# Rapid ethylene-triggered protein complex remodeling in dark grown hypocotyls

**DOI:** 10.1101/2025.05.15.654145

**Authors:** Youngwoo Lee, Hye Lin Park, Gyeong Mee Yoon, Daniel B Szymanski

**Author notes:** These authors contributed equally to this work (Youngwoo Lee, Hye Lin Park).

## Abstract

Protein-level cellular dynamics, including multimerization, play a crucial role in rapid adaptation of plants during developmental transitions and environmental stresses. The gaseous hormone ethylene is a key regulator of rapid cellular growth modulation. During soil emergence, etiolated seedlings undergo crucial morphological changes to their apical hooks and hypocotyls, with ethylene inhibiting hypocotyl axial elongation while promoting radial expansion. Ethylene triggers these growth responses within two hours; however, the protein machineries and cellular processes that mediate morphogenesis control are not known. Here, we used quantitative proteomics and co-fractionation mass spectrometry to test for rapid ethylene-dependent changes in protein abundance and protein complex composition. Protein multimerization responses were numerous and diverse. There were instances of protein complex assembly and disassembly, with varying degrees of completeness. Small-scale validation tests indicate that the identified proteins play a role in hypocotyl development and suggest that this novel approach to gene discovery identifies potential targets for ethylene-mediated growth regulation and enhanced seedling adaptability during early development.

## Introduction

Among many ethylene-regulated developmental processes (Abeles et al., 1992; Bleecker and Kende, 2000; Alonso and Ecker, 2001), ethylene modulation of hypocotyl development in darkness is a classic example of hormone-regulated developmental plasticity. It is thought that ethylene levels increase as the subterranean seedling encounters stiff physical barriers, and this leads to an altered growth habit that increases the likelihood of soil emergence (Von Arnim and Deng, 1996; Smalle et al., 1997). In dark-grown seedlings, ethylene induces biphasic hypocotyl growth inhibition (Phase I and II), with distinct mechanisms governing each phase over a two-hour timeline (Binder et al., 2004; Binder, 2007). Phase I begins rapidly, within 10-15 minutes of ethylene exposure, reducing the growth rate by ∼50 %. This initial suppression is mediated non-transcriptionally via Ethylene-insensitive 2 (EIN2), a positive regulator of ethylene signaling, which modulates cellular processes like microtubule stabilization and cell wall remodeling (Binder et al., 2004). Phase II follows as a sustained inhibition that initiates about 30 minutes after ethylene exposure. It requires the transcription factors EIN3/EIN3-like (EIL), which activate genes such as WAVE-DAMPENED-LIKE 5 (WDL5) to further suppress growth by enhancing microtubule stability (Sun et al., 2015). Ethylene primarily targets the epidermal cells of the hypocotyl (Smalle et al., 1997; Vandenbussche et al., 2007; Vandenbussche et al., 2012; Das et al., 2016; Seo and Yoon, 2019), where it suppresses elongation by disrupting cortical microtubule arrays and promoting microtubule stabilization, thus enforcing cell growth inhibition. Within two hours, growth is profoundly inhibited but not entirely halted, maintaining this low growth baseline. Despite this detailed characterization of the biphasic response in hypocotyl growth inhibition, the dynamic molecular mechanism governing protein abundance changes and complex formation such as multimerization during ethylene-mediated hypocotyl growth suppression remains poorly understood.

Ethylene signaling operates through several well-characterized signal transduction pathways. Upon binding to ethylene receptors at the ER, ethylene inactivates the receptors, releasing their inhibition of Ethylene-insensitive 2 (EIN2), which subsequently activates EIN3/EIN3-like (EIL) transcription factors. This triggers the expression of Ethylene Response Factor (ERF) transcription factors that regulate genes involved in growth suppression (Ju et al., 2012; Qiao et al., 2012; Wen et al., 2012). The downstream targets of ethylene that mediate growth modulation are not well characterized. At the cellular level, ethylene may inhibit axial cell elongation by reducing cell wall extensibility through altered transcription of several proteins, including expansins, xyloglucan endotransglucosylase/hydrolases (XTHs), and peroxidases (Van de Poel et al., 2015; Qin et al., 2019). Ethylene also modifies microtubule organization, which affects cellulose deposition patterns in the cell wall that could affect axial expansion (Van de Poel et al., 2015). Furthermore, ethylene exerts its negative impact on plant growth through crosstalk with other phytohormones. Ethylene enhances auxin activity through multiple mechanisms, upregulating PIN proteins and AUX1 to promote auxin transport and stimulating auxin biosynthesis (Ruzicka et al., 2007; Swarup et al., 2007; Liu et al., 2018), while simultaneously antagonizing gibberellin (GA) signaling by promoting DELLA protein accumulation and inducing GA catabolism via GA2ox (Achard et al., 2003; Schomburg et al., 2003; Yoo et al., 2009). The interaction with abscisic acid (ABA) adds another layer of complexity but is highly dependent on light conditions and involves complex hormonal crosstalk. In darkness, ABA has little to no effect on hypocotyl growth unless light signaling pathways are constitutively active, as in mutants like *cop1-4* or *pifq*, where ABA can then exert inhibitory effects via auxin transport and the upregulation of DELLA proteins that suppress GA signaling (Ghassemian et al., 2000; Liang et al., 2012; Peng et al., 2022). In contrast, in the presence of light, ethylene promotes hypocotyl elongation by facilitating COP1-mediated degradation of HY5 and activating PIF3-dependent auxin pathways (Yu et al., 2013).

Modern protein mass spectrometry offers a powerful systems-level approach for understanding proteome response to developmental, physiological, and environmental stimuli (Gingras et al., 2007; Nilsson et al., 2010; Aebersold and Mann, 2016; Lee et al., 2025). In ethylene signaling, proteomics has captured changes in protein abundance, localization, and post-translational modifications (PTMs). Gel-based proteomics (Prayitno et al., 2006; Zheng et al., 2013; Wang et al., 2015; Kim et al., 2018; Shin et al., 2020), stable and isobaric isotope labeling (Wang et al., 2015; Zhang et al., 2015; Du et al., 2016), and phosphoproteomics (Chen et al., 2011; Wang et al., 2015; Zhang et al., 2015) have been applied across plant species to characterize ethylene-responsive proteins and regulatory signaling networks. In climacteric fruits such as apples, kiwifruits, and bananas, ethylene treatment induces proteins involved in hormone biosynthesis, redox regulation, energy metabolism, and defense mechanisms (Zheng et al., 2013; Li et al., 2015; Du et al., 2016; Shin et al., 2020). In banana fruit, pretreatment with ethylene may enhance chilling and heat tolerance by upregulating proteins associated with ATP production, ROS scavenging, and protein turnover (Li et al., 2015; Du et al., 2016). In Arabidopsis, ethylene affects proteins involved in vesicle trafficking, oxidative stress, and hormone crosstalk, while repressing ribosomal proteins and brassinosteroid-regulated proteins (Chen et al., 2011). Phosphoproteomic studies using *ein2* mutants further revealed that ethylene signaling induces a general reduction in phosphorylation levels, with distinct motifs present in both ethylene-enhanced and -repressed phosphopeptides (Chen et al., 2011). Collectively, these studies underscore the versatility of ethylene signaling and highlight the critical roles of translational and post-translational regulation, including phosphorylation motifs and kinase activity, in orchestrating rapid and context-dependent proteomic responses.

Co-fractionation mass spectrometry (CFMS), a strategy that couples biochemical fractionation with mass spectrometry, enables the characterization of endogenous protein complex sizes (Kristensen et al., 2012) and organelle localizations (Andersen et al., 2003) without requiring transgenes or targeted enrichment. CFMS has been extensively utilized to determine the apparent masses and localization of protein complexes in diverse Arabidopsis tissues, including leaves (Aryal et al., 2014; Aryal et al., 2017; McBride et al., 2017) and roots (Gilbert and Schulze, 2019), as well as in organelles such as chloroplasts (Olinares et al., 2010) and mitochondria (Klodmann et al., 2010; Senkler et al., 2017). This approach has enabled the identification and characterization of numerous known and novel protein complexes across plant species (McBride et al., 2019; McWhite et al., 2020; Lee et al., 2021; Lee et al., 2025; Yang et al., 2025), and has been successfully used to monitor multimerization dynamics during diurnal cycles (Gorka et al., 2019), protein-ligand interactions (Veyel et al., 2017), and over evolutionary timescales (Lee and Szymanski, 2021). Reliable detection of multimerization variants is limited to moderately abundant proteins that can be reproducibly quantified using LC/MS and the subset of complexes that display significant size differences that can be detected using size exclusion chromatography (Lee and Szymanski, 2021). Alternative data-independent LC/MS acquisition and analysis strategies provide new possibilities for protein complex analysis that have the potential for greater coverage and reproducibility (Mallam et al., 2019; Heusel et al., 2020; Bludau et al., 2023). However, at present protein multimerization changes in response to plant hormone signaling have remained largely unexplored.

In this study, we performed proteomic analyses to investigate the protein-level responses underlying the short-term response of growth inhibition induced by ethylene treatment. By leveraging the etiolated hypocotyl growth inhibition system and careful dissection of the target organ, we analyzed the overall proteome-wide ethylene responses in both crude soluble and microsome-associated cellular fractions. Using CFMS and high-throughput statistical analyses, we profiled changes in protein multimerization states following ethylene exposure. Our data revealed proteins showing alterations in abundance, localization, and multimerization, many of which are associated with core cellular functions such as translation, vesicle trafficking, energy metabolism, and hormone-hormone signaling crosstalk. Notably, we observed a lack of correlation between ethylene-induced transcriptional changes and protein abundance alterations, highlighting the importance of translational and post-translational regulation, as well as protein-protein interactions, in ethylene-mediated growth inhibition. This work provides the first global view of ethylene-dependent multimerization variants and uncovers potential mechanisms of post-translational control during early hormone signaling. Together, our findings demonstrate the power of CFMS to uncover dynamic changes in protein complex assembly and offer a systems-level framework for dissecting ethylene signaling in plants.

## Results

### Exogenous ethylene inhibits hypocotyl cell growth

To recapitulate ethylene-mediated growth inhibition in response to ethylene exposure, we measured hypocotyl length of three-day-old, etiolated Arabidopsis seedlings exposed to ethylene for two hours (Figure 1A and B). Wild-type Col-0 seedlings showed significant hypocotyl growth inhibition and increased *Ethylene Response Factor 1* (*ERF1*) expression (Figure 1C), while ethylene-insensitive *ein2-5* mutant remained unresponsive (Figure 1B). Consistent with this, at the cellular level, ethylene treatment suppressed the elongation of cells below the cotyledon-hypocotyl junction in wild-type but not in *ein2-5* seedlings (Figure 1D and 1E). These cells typically cover the apical hook and double in length within a day in the dark in wild-type seedlings (Gendreau et al., 1997). Our results demonstrated that two hours of ethylene treatment were sufficient to inhibit hypocotyl elongation. We employed this ethylene treatment condition for subsequent quantitative proteomic analyses.

**Figure 1.**
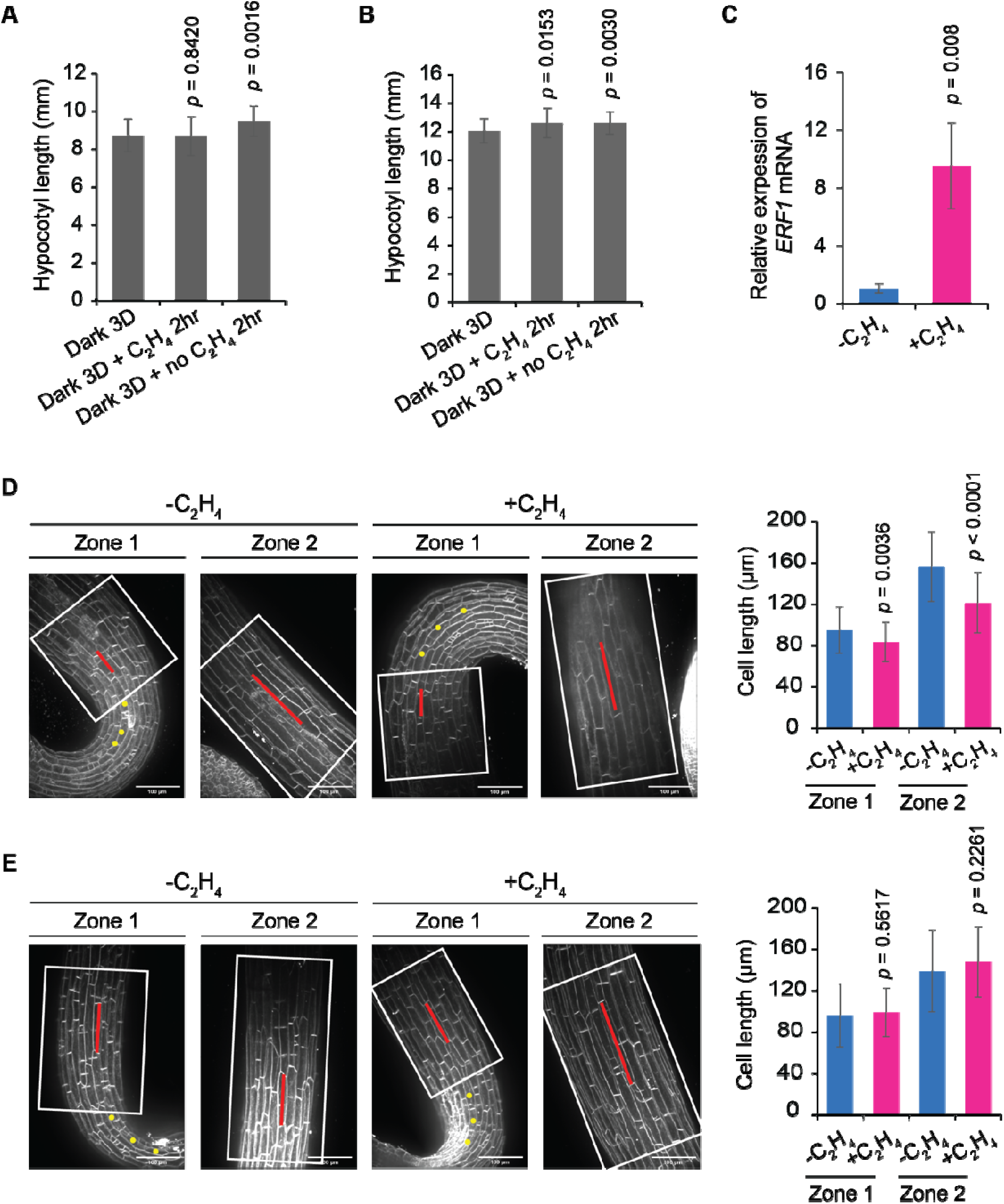
Ethylene treatment inhibits the growth of hypocotyl in WT but not in the *ein2-5* in the dark. **A-B,** Hypocotyl length of WT (**A**) and *ein2-5* (**B**). Three-day-old, etiolated seedlings (Dark 3D) were treated with or without 100 ppm ethylene gas for 2 hours. The graphs represent the mean of hypocotyl length and the error bars, SD (n ≥ 22 seedlings). Significance was determined by a two-tailed Student’s *t*-test to compare the hypocotyl length of seedlings with or without ethylene relative to Dark 3D. **C,** Quantitative gene expression analysis of ethylene response *ERF1* without or with ethylene treatment (100 ppm). Gene expression was normalized to an Actin control and presented relative to that without ethylene treatment. The error bar is SD (n = 3 biological replicates). Two-tailed Student’s *t-*test. **D-E,** Representative images for etiolated hypocotyls of WT (**D**) and *ein2-5* (**E**). The cell lengths of 4^th^-6^th^ (zone 1) and 7^th^-9^th^ (zone 2) cells from the apical hook were measured. The scale bars, 100 µm. The graph represents the average cell length of hypocotyls of WT and *ein2-5* after two hours with and without ethylene treatment. Yellow dots indicated the cells from the apical hook. Significance was determined by a two-tailed Student’s t-test, comparing the cell length of ethylene-treated to no ethylene-treated hypocotyls. The error bar is SD (n ≥ 3 seedlings and n ≥46 cells).

### Identification of soluble and membrane-associated proteins in etiolated hypocotyls

To analyze proteome-wide responses to ethylene during Arabidopsis seedling growth, we dissected hypocotyls from three-day-old, etiolated Col-0 and *ein2-5* mutant seedlings grown in the dark. For our quantitative analysis of protein abundance and crude subcellular localization, hypocotyls were fractionated by differential centrifugation into crude soluble (S200) and microsome-associated (P200) fractions (Figure 2A). The S200 contains soluble proteins from the apoplast, cytosol, and those from broken plastids (Aryal et al., 2014; McBride et al., 2019). The washed P200 fraction is highly enriched in true membrane-associated proteins (McBride et al., 2017). At a 1 % false discovery rate (FDR), 2,146 S200 and 1,079 P200 proteins were reliably detected in at least two out of three biological replicates (Supplemental Datasets S1, S2A, and S3A). We applied a label-free quantification (LFQ) normalization across all samples and defined reproducible proteins as those quantified in at least two out of three replicates. This filtering yielded 1,839 and 865 reproducible proteins from the S200 and P200 fractions, respectively, which were used for downstream analyses (Figure 2B and 2C; Supplemental Datasets S2B and S3B). The reproducibility among replicates and between treatments was high based on Pearson correlation coefficient (PCC) and hierarchical clustering analysis (Supplemental figure S1A and S1B). The third replicate of Col-0 +C_2_H_4_ S200 exhibited the lowest coverage and clustered separately due to a high number of missing values. Principal component analysis (PCA) clearly resolved genotypes, but the ethylene-treated groups were not resolved from controls, suggesting that ethylene does not have a clear global effect on protein abundance. (Supplemental figure S1C).

**Figure 2.**
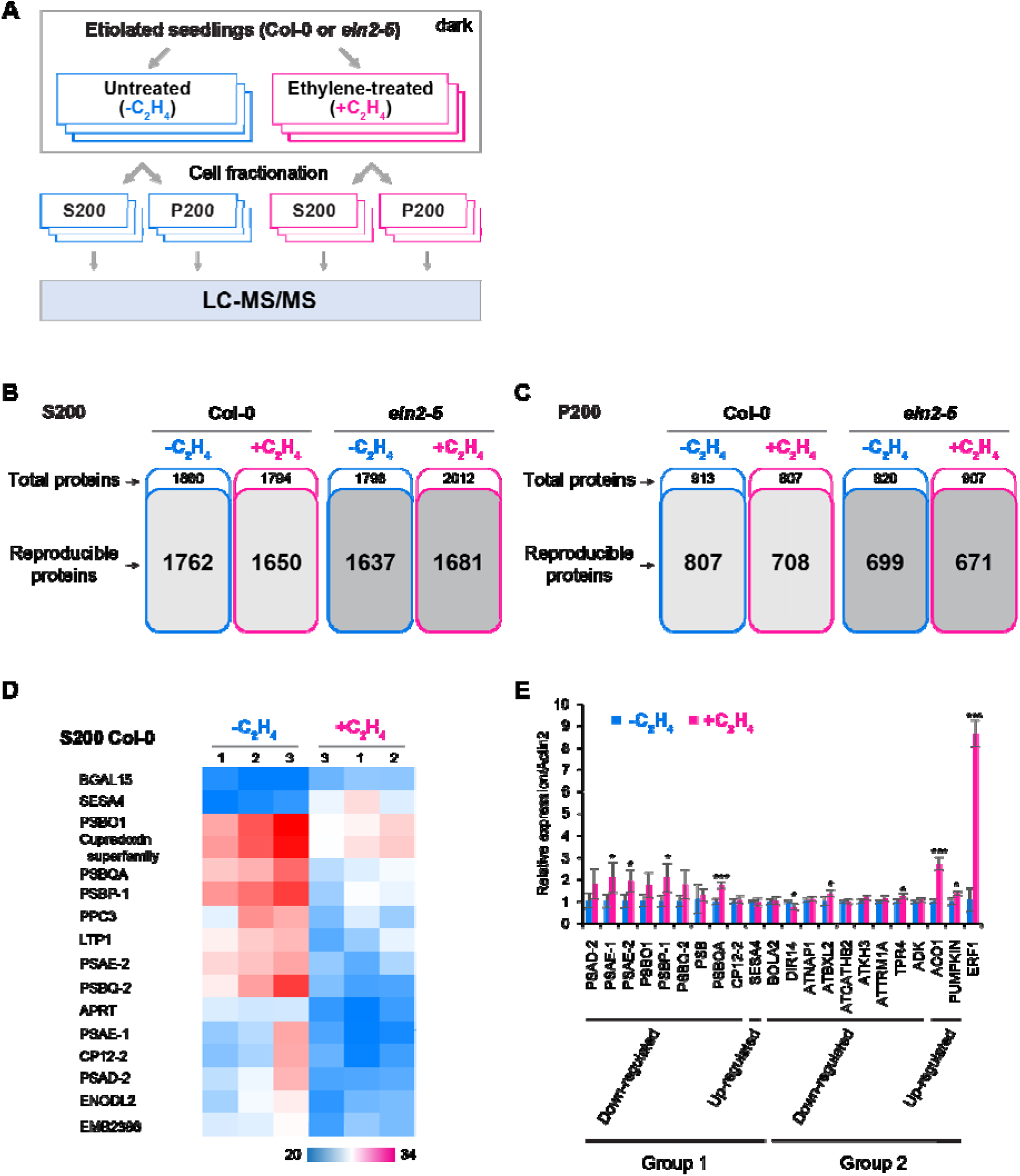
Quantified proteomes responding to the exogenous ethylene treatment. **A,** A workflow of comparative proteomics for elucidating the effects of ethylene on the distribution of proteins in different cellular fractions. The soluble (S200) and microsome-associated (P200) proteins were extracted from three-day-old, etiolated Col-0 and *ein2-5* hypocotyls treated with (+C_2_H_4_) or without (-C_2_H_4_) gaseous ethylene. Three independent samples were prepared for LC-MS/MS analysis. **B & C,** Proteome coverages of S200 (B) and P200 (C). Total proteins were defined as those identified in at least two out of three replicates. Reproducible proteins were defined as those quantified in at least two out of three replicates. The reproducible proteins in the bold text were used in the downstream analysis. **D,** A heatmap of Col-0 S200 proteins differentially responding to ethylene treatment, as determined by two-sample *t*-tests at 10 % FDR with a 4-fold change threshold. BGAL15: AT1G31740.1, SESA4: AT4G27170.1, PSBO1: AT5G66570.1, Cupredoxin superfamily AT1G20340.1, PSBQA: AT4G21280.1, PSBP-1: AT1G06680.1, PPC3: AT3G14940.1, LTP1: AT2G38540.1, PSAE-2: AT2G20260.1, PSBQ-2: AT4G05180.1, APRT: AT4G22570.1, PSAE-1: AT4G28750.1, CP12-2: AT3G62410.1, PSAD-2: AT1G03130.1, ENODL2: AT4G27520.1, EMB2386: AT1G02780.1. **E,** Relative expression of S200 proteins in Col-0 in response to ethylene treatment. Three-day-old dark-grown seedlings were treated with 100 ppm ethylene gas for 2 hours at room temperature in the dark prior to analysis. Gene expression was normalized to *Actin* control and presented relative to the untreated control. Expression of the ethylene-responsive gene *ERF1* was used as a positive control to confirm the effectiveness of ethylene treatment. Error bars represent standard deviation (SD) from three biological replicates (n = 3). Group1: differentially expressed proteins detected by statistics, Group2: differentially expressed proteins identified by count-based data filtering. BOLA2: AT5G09830.1, DIR14: AT4G11210.1), ATNAP1: AT4G26110.1, ATBXL2: AT1G02640.1, ATCATHB2: AT1G02305.1, ATKH3: AT1G33680.1, ATTRM1A: AT3G02320.1, TPR4: AT1G04530.1, ADK: AT2G37250.1, ACO1: AT2G19590.1, PUMPKIN: AT3G18680.1, ERF1: AT3G23240.1.

The breadth of metabolic pathways associated with the dataset was visualized by mapping the proteome of untreated dark-grown hypocotyls (Col-0-C_2_H_4_ S200) onto the Plant Metabolic Network (Supplemental figure S2). This metabolic map underscores that the dark-grown hypocotyl acts as metabolically active tissue, coordinating nitrogen assimilation (e.g., GS/GOGAT cycle), redox homeostasis (e.g., glutathione and thioredoxin systems), and amino acid metabolism (e.g., glutamate, aspartate, and branched-chain amino acids) to support rapid cell elongation (Bellegarde et al., 2019; Zayed et al., 2023; Zhu et al., 2025). The proteomics data are also consistent with catabolic reactions associated with mobilizing protein and lipids to enable new cell wall and membrane biogenesis (Kim et al., 2013; Fan et al., 2019; Xu and Fan, 2022). Gluconeogenesis, in particular, is essential for the seed-to-seedling transition, as it converts storage lipids into carbon and energy required for early development (Eastmond et al., 2015). The most highly abundant enzymes in the dataset map to central carbon metabolism (Supplemental figure S3), and the presence of highly abundant pyruvate orthophosphate dikinase and pyruvate carboxykinase (Supplemental Dataset S1E) may reflect gluconeogenesis in the cytosol. Our data provide a resource to analyze how ethylene modulates protein abundance in pathways associated with energy homeostasis and resource allocation in developing hypocotyls.

Next, we performed two-sample *t*-tests independently for each genotype and compared the statistical results to identify proteins that differed in response to ethylene treatment in Col-0 samples but not in *ein2-5* samples (Supplemental Datasets S2B and S3B). This analysis revealed 16 S200 proteins with at least 4-fold (or ± 2 log2 ratio of -C_2_H_4_ to +C_2_H_4_) abundance changes in response to ethylene in an EIN2-dependent manner (Figure 2D; Supplemental figure S1D). The 14 proteins exhibited reduced levels in the presence of ethylene, including extrinsic subunits of photosystem I and II. Additionally, AtPPC3 (AT3G14940.1) and LTP1 (AT2G38540.1) exhibited significant decreases after ethylene treatment. *AtPPC3* encodes a phosphoenolpyruvate carboxylase (PEPC) involved in oxaloacetate formation and is rapidly induced by nitrate treatment in roots (Vidal et al., 2013). In contrast, the abundance levels of β-galactosidase 15 (BGAL15; AT1G31740.1) and seed storage albumin 4 (SESA4; AT4G27170.1) were significantly increased following ethylene treatment. BGALs hydrolyze β-D-galactosides and are involved in flower senescence and fruit ripening (Ahn et al., 2007; Chandrasekar and van der Hoorn, 2016), and have been shown to regulate their own transcript levels during ripening (Alexander and Grierson, 2002; Moctezuma et al., 2003). SESA4 is important for nitrogen and sulfur provision during germination, and its transcript abundance decreases after brassinolide treatment in Arabidopsis seedlings (Goda et al., 2004). In contrast to the S200 dataset, *t*-tests within the P200 dataset did not reveal any proteins with significant ethylene-dependent changes exceeding the 4-fold difference (Supplemental figure S1D; Supplemental Dataset S3B).

To determine if observed protein level changes in response to ethylene were correlated with transcriptional alteration, we selected 21 high confidence protein hits from the S200 dataset based on their abundance and spectral counts (Supplemental Dataset S2C) and examined the transcript levels of the corresponding genes (Figure 2E). Seedling growth in darkness for three days with or without ethylene treatment was subjected to qPCR-based gene expression analysis. Unlike *ERF1* control, which showed an 8-fold increase in its transcript in response to ethylene, most of the selected genes showed no transcript changes, while a small subset exhibited concurrent changes at both transcript and protein abundance. ACC oxidase 1 (ACO1; AT2G19590.1) and an amino acid kinase family protein (PUMPKIN; AT3G18680.1), both putatively up-regulated in response to ethylene at the protein level, exhibited corresponding up-regulation at the transcript level. Conversely, the disease resistance-responsive (dirigent-like protein) family protein (DIR; AT4G11210.1), predicted as down-regulated in proteome data (Supplemental Dataset S2C), showed consistent down-regulation at the transcript level. These results suggest that ethylene predominantly regulates protein abundance through post-transcriptional mechanisms rather than transcriptional control during rapid hypocotyl growth inhibition.

### Multimerization changes in response to ethylene

To test for protein multimerization changes in response to exogenous ethylene treatment, we performed co-fractionation mass spectrometry (CFMS) on S200 fractions obtained from three-day-old, etiolated Col-0 hypocotyls, either treated with ethylene (SEC_Ethylene_) or left untreated (SEC_Untreated_) (Supplemental figure S4). Biological duplicates of SEC profiles are sufficient to gain reliable information on protein multimerization states (Aryal et al., 2014; Aryal et al., 2017; Gilbert and Schulze, 2019; McBride et al., 2019; Pang et al., 2020; Lee et al., 2021; Lee and Szymanski, 2021; Skinnider and Foster, 2021; Schlossarek et al., 2023). The vast majority of proteins identified in the SEC_Untreated_ and SEC_Ethylene_ samples were detected in both replicates, 2,908 and 2,692, respectively (Figure 3A and 3B; Supplemental Dataset S4). The replicate profiles demonstrated high reproducibility based on PCC values, which were maximal along the diagonal corresponding to identical column fractions in the replicates (Figure 3C and 3D). Most proteins had identical peak locations in the replicates, and approximately 95 % had peaks within a two-fraction shift between replicates for both SEC_Untreated_ and SEC_Ethylene_ (Figure 3E and 3F).

**Figure 3.**
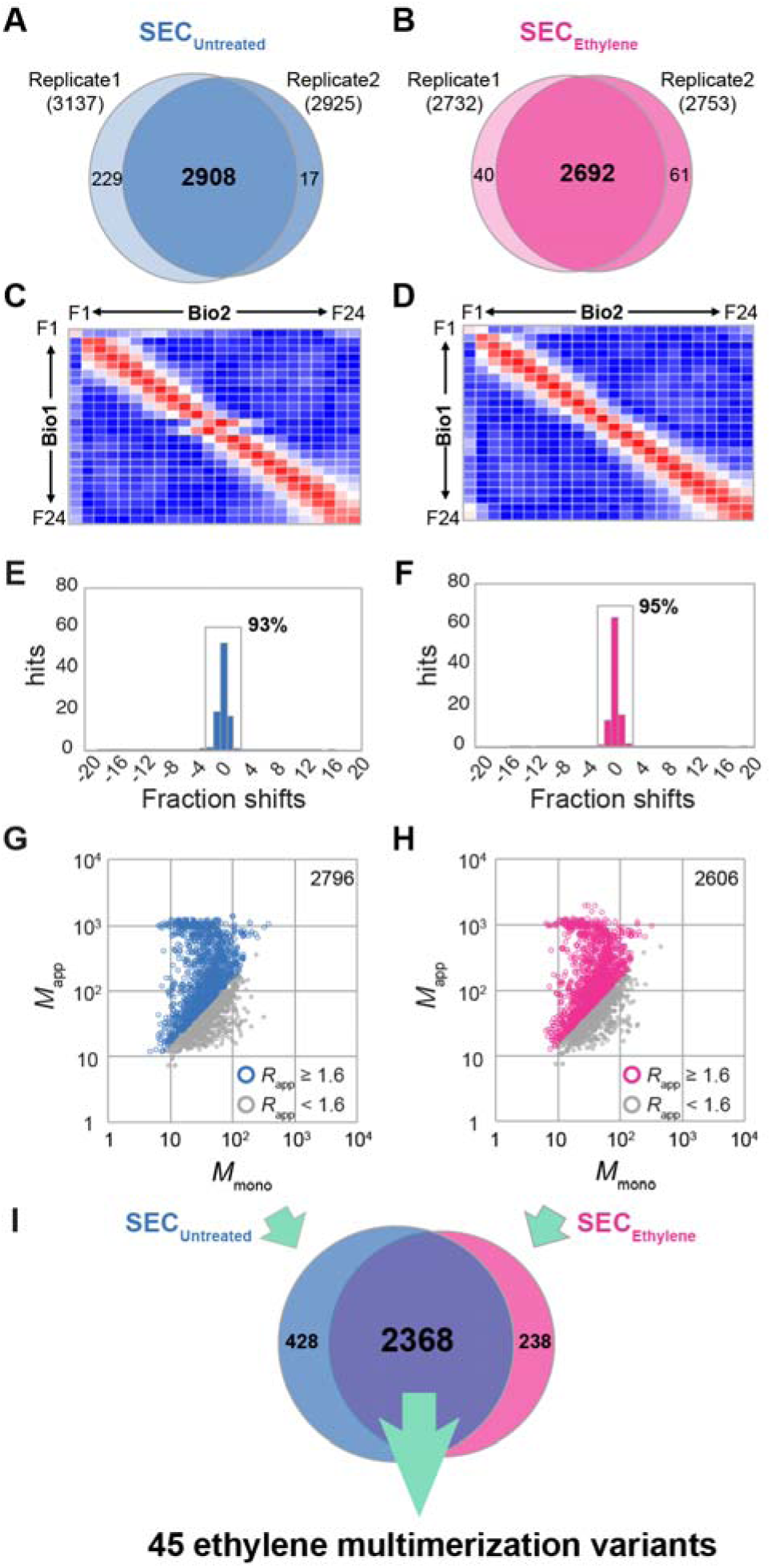
The CFMS pipeline allows analysis of protein multimerization responding to ethylene treatment. **A–F,** Reproducibility of the CFMS datasets. Protein overlaps between two biological replicates in the untreated control (A; SEC_Untreated_) and ethylene-treated (B; SEC_Ethylene_) datasets obtained from S200 fraction in Col-0 plants. Pearson correlation coefficients of protein abundances across fractions in the SEC_Untreated_ (C) and SEC_Ethylene_ (D). Column fraction shifts between the duplicates for all proteins in the SEC_Untreated_ (E) and SEC_Ethylene_ (F). Reproducible proteins present within a 2-fraction shift are boxed, with the ratio of reproducibility shown. **G–H,** Scatter plots of the *M*_mono_ and *M*_app_ of the reproducible proteins in the SEC_Untreated_ (G) and SEC_Ethylene_ (H). Open circles in grey are proteins with *R*_app_ < 1.6, while those in blue and pink have *R*_app_ ≥ 1.6. **I,** The test for ethylene multimerization variants. Numbers on the Venn diagram show reproducible proteins in the SEC_Untreated_ and SEC_Ethylene_.

Based on this two-fraction reproducibility threshold, we selected 2,796 SEC_Untreated_ and 2,606 SEC_Ethylene_ profiles for downstream analyses. To deconvolve proteins with multiple elution peaks, we applied a Gaussian peak detection algorithm (McBride et al., 2017). Proteins with multiple peaks were assigned with suffixes indicating peak numbers and are included in Supplemental Dataset S5A and S5B. When comparing apparent mass (*M*_app_) values to predicted monomeric mass (*M*_mono_) values of peaks, the distributions of points on both plots were skewed toward multimeric forms (Figure 2G and 2H), consistent with previous estimates for soluble proteins across various plant species and tissue types (Aryal et al., 2014; McBride et al., 2017; McBride et al., 2019; Lee et al., 2021; Lee and Szymanski, 2021; Lee et al., 2025). These results suggest that ethylene treatment does not induce global changes in protein multimerization. From both datasets, 2,368 proteins were reliably detected in both untreated control and ethylene-treated samples, while 428 and 238 proteins belonged only to SEC_Untreated_ and SEC_Ethylene_, respectively (Figure 3I).

To test for instances of ethylene-dependent changes in protein multimerization, we followed published methods (Lee and Szymanski, 2021) to conduct statistical tests of the *M*_app_ values for the 2,368 proteins (Figure 3I; Supplemental Data S6A). *ANOVA* tests were performed using a 5 % FDR threshold. Proteins flagged by this analysis were further filtered to remove potential false positives. Specifically, raw elution profiles were manually inspected to rule out cases where a minor or clearly co-eluting peak appeared in only one of the replicates. In addition, we identified two cases in which the means of the significantly different peaks were within two column fractions and flagged them as “less reliable” in column AB of the Supplemental Data S6B. Although alternative methods for profile comparisons might identify more multimerization variants (Mallam et al., 2019; Heusel et al., 2020; Bludau et al., 2023), our data filtering and analysis pipeline identified 45 highly reliable multimerization variants in response to ethylene (Supplemental Data S6B). Gene Ontology (GO) enrichment analysis at a 1 % false discovery rate (FDR) revealed “structural constituent of ribosome” and “response to cytokinin” as the most significantly enriched molecular function and biological process terms, respectively, among the ethylene-induced multimerization variants (Supplemental Data S7). Notably, eight ribosomal proteins showed altered multimerization states following ethylene treatment, suggesting that ribosomal protein complexes may be remodeled or that their assembly is dynamically regulated in response to ethylene signaling. These changes may be mediated by phosphorylation or other PTMs of ribosomal proteins or associated factors, potentially affecting ribosome biogenesis and function. These findings highlight that ethylene signaling extends beyond transcriptional regulation to influence translational control and ribosome assembly. Six other multimerization variants were associated with the “response to cytokinin” category, including Binding Partner of ACD11 1 (BPA1; AT5G16840.3) and Photosystem II Stability/Assembly Factor (HCF136; AT5G23120.1). BPA1 exhibited a shift from a monomeric peak with an apparent molecular mass of 31 kDa (*R*_app_ of 1.1) to a multimeric form with a *M*_app_ of 54.2 kDa (*R*_app_ of 2.0) following ethylene treatment, suggesting dimerization. BPA1 is a key regulator of reactive oxygen species (ROS)-mediated defense responses in plants (Li et al., 2019). Its dimeric form may reflect interaction with ACD11, potentially contributing to complex stabilization and proper regulation of programmed cell death. Conversely, HCF136 shifted from a dimeric form (*R*_app_ of 1.7) to a monomeric or proteolyzed form with a *M*_app_ of 27.1 kDa (*R*_app_ of 0.6) after ethylene exposure. HCF136 is essential for the stability and assembly of Photosystem II and selectively regulates PSII biogenesis, particularly under low-light conditions (Plücken et al., 2002). The observed decrease in abundance of PSII complex subunits (Figure 2D) may be associated with HCF136 degradation or disassembly under ethylene treatment. A summary of 26 multimerization variants identified at a stringent FDR (1 %) is provided in Table 1.

**Table 1.**
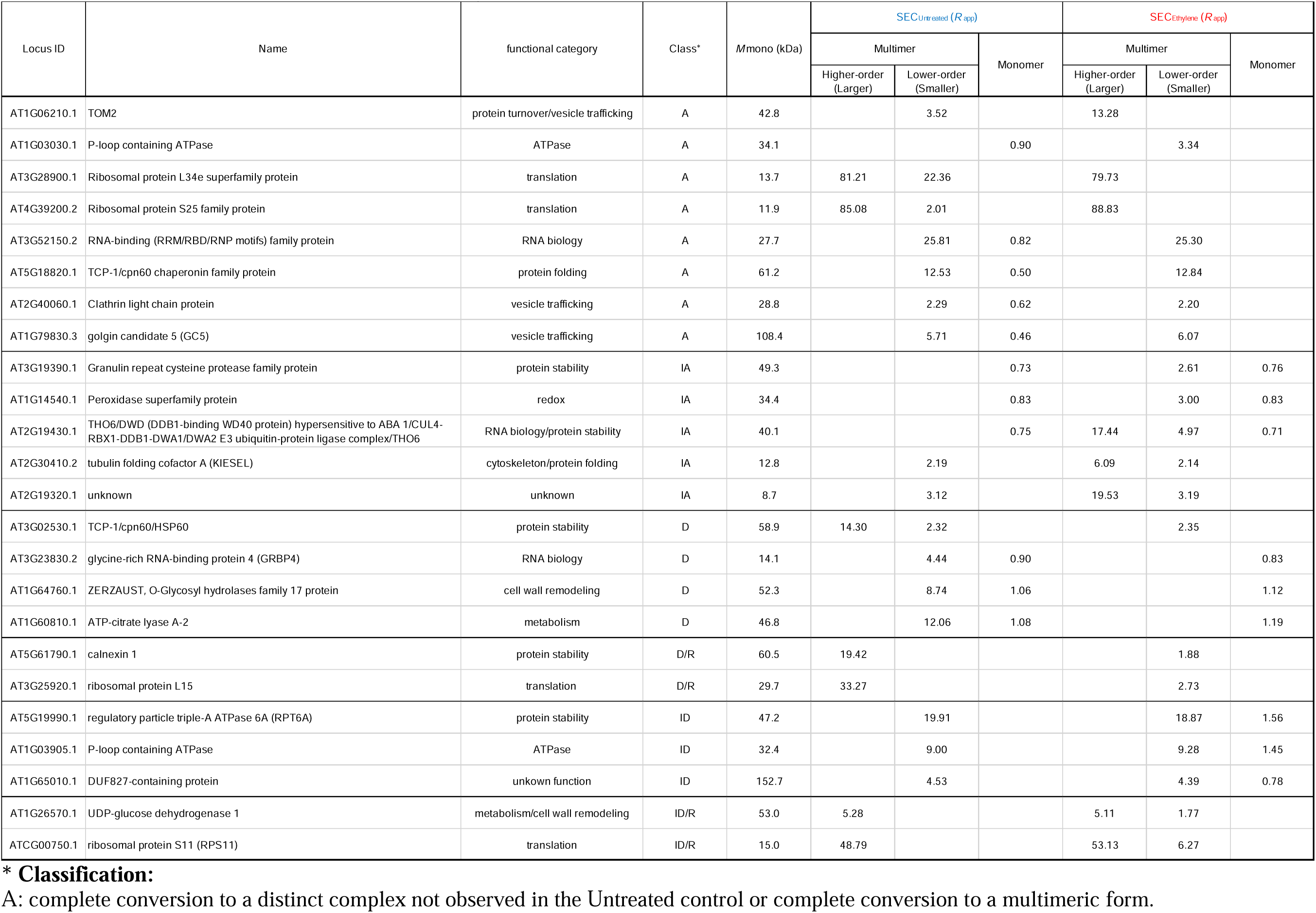
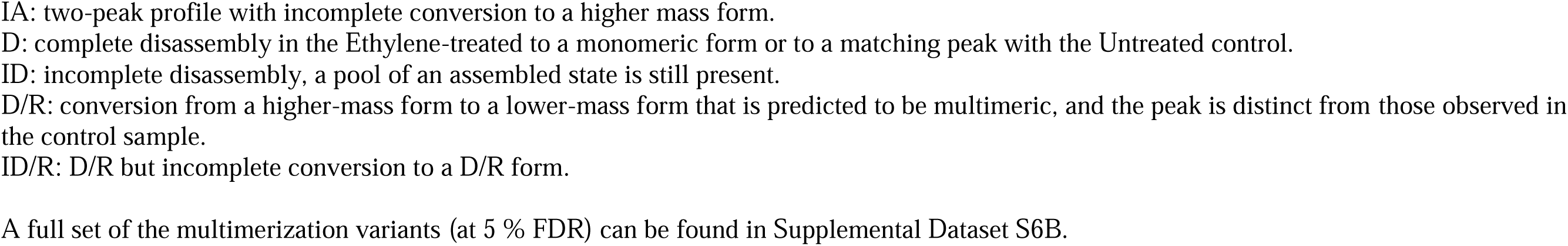
Multimerization variants after 2 hours of ethylene exposure.

| Locus ID | Name | functional category | Class* | Mmono (kDa) | SEC<sub>Untreated</sub> ($R_{app}$) | | SEC<sub>Ethylene</sub> ($R_{app}$) | | | |
| --- | --- | --- | --- | --- | --- | --- | --- | --- | --- | --- |
|  |  |  |  |  | Multimer |  | Monomer | Multimer |  | Monomer |
|  |  |  |  |  | Higher-order (Larger) | Lower-order (Smaller) |  | Higher-order (Larger) | Lower-order (Smaller) |  |
| AT1G06210.1 | TOM2 | protein turnover/vesicle trafficking | A | 42.8 |  | 3.52 |  | 13.28 |  |  |
| AT1G03030.1 | P-loop containing ATPase | ATPase | A | 34.1 |  |  | 0.90 |  | 3.34 |  |
| AT3G28900.1 | Ribosomal protein L34e superfamily protein | translation | A | 13.7 | 81.21 | 22.36 |  | 79.73 |  |  |
| AT4G39200.2 | Ribosomal protein S25 family protein | translation | A | 11.9 | 85.08 | 2.01 |  | 88.83 |  |  |
| AT3G52150.2 | RNA-binding (RRM/RBD/RNP motifs) family protein | RNA biology | A | 27.7 |  | 25.81 | 0.82 |  | 25.30 |  |
| AT5G18820.1 | TCP-1/cpn60 chaperonin family protein | protein folding | A | 61.2 |  | 12.53 | 0.50 |  | 12.84 |  |
| AT2G40060.1 | Clathrin light chain protein | vesicle trafficking | A | 28.8 |  | 2.29 | 0.62 |  | 2.20 |  |
| AT1G79830.3 | golgin candidate 5 (GC5) | vesicle trafficking | A | 108.4 |  | 5.71 | 0.46 |  | 6.07 |  |
| AT3G19390.1 | Granulin repeat cysteine protease family protein | protein stability | IA | 49.3 |  |  | 0.73 |  | 2.61 | 0.76 |
| AT1G14540.1 | Peroxidase superfamily protein | redox | IA | 34.4 |  |  | 0.83 |  | 3.00 | 0.83 |
| AT2G19430.1 | THO6/DWD (DDB1-binding WD40 protein) hypersensitive to ABA 1/CUL4-RBX1-DDB1-DWA1/DWA2 E3 ubiquitin-protein ligase complex/THO6 | RNA biology/protein stability | IA | 40.1 |  |  | 0.75 | 17.44 | 4.97 | 0.71 |
| AT2G30410.2 | tubulin folding cofactor A (KIESEL) | cytoskeleton/protein folding | IA | 12.8 |  | 2.19 |  | 6.09 | 2.14 |  |
| AT2G19320.1 | unknown | unknown | IA | 8.7 |  | 3.12 |  | 19.53 | 3.19 |  |
| AT3G02530.1 | TCP-1/cpn60/HSP60 | protein stability | D | 58.9 | 14.30 | 2.32 |  |  | 2.35 |  |
| AT3G23830.2 | glycine-rich RNA-binding protein 4 (GRBP4) | RNA biology | D | 14.1 |  | 4.44 | 0.90 |  |  | 0.83 |
| AT1G64760.1 | ZERZAUST, O-Glycosyl hydrolases family 17 protein | cell wall remodeling | D | 52.3 |  | 8.74 | 1.06 |  |  | 1.12 |
| AT1G60810.1 | ATP-citrate lyase A-2 | metabolism | D | 46.8 |  | 12.06 | 1.08 |  |  | 1.19 |
| AT5G61790.1 | calnexin 1 | protein stability | D/R | 60.5 | 19.42 |  |  |  | 1.88 |  |
| AT3G25920.1 | ribosomal protein L15 | translation | D/R | 29.7 | 33.27 |  |  |  | 2.73 |  |
| AT5G19990.1 | regulatory particle triple-A ATPase 6A (RPT6A) | protein stability | ID | 47.2 |  | 19.91 |  |  | 18.87 | 1.56 |
| AT1G03905.1 | P-loop containing ATPase | ATPase | ID | 32.4 |  | 9.00 |  |  | 9.28 | 1.45 |
| AT1G65010.1 | DUF827-containing protein | unkown function | ID | 152.7 |  | 4.53 |  |  | 4.39 | 0.78 |
| AT1G26570.1 | UDP-glucose dehydrogenase 1 | metabolism/cell wall remodeling | ID/R | 53.0 | 5.28 |  |  | 5.11 | 1.77 |  |
| ATCG00750.1 | ribosomal protein S11 (RPS11) | translation | ID/R | 15.0 | 48.79 |  |  | 53.13 | 6.27 |  |

**\* Classification:**
A: complete conversion to a distinct complex not observed in the Untreated control or complete conversion to a multimeric form.
IA: two-peak profile with incomplete conversion to a higher mass form.
D: complete disassembly in the Ethylene-treated to a monomeric form or to a matching peak with the Untreated control.
ID: incomplete disassembly, a pool of an assembled state is still present.
D/R: conversion from a higher-mass form to a lower-mass form that is predicted to be multimeric, and the peak is distinct from those observed in the control sample.
ID/R: D/R but incomplete conversion to a D/R form.
A full set of the multimerization variants (at 5 % FDR) can be found in Supplemental Dataset S6B.

We further classified the different types of multimerization behavior based on changes in apparent protein size and the partitioning of the protein pool between distinct forms, as indicated by multiple peaks in the SEC profiles (Table 1). In some instances, ethylene promoted conversion to a more highly assembled state with an increased apparent mass (classified as A or IA in Table 1). For example, the predicted CUL4 E3 ligase substrate receptor DWA1 (AT2G19430.1) assembled into two high-mass forms that were not present in the control (A). In other cases, such as the tubulin folding cofactor complex subunit KIESEL (AT2G30410.2), the increase in assembly was incomplete (IA) because in the presence of ethylene, the protein pool was partitioned between a distinct high-mass complex and a lower-mass complex that was also observed in the control. In contrast, ethylene also appeared to cause disassembly or rearrangement of protein from a higher-mass complex to a resolvable lower-mass form. This behavior was classified as disassembly (D) if the lower-mass form was consistent with the monomeric state, e.g., *R*_app_ < 1.6 as in GRBP4 (AT3G23830.2), or having a lower-mass peak matching a peak present in the control TCP-1 (AT3G02530.1). The dynamic was classified as disassembly/reassigned binding partner (D/R) if the lower-mass form was predicted to be multimeric, e.g., *R*_app_ > 1.6, and was unique to the ethylene-treated sample. The distinction is that the lower-mass complex could reflect partial disassembly or the repartitioning of the protein to a novel set of interactors. These *R*_app_ cut-offs are not foolproof metrics but use known information to predict the simplest explanations that underlie protein complex remodeling. D/R rearrangements were classified as incomplete (ID and ID/R) when the higher-mass peak persisted in the ethylene-treated samples, e.g., RPT6A (AT5G19990.1) and RPS11 (ATCG00750.1), respectively.

### Detailed analysis of selected ethylene-induced multimerization variants

We focused on a subset of interesting hits for deeper proteomic analyses and gene knockout experiments. DWD (DDB1-binding WD40 protein) hypersensitive to ABA 1 (DWA1 or THO6; AT2G19430.1) had a likely monomeric peak with a *M*_app_ of 30 kDa (*R*_app_ of 0.7) in both SEC_Untreated_ or SEC_Ethylene_ datasets (Figure 4A). Upon ethylene treatment, DWA1 displayed two additional higher mass peaks at 700 kDa (*R*_app_ of 17.4) and 200 kDa (*R*_app_ of 5.0), suggesting a transition to multimeric forms. Similarly, Aspartic proteinase A1 (APA1; AT1G11910.1) transitioned from a monomeric state (*R*_app_ of ∼0.7) to a higher-mass complex with a *R*_app_ of ∼12.1 in response to ethylene (Figure 4B). This enzyme is active as a monomer and contains a plant-specific insert (PSI) domain, which can influence enzymatic activity and protein interactions (Mazorra-Manzano et al., 2010). The PSI domain may mediate higher-order complexes with distinct (altered or reduced specificity) activity in response to ethylene signaling.

**Figure 4.**
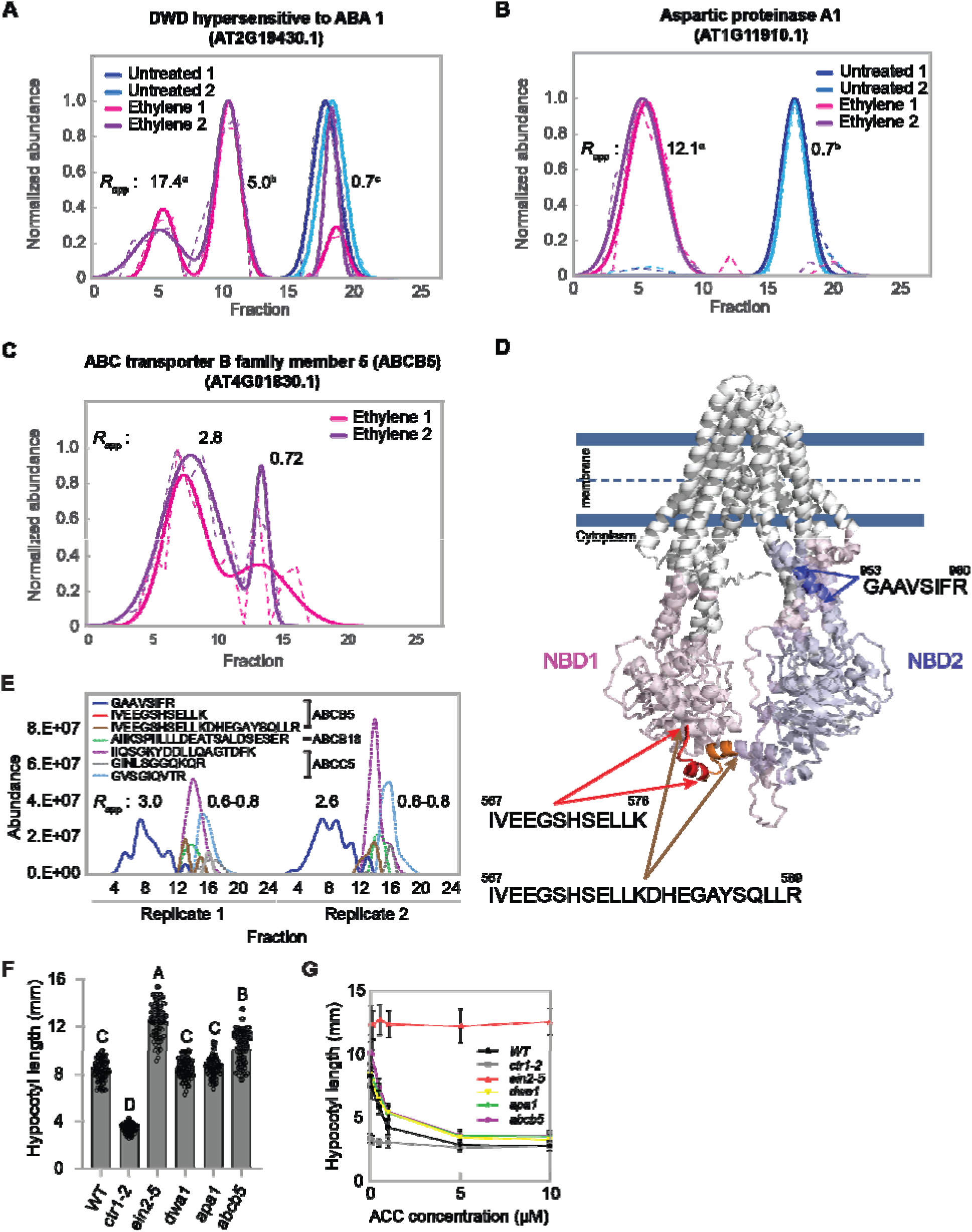
Protein multimerization variants in response to ethylene treatment. **A & B,** Elution profiles of proteins showing ethylene-dependent changes in multimerization states. **C,** Elution profiles of a protein exhibiting a putative ethylene-induced change in apparent localization. Elution profiles of proteins in SEC_Untreated_ (untreated control; blue and light blue) and SEC_Ethylene_ (ethylene-treated; pink and purple) datasets were visualized across two biological replicates. Dashed lines represent raw elution profiles, while solid lines indicate Gaussian-fitted profiles. *R*_app_ values are shown alongside each SEC peak, with statistical groupings used to highlight significant differences in protein multimerization states following 2 hours of ethylene treatment. **D,** Predicted molecular architecture of Arabidopsis ABCB5. The structural model of ABCB5 (Q9SYI3) was obtained from the AlphaFold Protein Structure Database (version updated November 2022) and visualized with the membrane. Three digested peptides identified in ethylene-treated hypocotyls are highlighted with their corresponding peptide sequences. Numbers above amino acid residues indicate their positions within the full-length ABCB5 protein sequence. NBD: nucleotide-binding domains. **E,** Peptide elution profiles of three ABC transporter proteins identified after ethylene treatment. Peptides derived from the same ABC transporter proteins are grouped using blankets. Peptide elution profiles were obtained from the SEC_Ethylene_ and reflect the multimerization states of the individual transporters. ABCB5: AT4G01830.1, ABCB18: AT3G28390.1, ABCC5: AT1G04120. **F**, The graph shows the quantification of hypocotyl lengths of seedlings grown on MS medium without ACC for 3 days. Statistical significance was determined using one-way *ANOVA* followed by Dunnett’s multiple comparisons test against the WT controls. Error bars, SD (n ≥ 70 seedlings). **G,** ACC dose-response curves for hypocotyl length of *dwa1*, *apa1*, and *abcb5* T-DNA mutant seedlings. Seedlings were grown in the media with different ACC concentrations indicated. *ctr1-2* and *ein2-5* were included as ethylene response controls. Error bars, SE (n ≥ 68 seedlings for each ACC concentration). One-way *ANOVA* with Dunnett’s test was performed to determine statistical significance.

Interestingly, ATP-binding cassette (ABC) transporter B family member 5 (ABCB5 or P-glycoprotein 5; AT4G01830.1) was classified as a potentially ethylene-induced protein complex since it was only detected after ethylene treatment (Figure 4C). ABCB5 exhibited multiple peaks with *R*_app_ values of 2.8 and 0.7 following ethylene treatment. The presence of ABC transporters in S200 proteomic experiments has long been somewhat of a mystery (Aryal et al., 2014; McBride et al., 2019). ABCB5 is a member of the broadly diversified plant ABC transporter family, which mediates the transport of various molecules, including auxin, across biological membranes (Sánchez-Fernández et al., 2001; Verrier et al., 2008). The ABCB transporters, including ABCB5, consist of four domains, two integral transmembrane domains (TMDs), each with six transmembrane helices, and two cytoplasmic nucleotide-binding domains (NBDs), arranged in the order TMD1-NBD1-TMD2-NBD2 (Figure 4D). In this study, three distinct ABCB5 peptides were identified, with two peptides located in NBD1 and one in NBD2. Peptides in NBD1 showed *R*_app_ values ranging from 0.6 to 0.8, while the peptide in NBD2 displayed *R*_app_ values between 2.6 and 3.0 (Figure 4E), suggesting proteolysis of the NBD1 and NBD2 regions in response to ethylene treatment. Peptides from two other ABC transporters, ABCB18 and ABCC5, identified in this study, also had *R*_app_ values of 0.6 to 0.8 and were all mapped to NBD1 sequences. Taken together, the proteolyzed NBD2 of ABCB5 may form a distinct protein complex in the cytoplasm during the ethylene response.

To test for a potential link between multimerization variability and hypocotyl development, we examined ethylene-dependent hypocotyl growth responses in *dwa1*, *apa1*, and *abcb5* single loss-of-function mutants (Figure 4F and G). The *dwa1* and *apa1* mutants showed comparable hypocotyl length to wild type when grown without ACC but exhibited reduced ethylene sensitivity with significantly longer hypocotyls when treated with higher ACC concentrations (Figure 4G). The *abcb5* mutant exhibited significantly longer hypocotyls compared to the wild type at various ACC concentrations but uniquely showed enhanced hypocotyl elongation without ethylene treatment (Figure 4F). Importantly, the proportional reduction in hypocotyl length in response to ethylene remained comparable between WT and *abcb5* mutants (Supplemental figure S5), suggesting that the longer hypocotyl phenotype may not be directly linked to ethylene sensitivity. These results suggest that DWA1 and APA1 contribute to ethylene-mediated hypocotyl growth regulation, whereas ABCB5 appears to have a broader function in modulating general hypocotyl development pathways, potentially including those regulated by auxin.

## Discussion

The analysis focused on the proteomic response of dissected hypocotyls after the seedlings were exposed to 100 ppm ethylene for 2 hours. A significant EIN2 and EIN3/EIL-dependent reduction in axial elongation was detectable in this time frame (Figure 1). The proteomic responses of dissected hypocotyls were analyzed following combinations of crude cell fractionation, SEC, an array of label-free quantitative proteomic, and statistical tests for proteins with altered properties in response to short-term ethylene treatment. Of the thousands of proteins in the soluble (S200) and crude microsomal (P200) proteins, only a small subset in the soluble displayed significant changes in abundance (Figure 2D; Supplemental Datasets S2 and S3) or multimerization (Figure 3; Supplemental Dataset S6). Collectively, this systems-level view indicates that ethylene affects very specific protein targets and unknown post-translational modifications may play an important role in orchestrating the transformation from axial to radial growth. Changes in protein complex composition are expected as they are frequently observed in the early steps of hormone-mediated signal transduction (Zeng et al., 2018). Post-translational modification, particularly phosphorylation, alters protein conformation and interaction capacity by introducing charge-mediated effects, inducing structural changes, or triggering allosteric signaling (Groban et al., 2006; Schonichen et al., 2013). These modifications can either stabilize or destabilize complexes depending on context, enabling precise control over signal transduction and stress responses. Our analyses, restricted to oxidation and N-terminal protein acetylation as the default variable modifications in MaxQuant. Further tests for a broader range of PTMs did not reveal any that correlated with the presence of a multimerization variant. The raw LC/MS files associated with these studies are available to enable additional analyses of PTMs and ethylene signaling. Protein cleavage serves as another critical mechanism by which it dismantles existing complexes or produces active protein fragments that initiate new functions (Berry et al., 2021; Zanotti et al., 2022). Additionally, subcellular redistribution redirects proteins to different cellular compartments, fundamentally changing their potential interaction partners (Park et al., 2023; Chien et al., 2024; Chien and Yoon, 2024; Park and Yoon, 2025).

The global multimerization response to plant hormone treatment has not been previously published. Here, we detected a diverse set of proteins with different types of multimerization changes. These changes are most likely to reflect non-covalent, reversible protein-protein interactions that are modulated during short-term ethylene treatment. The chromatographic and protein quantification methods are robust and have been used to quantify multimerization from a broad range of species and sample types (Andersen et al., 2003; Kristensen et al., 2012; Aryal et al., 2014; Aryal et al., 2017; McBride et al., 2017; Gilbert and Schulze, 2019; Gorka et al., 2019; McBride et al., 2019; McWhite et al., 2020; Pang et al., 2020; Lee et al., 2021; Lee and Szymanski, 2021; Lee et al., 2025). The protein complexes that are predicted to rearrange were selected after stringent filtering for reproducibility (Figure 3; Table 1; Supplemental Dataset S6A) and presence/absence scoring between treatments (Supplemental Dataset S6C).

The most reliable predictions of ethylene-dependent multimerization variants were those in which proteins were reproducibly detected in both ethylene-treated and control samples, and the apparent mass differences were significant at 1 % FDR. These SEC elution profiles were manually inspected to ensure there was no experimental evidence pointing to the existence of the alternative form in the counter sample (Table 1). There were 13 instances in which ethylene treatment led to the complete conversion of the soluble protein to a unique higher mass complex(es) not observed in the control (class A) or the incomplete conversion of a lower mass form in the control to a higher apparent mass species (class IA). These proteins included many proteins with known functions in translation, protein stability, RNA biology, and vesicle trafficking, and are strong candidates to mediate ethylene-dependent growth modulation. As one example, THO6/DWD (AT2G19430.1) appears to be a dual function protein. It is a subunit of the Arabidopsis THO/TREX mRNA export complex, which is thought to facilitate the transport of mRNA precursors and the biosynthesis of endogenous small interfering sRNAs in Arabidopsis (Yelina et al., 2010). THO6/DWD is also a substrate receptor of a CUL4-based E3 ubiquitin ligase targeting ABI5, and the mutant has been reported to be ABA hypersensitive (Lee et al., 2008; Lee et al., 2010). Based on the SEC profiles, we hypothesize that ethylene signaling promotes the assembly of monomeric THO6/DWD into two functionally distinct multimeric forms. Given that ABA is not actively involved in regulating hypocotyl elongation in darkness, the reported ABA hypersensitivity of *tho6/dwd* (Lee et al., 2008; Lee et al., 2010) may not directly account for the altered ethylene sensitivity we observed. More plausibly, DWA1 may have additional ABA-independent roles in ethylene signaling, such as mRNA export or sRNA biogenesis, affecting the stability or translation of ethylene signaling transcripts. Ethylene might play a role in this process by redirecting THO6/DWD into distinct protein complexes that function outside of ABA signaling.

Multimerization variability in conserved vesicle trafficking and proteolytic machineries. Of note, the TOM-like 2 ubiquitin receptor (TOM2; AT1G06210.1) binds to ubiquitinated auxin efflux carrier PIN2 and affects its trafficking and turnover in the lytic vacuole (Korbei et al., 2013). Mutation of TOM2 and other orthologs leads to ABA-hypersensitivity that is correlated with increased levels of the soluble ABA receptor, and its failure to traffic to the lytic vacuole (Mazorra-Manzano et al., 2010; Moulinier-Anzola et al., 2024). In our system, ethylene treatment promotes the assembly of a novel high apparent mass vacuolar aspartic proteinase A1 (APA1) protein complex (Shimaoka et al., 2004). The APA1 complexes in our S200 fraction likely originated from broken vacuoles, and we hypothesize that higher mass APA1 complexes reflect accelerated turnover of proteins trafficked to the vacuole (perhaps through TOM2-dependent trafficking). The mild ethylene insensitivity of *apa1* seedlings (Figure 4F) could reflect gene functions in the context of ABA signaling (Sebastian et al., 2020).

Decreased axial elongation has been correlated with observed ethylene-dependent reorientation of transverse cortical microtubule arrays and WAVE-DAMPENED2-LIKE proteins in the growth response of etiolated hypocotyls in many species, including Arabidopsis (Le et al., 2005; Sun et al., 2015). The tubulin-folding cofactor (TFC; AT2G30410.2) chaperone complex mediates α- and ß-tubulin heterodimer formation (Szymanski, 2002) and has been detected in leaves as *R*_app_ of ∼2.0 (Aryal et al., 2017; McBride et al., 2019; Lee and Szymanski, 2021). In etiolated hypocotyls, TFC-A assembles into a distinct higher mass complex that could alter chaperone activity and either the amount of tubulin heterodimers or the particular α- and ß-tubulin isoforms that are involved in microtubule responses to ethylene. We also detected protein complex rearrangements that are predicted to affect cell wall matrix properties. UDP-glucose dehydrogenase 1 (UGD1 at first mention; AT1G26570.1) generates UDP-glucuronic acid, an essential precursor for arabinose, xylose, galacturonic acid, and apiose sugars in a variety of matrix polysaccharides are required to assemble fully functional cell walls (Reboul et al., 2011). UGD1 was the only UDP-glucose dehydrogenase detected in our dataset with no significant change in abundance (Supplemental Datasets S2 and S3), and is expected to be active as a monomer (Campbell et al., 1997). Cell wall synthesis must persist to support radial expansion and shape change. It remains to be determined if the different UGD1 multimerization states affect activity and wall material properties in response to ethylene treatment.

There were also 11 examples in which ethylene promoted the disassembly or rearrangement of a protein into forms with a reduced apparent mass (Qiao et al., 2012; Wen et al., 2012). For the chaperones TCP-1 (AT3G02530.1) and calnexin (AT5G61790.1), as well as the proteasome regulatory particle 6A (AT5G19990.1), protein disassembly likely reflects a reduced activity and a fine-tuning of the cellular machineries that broadly influence the composition of the proteome. In ethylene signaling, protein complex remodeling and proteasome function are essential for rapid hypocotyl growth modulation. In the absence of ethylene, the receptor-CTR1 complex at the ER remains intact, keeping the signaling repressed (Ju et al., 2012). Upon ethylene perception, the complex disassembles, releasing EIN2, which activates the downstream ethylene signaling (Qiao et al., 2012; Wen et al., 2012). Simultaneously, ethylene promotes the degradation of EBF1/2, EIN3-targeting F-box proteins, leading to stabilization of EIN3, a key transcription factor required for ethylene-regulated growth responses (Guo and Ecker, 2003; Li et al., 2015; Merchante et al., 2015). The proteasome activity plays an integral role in these processes by regulating the turnover of key signaling components like EIN3 and EBF1/2, thus ensuring that ethylene response is quickly adjusted. Several recent studies also suggest that the proteasome itself may be dynamically regulated by post-translational modification, affecting its assembly and function (Kikuchi et al., 2010; Cui et al., 2014; Hirano et al., 2016; Marshall and Vierstra, 2019). This paper identifies dozens of additional proteins and protein complexes that are predicted to be targets of ethylene signaling.

ABCB5 is an interesting example, as it was reproducibly detected solely in the S200 fraction of ethylene-treated plants. Given the presence of multiple membrane spanning segments in the predicted structure of ABCB5 and the location of the peptides in the soluble domains, it is likely that these protein fragments are released by proteolysis (Figure 4D and 4E). The NBD2 domain mediates regulatory interaction with *cis-trans* peptidylprolyl isomerase Twisted Dwarf1 (Hao et al., 2020). It remains to be determined if the proteolytic release of NBD2 is of regulatory importance. We failed to observe a clear growth phenotype in seedlings that overexpressed soluble NBD1 or NBD2 domains, so the protein fragment does not appear to be sufficient to alter plant development. ABCB5 somehow limits hypocotyl elongation in the absence of ethylene and consistently showed similar reduction trends as WT in response to various ACC concentrations (Figure 4F and Supplemental figure S5), suggesting that ABCB5 is not likely to be involved in ethylene-regulated hypocotyl growth regulation. ABCB transporters have diverse hormone transport functions during plant development (Hu et al., 2025), and the precise role of ABCB5 remains to be determined.

## Conclusions and Future Perspectives

This paper describes a new quantitative proteomics pipeline and includes useful databases on the rapid systems-level responses of the etiolated hypocotyl proteome to ethylene. Reduced protein abundance appears to be focused on soluble proteins and may reflect a down-regulation of cellular machineries associated with rapid axial elongation. We also detected diverse types of multimerization variants that depict an active response that includes known and predicted ATPases, proteases, carbon flux into non-cellulosic polysaccharides (Reboul et al., 2011;Vaddepalli et al., 2017), vesicle trafficking, and cytoskeletal control. We predict that these multimerization variants reflect key points of developmental control in which post-translational modifications, altered subcellular localization, or proteolytic processing facilitate specific changes in protein-protein interactions and dramatic changes in growth habit. Further analyses are needed to validate their functional importance and determine how they operate in the context of ethylene signal transduction and developmental control.

## Material and Methods

### Plant materials and growth conditions

Arabidopsis Columbia (Col-0) and the ethylene-insensitive mutant, ein2-5, were used in this study (Alonso et al., 1999). All plants were sterilized and subsequently planted on the half-strengthen Murashige and Skoog (MS) media supplemented with 1 % sucrose and 0.8 % plant agar (pH 5.7). Following stratification and light activation, the plates were vertically positioned under a continuous dark chamber at 22°C for 3 days.

T-DNA insertion mutants, *apa1* (SALK_041027), *dwa1* (SALK_051022), and *abcb5* (CS1007014), were obtained from the Arabidopsis Biological Resource Center, and the ethylene signaling mutants, ctr1-2 (Kieber et al., 1993) and ein2-5 (Alonso et al., 1999), were used as a control.

### Ethylene treatment of etiolated seedlings

The etiolated seedlings were vertically placed in a modular incubator chamber (Billups-Rothenberg) for exogenous ethylene treatment in the dark. Ethylene gas (100 ppm) was introduced into the chamber with a flow rate of 20 cc/min for 5 min, after which the stopper was closed to maintain the ethylene-treated condition for 2 hours. Following the ethylene treatment, the root and apical hook were excised using a razor blade, and the hypocotyls were collected for RNA and protein extraction. All these procedures were conducted in a dark room illuminated with green light (GONHOM A19 Green Bulb E26 9W).

### Measurements of hypocotyl cell length

Etiolated hypocotyls were sampled after treatment with or without ethylene for 2 hours and stained with FM^TM^ 4-64 dye (Invitrogen, Waltham, MA, USA) following the manufacturer’s instruction. Zones were defined based on cell locations starting from the hook angle. The longitudinal cell growth was captured using an inverted confocal microscope (Zeiss, Jena, Germany). Cells from the 4^th^ to 6^th^ cells from the hook were designated as Zone 1, while those from the 7^th^ to 10^th^ cells were designated as Zone 2.

### Isolation of soluble (S200) and membrane-associated (P200) proteins

Hypocotyls were harvested from three-day-old, etiolated seedlings under a green light lamp (GONHOM A19 Green Bulb E26 9W) with an additional layer of green filter paper in a dark room. Proteins were extracted from 0.65 g of fresh hypocotyls as previously described (Aryal et al., 2014; McBride et al., 2017; McBride et al., 2019; Lee and Szymanski, 2021). In brief, the fresh tissues were ground in four volumes of a cold microsome isolation buffer (MIB) [50 mM HEPES/KOH (pH 7.5), 250 mM sorbitol, 50 mM KOAc, 2 mM Mg(OAc)_2_, 1 mM EDTA, 1 mM EGTA, 1 mM dithiothreitol (DTT), 2 mM PMSF and 1 % (v/v) protease inhibitor cocktail (160 mg/mL benzamidine-HCl, 100 mg/mL leupeptin, 12 mg/mL phenanthroline, 0.1 mg/mL aprotinin, and 0.1 mg/mL pepstatin A)] using a Polytron homogenizer (Brinkman Instruments, New York, NY, USA) with two 10-second grinding steps, interspersed with 1-minute rests. The homogenate was filtered through 4 layers of cheesecloth pre-soaked in cold MIB and then centrifuged on an Allegra X-30R centrifuge (Beckman Coulter Life Sciences, Indianapolis, IN, USA) at 1,000 × g for 10 minutes to remove debris. The supernatant was subsequently centrifuged at 200,000 × g for 20 minutes at 4°C using a Beckman Optima Ultracentrifuge (Beckman Coulter Life Sciences, Indianapolis, IN, USA) with a TLA110 rotor. The resulting supernatant was retained as the soluble fraction (S200) enriched in cytosolic proteins and soluble proteins released from broken chloroplasts.

To enrich a membrane-associated fraction (P200), the pellet obtained from the previous centrifugation was resuspended in cold MIB, incubated on ice for 10 minutes, and ultracentrifuged at 200,000 × g for 20 minutes at 4°C. This washing step was repeated twice. To solubilize P200 proteins, we applied a sodium cholate-based extraction method with the goal of profiling P200 protein complexes responsive to ethylene. The microsomal pellet was resuspended in 4 % cholate-containing HEPES buffer and incubated with gentle rotation for 1 hour at room temperature (Basu et al., 2008; McBride et al., 2017). Solubilized P200 proteins were enriched by ultracentrifugation at 200,000 × g for 20 minutes at 4°C.

For the comparative proteomics, three biological replicates were prepared for both S200 and P200 fractions obtained from Col-0 or *ein2-5* plants, either untreated (-C_2_H_4_) or treated with ethylene (+C_2_H_4_). Protein quantification was performed using a BCA assay kit according to the manufacturer’s protocol (Thermo Fisher Scientific Inc., Waltham, MA, USA). Protein yields per milligram of fresh hypocotyl were 0.55 µg/mg and 0.50 µg/mg for S200 and P200 samples, respectively.

### Size Exclusion Chromatography for Co-fractionation mass spectrometry (CFMS)

For CFMS analysis (Supplemental figure S4), approximately 400 µg of S200 proteins were obtained from about 0.65 g of etiolated Col-0 hypocotyls, either untreated (SEC_Untreated_) or treated with ethylene (SEC_Ethylene_), as described above. The S200 proteins were resolved over a Superdex Increase 200 10/300 GL column (GE Healthcare, Chicago, IL, USA) on an AKTA FPLC system (GE Healthcare, Chicago, IL, USA). The mobile phase consisted of 50 mM HEPES-KOH (pH 7.5), 100 mM NaCl, 10 mM MgCl_2_, 5 % glycerol, and 1 mM DTT, with a flow rate of 0.65 mL/min (Lee and Szymanski, 2021). Fractions of 500 µL were collected, totaling 24 fractions, including the first two void fractions. Proteins in each fraction were precipitated using a cold acetone method. Two biological replicates were prepared for each condition. Before sample separation, the column was calibrated using the Gel Filtration Standard Kit 1000 (MWGF1000, Sigma-Aldrich, St. Louis, MO, USA).

### LC-MS/MS sample preparation

Protein samples were digested using trypsin as described previously (McBride et al., 2019). Denatured protein samples in 8 M urea were reduced with 10 mM DTT for 45 minutes at 60°C, followed by alkylation with 20 mM iodoacetamide for 45 minutes at room temperature in the dark. The urea concentration was then diluted to below 1.5 M by adding 50 mM NH_4_HCO_3_ before trypsin digestion. Trypsin (Sigma-Aldrich, St. Louis, MO, USA) was added at an enzyme-to-protein ratio of 1:25, and digestion was performed at 37°C overnight. The reaction was stopped by adding trifluoroacetic acid. Digested peptides were purified using C18 Micro Spin Columns (74-4601, Harvard Apparatus, Holliston, MA, USA). Peptide concentrations were measured using a BCA assay (Thermo Fisher Scientific Inc.). For comparative proteomics, 1 μg of peptides was loaded onto the LC-MS/MS system. Peptide samples (SEC_Untreated_ and SEC_Ethylene_) from CFMS experiments were adjusted to equal volumes. The most concentrated sample had a peptide concentration of 1 μg/μL, and 1 μL of each sample was injected into the LC-MS/MS system.

### LC-MS/MS data acquisition

The S200 and P200 samples for comparative proteomics were analyzed as described previously (Lee et al., 2021). Peptide samples were analyzed using a reverse-phase LC-ESI-MS/MS system on a Dionex UltiMate 3000 RSLC nano system coupled with a Q-Exactive HF Hybrid Quadrupole-Orbitrap Mass Spectrometer (Thermo Fisher Scientific Inc., Waltham, MA, USA). Peptides were resolved over a 125-minute gradient at a flow rate of 300[nL/min. An MS survey scan was acquired in the 350–1,600 m/z range. MS/MS spectra were obtained by selecting the 20 most abundant precursor ions for sequencing using high-energy collisional dissociation (HCD) with a normalized collision energy of 27 %. A 15-second dynamic exclusion window was applied to minimize redundant sequencing of the same ion.

LC-MS/MS analysis for SEC samples was performed as described (Lee et al., 2025). Peptide samples were analyzed using a reverse-phase LC-ESI-MS/MS system on a Dionex UltiMate 3000 RSLC nano system coupled with an Orbitrap Fusion Lumos Tribrid Mass Spectrometer (Thermo Fisher Scientific Inc., Waltham, MA, USA). Peptides were first loaded onto a trap column (300 μm ID × 5 mm) packed with 5-μm, 100-Å PepMap C18 material and then separated on a reverse-phase analytical column (75 μm ID × 50 cm) packed with 2-μm, 100-Å PepMap C18 material (Thermo Fisher Scientific Inc.). Peptides were resolved at a flow rate of 200 nL/min over a 160-minute LC gradient. All data were acquired in positive ion mode in the Orbitrap mass analyzer using an HCD fragmentation scheme. The MS scan range was set to 375–1,500 m/z at a resolution of 120,000. The automatic gain control (AGC) target was set to standard, with a maximum injection time of 50 ms. A dynamic exclusion of 60 seconds and an intensity threshold of 5.0e3 were applied. MS data were acquired in data-dependent mode with a cycle time of 3 s per scan.

### Peptide Identification and Quantification

Protein identification and relative abundance quantification were performed using the Andromeda search engine within MaxQuant version 1.6.14.0 (Cox et al., 2014). MS/MS spectra were searched against the *Arabidopsis thaliana* TAIR10 protein database: Athaliana_167_TAIR10.protein.fa (Lamesch et al., 2012). For comparative proteomics, MS/MS spectra search for the S200 and P200 datasets was conducted independently. The search parameters were as follows: the “match between runs” function was enabled with a maximum matching time window of 0.7 min (default setting); one percent of the protein and peptide false discovery rate (FDR) was chosen; proteins identified by a single unique peptide were selected; label-free quantification (LFQ) was chosen; all other parameters were set to default. Peptide identification and quantification details are provided in Supplemental Dataset S1, while protein information for S200 and P200 is available in Supplemental Datasets S2 and S3, respectively. For CFMS analysis, searches were conducted with all fractions from biological duplicates in a single query, as previously described (Lee et al., 2025). The same search settings were applied, except that the LFQ method was not used. Peptide and protein details for CFMS are provided in Supplemental Dataset S4.

### Data filtering for protein identification in comparative proteomics (S200 and P200)

To identify differentially expressed proteins upon ethylene treatment, we applied a two-step filtering strategy to each dataset independently (Lee et al., 2025). First, for the count-based data filtering, proteins identified with MS/MS count(s) ≥ 1 in at least two out of three replicates were considered identified proteins (Supplemental Datasets S2A for S200 and S3A for P200). Among these, proteins with LFQ intensities greater than zero were classified as quantified proteins. Second, for protein intensity-based filtering, only quantified proteins with LFQ intensities greater than zero in at least two out of three replicates were retained for statistical analysis (Supplemental Datasets S2B and S3B).

### Determination of protein multimerization states

We applied an optimized Gaussian fitting algorithm to deconvolve the raw elution profiles into fitted peaks (McBride et al., 2017). The fitted peak locations were then converted into protein apparent mass values using the SEC column calibration curve described in the SEC method section. In cases where a protein elution profile could not be fitted to a Gaussian curve, the global maximum (*G*_max_) was replaced by the peak location. We selected peaks that were consistently present within a two-fraction shift between biological replicates for further analysis. A protein’s multimerization state (*R*_app_) was defined as the ratio of the *M*_app_ to the theoretical monomer mass (*M*_mono_) of the protein (Liu et al., 2008; Aryal et al., 2014; Lee and Szymanski, 2021). A threshold of *R*_app_ ≥ 1.6 was used to define proteins as part of a complex. Peaks with *M*_app_ values greater than 1,200 kDa were excluded to avoid those present in the first two void fractions from the SEC separations. Raw elution profiles along with Gaussian fitted profiles are available in Supplemental figure S6, and *M*_app_ and *R*_app_ values for reproducible proteins are provided in Supplemental Dataset S5.

### Detection of variations in protein abundance and multimerization states after ethylene treatment

In the comparative proteomics analysis, two-sample *t*-tests were performed between all quantified proteins, selected by protein intensity-based filtering, in the untreated (-C_2_H_4_) and ethylene-treated (+C_2_H_4_) datasets to identify differentially expressed proteins in response to ethylene treatment. LFQ intensity values were log2-transformed, and missing values were imputed by sampling from a normal distribution (width = 0.3, shift = 1.5) using Perseus version 2.0.7.0 (Tyanova et al., 2016). Statistically significant proteins were selected with a 10 % false discovery rate (FDR) and a 4-fold change threshold using Perseus’ built-in tool (see columns AS to AU and BB to BD in Supplemental Datasets S2B and S3B).

To find variants in protein multimerization, *ANOVA* and Tukey’s post hoc test were performed as described previously (Lee and Szymanski, 2021). The *ANOVA* tests were run against both *M*_app_ and *R*_app_ values for proteins present in both SEC_Ethylene_ and SEC_Untreated_ datasets. This multiple comparison test was set at a 5 % FDR to minimize likely false positives and negatives. Tukey’s post hoc test was then applied at a significance level of *p* < 0.05. Statistical results were manually verified by examining the raw elution profiles of the flagged proteins in both datasets to determine if significant differences were present, as indicated by the statistical tests (Supplemental Dataset S6).

To identify proteins that were “potentially degraded by ethylene” or “potentially induced or stabilized by ethylene” based on the SEC data, we identified proteins that had reproducible peaks in one condition and were completely absent from all column fractions from the counter sample (Figure 3I; Supplemental Dataset S6C). Unreliable proteins present in the counter sample were then classified as “Unreliable: present in its counterpart SEC sample” and annotated in column AL of Supplemental Dataset S6C. Additionally, if the putatively differentially expressed protein was present in the unfractionated S200 counter sample (Supplemental Dataset S2B), they were classified as “Unreliable: detected in the Ethylene-treated dataset” or “Unreliable: detected in the Untreated control dataset” and subsequently removed.

### Expression analysis of ethylene response genes by quantitative RT-PCR

Three-day-old, etiolated seedlings were treated with/without ethylene gas for 2 hours and then harvested hypocotyls under the green light condition, as aforementioned. Total RNA was extracted using RNeasy Plant Mini Kits (Qiagen, Hilden, Germany), and 1 μg of total RNA was used to synthesize cDNA with SuperScript II reverse transcriptase (Invitrogen, Waltham, MA, USA), following the manufacturers’ protocols. qRT-PCR was performed using PowerUP^TM^ SYBR^TM^ Green Master Mix (Applied Biosystems, Waltham, MA, USA). From the S200 dataset, 21 high-confidence protein hits were statistically selected based on their abundance (Group1; Figure 2D) and stringent count-based data filtering (Group2). Specifically, proteins were included if they were identified with MS/MS counts ≥ 1 in at least two out of three replicates and were not detected in any of the replicates from the counter sample. The selected proteins are summarized in Supplemental Dataset S2C, and the primers used for downstream validation are listed in Supplemental Table S1. Three biological replicates were analyzed, and the relative expression was normalized to *actin2* as an internal control. Statistical significance was determined using a two-tailed Student’s *t-*test.

### Gene ontology enrichment analysis

Gene Ontology (GO) enrichment analysis was performed using agriGO v2.0 Singular Enrichment Analysis (Tian et al., 2017). The enrichment was calculated using Fisher’s exact test against the TAIR genome locus (TAIR10_2017) background. The minimum number of mapping entries was 5, and Yekutieli (FDR under dependency) was used as a multi-test adjustment method. Significantly enriched GO terms in biological process (BP), GO molecular function (MF), and GO cellular component (CC) categories were reported at 5 % FDR.

### Statistical tests and data analysis

Statistical analyses were performed using R version 3.5.1 (R Core Team, 2018) on RStudio 1.1.463 (RStudio Team, 2018). Gaussian fitting was applied using MATLAB_R2021a, and data organization was done using Microsoft Excel (Office 365 for Mac).

## Data and materials availability

The MS raw files have been deposited into the ProteomeXchange Consortium via PRIDE under accession codes PXD061207 for the comparative proteomics and PXD061200 for the CFMS analysis. The Gaussian fitting code described in McBride *et al*. (2017) is available at (https://github.com/dlchenstat/Gaussian-fitting).

## Author contributions

All authors conceived and designed the study, performed data analyses, and wrote the paper. Y.L. and H.L.P. conducted the experiments. D.B.S. and G.M.Y. supervised this project.

## Acknowledgments

This material is based upon work supported by the National Science Foundation under Grant No. 1951819 to D.B.S. and Grant No. 2245525 to G.M.Y. Also, it is supported by a Purdue Center for Plant Biology seed grant to D.B.S. and G.M.Y. We thank Dr. Uma Aryal at the Purdue Proteomics Facility for running the LC/MS samples.

## Supplemental Datasets

Supplemental Dataset S1. Identified peptides and proteins in the S200 and P200 cellular fractions.

Supplemental Dataset S2. Soluble proteins (S200) and differentially expressed proteins upon ethylene treatment.

Supplemental Dataset S3. Microsome-associated proteins (P200) and differentially expressed proteins upon ethylene treatment.

Supplemental Dataset S4. Raw elution profiles of peptides and proteins in CFMS analysis.

Supplemental Dataset S5. Reproducible protein peaks in ethylene-untreated (SEC_Untreated_) and -treated (SEC_Ethylene_) CFMS datasets (Col-0 S200).

Supplemental Dataset S6. Protein multimerization variants affected by ethylene treatment (Col-0 S200).

Supplemental Dataset S7. Enriched GO terms of multimerization variants.

## Supplemental figures

Supplemental figure S1. Proteome changes in the soluble (S200) and microsome-associated (P200) cellular fractions of Col-0 and *ein2-5* etiolated plants as a response to ethylene treatment.

Supplemental figure S2. Enzyme coverages of metabolic pathways (top 80 % most abundant proteins of Col-0 S200 untreated control).

Supplemental figure S3. Enzyme coverages of metabolic pathways (top 5 % most abundant proteins of Col-0 S200 untreated control).

Supplemental figure S4. The CFMS pipeline to analyze Arabidopsis protein multimerization responses to gaseous ethylene treatment.

Supplemental figure S5. Normalized hypocotyl length of WT and *abcb5*.

Supplemental figure S6. Raw and Gaussian fitted protein elution profiles.

## Supplemental tables

Supplemental Table S1. Quantitative RT-PCR Primers.

## Notes

### Competing Interest Statement

The authors have declared no competing interest.

### Summary of Updates

Some texts have been revised for clarity

