## Supplemental figure S1 for "Rapid ethylene-triggered protein complex remodeling in dark grown hypocotyls"

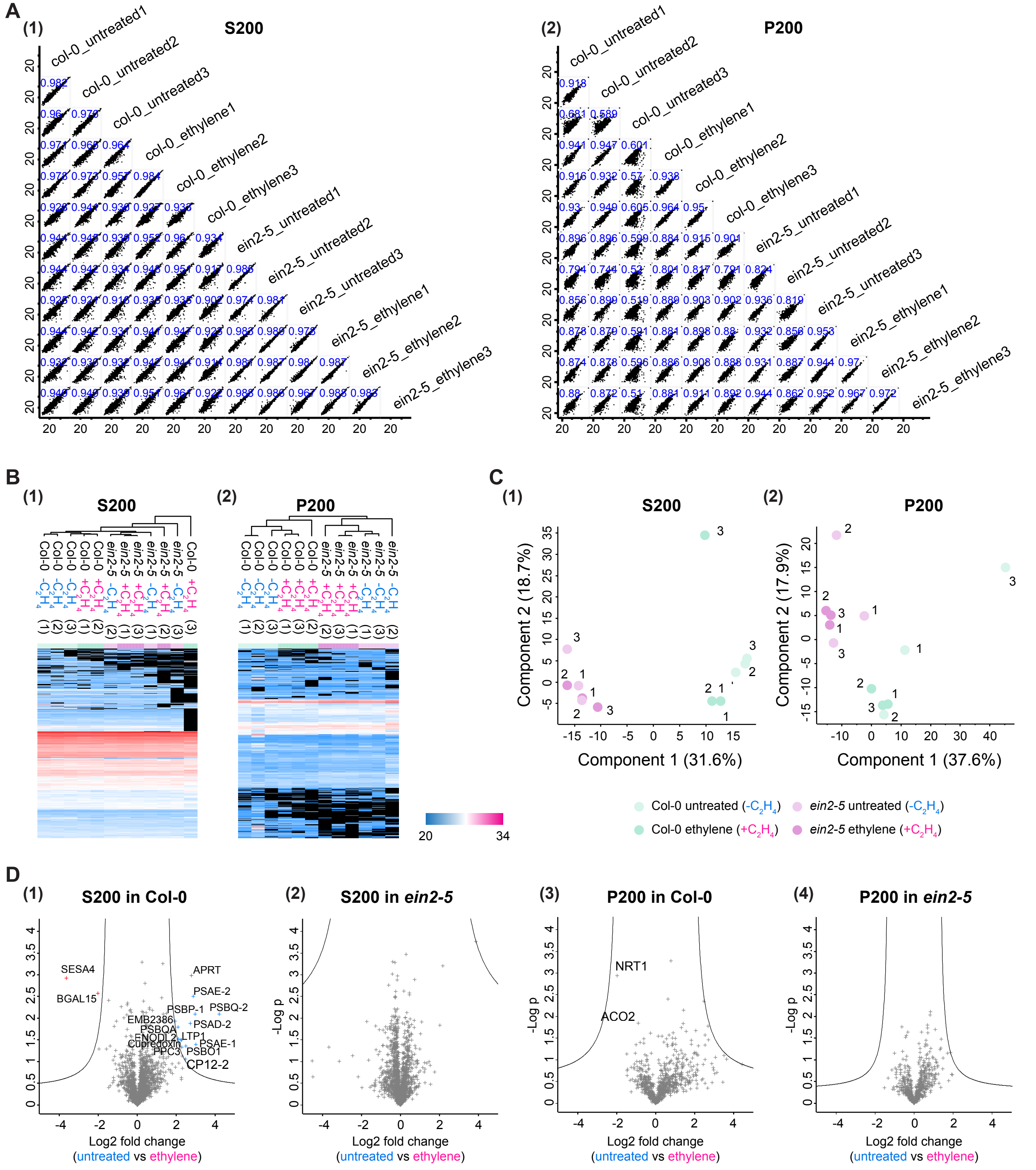


**Supplimental figure S1. Proteome changes in the soluble (S200) and microsome-associated (P200) cellular fractions of Col-0 and *ein2-5* etiolated plants as a response to ethylene treatment.**

**A,** *Pearson* correlation coefficient (PCC) of S200 (1) and P200 (2).

**B,** Hierarchical clustering and heatmap showing relative protein expression (log2-transformed LFQ protein intensities) of the differentially expressed proteins between the subsets. Columns correspond to the three replicates of the 4 subsets. Proteins that were not present in the datasets were rendered with black color.

**C,** Principal component analysis of S200 (1) and P200 (2) proteomes in Col-0 and *ein2-5* mutant plants with/without ethylene treatment. LFQ protein intensity values were log2-transformed, and then missing values were imputed by normal distribution (width = 0.3, shift = 1.5) using Perseus version 2.0.7.0.

**D,** Volcano plots depicting two-sample *t*-tests between all quantified proteins (0 values were imputed values) in untreated and ethylene-treated hypocotyls in S200 and P200 cellular fractions of Col-0 and *ein2-5* mutant plants. The lines indicate 10 % FDR and 2-fold changes (note: $Log_2$-transformed values).
