## Supplemental figure S2 for "Rapid ethylene-triggered protein complex remodeling in dark grown hypocotyls"

**Supplemental figure S2. Enzyme coverages of metabolic pathways (Col-0 S200 untreated control).**

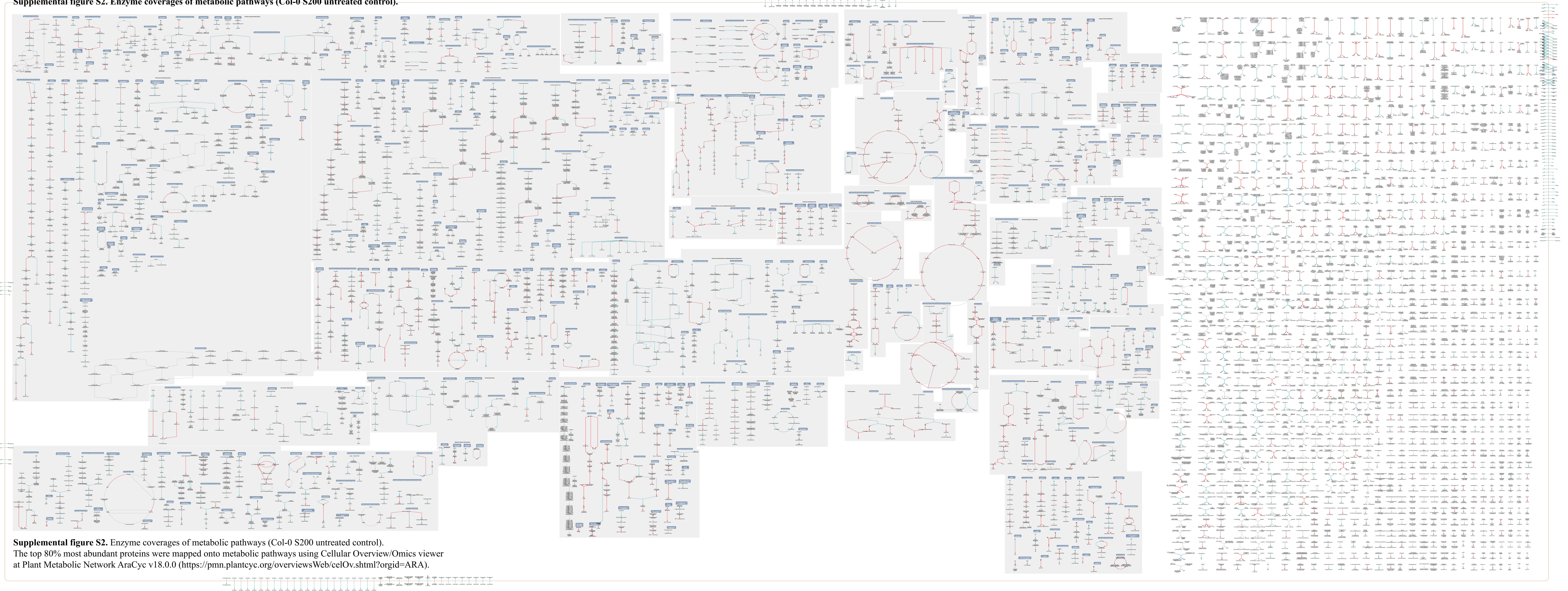

**Supplemental figure S2.** Enzyme coverages of metabolic pathways (Col-0 S200 untreated control).
The top 80% most abundant proteins were mapped onto metabolic pathways using Cellular Overview/Omics viewer at Plant Metabolic Network AraCyc v18.0.0 (https://pmn.plantcyc.org/overviewsWeb/celOv.shtml?orgid=ARA).
