## Supplemental figure S3 for "Rapid ethylene-triggered protein complex remodeling in dark grown hypocotyls"

**Supplemental figure S3. Enzyme coverages of metabolic pathways (top 5% most abundant proteins of Col-0 S200 untreated control).**

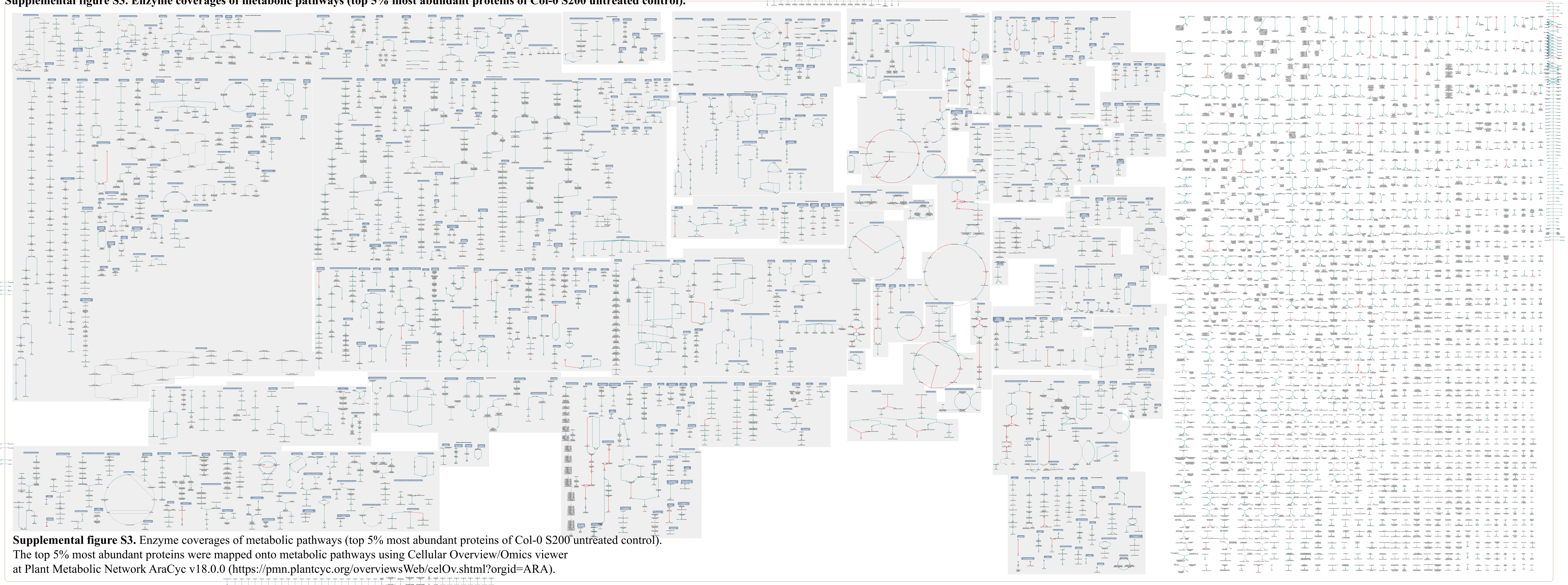

**Supplemental figure S3.** Enzyme coverages of metabolic pathways (top 5% most abundant proteins of Col-0 S200 untreated control). The top 5% most abundant proteins were mapped onto metabolic pathways using Cellular Overview/Omics viewer at Plant Metabolic Network AraCyc v18.0.0 (https://pmn.plantcyc.org/overviewsWeb/celOv.shtml?orgid=ARA).
