## Supplemental figure S4 for "Rapid ethylene-triggered protein complex remodeling in dark grown hypocotyls"

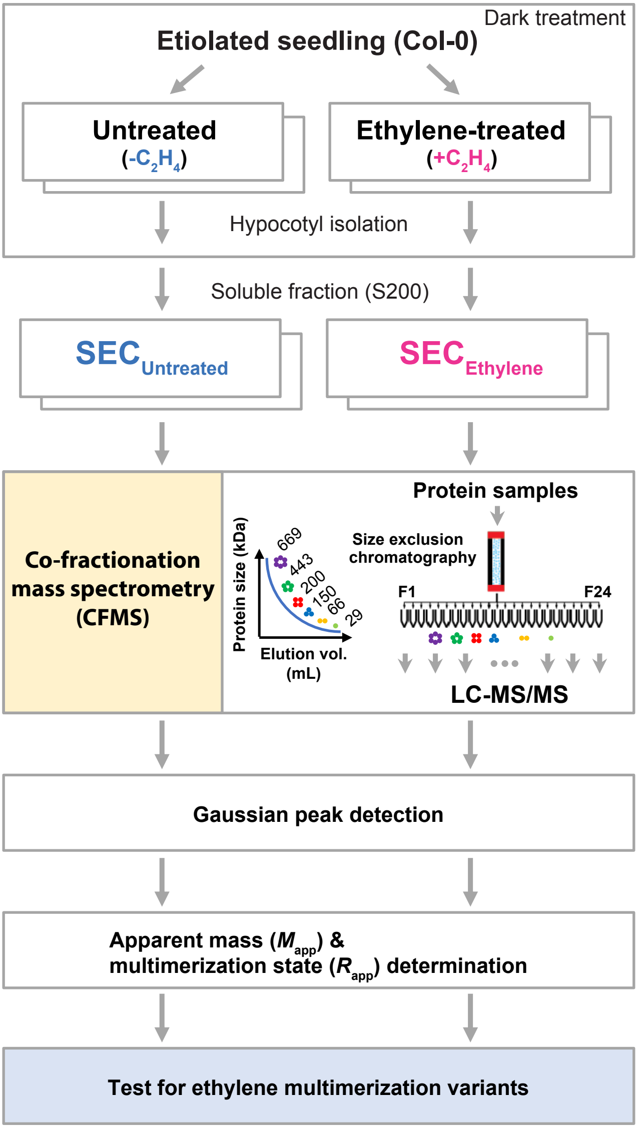


**Supplemental figure S4. The CFMS pipeline to analyze Arabidopsis protein multimerization responses to gaseous ethylene treatment.** Three-day-old, etiolated Arabidopsis seedlings are treated with exogenous ethylene for two hours. S200 protein complexes are prepared from hypocotyl tissues from the ethylene-treated and untreated etiolated seedlings under non-denaturing conditions. The endogenous protein complexes are fractionized in a size exclusion chromatography and then each fraction is analyzed by LC-MS/MS. The elution profiles are fitted to Gaussian peaks to determine proteins apparent mass ($M_{app}$) and multimerization state ($R_{app}$). *ANOVA* tests are used to detect multimerization variants that respond to the ethylene treatment.
