## Supplemental figure S5 for "Rapid ethylene-triggered protein complex remodeling in dark grown hypocotyls"

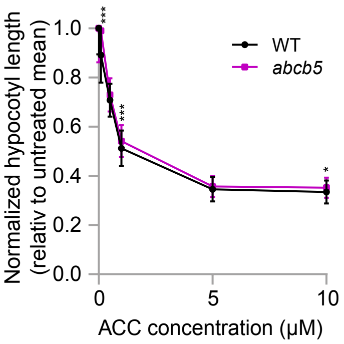


**Supplimental figure S5. Normalized hypocotyl length of WT and *abcb5*.** Three-day-old dark-grown WT and *abcb5* seedlings were grown on MS medium supplemented with varying concentrations of ACC. Hypocotyl lengths of individual seedlings were normalized to the mean length of untreated (0 µM ACC) seedlings of the corresponding genotype, which was set to 1. The graph shows the relative inhibition of hypocotyl elongation in response to ACC. Data represent mean ± SD. Statistical significance was determined using a two-tailed Student's t-test comparing *abcb5* to WT at each ACC concentration.
