## Supplemental figure S6 for "Rapid ethylene-triggered protein complex remodeling in dark grown hypocotyls"

### Supplemental figure S6. Raw and Gaussian fitted protein elution profiles (SEC)

### $SEC_{Untreated}$ replicate 1 elution profiles

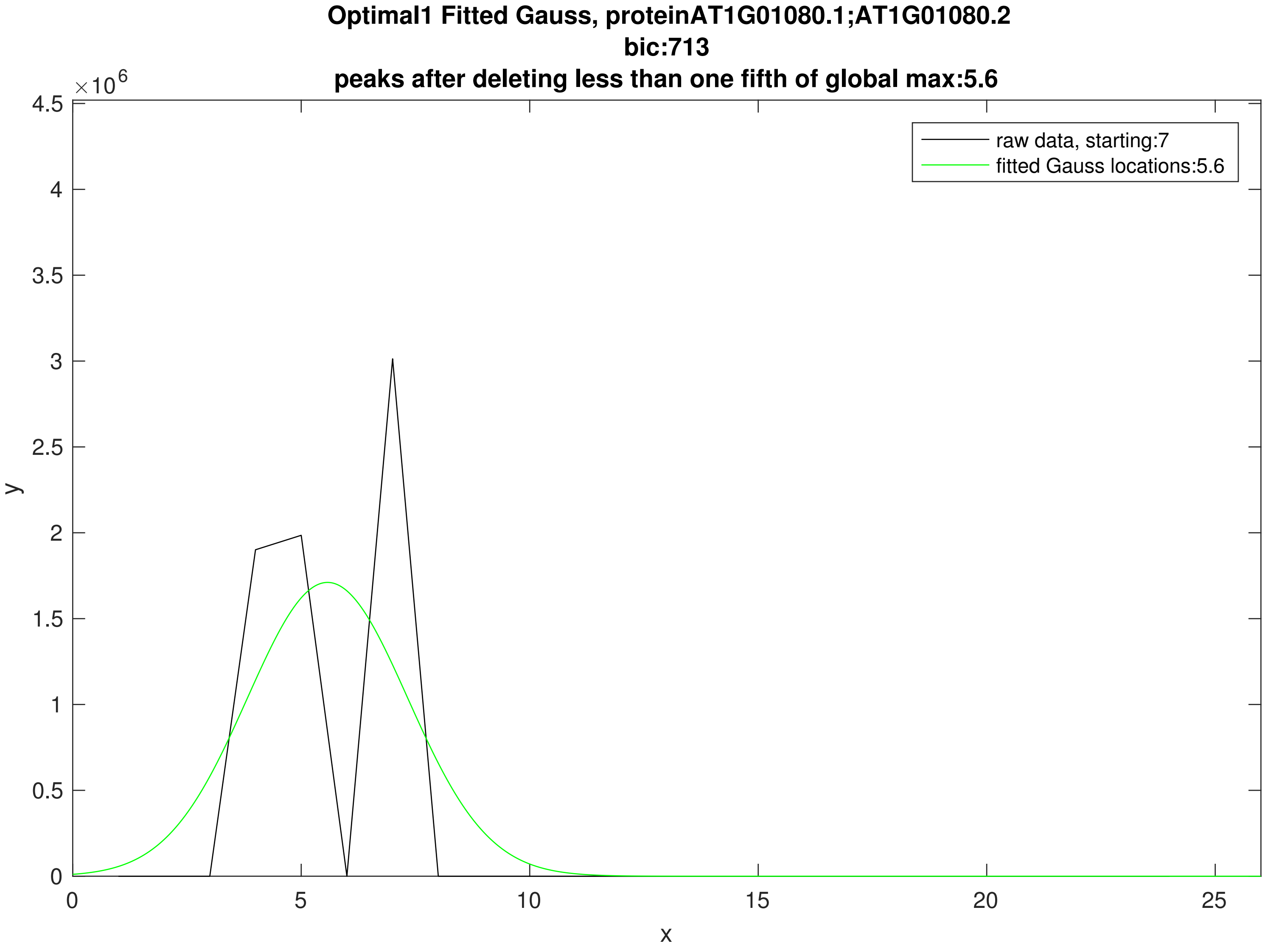
Optimal1 Fitted Gauss, proteinAT1G01080.1;AT1G01080.2
bic:713
peaks after deleting less than one fifth of global max:5.6
×10^6
4.5
4
3.5
3
2.5
2
1.5
1
0.5
0
y
0
5
10
15
20
25
x
raw data, starting:7
fitted Gauss locations:5.6

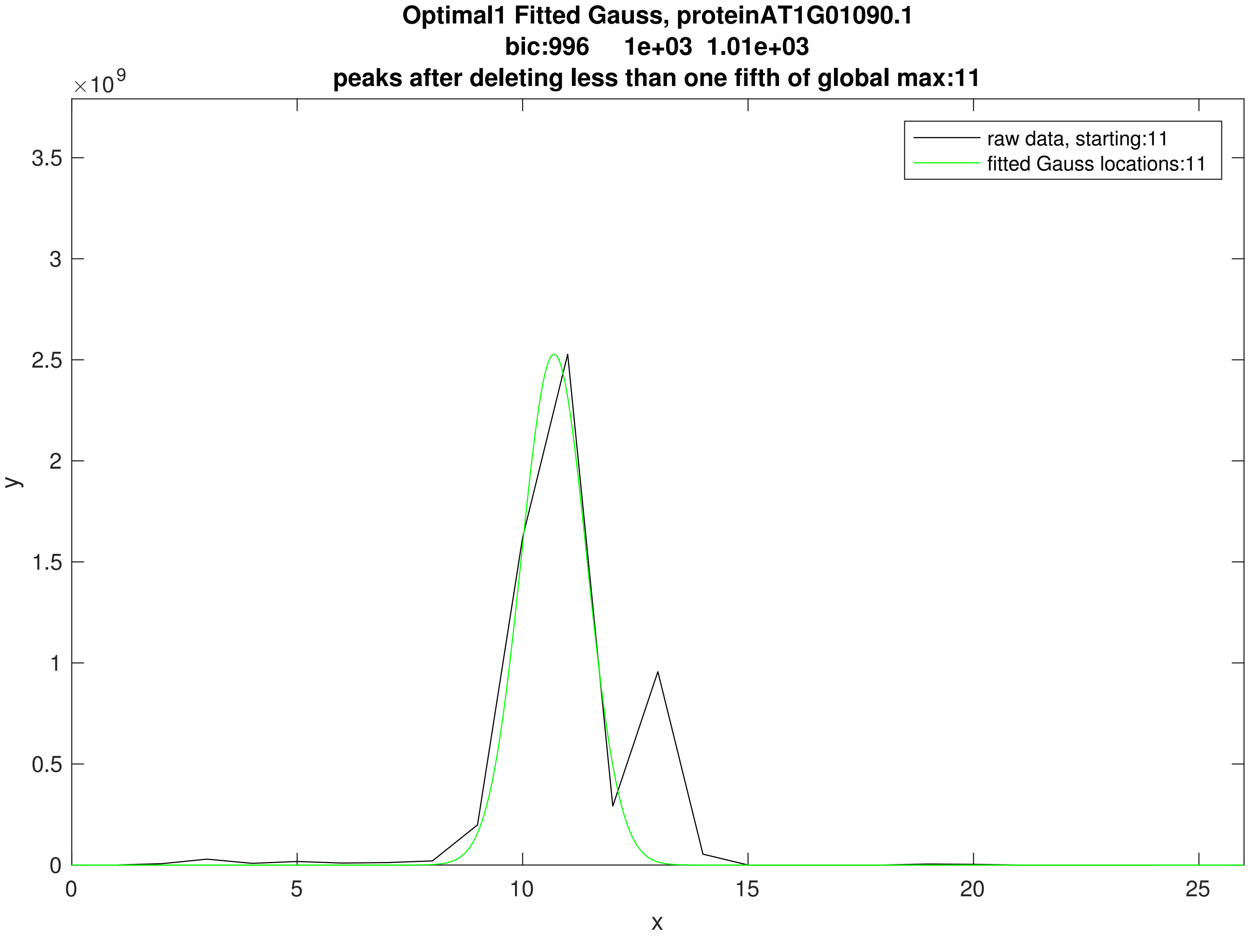
Optimal1 Fitted Gauss, proteinAT1G01090.1
bic:996 1e+03 1.01e+03
peaks after deleting less than one fifth of global max:11
×10^9
raw data, starting:11
fitted Gauss locations:11
3.5
3
2.5
2
1.5
1
0.5
0
y
0
5
10
15
20
25
x

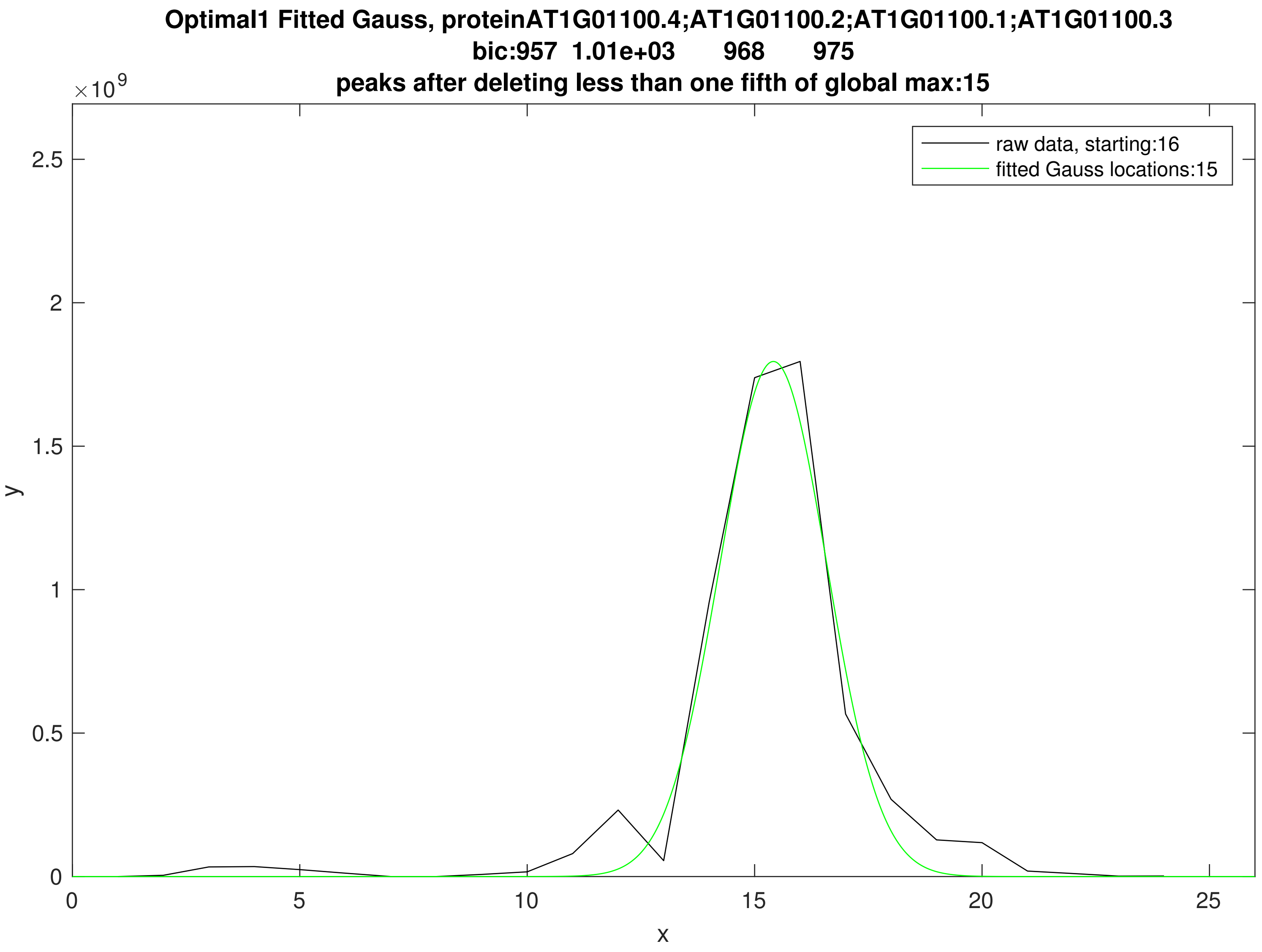
Optimal1 Fitted Gauss, proteinAT1G01100.4;AT1G01100.2;AT1G01100.1;AT1G01100.3
bic:957 1.01e+03 968 975
peaks after deleting less than one fifth of global max:15
×10^9
2.5
2
1.5
1
0.5
0
y
0
5
10
15
20
25
x
raw data, starting:16
fitted Gauss locations:15

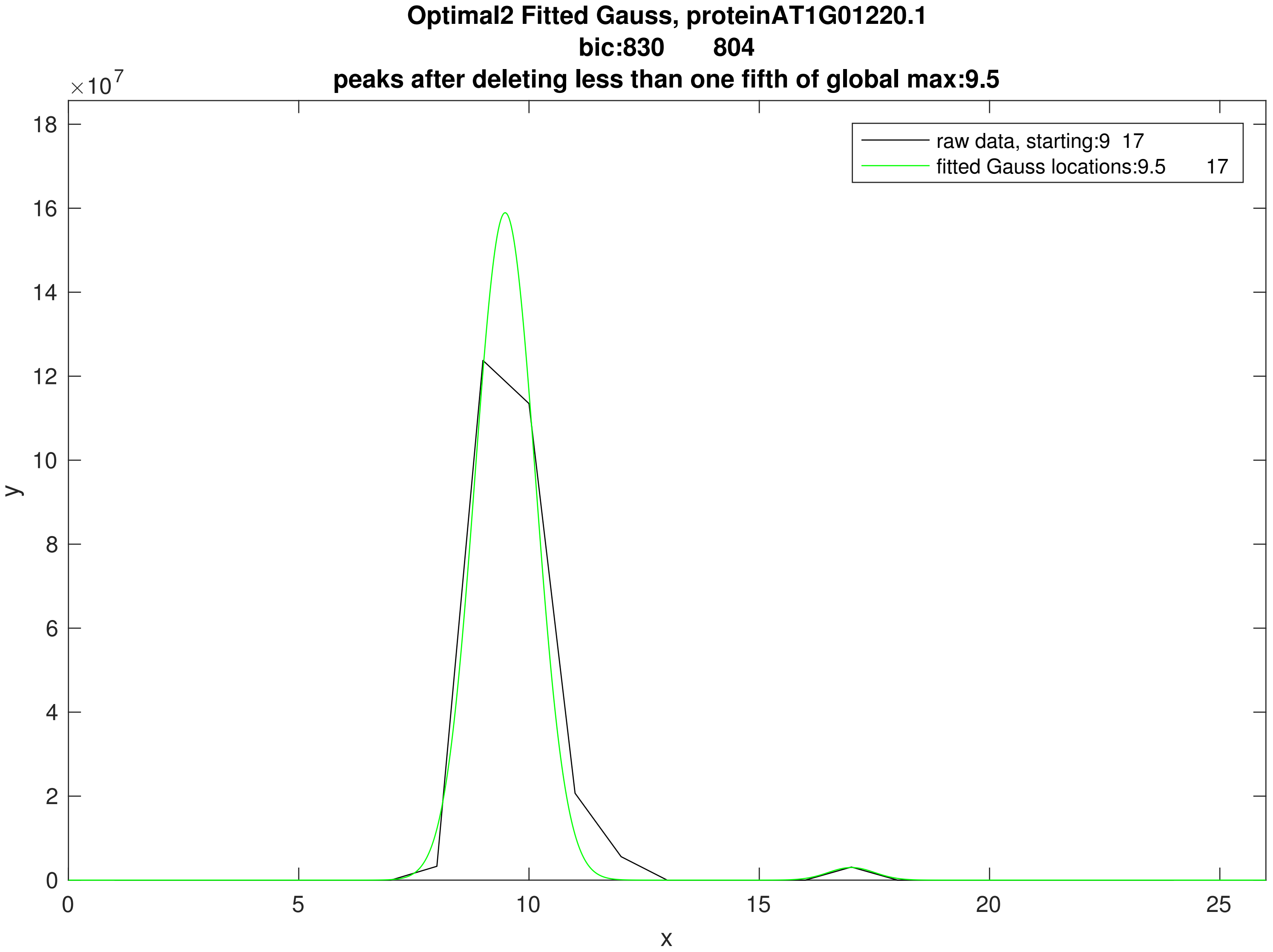
Optimal2 Fitted Gauss, proteinAT1G01220.1
bic:830 804
peaks after deleting less than one fifth of global max:9.5
×10^7
raw data, starting:9 17
fitted Gauss locations:9.5 17
y
x
0
2
4
6
8
10
12
14
16
18
0
5
10
15
20
25

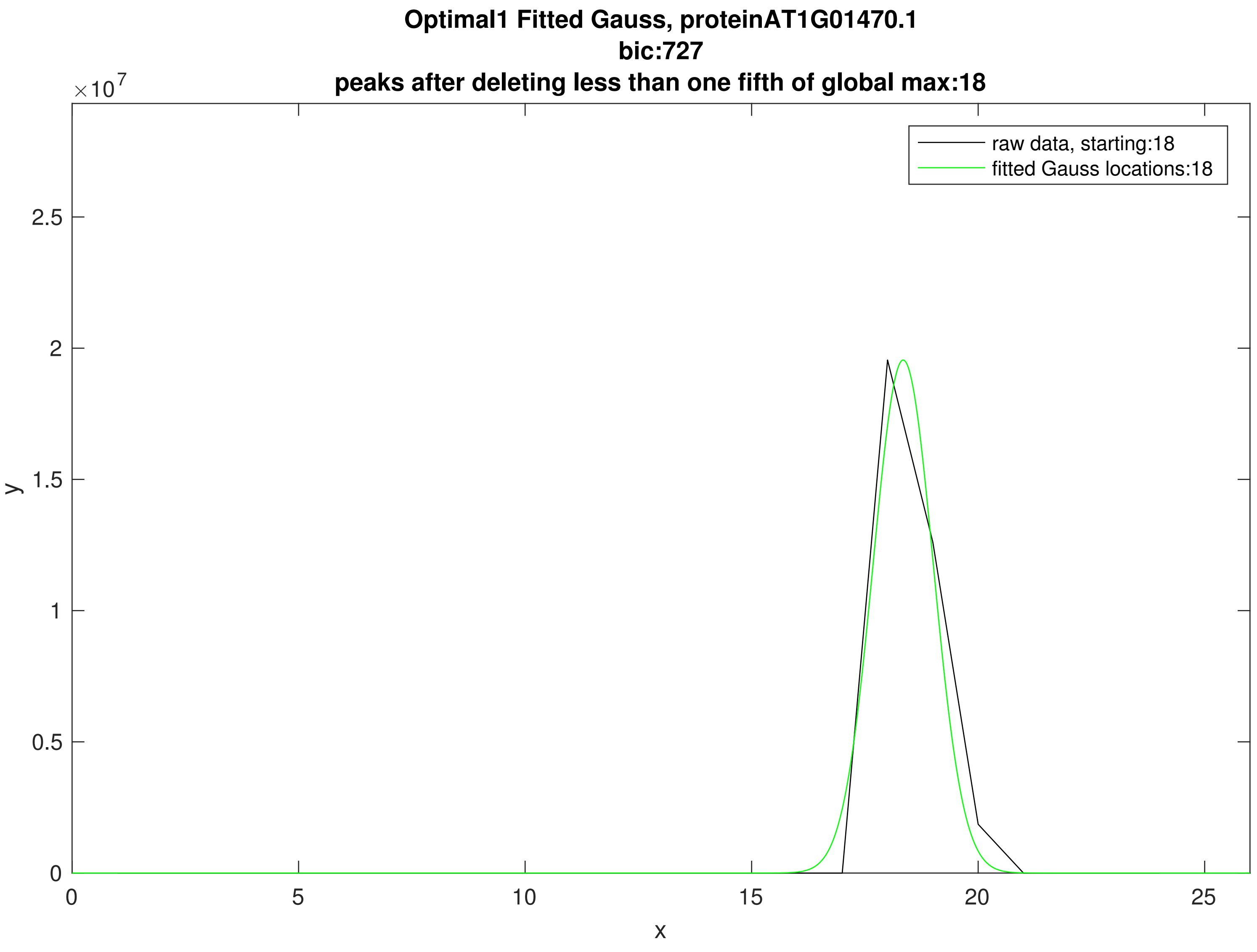
Optimal1 Fitted Gauss, proteinAT1G01470.1
bic:727
peaks after deleting less than one fifth of global max:18
×10^7
raw data, starting:18
fitted Gauss locations:18
y
x
0
0.5
1
1.5
2
2.5
0
5
10
15
20
25

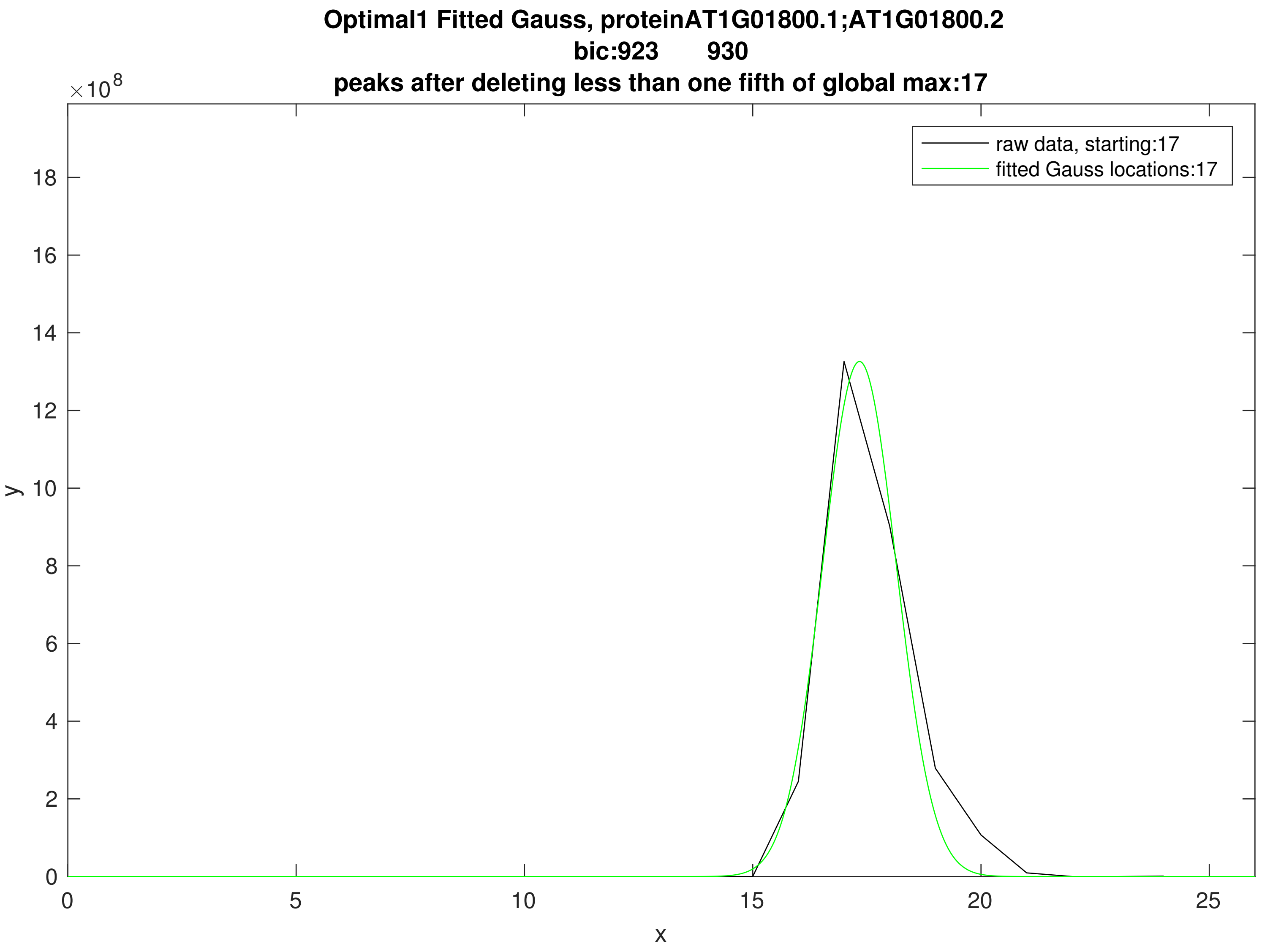

Optimal1 Fitted Gauss, proteinAT1G01800.1;AT1G01800.2
bic:923 930
peaks after deleting less than one fifth of global max:17
×10^8
raw data, starting:17
fitted Gauss locations:17
y
x

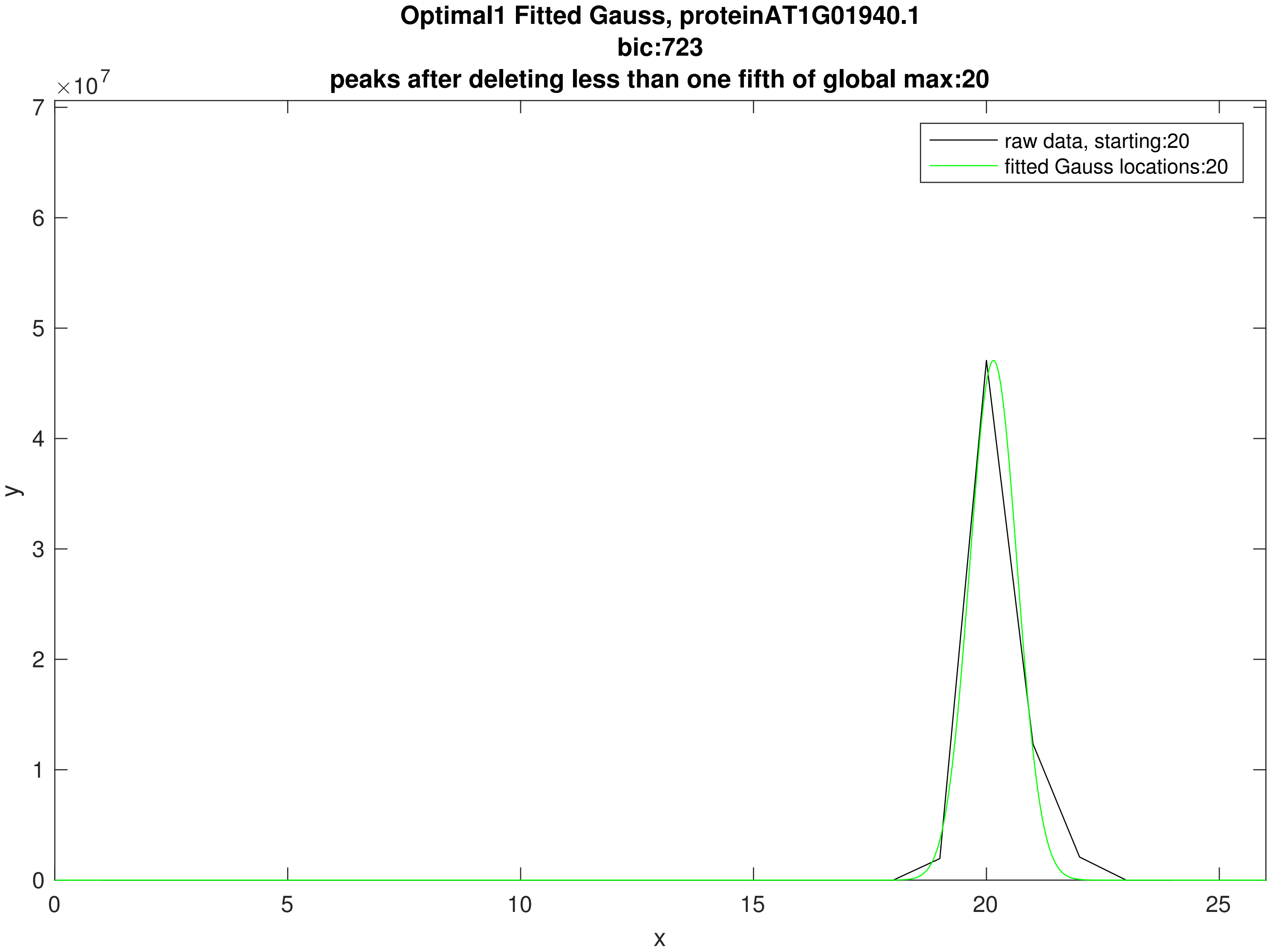
Optimal1 Fitted Gauss, proteinAT1G01940.1
bic:723
peaks after deleting less than one fifth of global max:20
×10^7
raw data, starting:20
fitted Gauss locations:20
y
x
0
1
2
3
4
5
6
7
0
5
10
15
20
25

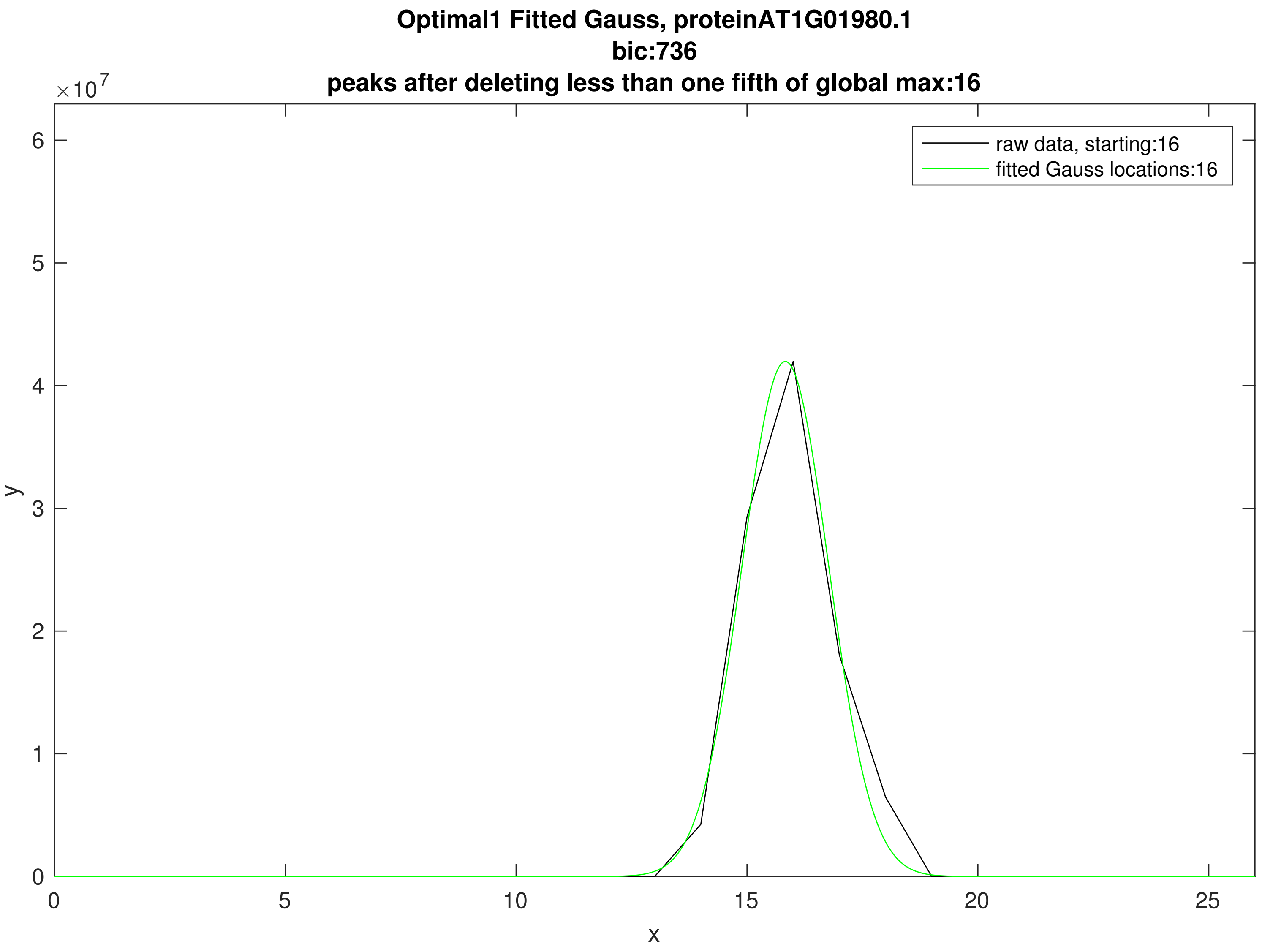
Optimal1 Fitted Gauss, proteinAT1G01980.1
bic:736
peaks after deleting less than one fifth of global max:16
×10^7
raw data, starting:16
fitted Gauss locations:16
y
x
0
1
2
3
4
5
6
0
5
10
15
20
25

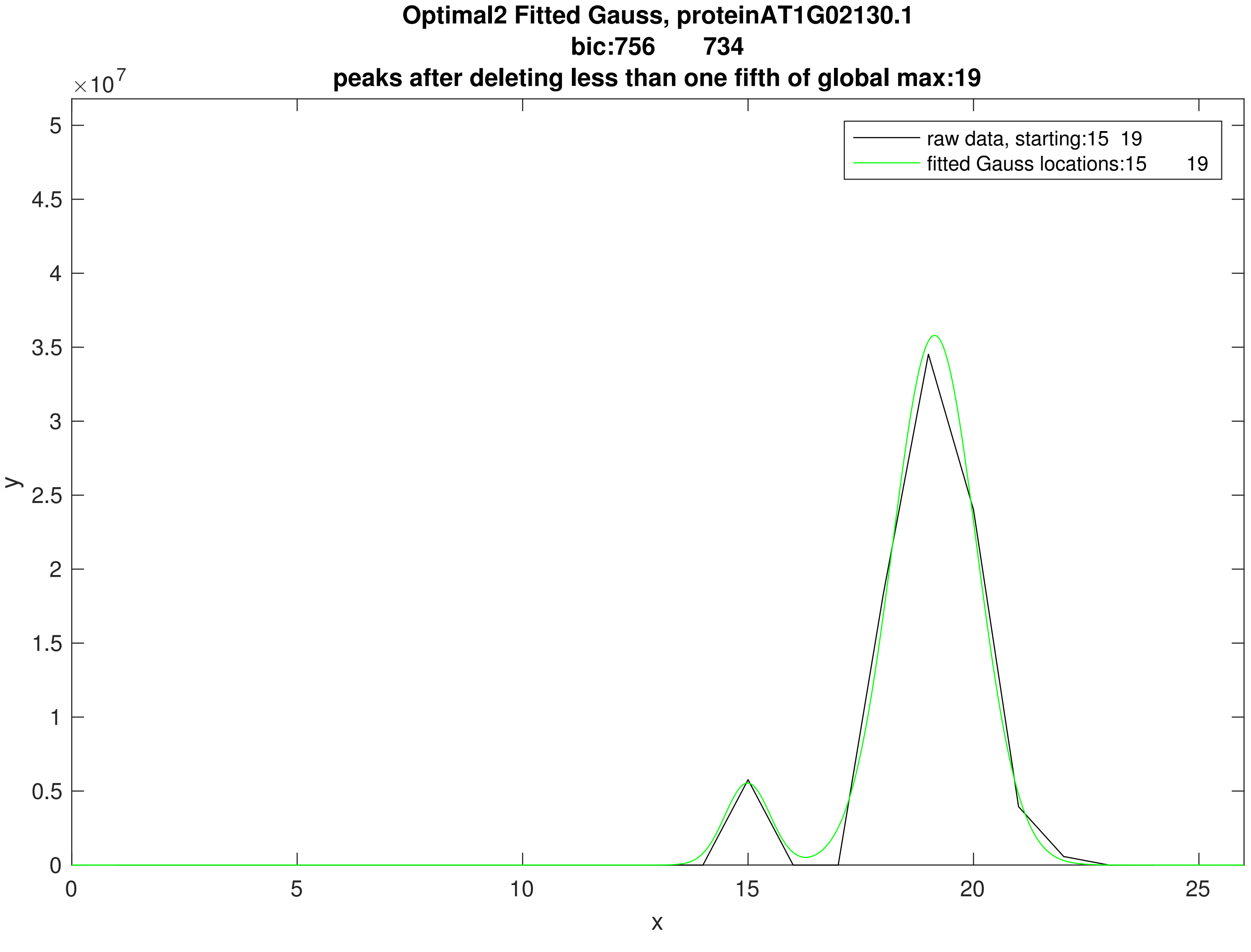
Optimal2 Fitted Gauss, proteinAT1G02130.1
bic:756 734
peaks after deleting less than one fifth of global max:19
×10^7
5
4.5
4
3.5
3
2.5
2
1.5
1
0.5
0
y
0
5
10
15
20
25
x
raw data, starting:15 19
fitted Gauss locations:15 19

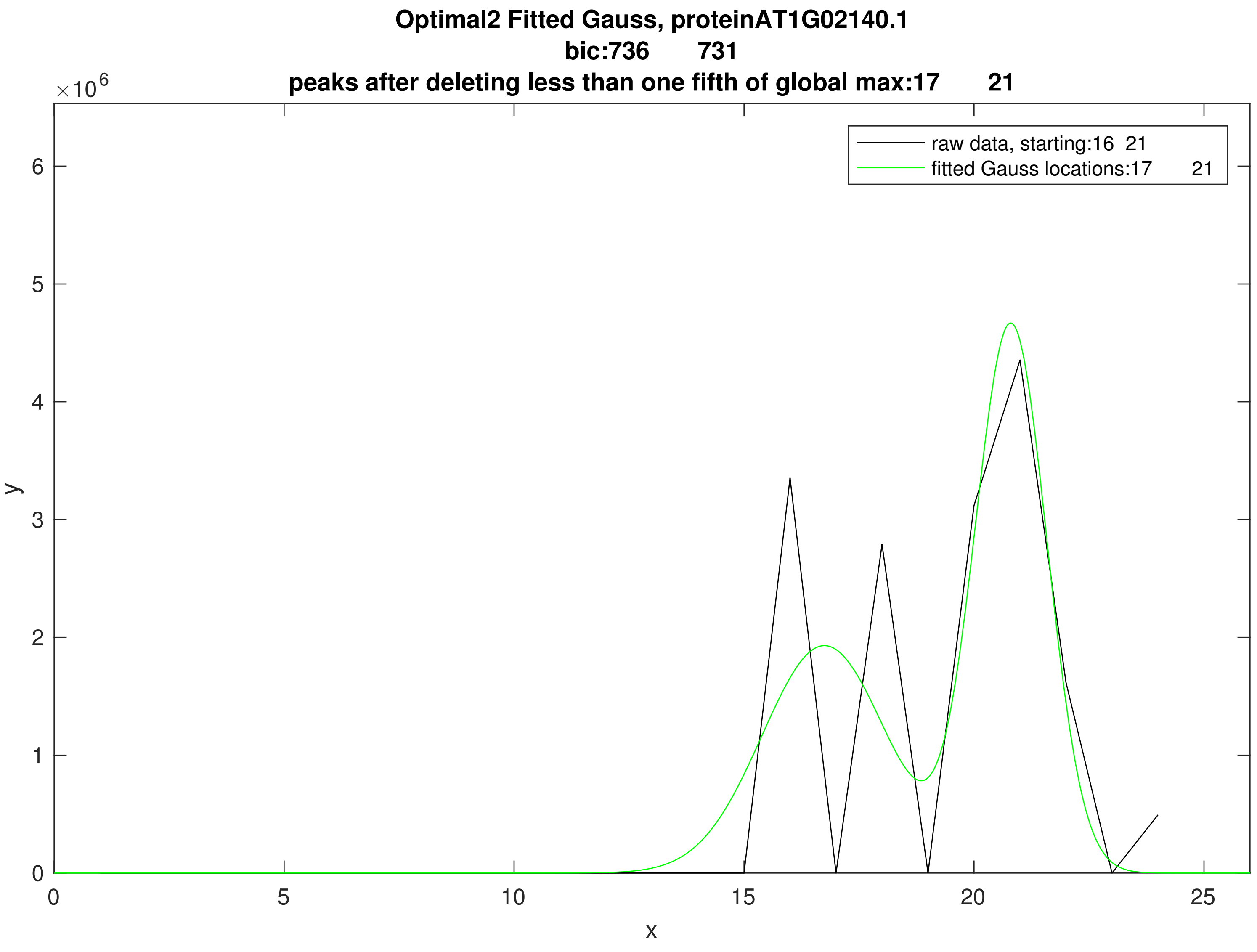
Optimal2 Fitted Gauss, proteinAT1G02140.1
bic:736 731
peaks after deleting less than one fifth of global max:17 21
×10^6
raw data, starting:16 21
fitted Gauss locations:17 21
y
x
0
1
2
3
4
5
6
0
5
10
15
20
25

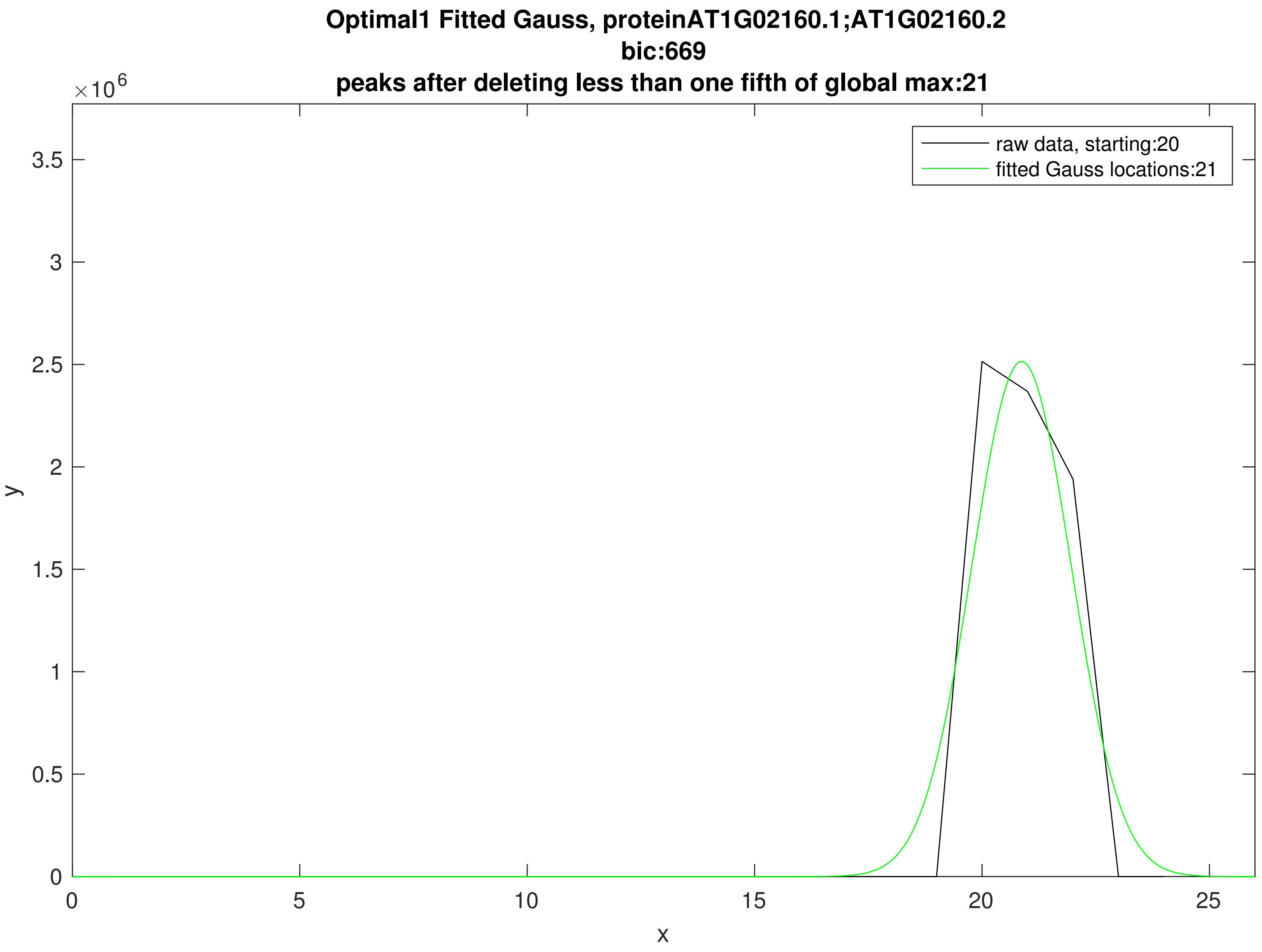

Optimal1 Fitted Gauss, proteinAT1G02160.1;AT1G02160.2
bic:669
peaks after deleting less than one fifth of global max:21
×10^6
3.5
3
2.5
2
1.5
1
0.5
0
y
0
5
10
15
20
25
x
raw data, starting:20
fitted Gauss locations:21

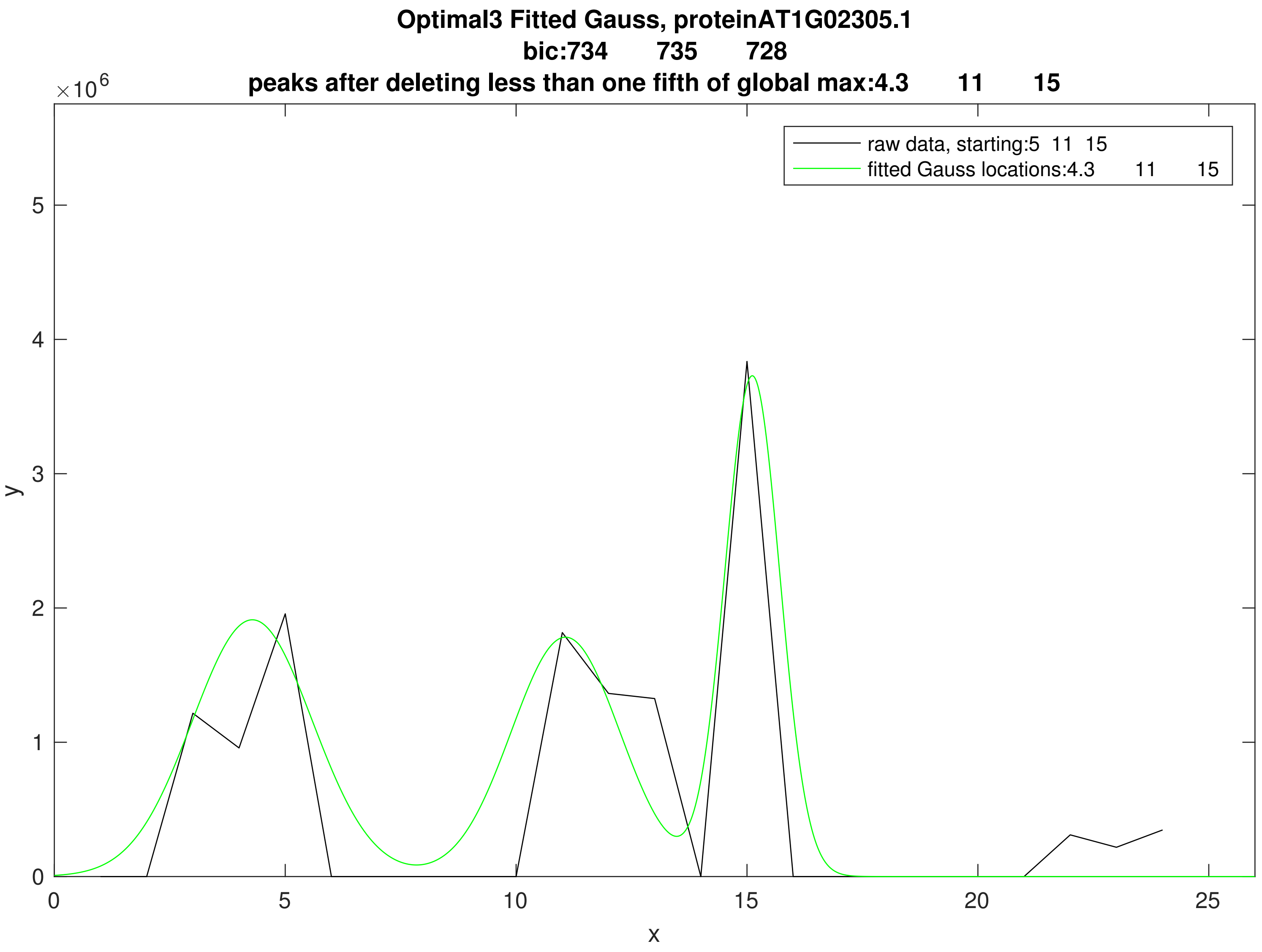

Optimal3 Fitted Gauss, proteinAT1G02305.1
bic:734 735 728
peaks after deleting less than one fifth of global max:4.3 11 15
×10^6
raw data, starting:5 11 15
fitted Gauss locations:4.3 11 15
y
x
0
1
2
3
4
5
0
5
10
15
20
25

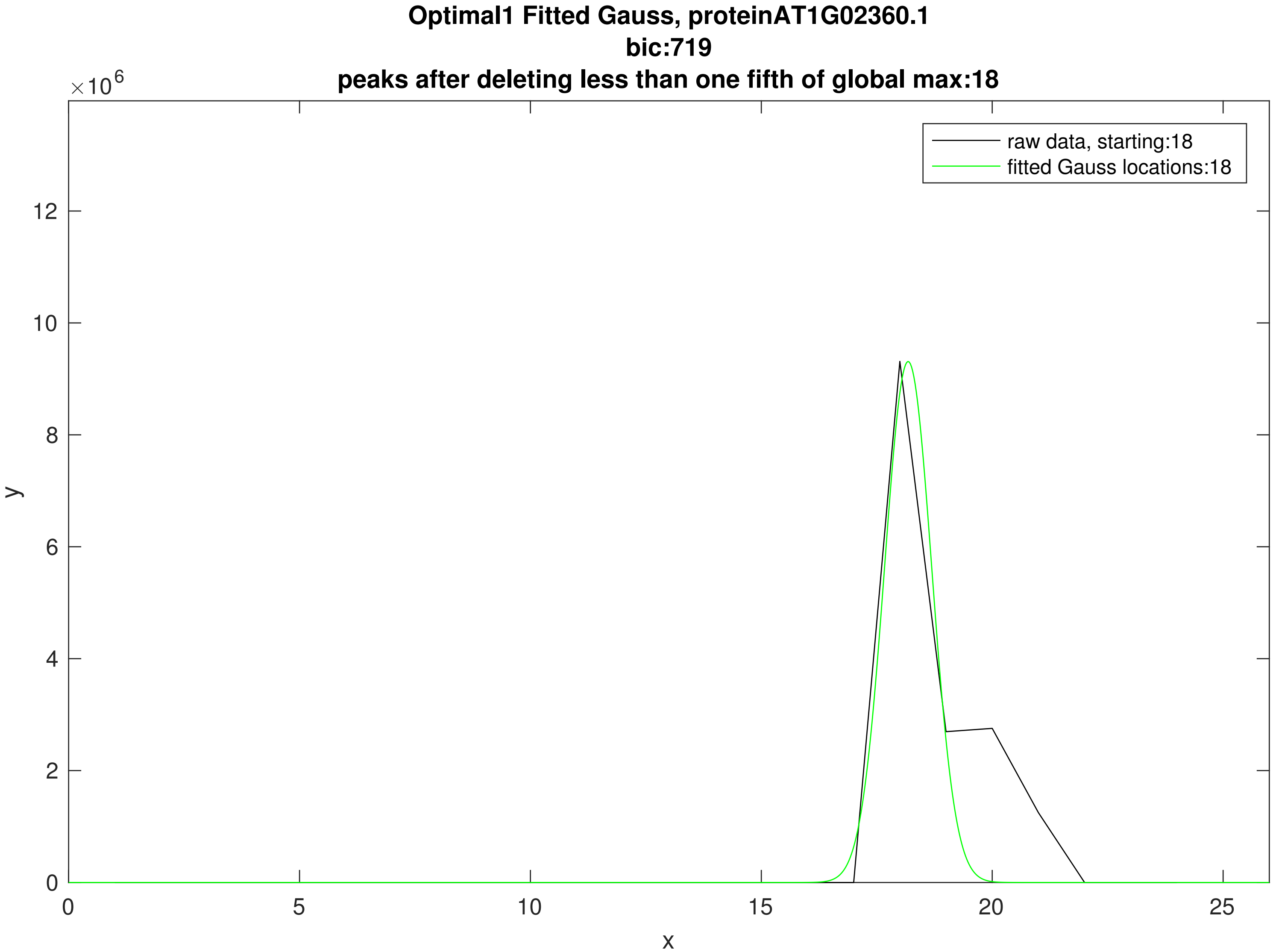

Optimal1 Fitted Gauss, proteinAT1G02360.1
bic:719
peaks after deleting less than one fifth of global max:18
×10^6
raw data, starting:18
fitted Gauss locations:18
12
10
8
6
4
2
0
y
0
5
10
15
20
25
x

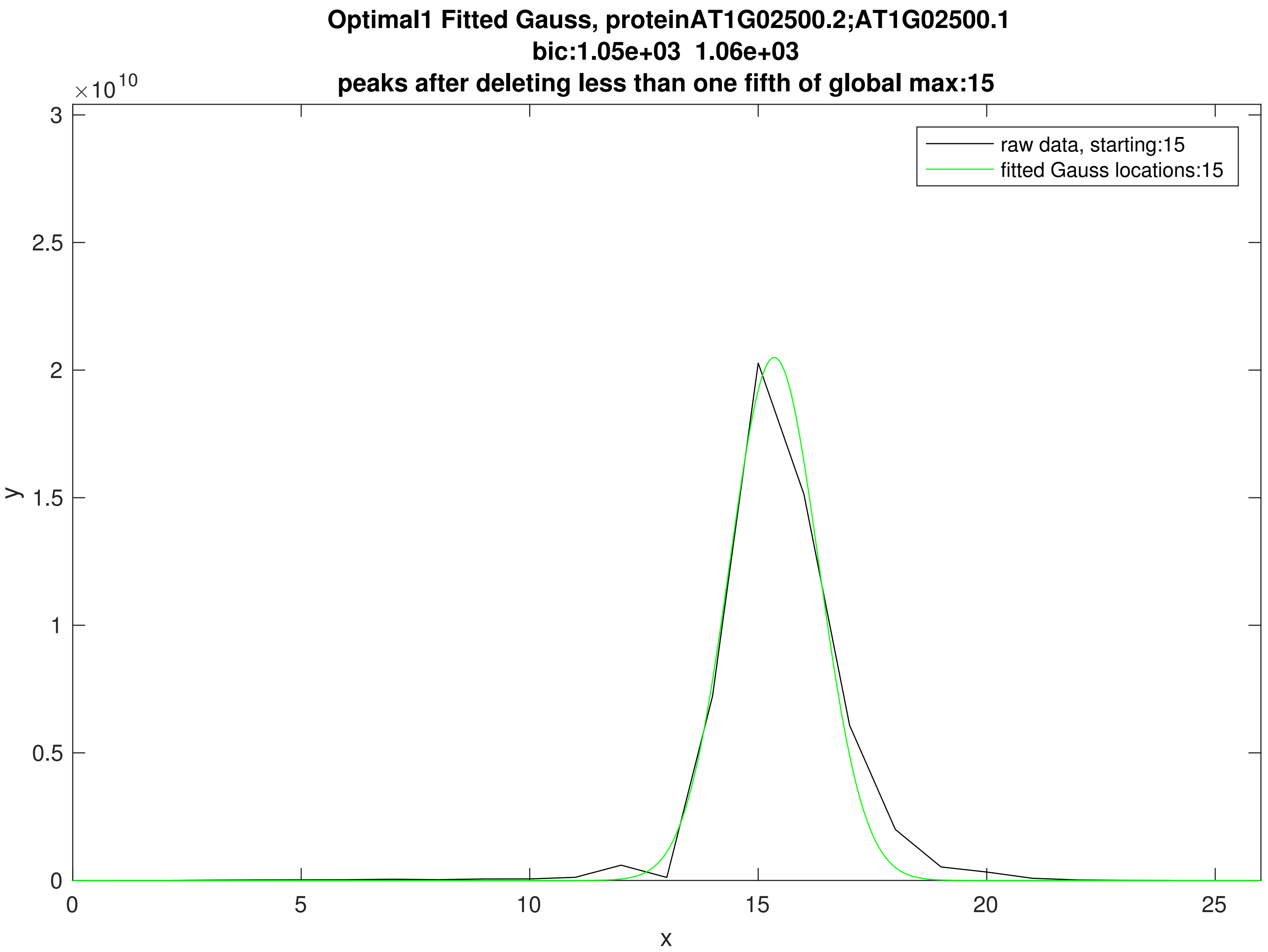
Optimal1 Fitted Gauss, proteinAT1G02500.2;AT1G02500.1
bic:1.05e+03  1.06e+03
peaks after deleting less than one fifth of global max:15
×10^10
3
2.5
2
1.5
1
0.5
0
y
0
5
10
15
20
25
x
raw data, starting:15
fitted Gauss locations:15

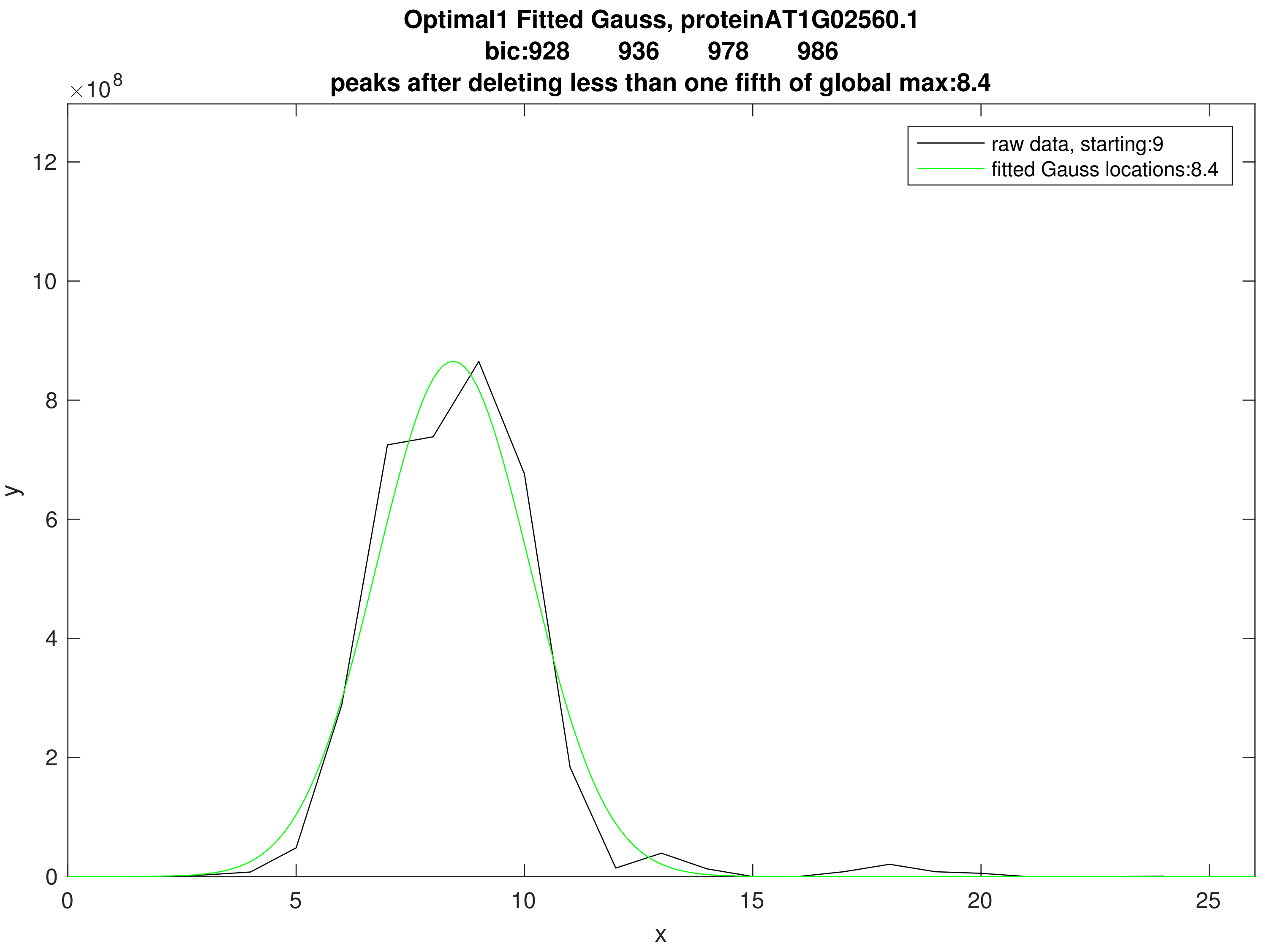

Optimal1 Fitted Gauss, proteinAT1G02560.1
bic:928 936 978 986
peaks after deleting less than one fifth of global max:8.4
×10^8
raw data, starting:9
fitted Gauss locations:8.4
y
x
0
2
4
6
8
10
12
0
5
10
15
20
25

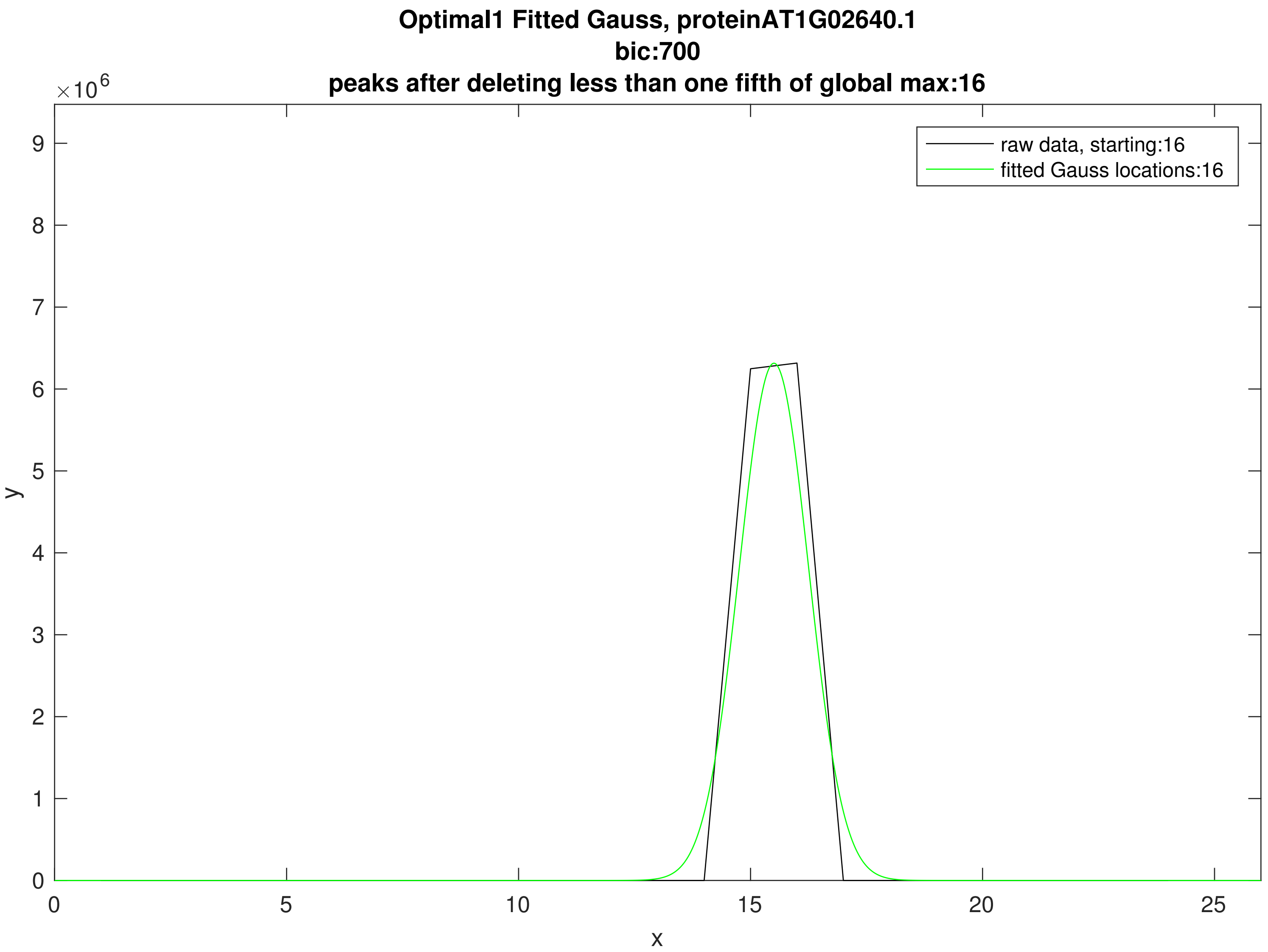

Optimal1 Fitted Gauss, proteinAT1G02640.1
bic:700
peaks after deleting less than one fifth of global max:16
×10^6
9
8
7
6
5
4
3
2
1
0
y
0
5
10
15
20
25
x
raw data, starting:16
fitted Gauss locations:16

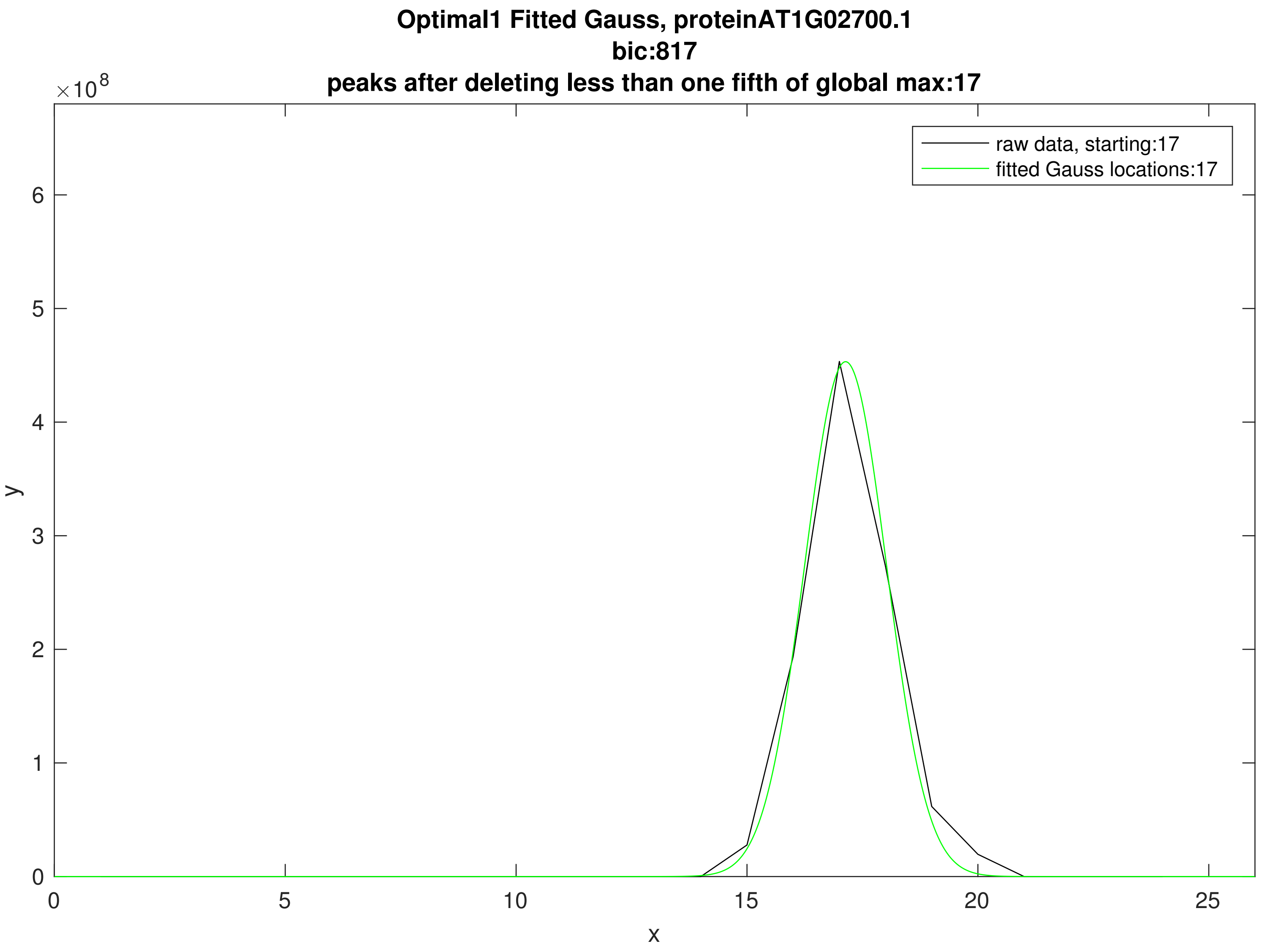
Optimal1 Fitted Gauss, proteinAT1G02700.1
bic:817
peaks after deleting less than one fifth of global max:17
×10^8
raw data, starting:17
fitted Gauss locations:17
y
x
0
1
2
3
4
5
6
0
5
10
15
20
25

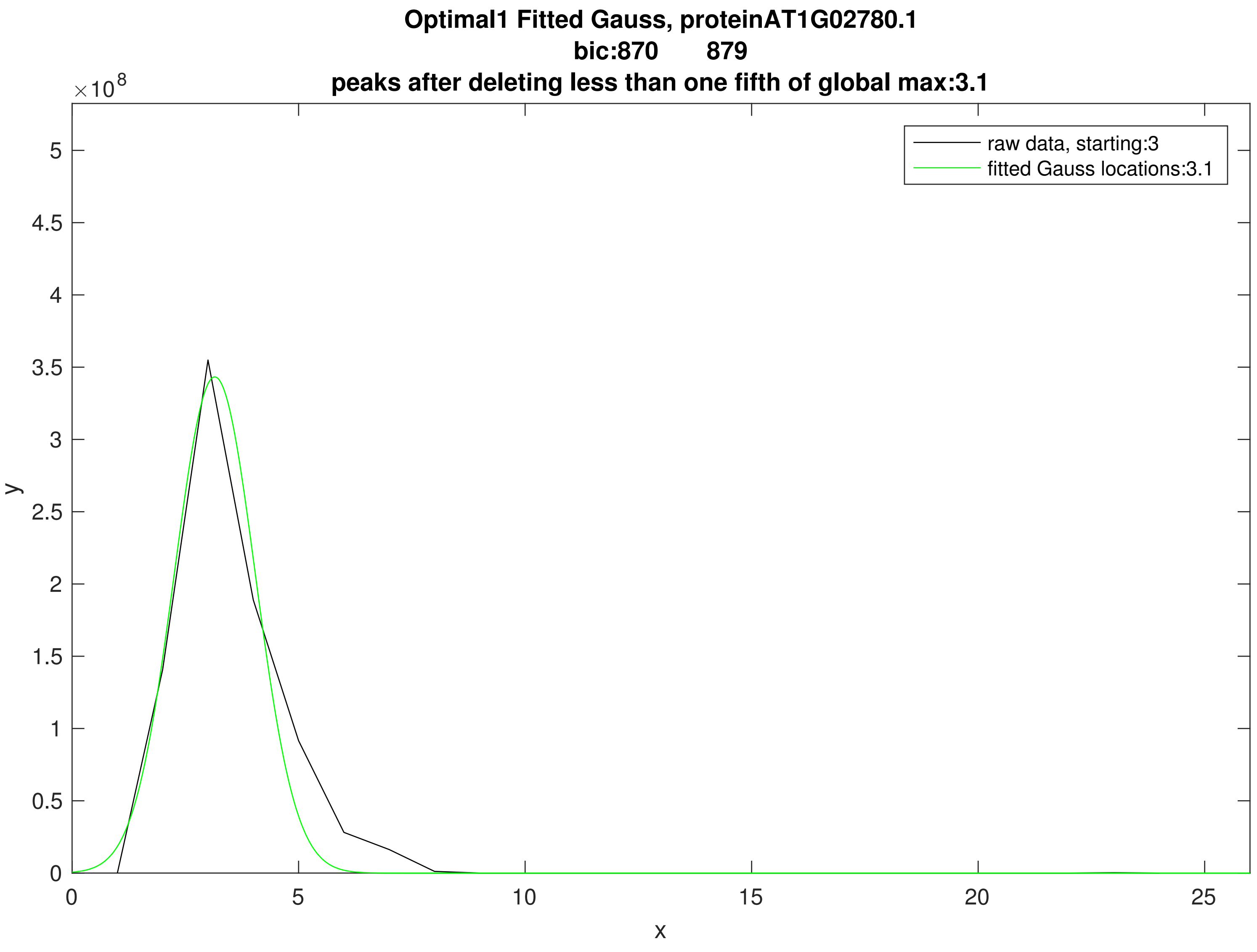

Optimal1 Fitted Gauss, proteinAT1G02780.1
bic:870 879
peaks after deleting less than one fifth of global max:3.1
×10^8
raw data, starting:3
fitted Gauss locations:3.1
y
x
0
0.5
1
1.5
2
2.5
3
3.5
4
4.5
5
0
5
10
15
20
25

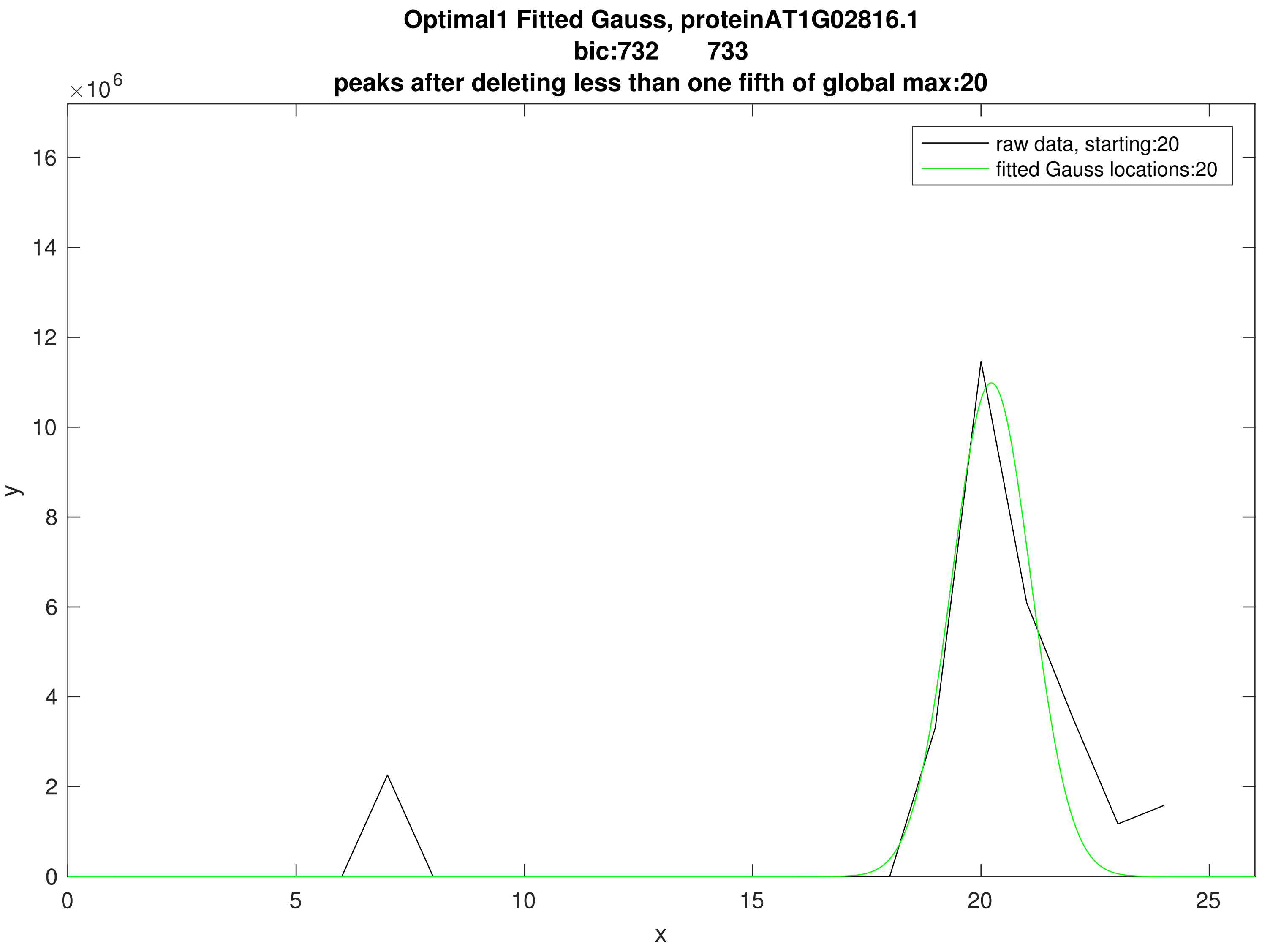
Optimal1 Fitted Gauss, proteinAT1G02816.1
bic:732 733
peaks after deleting less than one fifth of global max:20
×10^6
raw data, starting:20
fitted Gauss locations:20
y
x
0
2
4
6
8
10
12
14
16
0
5
10
15
20
25

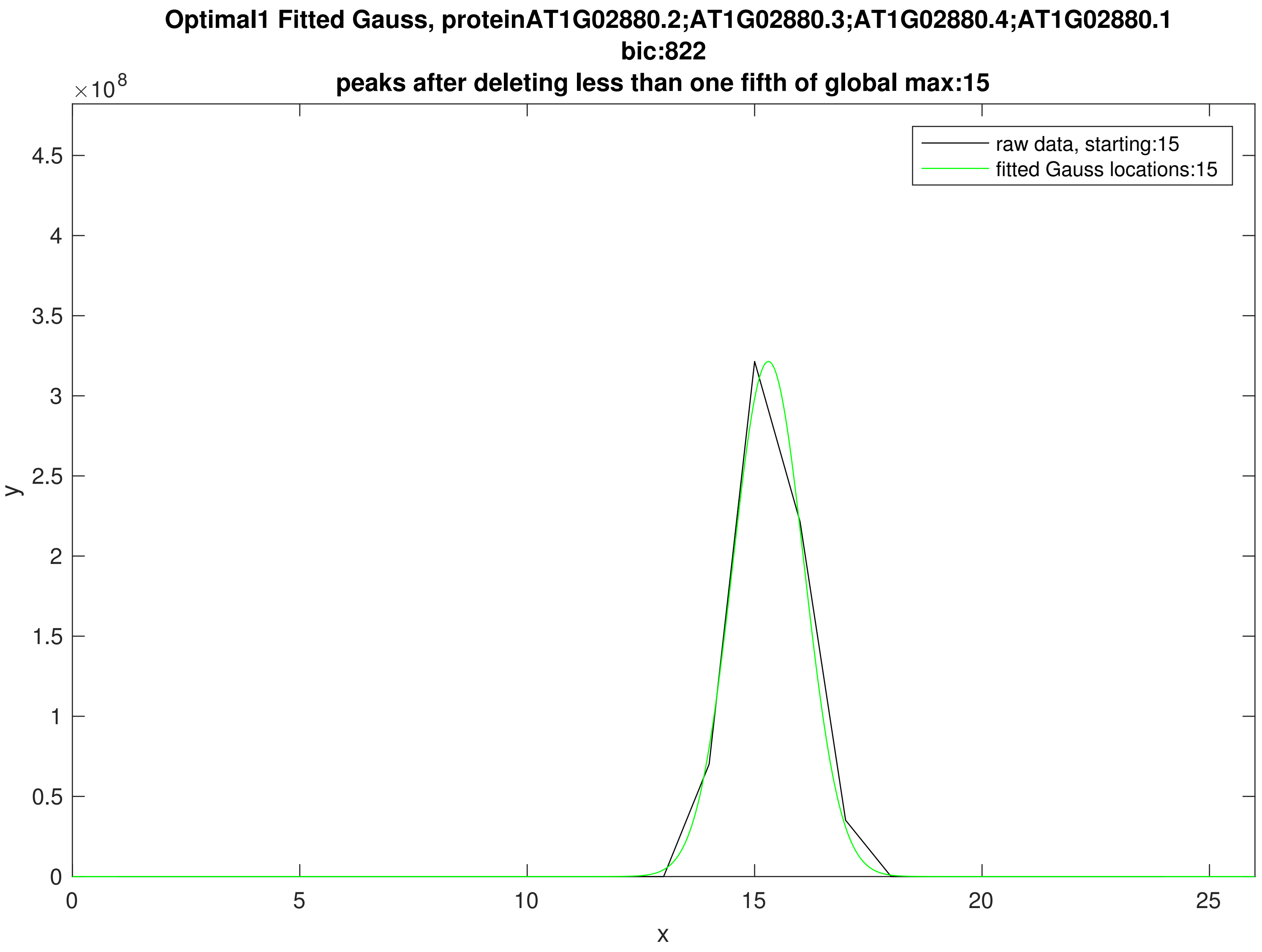

Optimal1 Fitted Gauss, proteinAT1G02880.2;AT1G02880.3;AT1G02880.4;AT1G02880.1
bic:822
peaks after deleting less than one fifth of global max:15
×10^8
raw data, starting:15
fitted Gauss locations:15
y
x
0
0.5
1
1.5
2
2.5
3
3.5
4
4.5
5
10
15
20
25

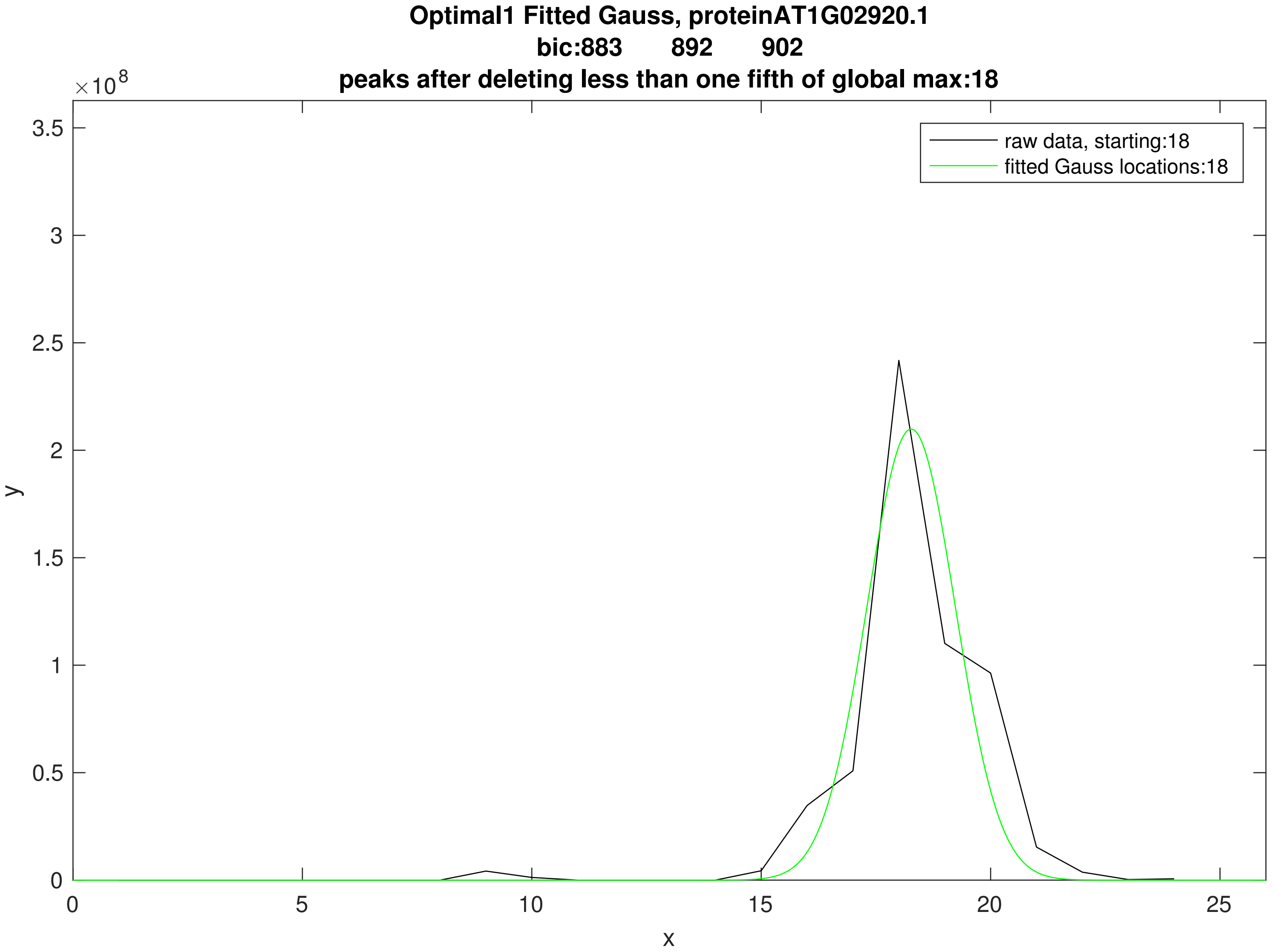

Optimal1 Fitted Gauss, proteinAT1G02920.1
bic:883 892 902
peaks after deleting less than one fifth of global max:18
×10^8
3.5
3
2.5
2
1.5
1
0.5
0
y
0
5
10
15
20
25
x
raw data, starting:18
fitted Gauss locations:18

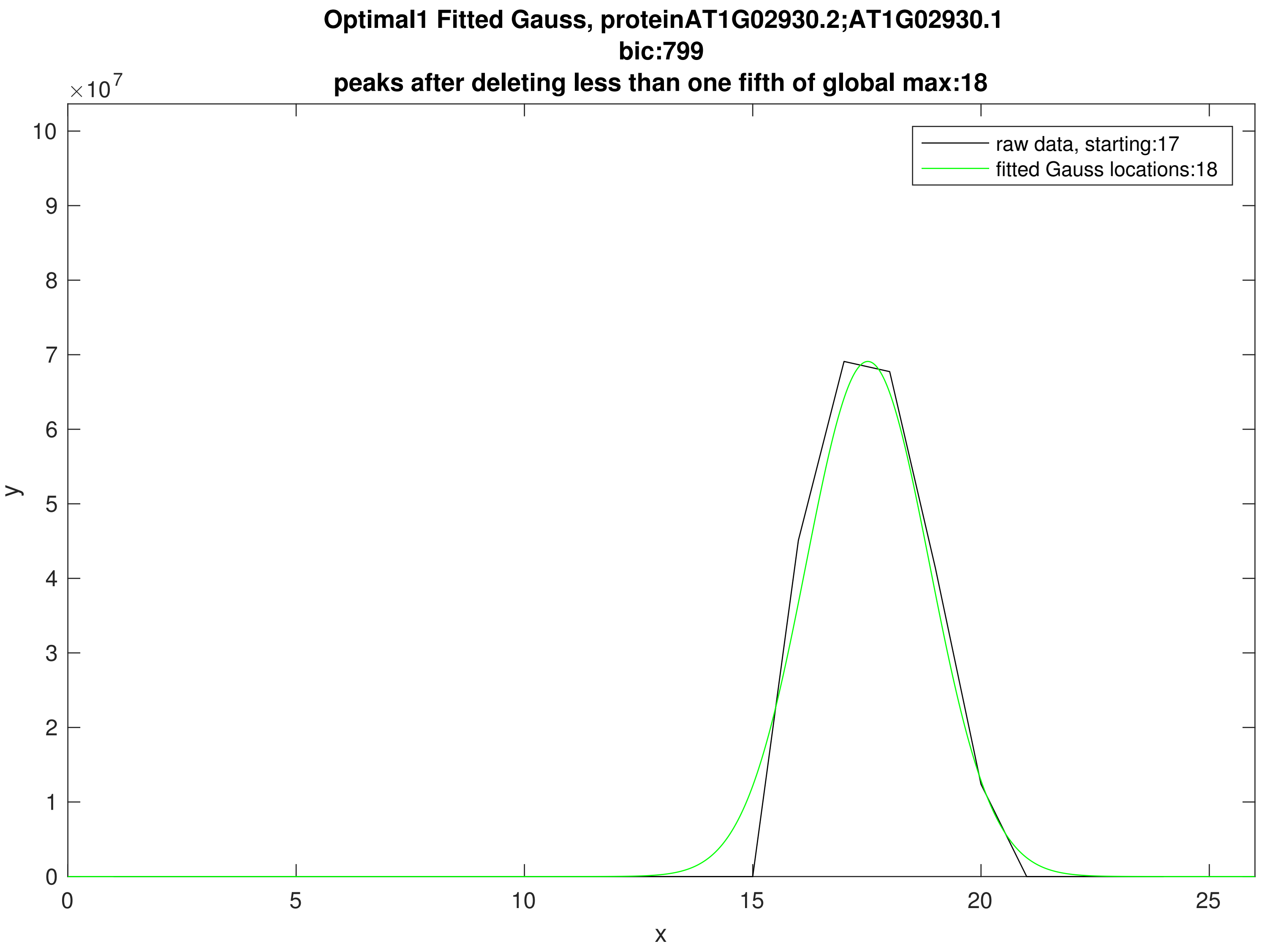
Optimal1 Fitted Gauss, proteinAT1G02930.2;AT1G02930.1
bic:799
peaks after deleting less than one fifth of global max:18
×10^7
raw data, starting:17
fitted Gauss locations:18
y
x
0
1
2
3
4
5
6
7
8
9
10
0
5
10
15
20
25

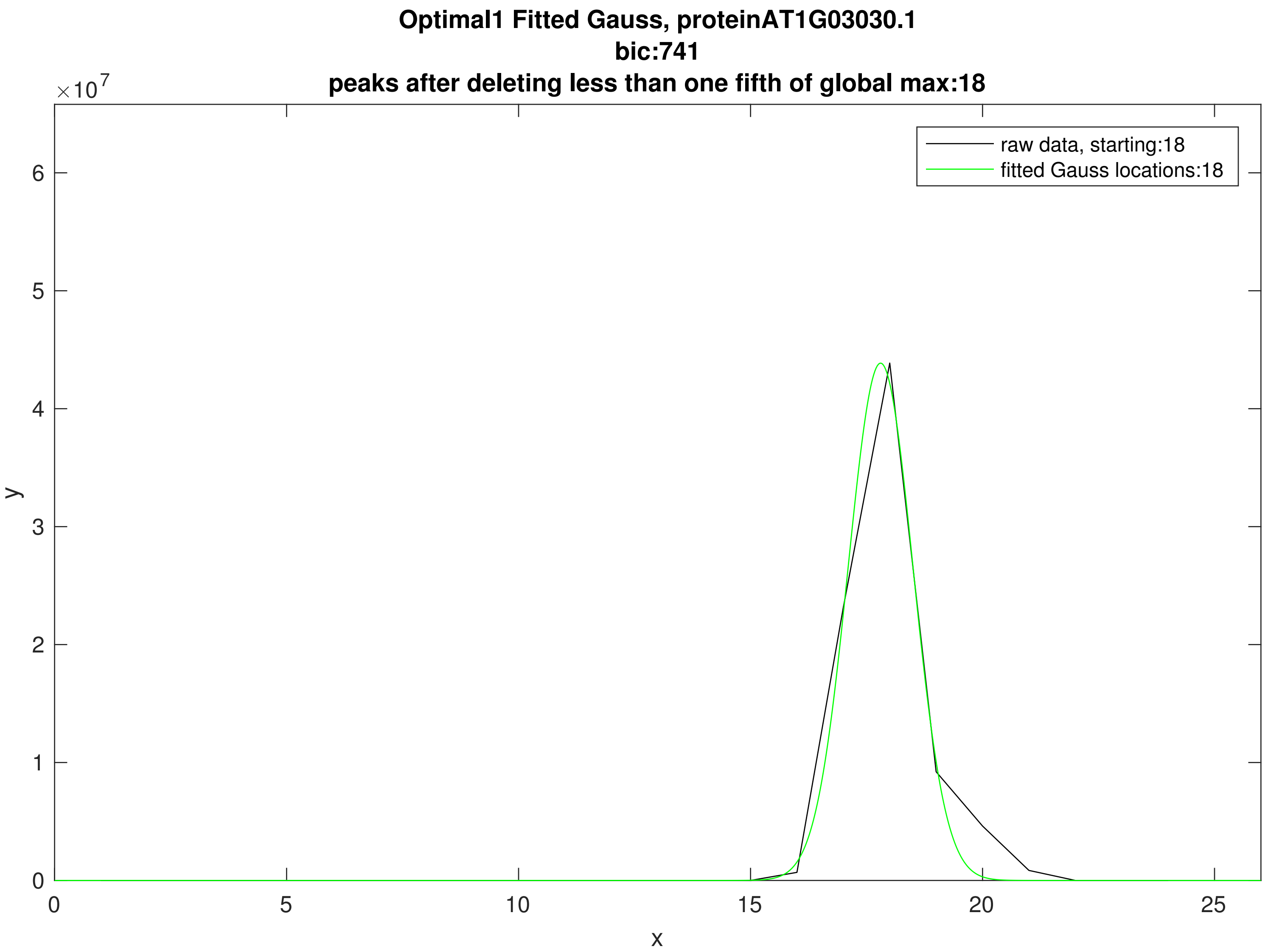
Optimal1 Fitted Gauss, proteinAT1G03030.1
bic:741
peaks after deleting less than one fifth of global max:18
×10^7
raw data, starting:18
fitted Gauss locations:18
y
x
0
1
2
3
4
5
6
0
5
10
15
20
25

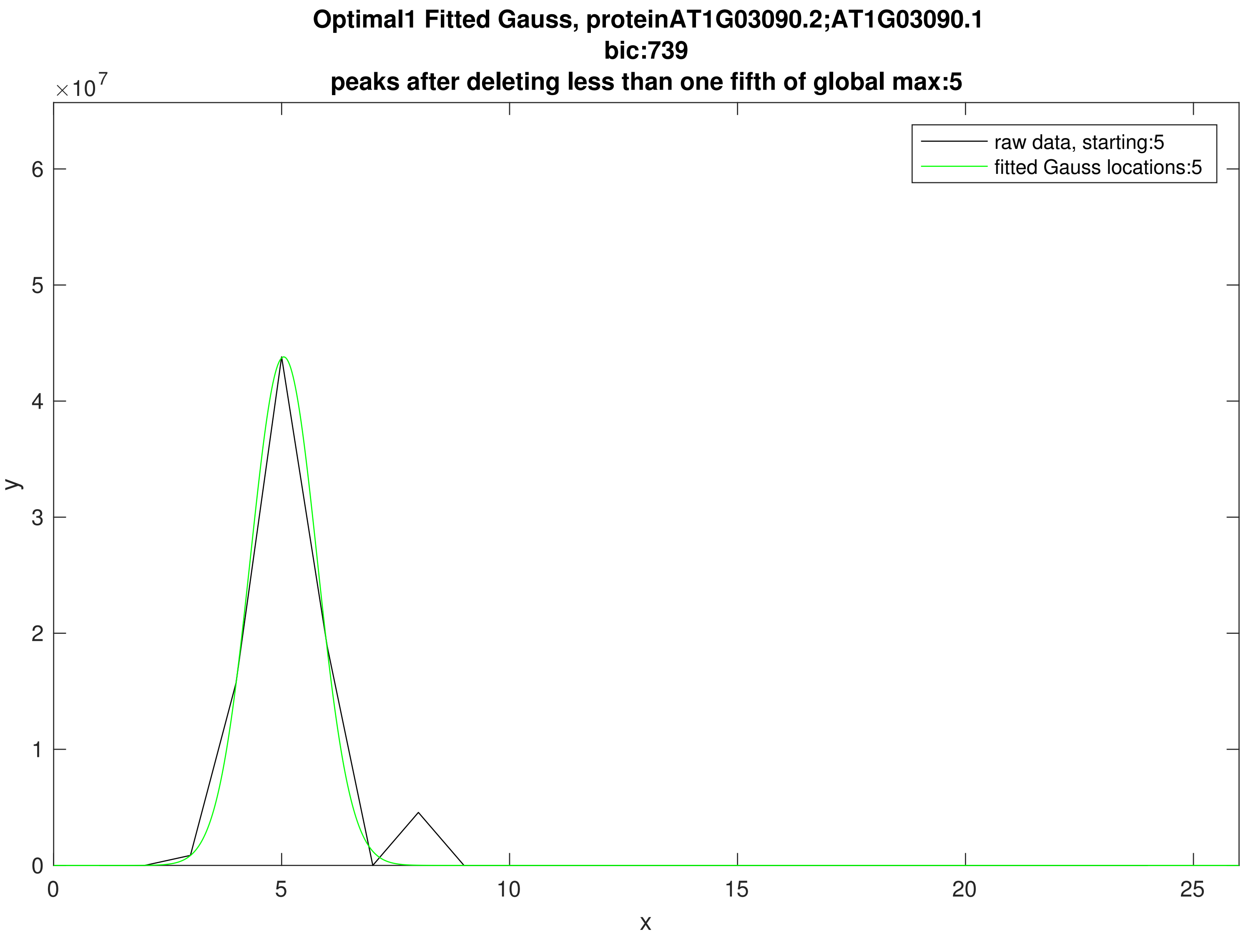

Optimal1 Fitted Gauss, proteinAT1G03090.2;AT1G03090.1
bic:739
peaks after deleting less than one fifth of global max:5
×10^7
raw data, starting:5
fitted Gauss locations:5
y
x
0
1
2
3
4
5
6
0
5
10
15
20
25

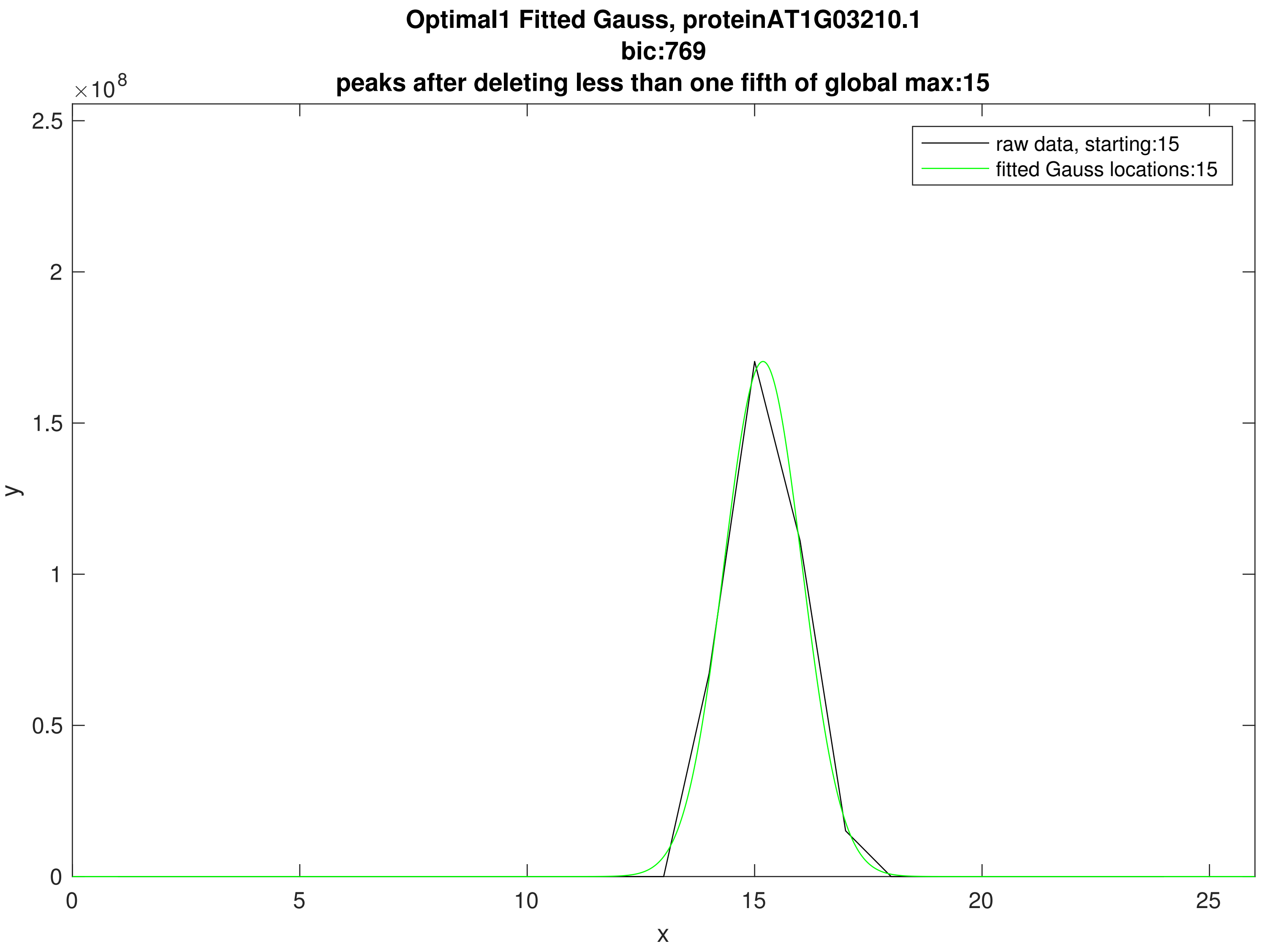

Optimal1 Fitted Gauss, proteinAT1G03210.1
bic:769
peaks after deleting less than one fifth of global max:15
×10^8
2.5
2
1.5
1
0.5
0
y
0
5
10
15
20
25
x
raw data, starting:15
fitted Gauss locations:15

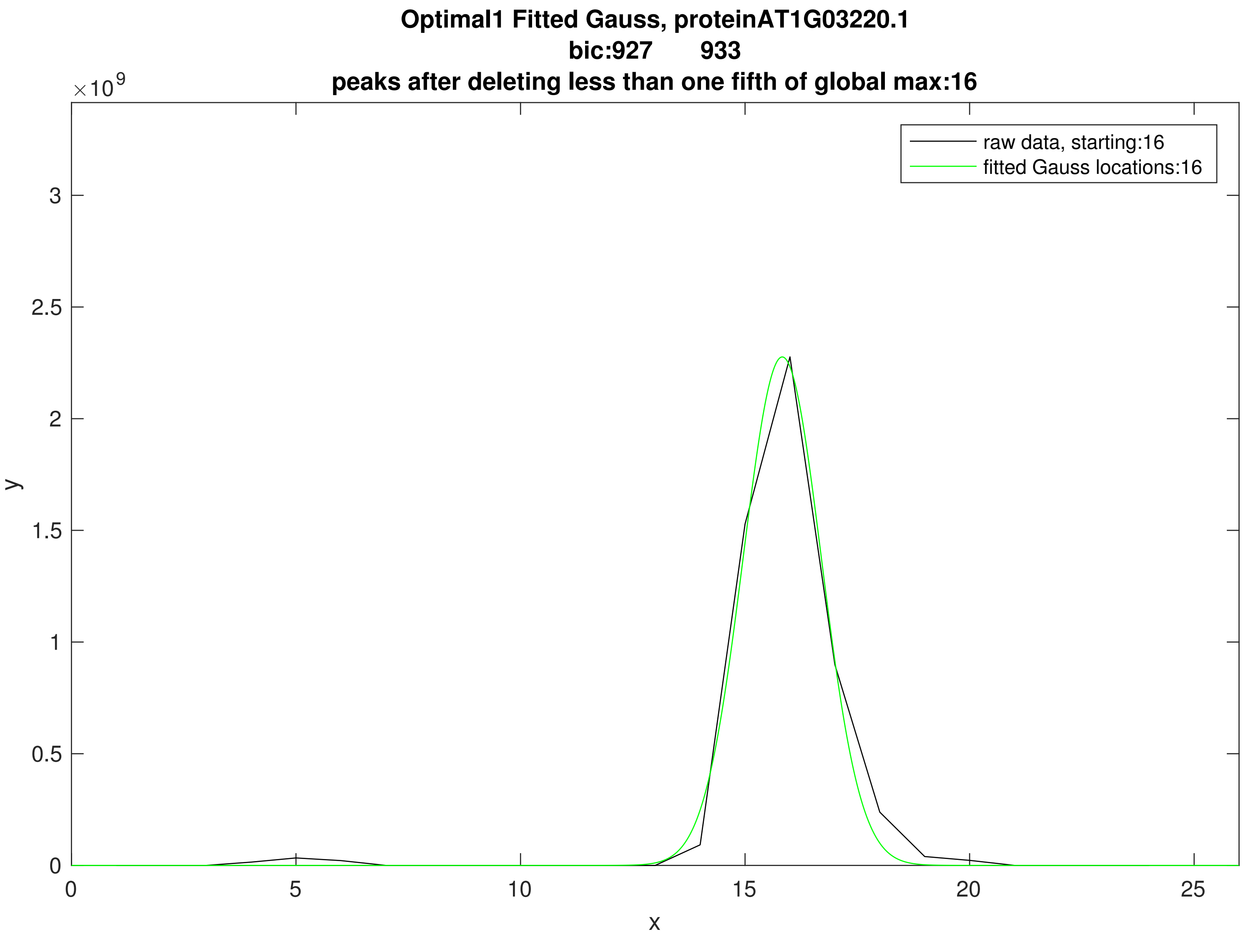
Optimal1 Fitted Gauss, proteinAT1G03220.1
bic:927 933
peaks after deleting less than one fifth of global max:16
×10^9
raw data, starting:16
fitted Gauss locations:16
y
x
0
0.5
1
1.5
2
2.5
3
0
5
10
15
20
25

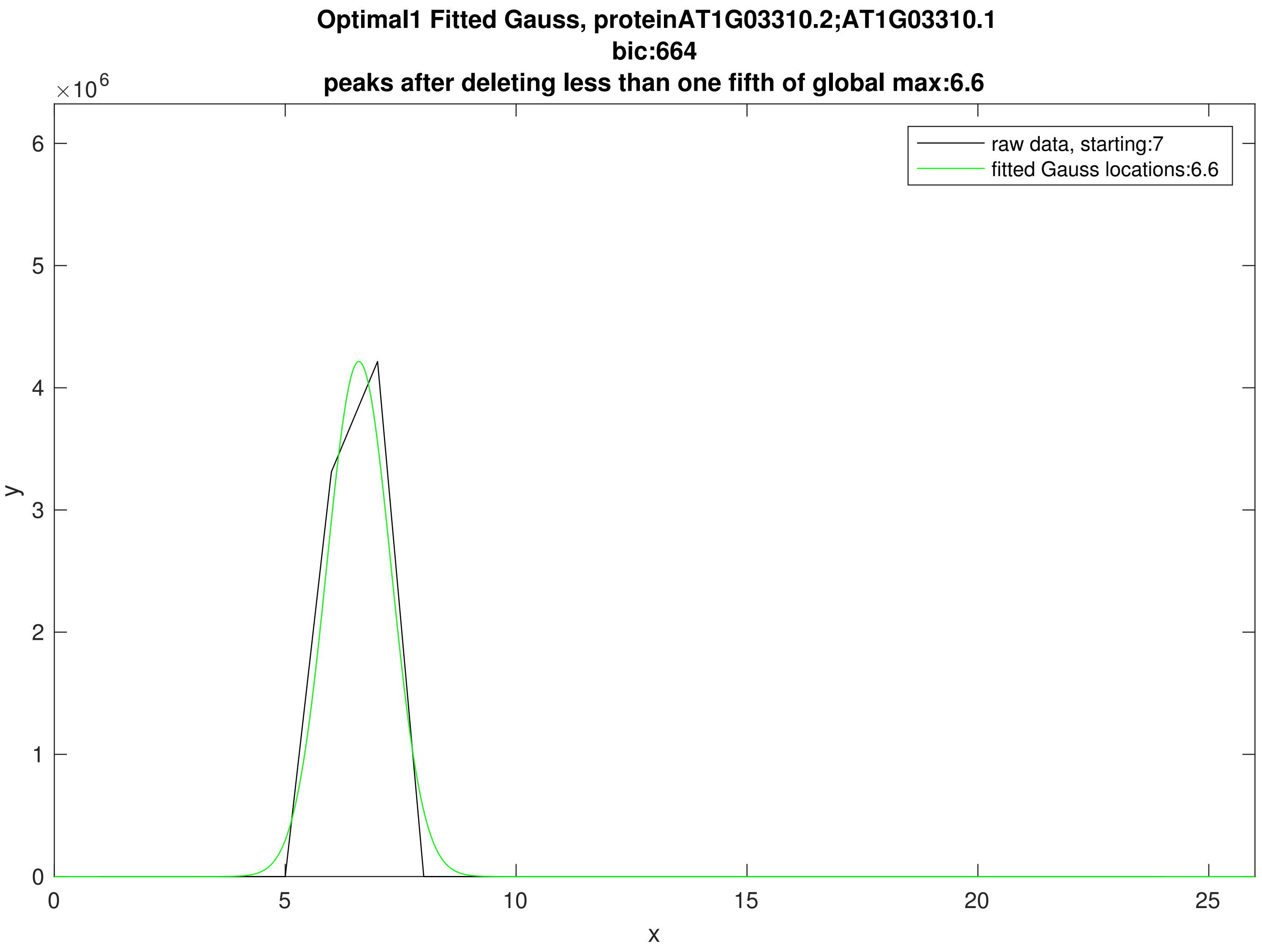

Optimal1 Fitted Gauss, proteinAT1G03310.2;AT1G03310.1
bic:664
peaks after deleting less than one fifth of global max:6.6
$\times 10^6$
raw data, starting:7
fitted Gauss locations:6.6
y
x
0
1
2
3
4
5
6
0
5
10
15
20
25

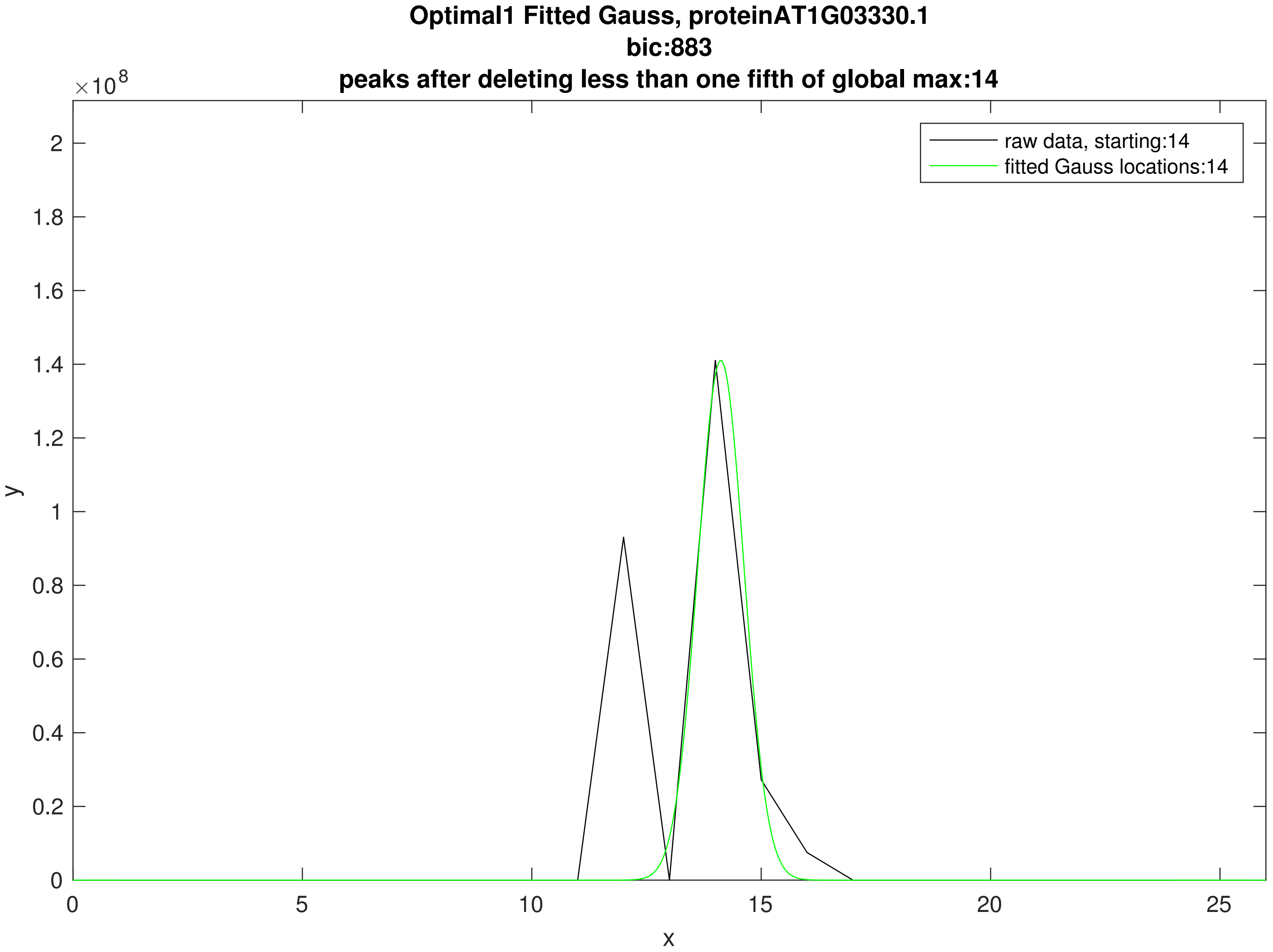
Optimal1 Fitted Gauss, proteinAT1G03330.1
bic:883
peaks after deleting less than one fifth of global max:14
×10^8
raw data, starting:14
fitted Gauss locations:14
y
x
0
0.2
0.4
0.6
0.8
1
1.2
1.4
1.6
1.8
2
5
10
15
20
25

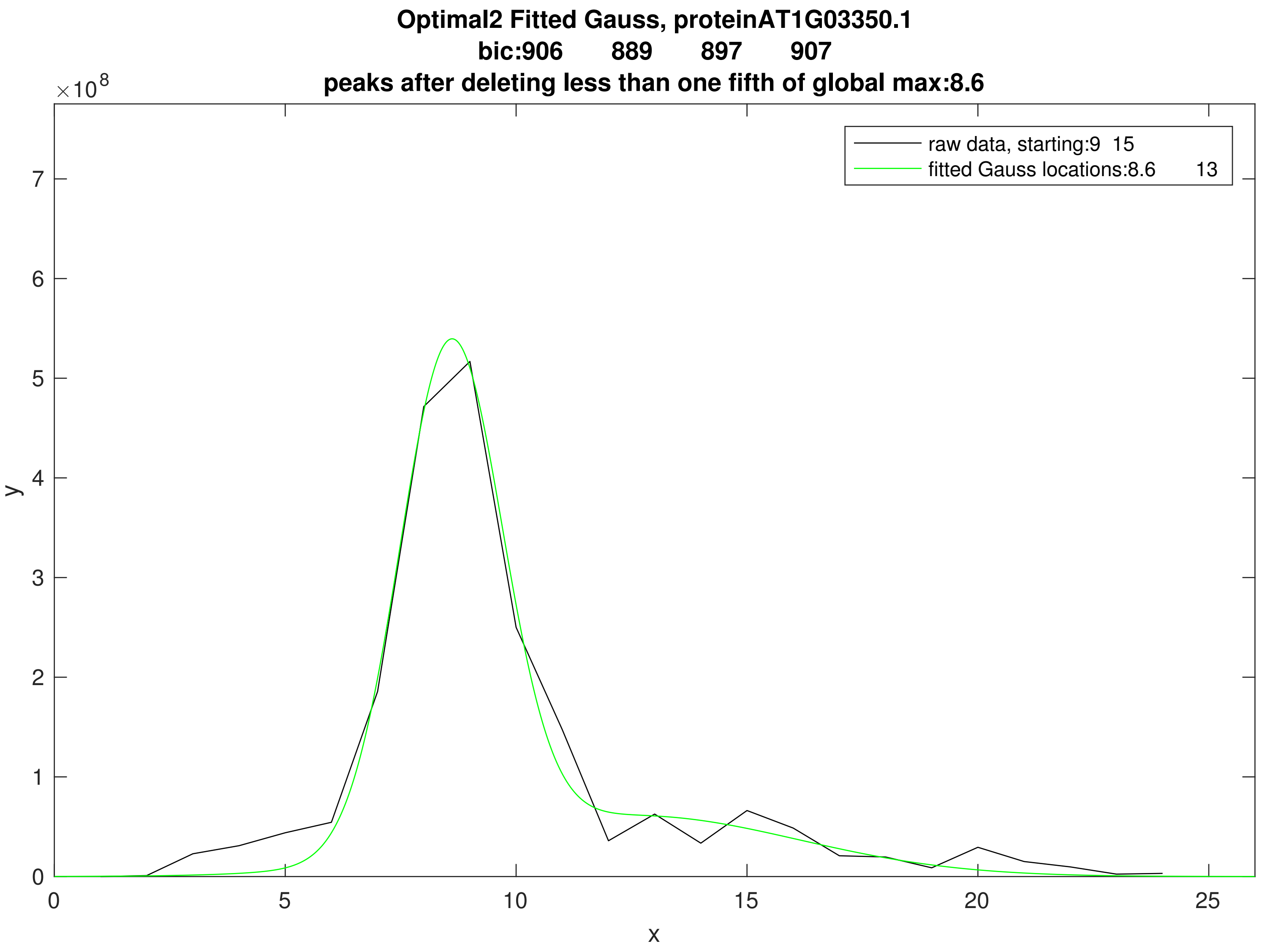
Optimal2 Fitted Gauss, proteinAT1G03350.1
bic:906 889 897 907
peaks after deleting less than one fifth of global max:8.6
×10^8
raw data, starting:9 15
fitted Gauss locations:8.6 13
y
x
0
1
2
3
4
5
6
7
0
5
10
15
20
25

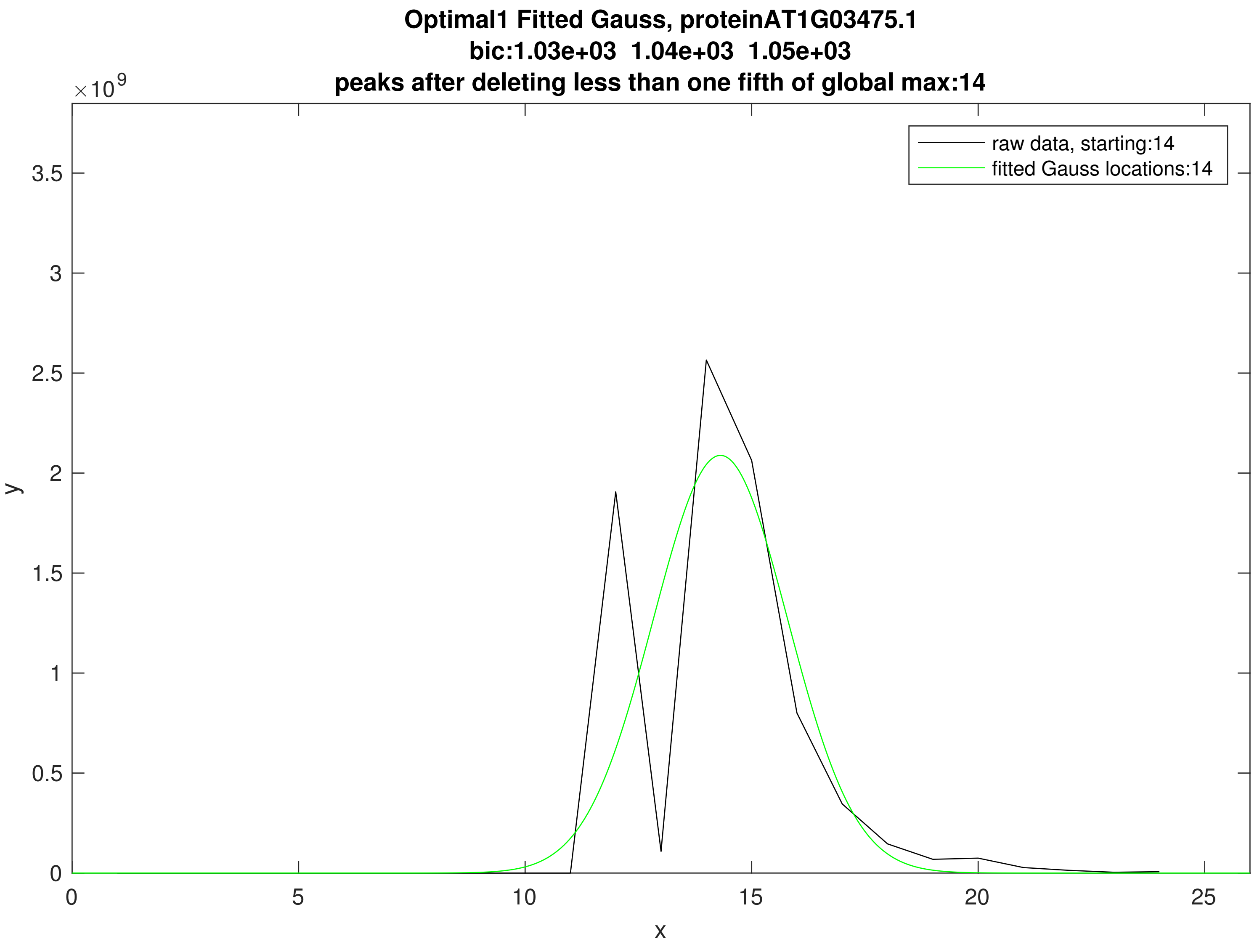

Optimal1 Fitted Gauss, proteinAT1G03475.1
bic:1.03e+03 1.04e+03 1.05e+03
peaks after deleting less than one fifth of global max:14
×10^9
raw data, starting:14
fitted Gauss locations:14
3.5
3
2.5
2
1.5
1
0.5
0
y
0
5
10
15
20
25
x

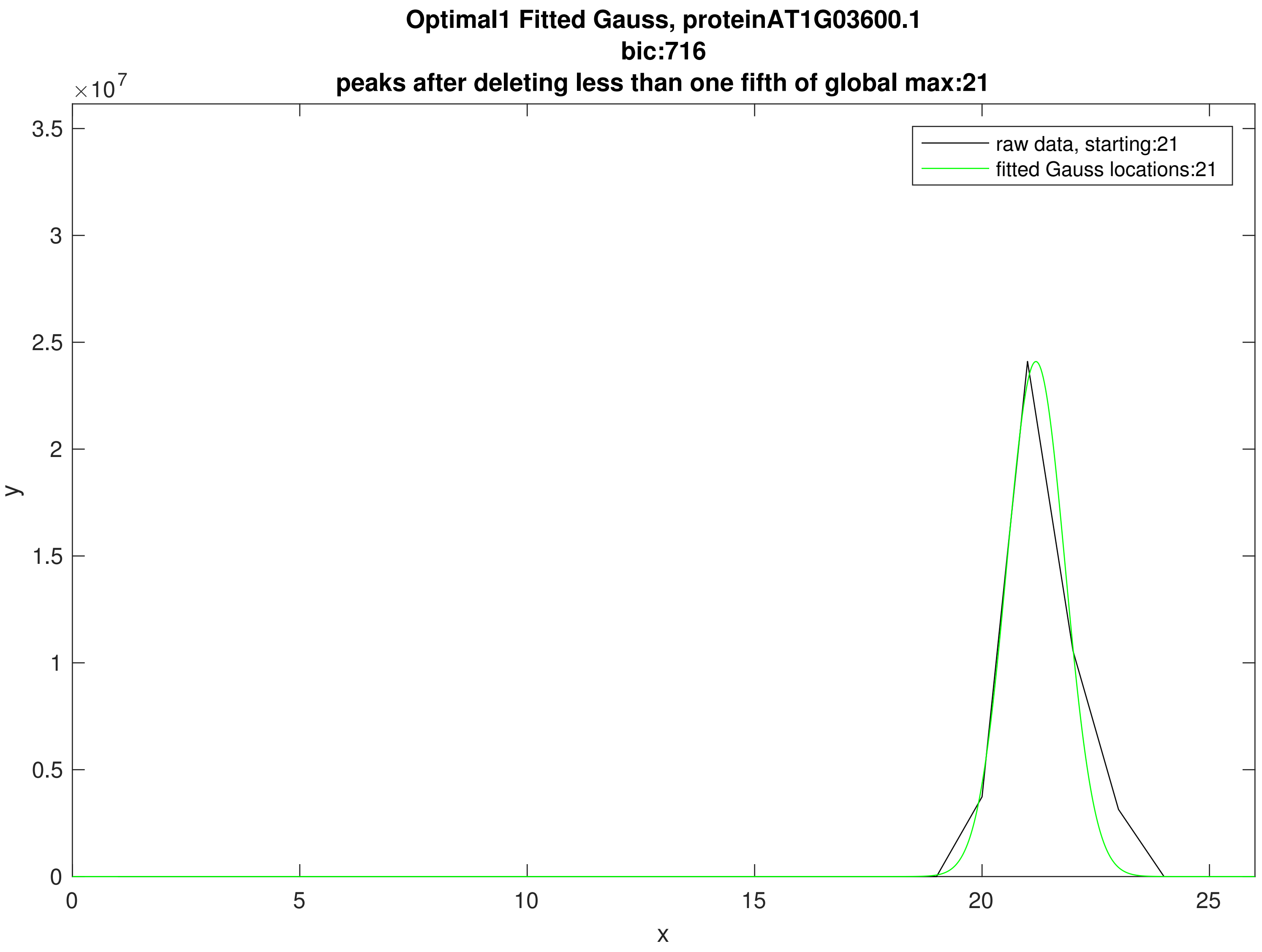
Optimal1 Fitted Gauss, proteinAT1G03600.1
bic:716
peaks after deleting less than one fifth of global max:21
×10^7
3.5
3
2.5
2
1.5
1
0.5
0
y
0
5
10
15
20
25
x
raw data, starting:21
fitted Gauss locations:21

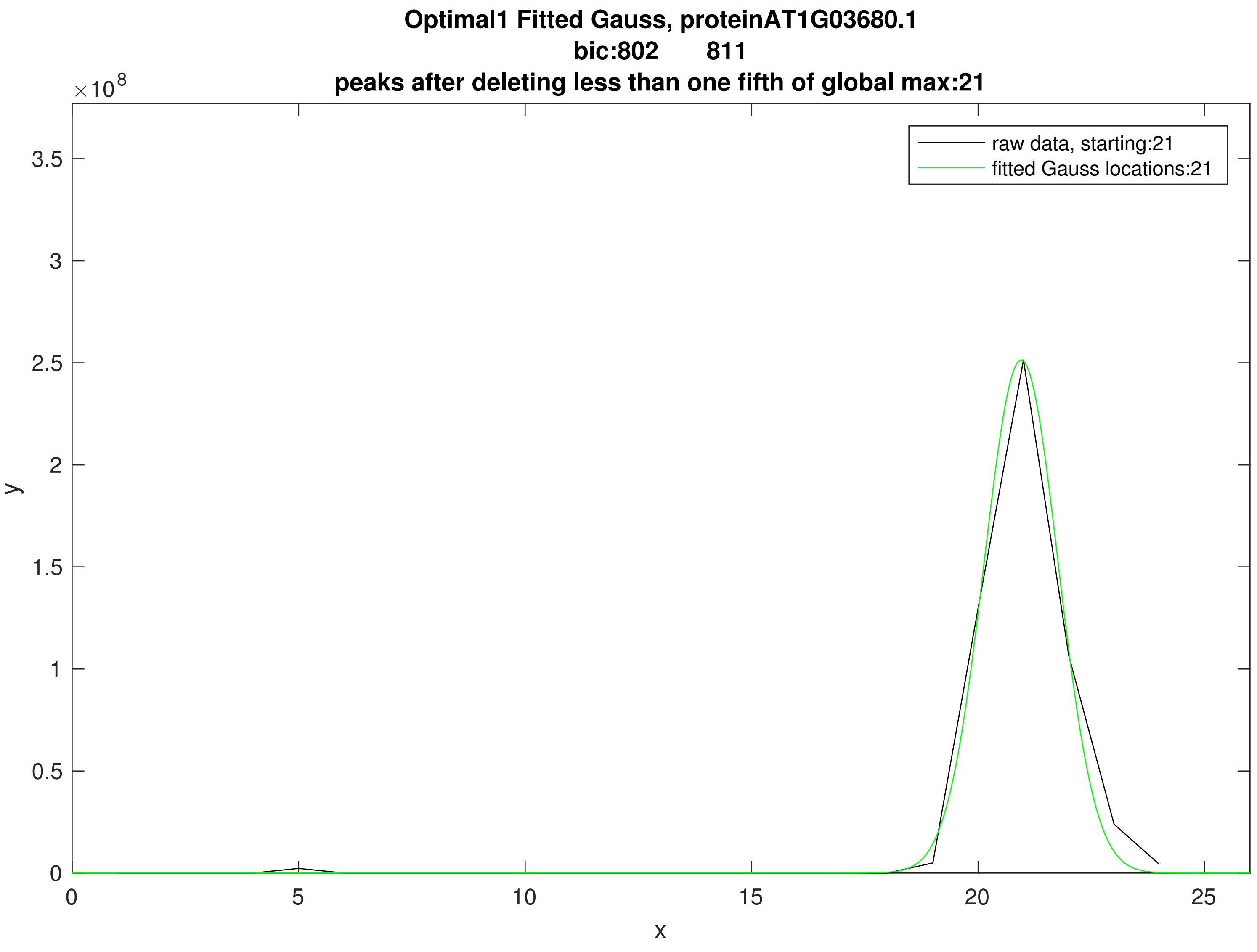
Optimal1 Fitted Gauss, proteinAT1G03680.1
bic:802 811
peaks after deleting less than one fifth of global max:21
×10^8
3.5
3
2.5
2
1.5
1
0.5
0
y
0
5
10
15
20
25
x
raw data, starting:21
fitted Gauss locations:21

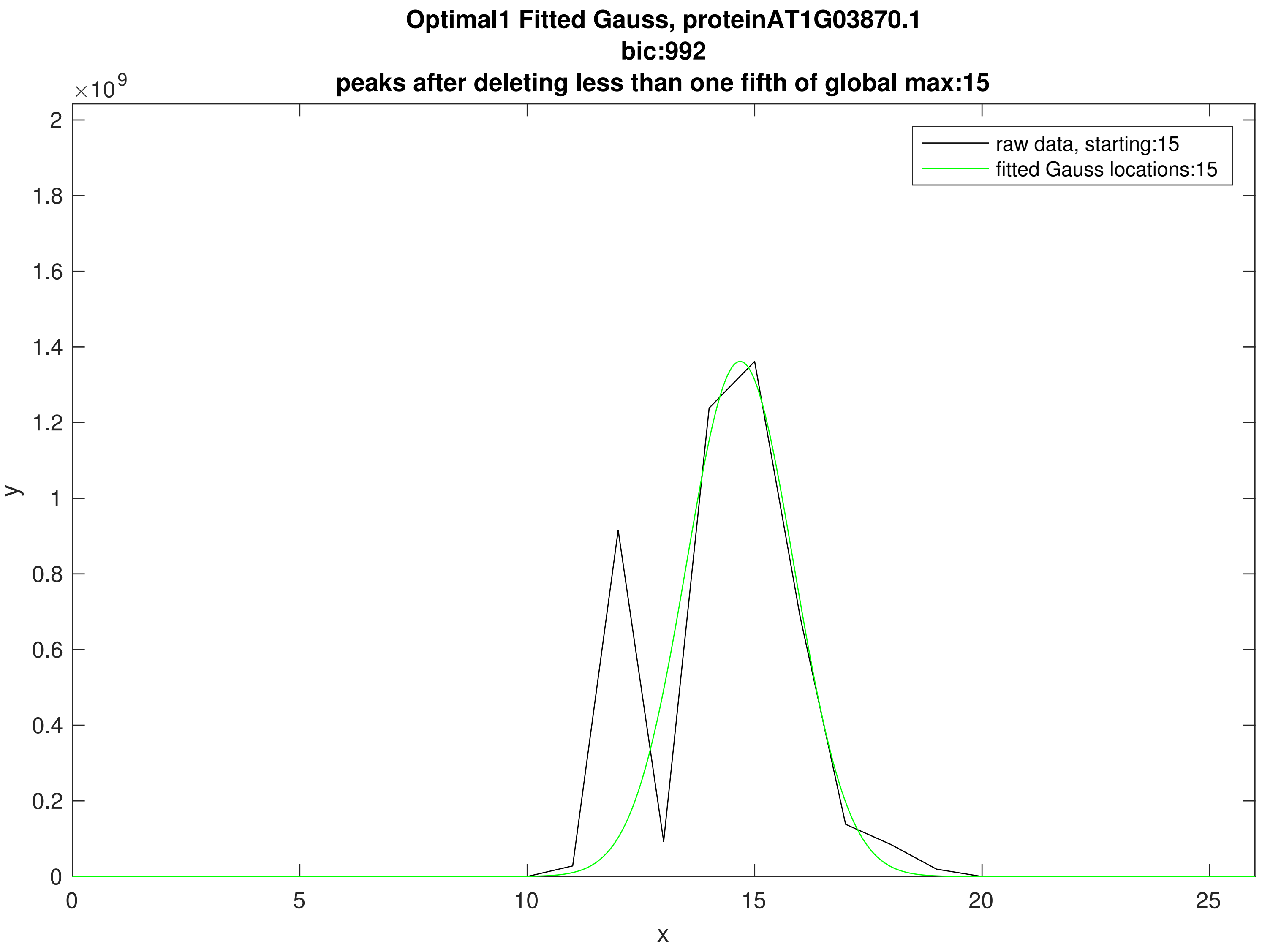
Optimal1 Fitted Gauss, proteinAT1G03870.1
bic:992
peaks after deleting less than one fifth of global max:15
×10^9
raw data, starting:15
fitted Gauss locations:15
y
x
0
0.2
0.4
0.6
0.8
1
1.2
1.4
1.6
1.8
2
0
5
10
15
20
25

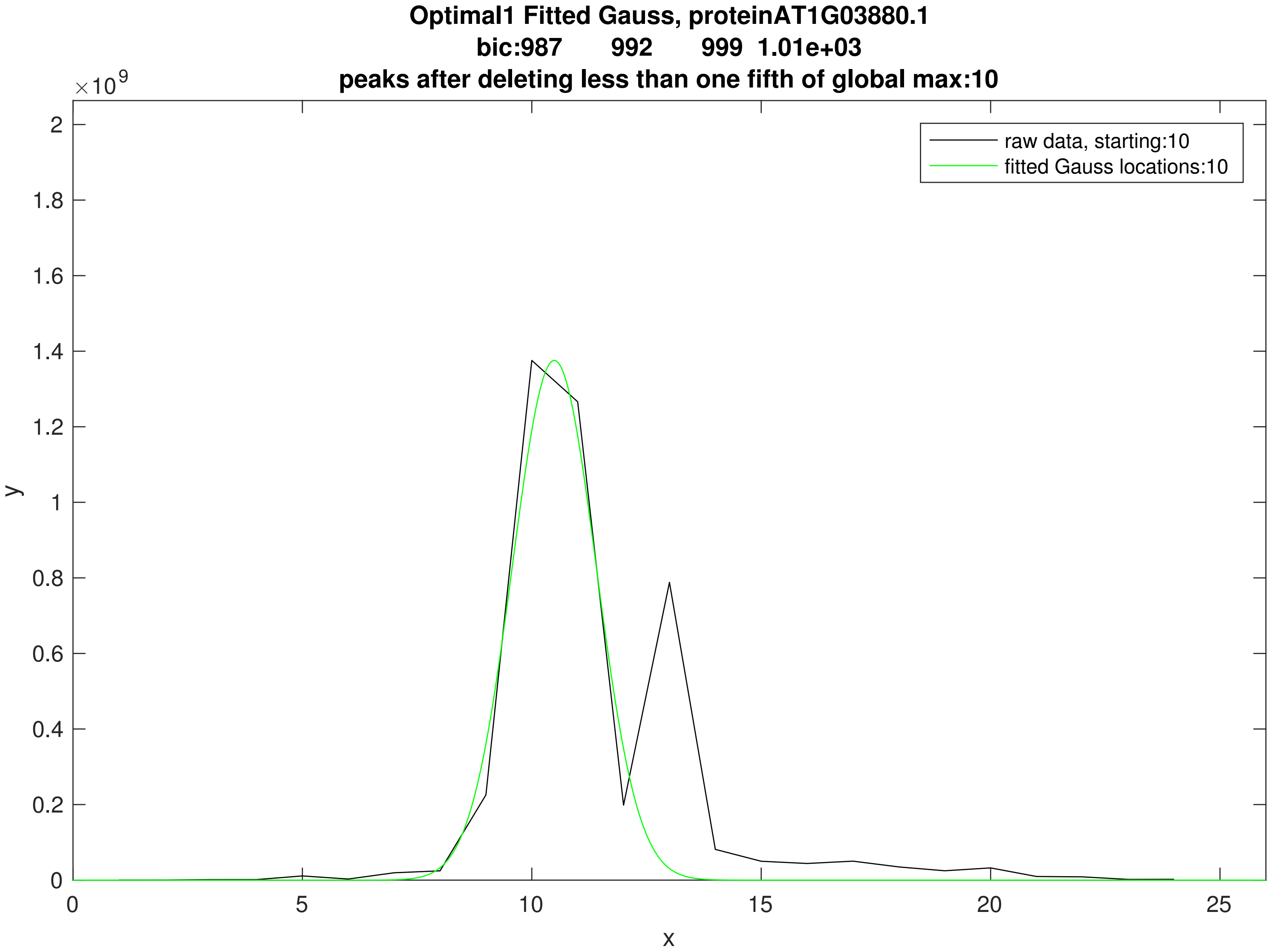

Optimal1 Fitted Gauss, proteinAT1G03880.1
bic:987 992 999 1.01e+03
peaks after deleting less than one fifth of global max:10
×10^9
raw data, starting:10
fitted Gauss locations:10
y
x
0
0.2
0.4
0.6
0.8
1
1.2
1.4
1.6
1.8
2
0
5
10
15
20
25

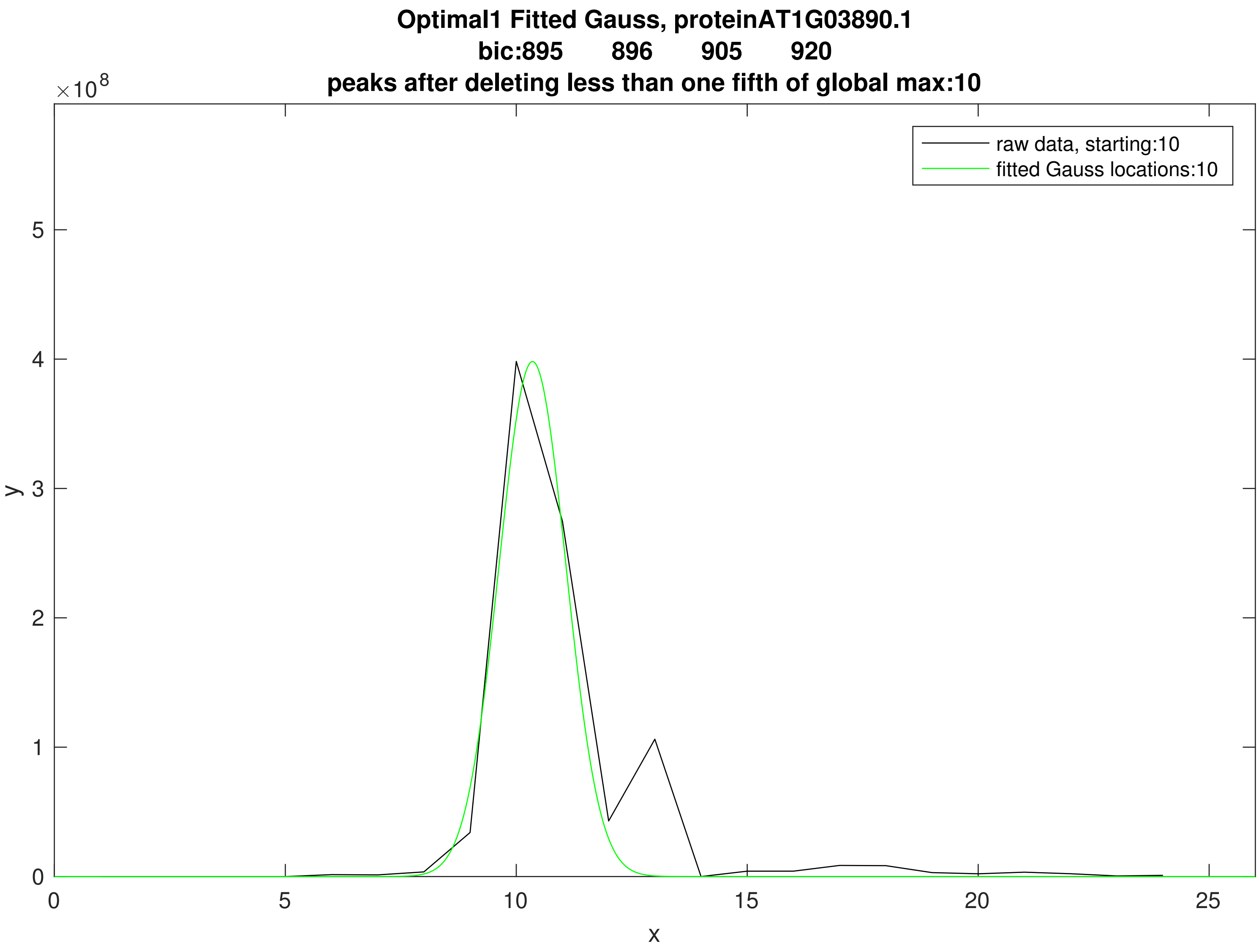

Optimal1 Fitted Gauss, proteinAT1G03890.1
bic:895 896 905 920
peaks after deleting less than one fifth of global max:10
×10^8
raw data, starting:10
fitted Gauss locations:10
y
x
0
1
2
3
4
5
10
15
20
25

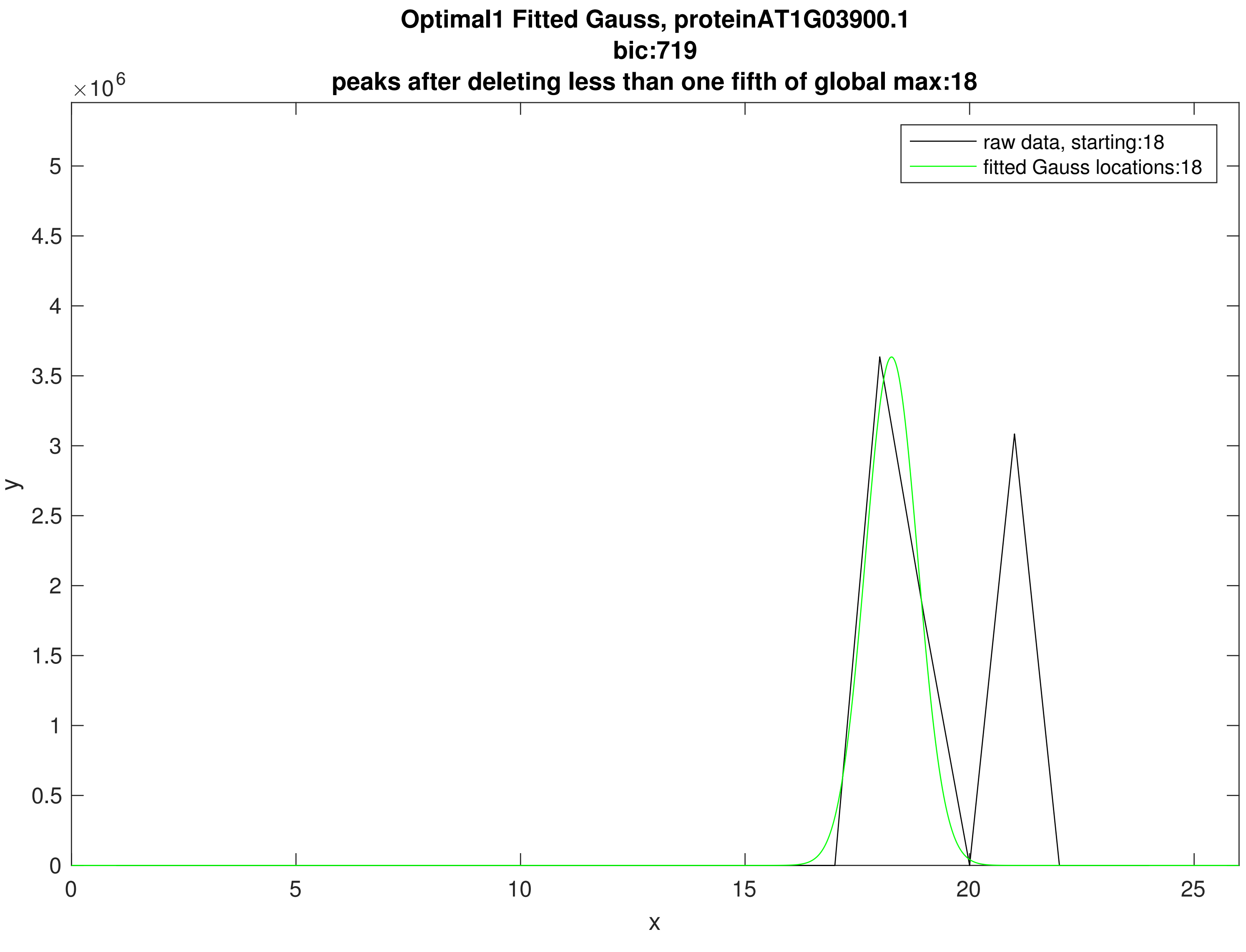
Optimal1 Fitted Gauss, proteinAT1G03900.1
bic:719
peaks after deleting less than one fifth of global max:18
×10^6
raw data, starting:18
fitted Gauss locations:18
y
x
0
0.5
1
1.5
2
2.5
3
3.5
4
4.5
5
0
5
10
15
20
25

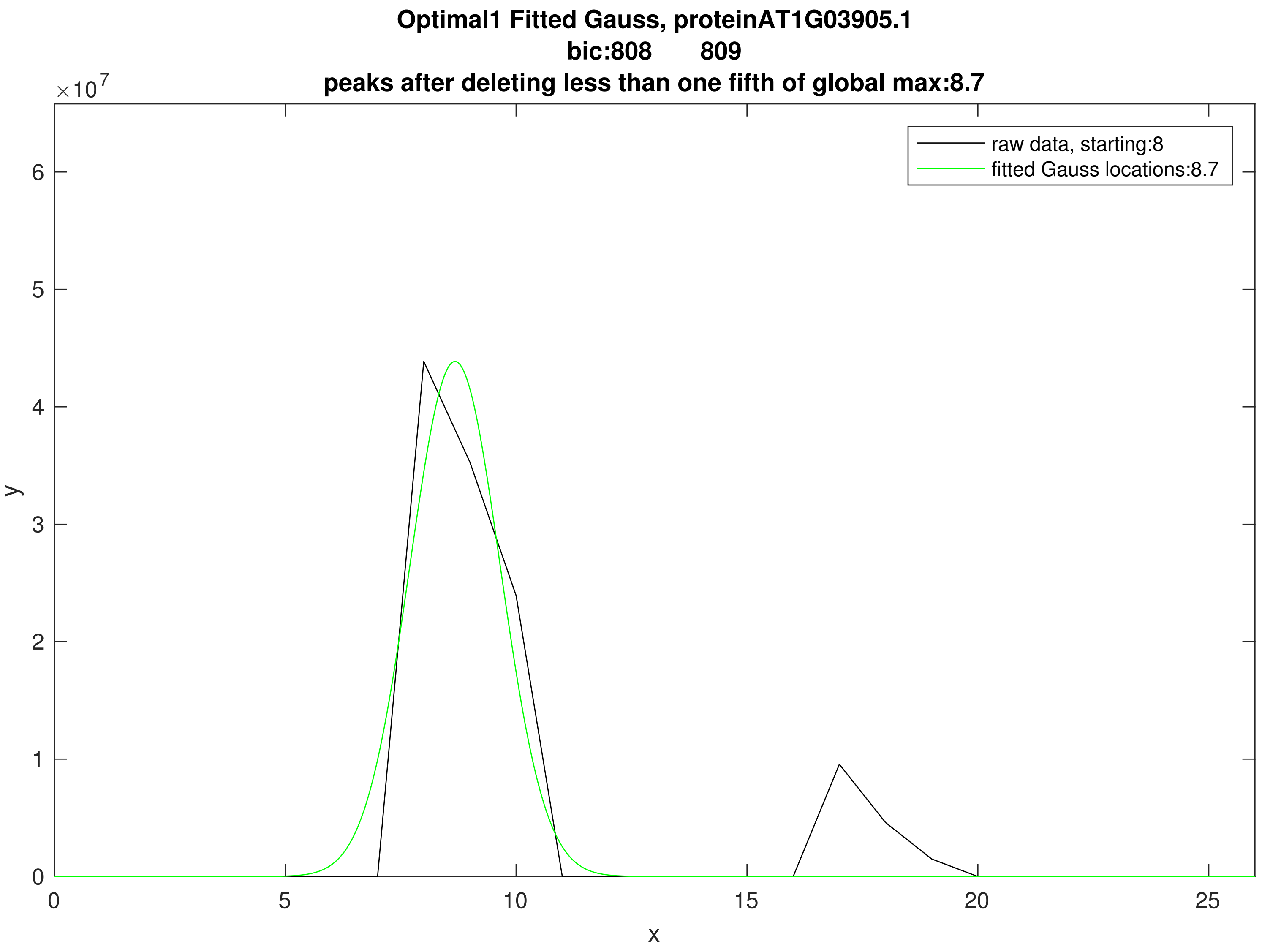

Optimal1 Fitted Gauss, proteinAT1G03905.1
bic:808 809
peaks after deleting less than one fifth of global max:8.7
×10^7
raw data, starting:8
fitted Gauss locations:8.7
y
x
0
1
2
3
4
5
6
0
5
10
15
20
25

**Optimal1 Fitted Gauss, proteinAT1G04080.2;AT1G04080.1;AT1G04080.3**
**bic:647**
**peaks after deleting less than one fifth of global max:6.7**

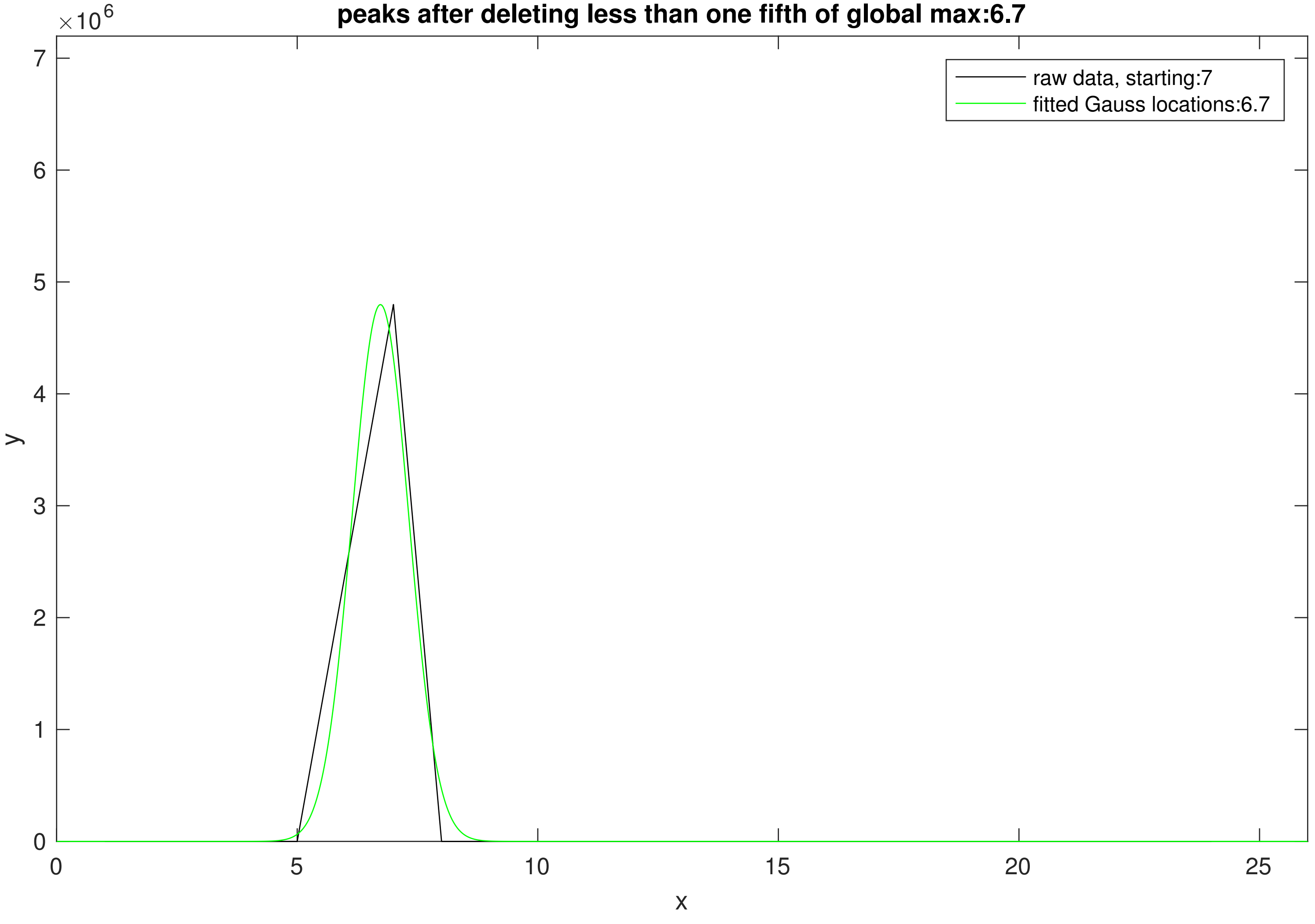

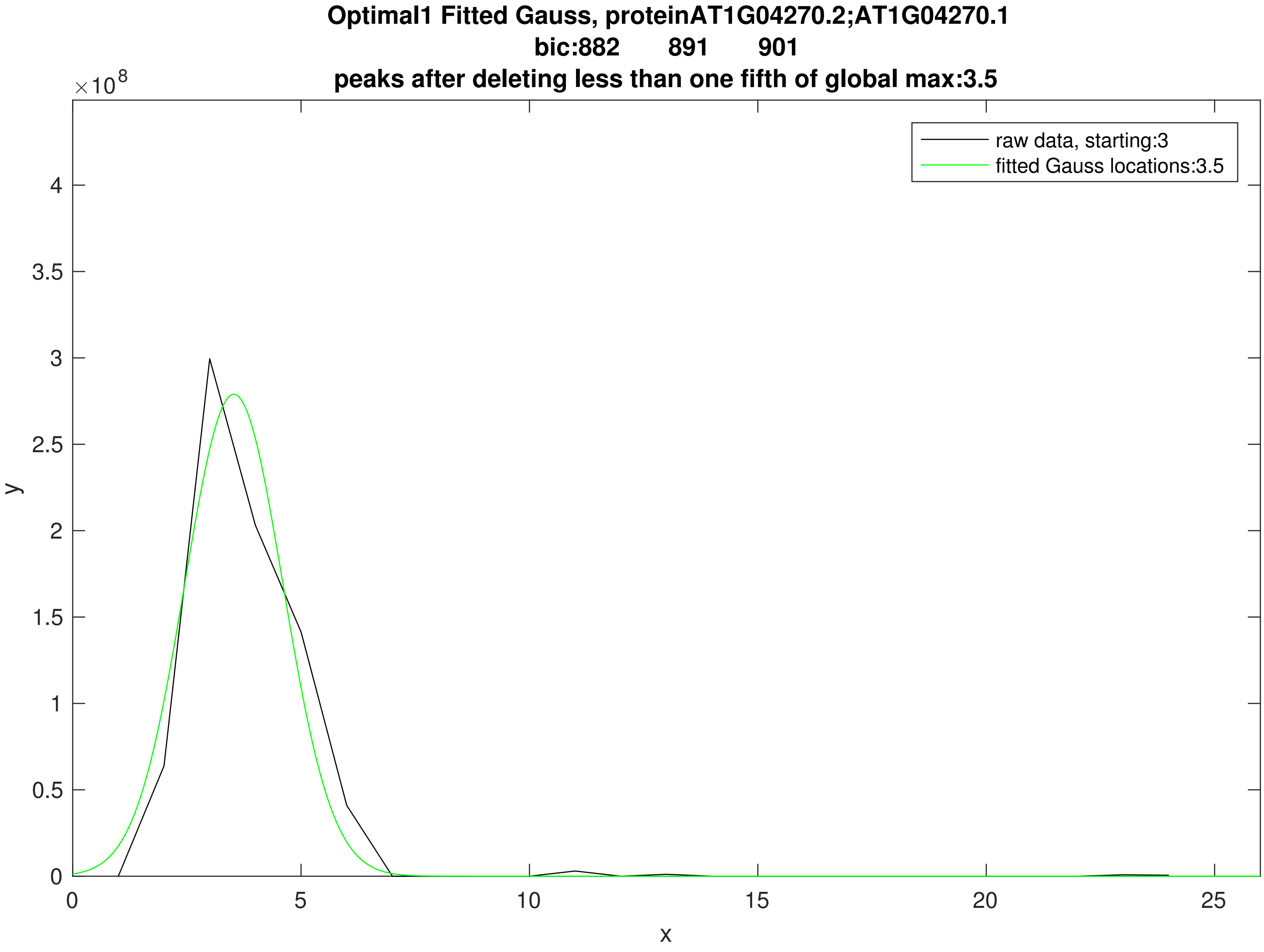
Optimal1 Fitted Gauss, proteinAT1G04270.2;AT1G04270.1
bic:882 891 901
peaks after deleting less than one fifth of global max:3.5
×10^8
raw data, starting:3
fitted Gauss locations:3.5
y
x
0
0.5
1
1.5
2
2.5
3
3.5
4
0
5
10
15
20
25

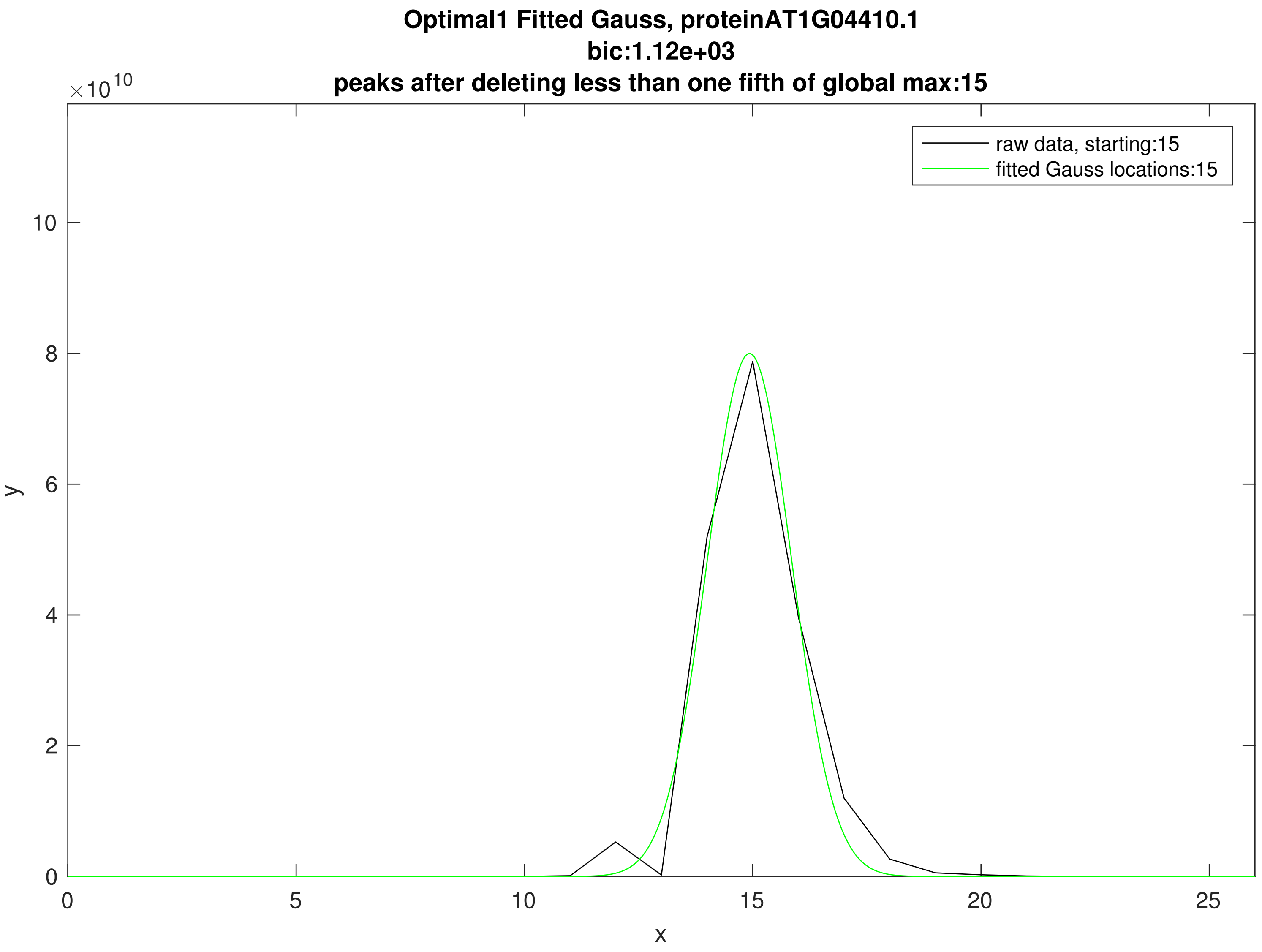
Optimal1 Fitted Gauss, proteinAT1G04410.1
bic:1.12e+03
peaks after deleting less than one fifth of global max:15
×10^10
raw data, starting:15
fitted Gauss locations:15
10
8
6
4
2
0
y
0
5
10
15
20
25
x

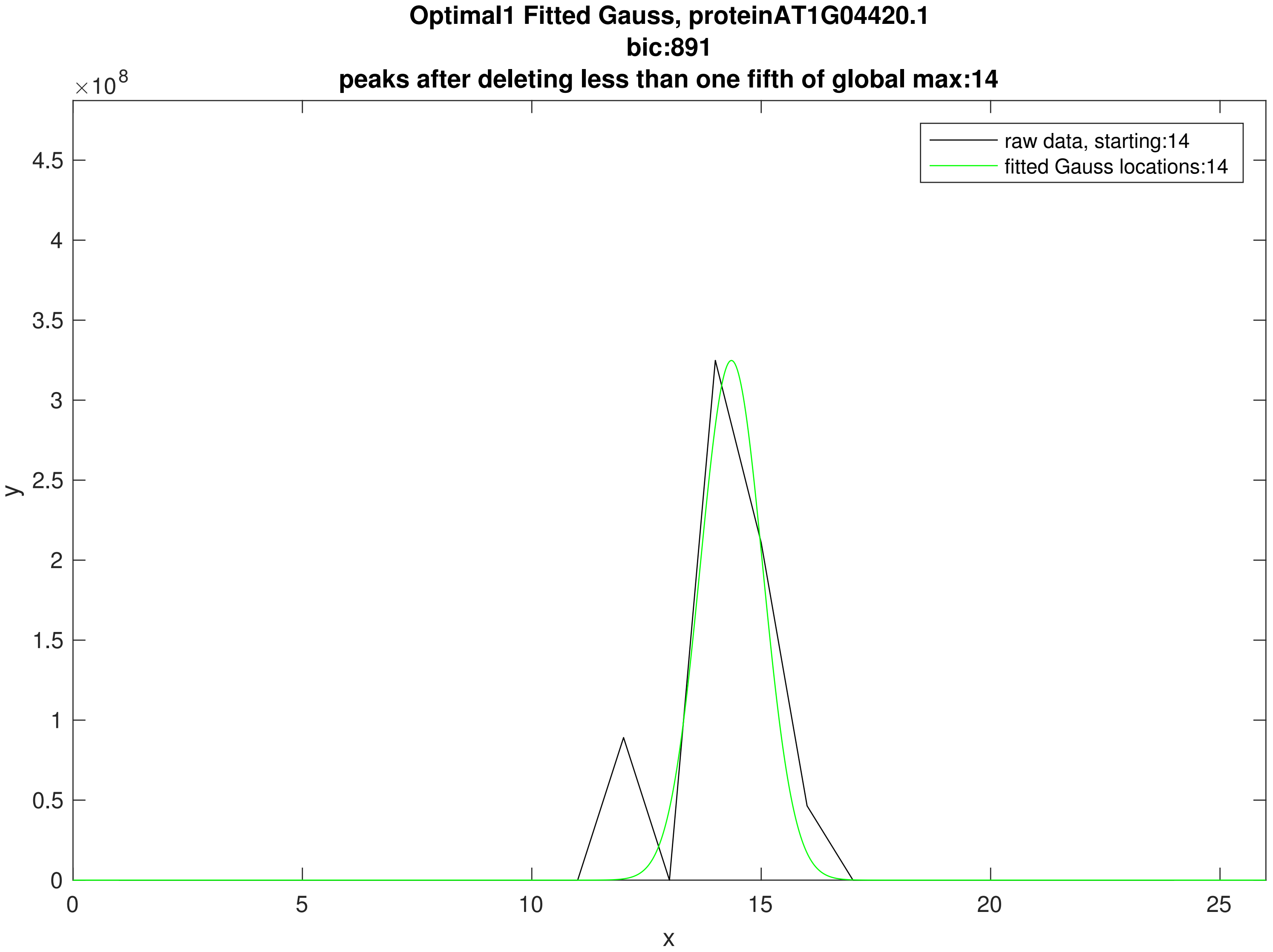

Optimal1 Fitted Gauss, proteinAT1G04420.1
bic:891
peaks after deleting less than one fifth of global max:14
×10^8
4.5
4
3.5
3
2.5
2
1.5
1
0.5
0
y
0
5
10
15
20
25
x
raw data, starting:14
fitted Gauss locations:14

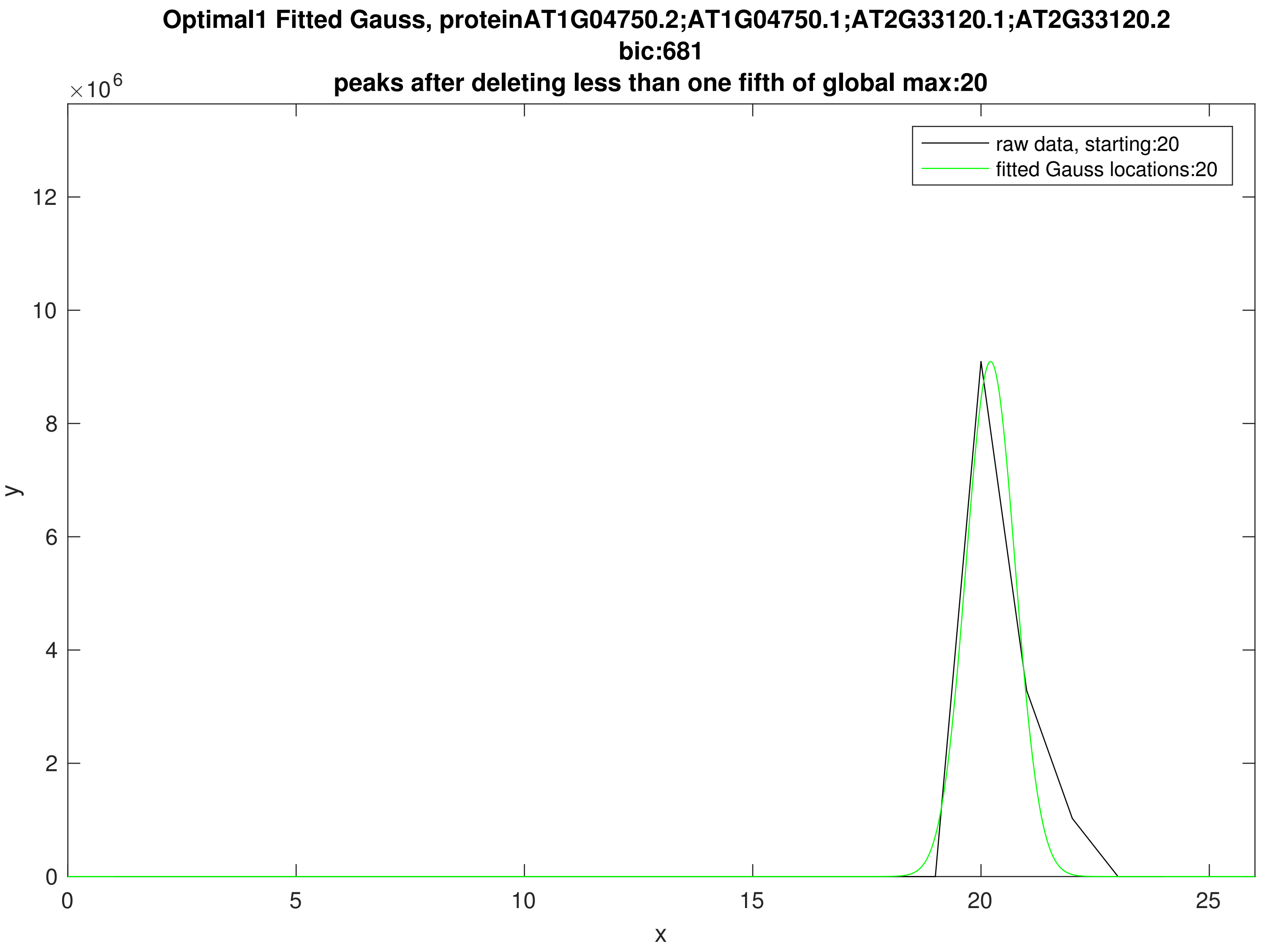

Optimal1 Fitted Gauss, proteinAT1G04750.2;AT1G04750.1;AT2G33120.1;AT2G33120.2
bic:681
peaks after deleting less than one fifth of global max:20
×10^6
raw data, starting:20
fitted Gauss locations:20
y
x
0
2
4
6
8
10
12
0
5
10
15
20
25

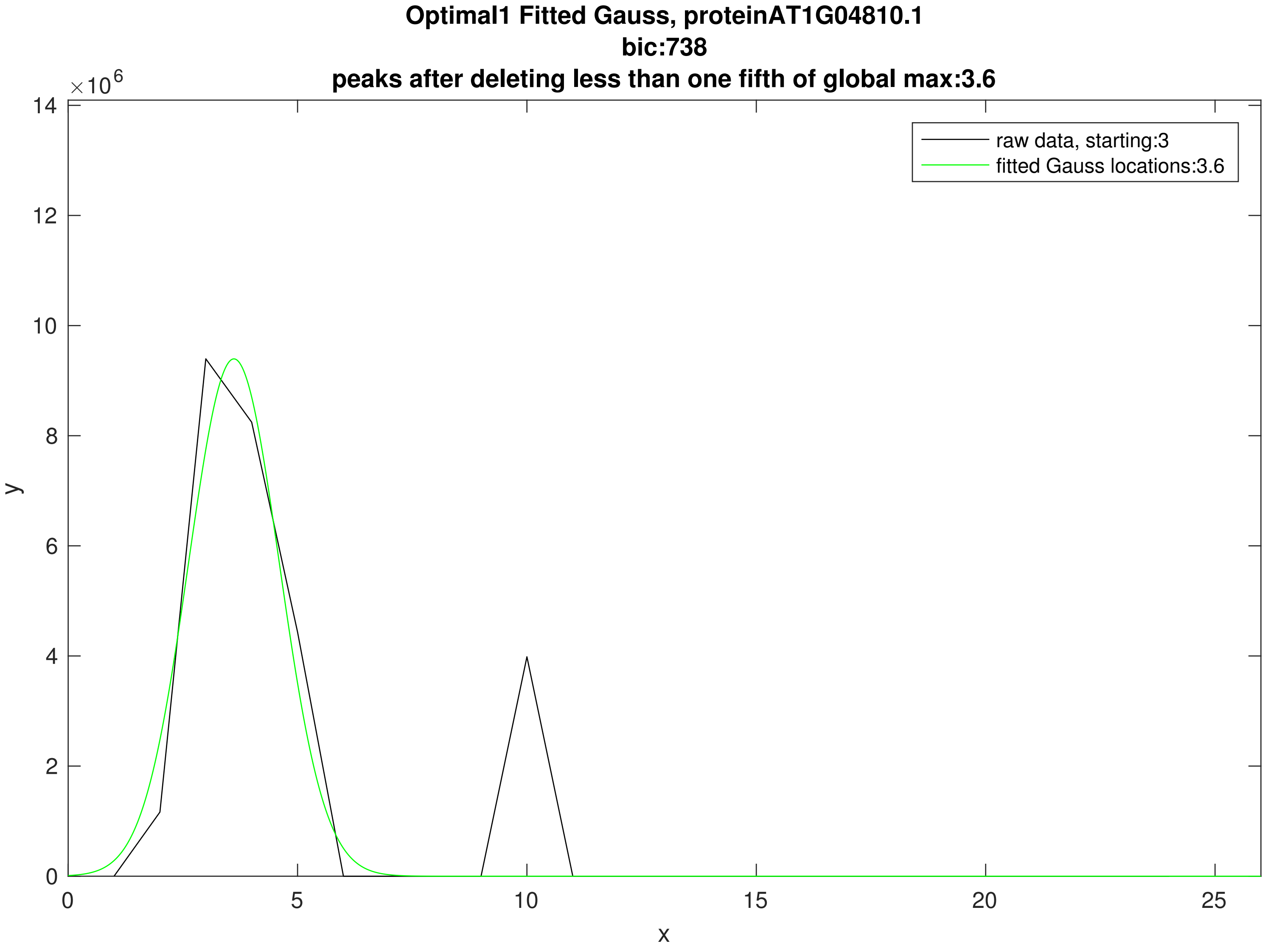
Optimal1 Fitted Gauss, proteinAT1G04810.1
bic:738
peaks after deleting less than one fifth of global max:3.6
×10^6
14
12
10
8
6
4
2
0
y
0
5
10
15
20
25
x
raw data, starting:3
fitted Gauss locations:3.6

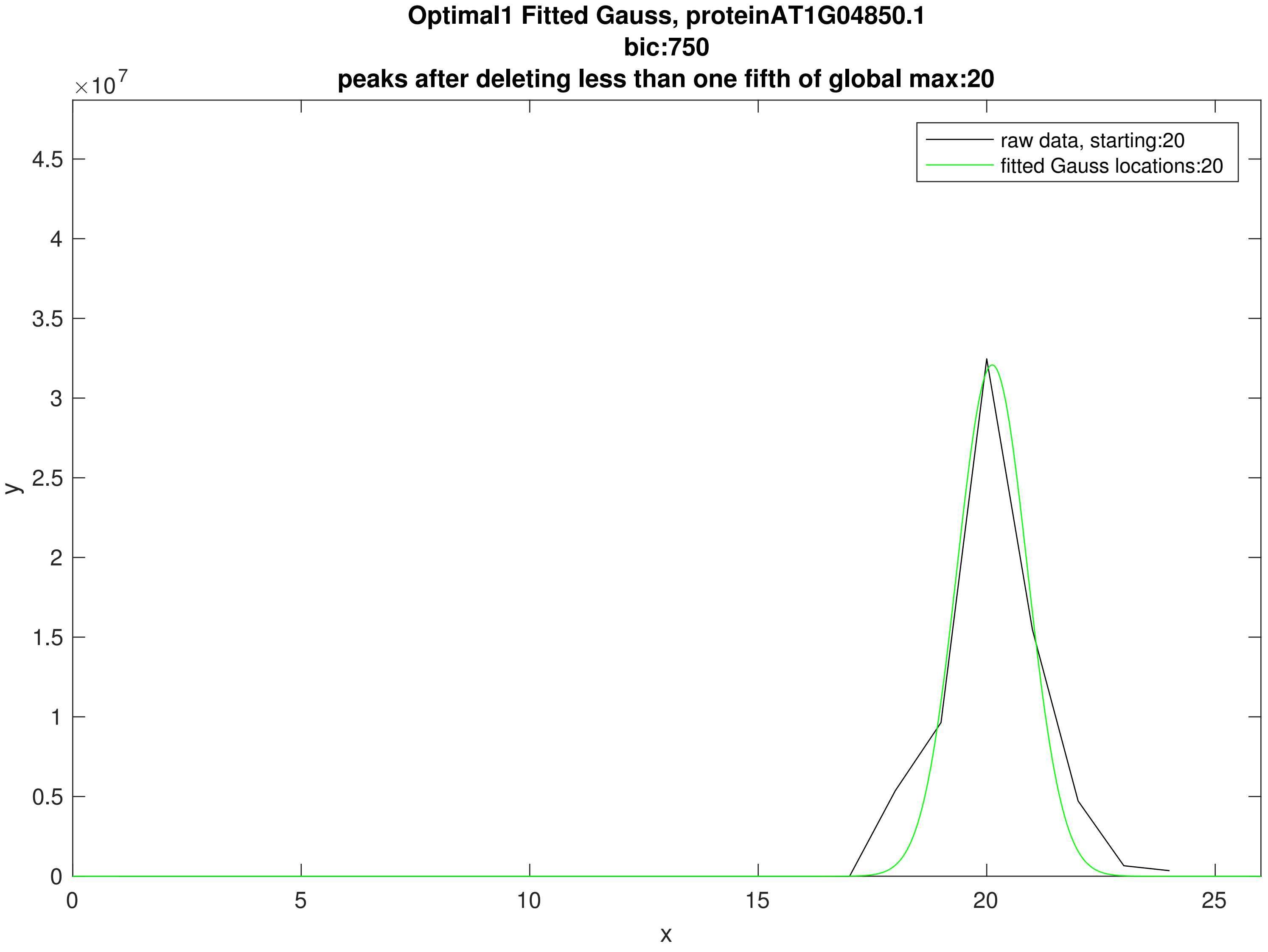
Optimal1 Fitted Gauss, proteinAT1G04850.1
bic:750
peaks after deleting less than one fifth of global max:20
×10^7
raw data, starting:20
fitted Gauss locations:20
y
x
0
0.5
1
1.5
2
2.5
3
3.5
4
4.5
5
10
15
20
25

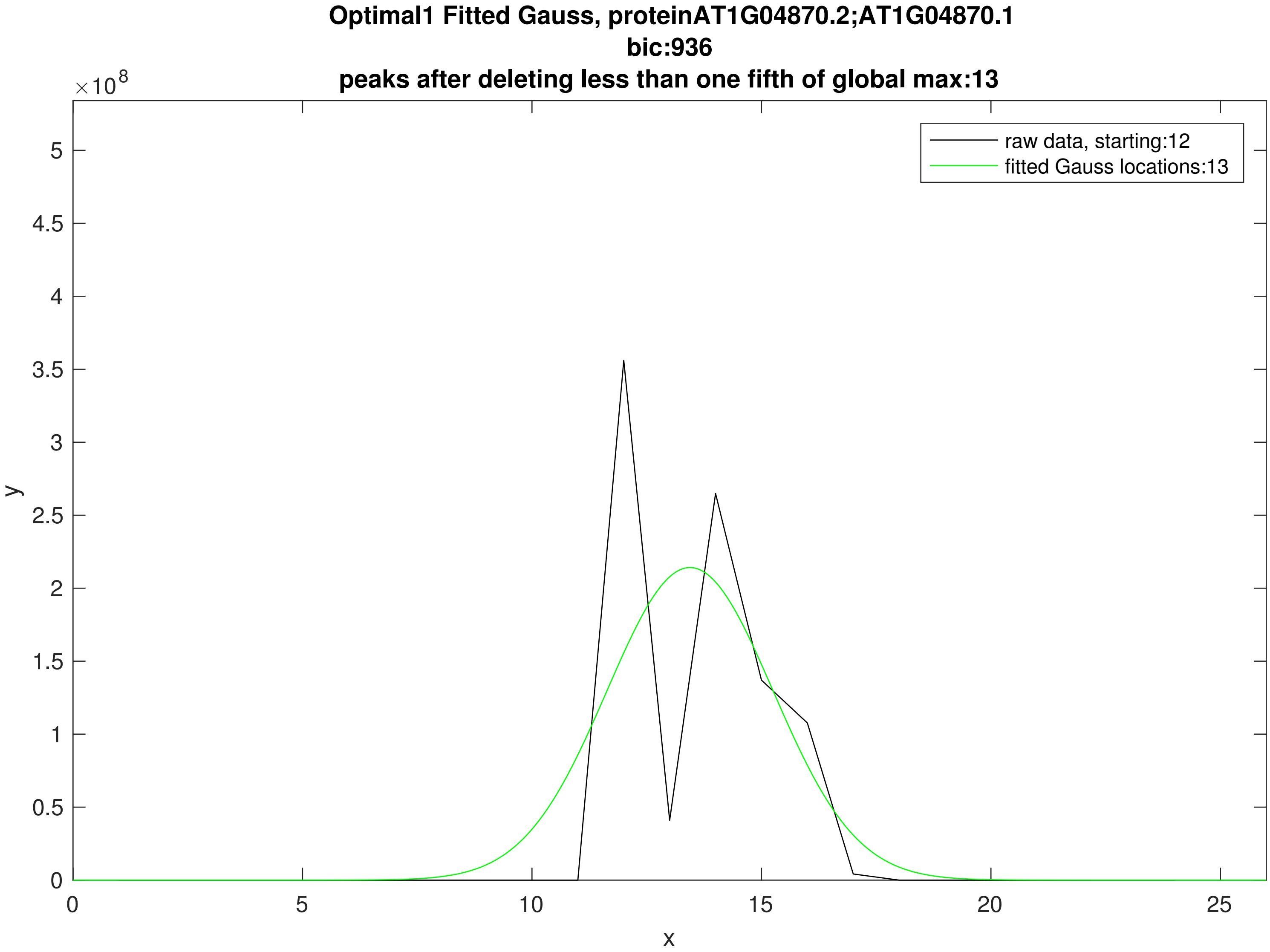

Optimal1 Fitted Gauss, proteinAT1G04870.2;AT1G04870.1
bic:936
peaks after deleting less than one fifth of global max:13
×10^8
raw data, starting:12
fitted Gauss locations:13
y
x
0
0.5
1
1.5
2
2.5
3
3.5
4
4.5
5
0
5
10
15
20
25

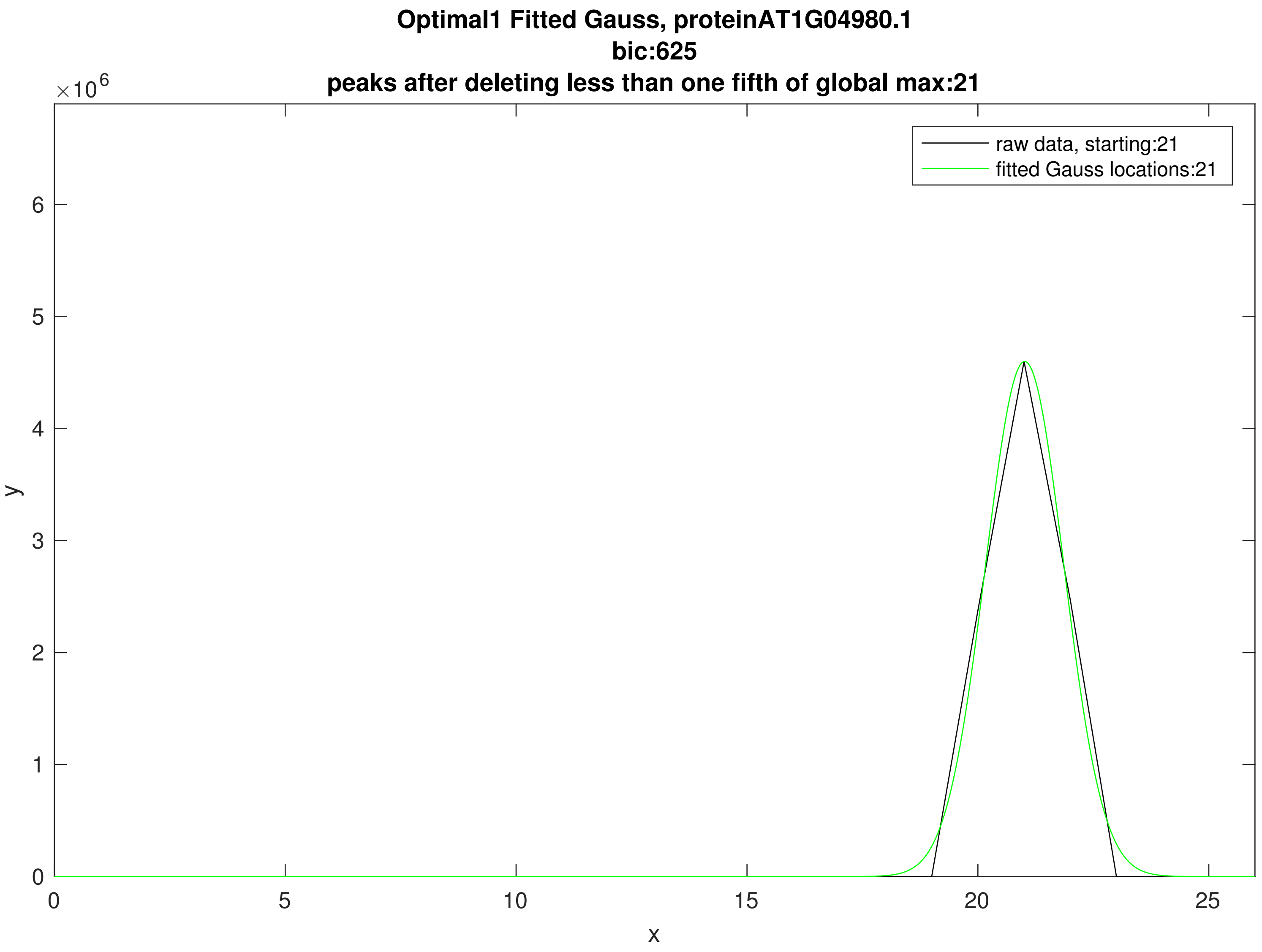

Optimal1 Fitted Gauss, proteinAT1G04980.1
bic:625
peaks after deleting less than one fifth of global max:21
×10^6
raw data, starting:21
fitted Gauss locations:21
y
x
0
1
2
3
4
5
6
0
5
10
15
20
25

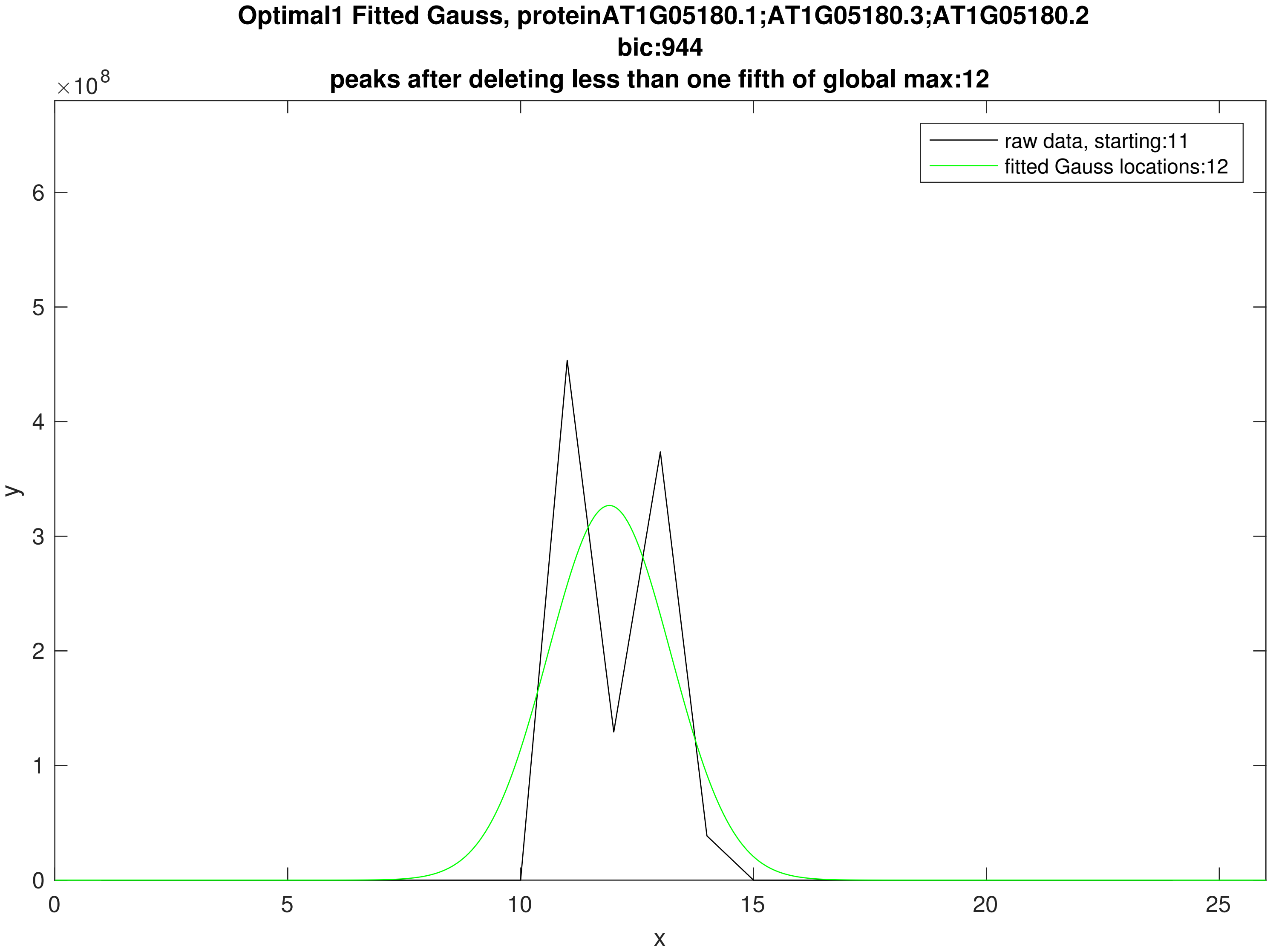

Optimal1 Fitted Gauss, proteinAT1G05180.1;AT1G05180.3;AT1G05180.2
bic:944
peaks after deleting less than one fifth of global max:12
×10^8
raw data, starting:11
fitted Gauss locations:12
y
x
0
1
2
3
4
5
6
0
5
10
15
20
25

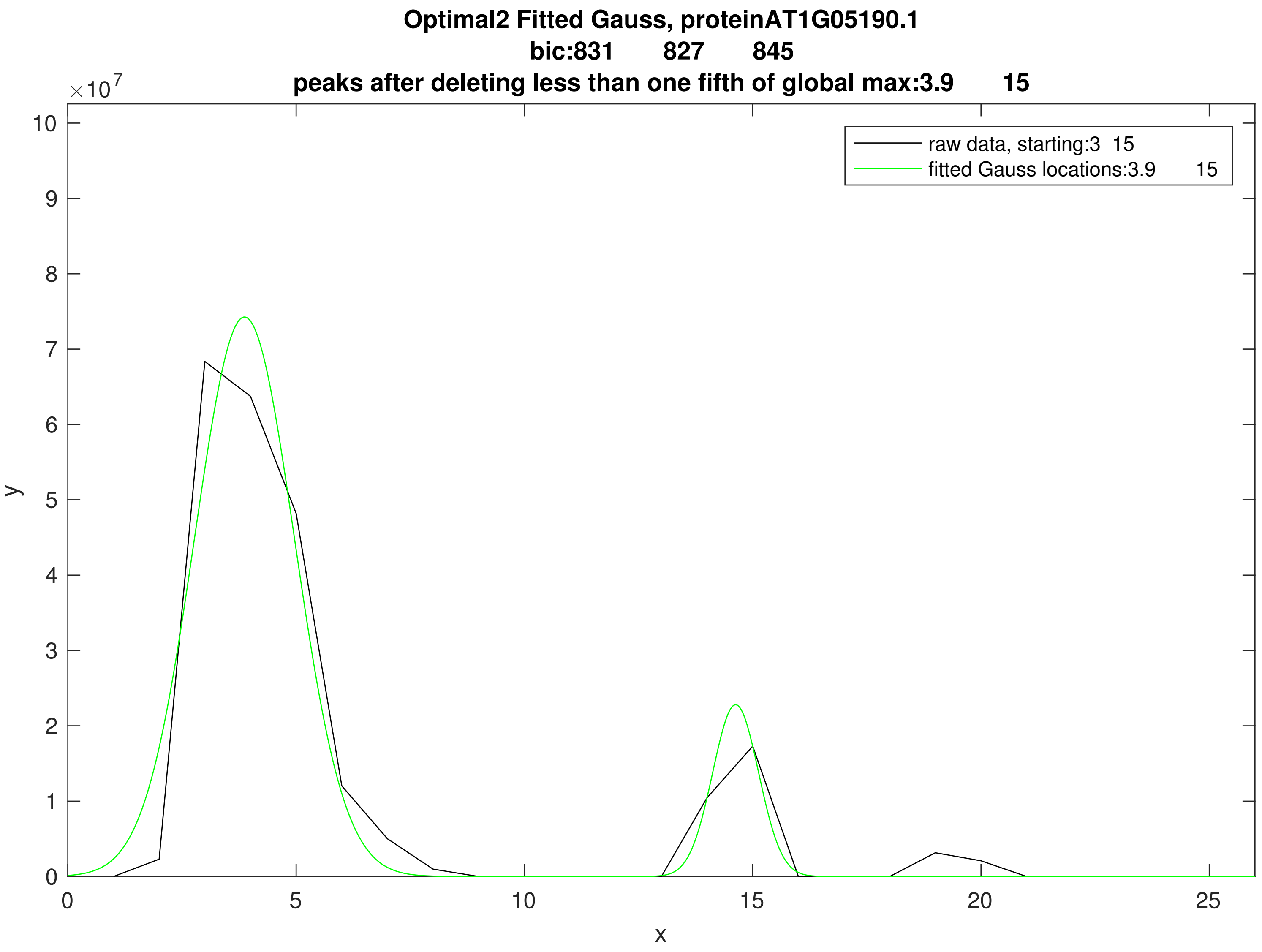

Optimal2 Fitted Gauss, proteinAT1G05190.1
bic:831 827 845
peaks after deleting less than one fifth of global max:3.9 15
×10^7
raw data, starting:3 15
fitted Gauss locations:3.9 15
y
x
0
1
2
3
4
5
6
7
8
9
10
0
5
10
15
20
25

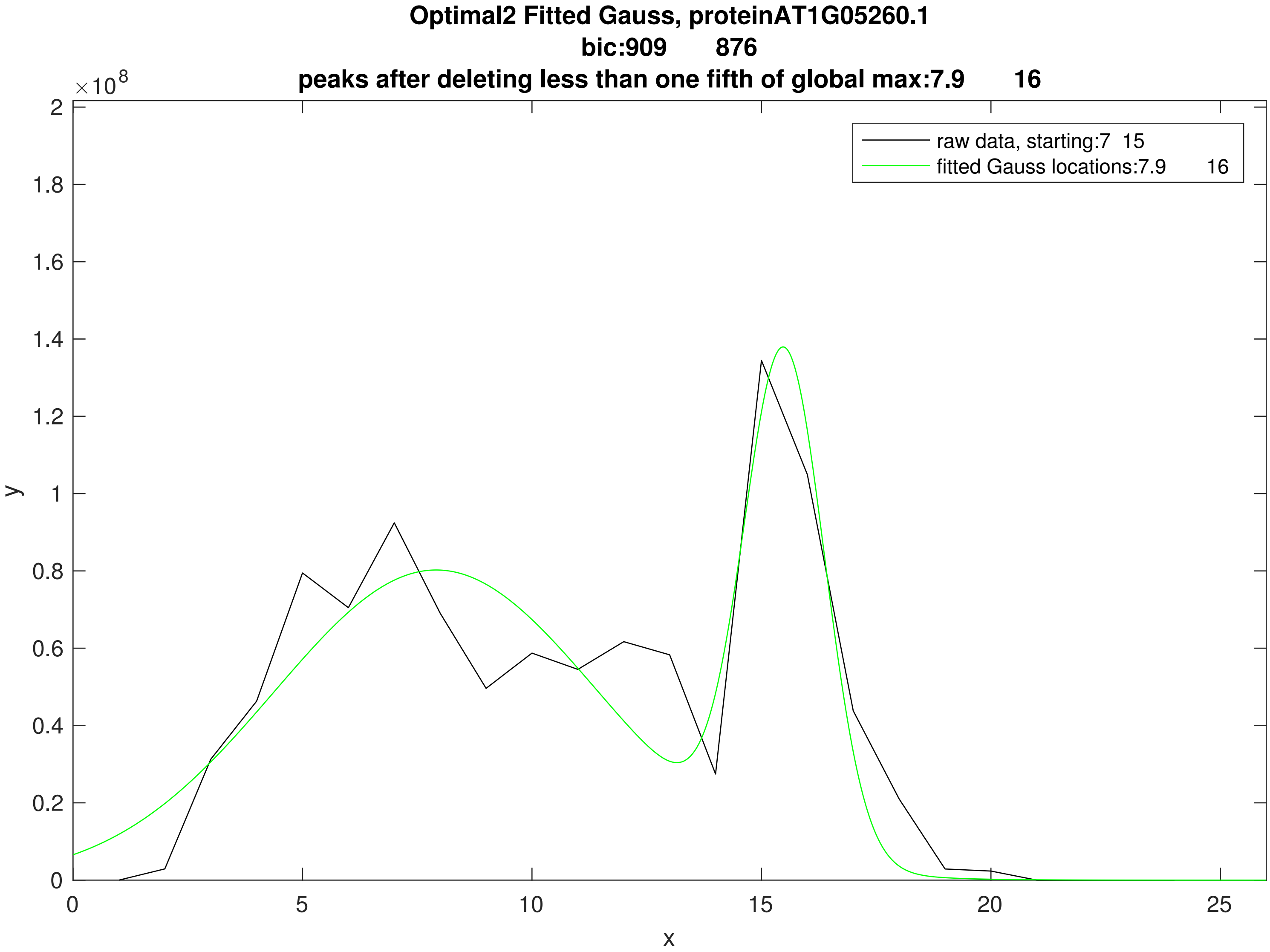

Optimal2 Fitted Gauss, proteinAT1G05260.1
bic:909 876
peaks after deleting less than one fifth of global max:7.9 16
×10^8
raw data, starting:7 15
fitted Gauss locations:7.9 16
y
x
0
0.2
0.4
0.6
0.8
1
1.2
1.4
1.6
1.8
2
5
10
15
20
25

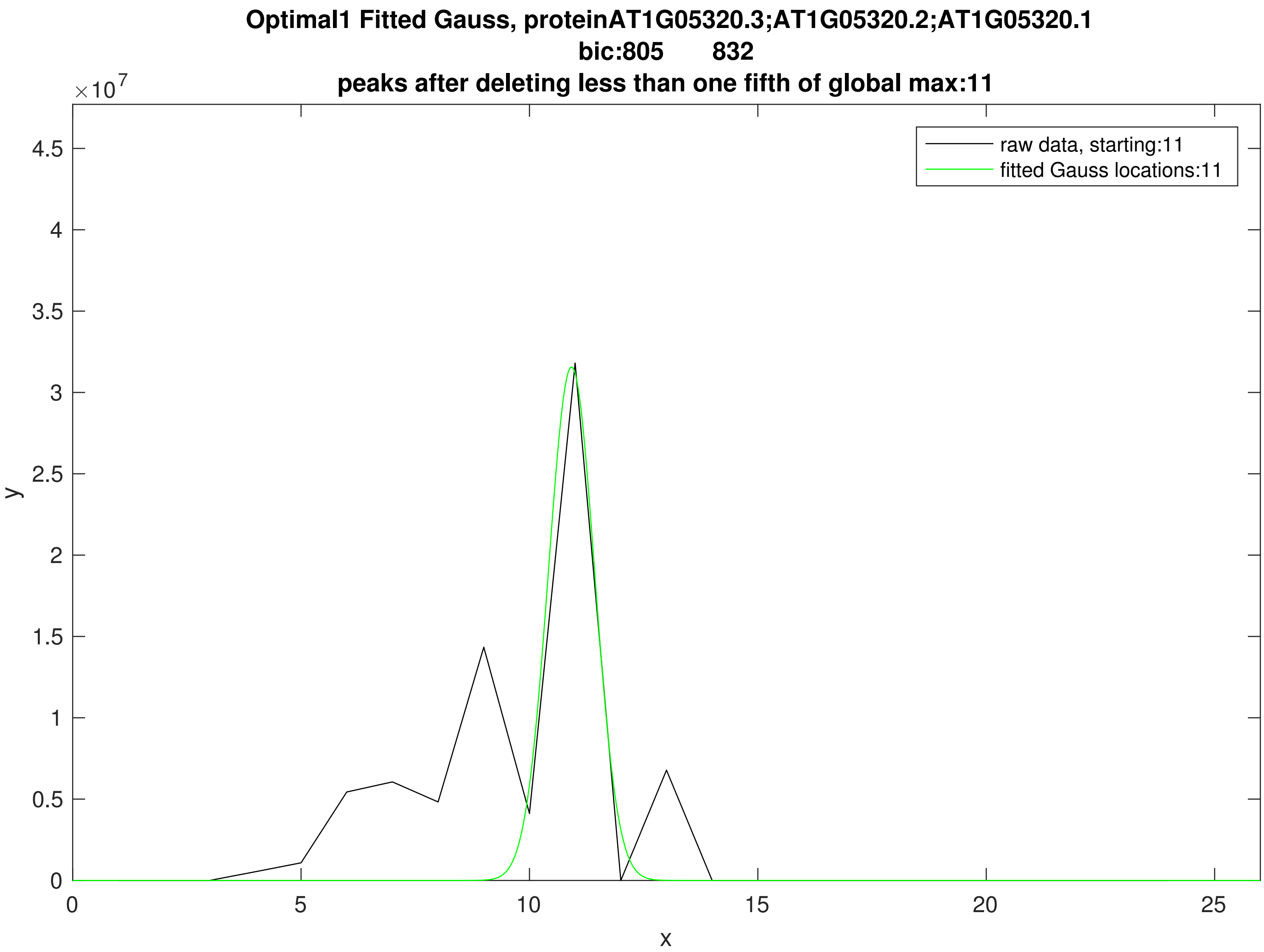

Optimal1 Fitted Gauss, proteinAT1G05320.3;AT1G05320.2;AT1G05320.1
bic:805 832
peaks after deleting less than one fifth of global max:11
×10^7
4.5
4
3.5
3
2.5
2
1.5
1
0.5
0
y
0
5
10
15
20
25
x
raw data, starting:11
fitted Gauss locations:11

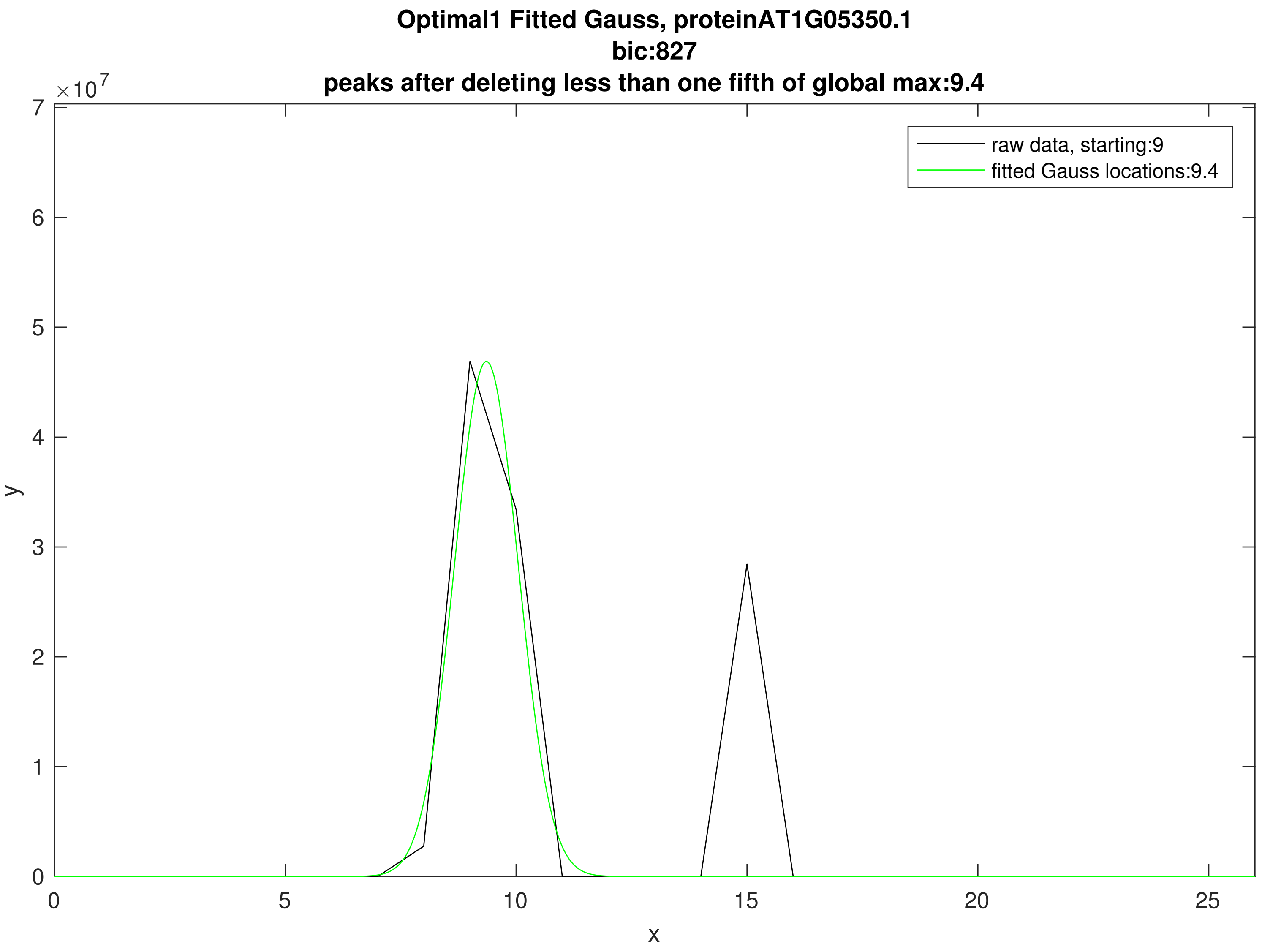

Optimal1 Fitted Gauss, proteinAT1G05350.1
bic:827
peaks after deleting less than one fifth of global max:9.4
×10^7
raw data, starting:9
fitted Gauss locations:9.4
y
x
0
1
2
3
4
5
6
7
0
5
10
15
20
25

#### Optimal1 Fitted Gauss, proteinAT1G05510.1
#### bic:1.01e+03  1.02e+03  1.03e+03
#### peaks after deleting less than one fifth of global max:15

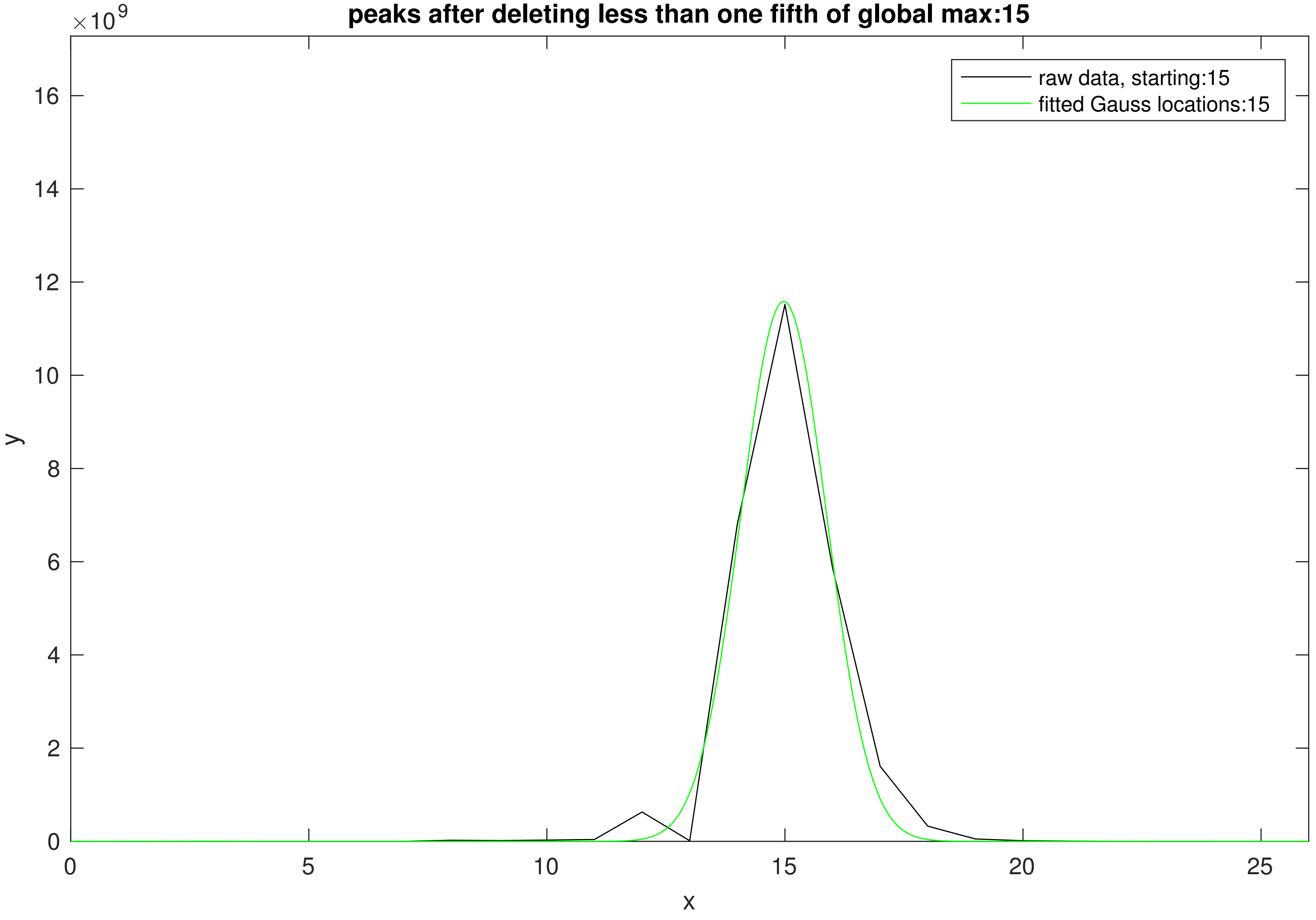

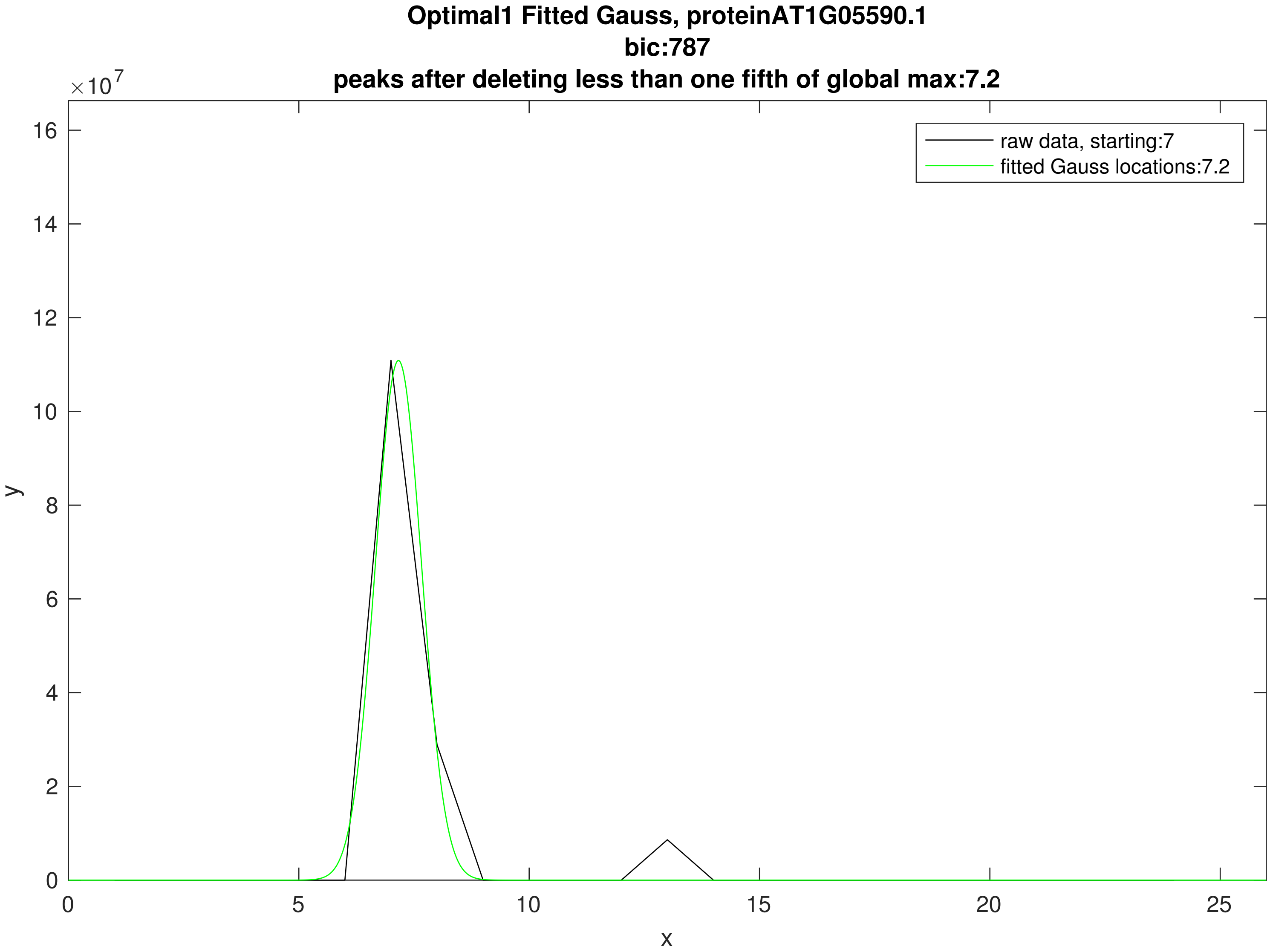
Optimal1 Fitted Gauss, proteinAT1G05590.1
bic:787
peaks after deleting less than one fifth of global max:7.2
×10^7
raw data, starting:7
fitted Gauss locations:7.2
y
x
0
2
4
6
8
10
12
14
16
0
5
10
15
20
25

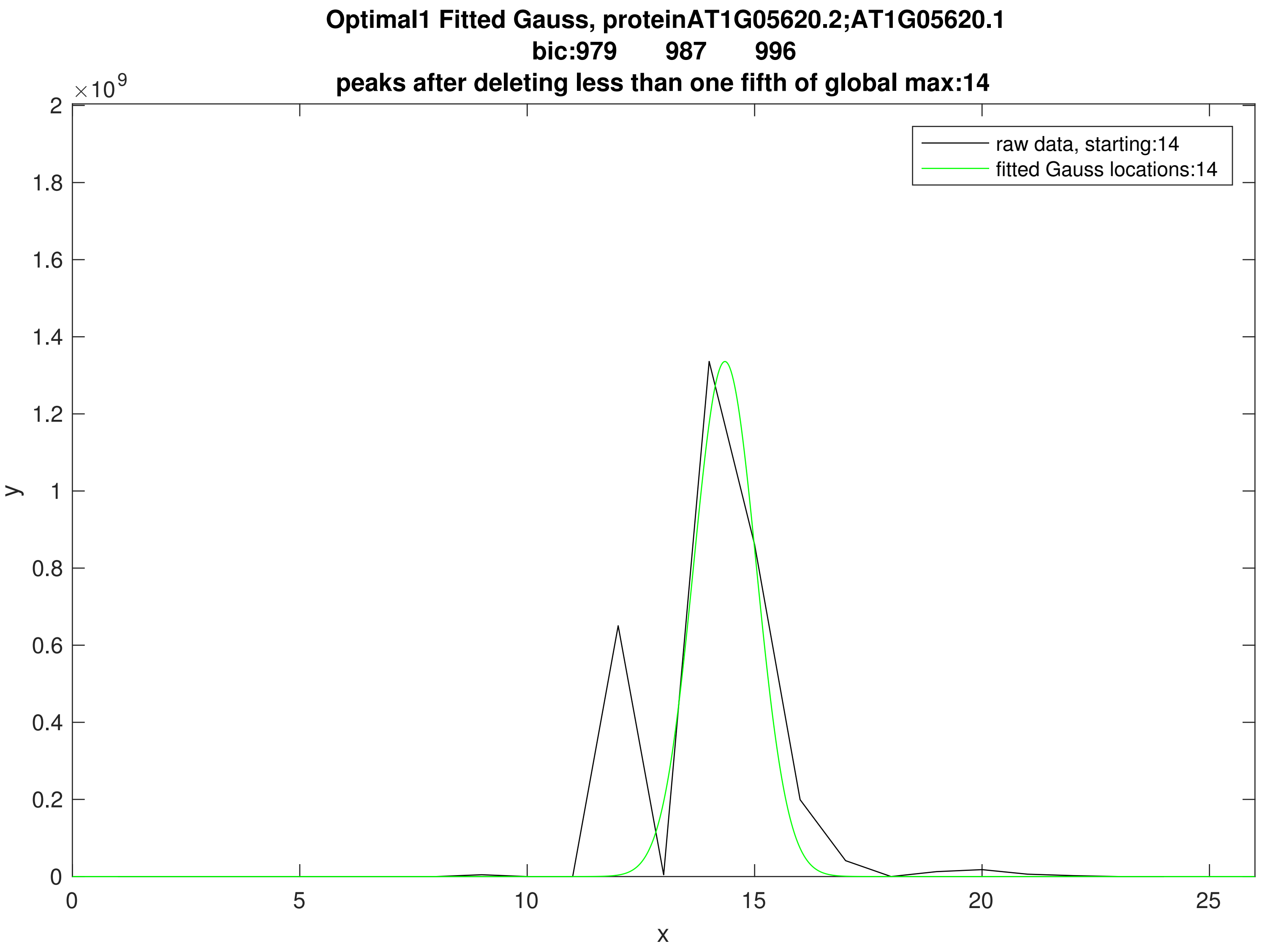

Optimal1 Fitted Gauss, proteinAT1G05620.2;AT1G05620.1
bic:979 987 996
peaks after deleting less than one fifth of global max:14
×10^9
2
1.8
1.6
1.4
1.2
1
0.8
0.6
0.4
0.2
0
y
0
5
10
15
20
25
x
raw data, starting:14
fitted Gauss locations:14

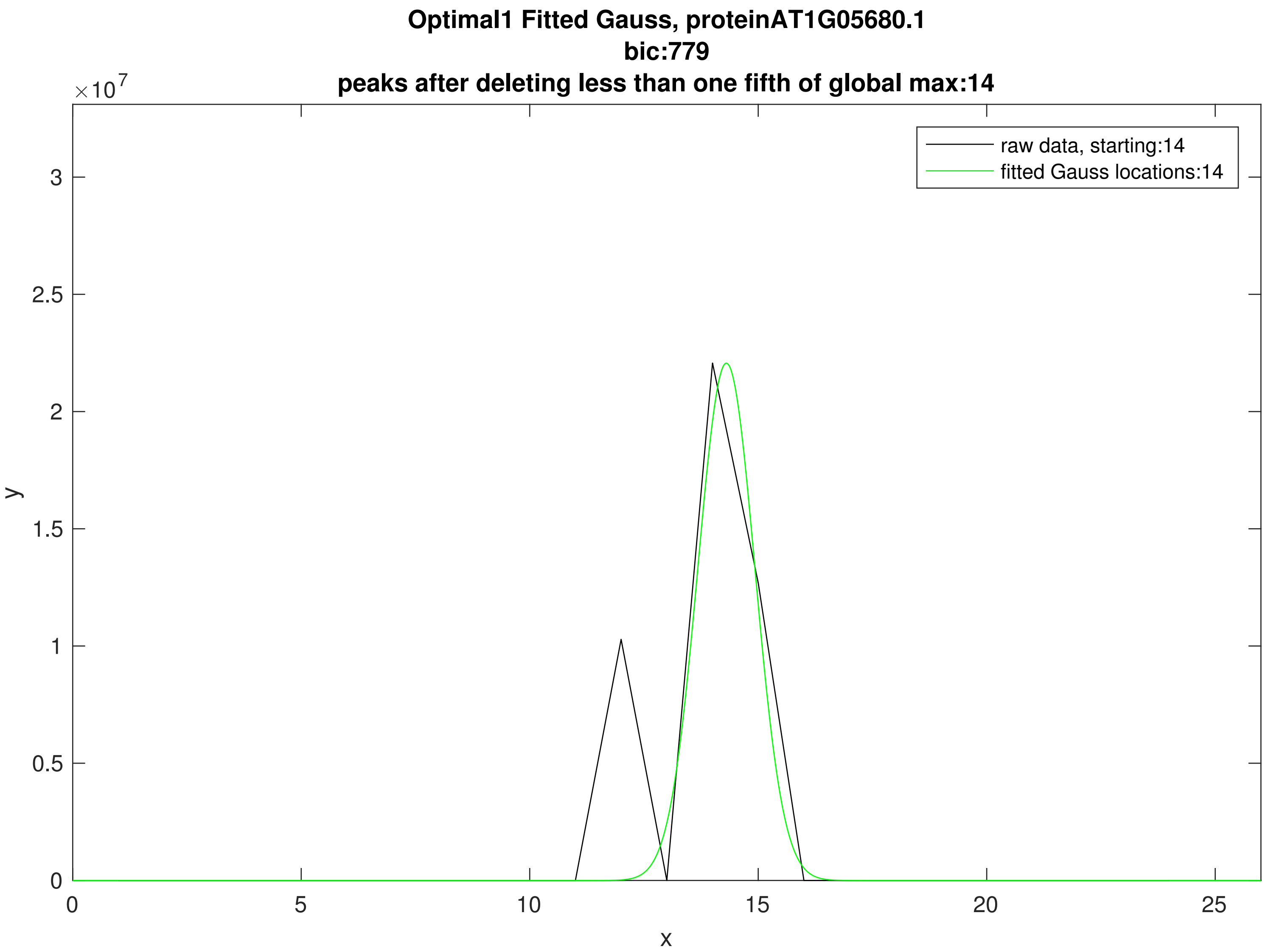
Optimal1 Fitted Gauss, proteinAT1G05680.1
bic:779
peaks after deleting less than one fifth of global max:14
×10^7
raw data, starting:14
fitted Gauss locations:14
y
x
0
0.5
1
1.5
2
2.5
3
0
5
10
15
20
25

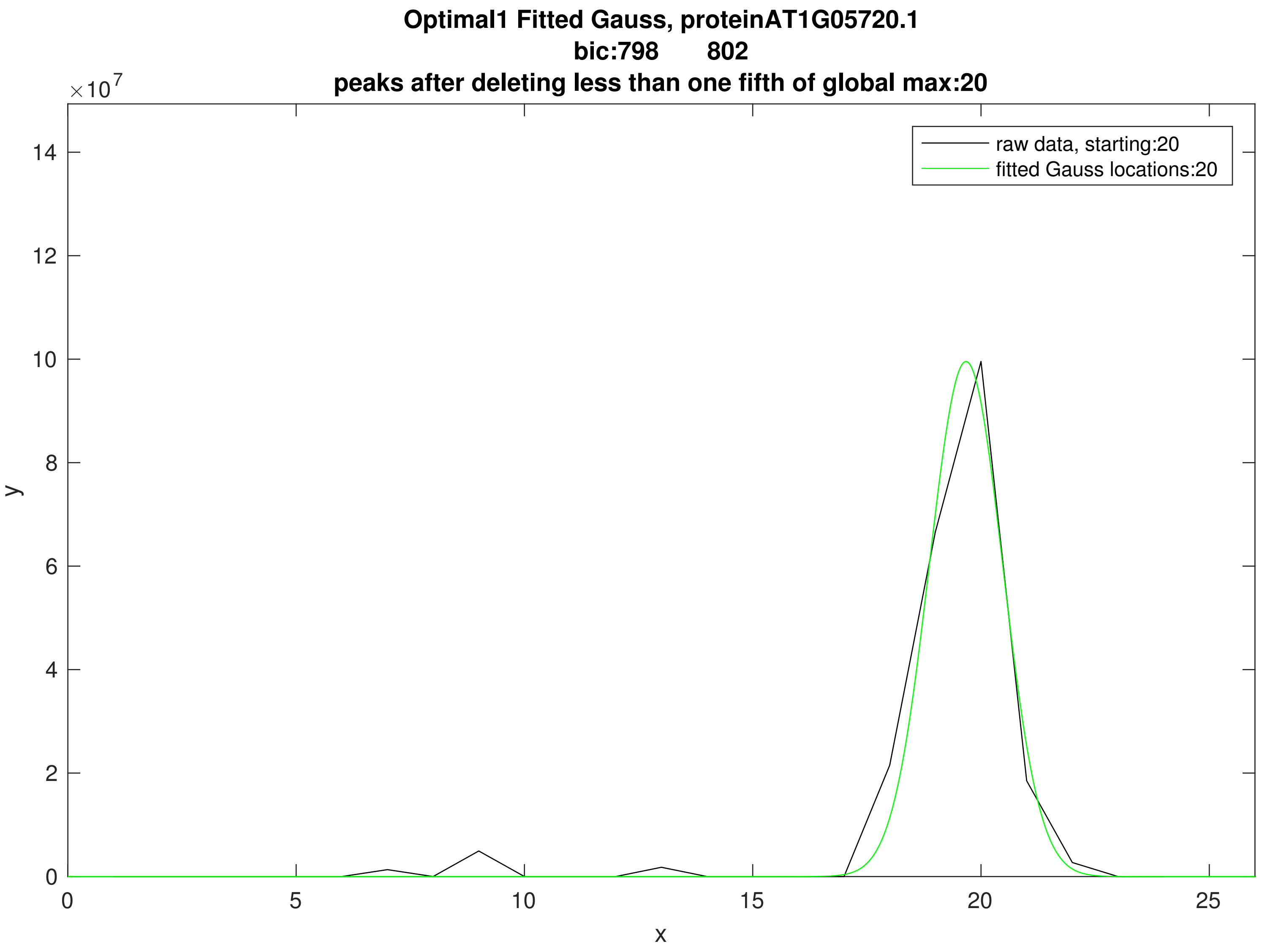
Optimal1 Fitted Gauss, proteinAT1G05720.1
bic:798 802
peaks after deleting less than one fifth of global max:20
$\times 10^7$
raw data, starting:20
fitted Gauss locations:20
y
x
0
2
4
6
8
10
12
14
0
5
10
15
20
25

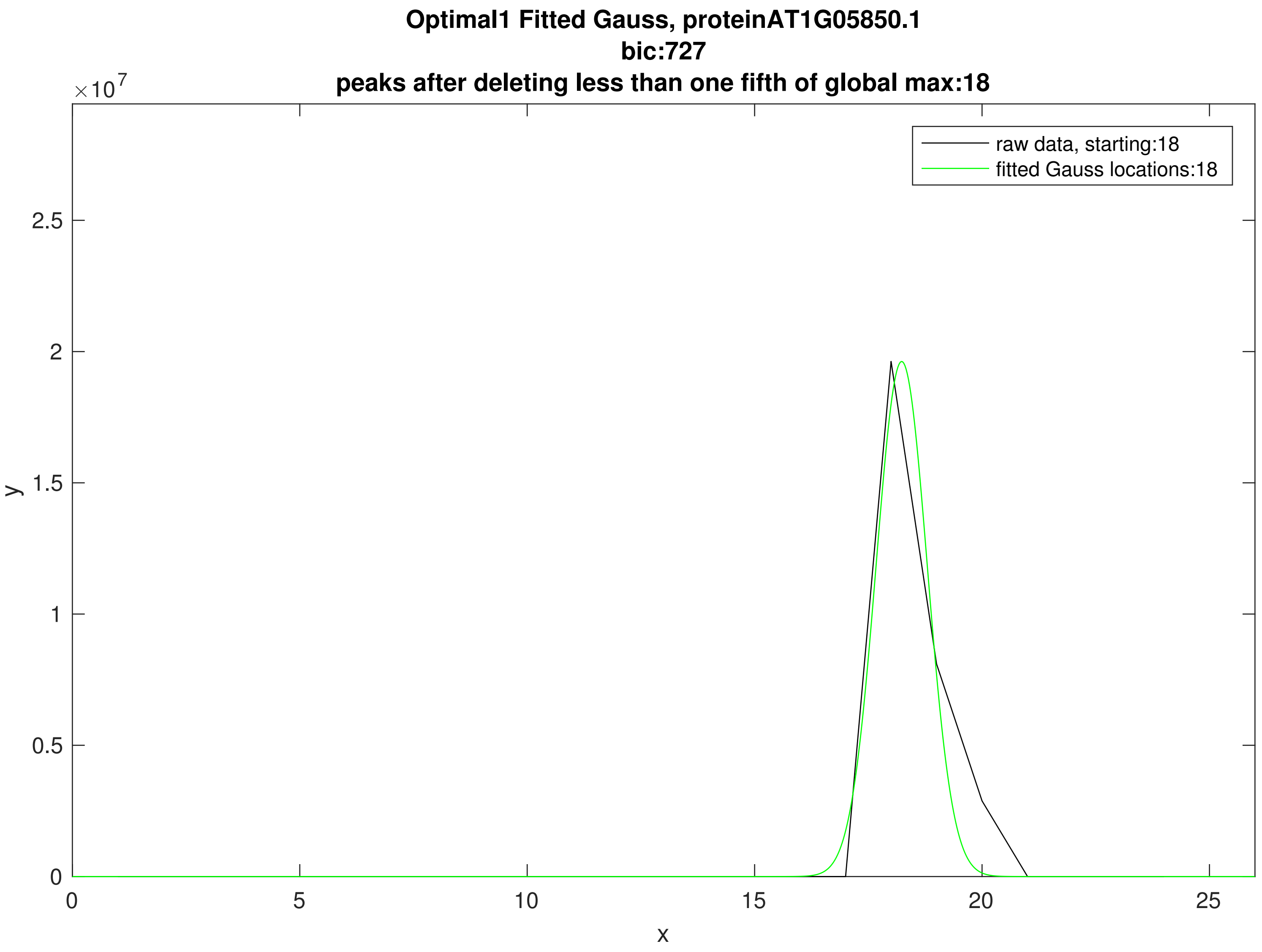
Optimal1 Fitted Gauss, proteinAT1G05850.1
bic:727
peaks after deleting less than one fifth of global max:18
×10^7
raw data, starting:18
fitted Gauss locations:18
y
x
0
0.5
1
1.5
2
2.5
0
5
10
15
20
25

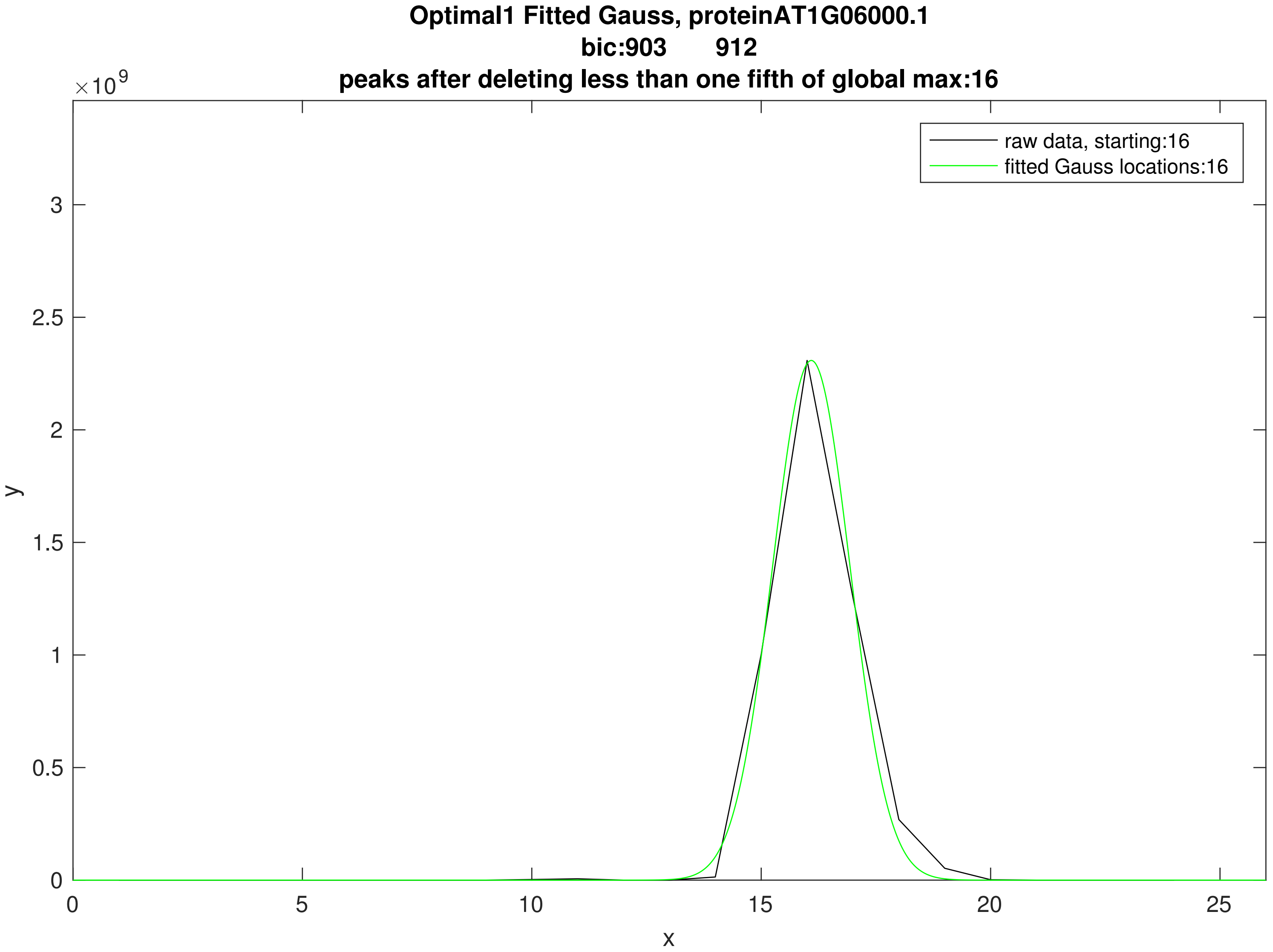
Optimal1 Fitted Gauss, proteinAT1G06000.1
bic:903 912
peaks after deleting less than one fifth of global max:16
×10^9
raw data, starting:16
fitted Gauss locations:16
y
x
0
0.5
1
1.5
2
2.5
3
0
5
10
15
20
25

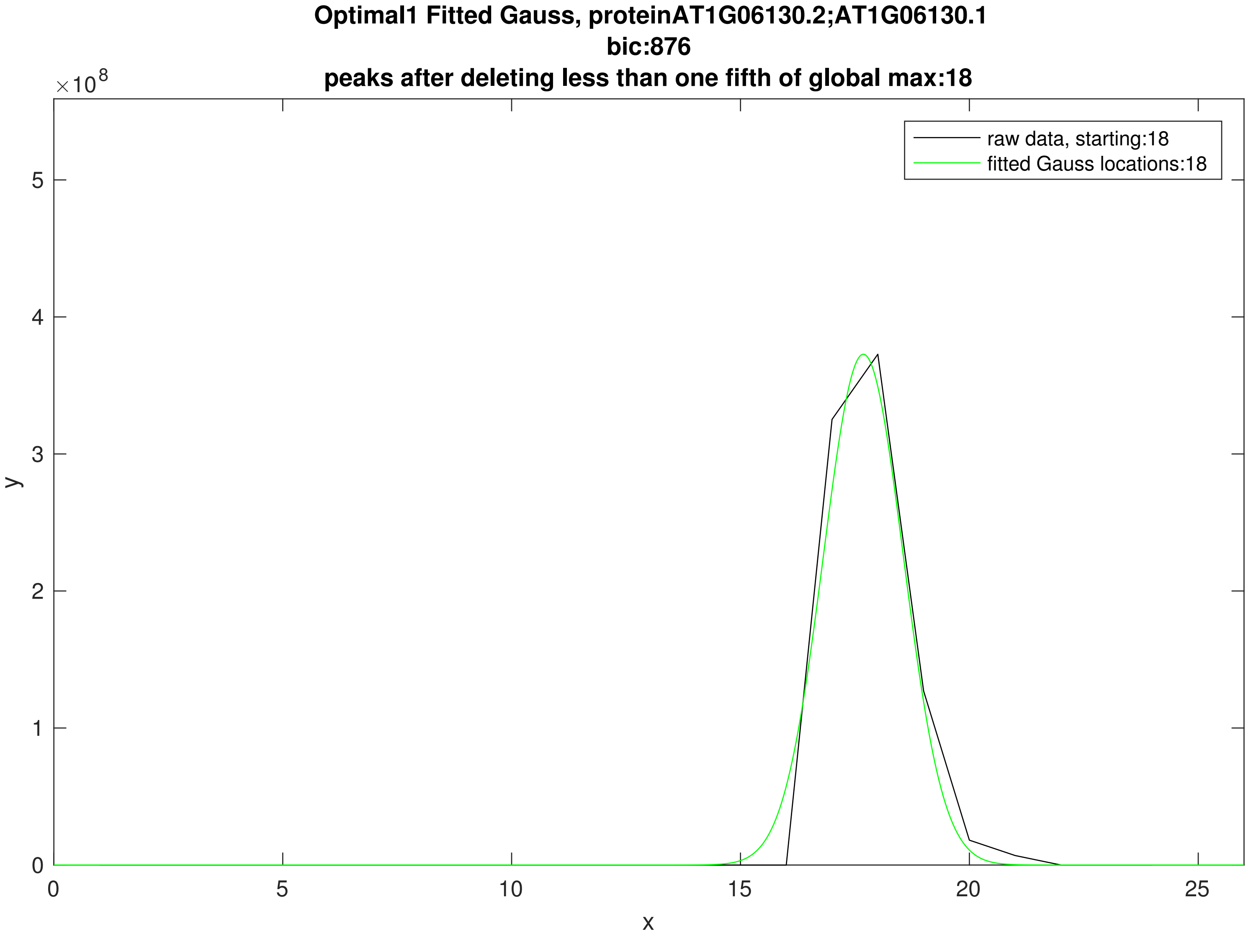

Optimal1 Fitted Gauss, proteinAT1G06130.2;AT1G06130.1
bic:876
peaks after deleting less than one fifth of global max:18
×10^8
5
4
3
2
1
0
y
0
5
10
15
20
25
x
raw data, starting:18
fitted Gauss locations:18

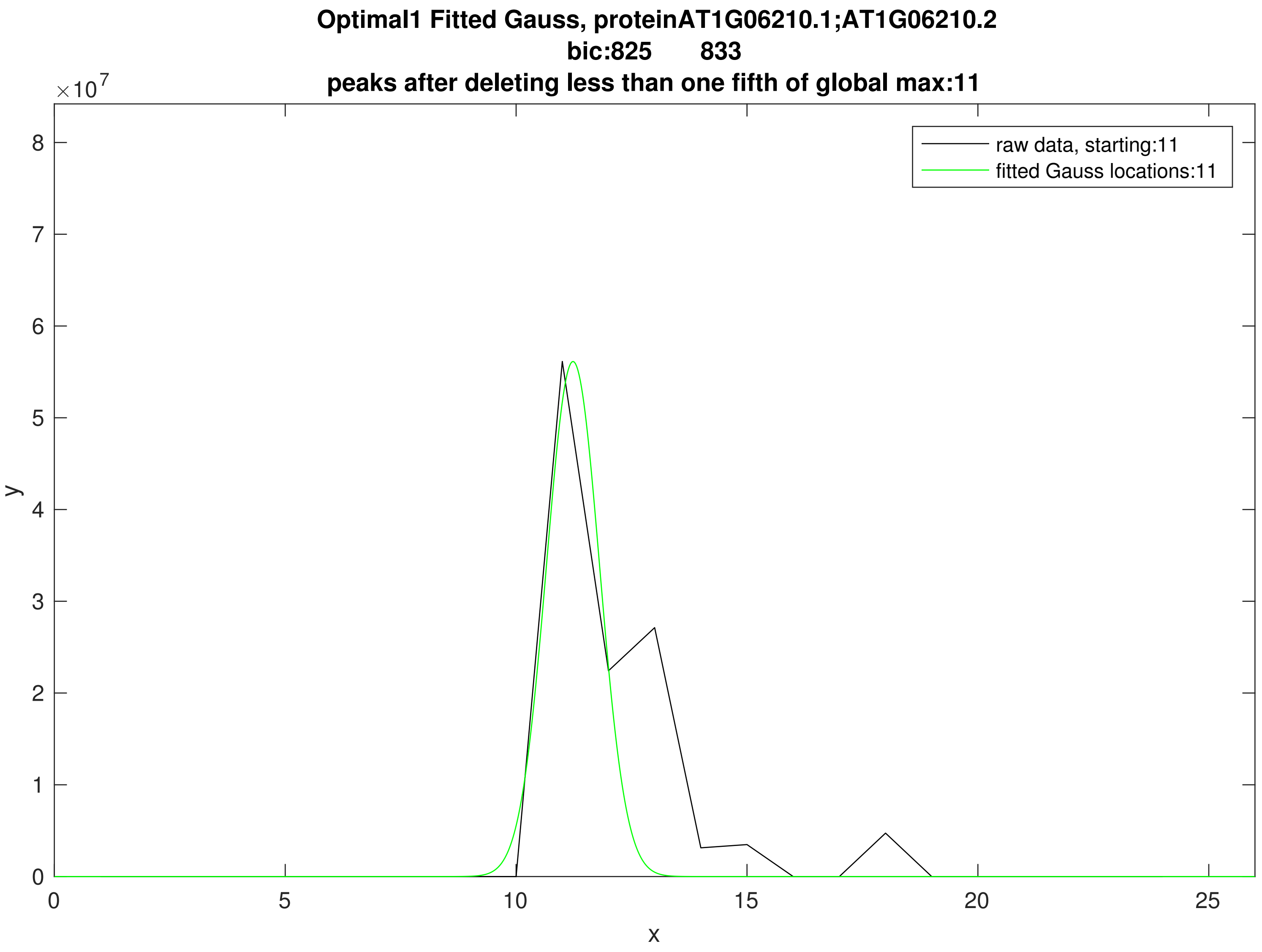
Optimal1 Fitted Gauss, proteinAT1G06210.1;AT1G06210.2
bic:825 833
peaks after deleting less than one fifth of global max:11
×10^7
raw data, starting:11
fitted Gauss locations:11
y
x
0
1
2
3
4
5
6
7
8
0
5
10
15
20
25

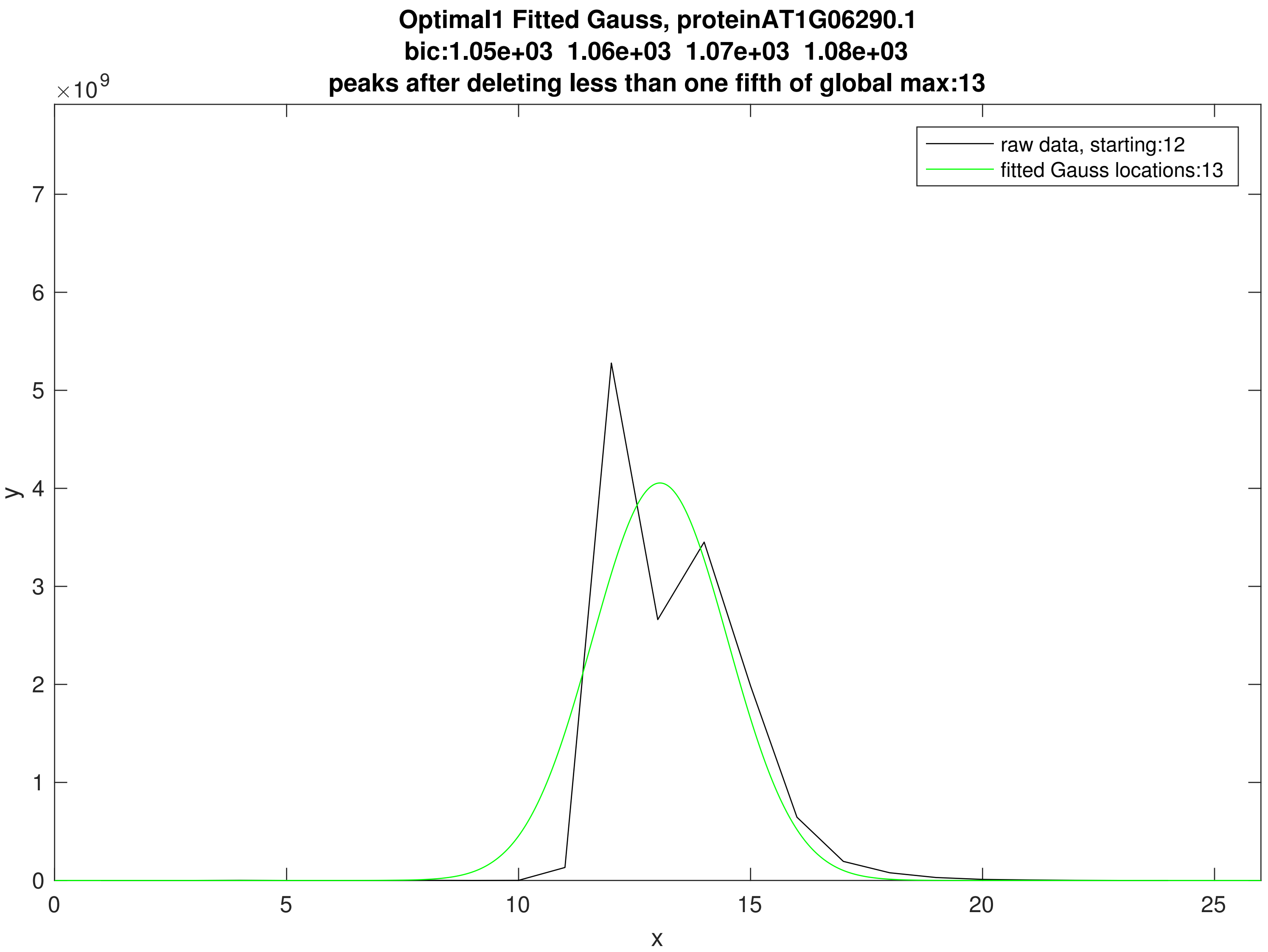
Optimal1 Fitted Gauss, proteinAT1G06290.1
bic:1.05e+03  1.06e+03  1.07e+03  1.08e+03
peaks after deleting less than one fifth of global max:13
×10^9
raw data, starting:12
fitted Gauss locations:13
y
x
0
1
2
3
4
5
6
7
0
5
10
15
20
25

Optimal1 Fitted Gauss, proteinAT1G06410.1
bic:719
peaks after deleting less than one fifth of global max:9.8

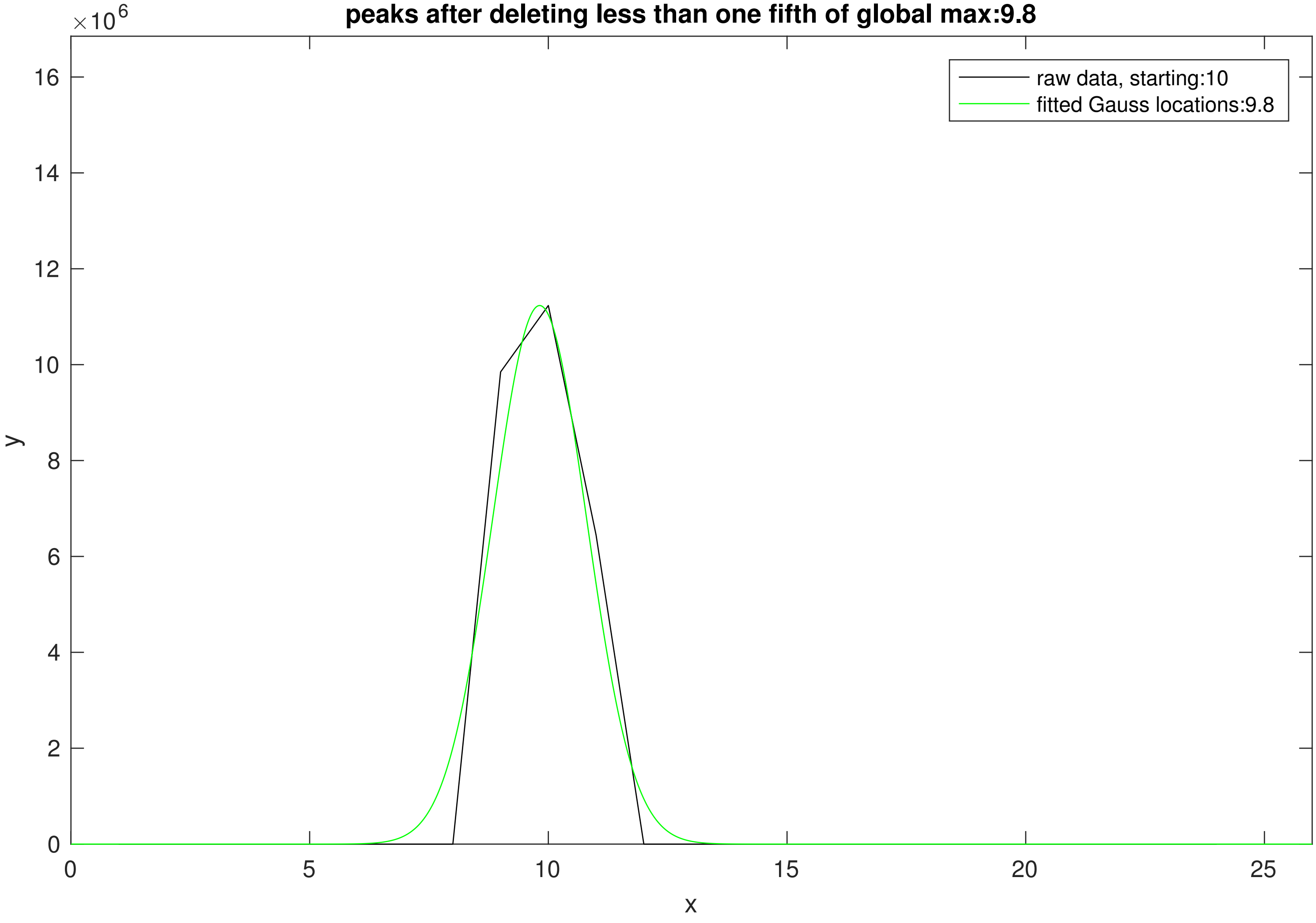

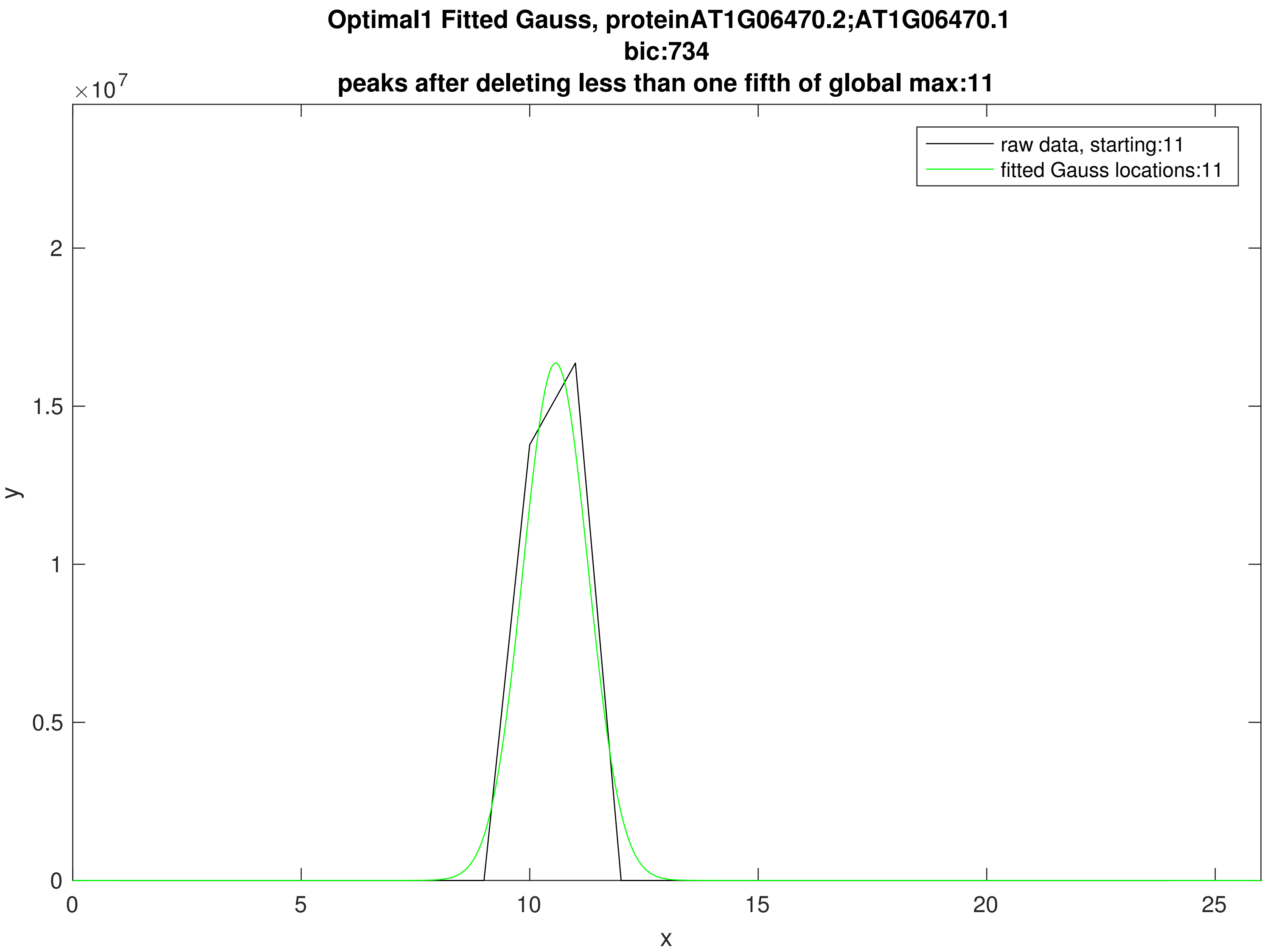

Optimal1 Fitted Gauss, proteinAT1G06470.2;AT1G06470.1
bic:734
peaks after deleting less than one fifth of global max:11
×10^7
raw data, starting:11
fitted Gauss locations:11
2
1.5
1
0.5
0
y
0
5
10
15
20
25
x

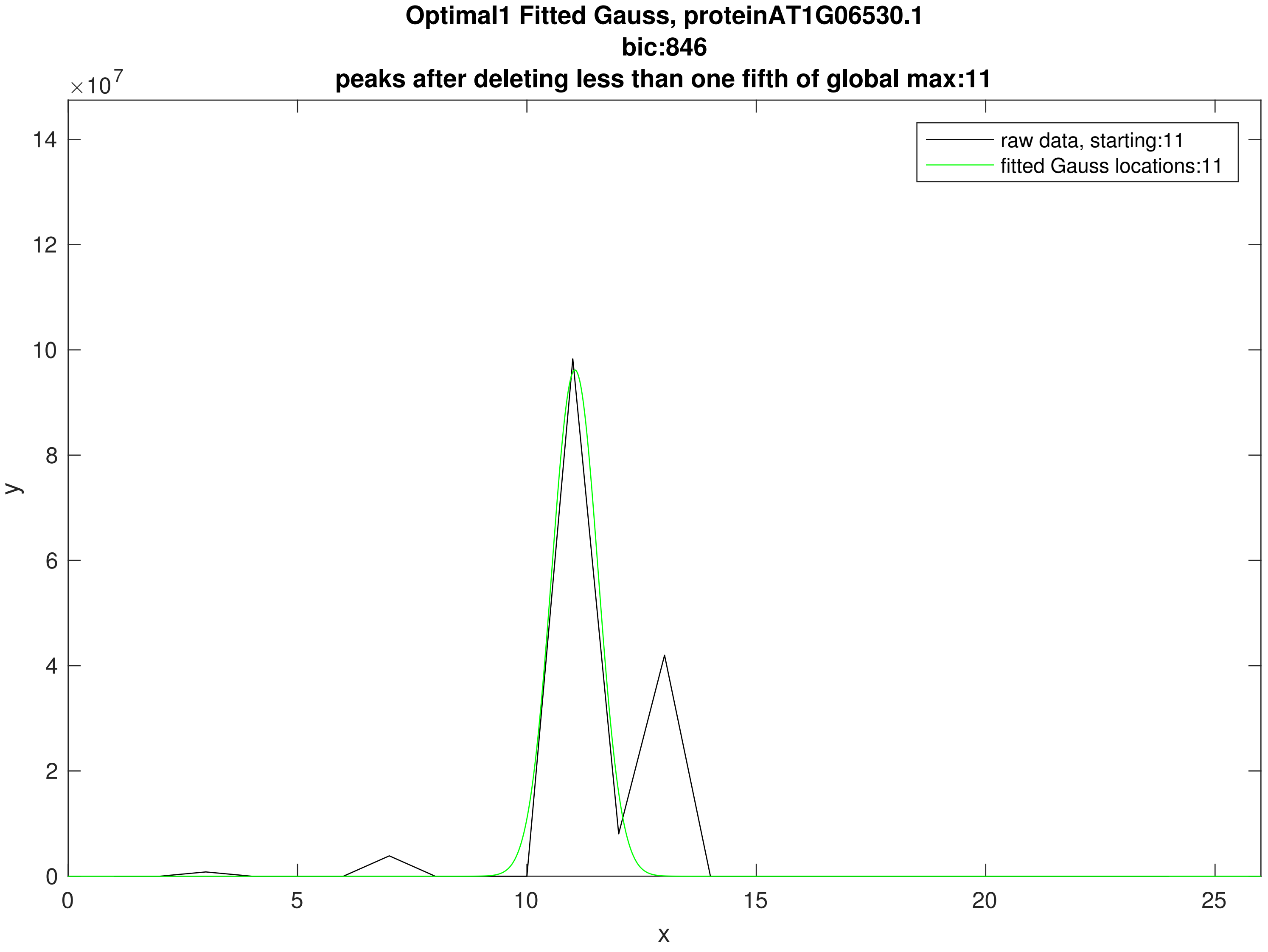
Optimal1 Fitted Gauss, proteinAT1G06530.1
bic:846
peaks after deleting less than one fifth of global max:11
×10^7
raw data, starting:11
fitted Gauss locations:11
y
x
0
2
4
6
8
10
12
14
0
5
10
15
20
25

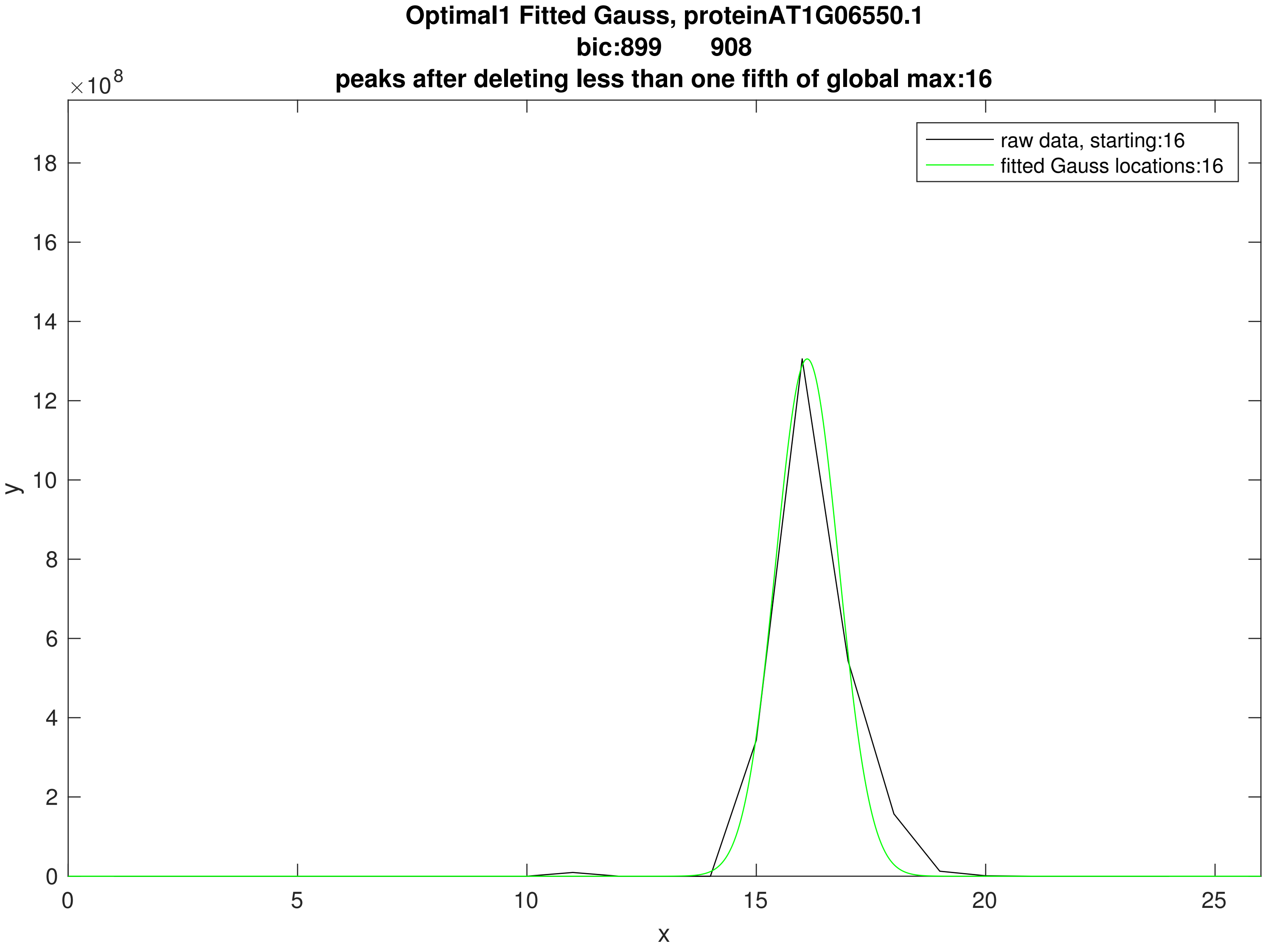
Optimal1 Fitted Gauss, proteinAT1G06550.1
bic:899 908
peaks after deleting less than one fifth of global max:16
×10^8
raw data, starting:16
fitted Gauss locations:16
y
x
0
2
4
6
8
10
12
14
16
18
0
5
10
15
20
25

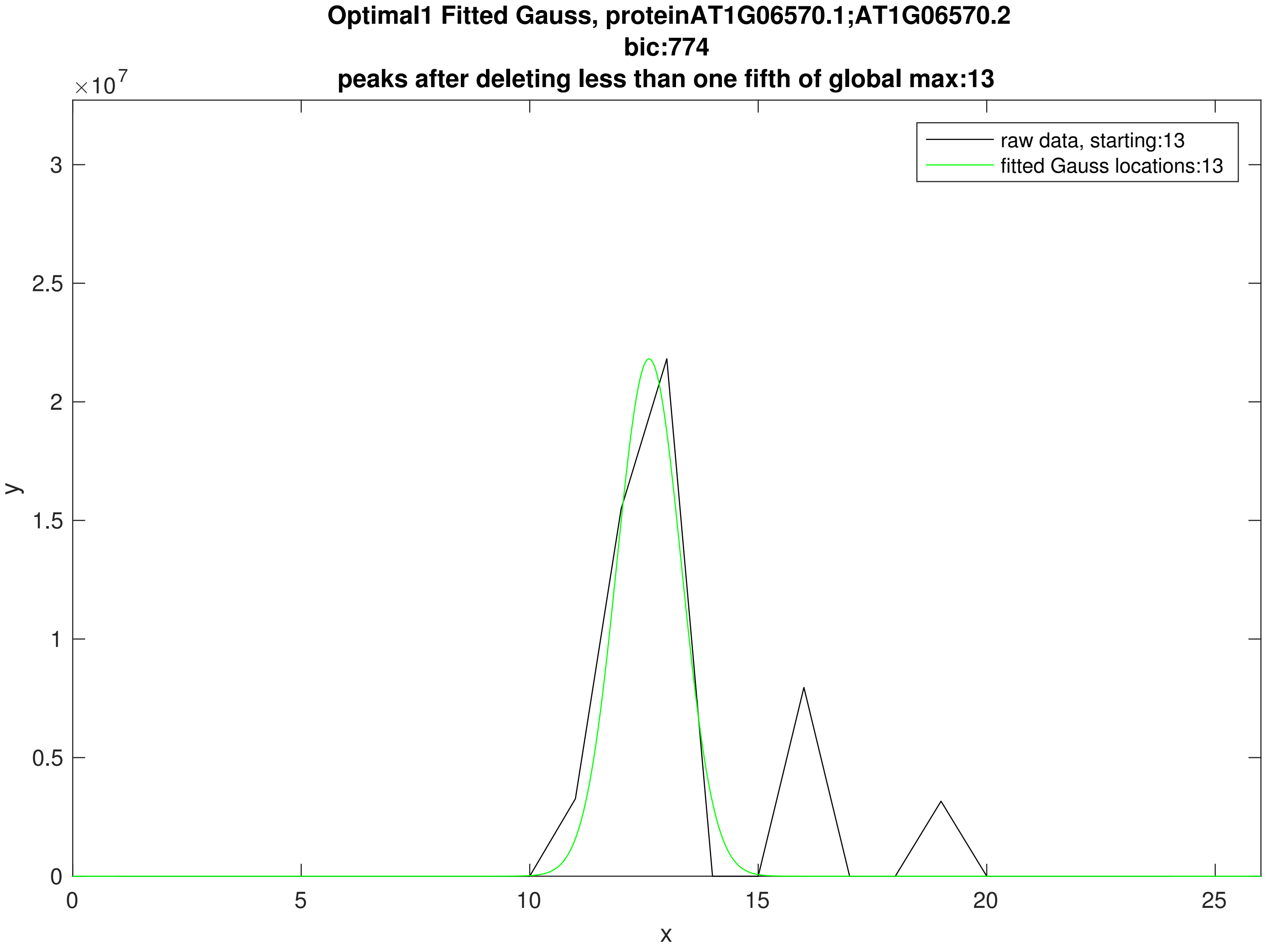
Optimal1 Fitted Gauss, proteinAT1G06570.1;AT1G06570.2
bic:774
peaks after deleting less than one fifth of global max:13
×10^7
raw data, starting:13
fitted Gauss locations:13
3
2.5
2
1.5
1
0.5
0
y
0
5
10
15
20
25
x

**Optimal1 Fitted Gauss, proteinAT1G06680.1;AT1G06680.2**
**bic:893 902**
**peaks after deleting less than one fifth of global max:19**

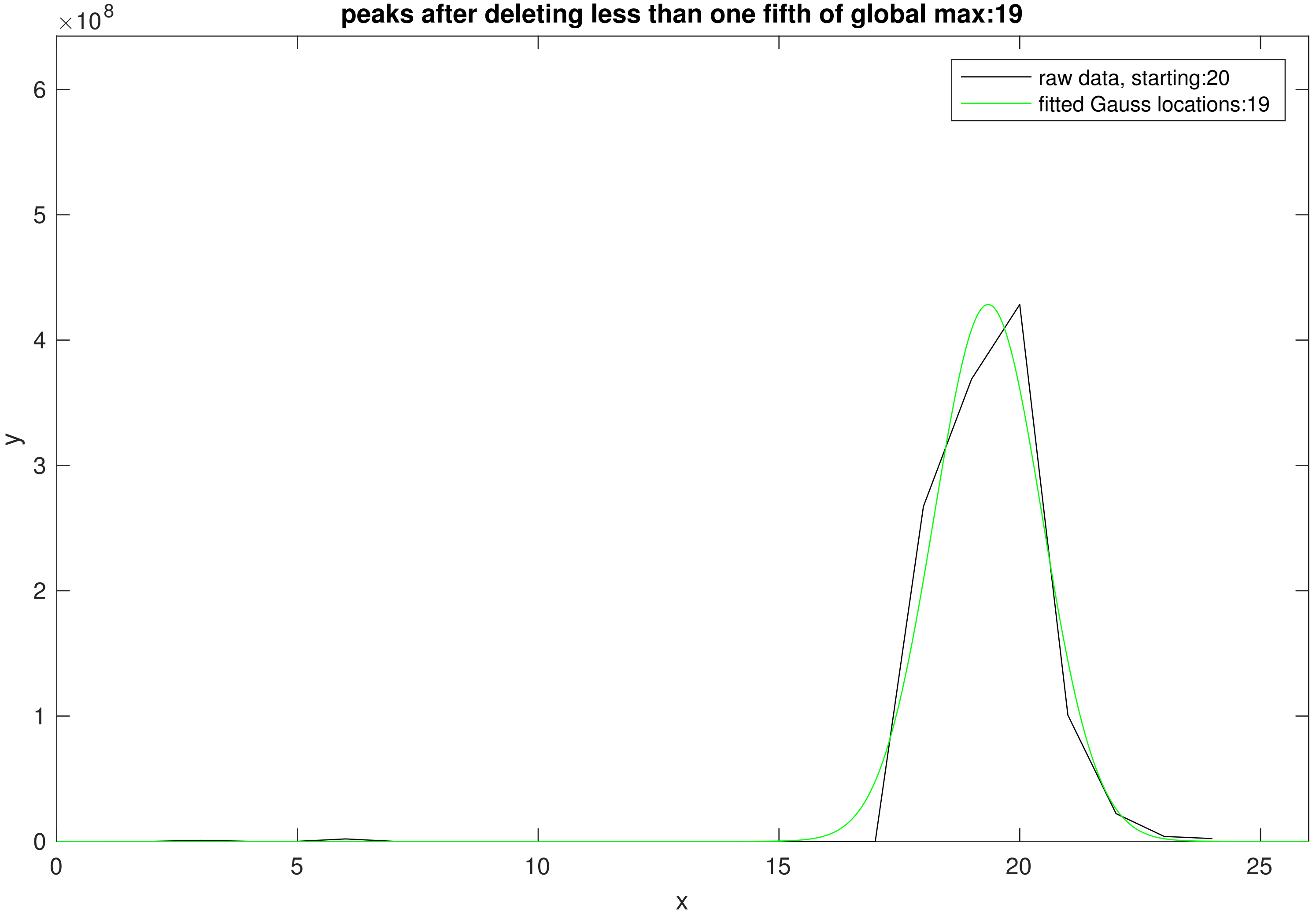

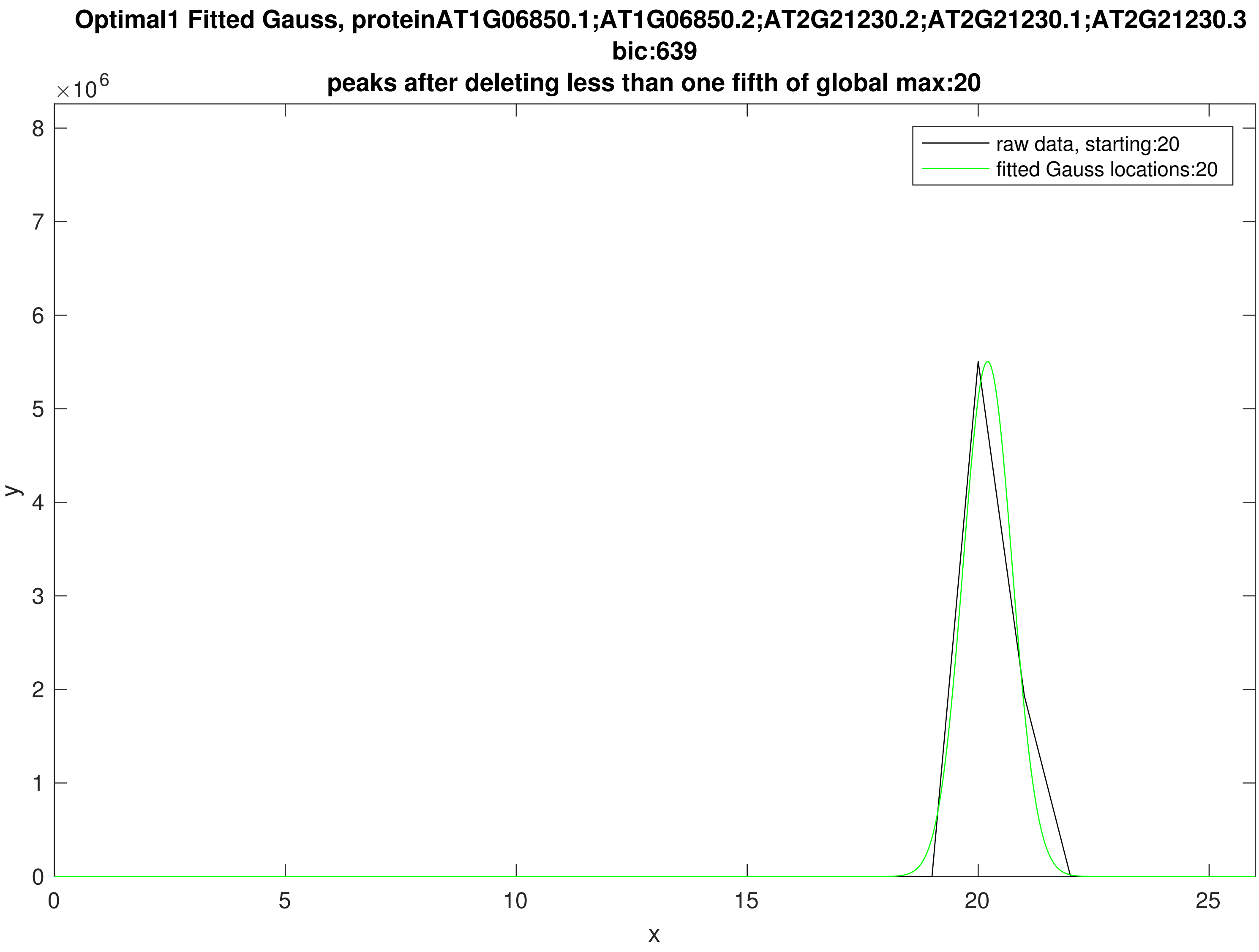
Optimal1 Fitted Gauss, proteinAT1G06850.1;AT1G06850.2;AT2G21230.2;AT2G21230.1;AT2G21230.3
bic:639
peaks after deleting less than one fifth of global max:20
×10^6
raw data, starting:20
fitted Gauss locations:20
y
x
0
1
2
3
4
5
6
7
8
0
5
10
15
20
25

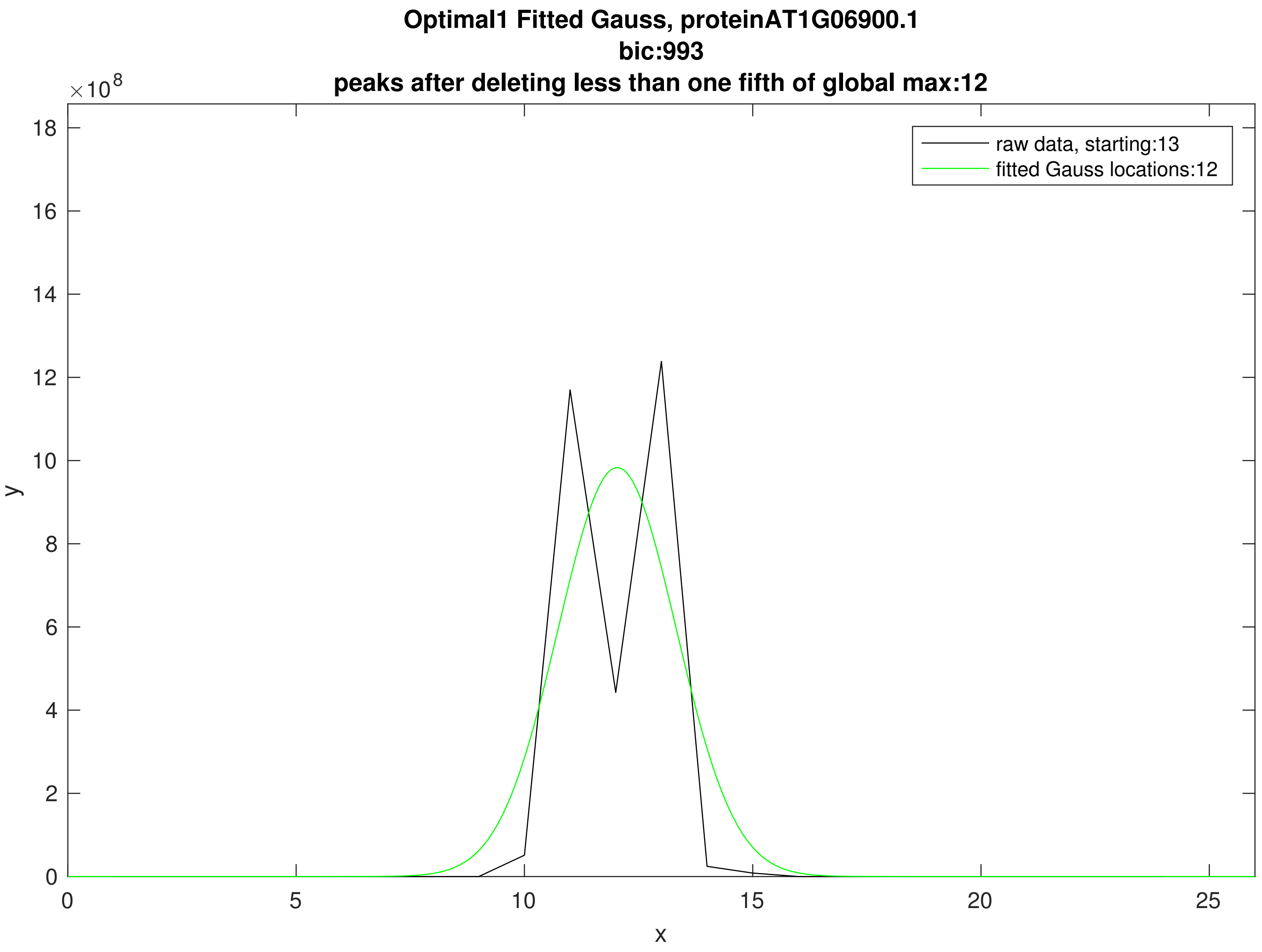
Optimal1 Fitted Gauss, proteinAT1G06900.1
bic:993
peaks after deleting less than one fifth of global max:12
×10^8
raw data, starting:13
fitted Gauss locations:12
18
16
14
12
10
8
6
4
2
0
y
0
5
10
15
20
25
x

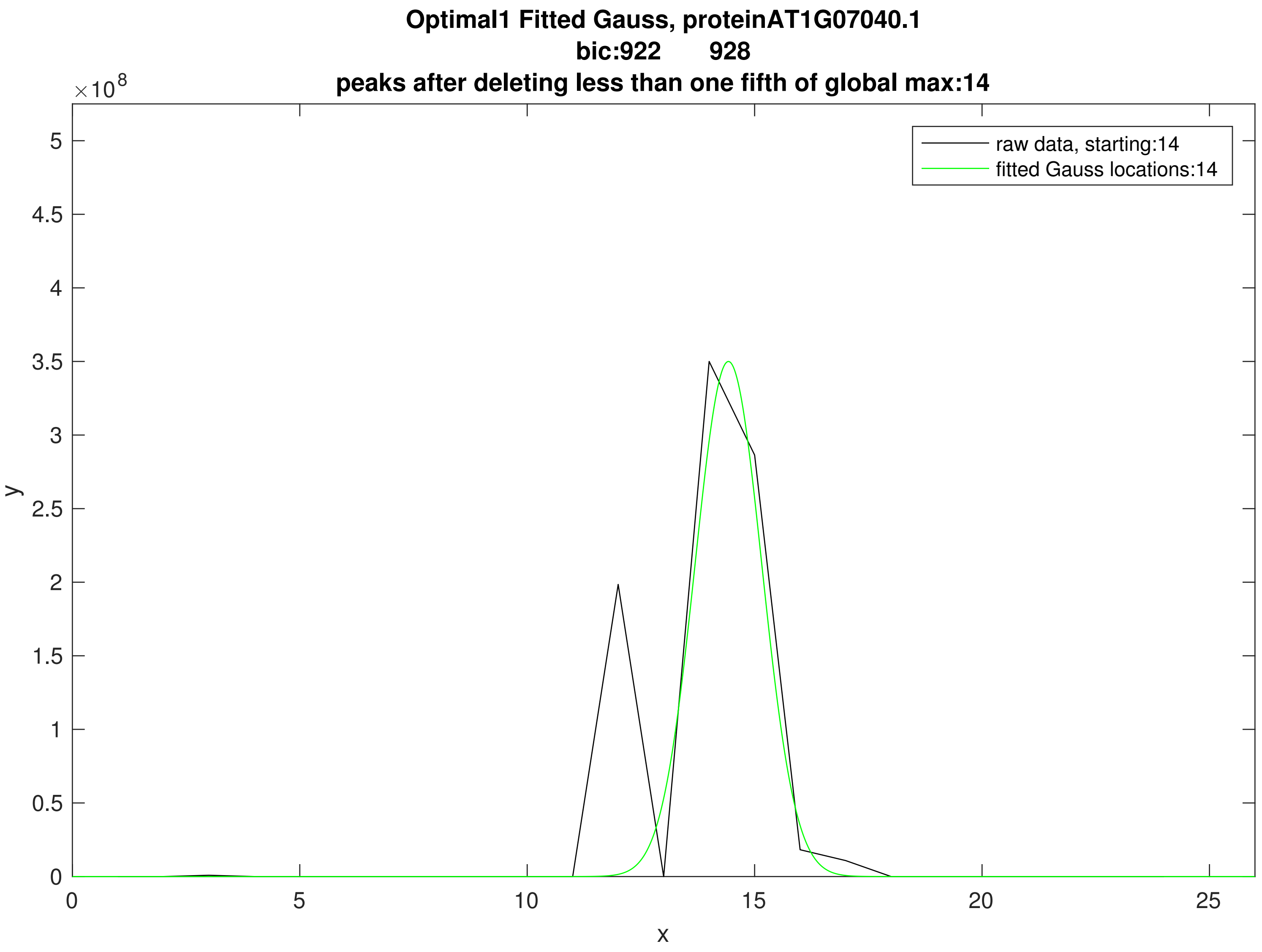
Optimal1 Fitted Gauss, proteinAT1G07040.1
bic:922 928
peaks after deleting less than one fifth of global max:14
×10^8
raw data, starting:14
fitted Gauss locations:14
y
x
0
0.5
1
1.5
2
2.5
3
3.5
4
4.5
5
0
5
10
15
20
25

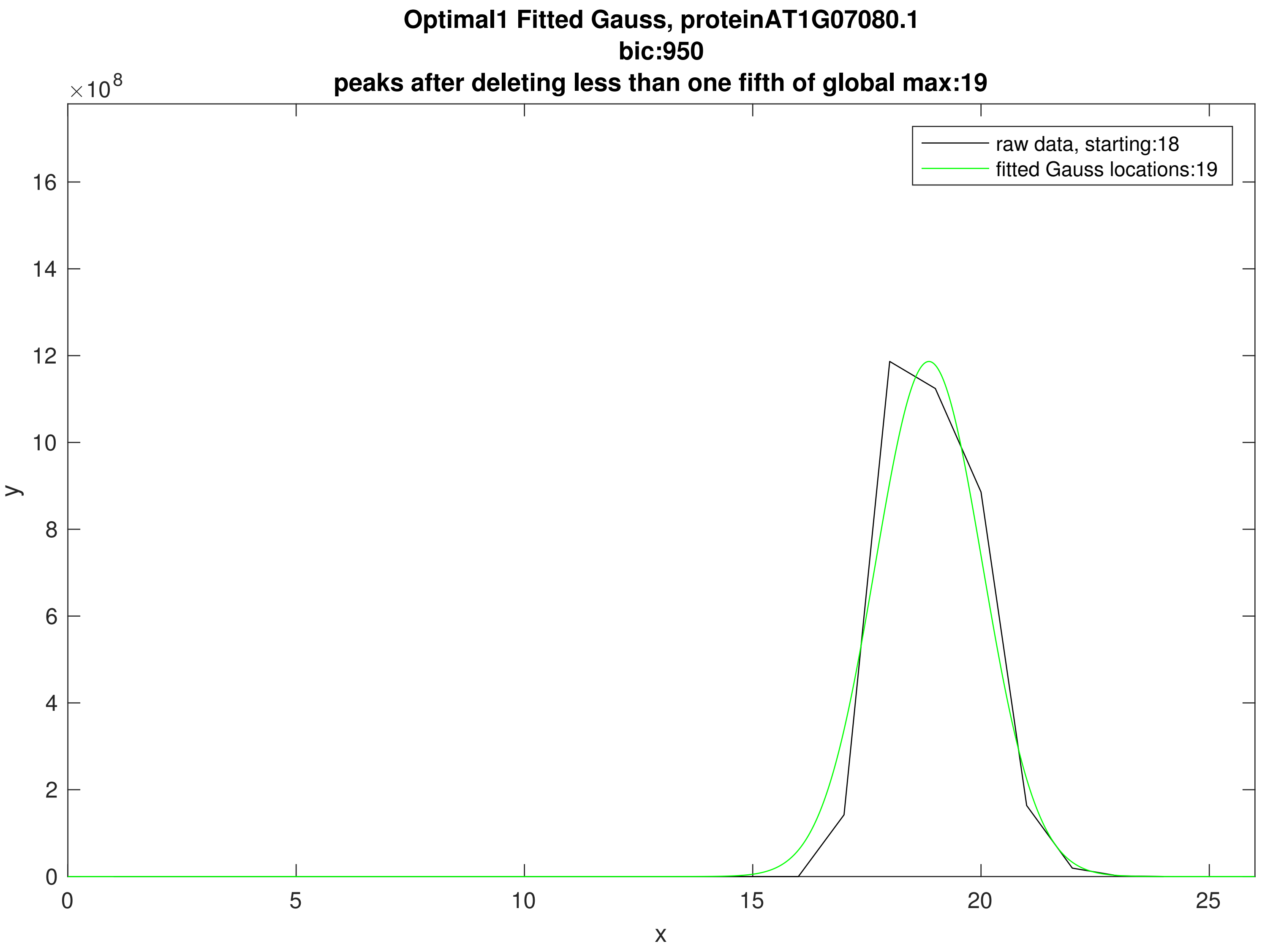

Optimal1 Fitted Gauss, proteinAT1G07080.1
bic:950
peaks after deleting less than one fifth of global max:19
×10^8
raw data, starting:18
fitted Gauss locations:19
16
14
12
10
8
6
4
2
0
y
0
5
10
15
20
25
x

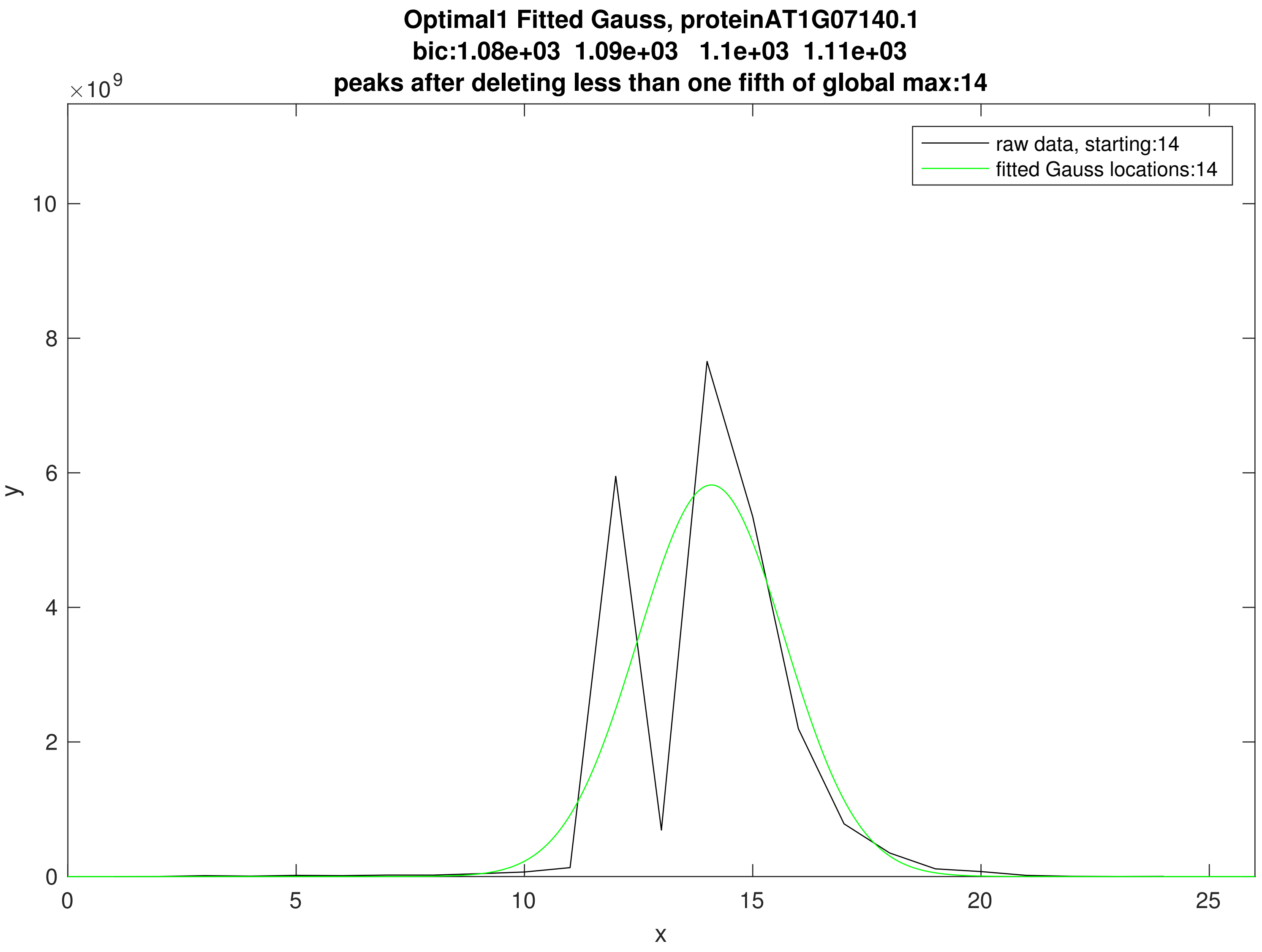
Optimal1 Fitted Gauss, proteinAT1G07140.1
bic:1.08e+03 1.09e+03 1.1e+03 1.11e+03
peaks after deleting less than one fifth of global max:14
×10^9
raw data, starting:14
fitted Gauss locations:14
y
x
0
2
4
6
8
10
0
5
10
15
20
25

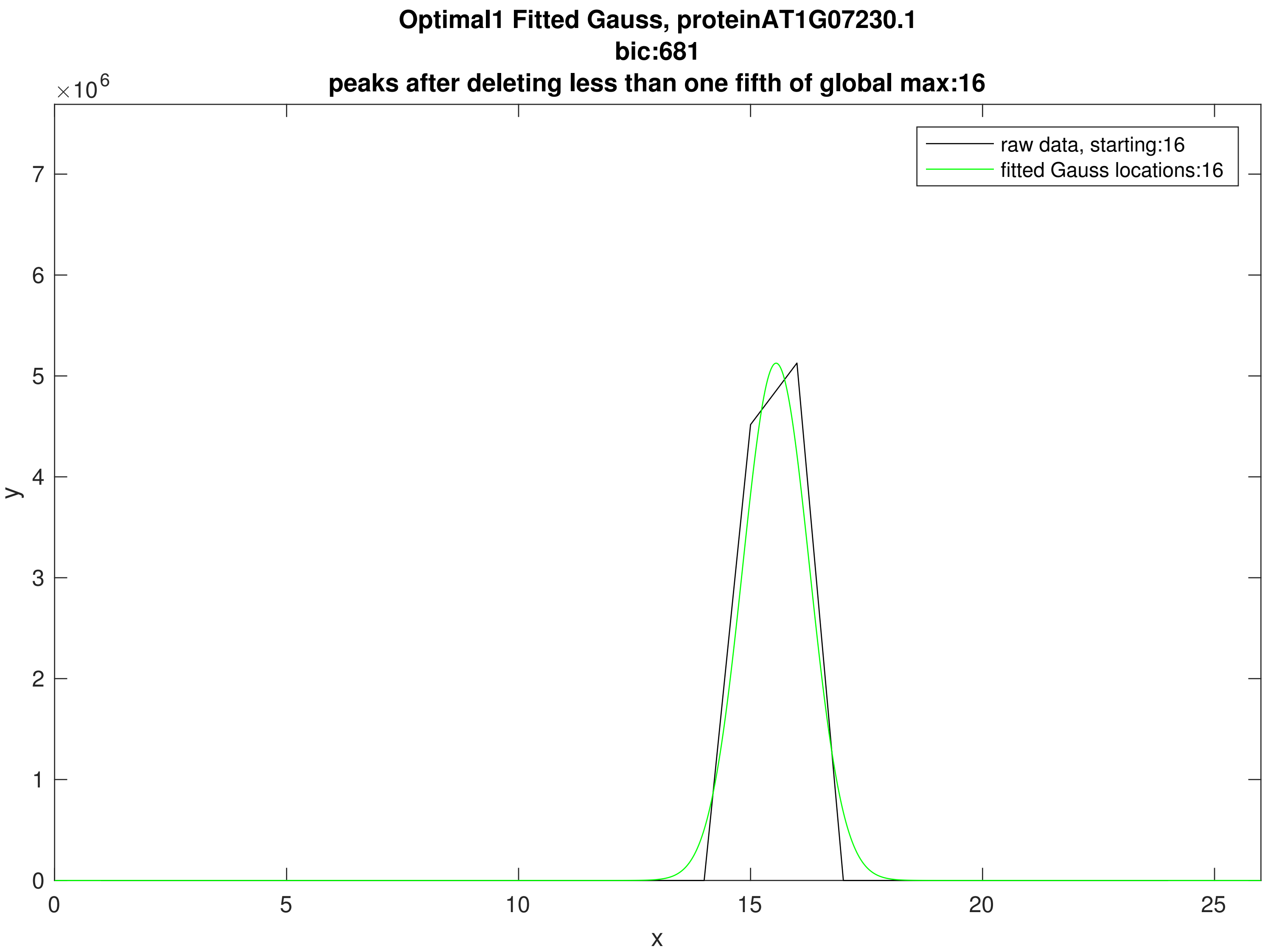

Optimal1 Fitted Gauss, proteinAT1G07230.1
bic:681
peaks after deleting less than one fifth of global max:16
×10^6
raw data, starting:16
fitted Gauss locations:16
7
6
5
4
3
2
1
0
y
0
5
10
15
20
25
x

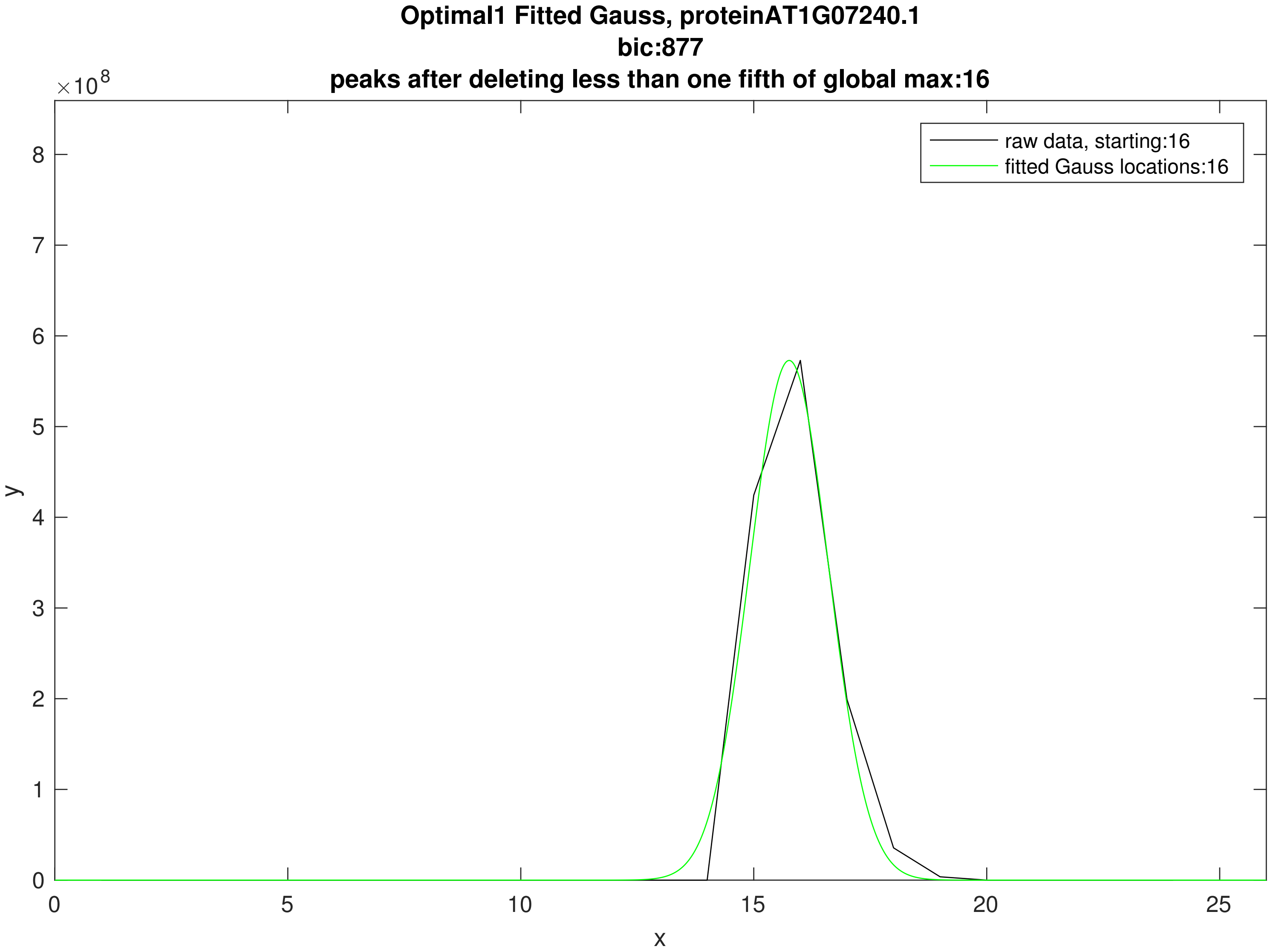

Optimal1 Fitted Gauss, proteinAT1G07240.1
bic:877
peaks after deleting less than one fifth of global max:16
×10^8
8
7
6
5
4
3
2
1
0
y
0
5
10
15
20
25
x
raw data, starting:16
fitted Gauss locations:16

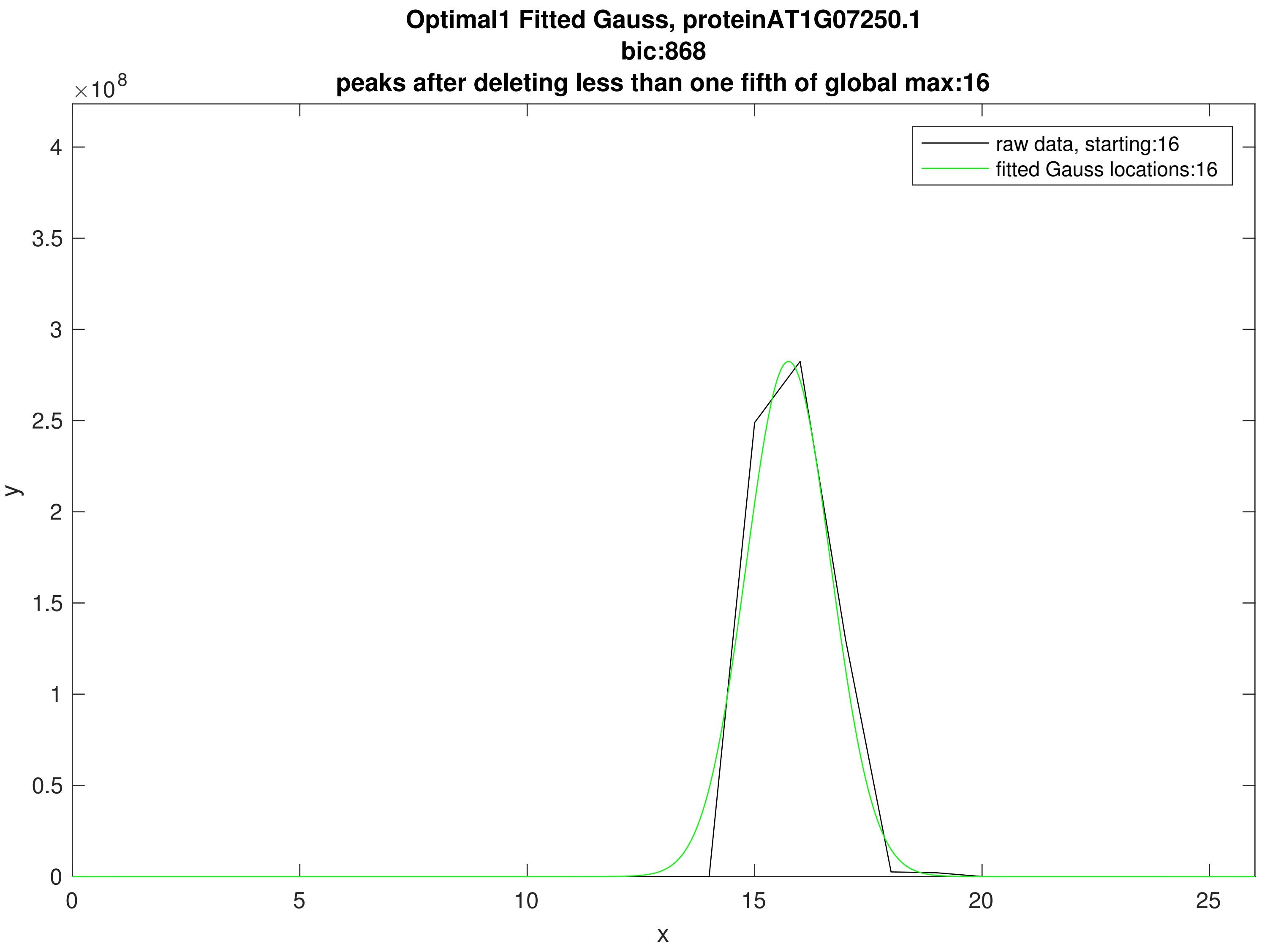
Optimal1 Fitted Gauss, proteinAT1G07250.1
bic:868
peaks after deleting less than one fifth of global max:16
×10^8
raw data, starting:16
fitted Gauss locations:16
y
x
0
0.5
1
1.5
2
2.5
3
3.5
4
0
5
10
15
20
25

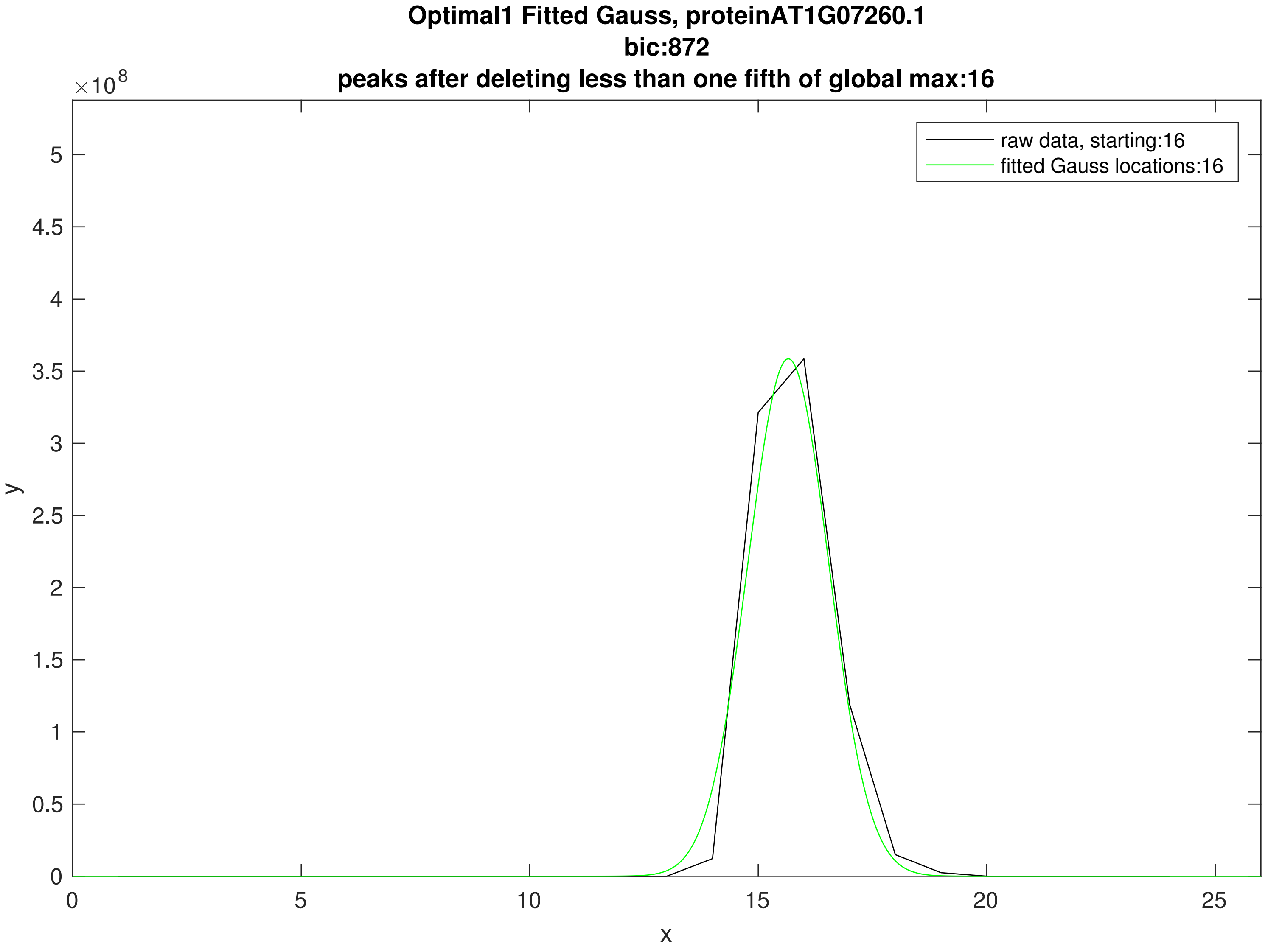

Optimal1 Fitted Gauss, proteinAT1G07260.1
bic:872
peaks after deleting less than one fifth of global max:16
×10^8
raw data, starting:16
fitted Gauss locations:16
y
x
0
0.5
1
1.5
2
2.5
3
3.5
4
4.5
5
0
5
10
15
20
25

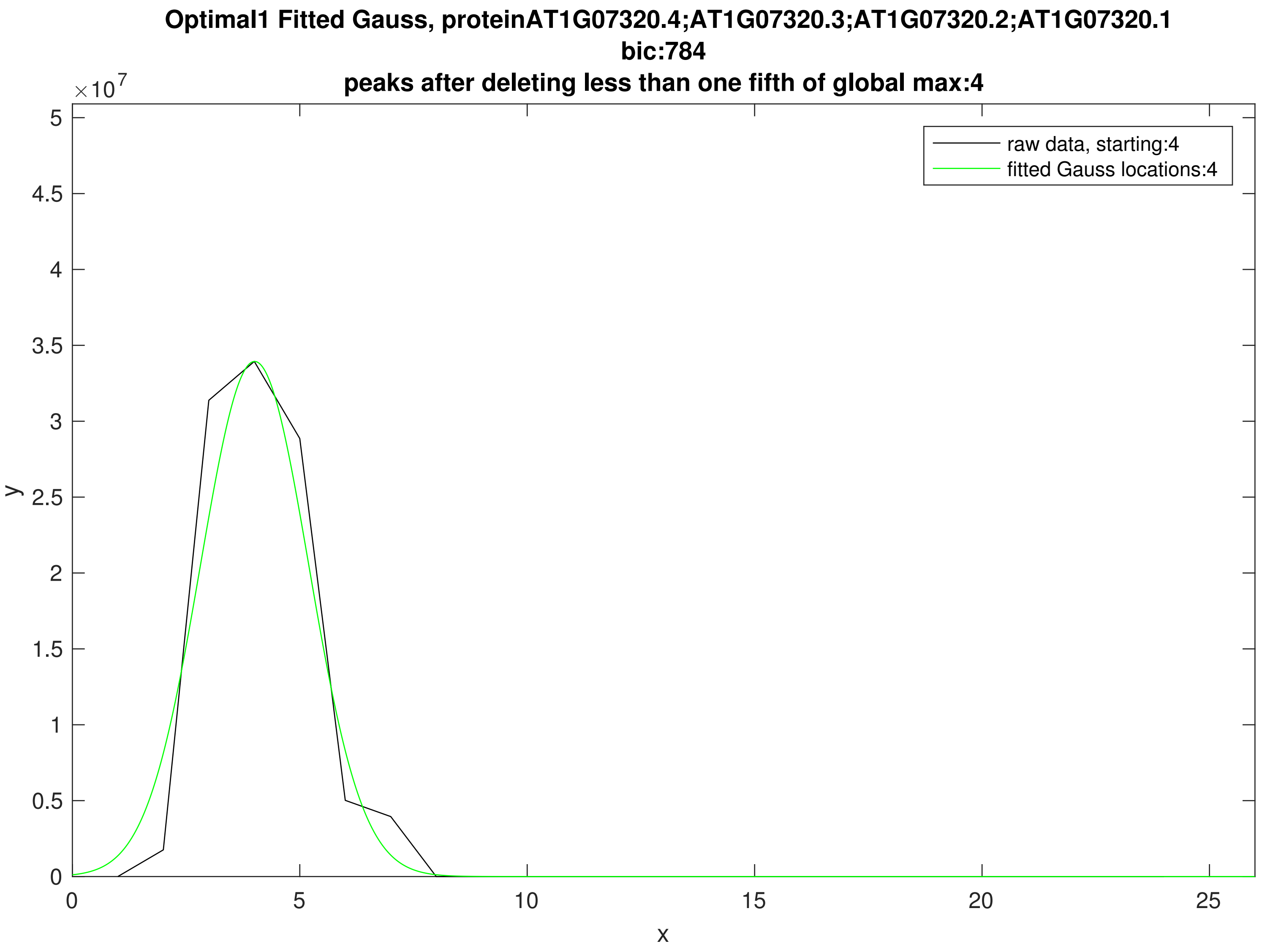

Optimal1 Fitted Gauss, proteinAT1G07320.4;AT1G07320.3;AT1G07320.2;AT1G07320.1
bic:784
peaks after deleting less than one fifth of global max:4
×10^7
raw data, starting:4
fitted Gauss locations:4
y
x
0
0.5
1
1.5
2
2.5
3
3.5
4
4.5
5
0
5
10
15
20
25

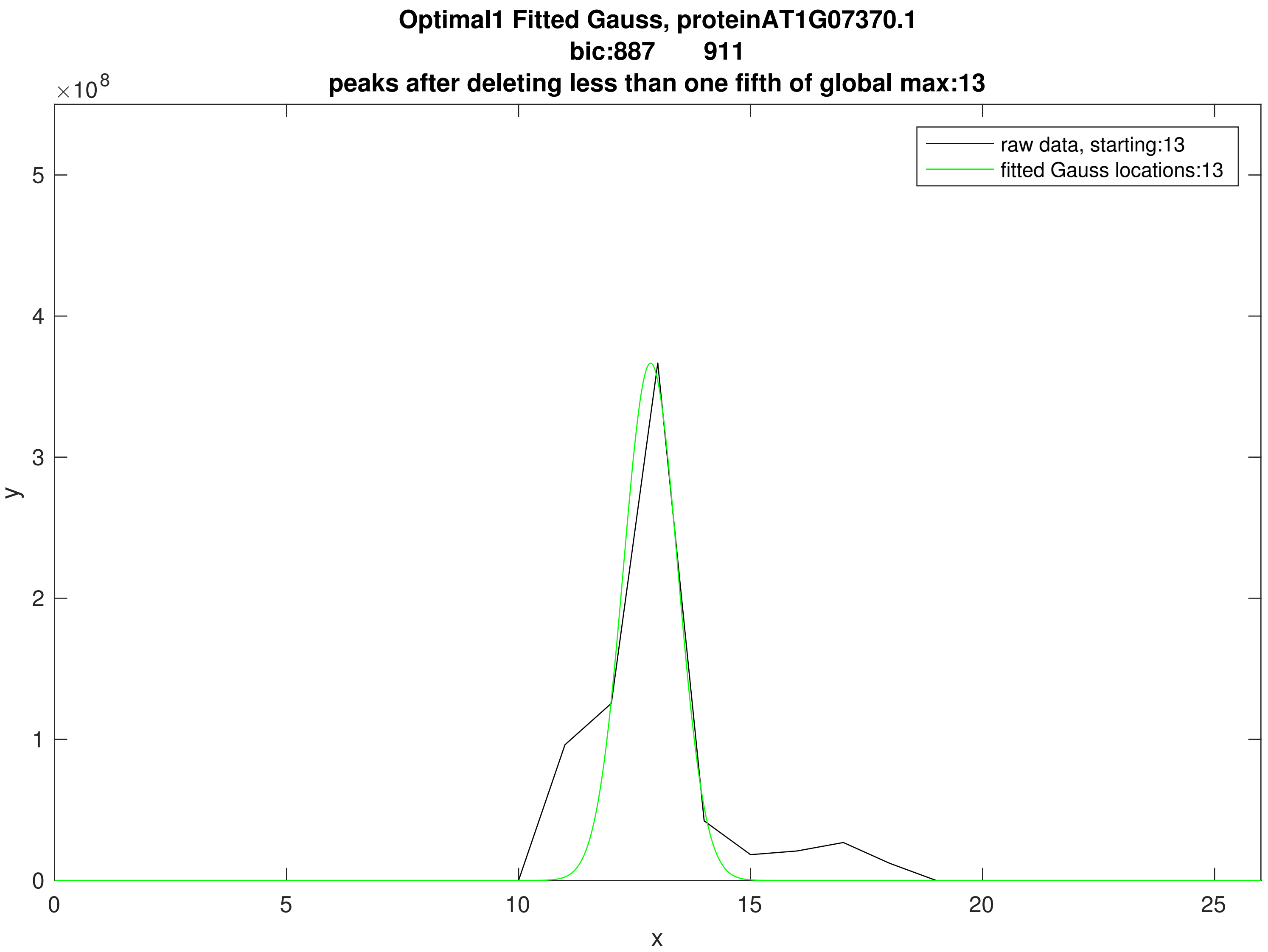

Optimal1 Fitted Gauss, proteinAT1G07370.1
bic:887 911
peaks after deleting less than one fifth of global max:13
×10^8
raw data, starting:13
fitted Gauss locations:13
y
x
0
1
2
3
4
5
0
5
10
15
20
25

#### Optimal1 Fitted Gauss, proteinAT1G07645.1
#### bic:958 960
#### peaks after deleting less than one fifth of global max:17

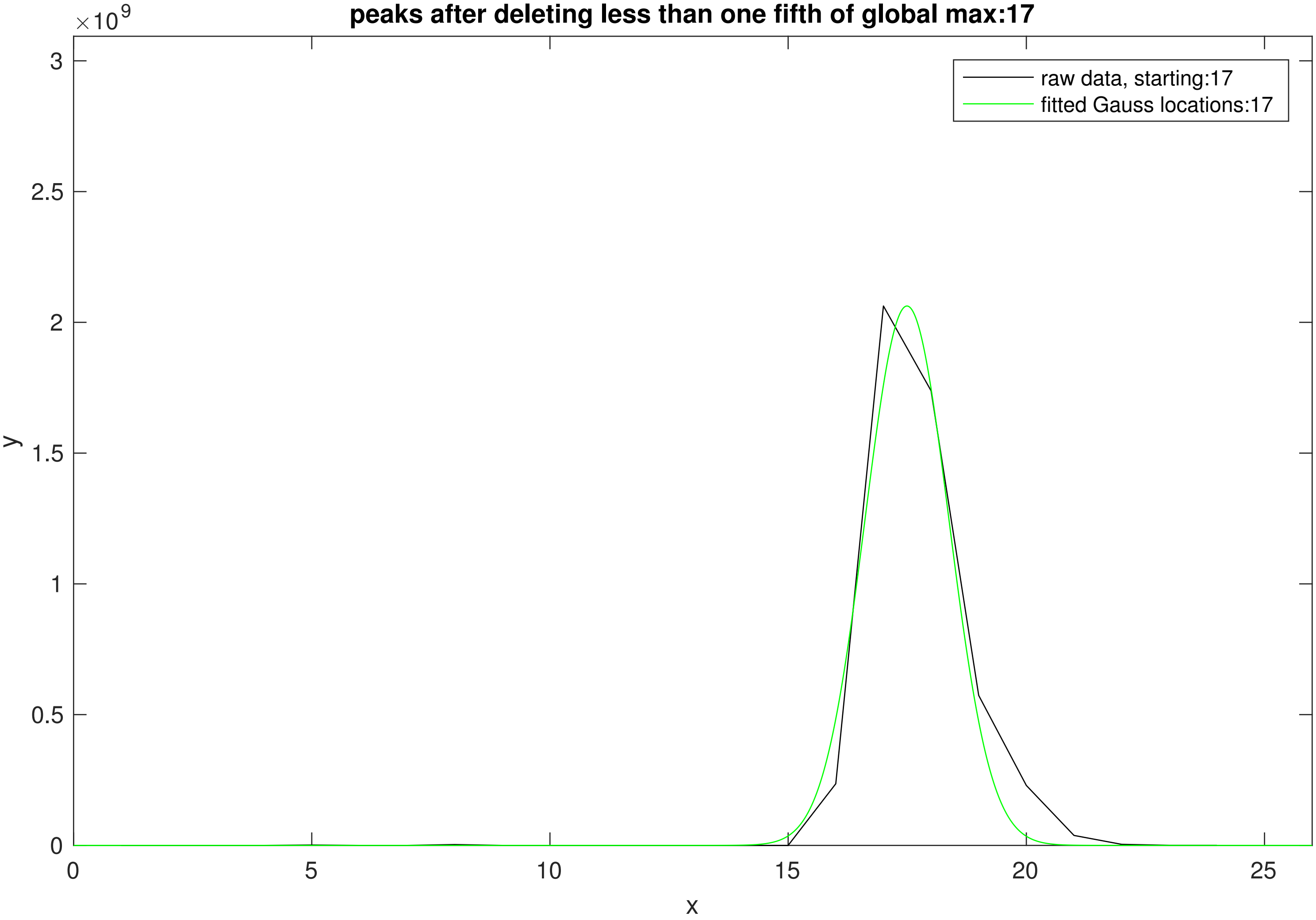

Optimal1 Fitted Gauss, proteinAT1G07700.2;AT1G07700.1;AT1G07700.3
bic:671
peaks after deleting less than one fifth of global max:18

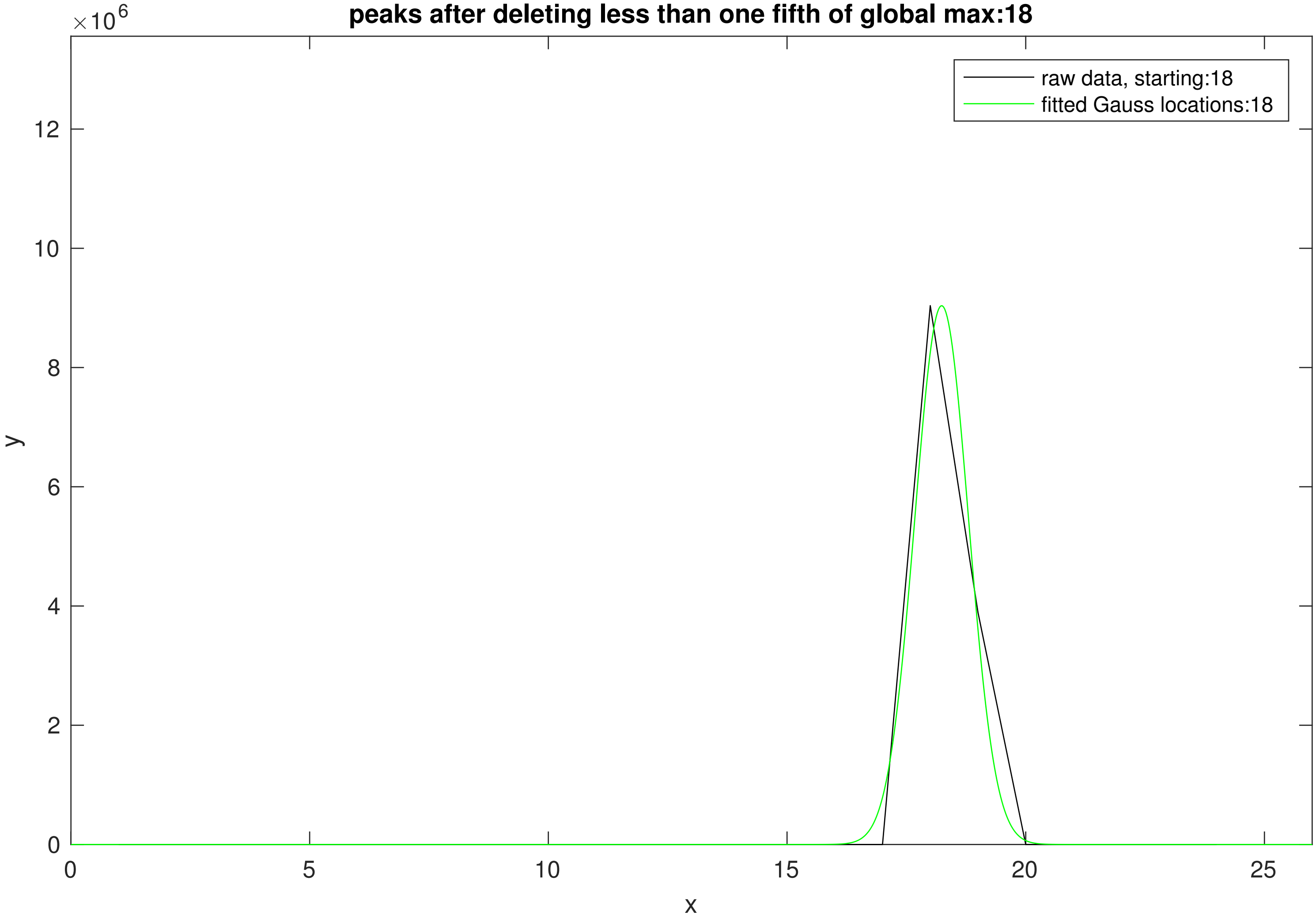

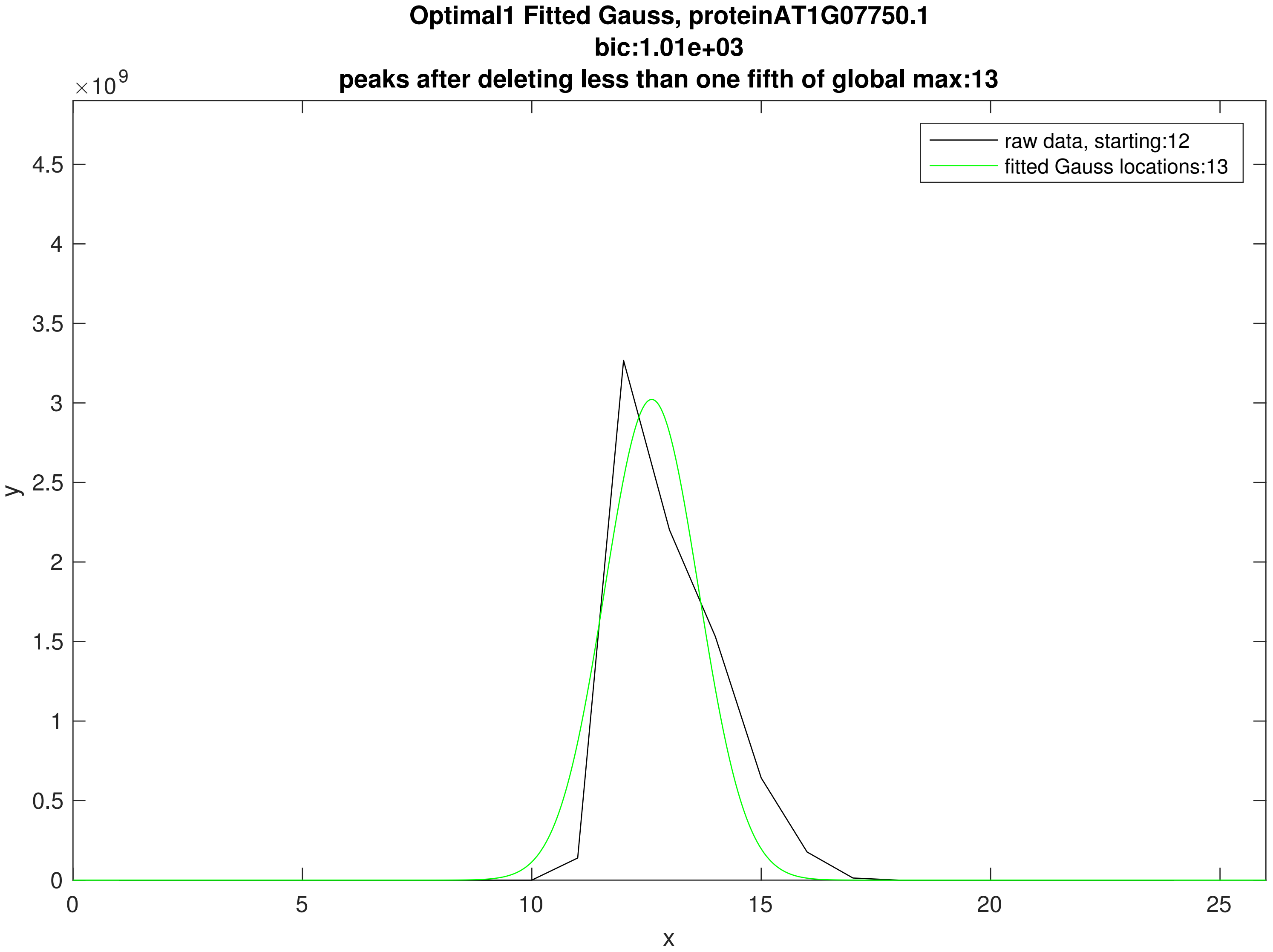

Optimal1 Fitted Gauss, proteinAT1G07750.1
bic:1.01e+03
peaks after deleting less than one fifth of global max:13
×10^9
raw data, starting:12
fitted Gauss locations:13
y
x

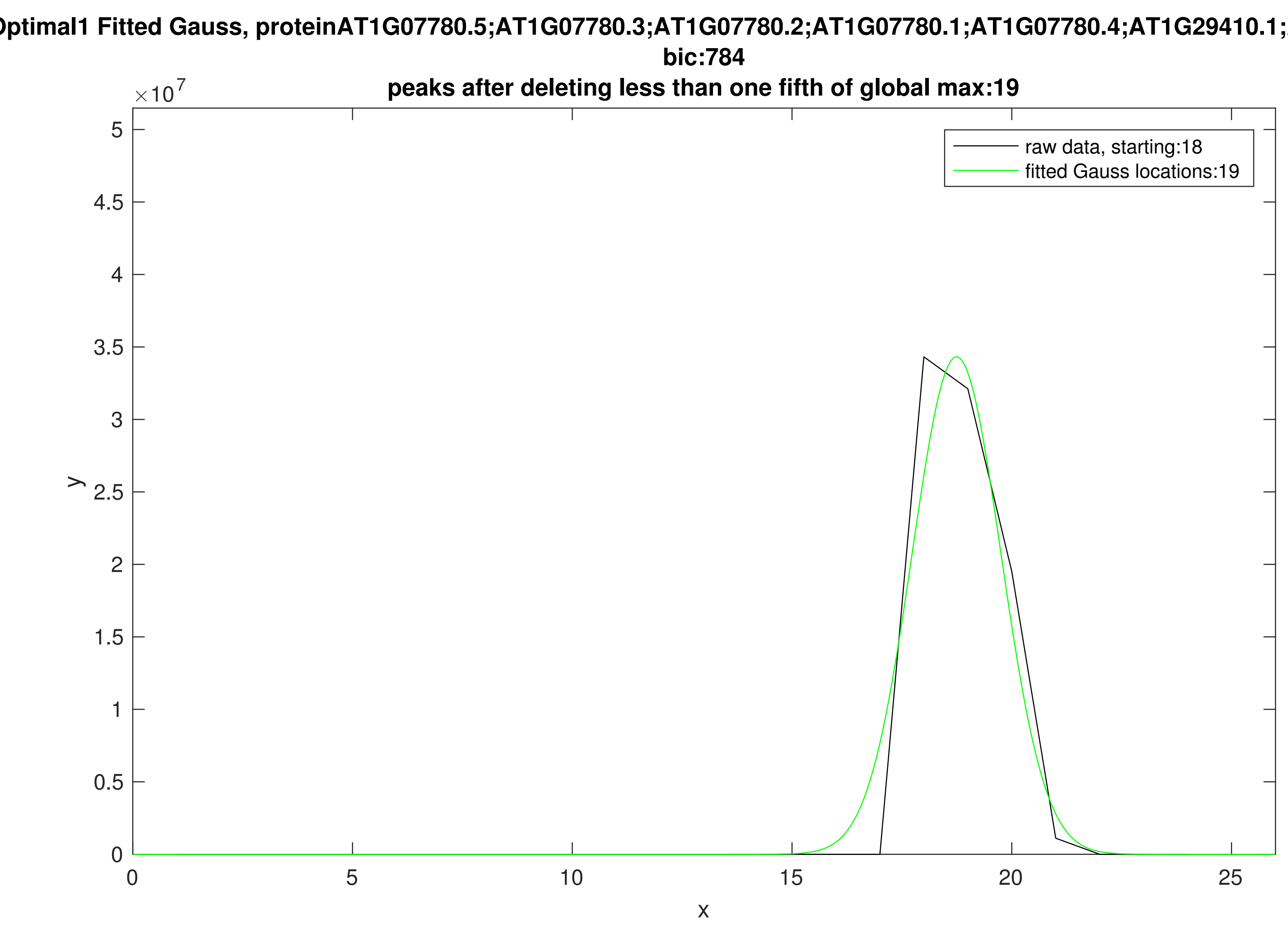
Optimal1 Fitted Gauss, proteinAT1G07780.5;AT1G07780.3;AT1G07780.2;AT1G07780.1;AT1G07780.4;AT1G29410.1;AT1
bic:784
peaks after deleting less than one fifth of global max:19
raw data, starting:18
fitted Gauss locations:19
×10^7
y
x
0
0.5
1
1.5
2
2.5
3
3.5
4
4.5
5
0
5
10
15
20
25

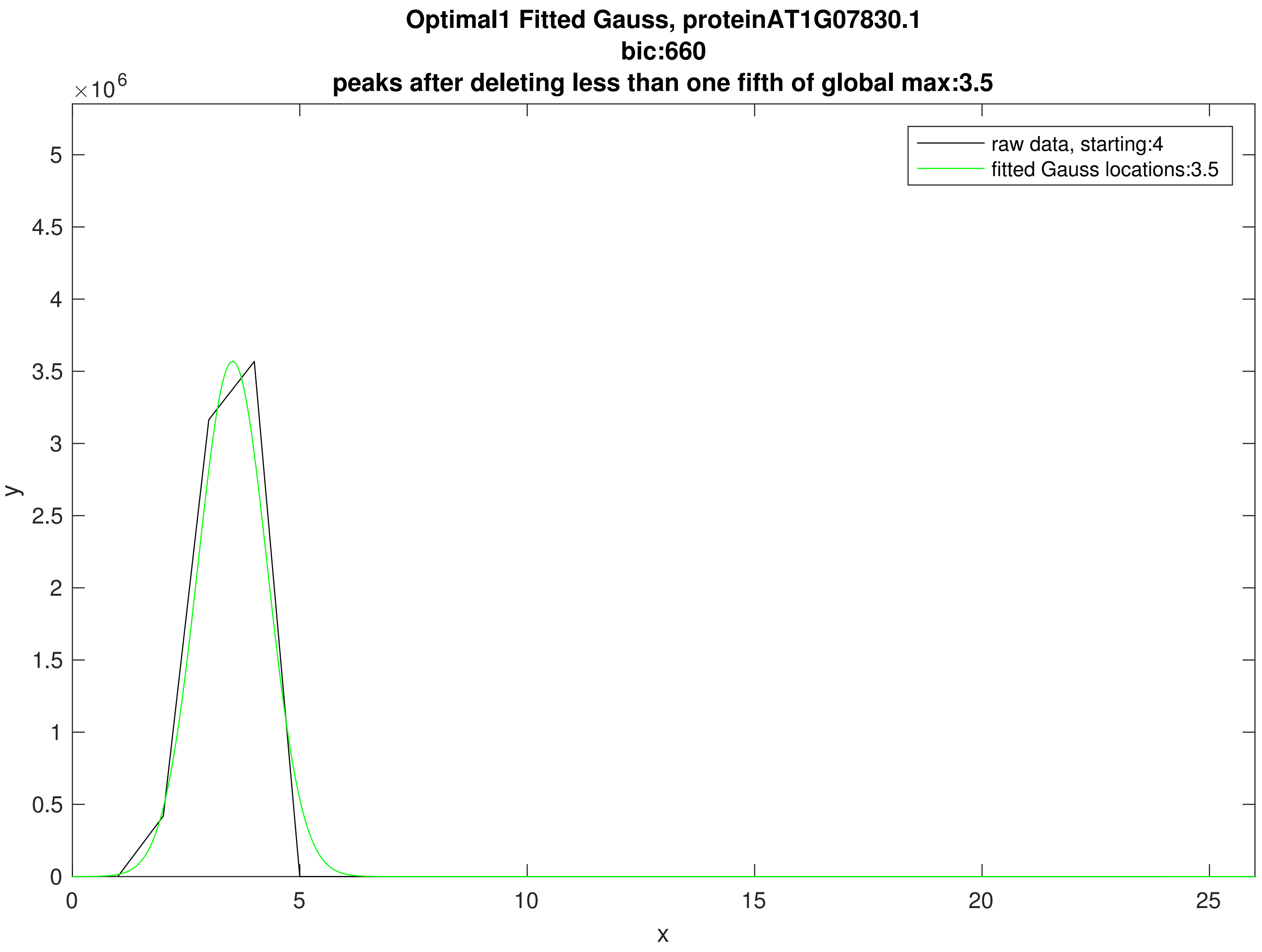

Optimal1 Fitted Gauss, proteinAT1G07830.1
bic:660
peaks after deleting less than one fifth of global max:3.5
×10^6
raw data, starting:4
fitted Gauss locations:3.5
y
x
0
0.5
1
1.5
2
2.5
3
3.5
4
4.5
5
10
15
20
25

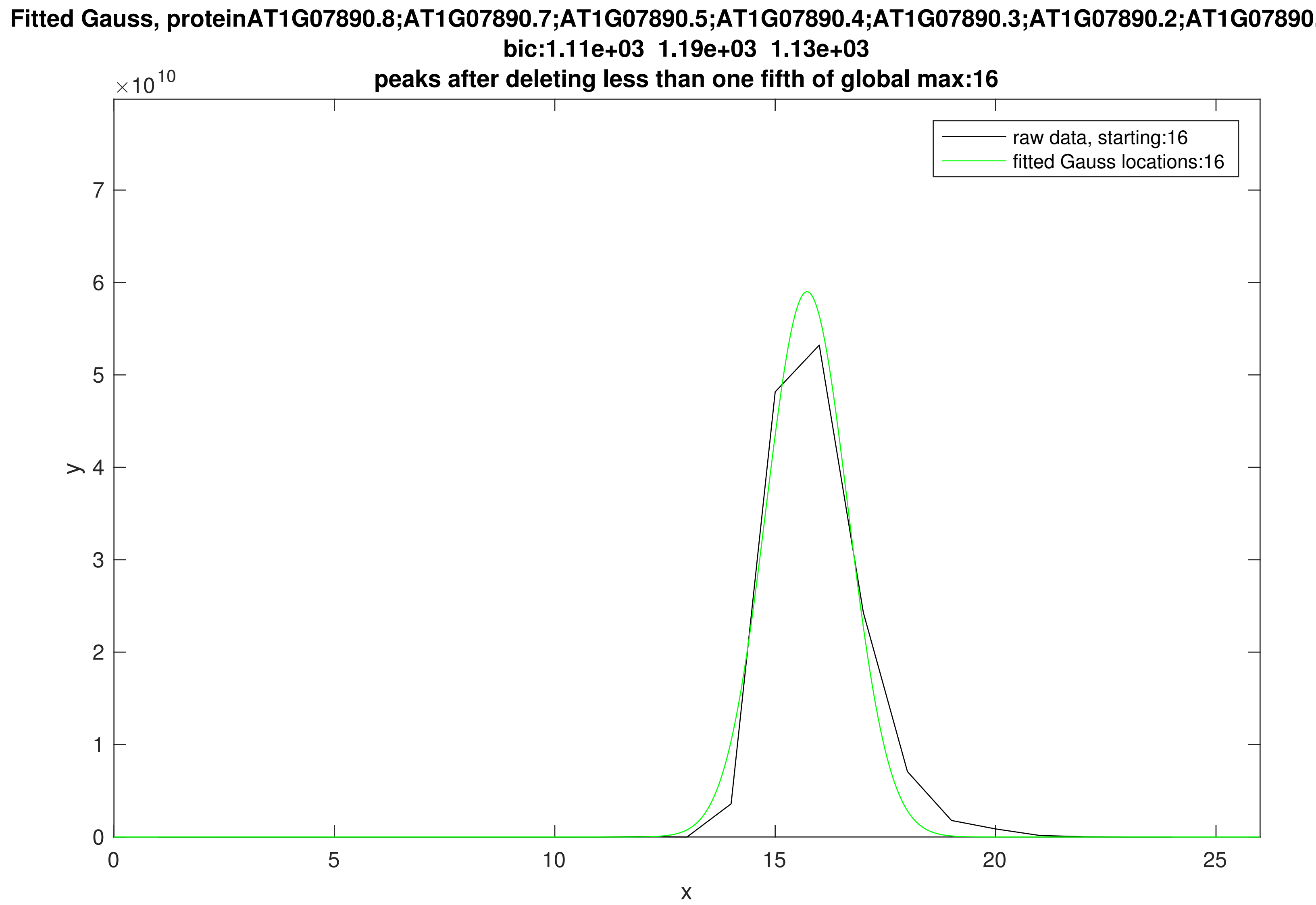
Fitted Gauss, proteinAT1G07890.8;AT1G07890.7;AT1G07890.5;AT1G07890.4;AT1G07890.3;AT1G07890.2;AT1G07890.
bic:1.11e+03 1.19e+03 1.13e+03
peaks after deleting less than one fifth of global max:16
×10^10
raw data, starting:16
fitted Gauss locations:16
y
x
0
1
2
3
4
5
6
7
0
5
10
15
20
25

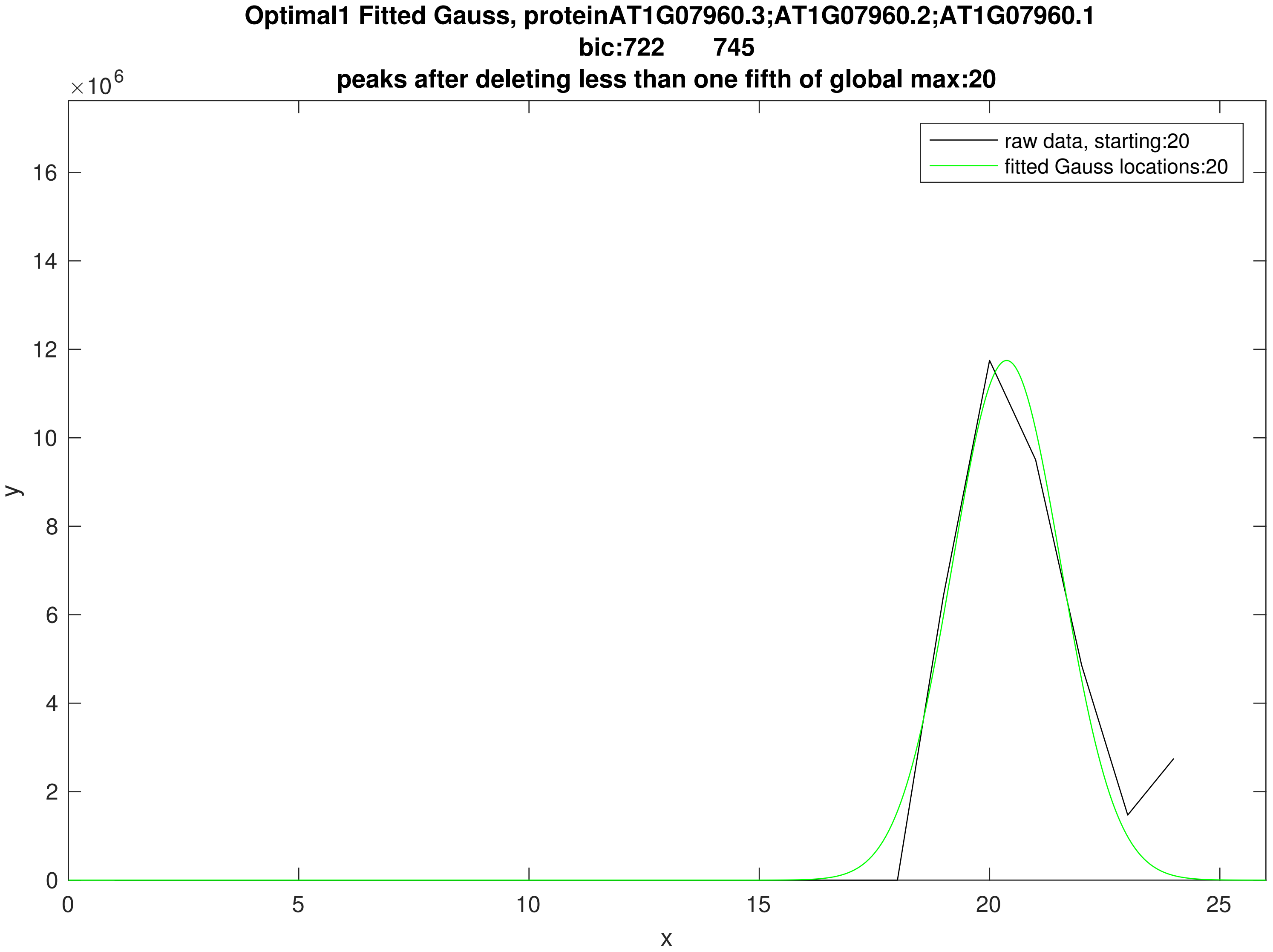

Optimal1 Fitted Gauss, proteinAT1G07960.3;AT1G07960.2;AT1G07960.1
bic:722 745
peaks after deleting less than one fifth of global max:20
×10^6
raw data, starting:20
fitted Gauss locations:20
y
x

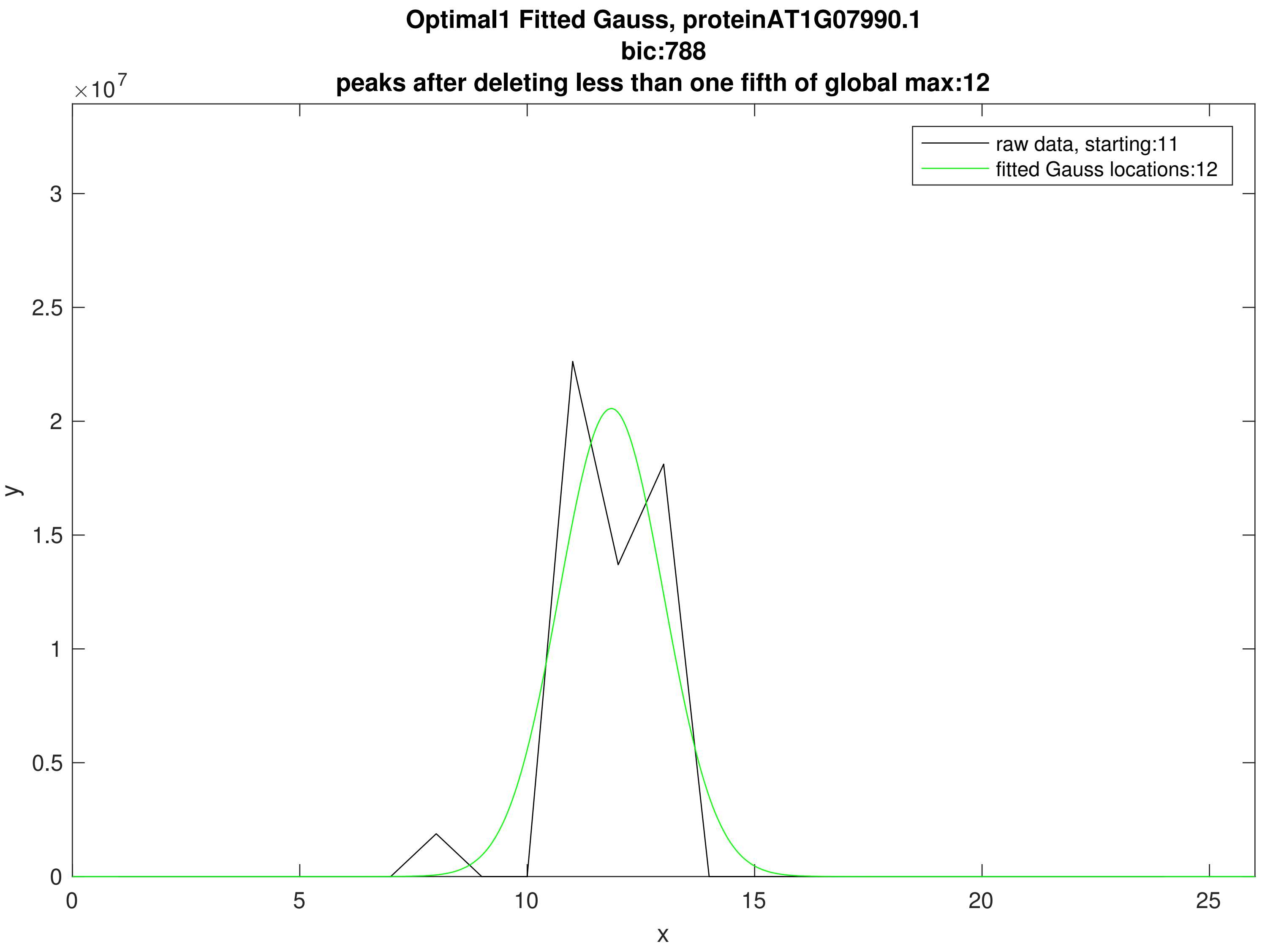
Optimal1 Fitted Gauss, proteinAT1G07990.1
bic:788
peaks after deleting less than one fifth of global max:12
×10^7
3
2.5
2
1.5
1
0.5
0
y
0
5
10
15
20
25
x
raw data, starting:11
fitted Gauss locations:12

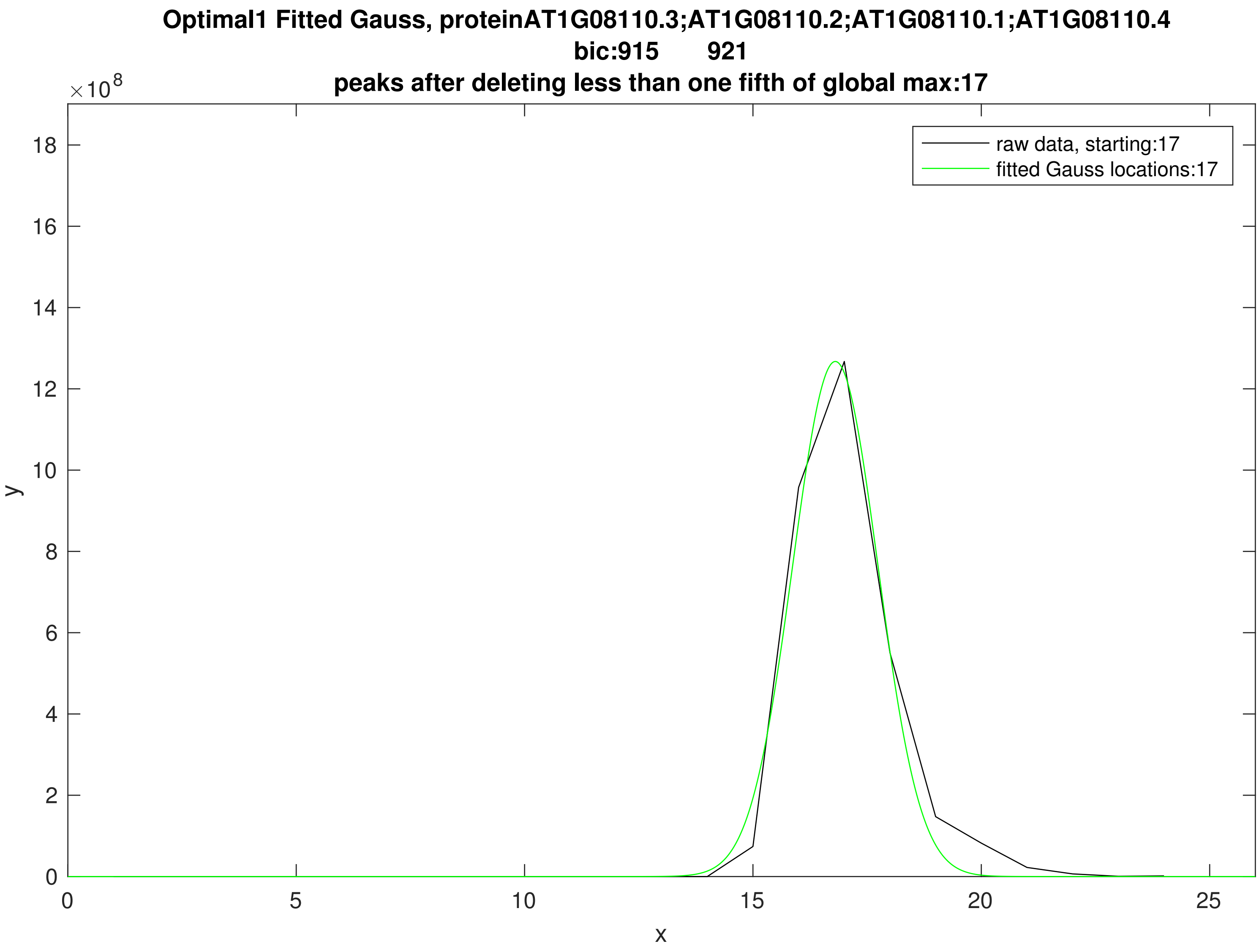

Optimal1 Fitted Gauss, proteinAT1G08110.3;AT1G08110.2;AT1G08110.1;AT1G08110.4
bic:915 921
peaks after deleting less than one fifth of global max:17
×10^8
raw data, starting:17
fitted Gauss locations:17
y
x
0
2
4
6
8
10
12
14
16
18
0
5
10
15
20
25

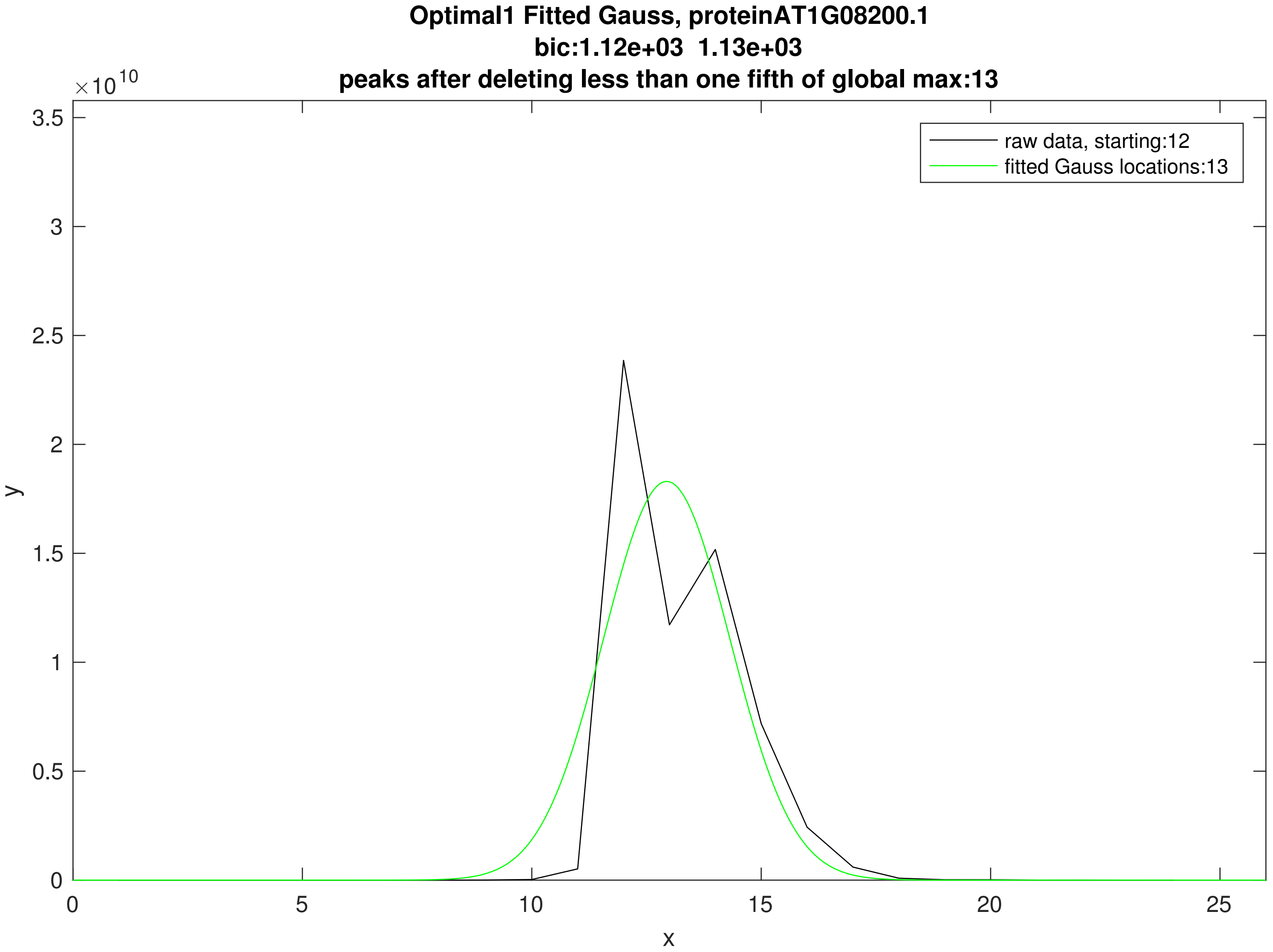

Optimal1 Fitted Gauss, proteinAT1G08200.1
bic:1.12e+03  1.13e+03
peaks after deleting less than one fifth of global max:13
×10^10
3.5
3
2.5
2
1.5
1
0.5
0
y
0
5
10
15
20
25
x
raw data, starting:12
fitted Gauss locations:13

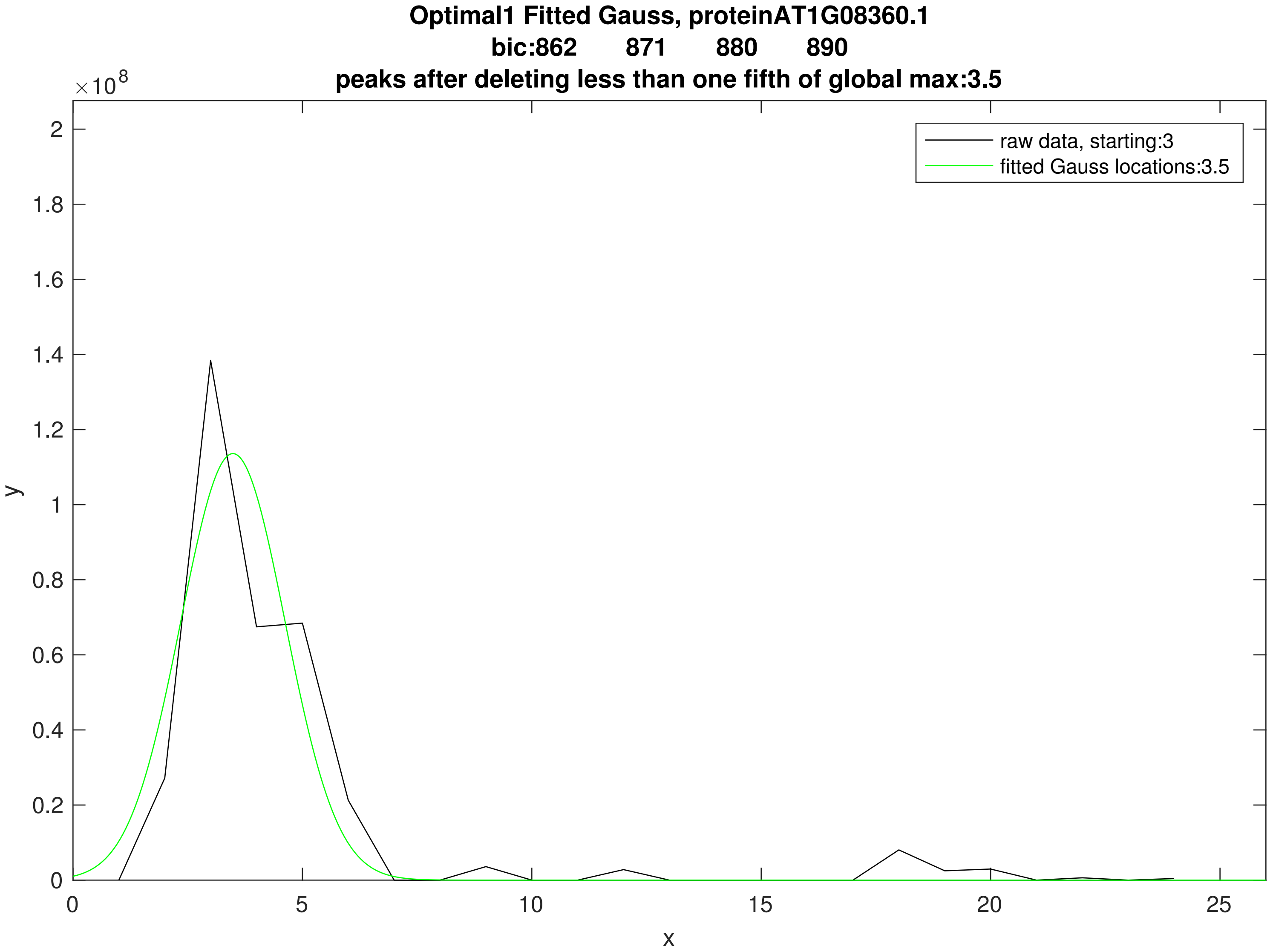

Optimal1 Fitted Gauss, proteinAT1G08360.1
bic:862 871 880 890
peaks after deleting less than one fifth of global max:3.5
×10^8
raw data, starting:3
fitted Gauss locations:3.5
y
x
0
0.2
0.4
0.6
0.8
1
1.2
1.4
1.6
1.8
2
0
5
10
15
20
25

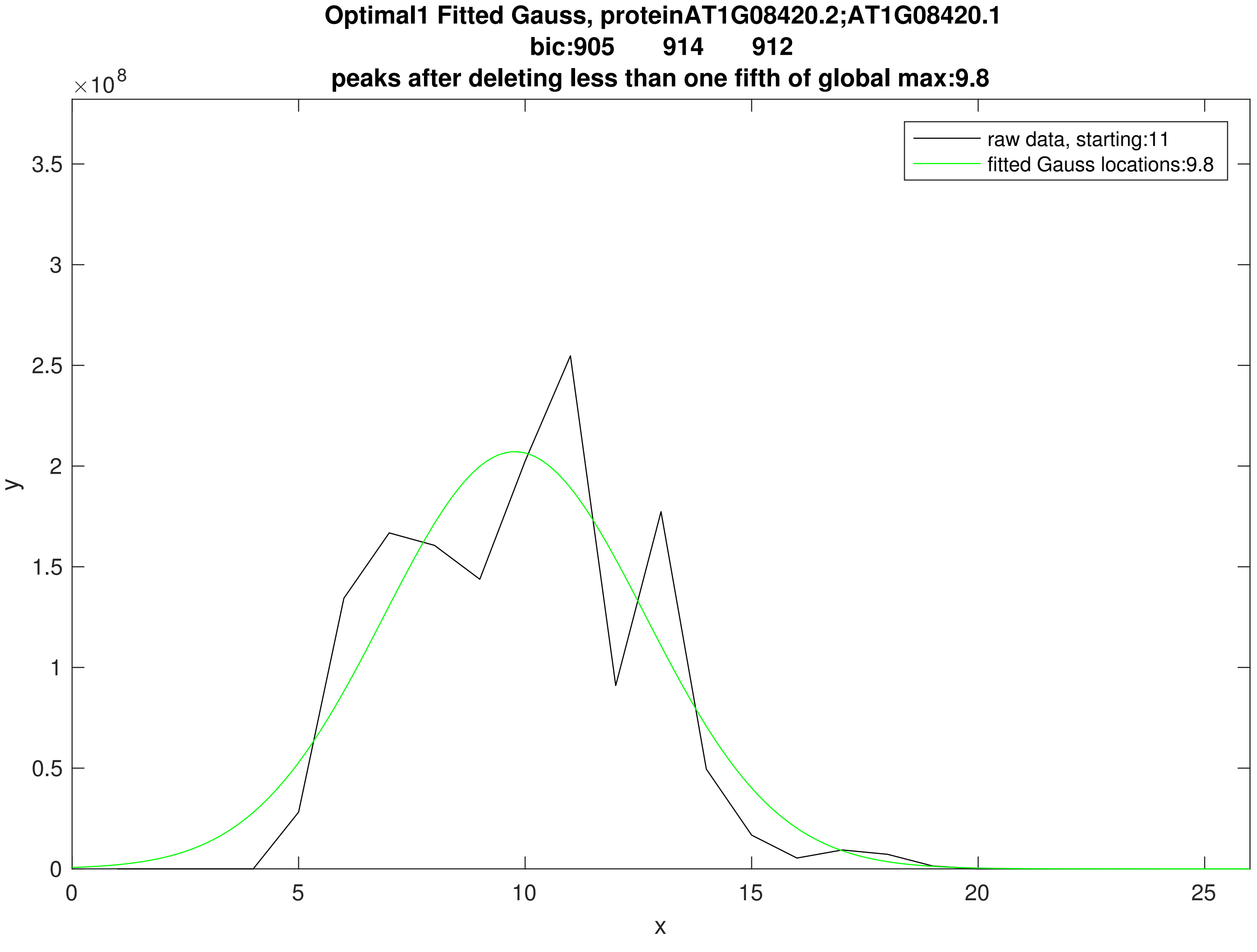

Optimal1 Fitted Gauss, proteinAT1G08420.2;AT1G08420.1
bic:905 914 912
peaks after deleting less than one fifth of global max:9.8
×10^8
raw data, starting:11
fitted Gauss locations:9.8
y
x
0
0.5
1
1.5
2
2.5
3
3.5
0
5
10
15
20
25

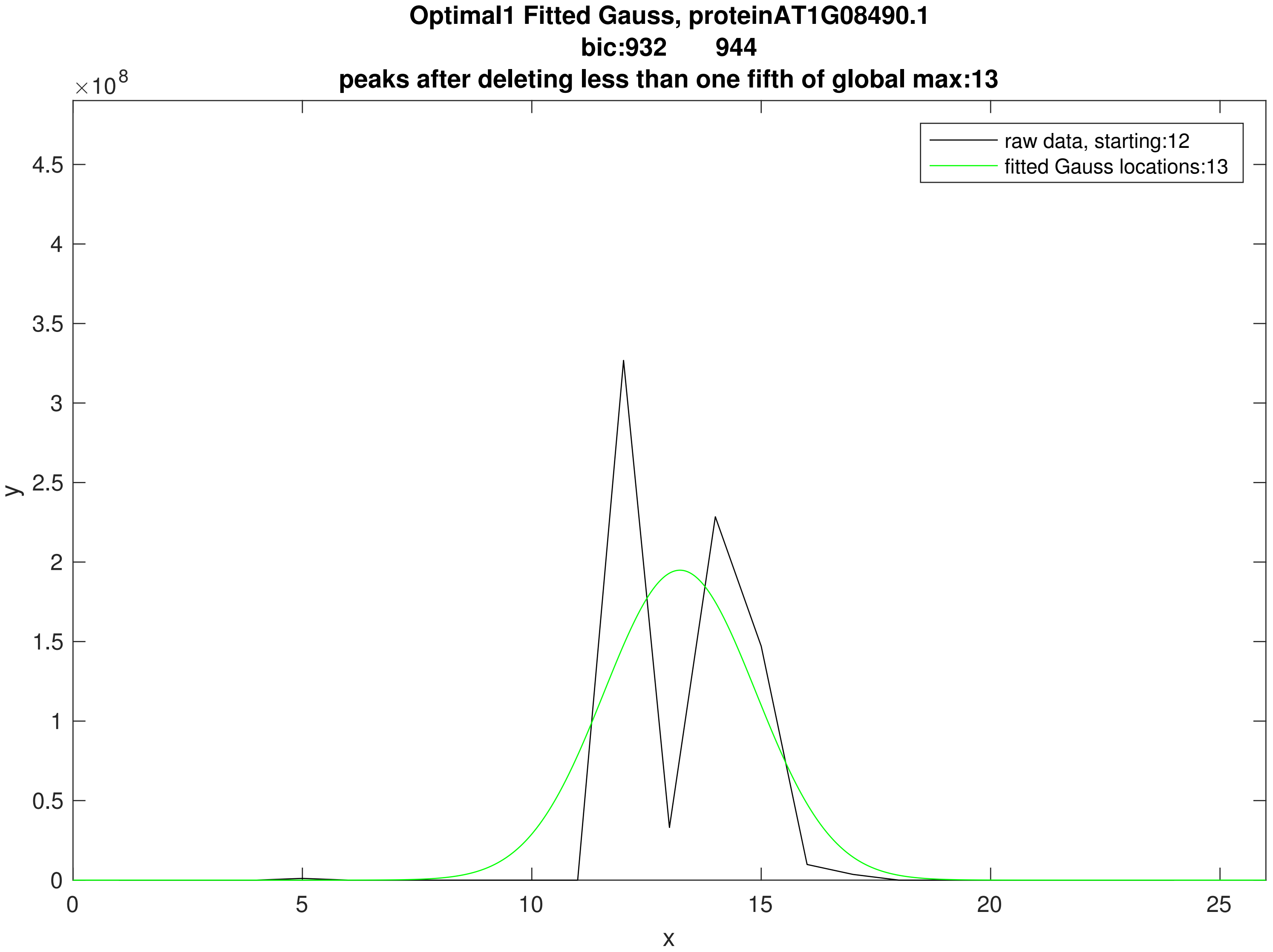
Optimal1 Fitted Gauss, proteinAT1G08490.1
bic:932 944
peaks after deleting less than one fifth of global max:13
×10^8
4.5
4
3.5
3
2.5
2
1.5
1
0.5
0
y
0
5
10
15
20
25
x
raw data, starting:12
fitted Gauss locations:13

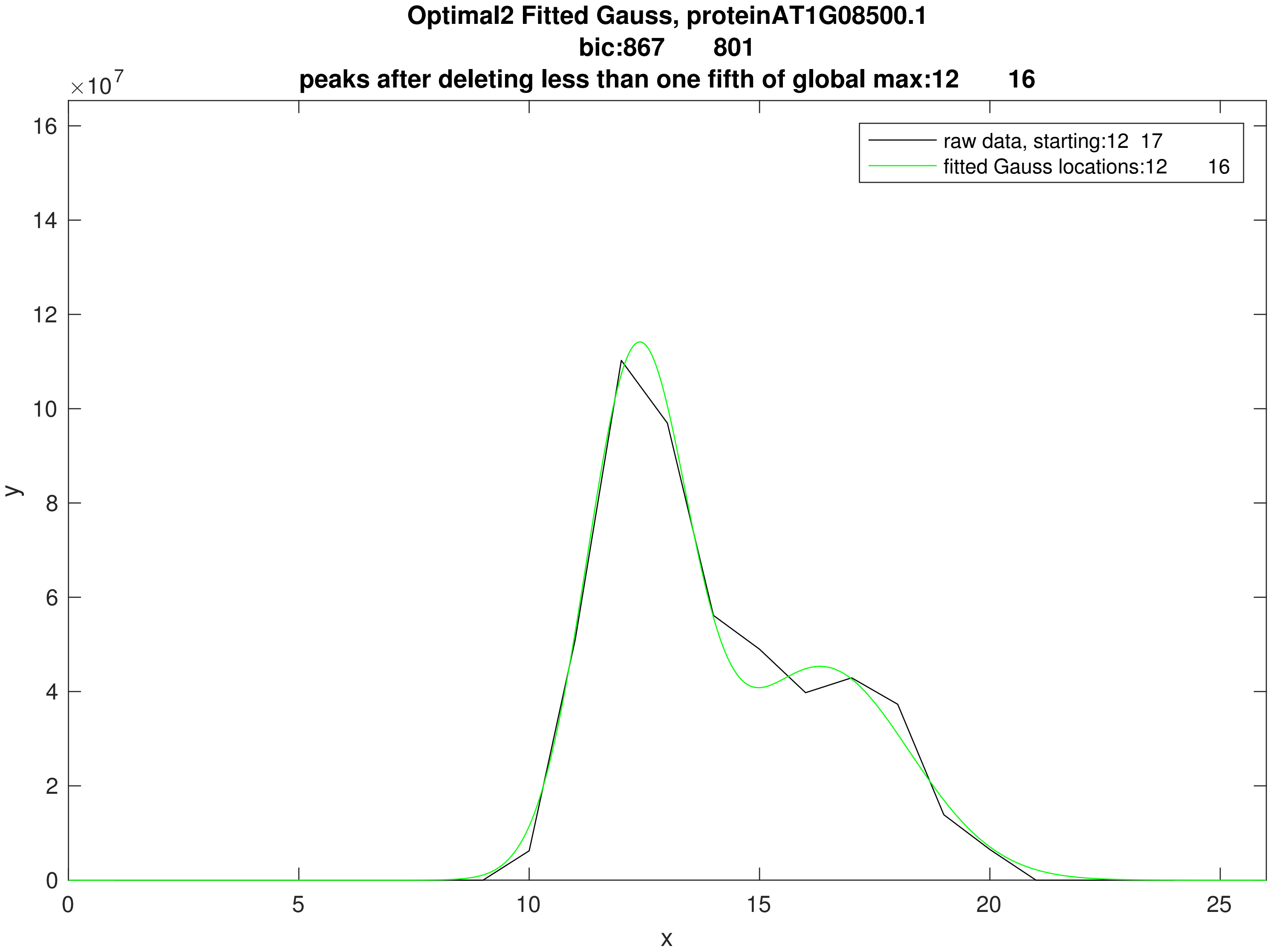
Optimal2 Fitted Gauss, proteinAT1G08500.1
bic:867 801
peaks after deleting less than one fifth of global max:12 16
×10^7
raw data, starting:12 17
fitted Gauss locations:12 16
y
x
0
2
4
6
8
10
12
14
16
0
5
10
15
20
25

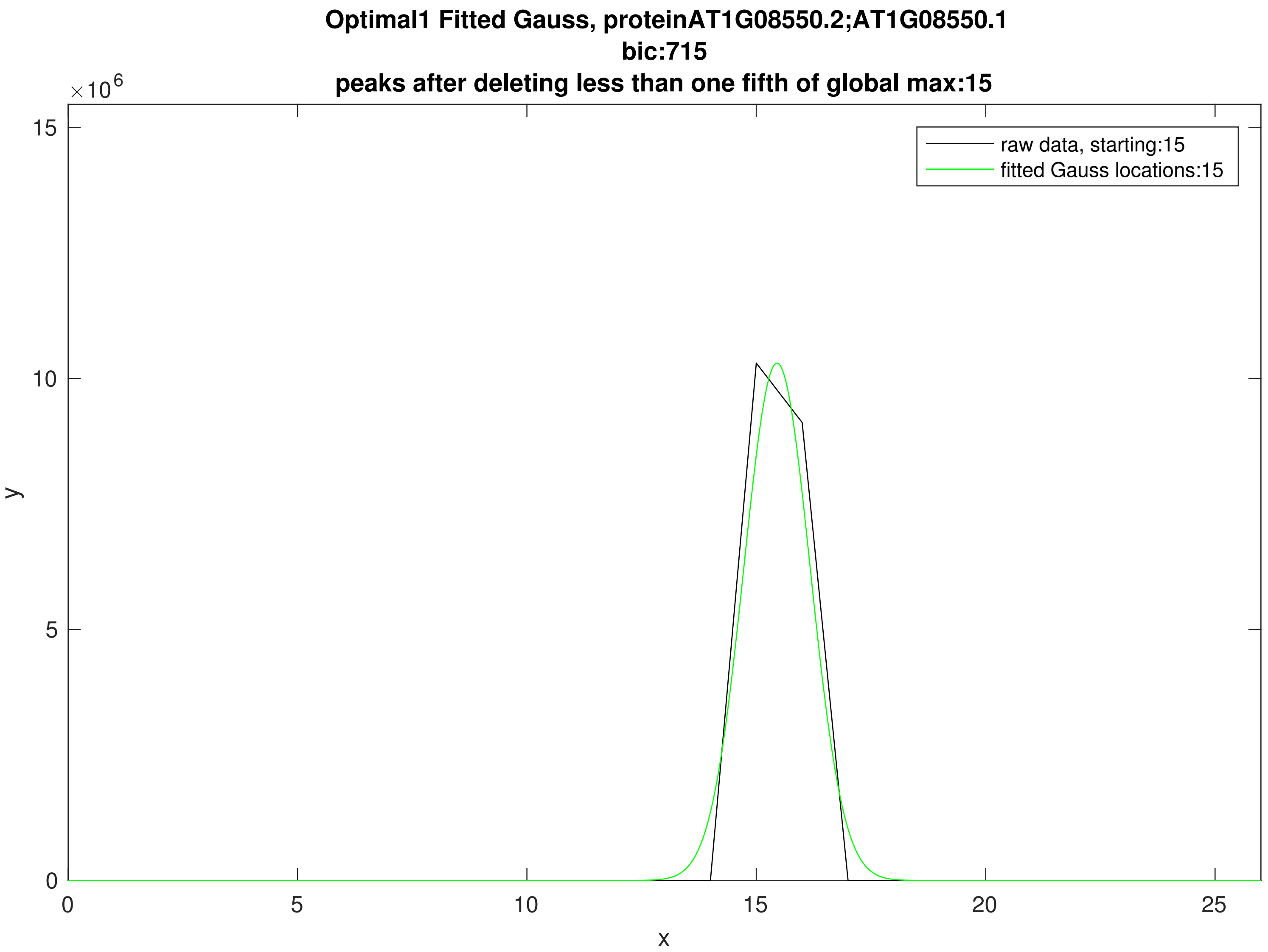

Optimal1 Fitted Gauss, proteinAT1G08550.2;AT1G08550.1
bic:715
peaks after deleting less than one fifth of global max:15
×10^6
raw data, starting:15
fitted Gauss locations:15
y
x
0
5
10
15
20
25

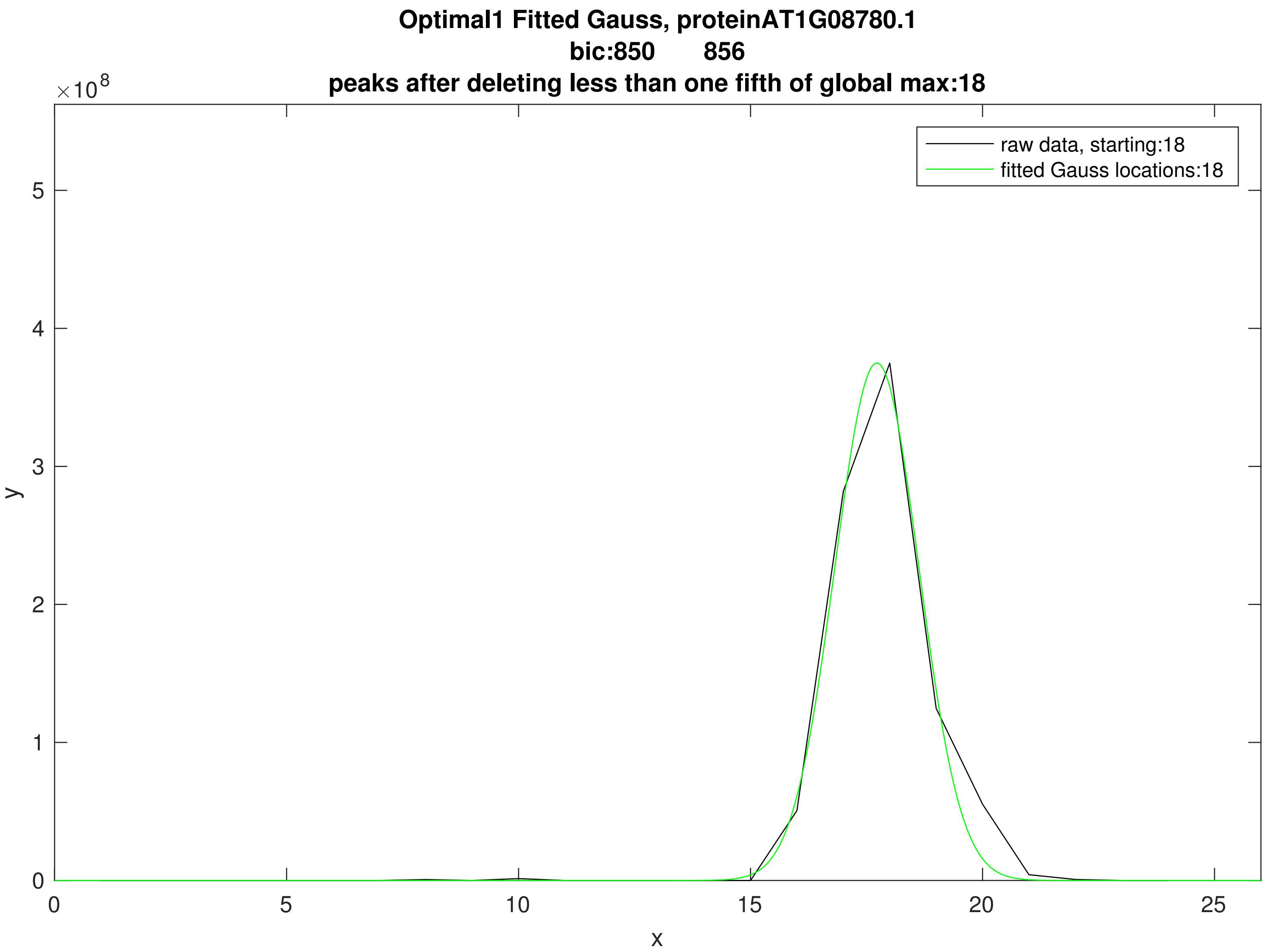

Optimal1 Fitted Gauss, proteinAT1G08780.1
bic:850 856
peaks after deleting less than one fifth of global max:18
×10^8
5
4
3
2
1
0
y
0
5
10
15
20
25
x
raw data, starting:18
fitted Gauss locations:18

Optimal1 Fitted Gauss, proteinAT1G08830.2;AT1G08830.1
bic:1.03e+03  1.04e+03
peaks after deleting less than one fifth of global max:17

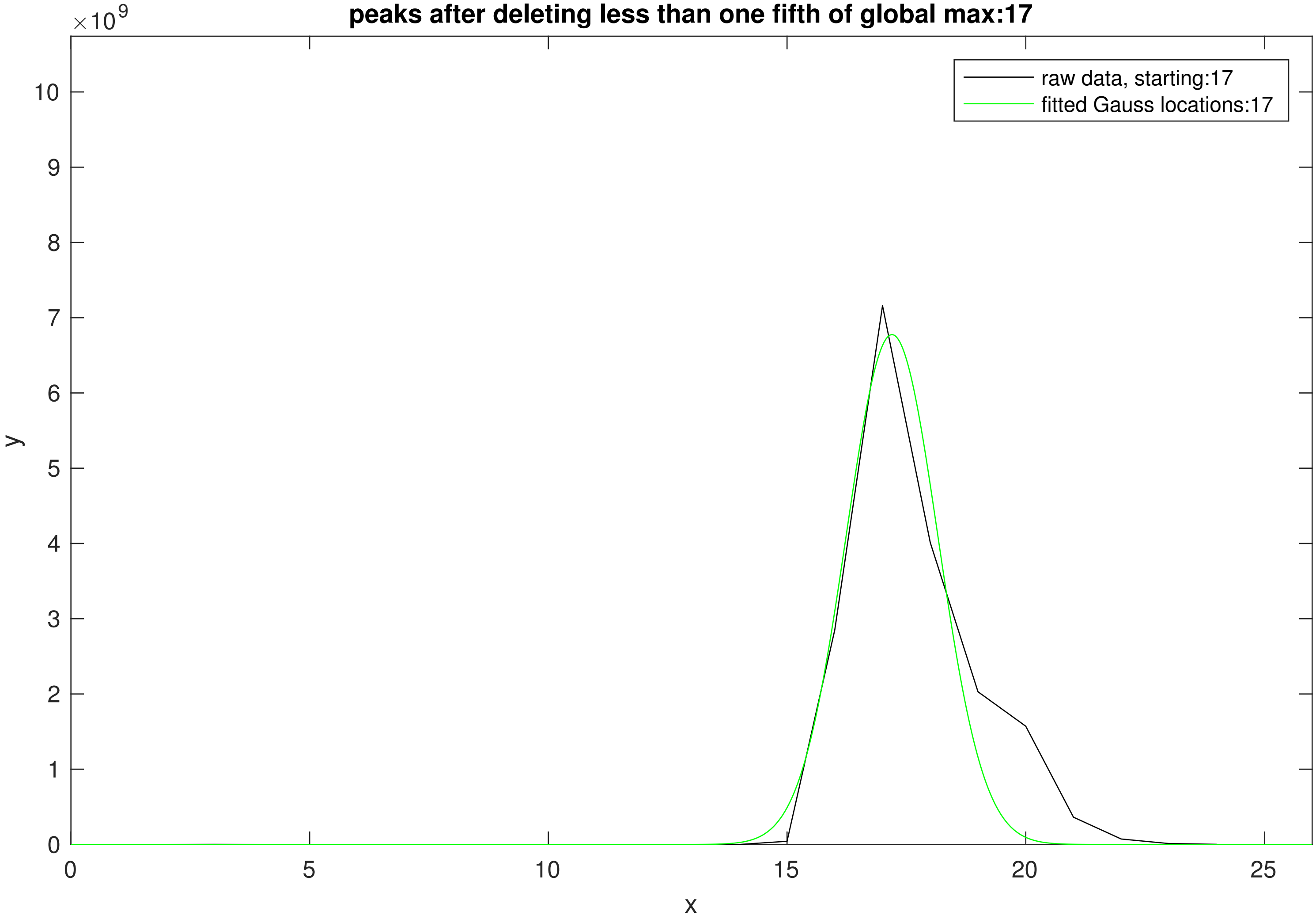

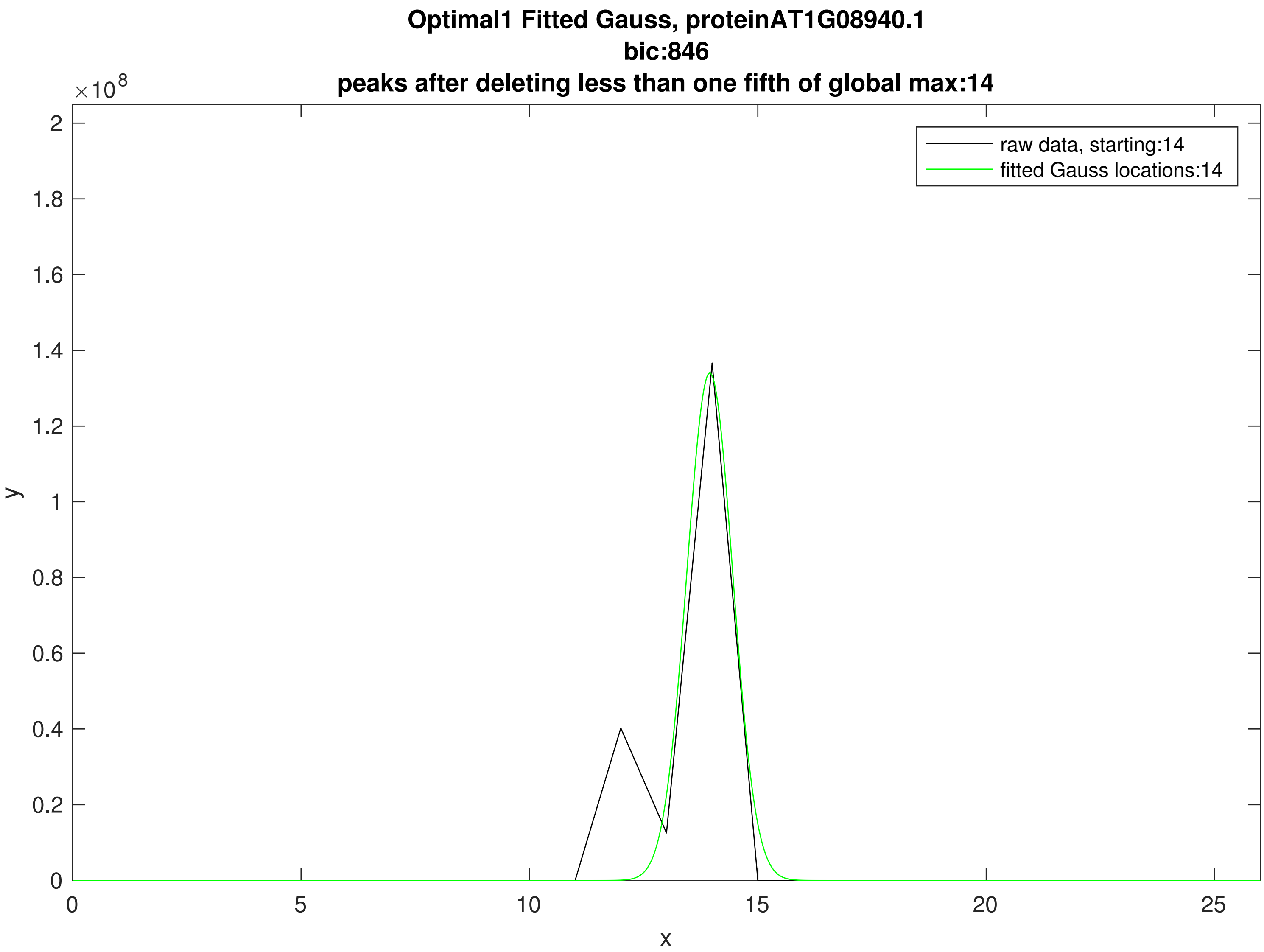

Optimal1 Fitted Gauss, proteinAT1G08940.1
bic:846
peaks after deleting less than one fifth of global max:14
×10^8
raw data, starting:14
fitted Gauss locations:14
y
x

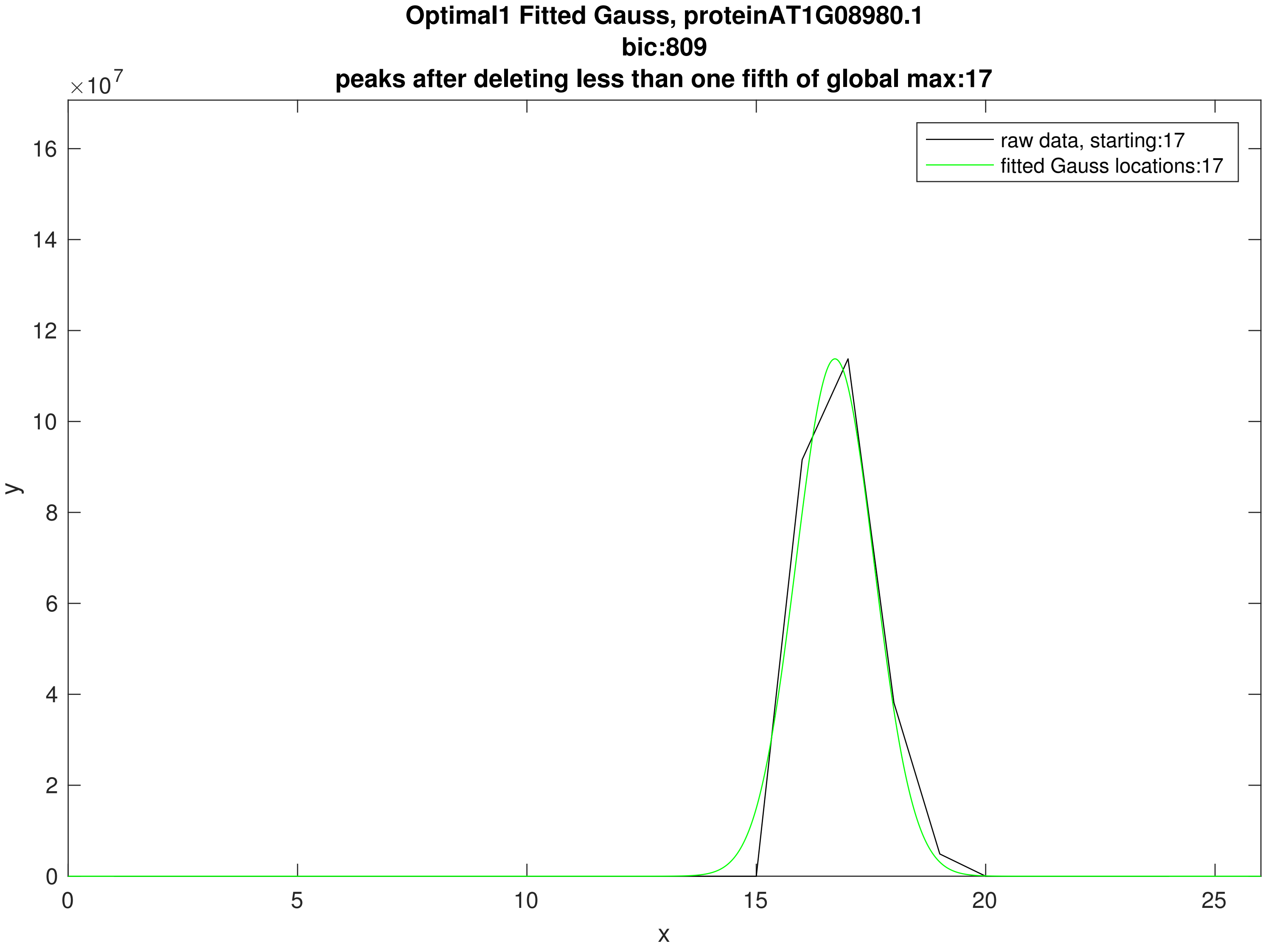

Optimal1 Fitted Gauss, proteinAT1G08980.1
bic:809
peaks after deleting less than one fifth of global max:17
×10^7
raw data, starting:17
fitted Gauss locations:17
y
x
0
2
4
6
8
10
12
14
16
0
5
10
15
20
25

**Optimal1 Fitted Gauss, proteinAT1G09010.1**
**bic:941 949**
**peaks after deleting less than one fifth of global max:14**

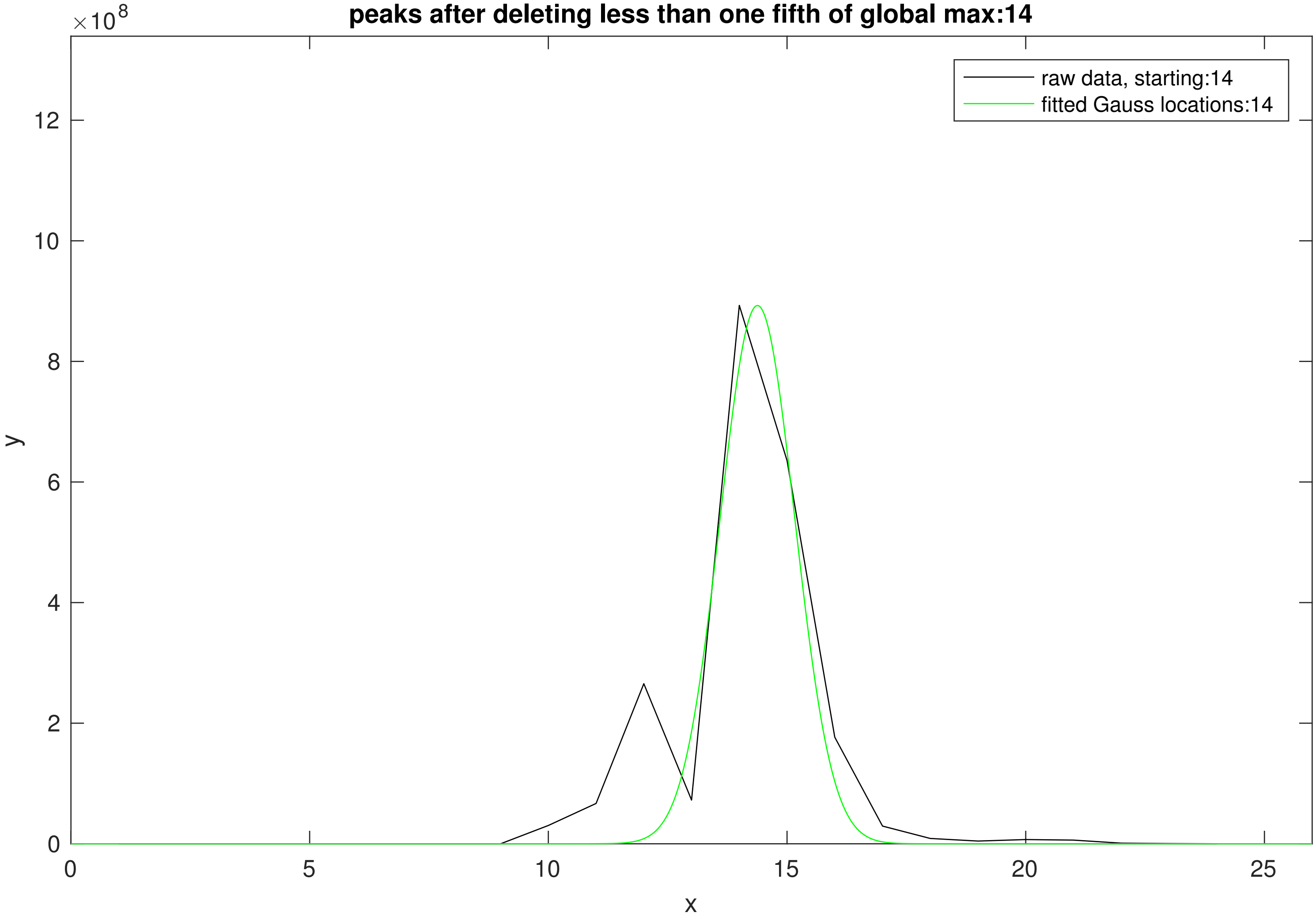

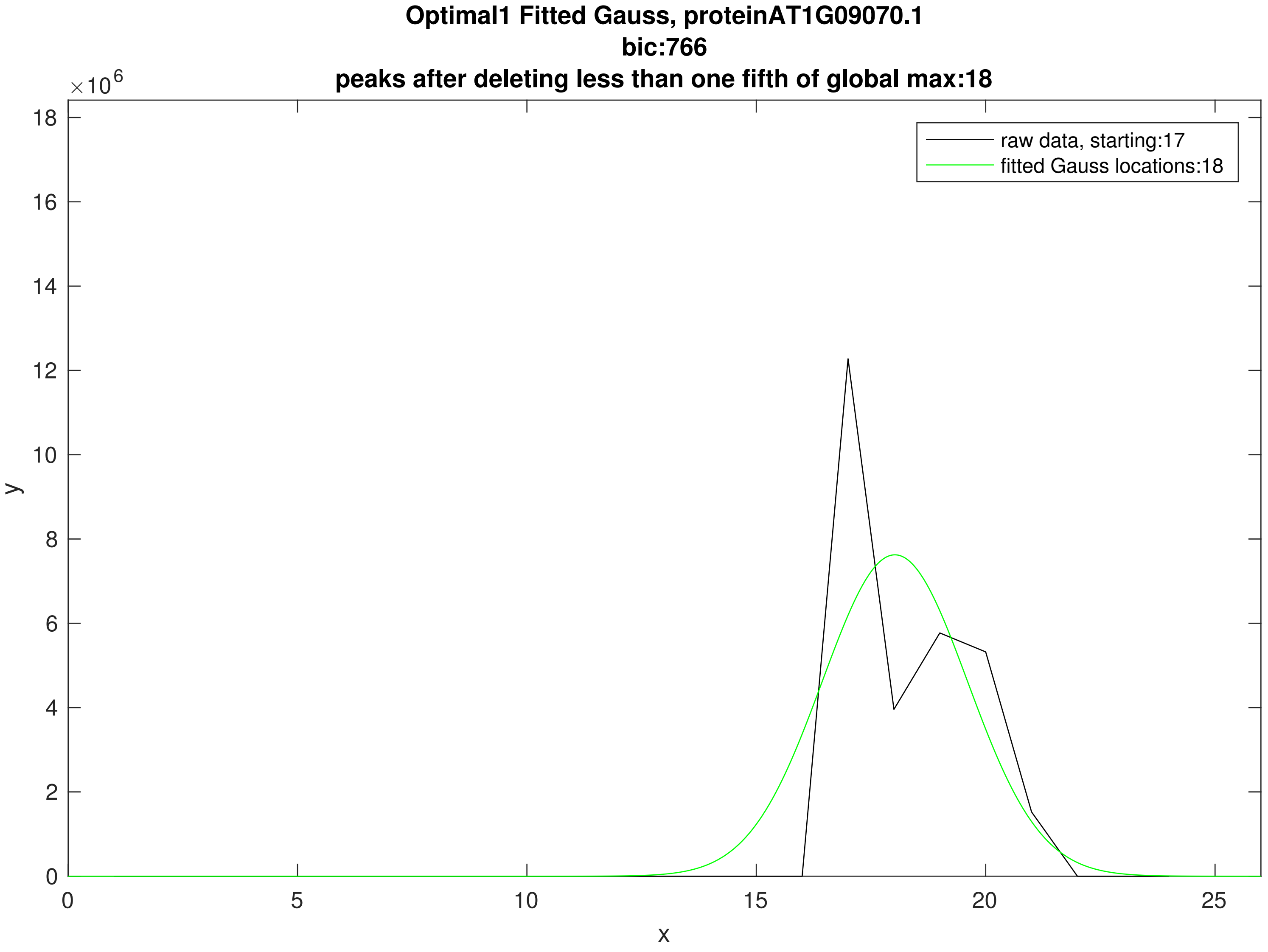

Optimal1 Fitted Gauss, proteinAT1G09070.1
bic:766
peaks after deleting less than one fifth of global max:18
×10^6
18
16
14
12
10
8
6
4
2
0
y
0
5
10
15
20
25
x
raw data, starting:17
fitted Gauss locations:18

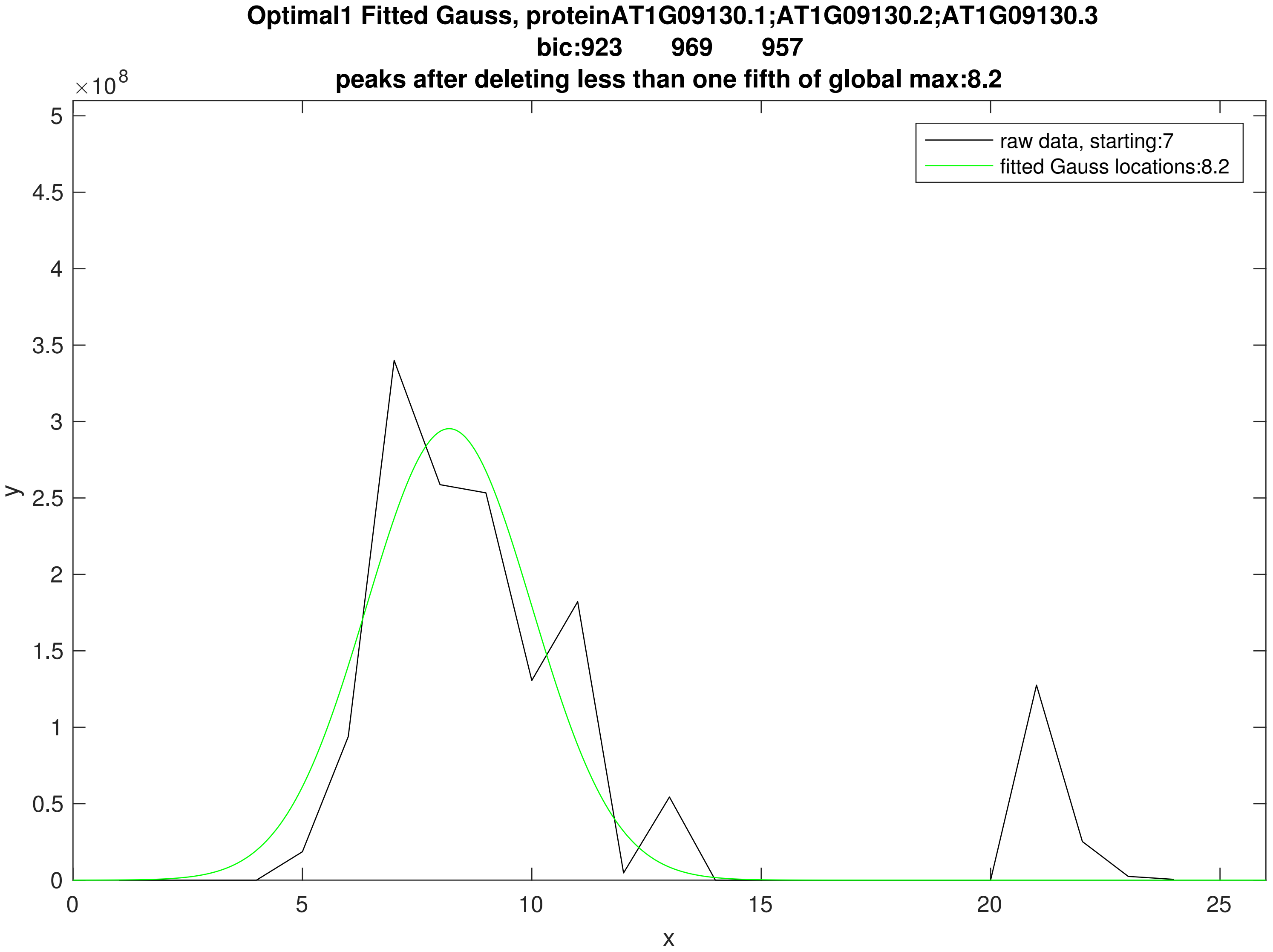

Optimal1 Fitted Gauss, proteinAT1G09130.1;AT1G09130.2;AT1G09130.3
bic:923 969 957
peaks after deleting less than one fifth of global max:8.2
×10^8
raw data, starting:7
fitted Gauss locations:8.2
y
x
0
0.5
1
1.5
2
2.5
3
3.5
4
4.5
5
0
5
10
15
20
25

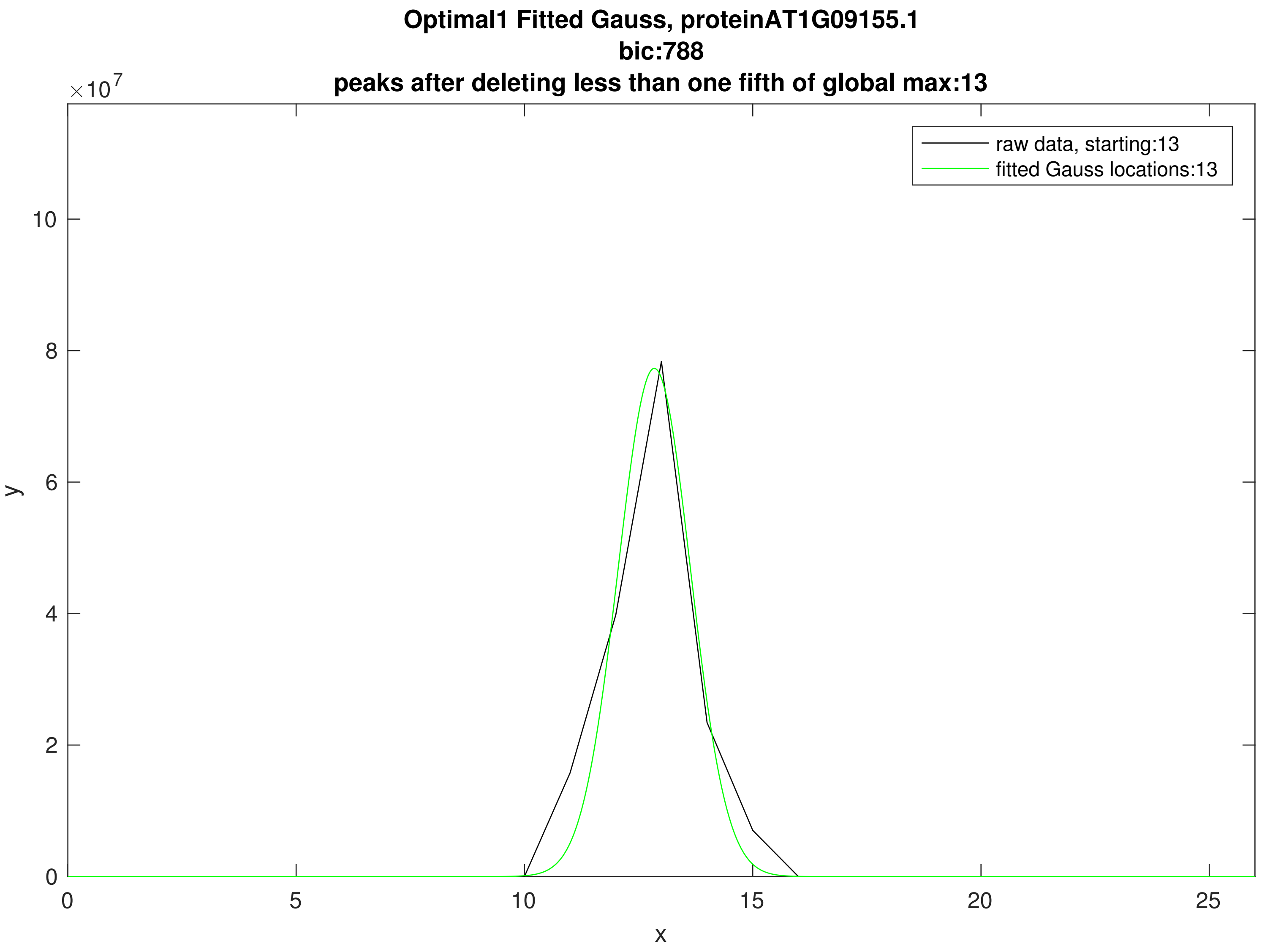

Optimal1 Fitted Gauss, proteinAT1G09155.1
bic:788
peaks after deleting less than one fifth of global max:13
×10^7
raw data, starting:13
fitted Gauss locations:13
y
x
0
2
4
6
8
10
5
15
20
25

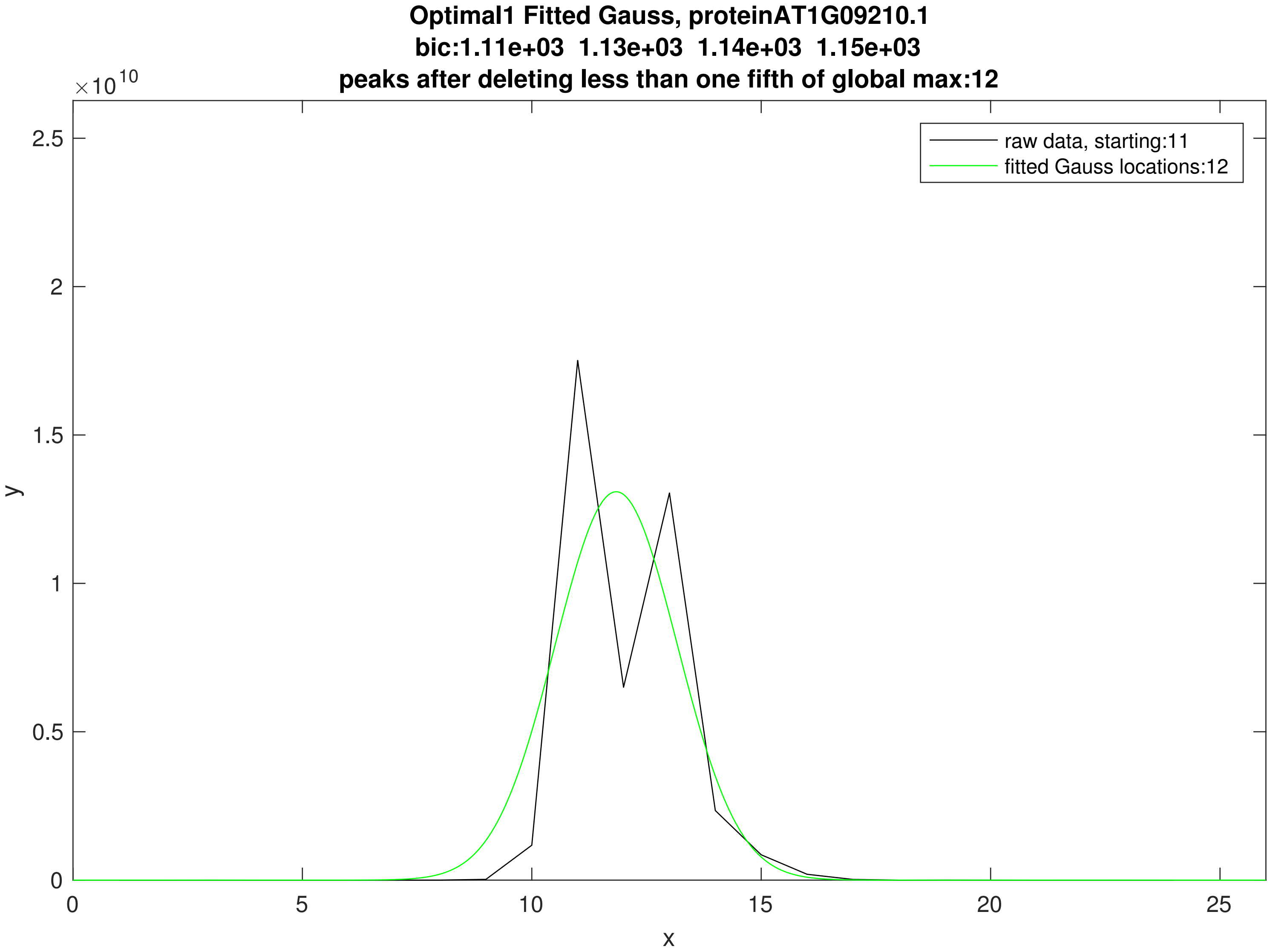

Optimal1 Fitted Gauss, proteinAT1G09210.1
bic:1.11e+03 1.13e+03 1.14e+03 1.15e+03
peaks after deleting less than one fifth of global max:12
×10^10
2.5
2
1.5
1
0.5
0
y
0
5
10
15
20
25
x
raw data, starting:11
fitted Gauss locations:12

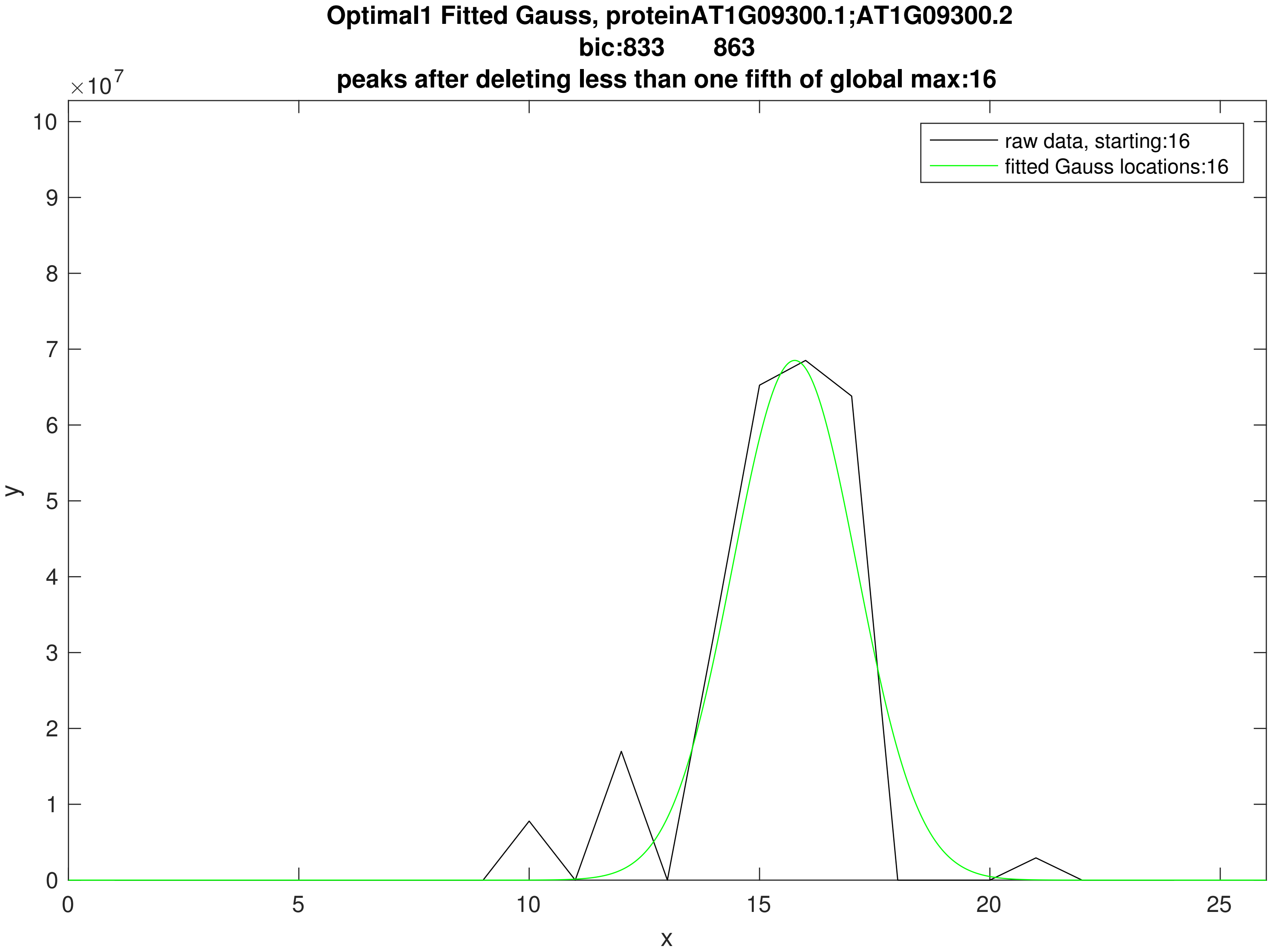

Optimal1 Fitted Gauss, proteinAT1G09300.1;AT1G09300.2
bic:833 863
peaks after deleting less than one fifth of global max:16
×10^7
raw data, starting:16
fitted Gauss locations:16
y
x
0
1
2
3
4
5
6
7
8
9
10
0
5
10
15
20
25

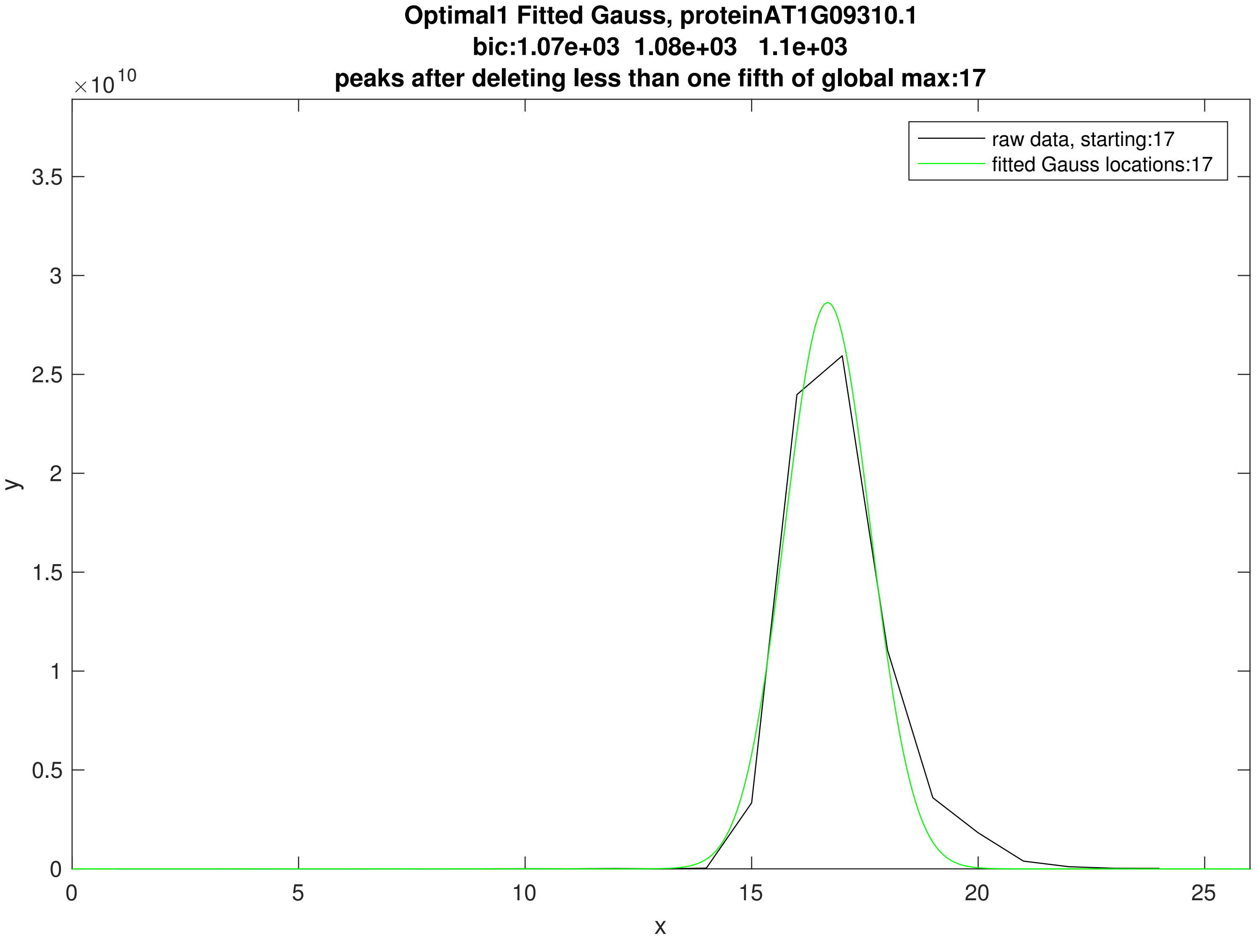

Optimal1 Fitted Gauss, proteinAT1G09310.1
bic:1.07e+03 1.08e+03 1.1e+03
peaks after deleting less than one fifth of global max:17
×10^10
raw data, starting:17
fitted Gauss locations:17
y
x
0
0.5
1
1.5
2
2.5
3
3.5
0
5
10
15
20
25

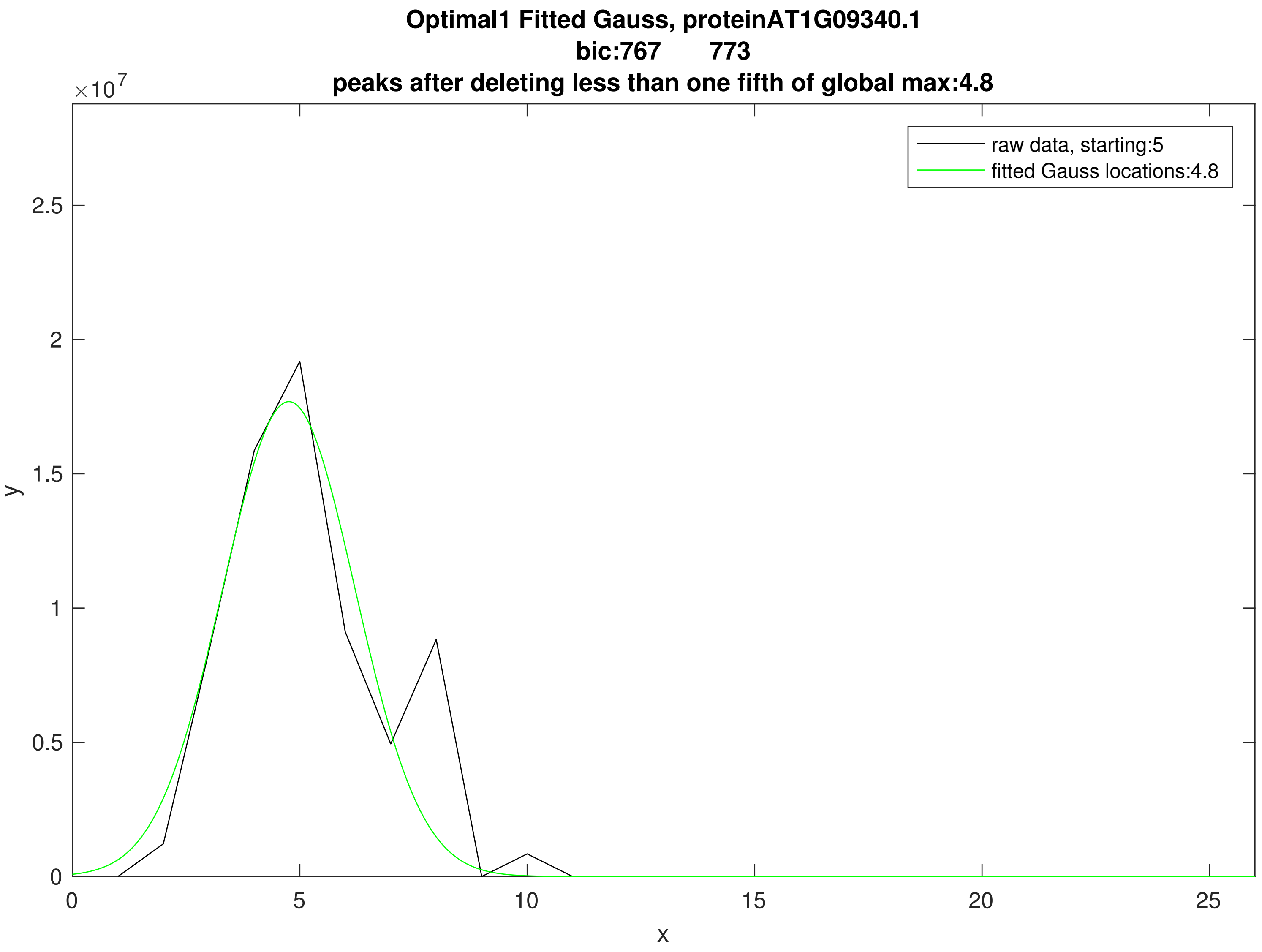

Optimal1 Fitted Gauss, proteinAT1G09340.1
bic:767 773
peaks after deleting less than one fifth of global max:4.8
×10^7
raw data, starting:5
fitted Gauss locations:4.8
y
x
0
0.5
1
1.5
2
2.5
0
5
10
15
20
25

**Optimal1 Fitted Gauss, proteinAT1G09430.1**
**bic:900 909**
**peaks after deleting less than one fifth of global max:15**

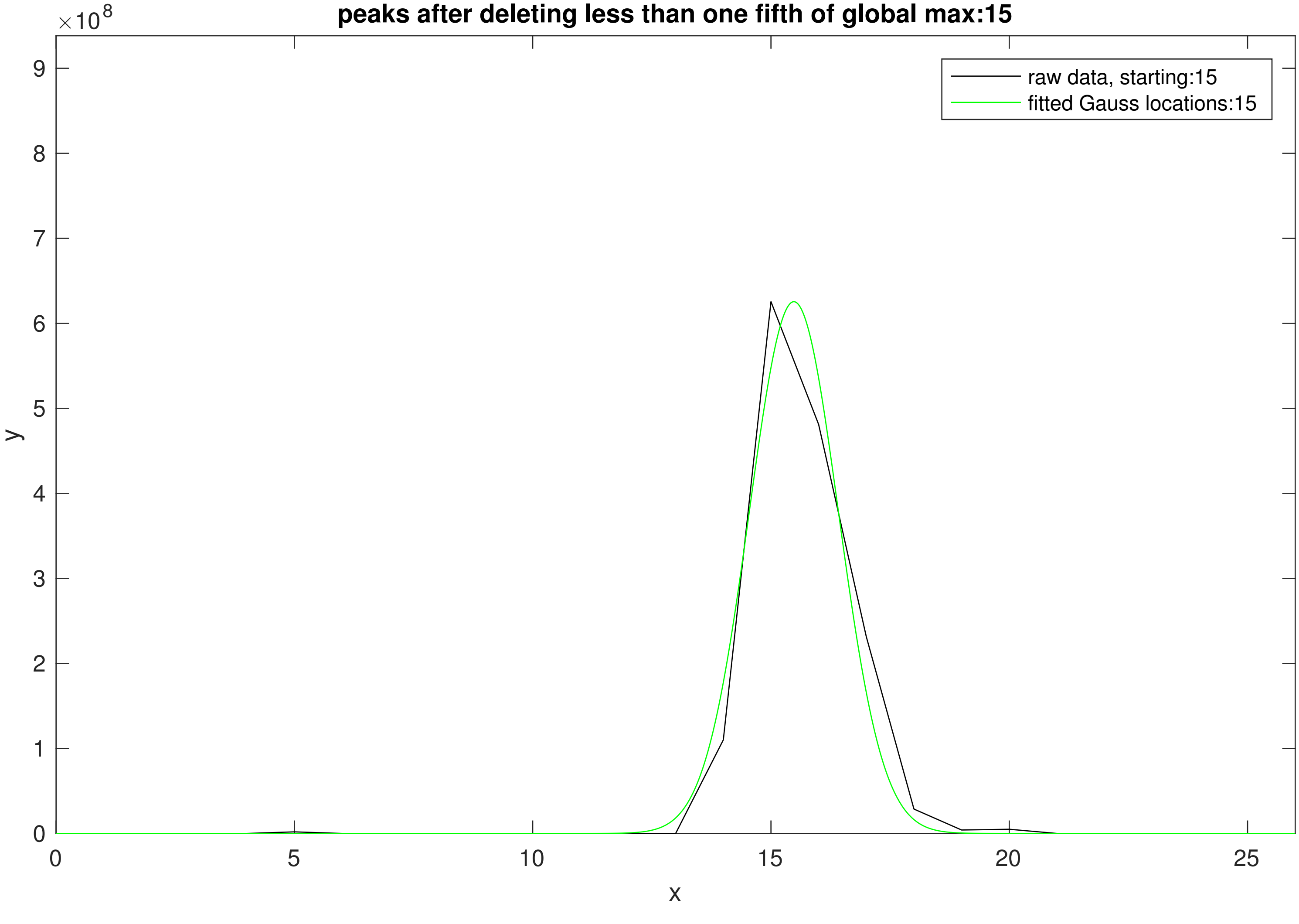

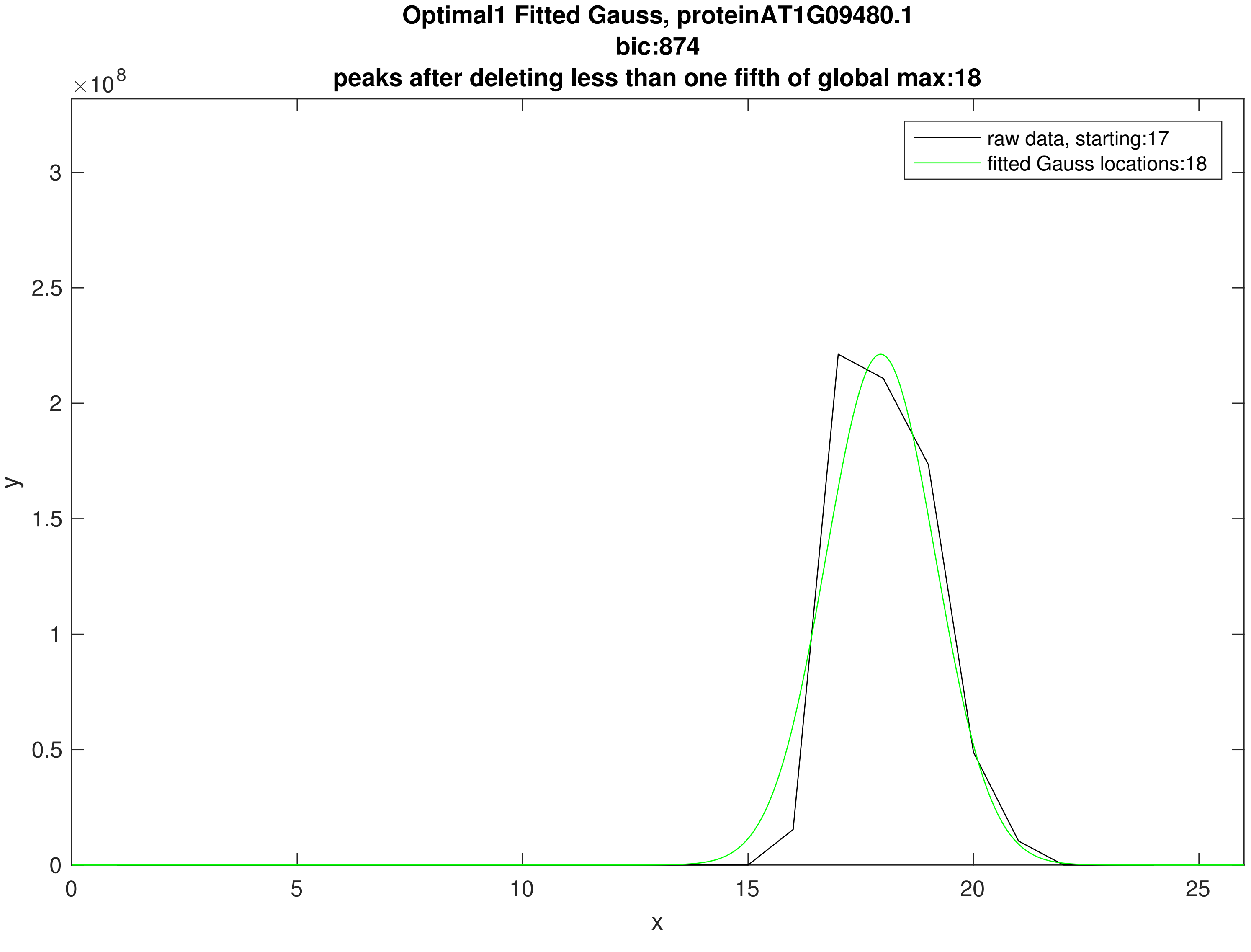
Optimal1 Fitted Gauss, proteinAT1G09480.1
bic:874
peaks after deleting less than one fifth of global max:18
×10^8
raw data, starting:17
fitted Gauss locations:18
y
x
0
0.5
1
1.5
2
2.5
3
0
5
10
15
20
25

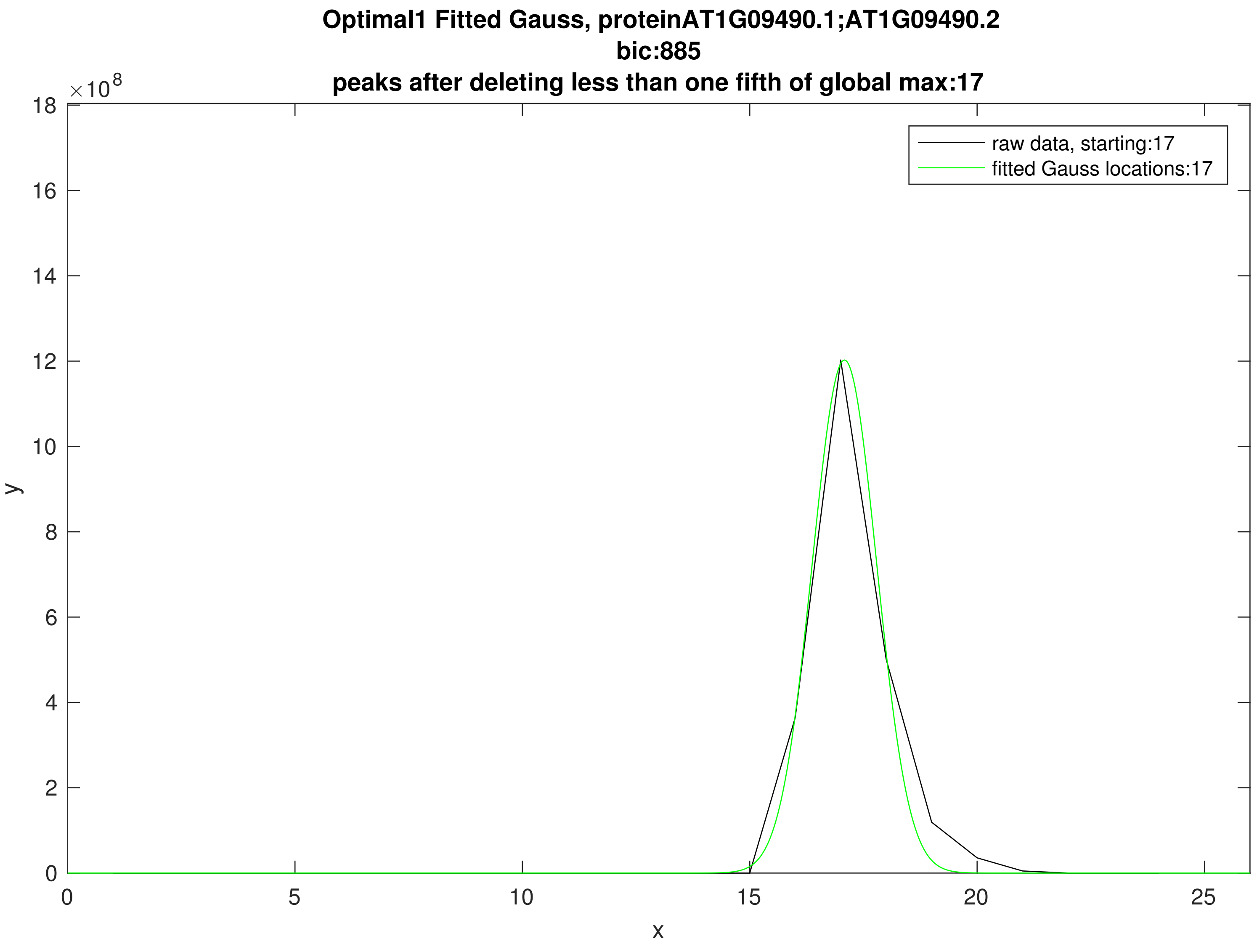

Optimal1 Fitted Gauss, proteinAT1G09490.1;AT1G09490.2
bic:885
peaks after deleting less than one fifth of global max:17
$\times 10^8$
18
16
14
12
10
8
6
4
2
0
y
0
5
10
15
20
25
x
raw data, starting:17
fitted Gauss locations:17

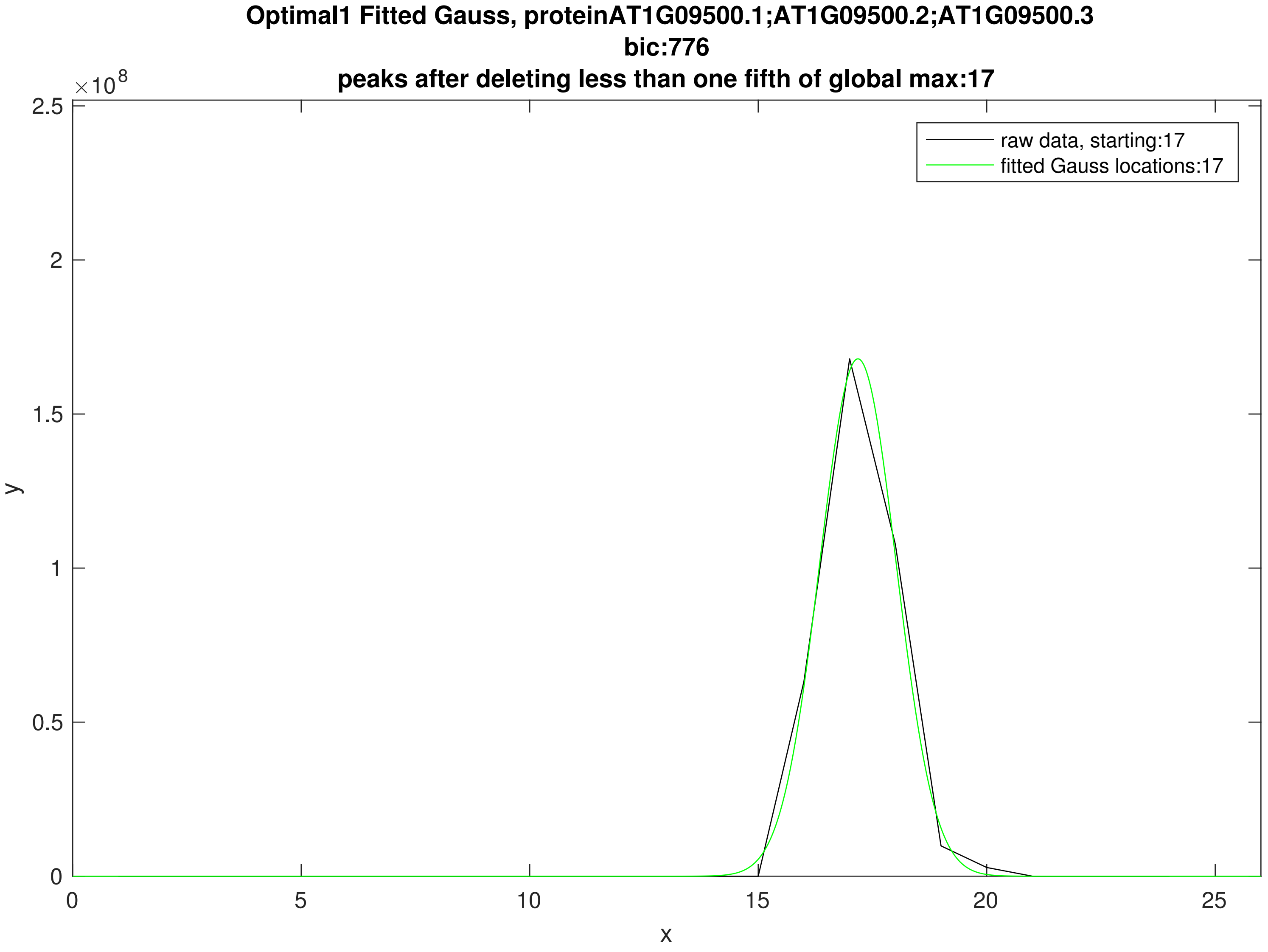
Optimal1 Fitted Gauss, proteinAT1G09500.1;AT1G09500.2;AT1G09500.3
bic:776
peaks after deleting less than one fifth of global max:17
×10^8
2.5
2
1.5
1
0.5
0
y
0
5
10
15
20
25
x
raw data, starting:17
fitted Gauss locations:17

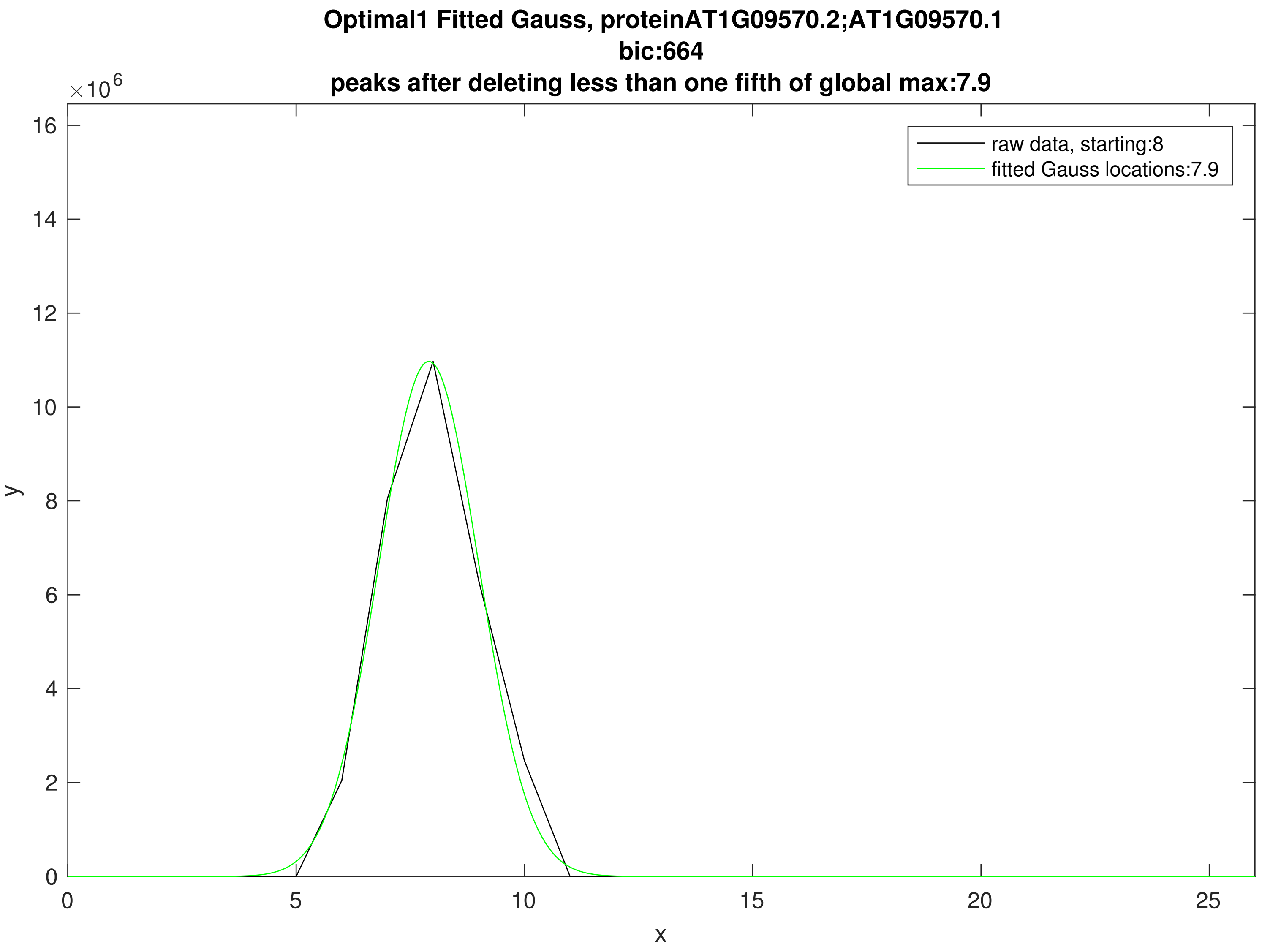

Optimal1 Fitted Gauss, proteinAT1G09570.2;AT1G09570.1
bic:664
peaks after deleting less than one fifth of global max:7.9
×10^6
raw data, starting:8
fitted Gauss locations:7.9
y
x
0
2
4
6
8
10
12
14
16
0
5
10
15
20
25

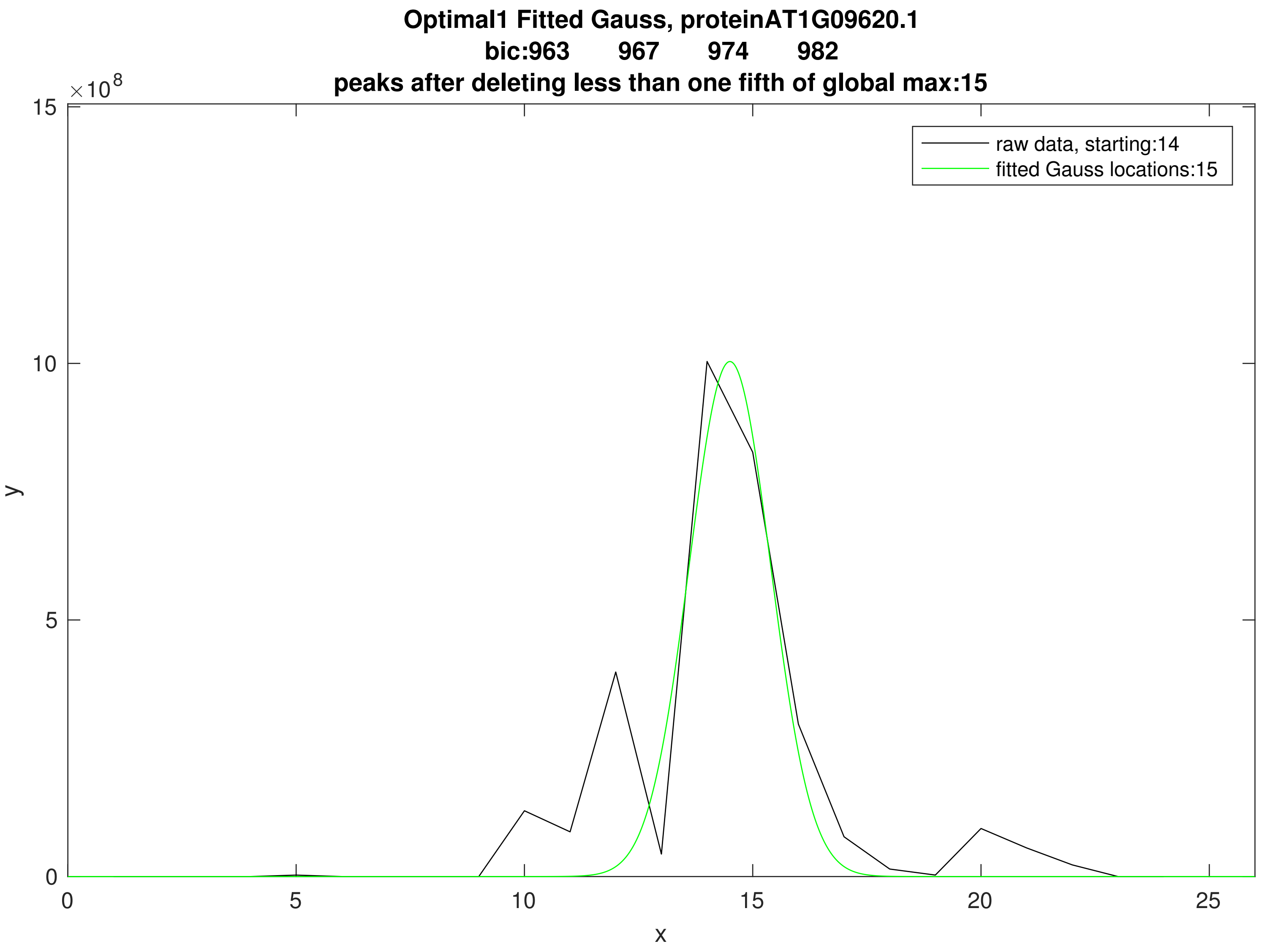

Optimal1 Fitted Gauss, proteinAT1G09620.1
bic:963 967 974 982
peaks after deleting less than one fifth of global max:15
×10^8
15
10
5
0
y
0
5
10
15
20
25
x
raw data, starting:14
fitted Gauss locations:15

**Optimal1 Fitted Gauss, proteinAT1G09640.1**
**bic:828**
**peaks after deleting less than one fifth of global max:4.9**

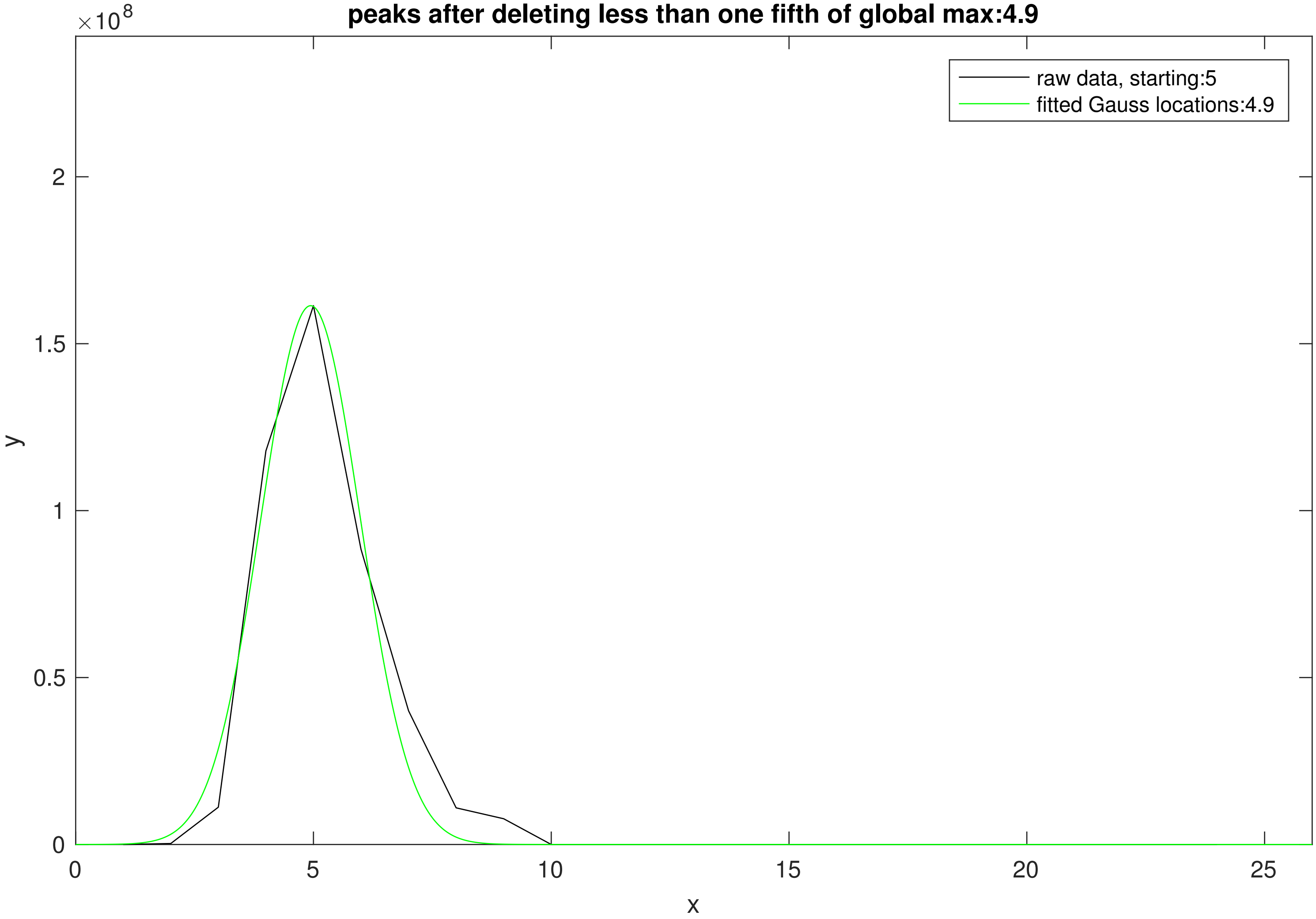

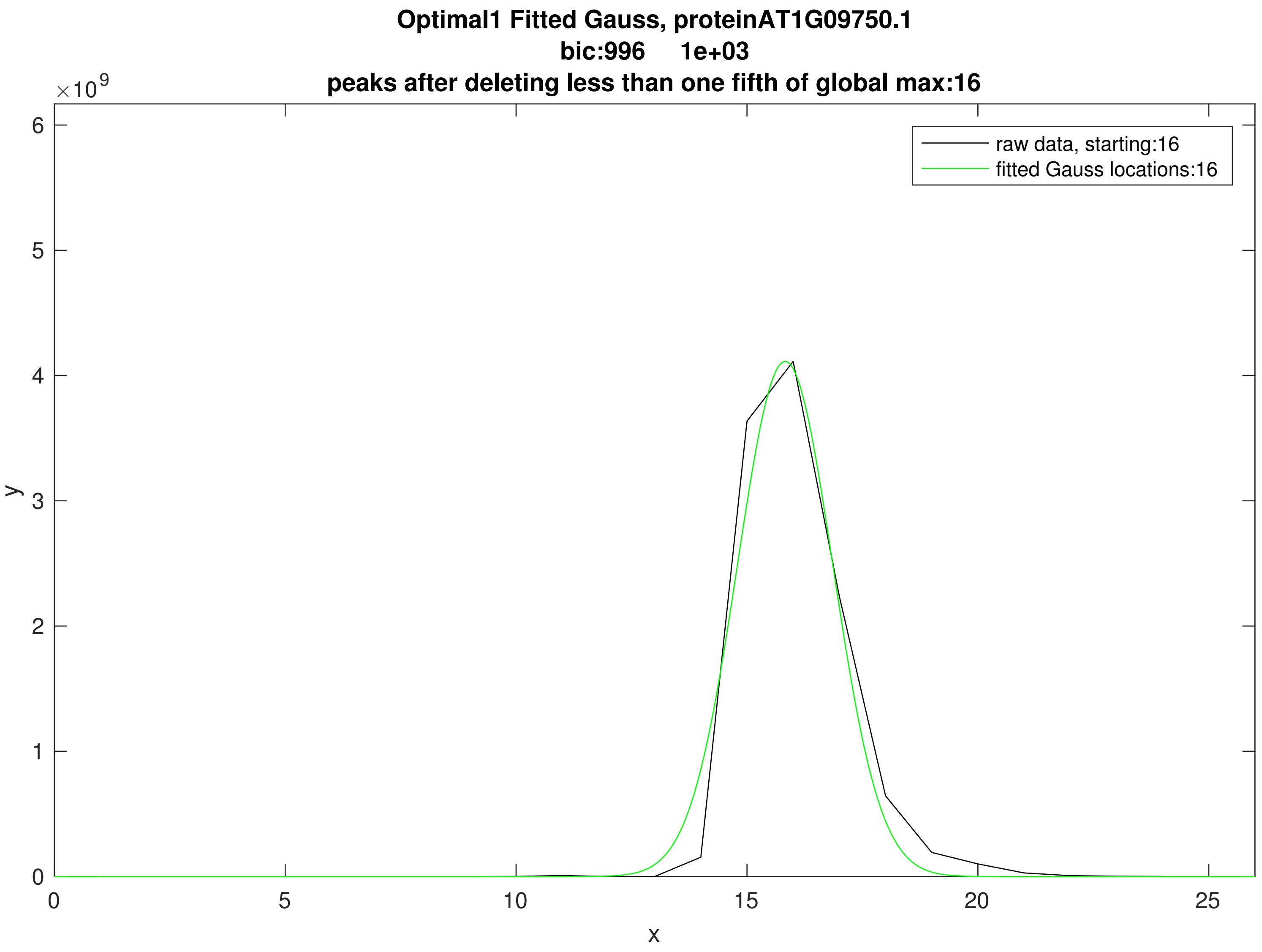
Optimal1 Fitted Gauss, proteinAT1G09750.1
bic:996 1e+03
peaks after deleting less than one fifth of global max:16
×10^9
6
5
4
3
2
1
0
y
0
5
10
15
20
25
x
raw data, starting:16
fitted Gauss locations:16

**Optimal1 Fitted Gauss, proteinAT1G09780.1**
**bic:1.08e+03  1.09e+03   1.1e+03**
**peaks after deleting less than one fifth of global max:15**

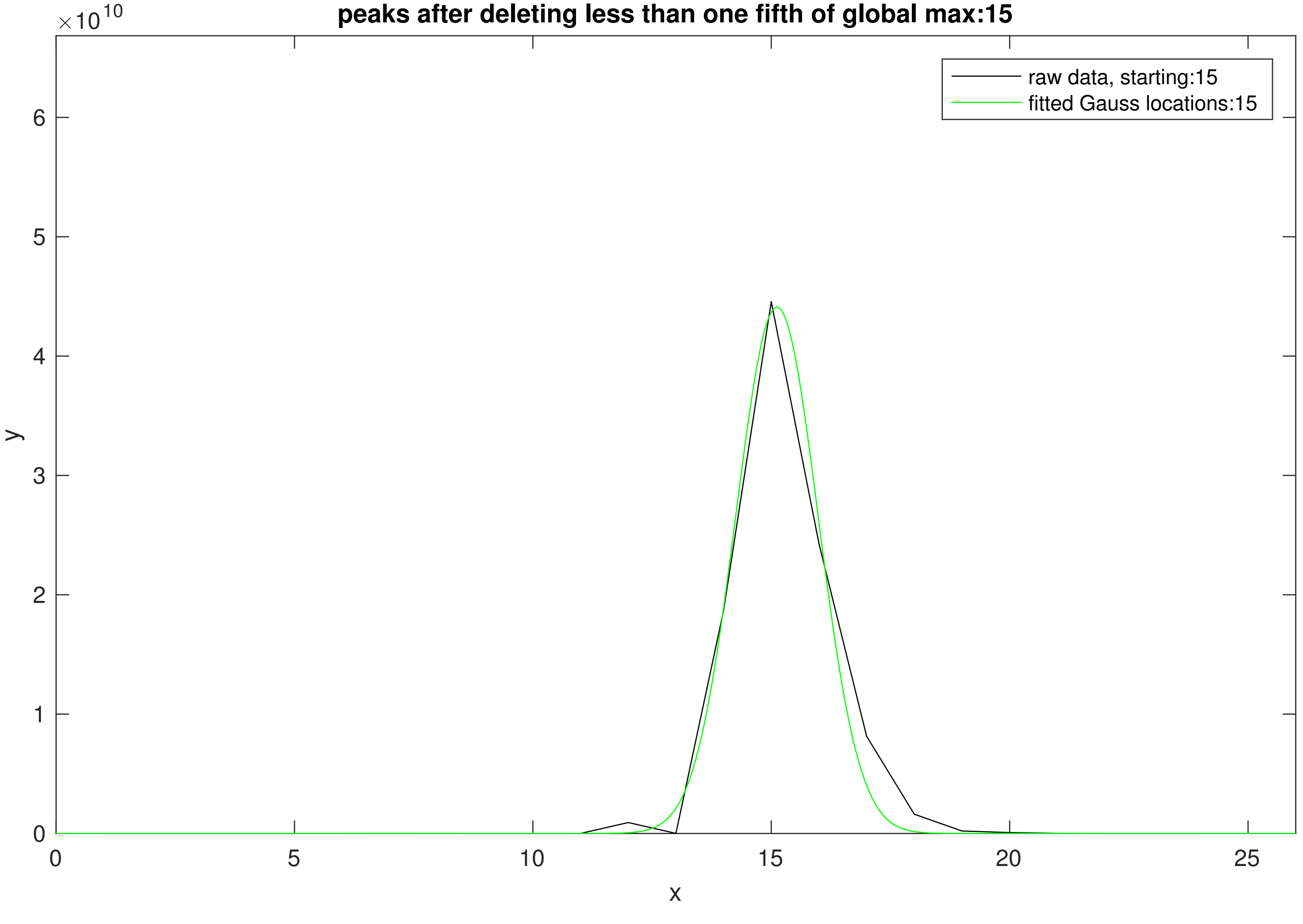

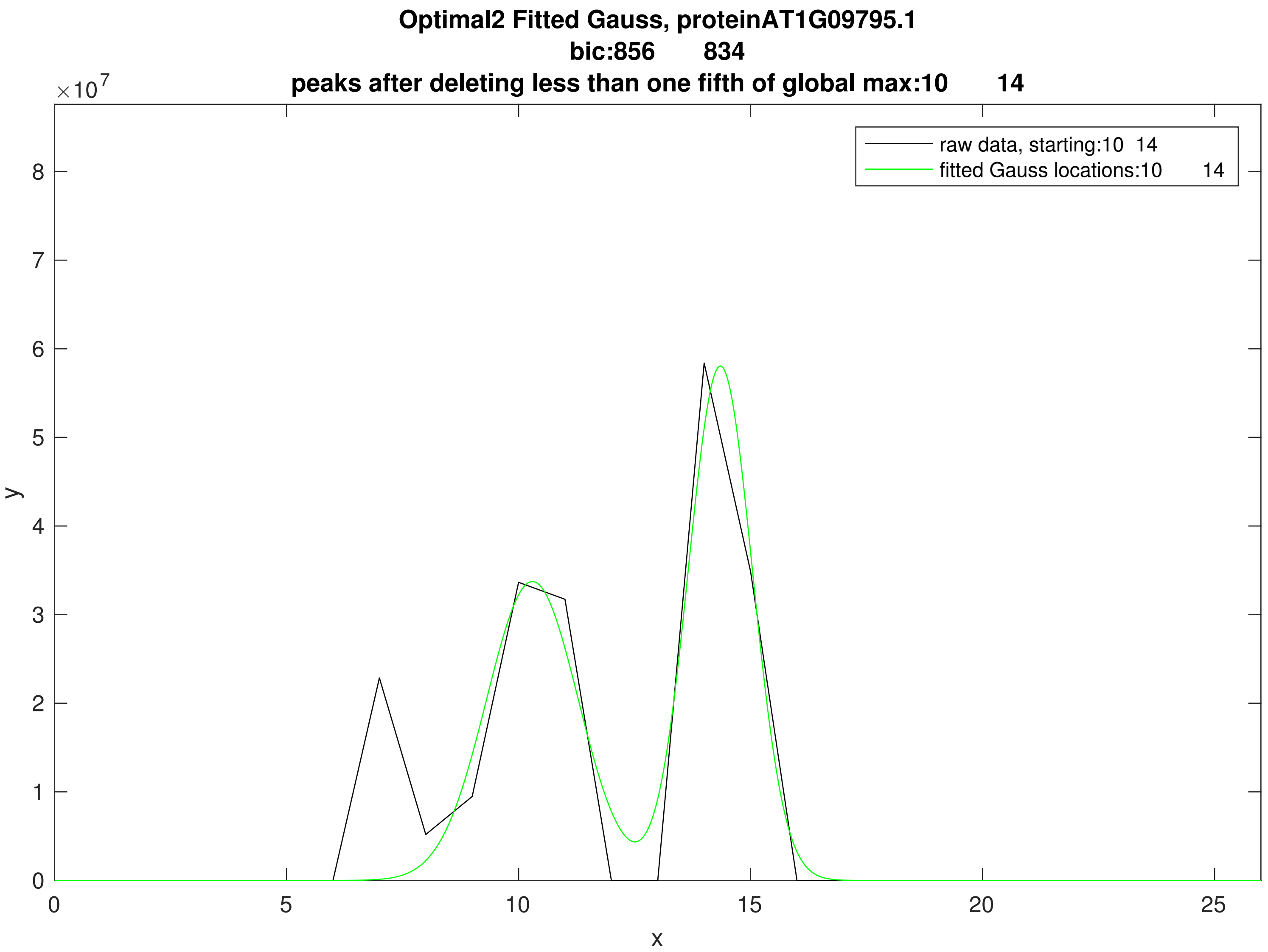
Optimal2 Fitted Gauss, proteinAT1G09795.1
bic:856 834
peaks after deleting less than one fifth of global max:10 14
×10^7
raw data, starting:10 14
fitted Gauss locations:10 14
y
x
0
1
2
3
4
5
6
7
8
0
5
10
15
20
25

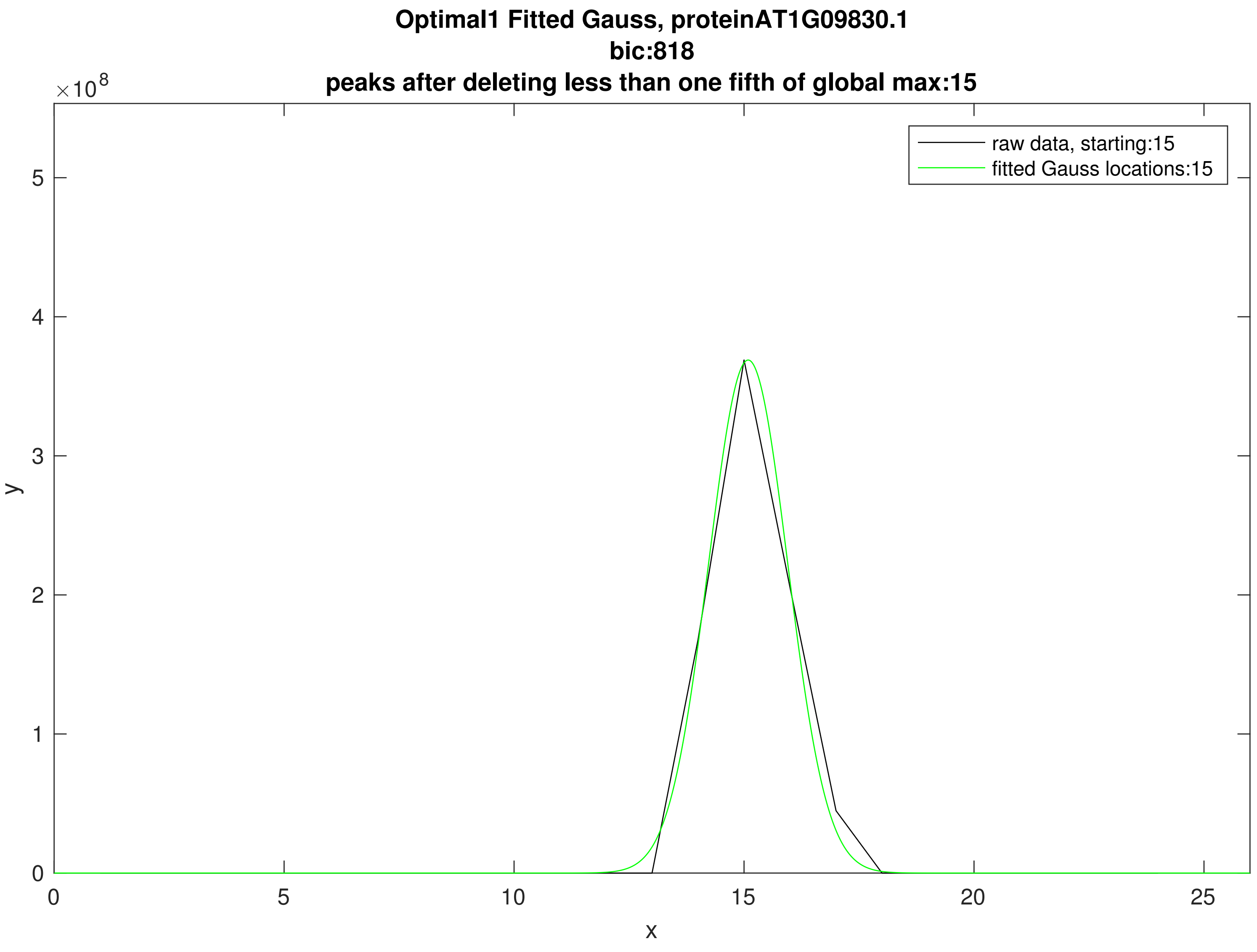
Optimal1 Fitted Gauss, proteinAT1G09830.1
bic:818
peaks after deleting less than one fifth of global max:15
×10^8
raw data, starting:15
fitted Gauss locations:15
y
x
0
1
2
3
4
5
0
5
10
15
20
25

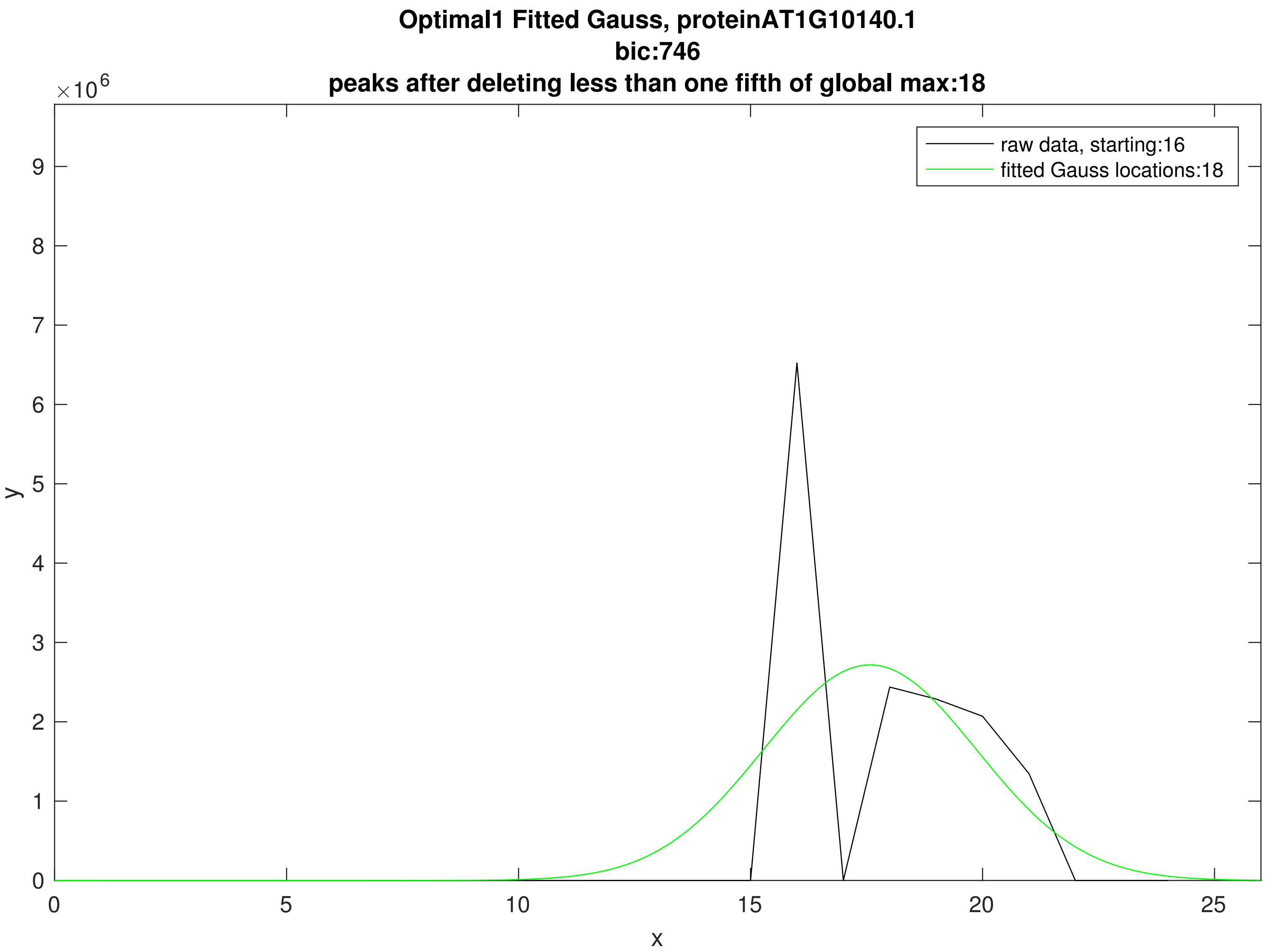

Optimal1 Fitted Gauss, proteinAT1G10140.1
bic:746
peaks after deleting less than one fifth of global max:18
×10^6
raw data, starting:16
fitted Gauss locations:18
y
x
0
1
2
3
4
5
6
7
8
9
0
5
10
15
20
25

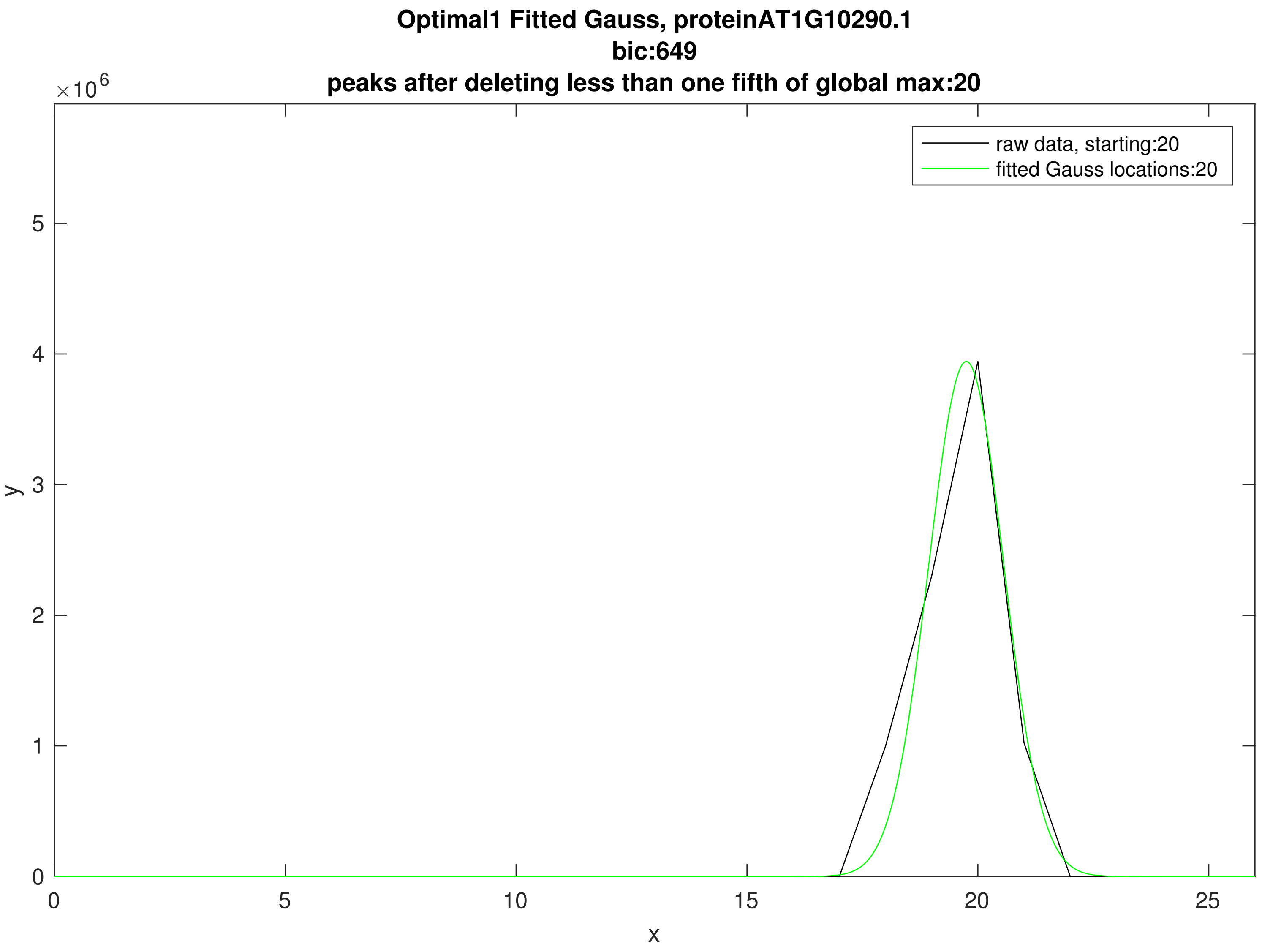

Optimal1 Fitted Gauss, proteinAT1G10290.1
bic:649
peaks after deleting less than one fifth of global max:20
×10^6
raw data, starting:20
fitted Gauss locations:20
y
x
0
1
2
3
4
5
0
5
10
15
20
25

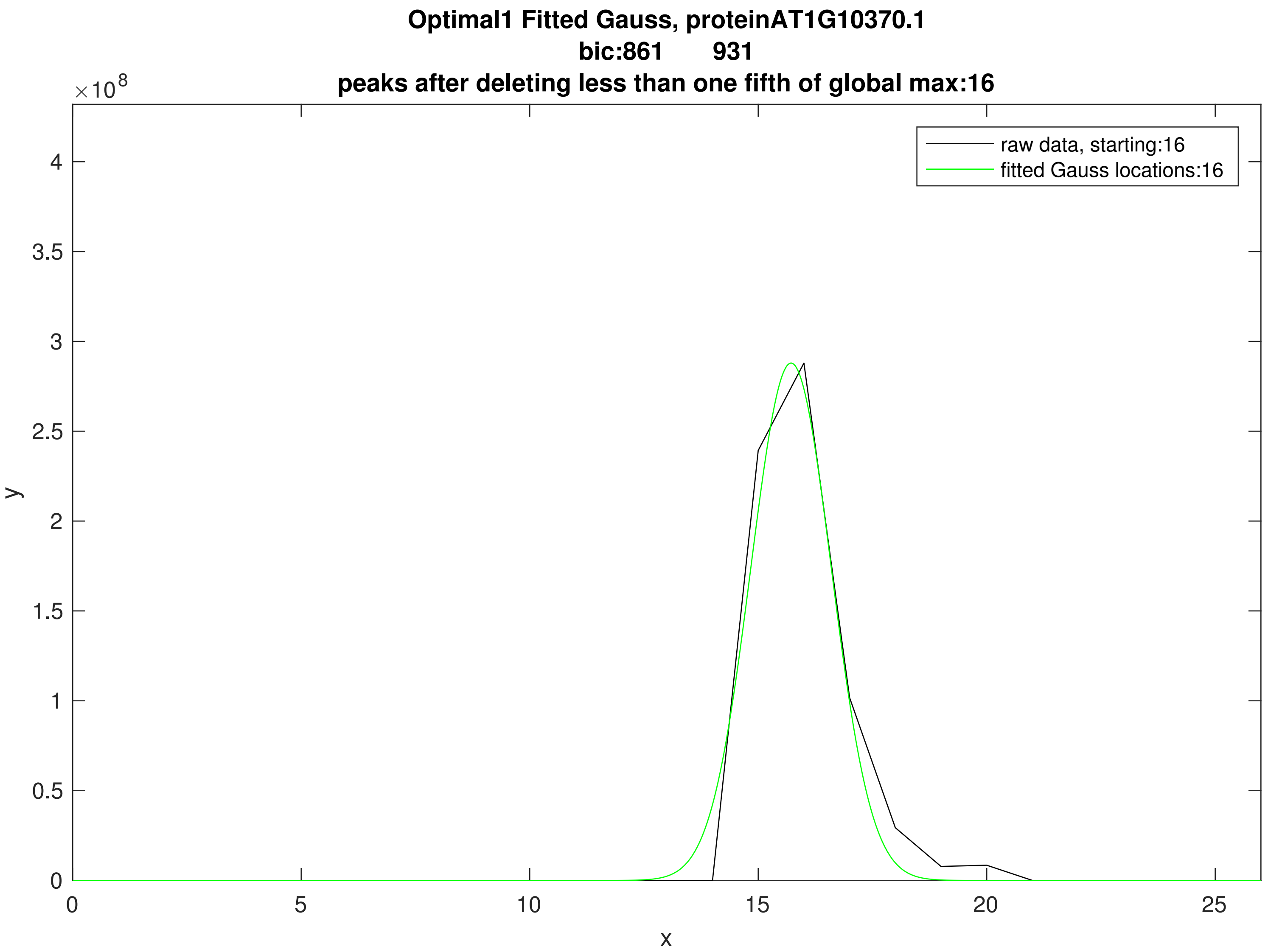

Optimal1 Fitted Gauss, proteinAT1G10370.1
bic:861 931
peaks after deleting less than one fifth of global max:16
×10^8
raw data, starting:16
fitted Gauss locations:16
y
x
0
0.5
1
1.5
2
2.5
3
3.5
4
0
5
10
15
20
25

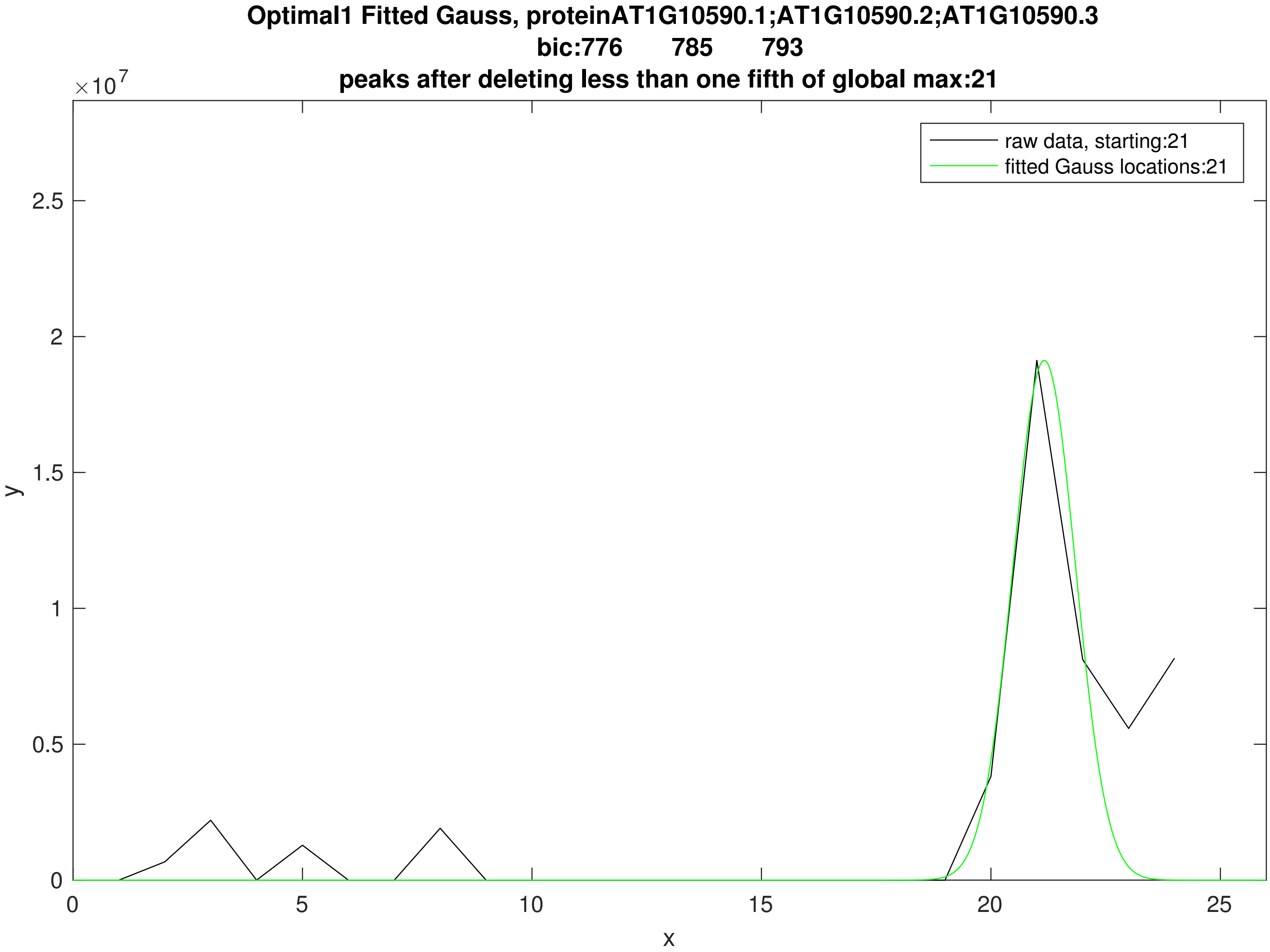

Optimal1 Fitted Gauss, proteinAT1G10590.1;AT1G10590.2;AT1G10590.3
bic:776 785 793
peaks after deleting less than one fifth of global max:21
×10^7
raw data, starting:21
fitted Gauss locations:21
y
x
0
0.5
1
1.5
2
2.5
0
5
10
15
20
25

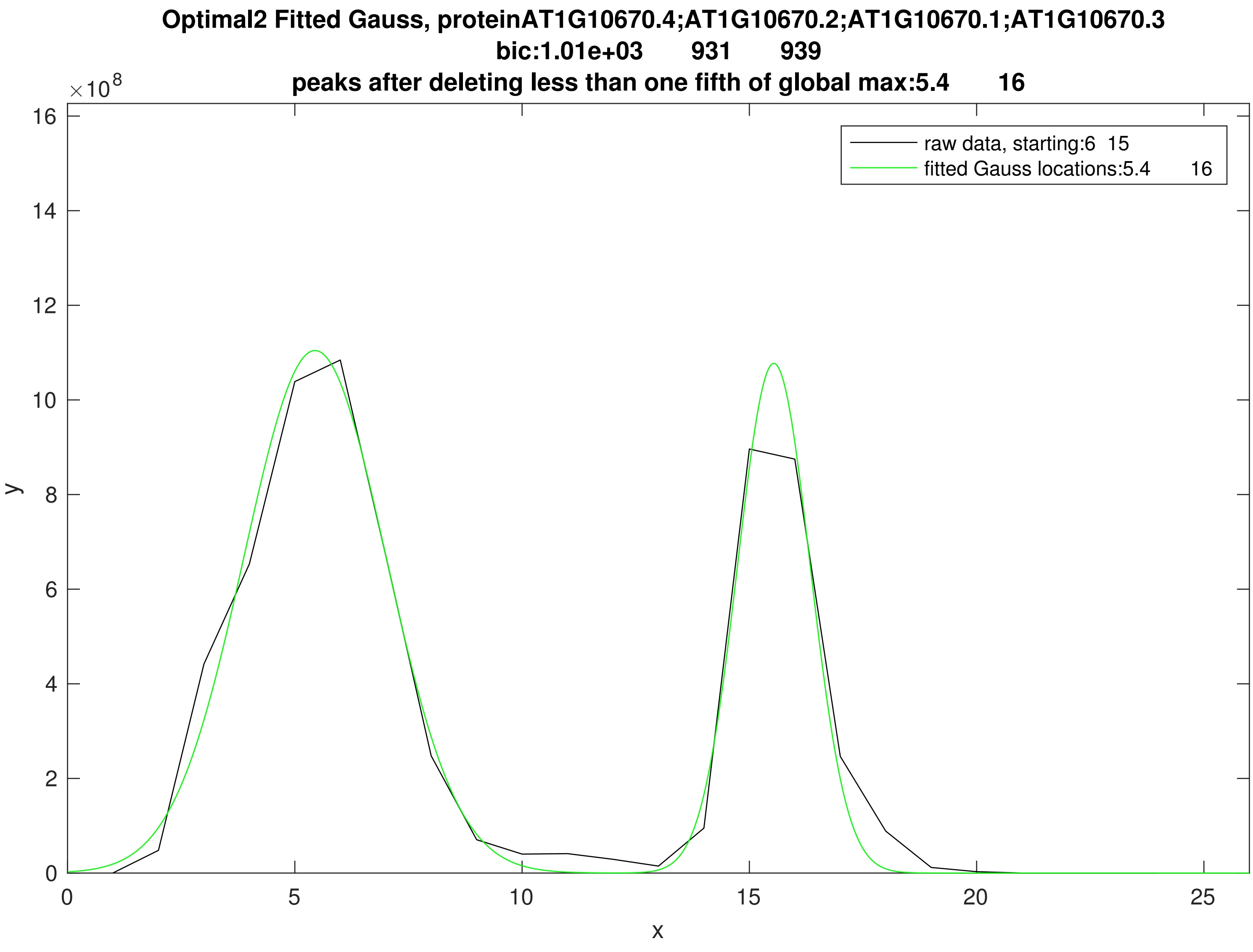
Optimal2 Fitted Gauss, proteinAT1G10670.4;AT1G10670.2;AT1G10670.1;AT1G10670.3
bic:1.01e+03 931 939
peaks after deleting less than one fifth of global max:5.4 16
raw data, starting:6 15
fitted Gauss locations:5.4 16
×10^8
y
x

**Optimal2 Fitted Gauss, proteinAT1G10700.1**
**bic:811 765**
**peaks after deleting less than one fifth of global max:4.1 8.7**

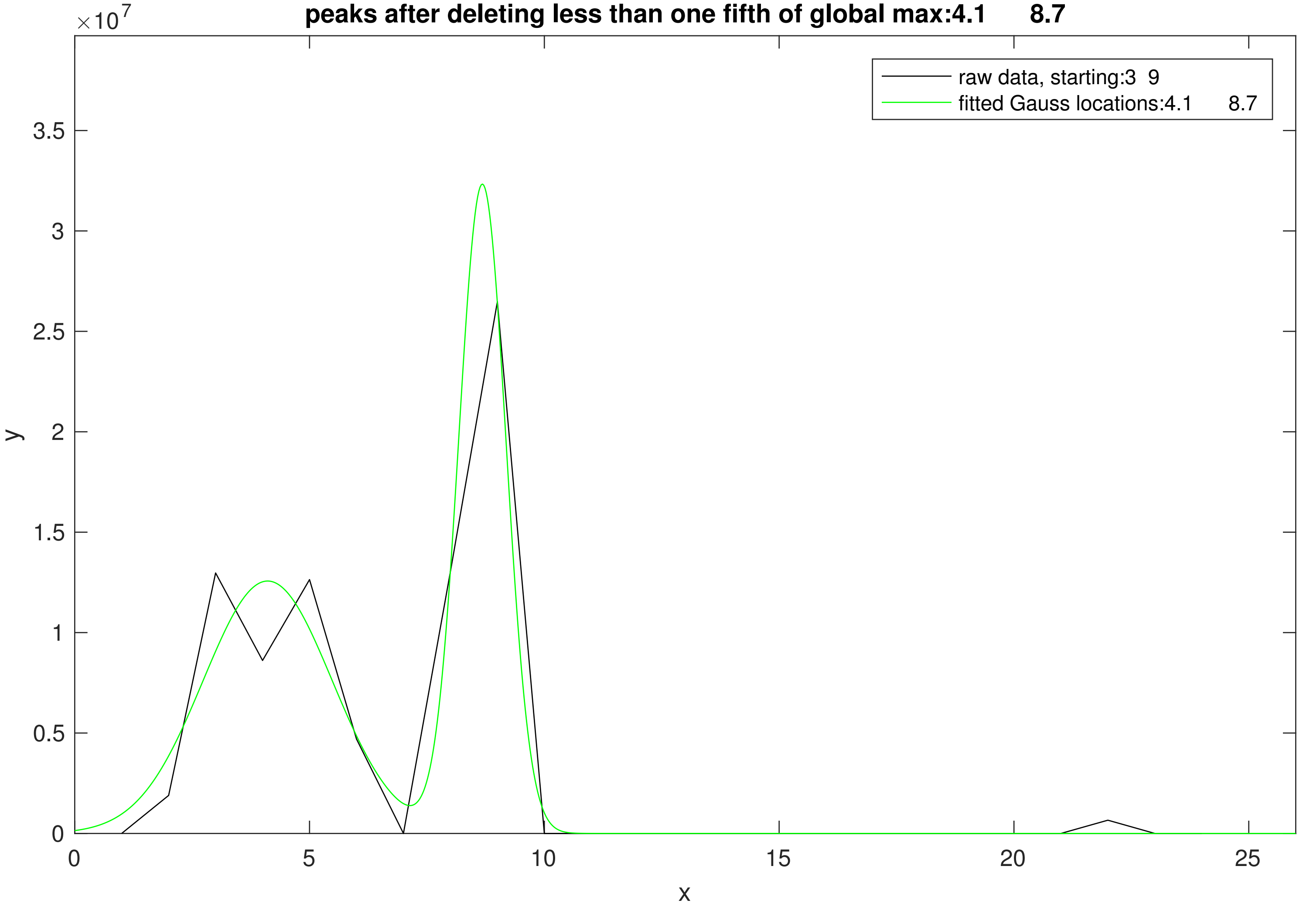

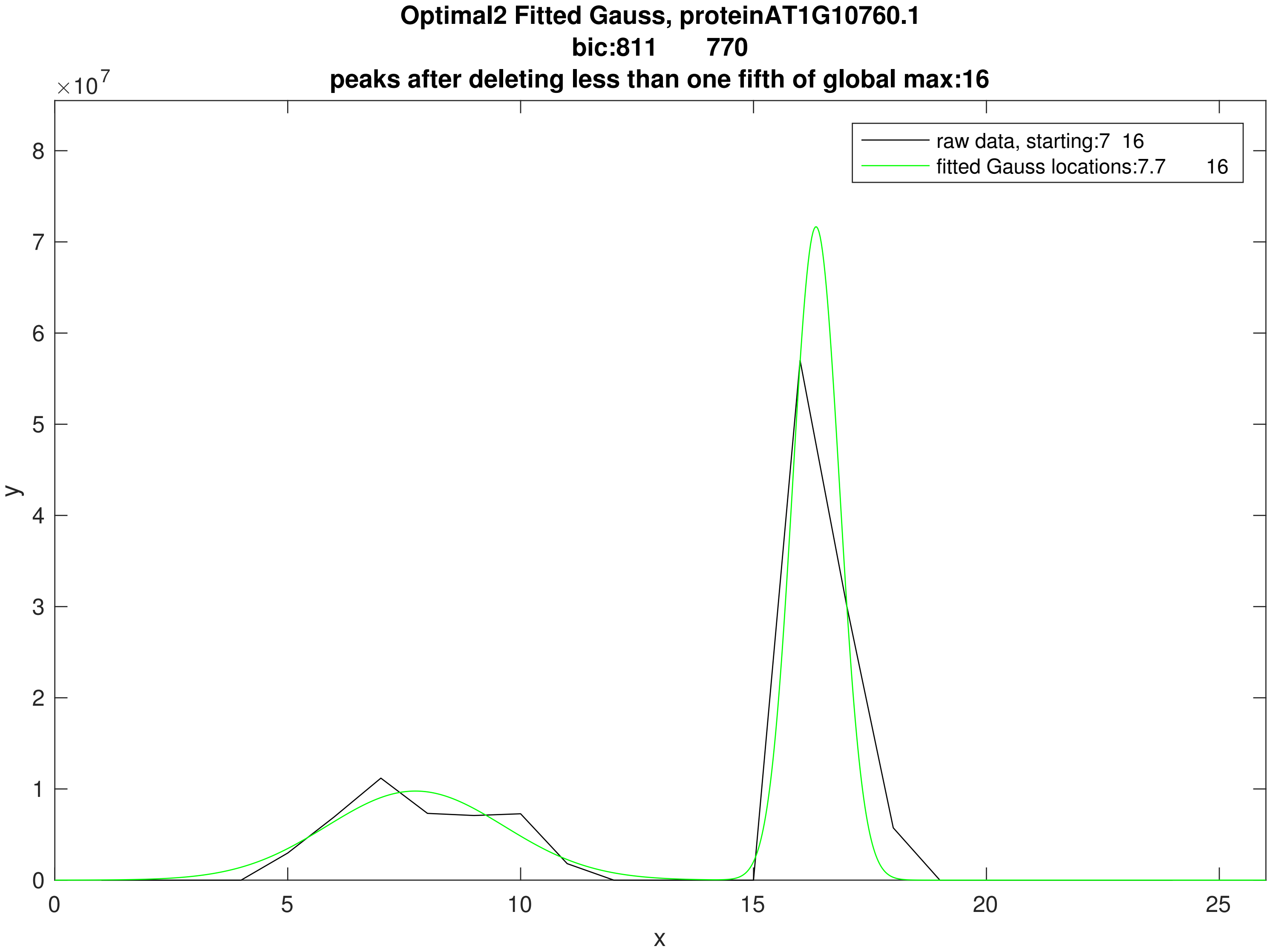
Optimal2 Fitted Gauss, proteinAT1G10760.1
bic:811 770
peaks after deleting less than one fifth of global max:16
×10^7
raw data, starting:7 16
fitted Gauss locations:7.7 16
y
x
0
1
2
3
4
5
6
7
8
0
5
10
15
20
25

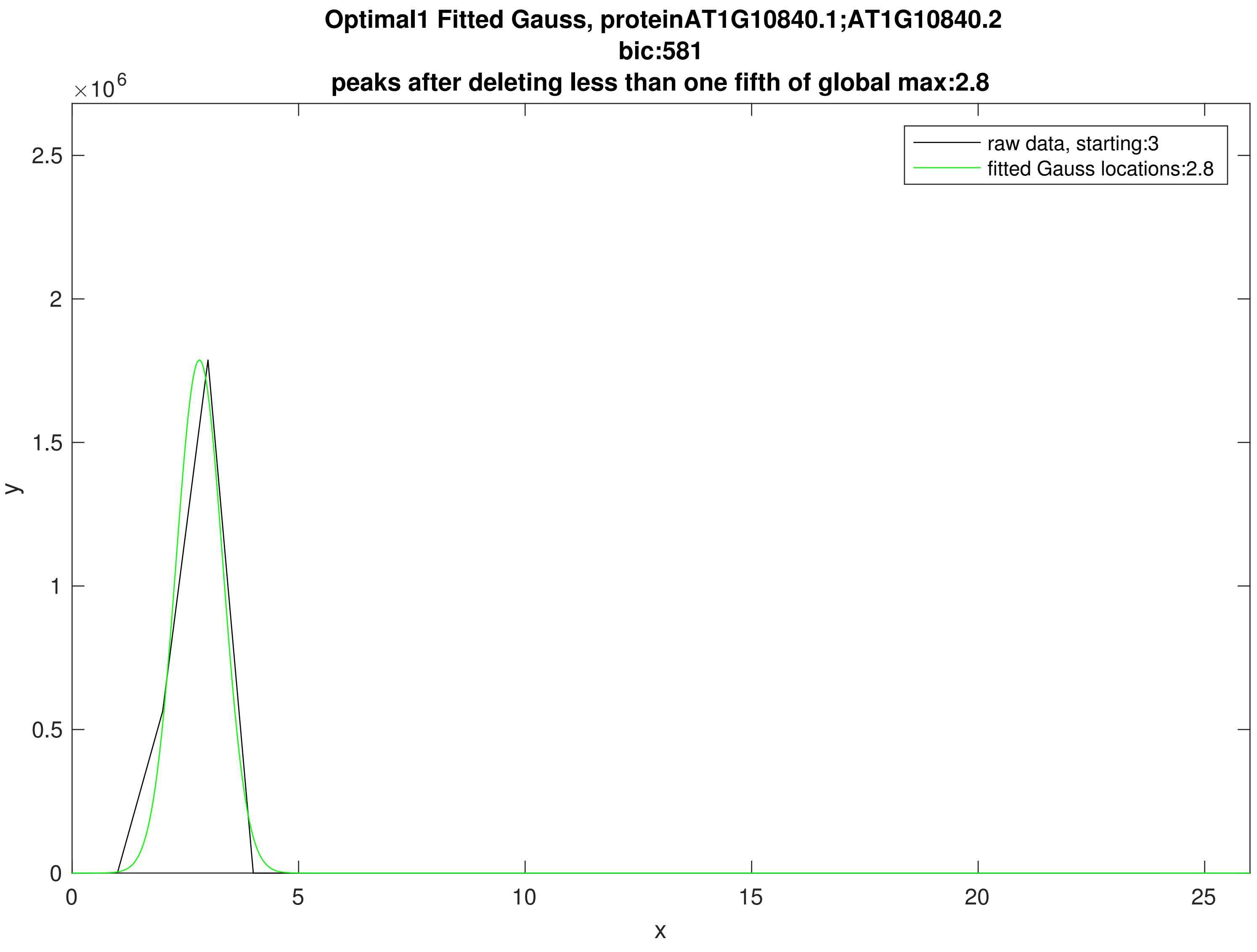
Optimal1 Fitted Gauss, proteinAT1G10840.1;AT1G10840.2
bic:581
peaks after deleting less than one fifth of global max:2.8
×10^6
raw data, starting:3
fitted Gauss locations:2.8
2.5
2
1.5
1
0.5
0
y
0
5
10
15
20
25
x

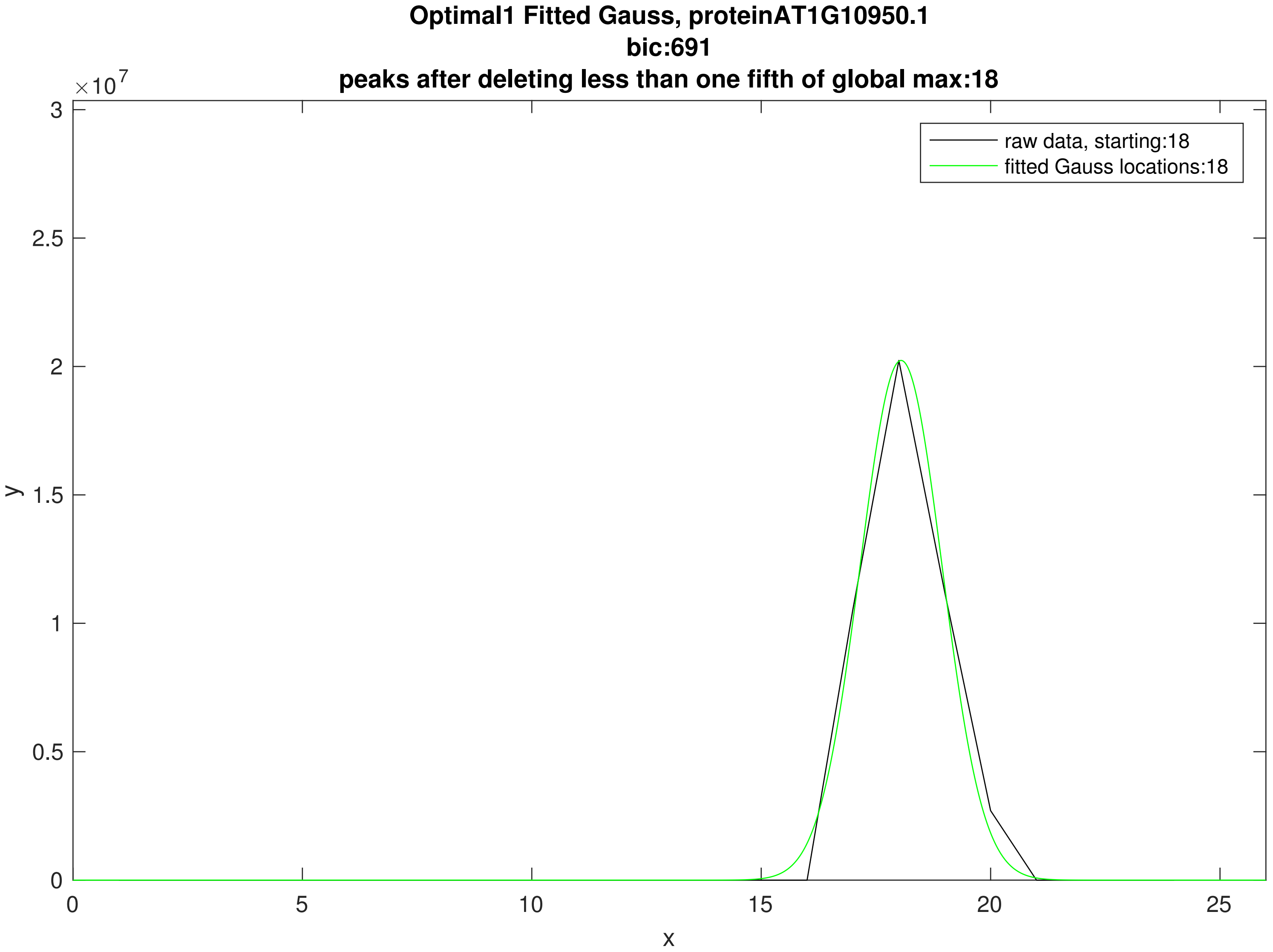
Optimal1 Fitted Gauss, proteinAT1G10950.1
bic:691
peaks after deleting less than one fifth of global max:18
×10^7
raw data, starting:18
fitted Gauss locations:18
y
x
0
0.5
1
1.5
2
2.5
3
0
5
10
15
20
25

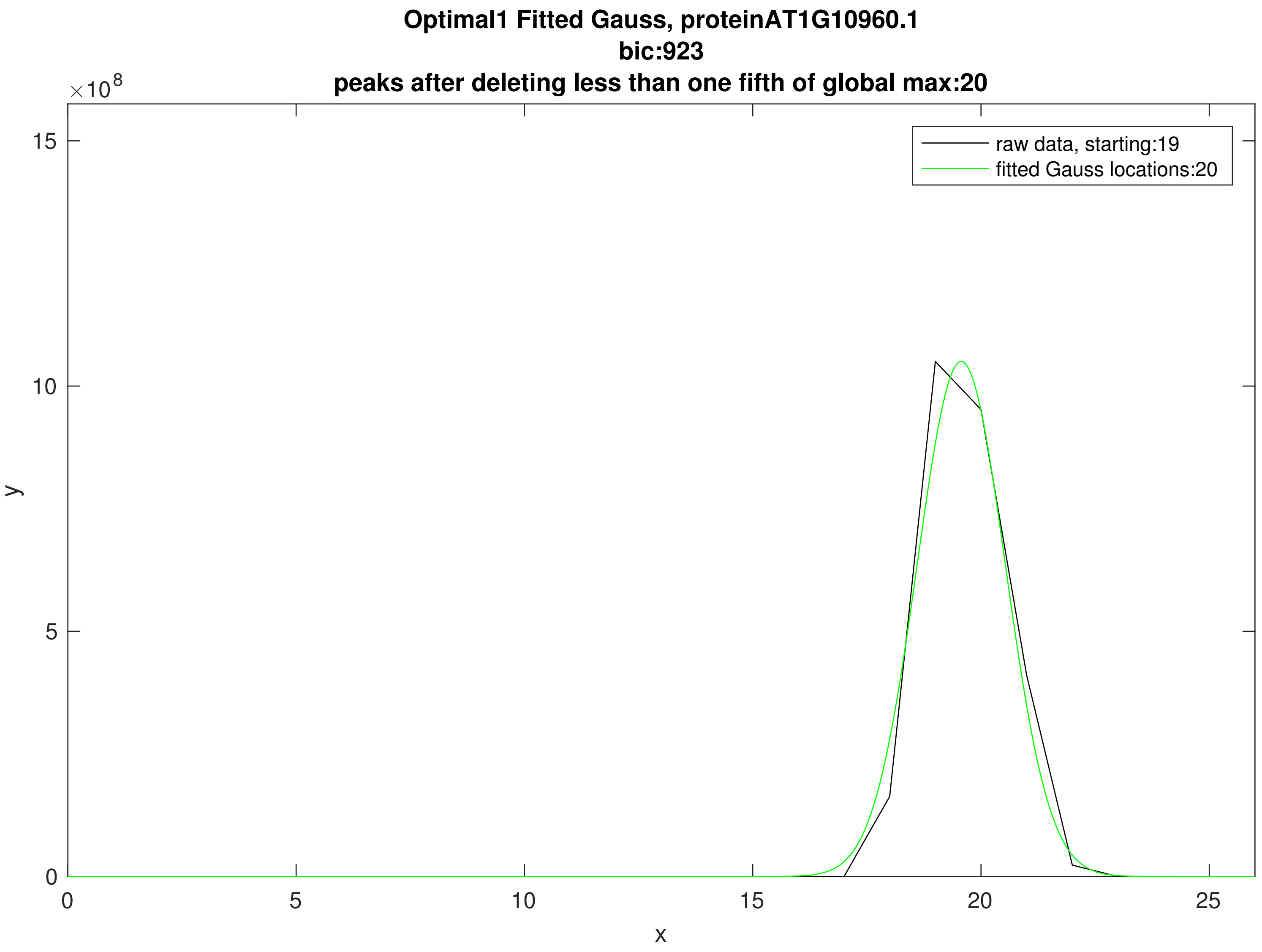

Optimal1 Fitted Gauss, proteinAT1G10960.1
bic:923
peaks after deleting less than one fifth of global max:20
×10^8
15
10
5
0
y
0
5
10
15
20
25
x
raw data, starting:19
fitted Gauss locations:20

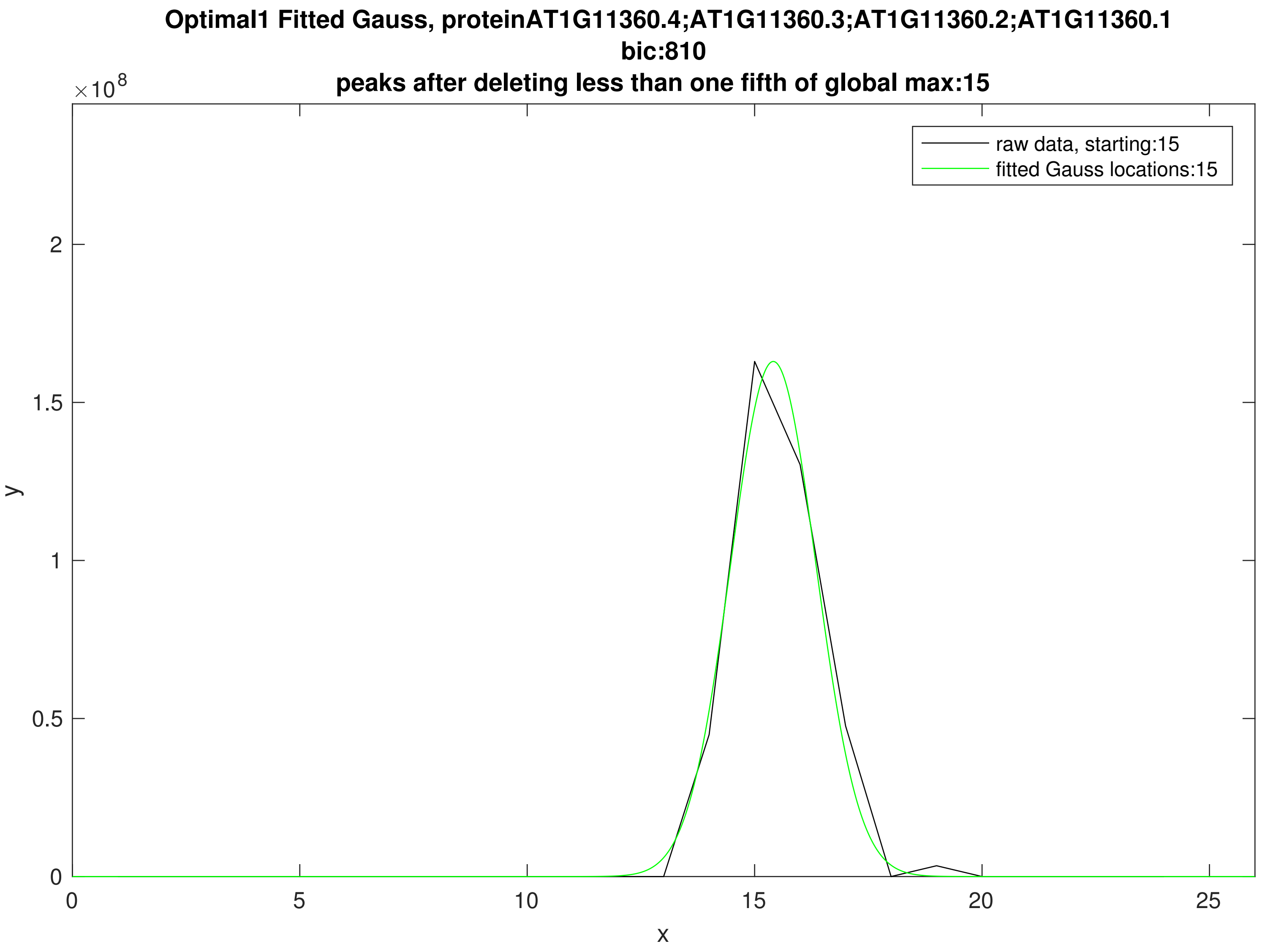
Optimal1 Fitted Gauss, proteinAT1G11360.4;AT1G11360.3;AT1G11360.2;AT1G11360.1
bic:810
peaks after deleting less than one fifth of global max:15
×10^8
raw data, starting:15
fitted Gauss locations:15
2
1.5
1
0.5
0
y
0
5
10
15
20
25
x

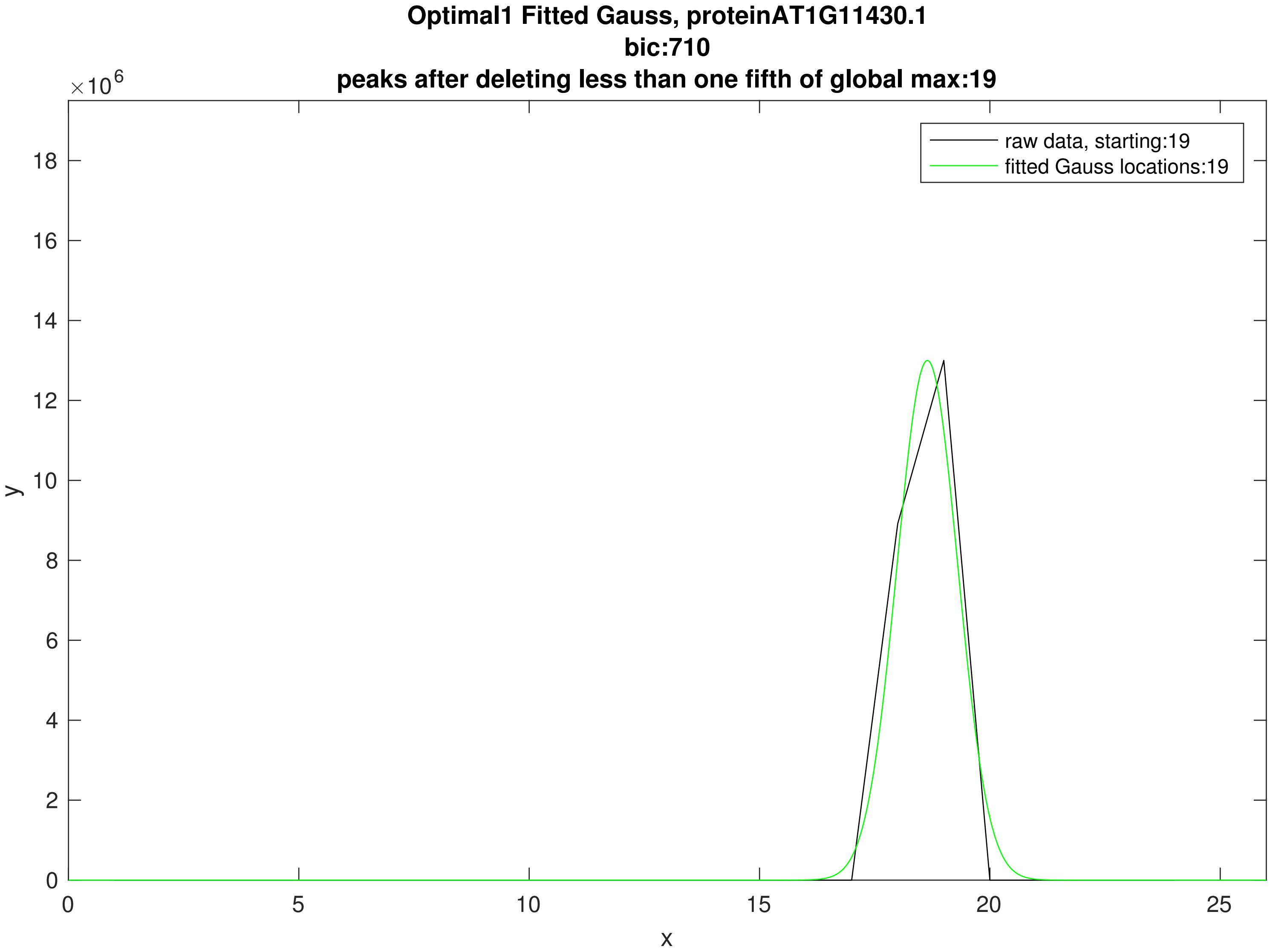
Optimal1 Fitted Gauss, proteinAT1G11430.1
bic:710
peaks after deleting less than one fifth of global max:19
×10^6
raw data, starting:19
fitted Gauss locations:19
y
x
0
2
4
6
8
10
12
14
16
18
0
5
10
15
20
25

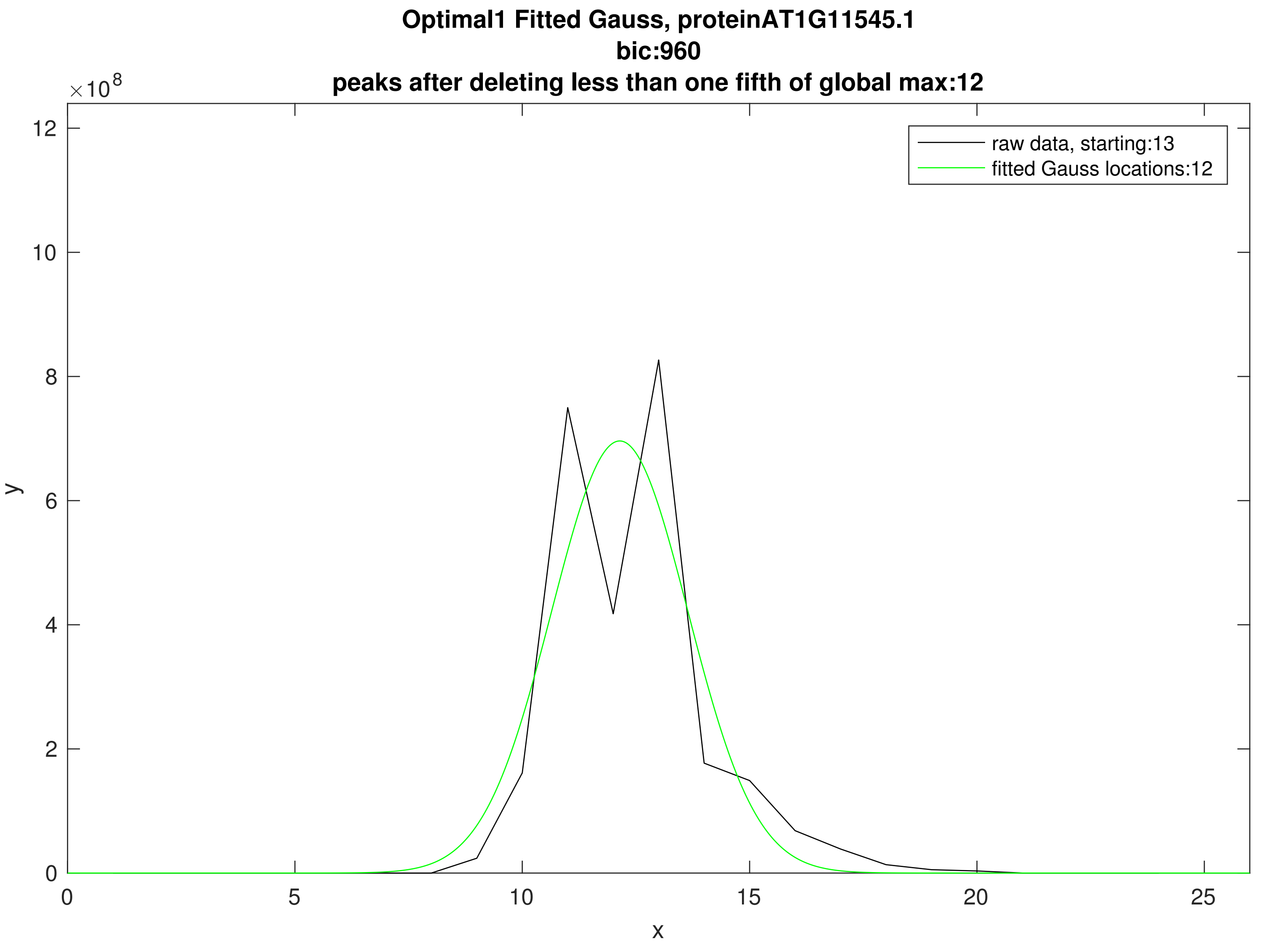
Optimal1 Fitted Gauss, proteinAT1G11545.1
bic:960
peaks after deleting less than one fifth of global max:12
×10^8
raw data, starting:13
fitted Gauss locations:12
y
x
0
2
4
6
8
10
12
0
5
10
15
20
25

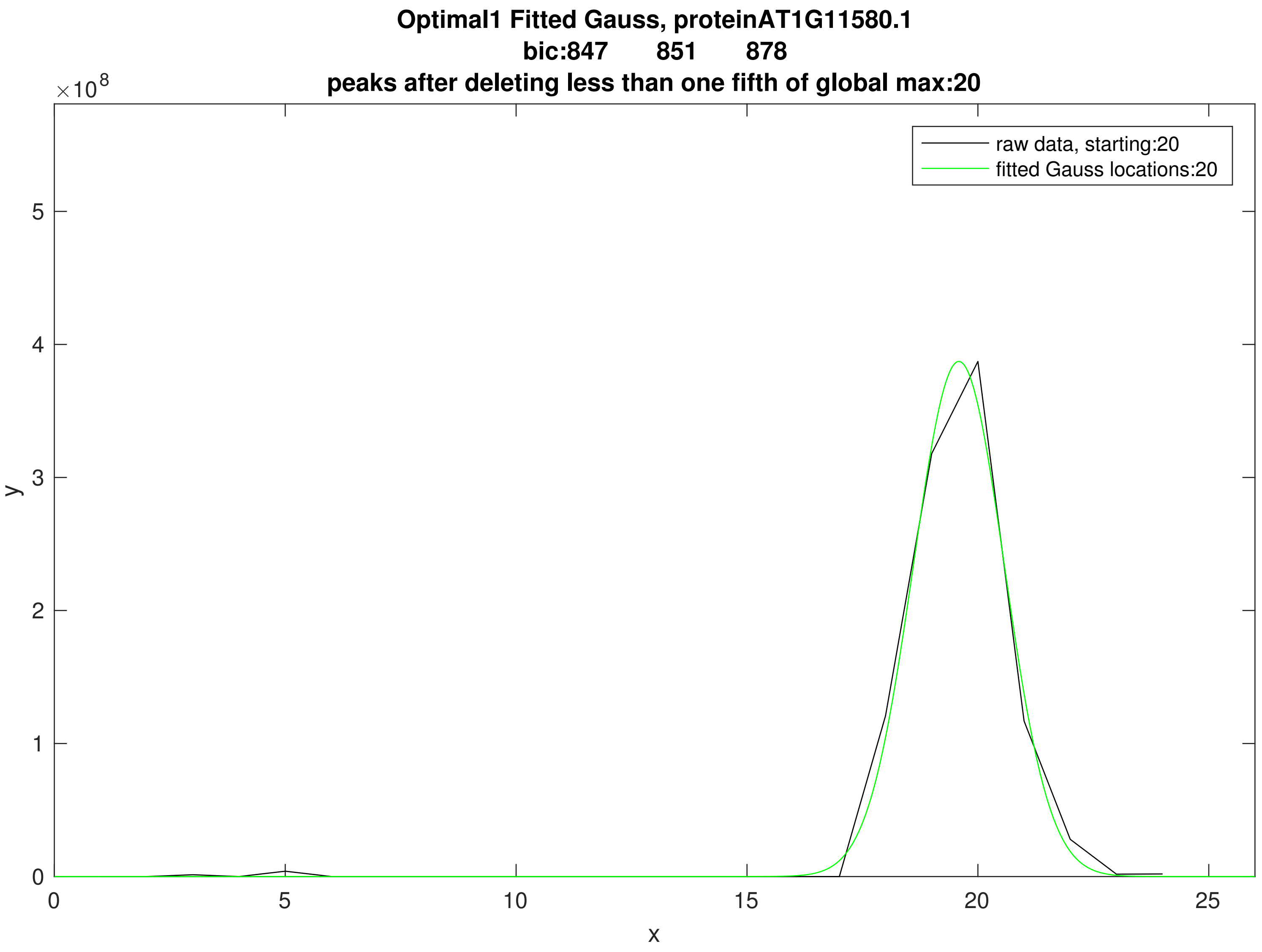

Optimal1 Fitted Gauss, proteinAT1G11580.1
bic:847 851 878
peaks after deleting less than one fifth of global max:20
×10^8
raw data, starting:20
fitted Gauss locations:20
5
4
3
2
1
0
y
0
5
10
15
20
25
x

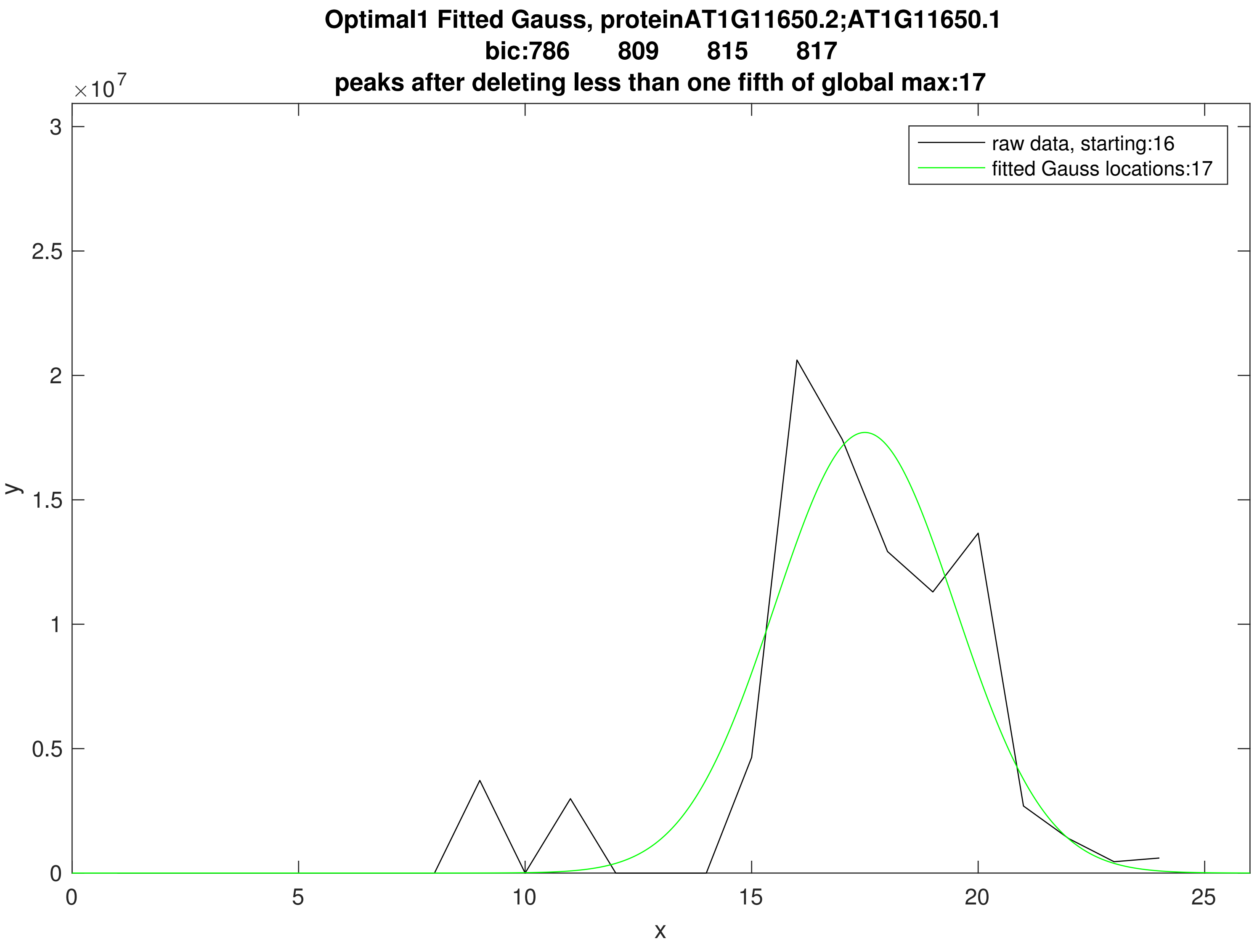
Optimal1 Fitted Gauss, proteinAT1G11650.2;AT1G11650.1
bic:786 809 815 817
peaks after deleting less than one fifth of global max:17
×10^7
raw data, starting:16
fitted Gauss locations:17
y
x
0
0.5
1
1.5
2
2.5
3
5
10
15
20
25

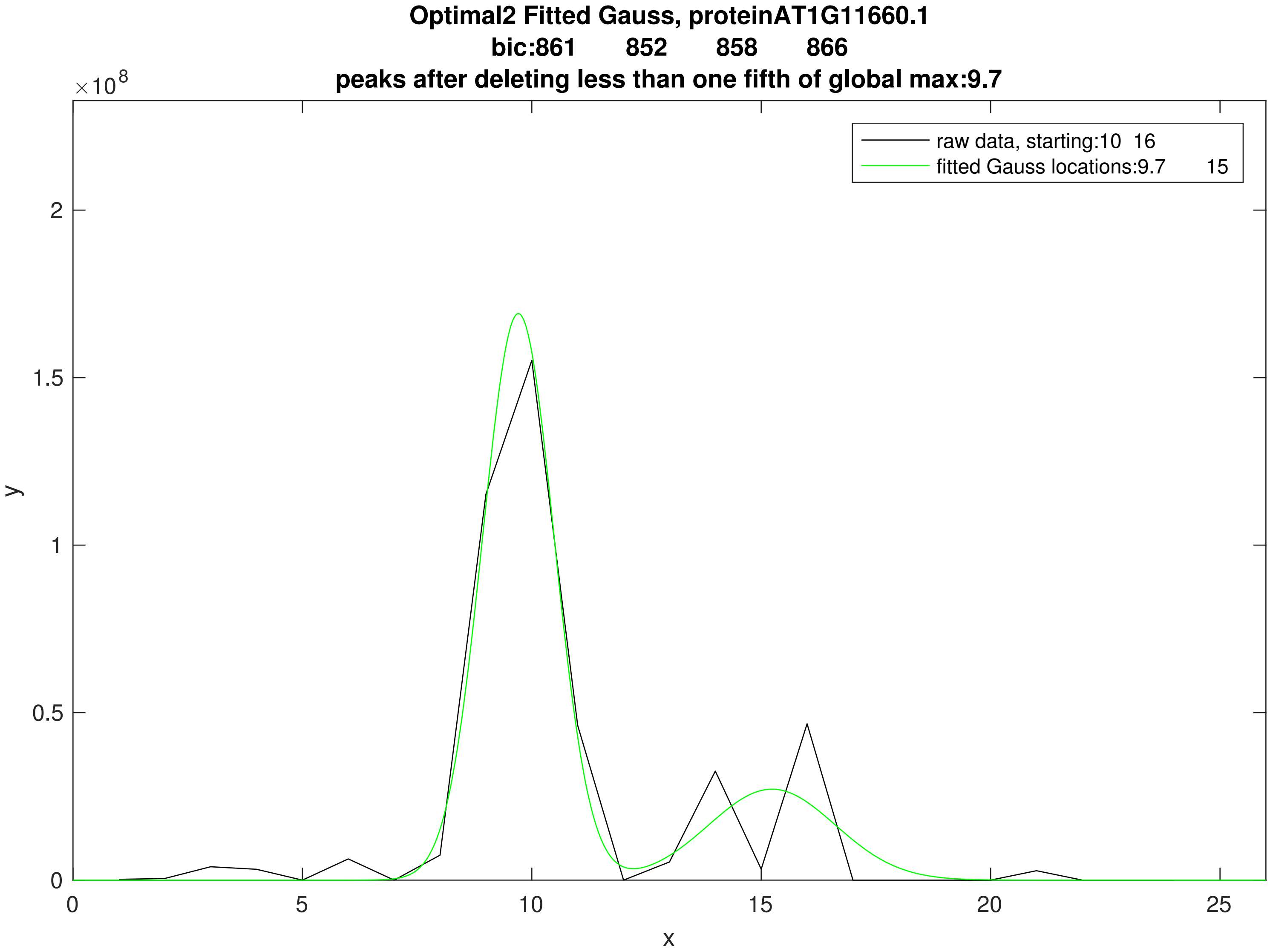

Optimal2 Fitted Gauss, proteinAT1G11660.1
bic:861 852 858 866
peaks after deleting less than one fifth of global max:9.7
raw data, starting:10 16
fitted Gauss locations:9.7 15
×10^8
y
x
0
0.5
1
1.5
2
5
10
15
20
25

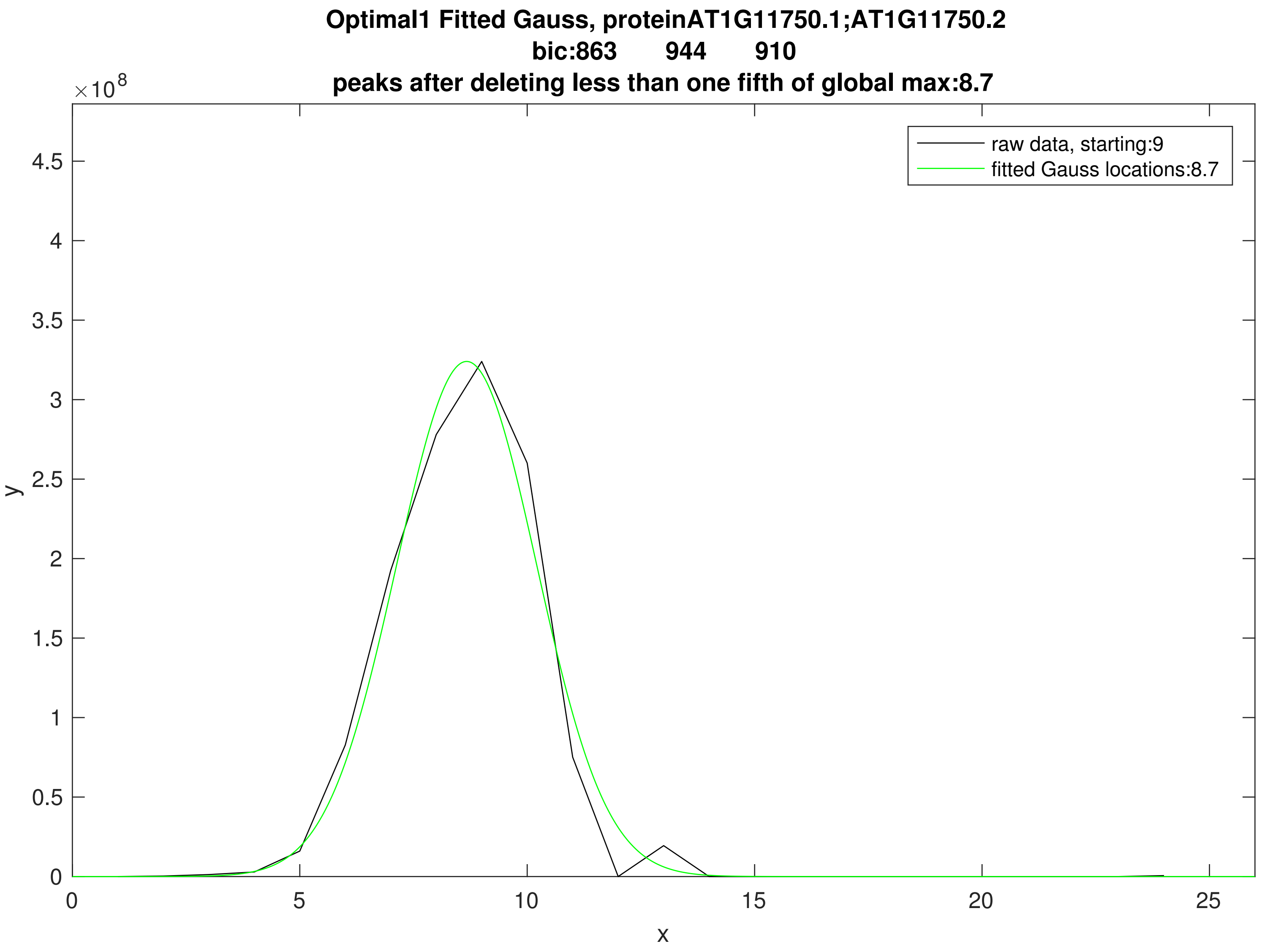

Optimal1 Fitted Gauss, proteinAT1G11750.1;AT1G11750.2
bic:863 944 910
peaks after deleting less than one fifth of global max:8.7
×10^8
raw data, starting:9
fitted Gauss locations:8.7
y
x
0
0.5
1
1.5
2
2.5
3
3.5
4
4.5
5
10
15
20
25

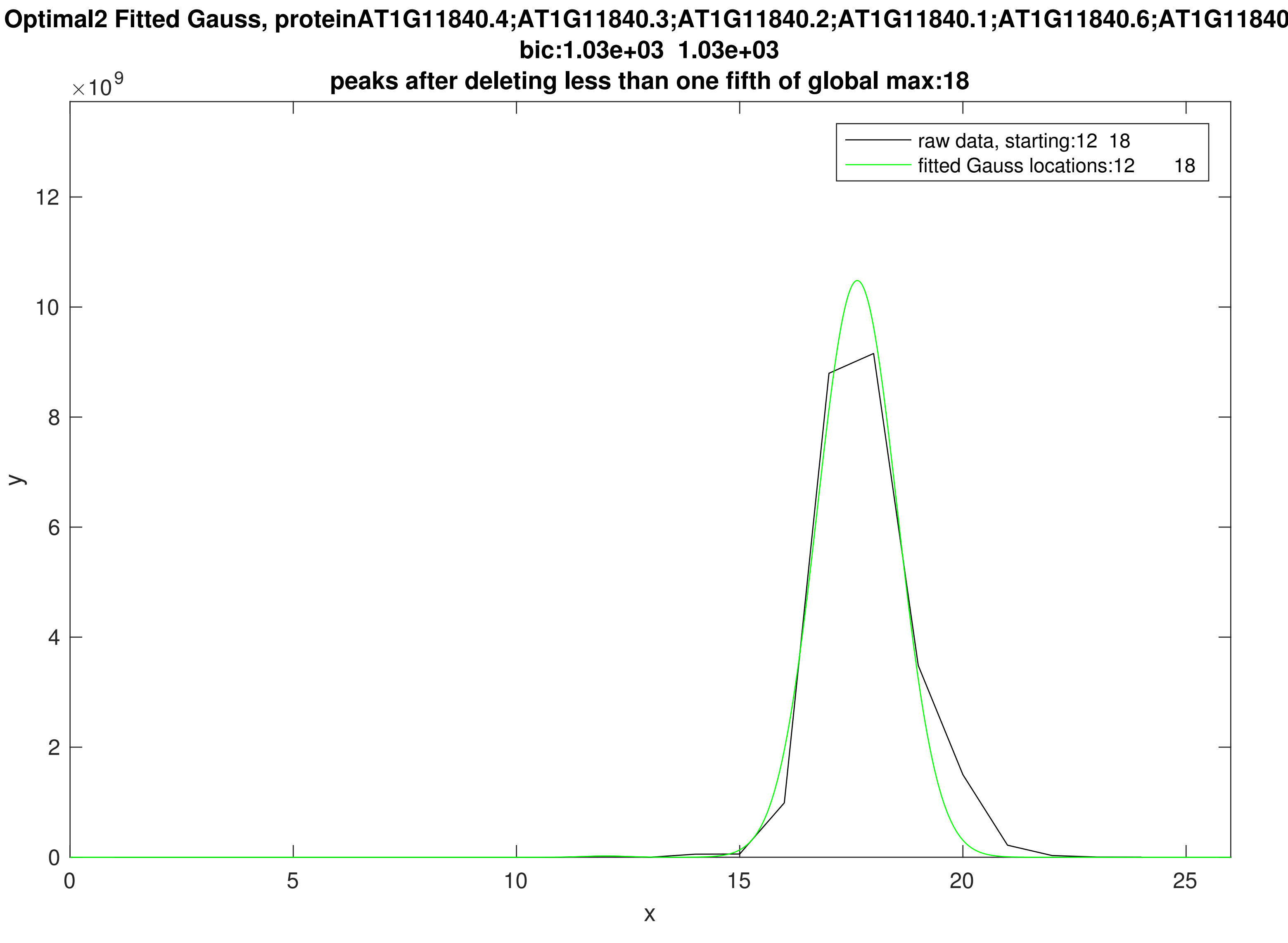

Optimal2 Fitted Gauss, proteinAT1G11840.4;AT1G11840.3;AT1G11840.2;AT1G11840.1;AT1G11840.6;AT1G11840
bic:1.03e+03 1.03e+03
peaks after deleting less than one fifth of global max:18
×10^9
raw data, starting:12 18
fitted Gauss locations:12 18
y
x
0
2
4
6
8
10
12
5
15
20
25

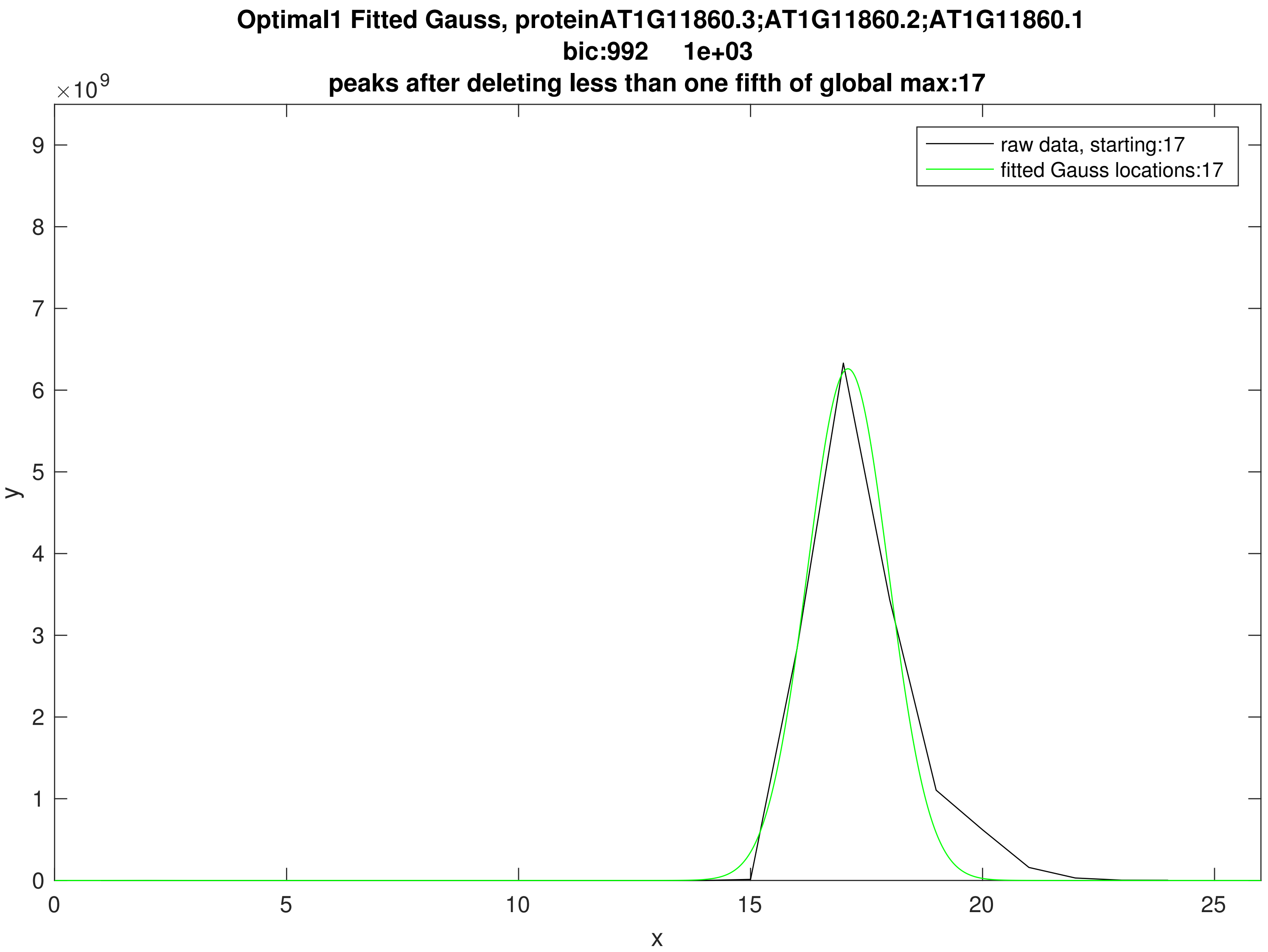
Optimal1 Fitted Gauss, proteinAT1G11860.3;AT1G11860.2;AT1G11860.1
bic:992     1e+03
peaks after deleting less than one fifth of global max:17
×10^9
raw data, starting:17
fitted Gauss locations:17
y
x
0
1
2
3
4
5
6
7
8
9
0
5
10
15
20
25

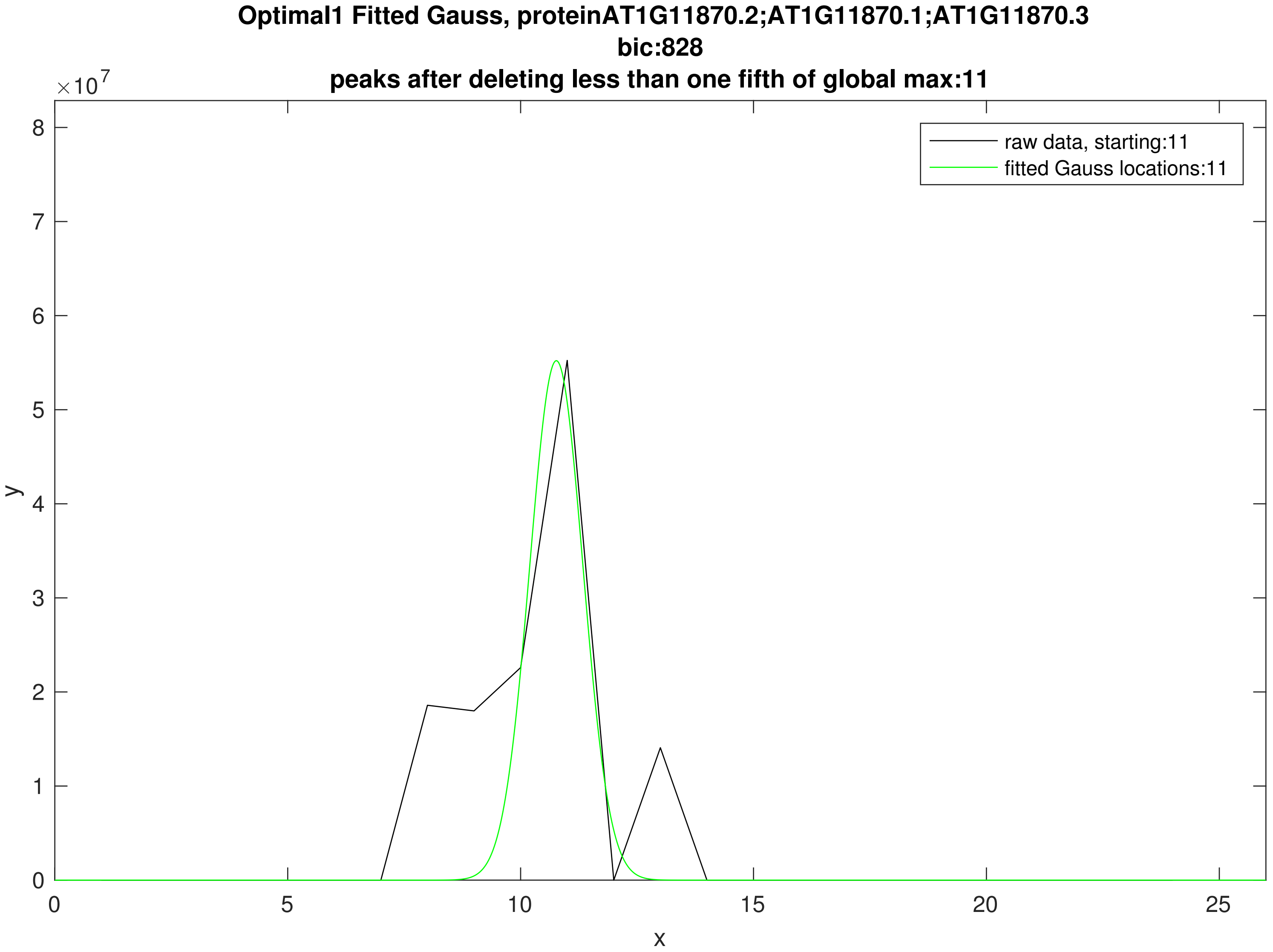

Optimal1 Fitted Gauss, proteinAT1G11870.2;AT1G11870.1;AT1G11870.3
bic:828
peaks after deleting less than one fifth of global max:11
×10^7
raw data, starting:11
fitted Gauss locations:11
y
x
0
1
2
3
4
5
6
7
8
5
10
15
20
25

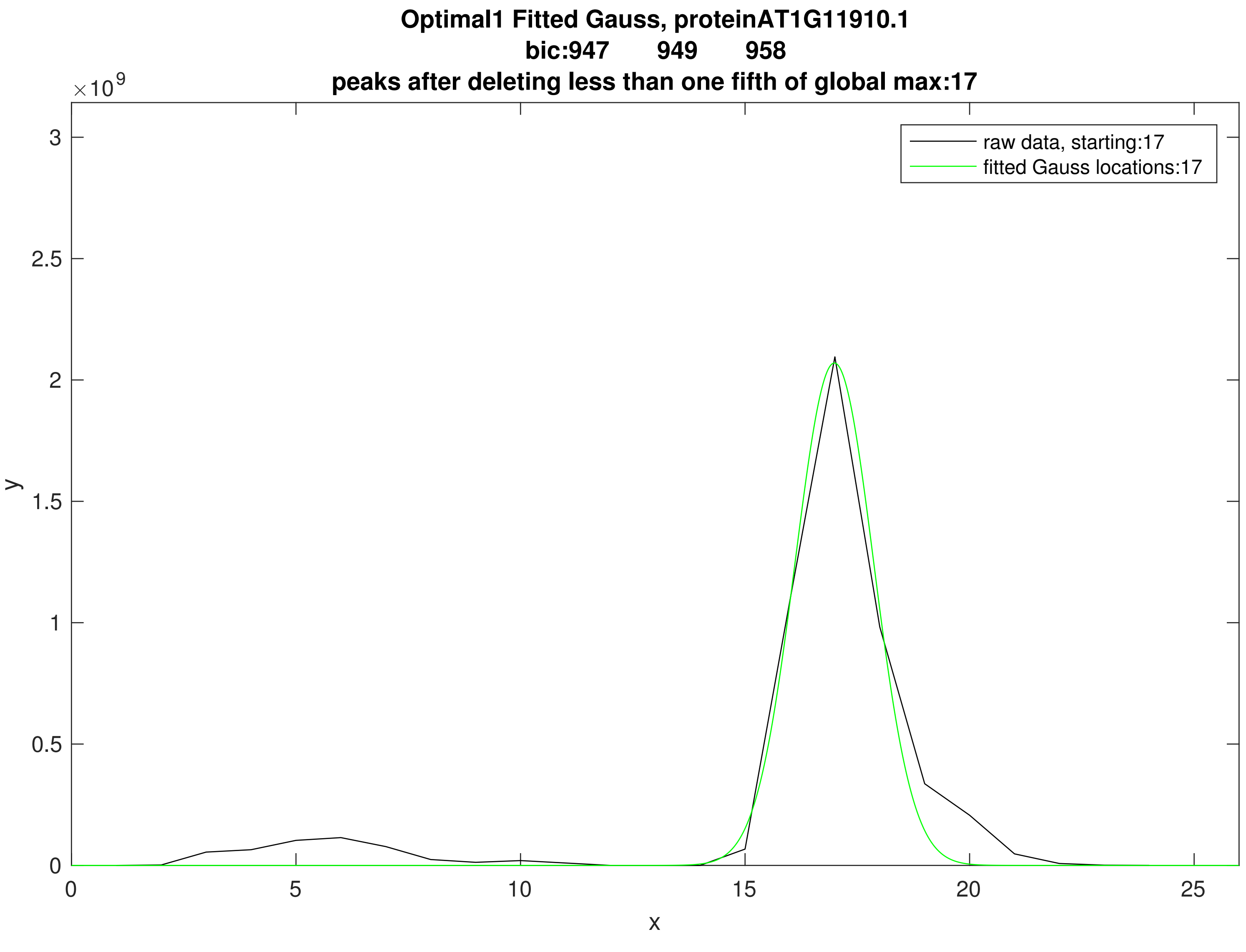
Optimal1 Fitted Gauss, proteinAT1G11910.1
bic:947 949 958
peaks after deleting less than one fifth of global max:17
×10^9
raw data, starting:17
fitted Gauss locations:17
y
x
0
0.5
1
1.5
2
2.5
3
0
5
10
15
20
25

**Optimal1 Fitted Gauss, proteinAT1G11930.2;AT1G11930.1**
**bic:794**
**peaks after deleting less than one fifth of global max:17**

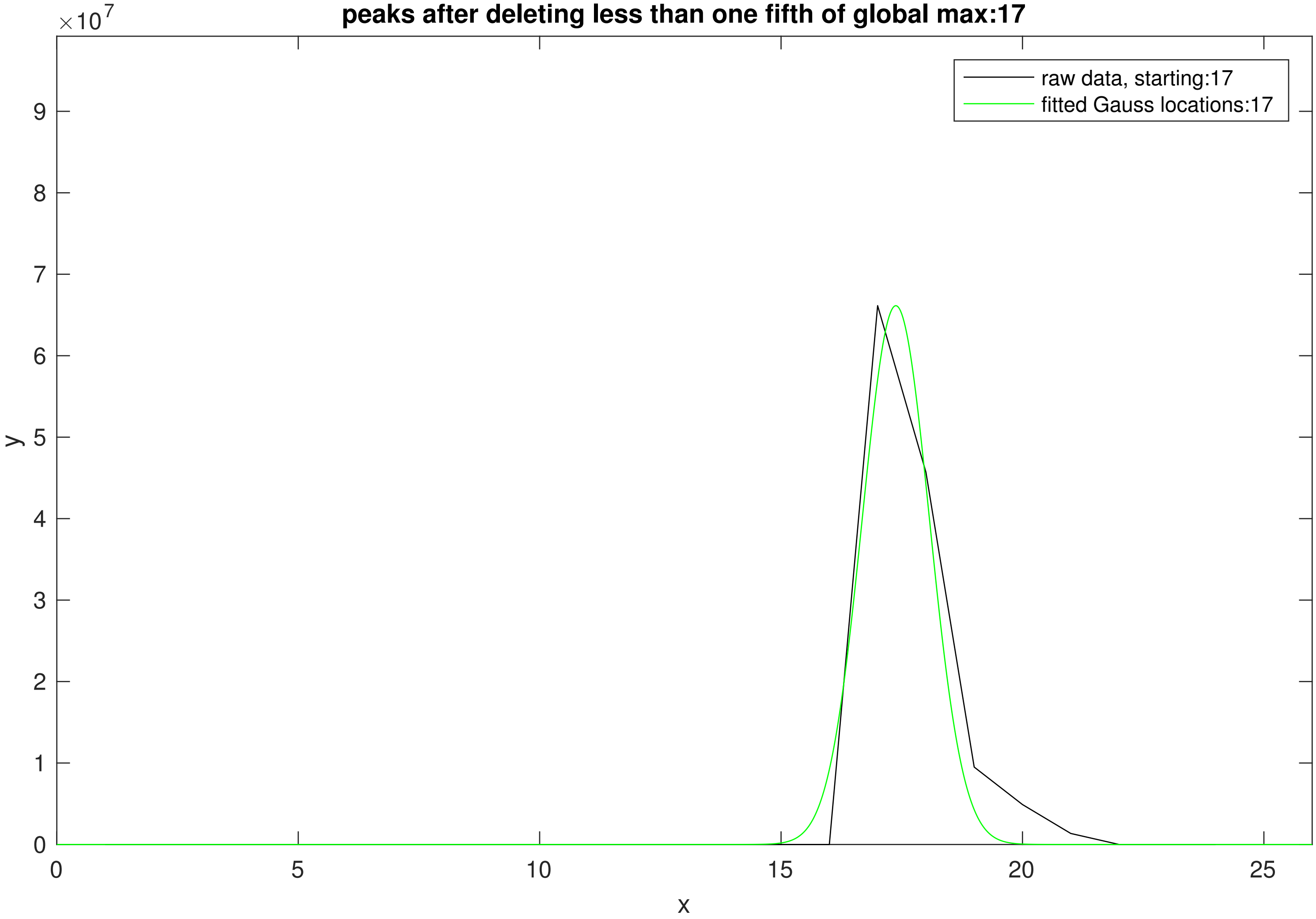

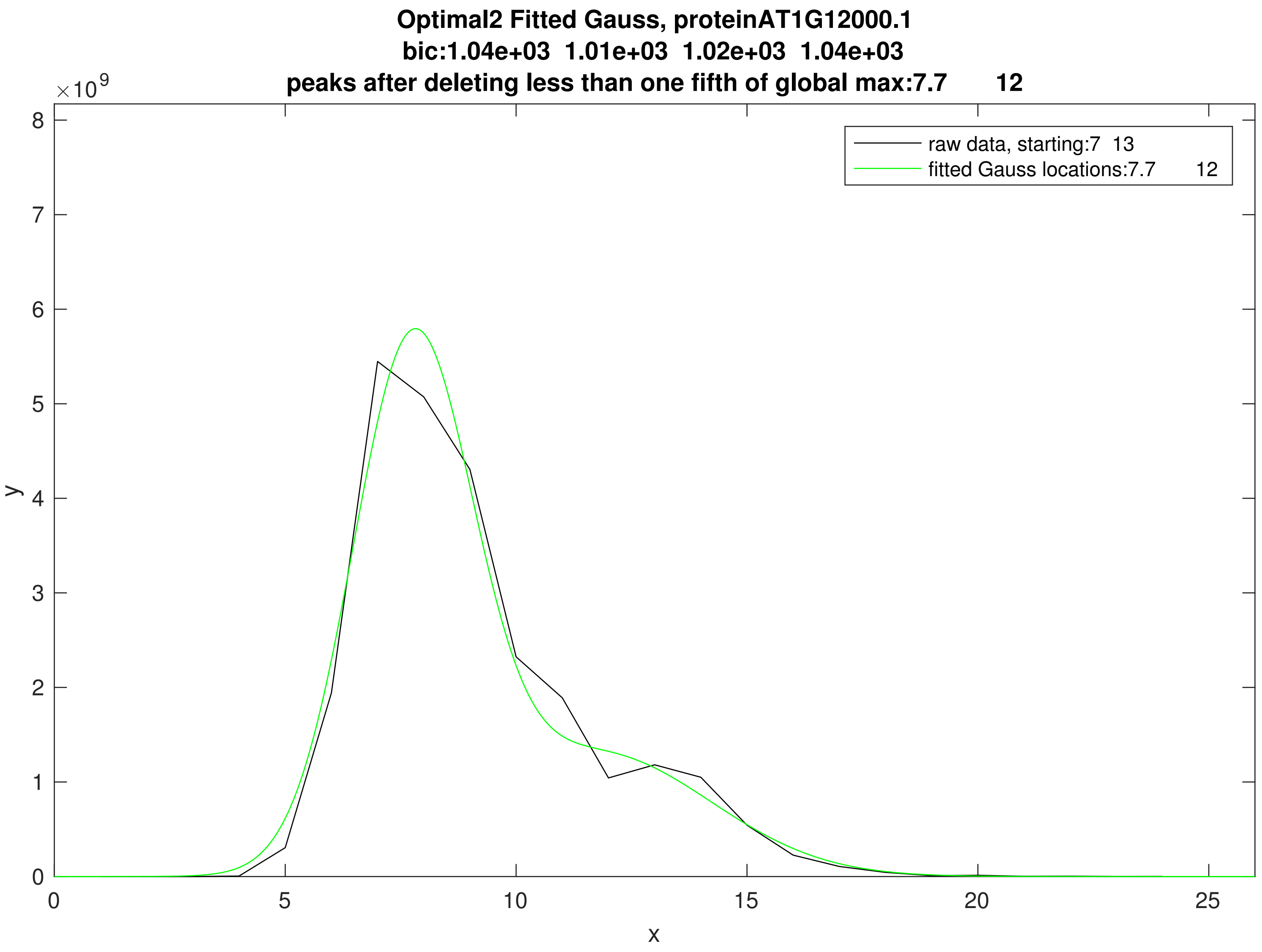
Optimal2 Fitted Gauss, proteinAT1G12000.1
bic:1.04e+03 1.01e+03 1.02e+03 1.04e+03
peaks after deleting less than one fifth of global max:7.7 12
×10^9
raw data, starting:7 13
fitted Gauss locations:7.7 12
y
x
0
1
2
3
4
5
6
7
8
0
5
10
15
20
25

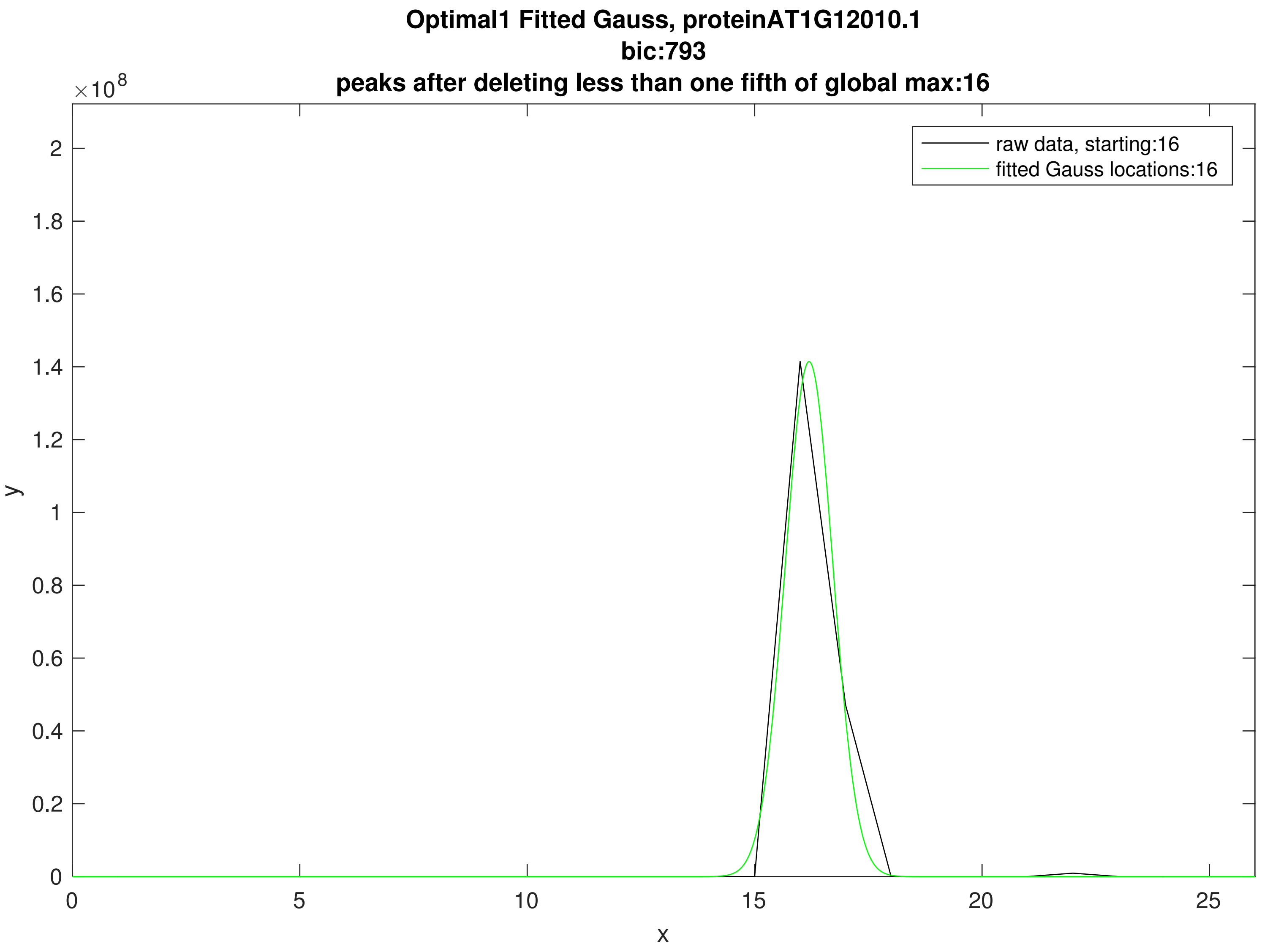

Optimal1 Fitted Gauss, proteinAT1G12010.1
bic:793
peaks after deleting less than one fifth of global max:16
×10^8
raw data, starting:16
fitted Gauss locations:16
y
x
0
0.2
0.4
0.6
0.8
1
1.2
1.4
1.6
1.8
2
0
5
10
15
20
25

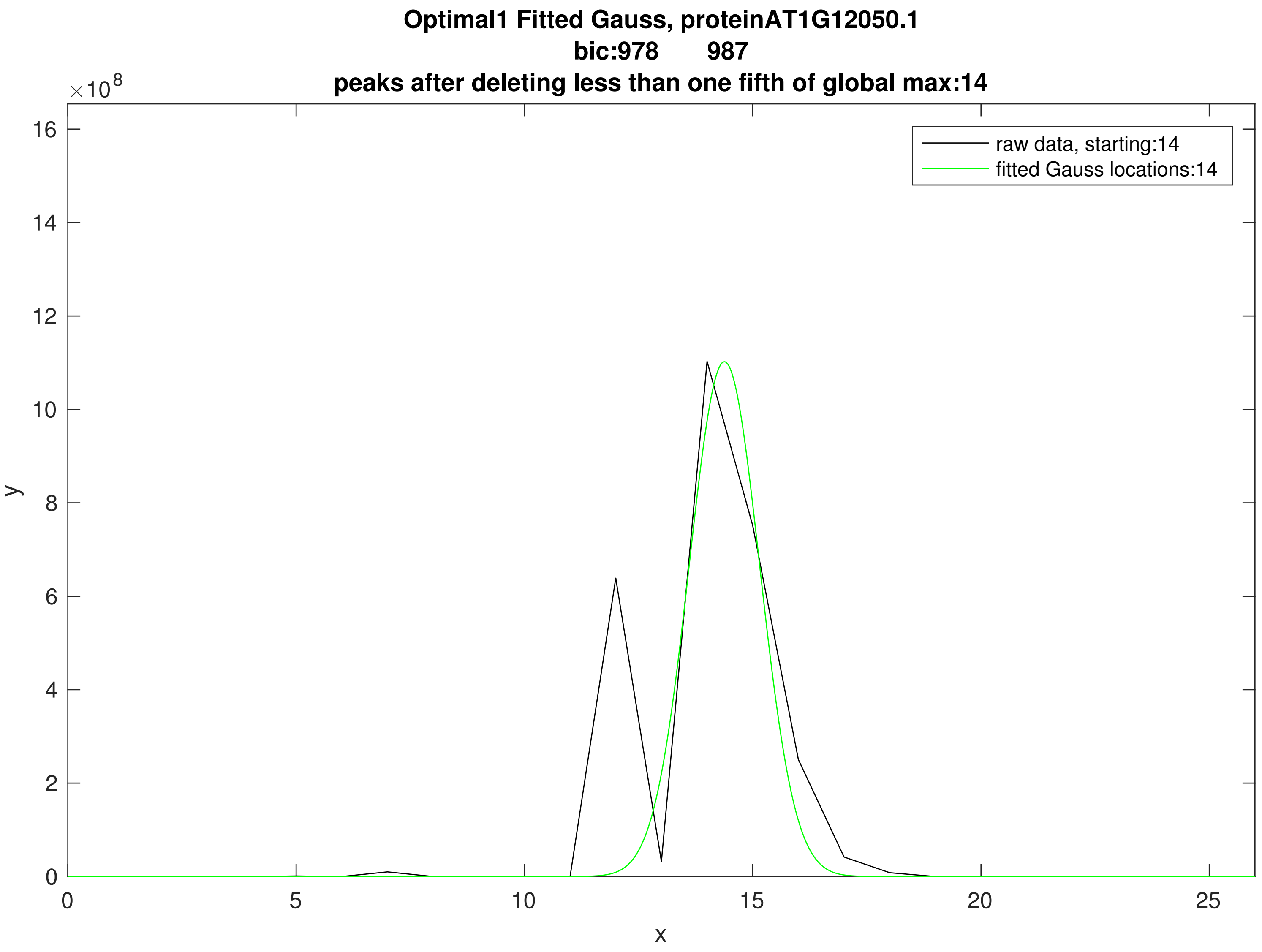
Optimal1 Fitted Gauss, proteinAT1G12050.1
bic:978 987
peaks after deleting less than one fifth of global max:14
×10^8
raw data, starting:14
fitted Gauss locations:14
y
x
0
2
4
6
8
10
12
14
16
0
5
10
15
20
25

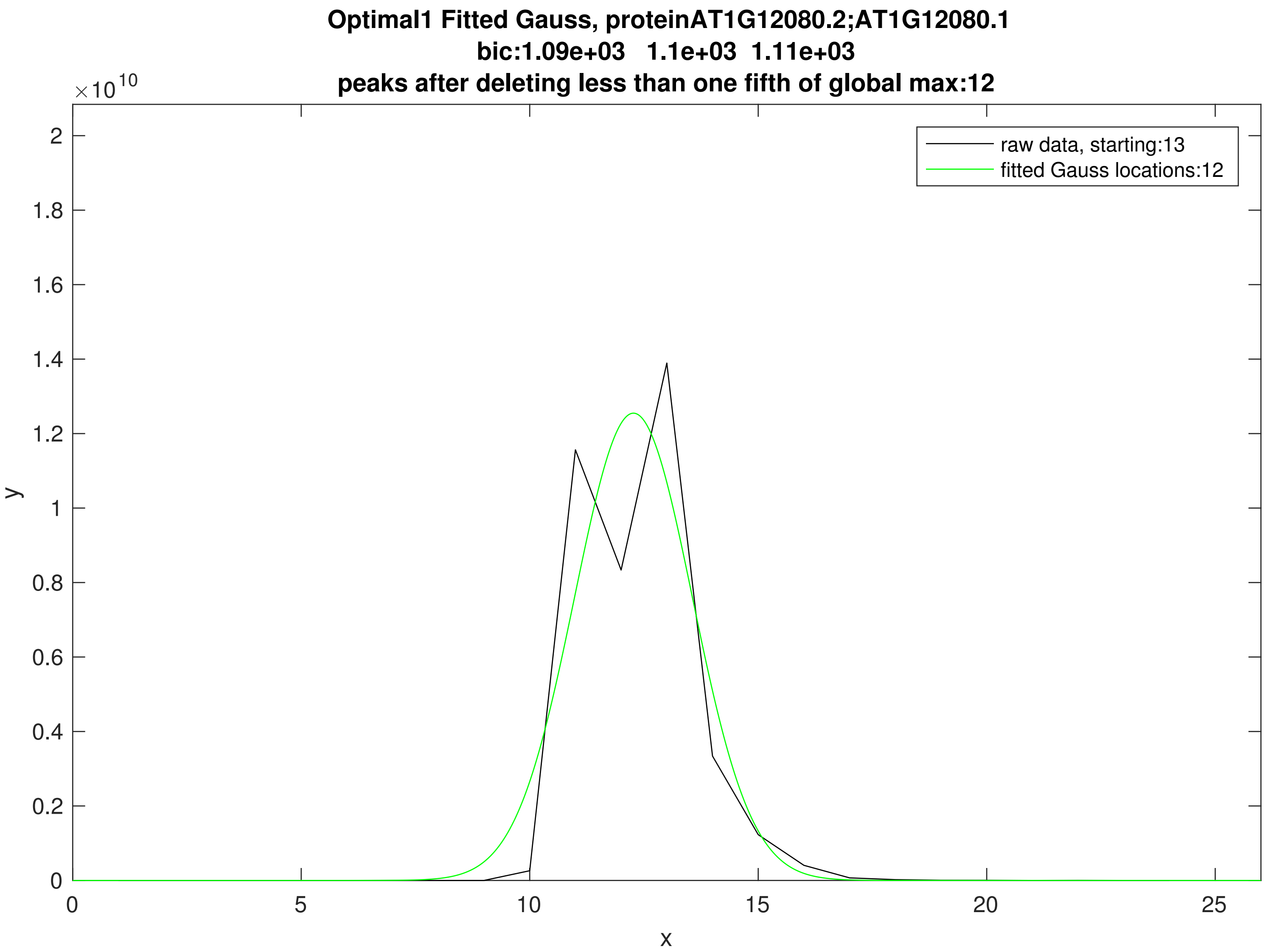

Optimal1 Fitted Gauss, proteinAT1G12080.2;AT1G12080.1
bic:1.09e+03 1.1e+03 1.11e+03
peaks after deleting less than one fifth of global max:12
×10^10
raw data, starting:13
fitted Gauss locations:12
y
x
0
0.2
0.4
0.6
0.8
1
1.2
1.4
1.6
1.8
2
0
5
10
15
20
25

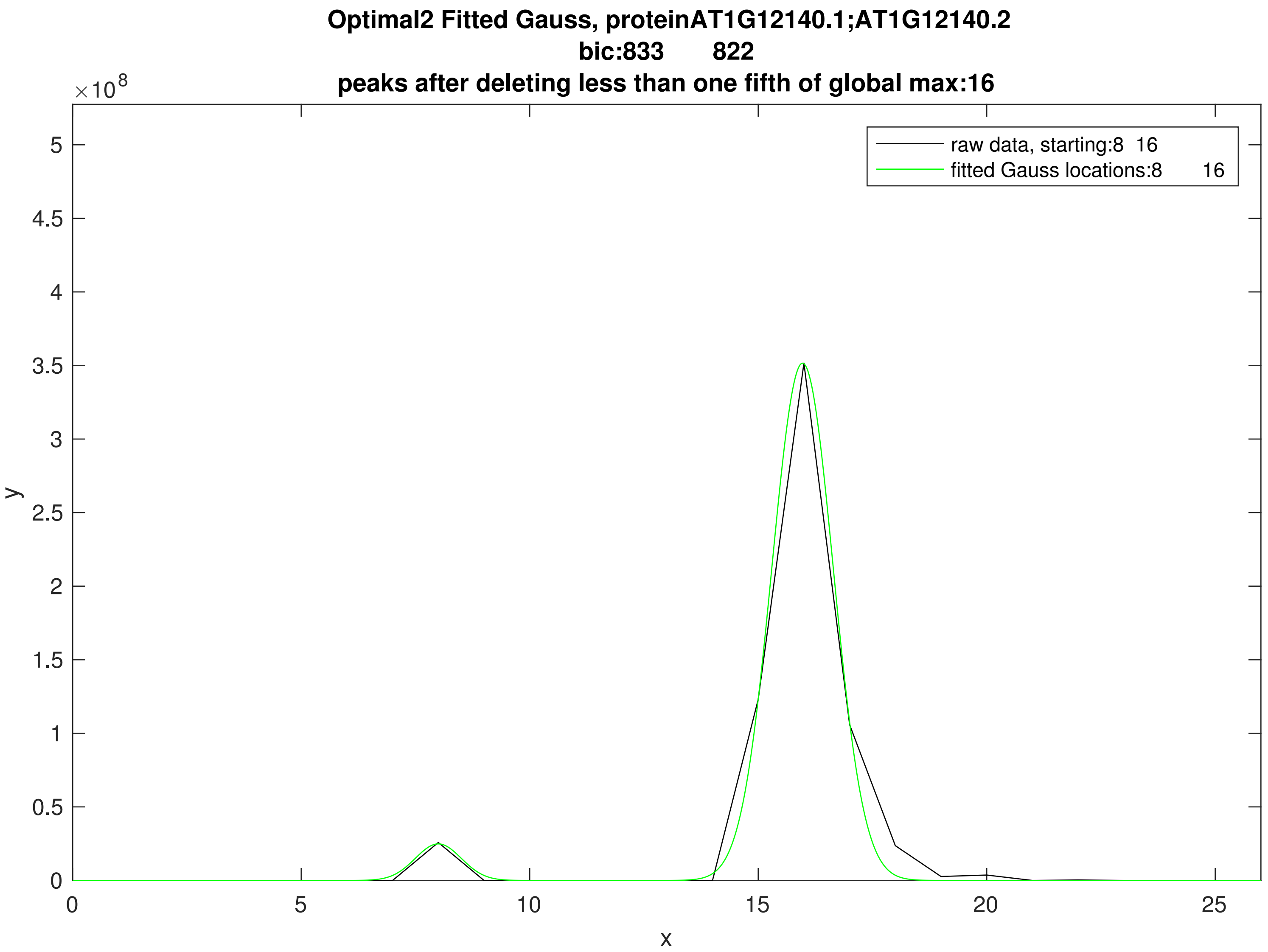
Optimal2 Fitted Gauss, proteinAT1G12140.1;AT1G12140.2
bic:833 822
peaks after deleting less than one fifth of global max:16
×10^8
raw data, starting:8 16
fitted Gauss locations:8 16
y
x
0
0.5
1
1.5
2
2.5
3
3.5
4
4.5
5
0
5
10
15
20
25

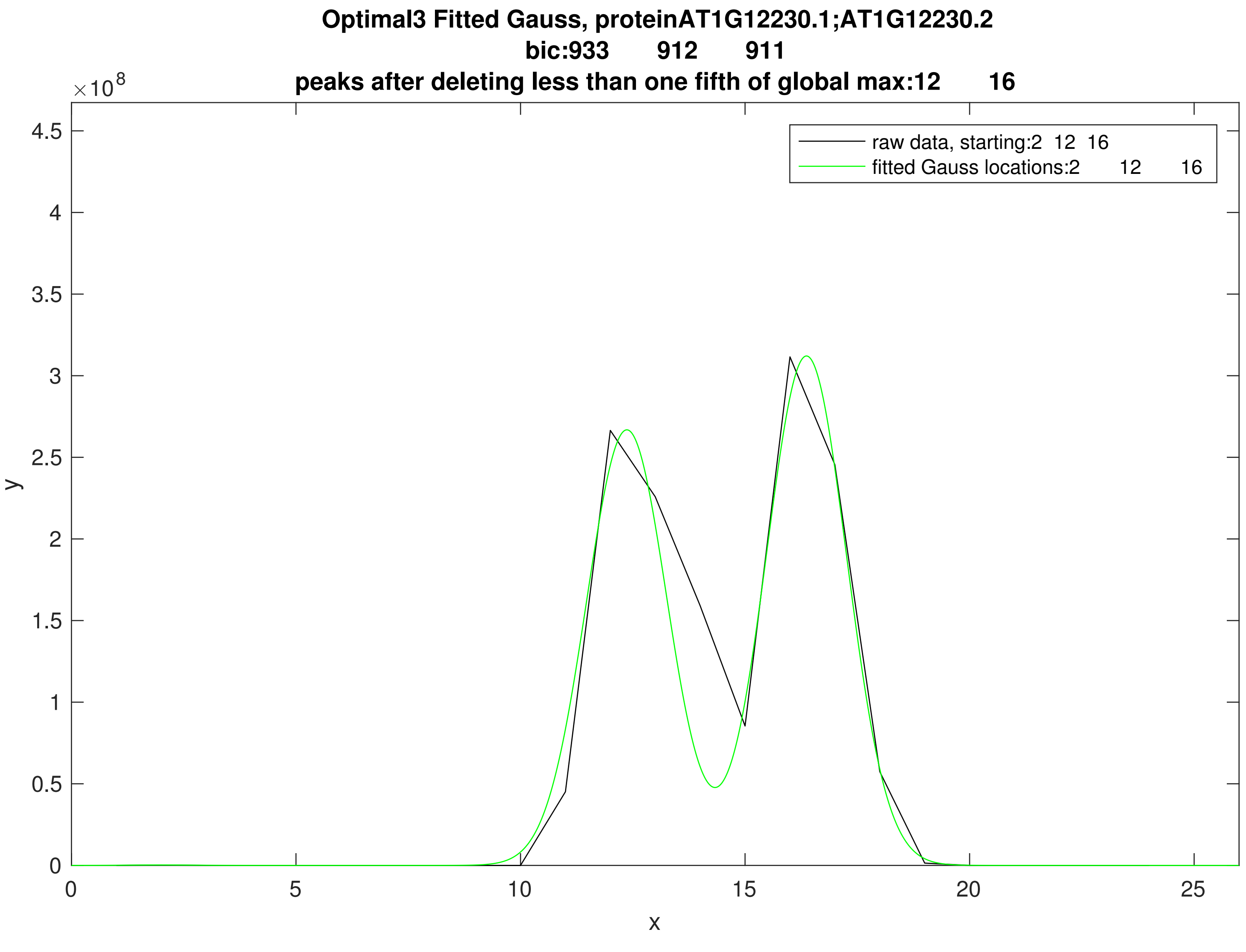

Optimal3 Fitted Gauss, proteinAT1G12230.1;AT1G12230.2
bic:933 912 911
peaks after deleting less than one fifth of global max:12 16
×10^8
raw data, starting:2 12 16
fitted Gauss locations:2 12 16
y
x
0
0.5
1
1.5
2
2.5
3
3.5
4
4.5
0
5
10
15
20
25

Optimal1 Fitted Gauss, proteinAT1G12240.1
bic:984 994 1e+03
peaks after deleting less than one fifth of global max:15

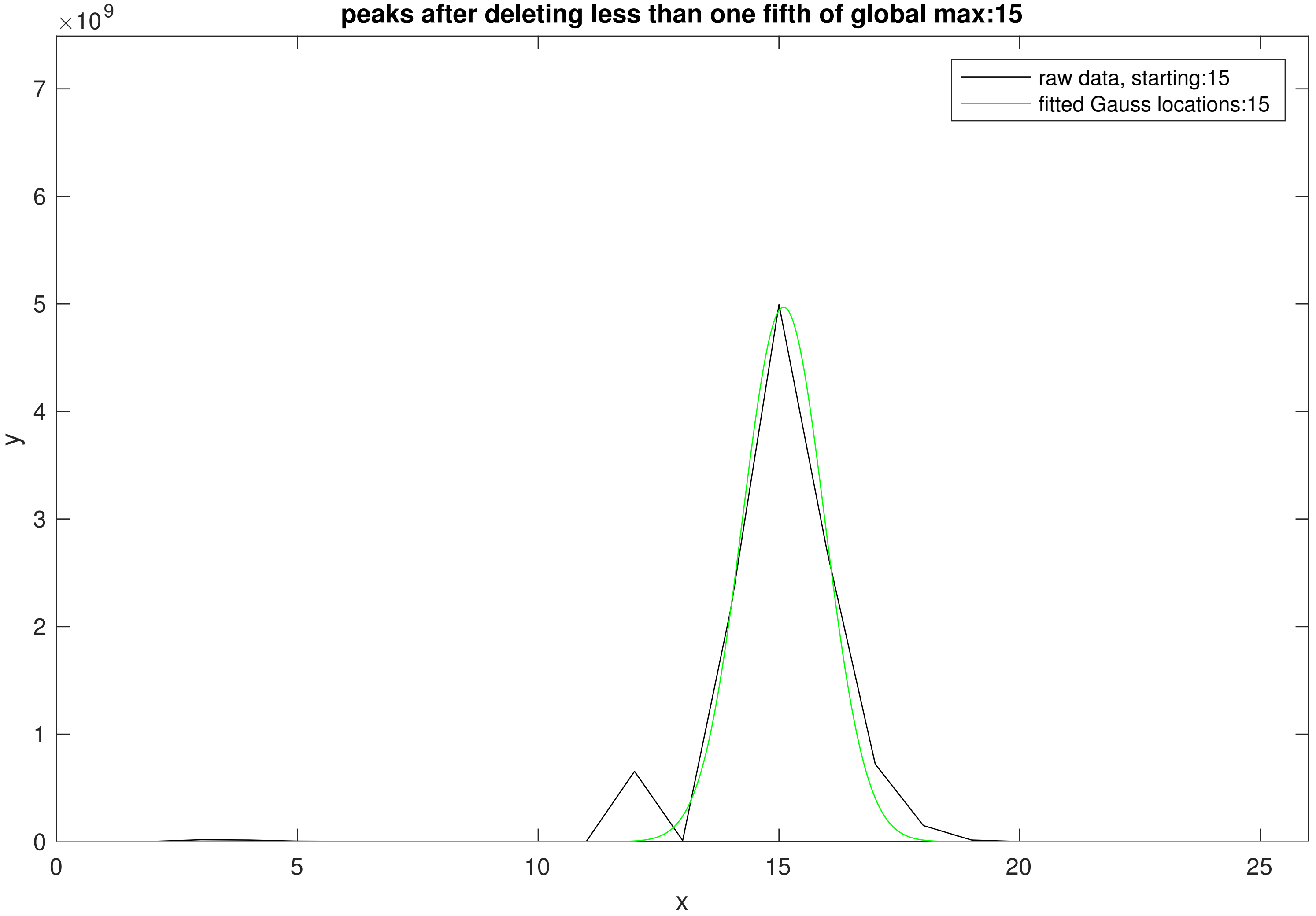

**Optimal1 Fitted Gauss, proteinAT1G12270.1**
**bic:886 887**
**peaks after deleting less than one fifth of global max:11**

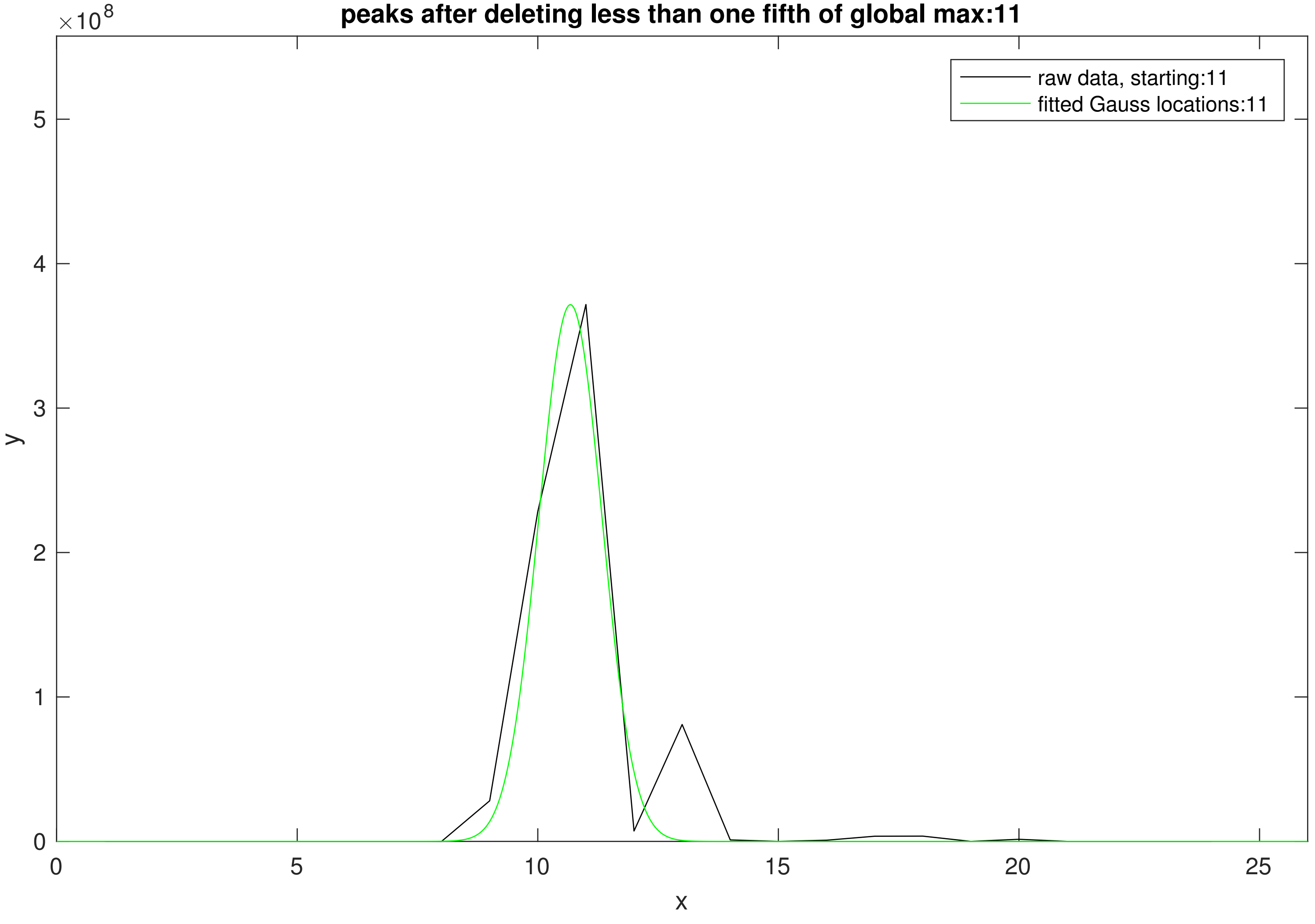

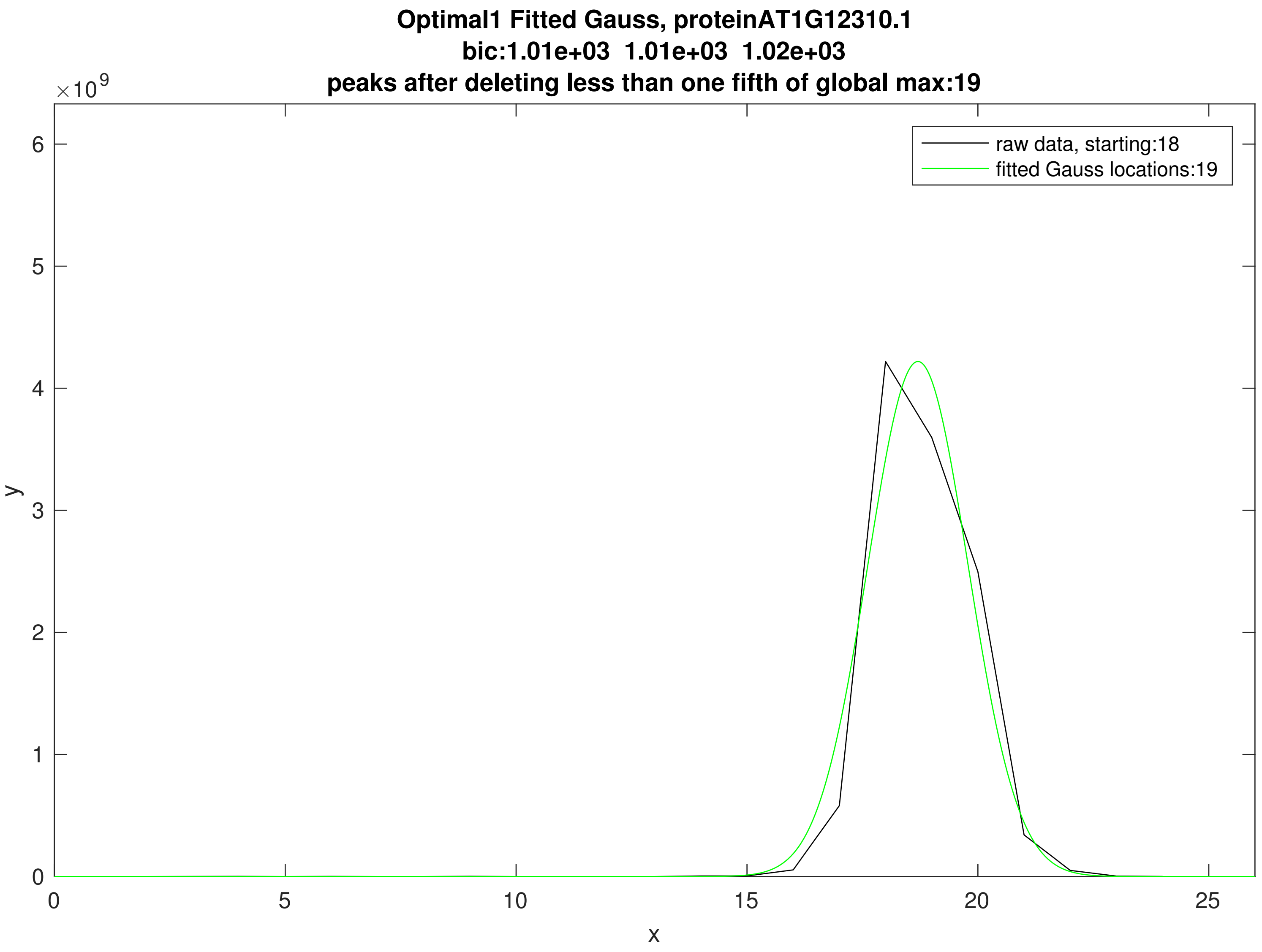
Optimal1 Fitted Gauss, proteinAT1G12310.1
bic:1.01e+03 1.01e+03 1.02e+03
peaks after deleting less than one fifth of global max:19
×10^9
raw data, starting:18
fitted Gauss locations:19
y
x
0
1
2
3
4
5
6
0
5
10
15
20
25

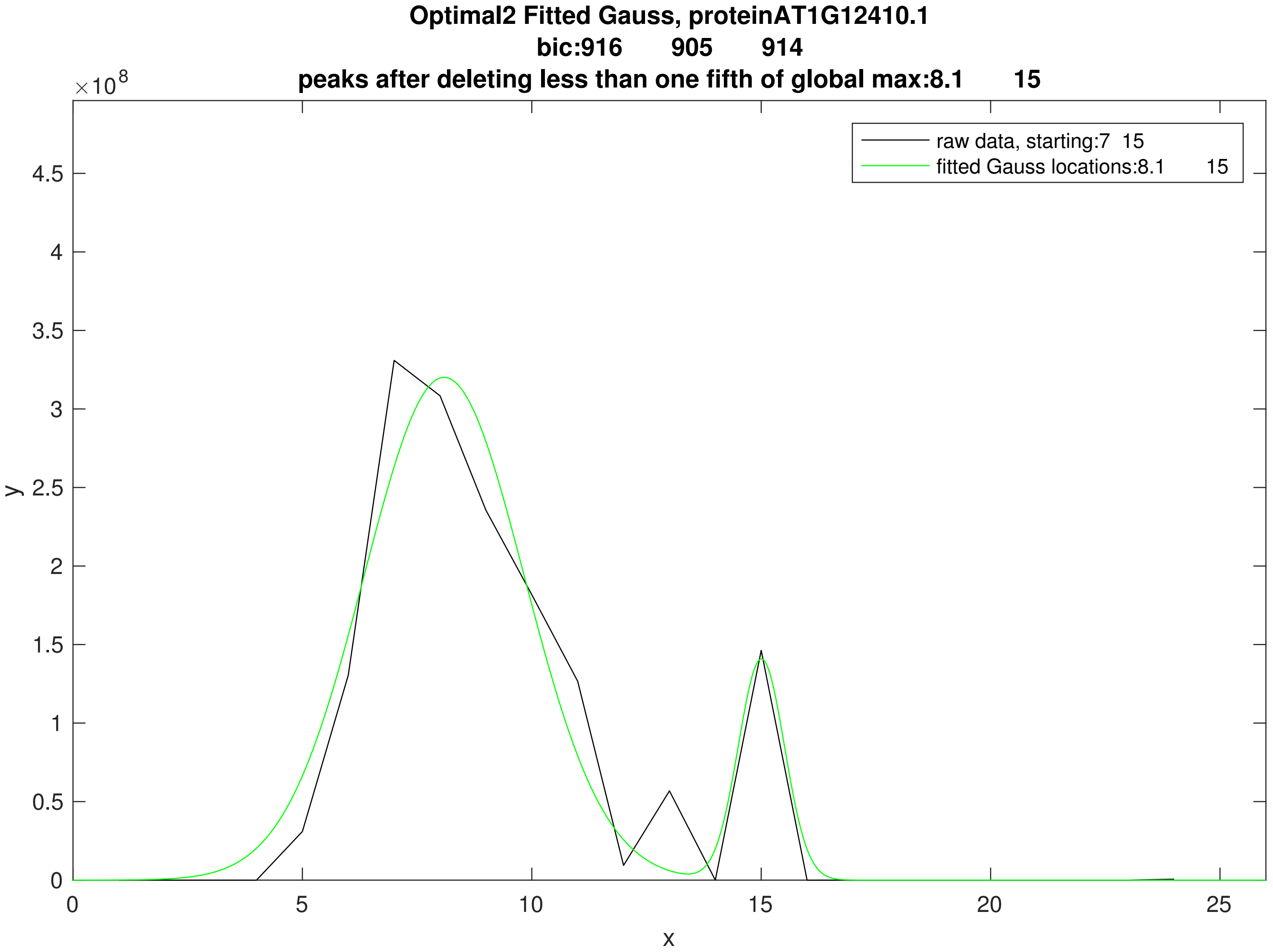
Optimal2 Fitted Gauss, proteinAT1G12410.1
bic:916 905 914
peaks after deleting less than one fifth of global max:8.1 15
×10^8
raw data, starting:7 15
fitted Gauss locations:8.1 15
y
x
0
0.5
1
1.5
2
2.5
3
3.5
4
4.5
5
10
15
20
25

#### Optimal1 Fitted Gauss, proteinAT1G12440.2;AT1G12440.1
#### bic:696
#### peaks after deleting less than one fifth of global max:18

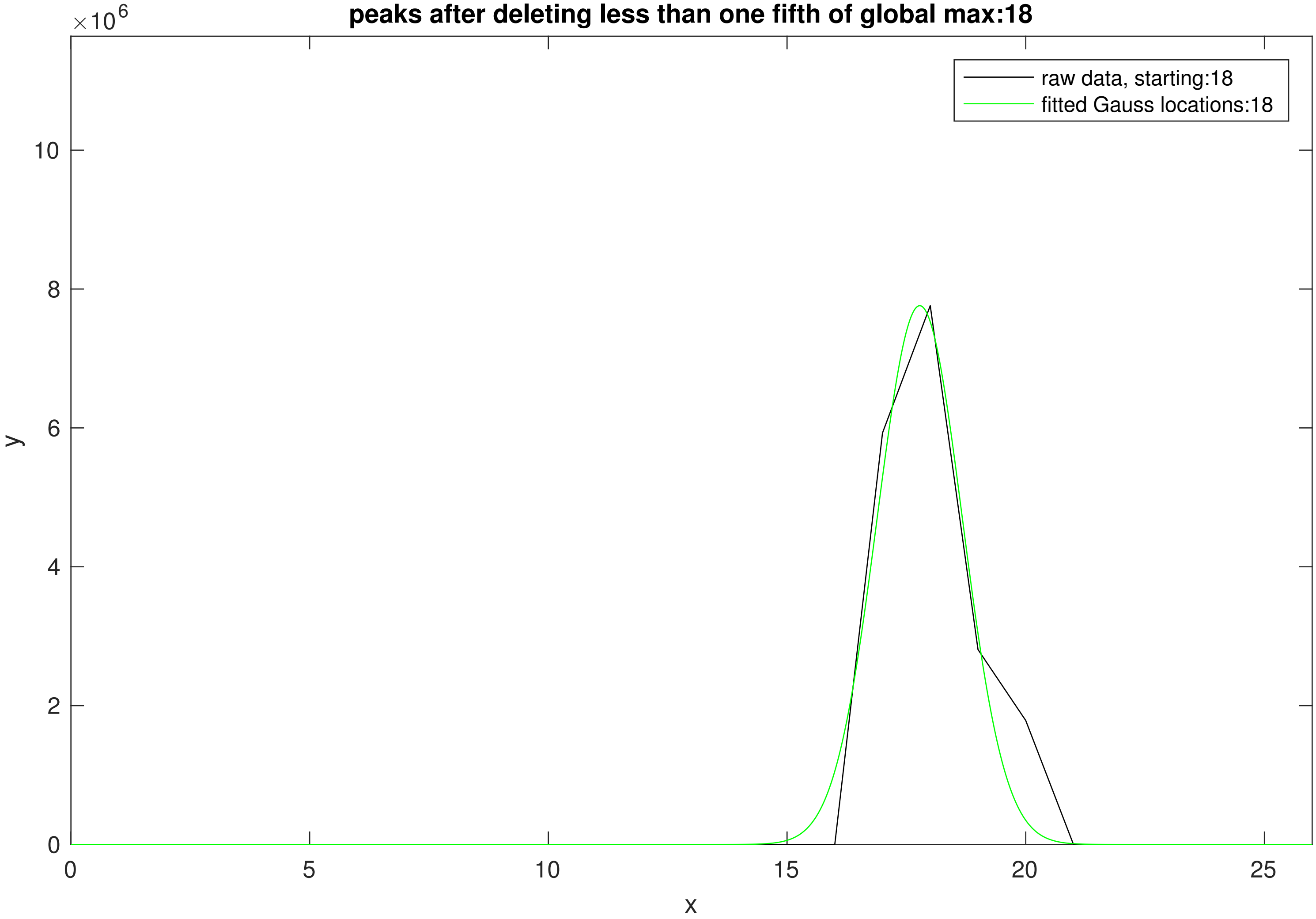

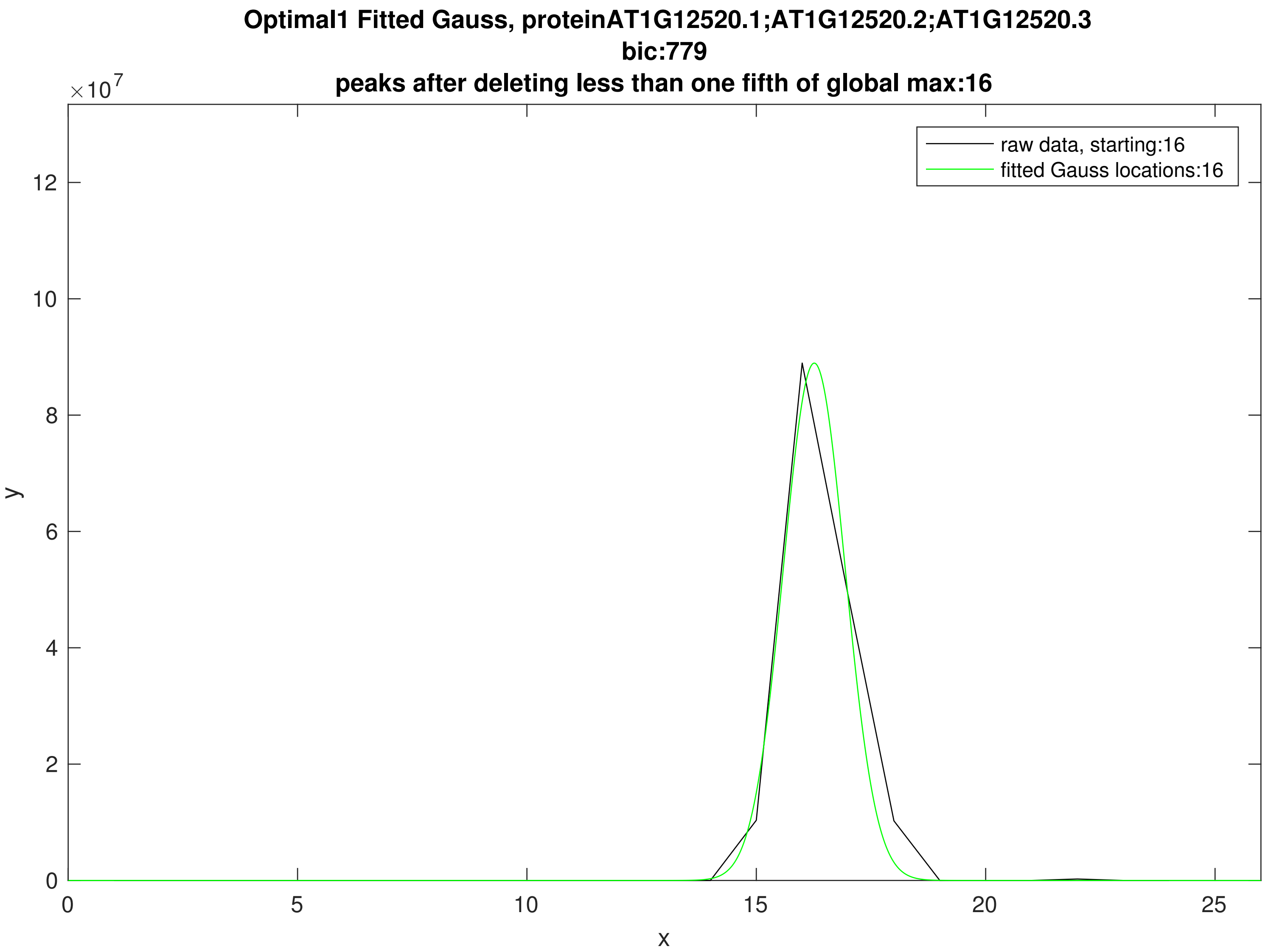
Optimal1 Fitted Gauss, proteinAT1G12520.1;AT1G12520.2;AT1G12520.3
bic:779
peaks after deleting less than one fifth of global max:16
×10^7
raw data, starting:16
fitted Gauss locations:16
y
x
0
2
4
6
8
10
12
0
5
10
15
20
25

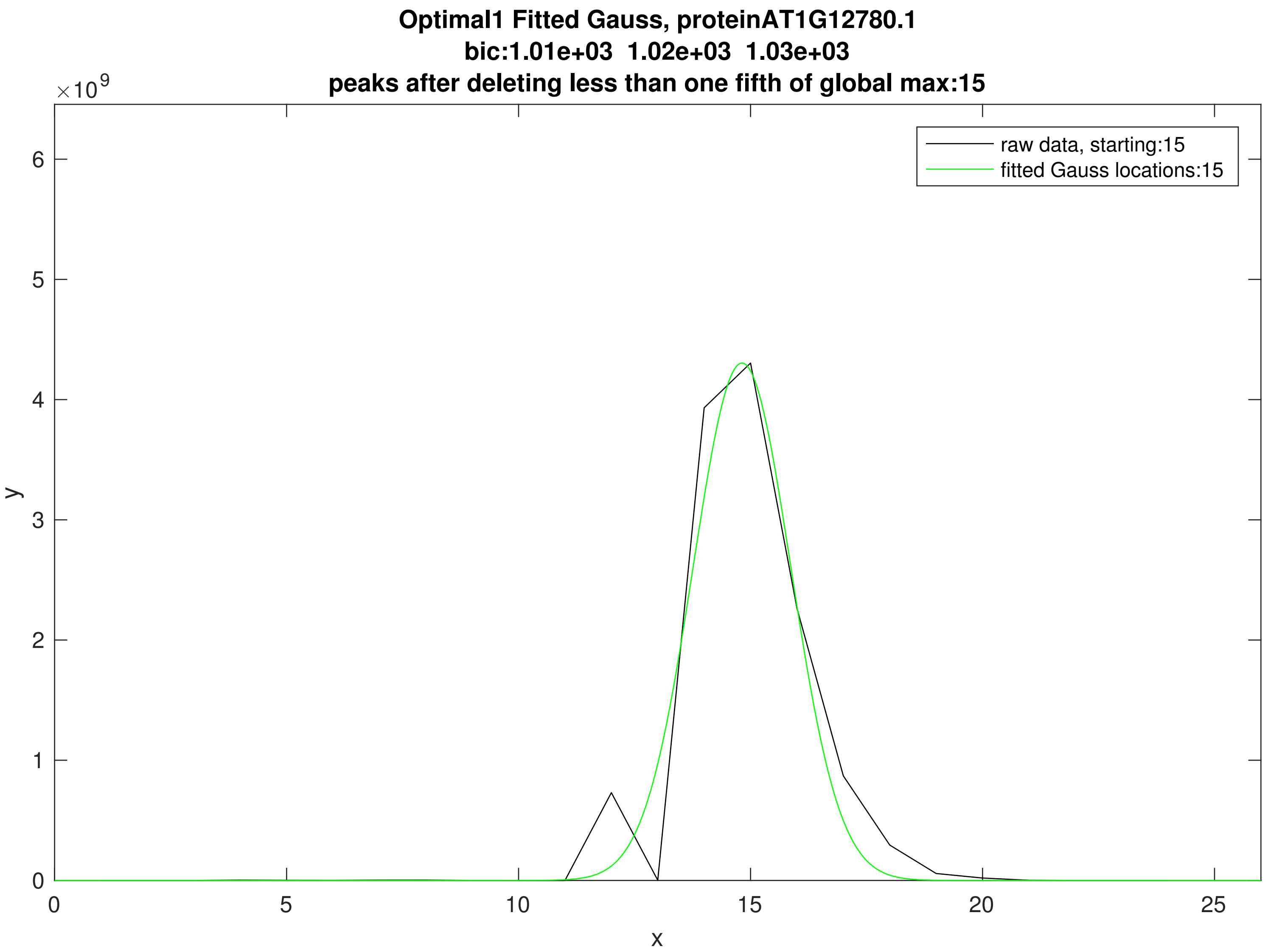
Optimal1 Fitted Gauss, proteinAT1G12780.1
bic:1.01e+03 1.02e+03 1.03e+03
peaks after deleting less than one fifth of global max:15
×10^9
raw data, starting:15
fitted Gauss locations:15
y
x
0
1
2
3
4
5
6
0
5
10
15
20
25

**Optimal1 Fitted Gauss, proteinAT1G12840.1**
**bic:801 805**
**peaks after deleting less than one fifth of global max:15**

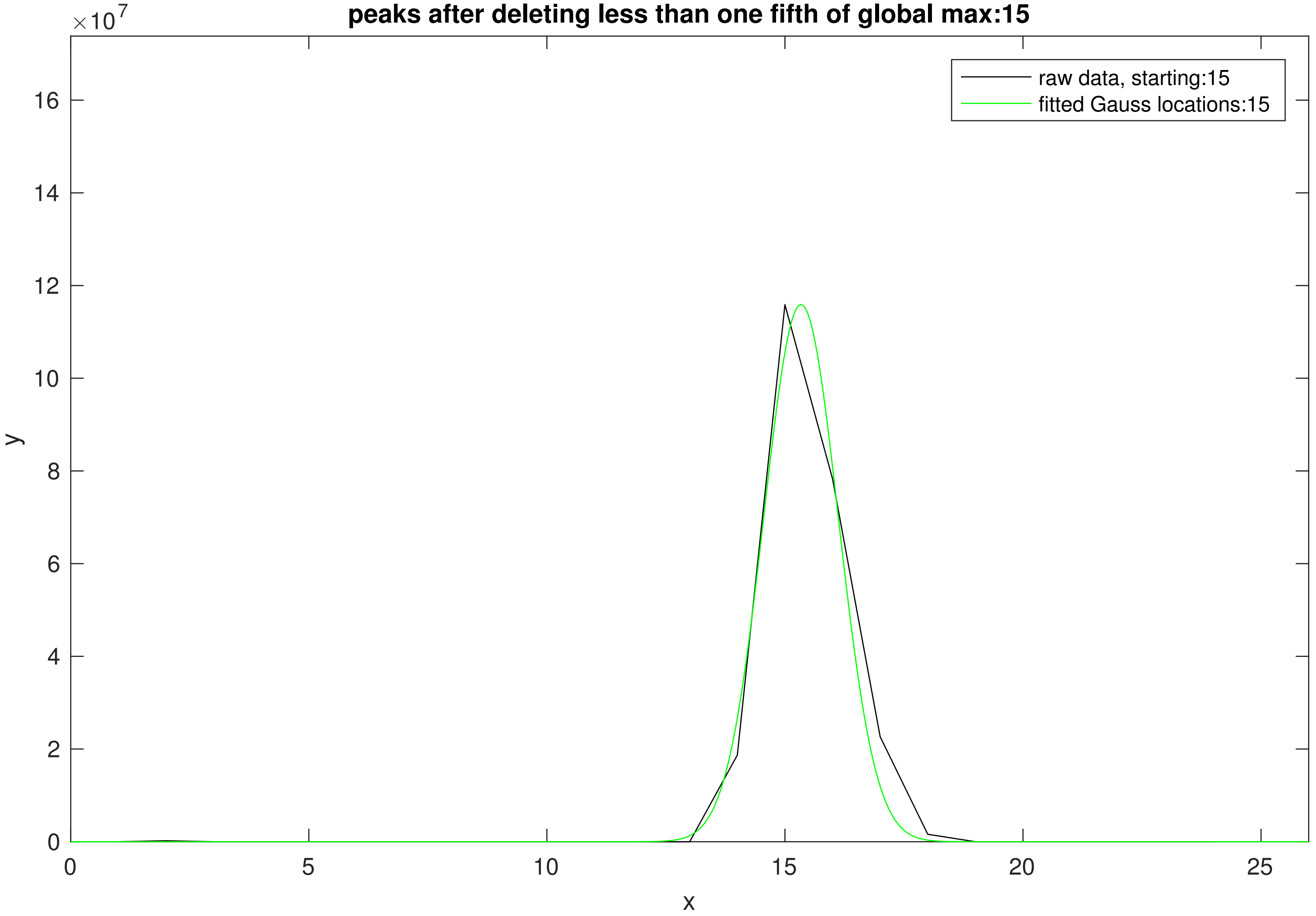

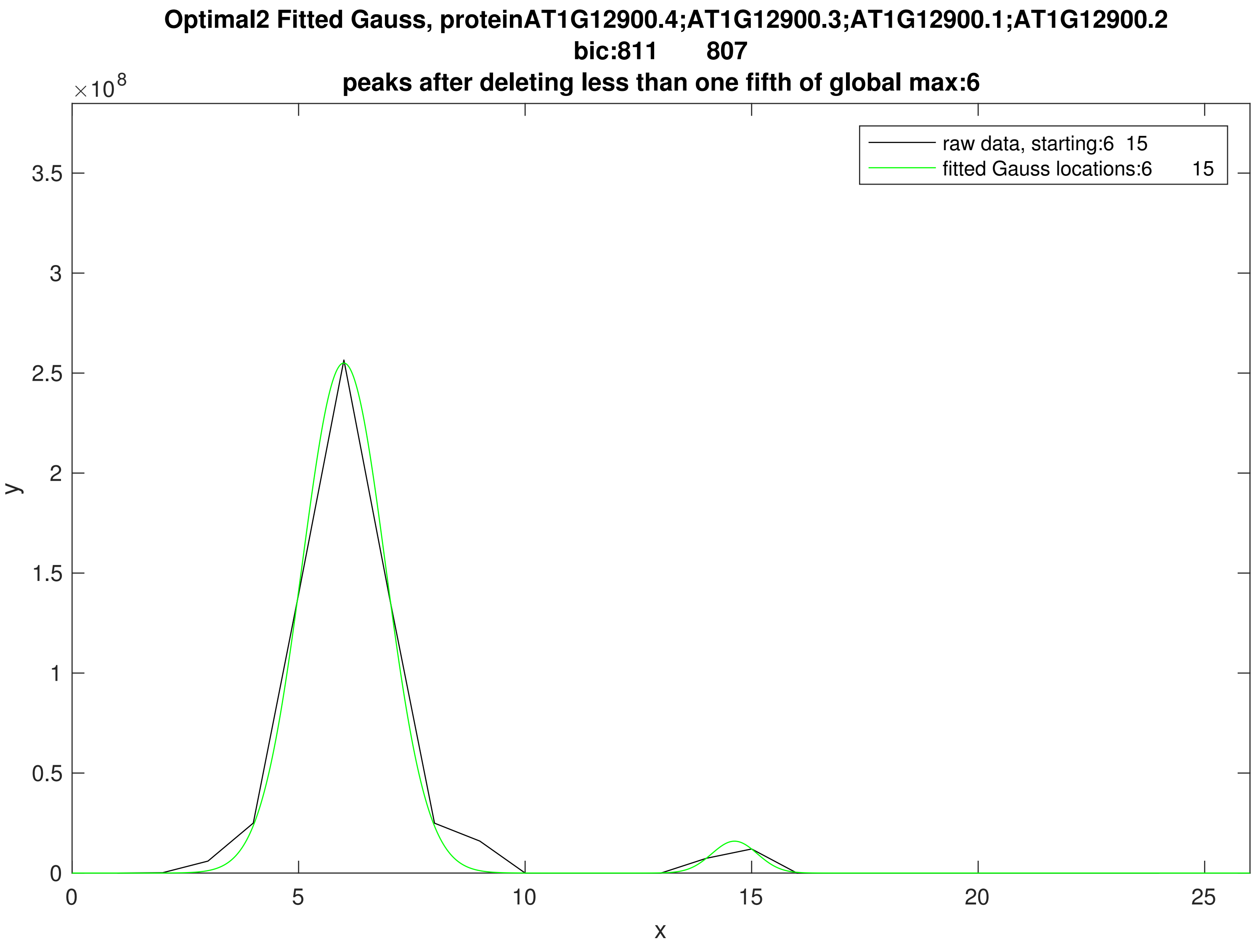
Optimal2 Fitted Gauss, proteinAT1G12900.4;AT1G12900.3;AT1G12900.1;AT1G12900.2
bic:811 807
peaks after deleting less than one fifth of global max:6
raw data, starting:6 15
fitted Gauss locations:6 15
$\times 10^8$
3.5
3
2.5
2
1.5
1
0.5
0
y
0
5
10
15
20
25
x

**Optimal1 Fitted Gauss, proteinAT1G12910.1**
**bic:738**
**peaks after deleting less than one fifth of global max:16**

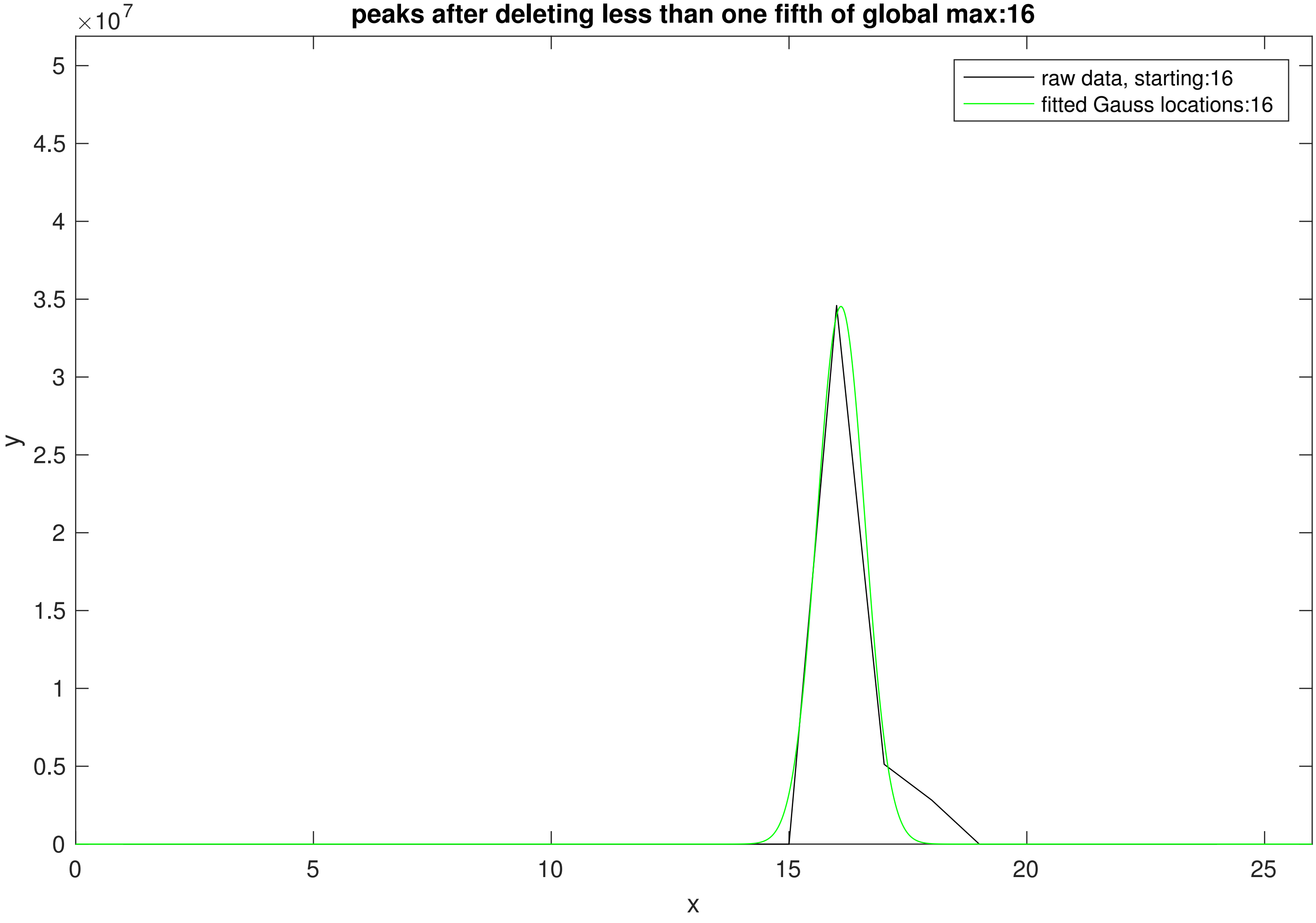

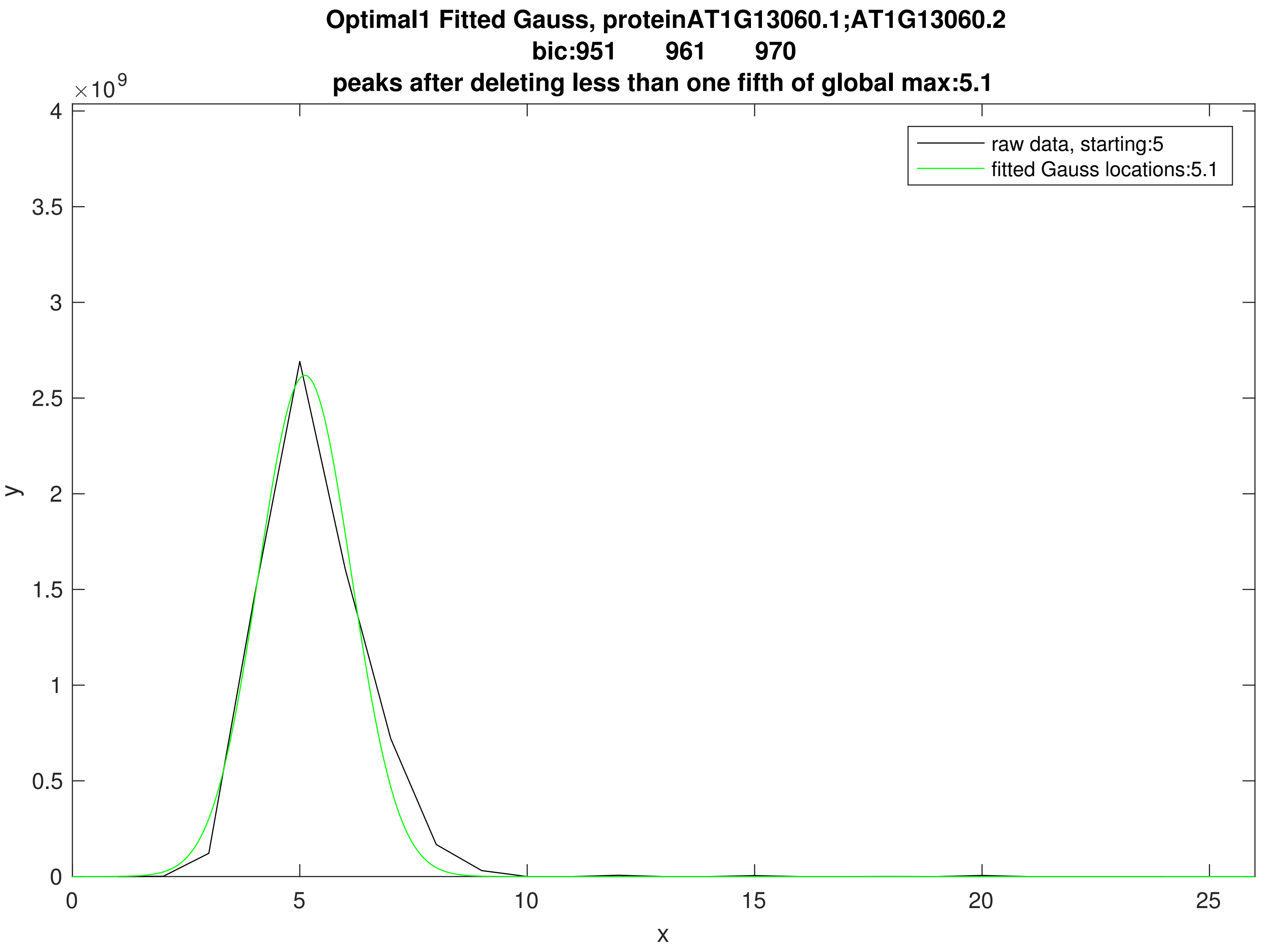
Optimal1 Fitted Gauss, proteinAT1G13060.1;AT1G13060.2
bic:951 961 970
peaks after deleting less than one fifth of global max:5.1
×10^9
4
3.5
3
2.5
2
1.5
1
0.5
0
y
0
5
10
15
20
25
x
raw data, starting:5
fitted Gauss locations:5.1

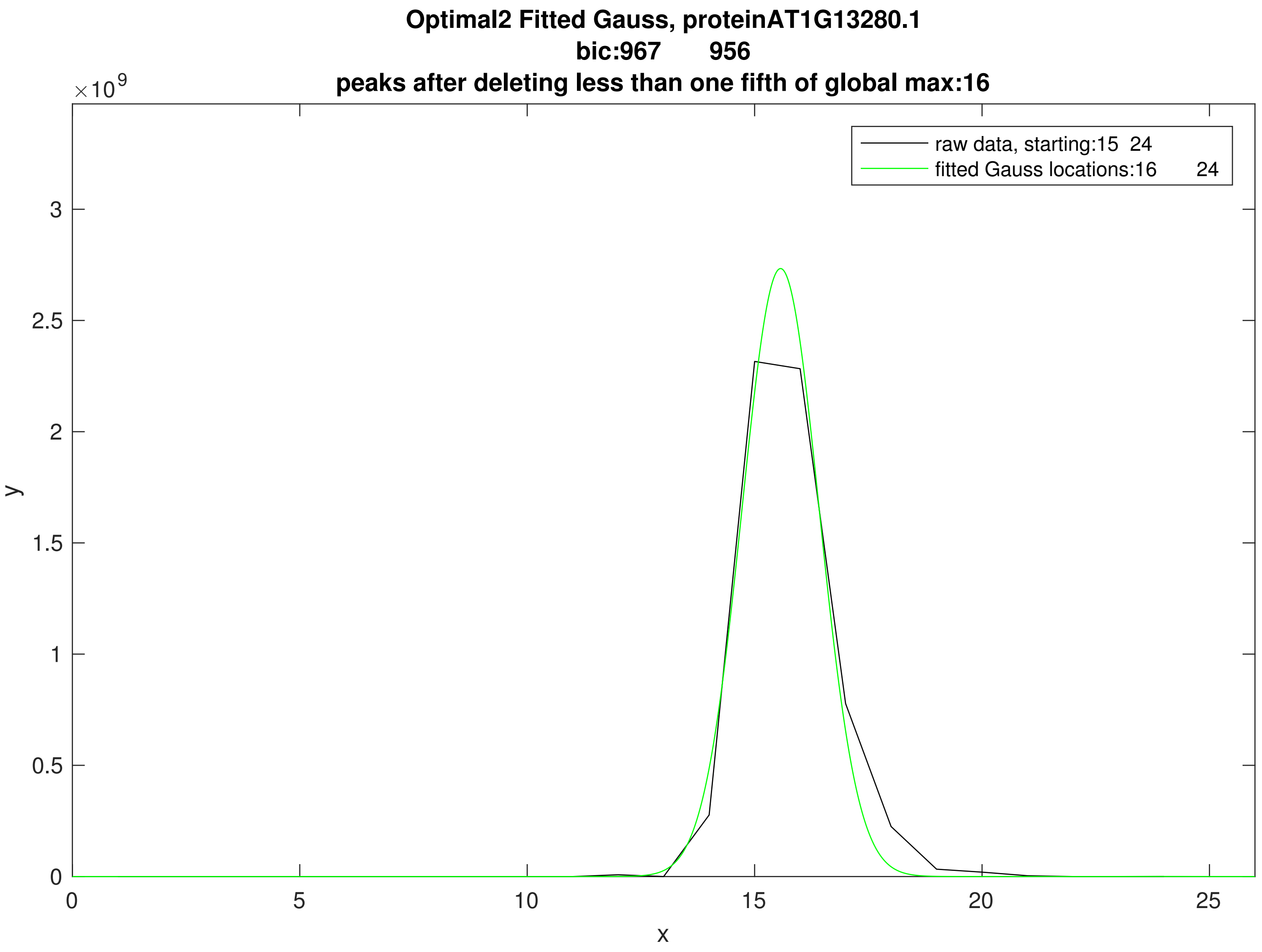

Optimal2 Fitted Gauss, proteinAT1G13280.1
bic:967 956
peaks after deleting less than one fifth of global max:16
×10^9
raw data, starting:15 24
fitted Gauss locations:16 24
3
2.5
2
1.5
1
0.5
0
y
0
5
10
15
20
25
x

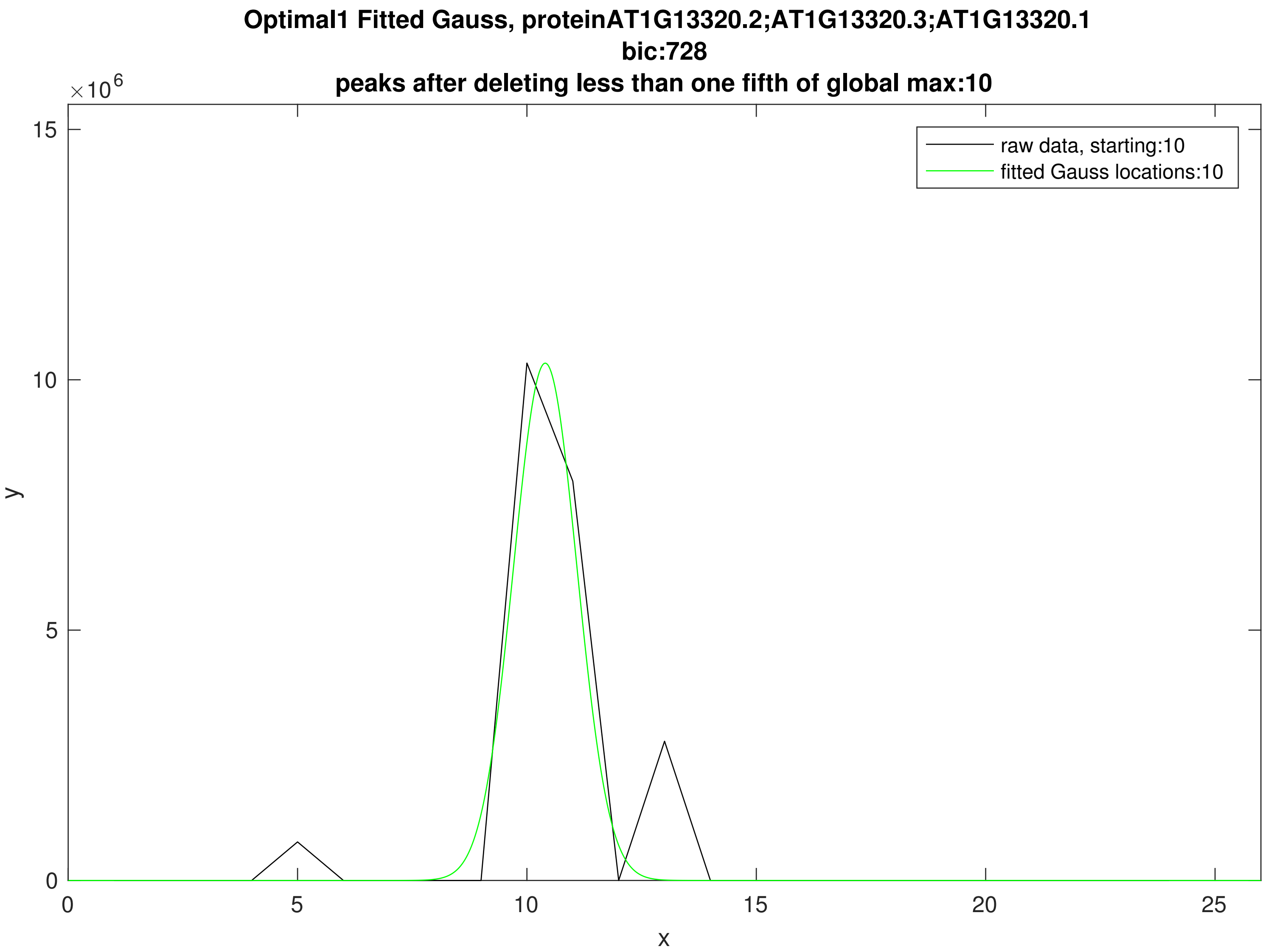

Optimal1 Fitted Gauss, proteinAT1G13320.2;AT1G13320.3;AT1G13320.1
bic:728
peaks after deleting less than one fifth of global max:10
×10^6
15
10
5
0
y
0
5
10
15
20
25
x
raw data, starting:10
fitted Gauss locations:10

**Optimal1 Fitted Gauss, proteinAT1G13440.1;AT1G13440.2**
**bic:1.02e+03  1.03e+03**
**peaks after deleting less than one fifth of global max:12**

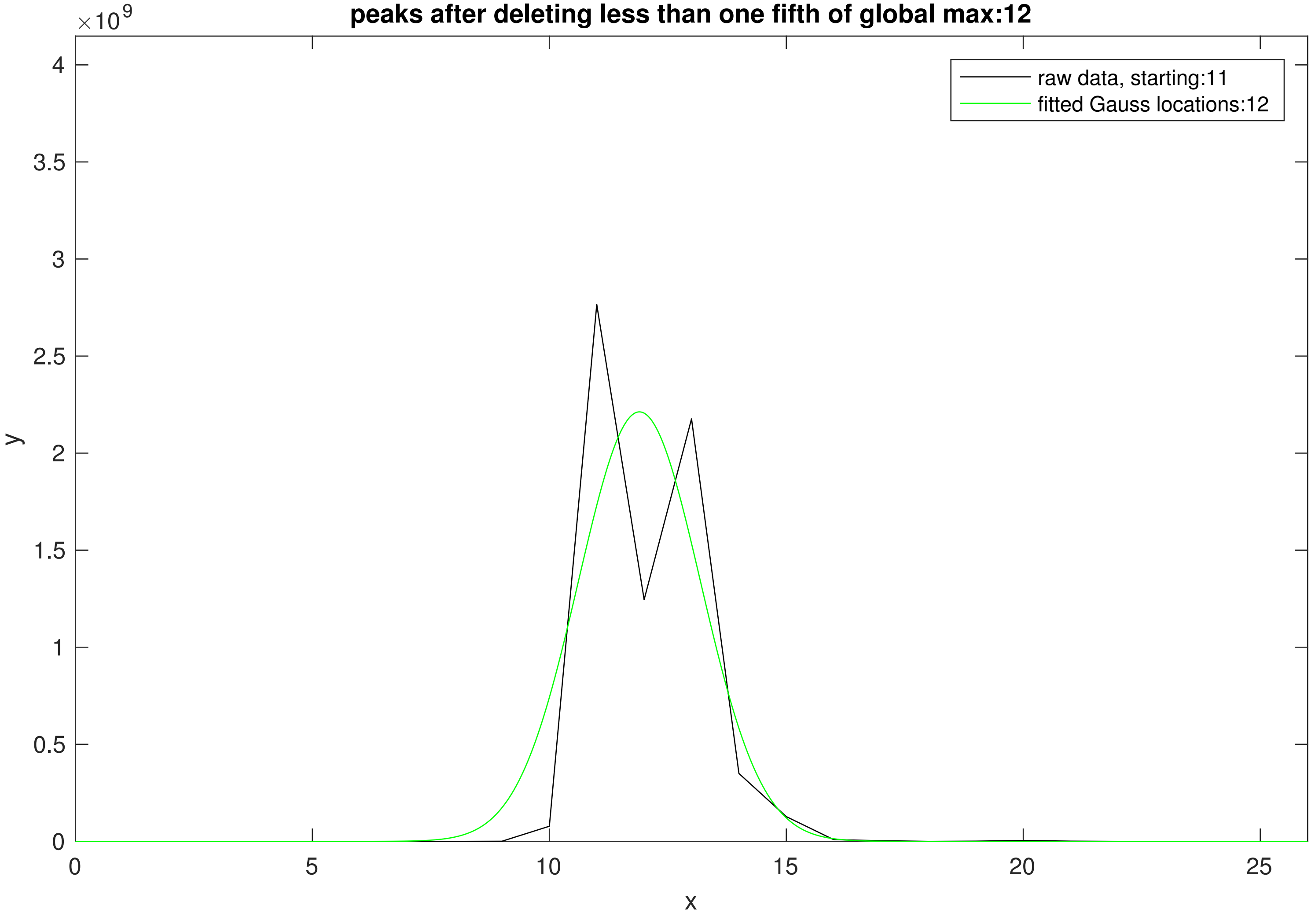

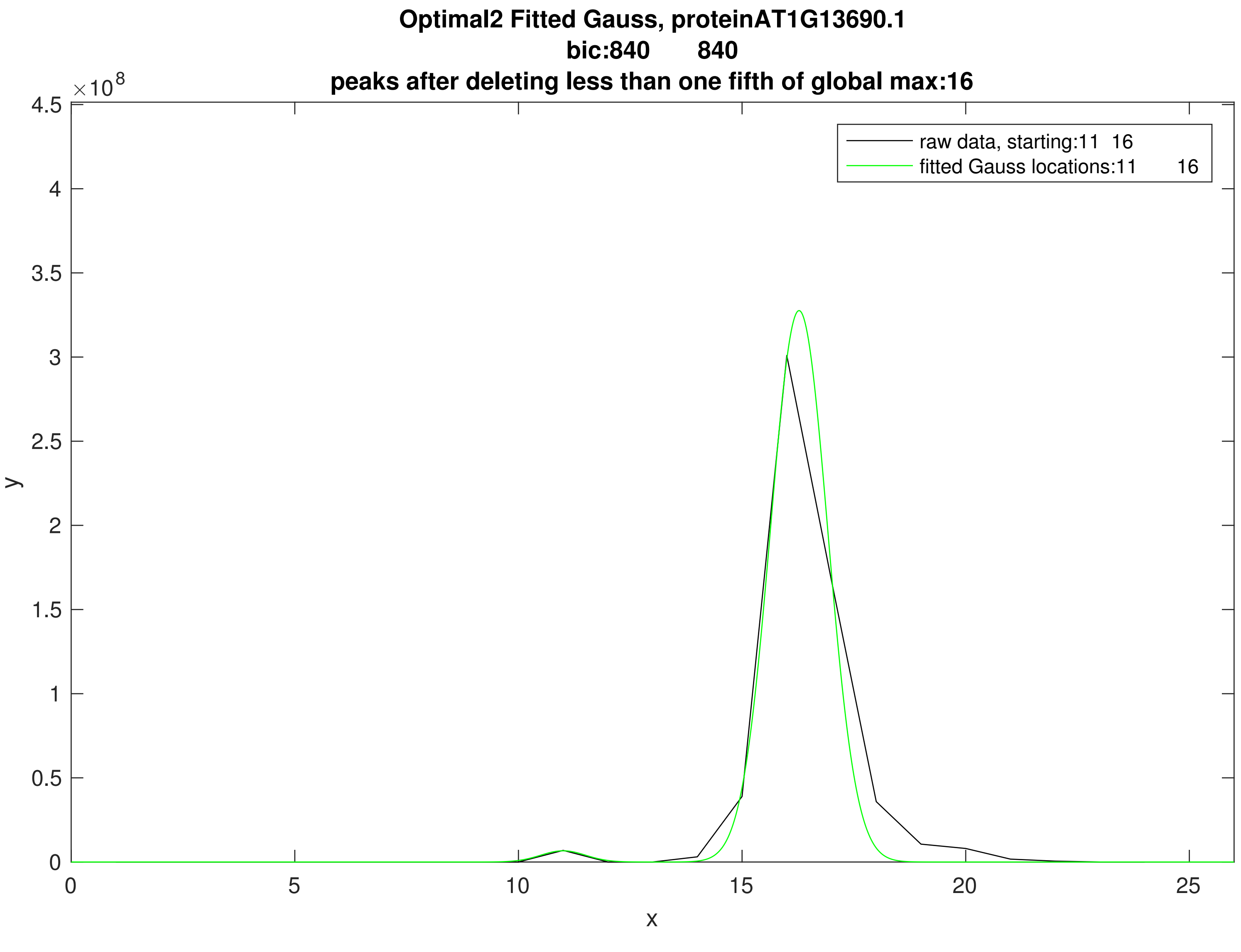

Optimal2 Fitted Gauss, proteinAT1G13690.1
bic:840 840
peaks after deleting less than one fifth of global max:16
×10^8
4.5
4
3.5
3
2.5
2
1.5
1
0.5
0
y
0
5
10
15
20
25
x
raw data, starting:11 16
fitted Gauss locations:11 16

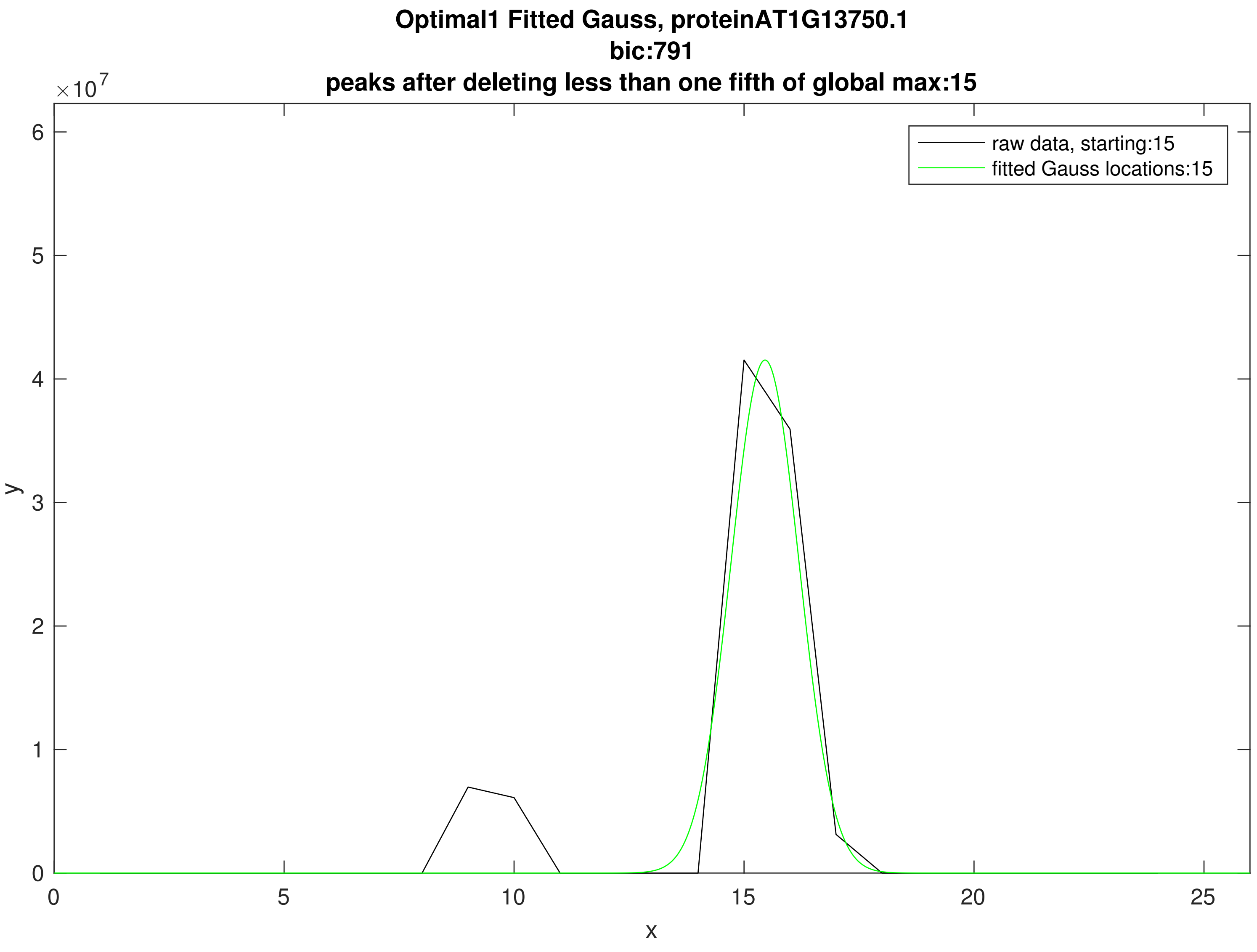

Optimal1 Fitted Gauss, proteinAT1G13750.1
bic:791
peaks after deleting less than one fifth of global max:15
×10^7
6
5
4
3
2
1
0
y
0
5
10
15
20
25
x
raw data, starting:15
fitted Gauss locations:15

**Optimal2 Fitted Gauss, proteinAT1G13900.1**
**bic:942 936 953 963**
**peaks after deleting less than one fifth of global max:11**

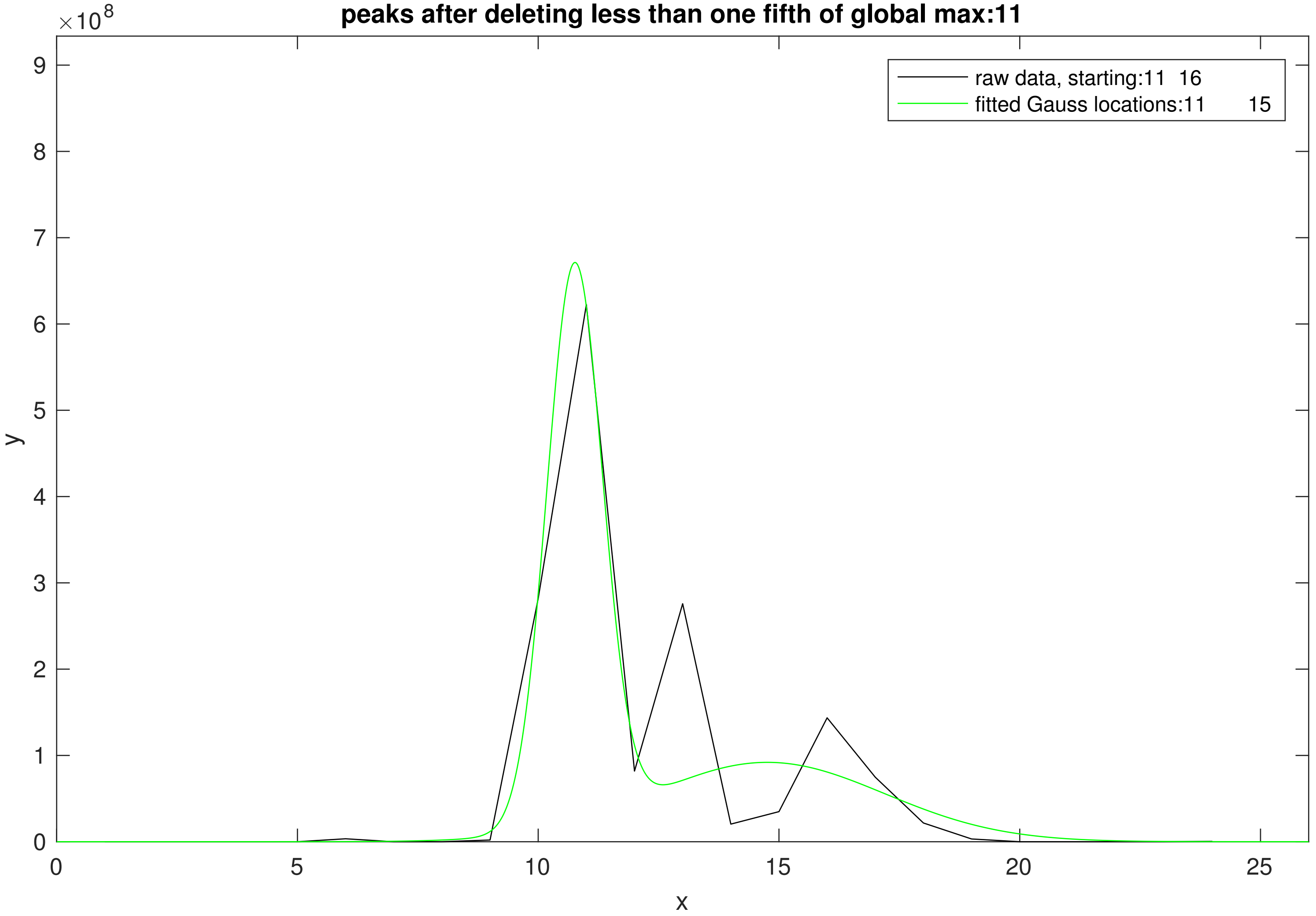

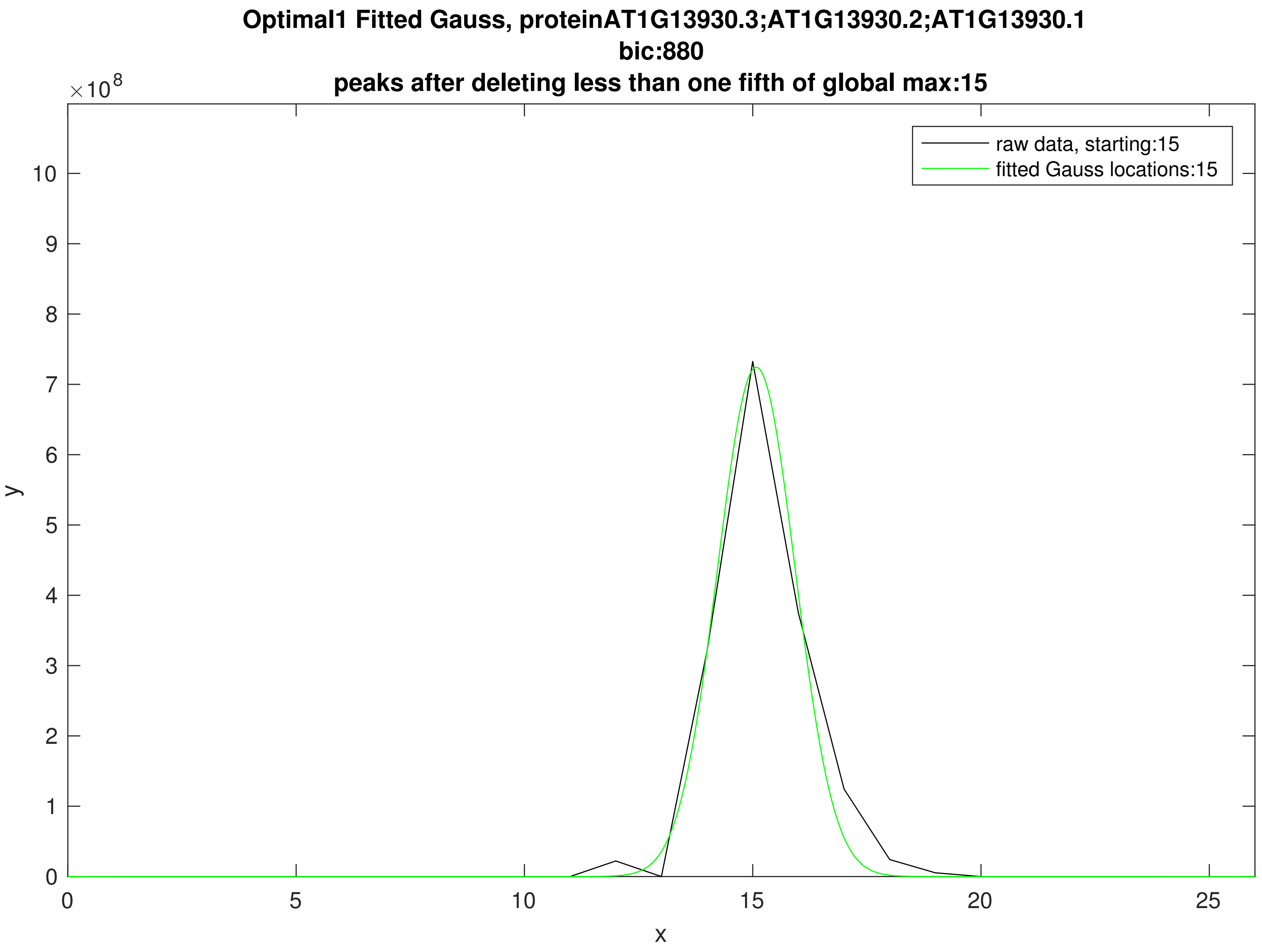

Optimal1 Fitted Gauss, proteinAT1G13930.3;AT1G13930.2;AT1G13930.1
bic:880
peaks after deleting less than one fifth of global max:15
$\times 10^8$
raw data, starting:15
fitted Gauss locations:15
y
x
0
1
2
3
4
5
6
7
8
9
10
0
5
10
15
20
25

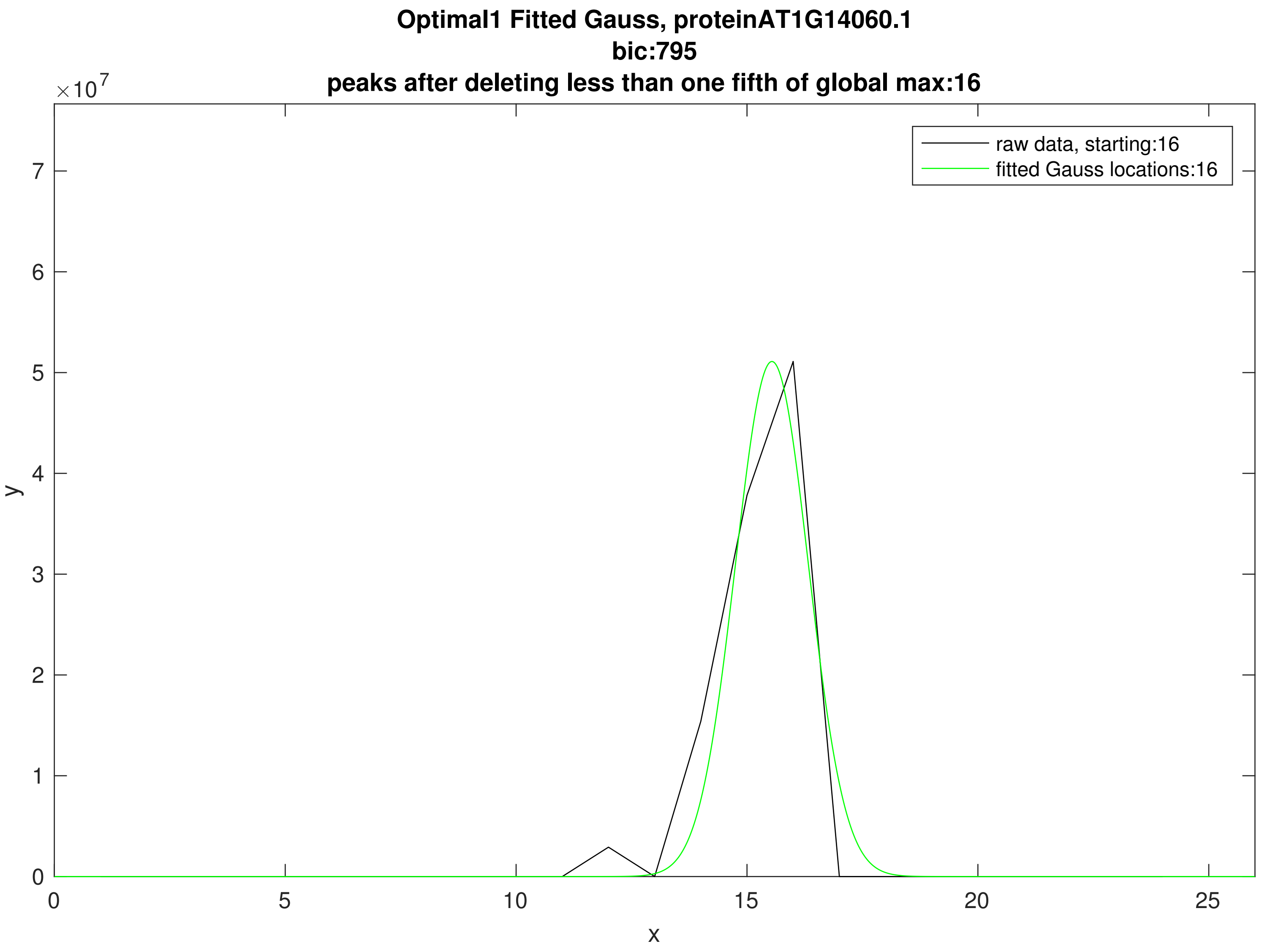

Optimal1 Fitted Gauss, proteinAT1G14060.1
bic:795
peaks after deleting less than one fifth of global max:16
×10^7
raw data, starting:16
fitted Gauss locations:16
y
x
0
1
2
3
4
5
6
7
0
5
10
15
20
25

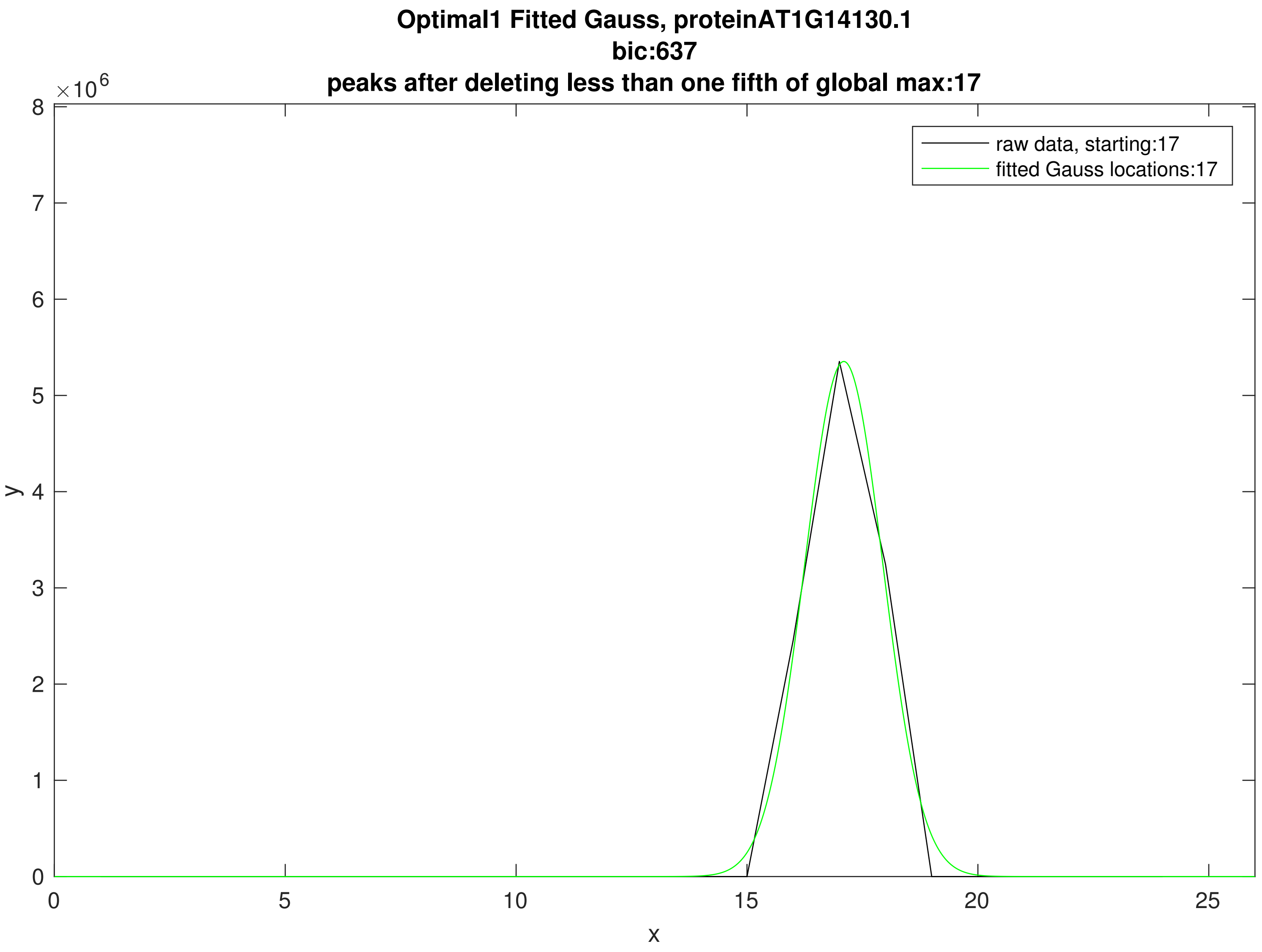

Optimal1 Fitted Gauss, proteinAT1G14130.1
bic:637
peaks after deleting less than one fifth of global max:17
$\times 10^6$
raw data, starting:17
fitted Gauss locations:17
y
x
0
1
2
3
4
5
6
7
8
0
5
10
15
20
25

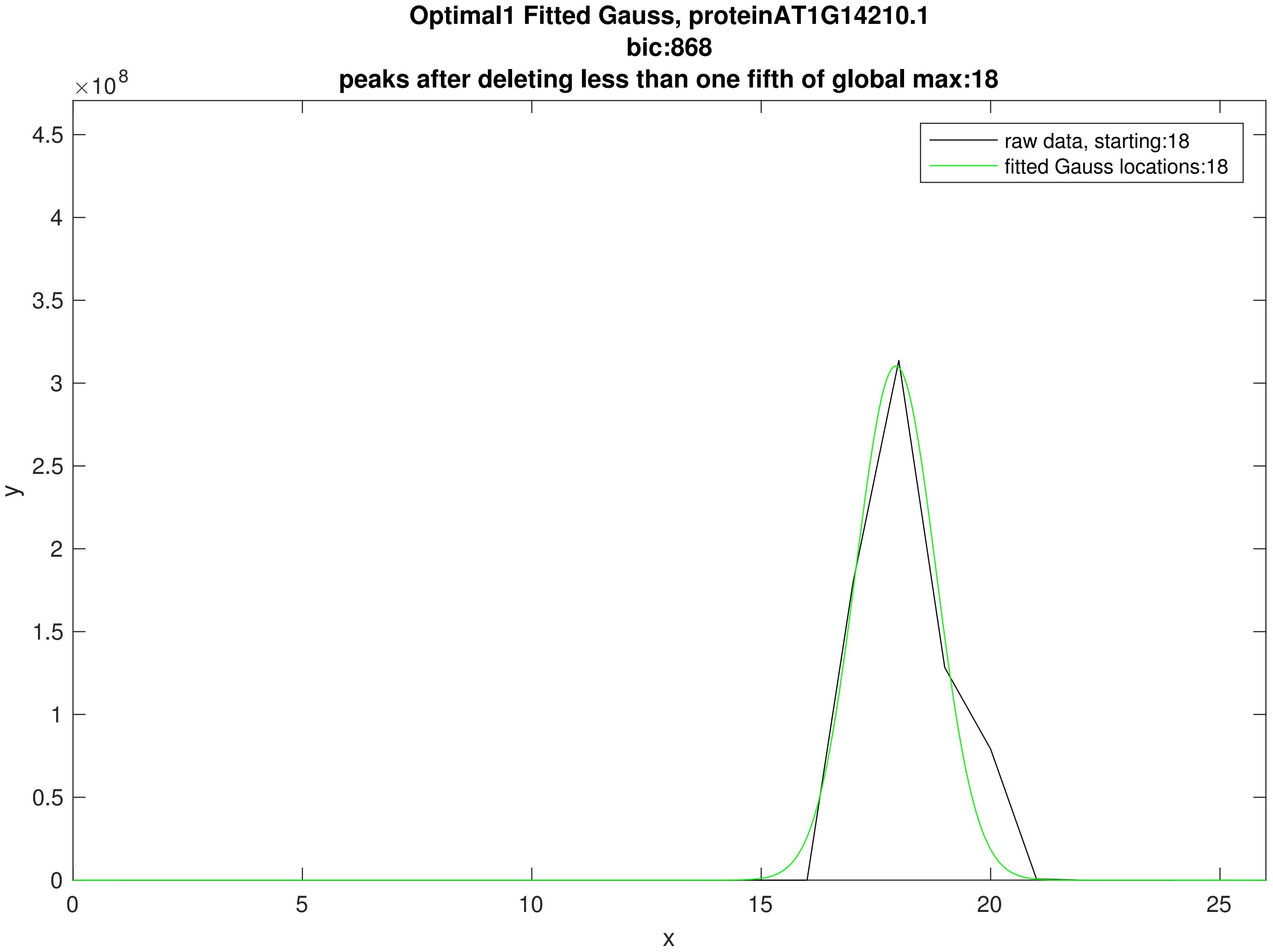
Optimal1 Fitted Gauss, proteinAT1G14210.1
bic:868
peaks after deleting less than one fifth of global max:18
×10^8
raw data, starting:18
fitted Gauss locations:18
y
x
0
0.5
1
1.5
2
2.5
3
3.5
4
4.5
5
10
15
20
25

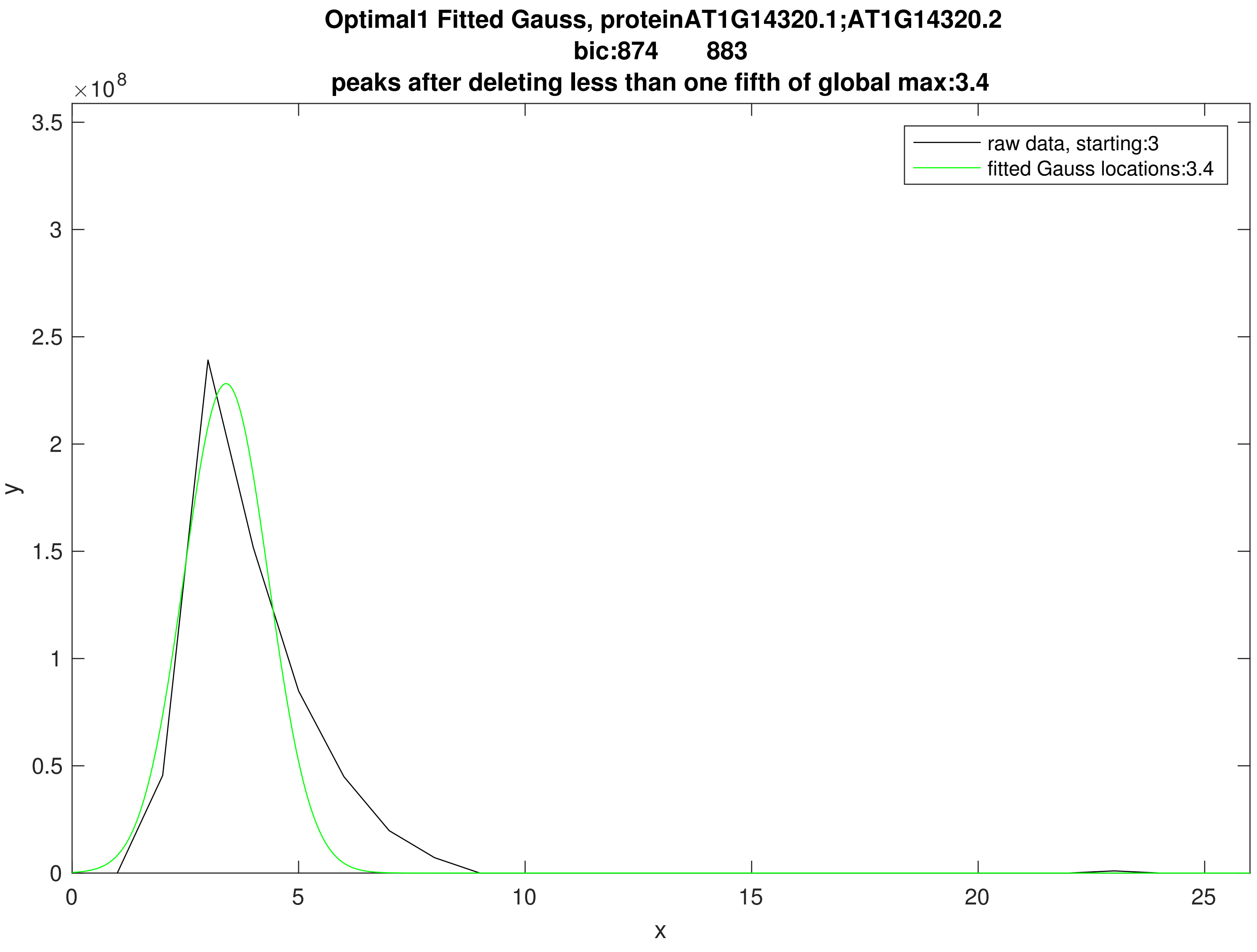

Optimal1 Fitted Gauss, proteinAT1G14320.1;AT1G14320.2
bic:874 883
peaks after deleting less than one fifth of global max:3.4
×10^8
raw data, starting:3
fitted Gauss locations:3.4
y
x
0
0.5
1
1.5
2
2.5
3
3.5
5
10
15
20
25

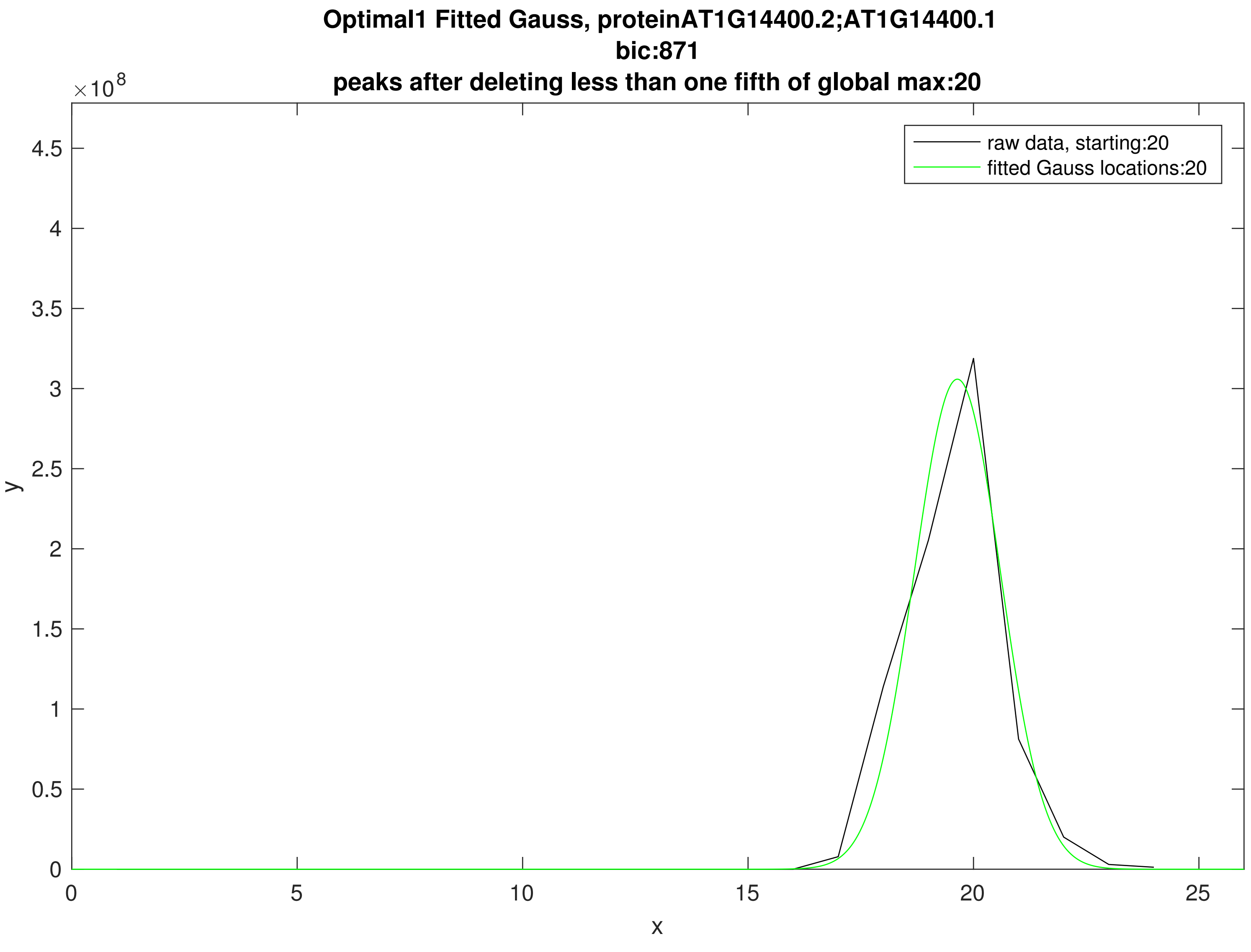

Optimal1 Fitted Gauss, proteinAT1G14400.2;AT1G14400.1
bic:871
peaks after deleting less than one fifth of global max:20
×10^8
raw data, starting:20
fitted Gauss locations:20
y
x
0
0.5
1
1.5
2
2.5
3
3.5
4
4.5
0
5
10
15
20
25

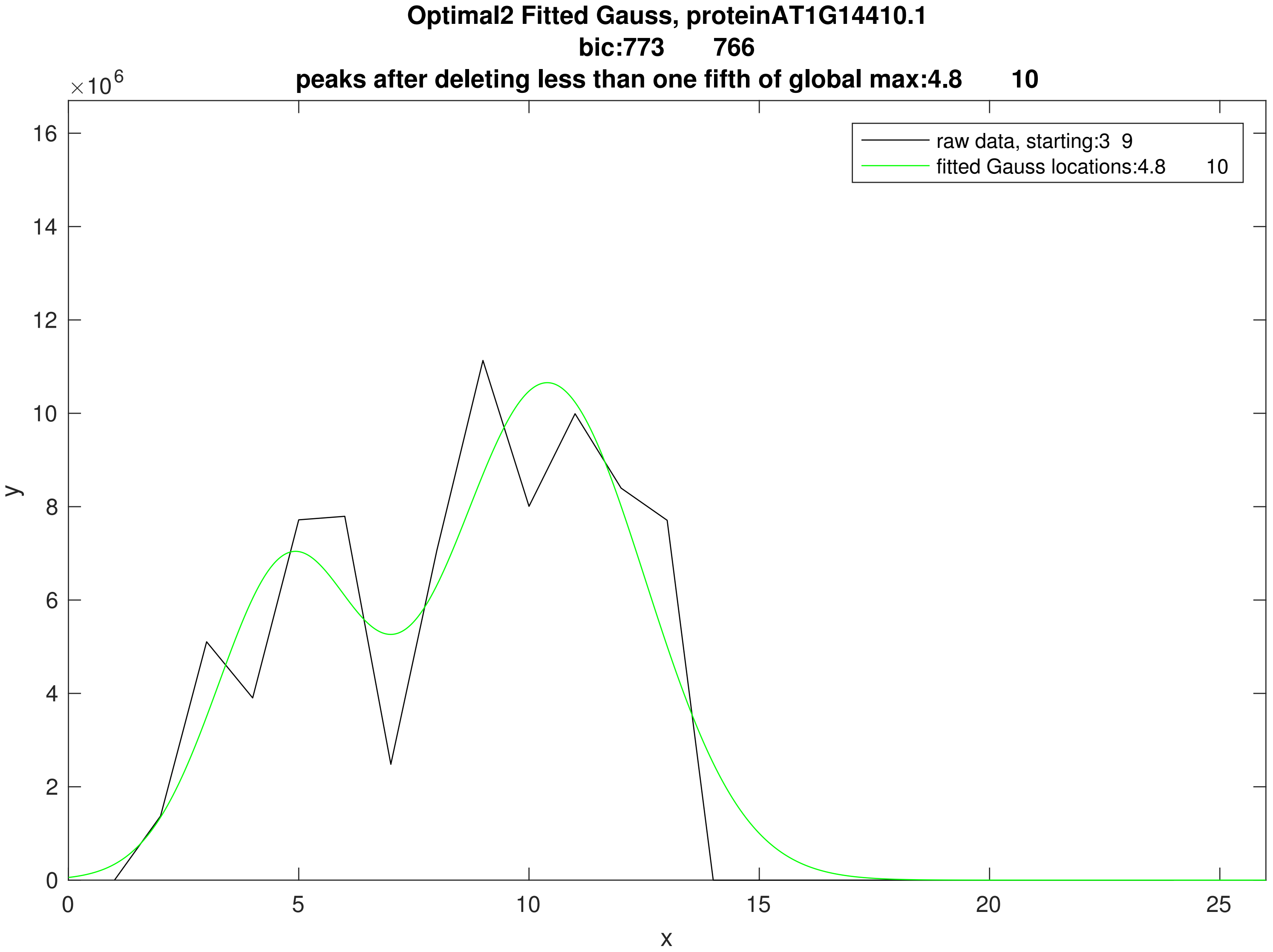
Optimal2 Fitted Gauss, proteinAT1G14410.1
bic:773 766
peaks after deleting less than one fifth of global max:4.8 10
×10^6
raw data, starting:3 9
fitted Gauss locations:4.8 10
y
x
0
2
4
6
8
10
12
14
16
0
5
10
15
20
25

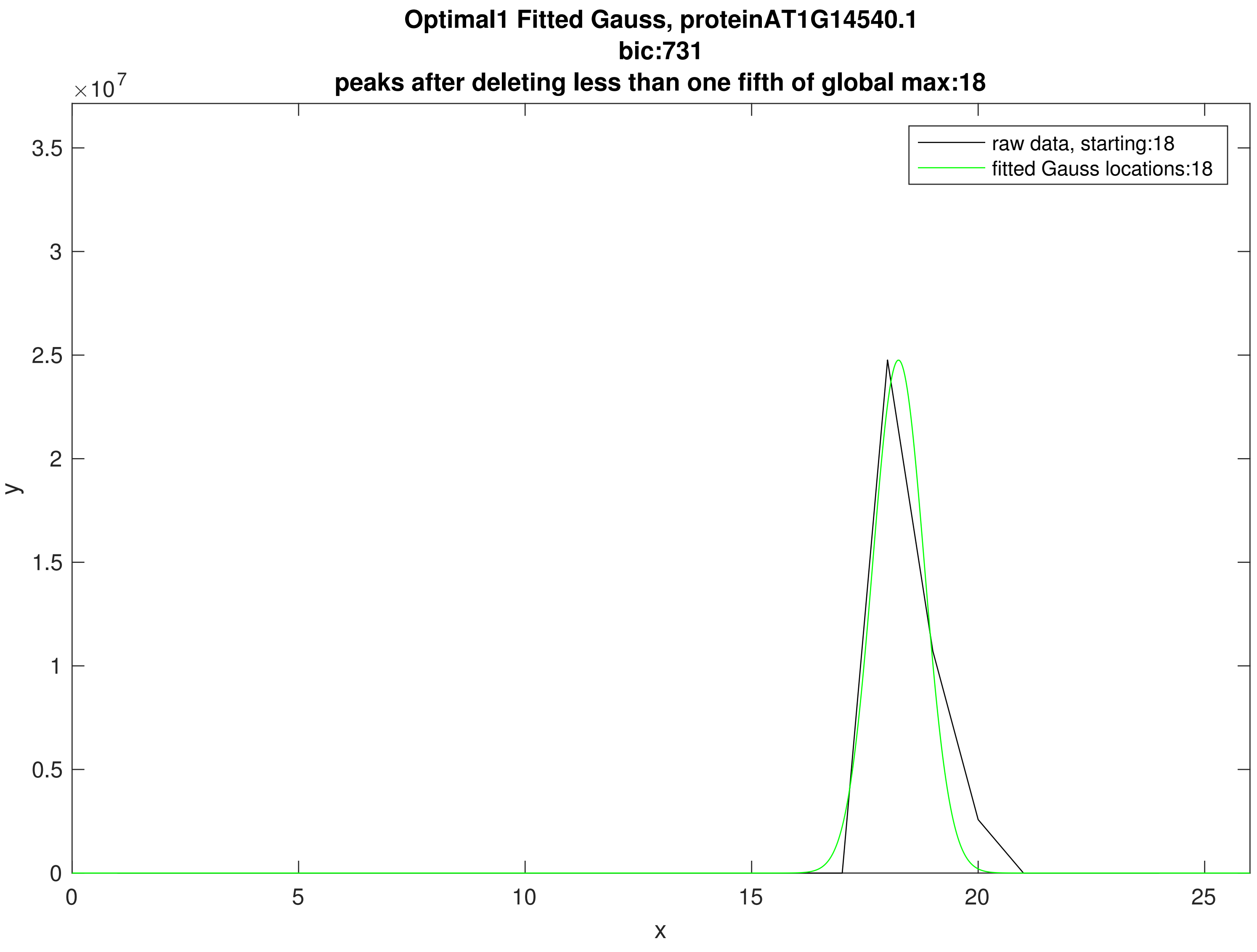
Optimal1 Fitted Gauss, proteinAT1G14540.1
bic:731
peaks after deleting less than one fifth of global max:18
×10^7
raw data, starting:18
fitted Gauss locations:18
y
x
0
0.5
1
1.5
2
2.5
3
3.5
0
5
10
15
20
25

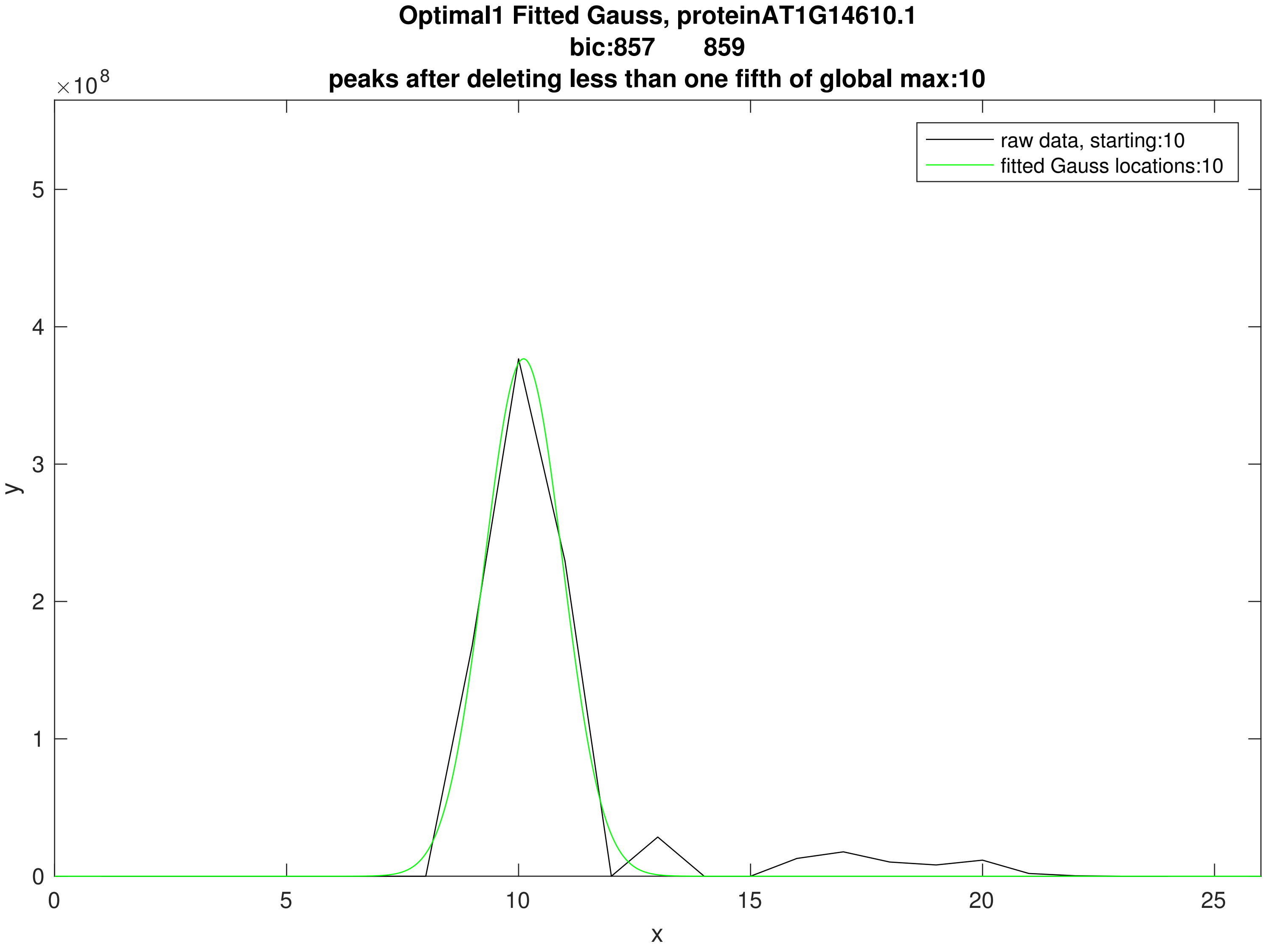

Optimal1 Fitted Gauss, proteinAT1G14610.1
bic:857 859
peaks after deleting less than one fifth of global max:10
×10^8
raw data, starting:10
fitted Gauss locations:10
5
4
3
2
1
0
y
0
5
10
15
20
25
x

Optimal1 Fitted Gauss, proteinAT1G14670.1;AT2G01970.1
bic:796 805
peaks after deleting less than one fifth of global max:18

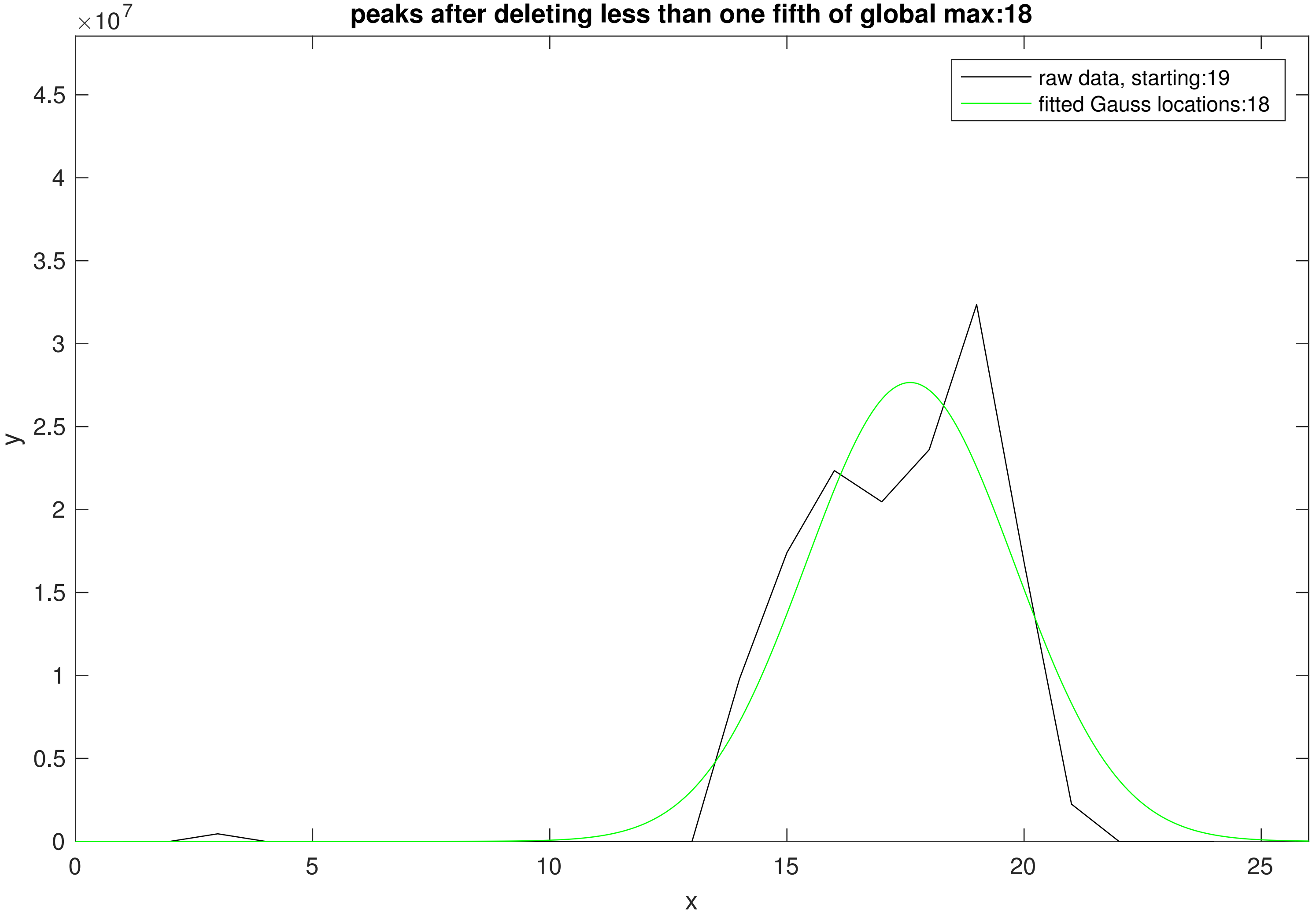

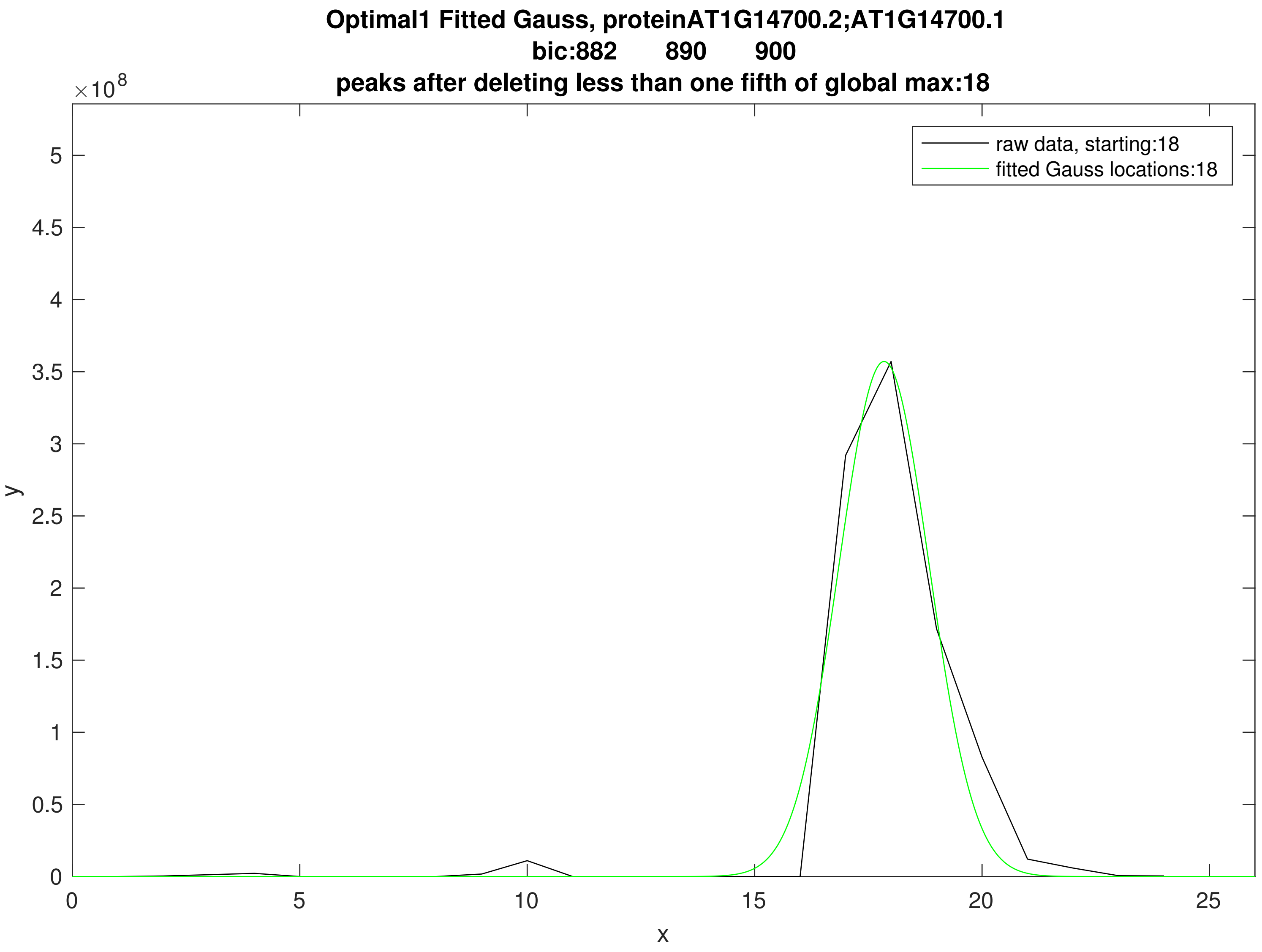
Optimal1 Fitted Gauss, proteinAT1G14700.2;AT1G14700.1
bic:882 890 900
peaks after deleting less than one fifth of global max:18
×10^8
5
4.5
4
3.5
3
2.5
2
1.5
1
0.5
0
y
0
5
10
15
20
25
x
raw data, starting:18
fitted Gauss locations:18

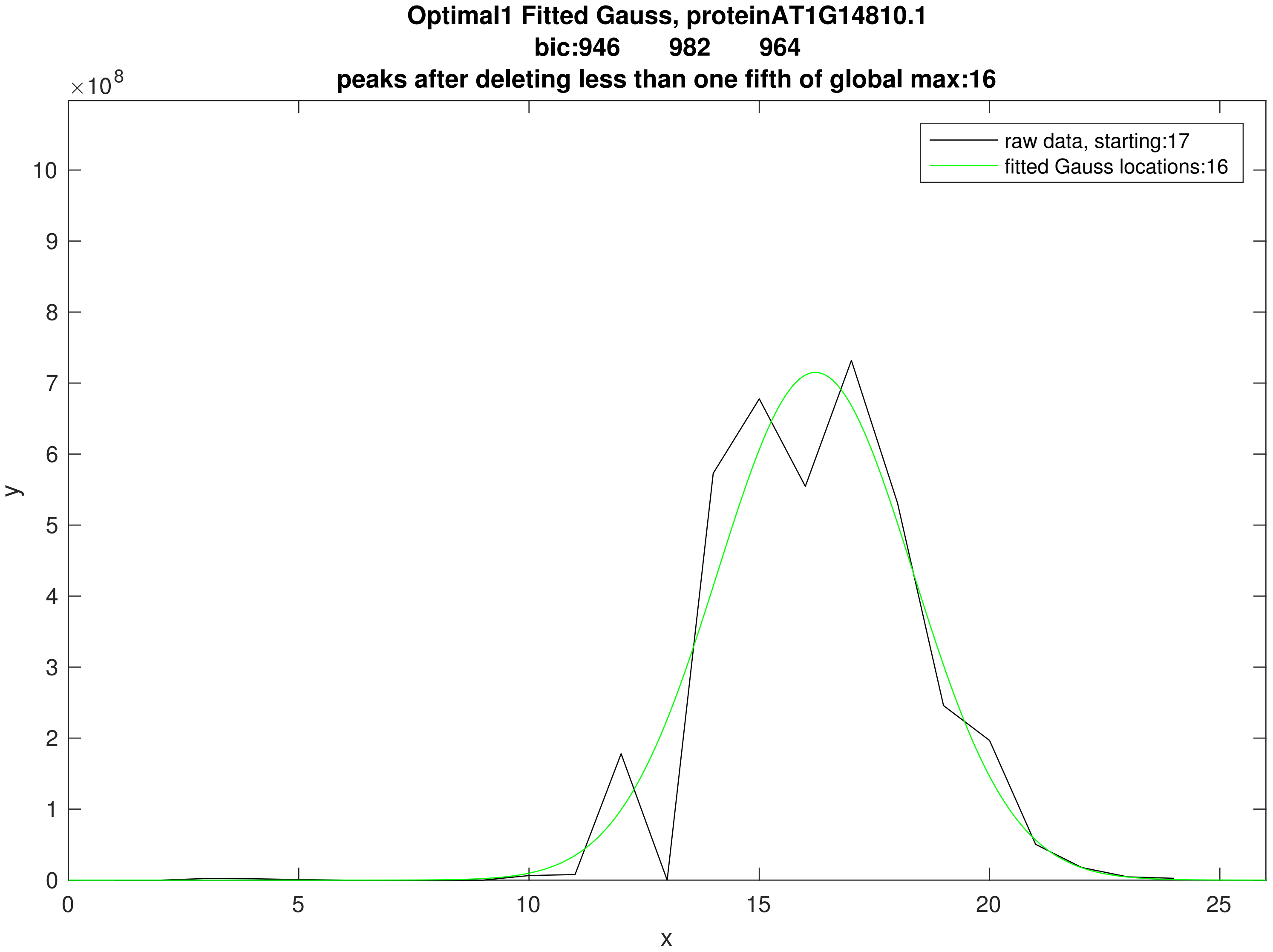

Optimal1 Fitted Gauss, proteinAT1G14810.1
bic:946 982 964
peaks after deleting less than one fifth of global max:16
×10^8
raw data, starting:17
fitted Gauss locations:16
y
x
0
1
2
3
4
5
6
7
8
9
10
0
5
10
15
20
25

**Optimal1 Fitted Gauss, proteinAT1G14930.1**
**bic:900 907**
**peaks after deleting less than one fifth of global max:20**

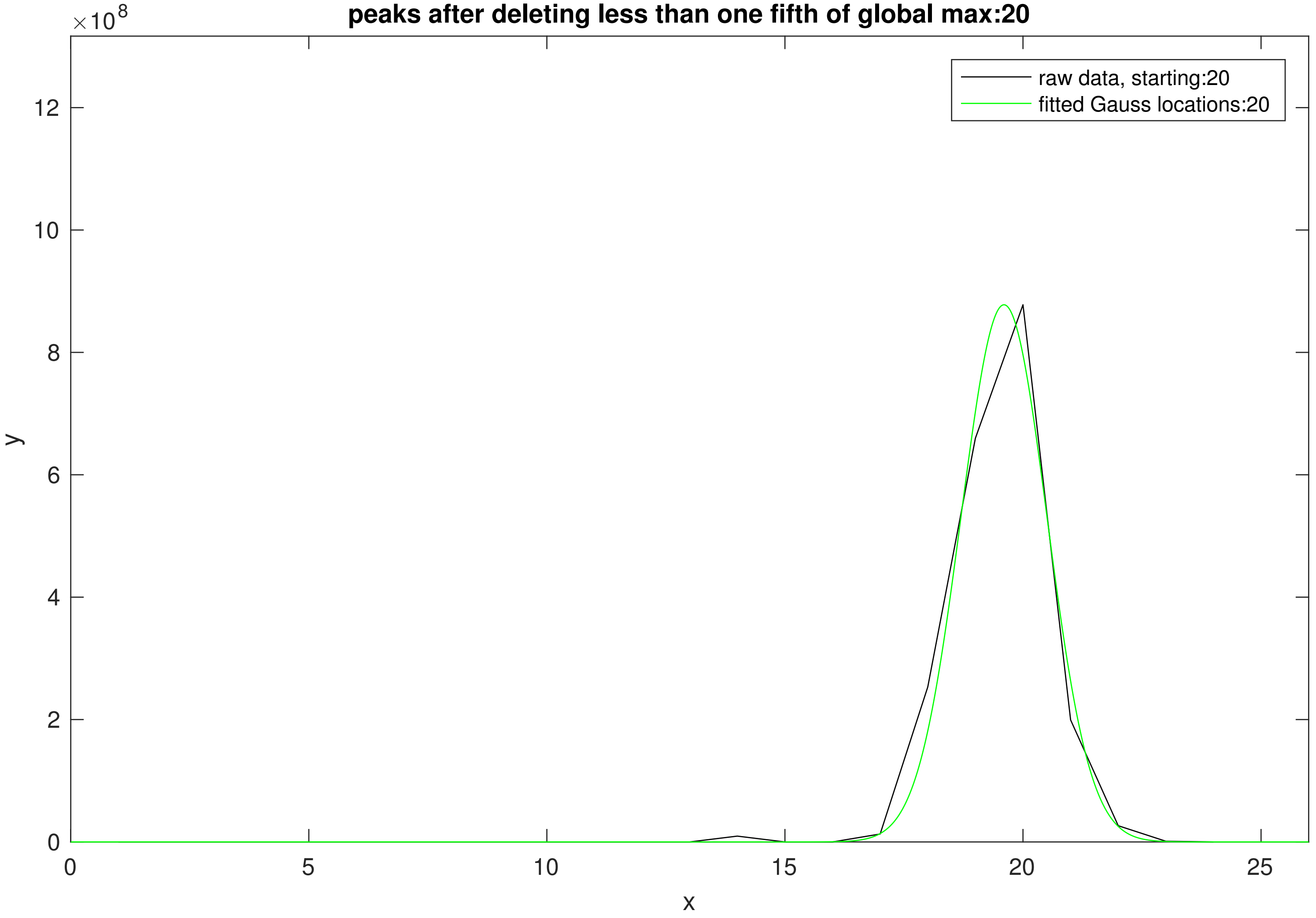

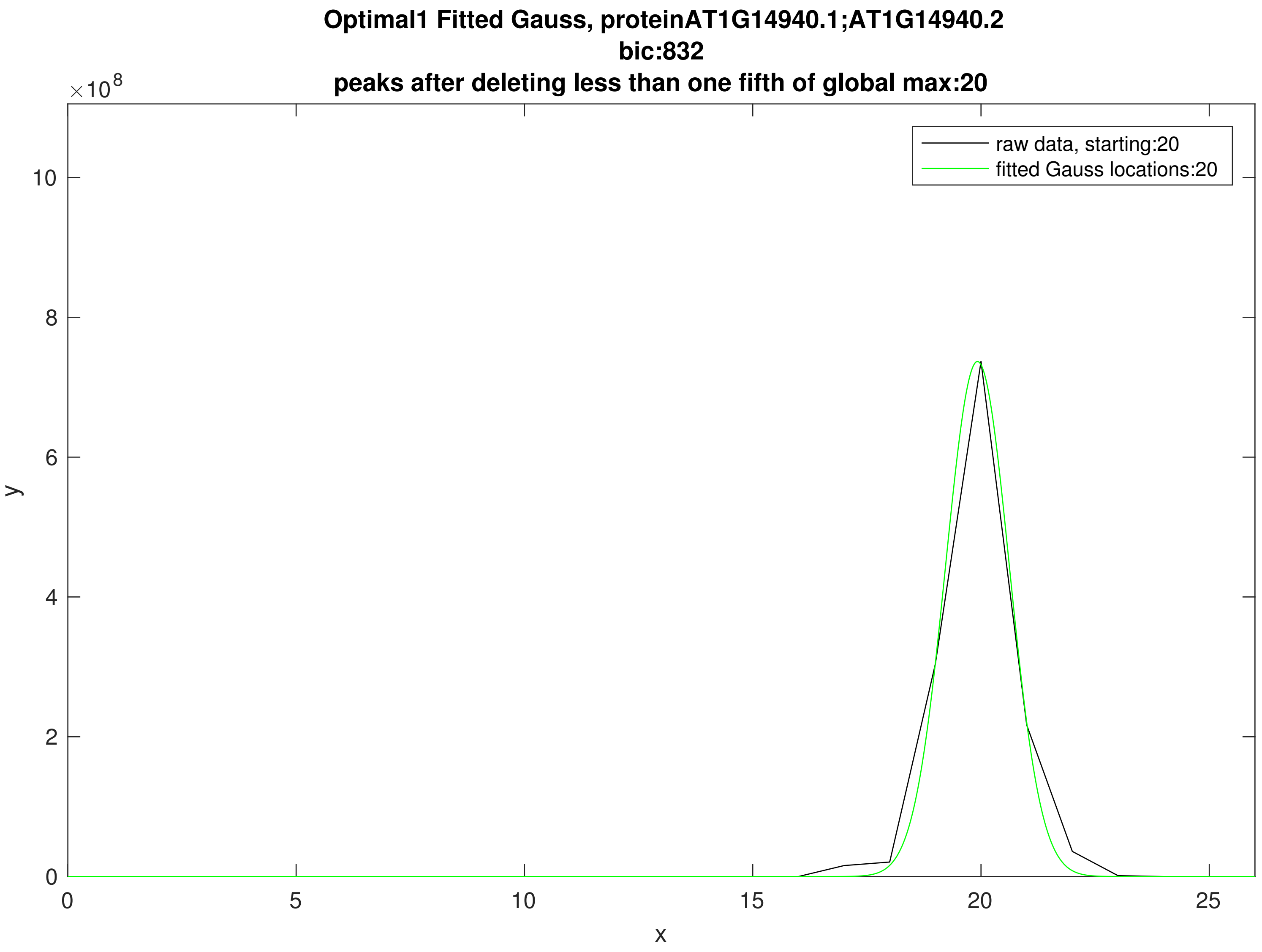
Optimal1 Fitted Gauss, proteinAT1G14940.1;AT1G14940.2
bic:832
peaks after deleting less than one fifth of global max:20
×10^8
raw data, starting:20
fitted Gauss locations:20
y
x
0
2
4
6
8
10
0
5
10
15
20
25

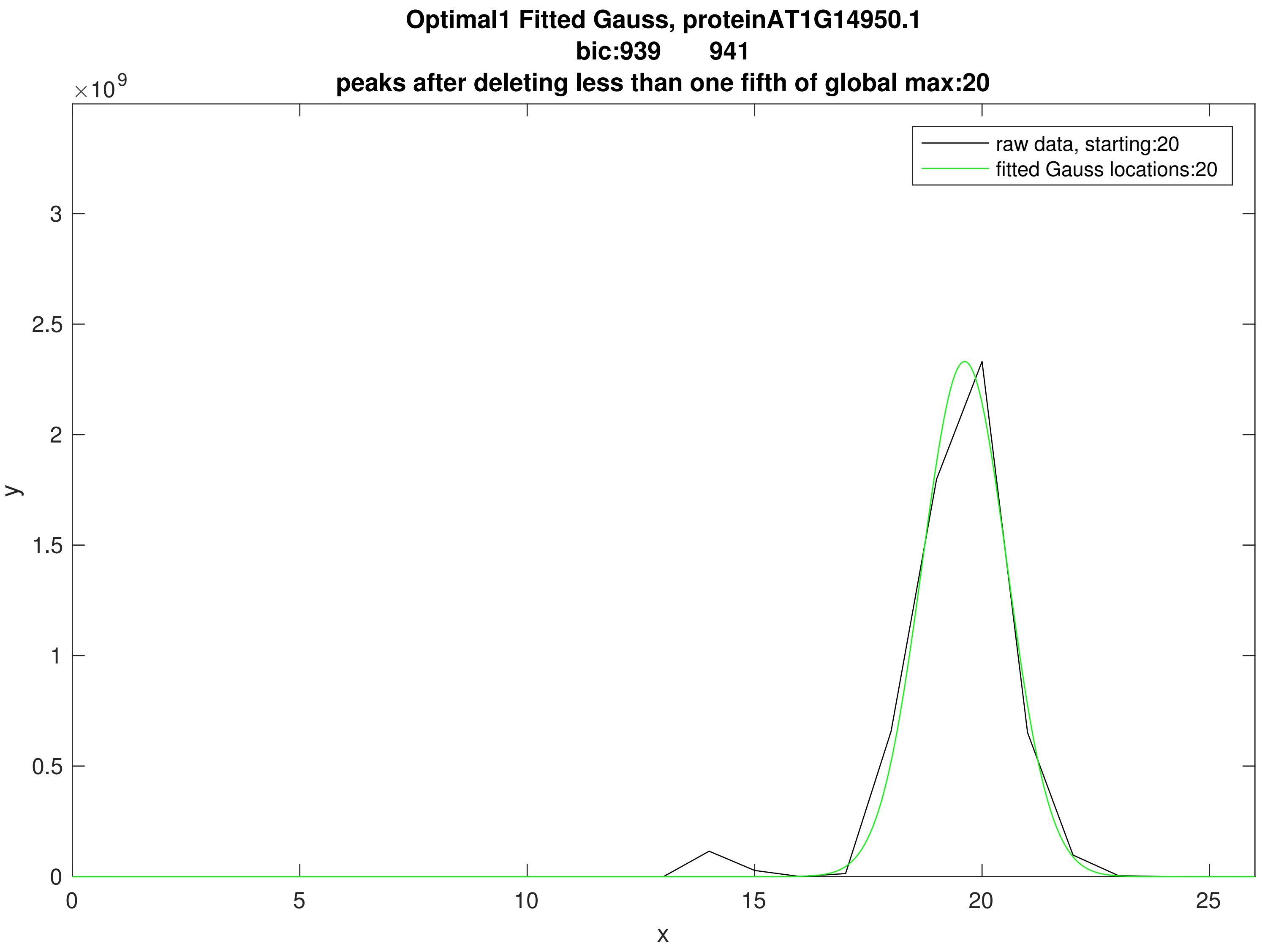

Optimal1 Fitted Gauss, proteinAT1G14950.1
bic:939 941
peaks after deleting less than one fifth of global max:20
×10^9
raw data, starting:20
fitted Gauss locations:20
y
x
0
0.5
1
1.5
2
2.5
3
0
5
10
15
20
25

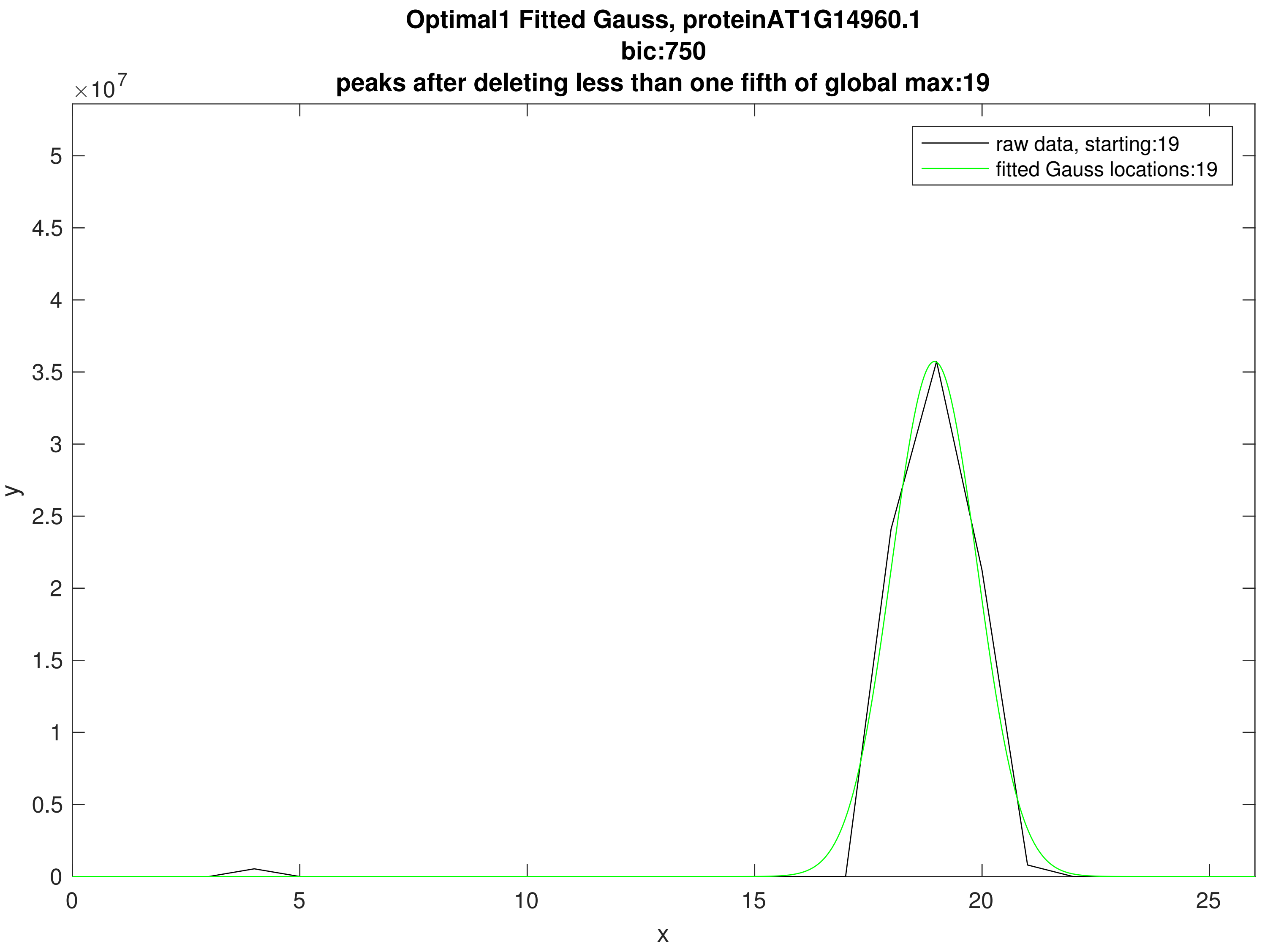
Optimal1 Fitted Gauss, proteinAT1G14960.1
bic:750
peaks after deleting less than one fifth of global max:19
×10^7
raw data, starting:19
fitted Gauss locations:19
y
x
0
0.5
1
1.5
2
2.5
3
3.5
4
4.5
5
0
5
10
15
20
25

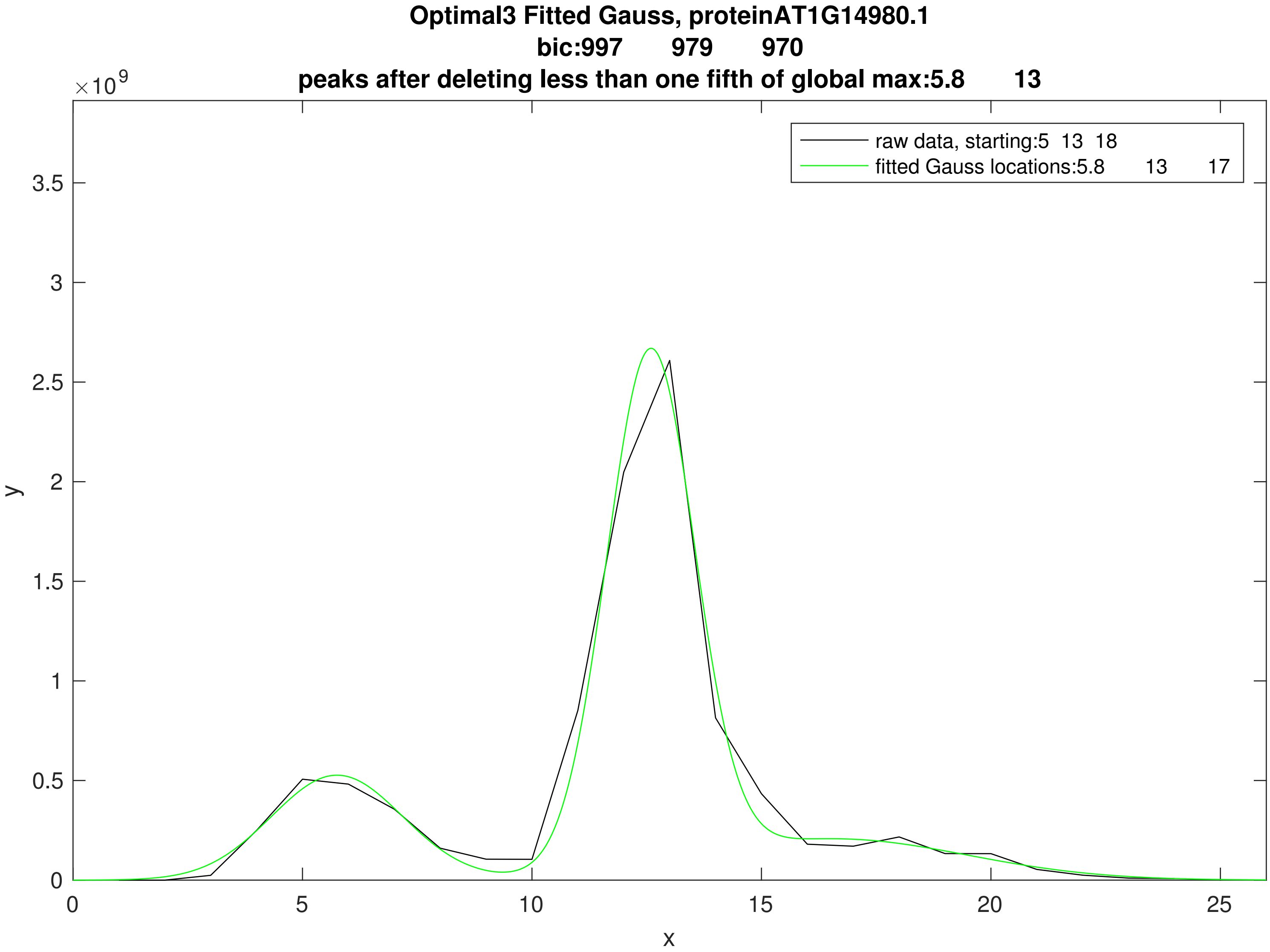

Optimal3 Fitted Gauss, proteinAT1G14980.1
bic:997 979 970
peaks after deleting less than one fifth of global max:5.8 13
×10^9
raw data, starting:5 13 18
fitted Gauss locations:5.8 13 17
3.5
3
2.5
2
1.5
1
0.5
0
y
0
5
10
15
20
25
x

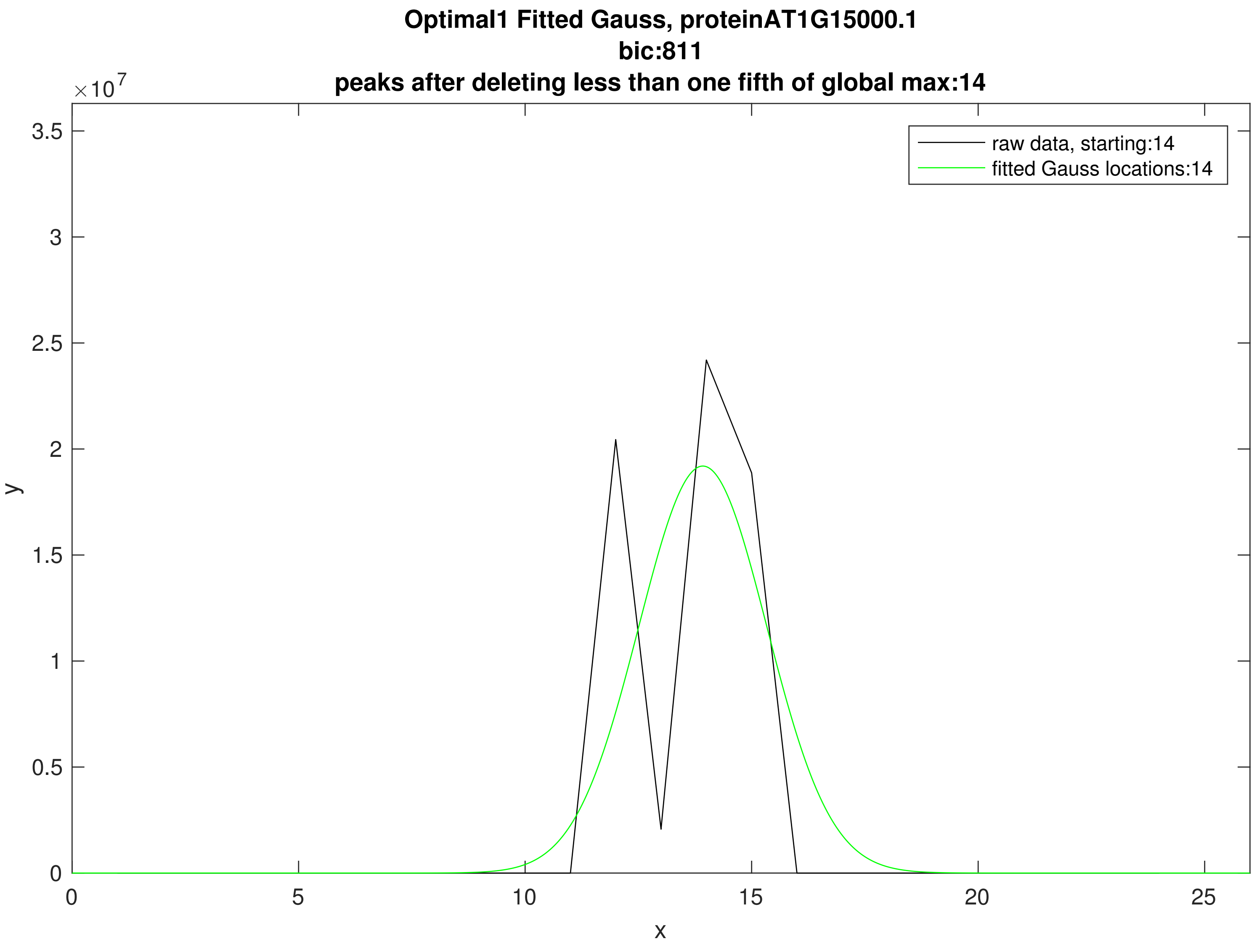
Optimal1 Fitted Gauss, proteinAT1G15000.1
bic:811
peaks after deleting less than one fifth of global max:14
×10^7
3.5
3
2.5
2
1.5
1
0.5
0
y
0
5
10
15
20
25
x
raw data, starting:14
fitted Gauss locations:14

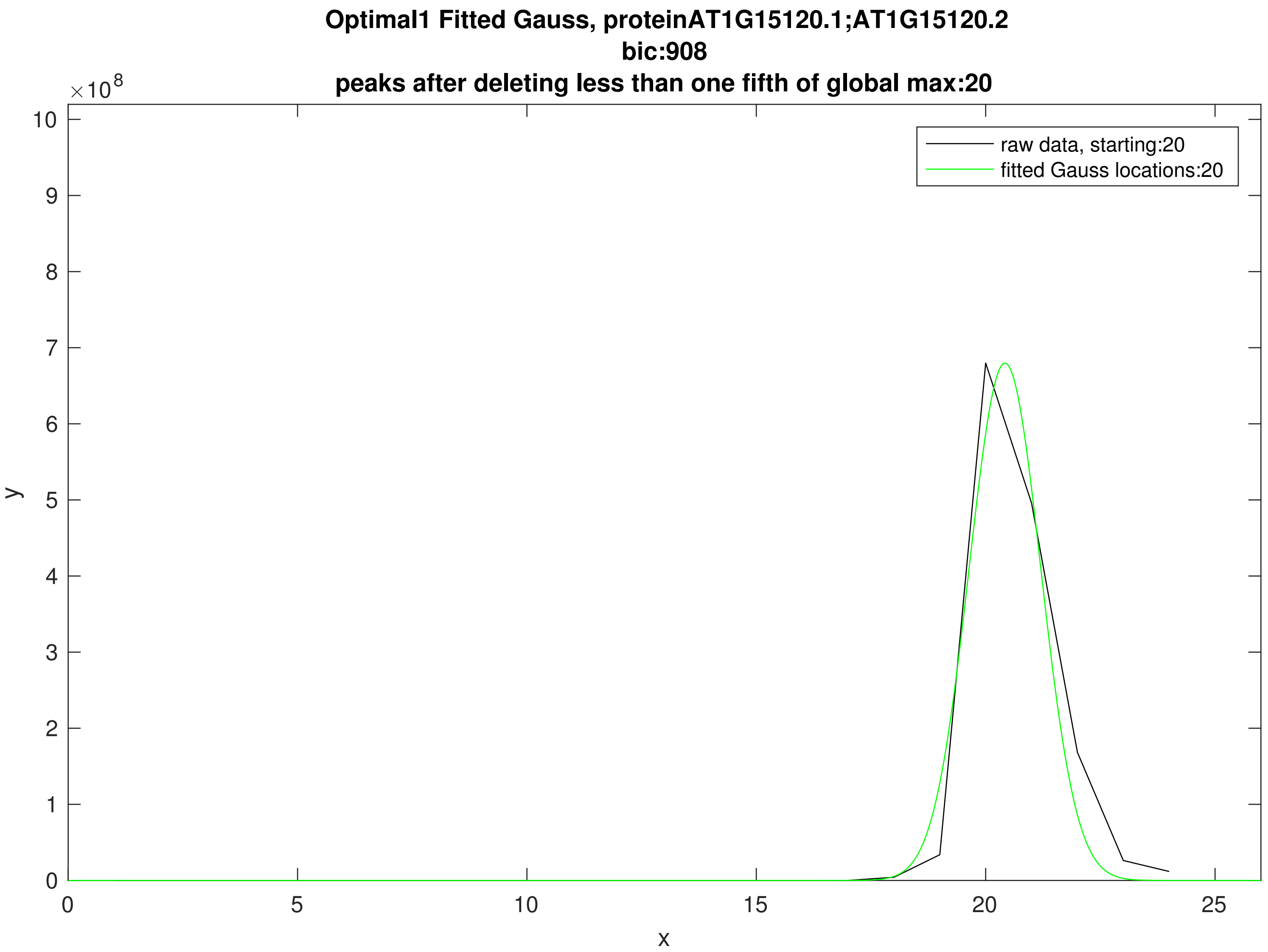
Optimal1 Fitted Gauss, proteinAT1G15120.1;AT1G15120.2
bic:908
peaks after deleting less than one fifth of global max:20
×10^8
raw data, starting:20
fitted Gauss locations:20
y
x
0
1
2
3
4
5
6
7
8
9
10
0
5
10
15
20
25

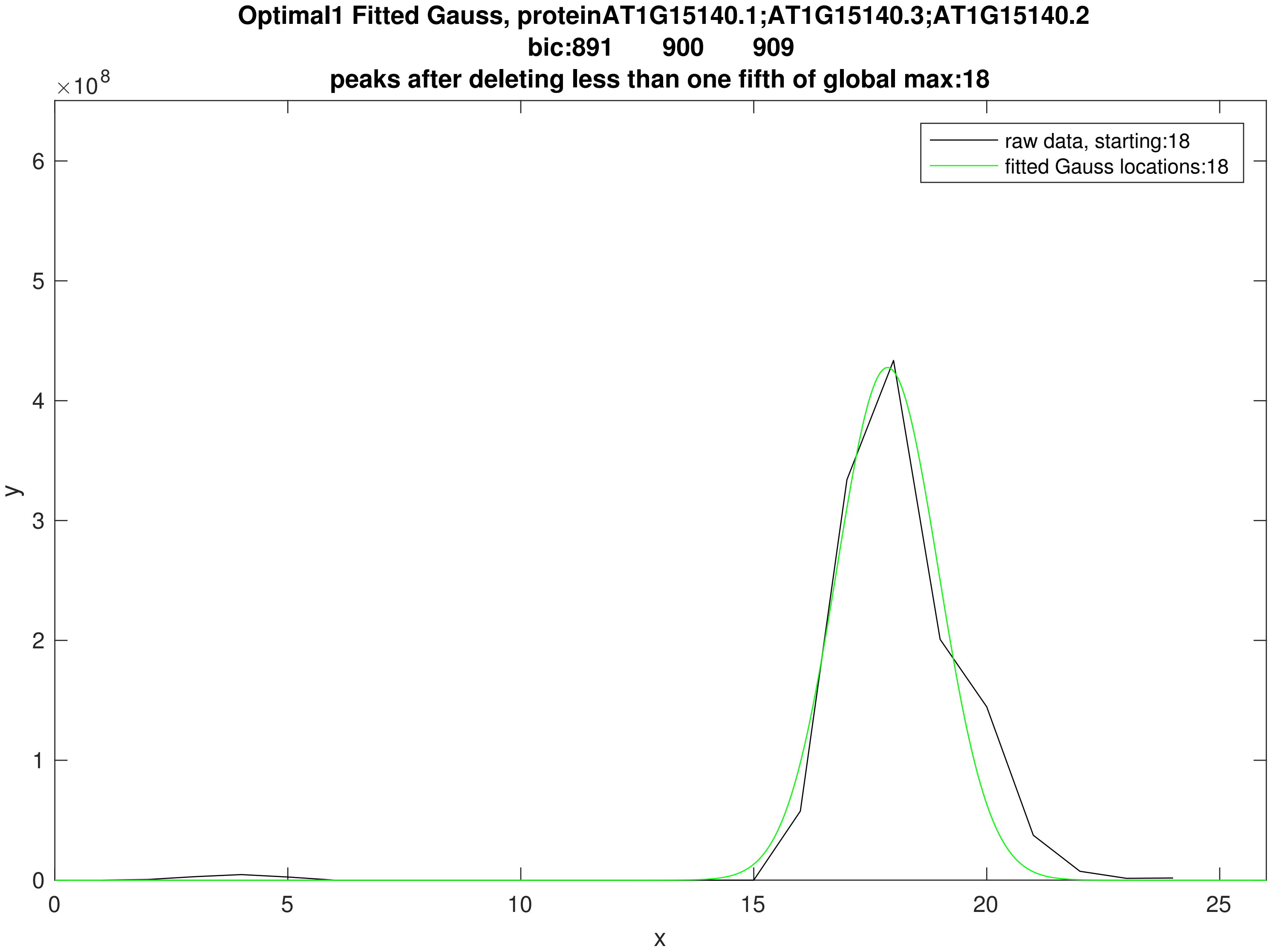

Optimal1 Fitted Gauss, proteinAT1G15140.1;AT1G15140.3;AT1G15140.2
bic:891 900 909
peaks after deleting less than one fifth of global max:18
×10^8
raw data, starting:18
fitted Gauss locations:18
y
x
0
1
2
3
4
5
6
0
5
10
15
20
25

Optimal1 Fitted Gauss, proteinAT1G15270.1
bic:779 811
peaks after deleting less than one fifth of global max:20

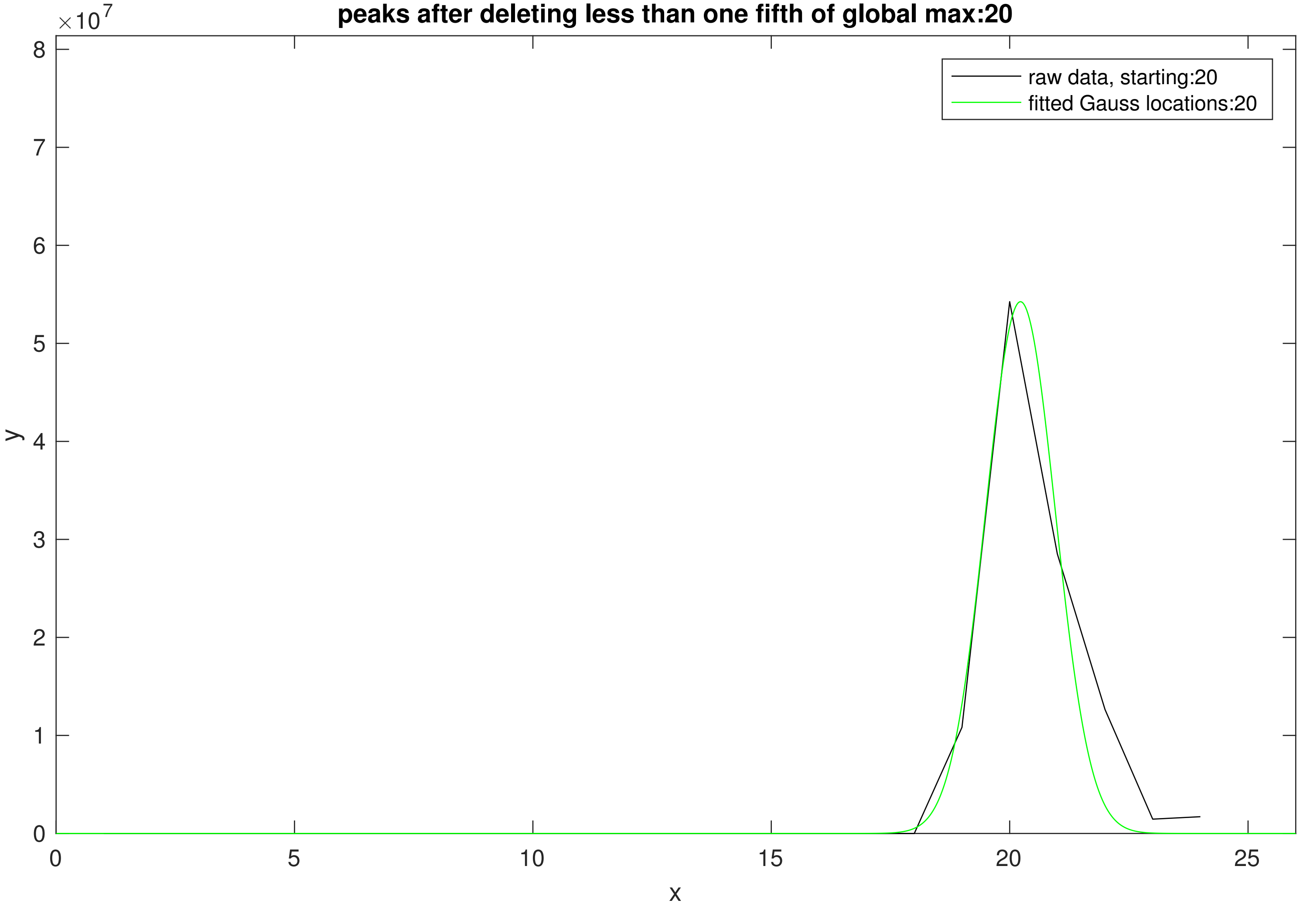

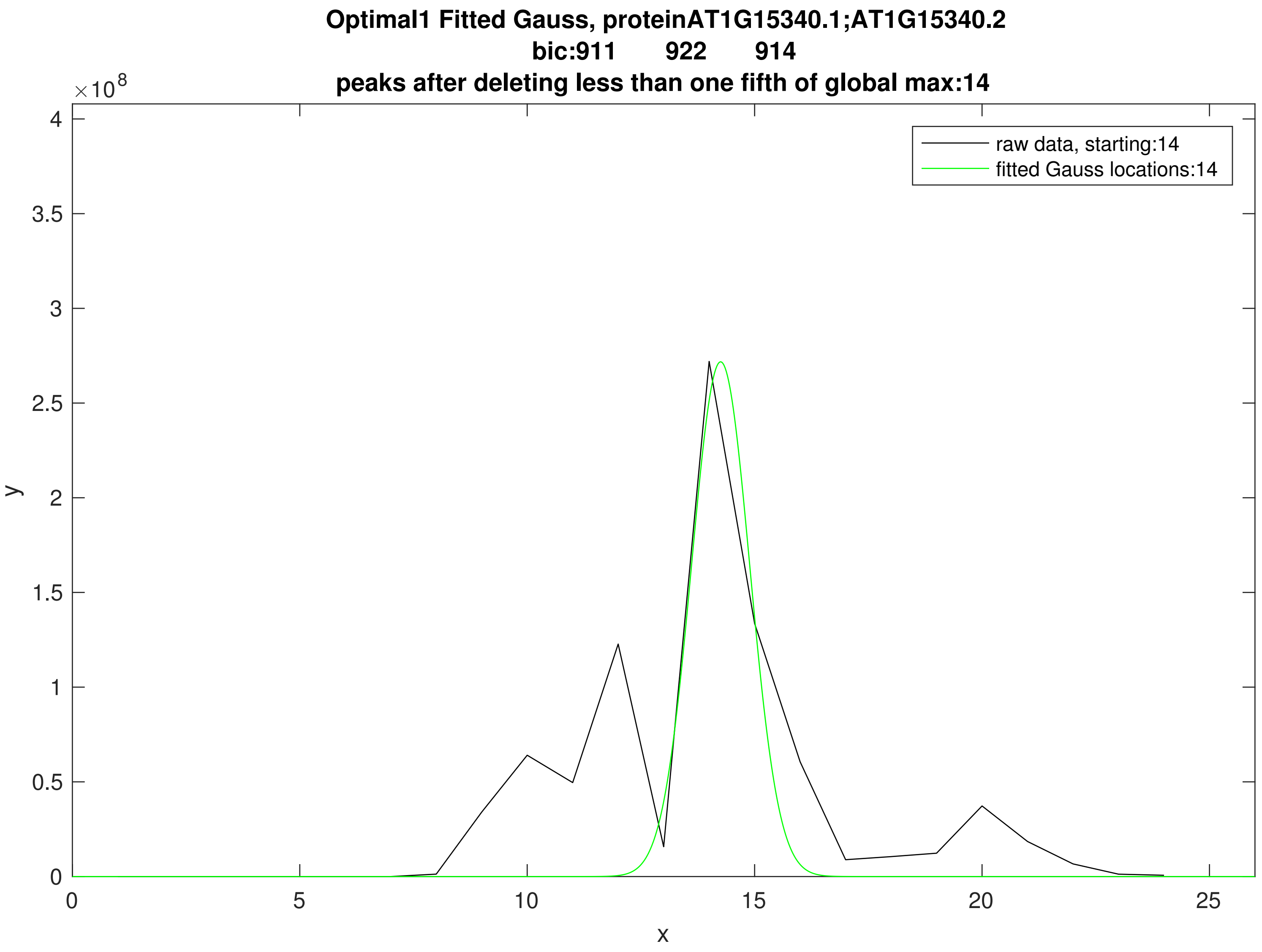
Optimal1 Fitted Gauss, proteinAT1G15340.1;AT1G15340.2
bic:911 922 914
peaks after deleting less than one fifth of global max:14
×10^8
raw data, starting:14
fitted Gauss locations:14
y
x
0
0.5
1
1.5
2
2.5
3
3.5
4
0
5
10
15
20
25

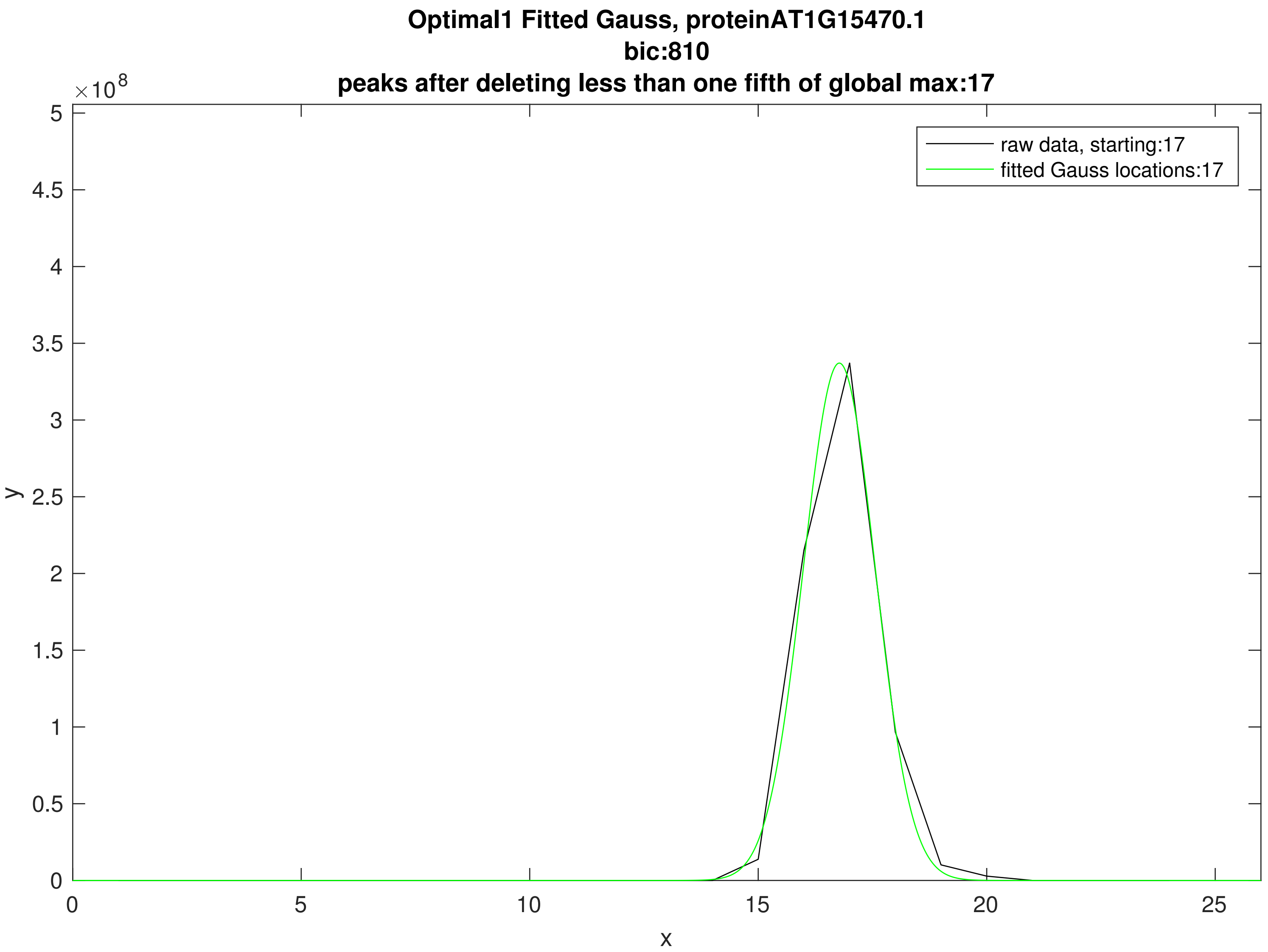

Optimal1 Fitted Gauss, proteinAT1G15470.1
bic:810
peaks after deleting less than one fifth of global max:17
×10^8
raw data, starting:17
fitted Gauss locations:17
y
x
0
0.5
1
1.5
2
2.5
3
3.5
4
4.5
5
0
5
10
15
20
25

**Optimal1 Fitted Gauss, proteinAT1G15790.2;AT1G15790.1**
**bic:711**
**peaks after deleting less than one fifth of global max:18**

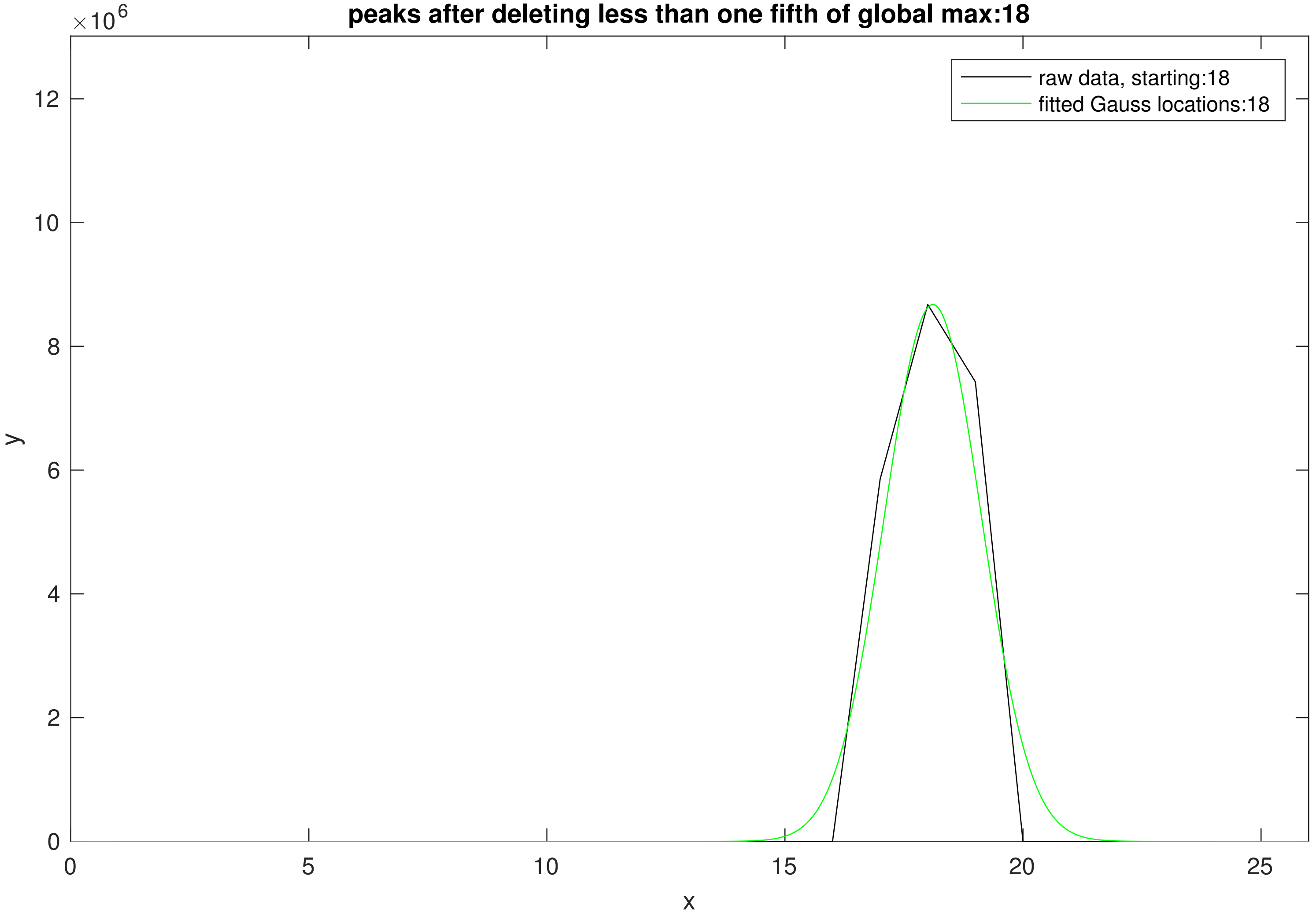

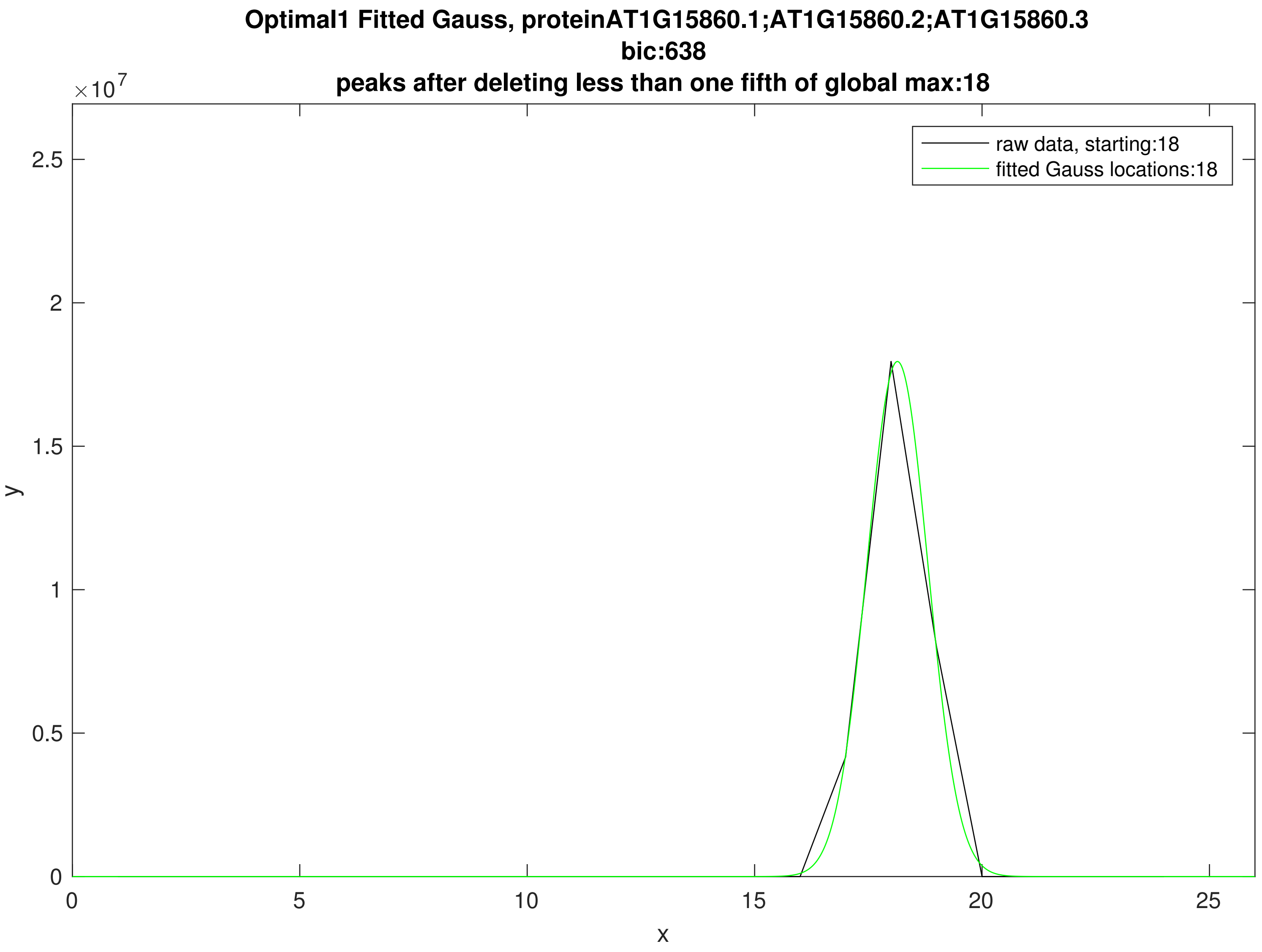

Optimal1 Fitted Gauss, proteinAT1G15860.1;AT1G15860.2;AT1G15860.3
bic:638
peaks after deleting less than one fifth of global max:18
×10^7
raw data, starting:18
fitted Gauss locations:18
2.5
2
1.5
1
0.5
0
y
0
5
10
15
20
25
x

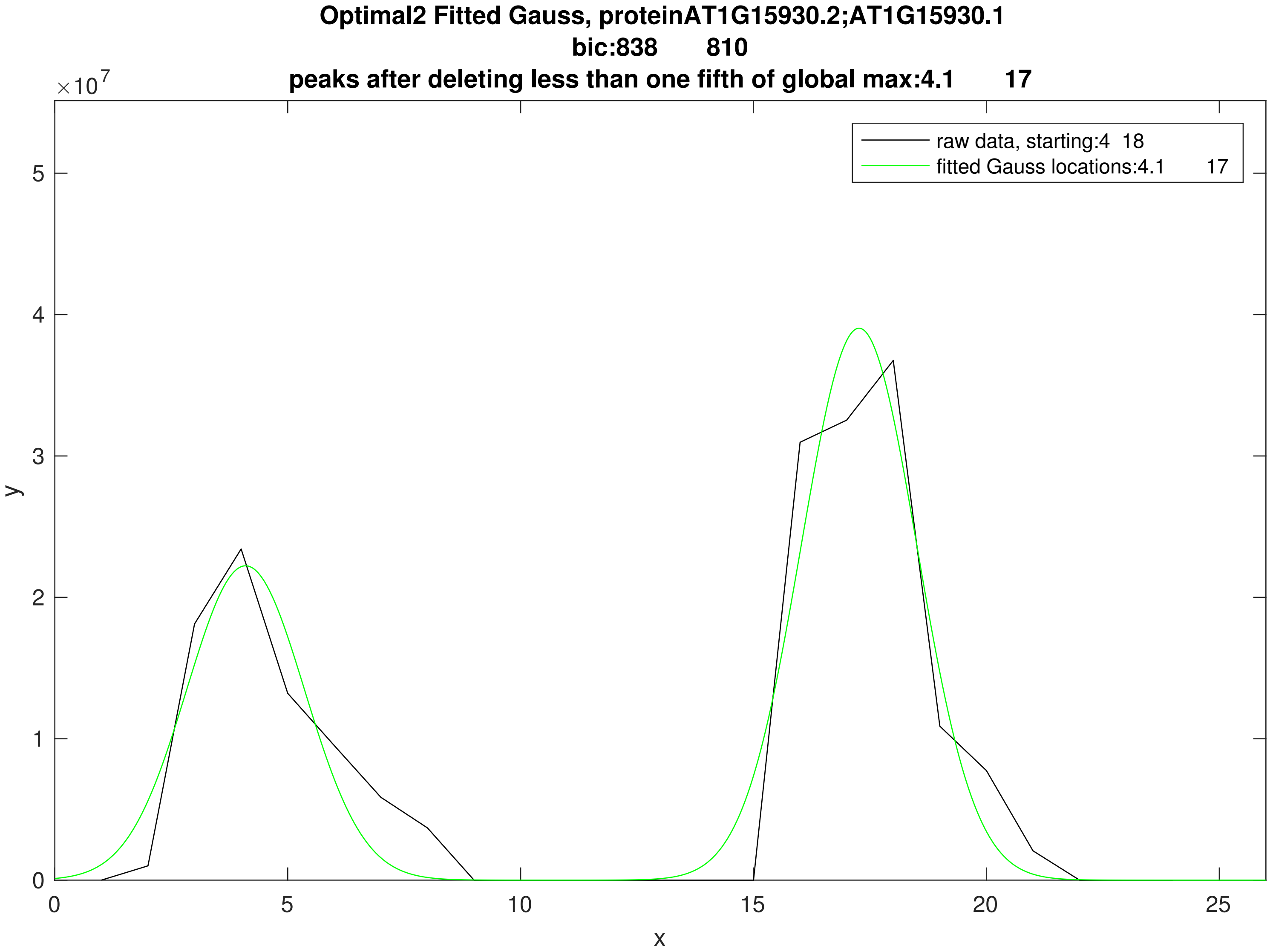

Optimal2 Fitted Gauss, proteinAT1G15930.2;AT1G15930.1
bic:838 810
peaks after deleting less than one fifth of global max:4.1 17
×10^7
raw data, starting:4 18
fitted Gauss locations:4.1 17
0
1
2
3
4
5
y
0
5
10
15
20
25
x

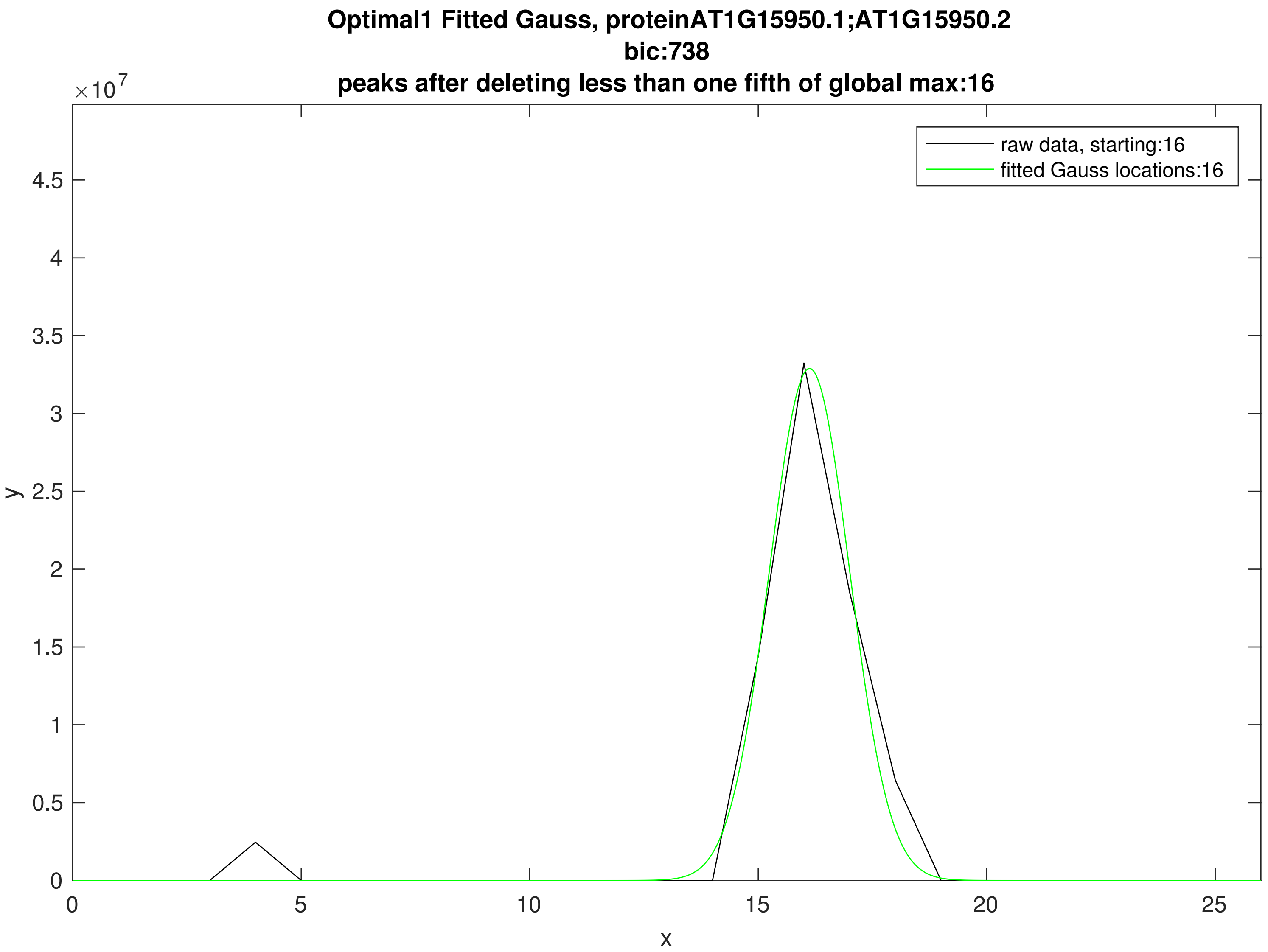
Optimal1 Fitted Gauss, proteinAT1G15950.1;AT1G15950.2
bic:738
peaks after deleting less than one fifth of global max:16
×10^7
raw data, starting:16
fitted Gauss locations:16
4.5
4
3.5
3
2.5
2
1.5
1
0.5
0
y
0
5
10
15
20
25
x

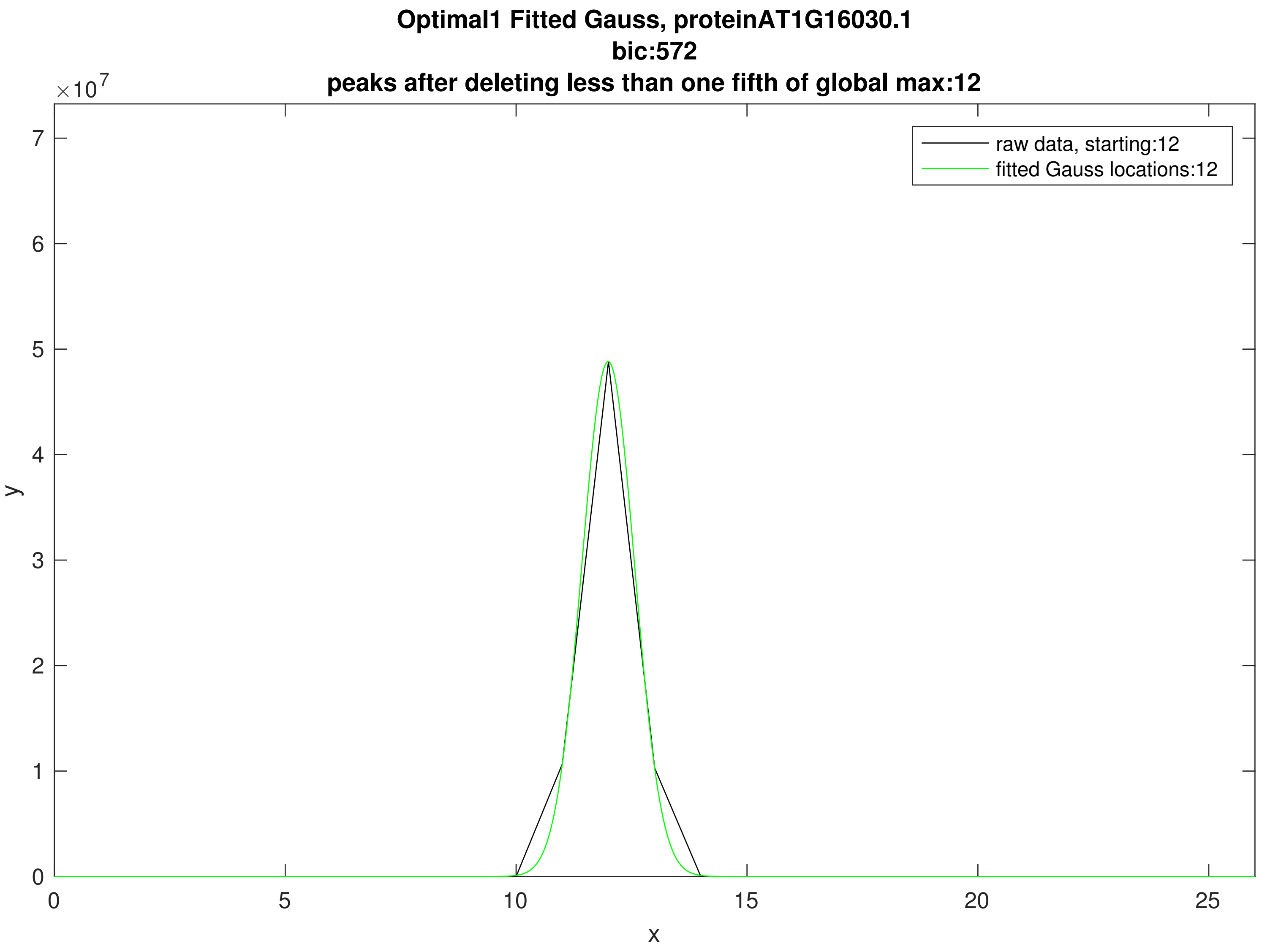

Optimal1 Fitted Gauss, proteinAT1G16030.1
bic:572
peaks after deleting less than one fifth of global max:12
×10^7
7
6
5
4
3
2
1
0
y
0
5
10
15
20
25
x
raw data, starting:12
fitted Gauss locations:12

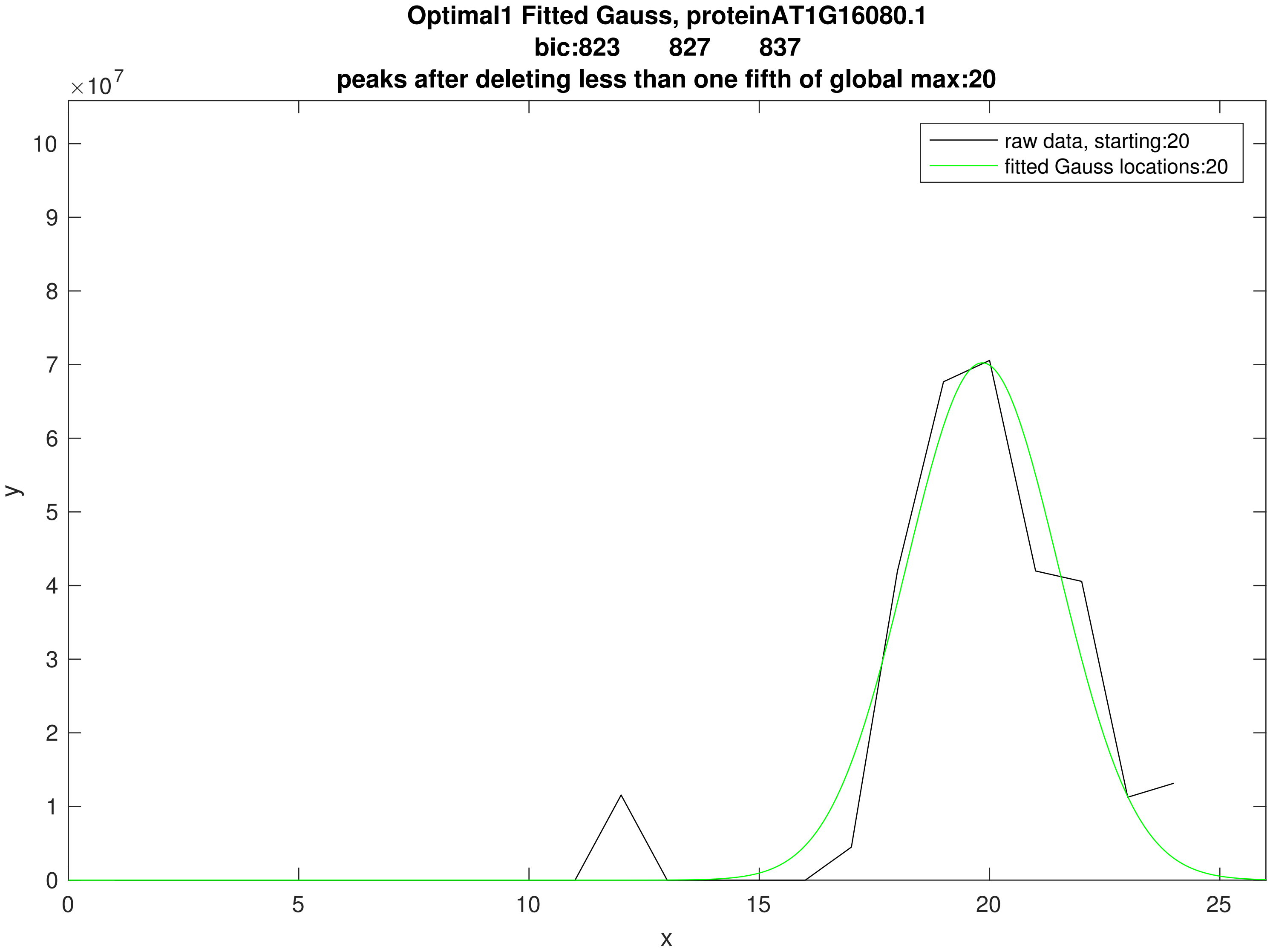
Optimal1 Fitted Gauss, proteinAT1G16080.1
bic:823 827 837
peaks after deleting less than one fifth of global max:20
×10^7
raw data, starting:20
fitted Gauss locations:20
y
x
0
1
2
3
4
5
6
7
8
9
10
0
5
10
15
20
25

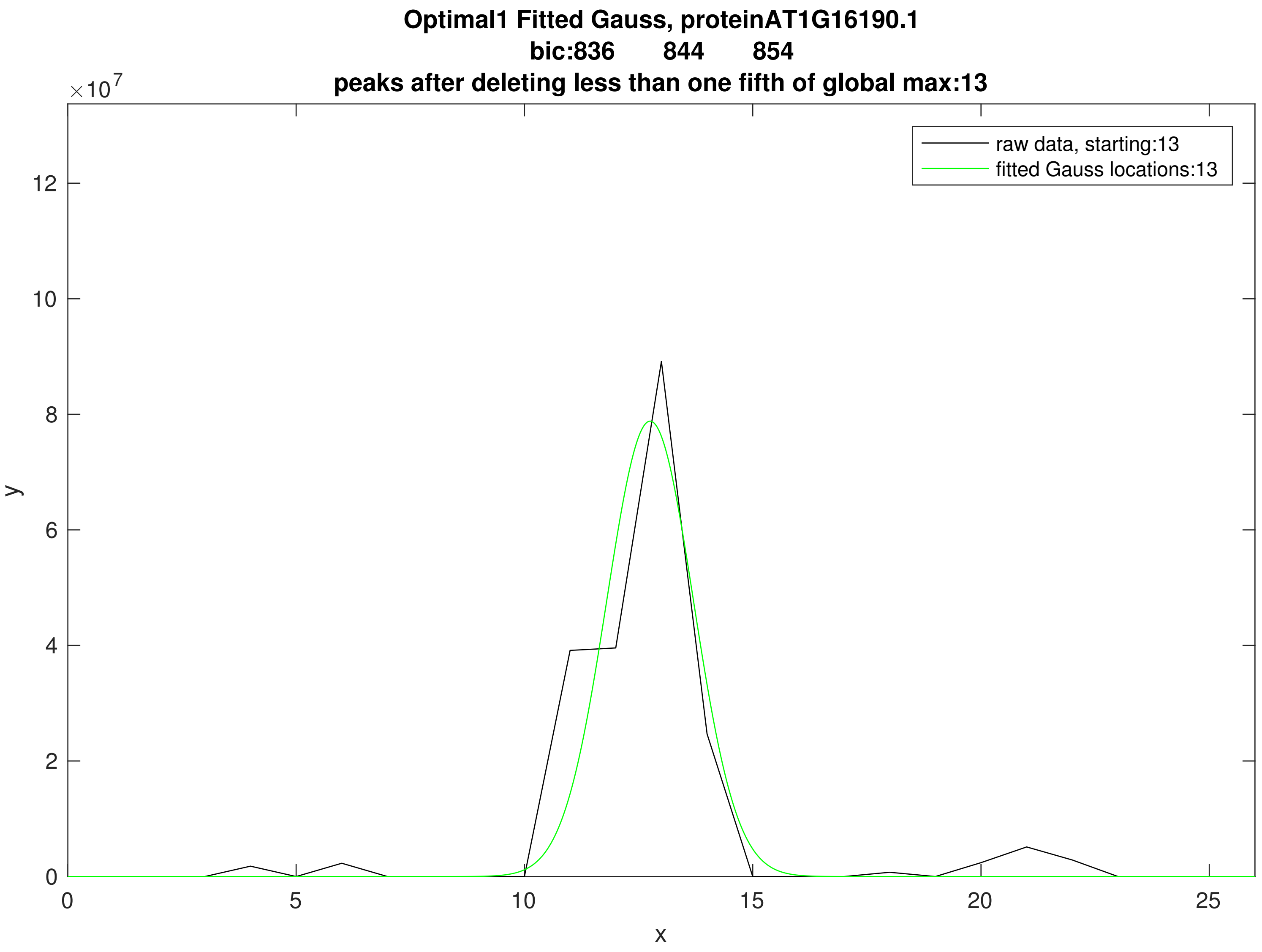
Optimal1 Fitted Gauss, proteinAT1G16190.1
bic:836 844 854
peaks after deleting less than one fifth of global max:13
×10^7
raw data, starting:13
fitted Gauss locations:13
y
x
0
2
4
6
8
10
12
0
5
10
15
20
25

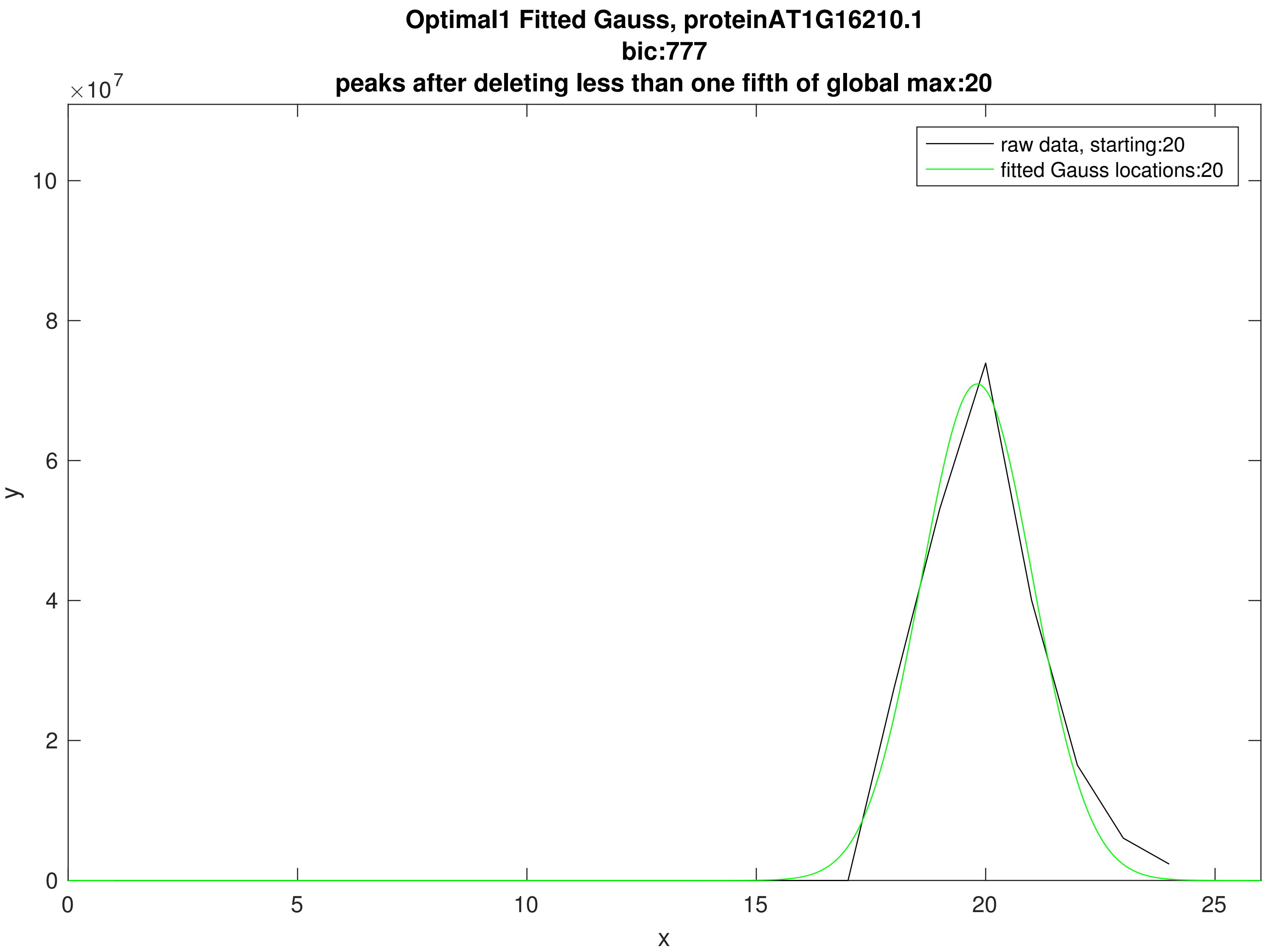

Optimal1 Fitted Gauss, proteinAT1G16210.1
bic:777
peaks after deleting less than one fifth of global max:20
×10^7
raw data, starting:20
fitted Gauss locations:20
y
x

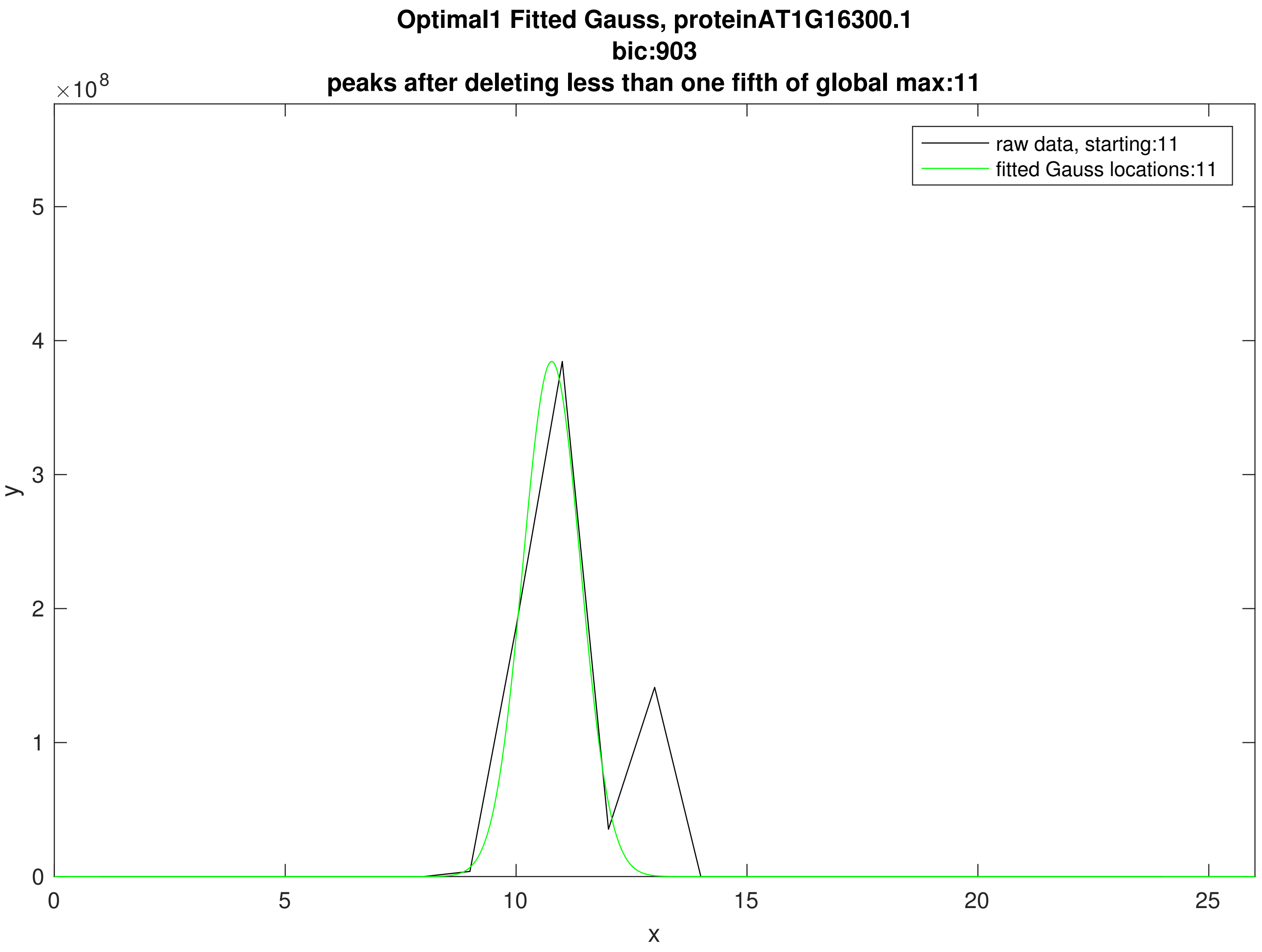
Optimal1 Fitted Gauss, proteinAT1G16300.1
bic:903
peaks after deleting less than one fifth of global max:11
×10^8
raw data, starting:11
fitted Gauss locations:11
y
x
0
1
2
3
4
5
0
5
10
15
20
25

**Optimal1 Fitted Gauss, proteinAT1G16340.3;AT1G16340.2;AT1G16340.1;AT1G16340.4**
**bic:816**
**peaks after deleting less than one fifth of global max:13**

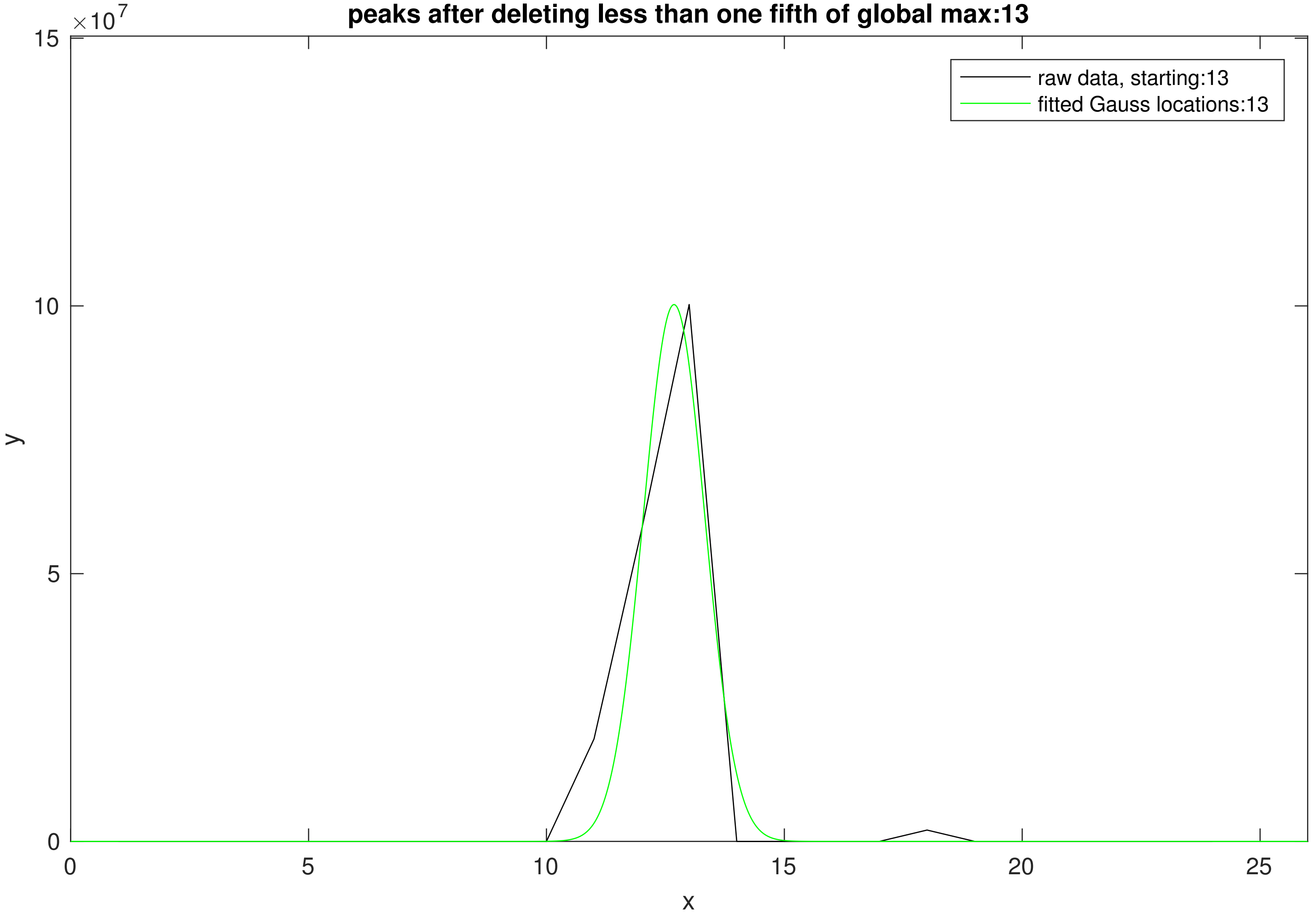

Optimal1 Fitted Gauss, proteinAT1G16350.1
bic:962  1.03e+03
peaks after deleting less than one fifth of global max:9.6

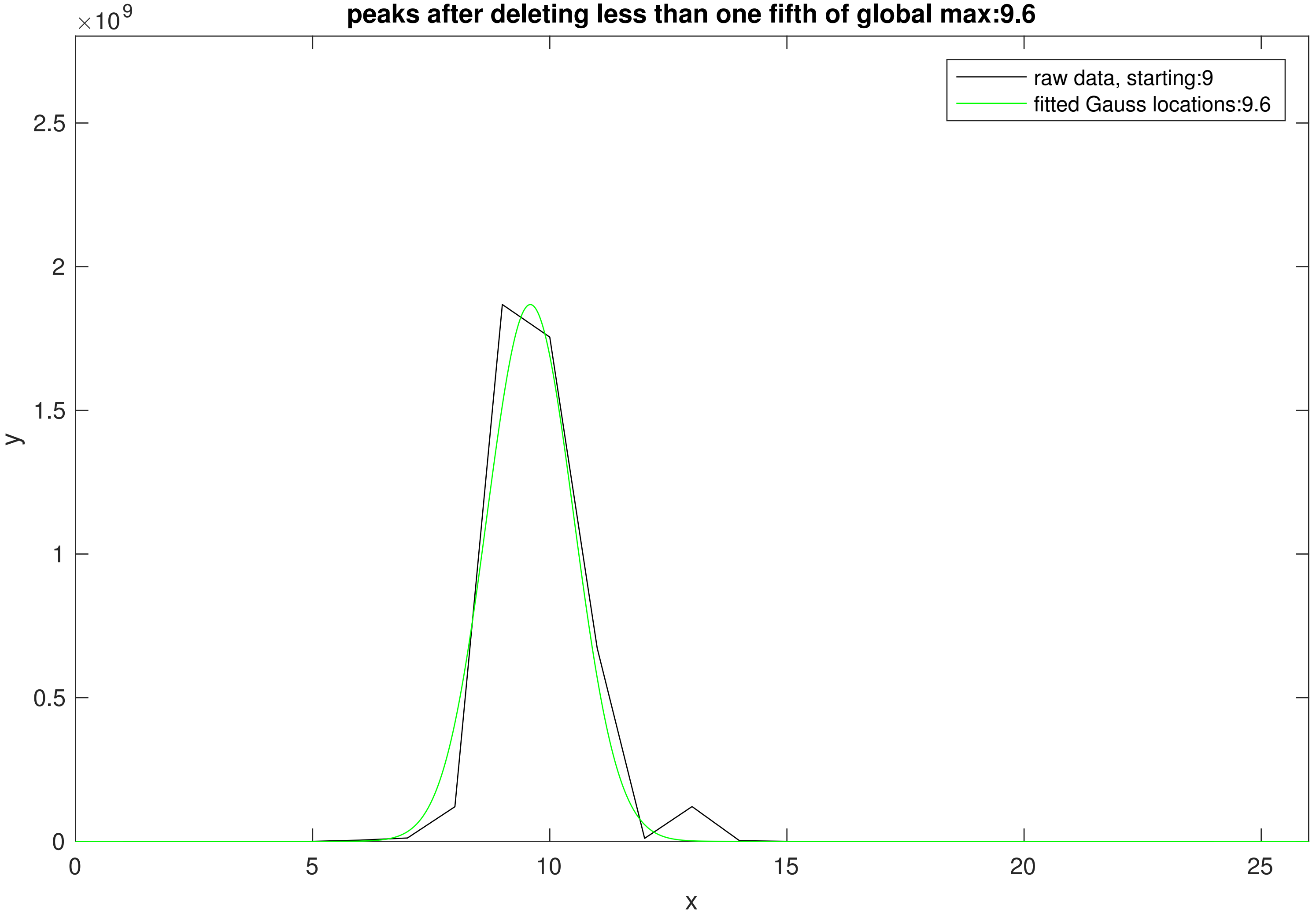

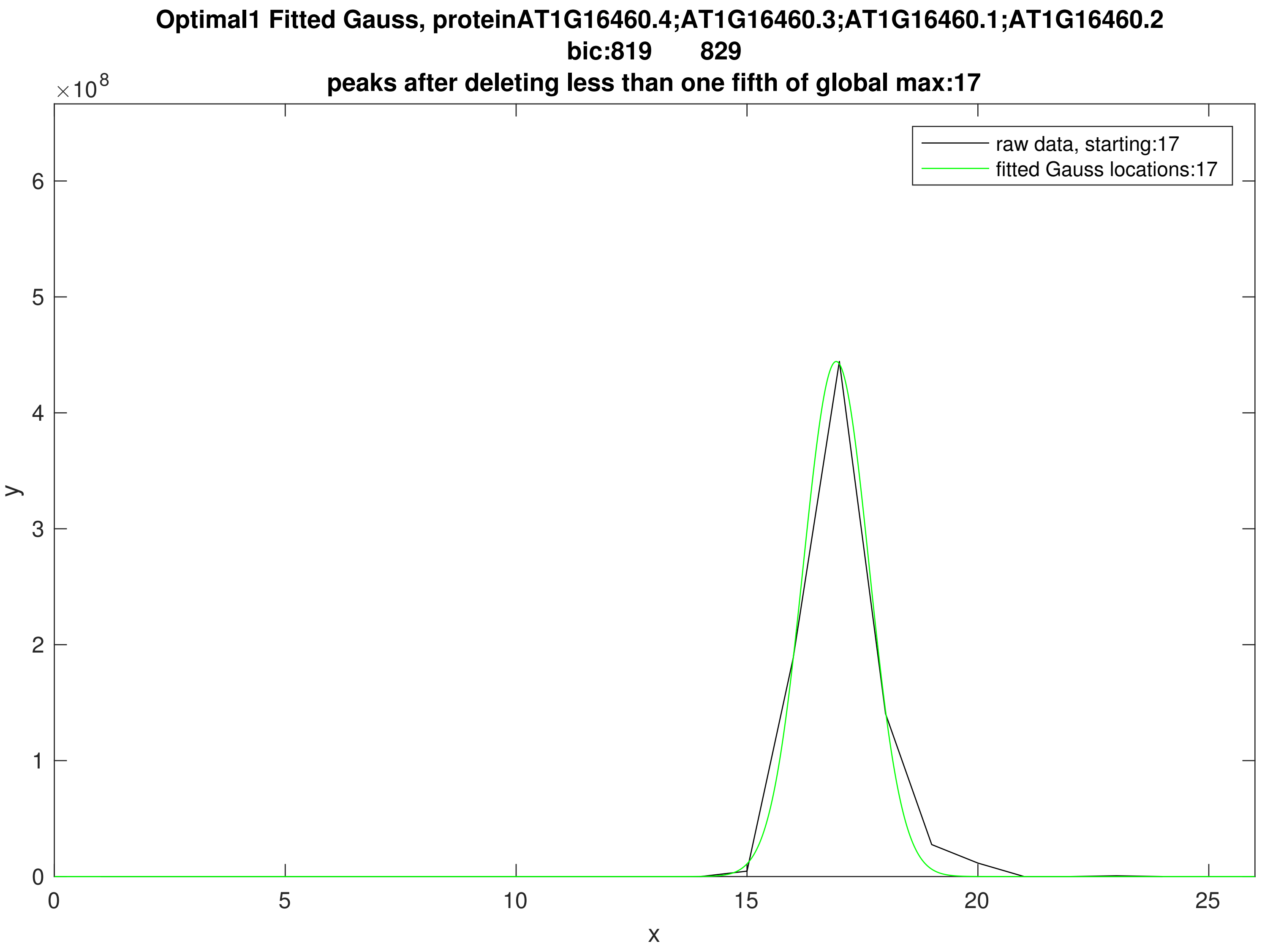

Optimal1 Fitted Gauss, proteinAT1G16460.4;AT1G16460.3;AT1G16460.1;AT1G16460.2
bic:819 829
peaks after deleting less than one fifth of global max:17
×10^8
raw data, starting:17
fitted Gauss locations:17
y
x
0
1
2
3
4
5
6
0
5
10
15
20
25

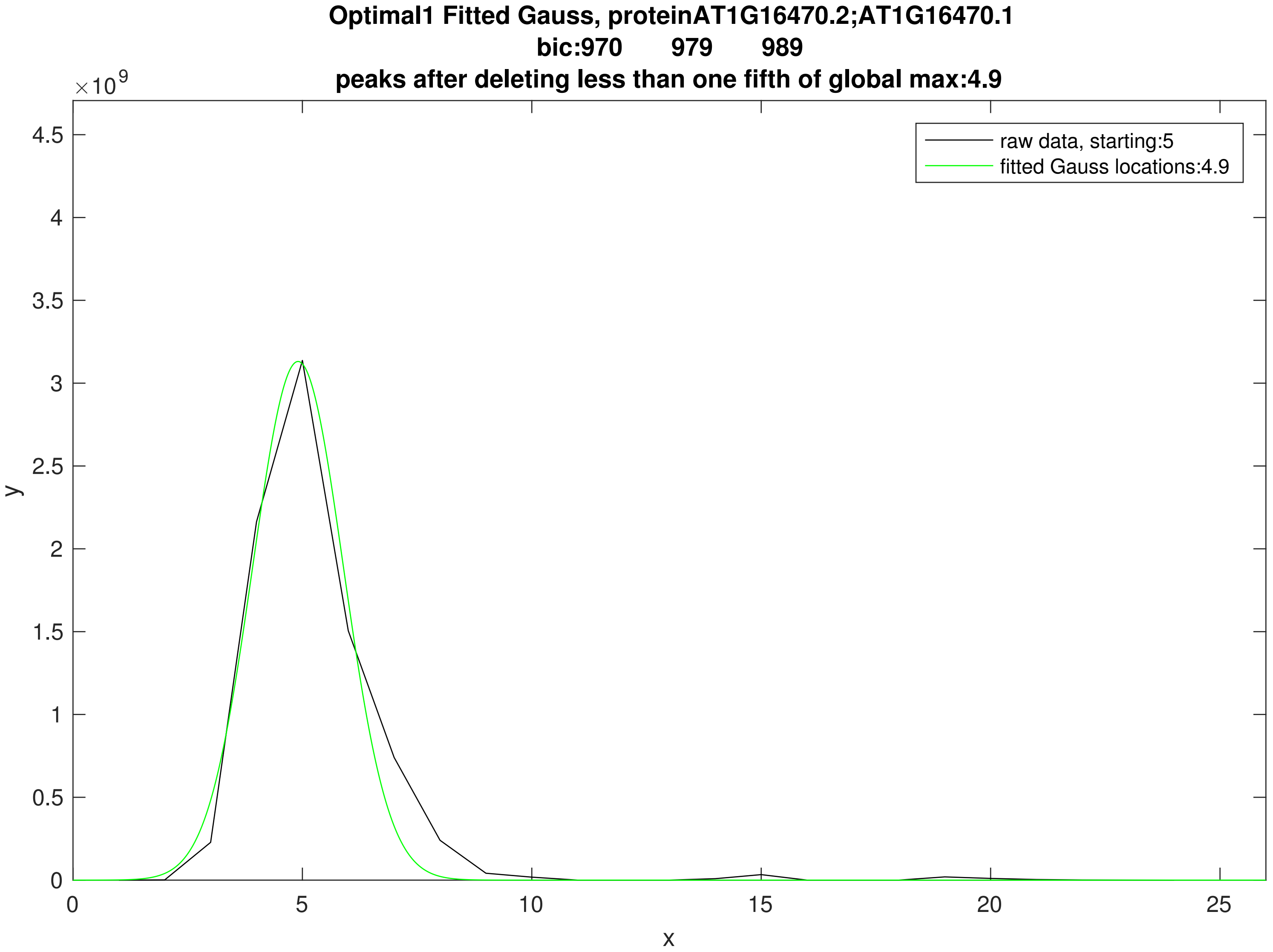

Optimal1 Fitted Gauss, proteinAT1G16470.2;AT1G16470.1
bic:970 979 989
peaks after deleting less than one fifth of global max:4.9
×10^9
raw data, starting:5
fitted Gauss locations:4.9
4.5
4
3.5
3
2.5
2
1.5
1
0.5
0
y
0
5
10
15
20
25
x

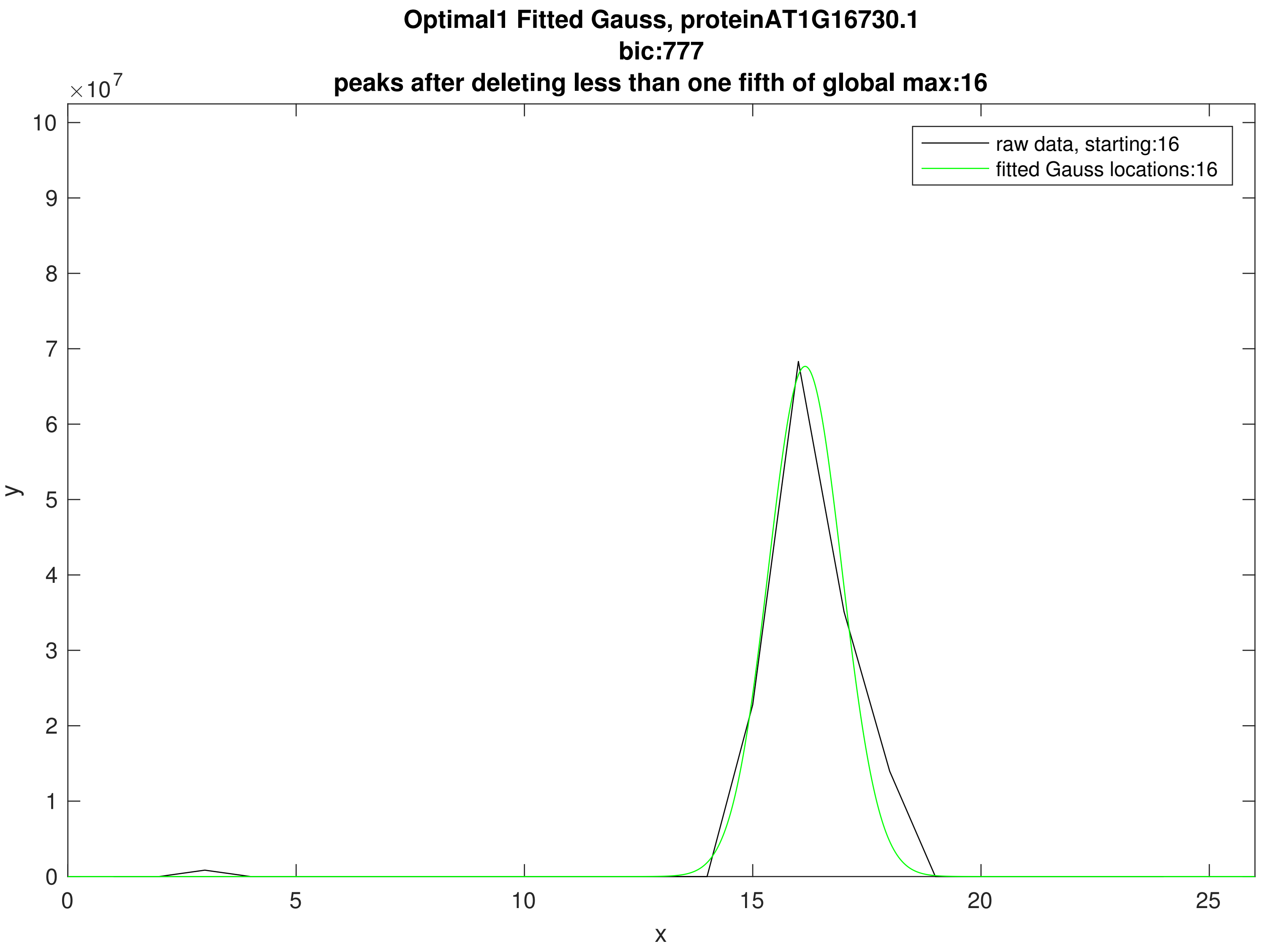

Optimal1 Fitted Gauss, proteinAT1G16730.1
bic:777
peaks after deleting less than one fifth of global max:16
×10^7
raw data, starting:16
fitted Gauss locations:16
y
x
0
1
2
3
4
5
6
7
8
9
10
0
5
10
15
20
25

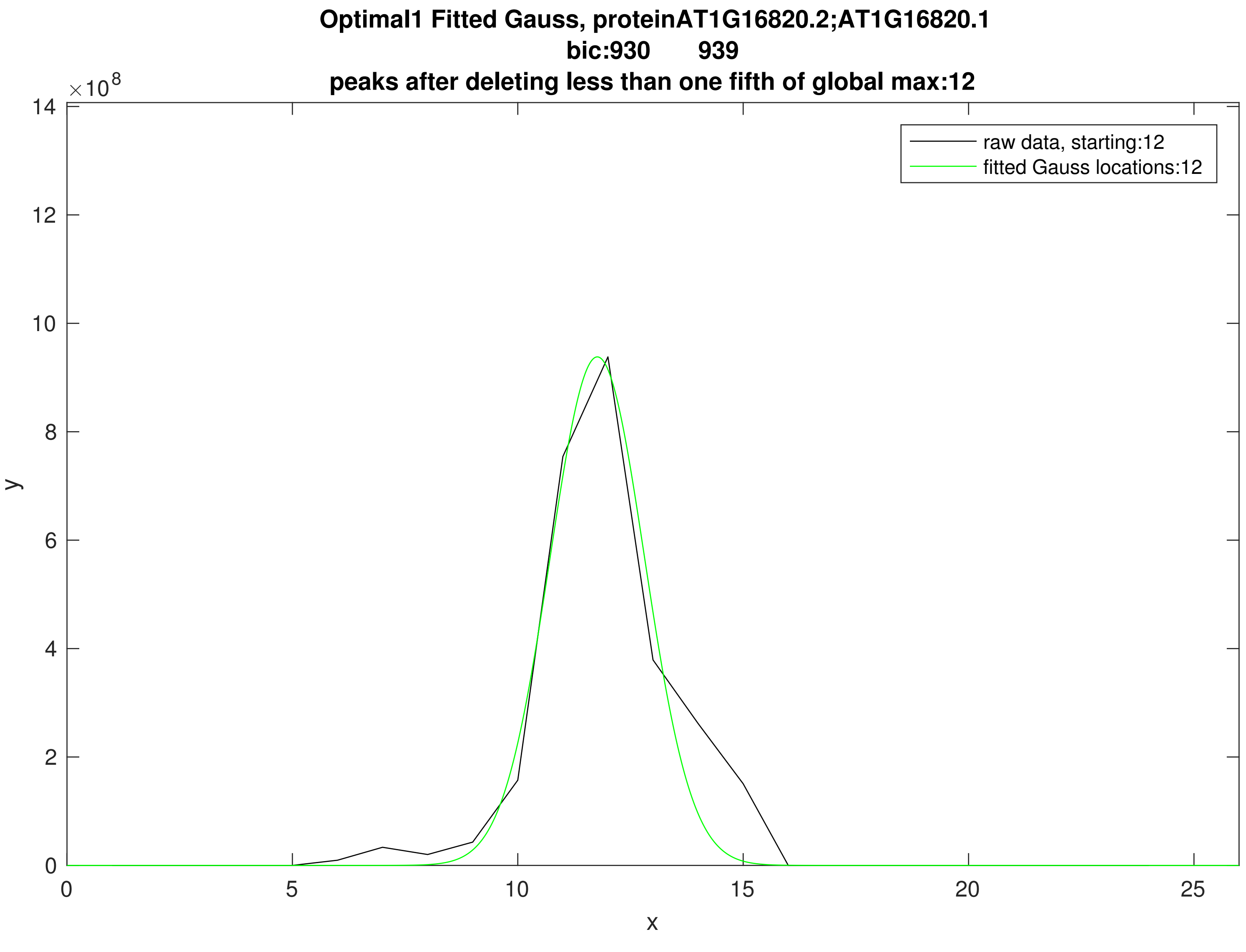

Optimal1 Fitted Gauss, proteinAT1G16820.2;AT1G16820.1
bic:930 939
peaks after deleting less than one fifth of global max:12
×10^8
14
12
10
8
6
4
2
0
y
0
5
10
15
20
25
x
raw data, starting:12
fitted Gauss locations:12

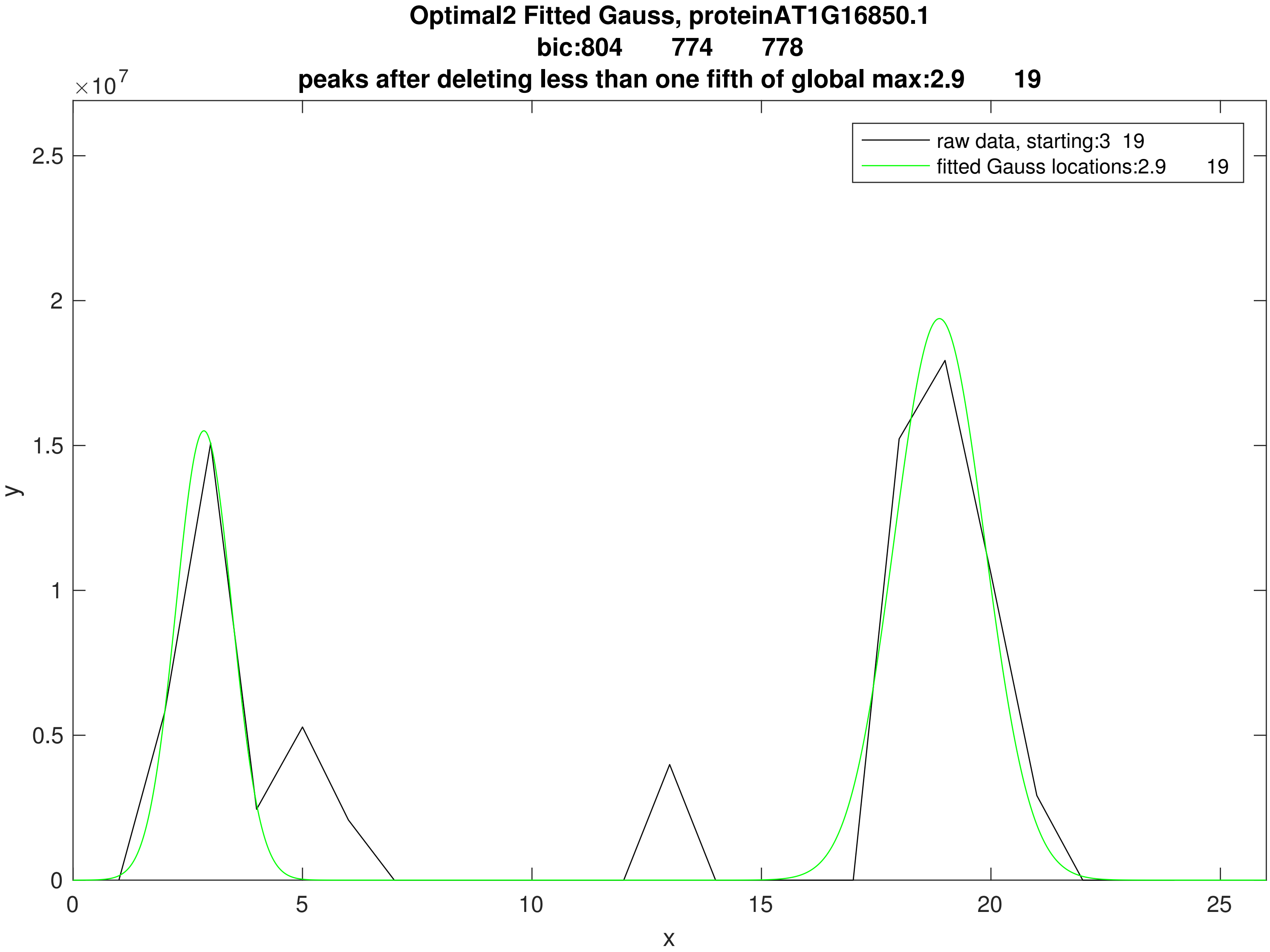

Optimal2 Fitted Gauss, proteinAT1G16850.1
bic:804 774 778
peaks after deleting less than one fifth of global max:2.9 19
×10^7
raw data, starting:3 19
fitted Gauss locations:2.9 19
2.5
2
1.5
1
0.5
0
y
0
5
10
15
20
25
x

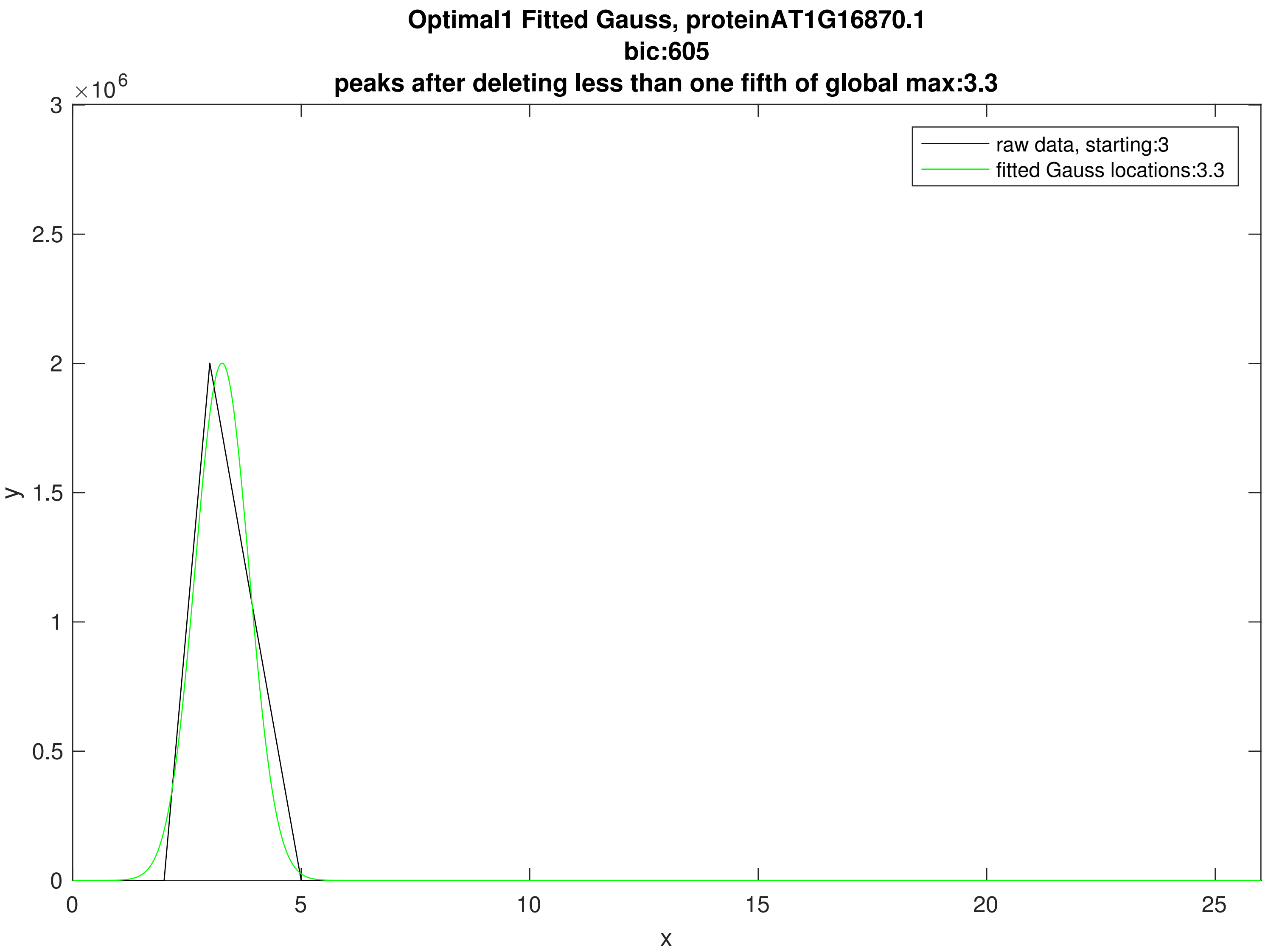

Optimal1 Fitted Gauss, proteinAT1G16870.1
bic:605
peaks after deleting less than one fifth of global max:3.3
×10^6
3
2.5
2
1.5
1
0.5
0
y
0
5
10
15
20
25
x
raw data, starting:3
fitted Gauss locations:3.3

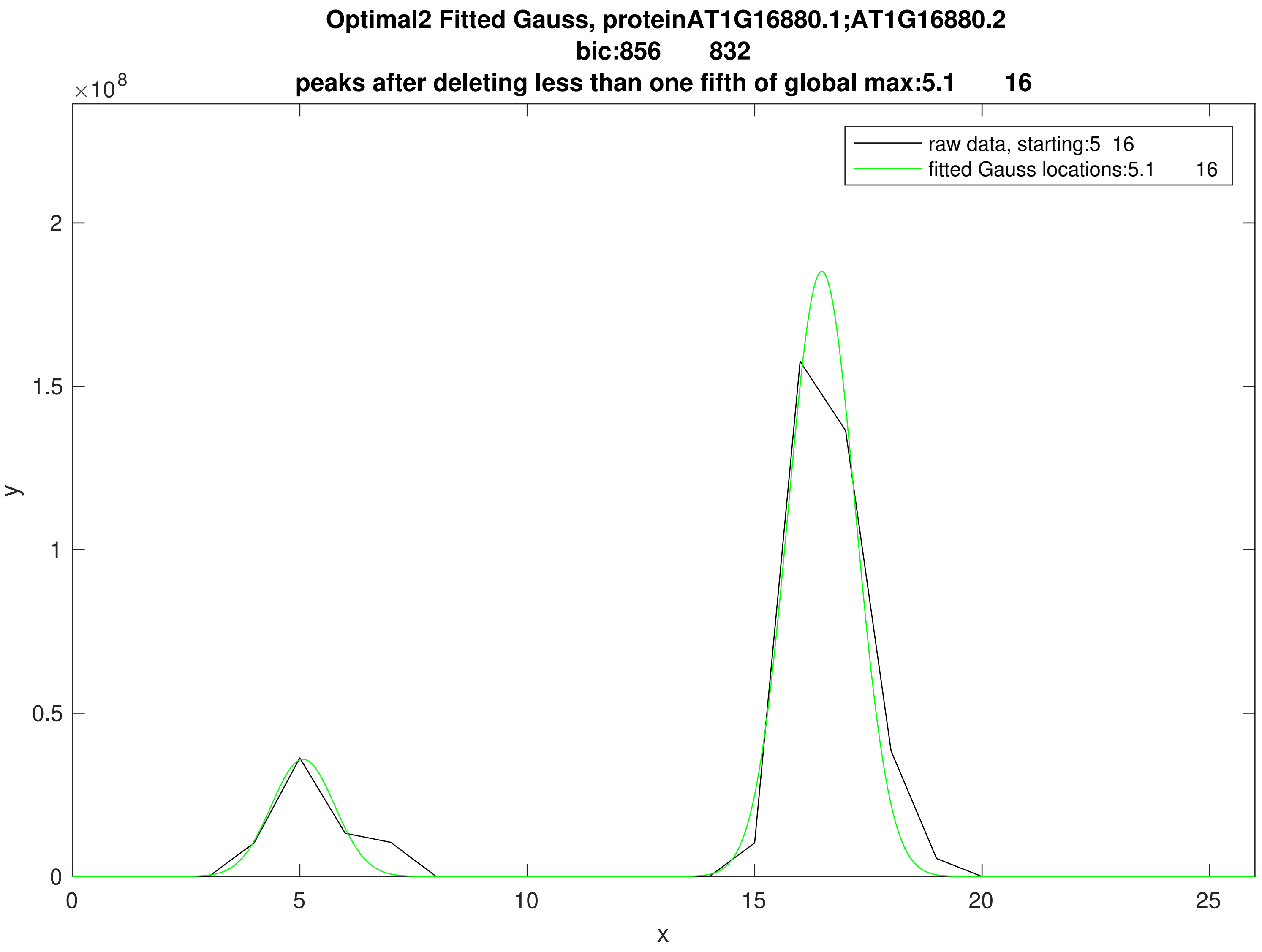

Optimal2 Fitted Gauss, proteinAT1G16880.1;AT1G16880.2
bic:856 832
peaks after deleting less than one fifth of global max:5.1 16
×10^8
raw data, starting:5 16
fitted Gauss locations:5.1 16
y
x
0
0.5
1
1.5
2
0
5
10
15
20
25

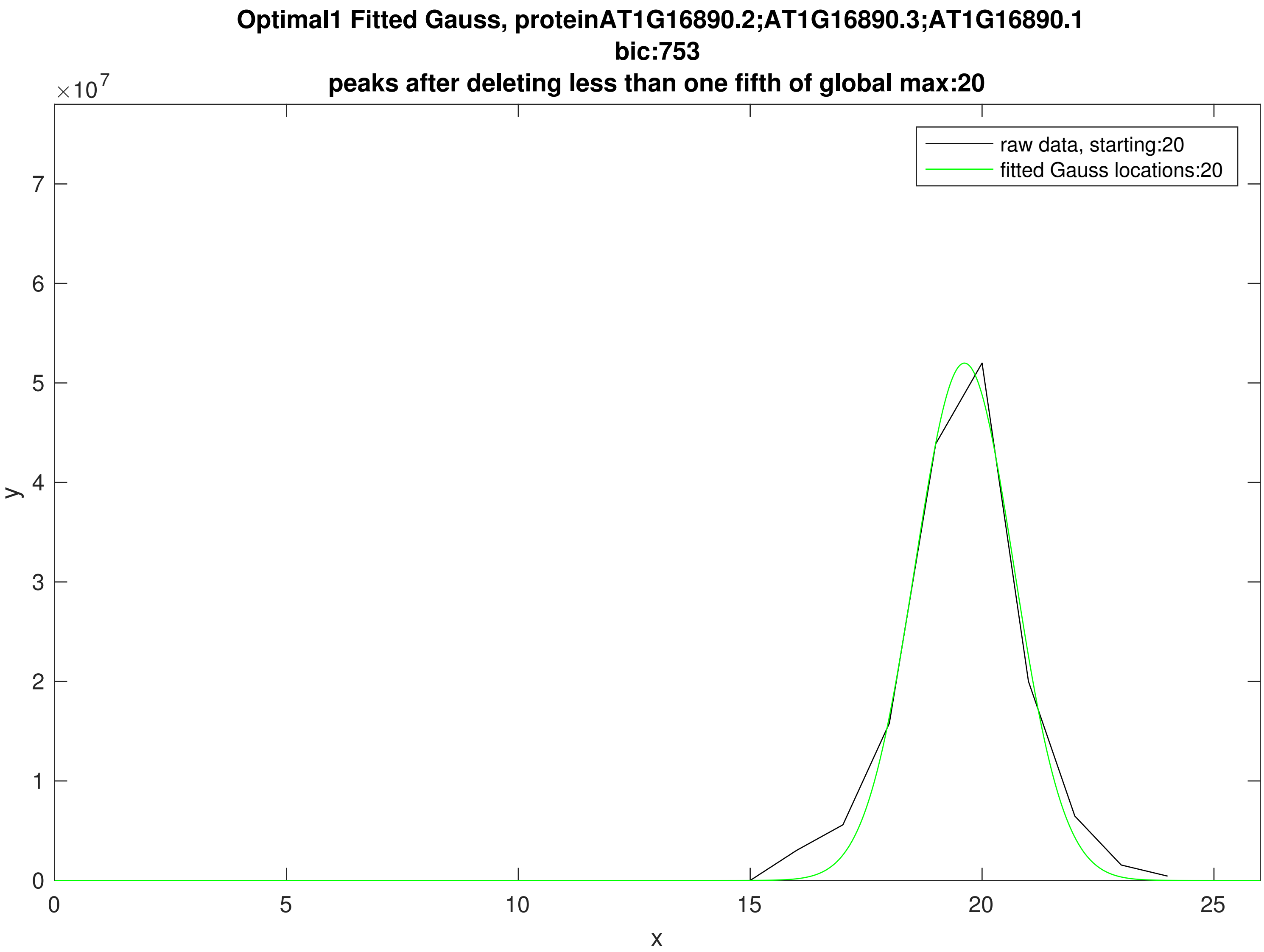
Optimal1 Fitted Gauss, proteinAT1G16890.2;AT1G16890.3;AT1G16890.1
bic:753
peaks after deleting less than one fifth of global max:20
×10^7
raw data, starting:20
fitted Gauss locations:20
y
x
0
1
2
3
4
5
6
7
0
5
10
15
20
25

#### Optimal1 Fitted Gauss, proteinAT1G16920.1;AT5G60860.1;AT3G15060.1
#### bic:759
#### peaks after deleting less than one fifth of global max:19

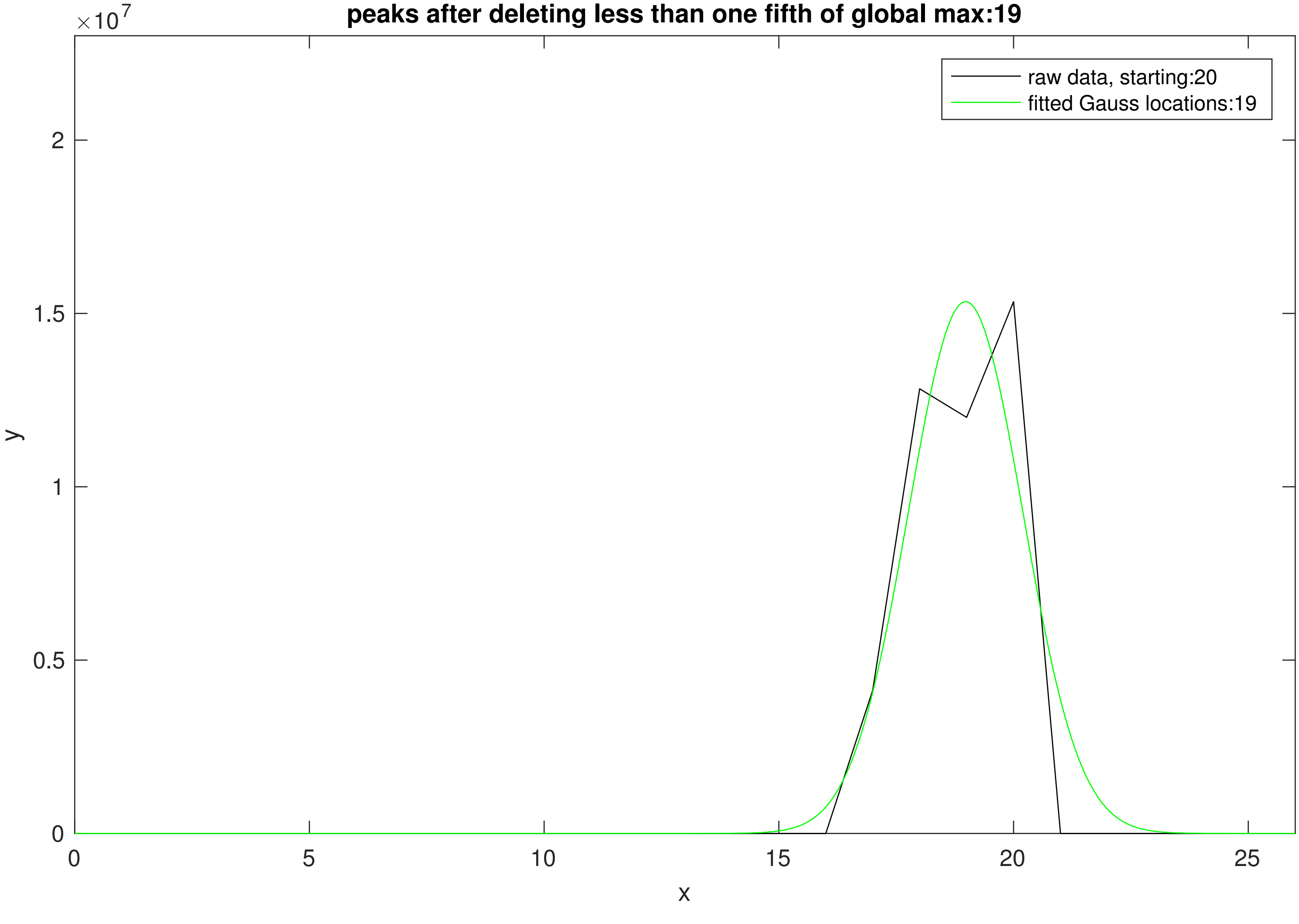

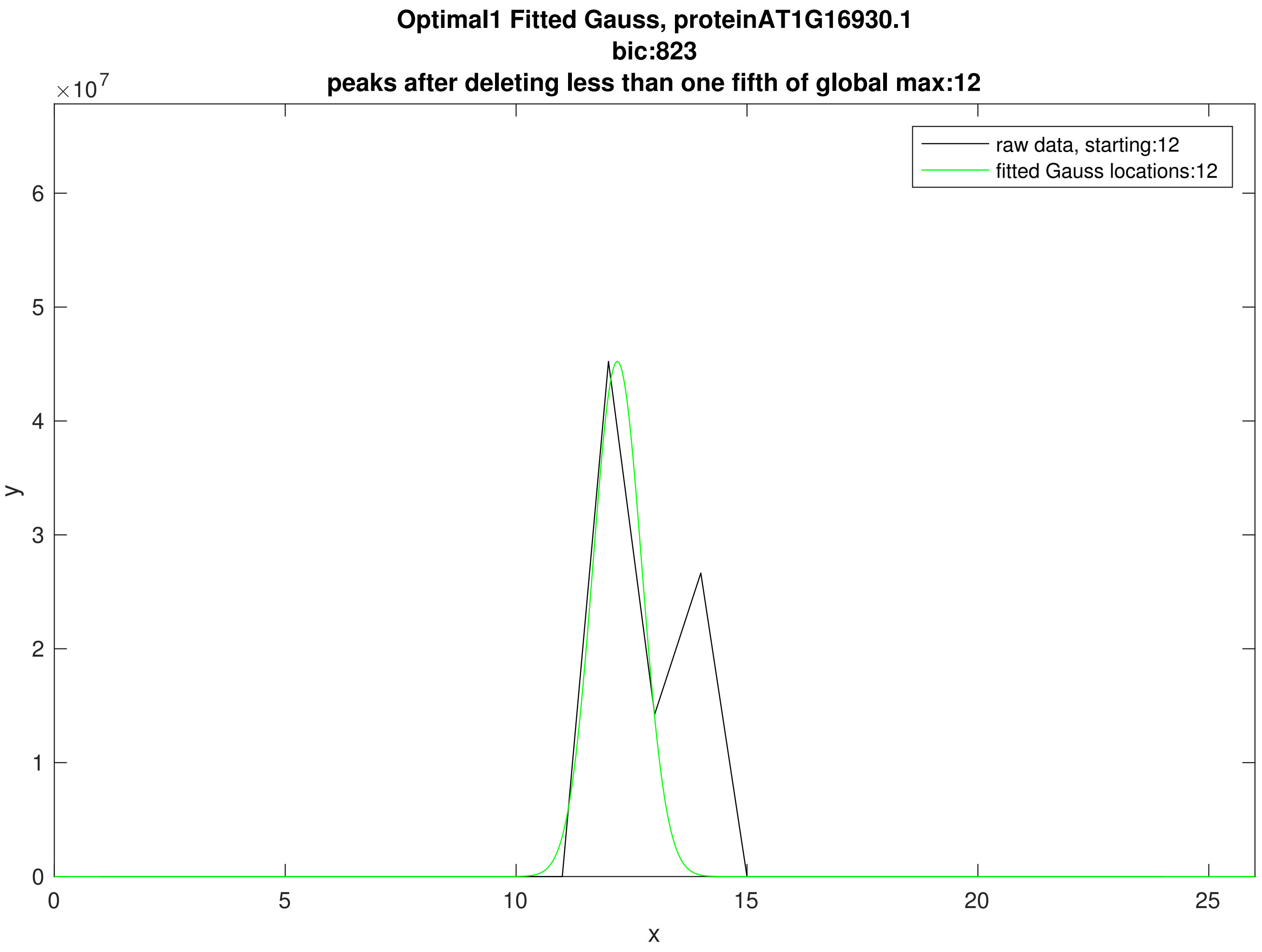

Optimal1 Fitted Gauss, proteinAT1G16930.1
bic:823
peaks after deleting less than one fifth of global max:12
×10^7
raw data, starting:12
fitted Gauss locations:12
y
x
0
1
2
3
4
5
6
0
5
10
15
20
25

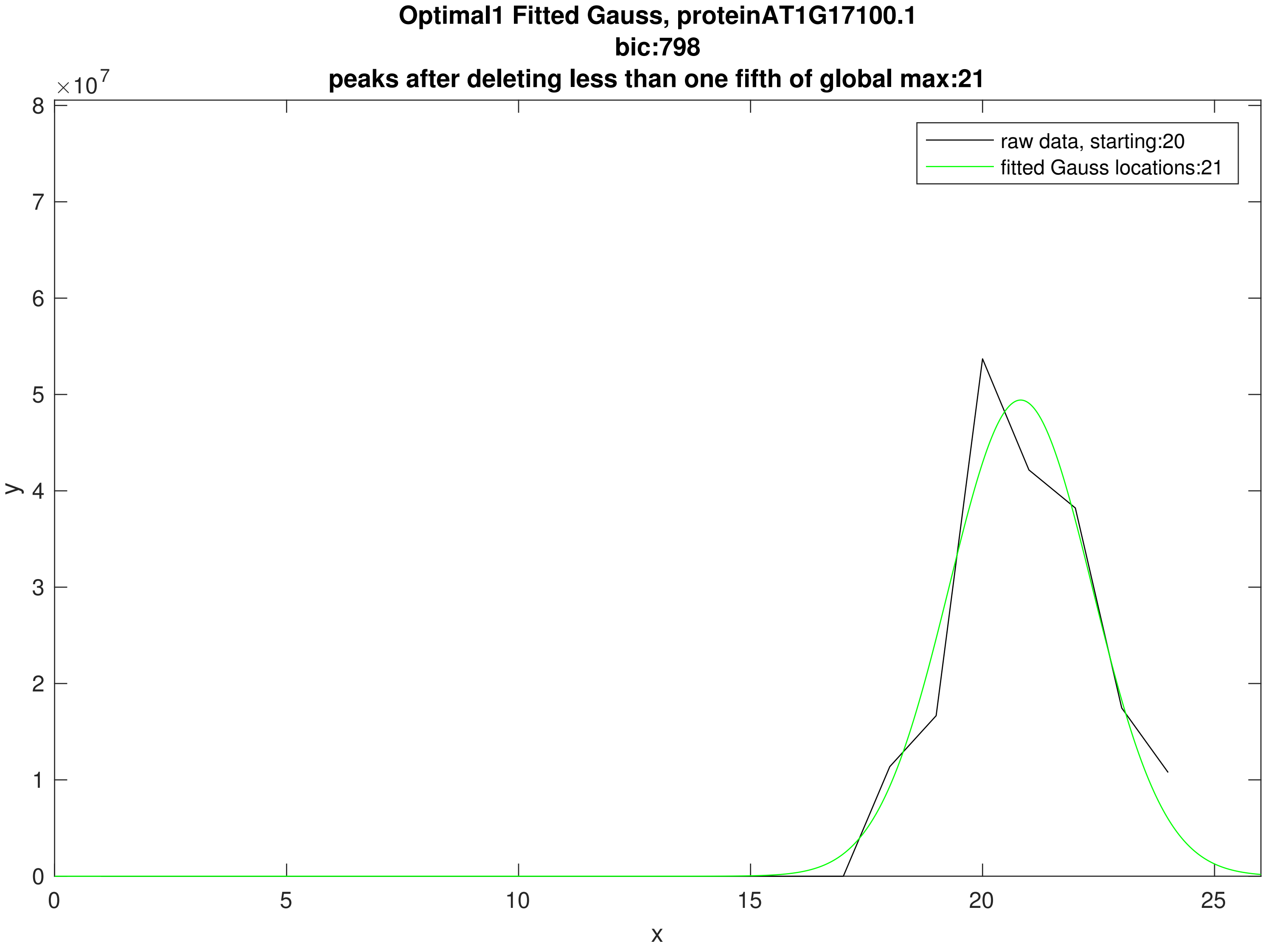
Optimal1 Fitted Gauss, proteinAT1G17100.1
bic:798
peaks after deleting less than one fifth of global max:21
×10^7
raw data, starting:20
fitted Gauss locations:21
y
x
0
1
2
3
4
5
6
7
8
0
5
10
15
20
25

Optimal1 Fitted Gauss, proteinAT1G17160.2;AT1G17160.1
bic:833
peaks after deleting less than one fifth of global max:14

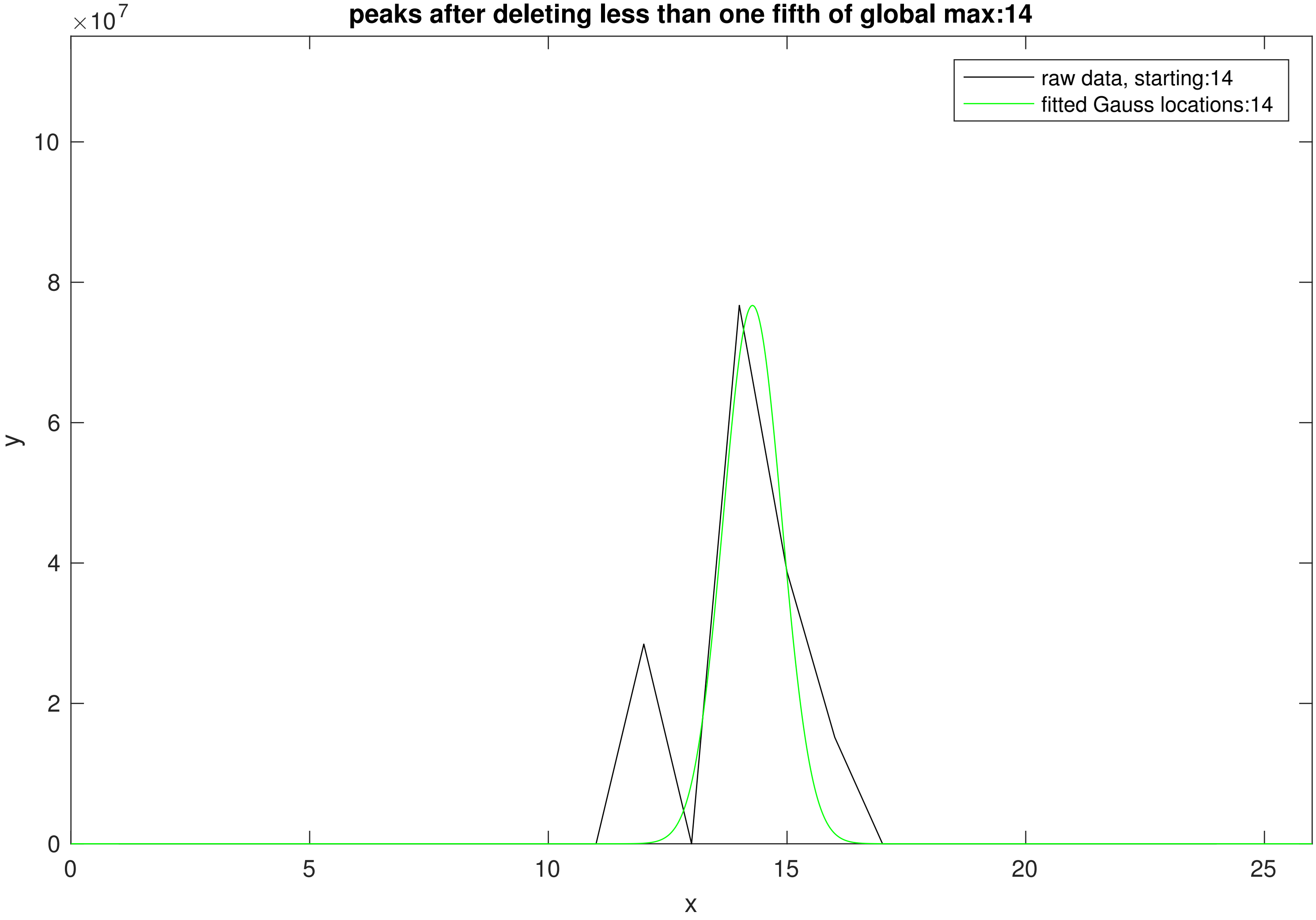

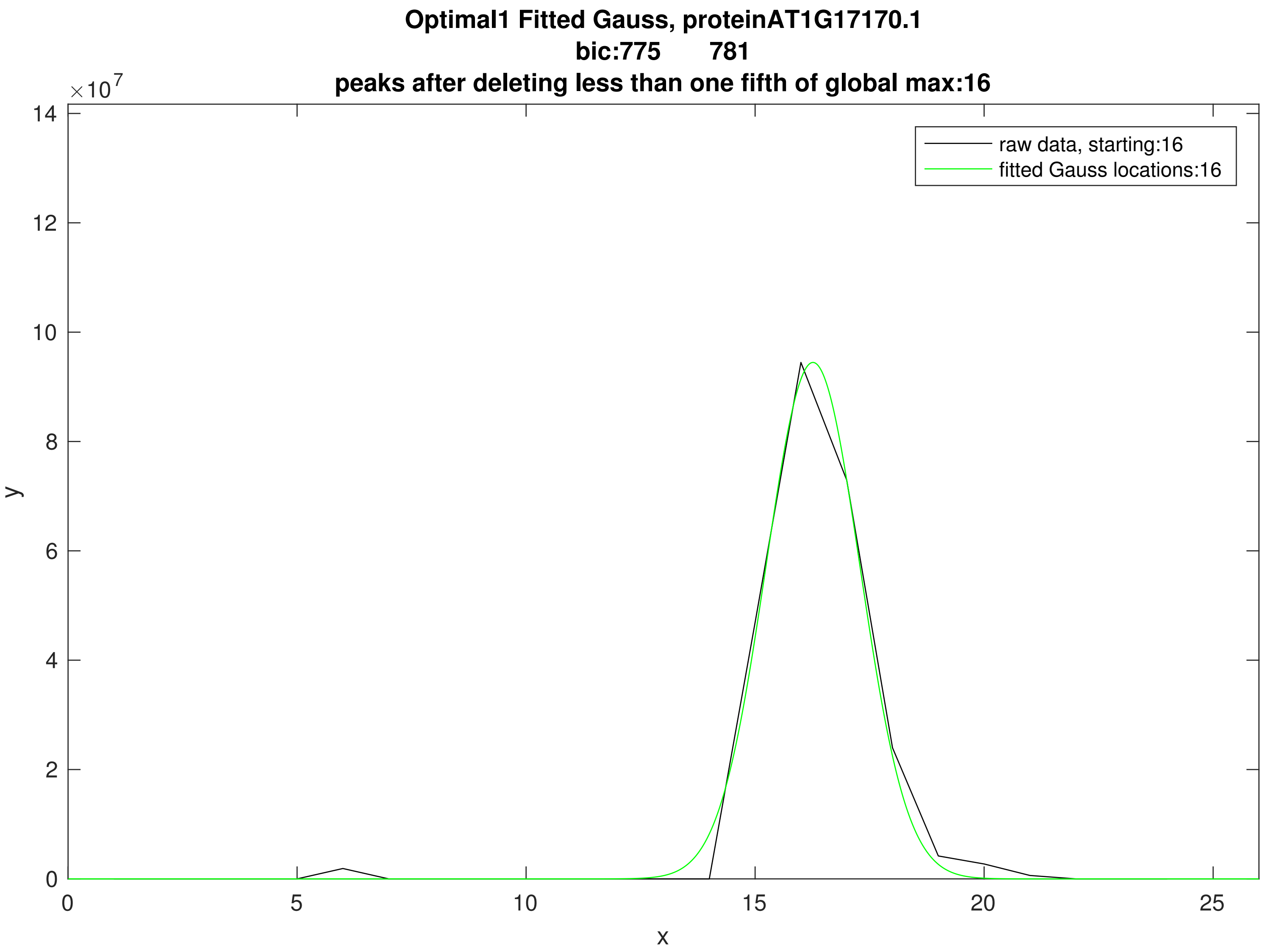

Optimal1 Fitted Gauss, proteinAT1G17170.1
bic:775 781
peaks after deleting less than one fifth of global max:16
×10^7
14
12
10
8
6
4
2
0
y
0
5
10
15
20
25
x
raw data, starting:16
fitted Gauss locations:16

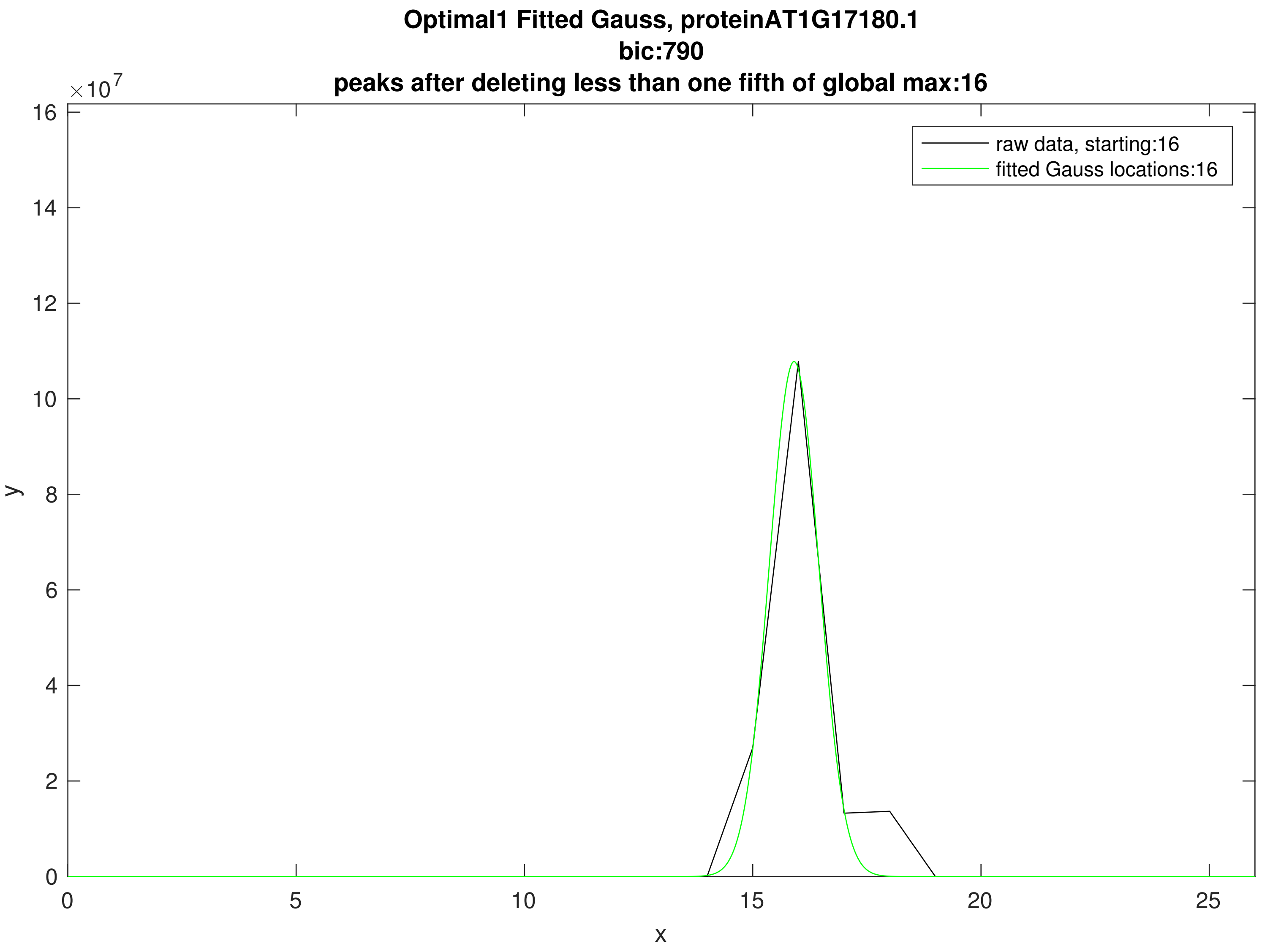
Optimal1 Fitted Gauss, proteinAT1G17180.1
bic:790
peaks after deleting less than one fifth of global max:16
×10^7
16
14
12
10
8
6
4
2
0
y
0
5
10
15
20
25
x
raw data, starting:16
fitted Gauss locations:16

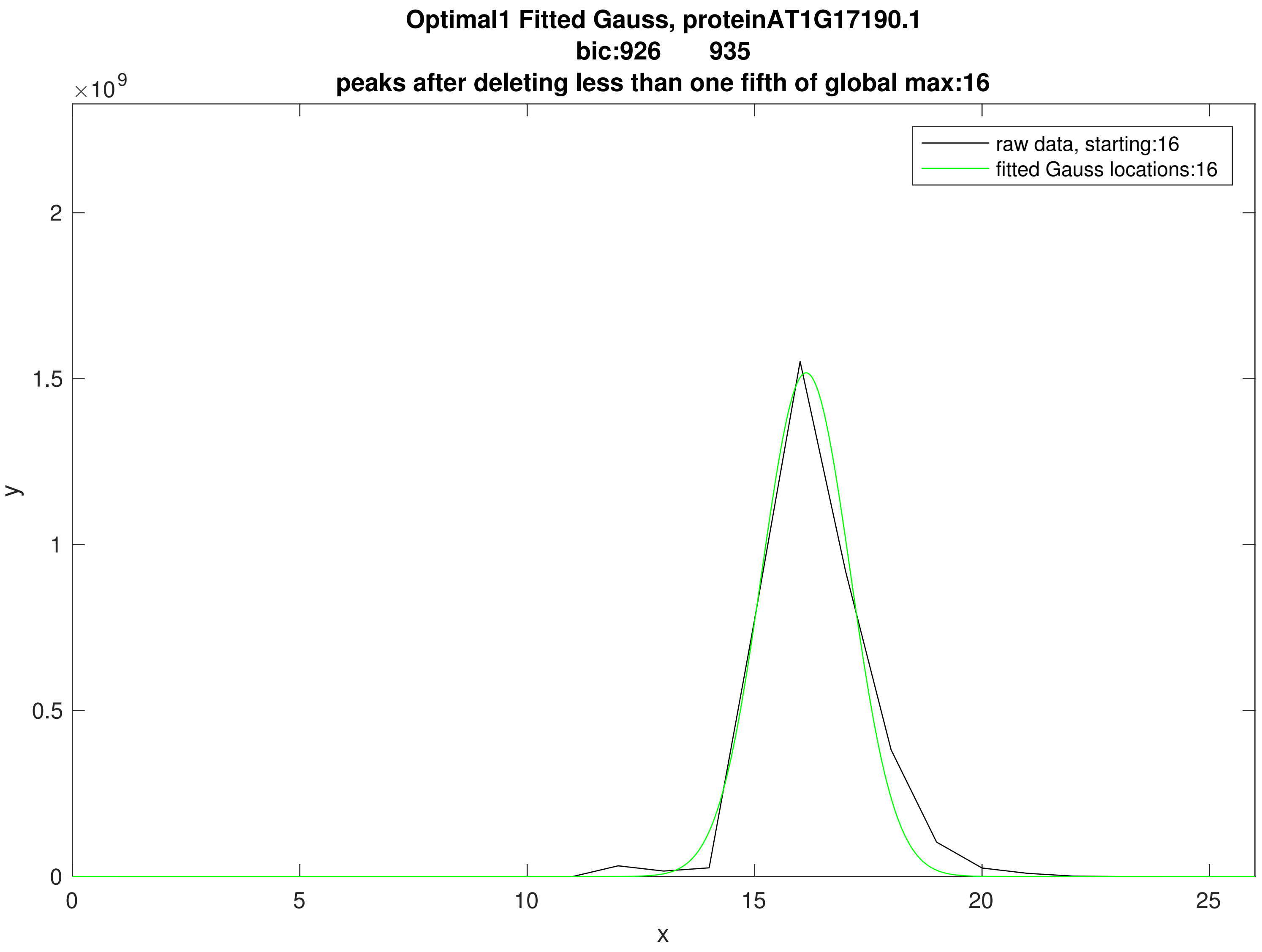
Optimal1 Fitted Gauss, proteinAT1G17190.1
bic:926 935
peaks after deleting less than one fifth of global max:16
×10^9
raw data, starting:16
fitted Gauss locations:16
y
x
0
0.5
1
1.5
2
0
5
10
15
20
25

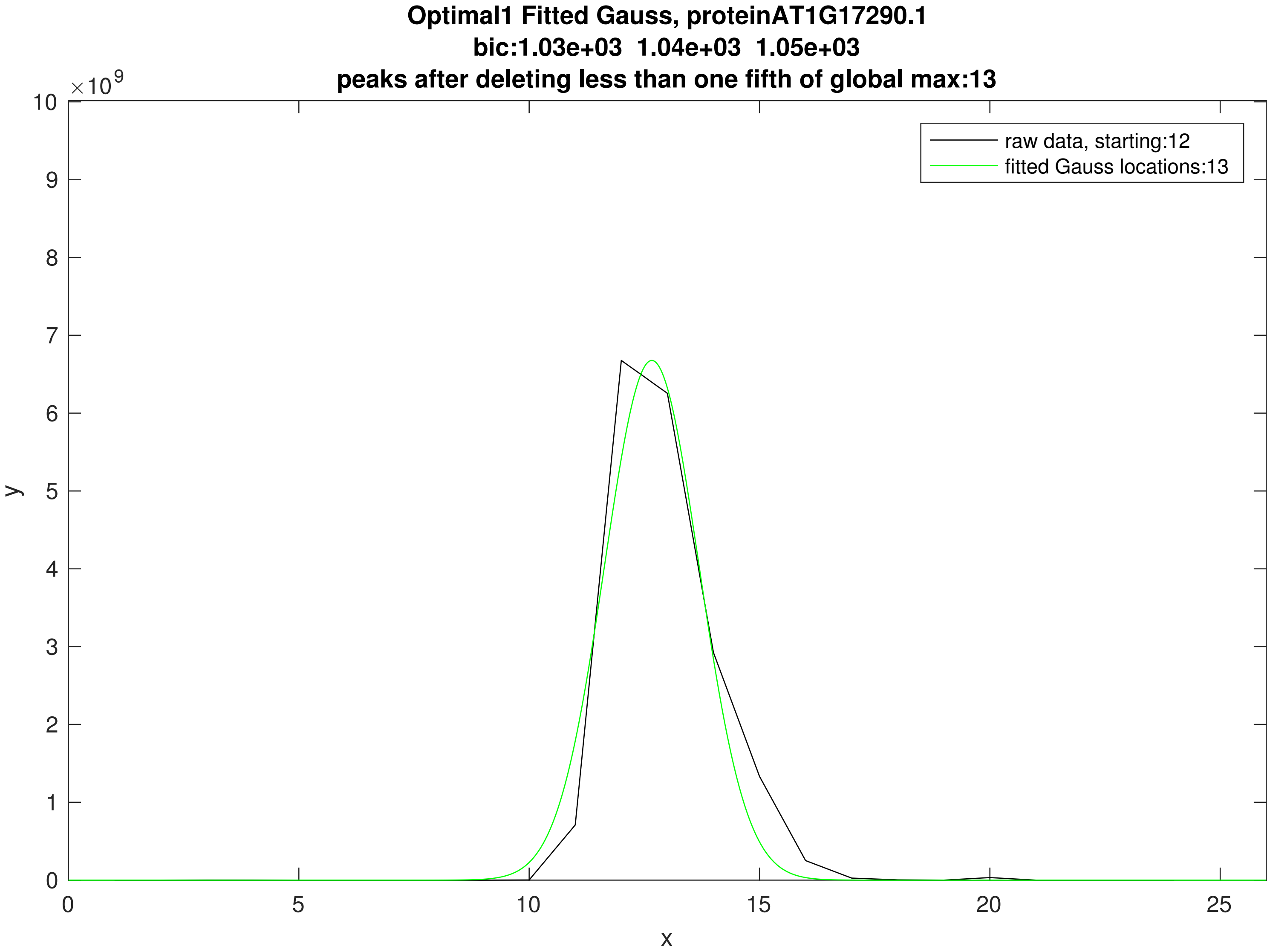

Optimal1 Fitted Gauss, proteinAT1G17290.1
bic:1.03e+03 1.04e+03 1.05e+03
peaks after deleting less than one fifth of global max:13
×10^9
10
9
8
7
6
5
4
3
2
1
0
y
0
5
10
15
20
25
x
raw data, starting:12
fitted Gauss locations:13

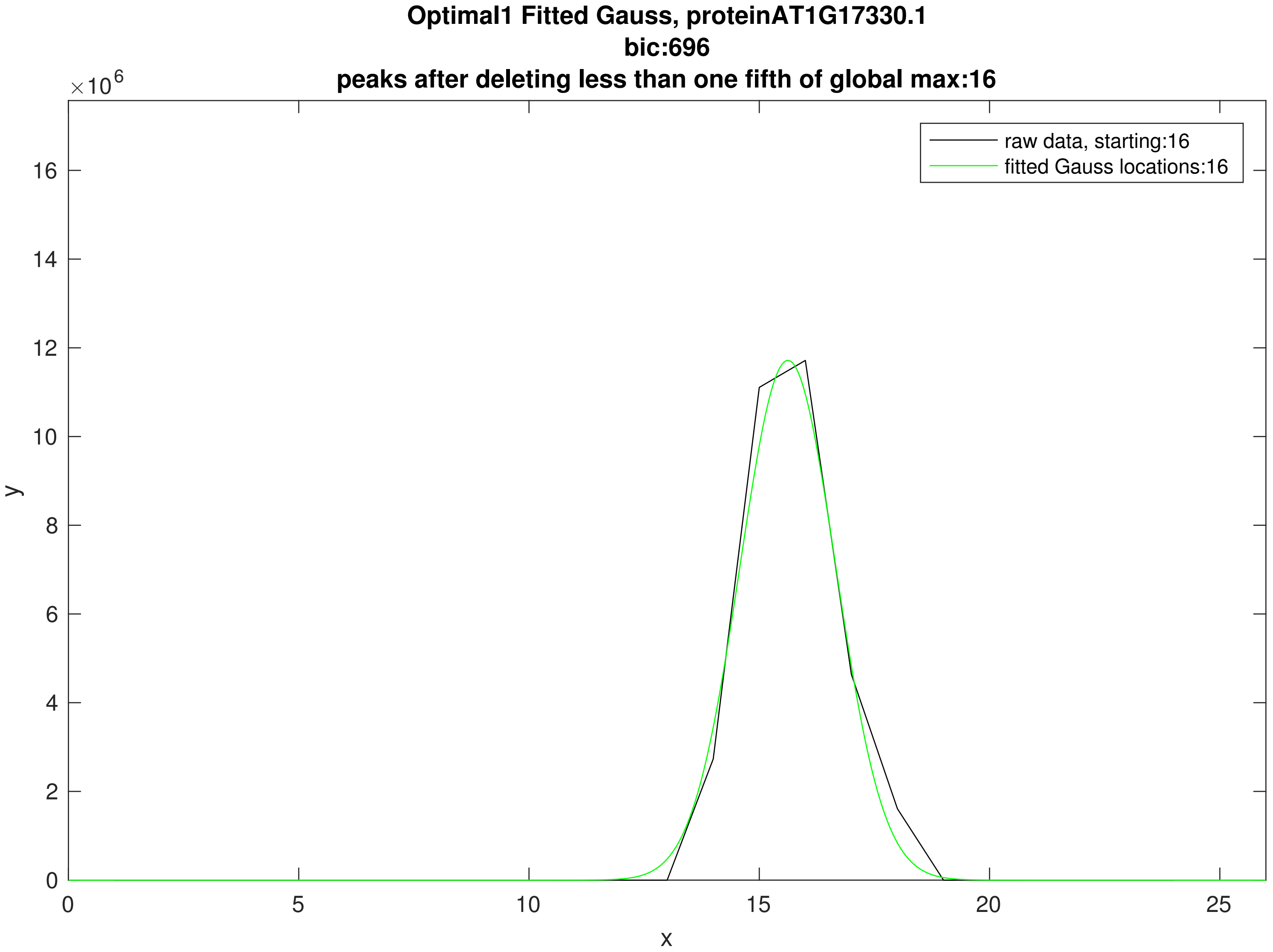

Optimal1 Fitted Gauss, proteinAT1G17330.1
bic:696
peaks after deleting less than one fifth of global max:16
$\times 10^6$
16
14
12
10
8
6
4
2
0
y
0
5
10
15
20
25
x
raw data, starting:16
fitted Gauss locations:16

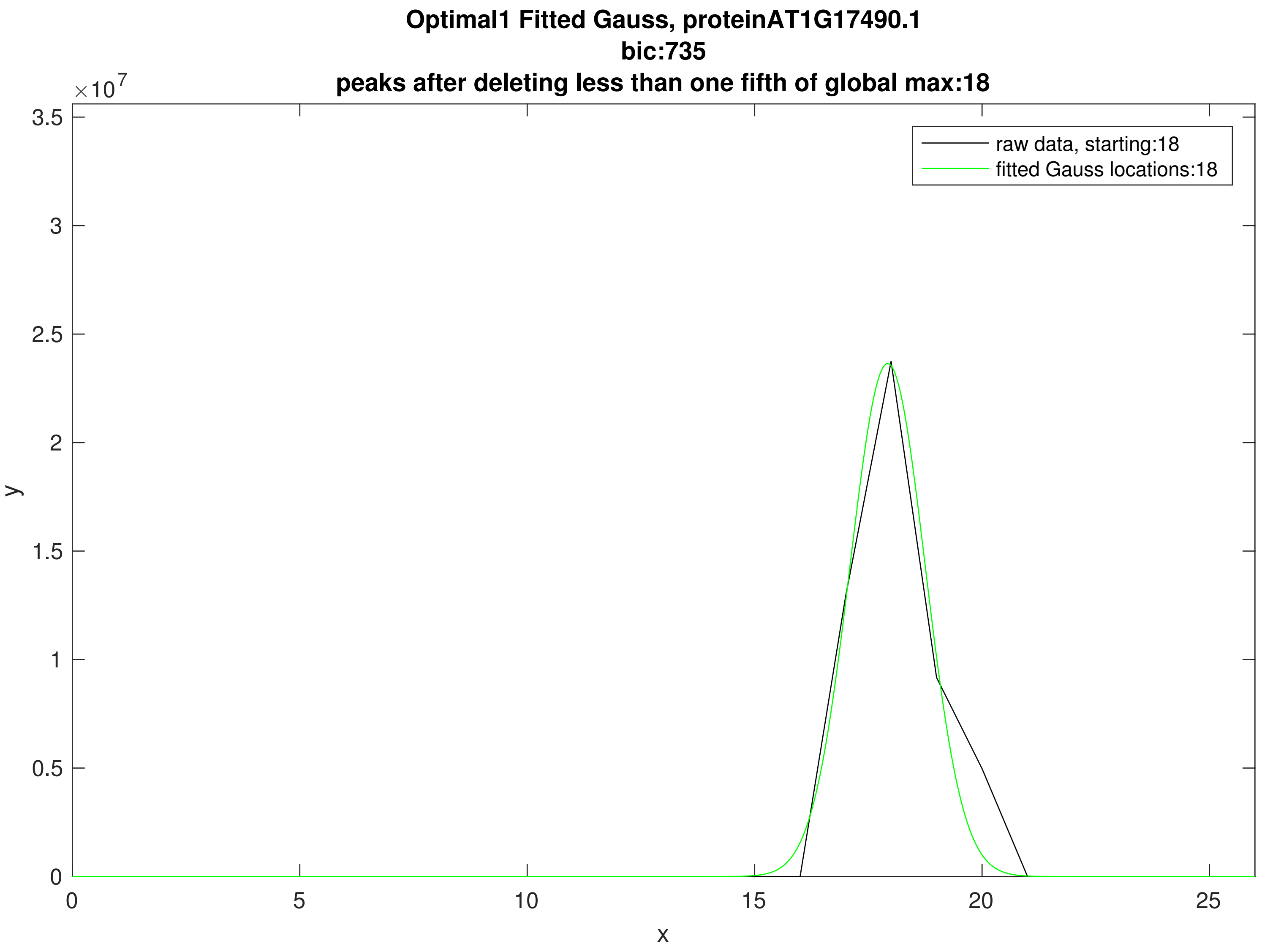
Optimal1 Fitted Gauss, proteinAT1G17490.1
bic:735
peaks after deleting less than one fifth of global max:18
×10^7
3.5
3
2.5
2
1.5
1
0.5
0
y
0
5
10
15
20
25
x
raw data, starting:18
fitted Gauss locations:18

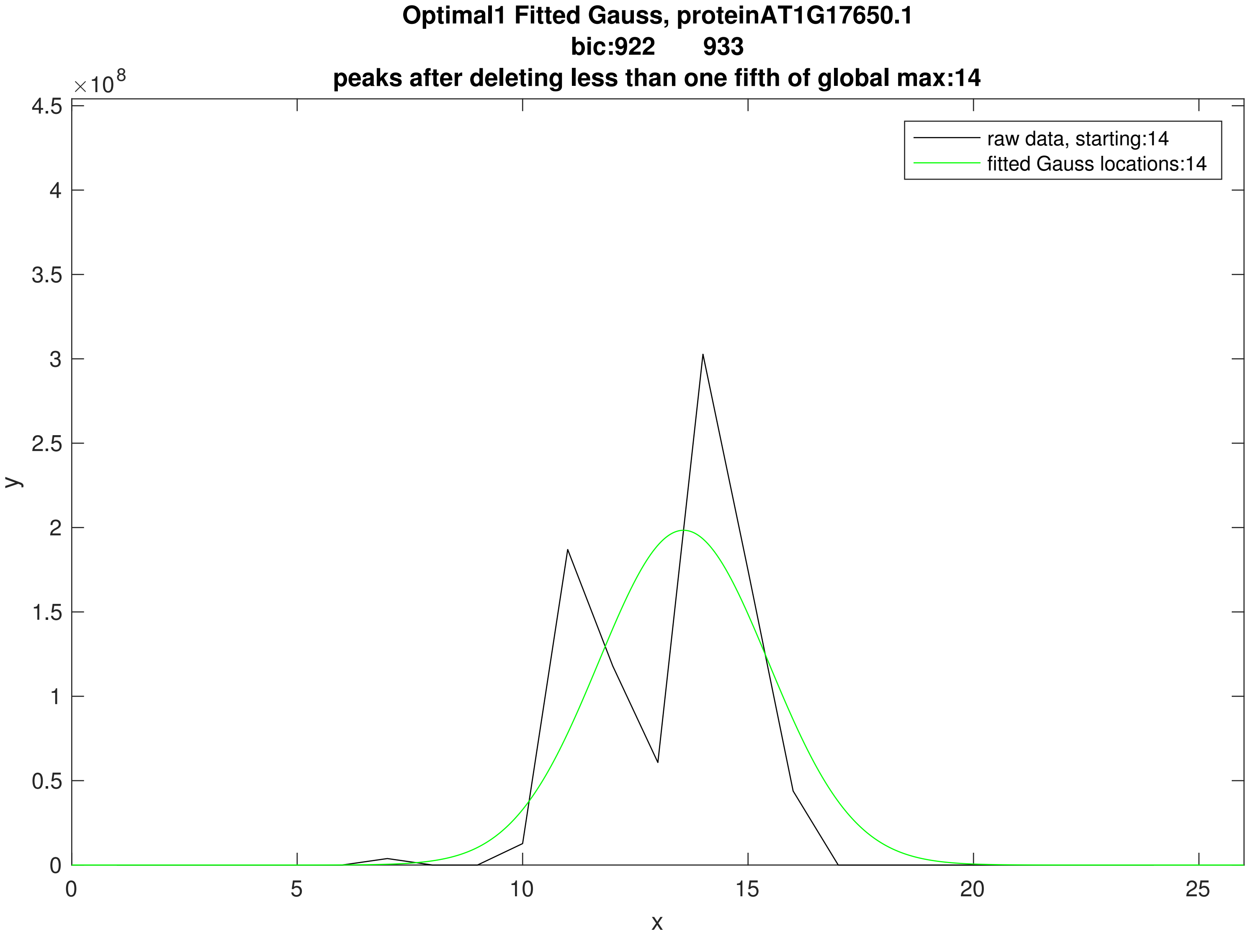
Optimal1 Fitted Gauss, proteinAT1G17650.1
bic:922 933
peaks after deleting less than one fifth of global max:14
×10^8
4.5
4
3.5
3
2.5
2
1.5
1
0.5
0
y
0
5
10
15
20
25
x
raw data, starting:14
fitted Gauss locations:14

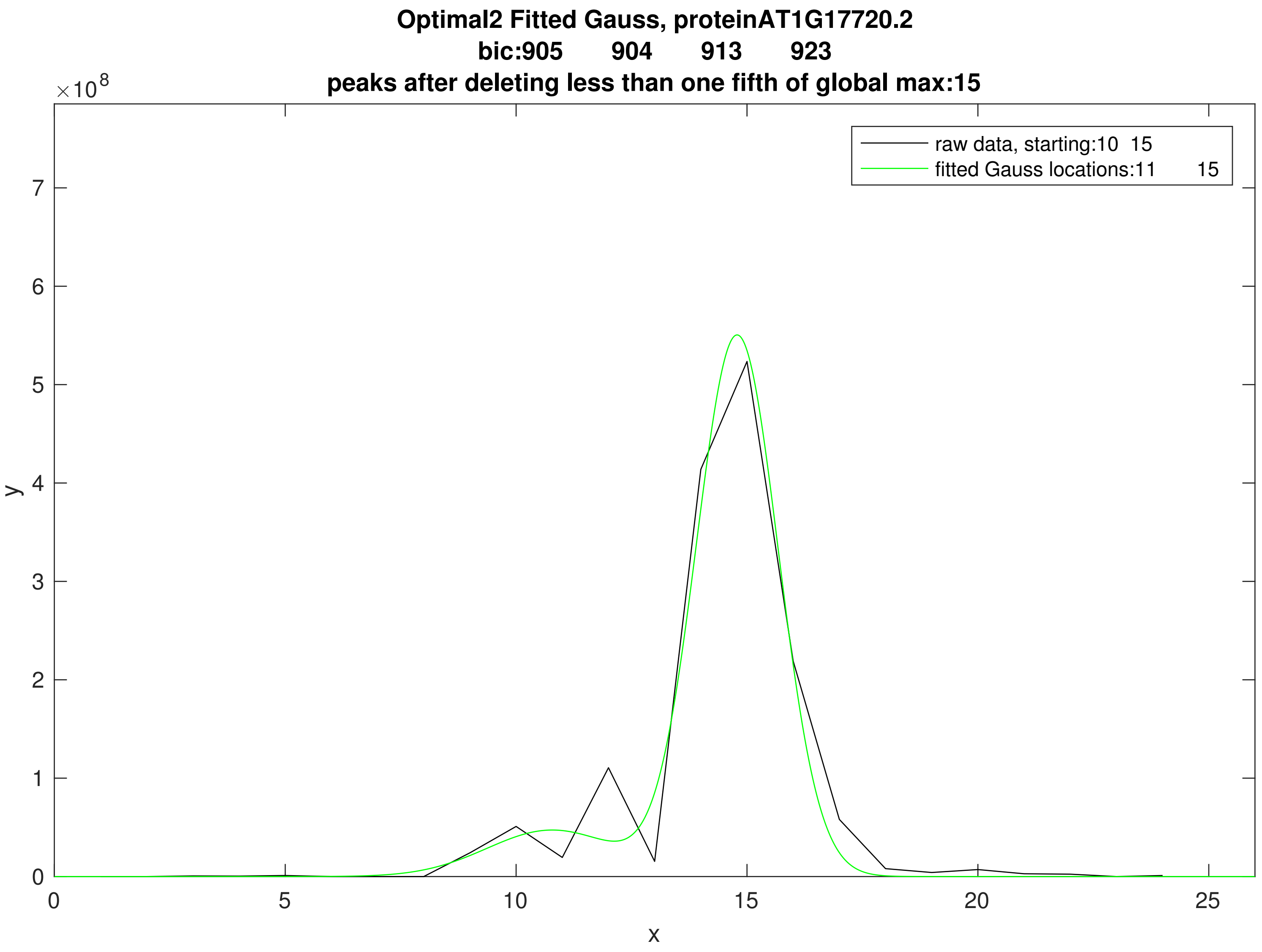
Optimal2 Fitted Gauss, proteinAT1G17720.2
bic:905 904 913 923
peaks after deleting less than one fifth of global max:15
raw data, starting:10 15
fitted Gauss locations:11 15
×10^8
y
x
0
1
2
3
4
5
6
7
0
5
10
15
20
25

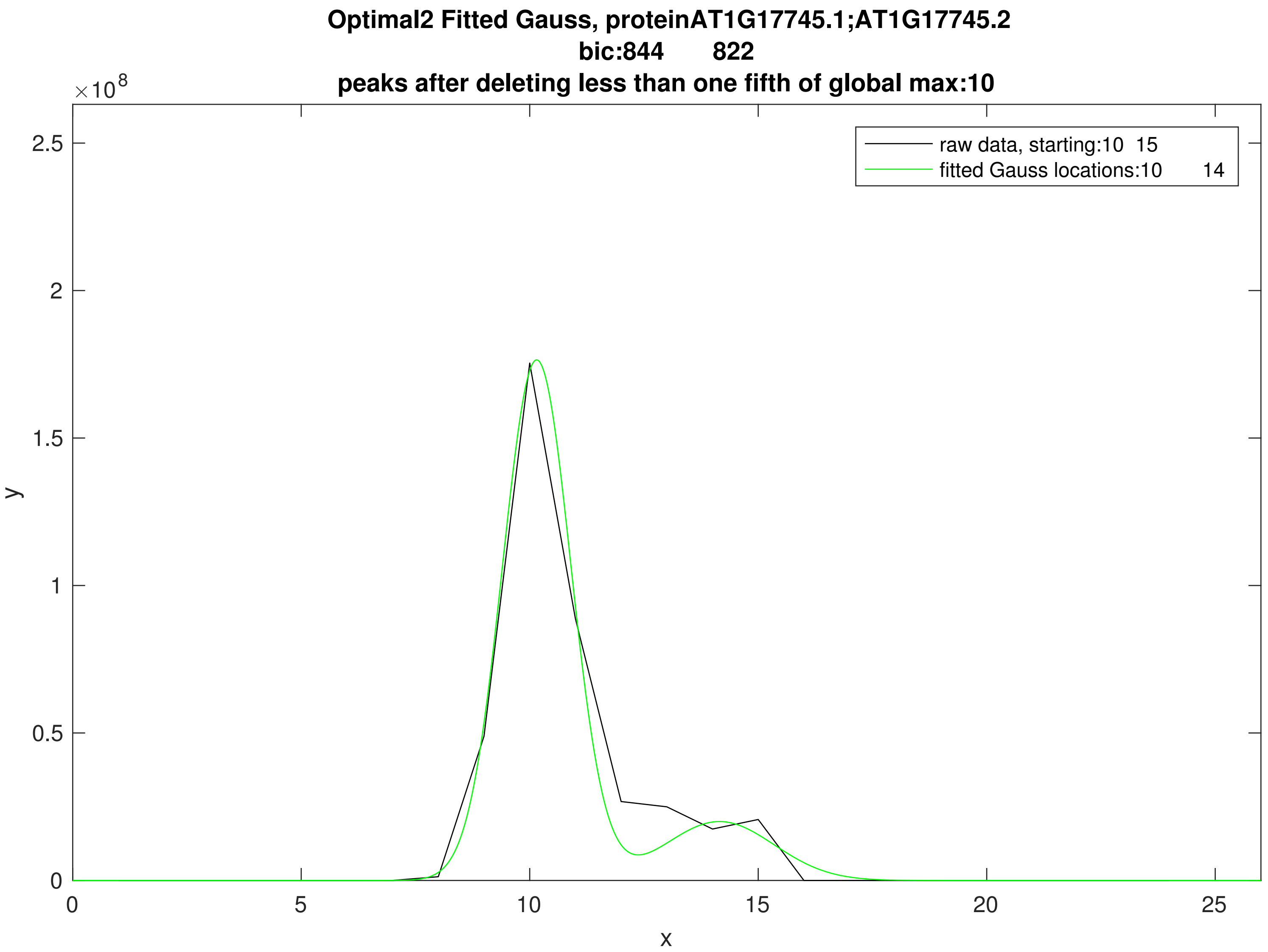
Optimal2 Fitted Gauss, proteinAT1G17745.1;AT1G17745.2
bic:844 822
peaks after deleting less than one fifth of global max:10
raw data, starting:10 15
fitted Gauss locations:10 14
×10^8
2.5
2
1.5
1
0.5
0
y
0
5
10
15
20
25
x

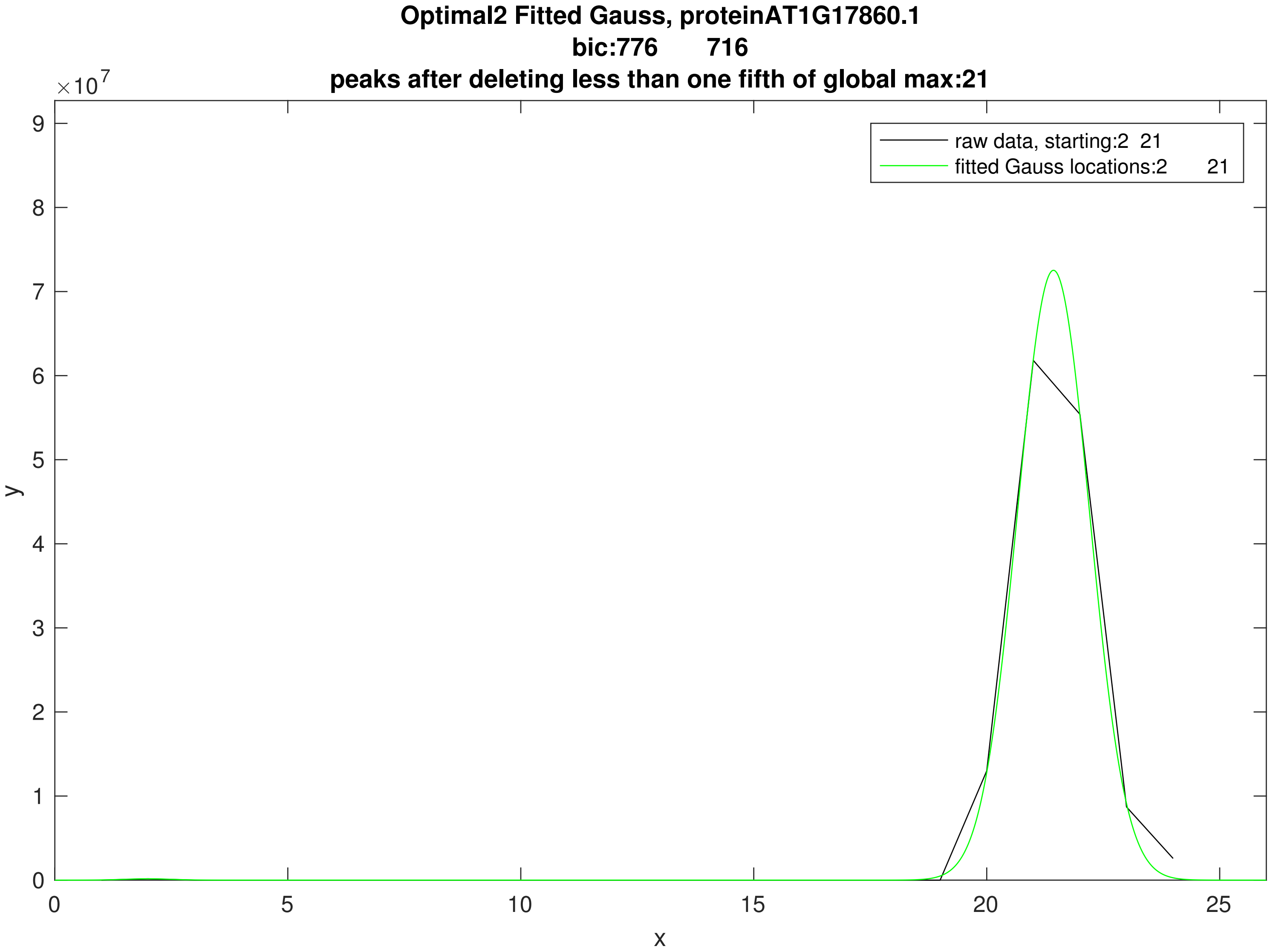
Optimal2 Fitted Gauss, proteinAT1G17860.1
bic:776 716
peaks after deleting less than one fifth of global max:21
×10^7
raw data, starting:2 21
fitted Gauss locations:2 21
y
x
0
1
2
3
4
5
6
7
8
9
0
5
10
15
20
25

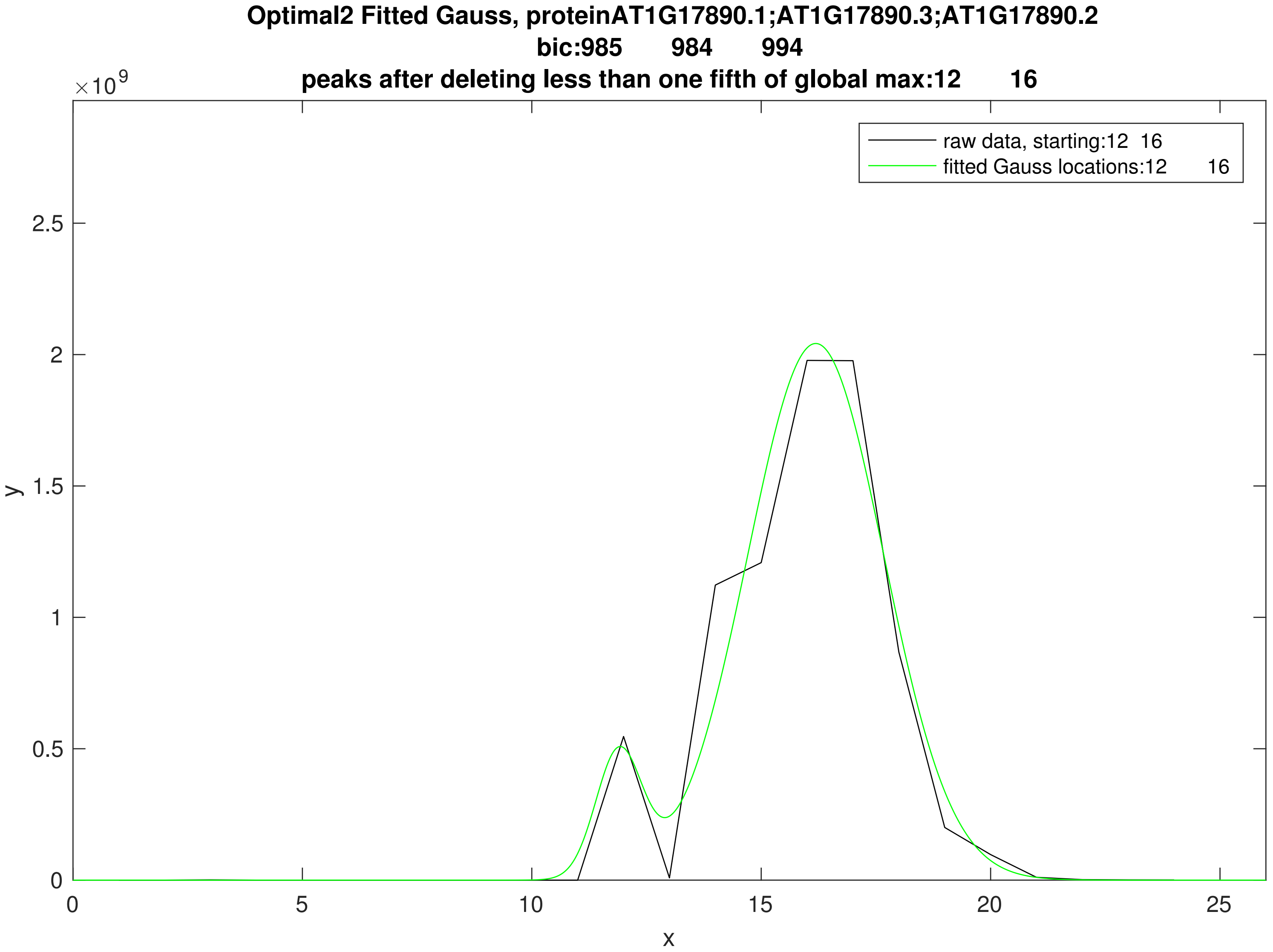
Optimal2 Fitted Gauss, proteinAT1G17890.1;AT1G17890.3;AT1G17890.2
bic:985 984 994
peaks after deleting less than one fifth of global max:12 16
raw data, starting:12 16
fitted Gauss locations:12 16
×10^9
y
x

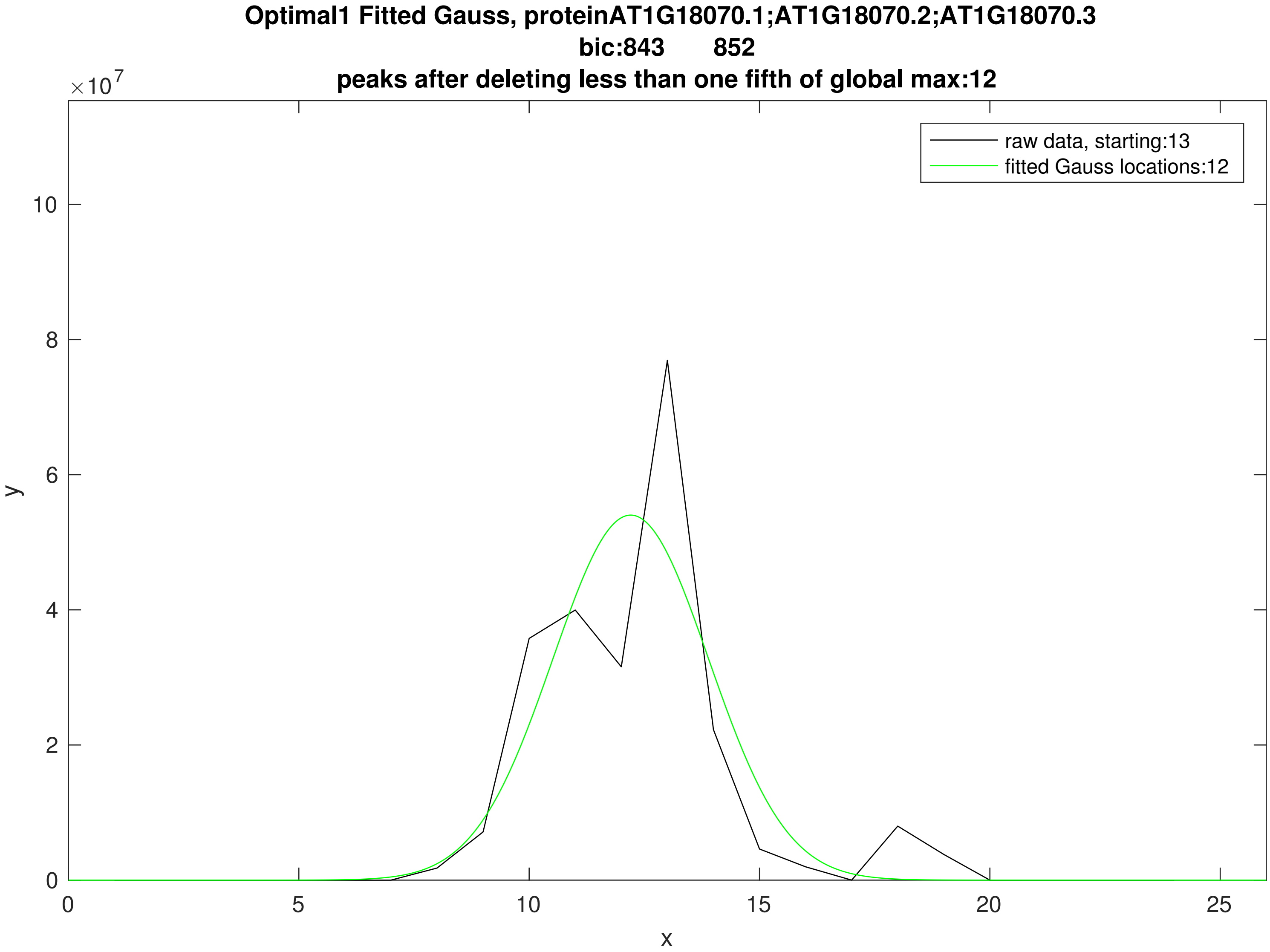

Optimal1 Fitted Gauss, proteinAT1G18070.1;AT1G18070.2;AT1G18070.3
bic:843 852
peaks after deleting less than one fifth of global max:12
×10^7
raw data, starting:13
fitted Gauss locations:12
10
8
6
4
2
0
y
0
5
10
15
20
25
x

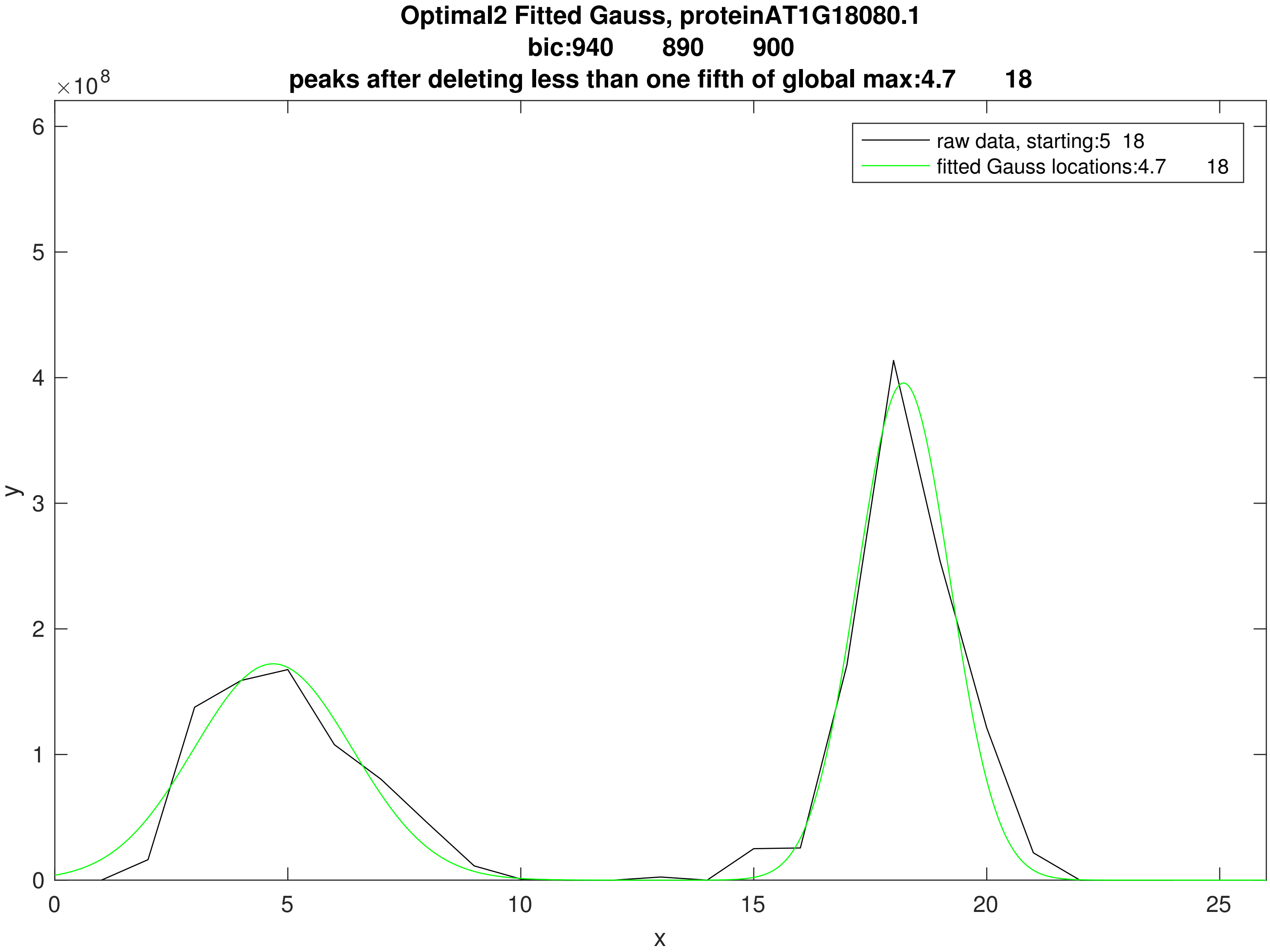
Optimal2 Fitted Gauss, proteinAT1G18080.1
bic:940 890 900
peaks after deleting less than one fifth of global max:4.7 18
×10^8
raw data, starting:5 18
fitted Gauss locations:4.7 18
y
x
0
1
2
3
4
5
6
0
5
10
15
20
25

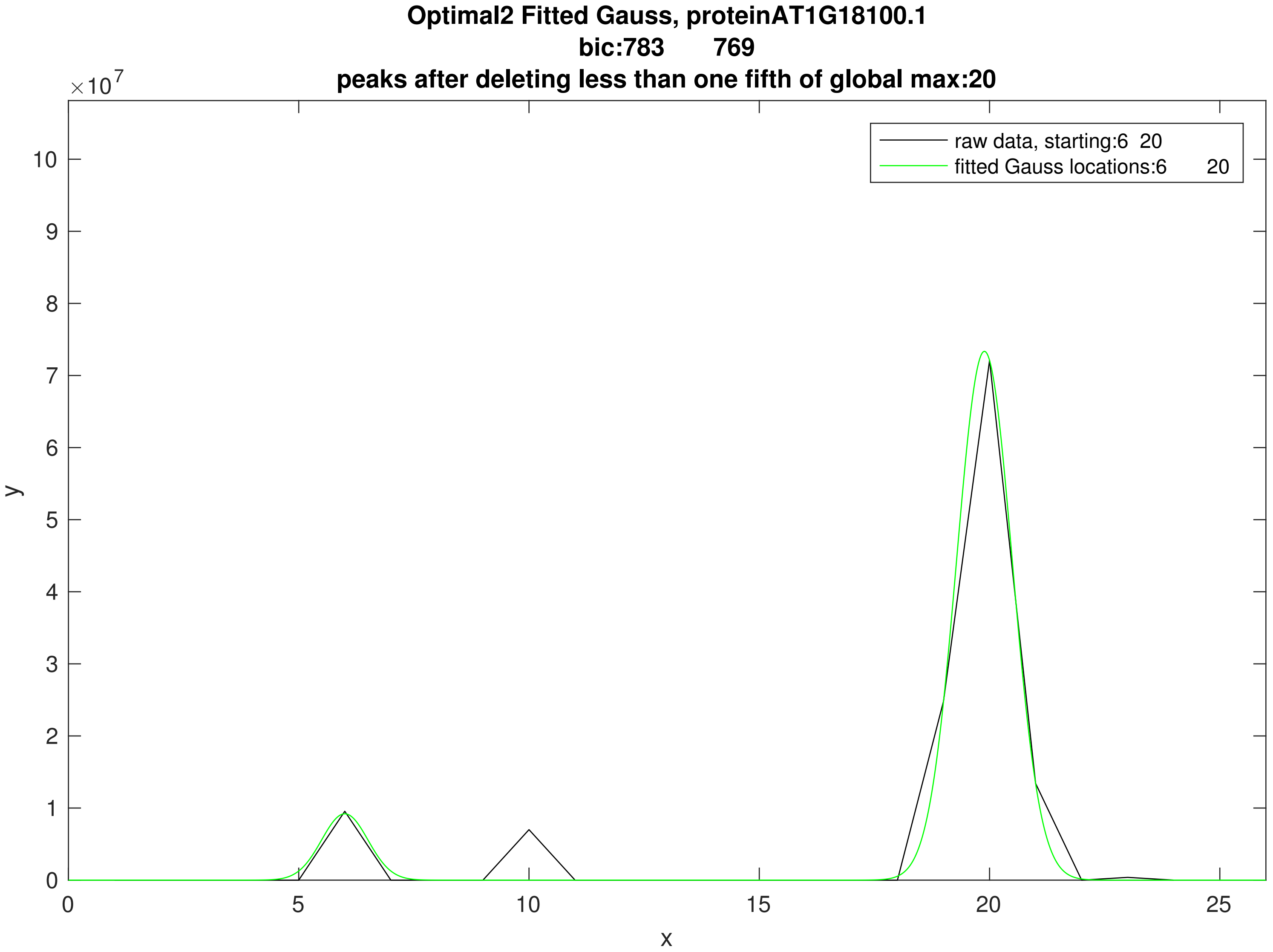

Optimal2 Fitted Gauss, proteinAT1G18100.1
bic:783 769
peaks after deleting less than one fifth of global max:20
×10^7
raw data, starting:6 20
fitted Gauss locations:6 20
y
x
0
1
2
3
4
5
6
7
8
9
10
0
5
10
15
20
25

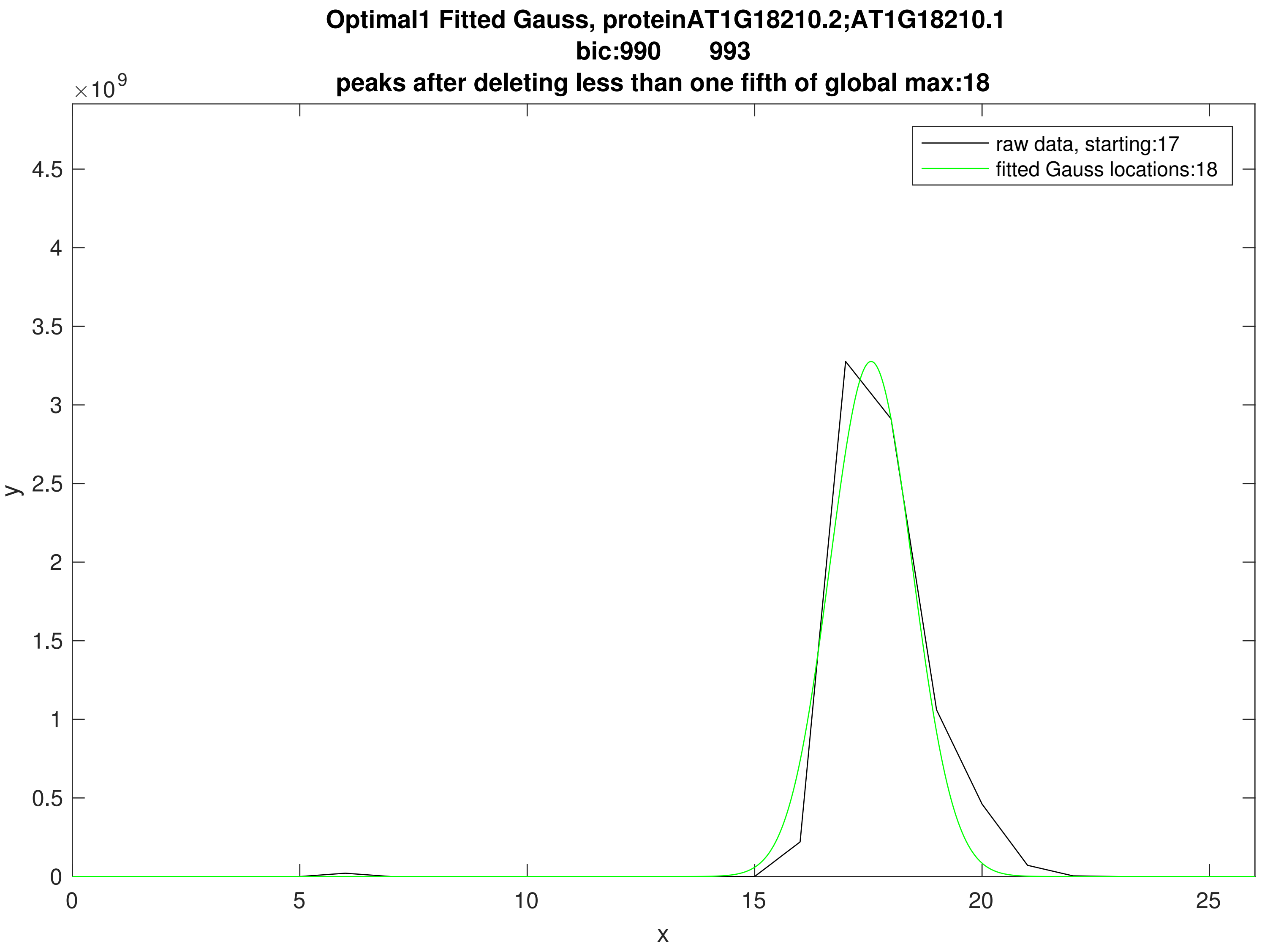

Optimal1 Fitted Gauss, proteinAT1G18210.2;AT1G18210.1
bic:990 993
peaks after deleting less than one fifth of global max:18
×10^9
raw data, starting:17
fitted Gauss locations:18
y
x
0
0.5
1
1.5
2
2.5
3
3.5
4
4.5
5
10
15
20
25

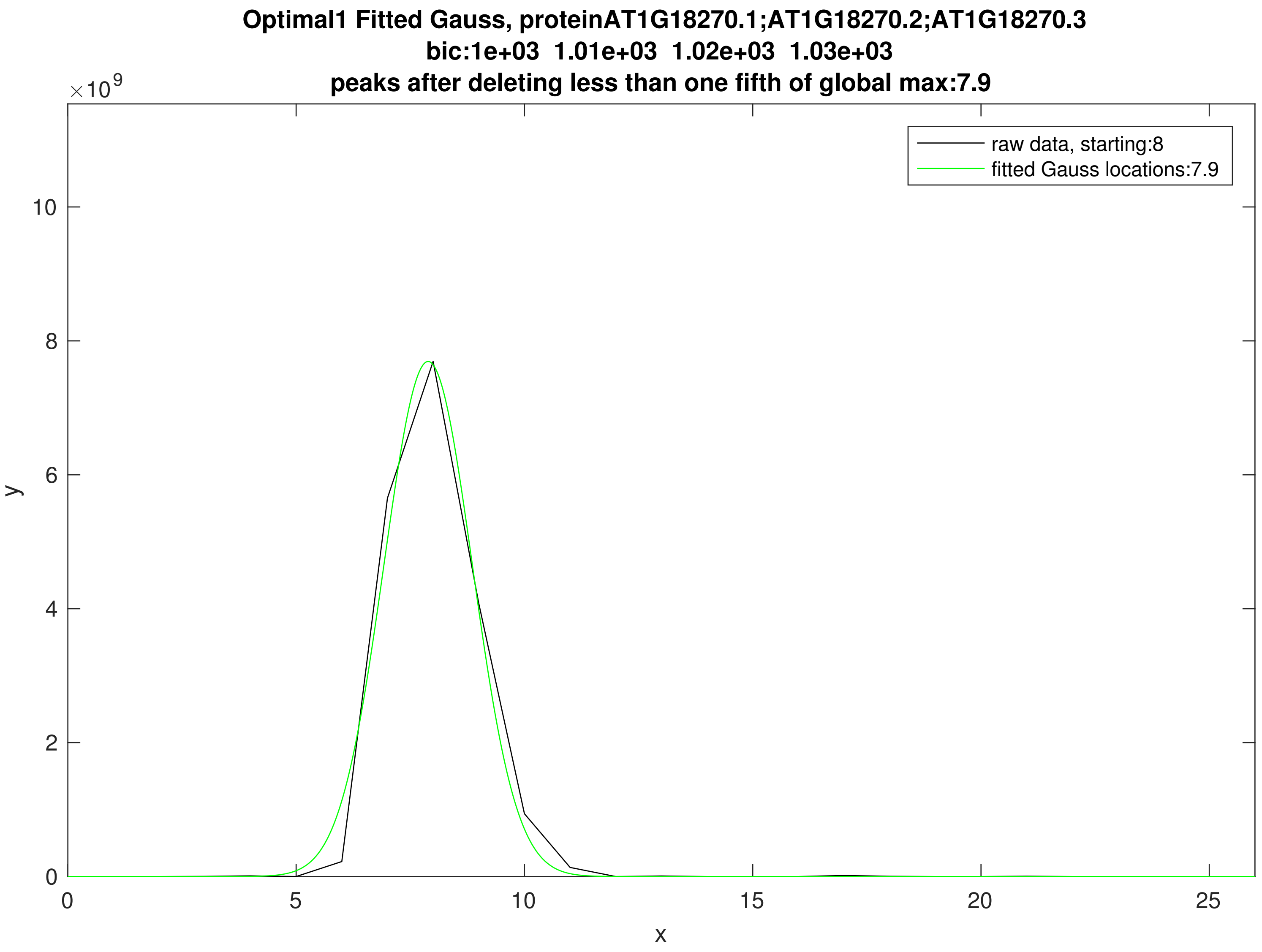

Optimal1 Fitted Gauss, proteinAT1G18270.1;AT1G18270.2;AT1G18270.3
bic:1e+03 1.01e+03 1.02e+03 1.03e+03
peaks after deleting less than one fifth of global max:7.9
×10^9
raw data, starting:8
fitted Gauss locations:7.9
10
8
6
4
2
0
y
0
5
10
15
20
25
x

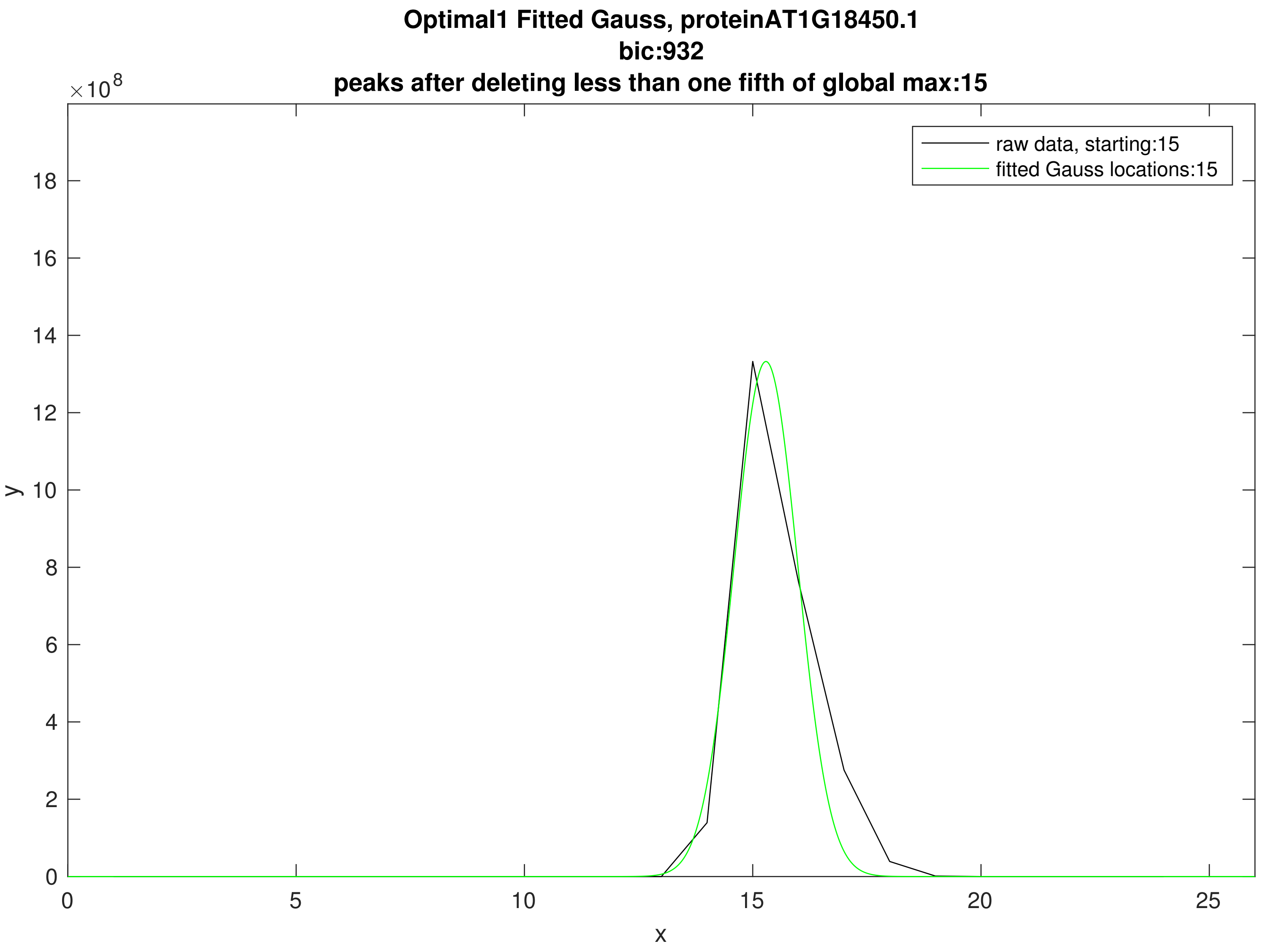

Optimal1 Fitted Gauss, proteinAT1G18450.1
bic:932
peaks after deleting less than one fifth of global max:15
×10^8
raw data, starting:15
fitted Gauss locations:15
y
x
0
2
4
6
8
10
12
14
16
18
0
5
10
15
20
25

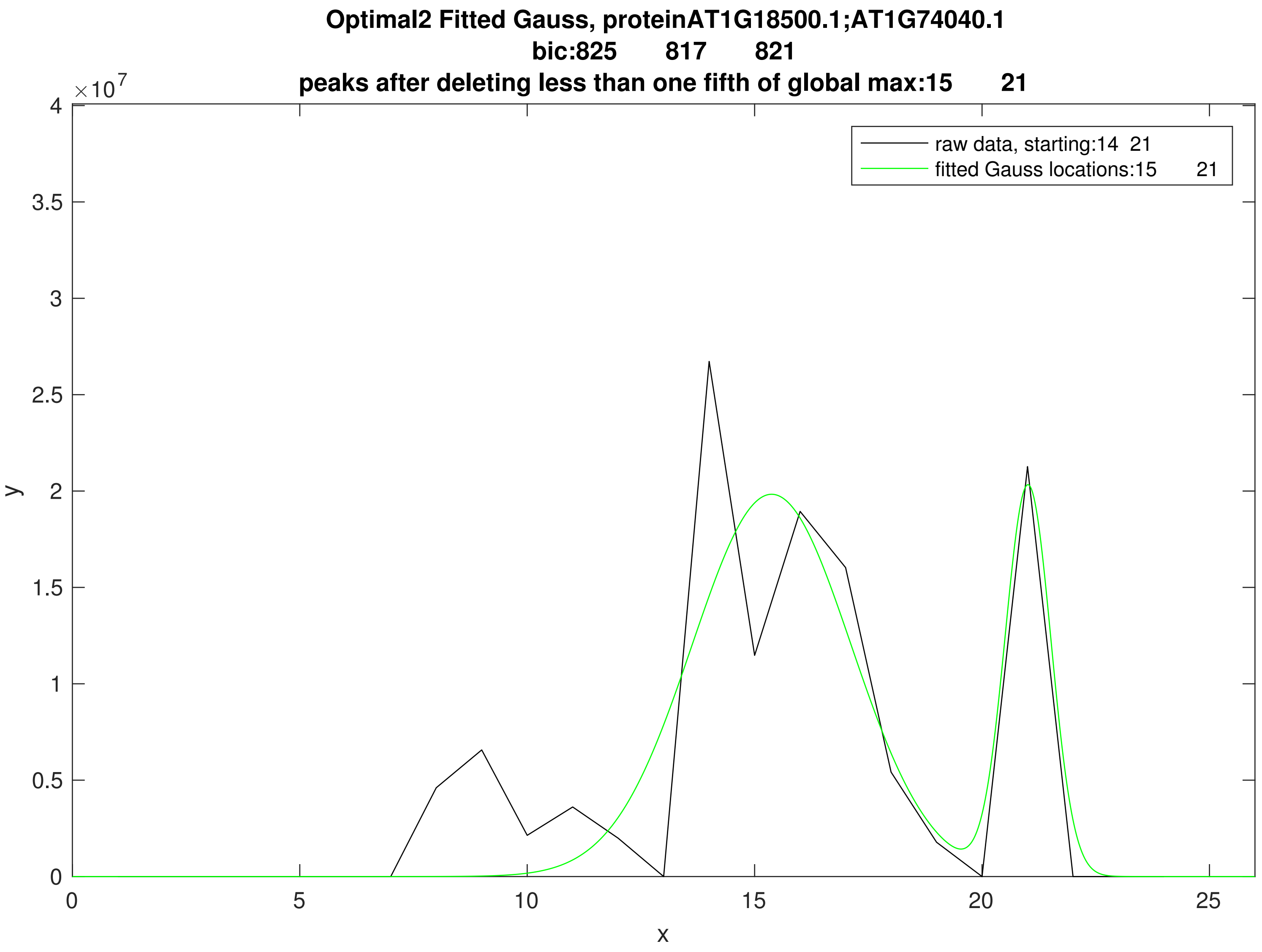
Optimal2 Fitted Gauss, proteinAT1G18500.1;AT1G74040.1
bic:825 817 821
peaks after deleting less than one fifth of global max:15 21
×10^7
raw data, starting:14 21
fitted Gauss locations:15 21
y
x
0
0.5
1
1.5
2
2.5
3
3.5
4
5
10
15
20
25

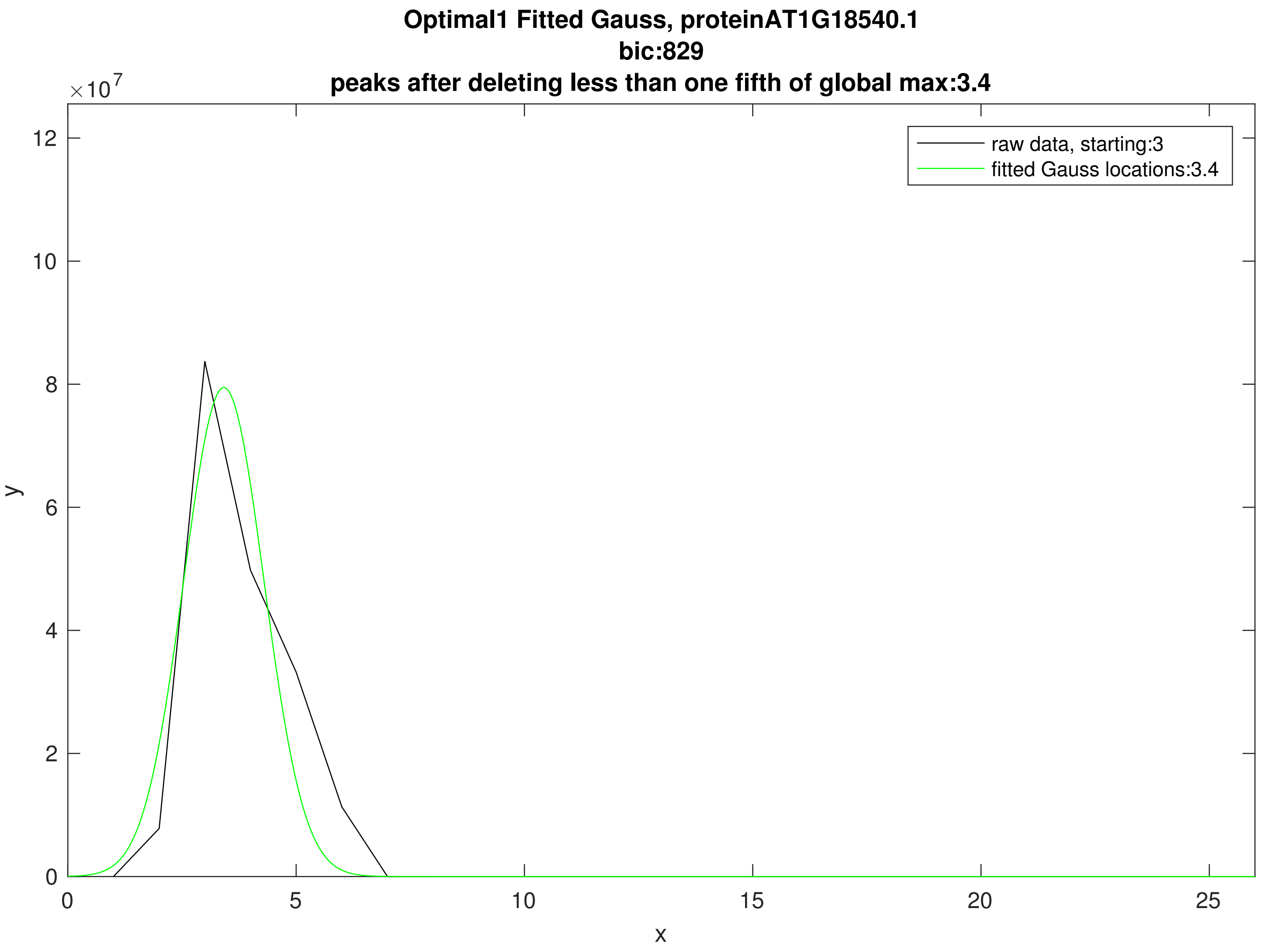

Optimal1 Fitted Gauss, proteinAT1G18540.1
bic:829
peaks after deleting less than one fifth of global max:3.4
×10^7
12
10
8
6
4
2
0
y
0
5
10
15
20
25
x
raw data, starting:3
fitted Gauss locations:3.4

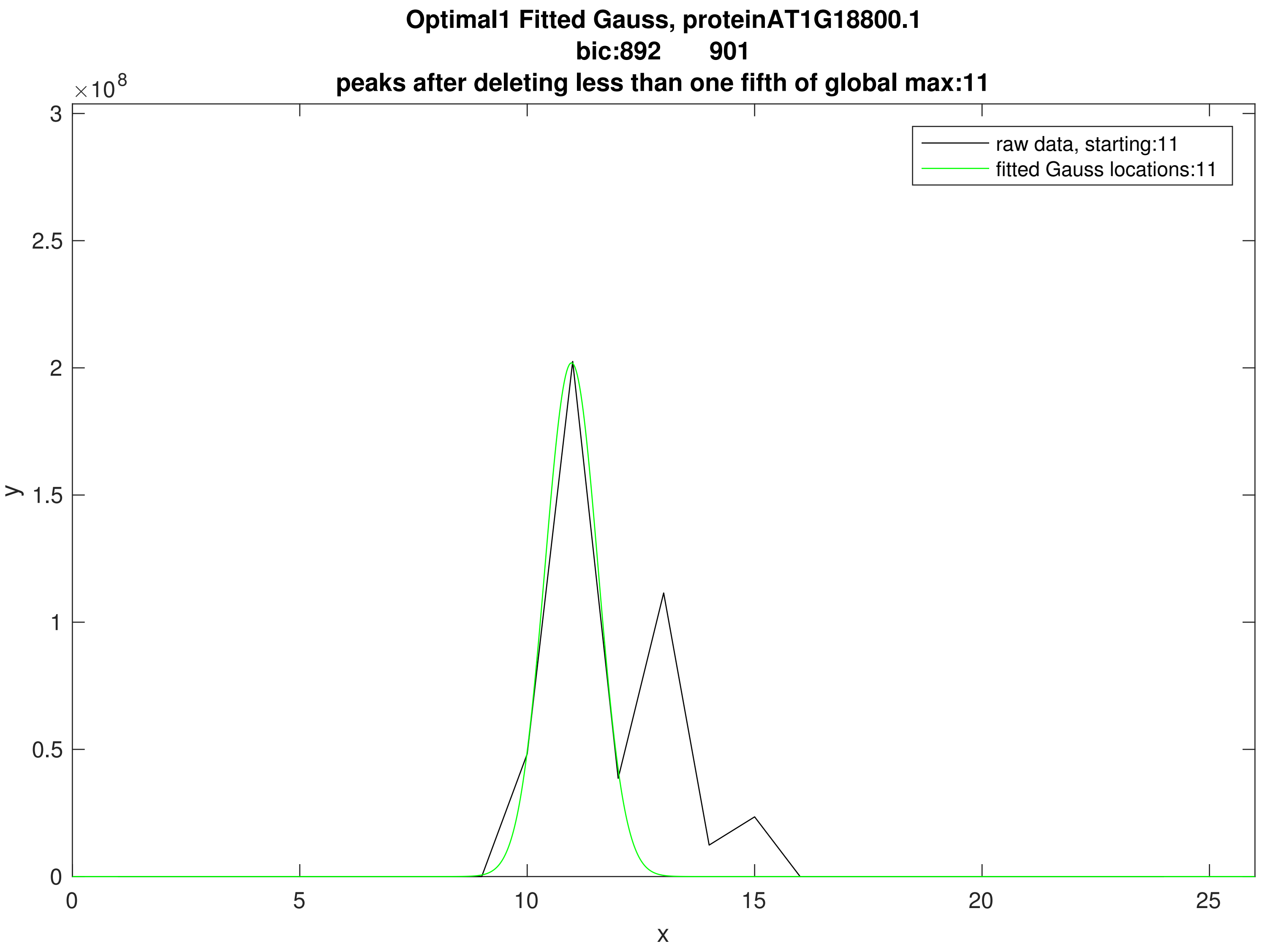
Optimal1 Fitted Gauss, proteinAT1G18800.1
bic:892 901
peaks after deleting less than one fifth of global max:11
×10^8
3
2.5
2
1.5
1
0.5
0
y
0
5
10
15
20
25
x
raw data, starting:11
fitted Gauss locations:11

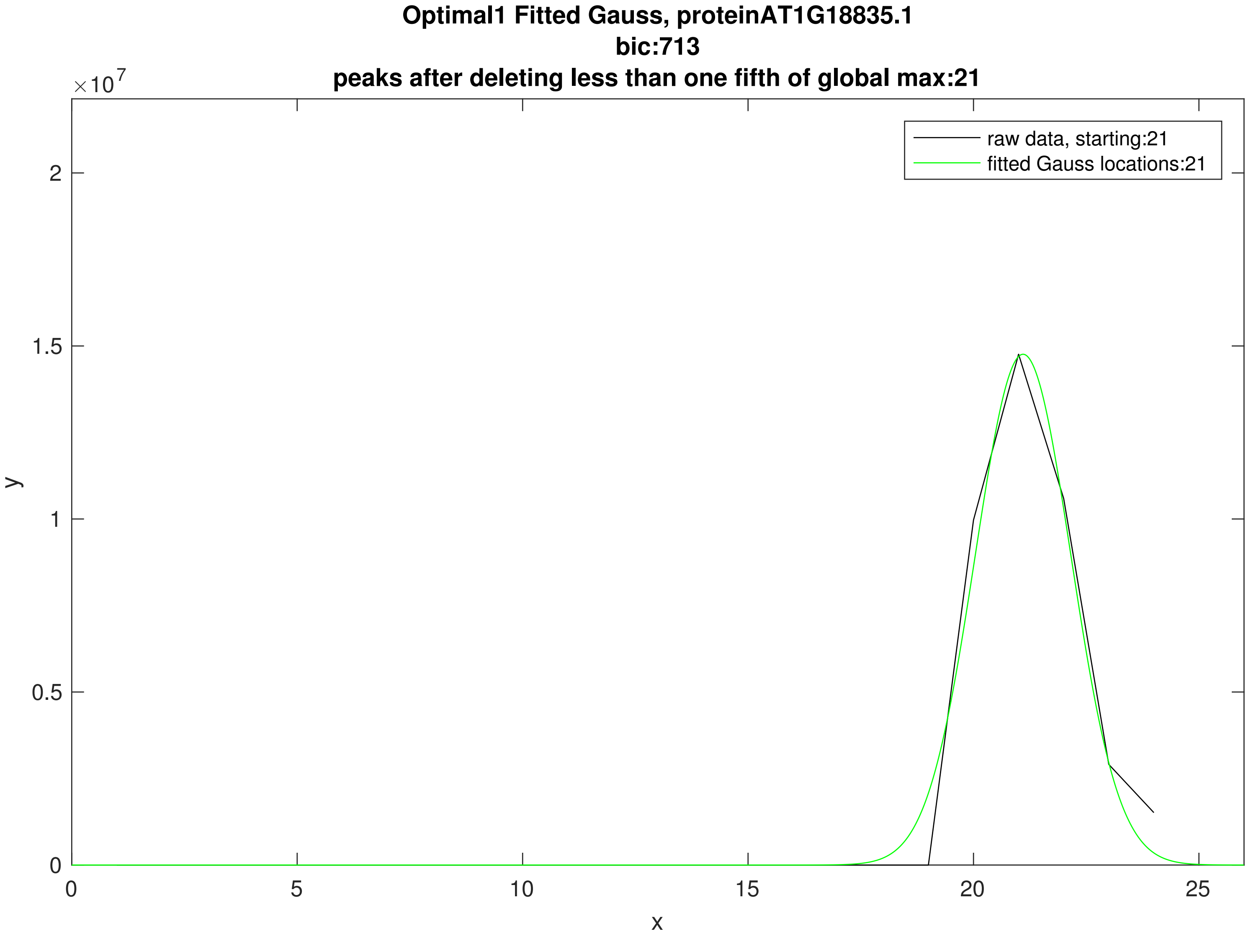

Optimal1 Fitted Gauss, proteinAT1G18835.1
bic:713
peaks after deleting less than one fifth of global max:21
×10^7
raw data, starting:21
fitted Gauss locations:21
y
x

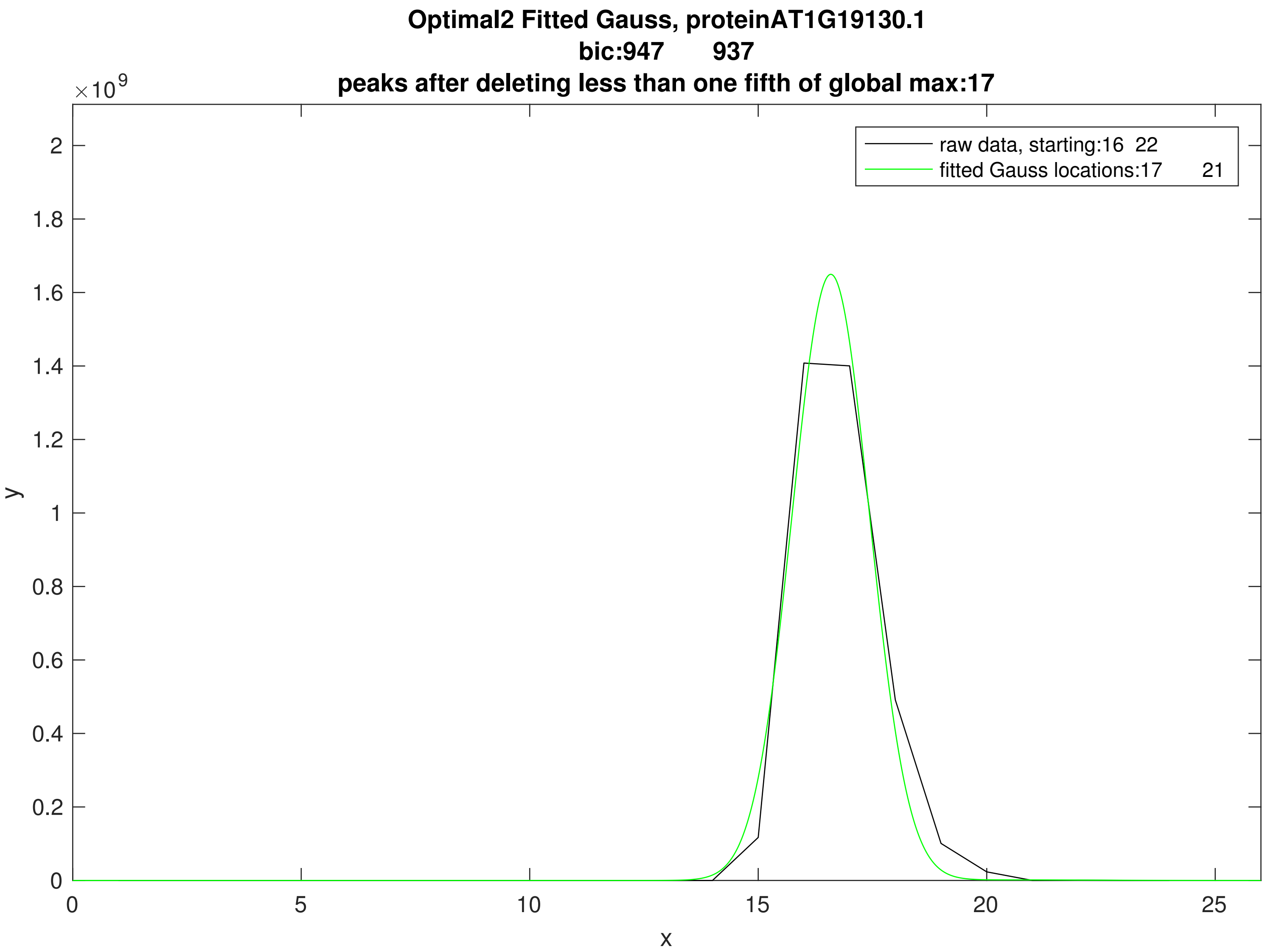
Optimal2 Fitted Gauss, proteinAT1G19130.1
bic:947 937
peaks after deleting less than one fifth of global max:17
×10^9
raw data, starting:16 22
fitted Gauss locations:17 21
y
x
0
0.2
0.4
0.6
0.8
1
1.2
1.4
1.6
1.8
2
0
5
10
15
20
25

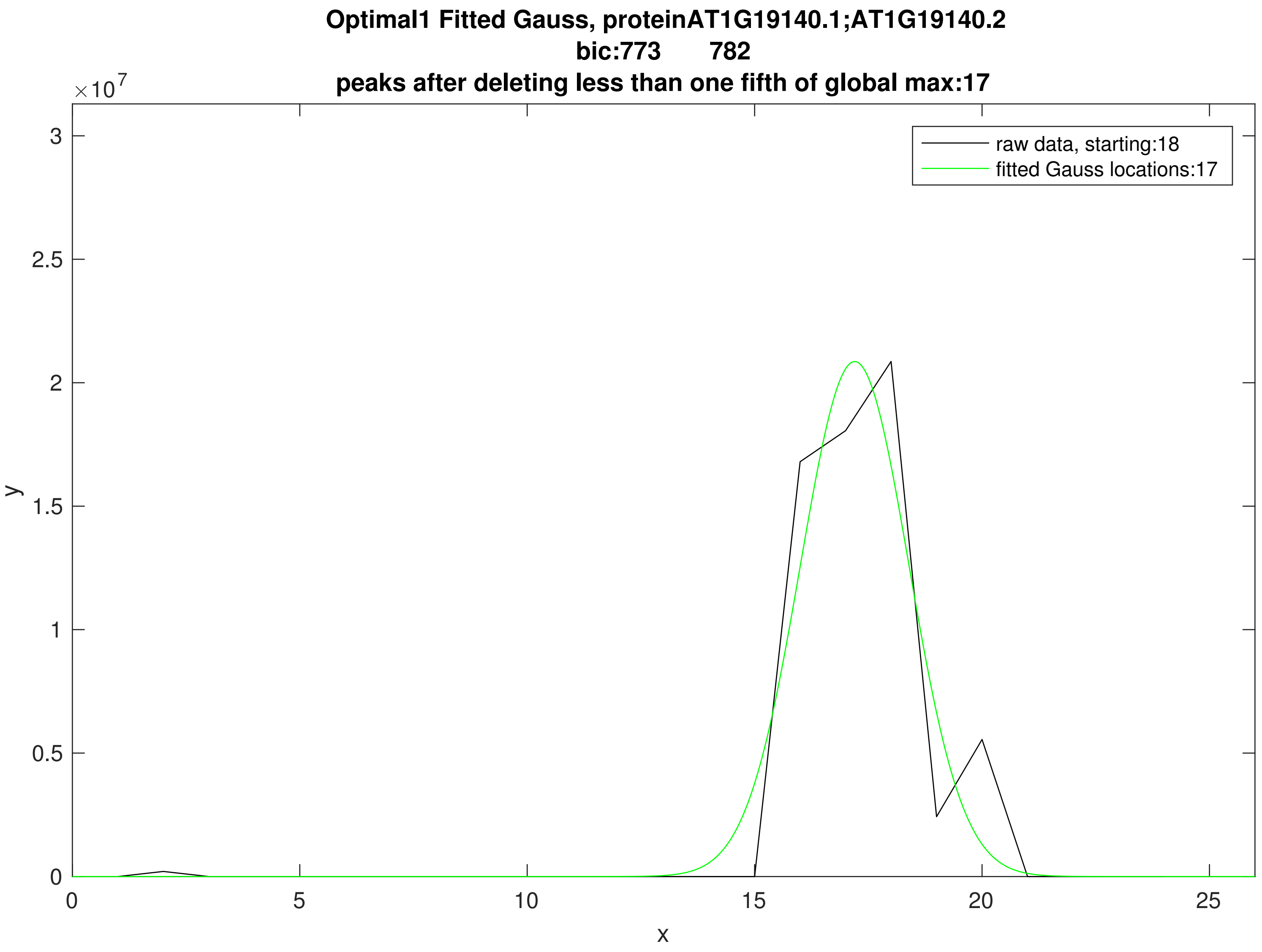

Optimal1 Fitted Gauss, proteinAT1G19140.1;AT1G19140.2
bic:773 782
peaks after deleting less than one fifth of global max:17
×10^7
raw data, starting:18
fitted Gauss locations:17
y
x
0
0.5
1
1.5
2
2.5
3
0
5
10
15
20
25

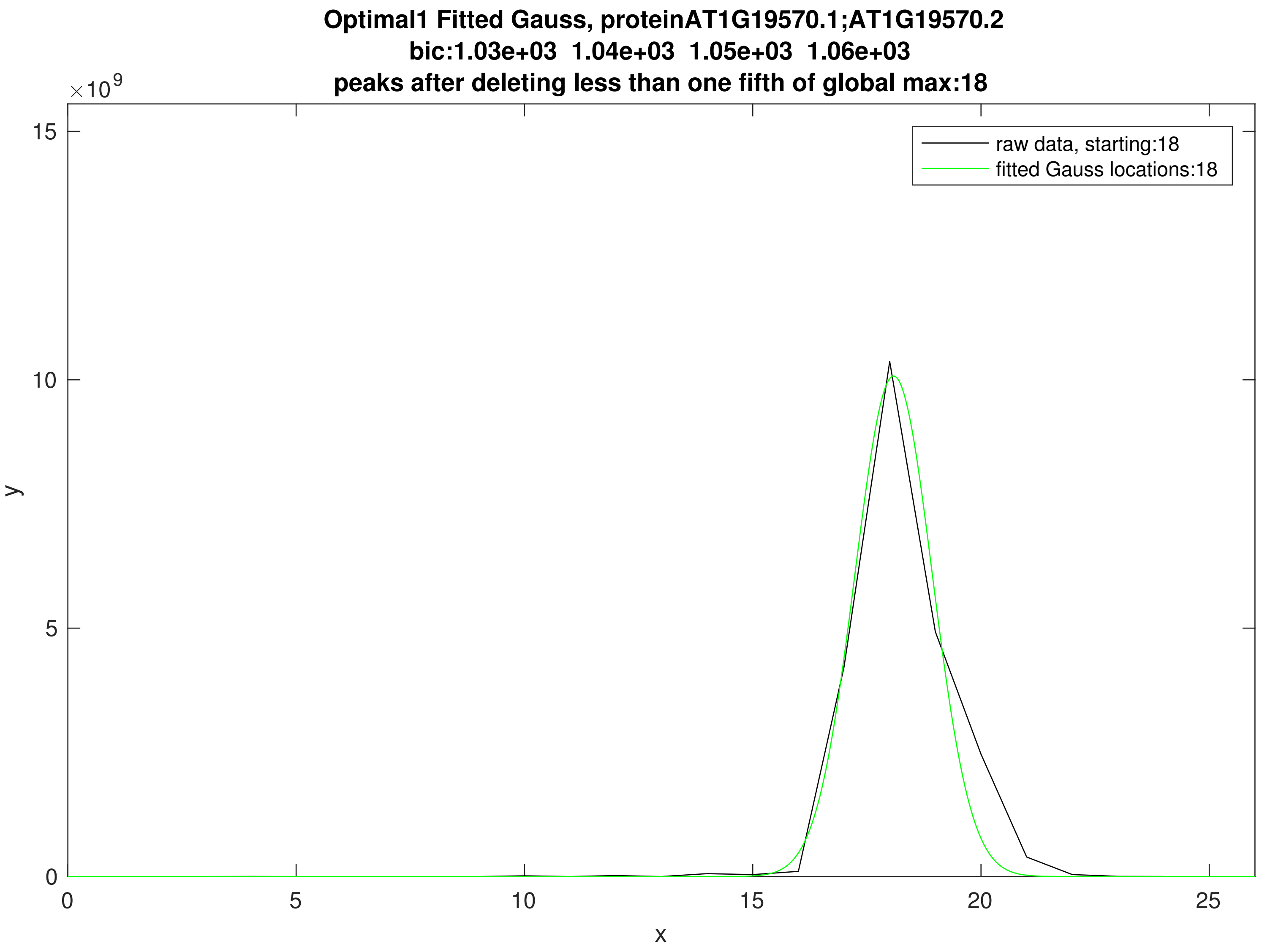

Optimal1 Fitted Gauss, proteinAT1G19570.1;AT1G19570.2
bic:1.03e+03 1.04e+03 1.05e+03 1.06e+03
peaks after deleting less than one fifth of global max:18
×10^9
raw data, starting:18
fitted Gauss locations:18
y
x
0
5
10
15
20
25

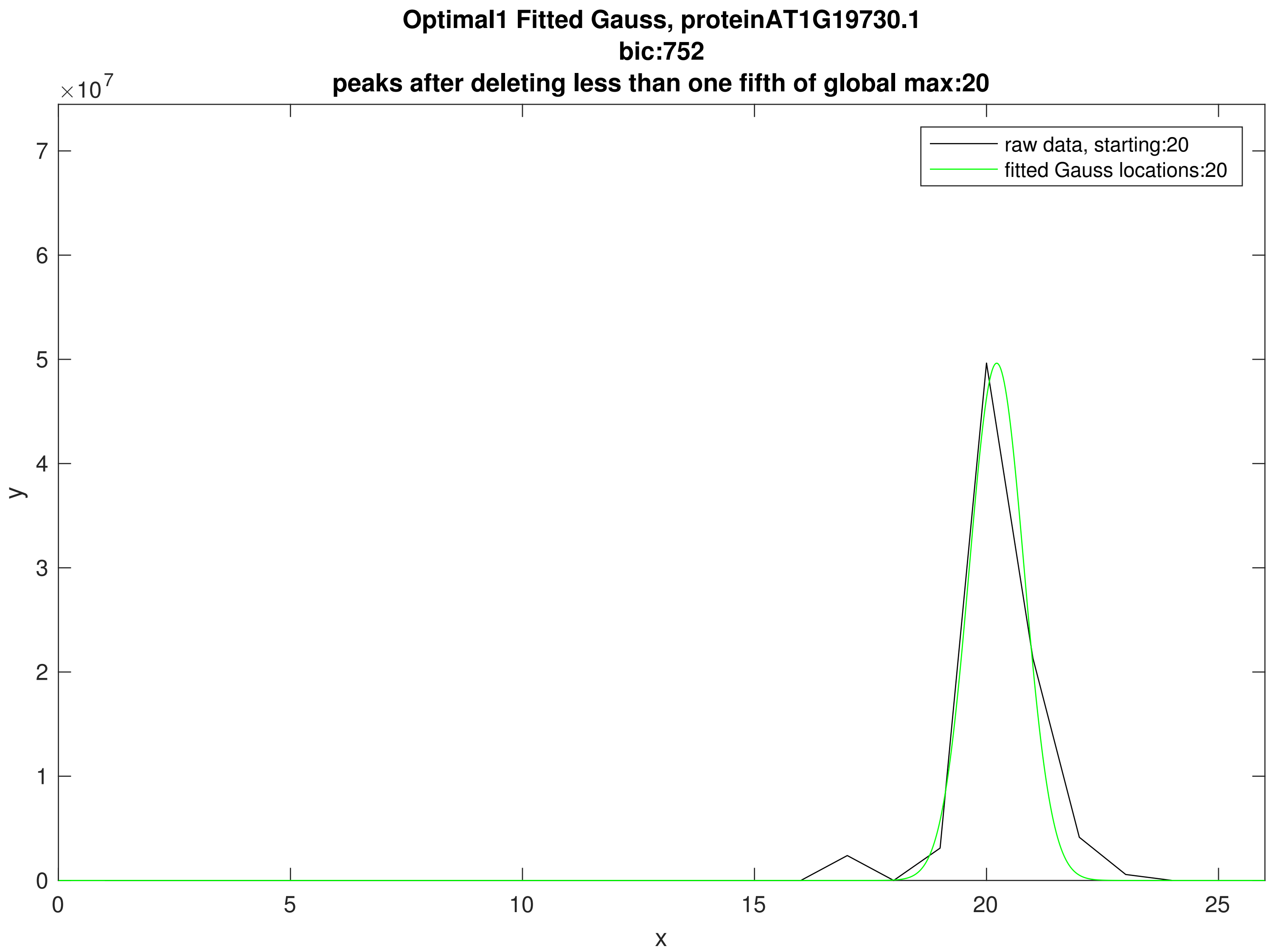

Optimal1 Fitted Gauss, proteinAT1G19730.1
bic:752
peaks after deleting less than one fifth of global max:20
×10^7
raw data, starting:20
fitted Gauss locations:20
y
x
0
1
2
3
4
5
6
7
0
5
10
15
20
25

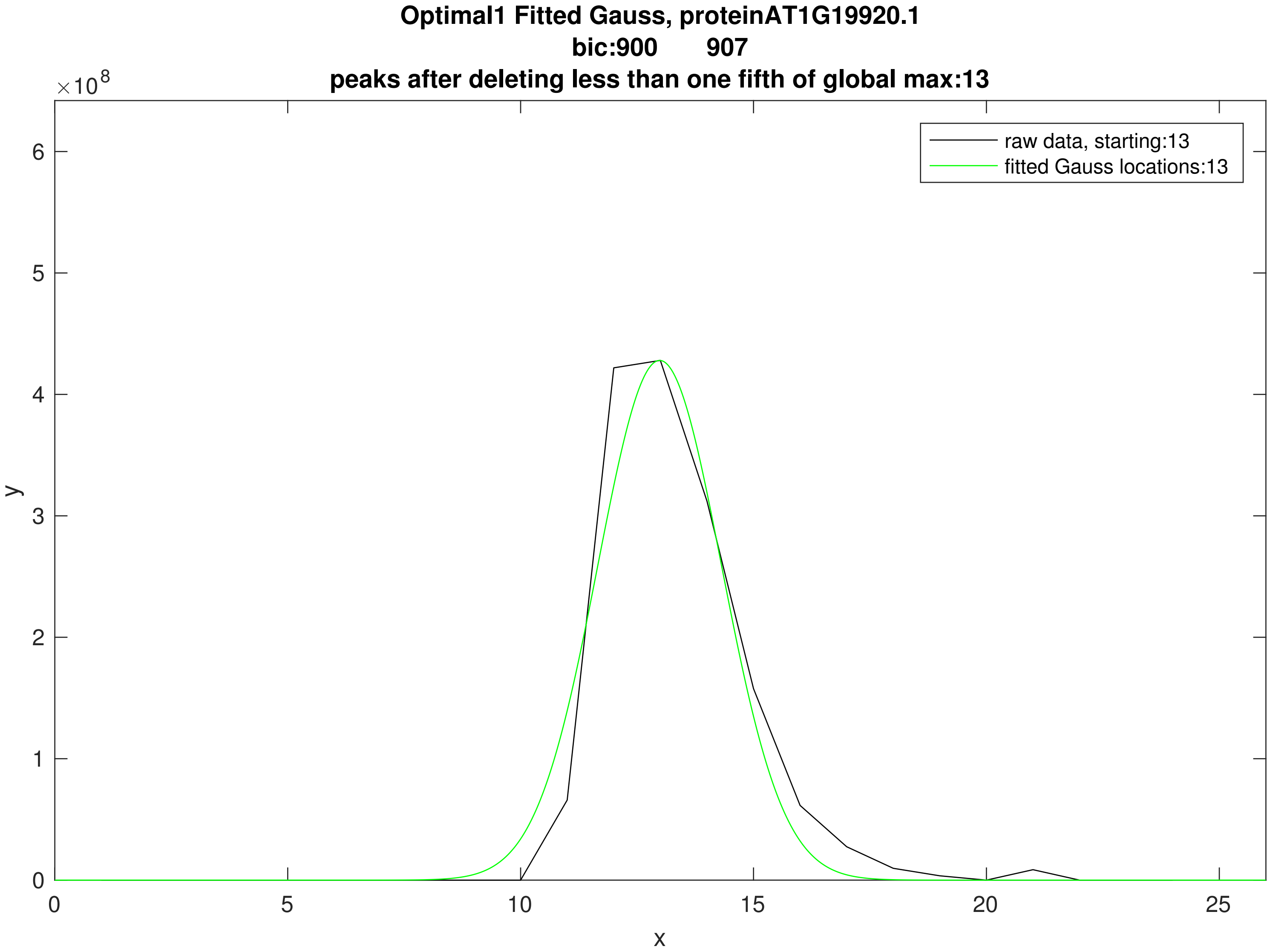

Optimal1 Fitted Gauss, proteinAT1G19920.1
bic:900 907
peaks after deleting less than one fifth of global max:13
×10^8
raw data, starting:13
fitted Gauss locations:13
y
x
0
1
2
3
4
5
6
0
5
10
15
20
25

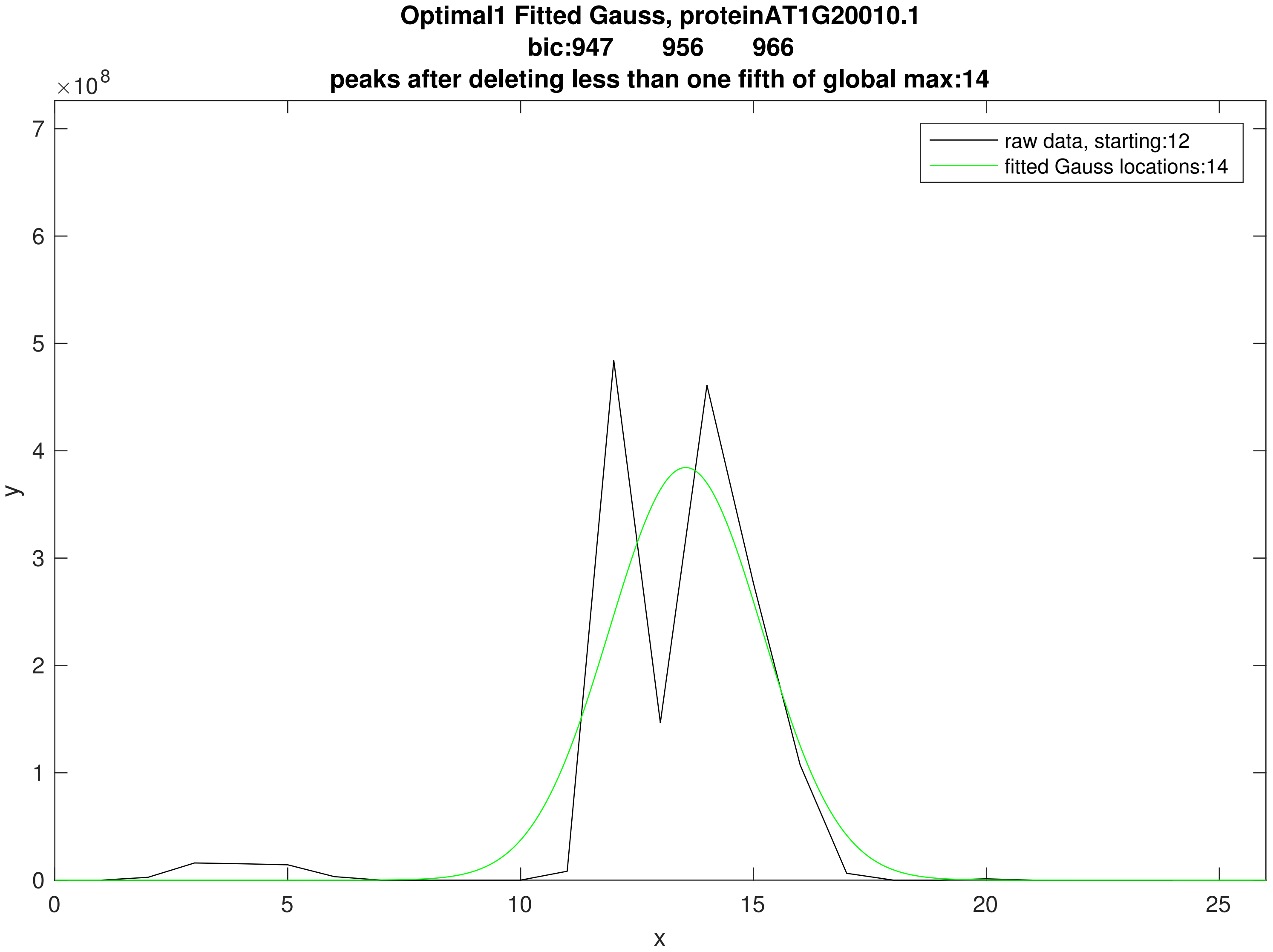

Optimal1 Fitted Gauss, proteinAT1G20010.1
bic:947 956 966
peaks after deleting less than one fifth of global max:14
×10^8
raw data, starting:12
fitted Gauss locations:14
y
x
0
1
2
3
4
5
6
7
0
5
10
15
20
25

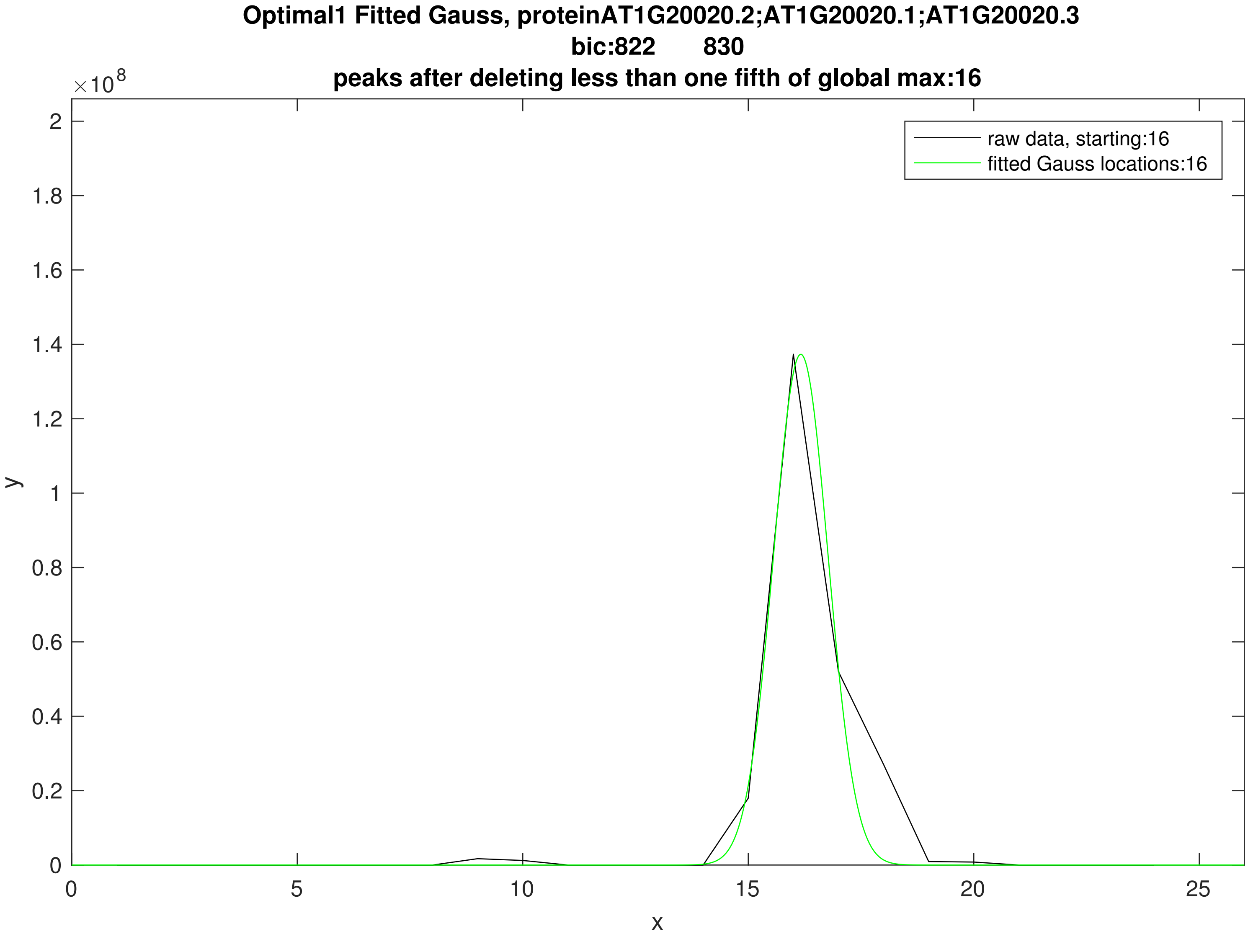

Optimal1 Fitted Gauss, proteinAT1G20020.2;AT1G20020.1;AT1G20020.3
bic:822 830
peaks after deleting less than one fifth of global max:16
×10^8
2
1.8
1.6
1.4
1.2
1
0.8
0.6
0.4
0.2
0
y
0
5
10
15
20
25
x
raw data, starting:16
fitted Gauss locations:16

**Optimal1 Fitted Gauss, proteinAT1G20160.2;AT1G20160.1**
**bic:918**
**peaks after deleting less than one fifth of global max:14**

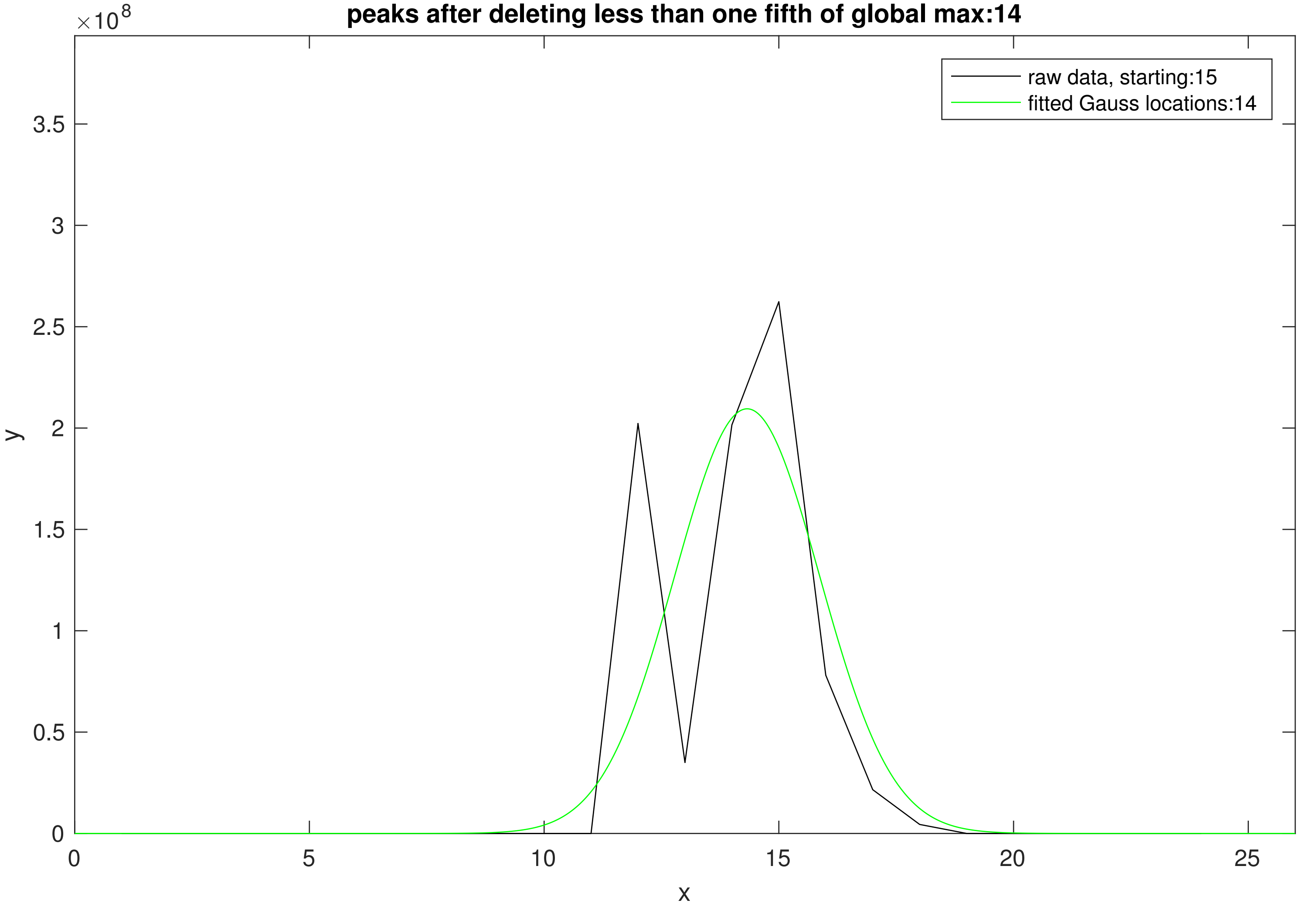

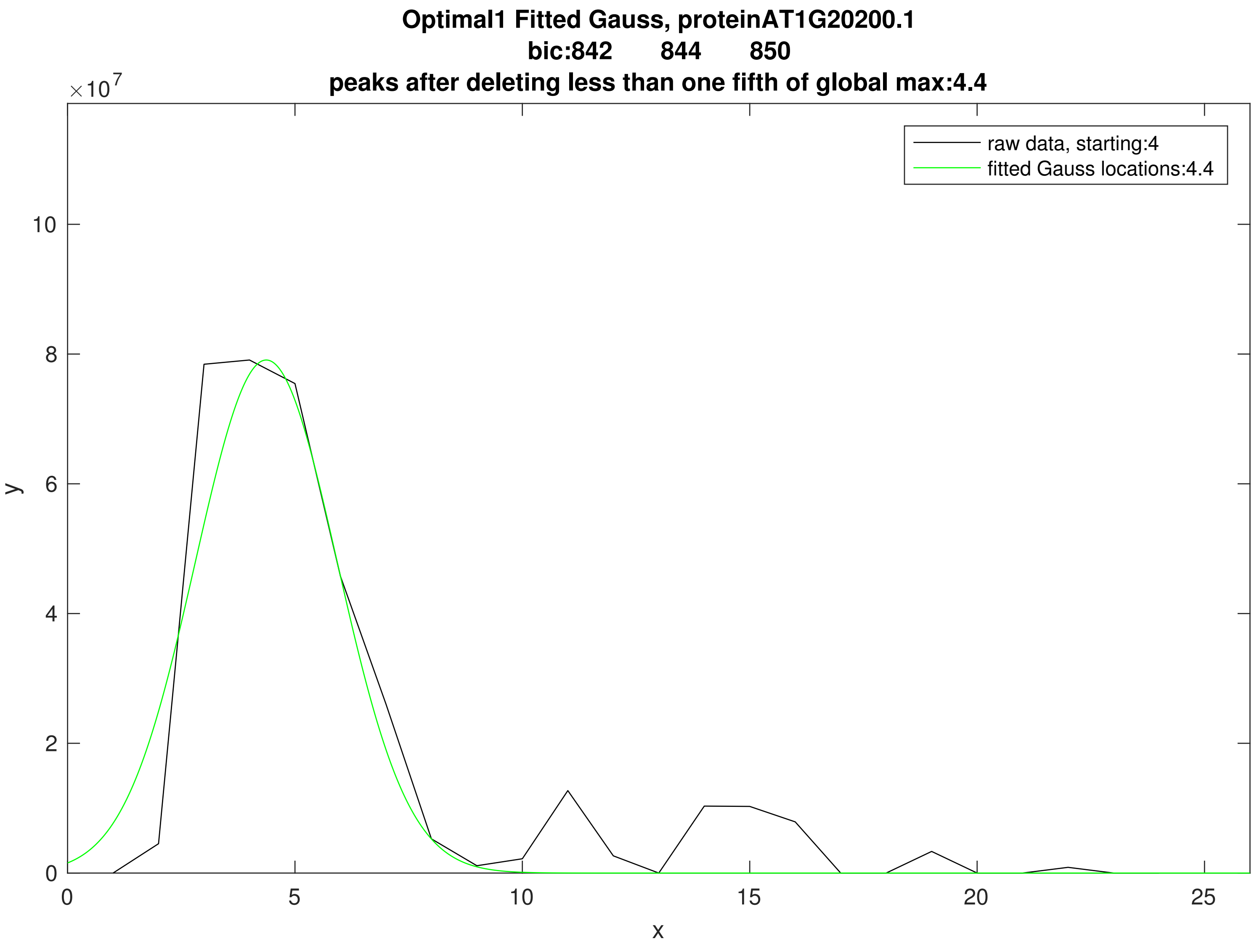

Optimal1 Fitted Gauss, proteinAT1G20200.1
bic:842 844 850
peaks after deleting less than one fifth of global max:4.4
×10^7
raw data, starting:4
fitted Gauss locations:4.4
10
8
6
4
2
0
y
0
5
10
15
20
25
x

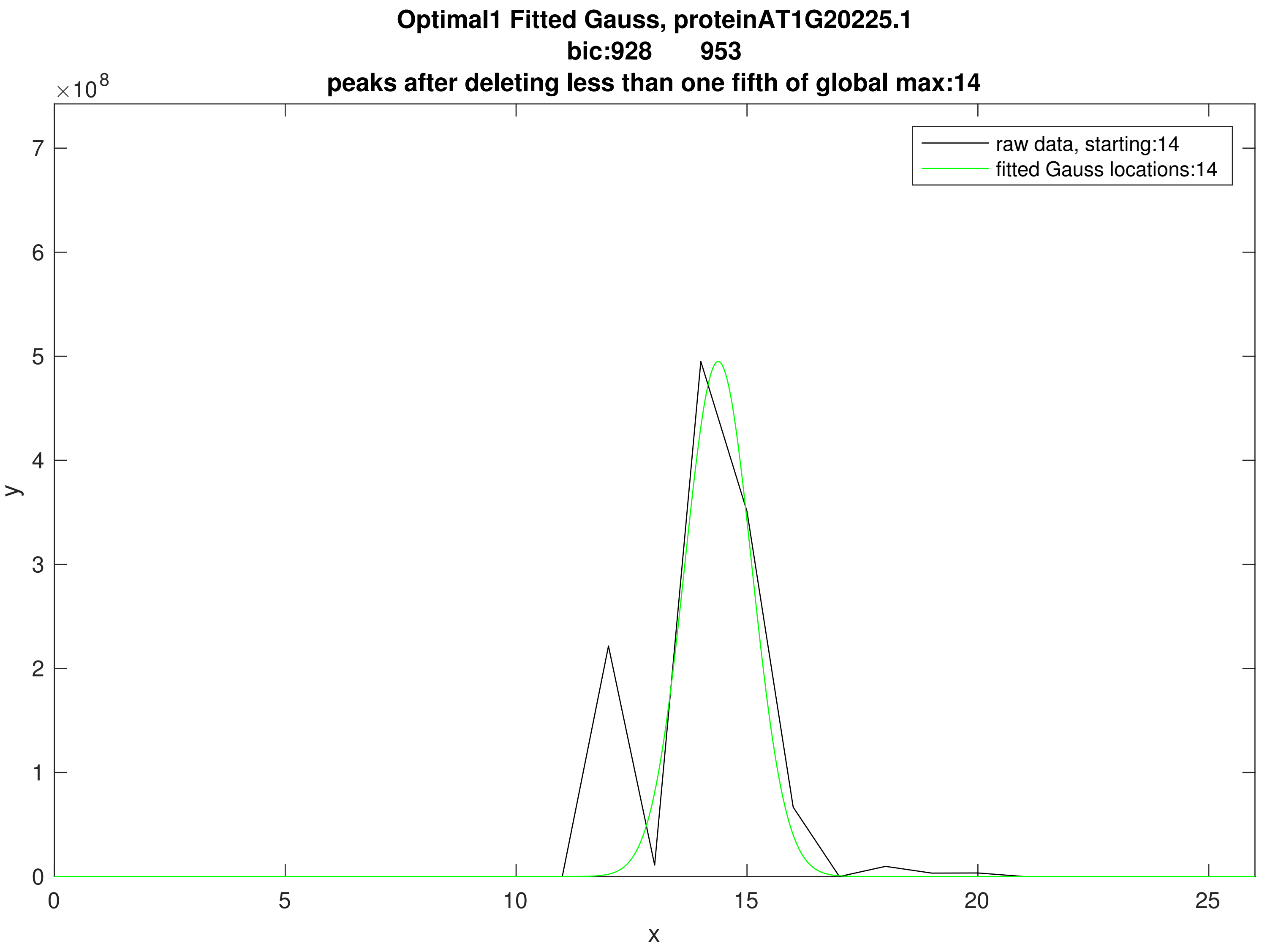

Optimal1 Fitted Gauss, proteinAT1G20225.1
bic:928 953
peaks after deleting less than one fifth of global max:14
×10^8
raw data, starting:14
fitted Gauss locations:14
y
x
0
1
2
3
4
5
6
7
0
5
10
15
20
25

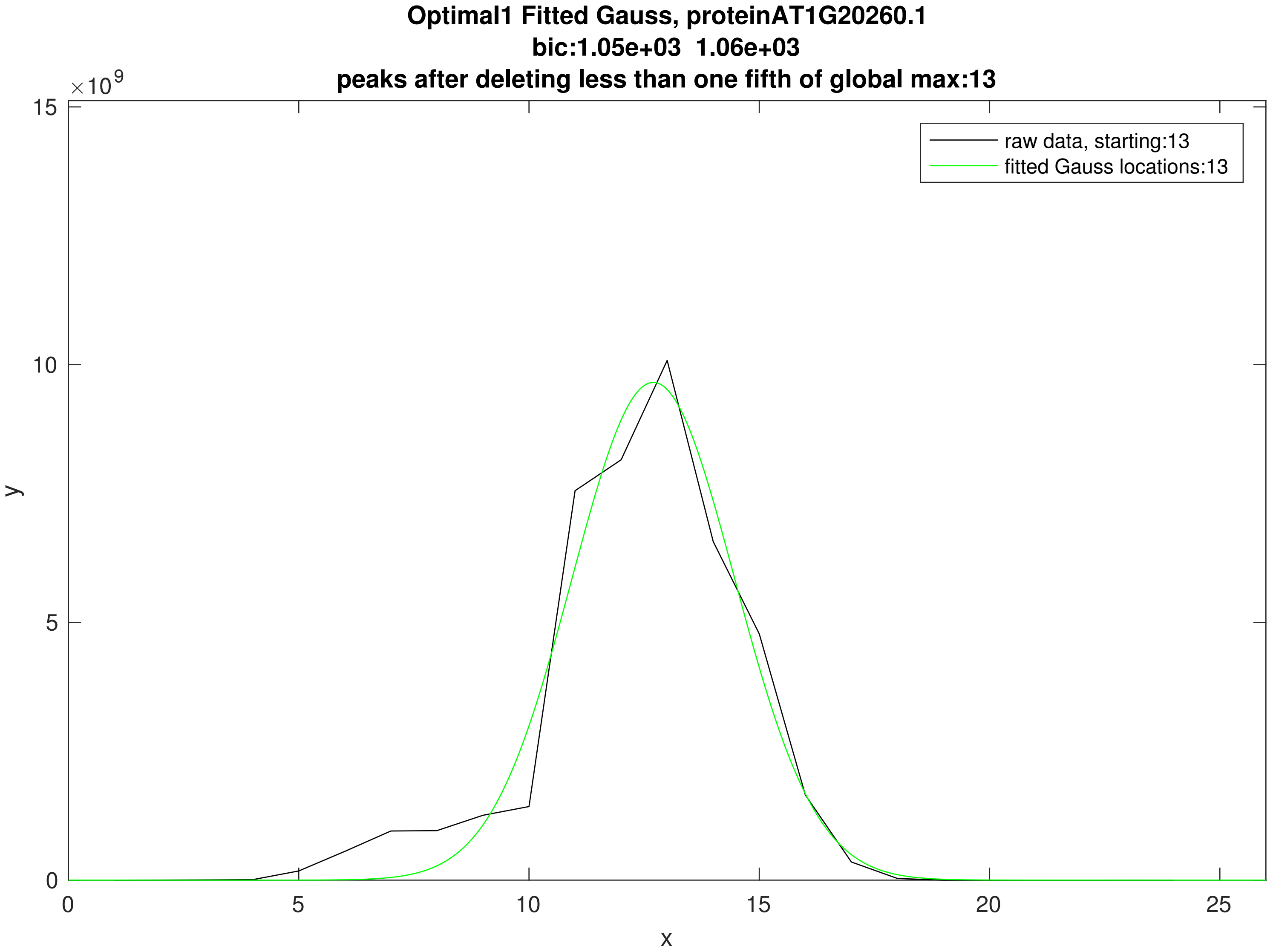
Optimal1 Fitted Gauss, proteinAT1G20260.1
bic:1.05e+03 1.06e+03
peaks after deleting less than one fifth of global max:13
×10^9
15
10
5
0
y
0
5
10
15
20
25
x
raw data, starting:13
fitted Gauss locations:13

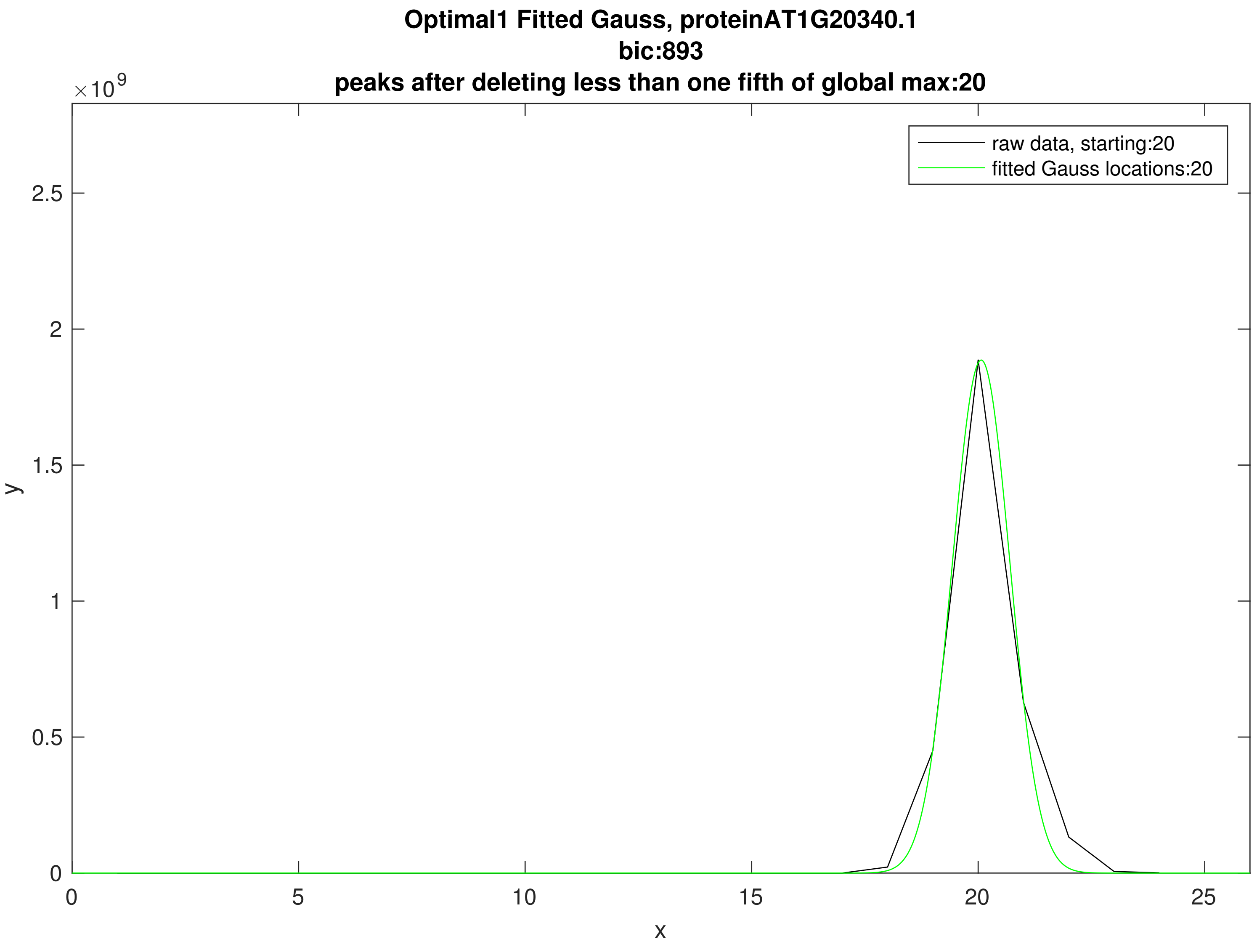
Optimal1 Fitted Gauss, proteinAT1G20340.1
bic:893
peaks after deleting less than one fifth of global max:20
×10^9
raw data, starting:20
fitted Gauss locations:20
y
x
0
0.5
1
1.5
2
2.5
0
5
10
15
20
25

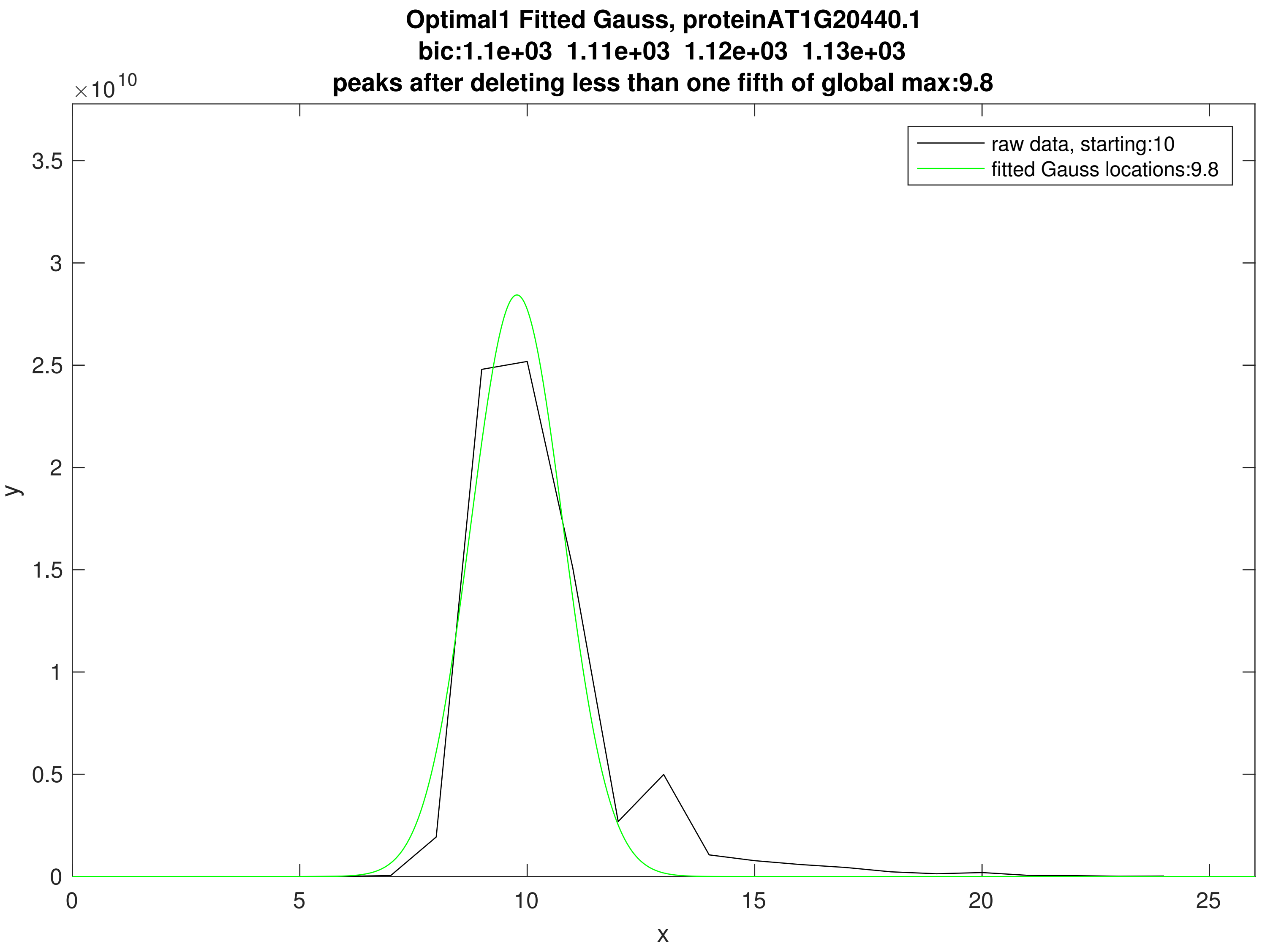

Optimal1 Fitted Gauss, proteinAT1G20440.1
bic:1.1e+03 1.11e+03 1.12e+03 1.13e+03
peaks after deleting less than one fifth of global max:9.8
×10^10
raw data, starting:10
fitted Gauss locations:9.8
y
x
0
0.5
1
1.5
2
2.5
3
3.5
5
10
15
20
25

**Optimal2 Fitted Gauss, proteinAT1G20450.2;AT1G20450.1**
**bic:998 990 994 1.01e+03**
**peaks after deleting less than one fifth of global max:10**

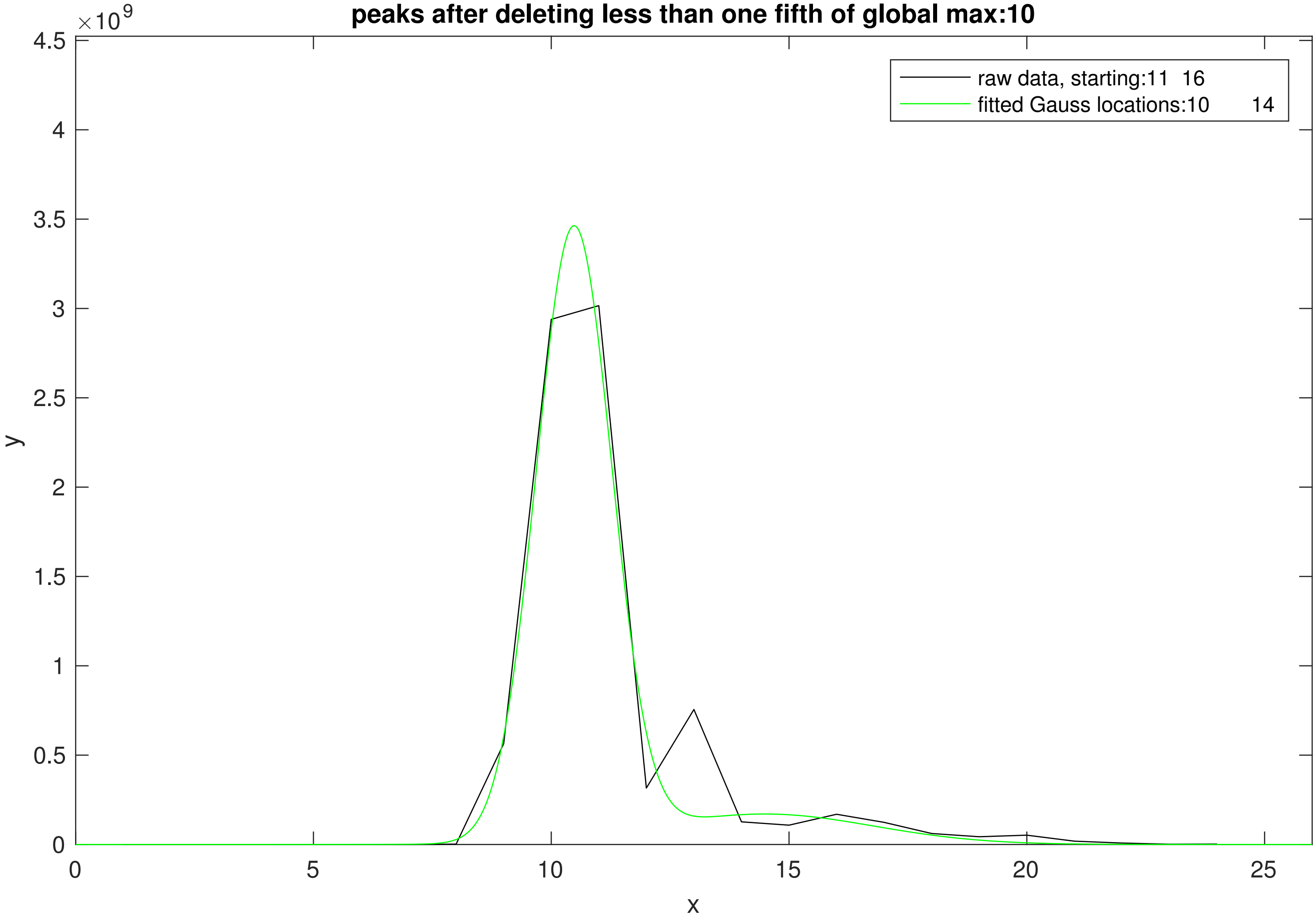

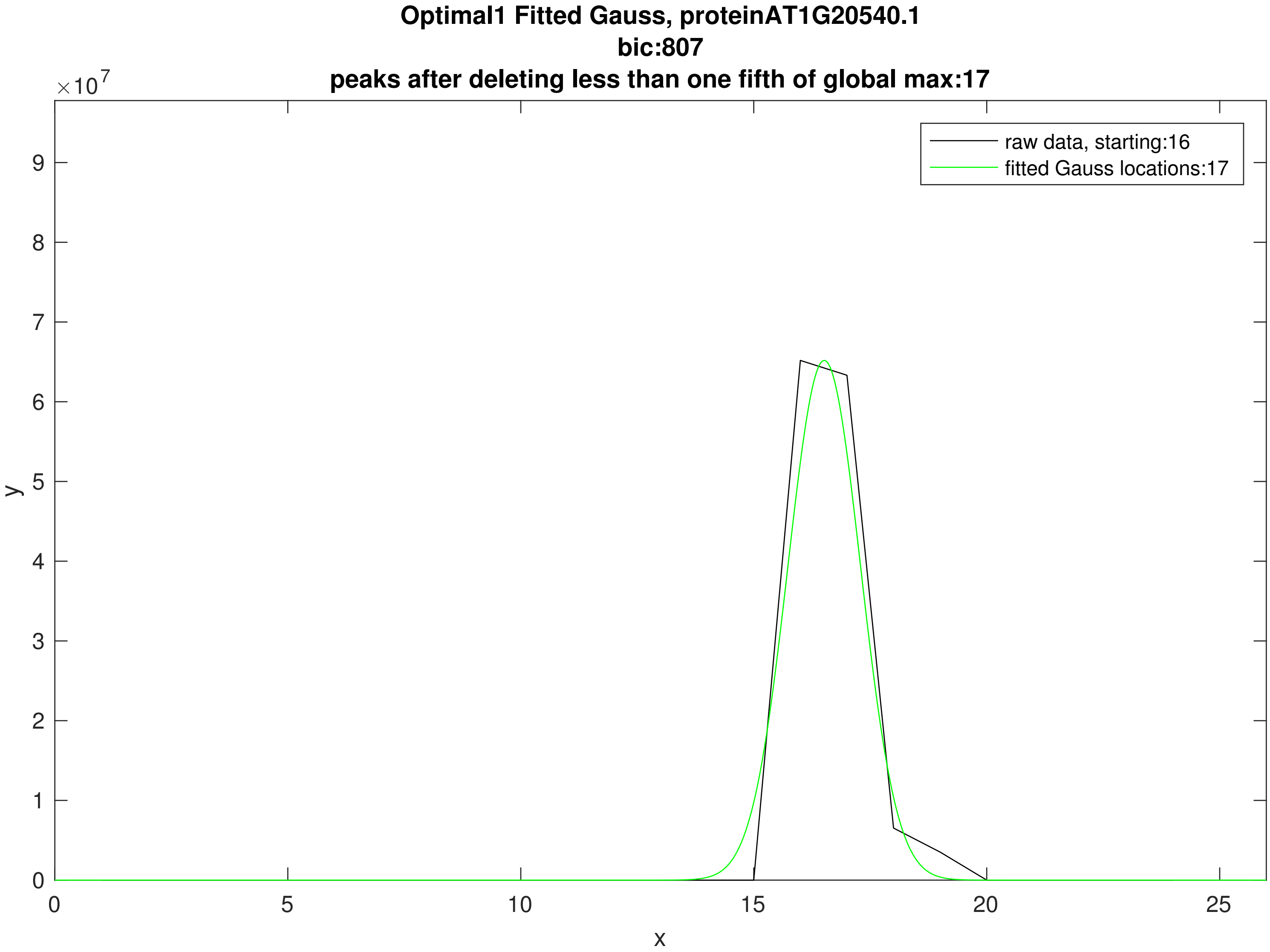
Optimal1 Fitted Gauss, proteinAT1G20540.1
bic:807
peaks after deleting less than one fifth of global max:17
×10^7
raw data, starting:16
fitted Gauss locations:17
y
x
0
1
2
3
4
5
6
7
8
9
0
5
10
15
20
25

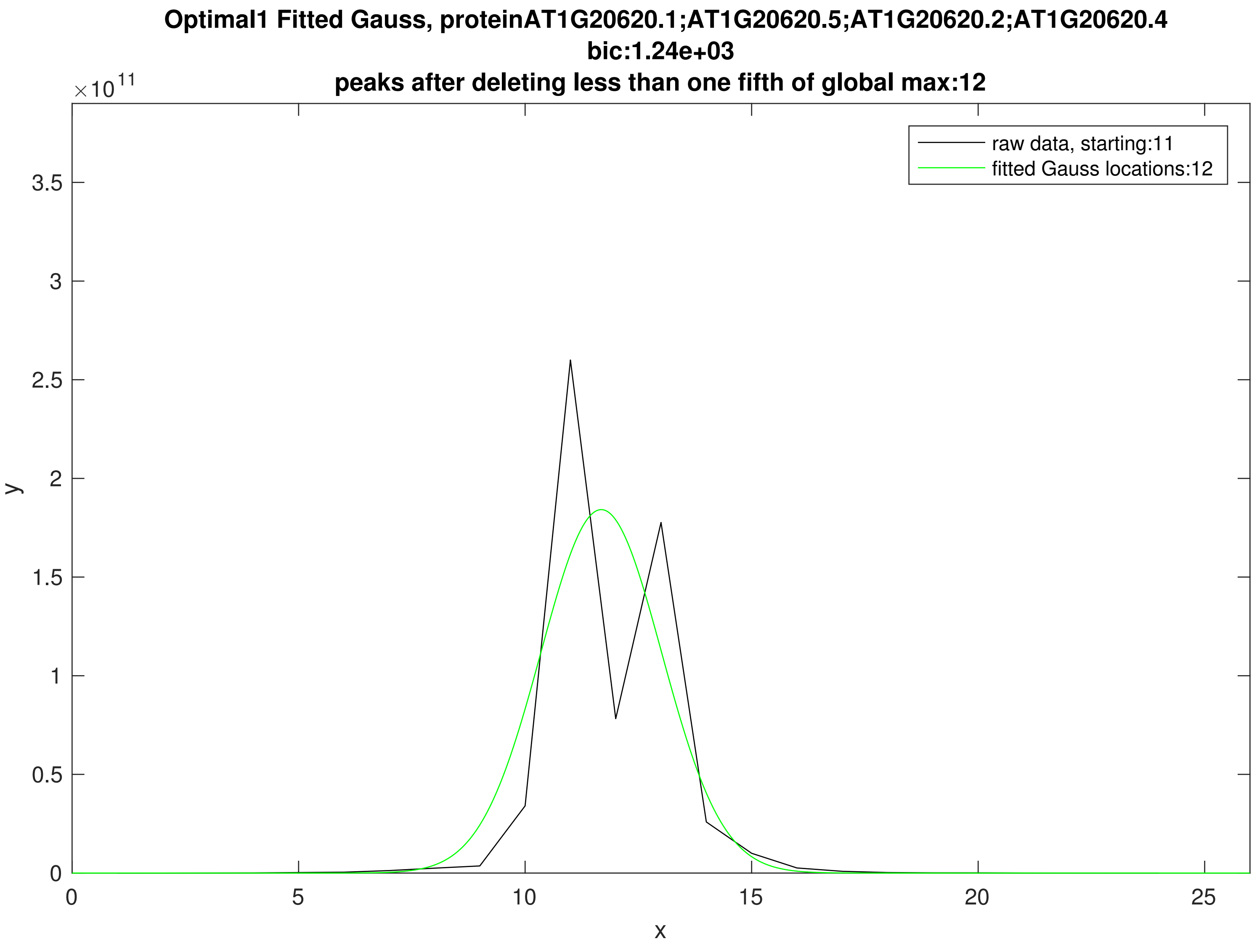
Optimal1 Fitted Gauss, proteinAT1G20620.1;AT1G20620.5;AT1G20620.2;AT1G20620.4
bic:1.24e+03
peaks after deleting less than one fifth of global max:12
×10^11
raw data, starting:11
fitted Gauss locations:12
y
x
0
0.5
1
1.5
2
2.5
3
3.5
0
5
10
15
20
25

**Optimal1 Fitted Gauss, proteinAT1G20630.1**
**bic:986 997 1.01e+03**
**peaks after deleting less than one fifth of global max:12**

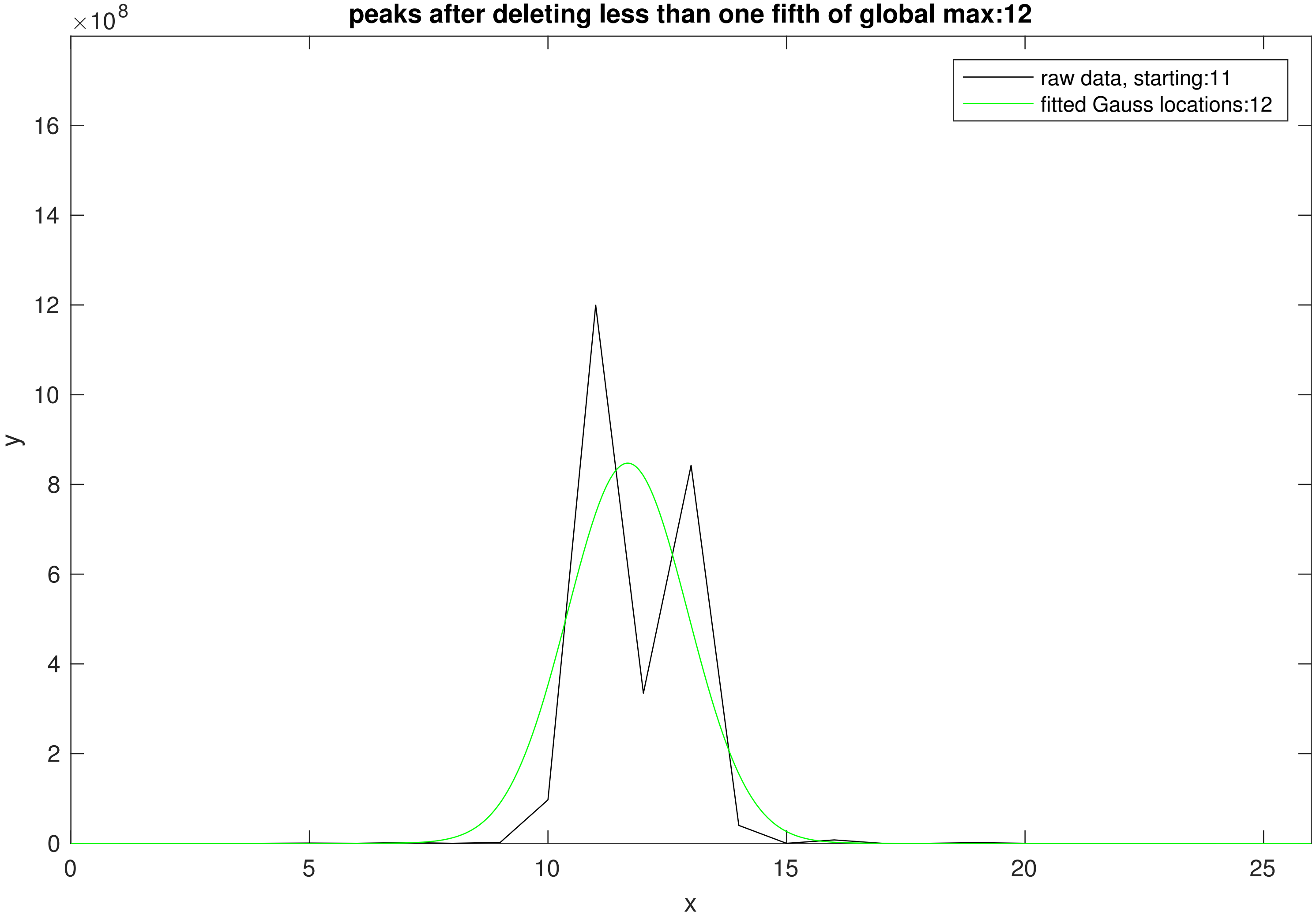

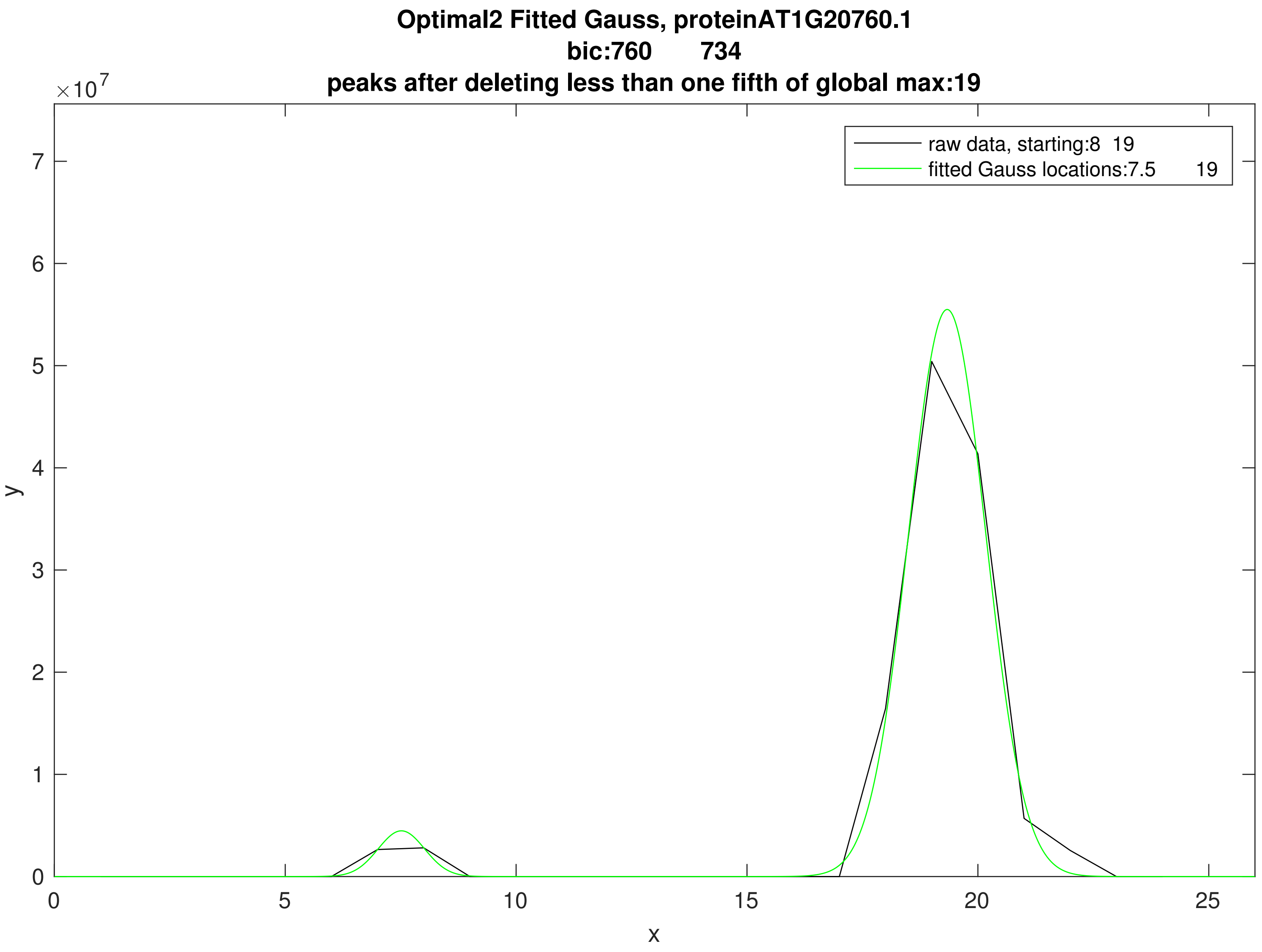

Optimal2 Fitted Gauss, proteinAT1G20760.1
bic:760 734
peaks after deleting less than one fifth of global max:19
×10^7
raw data, starting:8 19
fitted Gauss locations:7.5 19
y
x
0
1
2
3
4
5
6
7
0
5
10
15
20
25

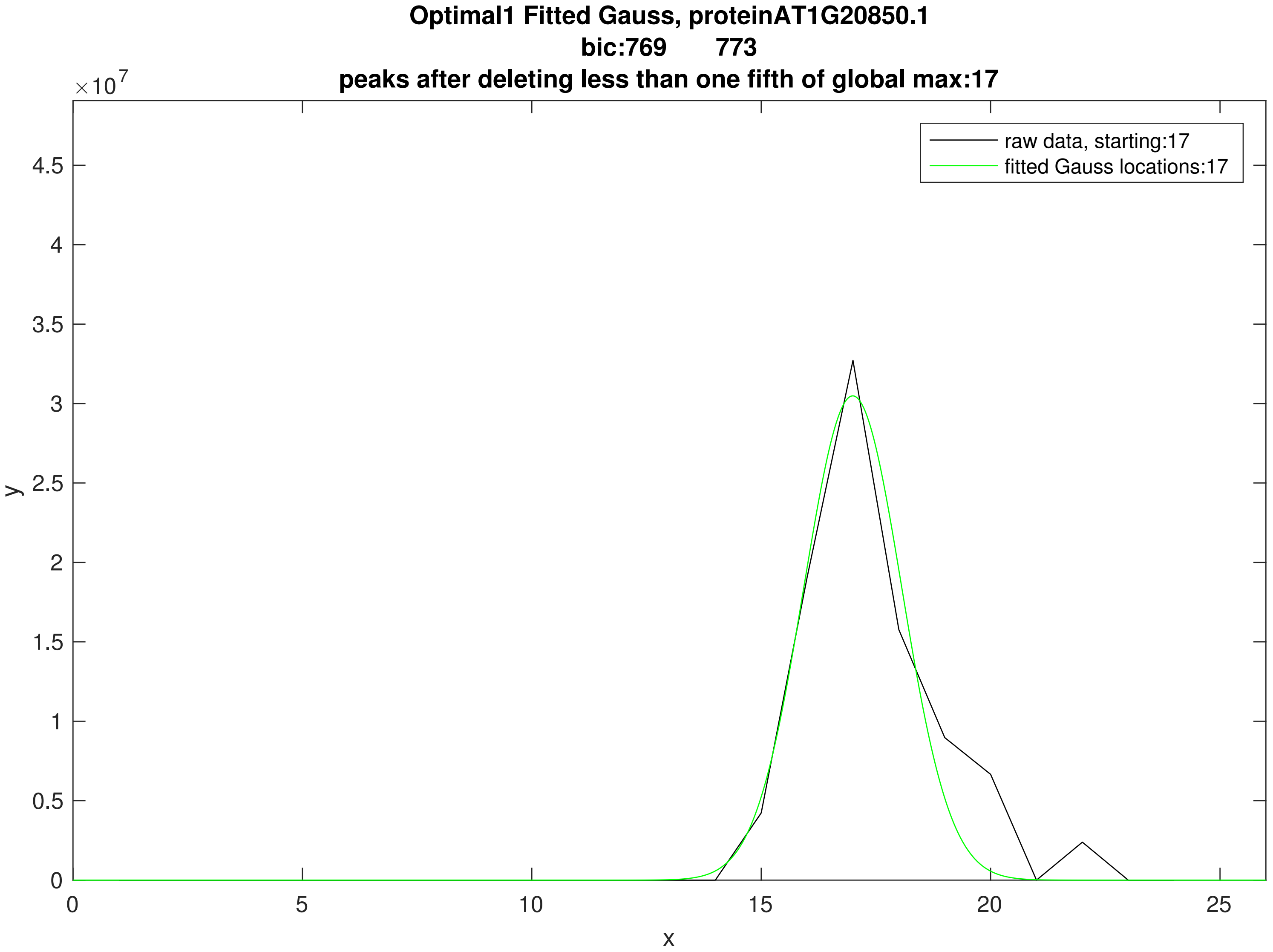

Optimal1 Fitted Gauss, proteinAT1G20850.1
bic:769 773
peaks after deleting less than one fifth of global max:17
×10^7
4.5
4
3.5
3
2.5
2
1.5
1
0.5
0
y
0
5
10
15
20
25
x
raw data, starting:17
fitted Gauss locations:17

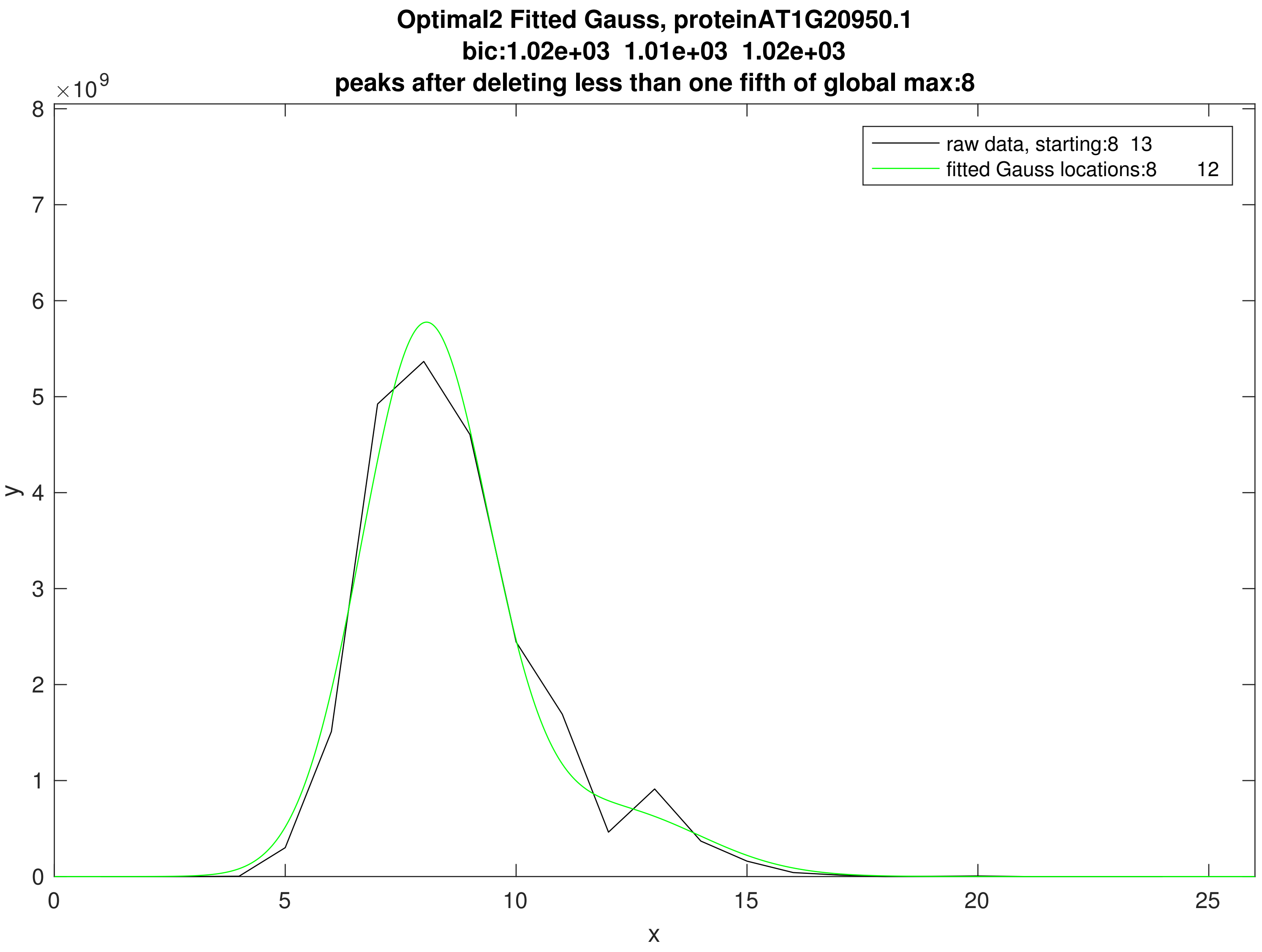

Optimal2 Fitted Gauss, proteinAT1G20950.1
bic:1.02e+03 1.01e+03 1.02e+03
peaks after deleting less than one fifth of global max:8
×10^9
raw data, starting:8 13
fitted Gauss locations:8 12
y
x
0
1
2
3
4
5
6
7
8
0
5
10
15
20
25

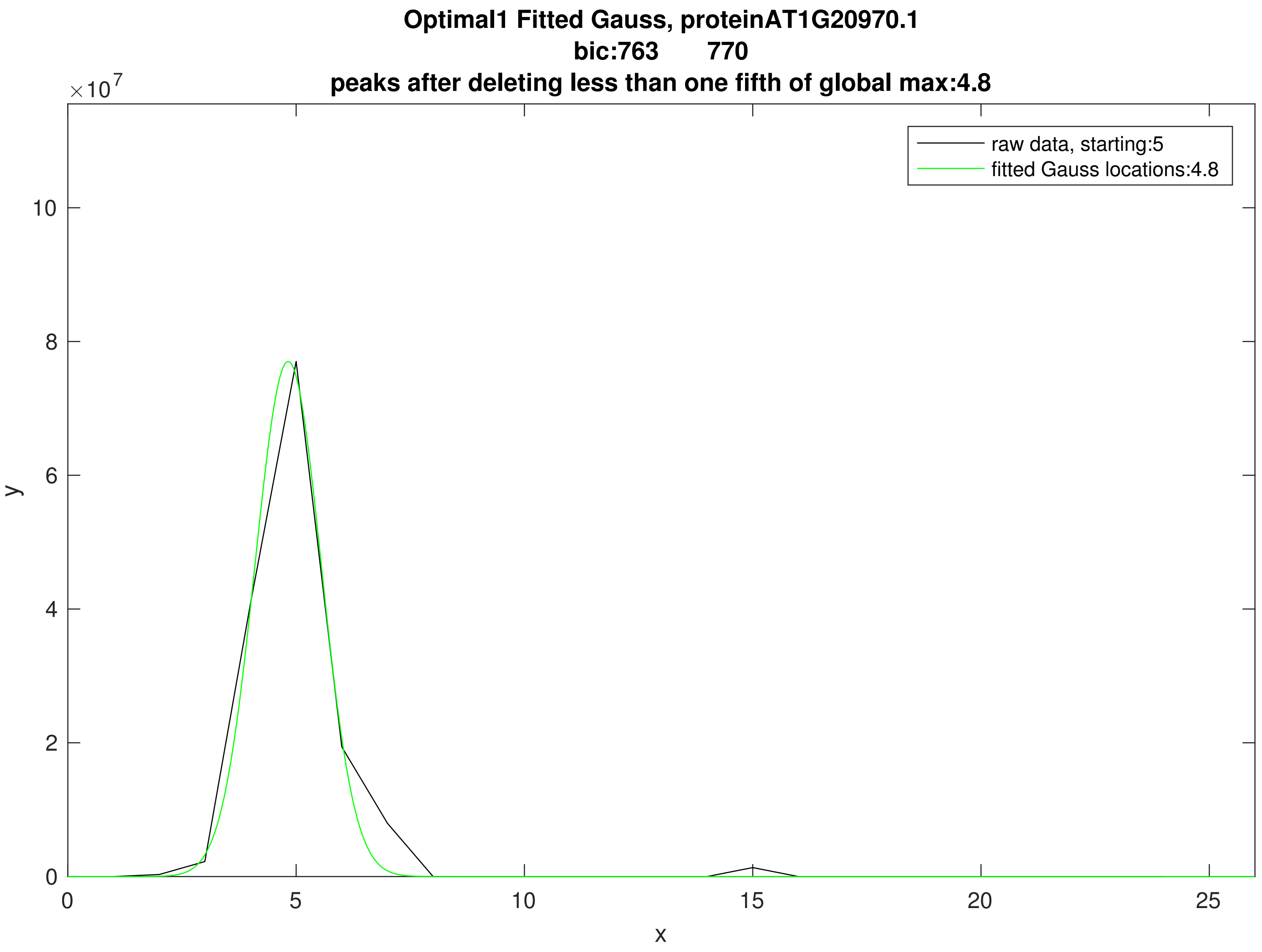
Optimal1 Fitted Gauss, proteinAT1G20970.1
bic:763 770
peaks after deleting less than one fifth of global max:4.8
×10^7
raw data, starting:5
fitted Gauss locations:4.8
y
x
0
2
4
6
8
10
0
5
10
15
20
25

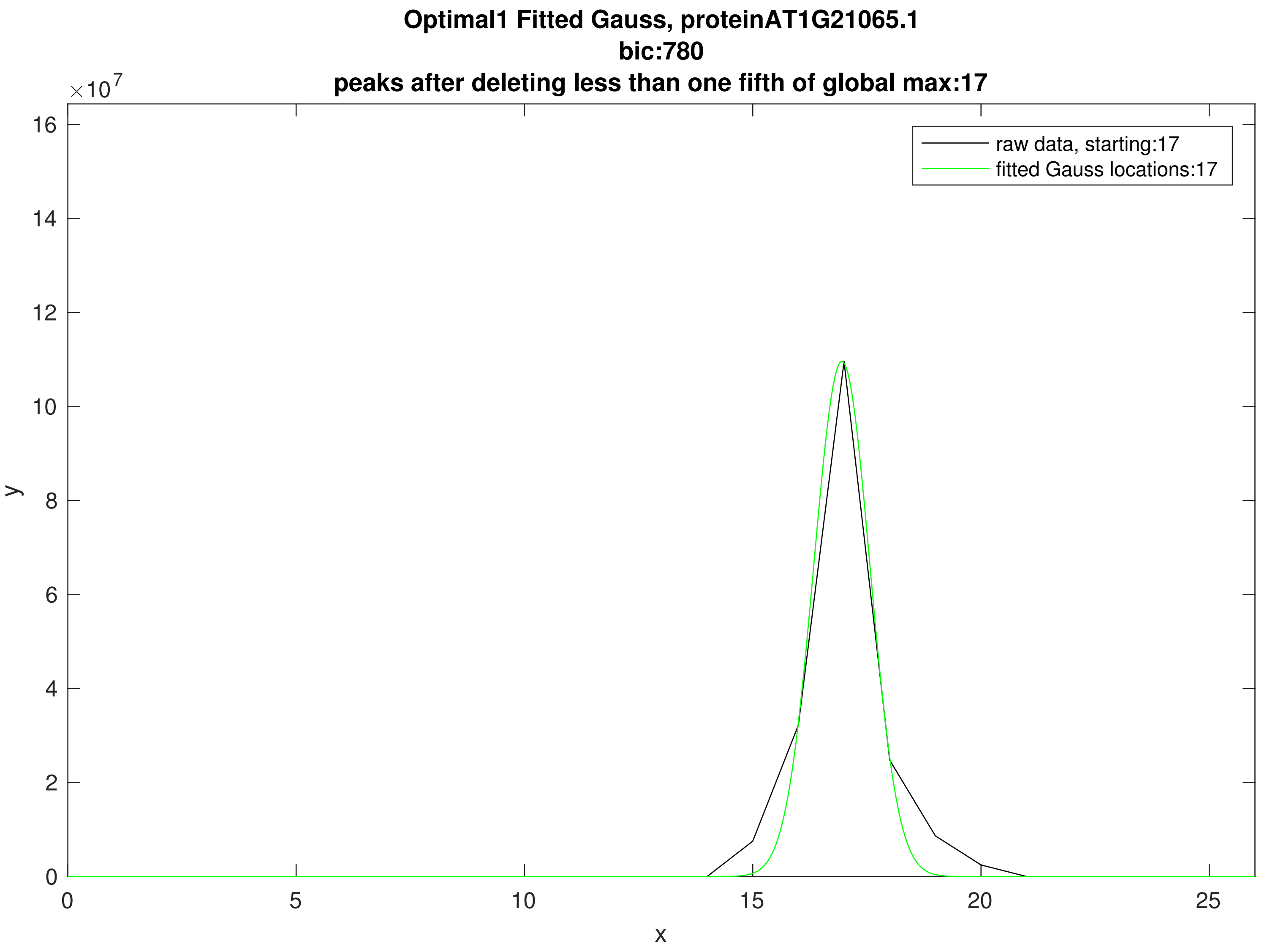
Optimal1 Fitted Gauss, proteinAT1G21065.1
bic:780
peaks after deleting less than one fifth of global max:17
×10^7
raw data, starting:17
fitted Gauss locations:17
y
x

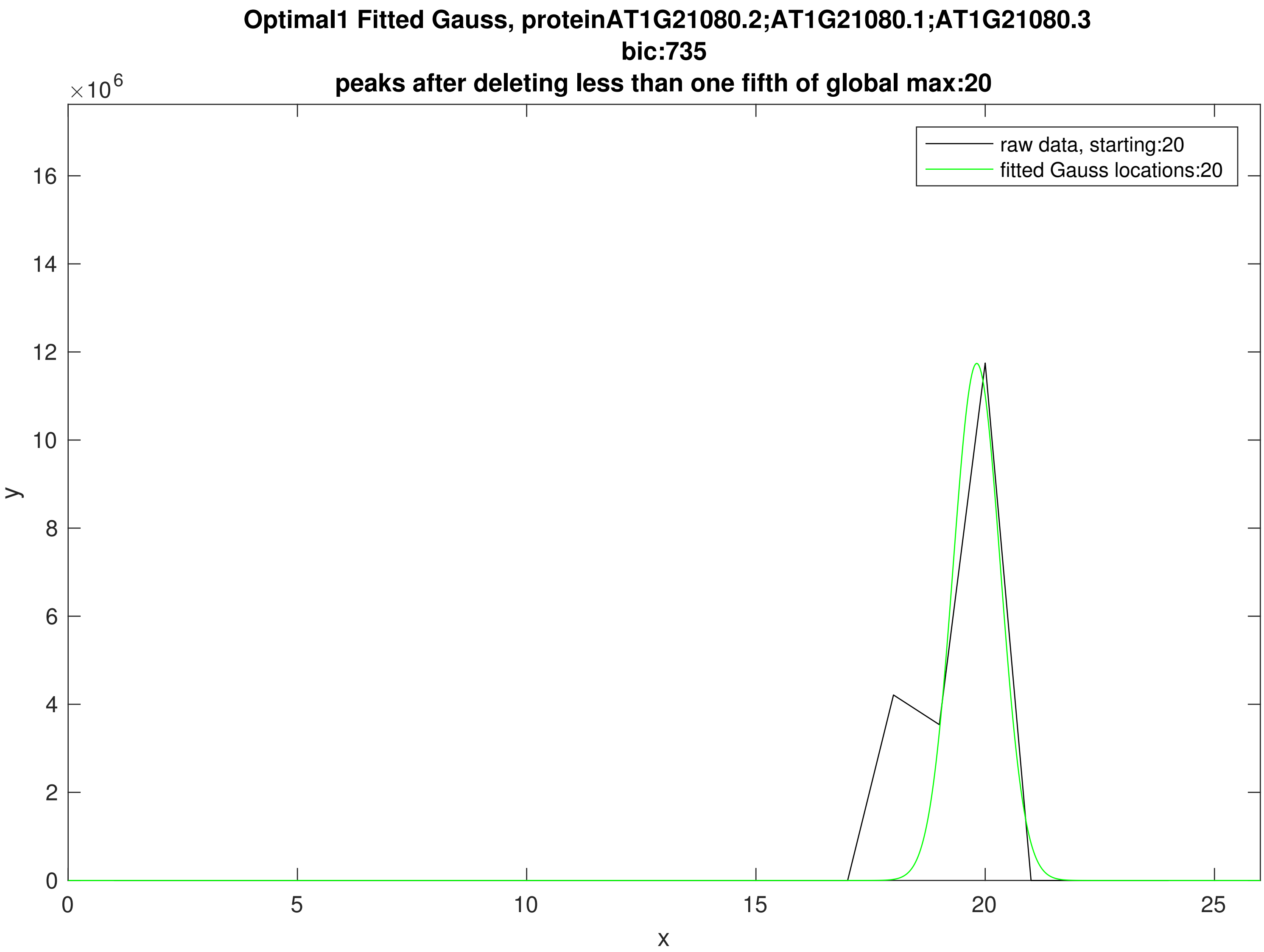

Optimal1 Fitted Gauss, proteinAT1G21080.2;AT1G21080.1;AT1G21080.3
bic:735
peaks after deleting less than one fifth of global max:20
×10^6
raw data, starting:20
fitted Gauss locations:20
y
x
0
2
4
6
8
10
12
14
16
0
5
10
15
20
25

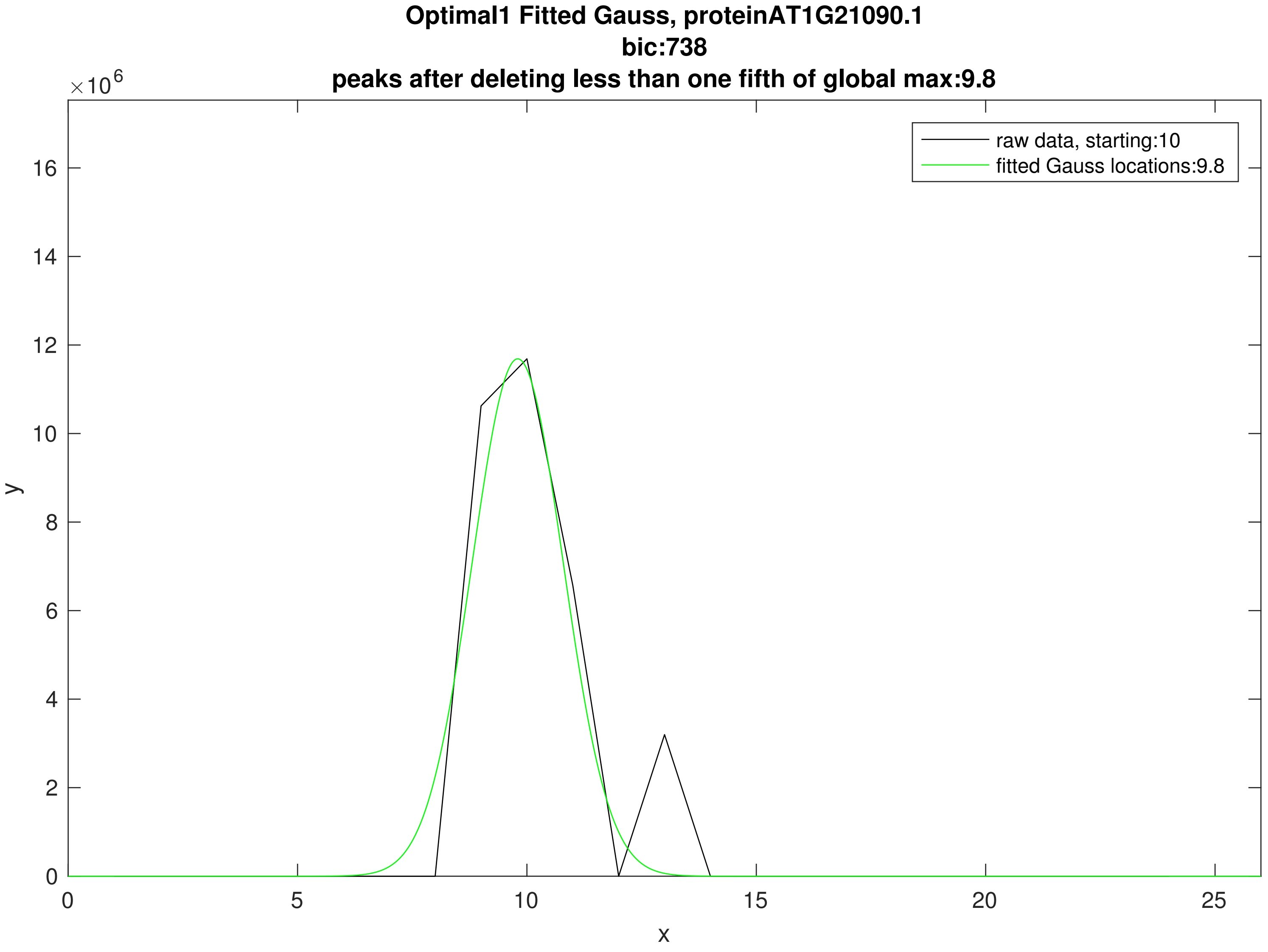

Optimal1 Fitted Gauss, proteinAT1G21090.1
bic:738
peaks after deleting less than one fifth of global max:9.8
×10^6
raw data, starting:10
fitted Gauss locations:9.8
16
14
12
10
8
6
4
2
0
y
0
5
10
15
20
25
x

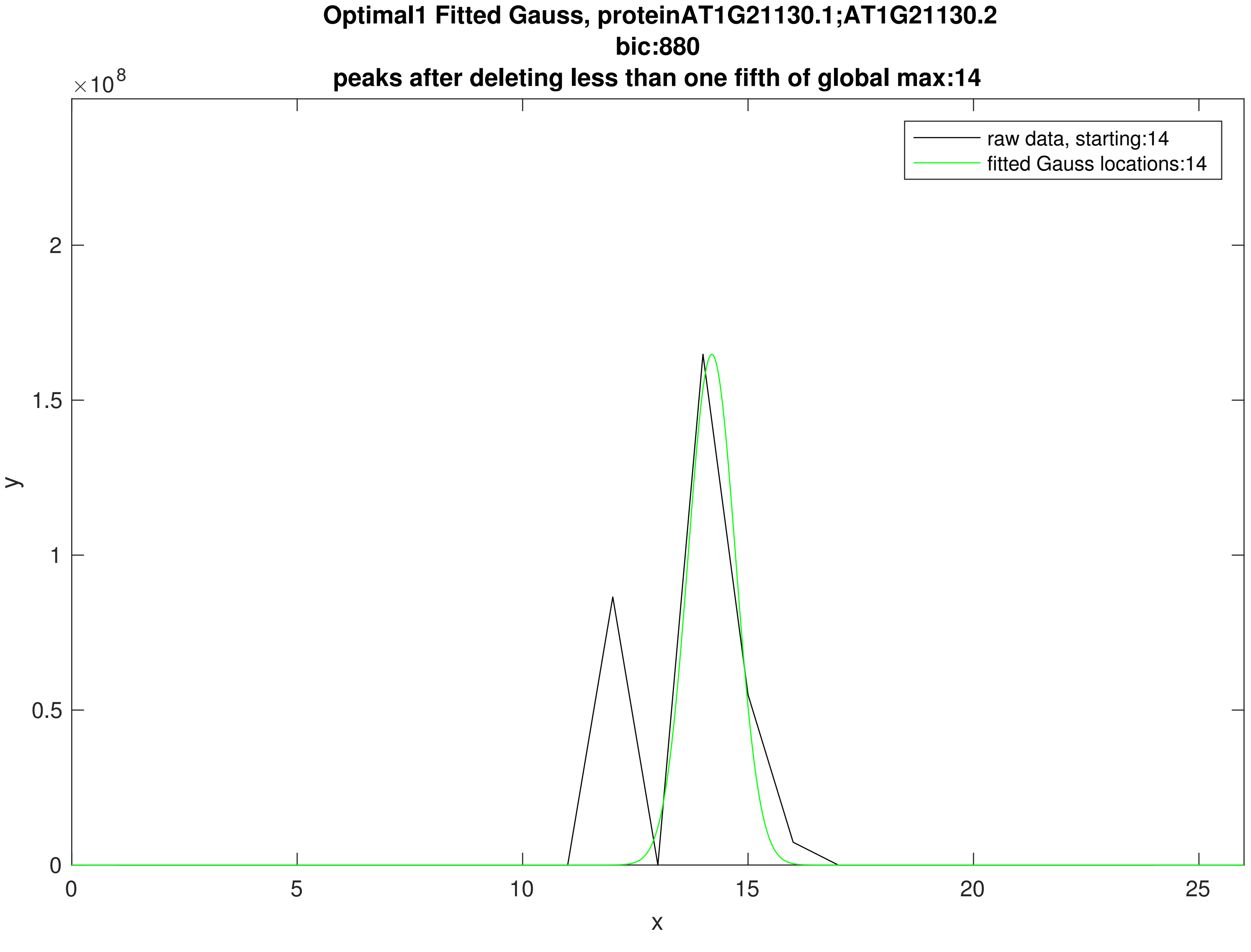
Optimal1 Fitted Gauss, proteinAT1G21130.1;AT1G21130.2
bic:880
peaks after deleting less than one fifth of global max:14
×10^8
raw data, starting:14
fitted Gauss locations:14
y
x
0
0.5
1
1.5
2
0
5
10
15
20
25

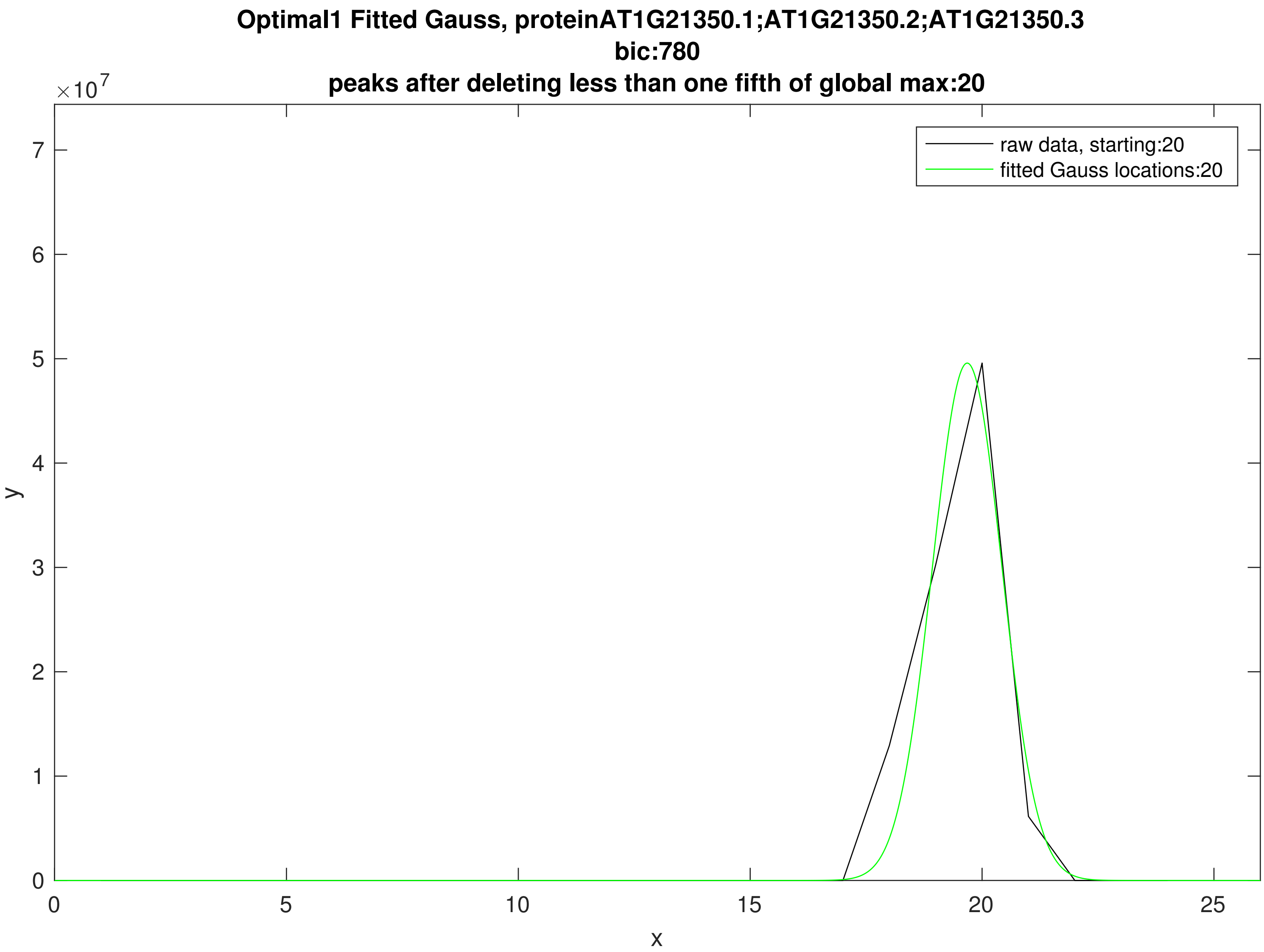

Optimal1 Fitted Gauss, proteinAT1G21350.1;AT1G21350.2;AT1G21350.3
bic:780
peaks after deleting less than one fifth of global max:20
×10^7
raw data, starting:20
fitted Gauss locations:20
y
x
0
1
2
3
4
5
6
7
0
5
10
15
20
25

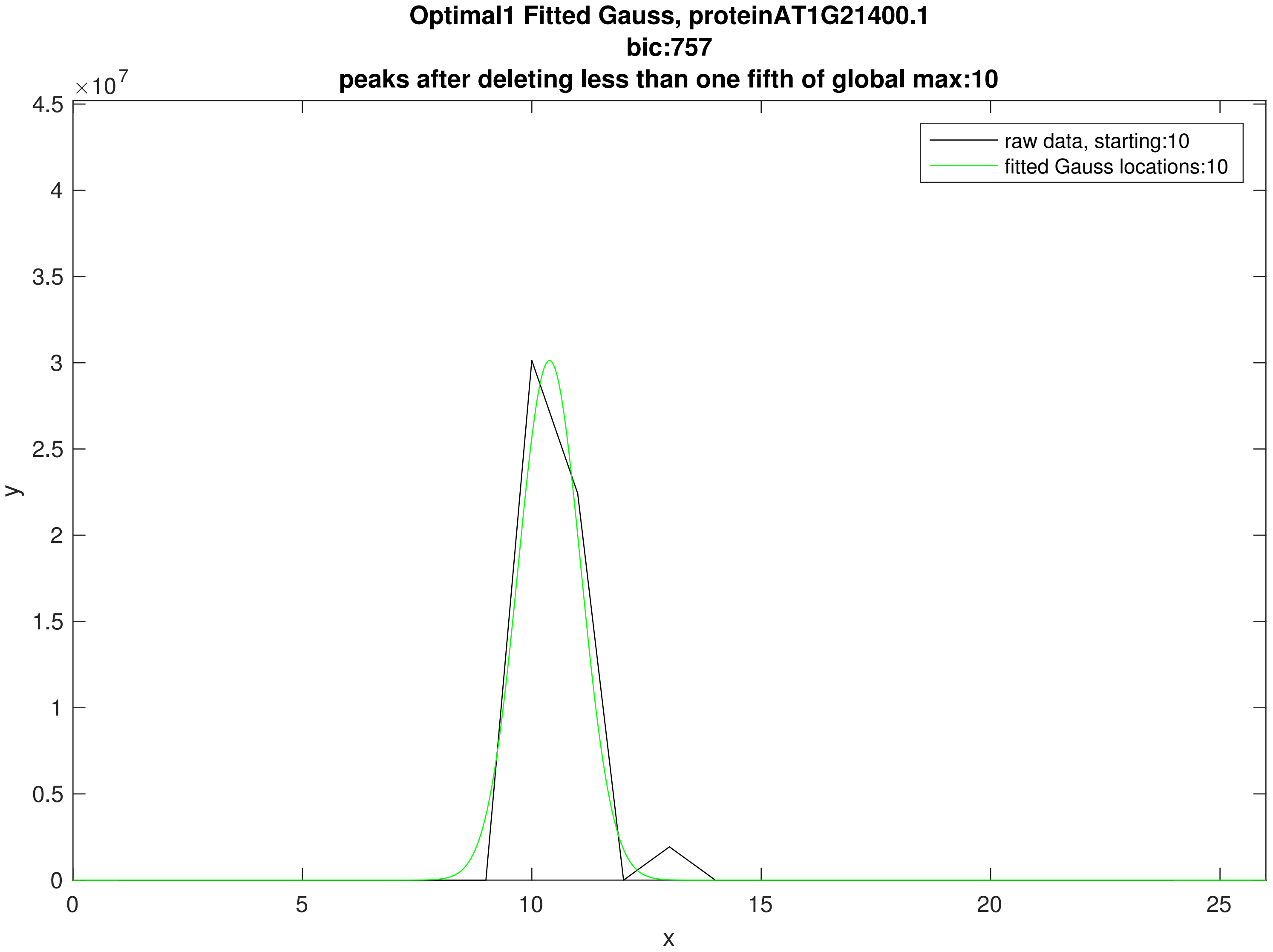

Optimal1 Fitted Gauss, proteinAT1G21400.1
bic:757
peaks after deleting less than one fifth of global max:10
×10^7
4.5
4
3.5
3
2.5
2
1.5
1
0.5
0
y
0
5
10
15
20
25
x
raw data, starting:10
fitted Gauss locations:10

**Optimal1 Fitted Gauss, proteinAT1G21440.1**
**bic:861**
**peaks after deleting less than one fifth of global max:14**

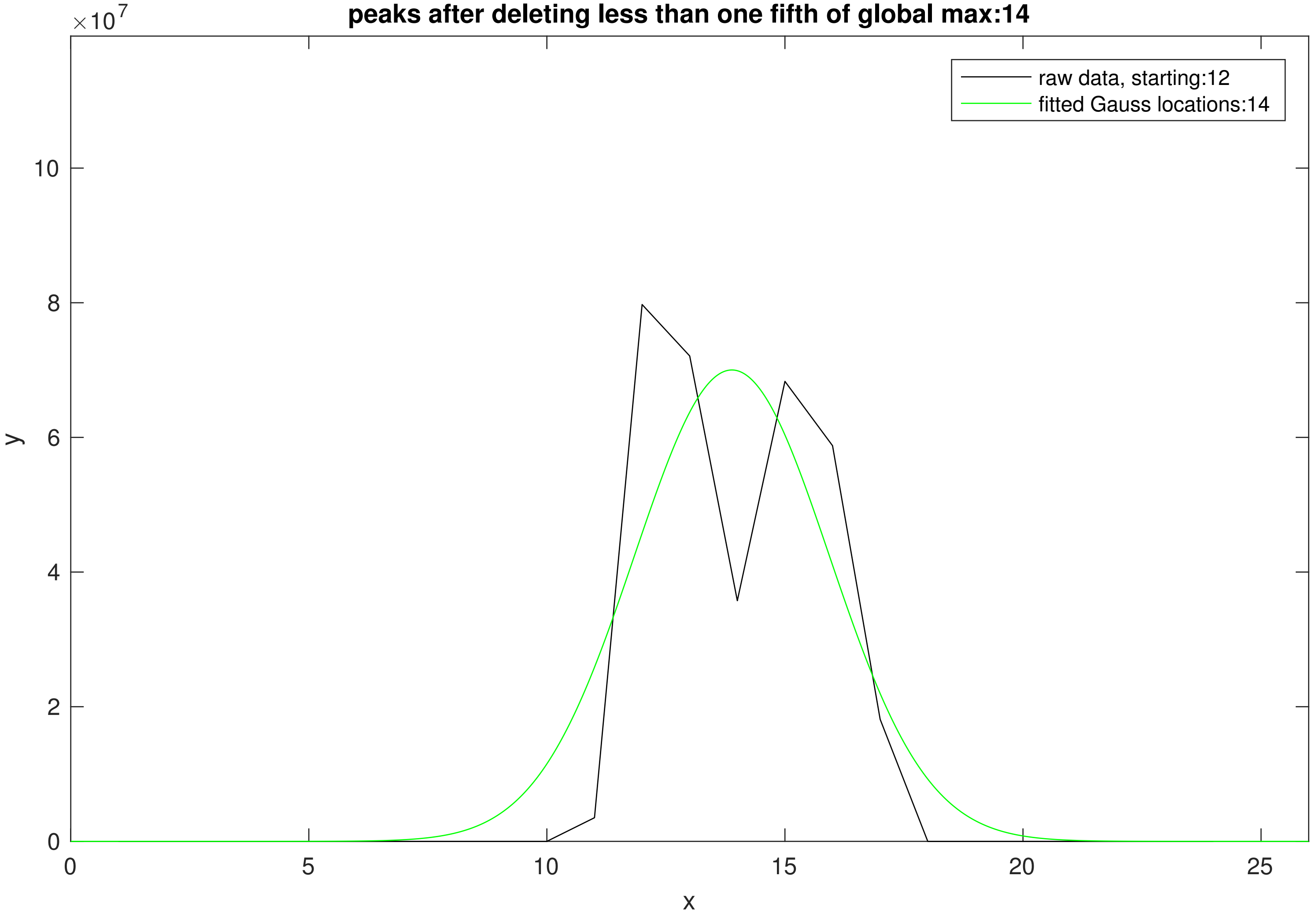

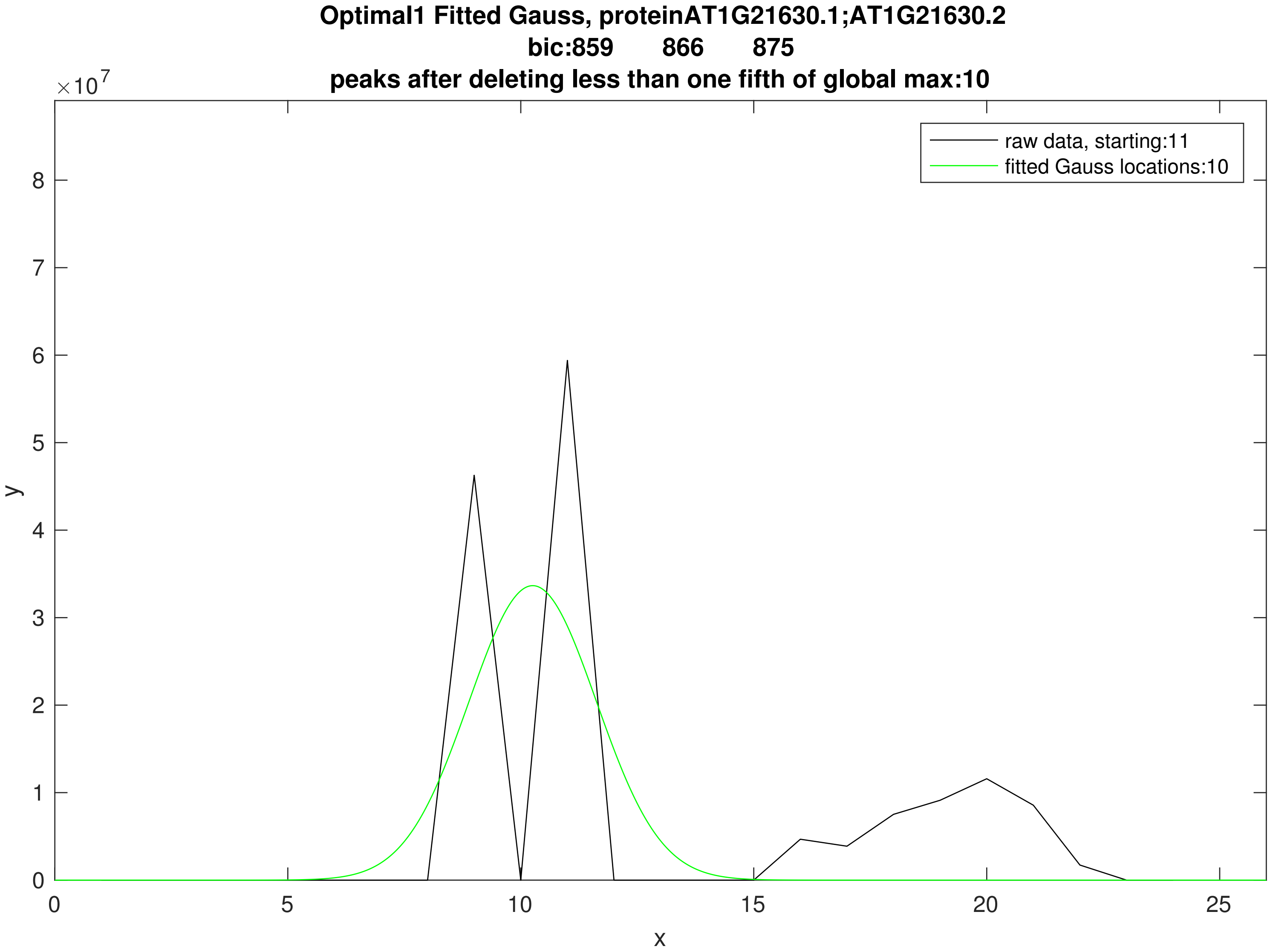

Optimal1 Fitted Gauss, proteinAT1G21630.1;AT1G21630.2
bic:859 866 875
peaks after deleting less than one fifth of global max:10
×10^7
raw data, starting:11
fitted Gauss locations:10
y
x
0
1
2
3
4
5
6
7
8
0
5
10
15
20
25

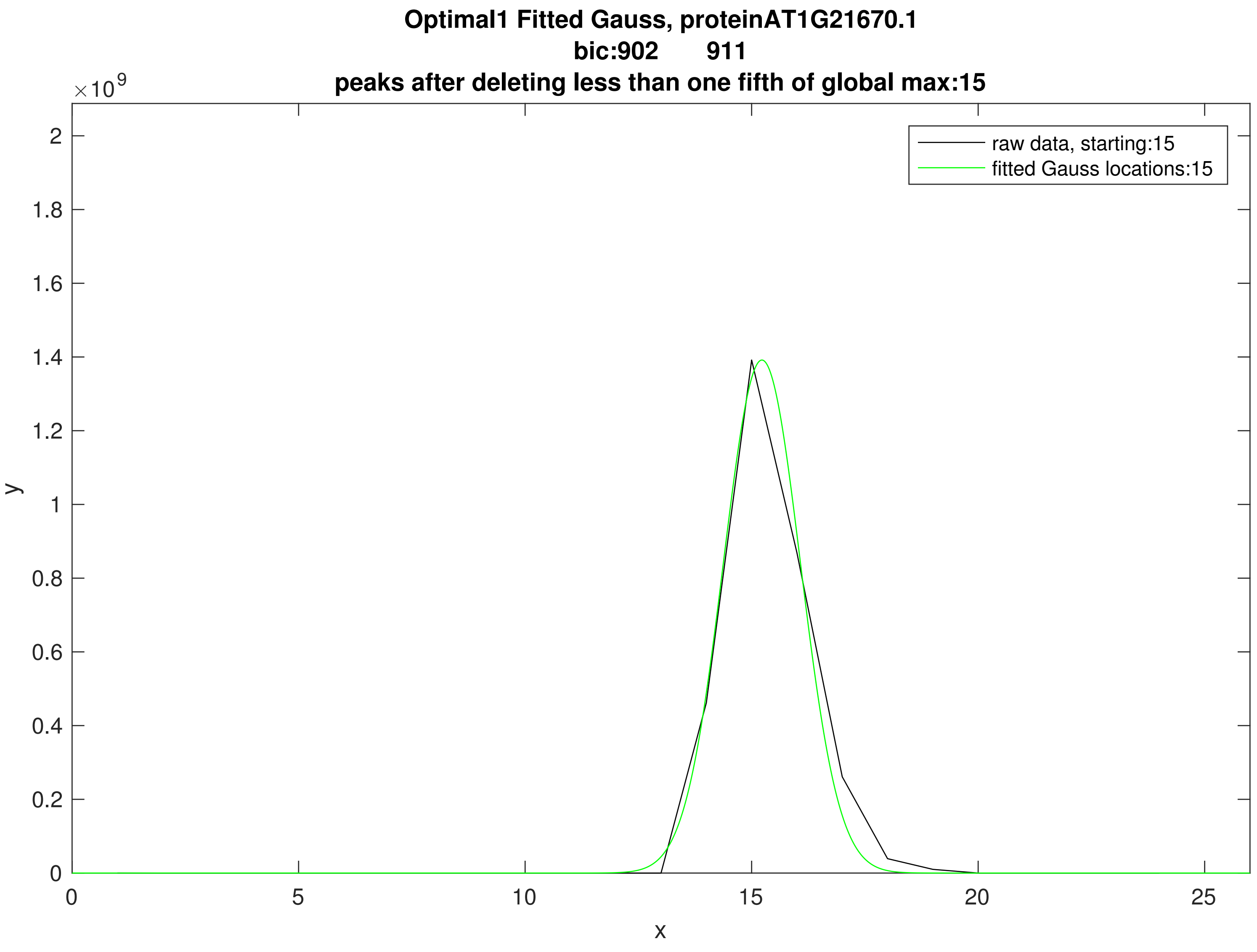

Optimal1 Fitted Gauss, proteinAT1G21670.1
bic:902 911
peaks after deleting less than one fifth of global max:15
×10^9
2
1.8
1.6
1.4
1.2
1
0.8
0.6
0.4
0.2
0
y
0
5
10
15
20
25
x
raw data, starting:15
fitted Gauss locations:15

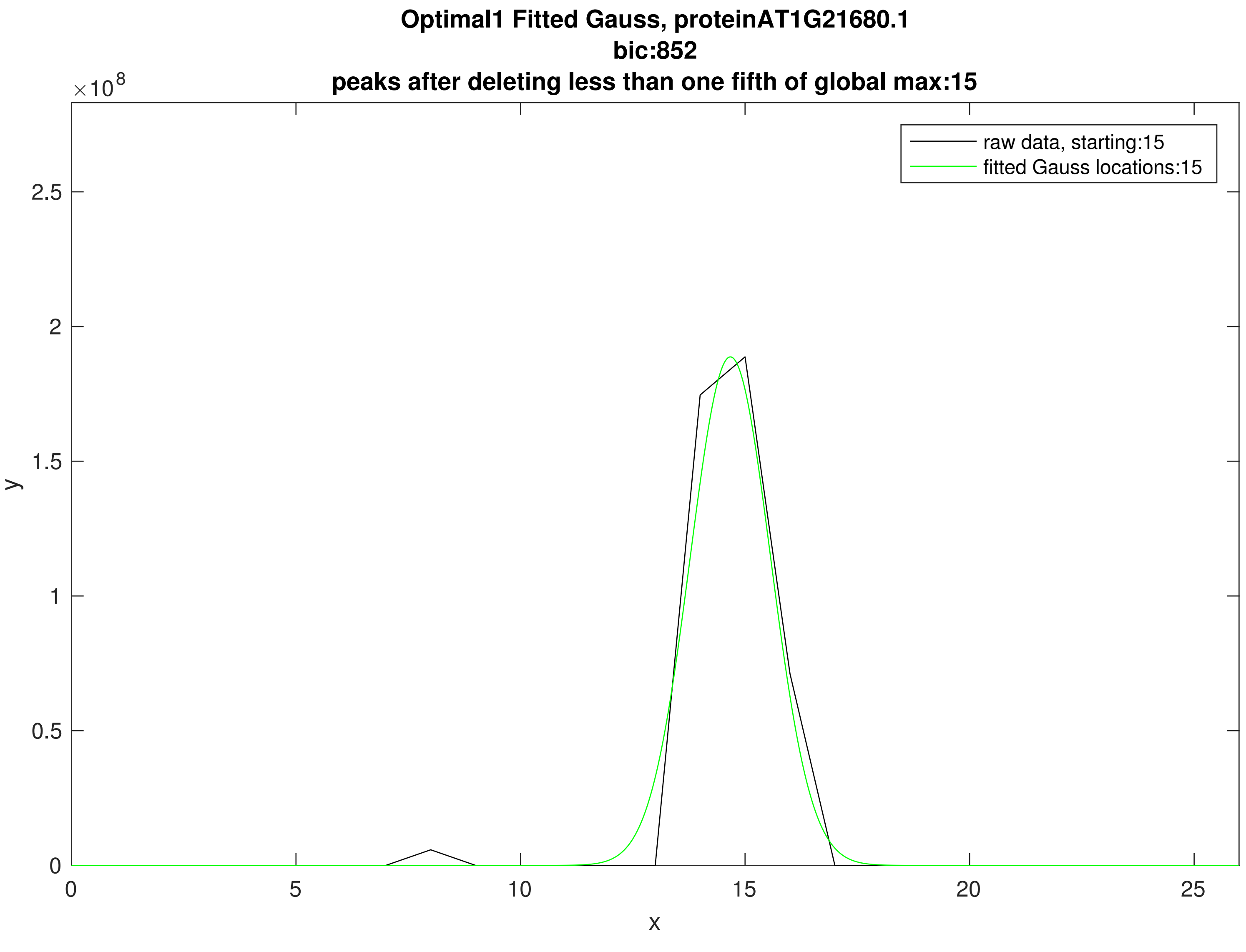
Optimal1 Fitted Gauss, proteinAT1G21680.1
bic:852
peaks after deleting less than one fifth of global max:15
×10^8
raw data, starting:15
fitted Gauss locations:15
y
x
0
0.5
1
1.5
2
2.5
0
5
10
15
20
25

**Optimal1 Fitted Gauss, proteinAT1G21720.1**
**bic:920 929**
**peaks after deleting less than one fifth of global max:5.4**

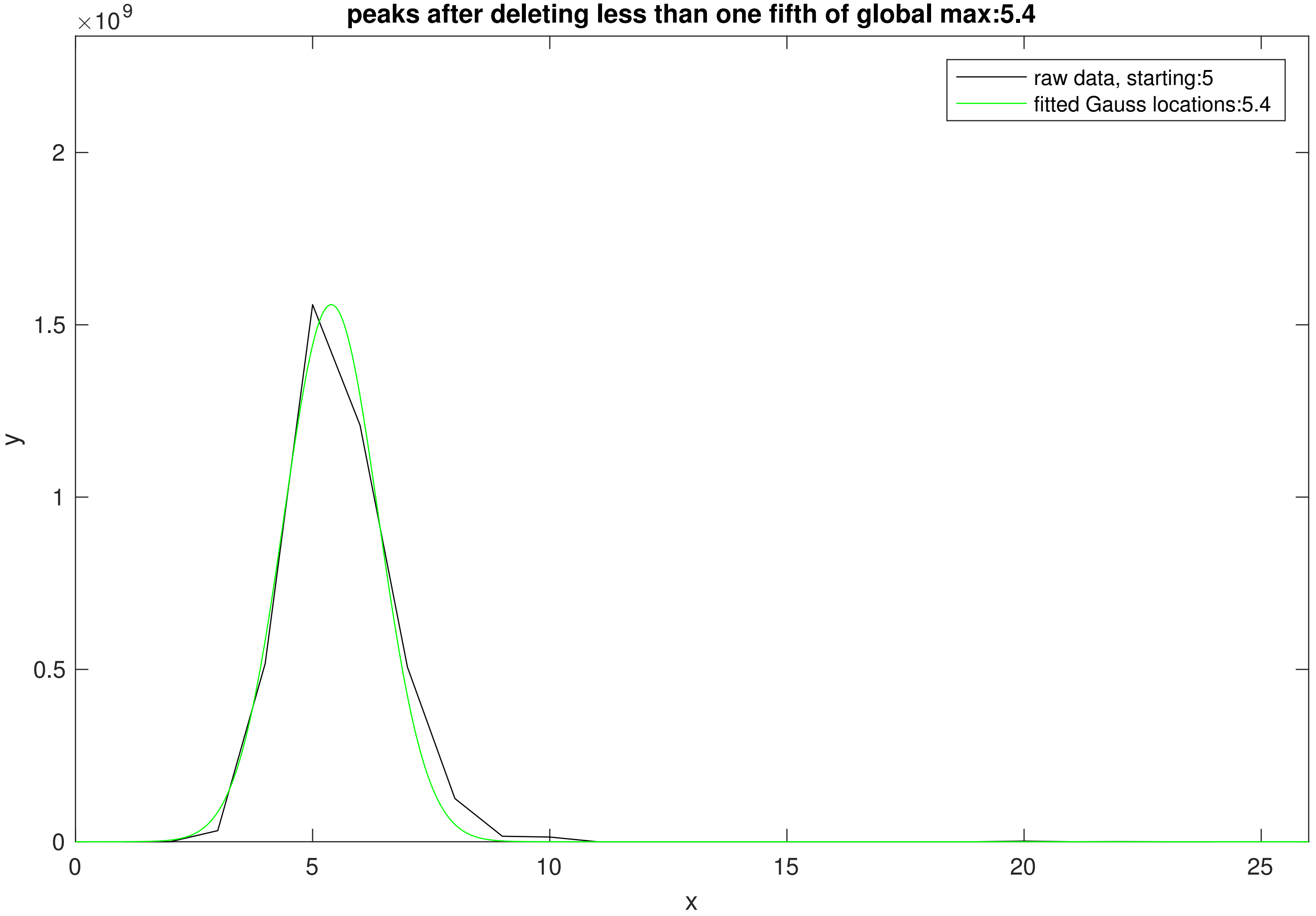

**Optimal1 Fitted Gauss, proteinAT1G21750.1;AT1G21750.2**
**bic:1.13e+03  1.15e+03  1.16e+03  1.17e+03**
**peaks after deleting less than one fifth of global max:13**

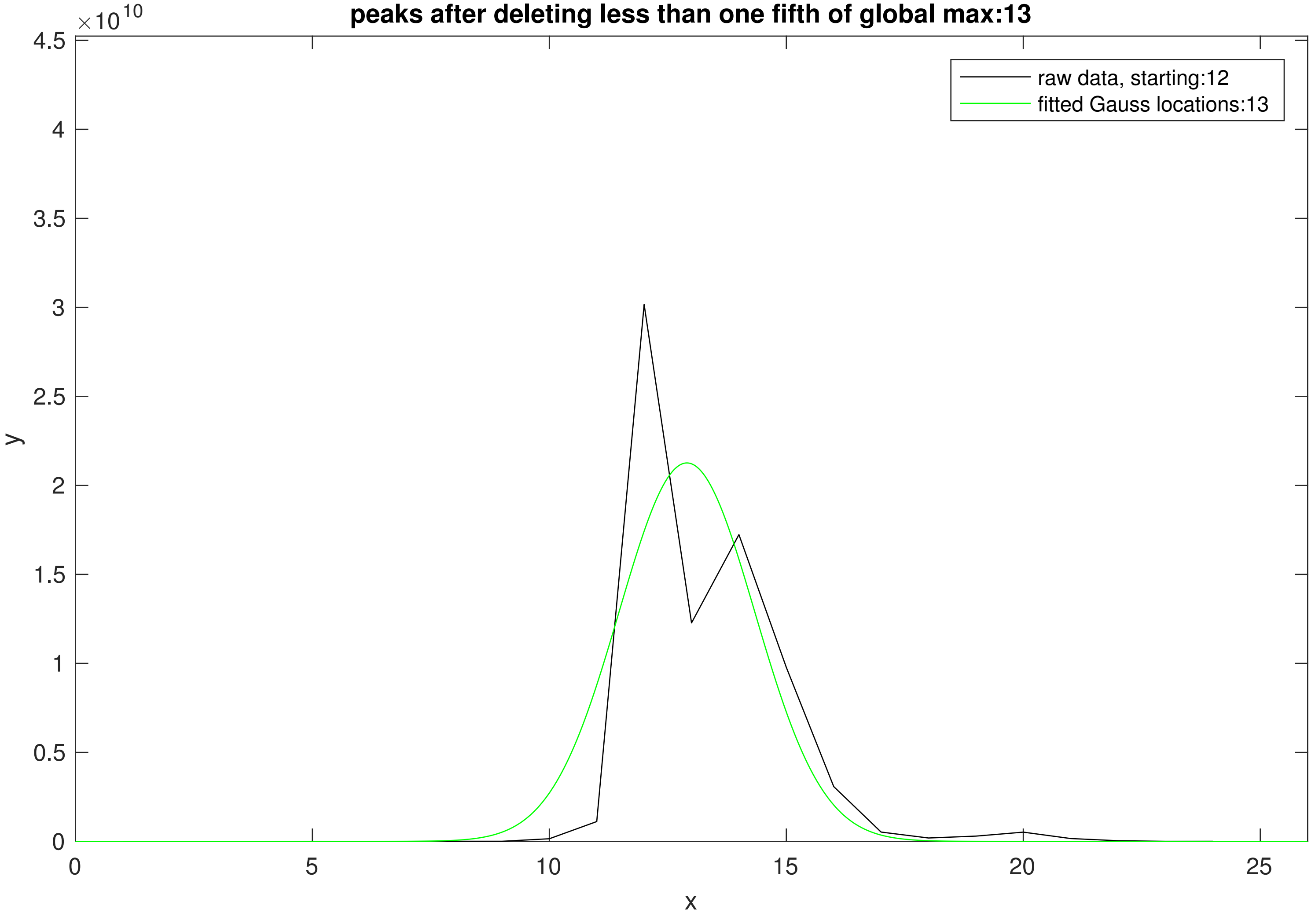

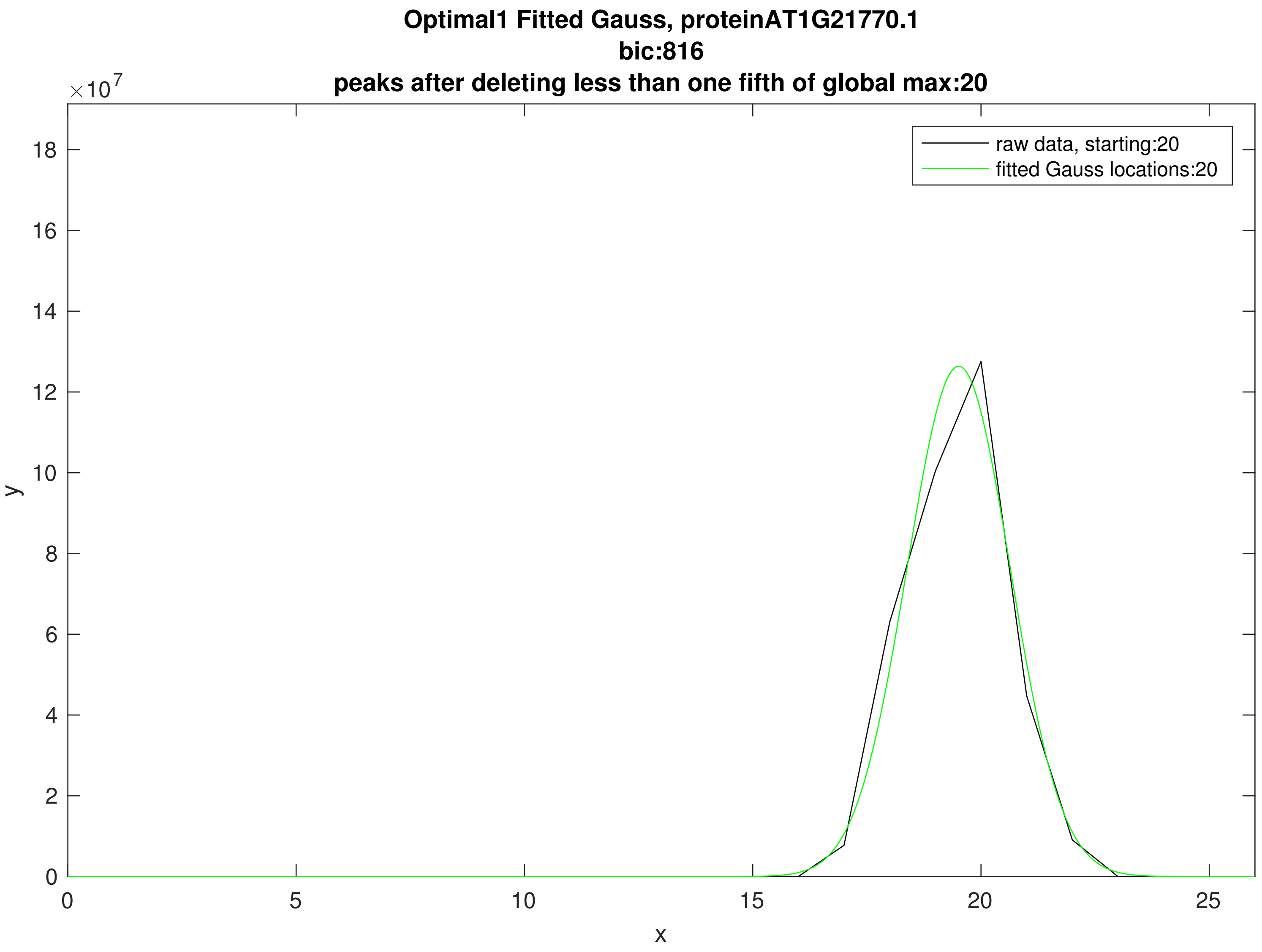

Optimal1 Fitted Gauss, proteinAT1G21770.1
bic:816
peaks after deleting less than one fifth of global max:20
×10^7
raw data, starting:20
fitted Gauss locations:20
y
x
0
2
4
6
8
10
12
14
16
18
0
5
10
15
20
25

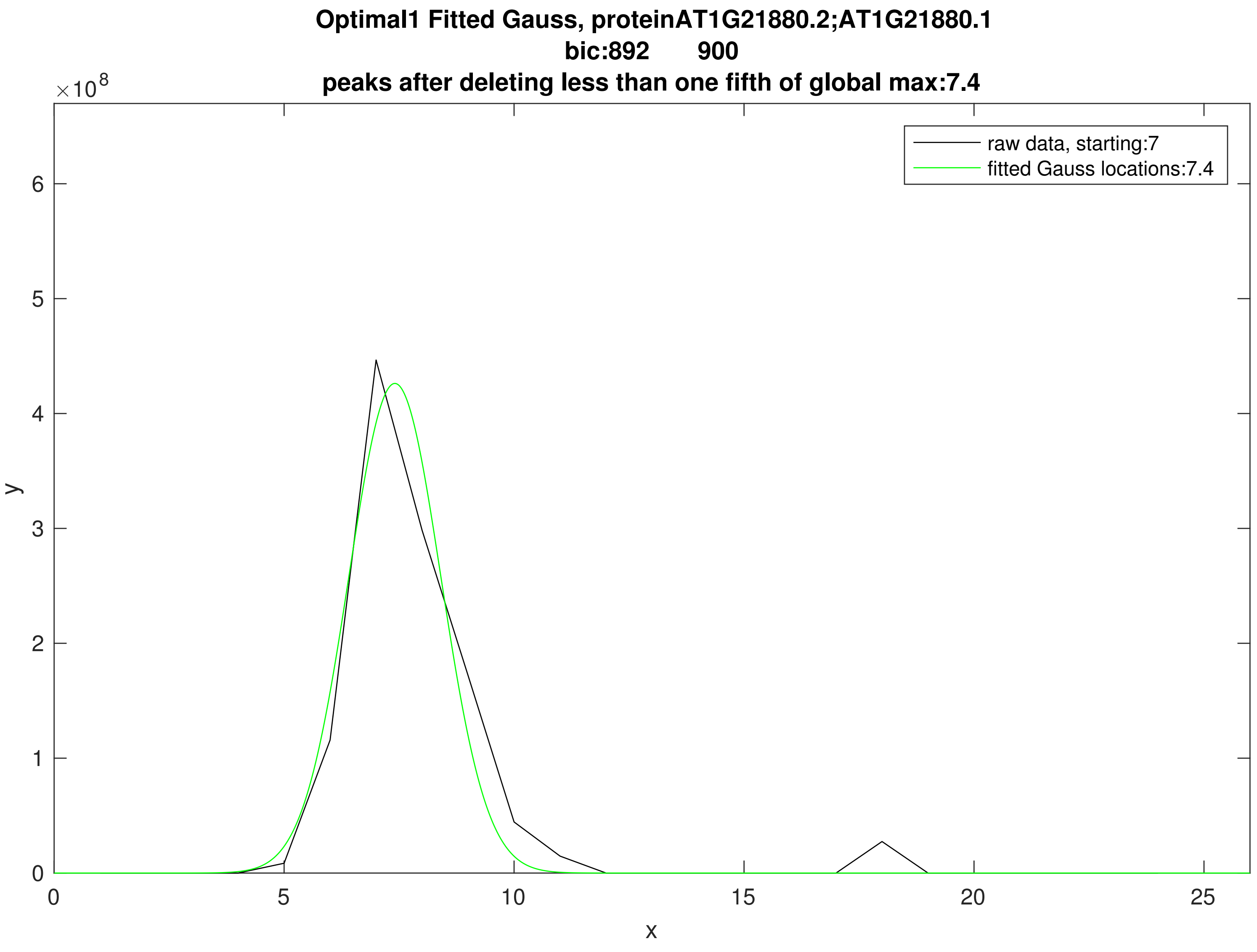
Optimal1 Fitted Gauss, proteinAT1G21880.2;AT1G21880.1
bic:892 900
peaks after deleting less than one fifth of global max:7.4
×10^8
raw data, starting:7
fitted Gauss locations:7.4
y
x
0
1
2
3
4
5
6
0
5
10
15
20
25

#### Optimal1 Fitted Gauss, proteinAT1G22020.1
#### bic:780
#### peaks after deleting less than one fifth of global max:11

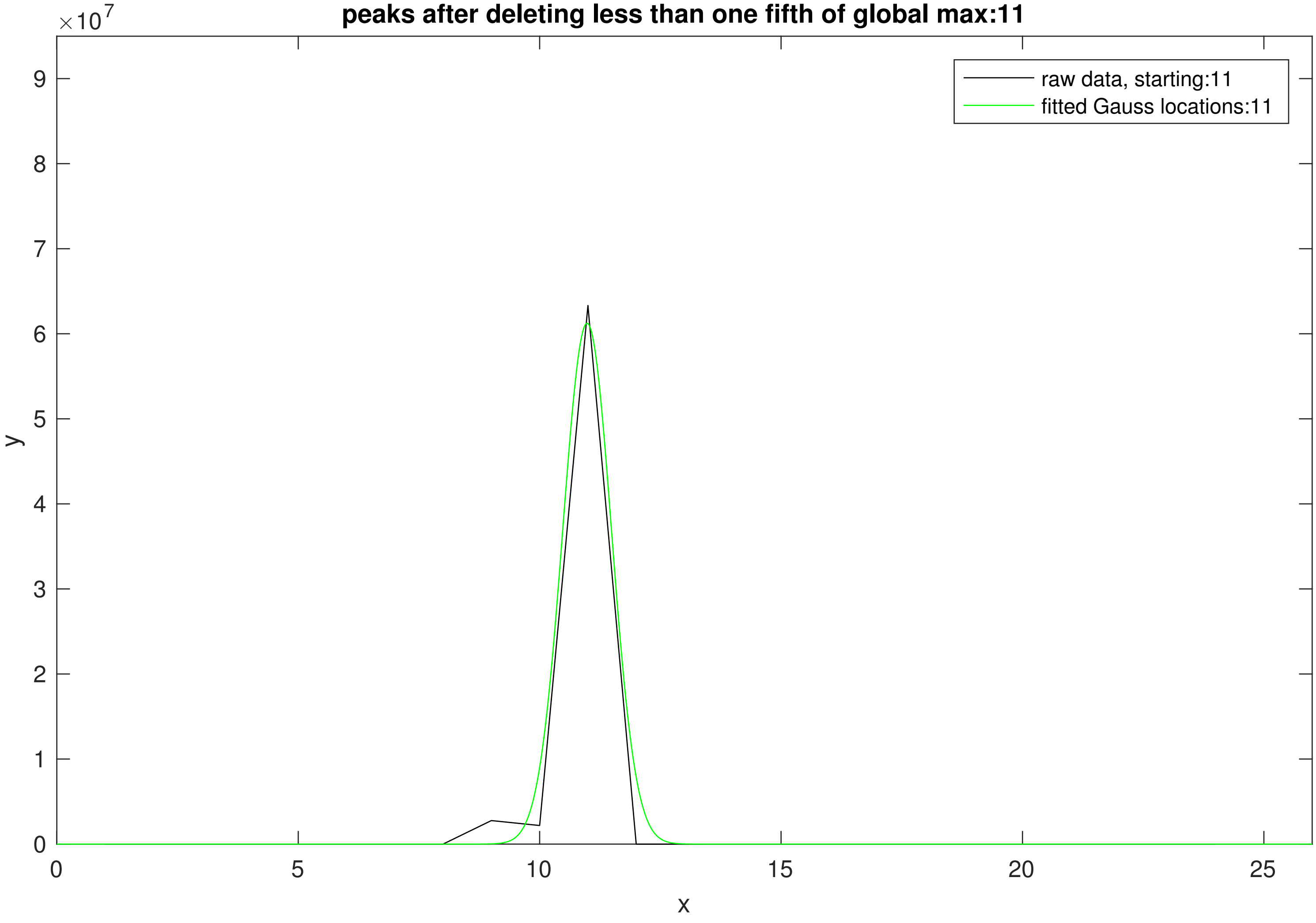

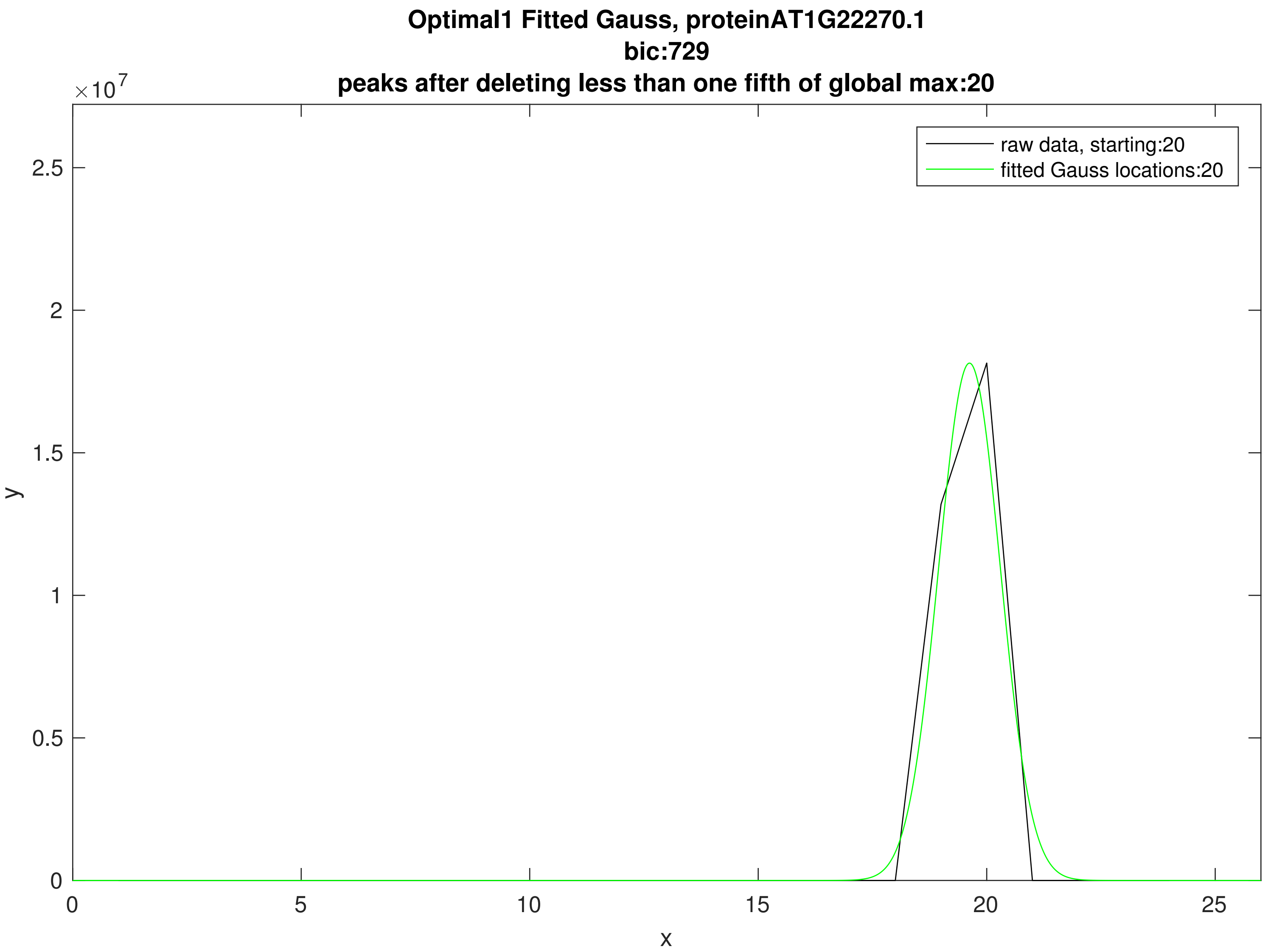

Optimal1 Fitted Gauss, proteinAT1G22270.1
bic:729
peaks after deleting less than one fifth of global max:20
×10^7
2.5
2
1.5
1
0.5
0
y
0
5
10
15
20
25
x
raw data, starting:20
fitted Gauss locations:20

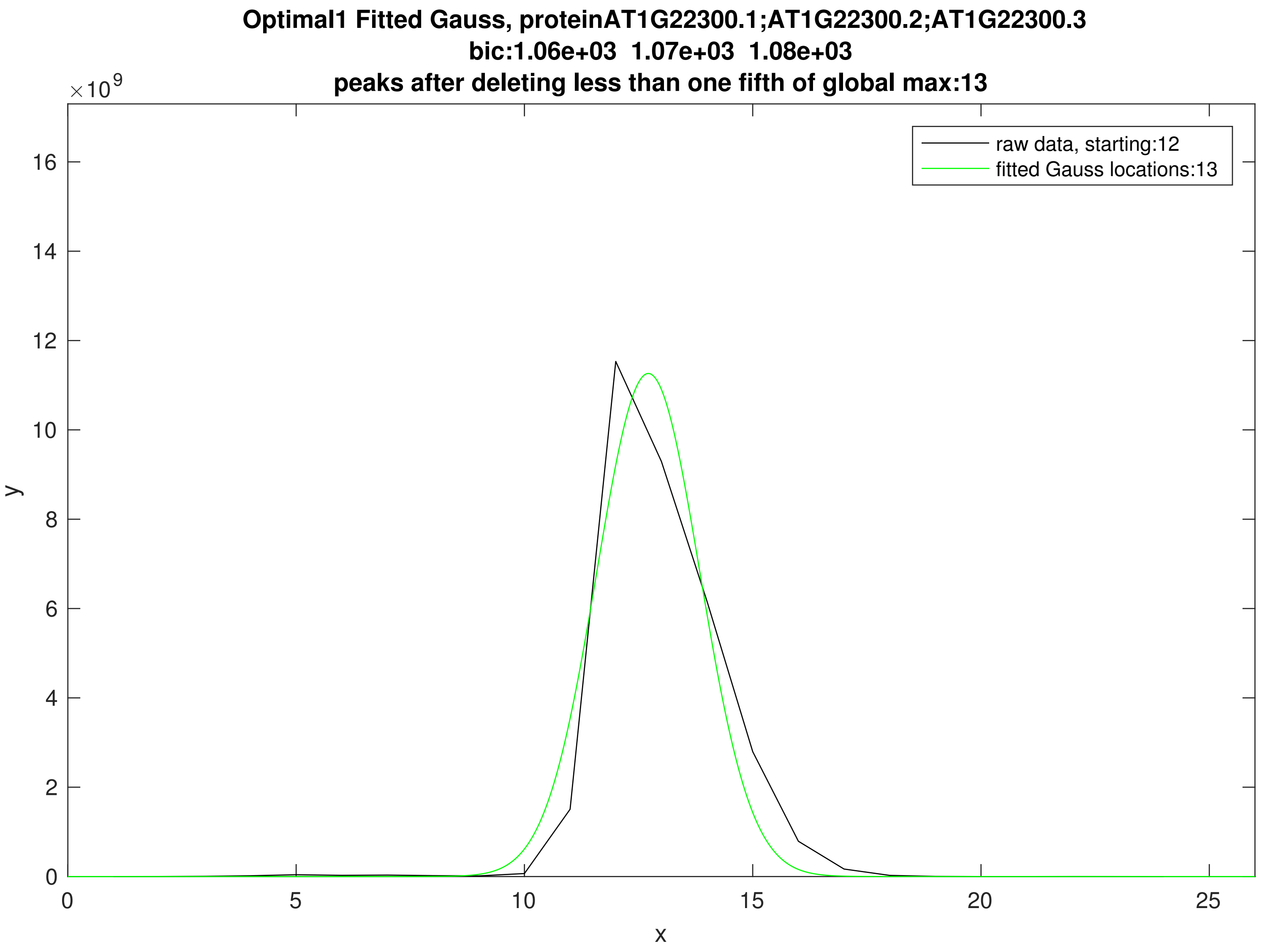
Optimal1 Fitted Gauss, proteinAT1G22300.1;AT1G22300.2;AT1G22300.3
bic:1.06e+03  1.07e+03  1.08e+03
peaks after deleting less than one fifth of global max:13
×10^9
raw data, starting:12
fitted Gauss locations:13
16
14
12
10
8
6
4
2
0
y
0
5
10
15
20
25
x

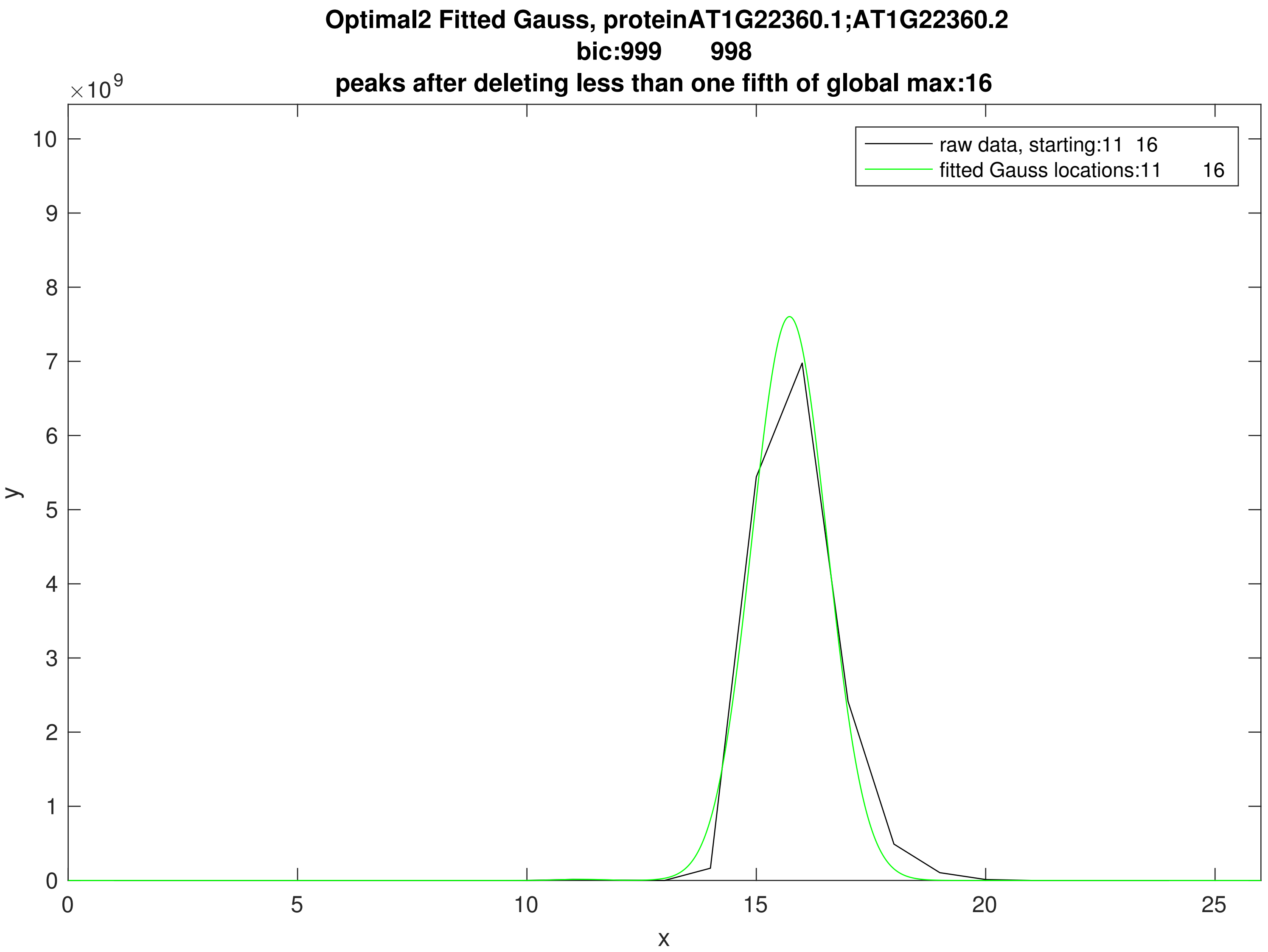
Optimal2 Fitted Gauss, proteinAT1G22360.1;AT1G22360.2
bic:999 998
peaks after deleting less than one fifth of global max:16
×10^9
raw data, starting:11 16
fitted Gauss locations:11 16
y
x
0
1
2
3
4
5
6
7
8
9
10
0
5
10
15
20
25

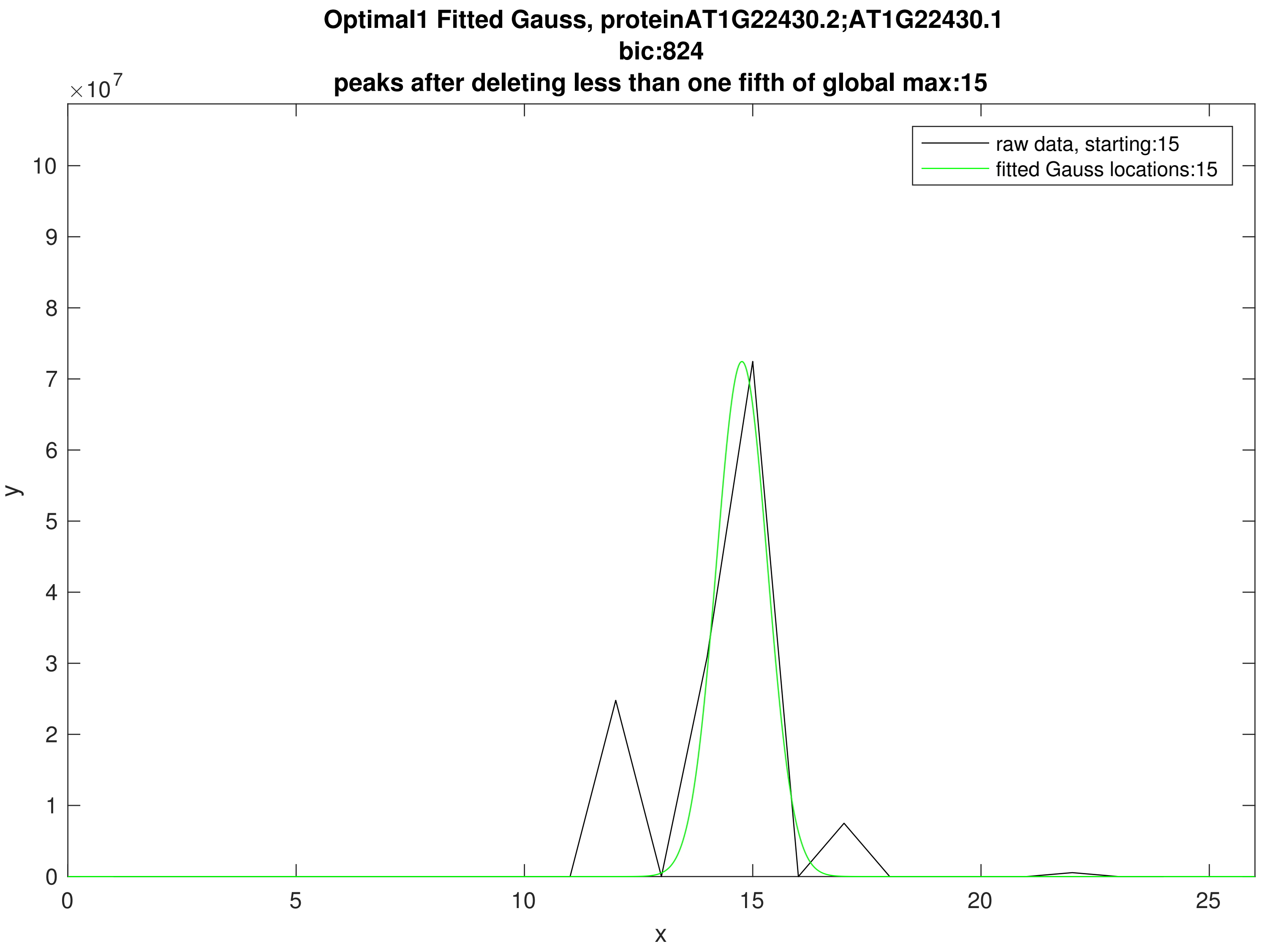
Optimal1 Fitted Gauss, proteinAT1G22430.2;AT1G22430.1
bic:824
peaks after deleting less than one fifth of global max:15
×10^7
raw data, starting:15
fitted Gauss locations:15
y
x
0
1
2
3
4
5
6
7
8
9
10
0
5
10
15
20
25

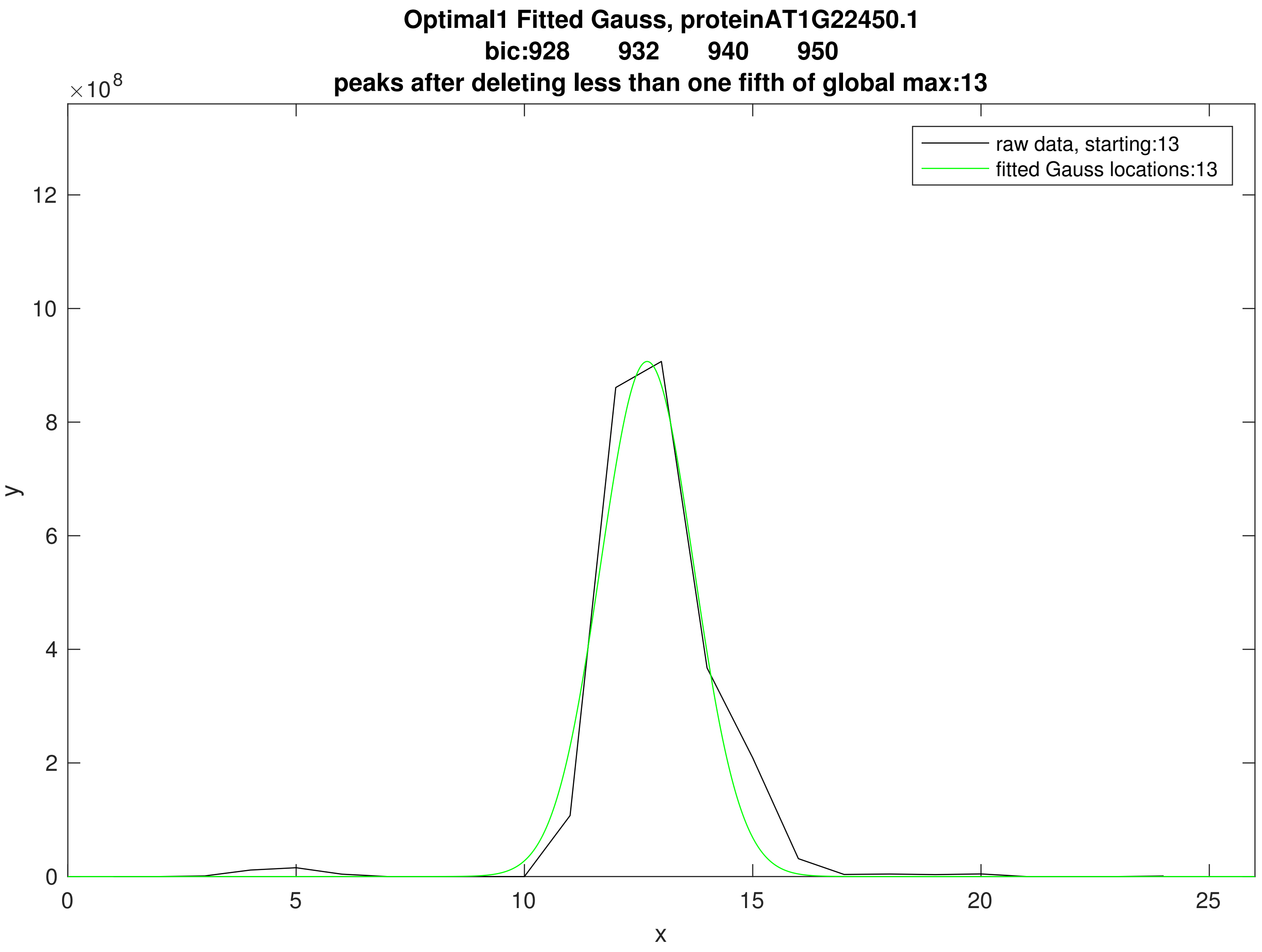

Optimal1 Fitted Gauss, proteinAT1G22450.1
bic:928 932 940 950
peaks after deleting less than one fifth of global max:13
×10^8
raw data, starting:13
fitted Gauss locations:13
12
10
8
6
4
2
0
y
0
5
10
15
20
25
x

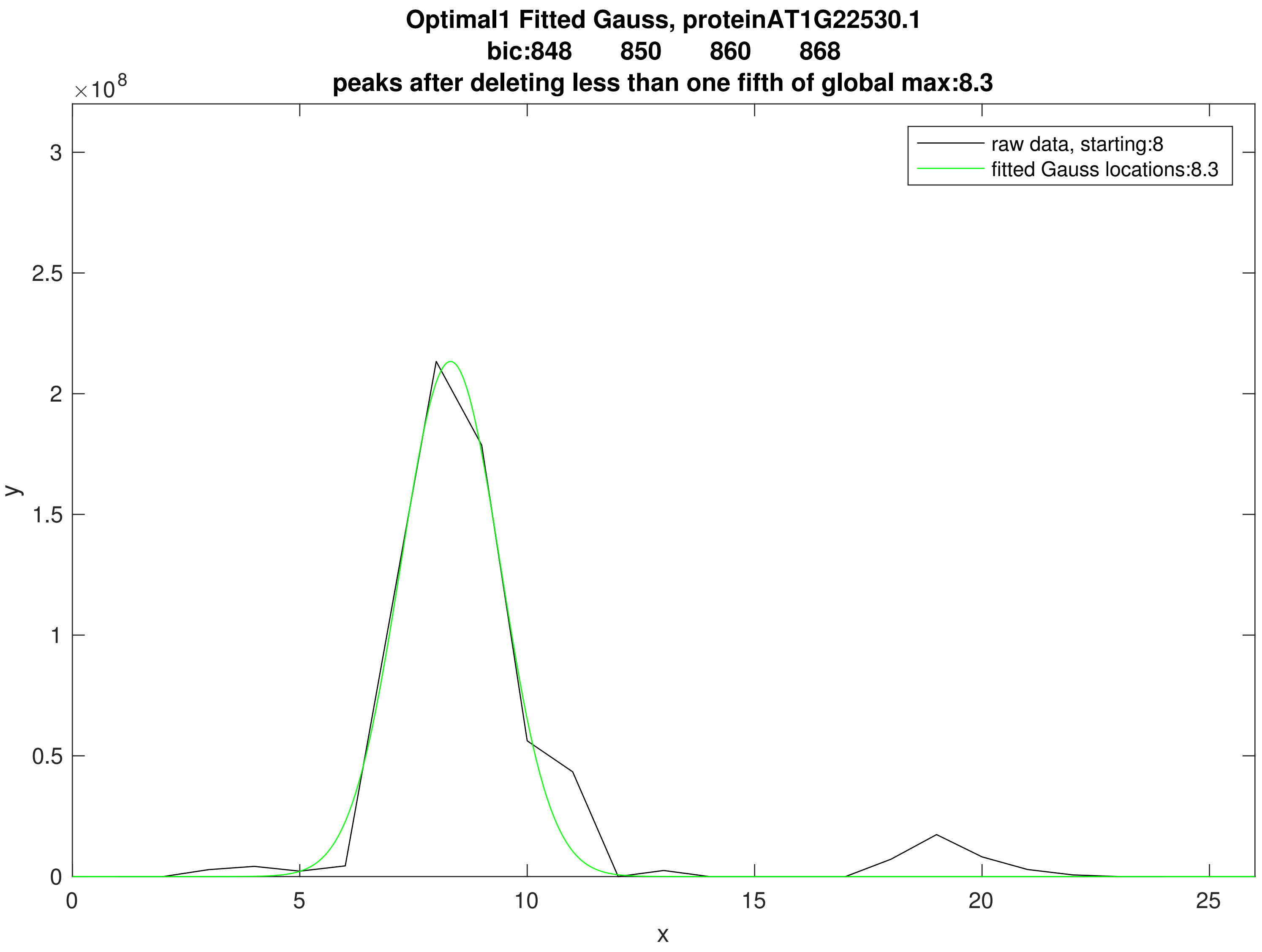
Optimal1 Fitted Gauss, proteinAT1G22530.1
bic:848 850 860 868
peaks after deleting less than one fifth of global max:8.3
×10^8
raw data, starting:8
fitted Gauss locations:8.3
3
2.5
2
1.5
1
0.5
0
y
0
5
10
15
20
25
x

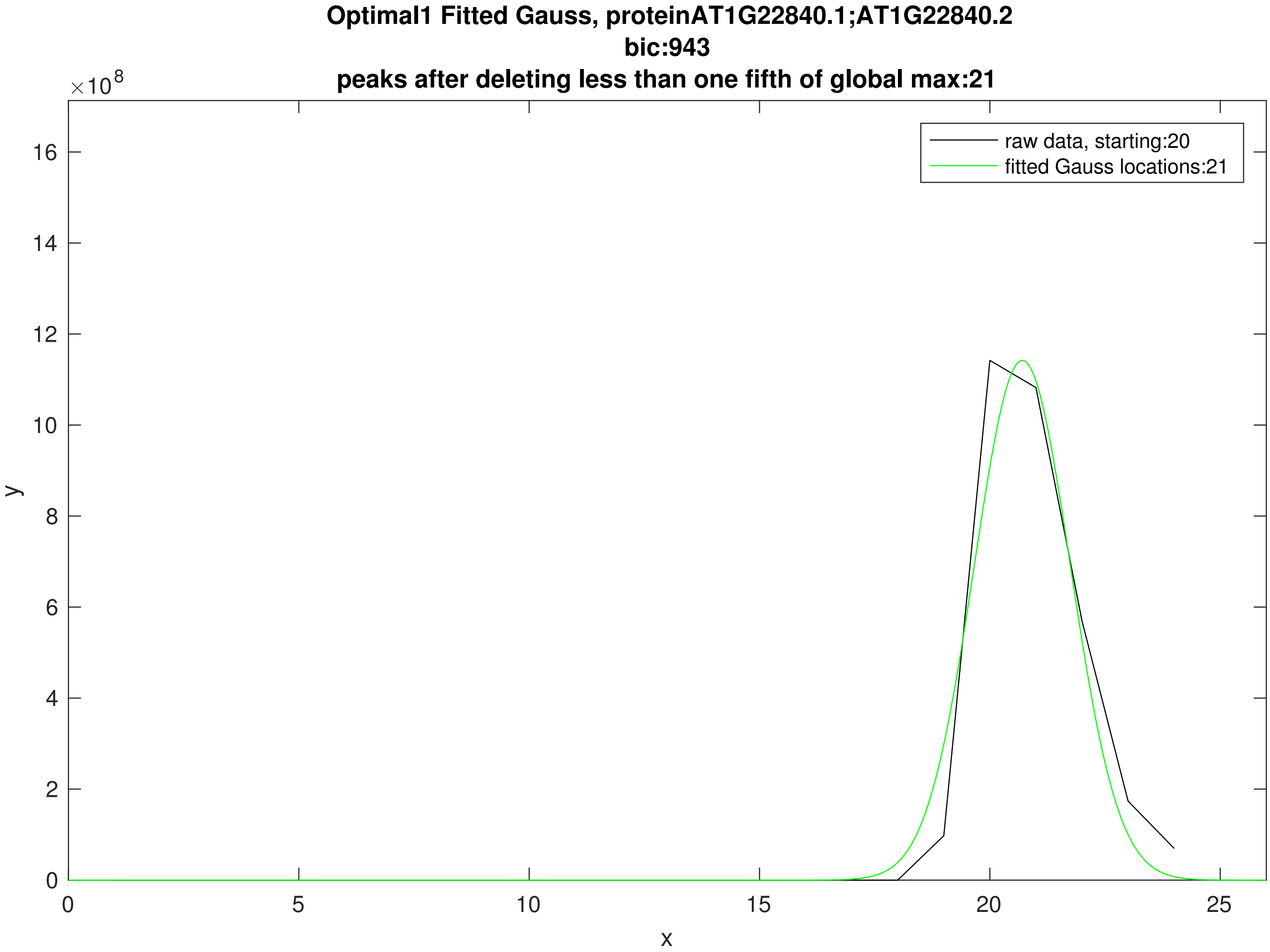
Optimal1 Fitted Gauss, proteinAT1G22840.1;AT1G22840.2
bic:943
peaks after deleting less than one fifth of global max:21
×10^8
raw data, starting:20
fitted Gauss locations:21
y
x
0
2
4
6
8
10
12
14
16
0
5
10
15
20
25

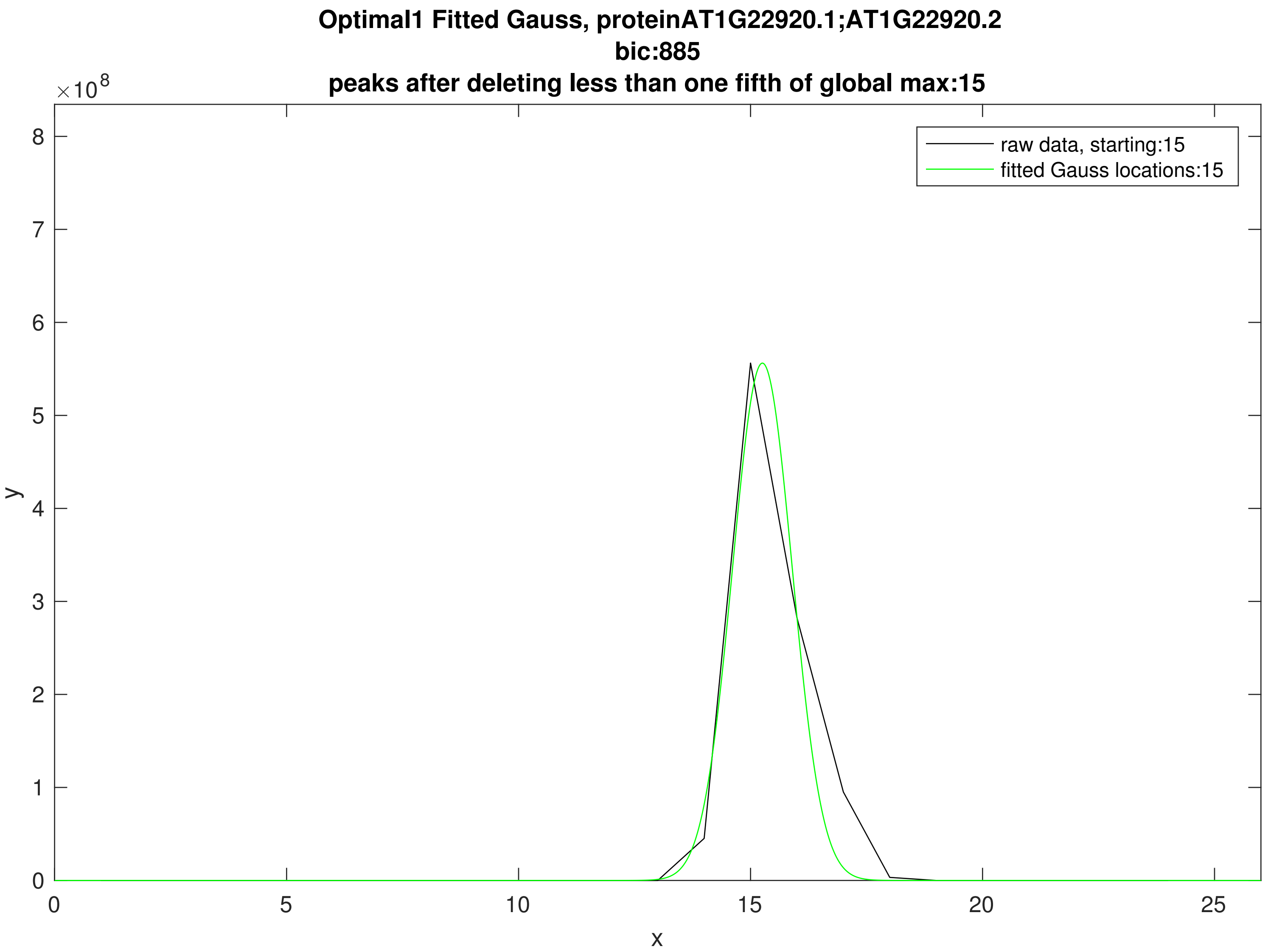

Optimal1 Fitted Gauss, proteinAT1G22920.1;AT1G22920.2
bic:885
peaks after deleting less than one fifth of global max:15
×10^8
raw data, starting:15
fitted Gauss locations:15
y
x
0
1
2
3
4
5
6
7
8
0
5
10
15
20
25

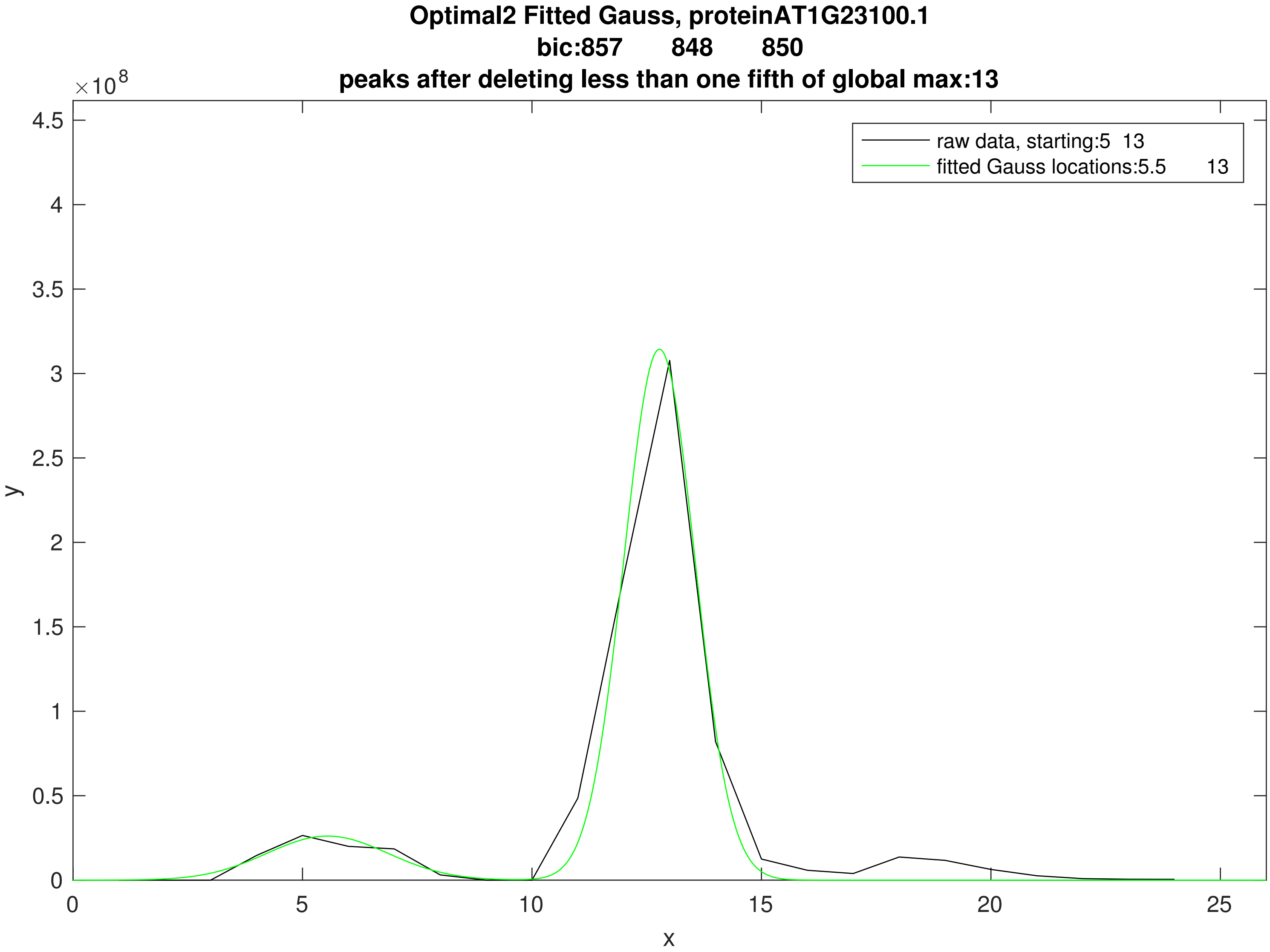

Optimal2 Fitted Gauss, proteinAT1G23100.1
bic:857 848 850
peaks after deleting less than one fifth of global max:13
×10^8
4.5
4
3.5
3
2.5
2
1.5
1
0.5
0
y
0
5
10
15
20
25
x
raw data, starting:5 13
fitted Gauss locations:5.5 13

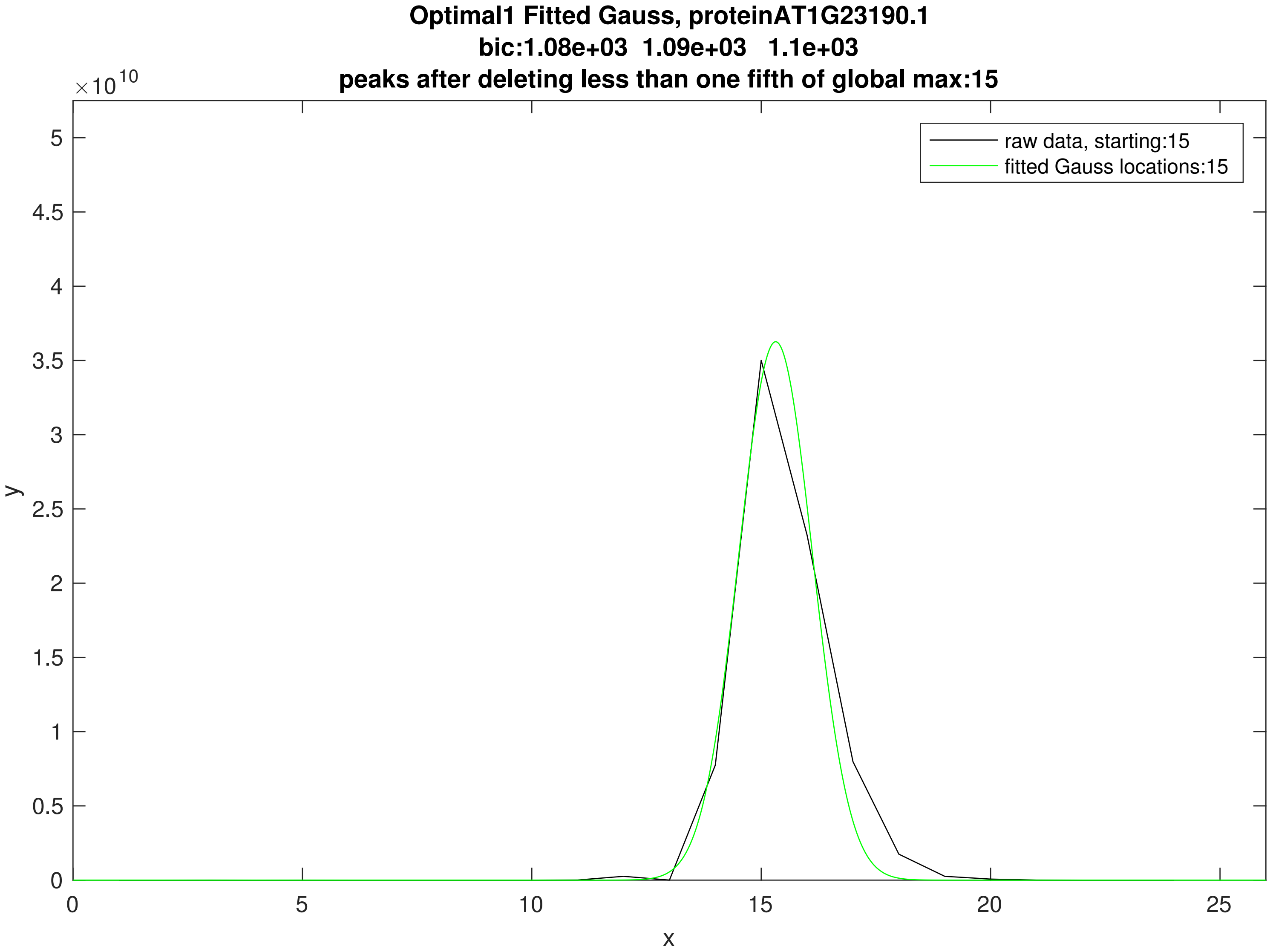

Optimal1 Fitted Gauss, proteinAT1G23190.1
bic:1.08e+03 1.09e+03 1.1e+03
peaks after deleting less than one fifth of global max:15
×10^10
5
4.5
4
3.5
3
2.5
2
1.5
1
0.5
0
y
0
5
10
15
20
25
x
raw data, starting:15
fitted Gauss locations:15

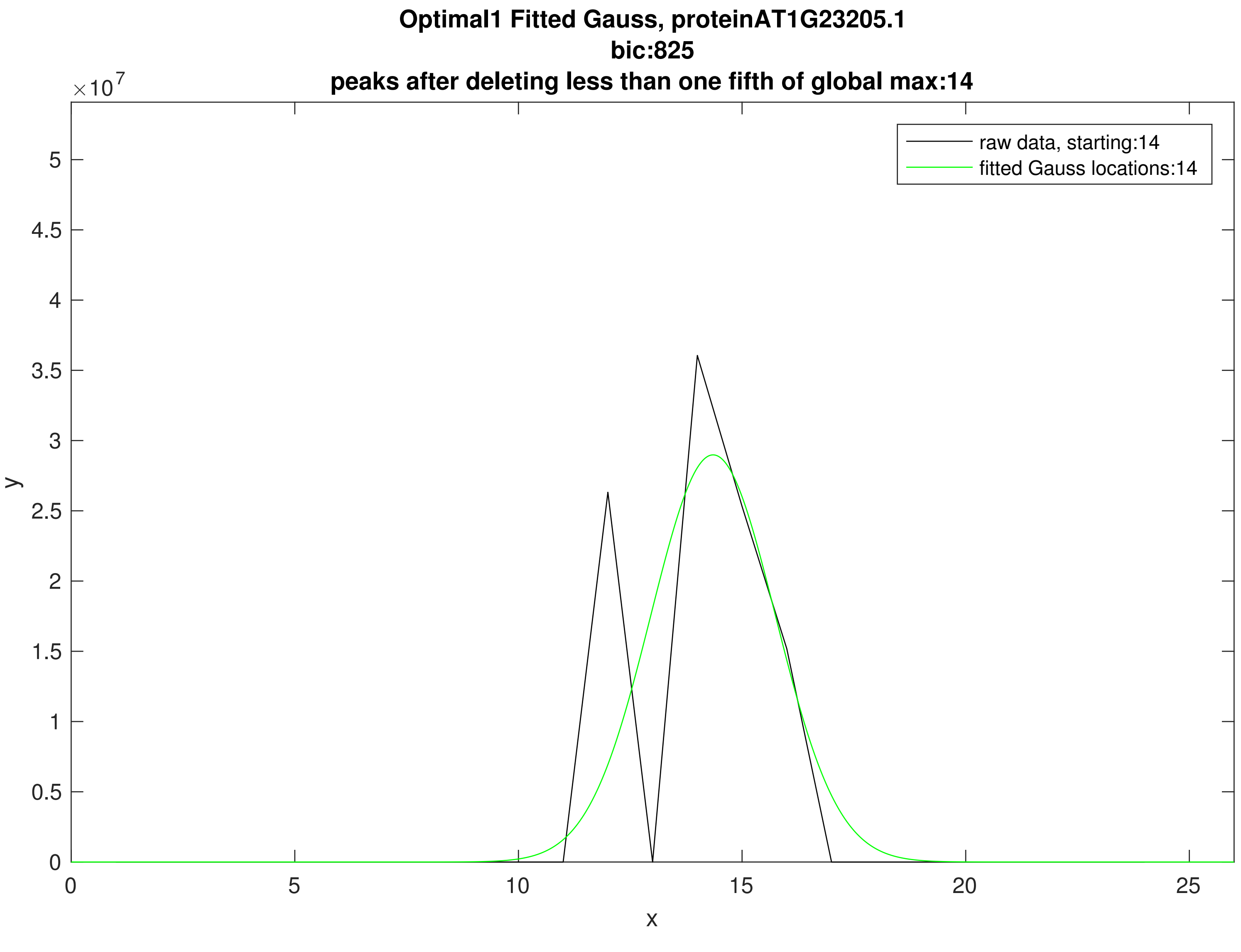

Optimal1 Fitted Gauss, proteinAT1G23205.1
bic:825
peaks after deleting less than one fifth of global max:14
×10^7
5
4.5
4
3.5
3
2.5
2
1.5
1
0.5
0
y
0
5
10
15
20
25
x
raw data, starting:14
fitted Gauss locations:14

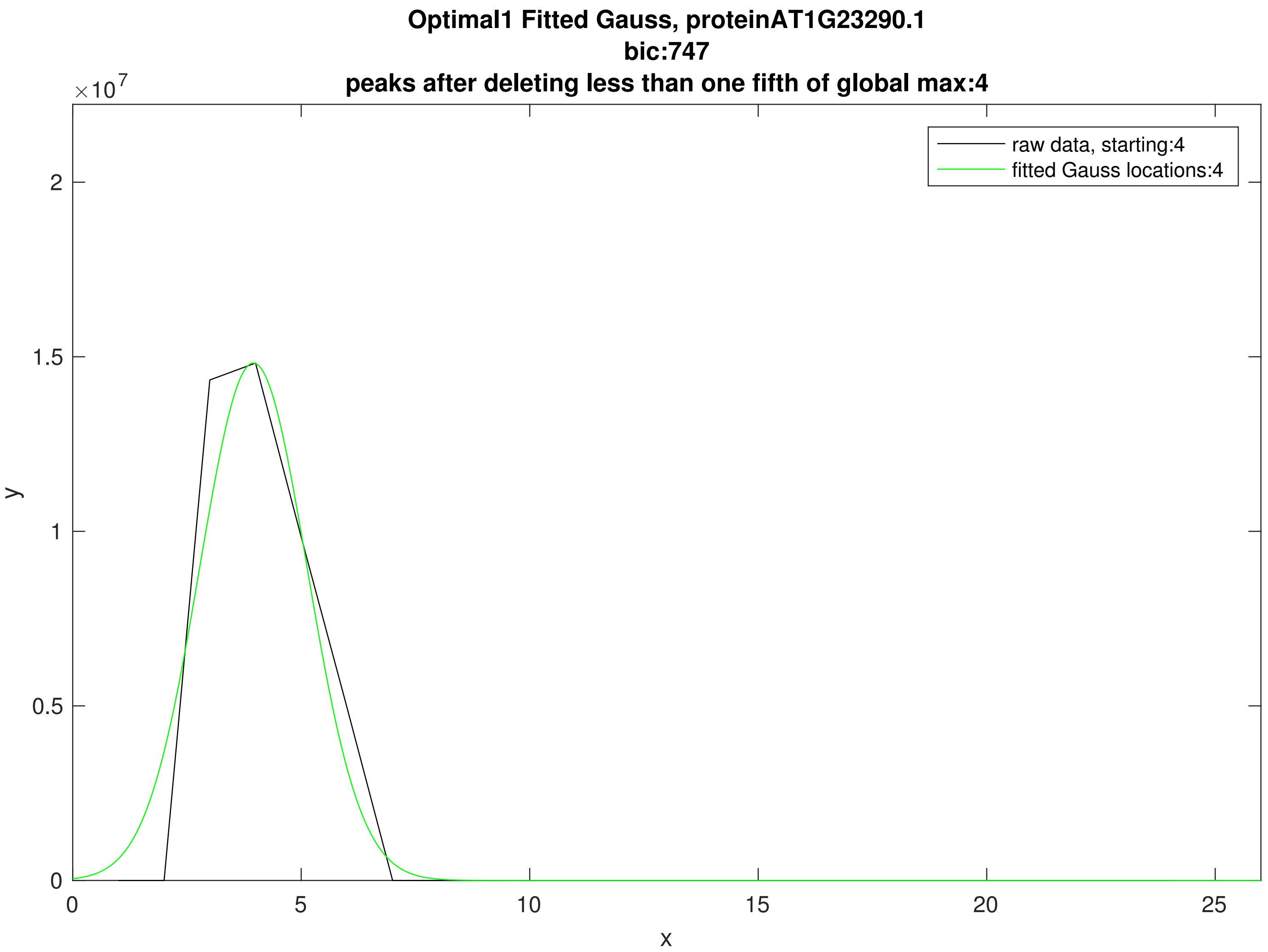

Optimal1 Fitted Gauss, proteinAT1G23290.1
bic:747
peaks after deleting less than one fifth of global max:4
×10^7
raw data, starting:4
fitted Gauss locations:4
2
1.5
1
0.5
0
y
0
5
10
15
20
25
x

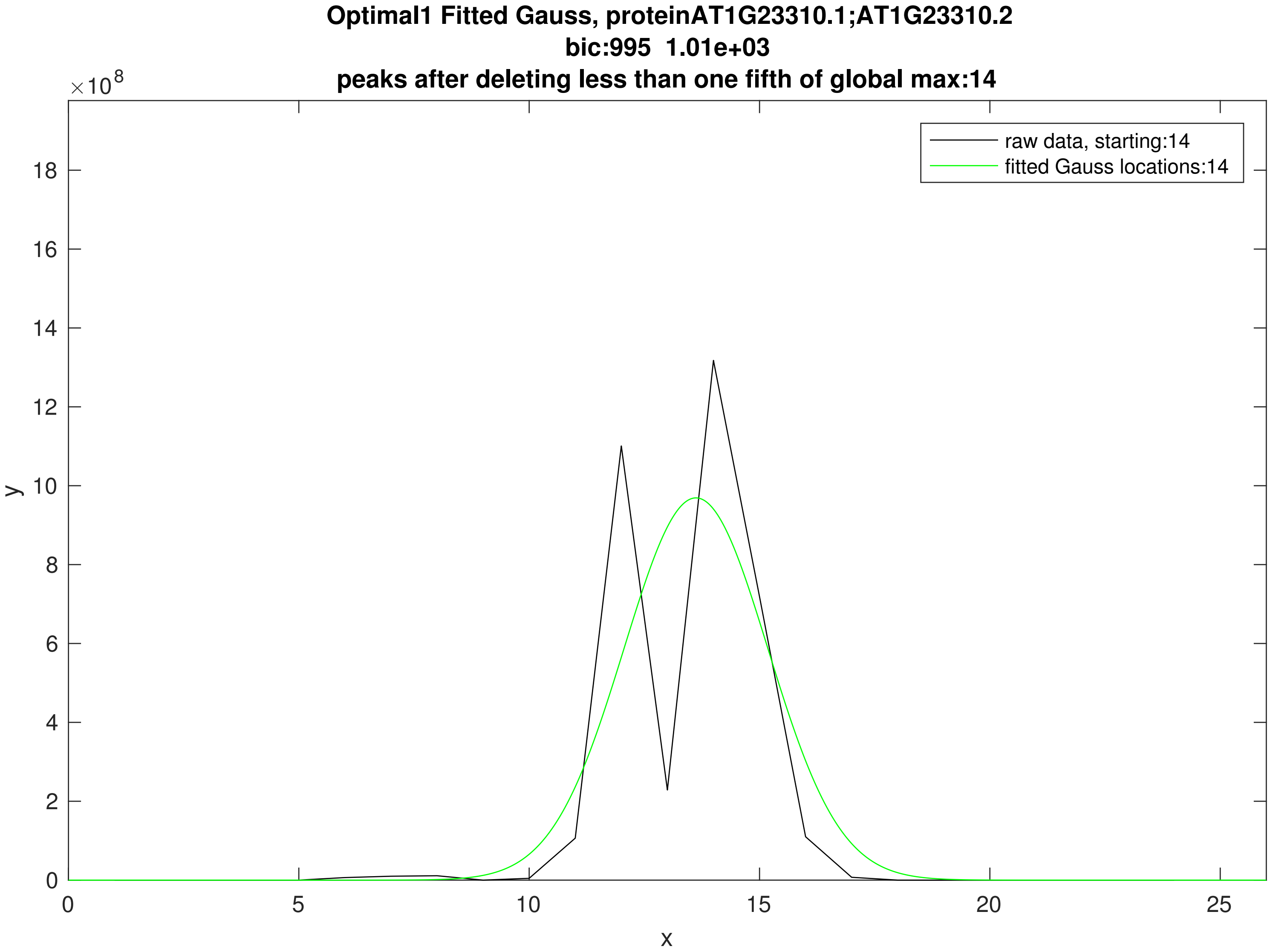

Optimal1 Fitted Gauss, proteinAT1G23310.1;AT1G23310.2
bic:995  1.01e+03
peaks after deleting less than one fifth of global max:14
×10^8
raw data, starting:14
fitted Gauss locations:14
y
x
0
2
4
6
8
10
12
14
16
18
0
5
10
15
20
25

Optimal1 Fitted Gauss, proteinAT1G23440.1
bic:904 906
peaks after deleting less than one fifth of global max:19

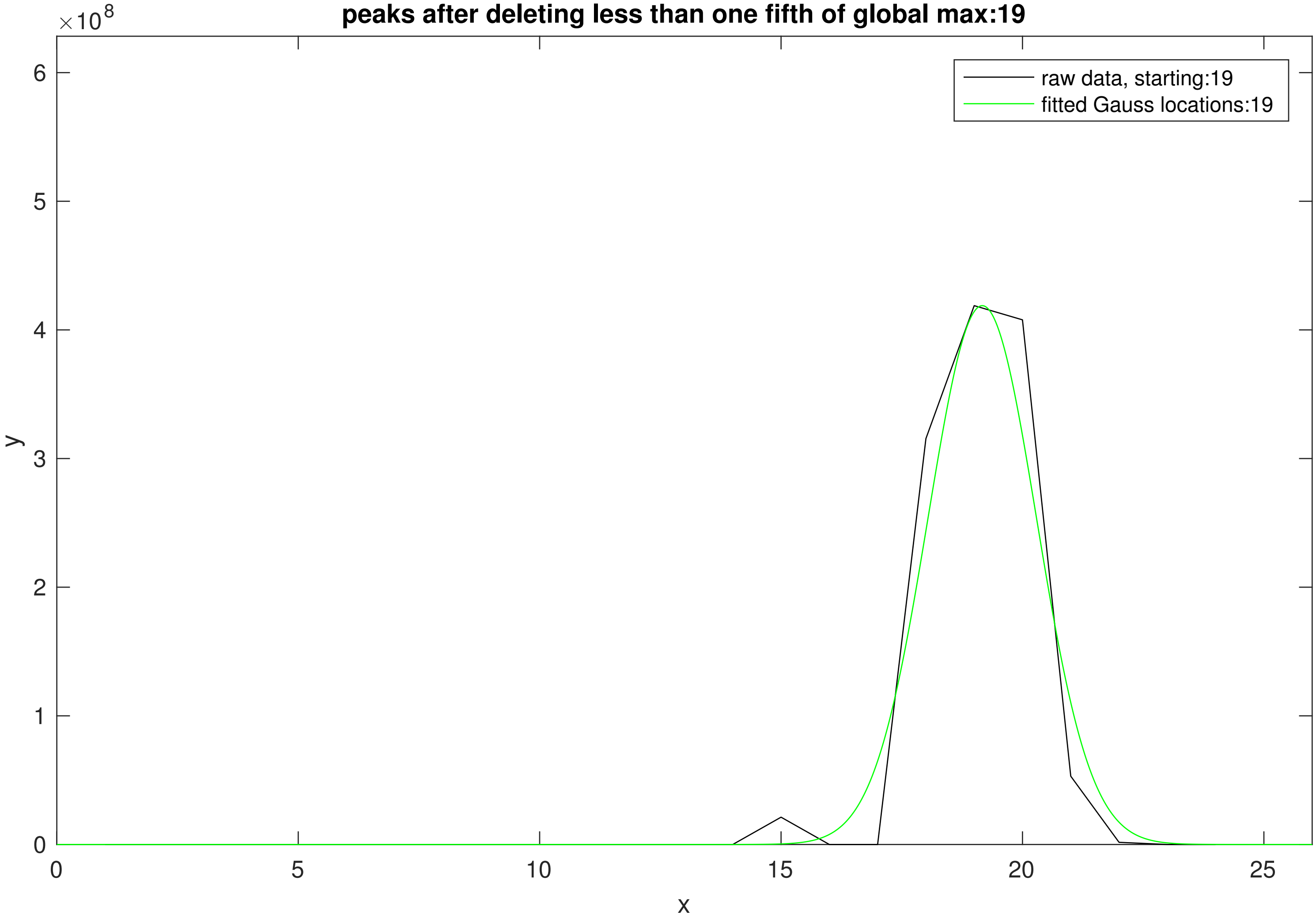

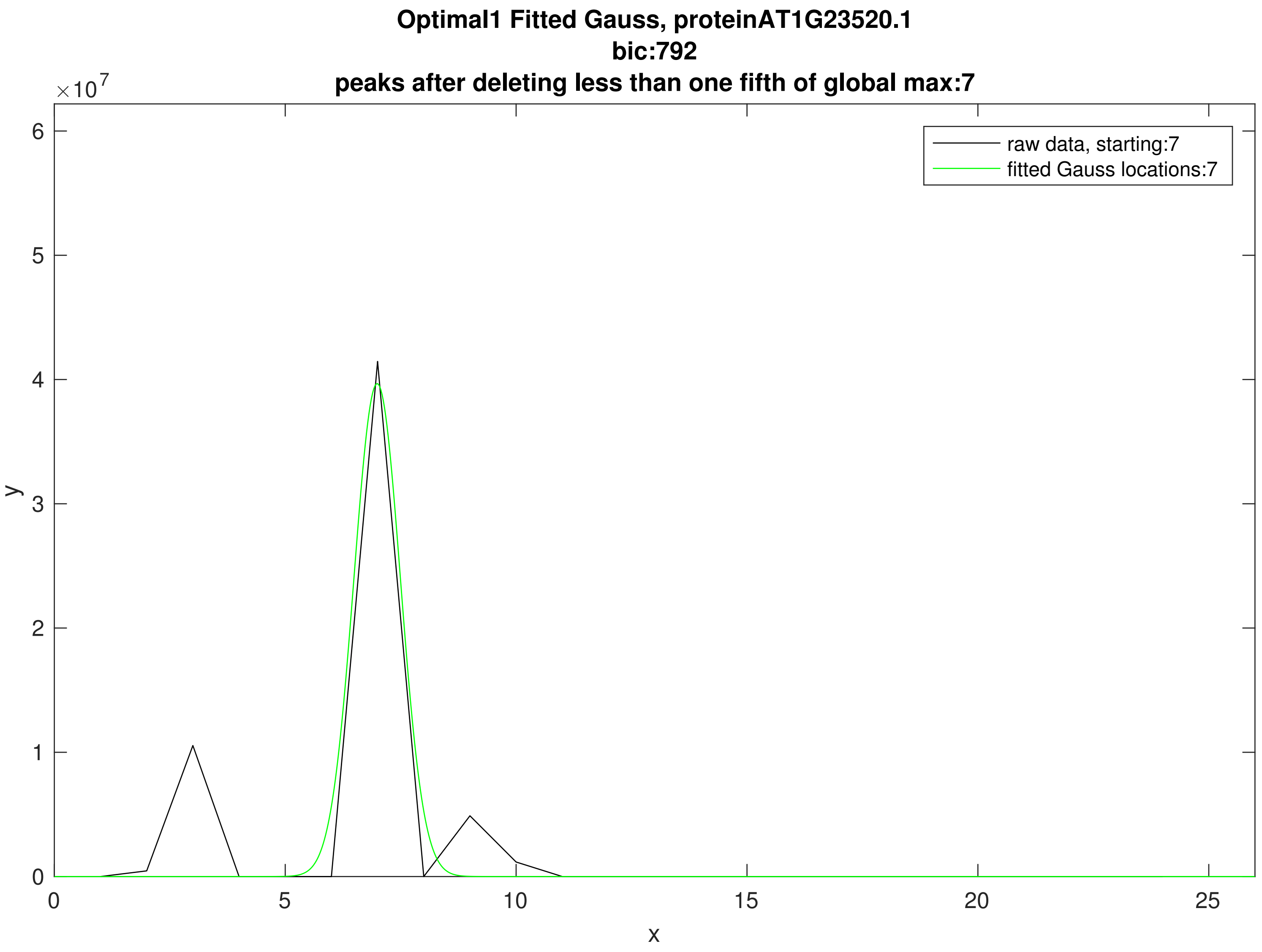

Optimal1 Fitted Gauss, proteinAT1G23520.1
bic:792
peaks after deleting less than one fifth of global max:7
×10^7
raw data, starting:7
fitted Gauss locations:7
y
x
0
1
2
3
4
5
6
0
5
10
15
20
25

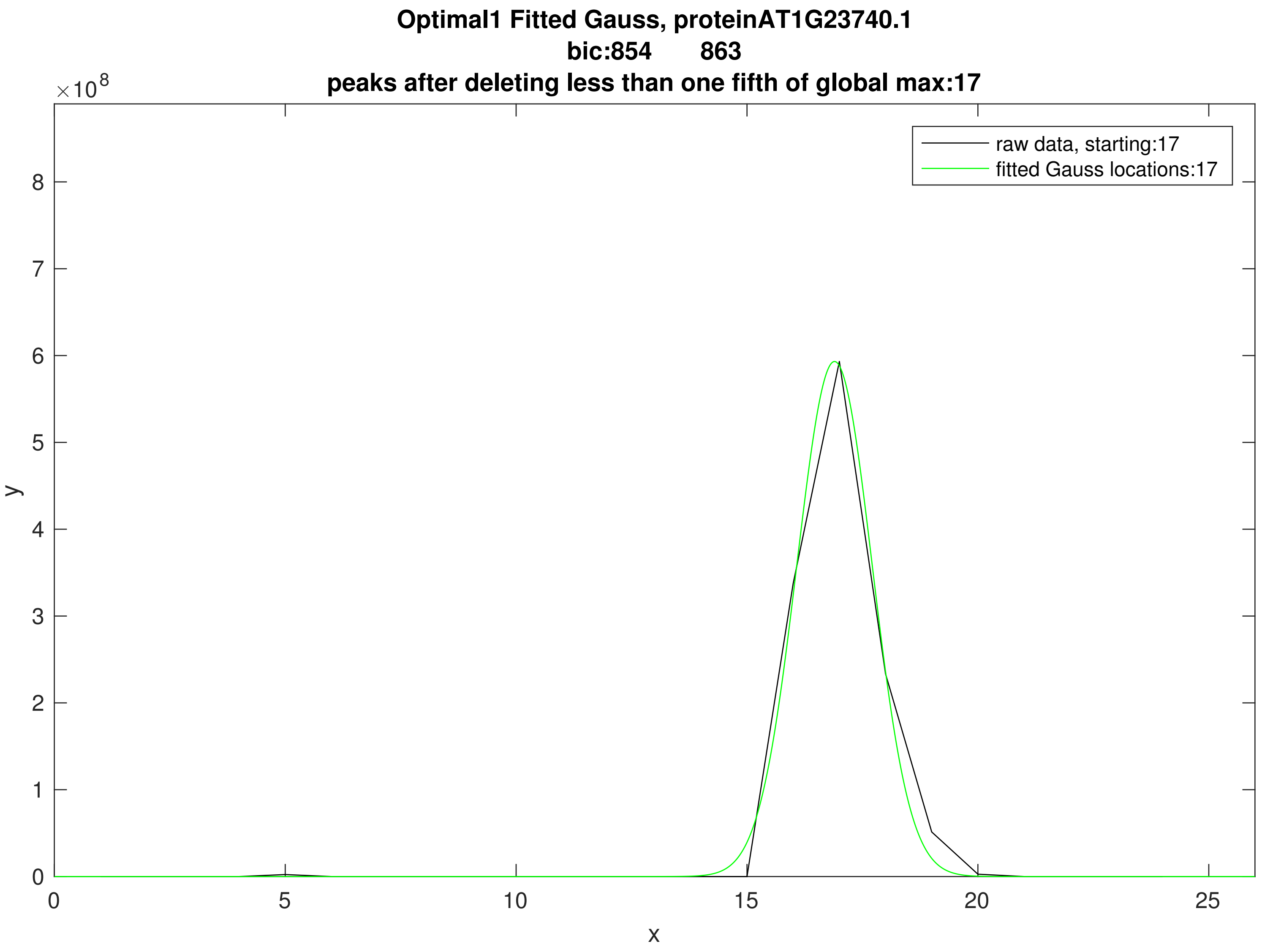
Optimal1 Fitted Gauss, proteinAT1G23740.1
bic:854 863
peaks after deleting less than one fifth of global max:17
×10^8
raw data, starting:17
fitted Gauss locations:17
y
x
0
1
2
3
4
5
6
7
8
0
5
10
15
20
25

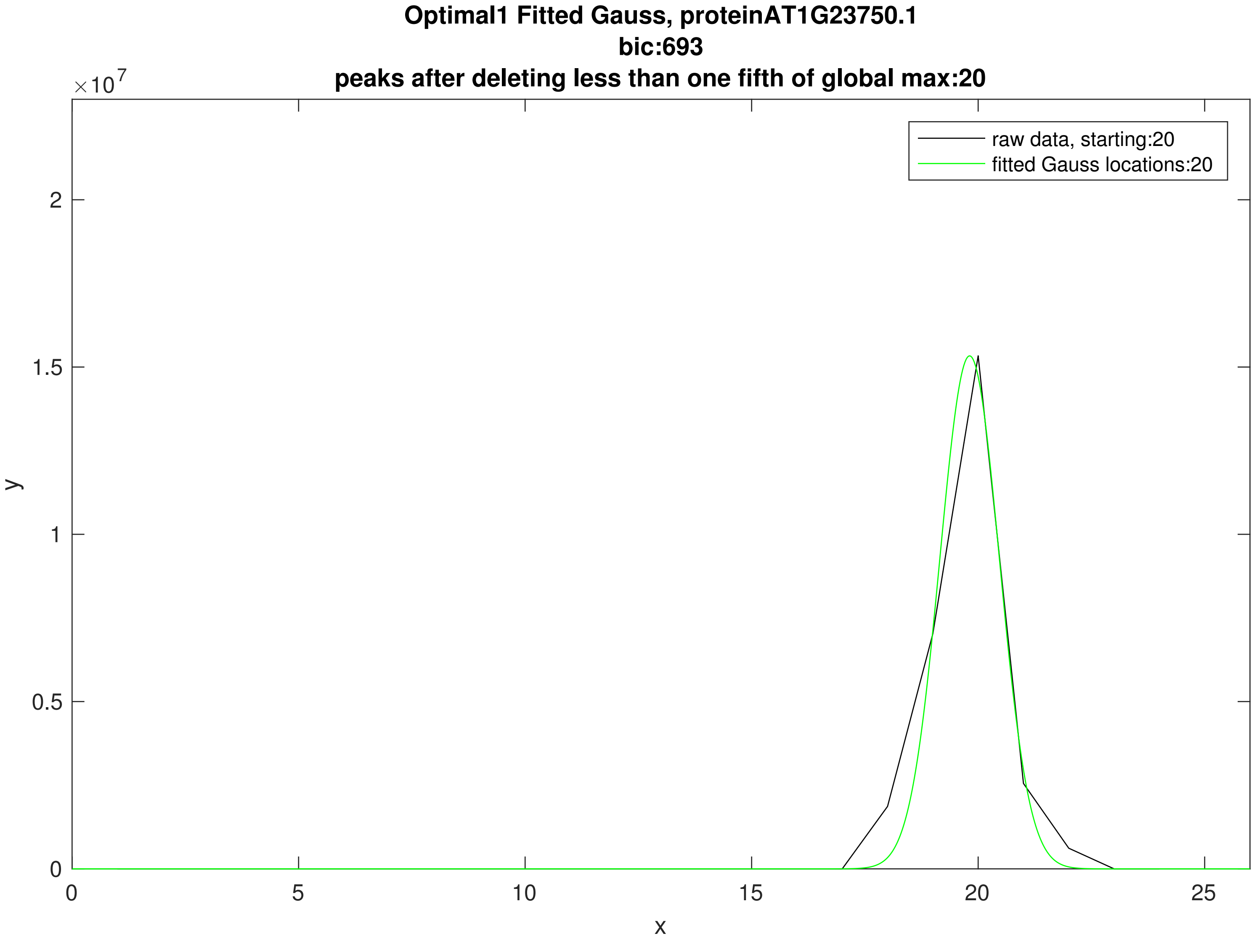

Optimal1 Fitted Gauss, proteinAT1G23750.1
bic:693
peaks after deleting less than one fifth of global max:20
×10^7
raw data, starting:20
fitted Gauss locations:20
2
1.5
1
0.5
0
y
0
5
10
15
20
25
x

**Optimal1 Fitted Gauss, proteinAT1G23800.1**
**bic:940 945**
**peaks after deleting less than one fifth of global max:10**

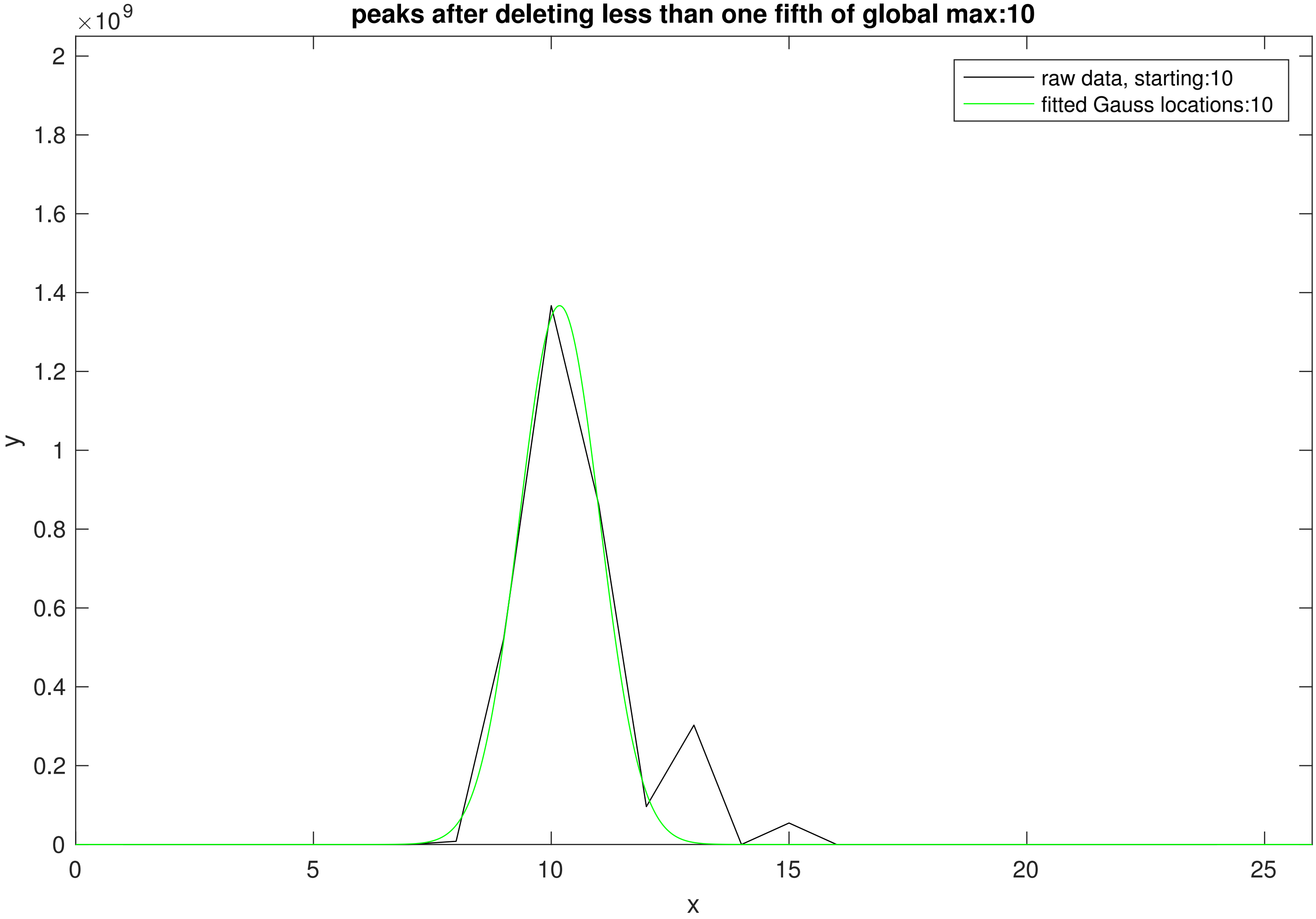

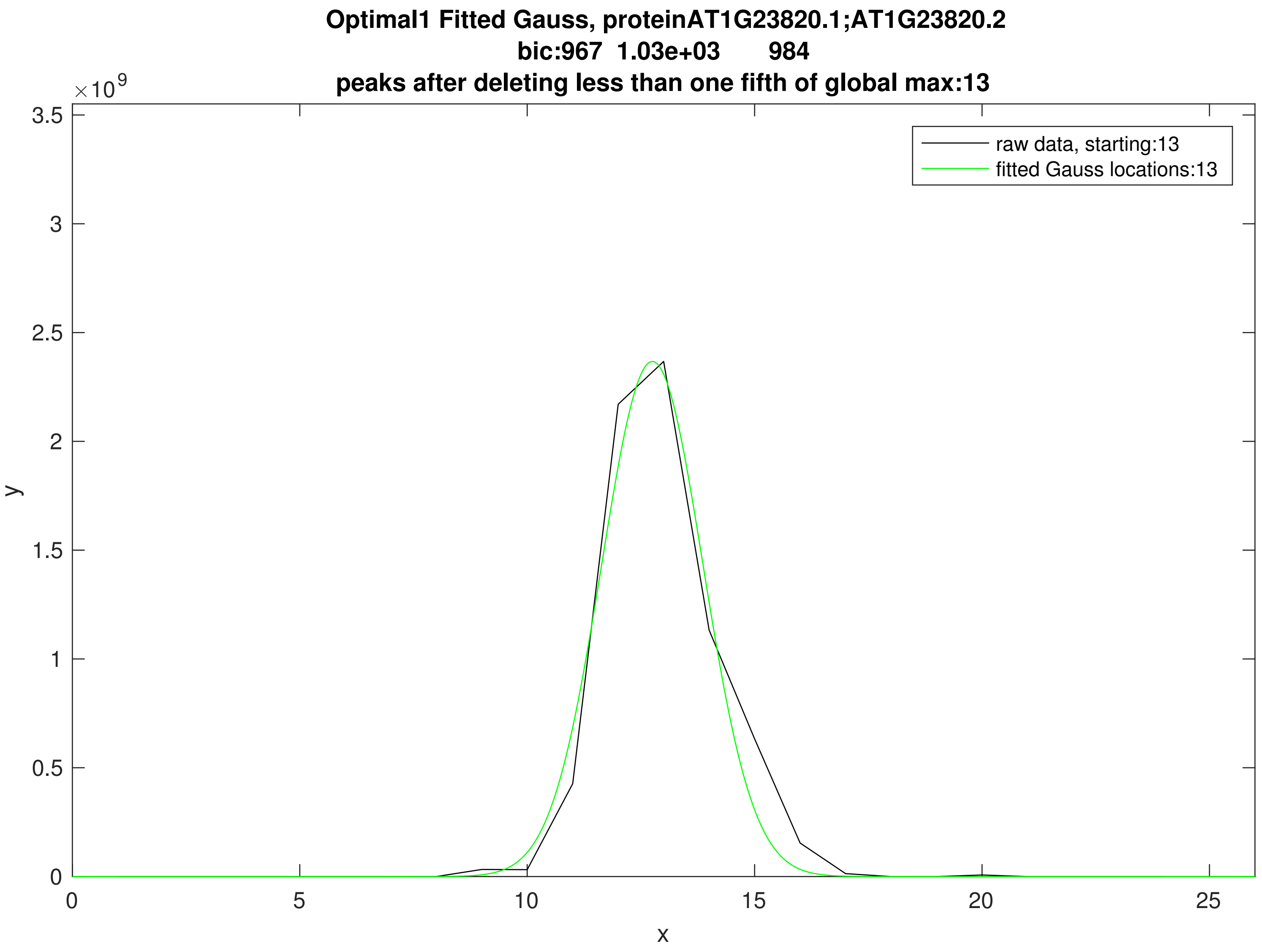

Optimal1 Fitted Gauss, proteinAT1G23820.1;AT1G23820.2
bic:967 1.03e+03 984
peaks after deleting less than one fifth of global max:13
×10^9
3.5
3
2.5
2
1.5
1
0.5
0
y
0
5
10
15
20
25
x
raw data, starting:13
fitted Gauss locations:13

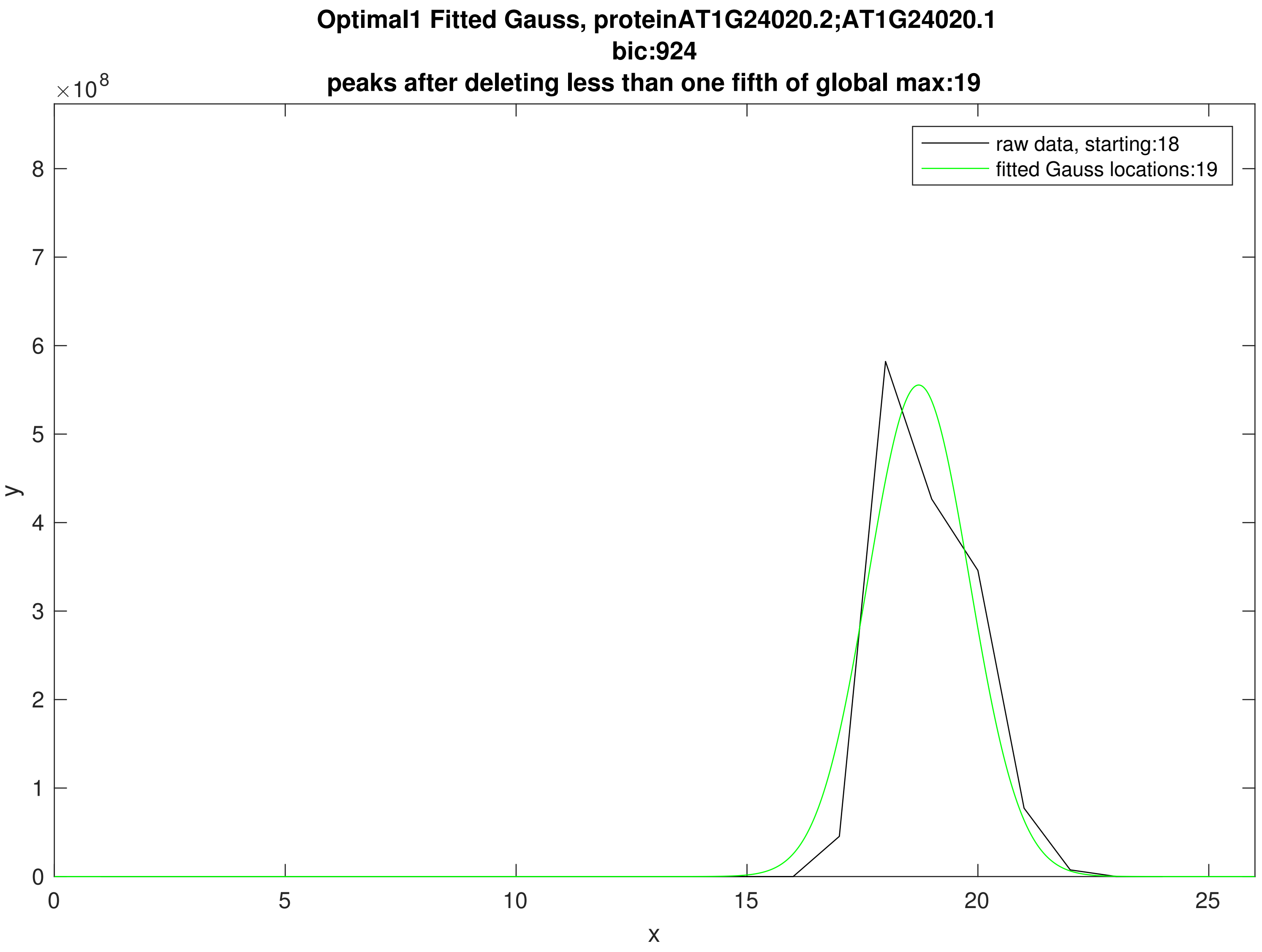
Optimal1 Fitted Gauss, proteinAT1G24020.2;AT1G24020.1
bic:924
peaks after deleting less than one fifth of global max:19
×10^8
raw data, starting:18
fitted Gauss locations:19
y
x
0
1
2
3
4
5
6
7
8
0
5
10
15
20
25

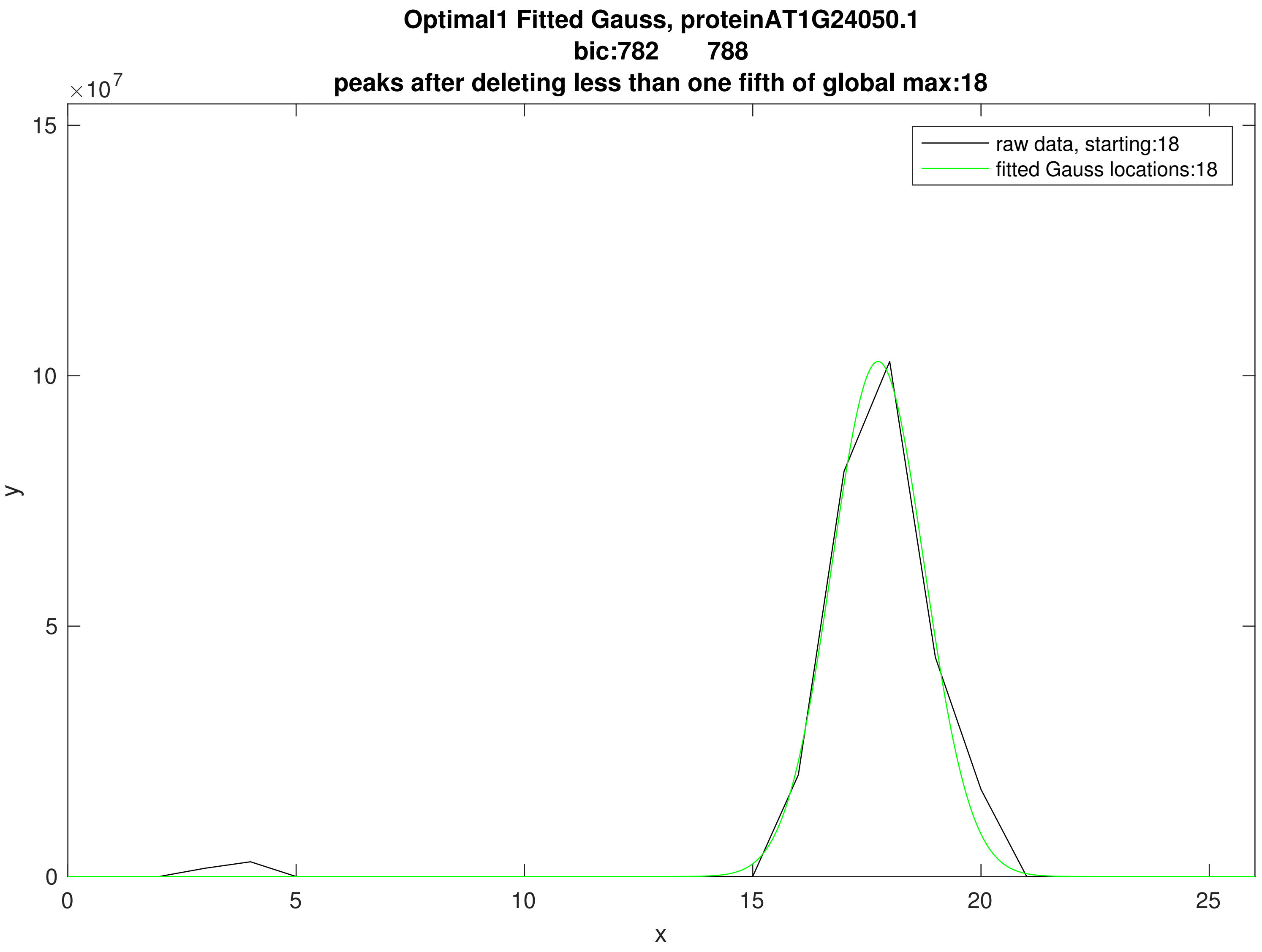
Optimal1 Fitted Gauss, proteinAT1G24050.1
bic:782 788
peaks after deleting less than one fifth of global max:18
×10^7
15
10
5
0
y
0
5
10
15
20
25
x
raw data, starting:18
fitted Gauss locations:18

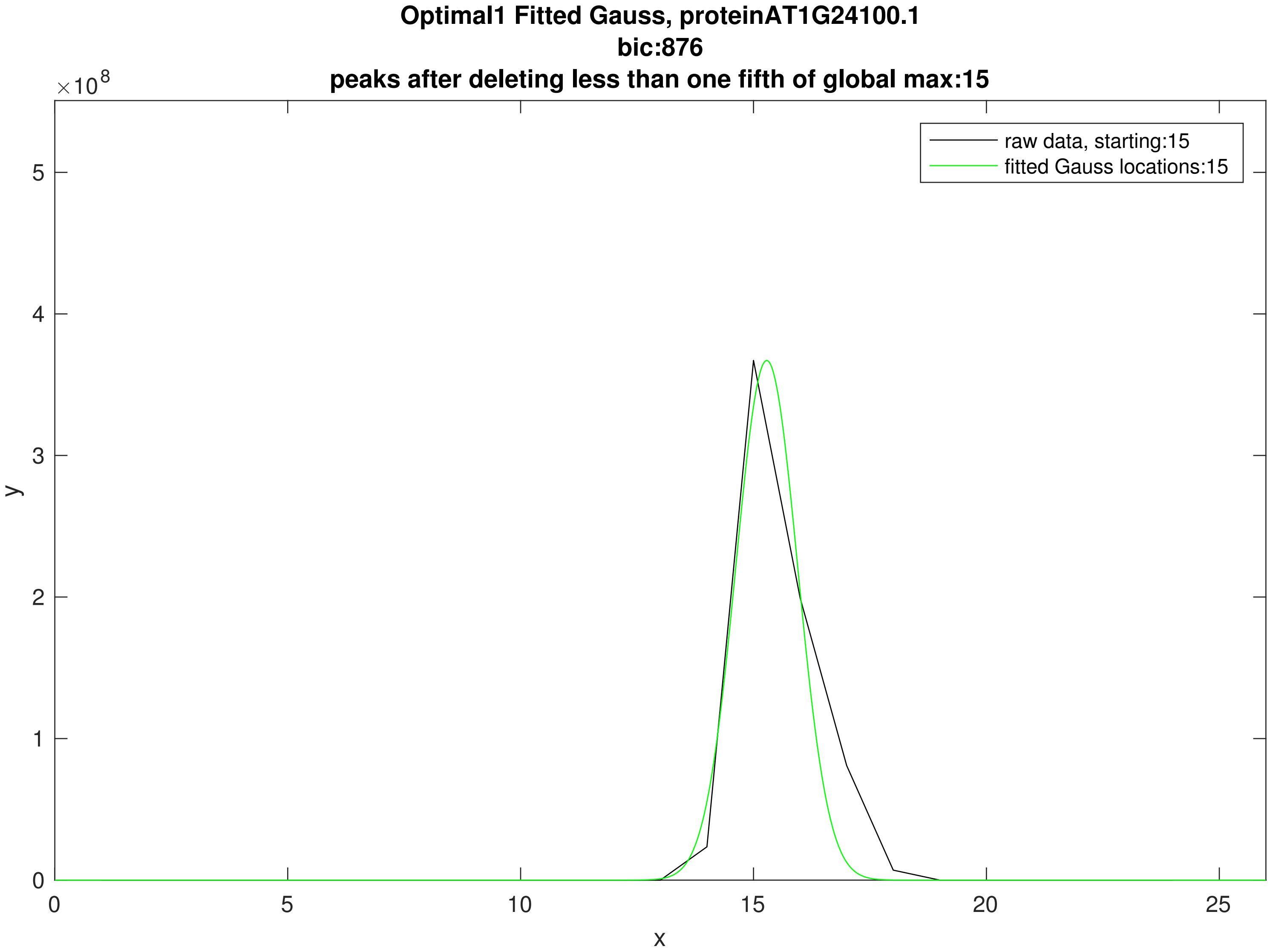

Optimal1 Fitted Gauss, proteinAT1G24100.1
bic:876
peaks after deleting less than one fifth of global max:15
×10^8
raw data, starting:15
fitted Gauss locations:15
y
x
0
1
2
3
4
5
0
5
10
15
20
25

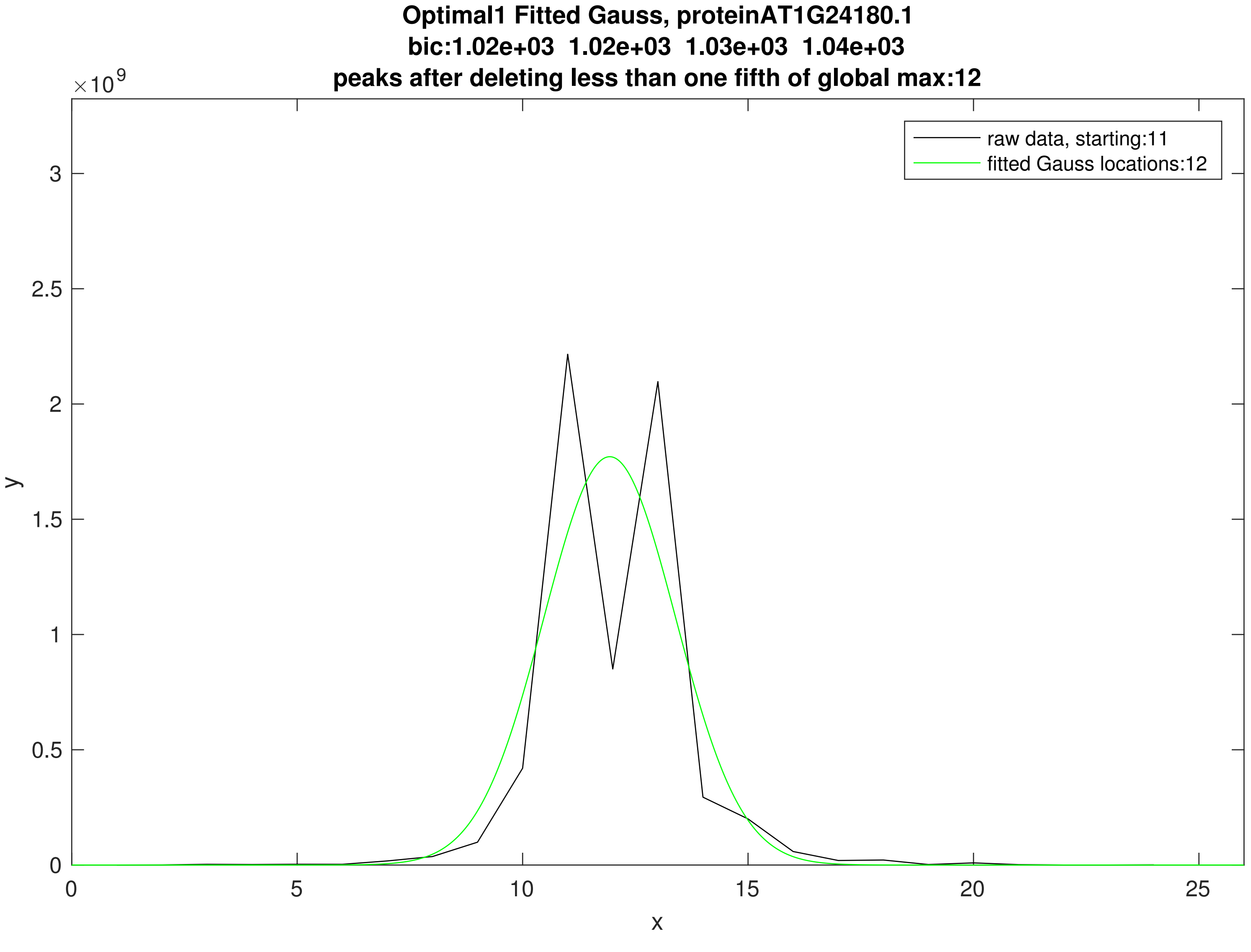

Optimal1 Fitted Gauss, proteinAT1G24180.1
bic:1.02e+03 1.02e+03 1.03e+03 1.04e+03
peaks after deleting less than one fifth of global max:12
×10^9
raw data, starting:11
fitted Gauss locations:12
3
2.5
2
1.5
1
0.5
0
y
0
5
10
15
20
25
x

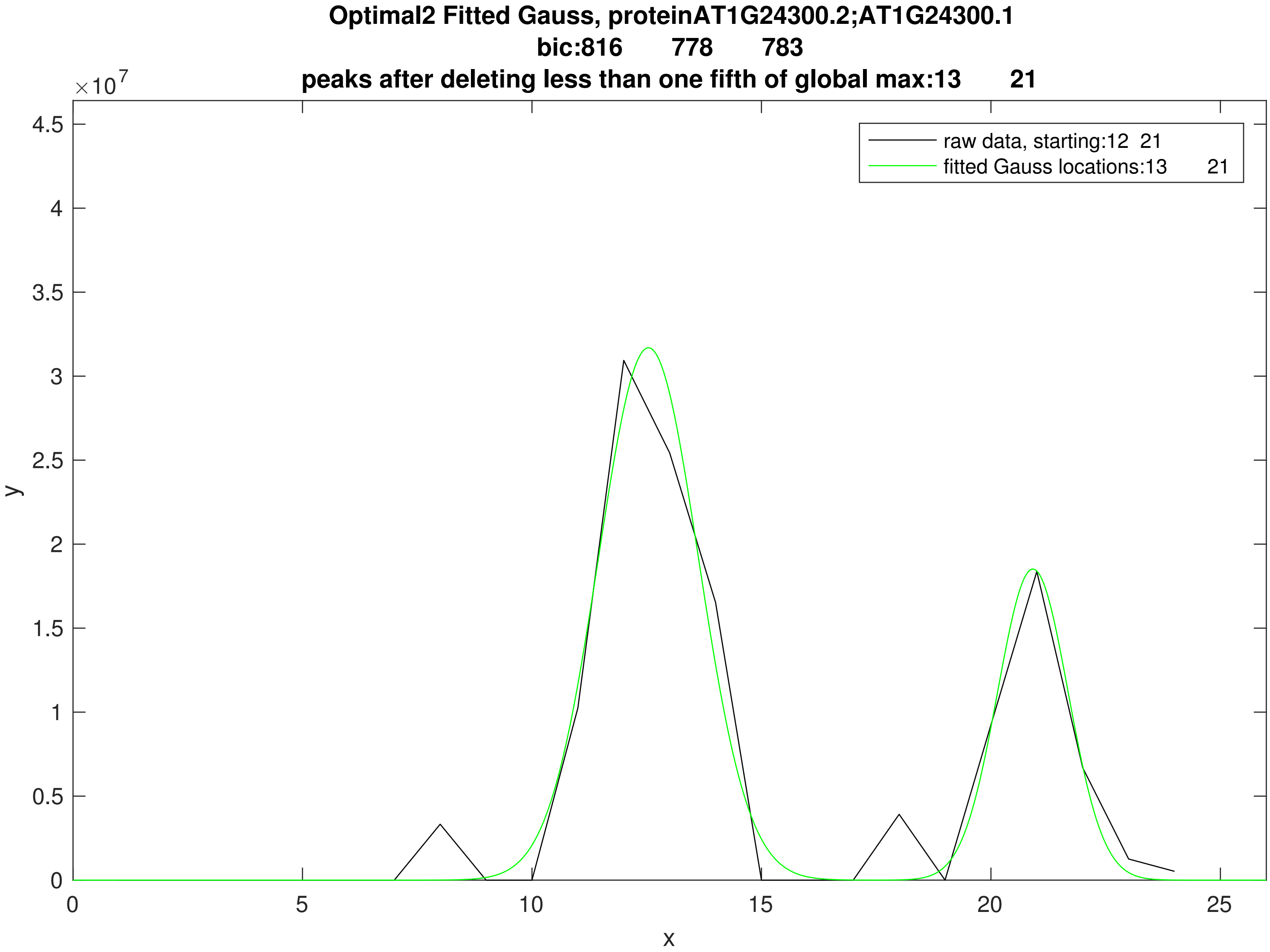

Optimal2 Fitted Gauss, proteinAT1G24300.2;AT1G24300.1
bic:816 778 783
peaks after deleting less than one fifth of global max:13 21
×10^7
raw data, starting:12 21
fitted Gauss locations:13 21
y
x
0
0.5
1
1.5
2
2.5
3
3.5
4
4.5
0
5
10
15
20
25

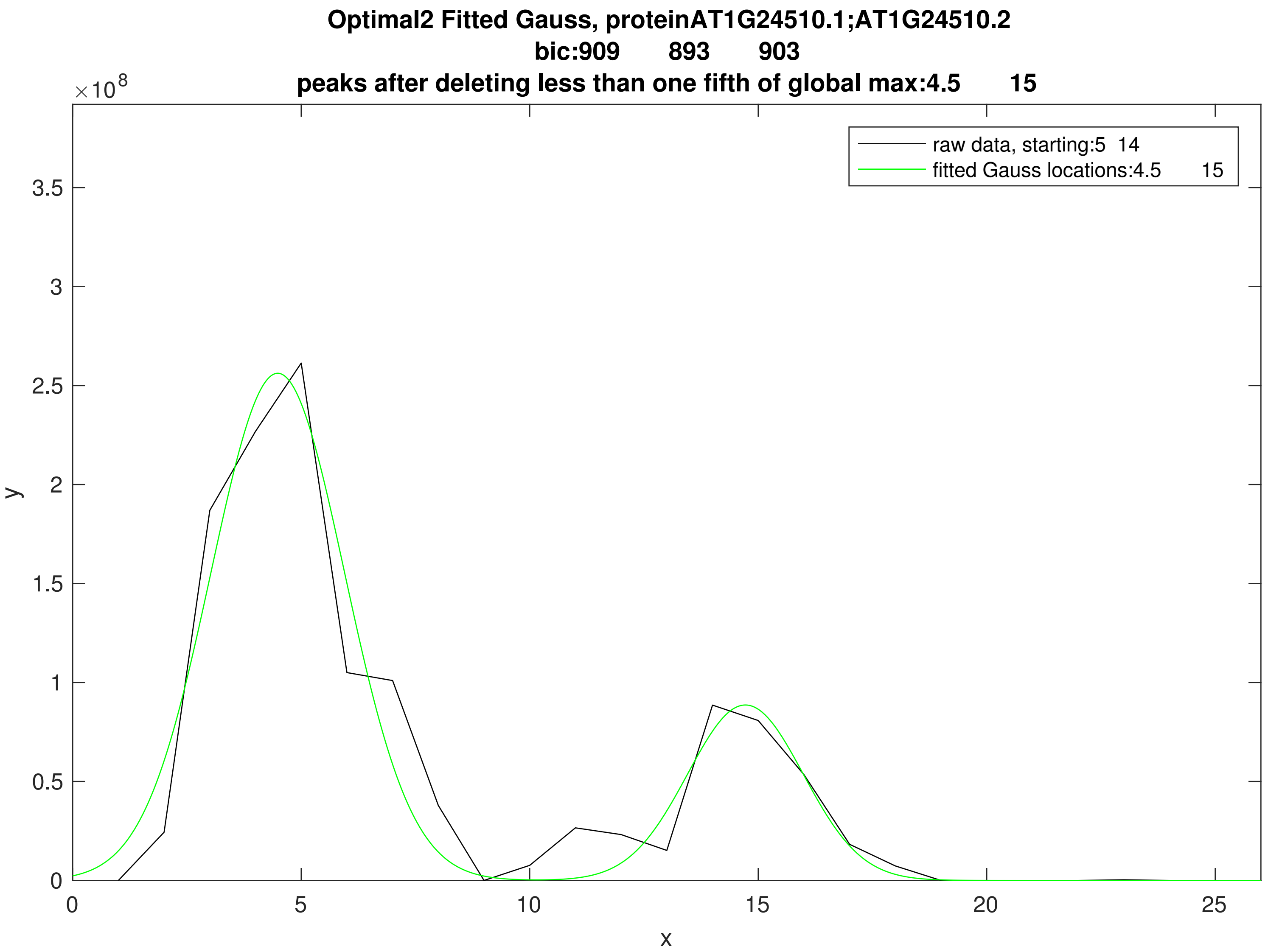

Optimal2 Fitted Gauss, proteinAT1G24510.1;AT1G24510.2
bic:909 893 903
peaks after deleting less than one fifth of global max:4.5 15
×10^8
raw data, starting:5 14
fitted Gauss locations:4.5 15
3.5
3
2.5
2
1.5
1
0.5
0
y
0
5
10
15
20
25
x

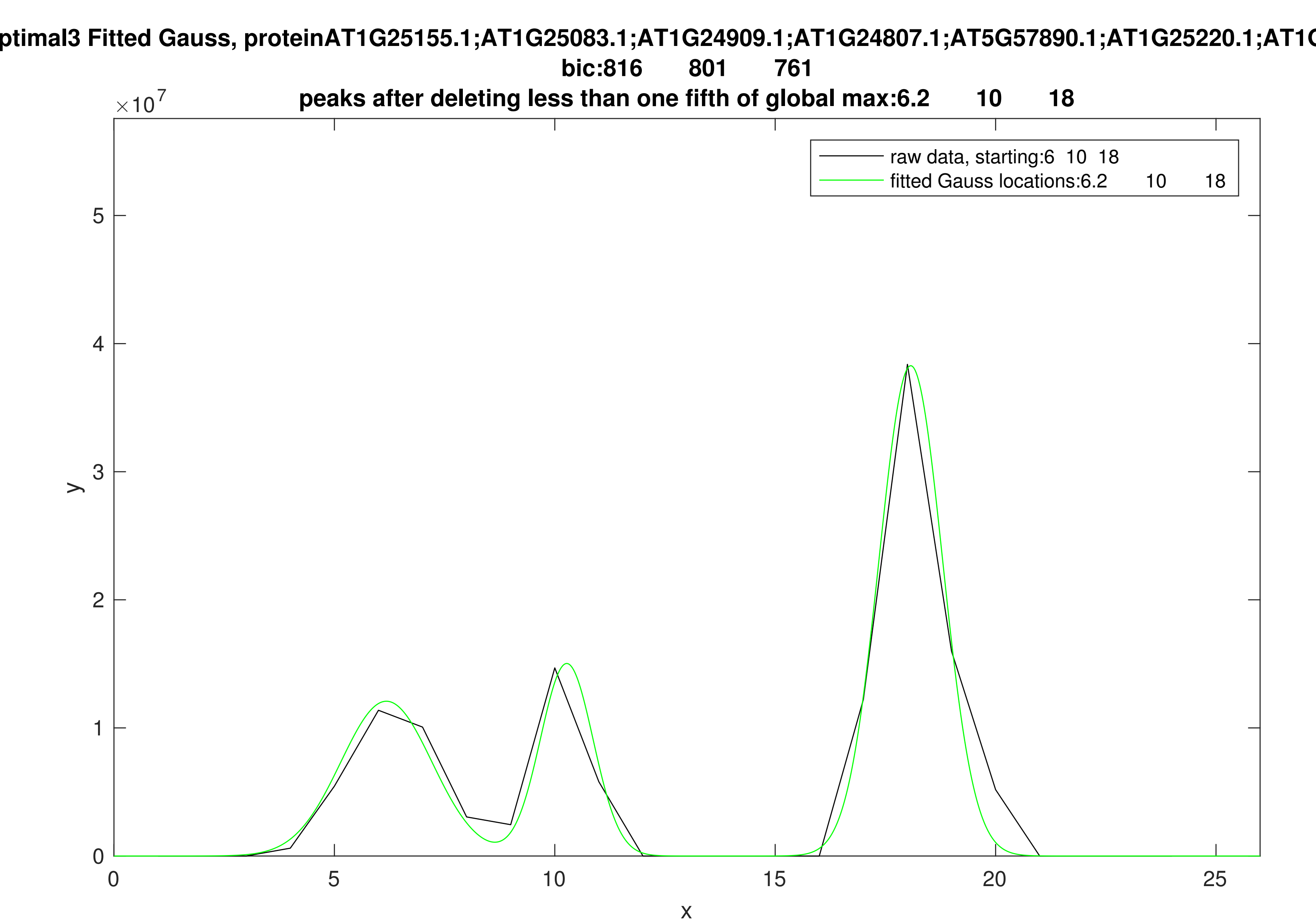

ptimal3 Fitted Gauss, proteinAT1G25155.1;AT1G25083.1;AT1G24909.1;AT1G24807.1;AT5G57890.1;AT1G25220.1;AT1G
bic:816 801 761
peaks after deleting less than one fifth of global max:6.2 10 18
×10^7
raw data, starting:6 10 18
fitted Gauss locations:6.2 10 18
y
x
0
1
2
3
4
5
0
5
10
15
20
25

**Optimal1 Fitted Gauss, proteinAT1G25211.1;AT1G25150.1;AT1G25055.1;AT1G24881.1;AT1G24800.1**
**bic:825**
**peaks after deleting less than one fifth of global max:14**

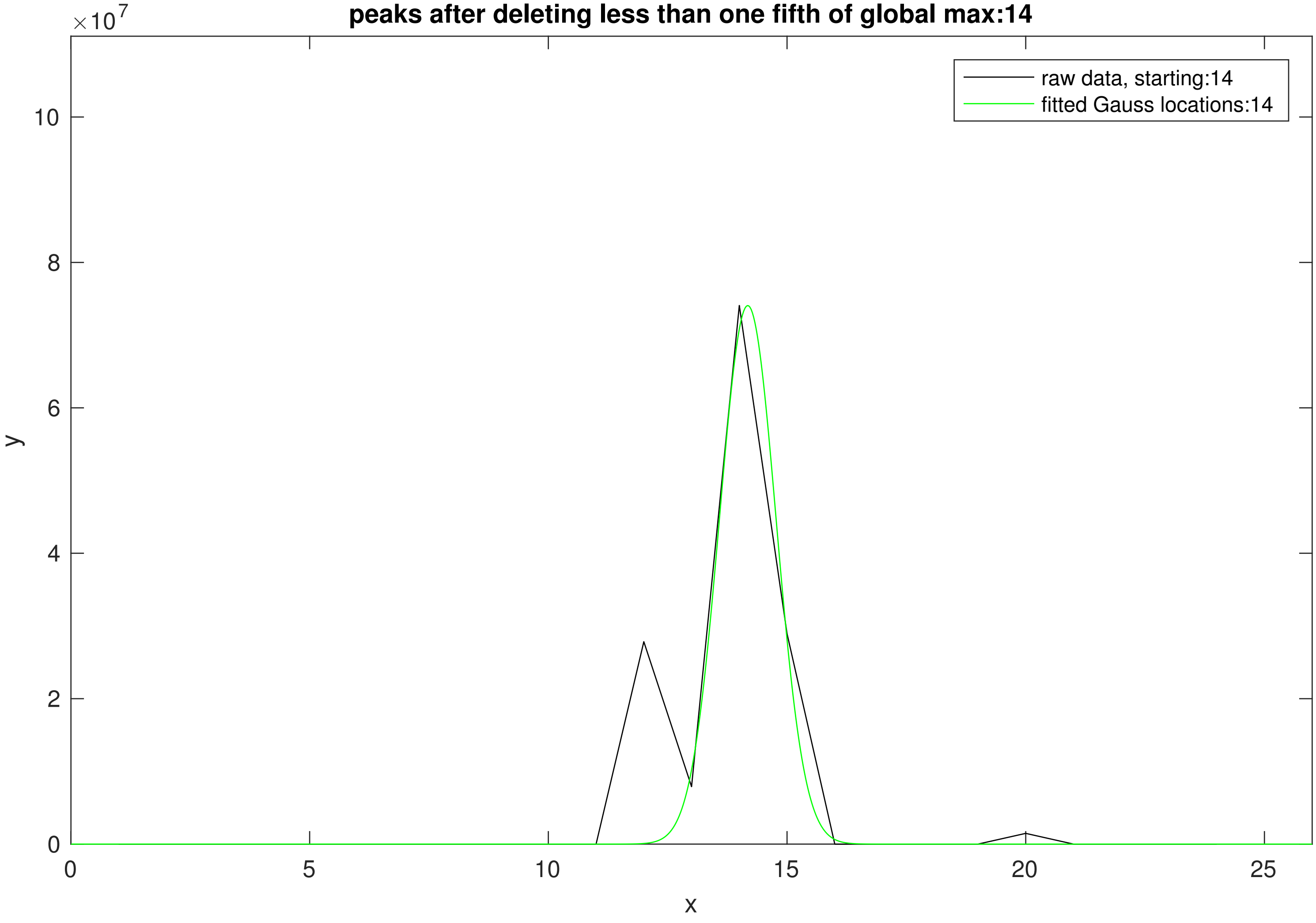

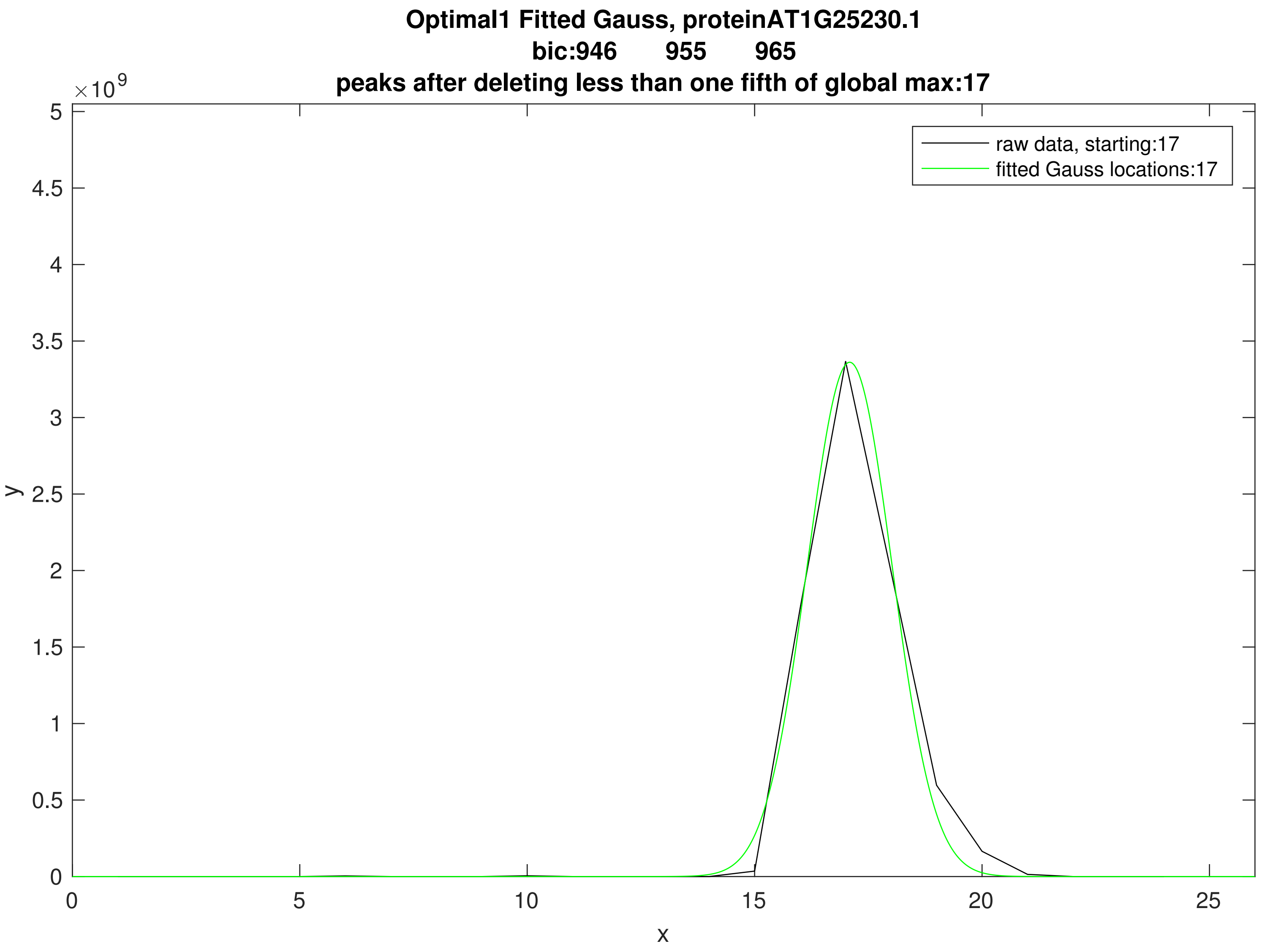

Optimal1 Fitted Gauss, proteinAT1G25230.1
bic:946 955 965
peaks after deleting less than one fifth of global max:17
×10^9
raw data, starting:17
fitted Gauss locations:17
y
x
0
0.5
1
1.5
2
2.5
3
3.5
4
4.5
5
0
5
10
15
20
25

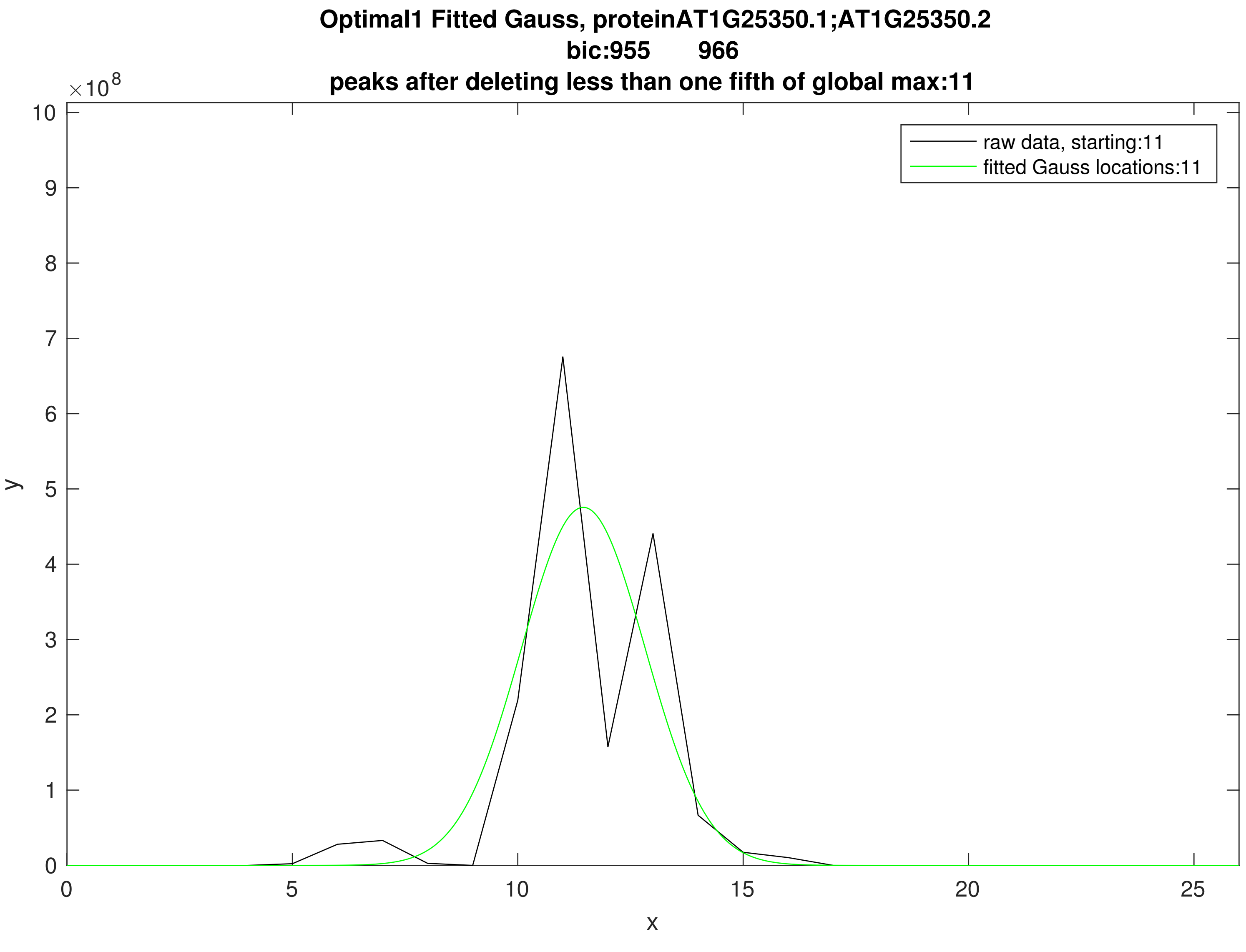

Optimal1 Fitted Gauss, proteinAT1G25350.1;AT1G25350.2
bic:955 966
peaks after deleting less than one fifth of global max:11
raw data, starting:11
fitted Gauss locations:11
y
x

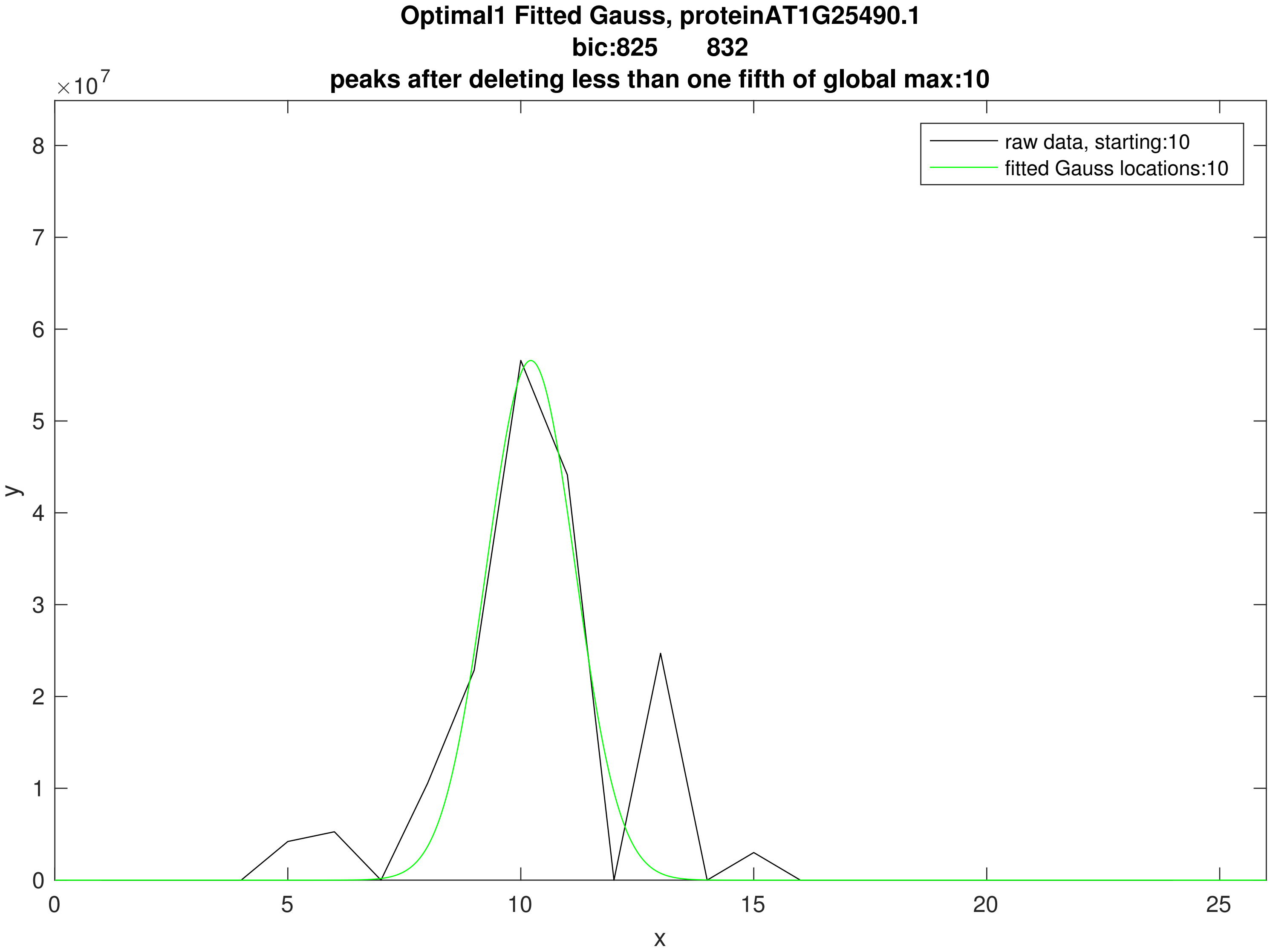

Optimal1 Fitted Gauss, proteinAT1G25490.1
bic:825 832
peaks after deleting less than one fifth of global max:10
×10^7
raw data, starting:10
fitted Gauss locations:10
y
x
0
1
2
3
4
5
6
7
8
0
5
10
15
20
25

Optimal1 Fitted Gauss, proteinAT1G25570.1
bic:808
peaks after deleting less than one fifth of global max:16

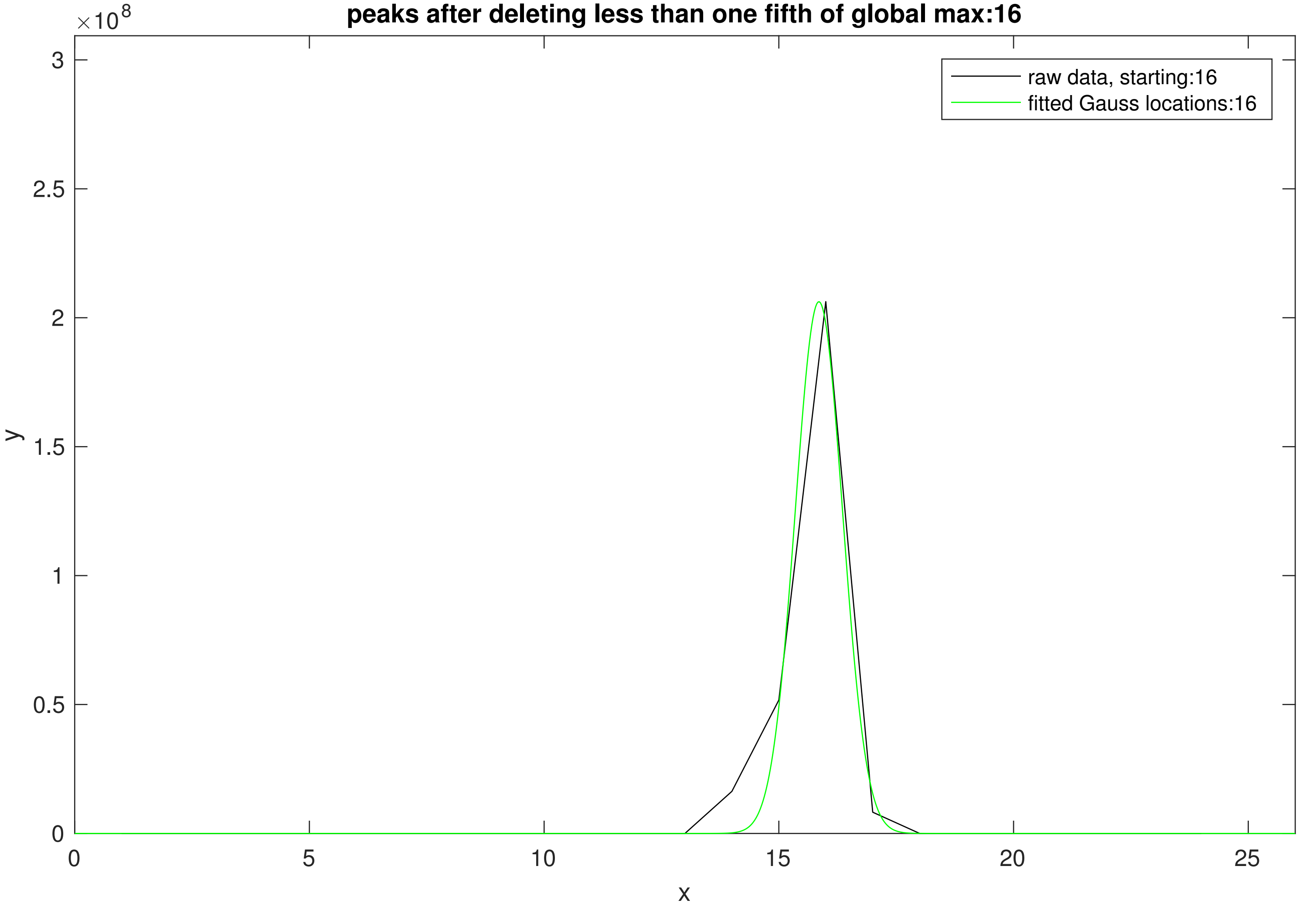

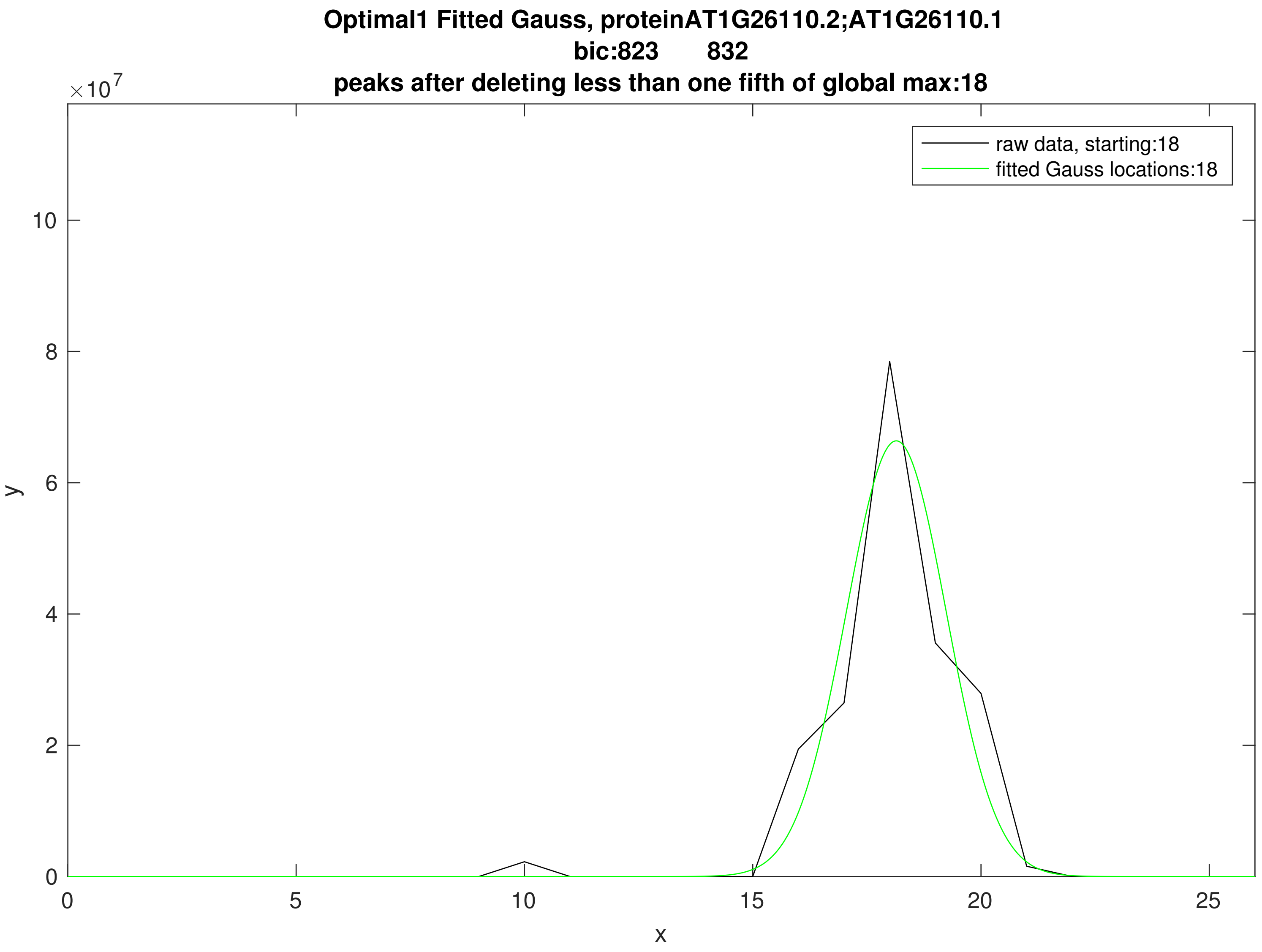

Optimal1 Fitted Gauss, proteinAT1G26110.2;AT1G26110.1
bic:823 832
peaks after deleting less than one fifth of global max:18
×10^7
raw data, starting:18
fitted Gauss locations:18
10
8
6
4
2
0
y
0
5
10
15
20
25
x

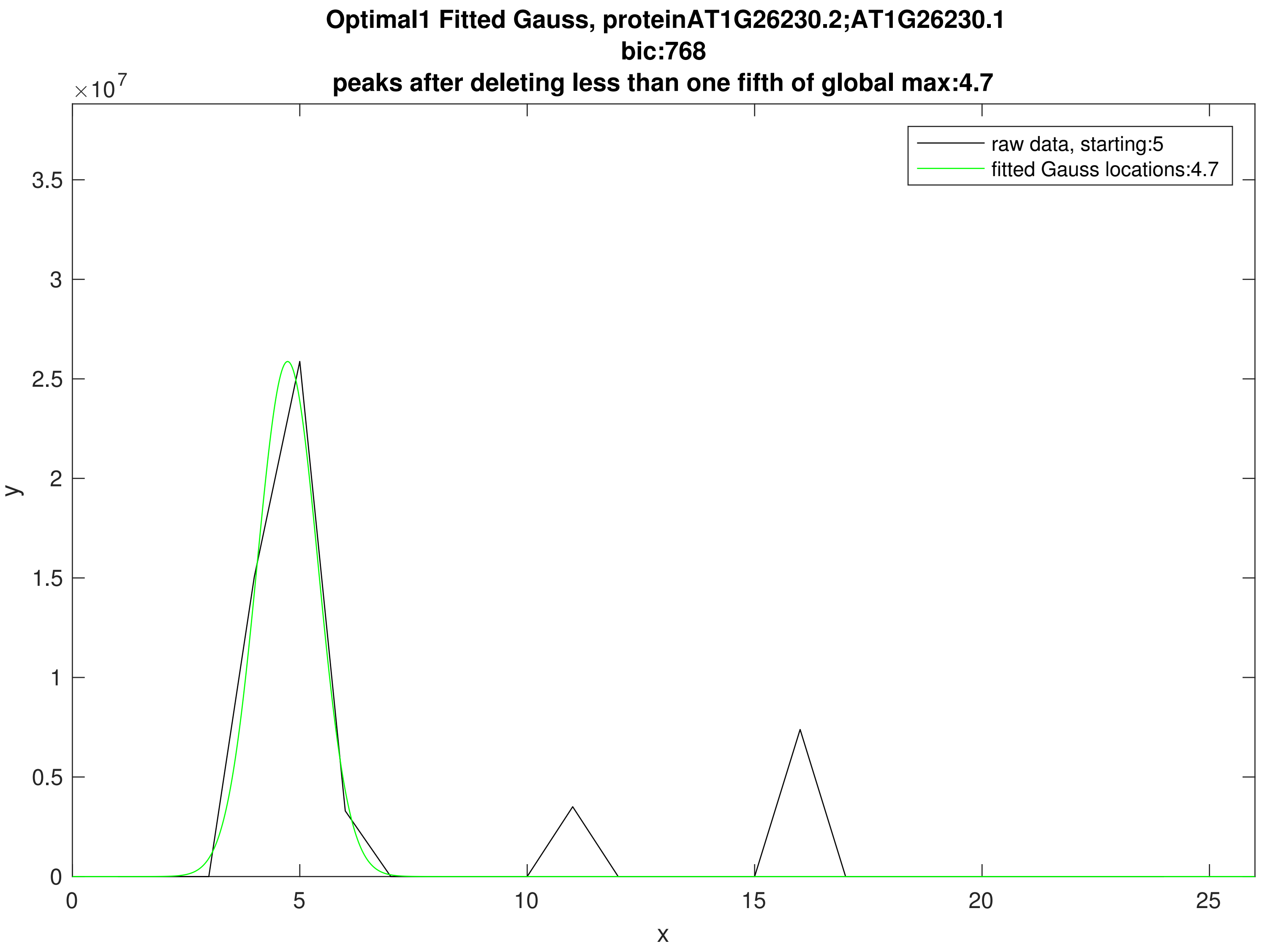

Optimal1 Fitted Gauss, proteinAT1G26230.2;AT1G26230.1
bic:768
peaks after deleting less than one fifth of global max:4.7
×10^7
raw data, starting:5
fitted Gauss locations:4.7
y
x
0
0.5
1
1.5
2
2.5
3
3.5
5
10
15
20
25

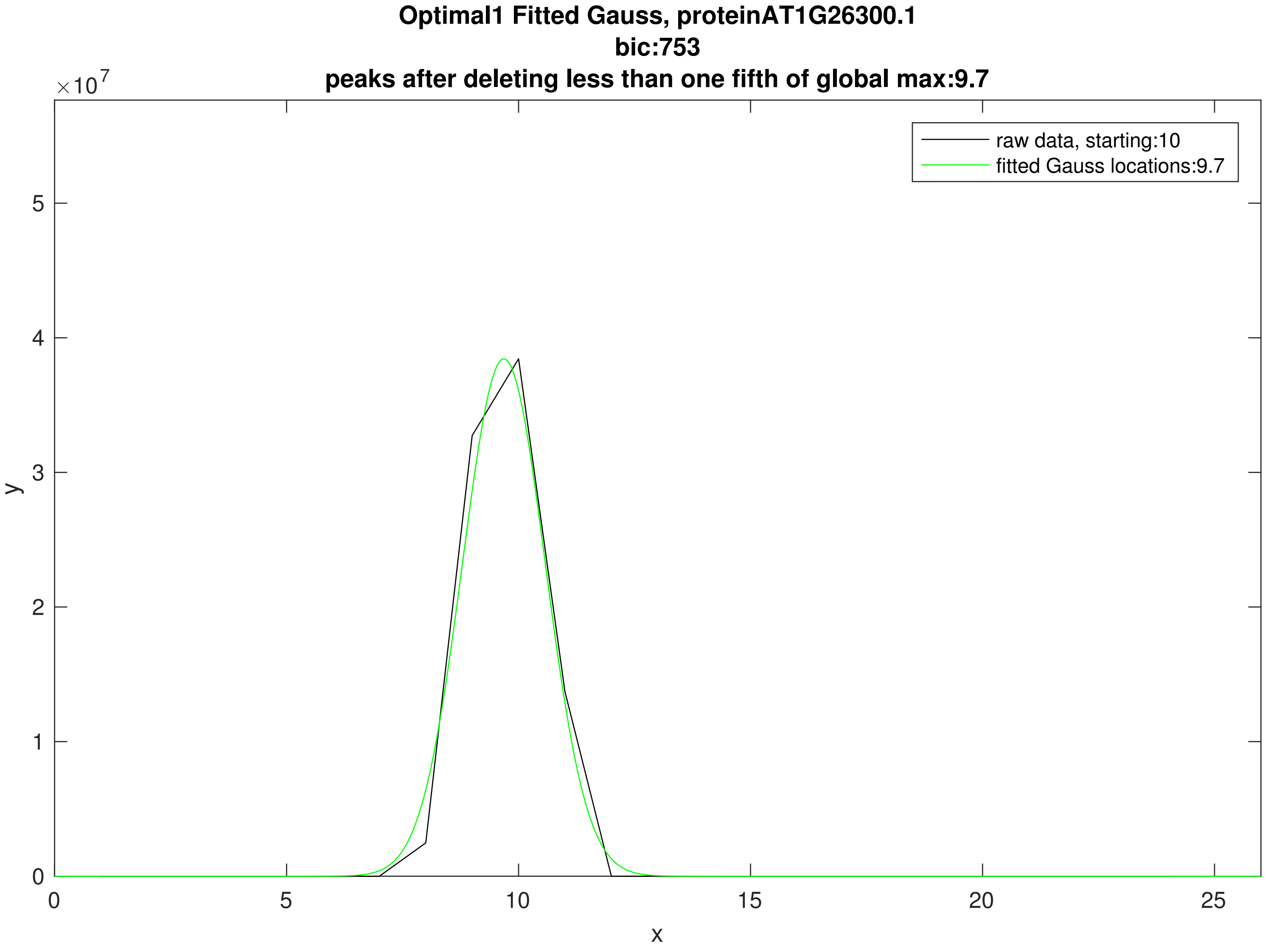

Optimal1 Fitted Gauss, proteinAT1G26300.1
bic:753
peaks after deleting less than one fifth of global max:9.7
×10^7
raw data, starting:10
fitted Gauss locations:9.7
y
x
0
1
2
3
4
5
0
5
10
15
20
25

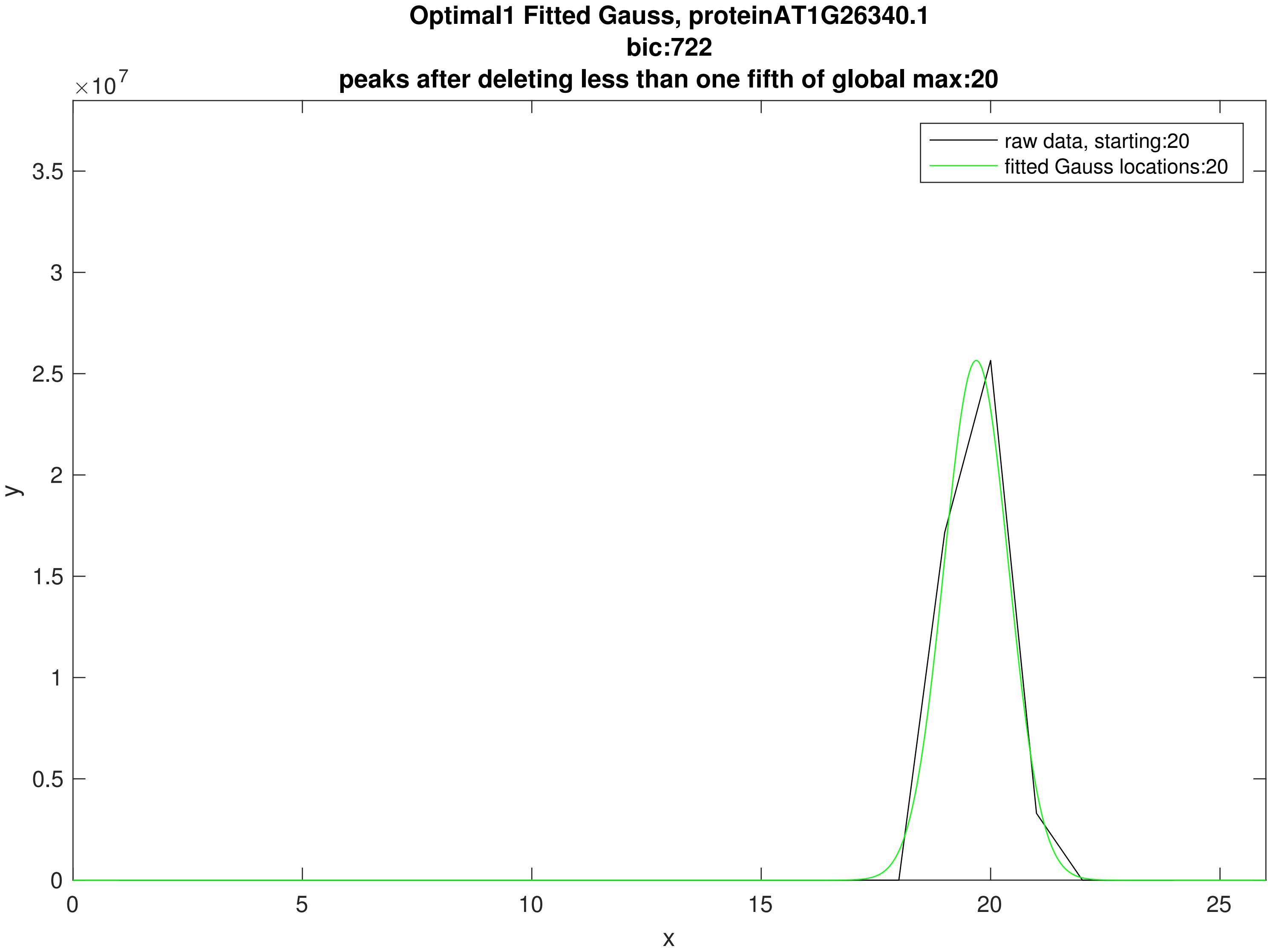
Optimal1 Fitted Gauss, proteinAT1G26340.1
bic:722
peaks after deleting less than one fifth of global max:20
×10^7
raw data, starting:20
fitted Gauss locations:20
y
x
0
0.5
1
1.5
2
2.5
3
3.5
0
5
10
15
20
25

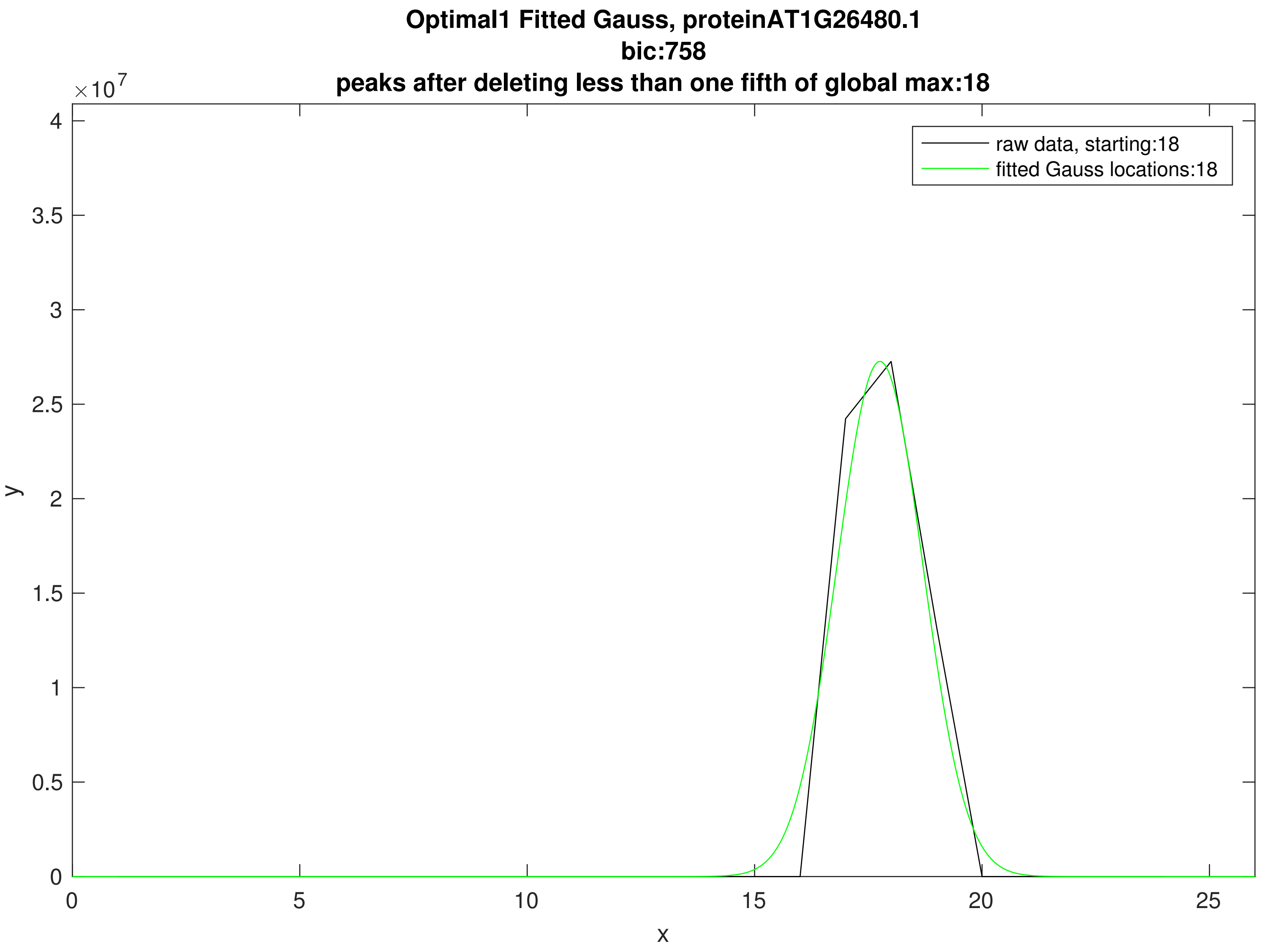
Optimal1 Fitted Gauss, proteinAT1G26480.1
bic:758
peaks after deleting less than one fifth of global max:18
×10^7
4
3.5
3
2.5
2
1.5
1
0.5
0
y
0
5
10
15
20
25
x
raw data, starting:18
fitted Gauss locations:18

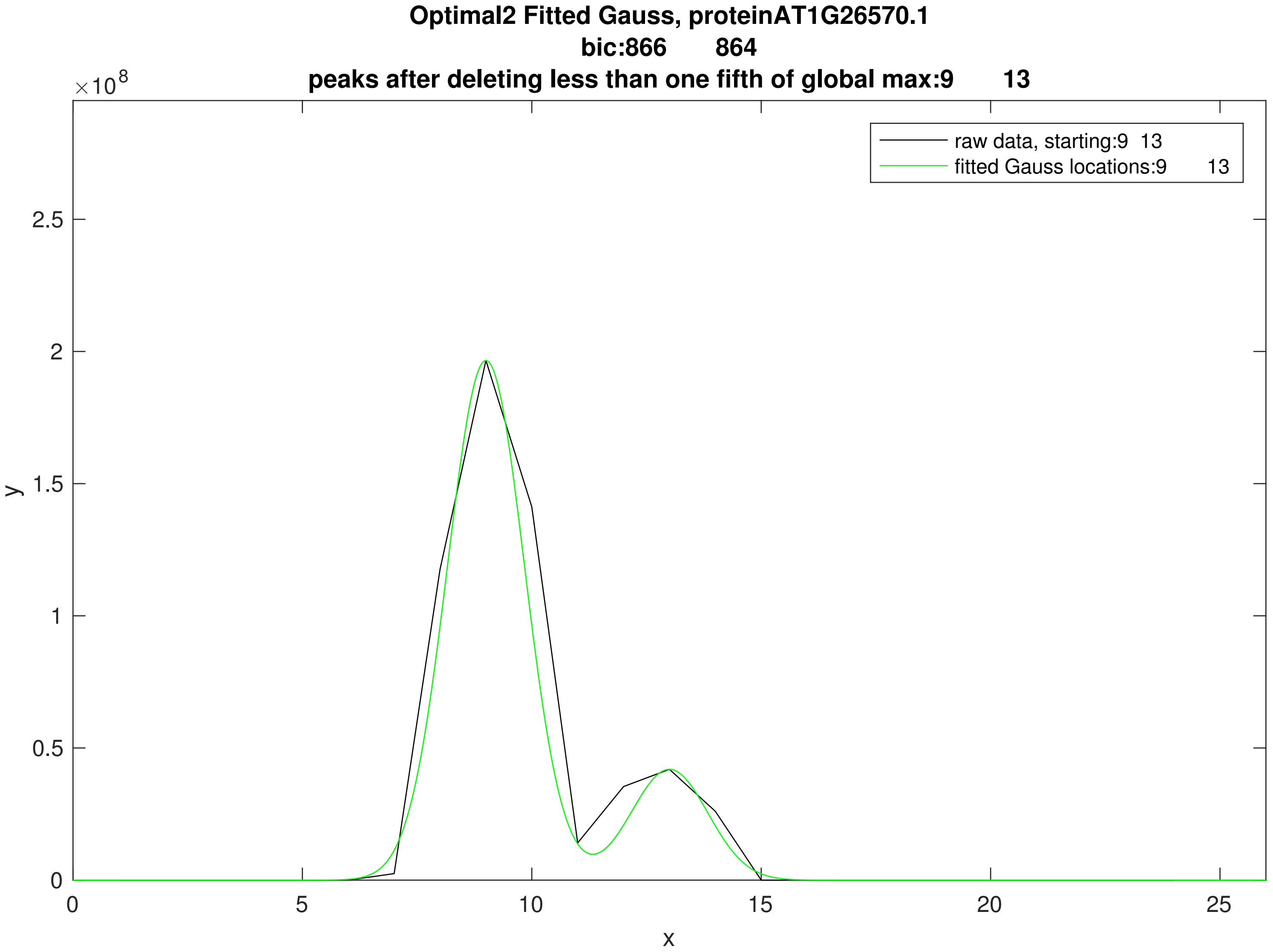

Optimal2 Fitted Gauss, proteinAT1G26570.1
bic:866 864
peaks after deleting less than one fifth of global max:9 13
×10^8
raw data, starting:9 13
fitted Gauss locations:9 13
2.5
2
1.5
1
0.5
0
y
0
5
10
15
20
25
x

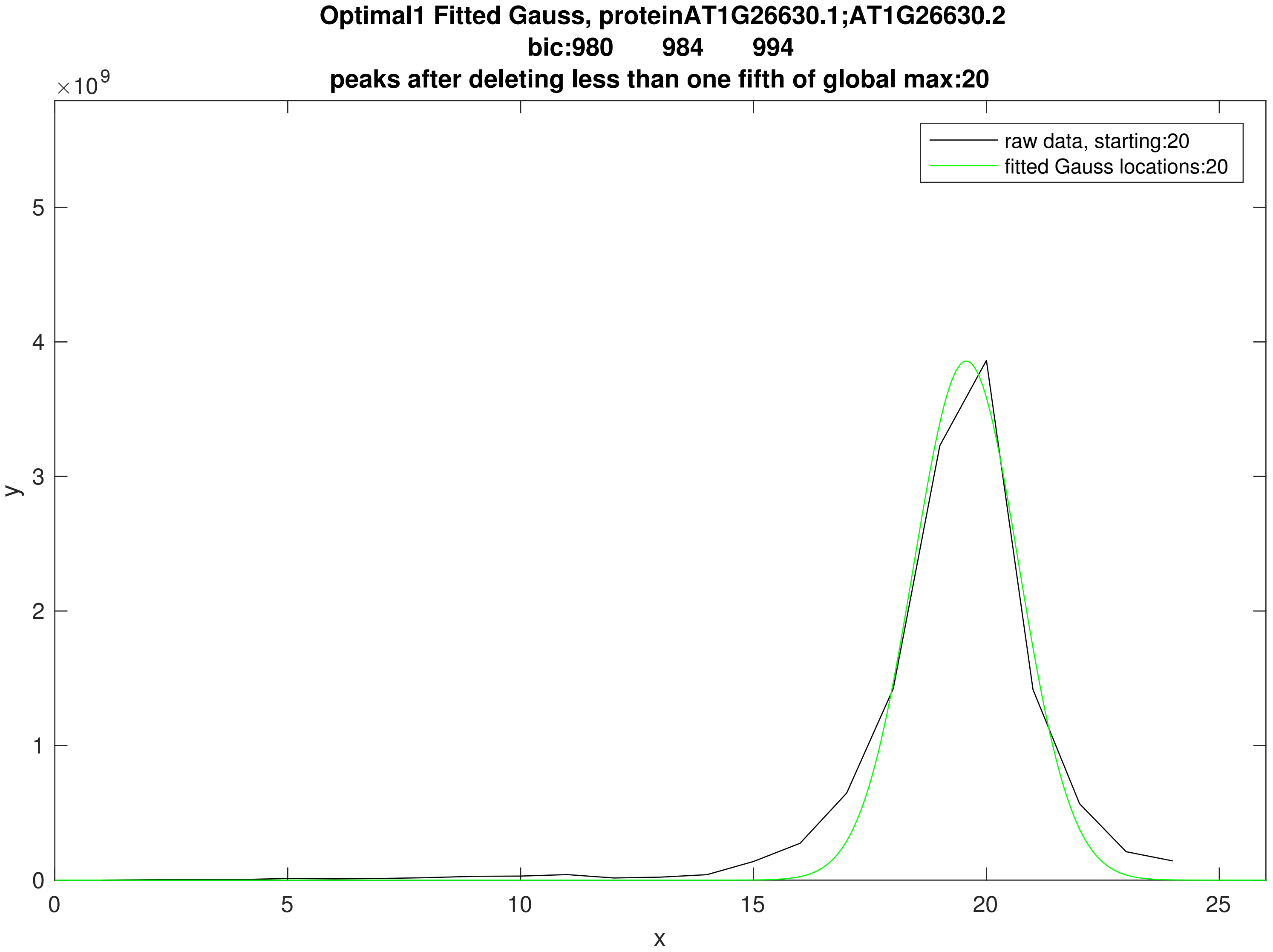
Optimal1 Fitted Gauss, proteinAT1G26630.1;AT1G26630.2
bic:980 984 994
peaks after deleting less than one fifth of global max:20
×10^9
raw data, starting:20
fitted Gauss locations:20
y
x
0
1
2
3
4
5
0
5
10
15
20
25

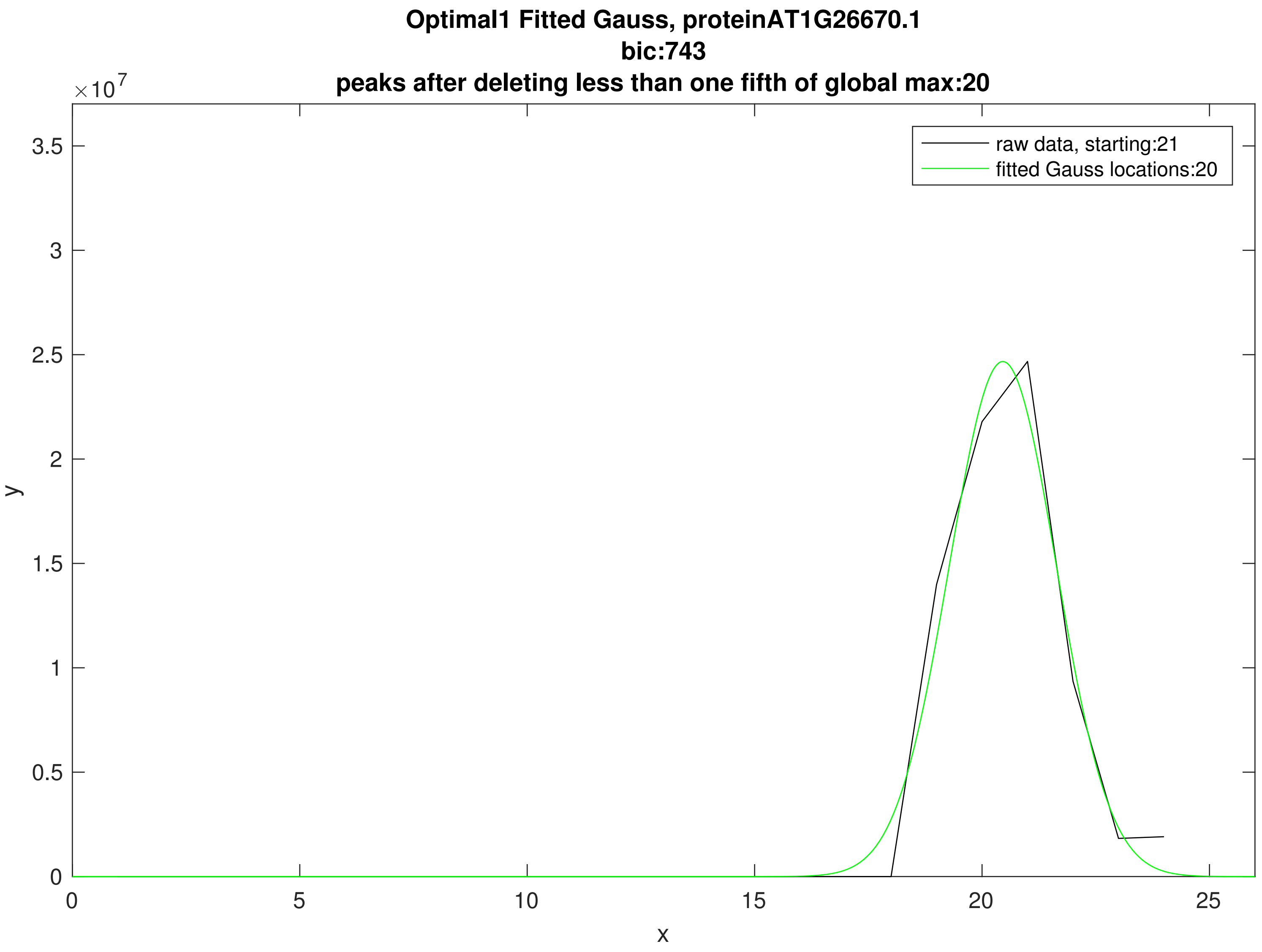
Optimal1 Fitted Gauss, proteinAT1G26670.1
bic:743
peaks after deleting less than one fifth of global max:20
×10^7
raw data, starting:21
fitted Gauss locations:20
3.5
3
2.5
2
1.5
1
0.5
0
y
0
5
10
15
20
25
x

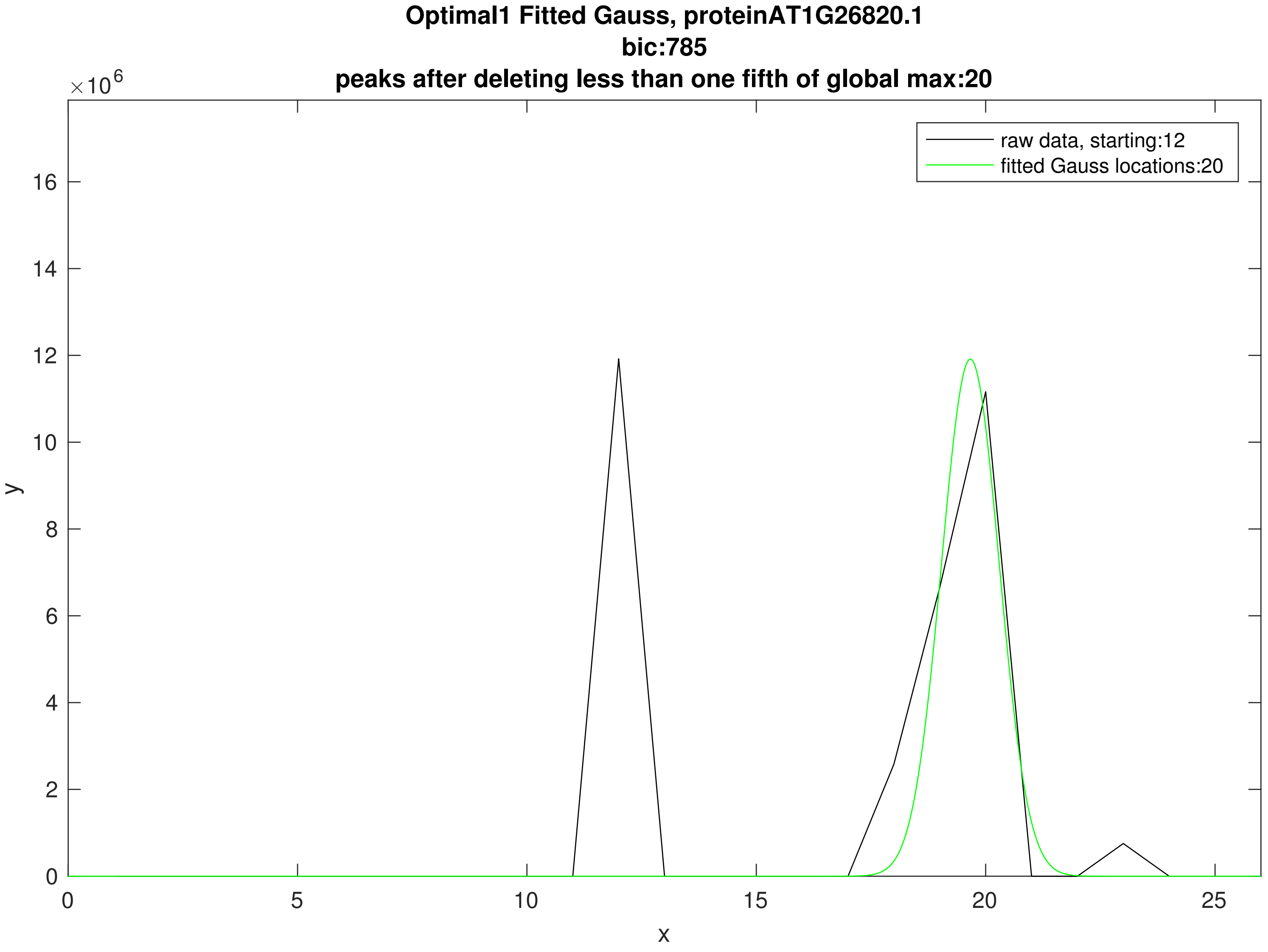

Optimal1 Fitted Gauss, proteinAT1G26820.1
bic:785
peaks after deleting less than one fifth of global max:20
×10^6
raw data, starting:12
fitted Gauss locations:20
y
x
0
2
4
6
8
10
12
14
16
5
15
20
25

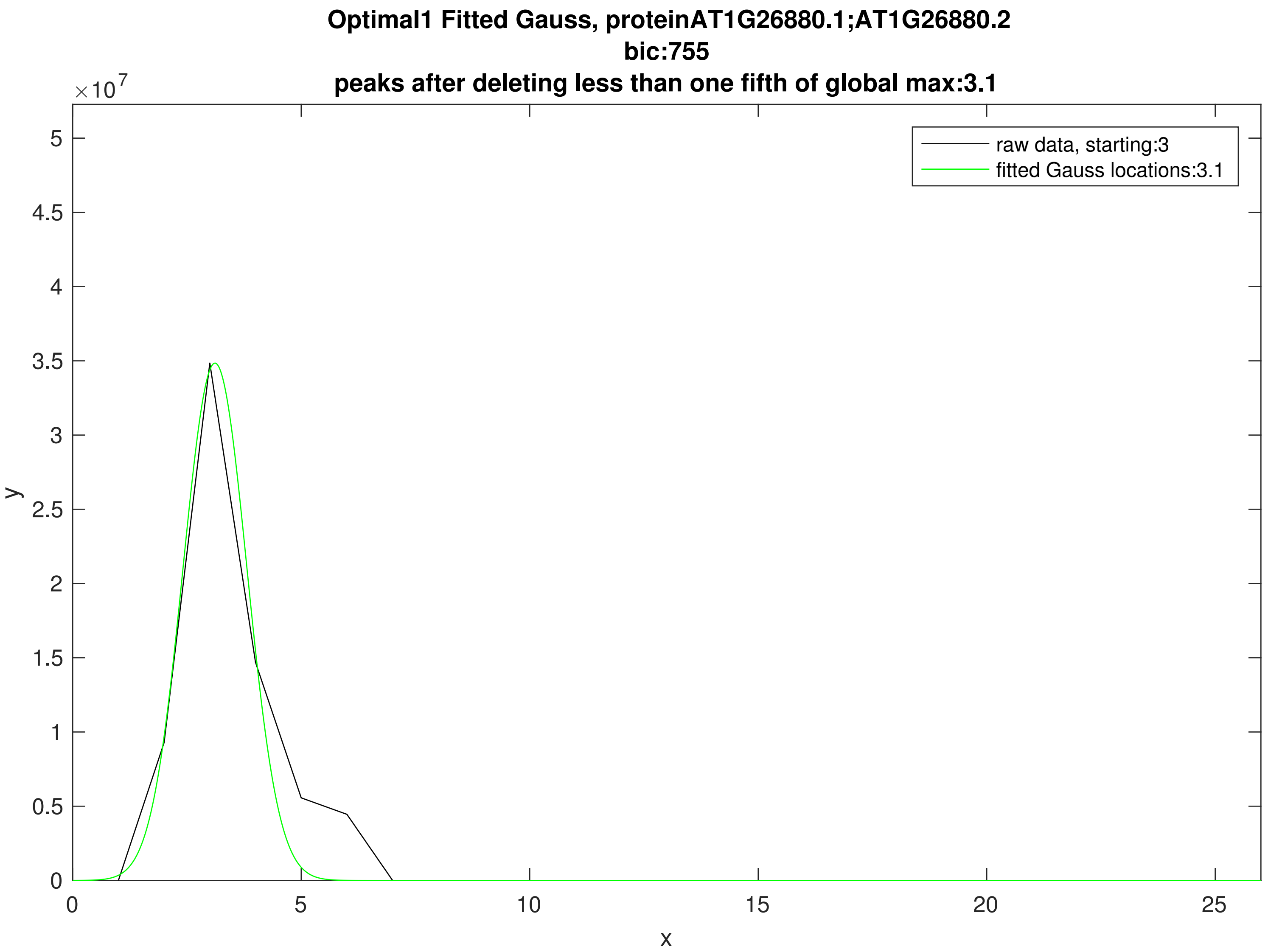

Optimal1 Fitted Gauss, proteinAT1G26880.1;AT1G26880.2
bic:755
peaks after deleting less than one fifth of global max:3.1
×10^7
raw data, starting:3
fitted Gauss locations:3.1
y
x
0
0.5
1
1.5
2
2.5
3
3.5
4
4.5
5
0
5
10
15
20
25

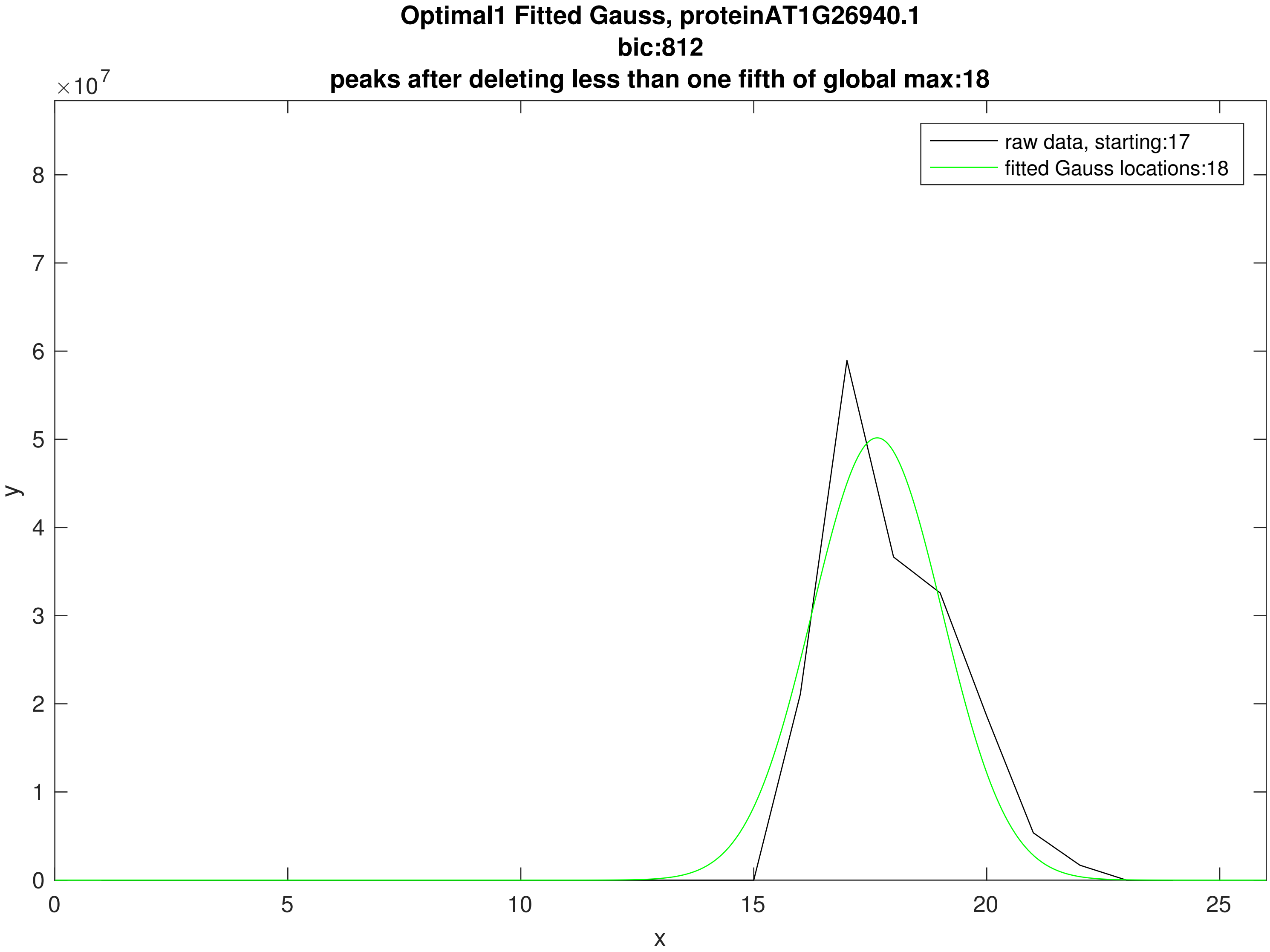

Optimal1 Fitted Gauss, proteinAT1G26940.1
bic:812
peaks after deleting less than one fifth of global max:18
×10^7
raw data, starting:17
fitted Gauss locations:18
8
7
6
5
4
3
2
1
0
y
0
5
10
15
20
25
x

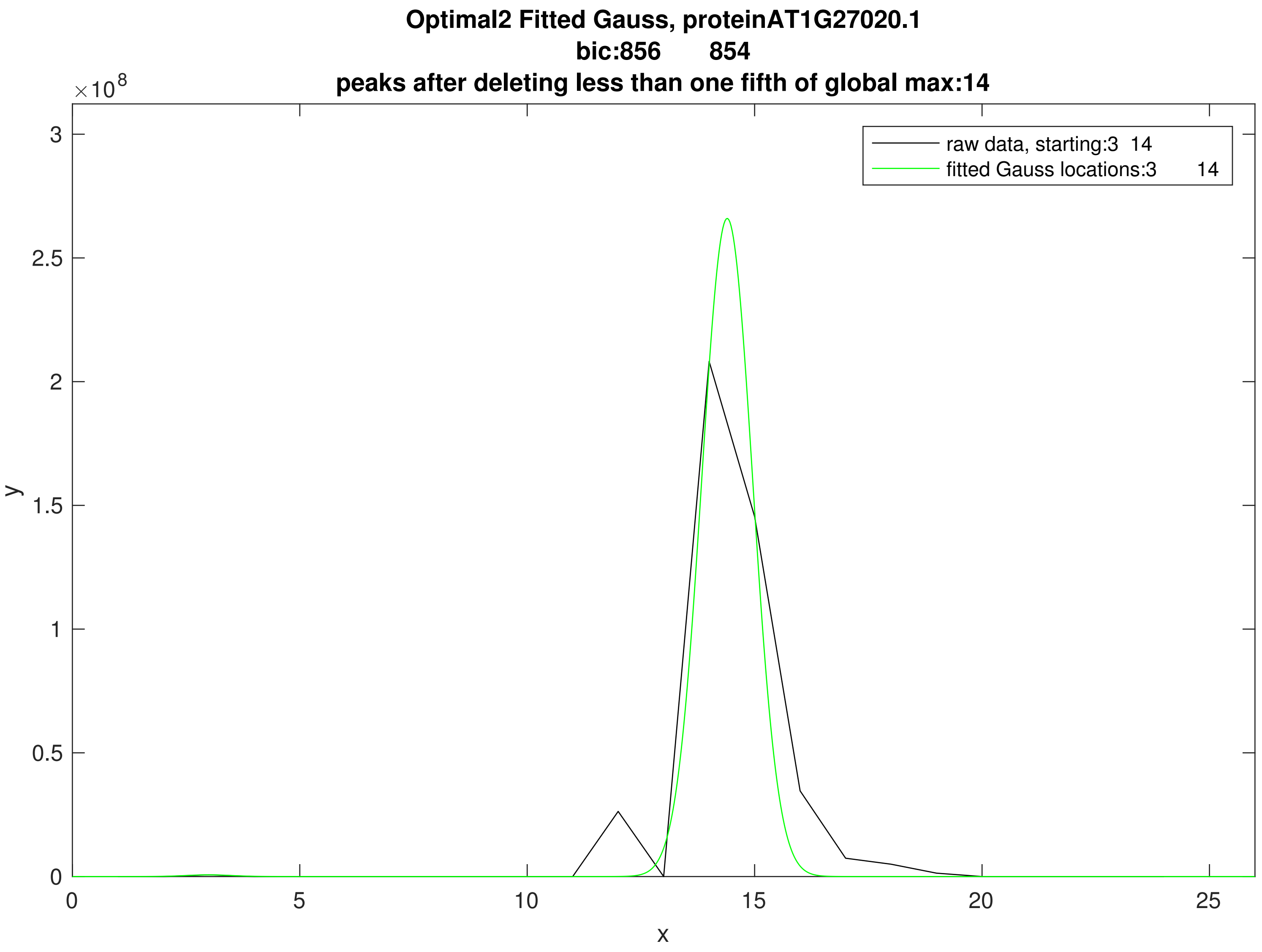

Optimal2 Fitted Gauss, proteinAT1G27020.1
bic:856 854
peaks after deleting less than one fifth of global max:14
×10^8
raw data, starting:3 14
fitted Gauss locations:3 14
y
x
0
0.5
1
1.5
2
2.5
3
0
5
10
15
20
25

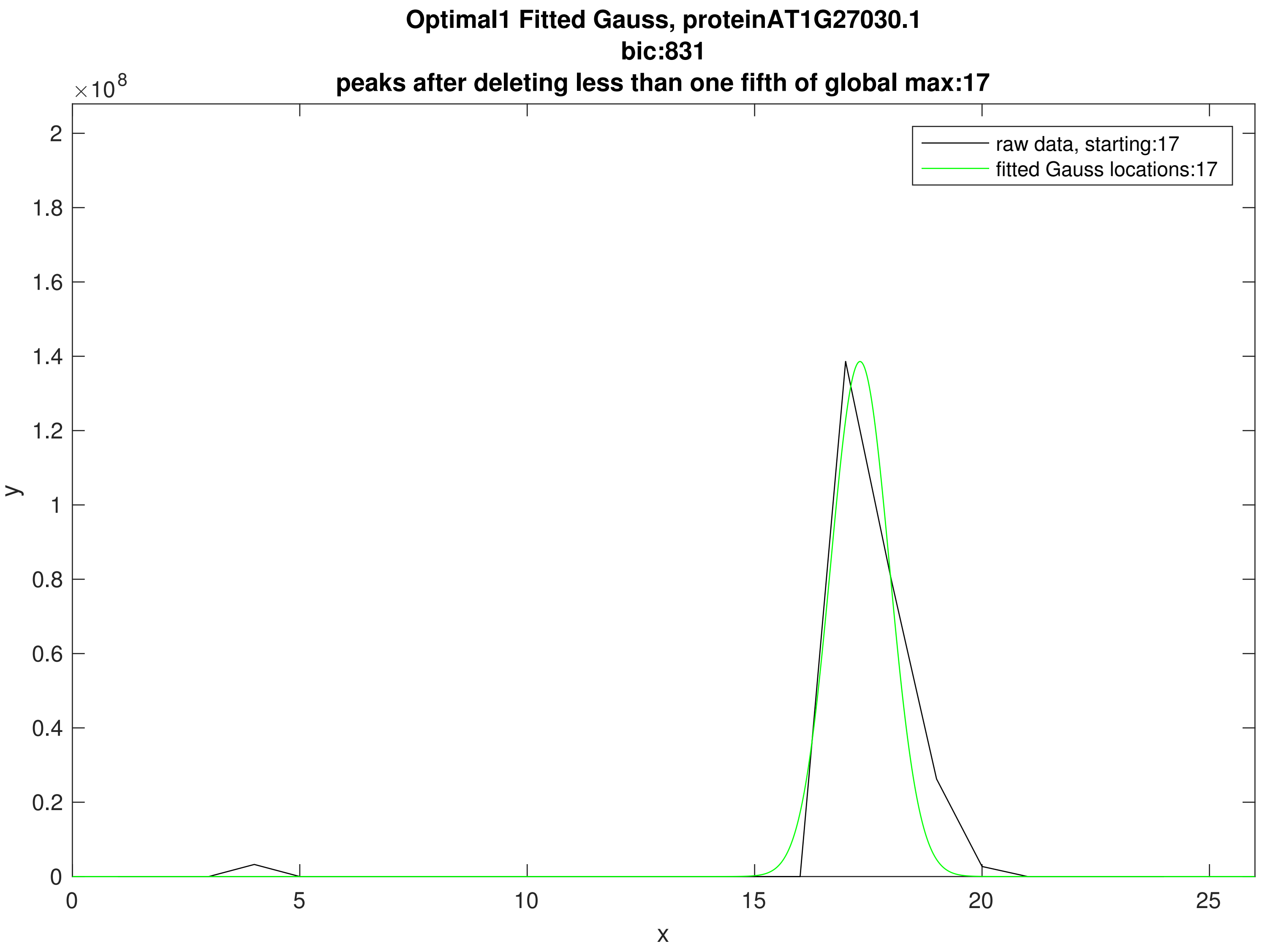
Optimal1 Fitted Gauss, proteinAT1G27030.1
bic:831
peaks after deleting less than one fifth of global max:17
×10^8
raw data, starting:17
fitted Gauss locations:17
y
x
0
0.2
0.4
0.6
0.8
1
1.2
1.4
1.6
1.8
2
5
10
15
20
25

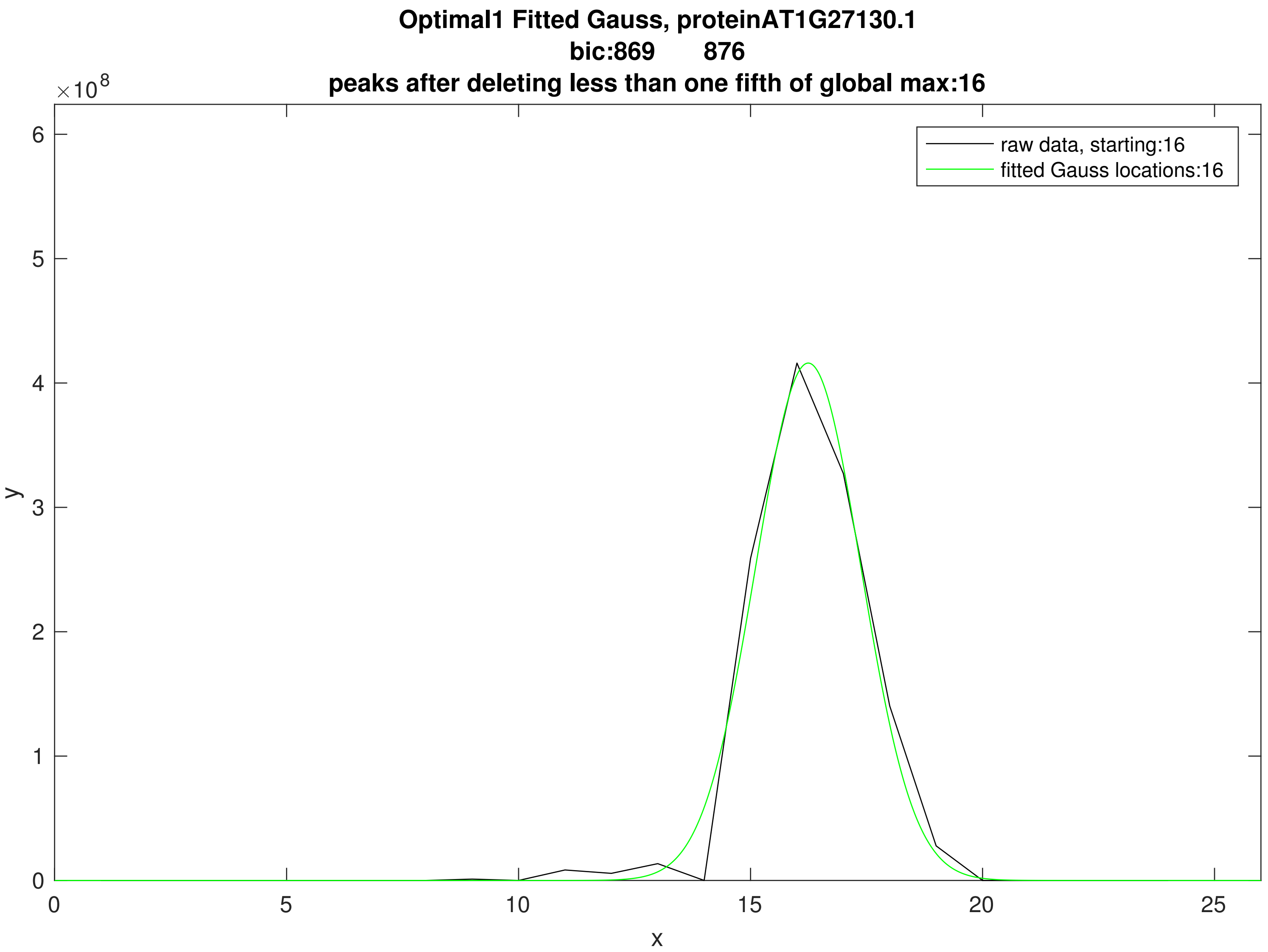
Optimal1 Fitted Gauss, proteinAT1G27130.1
bic:869 876
peaks after deleting less than one fifth of global max:16
×10^8
6
5
4
3
2
1
0
y
0
5
10
15
20
25
x
raw data, starting:16
fitted Gauss locations:16

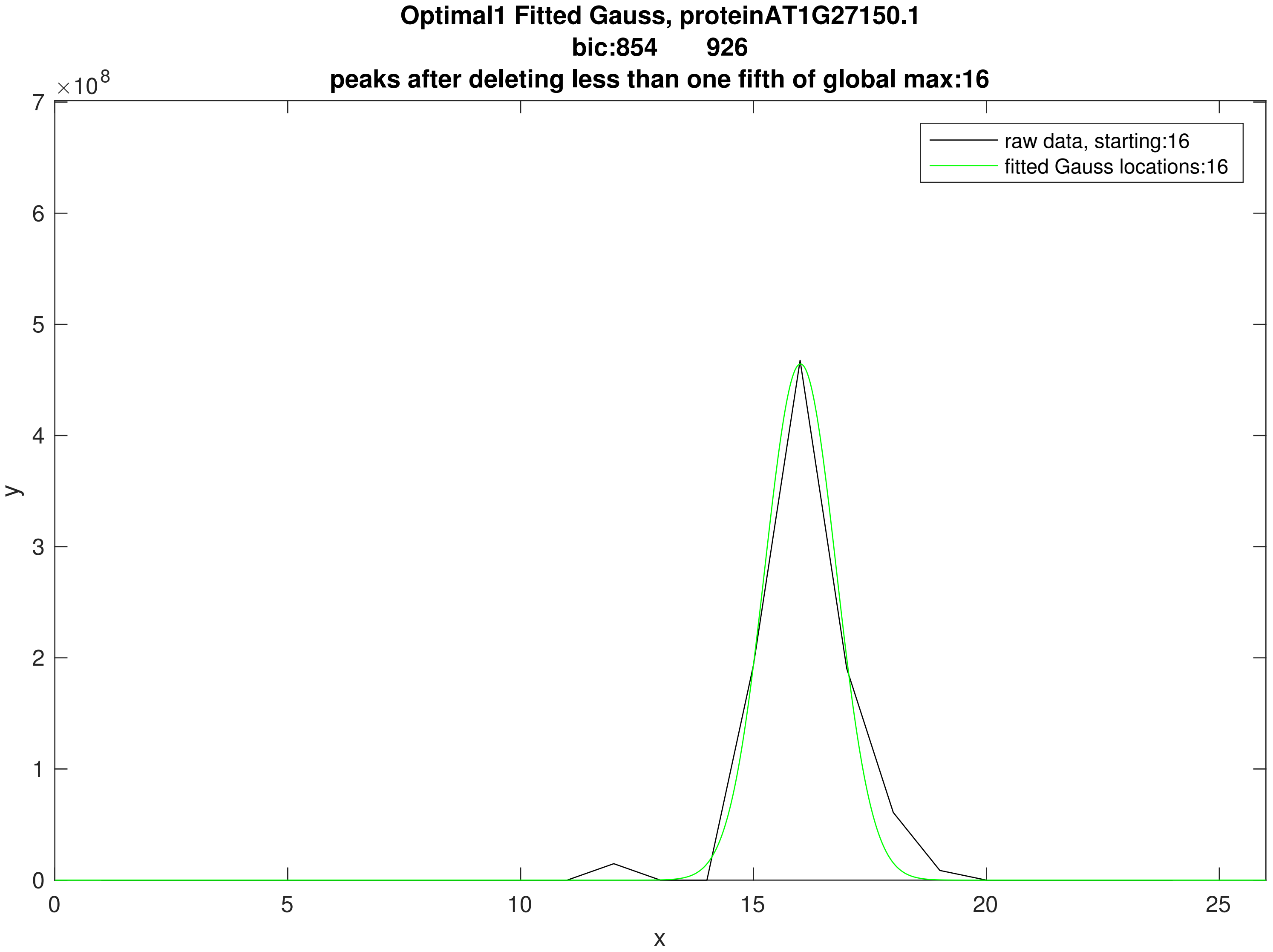
Optimal1 Fitted Gauss, proteinAT1G27150.1
bic:854 926
peaks after deleting less than one fifth of global max:16
×10^8
7
6
5
4
3
2
1
0
y
0
5
10
15
20
25
x
raw data, starting:16
fitted Gauss locations:16

#### Optimal1 Fitted Gauss, proteinAT1G27310.1
#### bic:879
#### peaks after deleting less than one fifth of global max:19

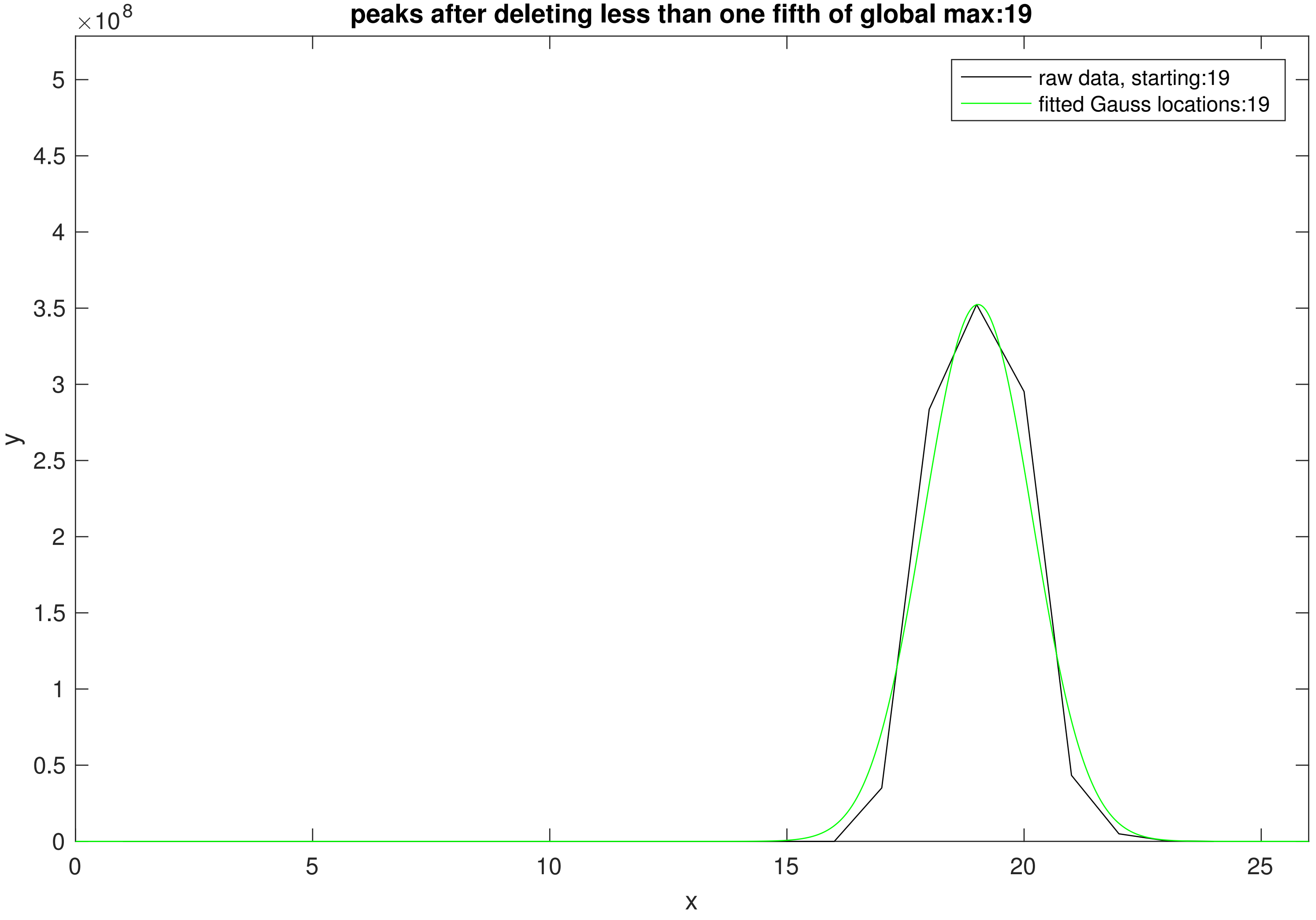

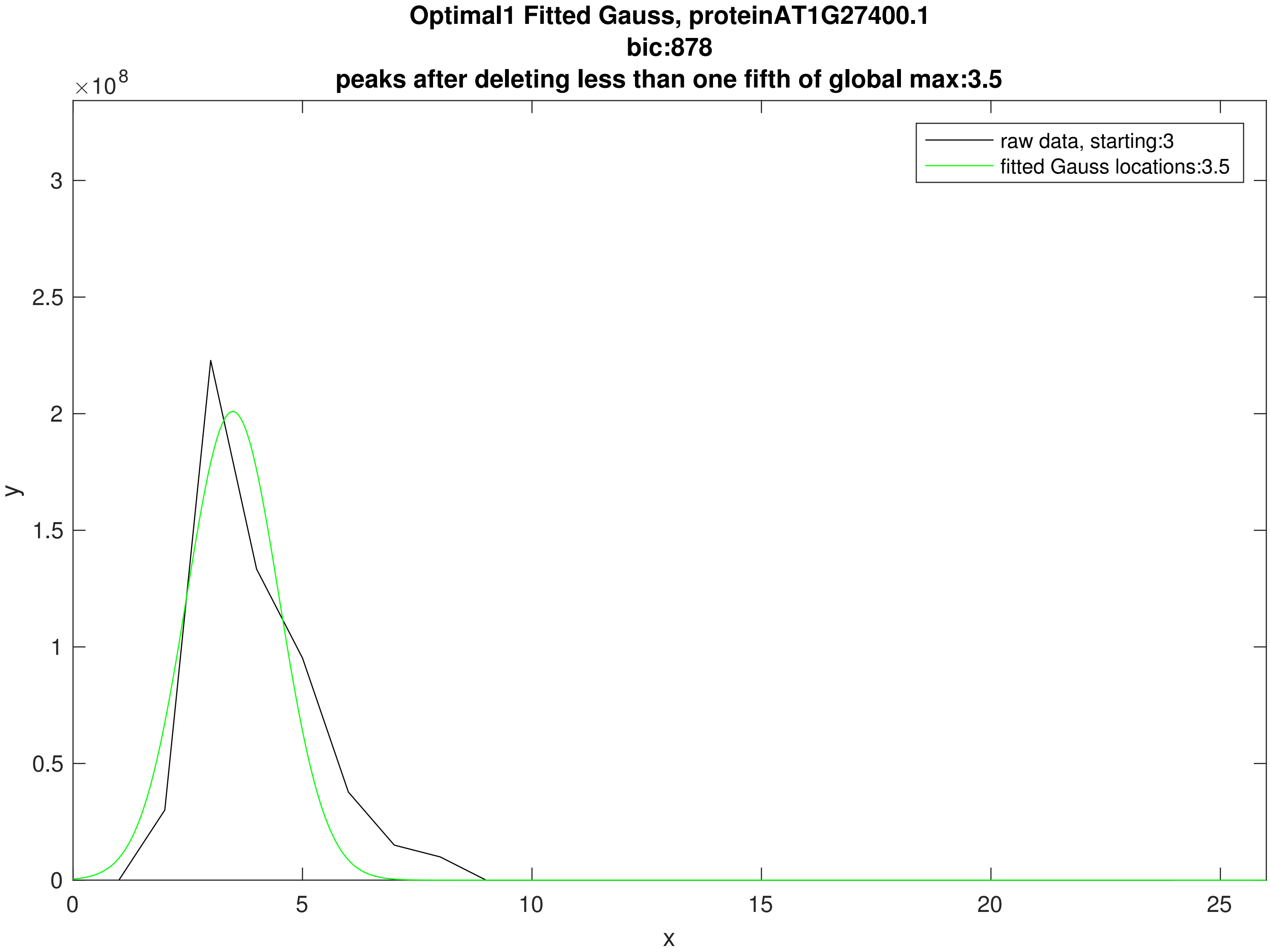
Optimal1 Fitted Gauss, proteinAT1G27400.1
bic:878
peaks after deleting less than one fifth of global max:3.5
×10^8
raw data, starting:3
fitted Gauss locations:3.5
y
x
0
0.5
1
1.5
2
2.5
3
0
5
10
15
20
25

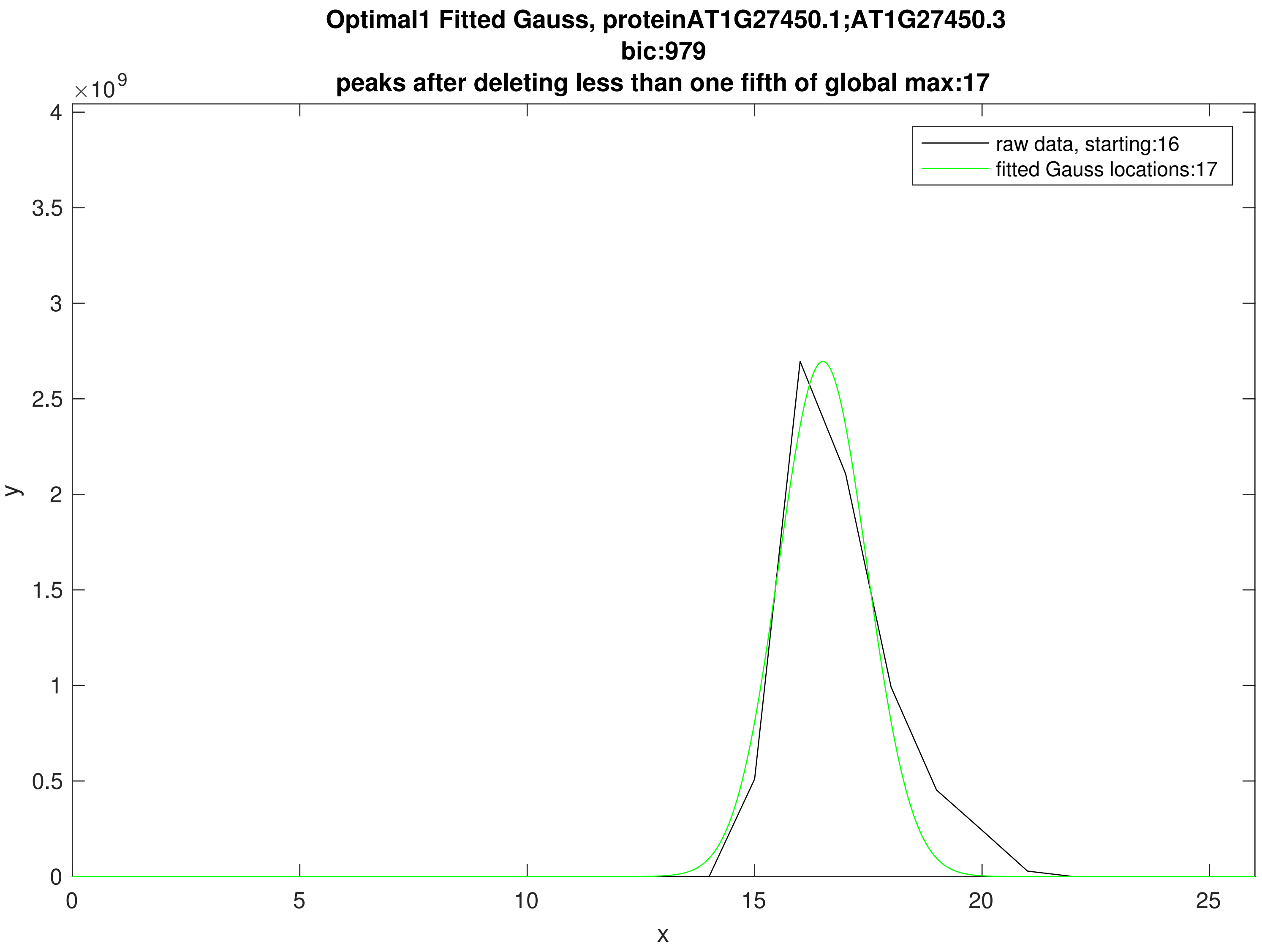
Optimal1 Fitted Gauss, proteinAT1G27450.1;AT1G27450.3
bic:979
peaks after deleting less than one fifth of global max:17
×10^9
raw data, starting:16
fitted Gauss locations:17
y
x
0
0.5
1
1.5
2
2.5
3
3.5
4
5
10
15
20
25

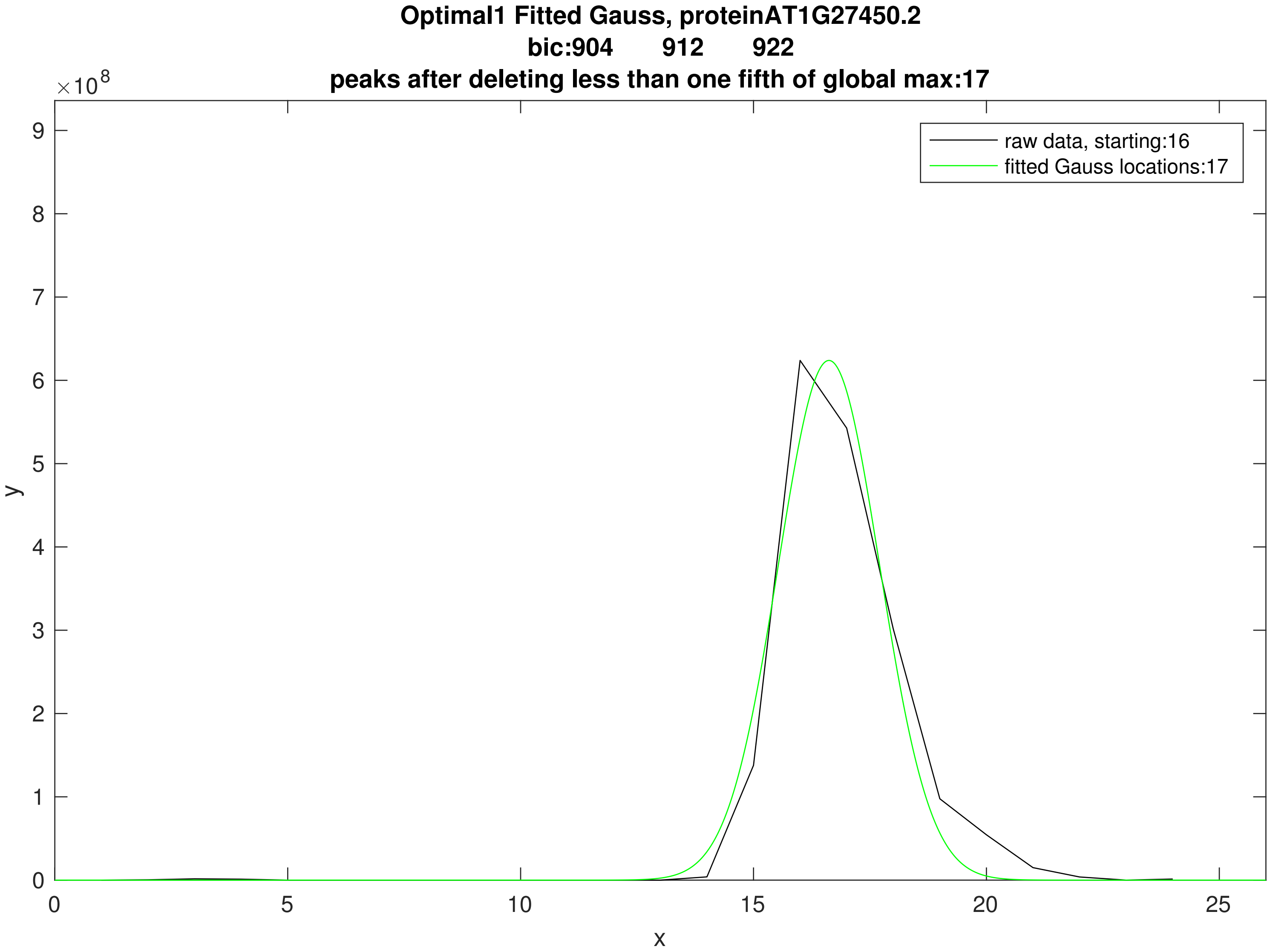
Optimal1 Fitted Gauss, proteinAT1G27450.2
bic:904 912 922
peaks after deleting less than one fifth of global max:17
×10^8
raw data, starting:16
fitted Gauss locations:17
y
x
0
1
2
3
4
5
6
7
8
9
0
5
10
15
20
25

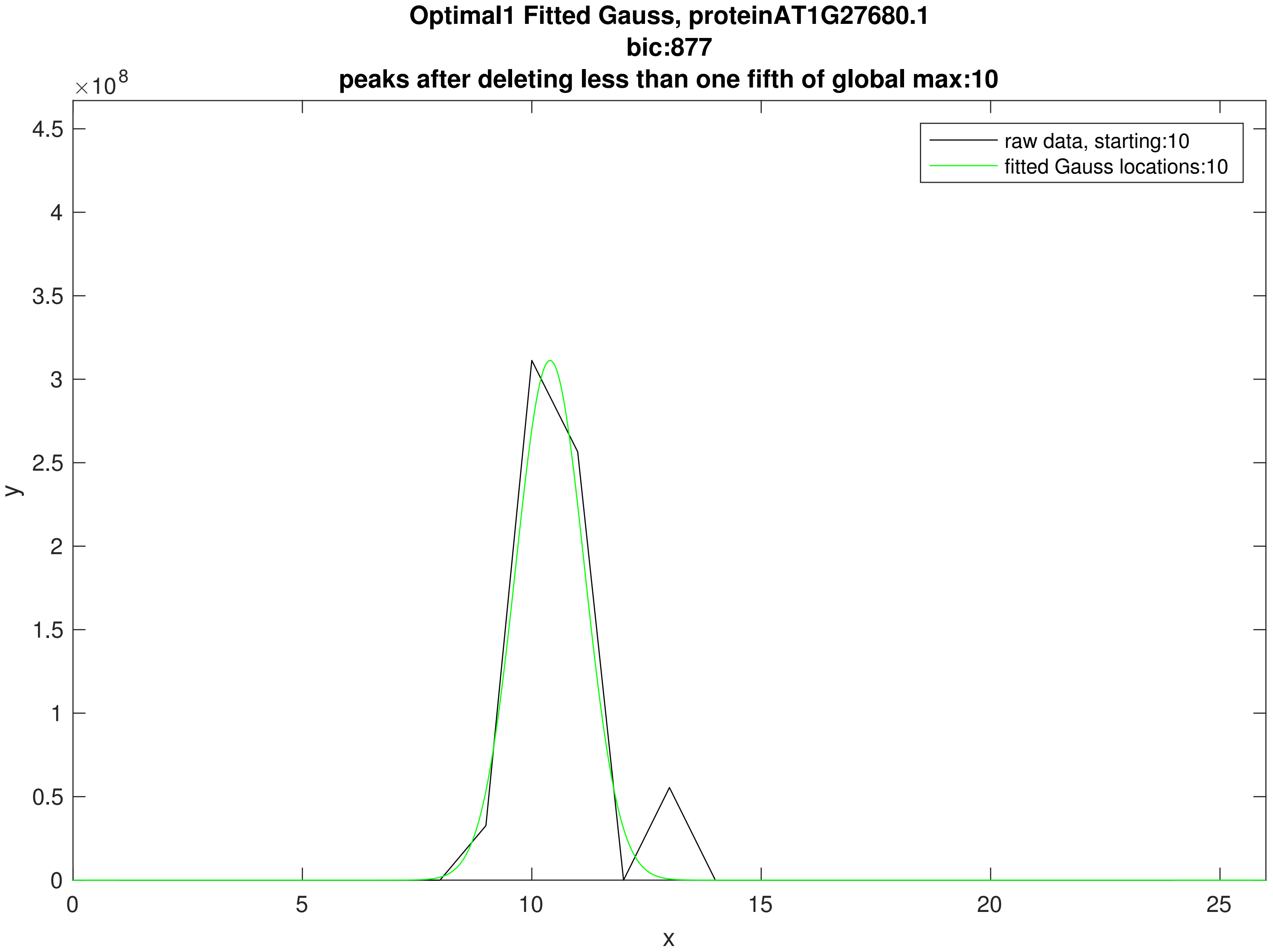

Optimal1 Fitted Gauss, proteinAT1G27680.1
bic:877
peaks after deleting less than one fifth of global max:10
×10^8
4.5
4
3.5
3
2.5
2
1.5
1
0.5
0
y
0
5
10
15
20
25
x
raw data, starting:10
fitted Gauss locations:10

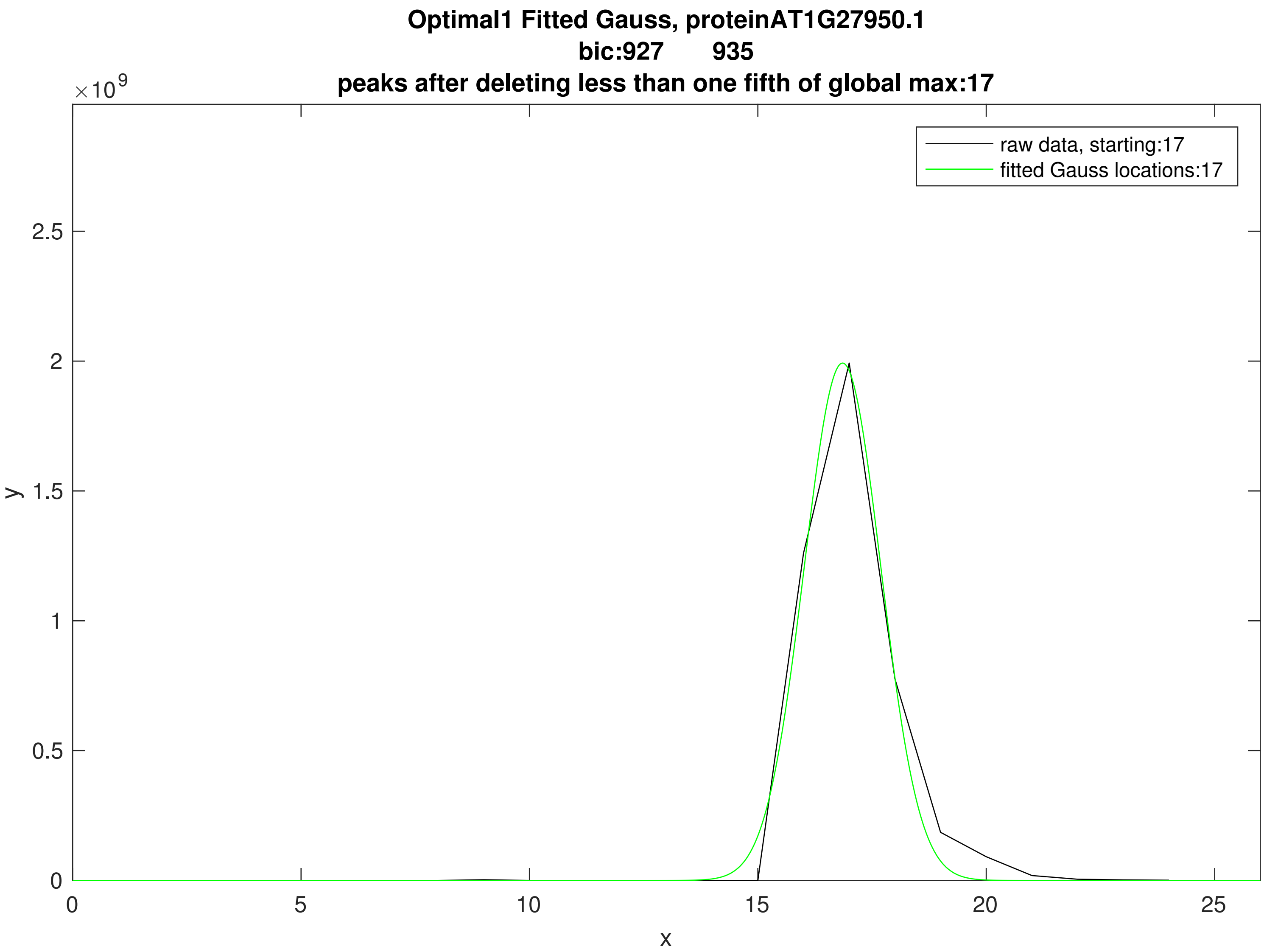

Optimal1 Fitted Gauss, proteinAT1G27950.1
bic:927 935
peaks after deleting less than one fifth of global max:17
×10^9
raw data, starting:17
fitted Gauss locations:17
2.5
2
1.5
1
0.5
0
y
0
5
10
15
20
25
x

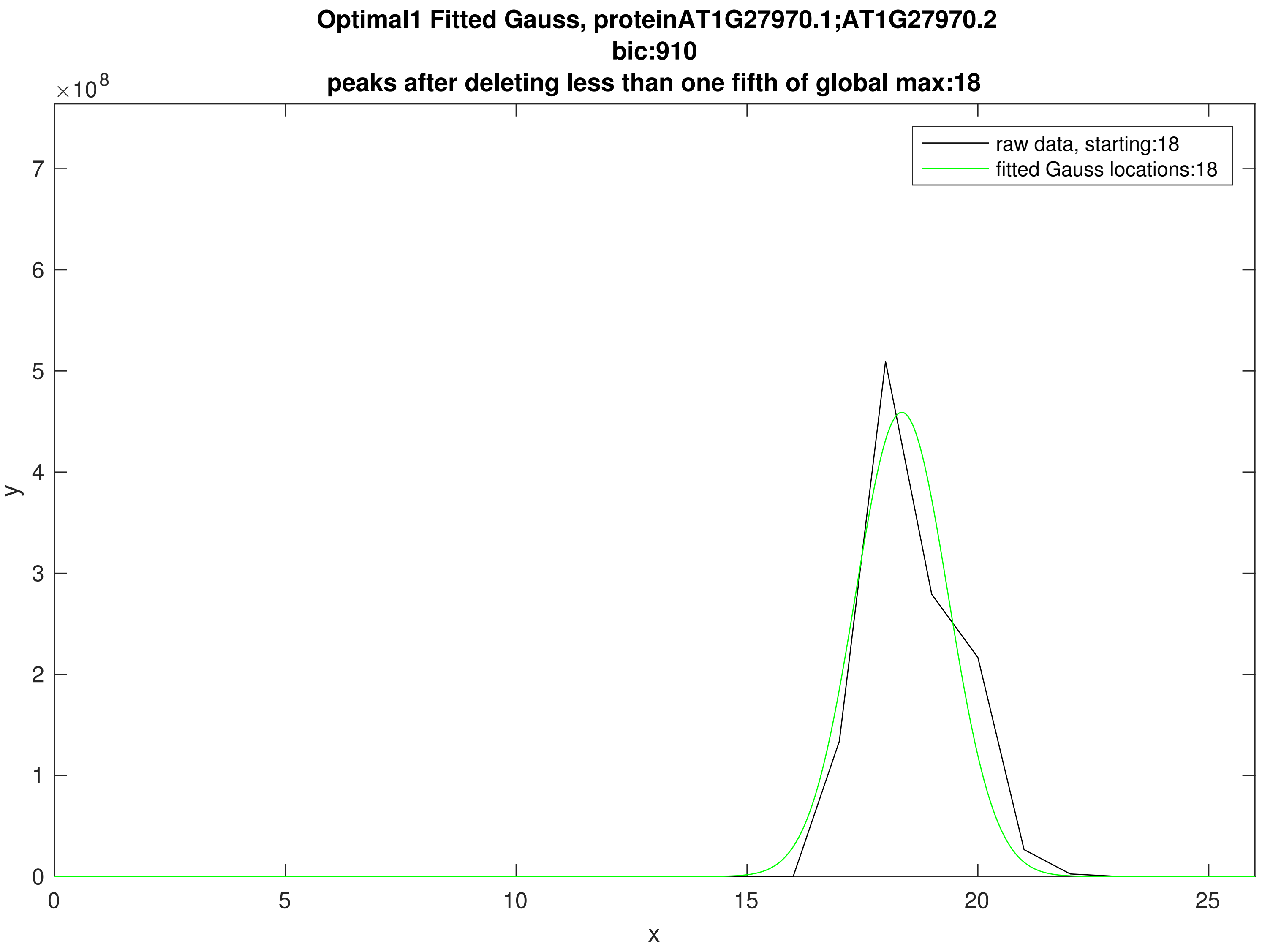
Optimal1 Fitted Gauss, proteinAT1G27970.1;AT1G27970.2
bic:910
peaks after deleting less than one fifth of global max:18
×10^8
raw data, starting:18
fitted Gauss locations:18
y
x
0
1
2
3
4
5
6
7
0
5
10
15
20
25

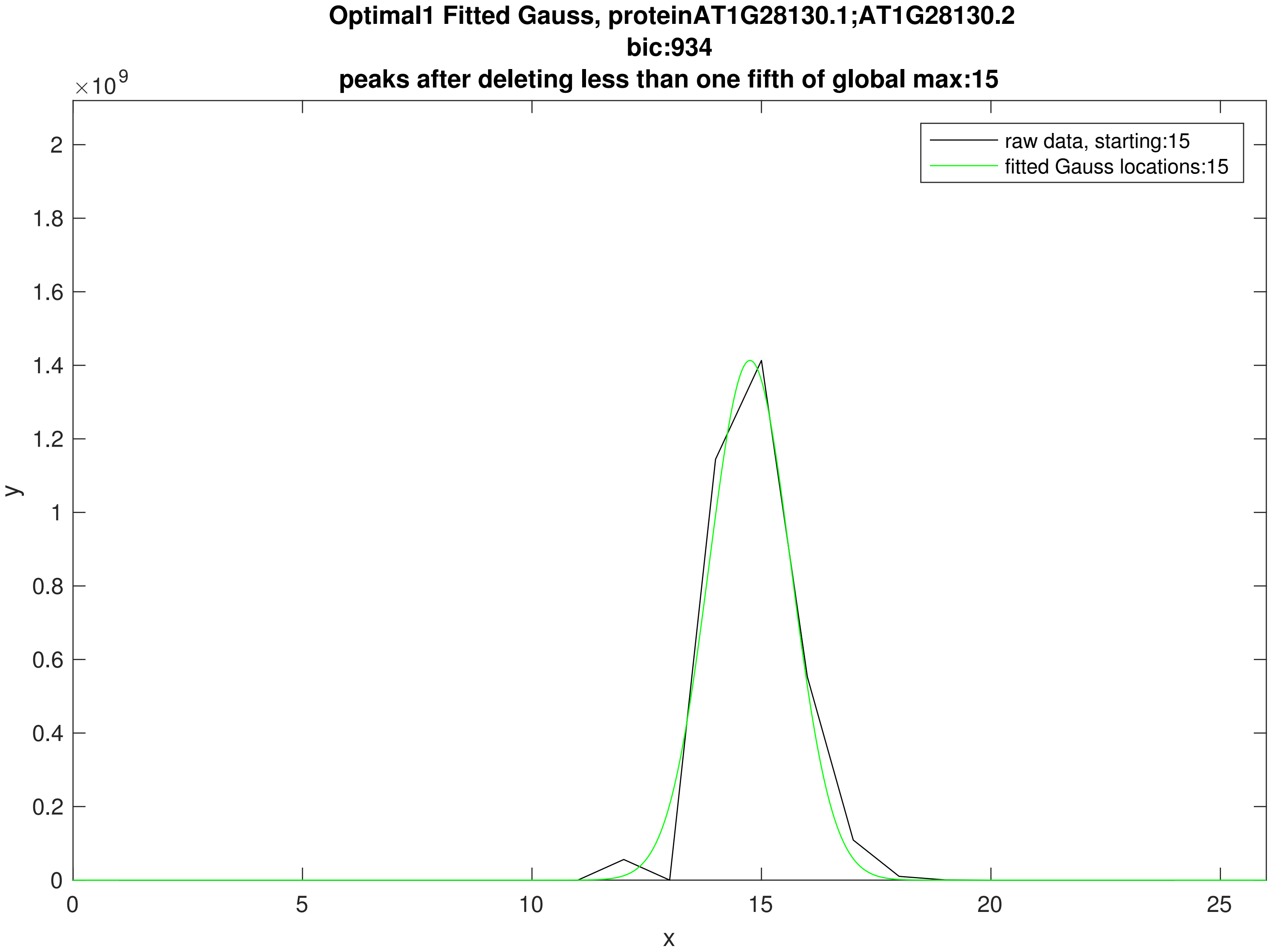

Optimal1 Fitted Gauss, proteinAT1G28130.1;AT1G28130.2
bic:934
peaks after deleting less than one fifth of global max:15
×10^9
raw data, starting:15
fitted Gauss locations:15
y
x
0
0.2
0.4
0.6
0.8
1
1.2
1.4
1.6
1.8
2
5
10
15
20
25

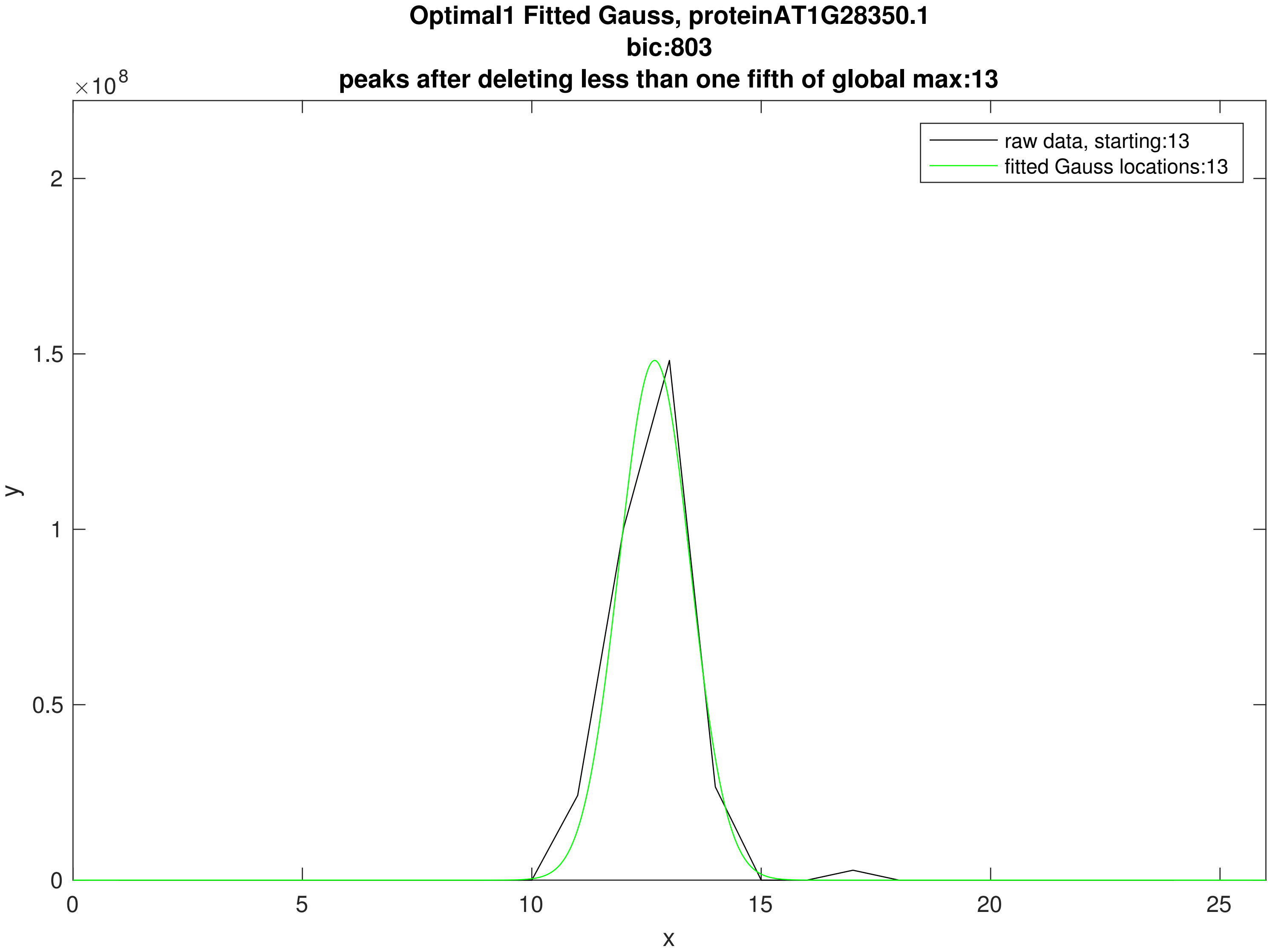

Optimal1 Fitted Gauss, proteinAT1G28350.1
bic:803
peaks after deleting less than one fifth of global max:13
×10^8
raw data, starting:13
fitted Gauss locations:13
y
x
0
0.5
1
1.5
2
0
5
10
15
20
25

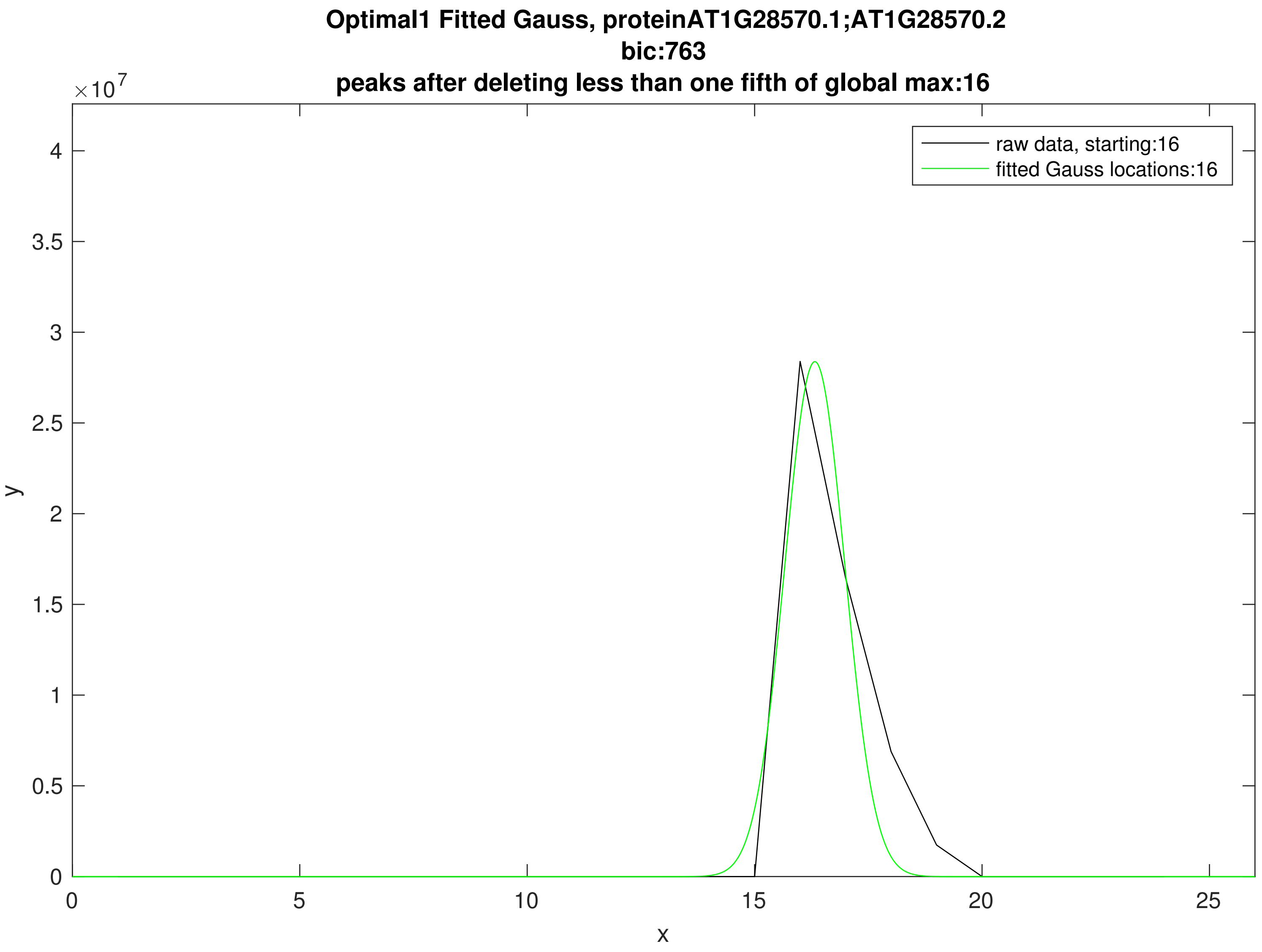

Optimal1 Fitted Gauss, proteinAT1G28570.1;AT1G28570.2
bic:763
peaks after deleting less than one fifth of global max:16
×10^7
raw data, starting:16
fitted Gauss locations:16
y
x
0
0.5
1
1.5
2
2.5
3
3.5
4
0
5
10
15
20
25

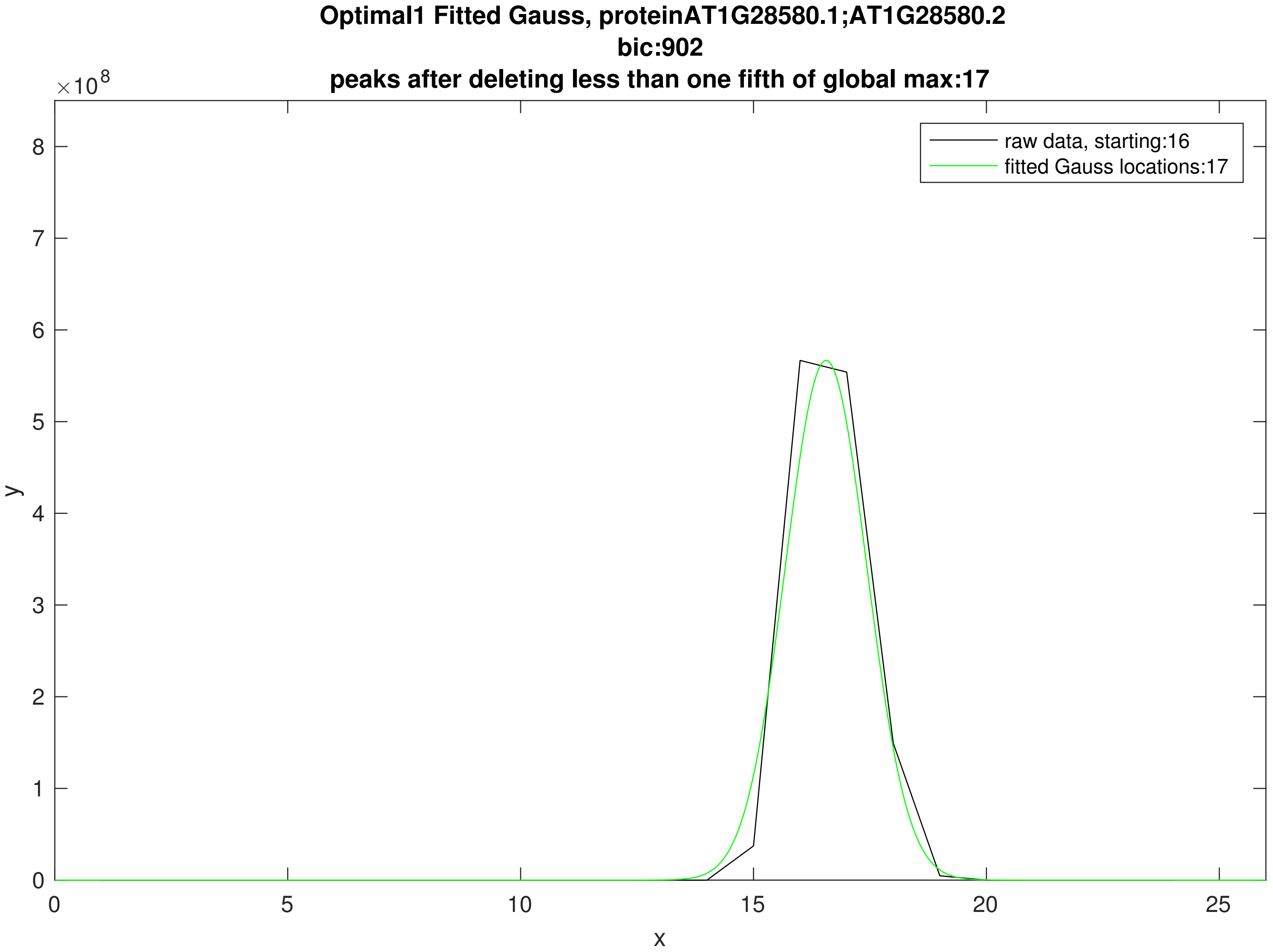

Optimal1 Fitted Gauss, proteinAT1G28580.1;AT1G28580.2
bic:902
peaks after deleting less than one fifth of global max:17
×10^8
raw data, starting:16
fitted Gauss locations:17
y
x
0
1
2
3
4
5
6
7
8
0
5
10
15
20
25

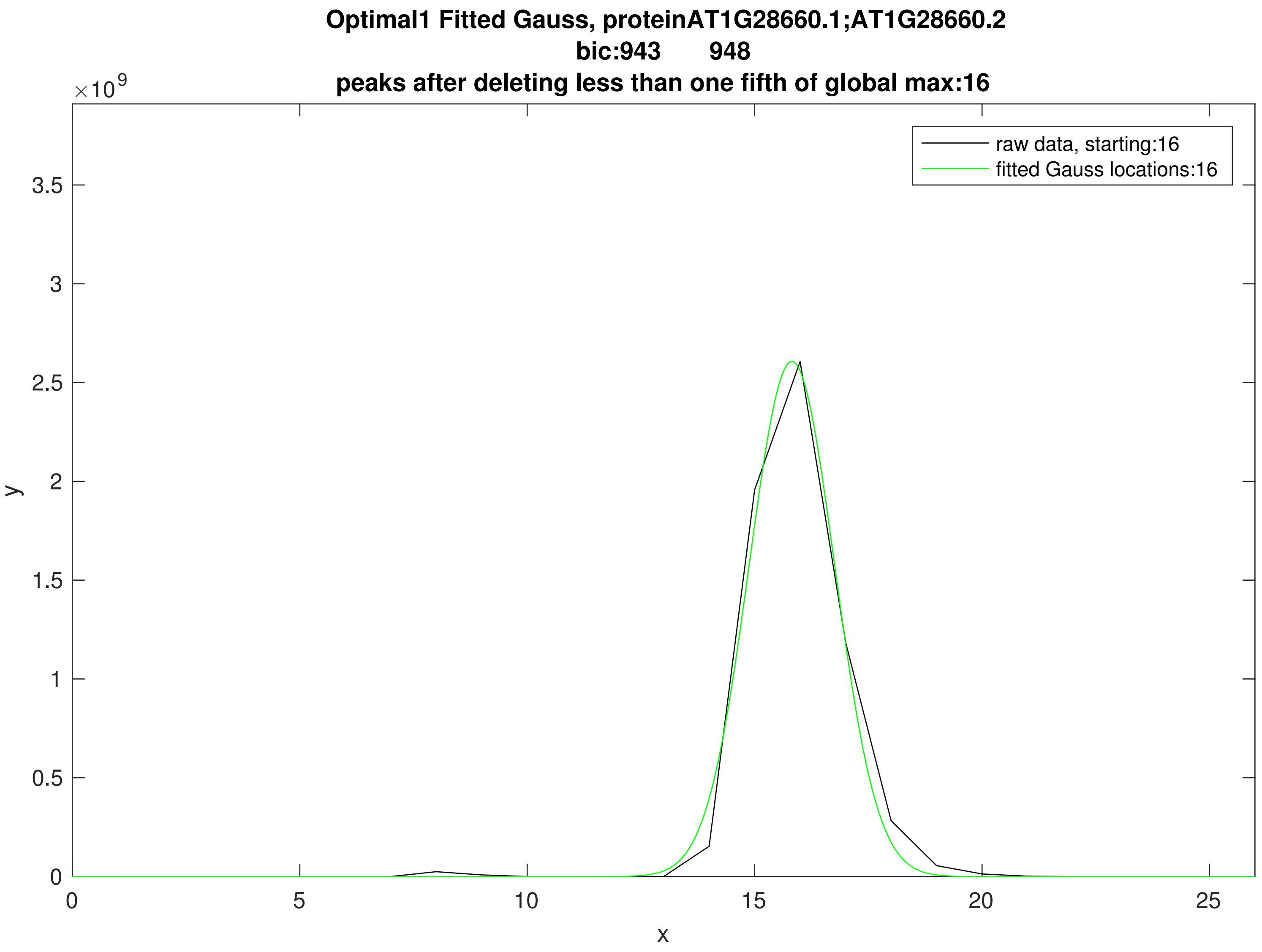
Optimal1 Fitted Gauss, proteinAT1G28660.1;AT1G28660.2
bic:943 948
peaks after deleting less than one fifth of global max:16
×10^9
3.5
3
2.5
2
1.5
1
0.5
0
y
0
5
10
15
20
25
x
raw data, starting:16
fitted Gauss locations:16

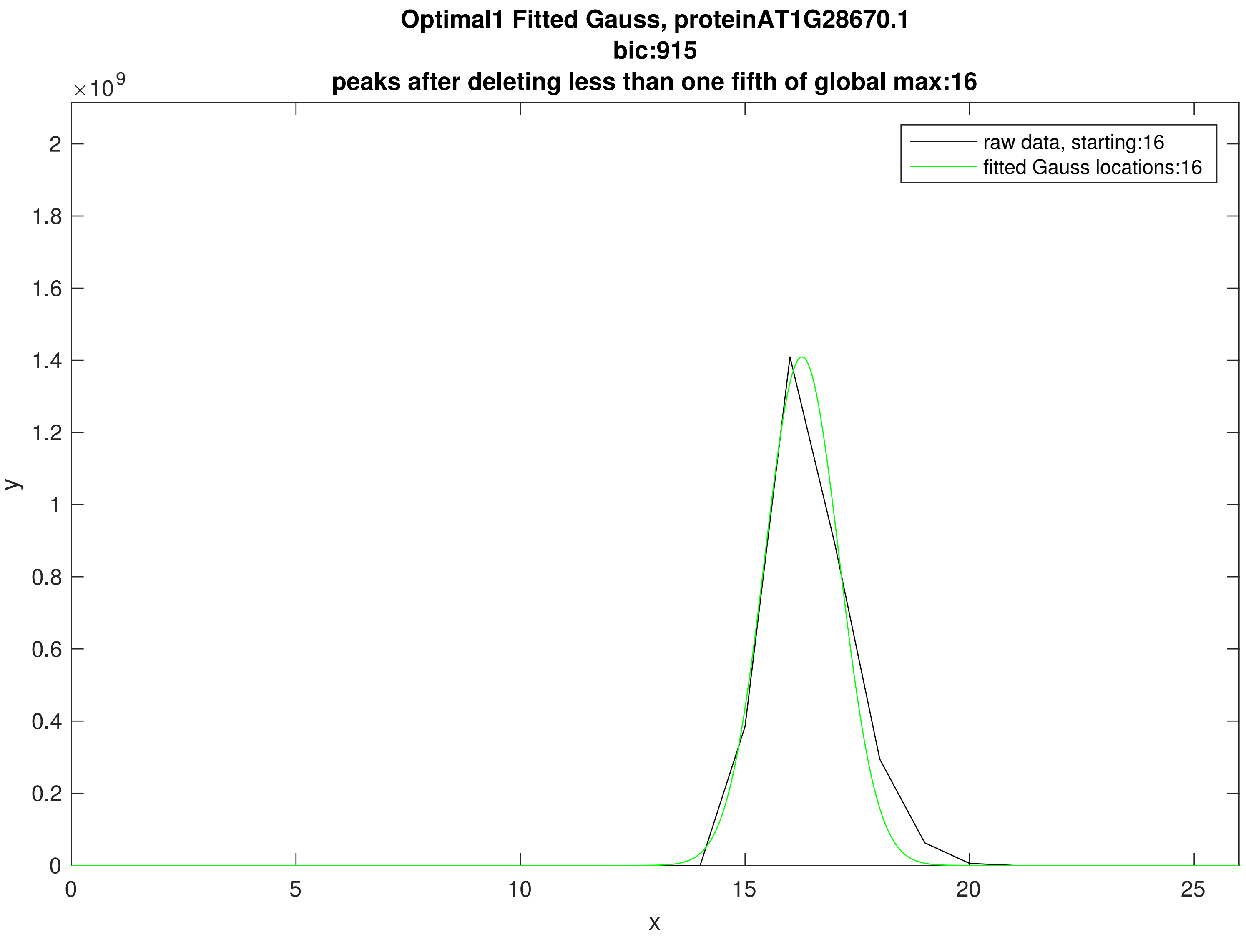

Optimal1 Fitted Gauss, proteinAT1G28670.1
bic:915
peaks after deleting less than one fifth of global max:16
×10^9
raw data, starting:16
fitted Gauss locations:16
y
x
0
0.2
0.4
0.6
0.8
1
1.2
1.4
1.6
1.8
2
0
5
10
15
20
25

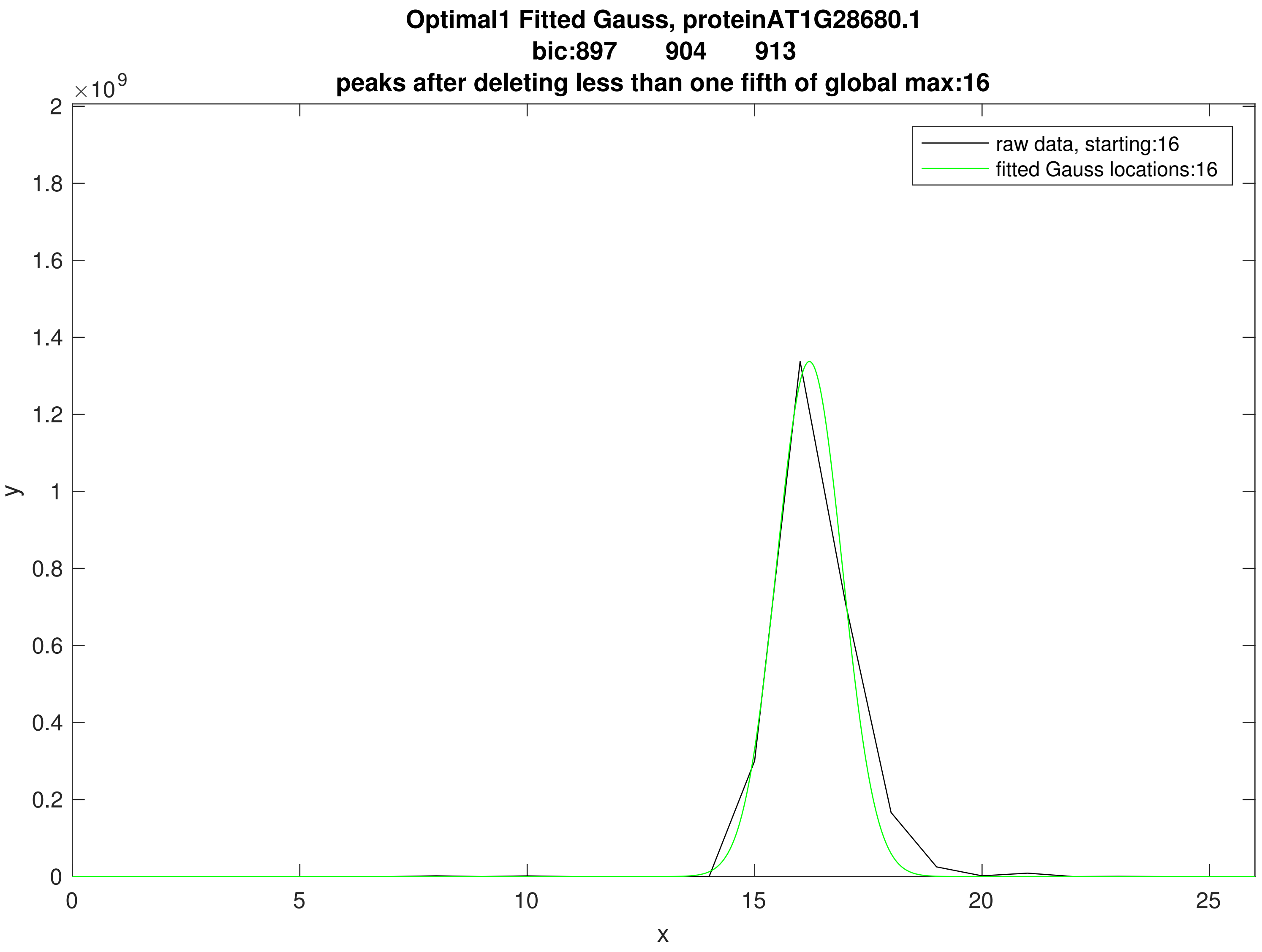

Optimal1 Fitted Gauss, proteinAT1G28680.1
bic:897 904 913
peaks after deleting less than one fifth of global max:16
×10^9
2
1.8
1.6
1.4
1.2
1
0.8
0.6
0.4
0.2
0
y
0
5
10
15
20
25
x
raw data, starting:16
fitted Gauss locations:16

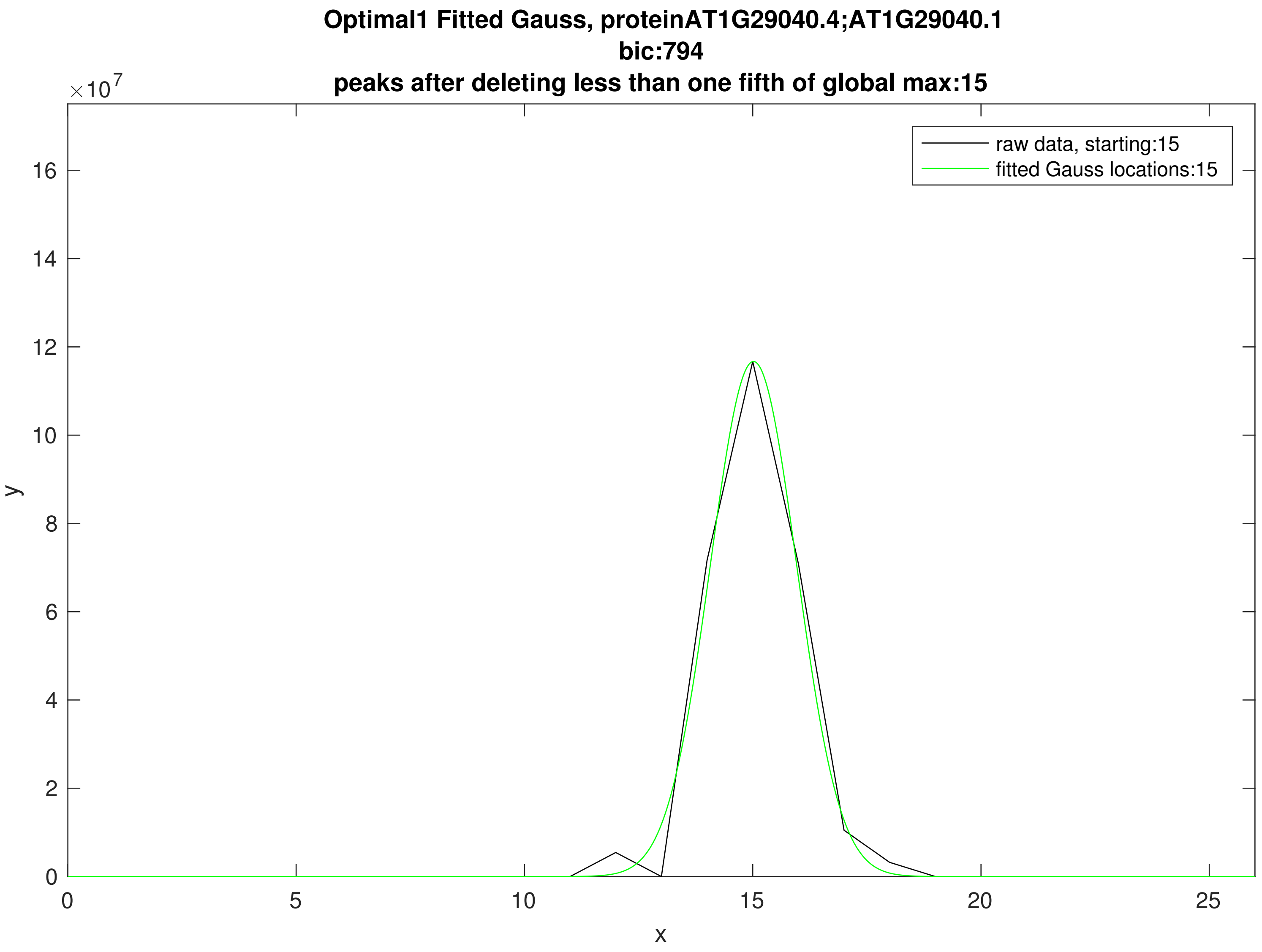

Optimal1 Fitted Gauss, proteinAT1G29040.4;AT1G29040.1
bic:794
peaks after deleting less than one fifth of global max:15
×10^7
raw data, starting:15
fitted Gauss locations:15
16
14
12
10
8
6
4
2
0
y
0
5
10
15
20
25
x

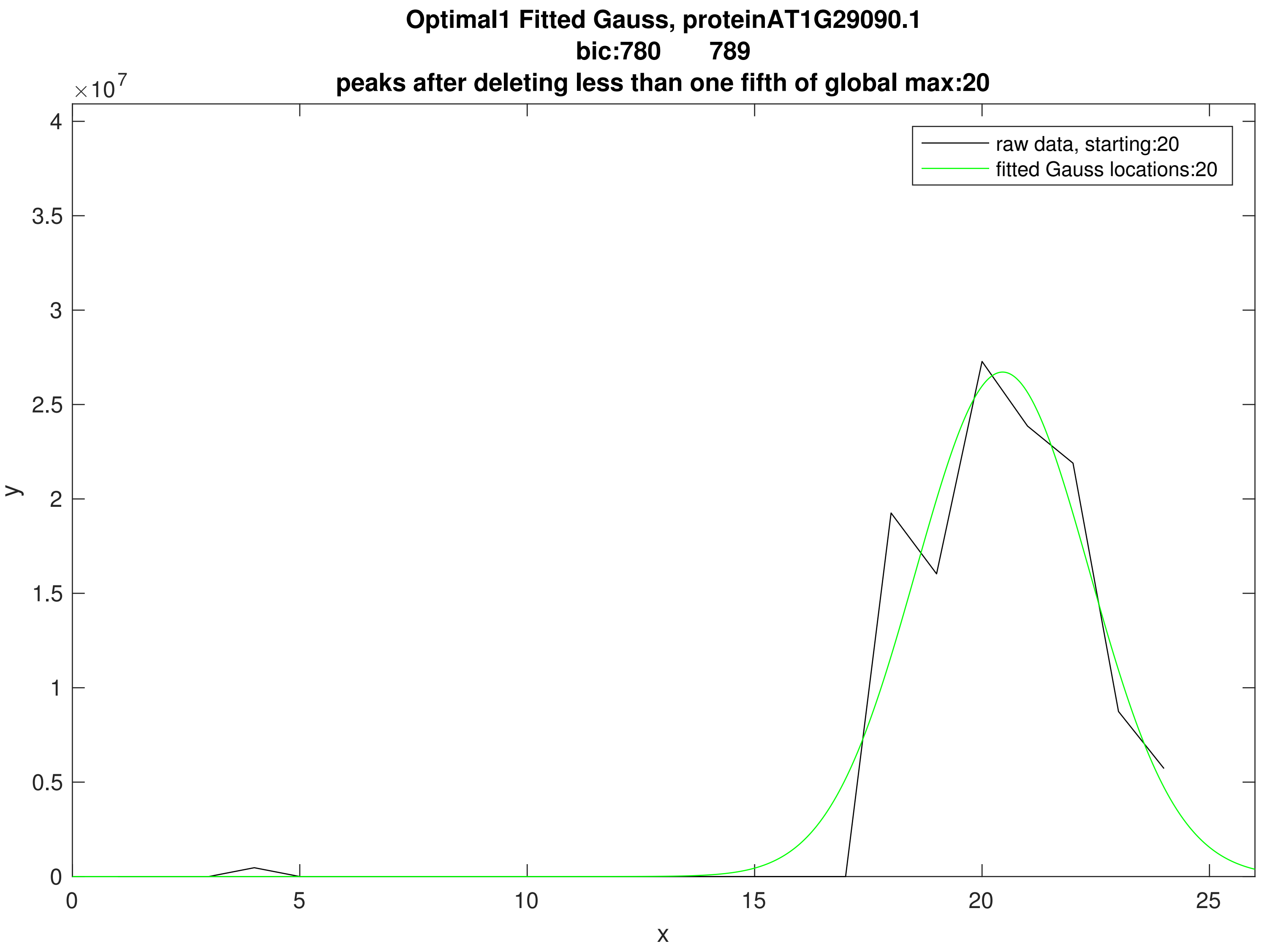
Optimal1 Fitted Gauss, proteinAT1G29090.1
bic:780 789
peaks after deleting less than one fifth of global max:20
×10^7
raw data, starting:20
fitted Gauss locations:20
y
x
0
0.5
1
1.5
2
2.5
3
3.5
4
0
5
10
15
20
25

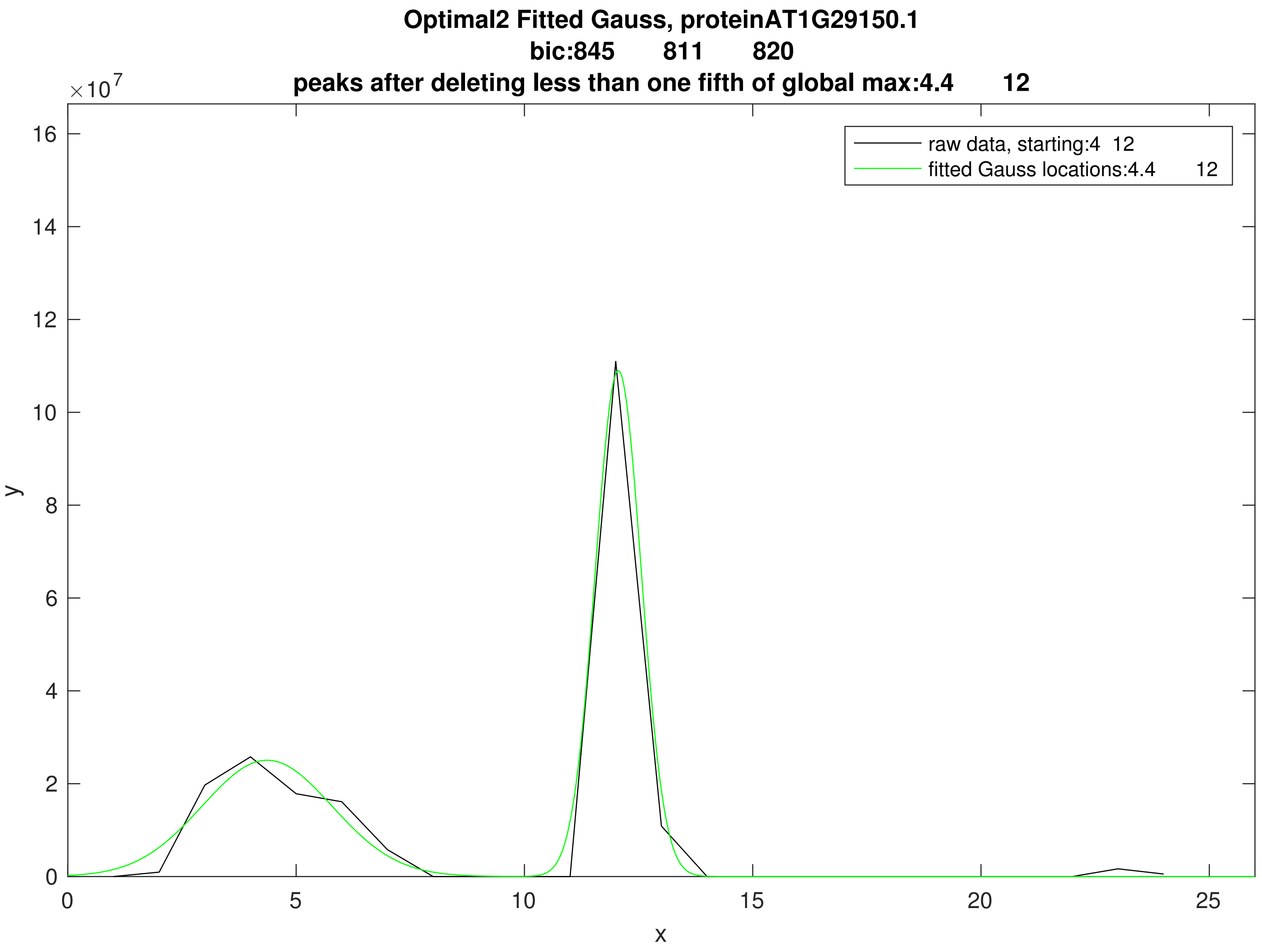
Optimal2 Fitted Gauss, proteinAT1G29150.1
bic:845 811 820
peaks after deleting less than one fifth of global max:4.4 12
×10^7
raw data, starting:4 12
fitted Gauss locations:4.4 12
y
x
0
2
4
6
8
10
12
14
16
0
5
10
15
20
25

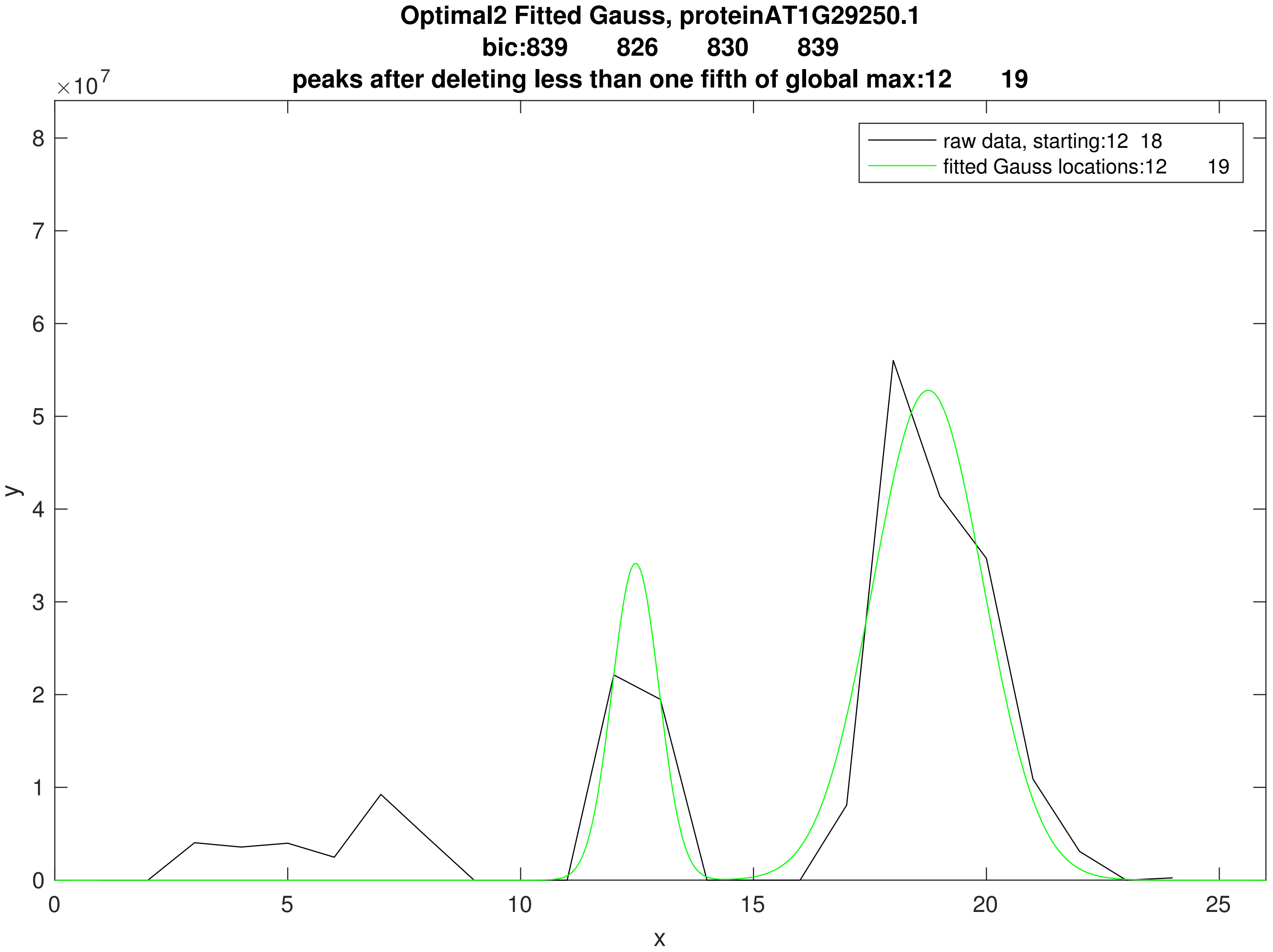

Optimal2 Fitted Gauss, proteinAT1G29250.1
bic:839 826 830 839
peaks after deleting less than one fifth of global max:12 19
×10^7
raw data, starting:12 18
fitted Gauss locations:12 19
y
x
0
1
2
3
4
5
6
7
8
0
5
10
15
20
25

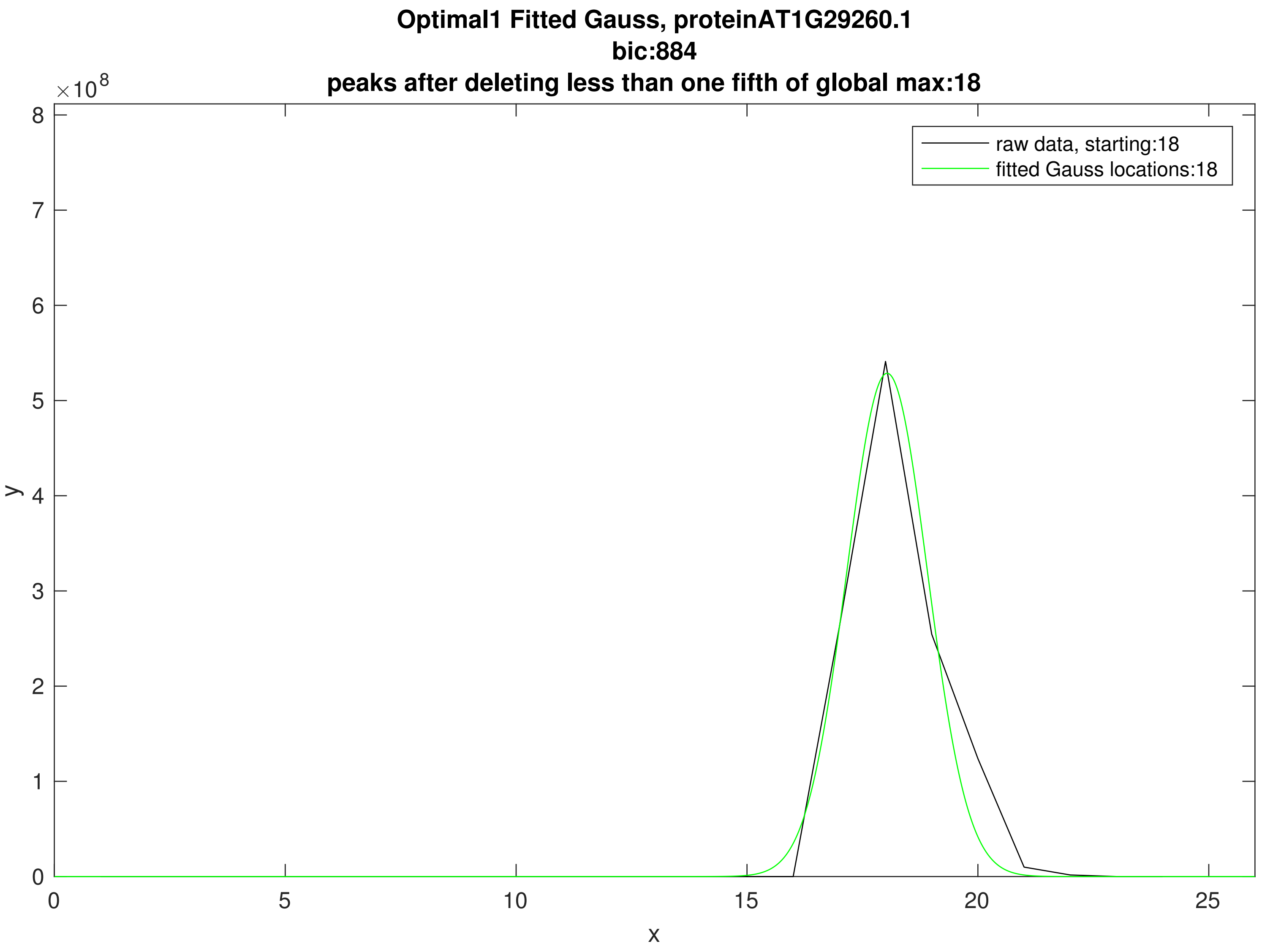
Optimal1 Fitted Gauss, proteinAT1G29260.1
bic:884
peaks after deleting less than one fifth of global max:18
×10^8
8
7
6
5
4
3
2
1
0
y
0
5
10
15
20
25
x
raw data, starting:18
fitted Gauss locations:18

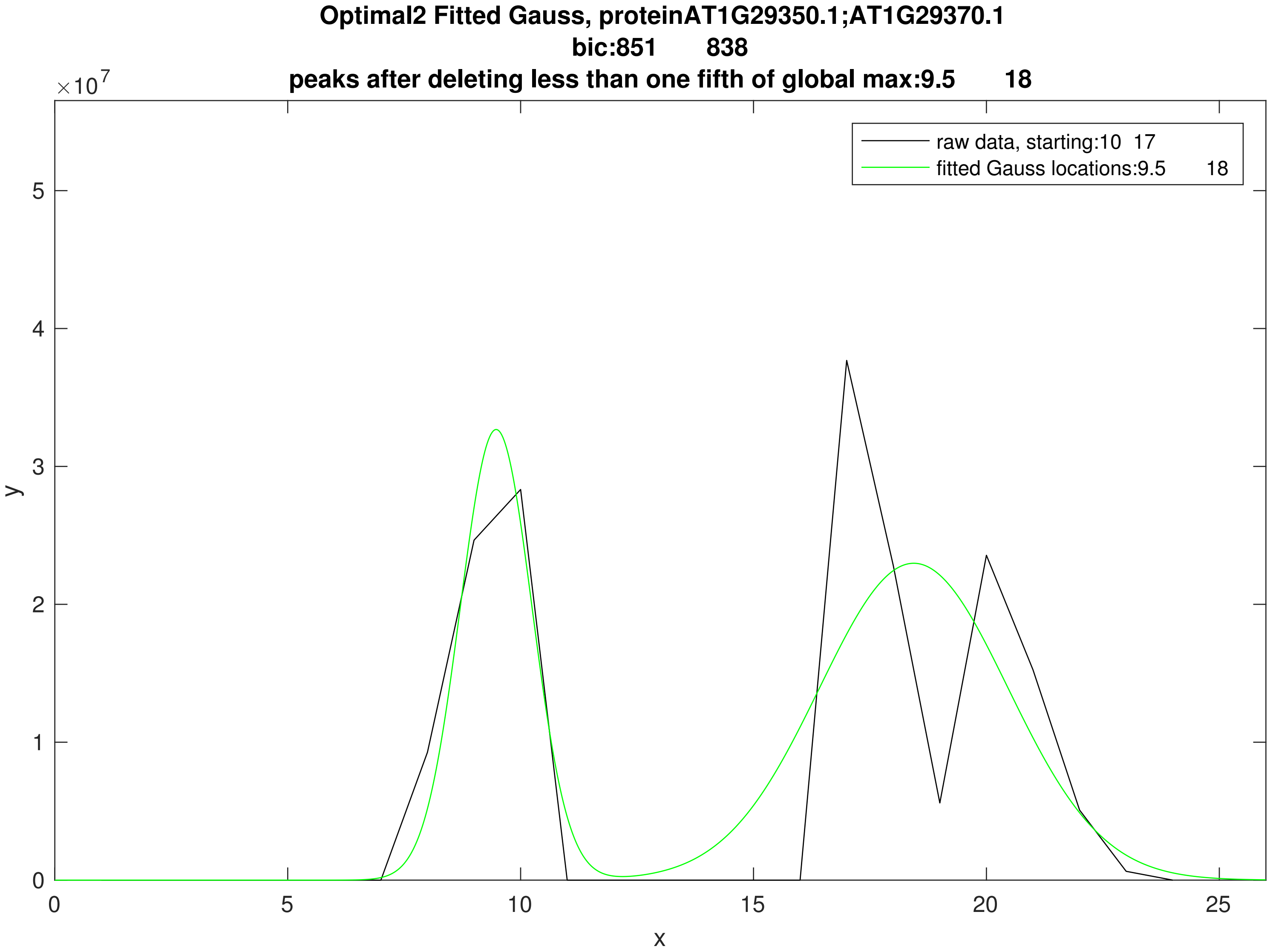

Optimal2 Fitted Gauss, proteinAT1G29350.1;AT1G29370.1
bic:851 838
peaks after deleting less than one fifth of global max:9.5 18
×10^7
raw data, starting:10 17
fitted Gauss locations:9.5 18
y
x
0
1
2
3
4
5
0
5
10
15
20
25

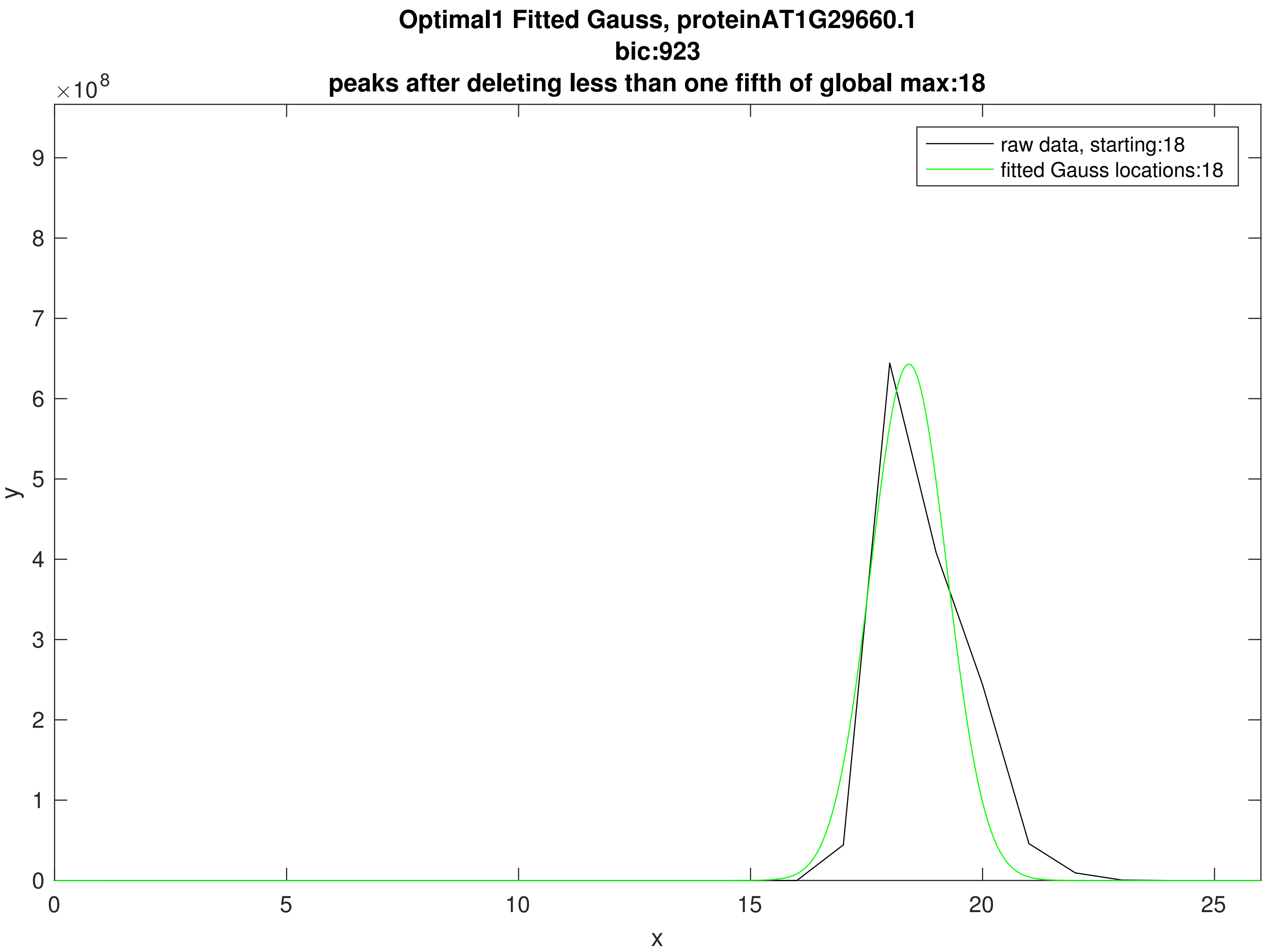

Optimal1 Fitted Gauss, proteinAT1G29660.1
bic:923
peaks after deleting less than one fifth of global max:18
×10^8
raw data, starting:18
fitted Gauss locations:18
y
x
0
1
2
3
4
5
6
7
8
9
0
5
10
15
20
25

**Optimal2 Fitted Gauss, proteinAT1G29670.1**
**bic:935 916**
**peaks after deleting less than one fifth of global max:16 22**

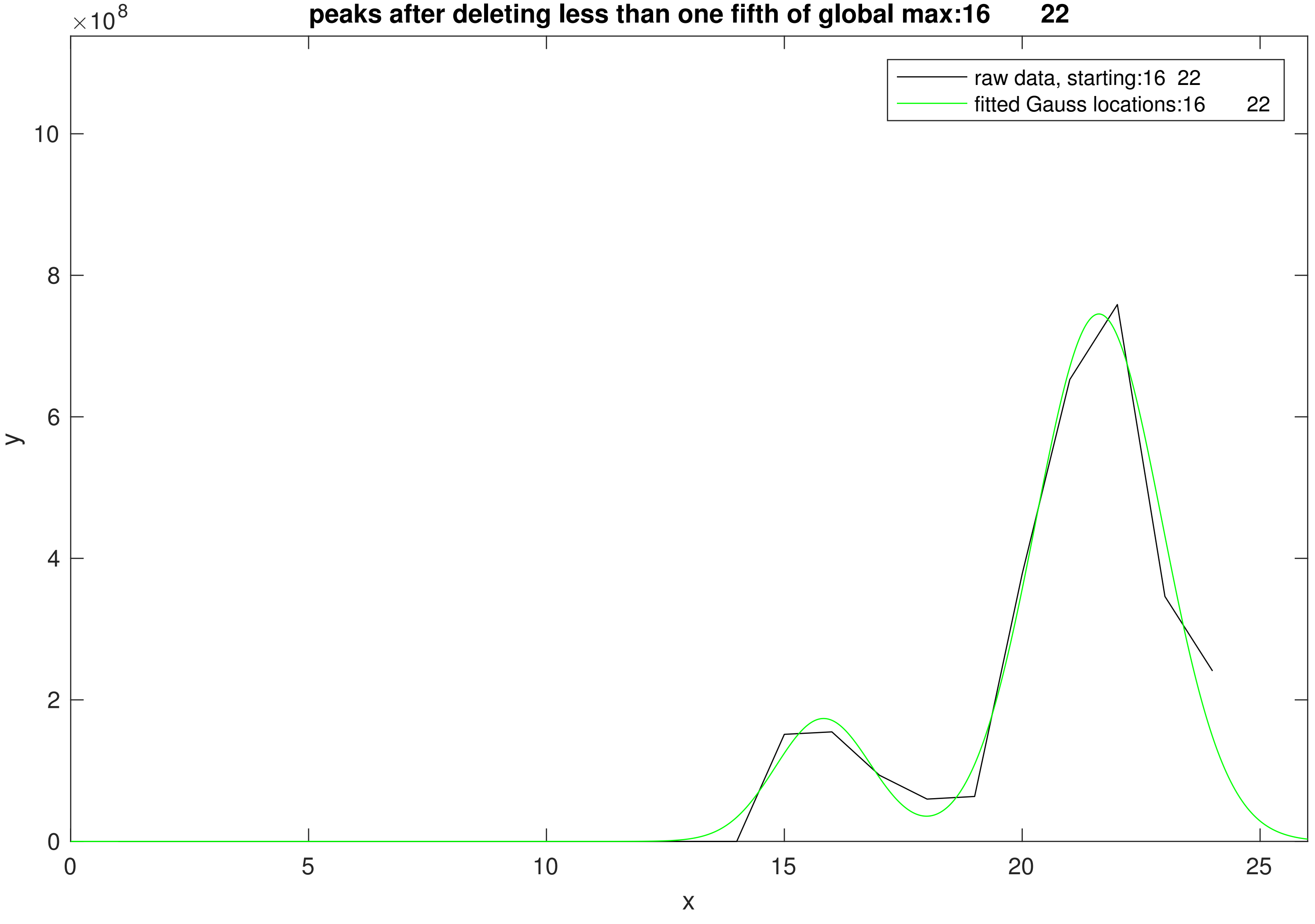

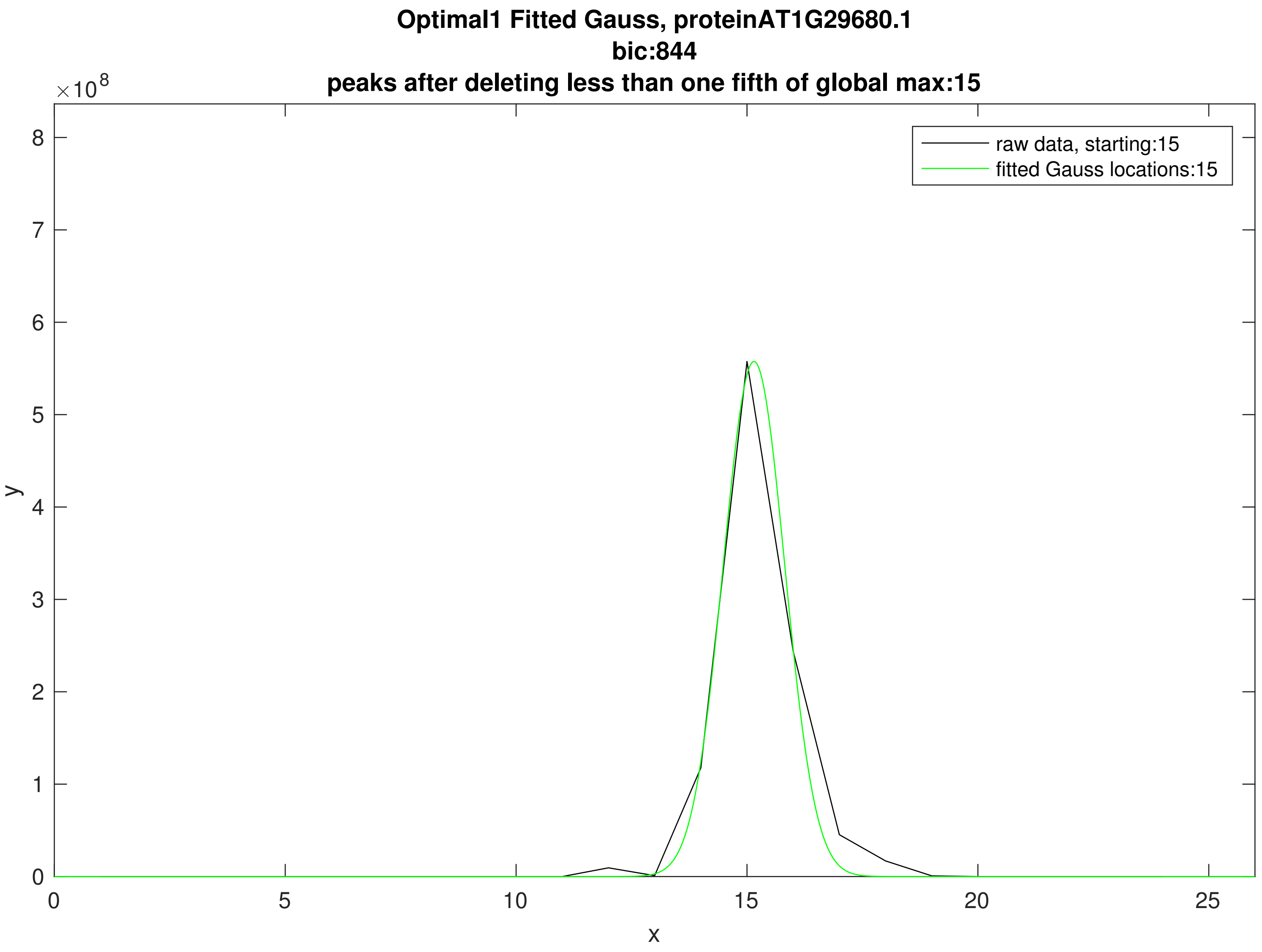

Optimal1 Fitted Gauss, proteinAT1G29680.1
bic:844
peaks after deleting less than one fifth of global max:15
×10^8
raw data, starting:15
fitted Gauss locations:15
y
x
0
1
2
3
4
5
6
7
8
0
5
10
15
20
25

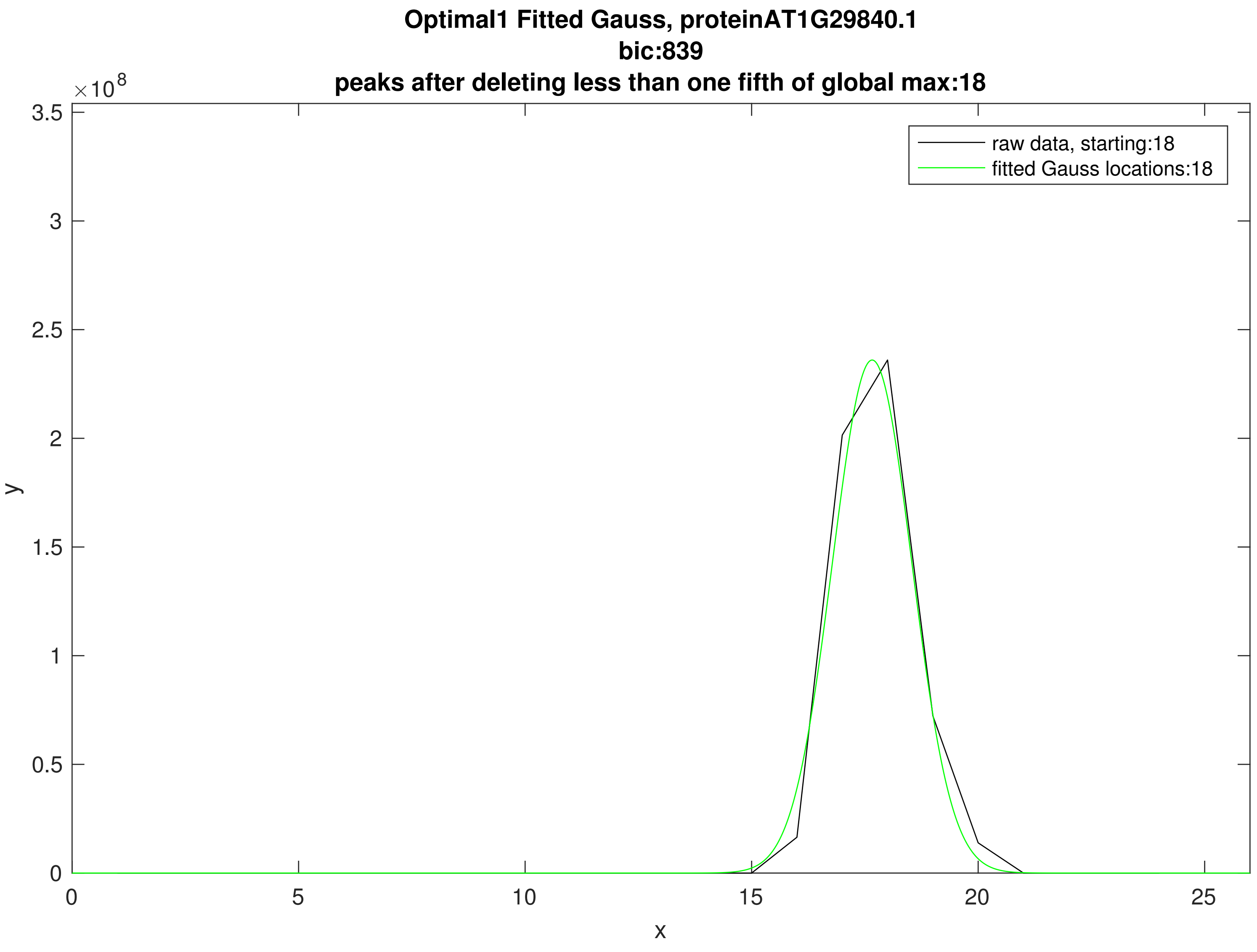
Optimal1 Fitted Gauss, proteinAT1G29840.1
bic:839
peaks after deleting less than one fifth of global max:18
raw data, starting:18
fitted Gauss locations:18
$\times 10^8$
y
x

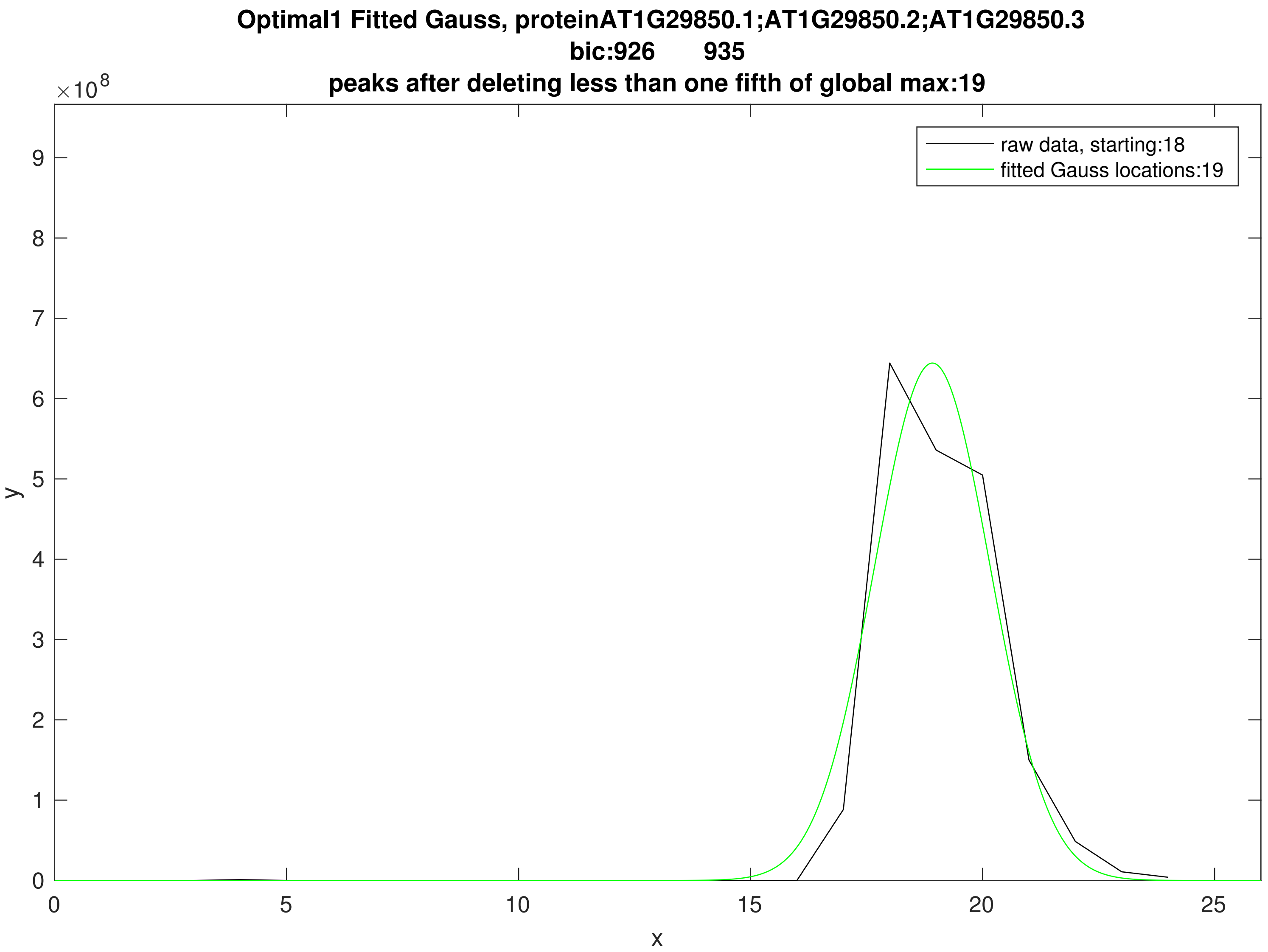

Optimal1 Fitted Gauss, proteinAT1G29850.1;AT1G29850.2;AT1G29850.3
bic:926 935
peaks after deleting less than one fifth of global max:19
×10^8
raw data, starting:18
fitted Gauss locations:19
y
x
0
1
2
3
4
5
6
7
8
9
0
5
10
15
20
25

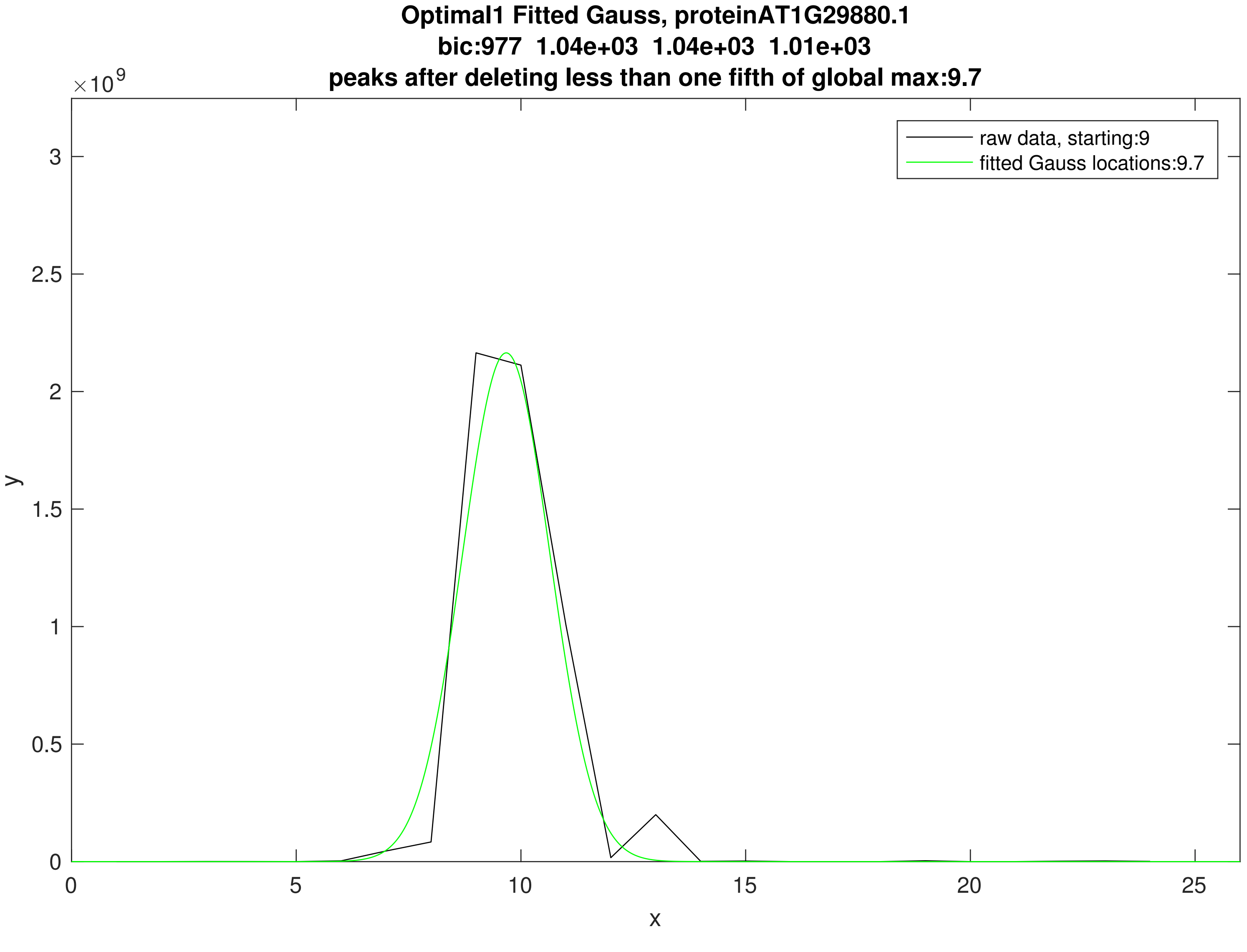

Optimal1 Fitted Gauss, proteinAT1G29880.1
bic:977 1.04e+03 1.04e+03 1.01e+03
peaks after deleting less than one fifth of global max:9.7
×10^9
raw data, starting:9
fitted Gauss locations:9.7
y
x
0
0.5
1
1.5
2
2.5
3
5
10
15
20
25

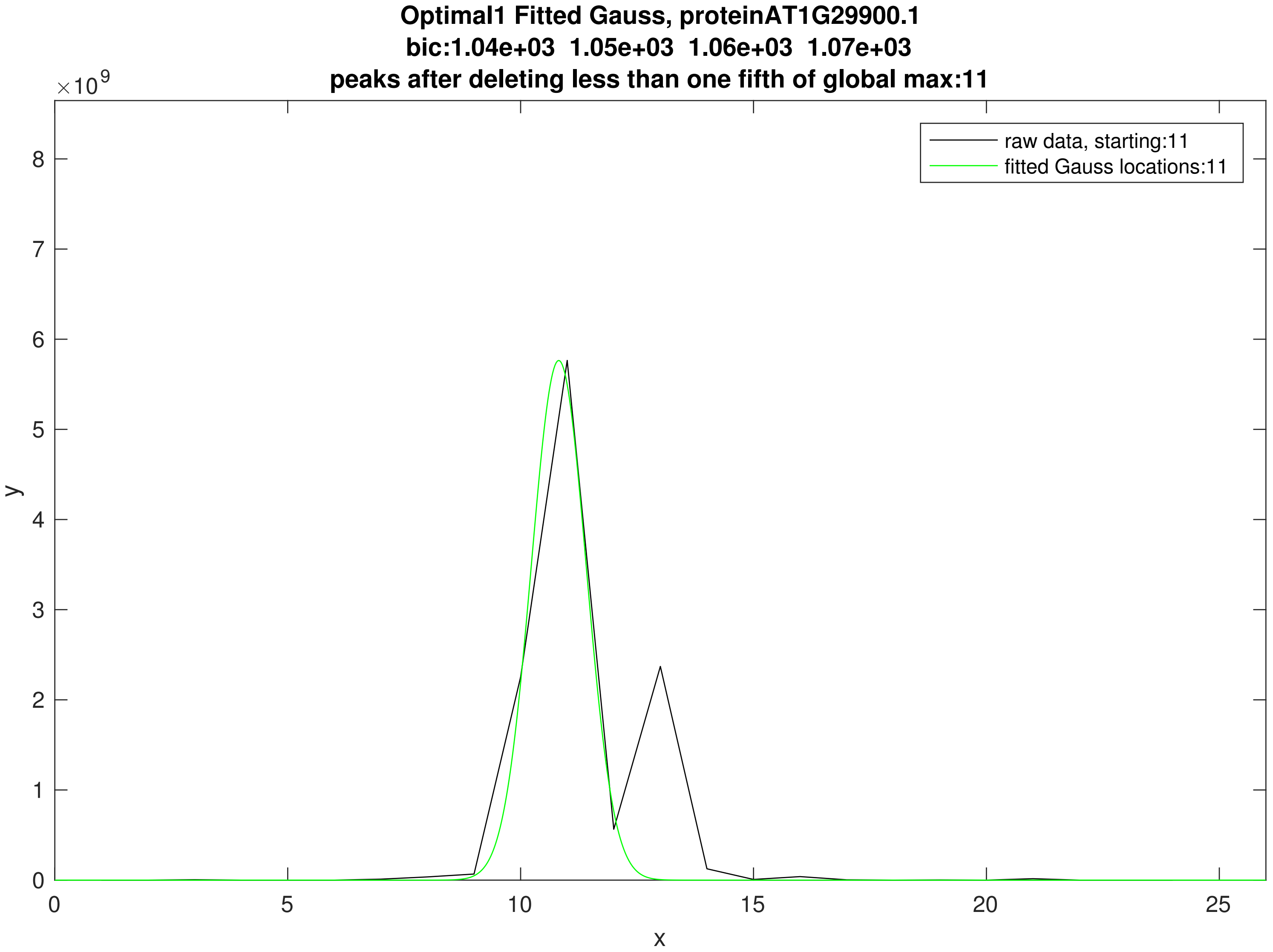

Optimal1 Fitted Gauss, proteinAT1G29900.1
bic:1.04e+03 1.05e+03 1.06e+03 1.07e+03
peaks after deleting less than one fifth of global max:11
×10^9
raw data, starting:11
fitted Gauss locations:11
y
x
0
1
2
3
4
5
6
7
8
0
5
10
15
20
25

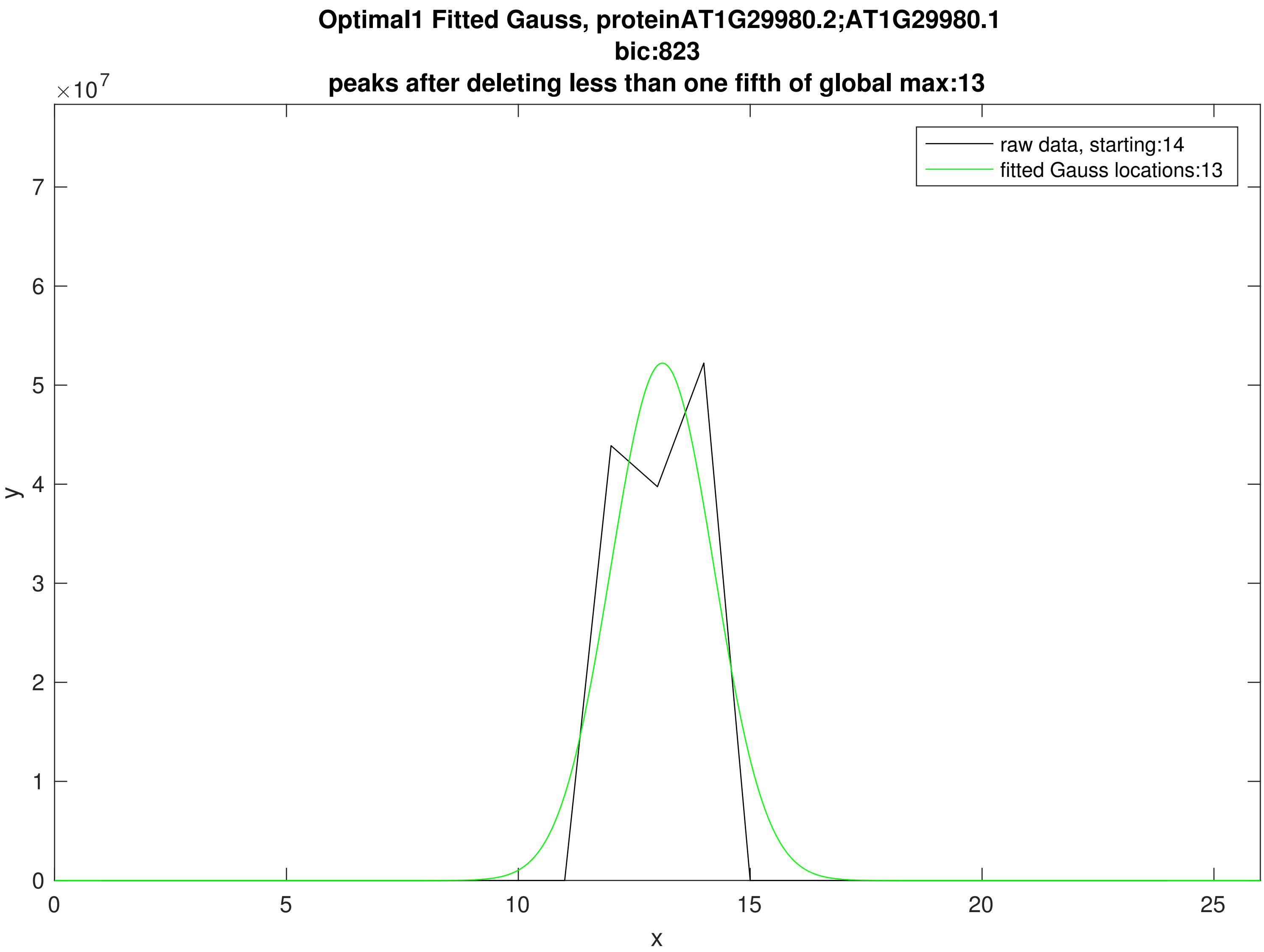

Optimal1 Fitted Gauss, proteinAT1G29980.2;AT1G29980.1
bic:823
peaks after deleting less than one fifth of global max:13
×10^7
raw data, starting:14
fitted Gauss locations:13
y
x
0
1
2
3
4
5
6
7
0
5
10
15
20
25

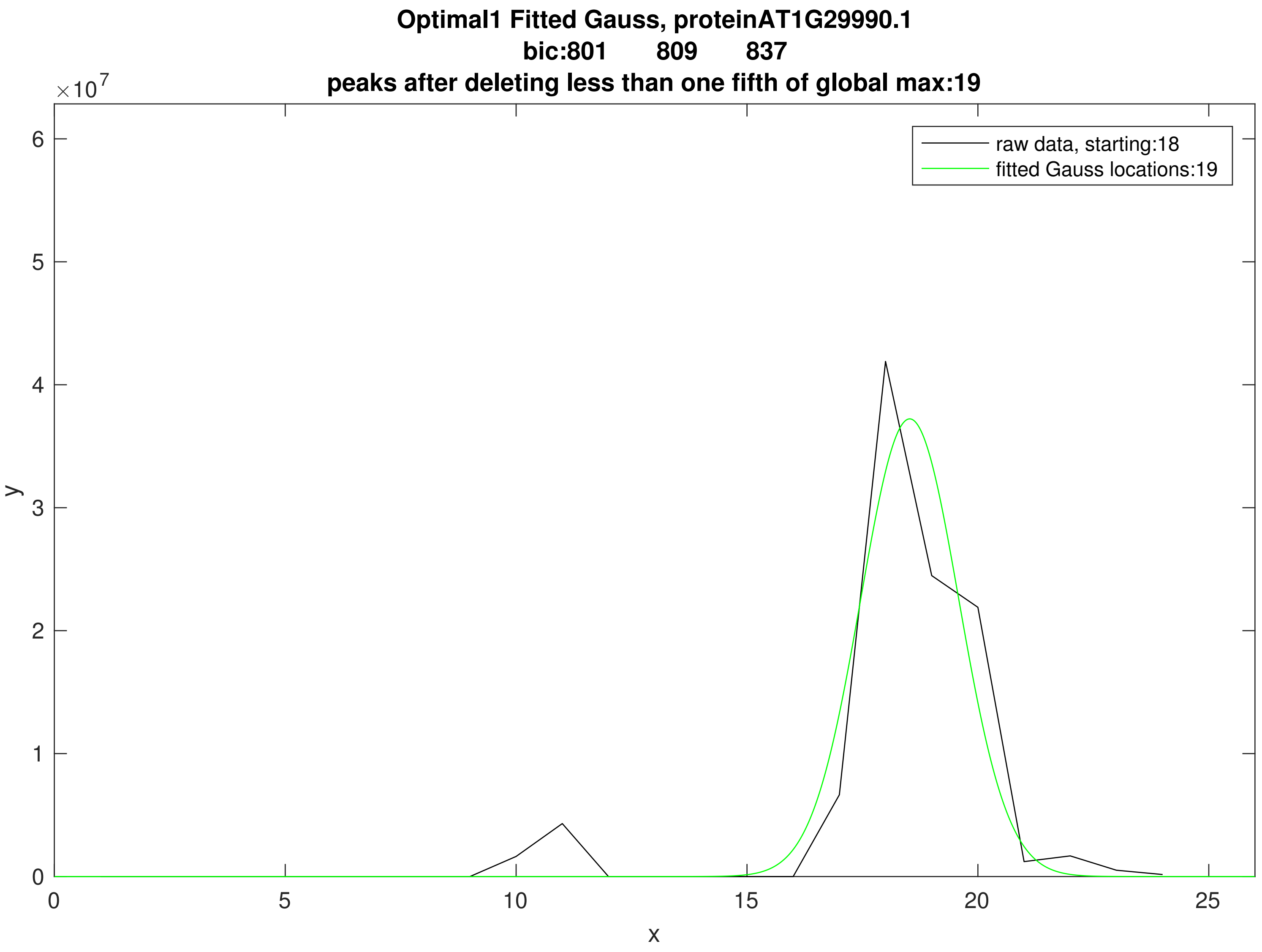
Optimal1 Fitted Gauss, proteinAT1G29990.1
bic:801 809 837
peaks after deleting less than one fifth of global max:19
×10^7
y
x
raw data, starting:18
fitted Gauss locations:19
0
1
2
3
4
5
6
0
5
10
15
20
25

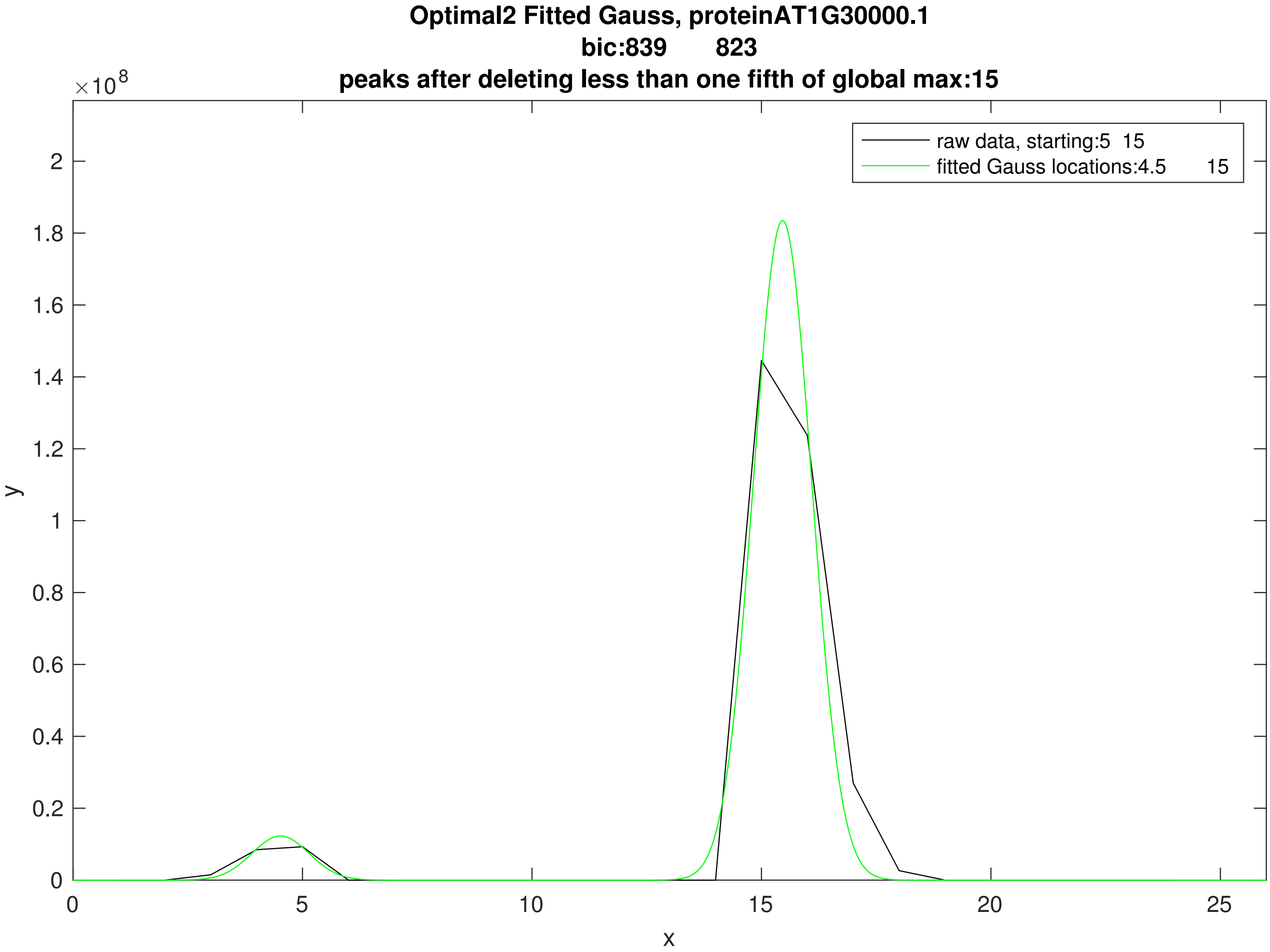

Optimal2 Fitted Gauss, proteinAT1G30000.1
bic:839 823
peaks after deleting less than one fifth of global max:15
×10^8
raw data, starting:5 15
fitted Gauss locations:4.5 15
y
x
0
0.2
0.4
0.6
0.8
1
1.2
1.4
1.6
1.8
2
5
10
15
20
25

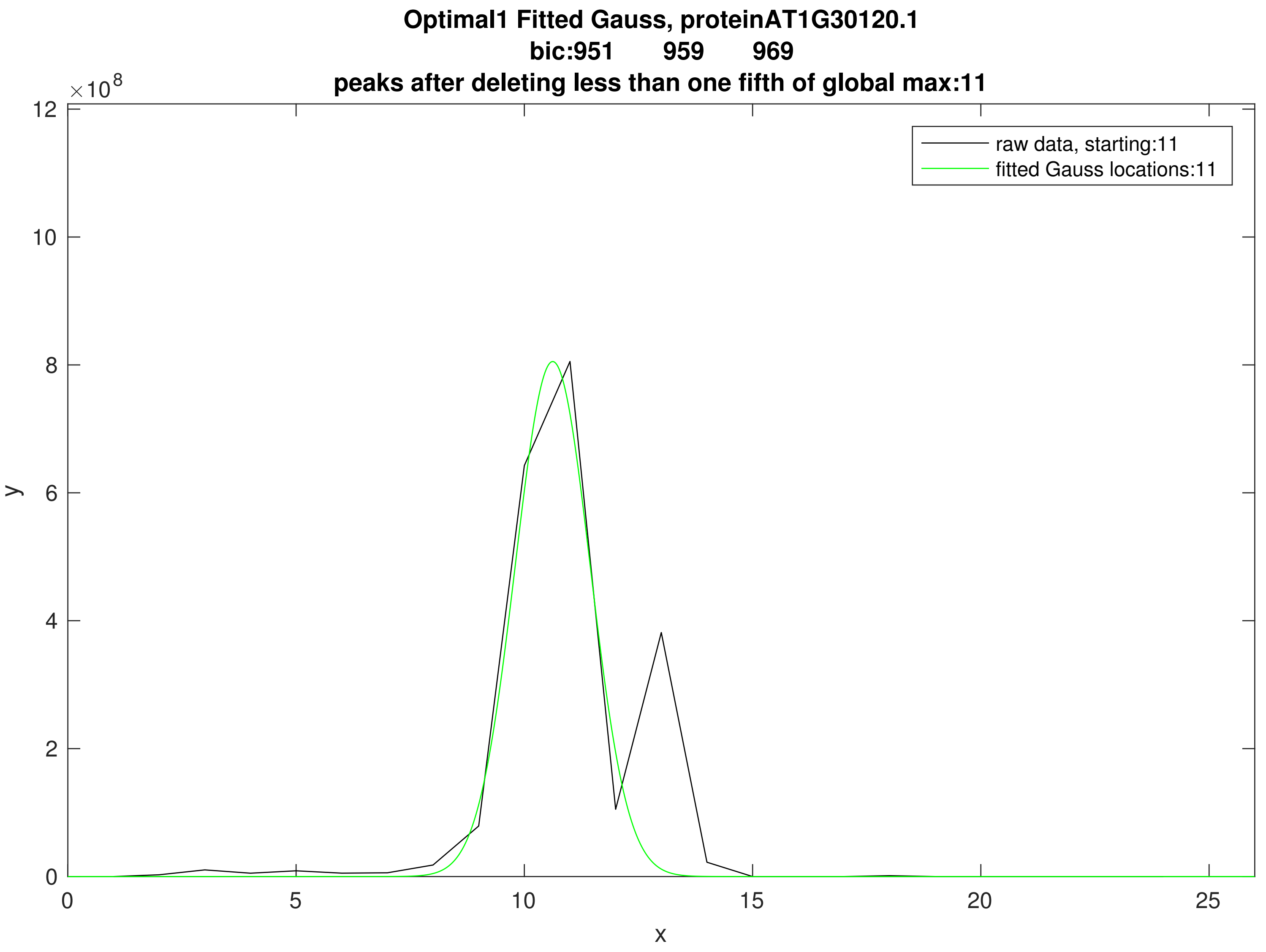

Optimal1 Fitted Gauss, proteinAT1G30120.1
bic:951 959 969
peaks after deleting less than one fifth of global max:11
×10^8
raw data, starting:11
fitted Gauss locations:11
y
x
0
2
4
6
8
10
12
0
5
10
15
20
25

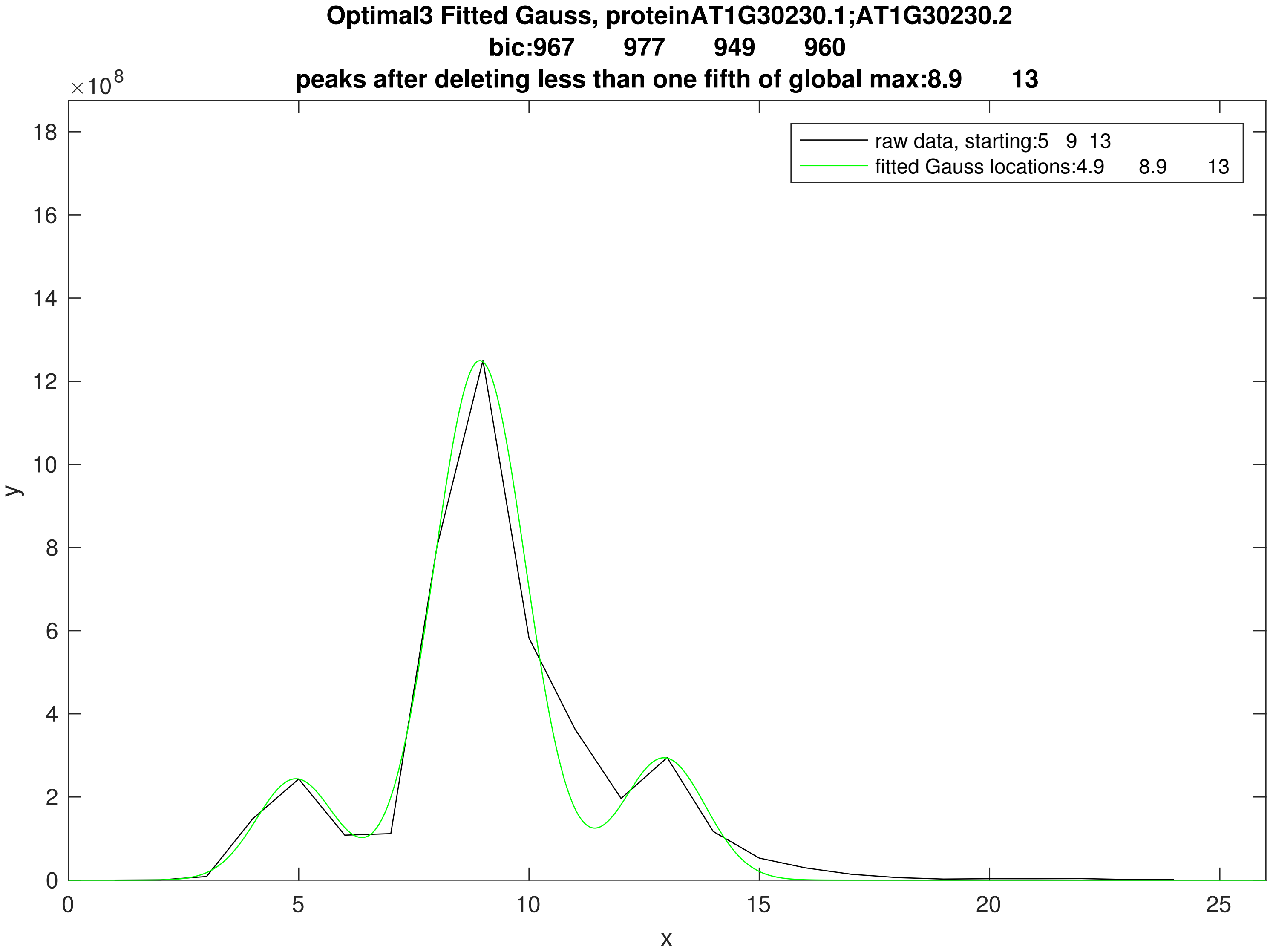

Optimal3 Fitted Gauss, proteinAT1G30230.1;AT1G30230.2
bic:967 977 949 960
peaks after deleting less than one fifth of global max:8.9 13
×10^8
raw data, starting:5 9 13
fitted Gauss locations:4.9 8.9 13
y
x
0
2
4
6
8
10
12
14
16
18
0
5
10
15
20
25

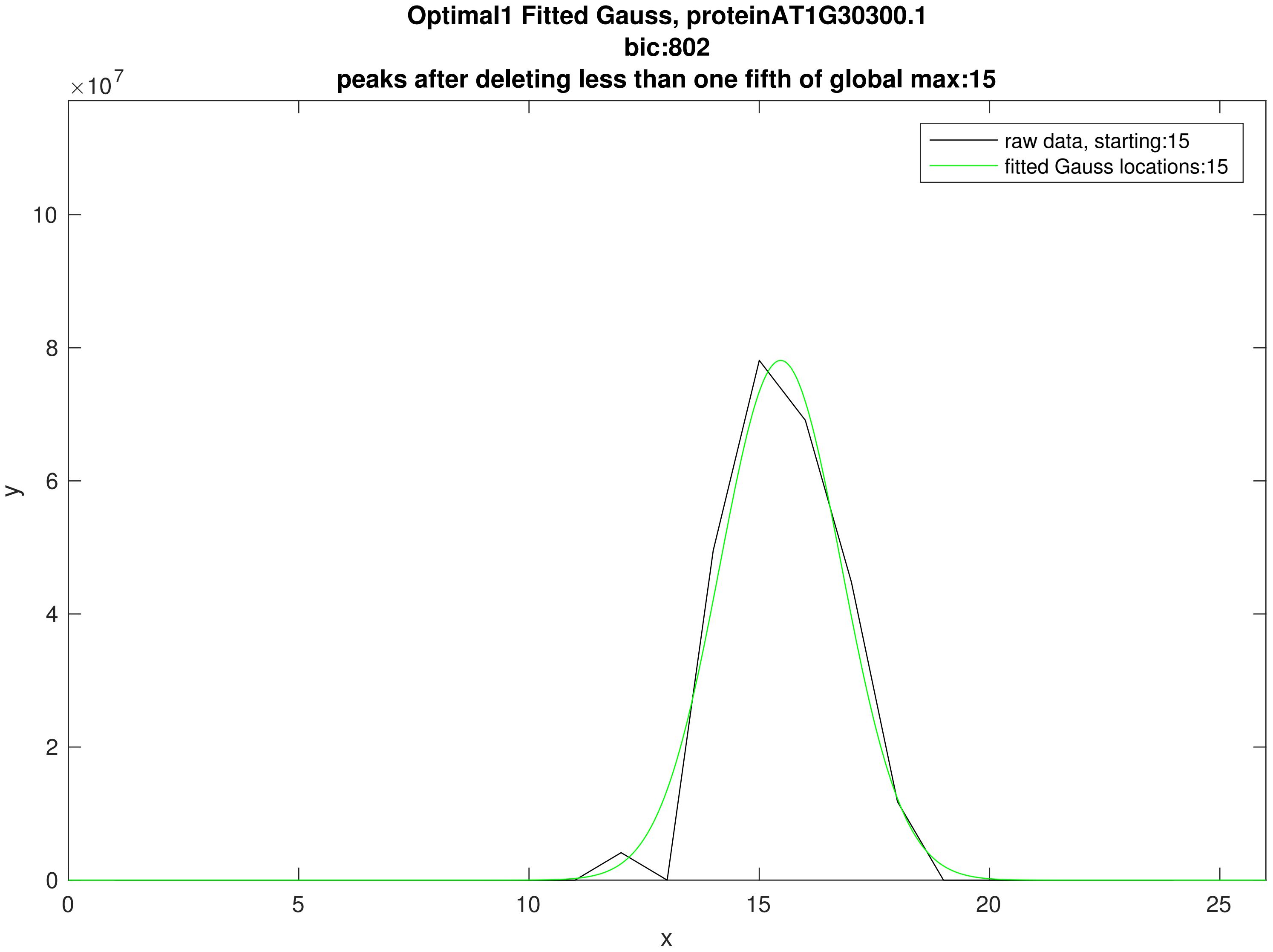
Optimal1 Fitted Gauss, proteinAT1G30300.1
bic:802
peaks after deleting less than one fifth of global max:15
×10^7
raw data, starting:15
fitted Gauss locations:15
y
0
2
4
6
8
10
x
0
5
10
15
20
25

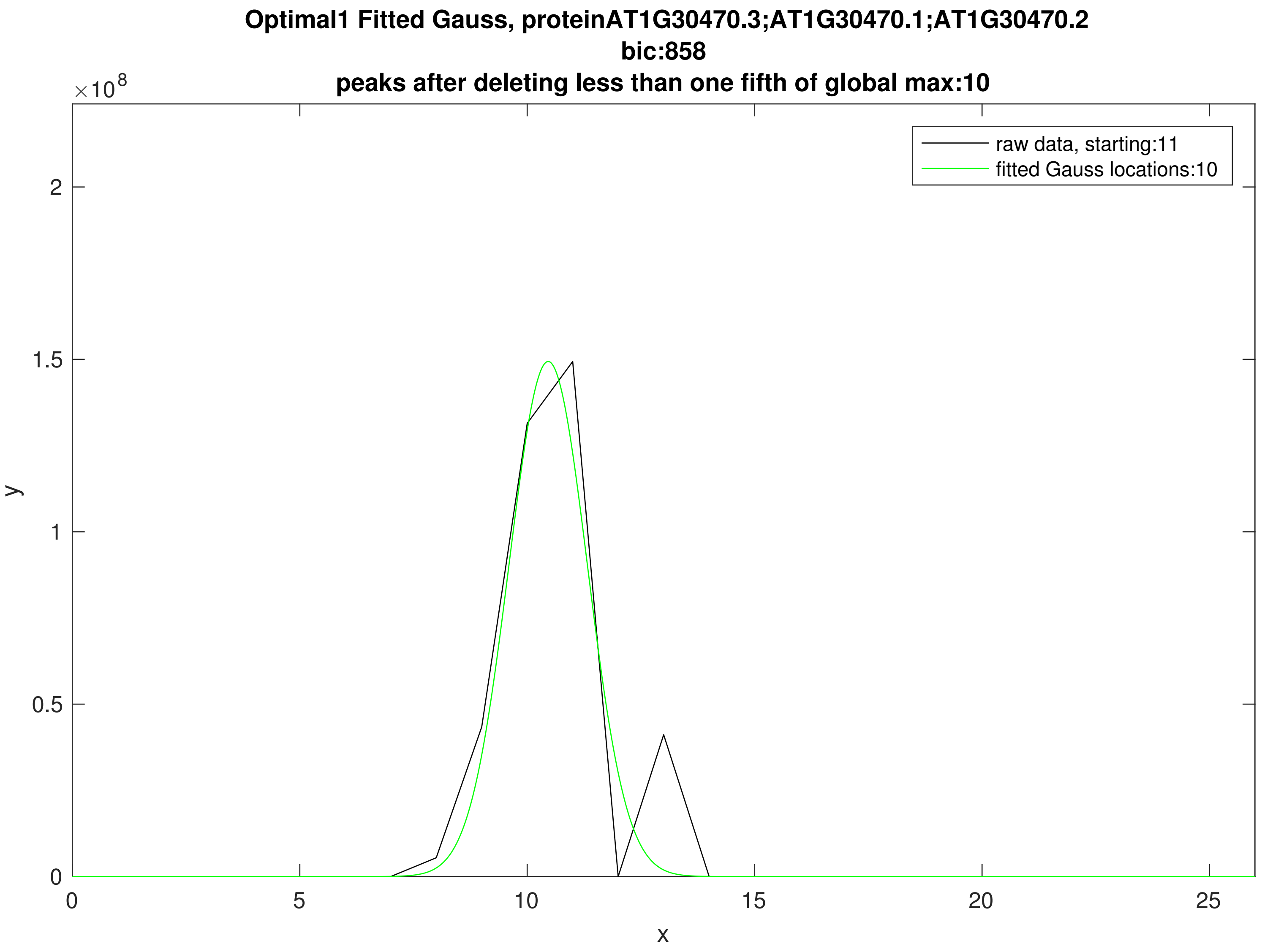
Optimal1 Fitted Gauss, proteinAT1G30470.3;AT1G30470.1;AT1G30470.2
bic:858
peaks after deleting less than one fifth of global max:10
×10^8
raw data, starting:11
fitted Gauss locations:10
y
x
0
0.5
1
1.5
2
0
5
10
15
20
25

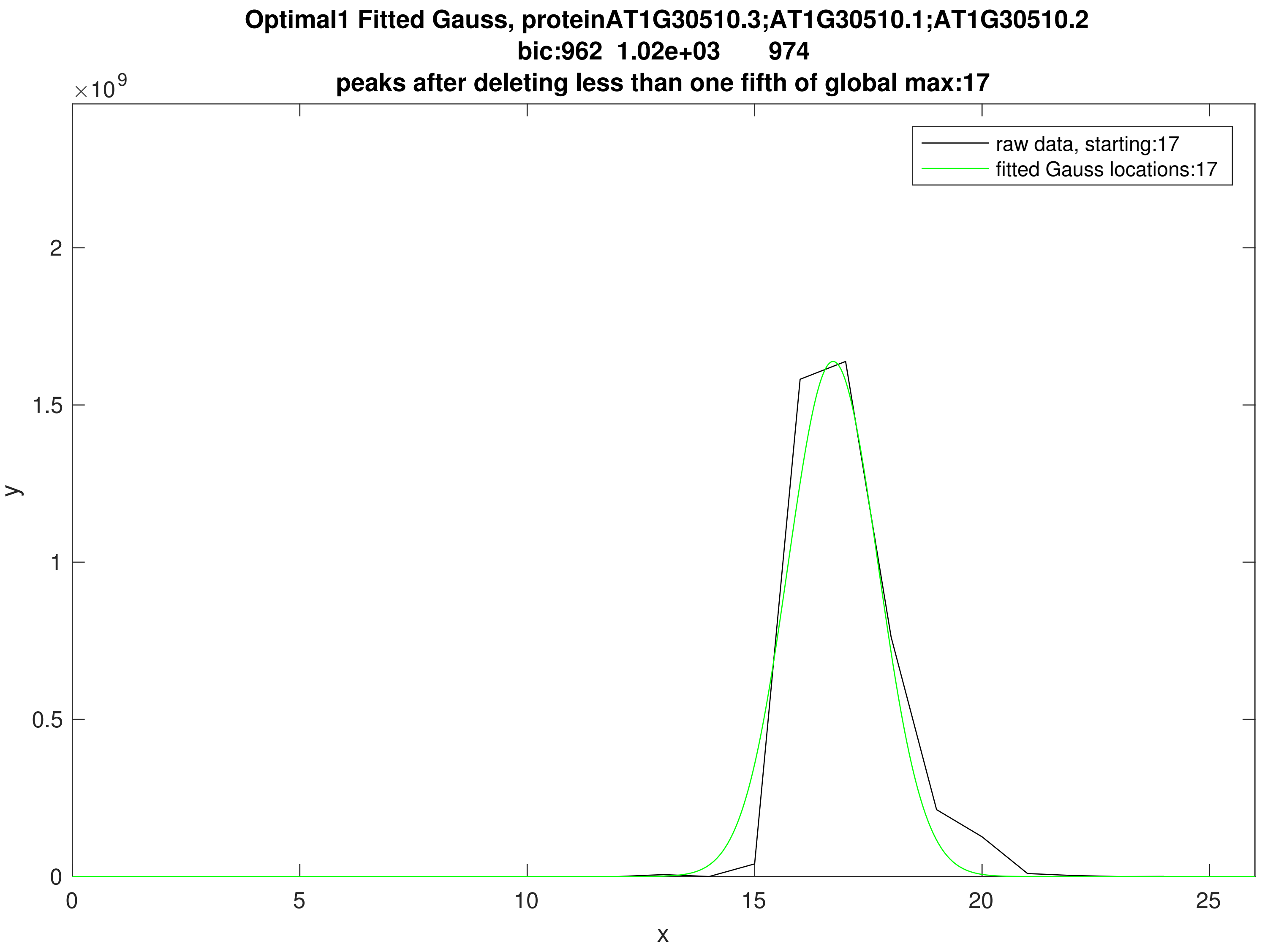

Optimal1 Fitted Gauss, proteinAT1G30510.3;AT1G30510.1;AT1G30510.2
bic:962 1.02e+03 974
peaks after deleting less than one fifth of global max:17
×10^9
raw data, starting:17
fitted Gauss locations:17
y
x
0
0.5
1
1.5
2
0
5
10
15
20
25

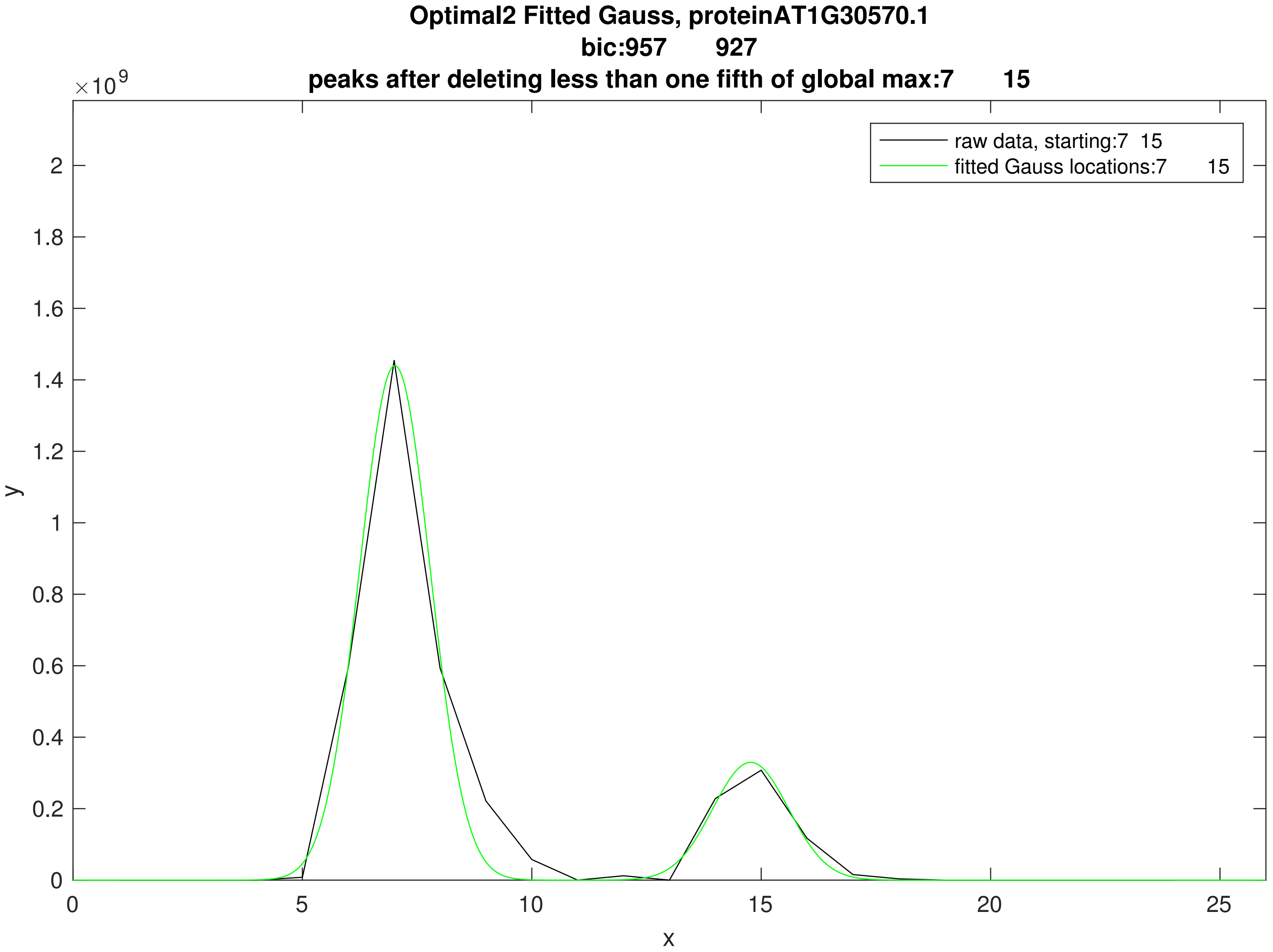
Optimal2 Fitted Gauss, proteinAT1G30570.1
bic:957 927
peaks after deleting less than one fifth of global max:7 15
×10^9
raw data, starting:7 15
fitted Gauss locations:7 15
y
x
0
0.2
0.4
0.6
0.8
1
1.2
1.4
1.6
1.8
2
5
10
15
20
25

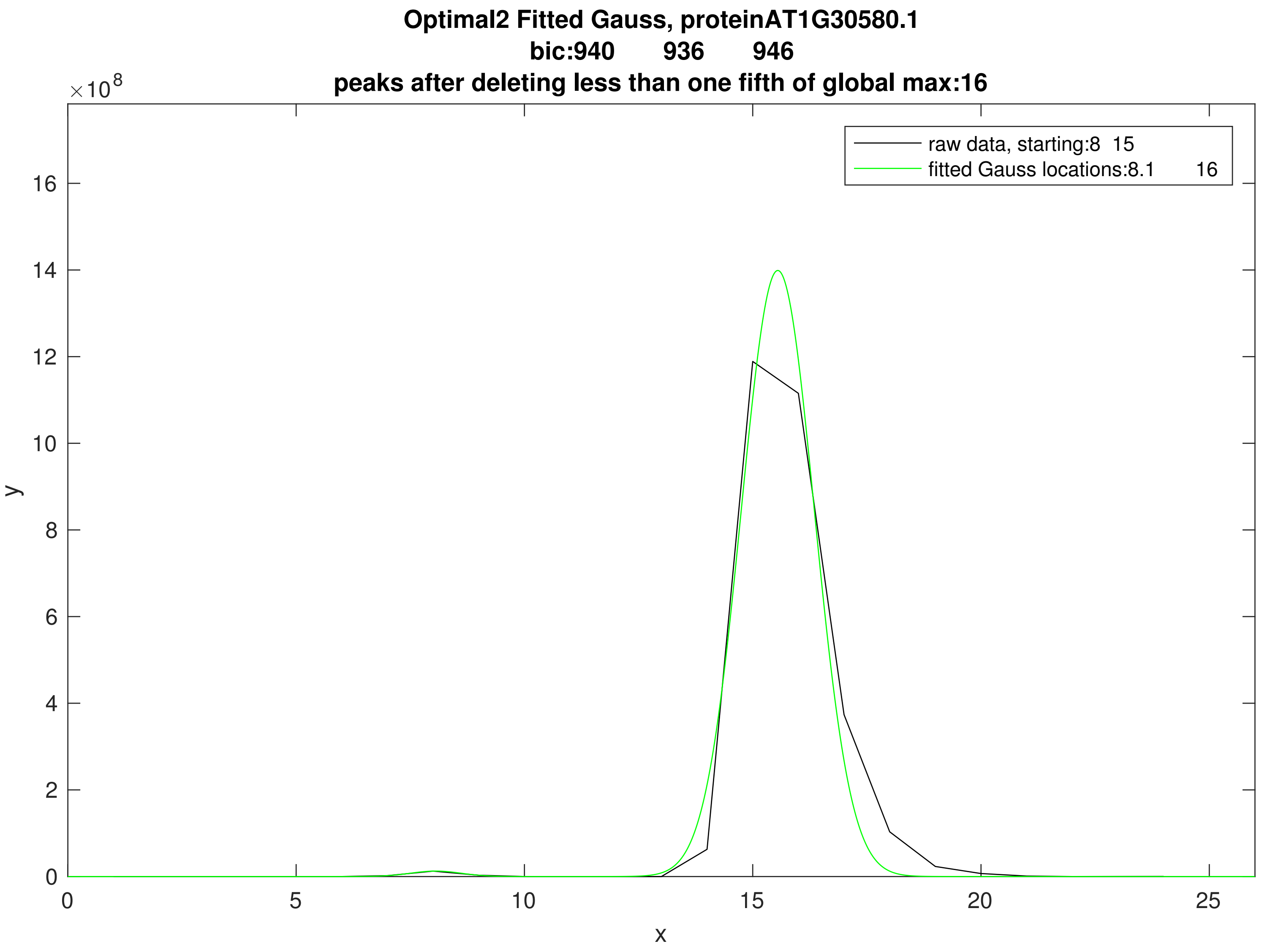

Optimal2 Fitted Gauss, proteinAT1G30580.1
bic:940 936 946
peaks after deleting less than one fifth of global max:16
×10^8
raw data, starting:8 15
fitted Gauss locations:8.1 16
16
14
12
10
8
6
4
2
0
y
0
5
10
15
20
25
x

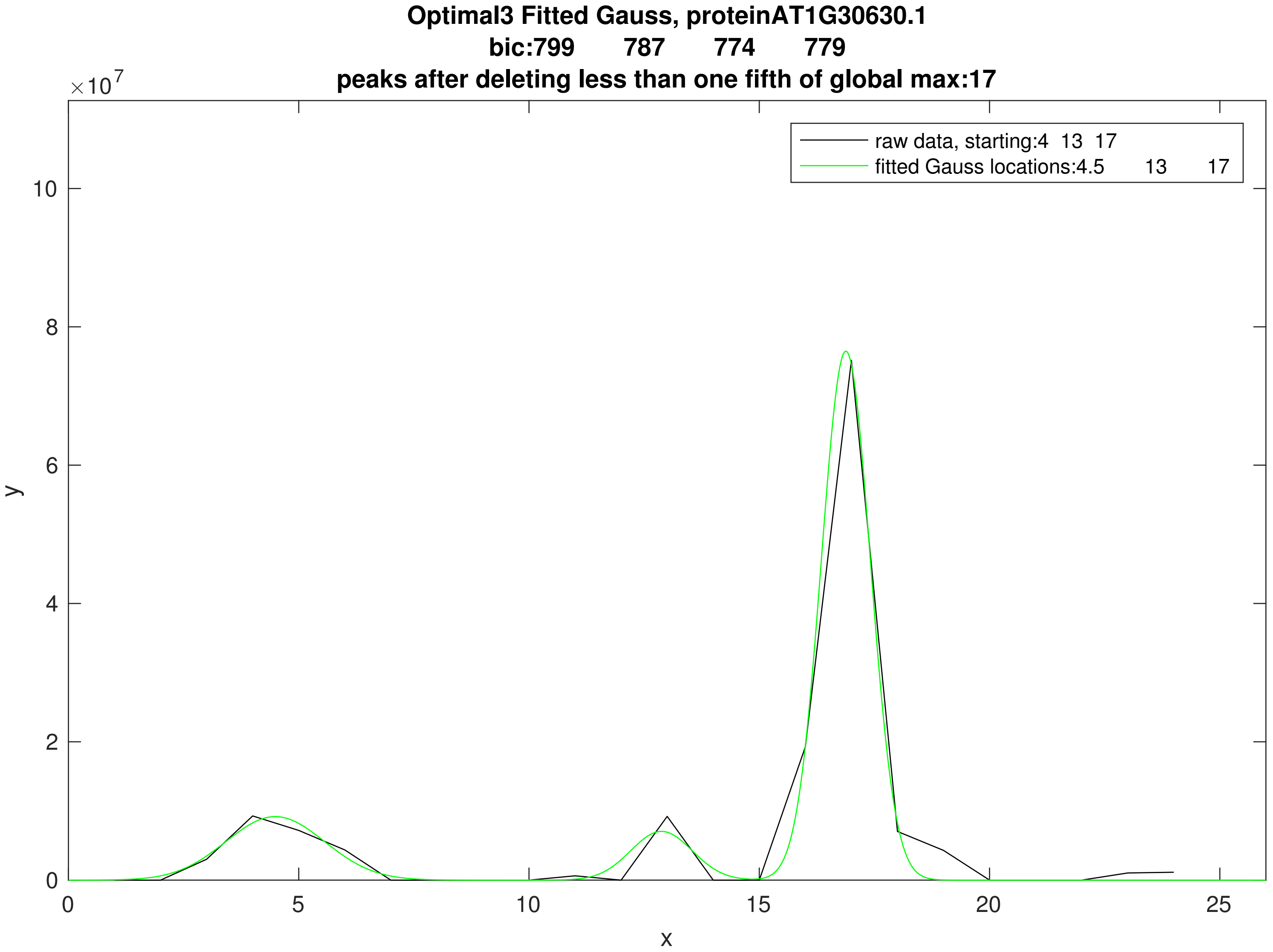
Optimal3 Fitted Gauss, proteinAT1G30630.1
bic:799 787 774 779
peaks after deleting less than one fifth of global max:17
×10^7
raw data, starting:4 13 17
fitted Gauss locations:4.5 13 17
10
8
6
4
2
0
y
0
5
10
15
20
25
x

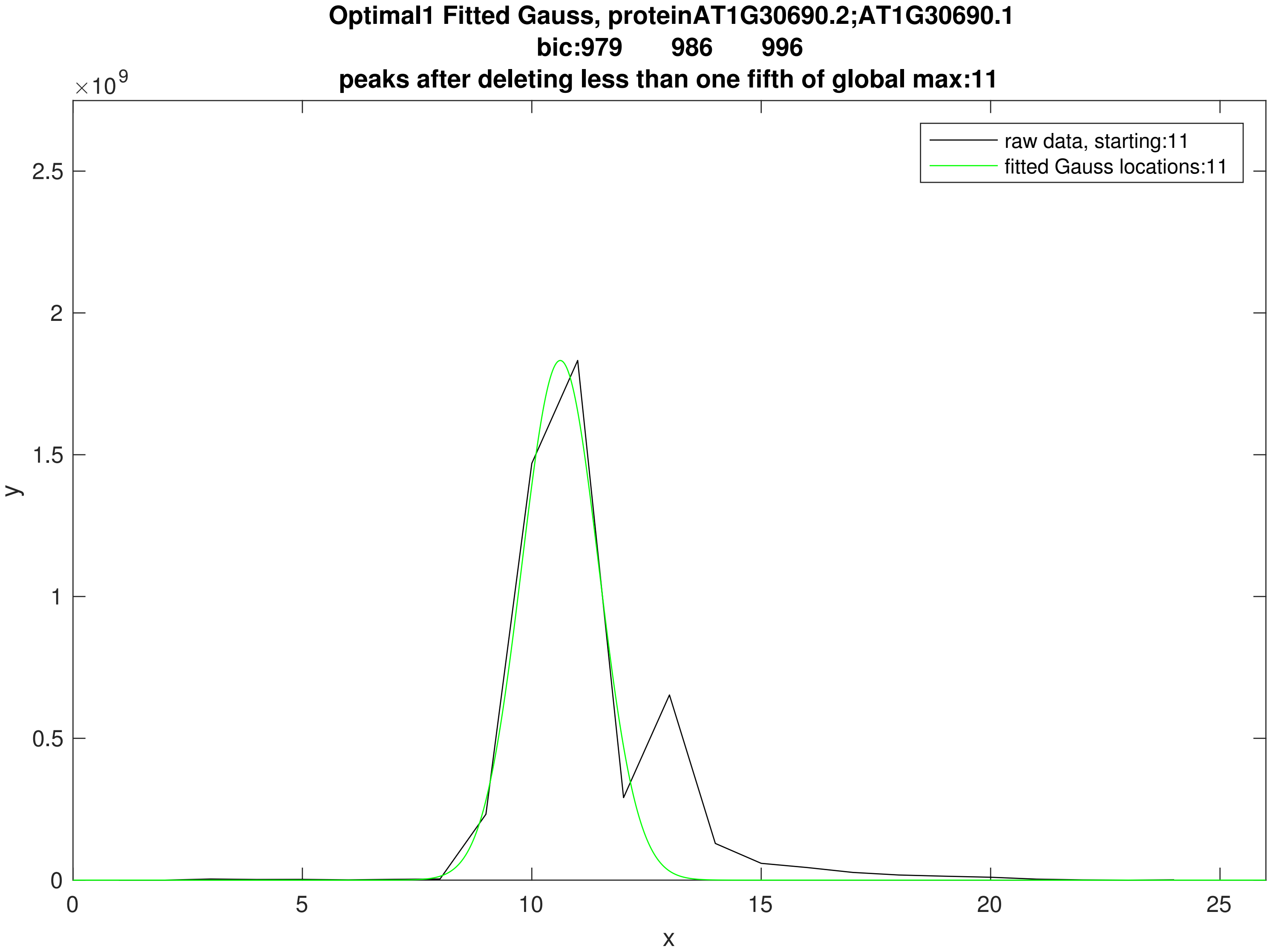

Optimal1 Fitted Gauss, proteinAT1G30690.2;AT1G30690.1
bic:979 986 996
peaks after deleting less than one fifth of global max:11
×10^9
2.5
2
1.5
1
0.5
0
y
0
5
10
15
20
25
x
raw data, starting:11
fitted Gauss locations:11

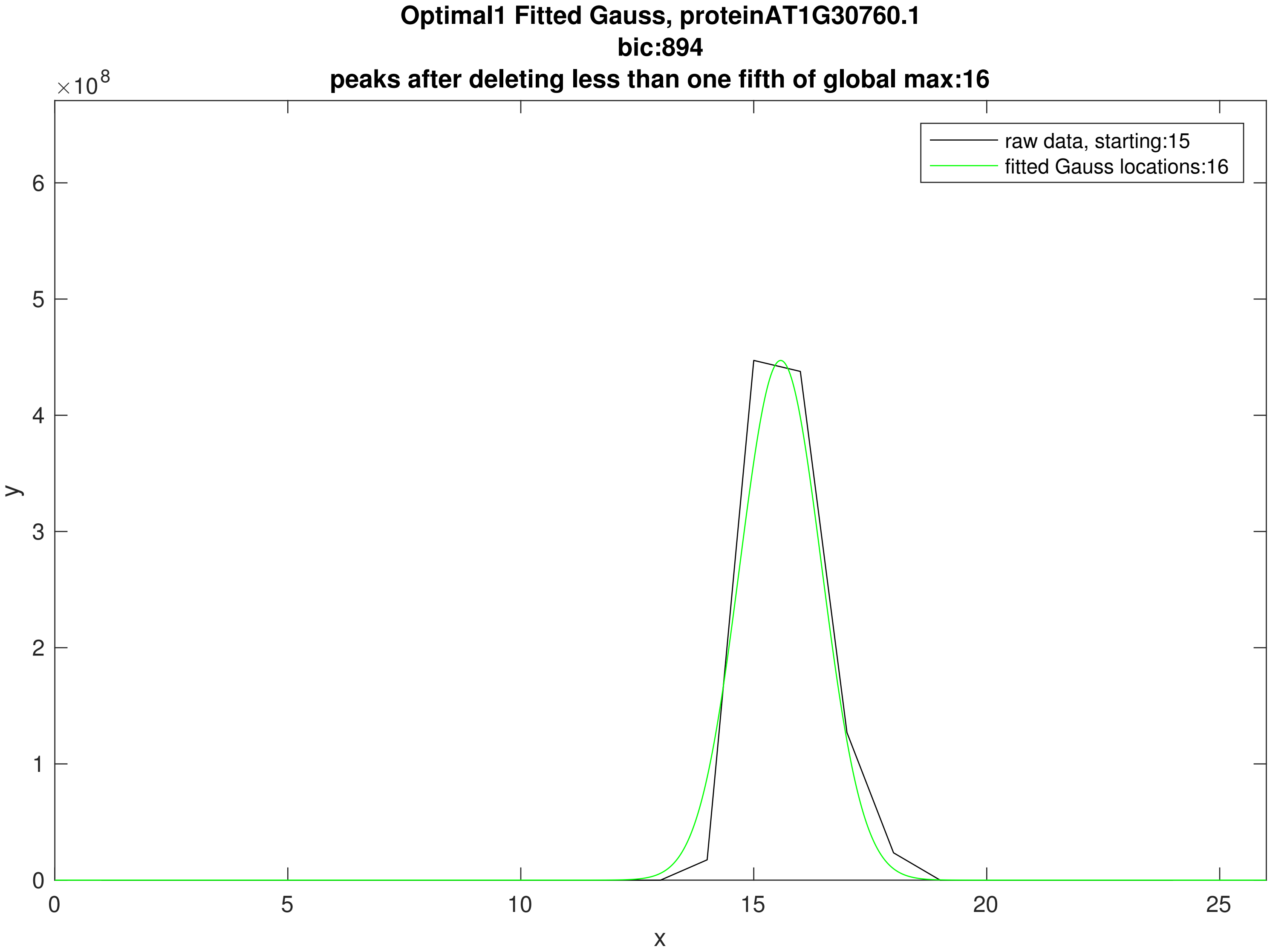

Optimal1 Fitted Gauss, proteinAT1G30760.1
bic:894
peaks after deleting less than one fifth of global max:16
×10^8
raw data, starting:15
fitted Gauss locations:16
y
x
0
1
2
3
4
5
6
0
5
10
15
20
25

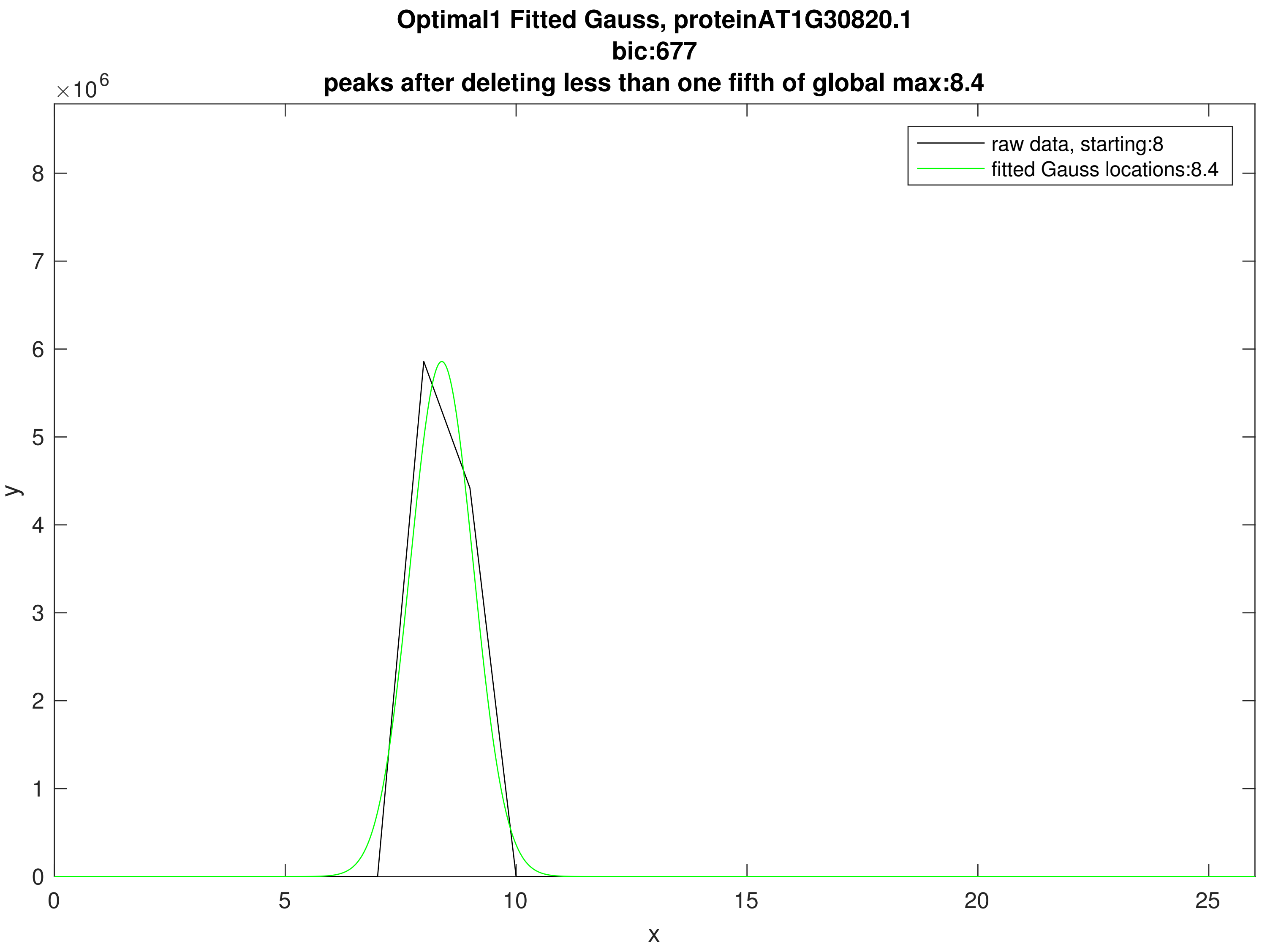

Optimal1 Fitted Gauss, proteinAT1G30820.1
bic:677
peaks after deleting less than one fifth of global max:8.4
×10^6
raw data, starting:8
fitted Gauss locations:8.4
y
x
0
1
2
3
4
5
6
7
8
0
5
10
15
20
25

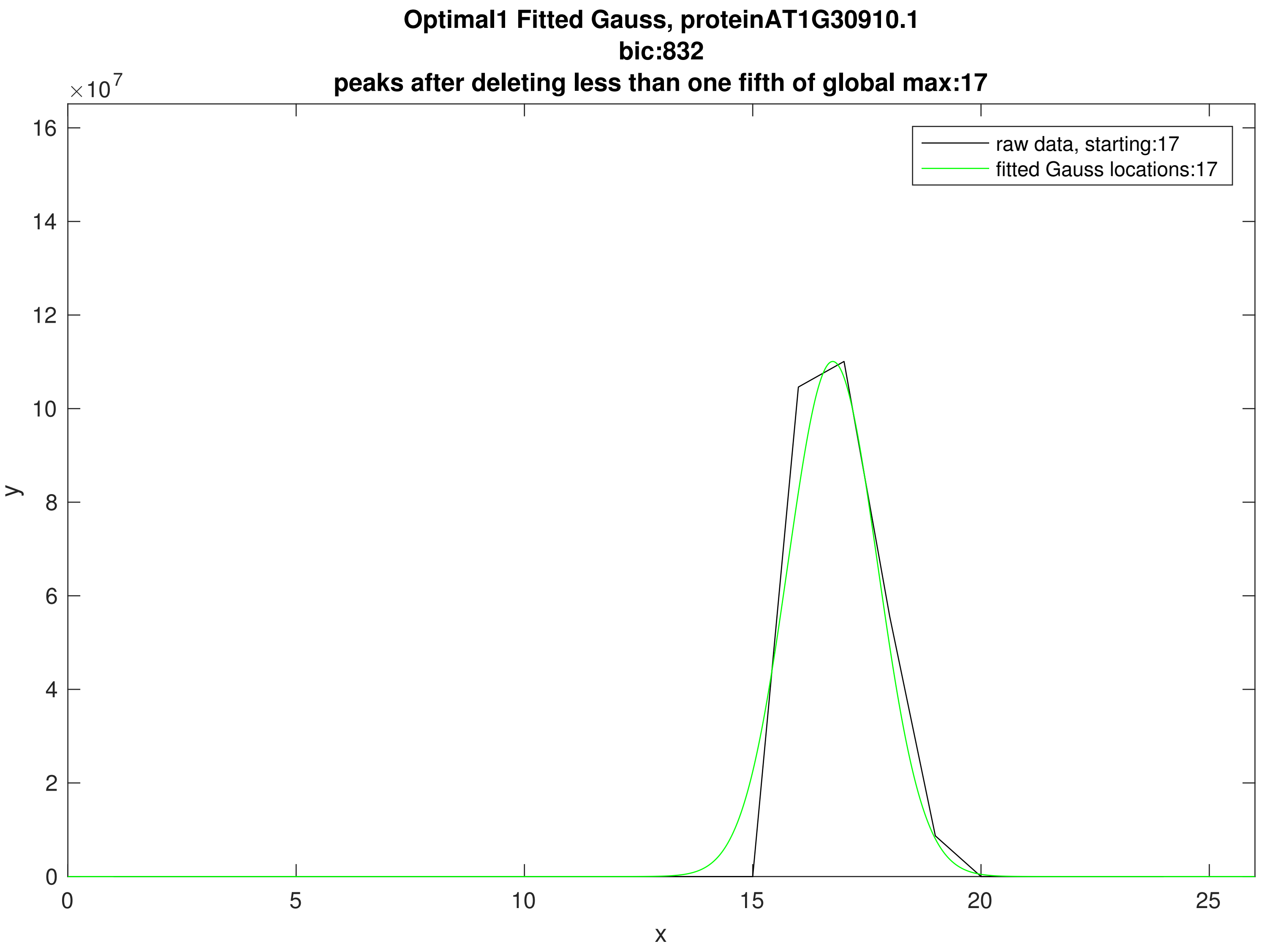
Optimal1 Fitted Gauss, proteinAT1G30910.1
bic:832
peaks after deleting less than one fifth of global max:17
×10^7
16
14
12
10
8
6
4
2
0
y
0
5
10
15
20
25
x
raw data, starting:17
fitted Gauss locations:17

**Optimal1 Fitted Gauss, proteinAT1G31070.2**
**bic:914 923**
**peaks after deleting less than one fifth of global max:15**

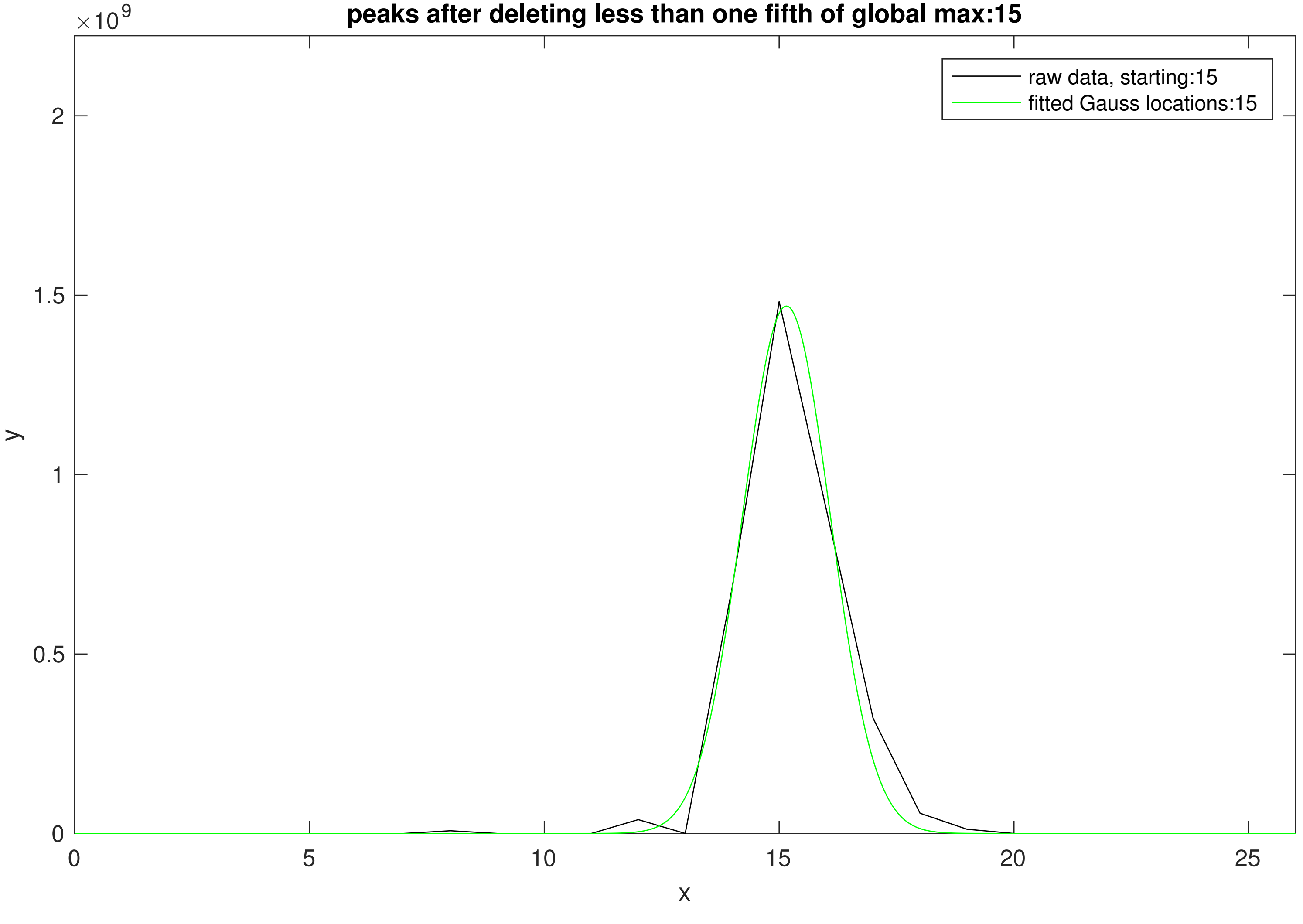

**Optimal1 Fitted Gauss, proteinAT1G31160.1**
**bic:814 823**
**peaks after deleting less than one fifth of global max:17**

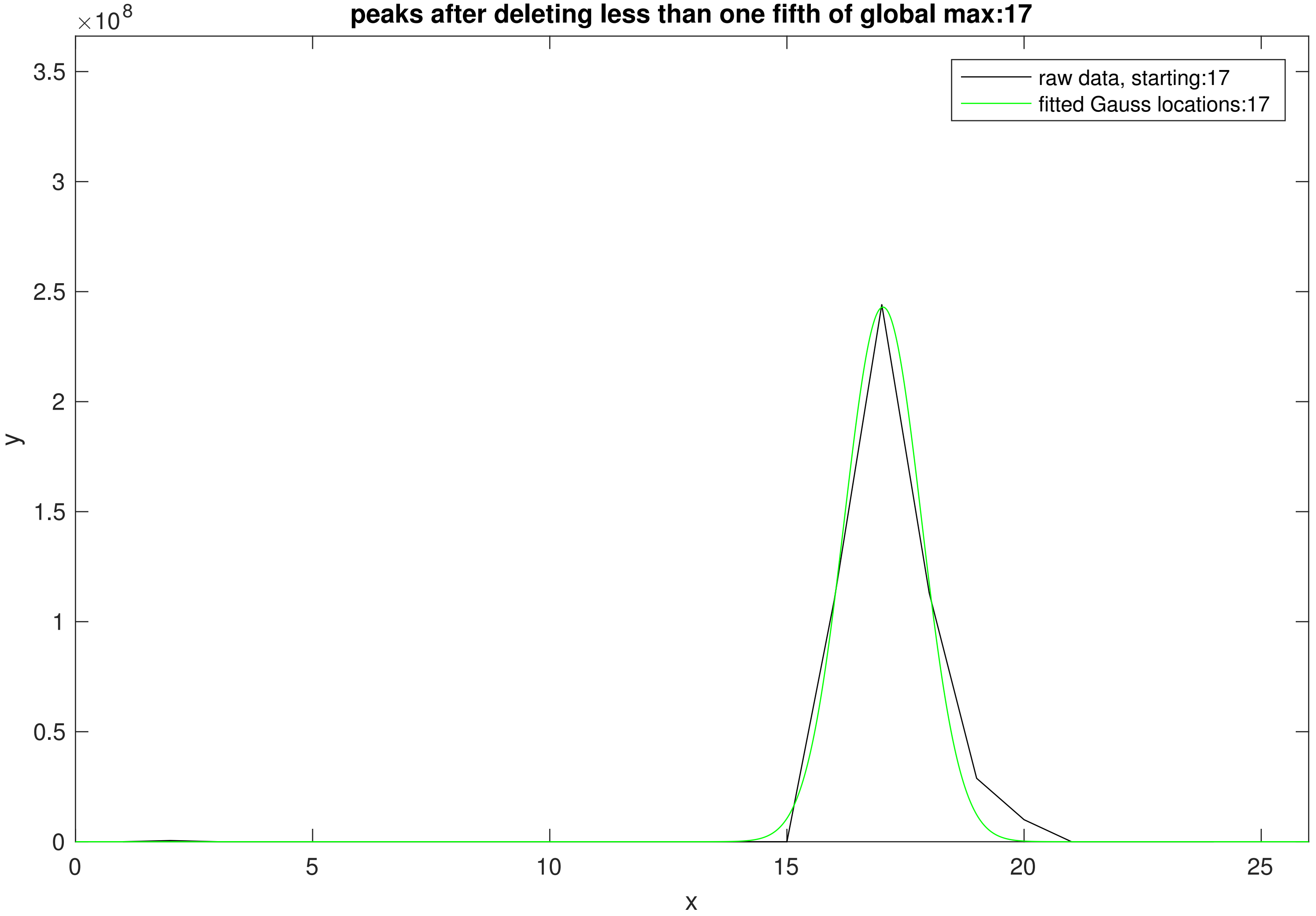

**Optimal1 Fitted Gauss, proteinAT1G31180.1**
**bic:1.01e+03  1.01e+03  1.02e+03**
**peaks after deleting less than one fifth of global max:14**

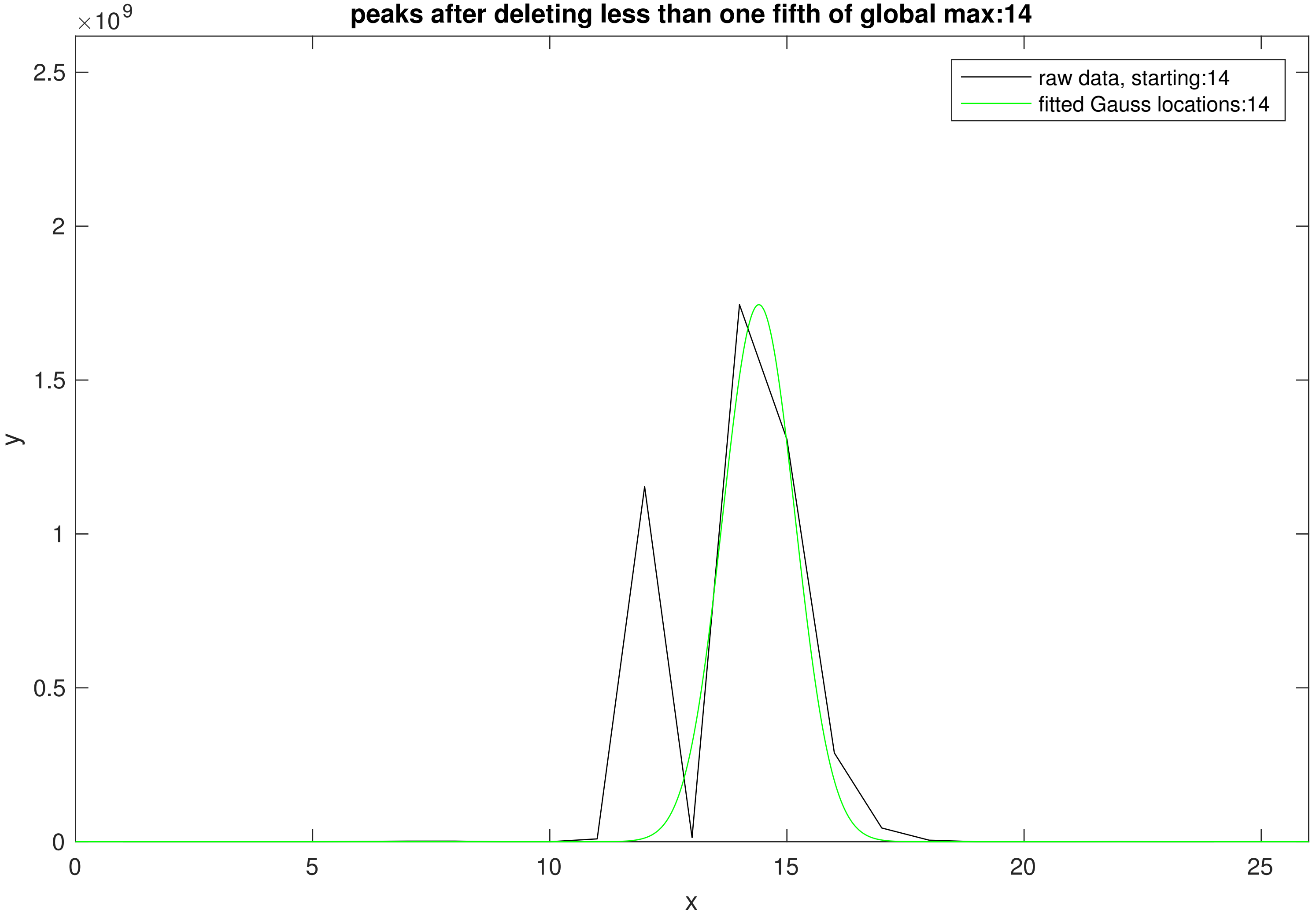

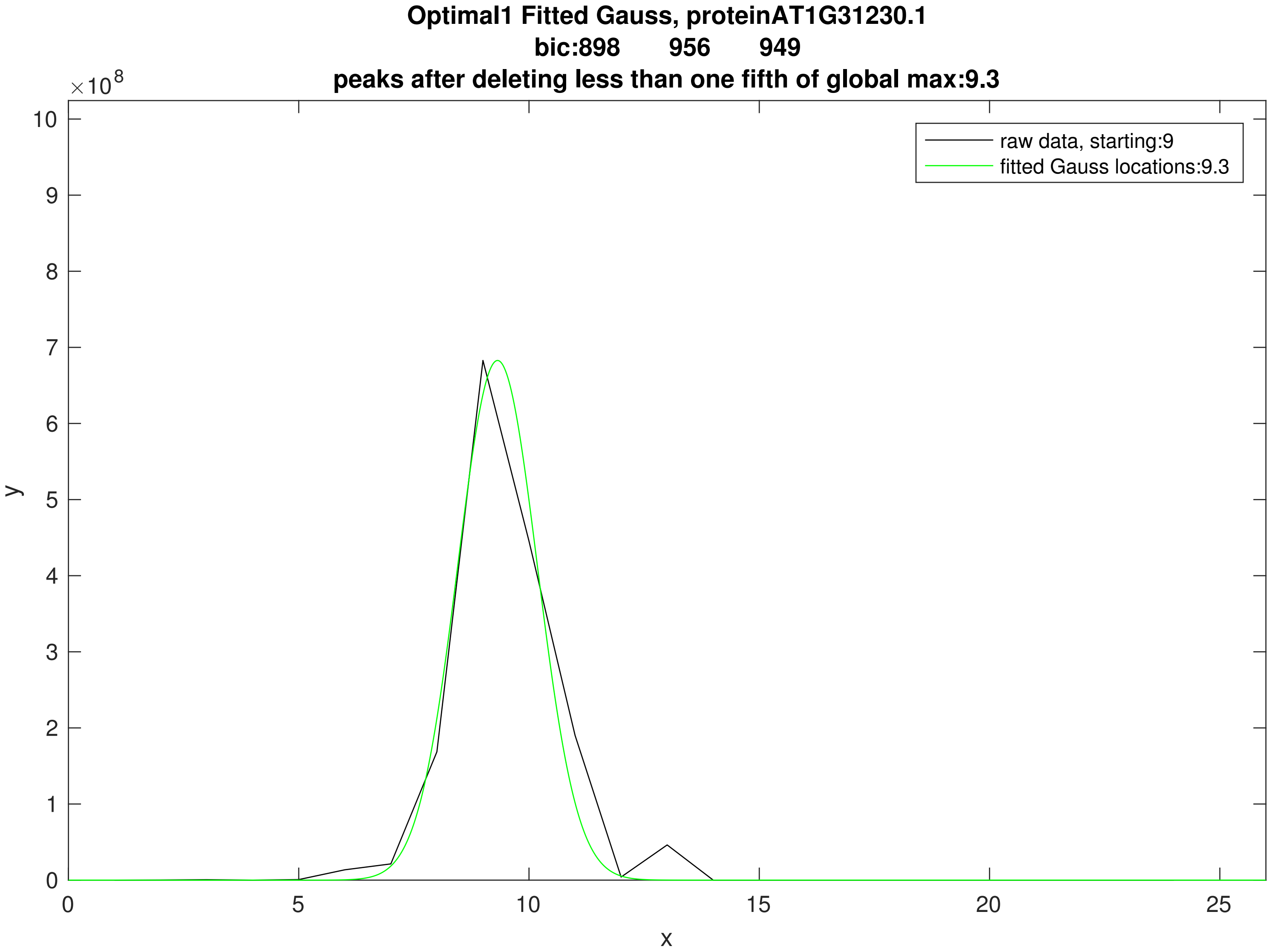
Optimal1 Fitted Gauss, proteinAT1G31230.1
bic:898 956 949
peaks after deleting less than one fifth of global max:9.3
×10^8
raw data, starting:9
fitted Gauss locations:9.3
y
x
0
1
2
3
4
5
6
7
8
9
10
15
20
25

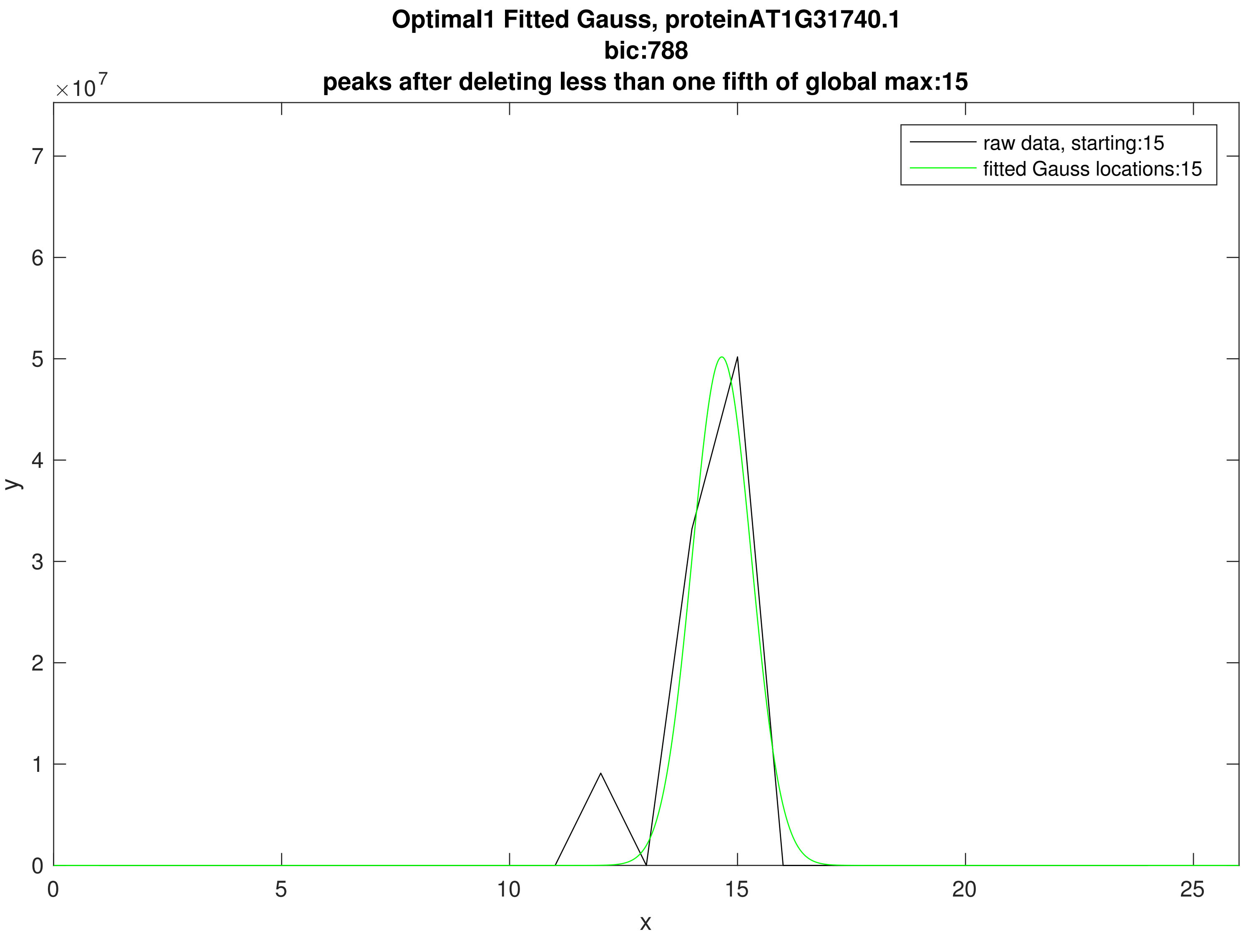
Optimal1 Fitted Gauss, proteinAT1G31740.1
bic:788
peaks after deleting less than one fifth of global max:15
×10^7
raw data, starting:15
fitted Gauss locations:15
y
x
0
1
2
3
4
5
6
7
0
5
10
15
20
25

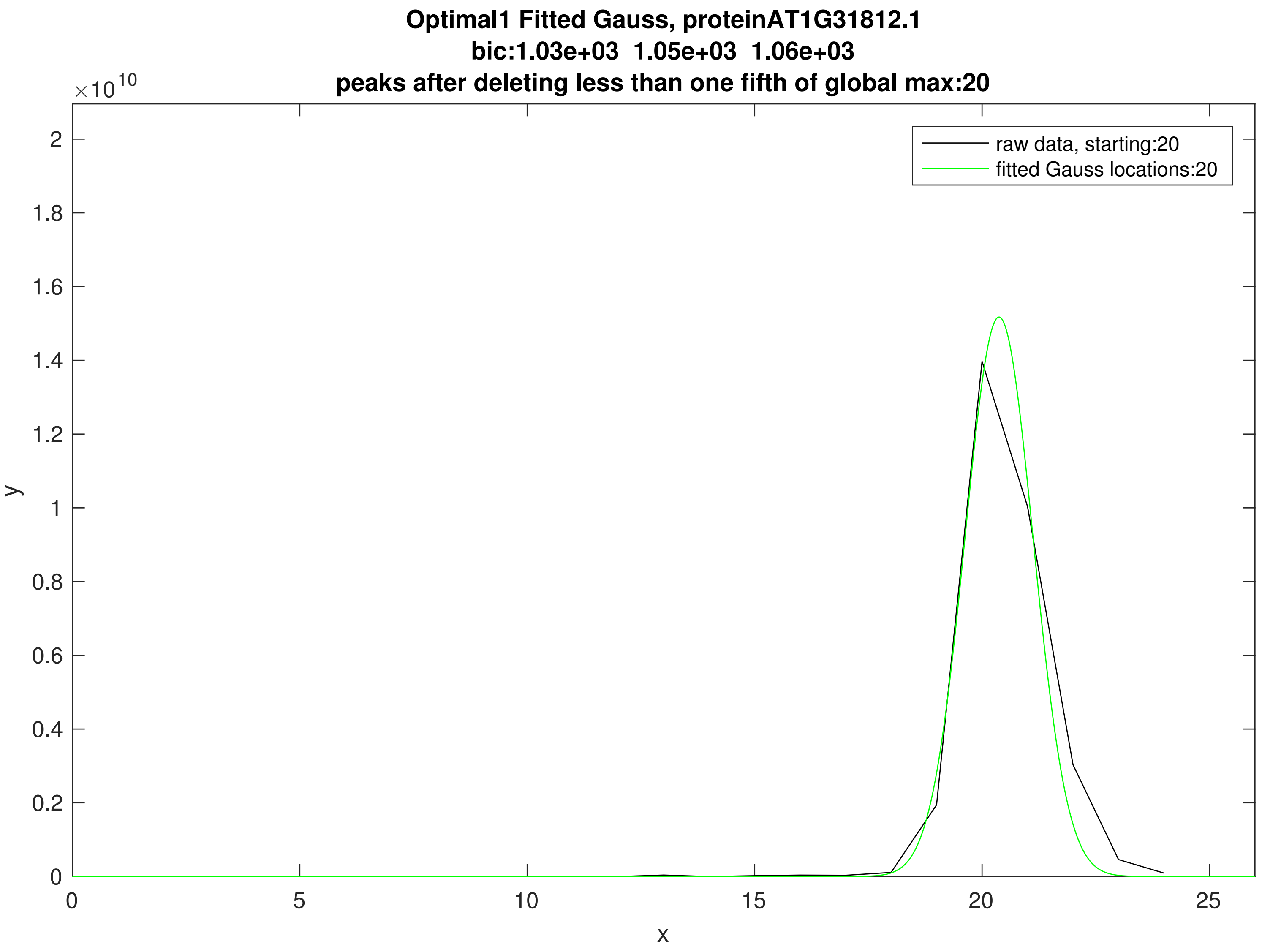
Optimal1 Fitted Gauss, proteinAT1G31812.1
bic:1.03e+03 1.05e+03 1.06e+03
peaks after deleting less than one fifth of global max:20
×10^10
raw data, starting:20
fitted Gauss locations:20
y
x
0
0.2
0.4
0.6
0.8
1
1.2
1.4
1.6
1.8
2
0
5
10
15
20
25

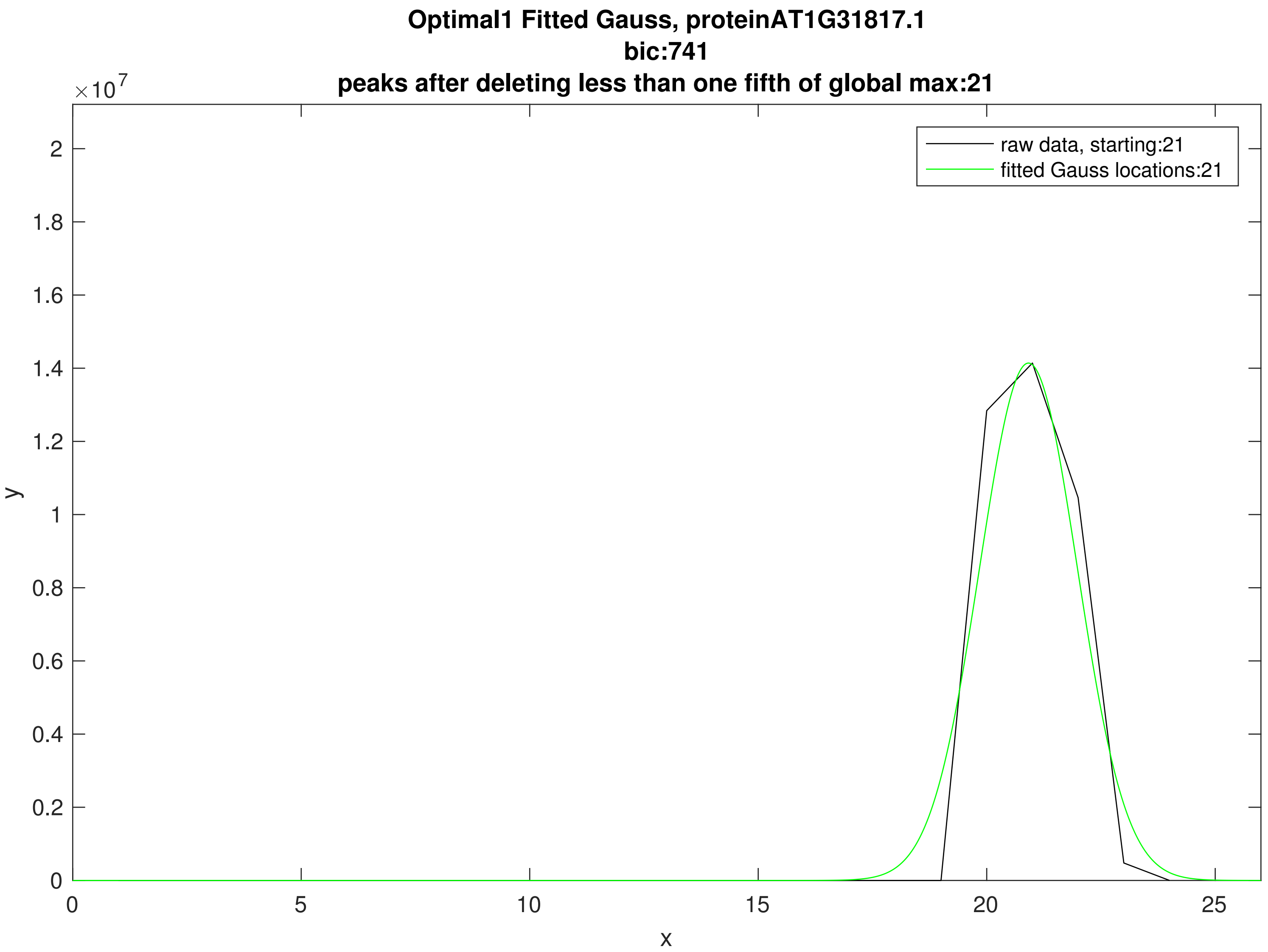

Optimal1 Fitted Gauss, proteinAT1G31817.1
bic:741
peaks after deleting less than one fifth of global max:21
×10^7
2
1.8
1.6
1.4
1.2
1
0.8
0.6
0.4
0.2
0
y
0
5
10
15
20
25
x
raw data, starting:21
fitted Gauss locations:21

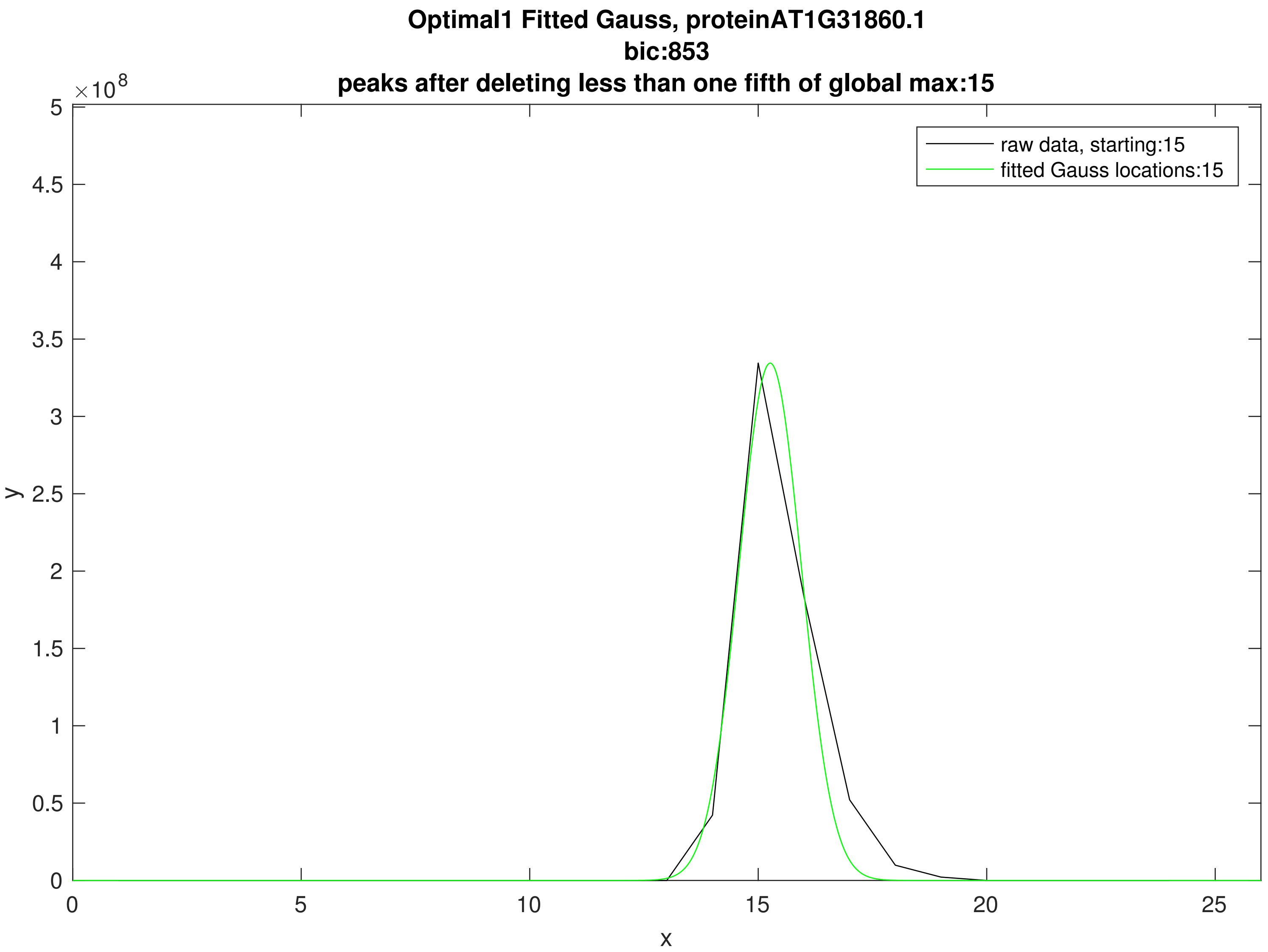
Optimal1 Fitted Gauss, proteinAT1G31860.1
bic:853
peaks after deleting less than one fifth of global max:15
×10^8
raw data, starting:15
fitted Gauss locations:15
y
x
0
0.5
1
1.5
2
2.5
3
3.5
4
4.5
5
0
5
10
15
20
25

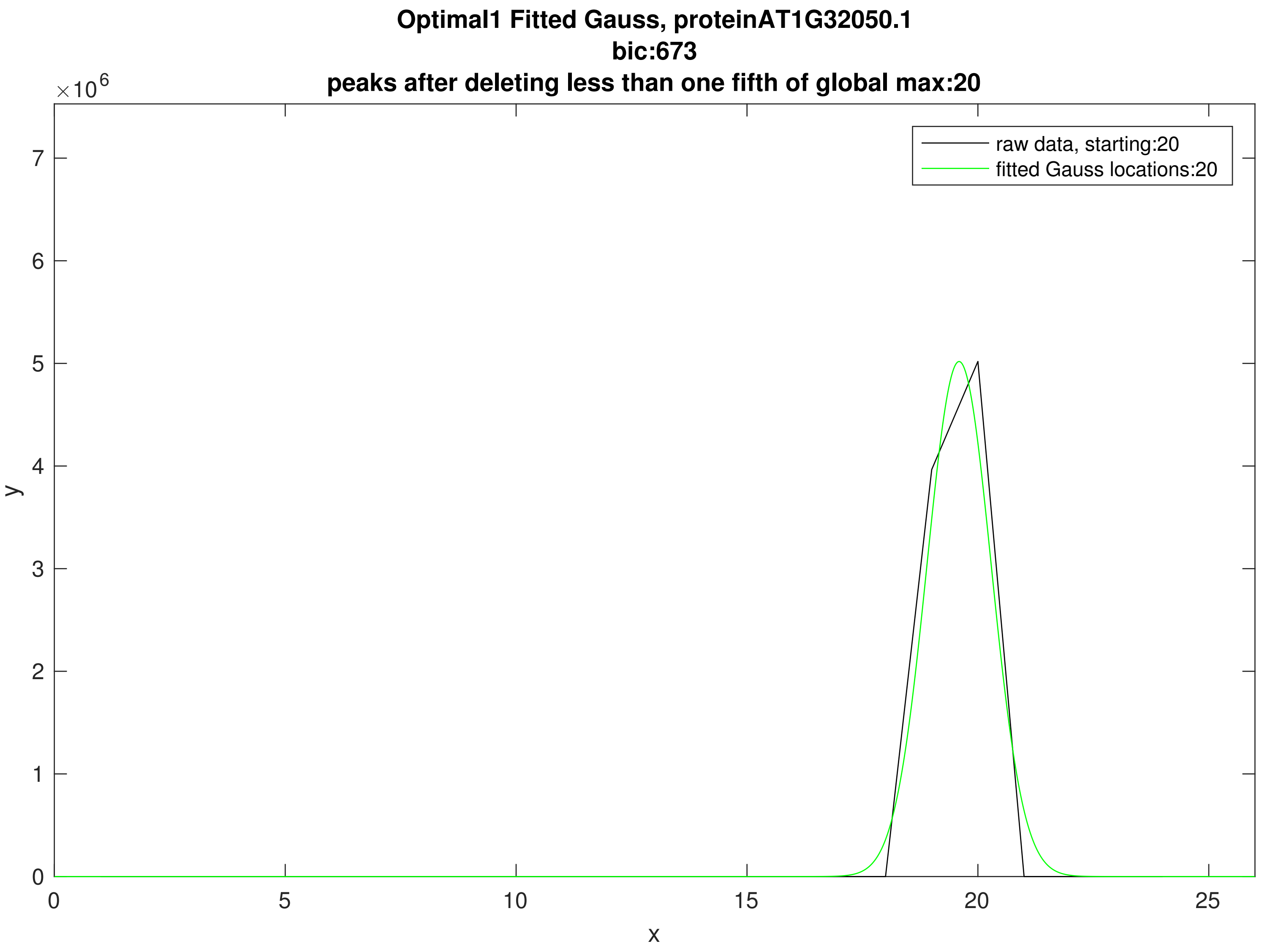
Optimal1 Fitted Gauss, proteinAT1G32050.1
bic:673
peaks after deleting less than one fifth of global max:20
×10^6
raw data, starting:20
fitted Gauss locations:20
y
x
0
1
2
3
4
5
6
7
0
5
10
15
20
25

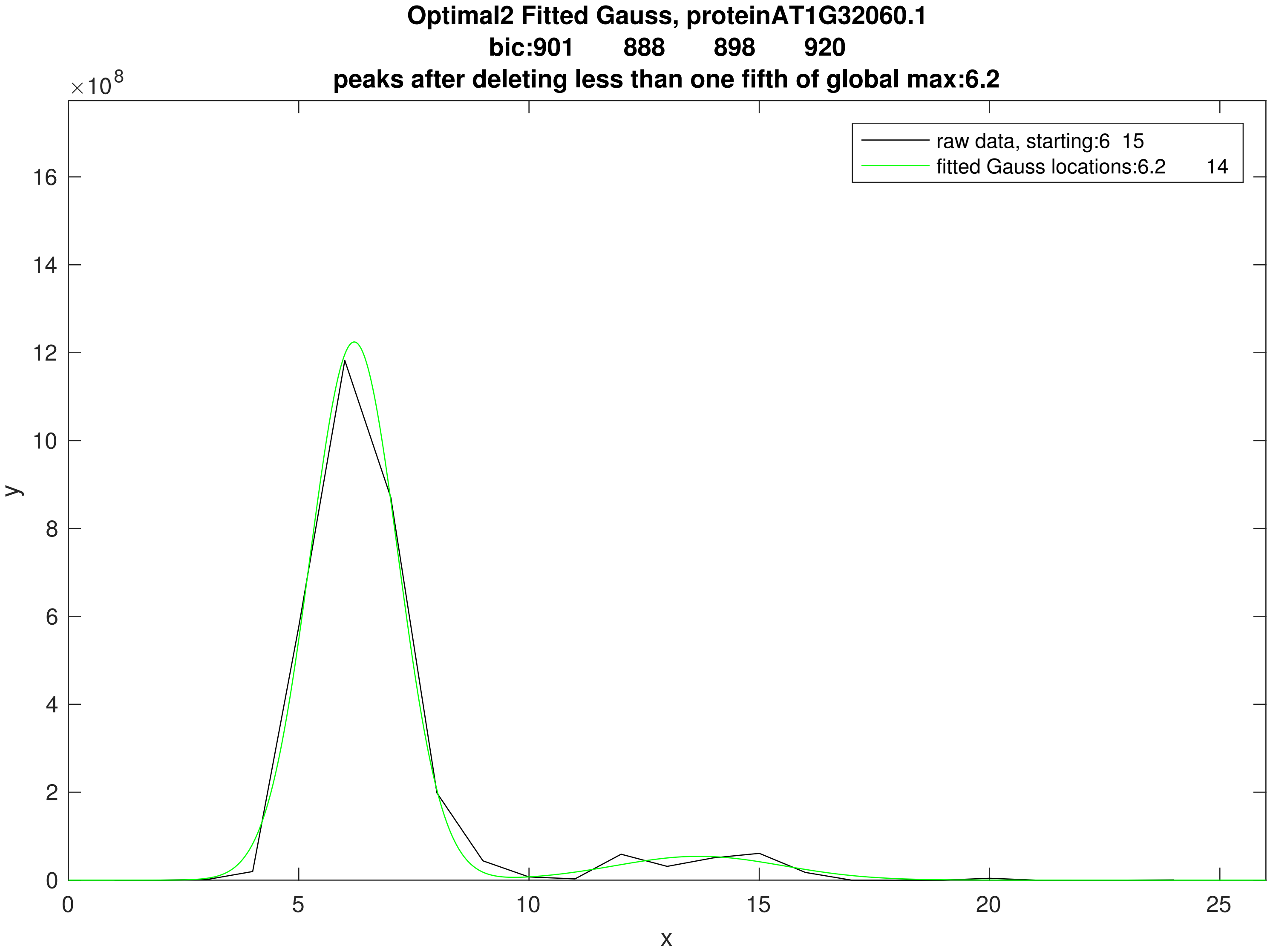

Optimal2 Fitted Gauss, proteinAT1G32060.1
bic:901 888 898 920
peaks after deleting less than one fifth of global max:6.2
×10^8
raw data, starting:6 15
fitted Gauss locations:6.2 14
y
x
0
2
4
6
8
10
12
14
16
0
5
10
15
20
25

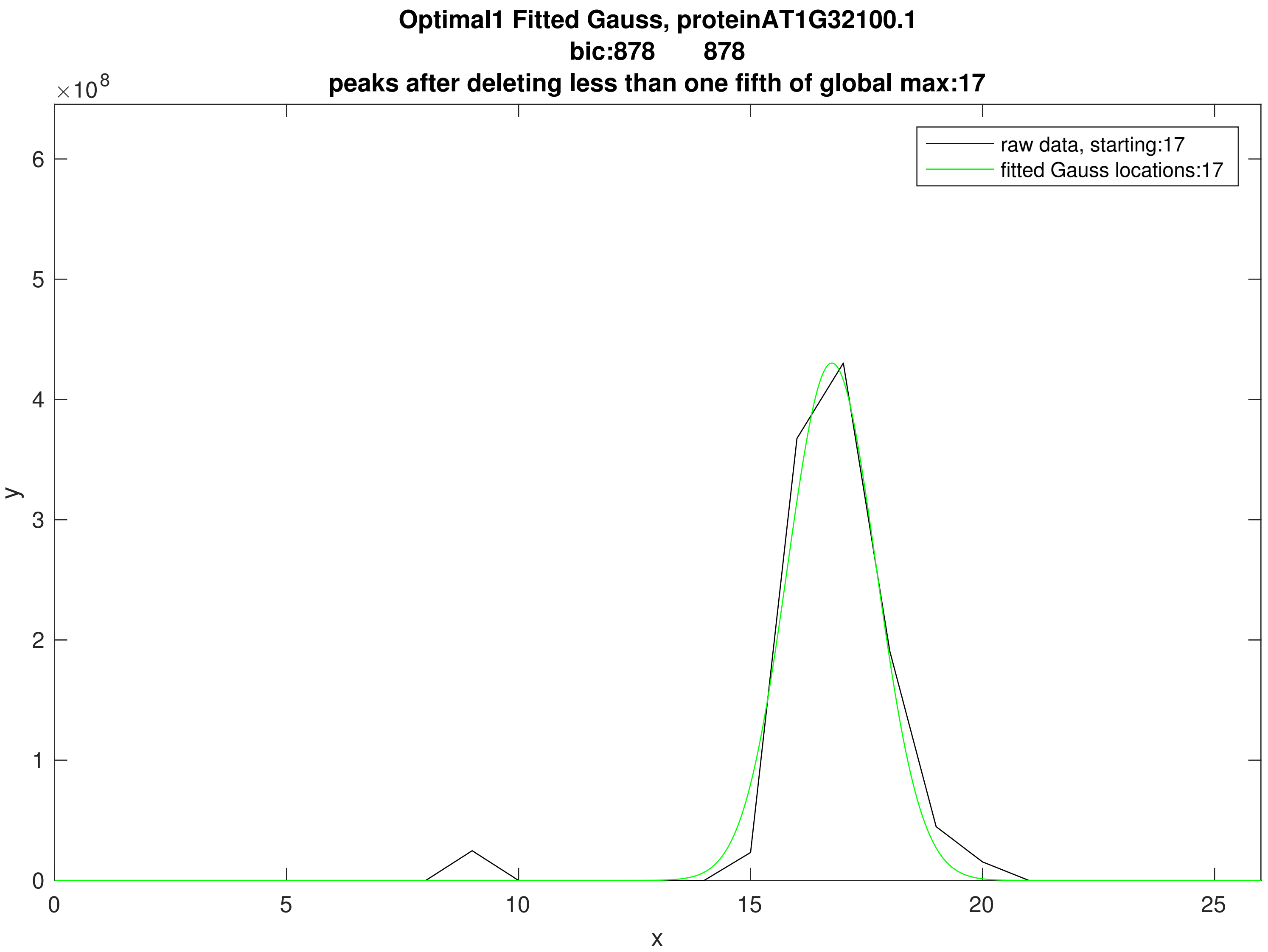

Optimal1 Fitted Gauss, proteinAT1G32100.1
bic:878 878
peaks after deleting less than one fifth of global max:17
×10^8
raw data, starting:17
fitted Gauss locations:17
y
x
0
1
2
3
4
5
6
0
5
10
15
20
25

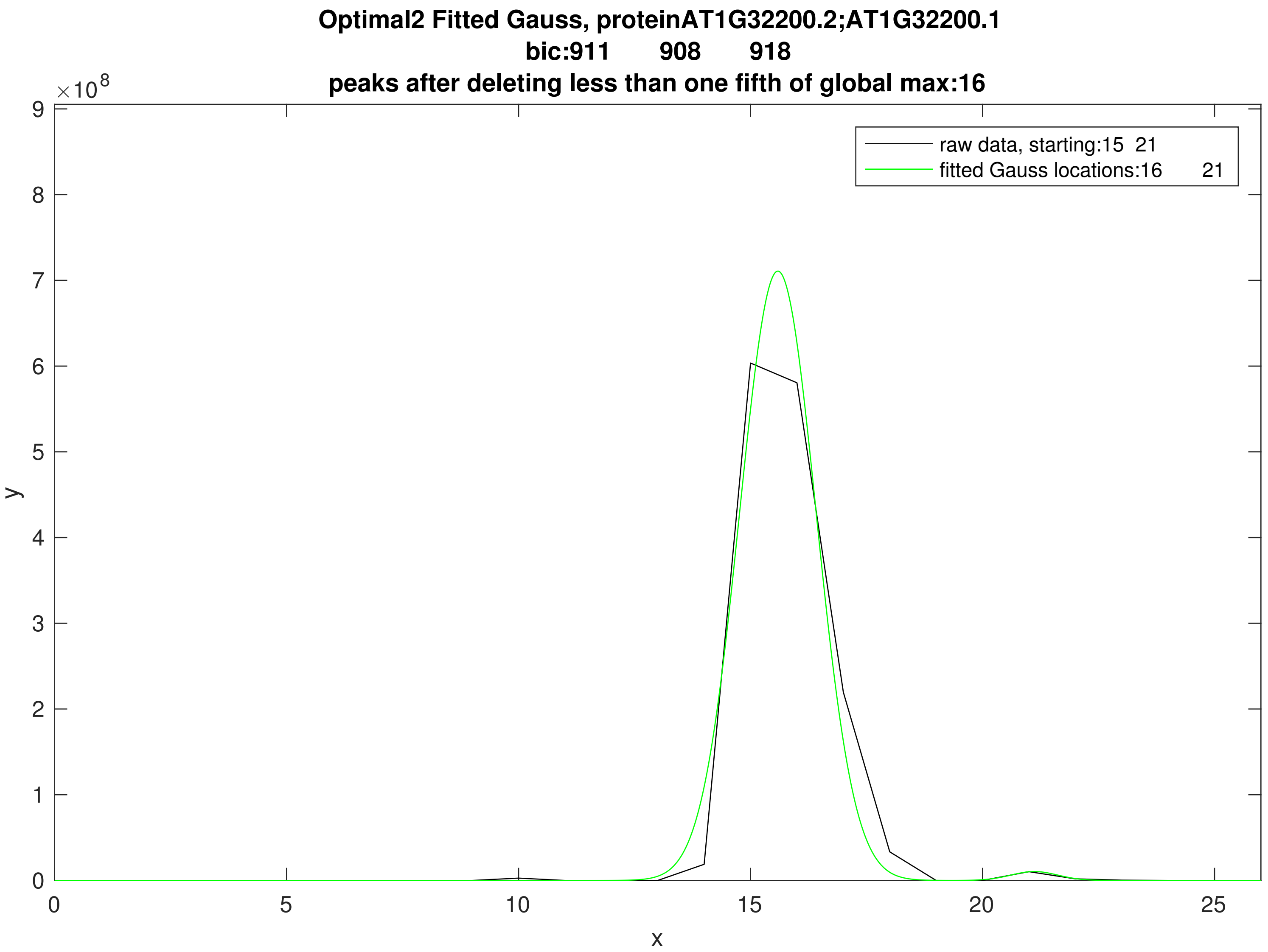

Optimal2 Fitted Gauss, proteinAT1G32200.2;AT1G32200.1
bic:911 908 918
peaks after deleting less than one fifth of global max:16
×10^8
raw data, starting:15 21
fitted Gauss locations:16 21
y
x
0
1
2
3
4
5
6
7
8
9
0
5
10
15
20
25

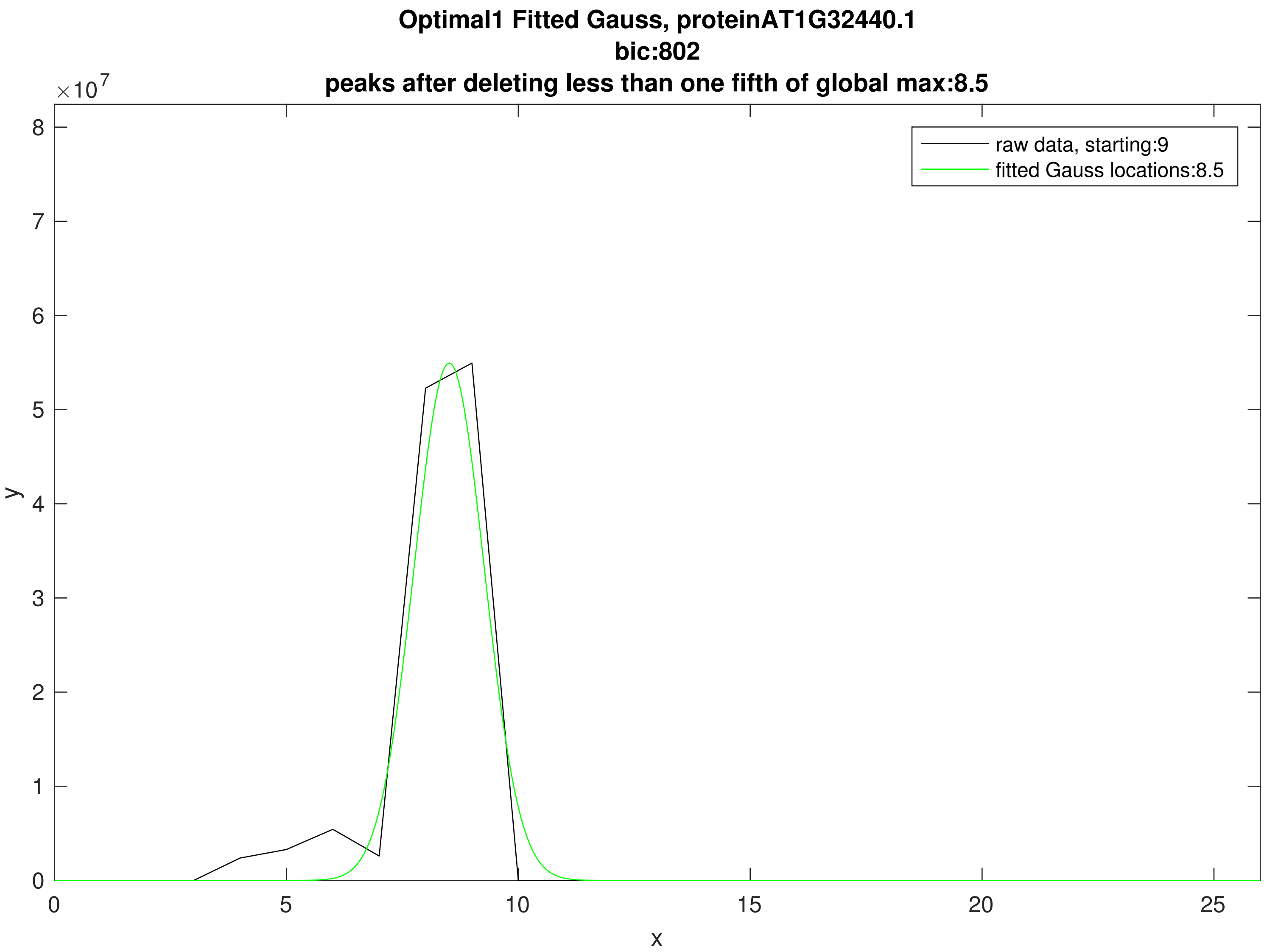

Optimal1 Fitted Gauss, proteinAT1G32440.1
bic:802
peaks after deleting less than one fifth of global max:8.5
×10^7
raw data, starting:9
fitted Gauss locations:8.5
y
x
0
1
2
3
4
5
6
7
8
0
5
10
15
20
25

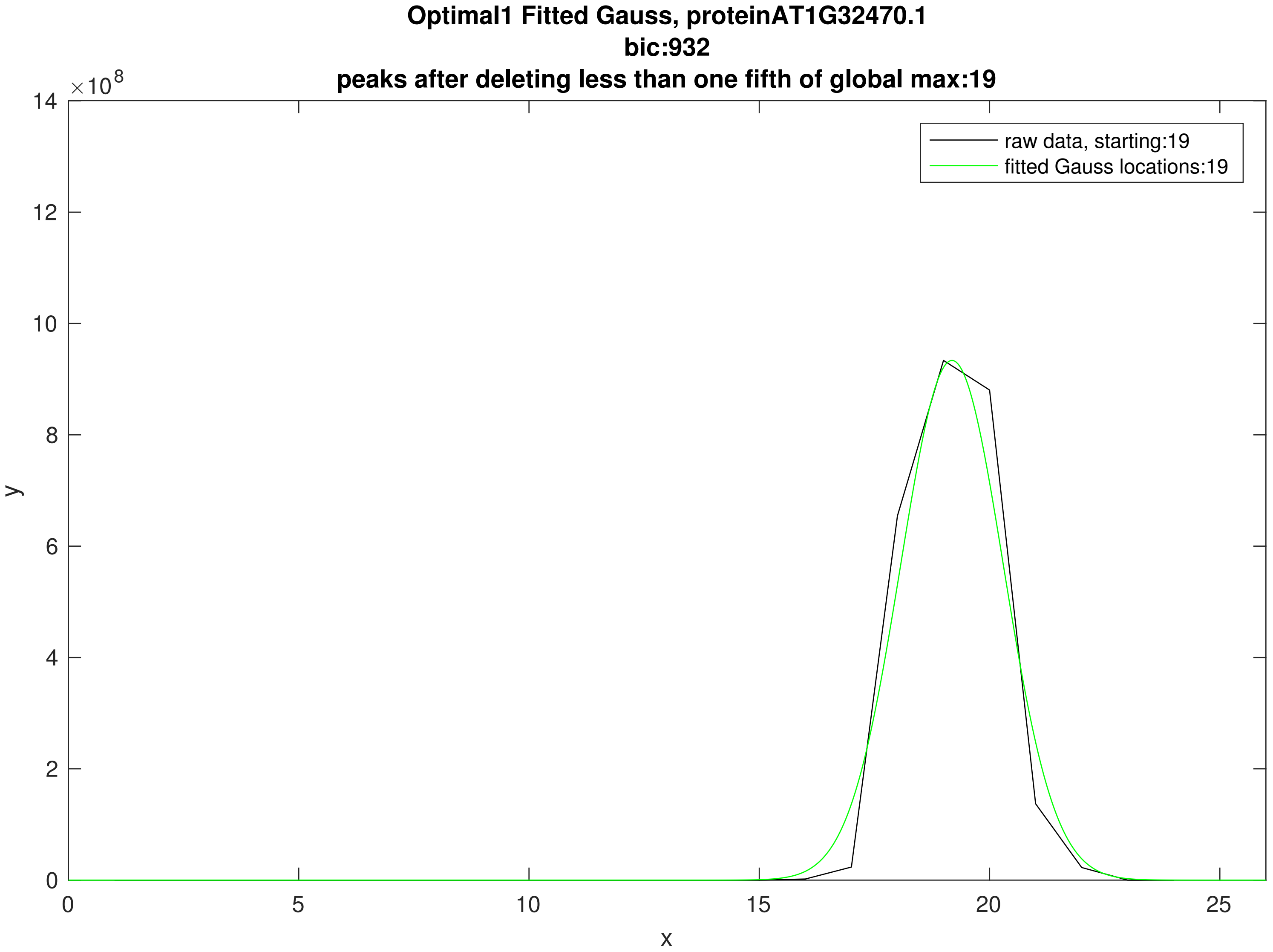
Optimal1 Fitted Gauss, proteinAT1G32470.1
bic:932
peaks after deleting less than one fifth of global max:19
×10^8
14
12
10
8
6
4
2
0
y
0
5
10
15
20
25
x
raw data, starting:19
fitted Gauss locations:19

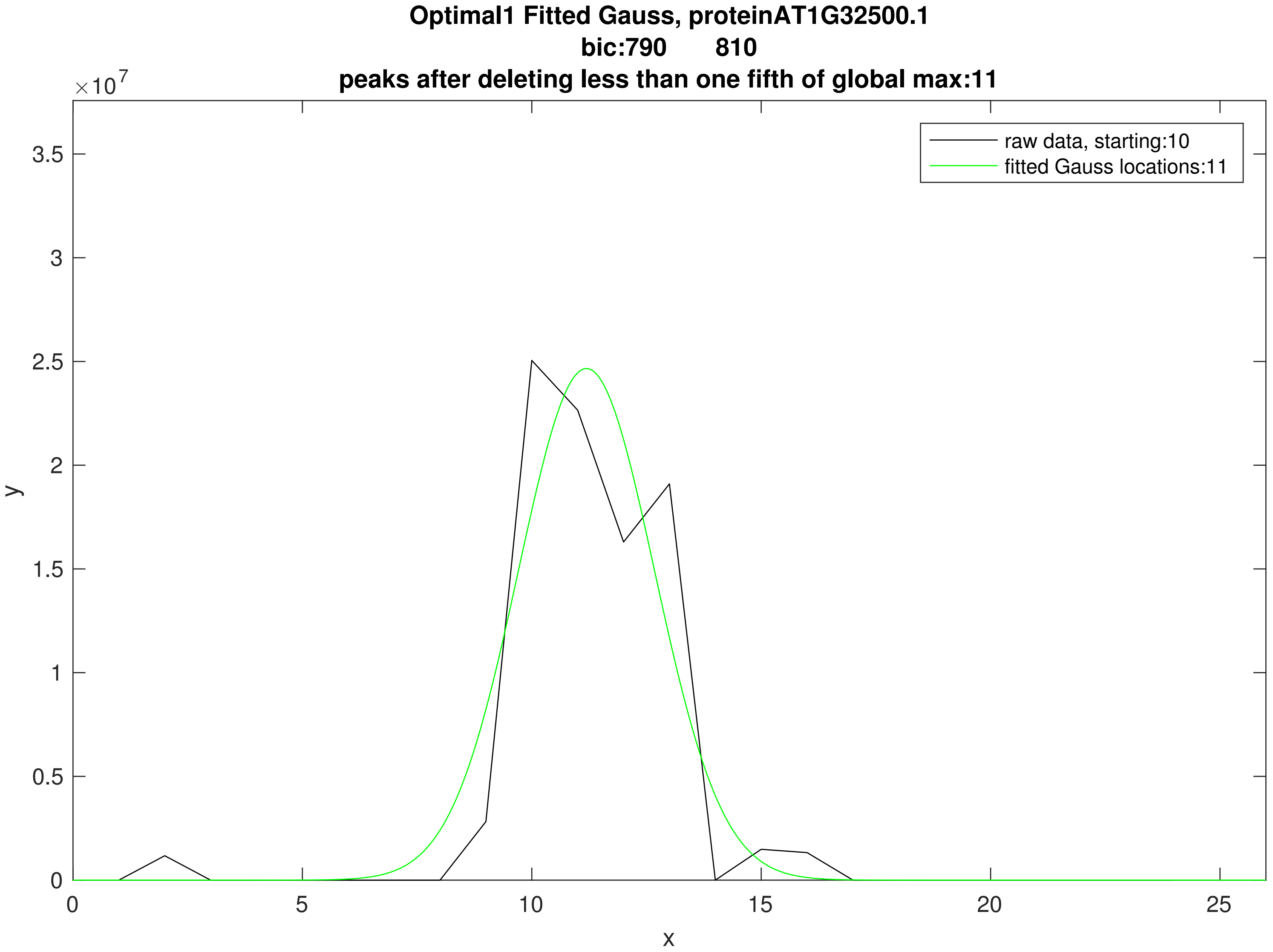

Optimal1 Fitted Gauss, proteinAT1G32500.1
bic:790 810
peaks after deleting less than one fifth of global max:11
×10^7
3.5
3
2.5
2
1.5
1
0.5
0
y
0
5
10
15
20
25
x
raw data, starting:10
fitted Gauss locations:11

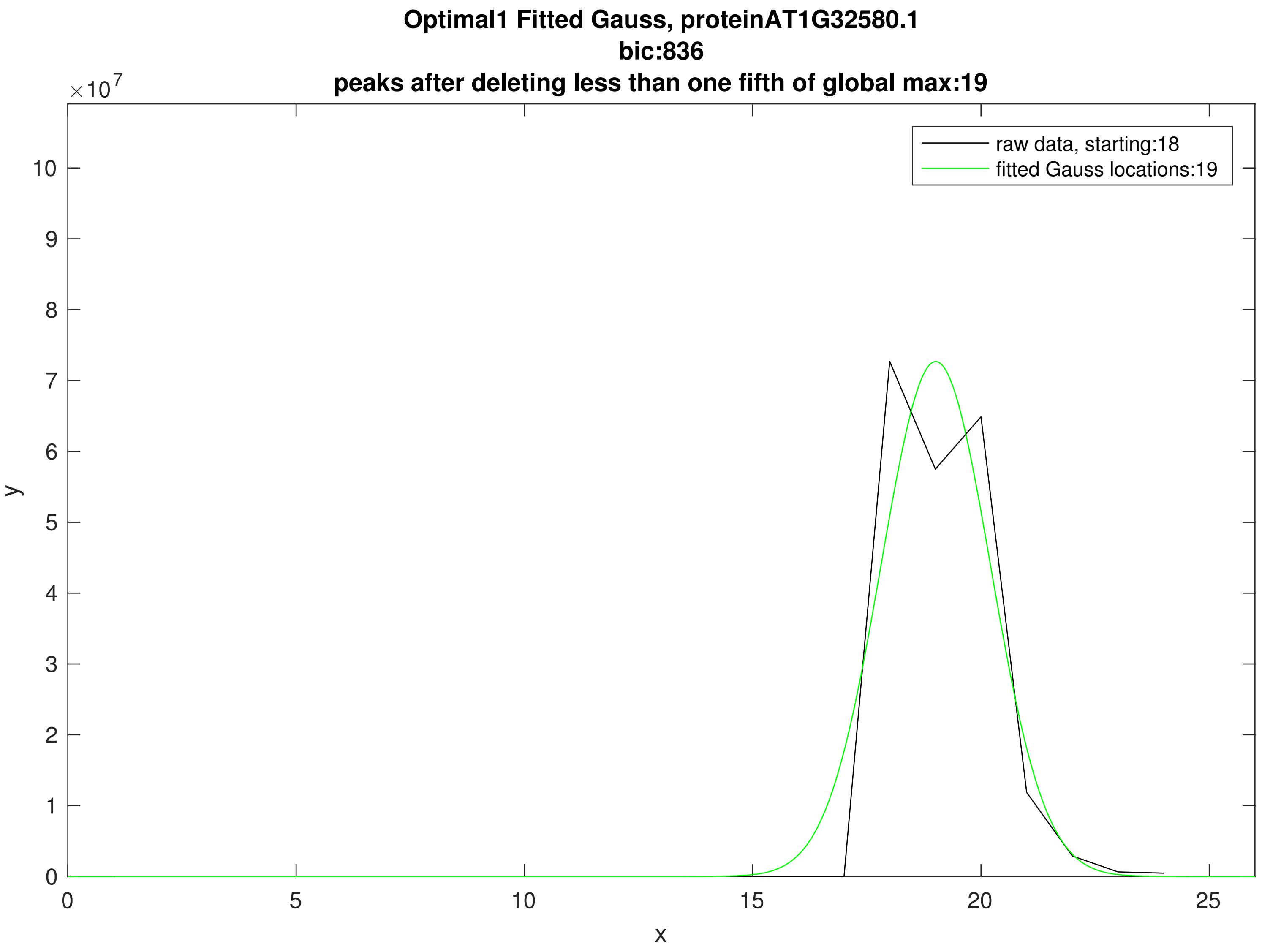

Optimal1 Fitted Gauss, proteinAT1G32580.1
bic:836
peaks after deleting less than one fifth of global max:19
×10^7
raw data, starting:18
fitted Gauss locations:19
y
x
0
1
2
3
4
5
6
7
8
9
10
0
5
10
15
20
25

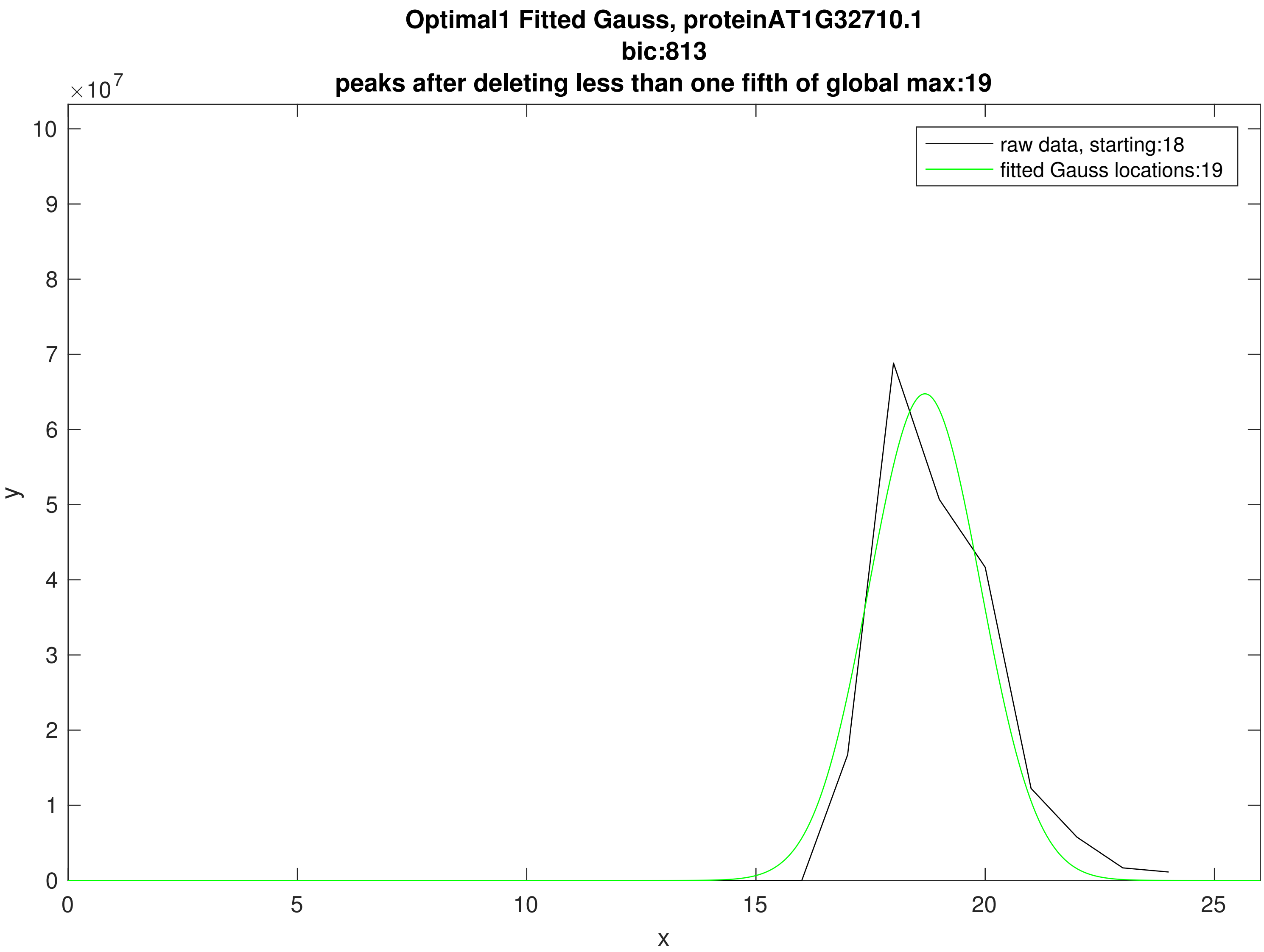

Optimal1 Fitted Gauss, proteinAT1G32710.1
bic:813
peaks after deleting less than one fifth of global max:19
×10^7
10
9
8
7
6
5
4
3
2
1
0
y
0
5
10
15
20
25
x
raw data, starting:18
fitted Gauss locations:19

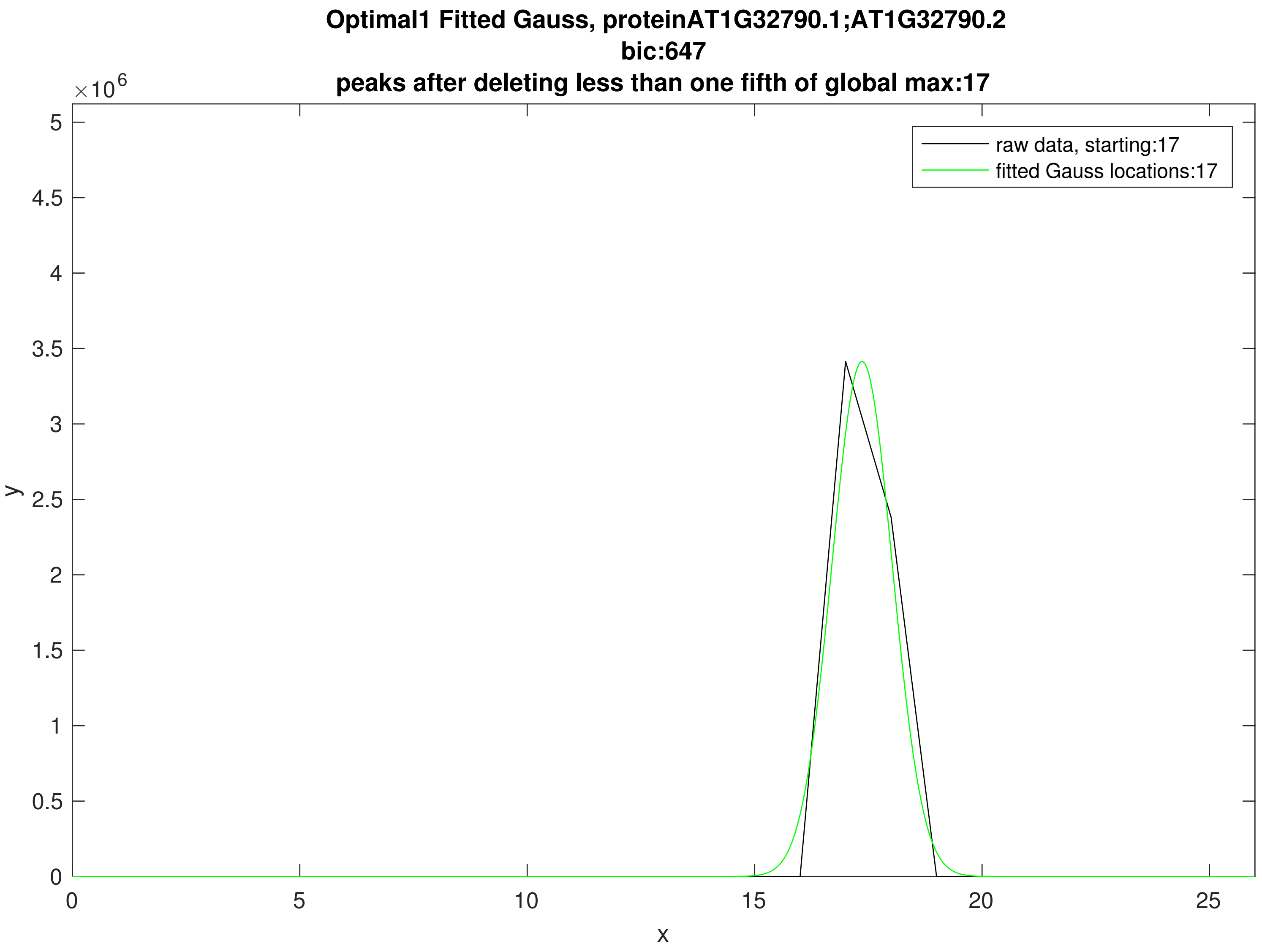
Optimal1 Fitted Gauss, proteinAT1G32790.1;AT1G32790.2
bic:647
peaks after deleting less than one fifth of global max:17
×10^6
raw data, starting:17
fitted Gauss locations:17
y
x
0
0.5
1
1.5
2
2.5
3
3.5
4
4.5
5
0
5
10
15
20
25

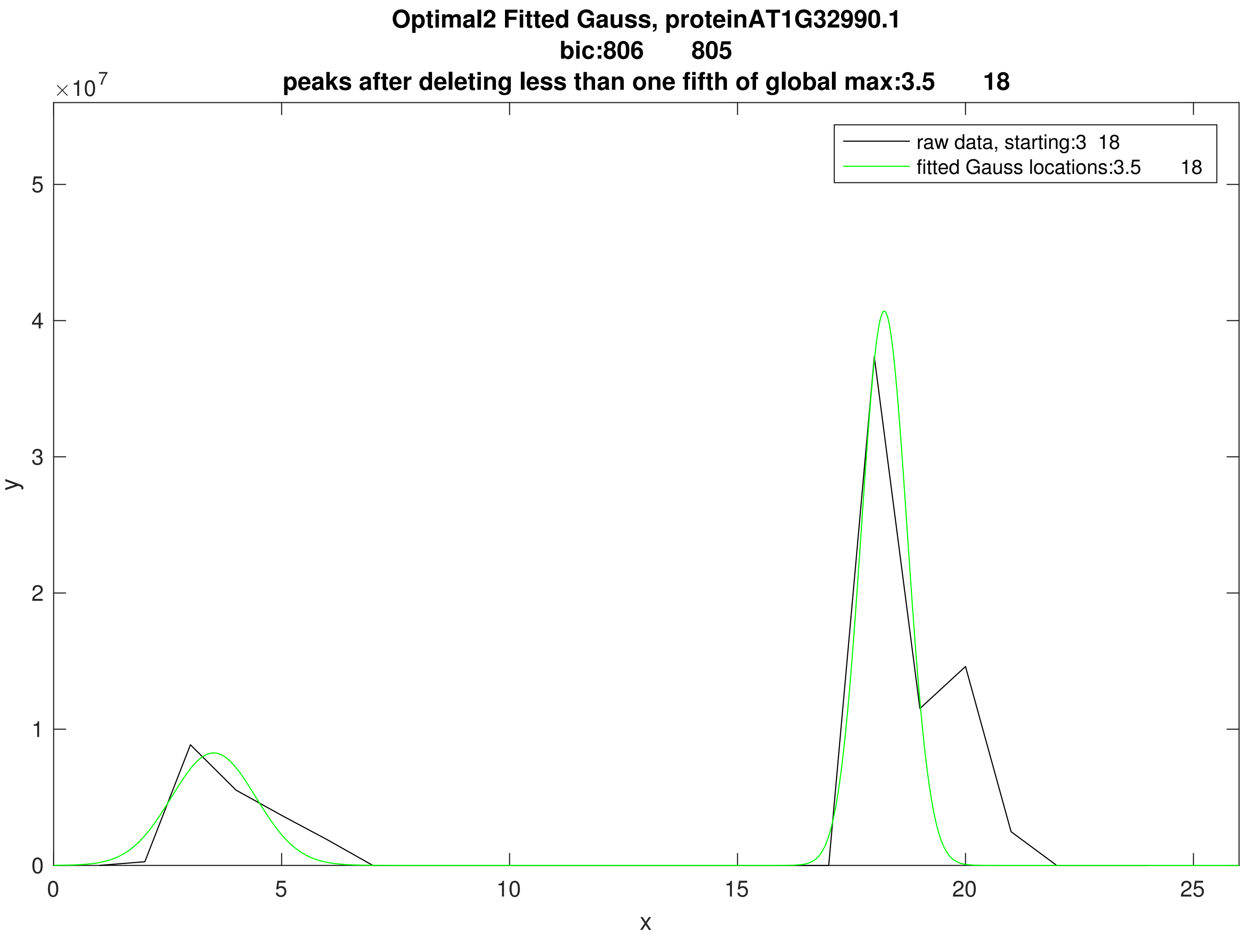

Optimal2 Fitted Gauss, proteinAT1G32990.1
bic:806 805
peaks after deleting less than one fifth of global max:3.5 18
×10^7
raw data, starting:3 18
fitted Gauss locations:3.5 18
y
x
0
1
2
3
4
5
0
5
10
15
20
25

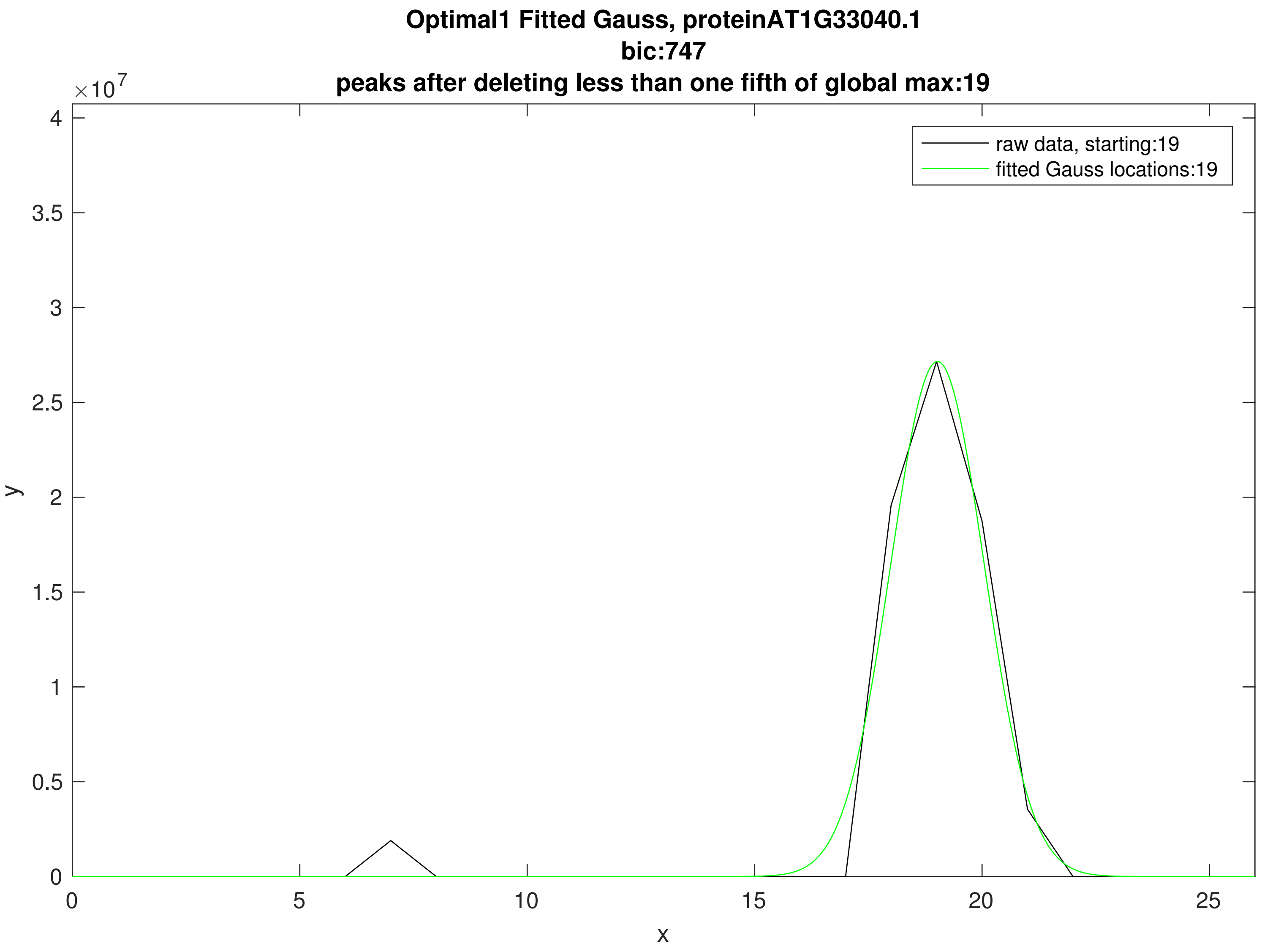
Optimal1 Fitted Gauss, proteinAT1G33040.1
bic:747
peaks after deleting less than one fifth of global max:19
×10^7
raw data, starting:19
fitted Gauss locations:19
y
x
0
0.5
1
1.5
2
2.5
3
3.5
4
5
10
15
20
25

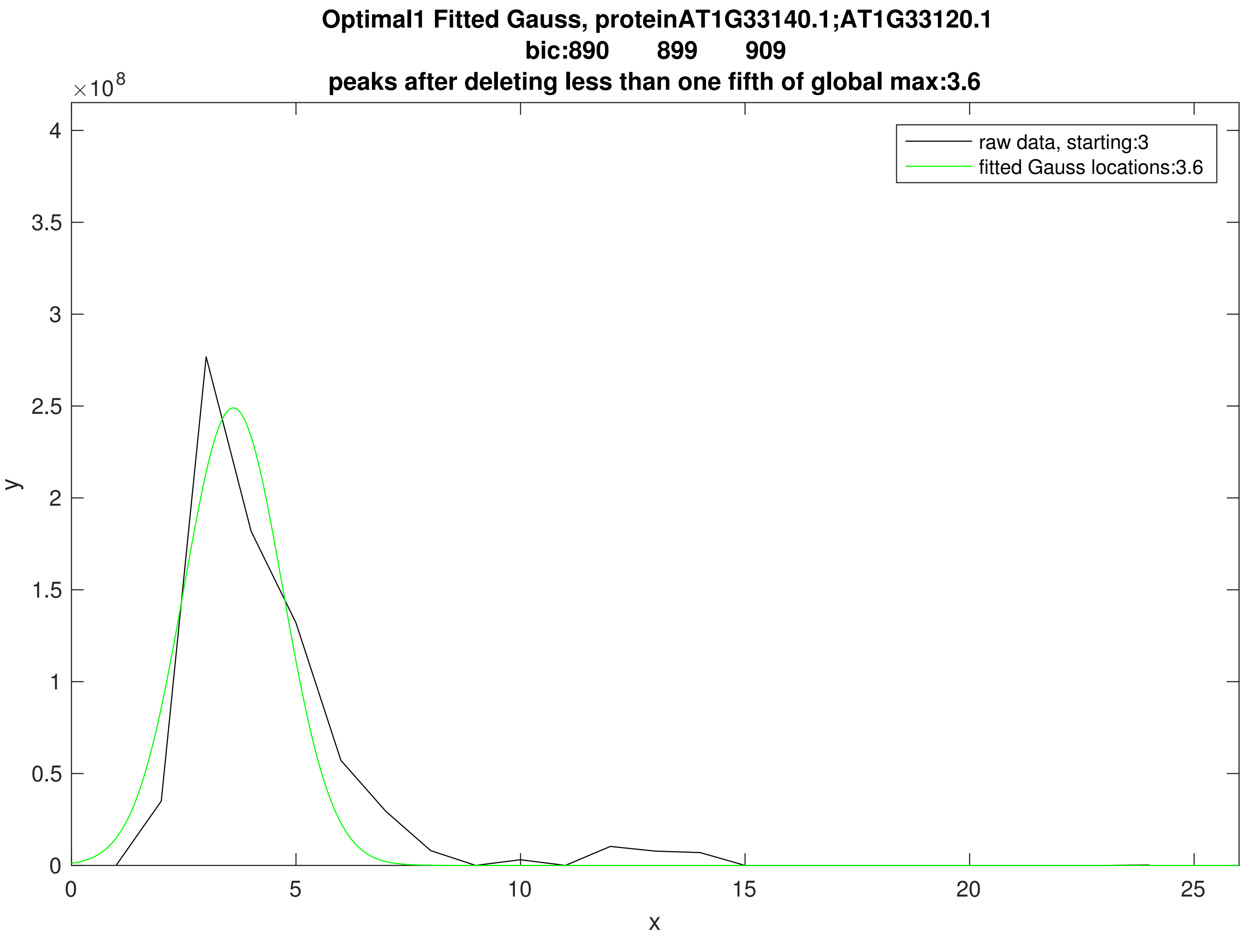
Optimal1 Fitted Gauss, proteinAT1G33140.1;AT1G33120.1
bic:890 899 909
peaks after deleting less than one fifth of global max:3.6
×10^8
raw data, starting:3
fitted Gauss locations:3.6
y
x
0
0.5
1
1.5
2
2.5
3
3.5
4
5
10
15
20
25

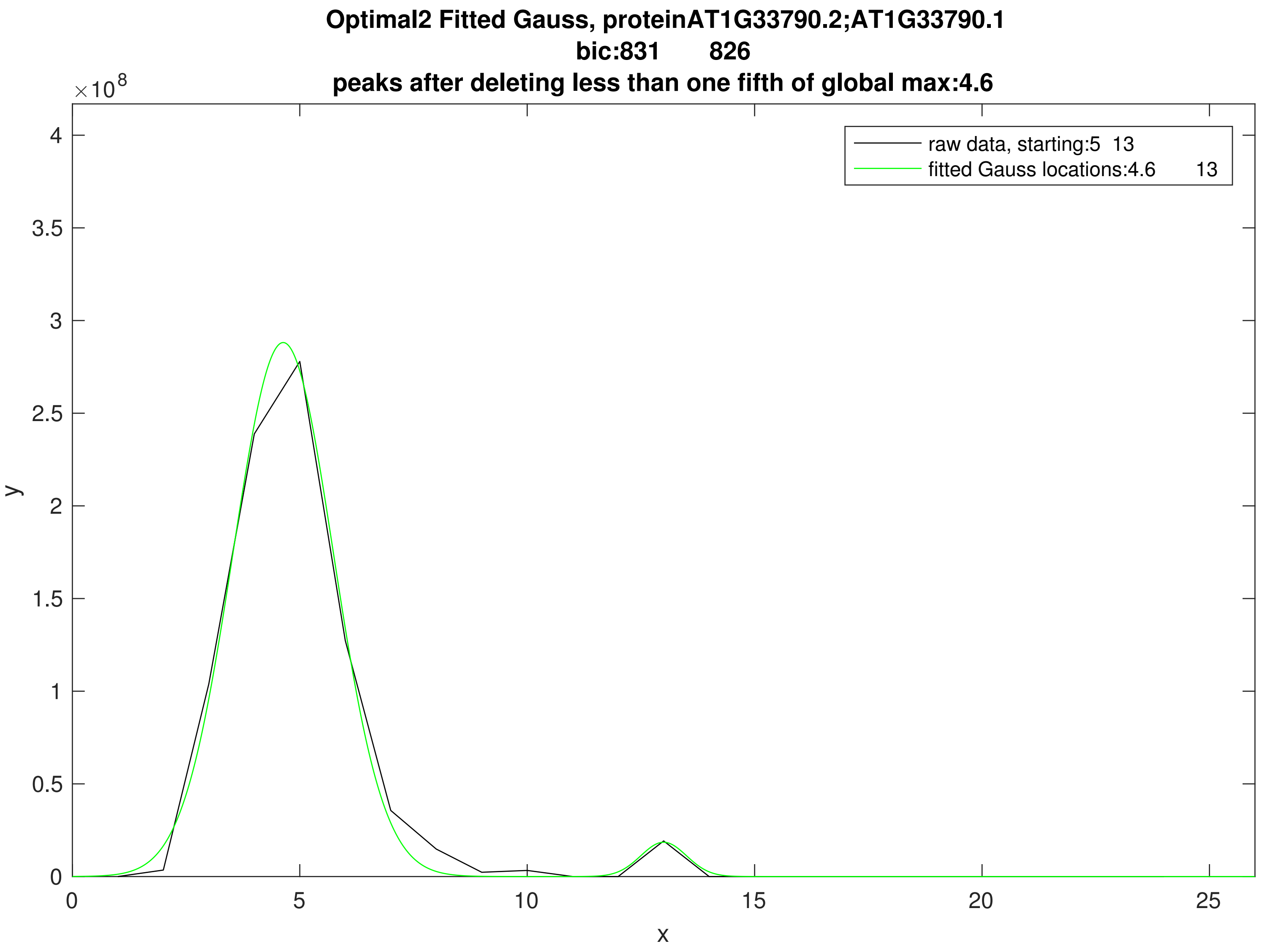

Optimal2 Fitted Gauss, proteinAT1G33790.2;AT1G33790.1
bic:831 826
peaks after deleting less than one fifth of global max:4.6
×10^8
4
3.5
3
2.5
2
1.5
1
0.5
0
y
0
5
10
15
20
25
x
raw data, starting:5 13
fitted Gauss locations:4.6 13

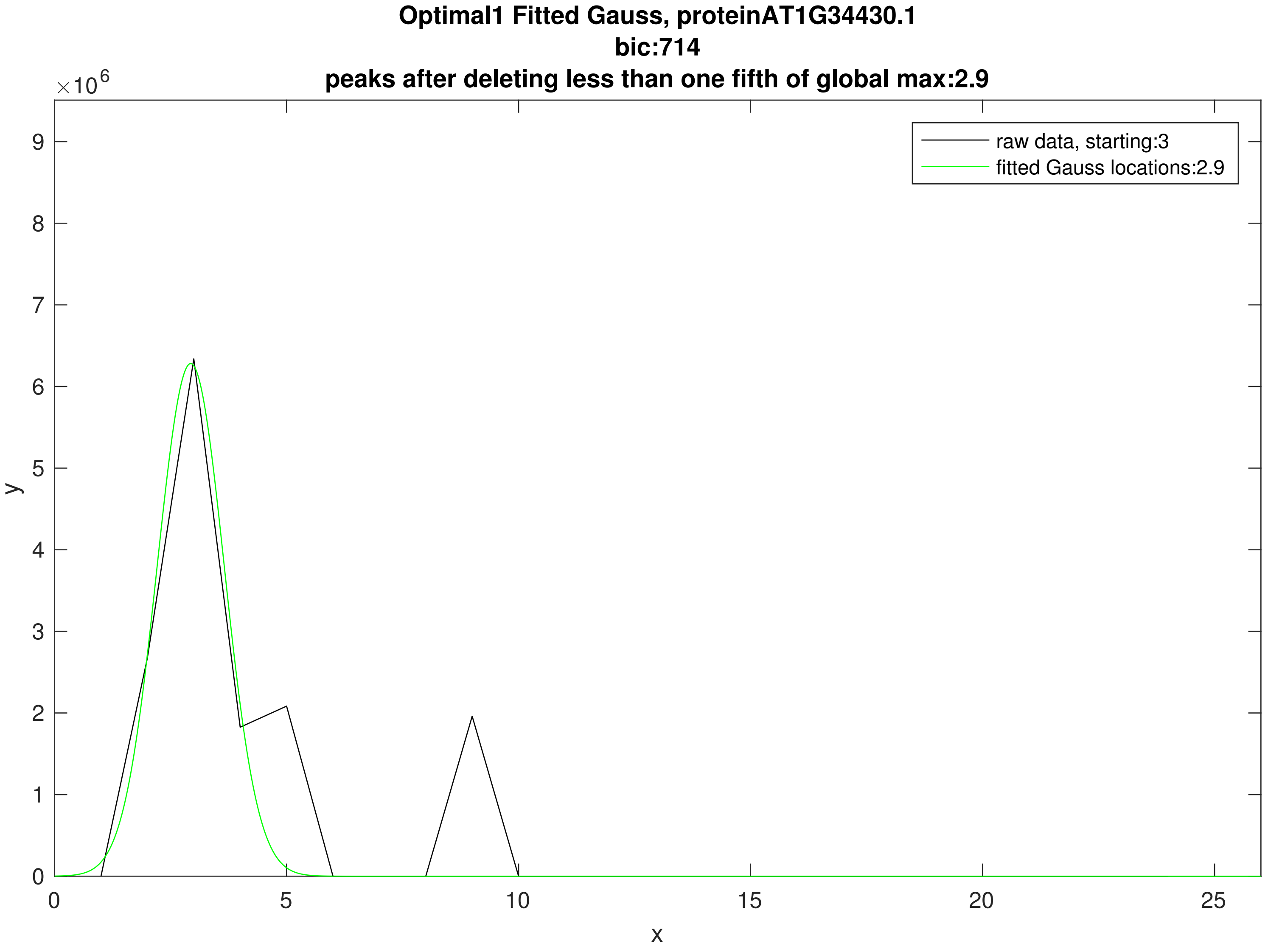

Optimal1 Fitted Gauss, proteinAT1G34430.1
bic:714
peaks after deleting less than one fifth of global max:2.9
×10^6
raw data, starting:3
fitted Gauss locations:2.9
y
x
0
1
2
3
4
5
6
7
8
9
0
5
10
15
20
25

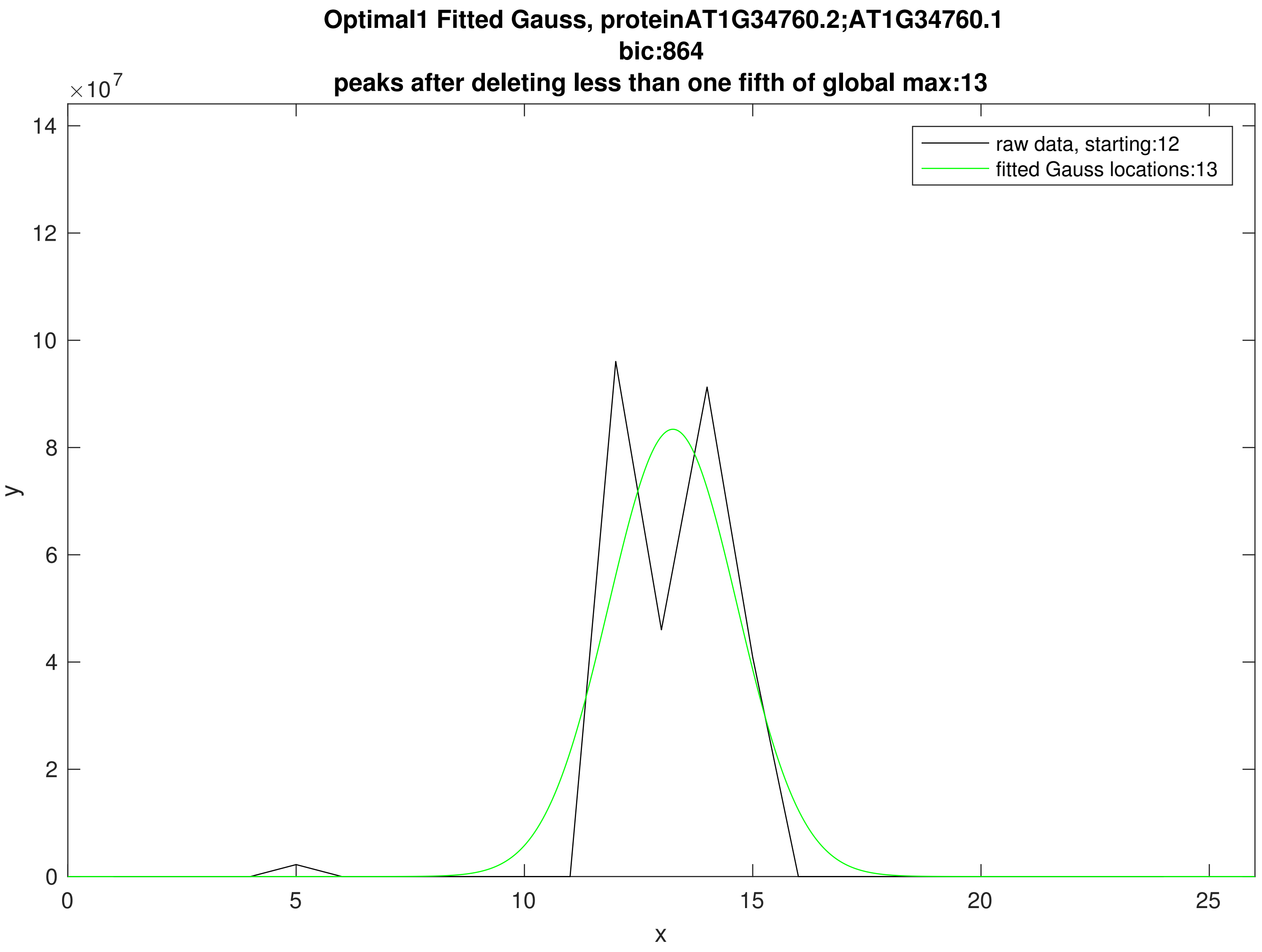
Optimal1 Fitted Gauss, proteinAT1G34760.2;AT1G34760.1
bic:864
peaks after deleting less than one fifth of global max:13
×10^7
14
12
10
8
6
4
2
0
y
0
5
10
15
20
25
x
raw data, starting:12
fitted Gauss locations:13

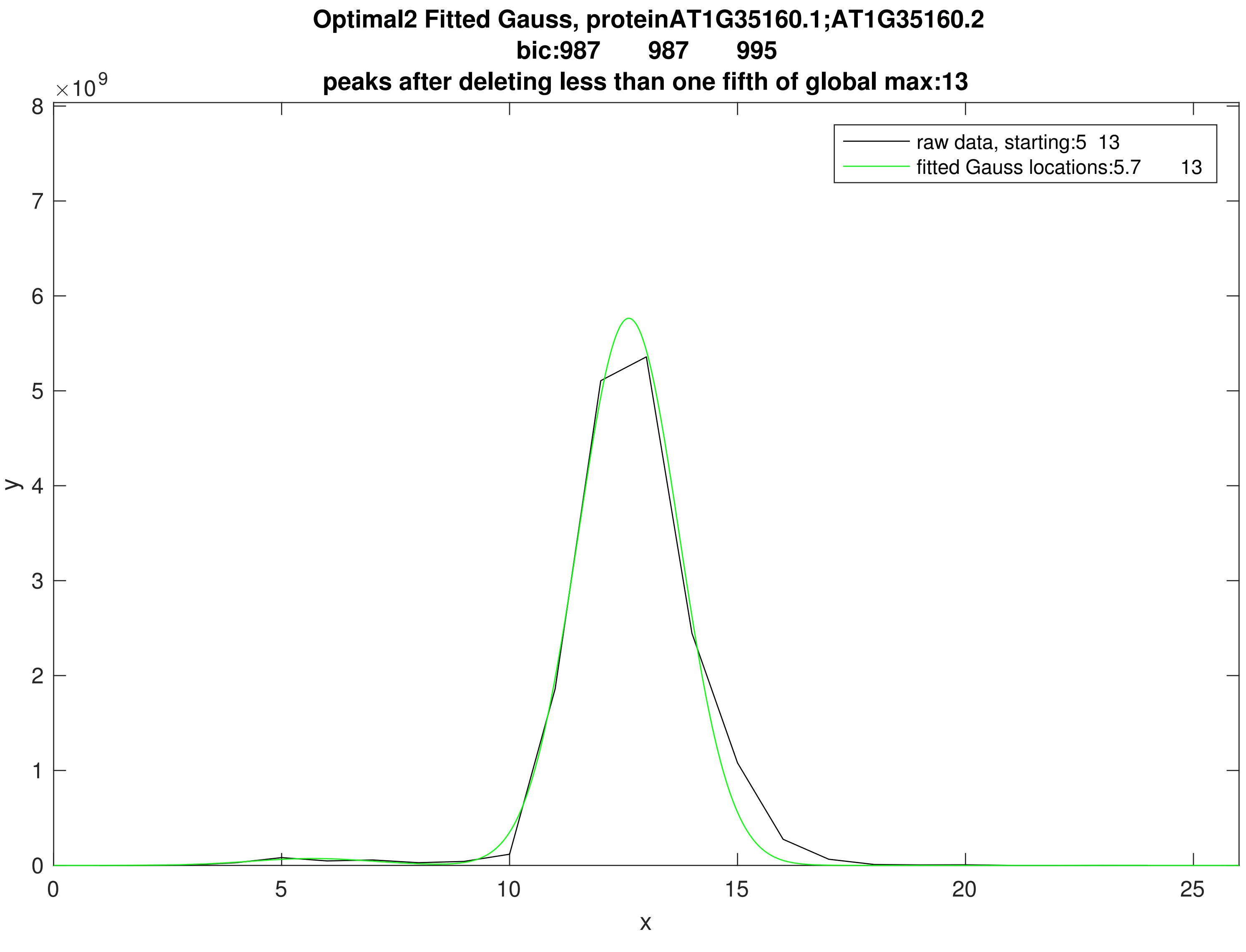
Optimal2 Fitted Gauss, proteinAT1G35160.1;AT1G35160.2
bic:987 987 995
peaks after deleting less than one fifth of global max:13
×10^9
raw data, starting:5 13
fitted Gauss locations:5.7 13
y
x
0
1
2
3
4
5
6
7
8
0
5
10
15
20
25

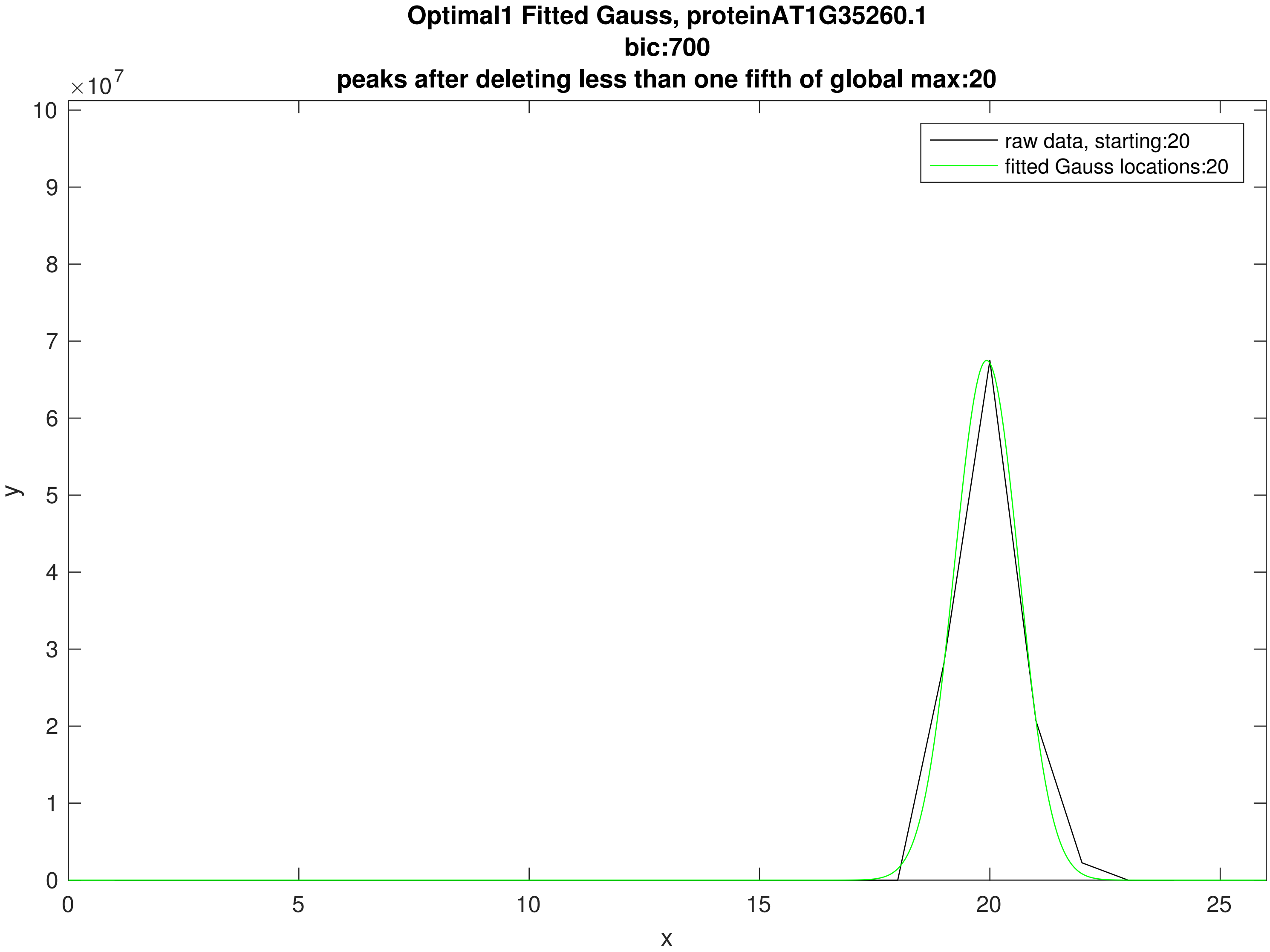

Optimal1 Fitted Gauss, proteinAT1G35260.1
bic:700
peaks after deleting less than one fifth of global max:20
×10^7
10
9
8
7
6
5
4
3
2
1
0
y
0
5
10
15
20
25
x
raw data, starting:20
fitted Gauss locations:20

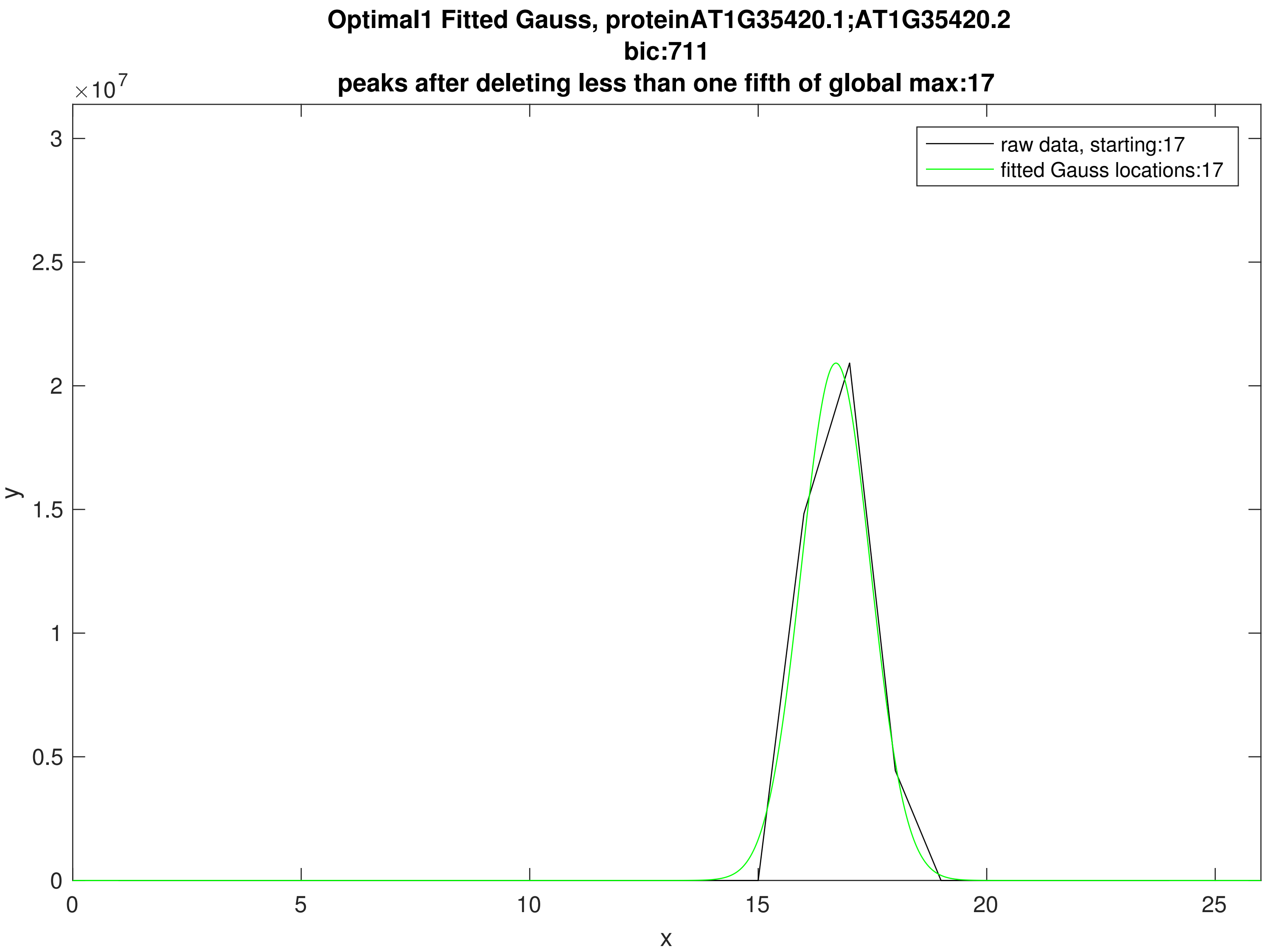
Optimal1 Fitted Gauss, proteinAT1G35420.1;AT1G35420.2
bic:711
peaks after deleting less than one fifth of global max:17
×10^7
3
2.5
2
1.5
1
0.5
0
y
0
5
10
15
20
25
x
raw data, starting:17
fitted Gauss locations:17

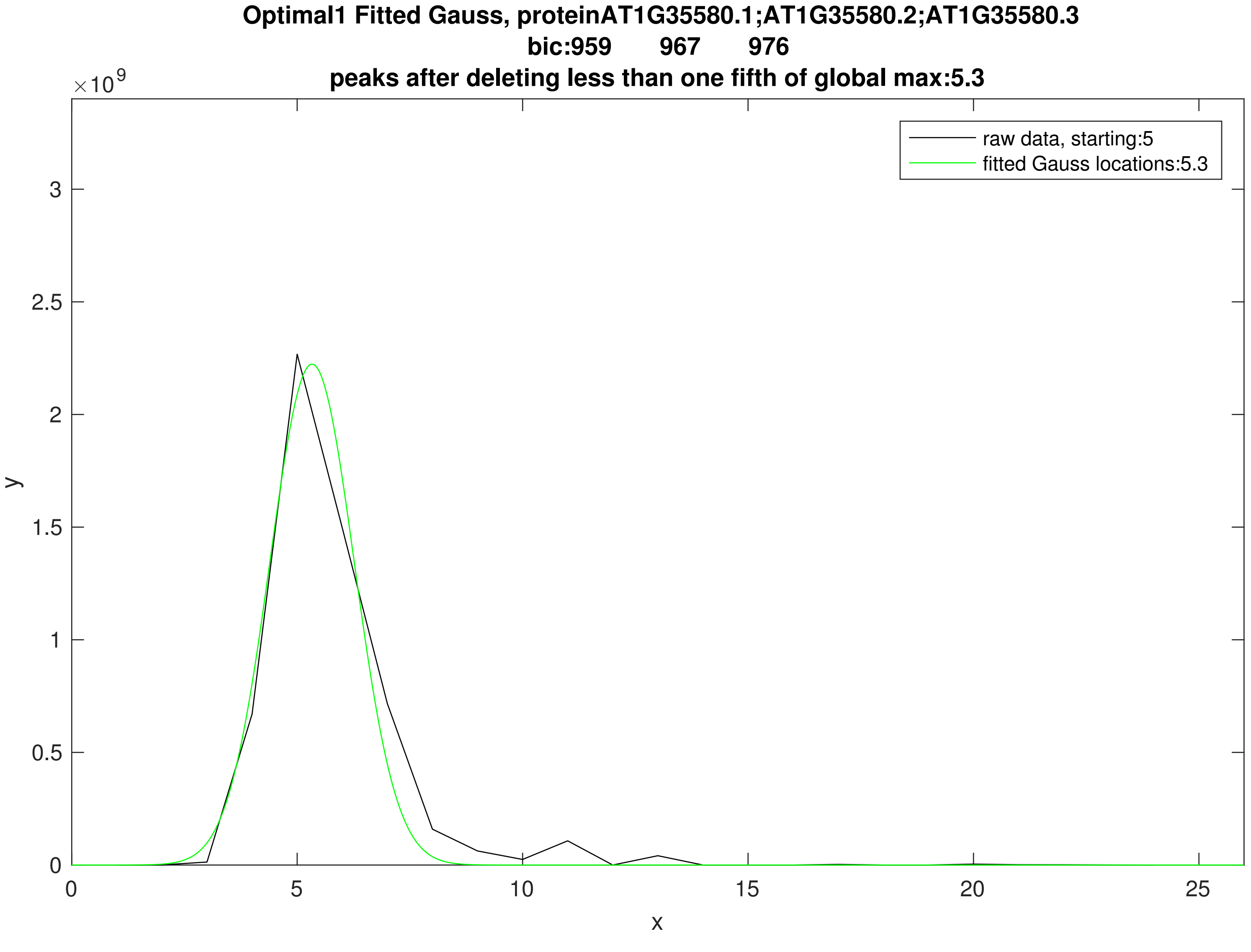

Optimal1 Fitted Gauss, proteinAT1G35580.1;AT1G35580.2;AT1G35580.3
bic:959 967 976
peaks after deleting less than one fifth of global max:5.3
×10^9
raw data, starting:5
fitted Gauss locations:5.3
y
x
0
0.5
1
1.5
2
2.5
3
5
10
15
20
25

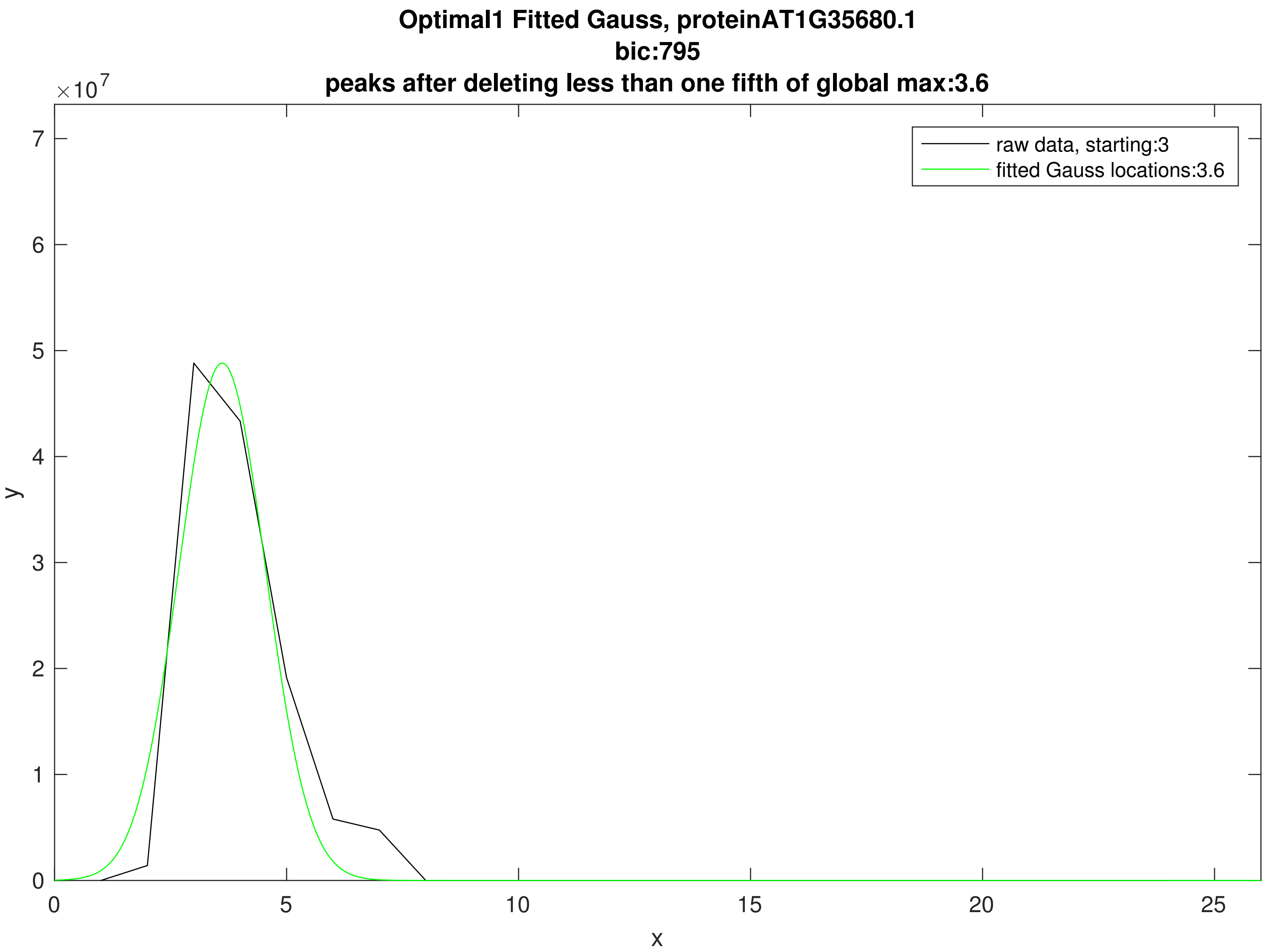

Optimal1 Fitted Gauss, proteinAT1G35680.1
bic:795
peaks after deleting less than one fifth of global max:3.6
×10^7
raw data, starting:3
fitted Gauss locations:3.6
y
x
0
1
2
3
4
5
6
7
0
5
10
15
20
25

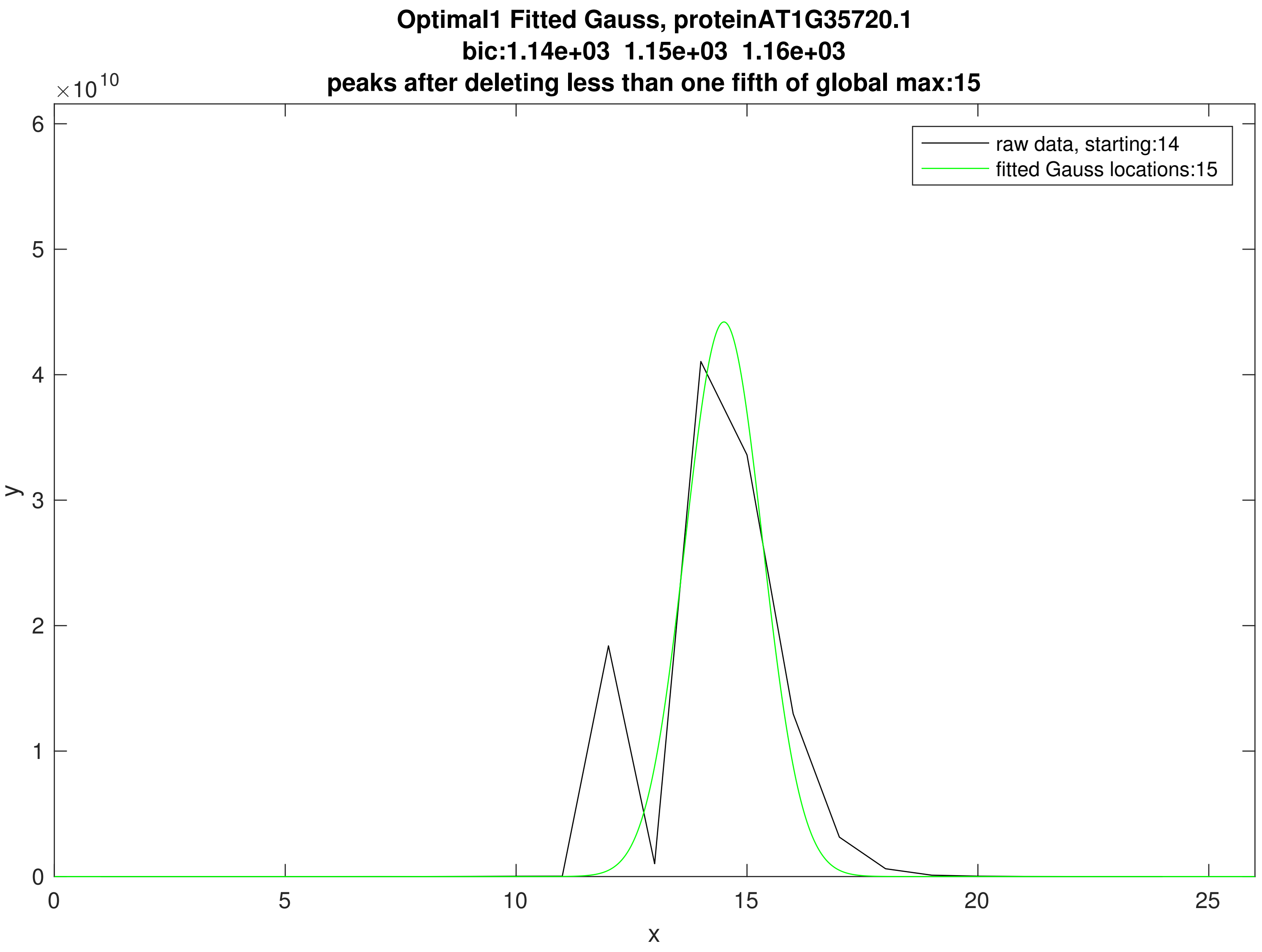
Optimal1 Fitted Gauss, proteinAT1G35720.1
bic:1.14e+03 1.15e+03 1.16e+03
peaks after deleting less than one fifth of global max:15
×10^10
raw data, starting:14
fitted Gauss locations:15
y
x
0
1
2
3
4
5
6
0
5
10
15
20
25

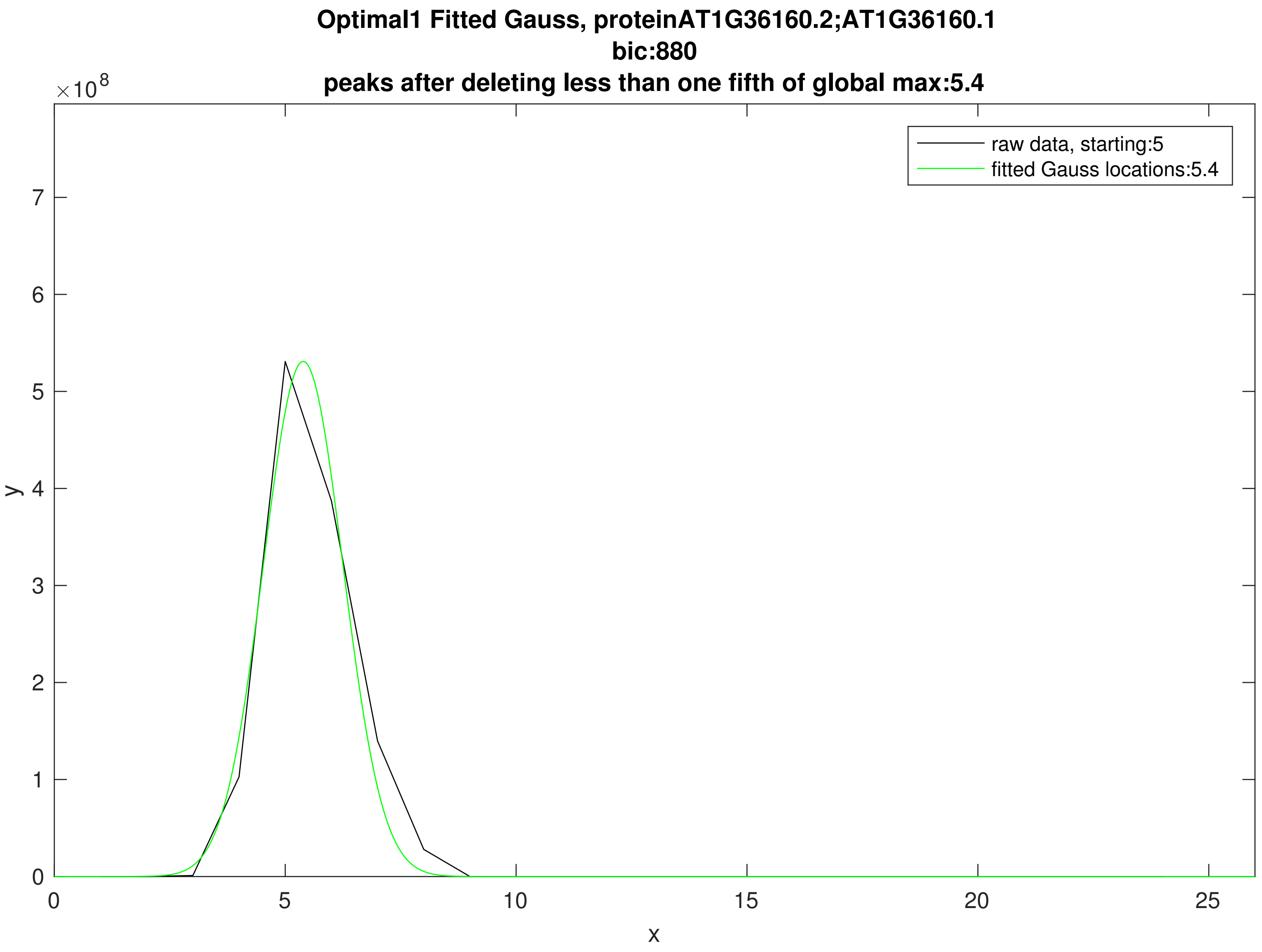

Optimal1 Fitted Gauss, proteinAT1G36160.2;AT1G36160.1
bic:880
peaks after deleting less than one fifth of global max:5.4
×10^8
raw data, starting:5
fitted Gauss locations:5.4
y
x
0
1
2
3
4
5
6
7
10
15
20
25

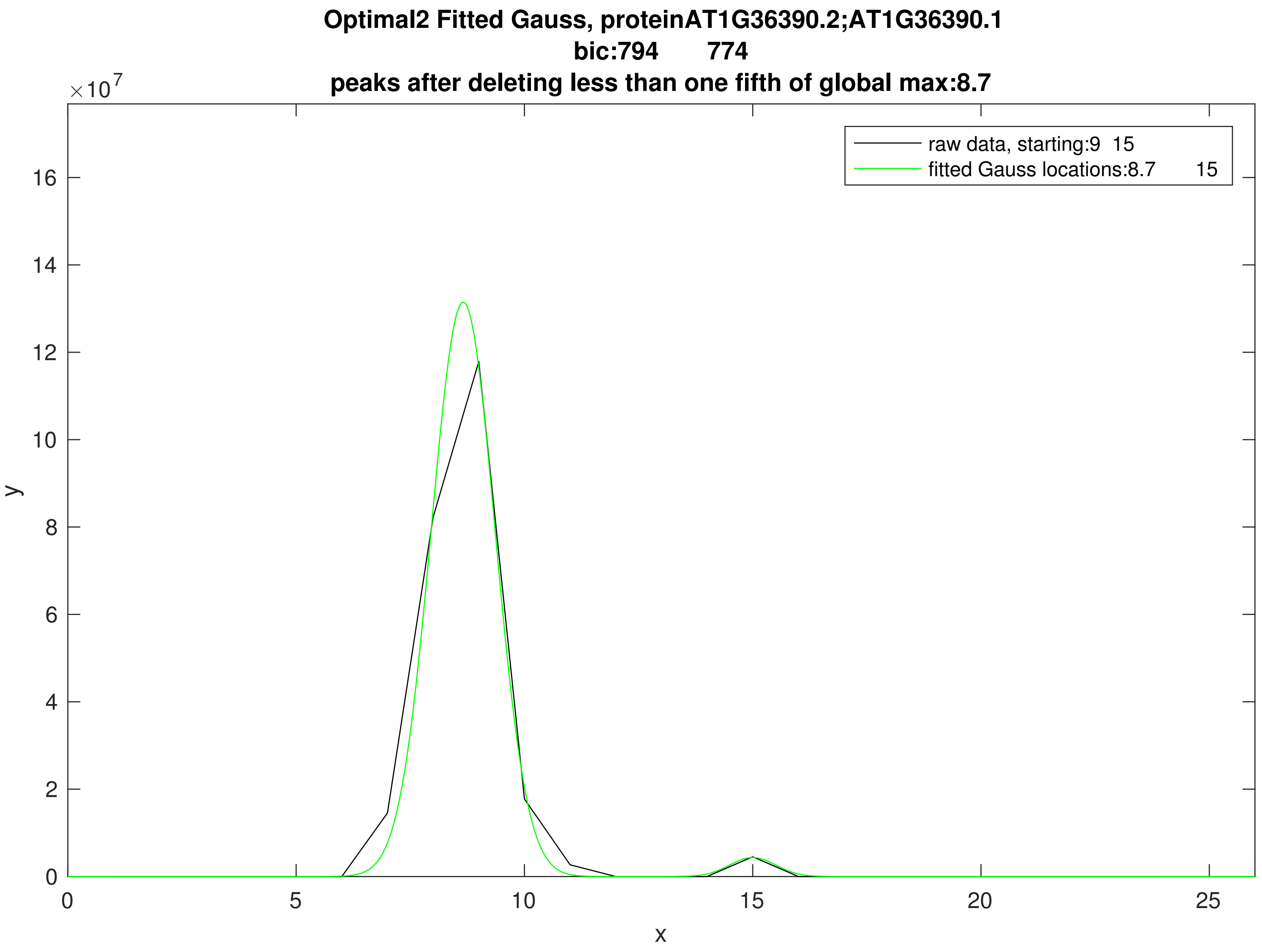
Optimal2 Fitted Gauss, proteinAT1G36390.2;AT1G36390.1
bic:794 774
peaks after deleting less than one fifth of global max:8.7
×10^7
raw data, starting:9 15
fitted Gauss locations:8.7 15
y
x
0
2
4
6
8
10
12
14
16
0
5
10
15
20
25

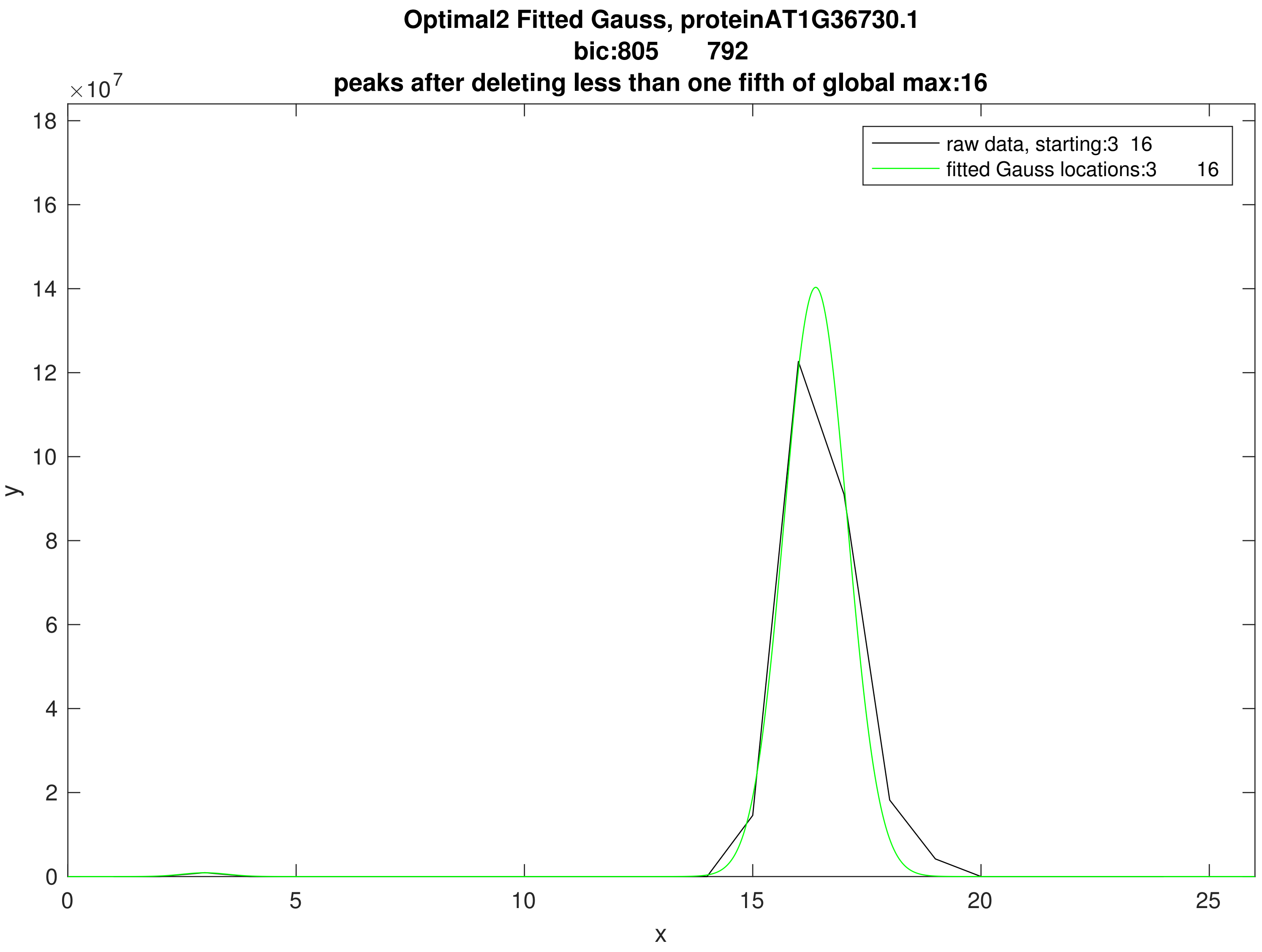

Optimal2 Fitted Gauss, proteinAT1G36730.1
bic:805 792
peaks after deleting less than one fifth of global max:16
×10^7
raw data, starting:3 16
fitted Gauss locations:3 16
y
x
0
2
4
6
8
10
12
14
16
18
5
15
20
25

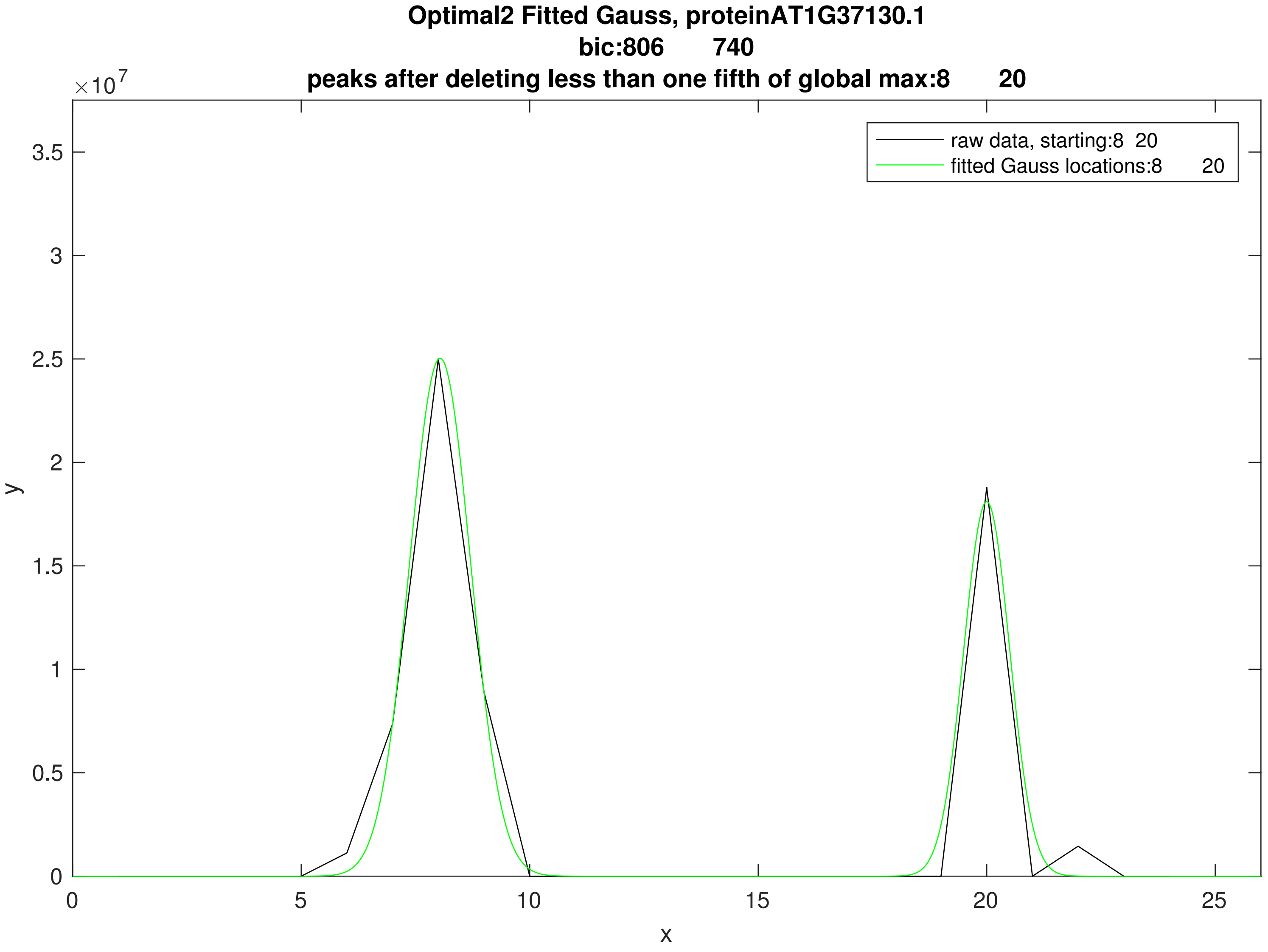
Optimal2 Fitted Gauss, proteinAT1G37130.1
bic:806 740
peaks after deleting less than one fifth of global max:8 20
×10^7
raw data, starting:8 20
fitted Gauss locations:8 20
3.5
3
2.5
2
1.5
1
0.5
0
y
0
5
10
15
20
25
x

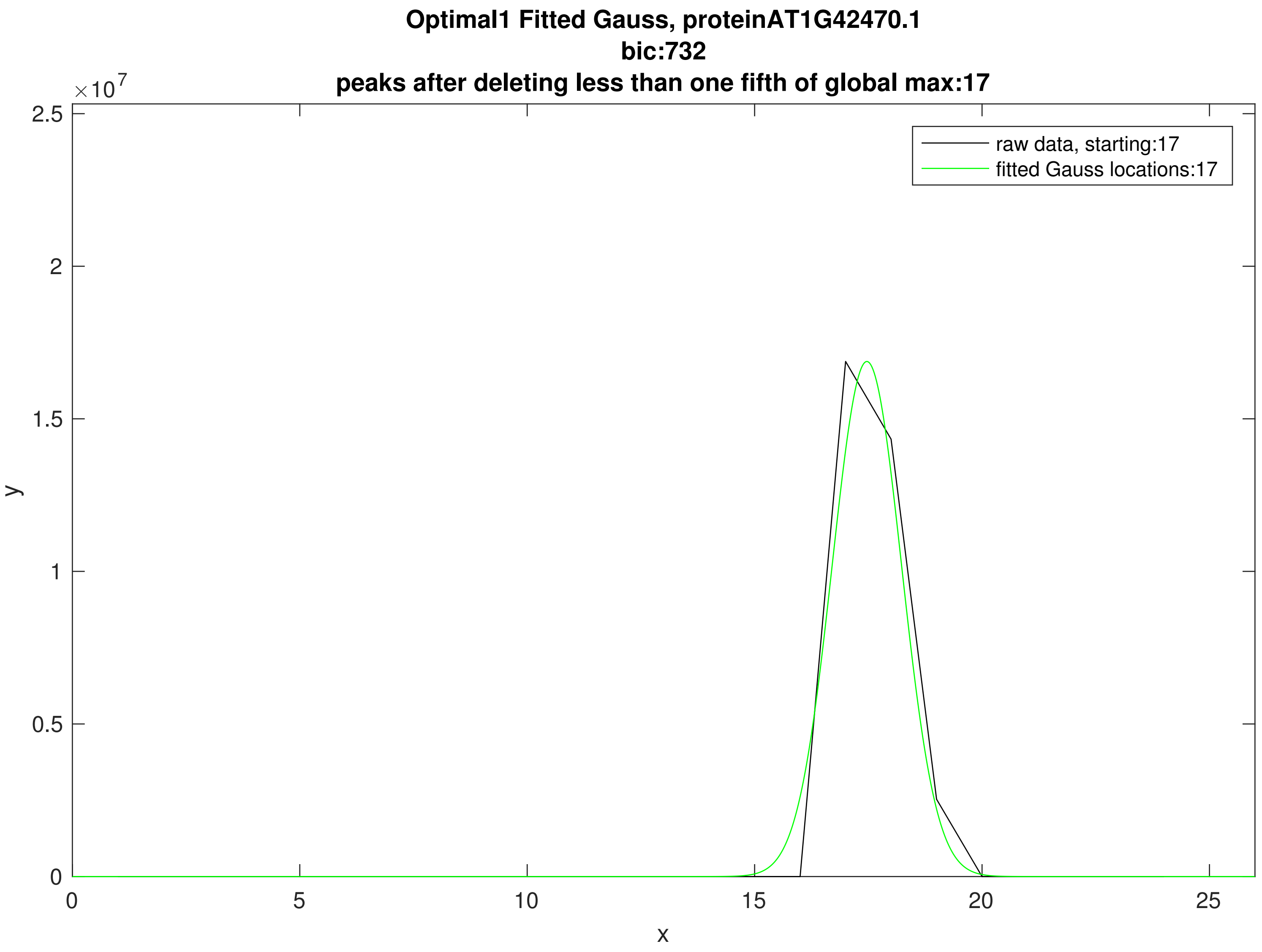
Optimal1 Fitted Gauss, proteinAT1G42470.1
bic:732
peaks after deleting less than one fifth of global max:17
×10^7
2.5
2
1.5
1
0.5
0
y
0
5
10
15
20
25
x
raw data, starting:17
fitted Gauss locations:17

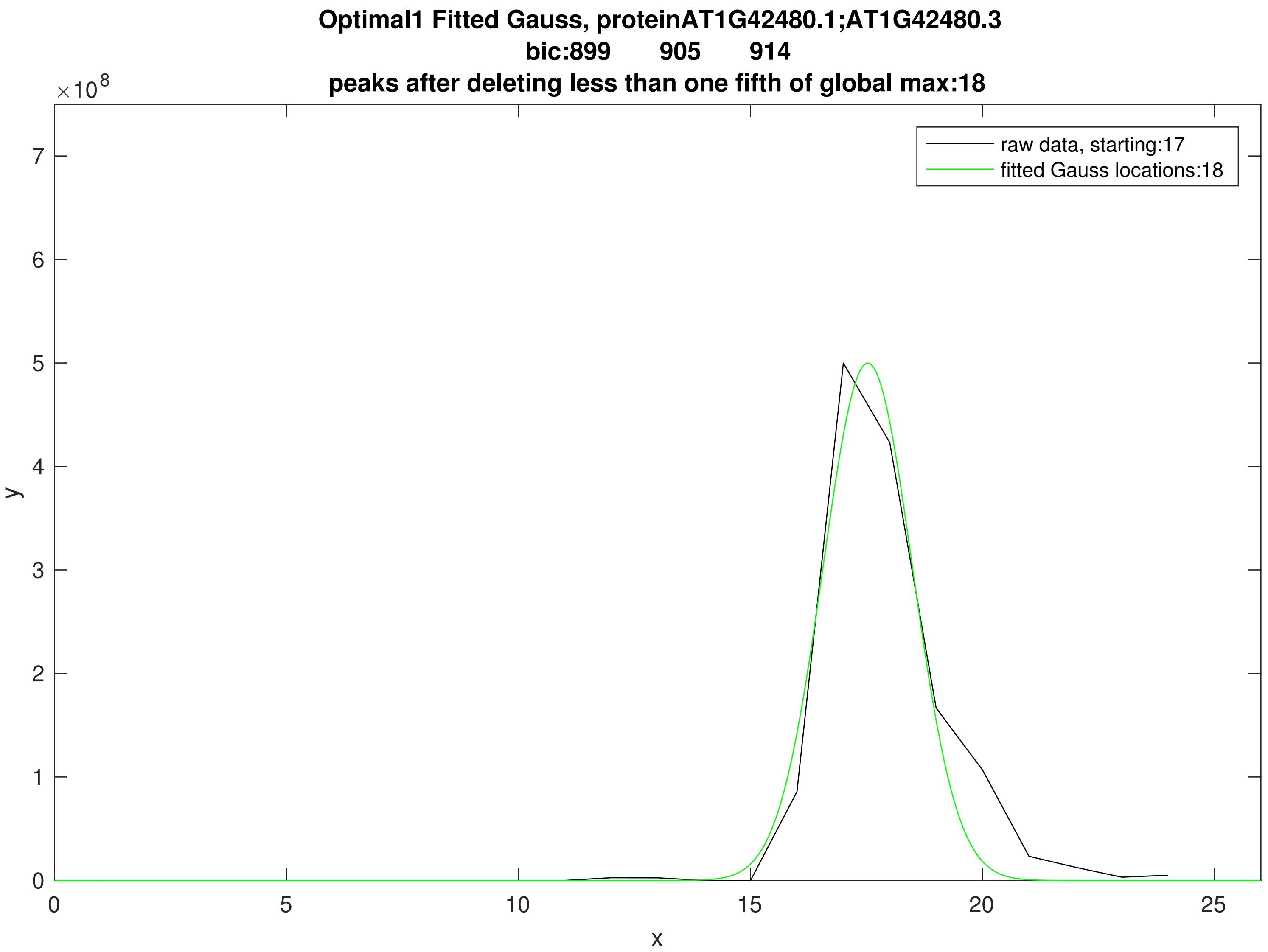

Optimal1 Fitted Gauss, proteinAT1G42480.1;AT1G42480.3
bic:899 905 914
peaks after deleting less than one fifth of global max:18
×10^8
raw data, starting:17
fitted Gauss locations:18
y
x
0
1
2
3
4
5
6
7
0
5
10
15
20
25

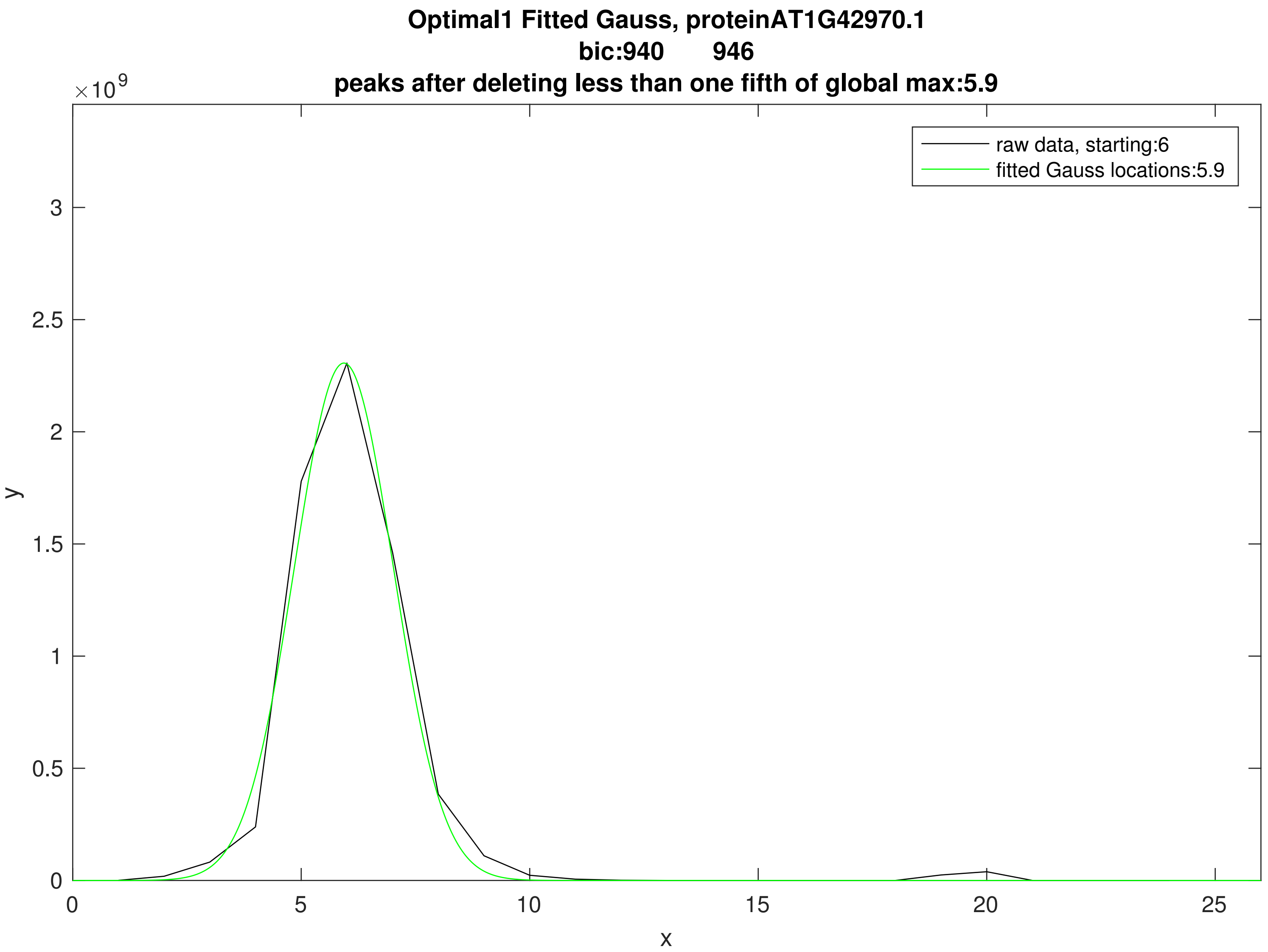

Optimal1 Fitted Gauss, proteinAT1G42970.1
bic:940 946
peaks after deleting less than one fifth of global max:5.9
×10^9
raw data, starting:6
fitted Gauss locations:5.9
3
2.5
2
1.5
1
0.5
0
y
0
5
10
15
20
25
x

l Gauss, proteinAT1G43170.9;AT1G43170.8;AT1G43170.7;AT1G43170.6;AT1G43170.5;AT1G43170.3;AT1G43170.2;AT1

bic:940

peaks after deleting less than one fifth of global max:3.5

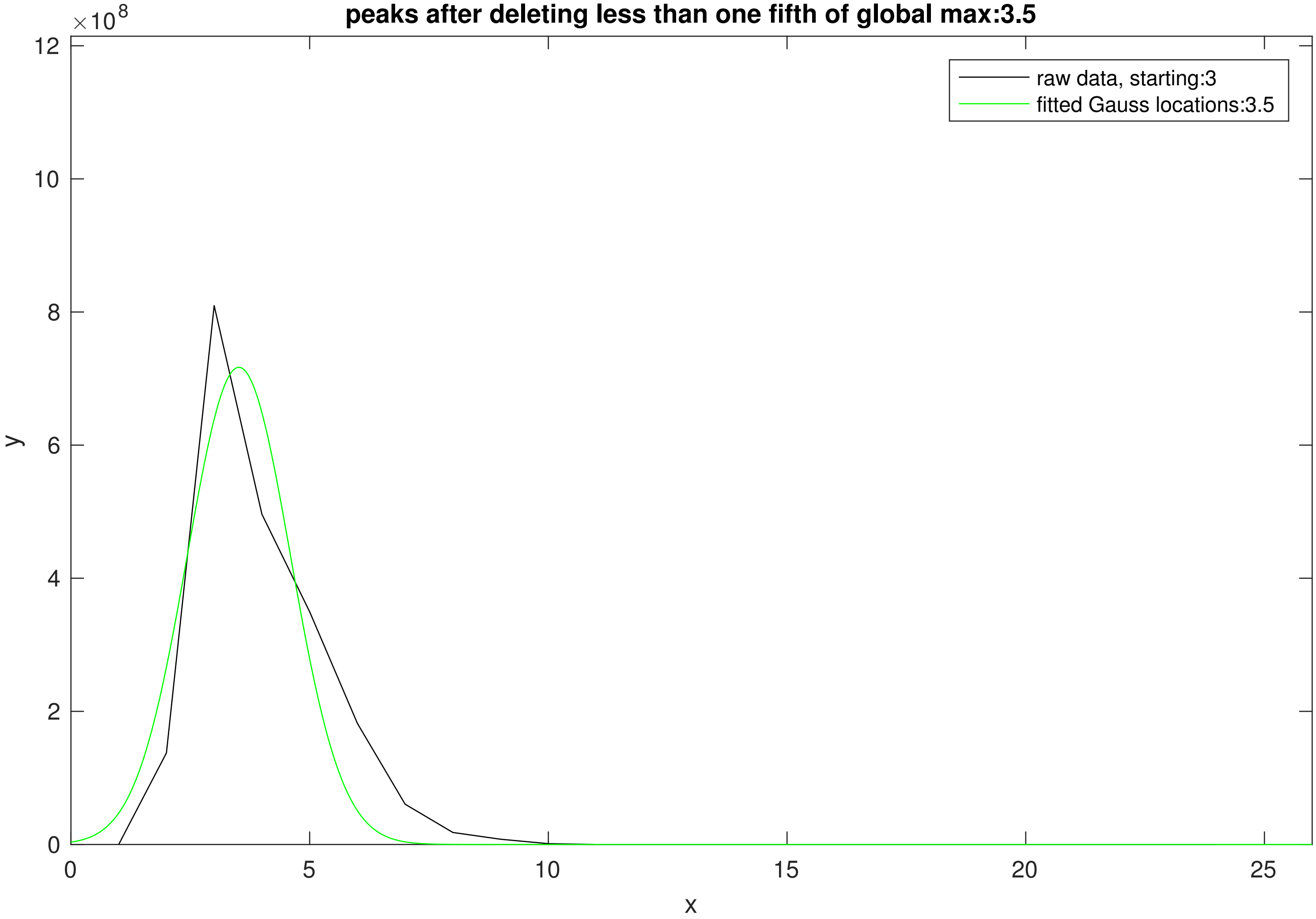

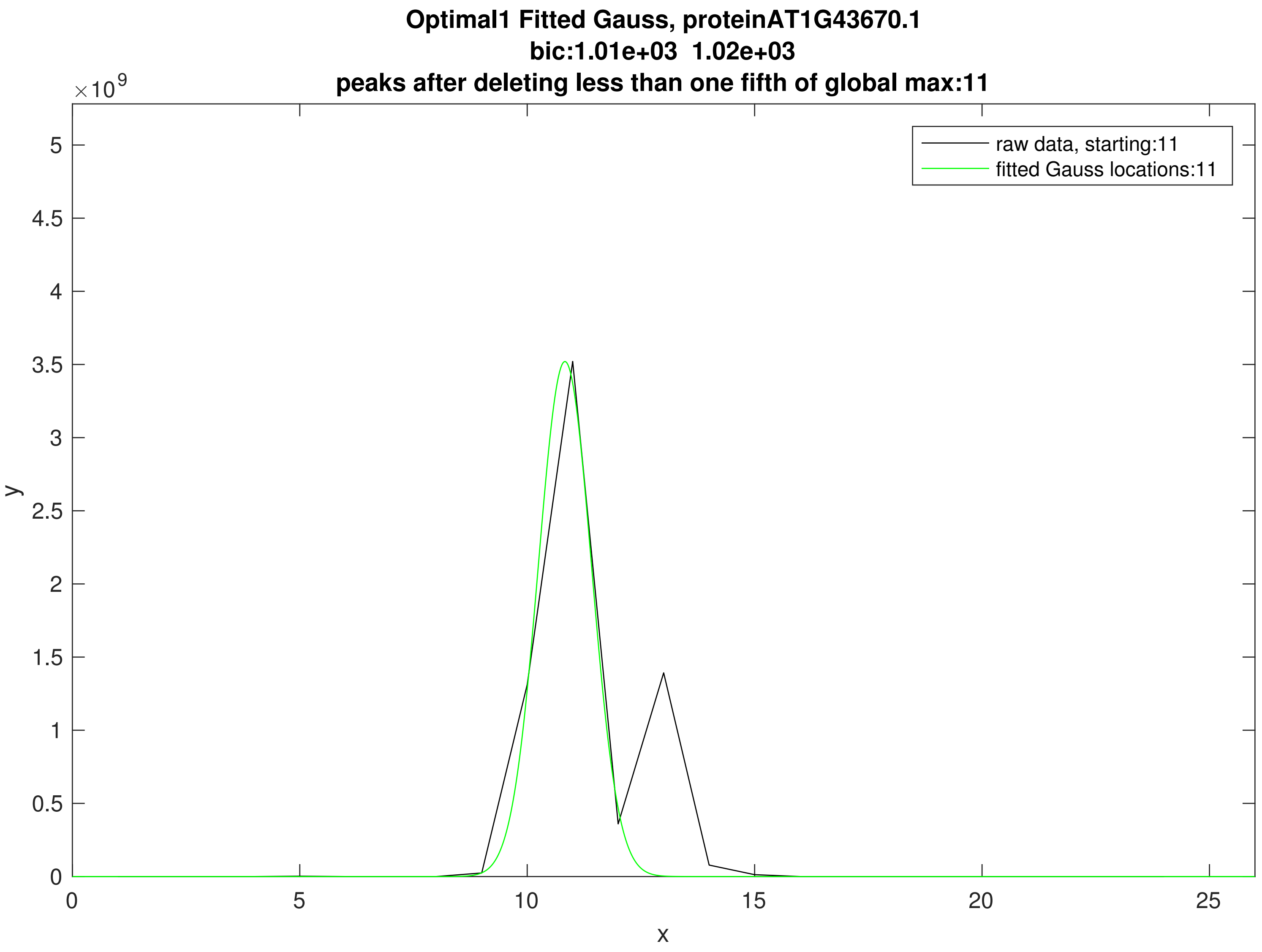
Optimal1 Fitted Gauss, proteinAT1G43670.1
bic:1.01e+03  1.02e+03
peaks after deleting less than one fifth of global max:11
×10^9
raw data, starting:11
fitted Gauss locations:11
y
x
0
0.5
1
1.5
2
2.5
3
3.5
4
4.5
5
0
5
10
15
20
25

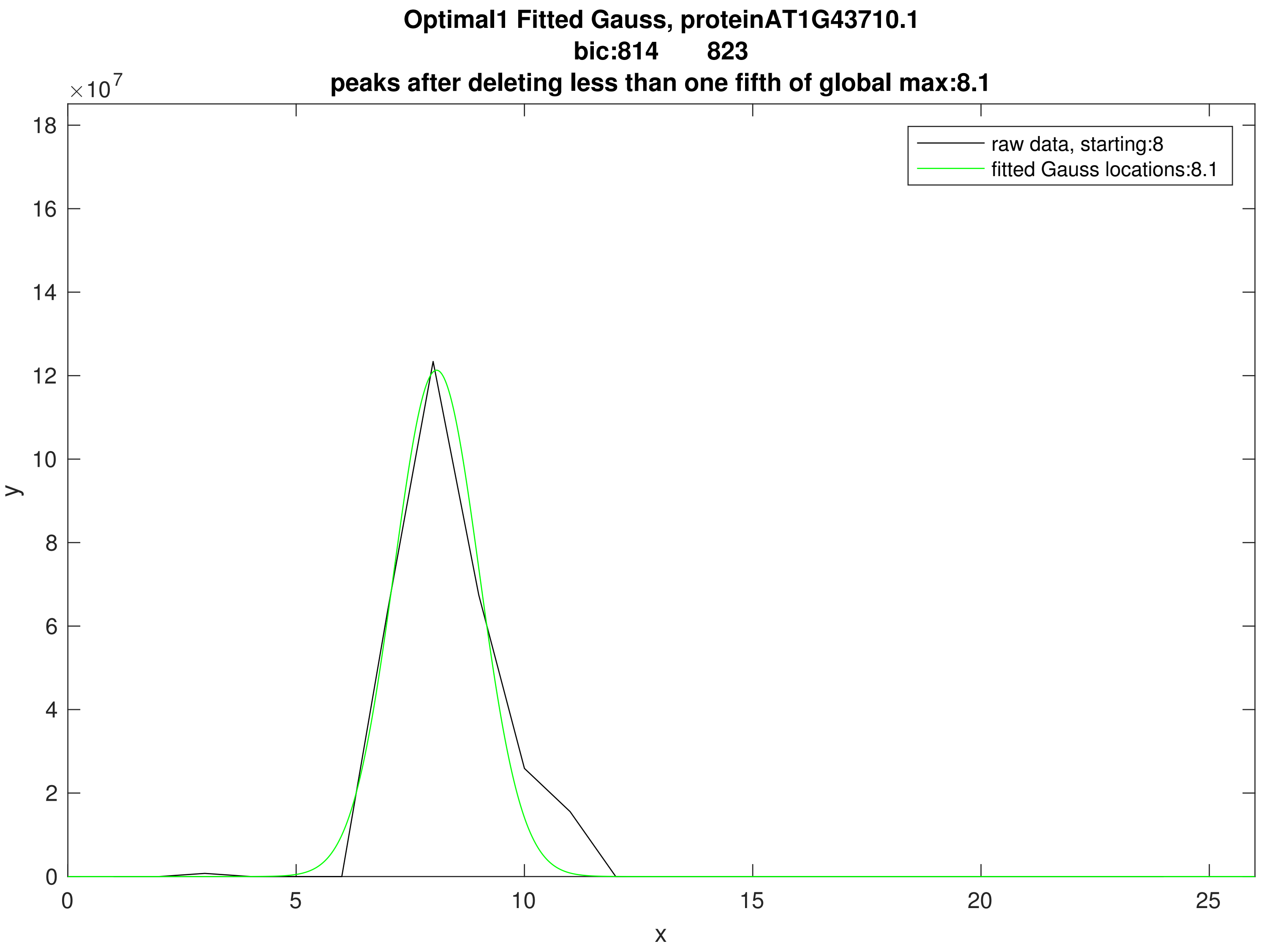
Optimal1 Fitted Gauss, proteinAT1G43710.1
bic:814 823
peaks after deleting less than one fifth of global max:8.1
×10^7
18
16
14
12
10
8
6
4
2
0
y
0
5
10
15
20
25
x
raw data, starting:8
fitted Gauss locations:8.1

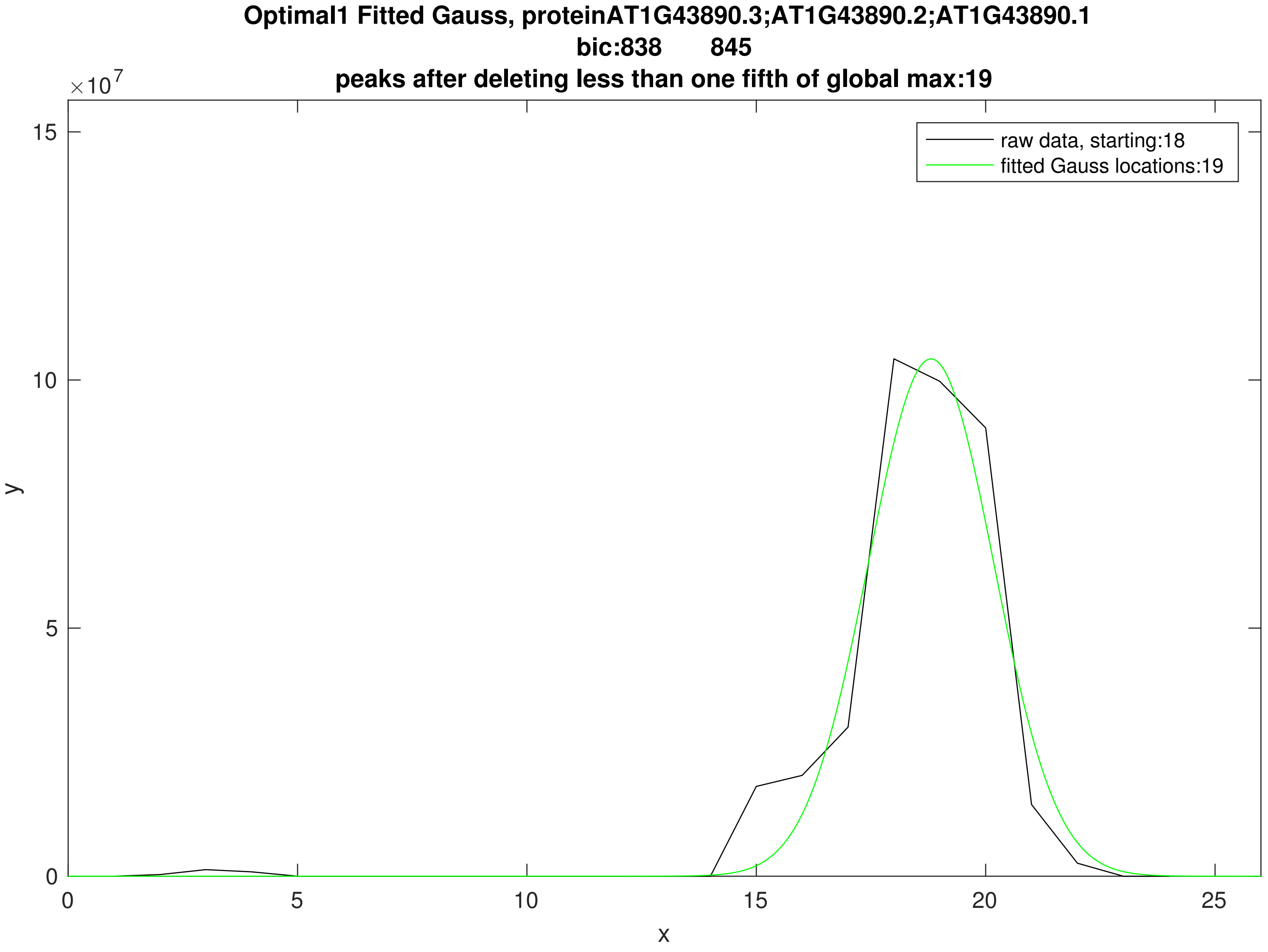

Optimal1 Fitted Gauss, proteinAT1G43890.3;AT1G43890.2;AT1G43890.1
bic:838 845
peaks after deleting less than one fifth of global max:19
×10^7
15
10
5
0
0
5
10
15
20
25
x
y
raw data, starting:18
fitted Gauss locations:19

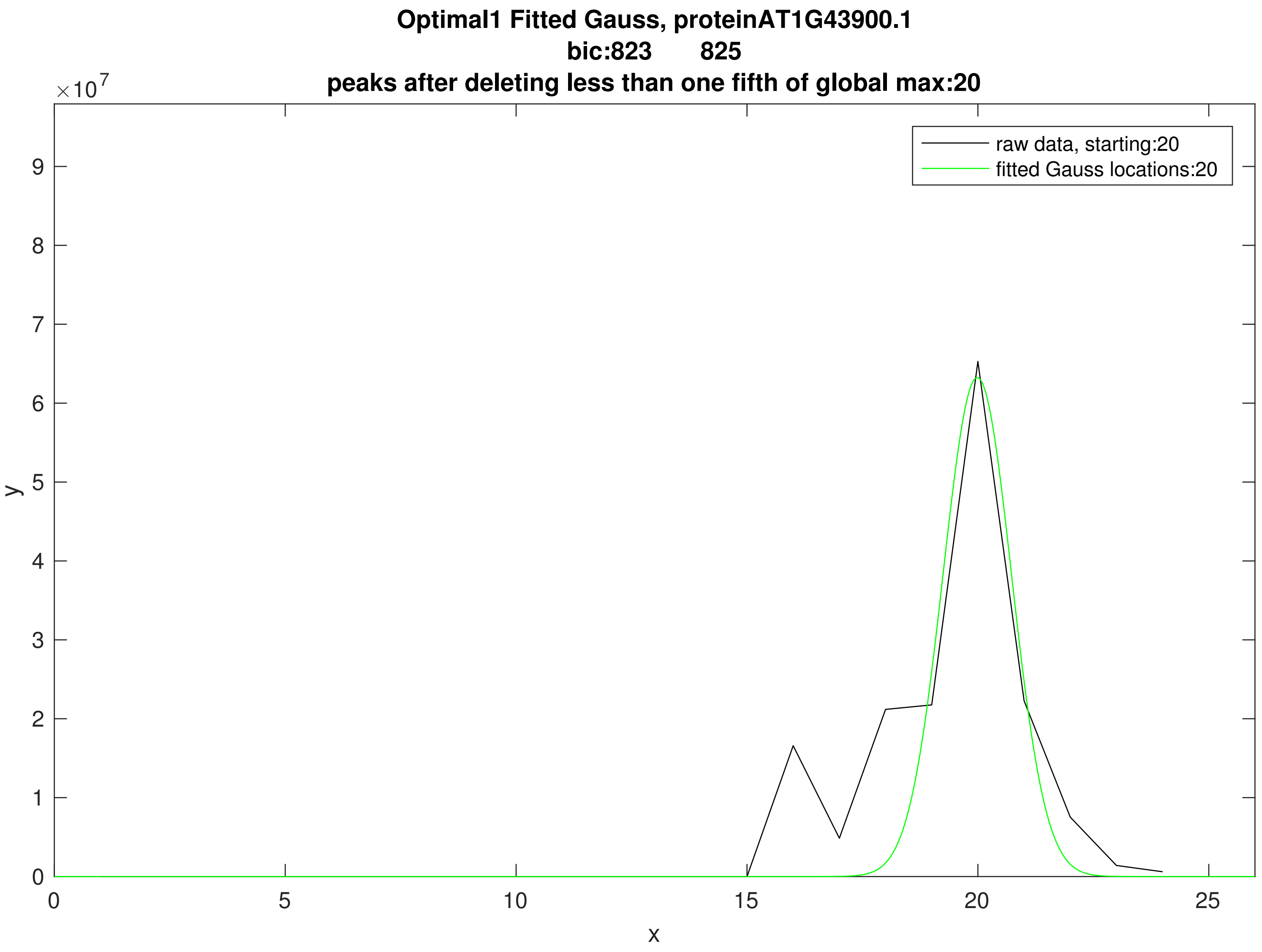

Optimal1 Fitted Gauss, proteinAT1G43900.1
bic:823 825
peaks after deleting less than one fifth of global max:20
×10^7
raw data, starting:20
fitted Gauss locations:20
y
x
0
1
2
3
4
5
6
7
8
9
0
5
10
15
20
25

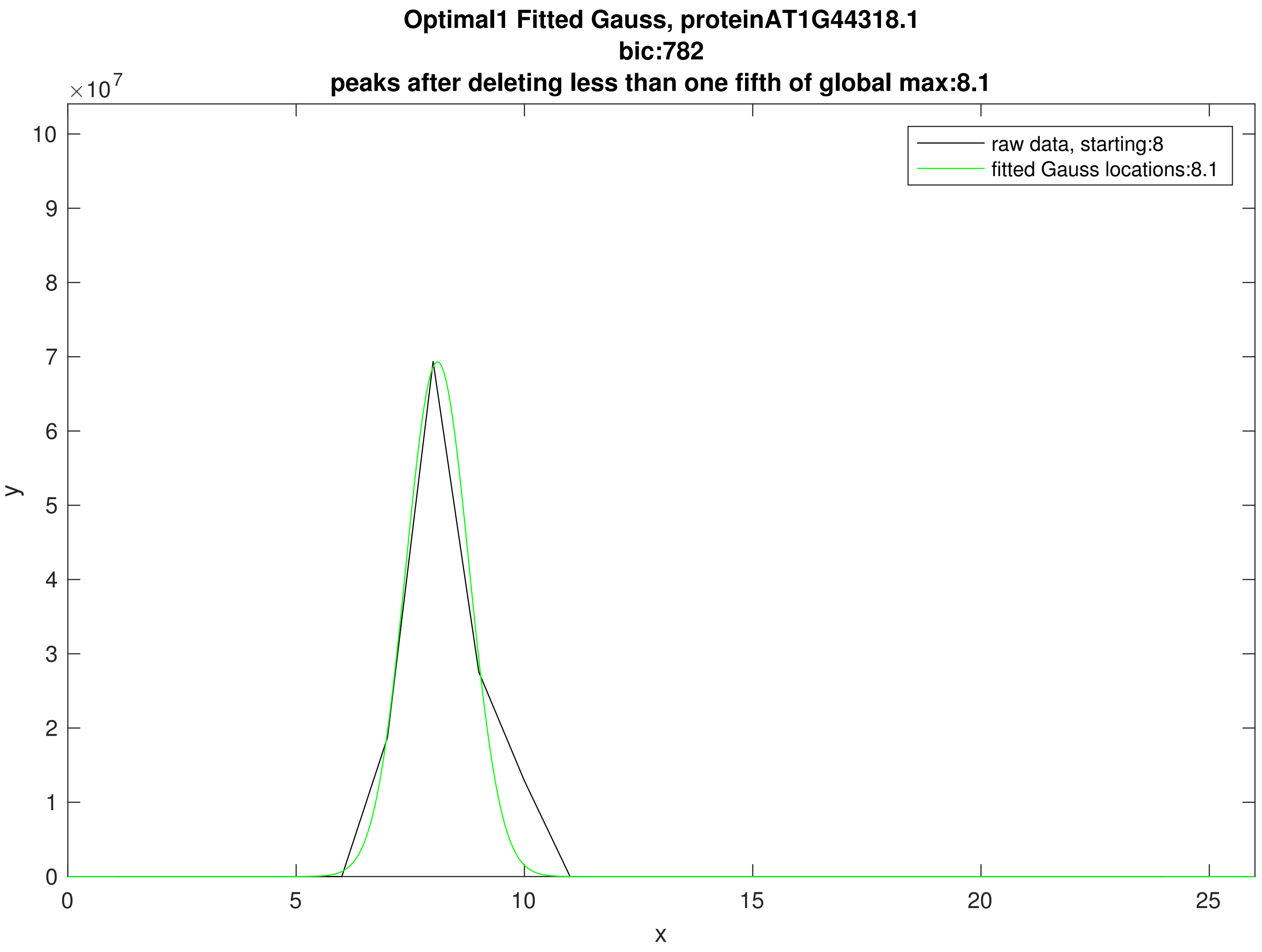
Optimal1 Fitted Gauss, proteinAT1G44318.1
bic:782
peaks after deleting less than one fifth of global max:8.1
×10^7
raw data, starting:8
fitted Gauss locations:8.1
y
x
0
1
2
3
4
5
6
7
8
9
10
0
5
10
15
20
25

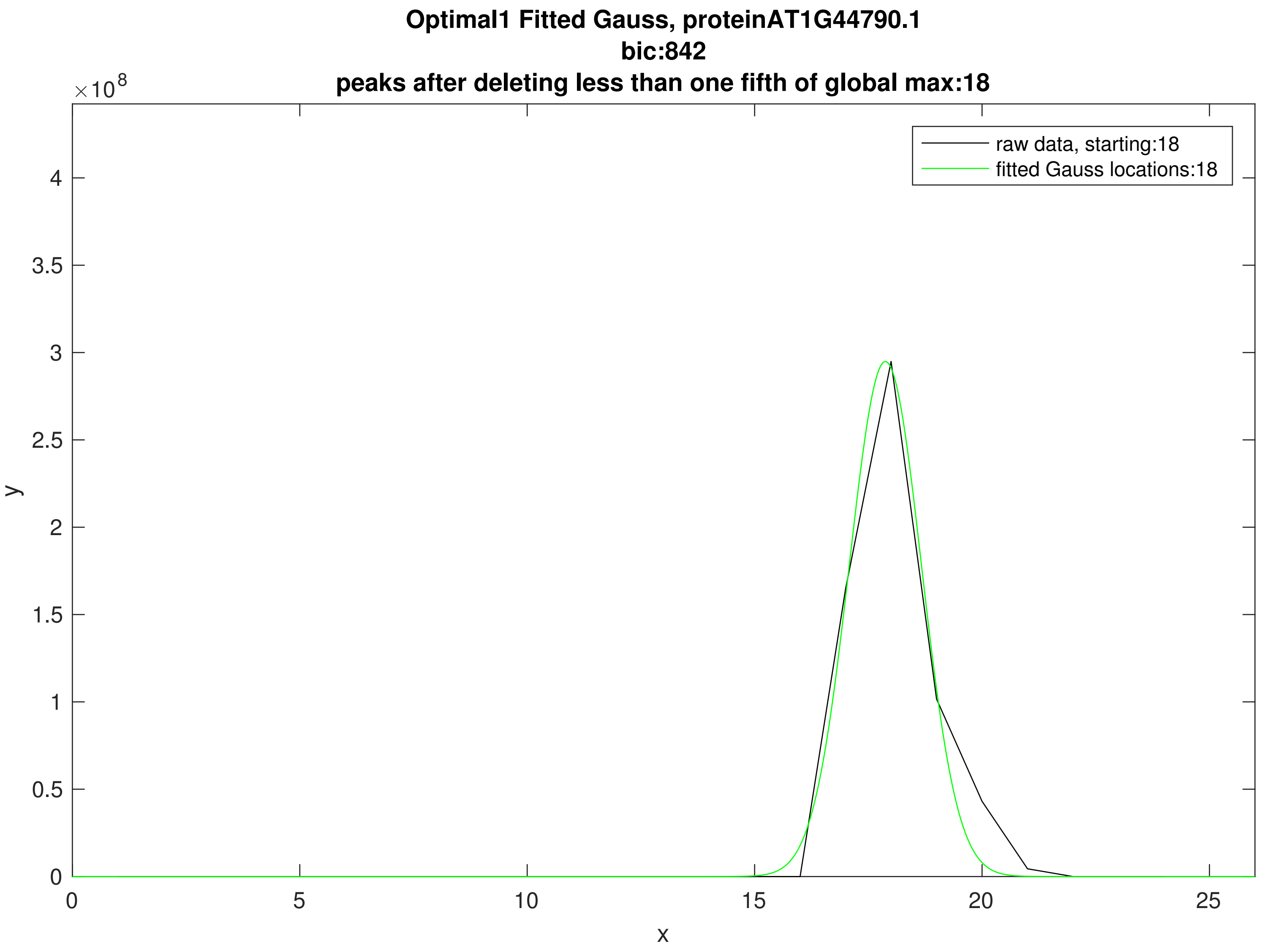
Optimal1 Fitted Gauss, proteinAT1G44790.1
bic:842
peaks after deleting less than one fifth of global max:18
×10^8
raw data, starting:18
fitted Gauss locations:18
y
x
0
0.5
1
1.5
2
2.5
3
3.5
4
0
5
10
15
20
25

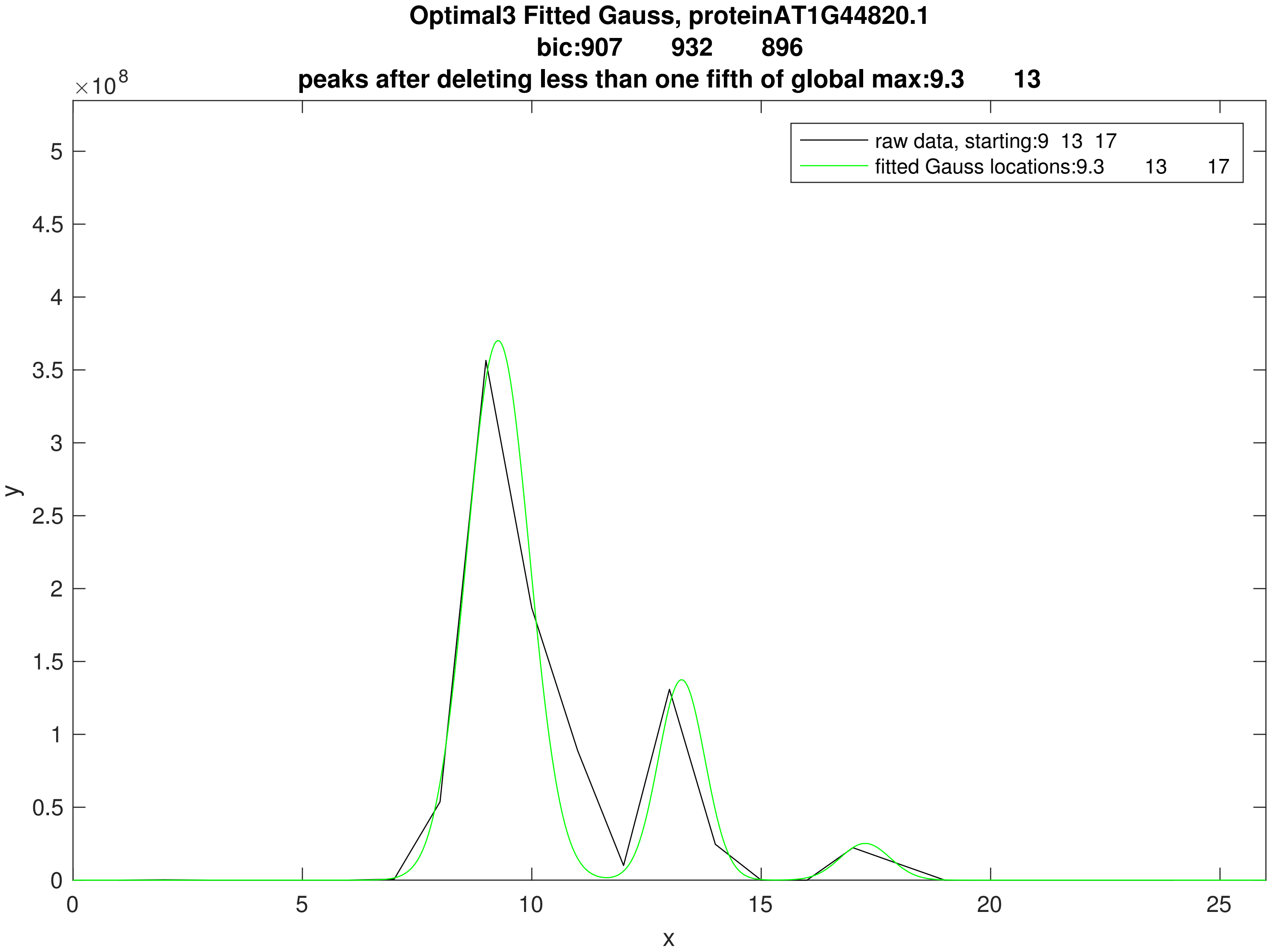
Optimal3 Fitted Gauss, proteinAT1G44820.1
bic:907 932 896
peaks after deleting less than one fifth of global max:9.3 13
$\times 10^8$
raw data, starting:9 13 17
fitted Gauss locations:9.3 13 17
y
x
0
0.5
1
1.5
2
2.5
3
3.5
4
4.5
5
0
5
10
15
20
25

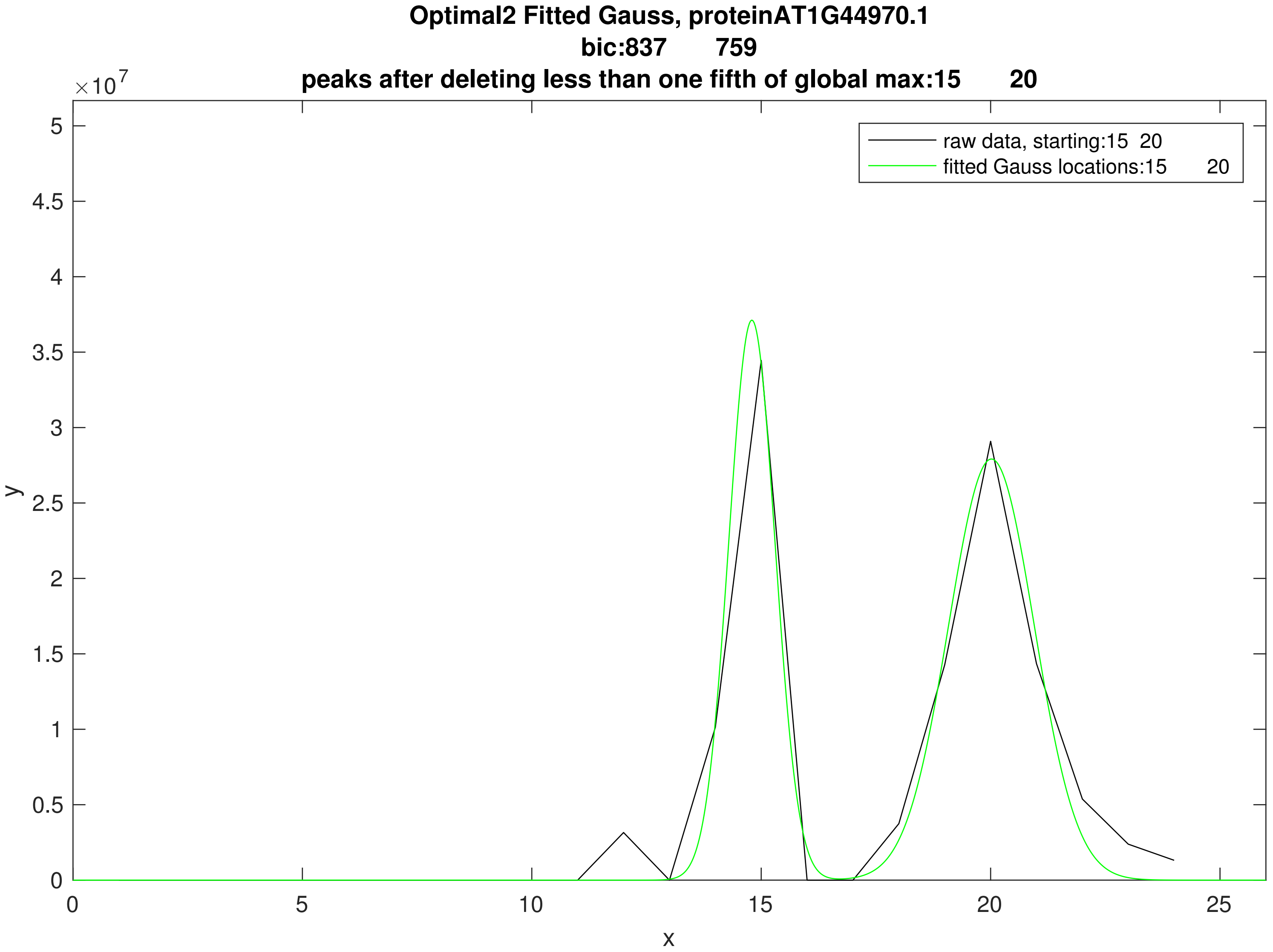

Optimal2 Fitted Gauss, proteinAT1G44970.1
bic:837 759
peaks after deleting less than one fifth of global max:15 20
×10^7
raw data, starting:15 20
fitted Gauss locations:15 20
y
x
0
0.5
1
1.5
2
2.5
3
3.5
4
4.5
5
0
5
10
15
20
25

**Optimal2 Fitted Gauss, proteinAT1G45130.1**
**bic:888 888 897**
**peaks after deleting less than one fifth of global max:17**

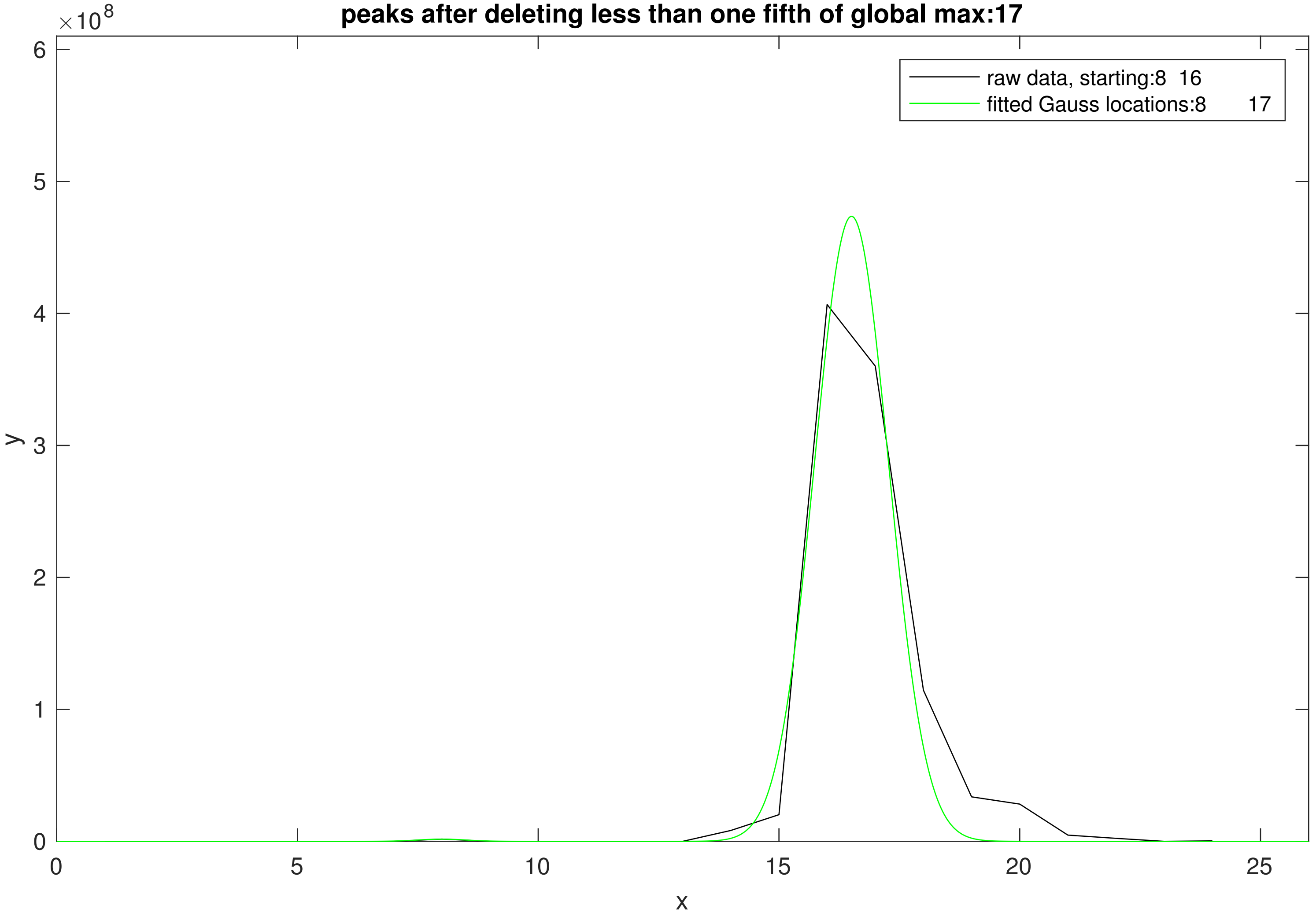

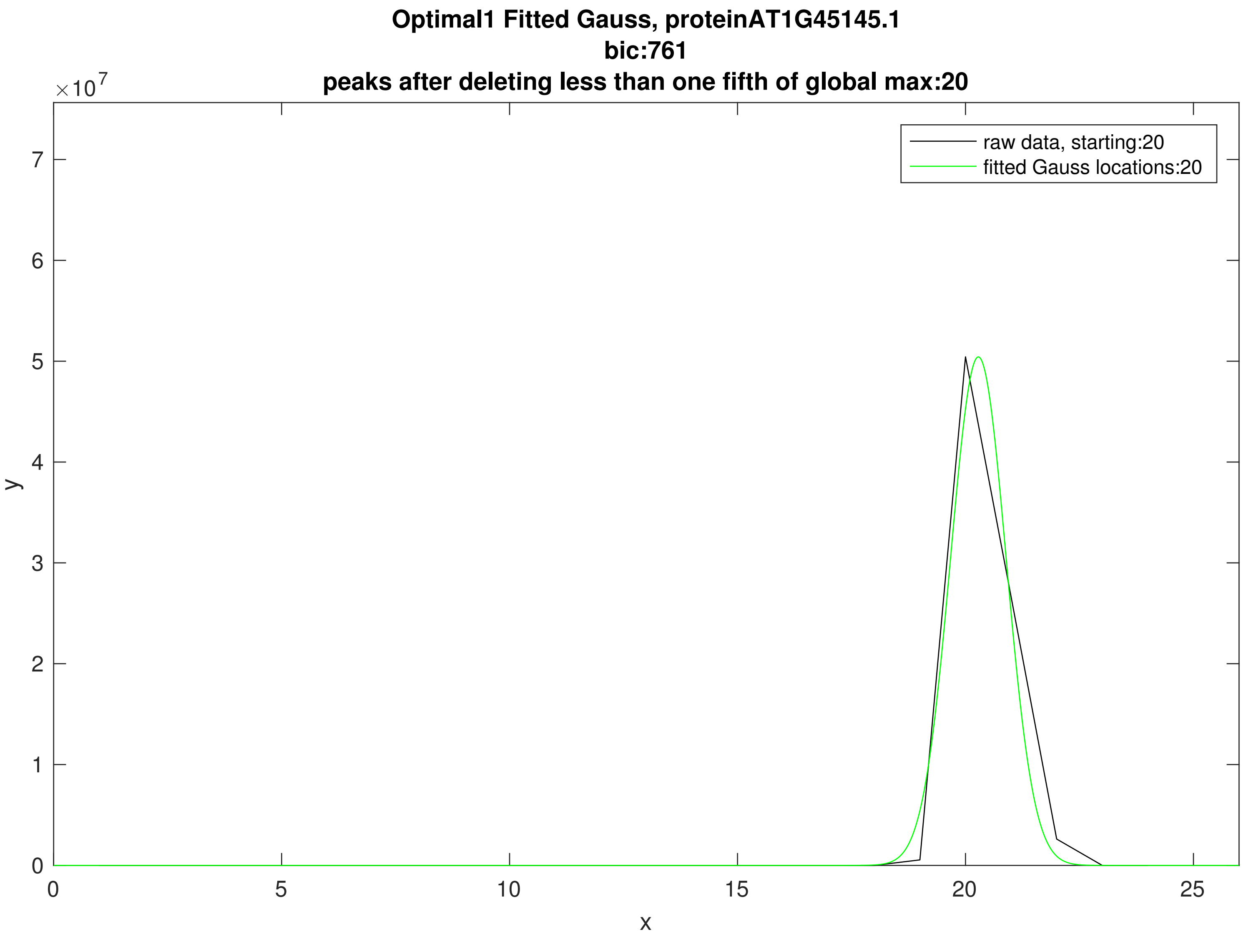

Optimal1 Fitted Gauss, proteinAT1G45145.1
bic:761
peaks after deleting less than one fifth of global max:20
×10^7
raw data, starting:20
fitted Gauss locations:20
y
x
0
1
2
3
4
5
6
7
0
5
10
15
20
25

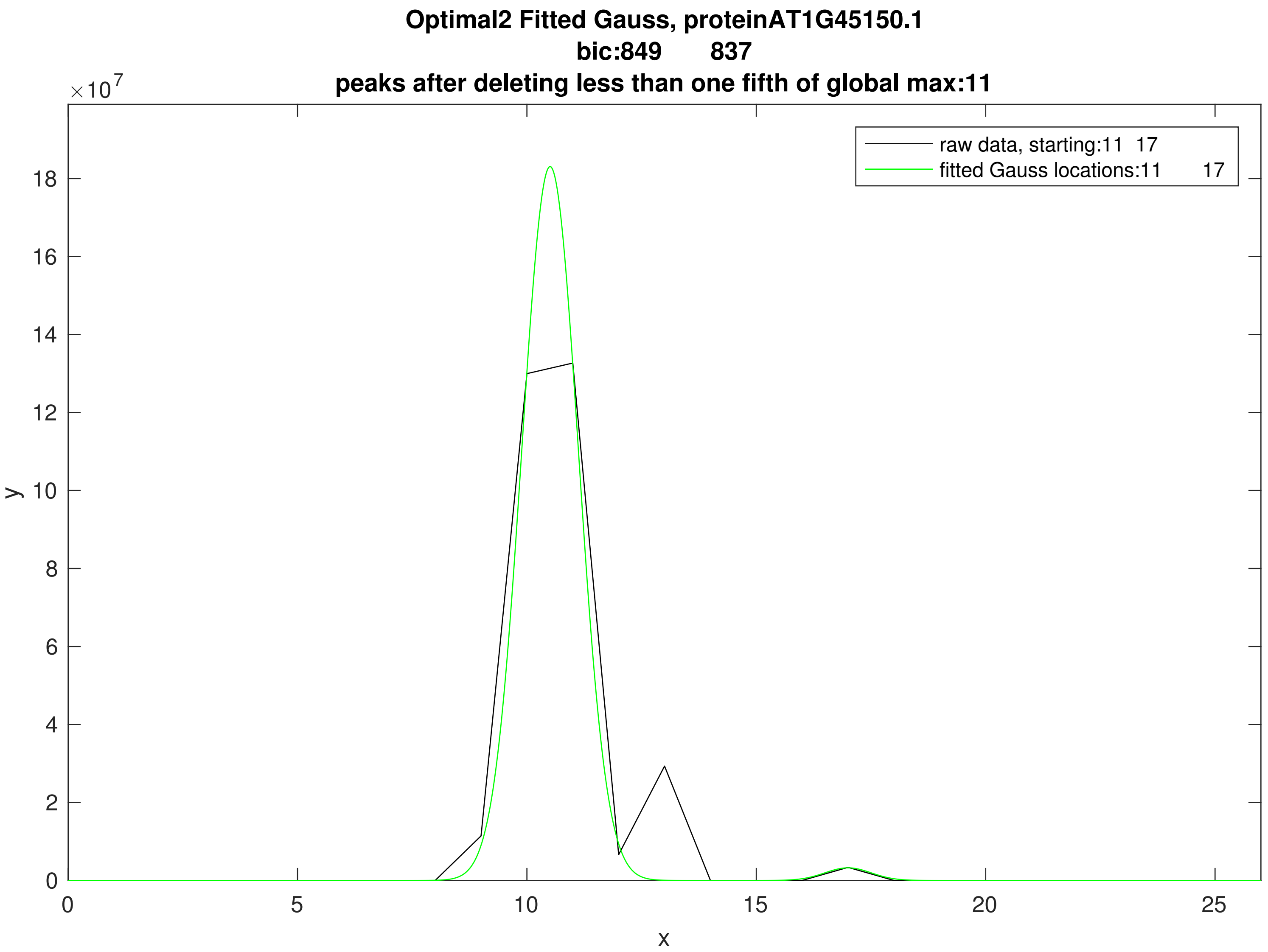
Optimal2 Fitted Gauss, proteinAT1G45150.1
bic:849 837
peaks after deleting less than one fifth of global max:11
×10^7
raw data, starting:11 17
fitted Gauss locations:11 17
y
x
0
2
4
6
8
10
12
14
16
18
0
5
10
15
20
25

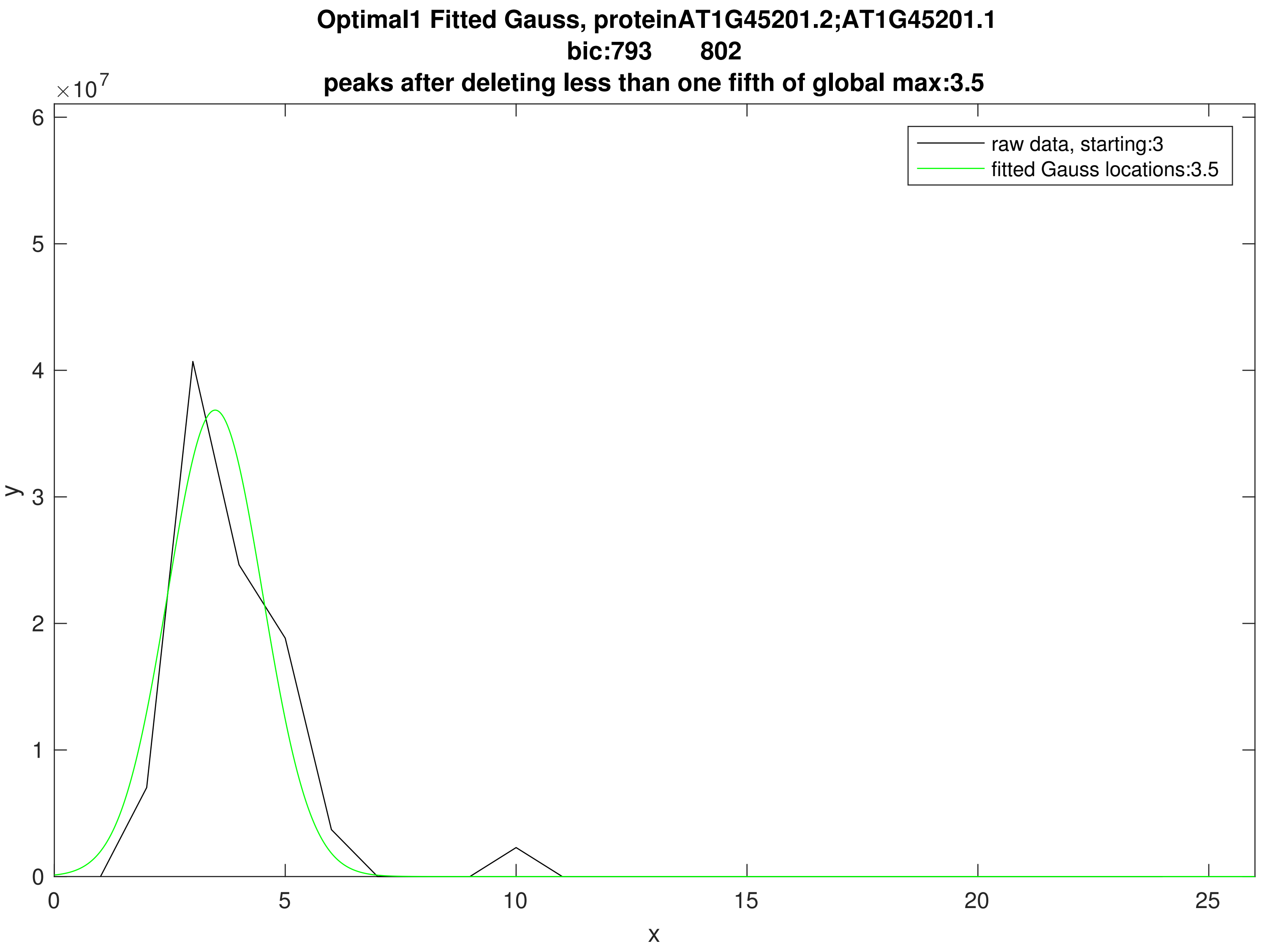
Optimal1 Fitted Gauss, proteinAT1G45201.2;AT1G45201.1
bic:793 802
peaks after deleting less than one fifth of global max:3.5
×10^7
raw data, starting:3
fitted Gauss locations:3.5
y
x
0
1
2
3
4
5
6
0
5
10
15
20
25

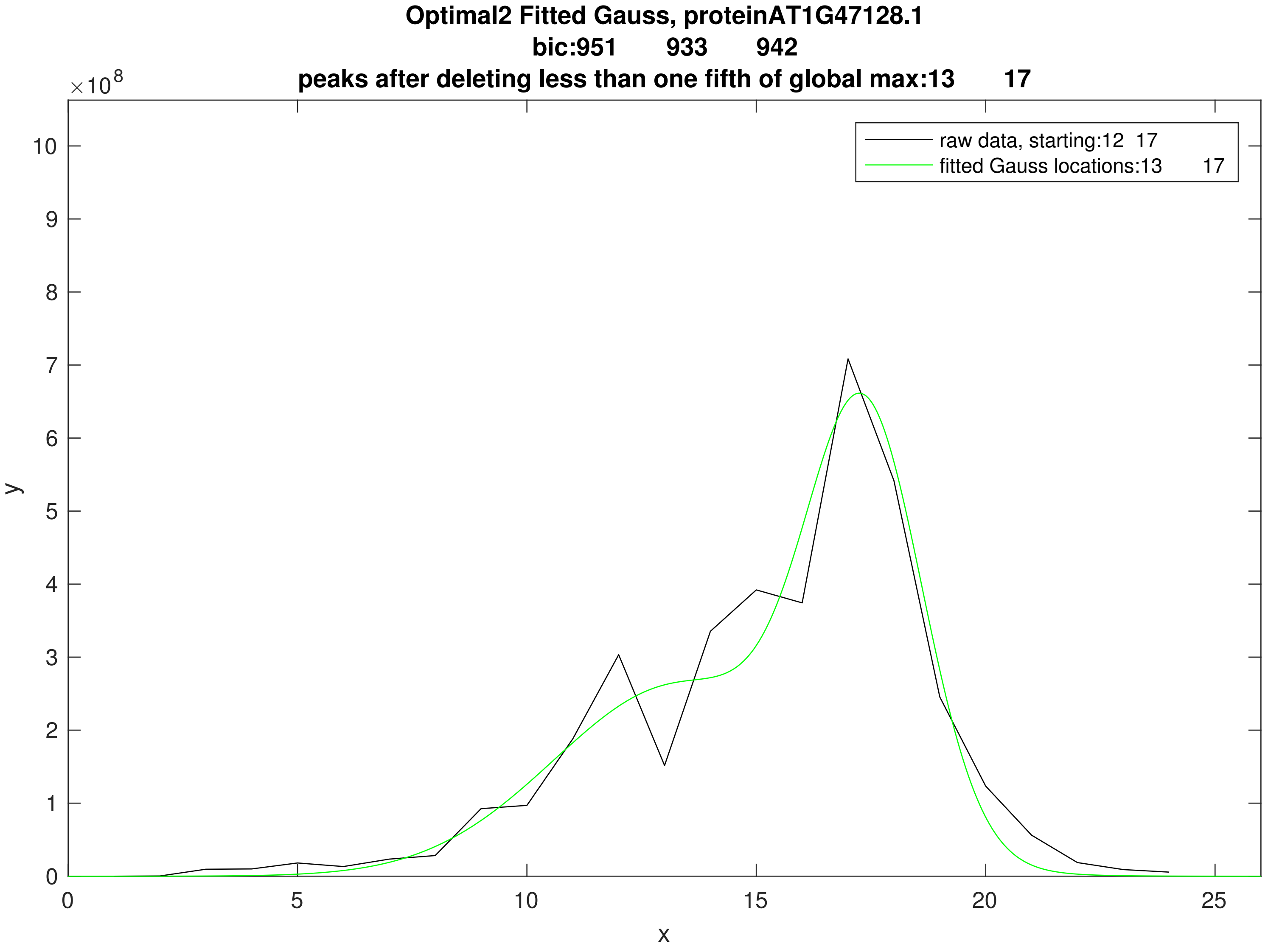
Optimal2 Fitted Gauss, proteinAT1G47128.1
bic:951 933 942
peaks after deleting less than one fifth of global max:13 17
×10^8
raw data, starting:12 17
fitted Gauss locations:13 17
y
x
0
1
2
3
4
5
6
7
8
9
10
0
5
10
15
20
25

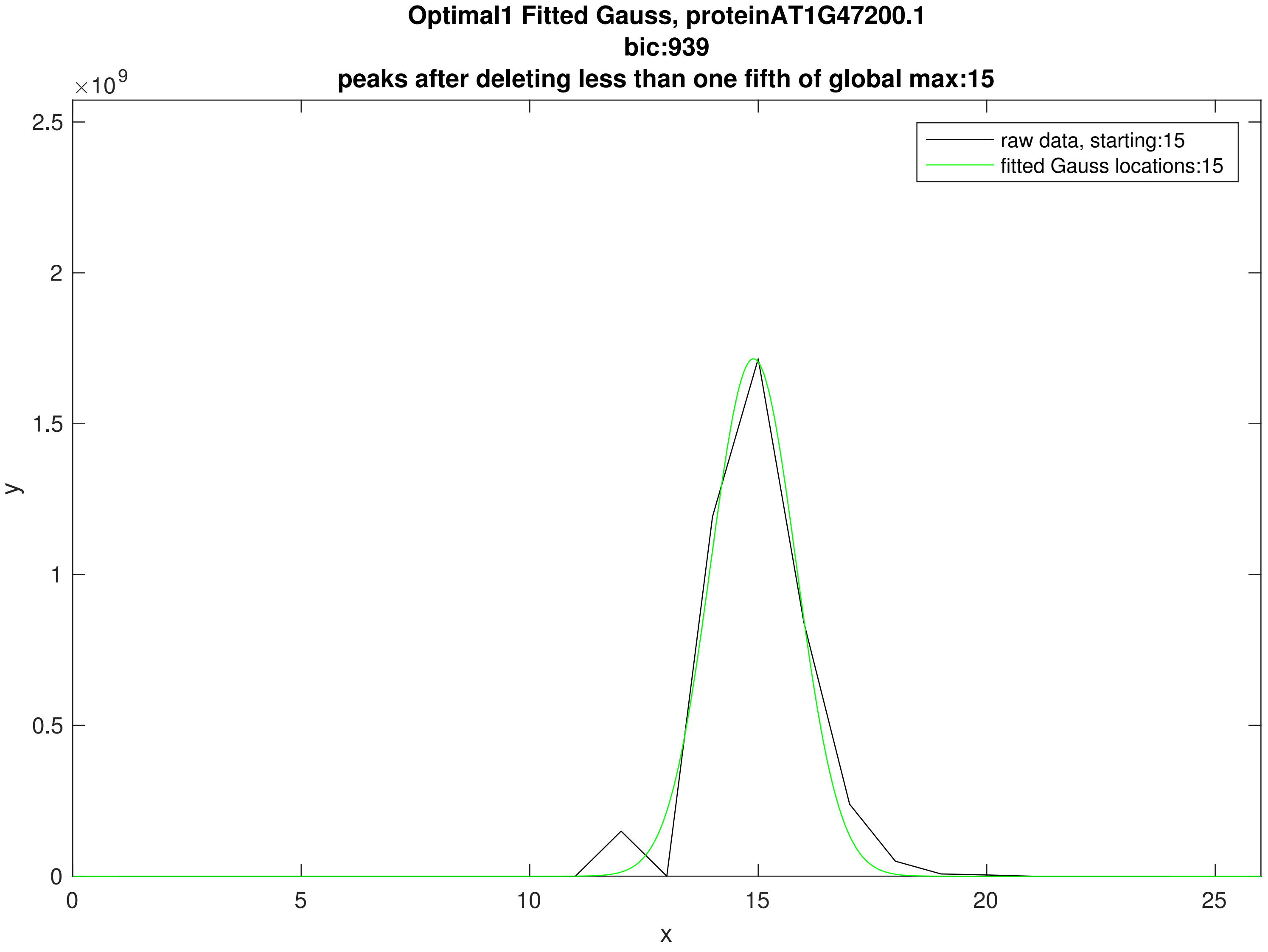

Optimal1 Fitted Gauss, proteinAT1G47200.1
bic:939
peaks after deleting less than one fifth of global max:15
×10^9
2.5
2
1.5
1
0.5
0
y
0
5
10
15
20
25
x
raw data, starting:15
fitted Gauss locations:15

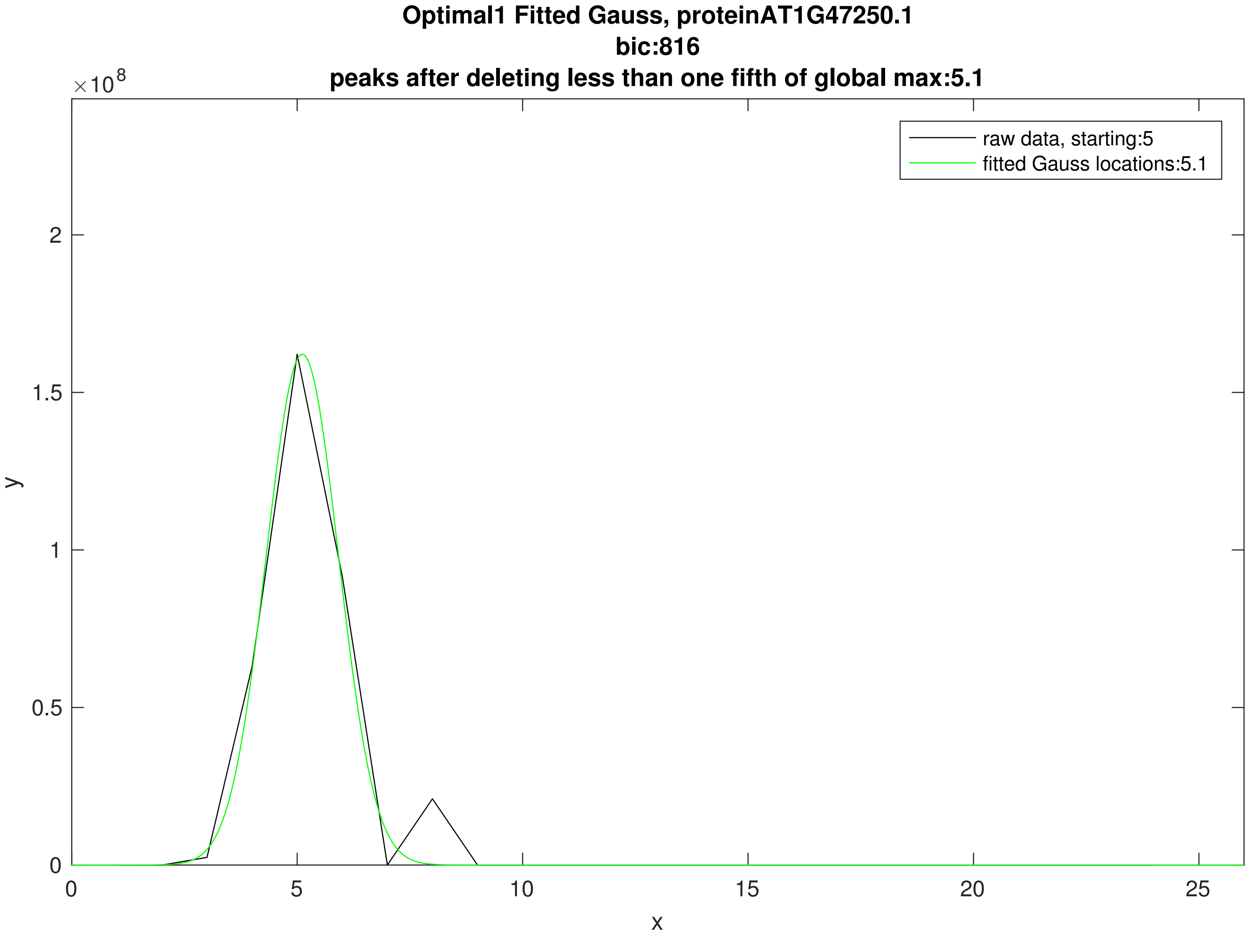

Optimal1 Fitted Gauss, proteinAT1G47250.1
bic:816
peaks after deleting less than one fifth of global max:5.1
×10^8
raw data, starting:5
fitted Gauss locations:5.1
2
1.5
1
0.5
0
y
0
5
10
15
20
25
x

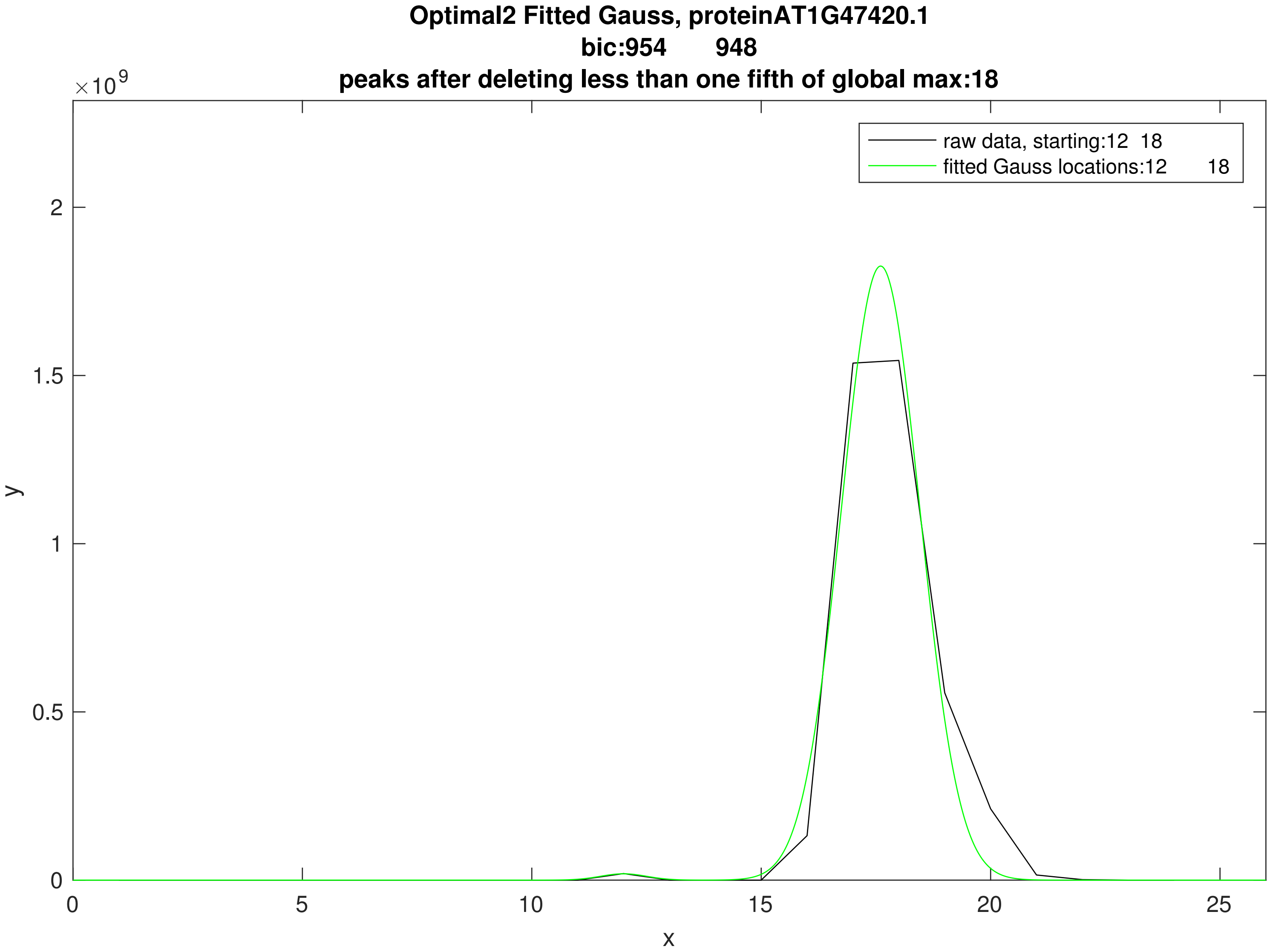
Optimal2 Fitted Gauss, proteinAT1G47420.1
bic:954 948
peaks after deleting less than one fifth of global max:18
×10^9
raw data, starting:12 18
fitted Gauss locations:12 18
y
x
0
0.5
1
1.5
2
5
10
15
20
25

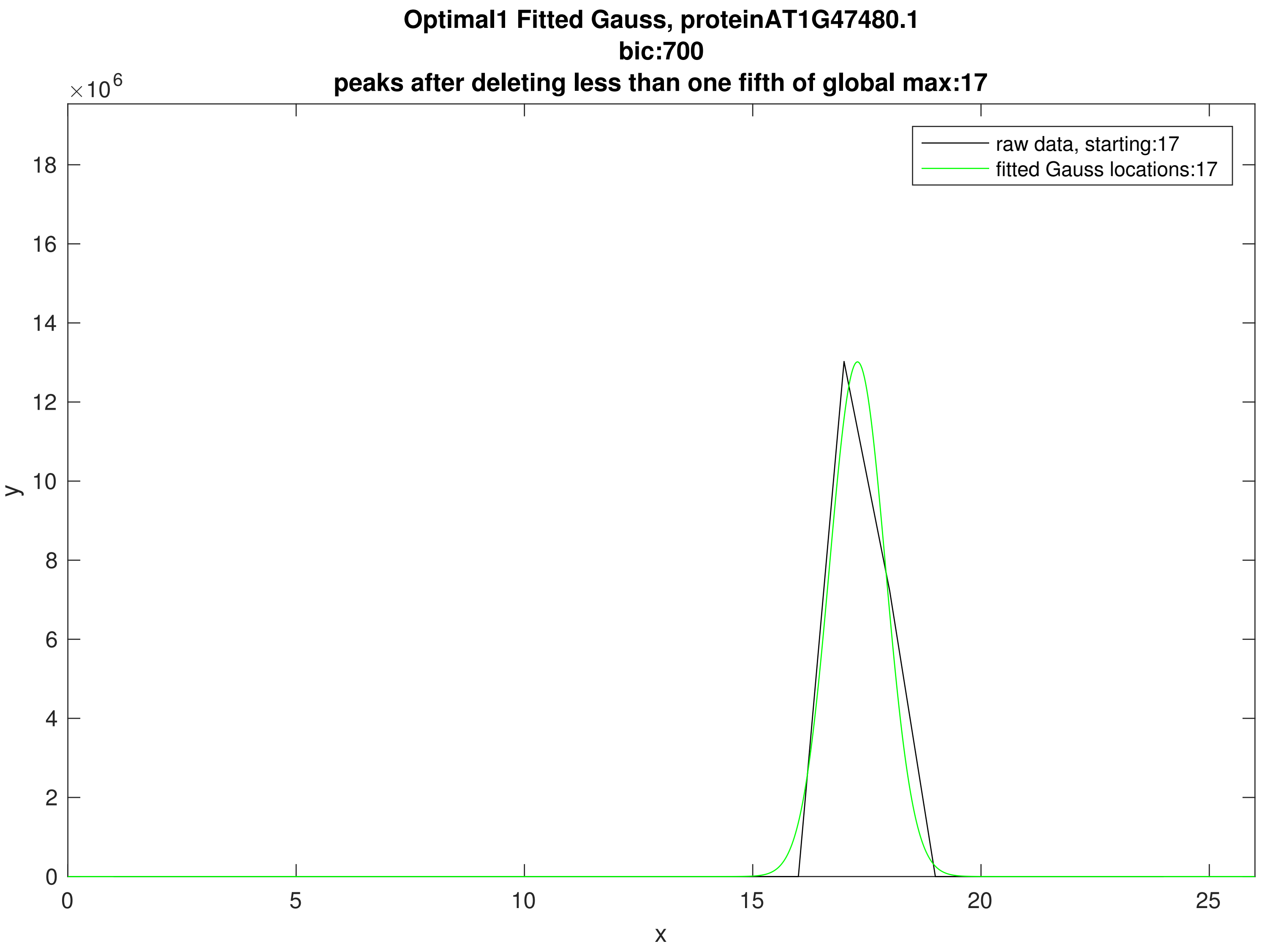
Optimal1 Fitted Gauss, proteinAT1G47480.1
bic:700
peaks after deleting less than one fifth of global max:17
×10^6
raw data, starting:17
fitted Gauss locations:17
y
x
0
2
4
6
8
10
12
14
16
18
0
5
10
15
20
25

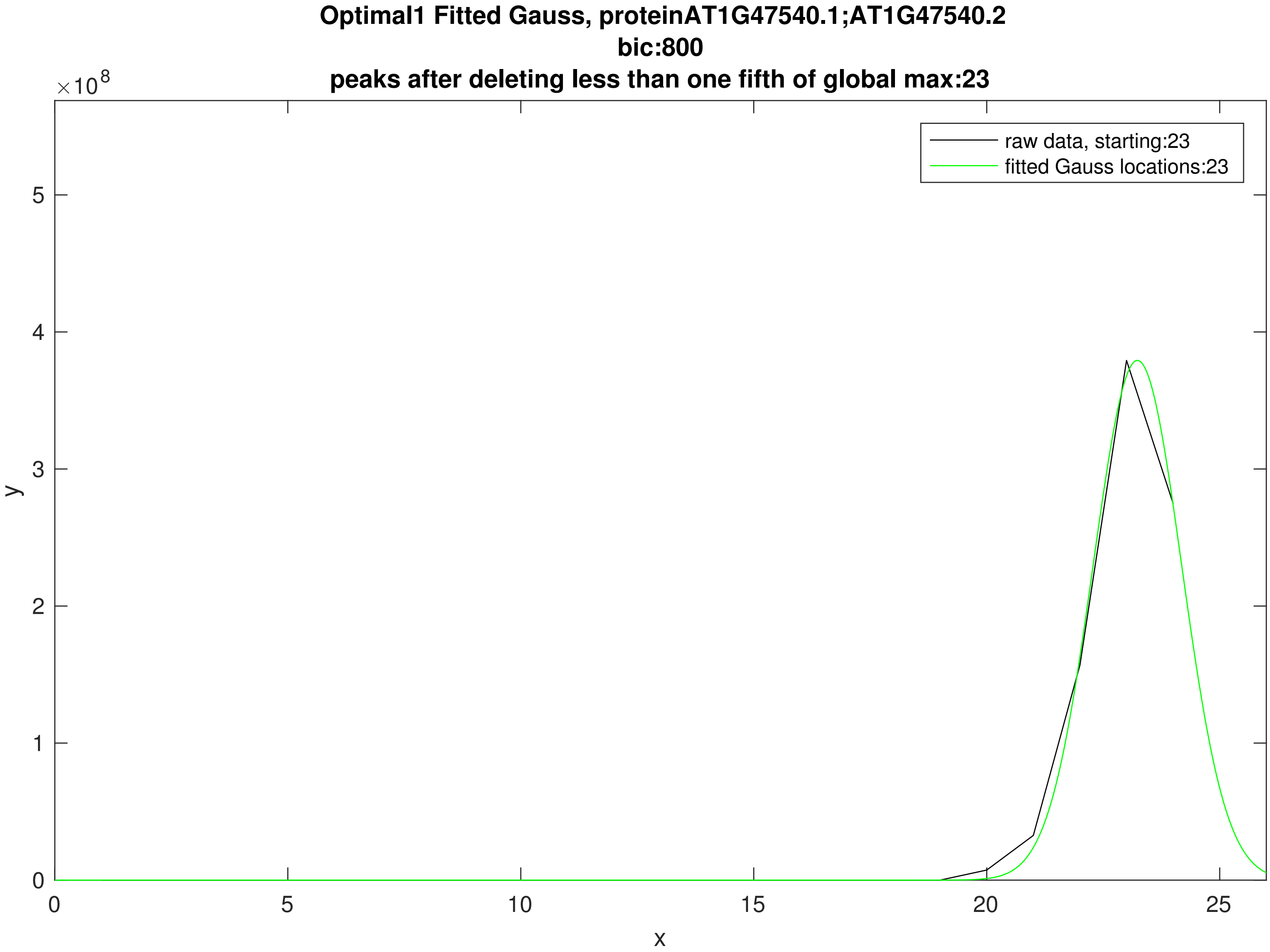

Optimal1 Fitted Gauss, proteinAT1G47540.1;AT1G47540.2
bic:800
peaks after deleting less than one fifth of global max:23
×10^8
raw data, starting:23
fitted Gauss locations:23
y
x
0
1
2
3
4
5
0
5
10
15
20
25

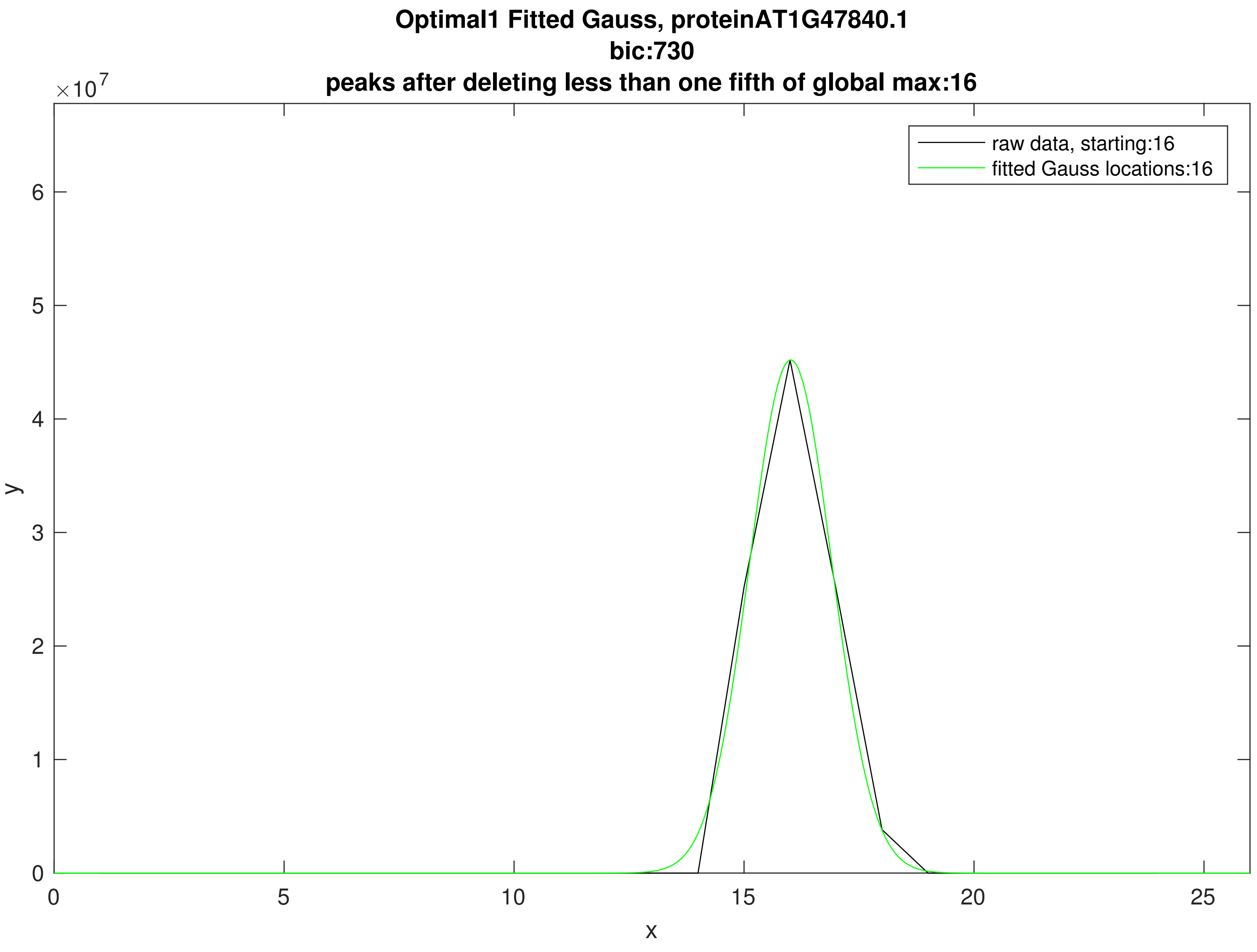
Optimal1 Fitted Gauss, proteinAT1G47840.1
bic:730
peaks after deleting less than one fifth of global max:16
×10^7
raw data, starting:16
fitted Gauss locations:16
y
x
0
1
2
3
4
5
6
0
5
10
15
20
25

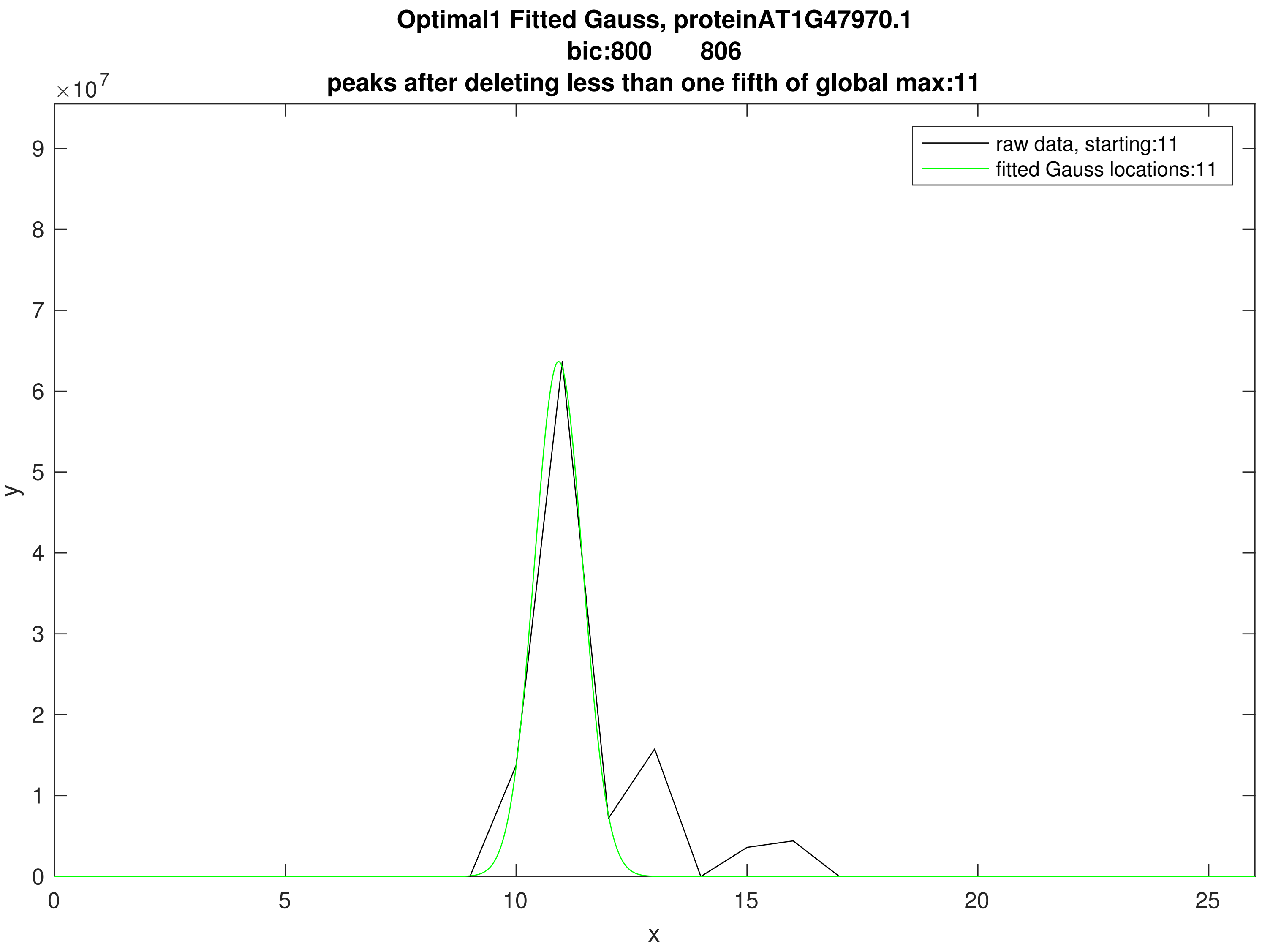

Optimal1 Fitted Gauss, proteinAT1G47970.1
bic:800 806
peaks after deleting less than one fifth of global max:11
×10^7
9
8
7
6
5
4
3
2
1
0
y
0
5
10
15
20
25
x
raw data, starting:11
fitted Gauss locations:11

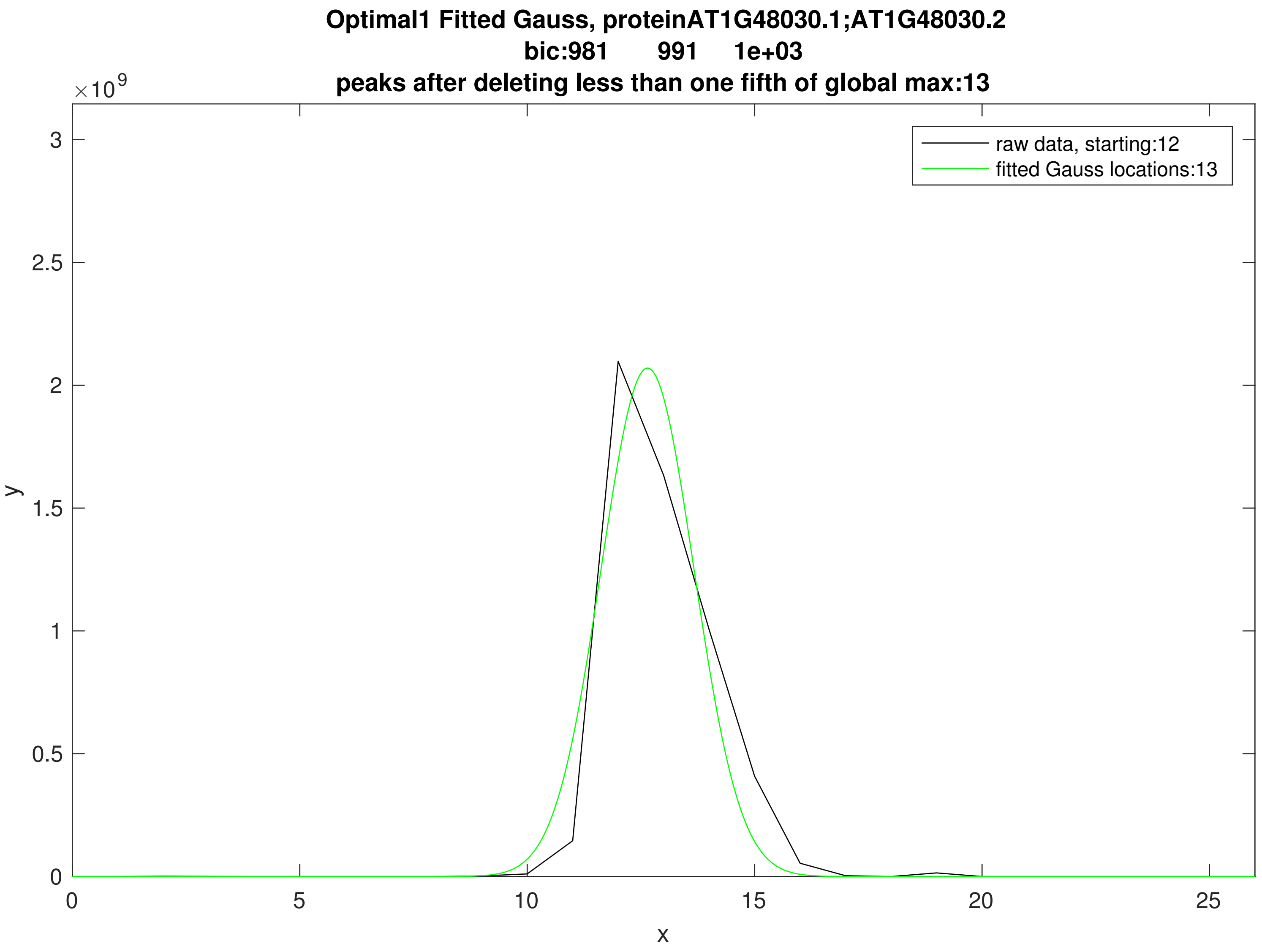

Optimal1 Fitted Gauss, proteinAT1G48030.1;AT1G48030.2
bic:981 991 1e+03
peaks after deleting less than one fifth of global max:13
×10^9
raw data, starting:12
fitted Gauss locations:13
y
x

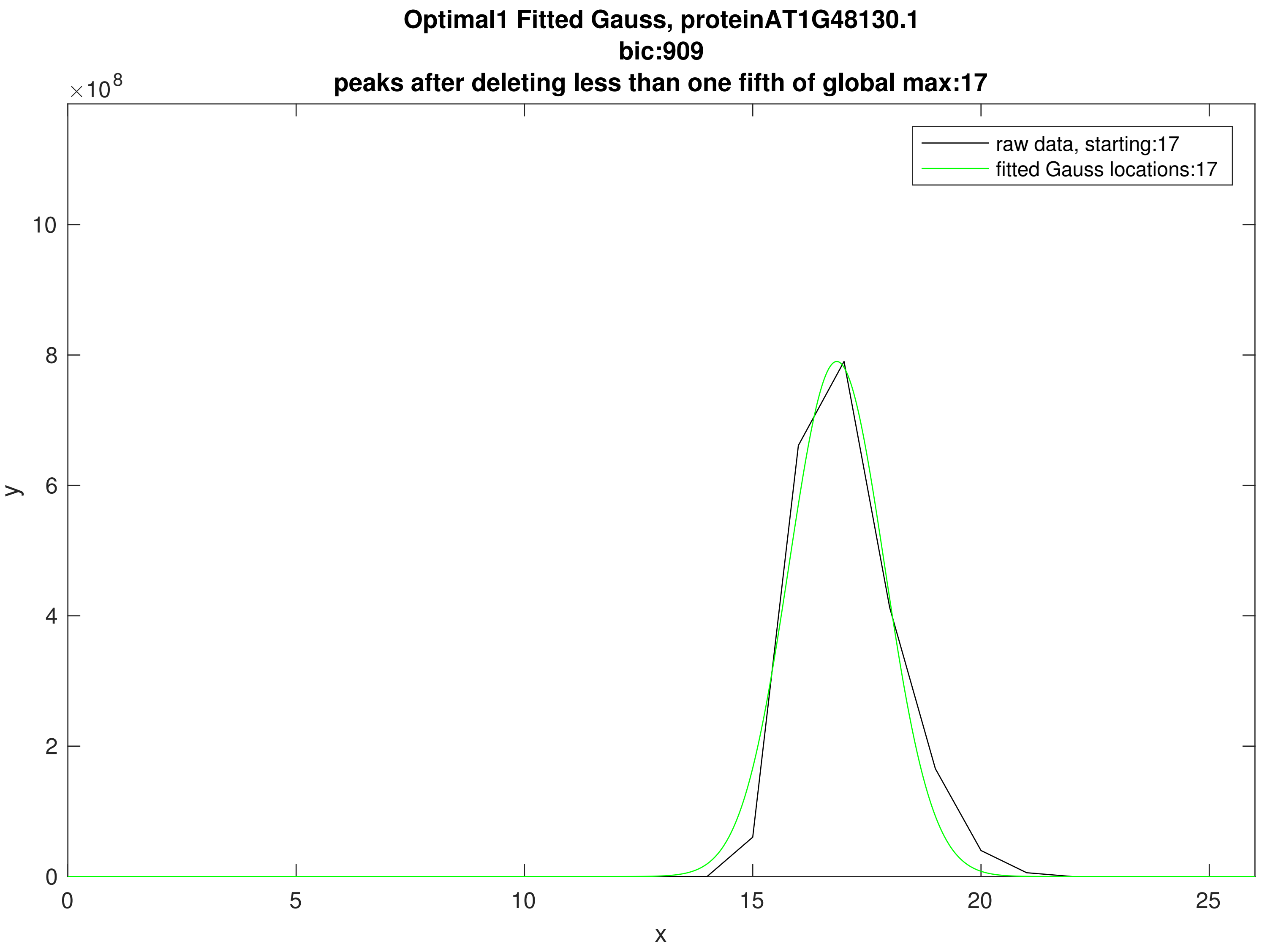
Optimal1 Fitted Gauss, proteinAT1G48130.1
bic:909
peaks after deleting less than one fifth of global max:17
×10^8
raw data, starting:17
fitted Gauss locations:17
y
x
0
2
4
6
8
10
0
5
10
15
20
25

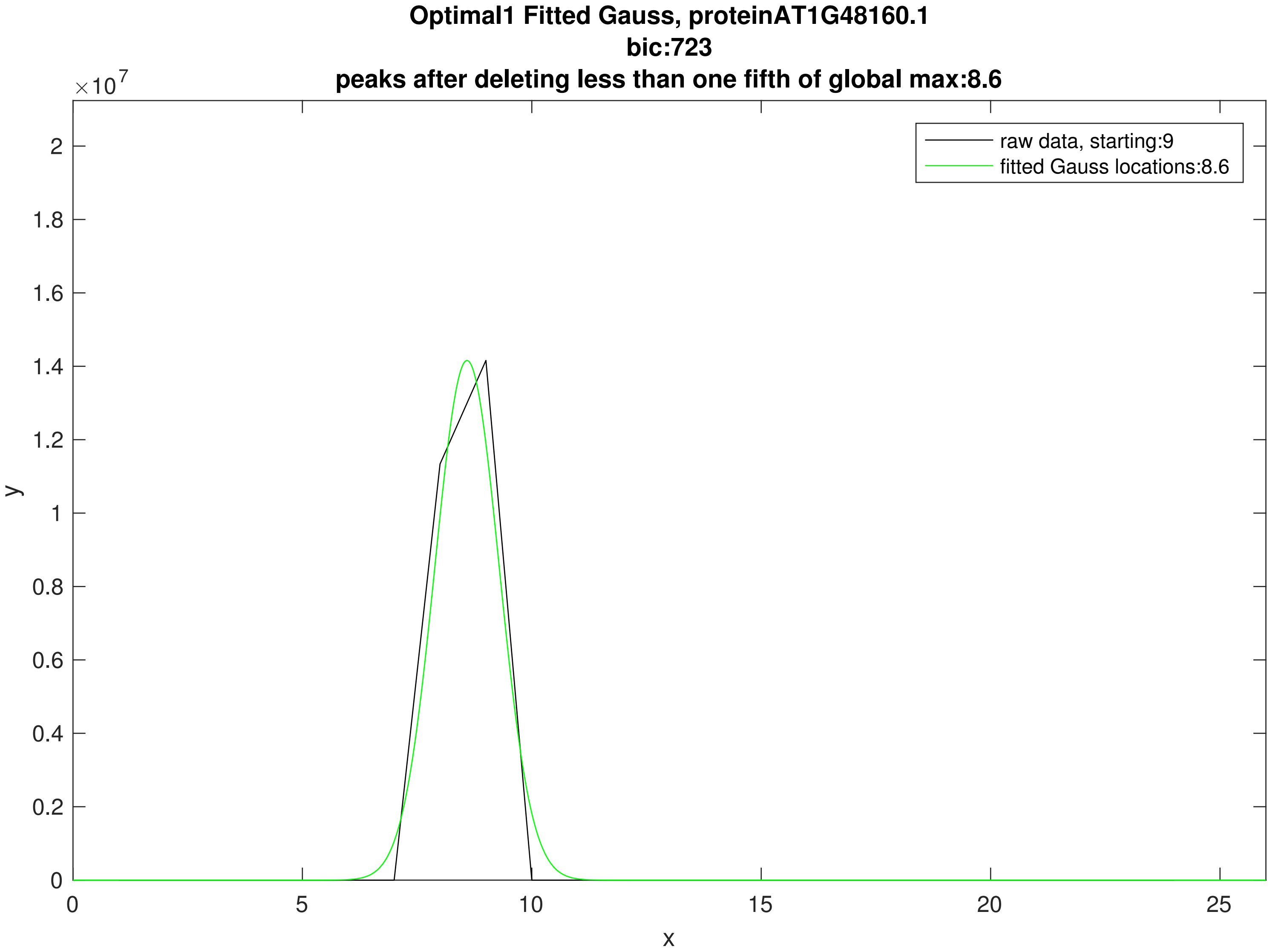
Optimal1 Fitted Gauss, proteinAT1G48160.1
bic:723
peaks after deleting less than one fifth of global max:8.6
×10^7
raw data, starting:9
fitted Gauss locations:8.6
y
x
0
0.2
0.4
0.6
0.8
1
1.2
1.4
1.6
1.8
2
5
10
15
20
25

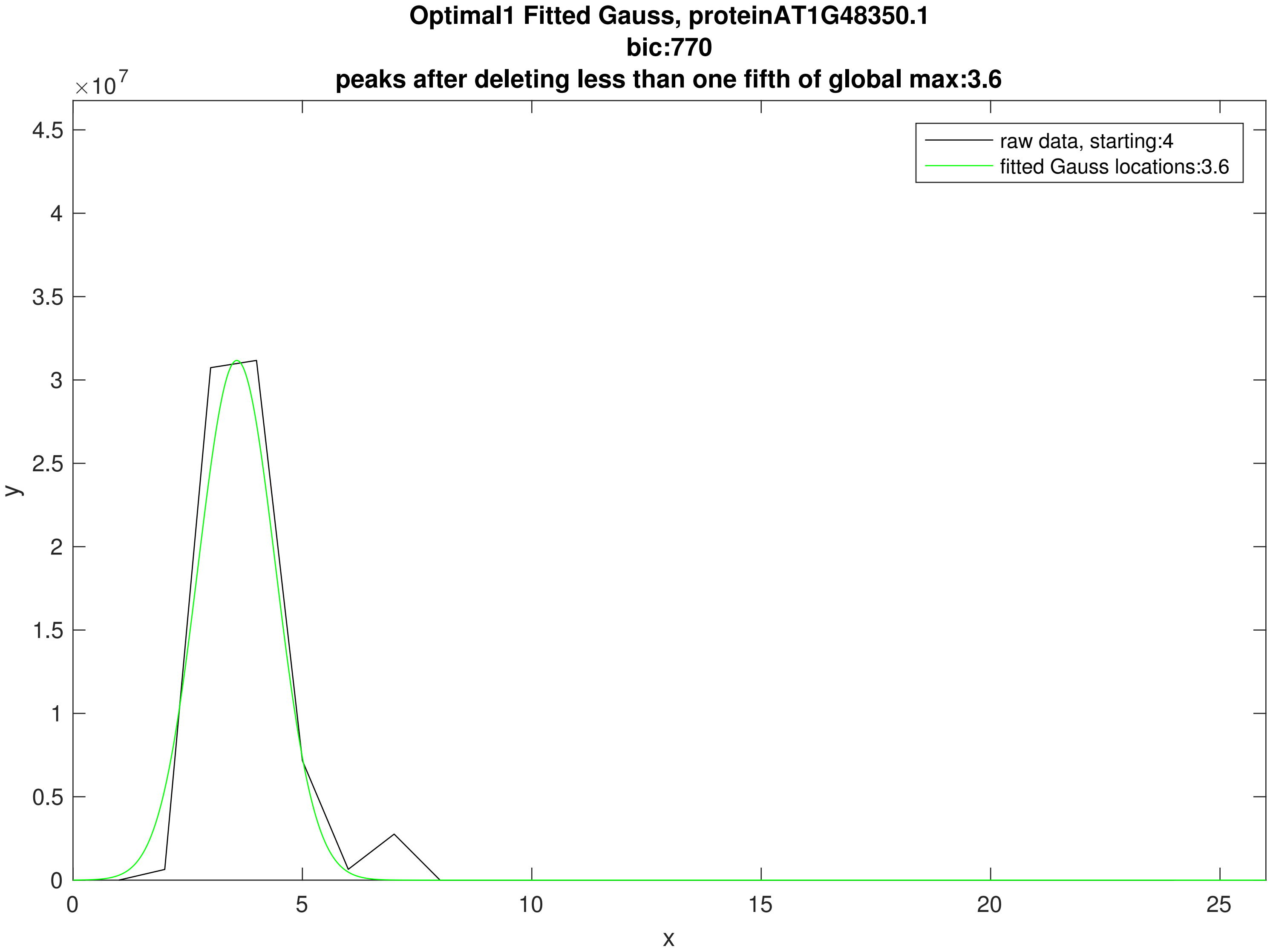

Optimal1 Fitted Gauss, proteinAT1G48350.1
bic:770
peaks after deleting less than one fifth of global max:3.6
×10^7
raw data, starting:4
fitted Gauss locations:3.6
y
x
0
0.5
1
1.5
2
2.5
3
3.5
4
4.5
0
5
10
15
20
25

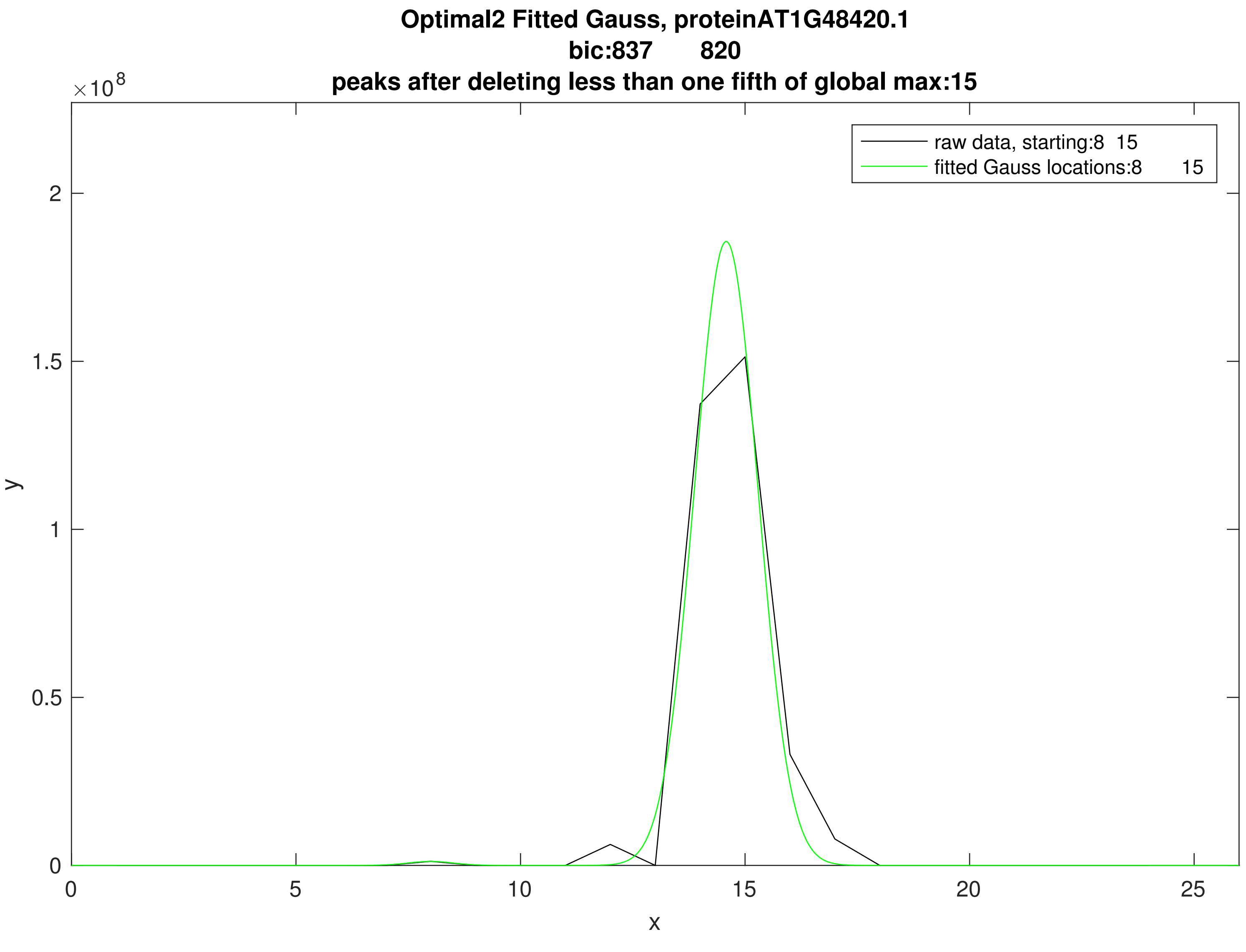

Optimal2 Fitted Gauss, proteinAT1G48420.1
bic:837 820
peaks after deleting less than one fifth of global max:15
×10^8
raw data, starting:8 15
fitted Gauss locations:8 15
y
x
0
0.5
1
1.5
2
0
5
10
15
20
25

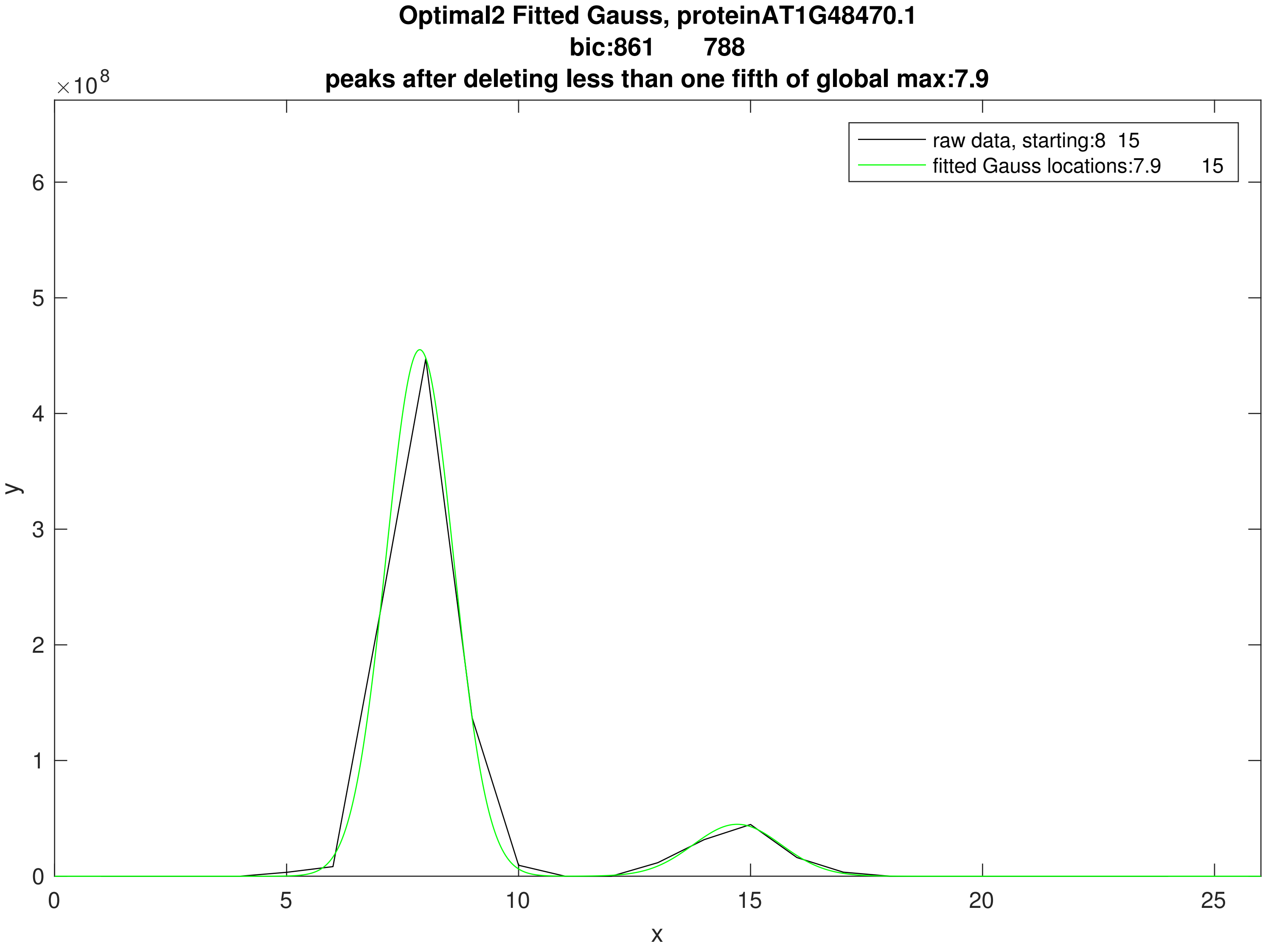

Optimal2 Fitted Gauss, proteinAT1G48470.1
bic:861 788
peaks after deleting less than one fifth of global max:7.9
×10^8
raw data, starting:8 15
fitted Gauss locations:7.9 15
y
x
0
1
2
3
4
5
6
0
5
10
15
20
25

**Optimal1 Fitted Gauss, proteinAT1G48520.1;AT1G48520.2;AT1G48520.3**
**bic:872 879**
**peaks after deleting less than one fifth of global max:11**

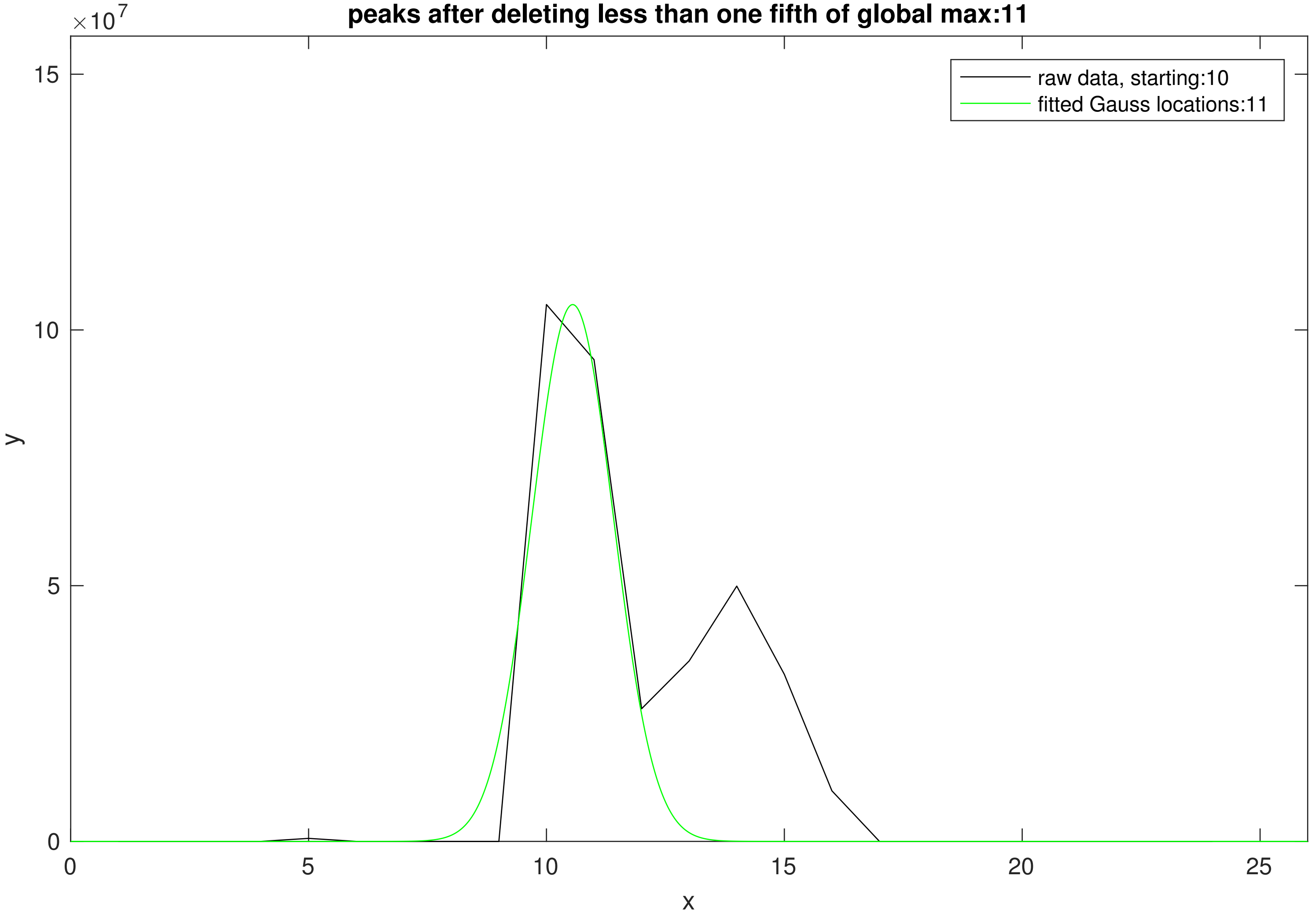

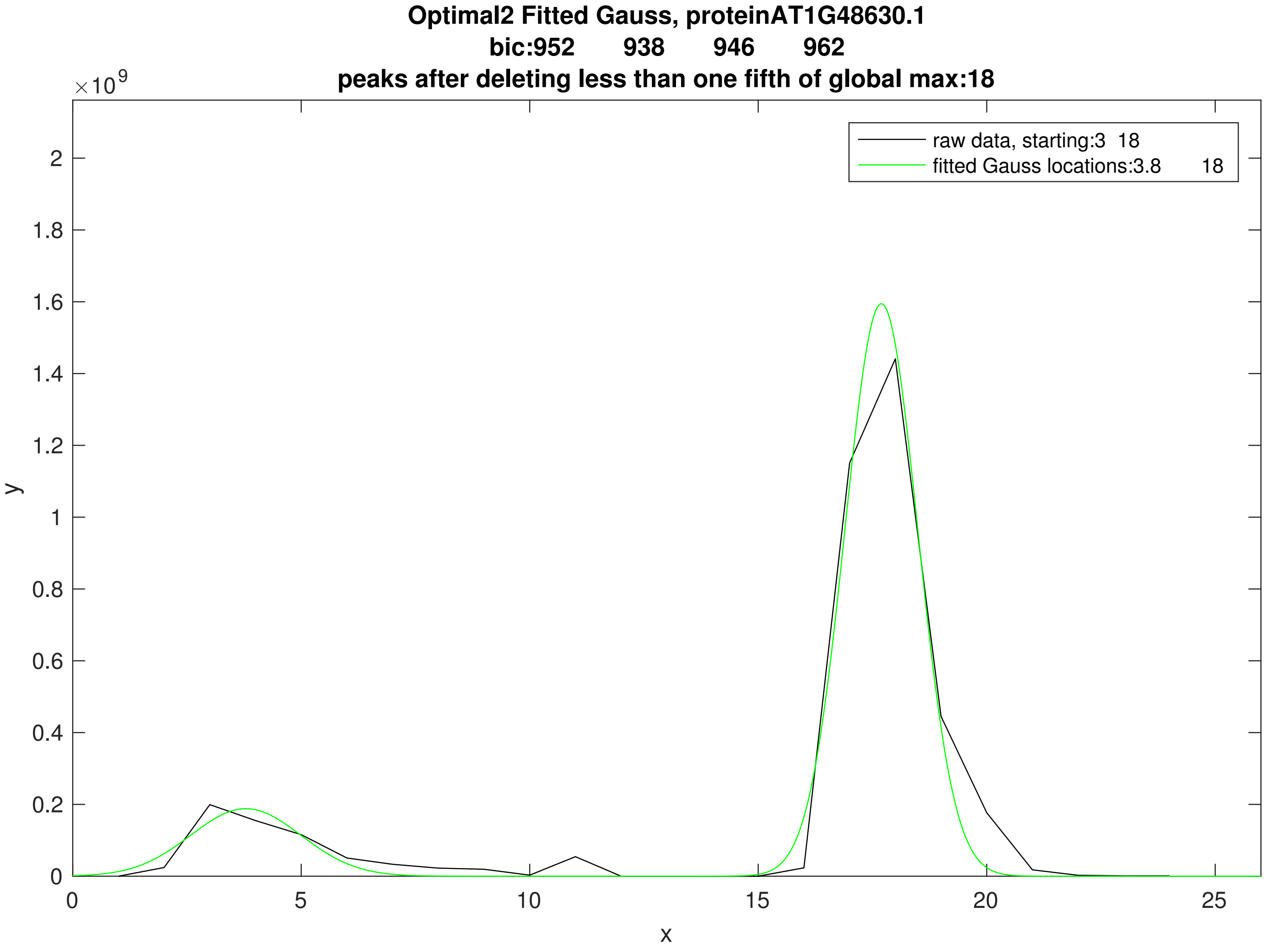

Optimal2 Fitted Gauss, proteinAT1G48630.1
bic:952 938 946 962
peaks after deleting less than one fifth of global max:18
×10^9
raw data, starting:3 18
fitted Gauss locations:3.8 18
2
1.8
1.6
1.4
1.2
1
0.8
0.6
0.4
0.2
0
y
0
5
10
15
20
25
x

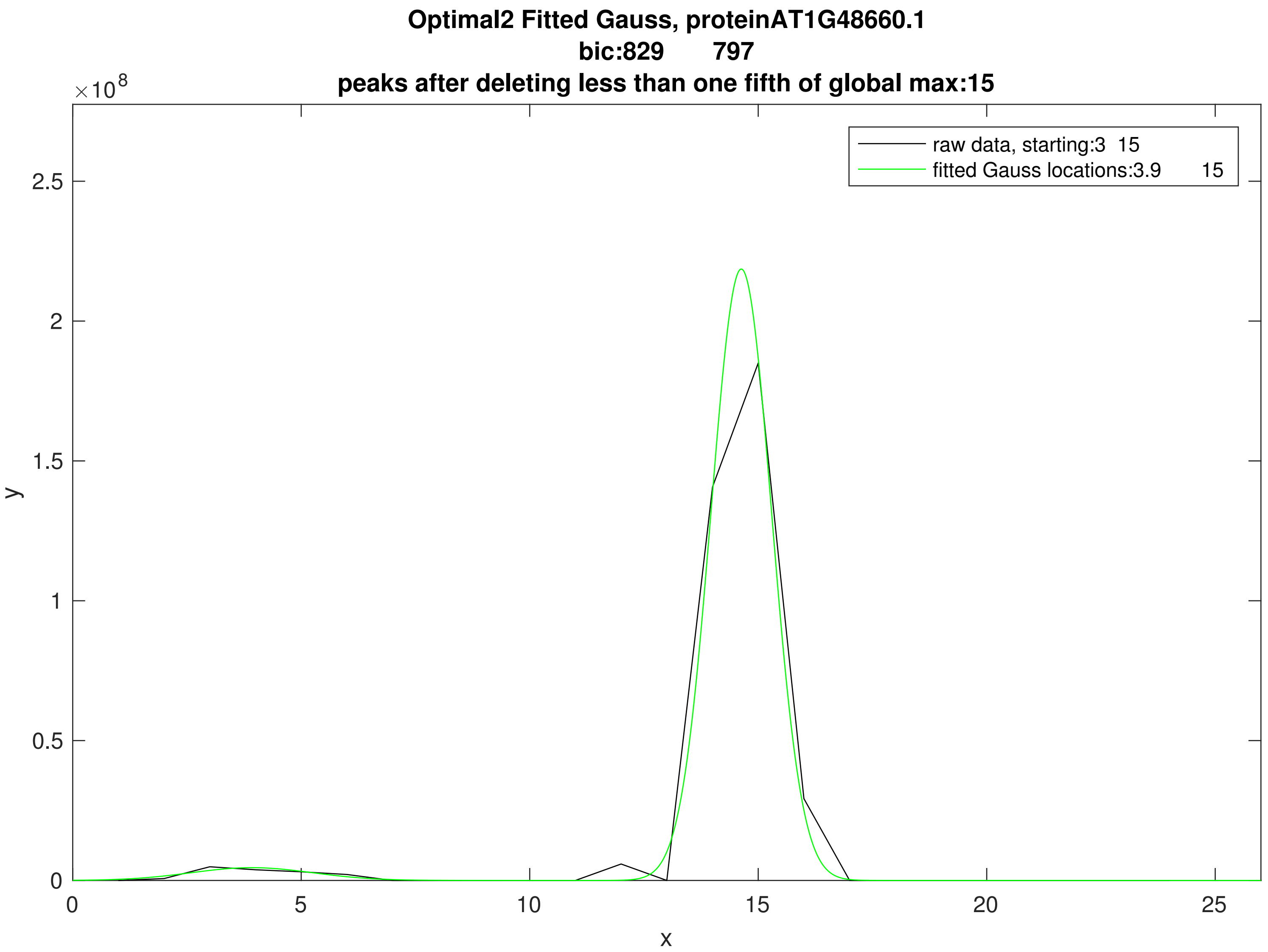
Optimal2 Fitted Gauss, proteinAT1G48660.1
bic:829 797
peaks after deleting less than one fifth of global max:15
raw data, starting:3 15
fitted Gauss locations:3.9 15
×10^8
2.5
2
1.5
1
0.5
0
y
0
5
10
15
20
25
x

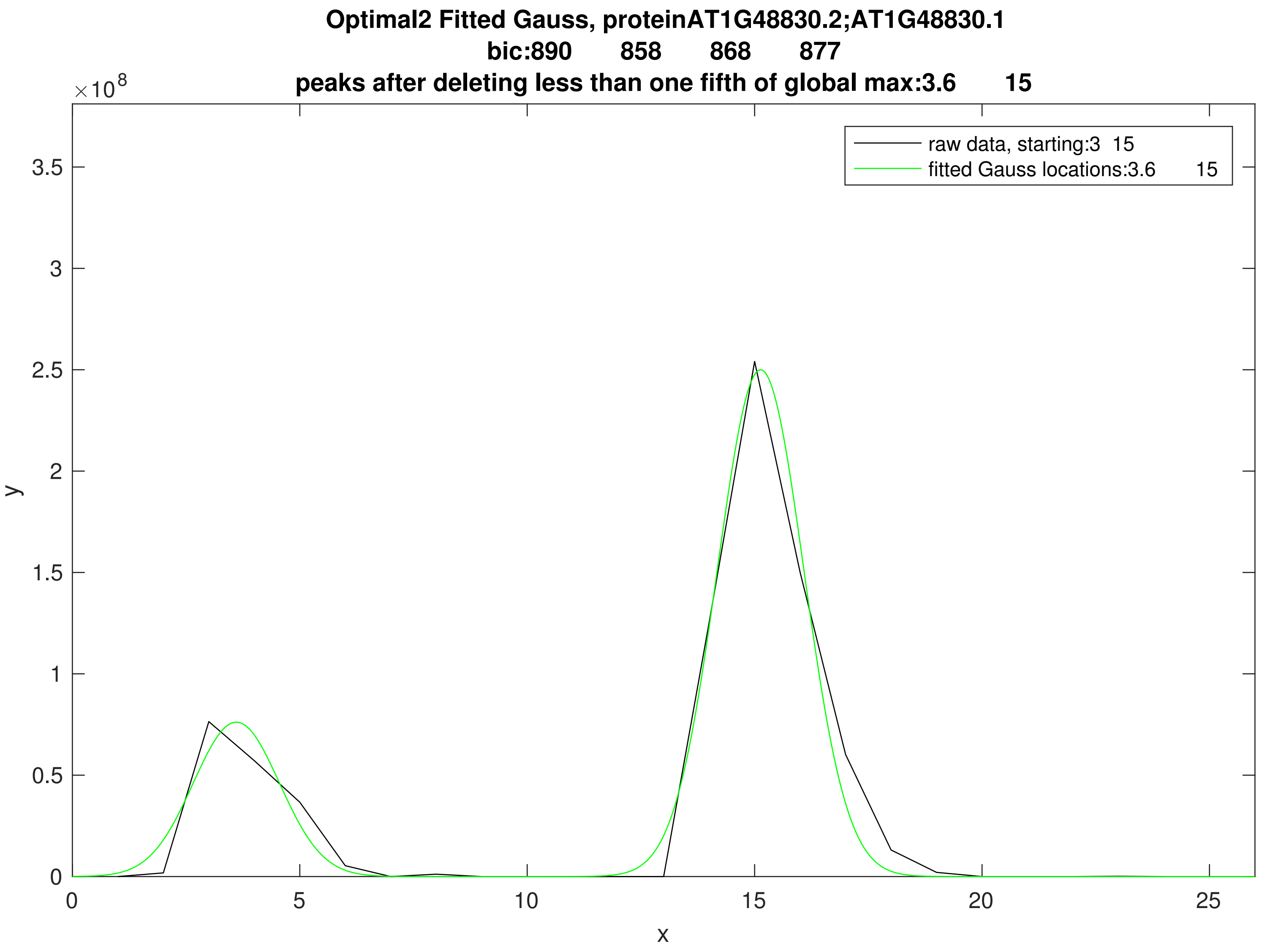
Optimal2 Fitted Gauss, proteinAT1G48830.2;AT1G48830.1
bic:890 858 868 877
peaks after deleting less than one fifth of global max:3.6 15
×10^8
3.5
3
2.5
2
1.5
1
0.5
0
y
0
5
10
15
20
25
x
raw data, starting:3 15
fitted Gauss locations:3.6 15

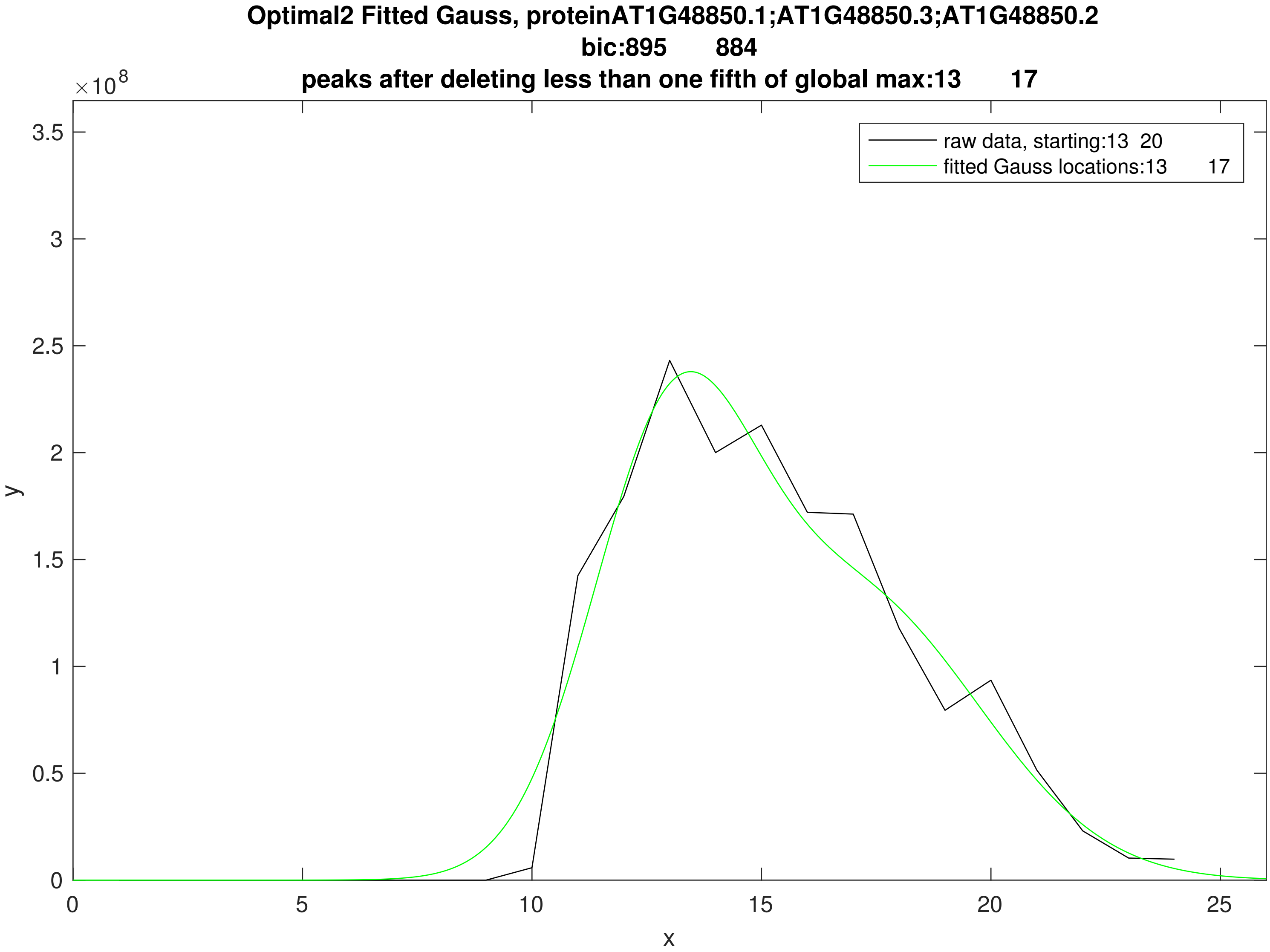

Optimal2 Fitted Gauss, proteinAT1G48850.1;AT1G48850.3;AT1G48850.2
bic:895 884
peaks after deleting less than one fifth of global max:13 17
×10^8
3.5
3
2.5
2
1.5
1
0.5
0
y
0
5
10
15
20
25
x
raw data, starting:13 20
fitted Gauss locations:13 17

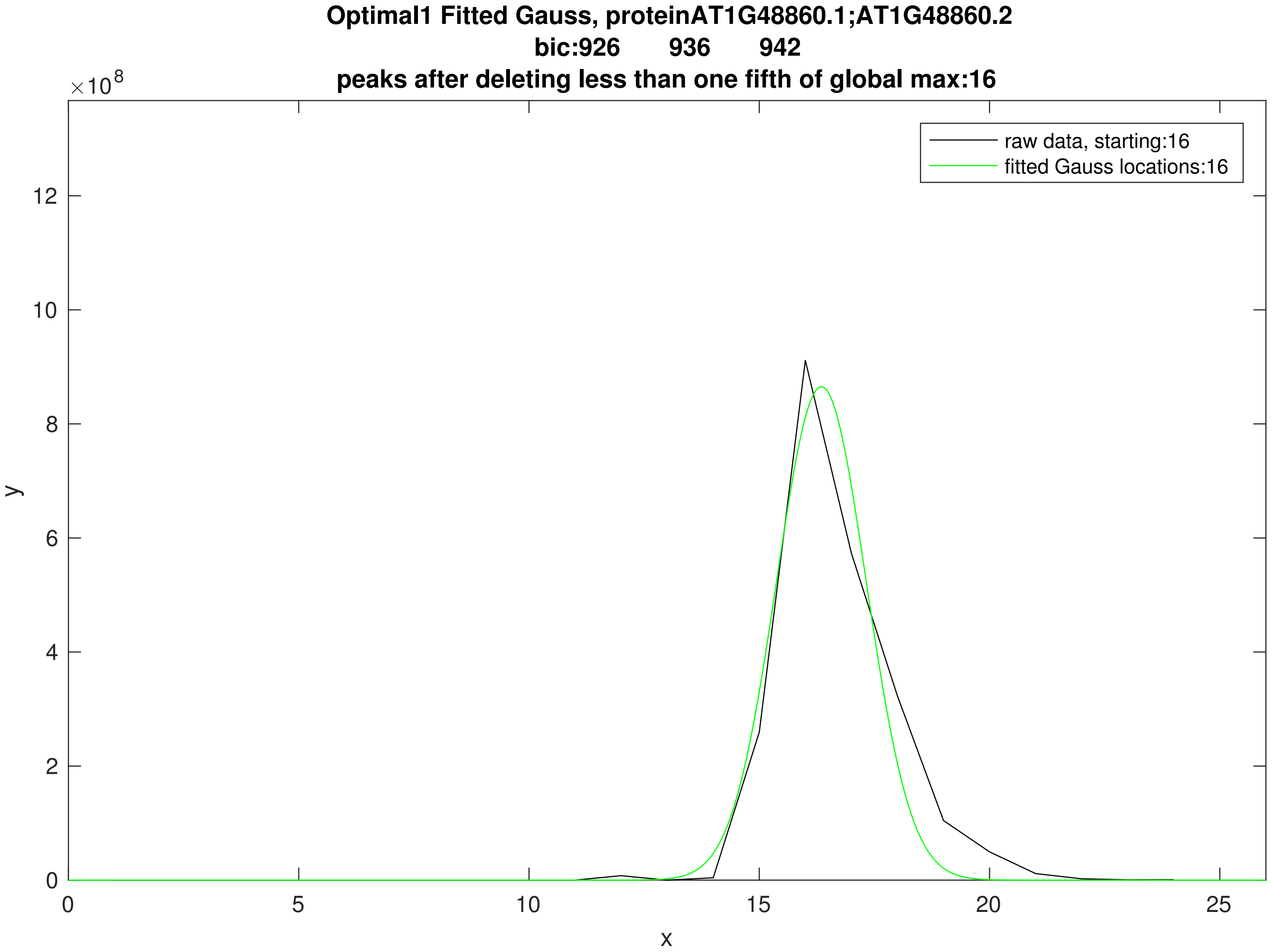

Optimal1 Fitted Gauss, proteinAT1G48860.1;AT1G48860.2
bic:926 936 942
peaks after deleting less than one fifth of global max:16
×10^8
raw data, starting:16
fitted Gauss locations:16
12
10
8
6
4
2
0
y
0
5
10
15
20
25
x

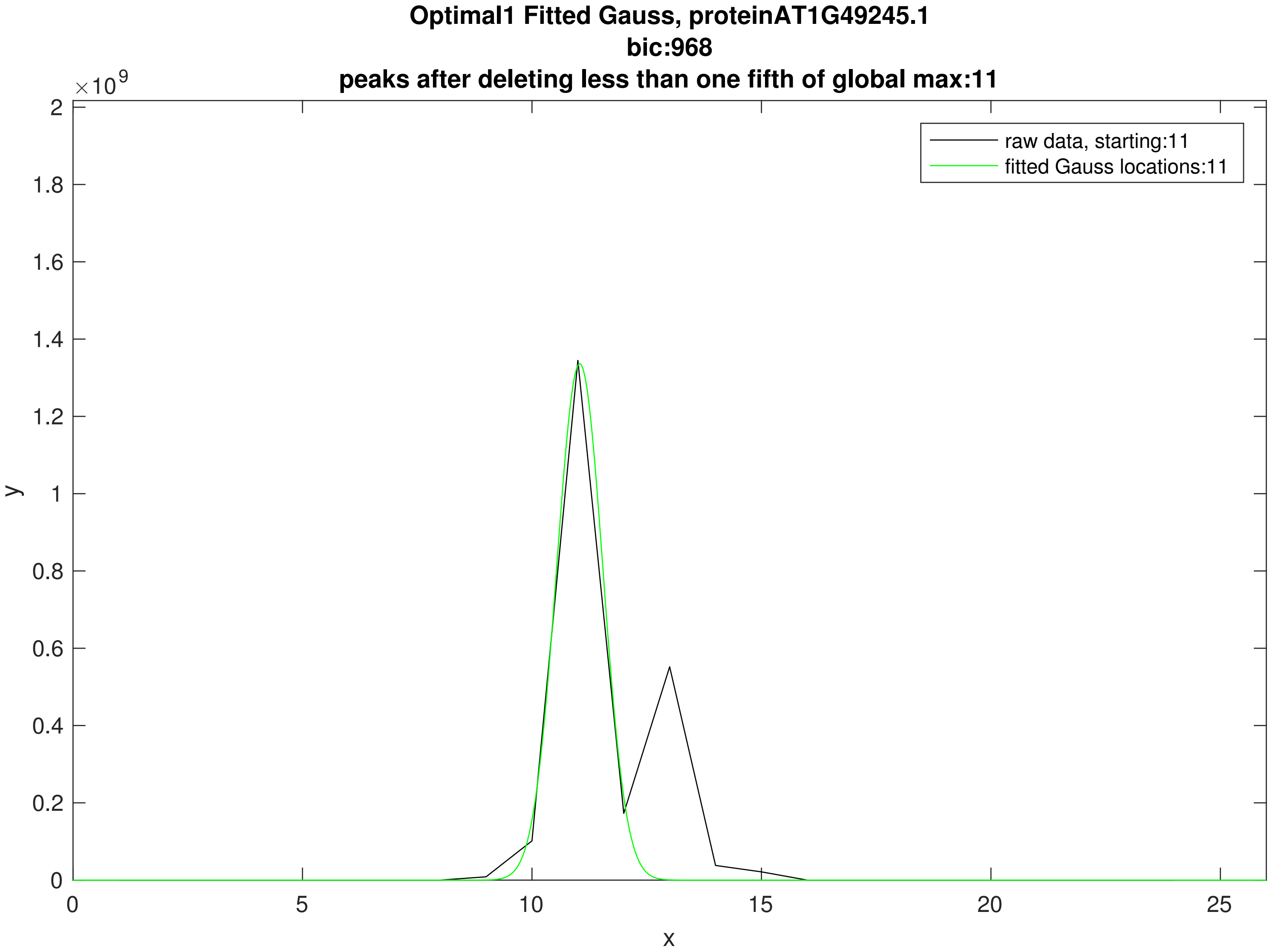
Optimal1 Fitted Gauss, proteinAT1G49245.1
bic:968
peaks after deleting less than one fifth of global max:11
×10^9
raw data, starting:11
fitted Gauss locations:11
y
x
0
0.2
0.4
0.6
0.8
1
1.2
1.4
1.6
1.8
2
5
10
15
20
25

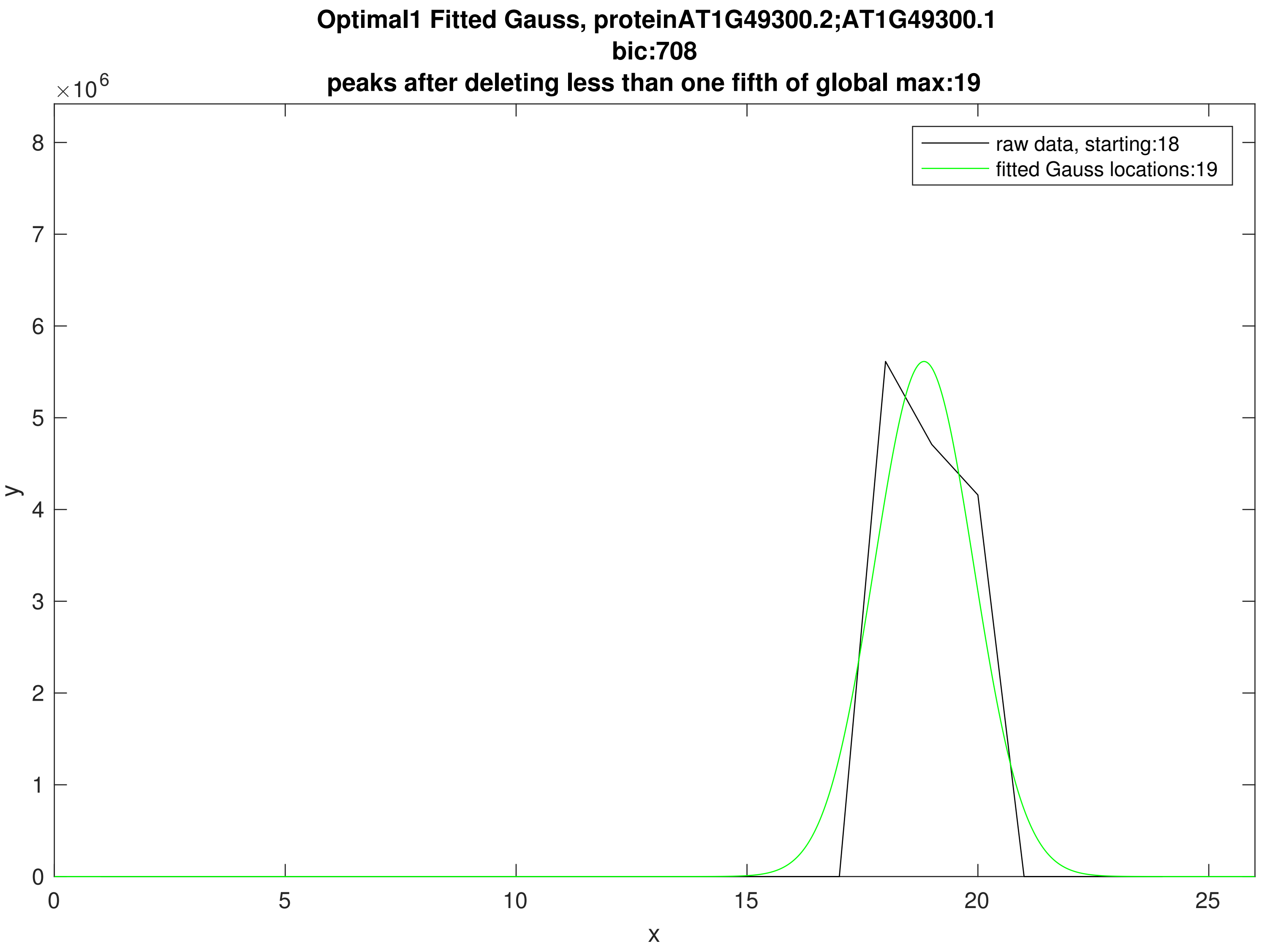

Optimal1 Fitted Gauss, proteinAT1G49300.2;AT1G49300.1
bic:708
peaks after deleting less than one fifth of global max:19
×10^6
raw data, starting:18
fitted Gauss locations:19
y
x
0
1
2
3
4
5
6
7
8
0
5
10
15
20
25

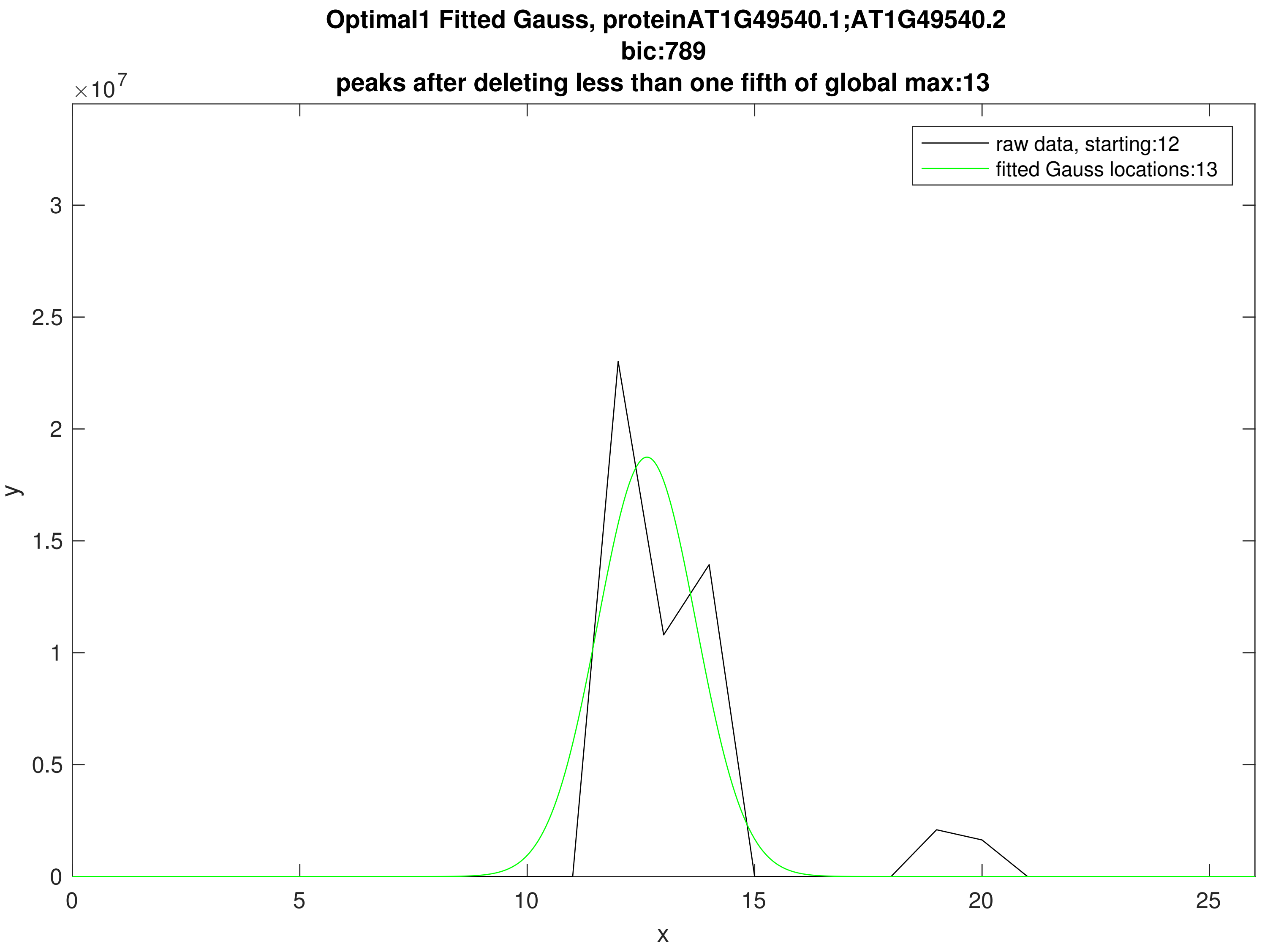

Optimal1 Fitted Gauss, proteinAT1G49540.1;AT1G49540.2
bic:789
peaks after deleting less than one fifth of global max:13
×10^7
raw data, starting:12
fitted Gauss locations:13
y
x
0
0.5
1
1.5
2
2.5
3
0
5
10
15
20
25

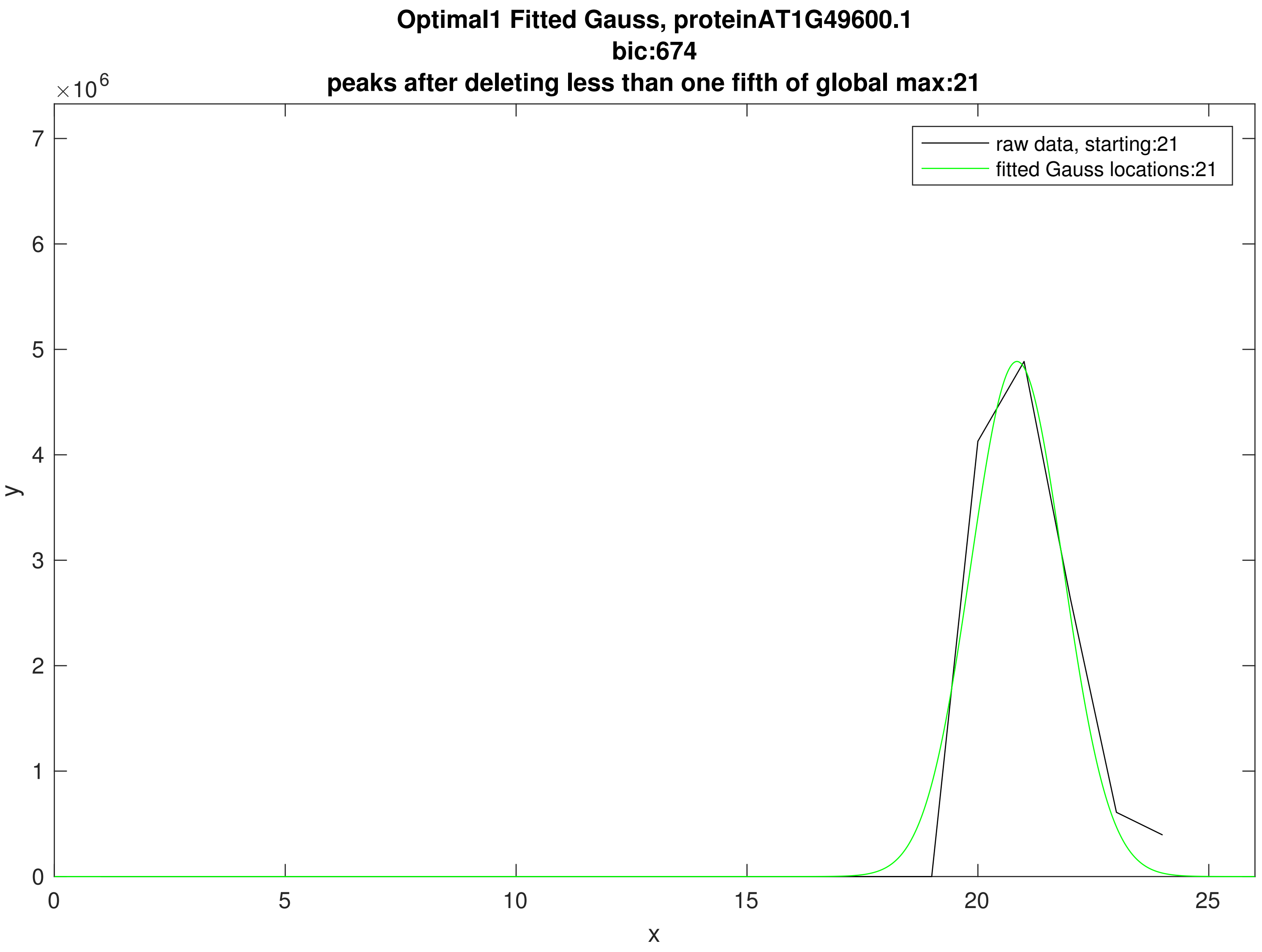

Optimal1 Fitted Gauss, proteinAT1G49600.1
bic:674
peaks after deleting less than one fifth of global max:21
×10^6
raw data, starting:21
fitted Gauss locations:21
y
x
0
1
2
3
4
5
6
7
0
5
10
15
20
25

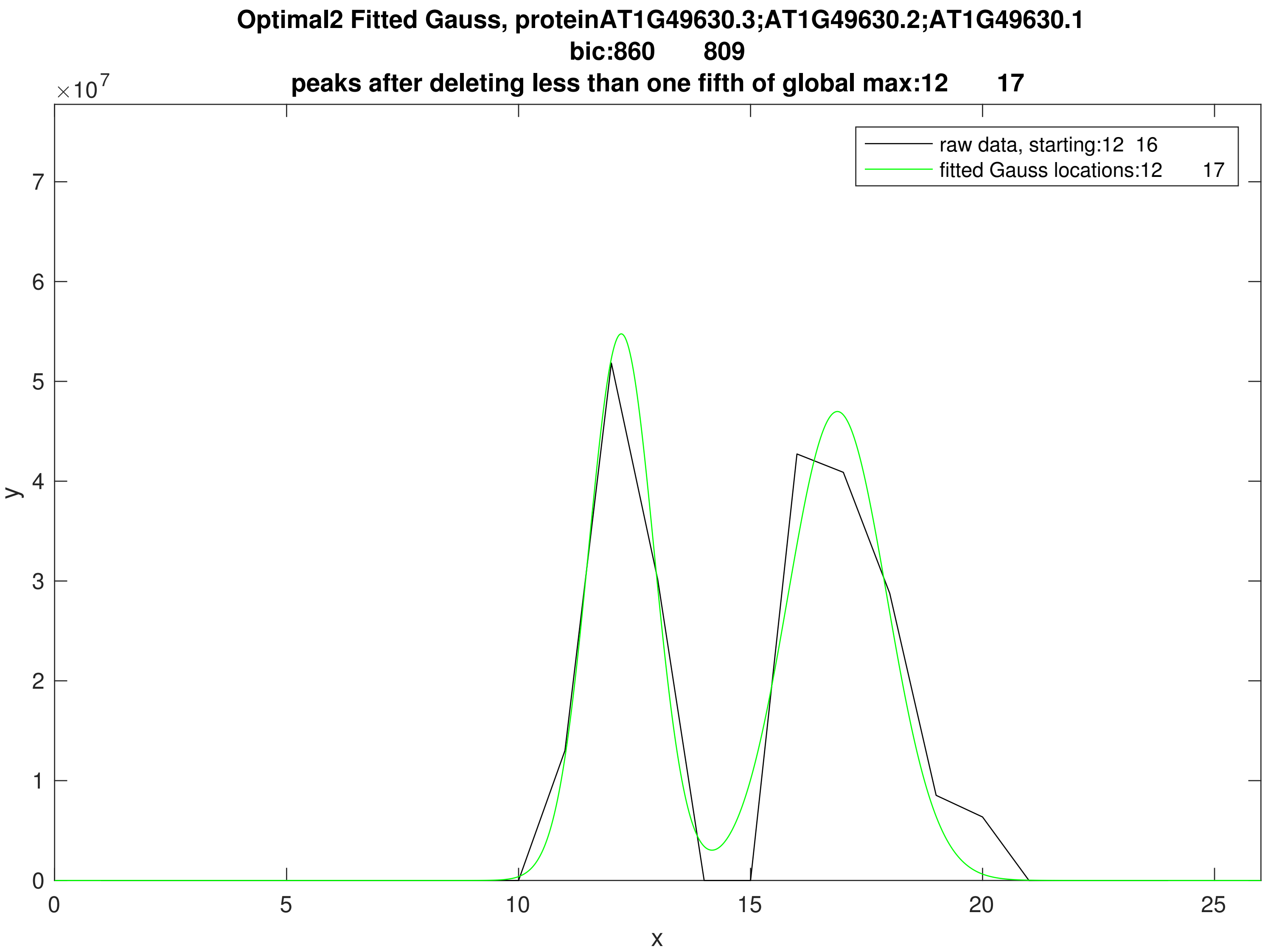
Optimal2 Fitted Gauss, proteinAT1G49630.3;AT1G49630.2;AT1G49630.1
bic:860 809
peaks after deleting less than one fifth of global max:12 17
×10^7
raw data, starting:12 16
fitted Gauss locations:12 17
y
x
0
1
2
3
4
5
6
7
0
5
10
15
20
25

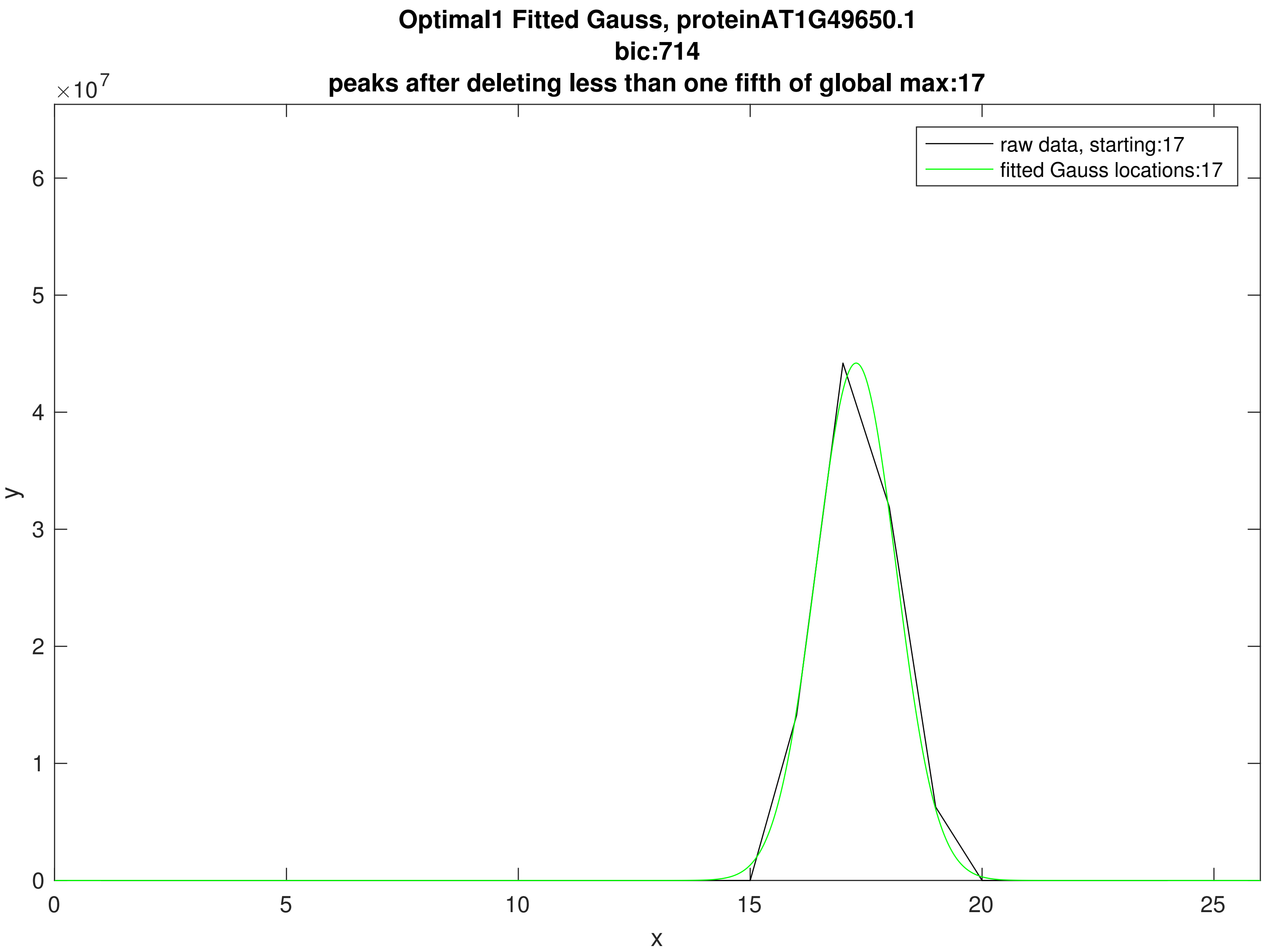

Optimal1 Fitted Gauss, proteinAT1G49650.1
bic:714
peaks after deleting less than one fifth of global max:17
×10^7
6
5
4
3
2
1
0
y
0
5
10
15
20
25
x
raw data, starting:17
fitted Gauss locations:17

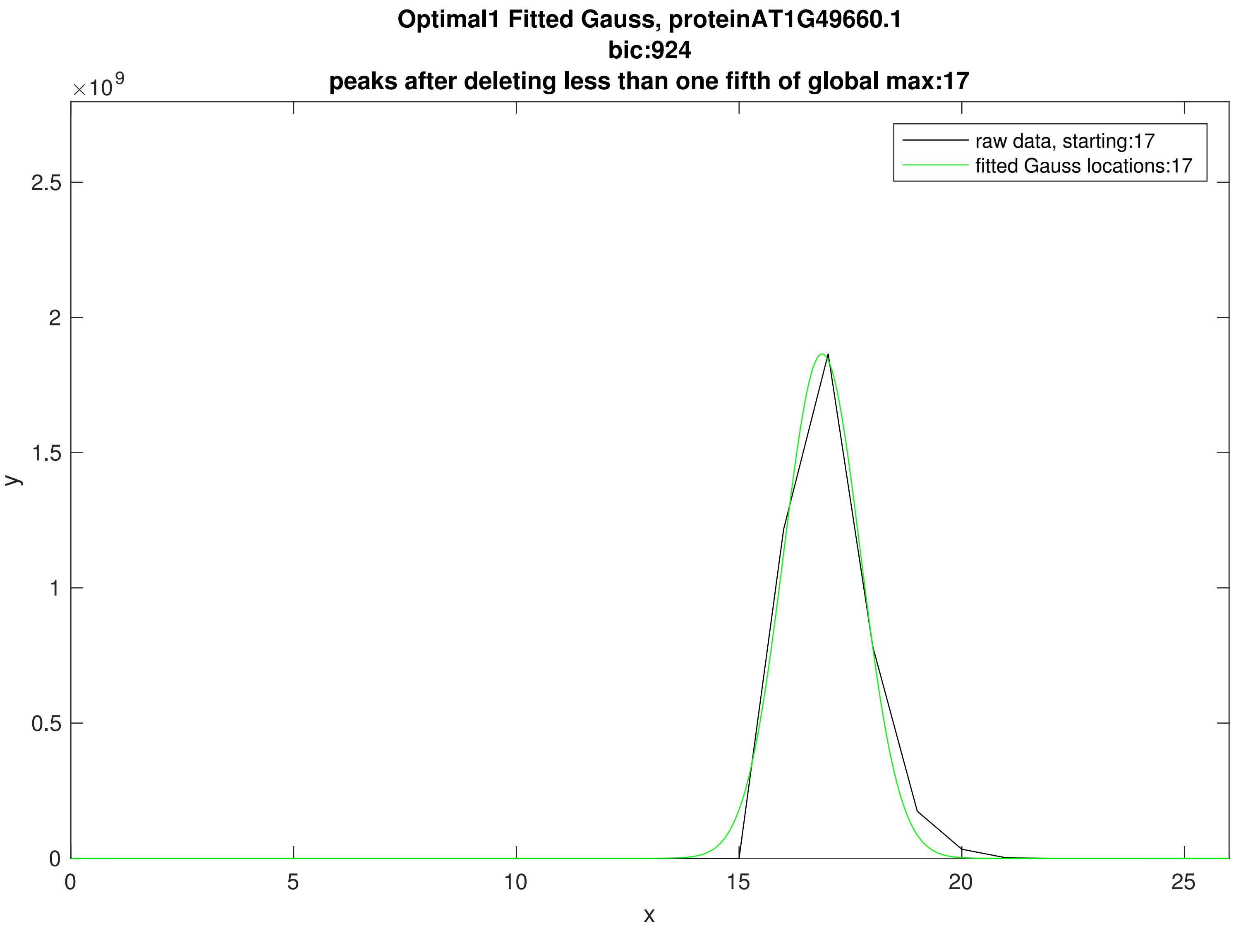

Optimal1 Fitted Gauss, proteinAT1G49660.1
bic:924
peaks after deleting less than one fifth of global max:17
×10^9
2.5
2
1.5
1
0.5
0
y
0
5
10
15
20
25
x
raw data, starting:17
fitted Gauss locations:17

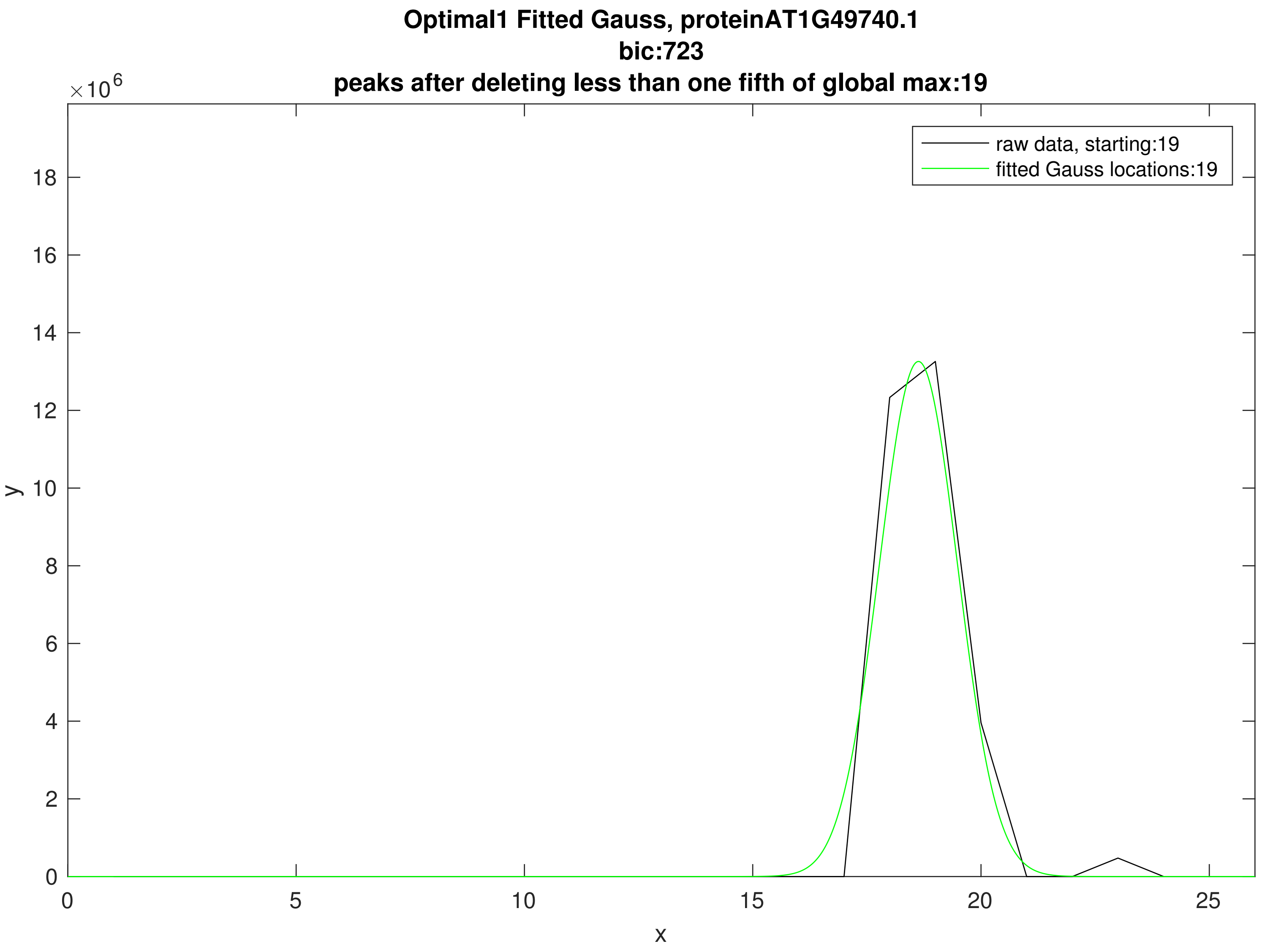

Optimal1 Fitted Gauss, proteinAT1G49740.1
bic:723
peaks after deleting less than one fifth of global max:19
×10^6
raw data, starting:19
fitted Gauss locations:19
y
x
0
2
4
6
8
10
12
14
16
18
0
5
10
15
20
25

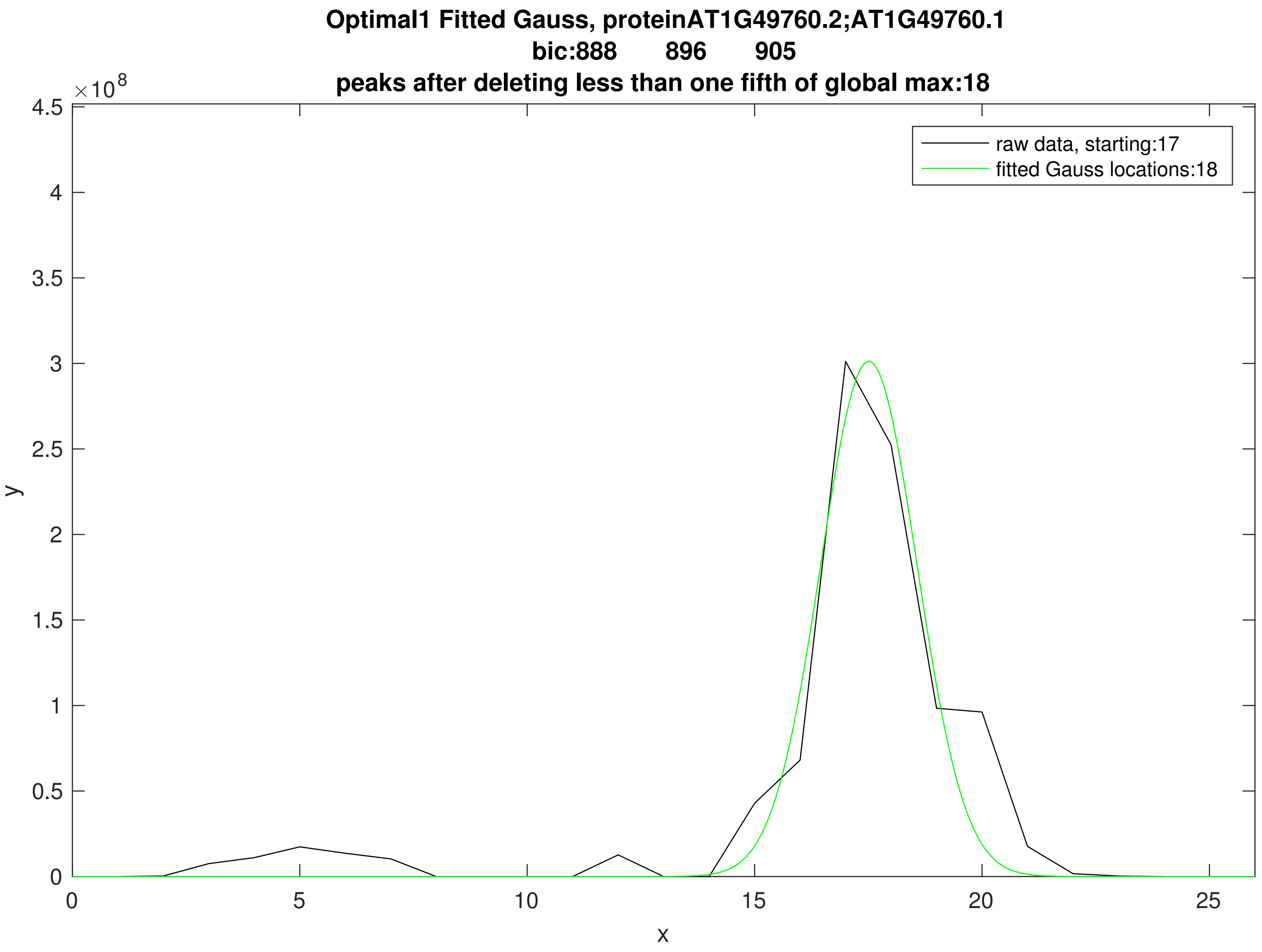

Optimal1 Fitted Gauss, proteinAT1G49760.2;AT1G49760.1
bic:888 896 905
peaks after deleting less than one fifth of global max:18
×10^8
4.5
4
3.5
3
2.5
2
1.5
1
0.5
0
y
0
5
10
15
20
25
x
raw data, starting:17
fitted Gauss locations:18

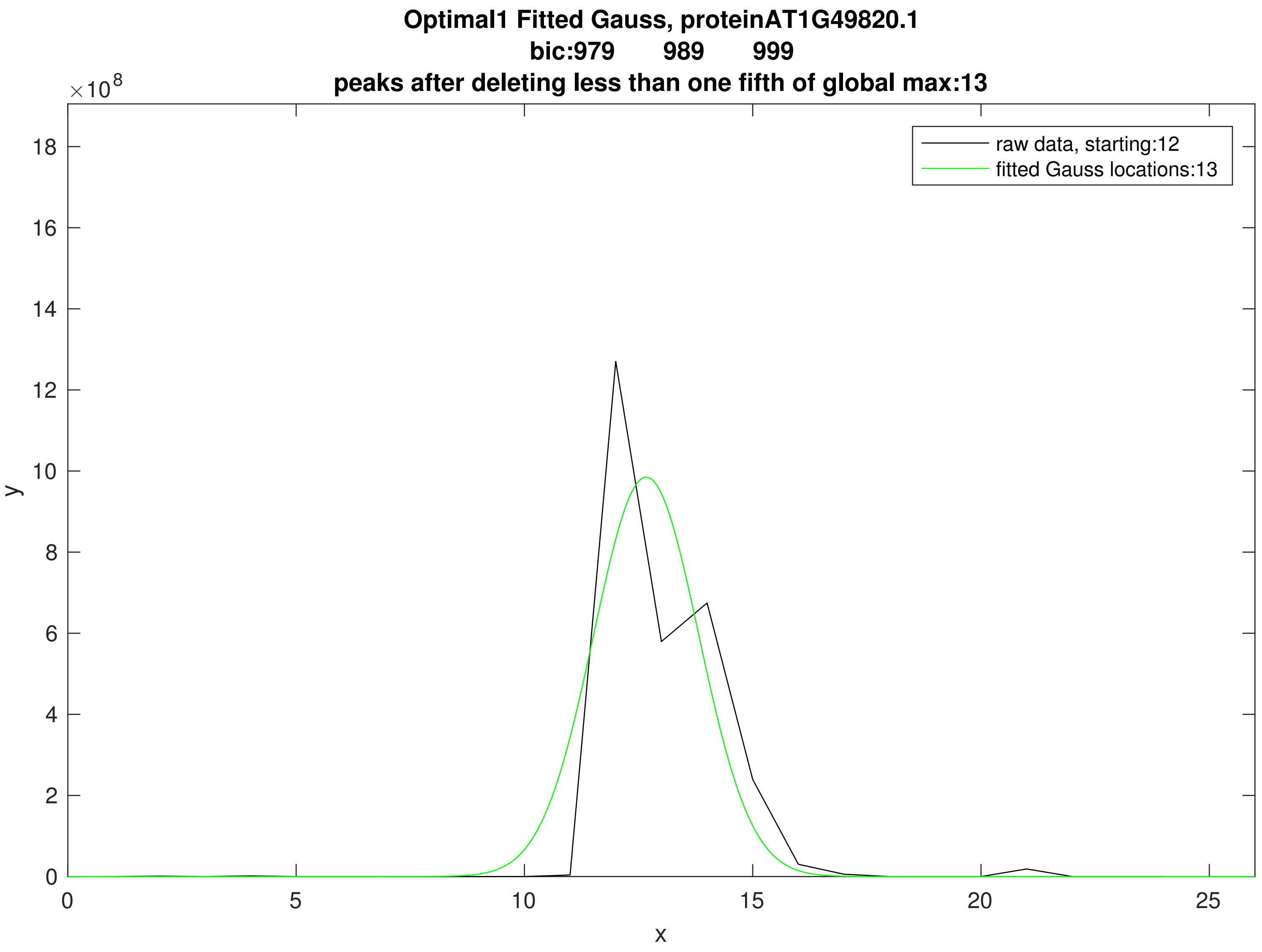
Optimal1 Fitted Gauss, proteinAT1G49820.1
bic:979 989 999
peaks after deleting less than one fifth of global max:13
×10^8
18
16
14
12
10
8
6
4
2
0
y
0
5
10
15
20
25
x
raw data, starting:12
fitted Gauss locations:13

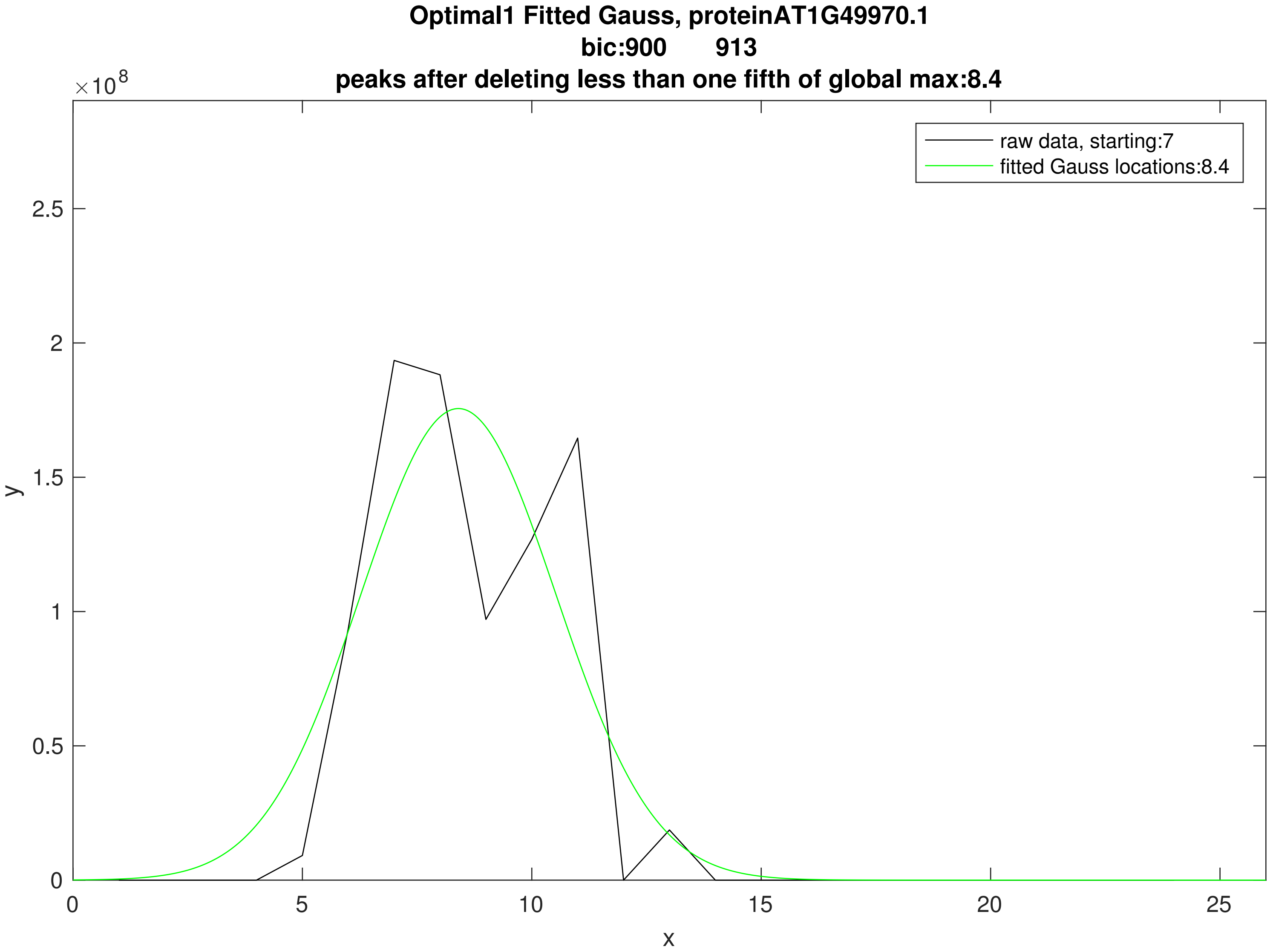
Optimal1 Fitted Gauss, proteinAT1G49970.1
bic:900 913
peaks after deleting less than one fifth of global max:8.4
×10^8
raw data, starting:7
fitted Gauss locations:8.4
y
x
0
0.5
1
1.5
2
2.5
0
5
10
15
20
25

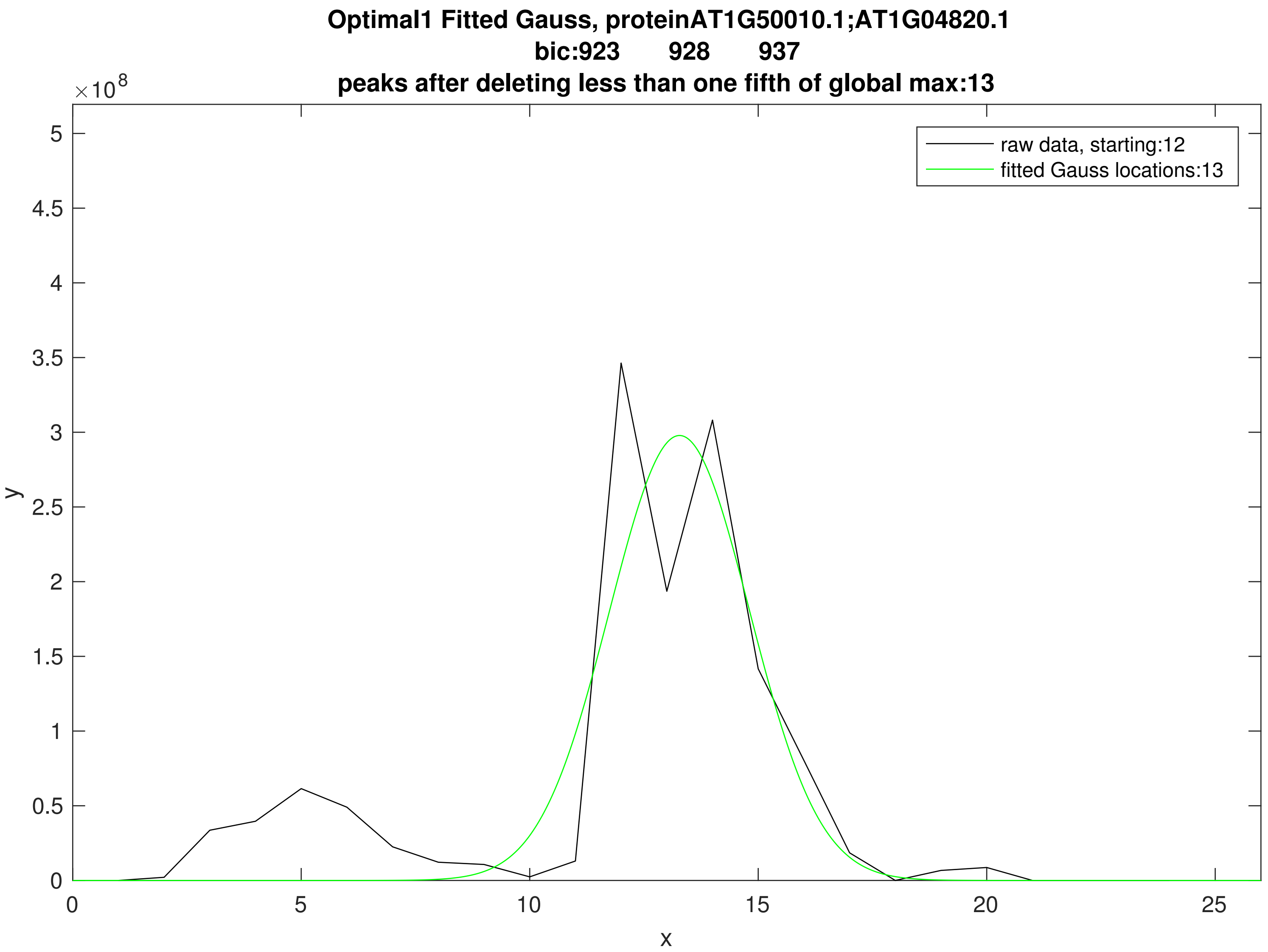
Optimal1 Fitted Gauss, proteinAT1G50010.1;AT1G04820.1
bic:923 928 937
peaks after deleting less than one fifth of global max:13
×10^8
raw data, starting:12
fitted Gauss locations:13
y
x
0
0.5
1
1.5
2
2.5
3
3.5
4
4.5
5
0
5
10
15
20
25

Optimal1 Fitted Gauss, proteinAT1G50110.1
bic:821
peaks after deleting less than one fifth of global max:15

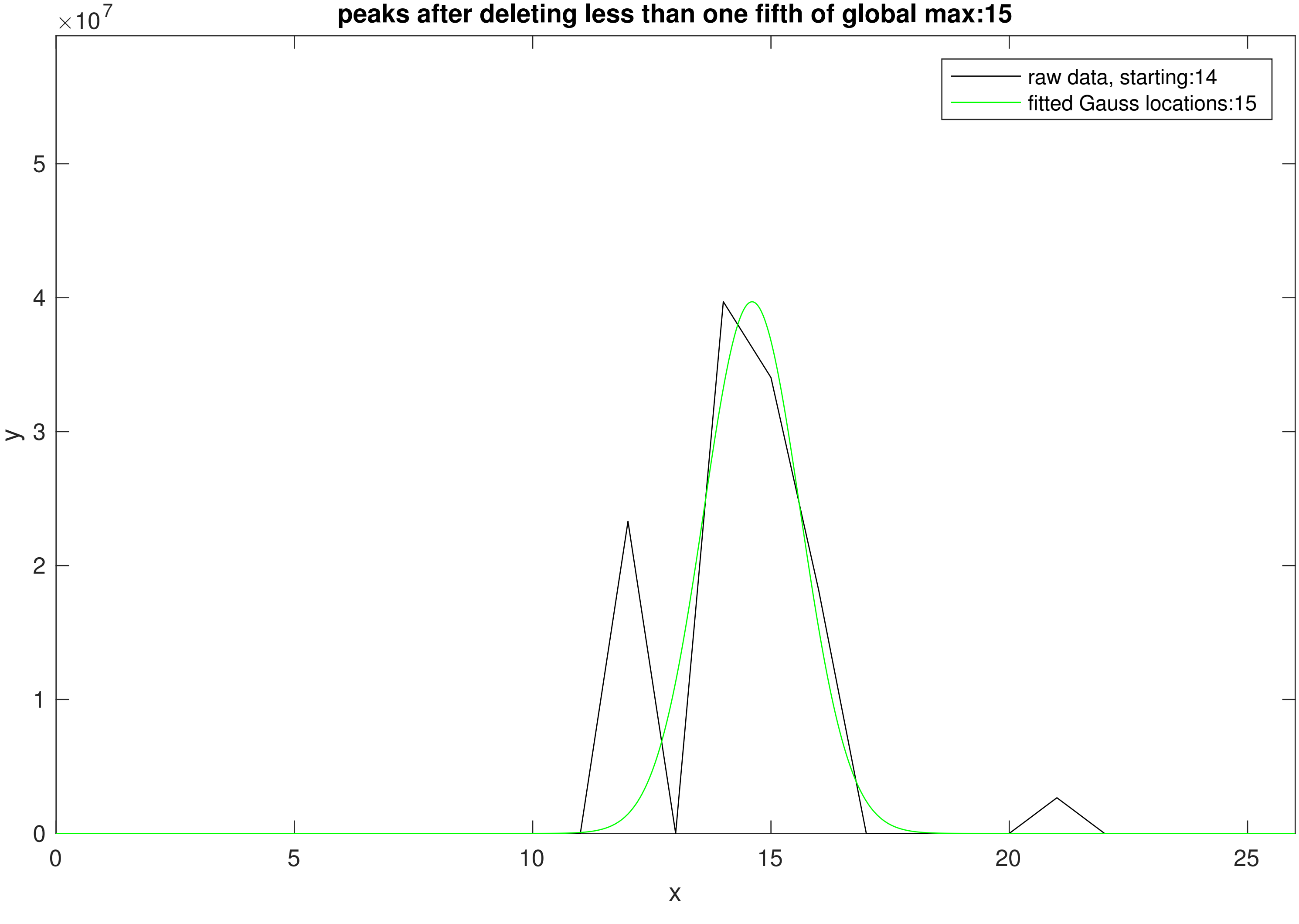

#### Optimal1 Fitted Gauss, proteinAT1G50200.1
#### bic:1.04e+03  1.05e+03  1.06e+03  1.07e+03
#### peaks after deleting less than one fifth of global max:11

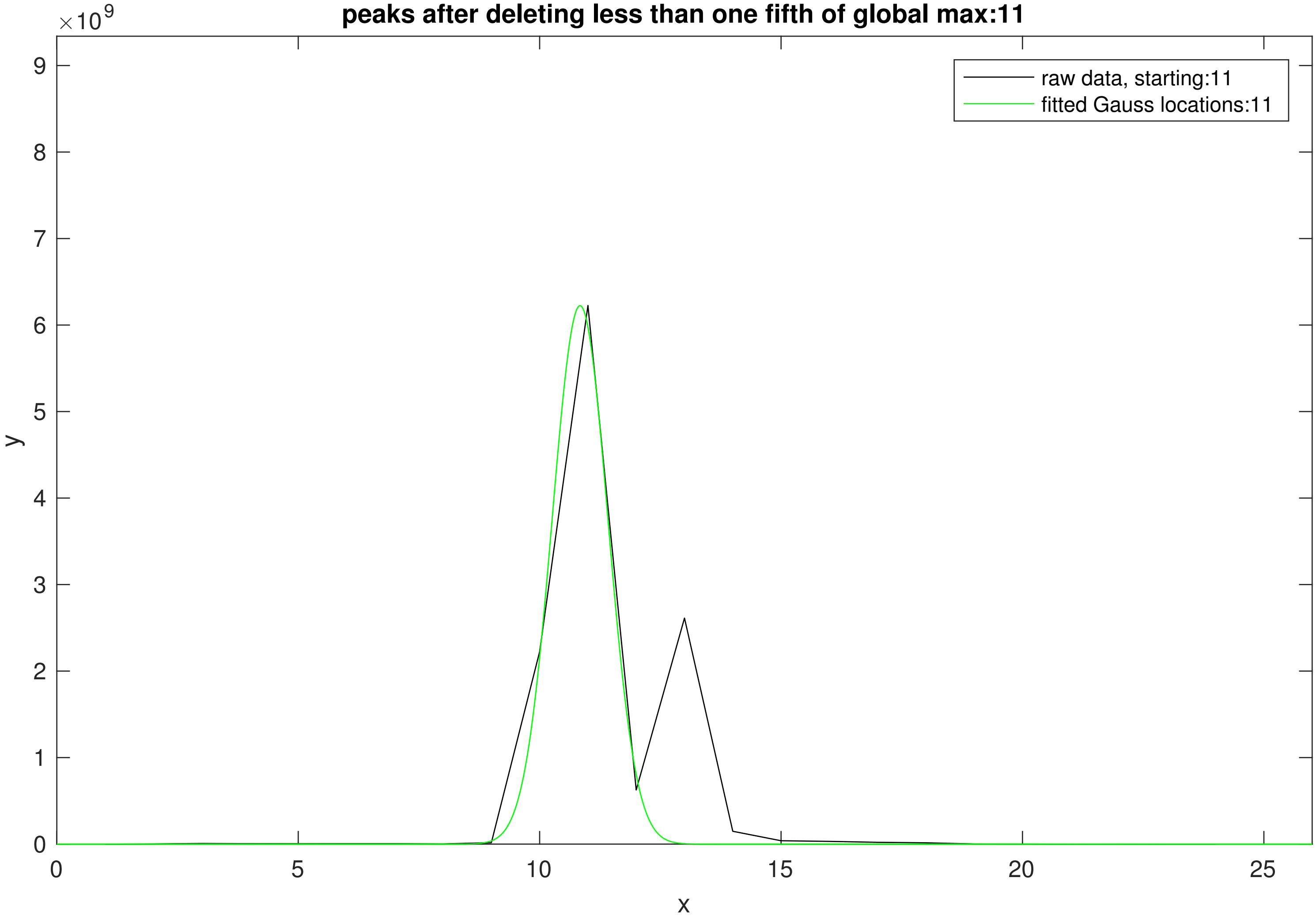

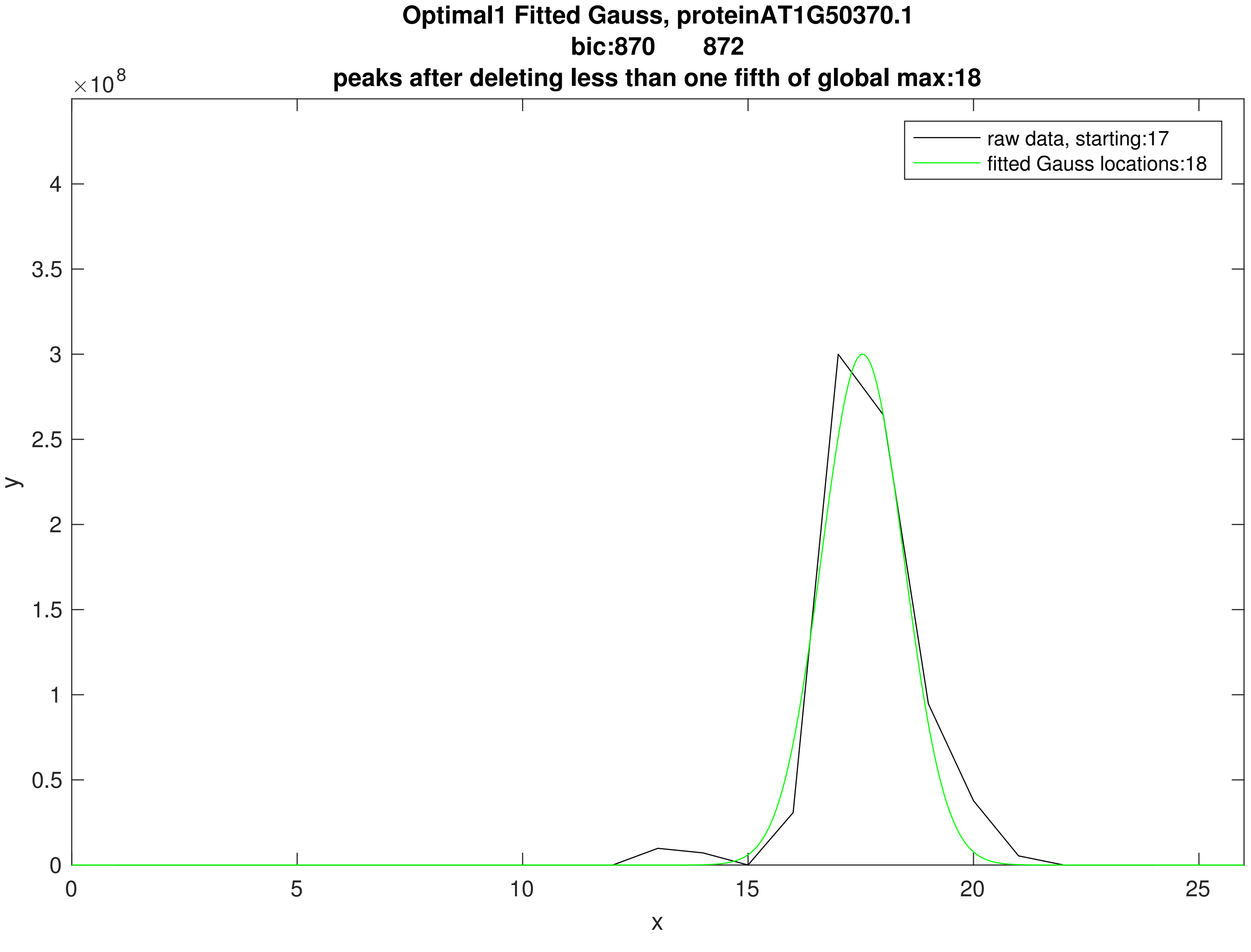

Optimal1 Fitted Gauss, proteinAT1G50370.1
bic:870 872
peaks after deleting less than one fifth of global max:18
×10^8
raw data, starting:17
fitted Gauss locations:18
4
3.5
3
2.5
2
1.5
1
0.5
0
y
0
5
10
15
20
25
x

**Optimal1 Fitted Gauss, proteinAT1G50380.1**
**bic:964**
**peaks after deleting less than one fifth of global max:15**

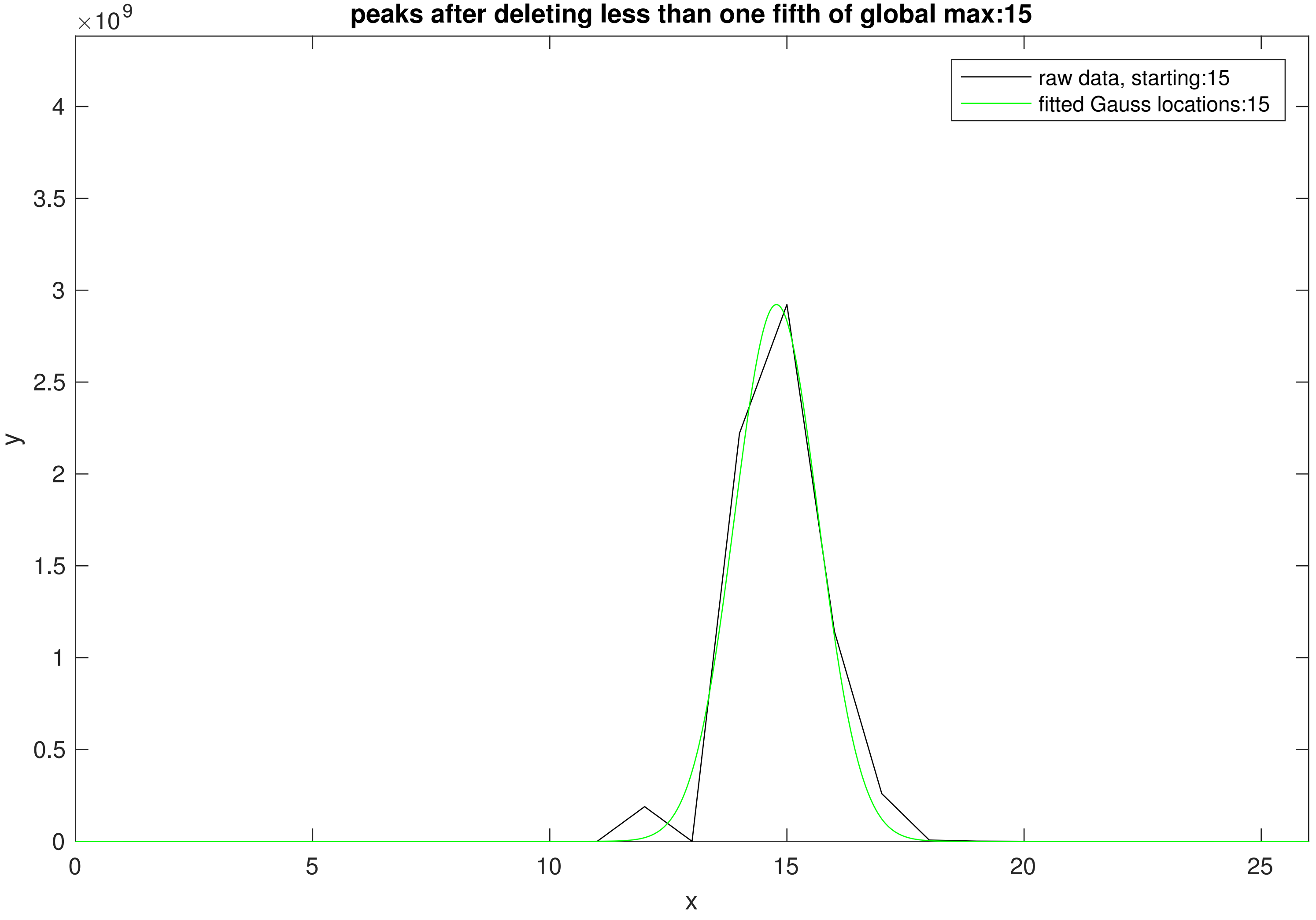

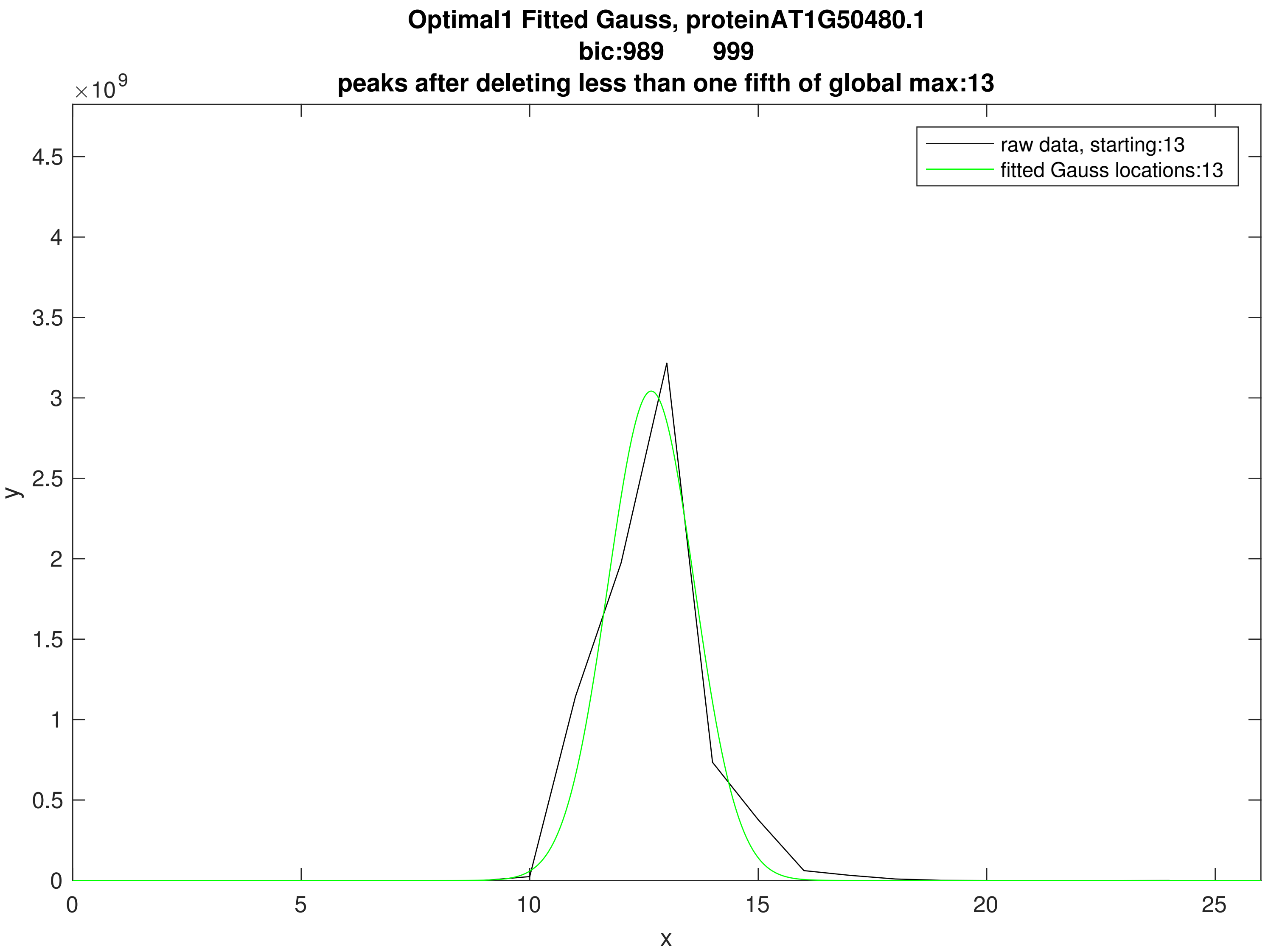

Optimal1 Fitted Gauss, proteinAT1G50480.1
bic:989 999
peaks after deleting less than one fifth of global max:13
×10^9
raw data, starting:13
fitted Gauss locations:13
y
x
0
0.5
1
1.5
2
2.5
3
3.5
4
4.5
5
10
15
20
25

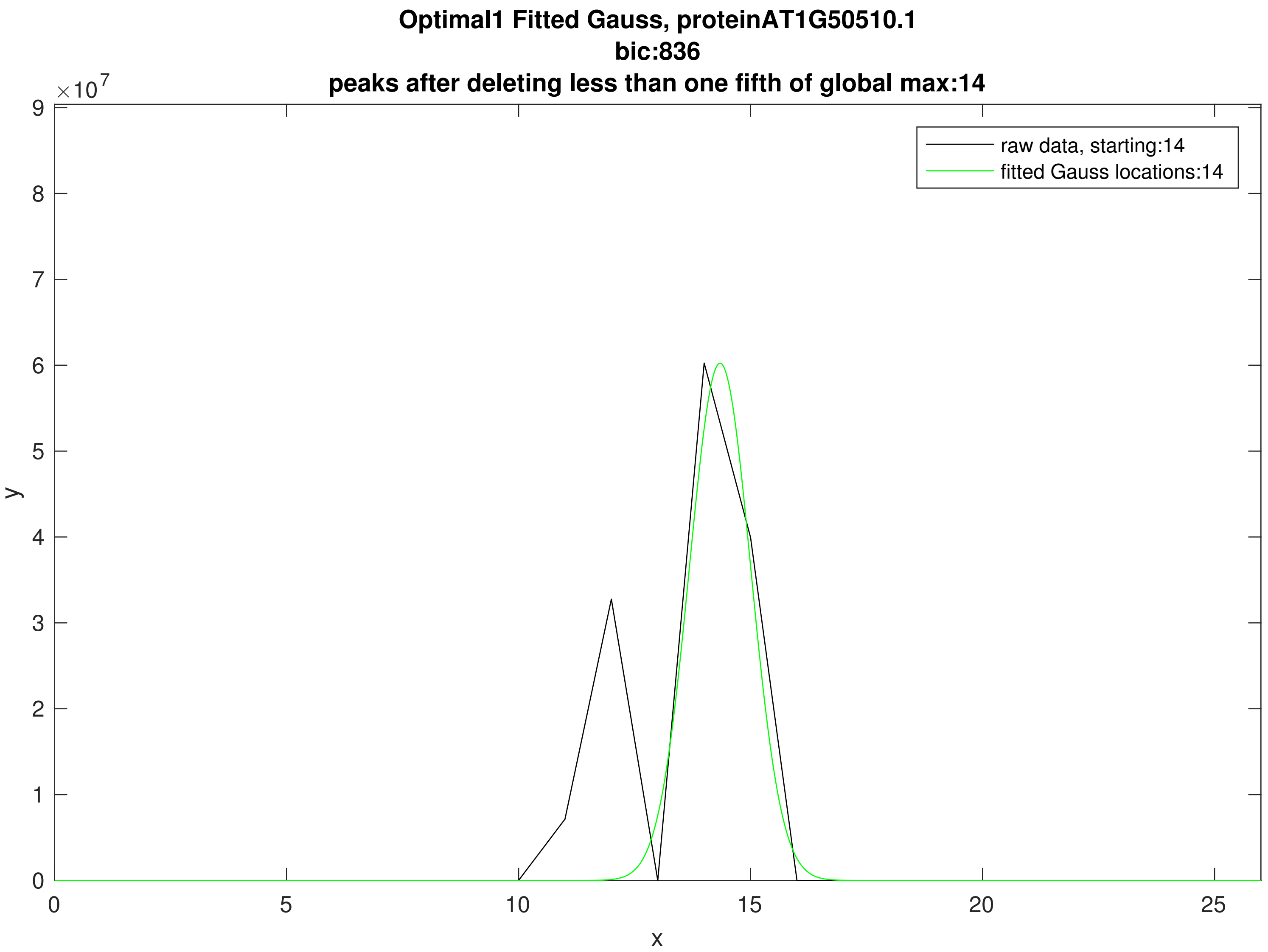

Optimal1 Fitted Gauss, proteinAT1G50510.1
bic:836
peaks after deleting less than one fifth of global max:14
×10^7
raw data, starting:14
fitted Gauss locations:14
y
x
0
5
10
15
20
25
0
1
2
3
4
5
6
7
8
9

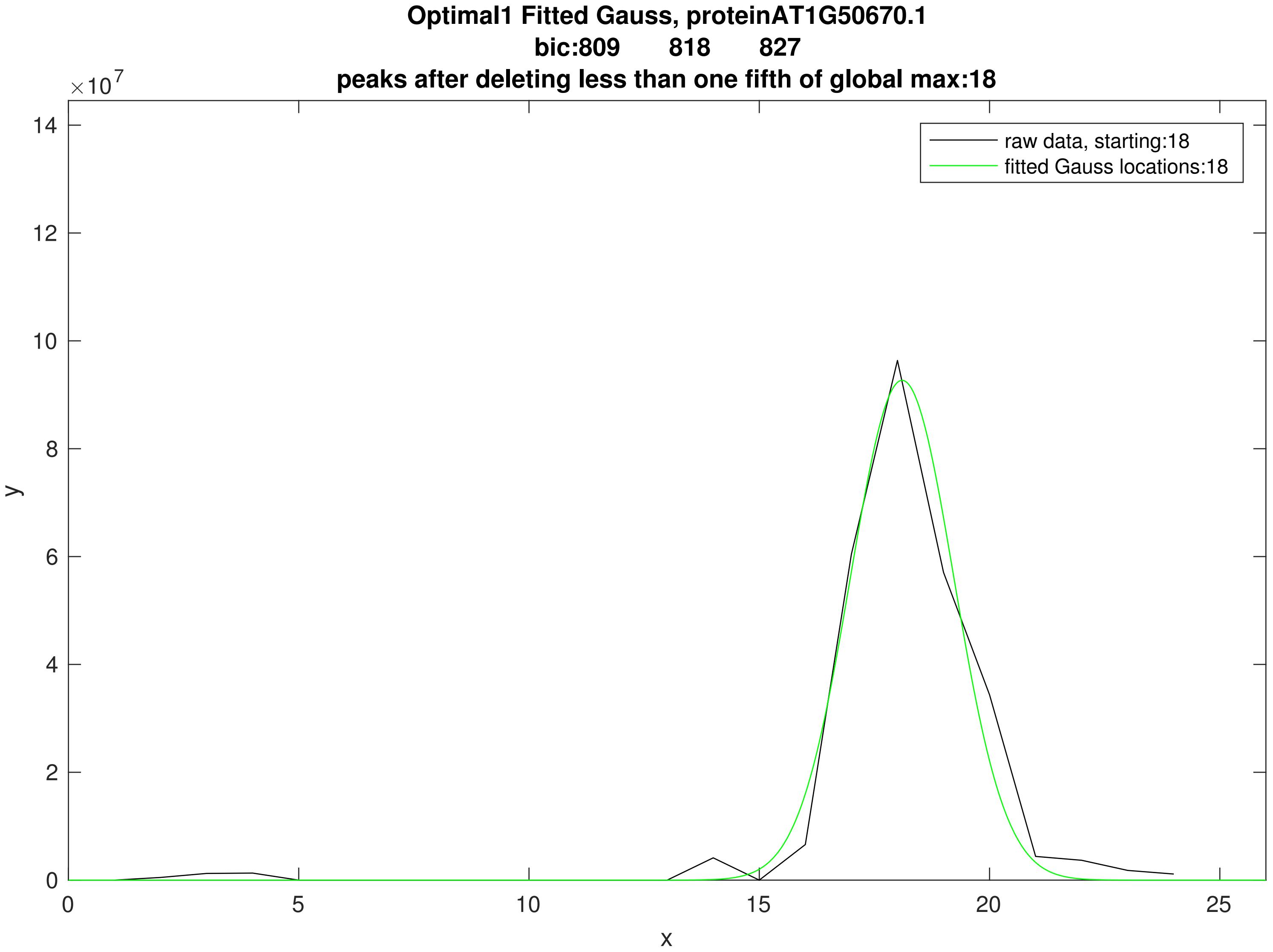
Optimal1 Fitted Gauss, proteinAT1G50670.1
bic:809 818 827
peaks after deleting less than one fifth of global max:18
×10^7
14
12
10
8
6
4
2
0
y
0
5
10
15
20
25
x
raw data, starting:18
fitted Gauss locations:18

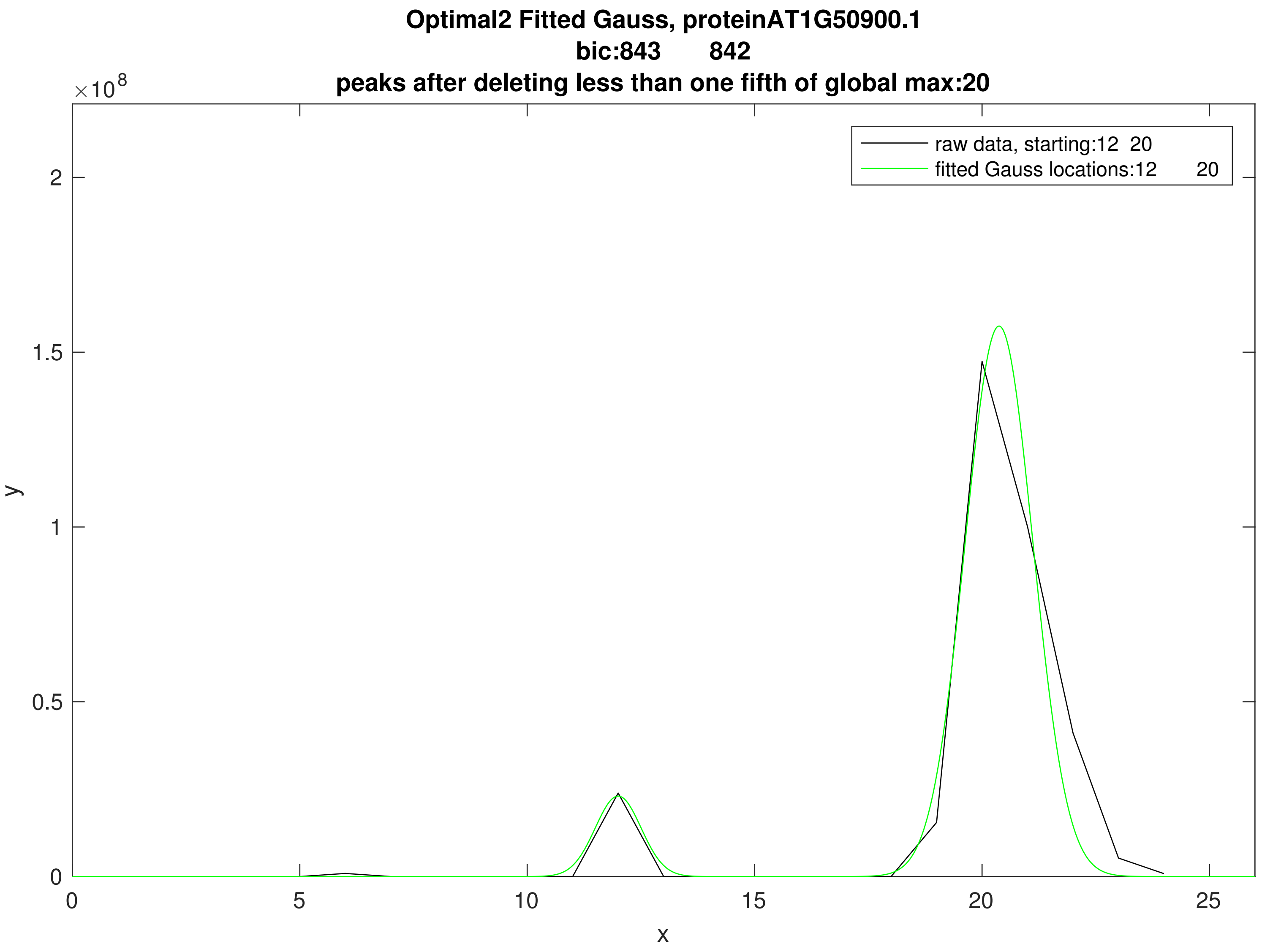

Optimal2 Fitted Gauss, proteinAT1G50900.1
bic:843 842
peaks after deleting less than one fifth of global max:20
×10^8
raw data, starting:12 20
fitted Gauss locations:12 20
y
x
0
0.5
1
1.5
2
0
5
10
15
20
25

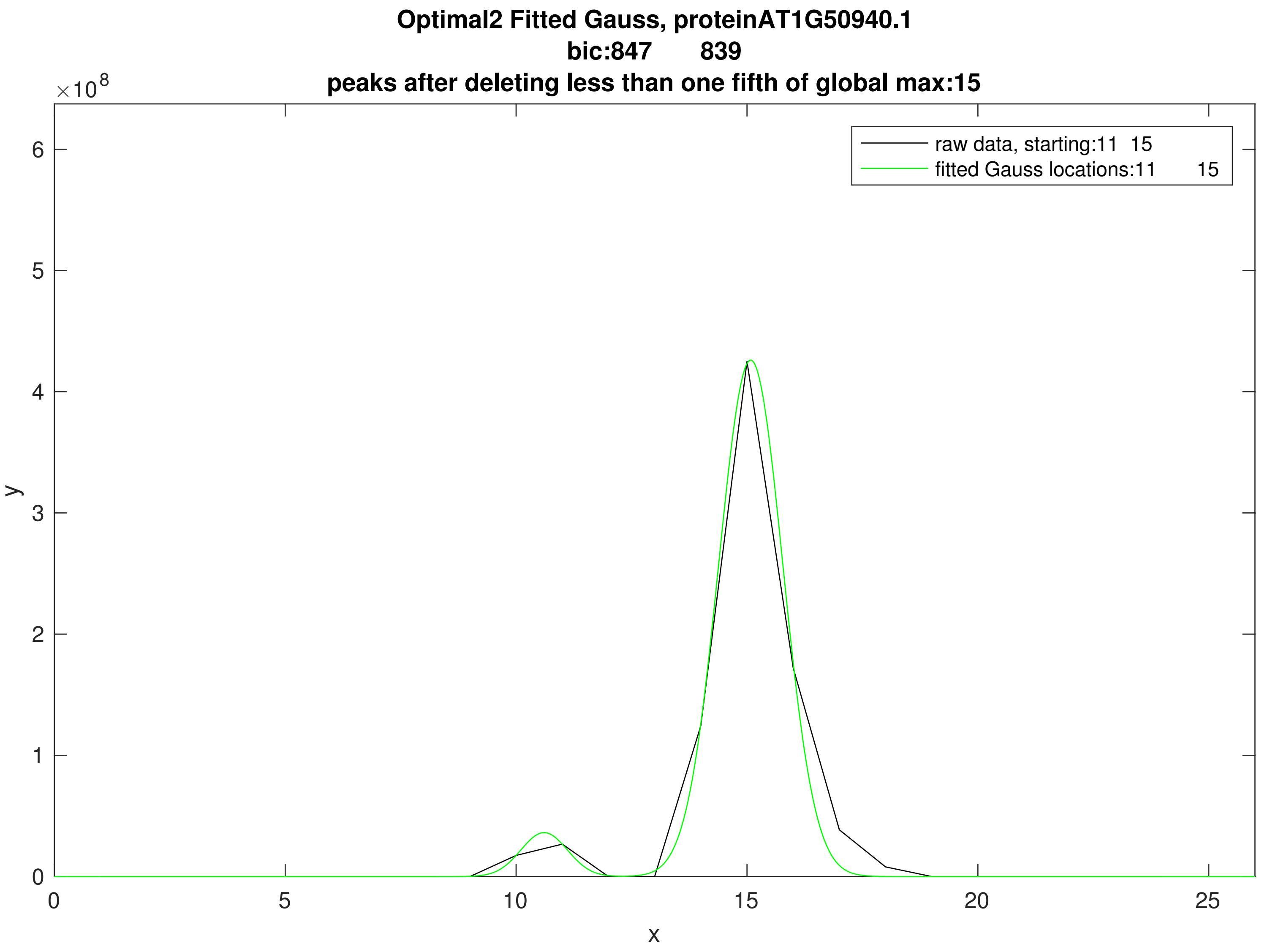
Optimal2 Fitted Gauss, proteinAT1G50940.1
bic:847 839
peaks after deleting less than one fifth of global max:15
×10^8
raw data, starting:11 15
fitted Gauss locations:11 15
y
x
0
1
2
3
4
5
6
0
5
10
15
20
25

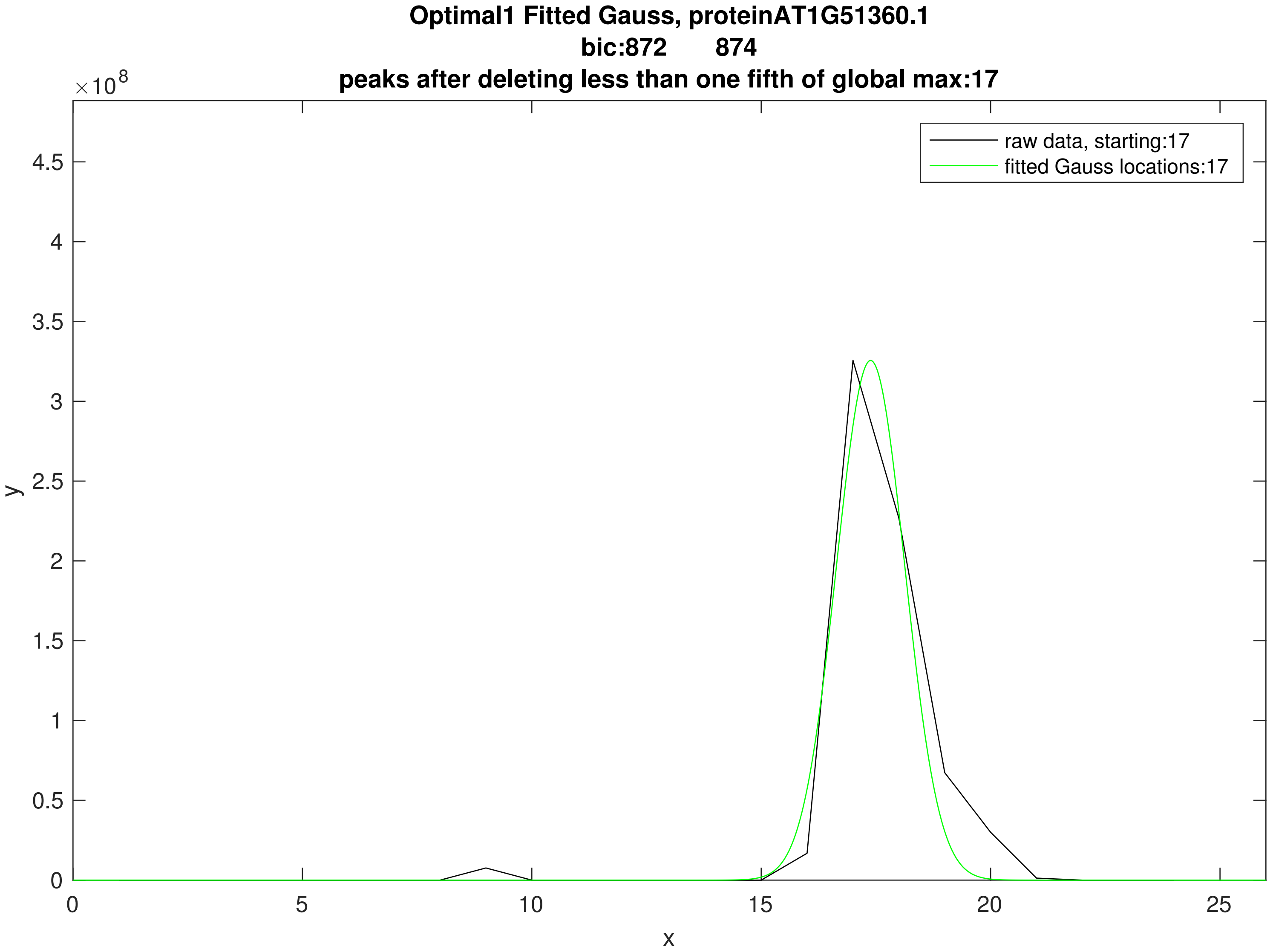

Optimal1 Fitted Gauss, proteinAT1G51360.1
bic:872 874
peaks after deleting less than one fifth of global max:17
×10^8
4.5
4
3.5
3
2.5
2
1.5
1
0.5
0
y
0
5
10
15
20
25
x
raw data, starting:17
fitted Gauss locations:17

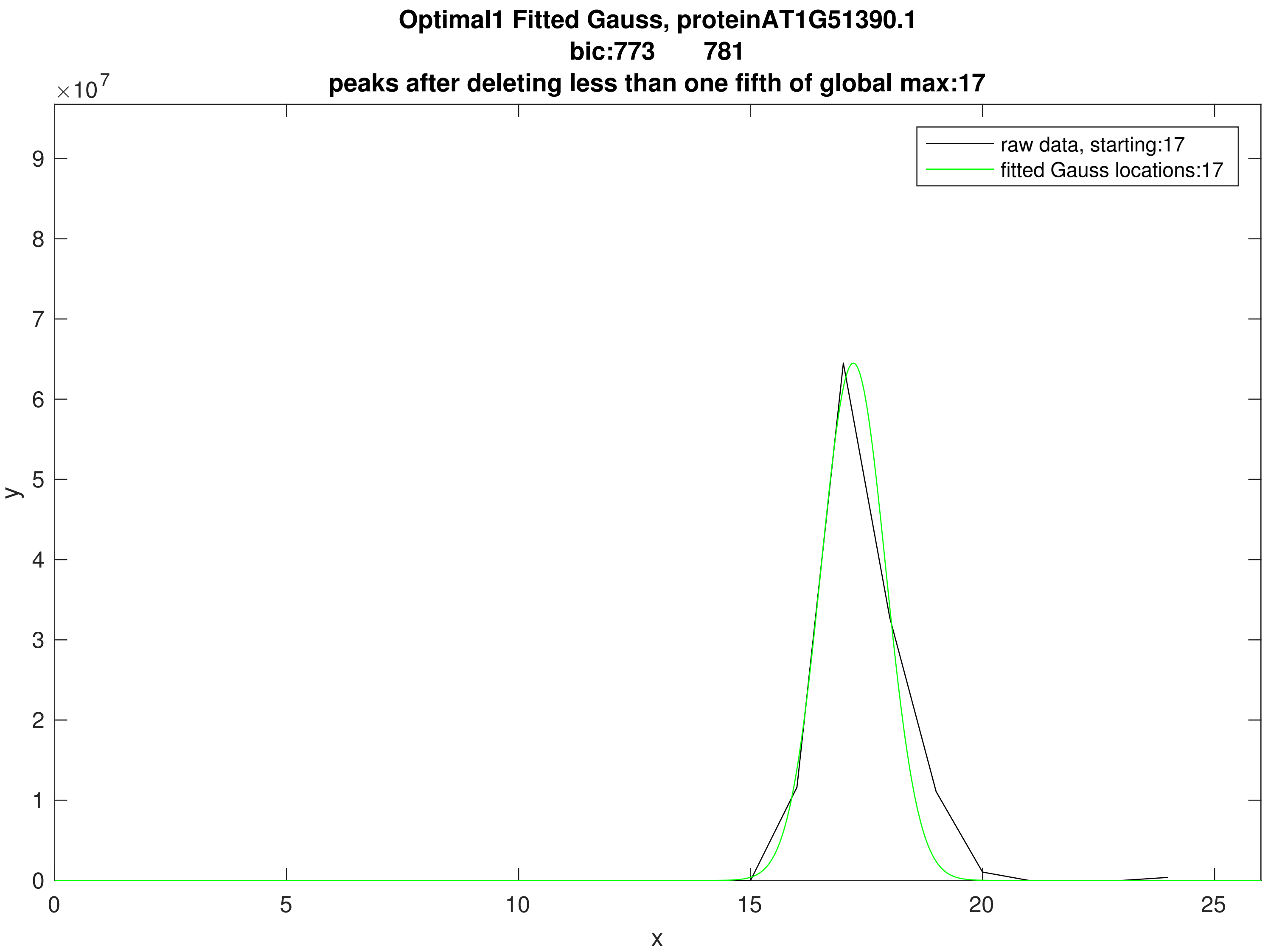

Optimal1 Fitted Gauss, proteinAT1G51390.1
bic:773 781
peaks after deleting less than one fifth of global max:17
×10^7
raw data, starting:17
fitted Gauss locations:17
y
x
0
1
2
3
4
5
6
7
8
9
0
5
10
15
20
25

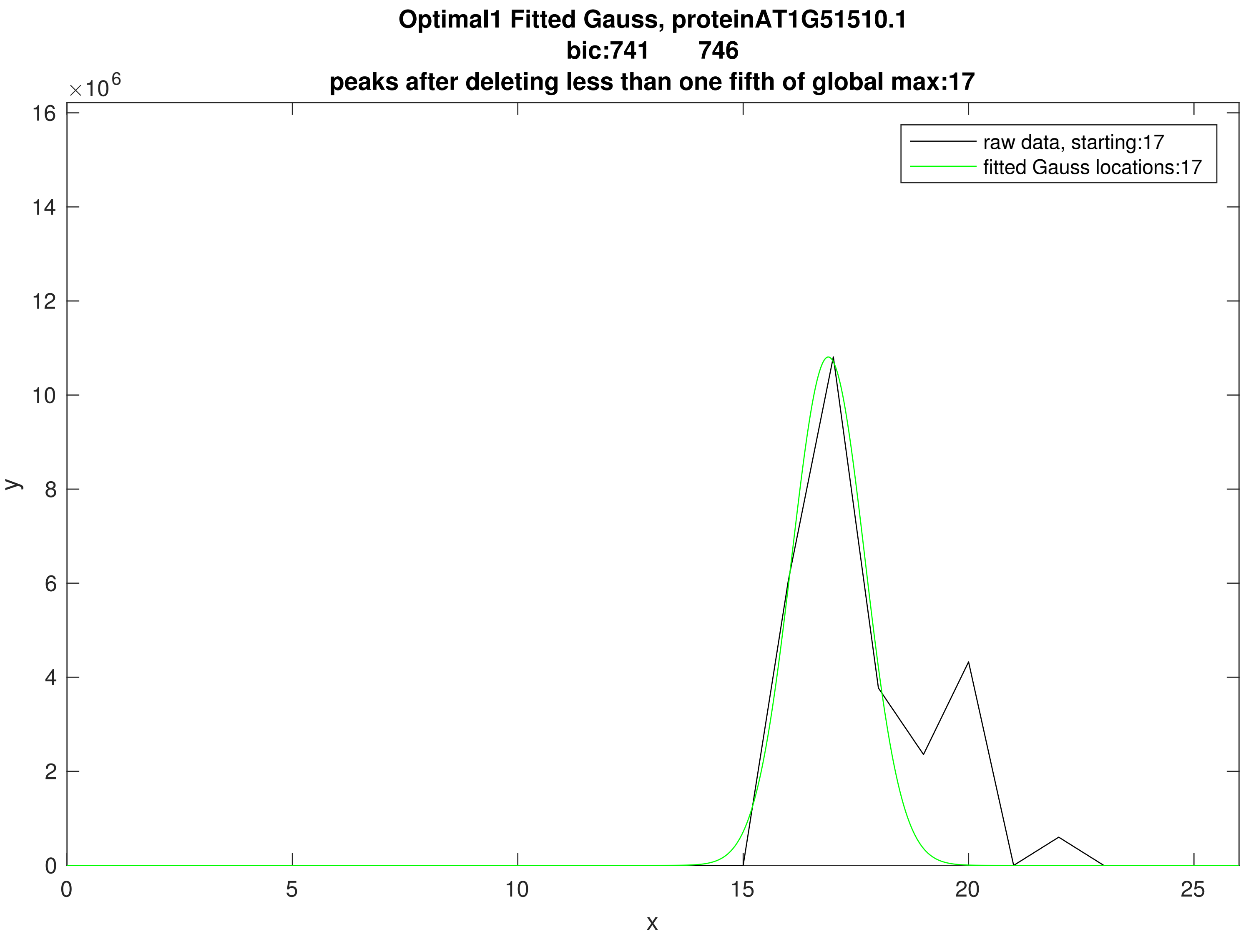

Optimal1 Fitted Gauss, proteinAT1G51510.1
bic:741 746
peaks after deleting less than one fifth of global max:17
×10^6
16
14
12
10
8
6
4
2
0
y
0
5
10
15
20
25
x
raw data, starting:17
fitted Gauss locations:17

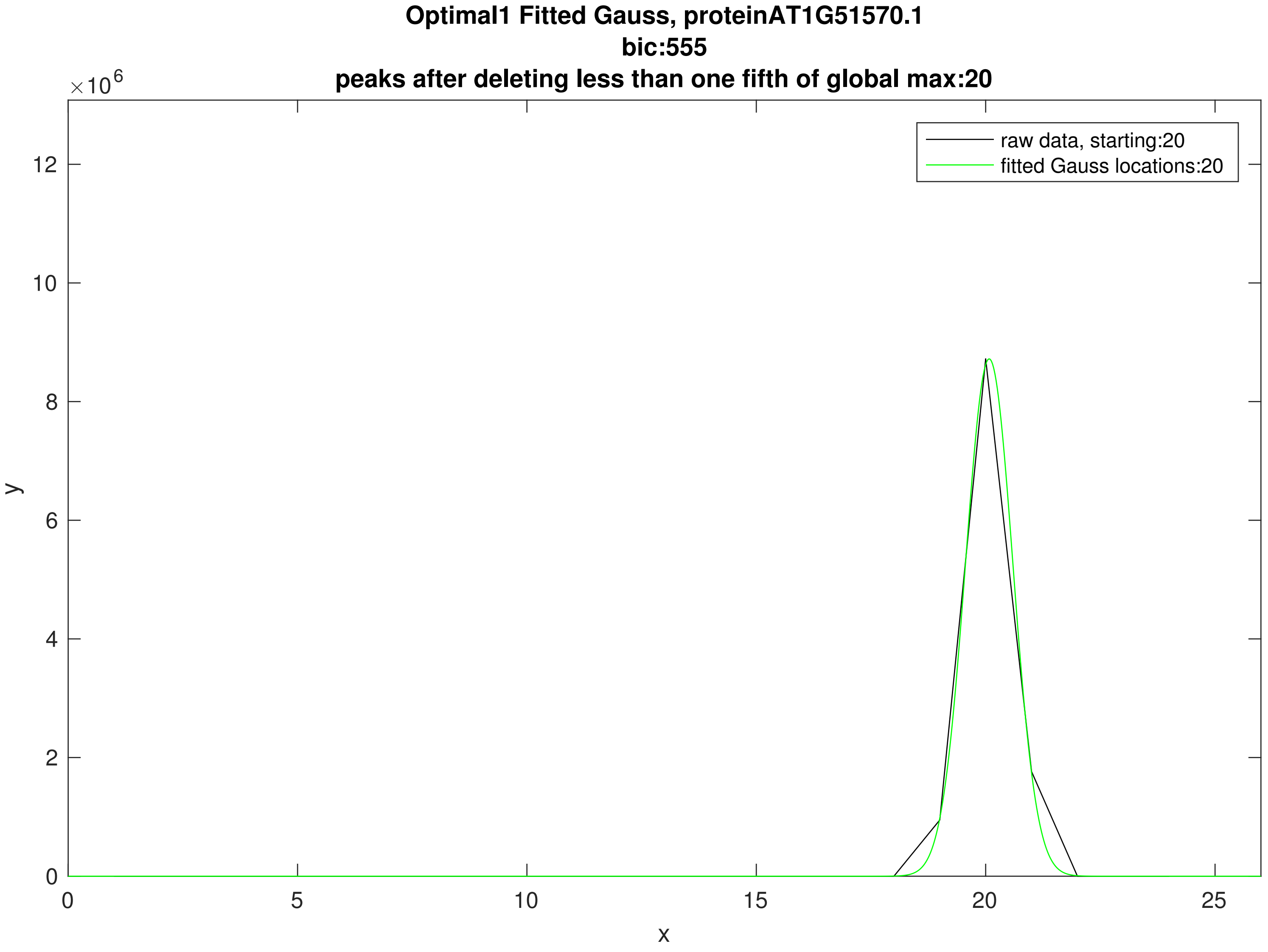

Optimal1 Fitted Gauss, proteinAT1G51570.1
bic:555
peaks after deleting less than one fifth of global max:20
×10^6
raw data, starting:20
fitted Gauss locations:20
y
x
0
2
4
6
8
10
12
0
5
10
15
20
25

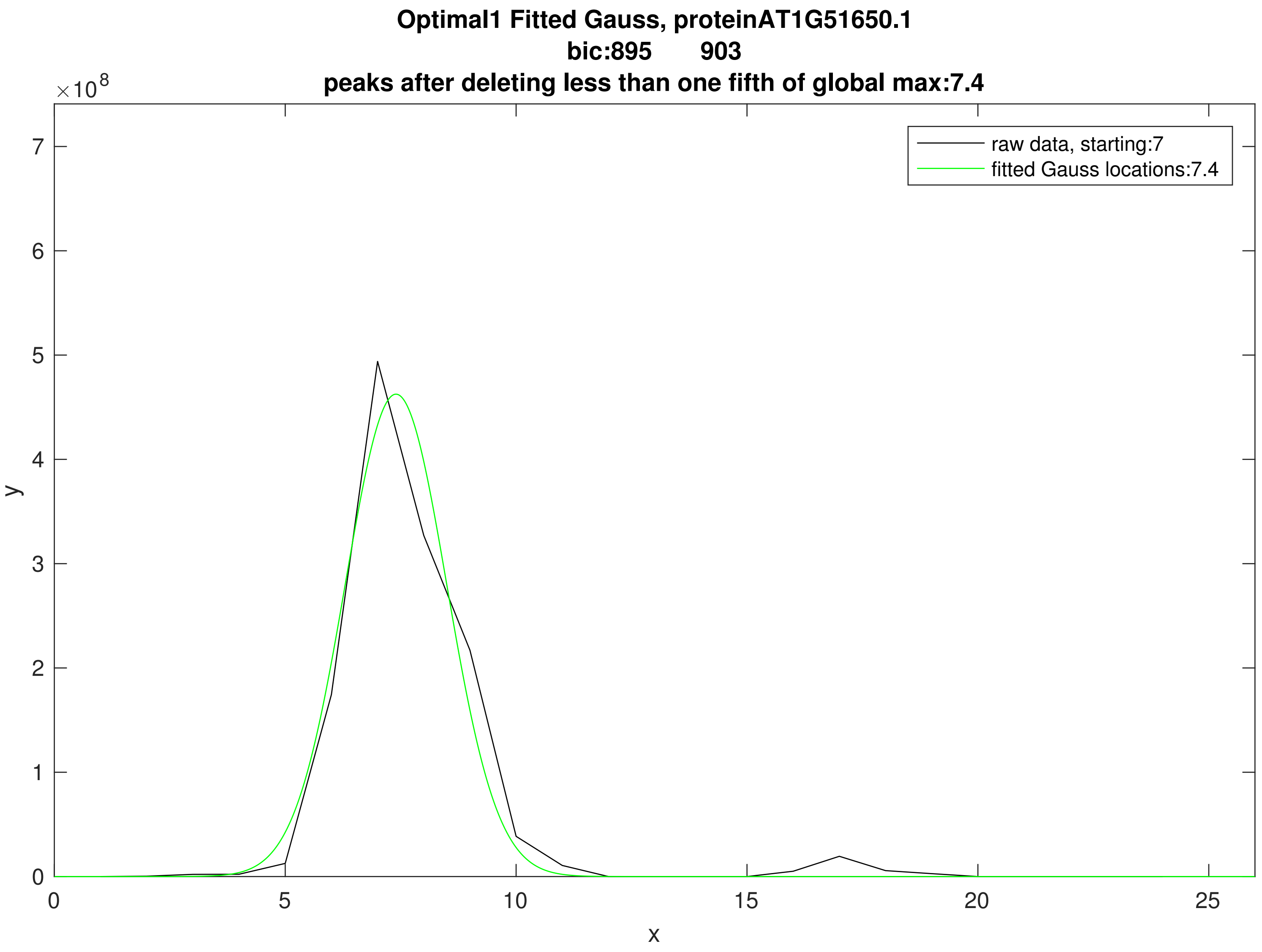
Optimal1 Fitted Gauss, proteinAT1G51650.1
bic:895 903
peaks after deleting less than one fifth of global max:7.4
×10^8
raw data, starting:7
fitted Gauss locations:7.4
y
x
0
1
2
3
4
5
6
7
10
15
20
25

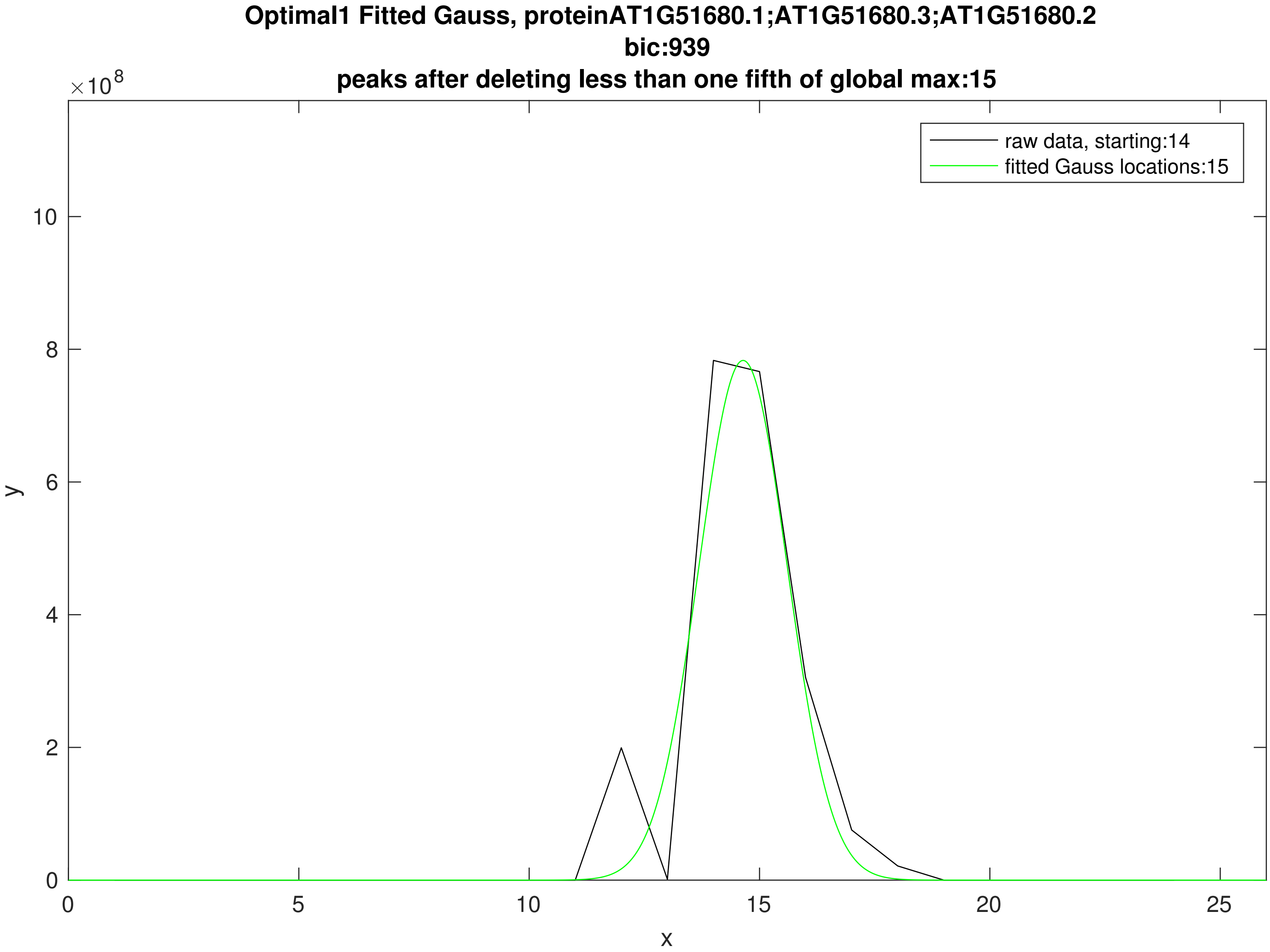
Optimal1 Fitted Gauss, proteinAT1G51680.1;AT1G51680.3;AT1G51680.2
bic:939
peaks after deleting less than one fifth of global max:15
×10^8
raw data, starting:14
fitted Gauss locations:15
y
x
0
2
4
6
8
10
5
15
20
25

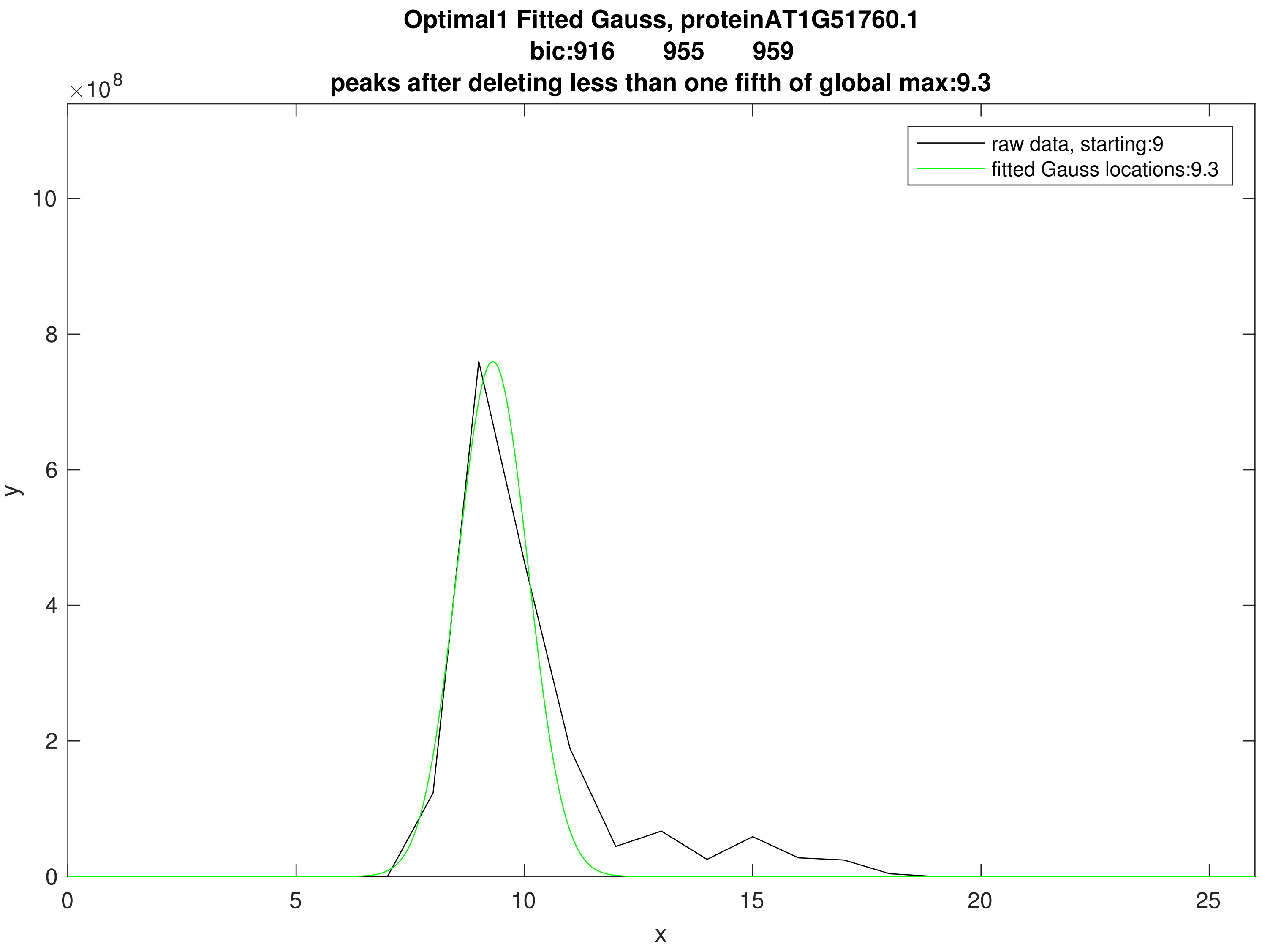
Optimal1 Fitted Gauss, proteinAT1G51760.1
bic:916 955 959
peaks after deleting less than one fifth of global max:9.3
×10^8
raw data, starting:9
fitted Gauss locations:9.3
y
x
0
2
4
6
8
10
0
5
10
15
20
25

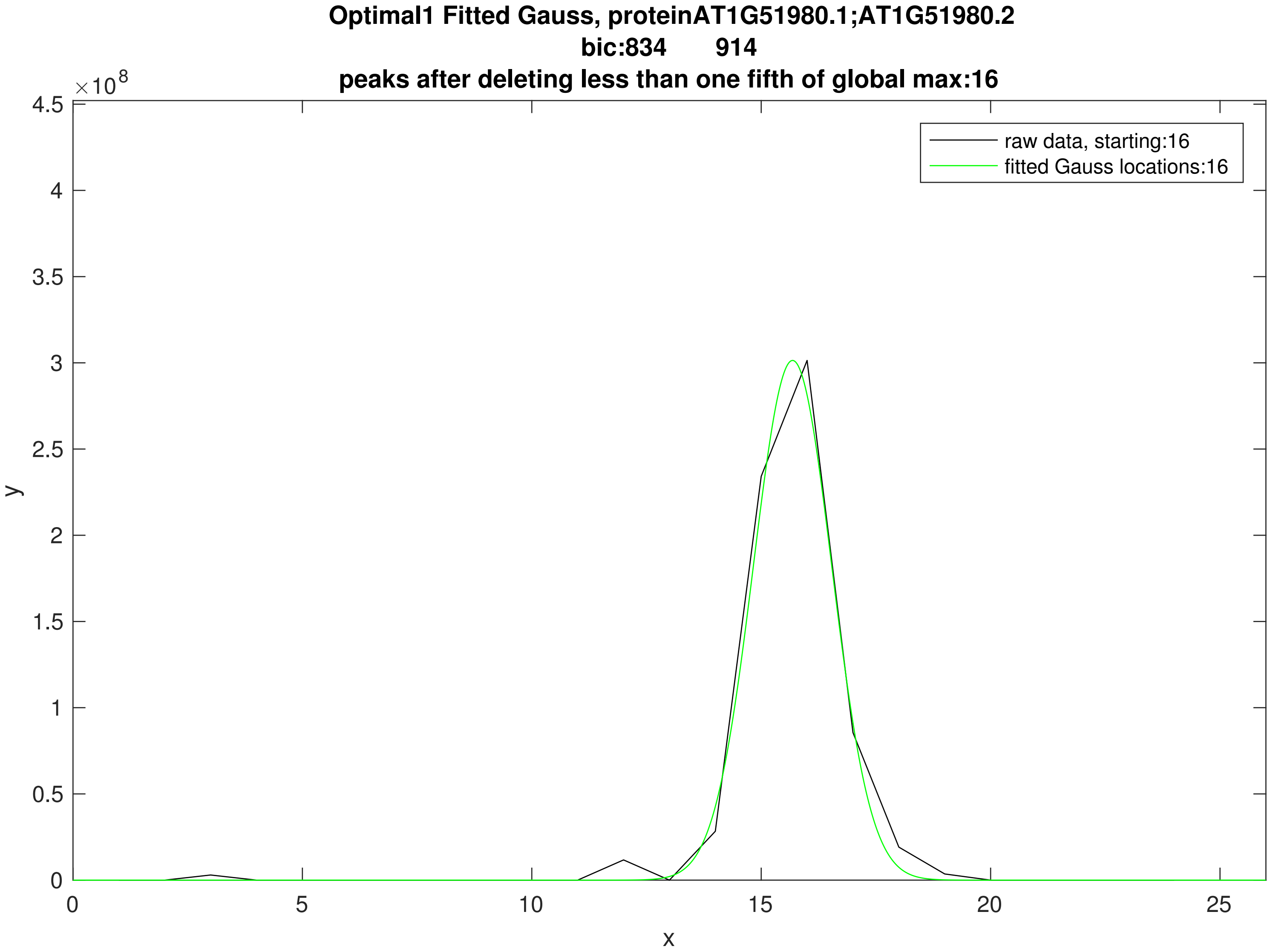

Optimal1 Fitted Gauss, proteinAT1G51980.1;AT1G51980.2
bic:834 914
peaks after deleting less than one fifth of global max:16
×10^8
4.5
4
3.5
3
2.5
2
1.5
1
0.5
0
y
0
5
10
15
20
25
x
raw data, starting:16
fitted Gauss locations:16

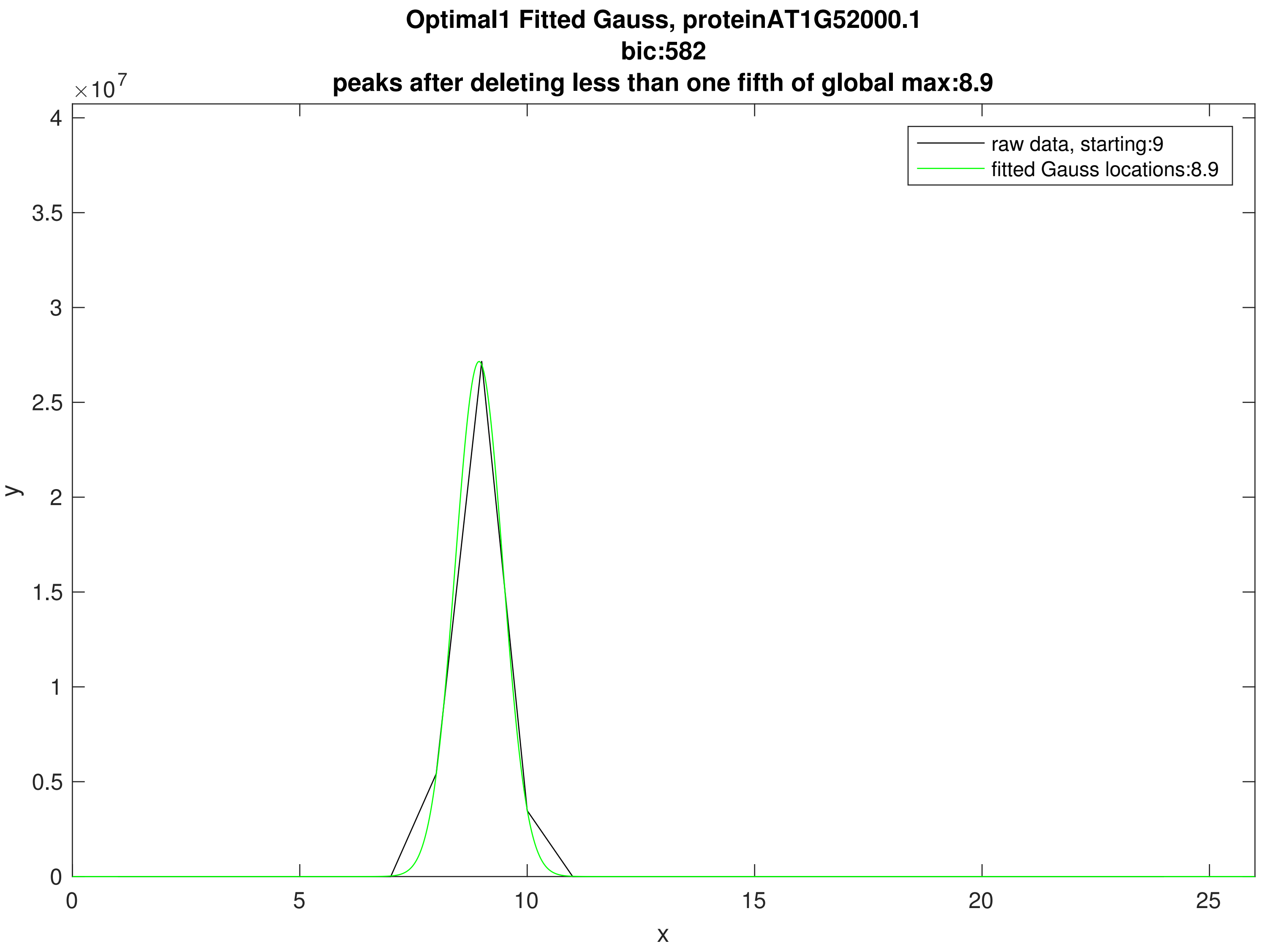

Optimal1 Fitted Gauss, proteinAT1G52000.1
bic:582
peaks after deleting less than one fifth of global max:8.9
×10^7
raw data, starting:9
fitted Gauss locations:8.9
y
x
0
0.5
1
1.5
2
2.5
3
3.5
4
5
10
15
20
25

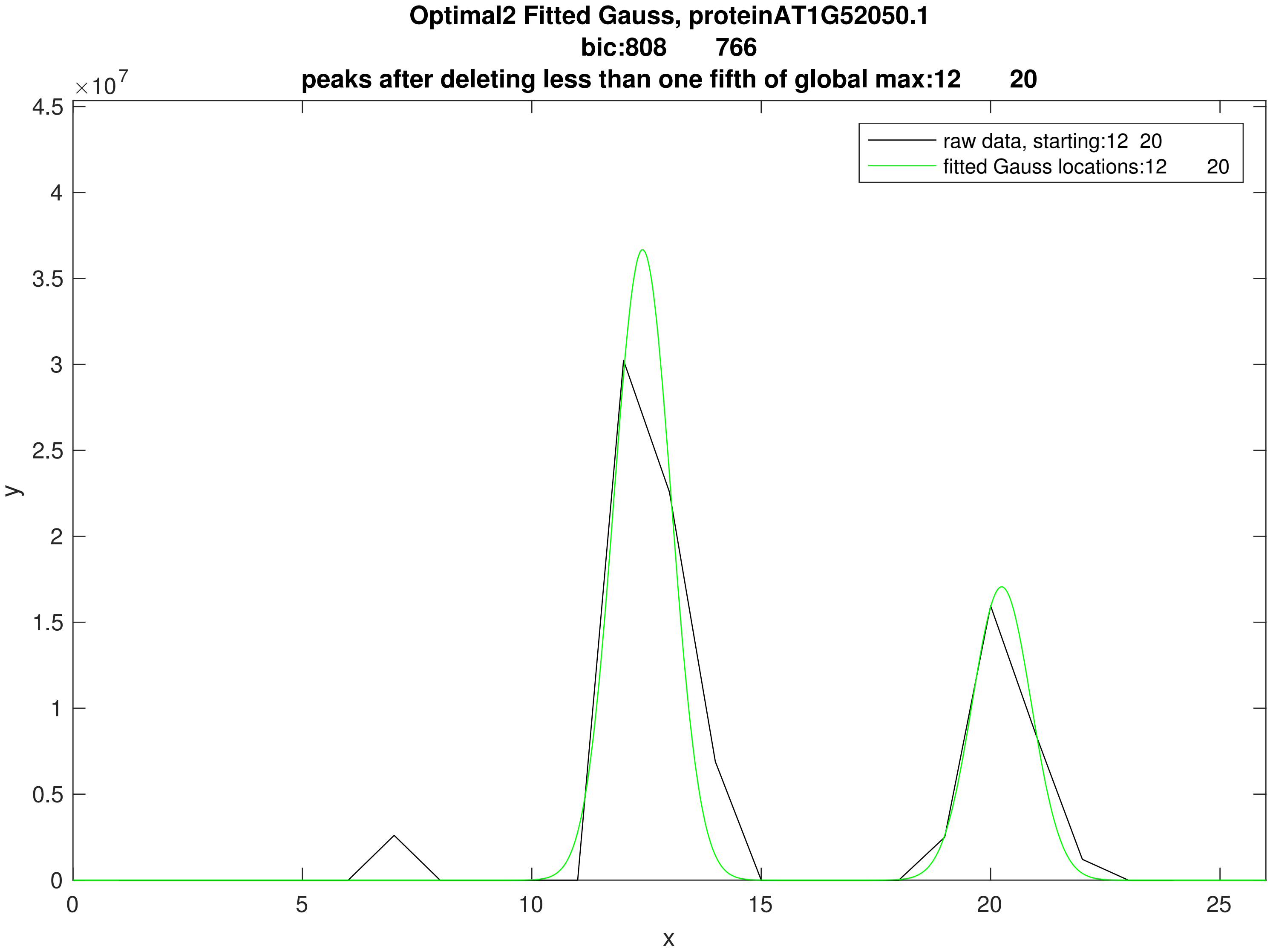

Optimal2 Fitted Gauss, proteinAT1G52050.1
bic:808 766
peaks after deleting less than one fifth of global max:12 20
raw data, starting:12 20
fitted Gauss locations:12 20
×10^7
x
y

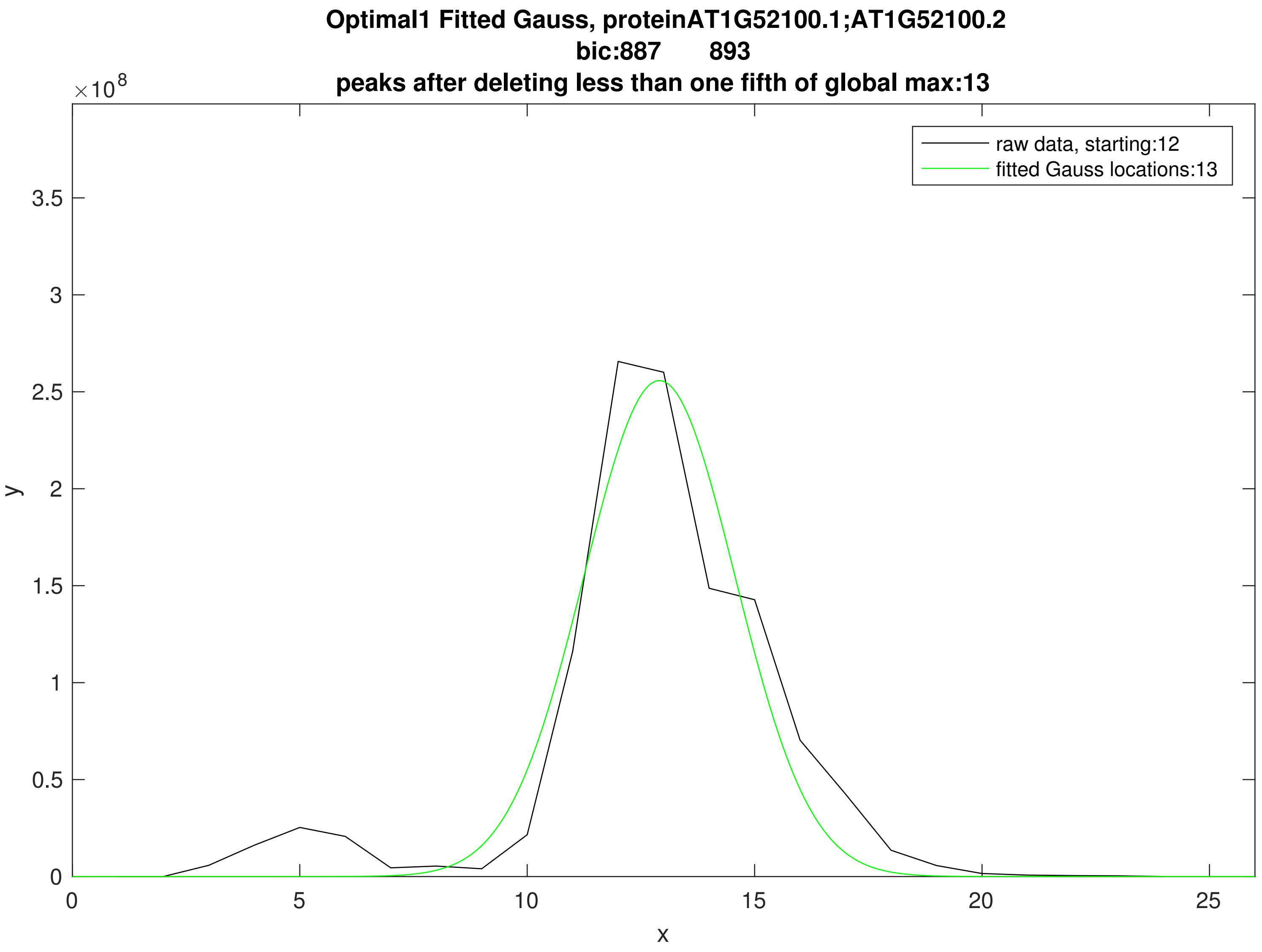
Optimal1 Fitted Gauss, proteinAT1G52100.1;AT1G52100.2
bic:887 893
peaks after deleting less than one fifth of global max:13
×10^8
raw data, starting:12
fitted Gauss locations:13
y
x
0
0.5
1
1.5
2
2.5
3
3.5
0
5
10
15
20
25

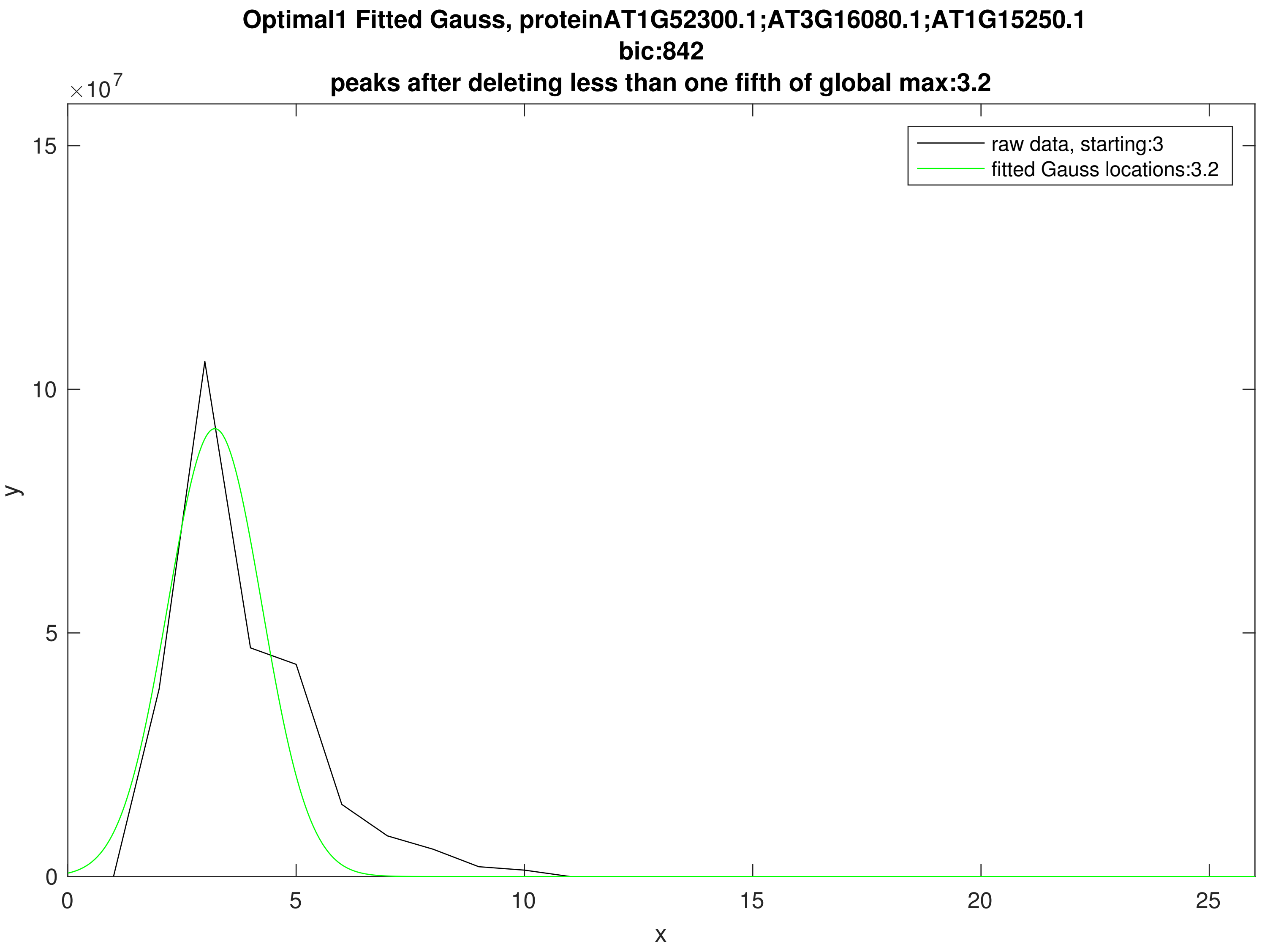
Optimal1 Fitted Gauss, proteinAT1G52300.1;AT3G16080.1;AT1G15250.1
bic:842
peaks after deleting less than one fifth of global max:3.2
$\times 10^7$
15
10
5
0
y
0
5
10
15
20
25
x
raw data, starting:3
fitted Gauss locations:3.2

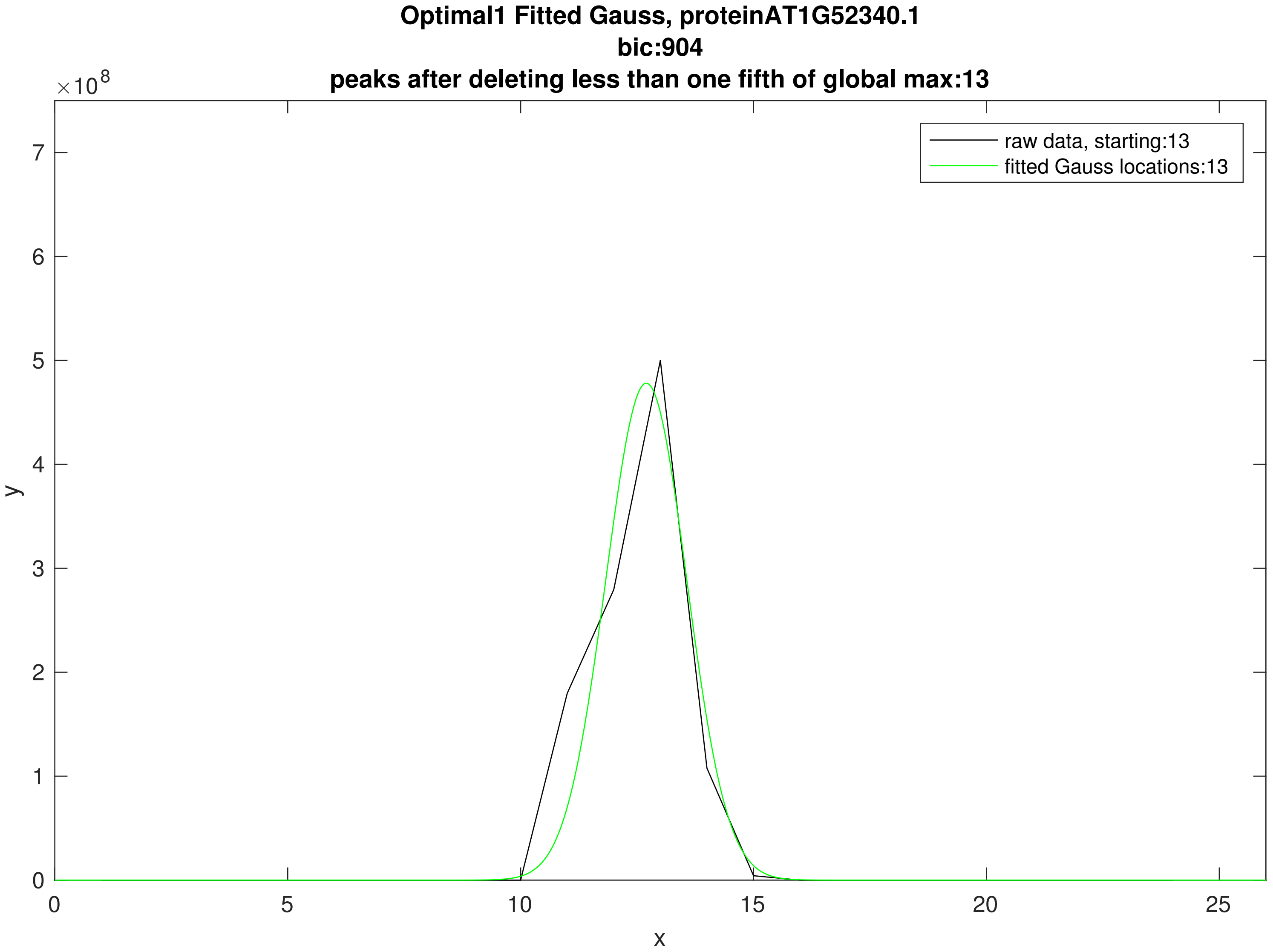

Optimal1 Fitted Gauss, proteinAT1G52340.1
bic:904
peaks after deleting less than one fifth of global max:13
×10^8
raw data, starting:13
fitted Gauss locations:13
y
x
0
1
2
3
4
5
6
7
0
5
10
15
20
25

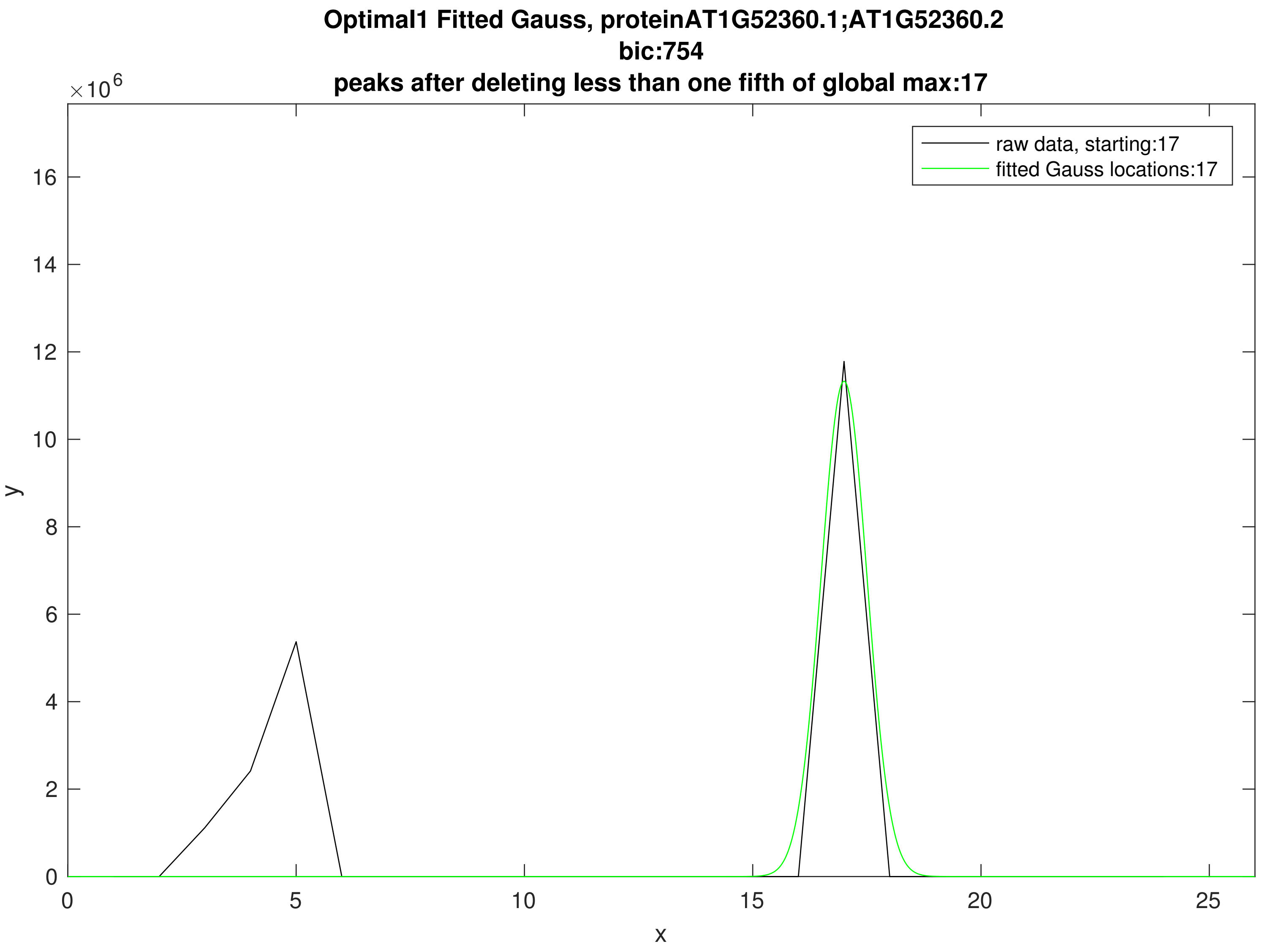

Optimal1 Fitted Gauss, proteinAT1G52360.1;AT1G52360.2
bic:754
peaks after deleting less than one fifth of global max:17
×10^6
raw data, starting:17
fitted Gauss locations:17
y
x
0
2
4
6
8
10
12
14
16
0
5
10
15
20
25

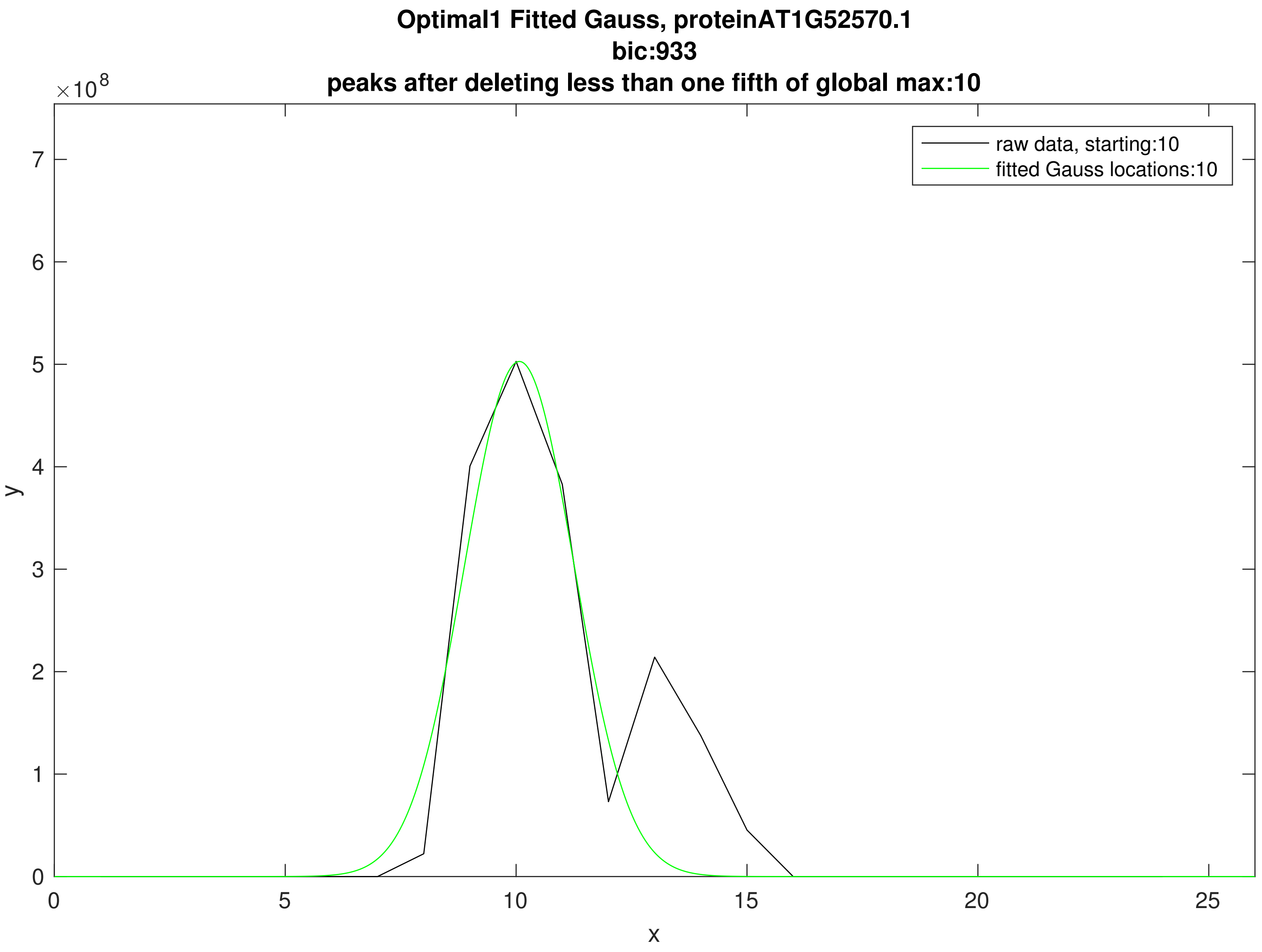

Optimal1 Fitted Gauss, proteinAT1G52570.1
bic:933
peaks after deleting less than one fifth of global max:10
×10^8
raw data, starting:10
fitted Gauss locations:10
y
x
0
1
2
3
4
5
6
7
0
5
10
15
20
25

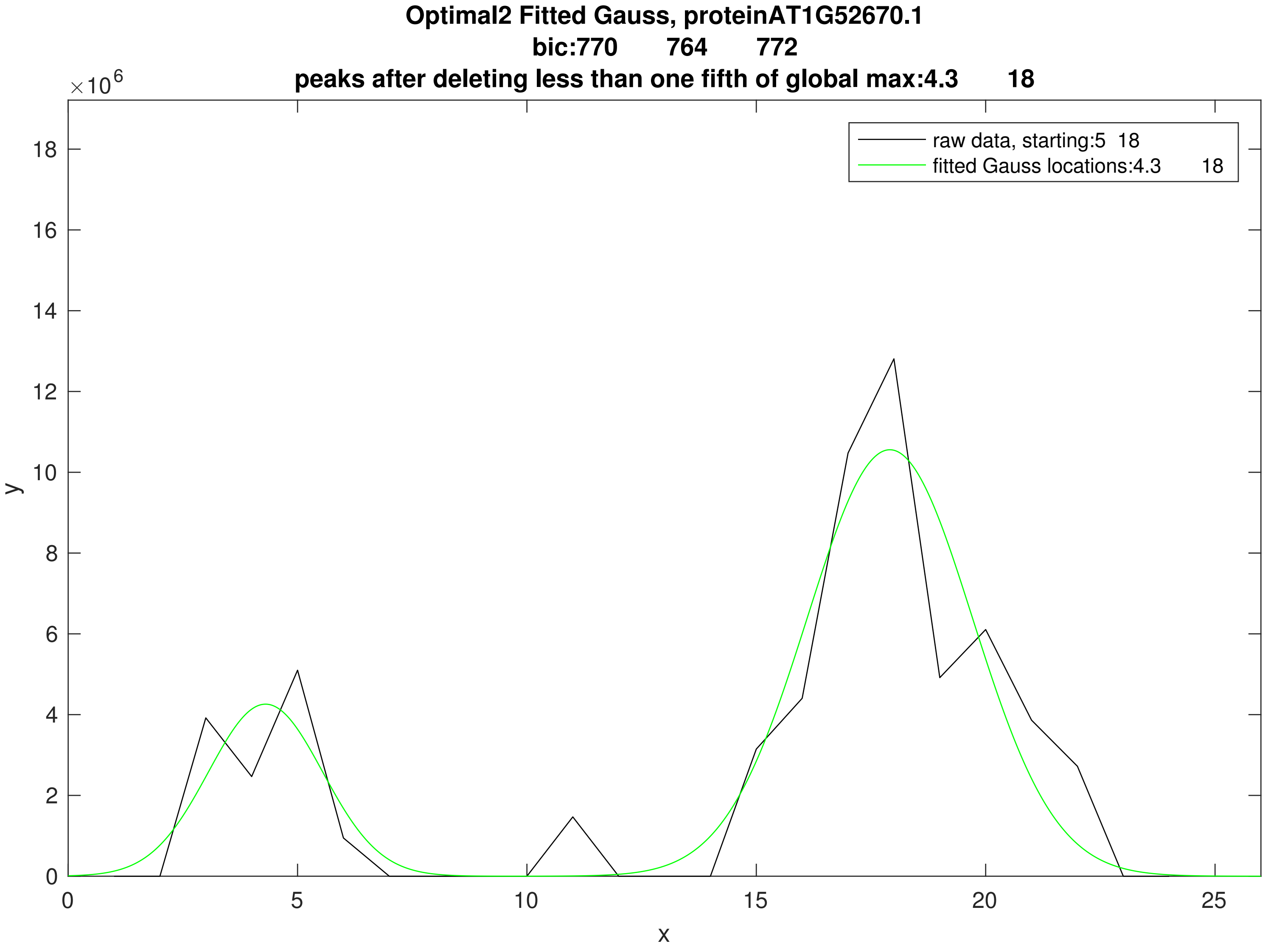

Optimal2 Fitted Gauss, proteinAT1G52670.1
bic:770 764 772
peaks after deleting less than one fifth of global max:4.3 18
×10^6
raw data, starting:5 18
fitted Gauss locations:4.3 18
y
x
0
2
4
6
8
10
12
14
16
18
0
5
10
15
20
25

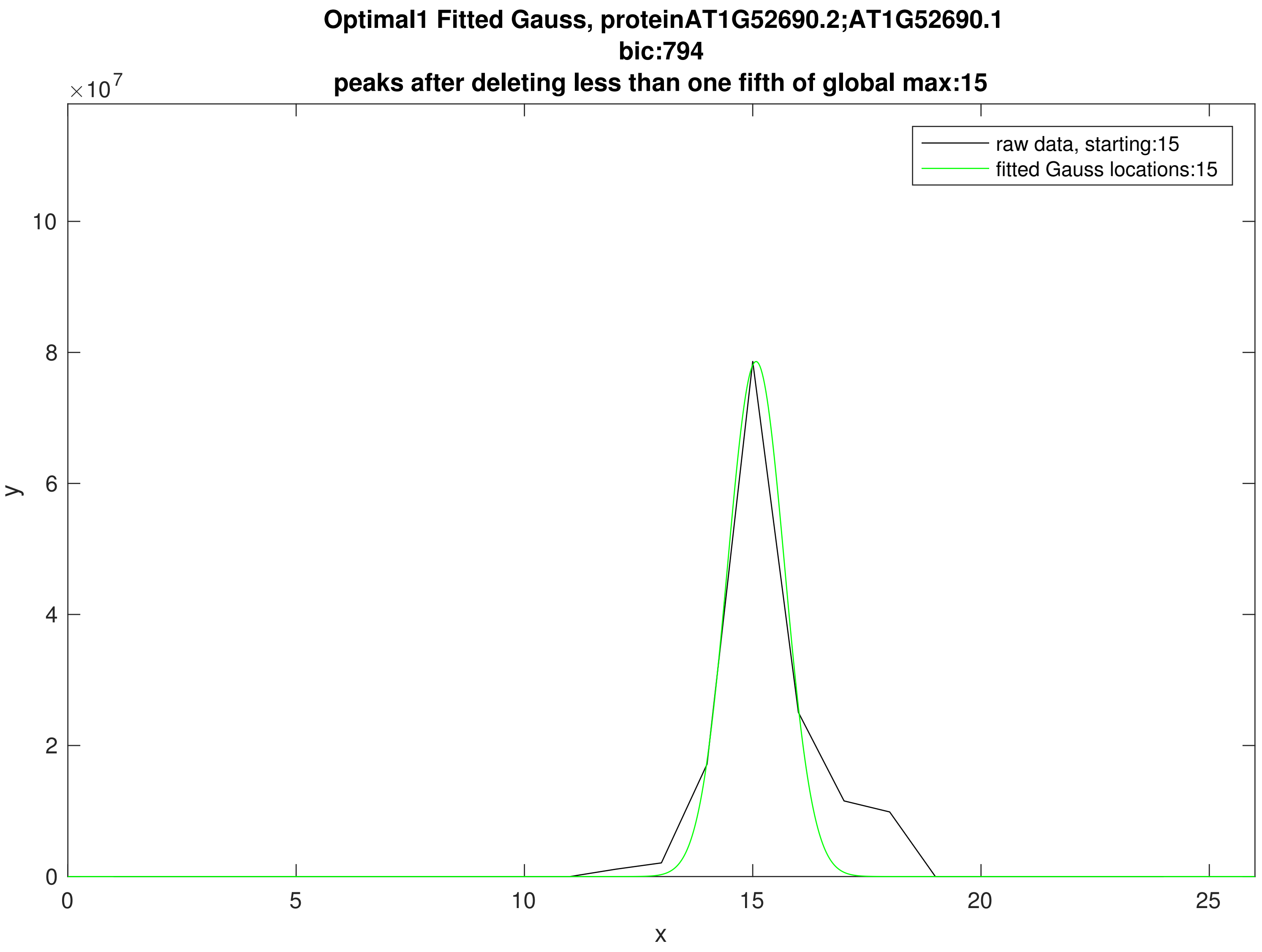

Optimal1 Fitted Gauss, proteinAT1G52690.2;AT1G52690.1
bic:794
peaks after deleting less than one fifth of global max:15
×10^7
raw data, starting:15
fitted Gauss locations:15
10
8
6
4
2
0
y
0
5
10
15
20
25
x

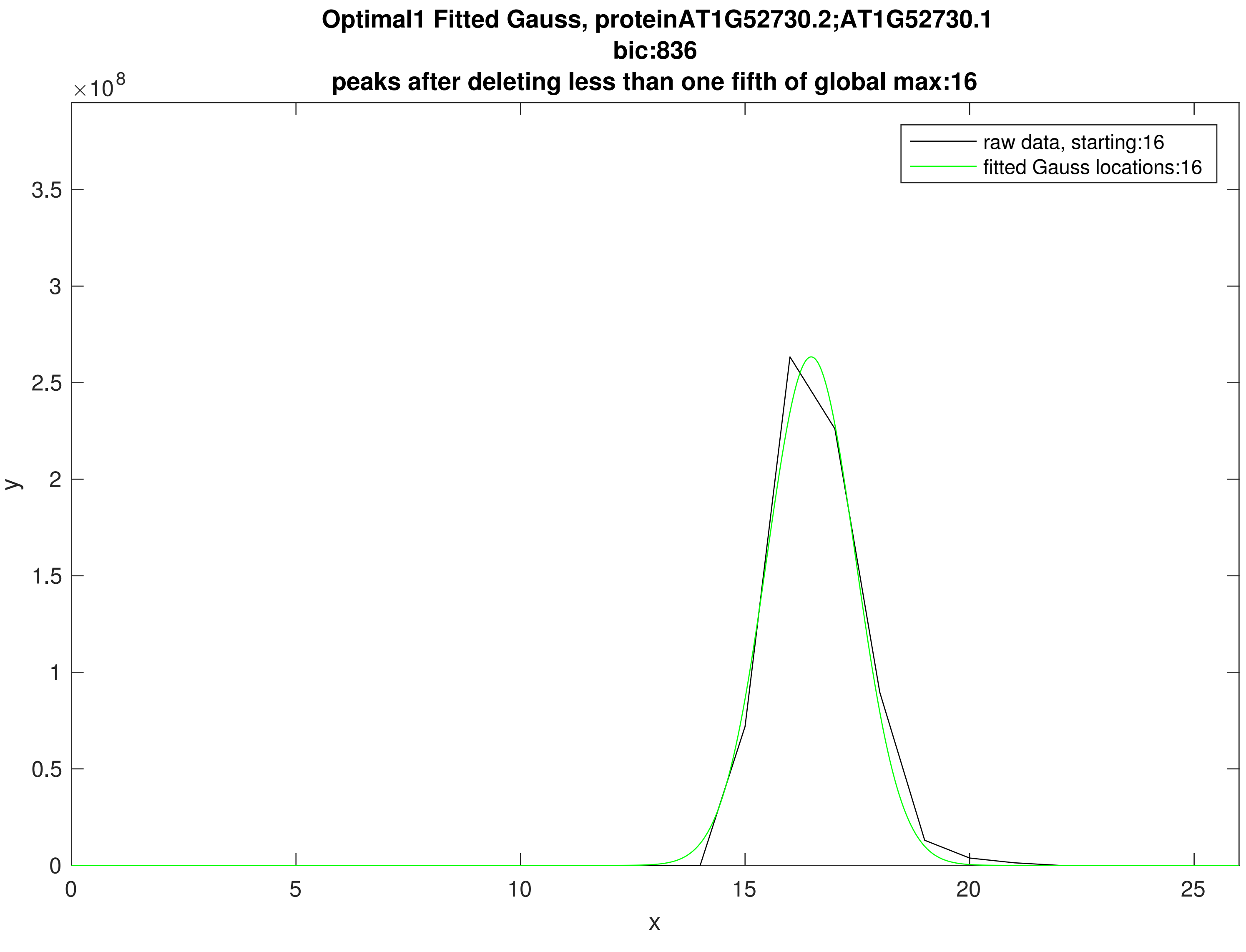

Optimal1 Fitted Gauss, proteinAT1G52730.2;AT1G52730.1
bic:836
peaks after deleting less than one fifth of global max:16
×10^8
raw data, starting:16
fitted Gauss locations:16
3.5
3
2.5
2
1.5
1
0.5
0
y
0
5
10
15
20
25
x

Optimal1 Fitted Gauss, proteinAT1G53030.1
bic:664
peaks after deleting less than one fifth of global max:20

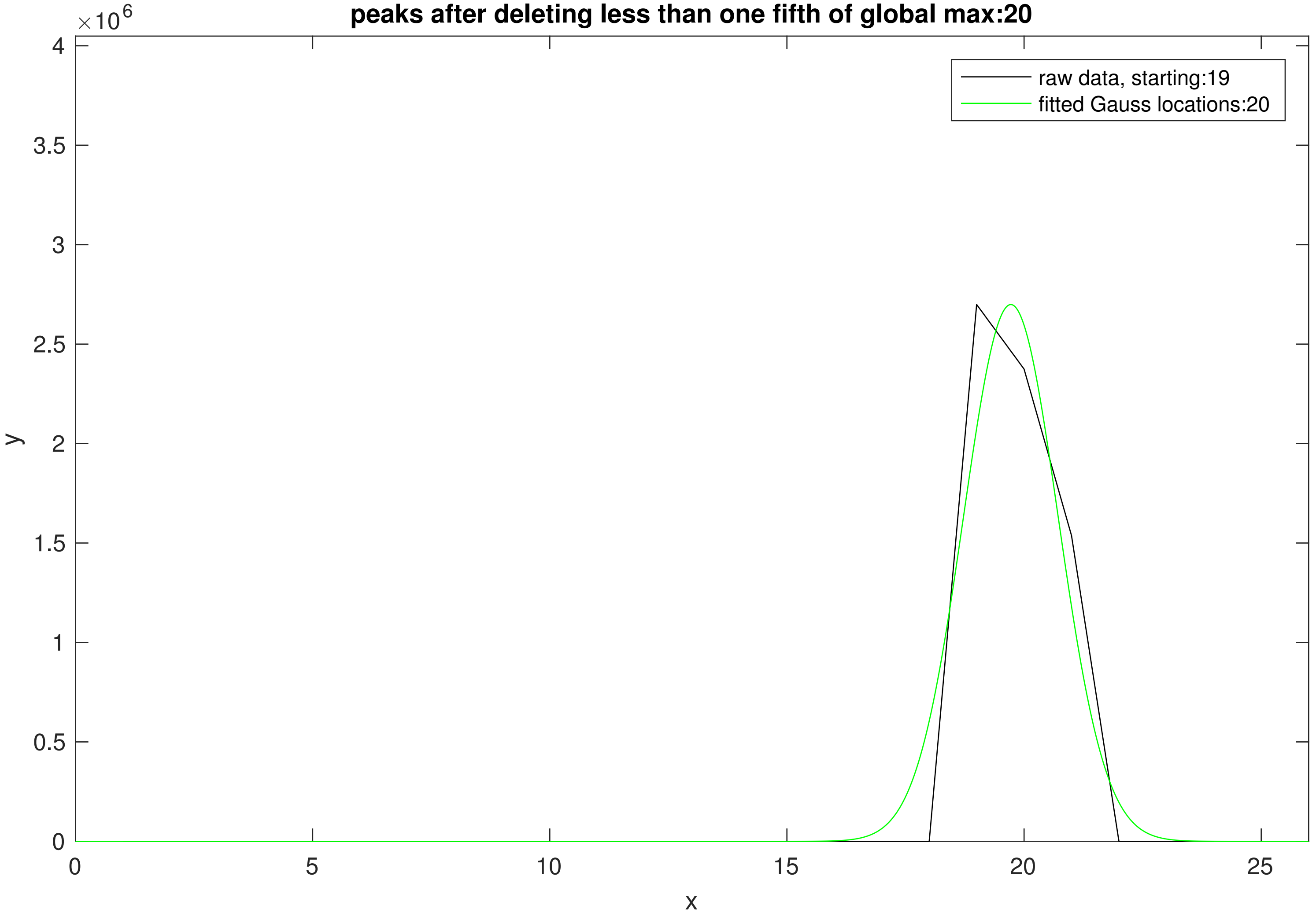

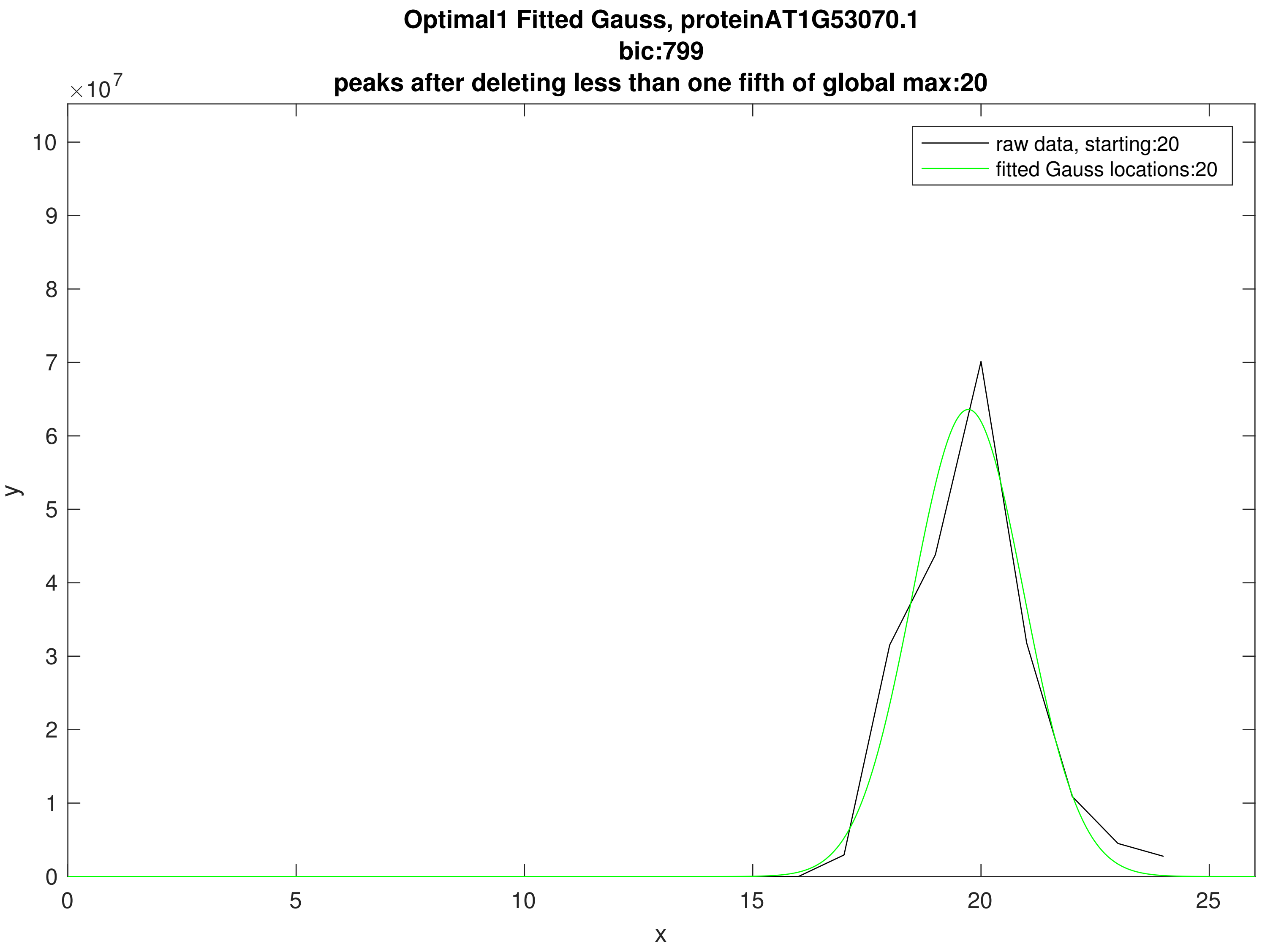

Optimal1 Fitted Gauss, proteinAT1G53070.1
bic:799
peaks after deleting less than one fifth of global max:20
×10^7
10
9
8
7
6
5
4
3
2
1
0
y
0
5
10
15
20
25
x
raw data, starting:20
fitted Gauss locations:20

Optimal1 Fitted Gauss, proteinAT1G53240.1
bic:1.08e+03  1.09e+03
peaks after deleting less than one fifth of global max:16

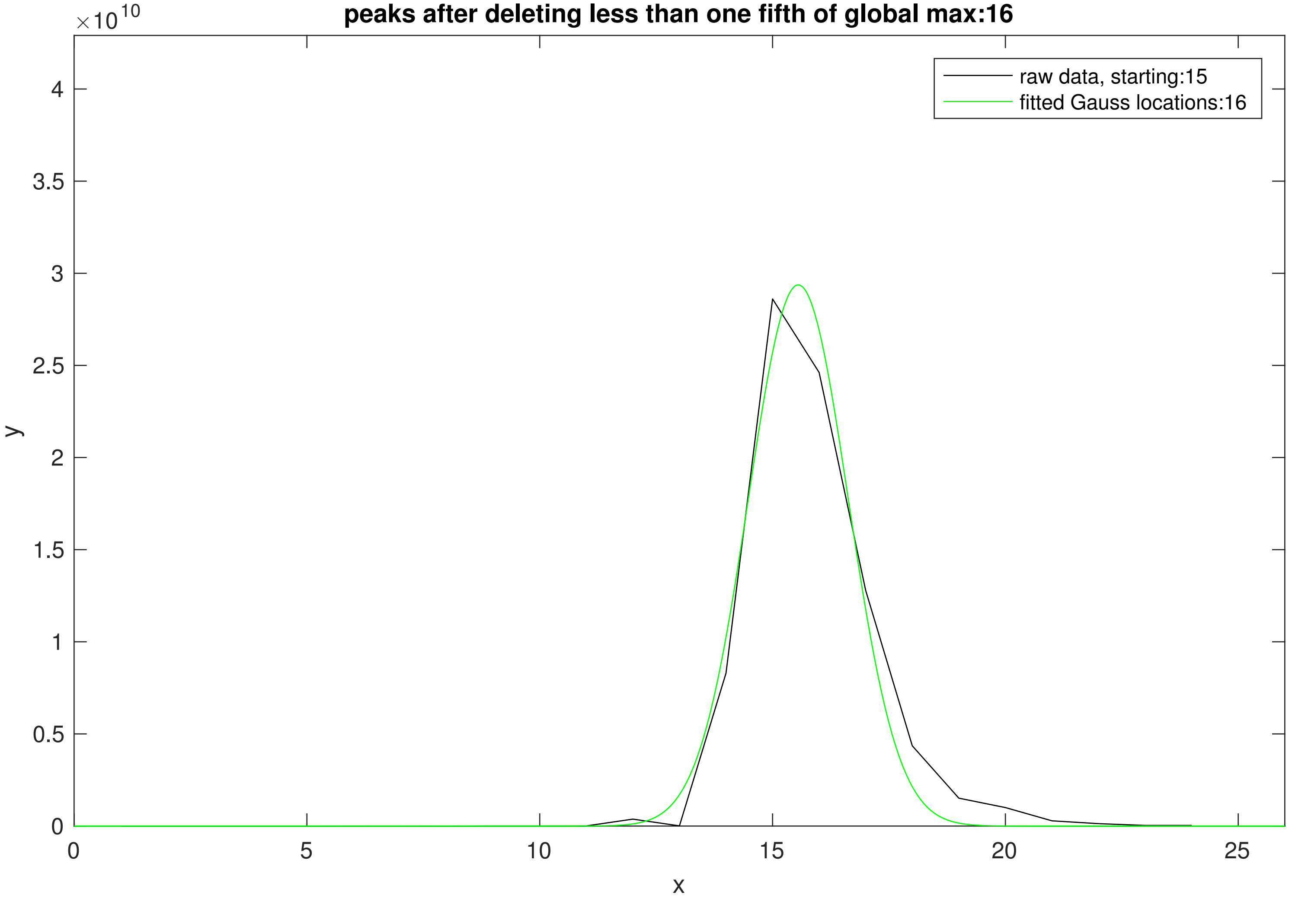

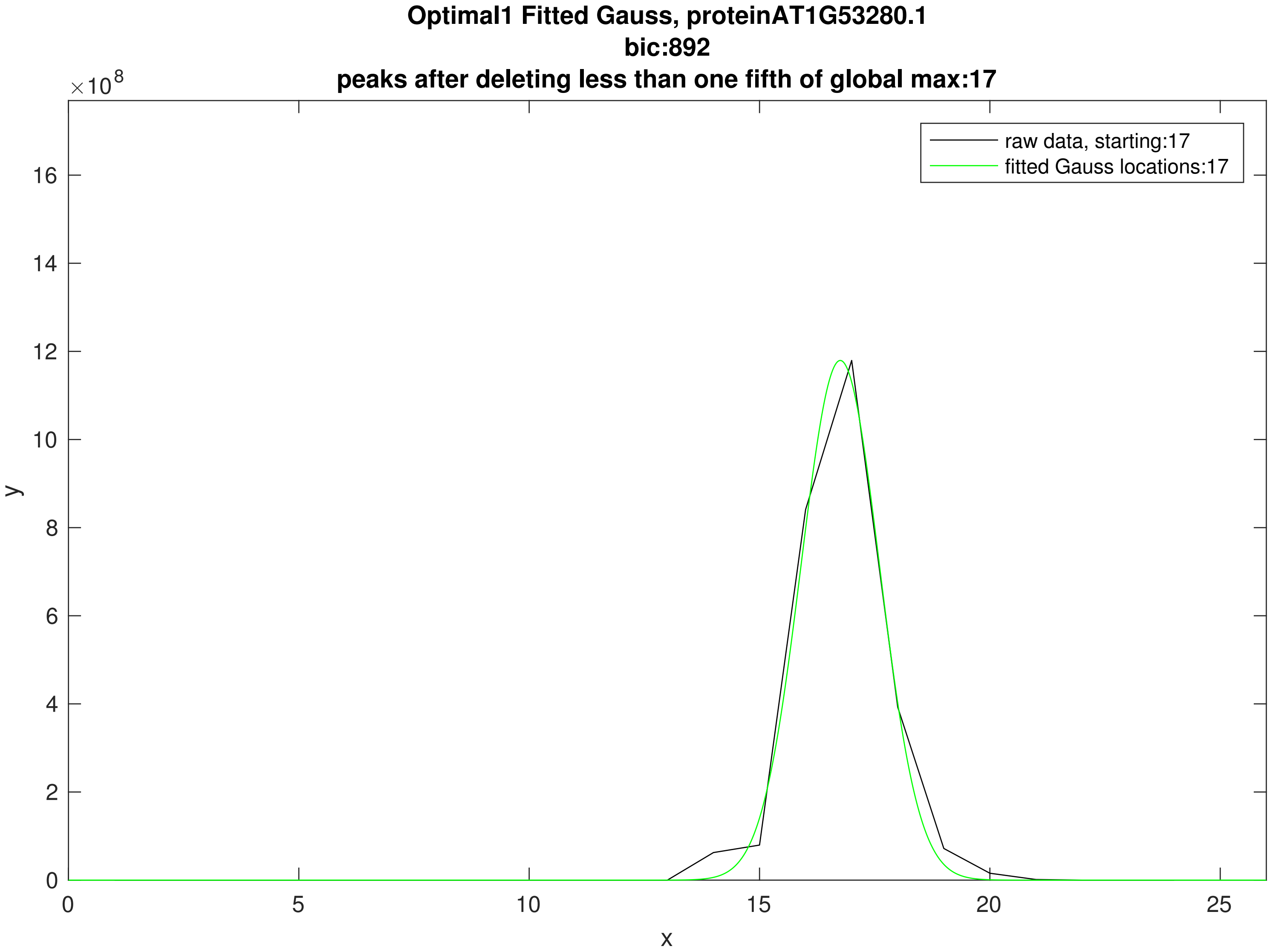

Optimal1 Fitted Gauss, proteinAT1G53280.1
bic:892
peaks after deleting less than one fifth of global max:17
×10^8
16
14
12
10
8
6
4
2
0
y
0
5
10
15
20
25
x
raw data, starting:17
fitted Gauss locations:17

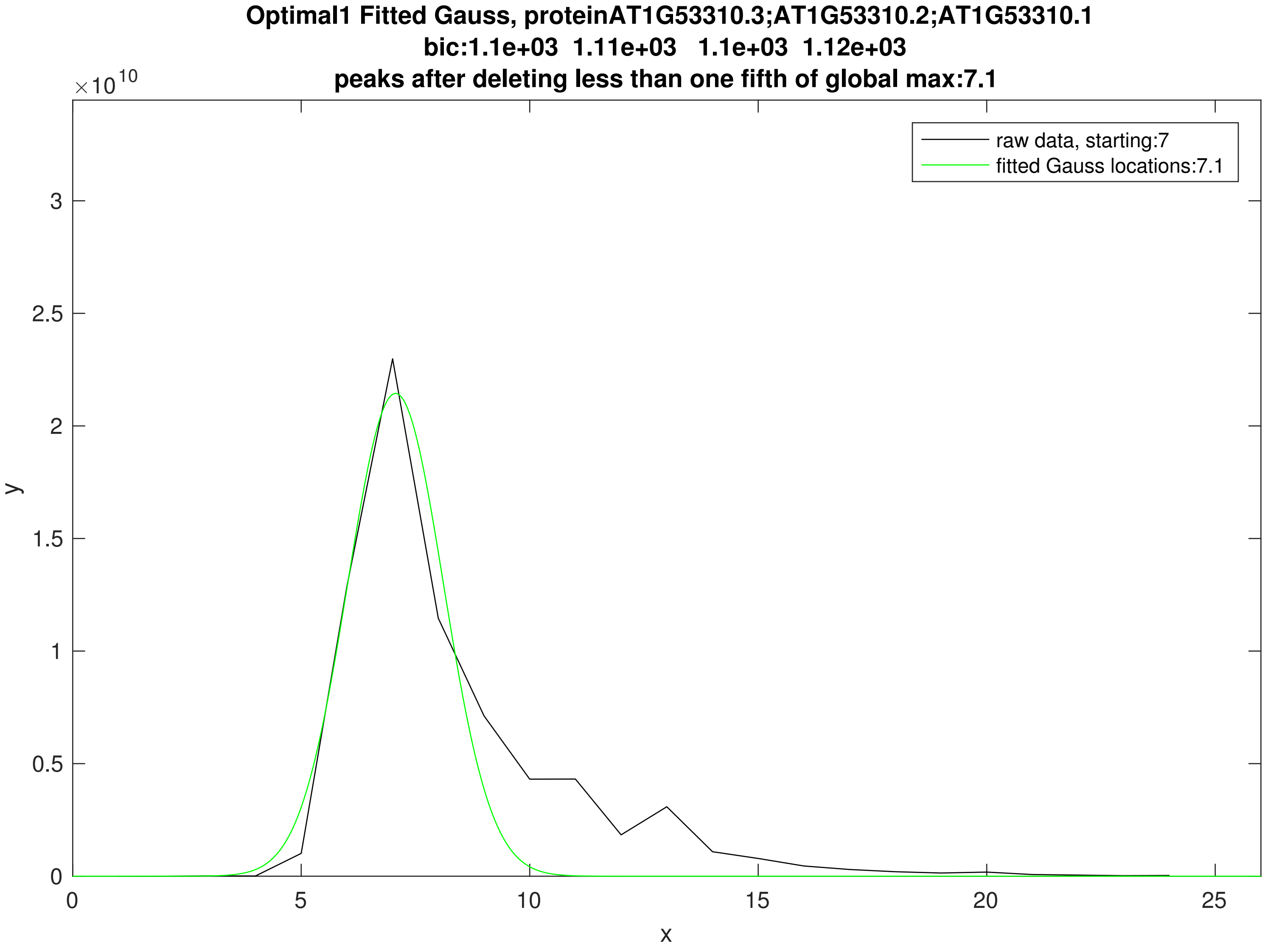

Optimal1 Fitted Gauss, proteinAT1G53310.3;AT1G53310.2;AT1G53310.1
bic:1.1e+03 1.11e+03 1.1e+03 1.12e+03
peaks after deleting less than one fifth of global max:7.1
×10^10
raw data, starting:7
fitted Gauss locations:7.1
y
x
0
0.5
1
1.5
2
2.5
3
0
5
10
15
20
25

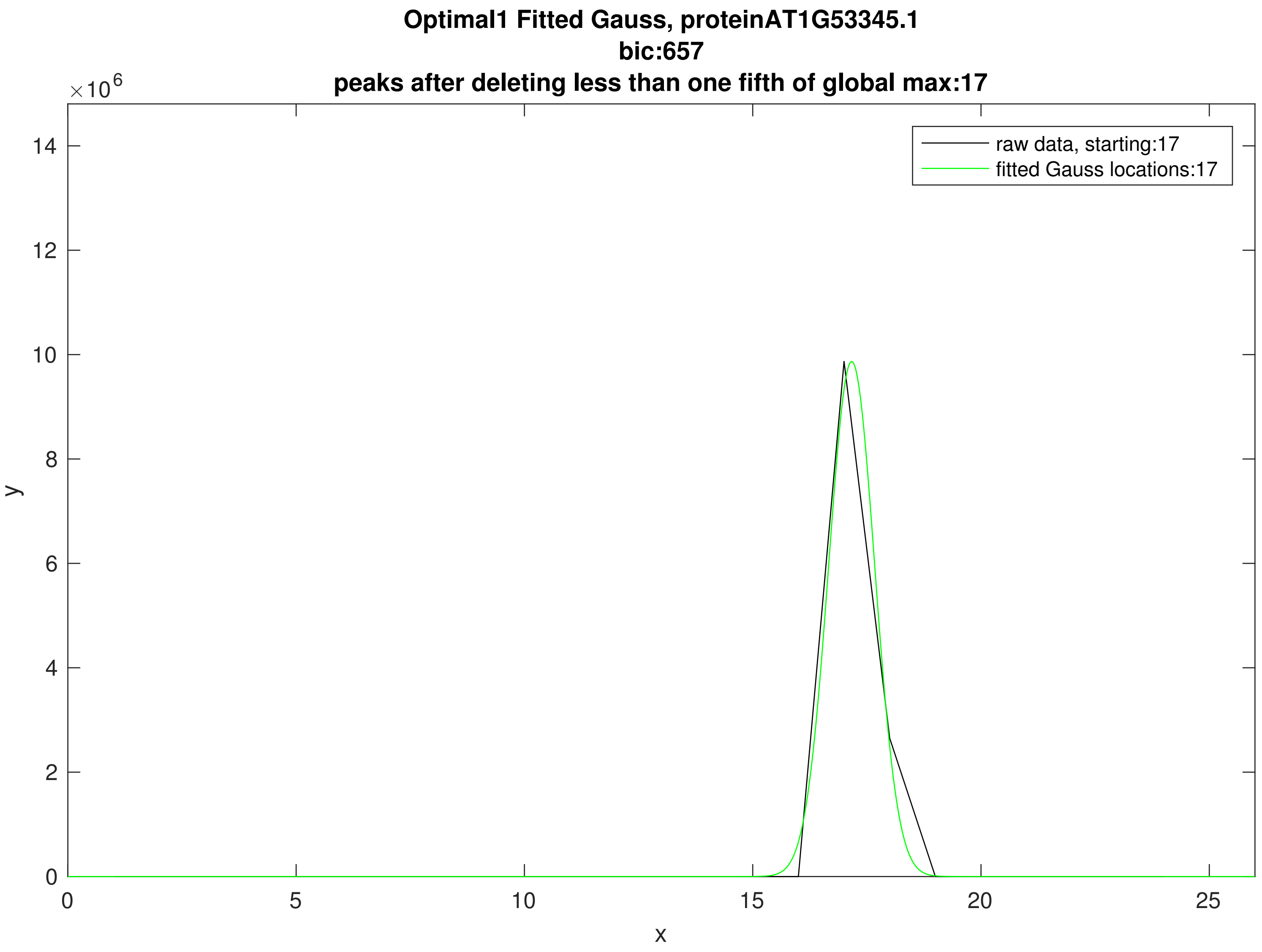

Optimal1 Fitted Gauss, proteinAT1G53345.1
bic:657
peaks after deleting less than one fifth of global max:17
×10^6
raw data, starting:17
fitted Gauss locations:17
y
x
0
2
4
6
8
10
12
14
0
5
10
15
20
25

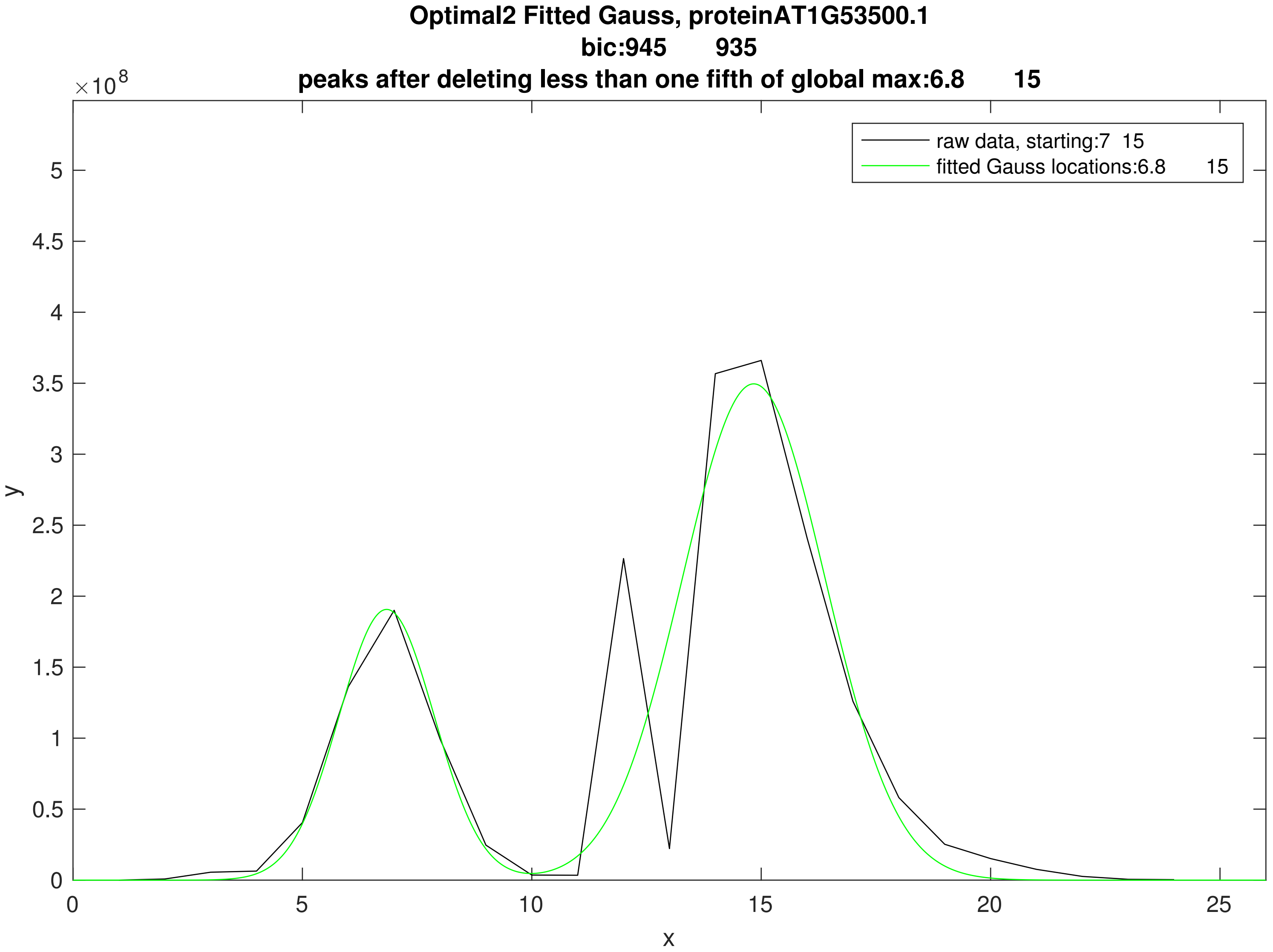
Optimal2 Fitted Gauss, proteinAT1G53500.1
bic:945 935
peaks after deleting less than one fifth of global max:6.8 15
×10^8
raw data, starting:7 15
fitted Gauss locations:6.8 15
y
x
0
0.5
1
1.5
2
2.5
3
3.5
4
4.5
5
0
5
10
15
20
25

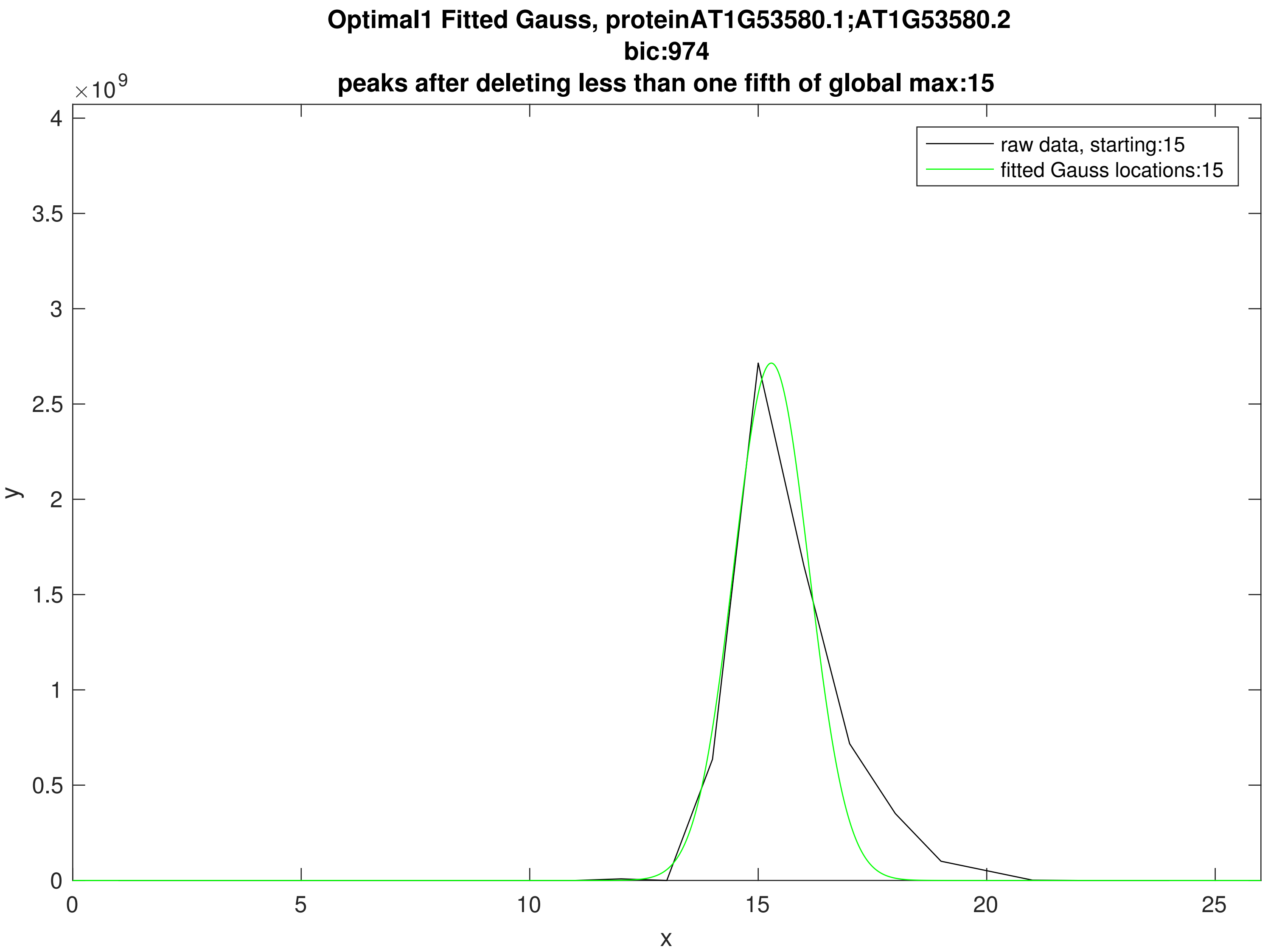

Optimal1 Fitted Gauss, proteinAT1G53580.1;AT1G53580.2
bic:974
peaks after deleting less than one fifth of global max:15
×10^9
4
3.5
3
2.5
2
1.5
1
0.5
0
y
0
5
10
15
20
25
x
raw data, starting:15
fitted Gauss locations:15

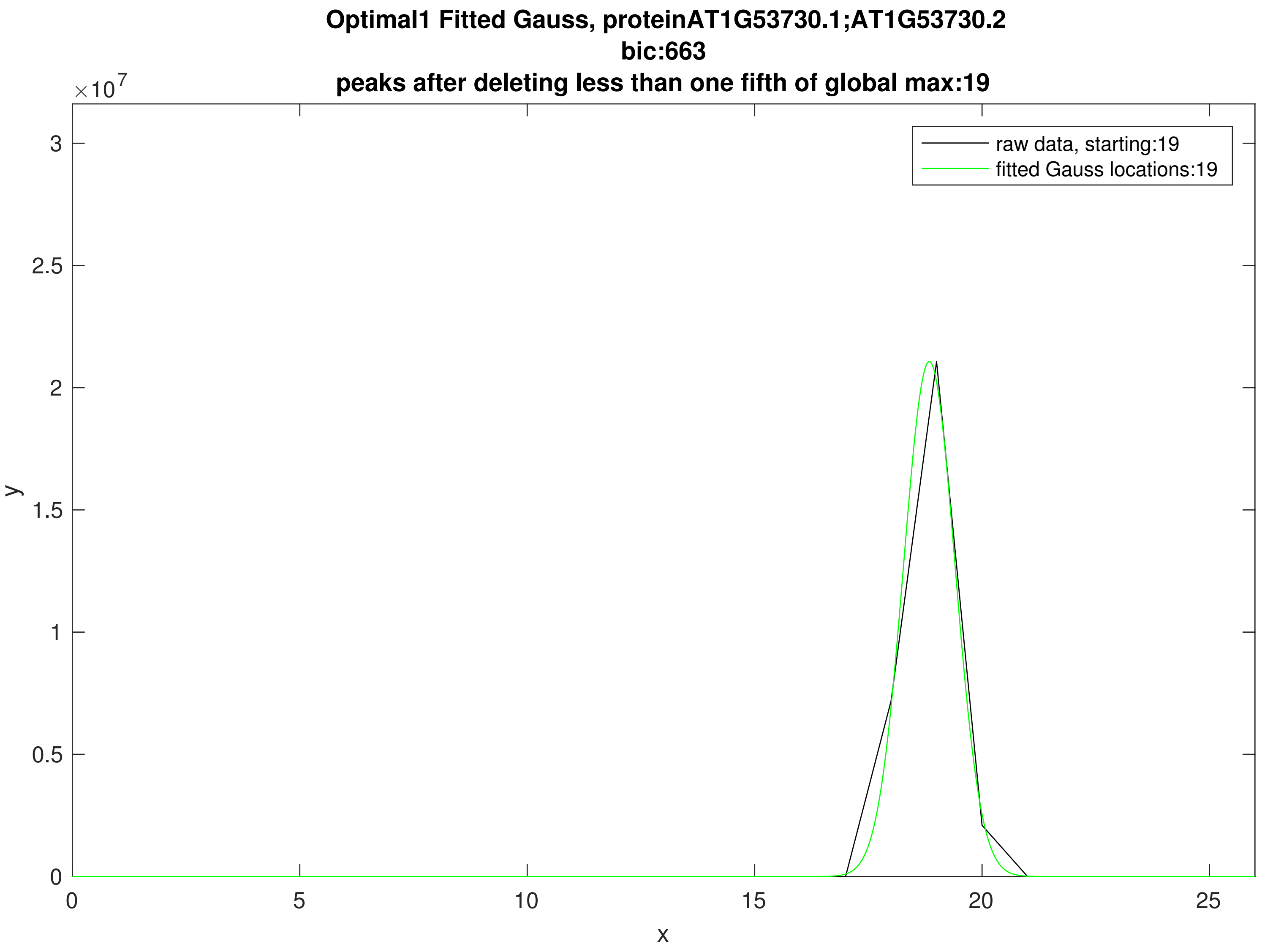

Optimal1 Fitted Gauss, proteinAT1G53730.1;AT1G53730.2
bic:663
peaks after deleting less than one fifth of global max:19
×10^7
3
2.5
2
1.5
1
0.5
0
y
0
5
10
15
20
25
x
raw data, starting:19
fitted Gauss locations:19

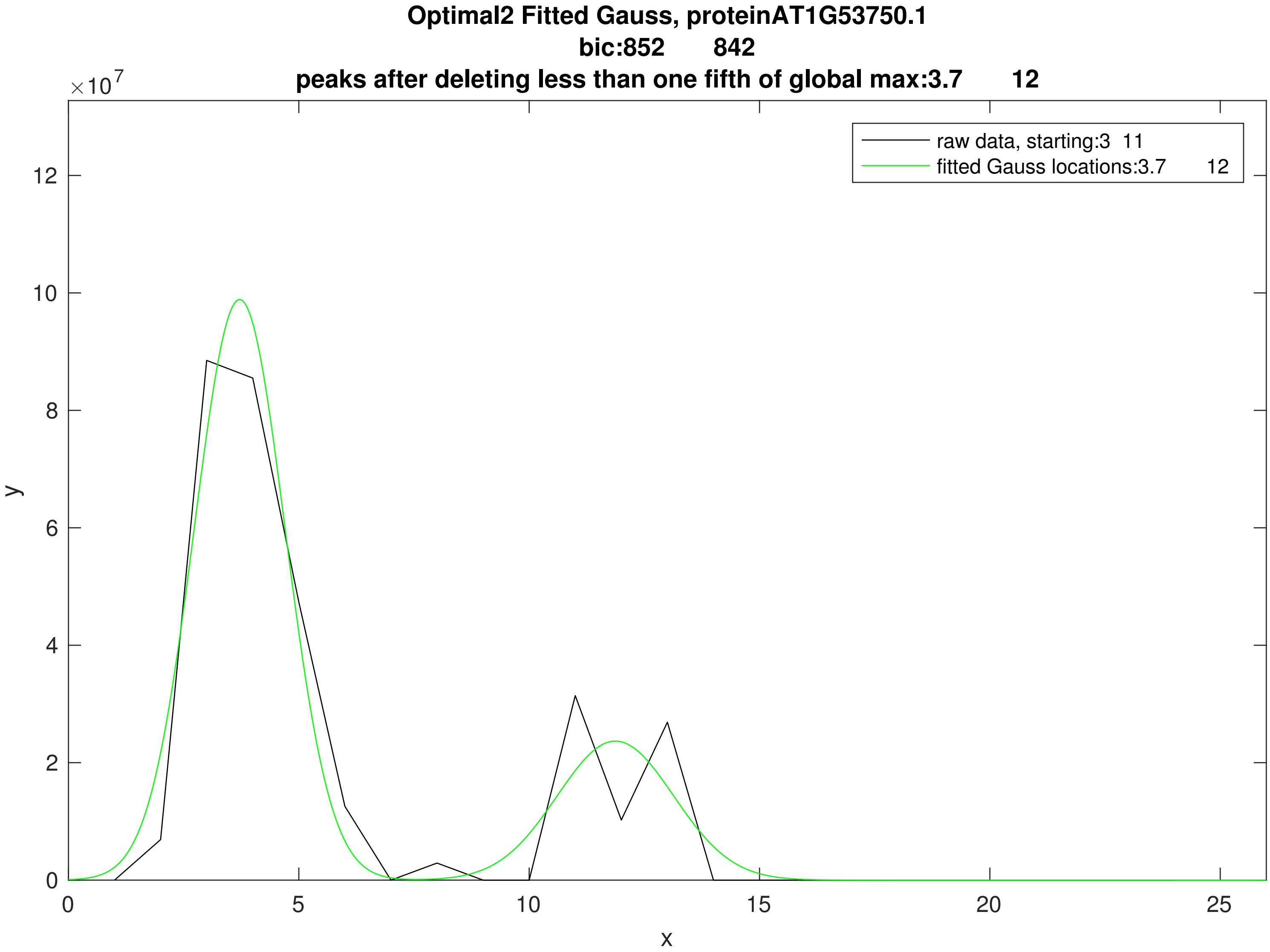

Optimal2 Fitted Gauss, proteinAT1G53750.1
bic:852 842
peaks after deleting less than one fifth of global max:3.7 12
×10^7
raw data, starting:3 11
fitted Gauss locations:3.7 12
y
x
0
2
4
6
8
10
12
0
5
10
15
20
25

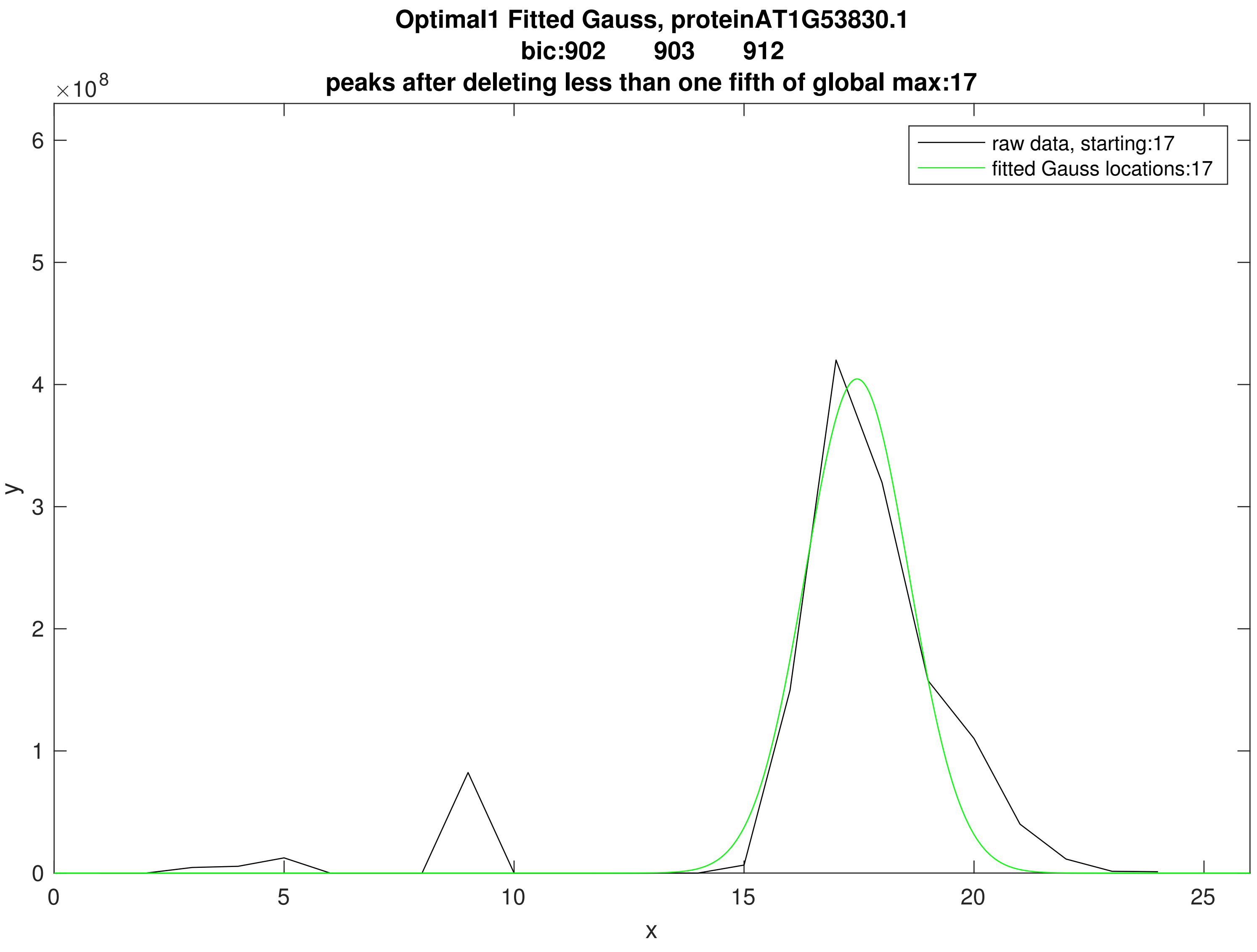

Optimal1 Fitted Gauss, proteinAT1G53830.1
bic:902 903 912
peaks after deleting less than one fifth of global max:17
×10^8
raw data, starting:17
fitted Gauss locations:17
y
x
0
1
2
3
4
5
6
0
5
10
15
20
25

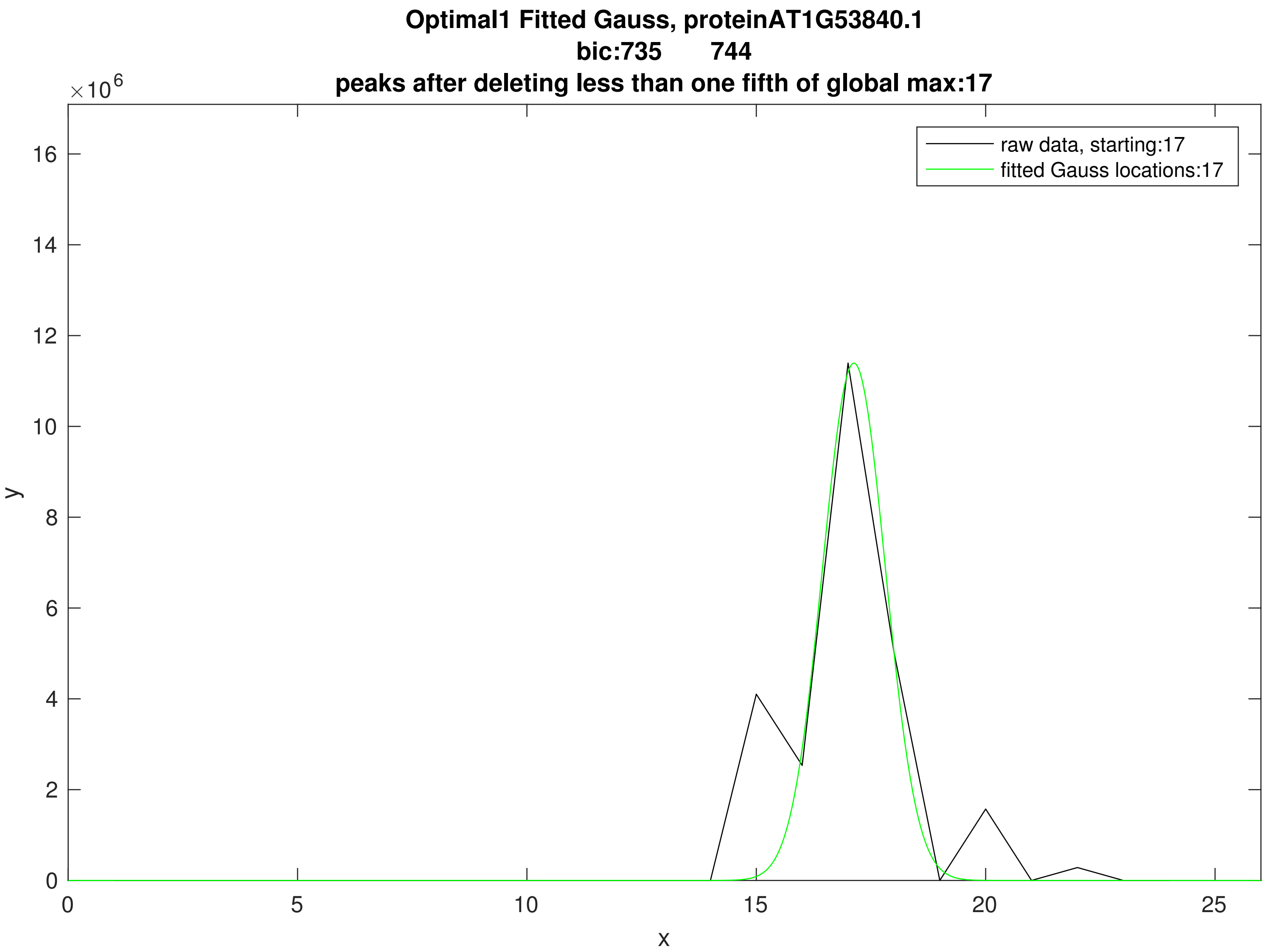
Optimal1 Fitted Gauss, proteinAT1G53840.1
bic:735 744
peaks after deleting less than one fifth of global max:17
×10^6
raw data, starting:17
fitted Gauss locations:17
y
x
0
2
4
6
8
10
12
14
16
0
5
10
15
20
25

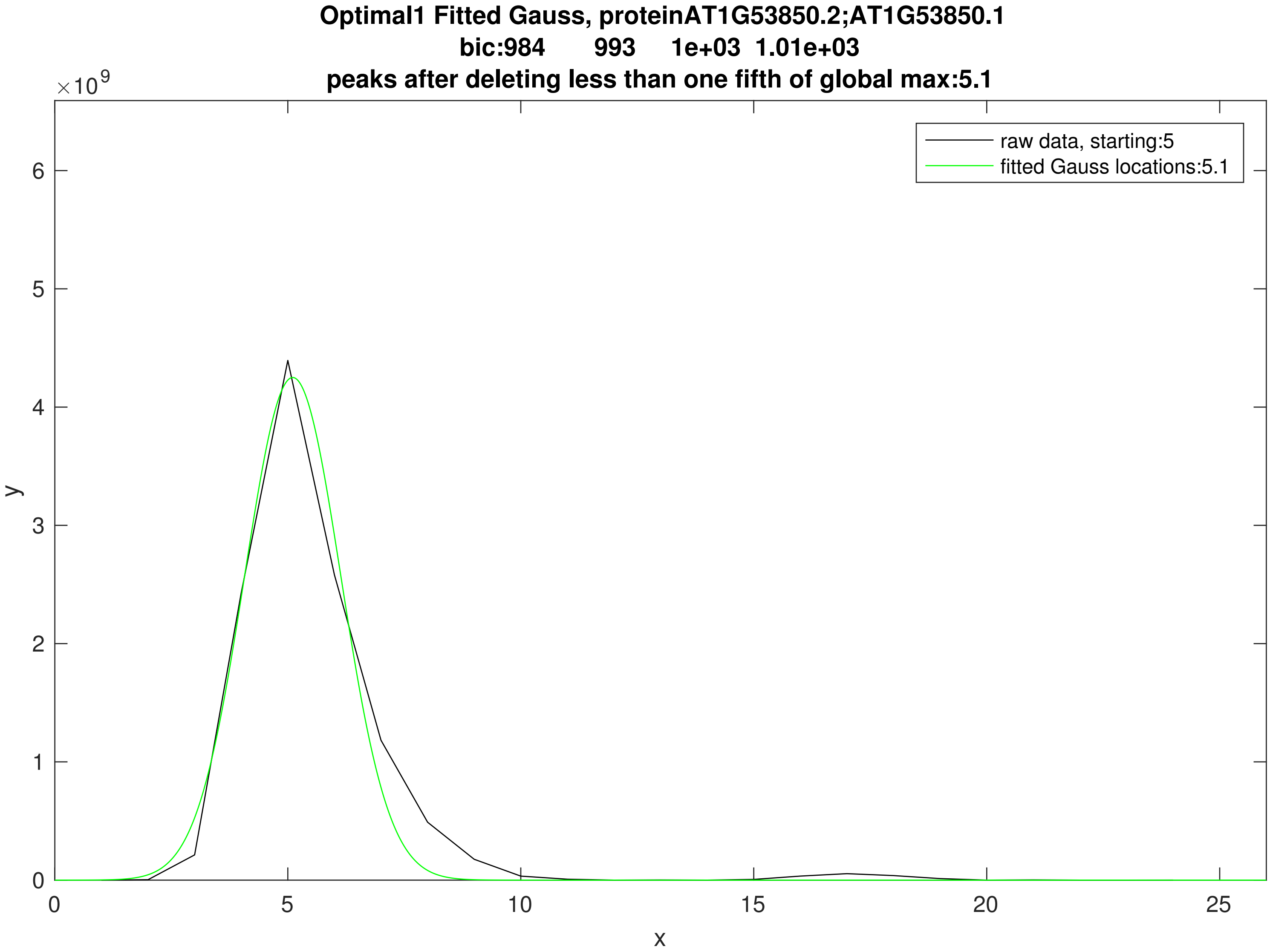

Optimal1 Fitted Gauss, proteinAT1G53850.2;AT1G53850.1
bic:984 993 1e+03 1.01e+03
peaks after deleting less than one fifth of global max:5.1
×10^9
raw data, starting:5
fitted Gauss locations:5.1
y
x
0
1
2
3
4
5
6
0
5
10
15
20
25

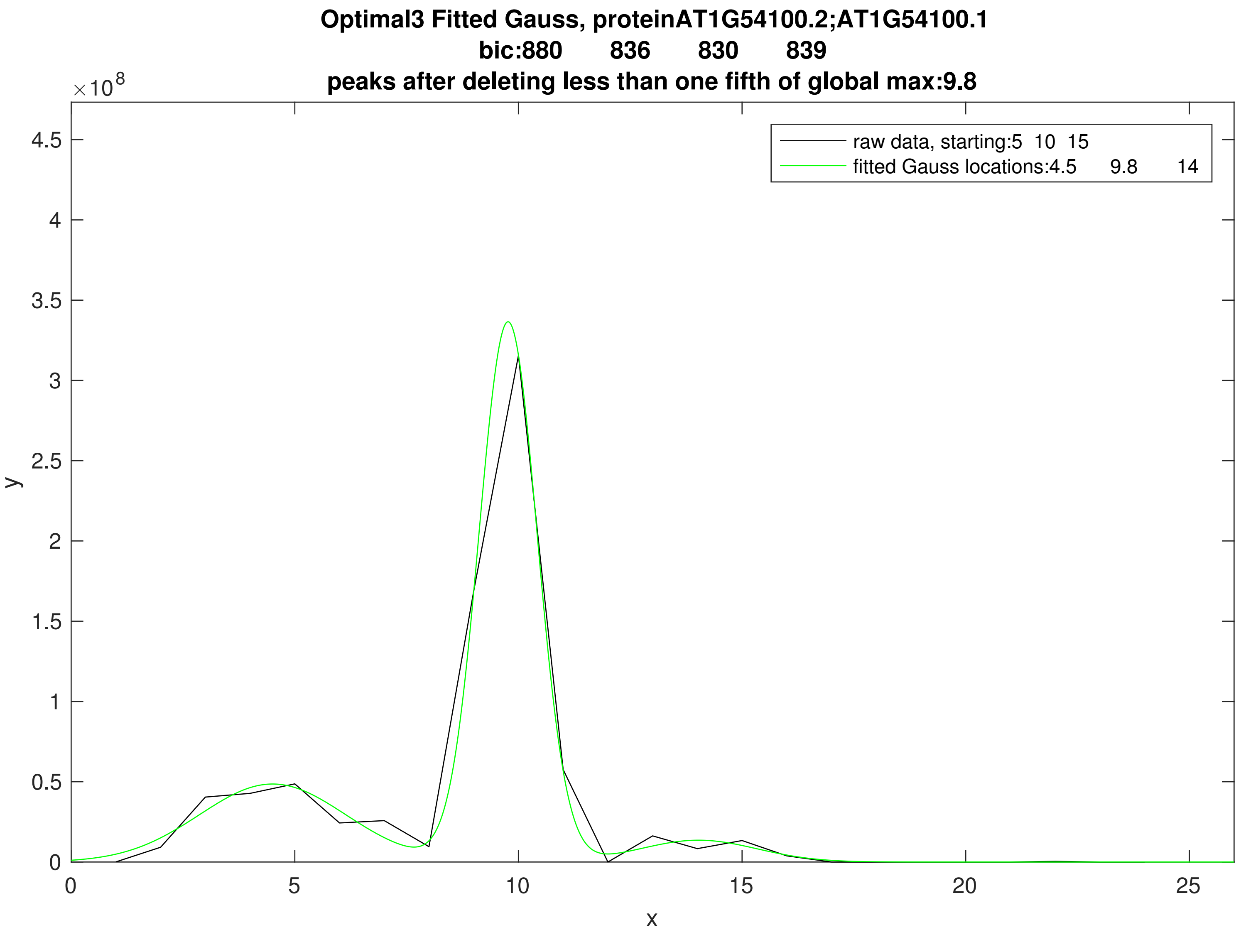

Optimal3 Fitted Gauss, proteinAT1G54100.2;AT1G54100.1
bic:880 836 830 839
peaks after deleting less than one fifth of global max:9.8
×10^8
raw data, starting:5 10 15
fitted Gauss locations:4.5 9.8 14
y
x
0
0.5
1
1.5
2
2.5
3
3.5
4
4.5
5
10
15
20
25

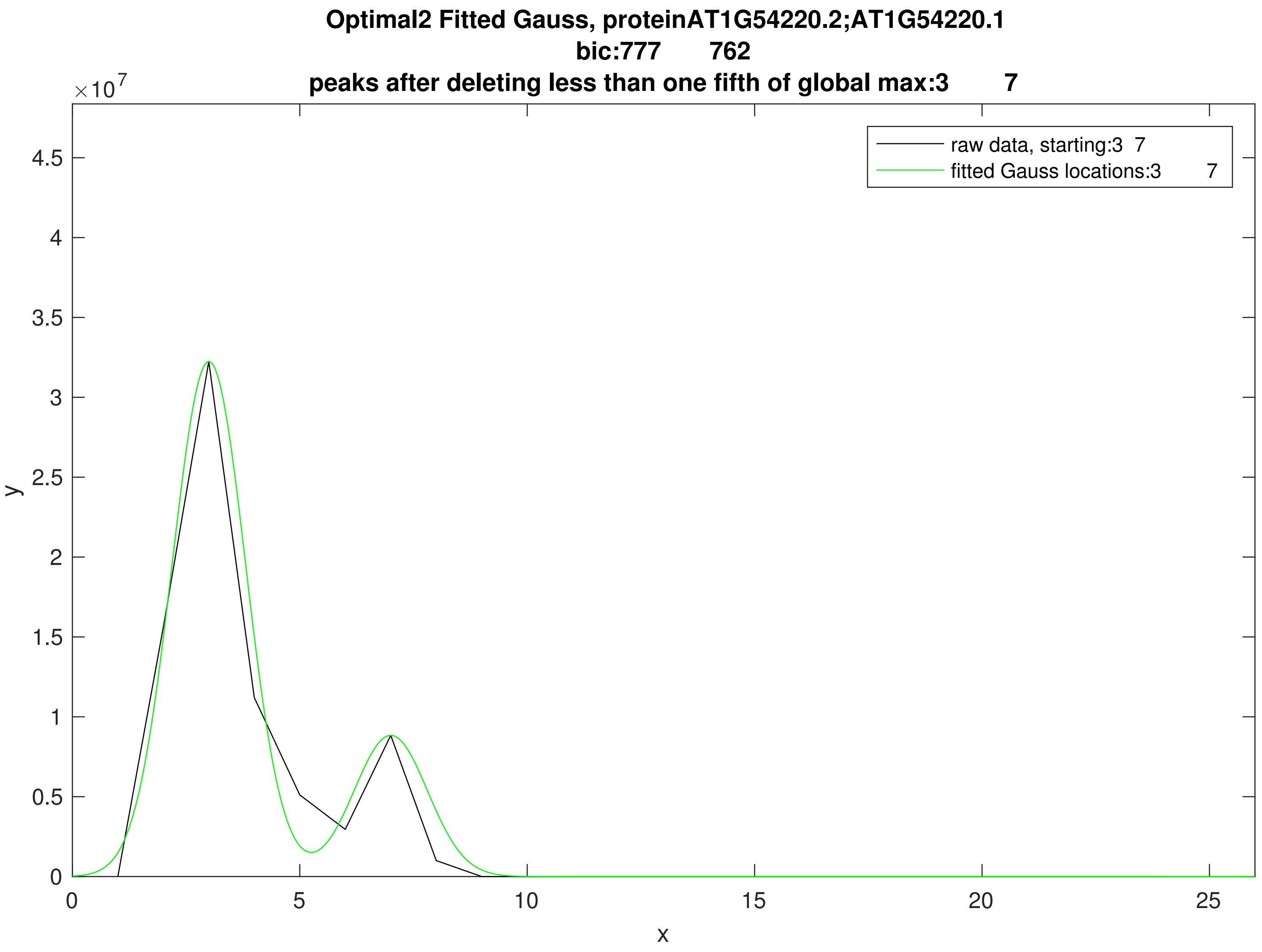

Optimal2 Fitted Gauss, proteinAT1G54220.2;AT1G54220.1
bic:777 762
peaks after deleting less than one fifth of global max:3 7
×10^7
raw data, starting:3 7
fitted Gauss locations:3 7
y
x
0
0.5
1
1.5
2
2.5
3
3.5
4
4.5
5
10
15
20
25

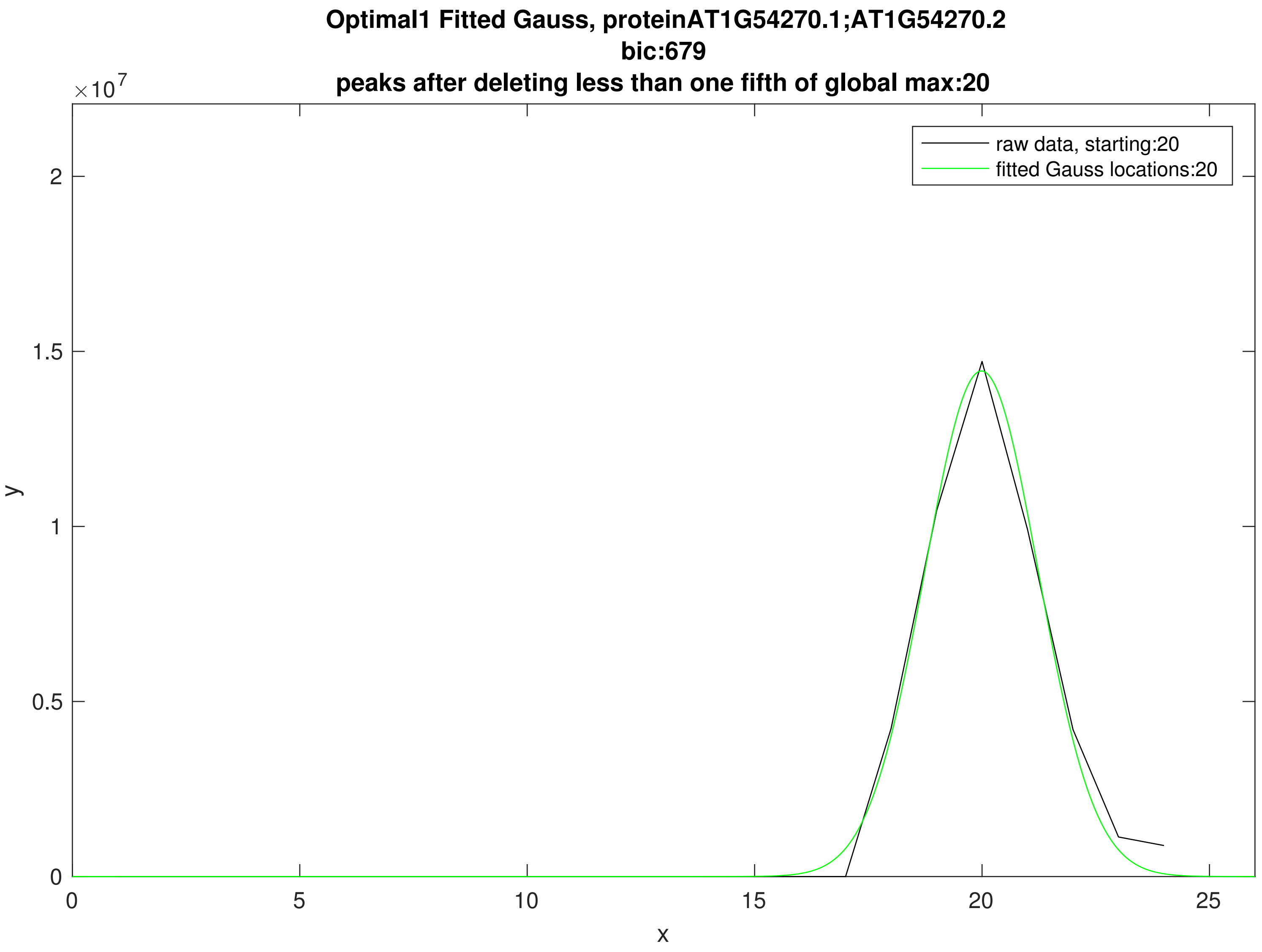
Optimal1 Fitted Gauss, proteinAT1G54270.1;AT1G54270.2
bic:679
peaks after deleting less than one fifth of global max:20
×10^7
raw data, starting:20
fitted Gauss locations:20
2
1.5
1
0.5
0
y
0
5
10
15
20
25
x

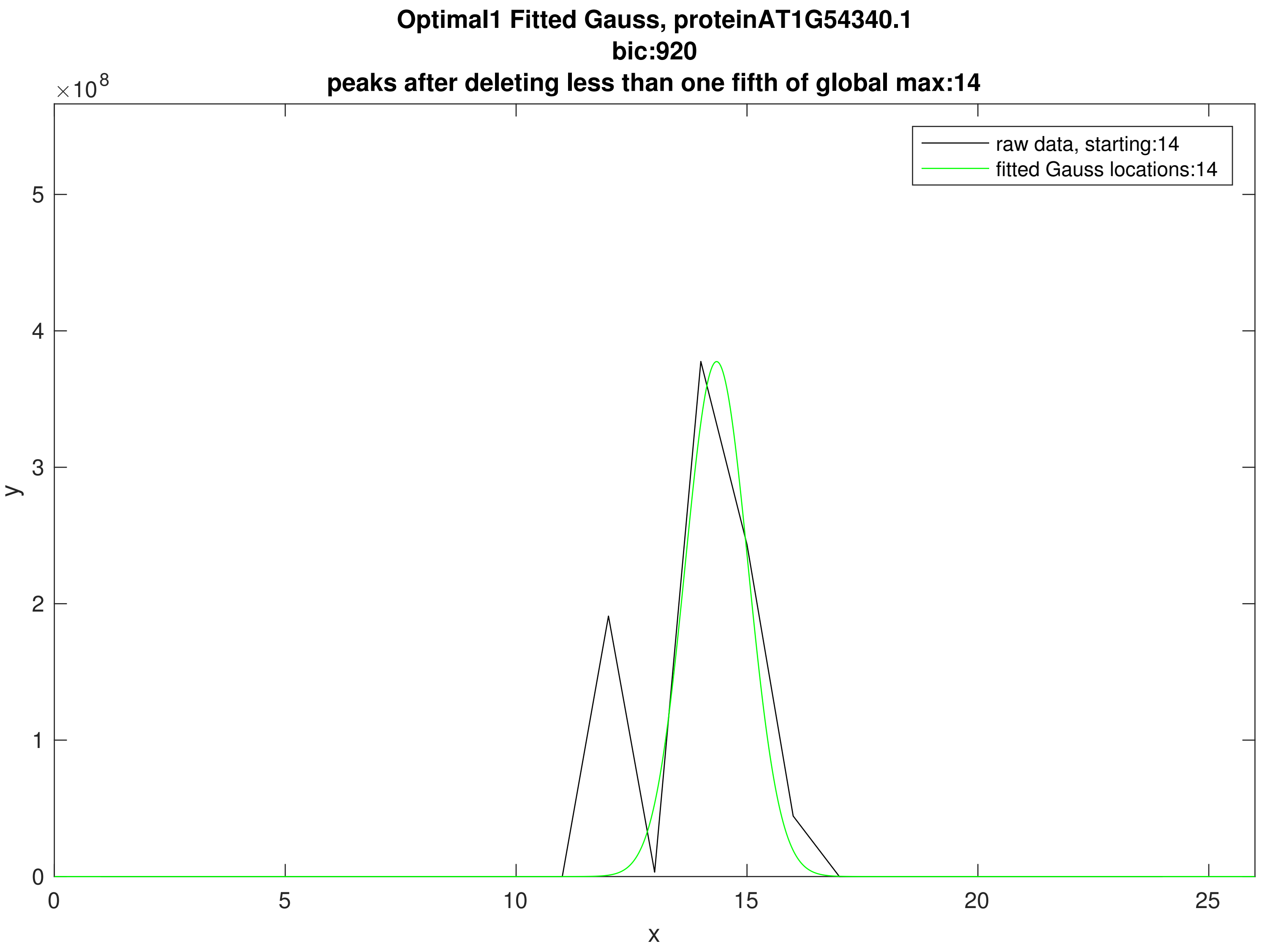
Optimal1 Fitted Gauss, proteinAT1G54340.1
bic:920
peaks after deleting less than one fifth of global max:14
×10^8
raw data, starting:14
fitted Gauss locations:14
y
x
0
1
2
3
4
5
0
5
10
15
20
25

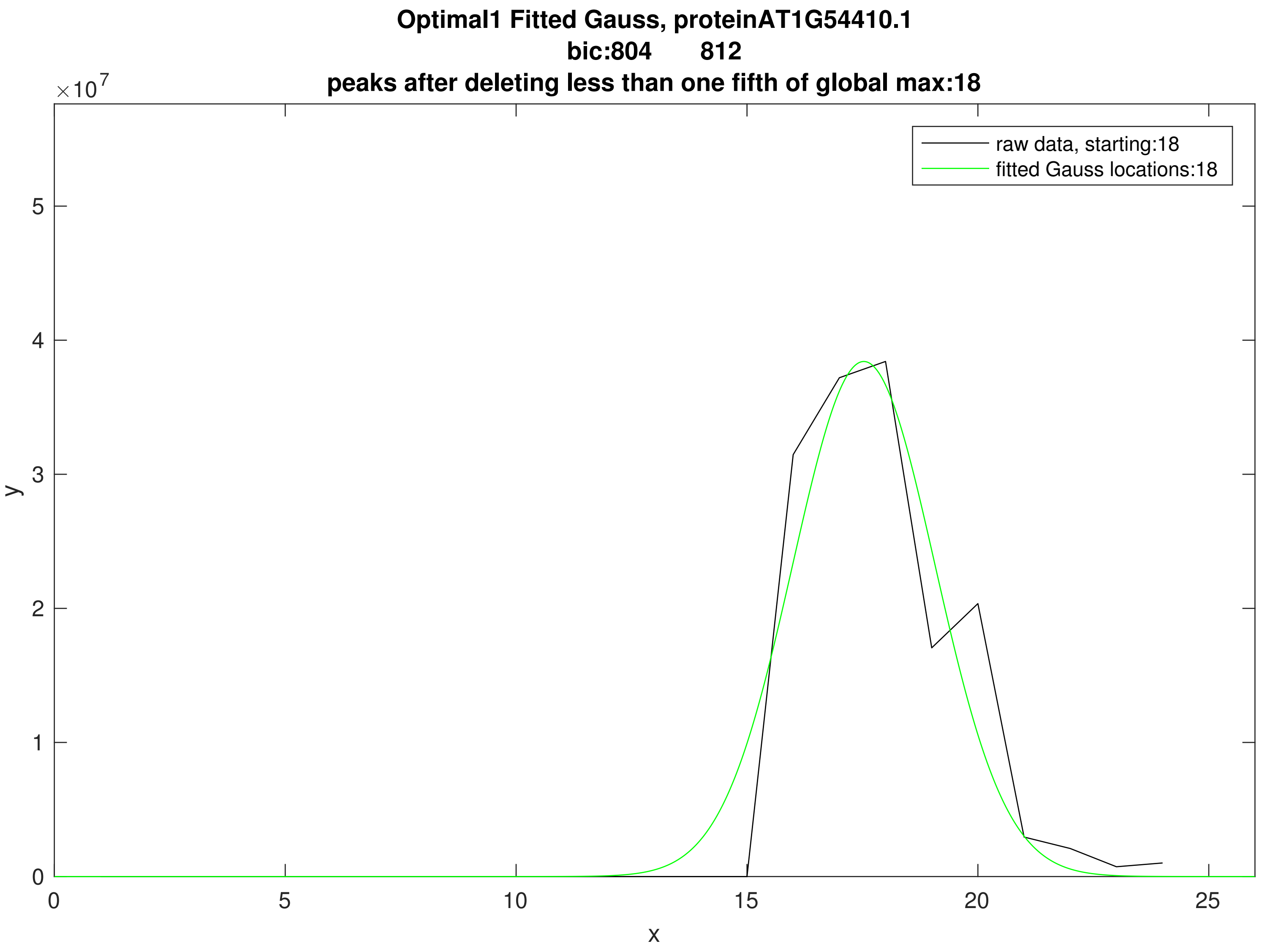

Optimal1 Fitted Gauss, proteinAT1G54410.1
bic:804 812
peaks after deleting less than one fifth of global max:18
×10^7
raw data, starting:18
fitted Gauss locations:18
y
x
0
1
2
3
4
5
0
5
10
15
20
25

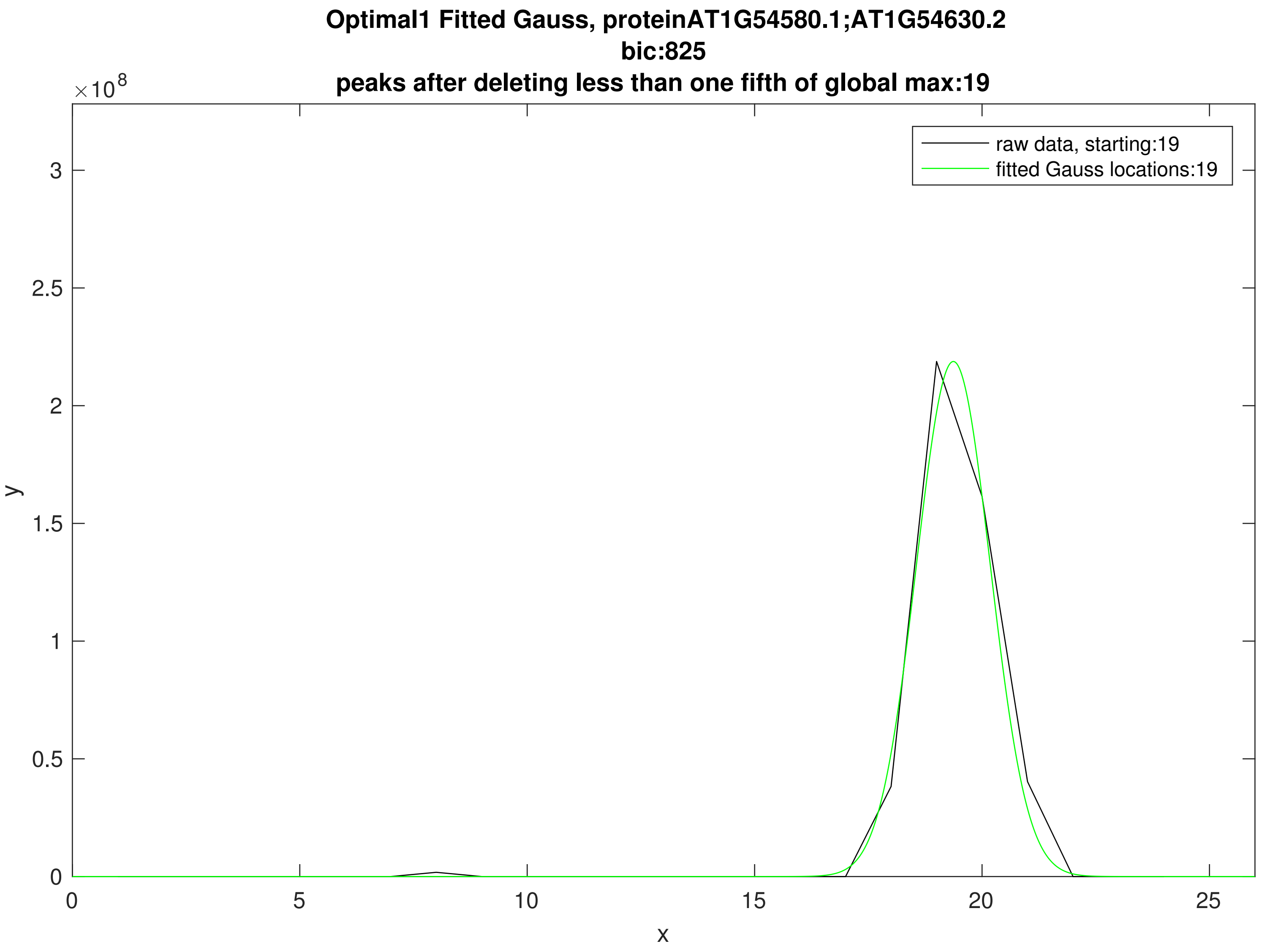
Optimal1 Fitted Gauss, proteinAT1G54580.1;AT1G54630.2
bic:825
peaks after deleting less than one fifth of global max:19
×10^8
raw data, starting:19
fitted Gauss locations:19
y
x
0
0.5
1
1.5
2
2.5
3
0
5
10
15
20
25

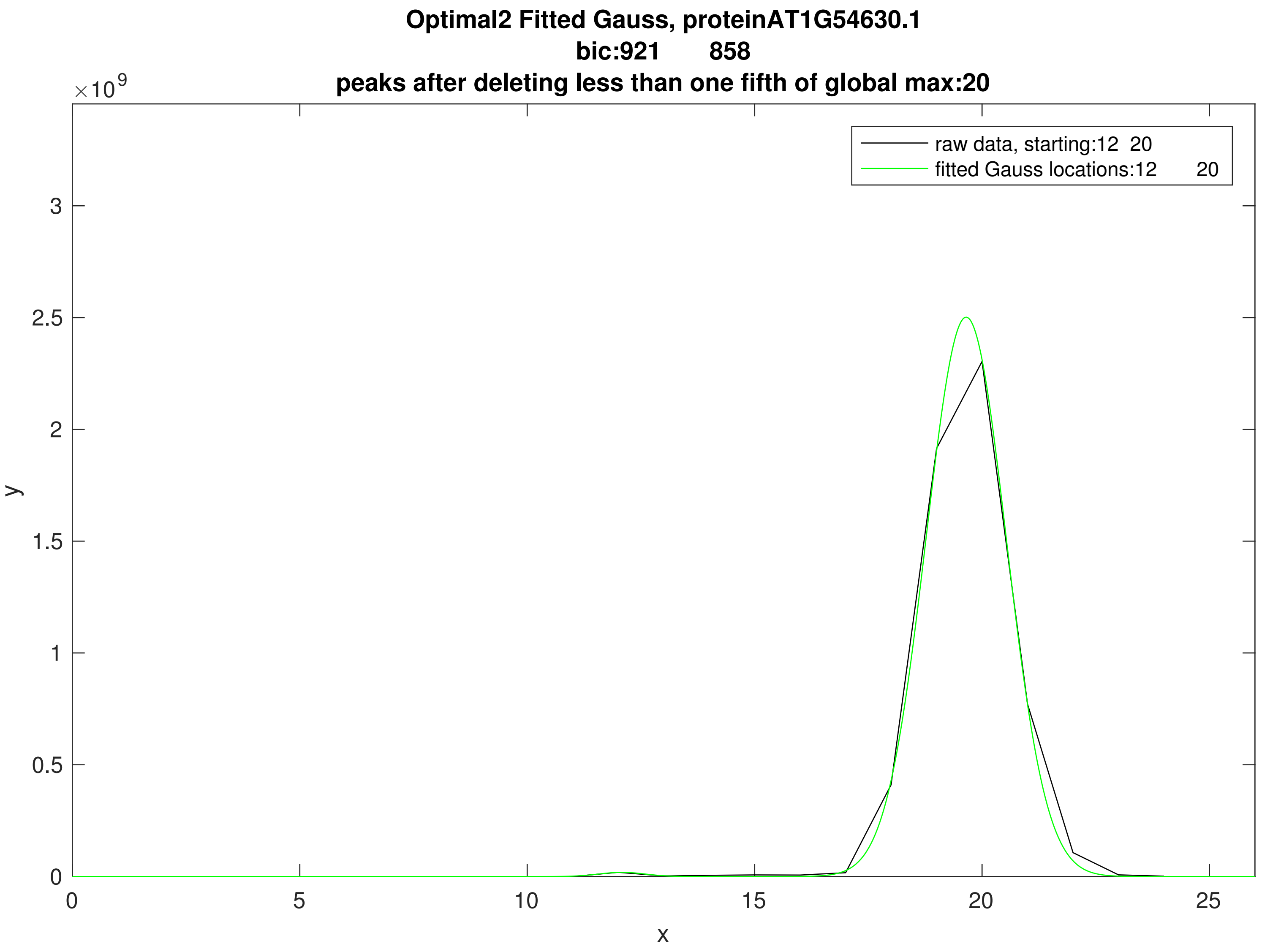

Optimal2 Fitted Gauss, proteinAT1G54630.1
bic:921 858
peaks after deleting less than one fifth of global max:20
×10^9
raw data, starting:12 20
fitted Gauss locations:12 20
3
2.5
2
1.5
1
0.5
0
y
0
5
10
15
20
25
x

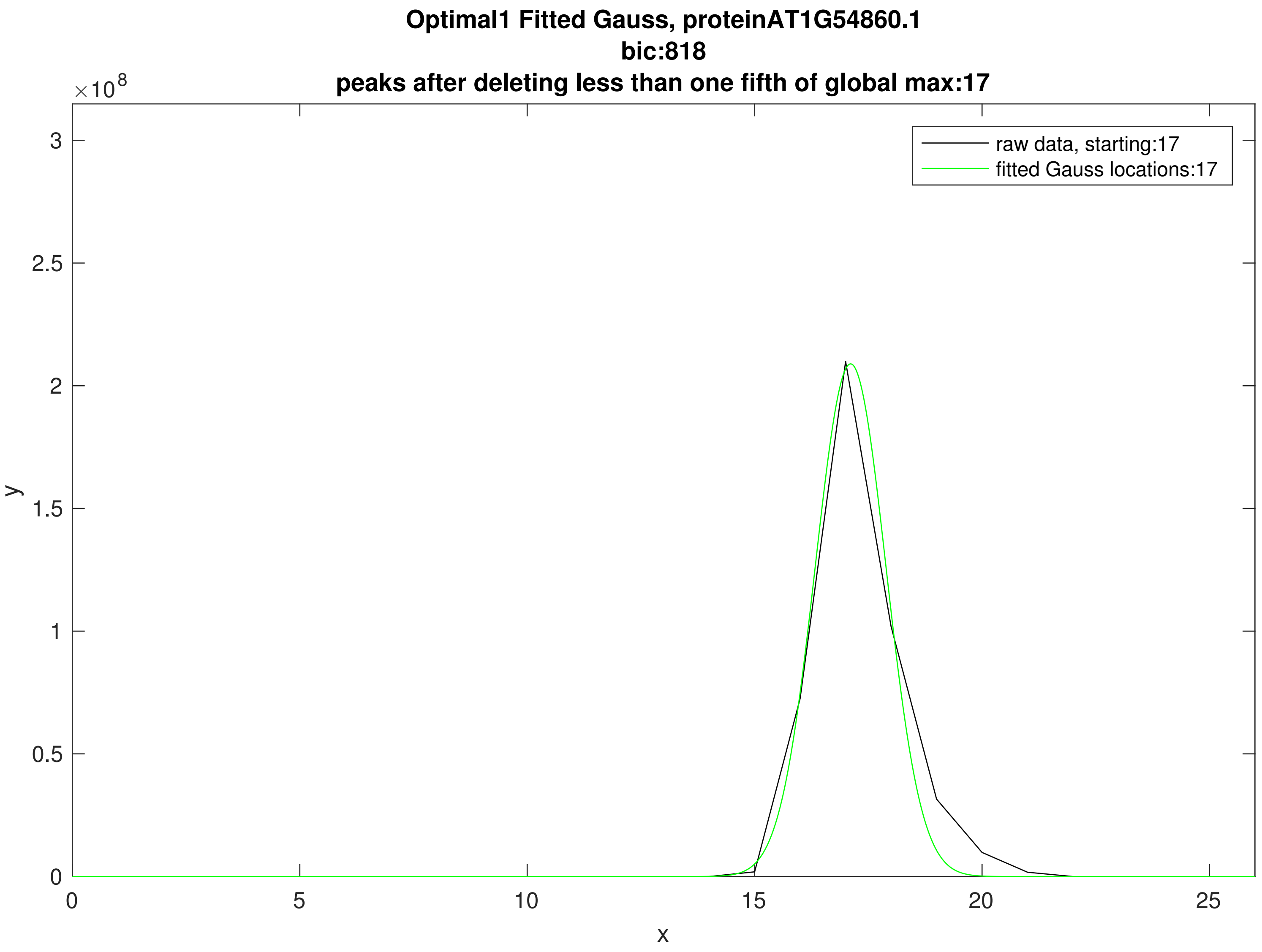

Optimal1 Fitted Gauss, proteinAT1G54860.1
bic:818
peaks after deleting less than one fifth of global max:17
×10^8
raw data, starting:17
fitted Gauss locations:17
y
x
0
0.5
1
1.5
2
2.5
3
0
5
10
15
20
25

**Optimal1 Fitted Gauss, proteinAT1G54870.1**
**bic:1.06e+03  1.07e+03**
**peaks after deleting less than one fifth of global max:12**

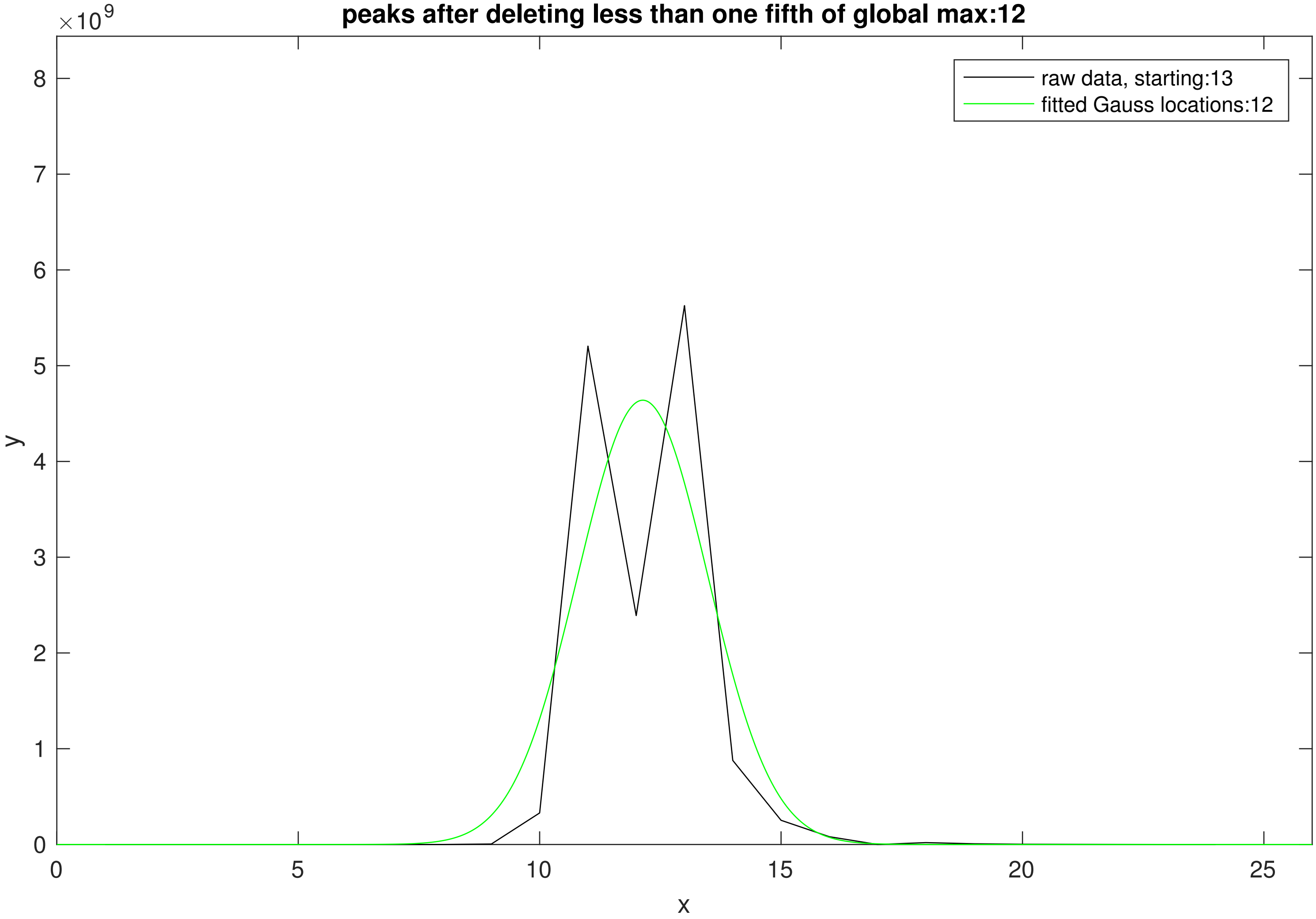

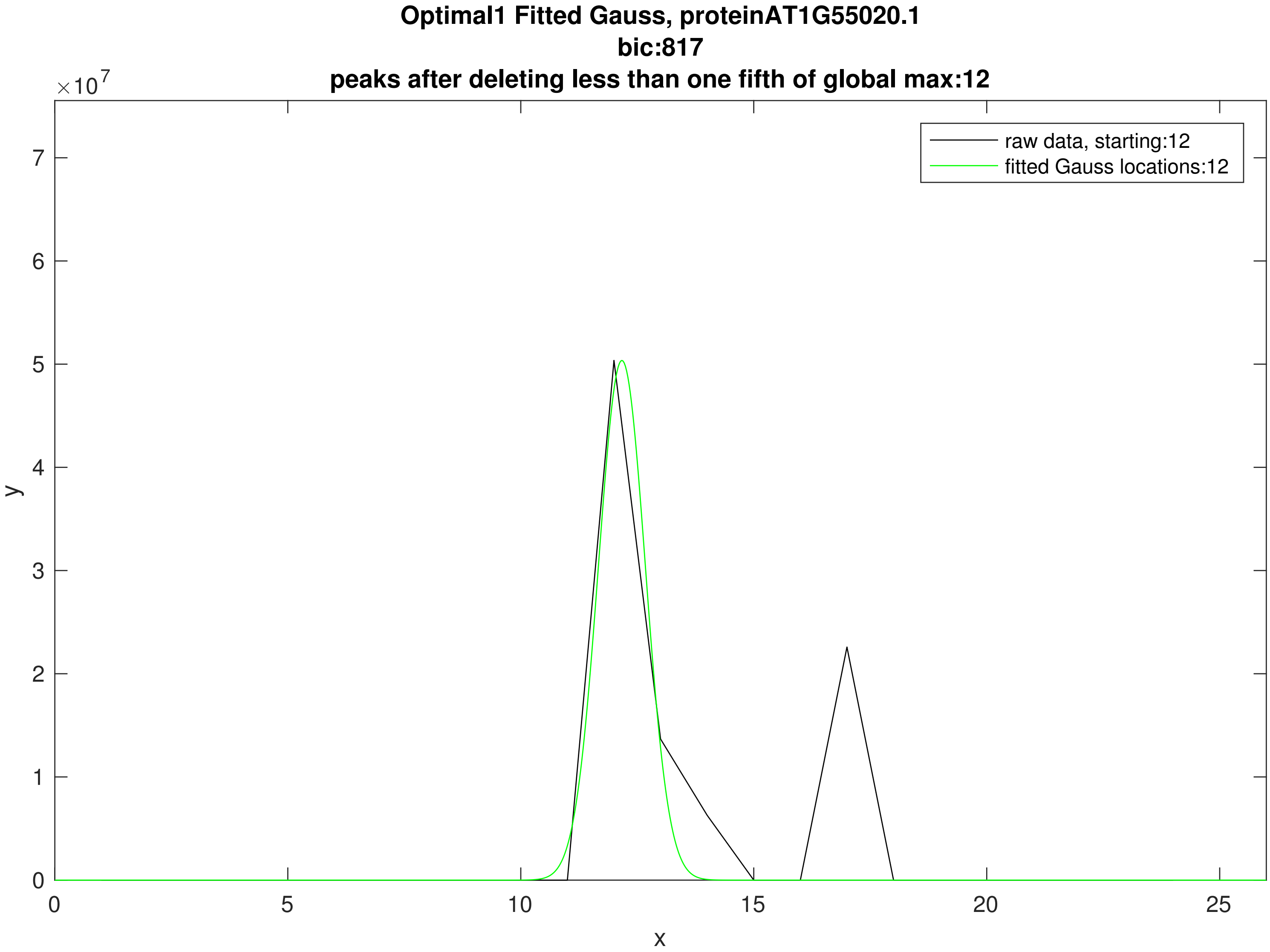

Optimal1 Fitted Gauss, proteinAT1G55020.1
bic:817
peaks after deleting less than one fifth of global max:12
×10^7
raw data, starting:12
fitted Gauss locations:12
y
x
0
1
2
3
4
5
6
7
0
5
10
15
20
25

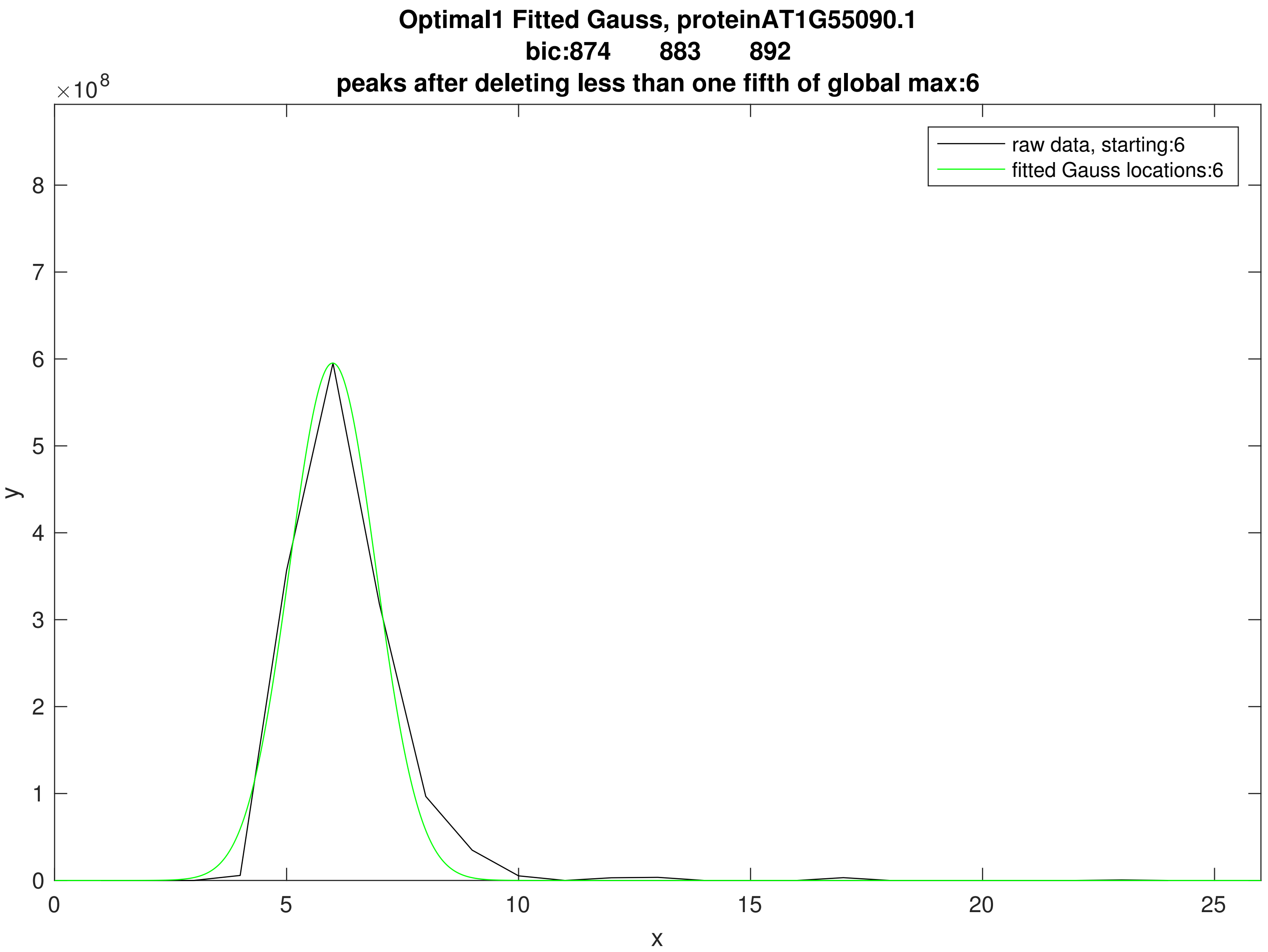
Optimal1 Fitted Gauss, proteinAT1G55090.1
bic:874 883 892
peaks after deleting less than one fifth of global max:6
×10^8
raw data, starting:6
fitted Gauss locations:6
y
x
0
1
2
3
4
5
6
7
8
0
5
10
15
20
25

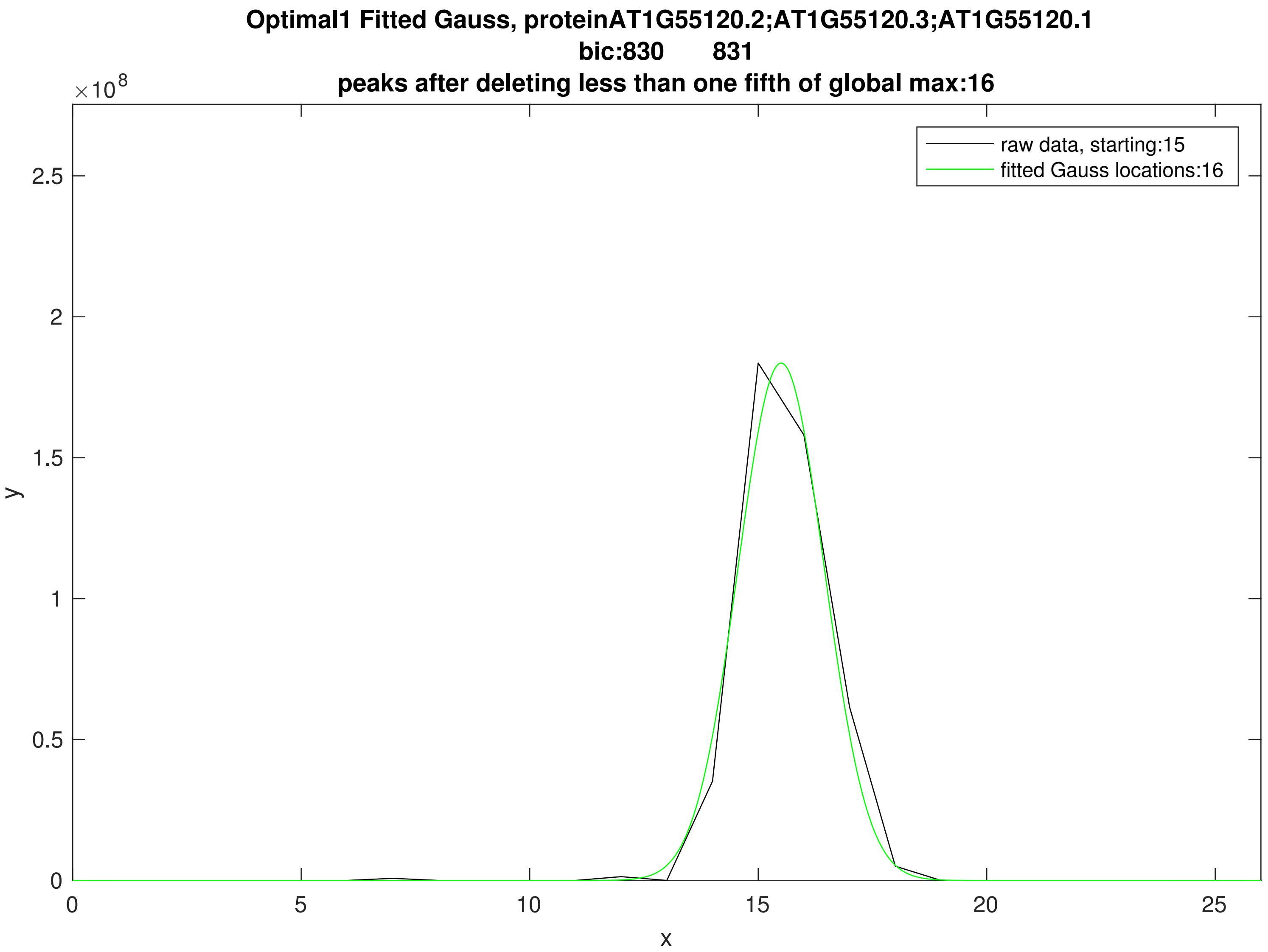

Optimal1 Fitted Gauss, proteinAT1G55120.2;AT1G55120.3;AT1G55120.1
bic:830 831
peaks after deleting less than one fifth of global max:16
×10^8
2.5
2
1.5
1
0.5
0
y
0
5
10
15
20
25
x
raw data, starting:15
fitted Gauss locations:16

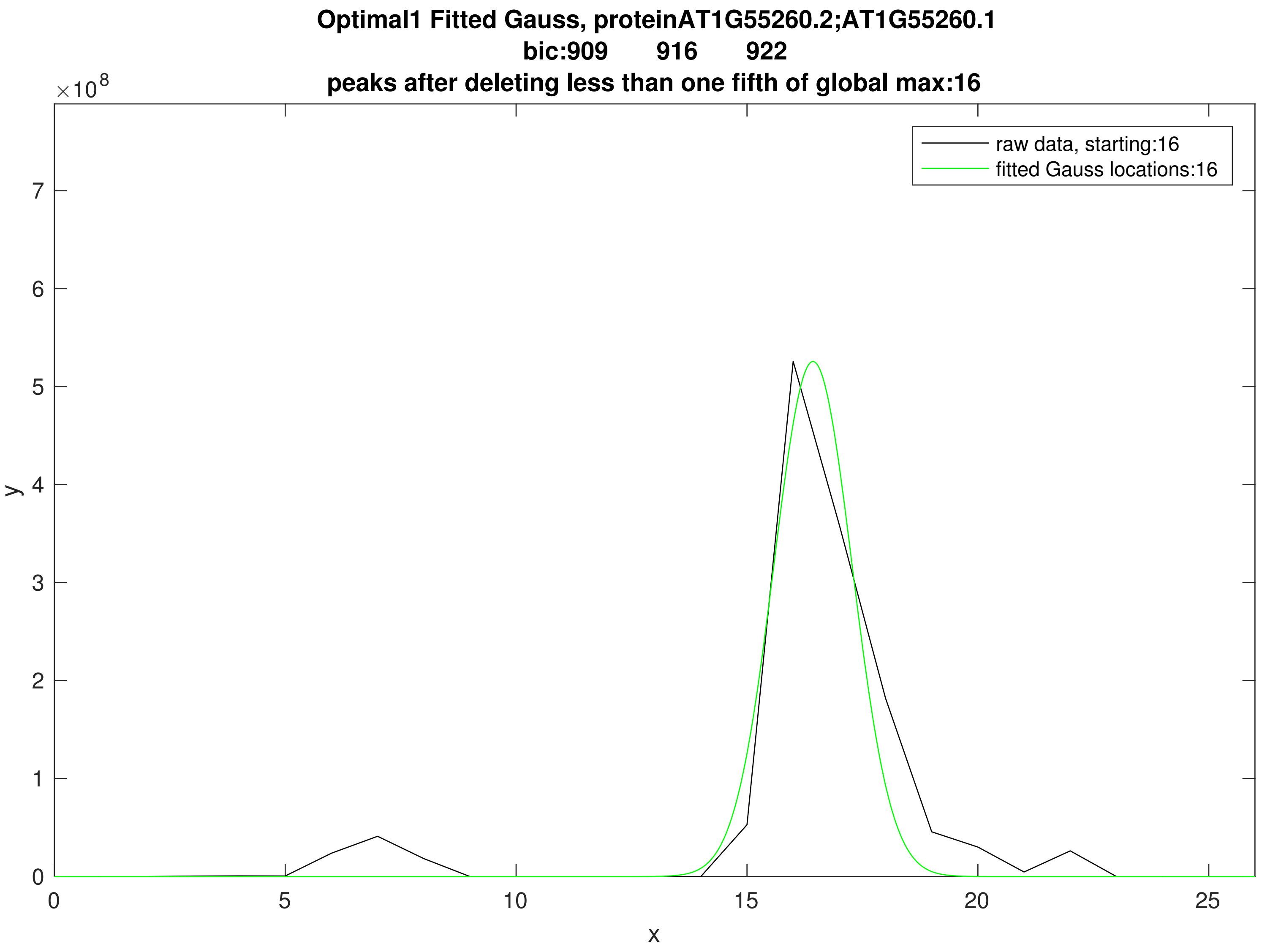

Optimal1 Fitted Gauss, proteinAT1G55260.2;AT1G55260.1
bic:909 916 922
peaks after deleting less than one fifth of global max:16
×10^8
raw data, starting:16
fitted Gauss locations:16
y
x
0
1
2
3
4
5
6
7
0
5
10
15
20
25

#### Optimal1 Fitted Gauss, proteinAT1G55265.1
#### bic:682
#### peaks after deleting less than one fifth of global max:22

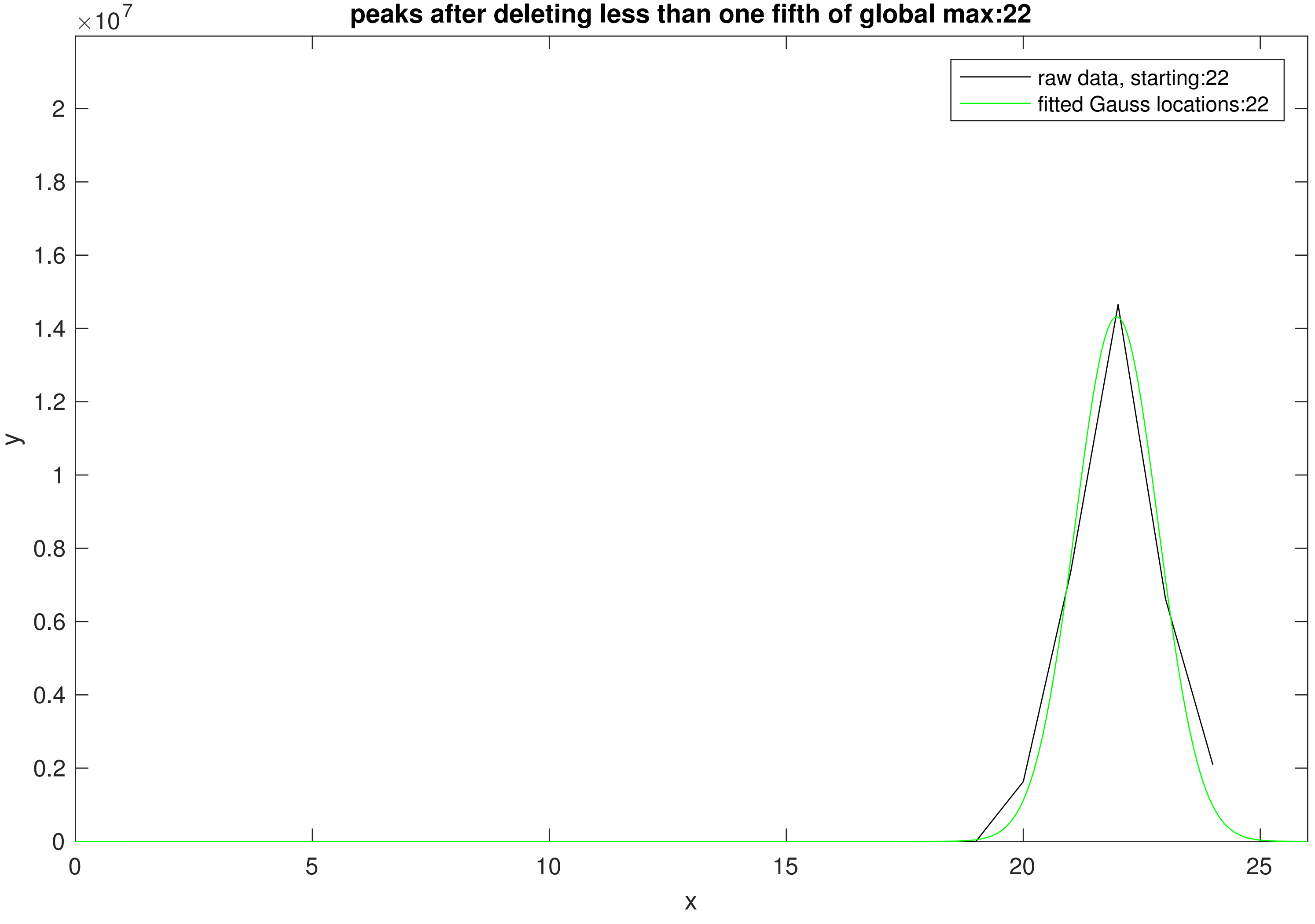

#### Optimal1 Fitted Gauss, proteinAT1G55490.2;AT1G55490.1
#### bic:1.04e+03 1.04e+03 1.05e+03 1.06e+03
#### peaks after deleting less than one fifth of global max:5

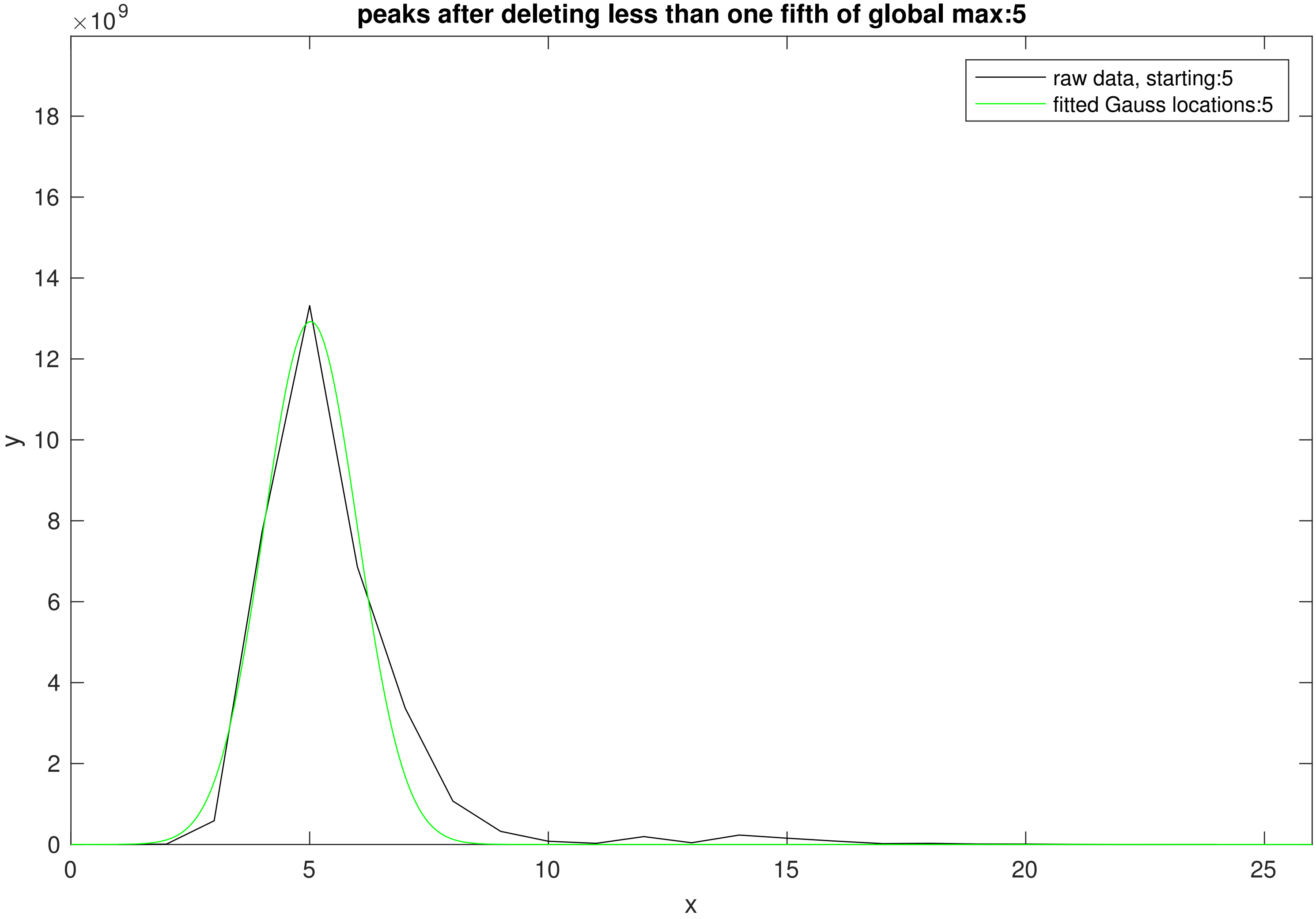

#### Optimal1 Fitted Gauss, proteinAT1G55570.1
#### bic:888
#### peaks after deleting less than one fifth of global max:15

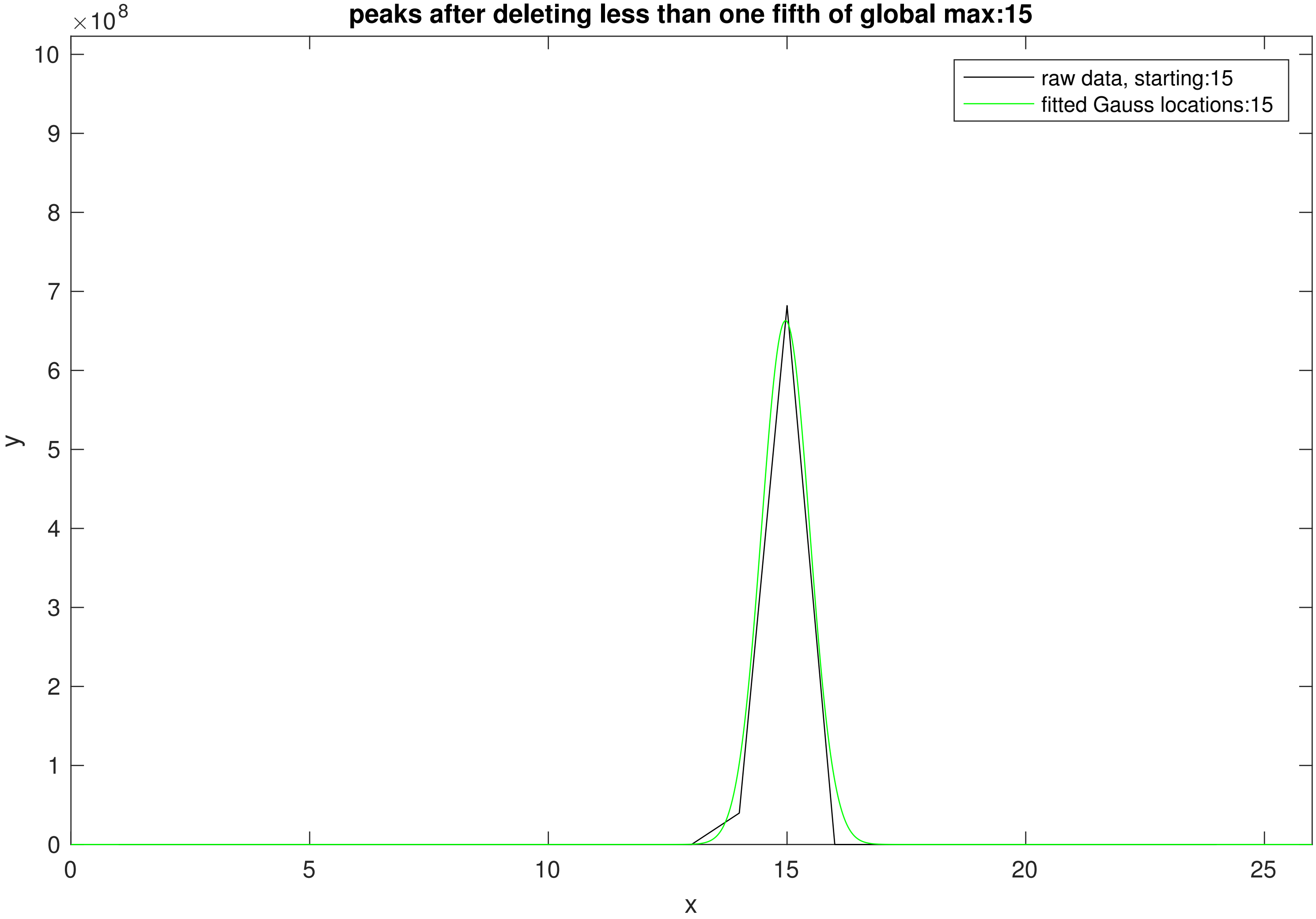

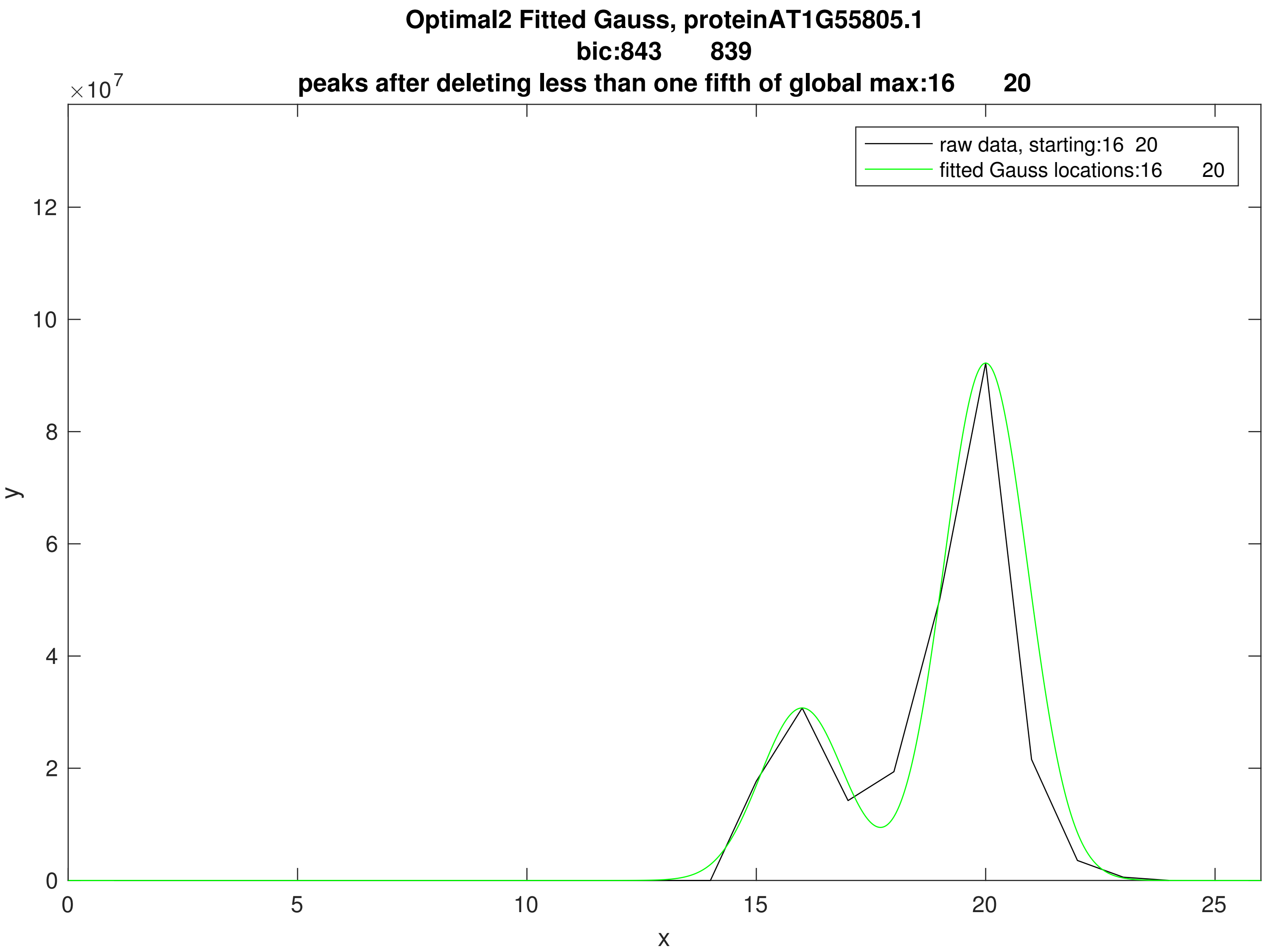
Optimal2 Fitted Gauss, proteinAT1G55805.1
bic:843 839
peaks after deleting less than one fifth of global max:16 20
×10^7
raw data, starting:16 20
fitted Gauss locations:16 20
y
x
0
2
4
6
8
10
12
0
5
10
15
20
25

**Optimal1 Fitted Gauss, proteinAT1G56070.1**
**bic:1.11e+03  1.12e+03  1.14e+03  1.15e+03**
**peaks after deleting less than one fifth of global max:13**

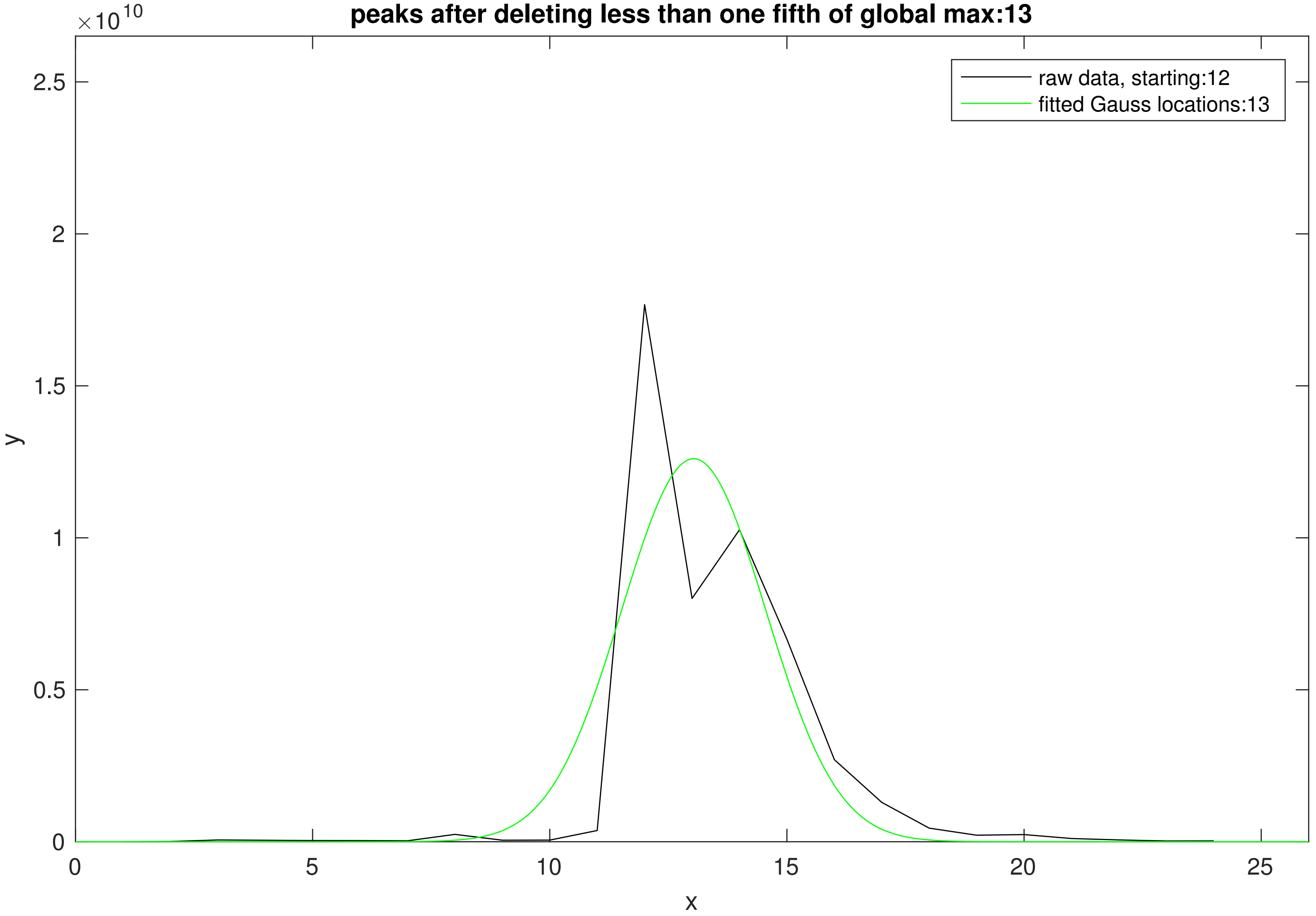

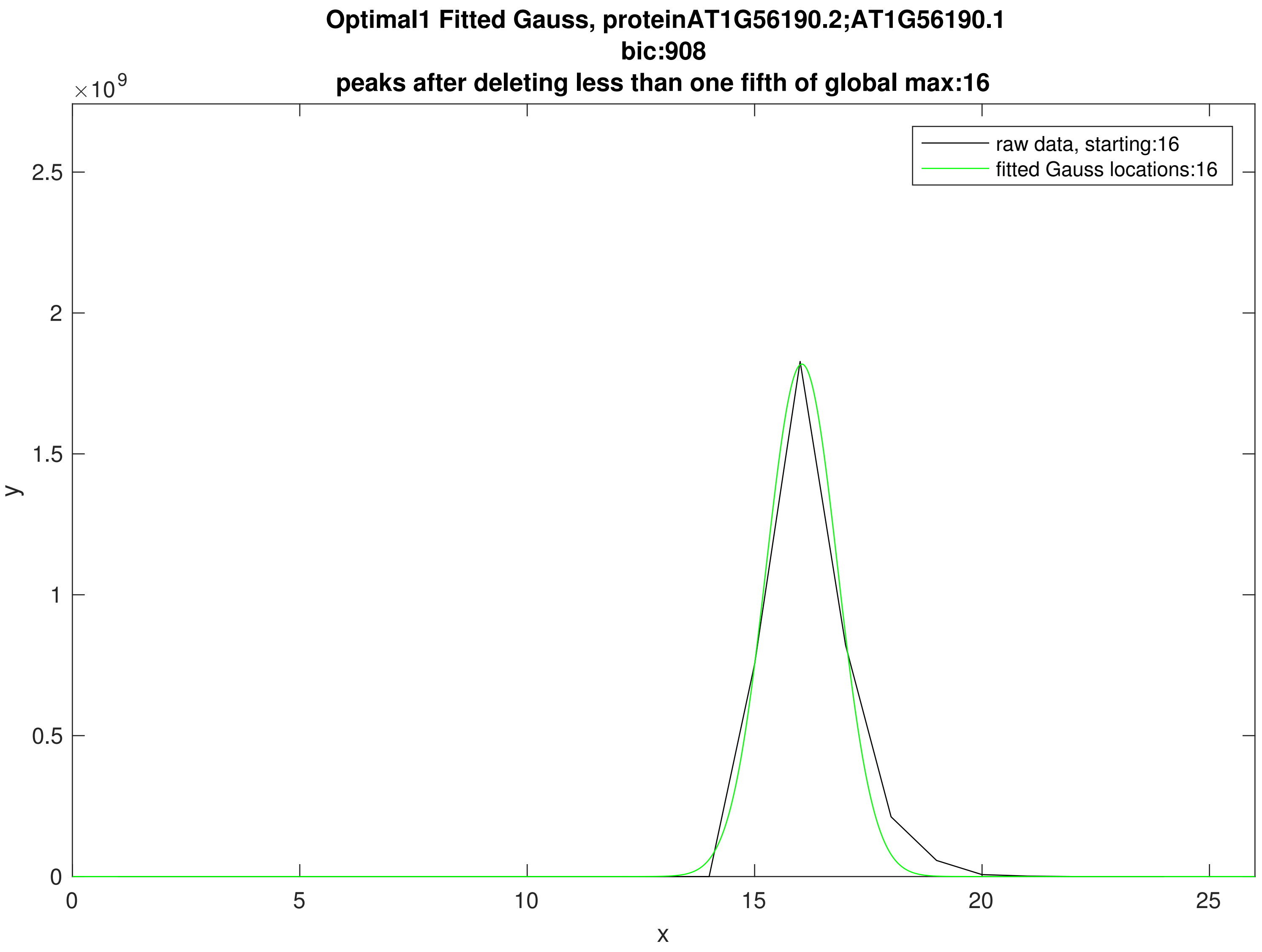

Optimal1 Fitted Gauss, proteinAT1G56190.2;AT1G56190.1
bic:908
peaks after deleting less than one fifth of global max:16
×10^9
raw data, starting:16
fitted Gauss locations:16
2.5
2
1.5
1
0.5
0
y
0
5
10
15
20
25
x

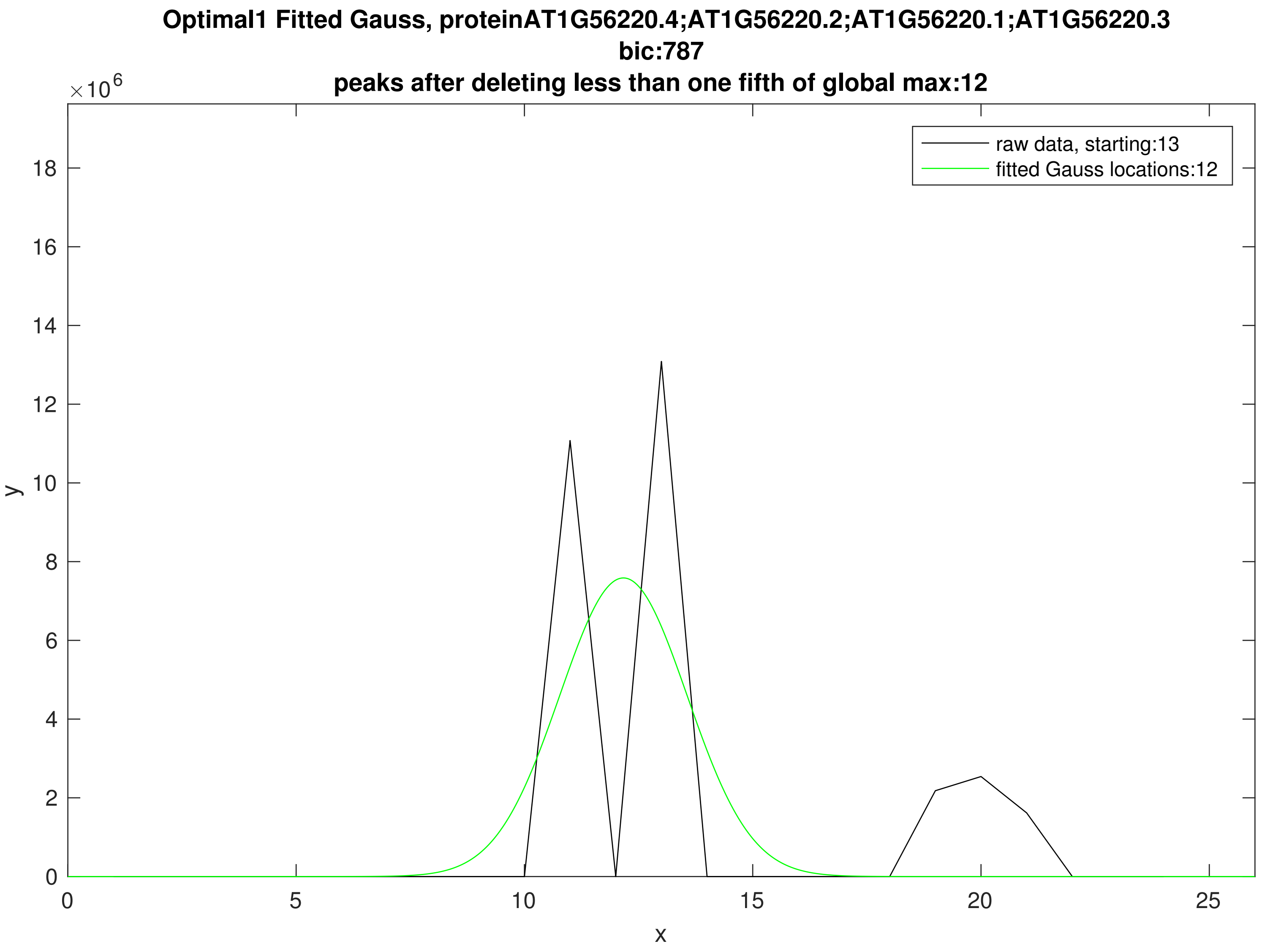
Optimal1 Fitted Gauss, proteinAT1G56220.4;AT1G56220.2;AT1G56220.1;AT1G56220.3
bic:787
peaks after deleting less than one fifth of global max:12
×10^6
raw data, starting:13
fitted Gauss locations:12
y
x
0
2
4
6
8
10
12
14
16
18
0
5
10
15
20
25

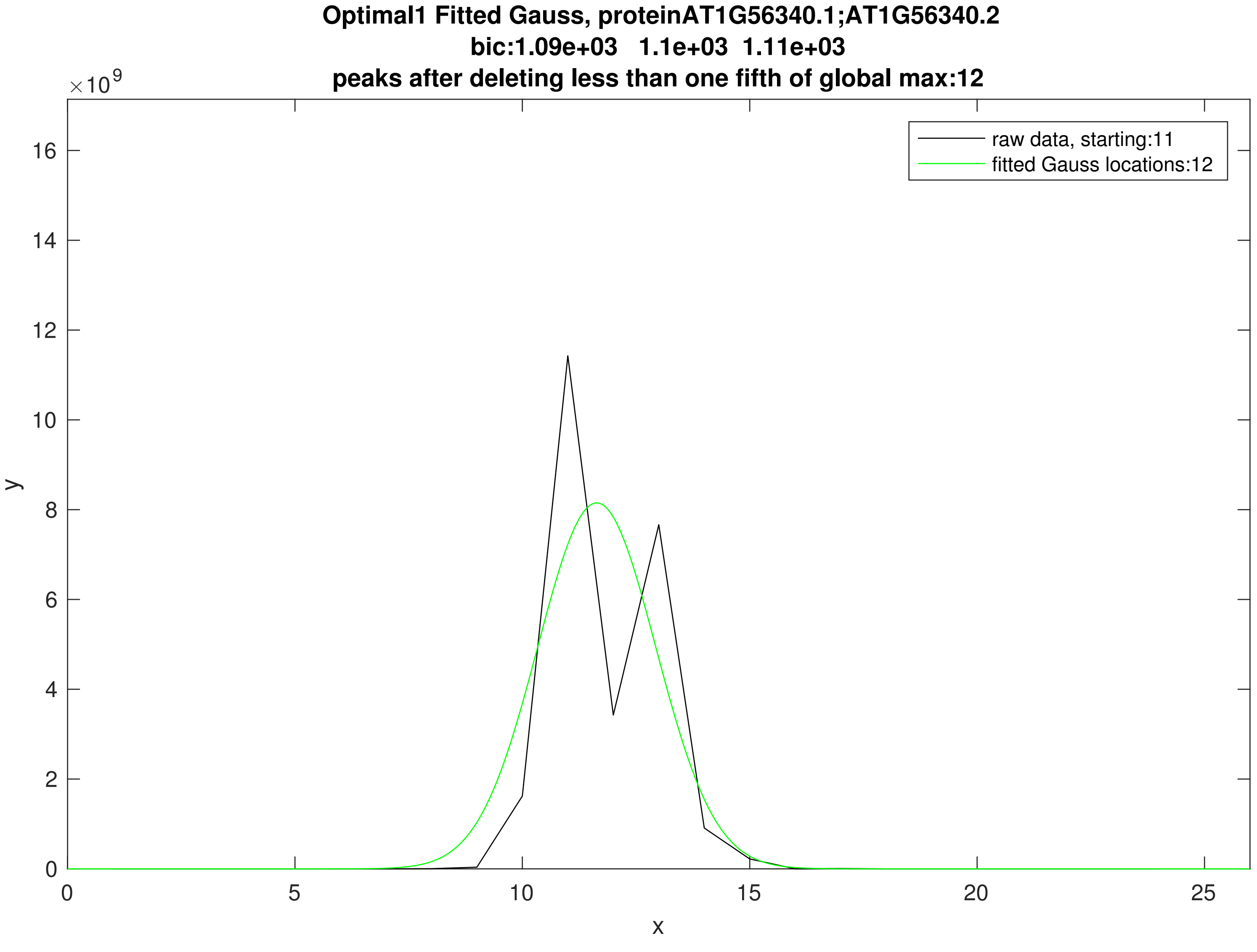

Optimal1 Fitted Gauss, proteinAT1G56340.1;AT1G56340.2
bic:1.09e+03 1.1e+03 1.11e+03
peaks after deleting less than one fifth of global max:12
×10^9
raw data, starting:11
fitted Gauss locations:12
16
14
12
10
8
6
4
2
0
y
0
5
10
15
20
25
x

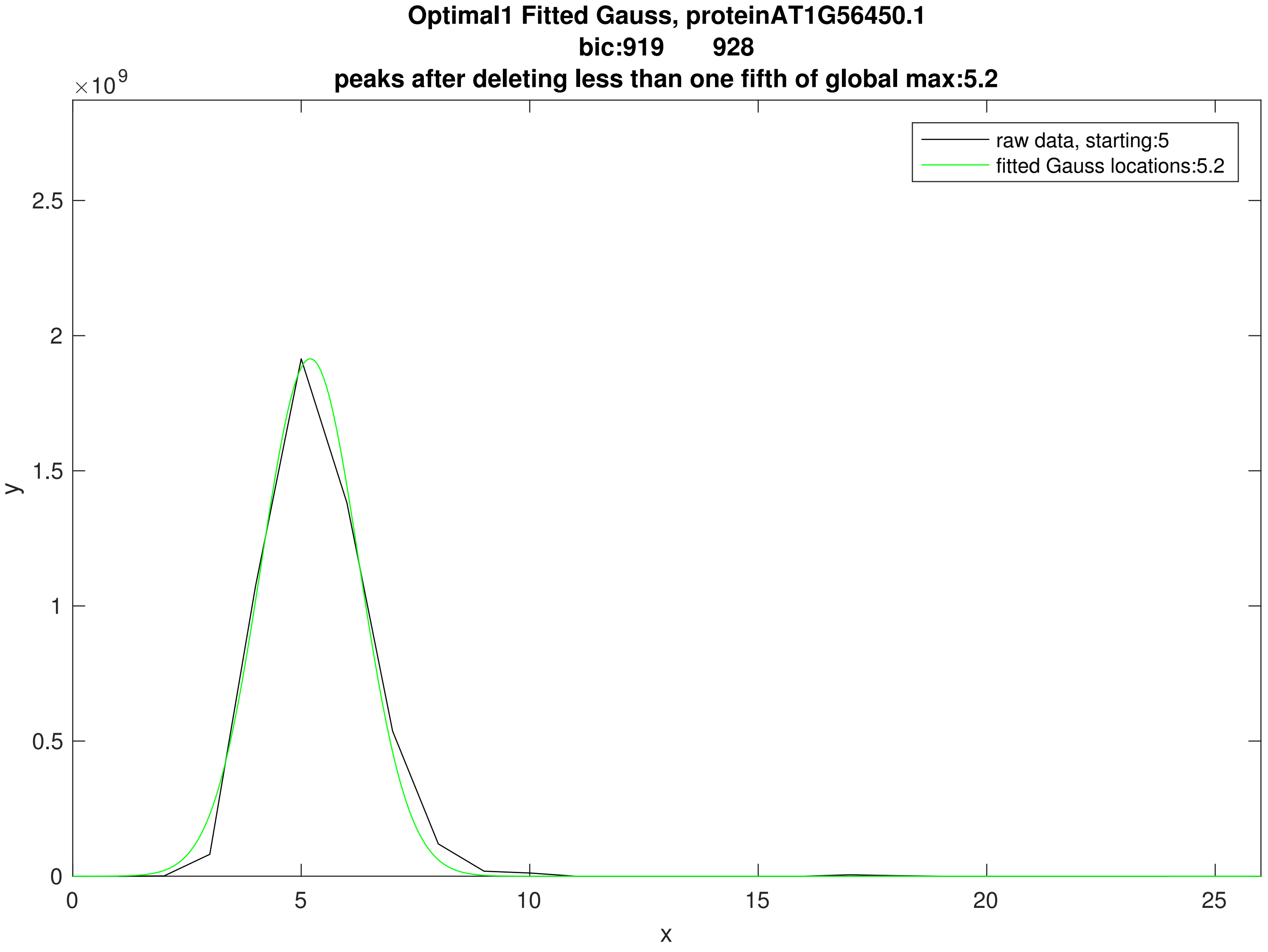
Optimal1 Fitted Gauss, proteinAT1G56450.1
bic:919 928
peaks after deleting less than one fifth of global max:5.2
raw data, starting:5
fitted Gauss locations:5.2
×10^9
2.5
2
1.5
1
0.5
0
y
0
5
10
15
20
25
x

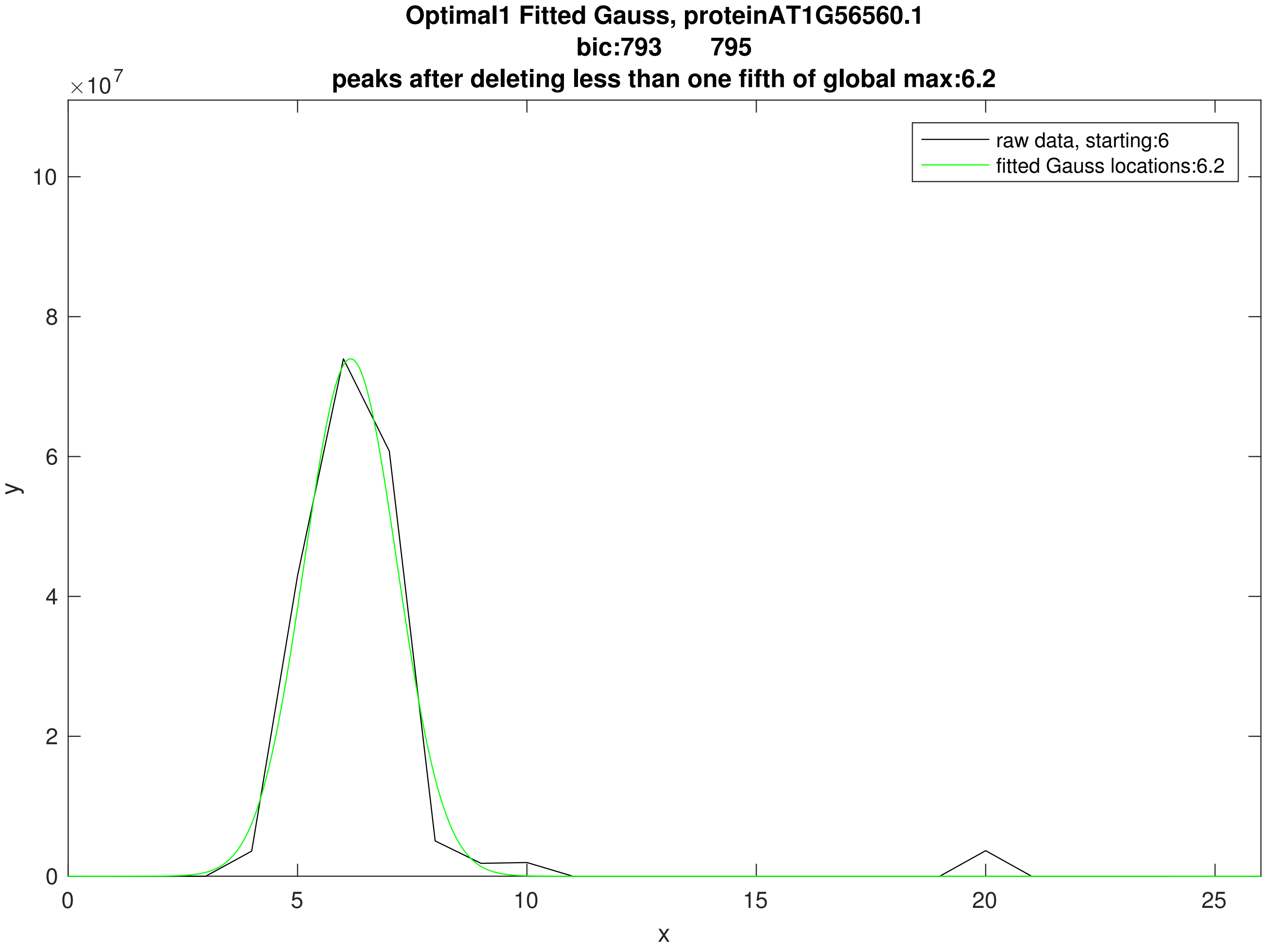

Optimal1 Fitted Gauss, proteinAT1G56560.1
bic:793 795
peaks after deleting less than one fifth of global max:6.2
×10^7
raw data, starting:6
fitted Gauss locations:6.2
y
x
0
2
4
6
8
10
0
5
10
15
20
25

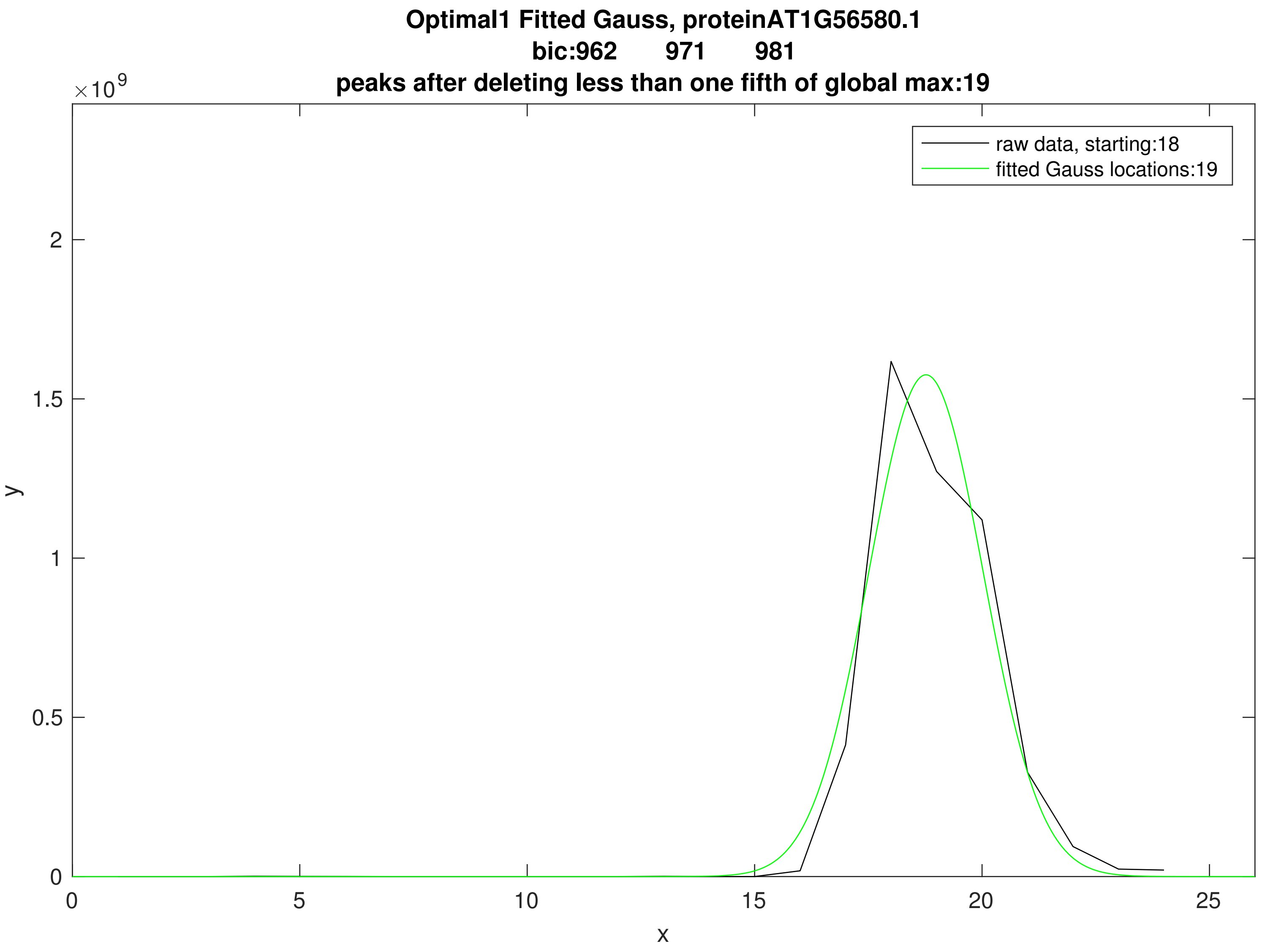
Optimal1 Fitted Gauss, proteinAT1G56580.1
bic:962 971 981
peaks after deleting less than one fifth of global max:19
×10^9
raw data, starting:18
fitted Gauss locations:19
y
x
0
0.5
1
1.5
2
0
5
10
15
20
25

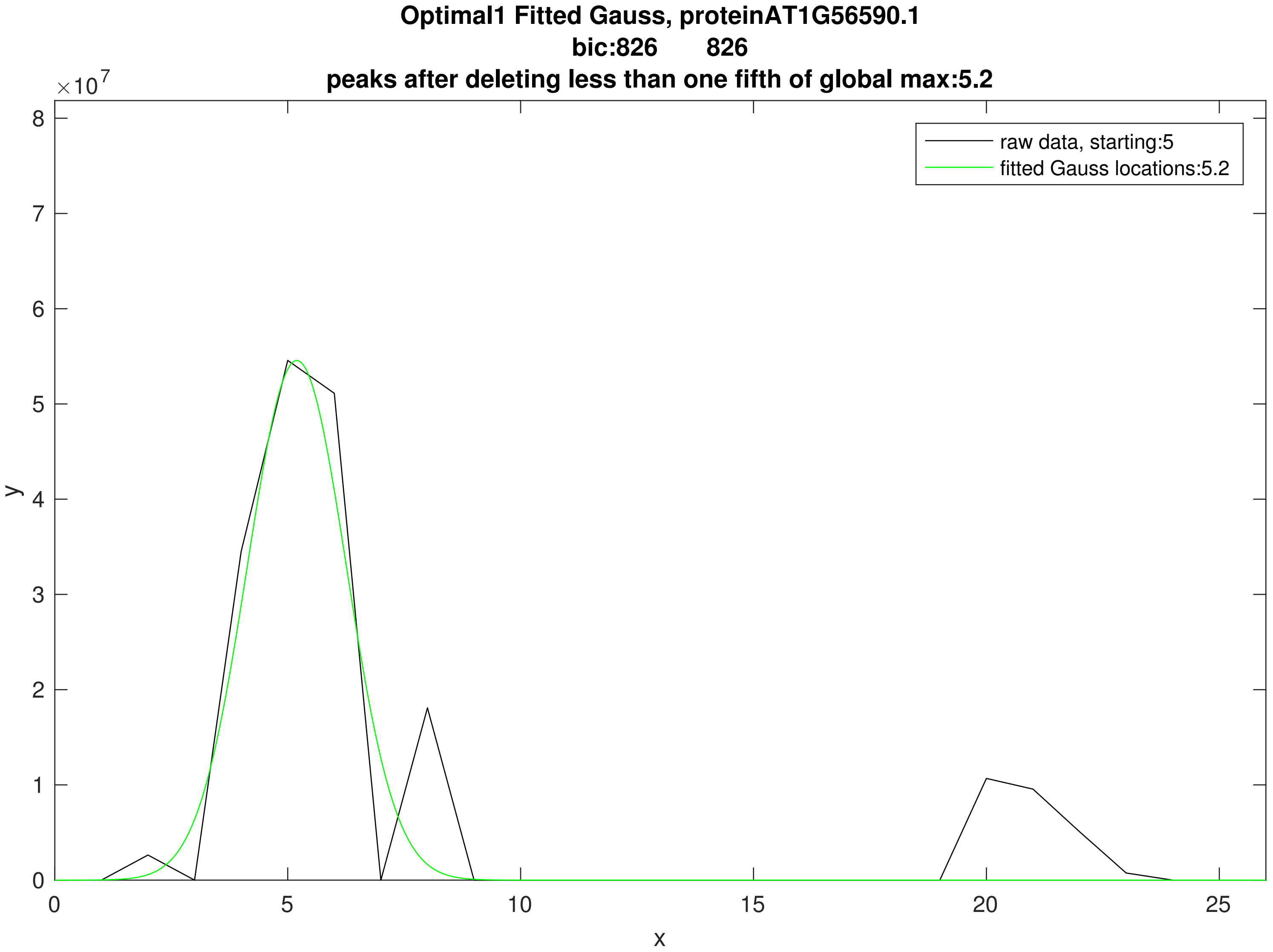

Optimal1 Fitted Gauss, proteinAT1G56590.1
bic:826 826
peaks after deleting less than one fifth of global max:5.2
×10^7
raw data, starting:5
fitted Gauss locations:5.2
y
x
0
1
2
3
4
5
6
7
8
0
5
10
15
20
25

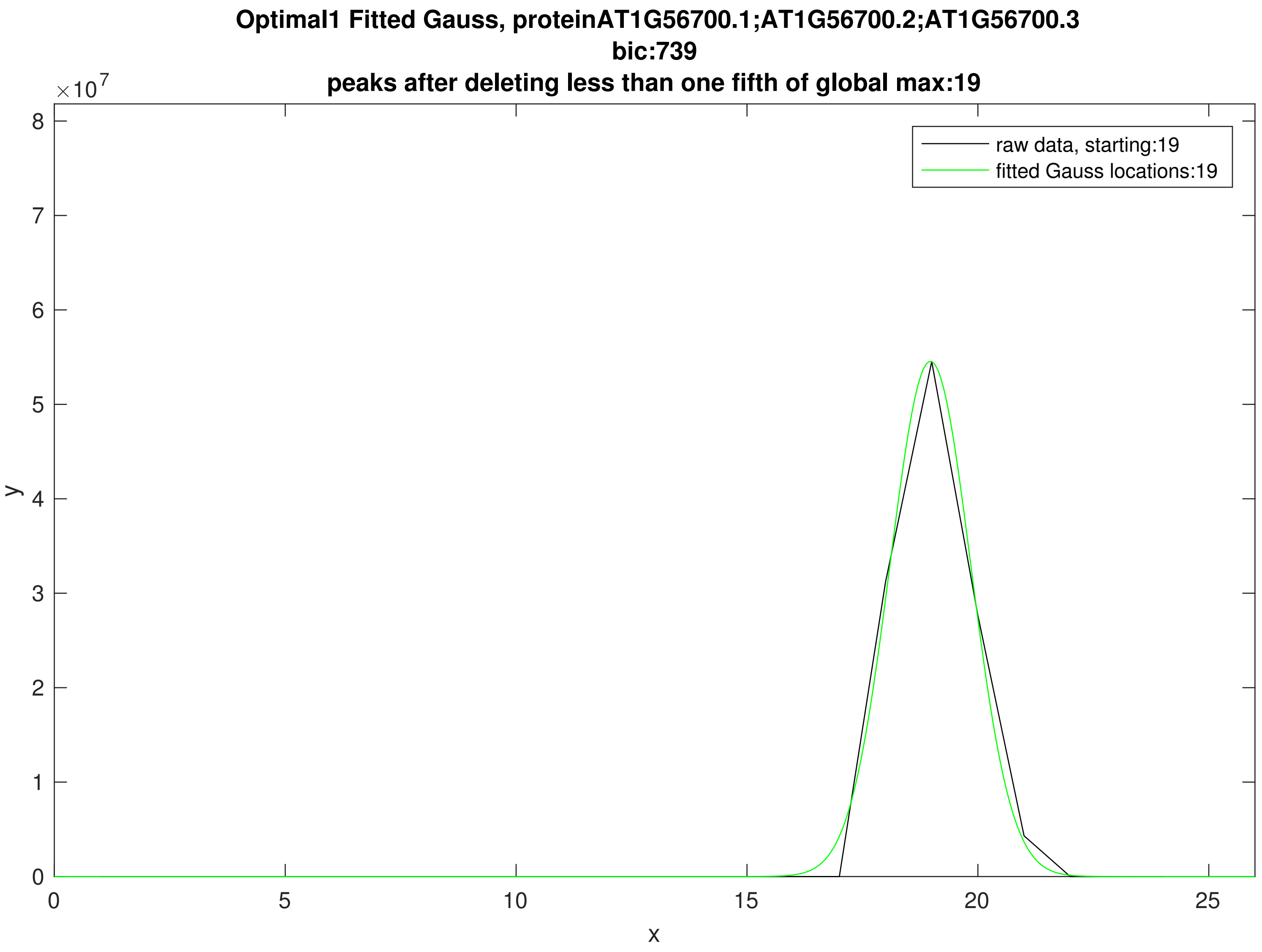
Optimal1 Fitted Gauss, proteinAT1G56700.1;AT1G56700.2;AT1G56700.3
bic:739
peaks after deleting less than one fifth of global max:19
×10^7
8
7
6
5
4
3
2
1
0
y
0
5
10
15
20
25
x
raw data, starting:19
fitted Gauss locations:19

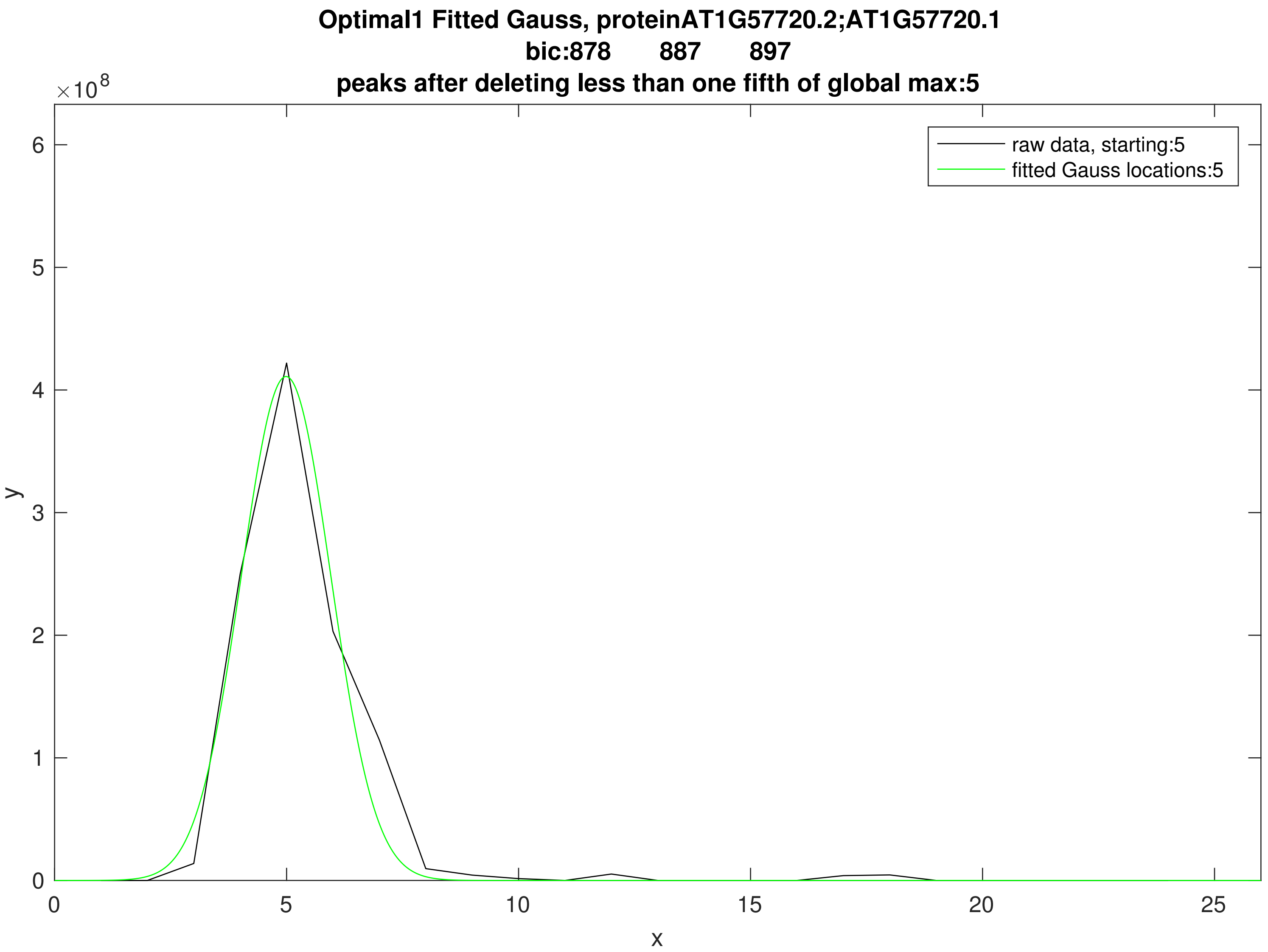

Optimal1 Fitted Gauss, proteinAT1G57720.2;AT1G57720.1
bic:878 887 897
peaks after deleting less than one fifth of global max:5
×10^8
raw data, starting:5
fitted Gauss locations:5
y
x
0
1
2
3
4
5
6
10
15
20
25

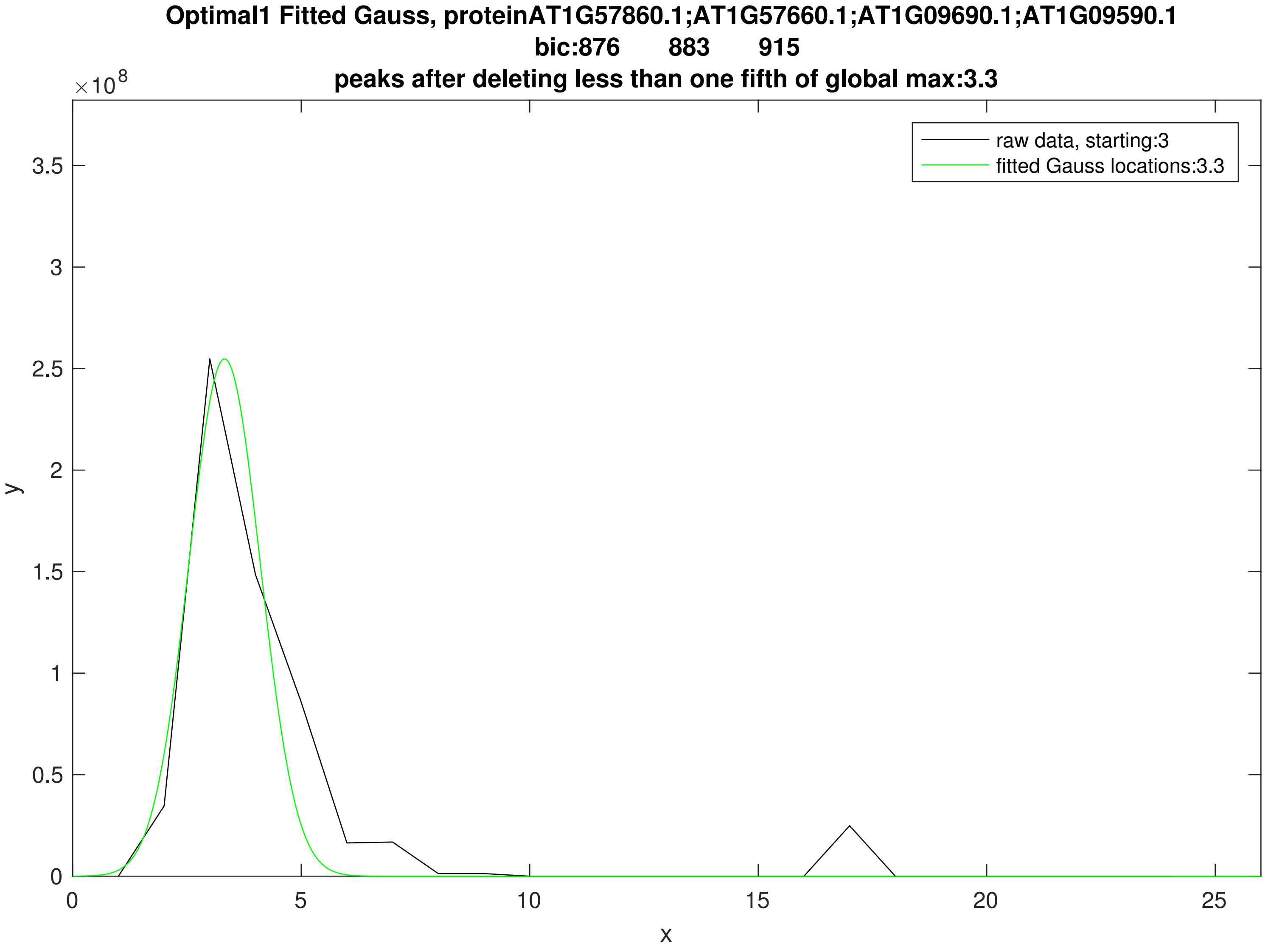

Optimal1 Fitted Gauss, proteinAT1G57860.1;AT1G57660.1;AT1G09690.1;AT1G09590.1
bic:876 883 915
peaks after deleting less than one fifth of global max:3.3
×10^8
raw data, starting:3
fitted Gauss locations:3.3
y
x
0
0.5
1
1.5
2
2.5
3
3.5
0
5
10
15
20
25

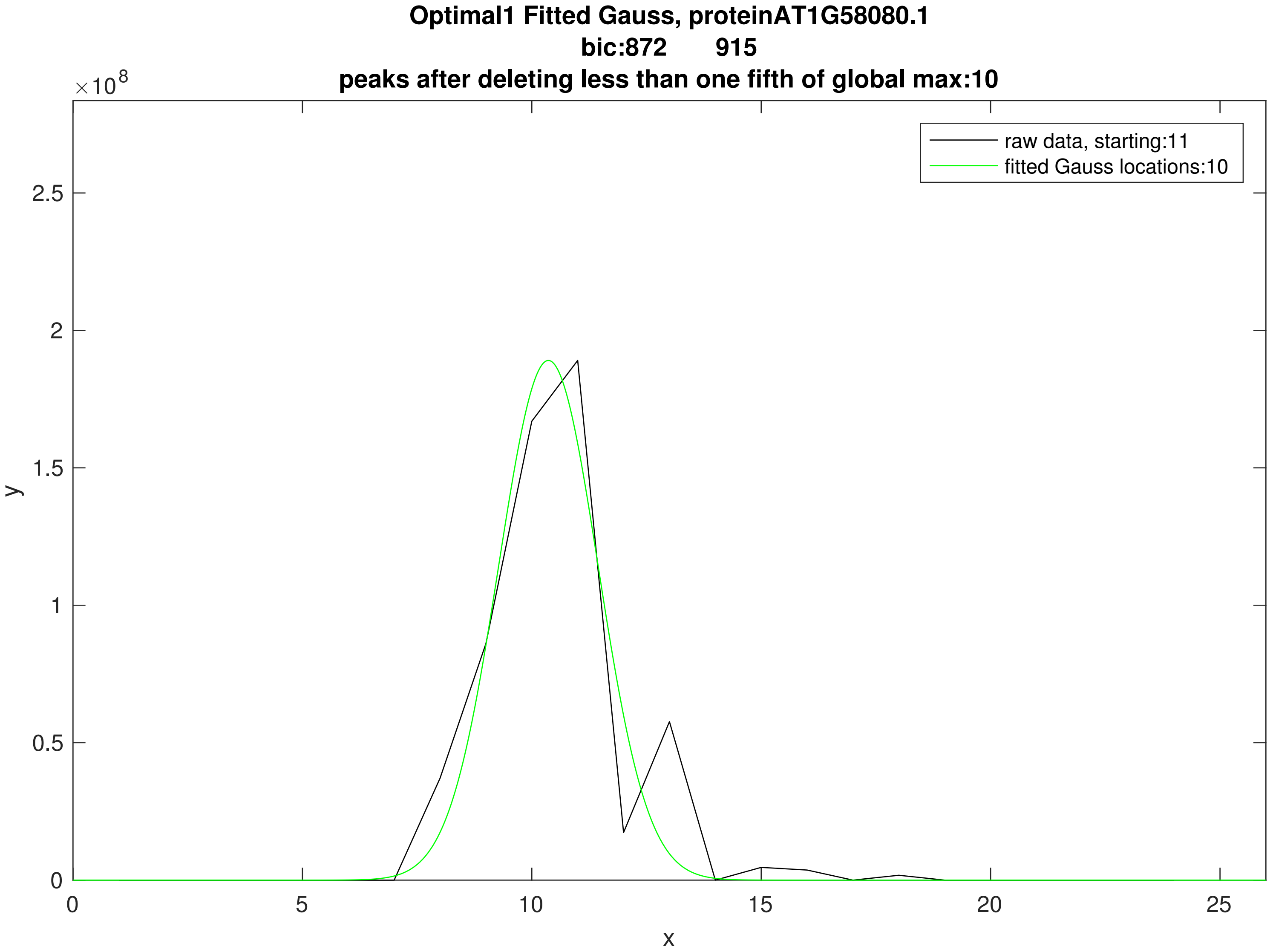
Optimal1 Fitted Gauss, proteinAT1G58080.1
bic:872 915
peaks after deleting less than one fifth of global max:10
×10^8
raw data, starting:11
fitted Gauss locations:10
y
x
0
0.5
1
1.5
2
2.5
0
5
10
15
20
25

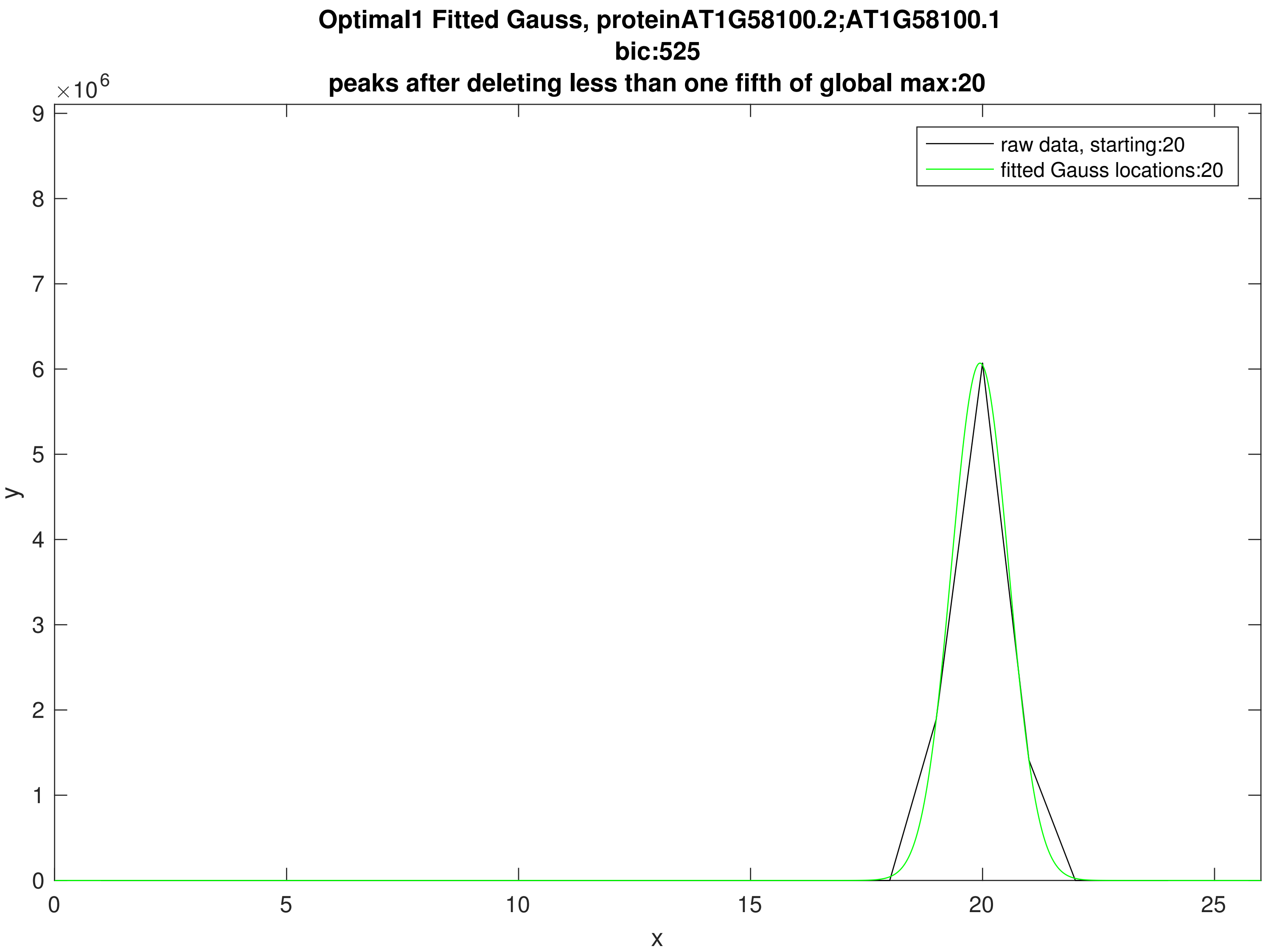

Optimal1 Fitted Gauss, proteinAT1G58100.2;AT1G58100.1
bic:525
peaks after deleting less than one fifth of global max:20
×10^6
raw data, starting:20
fitted Gauss locations:20
y
x
0
1
2
3
4
5
6
7
8
9
0
5
10
15
20
25

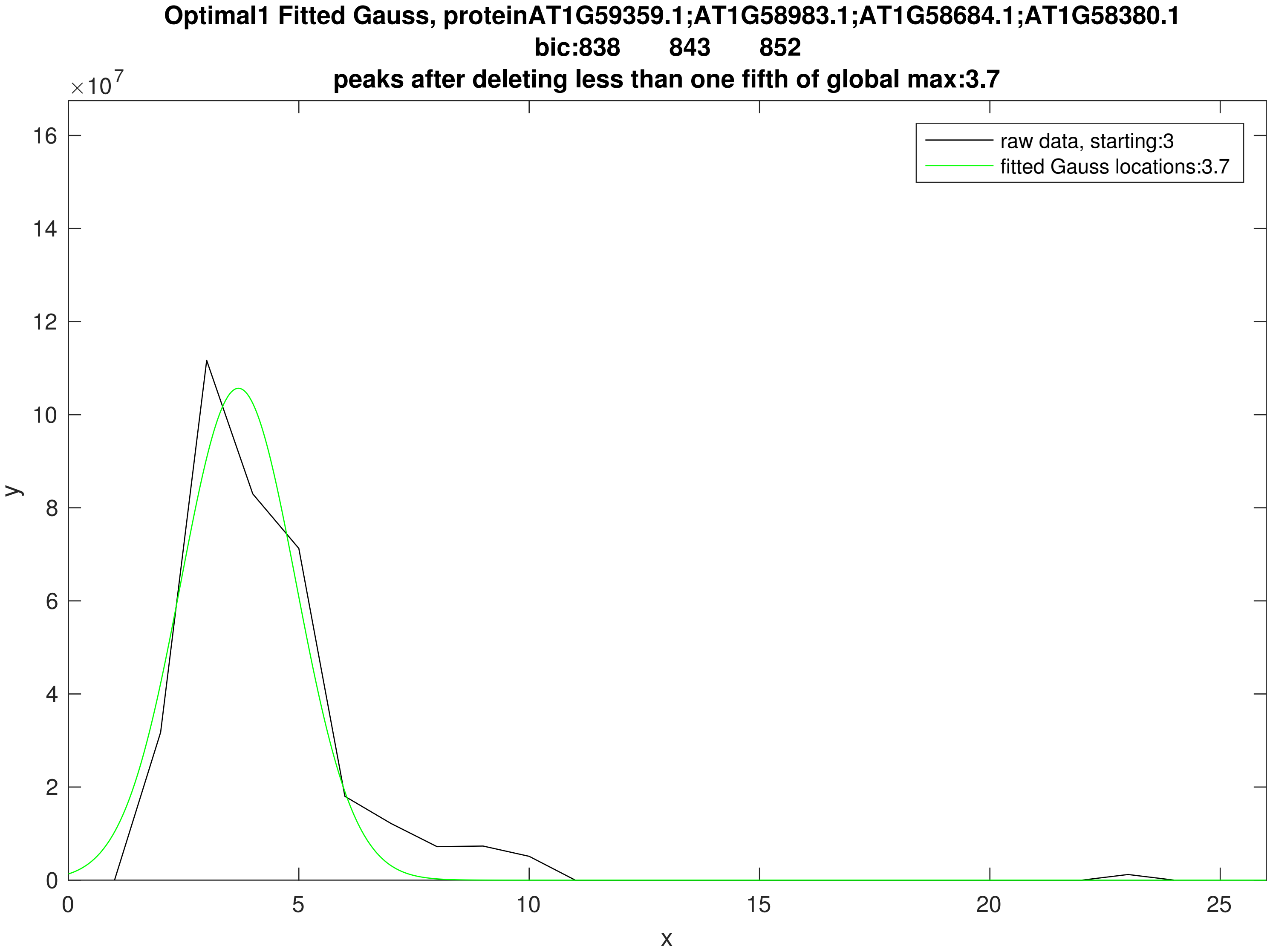
Optimal1 Fitted Gauss, proteinAT1G59359.1;AT1G58983.1;AT1G58684.1;AT1G58380.1
bic:838 843 852
peaks after deleting less than one fifth of global max:3.7
×10^7
raw data, starting:3
fitted Gauss locations:3.7
y
x
0
2
4
6
8
10
12
14
16
0
5
10
15
20
25

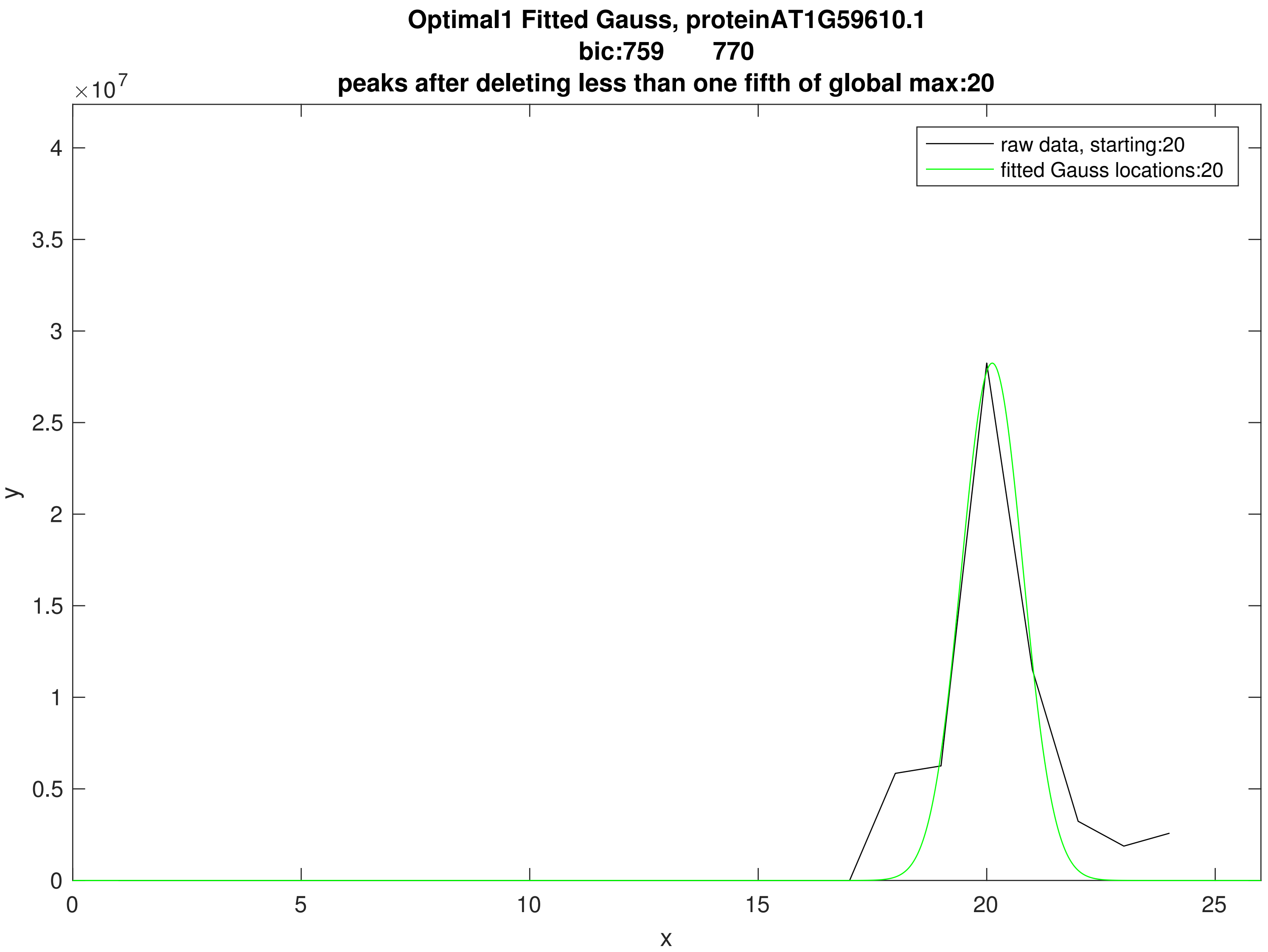
Optimal1 Fitted Gauss, proteinAT1G59610.1
bic:759 770
peaks after deleting less than one fifth of global max:20
×10^7
raw data, starting:20
fitted Gauss locations:20
y
x
0
0.5
1
1.5
2
2.5
3
3.5
4
0
5
10
15
20
25

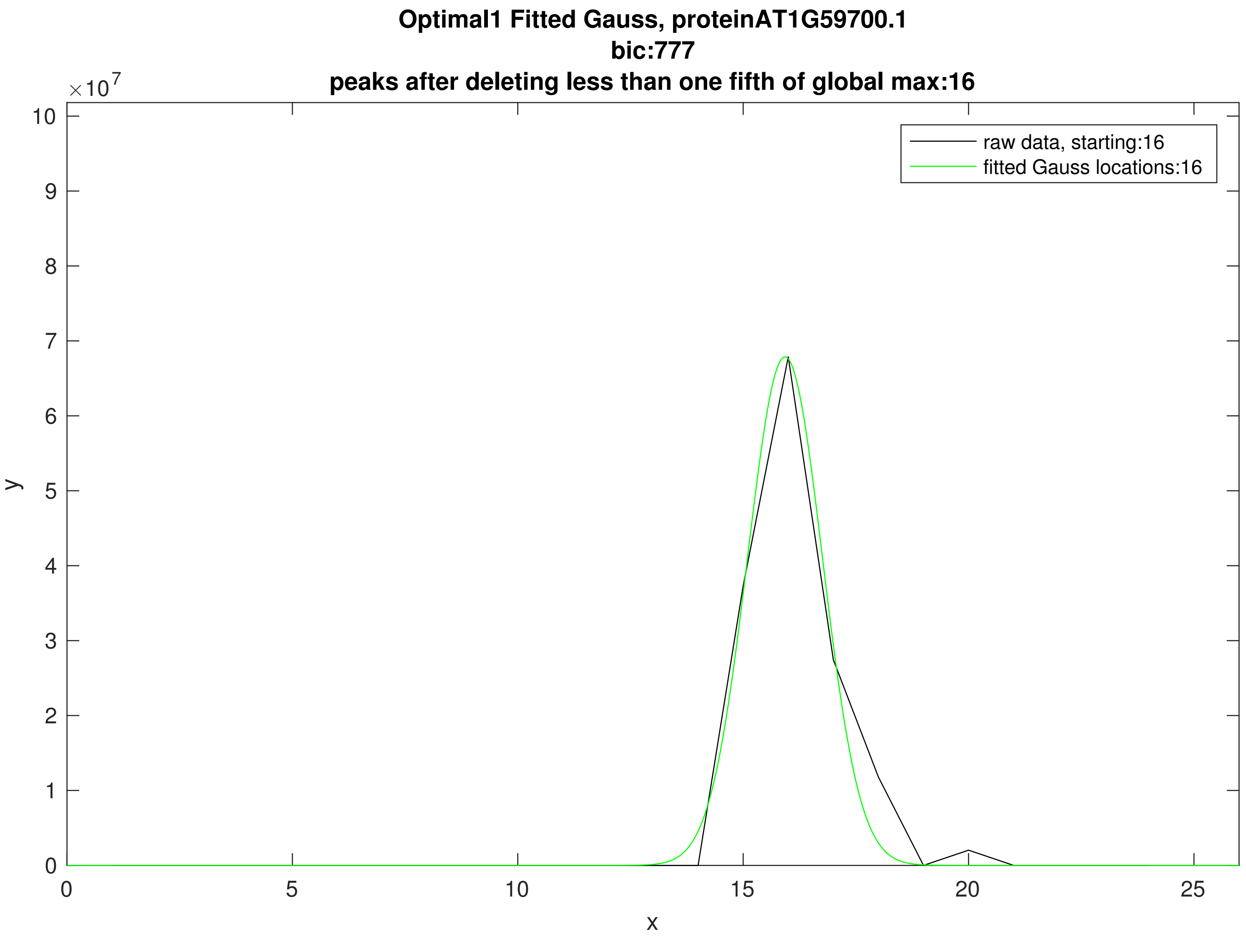
Optimal1 Fitted Gauss, proteinAT1G59700.1
bic:777
peaks after deleting less than one fifth of global max:16
×10^7
raw data, starting:16
fitted Gauss locations:16
y
x
0
1
2
3
4
5
6
7
8
9
10
0
5
10
15
20
25

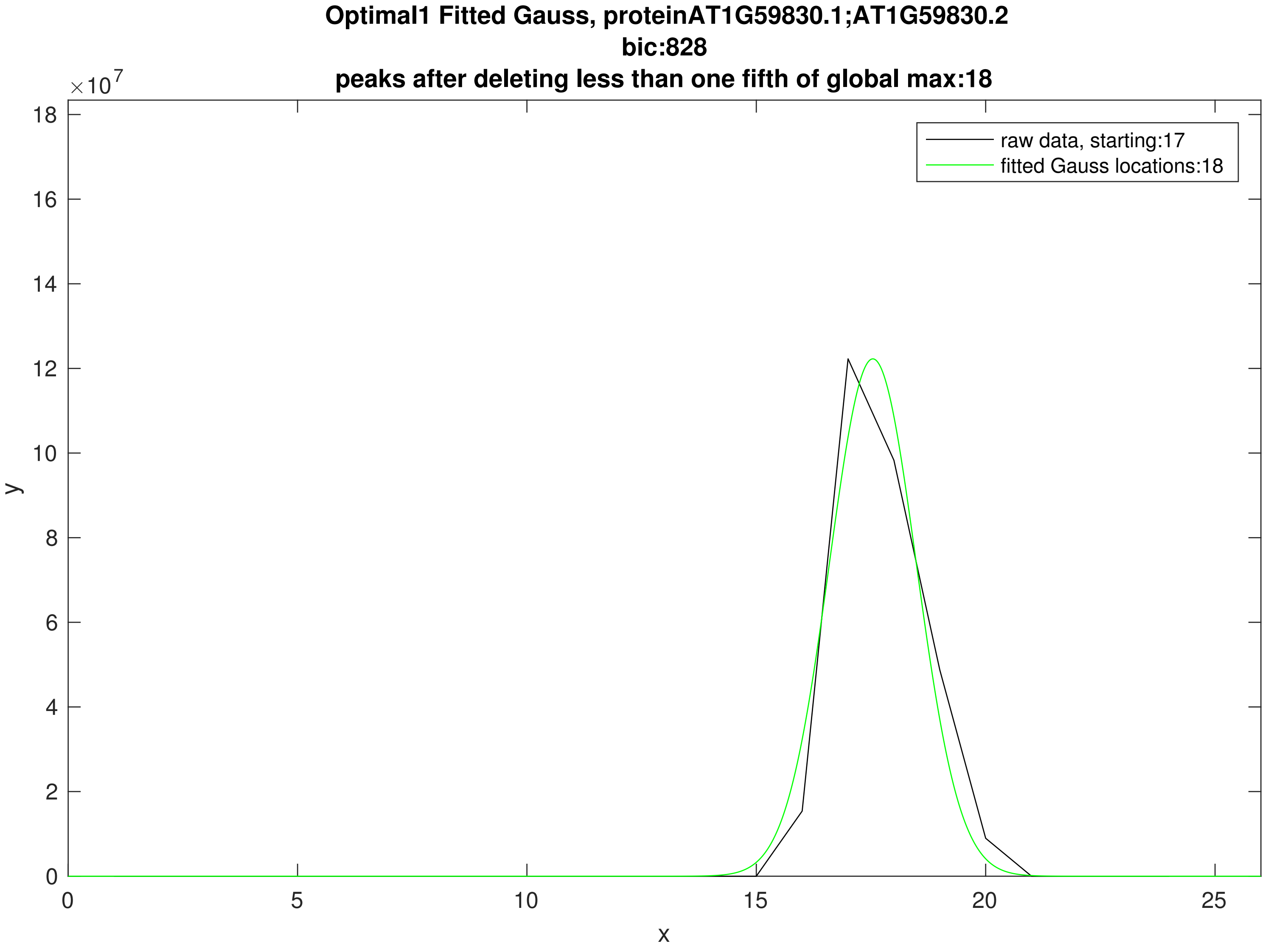

Optimal1 Fitted Gauss, proteinAT1G59830.1;AT1G59830.2
bic:828
peaks after deleting less than one fifth of global max:18
×10^7
raw data, starting:17
fitted Gauss locations:18
y
x
0
2
4
6
8
10
12
14
16
18
0
5
10
15
20
25

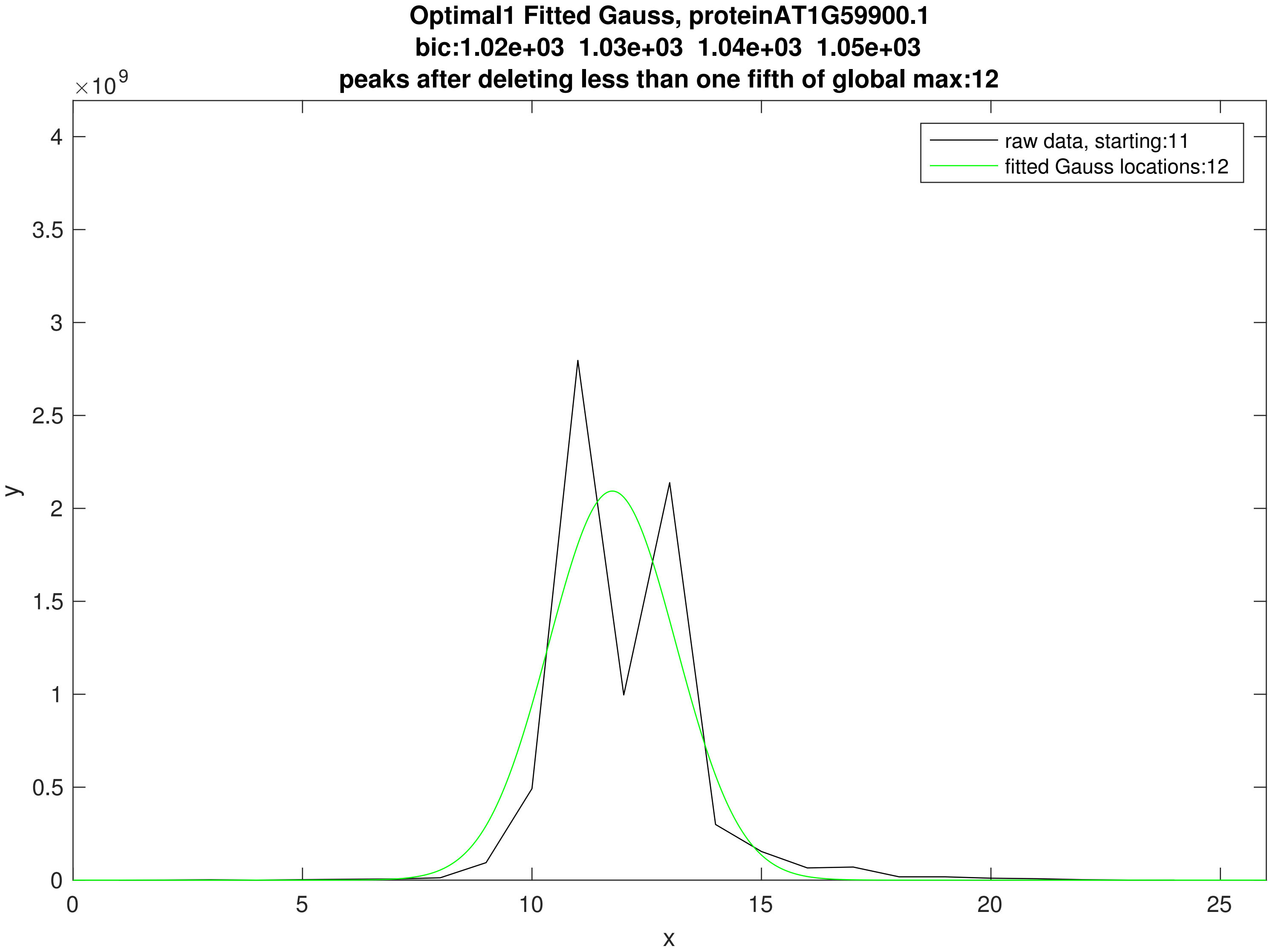
Optimal1 Fitted Gauss, proteinAT1G59900.1
bic:1.02e+03  1.03e+03  1.04e+03  1.05e+03
peaks after deleting less than one fifth of global max:12
×10^9
raw data, starting:11
fitted Gauss locations:12
y
x
0
0.5
1
1.5
2
2.5
3
3.5
4
0
5
10
15
20
25

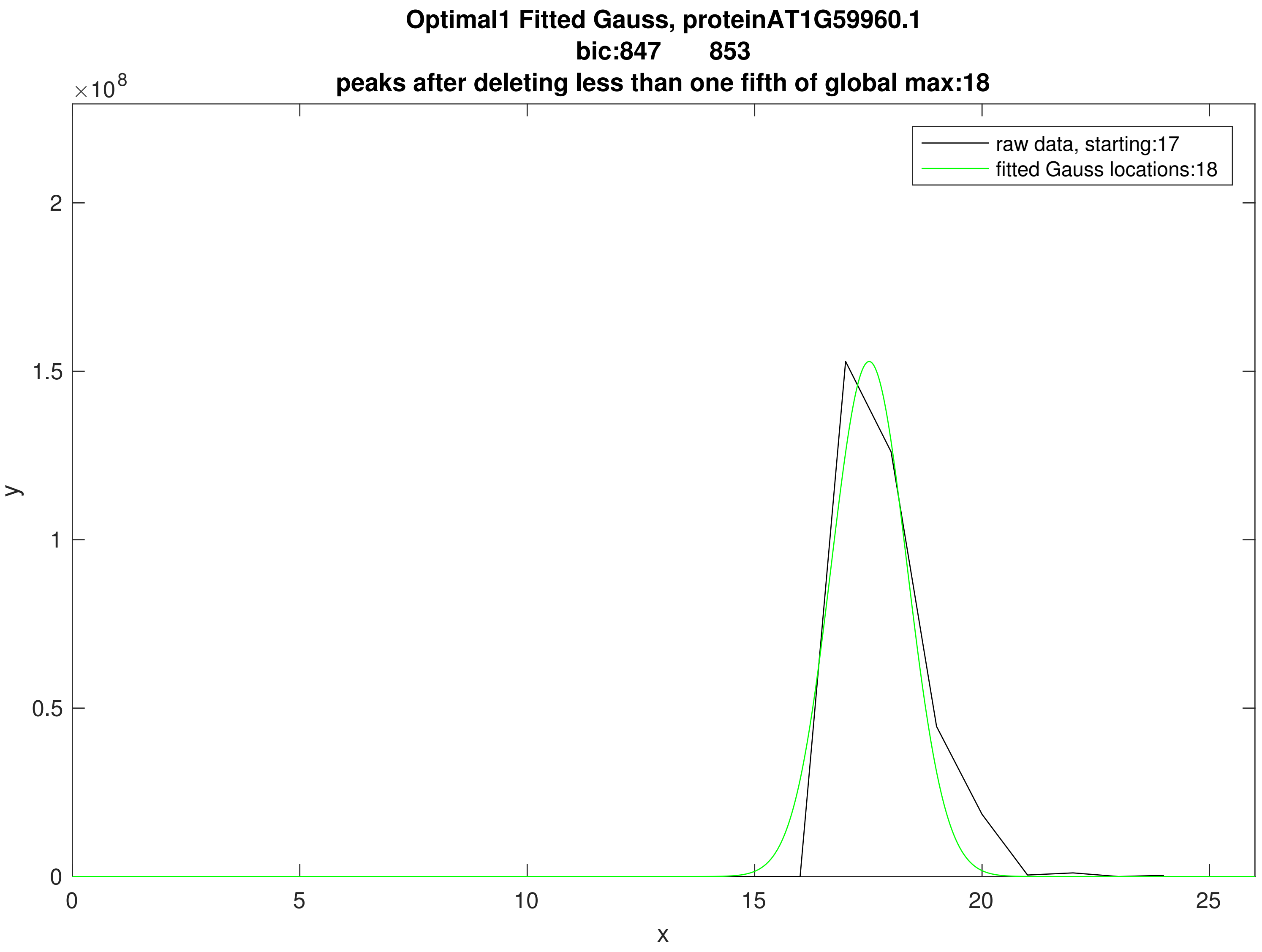

Optimal1 Fitted Gauss, proteinAT1G59960.1
bic:847 853
peaks after deleting less than one fifth of global max:18
×10^8
raw data, starting:17
fitted Gauss locations:18
y
x
0
0.5
1
1.5
2
0
5
10
15
20
25

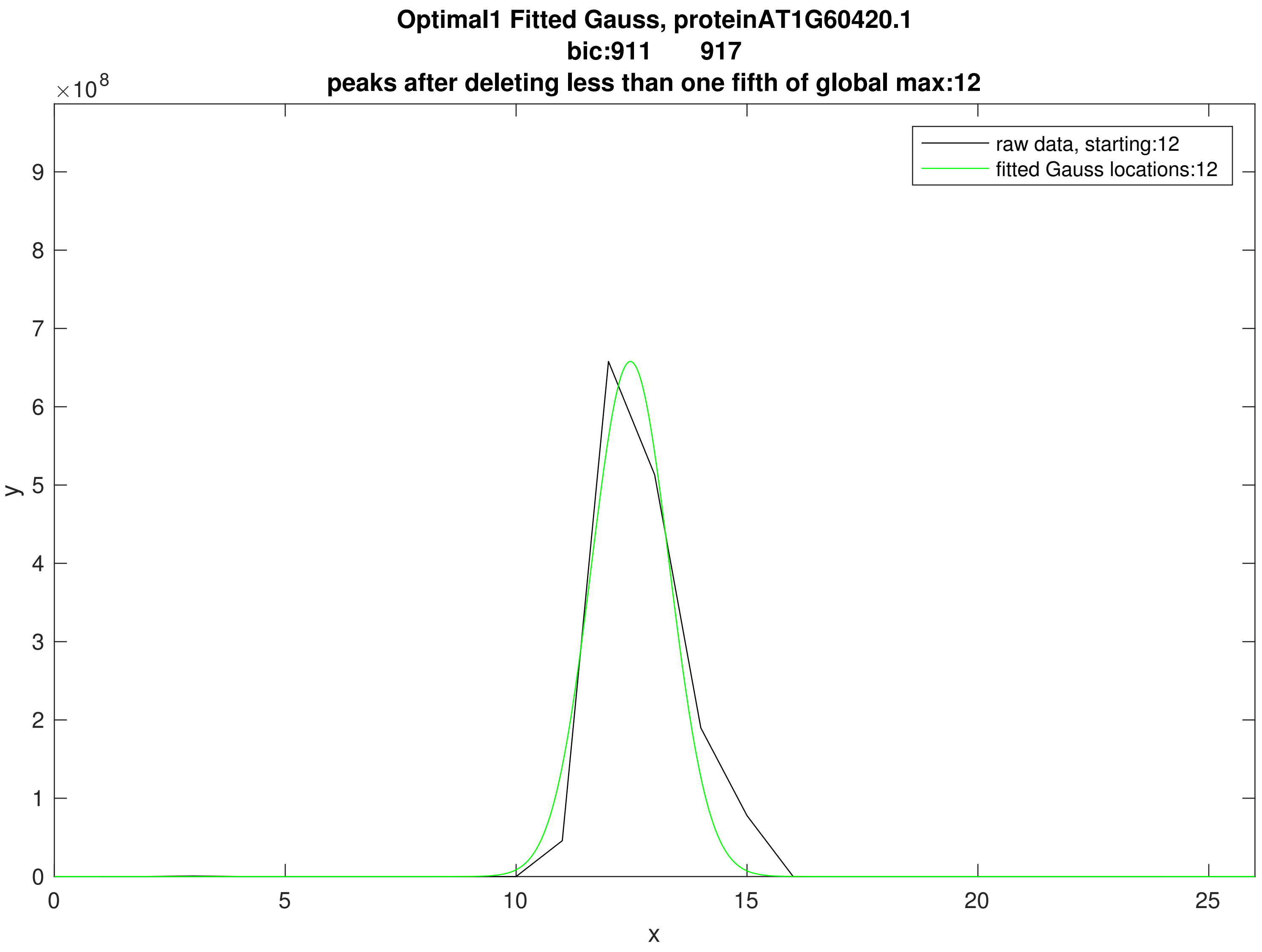
Optimal1 Fitted Gauss, proteinAT1G60420.1
bic:911 917
peaks after deleting less than one fifth of global max:12
×10^8
raw data, starting:12
fitted Gauss locations:12
y
x
0
1
2
3
4
5
6
7
8
9
0
5
10
15
20
25

Optimal1 Fitted Gauss, proteinAT1G60660.1
bic:696
peaks after deleting less than one fifth of global max:20

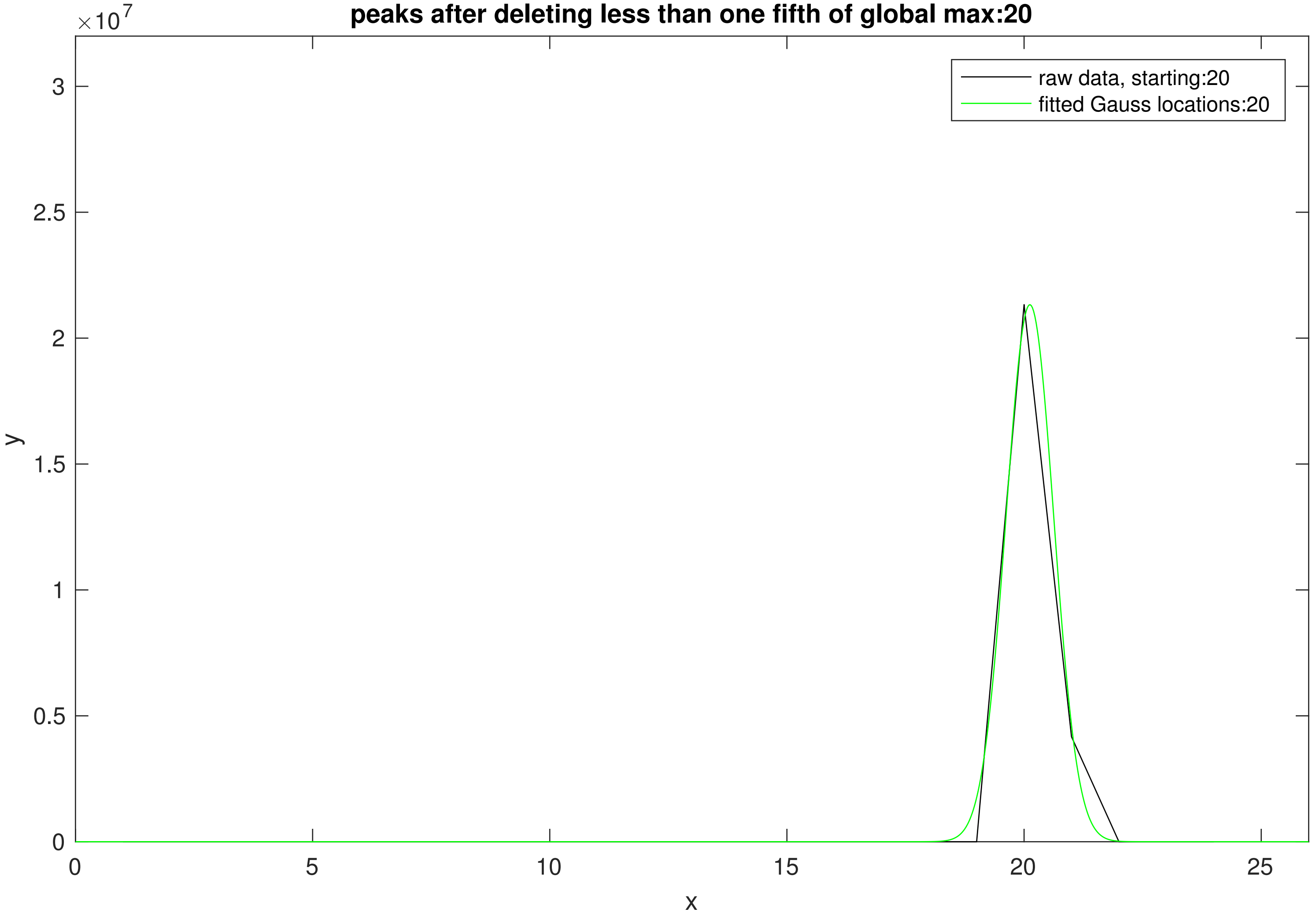

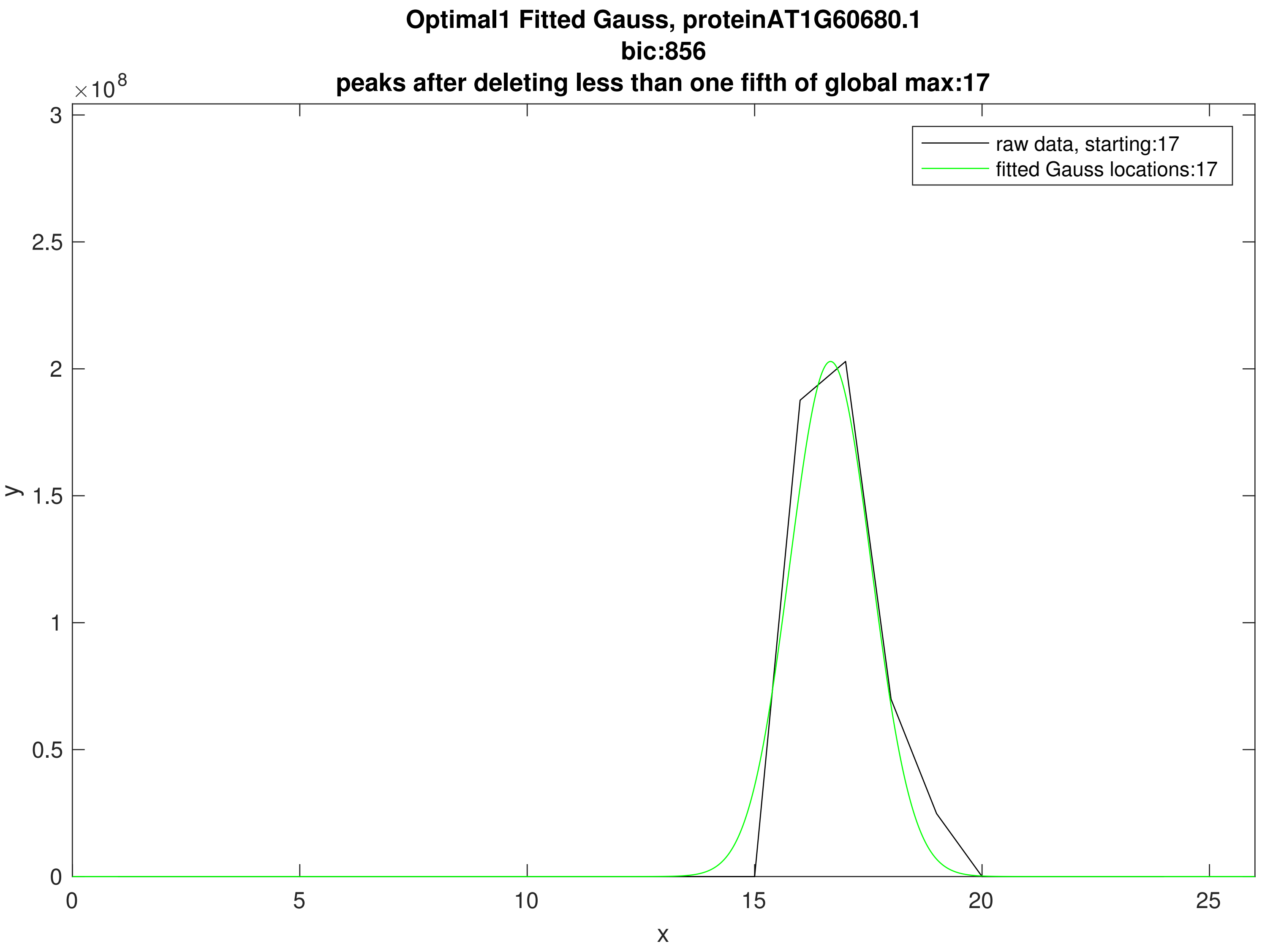

Optimal1 Fitted Gauss, proteinAT1G60680.1
bic:856
peaks after deleting less than one fifth of global max:17
×10^8
raw data, starting:17
fitted Gauss locations:17
y
x
0
0.5
1
1.5
2
2.5
3
0
5
10
15
20
25

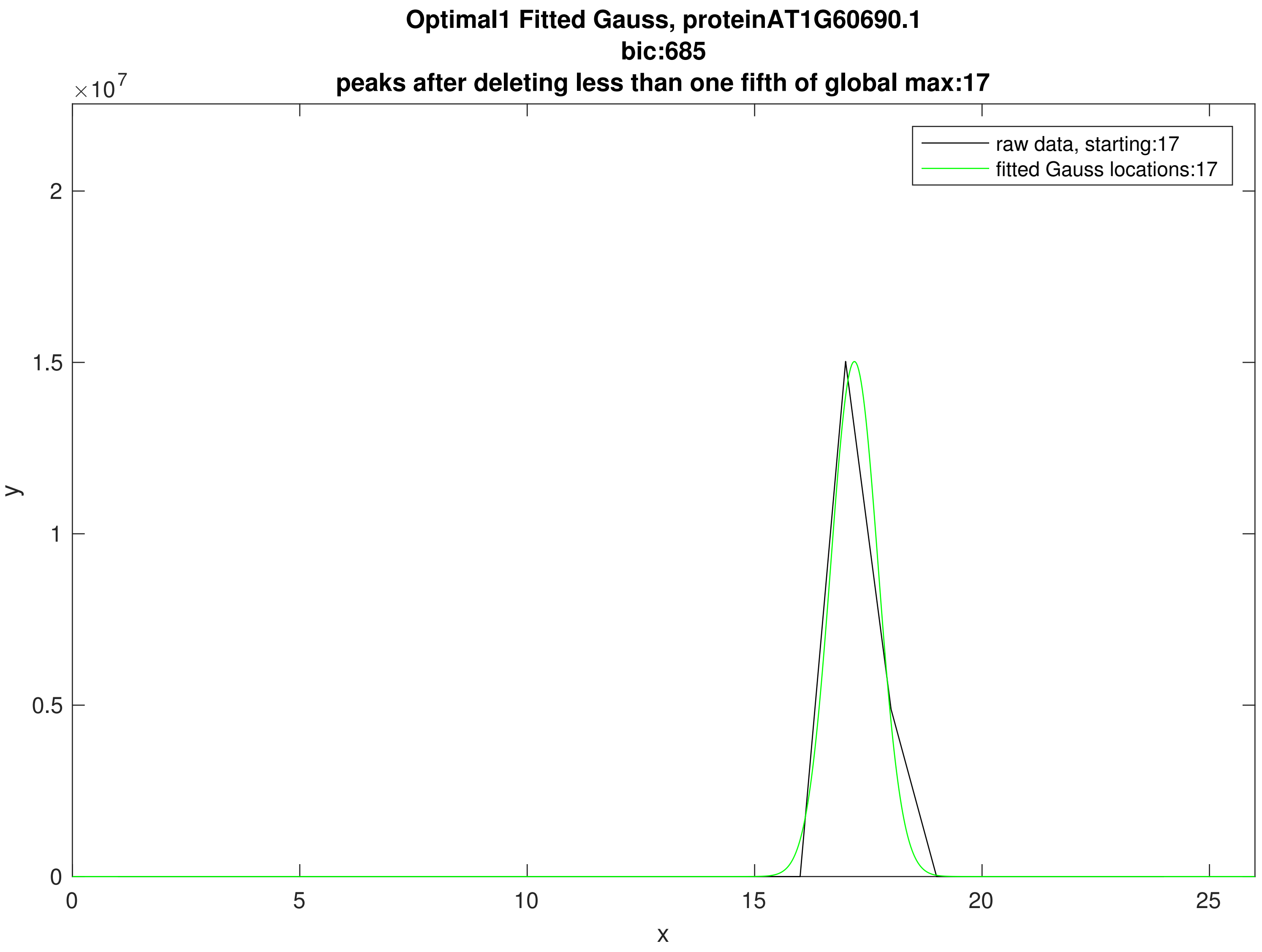
Optimal1 Fitted Gauss, proteinAT1G60690.1
bic:685
peaks after deleting less than one fifth of global max:17
×10^7
raw data, starting:17
fitted Gauss locations:17
y
x
0
0.5
1
1.5
2
5
10
15
20
25

**Optimal2 Fitted Gauss, proteinAT1G60710.1**
**bic:952 950 964**
**peaks after deleting less than one fifth of global max:16**

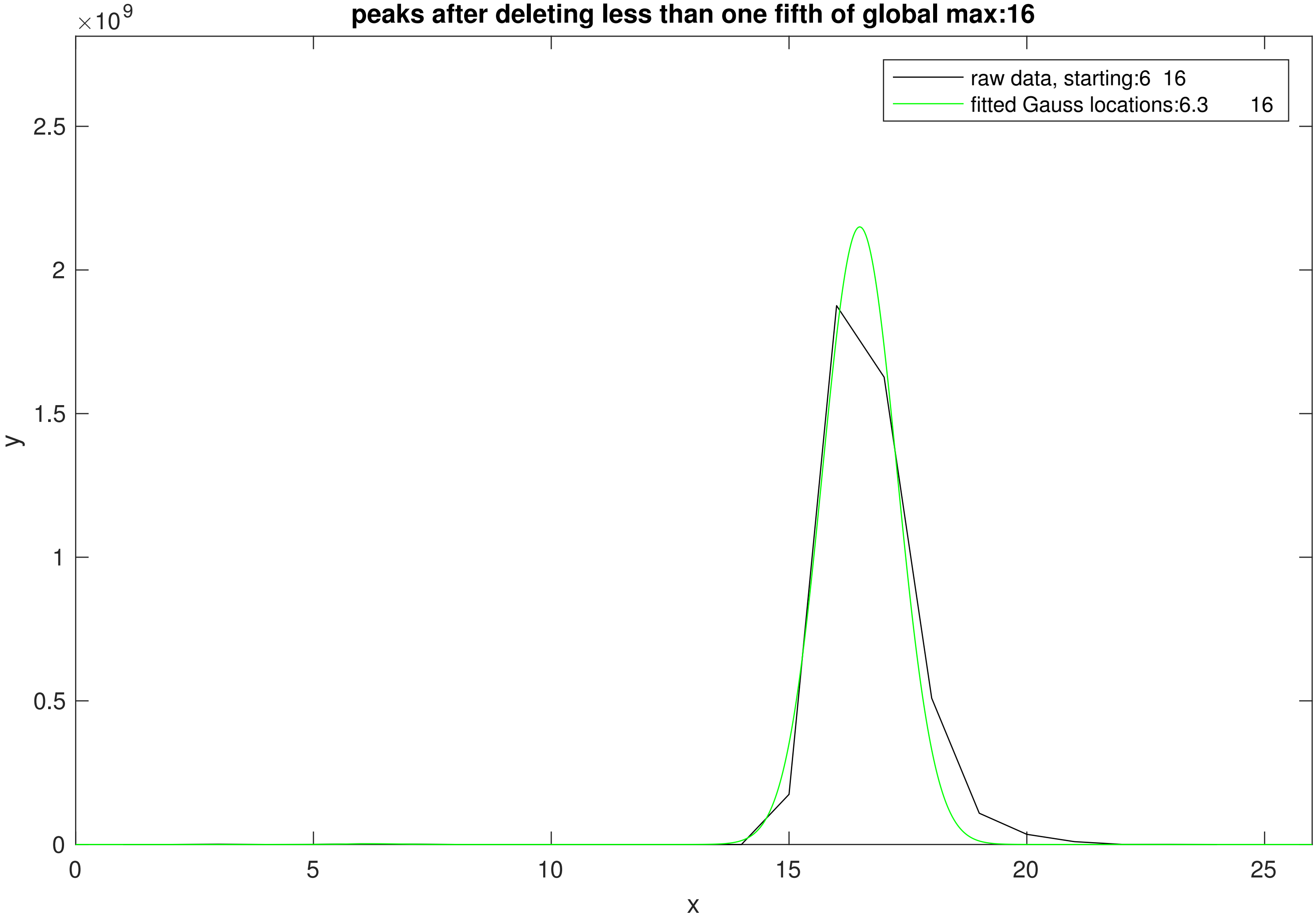

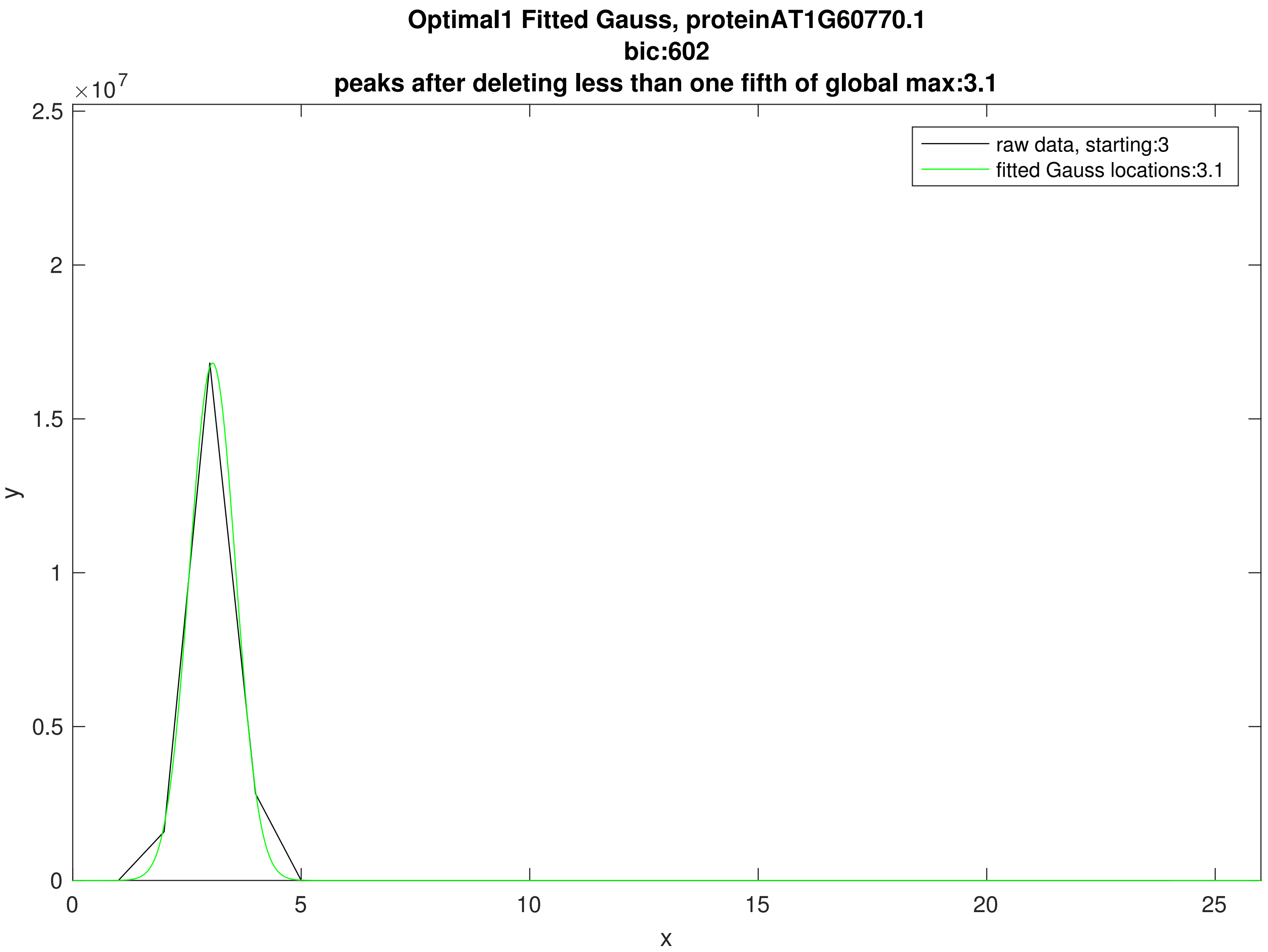

Optimal1 Fitted Gauss, proteinAT1G60770.1
bic:602
peaks after deleting less than one fifth of global max:3.1
×10^7
2.5
2
1.5
1
0.5
0
y
0
5
10
15
20
25
x
raw data, starting:3
fitted Gauss locations:3.1

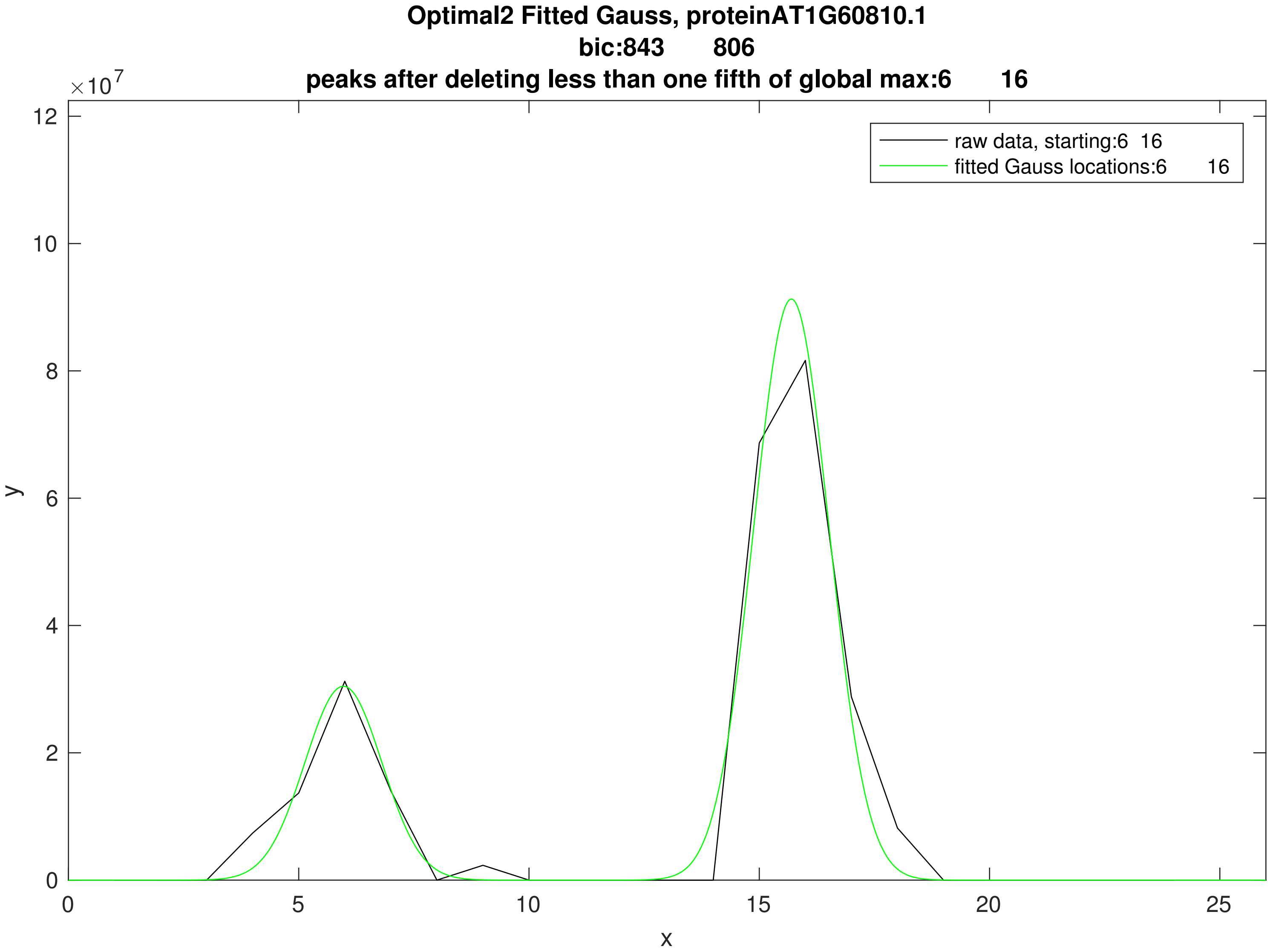

Optimal2 Fitted Gauss, proteinAT1G60810.1
bic:843 806
peaks after deleting less than one fifth of global max:6 16
×10^7
raw data, starting:6 16
fitted Gauss locations:6 16
y
x
0
2
4
6
8
10
12
5
15
20
25

**Optimal1 Fitted Gauss, proteinAT1G60950.1**
**bic:900 901**
**peaks after deleting less than one fifth of global max:19**

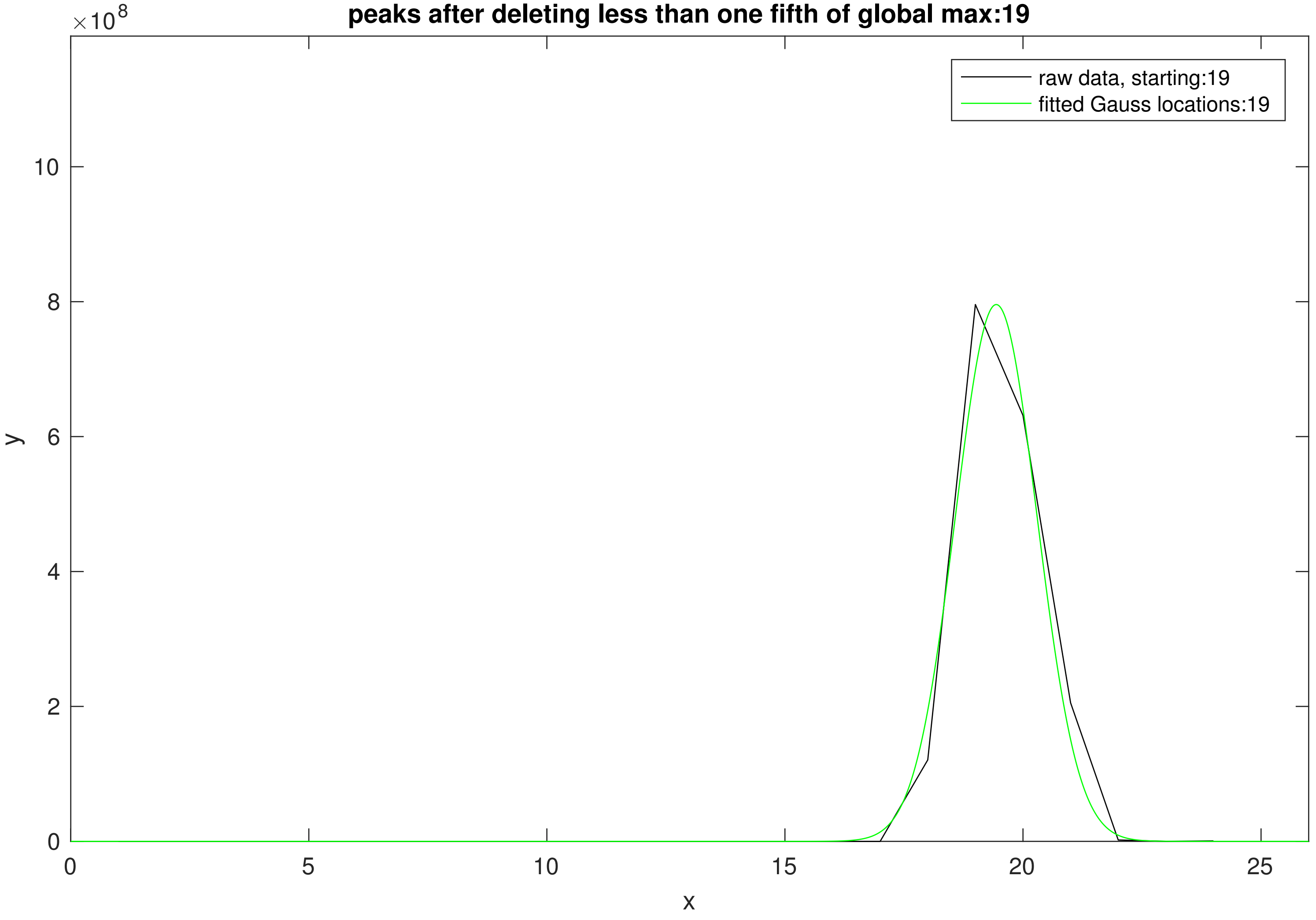

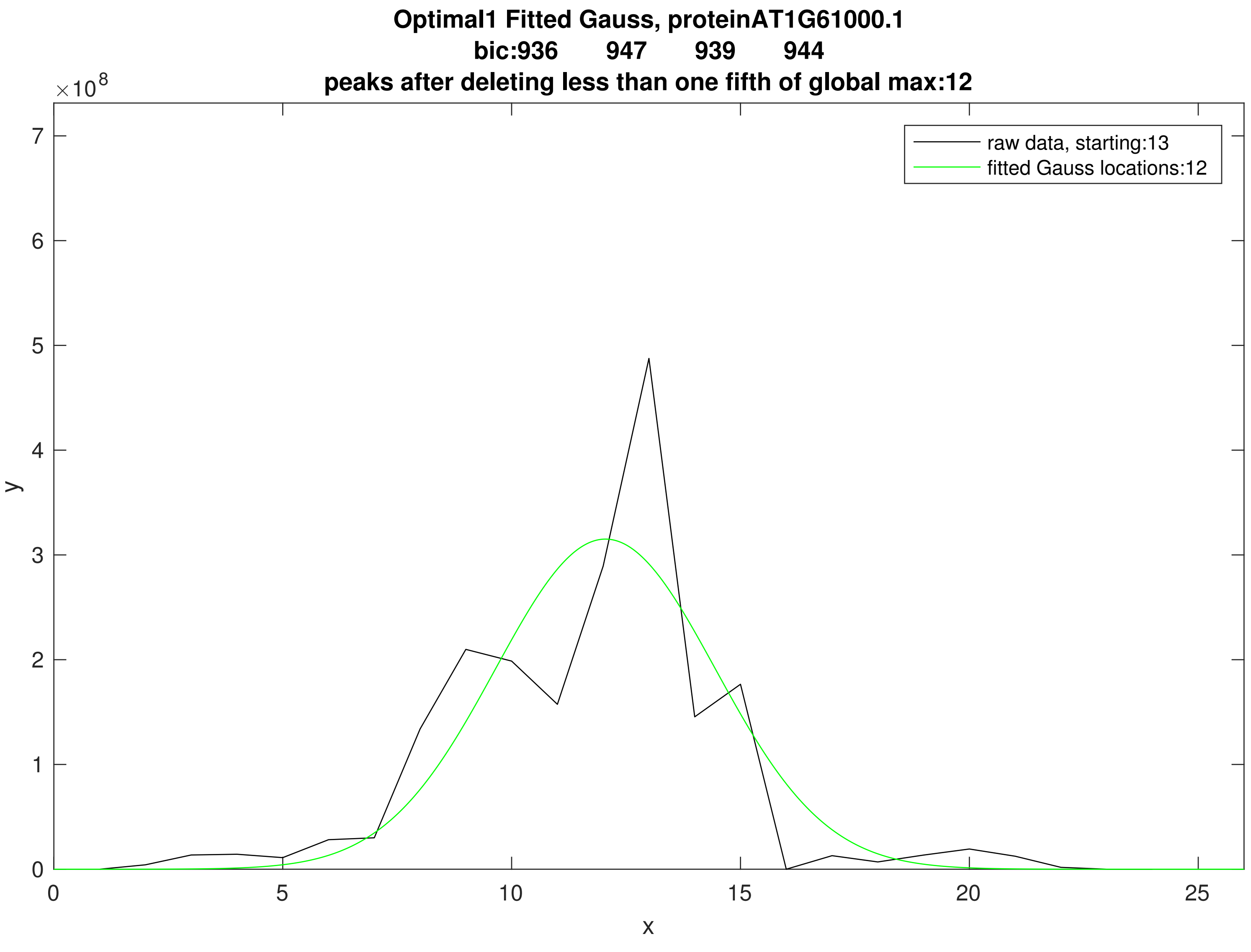
Optimal1 Fitted Gauss, proteinAT1G61000.1
bic:936 947 939 944
peaks after deleting less than one fifth of global max:12
×10^8
raw data, starting:13
fitted Gauss locations:12
y
x
0
1
2
3
4
5
6
7
0
5
10
15
20
25

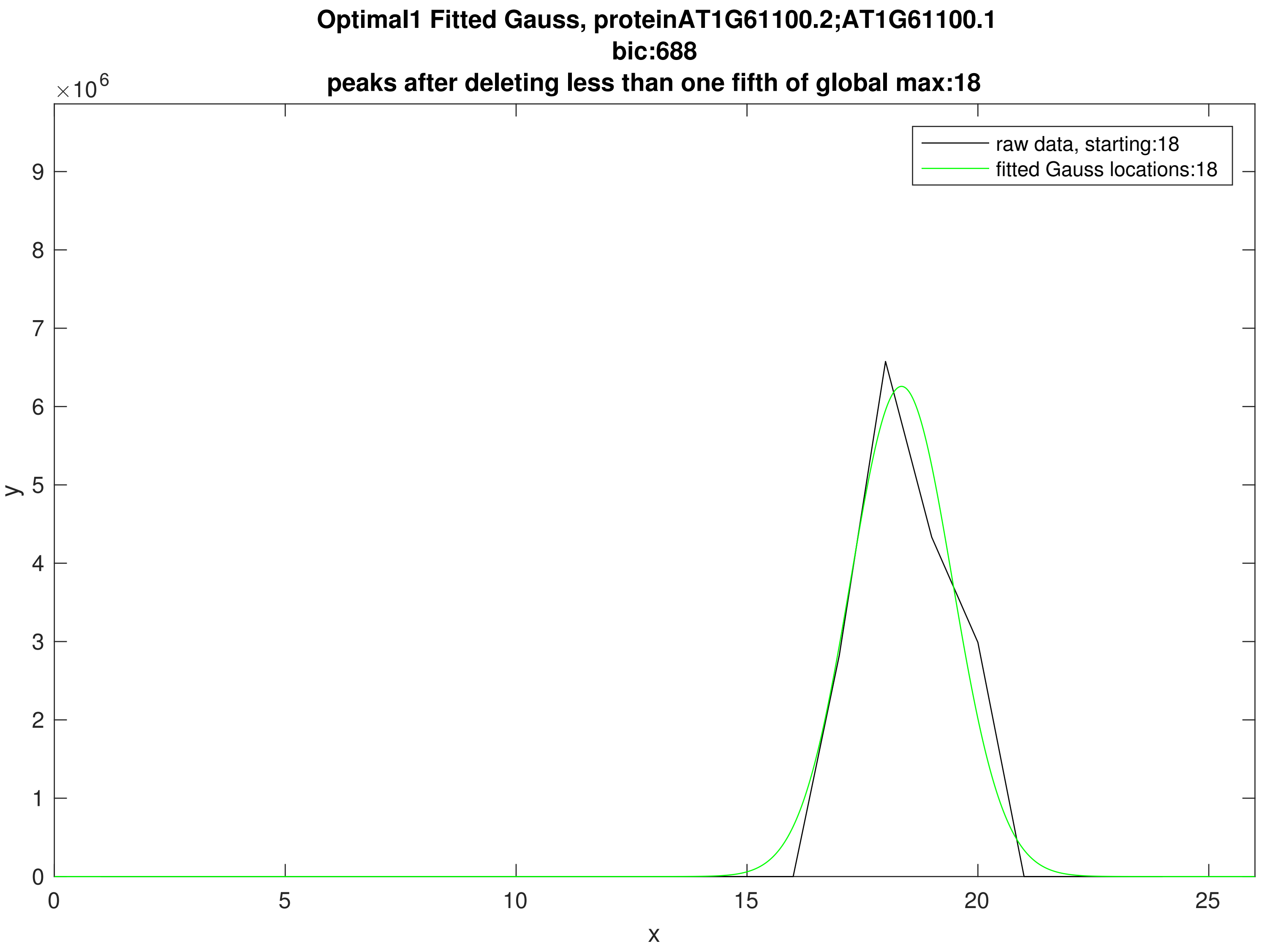
Optimal1 Fitted Gauss, proteinAT1G61100.2;AT1G61100.1
bic:688
peaks after deleting less than one fifth of global max:18
×10^6
raw data, starting:18
fitted Gauss locations:18
y
x
0
1
2
3
4
5
6
7
8
9
0
5
10
15
20
25

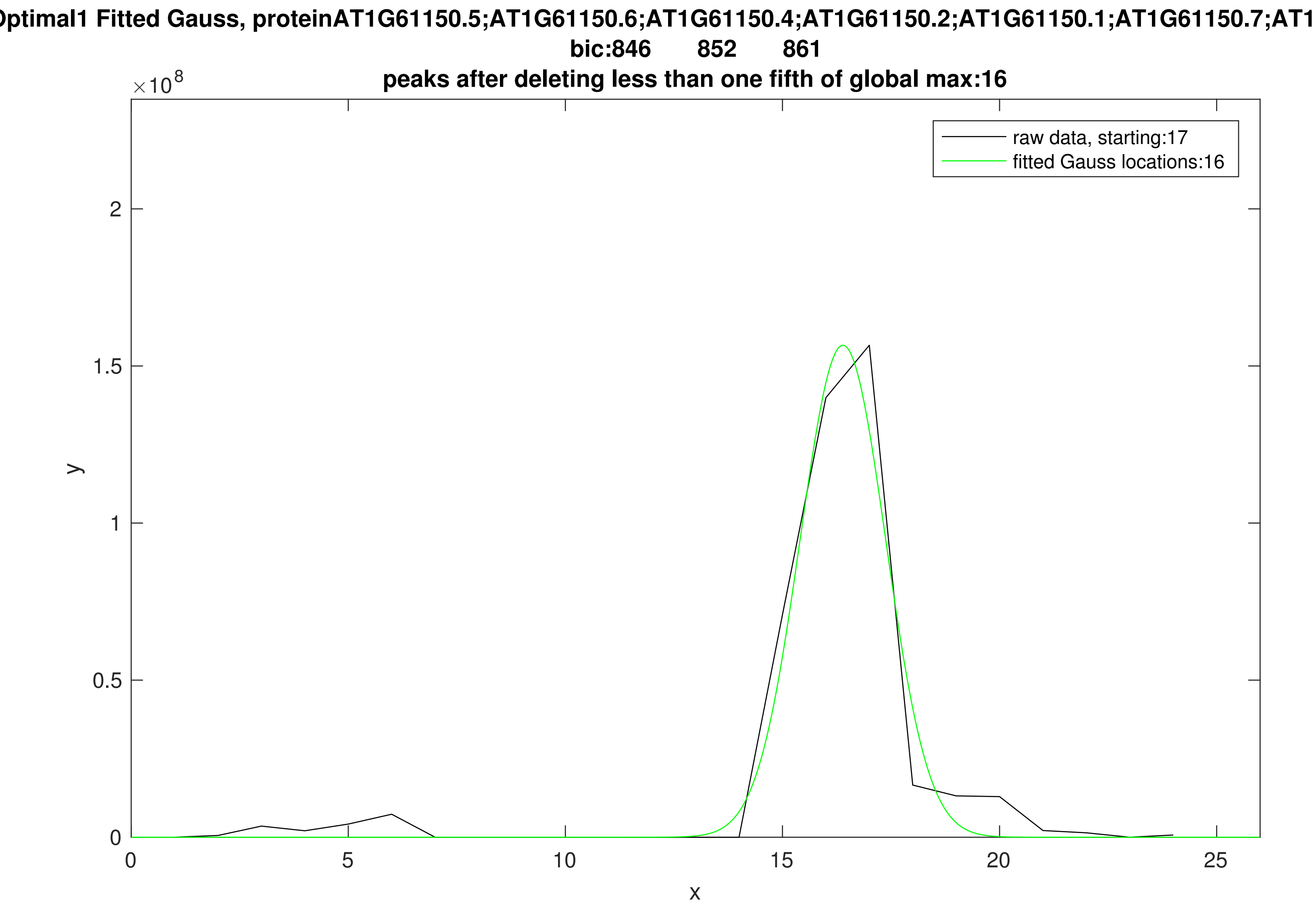
Optimal1 Fitted Gauss, proteinAT1G61150.5;AT1G61150.6;AT1G61150.4;AT1G61150.2;AT1G61150.1;AT1G61150.7;AT1
bic:846 852 861
peaks after deleting less than one fifth of global max:16
$\times 10^8$
raw data, starting:17
fitted Gauss locations:16
y
x
0
0.5
1
1.5
2
0
5
10
15
20
25

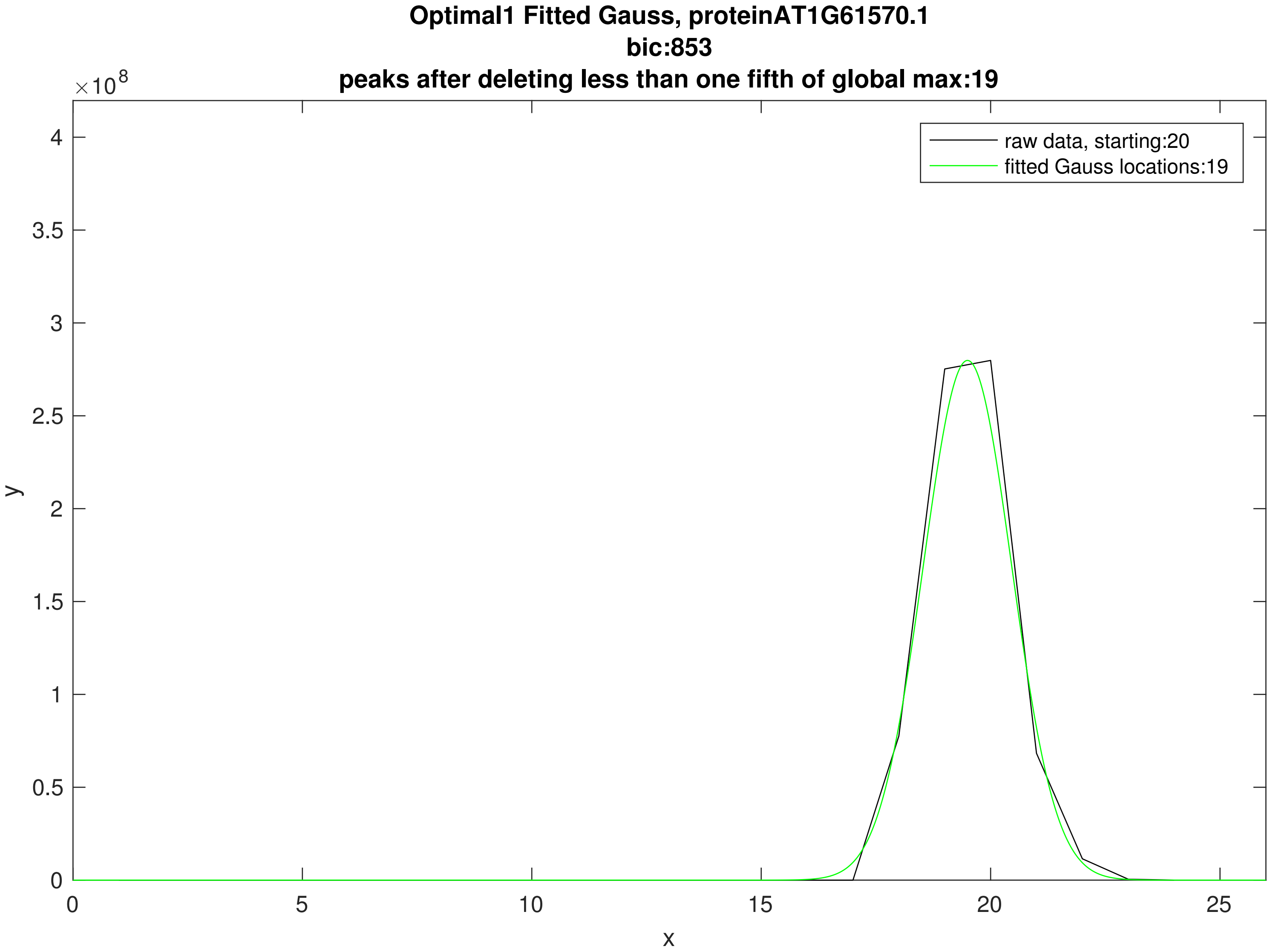

Optimal1 Fitted Gauss, proteinAT1G61570.1
bic:853
peaks after deleting less than one fifth of global max:19
×10^8
raw data, starting:20
fitted Gauss locations:19
y
x
0
0.5
1
1.5
2
2.5
3
3.5
4
0
5
10
15
20
25

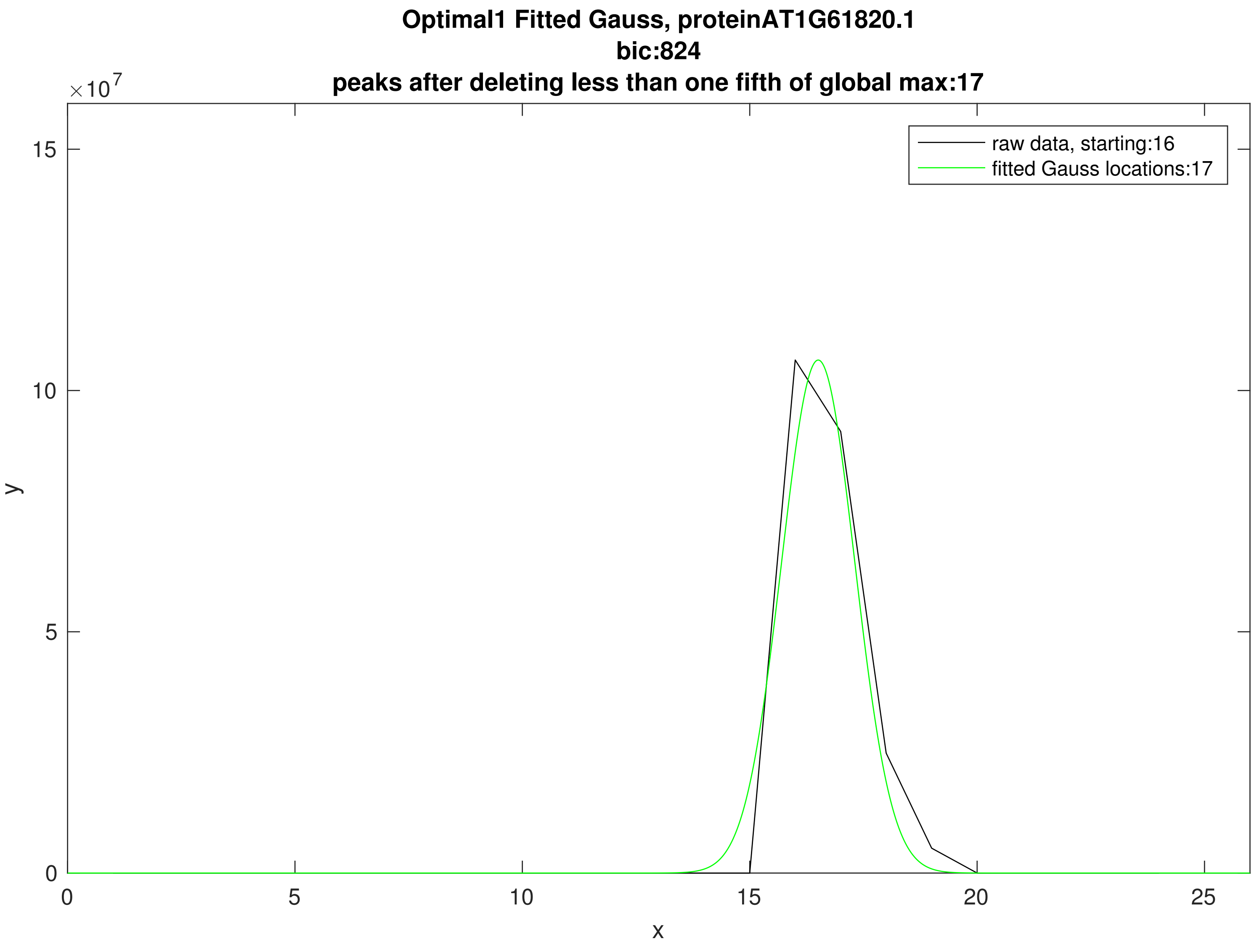
Optimal1 Fitted Gauss, proteinAT1G61820.1
bic:824
peaks after deleting less than one fifth of global max:17
×10^7
15
10
5
0
y
0
5
10
15
20
25
x
raw data, starting:16
fitted Gauss locations:17

#### Optimal1 Fitted Gauss, proteinAT1G61900.2;AT1G61900.3;AT1G61900.1
#### bic:912
#### peaks after deleting less than one fifth of global max:11

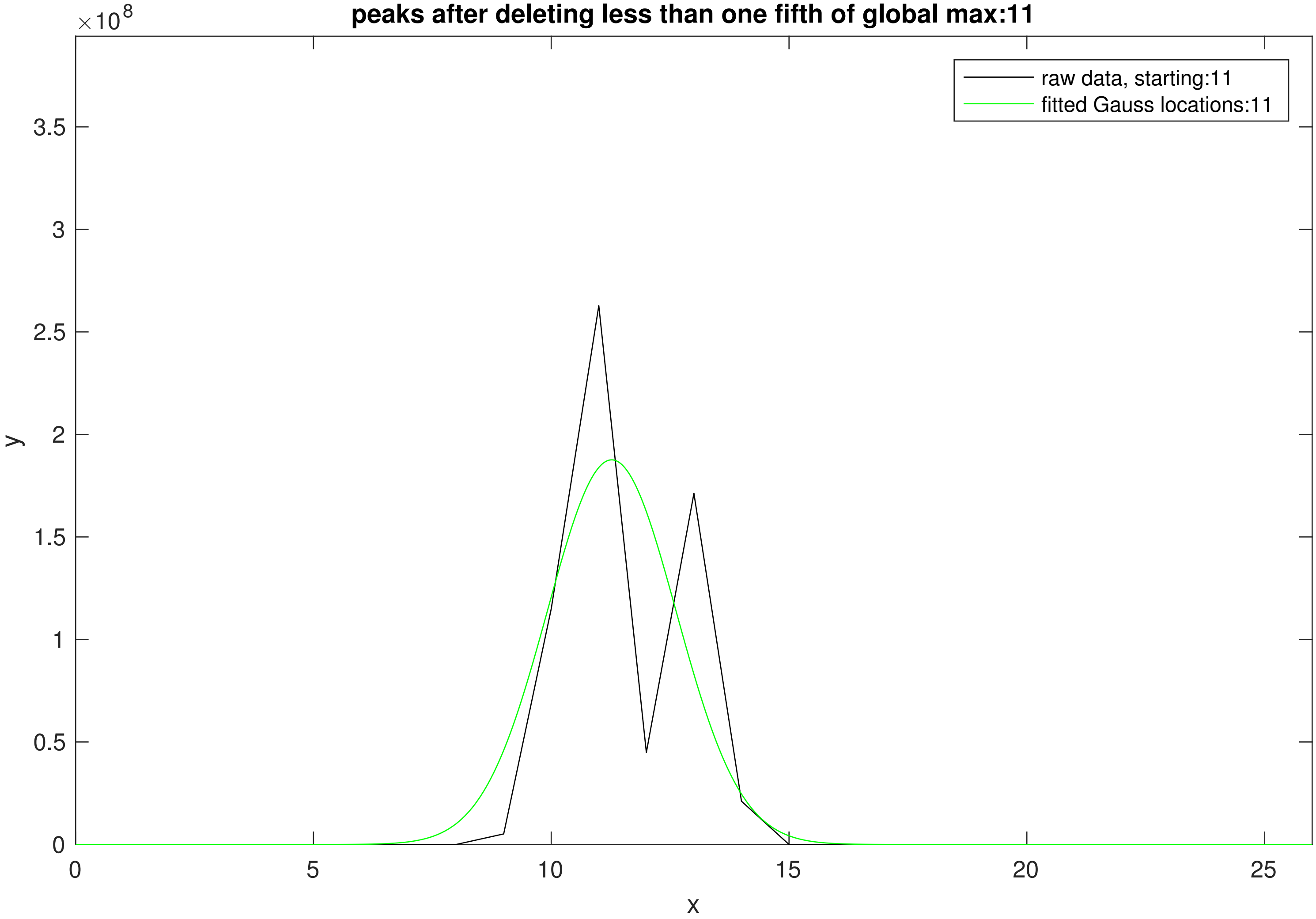

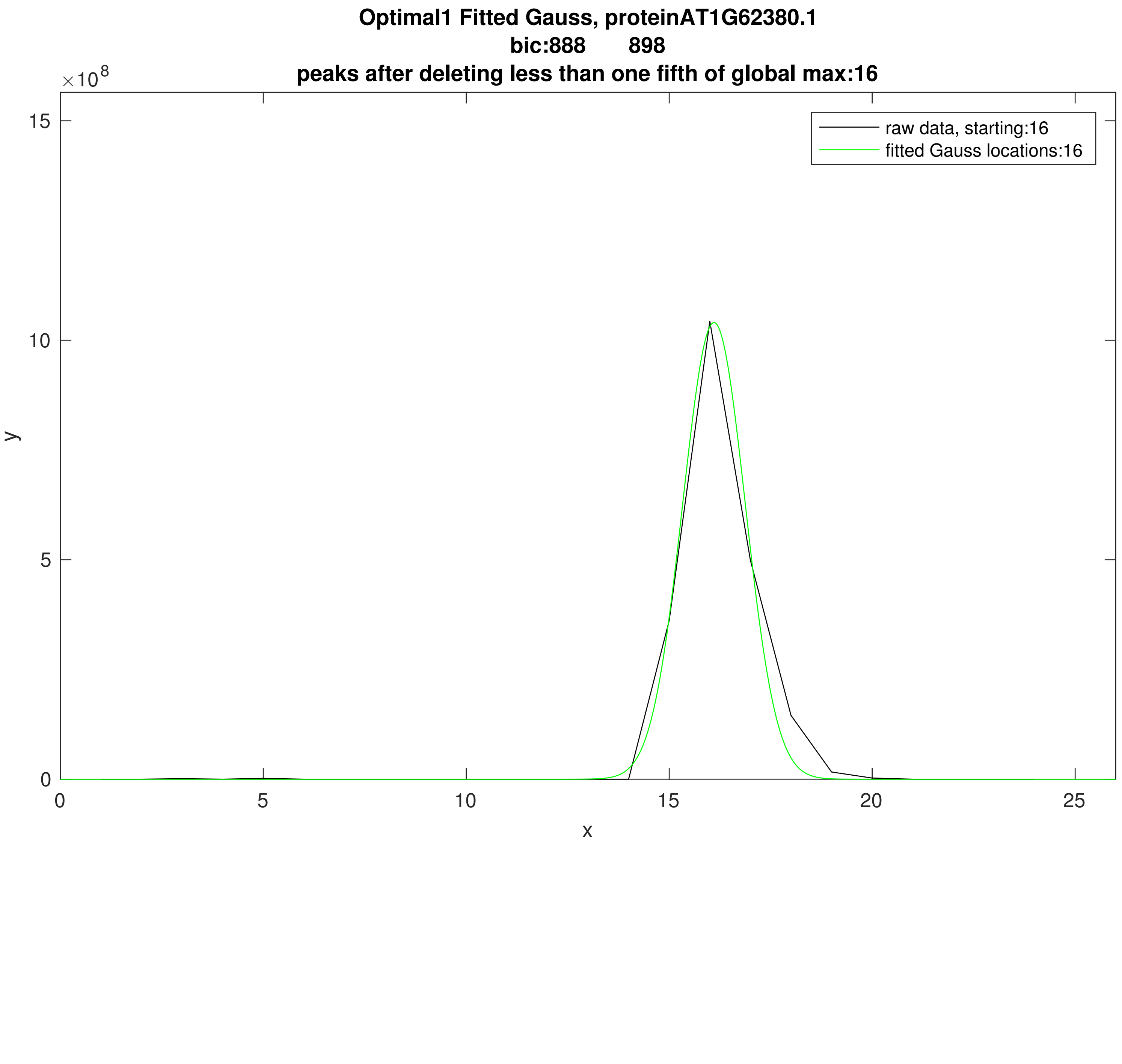

Optimal1 Fitted Gauss, proteinAT1G62380.1
bic:888 898
peaks after deleting less than one fifth of global max:16
×10^8
15
10
5
0
y
0
5
10
15
20
25
x
raw data, starting:16
fitted Gauss locations:16

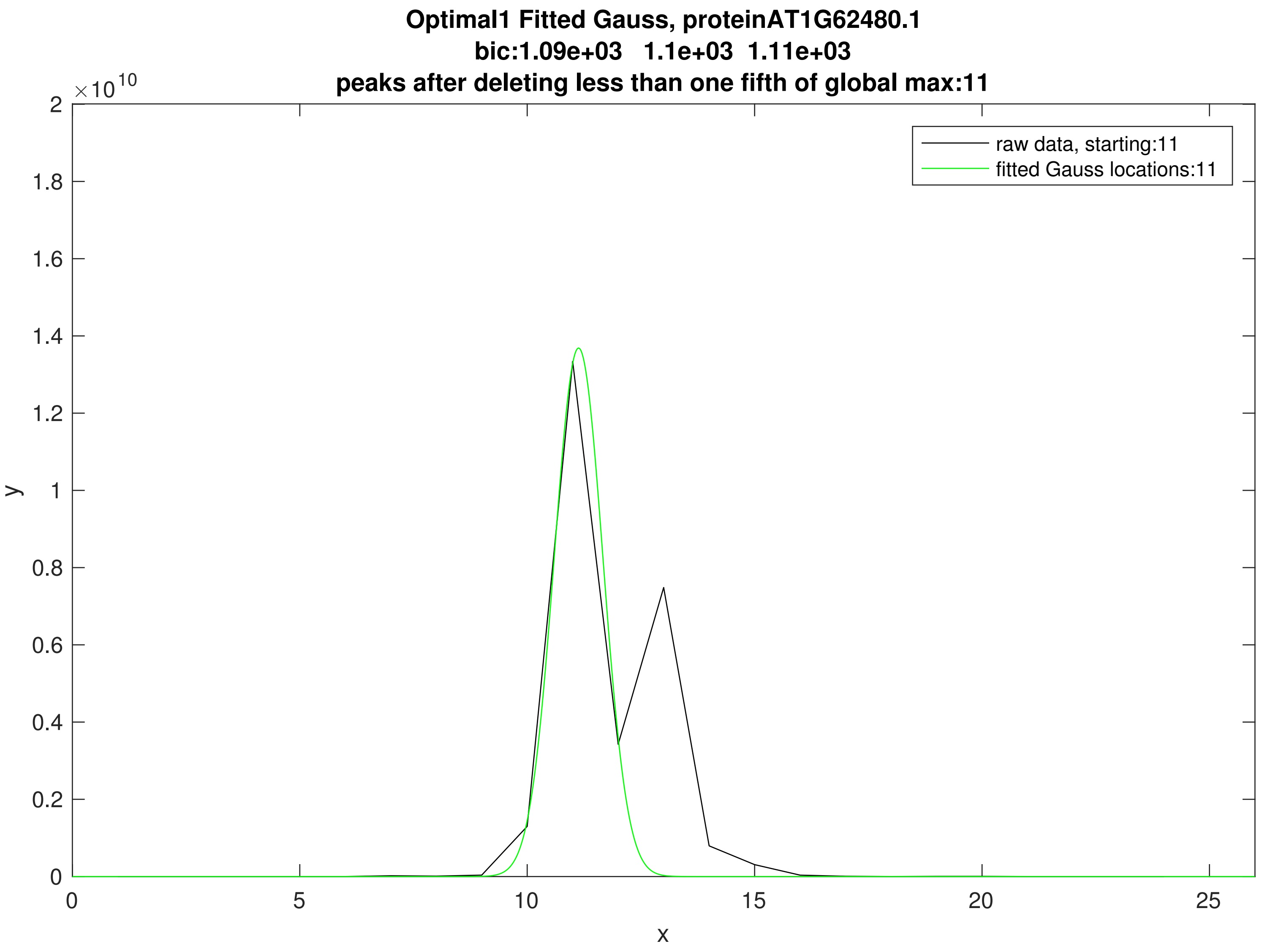

Optimal1 Fitted Gauss, proteinAT1G62480.1
bic:1.09e+03 1.1e+03 1.11e+03
peaks after deleting less than one fifth of global max:11
×10^10
raw data, starting:11
fitted Gauss locations:11
y
x

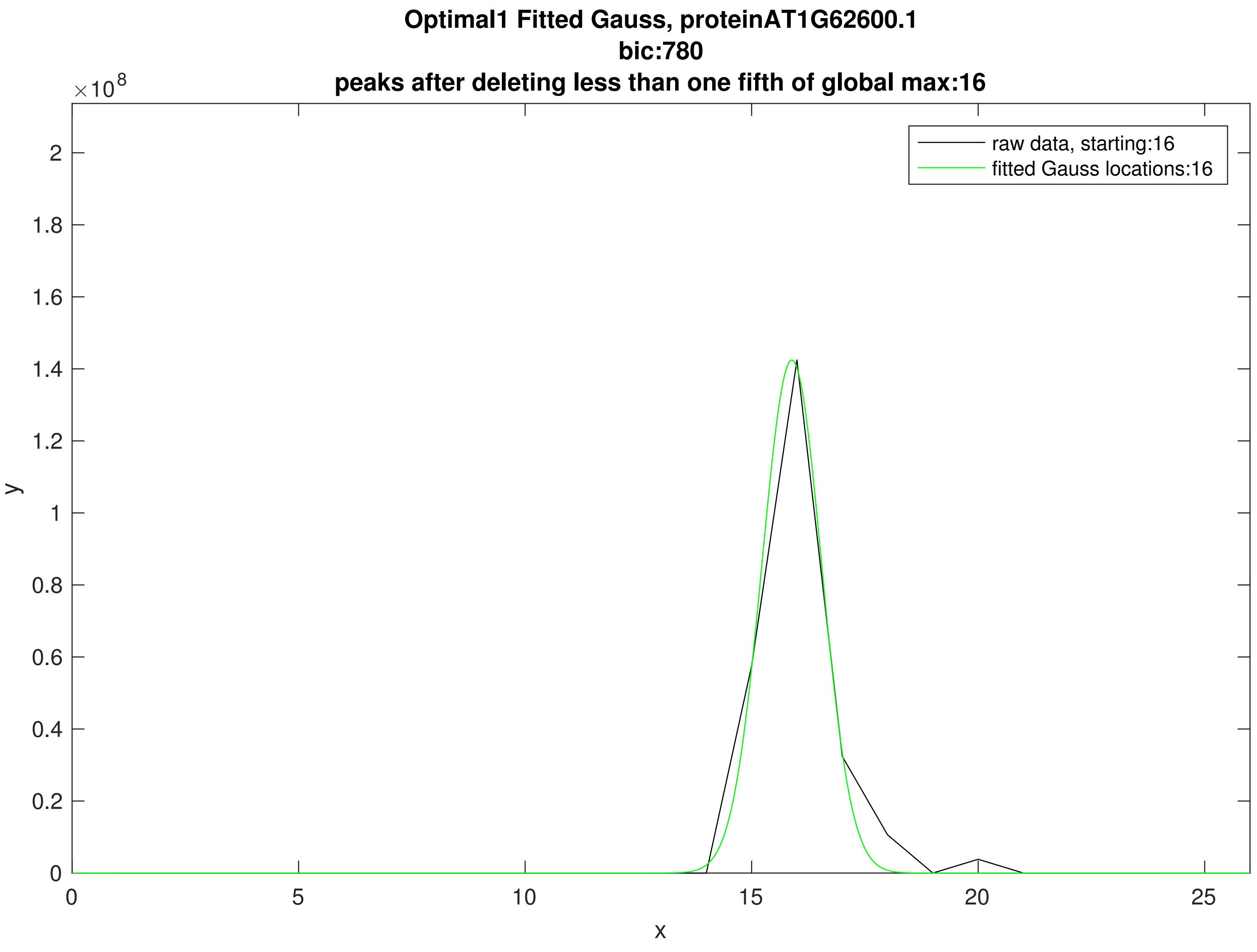
Optimal1 Fitted Gauss, proteinAT1G62600.1
bic:780
peaks after deleting less than one fifth of global max:16
×10^8
raw data, starting:16
fitted Gauss locations:16
y
x
0
0.2
0.4
0.6
0.8
1
1.2
1.4
1.6
1.8
2
5
10
15
20
25

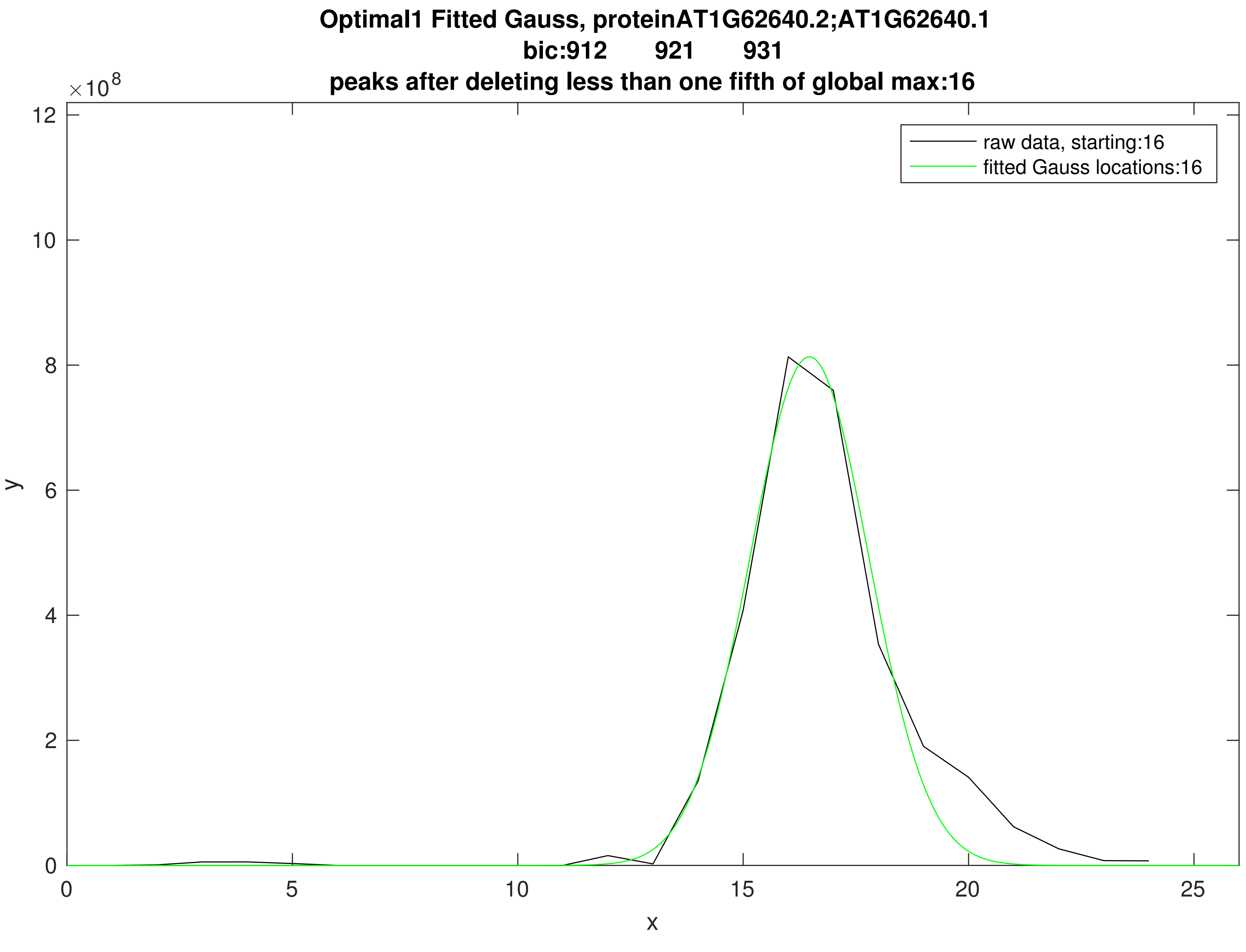
Optimal1 Fitted Gauss, proteinAT1G62640.2;AT1G62640.1
bic:912 921 931
peaks after deleting less than one fifth of global max:16
×10^8
raw data, starting:16
fitted Gauss locations:16
y
x
0
2
4
6
8
10
12
0
5
10
15
20
25

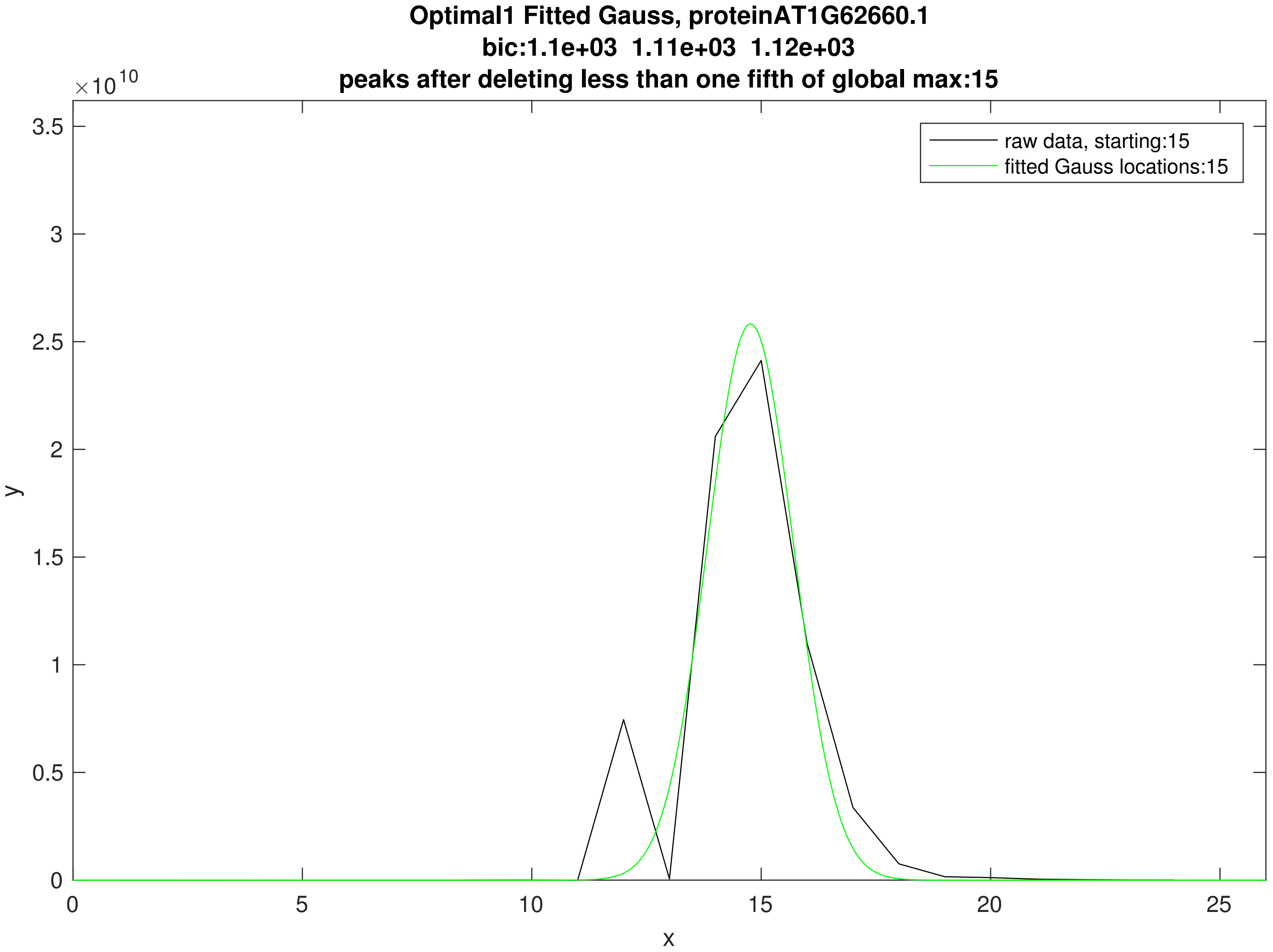

Optimal1 Fitted Gauss, proteinAT1G62660.1
bic:1.1e+03 1.11e+03 1.12e+03
peaks after deleting less than one fifth of global max:15
×10^10
raw data, starting:15
fitted Gauss locations:15
y
x
0
0.5
1
1.5
2
2.5
3
3.5
0
5
10
15
20
25

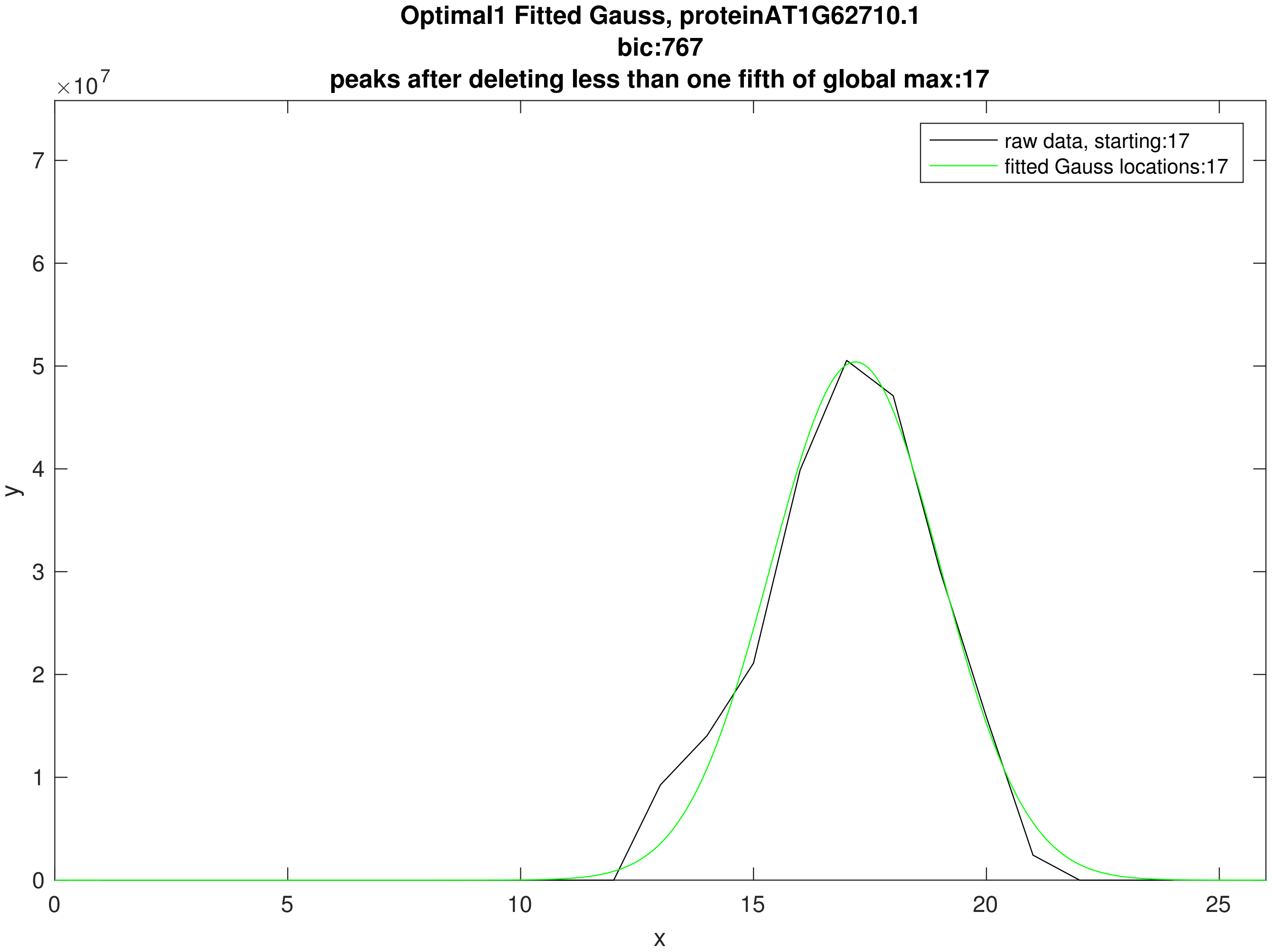
Optimal1 Fitted Gauss, proteinAT1G62710.1
bic:767
peaks after deleting less than one fifth of global max:17
×10^7
raw data, starting:17
fitted Gauss locations:17
y
x
0
1
2
3
4
5
6
7
0
5
10
15
20
25

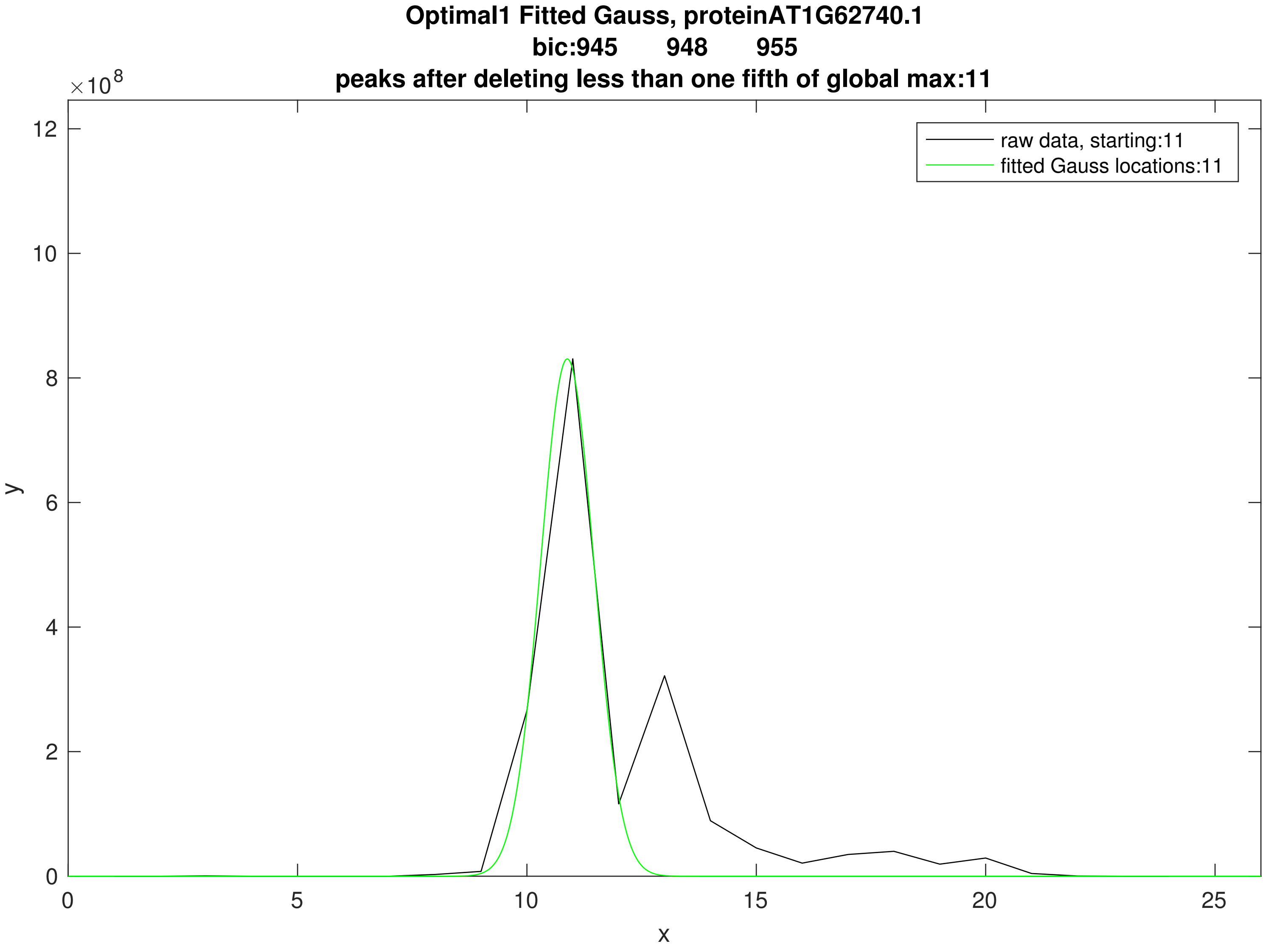

Optimal1 Fitted Gauss, proteinAT1G62740.1
bic:945 948 955
peaks after deleting less than one fifth of global max:11
×10^8
12
10
8
6
4
2
0
y
0
5
10
15
20
25
x
raw data, starting:11
fitted Gauss locations:11

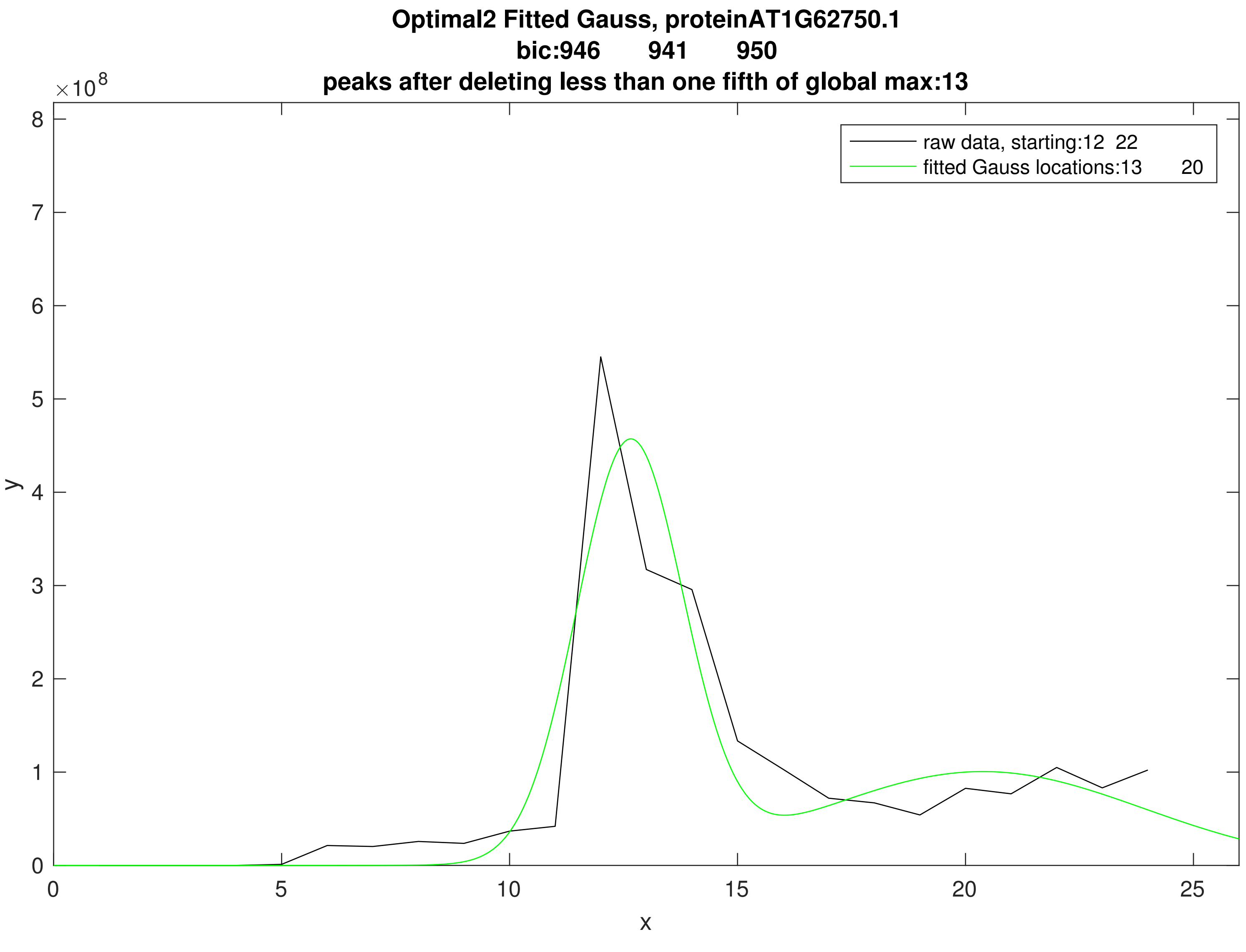

Optimal2 Fitted Gauss, proteinAT1G62750.1
bic:946 941 950
peaks after deleting less than one fifth of global max:13
×10^8
y
x
raw data, starting:12 22
fitted Gauss locations:13 20
0
1
2
3
4
5
6
7
8
0
5
10
15
20
25

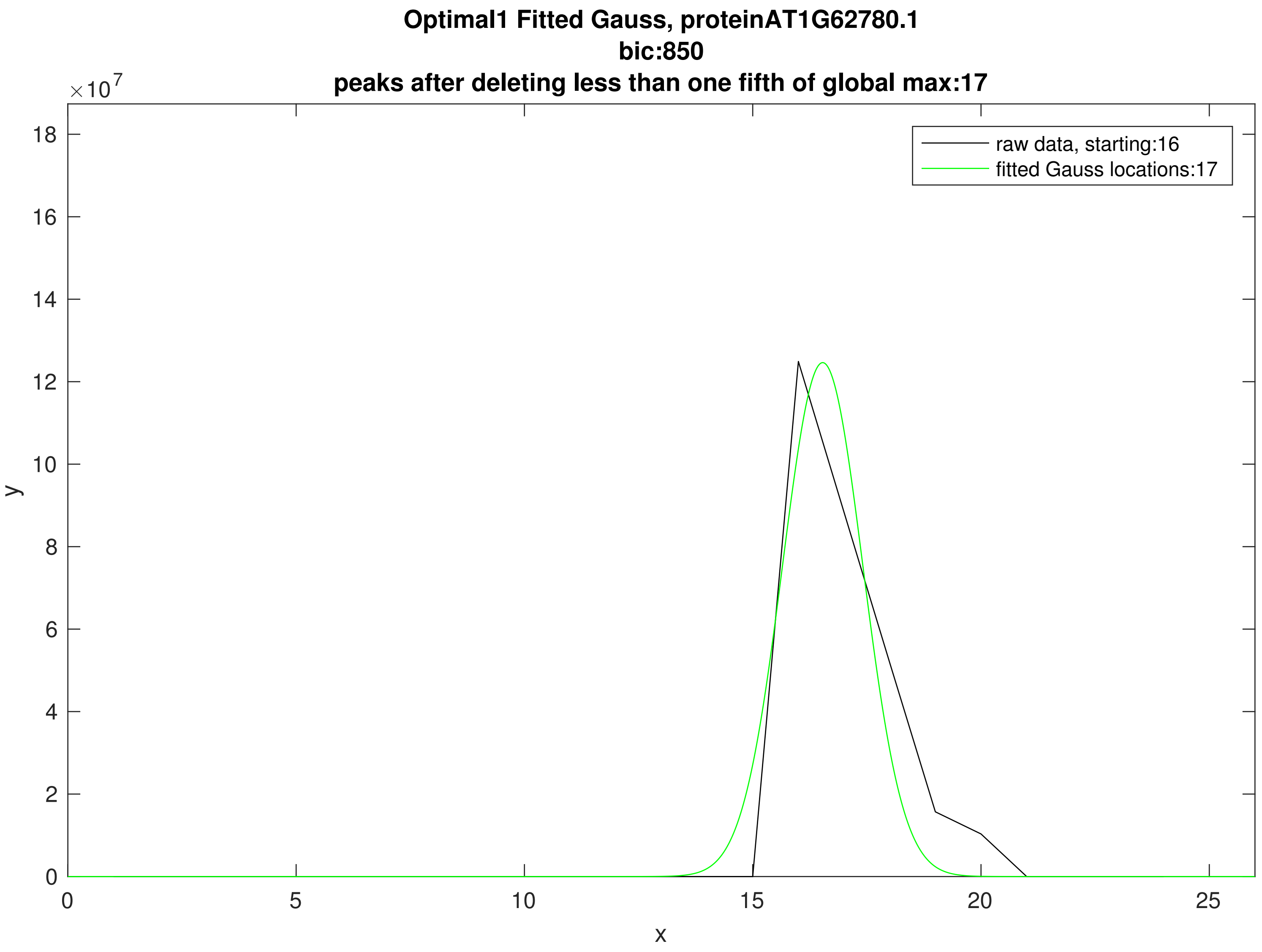

Optimal1 Fitted Gauss, proteinAT1G62780.1
bic:850
peaks after deleting less than one fifth of global max:17
×10^7
raw data, starting:16
fitted Gauss locations:17
y
x
0
2
4
6
8
10
12
14
16
18
0
5
10
15
20
25

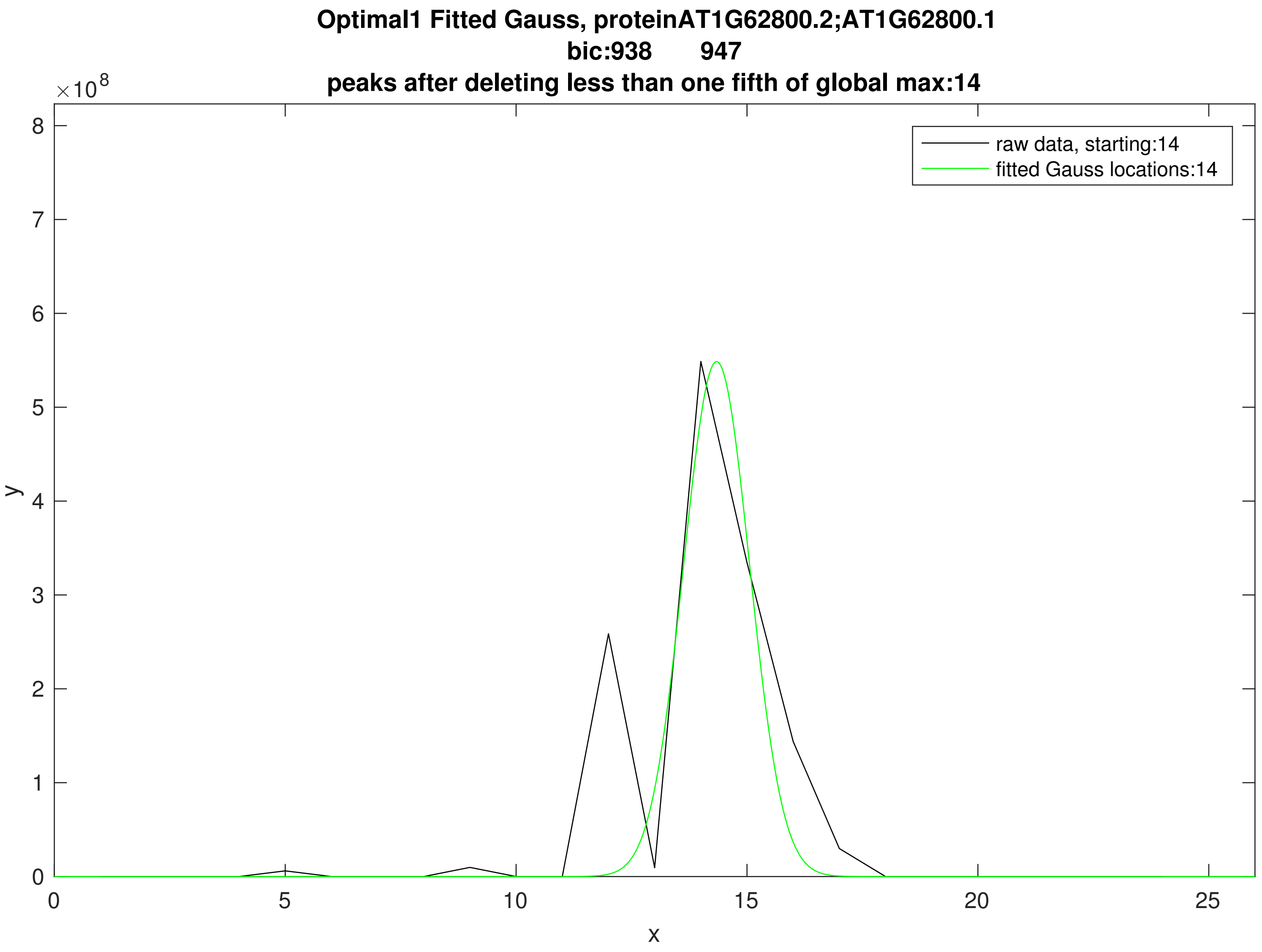

Optimal1 Fitted Gauss, proteinAT1G62800.2;AT1G62800.1
bic:938 947
peaks after deleting less than one fifth of global max:14
×10^8
raw data, starting:14
fitted Gauss locations:14
y
x
0
1
2
3
4
5
6
7
8
0
5
10
15
20
25

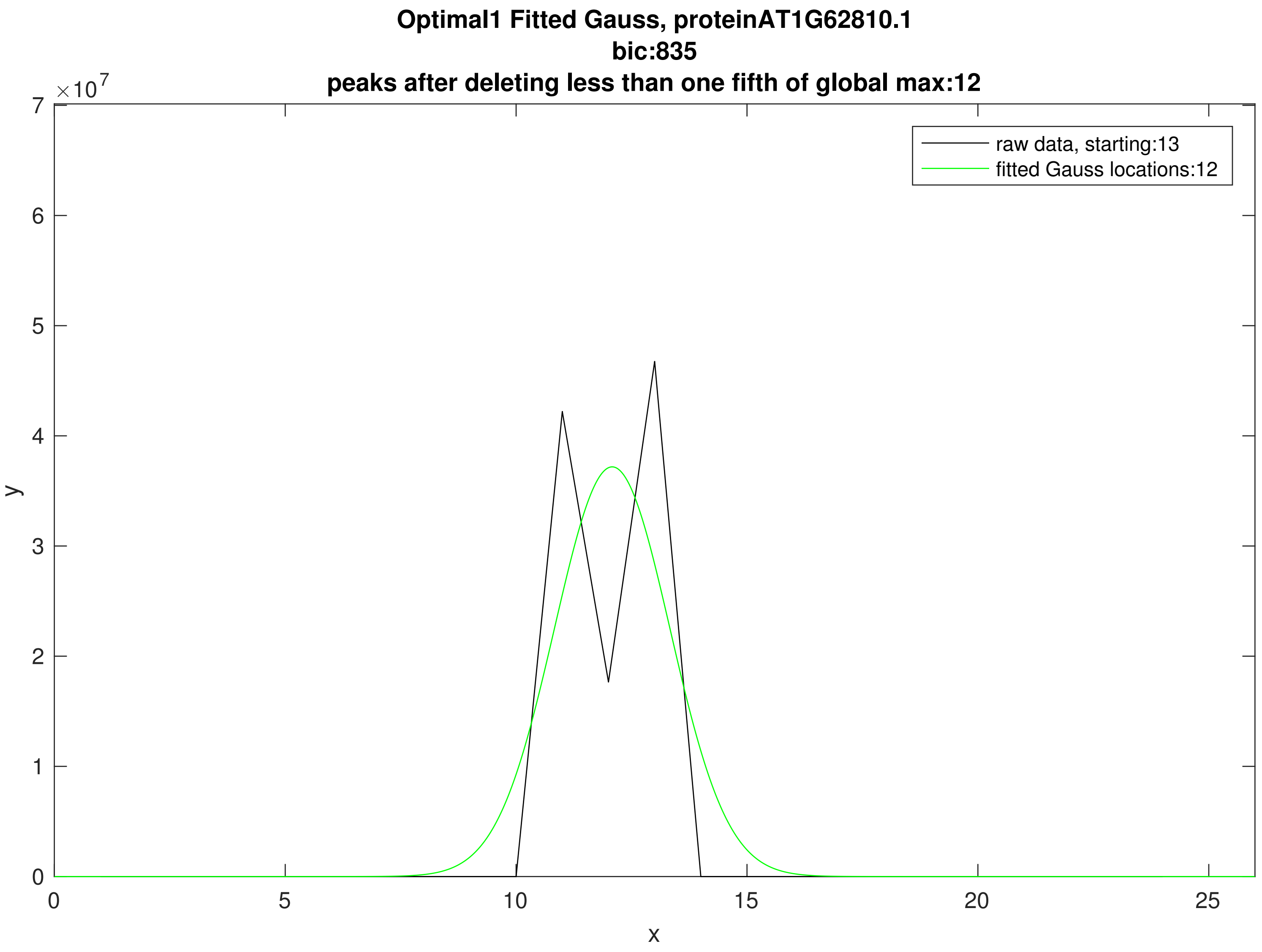
Optimal1 Fitted Gauss, proteinAT1G62810.1
bic:835
peaks after deleting less than one fifth of global max:12
raw data, starting:13
fitted Gauss locations:12
×10^7
y
x

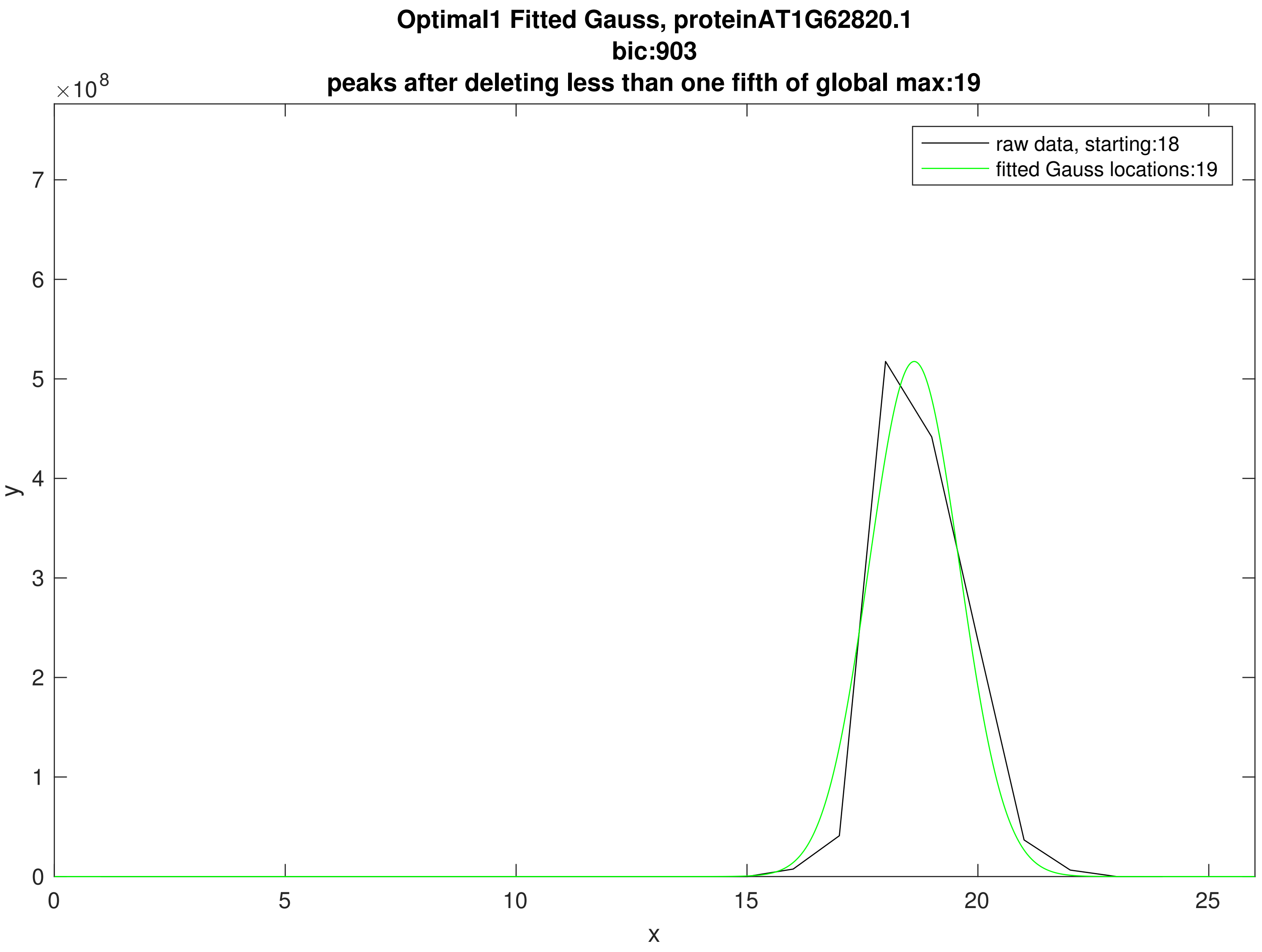

Optimal1 Fitted Gauss, proteinAT1G62820.1
bic:903
peaks after deleting less than one fifth of global max:19
×10^8
raw data, starting:18
fitted Gauss locations:19
y
x
0
1
2
3
4
5
6
7
0
5
10
15
20
25

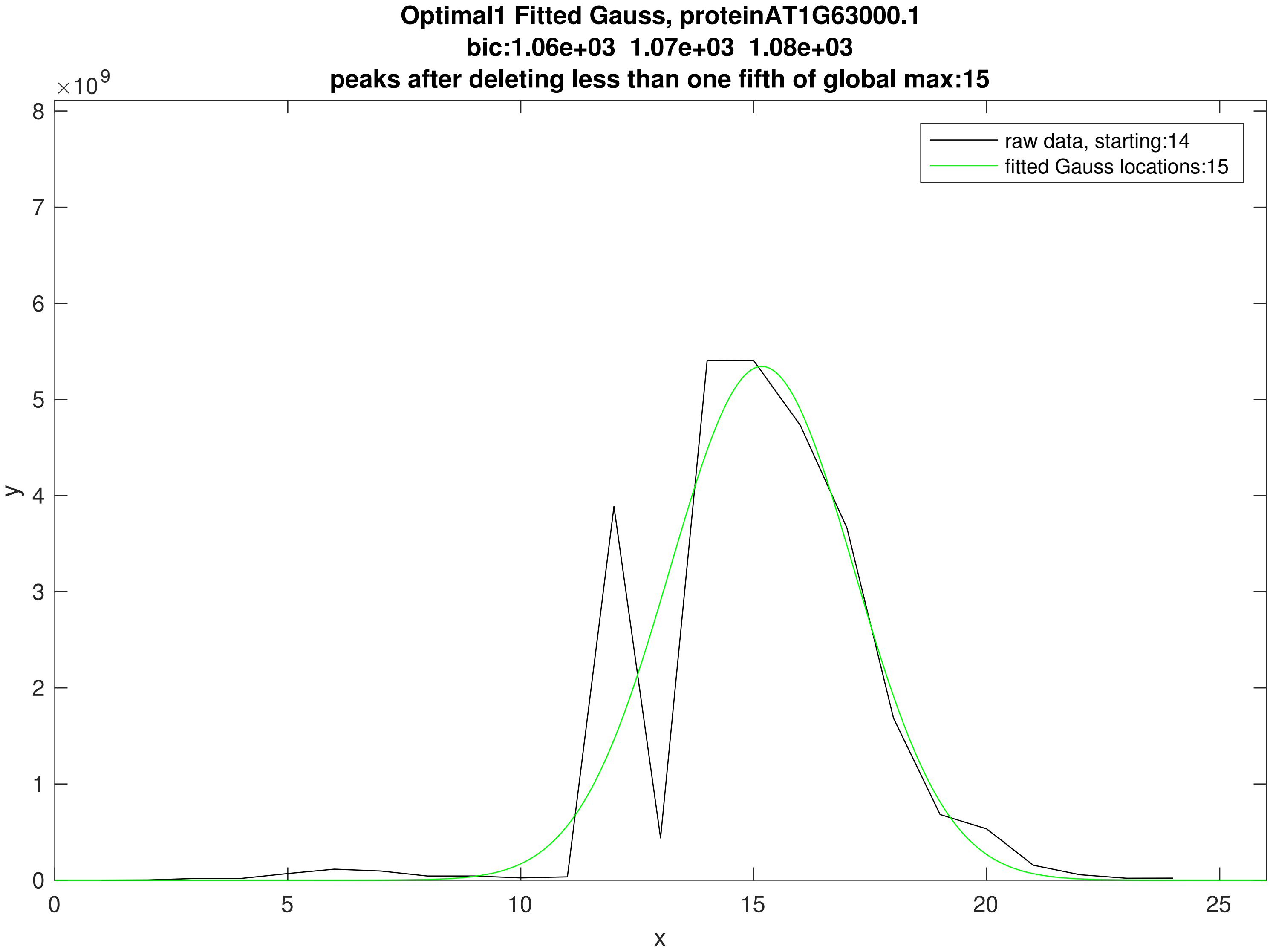

Optimal1 Fitted Gauss, proteinAT1G63000.1
bic:1.06e+03 1.07e+03 1.08e+03
peaks after deleting less than one fifth of global max:15
×10^9
raw data, starting:14
fitted Gauss locations:15
y
x
0
1
2
3
4
5
6
7
8
0
5
10
15
20
25

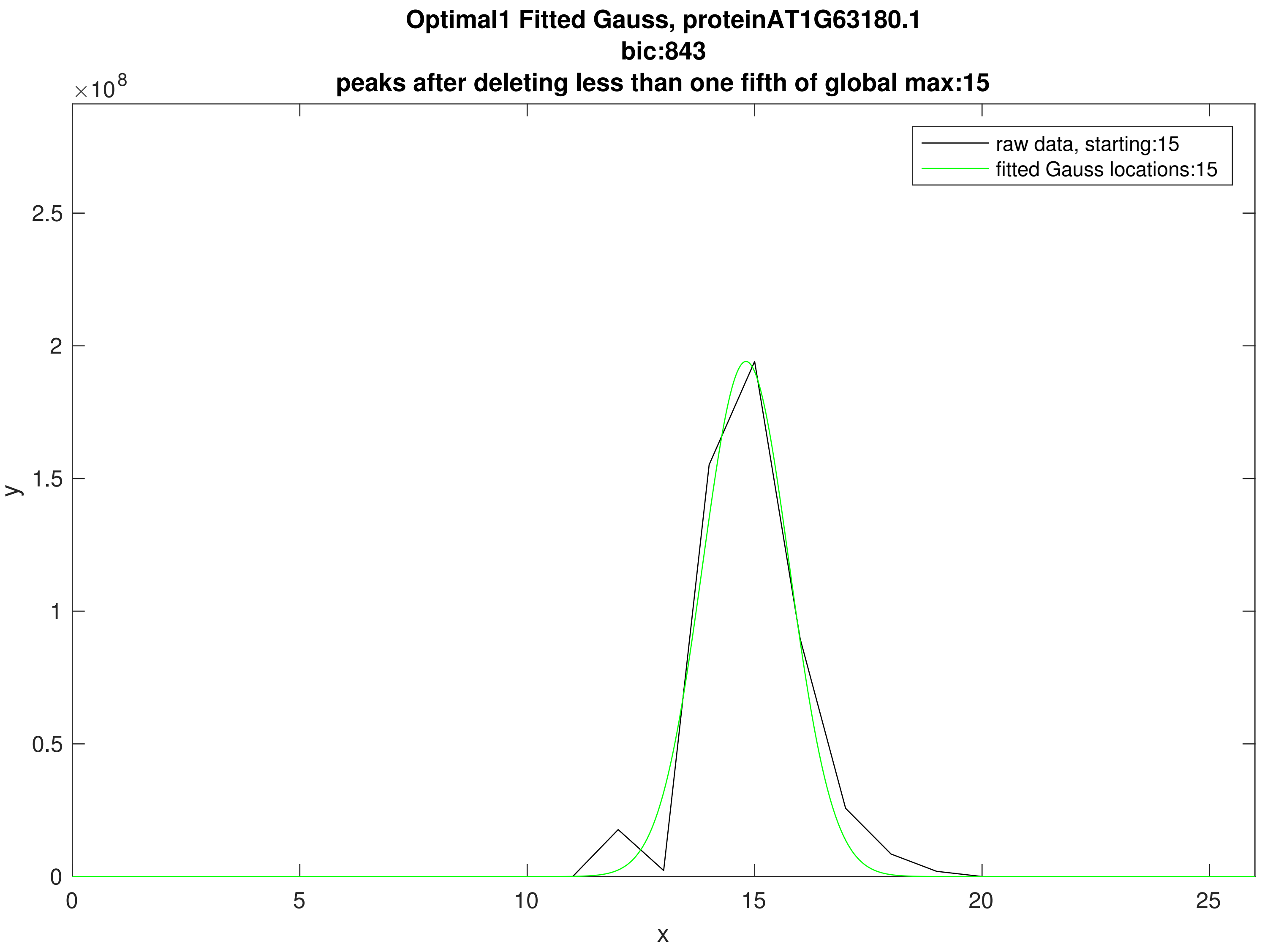

Optimal1 Fitted Gauss, proteinAT1G63180.1
bic:843
peaks after deleting less than one fifth of global max:15
×10^8
raw data, starting:15
fitted Gauss locations:15
y
x
0
0.5
1
1.5
2
2.5
0
5
10
15
20
25

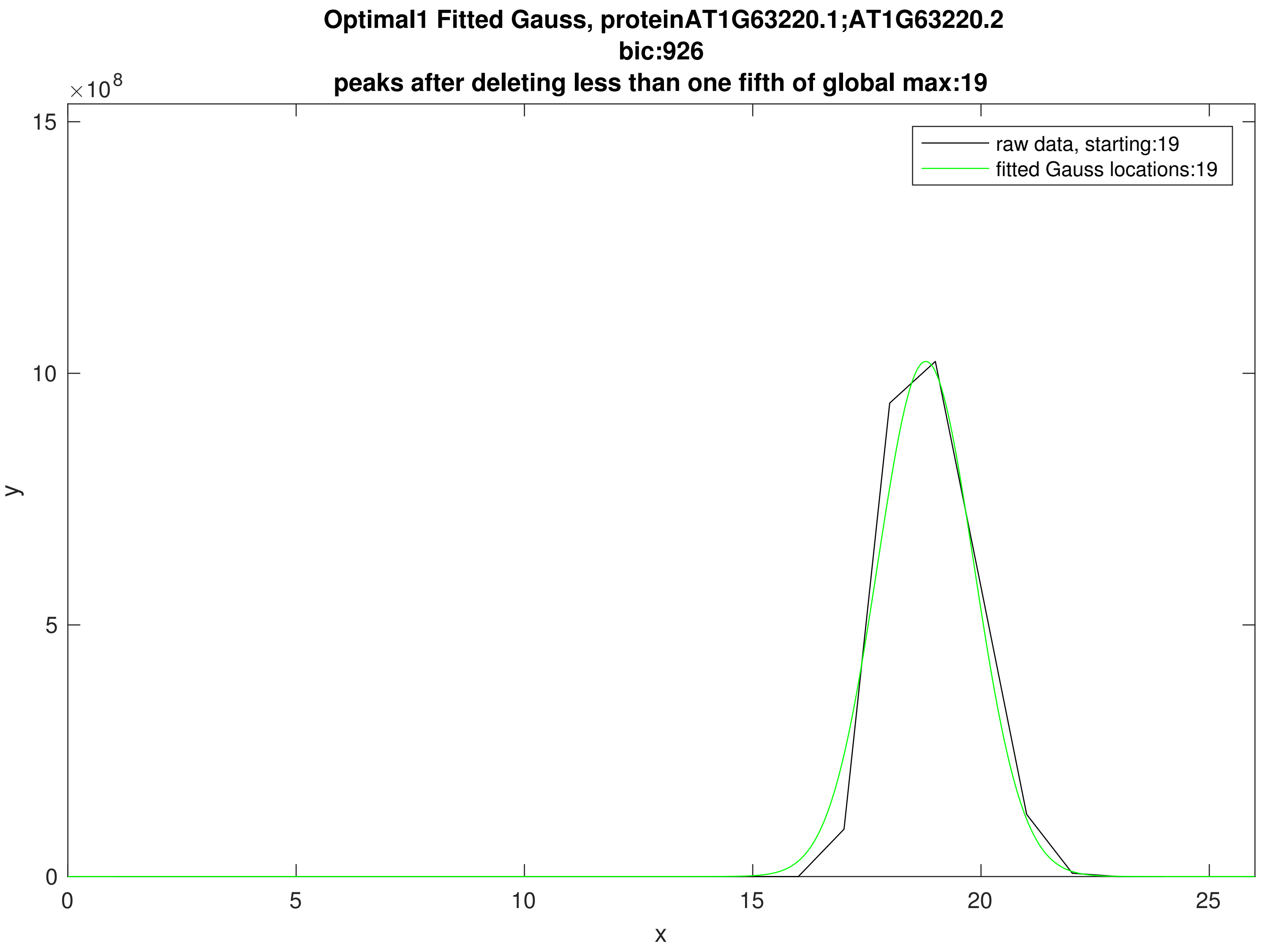

Optimal1 Fitted Gauss, proteinAT1G63220.1;AT1G63220.2
bic:926
peaks after deleting less than one fifth of global max:19
×10^8
15
10
5
0
y
0
5
10
15
20
25
x
raw data, starting:19
fitted Gauss locations:19

**Optimal1 Fitted Gauss, proteinAT1G63290.1**
**bic:836**
**peaks after deleting less than one fifth of global max:15**

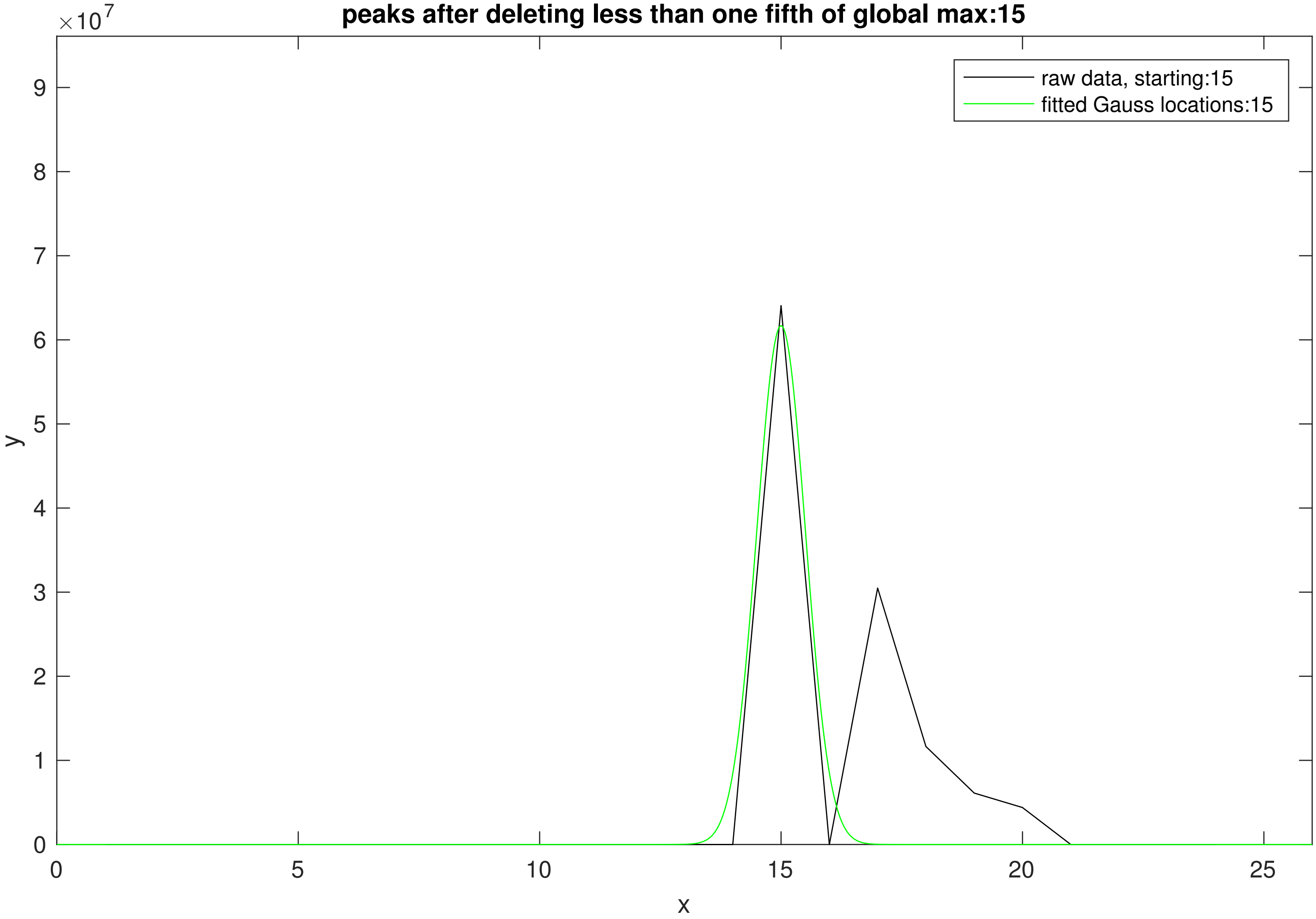

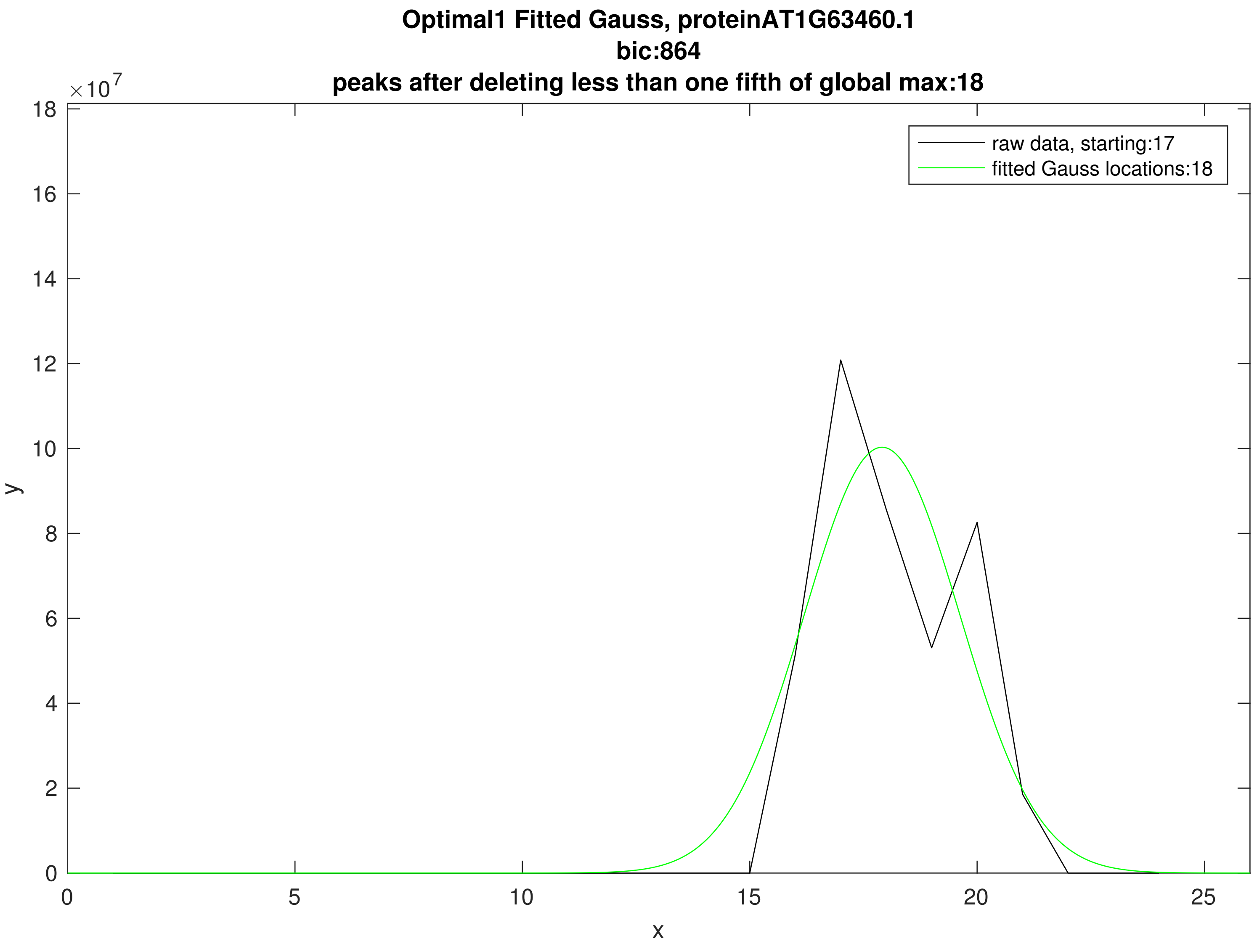

Optimal1 Fitted Gauss, proteinAT1G63460.1
bic:864
peaks after deleting less than one fifth of global max:18
×10^7
raw data, starting:17
fitted Gauss locations:18
y
x
0
2
4
6
8
10
12
14
16
18
0
5
10
15
20
25

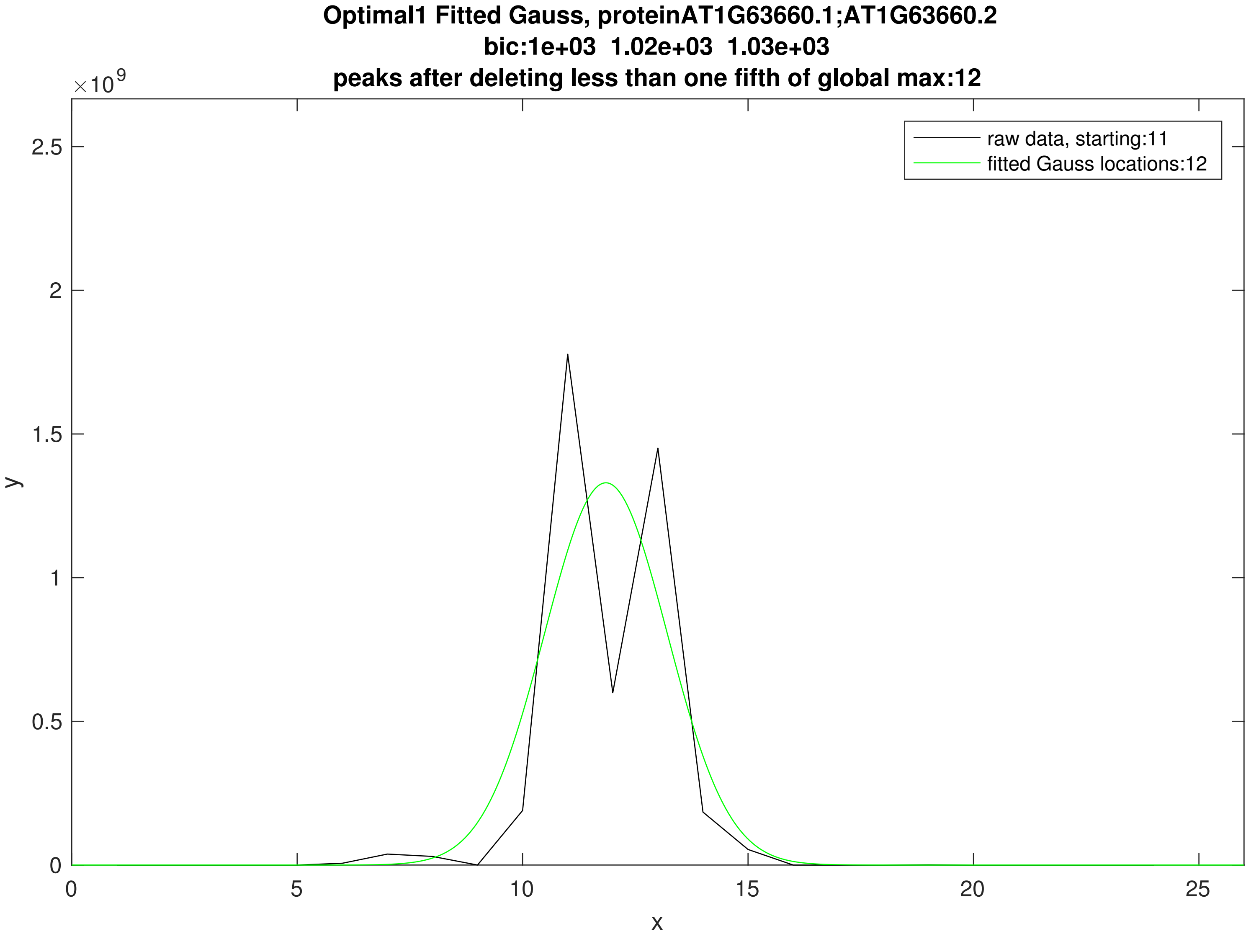
Optimal1 Fitted Gauss, proteinAT1G63660.1;AT1G63660.2
bic:1e+03  1.02e+03  1.03e+03
peaks after deleting less than one fifth of global max:12
×10^9
raw data, starting:11
fitted Gauss locations:12
y
x
0
0.5
1
1.5
2
2.5
5
10
15
20
25

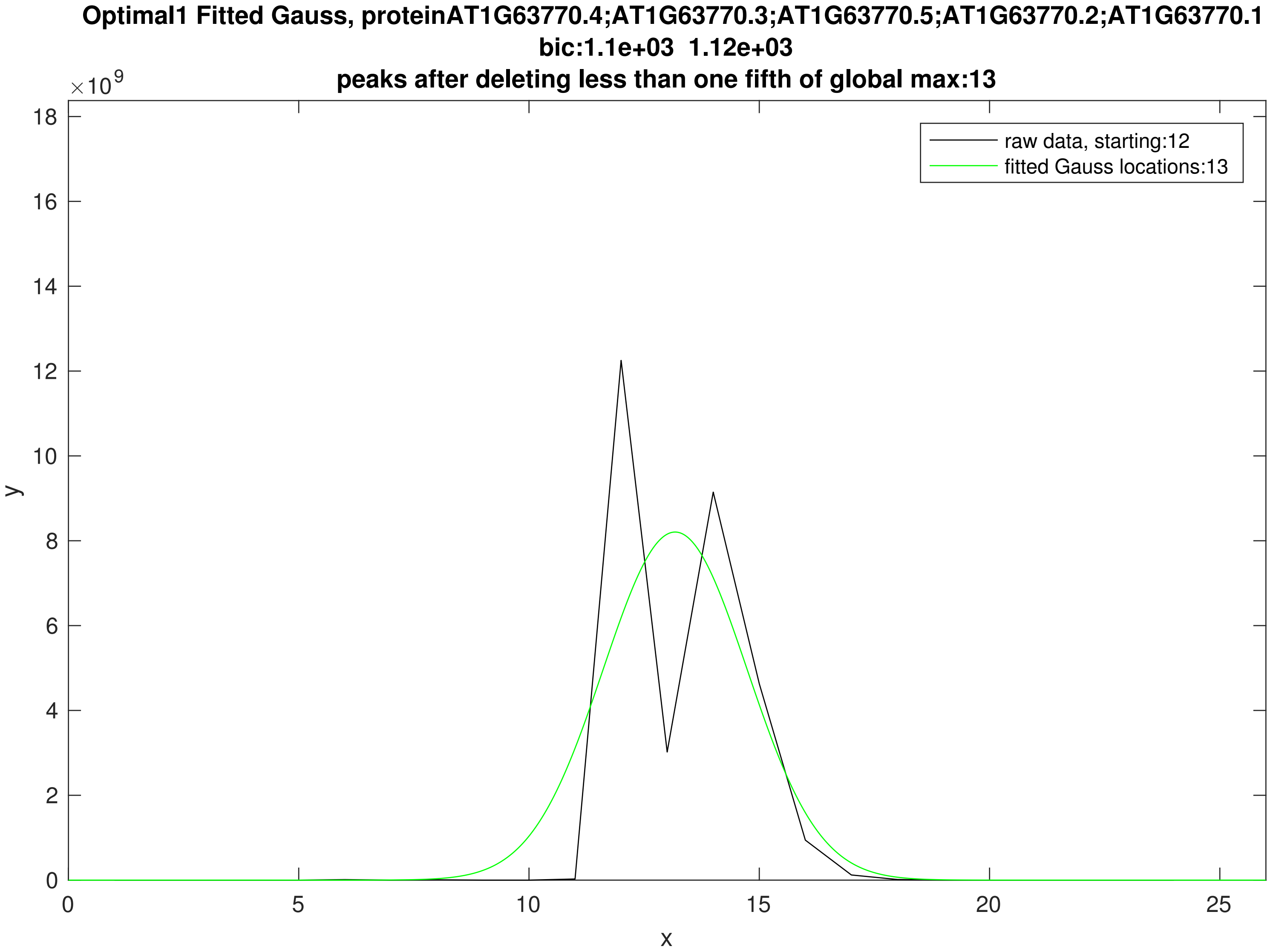
Optimal1 Fitted Gauss, proteinAT1G63770.4;AT1G63770.3;AT1G63770.5;AT1G63770.2;AT1G63770.1
bic:1.1e+03  1.12e+03
peaks after deleting less than one fifth of global max:13
×10^9
raw data, starting:12
fitted Gauss locations:13
y
x
0
2
4
6
8
10
12
14
16
18
0
5
10
15
20
25

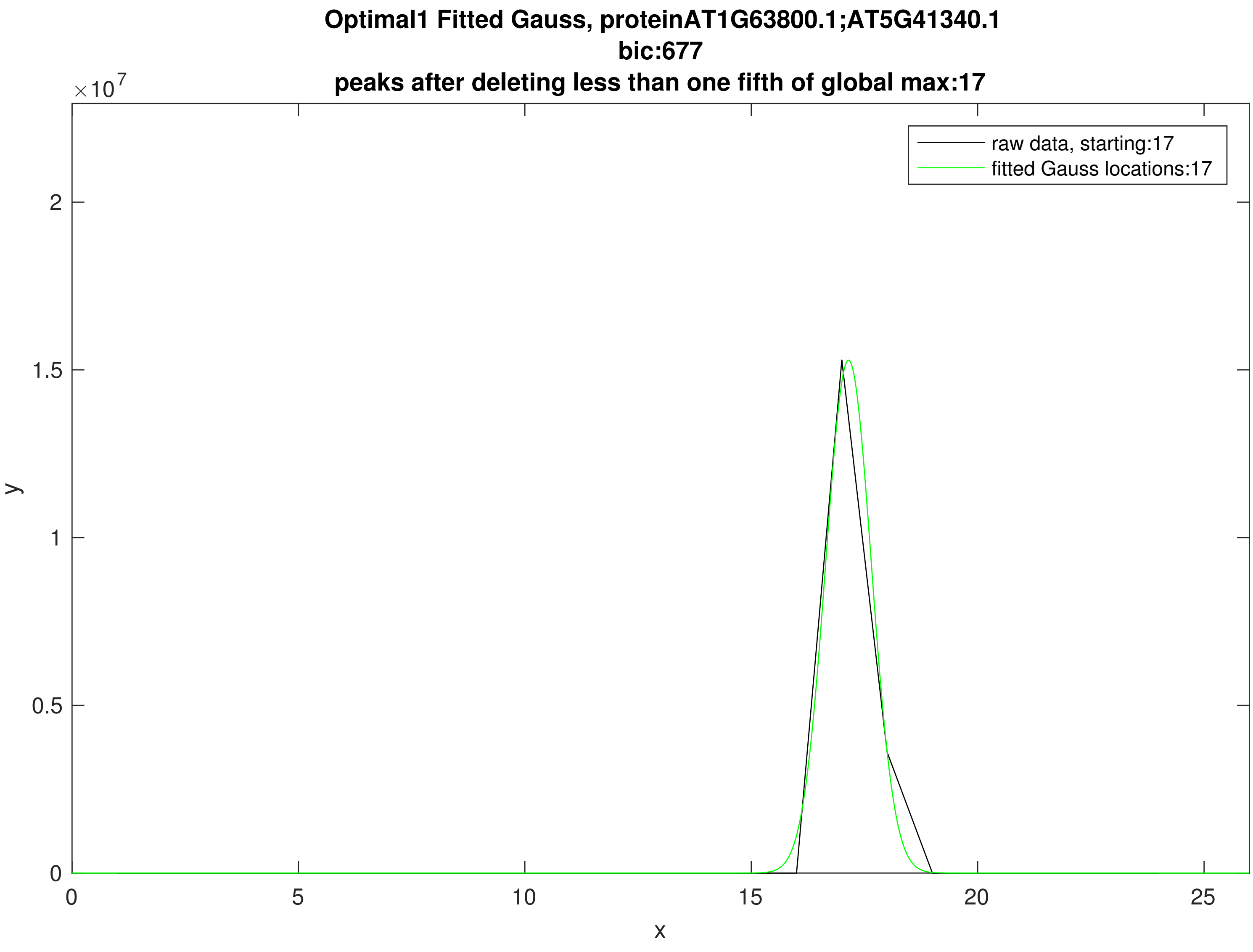

Optimal1 Fitted Gauss, proteinAT1G63800.1;AT5G41340.1
bic:677
peaks after deleting less than one fifth of global max:17
×10^7
raw data, starting:17
fitted Gauss locations:17
y
x
0
0.5
1
1.5
2
5
10
15
20
25

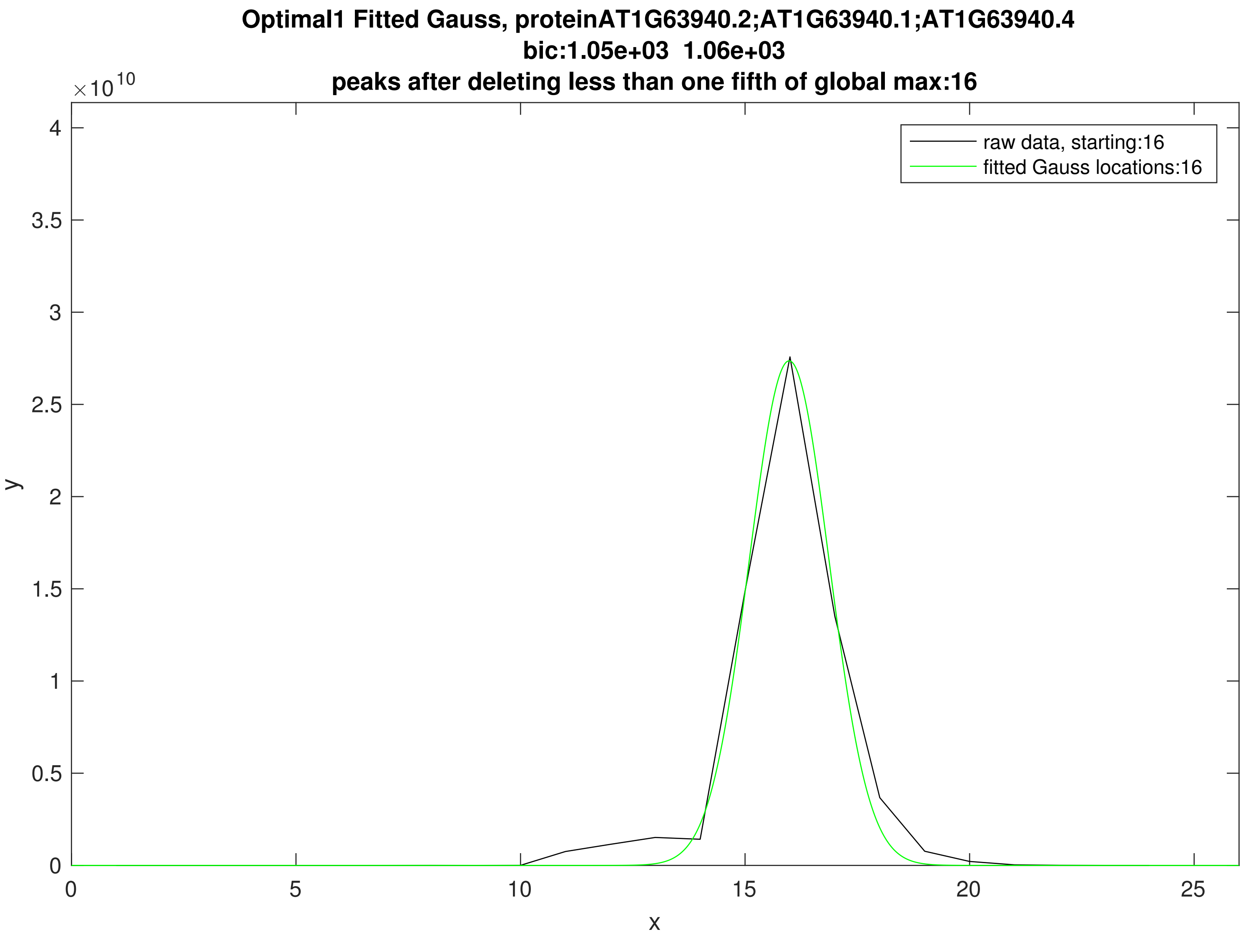
Optimal1 Fitted Gauss, proteinAT1G63940.2;AT1G63940.1;AT1G63940.4
bic:1.05e+03  1.06e+03
peaks after deleting less than one fifth of global max:16
×10^10
4
3.5
3
2.5
2
1.5
1
0.5
0
y
0
5
10
15
20
25
x
raw data, starting:16
fitted Gauss locations:16

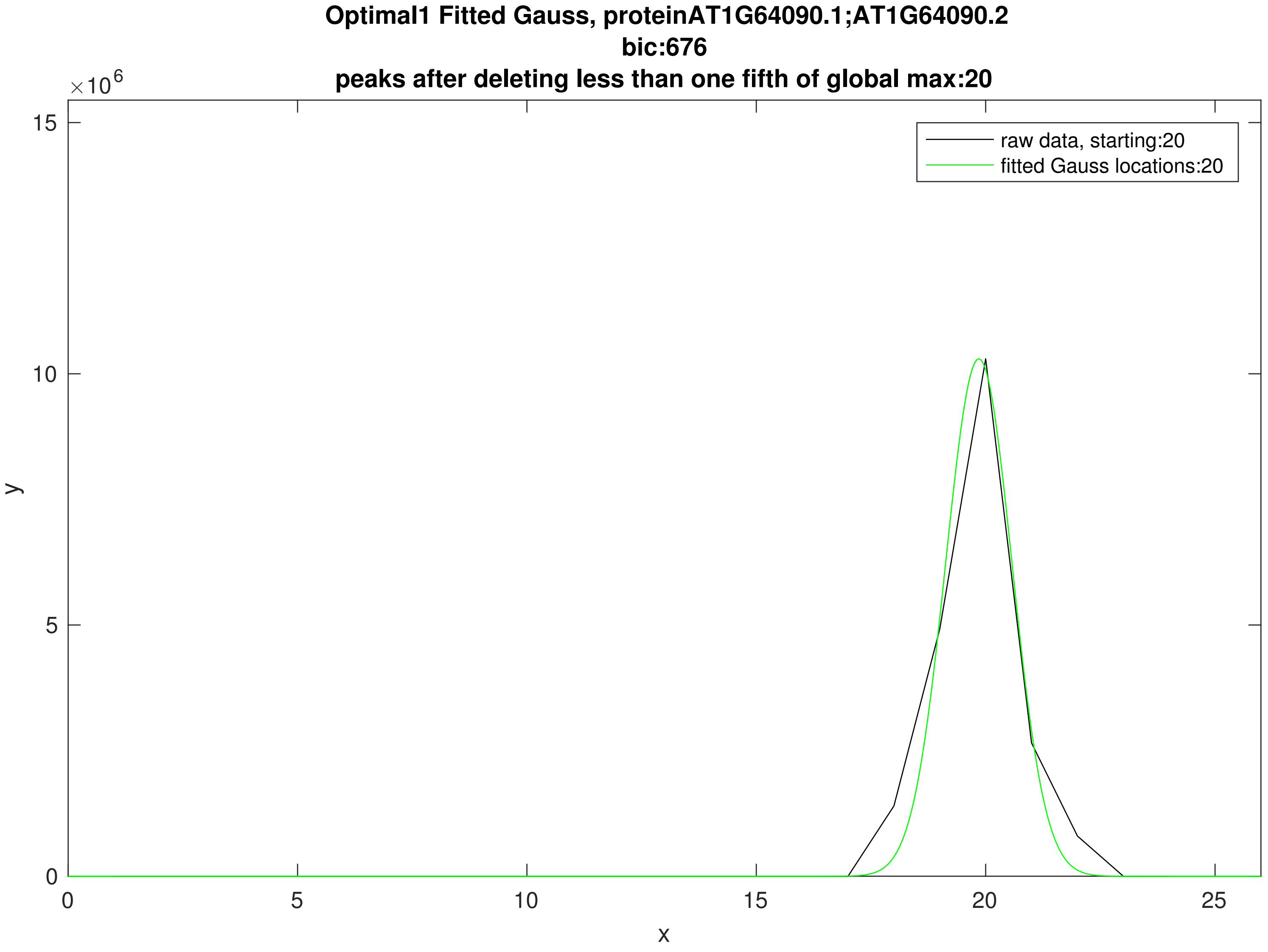

Optimal1 Fitted Gauss, proteinAT1G64090.1;AT1G64090.2
bic:676
peaks after deleting less than one fifth of global max:20
×10^6
15
10
5
0
y
0
5
10
15
20
25
x
raw data, starting:20
fitted Gauss locations:20

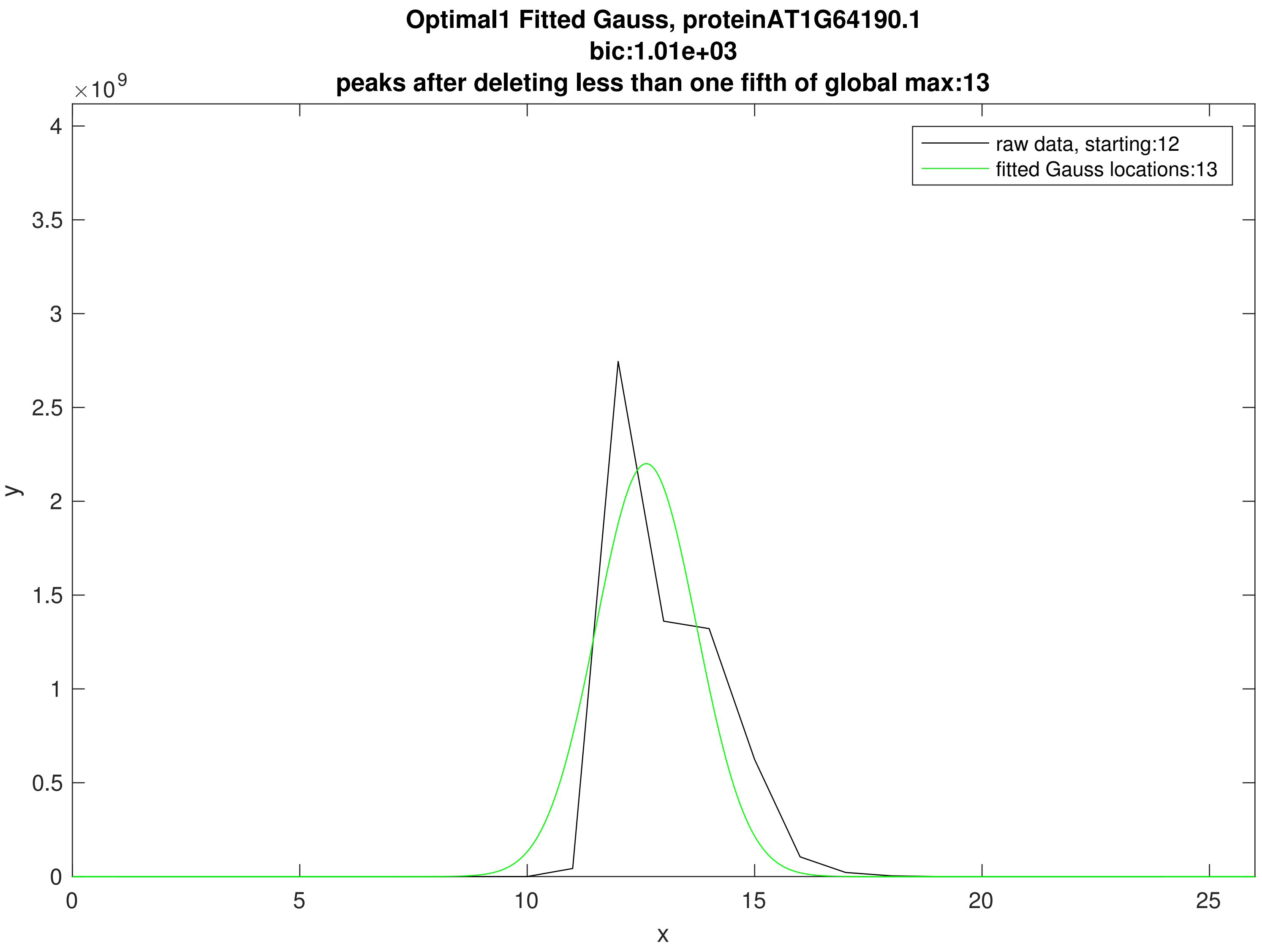
Optimal1 Fitted Gauss, proteinAT1G64190.1
bic:1.01e+03
peaks after deleting less than one fifth of global max:13
×10^9
raw data, starting:12
fitted Gauss locations:13
y
x
0
0.5
1
1.5
2
2.5
3
3.5
4
0
5
10
15
20
25

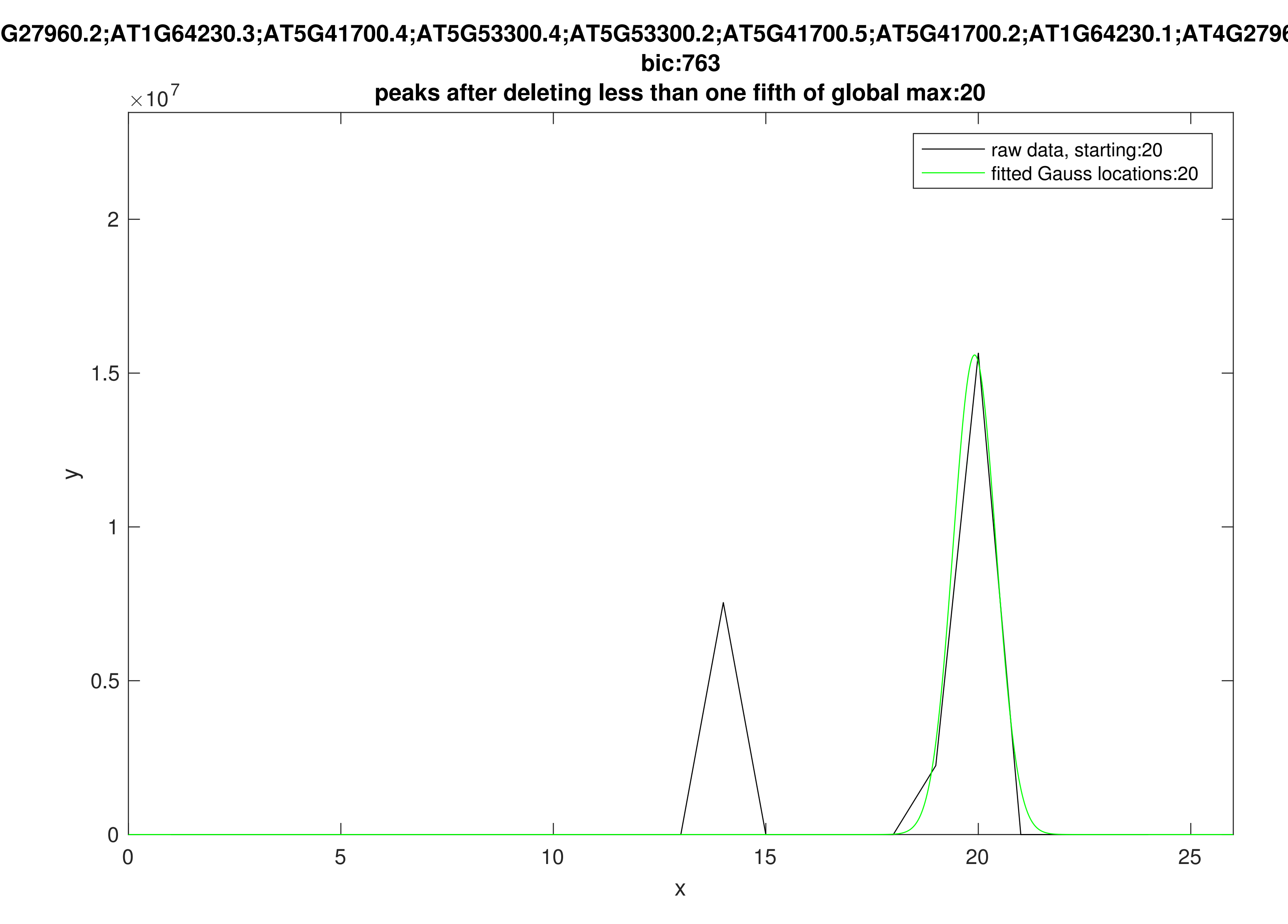

G27960.2;AT1G64230.3;AT5G41700.4;AT5G53300.4;AT5G53300.2;AT5G41700.5;AT5G41700.2;AT1G64230.1;AT4G2796
bic:763
peaks after deleting less than one fifth of global max:20
×10^7
raw data, starting:20
fitted Gauss locations:20
y
x
0
0.5
1
1.5
2
5
10
15
20
25

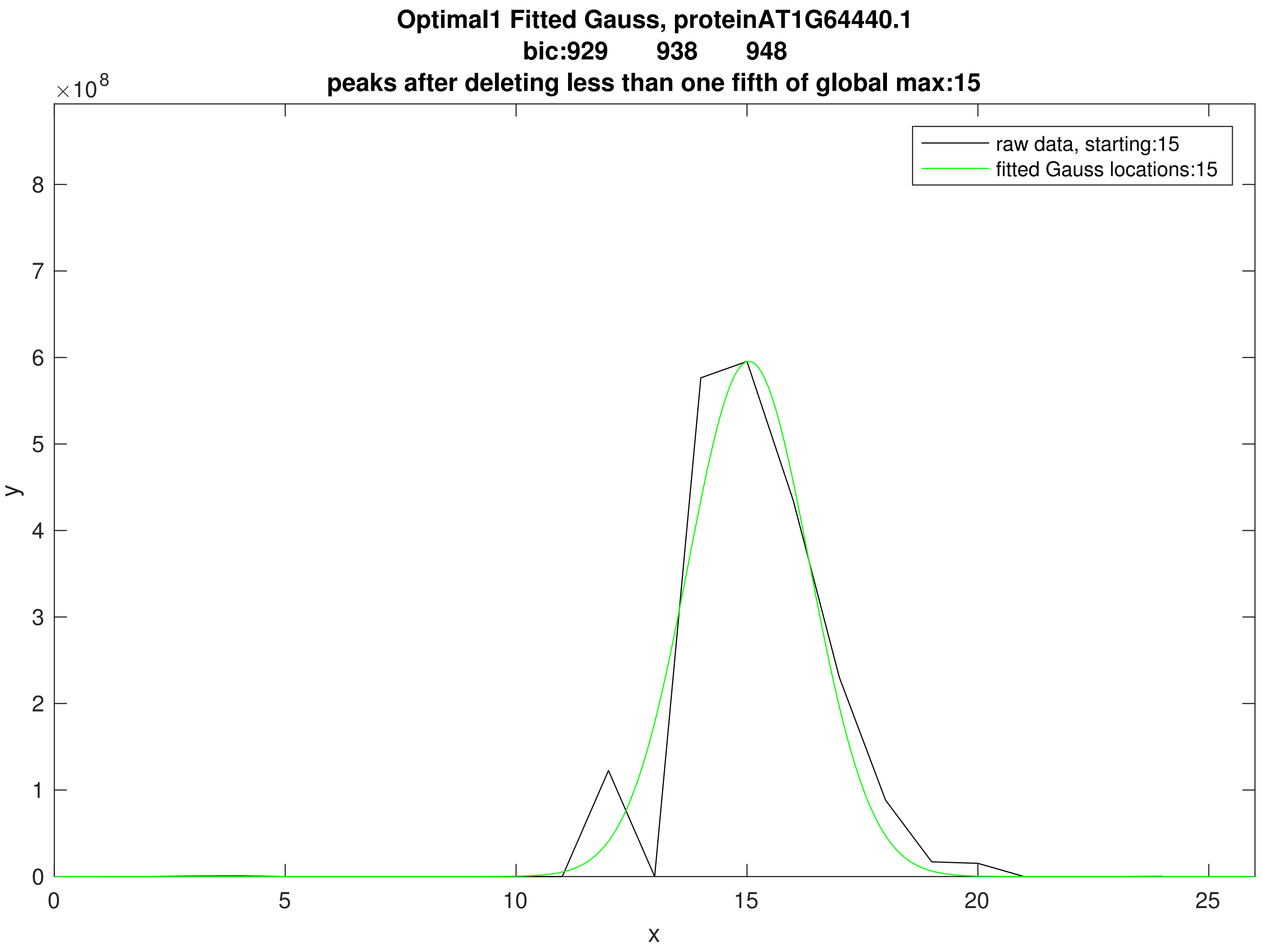

Optimal1 Fitted Gauss, proteinAT1G64440.1
bic:929 938 948
peaks after deleting less than one fifth of global max:15
×10^8
raw data, starting:15
fitted Gauss locations:15
y
x
0
1
2
3
4
5
6
7
8
0
5
10
15
20
25

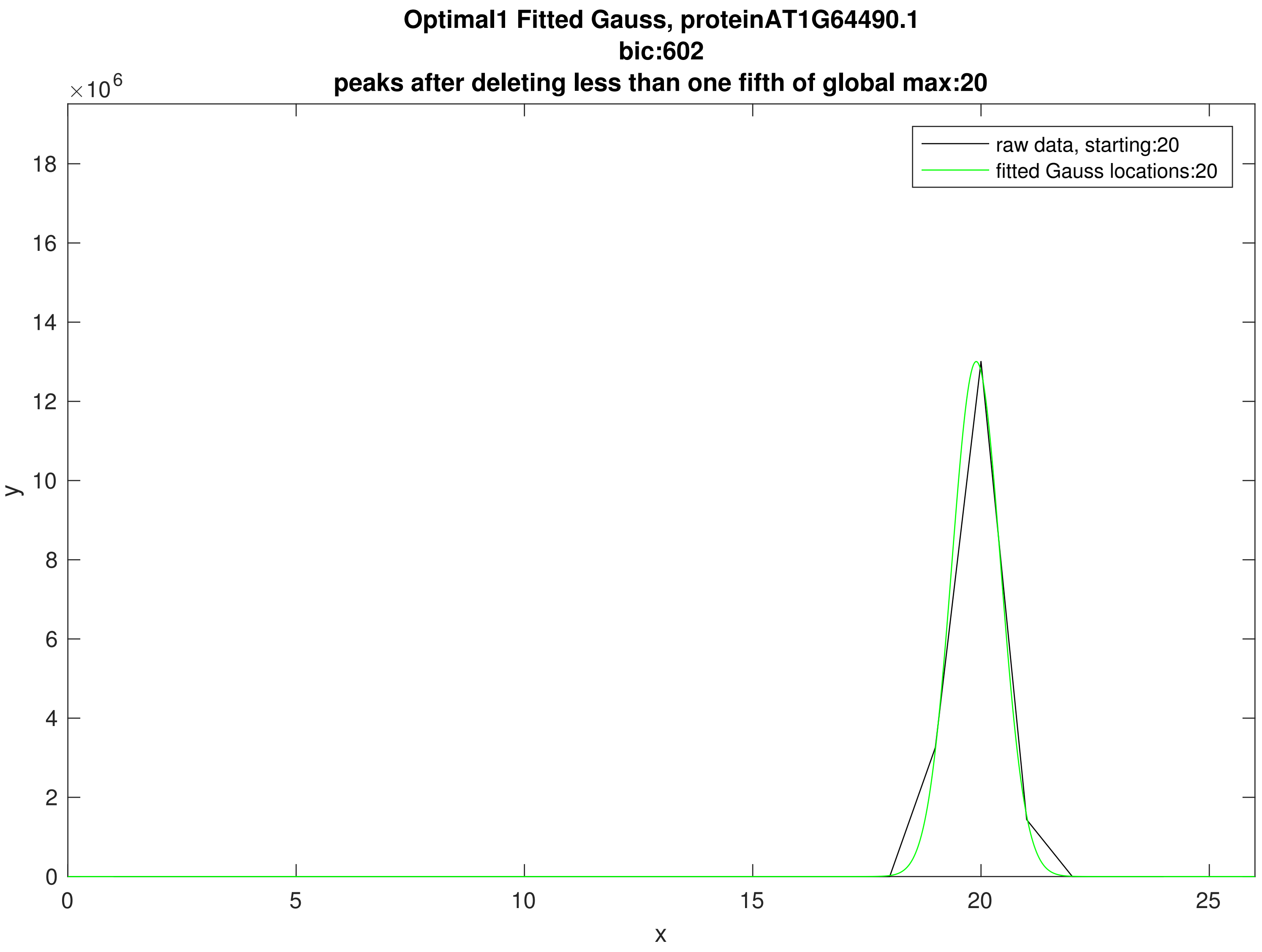

Optimal1 Fitted Gauss, proteinAT1G64490.1
bic:602
peaks after deleting less than one fifth of global max:20
×10^6
raw data, starting:20
fitted Gauss locations:20
y
x
0
2
4
6
8
10
12
14
16
18
0
5
10
15
20
25

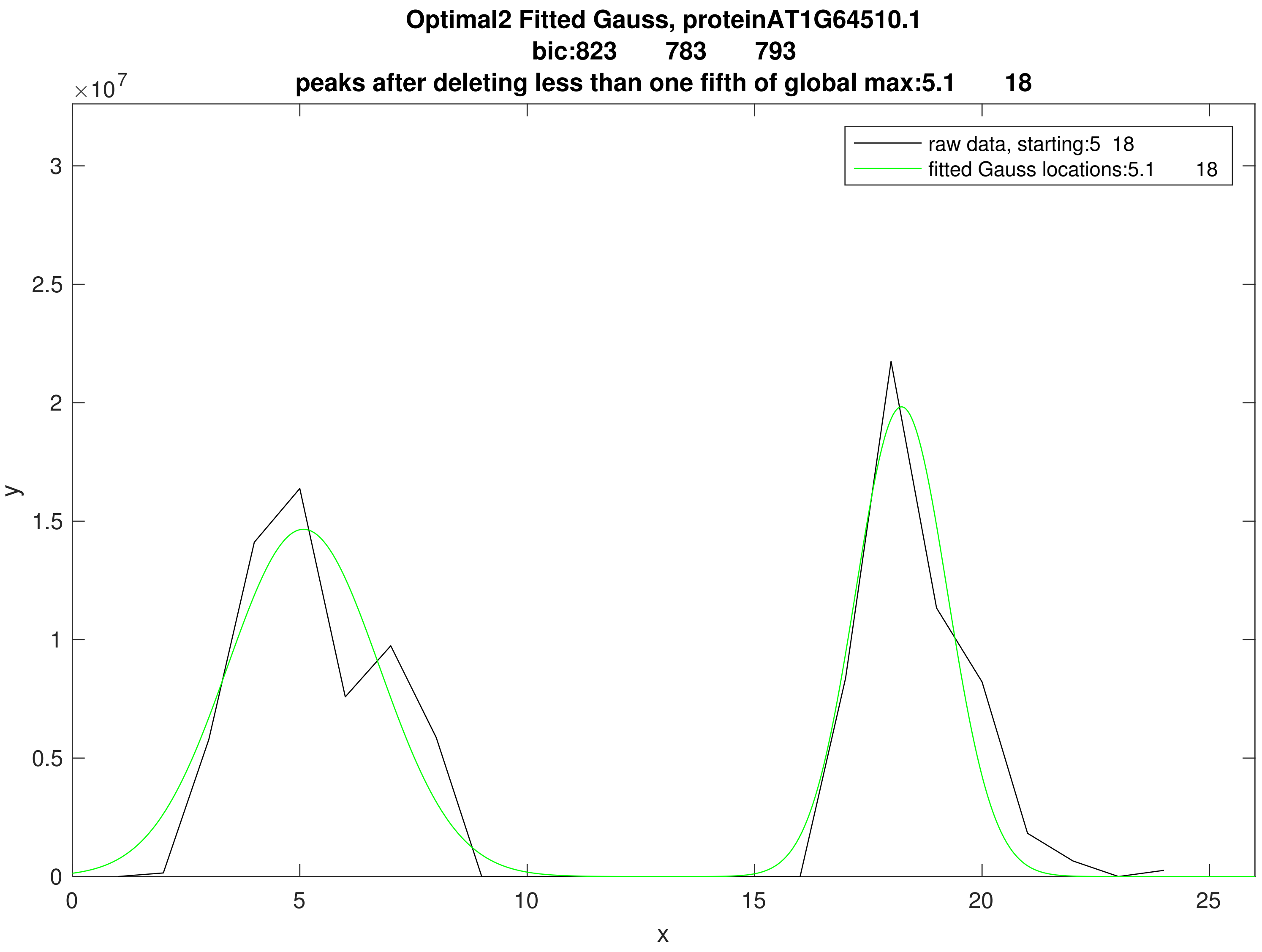
Optimal2 Fitted Gauss, proteinAT1G64510.1
bic:823 783 793
peaks after deleting less than one fifth of global max:5.1 18
raw data, starting:5 18
fitted Gauss locations:5.1 18
×10^7
y
x

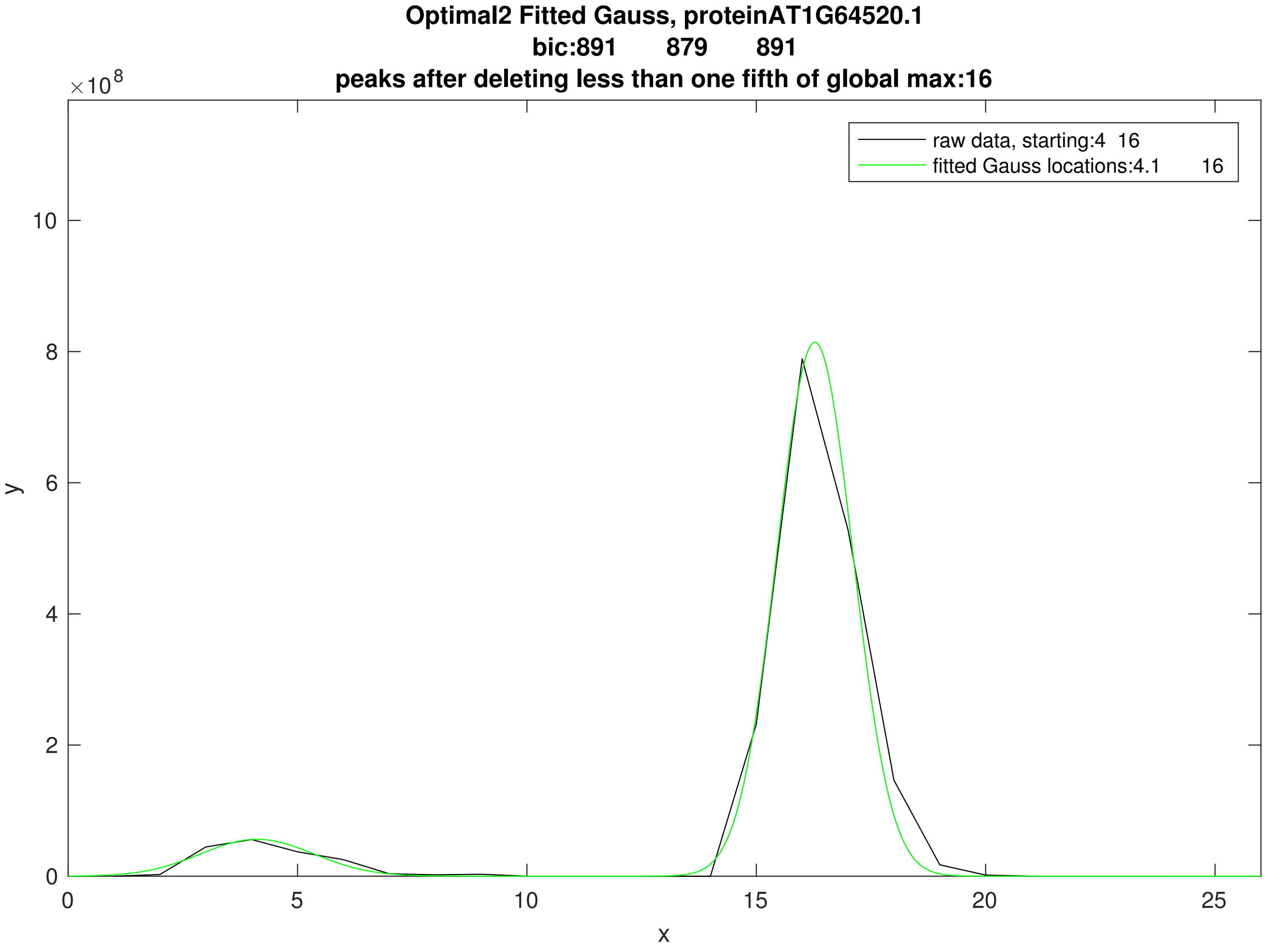

Optimal2 Fitted Gauss, proteinAT1G64520.1
bic:891 879 891
peaks after deleting less than one fifth of global max:16
×10^8
raw data, starting:4 16
fitted Gauss locations:4.1 16
y
x
0
2
4
6
8
10
0
5
10
15
20
25

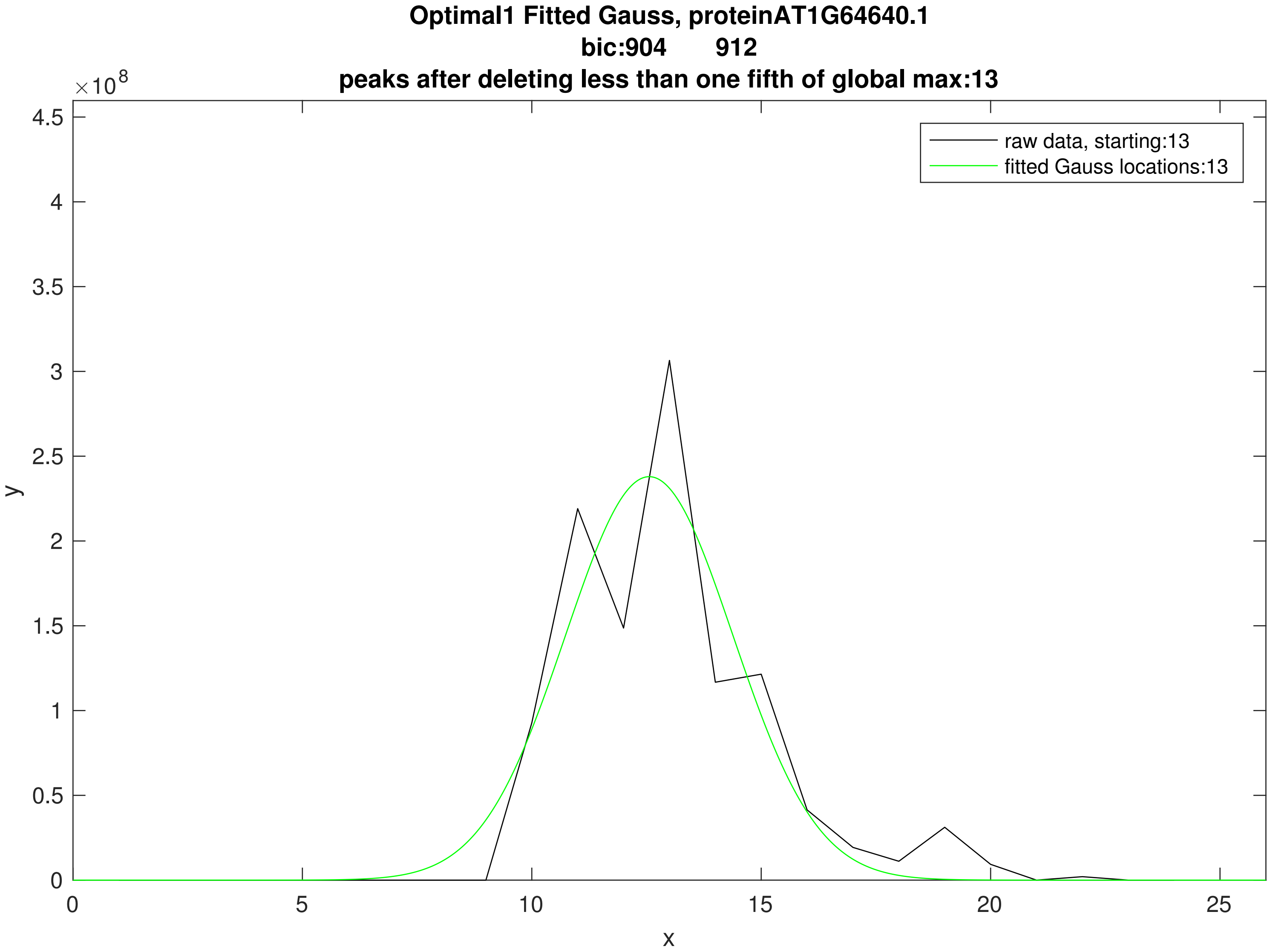

Optimal1 Fitted Gauss, proteinAT1G64640.1
bic:904 912
peaks after deleting less than one fifth of global max:13
×10^8
4.5
4
3.5
3
2.5
2
1.5
1
0.5
0
y
0
5
10
15
20
25
x
raw data, starting:13
fitted Gauss locations:13

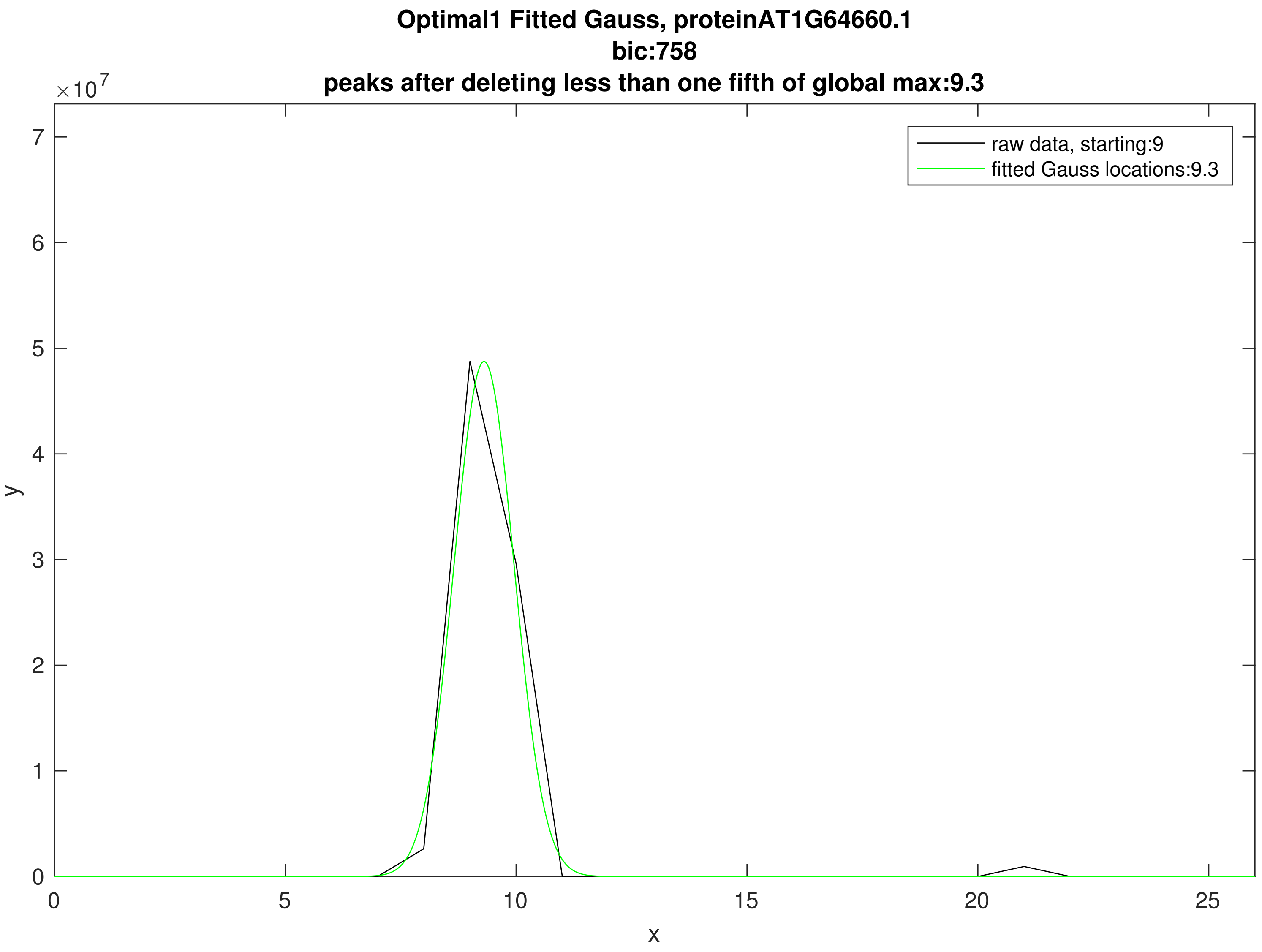

Optimal1 Fitted Gauss, proteinAT1G64660.1
bic:758
peaks after deleting less than one fifth of global max:9.3
×10^7
7
6
5
4
3
2
1
0
y
0
5
10
15
20
25
x
raw data, starting:9
fitted Gauss locations:9.3

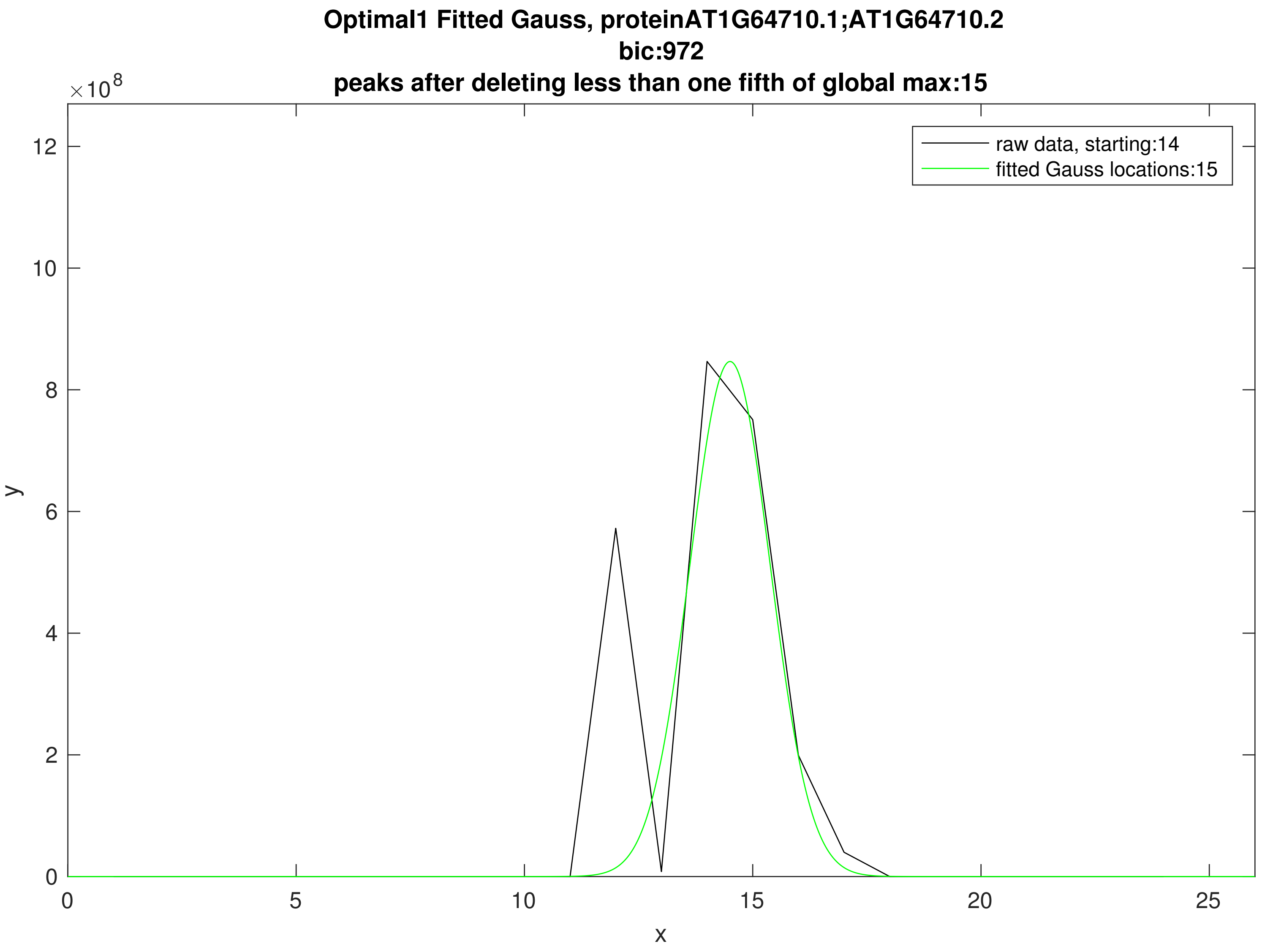
Optimal1 Fitted Gauss, proteinAT1G64710.1;AT1G64710.2
bic:972
peaks after deleting less than one fifth of global max:15
×10^8
raw data, starting:14
fitted Gauss locations:15
y
x
0
2
4
6
8
10
12
0
5
10
15
20
25

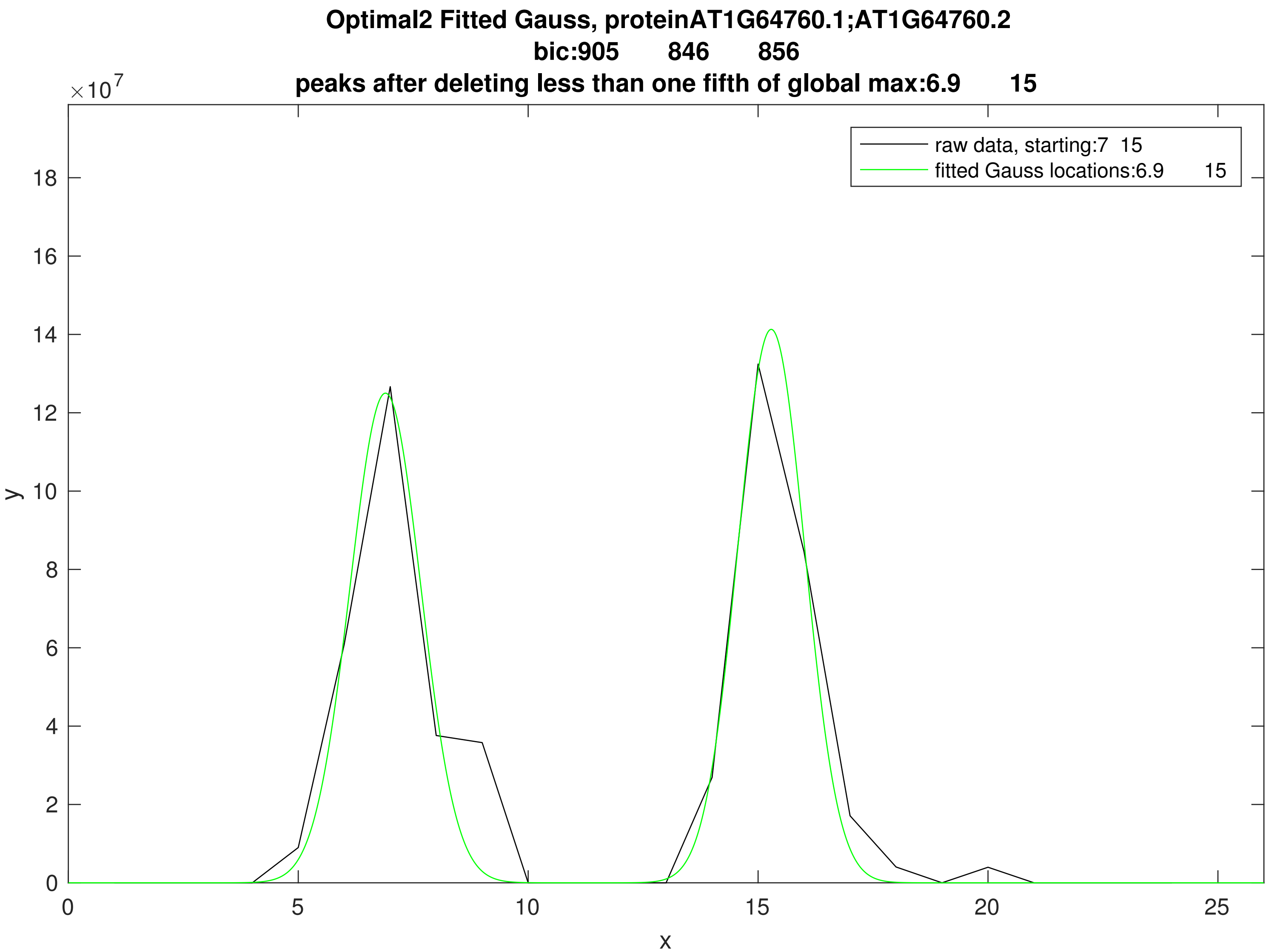

Optimal2 Fitted Gauss, proteinAT1G64760.1;AT1G64760.2
bic:905 846 856
peaks after deleting less than one fifth of global max:6.9 15
×10^7
raw data, starting:7 15
fitted Gauss locations:6.9 15
y
x
0
2
4
6
8
10
12
14
16
18
0
5
10
15
20
25

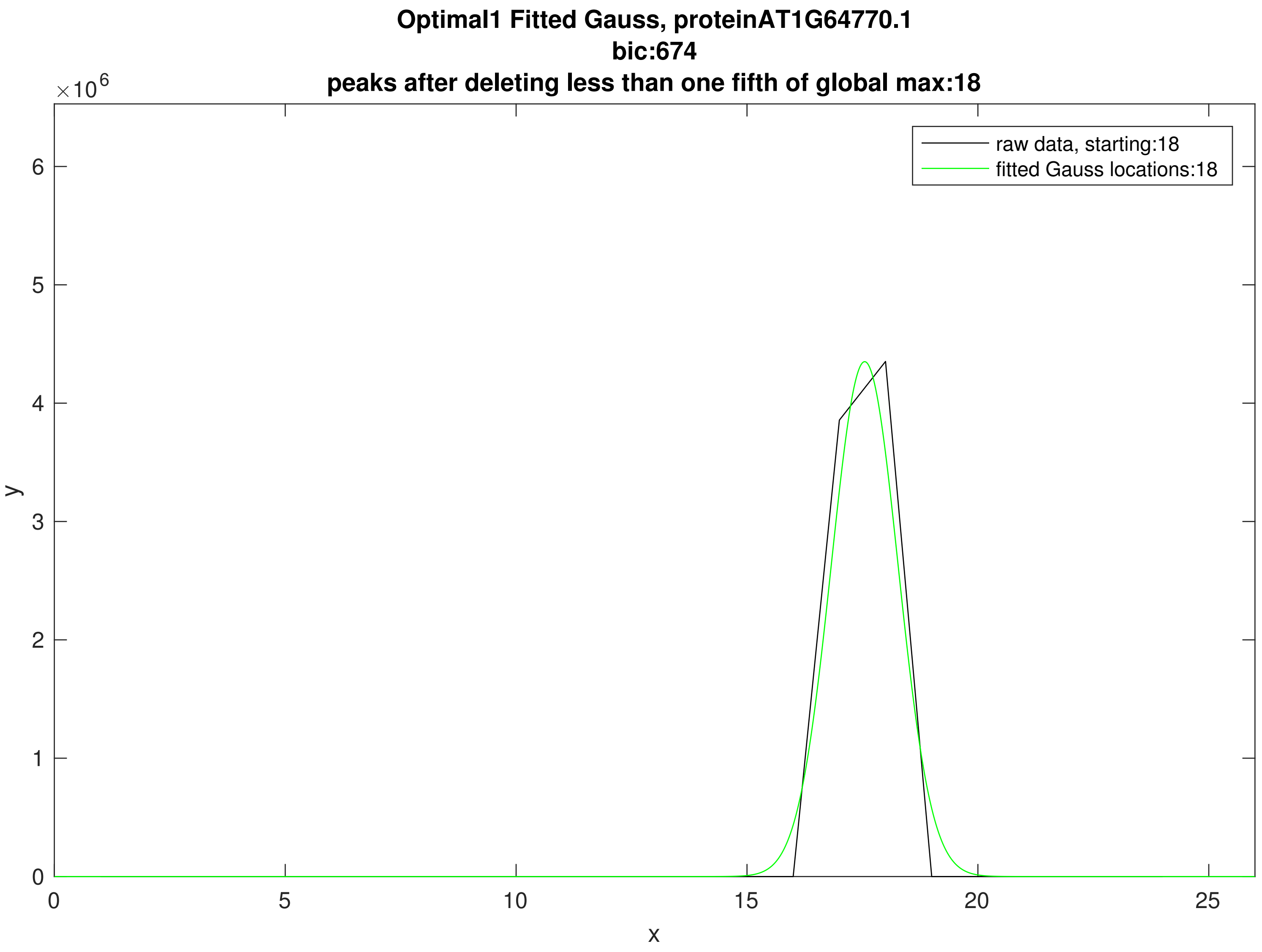

Optimal1 Fitted Gauss, proteinAT1G64770.1
bic:674
peaks after deleting less than one fifth of global max:18
×10^6
raw data, starting:18
fitted Gauss locations:18
y
x
0
1
2
3
4
5
6
0
5
10
15
20
25

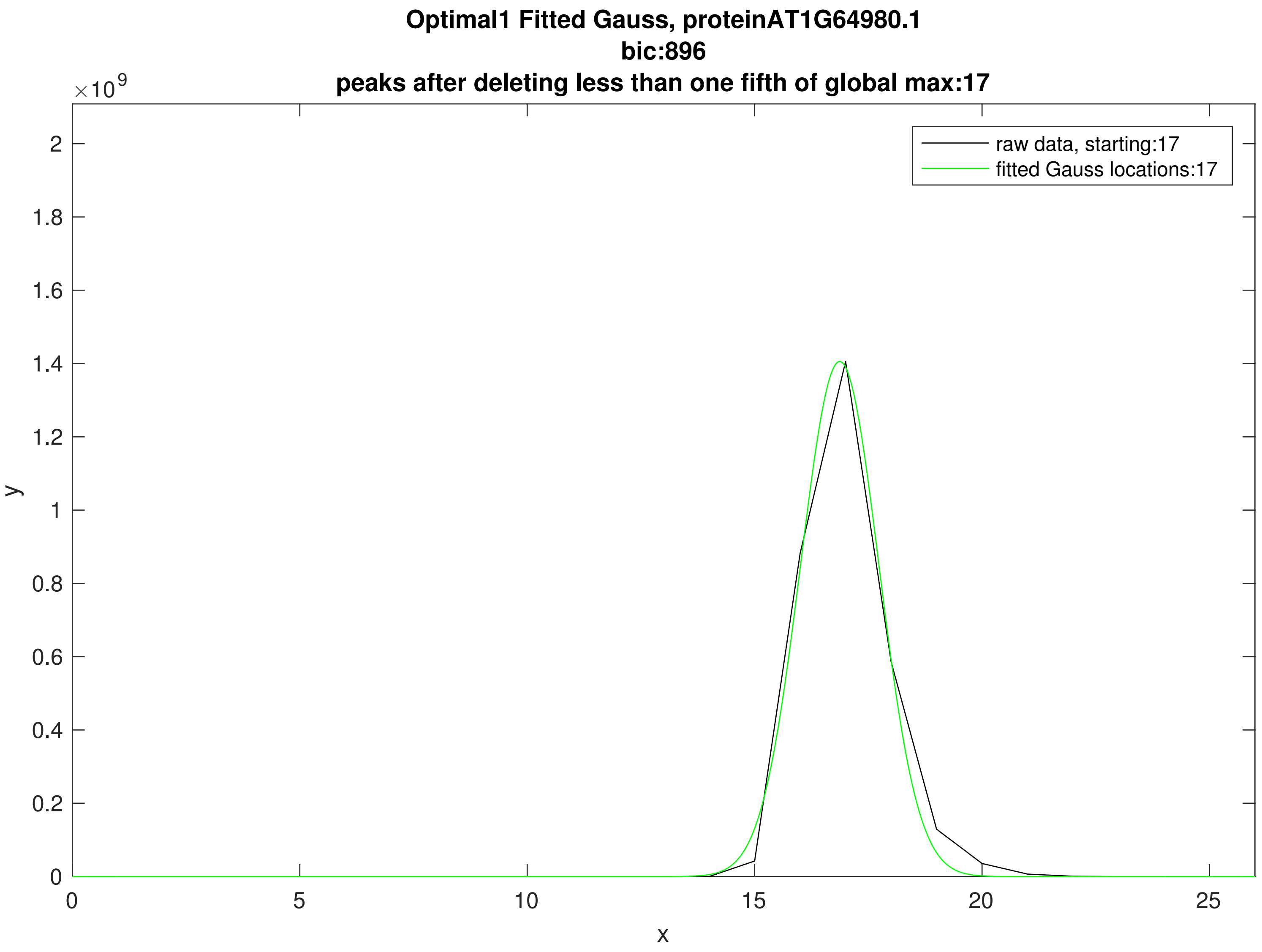

Optimal1 Fitted Gauss, proteinAT1G64980.1
bic:896
peaks after deleting less than one fifth of global max:17
×10^9
raw data, starting:17
fitted Gauss locations:17
y
x
0
0.2
0.4
0.6
0.8
1
1.2
1.4
1.6
1.8
2
0
5
10
15
20
25

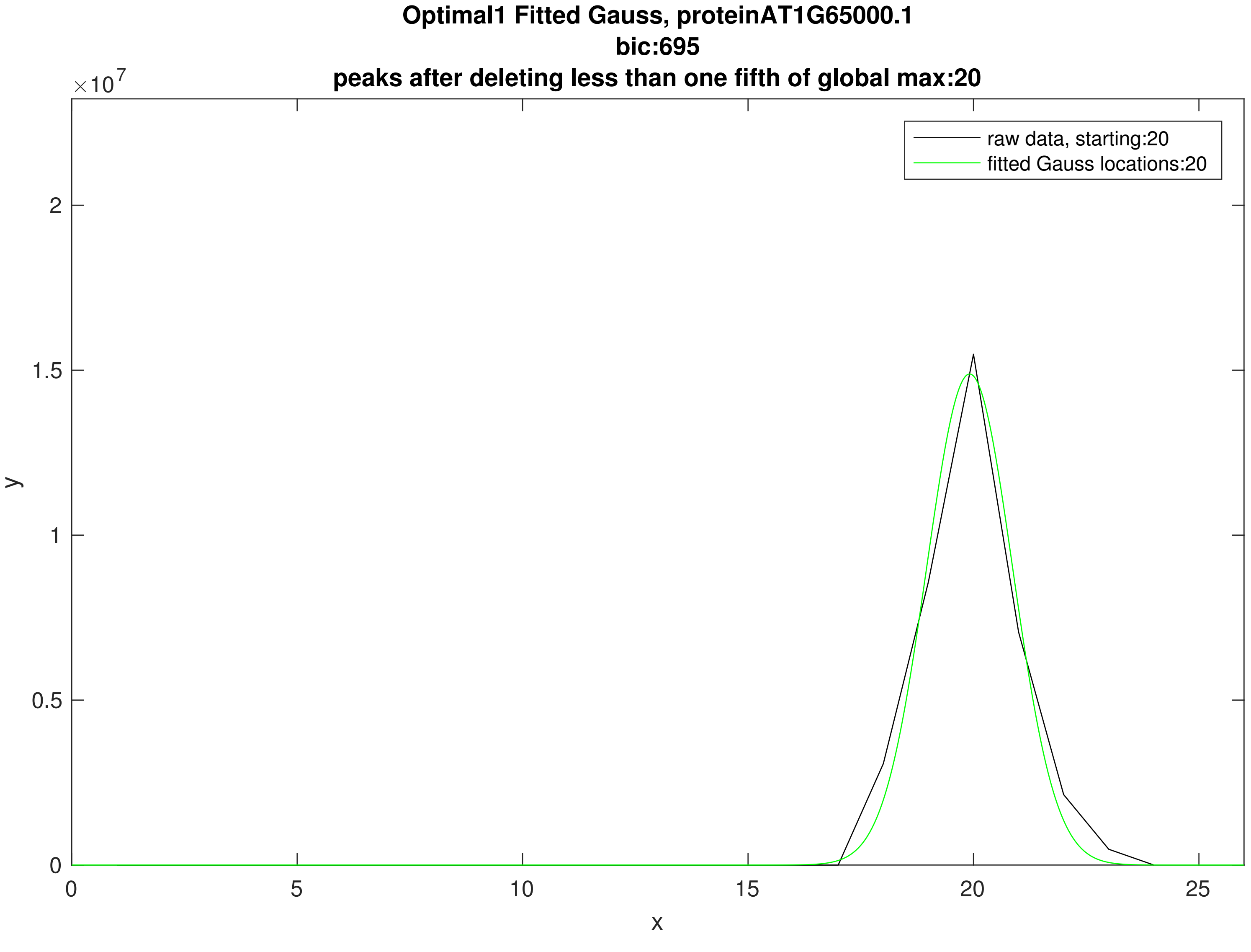
Optimal1 Fitted Gauss, proteinAT1G65000.1
bic:695
peaks after deleting less than one fifth of global max:20
×10^7
raw data, starting:20
fitted Gauss locations:20
y
x
0
0.5
1
1.5
2
0
5
10
15
20
25

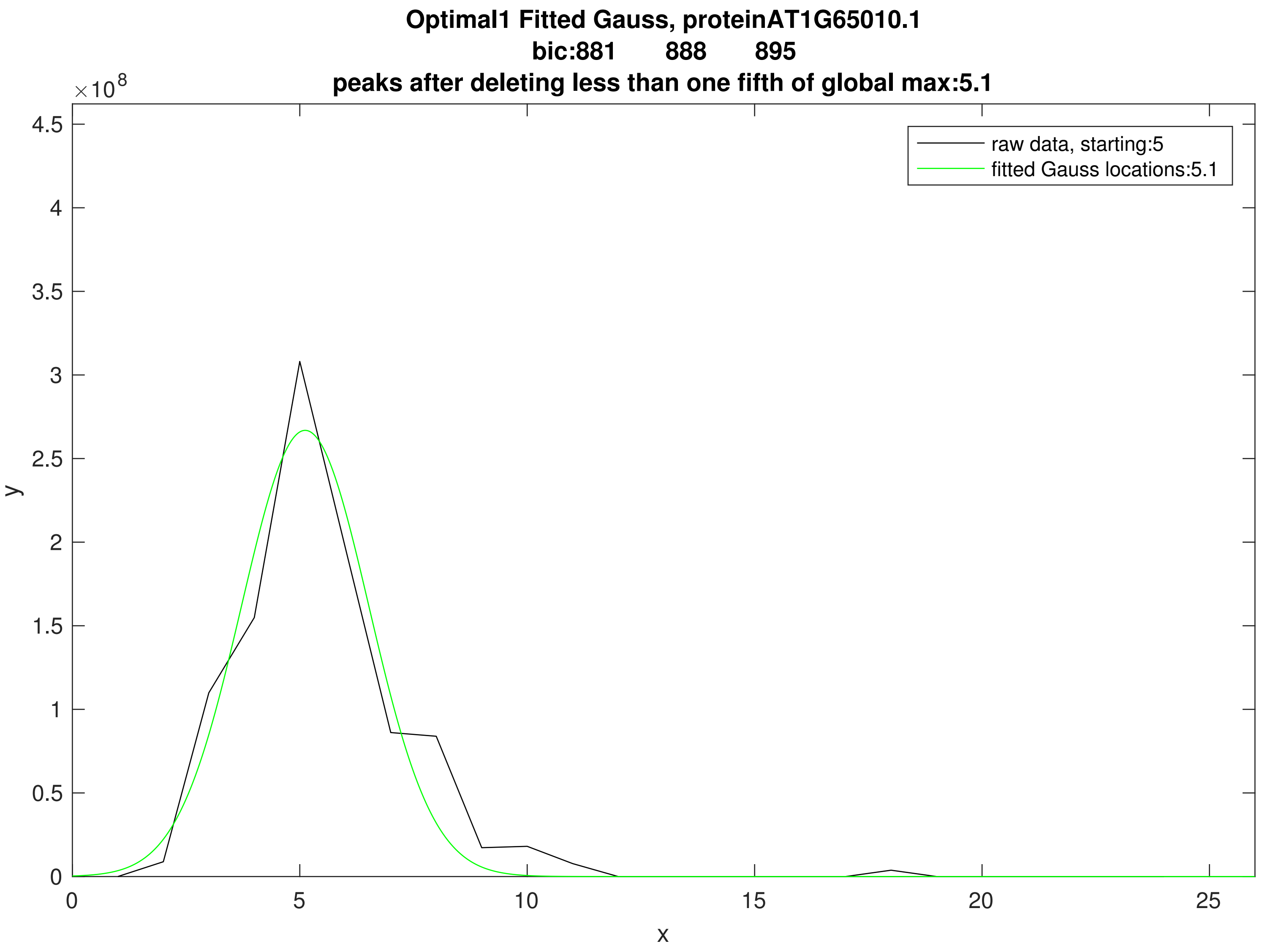
Optimal1 Fitted Gauss, proteinAT1G65010.1
bic:881 888 895
peaks after deleting less than one fifth of global max:5.1
×10^8
4.5
4
3.5
3
2.5
2
1.5
1
0.5
0
y
0
5
10
15
20
25
x
raw data, starting:5
fitted Gauss locations:5.1

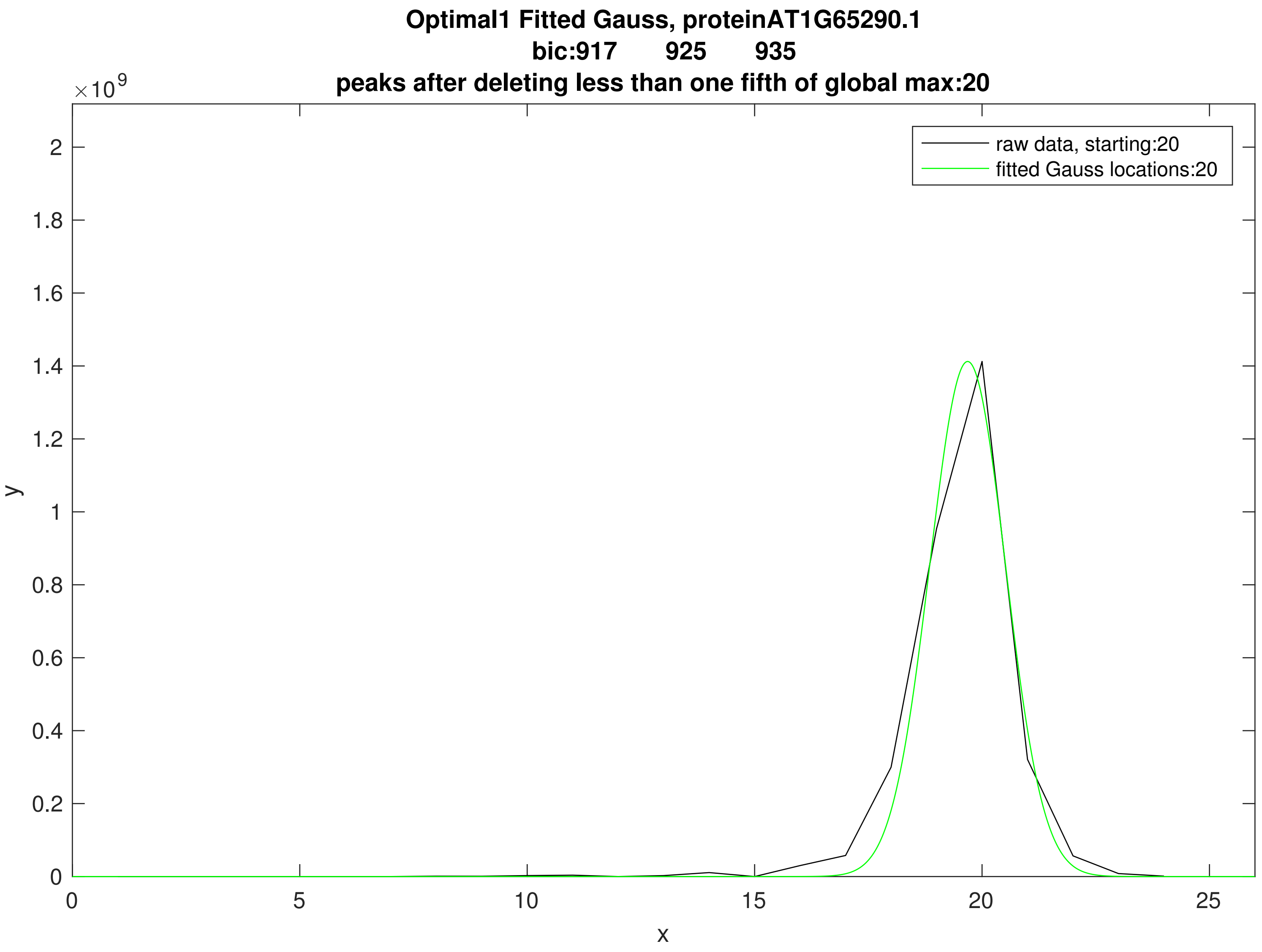
Optimal1 Fitted Gauss, proteinAT1G65290.1
bic:917 925 935
peaks after deleting less than one fifth of global max:20
×10^9
raw data, starting:20
fitted Gauss locations:20
y
x
0
0.2
0.4
0.6
0.8
1
1.2
1.4
1.6
1.8
2
0
5
10
15
20
25

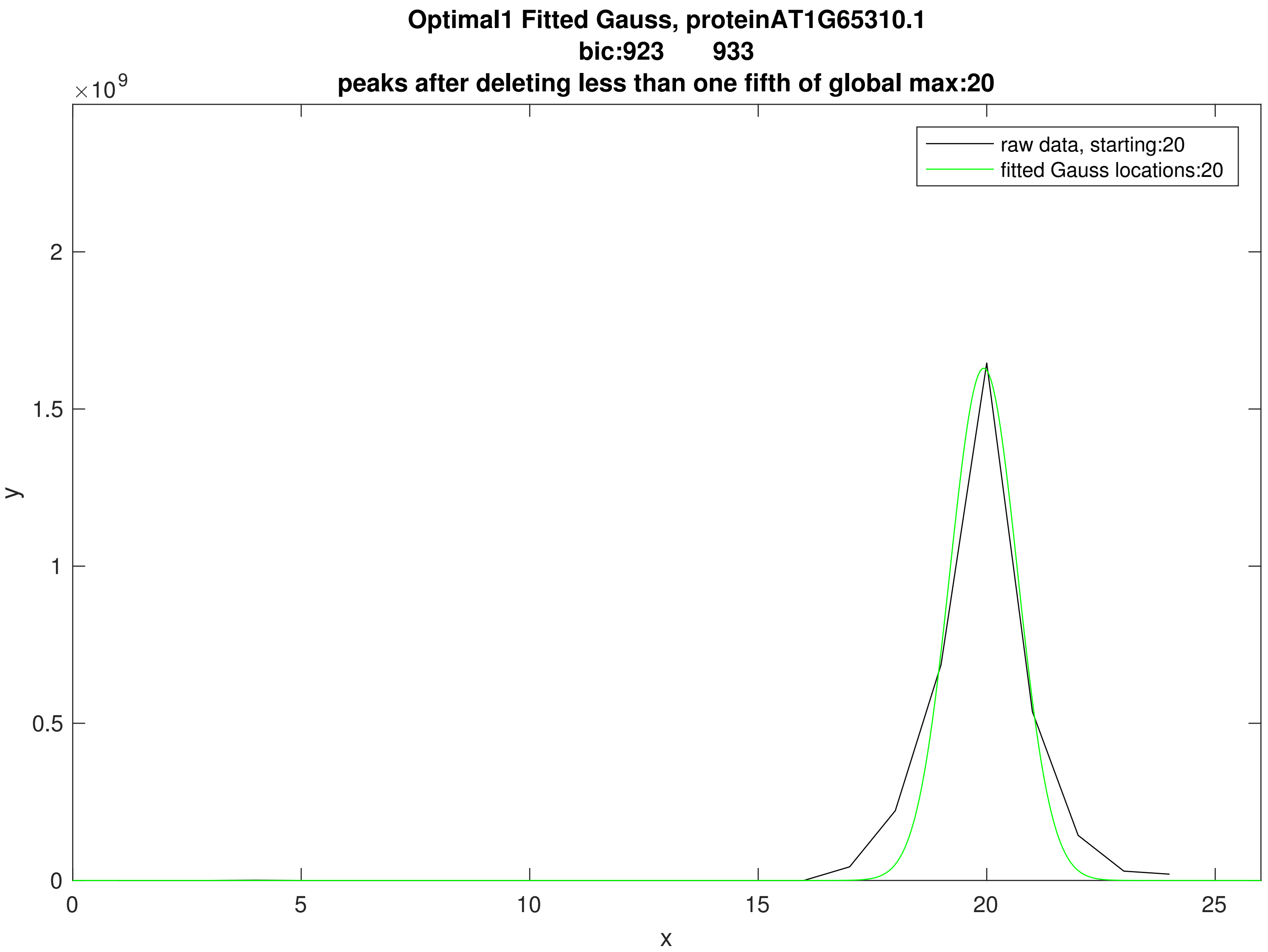

Optimal1 Fitted Gauss, proteinAT1G65310.1
bic:923 933
peaks after deleting less than one fifth of global max:20
×10^9
raw data, starting:20
fitted Gauss locations:20
y
x
0
0.5
1
1.5
2
0
5
10
15
20
25

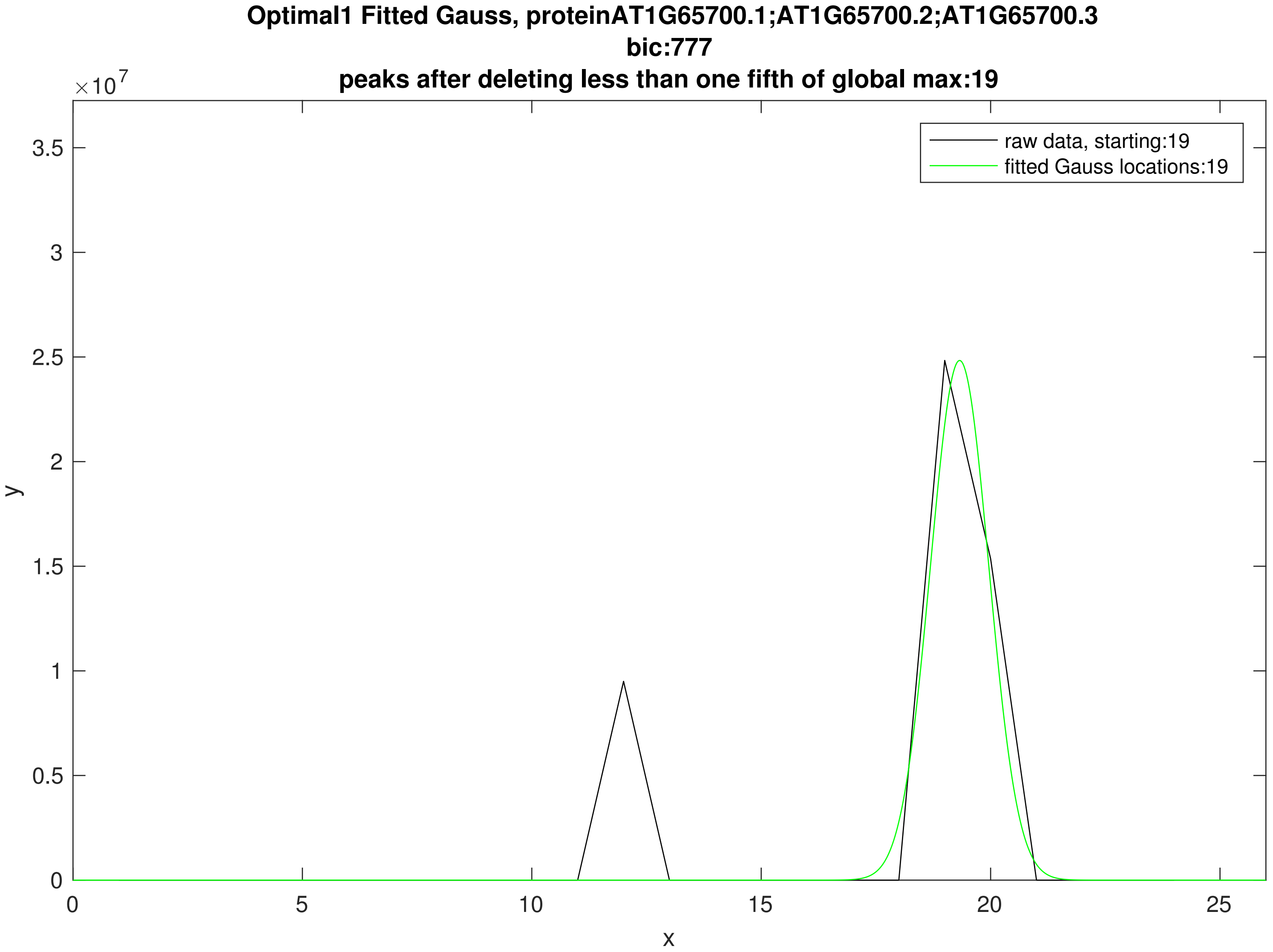

Optimal1 Fitted Gauss, proteinAT1G65700.1;AT1G65700.2;AT1G65700.3
bic:777
peaks after deleting less than one fifth of global max:19
×10^7
3.5
3
2.5
2
1.5
1
0.5
0
y
0
5
10
15
20
25
x
raw data, starting:19
fitted Gauss locations:19

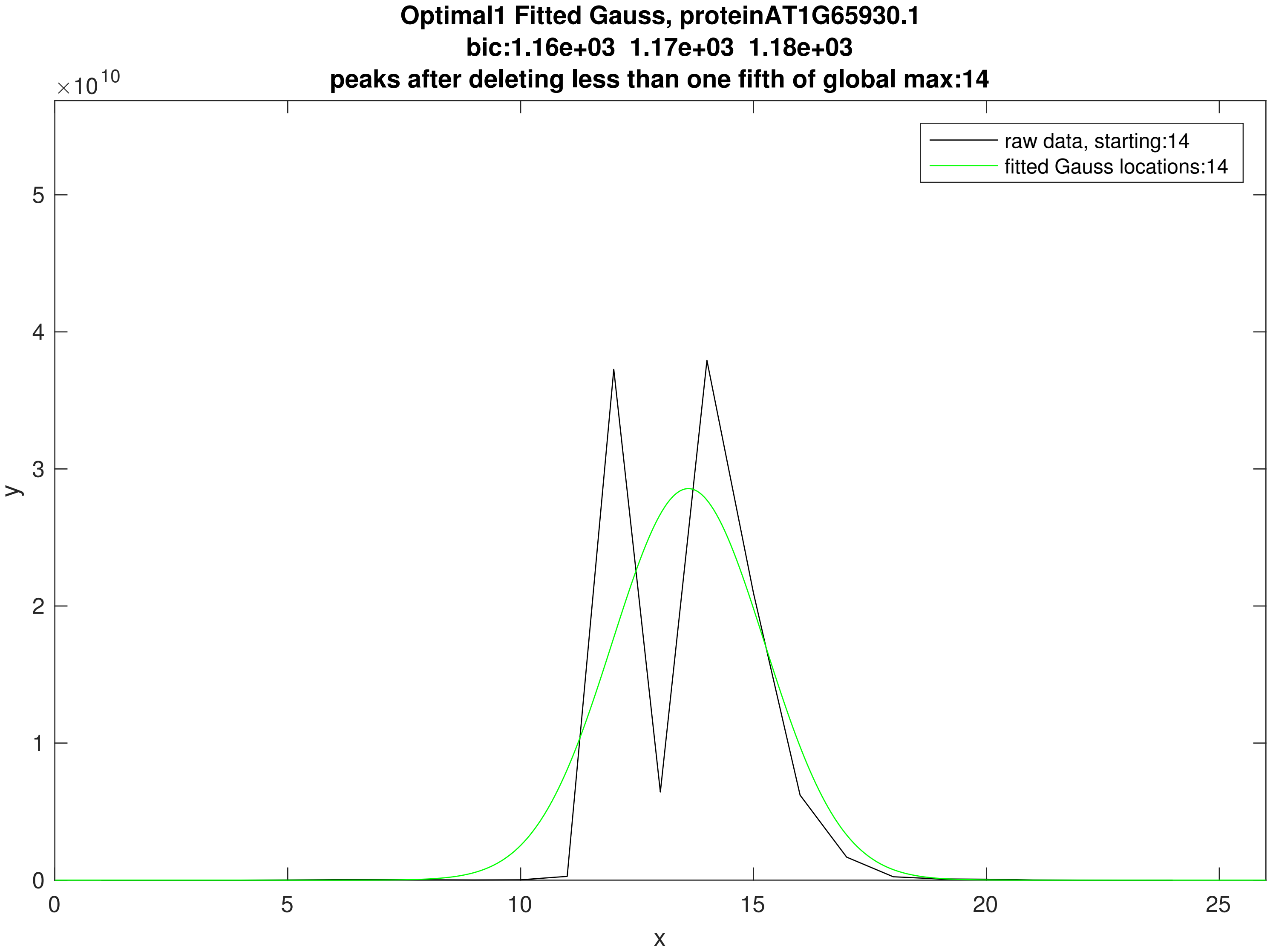

Optimal1 Fitted Gauss, proteinAT1G65930.1
bic:1.16e+03 1.17e+03 1.18e+03
peaks after deleting less than one fifth of global max:14
×10^10
raw data, starting:14
fitted Gauss locations:14
y
x
0
1
2
3
4
5
0
5
10
15
20
25

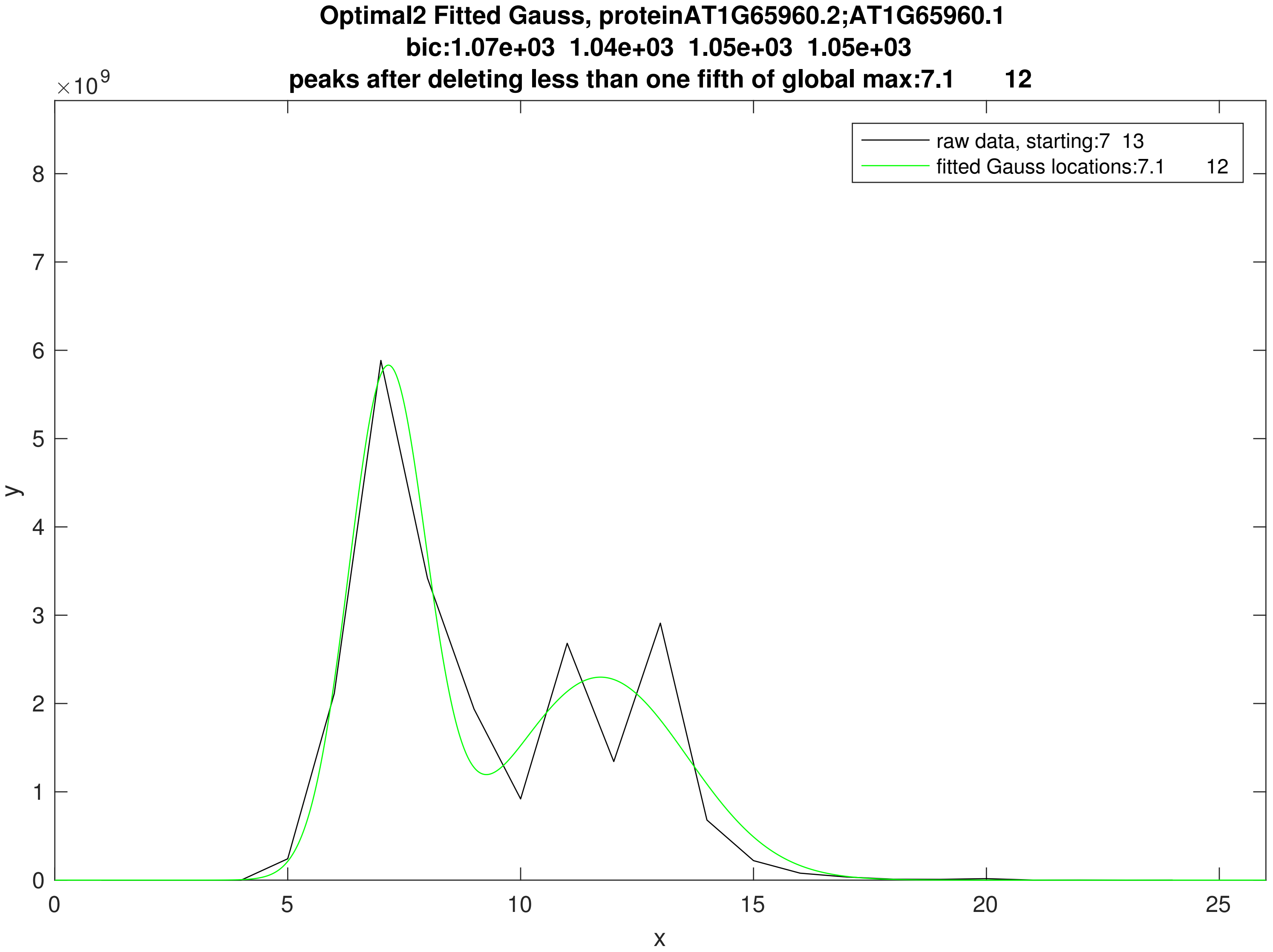

Optimal2 Fitted Gauss, proteinAT1G65960.2;AT1G65960.1
bic:1.07e+03 1.04e+03 1.05e+03 1.05e+03
peaks after deleting less than one fifth of global max:7.1 12
×10^9
raw data, starting:7 13
fitted Gauss locations:7.1 12
y
x

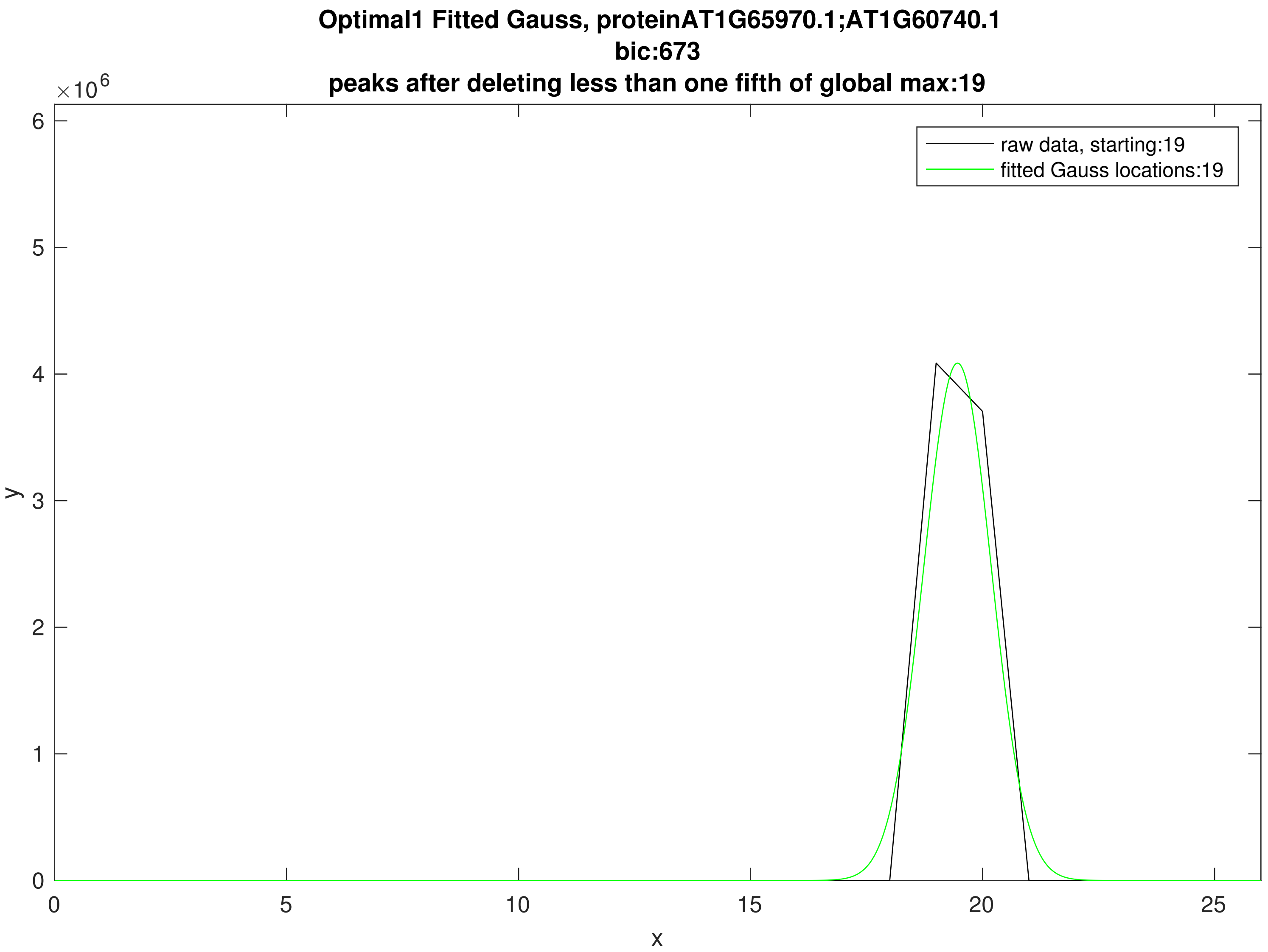
Optimal1 Fitted Gauss, proteinAT1G65970.1;AT1G60740.1
bic:673
peaks after deleting less than one fifth of global max:19
×10^6
raw data, starting:19
fitted Gauss locations:19
y
x
0
1
2
3
4
5
6
0
5
10
15
20
25

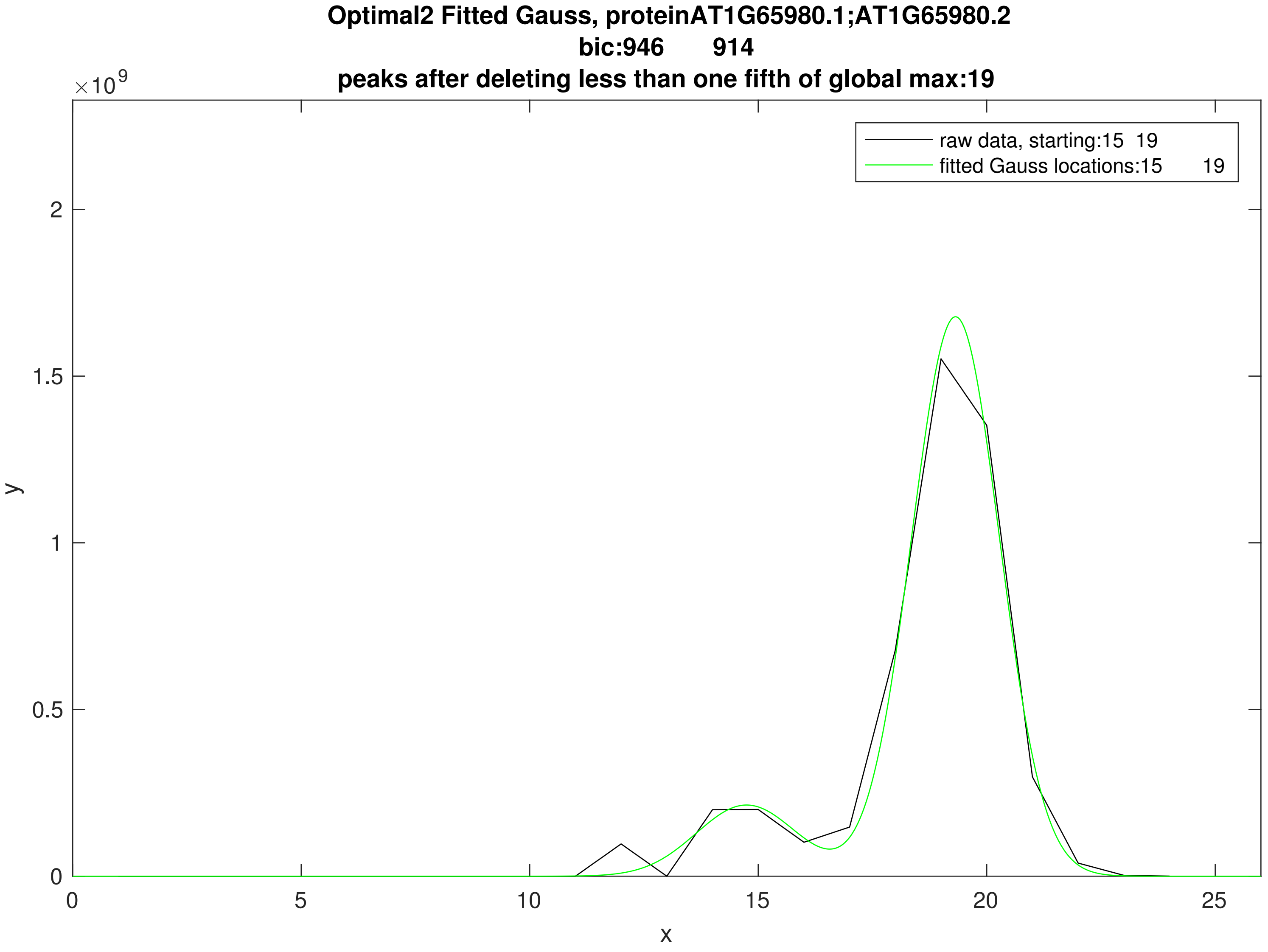

Optimal2 Fitted Gauss, proteinAT1G65980.1;AT1G65980.2
bic:946 914
peaks after deleting less than one fifth of global max:19
×10^9
raw data, starting:15 19
fitted Gauss locations:15 19
y
x
0
0.5
1
1.5
2
0
5
10
15
20
25

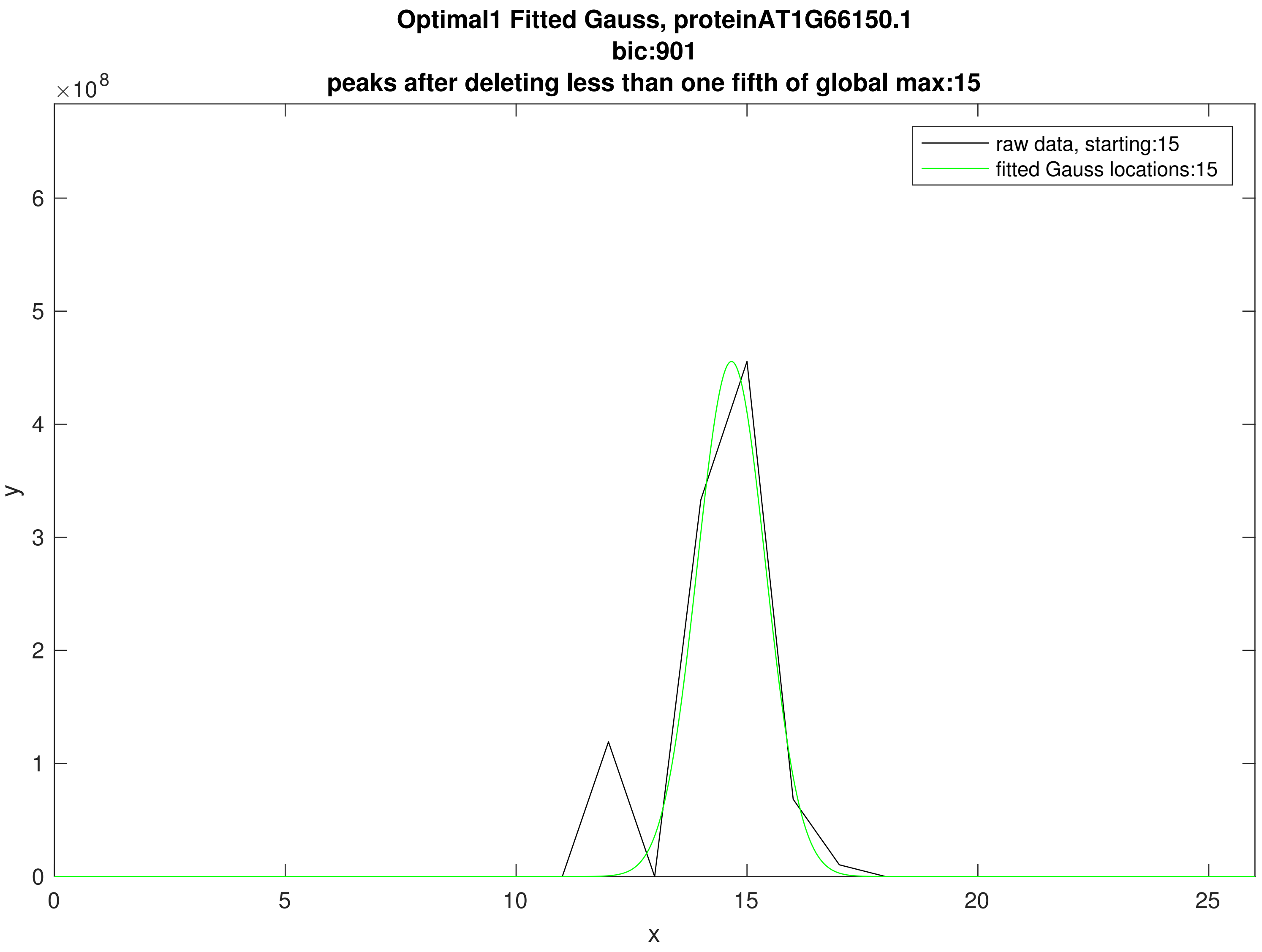

Optimal1 Fitted Gauss, proteinAT1G66150.1
bic:901
peaks after deleting less than one fifth of global max:15
×10^8
raw data, starting:15
fitted Gauss locations:15
y
x
0
1
2
3
4
5
6
0
5
10
15
20
25

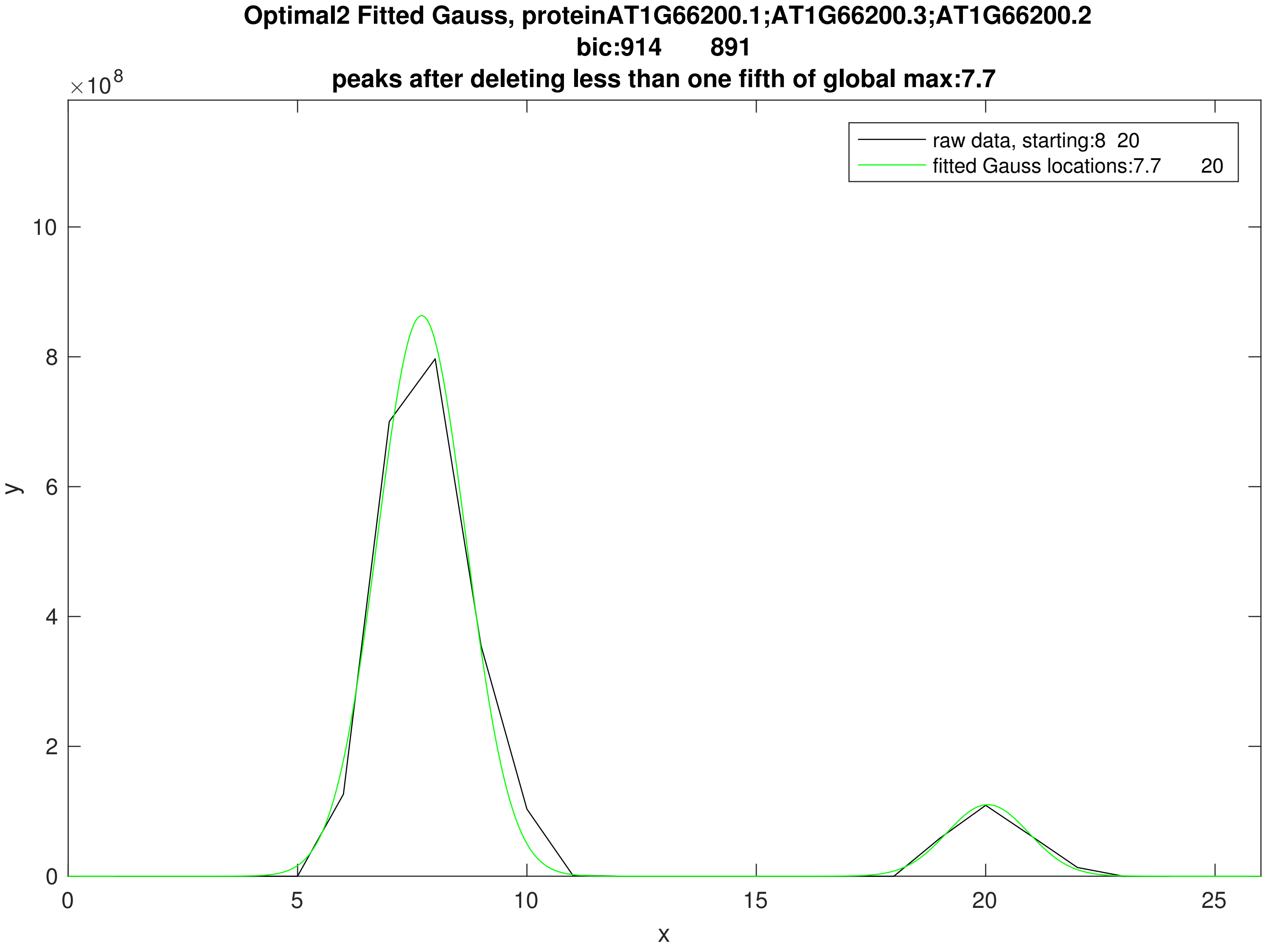

Optimal2 Fitted Gauss, proteinAT1G66200.1;AT1G66200.3;AT1G66200.2
bic:914 891
peaks after deleting less than one fifth of global max:7.7
×10^8
raw data, starting:8 20
fitted Gauss locations:7.7 20
y
x
0
2
4
6
8
10
5
15
20
25

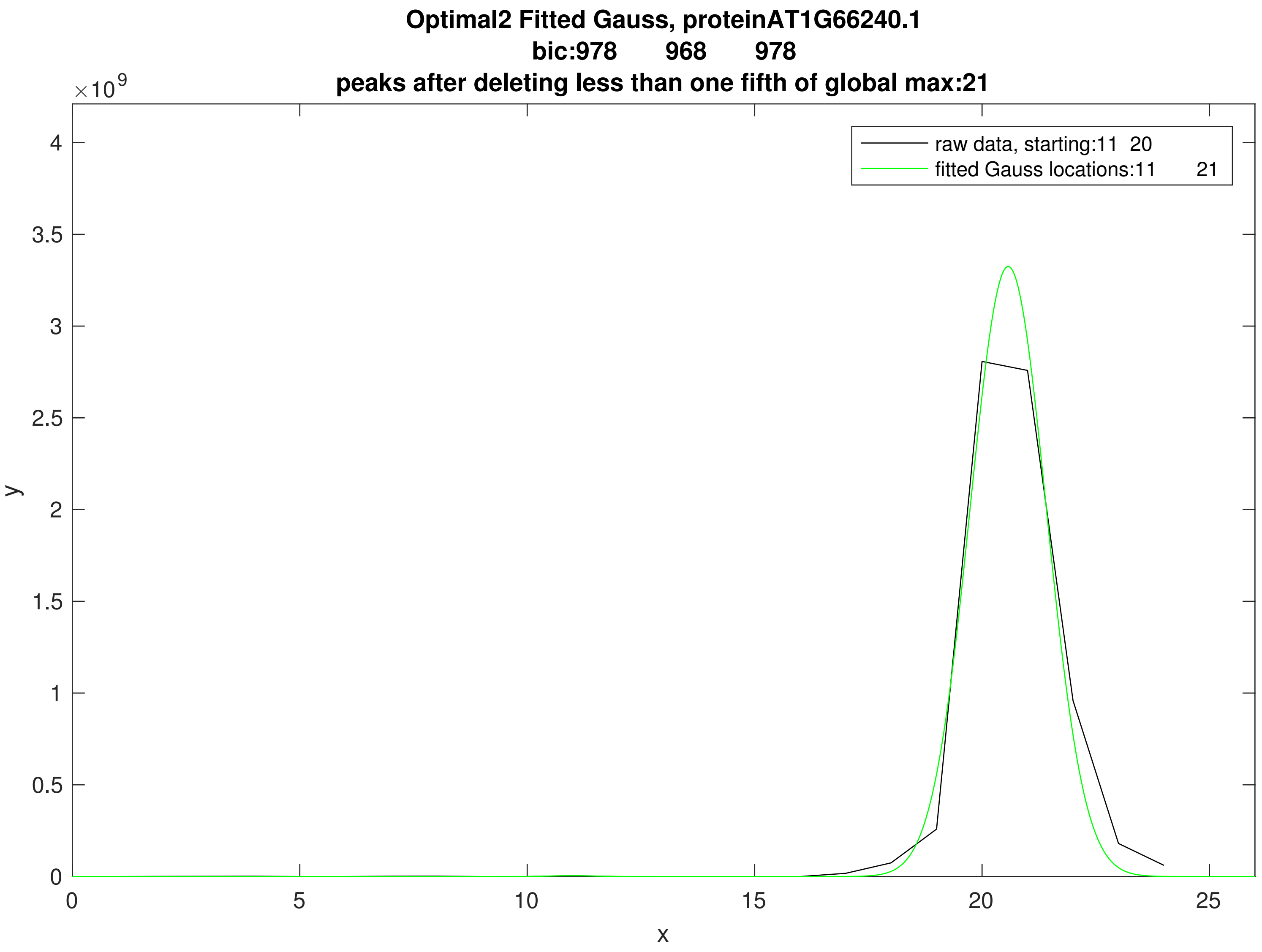
Optimal2 Fitted Gauss, proteinAT1G66240.1
bic:978 968 978
peaks after deleting less than one fifth of global max:21
×10^9
4
3.5
3
2.5
2
1.5
1
0.5
0
y
0
5
10
15
20
25
x
raw data, starting:11 20
fitted Gauss locations:11 21

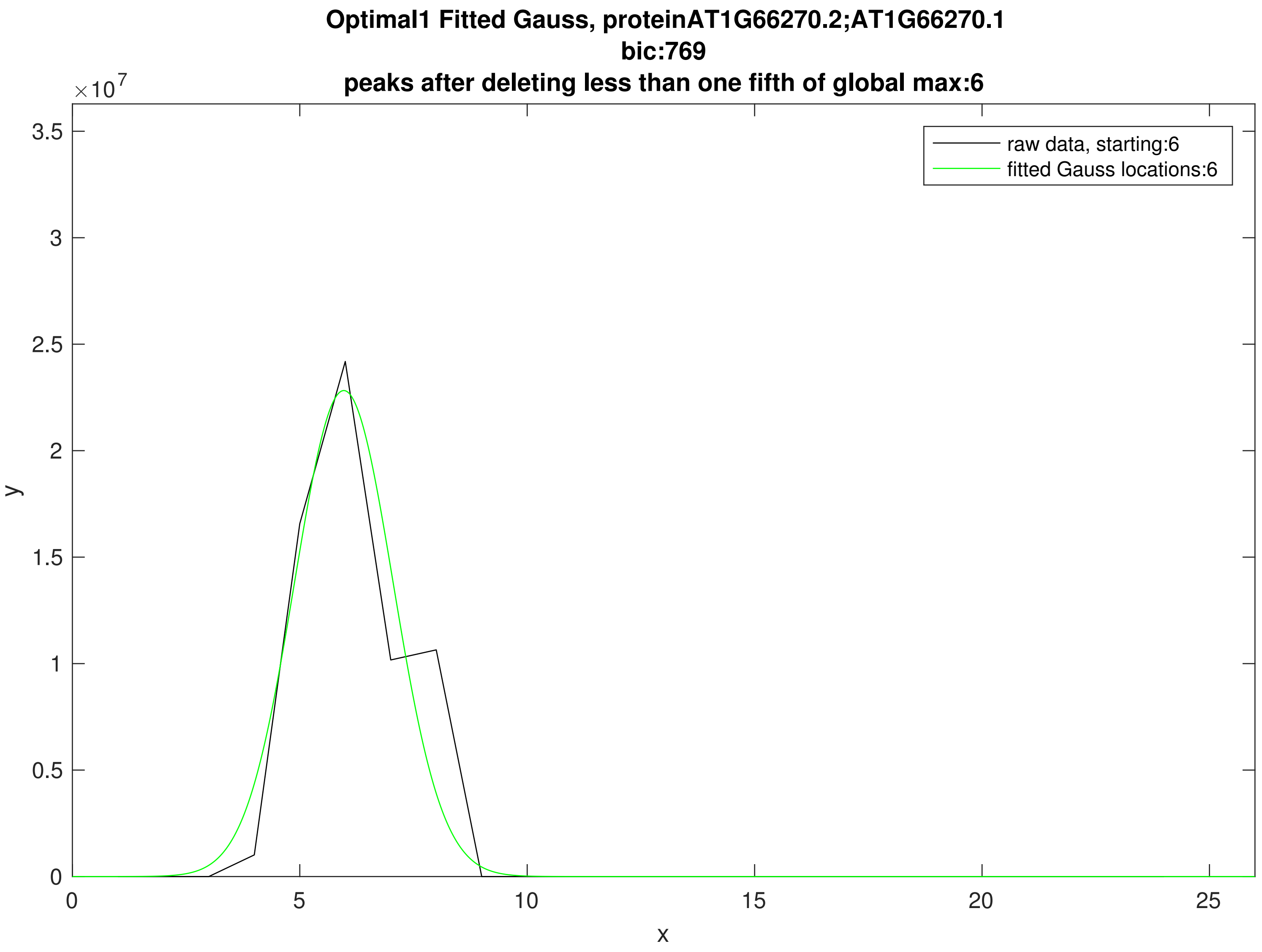

Optimal1 Fitted Gauss, proteinAT1G66270.2;AT1G66270.1
bic:769
peaks after deleting less than one fifth of global max:6
×10^7
raw data, starting:6
fitted Gauss locations:6
y
x
0
0.5
1
1.5
2
2.5
3
3.5
0
5
10
15
20
25

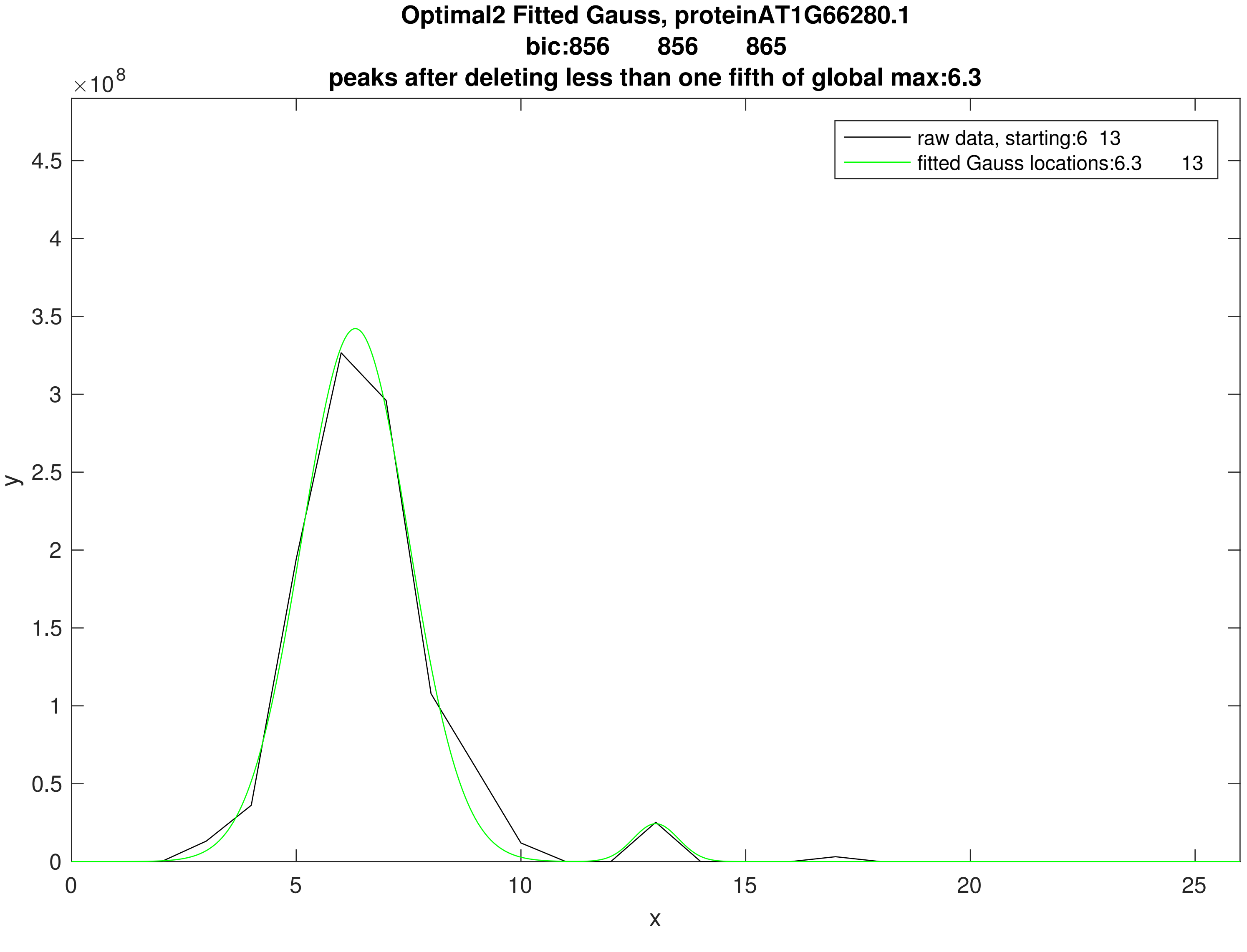

Optimal2 Fitted Gauss, proteinAT1G66280.1
bic:856 856 865
peaks after deleting less than one fifth of global max:6.3
×10^8
raw data, starting:6 13
fitted Gauss locations:6.3 13
y
x
0
0.5
1
1.5
2
2.5
3
3.5
4
4.5
5
10
15
20
25

**Optimal1 Fitted Gauss, proteinAT1G66400.1**
**bic:703**
**peaks after deleting less than one fifth of global max:19**

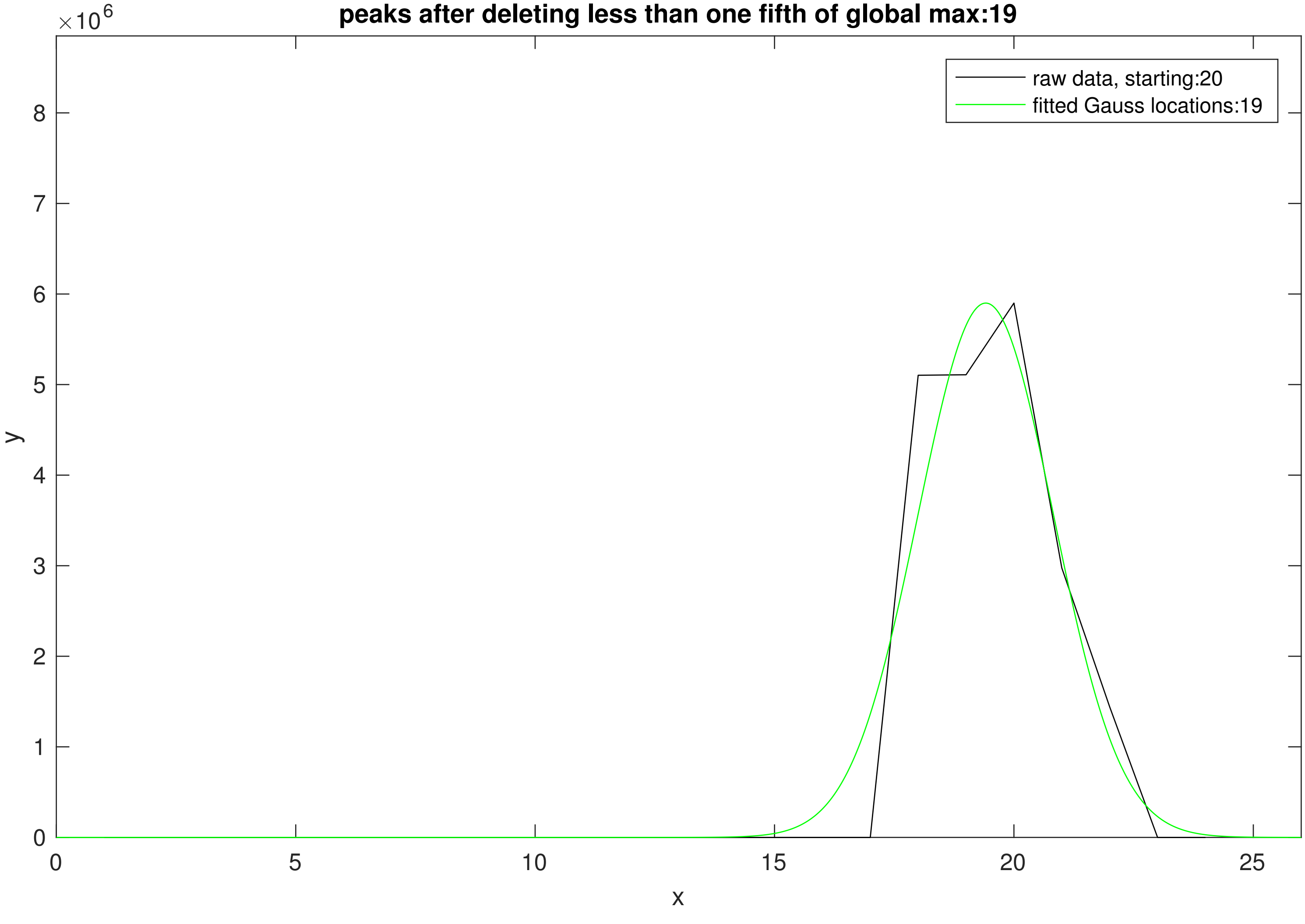

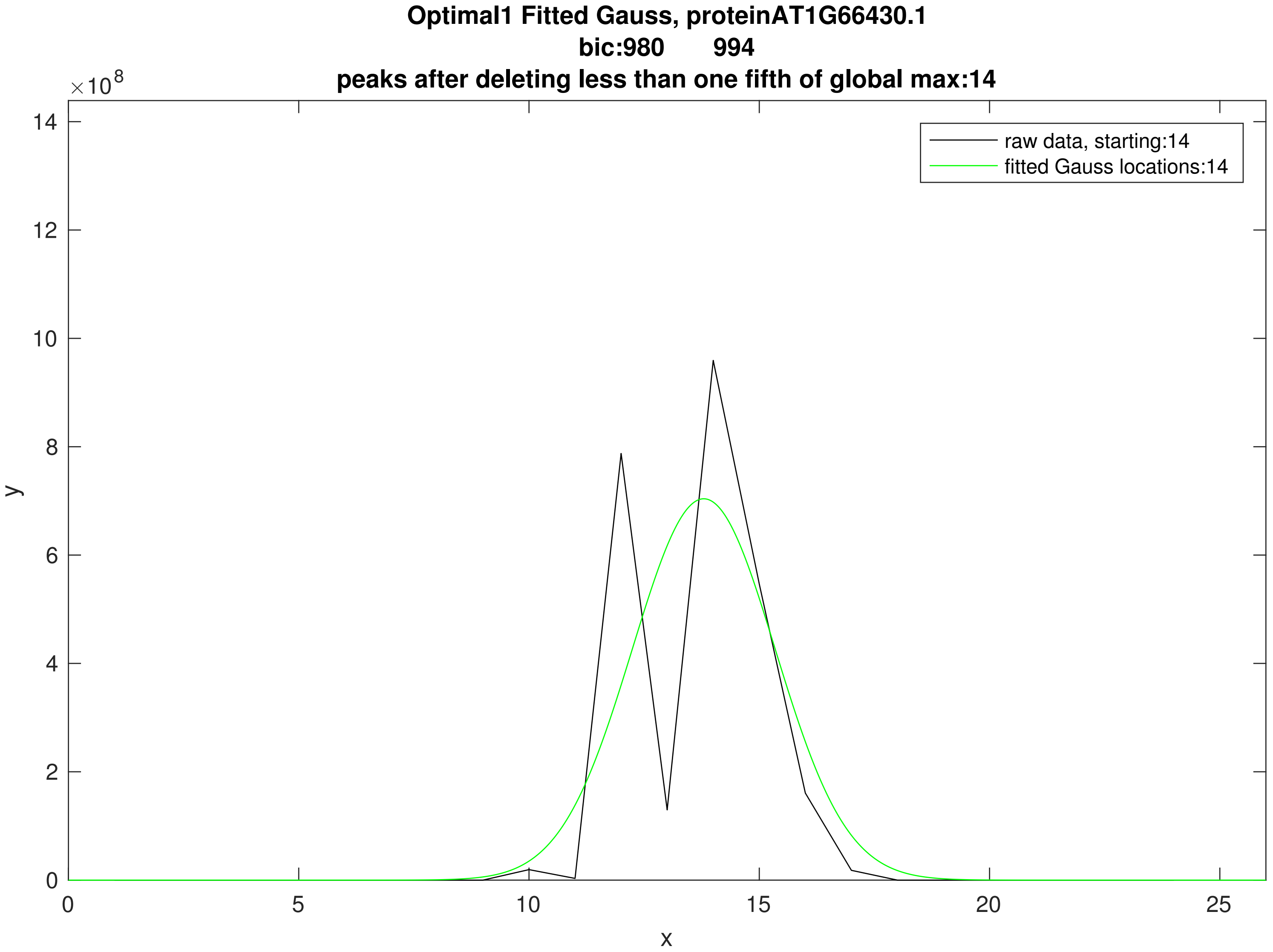

Optimal1 Fitted Gauss, proteinAT1G66430.1
bic:980 994
peaks after deleting less than one fifth of global max:14
×10^8
raw data, starting:14
fitted Gauss locations:14
y
x
0
2
4
6
8
10
12
14
0
5
10
15
20
25

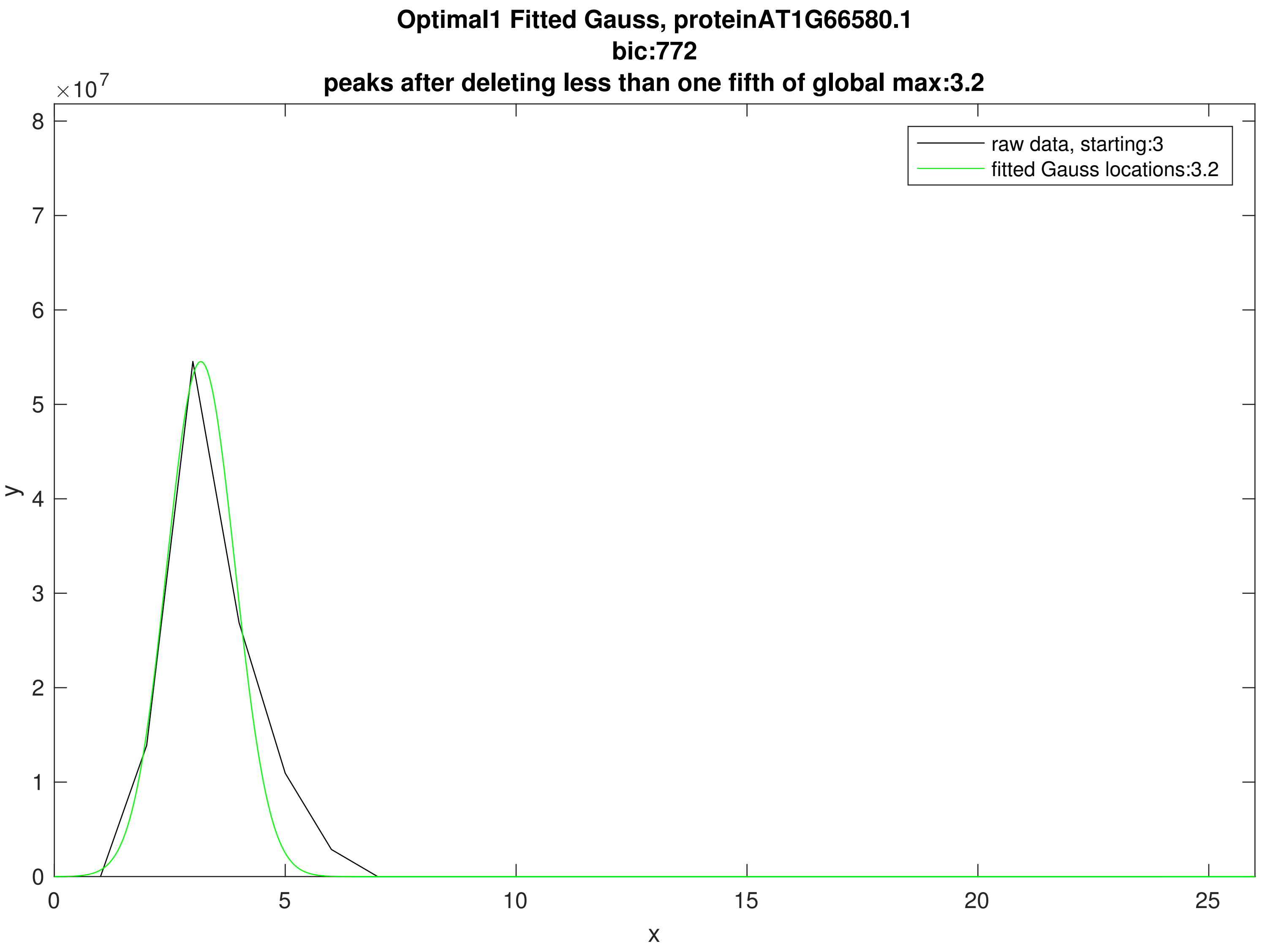

Optimal1 Fitted Gauss, proteinAT1G66580.1
bic:772
peaks after deleting less than one fifth of global max:3.2
×10^7
raw data, starting:3
fitted Gauss locations:3.2
y
x
0
1
2
3
4
5
6
7
8
0
5
10
15
20
25

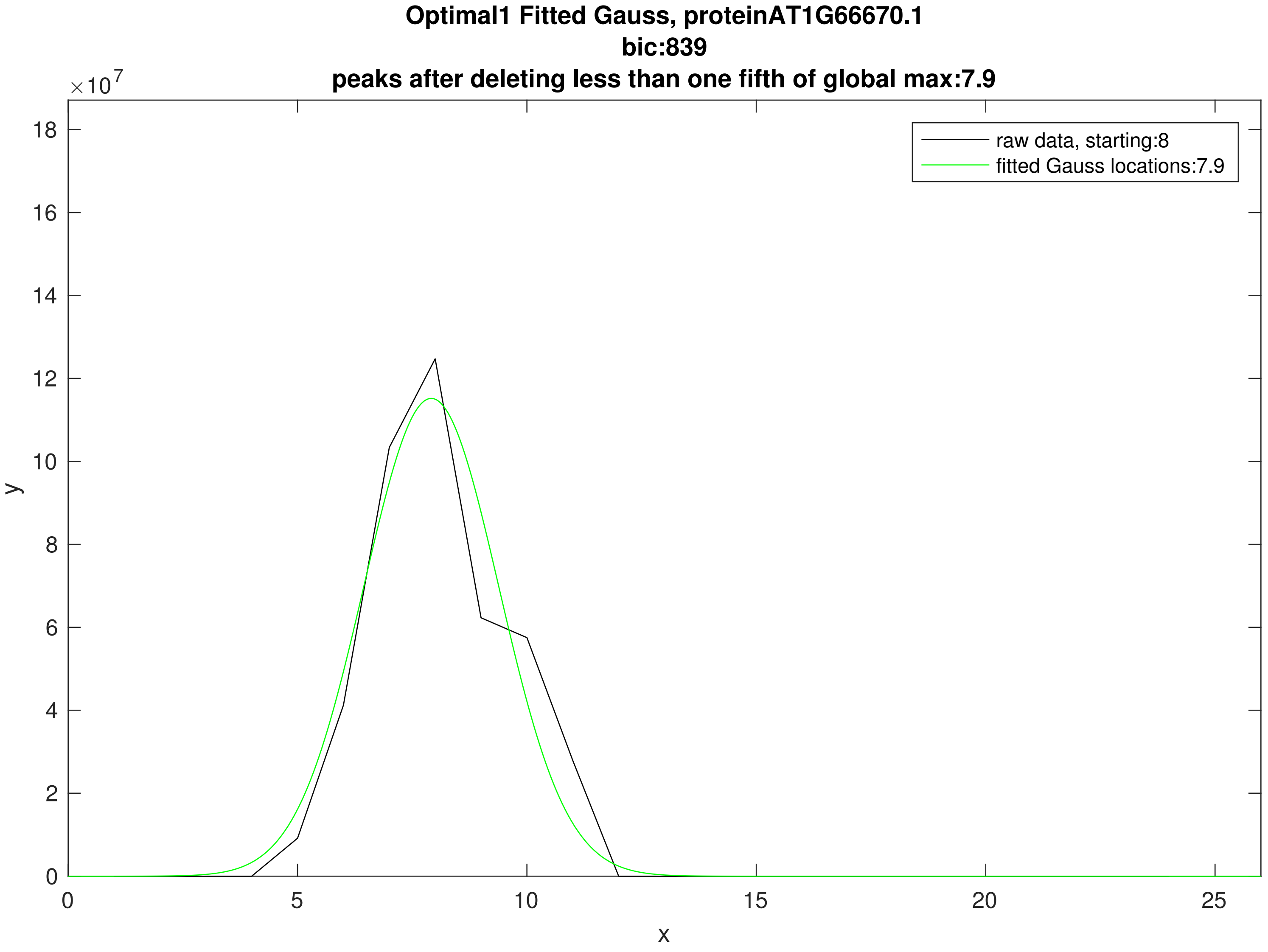

Optimal1 Fitted Gauss, proteinAT1G66670.1
bic:839
peaks after deleting less than one fifth of global max:7.9
×10^7
18
16
14
12
10
8
6
4
2
0
y
0
5
10
15
20
25
x
raw data, starting:8
fitted Gauss locations:7.9

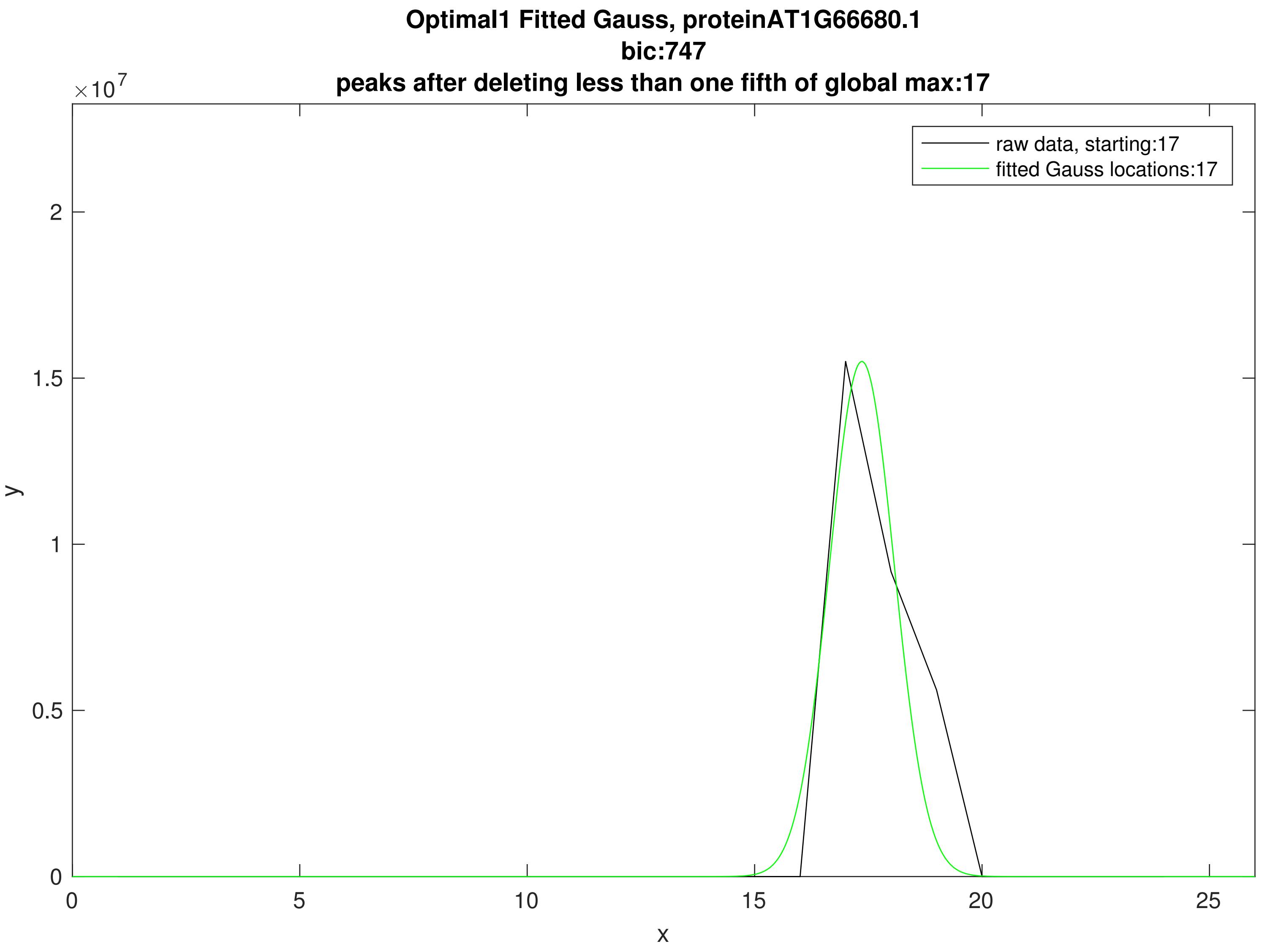

Optimal1 Fitted Gauss, proteinAT1G66680.1
bic:747
peaks after deleting less than one fifth of global max:17
×10^7
raw data, starting:17
fitted Gauss locations:17
2
1.5
1
0.5
0
y
0
5
10
15
20
25
x

Optimal1 Fitted Gauss, proteinAT1G67090.1
bic:1.02e+03  1.03e+03  1.04e+03
peaks after deleting less than one fifth of global max:6.8

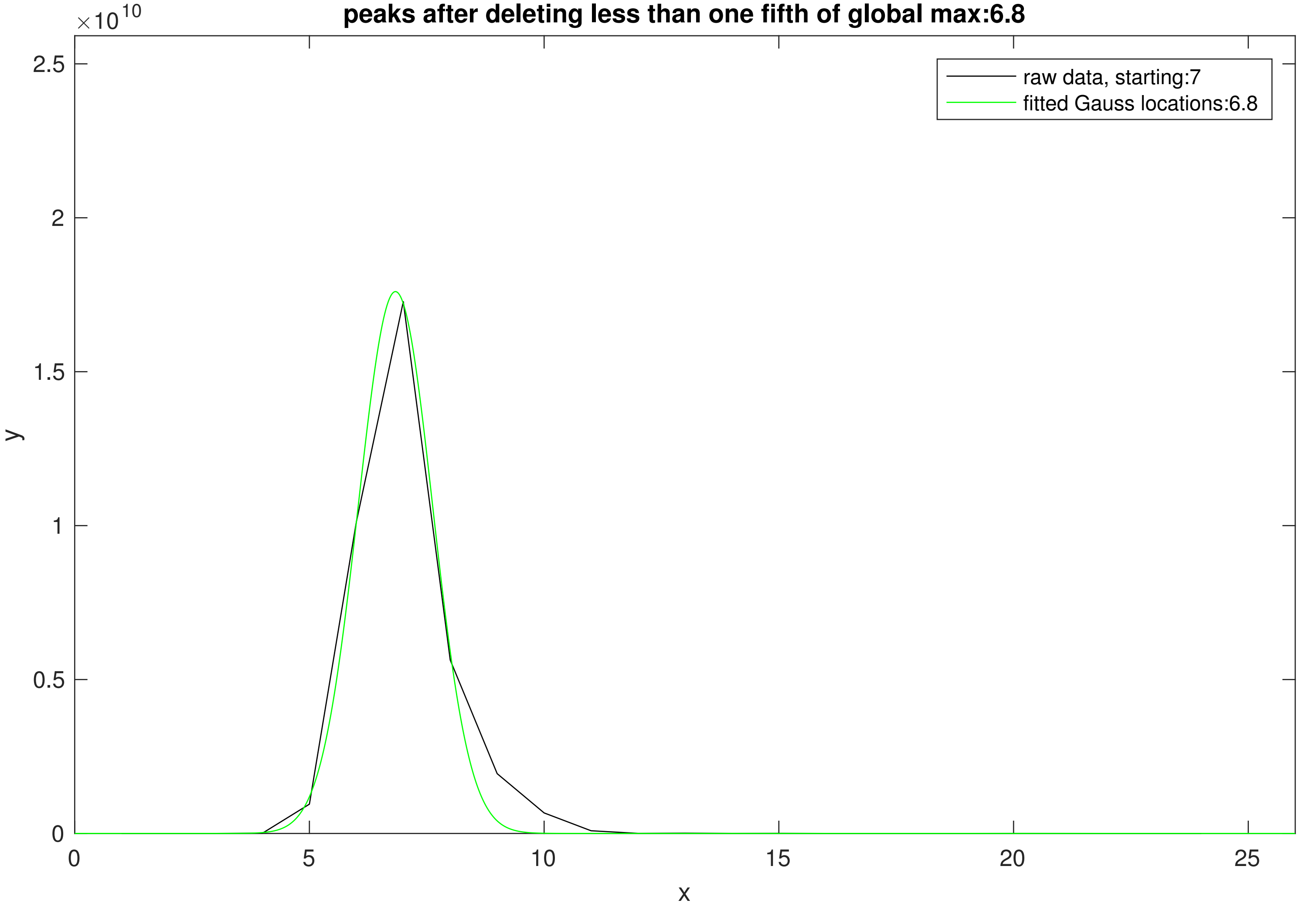

**Optimal1 Fitted Gauss, proteinAT1G67148.1**
**bic:635**
**peaks after deleting less than one fifth of global max:22**

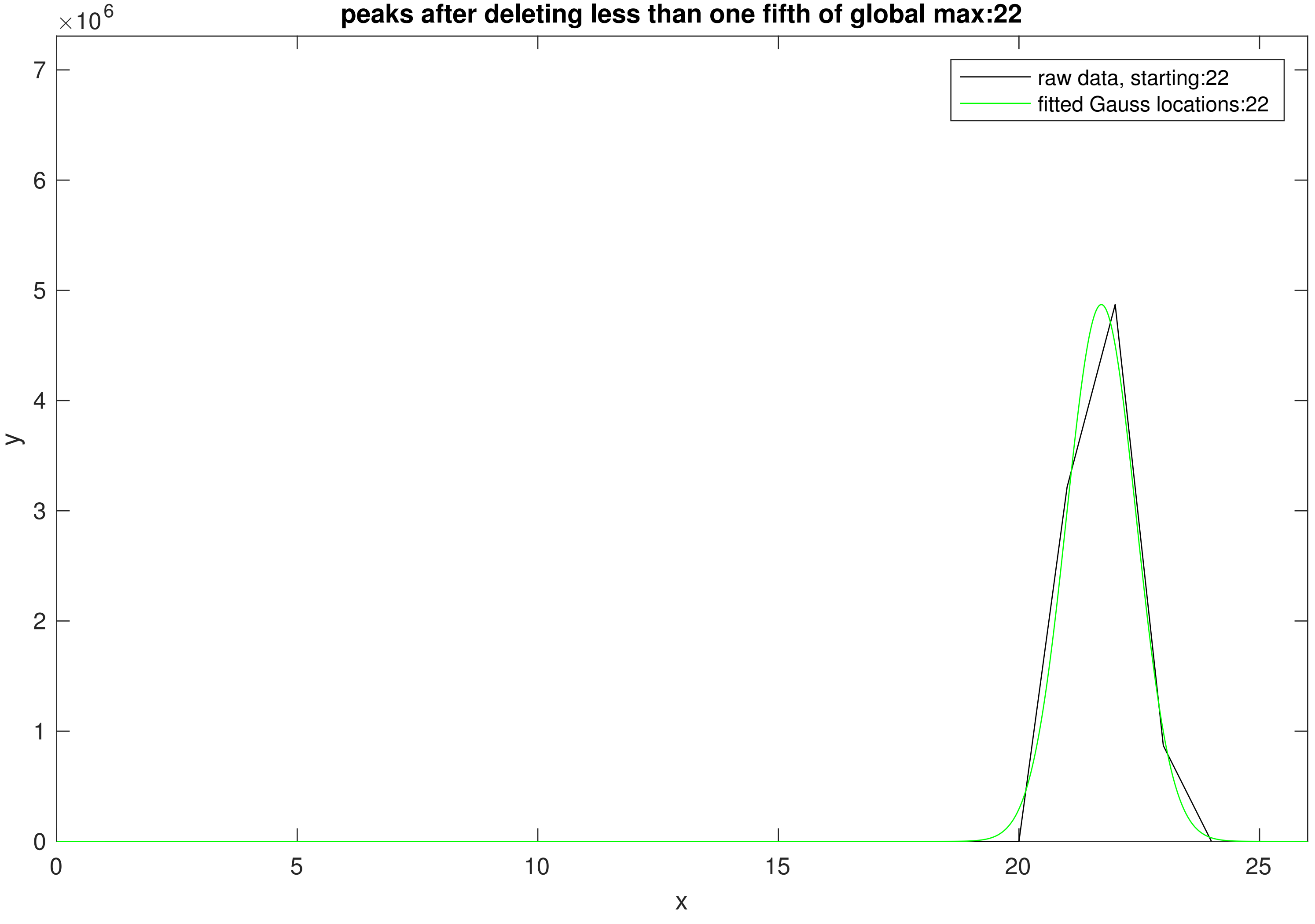

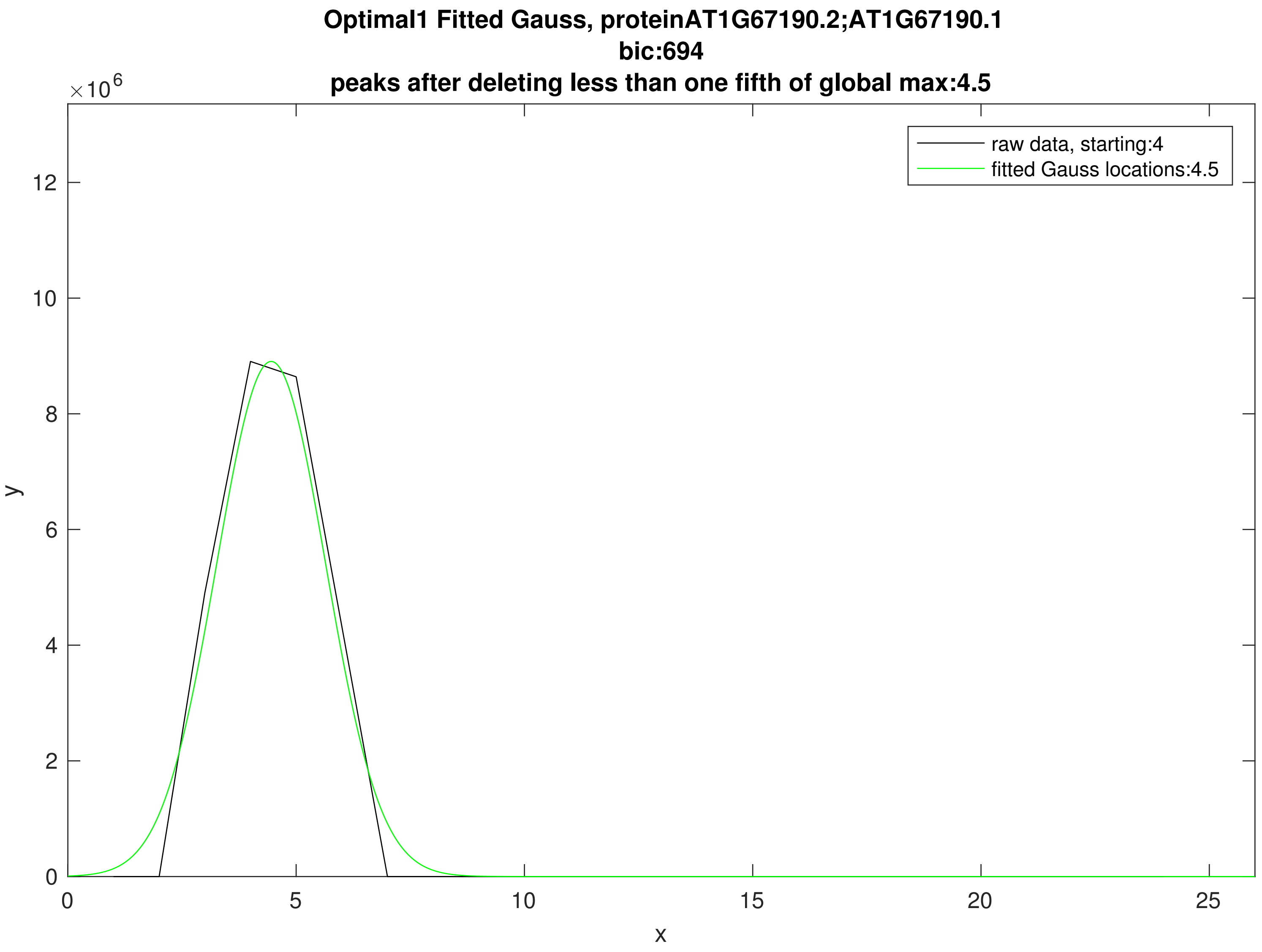

Optimal1 Fitted Gauss, proteinAT1G67190.2;AT1G67190.1
bic:694
peaks after deleting less than one fifth of global max:4.5
×10^6
raw data, starting:4
fitted Gauss locations:4.5
y
x
0
2
4
6
8
10
12
0
5
10
15
20
25

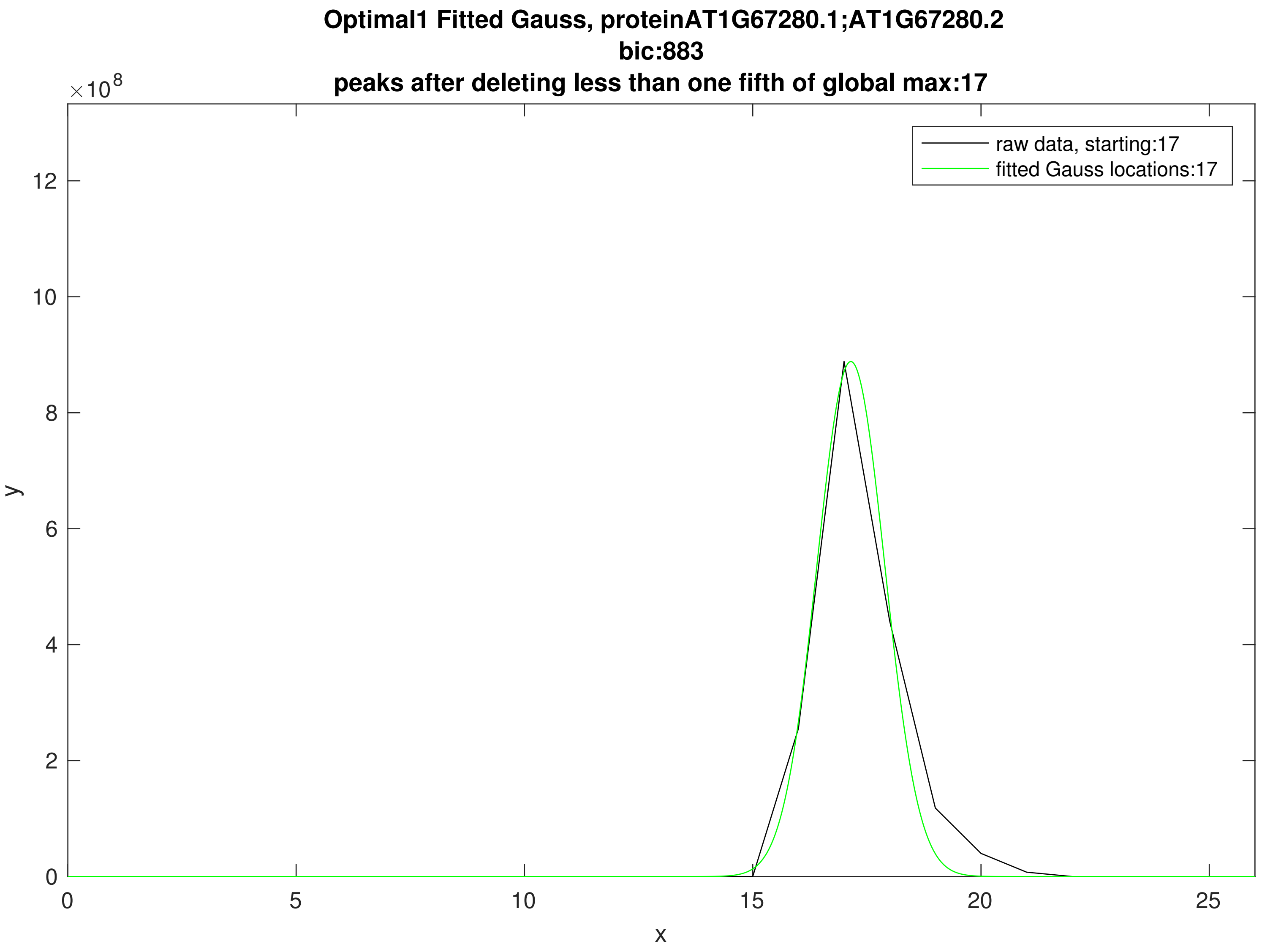

Optimal1 Fitted Gauss, proteinAT1G67280.1;AT1G67280.2
bic:883
peaks after deleting less than one fifth of global max:17
×10^8
raw data, starting:17
fitted Gauss locations:17
y
x
0
2
4
6
8
10
12
5
15
20
25

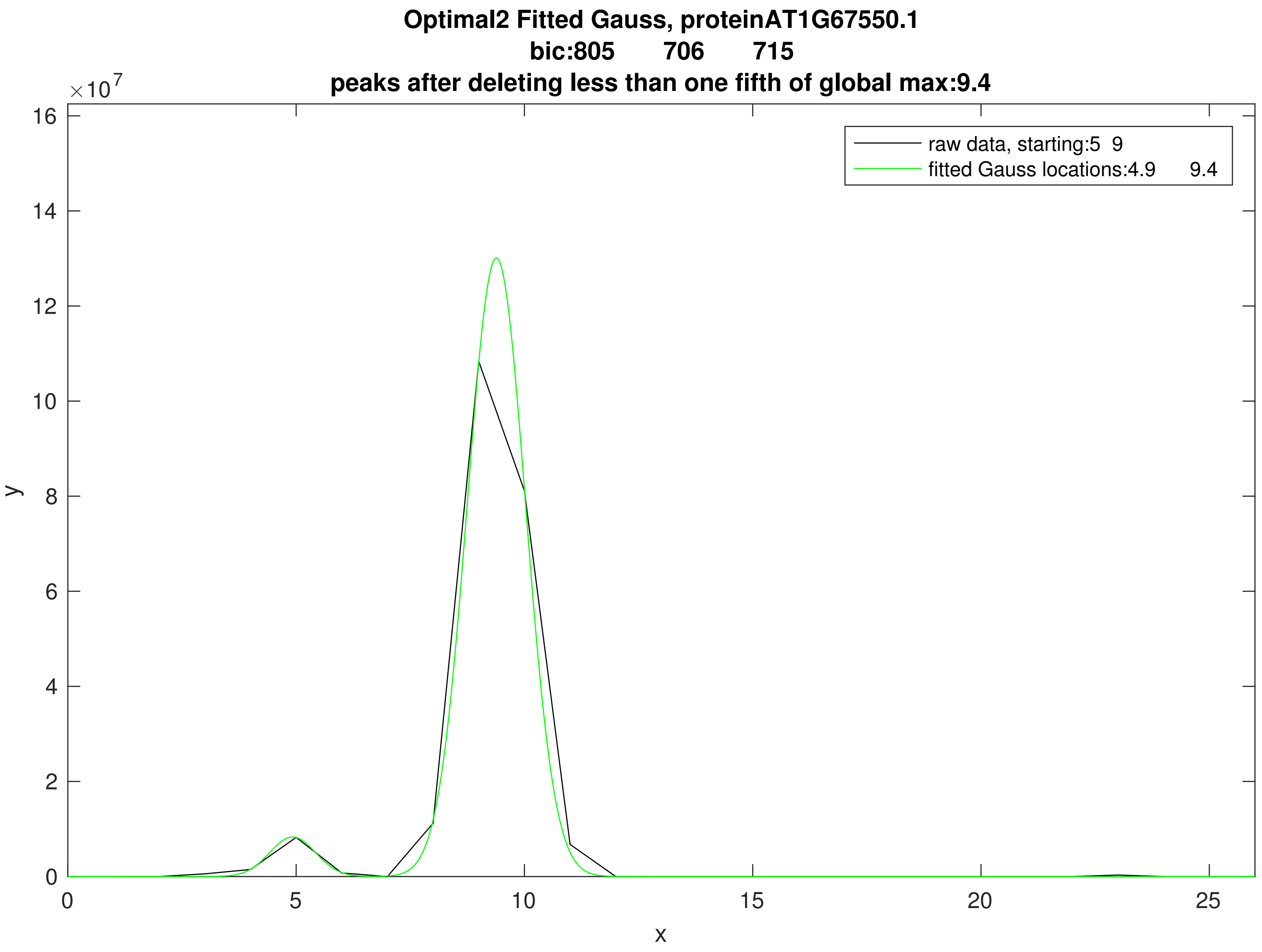
Optimal2 Fitted Gauss, proteinAT1G67550.1
bic:805 706 715
peaks after deleting less than one fifth of global max:9.4
×10^7
raw data, starting:5 9
fitted Gauss locations:4.9 9.4
y
x
0
2
4
6
8
10
12
14
16
0
5
10
15
20
25

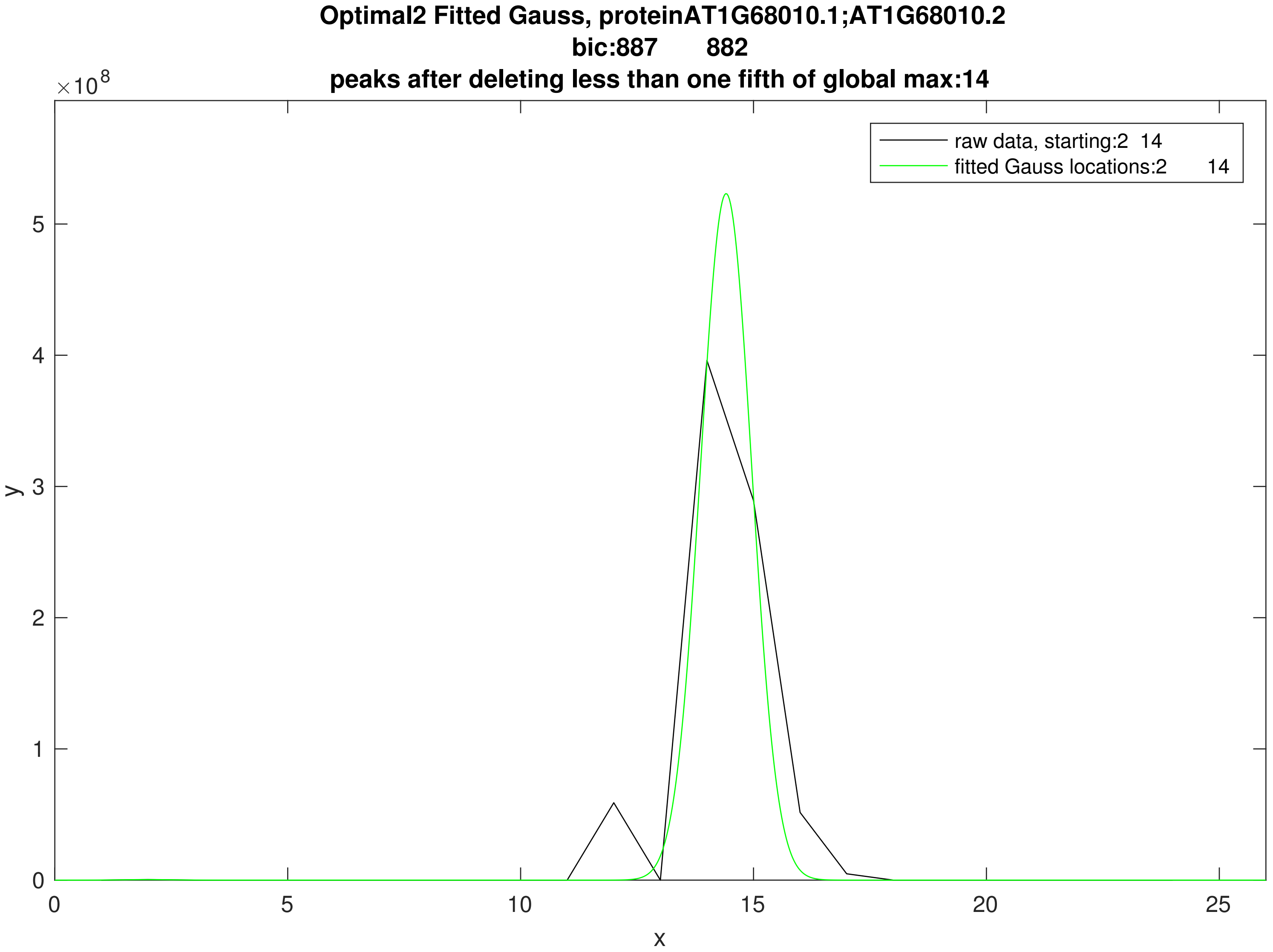
Optimal2 Fitted Gauss, proteinAT1G68010.1;AT1G68010.2
bic:887 882
peaks after deleting less than one fifth of global max:14
×10^8
raw data, starting:2 14
fitted Gauss locations:2 14
y
x
0
1
2
3
4
5
10
15
20
25

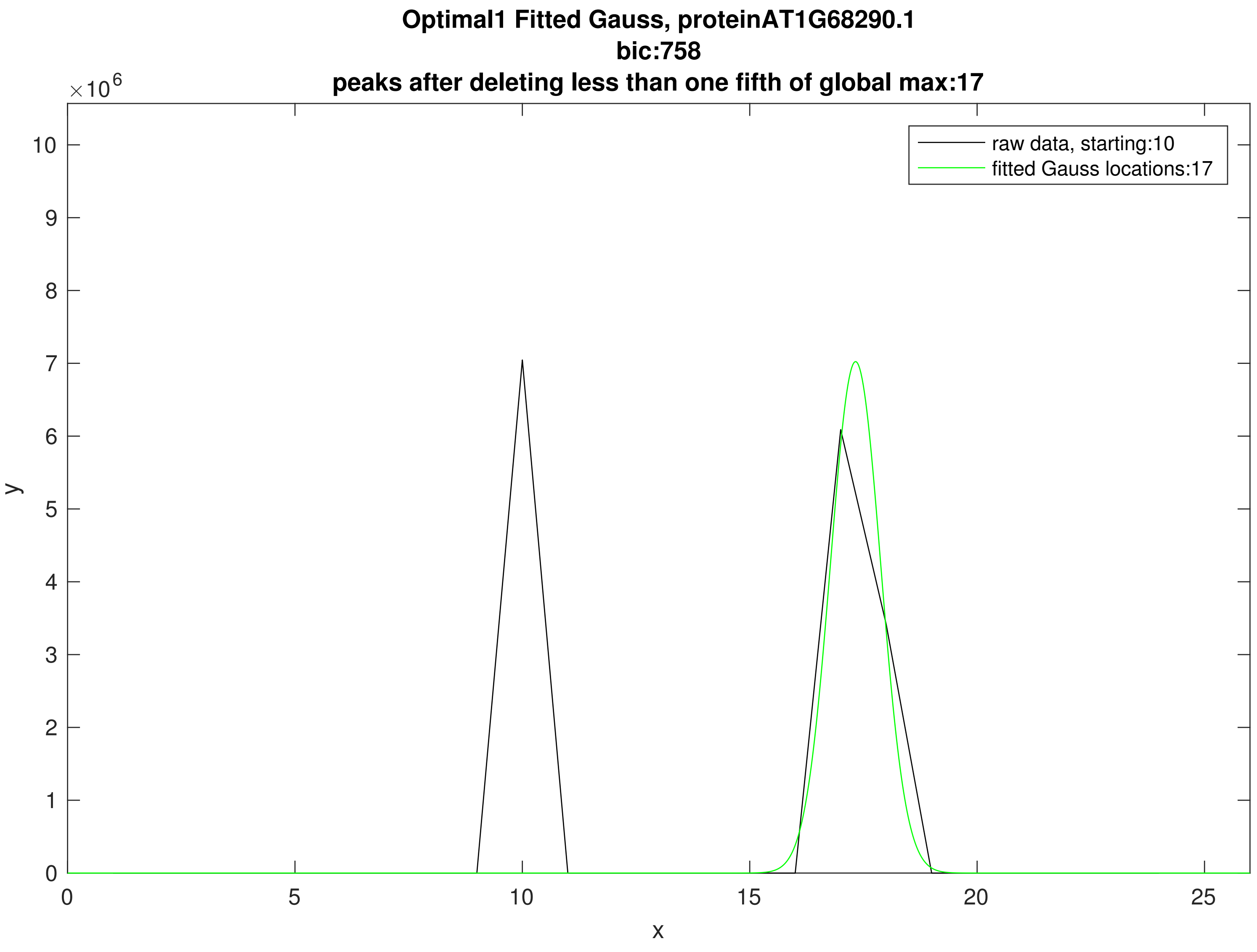
Optimal1 Fitted Gauss, proteinAT1G68290.1
bic:758
peaks after deleting less than one fifth of global max:17
×10^6
raw data, starting:10
fitted Gauss locations:17
y
x
0
1
2
3
4
5
6
7
8
9
10
0
5
10
15
20
25

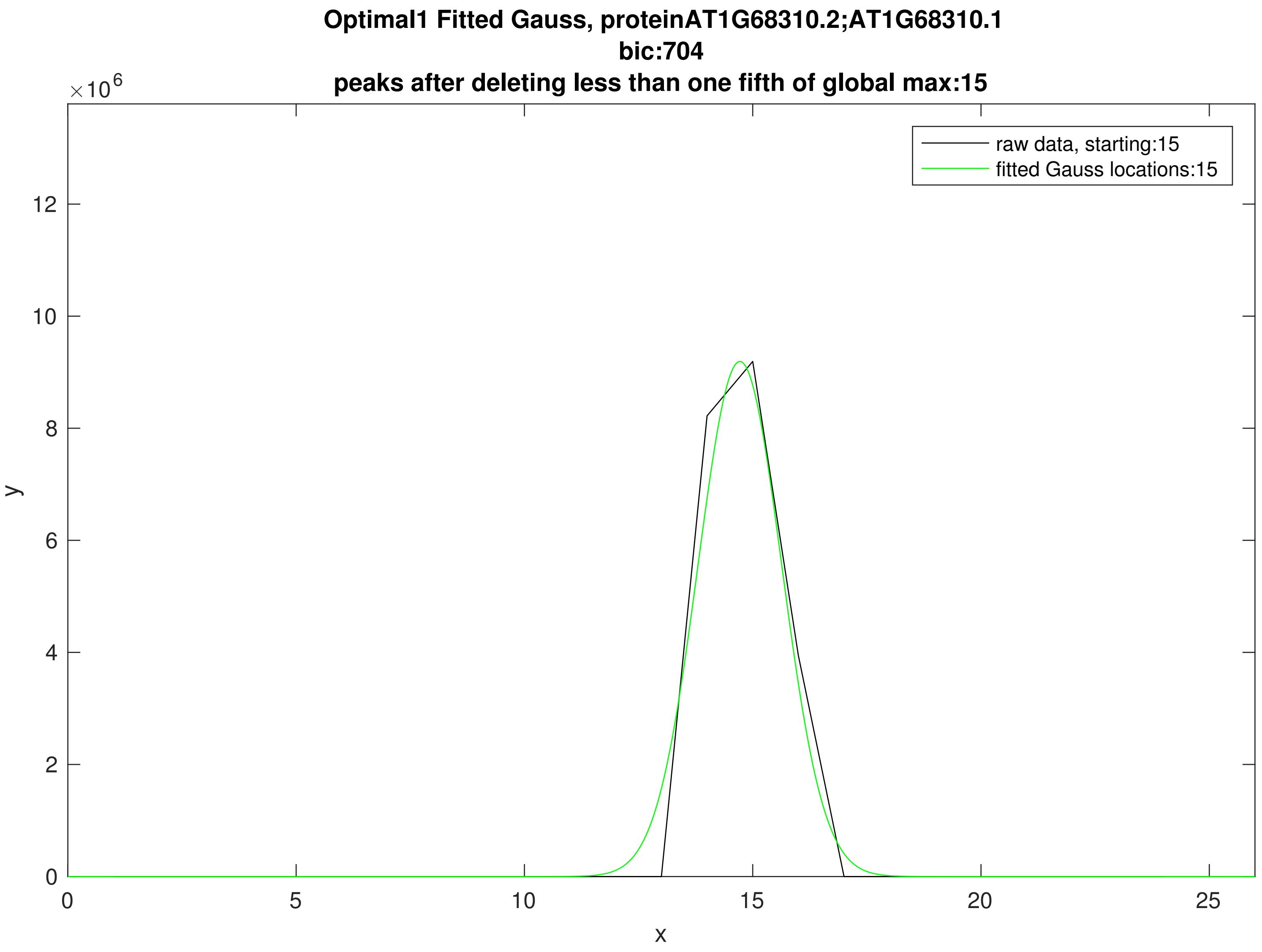

Optimal1 Fitted Gauss, proteinAT1G68310.2;AT1G68310.1
bic:704
peaks after deleting less than one fifth of global max:15
×10^6
raw data, starting:15
fitted Gauss locations:15
y
x
0
2
4
6
8
10
12
0
5
10
15
20
25

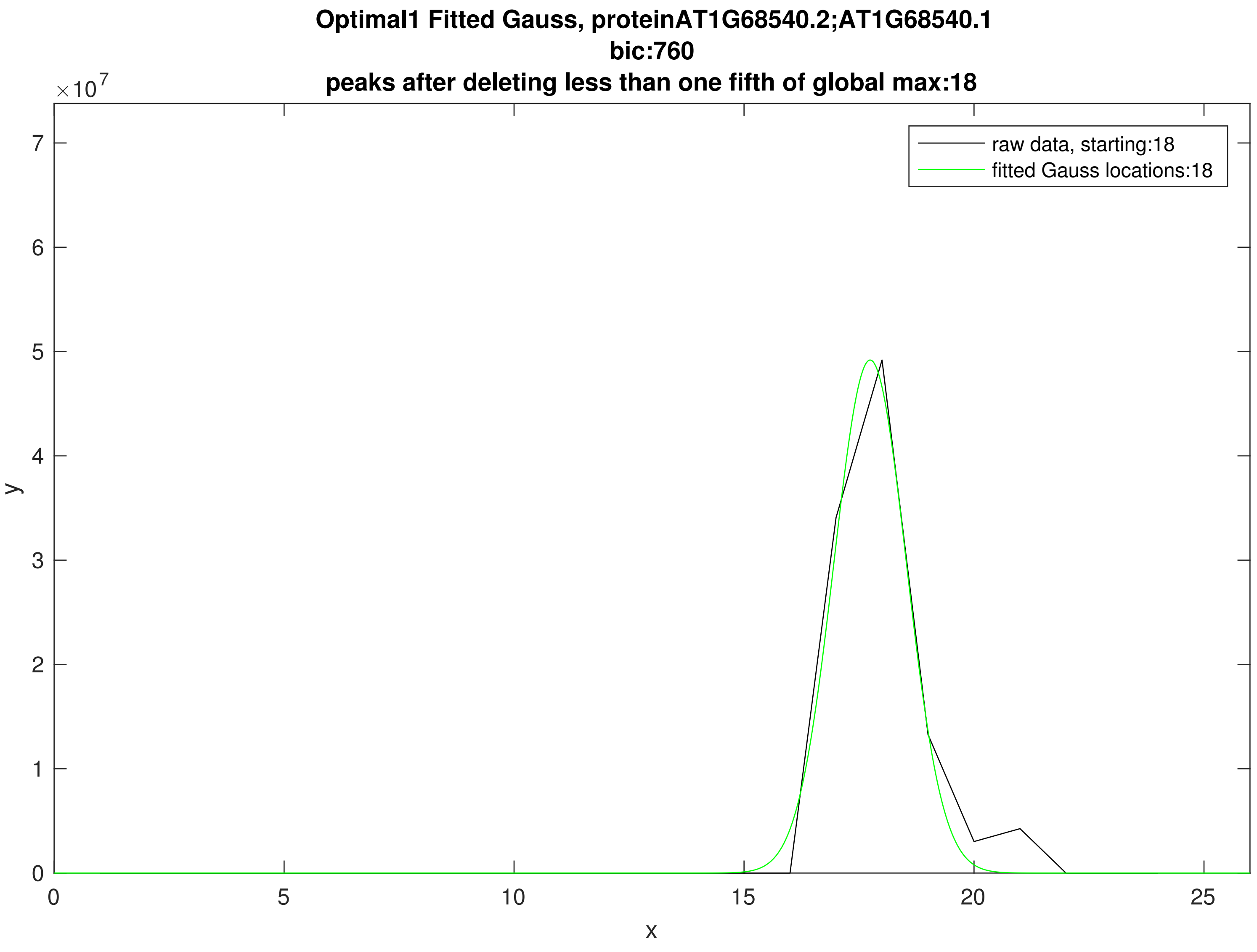

Optimal1 Fitted Gauss, proteinAT1G68540.2;AT1G68540.1
bic:760
peaks after deleting less than one fifth of global max:18
×10^7
raw data, starting:18
fitted Gauss locations:18
y
x
0
1
2
3
4
5
6
7
0
5
10
15
20
25

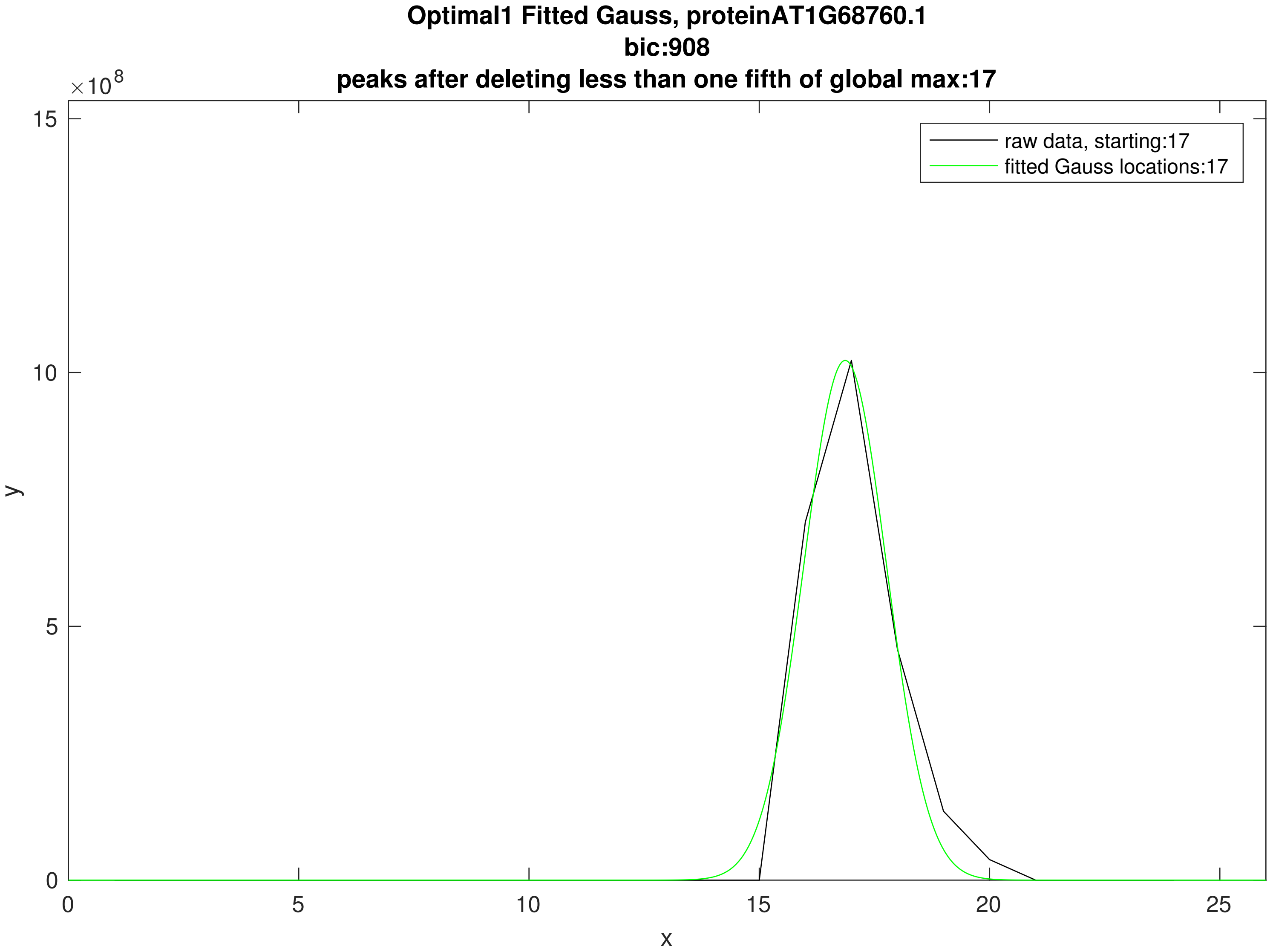
Optimal1 Fitted Gauss, proteinAT1G68760.1
bic:908
peaks after deleting less than one fifth of global max:17
×10^8
15
10
5
0
y
0
5
10
15
20
25
x
raw data, starting:17
fitted Gauss locations:17

**Optimal1 Fitted Gauss, proteinAT1G69295.1;AT1G26450.1**
**bic:802**
**peaks after deleting less than one fifth of global max:17**

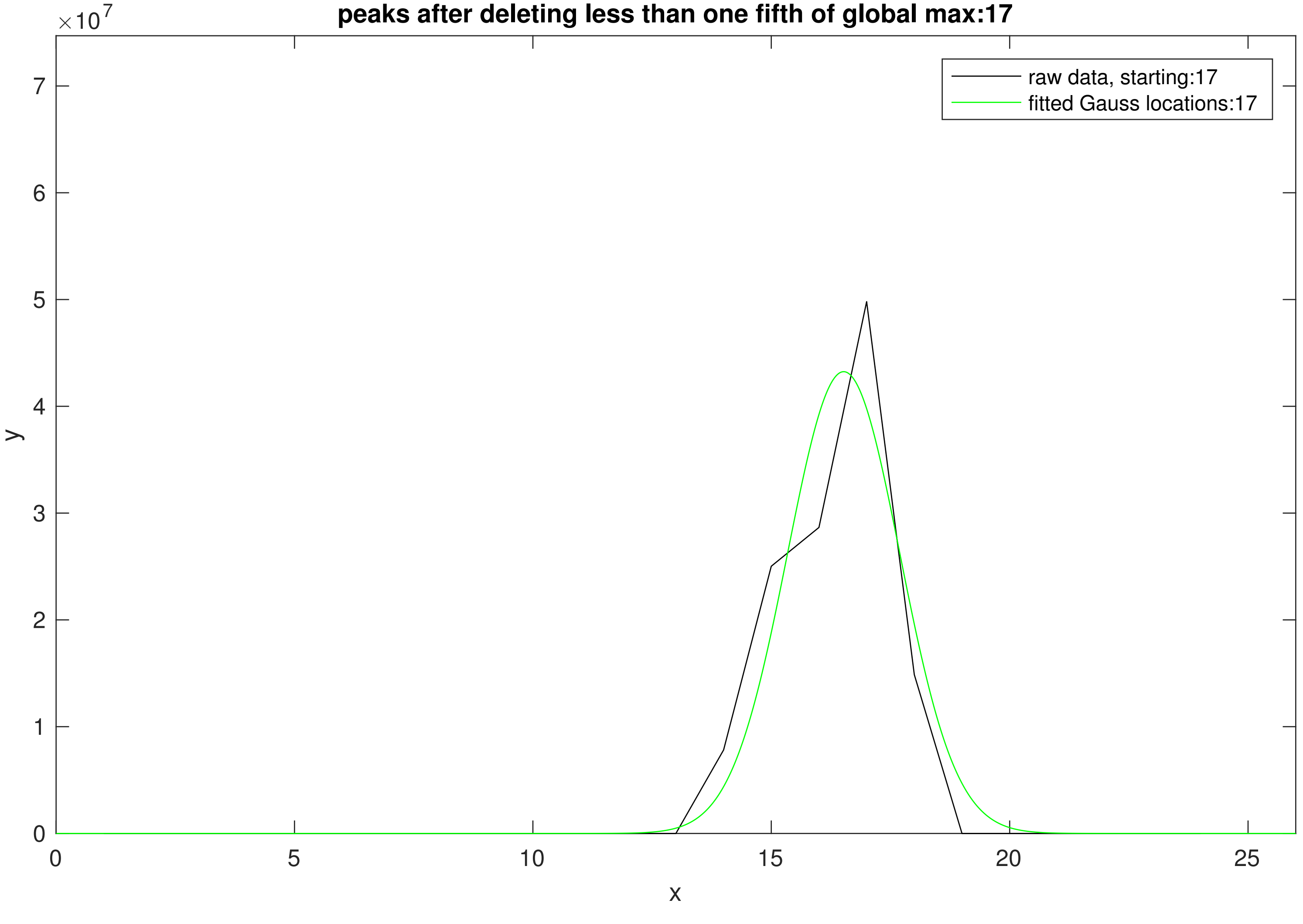

Optimal1 Fitted Gauss, proteinAT1G69340.1
bic:718 777
peaks after deleting less than one fifth of global max:16

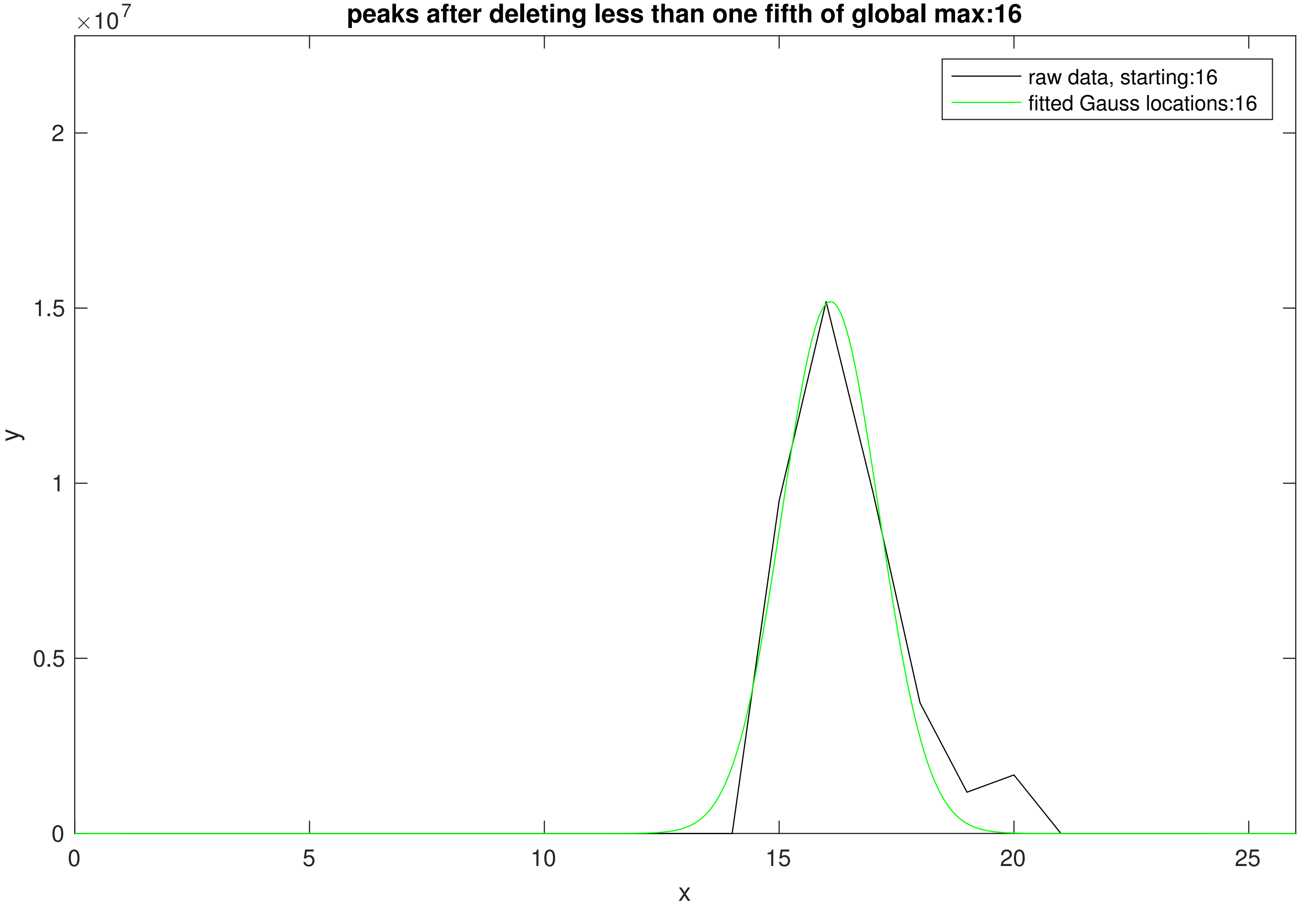

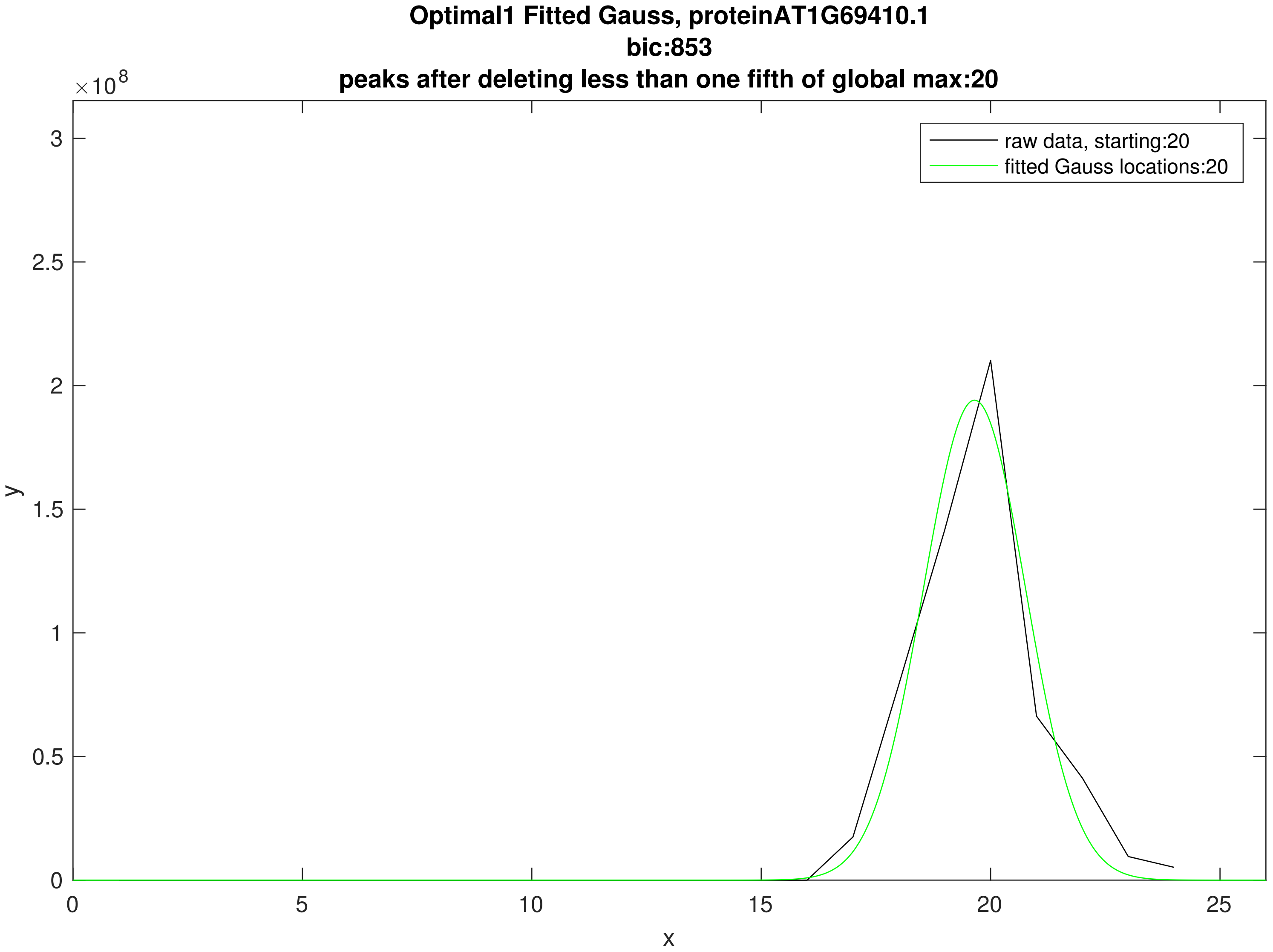
Optimal1 Fitted Gauss, proteinAT1G69410.1
bic:853
peaks after deleting less than one fifth of global max:20
×10^8
raw data, starting:20
fitted Gauss locations:20
y
x
0
0.5
1
1.5
2
2.5
3
0
5
10
15
20
25

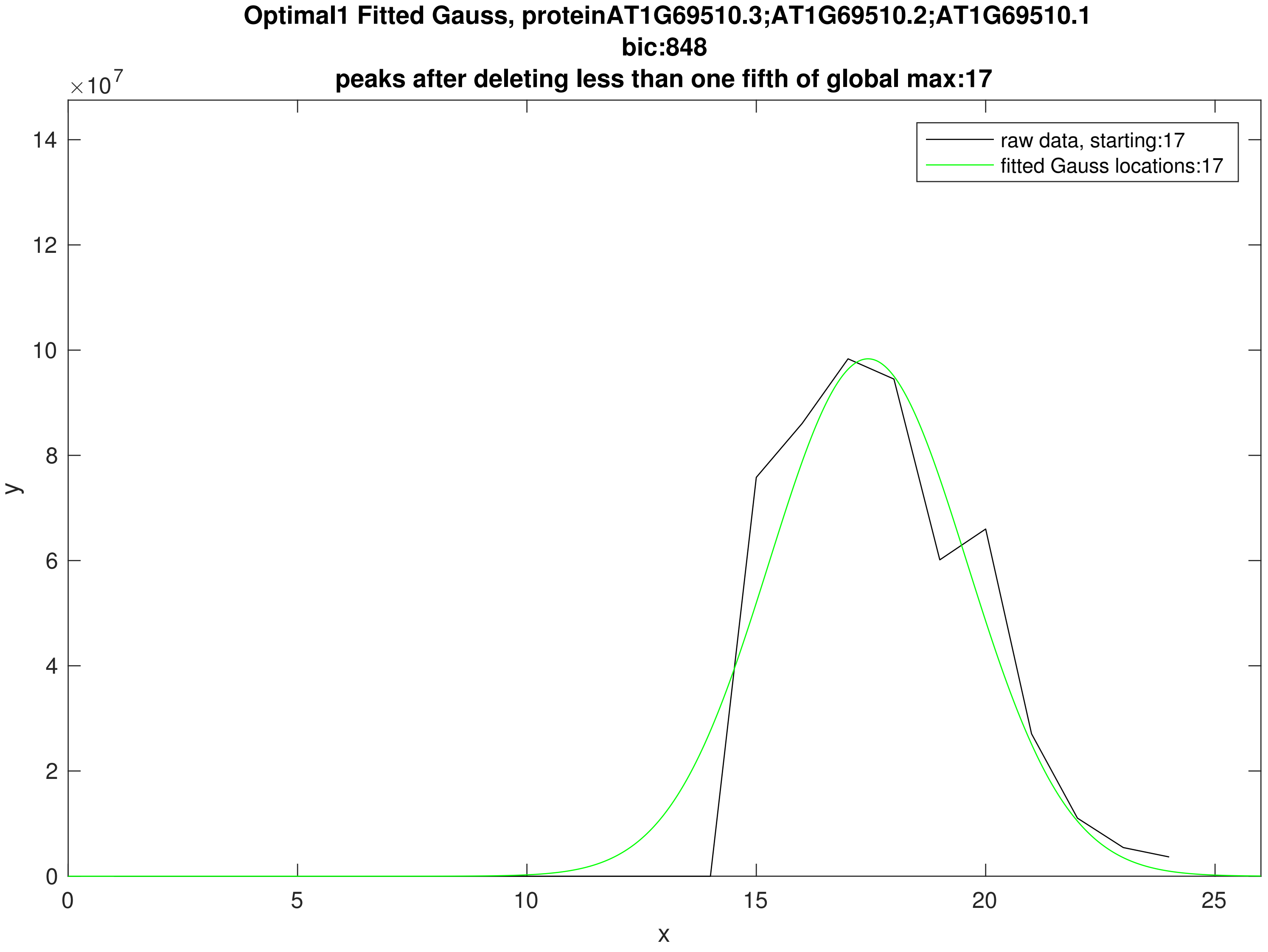

Optimal1 Fitted Gauss, proteinAT1G69510.3;AT1G69510.2;AT1G69510.1
bic:848
peaks after deleting less than one fifth of global max:17
×10^7
raw data, starting:17
fitted Gauss locations:17
y
x
0
2
4
6
8
10
12
14
0
5
10
15
20
25

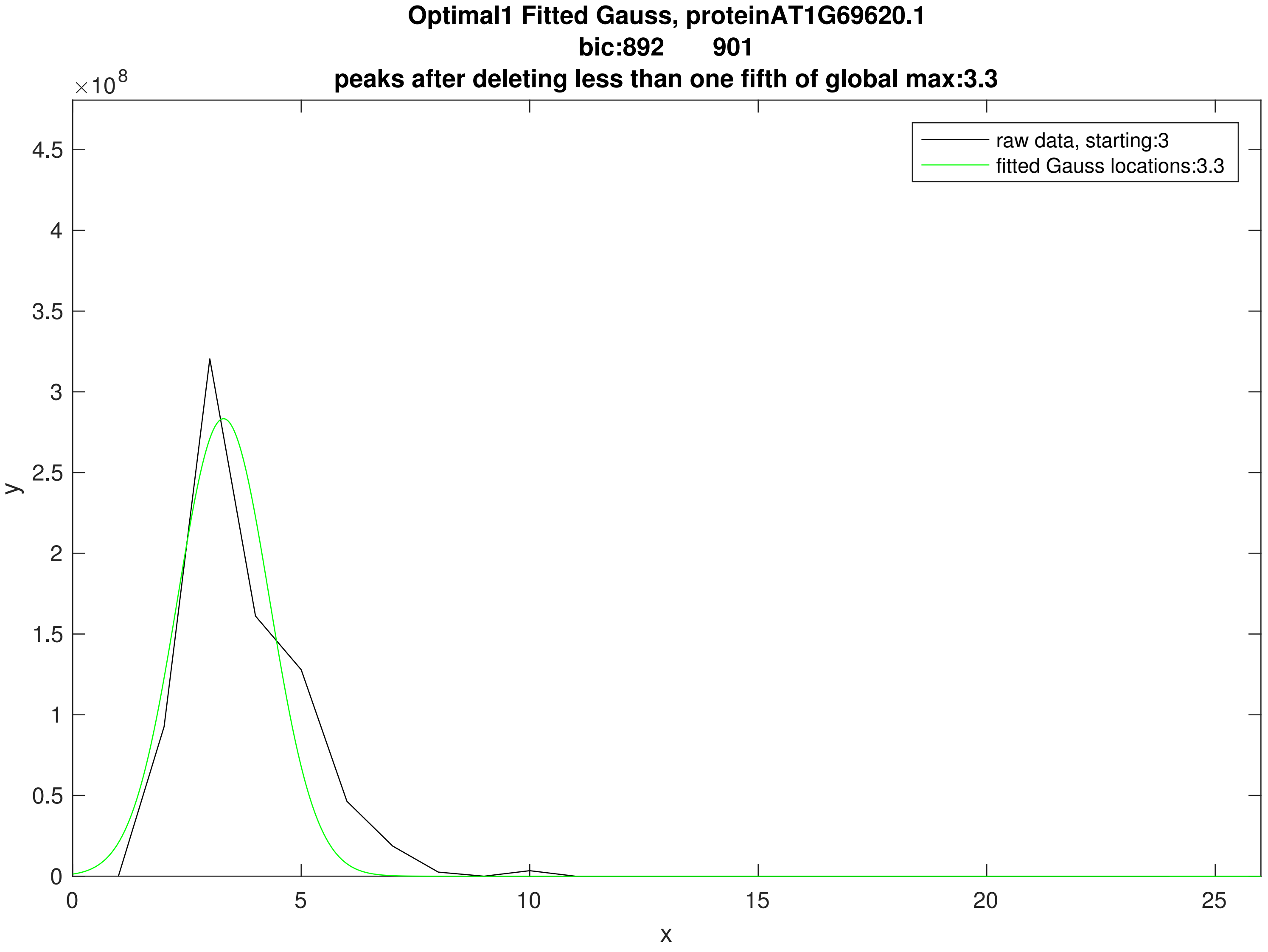

Optimal1 Fitted Gauss, proteinAT1G69620.1
bic:892 901
peaks after deleting less than one fifth of global max:3.3
×10^8
raw data, starting:3
fitted Gauss locations:3.3
y
x
0
0.5
1
1.5
2
2.5
3
3.5
4
4.5
0
5
10
15
20
25

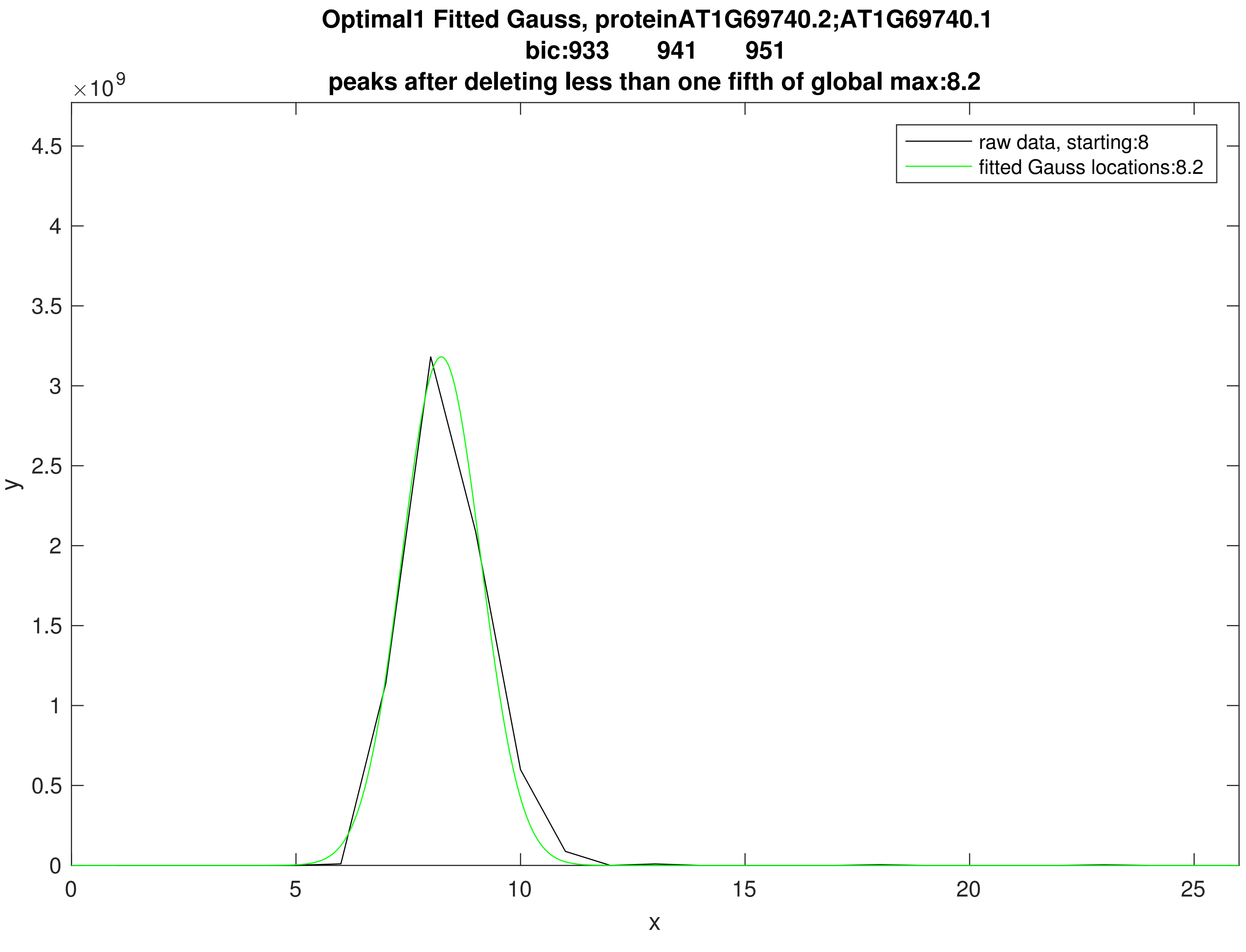
Optimal1 Fitted Gauss, proteinAT1G69740.2;AT1G69740.1
bic:933 941 951
peaks after deleting less than one fifth of global max:8.2
×10^9
4.5
4
3.5
3
2.5
2
1.5
1
0.5
0
y
0
5
10
15
20
25
x
raw data, starting:8
fitted Gauss locations:8.2

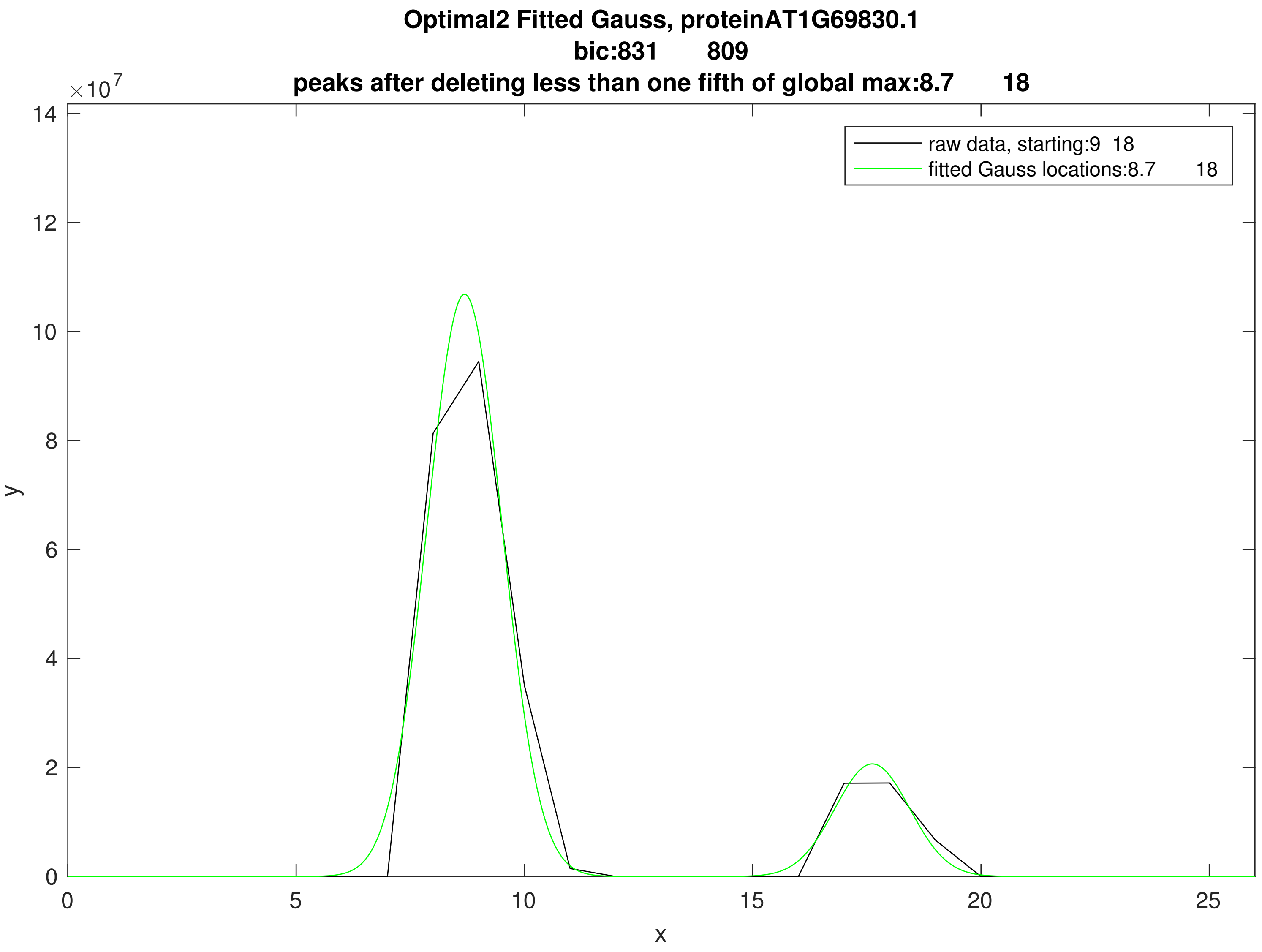
Optimal2 Fitted Gauss, proteinAT1G69830.1
bic:831 809
peaks after deleting less than one fifth of global max:8.7 18
×10^7
14
12
10
8
6
4
2
0
y
0
5
10
15
20
25
x
raw data, starting:9 18
fitted Gauss locations:8.7 18

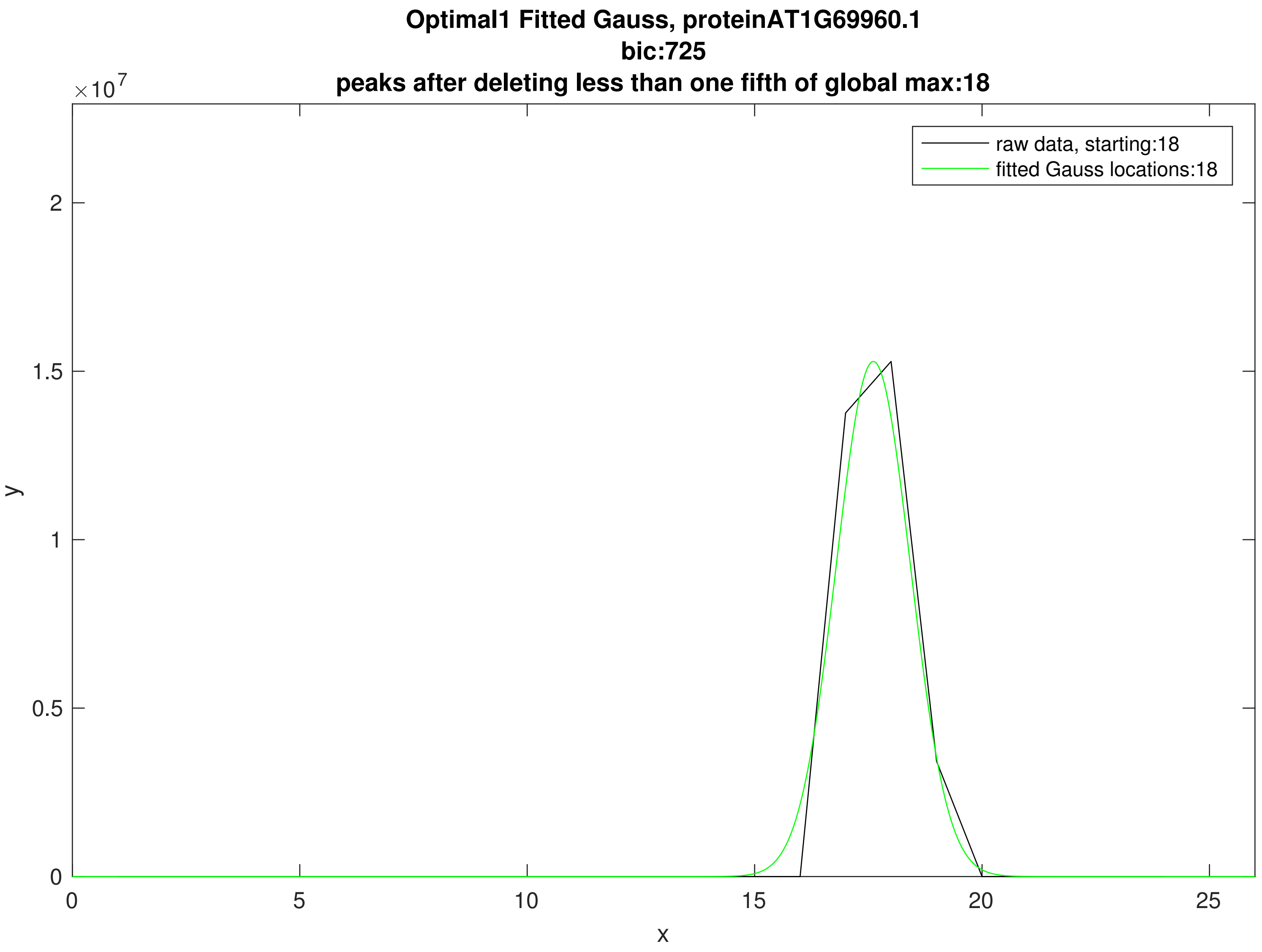

Optimal1 Fitted Gauss, proteinAT1G69960.1
bic:725
peaks after deleting less than one fifth of global max:18
×10^7
raw data, starting:18
fitted Gauss locations:18
2
1.5
1
0.5
0
y
0
5
10
15
20
25
x

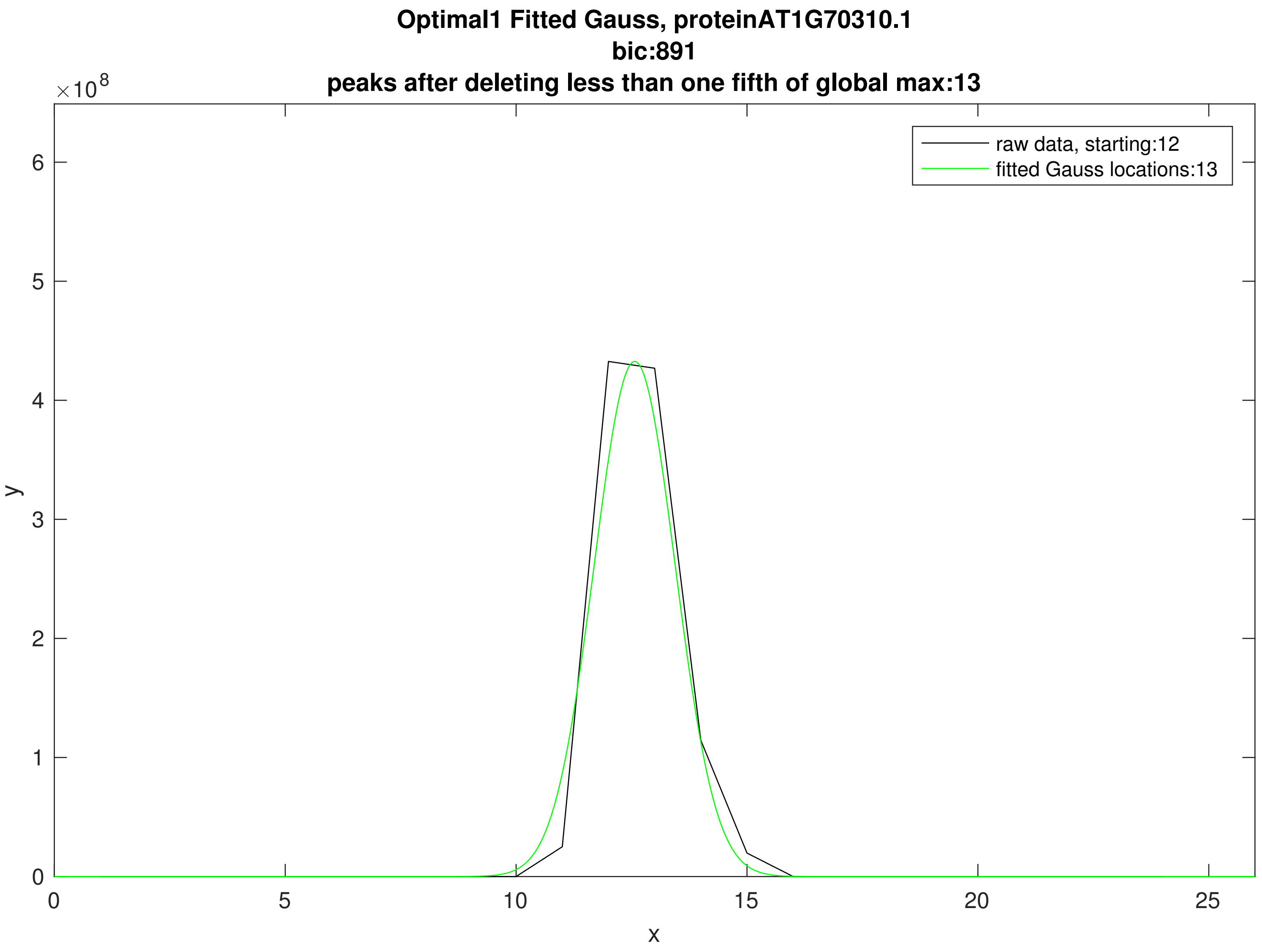

Optimal1 Fitted Gauss, proteinAT1G70310.1
bic:891
peaks after deleting less than one fifth of global max:13
×10^8
raw data, starting:12
fitted Gauss locations:13
y
x
0
1
2
3
4
5
6
0
5
10
15
20
25

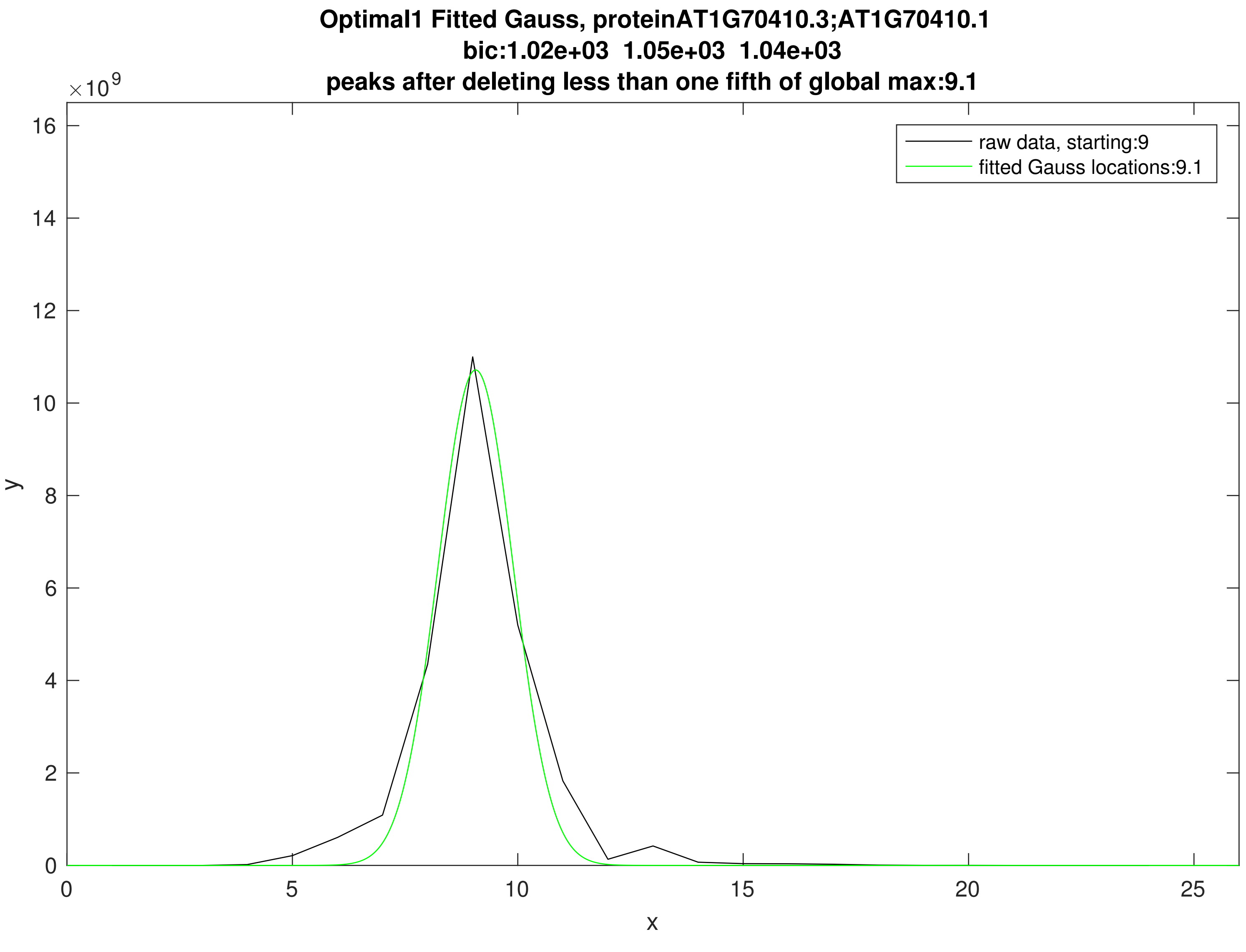

Optimal1 Fitted Gauss, proteinAT1G70410.3;AT1G70410.1
bic:1.02e+03  1.05e+03  1.04e+03
peaks after deleting less than one fifth of global max:9.1
×10^9
raw data, starting:9
fitted Gauss locations:9.1
y
x
0
2
4
6
8
10
12
14
16
0
5
10
15
20
25

**Optimal1 Fitted Gauss, proteinAT1G70520.1**
**bic:745**
**peaks after deleting less than one fifth of global max:17**

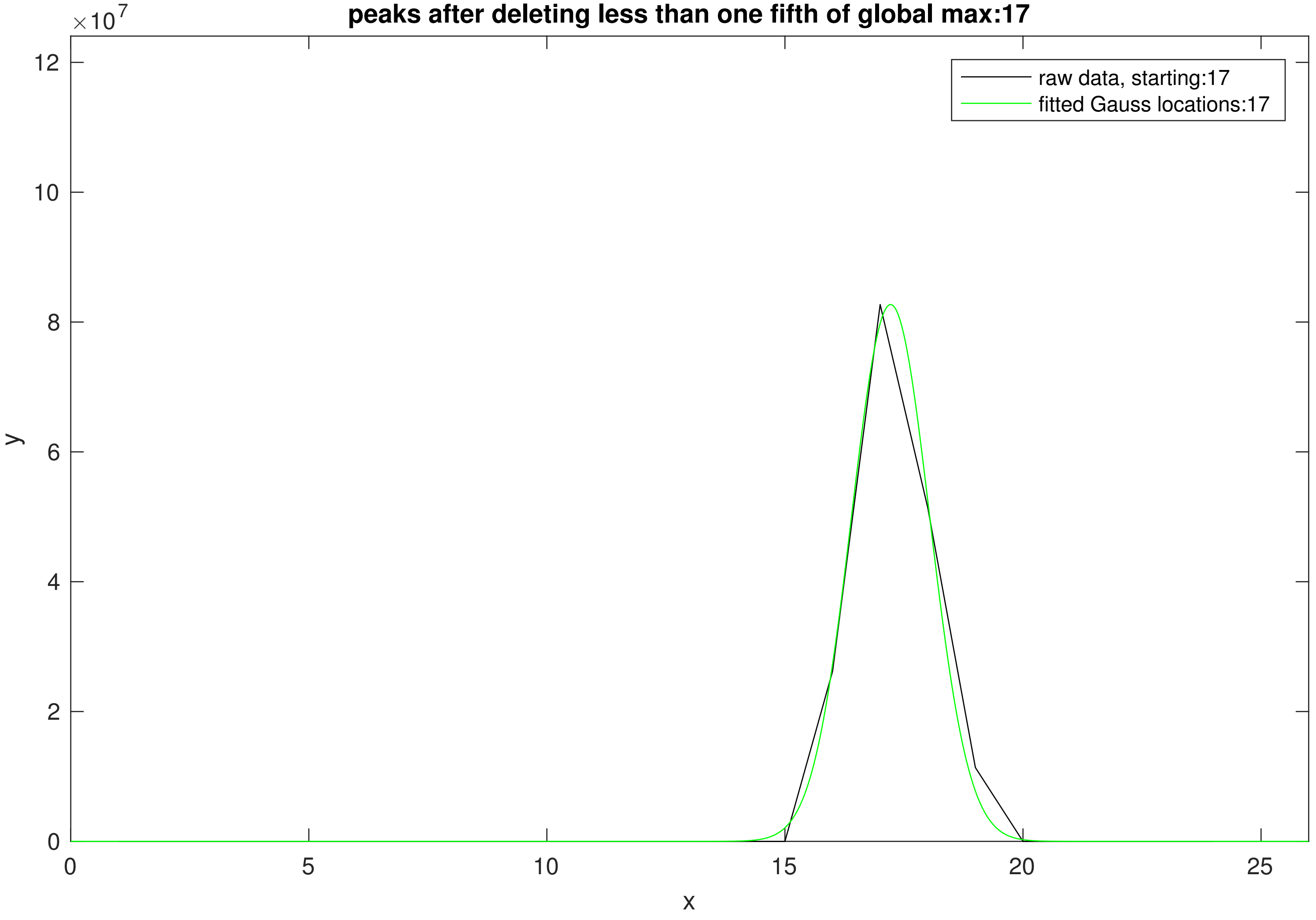

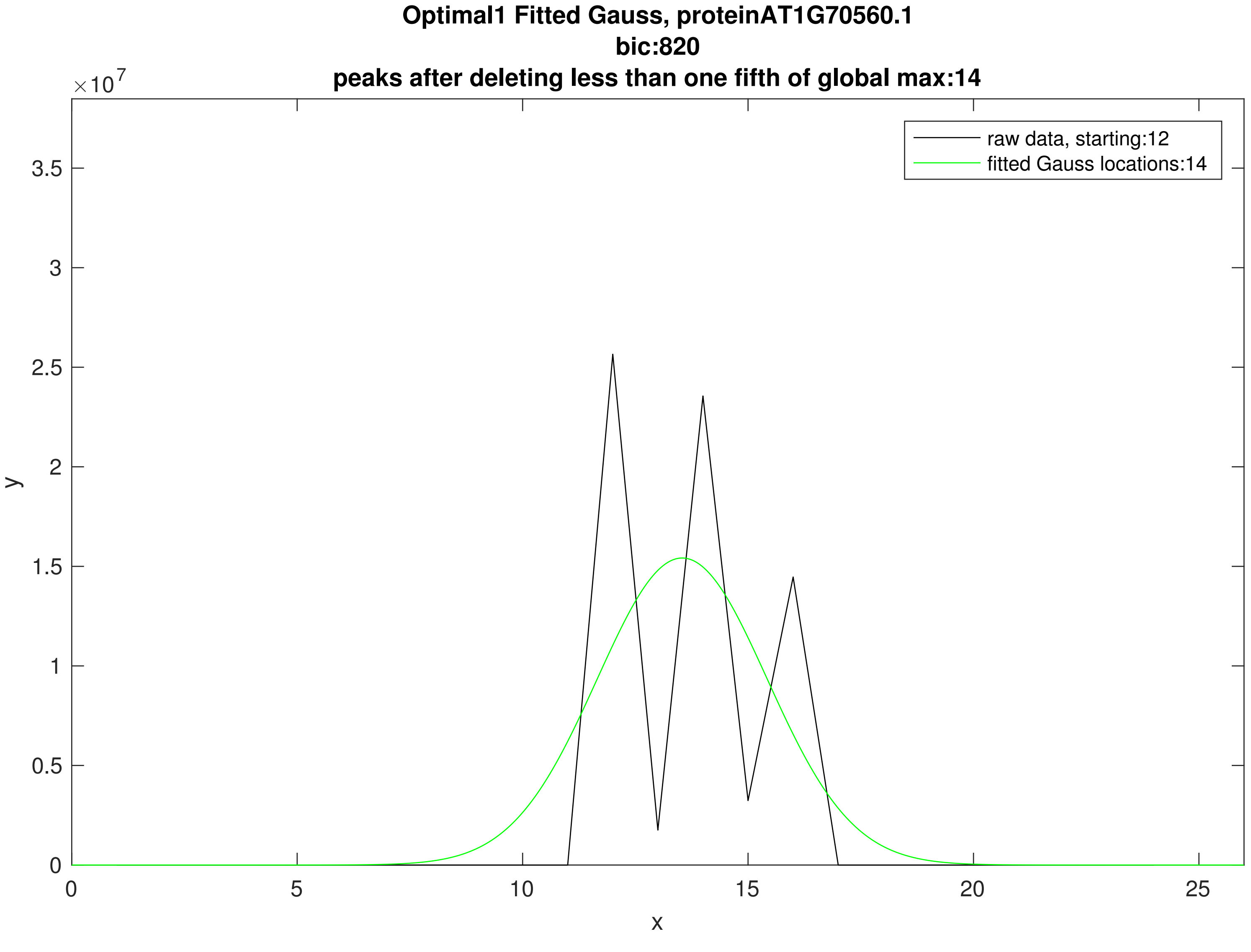

Optimal1 Fitted Gauss, proteinAT1G70560.1
bic:820
peaks after deleting less than one fifth of global max:14
×10^7
raw data, starting:12
fitted Gauss locations:14
3.5
3
2.5
2
1.5
1
0.5
0
y
0
5
10
15
20
25
x

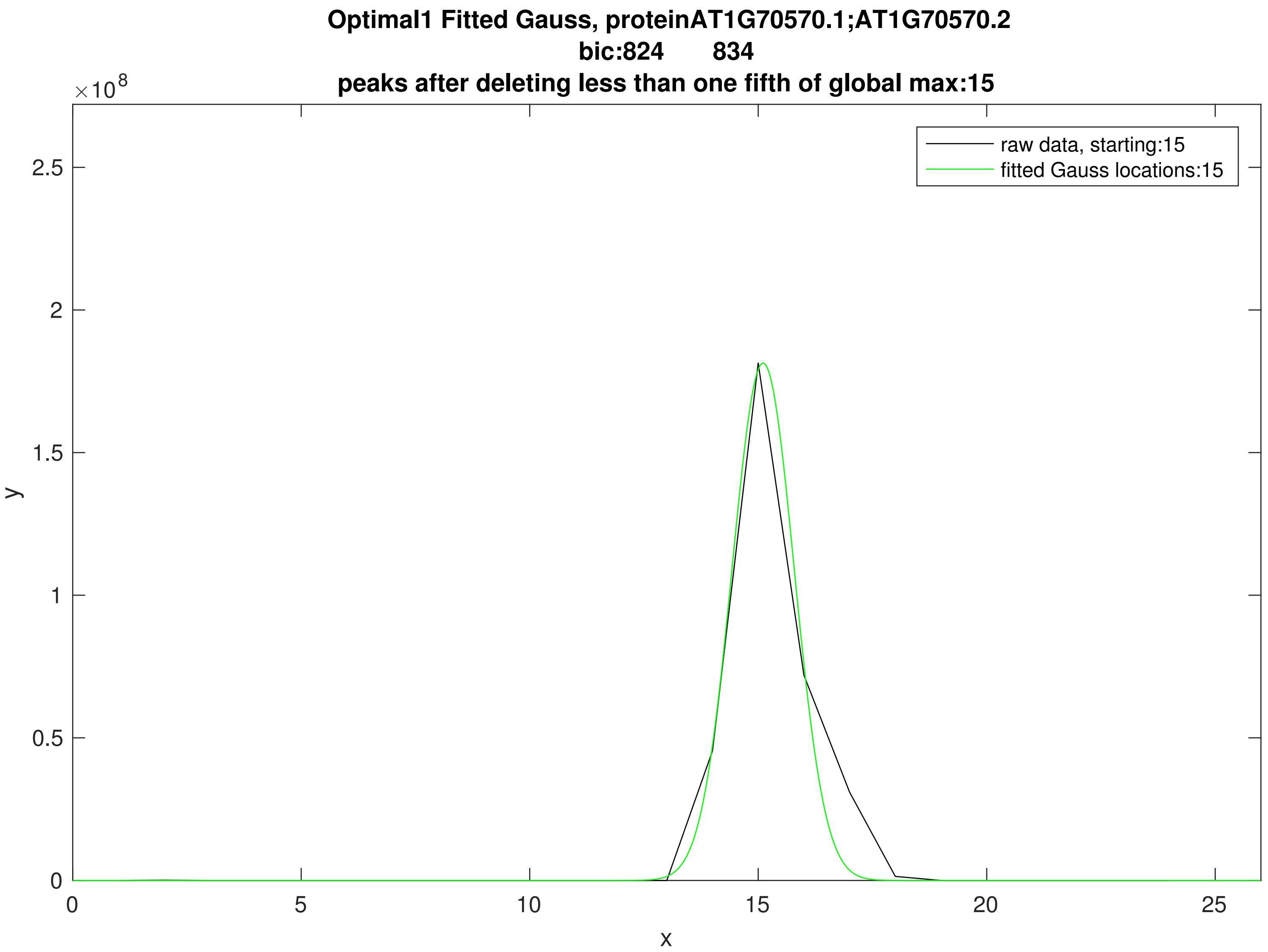
Optimal1 Fitted Gauss, proteinAT1G70570.1;AT1G70570.2
bic:824 834
peaks after deleting less than one fifth of global max:15
×10^8
2.5
2
1.5
1
0.5
0
y
0
5
10
15
20
25
x
raw data, starting:15
fitted Gauss locations:15

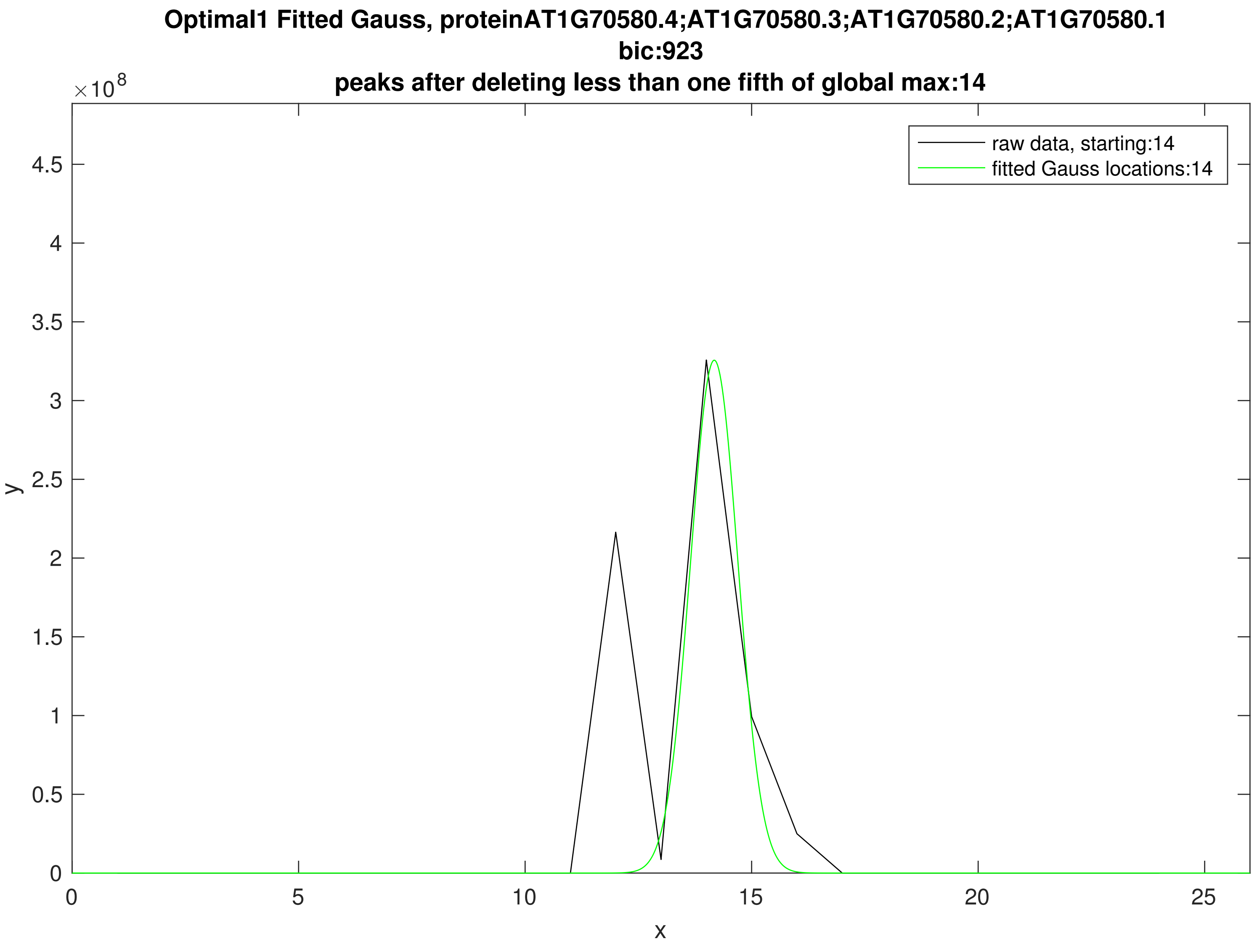

Optimal1 Fitted Gauss, proteinAT1G70580.4;AT1G70580.3;AT1G70580.2;AT1G70580.1
bic:923
peaks after deleting less than one fifth of global max:14
×10^8
4.5
4
3.5
3
2.5
2
1.5
1
0.5
0
y
0
5
10
15
20
25
x
raw data, starting:14
fitted Gauss locations:14

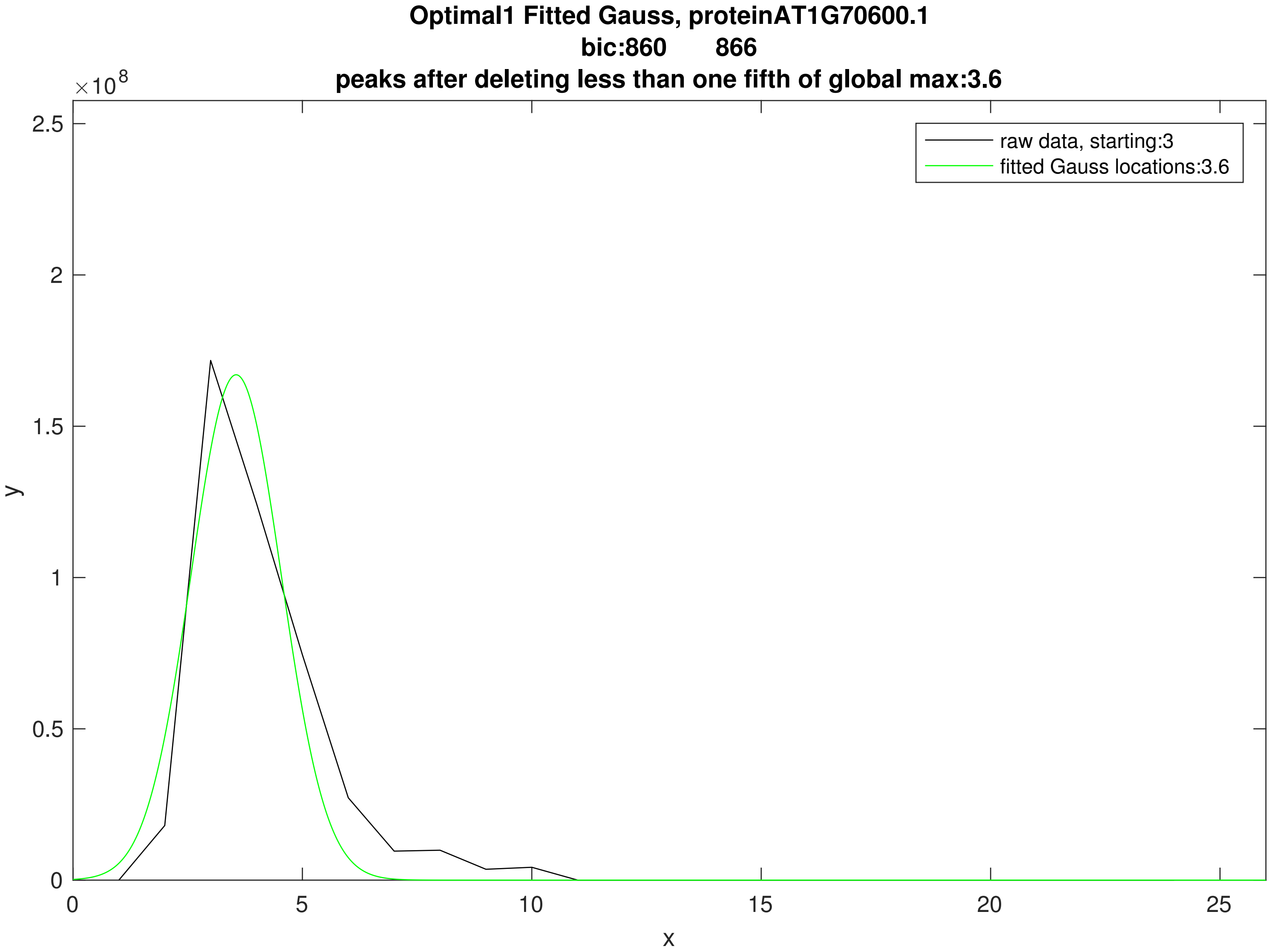

Optimal1 Fitted Gauss, proteinAT1G70600.1
bic:860 866
peaks after deleting less than one fifth of global max:3.6
×10^8
raw data, starting:3
fitted Gauss locations:3.6
y
x
0
0.5
1
1.5
2
2.5
5
10
15
20
25

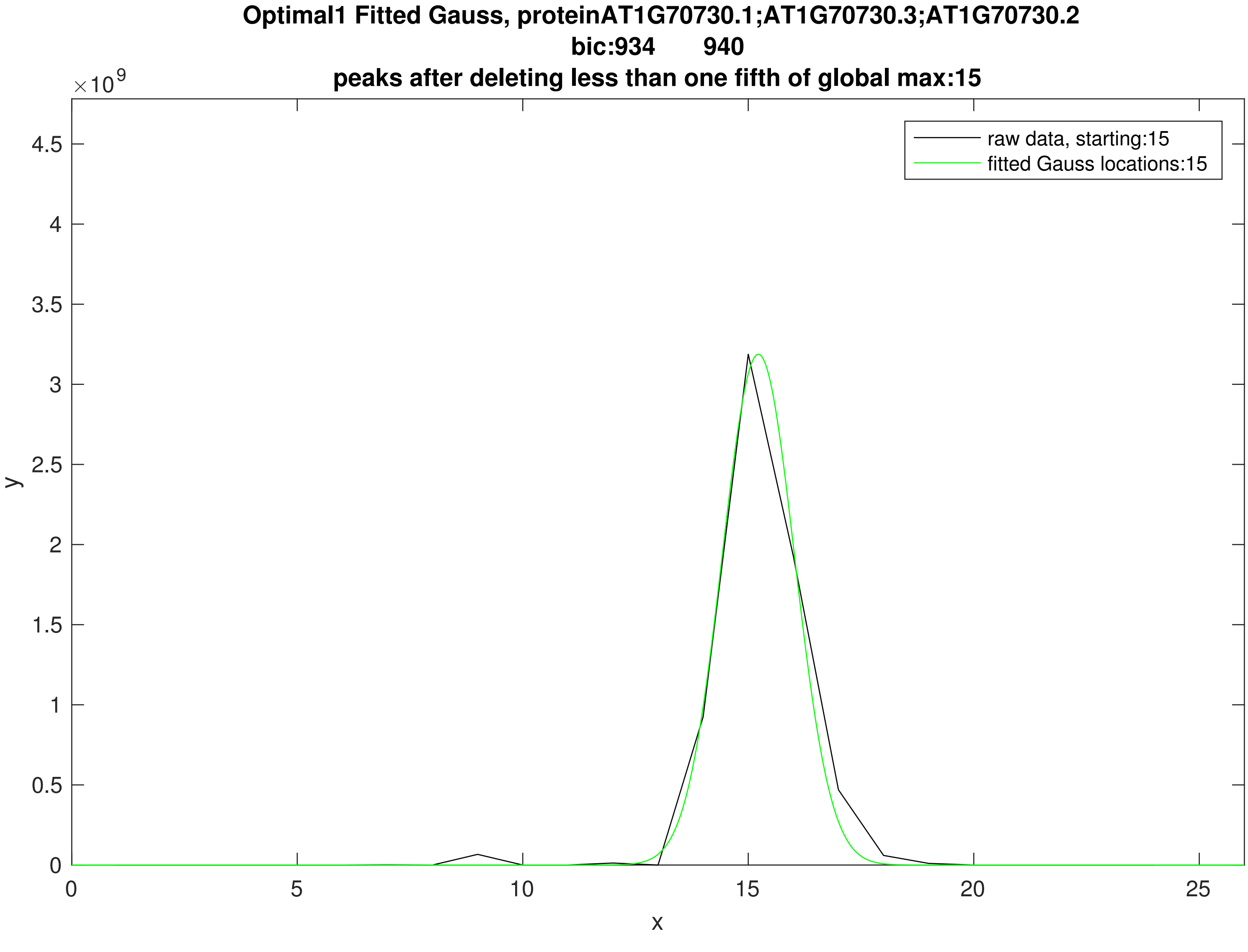
Optimal1 Fitted Gauss, proteinAT1G70730.1;AT1G70730.3;AT1G70730.2
bic:934 940
peaks after deleting less than one fifth of global max:15
×10^9
raw data, starting:15
fitted Gauss locations:15
y
x
0
0.5
1
1.5
2
2.5
3
3.5
4
4.5
5
10
15
20
25

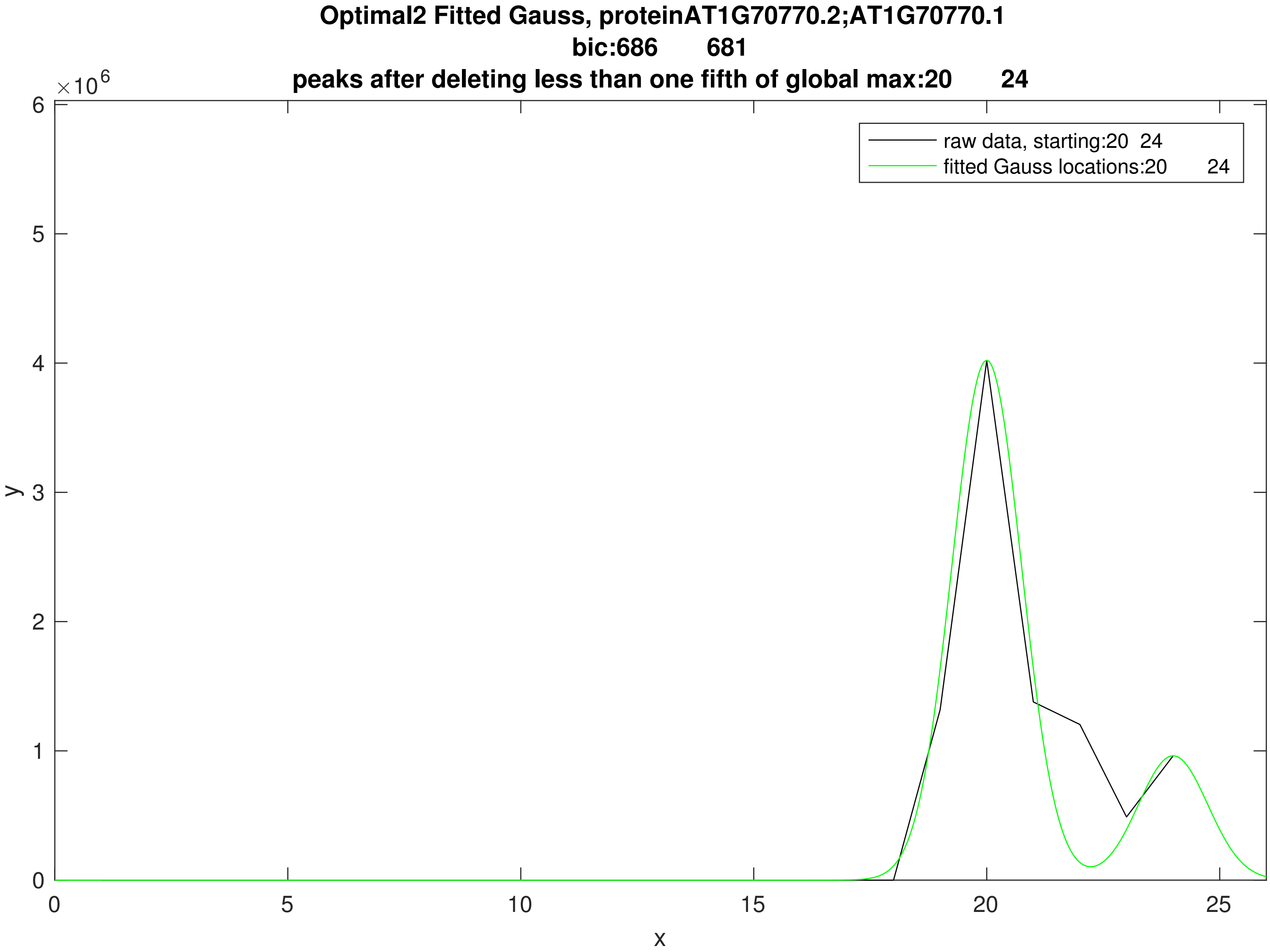
Optimal2 Fitted Gauss, proteinAT1G70770.2;AT1G70770.1
bic:686 681
peaks after deleting less than one fifth of global max:20 24
raw data, starting:20 24
fitted Gauss locations:20 24
y
x
×10^6
0
1
2
3
4
5
6
0
5
10
15
20
25

**Optimal1 Fitted Gauss, proteinAT1G70810.1**
**bic:760**
**peaks after deleting less than one fifth of global max:19**

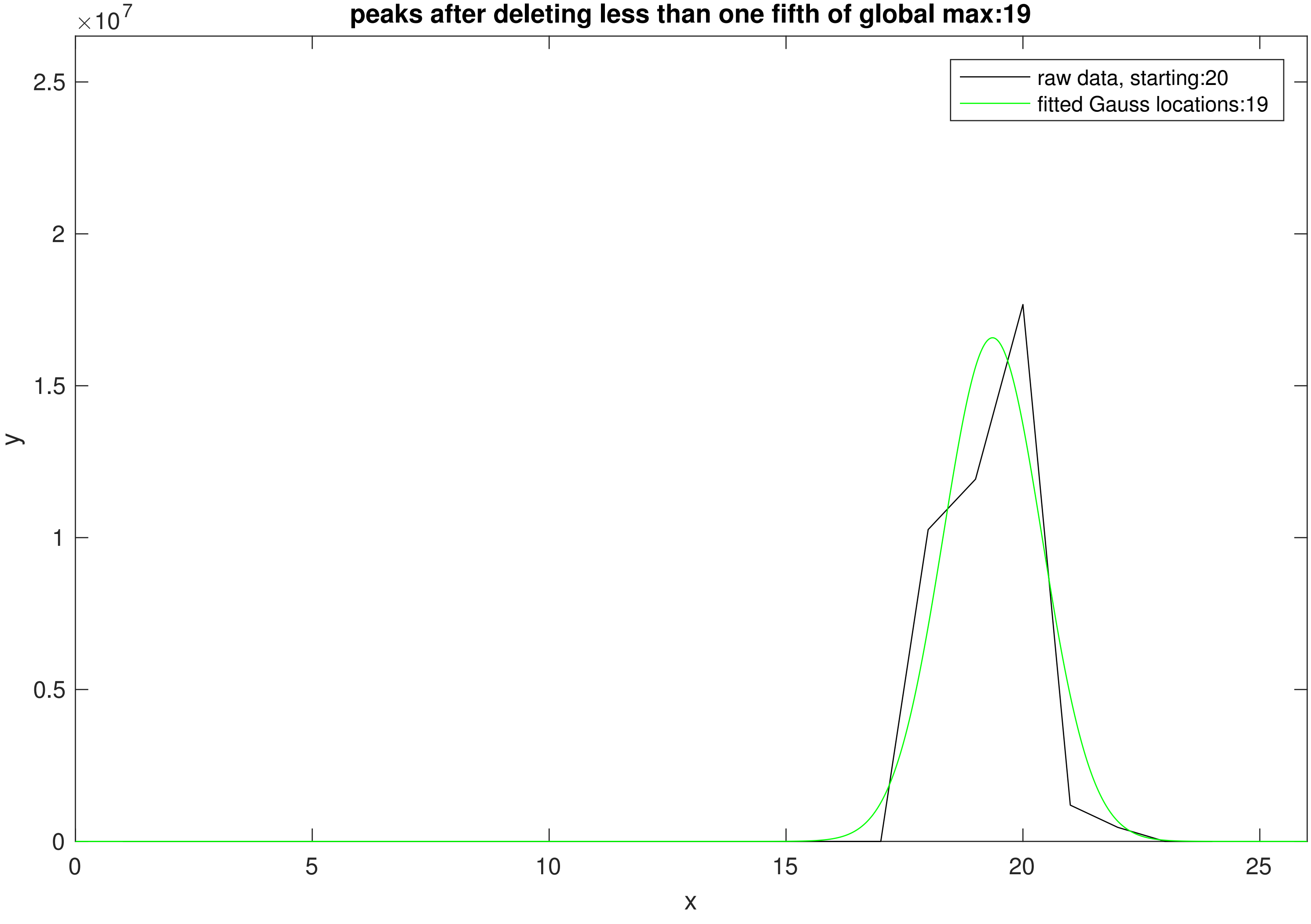

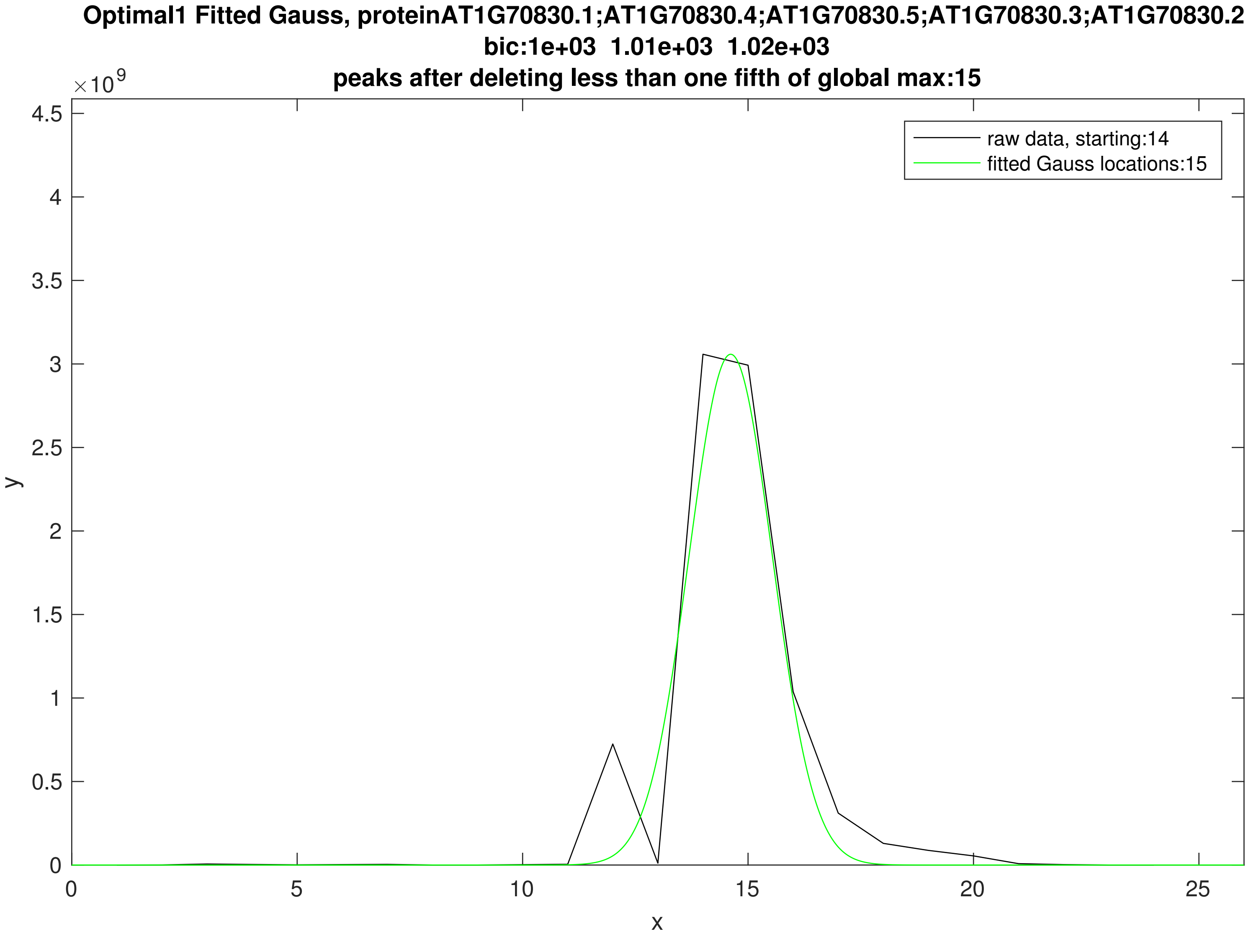

Optimal1 Fitted Gauss, proteinAT1G70830.1;AT1G70830.4;AT1G70830.5;AT1G70830.3;AT1G70830.2
bic:1e+03 1.01e+03 1.02e+03
peaks after deleting less than one fifth of global max:15
×10^9
4.5
4
3.5
3
2.5
2
1.5
1
0.5
0
y
0
5
10
15
20
25
x
raw data, starting:14
fitted Gauss locations:15

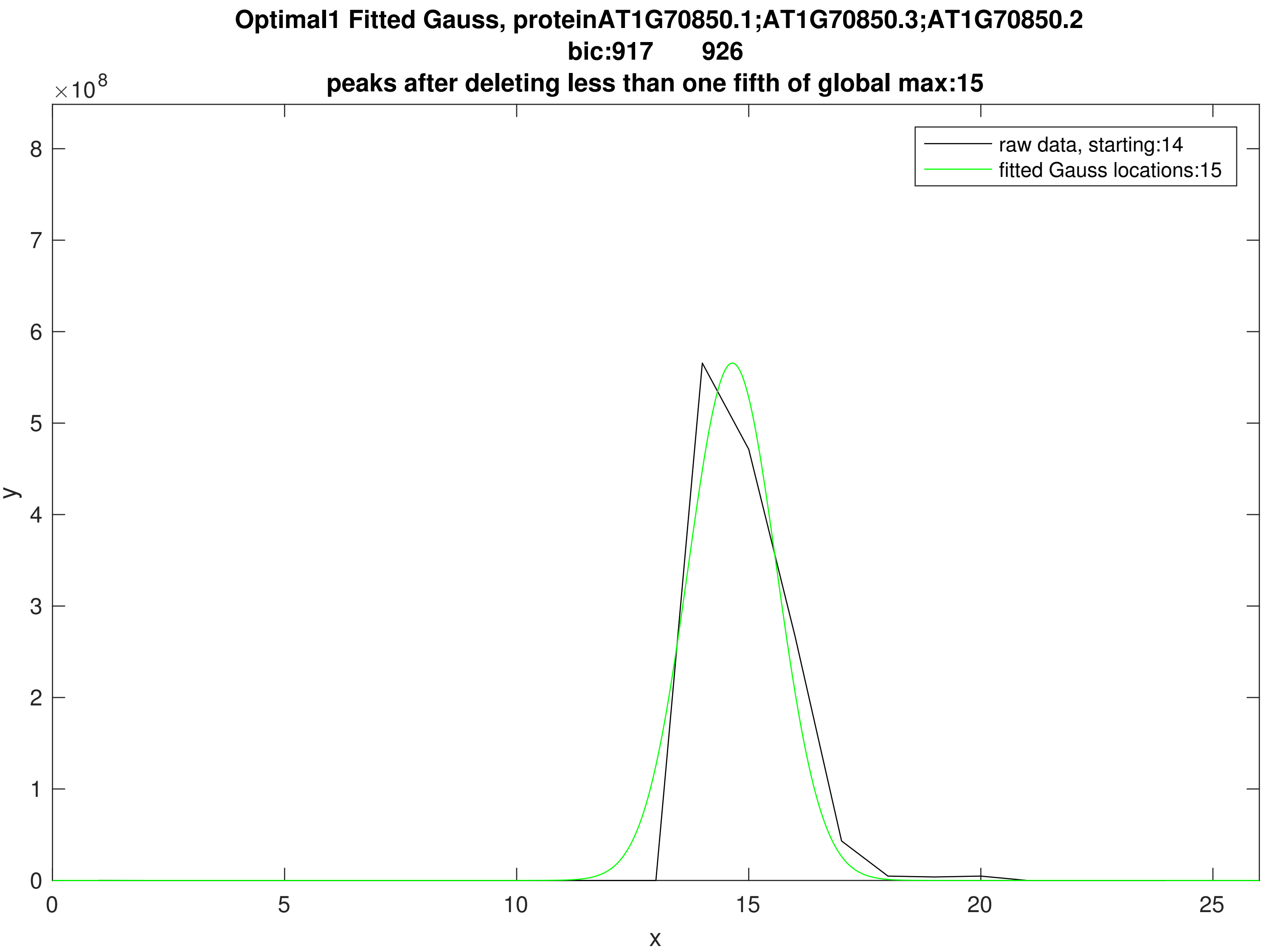

Optimal1 Fitted Gauss, proteinAT1G70850.1;AT1G70850.3;AT1G70850.2
bic:917 926
peaks after deleting less than one fifth of global max:15
×10^8
raw data, starting:14
fitted Gauss locations:15
y
x
0
1
2
3
4
5
6
7
8
0
5
10
15
20
25

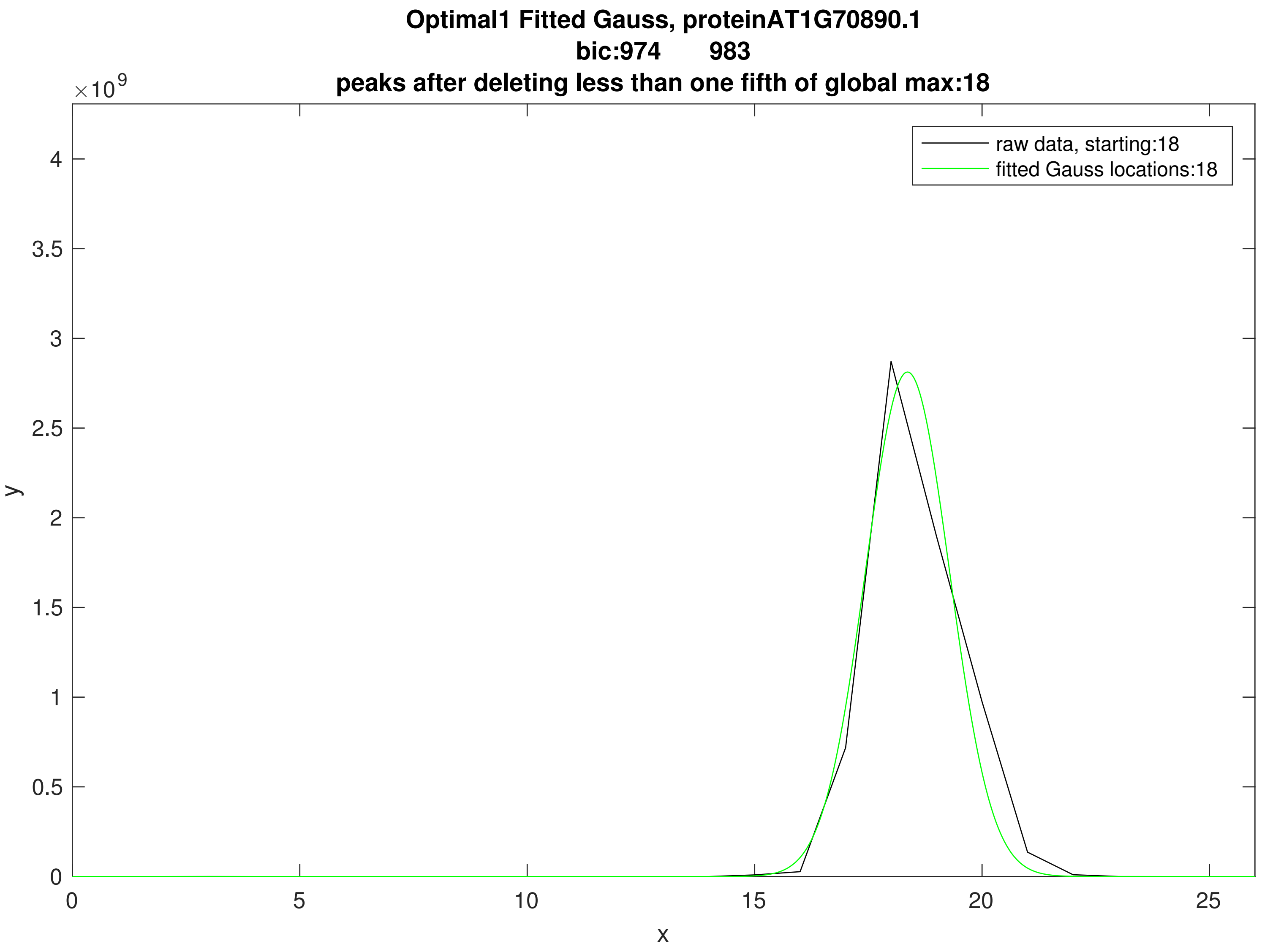

Optimal1 Fitted Gauss, proteinAT1G70890.1
bic:974 983
peaks after deleting less than one fifth of global max:18
×10^9
raw data, starting:18
fitted Gauss locations:18
y
x
0
0.5
1
1.5
2
2.5
3
3.5
4
0
5
10
15
20
25

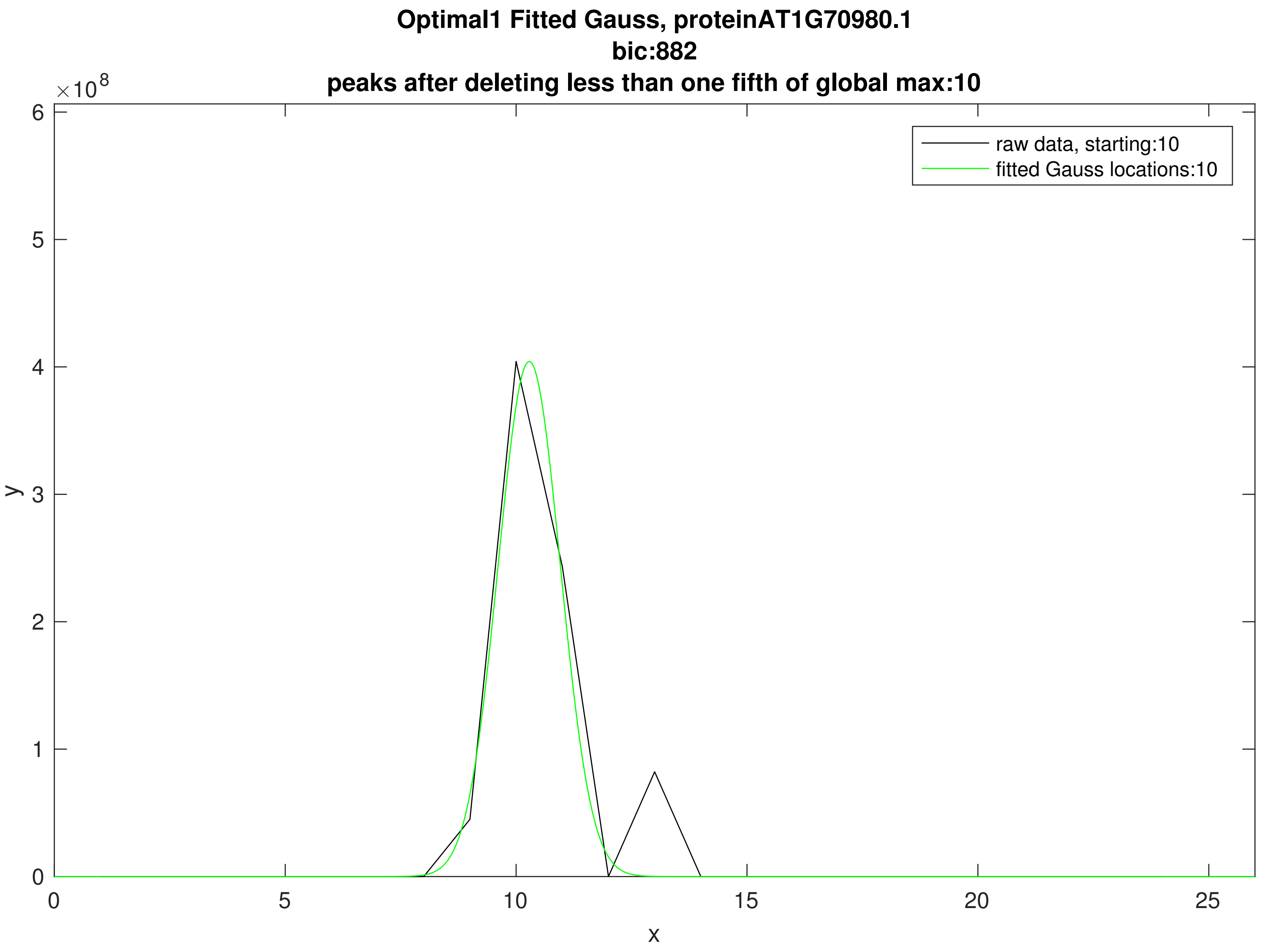

Optimal1 Fitted Gauss, proteinAT1G70980.1
bic:882
peaks after deleting less than one fifth of global max:10
×10^8
6
5
4
3
2
1
0
y
0
5
10
15
20
25
x
raw data, starting:10
fitted Gauss locations:10

Optimal1 Fitted Gauss, proteinAT1G71310.1;AT1G71310.2;AT1G71310.3
bic:825 835
peaks after deleting less than one fifth of global max:11

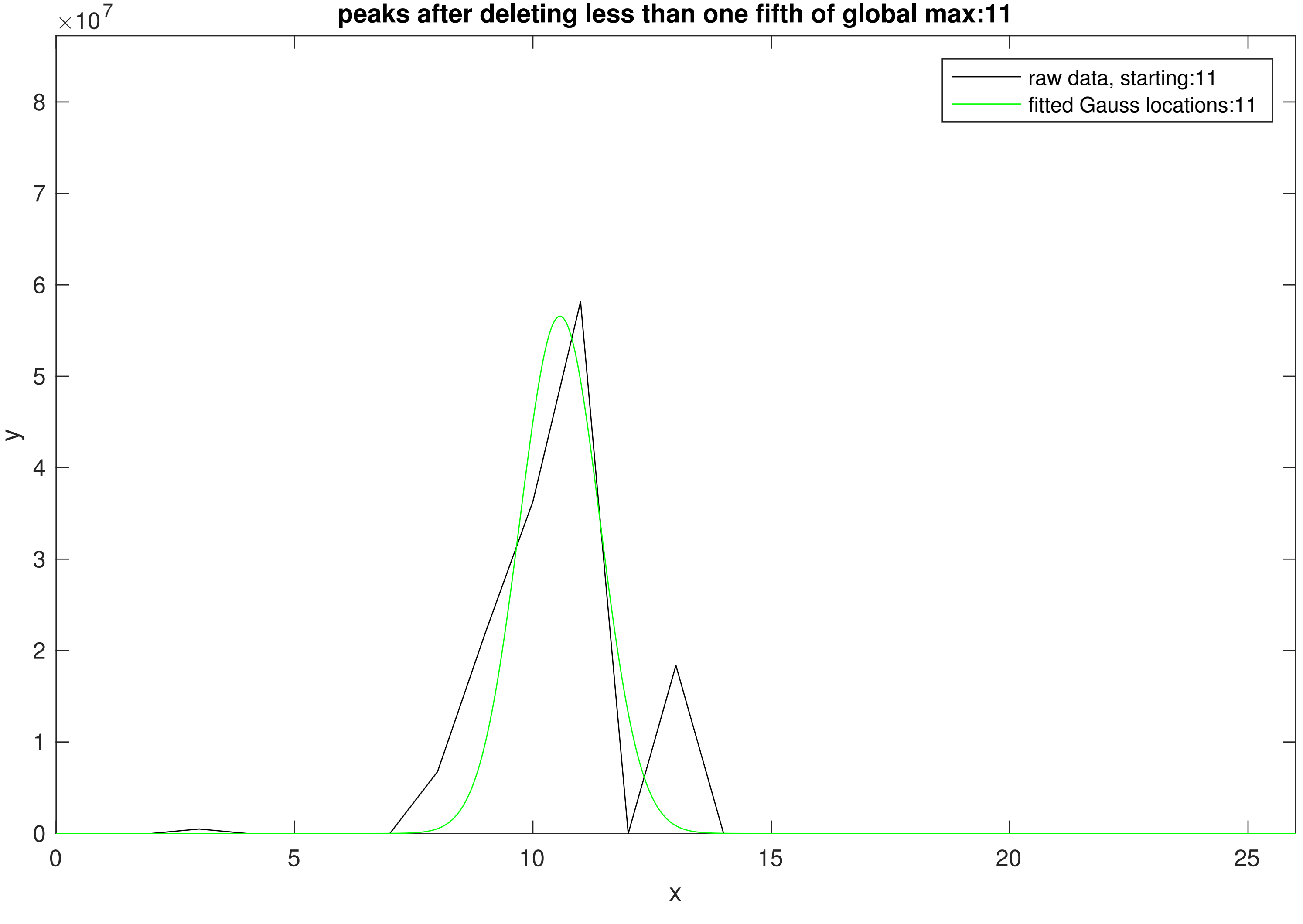

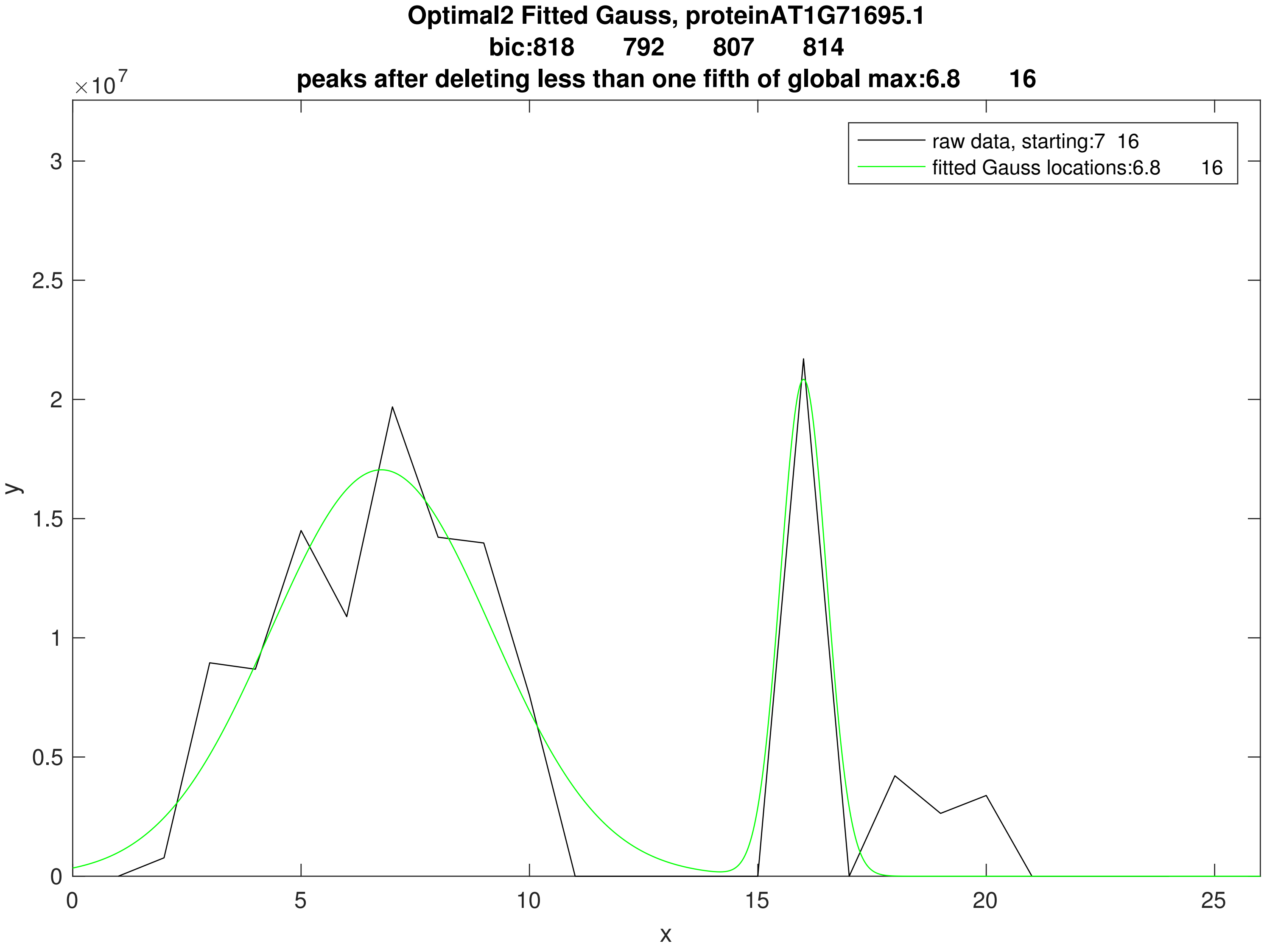
Optimal2 Fitted Gauss, proteinAT1G71695.1
bic:818 792 807 814
peaks after deleting less than one fifth of global max:6.8 16
×10^7
raw data, starting:7 16
fitted Gauss locations:6.8 16
y
x
0
0.5
1
1.5
2
2.5
3
5
10
15
20
25

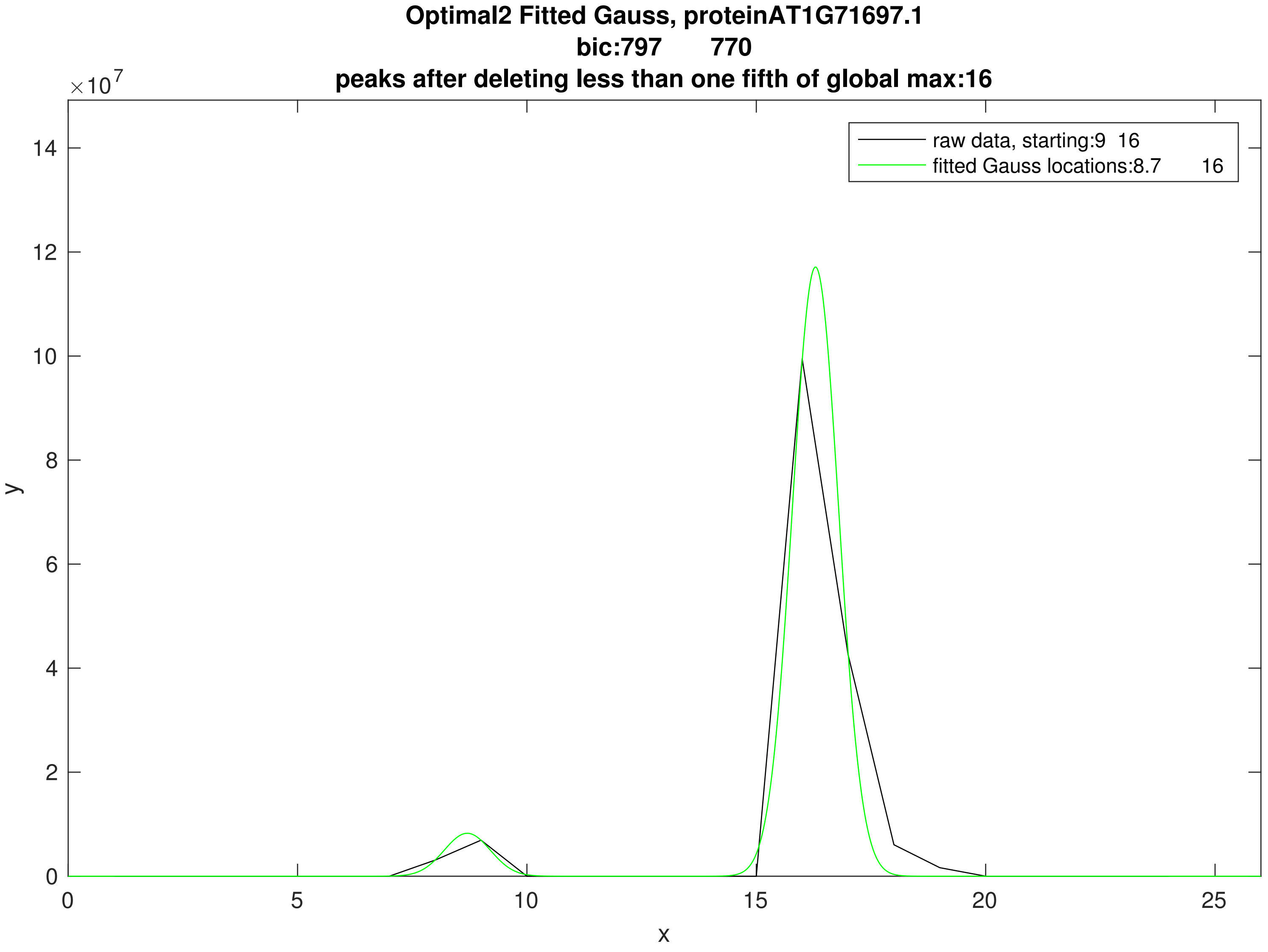

Optimal2 Fitted Gauss, proteinAT1G71697.1
bic:797 770
peaks after deleting less than one fifth of global max:16
×10^7
raw data, starting:9 16
fitted Gauss locations:8.7 16
y
x
0
2
4
6
8
10
12
14
0
5
10
15
20
25

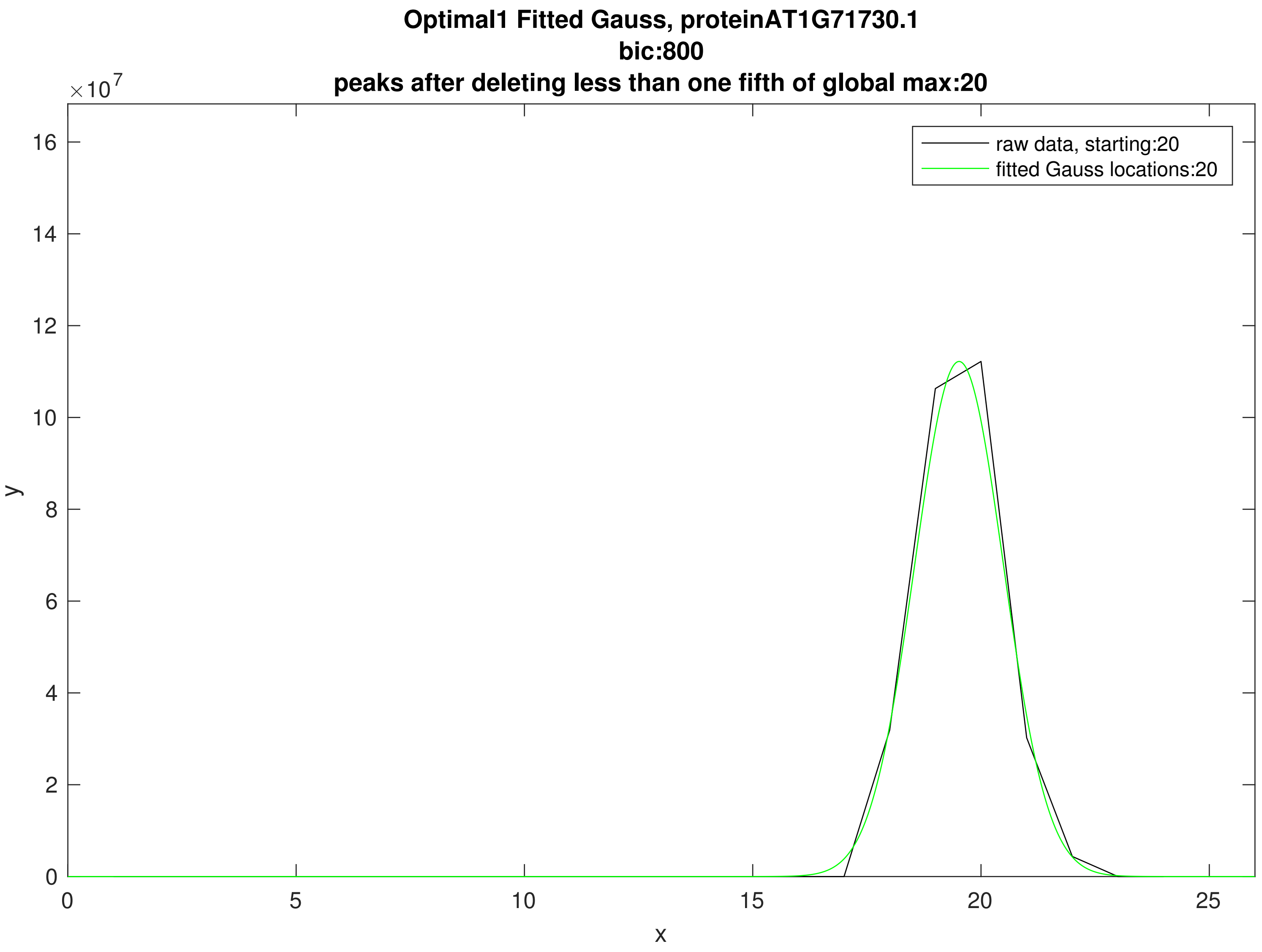

Optimal1 Fitted Gauss, proteinAT1G71730.1
bic:800
peaks after deleting less than one fifth of global max:20
×10^7
16
14
12
10
8
6
4
2
0
y
0
5
10
15
20
25
x
raw data, starting:20
fitted Gauss locations:20

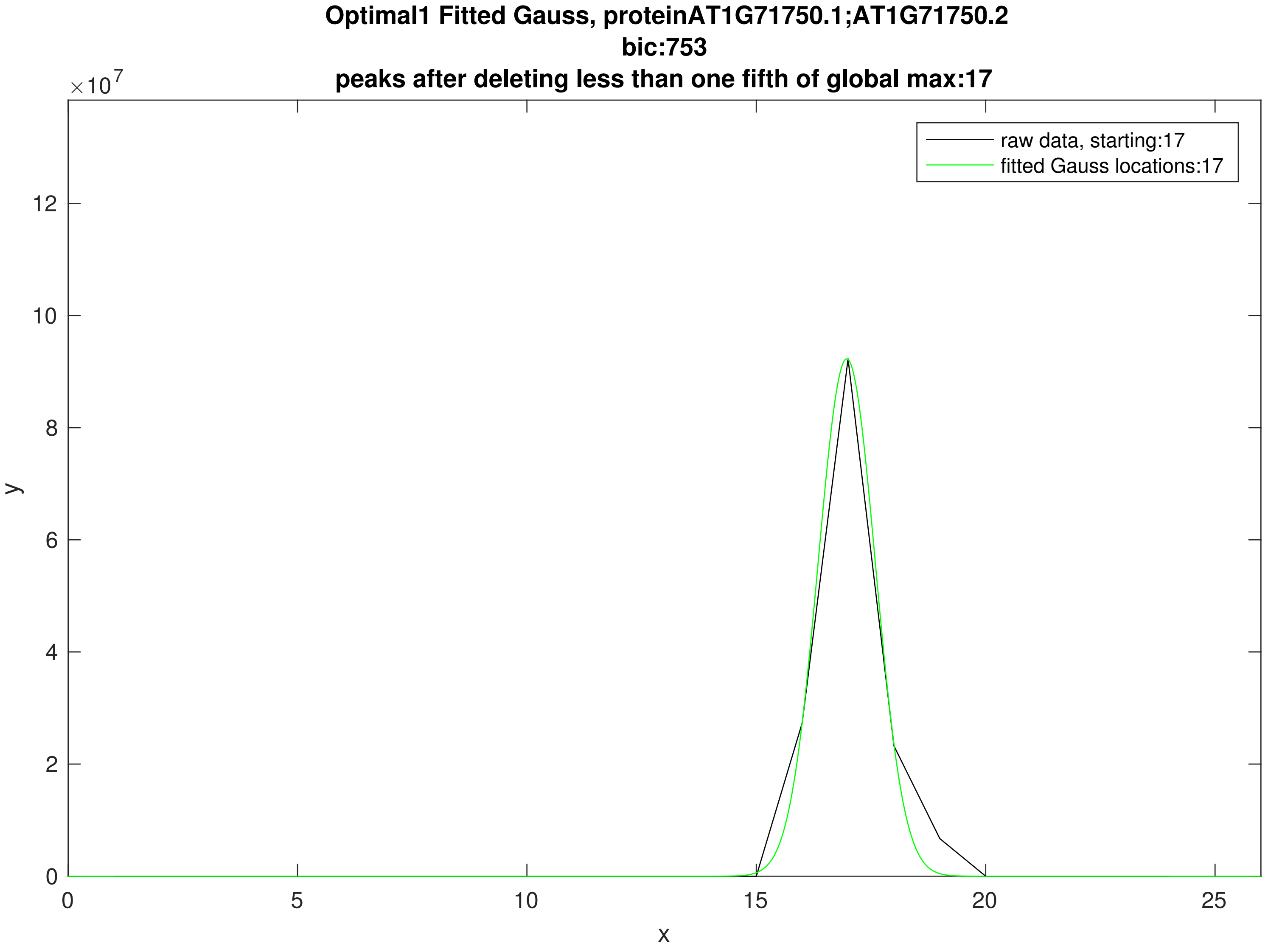

Optimal1 Fitted Gauss, proteinAT1G71750.1;AT1G71750.2
bic:753
peaks after deleting less than one fifth of global max:17
×10^7
raw data, starting:17
fitted Gauss locations:17
y
x
0
2
4
6
8
10
12
0
5
10
15
20
25

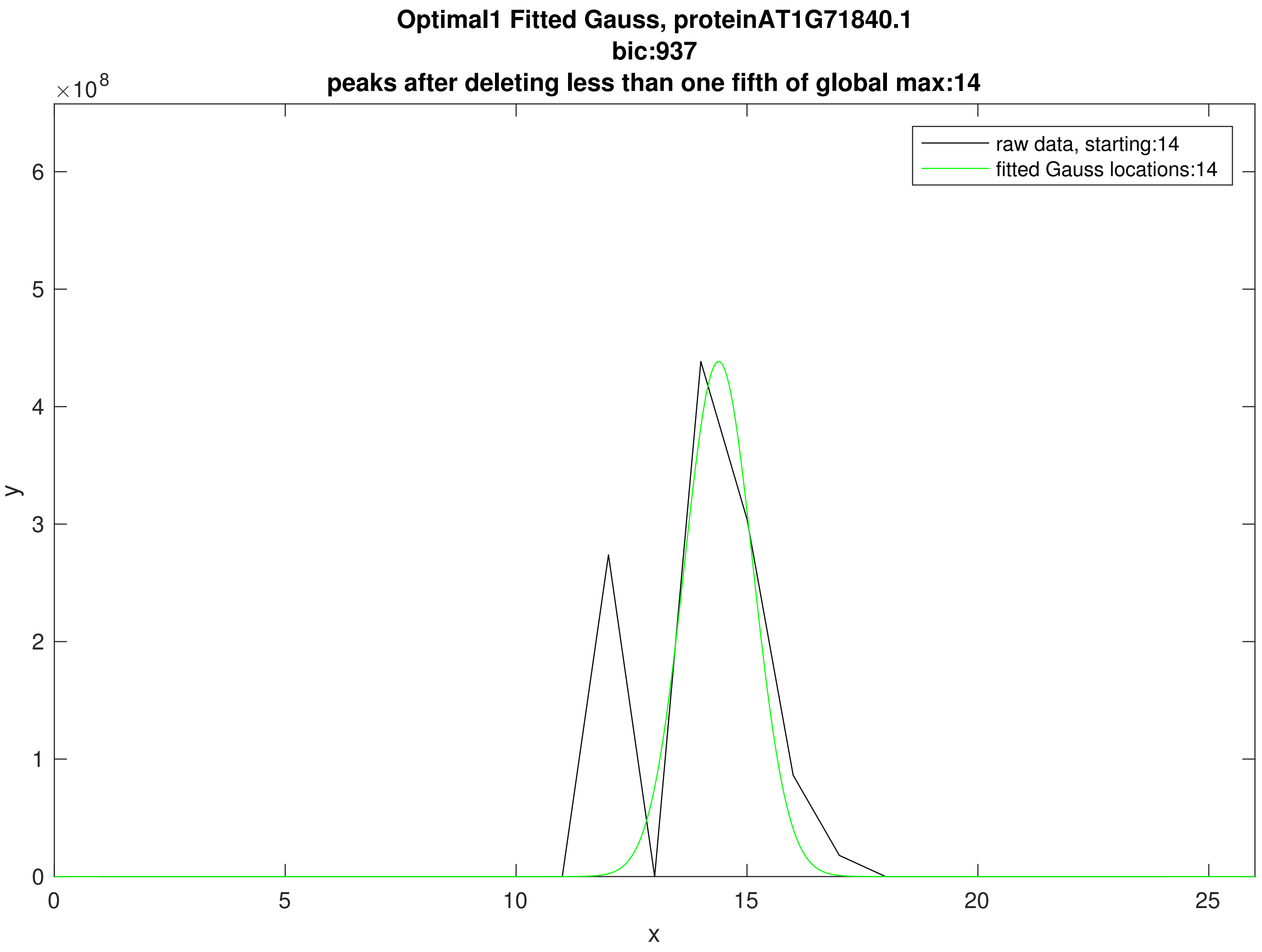

Optimal1 Fitted Gauss, proteinAT1G71840.1
bic:937
peaks after deleting less than one fifth of global max:14
×10^8
raw data, starting:14
fitted Gauss locations:14
y
x
0
1
2
3
4
5
6
0
5
10
15
20
25

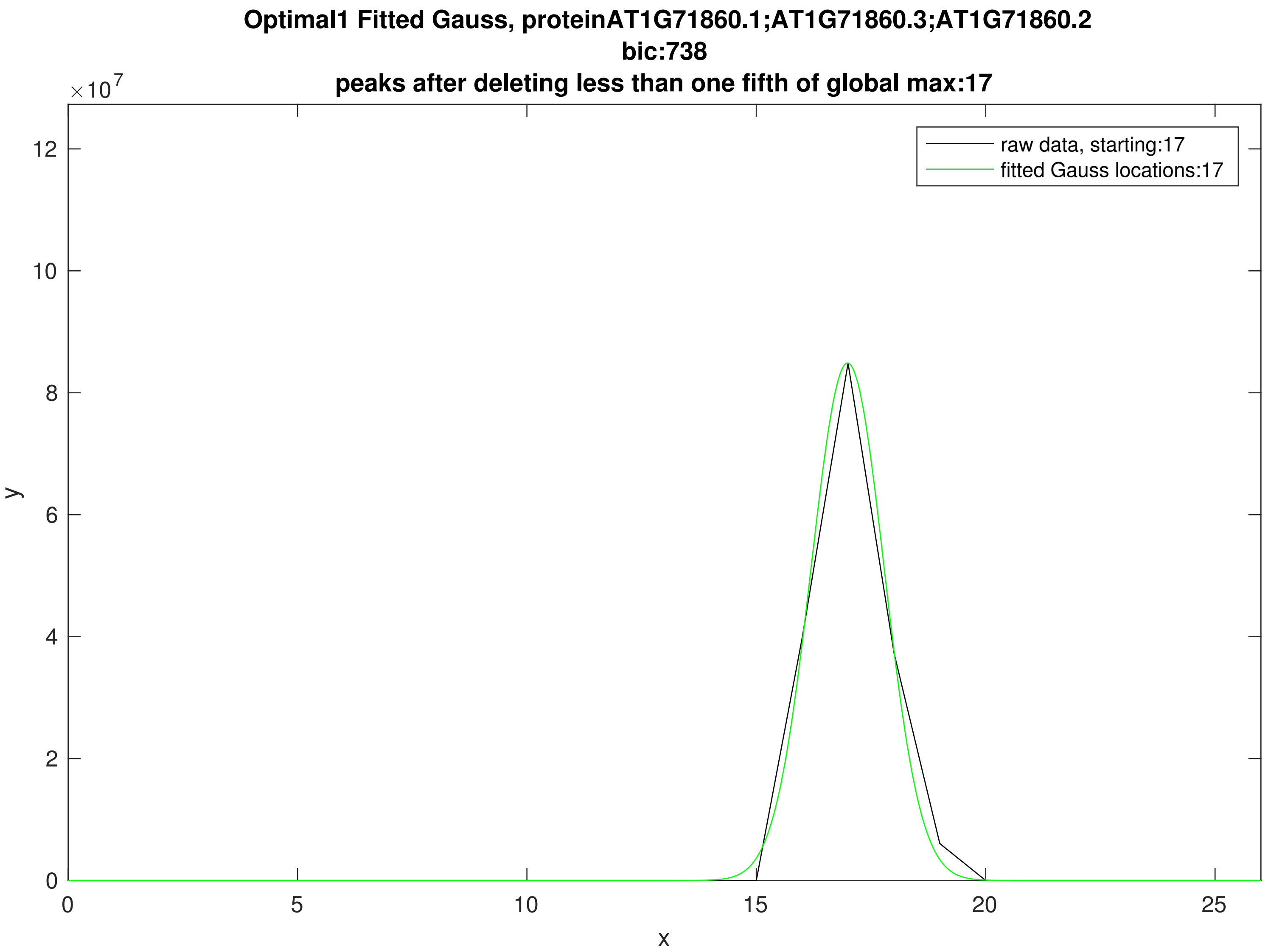

Optimal1 Fitted Gauss, proteinAT1G71860.1;AT1G71860.3;AT1G71860.2
bic:738
peaks after deleting less than one fifth of global max:17
×10^7
12
10
8
6
4
2
0
y
0
5
10
15
20
25
x
raw data, starting:17
fitted Gauss locations:17

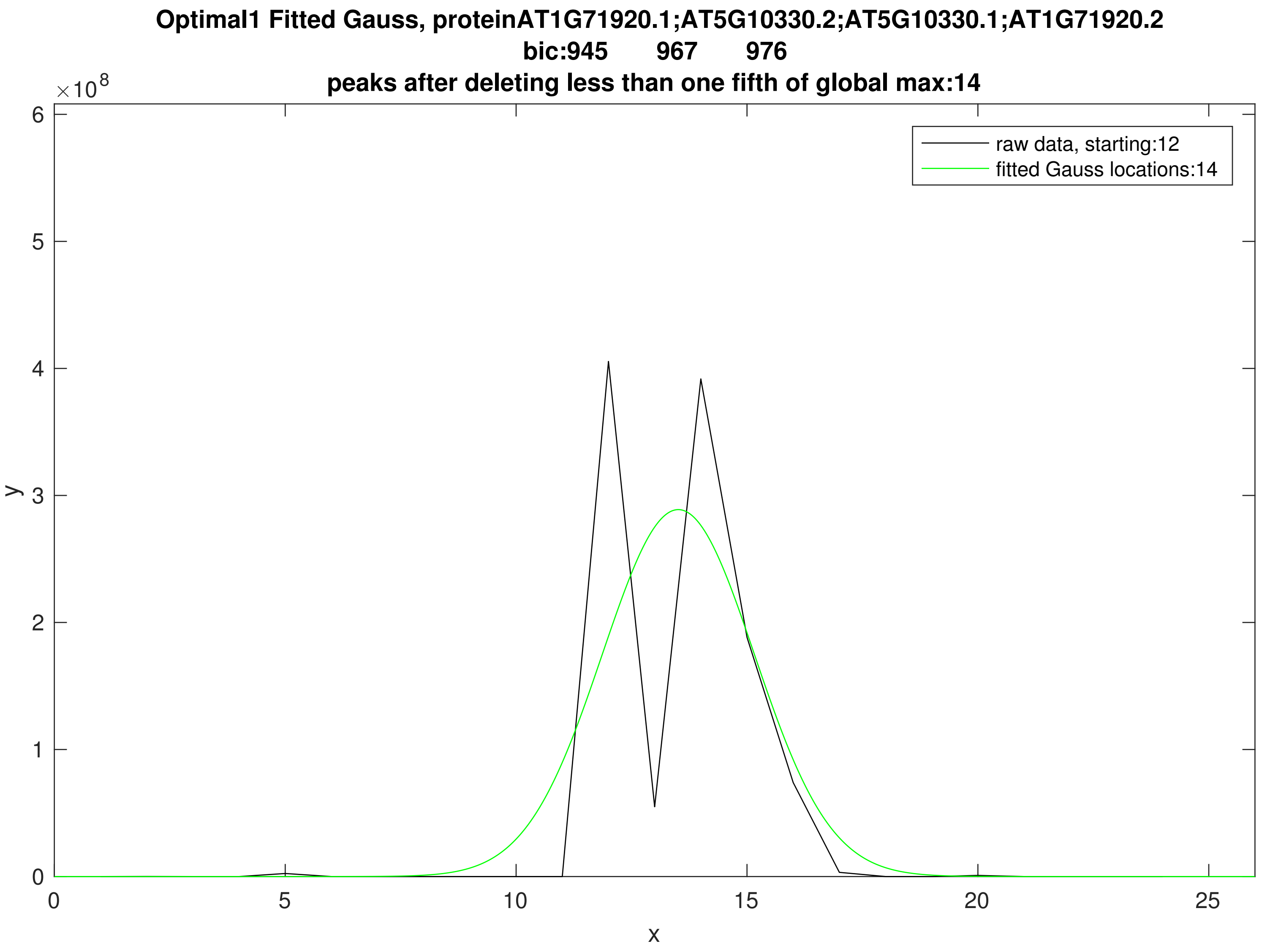

Optimal1 Fitted Gauss, proteinAT1G71920.1;AT5G10330.2;AT5G10330.1;AT1G71920.2
bic:945 967 976
peaks after deleting less than one fifth of global max:14
×10^8
6
5
4
3
2
1
0
y
0
5
10
15
20
25
x
raw data, starting:12
fitted Gauss locations:14

**Optimal1 Fitted Gauss, proteinAT1G71950.1**
**bic:966**
**peaks after deleting less than one fifth of global max:13**

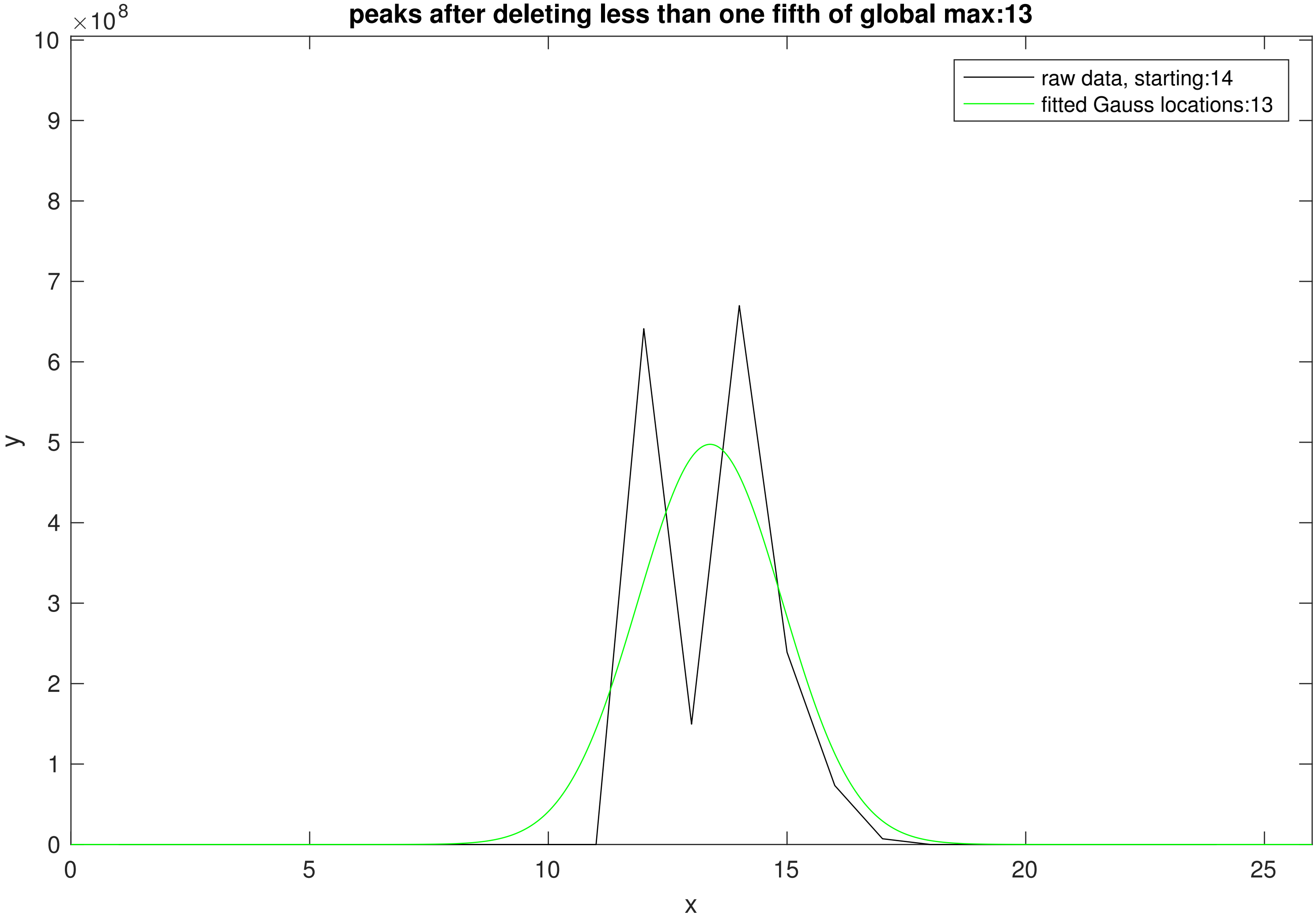

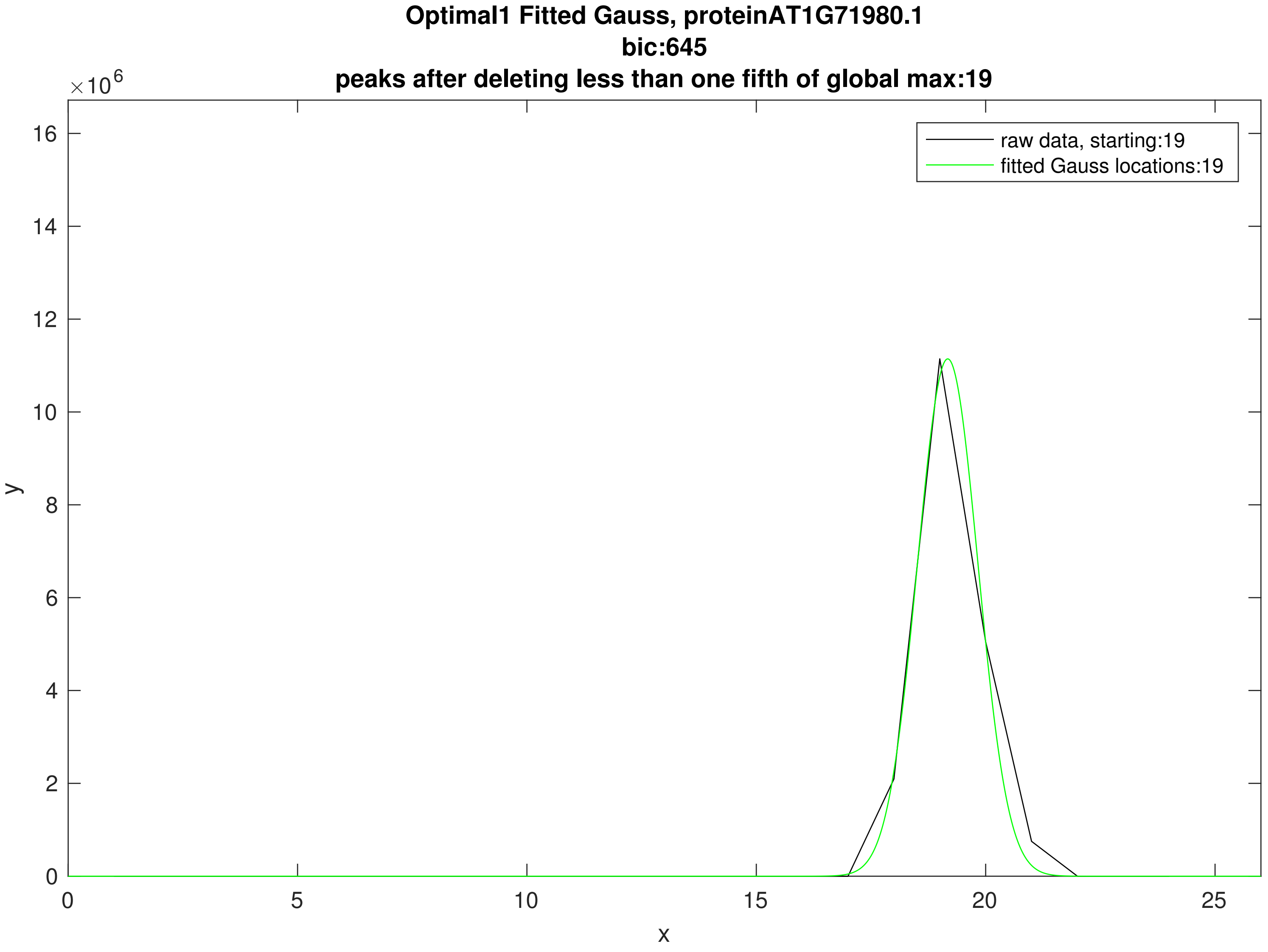

Optimal1 Fitted Gauss, proteinAT1G71980.1
bic:645
peaks after deleting less than one fifth of global max:19
×10^6
16
14
12
10
8
6
4
2
0
y
0
5
10
15
20
25
x
raw data, starting:19
fitted Gauss locations:19

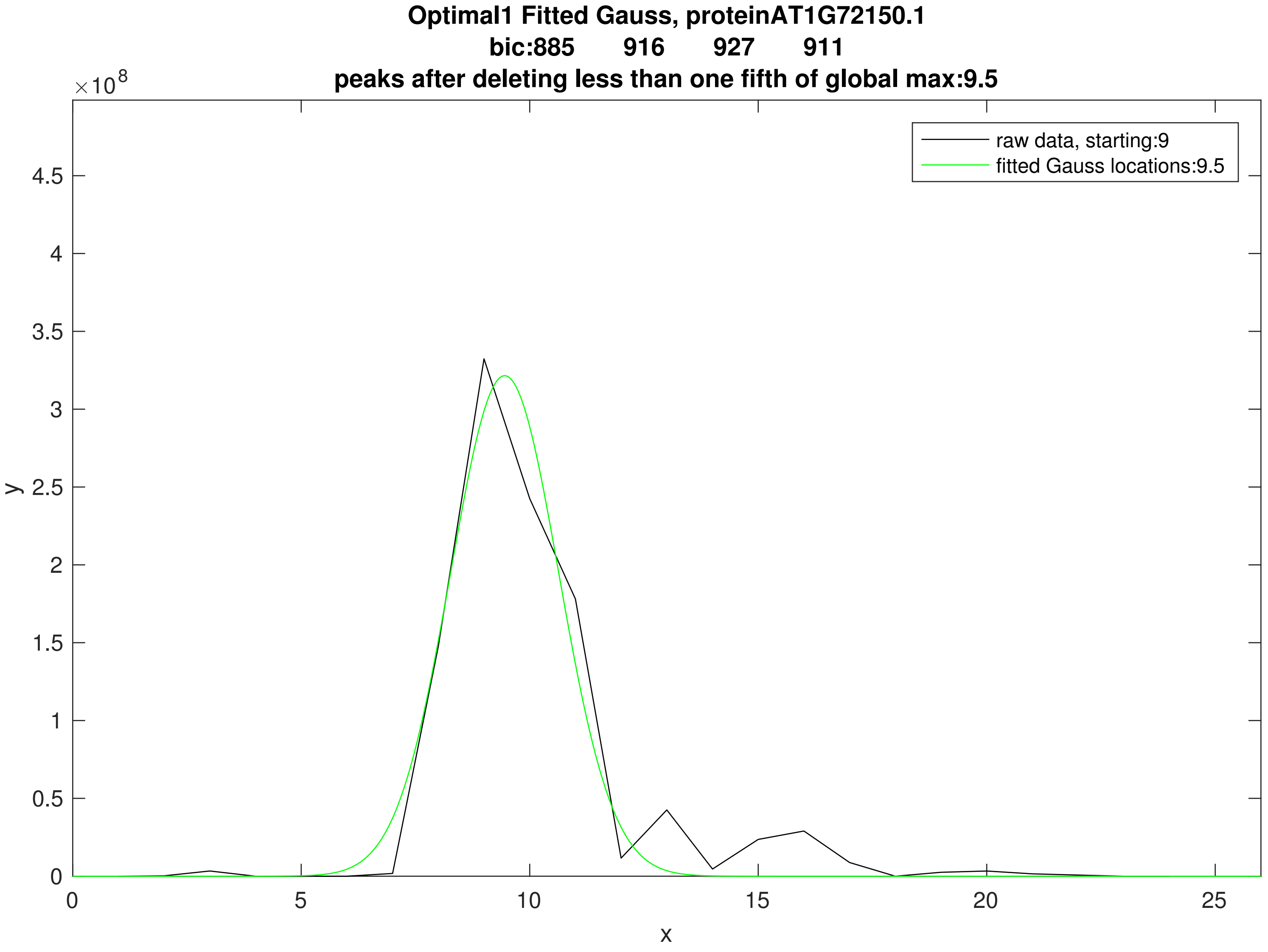
Optimal1 Fitted Gauss, proteinAT1G72150.1
bic:885 916 927 911
peaks after deleting less than one fifth of global max:9.5
×10^8
raw data, starting:9
fitted Gauss locations:9.5
y
x
0
0.5
1
1.5
2
2.5
3
3.5
4
4.5
5
10
15
20
25

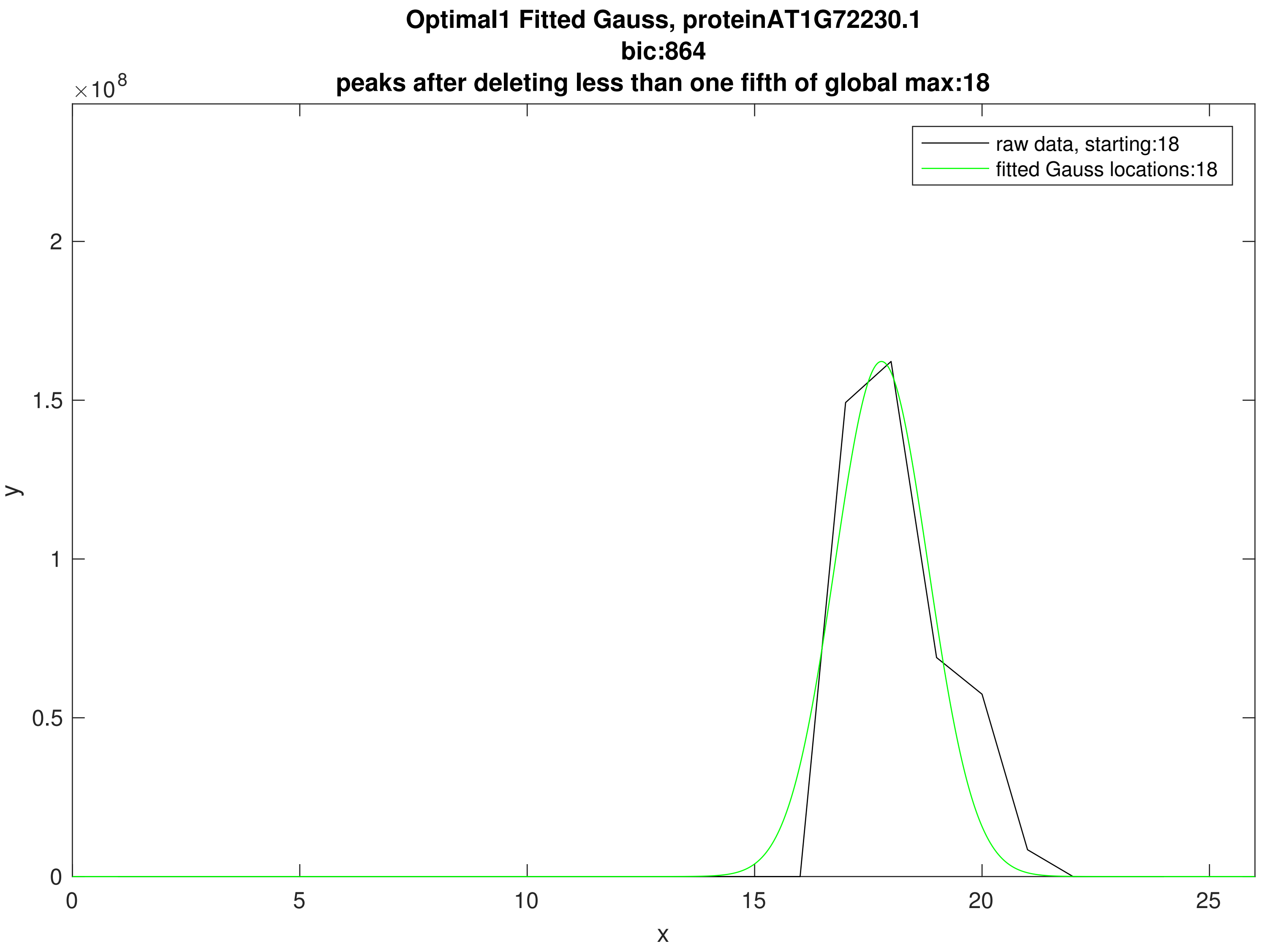
Optimal1 Fitted Gauss, proteinAT1G72230.1
bic:864
peaks after deleting less than one fifth of global max:18
×10^8
raw data, starting:18
fitted Gauss locations:18
2
1.5
1
0.5
0
y
0
5
10
15
20
25
x

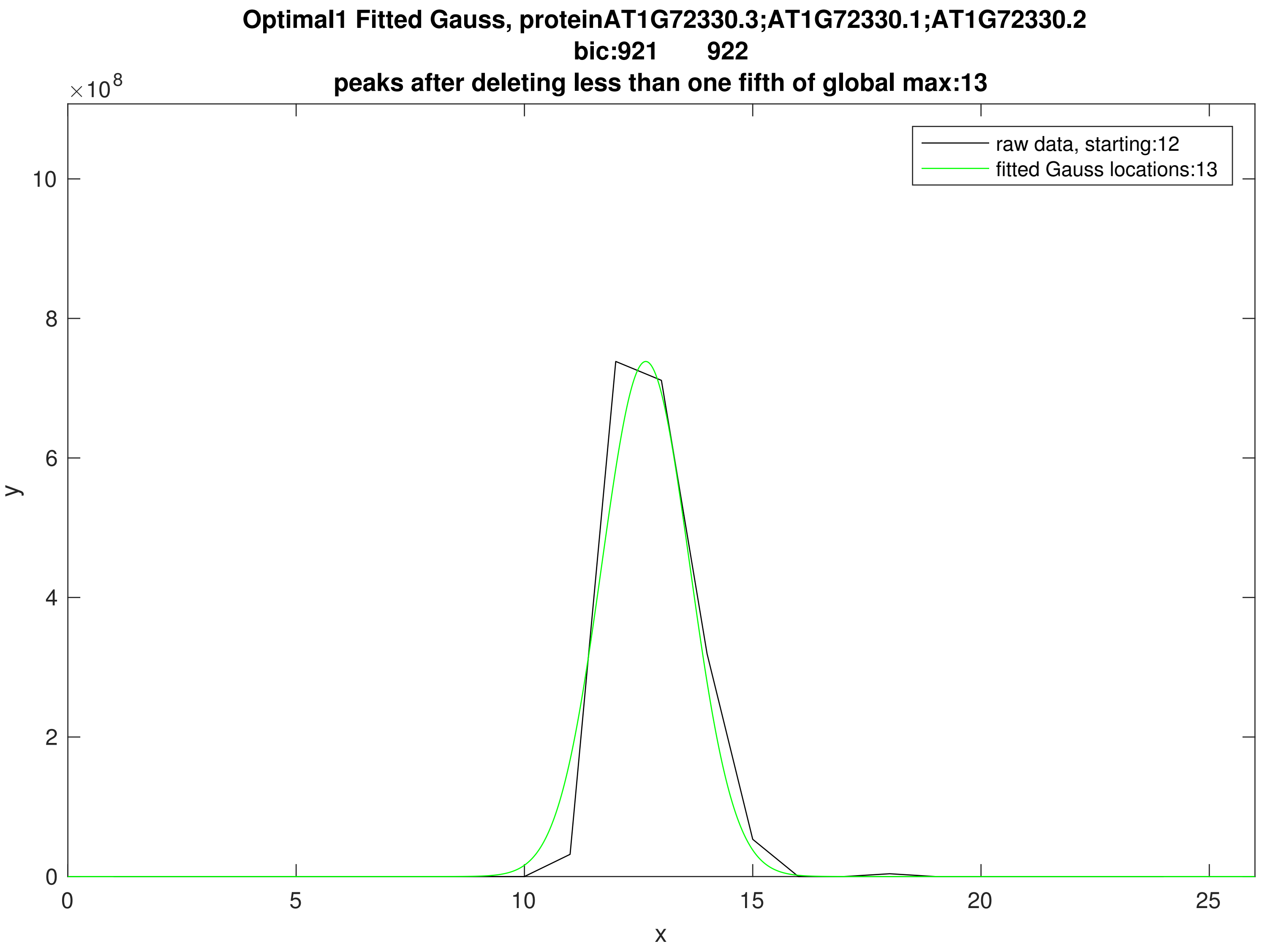
Optimal1 Fitted Gauss, proteinAT1G72330.3;AT1G72330.1;AT1G72330.2
bic:921 922
peaks after deleting less than one fifth of global max:13
×10^8
raw data, starting:12
fitted Gauss locations:13
y
x
0
2
4
6
8
10
5
15
20
25

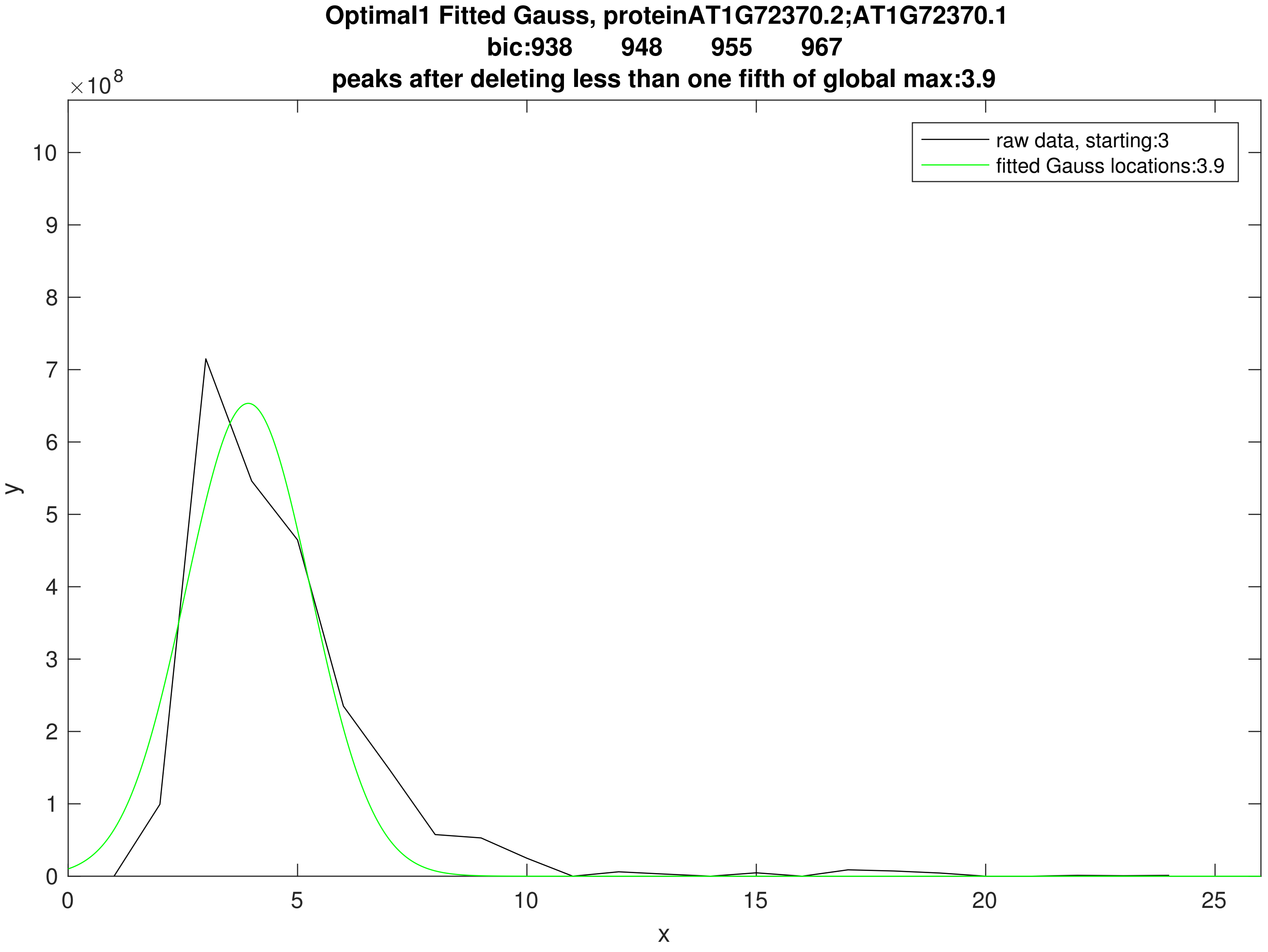
Optimal1 Fitted Gauss, proteinAT1G72370.2;AT1G72370.1
bic:938 948 955 967
peaks after deleting less than one fifth of global max:3.9
×10^8
raw data, starting:3
fitted Gauss locations:3.9
y
x
0
1
2
3
4
5
6
7
8
9
10
0
5
10
15
20
25

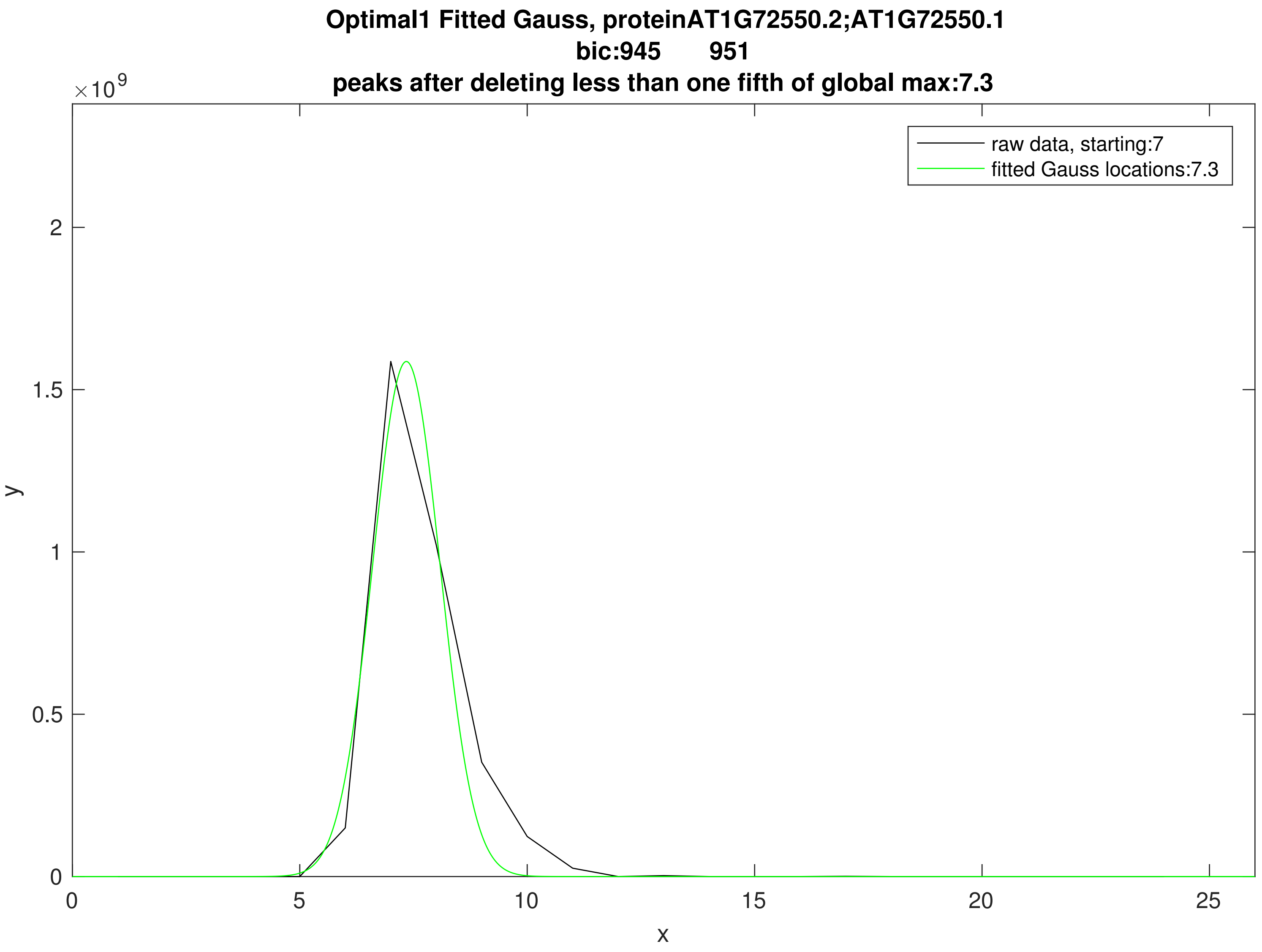

Optimal1 Fitted Gauss, proteinAT1G72550.2;AT1G72550.1
bic:945 951
peaks after deleting less than one fifth of global max:7.3
×10^9
raw data, starting:7
fitted Gauss locations:7.3
y
x
0
0.5
1
1.5
2
5
10
15
20
25

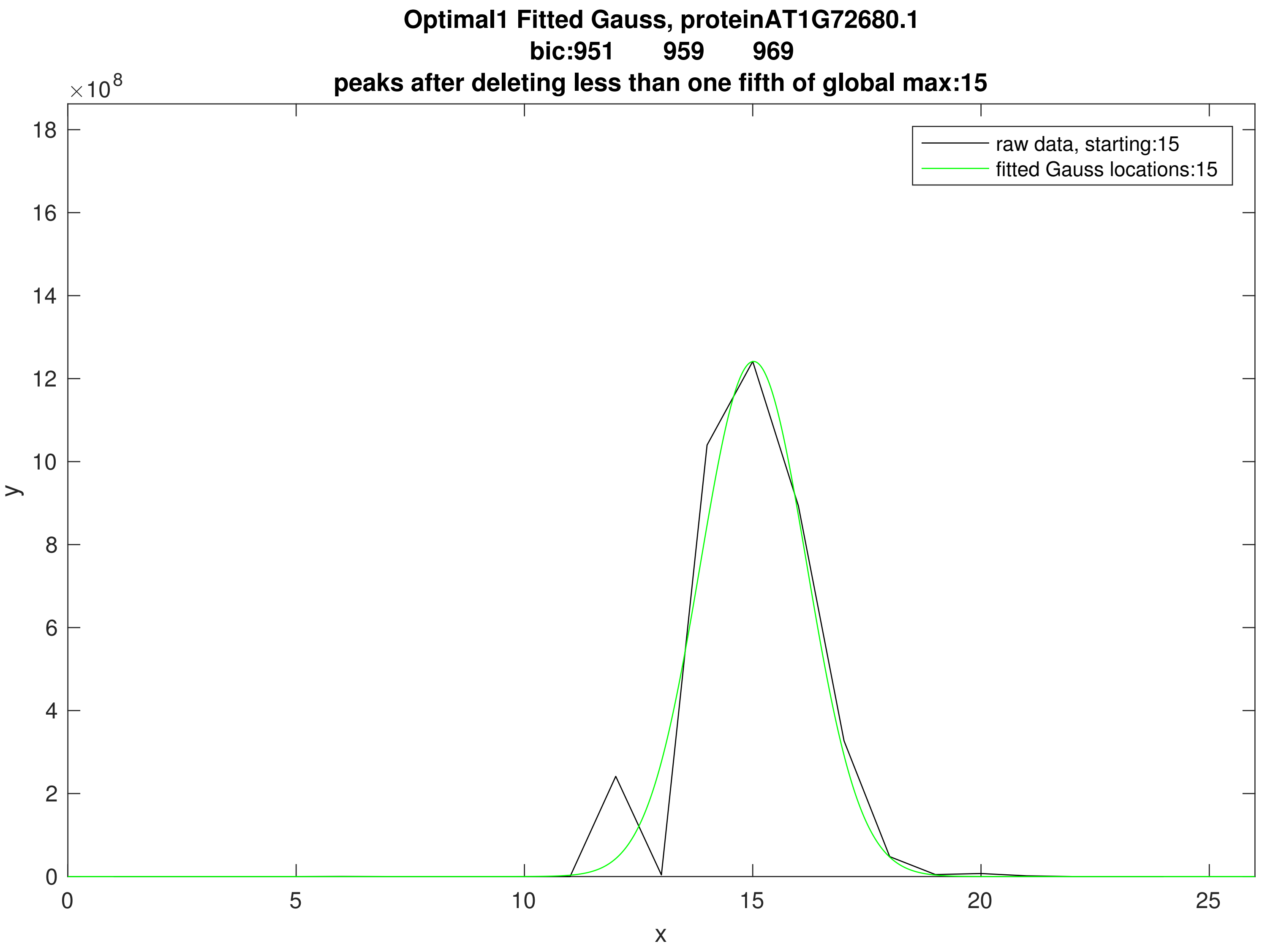

Optimal1 Fitted Gauss, proteinAT1G72680.1
bic:951 959 969
peaks after deleting less than one fifth of global max:15
×10^8
18
16
14
12
10
8
6
4
2
0
y
0
5
10
15
20
25
x
raw data, starting:15
fitted Gauss locations:15

#### Optimal1 Fitted Gauss, proteinAT1G72880.2;AT1G72880.1
#### bic:844
#### peaks after deleting less than one fifth of global max:11

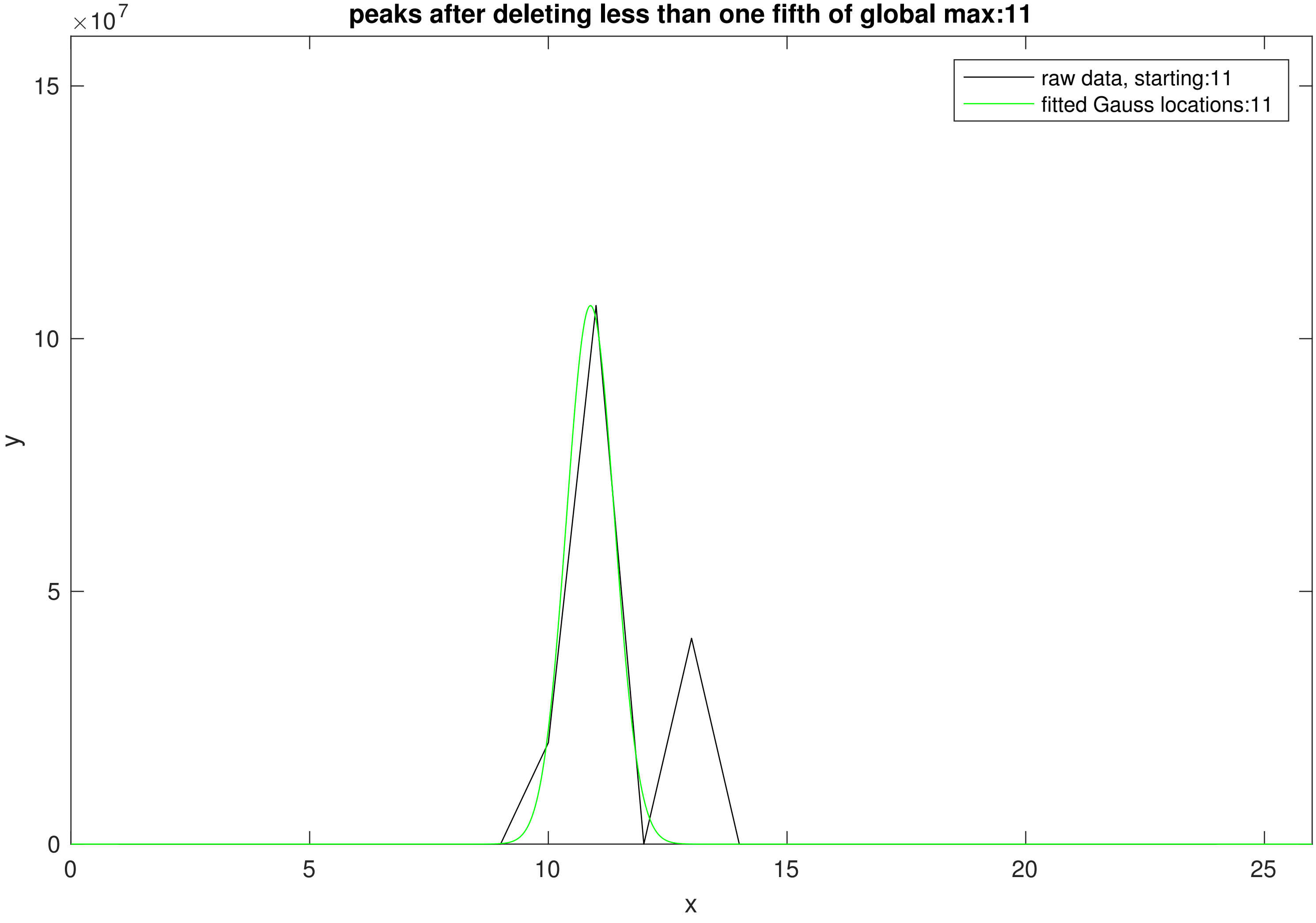

**Optimal1 Fitted Gauss, proteinAT1G73080.1**
**bic:727**
**peaks after deleting less than one fifth of global max:11**

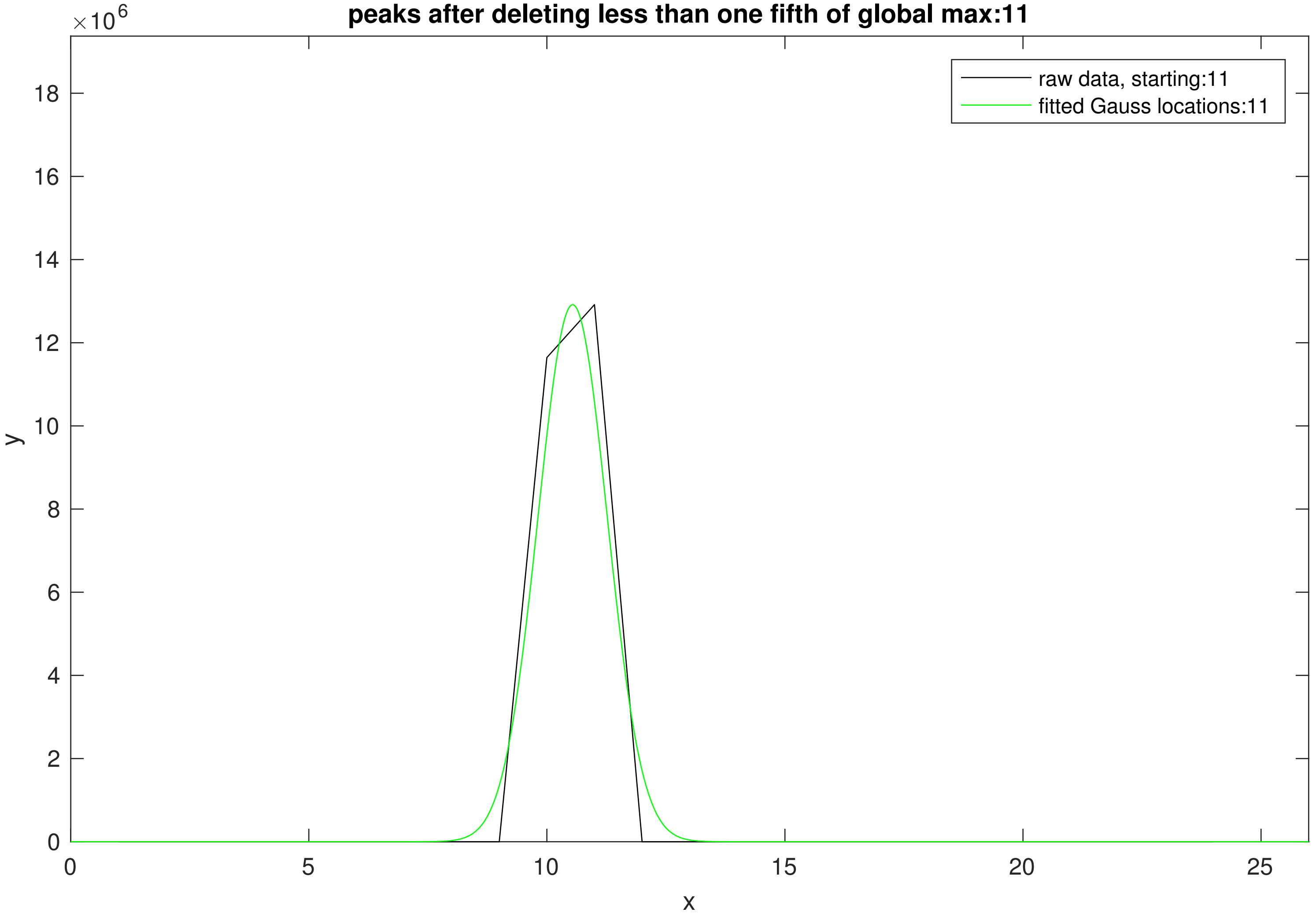

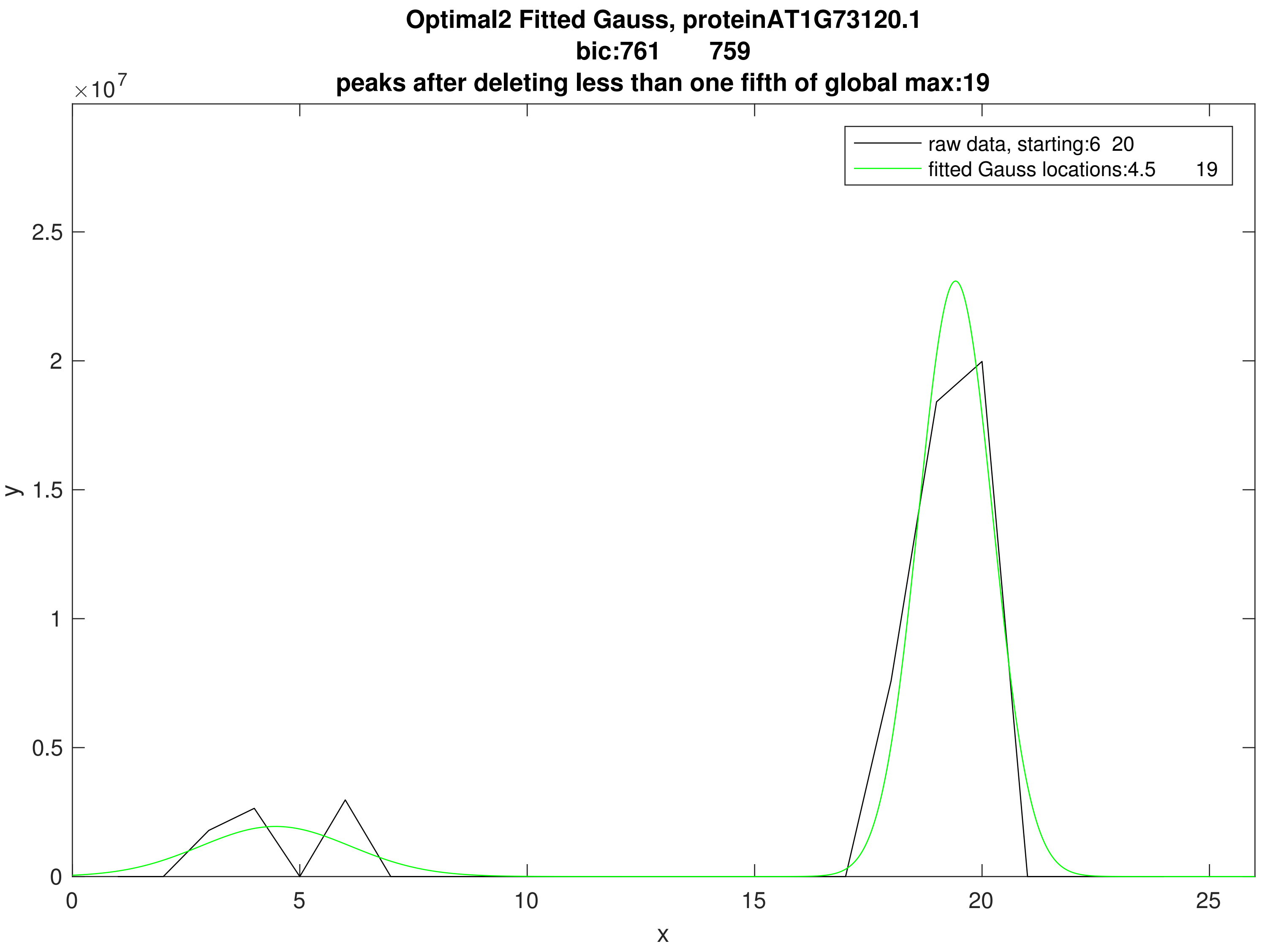

Optimal2 Fitted Gauss, proteinAT1G73120.1
bic:761 759
peaks after deleting less than one fifth of global max:19
×10^7
raw data, starting:6 20
fitted Gauss locations:4.5 19
y
x
0
0.5
1
1.5
2
2.5
5
10
15
20
25

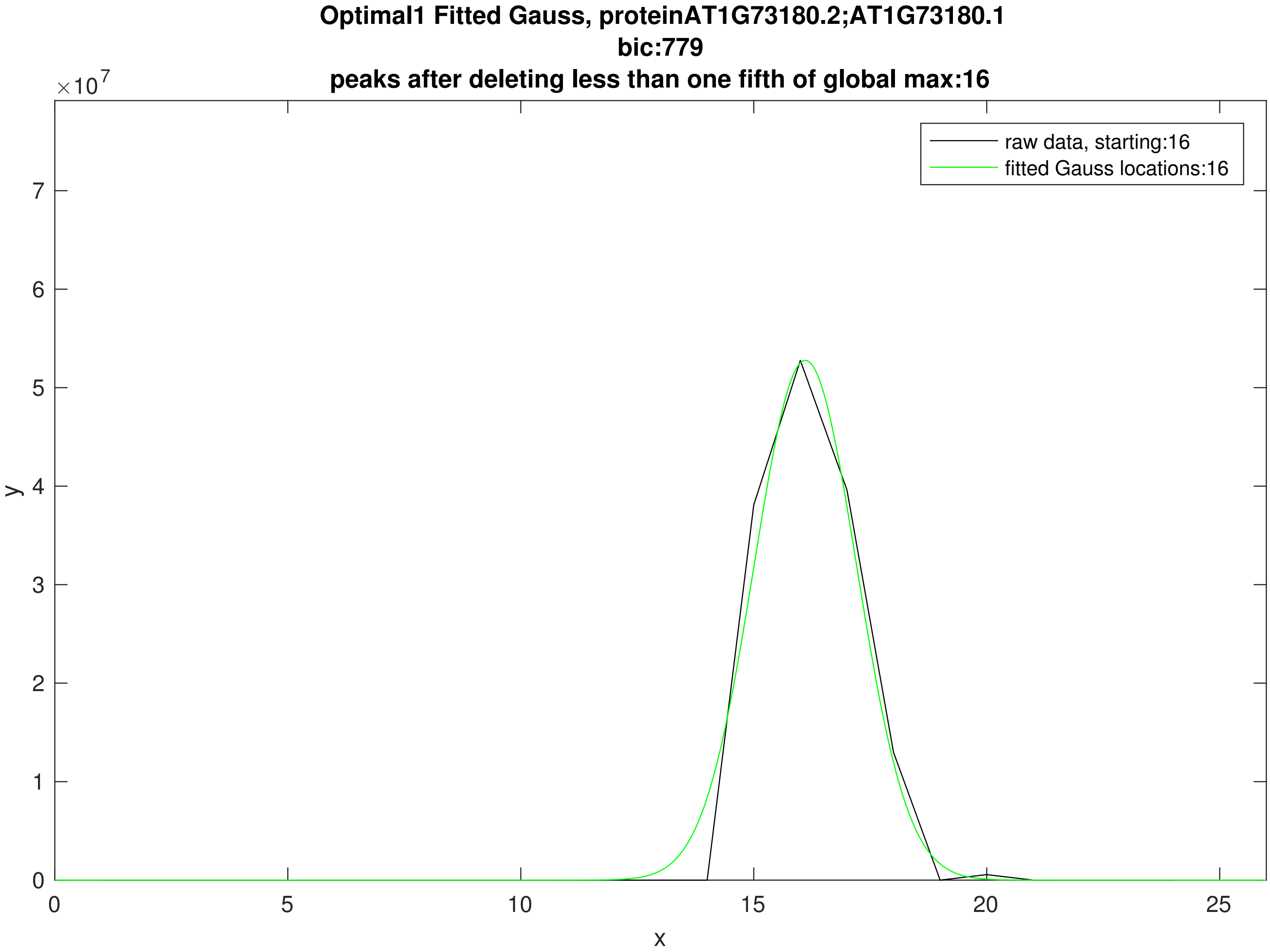

Optimal1 Fitted Gauss, proteinAT1G73180.2;AT1G73180.1
bic:779
peaks after deleting less than one fifth of global max:16
×10^7
raw data, starting:16
fitted Gauss locations:16
y
x
0
1
2
3
4
5
6
7
0
5
10
15
20
25

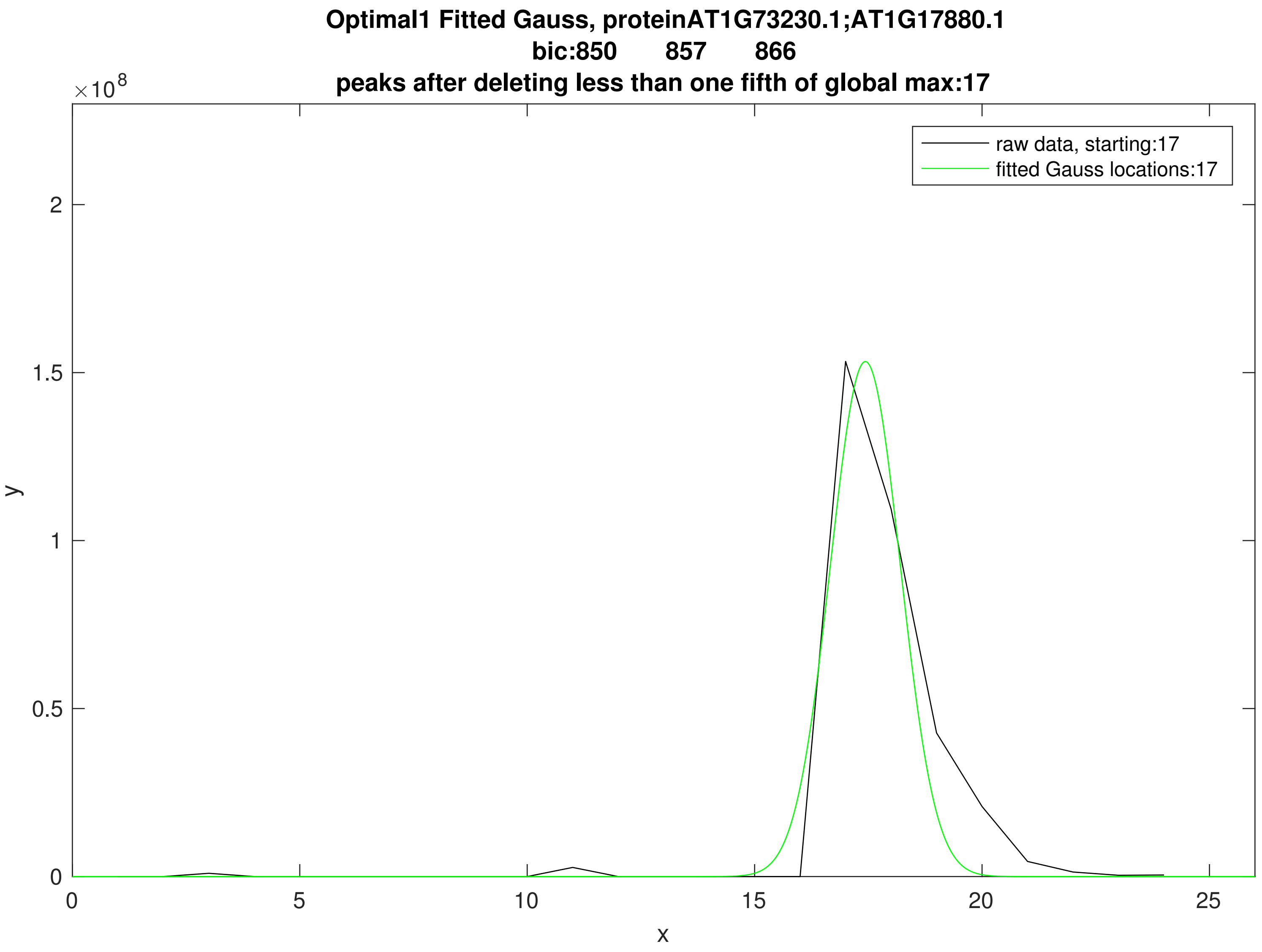

Optimal1 Fitted Gauss, proteinAT1G73230.1;AT1G17880.1
bic:850 857 866
peaks after deleting less than one fifth of global max:17
×10^8
raw data, starting:17
fitted Gauss locations:17
y
x
0
0.5
1
1.5
2
0
5
10
15
20
25

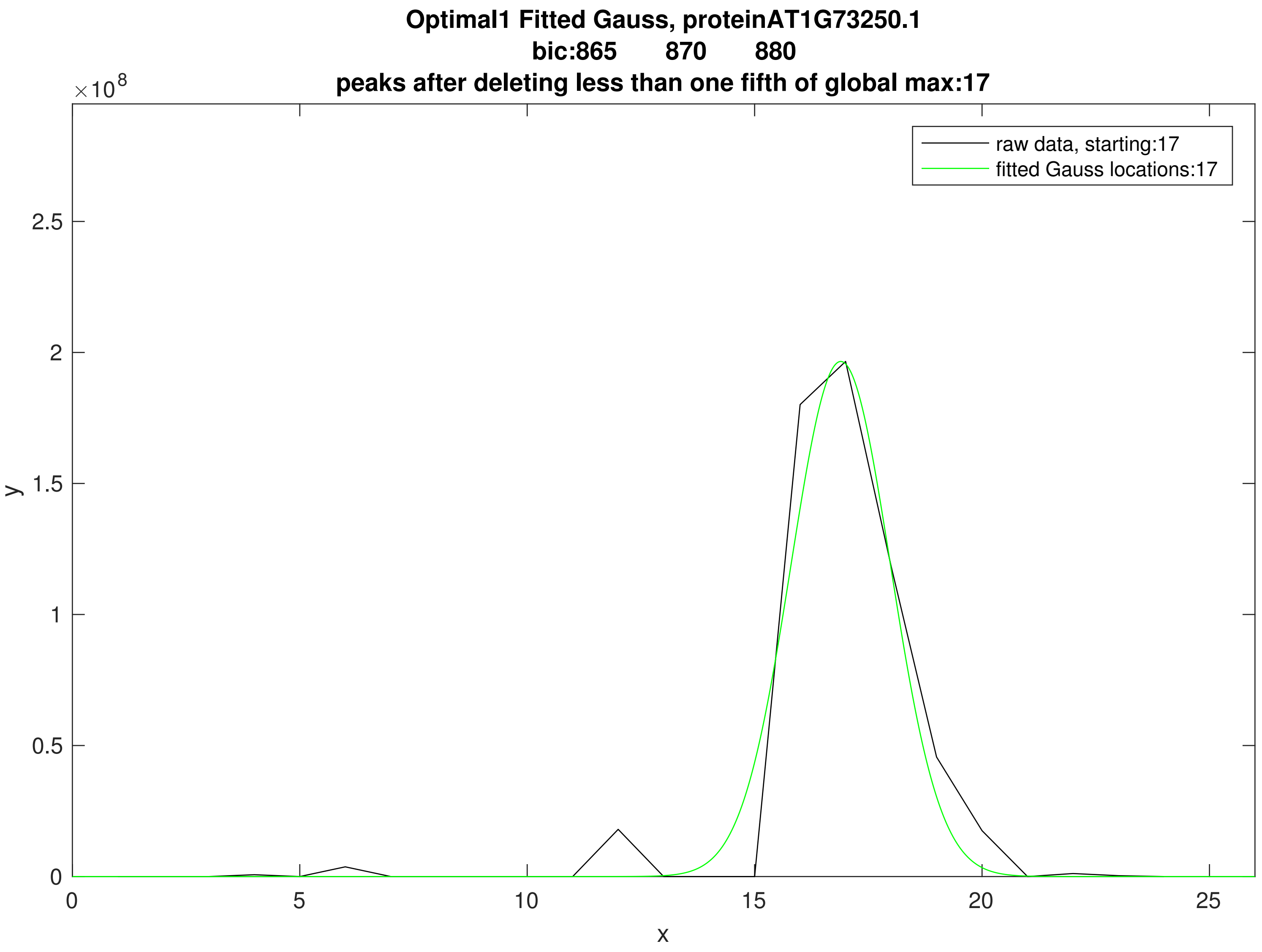

Optimal1 Fitted Gauss, proteinAT1G73250.1
bic:865 870 880
peaks after deleting less than one fifth of global max:17
×10^8
raw data, starting:17
fitted Gauss locations:17
y
x
0
0.5
1
1.5
2
2.5
0
5
10
15
20
25

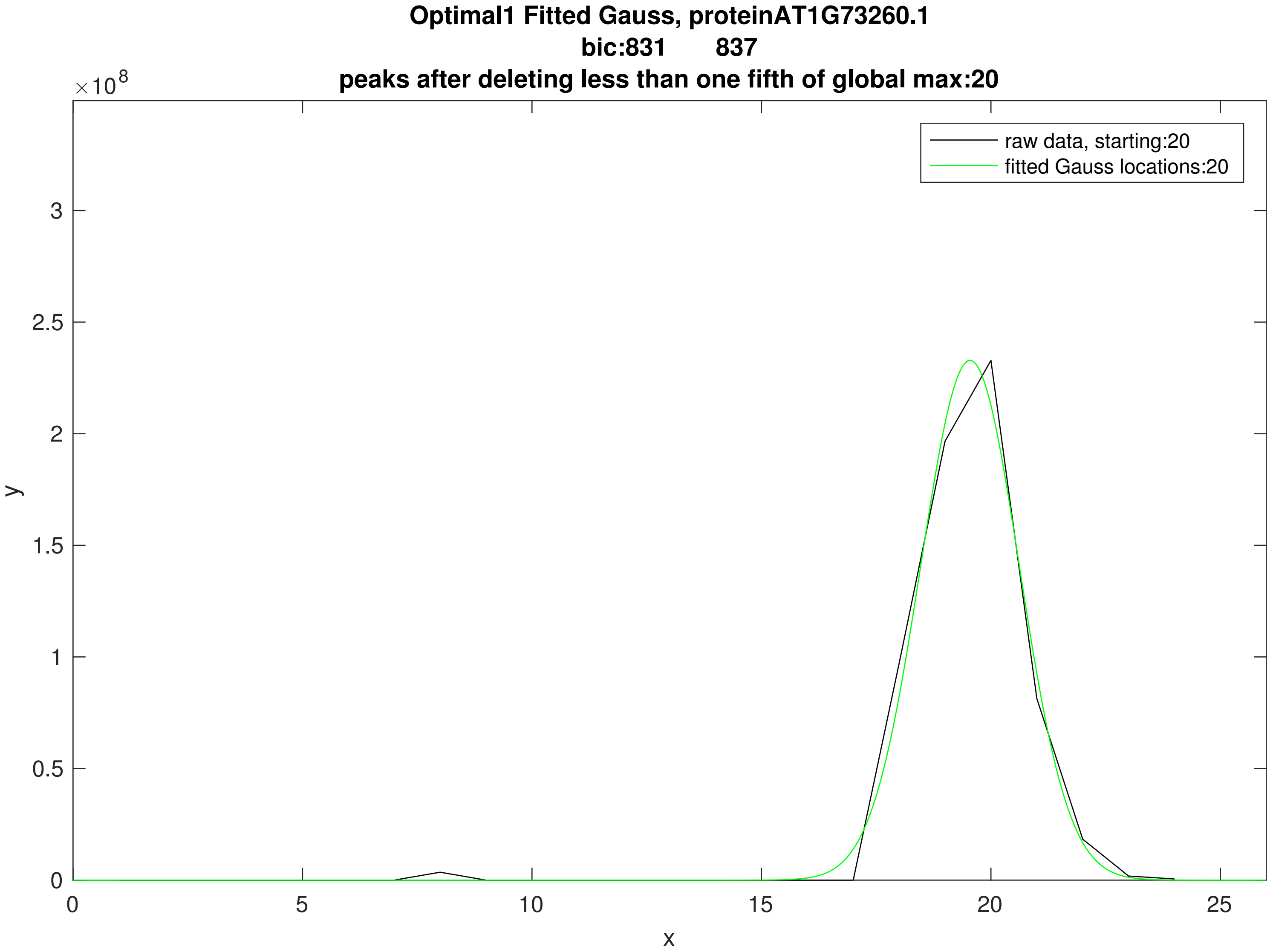

Optimal1 Fitted Gauss, proteinAT1G73260.1
bic:831 837
peaks after deleting less than one fifth of global max:20
×10^8
raw data, starting:20
fitted Gauss locations:20
3
2.5
2
1.5
1
0.5
0
y
0
5
10
15
20
25
x

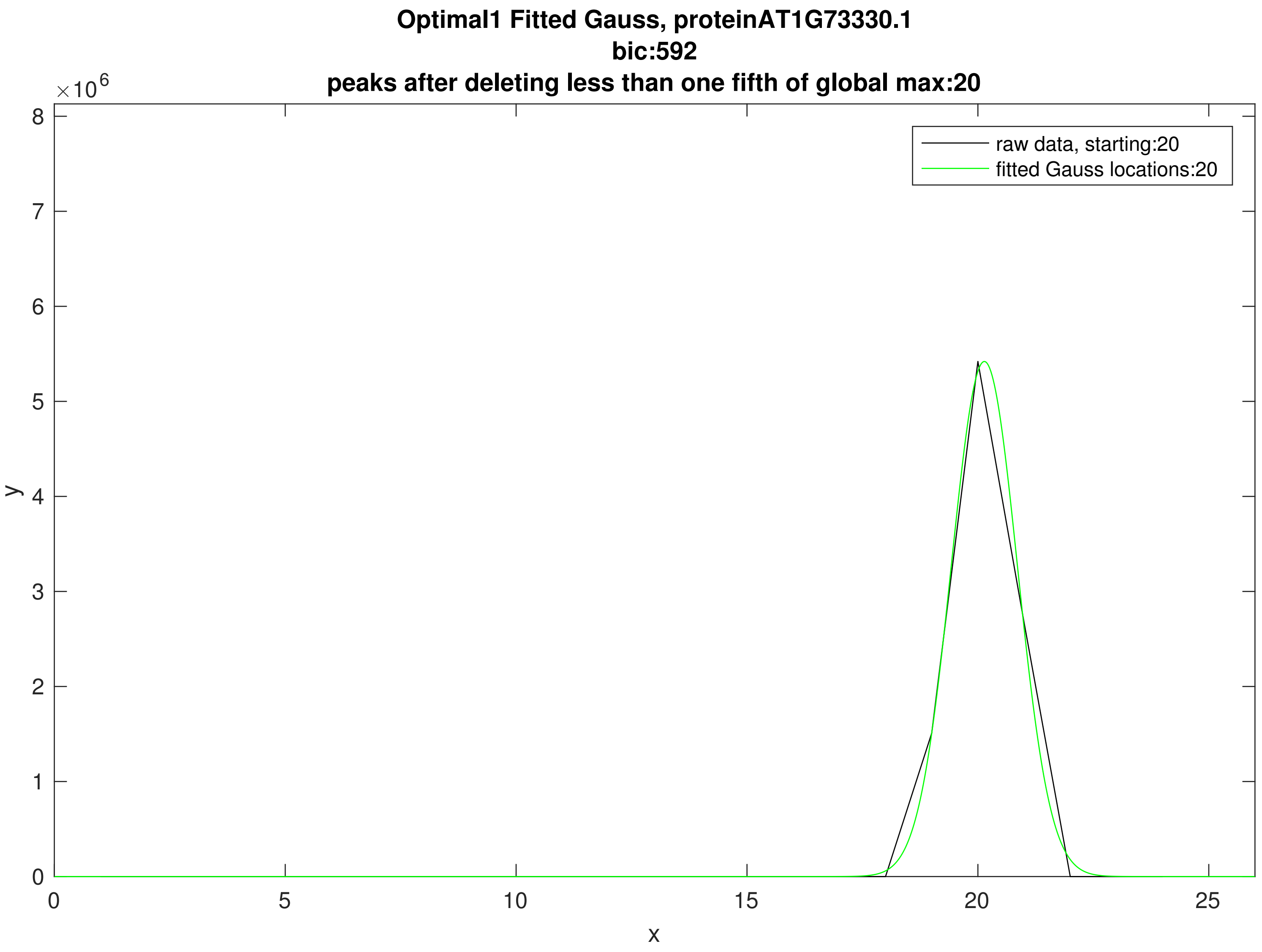
Optimal1 Fitted Gauss, proteinAT1G73330.1
bic:592
peaks after deleting less than one fifth of global max:20
×10^6
raw data, starting:20
fitted Gauss locations:20
y
x
0
1
2
3
4
5
6
7
8
0
5
10
15
20
25

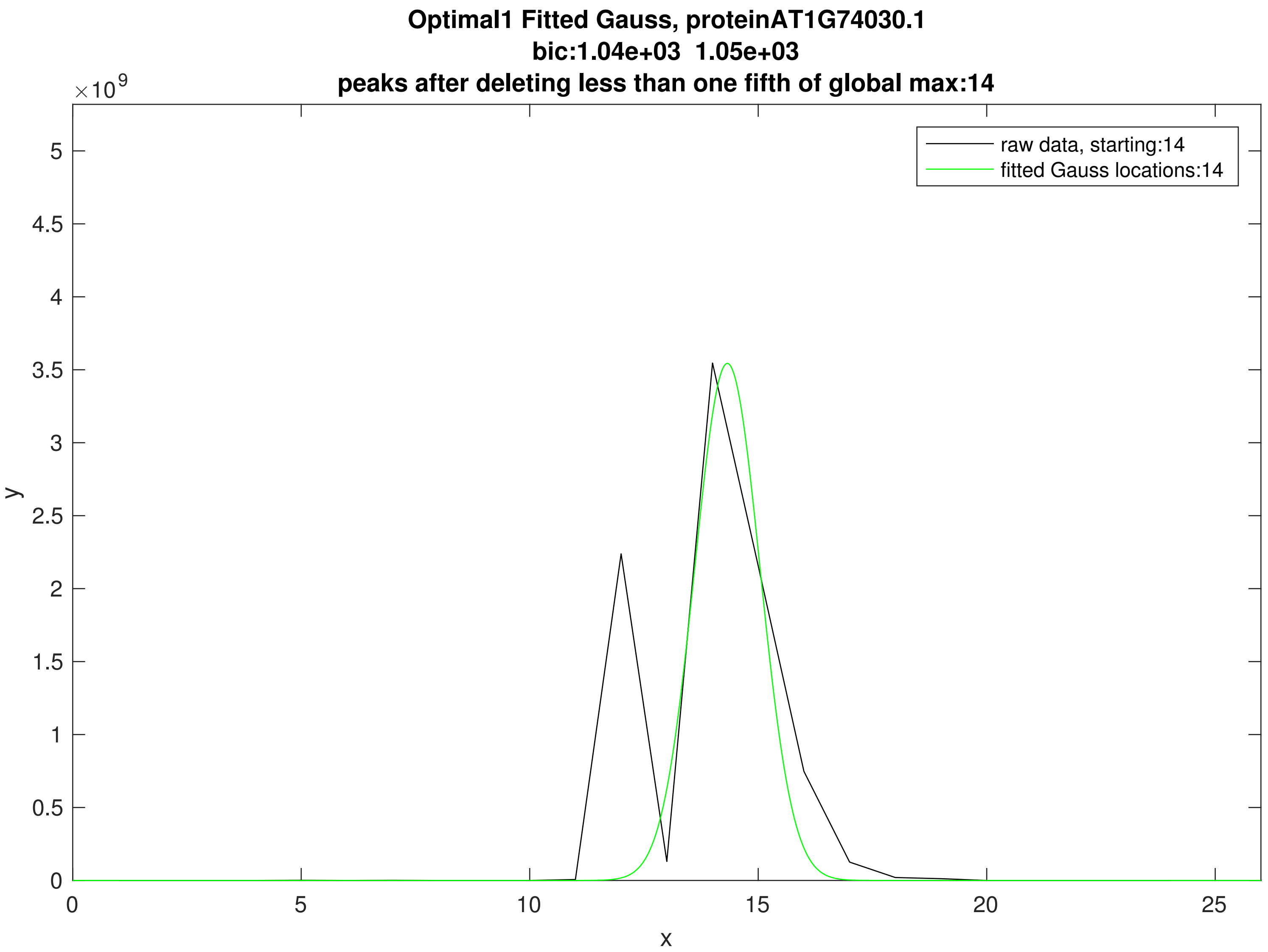

Optimal1 Fitted Gauss, proteinAT1G74030.1
bic:1.04e+03  1.05e+03
peaks after deleting less than one fifth of global max:14
×10^9
5
4.5
4
3.5
3
2.5
2
1.5
1
0.5
0
y
0
5
10
15
20
25
x
raw data, starting:14
fitted Gauss locations:14

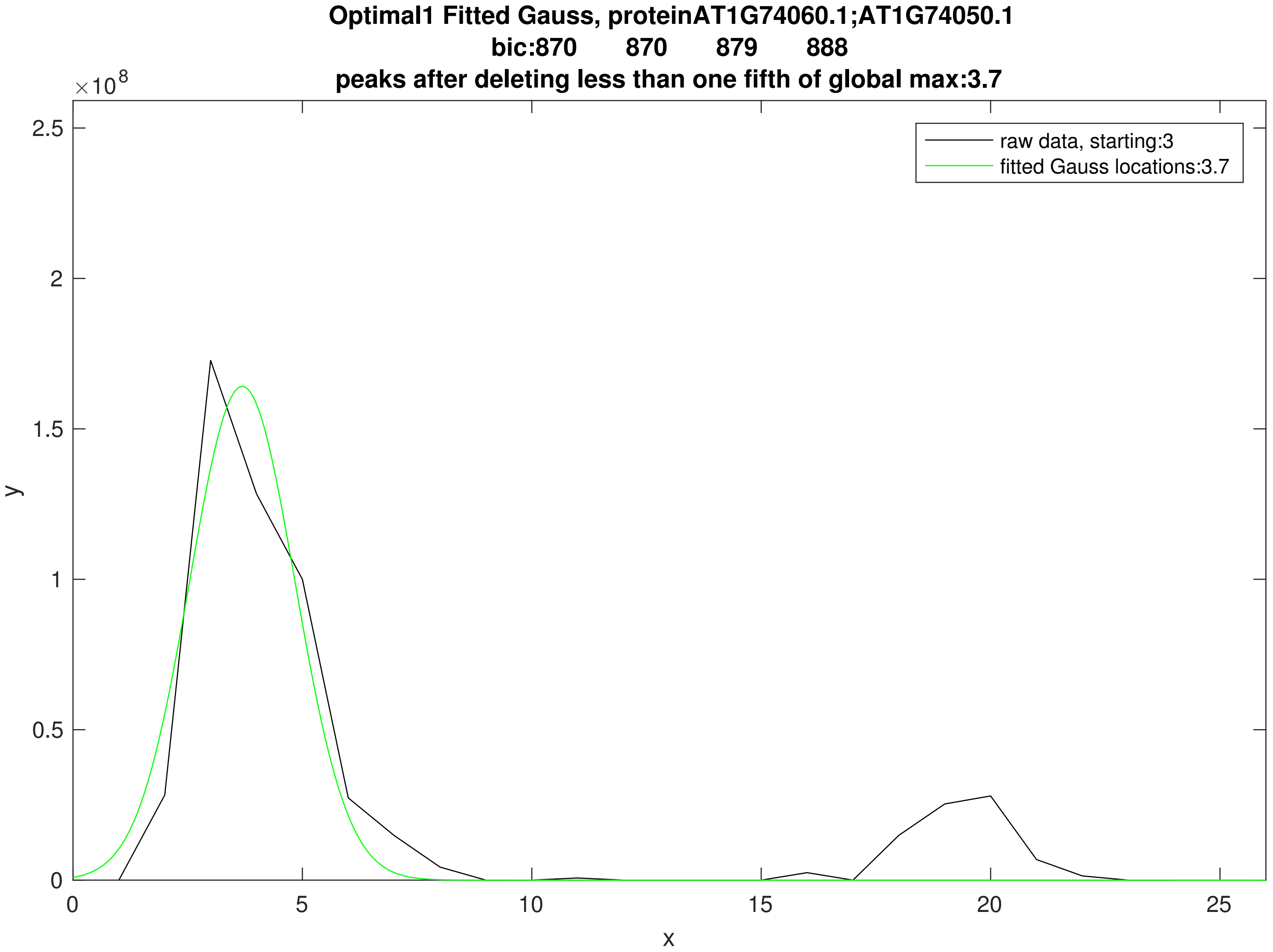

Optimal1 Fitted Gauss, proteinAT1G74060.1;AT1G74050.1
bic:870 870 879 888
peaks after deleting less than one fifth of global max:3.7
×10^8
raw data, starting:3
fitted Gauss locations:3.7
y
x
0
0.5
1
1.5
2
2.5
0
5
10
15
20
25

**Optimal1 Fitted Gauss, proteinAT1G74100.1**
**bic:769**
**peaks after deleting less than one fifth of global max:17**

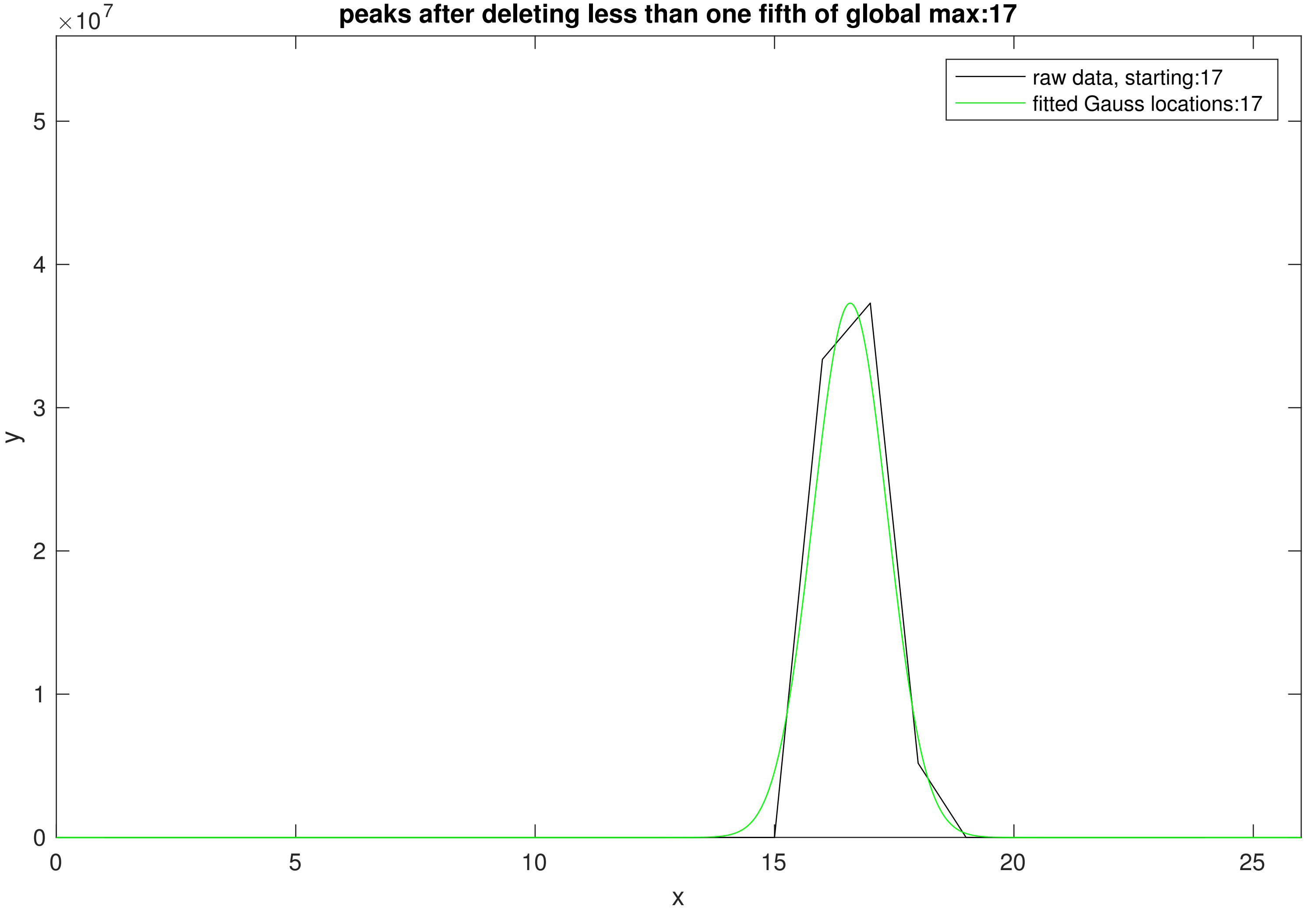

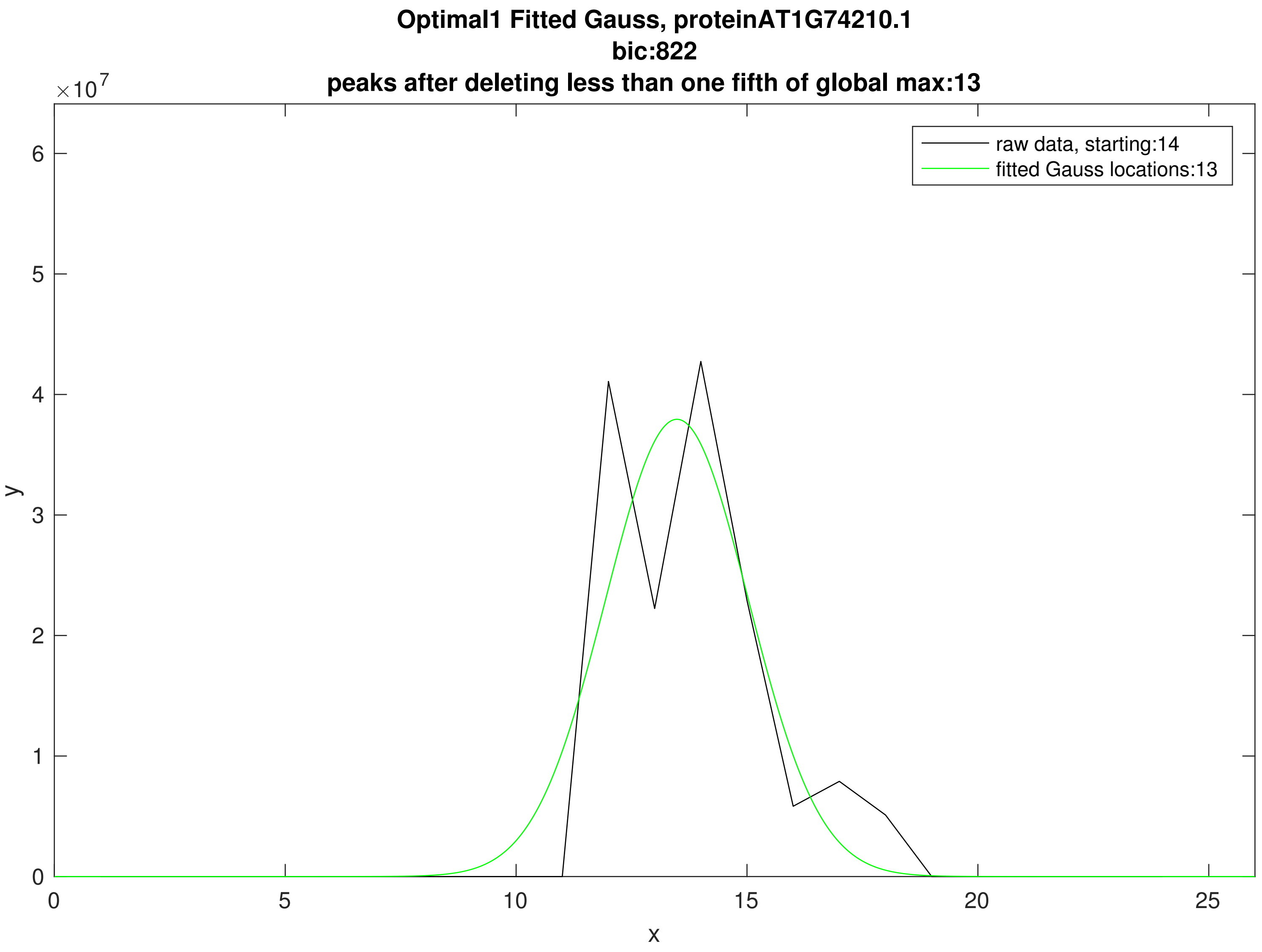

Optimal1 Fitted Gauss, proteinAT1G74210.1
bic:822
peaks after deleting less than one fifth of global max:13
×10^7
6
5
4
3
2
1
0
y
0
5
10
15
20
25
x
raw data, starting:14
fitted Gauss locations:13

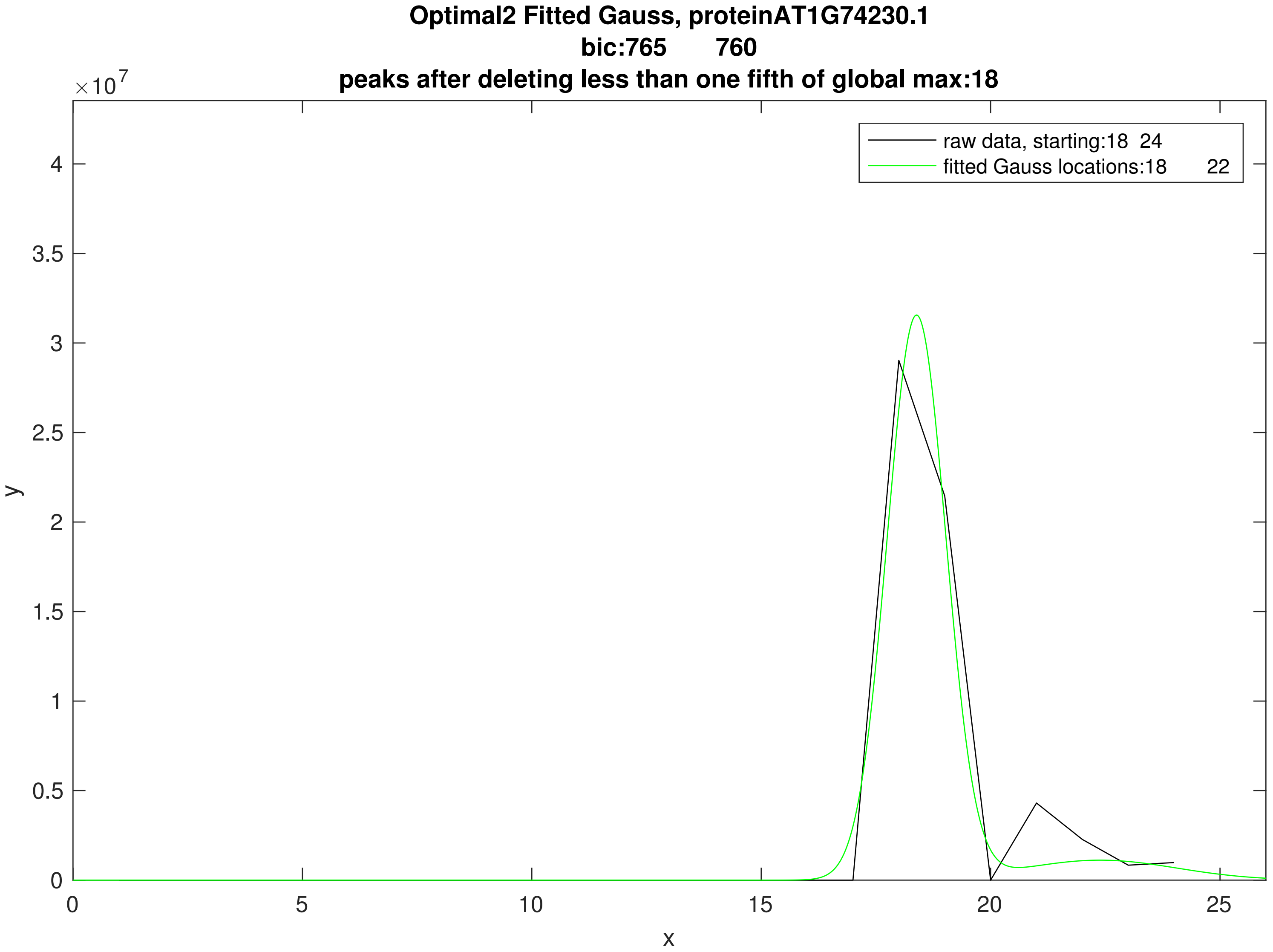

Optimal2 Fitted Gauss, proteinAT1G74230.1
bic:765 760
peaks after deleting less than one fifth of global max:18
×10^7
raw data, starting:18 24
fitted Gauss locations:18 22
4
3.5
3
2.5
2
1.5
1
0.5
0
y
0
5
10
15
20
25
x

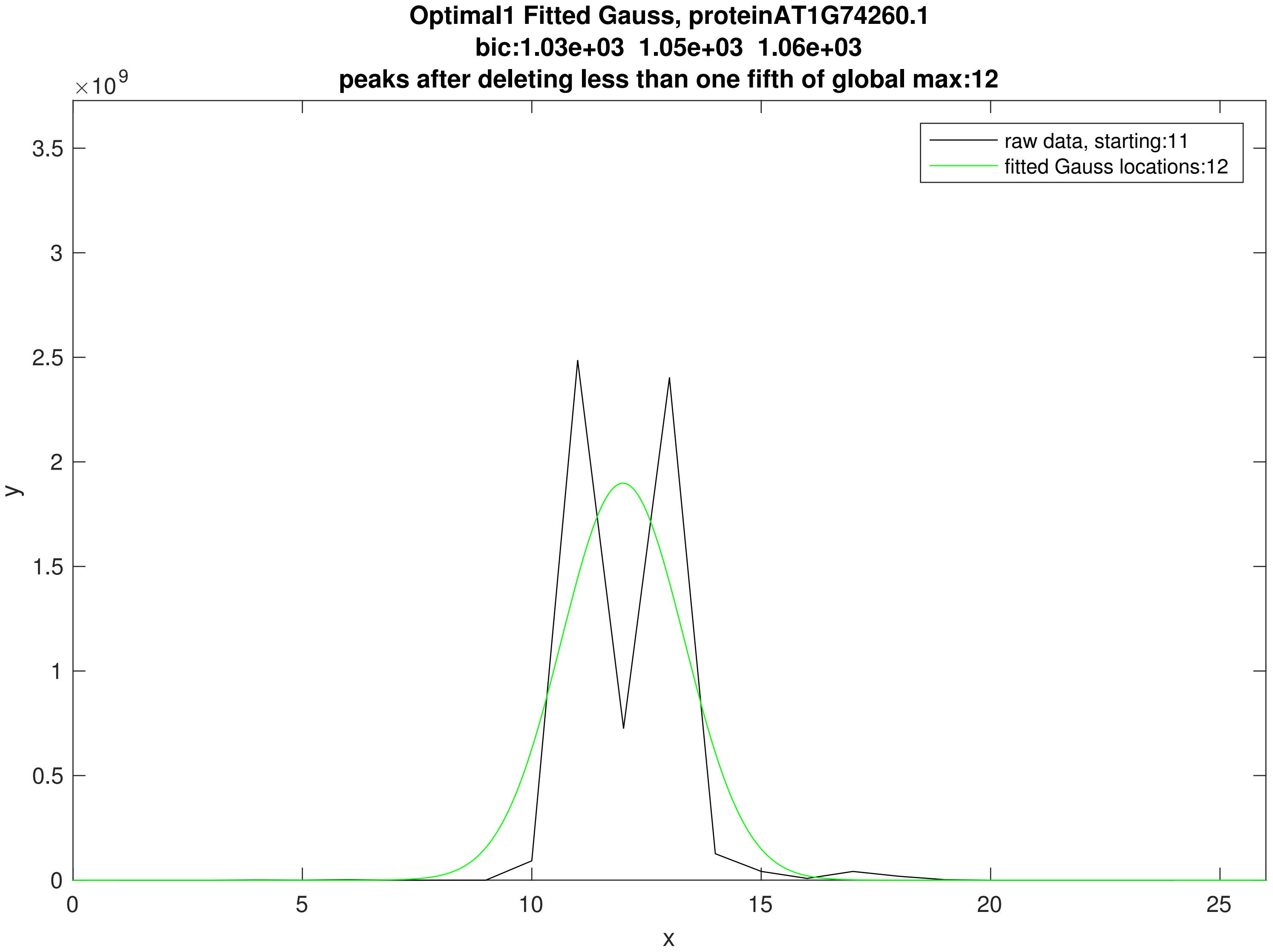
Optimal1 Fitted Gauss, proteinAT1G74260.1
bic:1.03e+03  1.05e+03  1.06e+03
peaks after deleting less than one fifth of global max:12
×10^9
3.5
3
2.5
2
1.5
1
0.5
0
y
0
5
10
15
20
25
x
raw data, starting:11
fitted Gauss locations:12

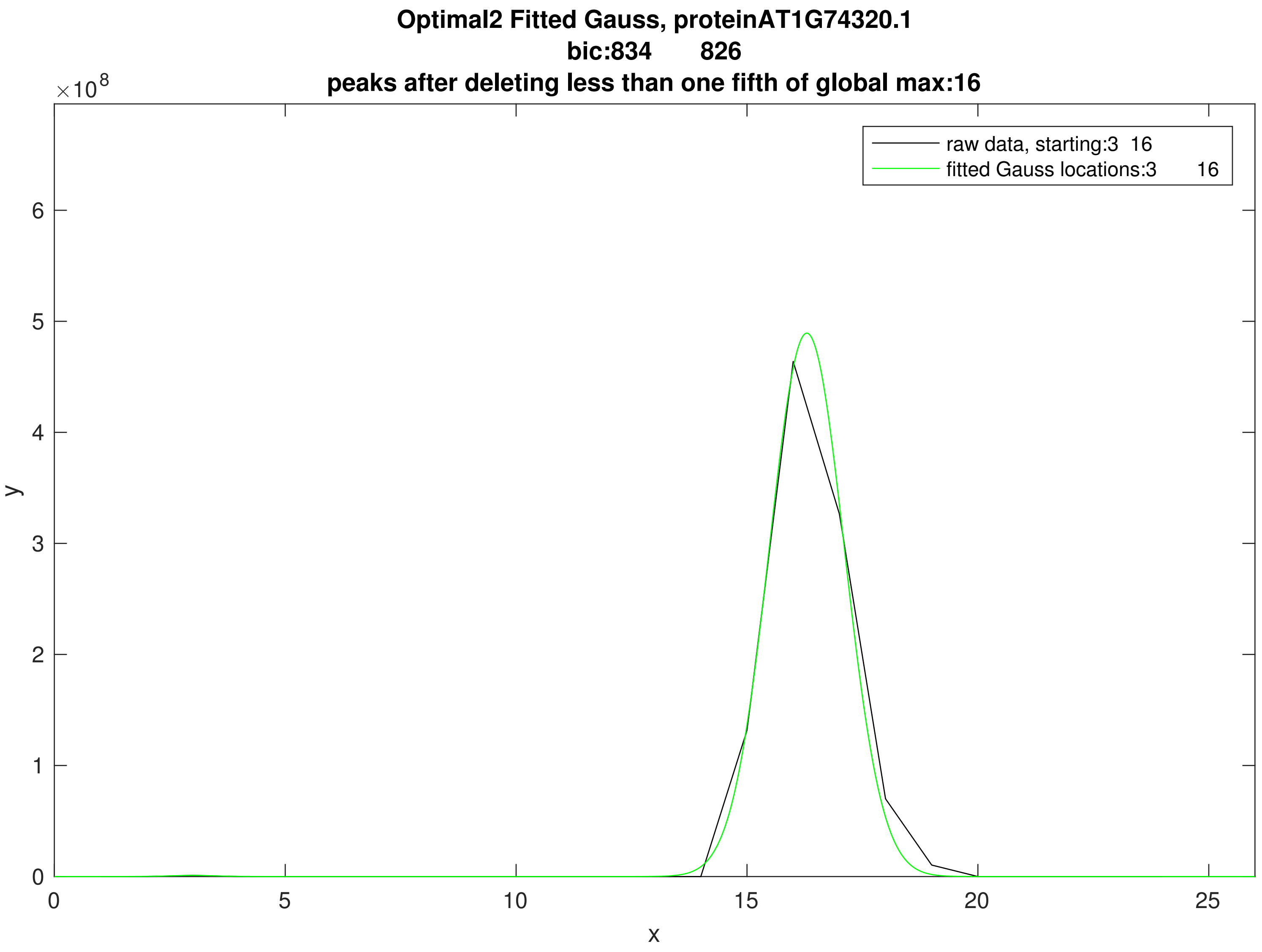

Optimal2 Fitted Gauss, proteinAT1G74320.1
bic:834 826
peaks after deleting less than one fifth of global max:16
×10^8
raw data, starting:3 16
fitted Gauss locations:3 16
y
x
0
1
2
3
4
5
6
0
5
10
15
20
25

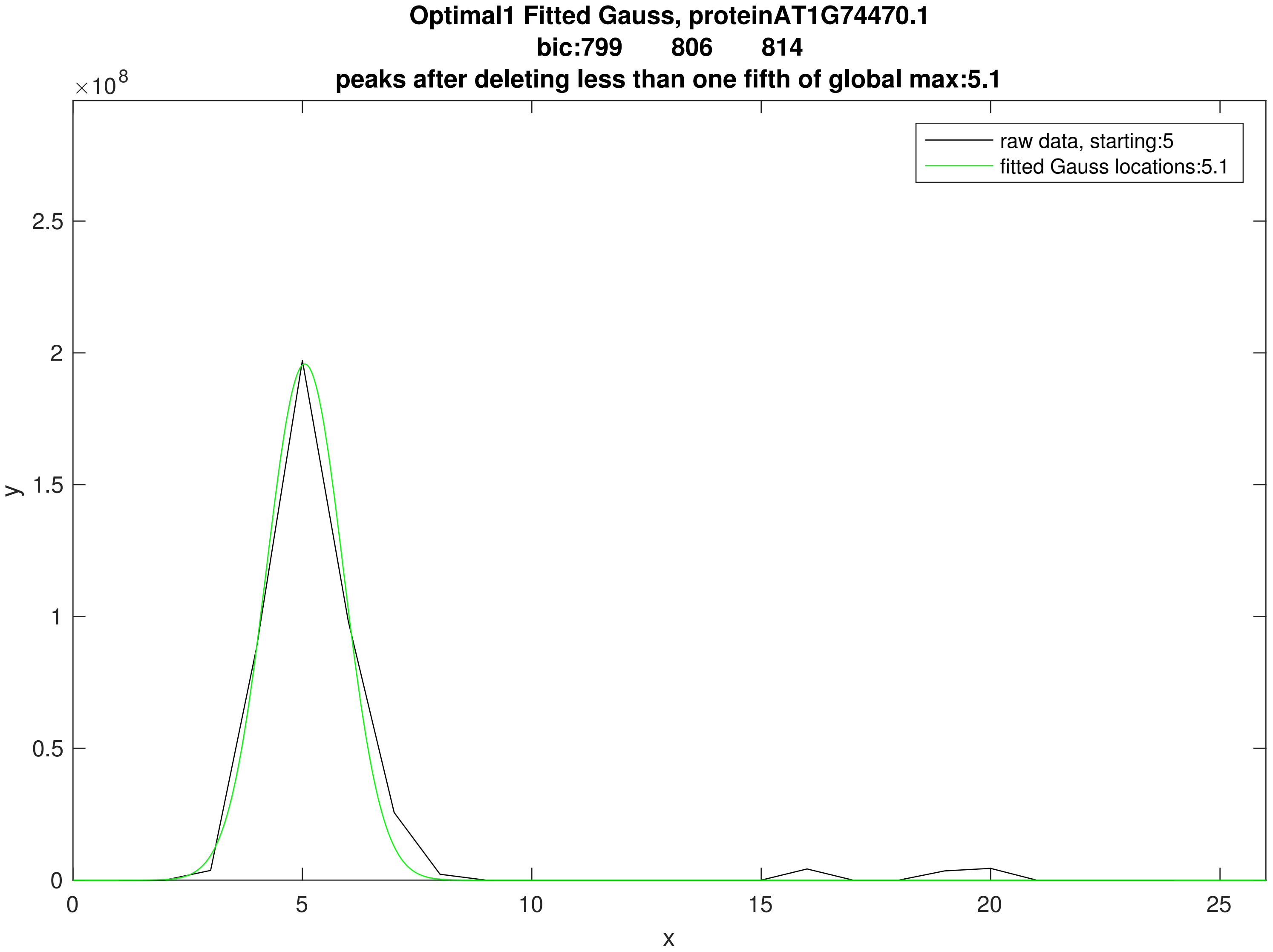

Optimal1 Fitted Gauss, proteinAT1G74470.1
bic:799 806 814
peaks after deleting less than one fifth of global max:5.1
×10^8
raw data, starting:5
fitted Gauss locations:5.1
y
x
0
0.5
1
1.5
2
2.5
0
5
10
15
20
25

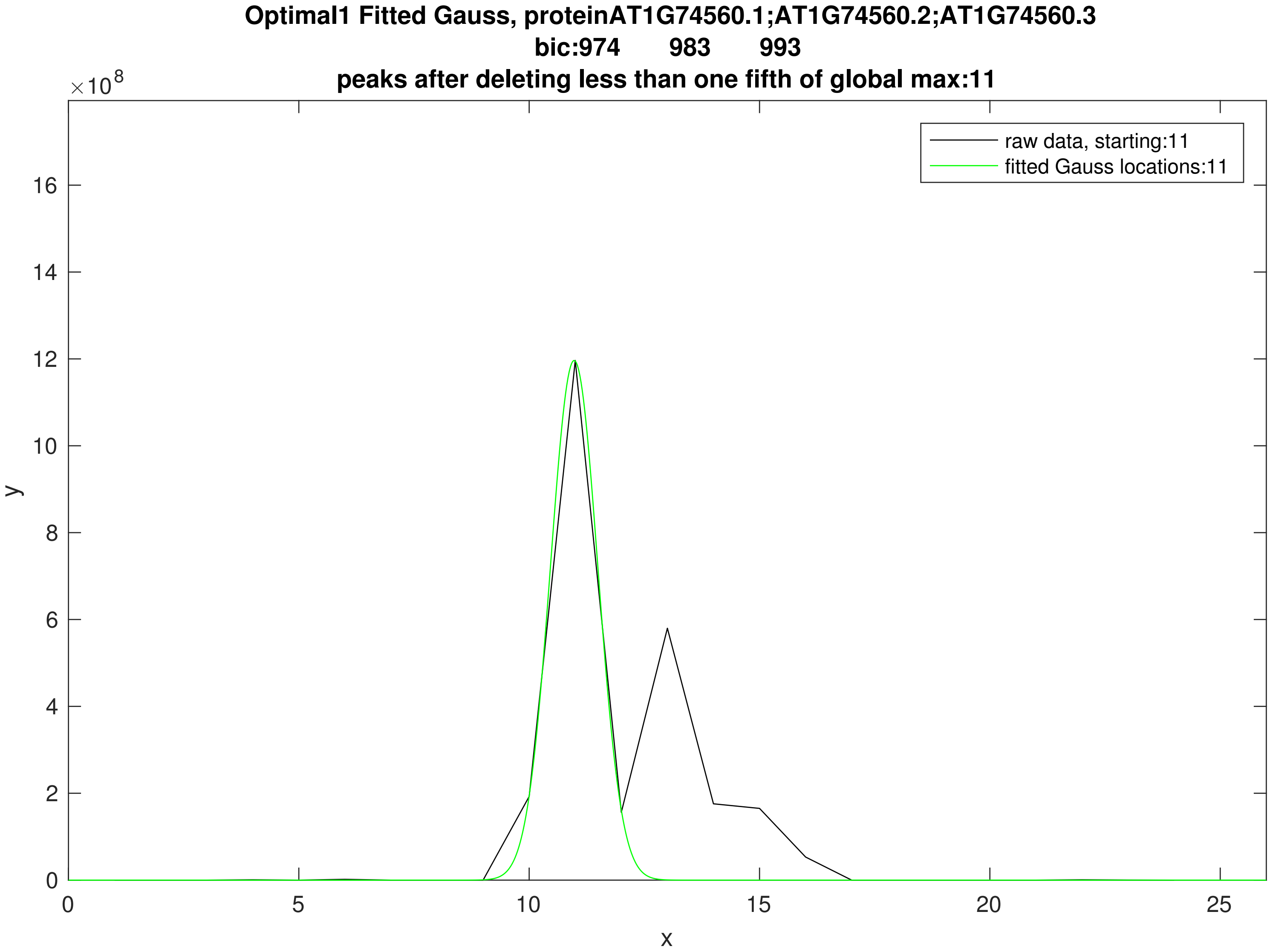

Optimal1 Fitted Gauss, proteinAT1G74560.1;AT1G74560.2;AT1G74560.3
bic:974 983 993
peaks after deleting less than one fifth of global max:11
×10^8
raw data, starting:11
fitted Gauss locations:11
y
x
0
2
4
6
8
10
12
14
16
0
5
10
15
20
25

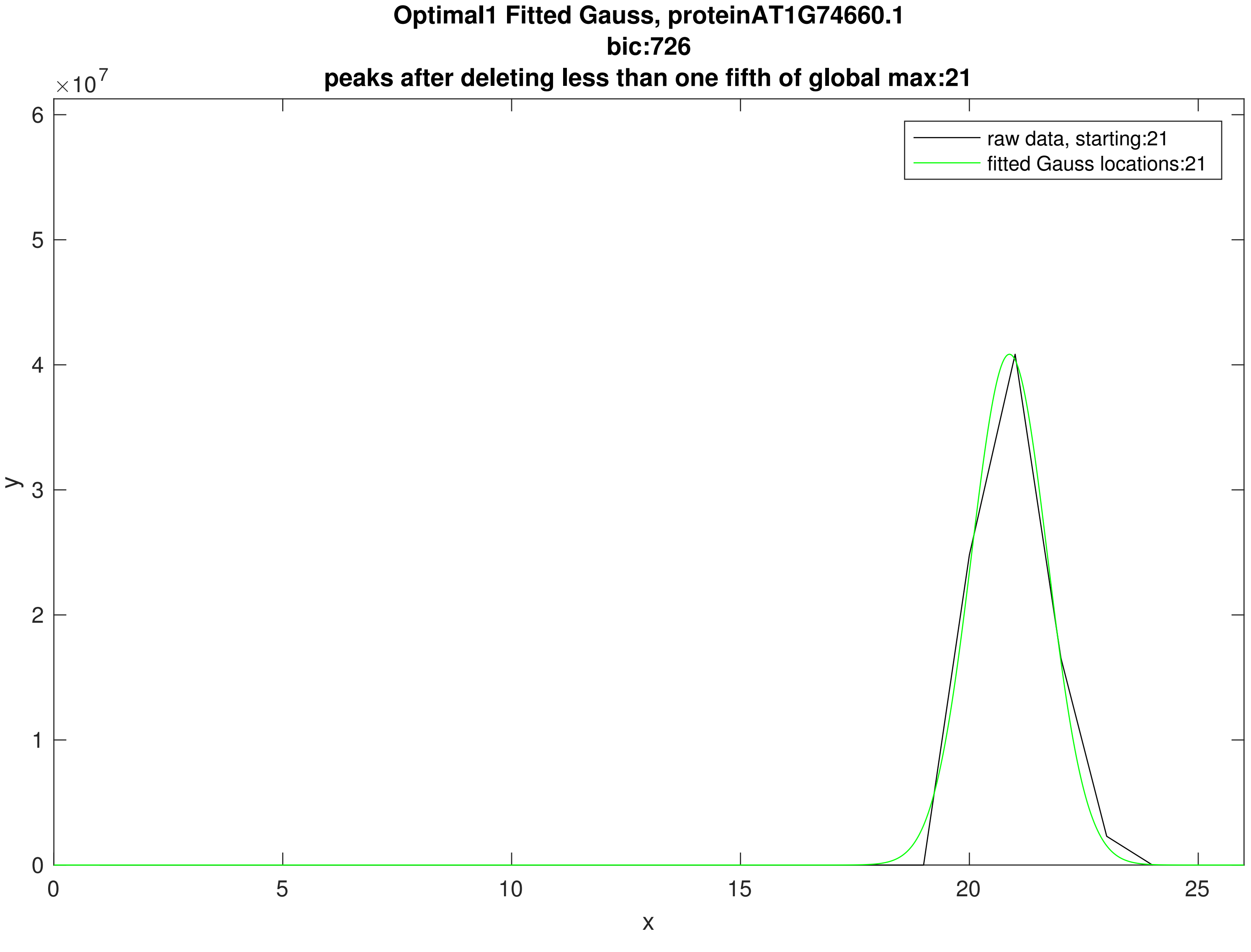

Optimal1 Fitted Gauss, proteinAT1G74660.1
bic:726
peaks after deleting less than one fifth of global max:21
×10^7
6
5
4
3
2
1
0
y
0
5
10
15
20
25
x
raw data, starting:21
fitted Gauss locations:21

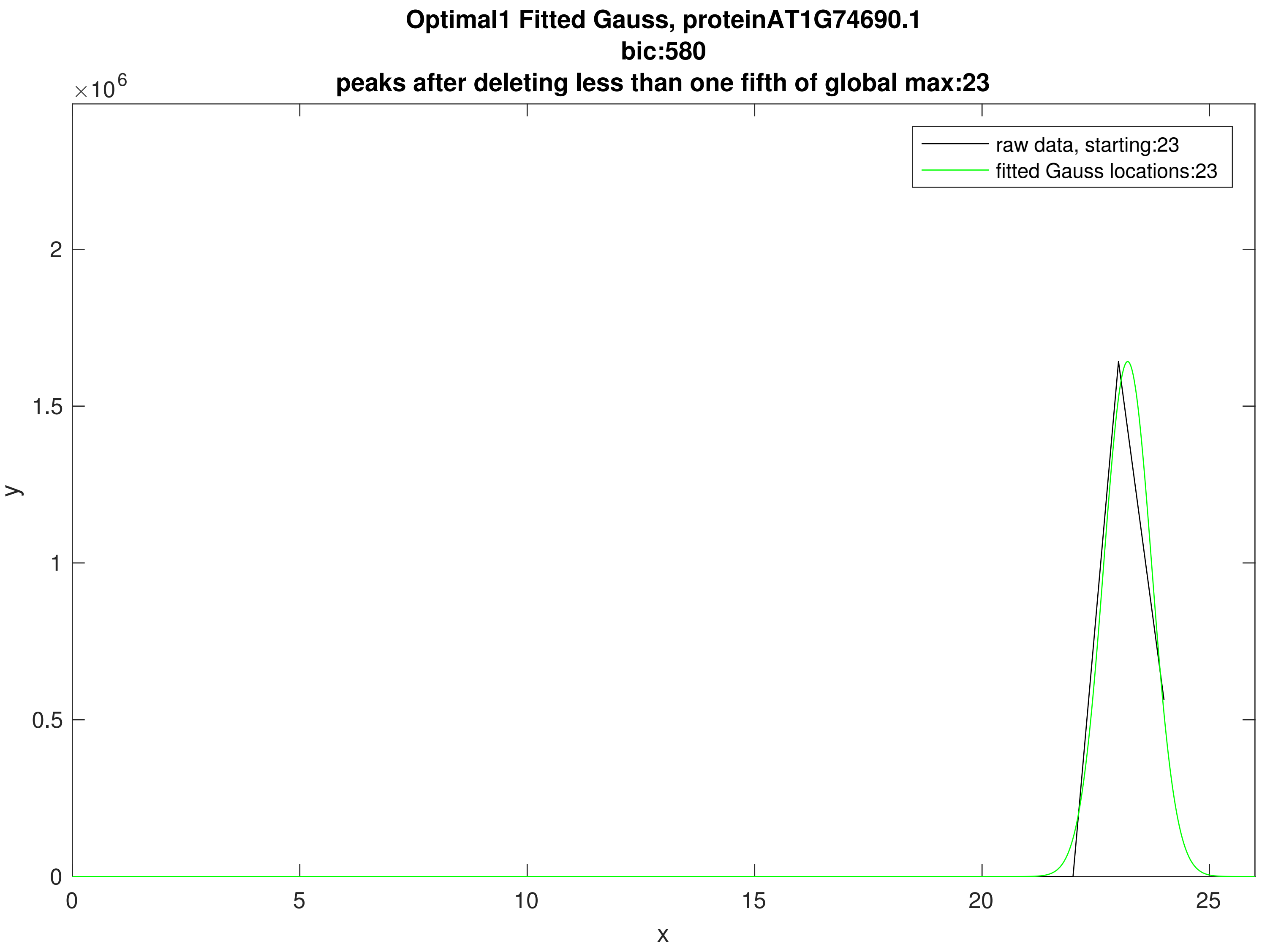

Optimal1 Fitted Gauss, proteinAT1G74690.1
bic:580
peaks after deleting less than one fifth of global max:23
$\times 10^6$
raw data, starting:23
fitted Gauss locations:23
y
x
0
0.5
1
1.5
2
5
10
15
20
25

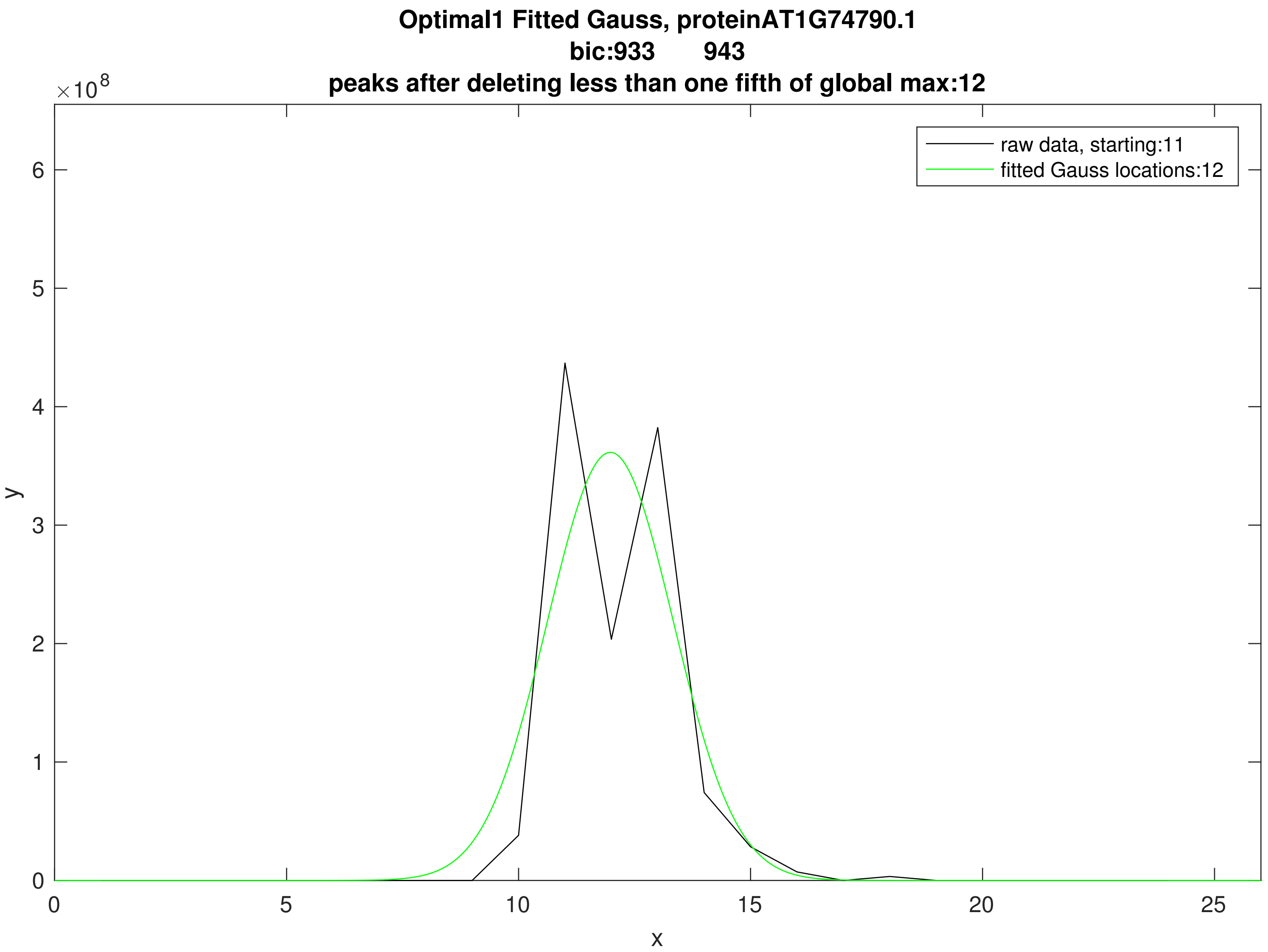

Optimal1 Fitted Gauss, proteinAT1G74790.1
bic:933 943
peaks after deleting less than one fifth of global max:12
×10^8
raw data, starting:11
fitted Gauss locations:12
y
x
0
1
2
3
4
5
6
0
5
10
15
20
25

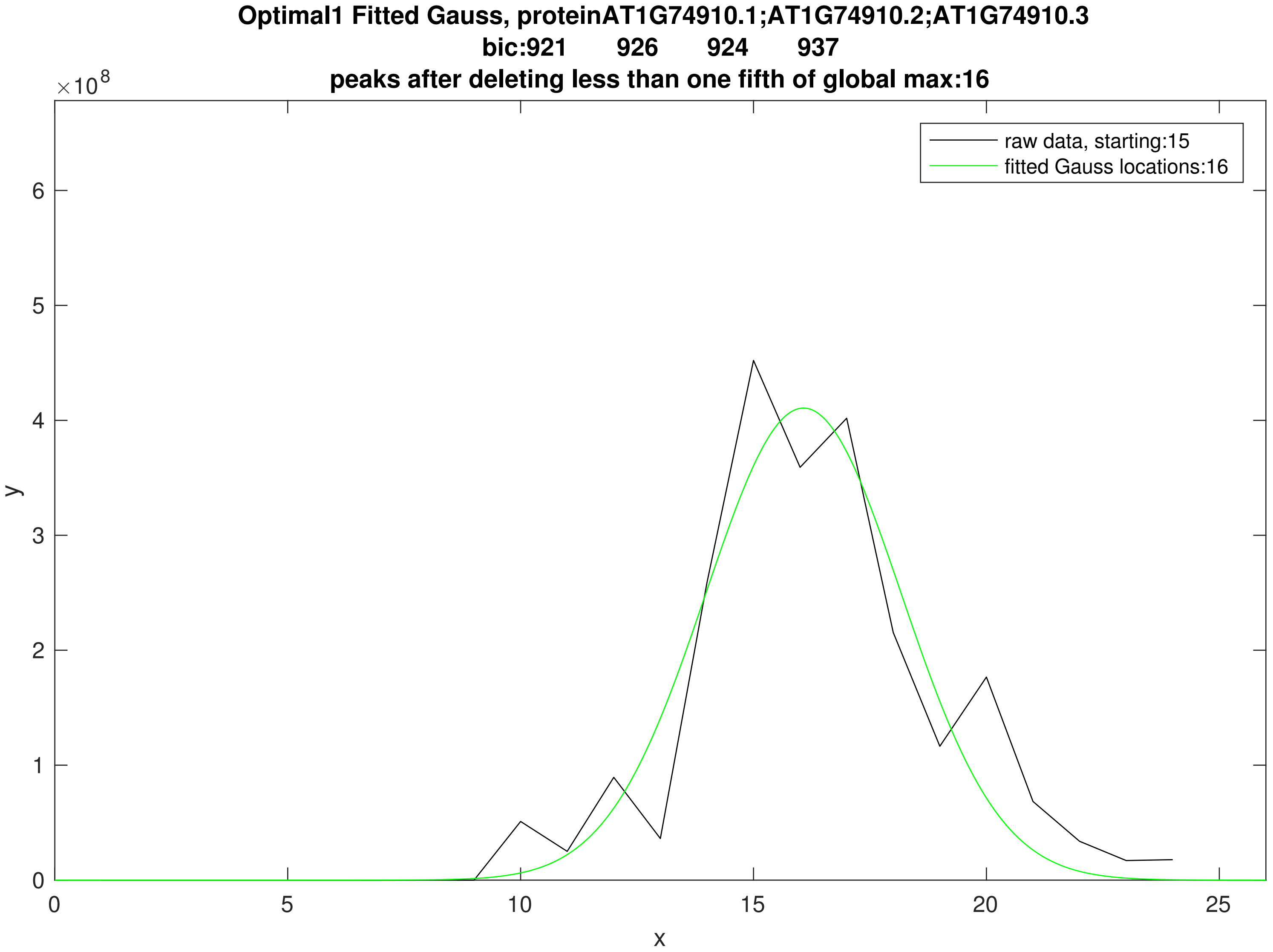

Optimal1 Fitted Gauss, proteinAT1G74910.1;AT1G74910.2;AT1G74910.3
bic:921 926 924 937
peaks after deleting less than one fifth of global max:16
×10^8
raw data, starting:15
fitted Gauss locations:16
y
x
0
1
2
3
4
5
6
0
5
10
15
20
25

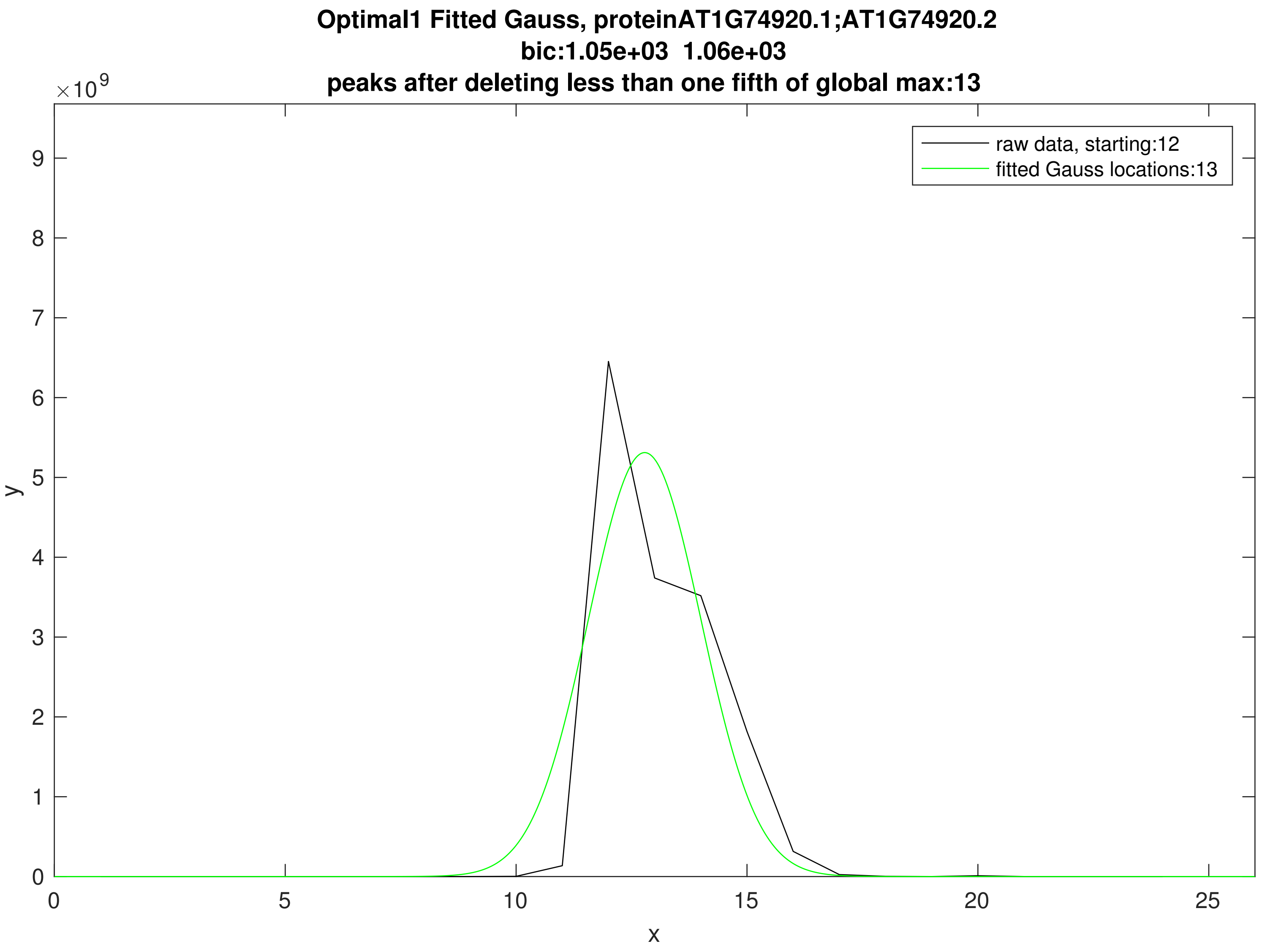

Optimal1 Fitted Gauss, proteinAT1G74920.1;AT1G74920.2
bic:1.05e+03 1.06e+03
peaks after deleting less than one fifth of global max:13
×10^9
raw data, starting:12
fitted Gauss locations:13
y
x
0
1
2
3
4
5
6
7
8
9
0
5
10
15
20
25

Optimal1 Fitted Gauss, proteinAT1G74960.3;AT1G74960.2;AT1G74960.1
bic:912
peaks after deleting less than one fifth of global max:15

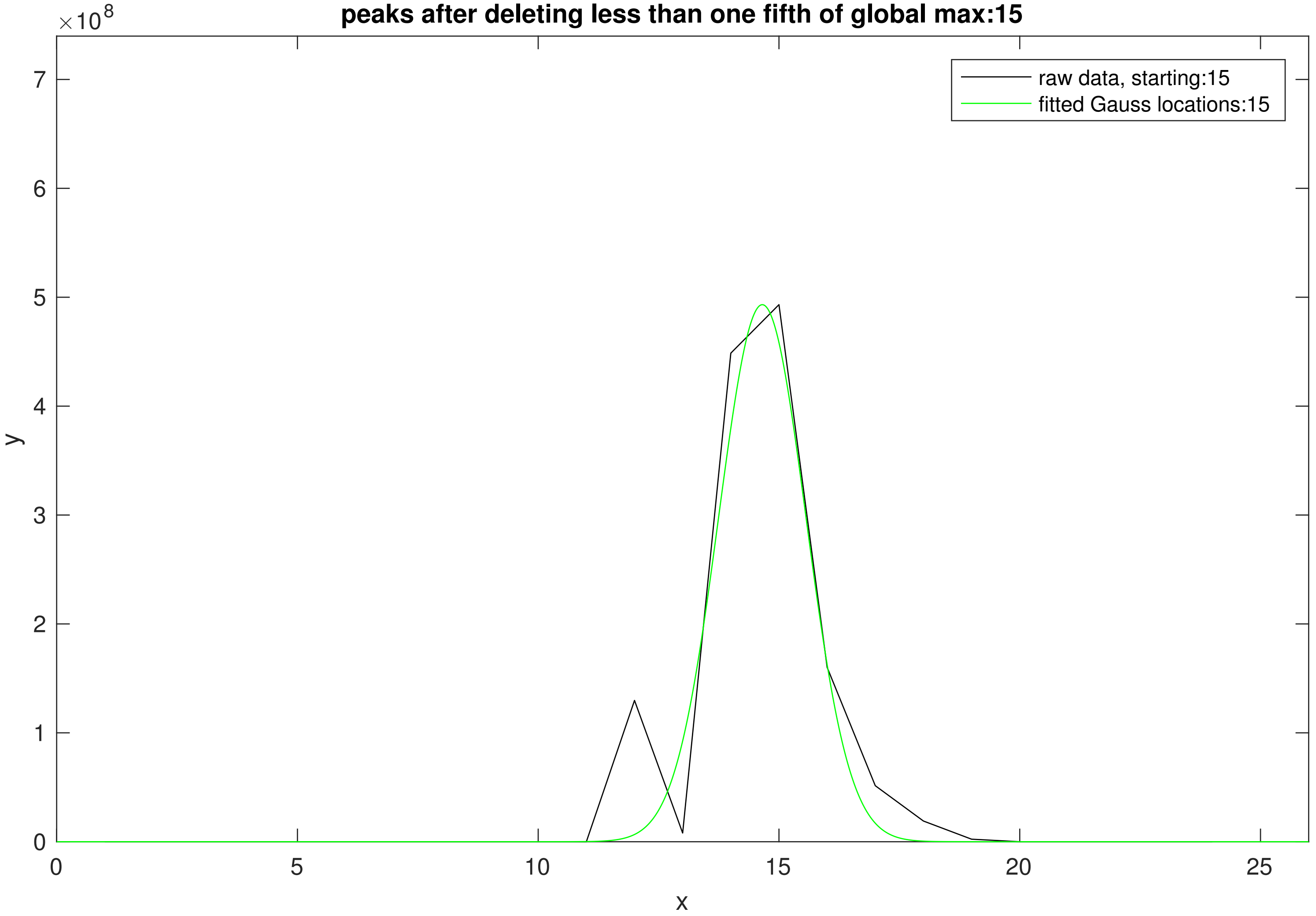

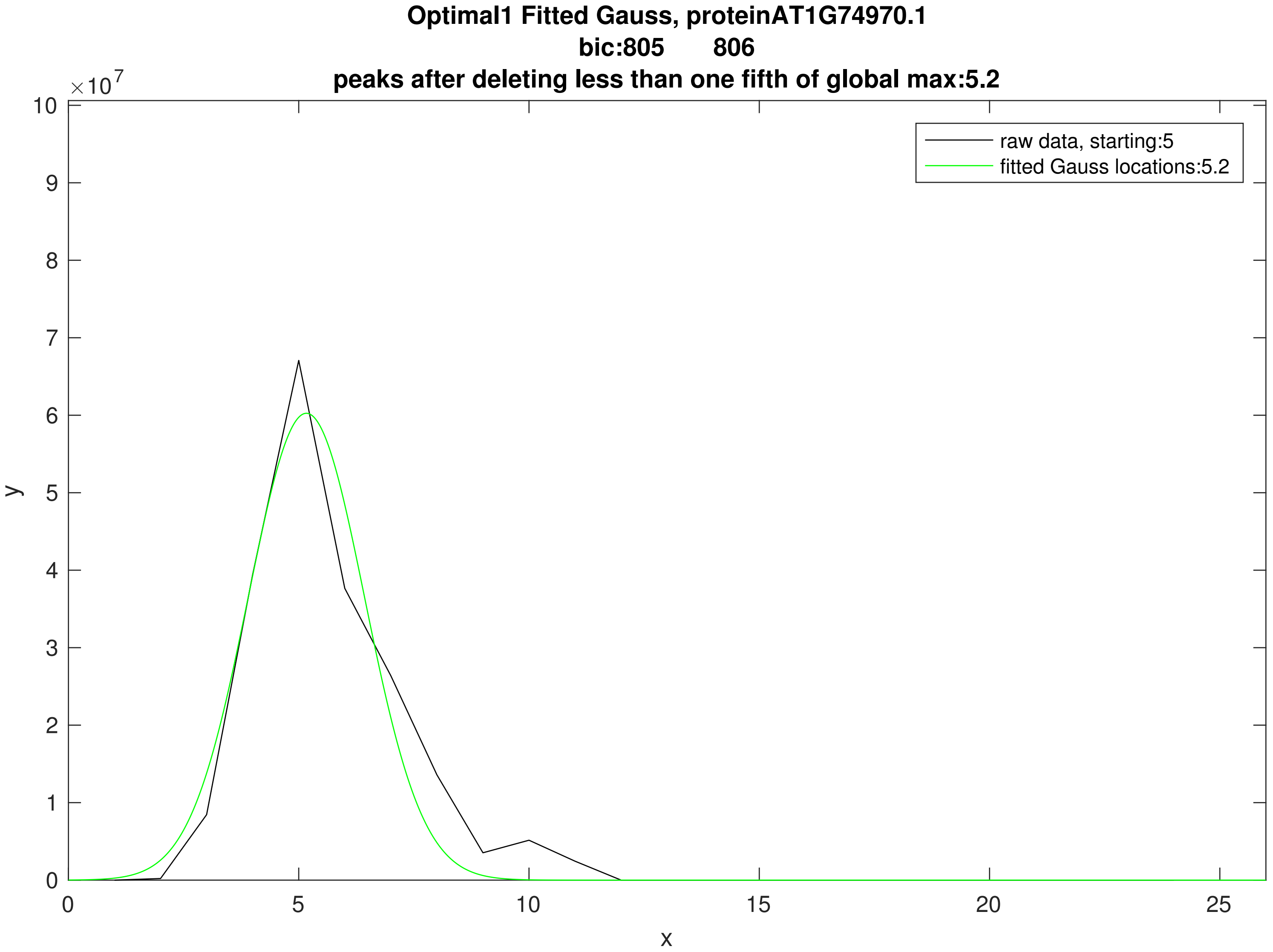
Optimal1 Fitted Gauss, proteinAT1G74970.1
bic:805 806
peaks after deleting less than one fifth of global max:5.2
×10^7
10
9
8
7
6
5
4
3
2
1
0
y
0
5
10
15
20
25
x
raw data, starting:5
fitted Gauss locations:5.2

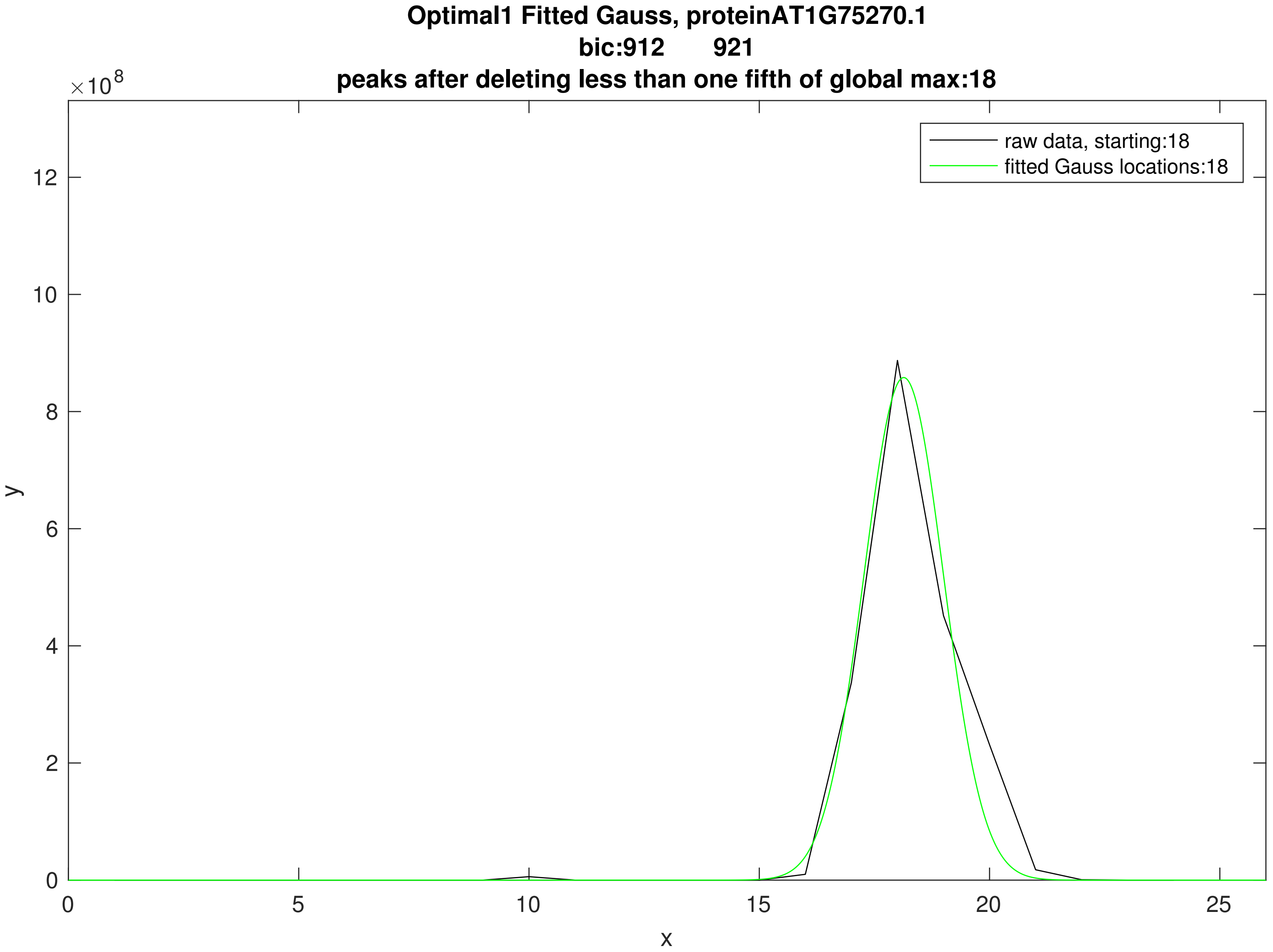
Optimal1 Fitted Gauss, proteinAT1G75270.1
bic:912 921
peaks after deleting less than one fifth of global max:18
×10^8
raw data, starting:18
fitted Gauss locations:18
y
x
0
2
4
6
8
10
12
0
5
10
15
20
25

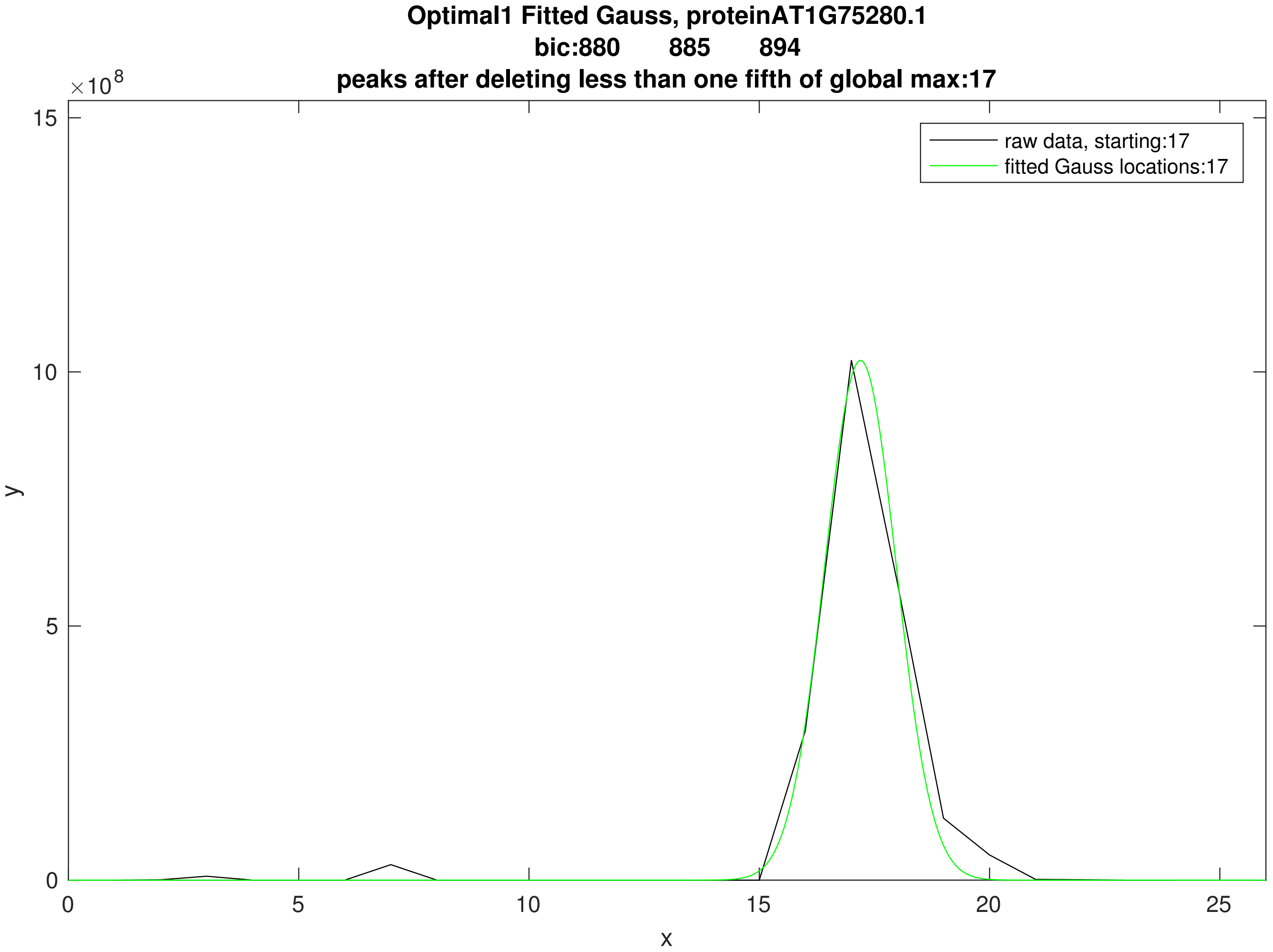

Optimal1 Fitted Gauss, proteinAT1G75280.1
bic:880 885 894
peaks after deleting less than one fifth of global max:17
×10^8
15
10
5
0
y
0
5
10
15
20
25
x
raw data, starting:17
fitted Gauss locations:17

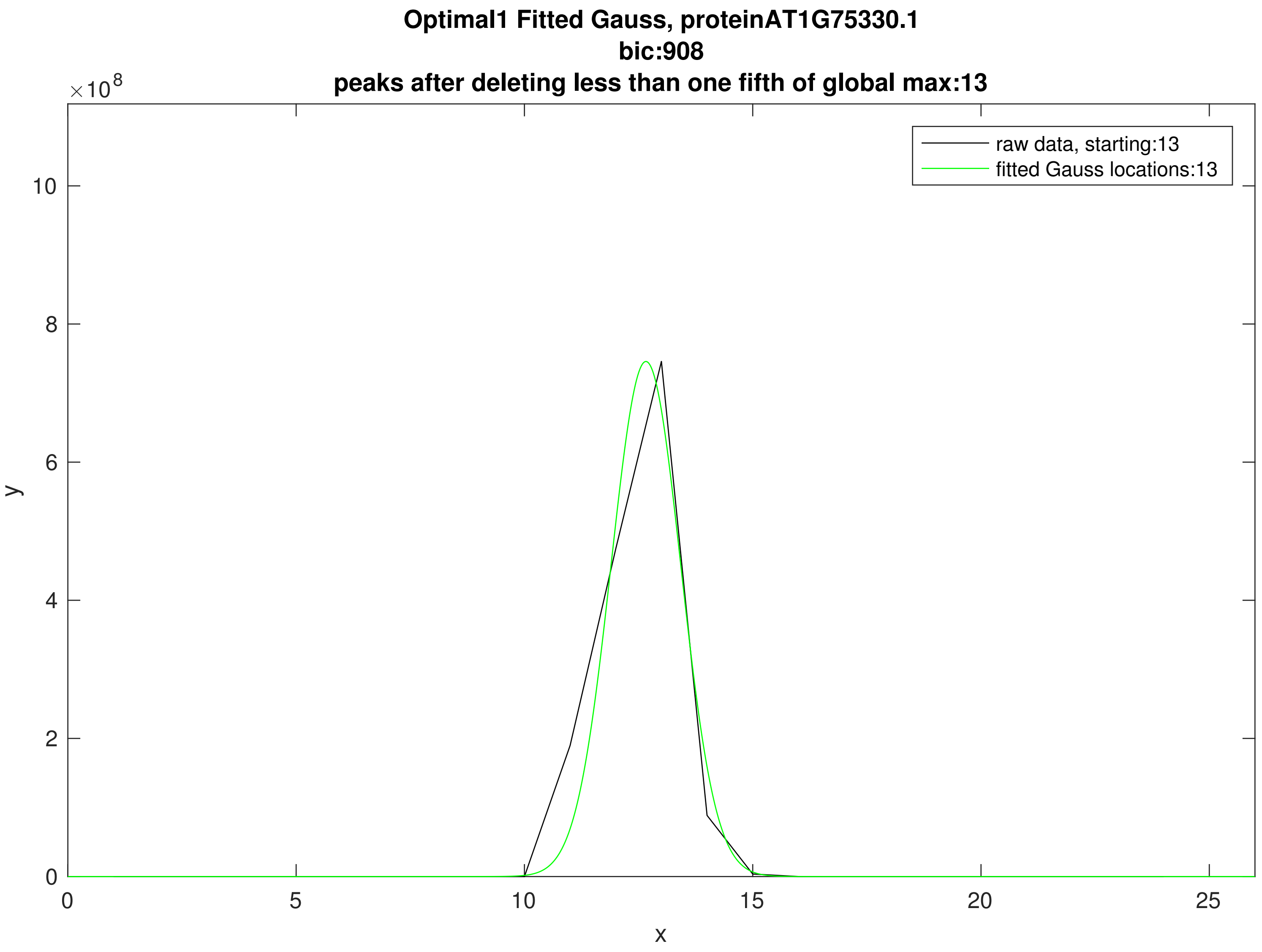
Optimal1 Fitted Gauss, proteinAT1G75330.1
bic:908
peaks after deleting less than one fifth of global max:13
$\times 10^8$
raw data, starting:13
fitted Gauss locations:13
y
x
0
2
4
6
8
10
0
5
10
15
20
25

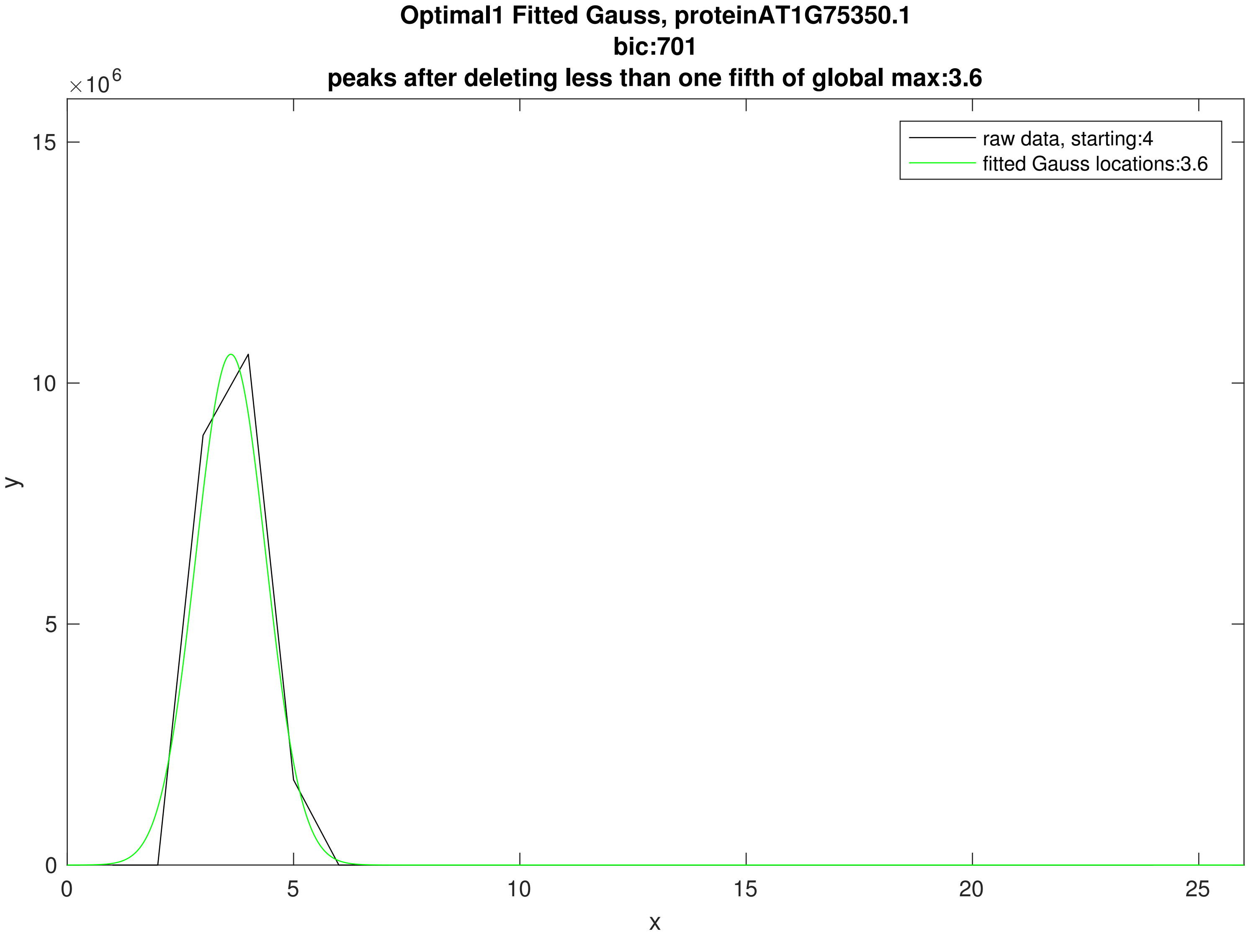
Optimal1 Fitted Gauss, proteinAT1G75350.1
bic:701
peaks after deleting less than one fifth of global max:3.6
×10^6
raw data, starting:4
fitted Gauss locations:3.6
y
x
0
5
10
15
20
25

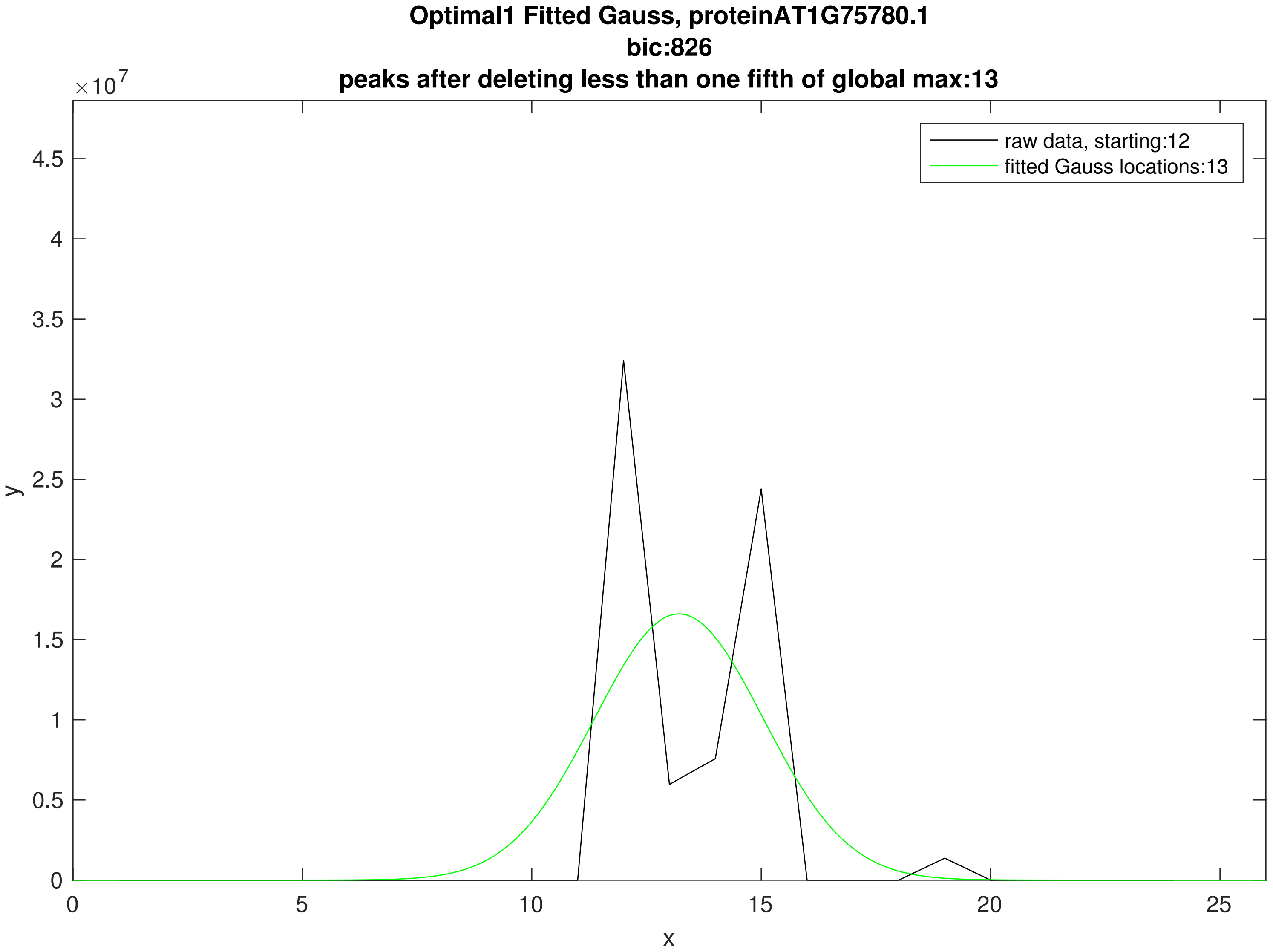
Optimal1 Fitted Gauss, proteinAT1G75780.1
bic:826
peaks after deleting less than one fifth of global max:13
×10^7
raw data, starting:12
fitted Gauss locations:13
4.5
4
3.5
3
2.5
2
1.5
1
0.5
0
y
0
5
10
15
20
25
x

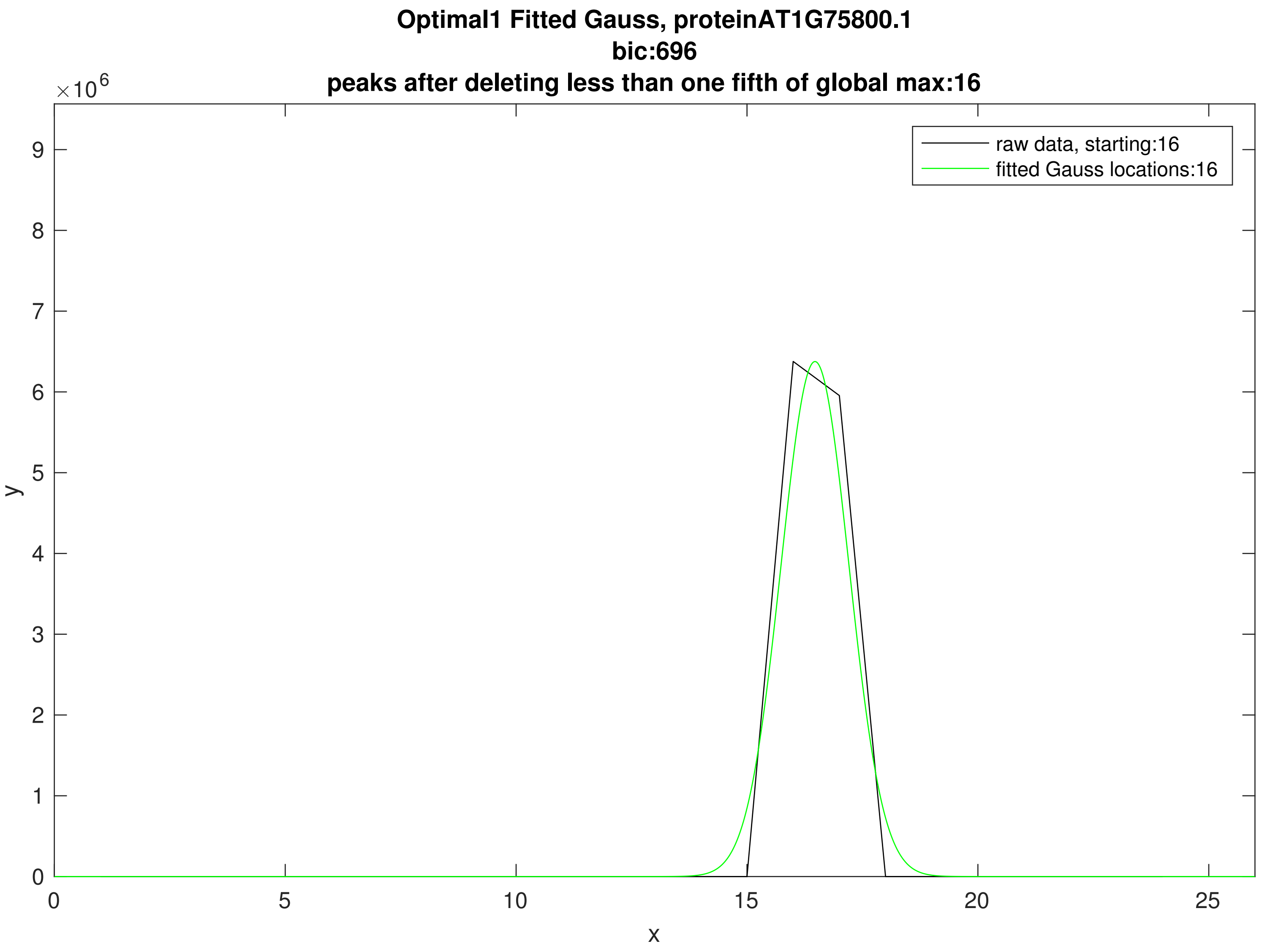

Optimal1 Fitted Gauss, proteinAT1G75800.1
bic:696
peaks after deleting less than one fifth of global max:16
×10^6
raw data, starting:16
fitted Gauss locations:16
y
x
0
5
10
15
20
25
1
2
3
4
5
6
7
8
9

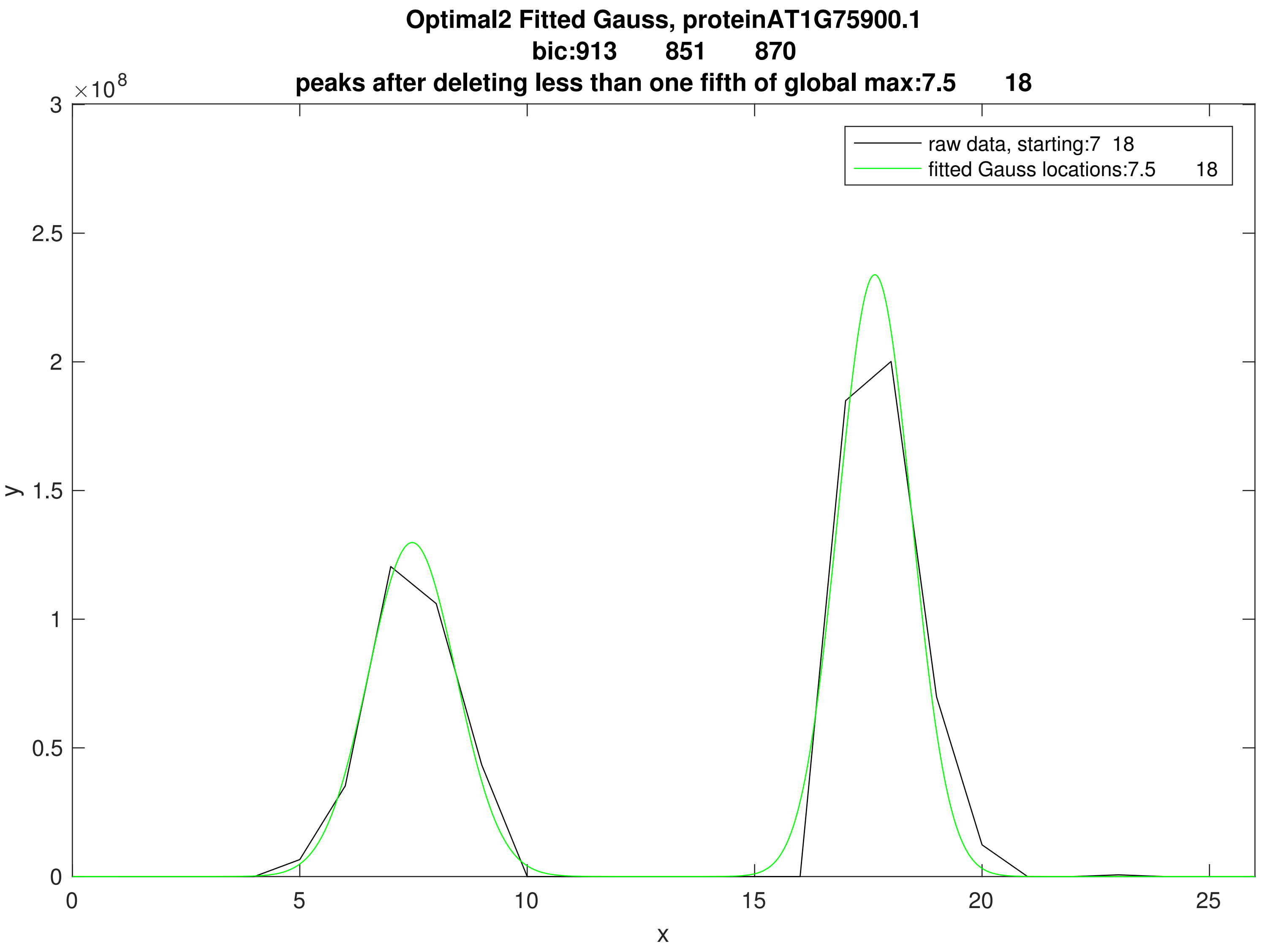

Optimal2 Fitted Gauss, proteinAT1G75900.1
bic:913 851 870
peaks after deleting less than one fifth of global max:7.5 18
×10^8
raw data, starting:7 18
fitted Gauss locations:7.5 18
y
x
0
0.5
1
1.5
2
2.5
3
5
10
15
20
25

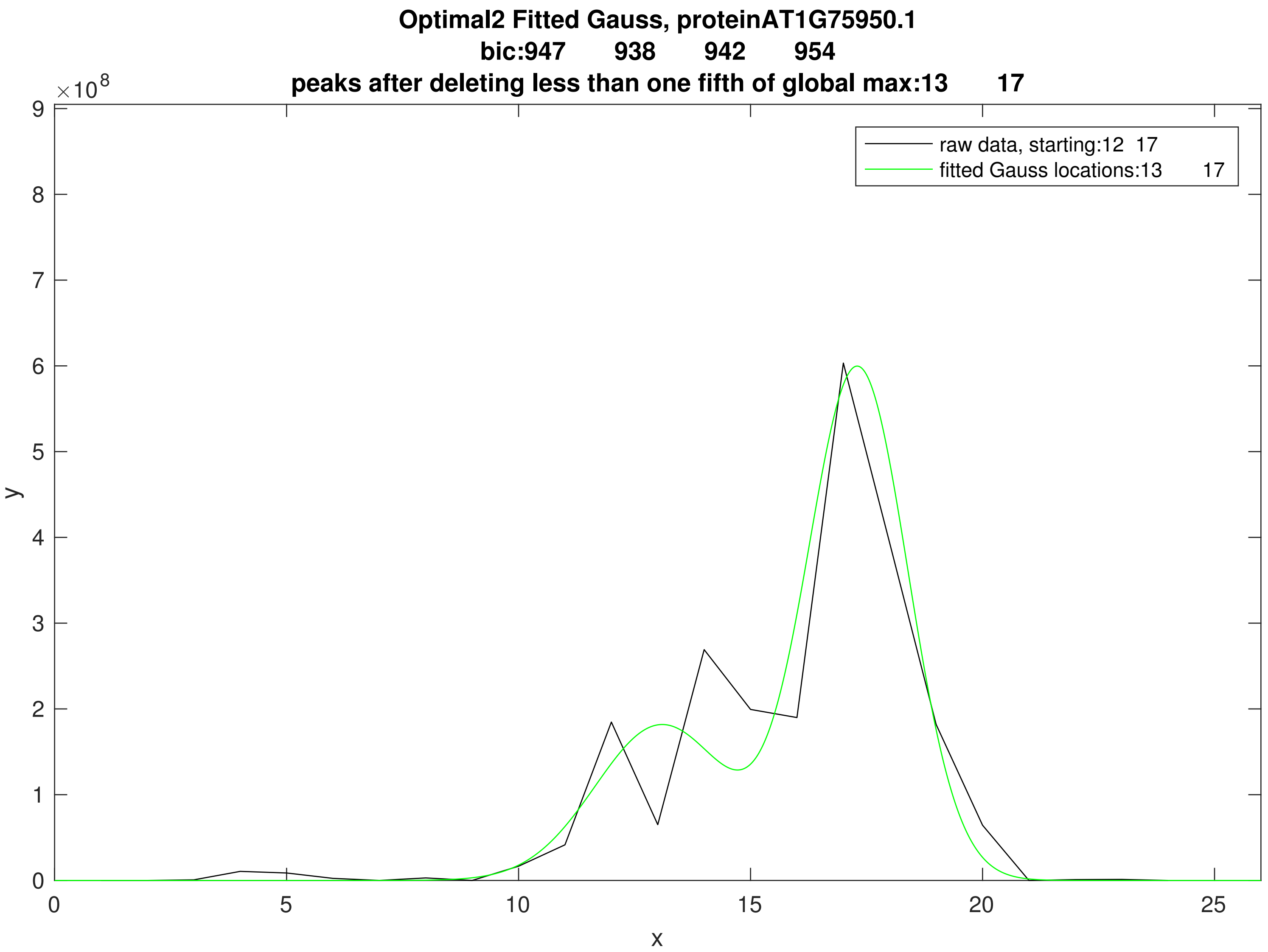

Optimal2 Fitted Gauss, proteinAT1G75950.1
bic:947 938 942 954
peaks after deleting less than one fifth of global max:13 17
×10^8
raw data, starting:12 17
fitted Gauss locations:13 17
y
x

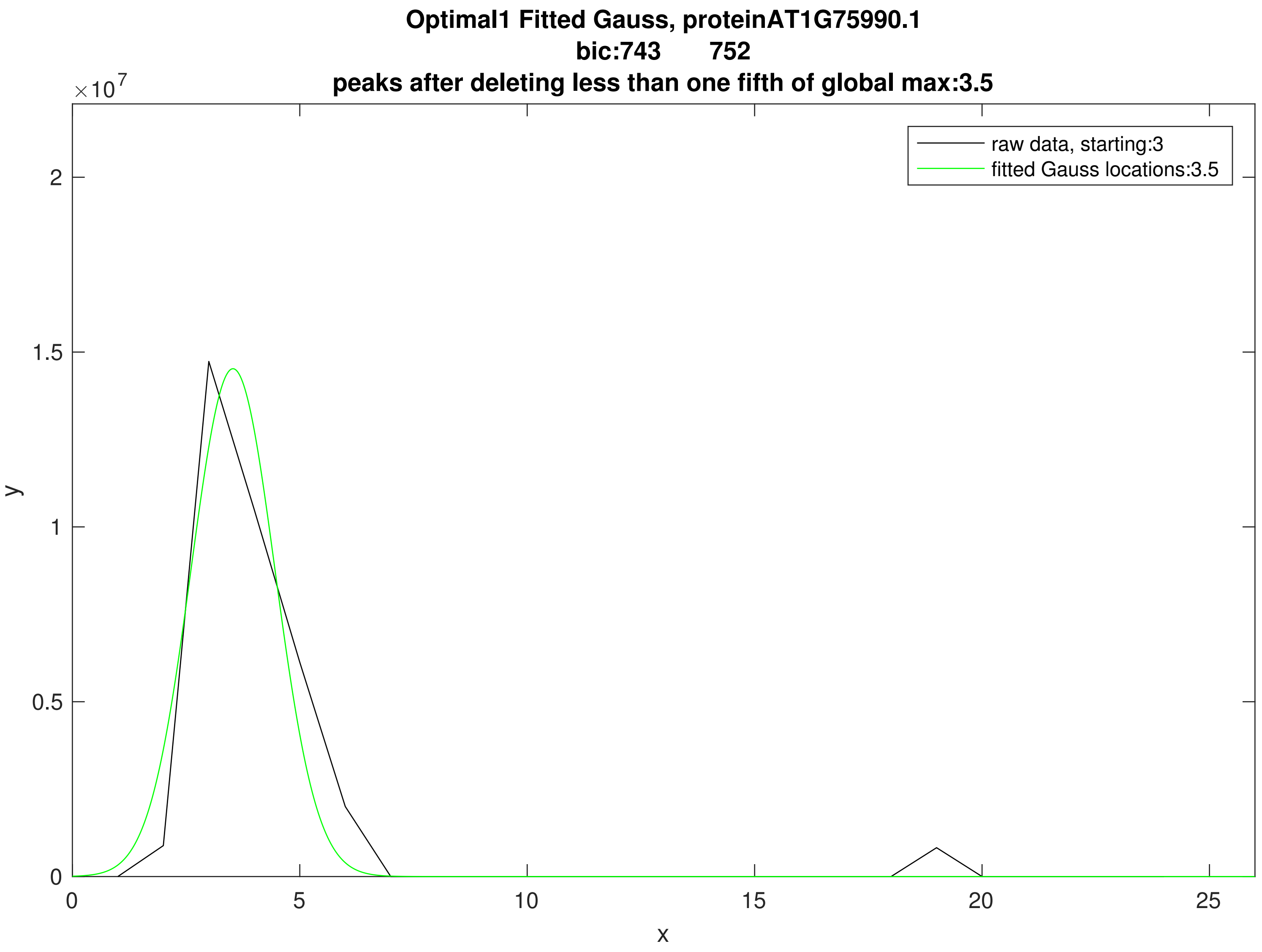
Optimal1 Fitted Gauss, proteinAT1G75990.1
bic:743 752
peaks after deleting less than one fifth of global max:3.5
×10^7
raw data, starting:3
fitted Gauss locations:3.5
y
x
0
0.5
1
1.5
2
5
10
15
20
25

**Optimal2 Fitted Gauss, proteinAT1G76010.1**
**bic:753 746**
**peaks after deleting less than one fifth of global max:4.8 11**

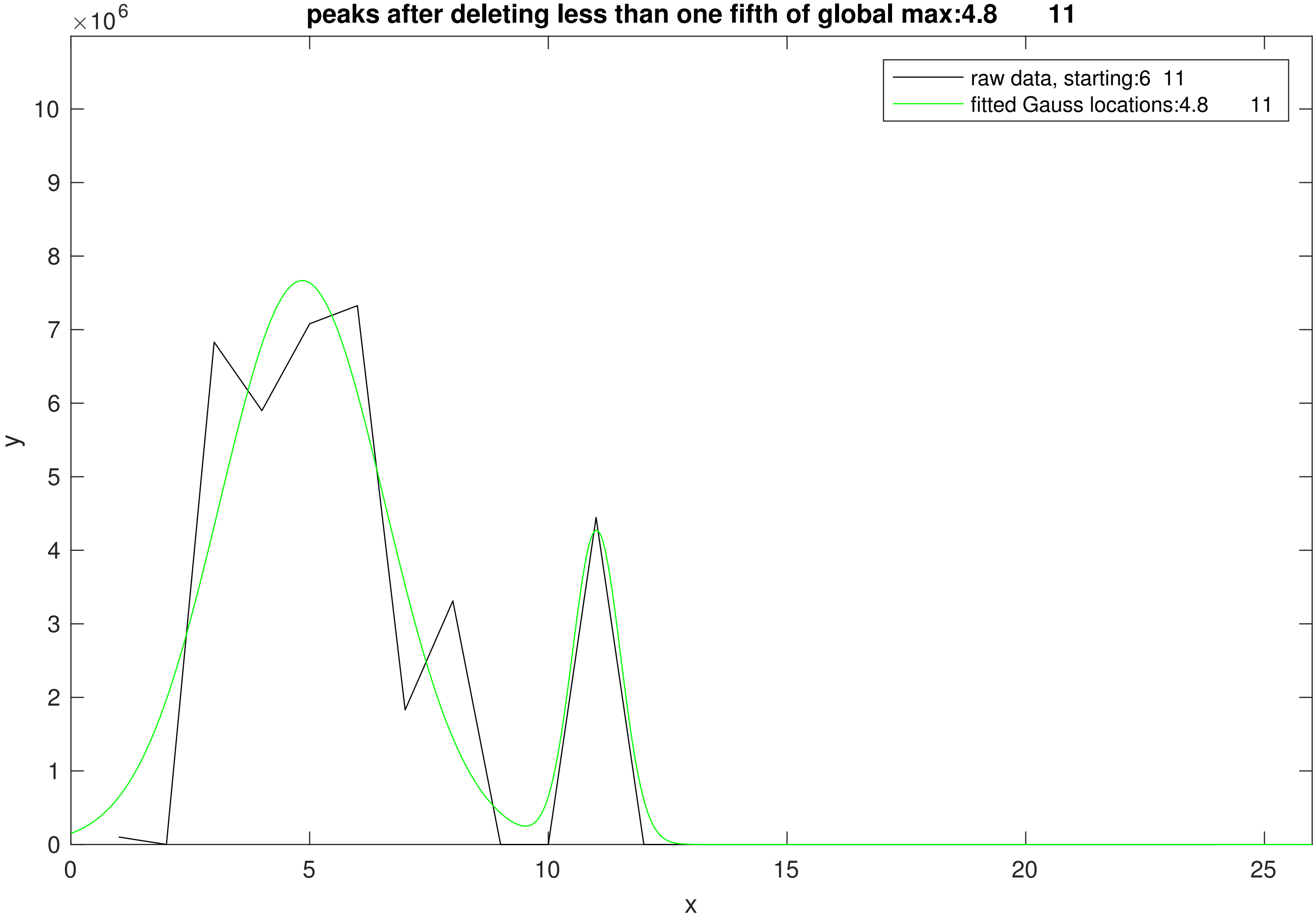

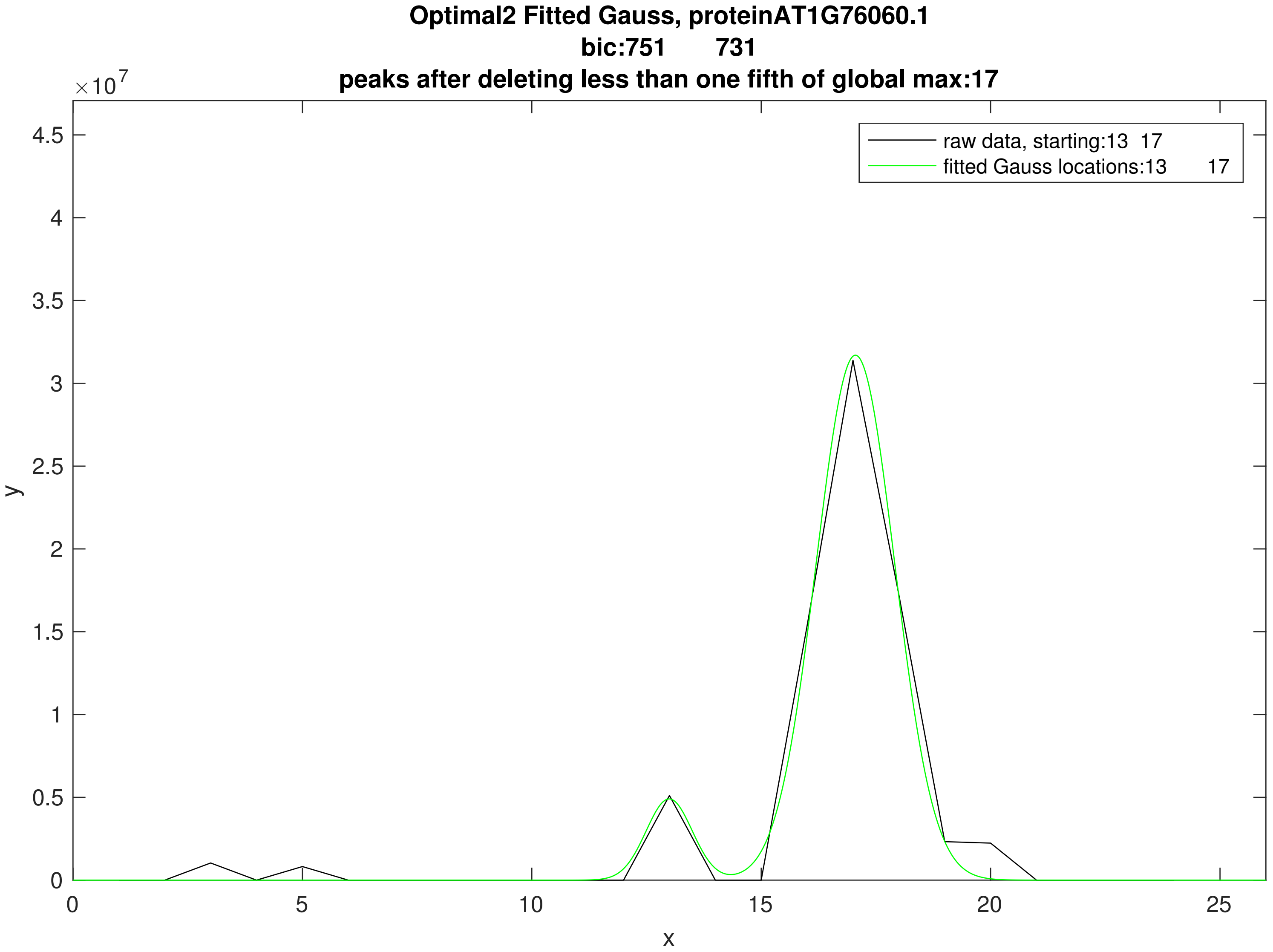
Optimal2 Fitted Gauss, proteinAT1G76060.1
bic:751 731
peaks after deleting less than one fifth of global max:17
×10^7
raw data, starting:13 17
fitted Gauss locations:13 17
y
x
0
0.5
1
1.5
2
2.5
3
3.5
4
4.5
0
5
10
15
20
25

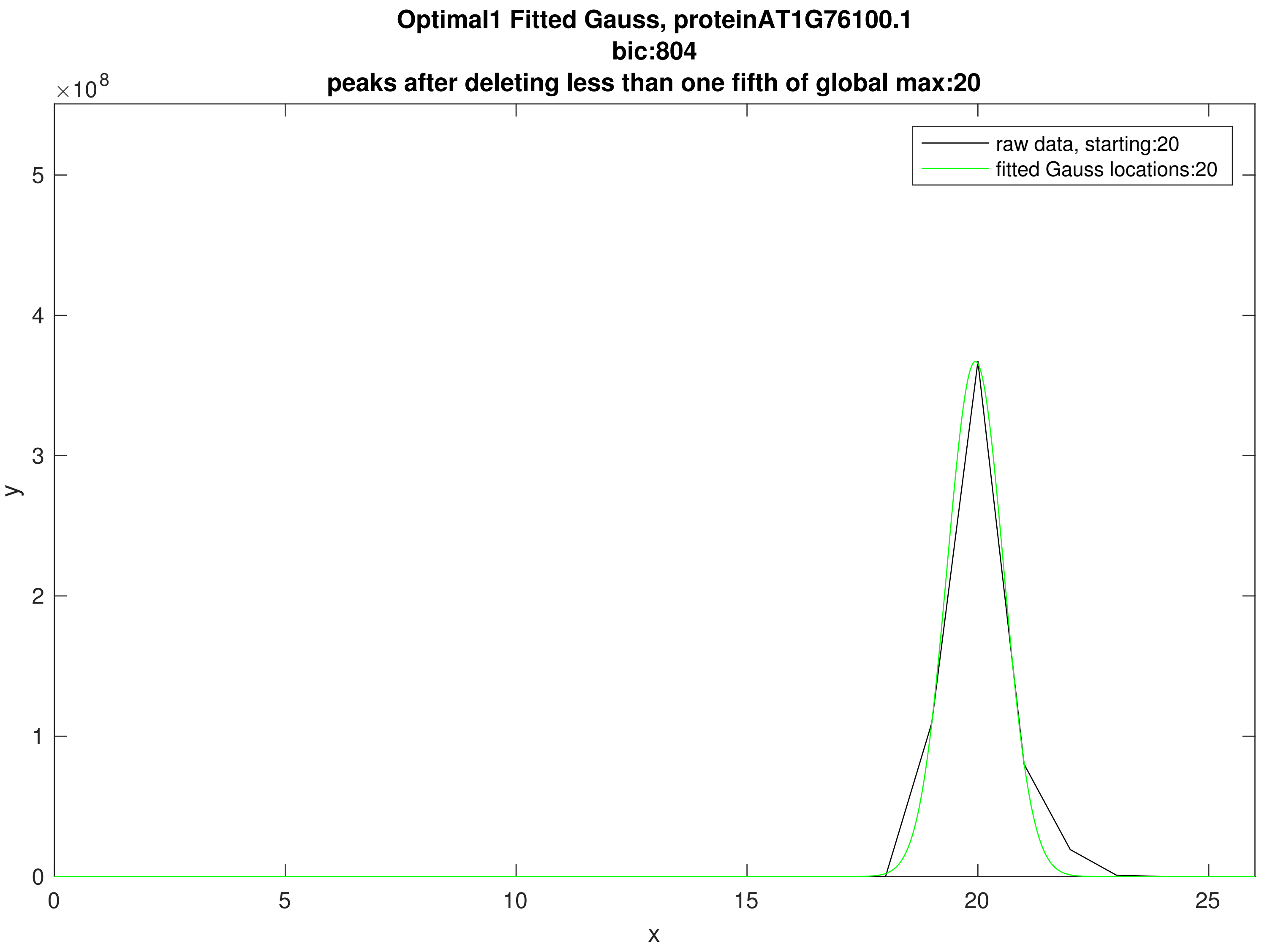

Optimal1 Fitted Gauss, proteinAT1G76100.1
bic:804
peaks after deleting less than one fifth of global max:20
$\times 10^8$
raw data, starting:20
fitted Gauss locations:20
y
x
0
1
2
3
4
5
0
5
10
15
20
25

Optimal1 Fitted Gauss, proteinAT1G76140.1;AT1G76140.2
bic:1.01e+03
peaks after deleting less than one fifth of global max:14

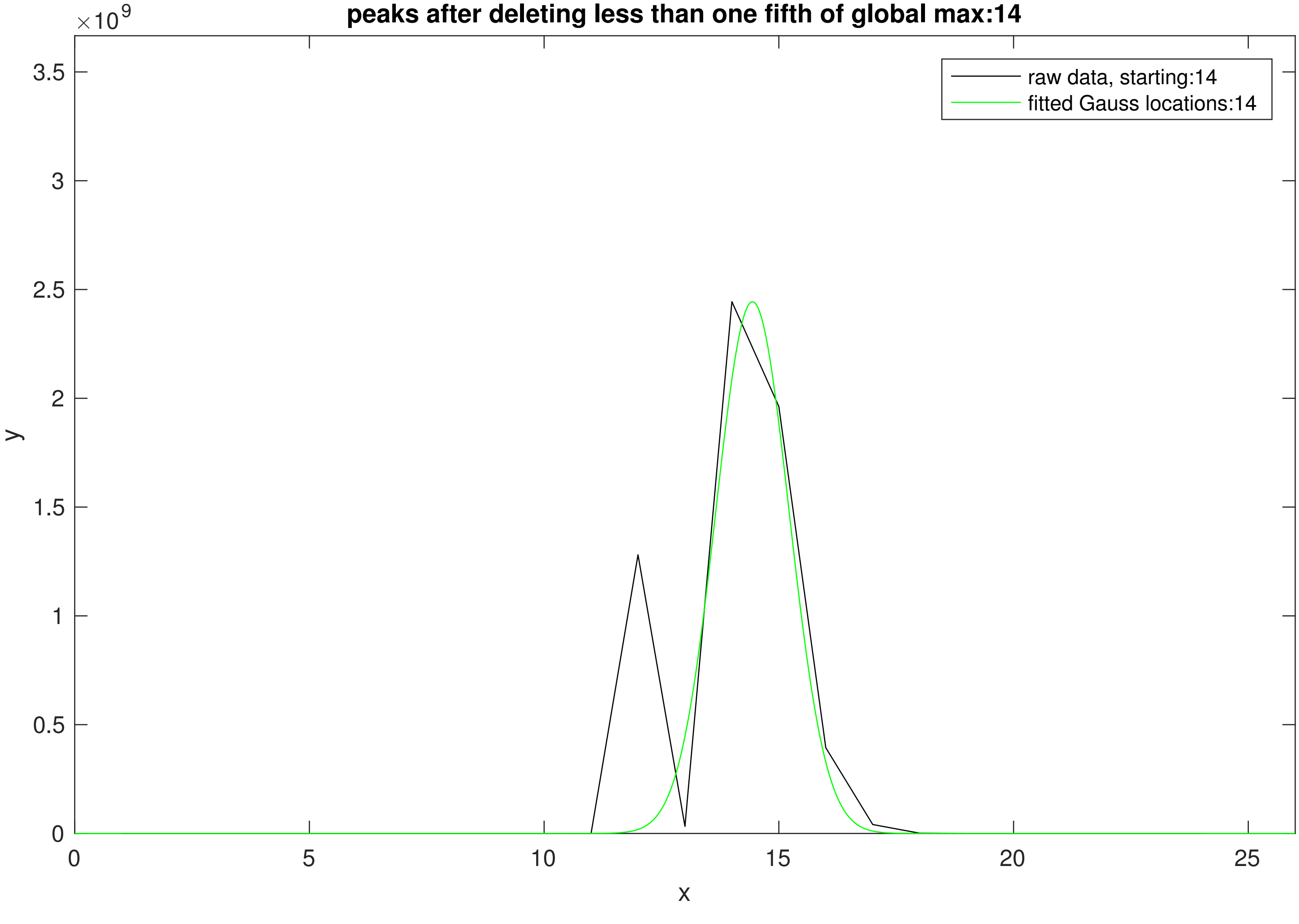

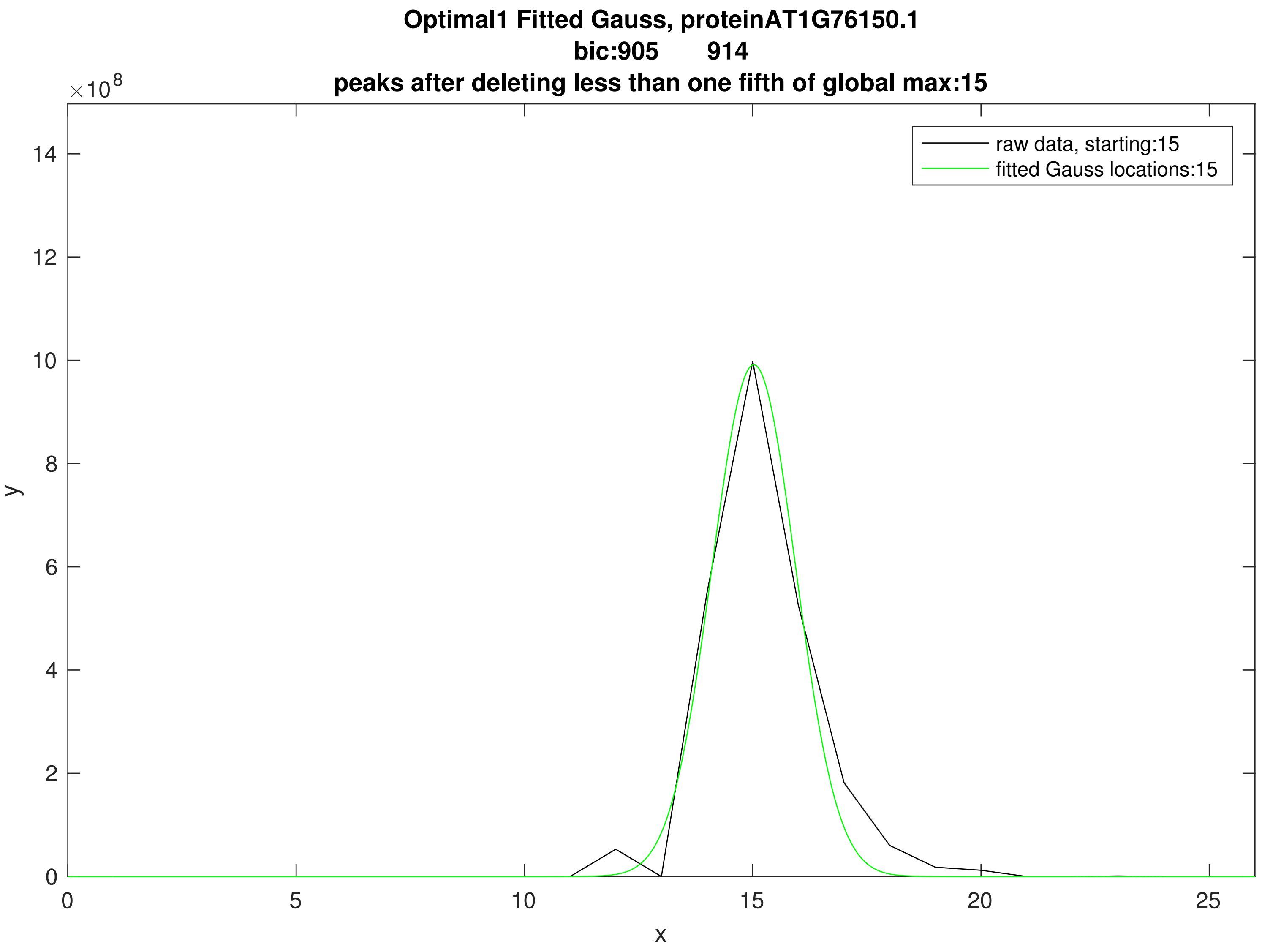

Optimal1 Fitted Gauss, proteinAT1G76150.1
bic:905 914
peaks after deleting less than one fifth of global max:15
×10^8
raw data, starting:15
fitted Gauss locations:15
y
x
0
2
4
6
8
10
12
14
5
15
20
25

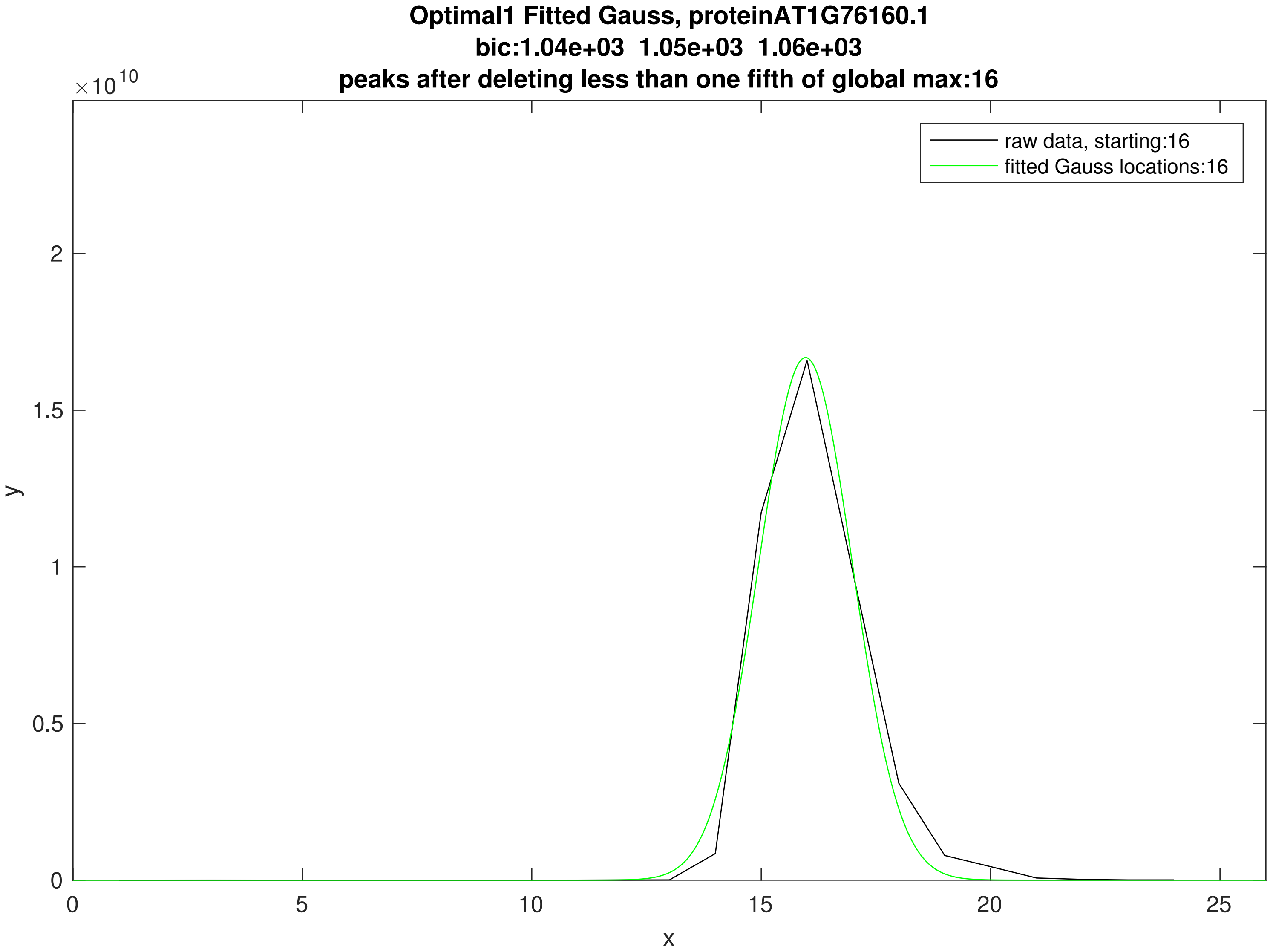

Optimal1 Fitted Gauss, proteinAT1G76160.1
bic:1.04e+03  1.05e+03  1.06e+03
peaks after deleting less than one fifth of global max:16
×10^10
raw data, starting:16
fitted Gauss locations:16
y
x
0
0.5
1
1.5
2
0
5
10
15
20
25

**Optimal1 Fitted Gauss, proteinAT1G76180.2;AT1G76180.1**
**bic:1.09e+03   1.1e+03   1.1e+03  1.11e+03**
**peaks after deleting less than one fifth of global max:13**

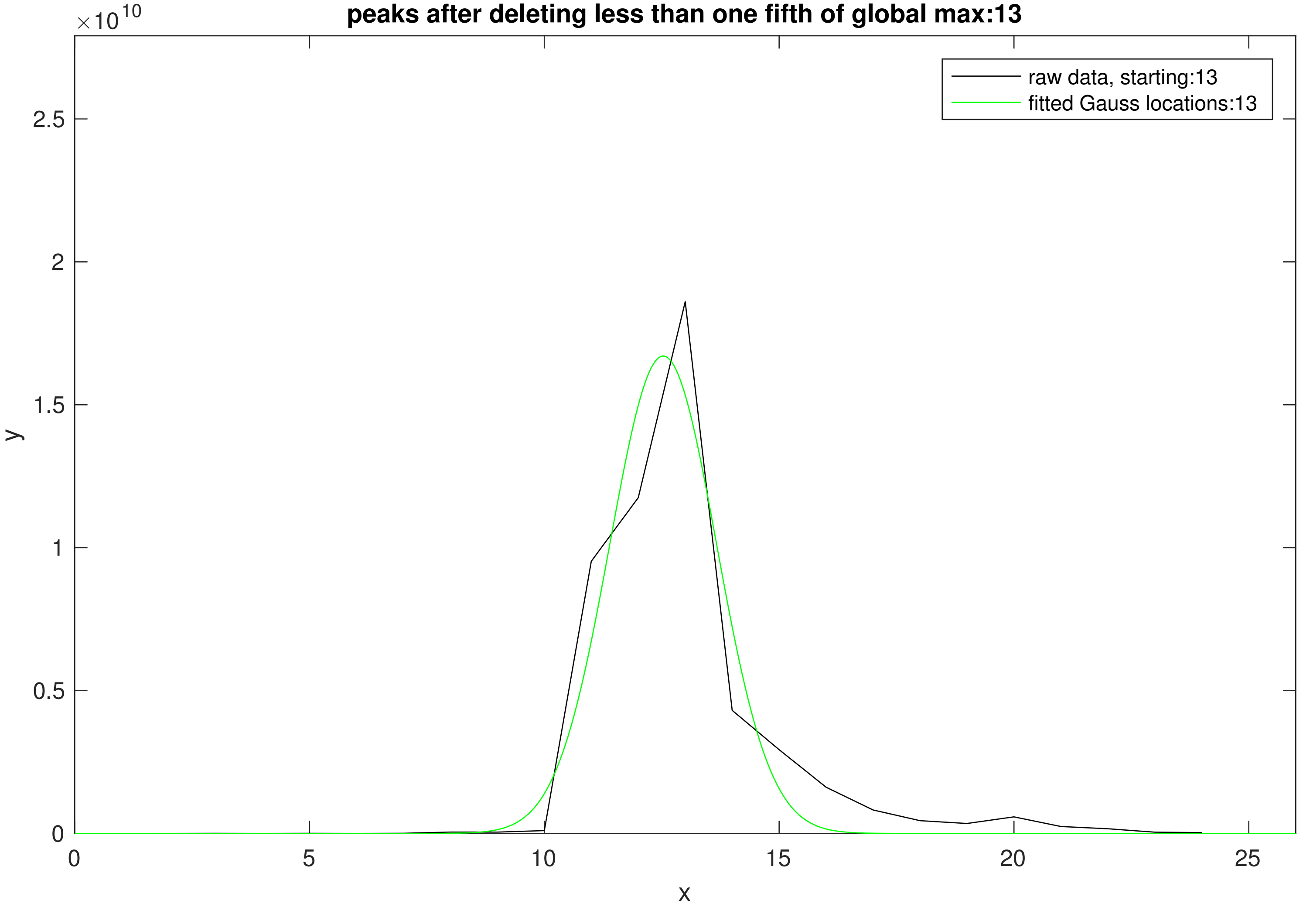

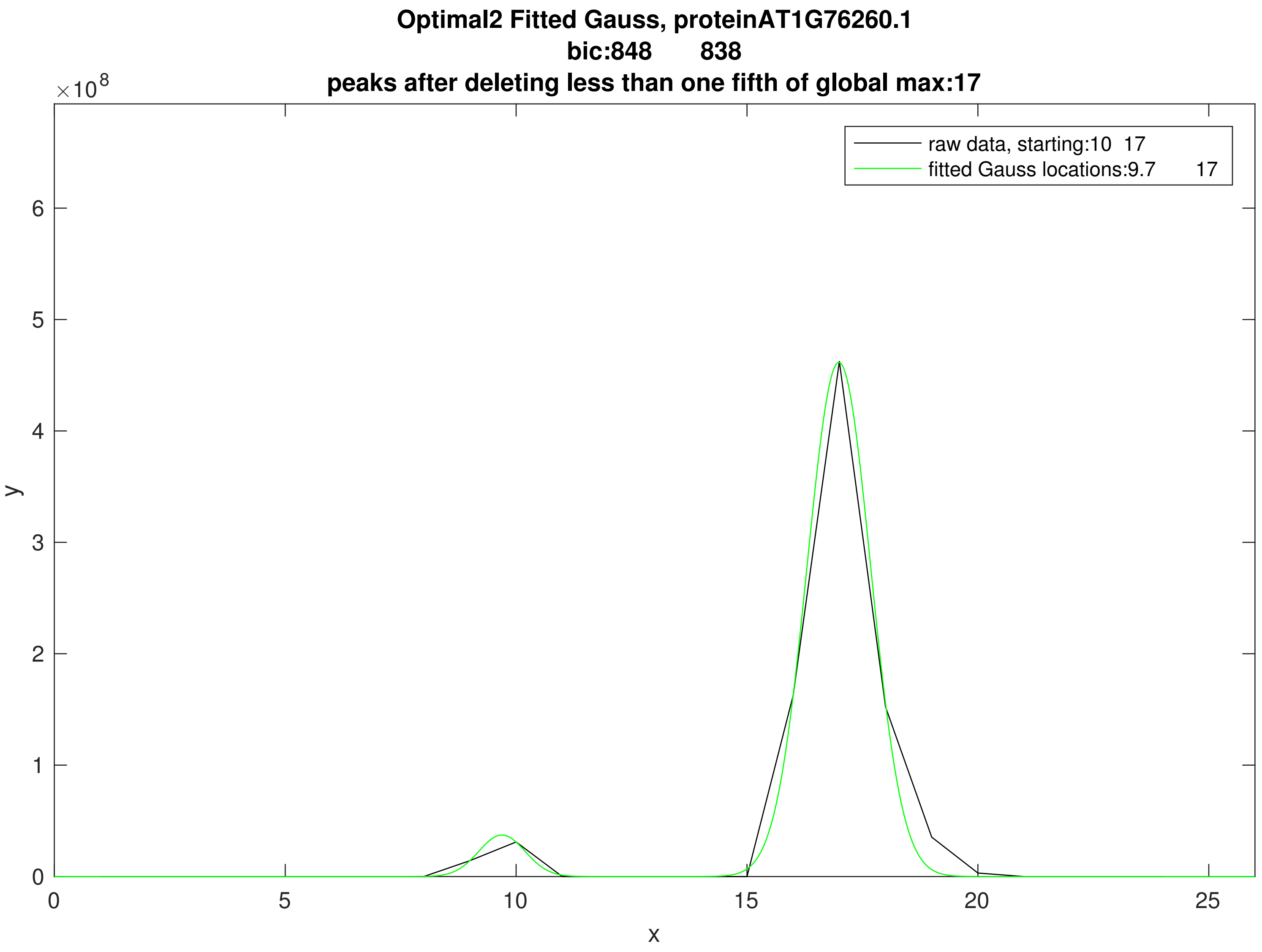
Optimal2 Fitted Gauss, proteinAT1G76260.1
bic:848 838
peaks after deleting less than one fifth of global max:17
×10^8
raw data, starting:10 17
fitted Gauss locations:9.7 17
y
x
0
1
2
3
4
5
6
0
5
10
15
20
25

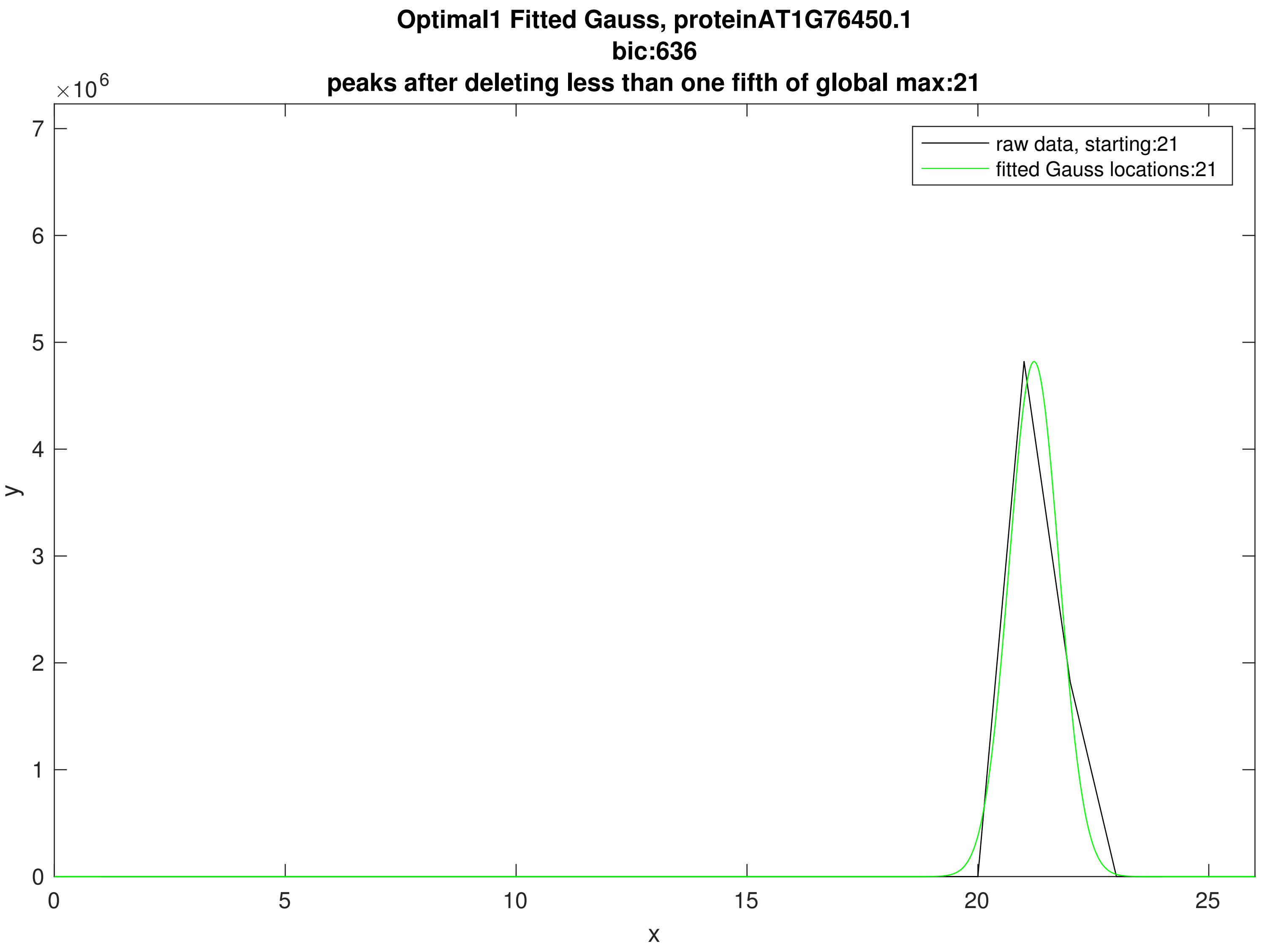

Optimal1 Fitted Gauss, proteinAT1G76450.1
bic:636
peaks after deleting less than one fifth of global max:21
×10^6
raw data, starting:21
fitted Gauss locations:21
y
x
0
1
2
3
4
5
6
7
0
5
10
15
20
25

**Optimal1 Fitted Gauss, proteinAT1G76550.1**
**bic:920 927 938**
**peaks after deleting less than one fifth of global max:7.6**

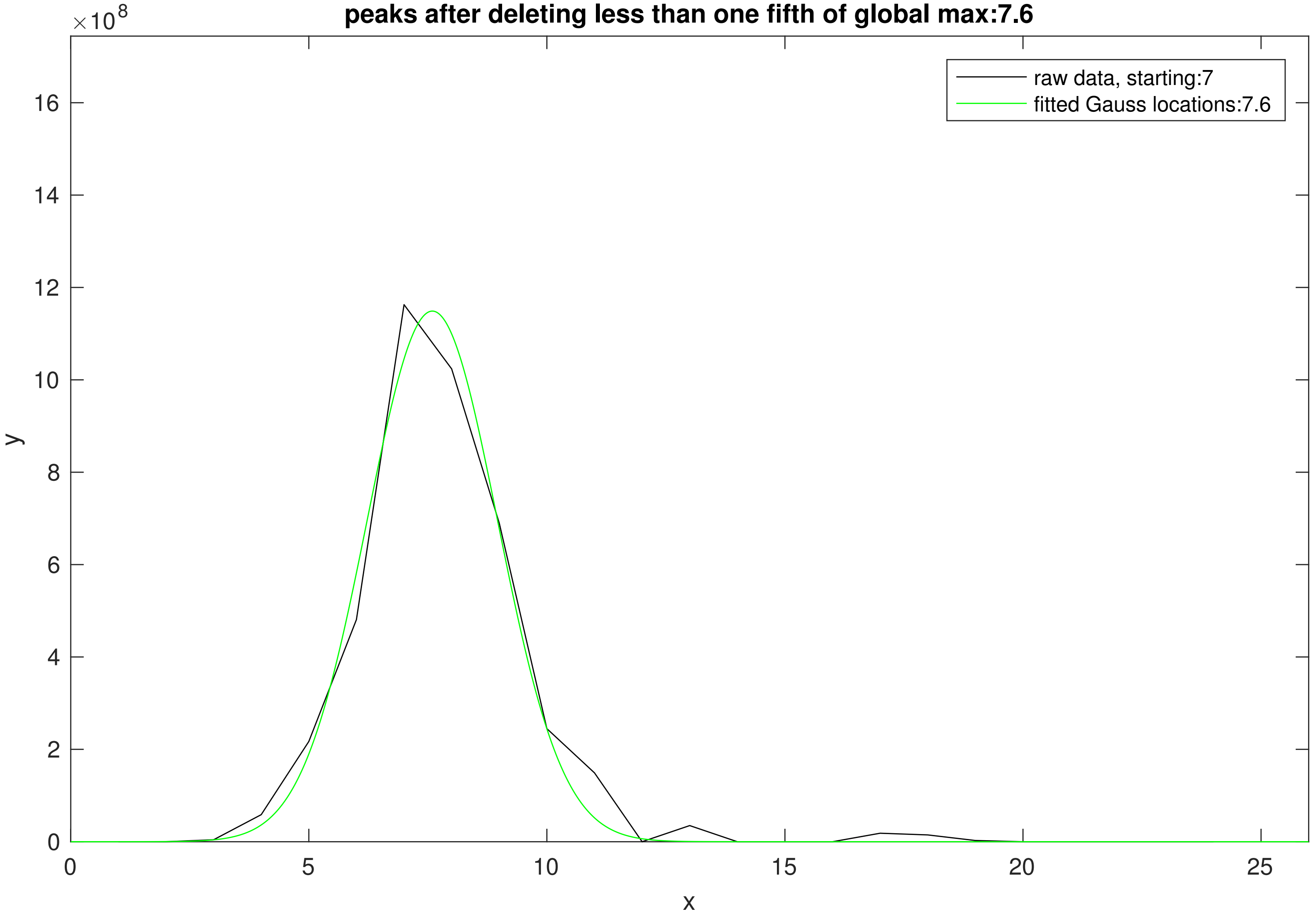

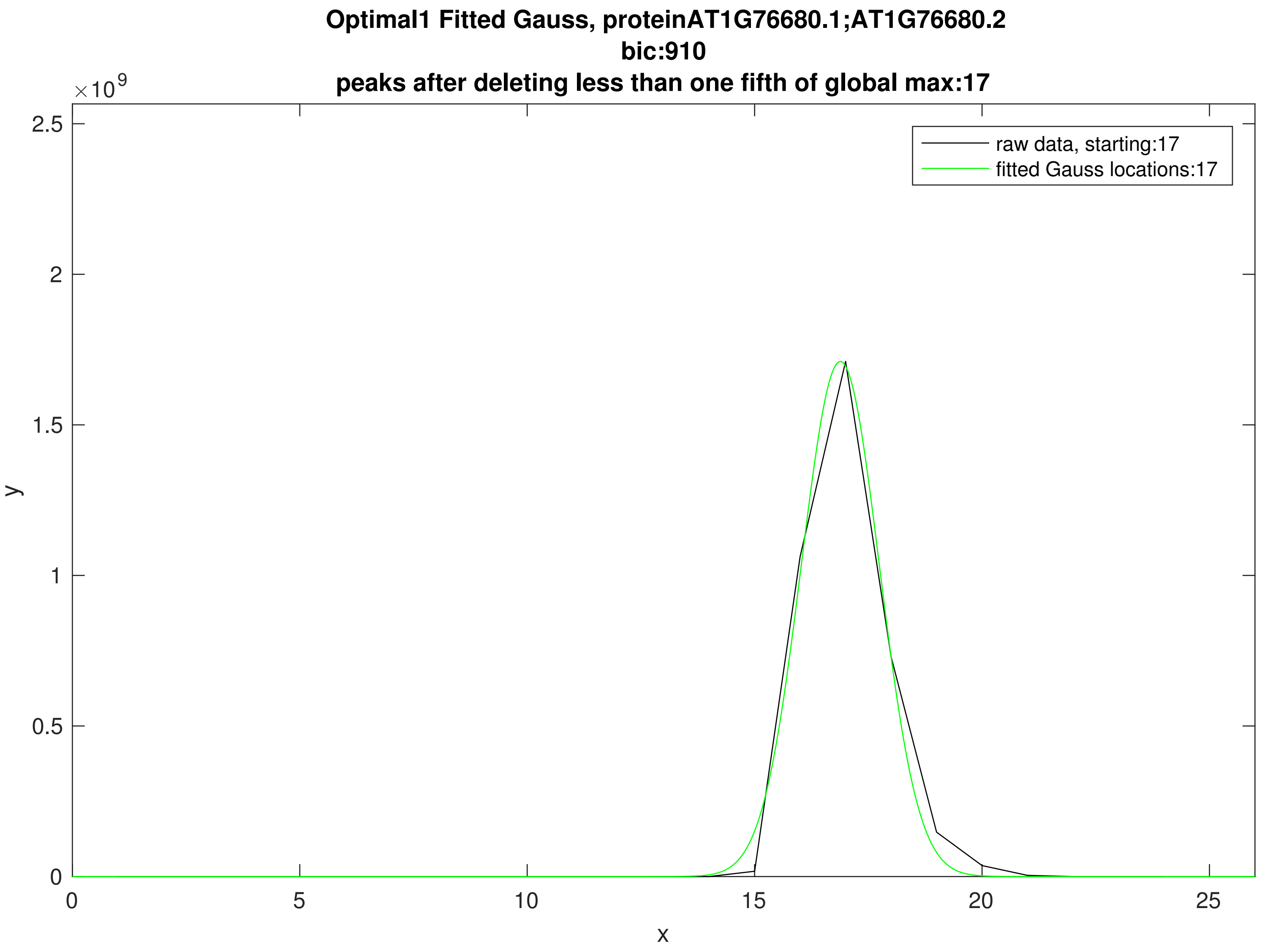

Optimal1 Fitted Gauss, proteinAT1G76680.1;AT1G76680.2
bic:910
peaks after deleting less than one fifth of global max:17
×10^9
2.5
2
1.5
1
0.5
0
y
0
5
10
15
20
25
x
raw data, starting:17
fitted Gauss locations:17

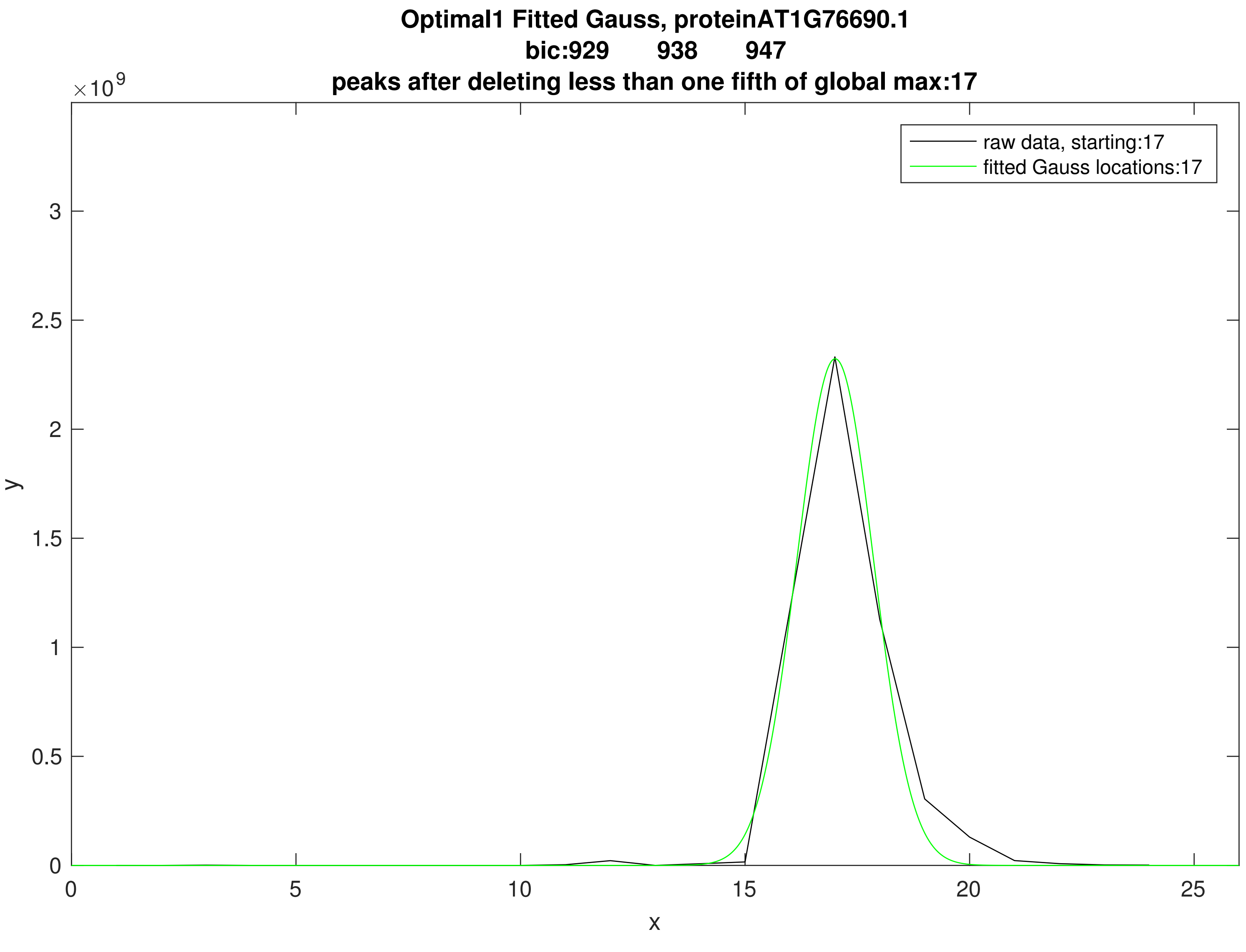

Optimal1 Fitted Gauss, proteinAT1G76690.1
bic:929 938 947
peaks after deleting less than one fifth of global max:17
×10^9
raw data, starting:17
fitted Gauss locations:17
y
x
0
0.5
1
1.5
2
2.5
3
0
5
10
15
20
25

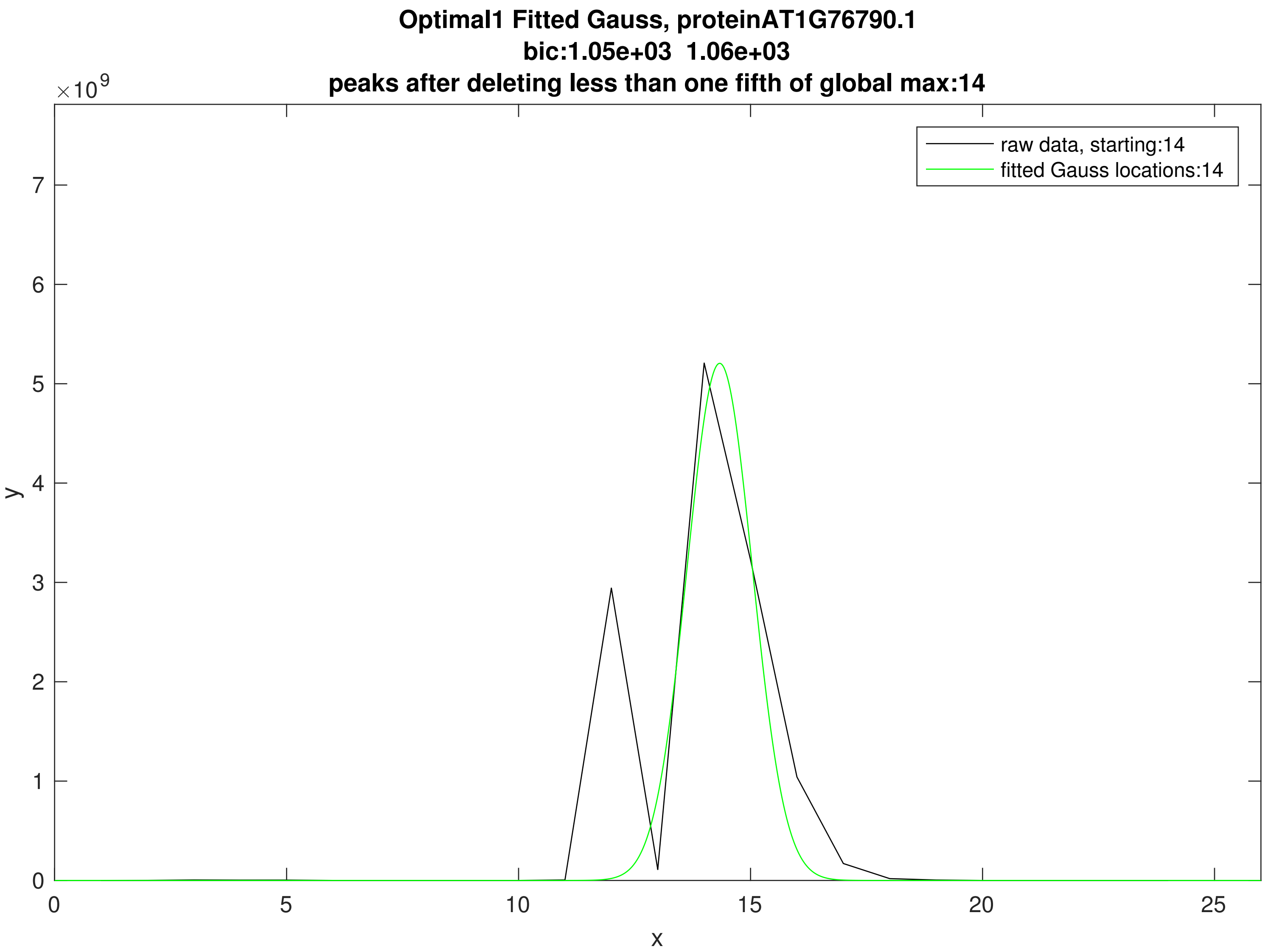

Optimal1 Fitted Gauss, proteinAT1G76790.1
bic:1.05e+03 1.06e+03
peaks after deleting less than one fifth of global max:14
×10^9
raw data, starting:14
fitted Gauss locations:14
y
x
0
1
2
3
4
5
6
7
0
5
10
15
20
25

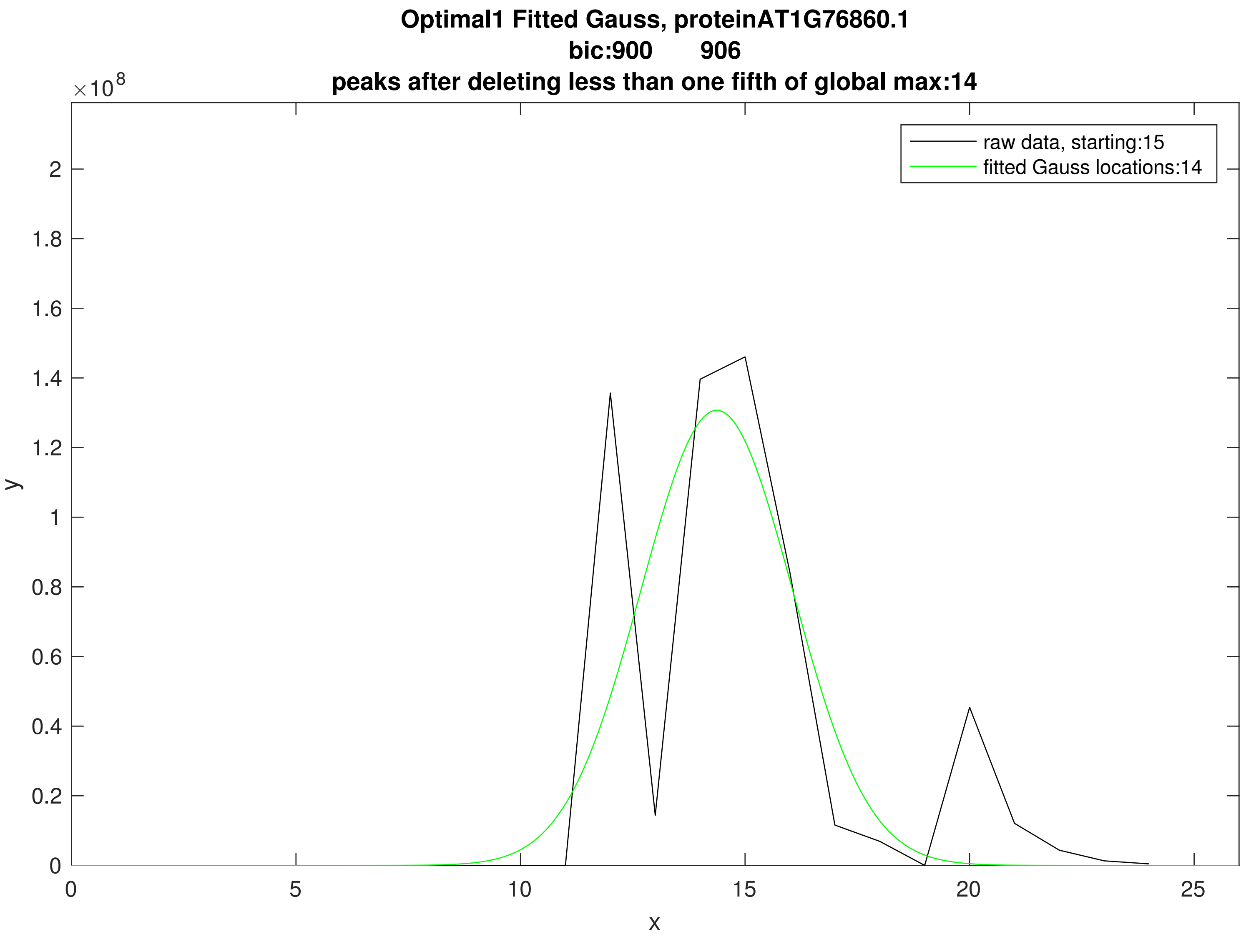
Optimal1 Fitted Gauss, proteinAT1G76860.1
bic:900 906
peaks after deleting less than one fifth of global max:14
×10^8
raw data, starting:15
fitted Gauss locations:14
y
x
0
0.2
0.4
0.6
0.8
1
1.2
1.4
1.6
1.8
2
5
10
15
20
25

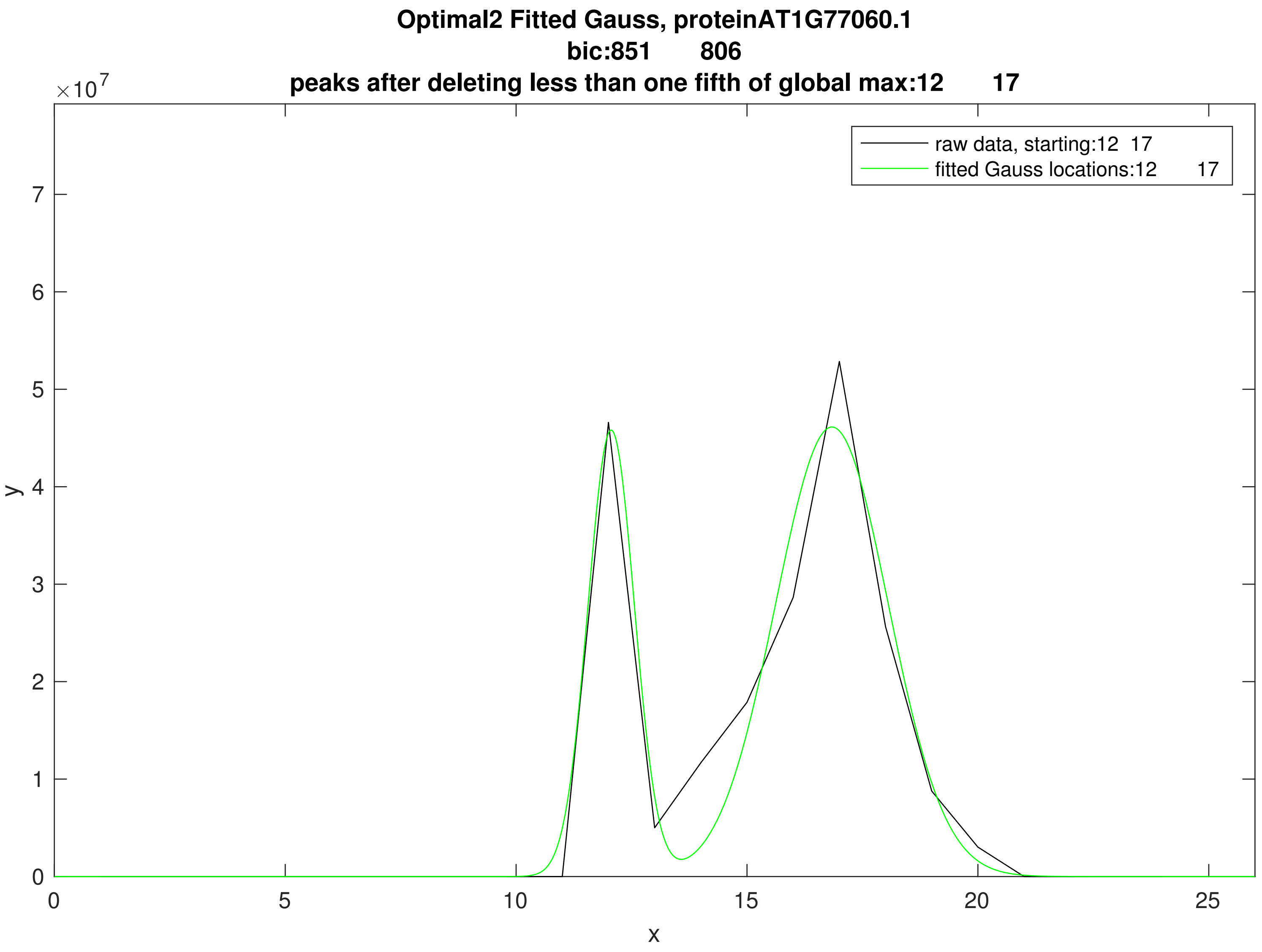

Optimal2 Fitted Gauss, proteinAT1G77060.1
bic:851 806
peaks after deleting less than one fifth of global max:12 17
×10^7
raw data, starting:12 17
fitted Gauss locations:12 17
y
x
0
1
2
3
4
5
6
7
0
5
10
15
20
25

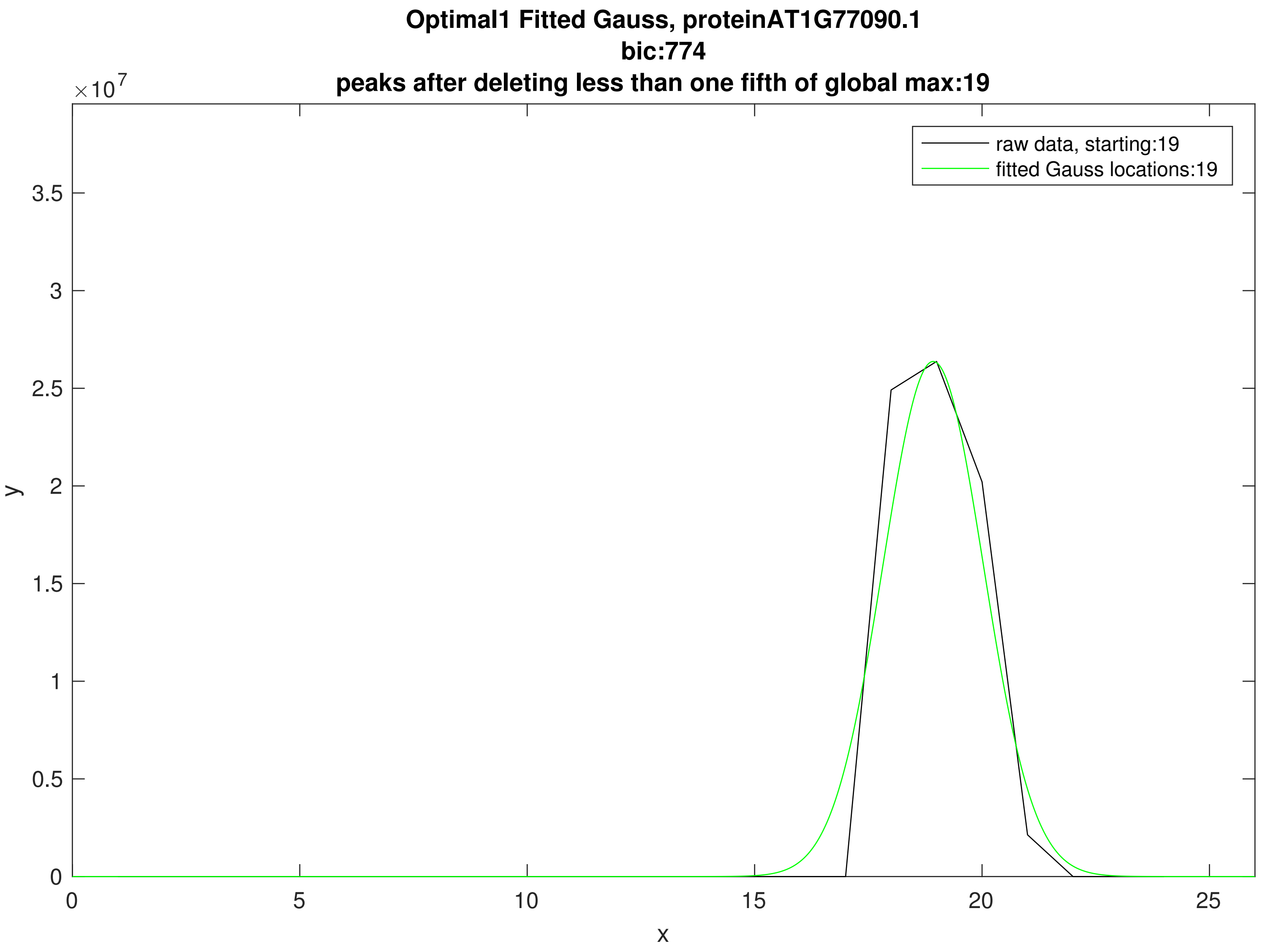

Optimal1 Fitted Gauss, proteinAT1G77090.1
bic:774
peaks after deleting less than one fifth of global max:19
×10^7
raw data, starting:19
fitted Gauss locations:19
3.5
3
2.5
2
1.5
1
0.5
0
y
0
5
10
15
20
25
x

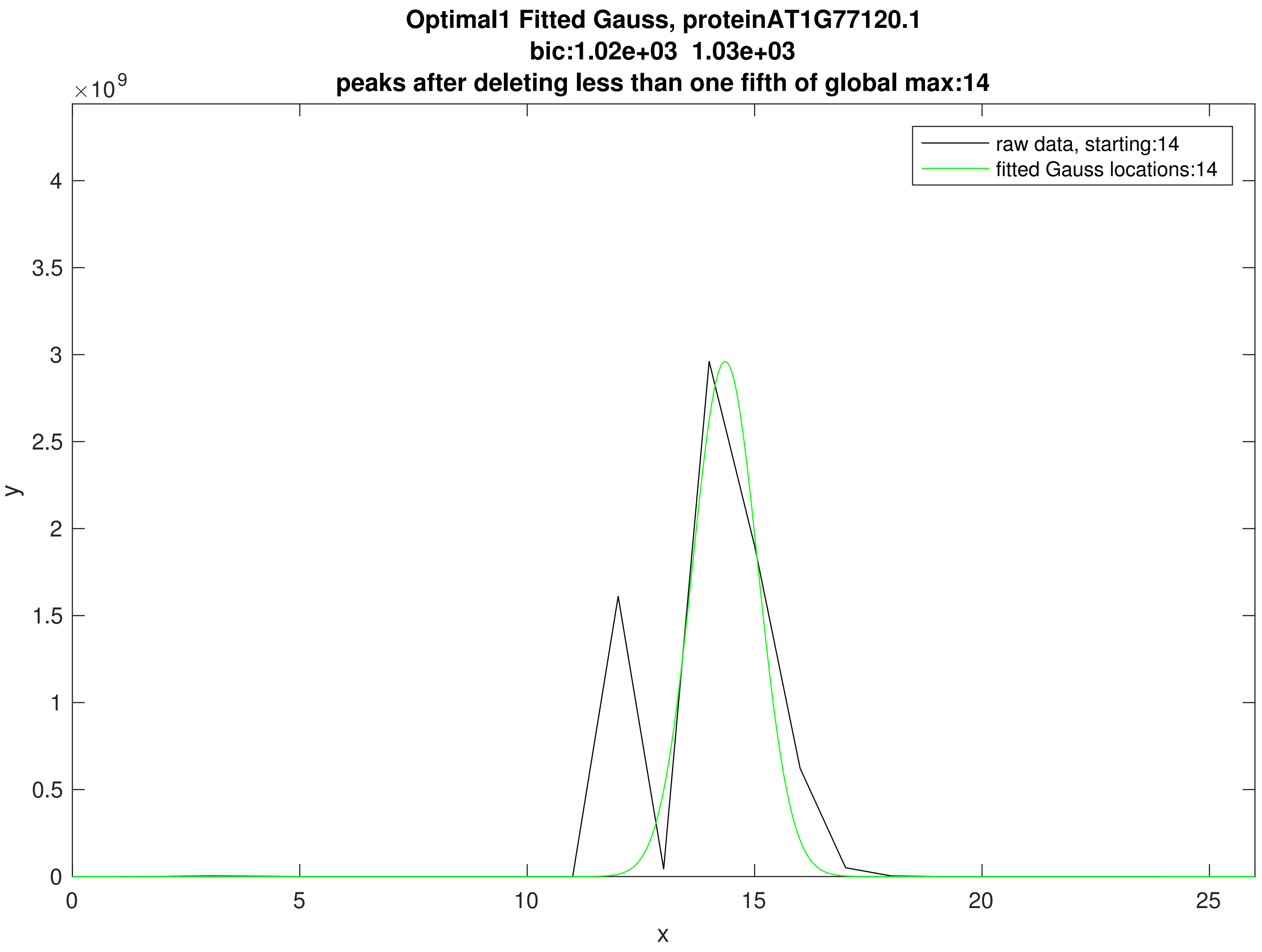

Optimal1 Fitted Gauss, proteinAT1G77120.1
bic:1.02e+03  1.03e+03
peaks after deleting less than one fifth of global max:14
×10^9
raw data, starting:14
fitted Gauss locations:14
4
3.5
3
2.5
2
1.5
1
0.5
0
y
0
5
10
15
20
25
x

**Optimal1 Fitted Gauss, proteinAT1G77330.1**
**bic:1.02e+03  1.03e+03  1.05e+03**
**peaks after deleting less than one fifth of global max:17**

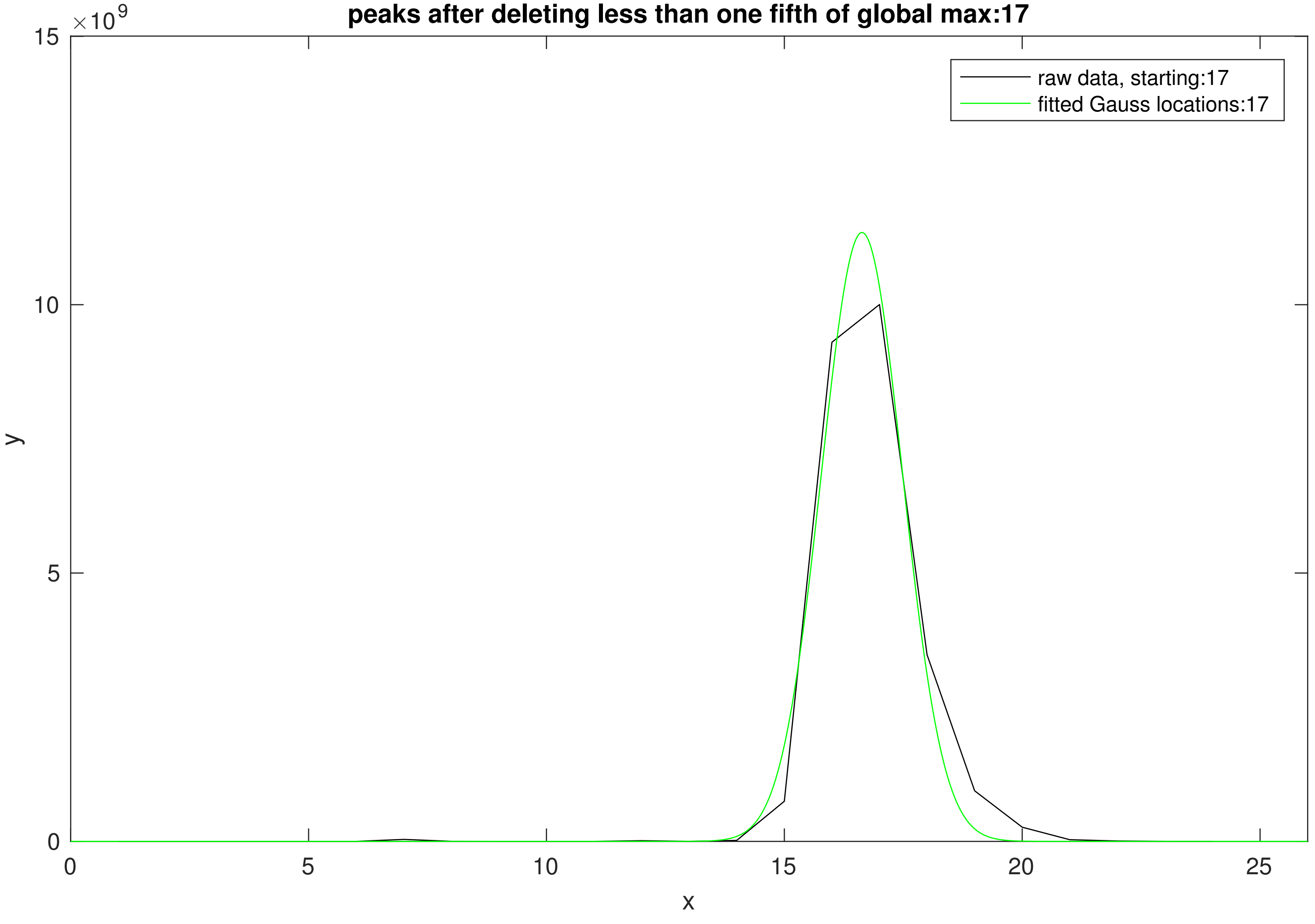

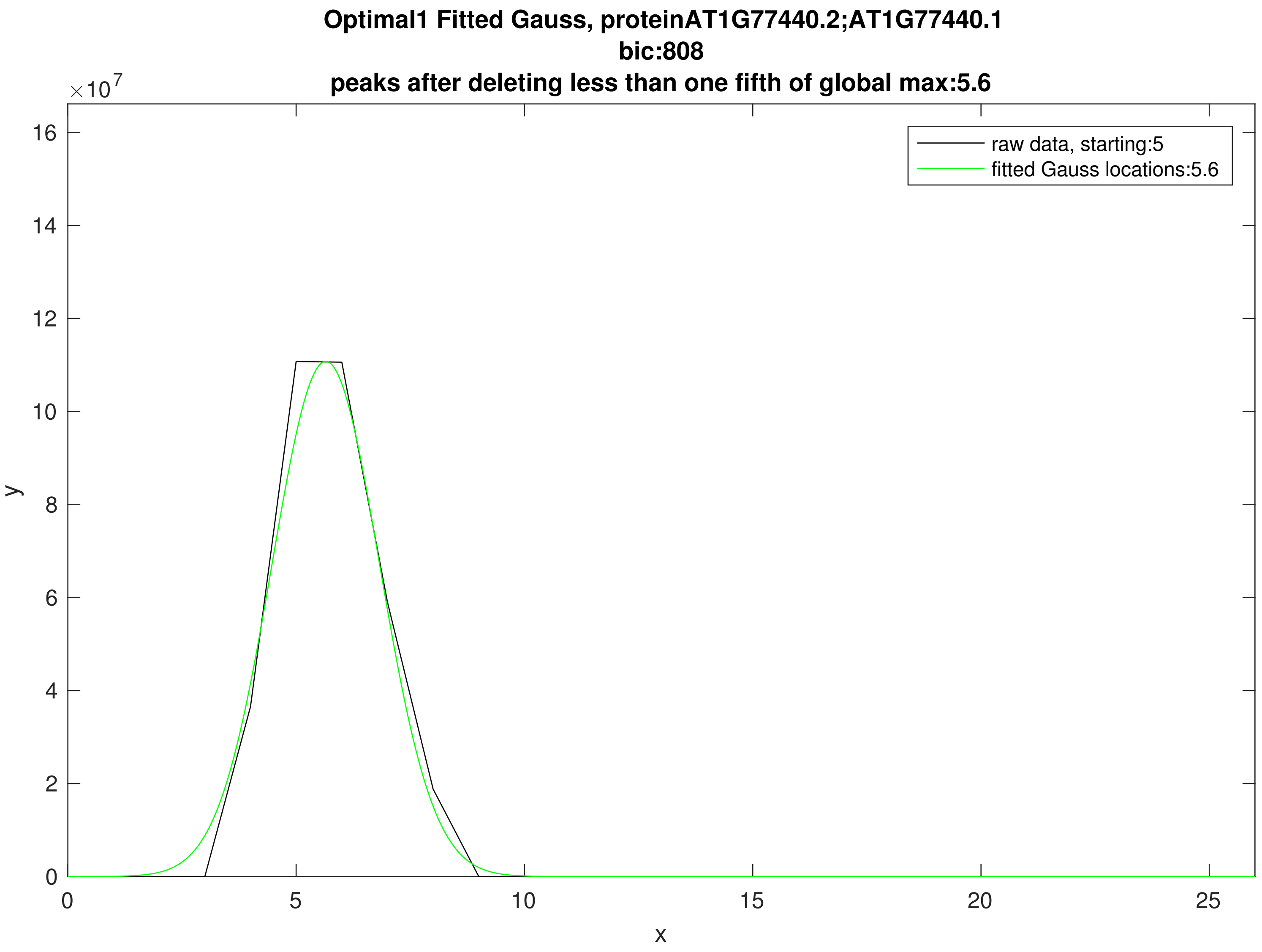

Optimal1 Fitted Gauss, proteinAT1G77440.2;AT1G77440.1
bic:808
peaks after deleting less than one fifth of global max:5.6
×10^7
16
14
12
10
8
6
4
2
0
y
0
5
10
15
20
25
x
raw data, starting:5
fitted Gauss locations:5.6

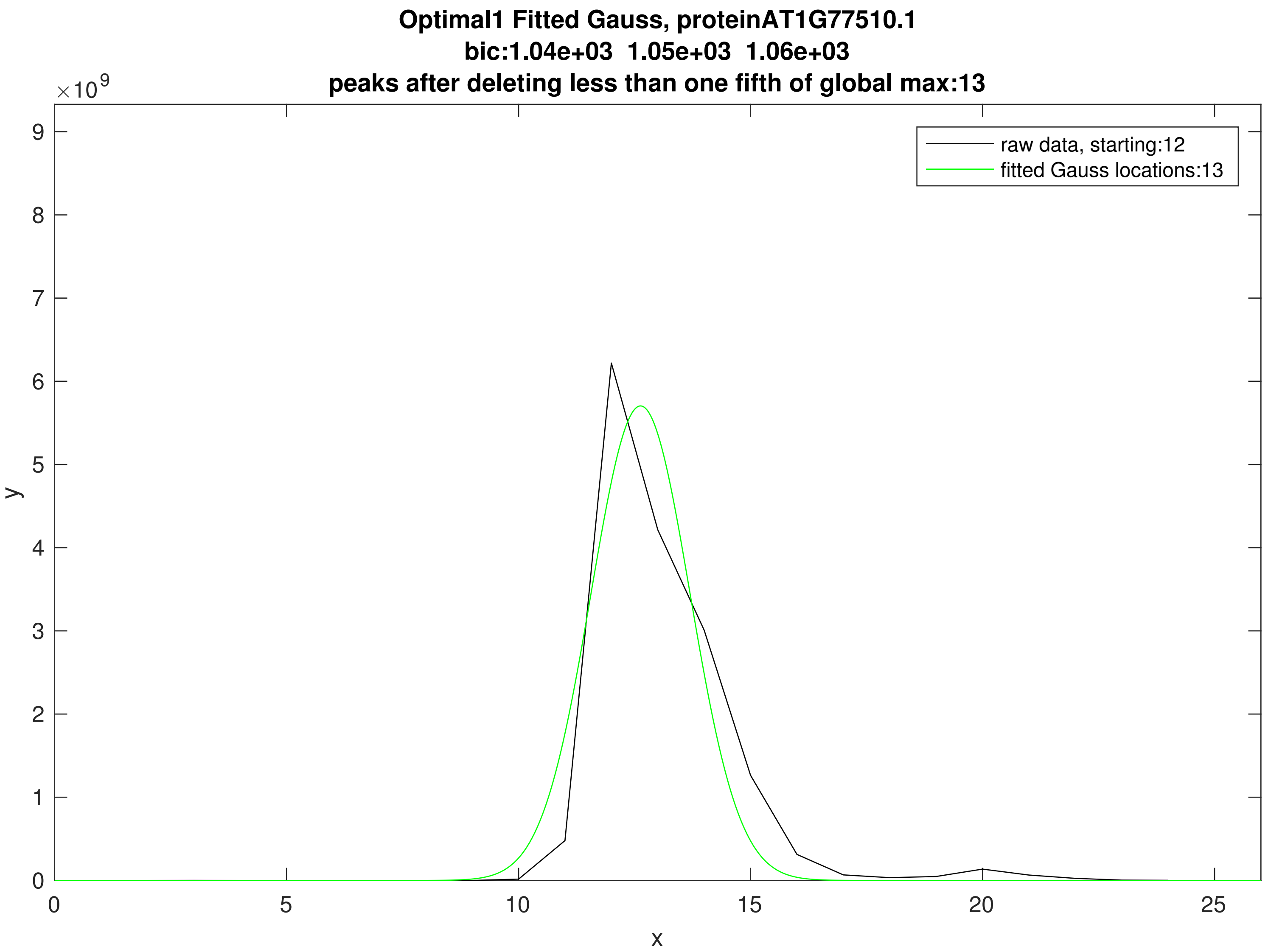

Optimal1 Fitted Gauss, proteinAT1G77510.1
bic:1.04e+03 1.05e+03 1.06e+03
peaks after deleting less than one fifth of global max:13
×10^9
raw data, starting:12
fitted Gauss locations:13
y
x
0
1
2
3
4
5
6
7
8
9
0
5
10
15
20
25

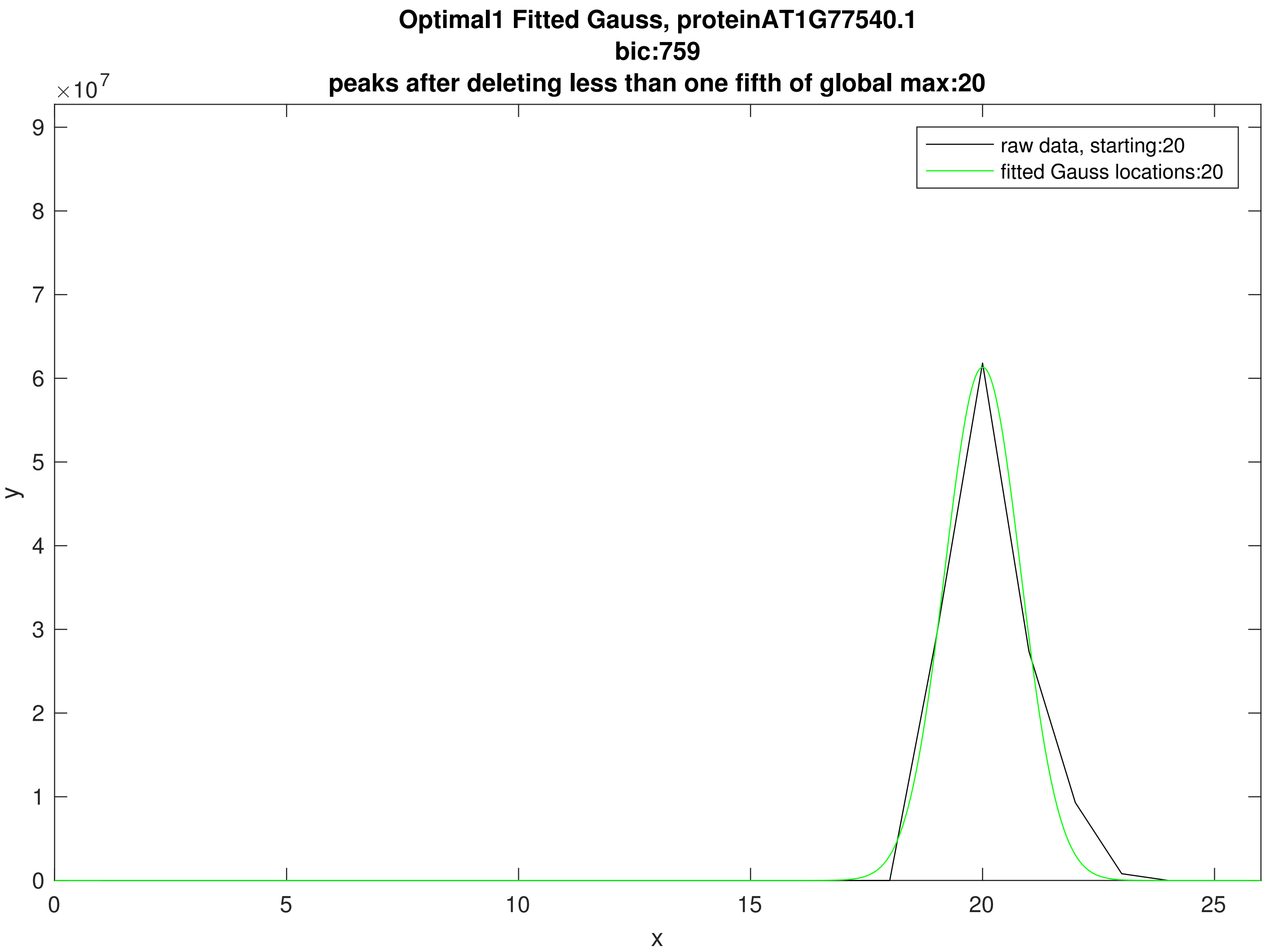

Optimal1 Fitted Gauss, proteinAT1G77540.1
bic:759
peaks after deleting less than one fifth of global max:20
×10^7
9
8
7
6
5
4
3
2
1
0
y
raw data, starting:20
fitted Gauss locations:20
0
5
10
15
20
25
x

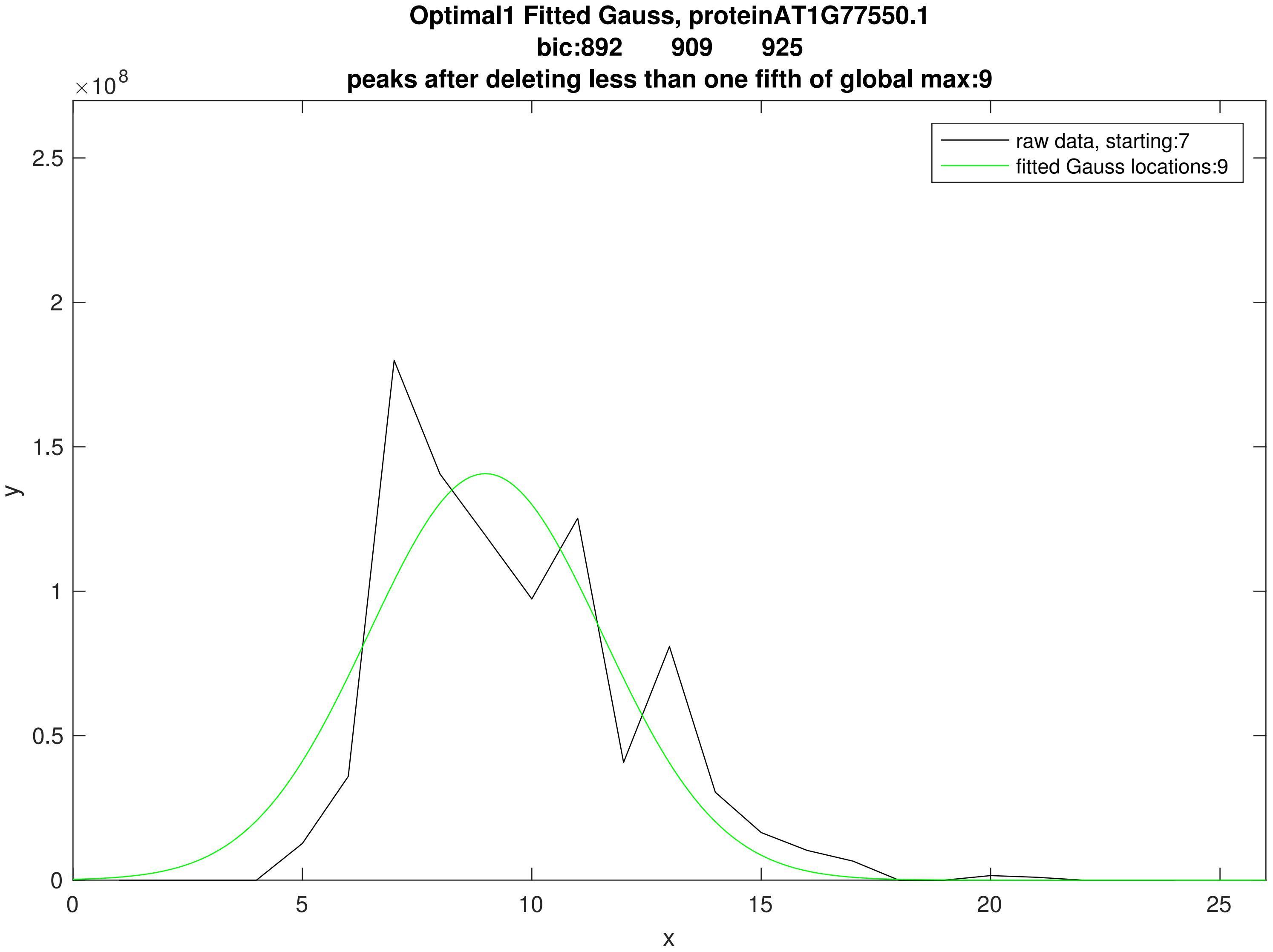

Optimal1 Fitted Gauss, proteinAT1G77550.1
bic:892 909 925
peaks after deleting less than one fifth of global max:9
×10^8
raw data, starting:7
fitted Gauss locations:9
y
x
0
0.5
1
1.5
2
2.5
5
10
15
20
25

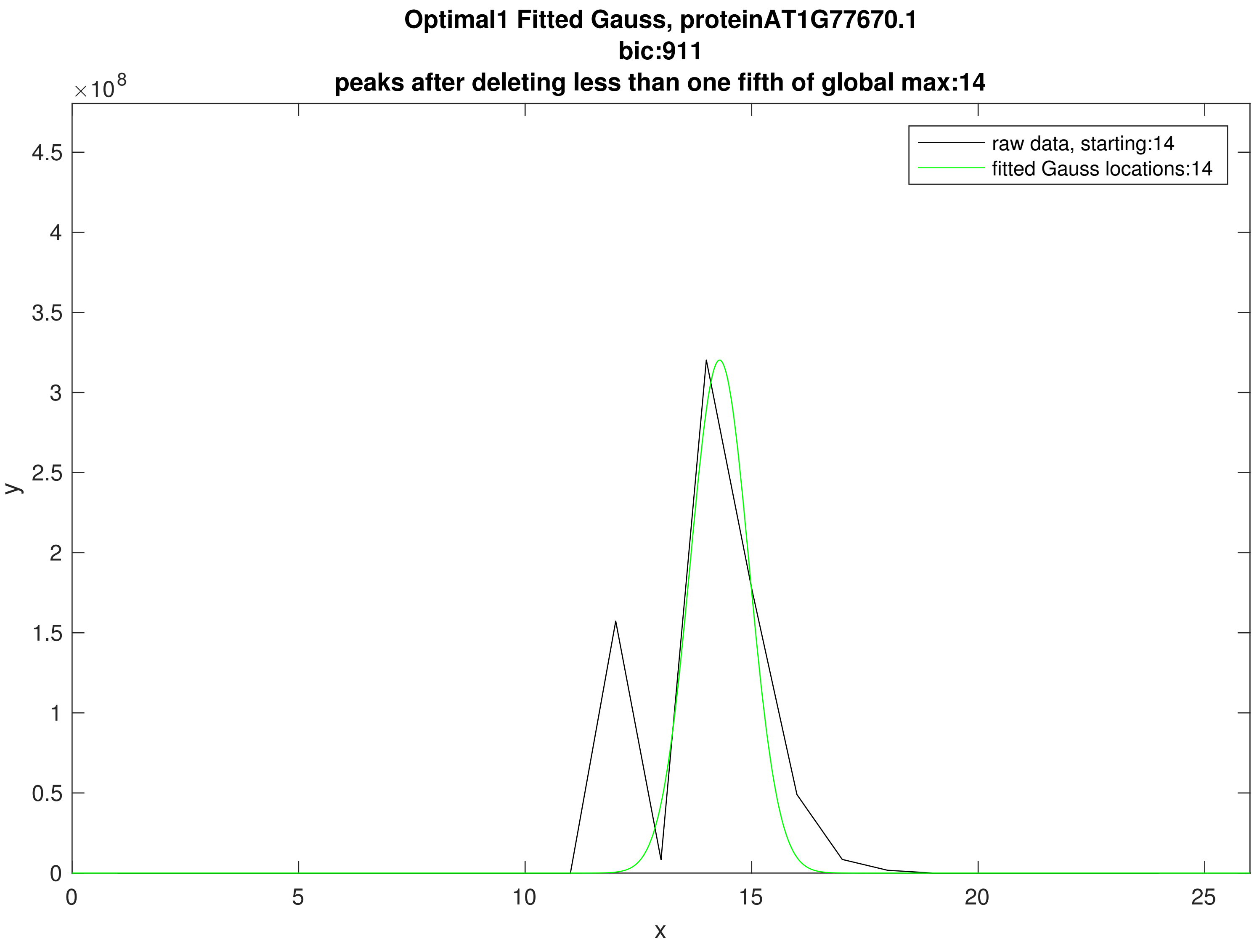

Optimal1 Fitted Gauss, proteinAT1G77670.1
bic:911
peaks after deleting less than one fifth of global max:14
×10^8
4.5
4
3.5
3
2.5
2
1.5
1
0.5
0
y
0
5
10
15
20
25
x
raw data, starting:14
fitted Gauss locations:14

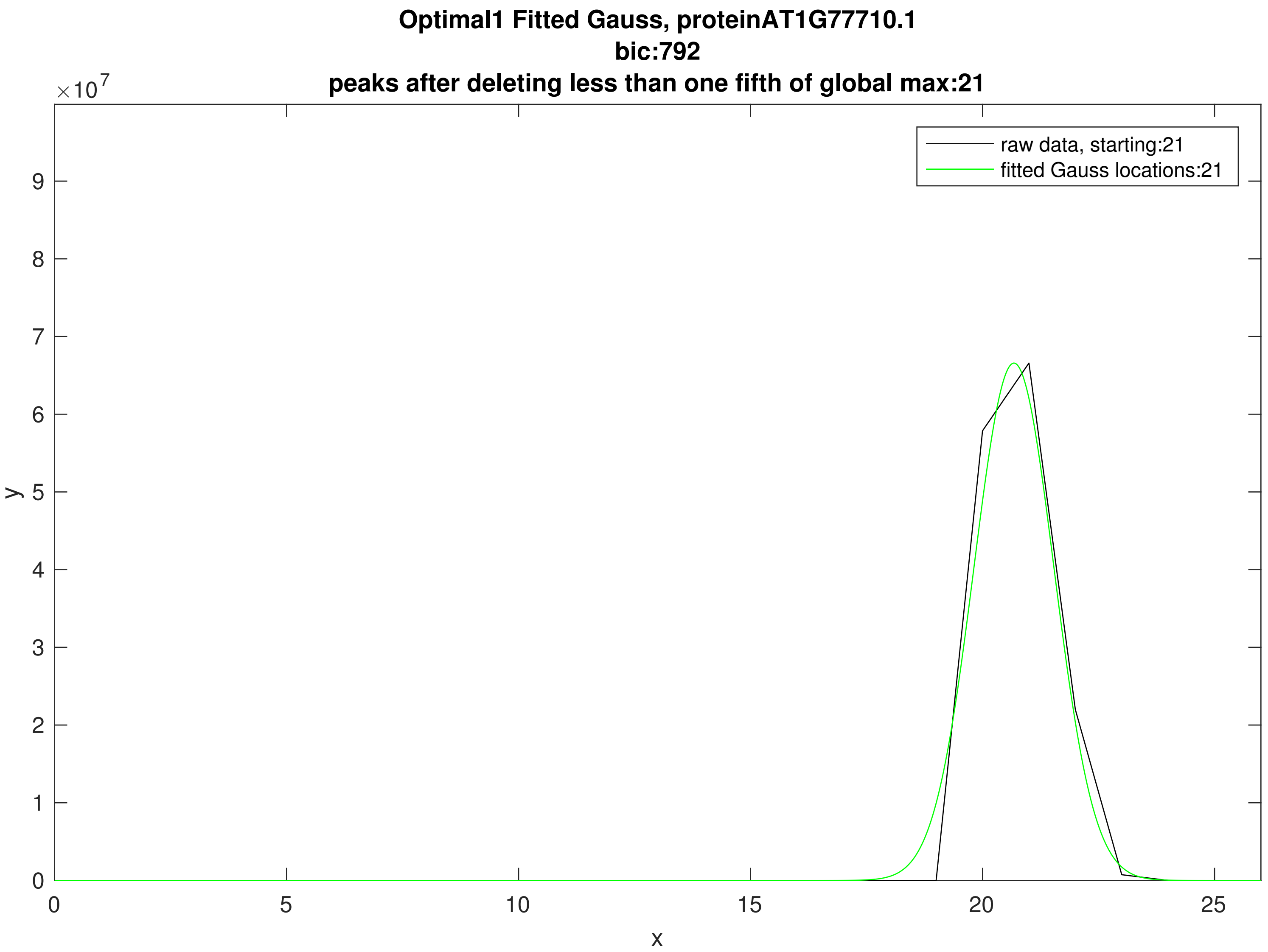
Optimal1 Fitted Gauss, proteinAT1G77710.1
bic:792
peaks after deleting less than one fifth of global max:21
×10^7
raw data, starting:21
fitted Gauss locations:21
y
x
0
1
2
3
4
5
6
7
8
9
0
5
10
15
20
25

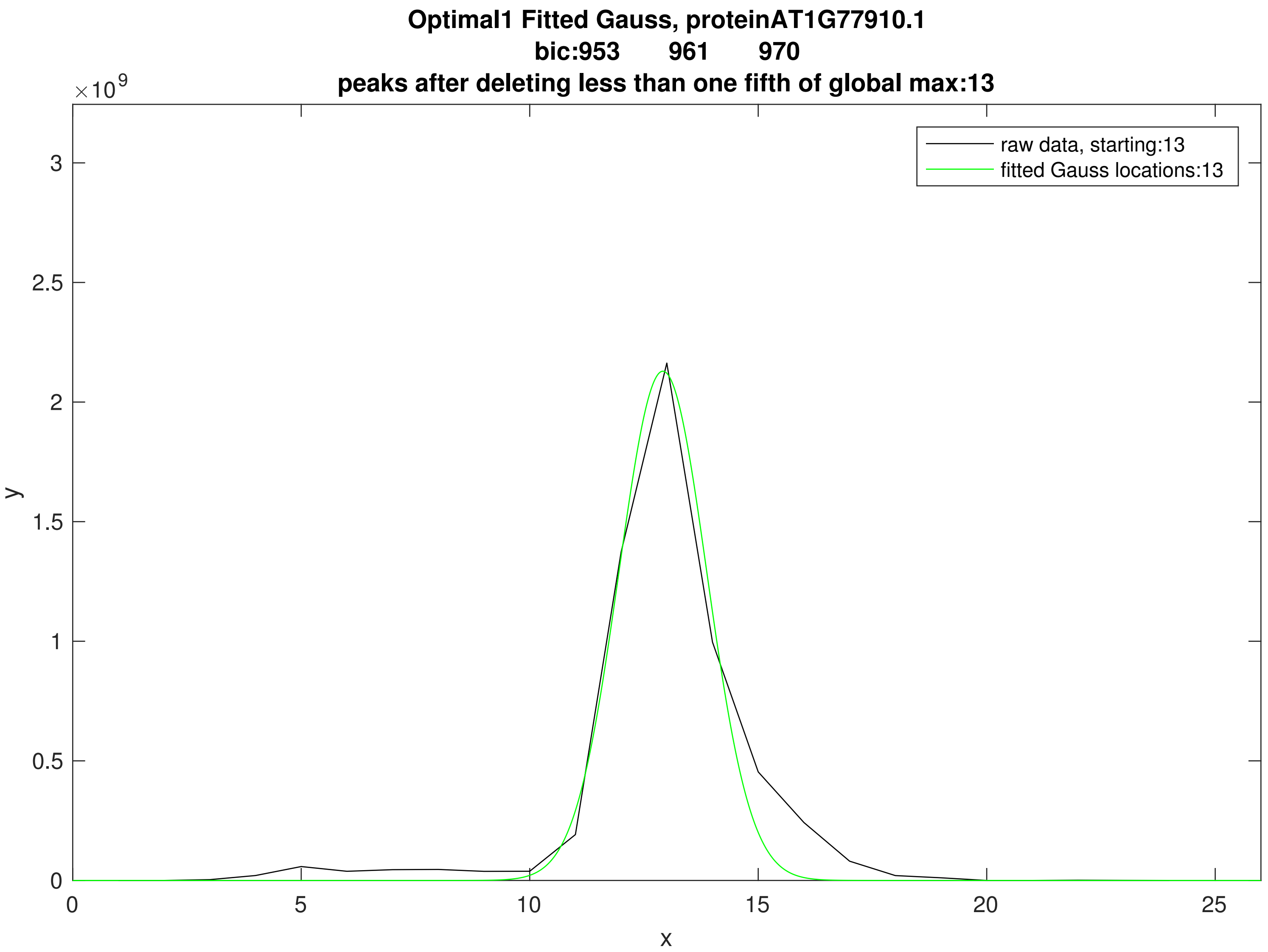
Optimal1 Fitted Gauss, proteinAT1G77910.1
bic:953 961 970
peaks after deleting less than one fifth of global max:13
×10^9
raw data, starting:13
fitted Gauss locations:13
y
x
0
0.5
1
1.5
2
2.5
3
0
5
10
15
20
25

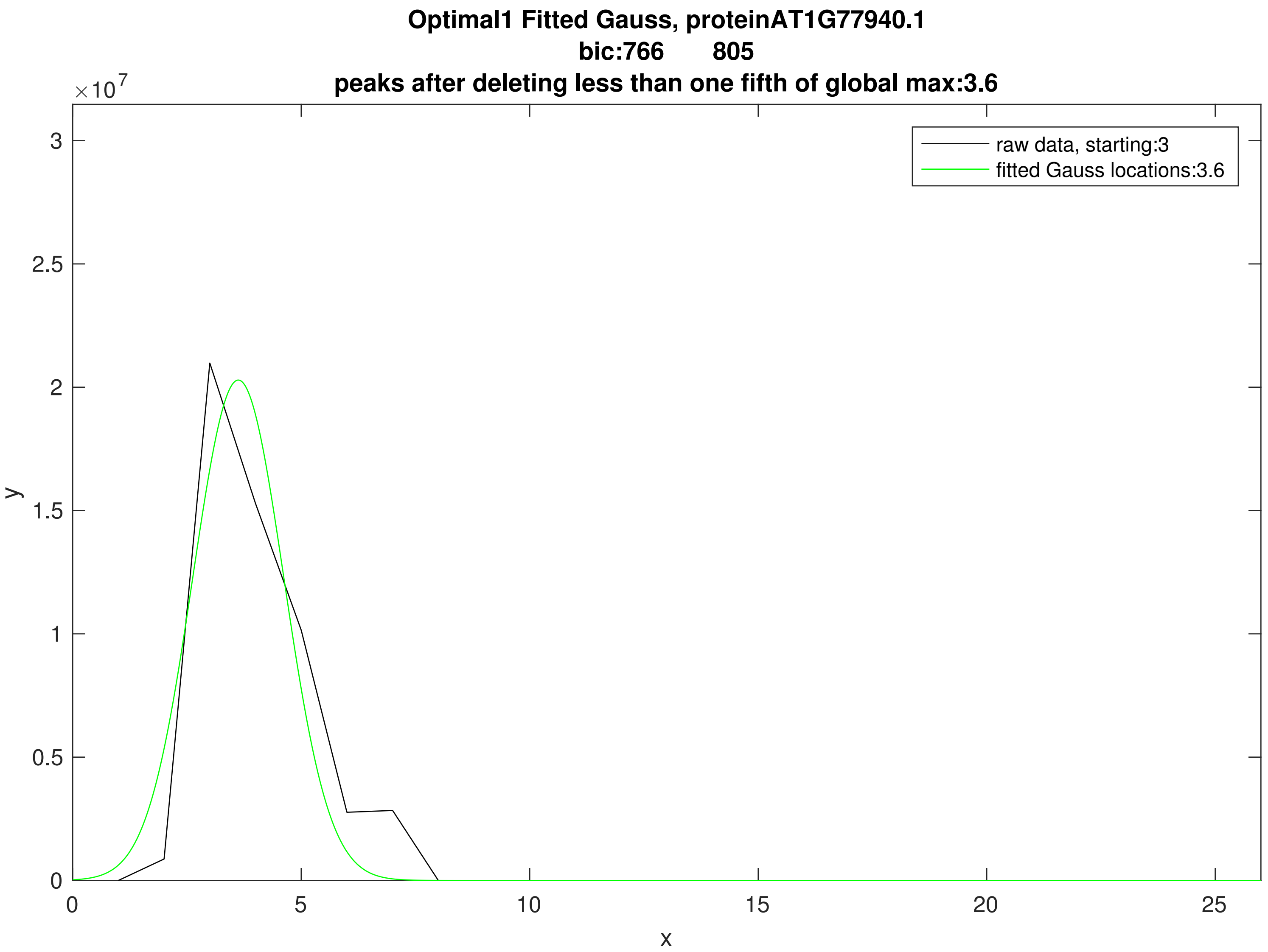
Optimal1 Fitted Gauss, proteinAT1G77940.1
bic:766 805
peaks after deleting less than one fifth of global max:3.6
×10^7
raw data, starting:3
fitted Gauss locations:3.6
y
x
0
0.5
1
1.5
2
2.5
3
0
5
10
15
20
25

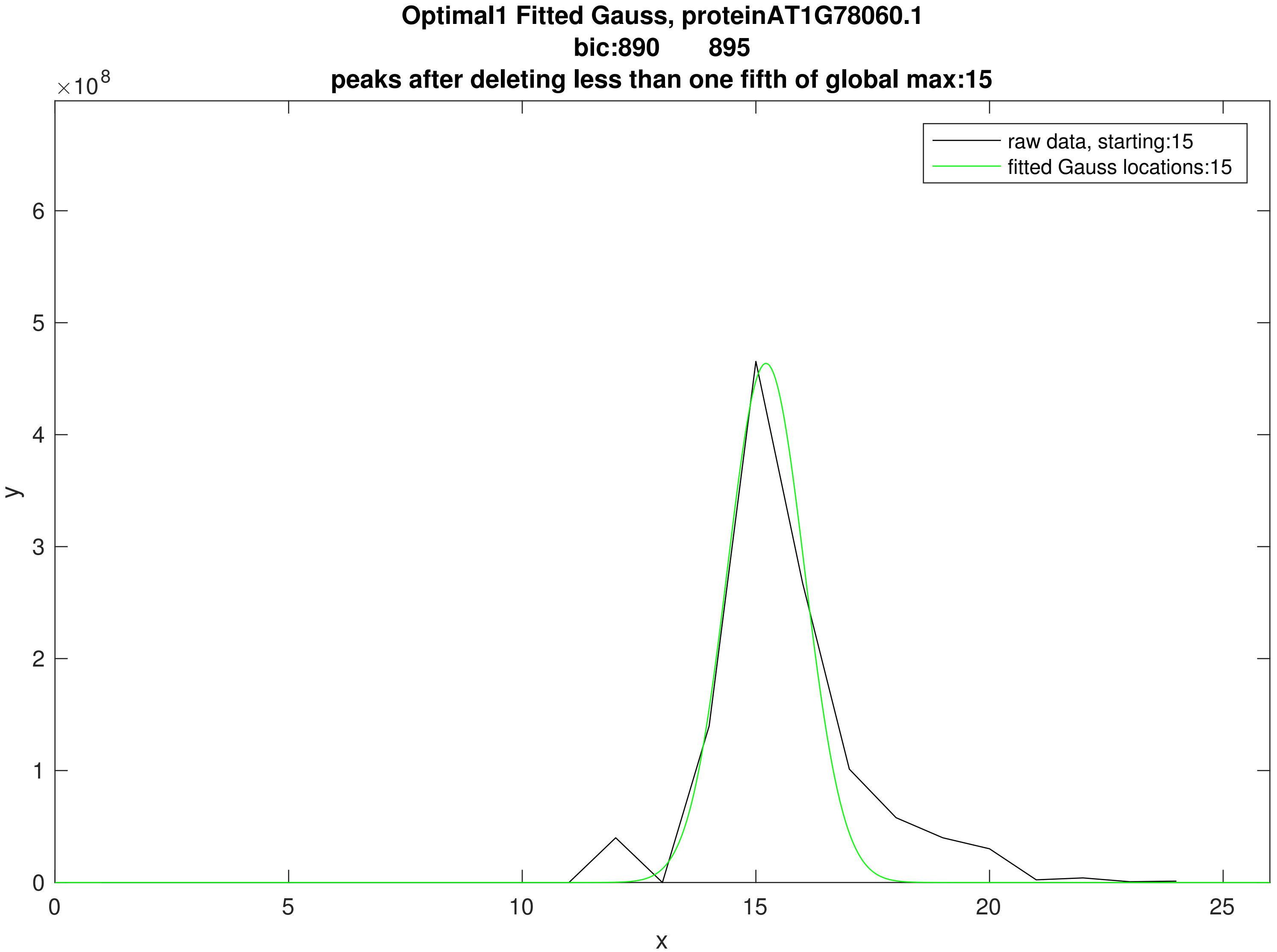

Optimal1 Fitted Gauss, proteinAT1G78060.1
bic:890 895
peaks after deleting less than one fifth of global max:15
×10^8
raw data, starting:15
fitted Gauss locations:15
y
x
0
1
2
3
4
5
6
0
5
10
15
20
25

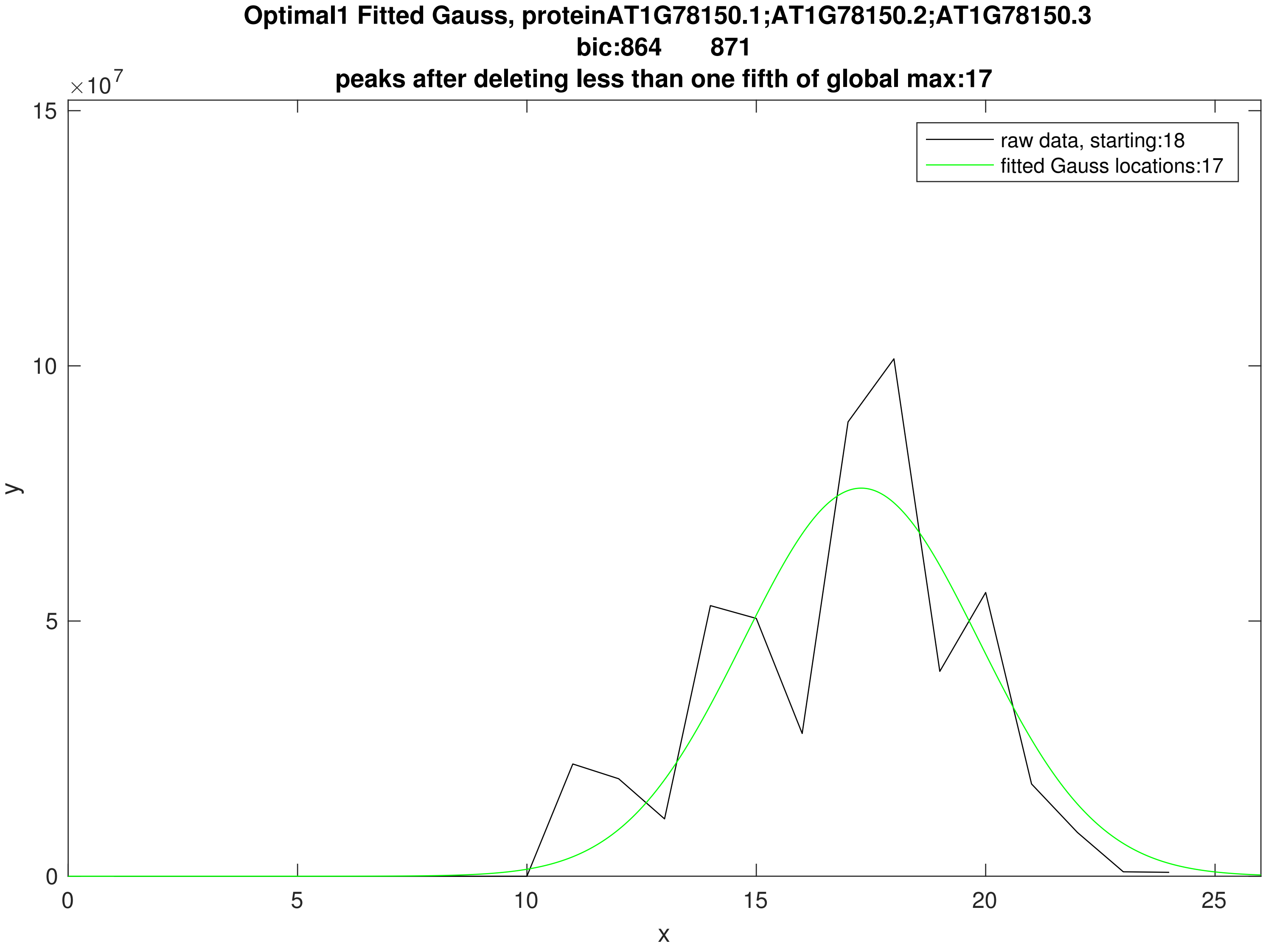
Optimal1 Fitted Gauss, proteinAT1G78150.1;AT1G78150.2;AT1G78150.3
bic:864 871
peaks after deleting less than one fifth of global max:17
×10^7
15
10
5
0
y
0
5
10
15
20
25
x
raw data, starting:18
fitted Gauss locations:17

Optimal1 Fitted Gauss, proteinAT1G78260.2;AT1G78260.1;AT1G22330.1
bic:723
peaks after deleting less than one fifth of global max:21

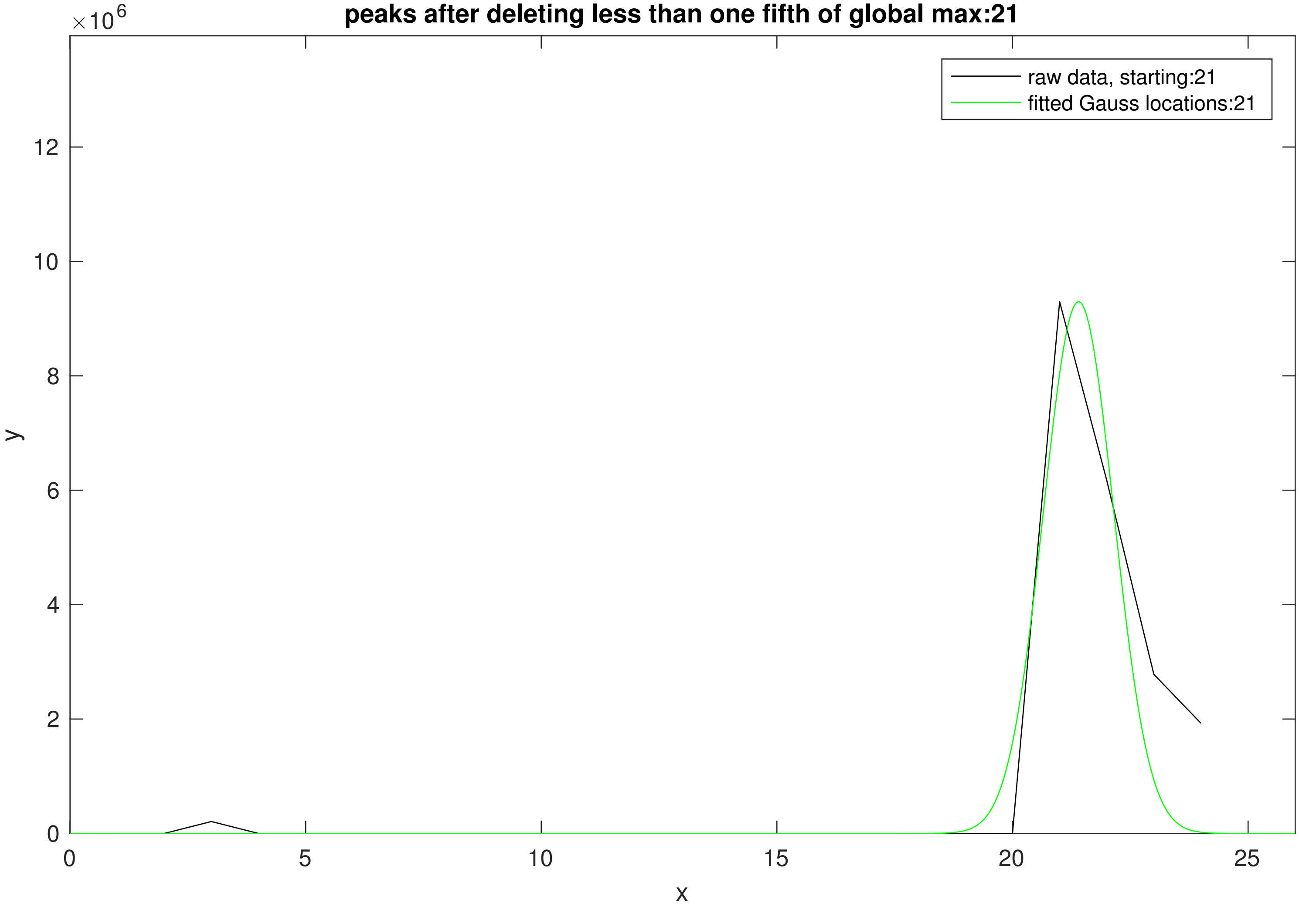

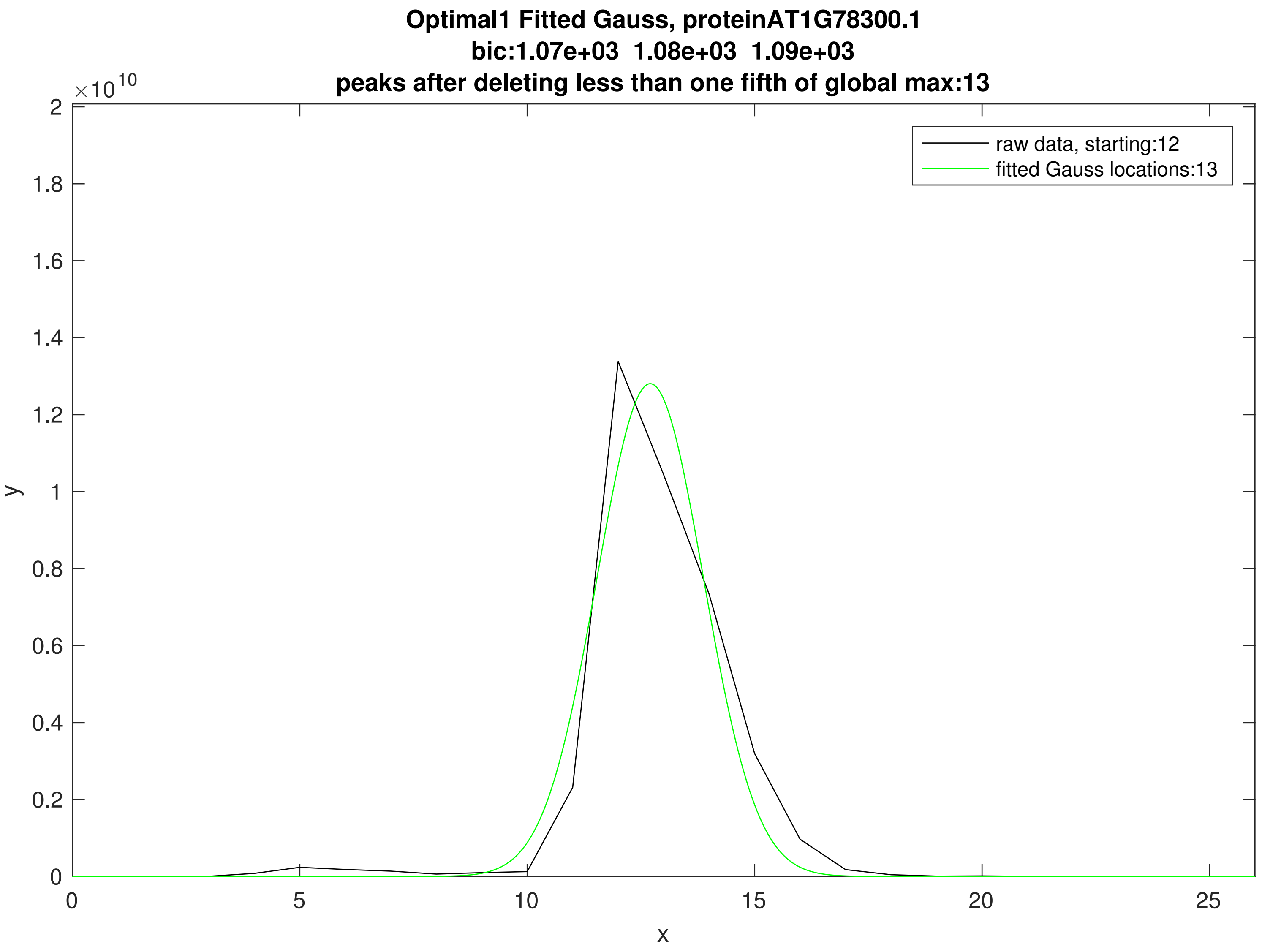

Optimal1 Fitted Gauss, proteinAT1G78300.1
bic:1.07e+03  1.08e+03  1.09e+03
peaks after deleting less than one fifth of global max:13
×10^10
raw data, starting:12
fitted Gauss locations:13
y
x
0
0.2
0.4
0.6
0.8
1
1.2
1.4
1.6
1.8
2
5
10
15
20
25

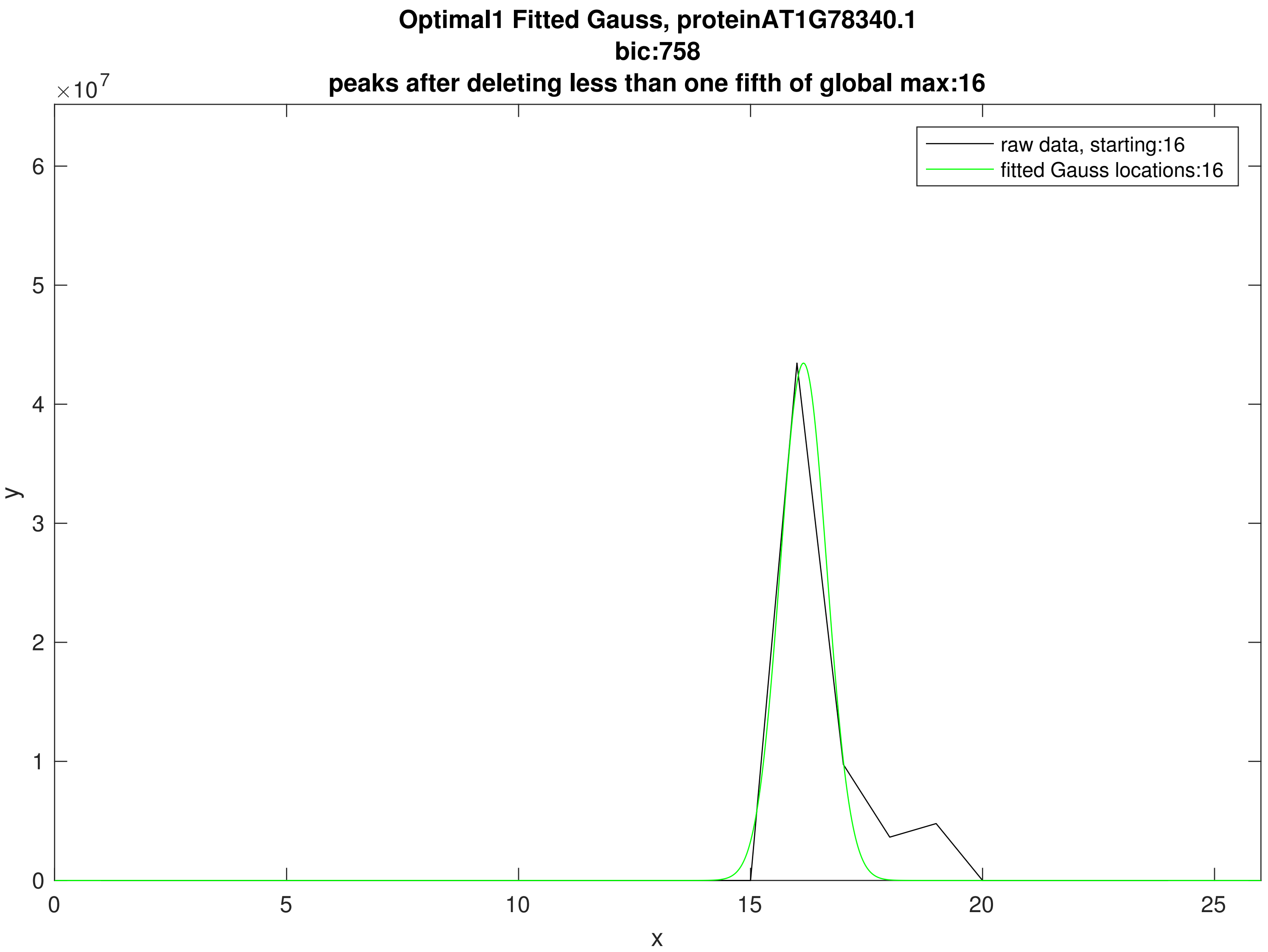
Optimal1 Fitted Gauss, proteinAT1G78340.1
bic:758
peaks after deleting less than one fifth of global max:16
×10^7
raw data, starting:16
fitted Gauss locations:16
y
x
0
1
2
3
4
5
6
0
5
10
15
20
25

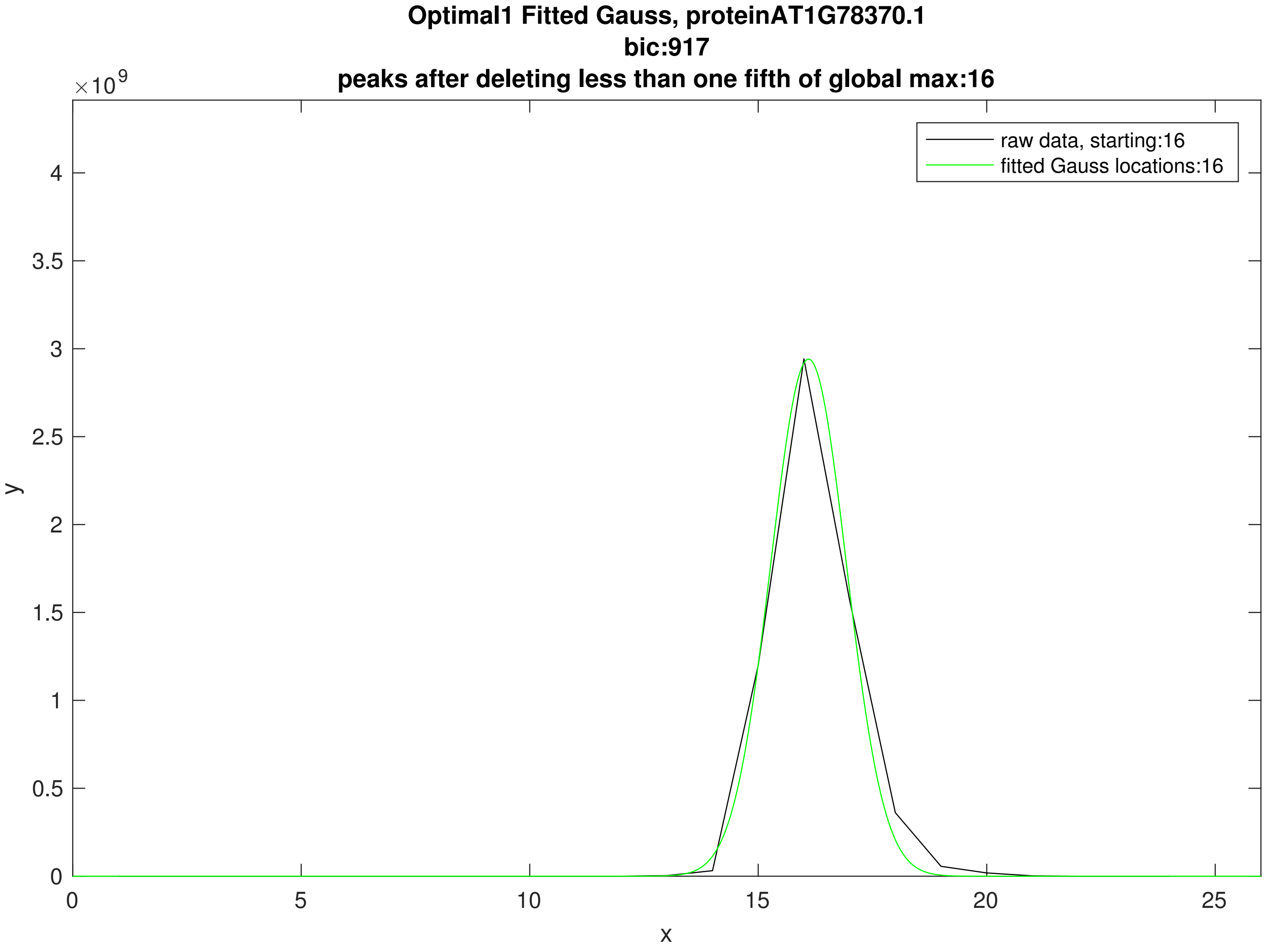
Optimal1 Fitted Gauss, proteinAT1G78370.1
bic:917
peaks after deleting less than one fifth of global max:16
×10^9
raw data, starting:16
fitted Gauss locations:16
y
x
0
0.5
1
1.5
2
2.5
3
3.5
4
5
10
15
20
25

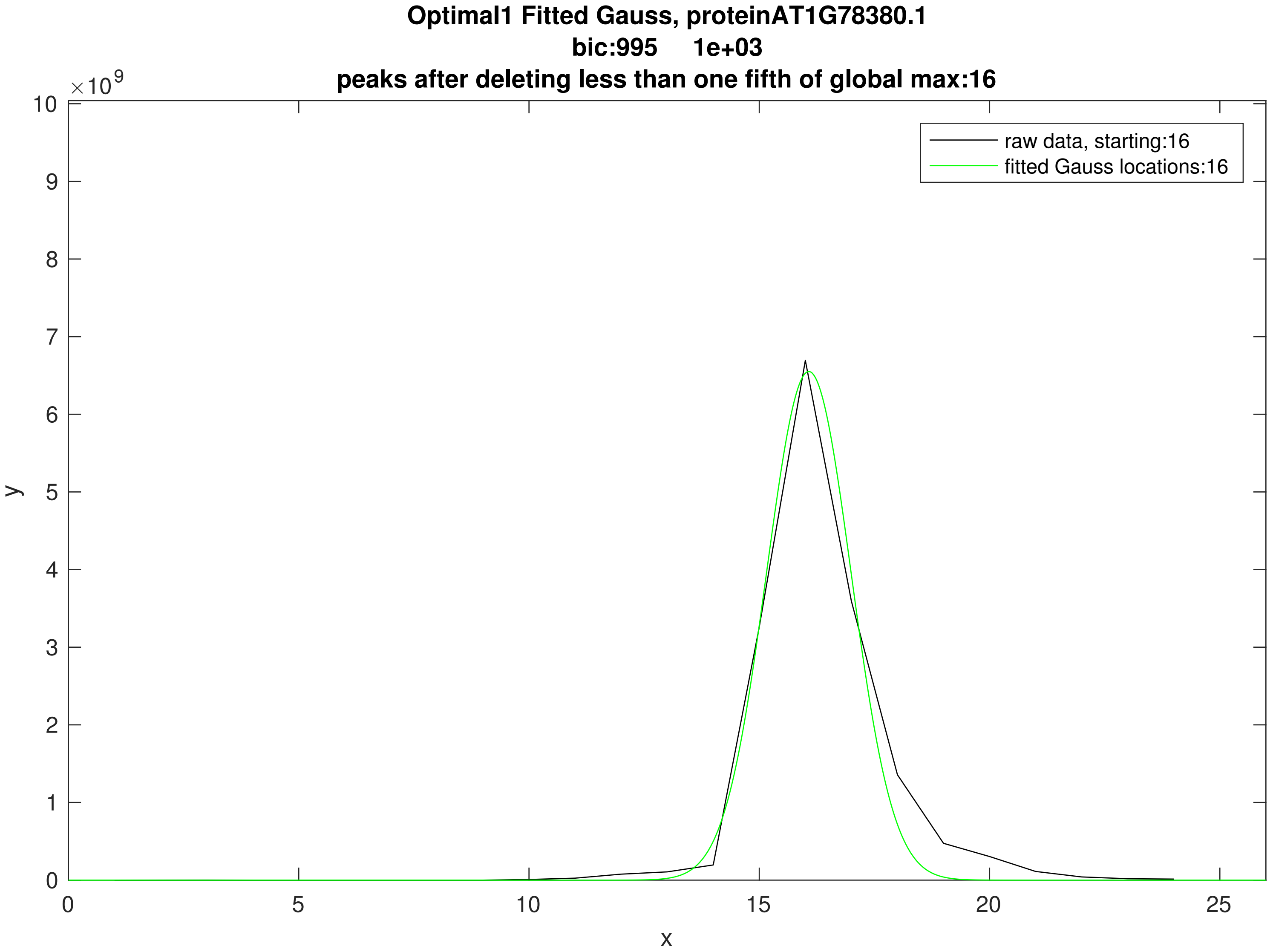
Optimal1 Fitted Gauss, proteinAT1G78380.1
bic:995 1e+03
peaks after deleting less than one fifth of global max:16
×10^9
10
9
8
7
6
5
4
3
2
1
0
y
0
5
10
15
20
25
x
raw data, starting:16
fitted Gauss locations:16

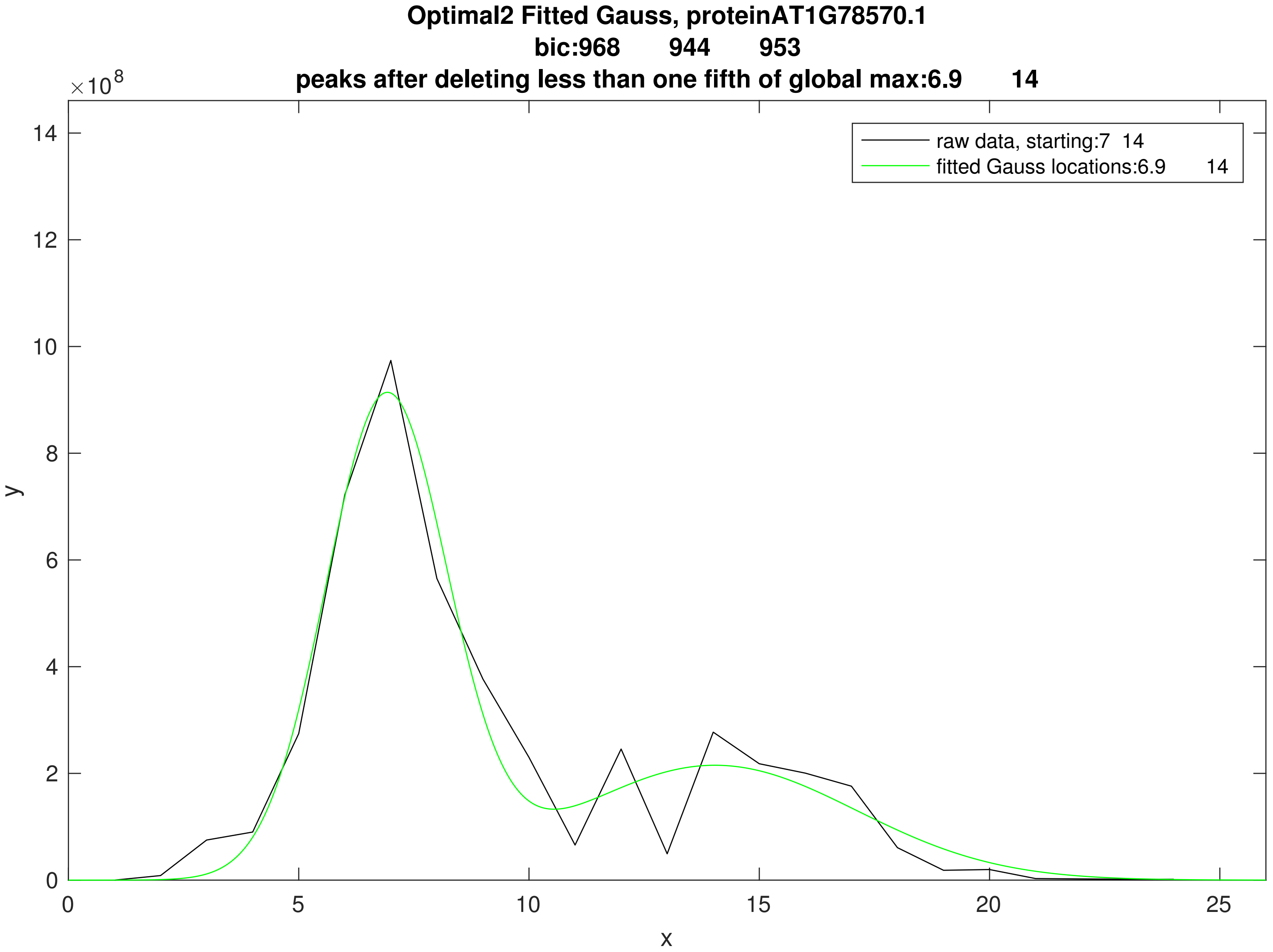

Optimal2 Fitted Gauss, proteinAT1G78570.1
bic:968 944 953
peaks after deleting less than one fifth of global max:6.9 14
×10^8
raw data, starting:7 14
fitted Gauss locations:6.9 14
y
x
0
2
4
6
8
10
12
14
0
5
10
15
20
25

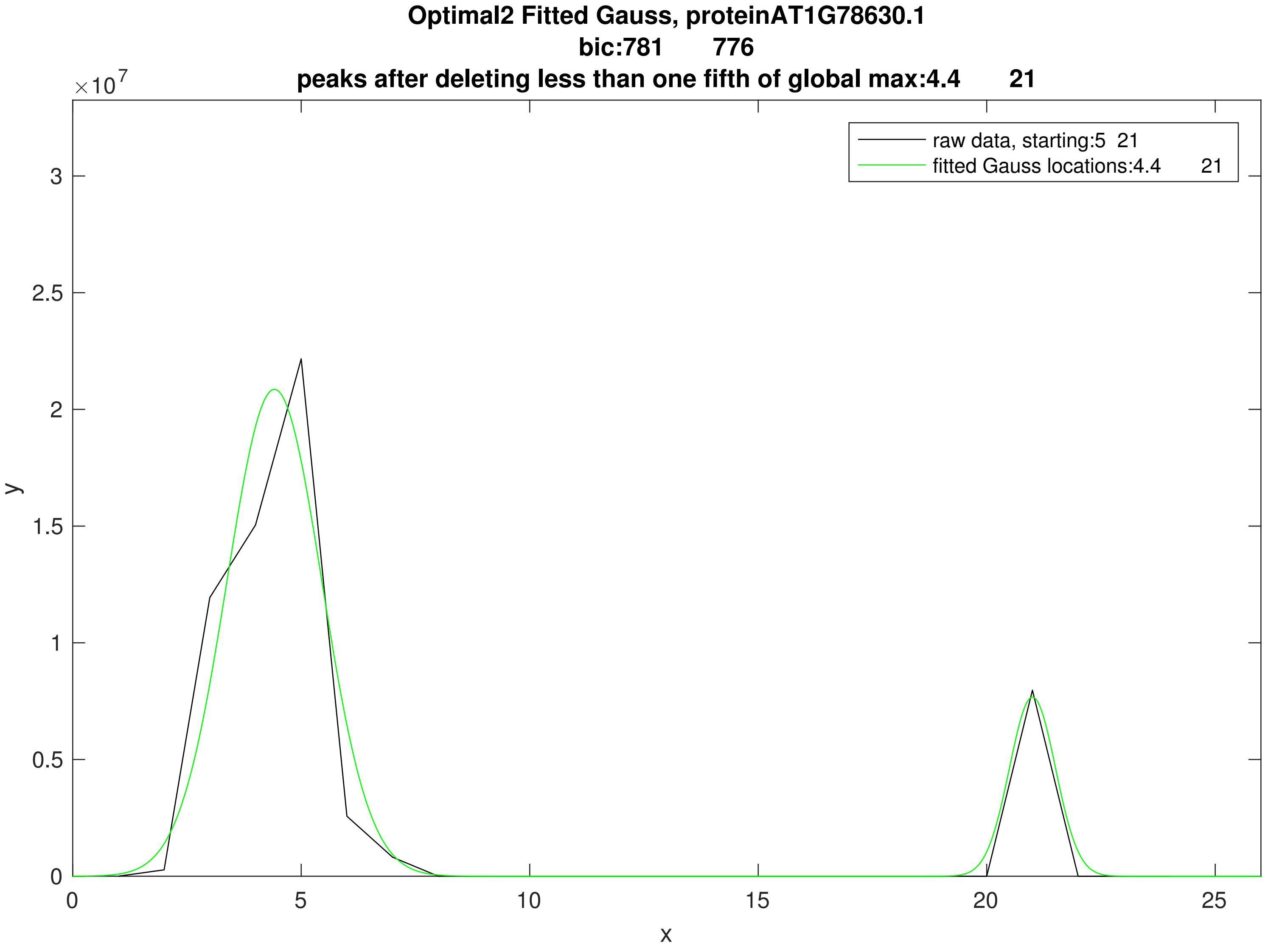
Optimal2 Fitted Gauss, proteinAT1G78630.1
bic:781 776
peaks after deleting less than one fifth of global max:4.4 21
×10^7
raw data, starting:5 21
fitted Gauss locations:4.4 21
y
x
0
0.5
1
1.5
2
2.5
3
5
10
15
20
25

Optimal1 Fitted Gauss, proteinAT1G78660.2;AT1G78660.3;AT1G78660.1
bic:877 878
peaks after deleting less than one fifth of global max:15

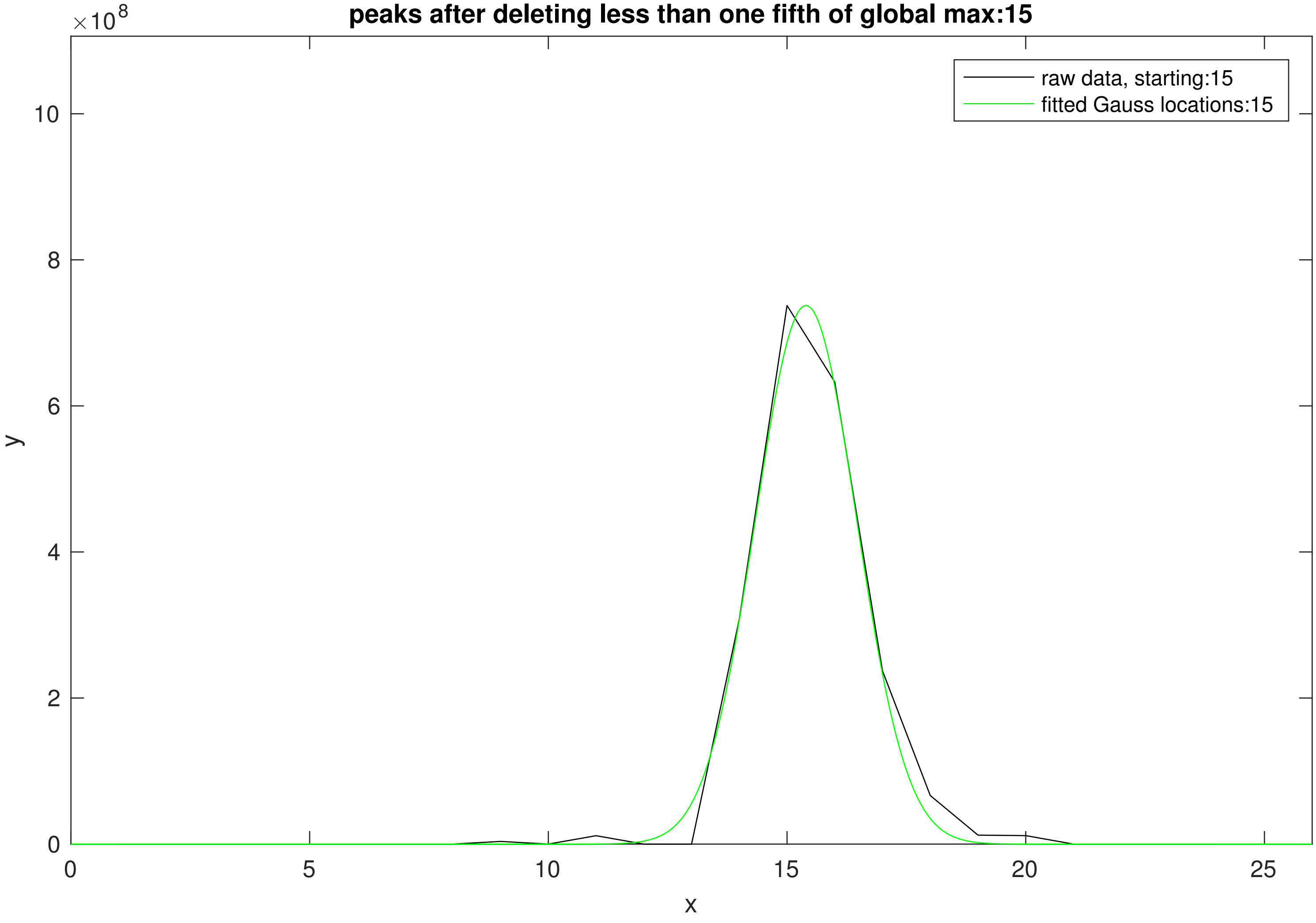

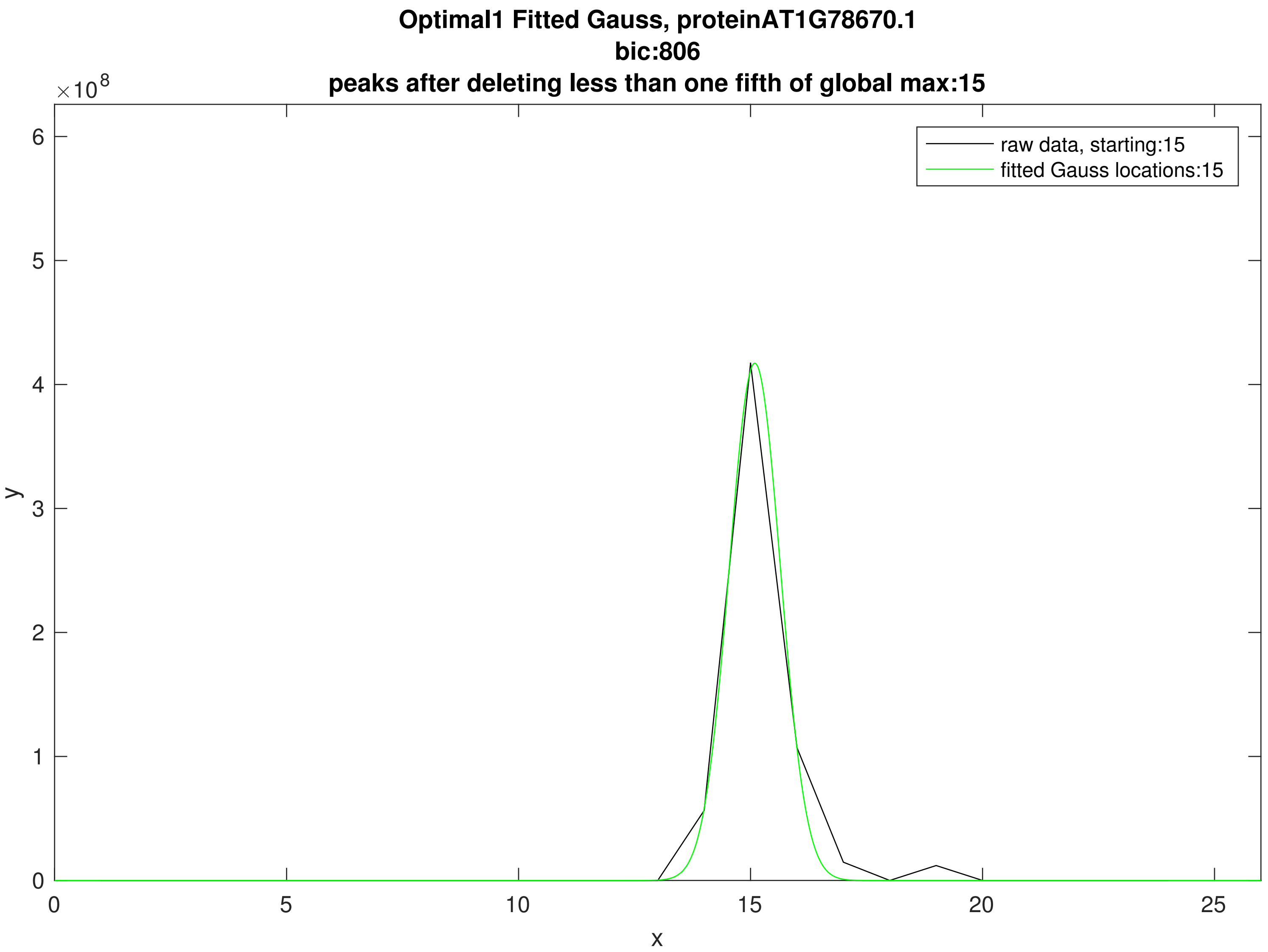
Optimal1 Fitted Gauss, proteinAT1G78670.1
bic:806
peaks after deleting less than one fifth of global max:15
×10^8
6
5
4
3
2
1
0
y
0
5
10
15
20
25
x
raw data, starting:15
fitted Gauss locations:15

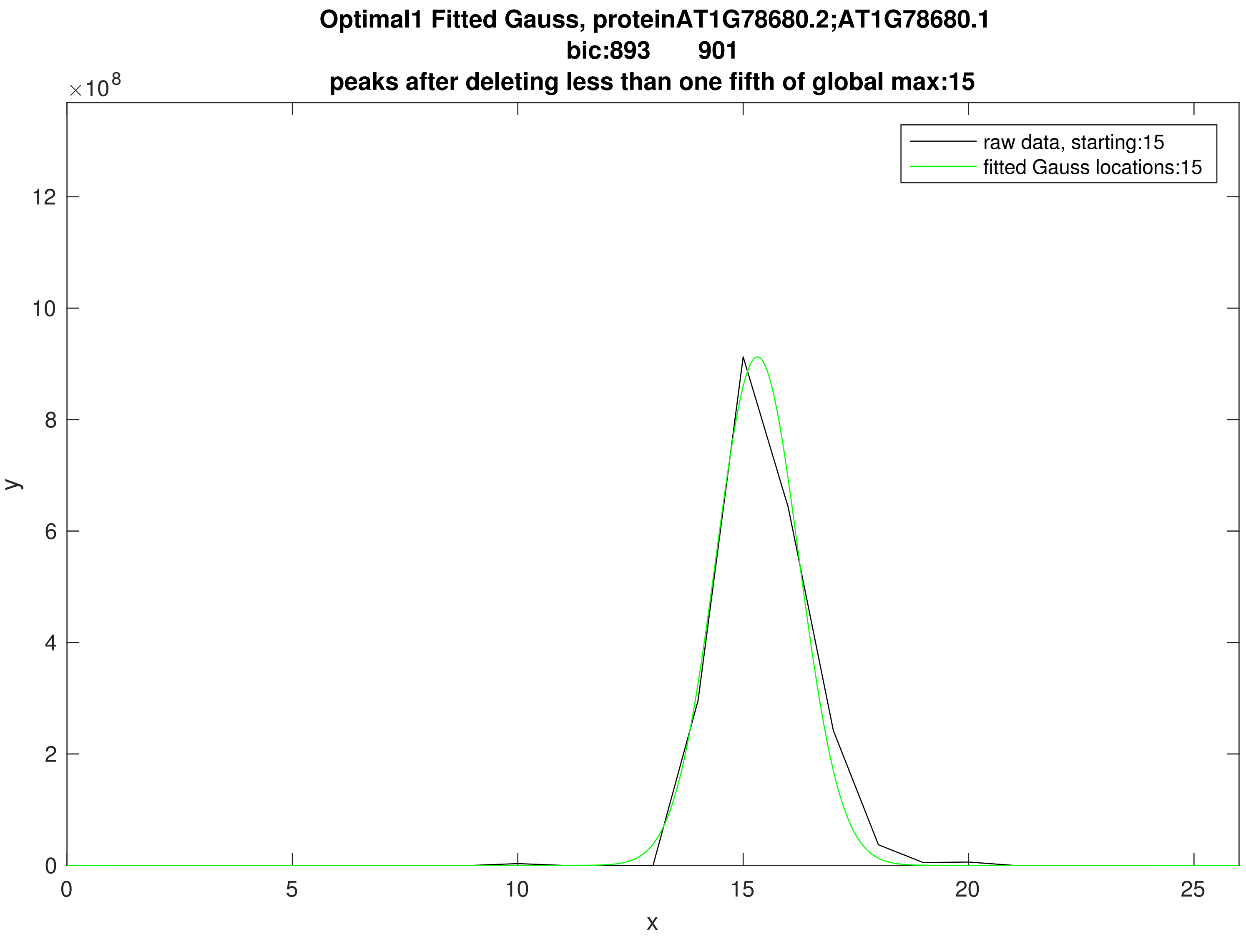
Optimal1 Fitted Gauss, proteinAT1G78680.2;AT1G78680.1
bic:893 901
peaks after deleting less than one fifth of global max:15
×10^8
raw data, starting:15
fitted Gauss locations:15
12
10
8
6
4
2
0
y
0
5
10
15
20
25
x

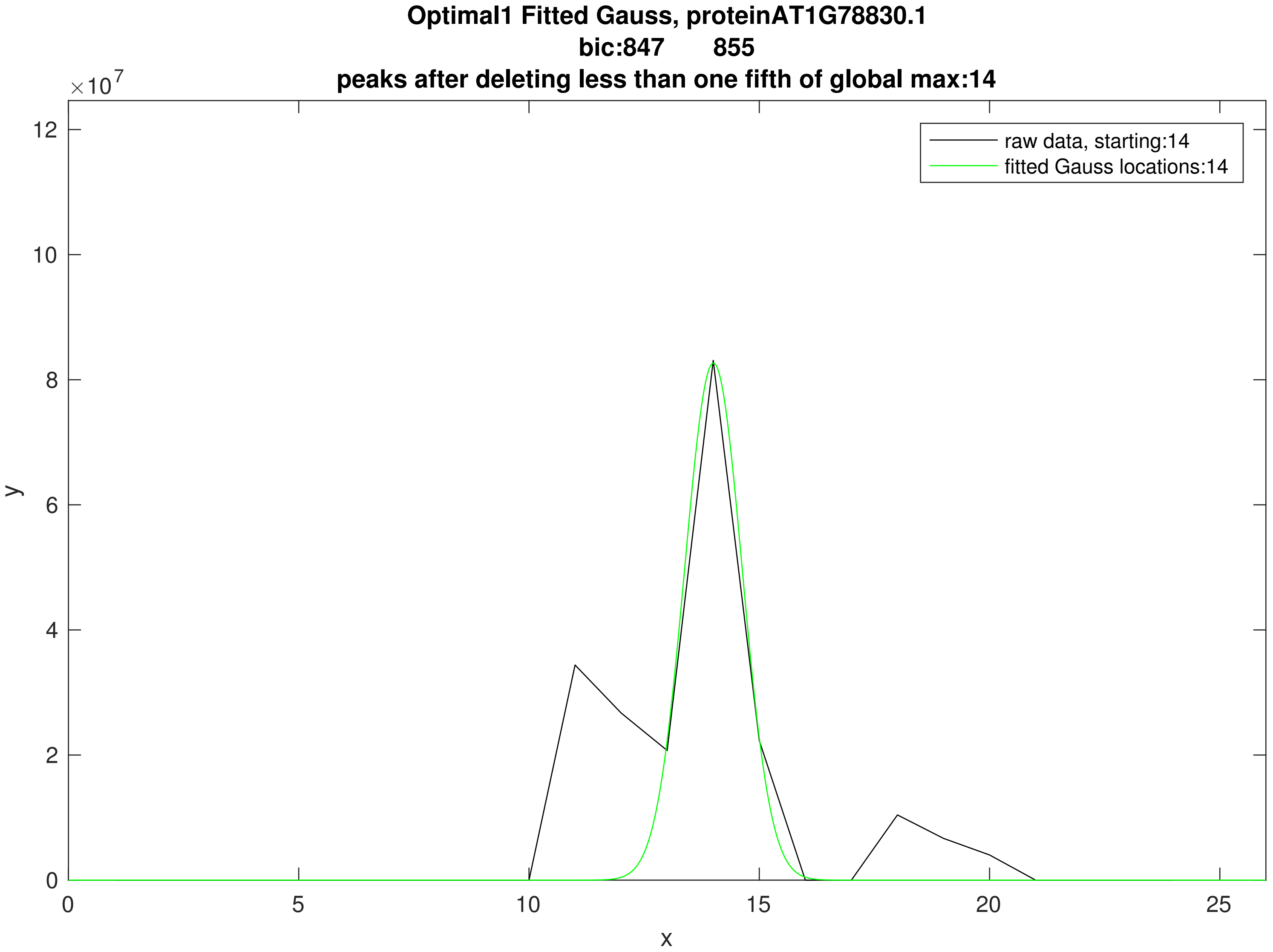
Optimal1 Fitted Gauss, proteinAT1G78830.1
bic:847 855
peaks after deleting less than one fifth of global max:14
×10^7
raw data, starting:14
fitted Gauss locations:14
y
x
0
2
4
6
8
10
12
5
15
20
25

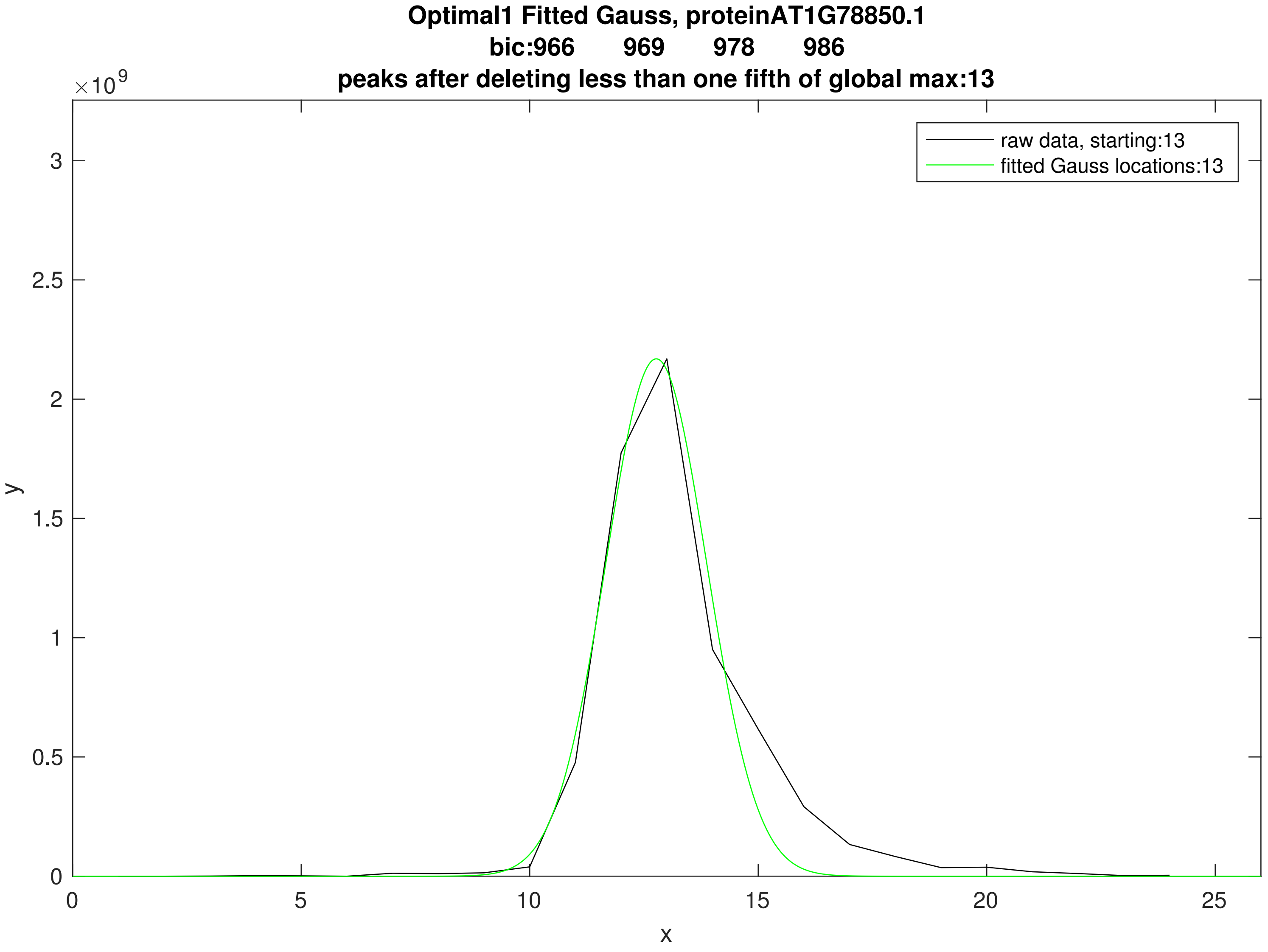
Optimal1 Fitted Gauss, proteinAT1G78850.1
bic:966 969 978 986
peaks after deleting less than one fifth of global max:13
×10^9
3
2.5
2
1.5
1
0.5
0
y
0
5
10
15
20
25
x
raw data, starting:13
fitted Gauss locations:13

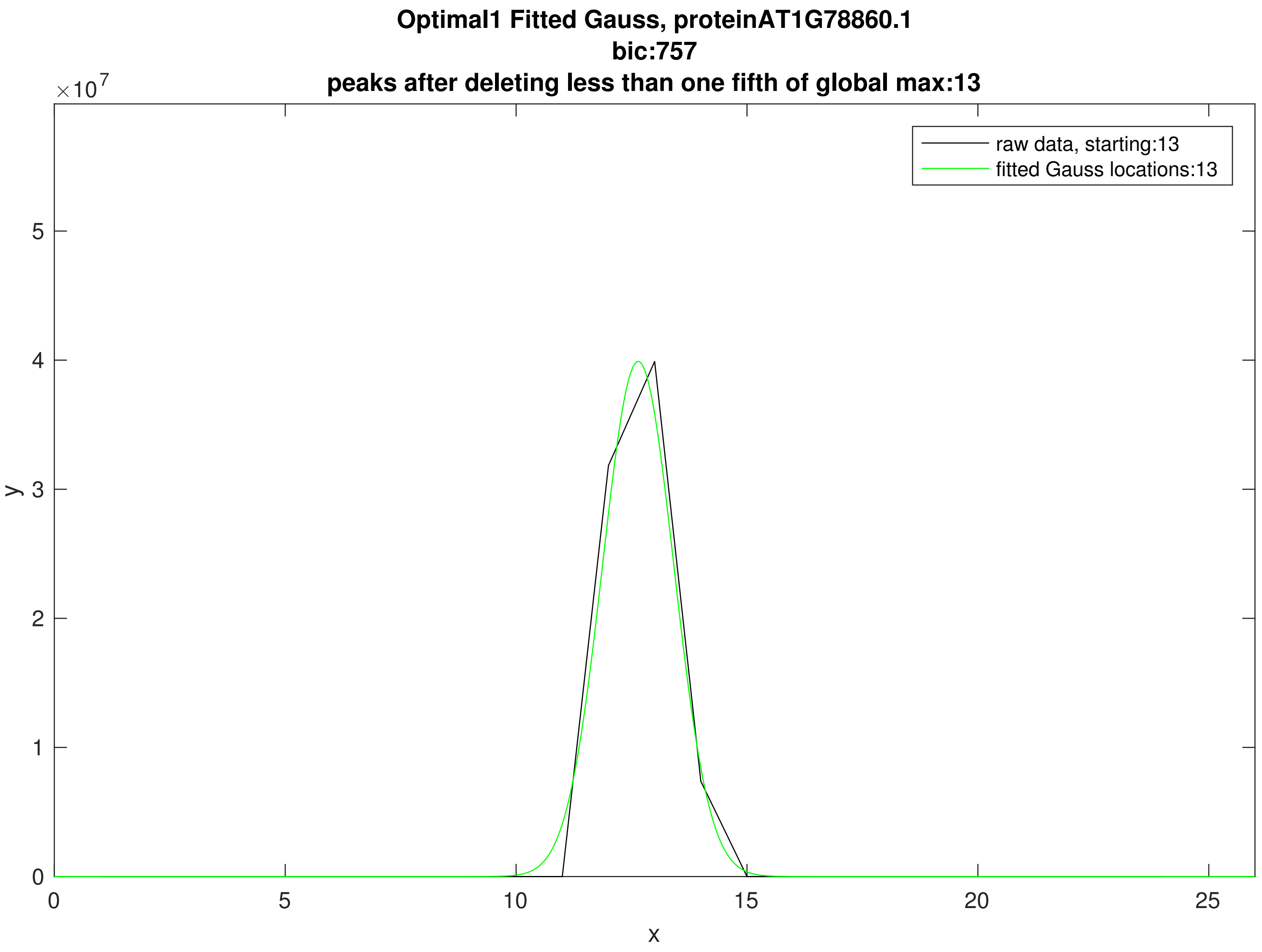
Optimal1 Fitted Gauss, proteinAT1G78860.1
bic:757
peaks after deleting less than one fifth of global max:13
$\times 10^7$
raw data, starting:13
fitted Gauss locations:13
y
x
0
1
2
3
4
5
0
5
10
15
20
25

**Optimal1 Fitted Gauss, proteinAT1G78870.1;AT1G78870.2;AT1G78870.3**
**bic:698**
**peaks after deleting less than one fifth of global max:20**

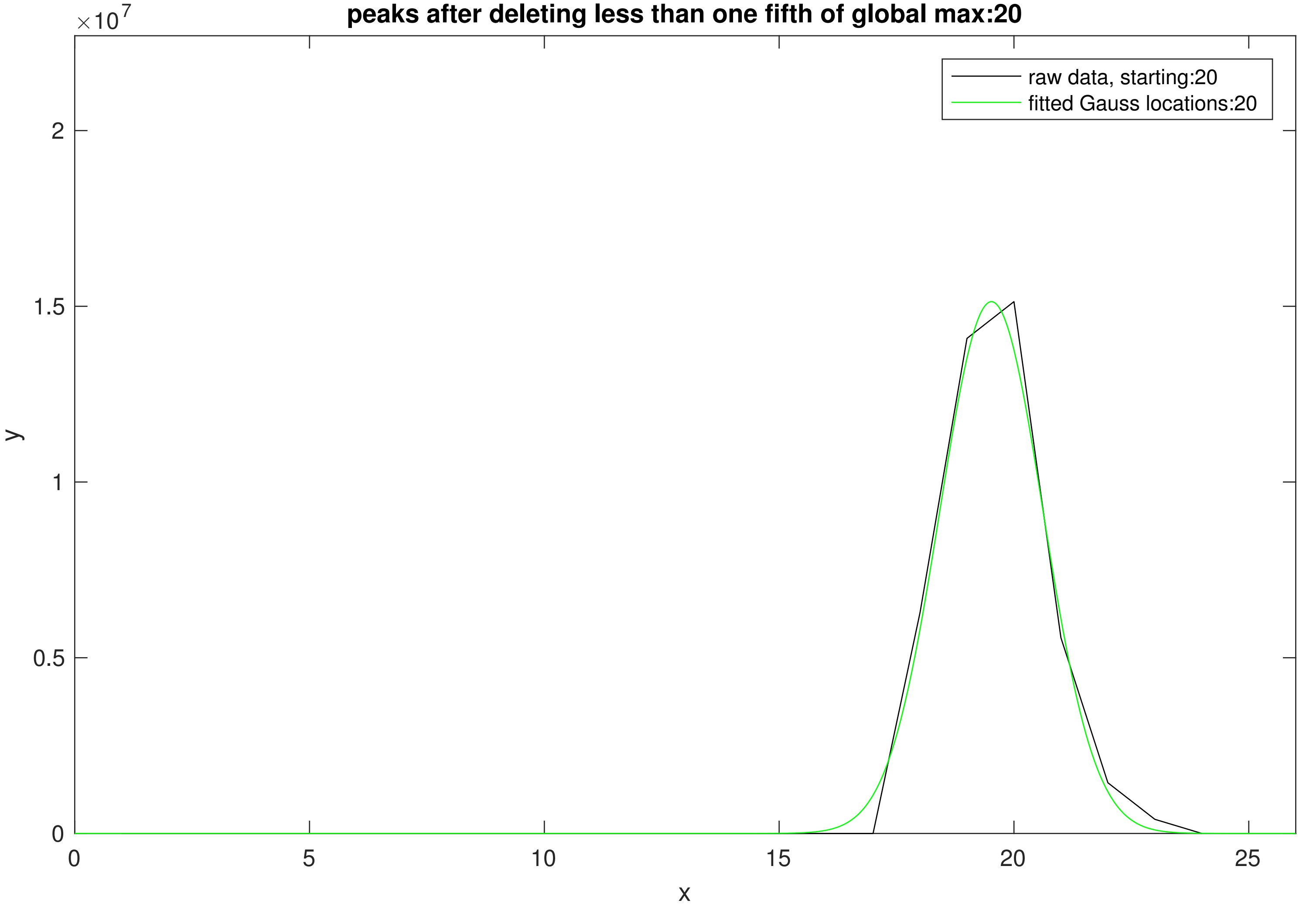

Optimal1 Fitted Gauss, proteinAT1G78900.2;AT1G78900.1
bic:1.06e+03  1.07e+03
peaks after deleting less than one fifth of global max:12

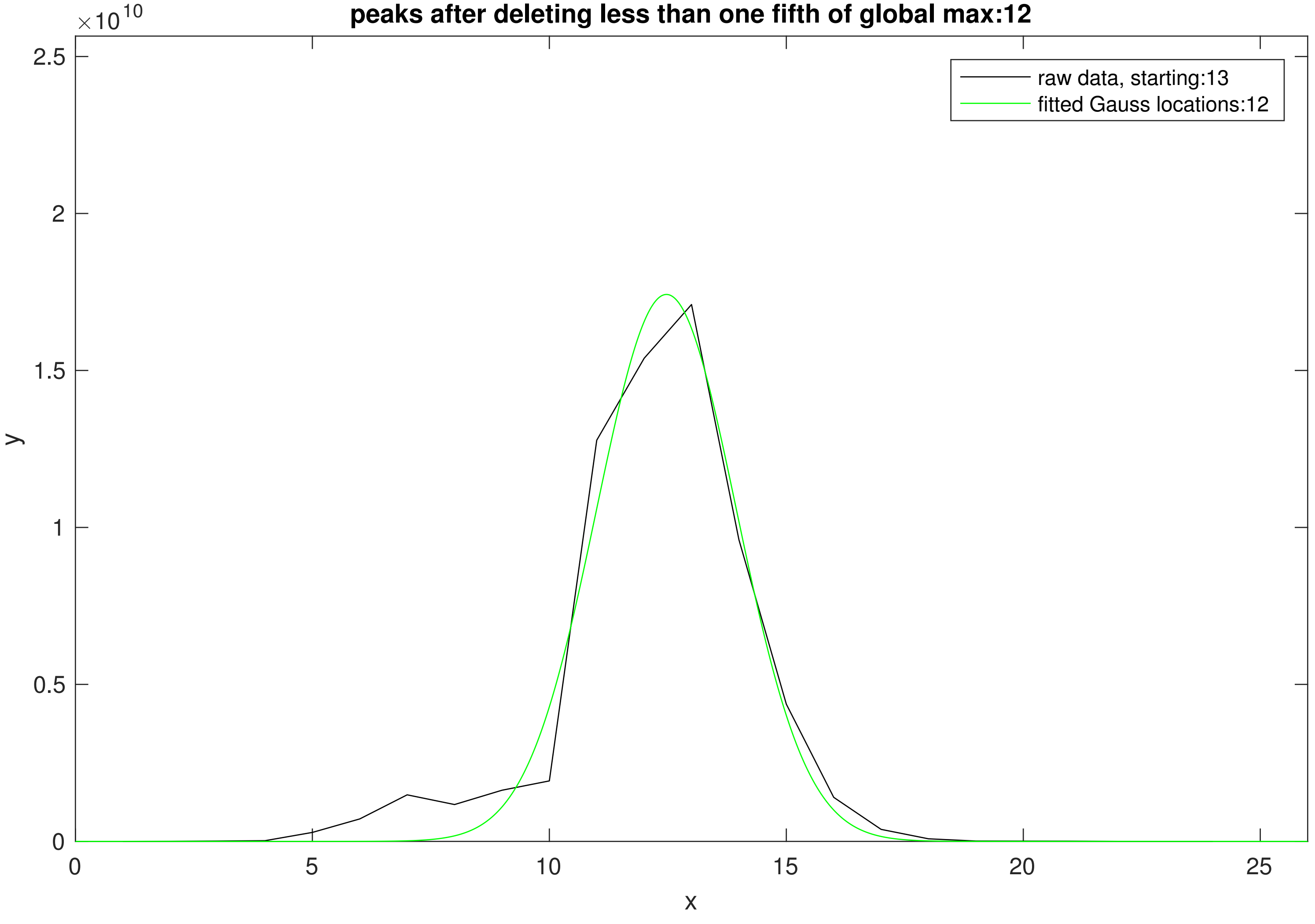

Optimal1 Fitted Gauss, proteinAT1G79210.3;AT1G79210.2;AT1G79210.1
bic:844
peaks after deleting less than one fifth of global max:5

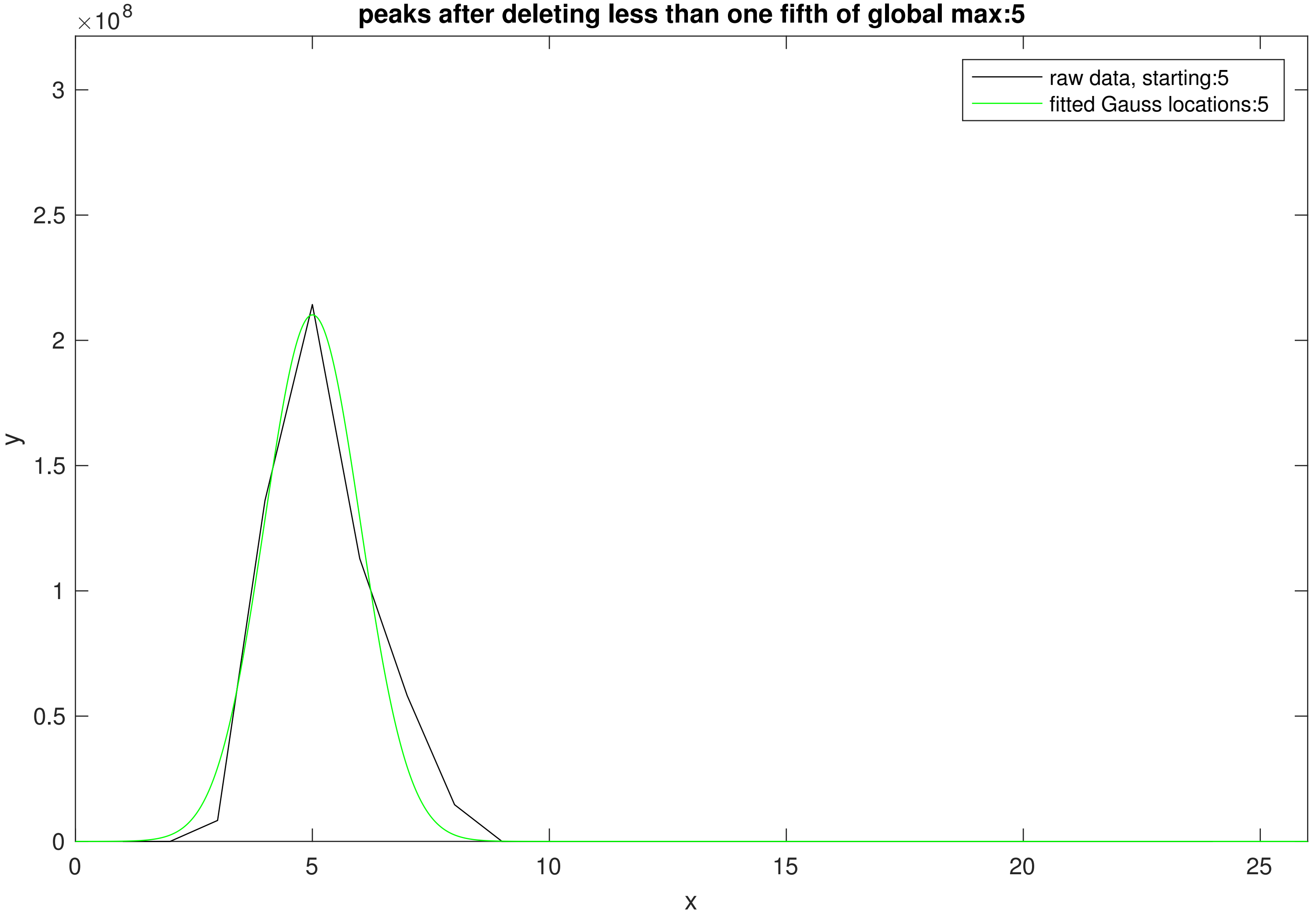

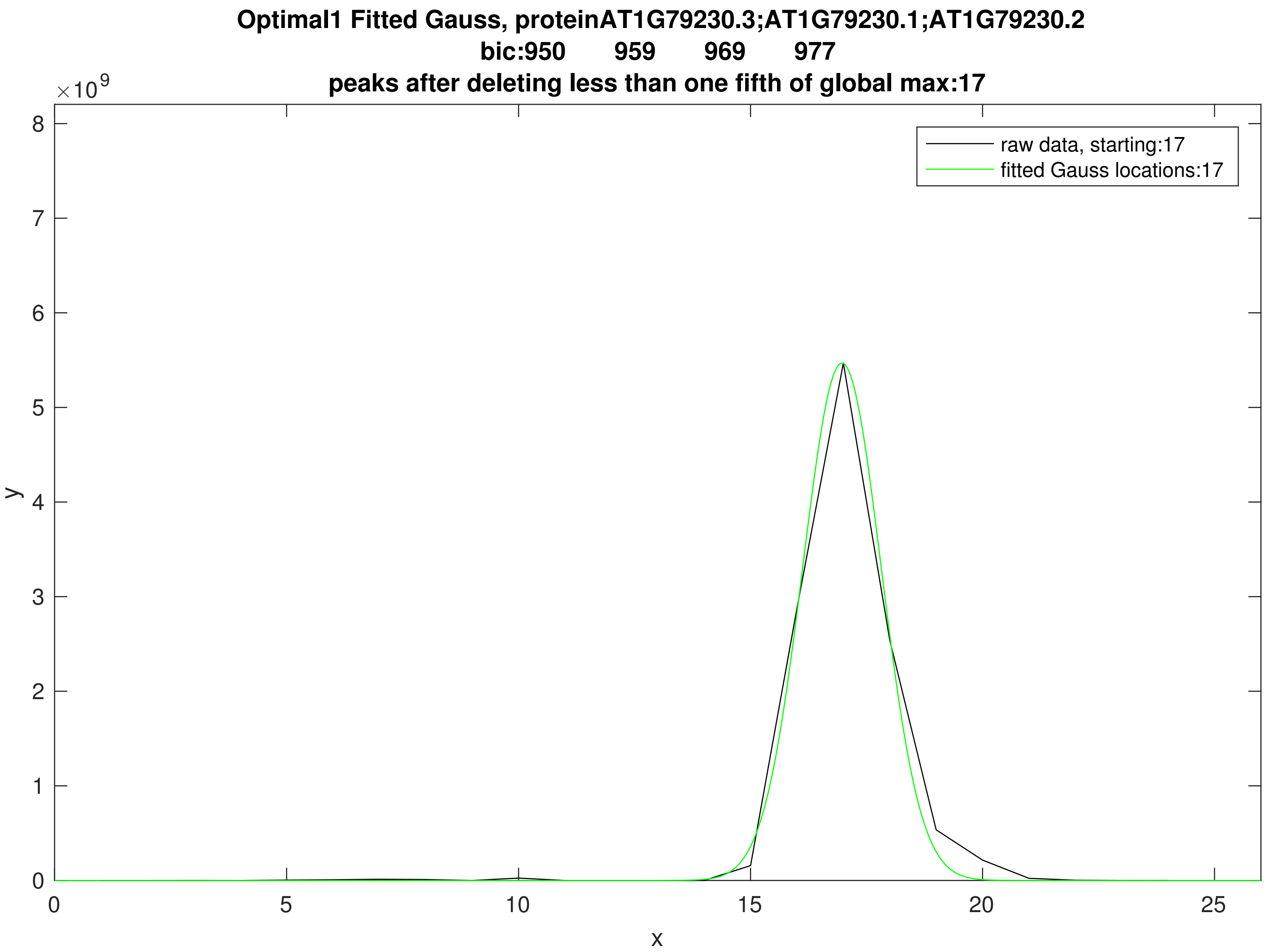

Optimal1 Fitted Gauss, proteinAT1G79230.3;AT1G79230.1;AT1G79230.2
bic:950 959 969 977
peaks after deleting less than one fifth of global max:17
×10^9
raw data, starting:17
fitted Gauss locations:17
y
x

**Optimal1 Fitted Gauss, proteinAT1G79260.1**
**bic:834 837 847**
**peaks after deleting less than one fifth of global max:17**

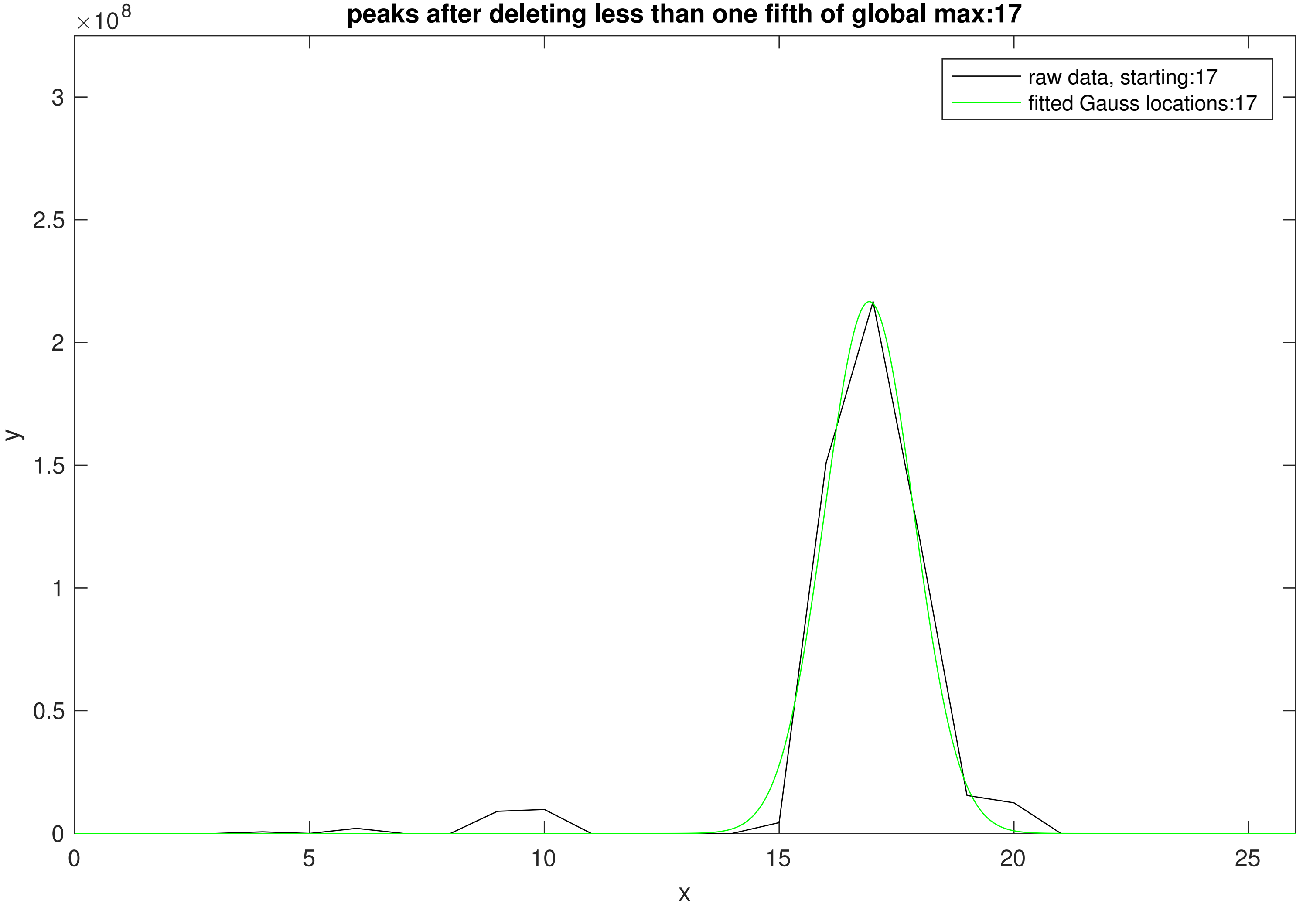

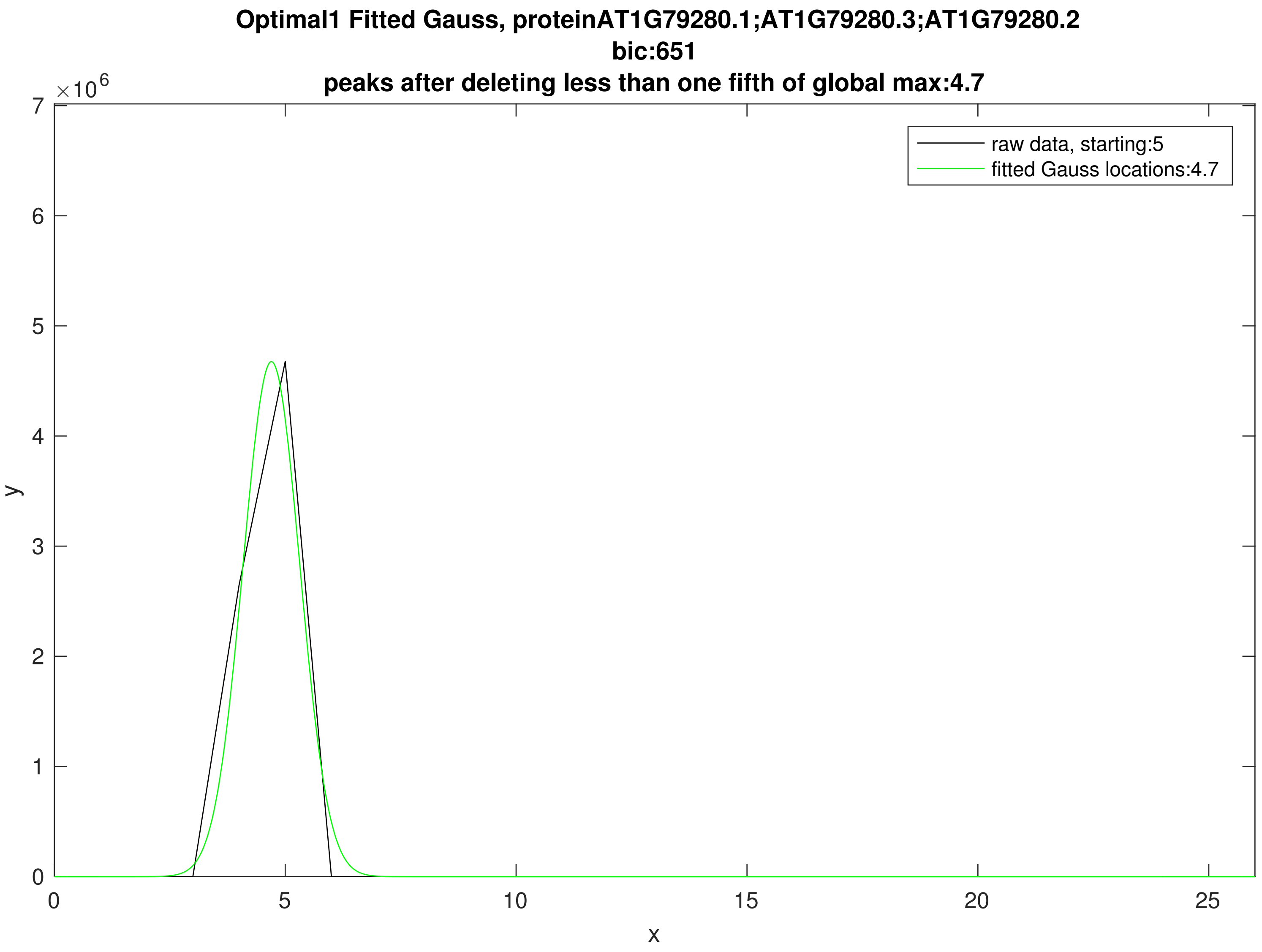
Optimal1 Fitted Gauss, proteinAT1G79280.1;AT1G79280.3;AT1G79280.2
bic:651
peaks after deleting less than one fifth of global max:4.7
$\times 10^6$
raw data, starting:5
fitted Gauss locations:4.7
y
x
0
1
2
3
4
5
6
7
0
5
10
15
20
25

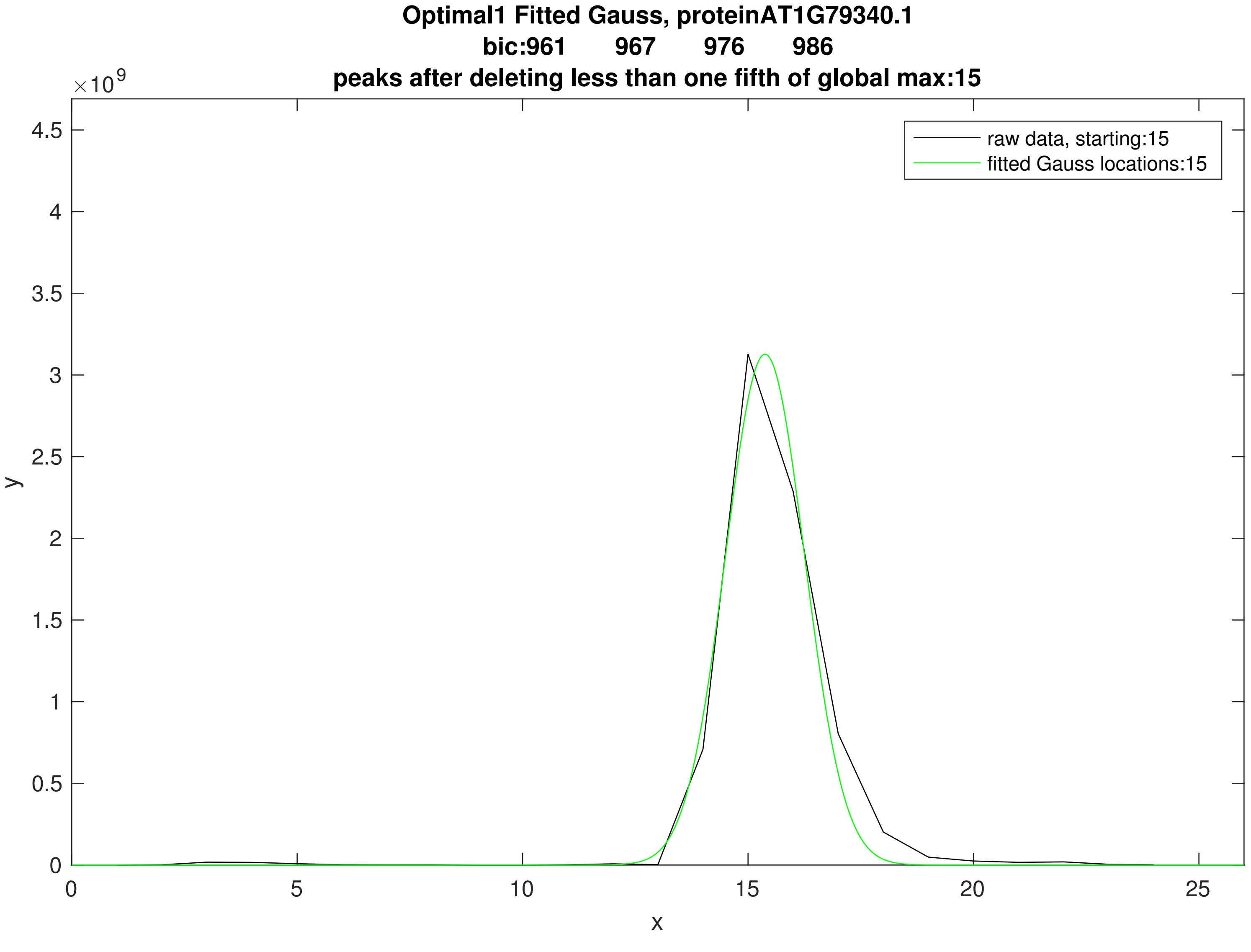
Optimal1 Fitted Gauss, proteinAT1G79340.1
bic:961 967 976 986
peaks after deleting less than one fifth of global max:15
×10^9
raw data, starting:15
fitted Gauss locations:15
4.5
4
3.5
3
2.5
2
1.5
1
0.5
0
y
0
5
10
15
20
25
x

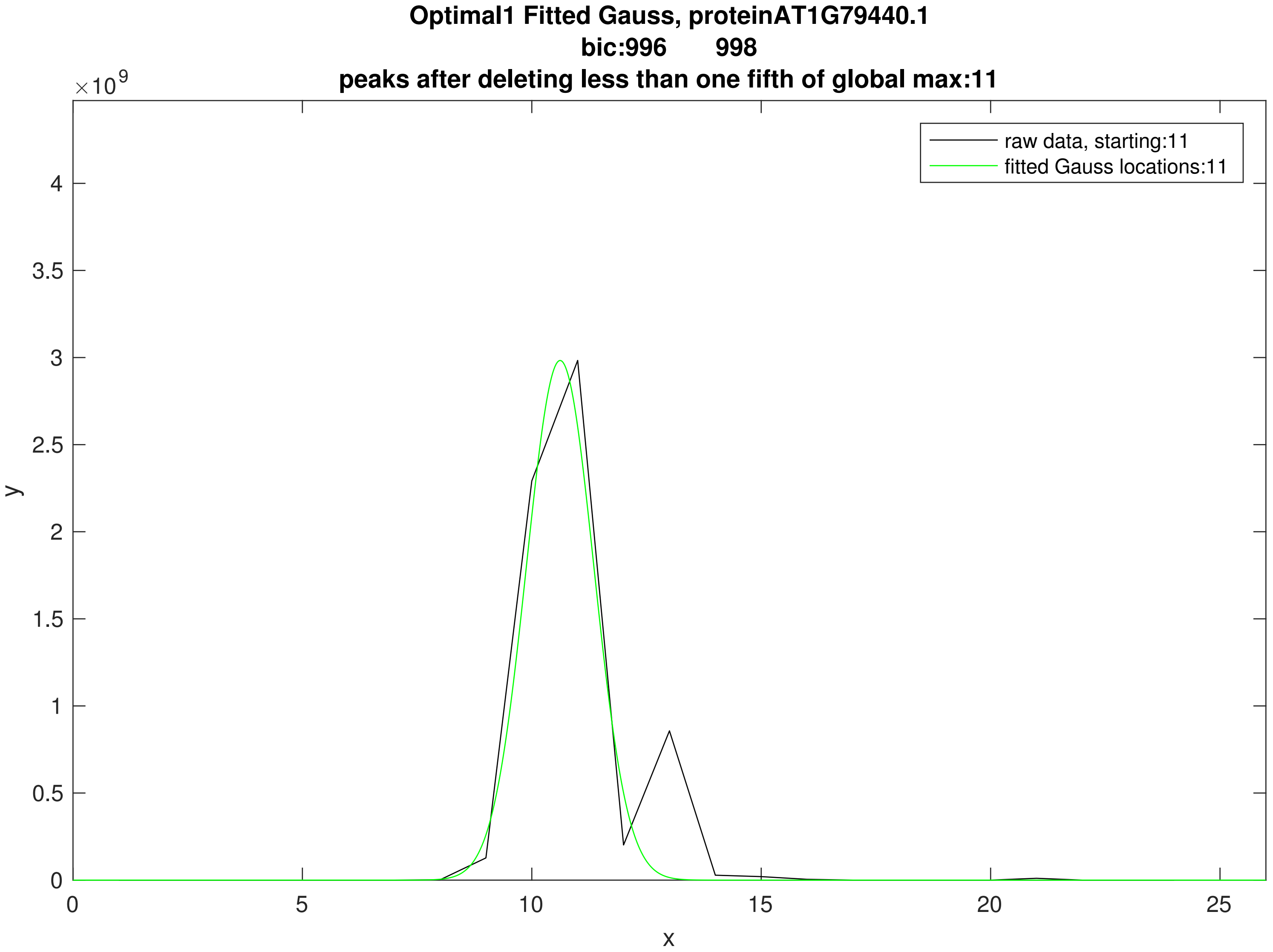

Optimal1 Fitted Gauss, proteinAT1G79440.1
bic:996 998
peaks after deleting less than one fifth of global max:11
×10^9
raw data, starting:11
fitted Gauss locations:11
y
x
0
0.5
1
1.5
2
2.5
3
3.5
4
0
5
10
15
20
25

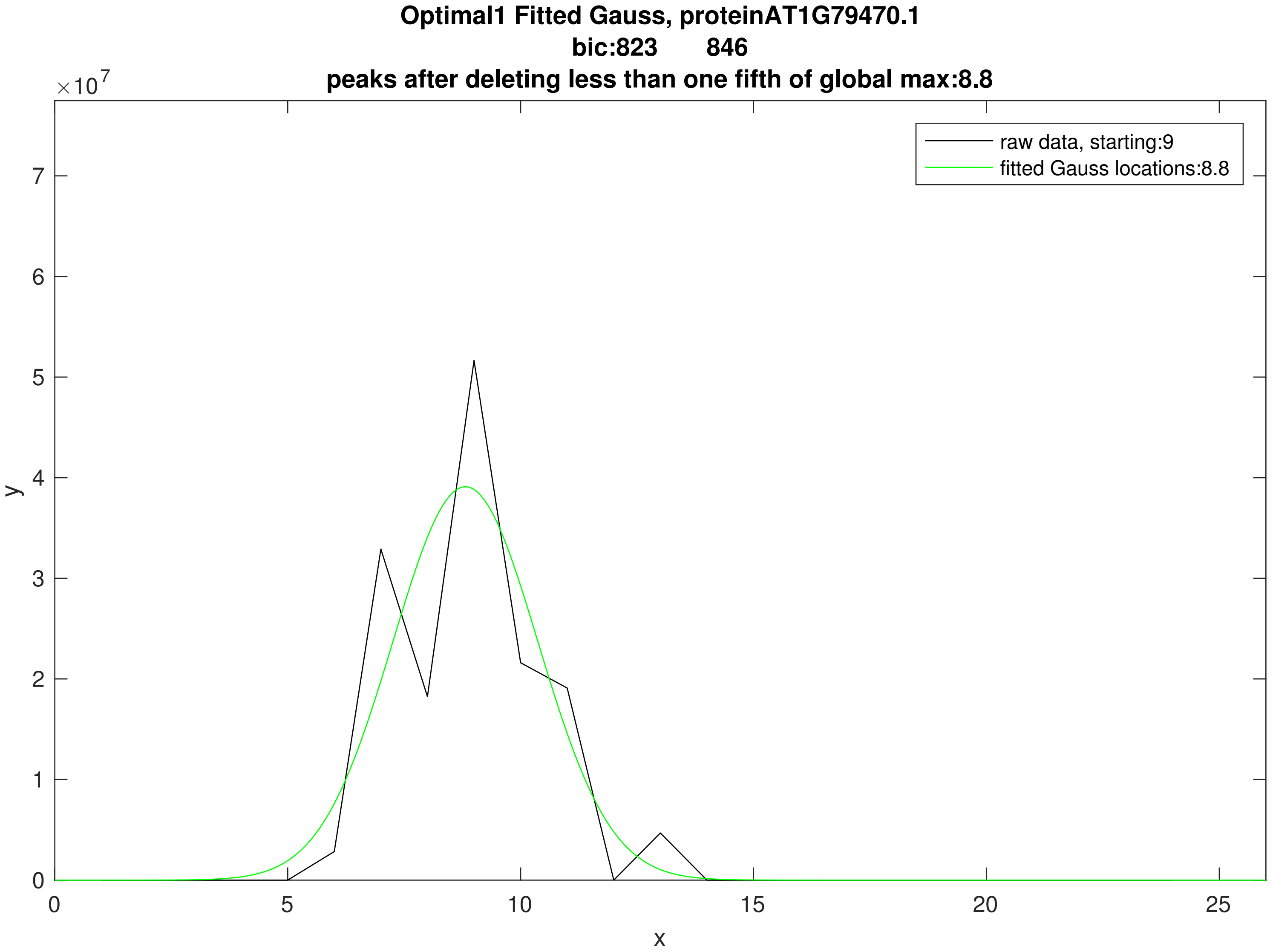

Optimal1 Fitted Gauss, proteinAT1G79470.1
bic:823 846
peaks after deleting less than one fifth of global max:8.8
×10^7
raw data, starting:9
fitted Gauss locations:8.8
y
x
0
1
2
3
4
5
6
7
0
5
10
15
20
25

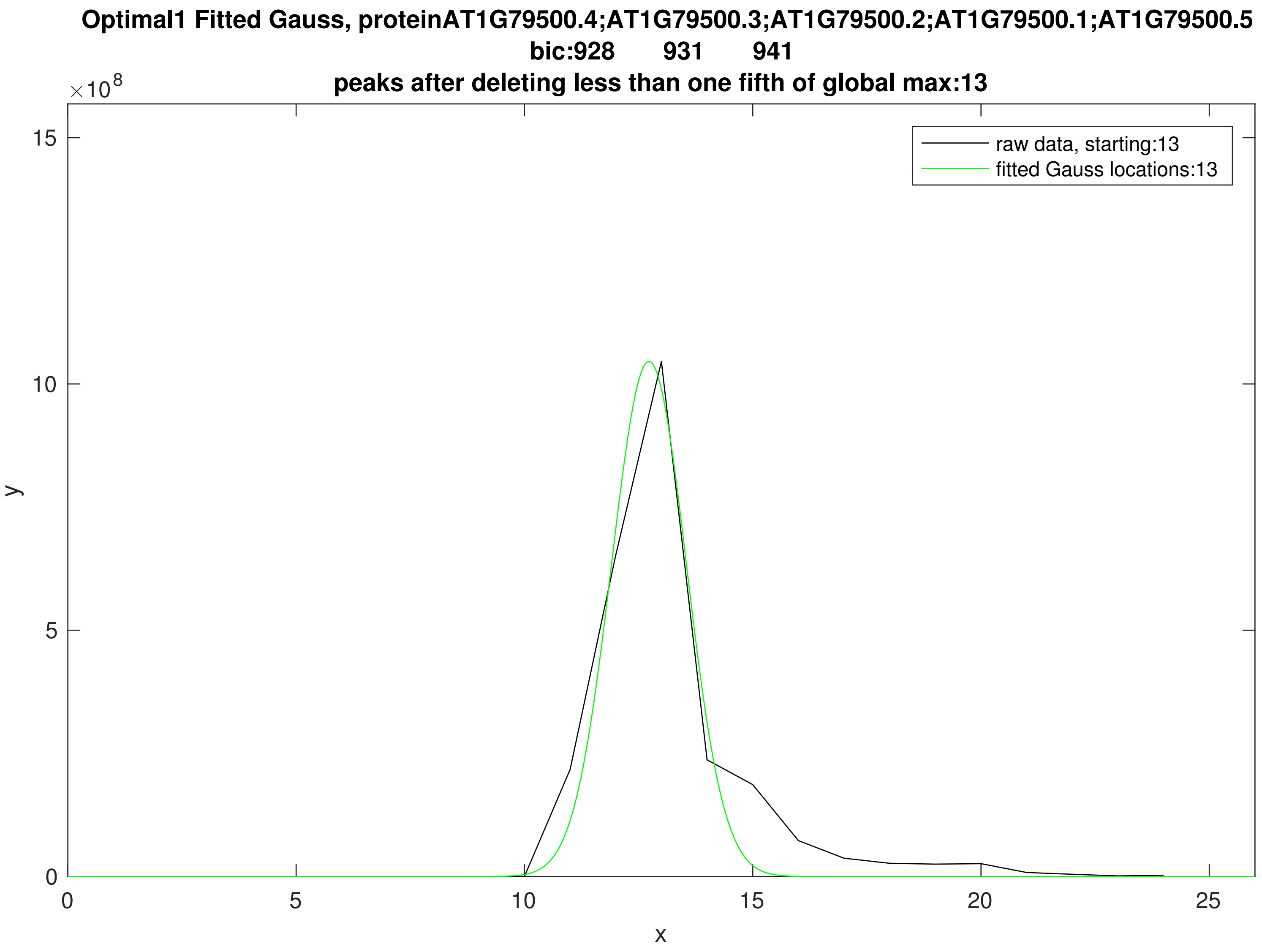

Optimal1 Fitted Gauss, proteinAT1G79500.4;AT1G79500.3;AT1G79500.2;AT1G79500.1;AT1G79500.5
bic:928 931 941
peaks after deleting less than one fifth of global max:13
×10^8
15
10
5
0
y
0
5
10
15
20
25
x
raw data, starting:13
fitted Gauss locations:13

**Optimal1 Fitted Gauss, proteinAT1G79530.1**
**bic:1.05e+03  1.06e+03  1.07e+03**
**peaks after deleting less than one fifth of global max:11**

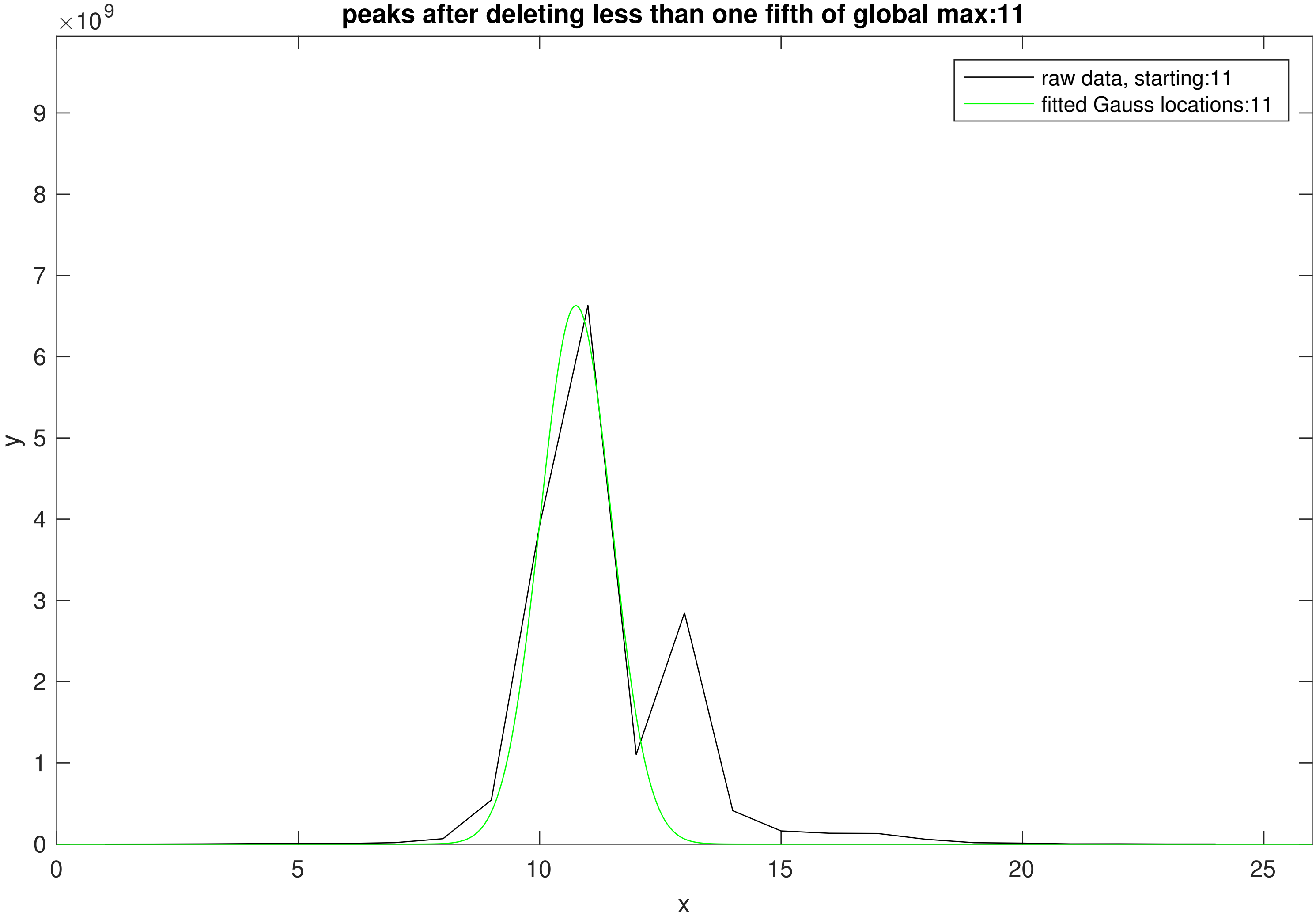

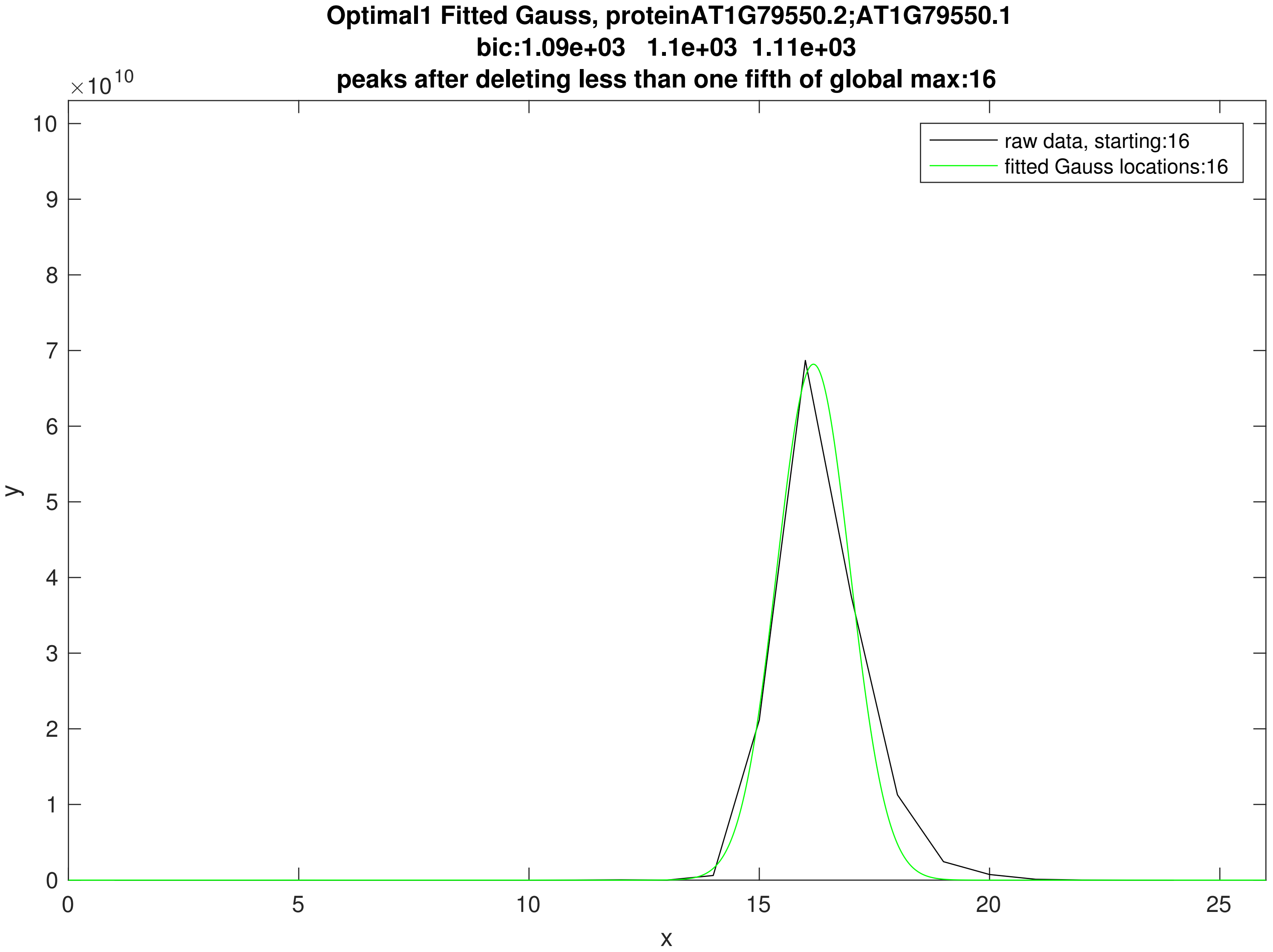

Optimal1 Fitted Gauss, proteinAT1G79550.2;AT1G79550.1
bic:1.09e+03 1.1e+03 1.11e+03
peaks after deleting less than one fifth of global max:16
×10^10
raw data, starting:16
fitted Gauss locations:16
y
x
0
1
2
3
4
5
6
7
8
9
10
0
5
10
15
20
25

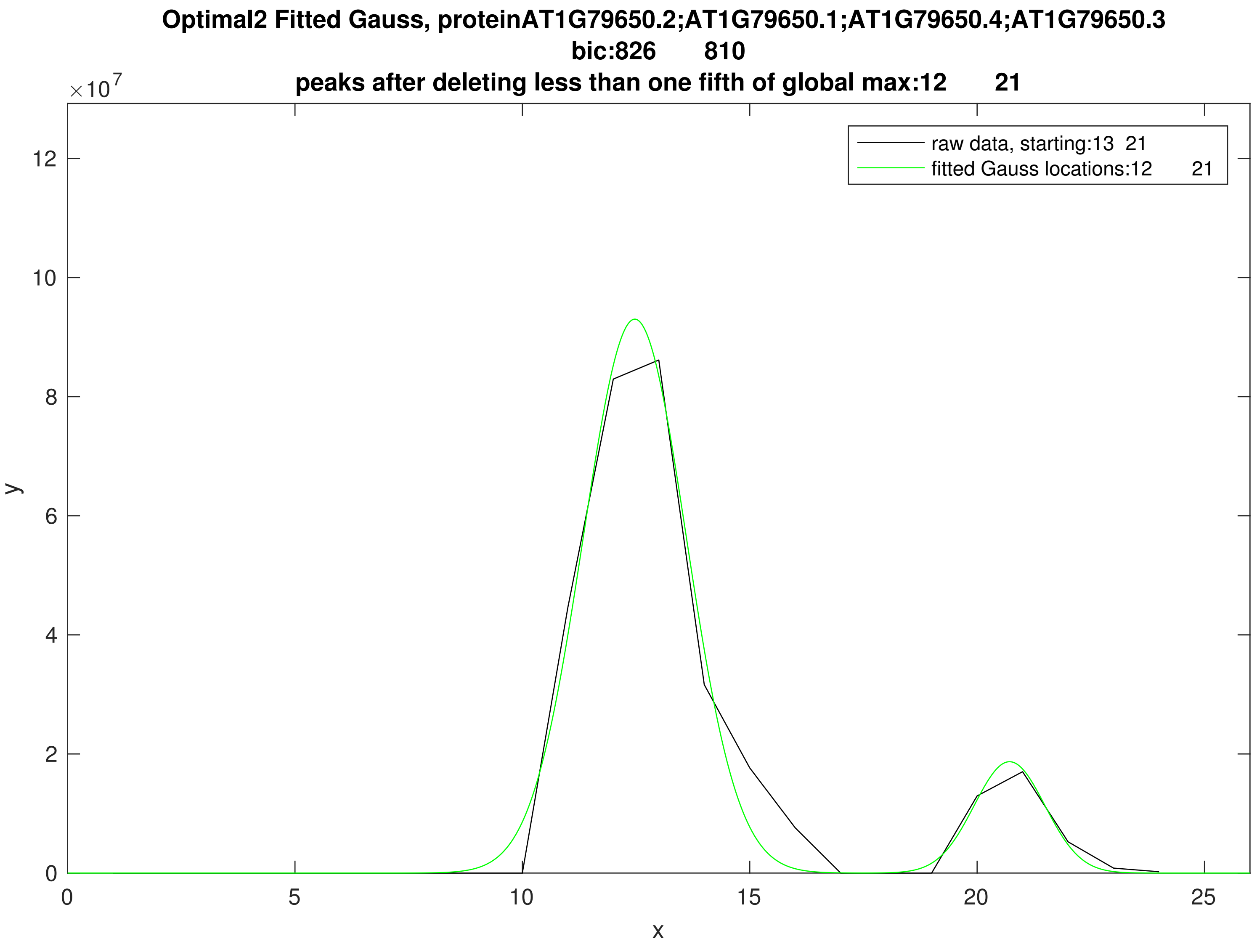

Optimal2 Fitted Gauss, proteinAT1G79650.2;AT1G79650.1;AT1G79650.4;AT1G79650.3
bic:826 810
peaks after deleting less than one fifth of global max:12 21
×10^7
raw data, starting:13 21
fitted Gauss locations:12 21
y
x
0
2
4
6
8
10
12
0
5
10
15
20
25

**Optimal1 Fitted Gauss, proteinAT1G79690.1**
**bic:1.06e+03  1.08e+03**
**peaks after deleting less than one fifth of global max:13**

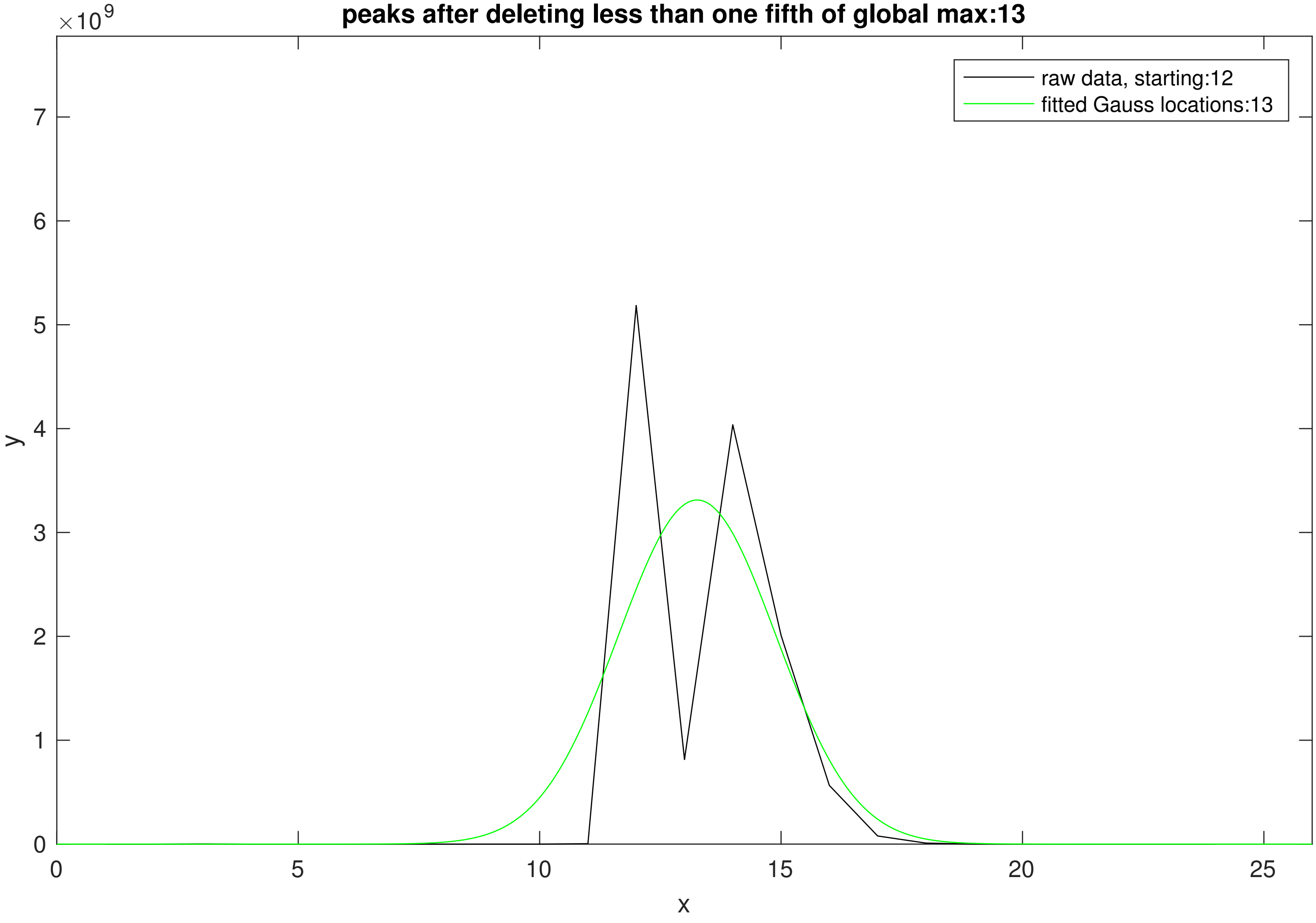

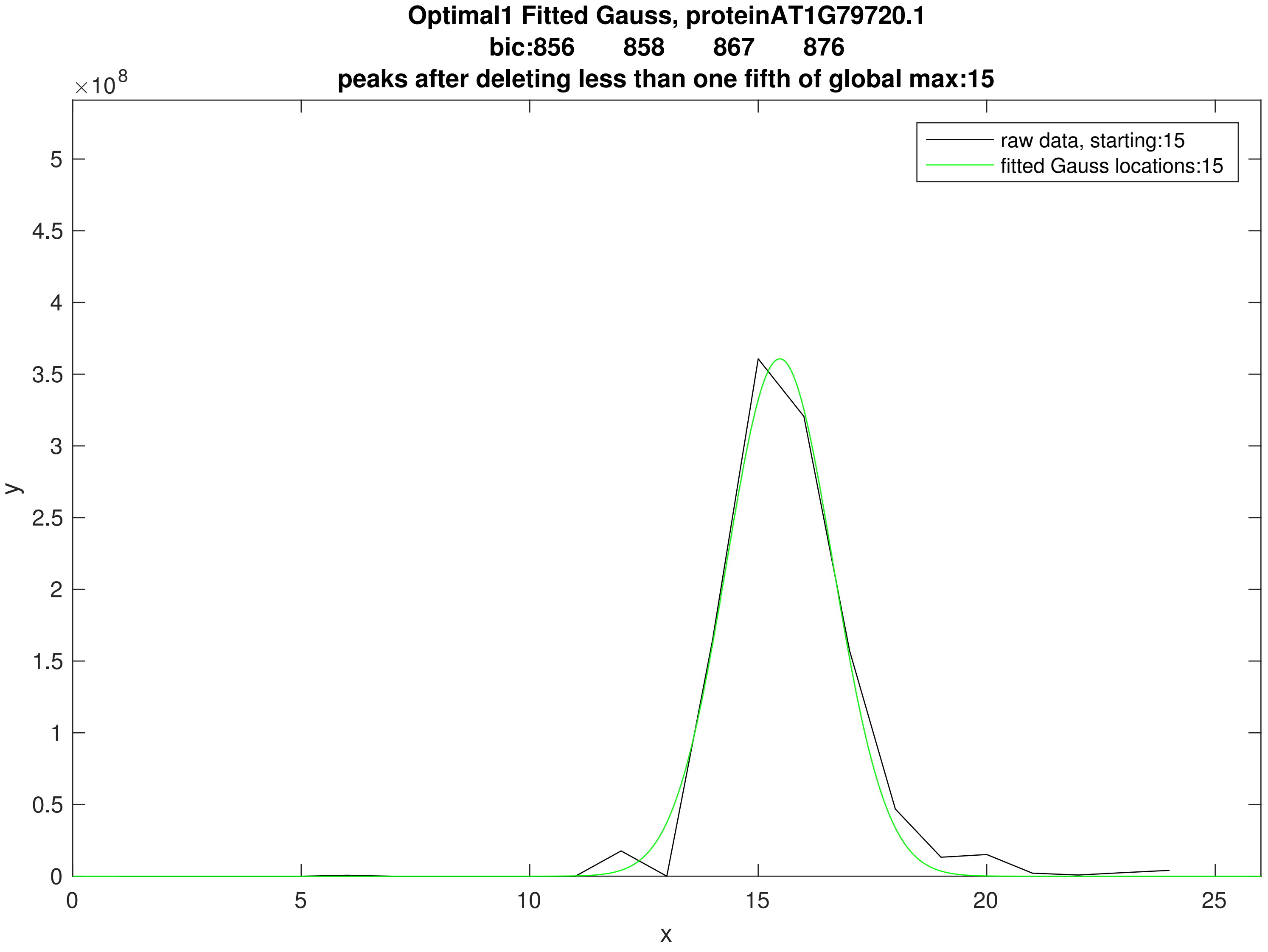

Optimal1 Fitted Gauss, proteinAT1G79720.1
bic:856 858 867 876
peaks after deleting less than one fifth of global max:15
×10^8
raw data, starting:15
fitted Gauss locations:15
y
x
0
0.5
1
1.5
2
2.5
3
3.5
4
4.5
5
0
5
10
15
20
25

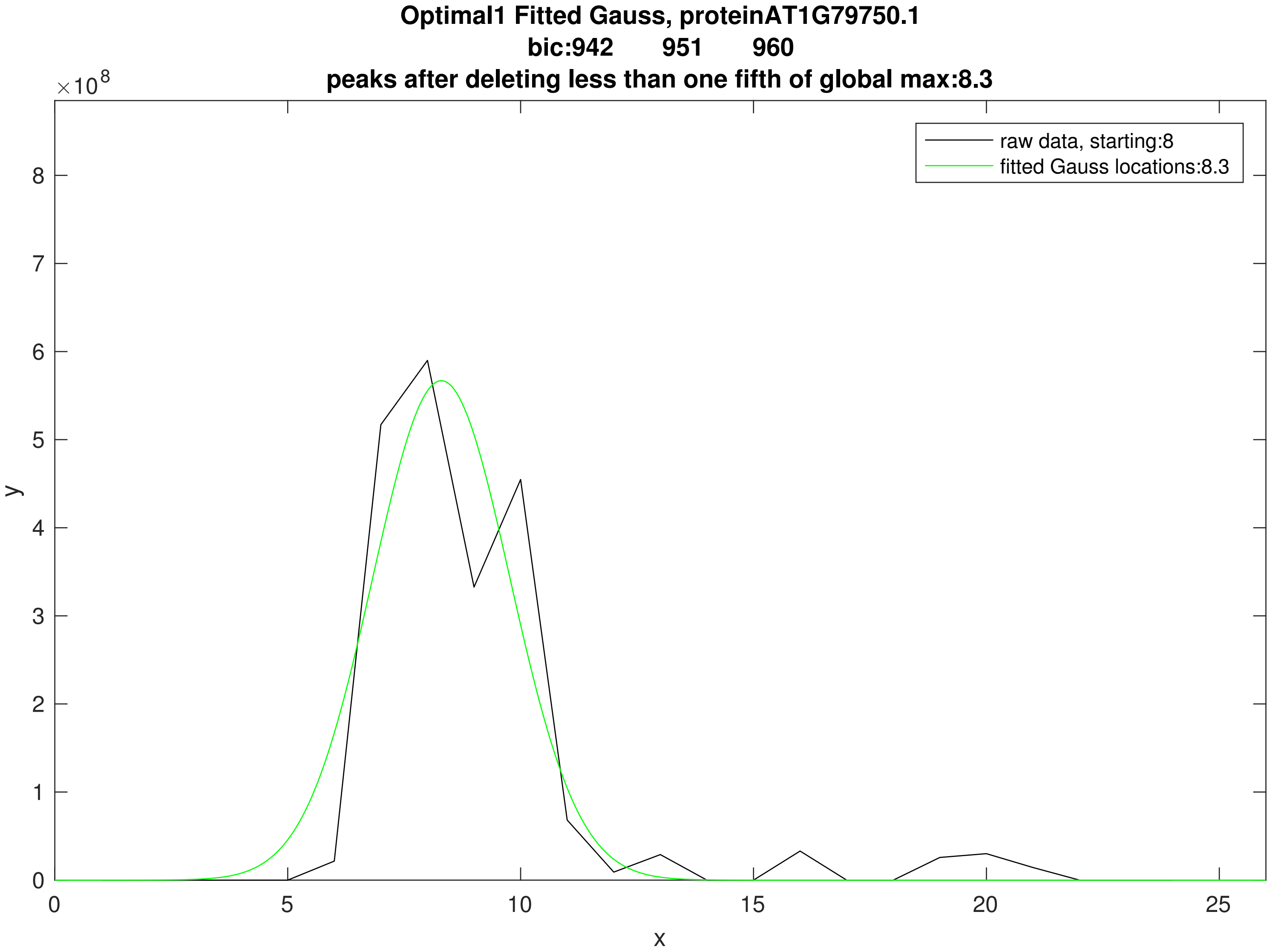
Optimal1 Fitted Gauss, proteinAT1G79750.1
bic:942 951 960
peaks after deleting less than one fifth of global max:8.3
×10^8
raw data, starting:8
fitted Gauss locations:8.3
y
x

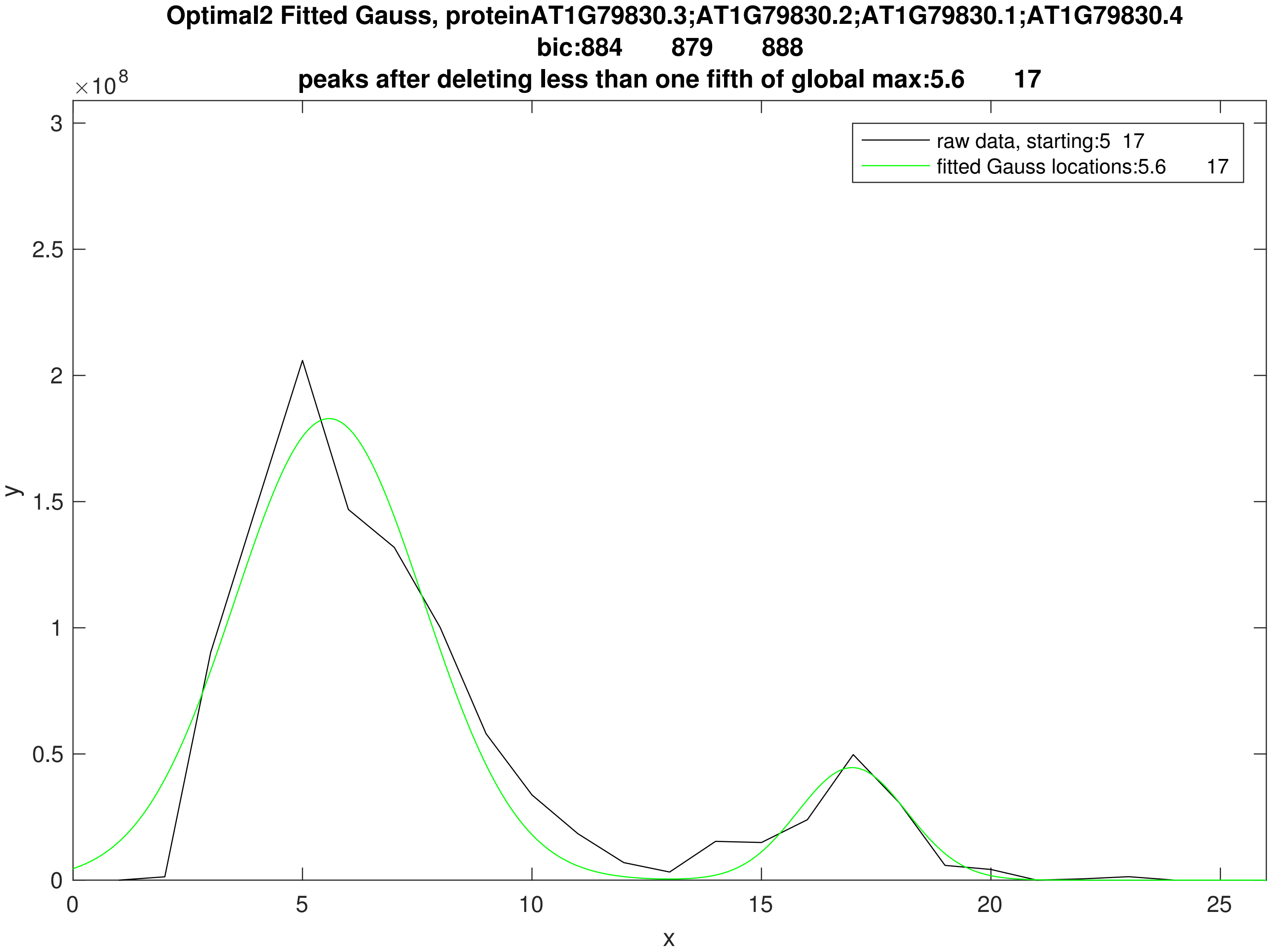
Optimal2 Fitted Gauss, proteinAT1G79830.3;AT1G79830.2;AT1G79830.1;AT1G79830.4
bic:884 879 888
peaks after deleting less than one fifth of global max:5.6 17
×10^8
raw data, starting:5 17
fitted Gauss locations:5.6 17
y
x
0
0.5
1
1.5
2
2.5
3
0
5
10
15
20
25

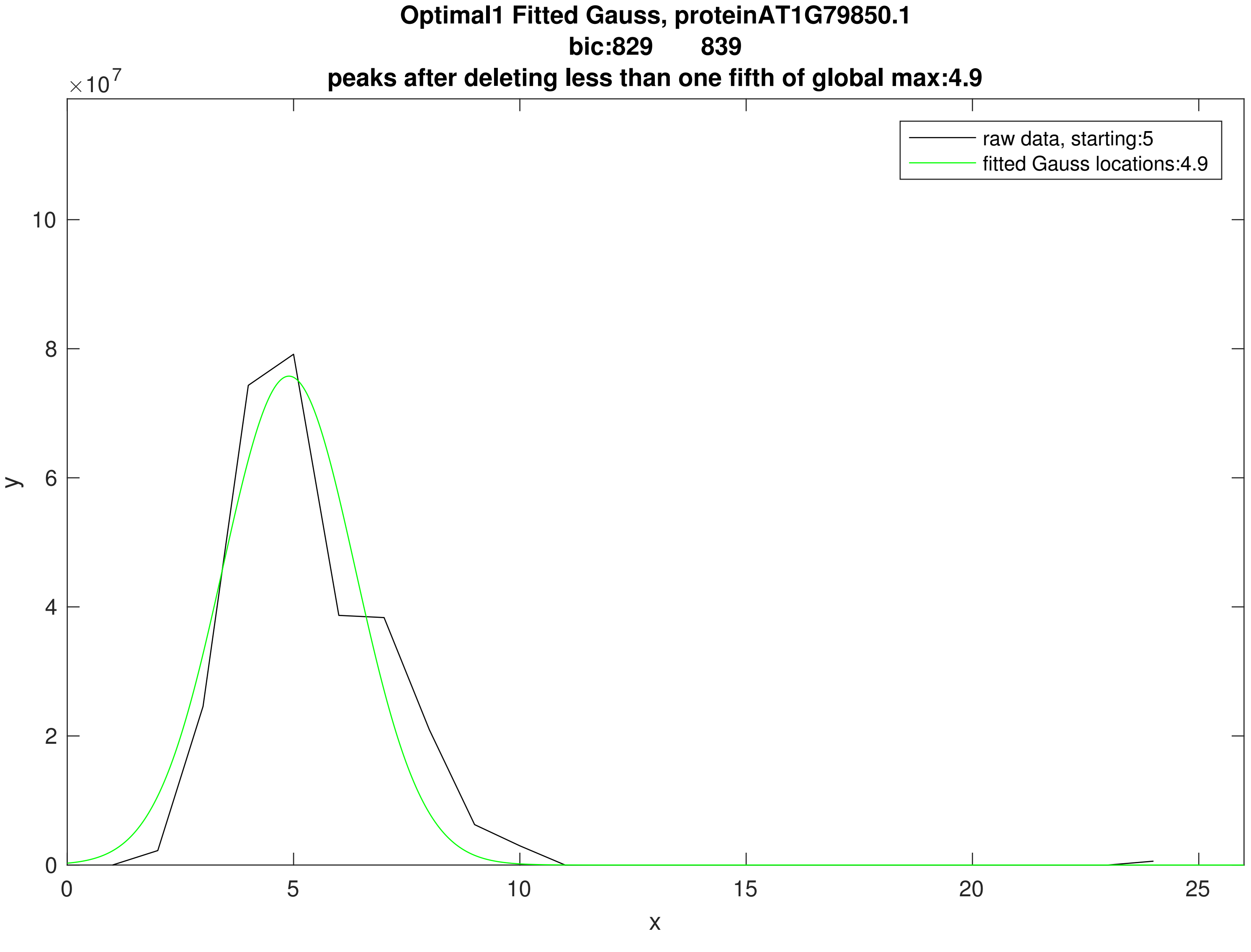
Optimal1 Fitted Gauss, proteinAT1G79850.1
bic:829 839
peaks after deleting less than one fifth of global max:4.9
×10^7
raw data, starting:5
fitted Gauss locations:4.9
y
x
0
2
4
6
8
10
0
5
10
15
20
25

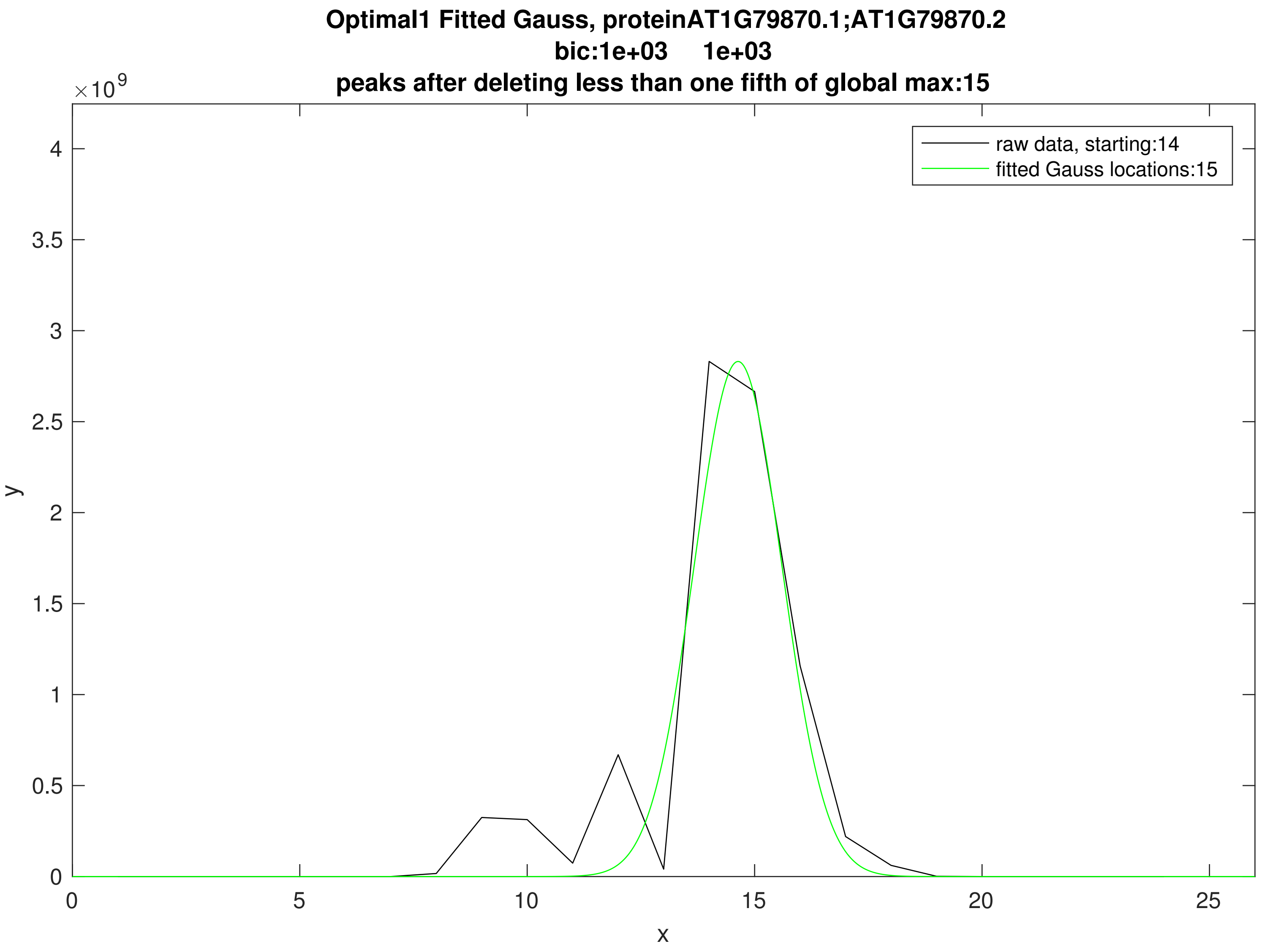
Optimal1 Fitted Gauss, proteinAT1G79870.1;AT1G79870.2
bic:1e+03 1e+03
peaks after deleting less than one fifth of global max:15
×10^9
raw data, starting:14
fitted Gauss locations:15
y
x
0
0.5
1
1.5
2
2.5
3
3.5
4
5
10
15
20
25

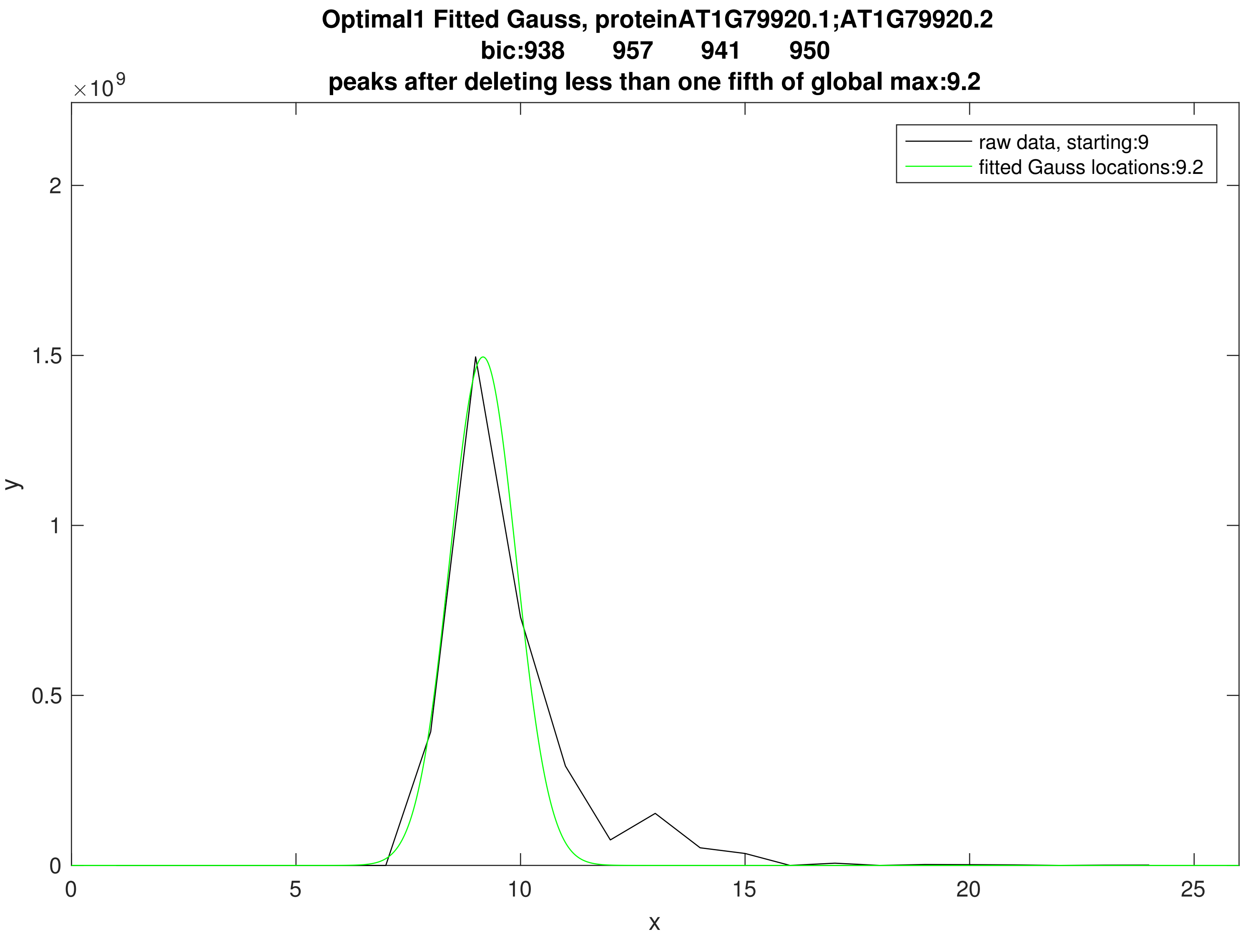
Optimal1 Fitted Gauss, proteinAT1G79920.1;AT1G79920.2
bic:938 957 941 950
peaks after deleting less than one fifth of global max:9.2
×10^9
raw data, starting:9
fitted Gauss locations:9.2
y
x
0
0.5
1
1.5
2
0
5
10
15
20
25

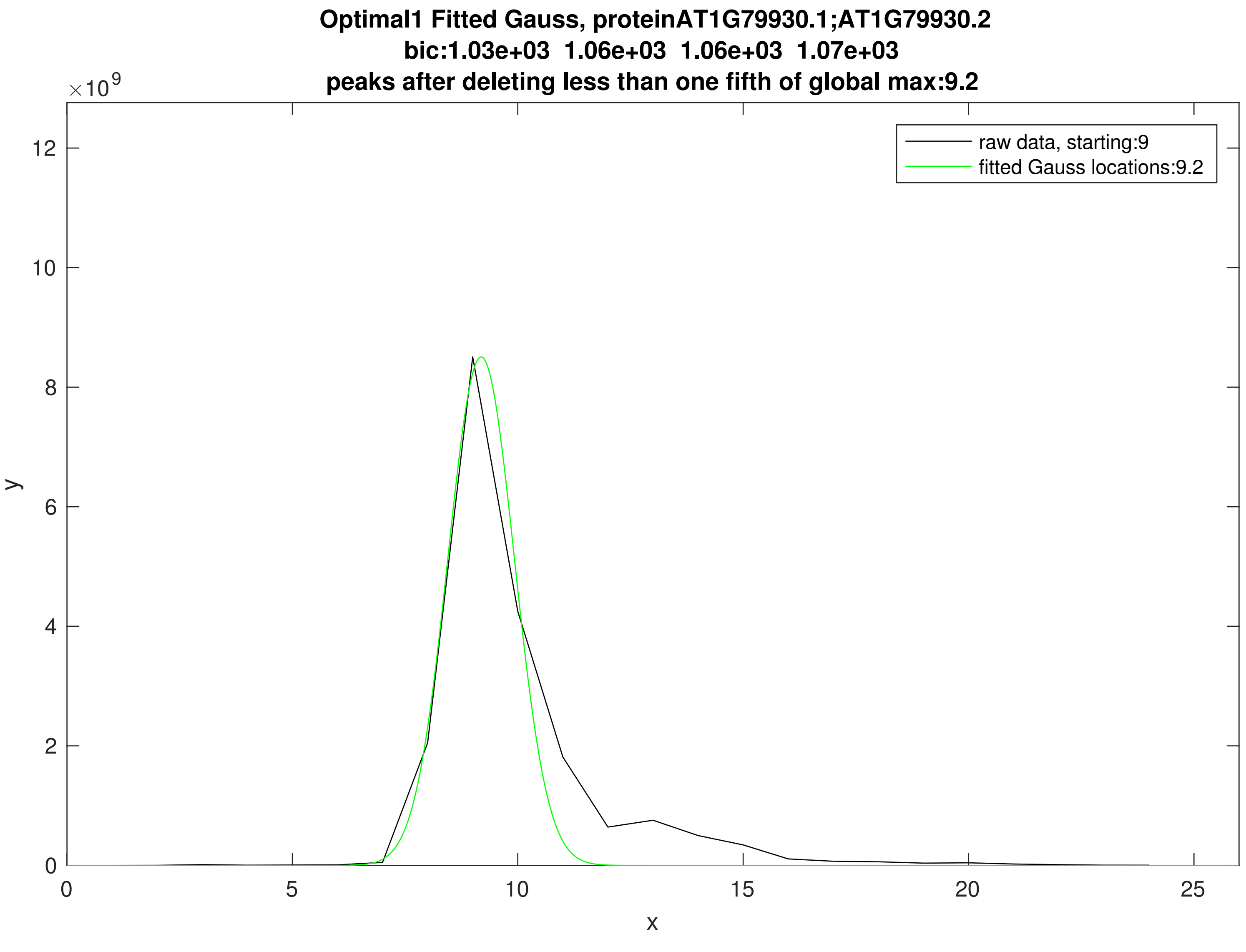
Optimal1 Fitted Gauss, proteinAT1G79930.1;AT1G79930.2
bic:1.03e+03 1.06e+03 1.06e+03 1.07e+03
peaks after deleting less than one fifth of global max:9.2
×10^9
raw data, starting:9
fitted Gauss locations:9.2
y
x
0
2
4
6
8
10
12
0
5
10
15
20
25

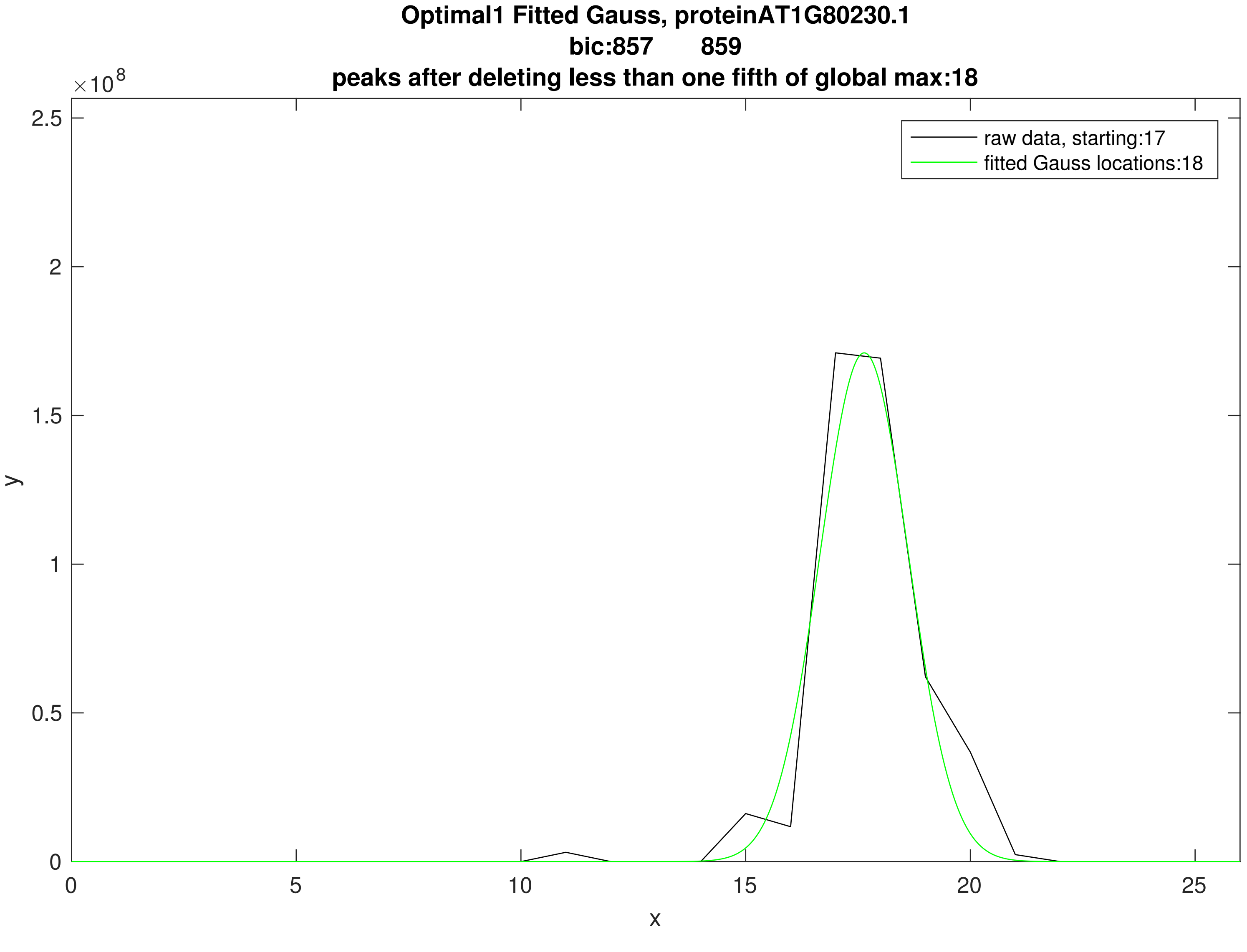
Optimal1 Fitted Gauss, proteinAT1G80230.1
bic:857 859
peaks after deleting less than one fifth of global max:18
×10^8
2.5
2
1.5
1
0.5
0
y
0
5
10
15
20
25
x
raw data, starting:17
fitted Gauss locations:18

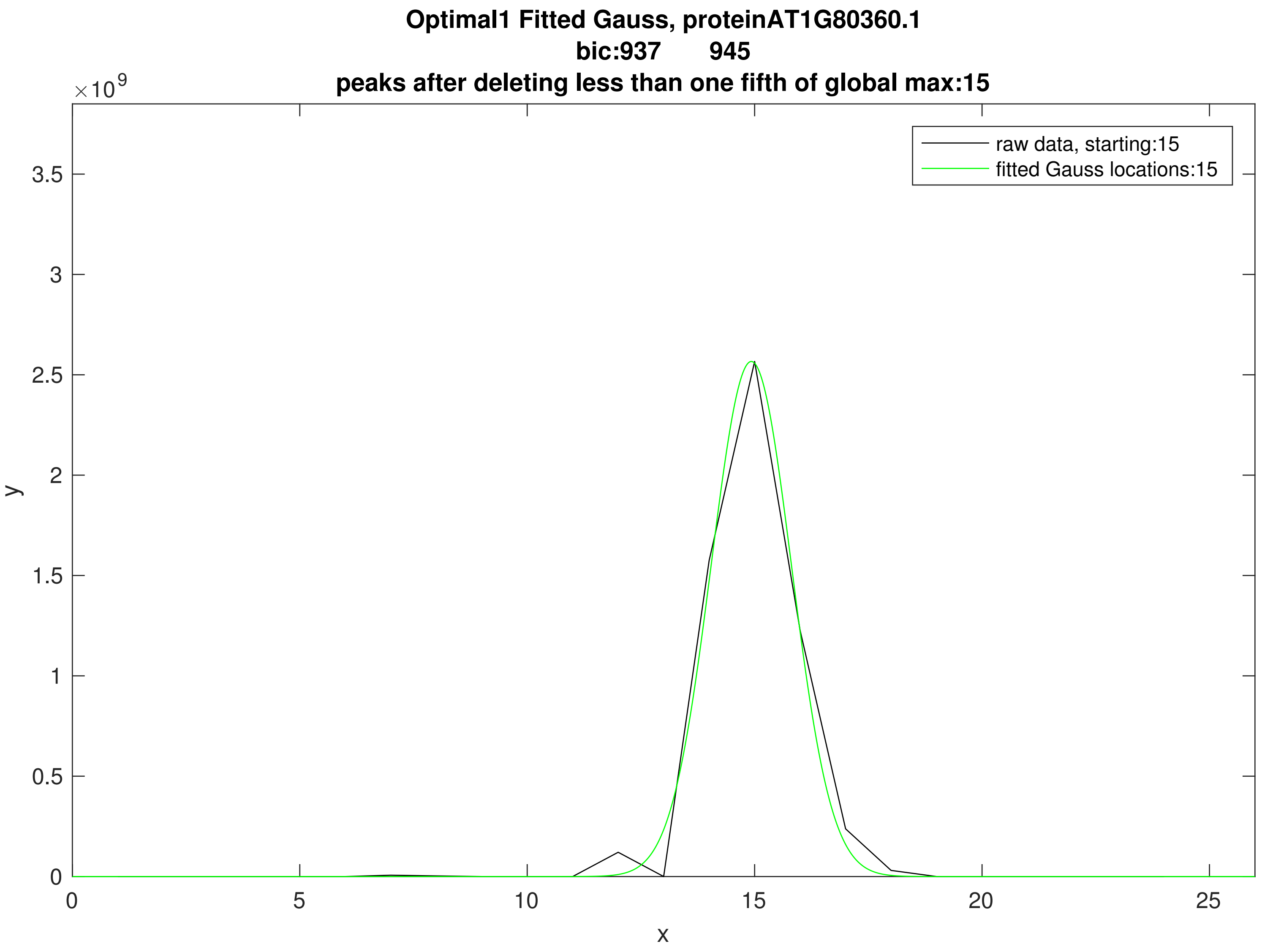

Optimal1 Fitted Gauss, proteinAT1G80360.1
bic:937 945
peaks after deleting less than one fifth of global max:15
×10^9
raw data, starting:15
fitted Gauss locations:15
y
x
0
0.5
1
1.5
2
2.5
3
3.5
5
10
15
20
25

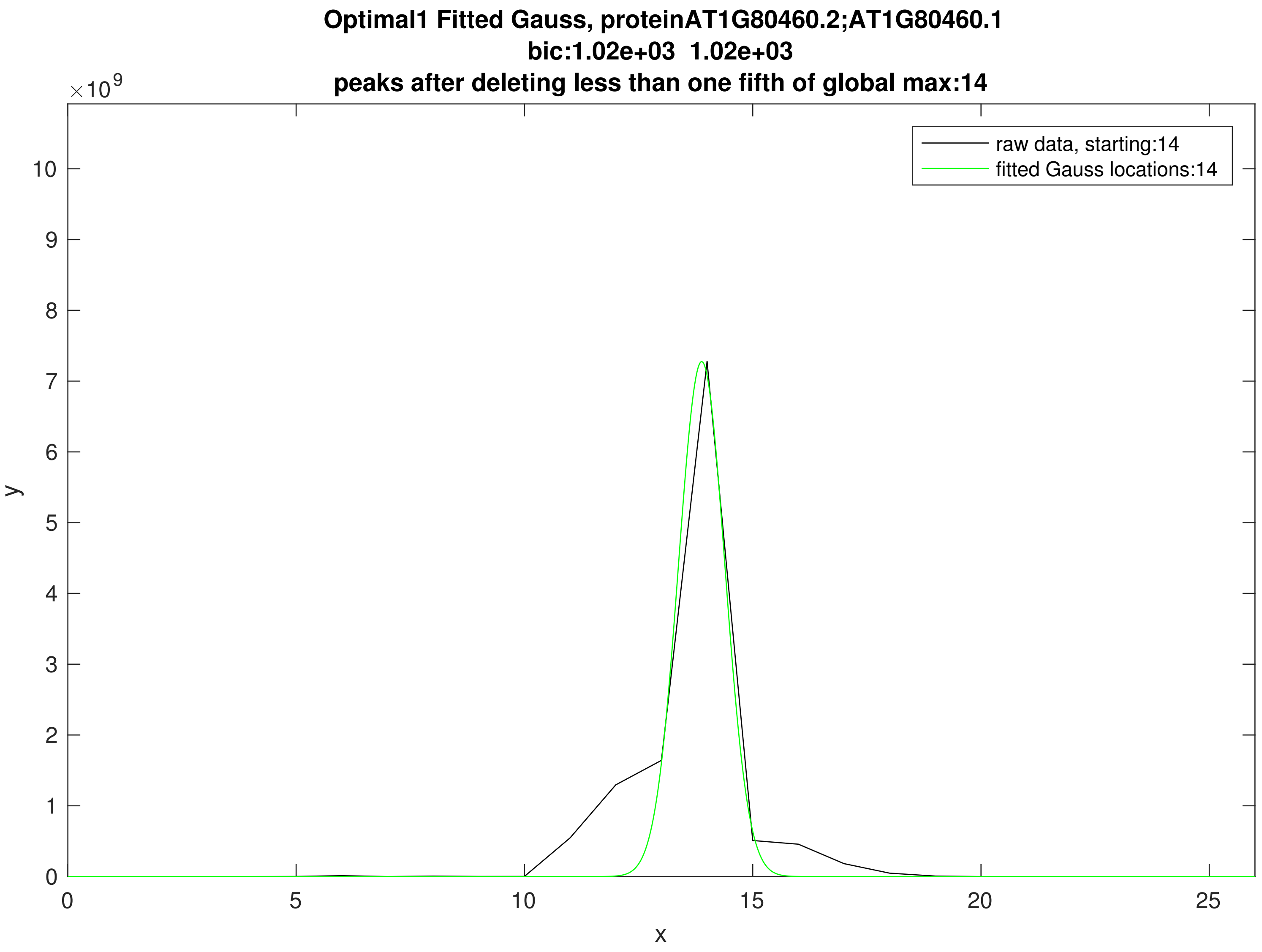

Optimal1 Fitted Gauss, proteinAT1G80460.2;AT1G80460.1
bic:1.02e+03  1.02e+03
peaks after deleting less than one fifth of global max:14
×10^9
raw data, starting:14
fitted Gauss locations:14
y
x
0
1
2
3
4
5
6
7
8
9
10
15
20
25

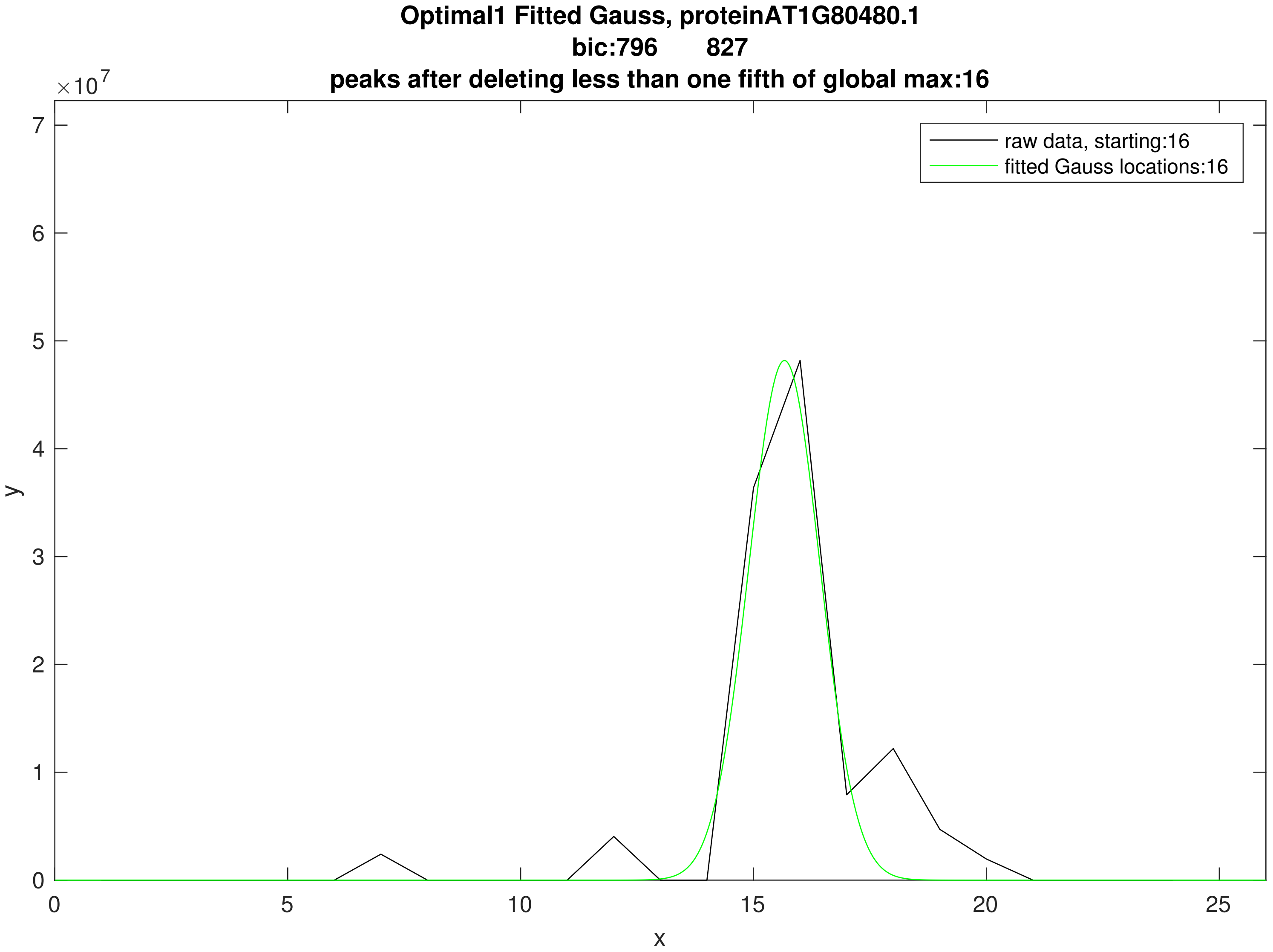
Optimal1 Fitted Gauss, proteinAT1G80480.1
bic:796 827
peaks after deleting less than one fifth of global max:16
×10^7
raw data, starting:16
fitted Gauss locations:16
y
x
0
1
2
3
4
5
6
7
0
5
10
15
20
25

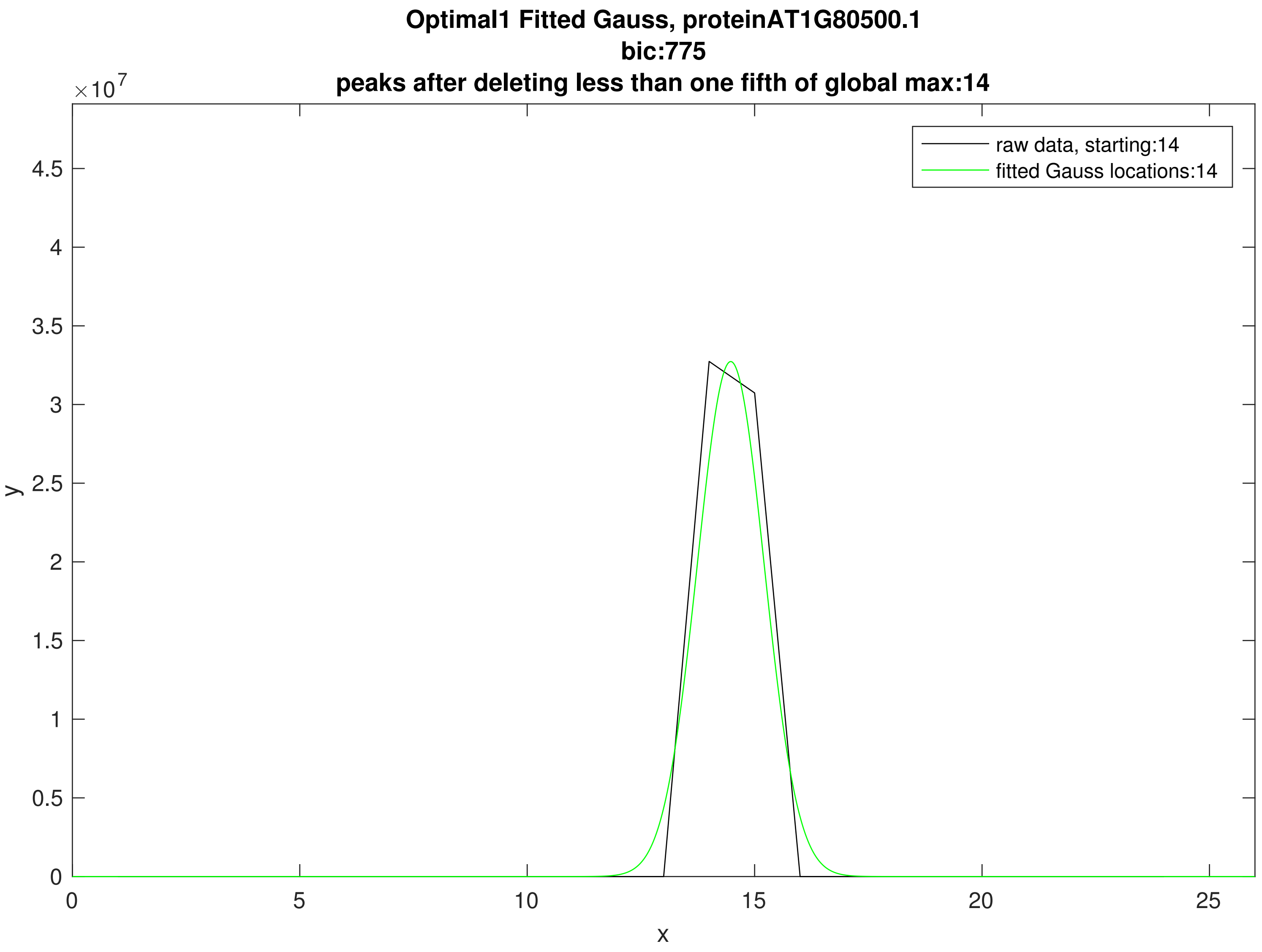

Optimal1 Fitted Gauss, proteinAT1G80500.1
bic:775
peaks after deleting less than one fifth of global max:14
×10^7
4.5
4
3.5
3
2.5
2
1.5
1
0.5
0
y
0
5
10
15
20
25
x
raw data, starting:14
fitted Gauss locations:14

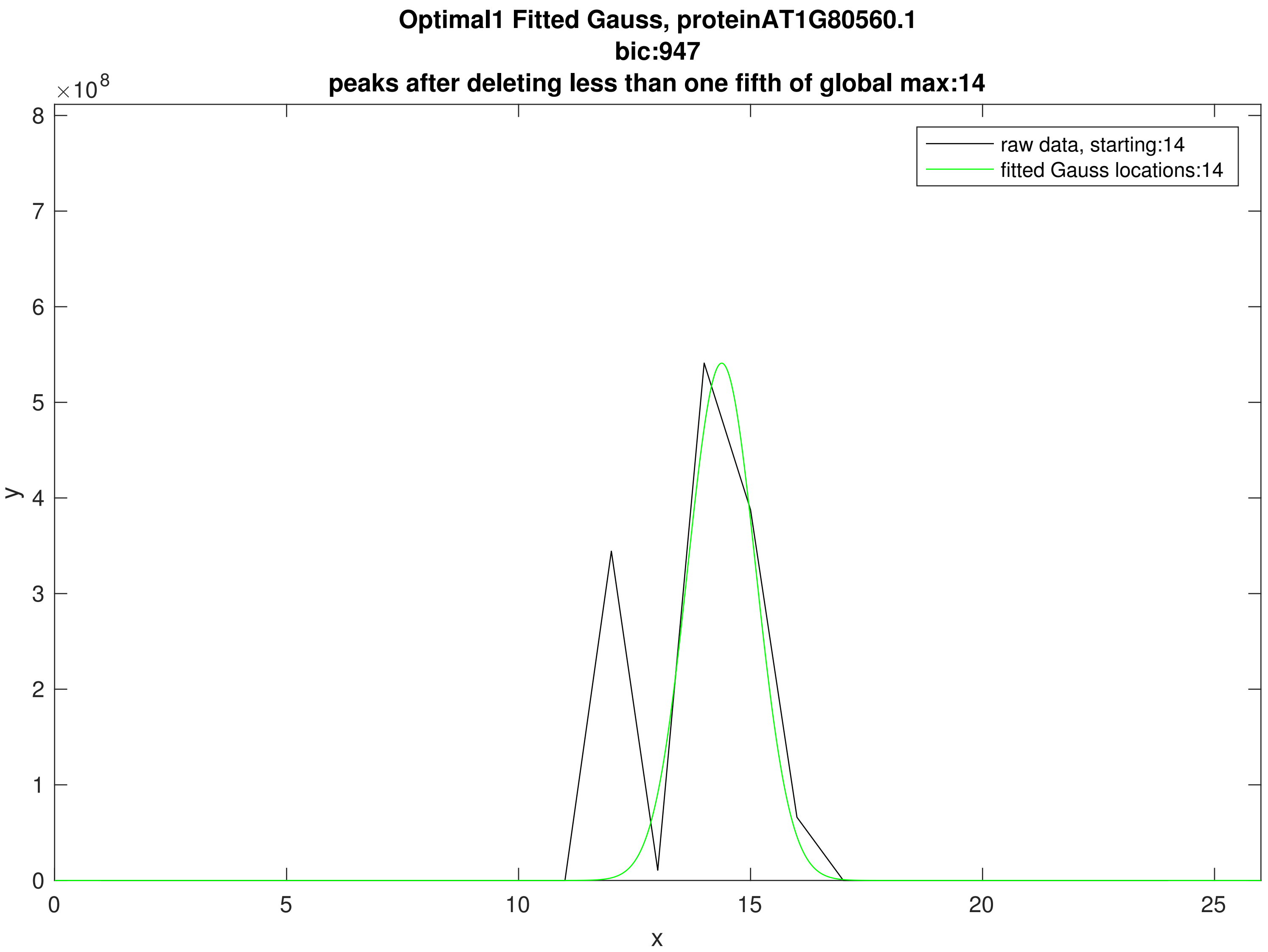

Optimal1 Fitted Gauss, proteinAT1G80560.1
bic:947
peaks after deleting less than one fifth of global max:14
×10^8
raw data, starting:14
fitted Gauss locations:14
y
x
0
1
2
3
4
5
6
7
8
0
5
10
15
20
25

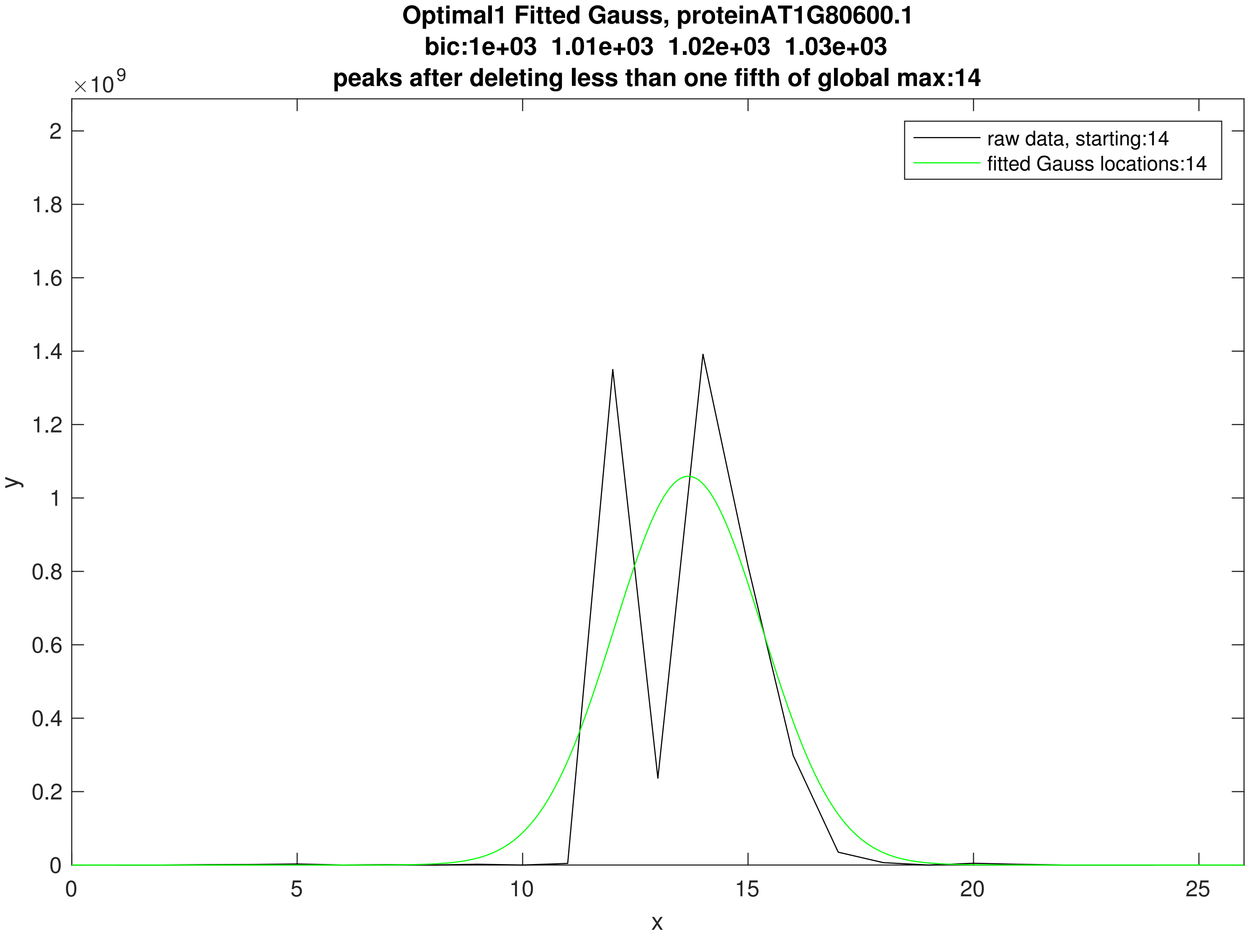
Optimal1 Fitted Gauss, proteinAT1G80600.1
bic:1e+03  1.01e+03  1.02e+03  1.03e+03
peaks after deleting less than one fifth of global max:14
×10^9
raw data, starting:14
fitted Gauss locations:14
y
x
0
0.2
0.4
0.6
0.8
1
1.2
1.4
1.6
1.8
2
5
10
15
20
25

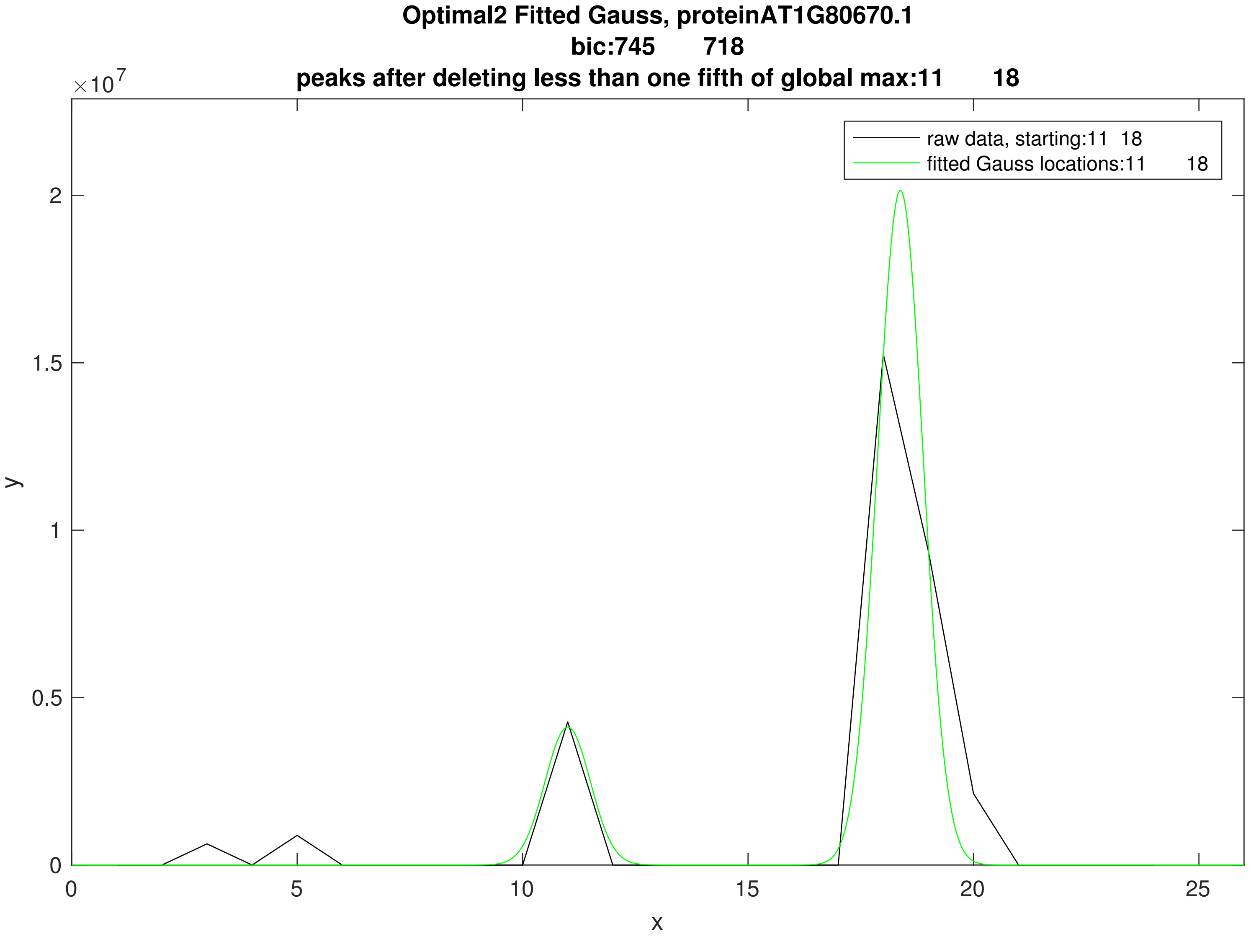

Optimal2 Fitted Gauss, proteinAT1G80670.1
bic:745 718
peaks after deleting less than one fifth of global max:11 18
×10^7
raw data, starting:11 18
fitted Gauss locations:11 18
y
x
0
0.5
1
1.5
2
5
10
15
20
25

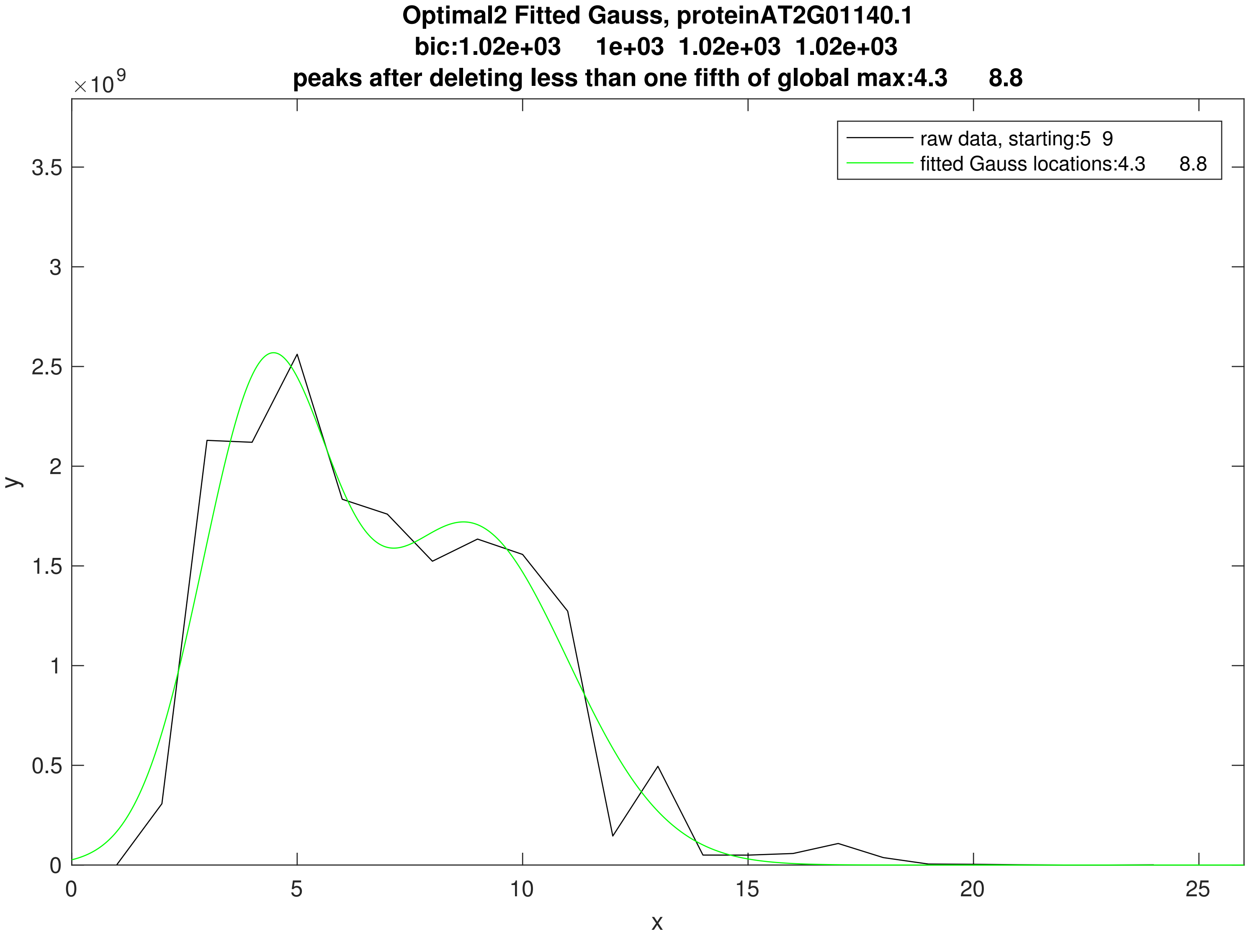
Optimal2 Fitted Gauss, proteinAT2G01140.1
bic:1.02e+03 1e+03 1.02e+03 1.02e+03
peaks after deleting less than one fifth of global max:4.3 8.8
×10^9
raw data, starting:5 9
fitted Gauss locations:4.3 8.8
y
x
0
0.5
1
1.5
2
2.5
3
3.5
0
5
10
15
20
25

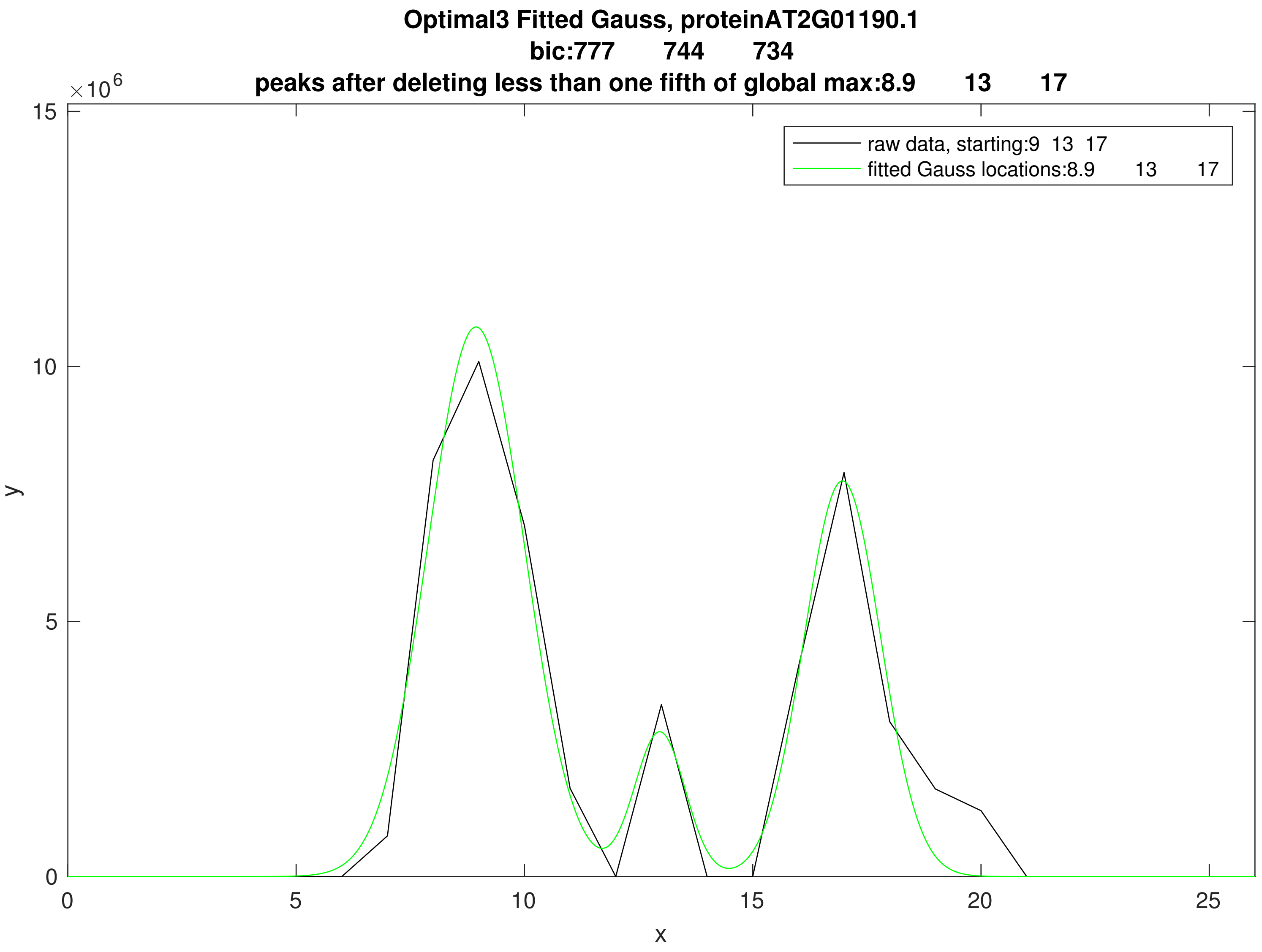
Optimal3 Fitted Gauss, proteinAT2G01190.1
bic:777 744 734
peaks after deleting less than one fifth of global max:8.9 13 17
×10^6
raw data, starting:9 13 17
fitted Gauss locations:8.9 13 17
y
x
0
5
10
15
20
25

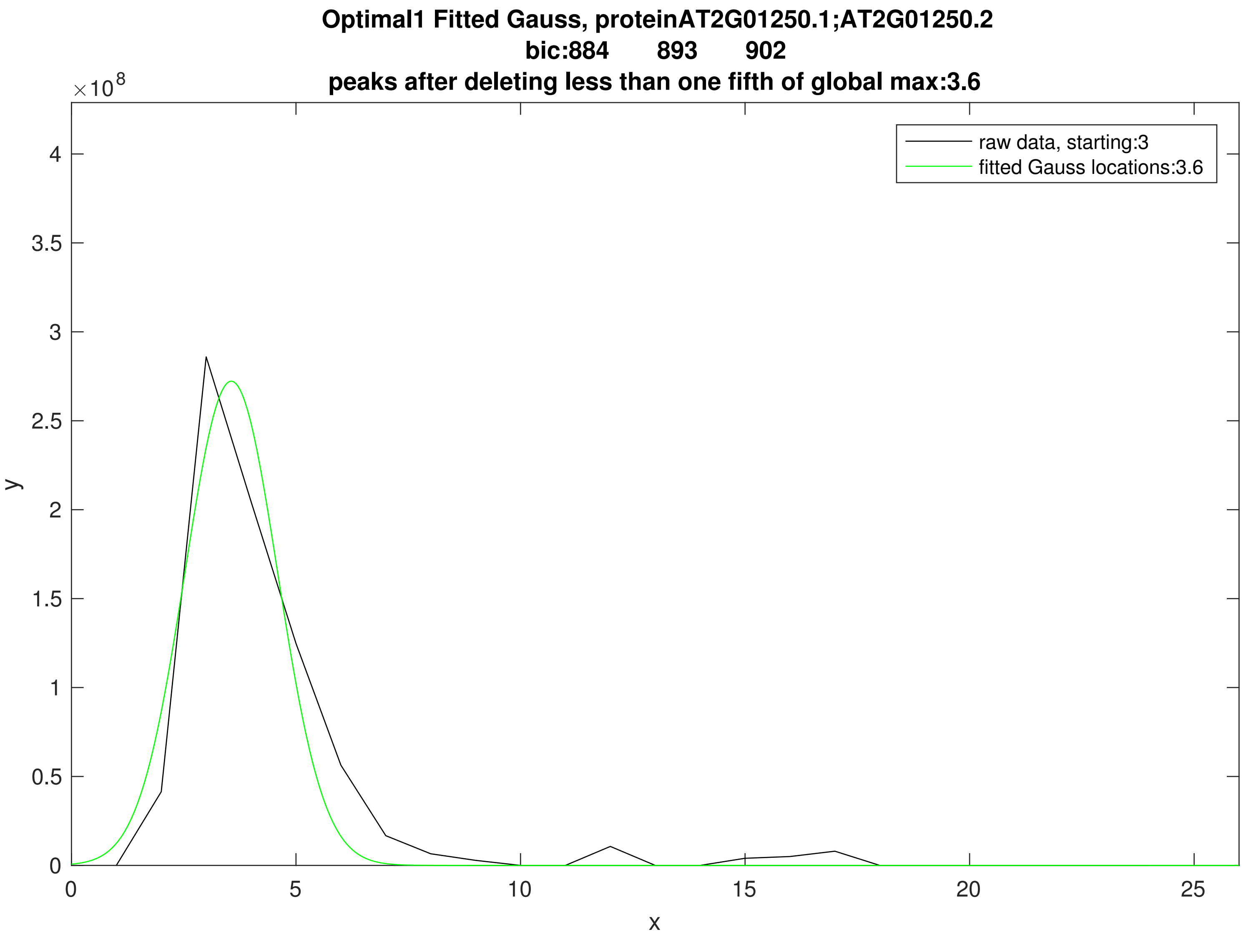

Optimal1 Fitted Gauss, proteinAT2G01250.1;AT2G01250.2
bic:884 893 902
peaks after deleting less than one fifth of global max:3.6
×10^8
raw data, starting:3
fitted Gauss locations:3.6
y
x
0
0.5
1
1.5
2
2.5
3
3.5
4
5
10
15
20
25

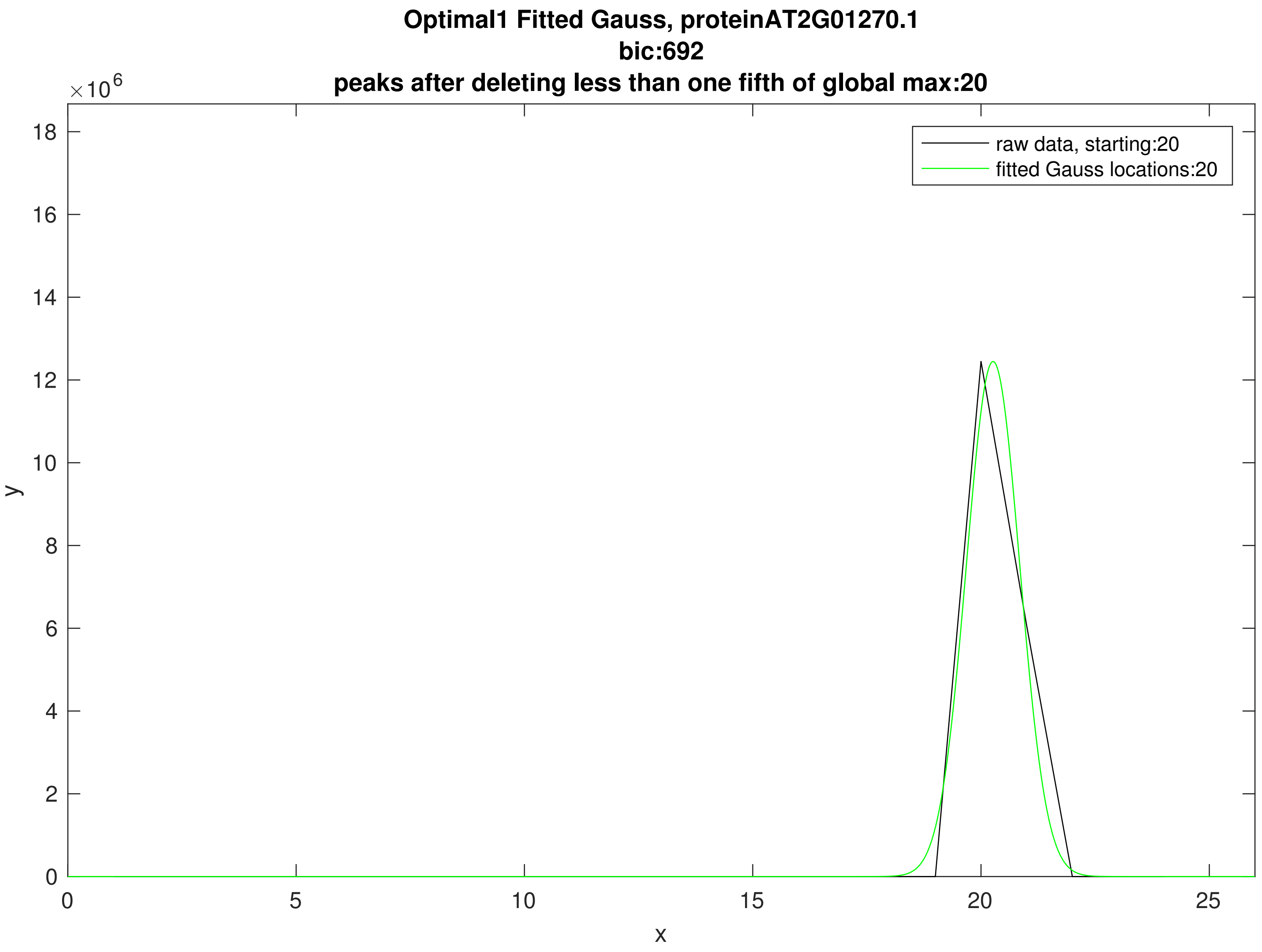
Optimal1 Fitted Gauss, proteinAT2G01270.1
bic:692
peaks after deleting less than one fifth of global max:20
×10^6
raw data, starting:20
fitted Gauss locations:20
y
x
0
2
4
6
8
10
12
14
16
18
0
5
10
15
20
25

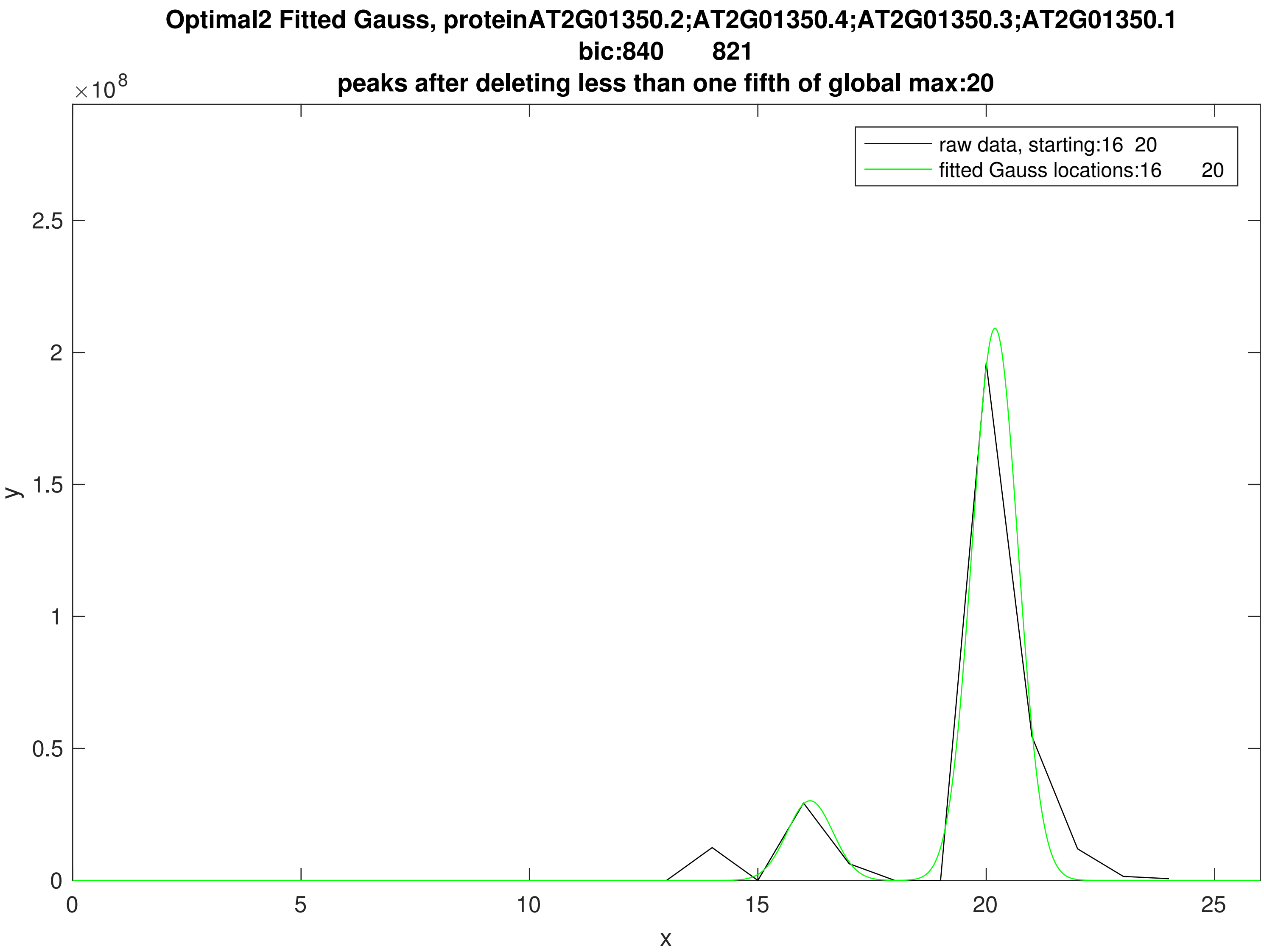
Optimal2 Fitted Gauss, proteinAT2G01350.2;AT2G01350.4;AT2G01350.3;AT2G01350.1
bic:840 821
peaks after deleting less than one fifth of global max:20
×10^8
raw data, starting:16 20
fitted Gauss locations:16 20
y
x
0
0.5
1
1.5
2
2.5
0
5
10
15
20
25

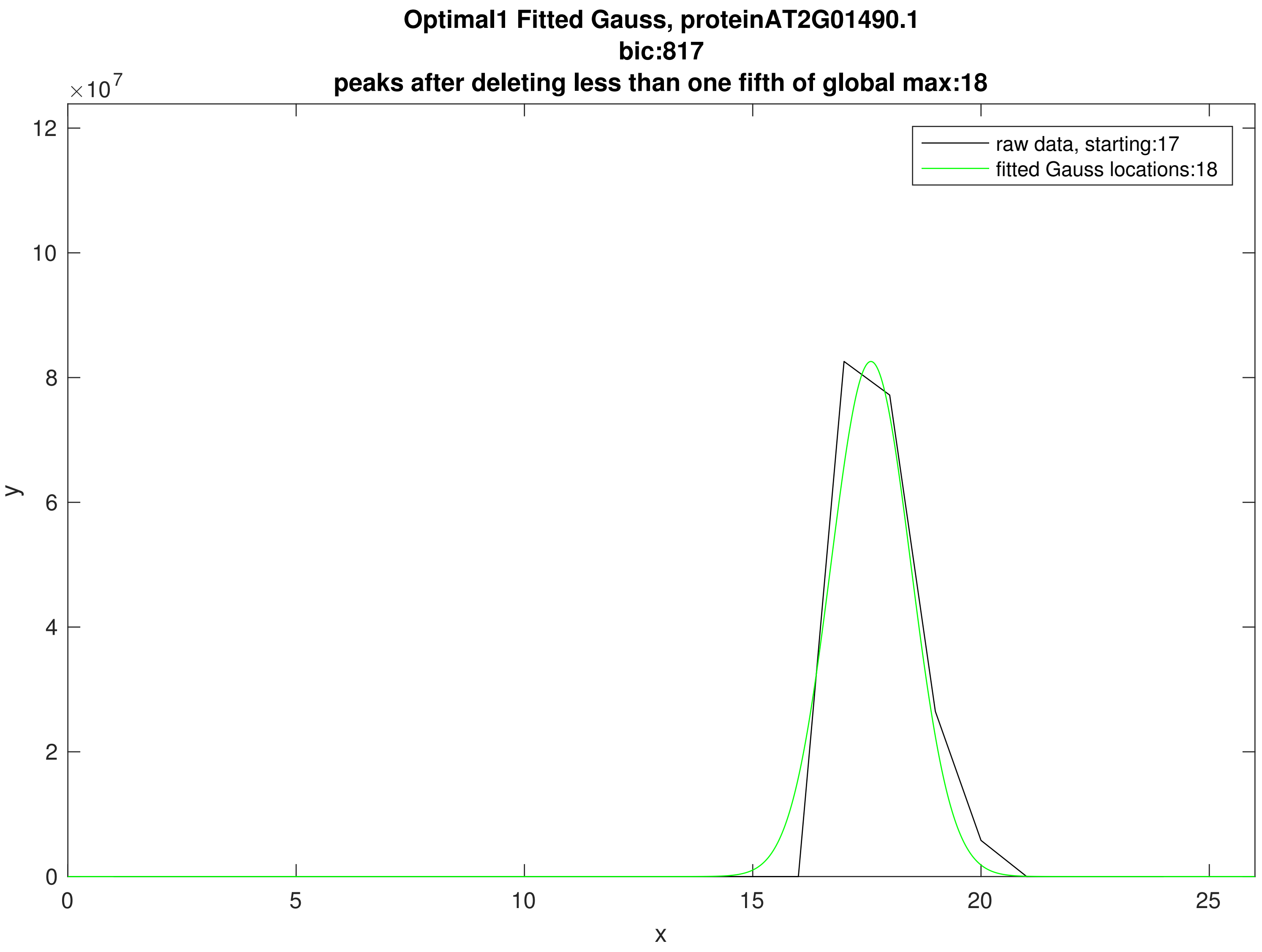

Optimal1 Fitted Gauss, proteinAT2G01490.1
bic:817
peaks after deleting less than one fifth of global max:18
×10^7
raw data, starting:17
fitted Gauss locations:18
y
x
0
2
4
6
8
10
12
0
5
10
15
20
25

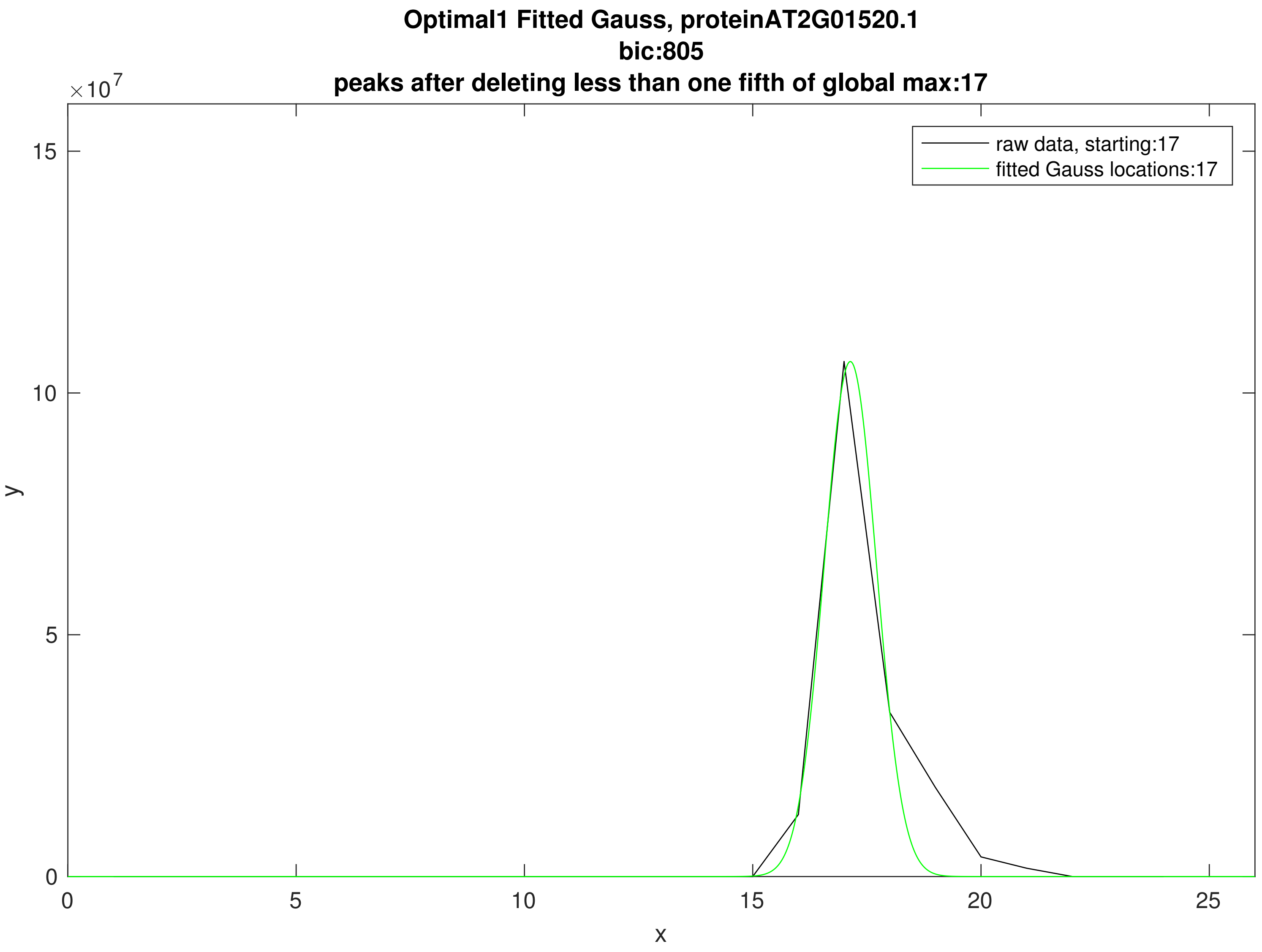

Optimal1 Fitted Gauss, proteinAT2G01520.1
bic:805
peaks after deleting less than one fifth of global max:17
×10^7
15
10
5
0
y
0
5
10
15
20
25
x
raw data, starting:17
fitted Gauss locations:17

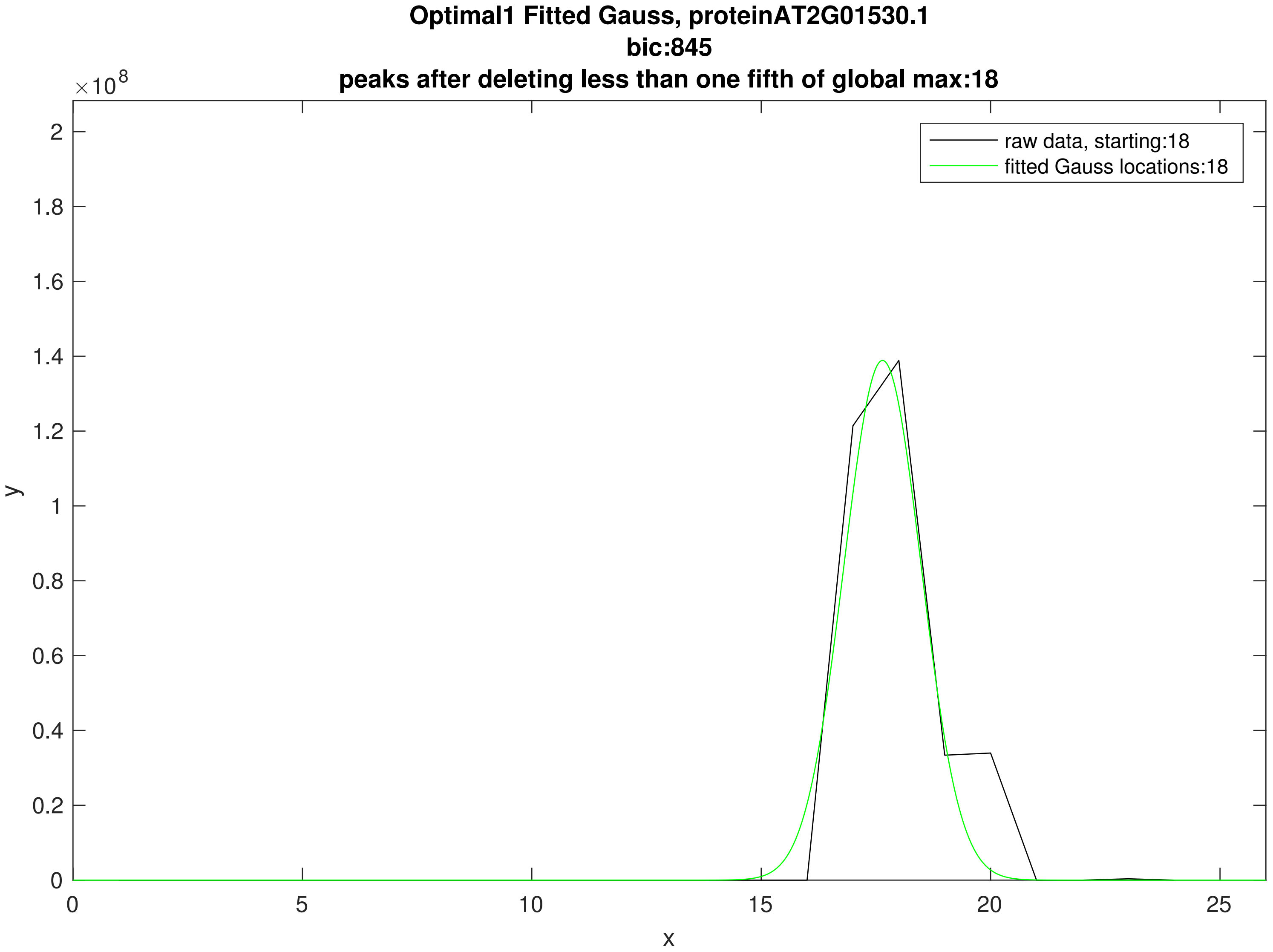
Optimal1 Fitted Gauss, proteinAT2G01530.1
bic:845
peaks after deleting less than one fifth of global max:18
×10^8
raw data, starting:18
fitted Gauss locations:18
y
x
0
0.2
0.4
0.6
0.8
1
1.2
1.4
1.6
1.8
2
0
5
10
15
20
25

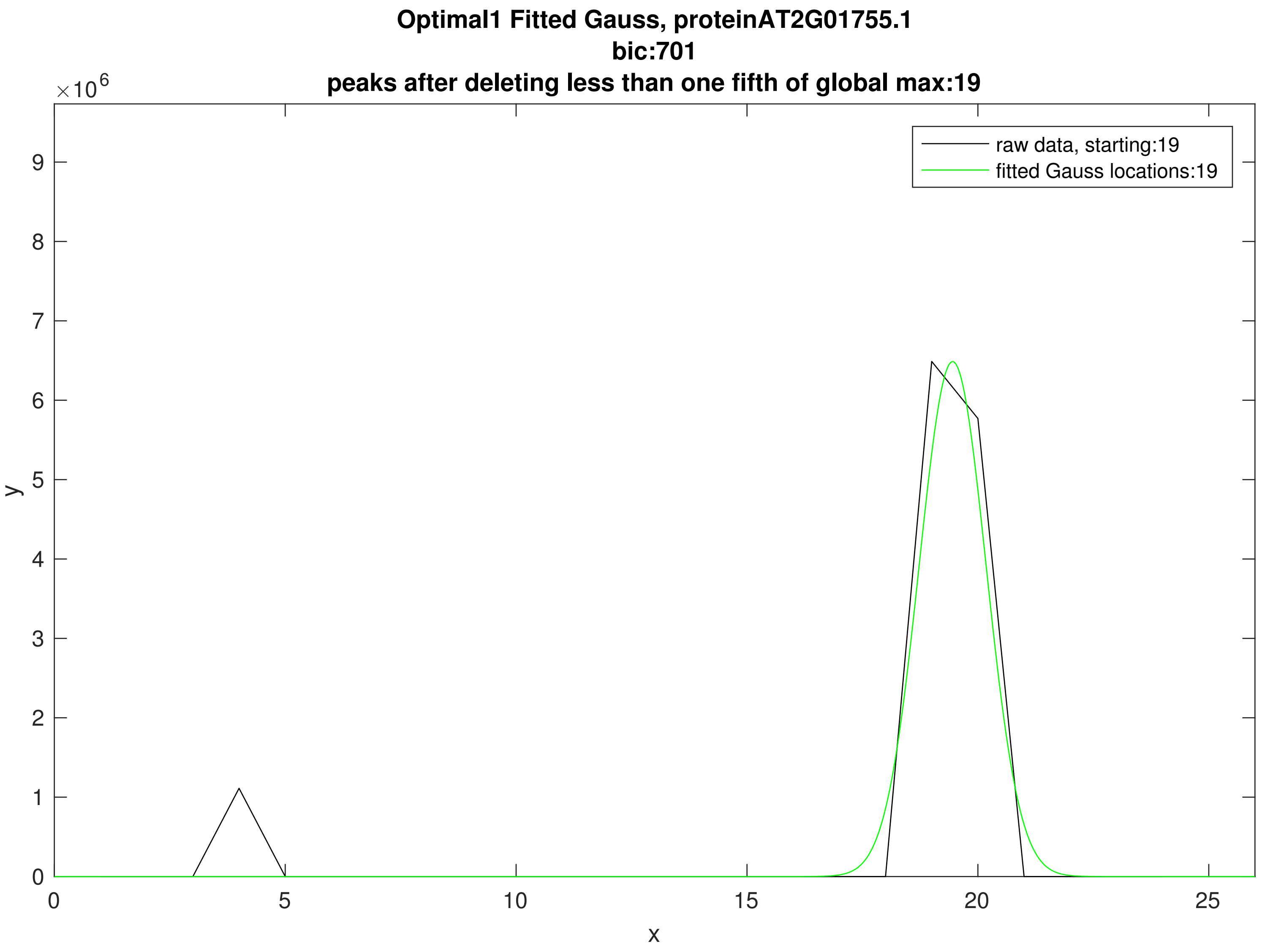

Optimal1 Fitted Gauss, proteinAT2G01755.1
bic:701
peaks after deleting less than one fifth of global max:19
×10^6
raw data, starting:19
fitted Gauss locations:19
y
x
0
1
2
3
4
5
6
7
8
9
0
5
10
15
20
25

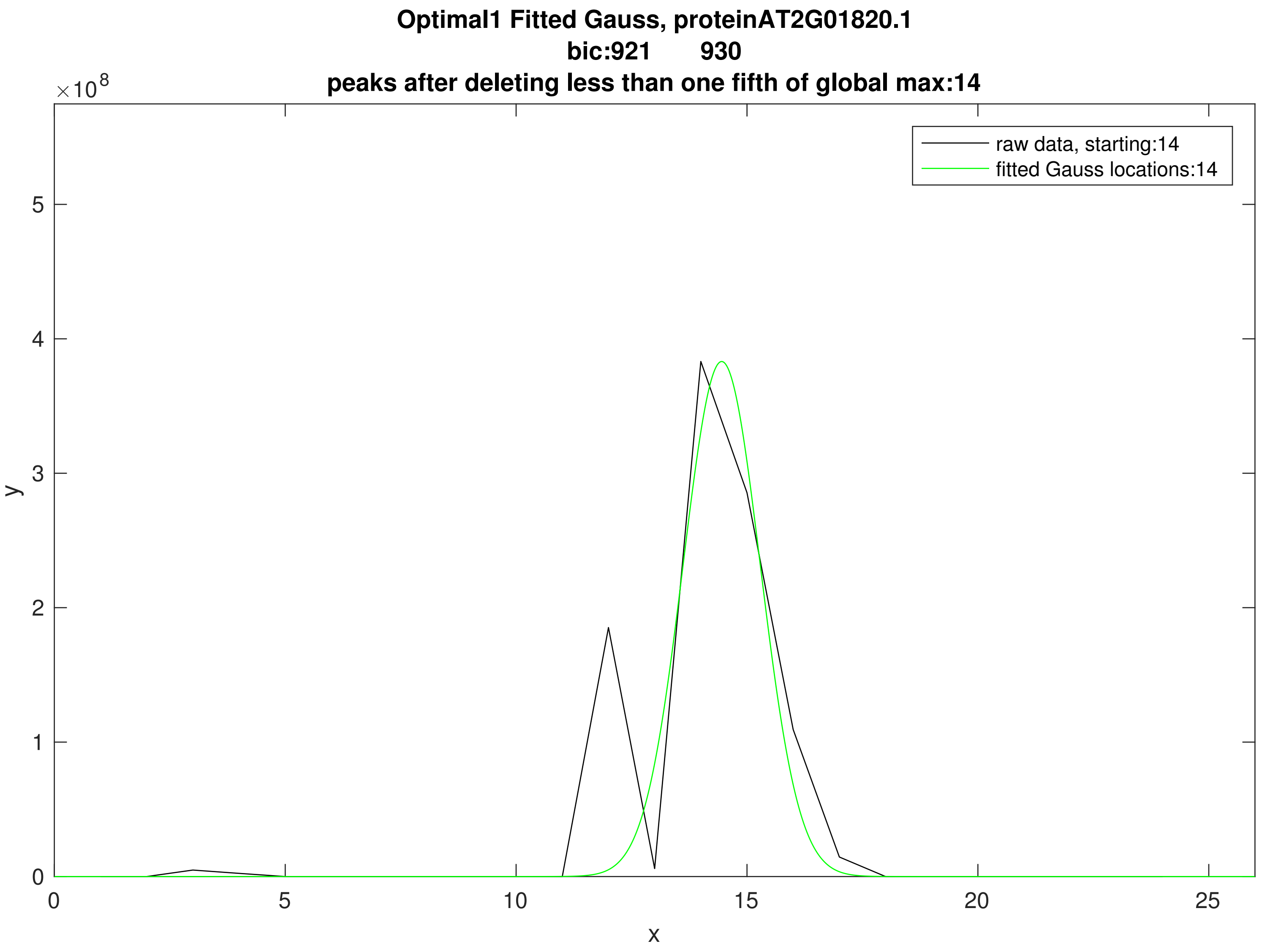

Optimal1 Fitted Gauss, proteinAT2G01820.1
bic:921 930
peaks after deleting less than one fifth of global max:14
×10^8
raw data, starting:14
fitted Gauss locations:14
y
x
0
1
2
3
4
5
0
5
10
15
20
25

**Optimal1 Fitted Gauss, proteinAT2G01890.1;AT2G01890.2**
**bic:823**
**peaks after deleting less than one fifth of global max:18**

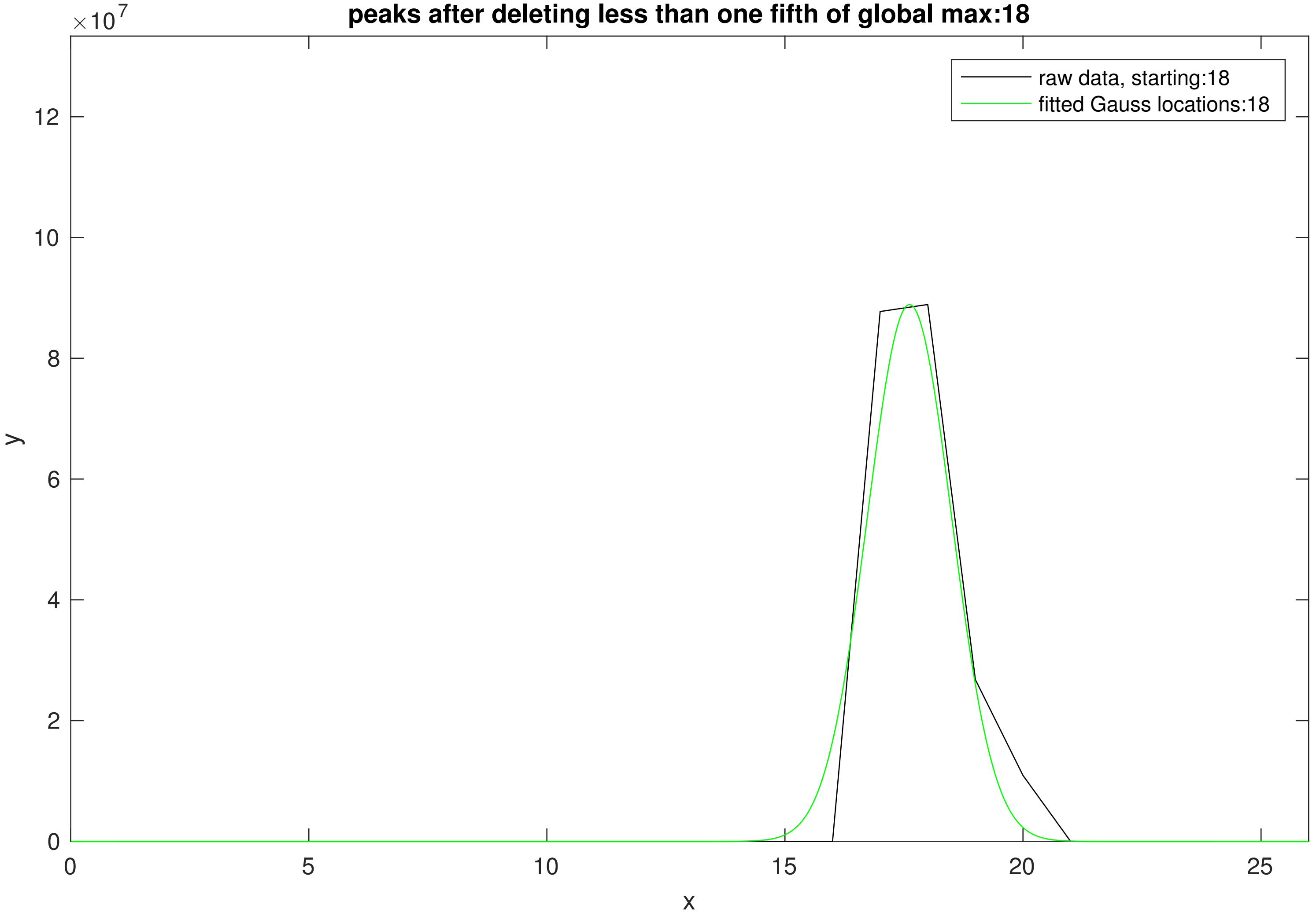

**Optimal1 Fitted Gauss, proteinAT2G02050.1**
**bic:768**
**peaks after deleting less than one fifth of global max:20**

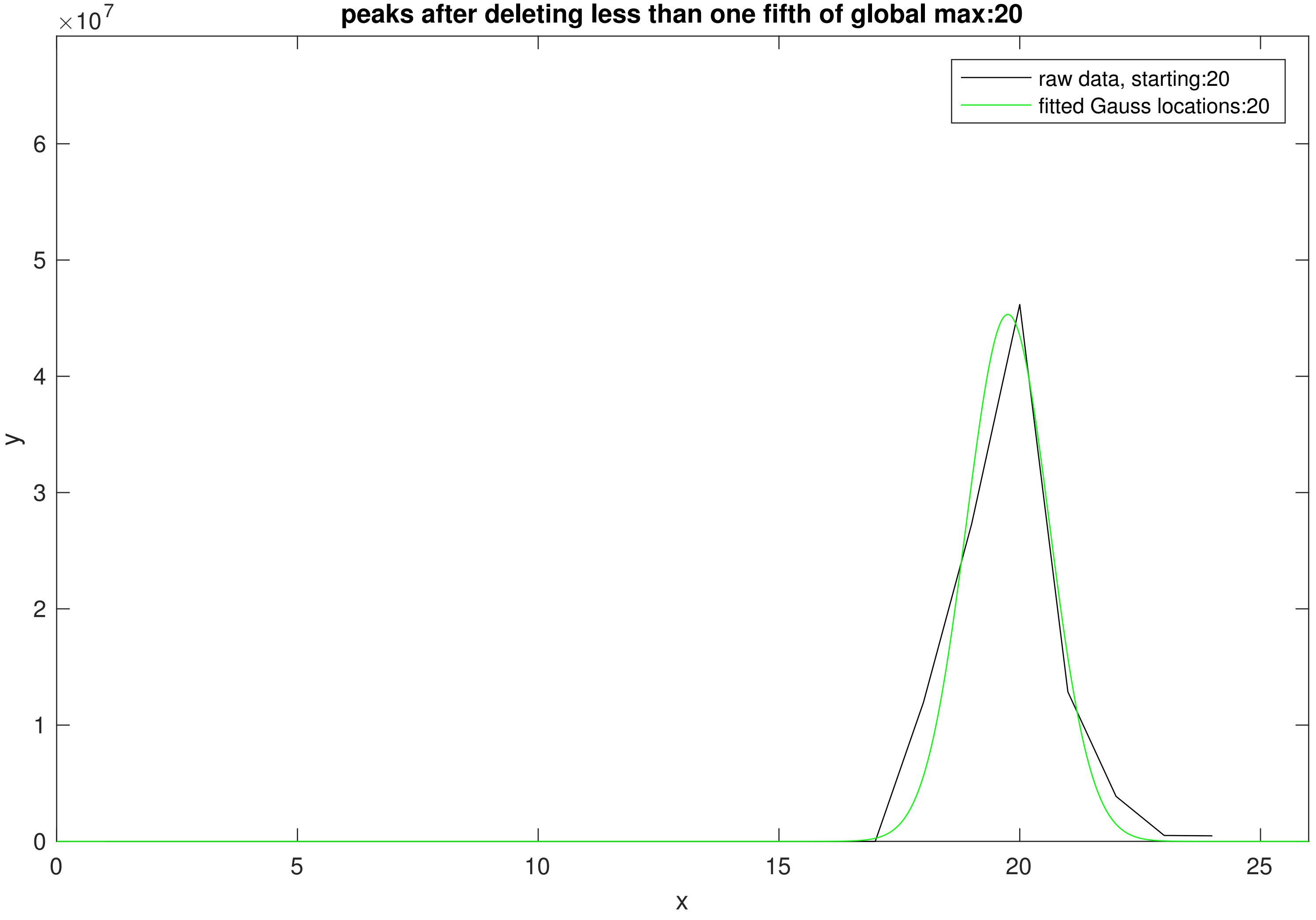

**Optimal1 Fitted Gauss, proteinAT2G02390.1;AT2G02390.2;AT2G02390.3**
**bic:868 876 886**
**peaks after deleting less than one fifth of global max:15**

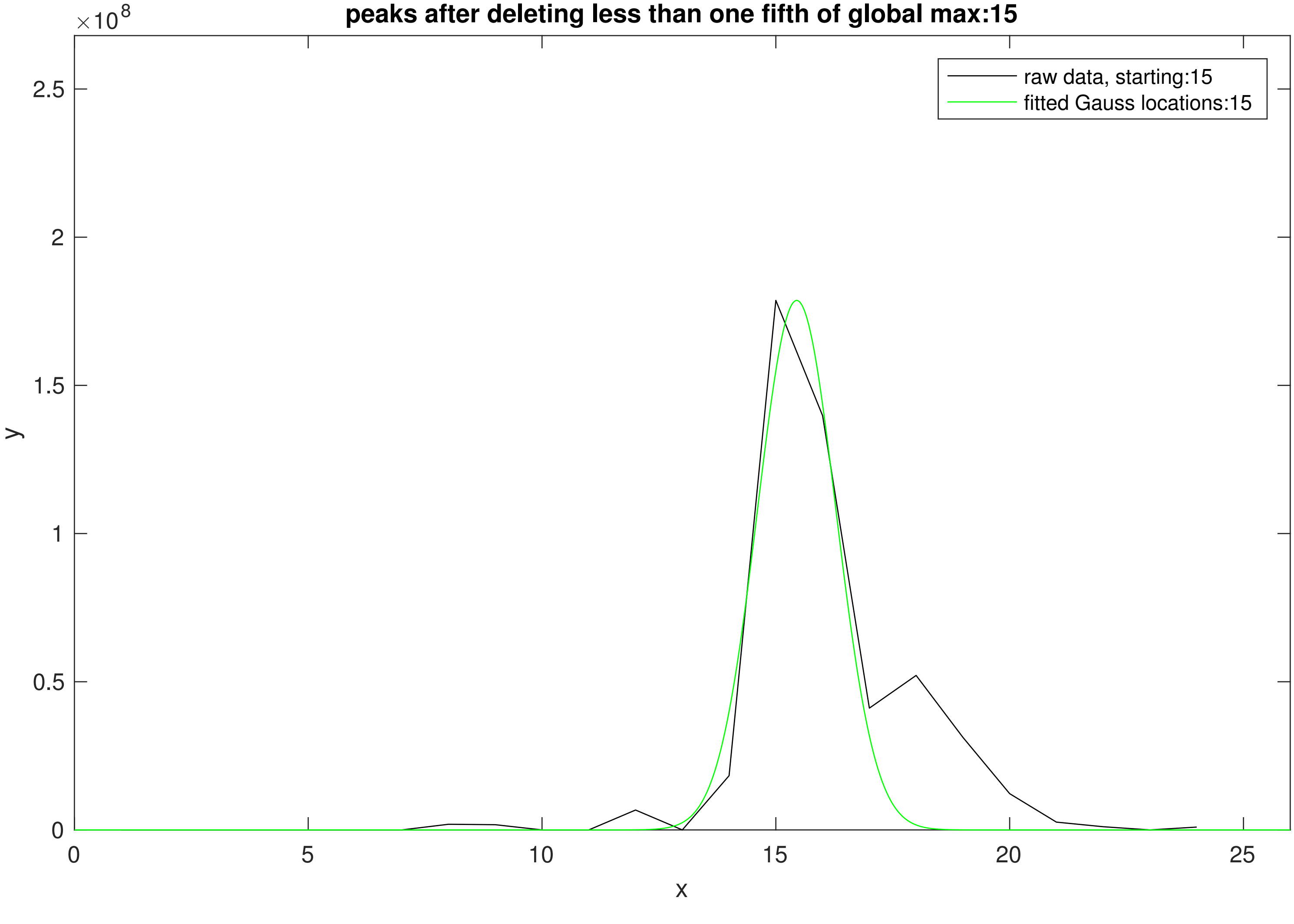

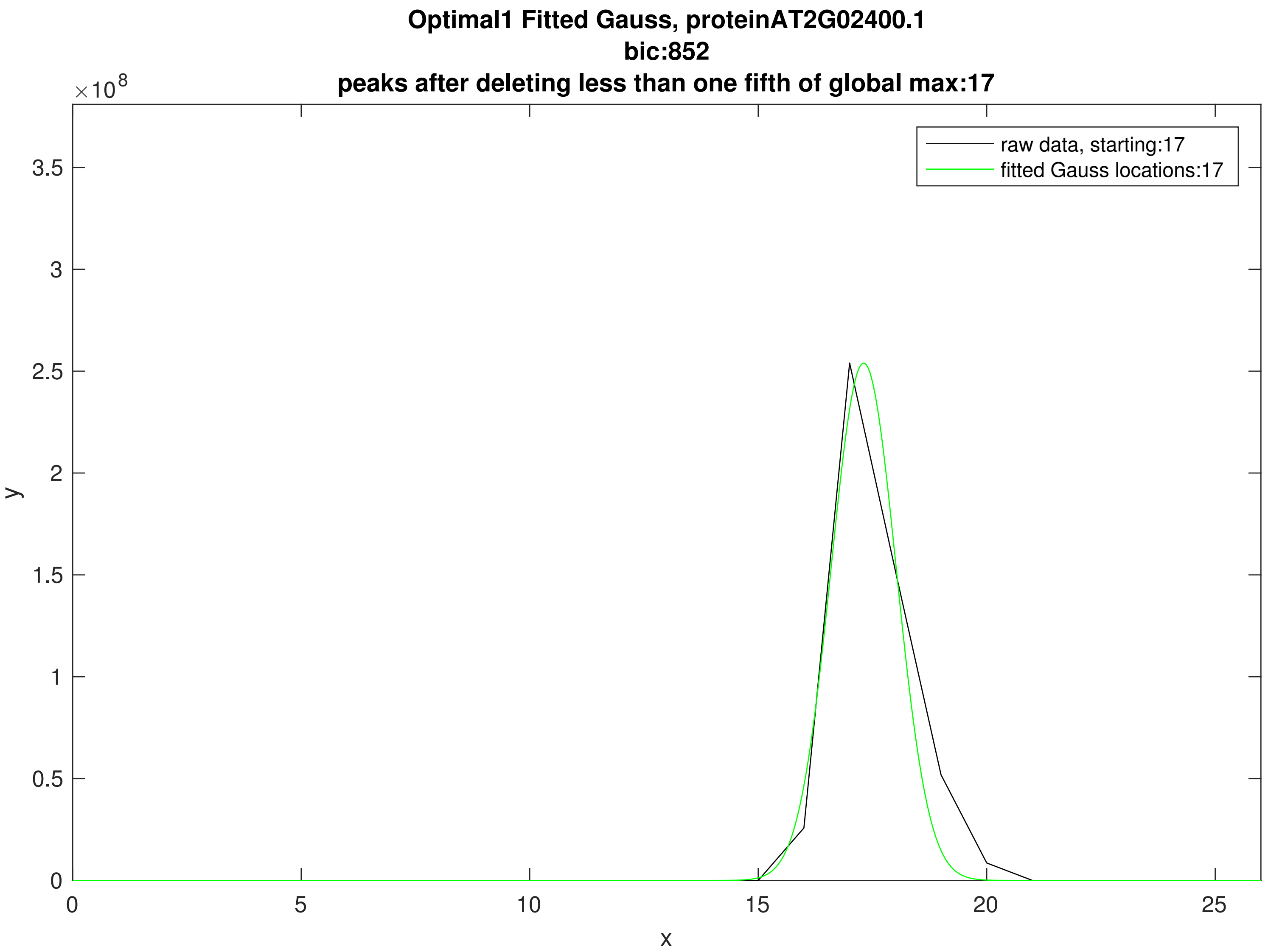

Optimal1 Fitted Gauss, proteinAT2G02400.1
bic:852
peaks after deleting less than one fifth of global max:17
×10^8
3.5
3
2.5
2
1.5
1
0.5
0
y
0
5
10
15
20
25
x
raw data, starting:17
fitted Gauss locations:17

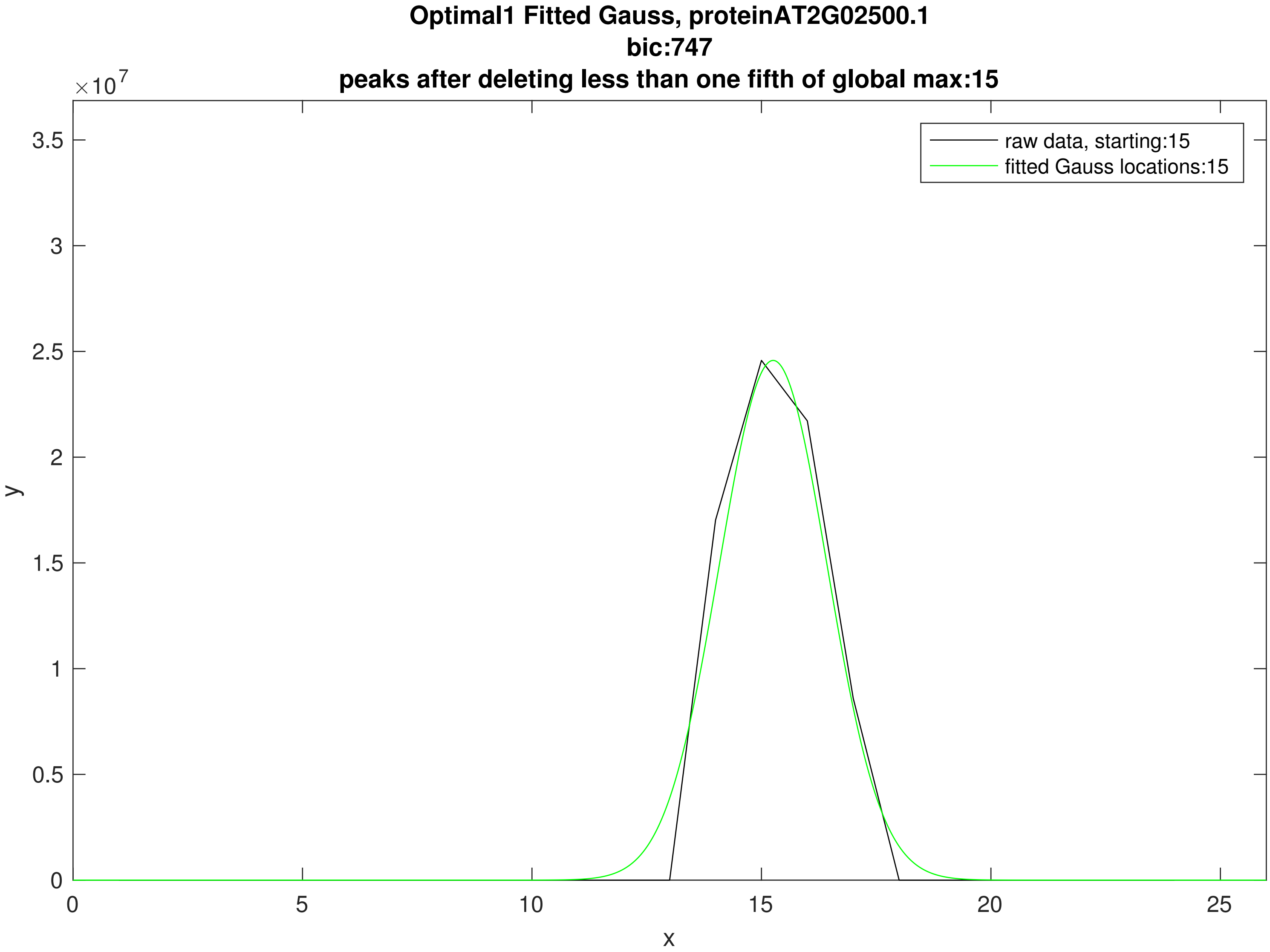

Optimal1 Fitted Gauss, proteinAT2G02500.1
bic:747
peaks after deleting less than one fifth of global max:15
×10^7
3.5
3
2.5
2
1.5
1
0.5
0
y
0
5
10
15
20
25
x
raw data, starting:15
fitted Gauss locations:15

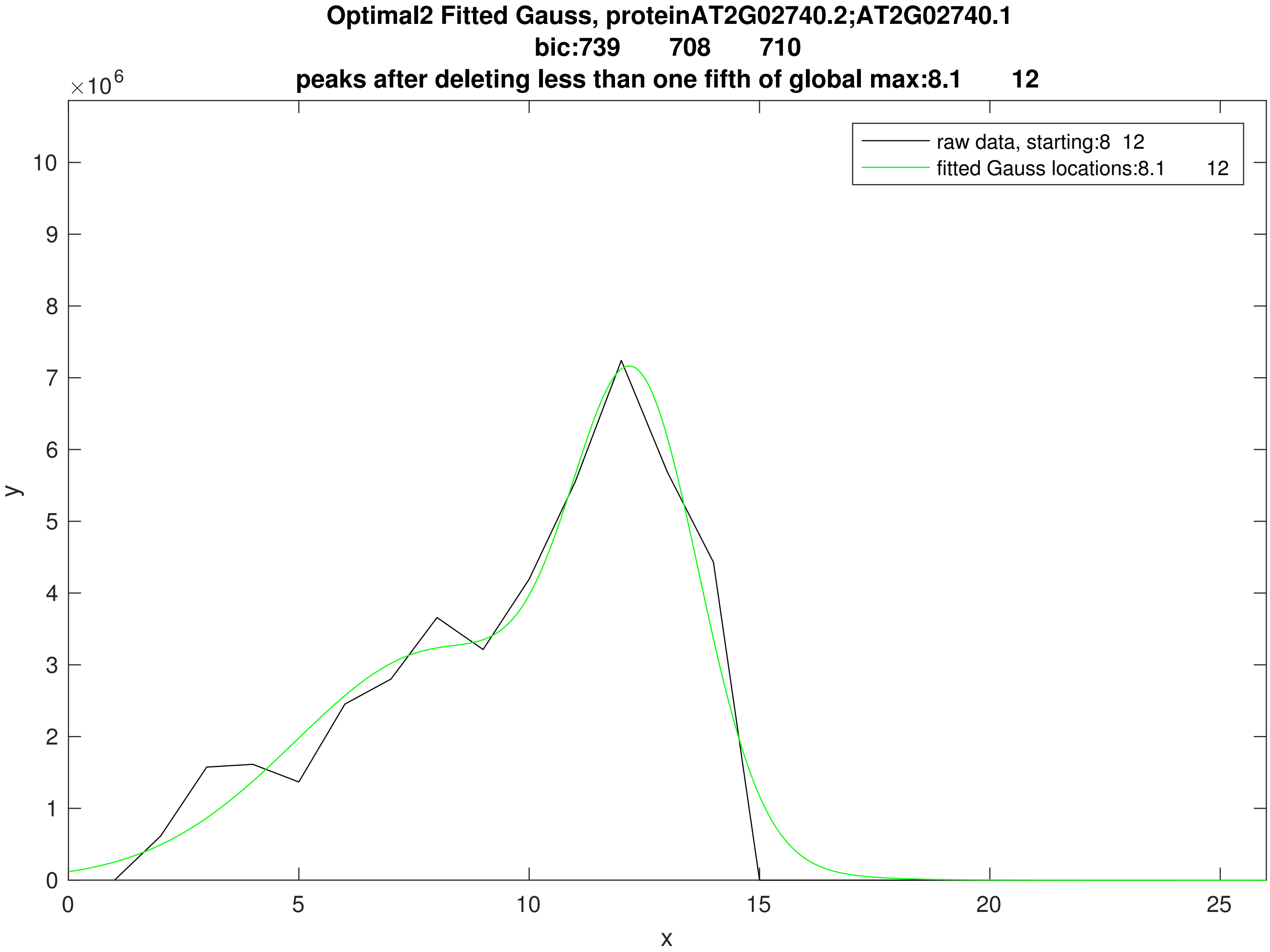

Optimal2 Fitted Gauss, proteinAT2G02740.2;AT2G02740.1
bic:739 708 710
peaks after deleting less than one fifth of global max:8.1 12
×10^6
raw data, starting:8 12
fitted Gauss locations:8.1 12
y
x
0
1
2
3
4
5
6
7
8
9
10
0
5
10
15
20
25

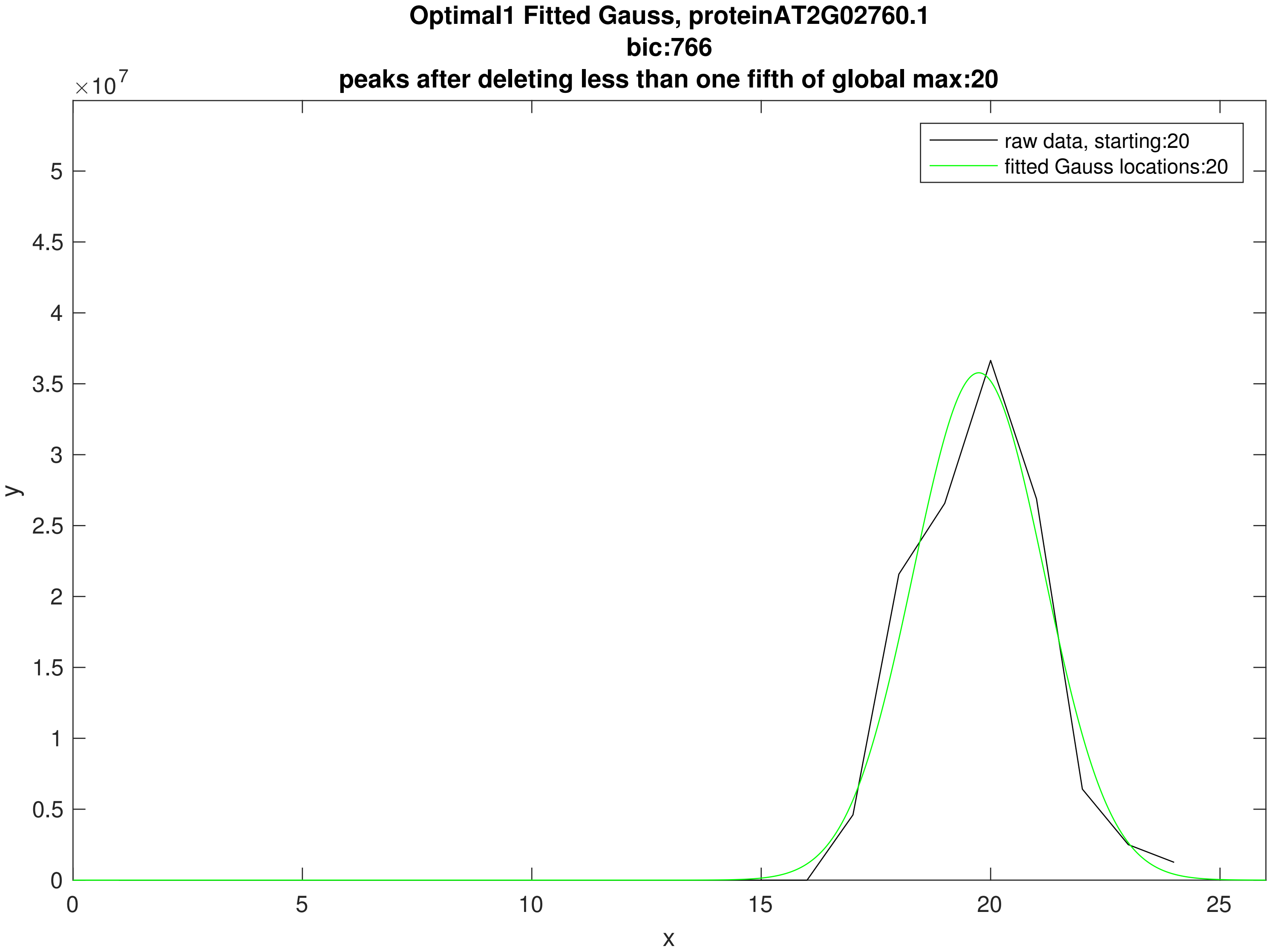
Optimal1 Fitted Gauss, proteinAT2G02760.1
bic:766
peaks after deleting less than one fifth of global max:20
×10^7
raw data, starting:20
fitted Gauss locations:20
y
x
0
0.5
1
1.5
2
2.5
3
3.5
4
4.5
5
0
5
10
15
20
25

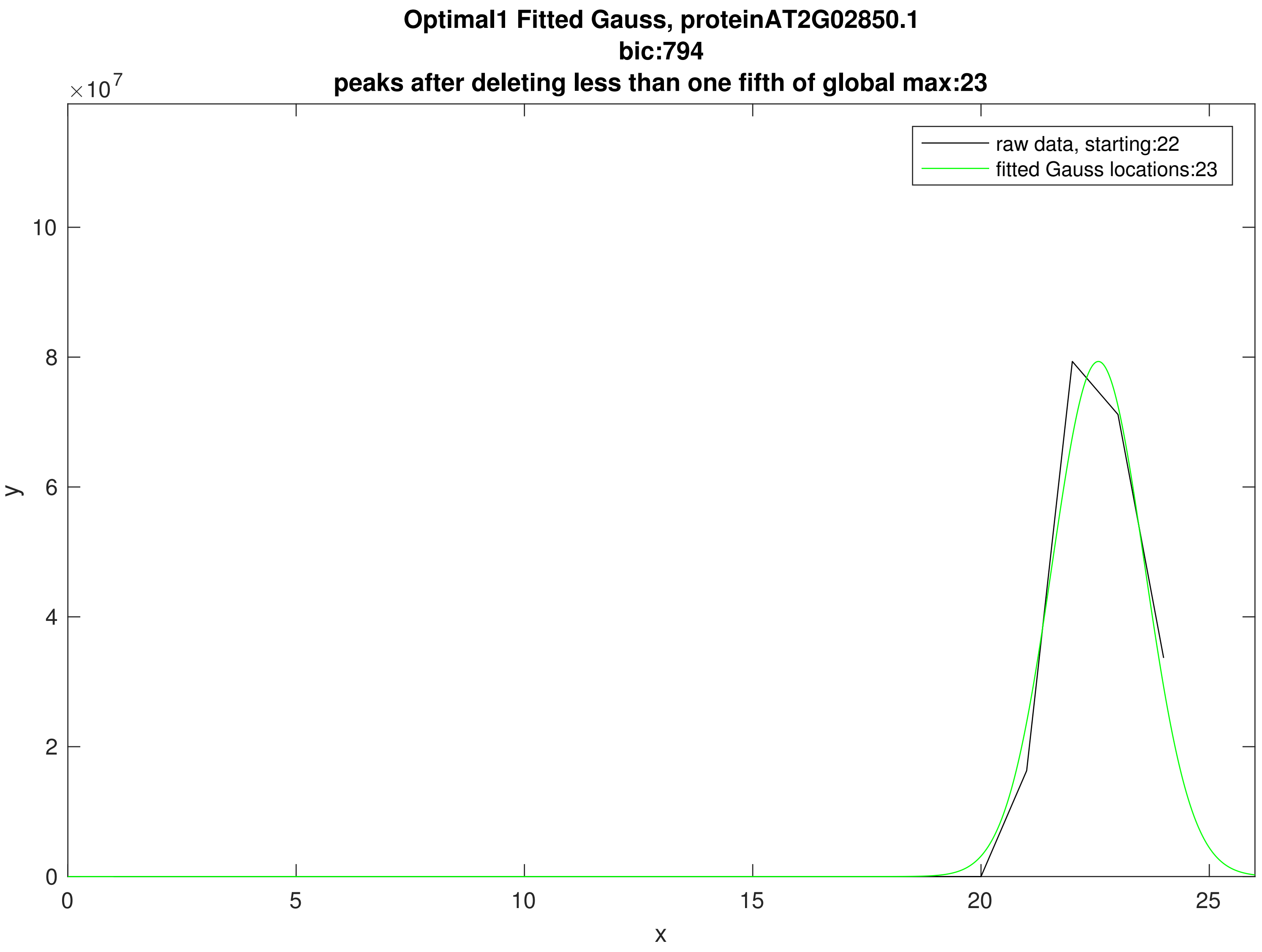

Optimal1 Fitted Gauss, proteinAT2G02850.1
bic:794
peaks after deleting less than one fifth of global max:23
×10^7
raw data, starting:22
fitted Gauss locations:23
y
x
0
2
4
6
8
10
5
15
20
25

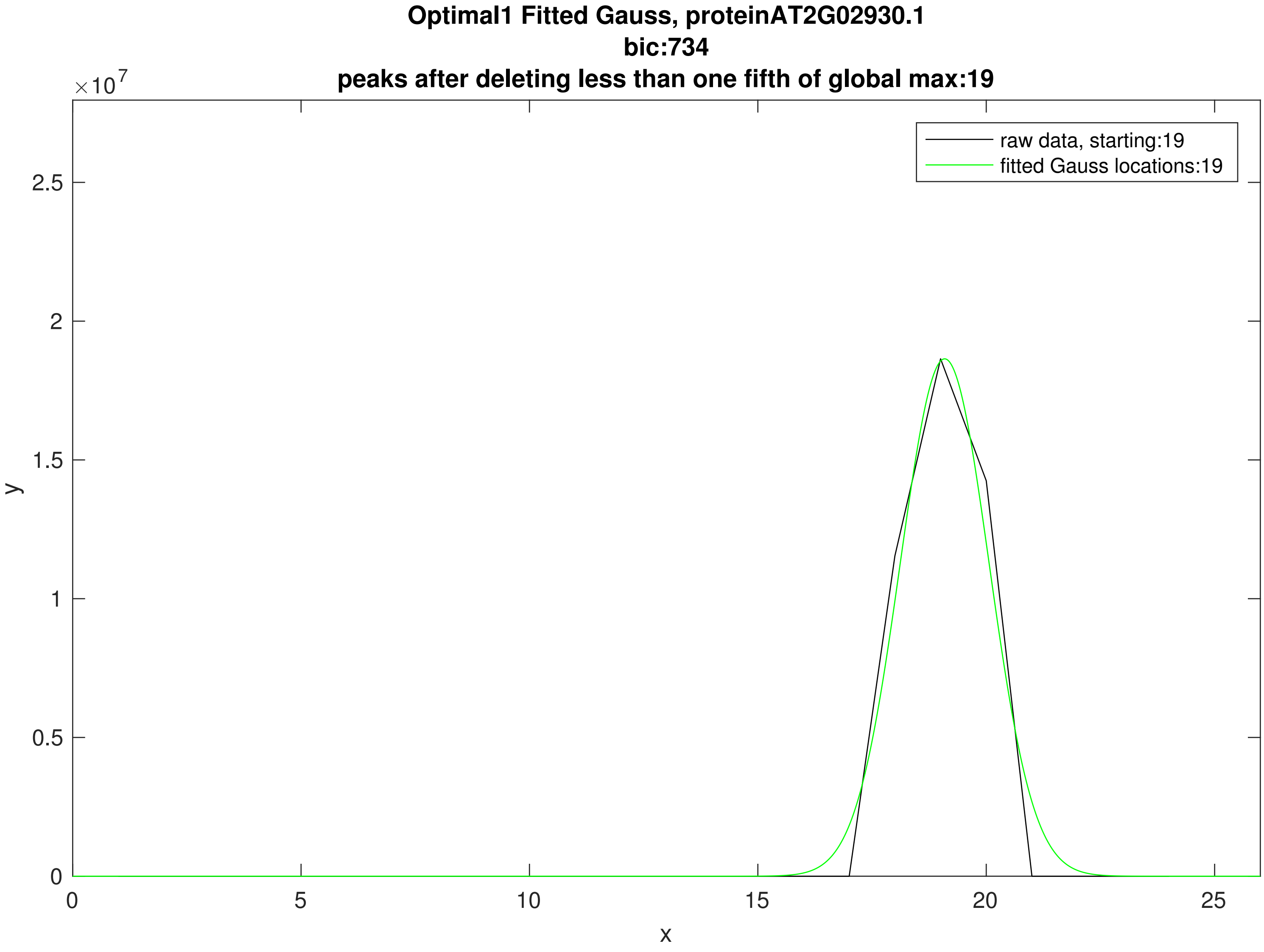

Optimal1 Fitted Gauss, proteinAT2G02930.1
bic:734
peaks after deleting less than one fifth of global max:19
×10^7
raw data, starting:19
fitted Gauss locations:19
2.5
2
1.5
1
0.5
0
y
0
5
10
15
20
25
x

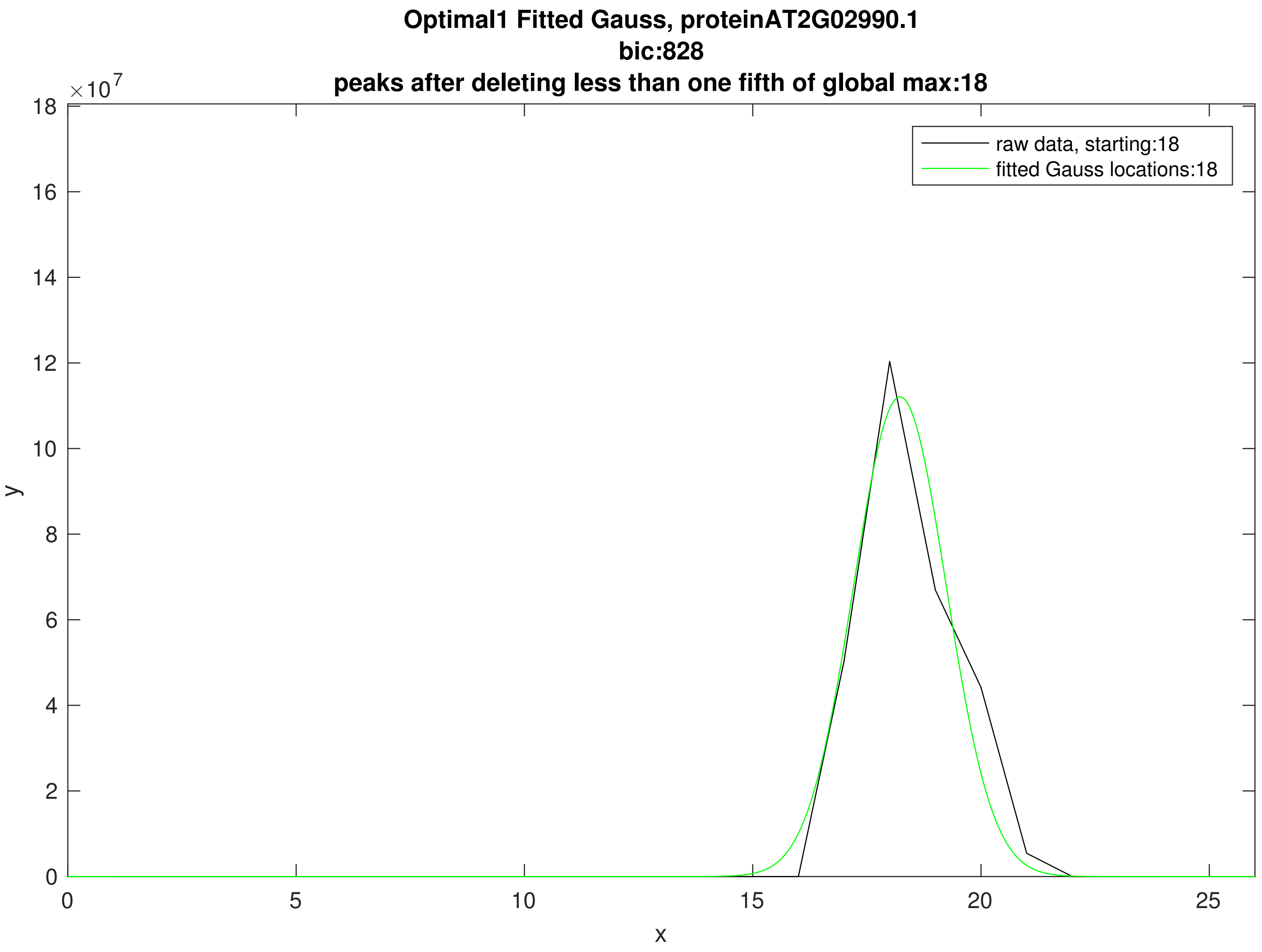
Optimal1 Fitted Gauss, proteinAT2G02990.1
bic:828
peaks after deleting less than one fifth of global max:18
$\times 10^7$
raw data, starting:18
fitted Gauss locations:18
y
x
0
2
4
6
8
10
12
14
16
18
0
5
10
15
20
25

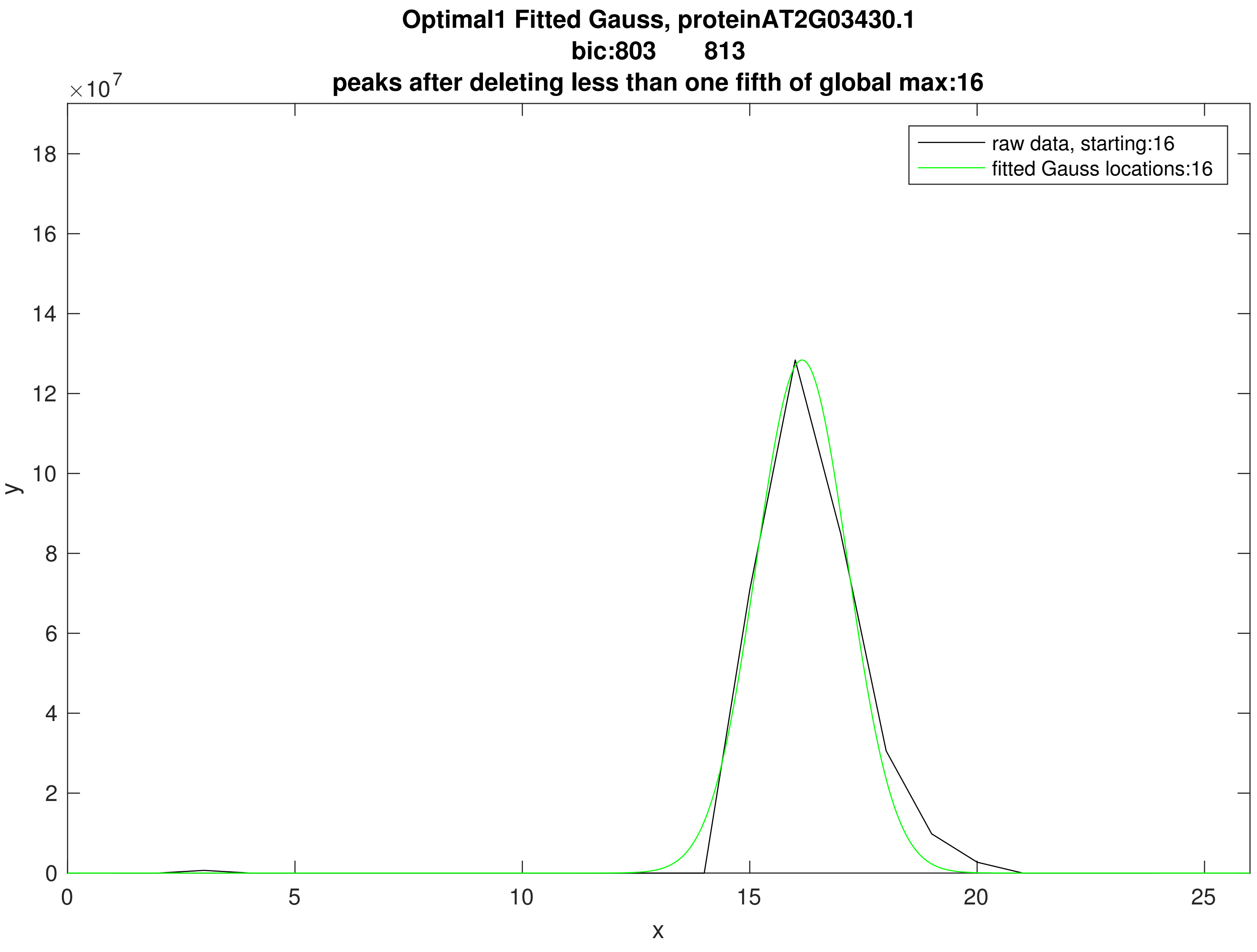
Optimal1 Fitted Gauss, proteinAT2G03430.1
bic:803 813
peaks after deleting less than one fifth of global max:16
×10^7
raw data, starting:16
fitted Gauss locations:16
y
x
0
2
4
6
8
10
12
14
16
18
0
5
10
15
20
25

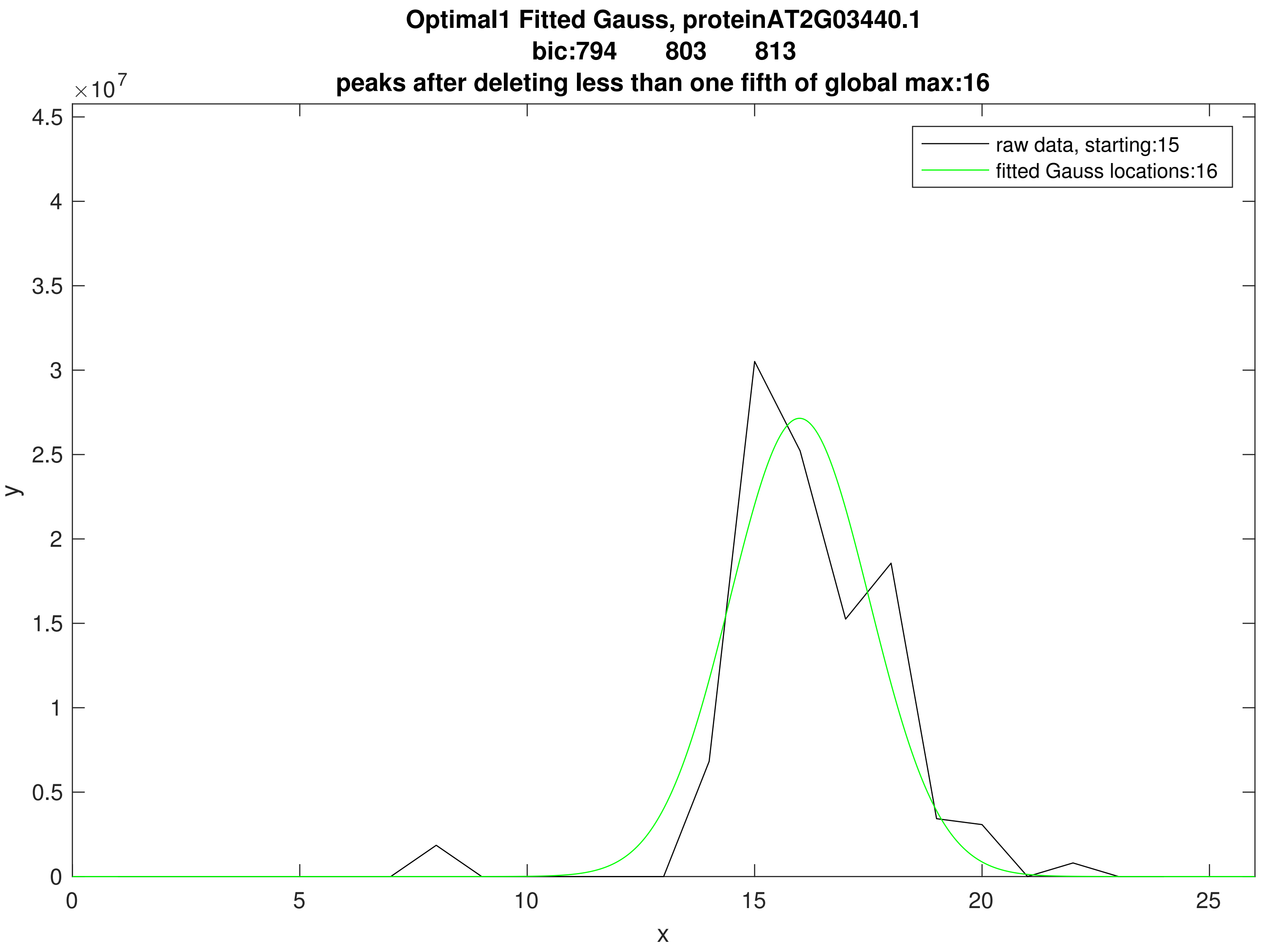
Optimal1 Fitted Gauss, proteinAT2G03440.1
bic:794 803 813
peaks after deleting less than one fifth of global max:16
×10^7
raw data, starting:15
fitted Gauss locations:16
y
x
0
0.5
1
1.5
2
2.5
3
3.5
4
4.5
5
10
15
20
25

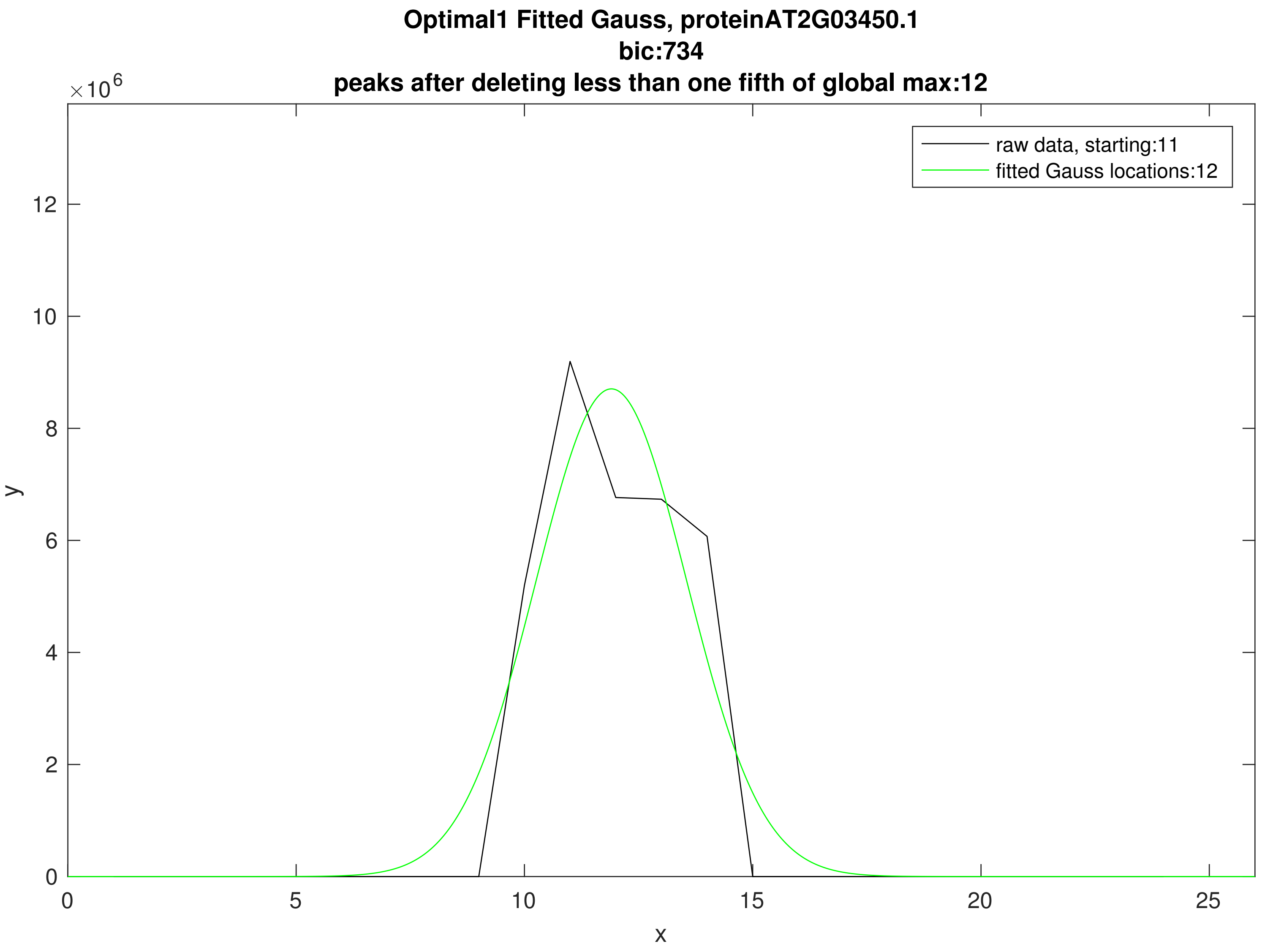
Optimal1 Fitted Gauss, proteinAT2G03450.1
bic:734
peaks after deleting less than one fifth of global max:12
×10^6
raw data, starting:11
fitted Gauss locations:12
y
x
0
2
4
6
8
10
12
0
5
10
15
20
25

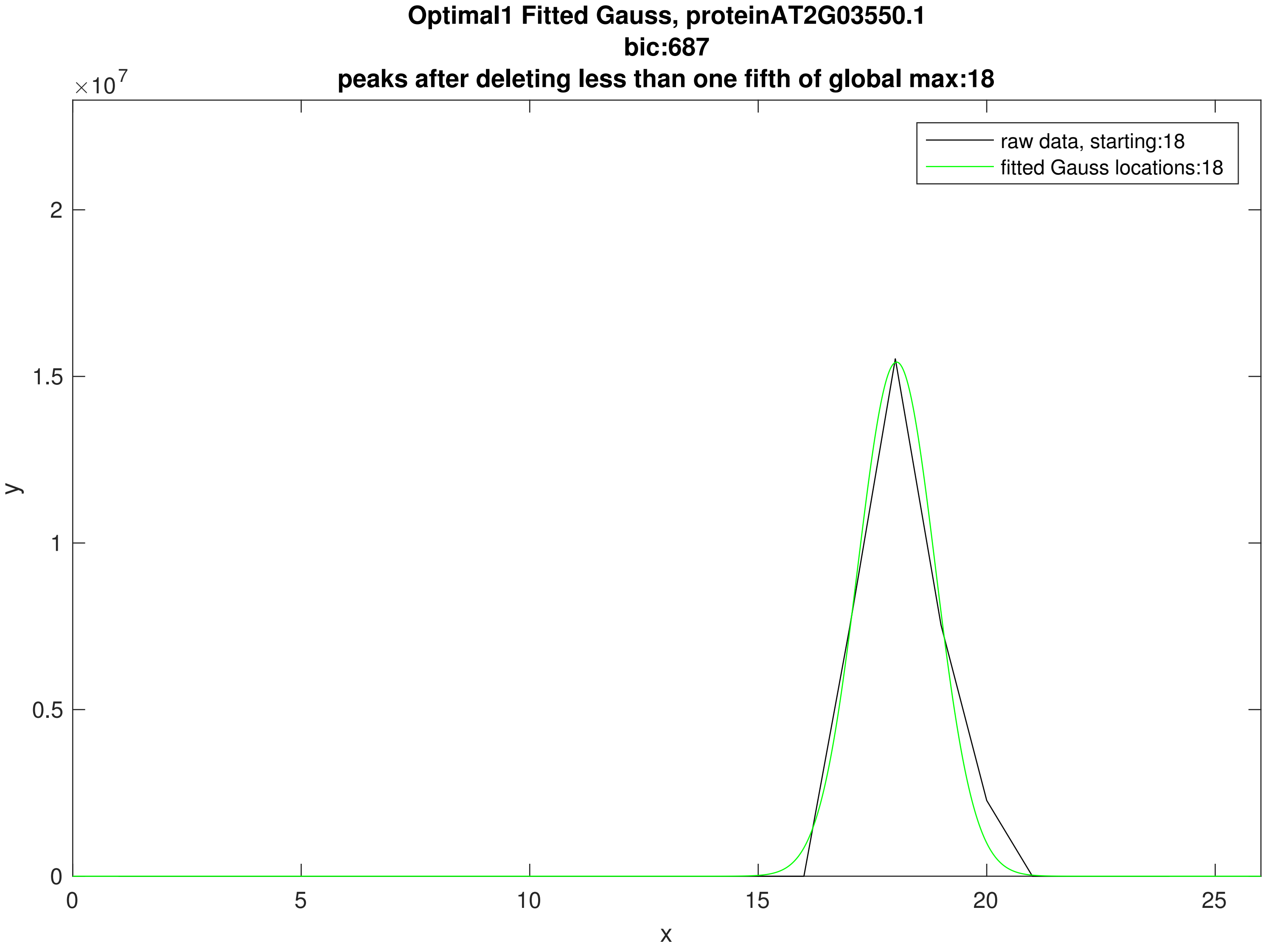
Optimal1 Fitted Gauss, proteinAT2G03550.1
bic:687
peaks after deleting less than one fifth of global max:18
×10^7
raw data, starting:18
fitted Gauss locations:18
y
x
0
0.5
1
1.5
2
0
5
10
15
20
25

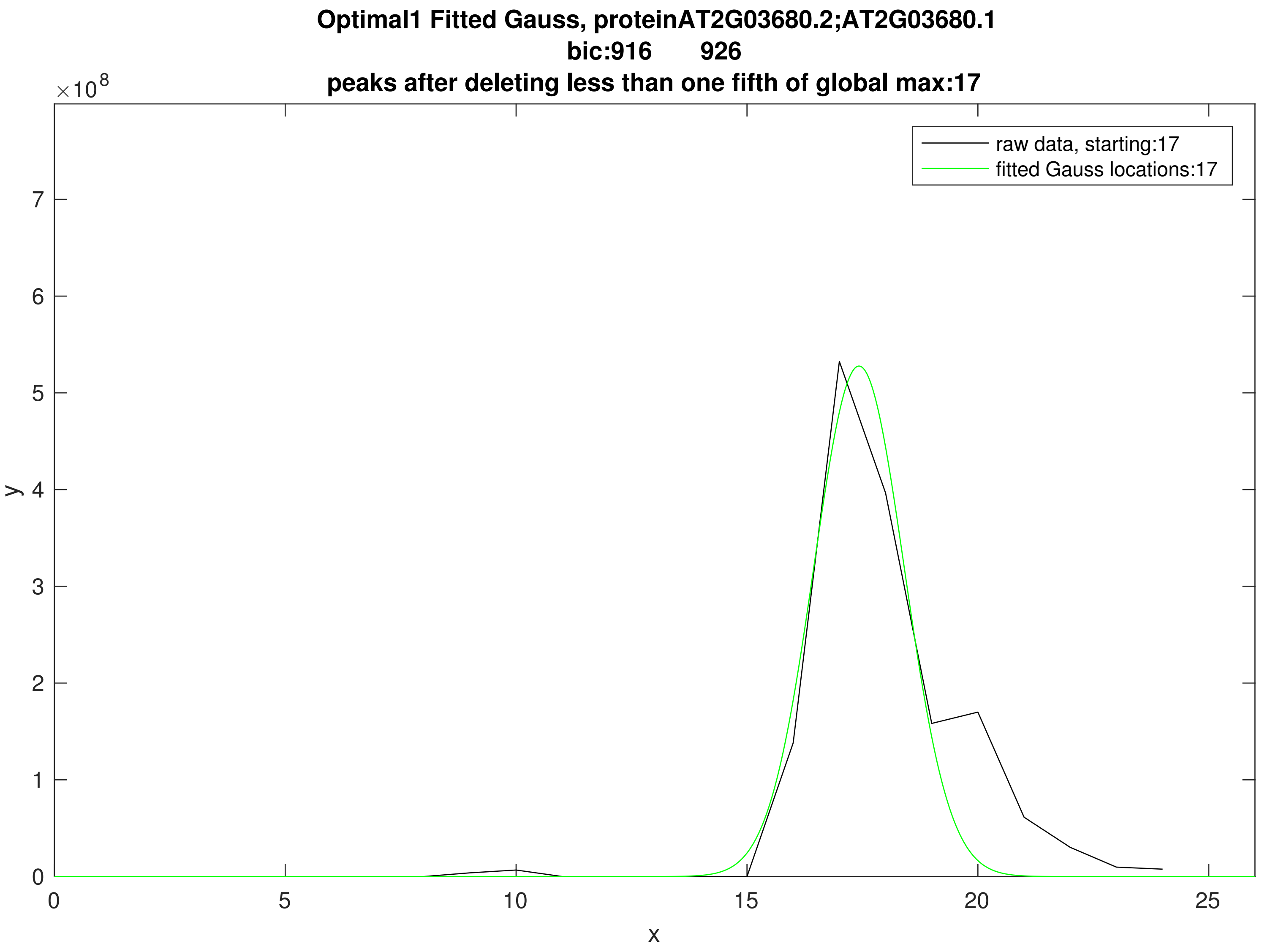
Optimal1 Fitted Gauss, proteinAT2G03680.2;AT2G03680.1
bic:916 926
peaks after deleting less than one fifth of global max:17
×10^8
raw data, starting:17
fitted Gauss locations:17
y
x
0
1
2
3
4
5
6
7
0
5
10
15
20
25

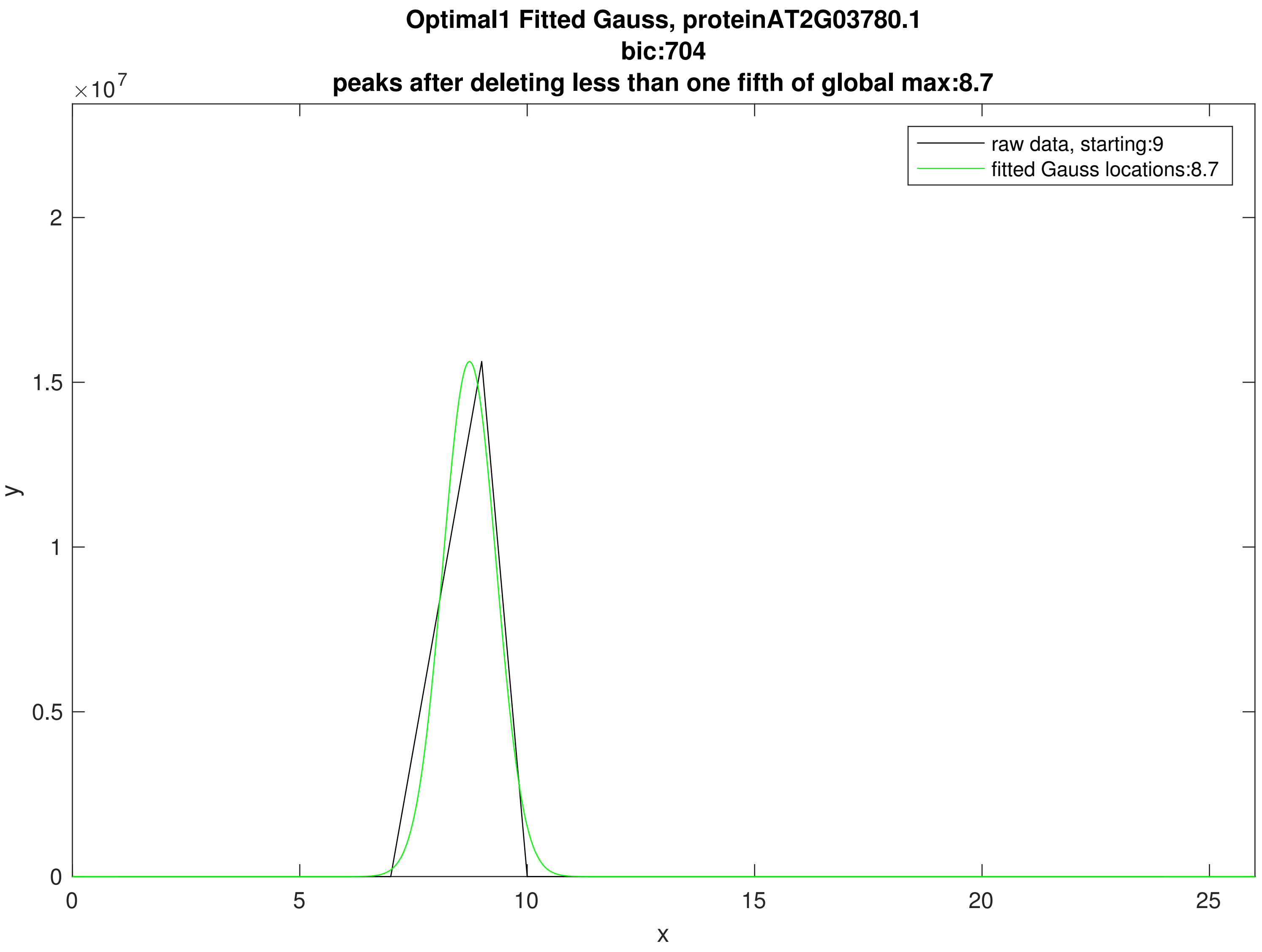

Optimal1 Fitted Gauss, proteinAT2G03780.1
bic:704
peaks after deleting less than one fifth of global max:8.7
×10^7
raw data, starting:9
fitted Gauss locations:8.7
2
1.5
1
0.5
0
y
0
5
10
15
20
25
x

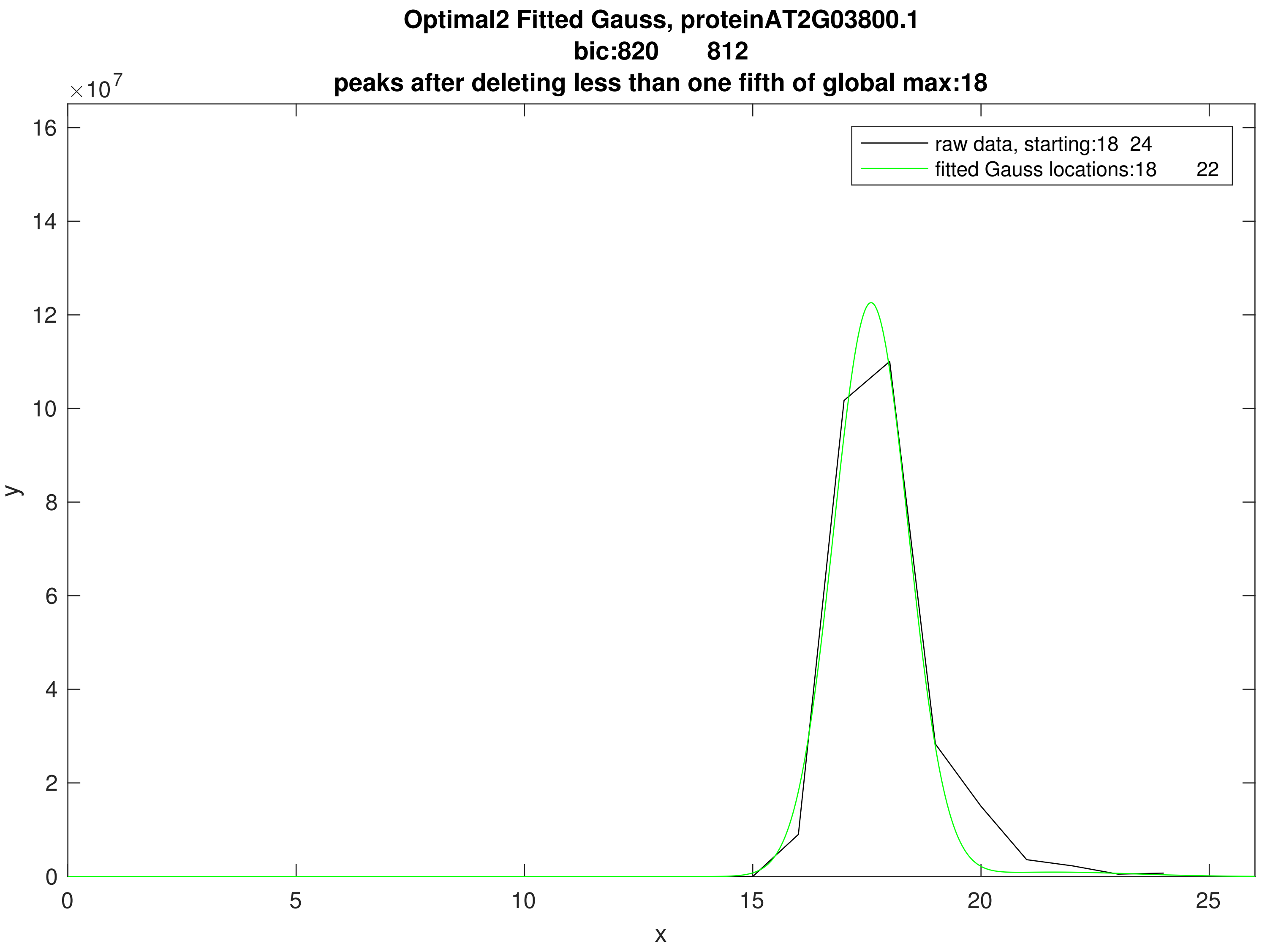

Optimal2 Fitted Gauss, proteinAT2G03800.1
bic:820 812
peaks after deleting less than one fifth of global max:18
×10^7
raw data, starting:18 24
fitted Gauss locations:18 22
y
x
0
2
4
6
8
10
12
14
16
5
15
20
25

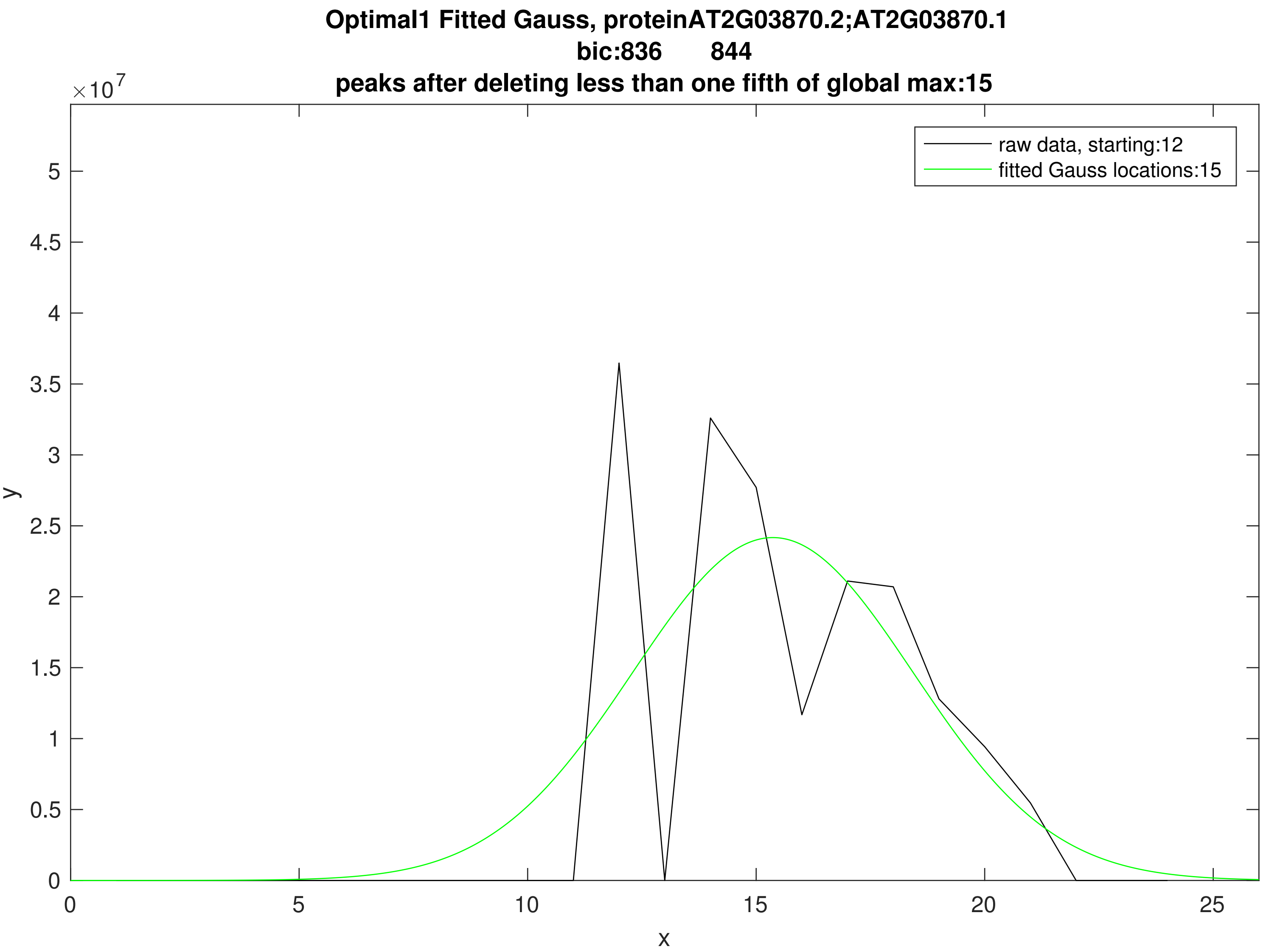

Optimal1 Fitted Gauss, proteinAT2G03870.2;AT2G03870.1
bic:836 844
peaks after deleting less than one fifth of global max:15
×10^7
raw data, starting:12
fitted Gauss locations:15
y
x
0
0.5
1
1.5
2
2.5
3
3.5
4
4.5
5
0
5
10
15
20
25

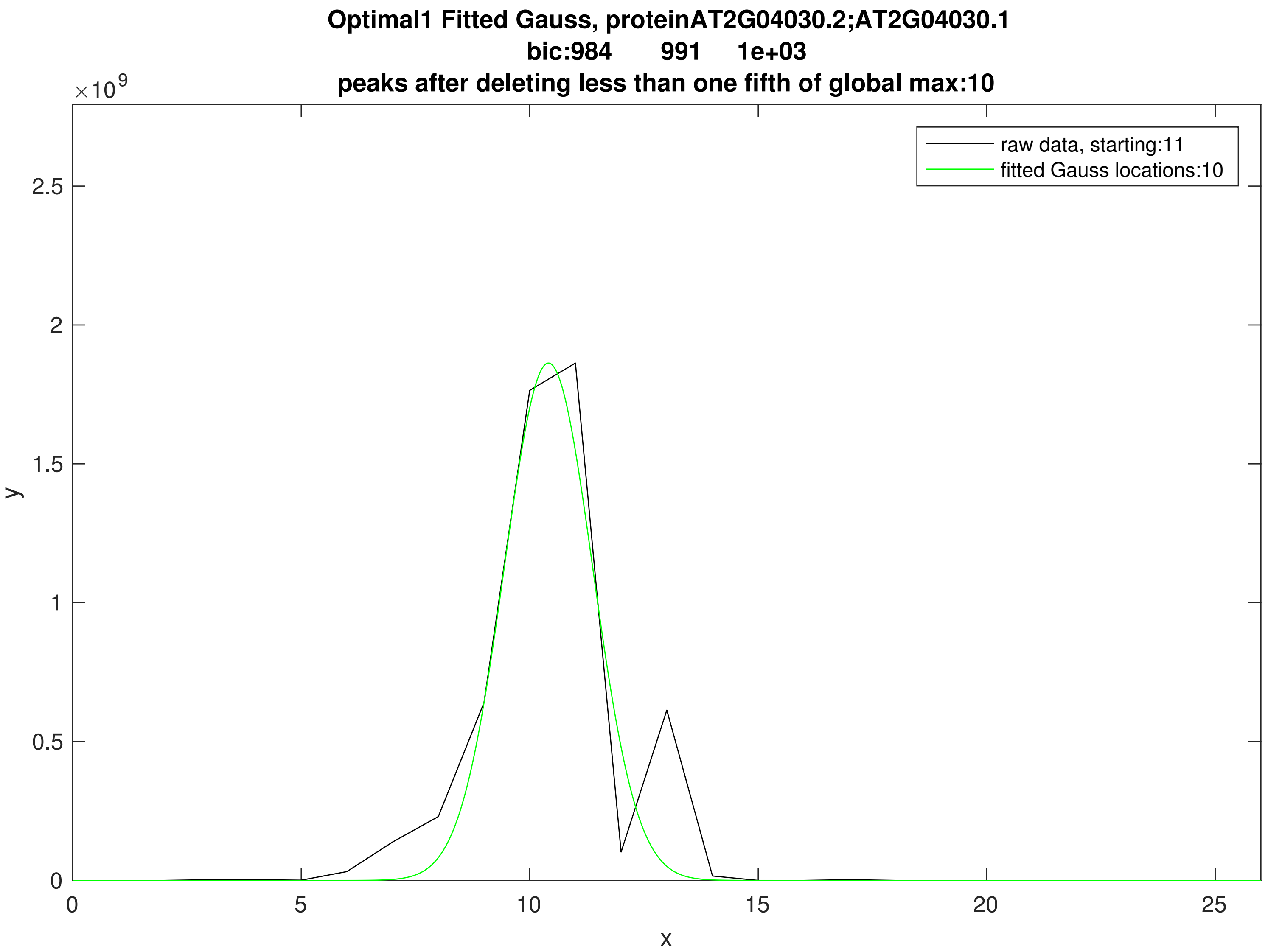

Optimal1 Fitted Gauss, proteinAT2G04030.2;AT2G04030.1
bic:984 991 1e+03
peaks after deleting less than one fifth of global max:10
×10^9
raw data, starting:11
fitted Gauss locations:10
2.5
2
1.5
1
0.5
0
y
0
5
10
15
20
25
x

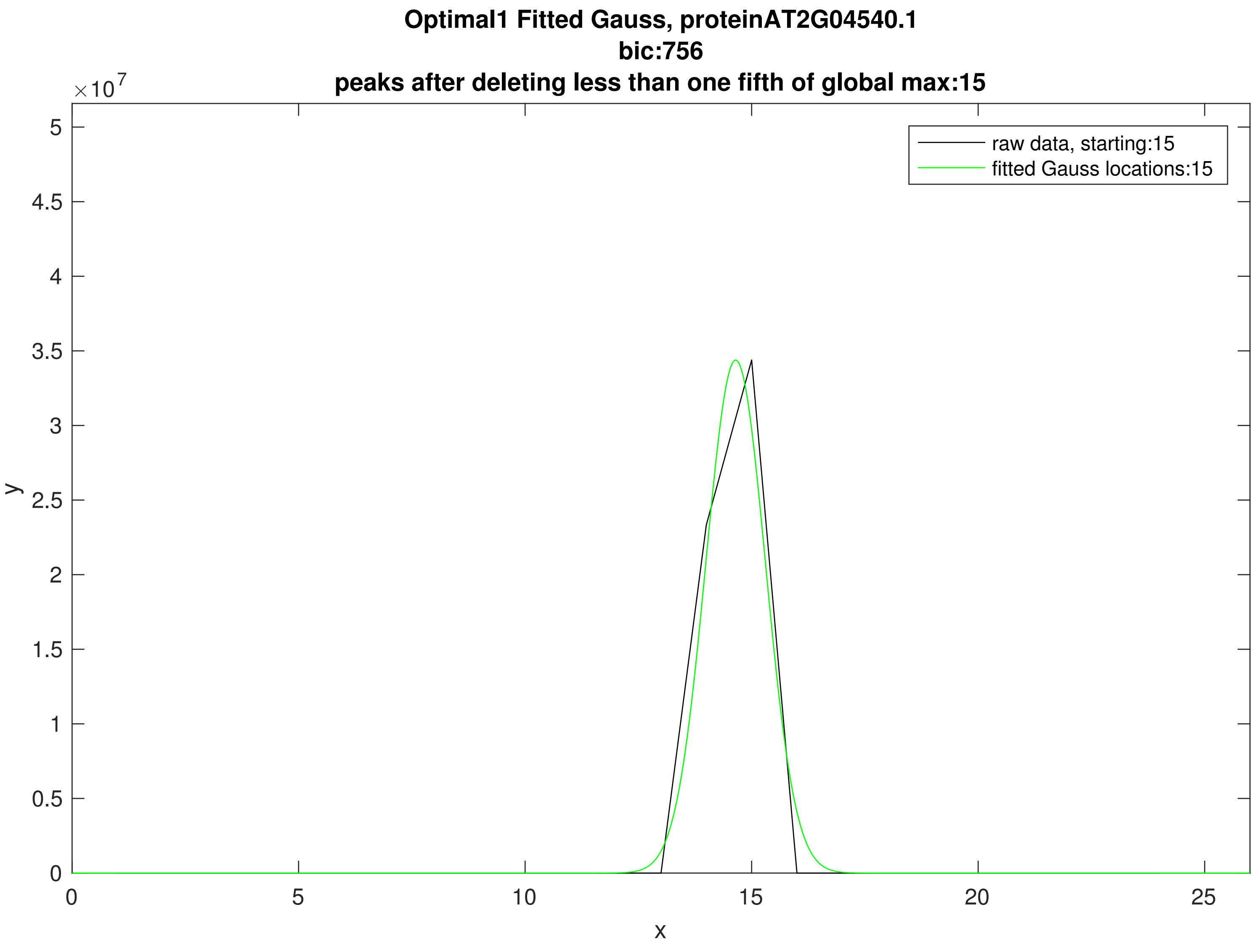
Optimal1 Fitted Gauss, proteinAT2G04540.1
bic:756
peaks after deleting less than one fifth of global max:15
×10^7
5
4.5
4
3.5
3
2.5
2
1.5
1
0.5
0
y
0
5
10
15
20
25
x
raw data, starting:15
fitted Gauss locations:15

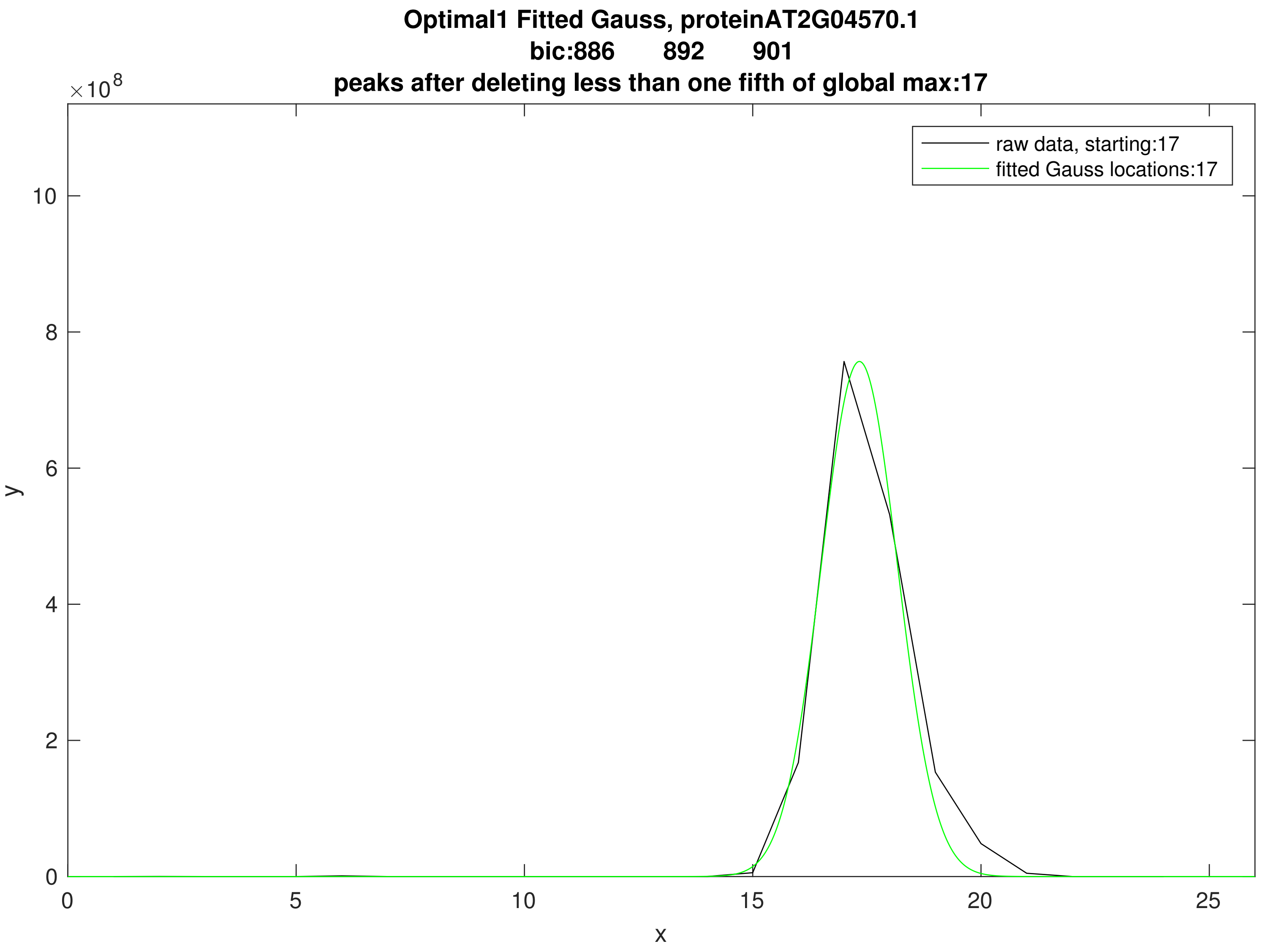
Optimal1 Fitted Gauss, proteinAT2G04570.1
bic:886 892 901
peaks after deleting less than one fifth of global max:17
×10^8
raw data, starting:17
fitted Gauss locations:17
10
8
6
4
2
0
y
0
5
10
15
20
25
x

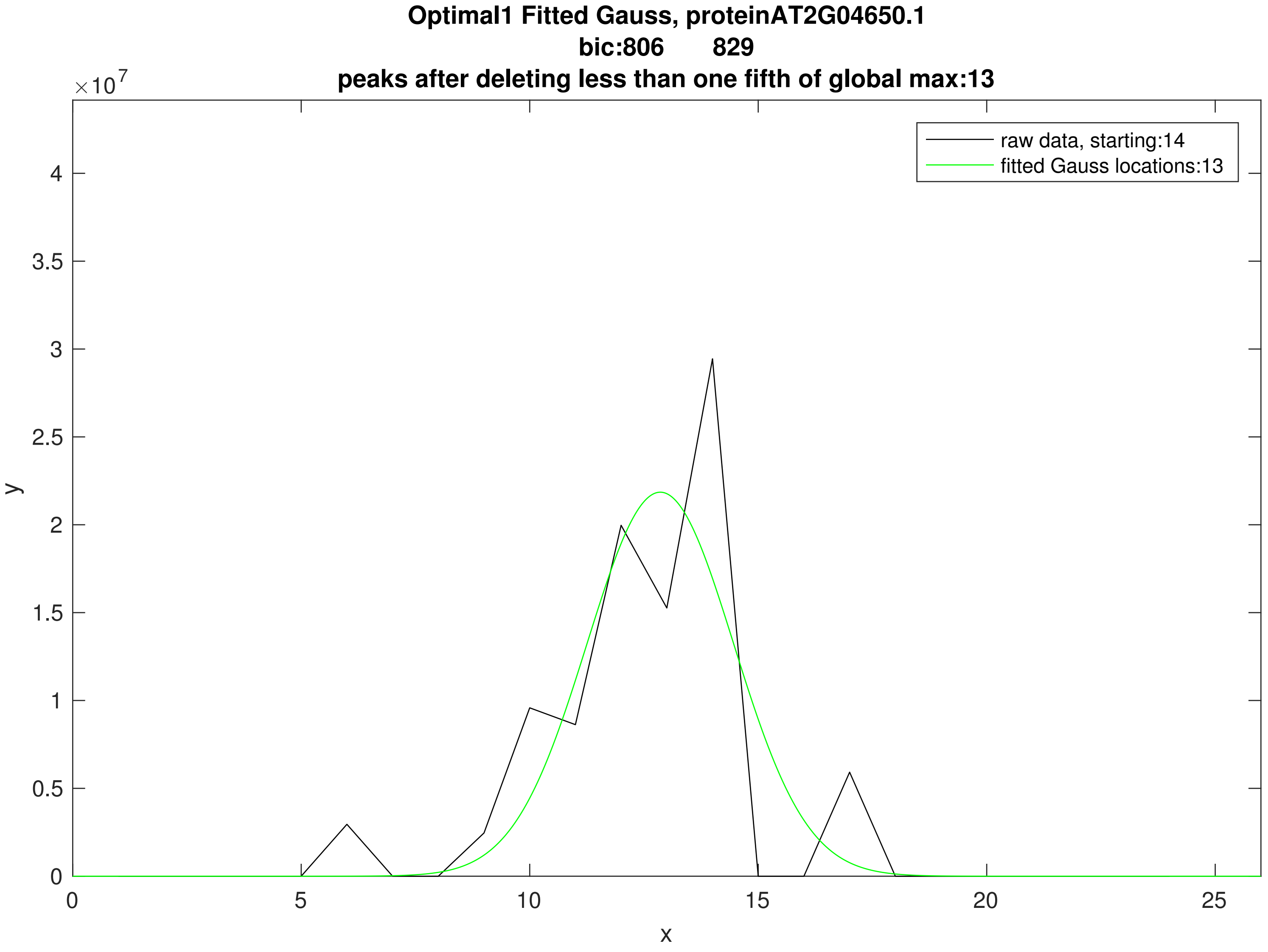

Optimal1 Fitted Gauss, proteinAT2G04650.1
bic:806 829
peaks after deleting less than one fifth of global max:13
×10^7
raw data, starting:14
fitted Gauss locations:13
y
x
0
0.5
1
1.5
2
2.5
3
3.5
4
5
10
15
20
25

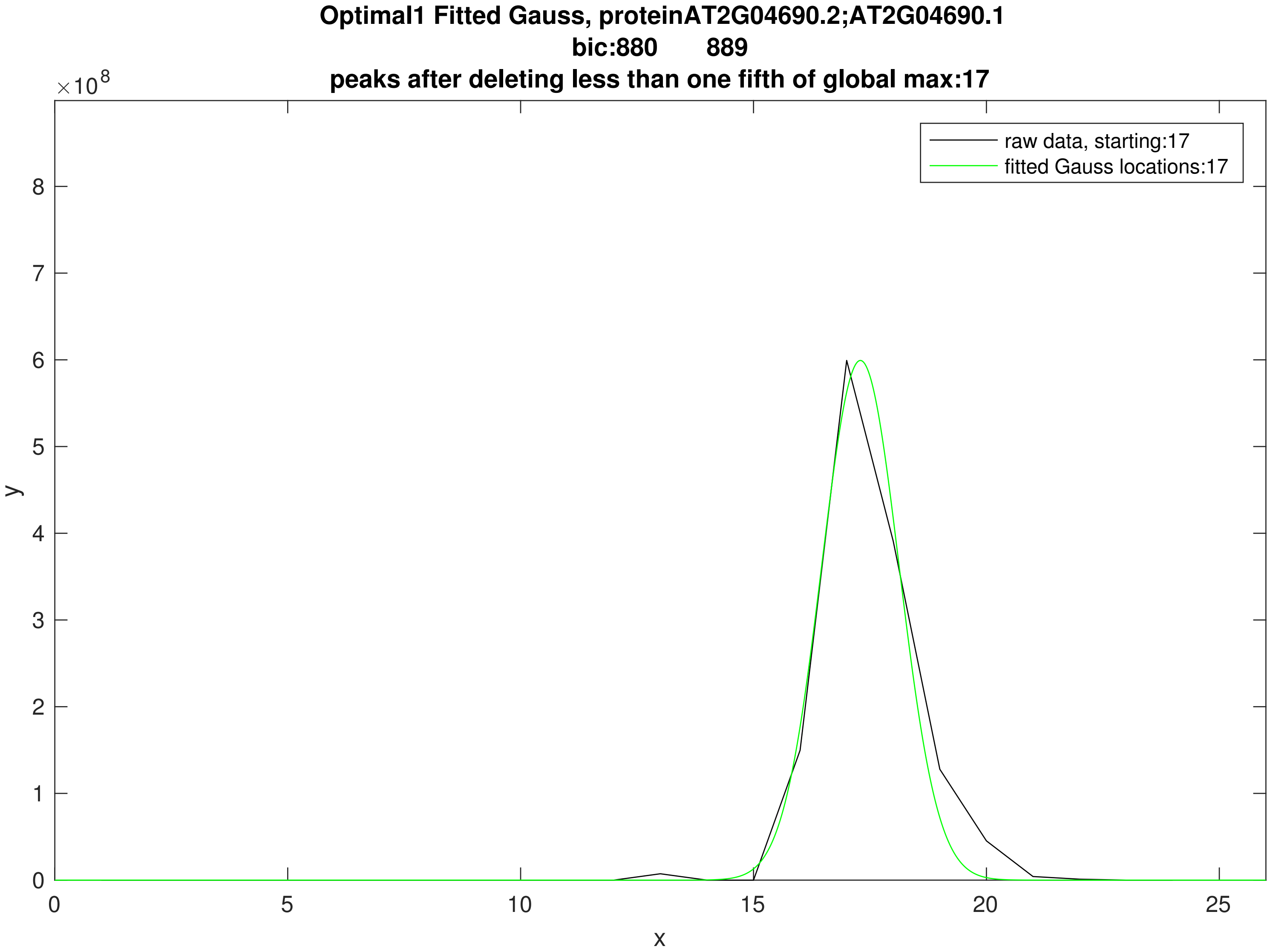
Optimal1 Fitted Gauss, proteinAT2G04690.2;AT2G04690.1
bic:880 889
peaks after deleting less than one fifth of global max:17
×10^8
raw data, starting:17
fitted Gauss locations:17
y
x
0
1
2
3
4
5
6
7
8
0
5
10
15
20
25

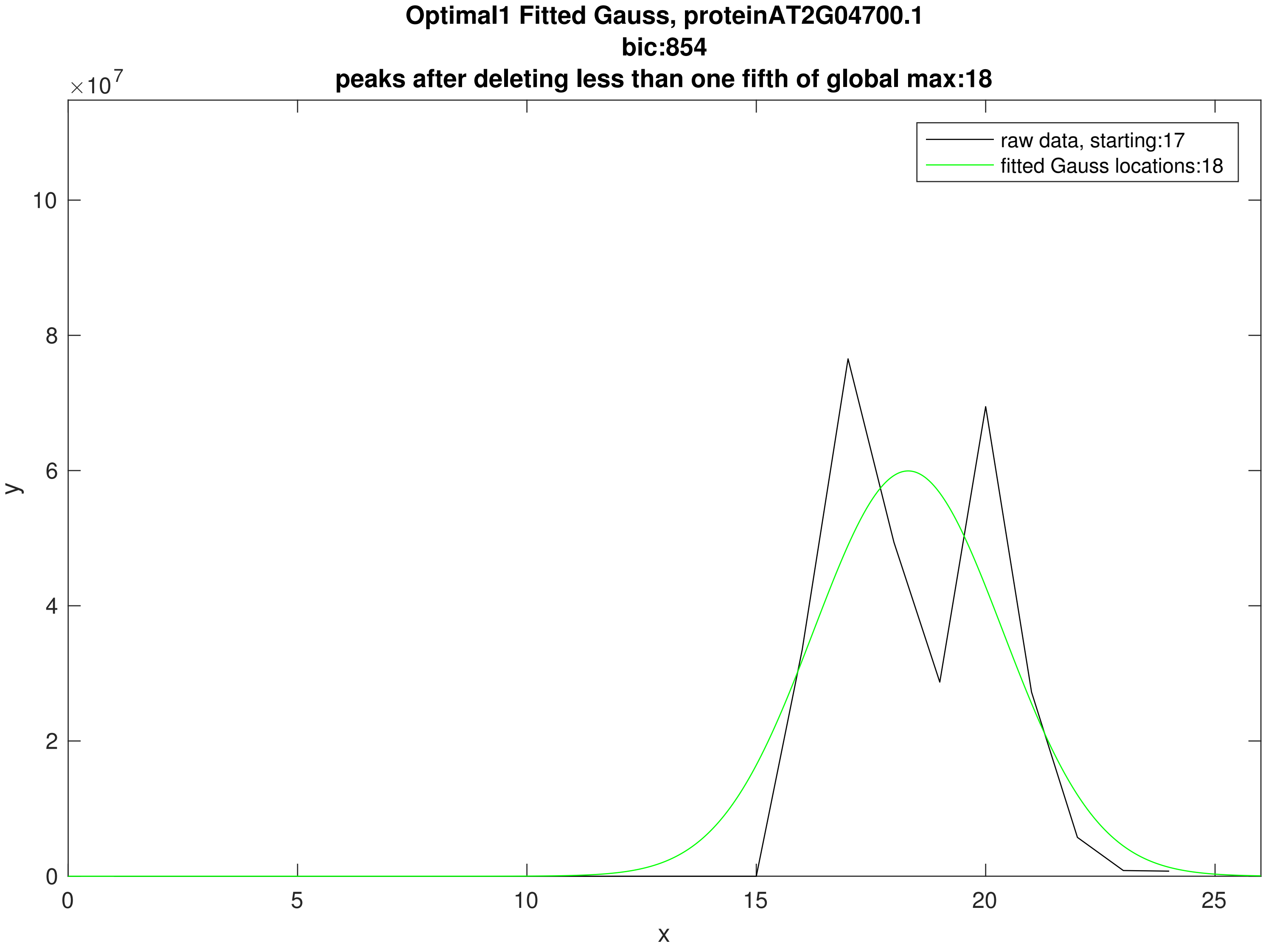
Optimal1 Fitted Gauss, proteinAT2G04700.1
bic:854
peaks after deleting less than one fifth of global max:18
×10^7
raw data, starting:17
fitted Gauss locations:18
y
x
0
2
4
6
8
10
0
5
10
15
20
25

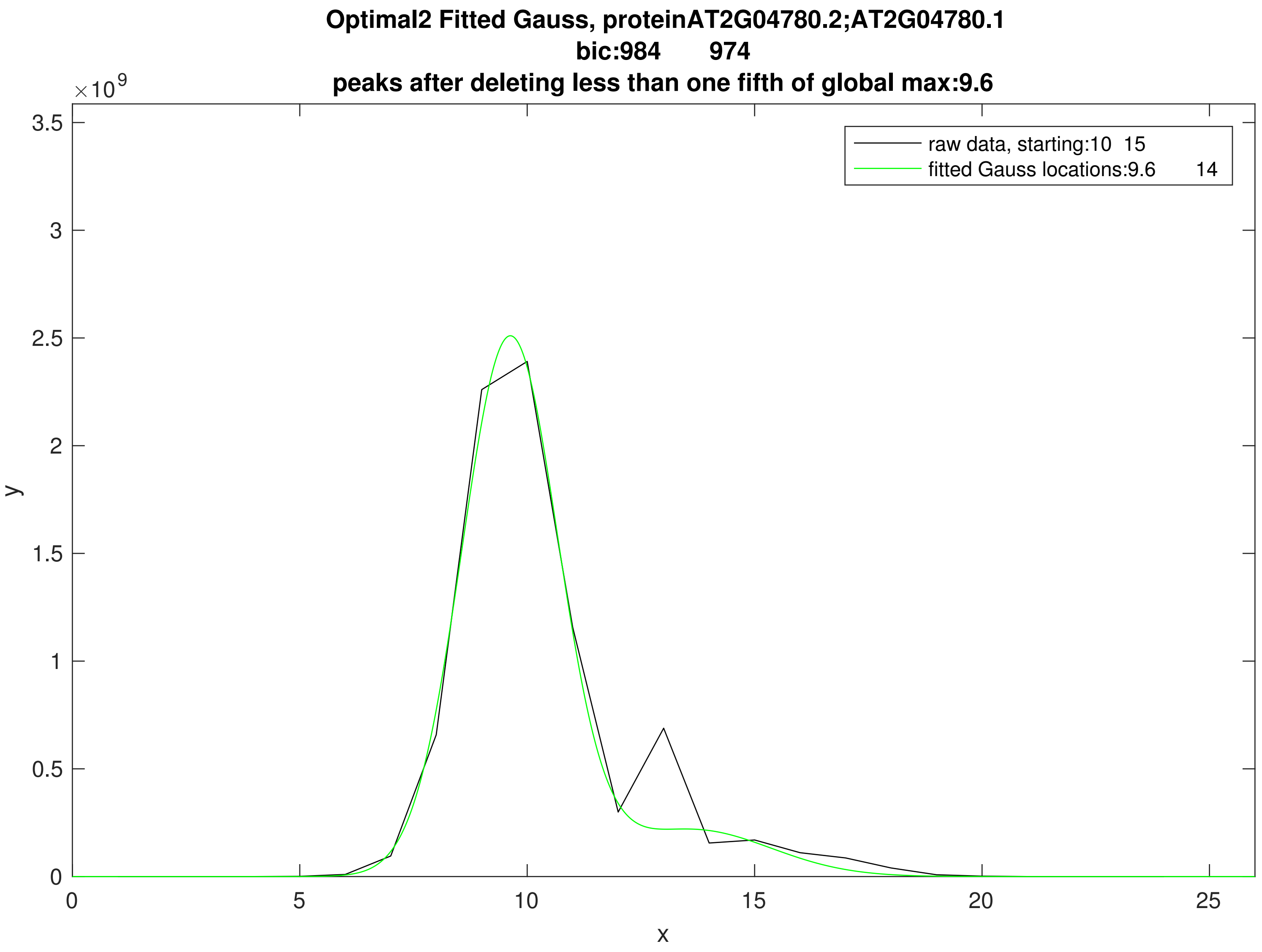
Optimal2 Fitted Gauss, proteinAT2G04780.2;AT2G04780.1
bic:984 974
peaks after deleting less than one fifth of global max:9.6
×10^9
3.5
3
2.5
2
1.5
1
0.5
0
y
0
5
10
15
20
25
x
raw data, starting:10 15
fitted Gauss locations:9.6 14

**Optimal1 Fitted Gauss, proteinAT2G04845.1;AT2G04845.2**
**bic:684**
**peaks after deleting less than one fifth of global max:18**

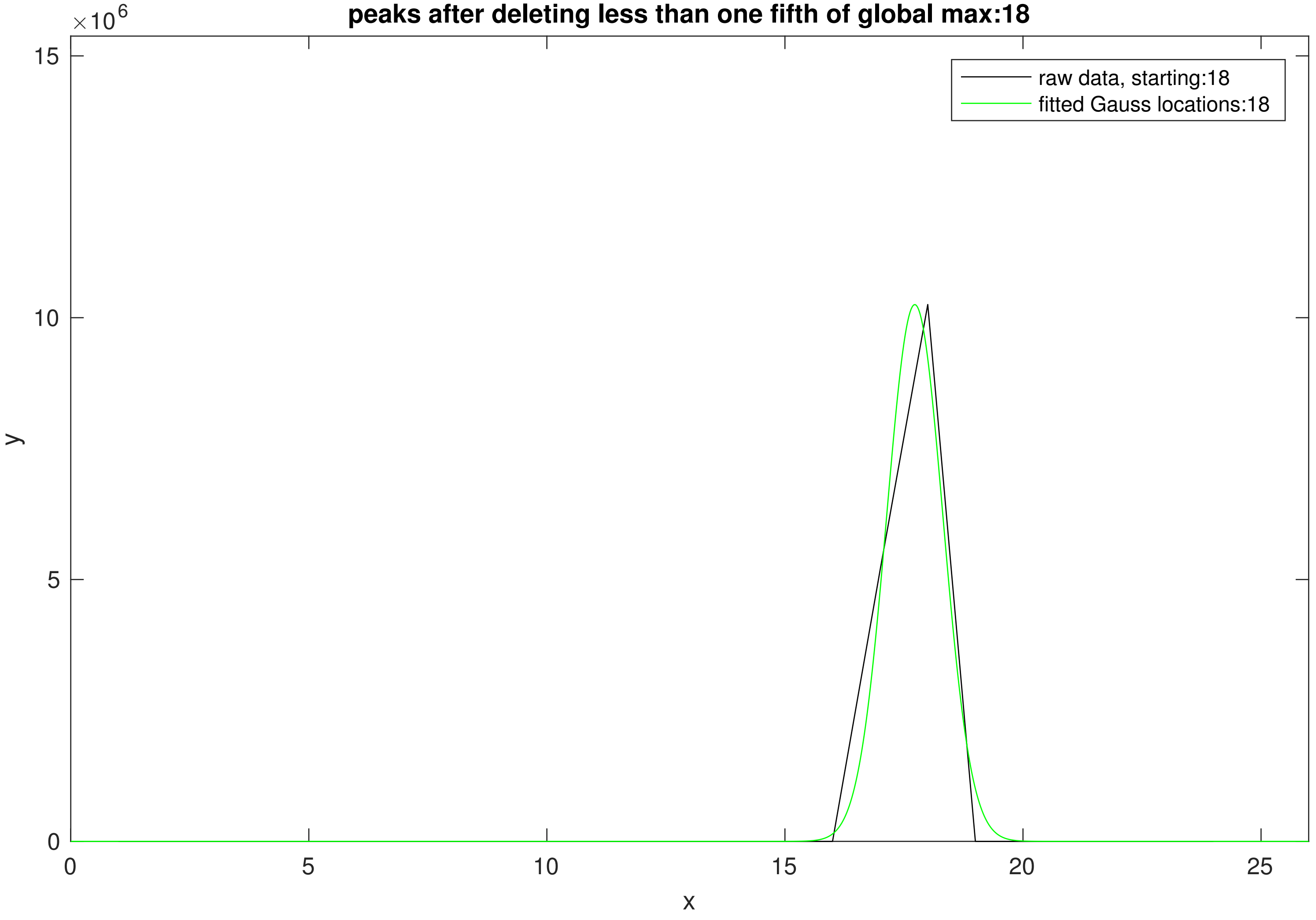

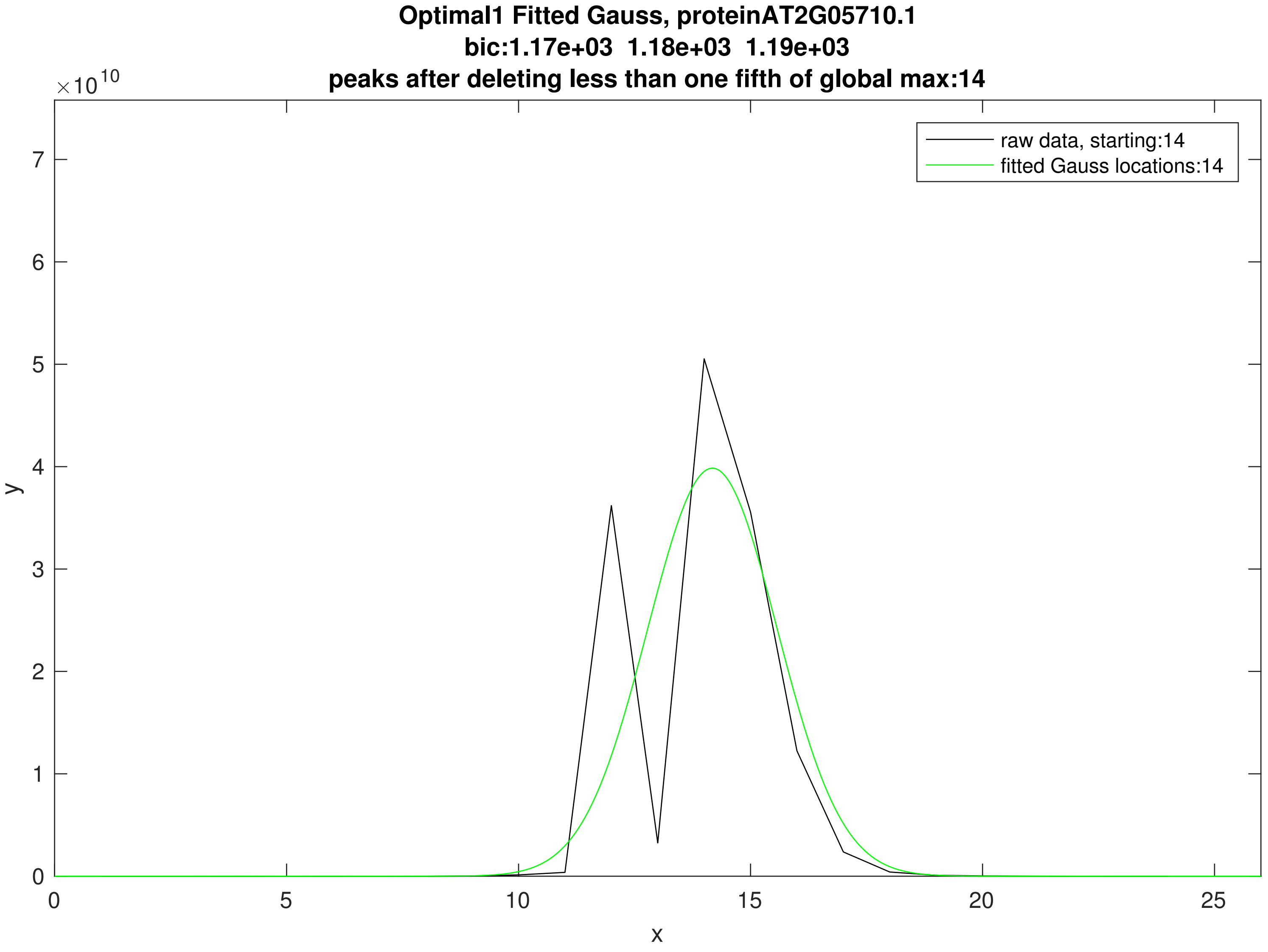

Optimal1 Fitted Gauss, proteinAT2G05710.1
bic:1.17e+03 1.18e+03 1.19e+03
peaks after deleting less than one fifth of global max:14
×10^10
raw data, starting:14
fitted Gauss locations:14
y
x
0
1
2
3
4
5
6
7
0
5
10
15
20
25

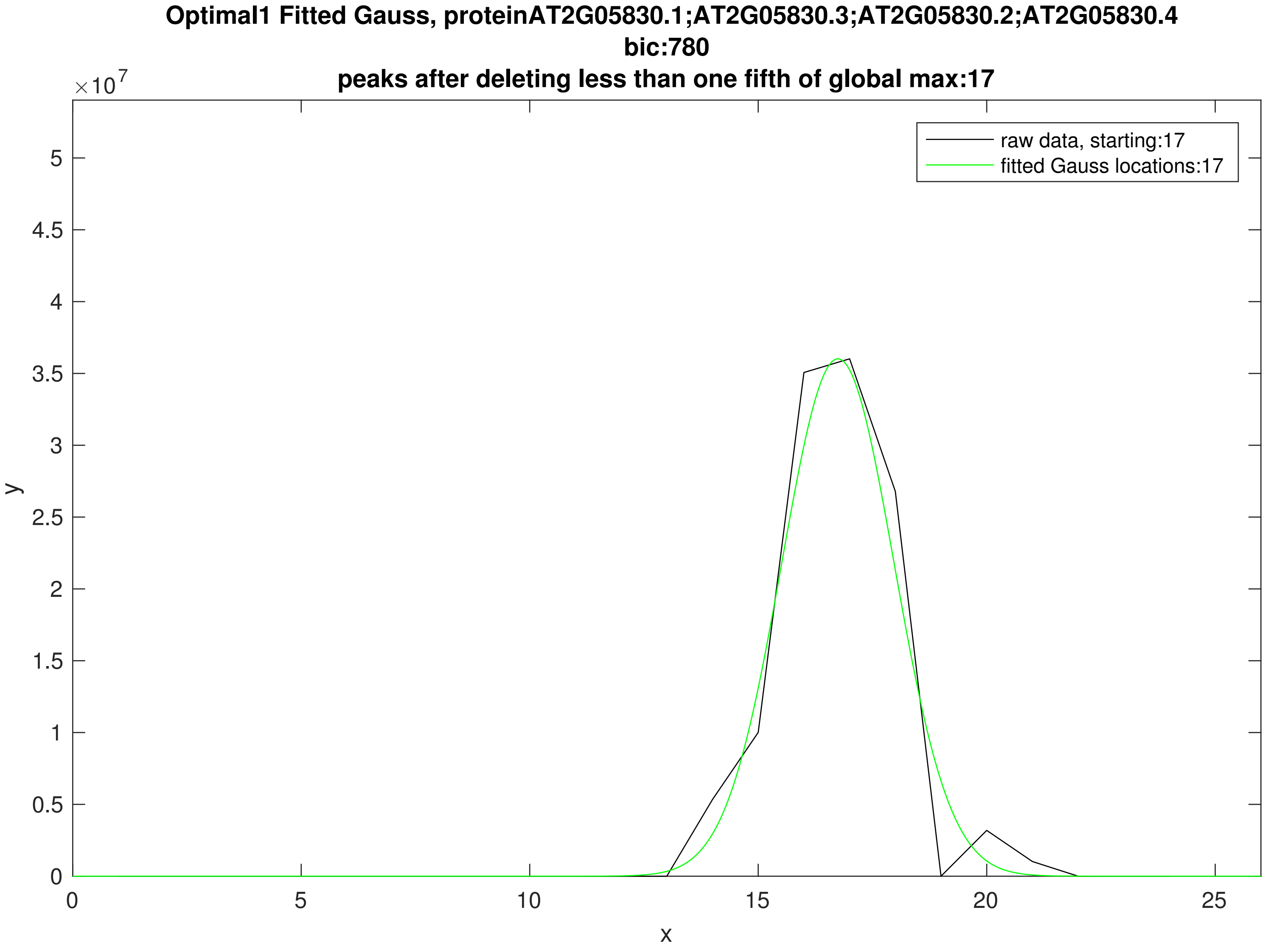

Optimal1 Fitted Gauss, proteinAT2G05830.1;AT2G05830.3;AT2G05830.2;AT2G05830.4
bic:780
peaks after deleting less than one fifth of global max:17
×10^7
raw data, starting:17
fitted Gauss locations:17
y
x
0
0.5
1
1.5
2
2.5
3
3.5
4
4.5
5
0
5
10
15
20
25

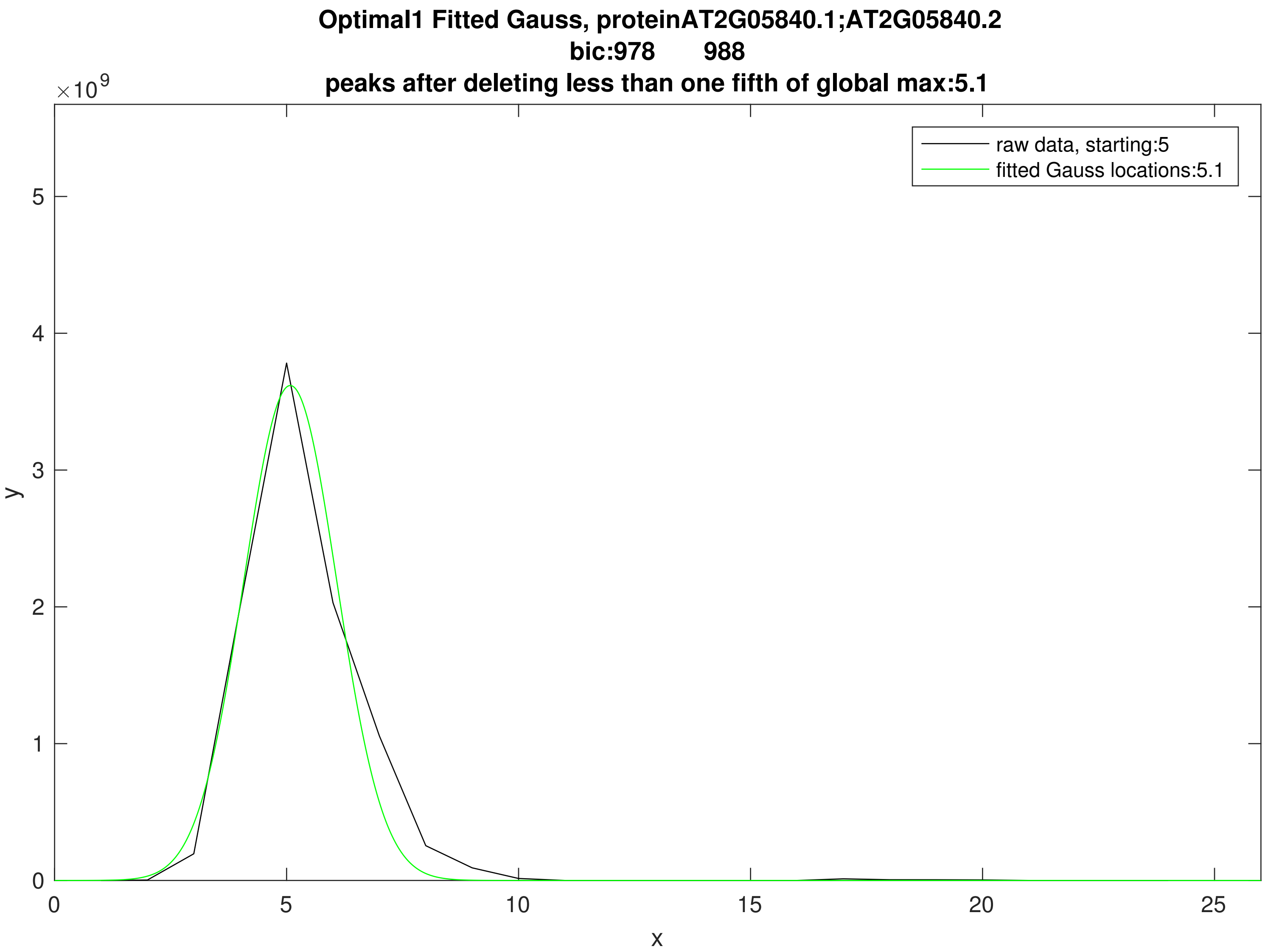

Optimal1 Fitted Gauss, proteinAT2G05840.1;AT2G05840.2
bic:978 988
peaks after deleting less than one fifth of global max:5.1
×10^9
raw data, starting:5
fitted Gauss locations:5.1
y
x
0
1
2
3
4
5
0
5
10
15
20
25

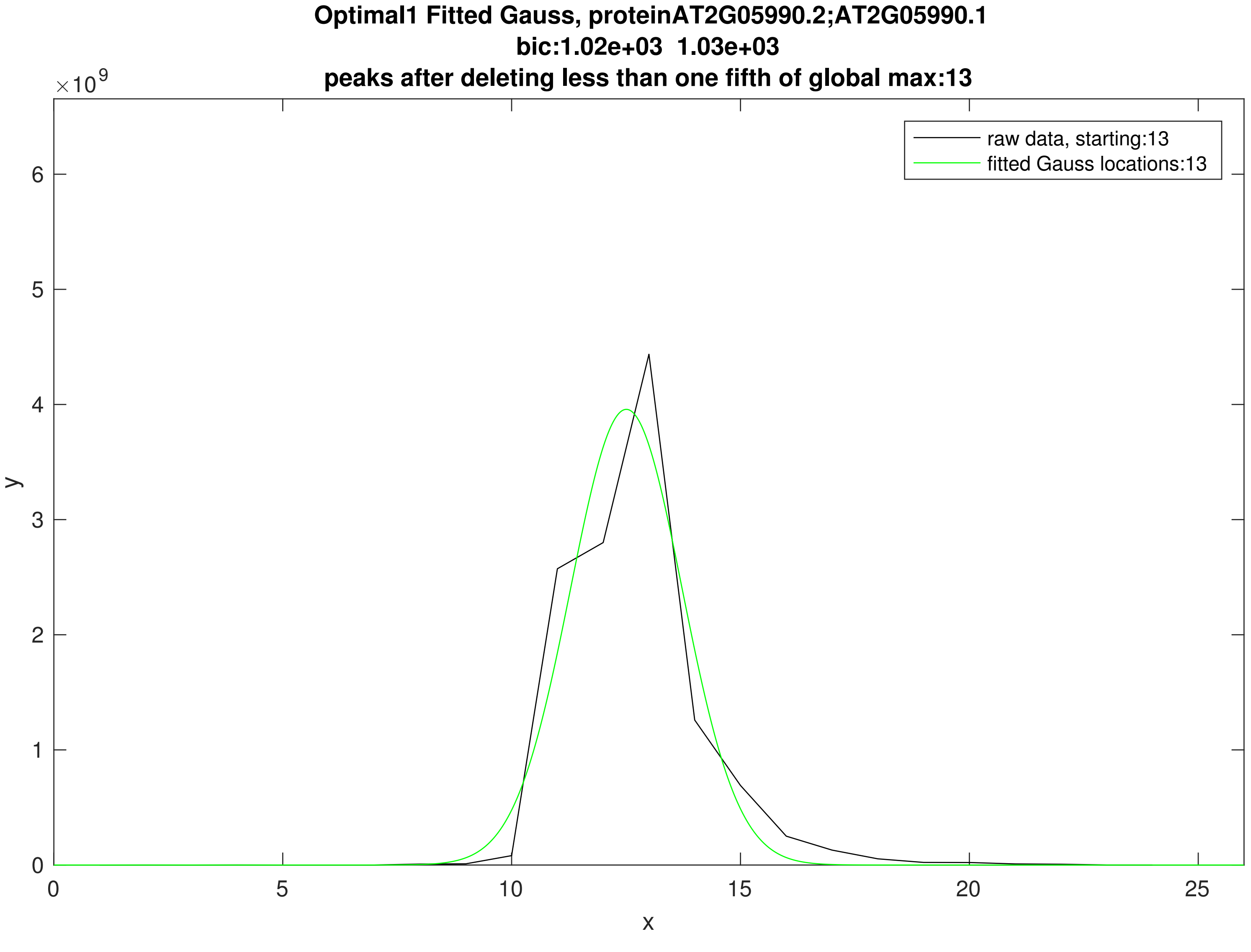
Optimal1 Fitted Gauss, proteinAT2G05990.2;AT2G05990.1
bic:1.02e+03  1.03e+03
peaks after deleting less than one fifth of global max:13
×10^9
raw data, starting:13
fitted Gauss locations:13
y
x
0
1
2
3
4
5
6
0
5
10
15
20
25

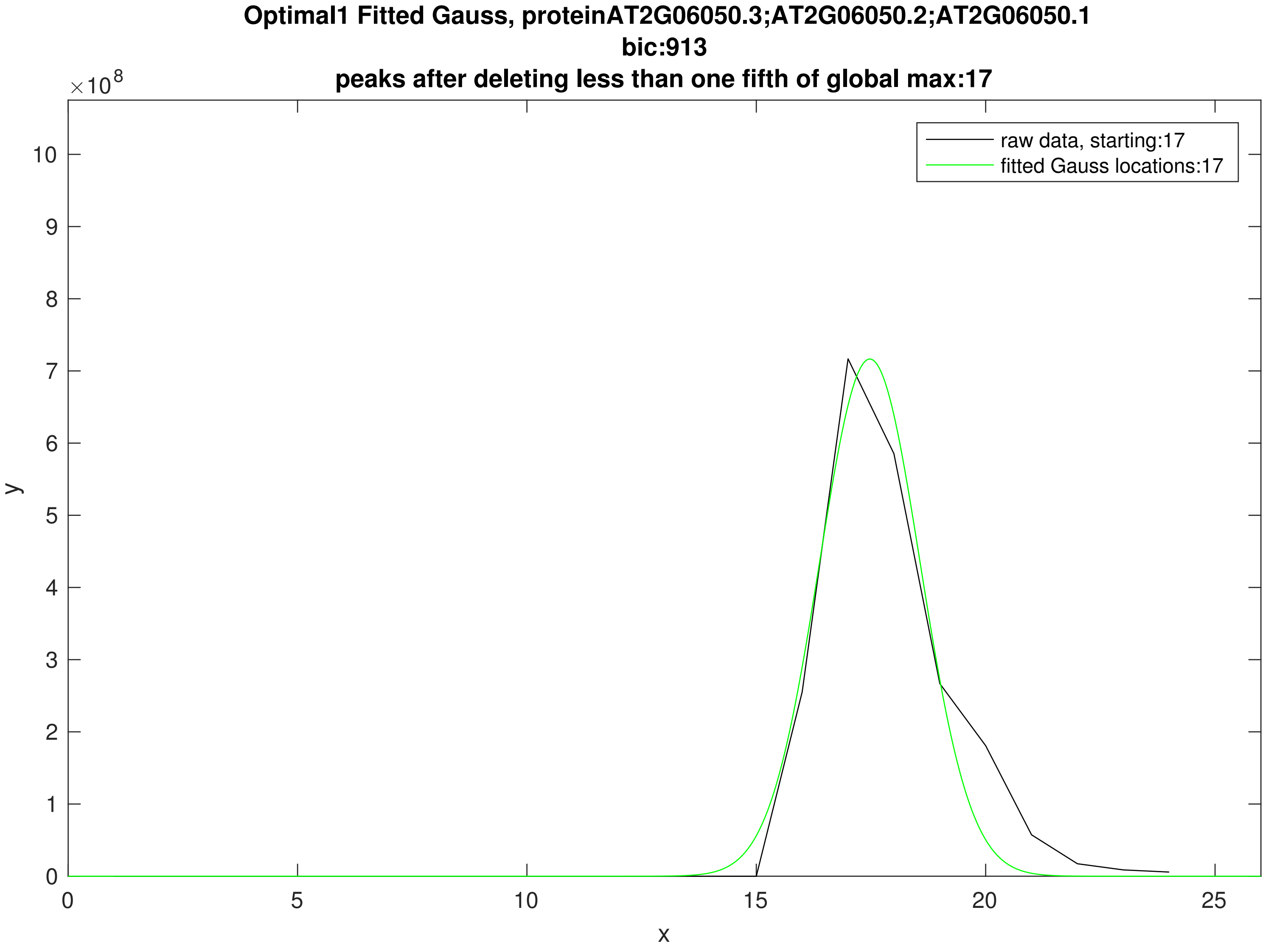

Optimal1 Fitted Gauss, proteinAT2G06050.3;AT2G06050.2;AT2G06050.1
bic:913
peaks after deleting less than one fifth of global max:17
×10^8
raw data, starting:17
fitted Gauss locations:17
y
x
0
1
2
3
4
5
6
7
8
9
10
0
5
10
15
20
25

**Optimal2 Fitted Gauss, proteinAT2G06850.1**
**bic:760 670**
**peaks after deleting less than one fifth of global max:3.4 12**

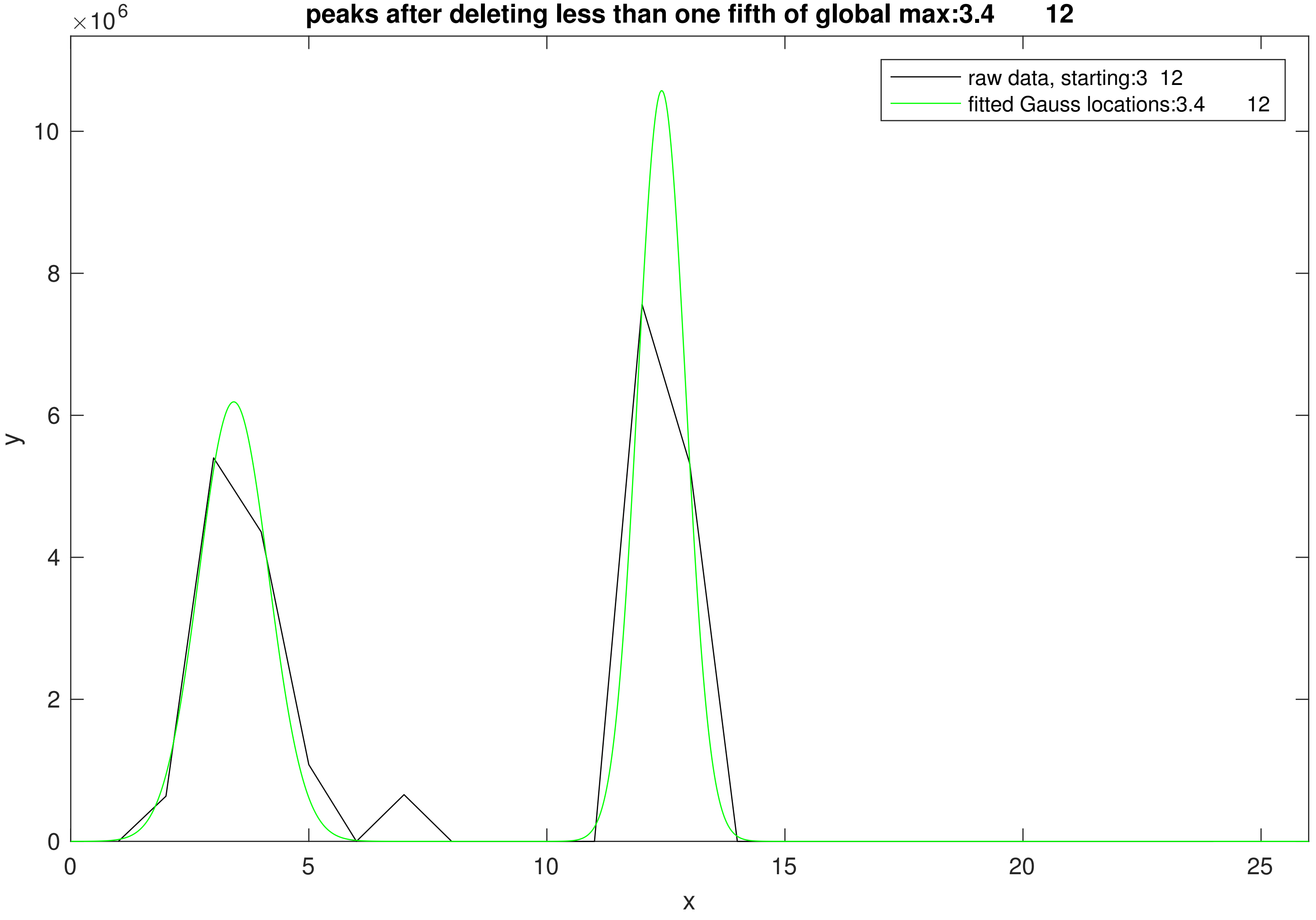

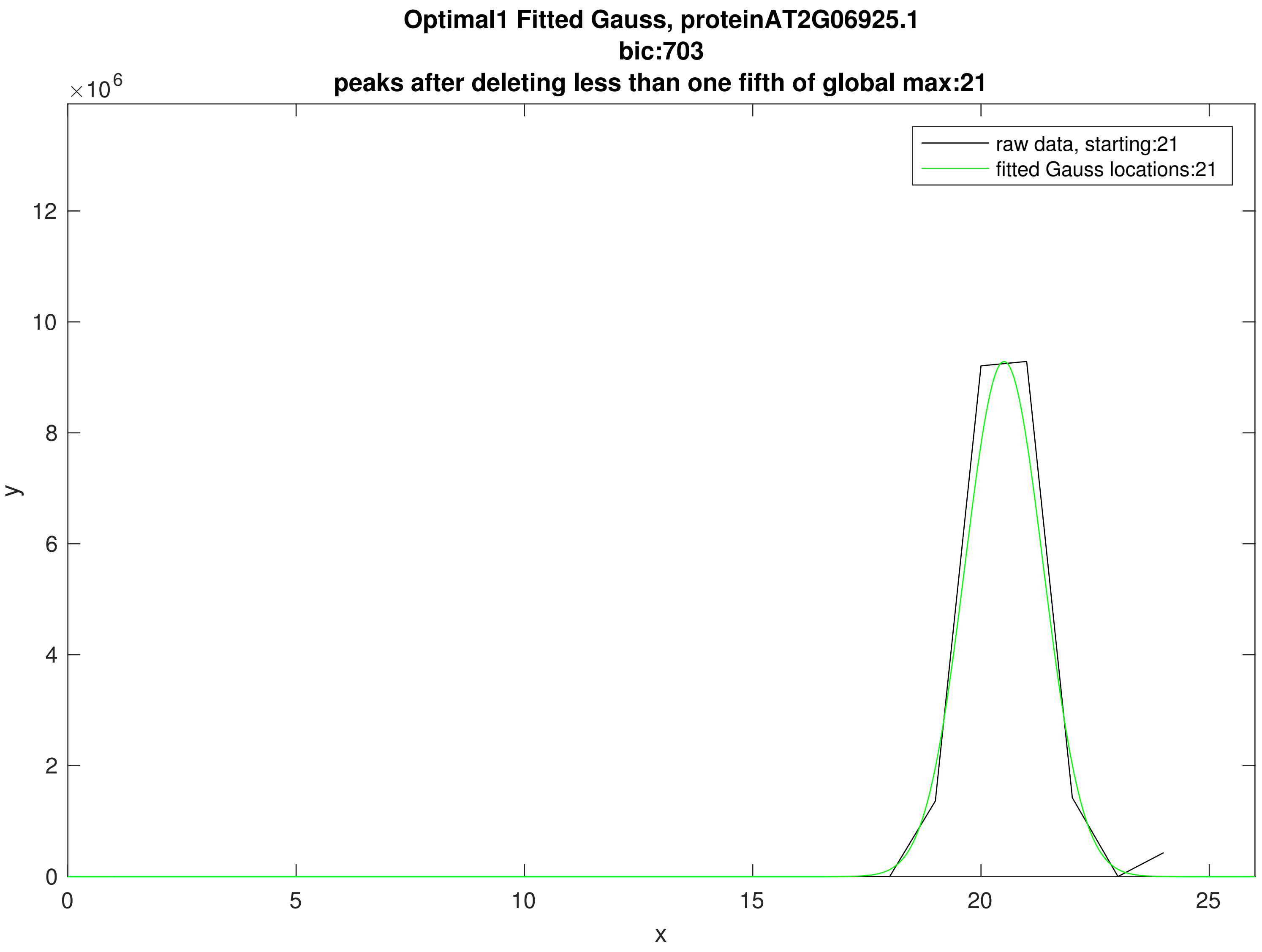

Optimal1 Fitted Gauss, proteinAT2G06925.1
bic:703
peaks after deleting less than one fifth of global max:21
×10^6
raw data, starting:21
fitted Gauss locations:21
12
10
8
6
4
2
0
y
0
5
10
15
20
25
x

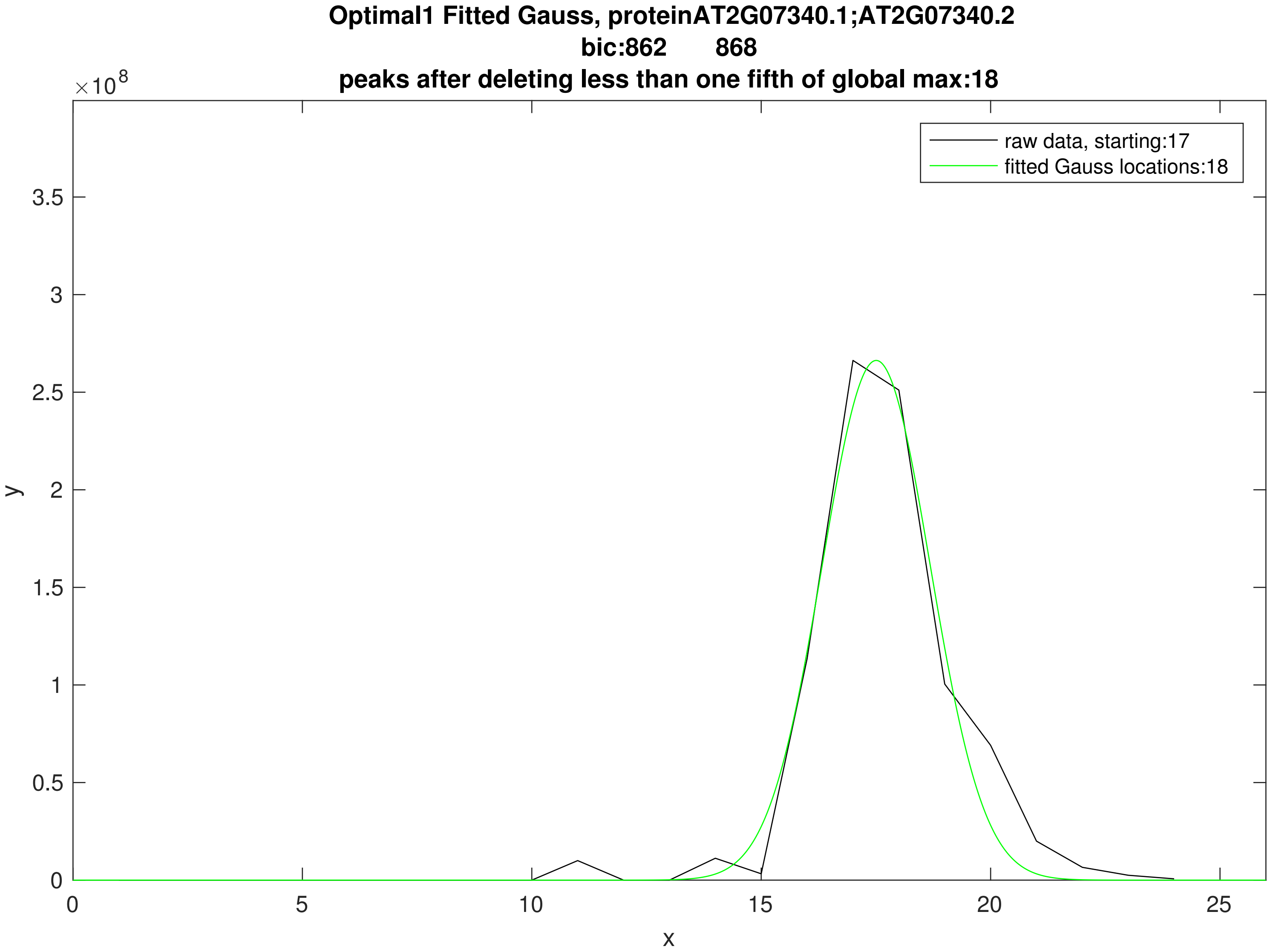

Optimal1 Fitted Gauss, proteinAT2G07340.1;AT2G07340.2
bic:862 868
peaks after deleting less than one fifth of global max:18
×10^8
raw data, starting:17
fitted Gauss locations:18
y
x
0
0.5
1
1.5
2
2.5
3
3.5
0
5
10
15
20
25

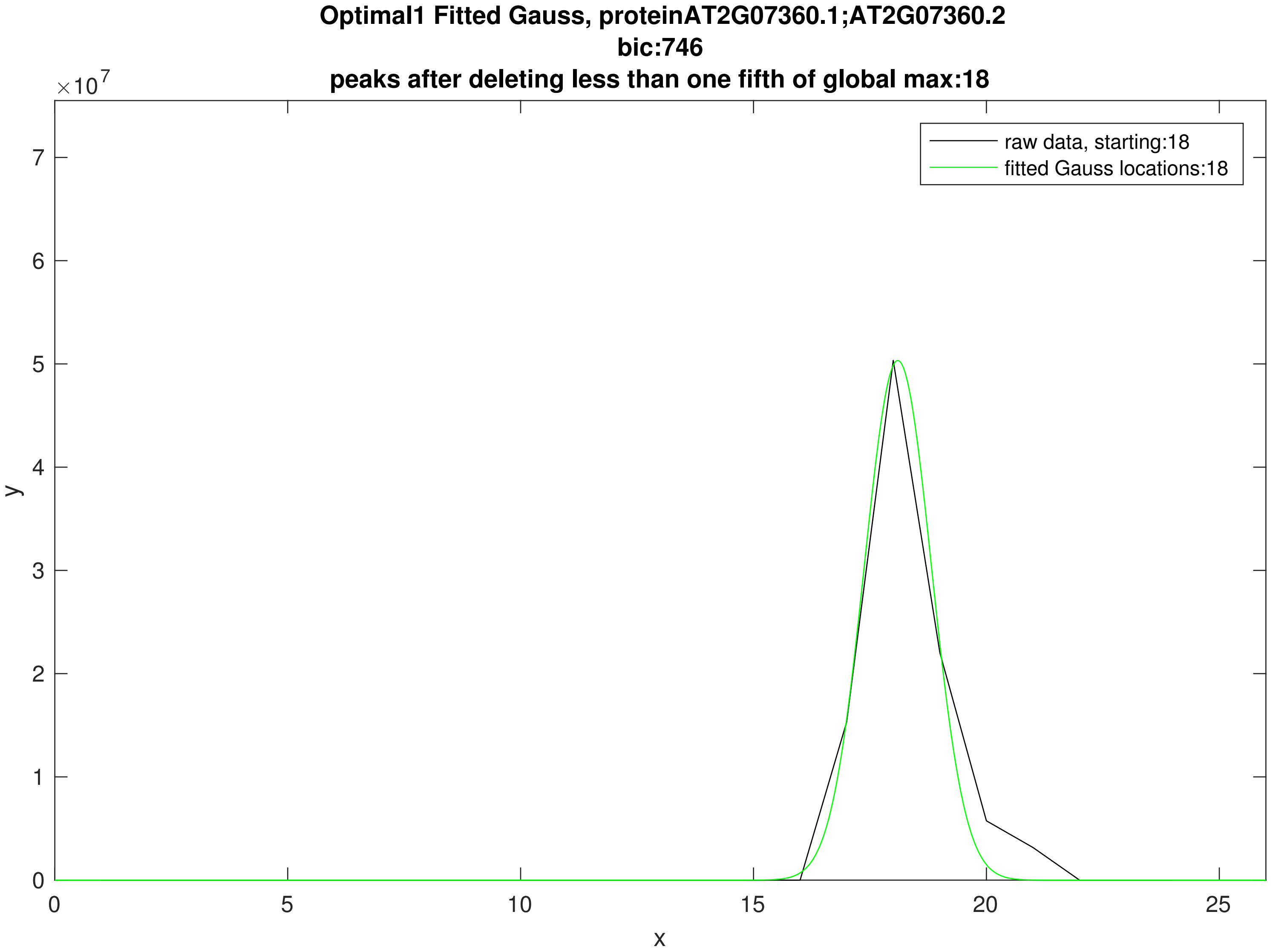

Optimal1 Fitted Gauss, proteinAT2G07360.1;AT2G07360.2
bic:746
peaks after deleting less than one fifth of global max:18
×10^7
raw data, starting:18
fitted Gauss locations:18
y
x
0
1
2
3
4
5
6
7
0
5
10
15
20
25

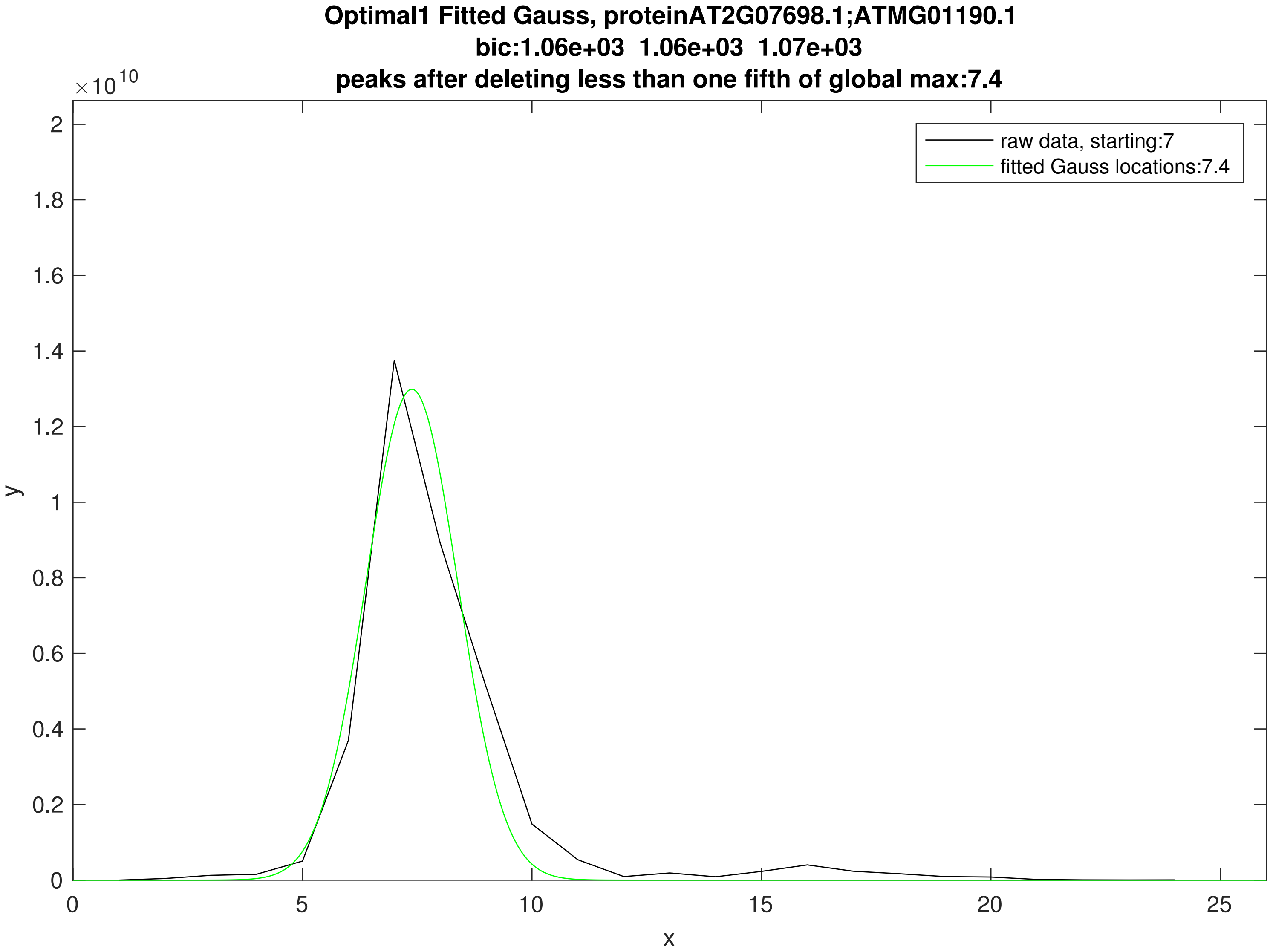

Optimal1 Fitted Gauss, proteinAT2G07698.1;ATMG01190.1
bic:1.06e+03  1.06e+03  1.07e+03
peaks after deleting less than one fifth of global max:7.4
×10^10
raw data, starting:7
fitted Gauss locations:7.4
y
x
0
0.2
0.4
0.6
0.8
1
1.2
1.4
1.6
1.8
2
5
10
15
20
25

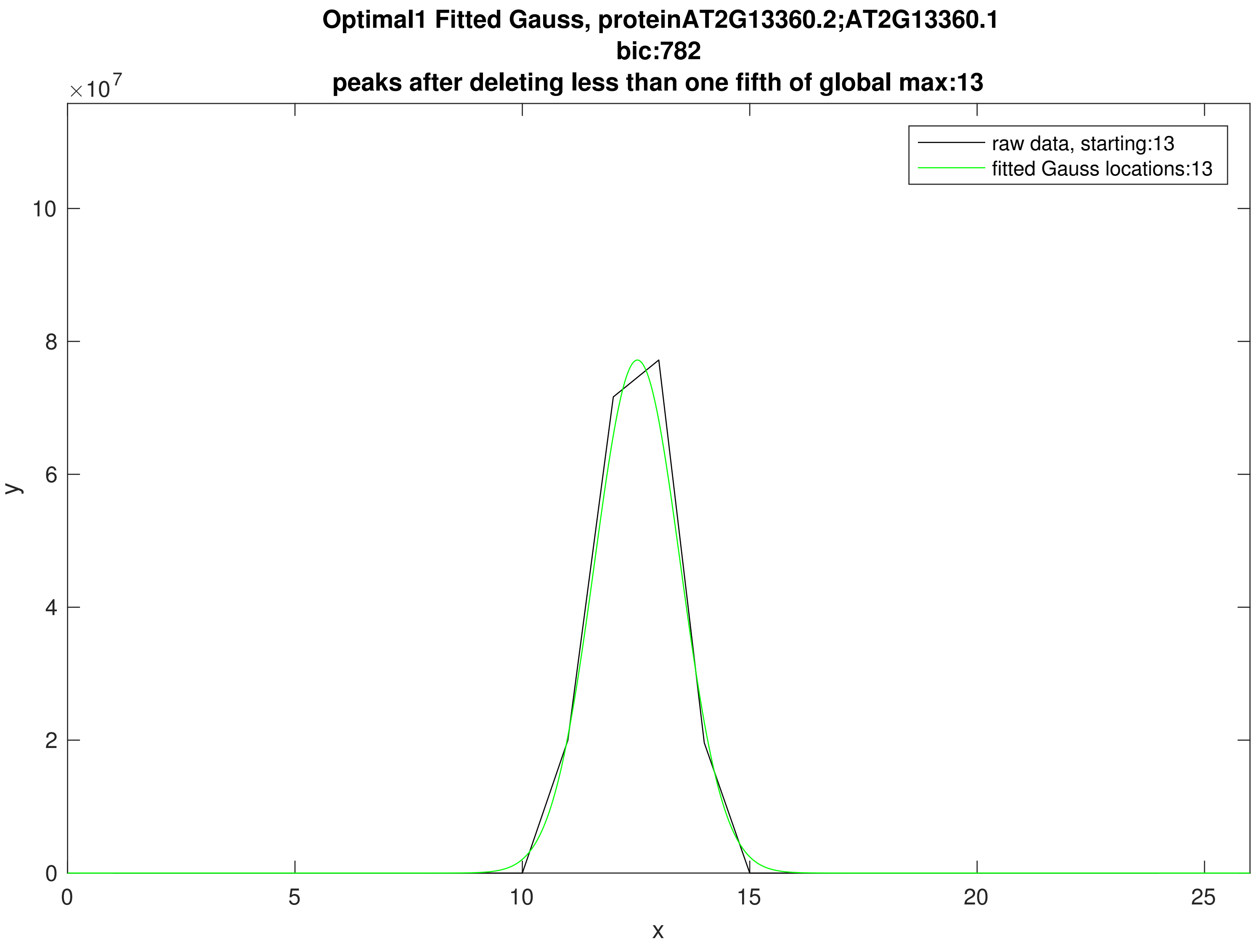
Optimal1 Fitted Gauss, proteinAT2G13360.2;AT2G13360.1
bic:782
peaks after deleting less than one fifth of global max:13
×10^7
raw data, starting:13
fitted Gauss locations:13
y
x
0
2
4
6
8
10
0
5
10
15
20
25

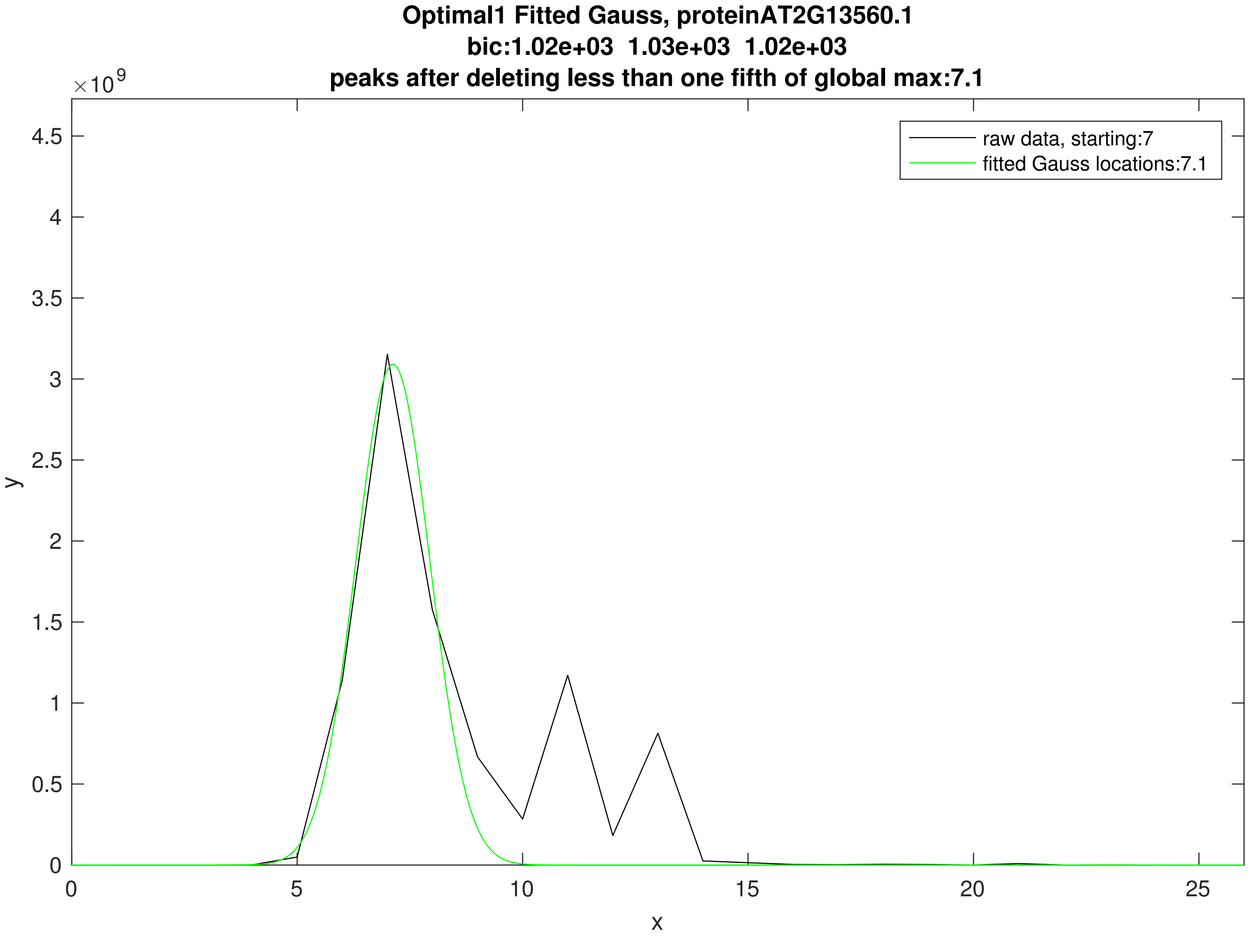
Optimal1 Fitted Gauss, proteinAT2G13560.1
bic:1.02e+03  1.03e+03  1.02e+03
peaks after deleting less than one fifth of global max:7.1
×10^9
4.5
4
3.5
3
2.5
2
1.5
1
0.5
0
y
0
5
10
15
20
25
x
raw data, starting:7
fitted Gauss locations:7.1

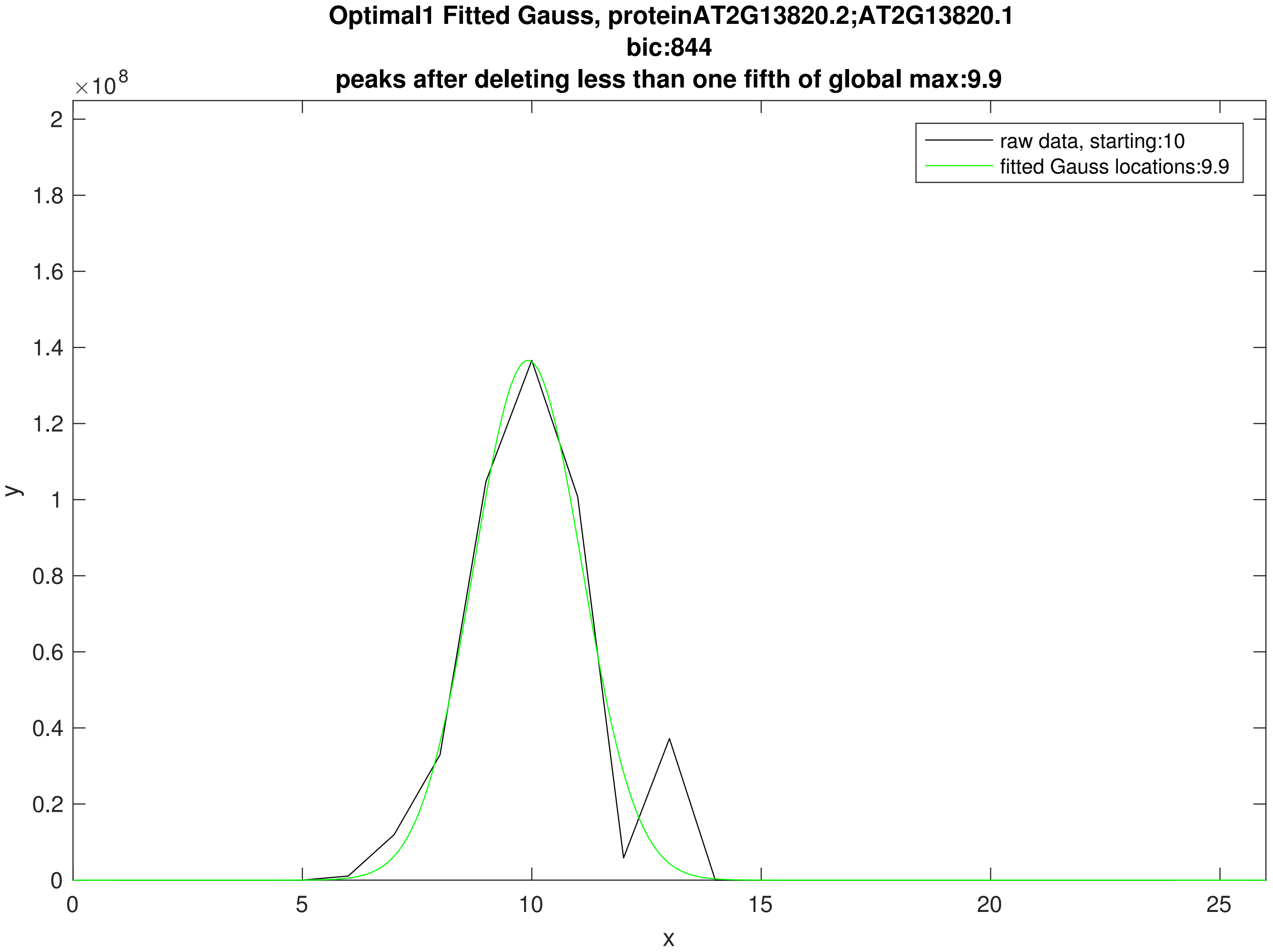

Optimal1 Fitted Gauss, proteinAT2G13820.2;AT2G13820.1
bic:844
peaks after deleting less than one fifth of global max:9.9
×10^8
raw data, starting:10
fitted Gauss locations:9.9
y
x
0
0.2
0.4
0.6
0.8
1
1.2
1.4
1.6
1.8
2
5
10
15
20
25

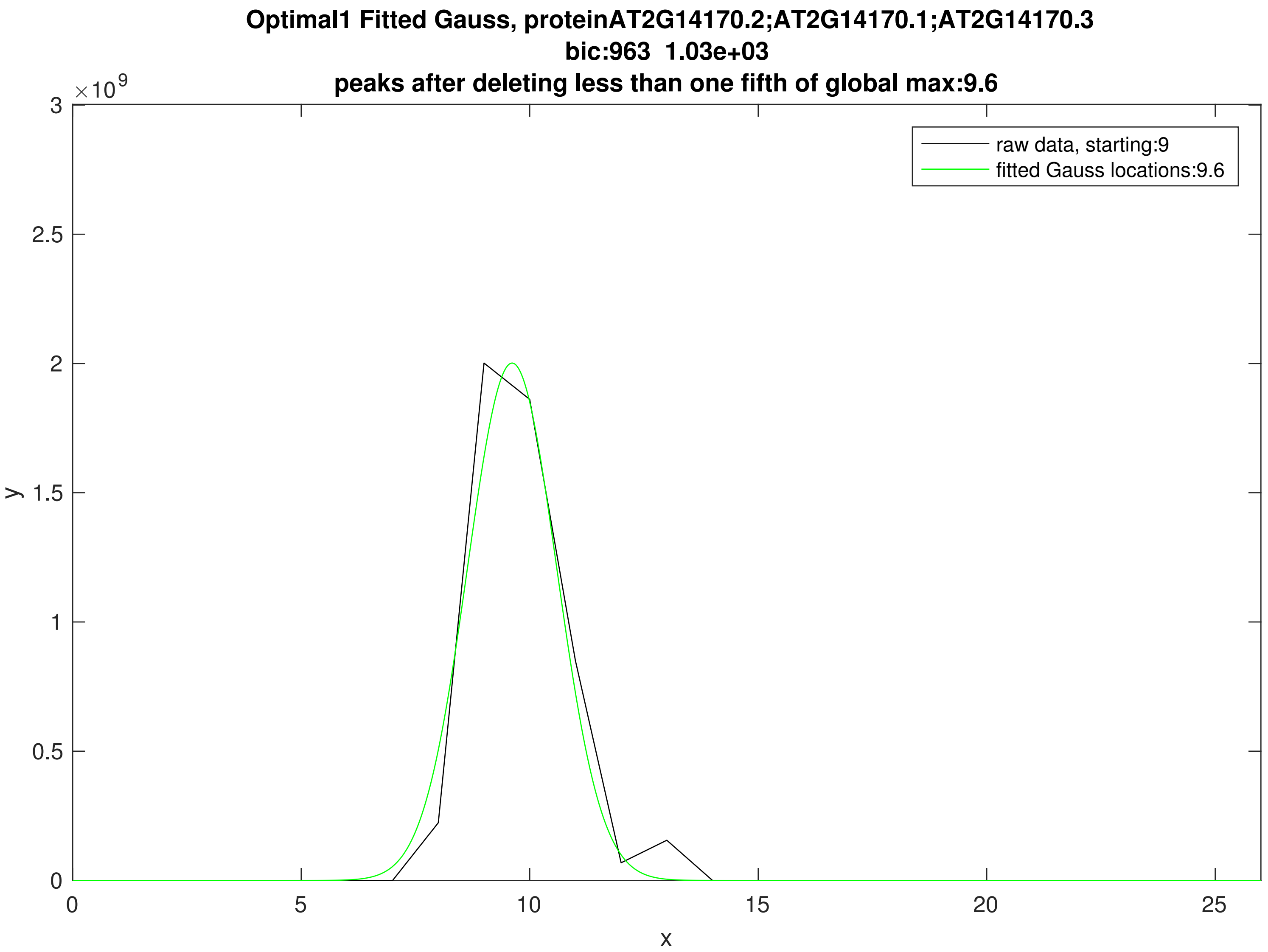

Optimal1 Fitted Gauss, proteinAT2G14170.2;AT2G14170.1;AT2G14170.3
bic:963 1.03e+03
peaks after deleting less than one fifth of global max:9.6
×10^9
3
2.5
2
1.5
1
0.5
0
y
0
5
10
15
20
25
x
raw data, starting:9
fitted Gauss locations:9.6

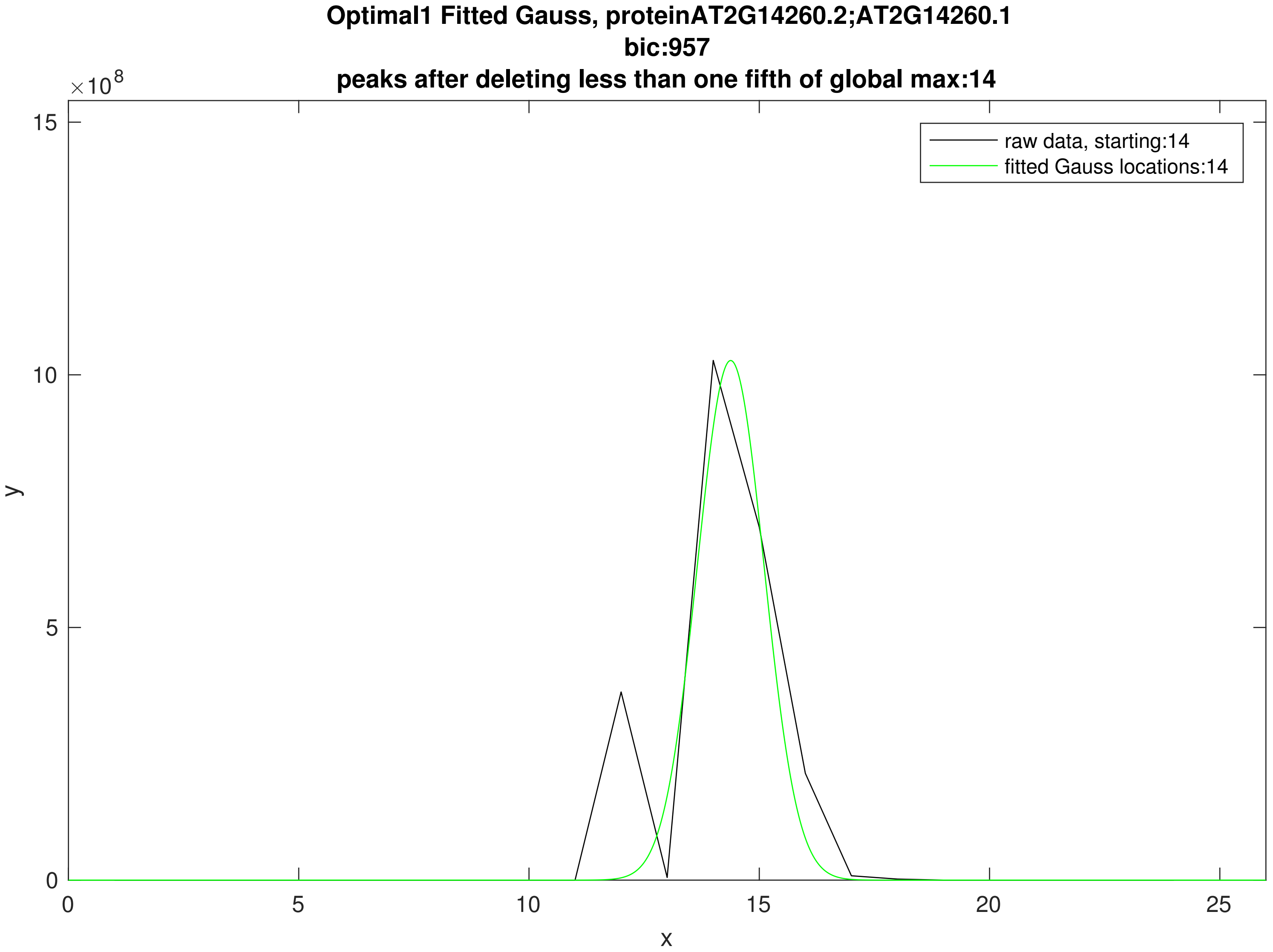
Optimal1 Fitted Gauss, proteinAT2G14260.2;AT2G14260.1
bic:957
peaks after deleting less than one fifth of global max:14
×10^8
raw data, starting:14
fitted Gauss locations:14
y
x
0
5
10
15
20
25

**Optimal1 Fitted Gauss, proteinAT2G14285.1;AT4G30220.1;AT4G30220.2**
**bic:701**
**peaks after deleting less than one fifth of global max:21**

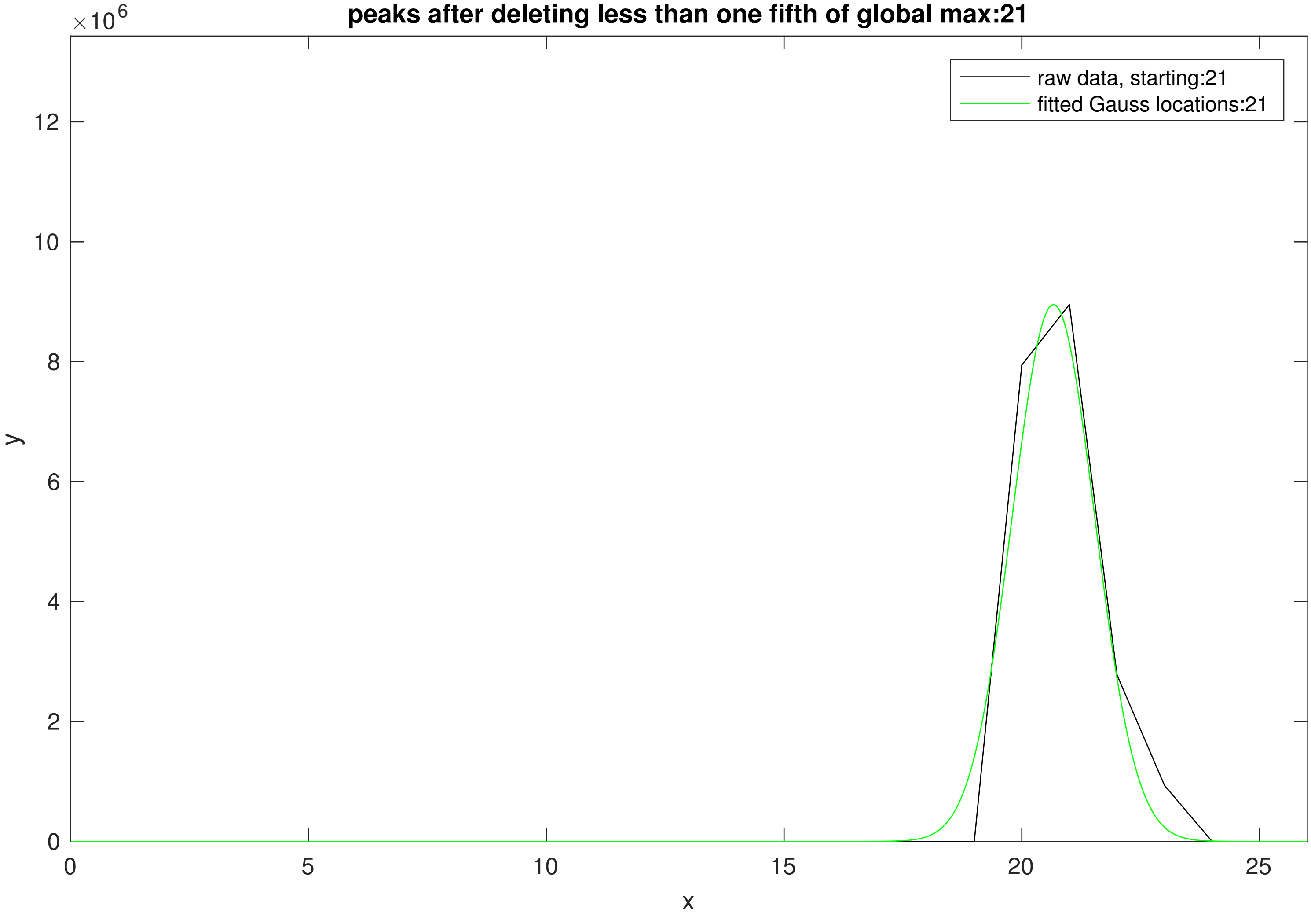

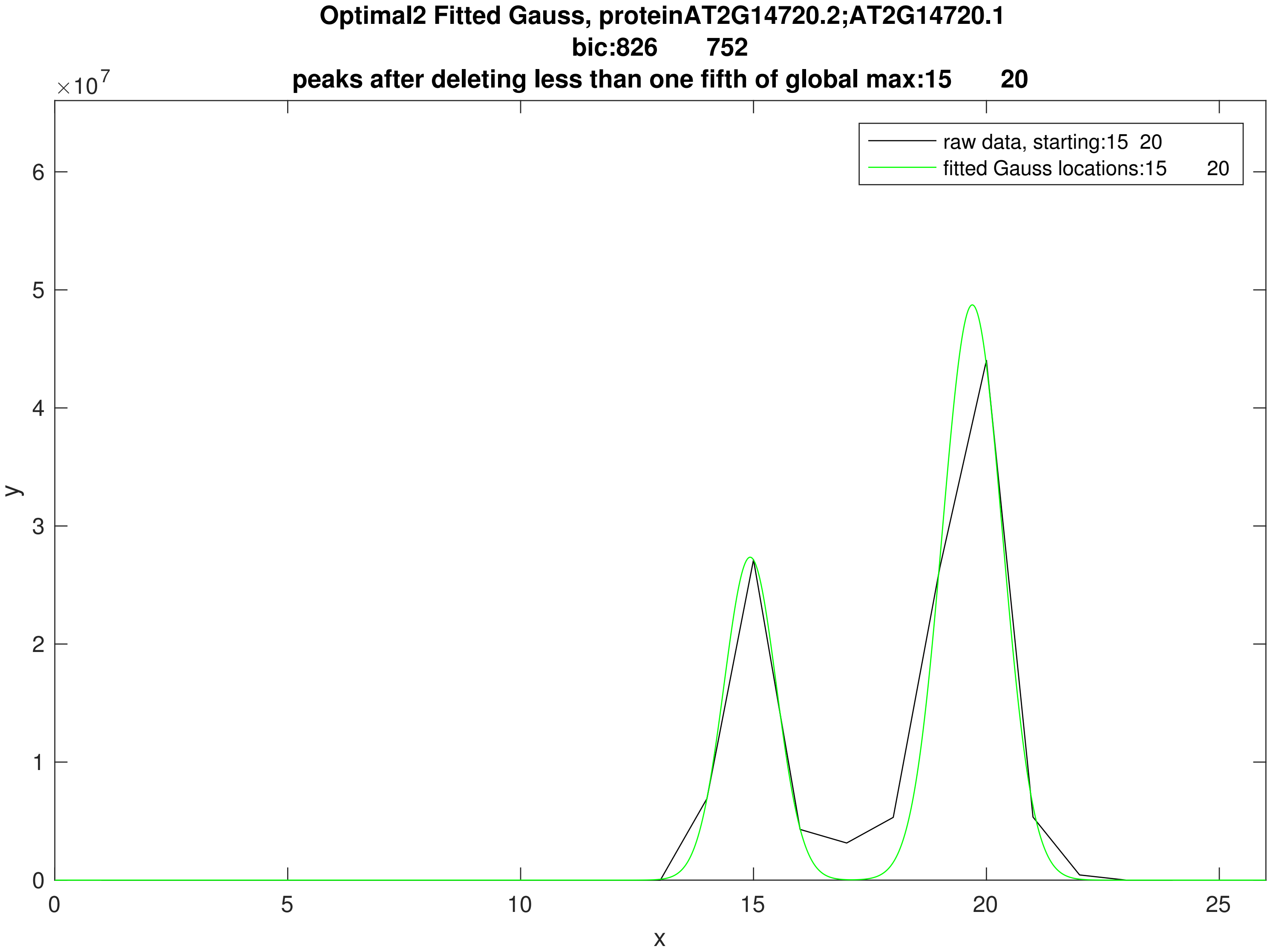
Optimal2 Fitted Gauss, proteinAT2G14720.2;AT2G14720.1
bic:826 752
peaks after deleting less than one fifth of global max:15 20
×10^7
raw data, starting:15 20
fitted Gauss locations:15 20
y
x
0
1
2
3
4
5
6
0
5
10
15
20
25

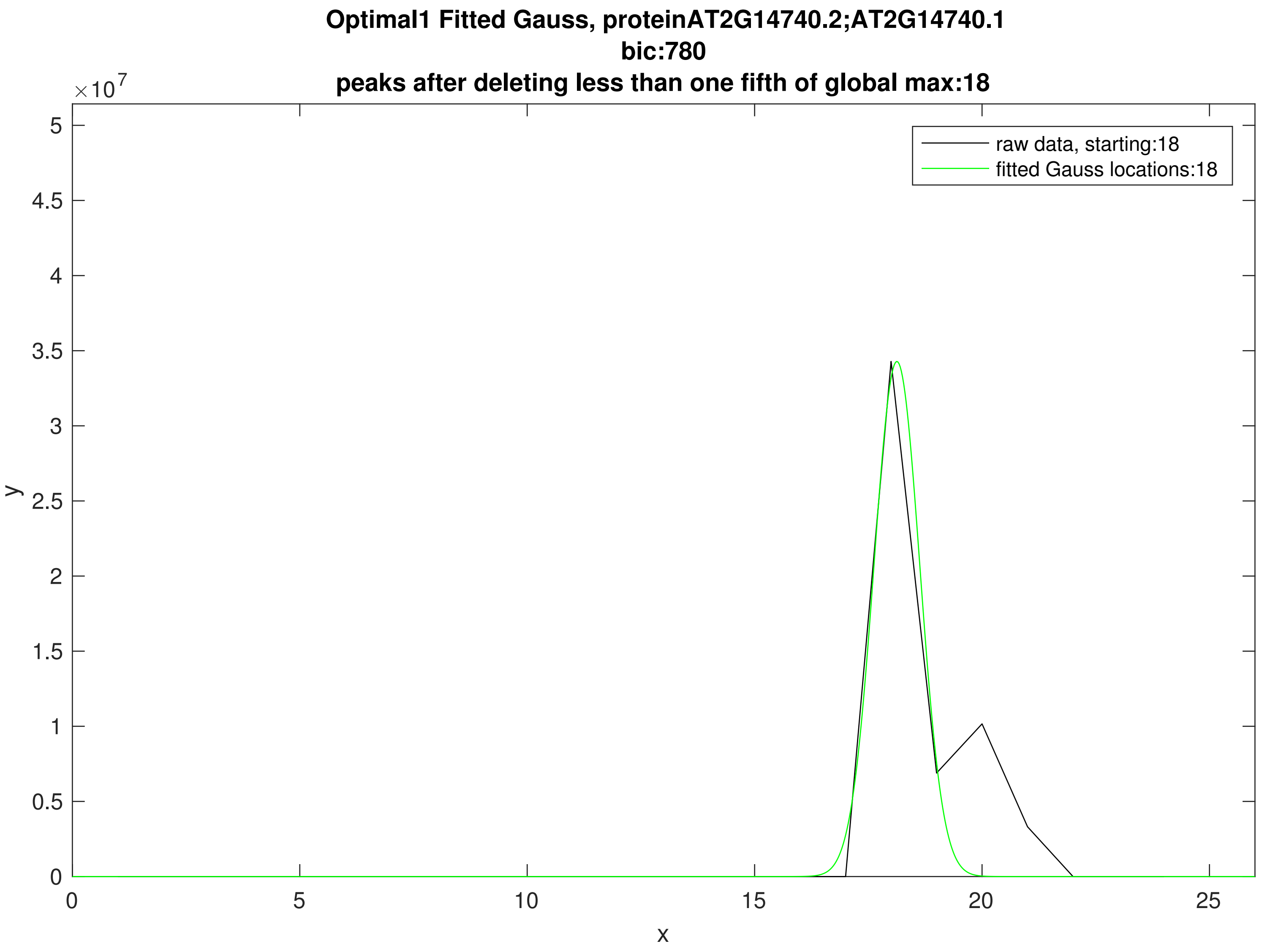
Optimal1 Fitted Gauss, proteinAT2G14740.2;AT2G14740.1
bic:780
peaks after deleting less than one fifth of global max:18
×10^7
raw data, starting:18
fitted Gauss locations:18
y
x
0
0.5
1
1.5
2
2.5
3
3.5
4
4.5
5
0
5
10
15
20
25

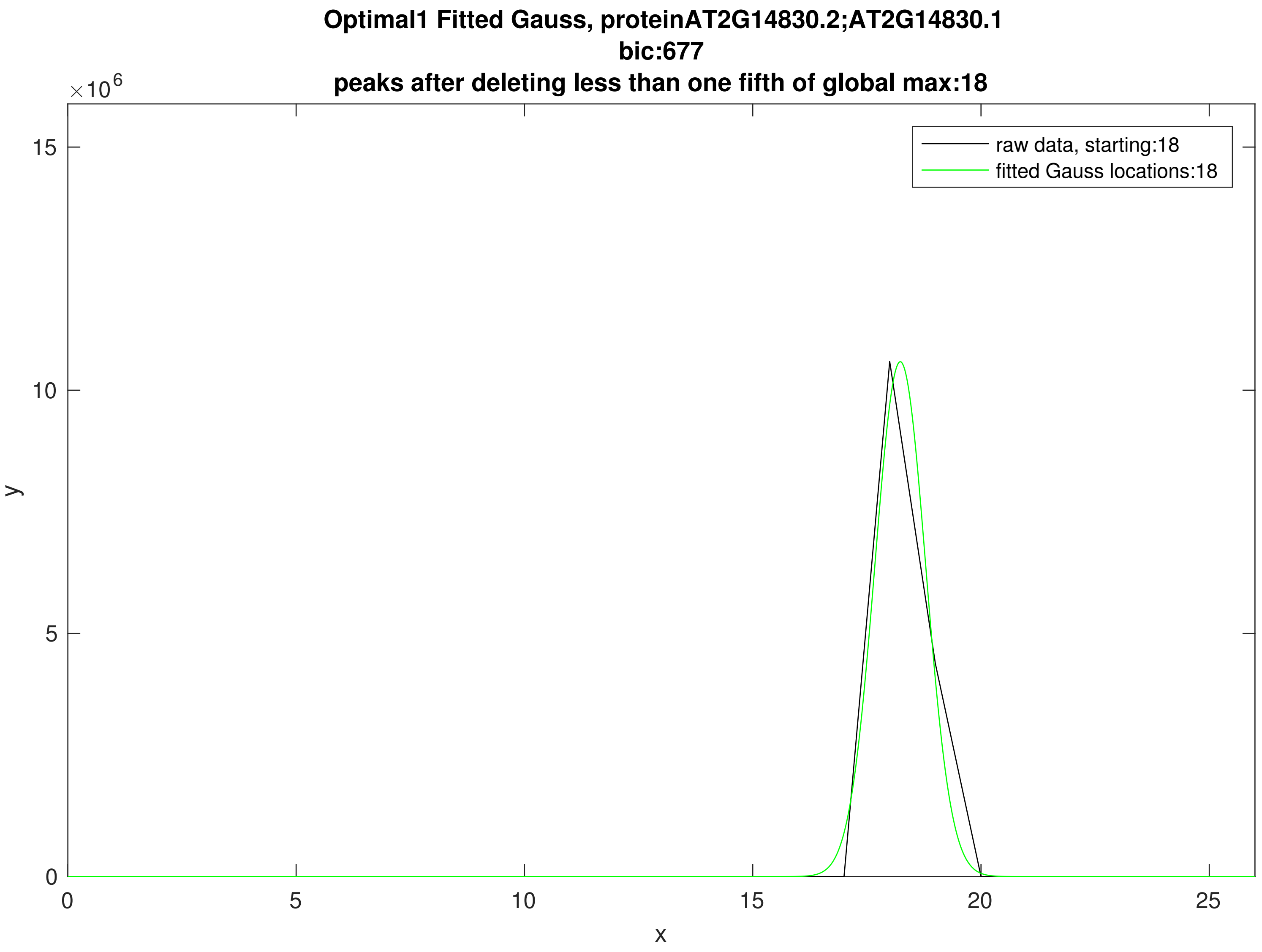
Optimal1 Fitted Gauss, proteinAT2G14830.2;AT2G14830.1
bic:677
peaks after deleting less than one fifth of global max:18
×10^6
raw data, starting:18
fitted Gauss locations:18
y
x
0
5
10
15
20
25

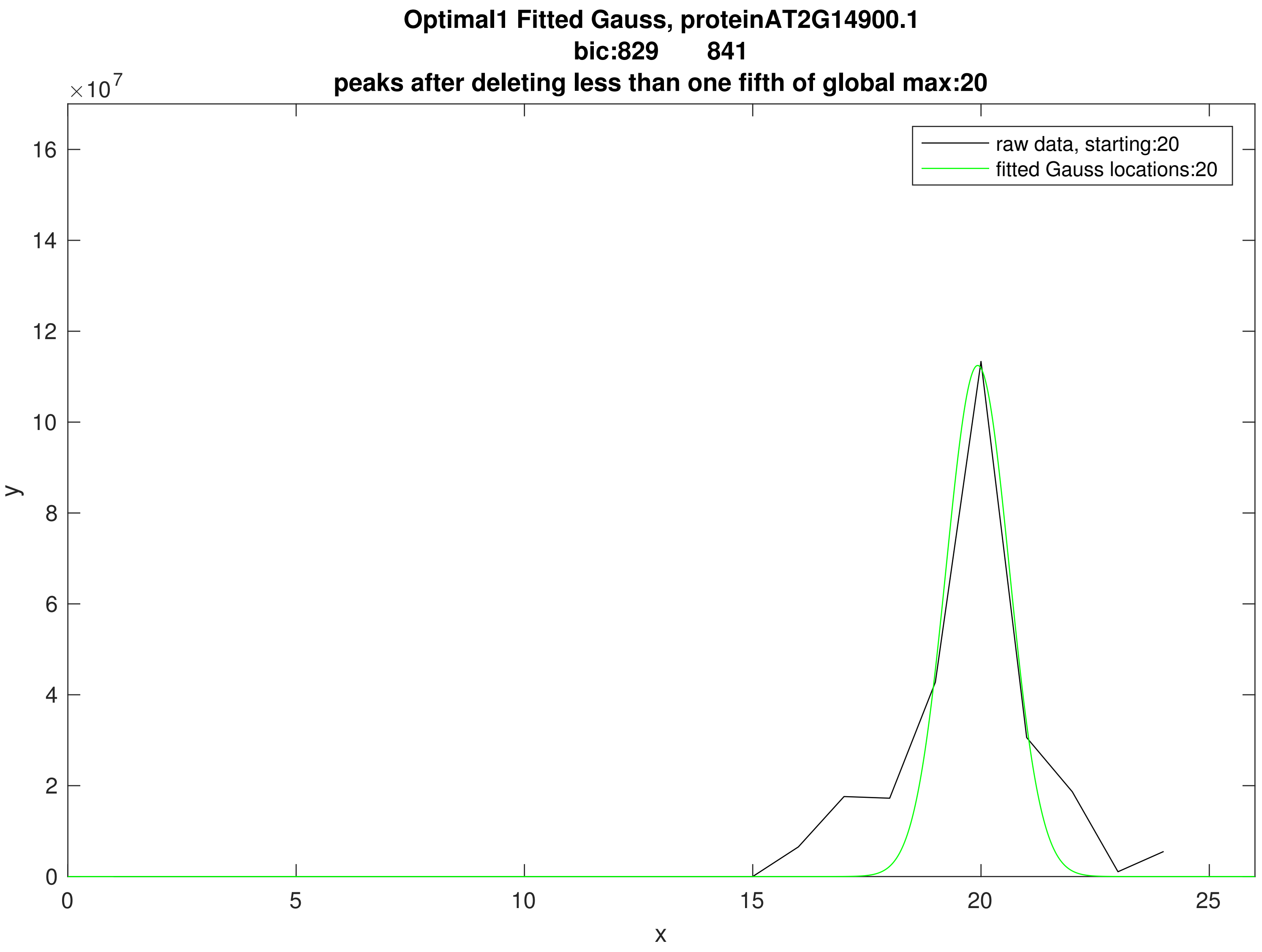

Optimal1 Fitted Gauss, proteinAT2G14900.1
bic:829 841
peaks after deleting less than one fifth of global max:20
×10^7
16
14
12
10
8
6
4
2
0
y
0
5
10
15
20
25
x
raw data, starting:20
fitted Gauss locations:20

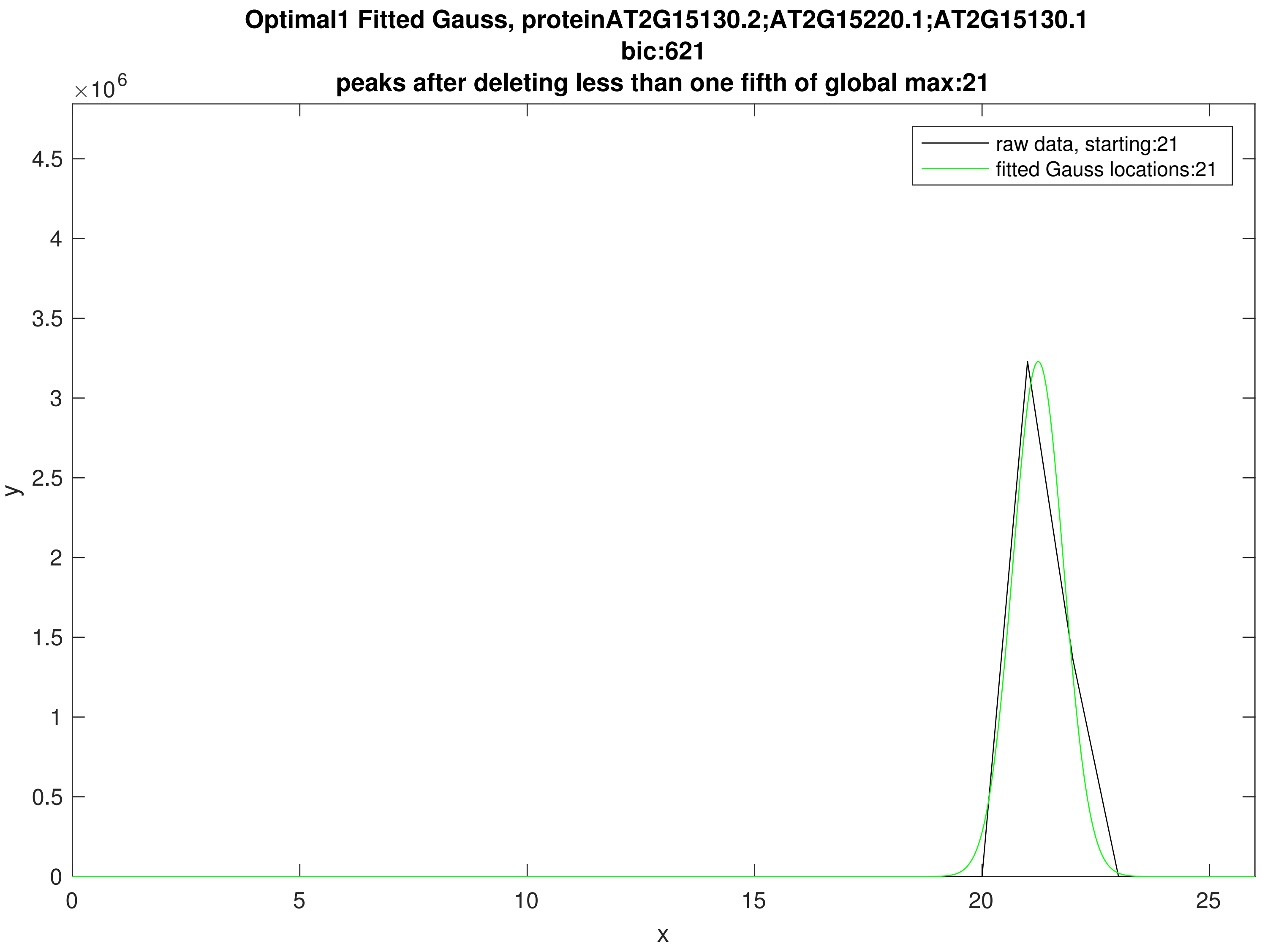
Optimal1 Fitted Gauss, proteinAT2G15130.2;AT2G15220.1;AT2G15130.1
bic:621
peaks after deleting less than one fifth of global max:21
×10^6
4.5
4
3.5
3
2.5
2
1.5
1
0.5
0
y
0
5
10
15
20
25
x
raw data, starting:21
fitted Gauss locations:21

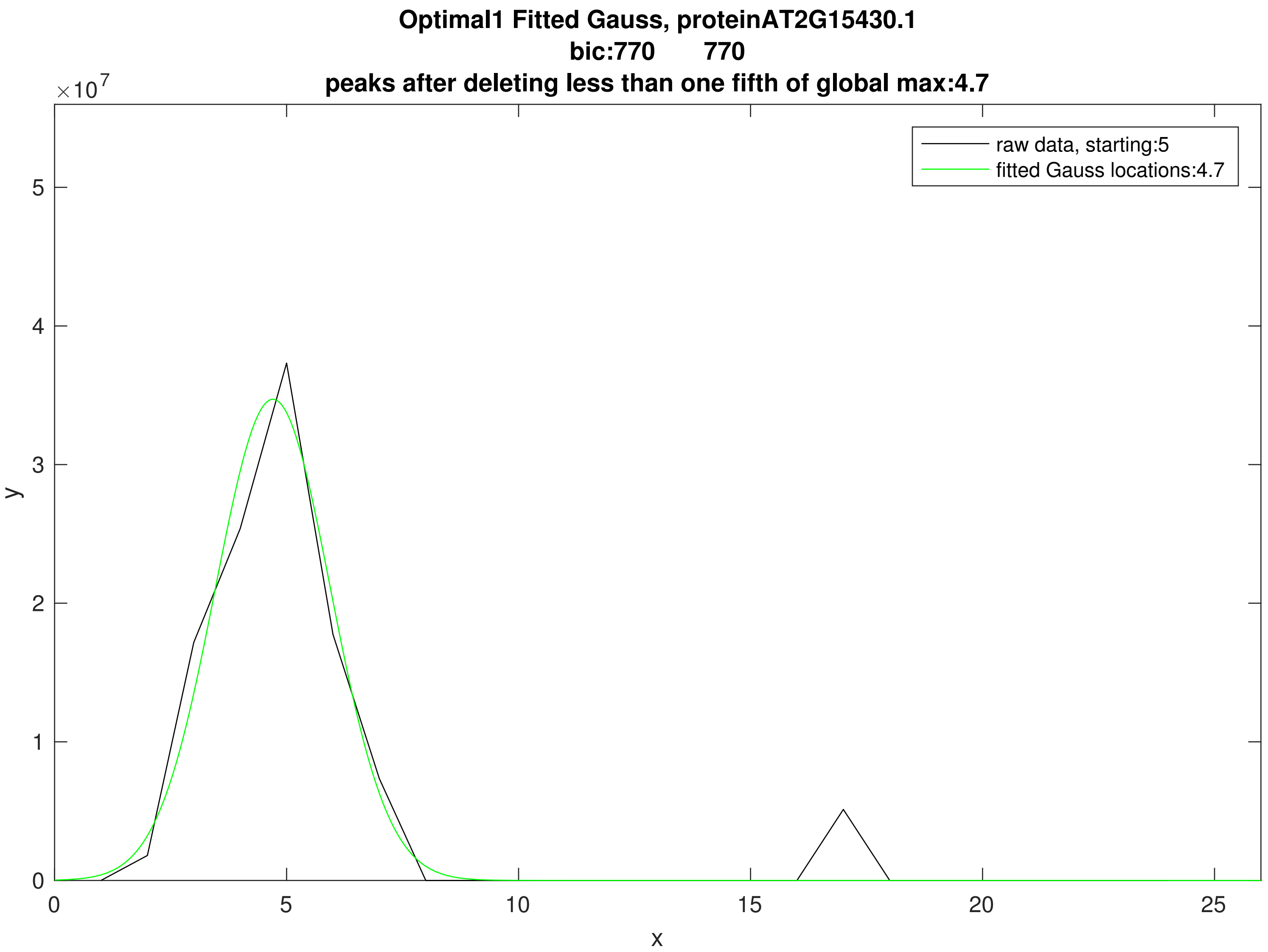

Optimal1 Fitted Gauss, proteinAT2G15430.1
bic:770 770
peaks after deleting less than one fifth of global max:4.7
×10^7
raw data, starting:5
fitted Gauss locations:4.7
5
4
3
2
1
0
y
0
5
10
15
20
25
x

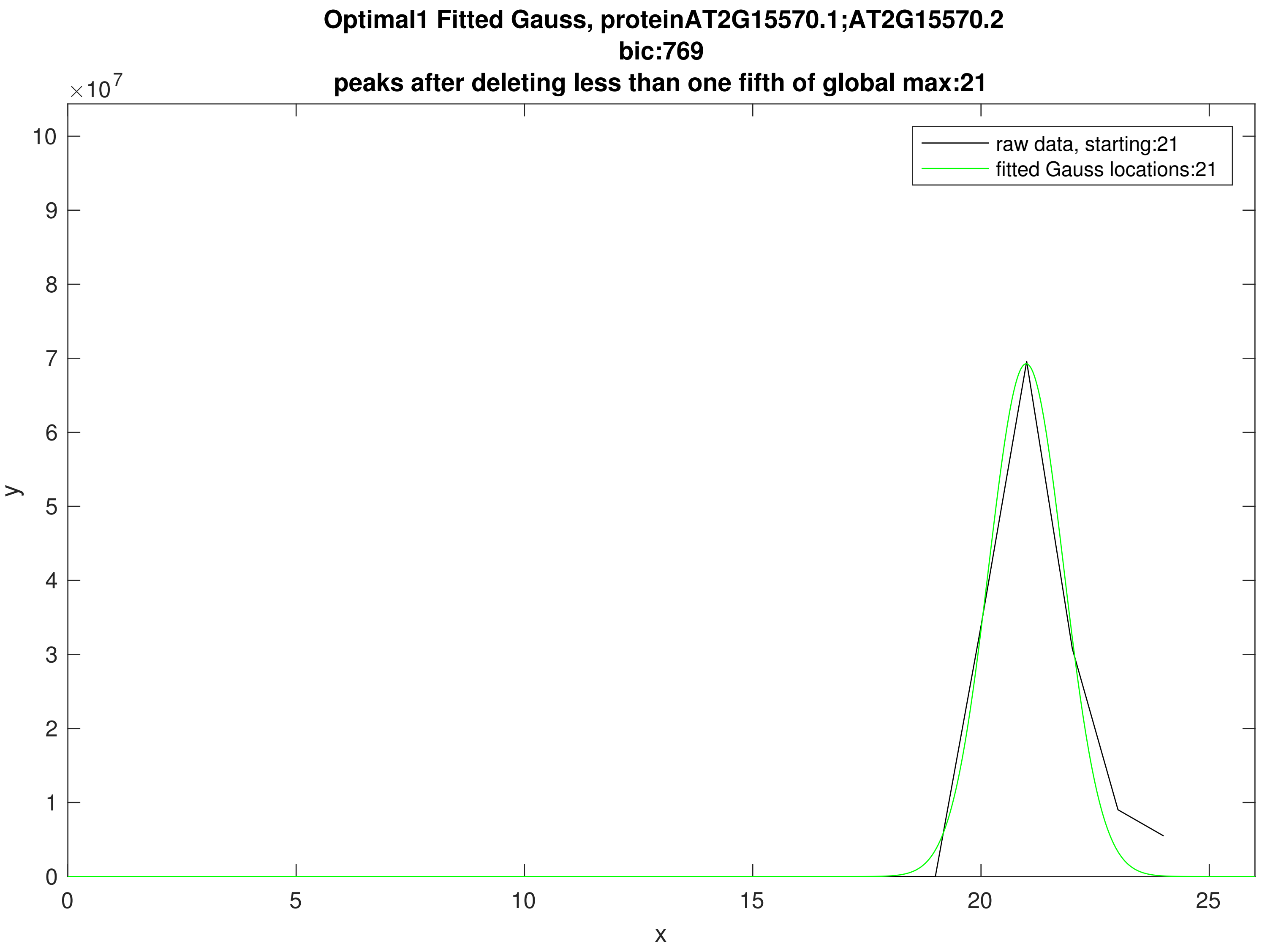
Optimal1 Fitted Gauss, proteinAT2G15570.1;AT2G15570.2
bic:769
peaks after deleting less than one fifth of global max:21
×10^7
raw data, starting:21
fitted Gauss locations:21
y
x
0
1
2
3
4
5
6
7
8
9
10
0
5
10
15
20
25

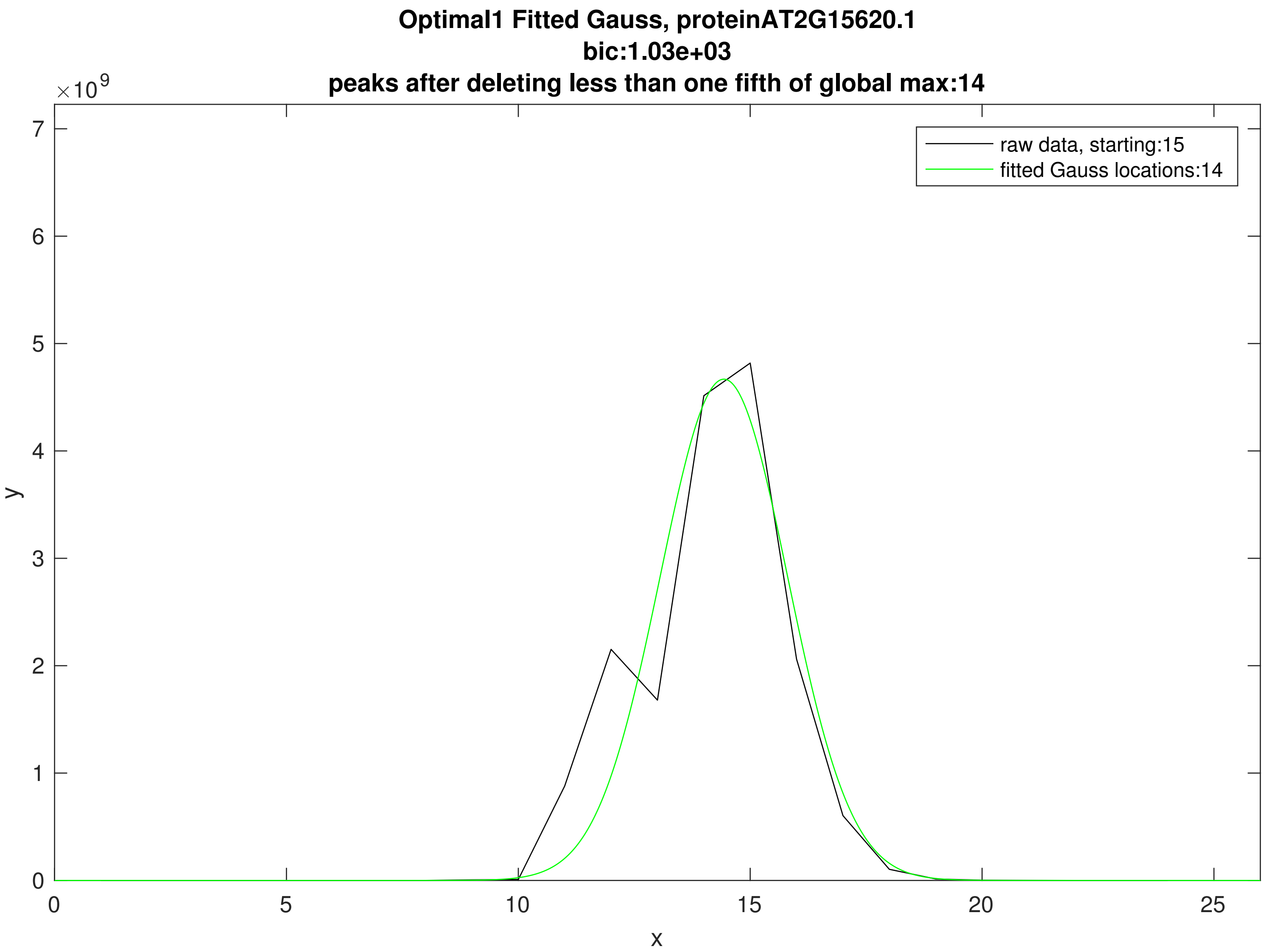

Optimal1 Fitted Gauss, proteinAT2G15620.1
bic:1.03e+03
peaks after deleting less than one fifth of global max:14
×10^9
7
6
5
4
3
2
1
0
y
0
5
10
15
20
25
x
raw data, starting:15
fitted Gauss locations:14

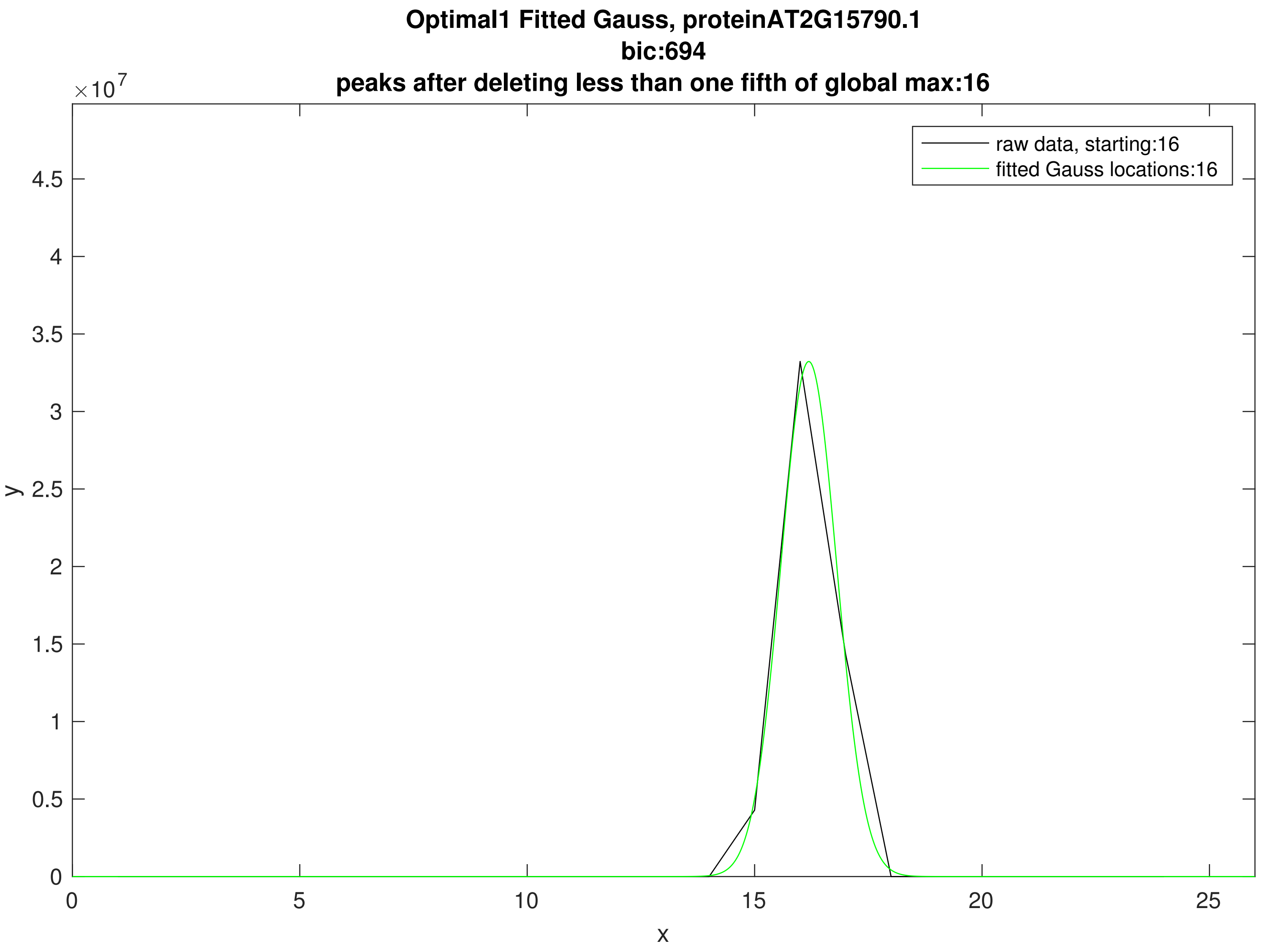

Optimal1 Fitted Gauss, proteinAT2G15790.1
bic:694
peaks after deleting less than one fifth of global max:16
×10^7
4.5
4
3.5
3
2.5
2
1.5
1
0.5
0
y
0
5
10
15
20
25
x
raw data, starting:16
fitted Gauss locations:16

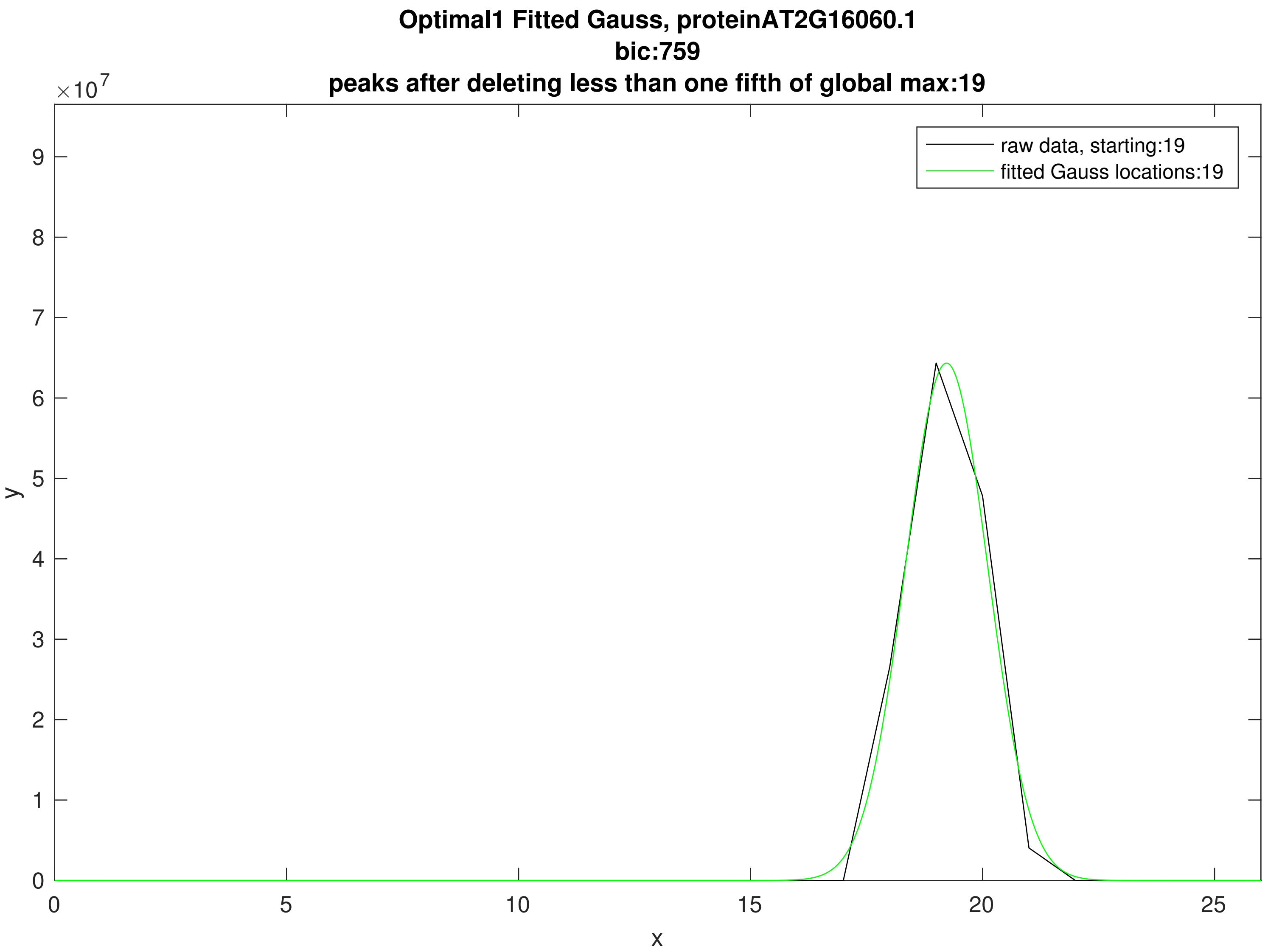

Optimal1 Fitted Gauss, proteinAT2G16060.1
bic:759
peaks after deleting less than one fifth of global max:19
×10^7
raw data, starting:19
fitted Gauss locations:19
y
x
0
1
2
3
4
5
6
7
8
9
0
5
10
15
20
25

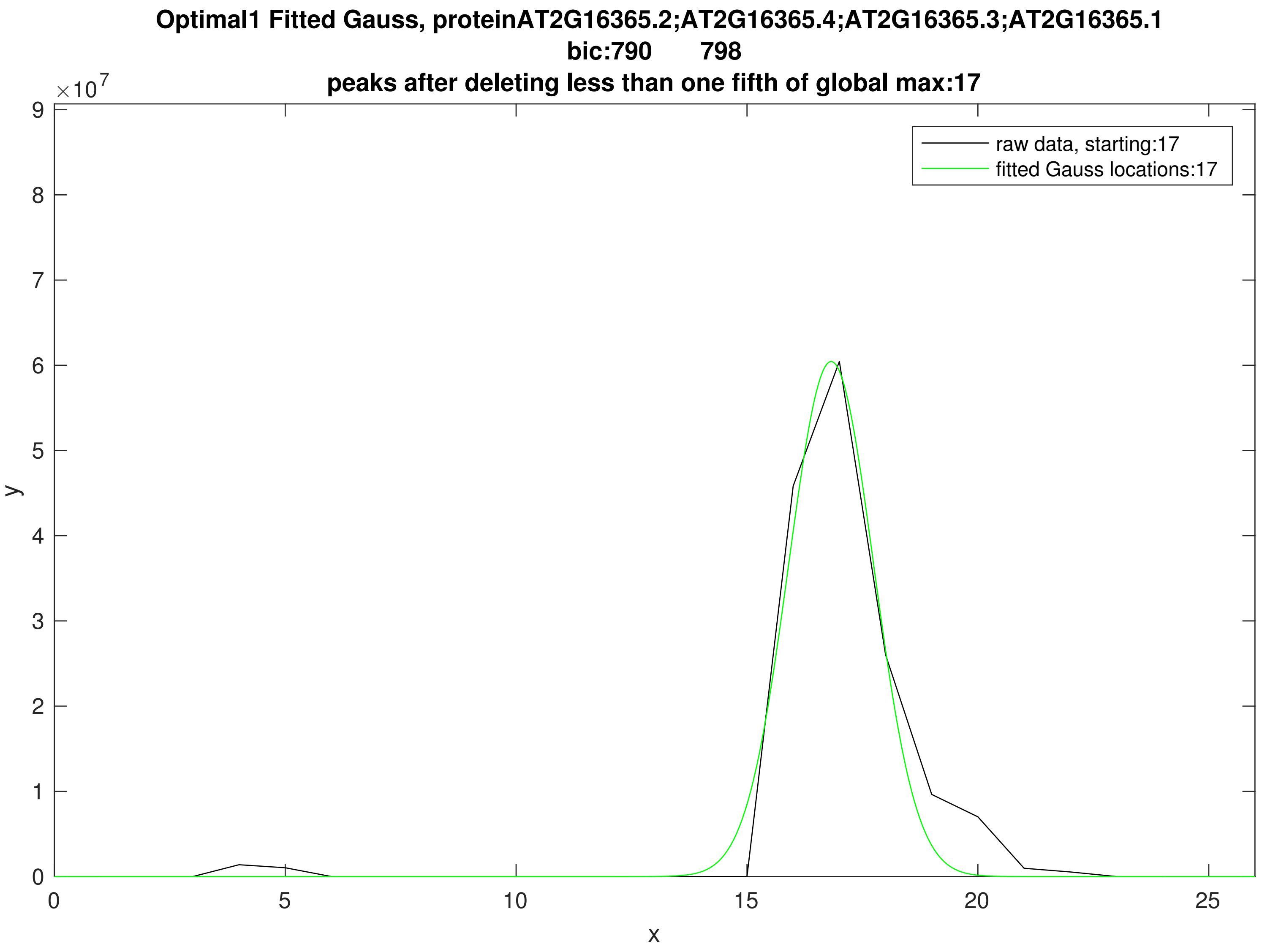
Optimal1 Fitted Gauss, proteinAT2G16365.2;AT2G16365.4;AT2G16365.3;AT2G16365.1
bic:790 798
peaks after deleting less than one fifth of global max:17
×10^7
raw data, starting:17
fitted Gauss locations:17
y
x

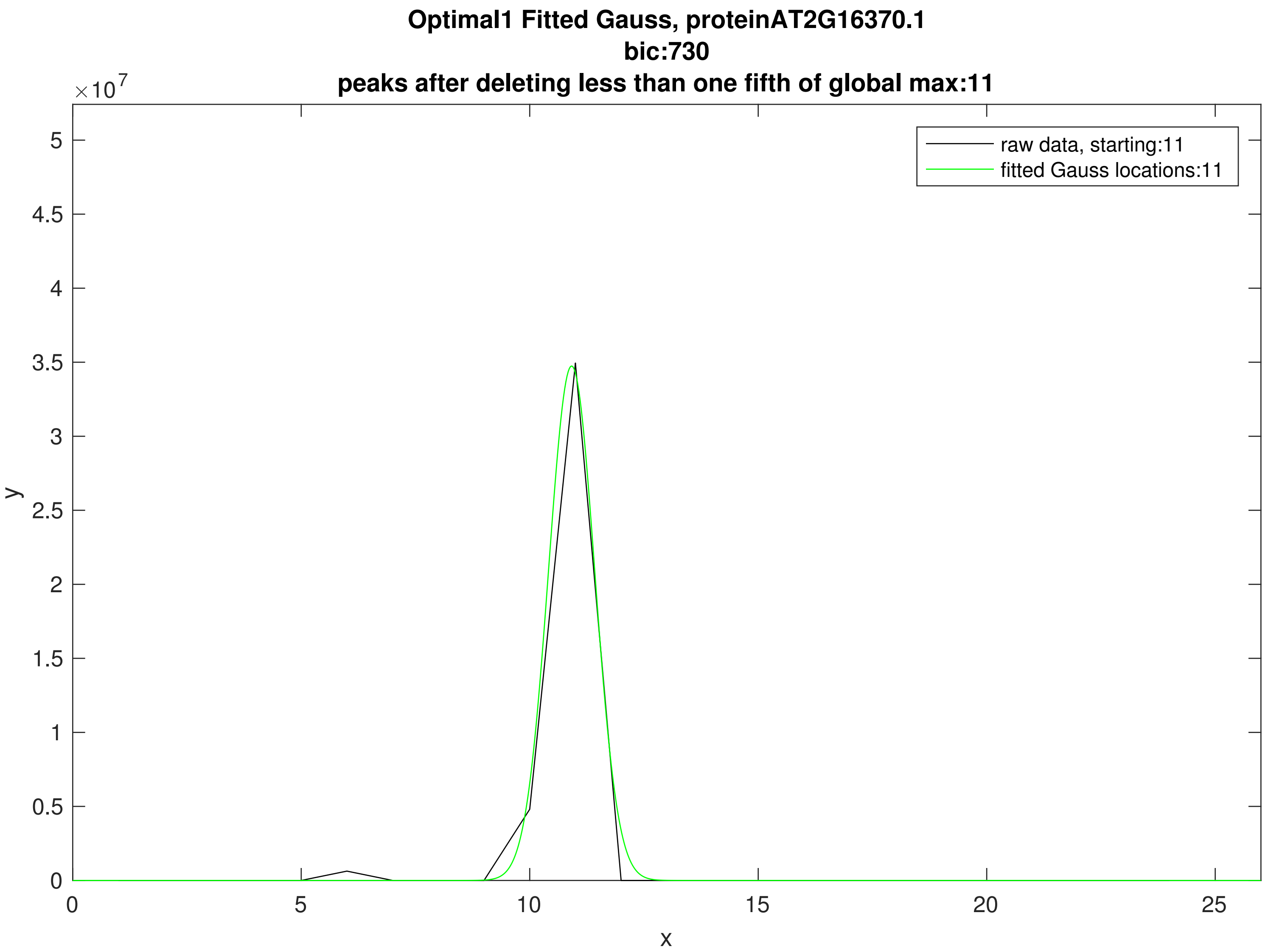

Optimal1 Fitted Gauss, proteinAT2G16370.1
bic:730
peaks after deleting less than one fifth of global max:11
×10^7
raw data, starting:11
fitted Gauss locations:11
y
x
0
0.5
1
1.5
2
2.5
3
3.5
4
4.5
5
0
5
10
15
20
25

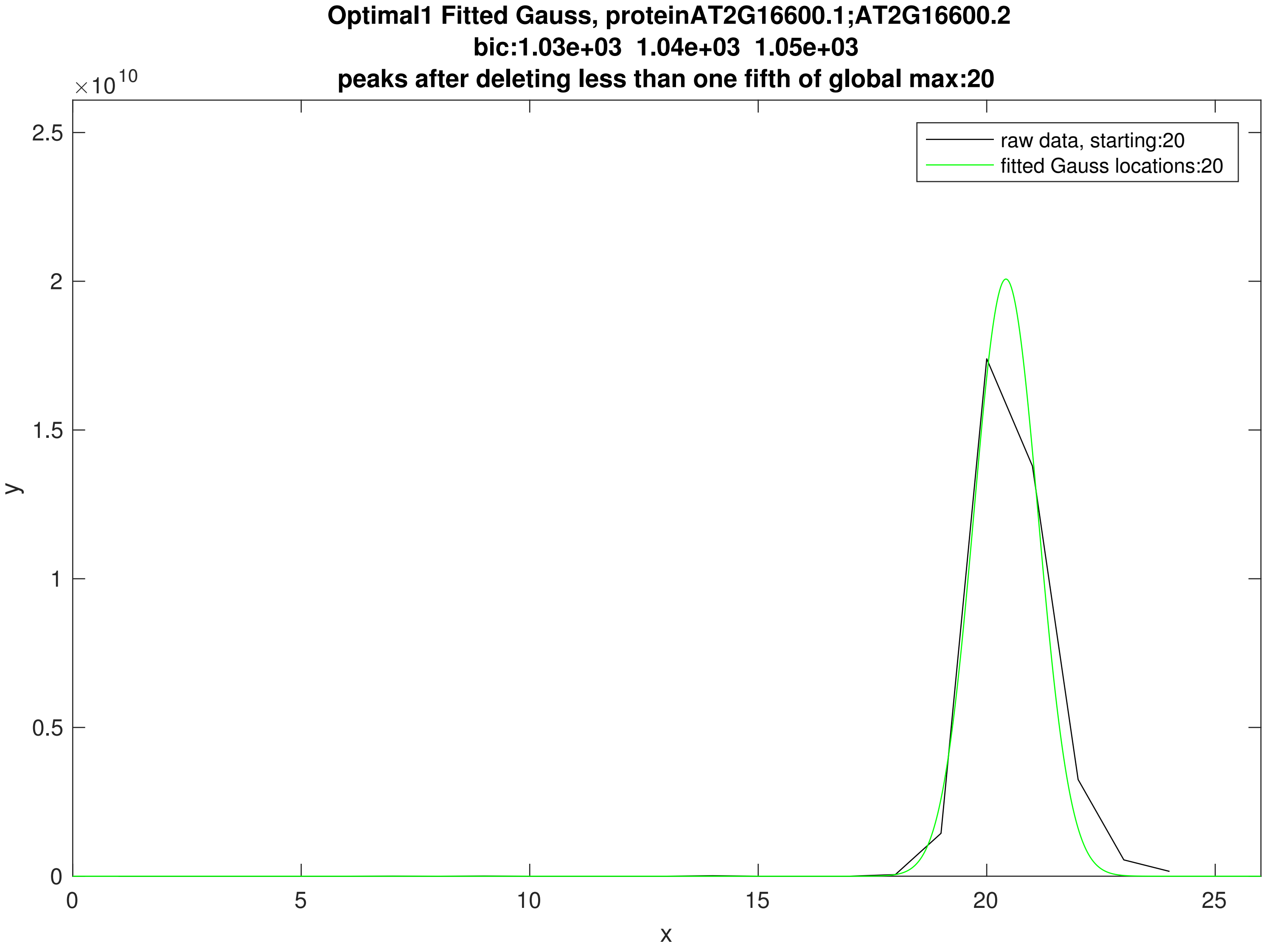
Optimal1 Fitted Gauss, proteinAT2G16600.1;AT2G16600.2
bic:1.03e+03  1.04e+03  1.05e+03
peaks after deleting less than one fifth of global max:20
×10^10
raw data, starting:20
fitted Gauss locations:20
y
x
0
0.5
1
1.5
2
2.5
5
10
15
20
25

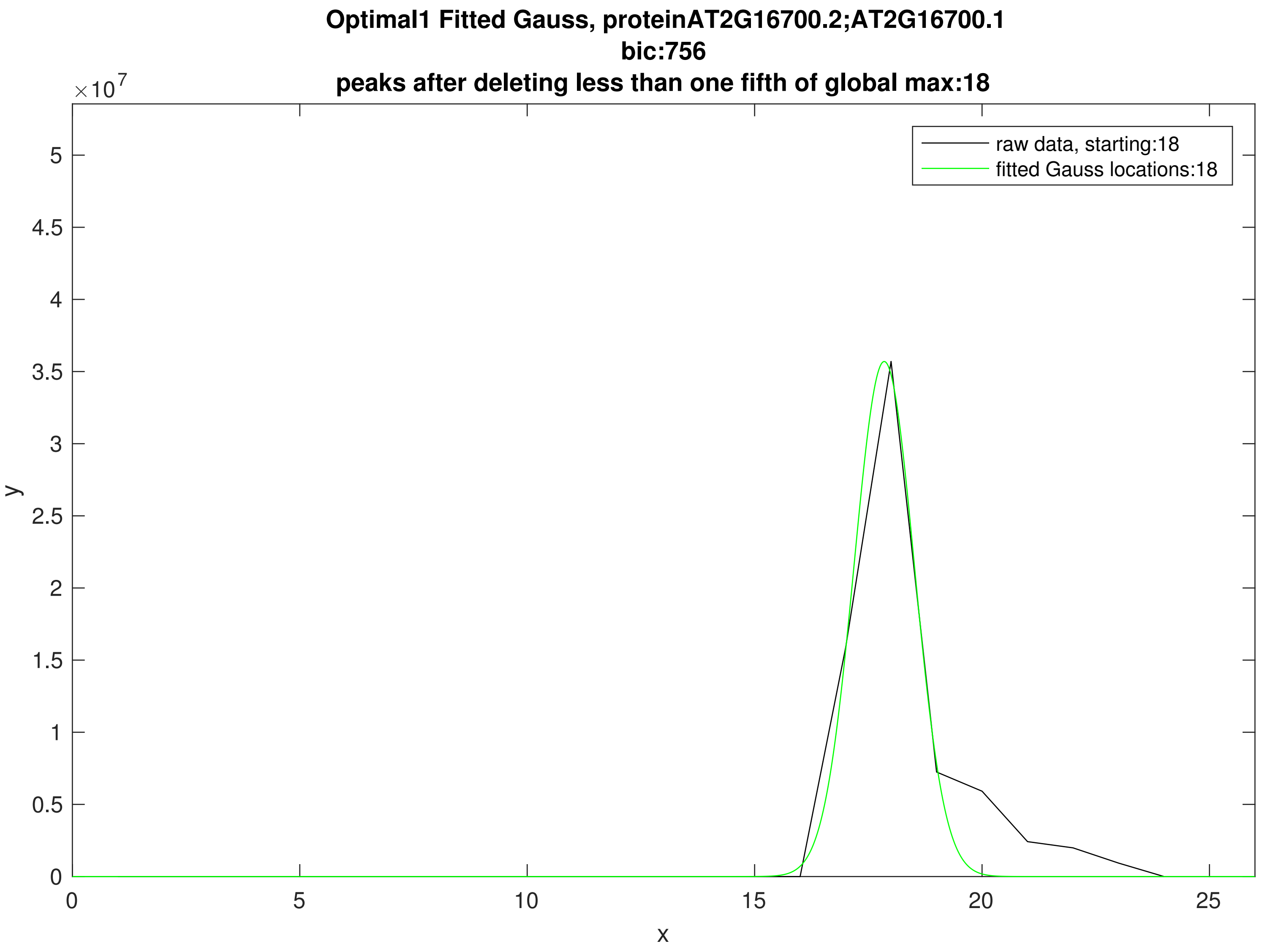

Optimal1 Fitted Gauss, proteinAT2G16700.2;AT2G16700.1
bic:756
peaks after deleting less than one fifth of global max:18
×10^7
raw data, starting:18
fitted Gauss locations:18
y
x
0
0.5
1
1.5
2
2.5
3
3.5
4
4.5
5
0
5
10
15
20
25

Optimal1 Fitted Gauss, proteinAT2G16780.1
bic:898 900
peaks after deleting less than one fifth of global max:15

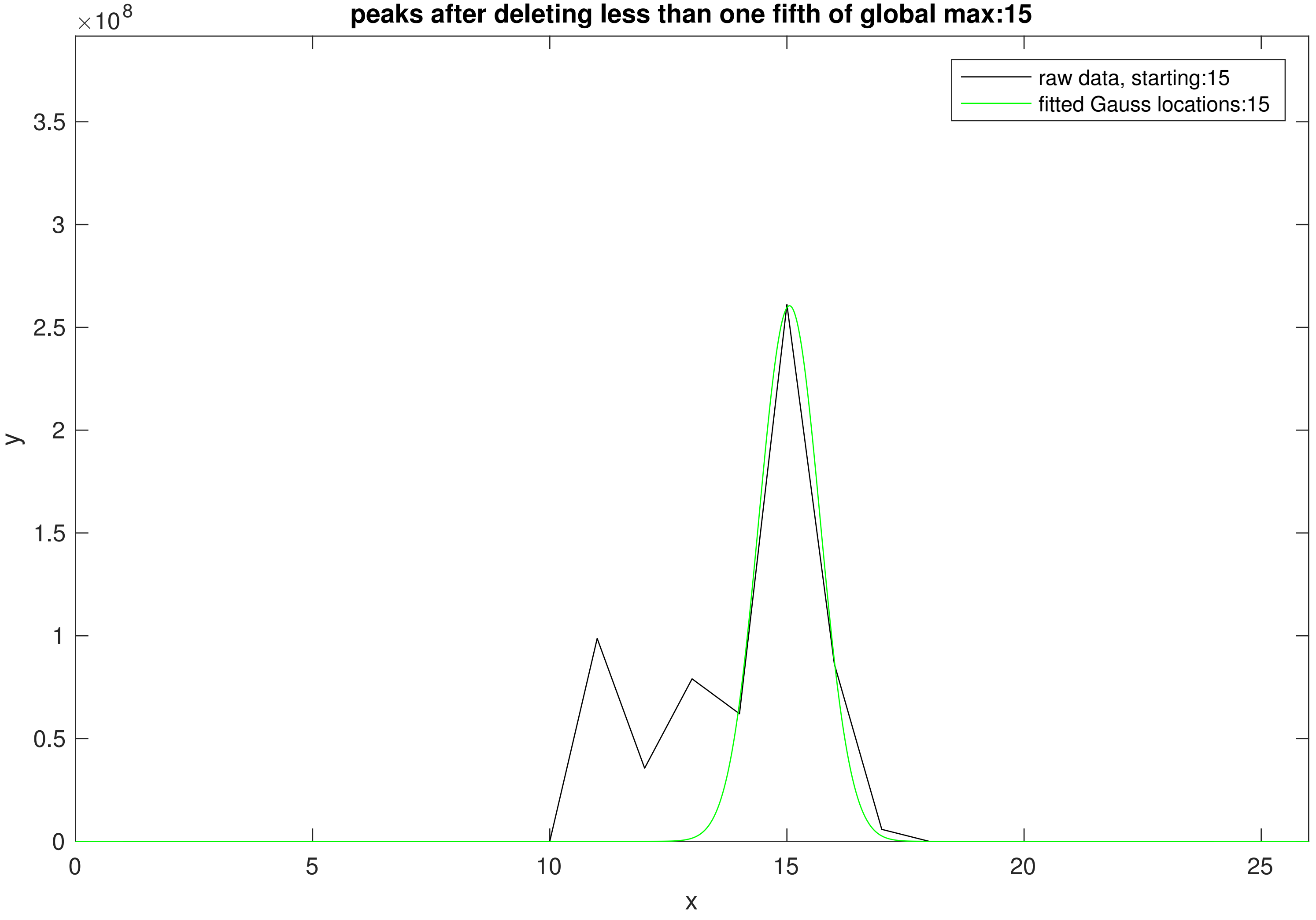

**Optimal1 Fitted Gauss, proteinAT2G16940.1;AT2G16940.3;AT2G16940.2**
**bic:679**
**peaks after deleting less than one fifth of global max:19**

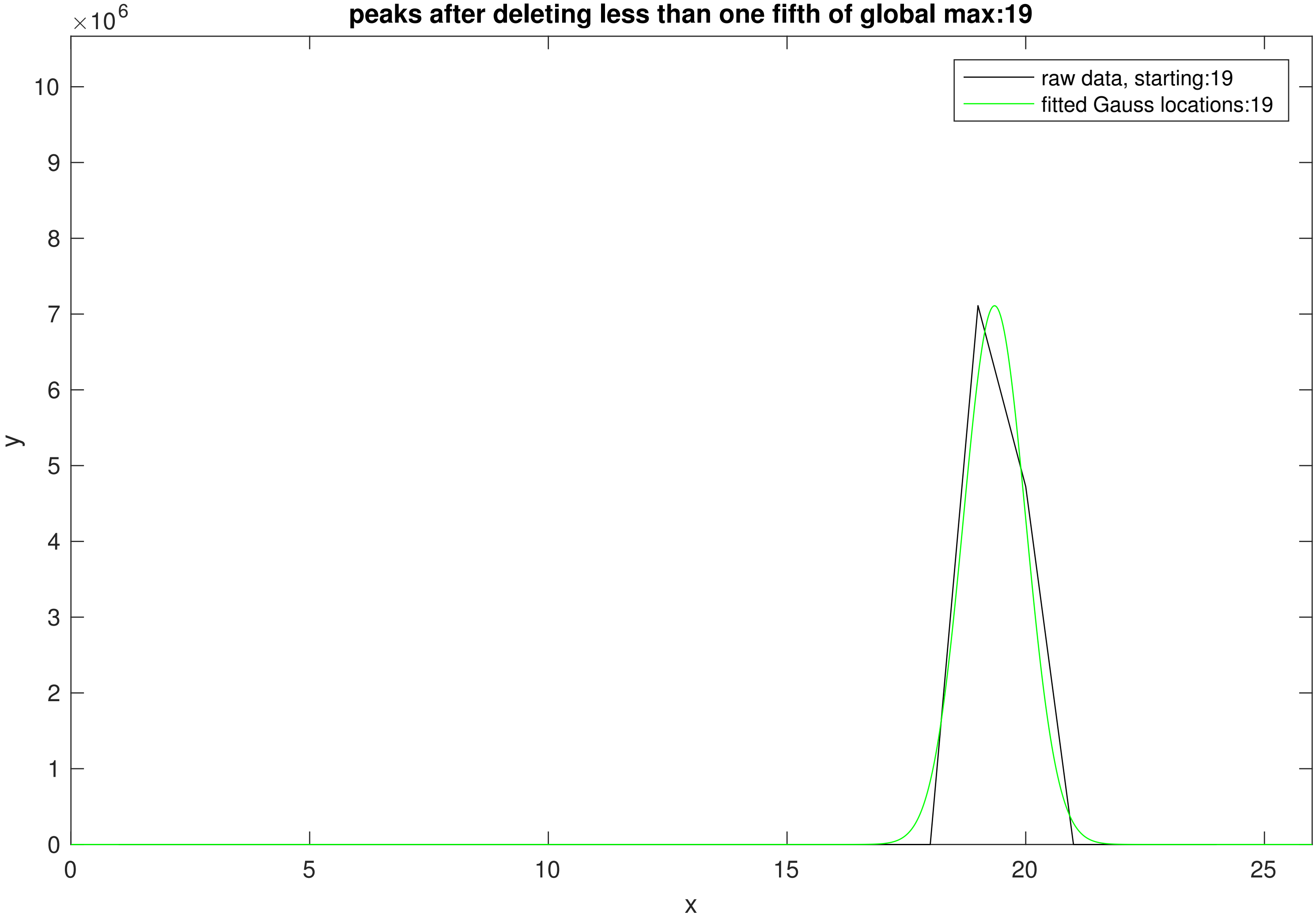

Optimal1 Fitted Gauss, proteinAT2G16950.2;AT2G16950.1
bic:696
peaks after deleting less than one fifth of global max:13

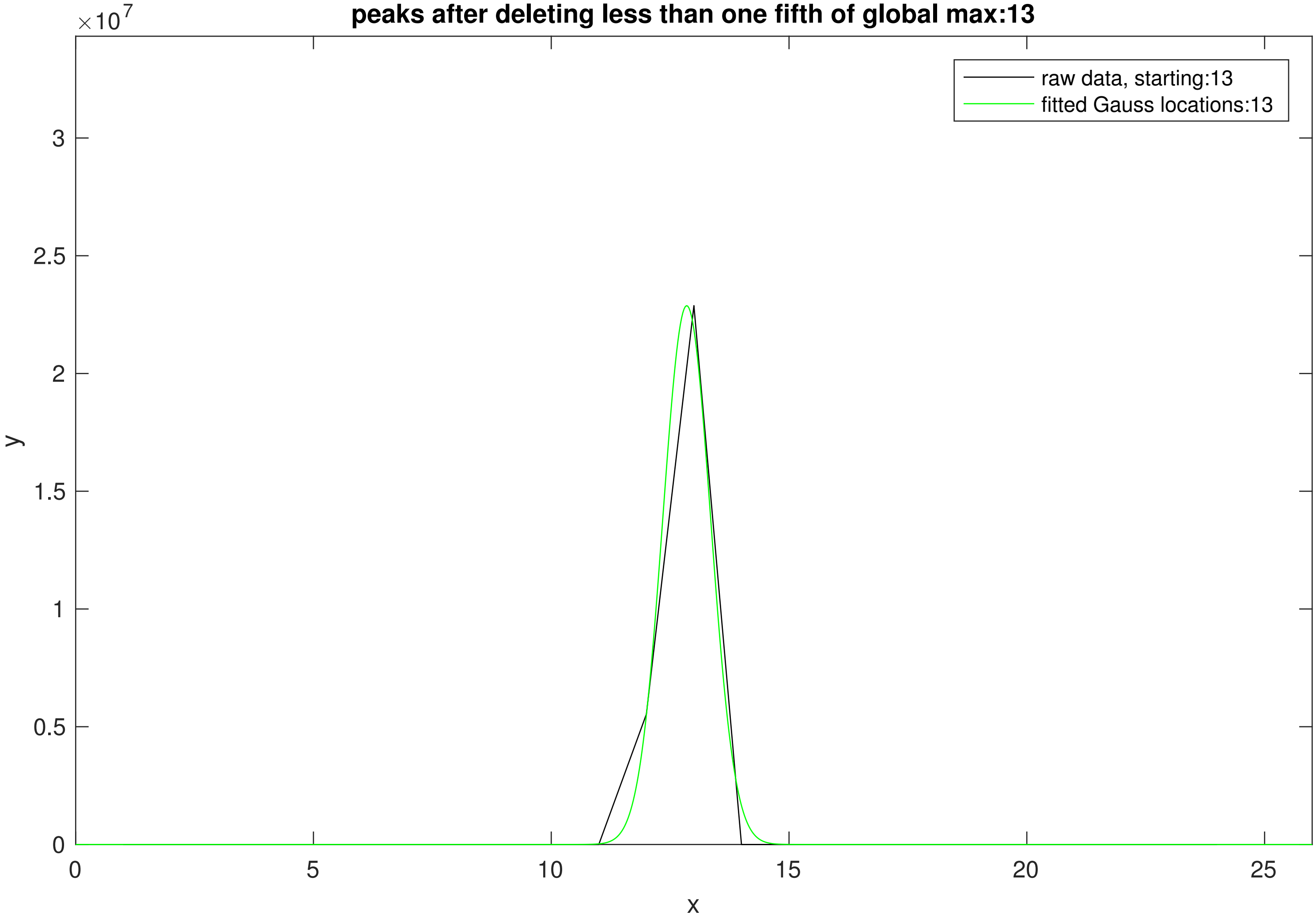

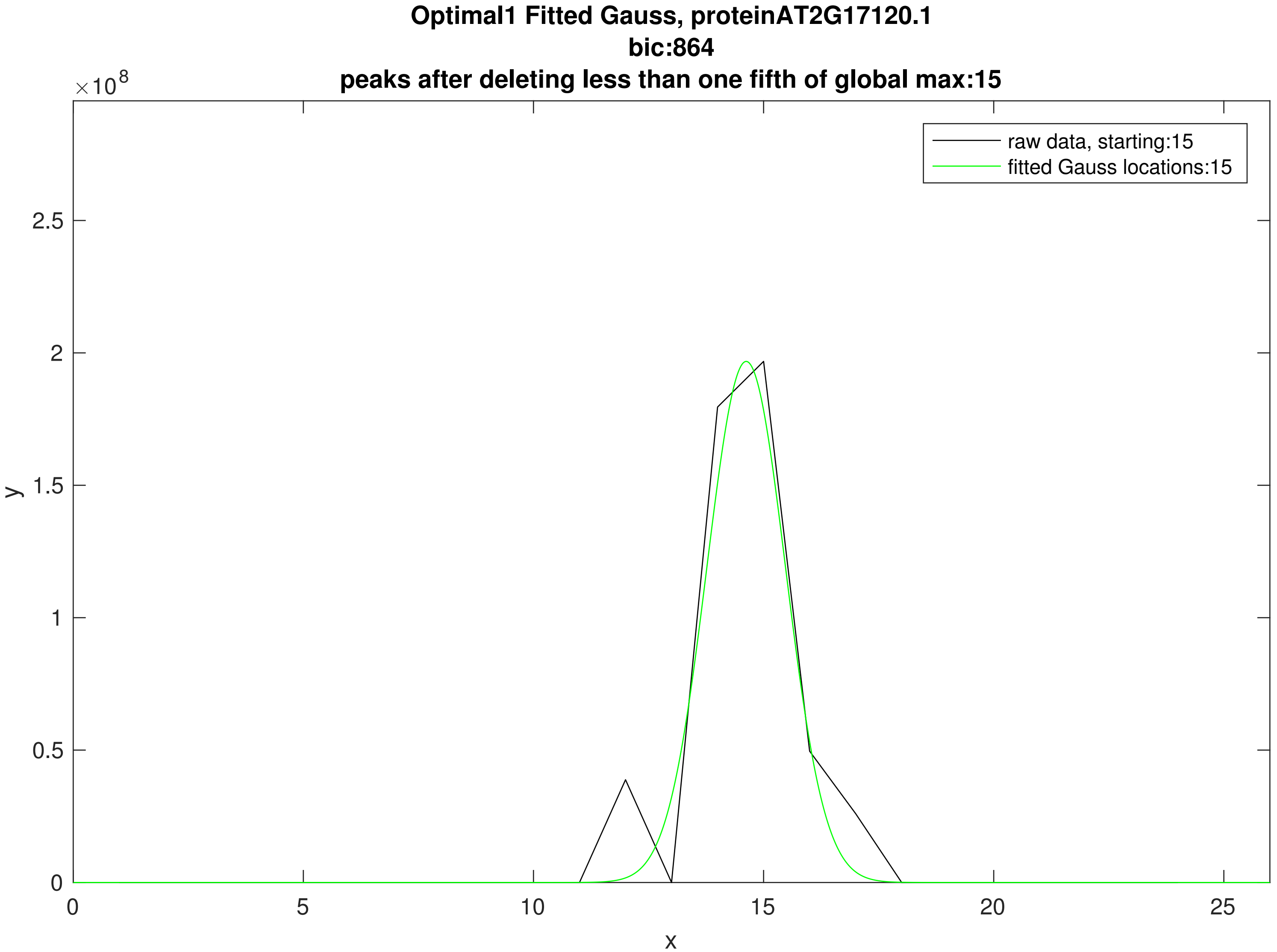
Optimal1 Fitted Gauss, proteinAT2G17120.1
bic:864
peaks after deleting less than one fifth of global max:15
×10^8
raw data, starting:15
fitted Gauss locations:15
2.5
2
1.5
1
0.5
0
y
0
5
10
15
20
25
x

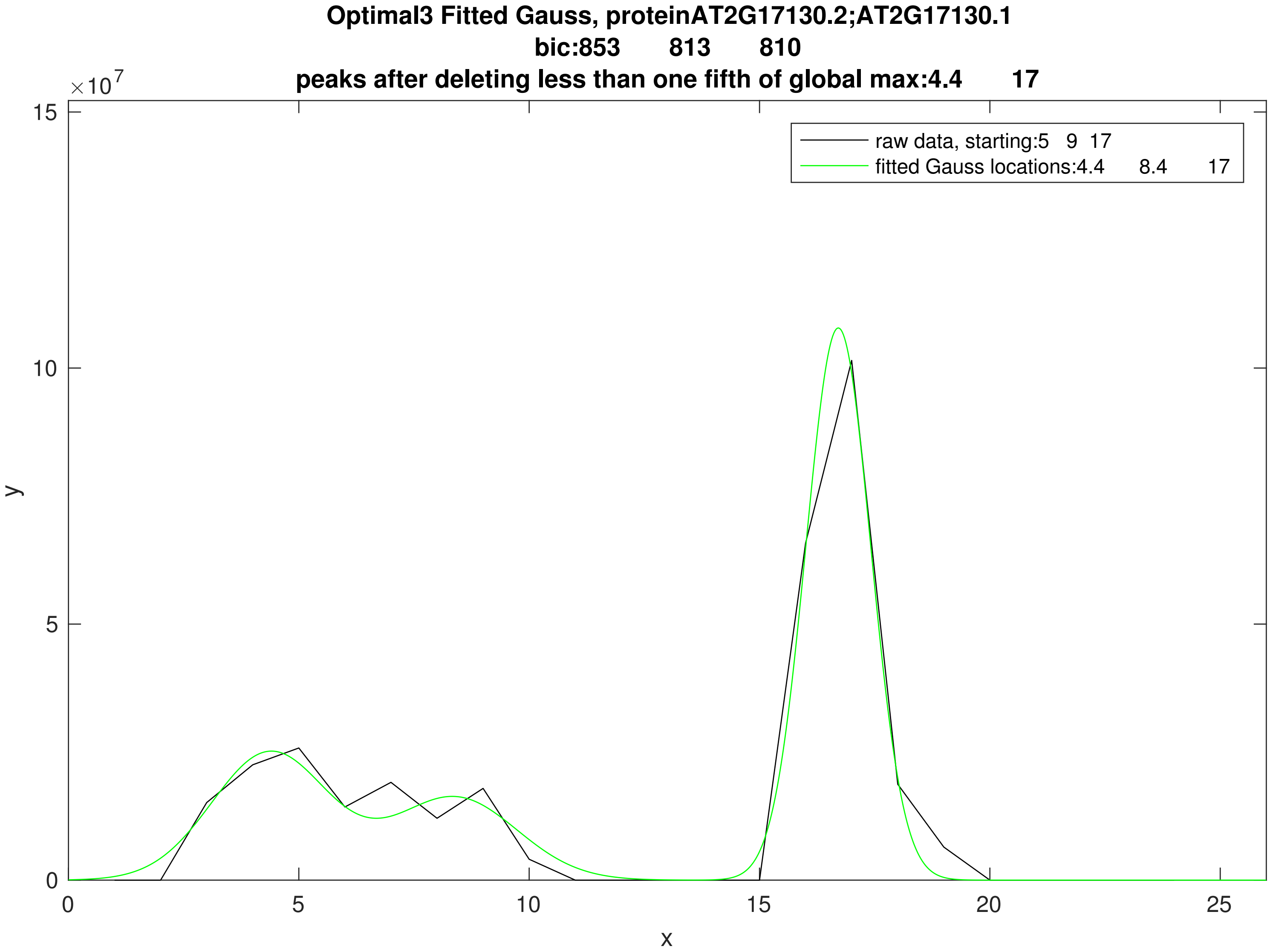

Optimal3 Fitted Gauss, proteinAT2G17130.2;AT2G17130.1
bic:853 813 810
peaks after deleting less than one fifth of global max:4.4 17
×10^7
15
10
5
0
y
0
5
10
15
20
25
x
raw data, starting:5 9 17
fitted Gauss locations:4.4 8.4 17

#### Optimal1 Fitted Gauss, proteinAT2G17190.1
#### bic:759
#### peaks after deleting less than one fifth of global max:19

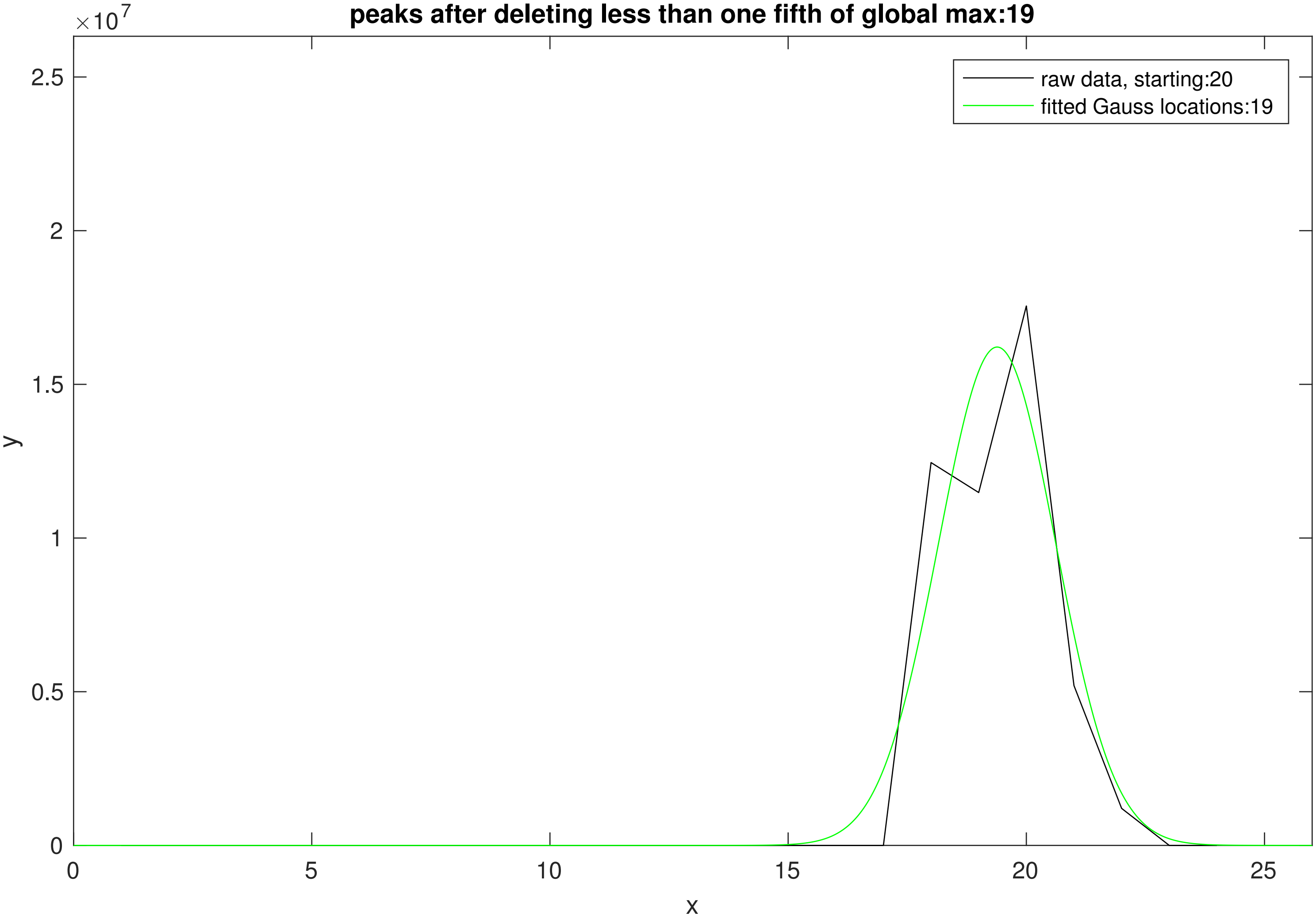

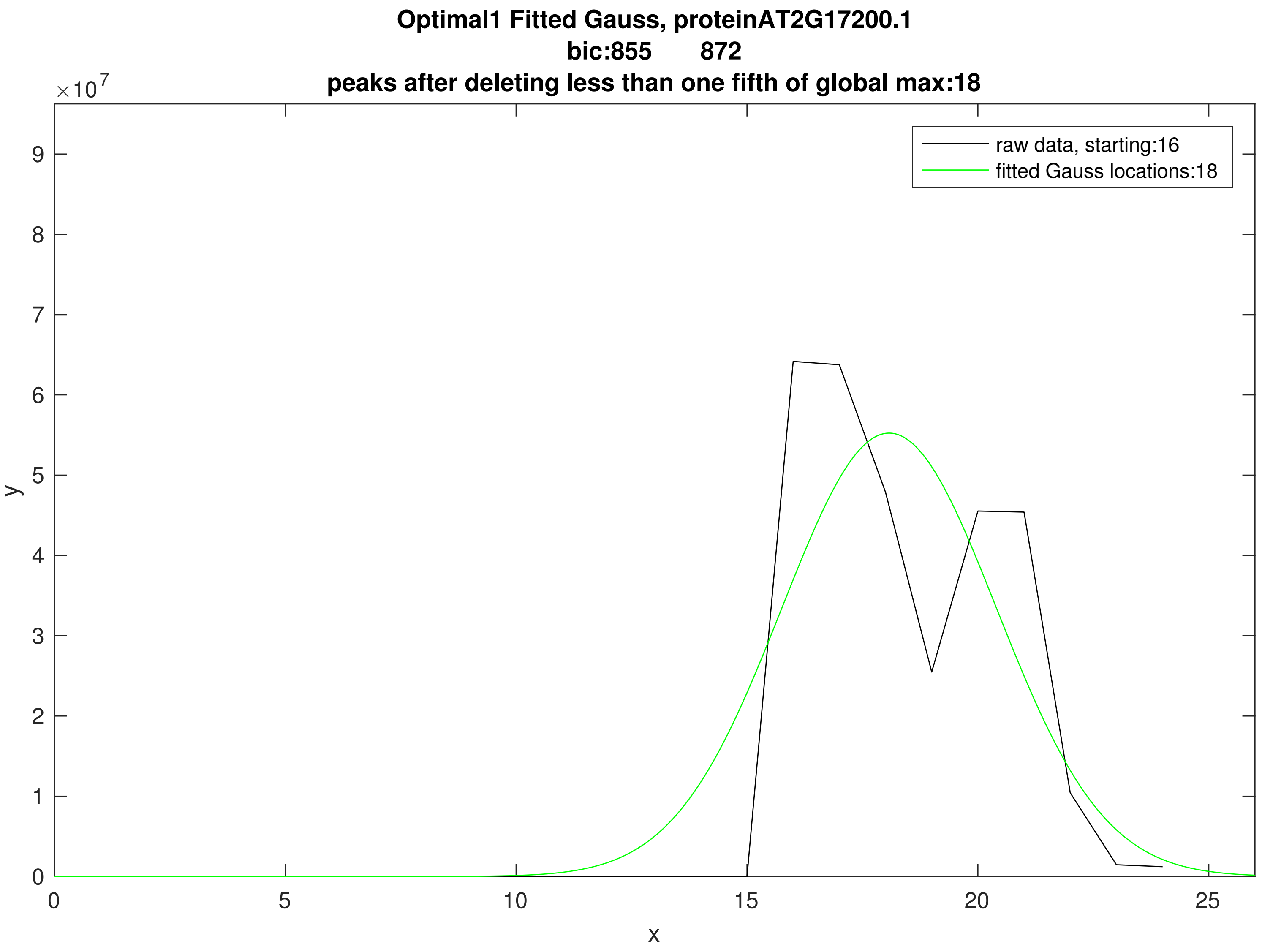

Optimal1 Fitted Gauss, proteinAT2G17200.1
bic:855 872
peaks after deleting less than one fifth of global max:18
×10^7
y
x
raw data, starting:16
fitted Gauss locations:18

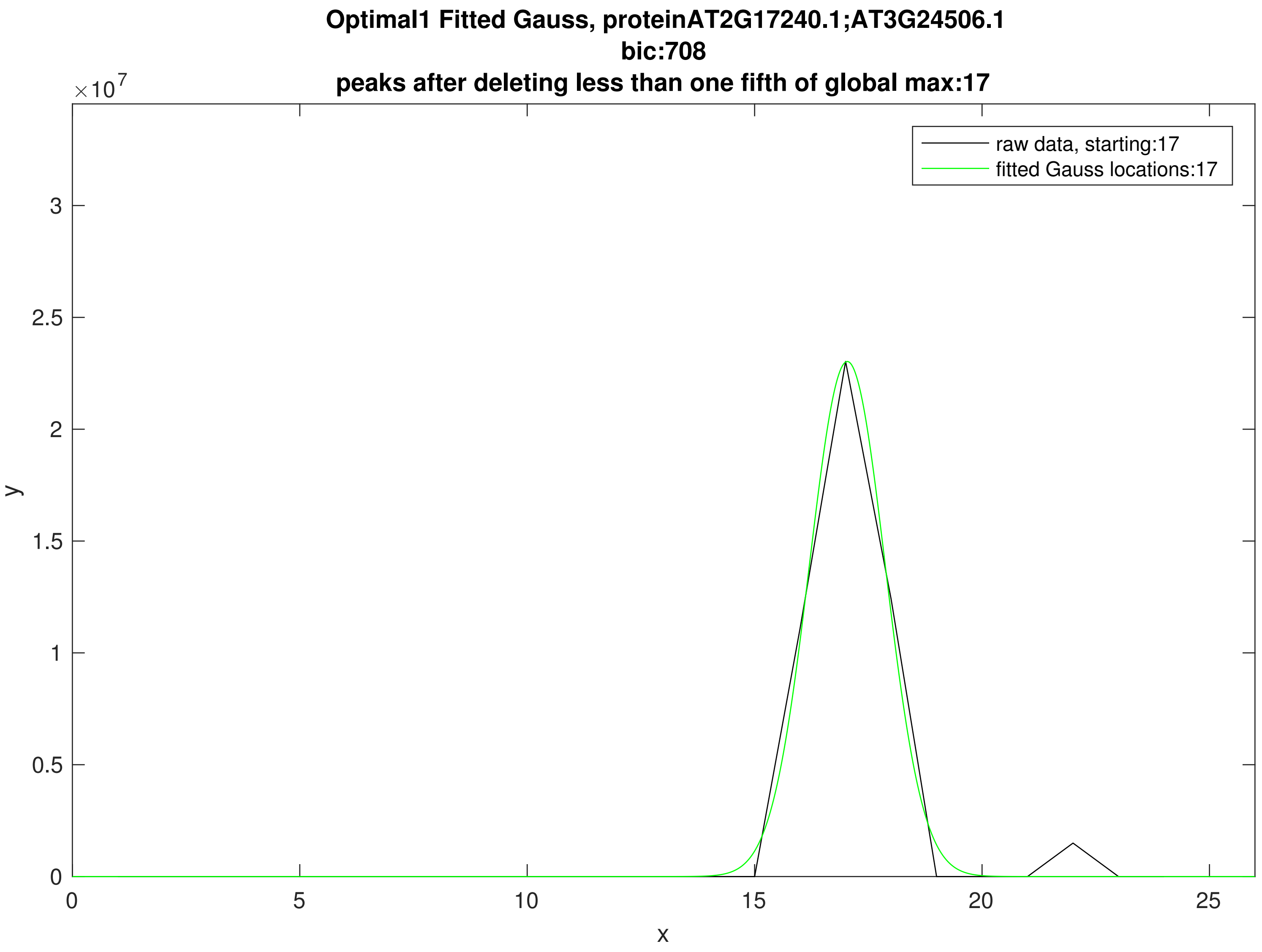

Optimal1 Fitted Gauss, proteinAT2G17240.1;AT3G24506.1
bic:708
peaks after deleting less than one fifth of global max:17
×10^7
raw data, starting:17
fitted Gauss locations:17
y
x
0
0.5
1
1.5
2
2.5
3
0
5
10
15
20
25

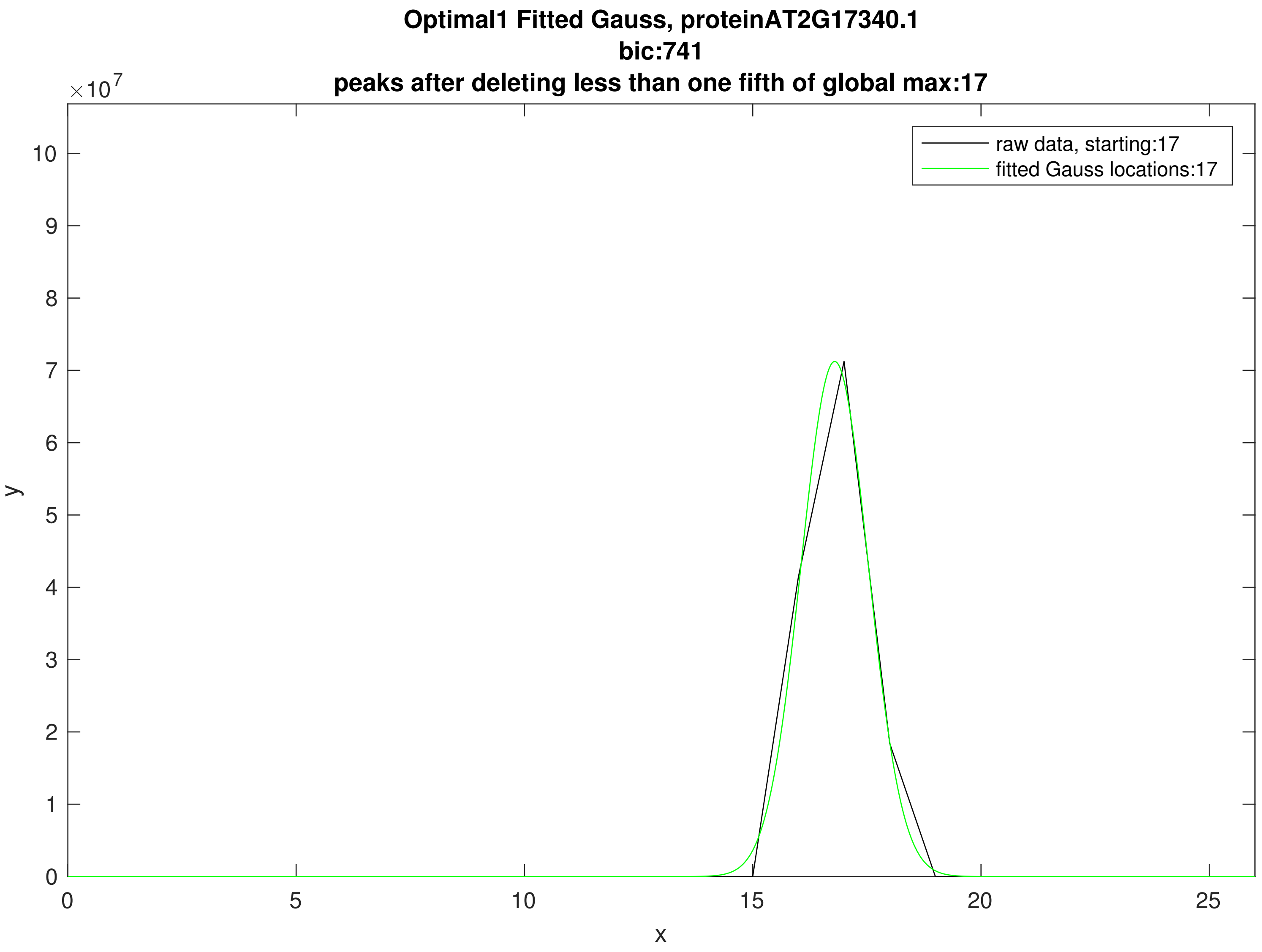
Optimal1 Fitted Gauss, proteinAT2G17340.1
bic:741
peaks after deleting less than one fifth of global max:17
×10^7
raw data, starting:17
fitted Gauss locations:17
y
x
0
1
2
3
4
5
6
7
8
9
10
0
5
10
15
20
25

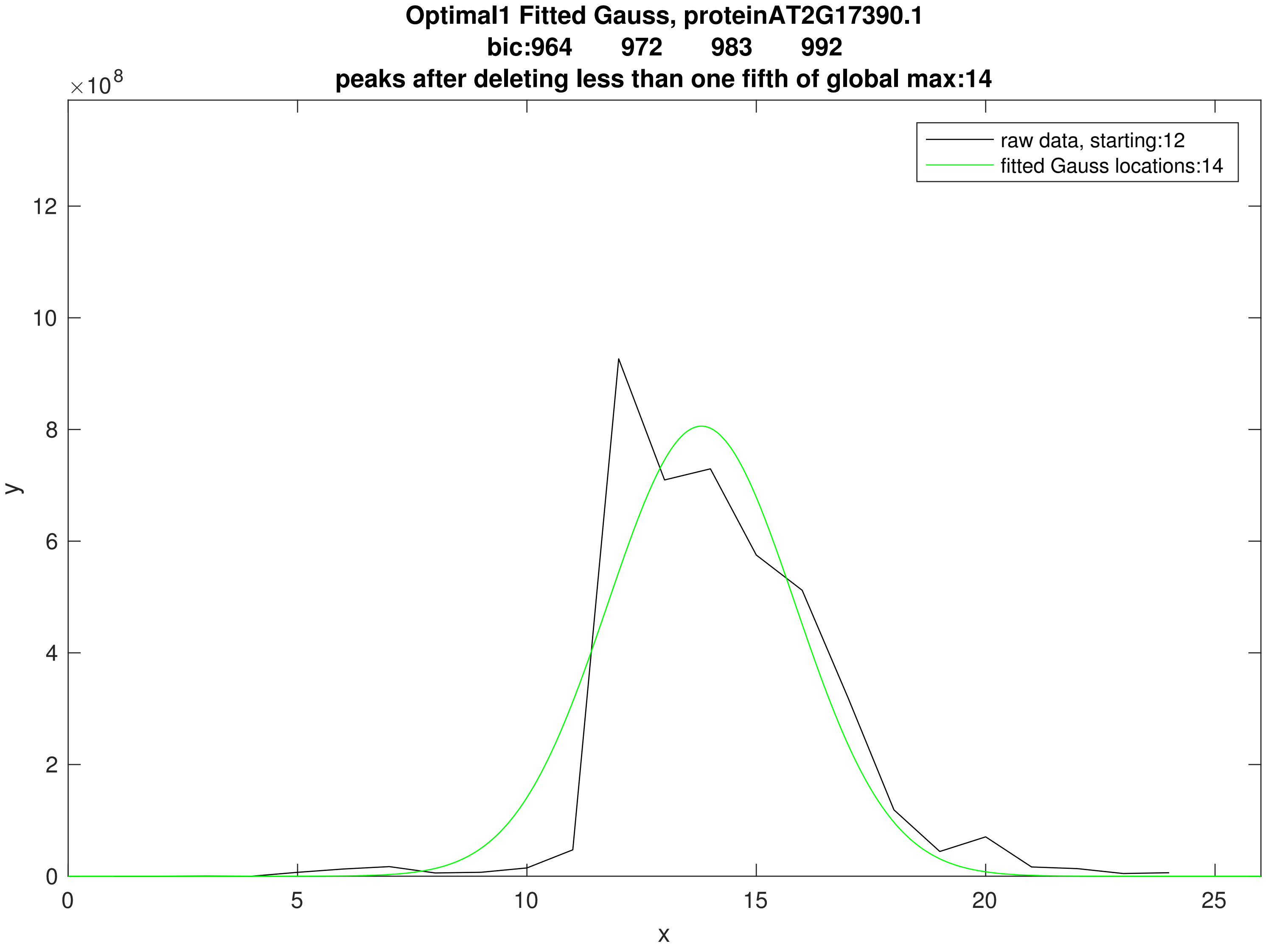

Optimal1 Fitted Gauss, proteinAT2G17390.1
bic:964 972 983 992
peaks after deleting less than one fifth of global max:14
×10^8
raw data, starting:12
fitted Gauss locations:14
y
x
0
2
4
6
8
10
12
0
5
10
15
20
25

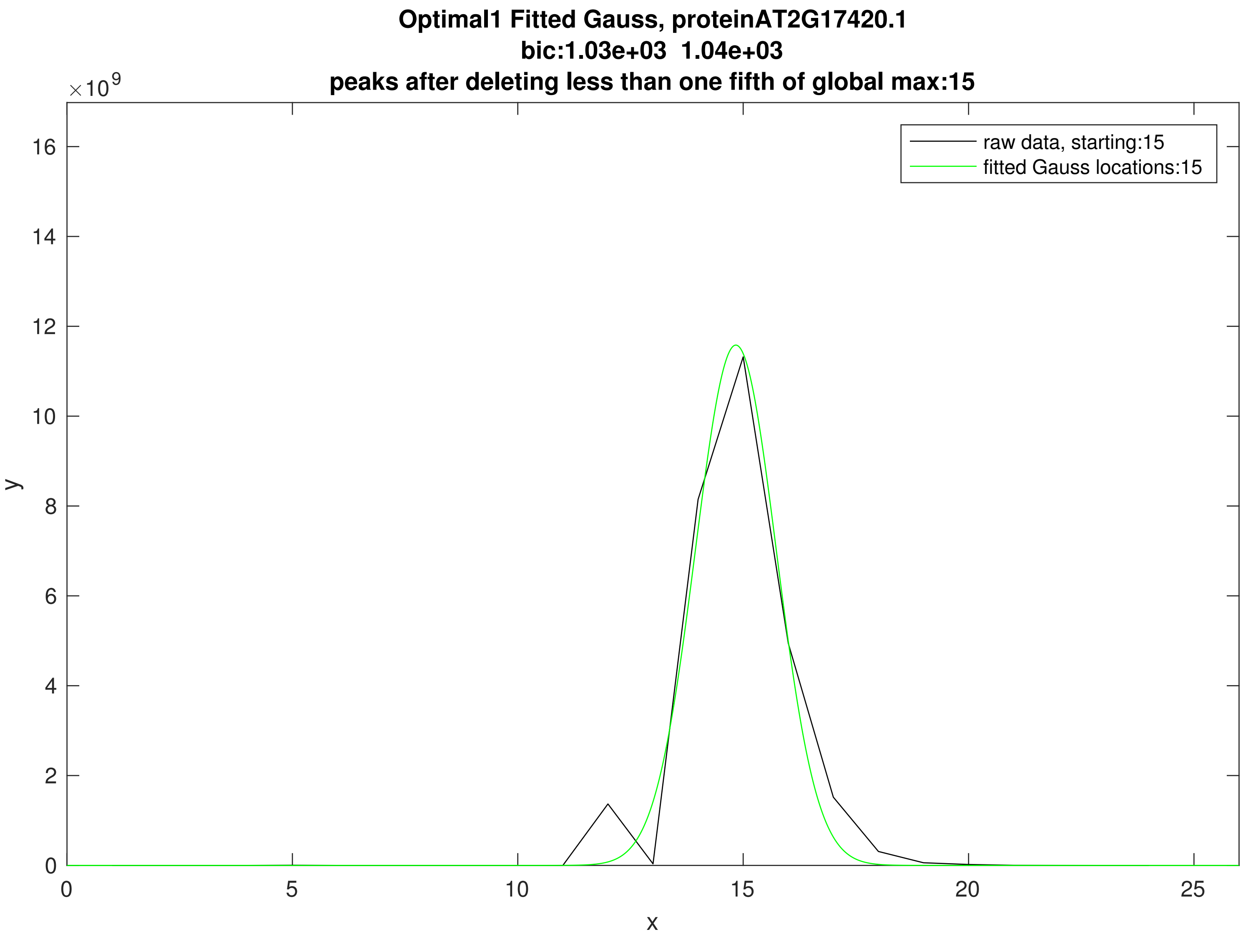

Optimal1 Fitted Gauss, proteinAT2G17420.1
bic:1.03e+03  1.04e+03
peaks after deleting less than one fifth of global max:15
×10^9
raw data, starting:15
fitted Gauss locations:15
y
x
0
2
4
6
8
10
12
14
16
0
5
10
15
20
25

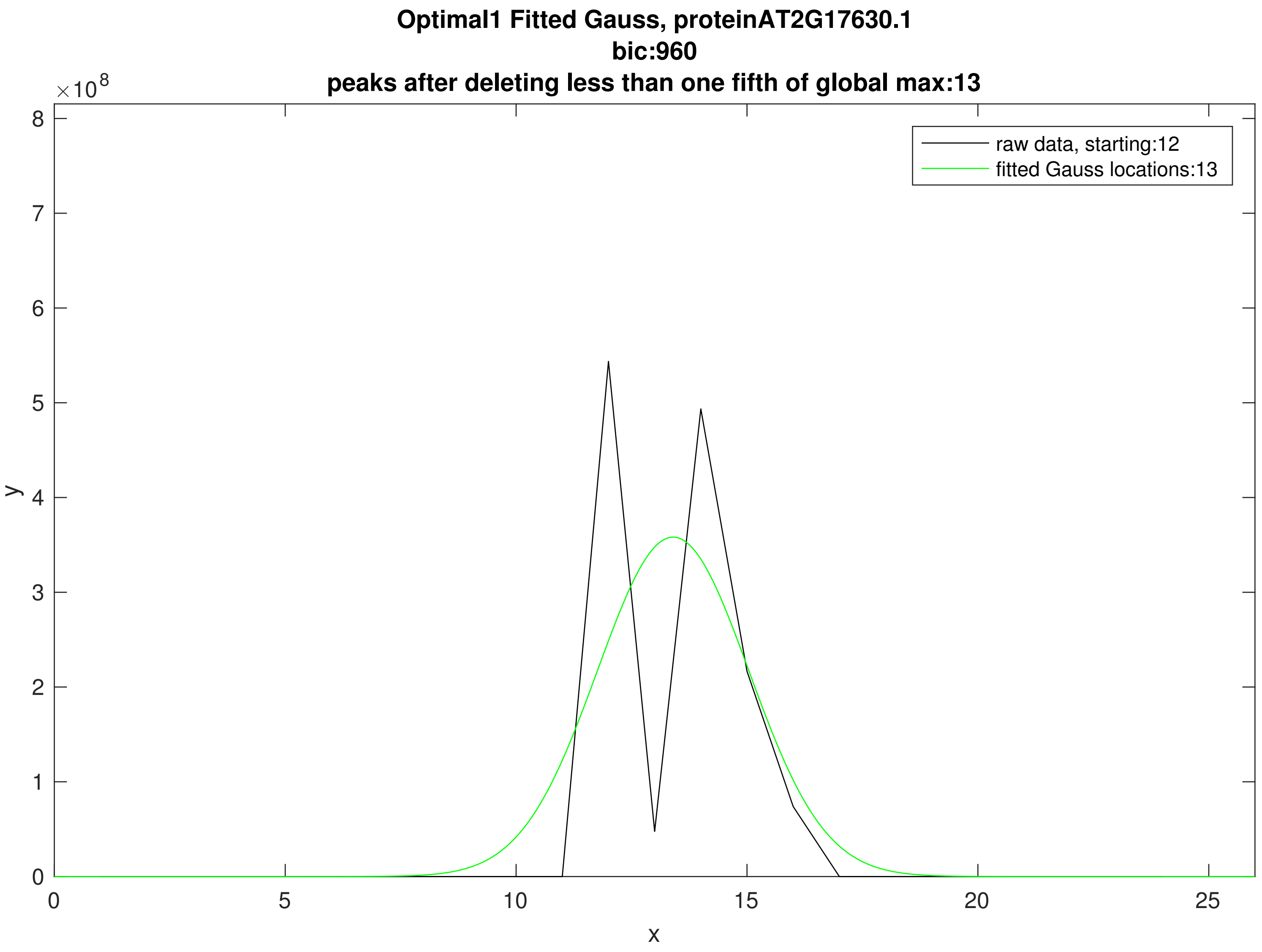

Optimal1 Fitted Gauss, proteinAT2G17630.1
bic:960
peaks after deleting less than one fifth of global max:13
×10^8
raw data, starting:12
fitted Gauss locations:13
y
x
0
1
2
3
4
5
6
7
8
0
5
10
15
20
25

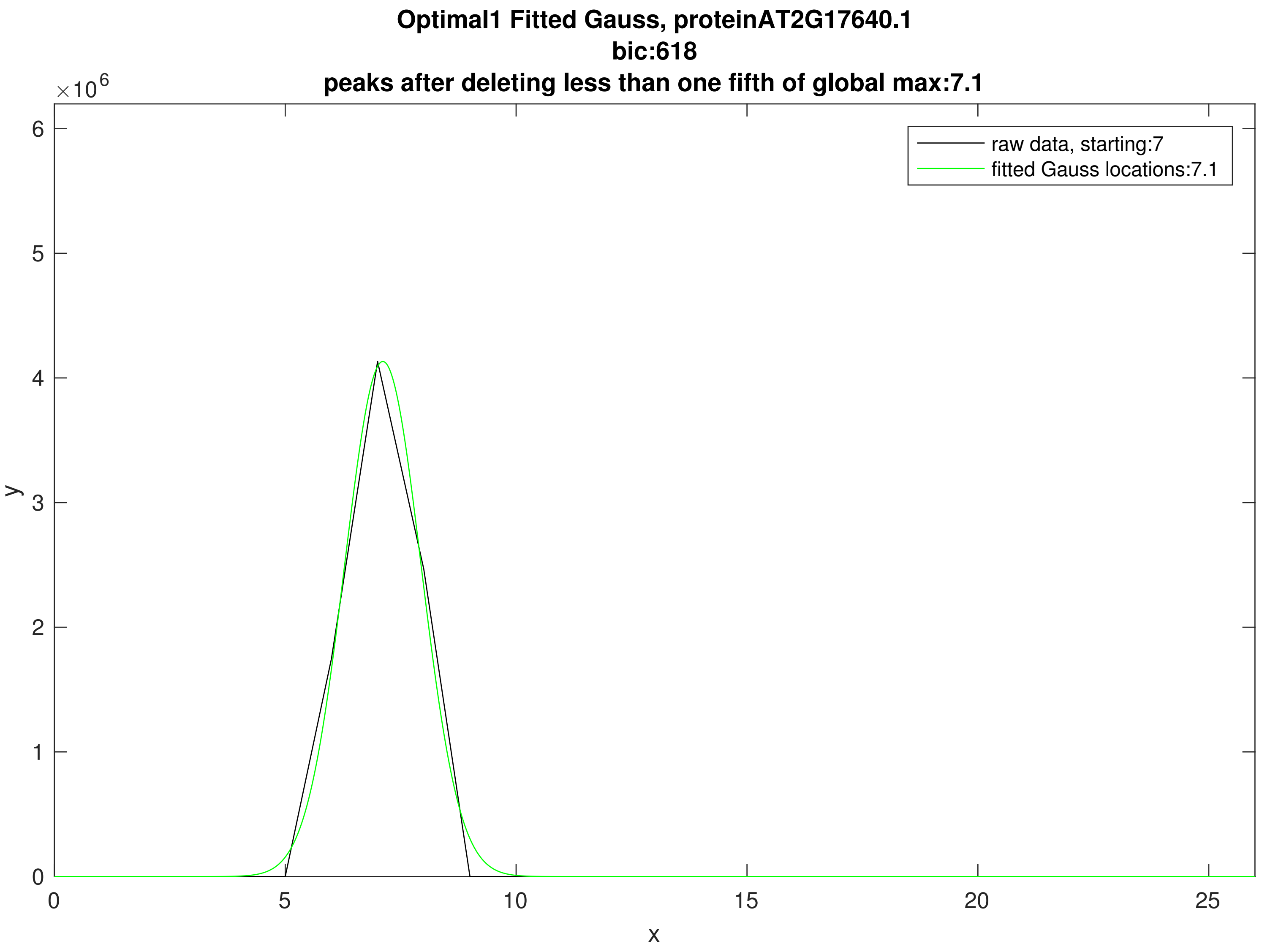
Optimal1 Fitted Gauss, proteinAT2G17640.1
bic:618
peaks after deleting less than one fifth of global max:7.1
×10^6
raw data, starting:7
fitted Gauss locations:7.1
y
x
0
1
2
3
4
5
6
0
5
10
15
20
25

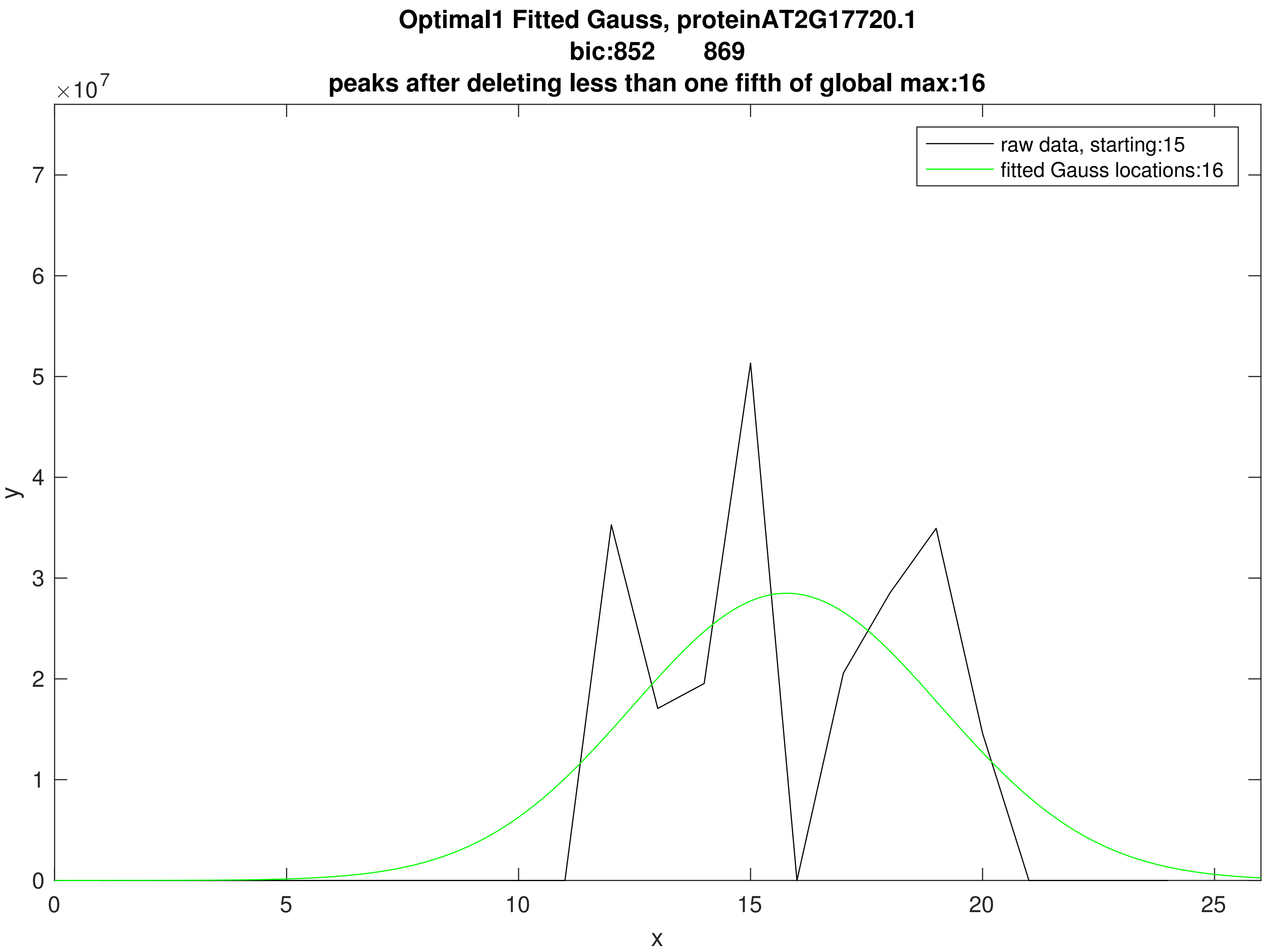
Optimal1 Fitted Gauss, proteinAT2G17720.1
bic:852 869
peaks after deleting less than one fifth of global max:16
×10^7
y
x
0
1
2
3
4
5
6
7
0
5
10
15
20
25
raw data, starting:15
fitted Gauss locations:16

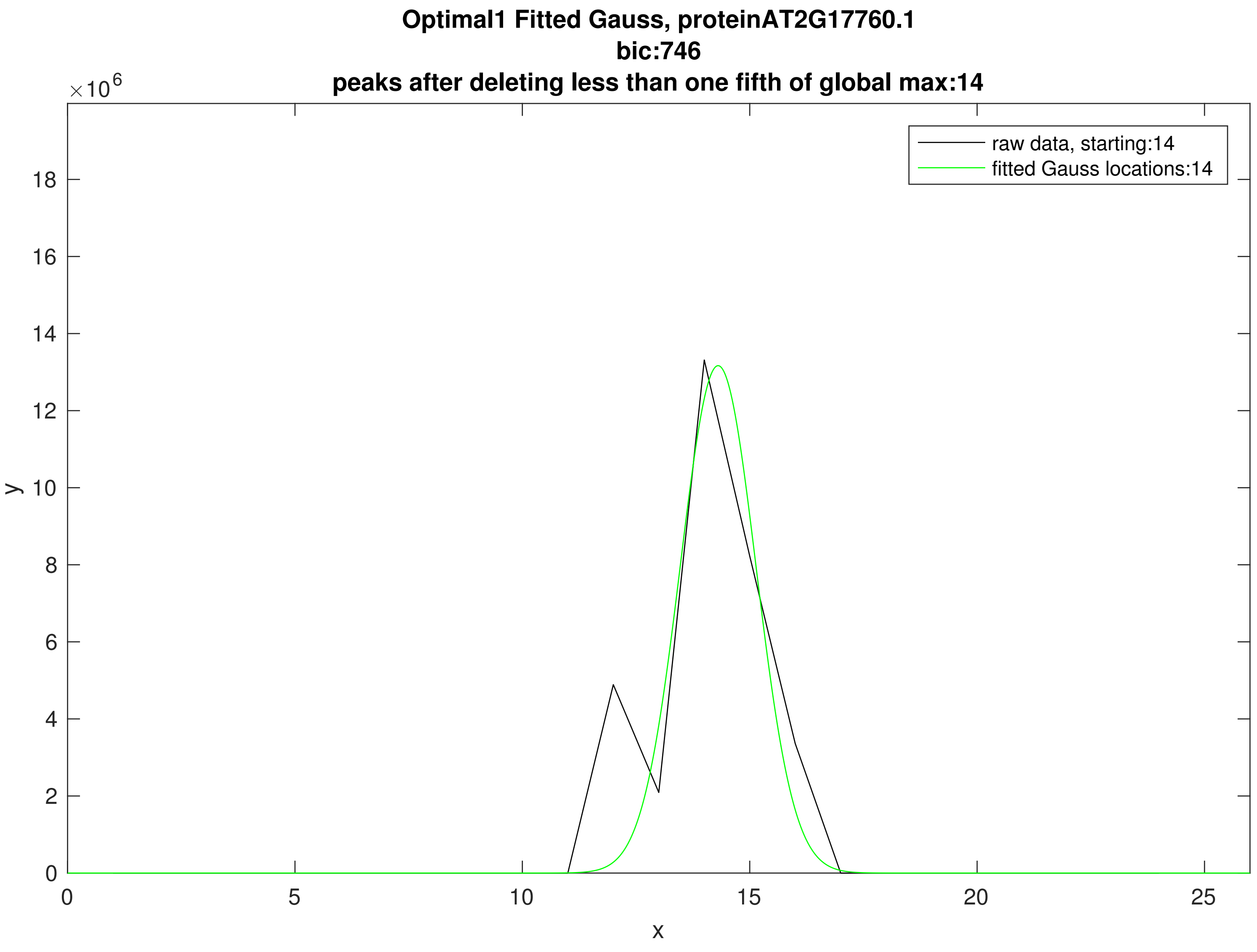

Optimal1 Fitted Gauss, proteinAT2G17760.1
bic:746
peaks after deleting less than one fifth of global max:14
×10^6
raw data, starting:14
fitted Gauss locations:14
y
x
0
2
4
6
8
10
12
14
16
18
0
5
10
15
20
25

**Optimal1 Fitted Gauss, proteinAT2G17870.1**
**bic:794 796 804 813**
**peaks after deleting less than one fifth of global max:17**

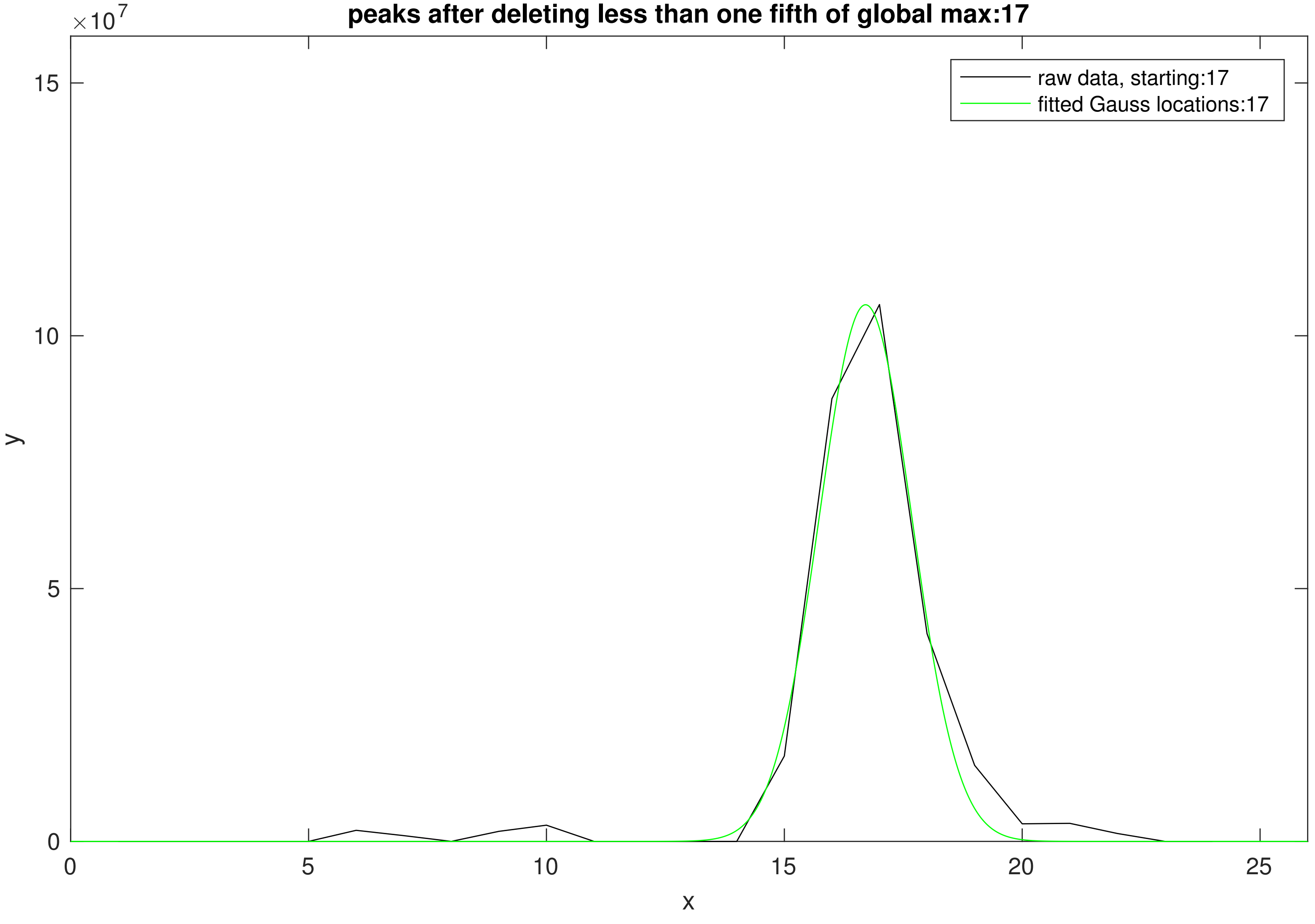

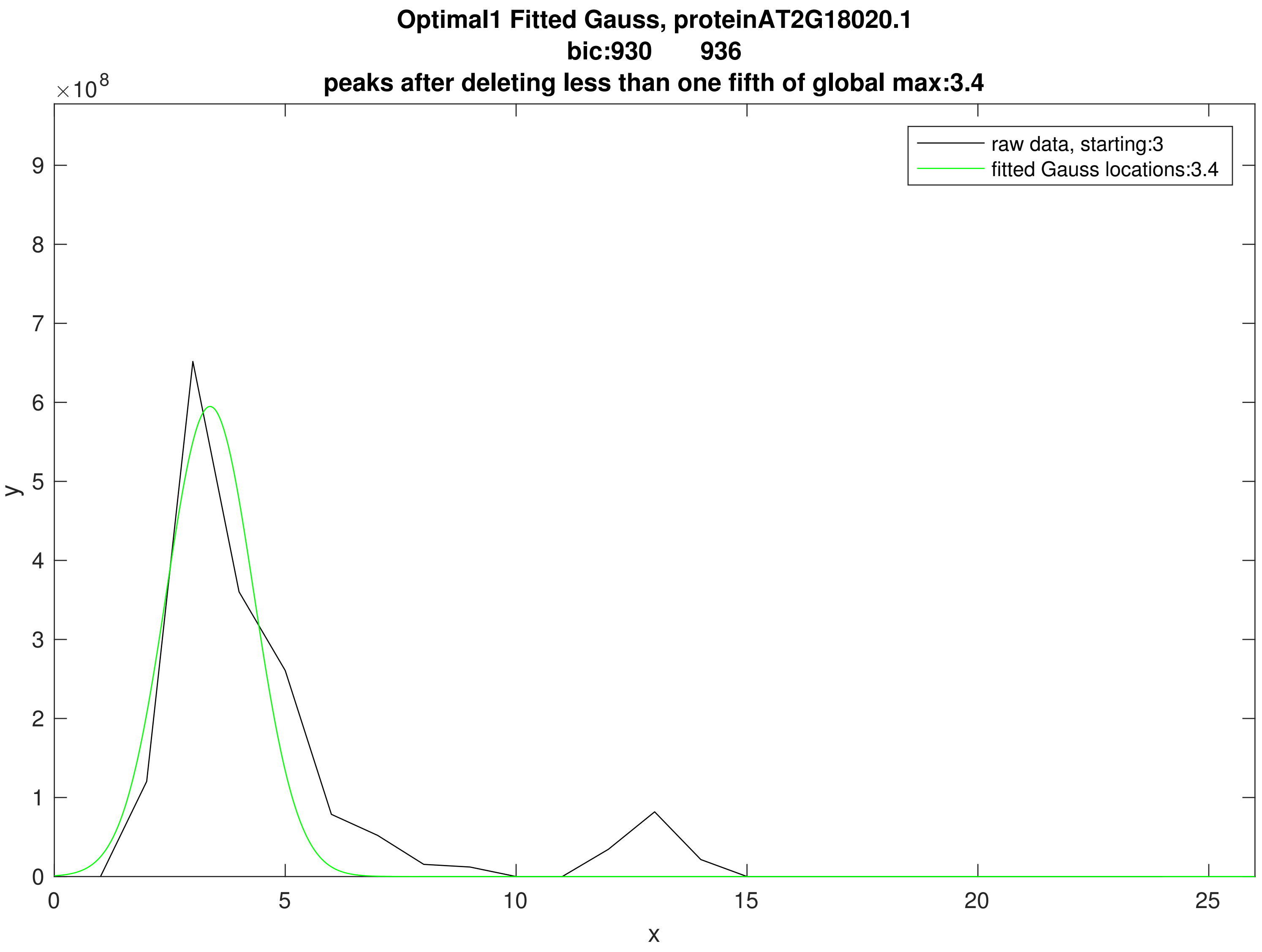
Optimal1 Fitted Gauss, proteinAT2G18020.1
bic:930 936
peaks after deleting less than one fifth of global max:3.4
×10^8
raw data, starting:3
fitted Gauss locations:3.4
y
x
0
1
2
3
4
5
6
7
8
9
0
5
10
15
20
25

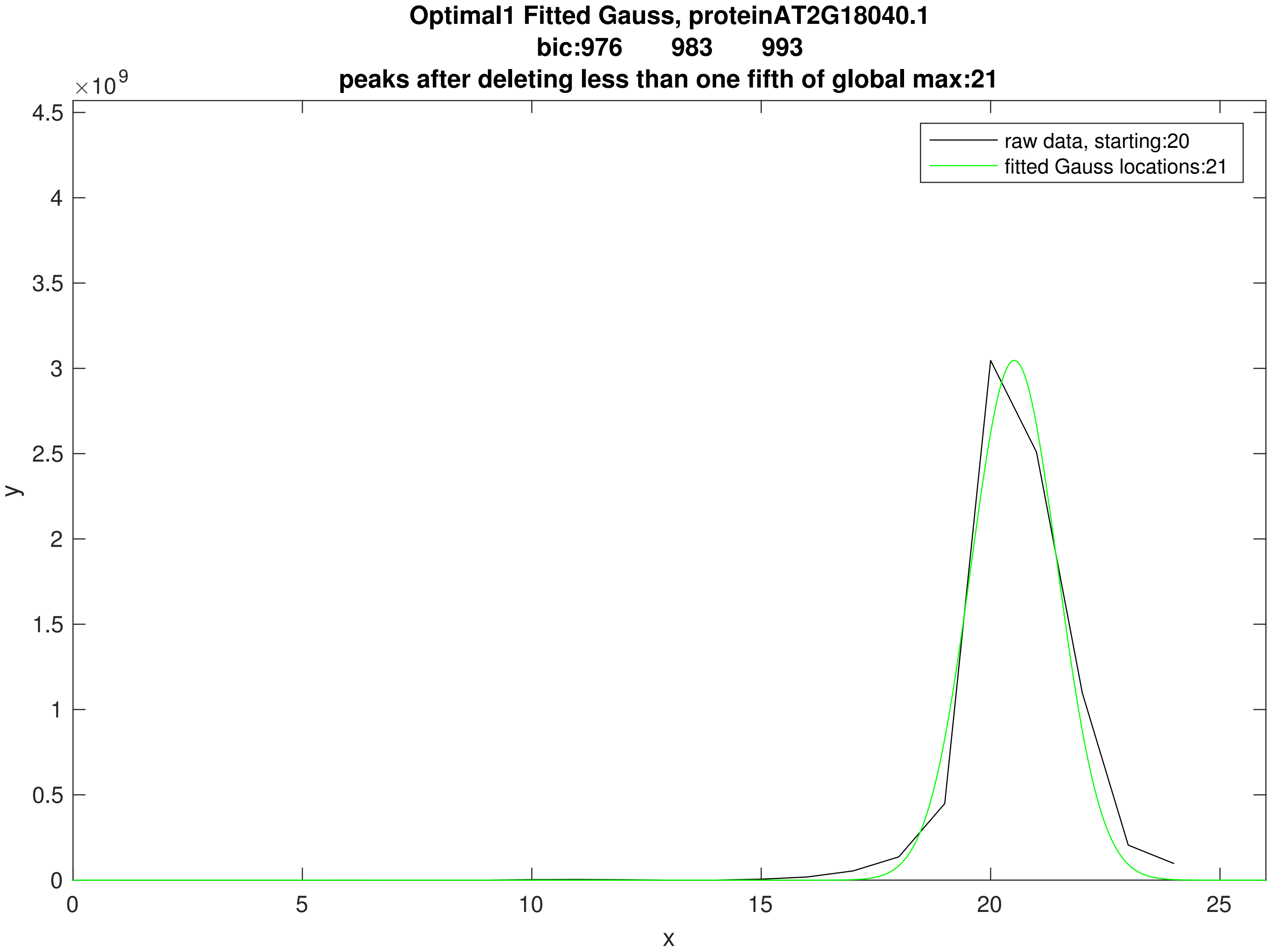
Optimal1 Fitted Gauss, proteinAT2G18040.1
bic:976 983 993
peaks after deleting less than one fifth of global max:21
×10^9
raw data, starting:20
fitted Gauss locations:21
y
x
0
0.5
1
1.5
2
2.5
3
3.5
4
4.5
0
5
10
15
20
25

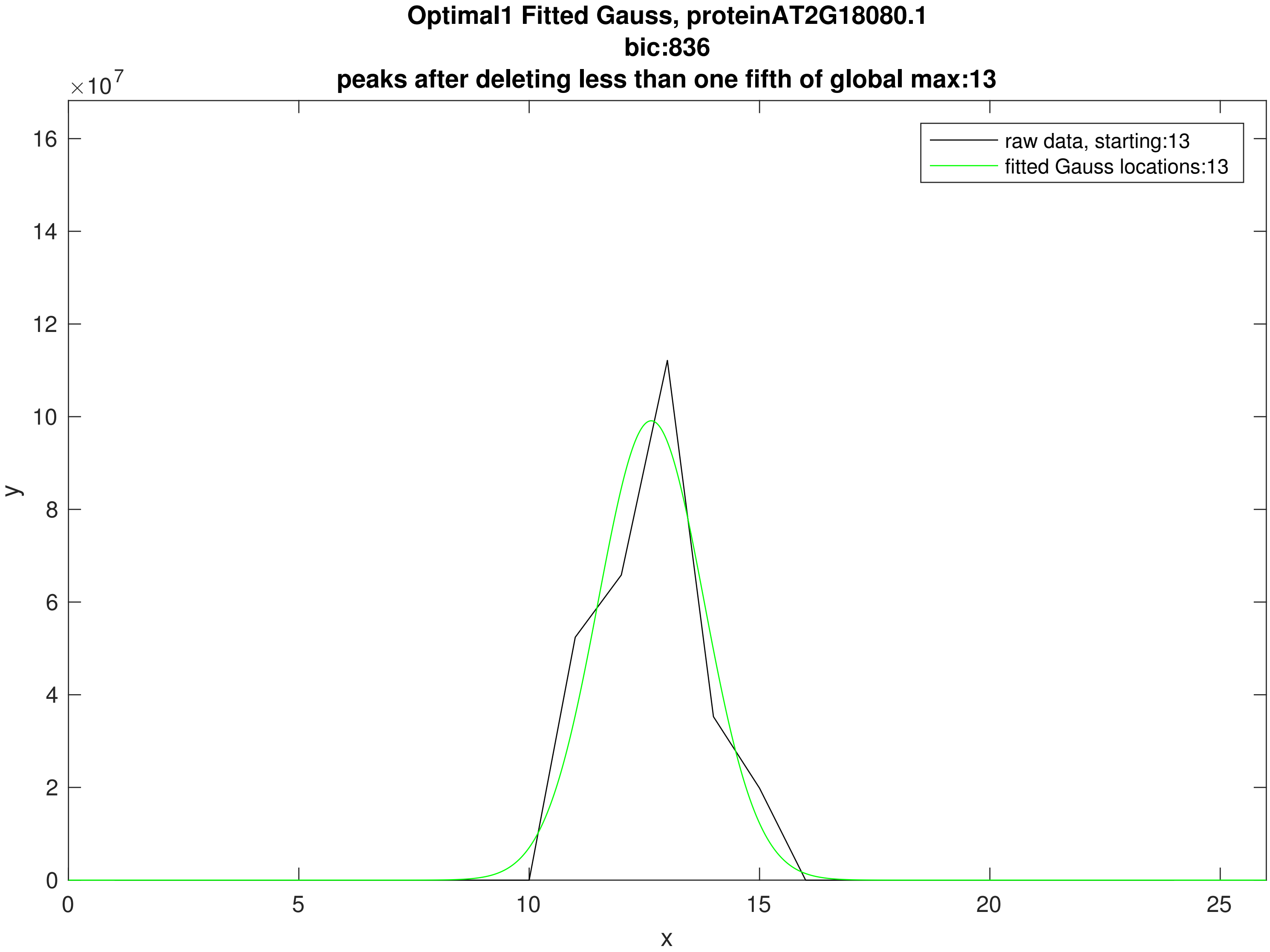
Optimal1 Fitted Gauss, proteinAT2G18080.1
bic:836
peaks after deleting less than one fifth of global max:13
×10^7
raw data, starting:13
fitted Gauss locations:13
y
x
0
2
4
6
8
10
12
14
16
0
5
10
15
20
25

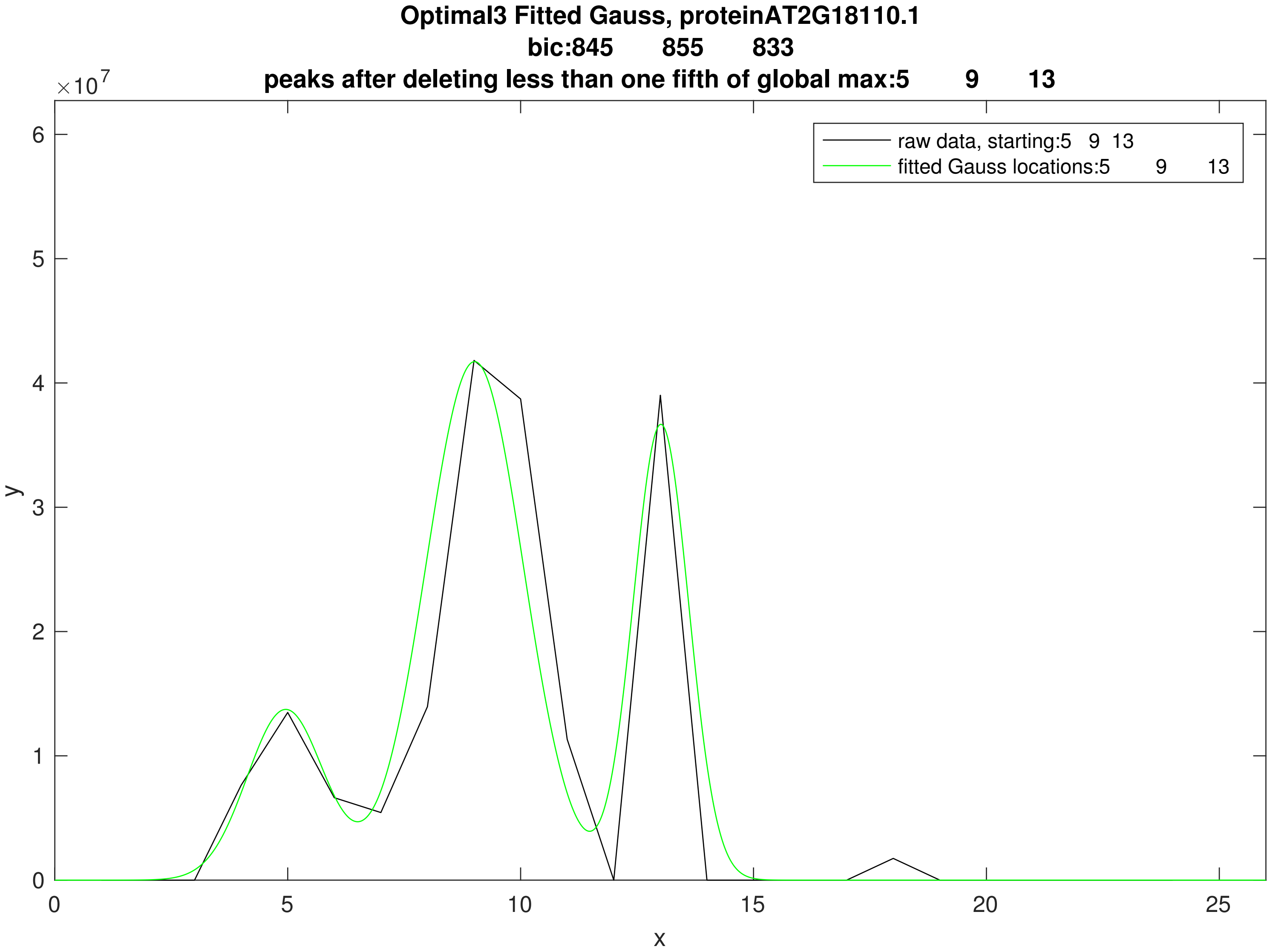
Optimal3 Fitted Gauss, proteinAT2G18110.1
bic:845 855 833
peaks after deleting less than one fifth of global max:5 9 13
×10^7
raw data, starting:5 9 13
fitted Gauss locations:5 9 13
y
x
0
1
2
3
4
5
6
0
5
10
15
20
25

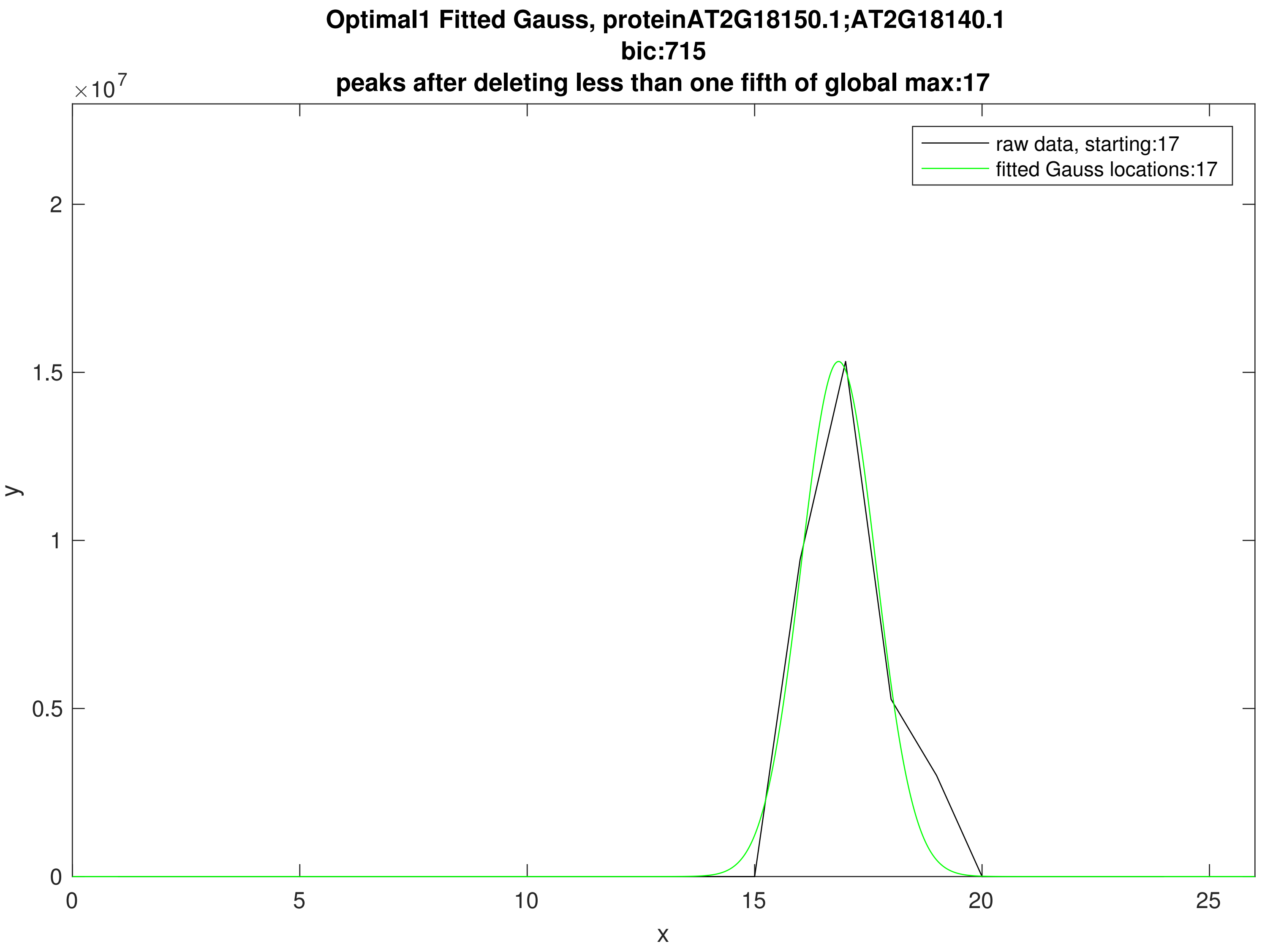

Optimal1 Fitted Gauss, proteinAT2G18150.1;AT2G18140.1
bic:715
peaks after deleting less than one fifth of global max:17
×10^7
raw data, starting:17
fitted Gauss locations:17
2
1.5
1
0.5
0
y
0
5
10
15
20
25
x

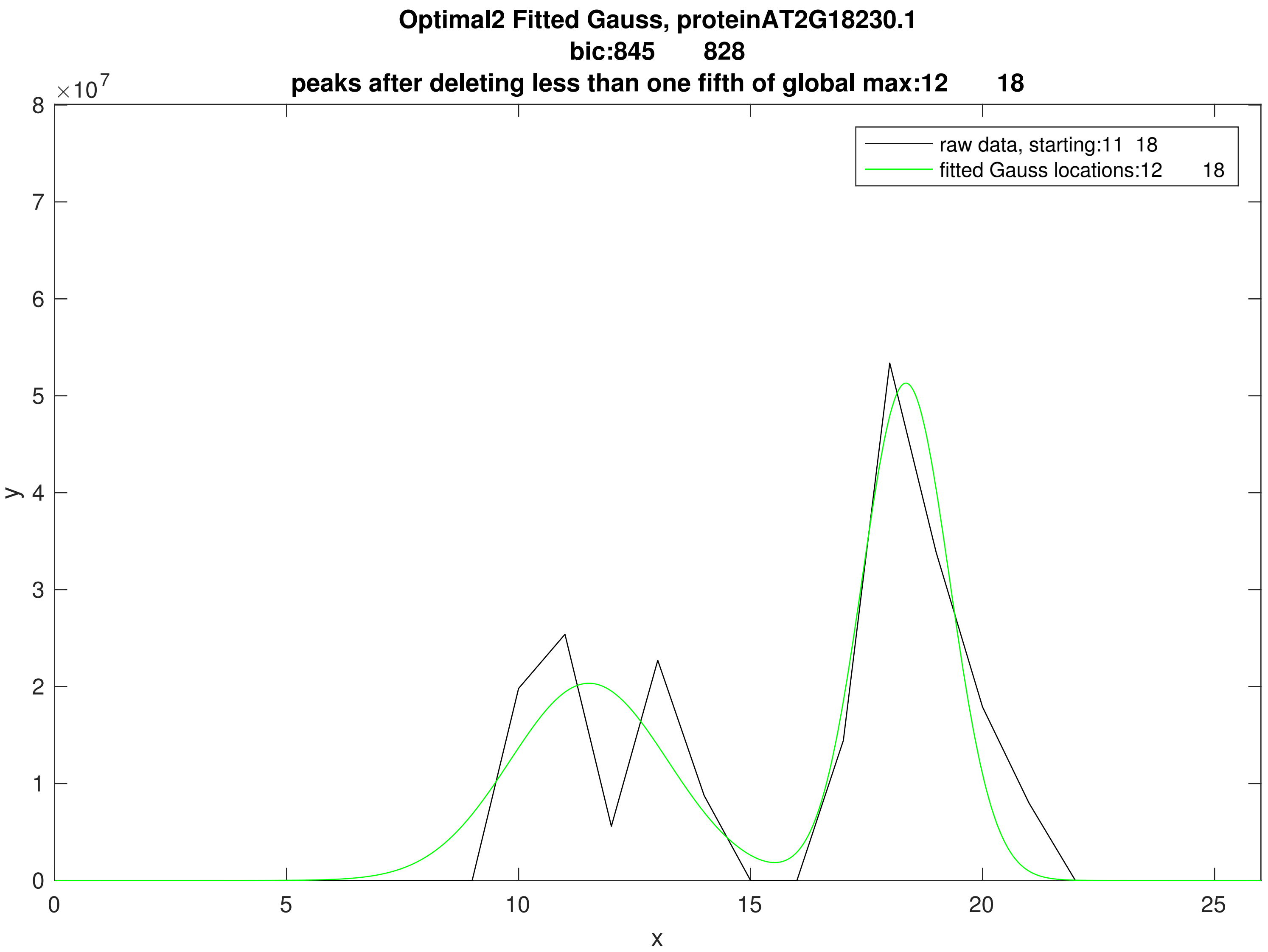

Optimal2 Fitted Gauss, proteinAT2G18230.1
bic:845 828
peaks after deleting less than one fifth of global max:12 18
×10^7
raw data, starting:11 18
fitted Gauss locations:12 18
y
x
0
1
2
3
4
5
6
7
8
0
5
10
15
20
25

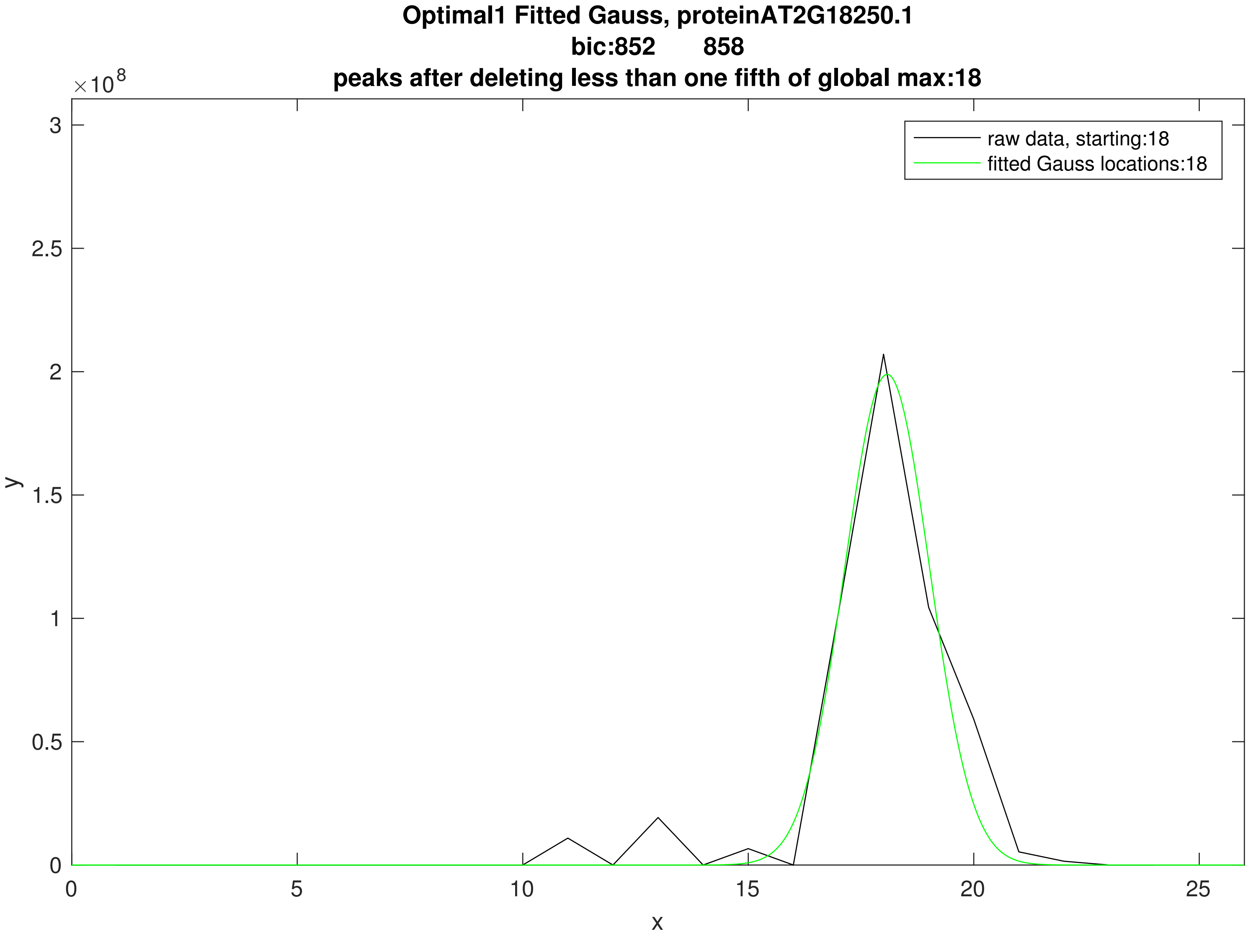

Optimal1 Fitted Gauss, proteinAT2G18250.1
bic:852 858
peaks after deleting less than one fifth of global max:18
×10^8
3
2.5
2
1.5
1
0.5
0
y
0
5
10
15
20
25
x
raw data, starting:18
fitted Gauss locations:18

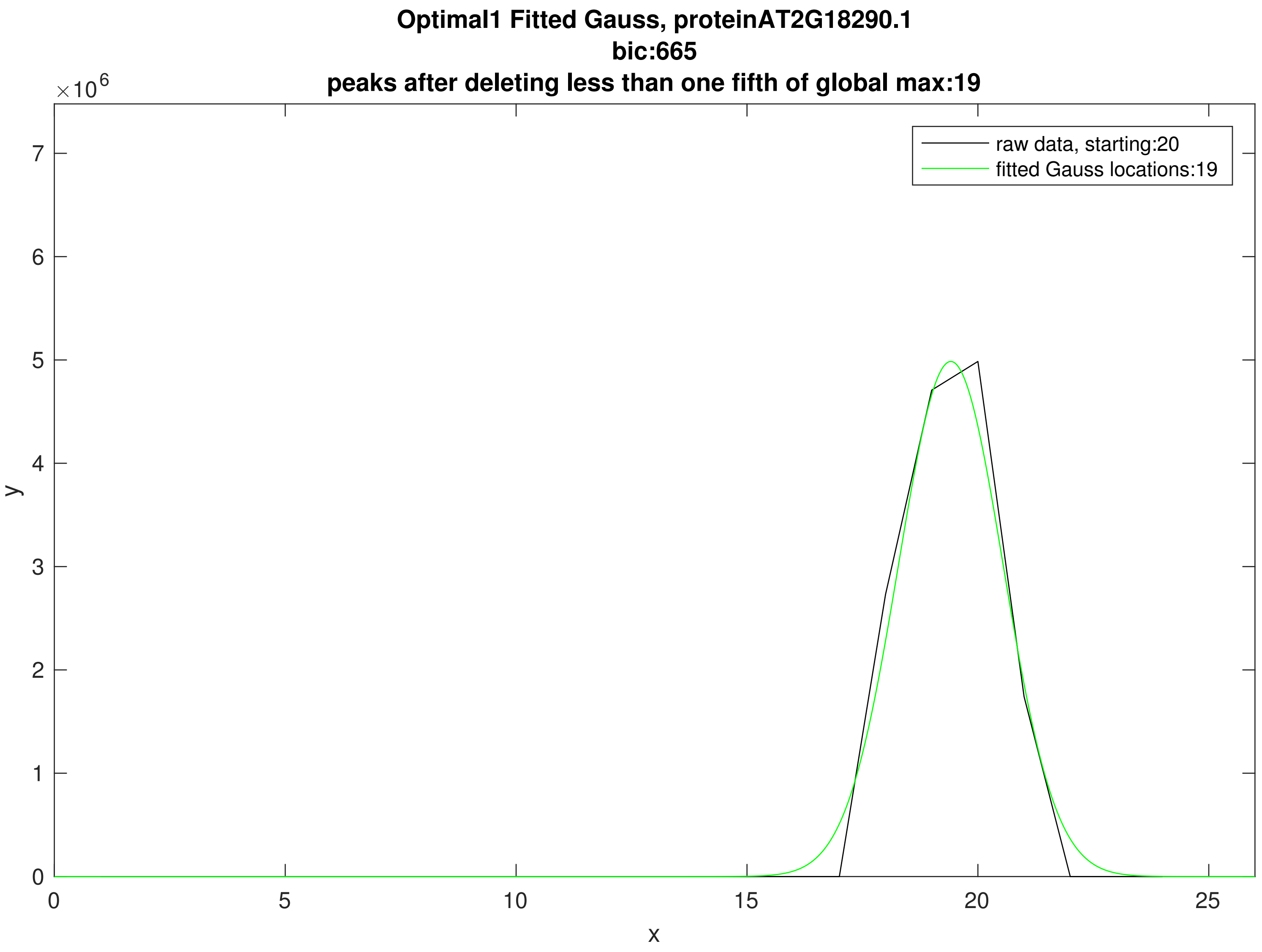

Optimal1 Fitted Gauss, proteinAT2G18290.1
bic:665
peaks after deleting less than one fifth of global max:19
×10^6
raw data, starting:20
fitted Gauss locations:19
y
x
0
1
2
3
4
5
6
7
0
5
10
15
20
25

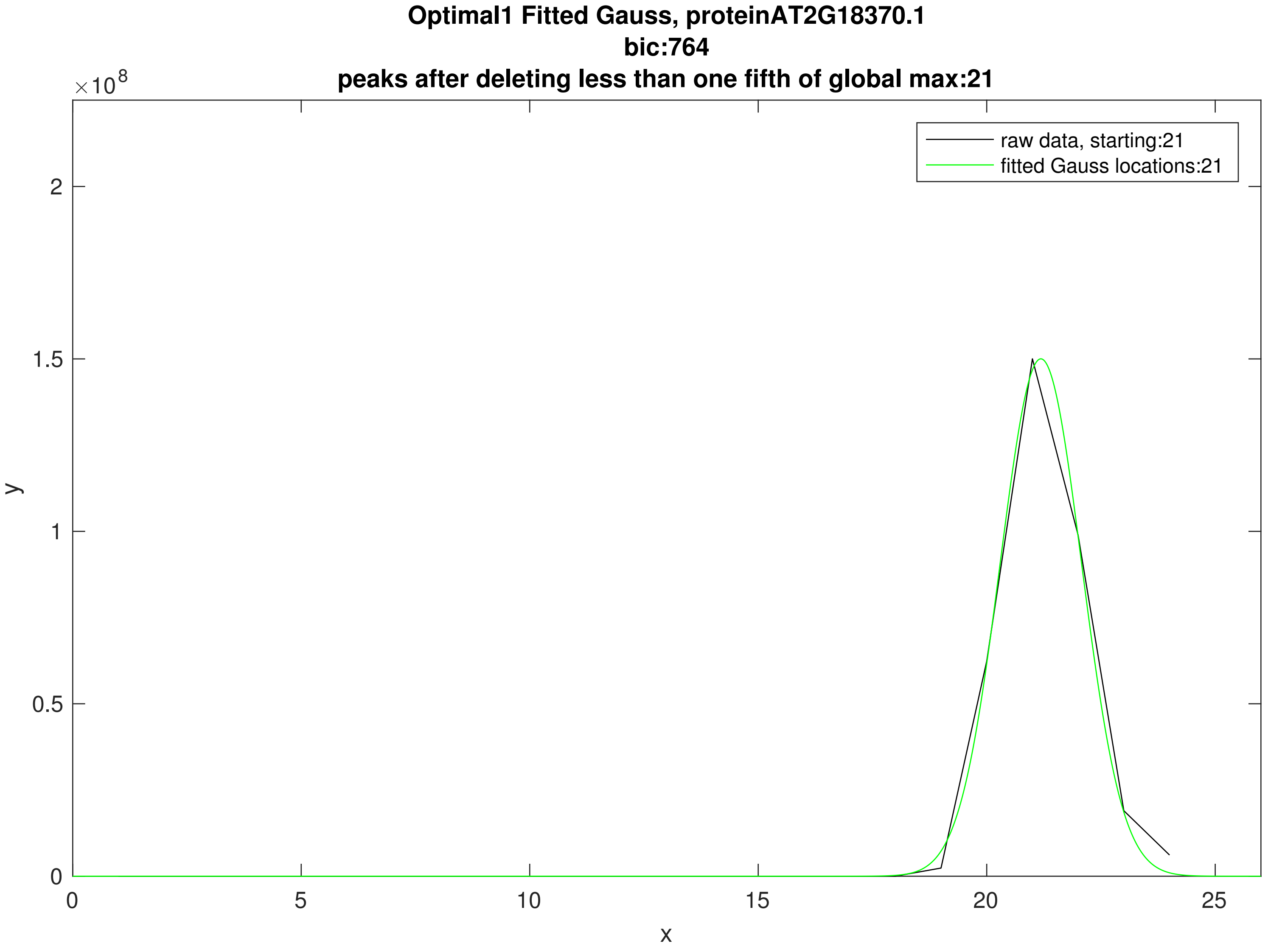

Optimal1 Fitted Gauss, proteinAT2G18370.1
bic:764
peaks after deleting less than one fifth of global max:21
×10^8
2
1.5
1
0.5
0
y
0
5
10
15
20
25
x
raw data, starting:21
fitted Gauss locations:21

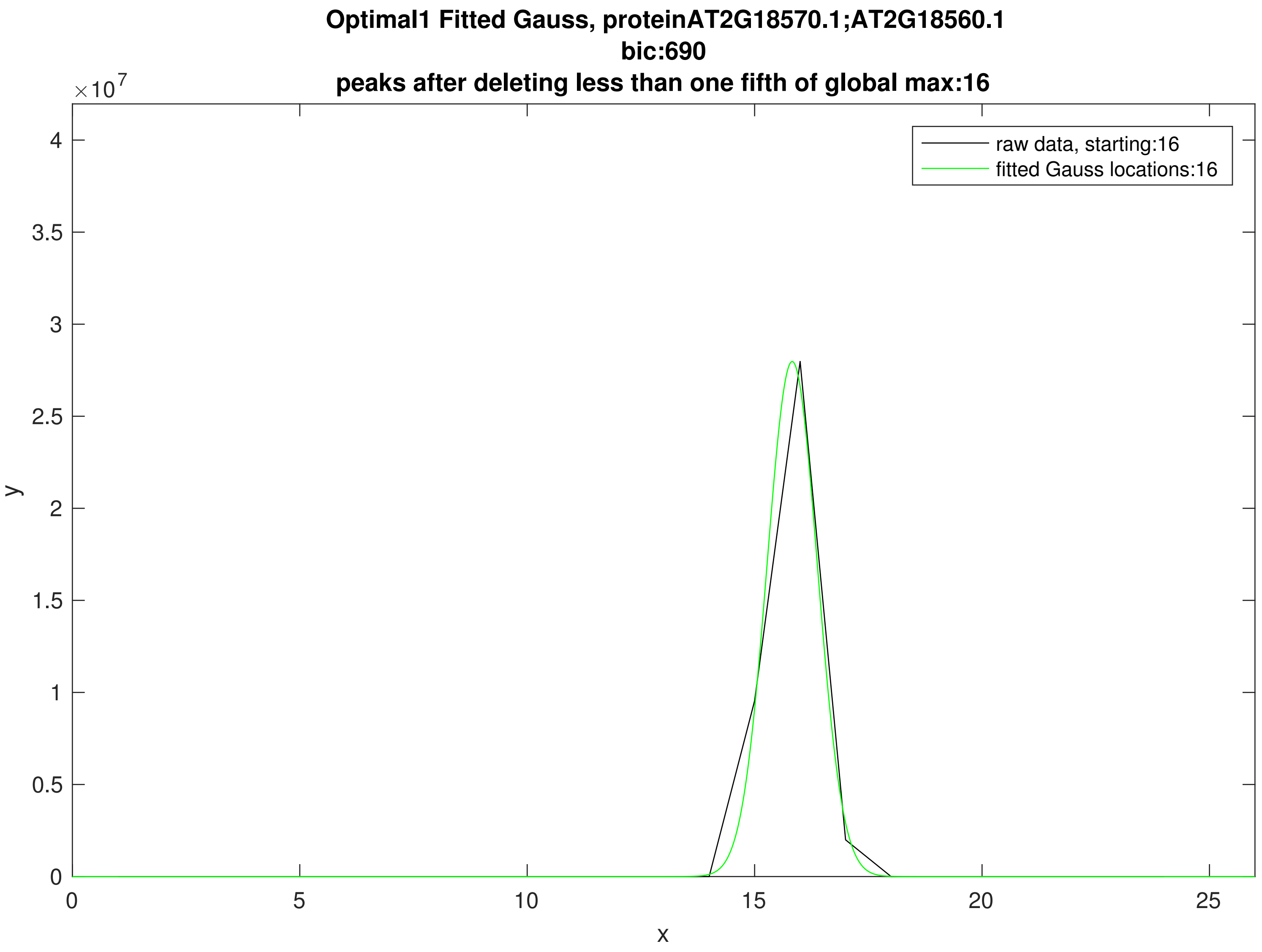

Optimal1 Fitted Gauss, proteinAT2G18570.1;AT2G18560.1
bic:690
peaks after deleting less than one fifth of global max:16
×10^7
raw data, starting:16
fitted Gauss locations:16
y
x
0
0.5
1
1.5
2
2.5
3
3.5
4
5
10
15
20
25

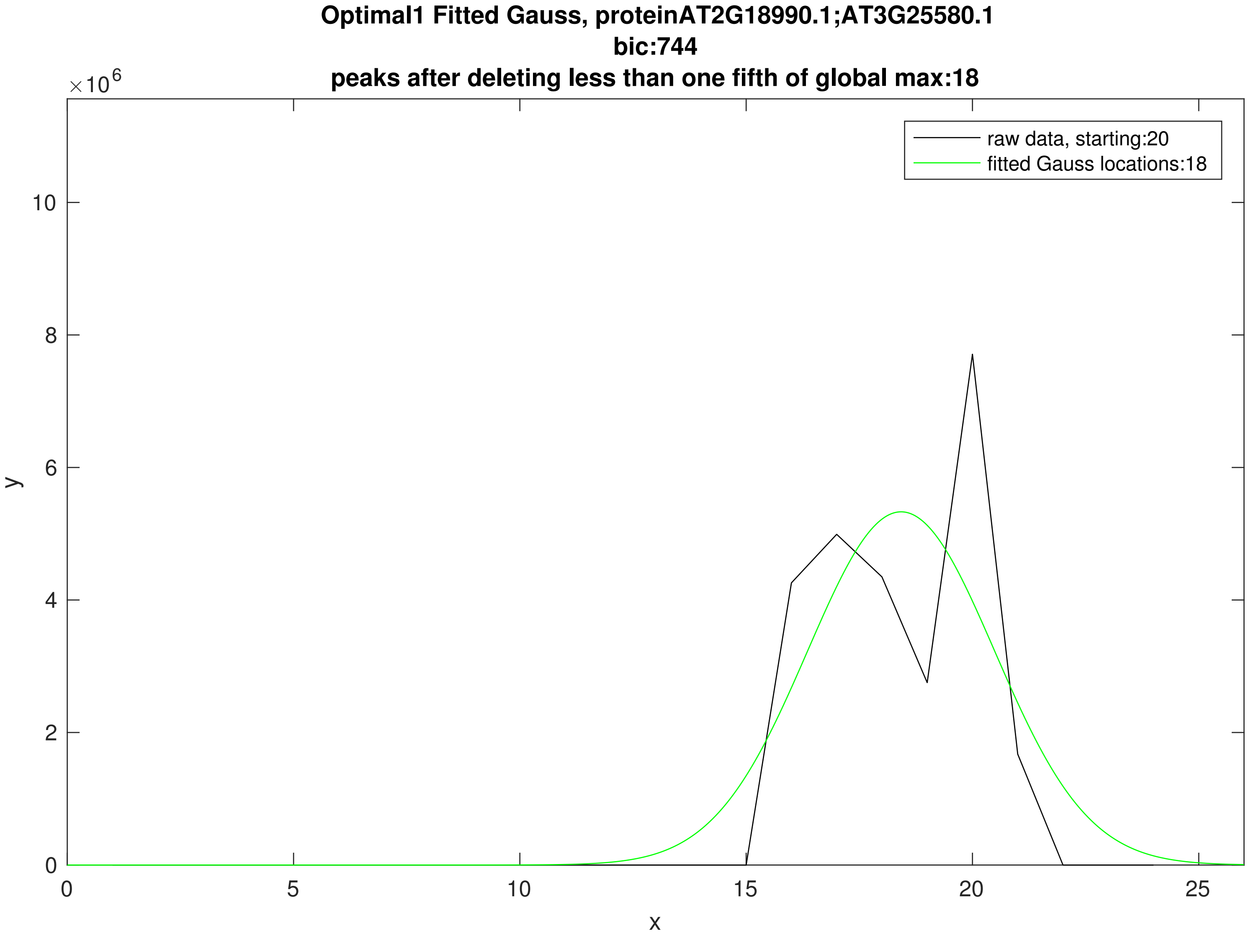

Optimal1 Fitted Gauss, proteinAT2G18990.1;AT3G25580.1
bic:744
peaks after deleting less than one fifth of global max:18
×10^6
raw data, starting:20
fitted Gauss locations:18
y
x
0
2
4
6
8
10
0
5
10
15
20
25

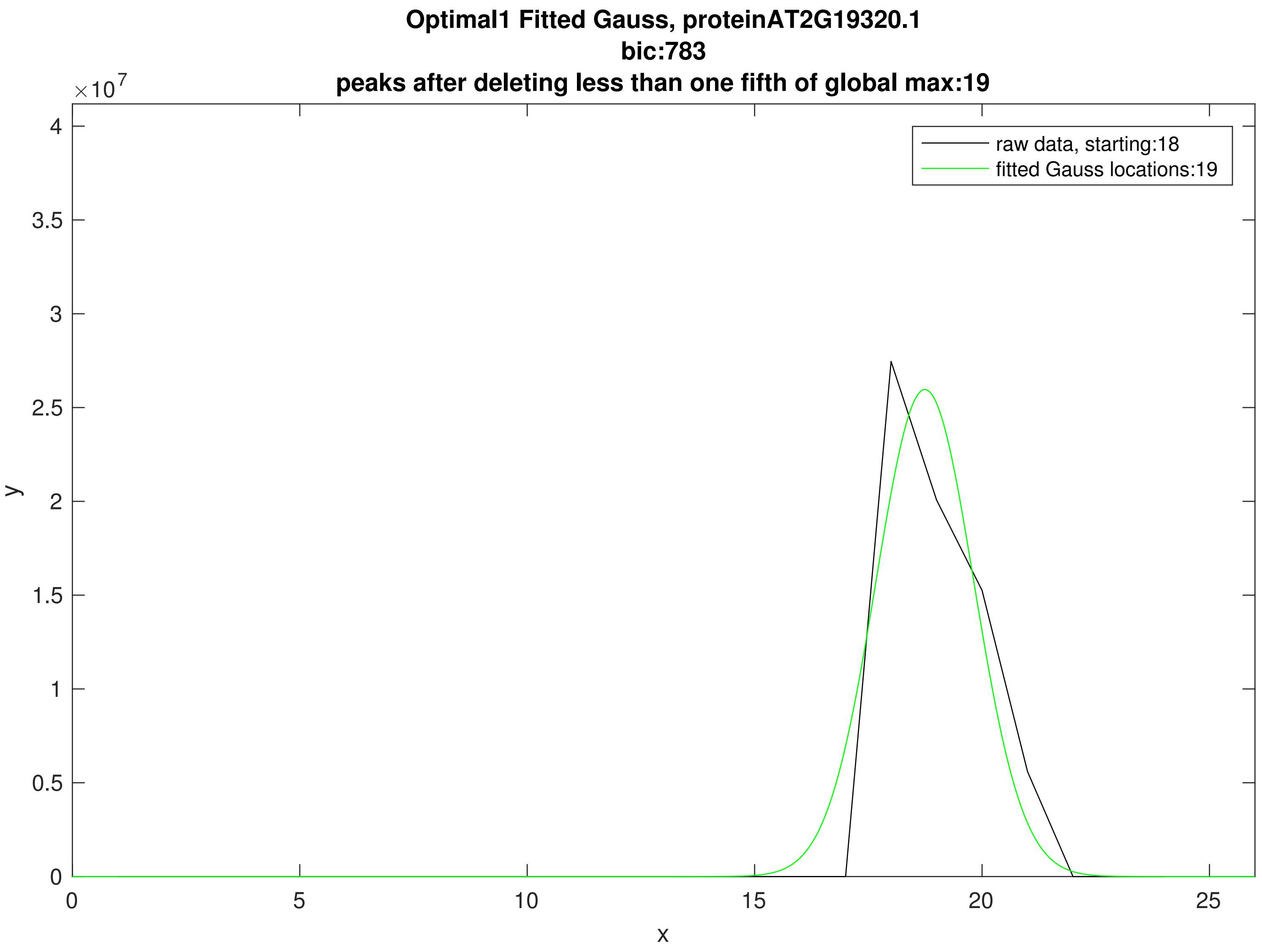

Optimal1 Fitted Gauss, proteinAT2G19320.1
bic:783
peaks after deleting less than one fifth of global max:19
raw data, starting:18
fitted Gauss locations:19
$\times 10^7$
4
3.5
3
2.5
2
1.5
1
0.5
0
y
0
5
10
15
20
25
x

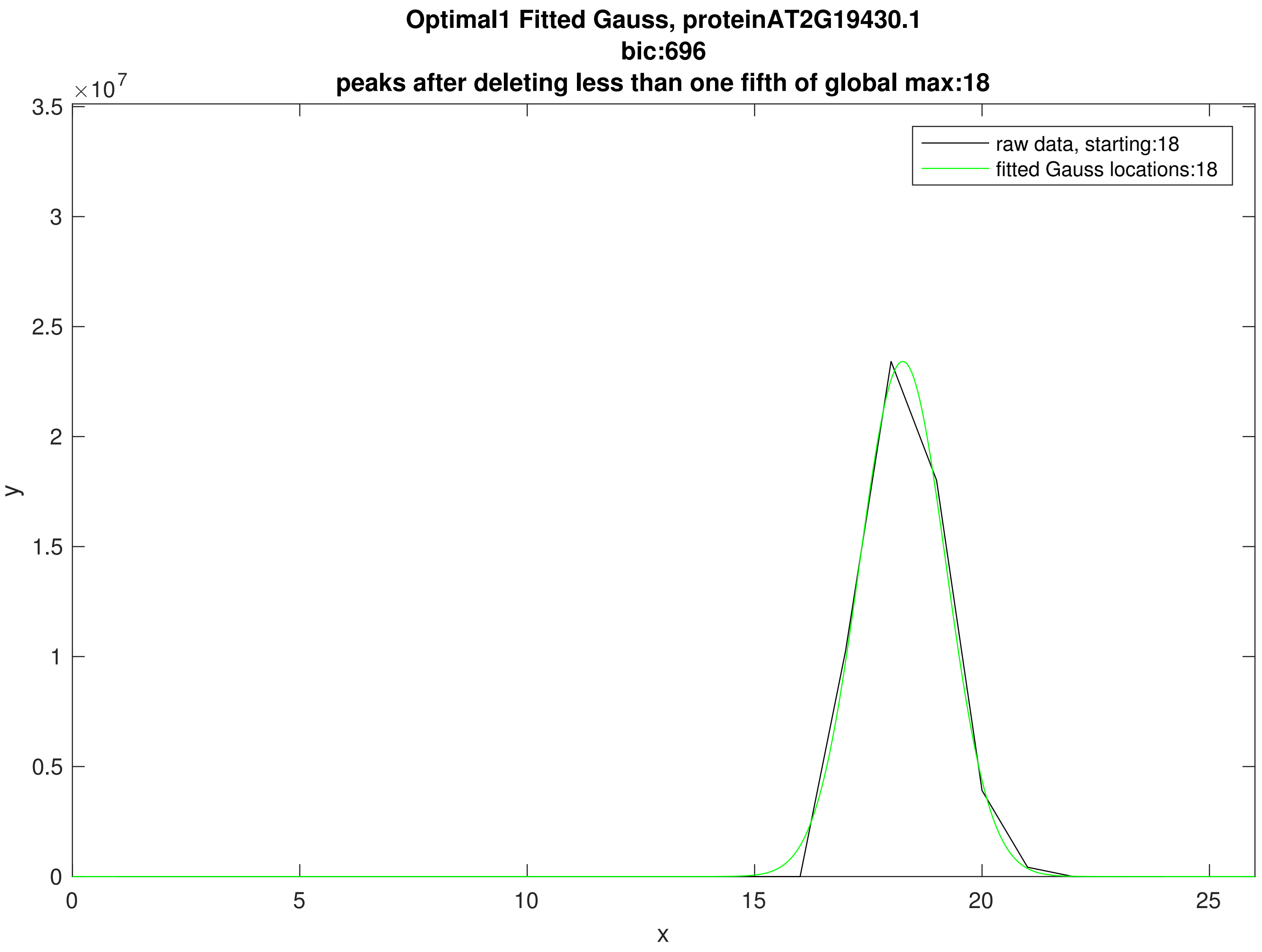
Optimal1 Fitted Gauss, proteinAT2G19430.1
bic:696
peaks after deleting less than one fifth of global max:18
×10^7
3.5
3
2.5
2
1.5
1
0.5
0
y
0
5
10
15
20
25
x
raw data, starting:18
fitted Gauss locations:18

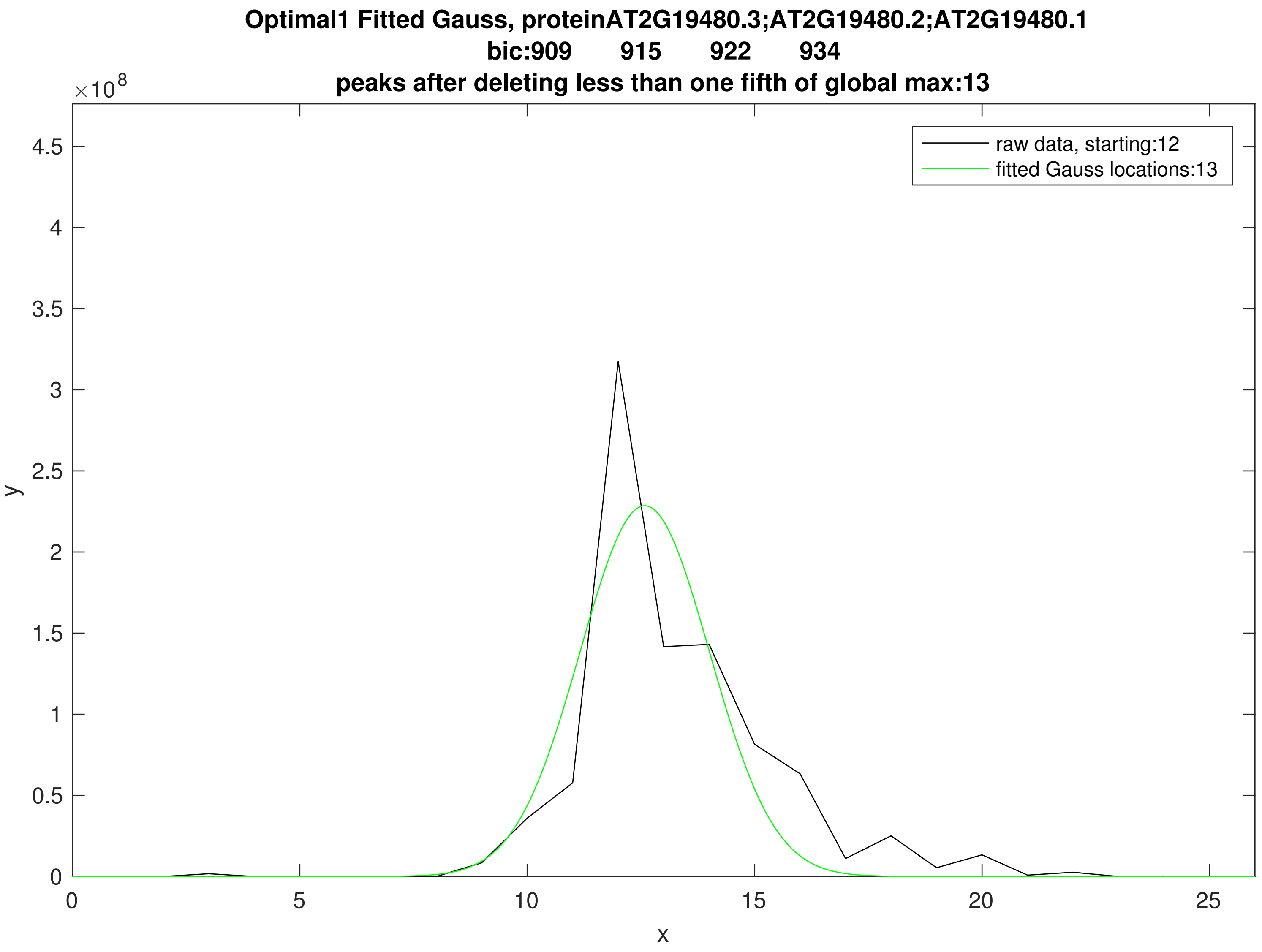

Optimal1 Fitted Gauss, proteinAT2G19480.3;AT2G19480.2;AT2G19480.1
bic:909 915 922 934
peaks after deleting less than one fifth of global max:13
×10^8
raw data, starting:12
fitted Gauss locations:13
y
x
0
0.5
1
1.5
2
2.5
3
3.5
4
4.5
0
5
10
15
20
25

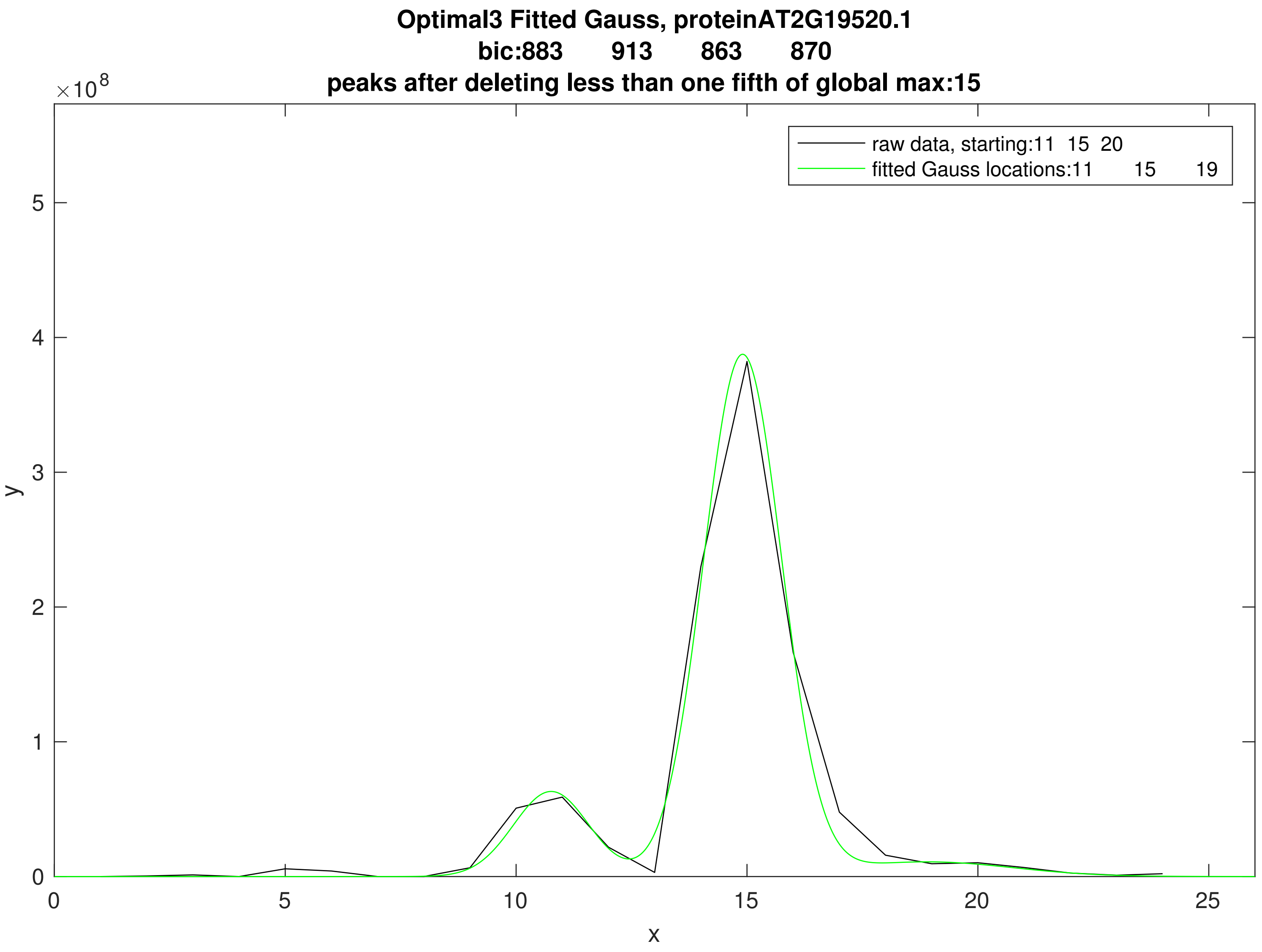

Optimal3 Fitted Gauss, proteinAT2G19520.1
bic:883 913 863 870
peaks after deleting less than one fifth of global max:15
×10^8
raw data, starting:11 15 20
fitted Gauss locations:11 15 19
y
x
0
1
2
3
4
5
0
5
10
15
20
25

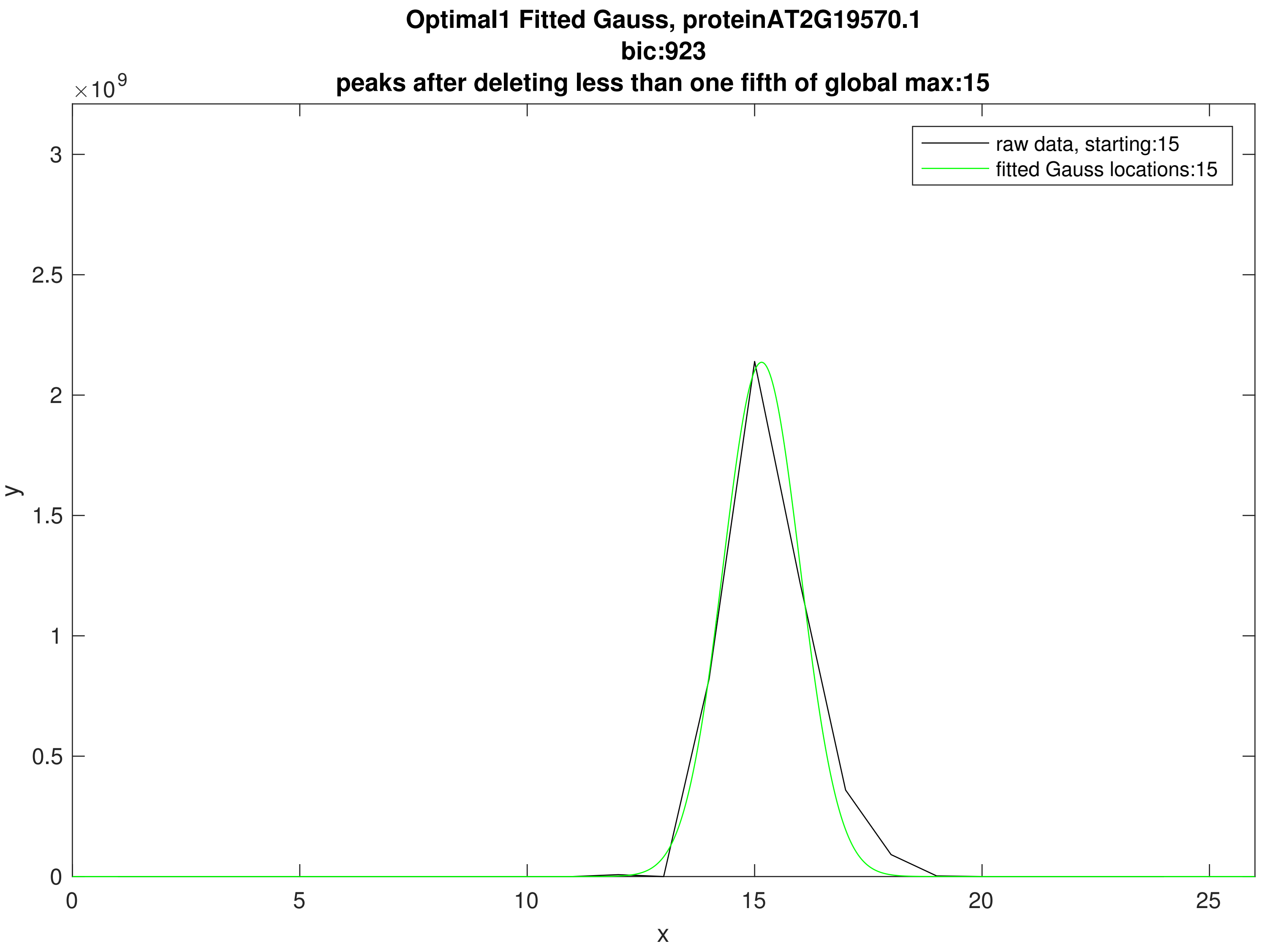

Optimal1 Fitted Gauss, proteinAT2G19570.1
bic:923
peaks after deleting less than one fifth of global max:15
×10^9
3
2.5
2
1.5
1
0.5
0
y
0
5
10
15
20
25
x
raw data, starting:15
fitted Gauss locations:15

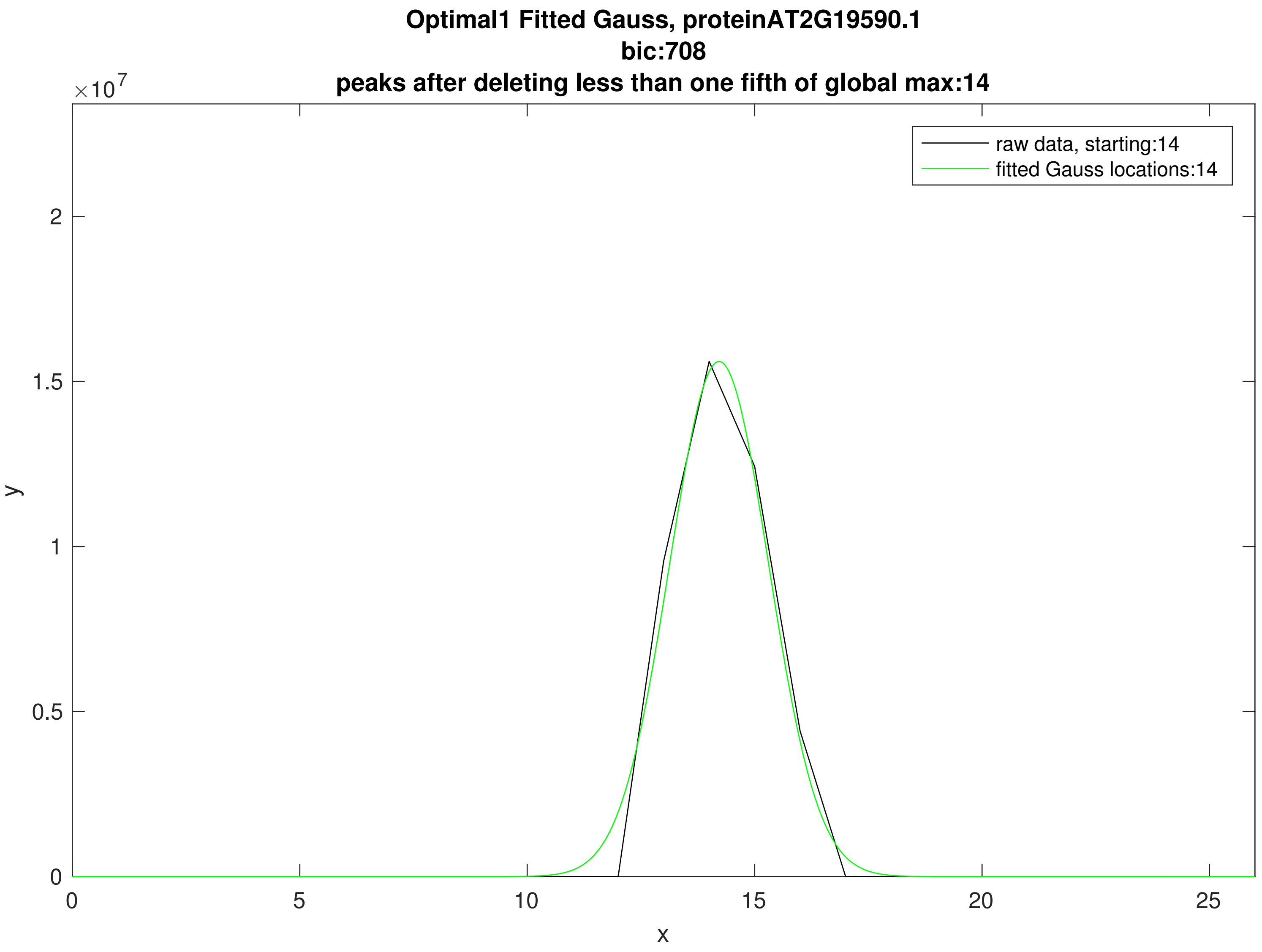

Optimal1 Fitted Gauss, proteinAT2G19590.1
bic:708
peaks after deleting less than one fifth of global max:14
×10^7
raw data, starting:14
fitted Gauss locations:14
2
1.5
1
0.5
0
y
0
5
10
15
20
25
x

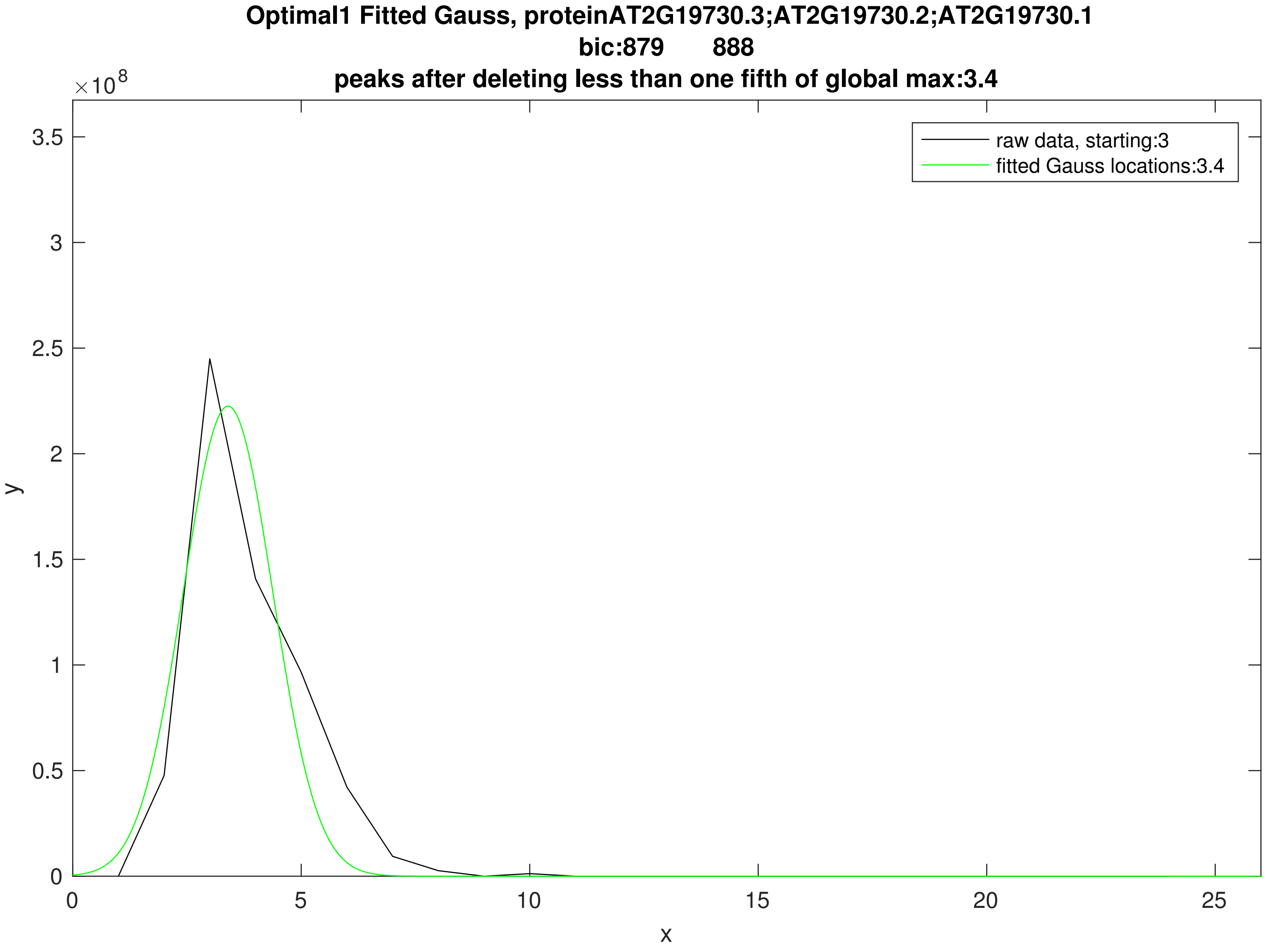

Optimal1 Fitted Gauss, proteinAT2G19730.3;AT2G19730.2;AT2G19730.1
bic:879 888
peaks after deleting less than one fifth of global max:3.4
×10^8
raw data, starting:3
fitted Gauss locations:3.4
y
x
0
0.5
1
1.5
2
2.5
3
3.5
5
10
15
20
25

Optimal1 Fitted Gauss, proteinAT2G19740.1;AT5G56710.2;AT5G56710.1;AT4G26230.1
bic:810
peaks after deleting less than one fifth of global max:3.5

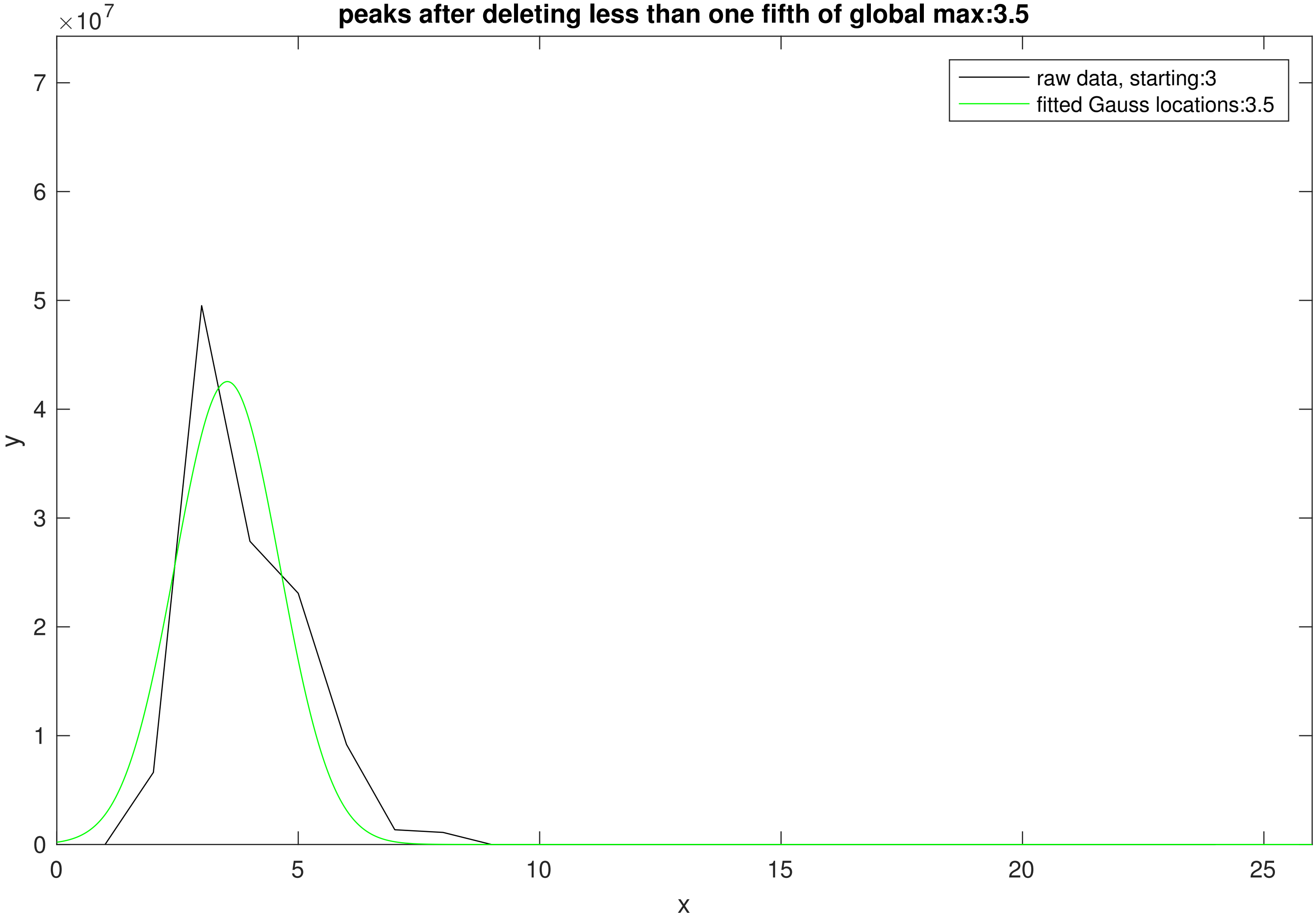

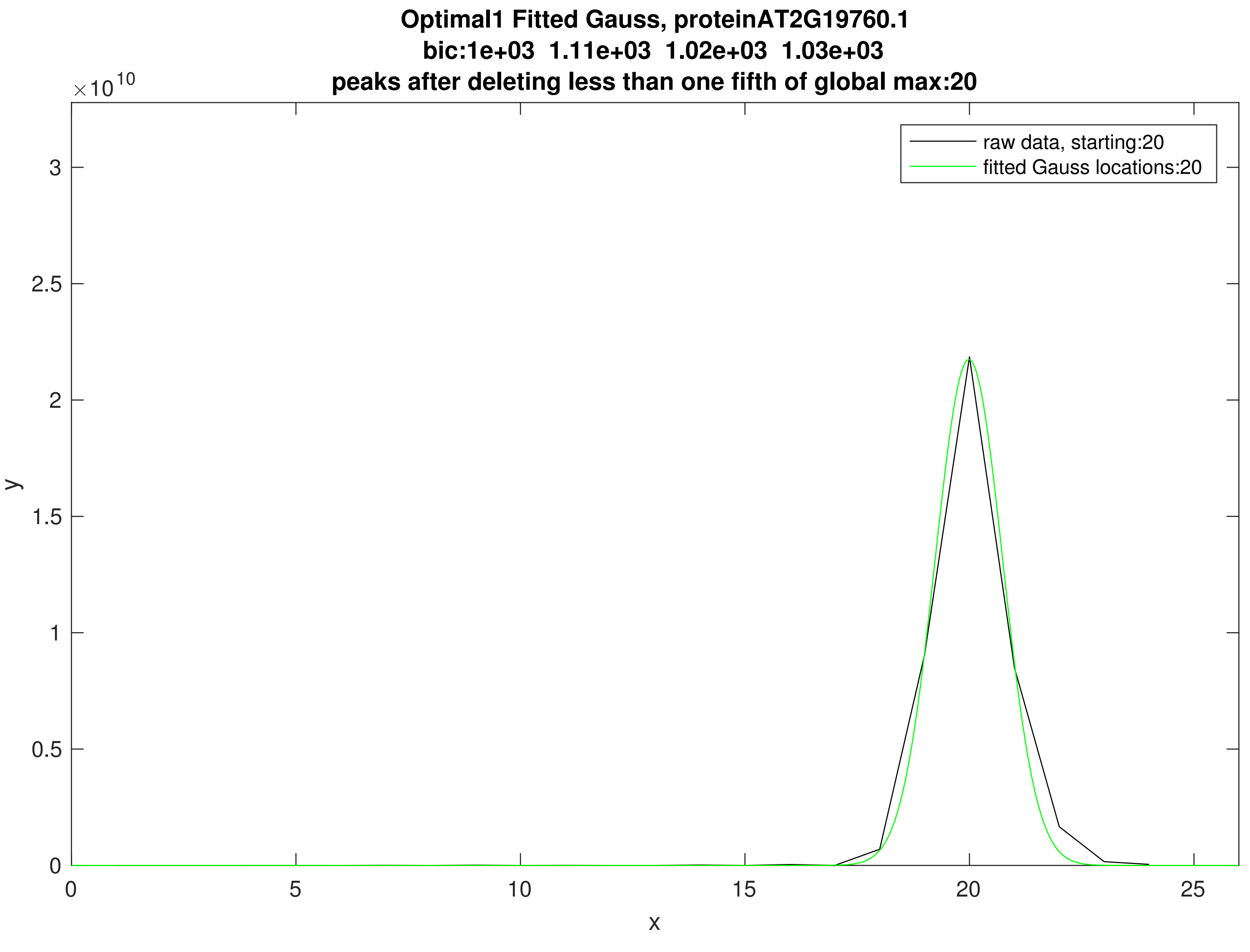

Optimal1 Fitted Gauss, proteinAT2G19760.1
bic:1e+03 1.11e+03 1.02e+03 1.03e+03
peaks after deleting less than one fifth of global max:20
×10^10
raw data, starting:20
fitted Gauss locations:20
y
x
0
0.5
1
1.5
2
2.5
3
5
10
15
20
25

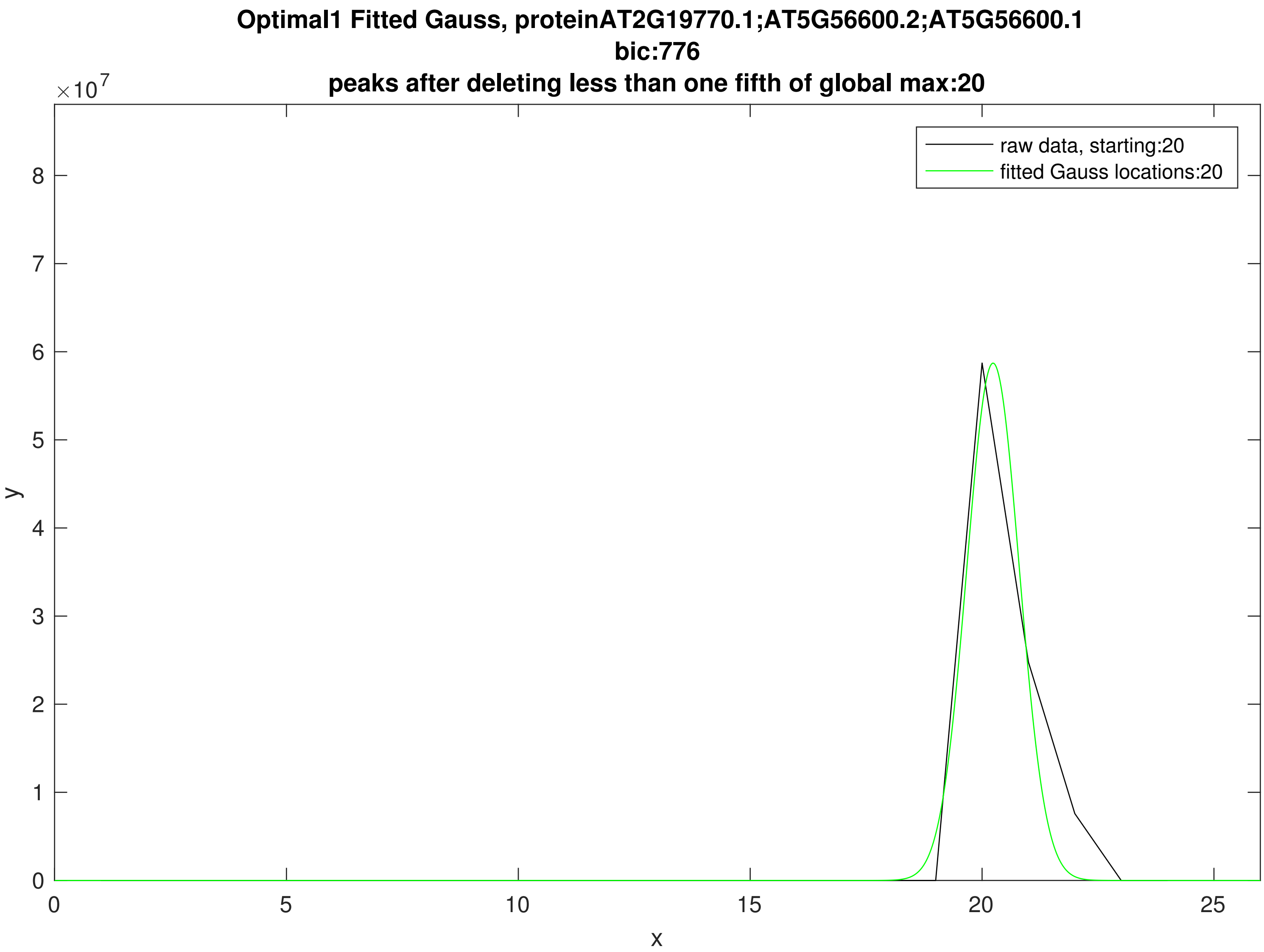
Optimal1 Fitted Gauss, proteinAT2G19770.1;AT5G56600.2;AT5G56600.1
bic:776
peaks after deleting less than one fifth of global max:20
×10^7
raw data, starting:20
fitted Gauss locations:20
y
x
0
1
2
3
4
5
6
7
8
0
5
10
15
20
25

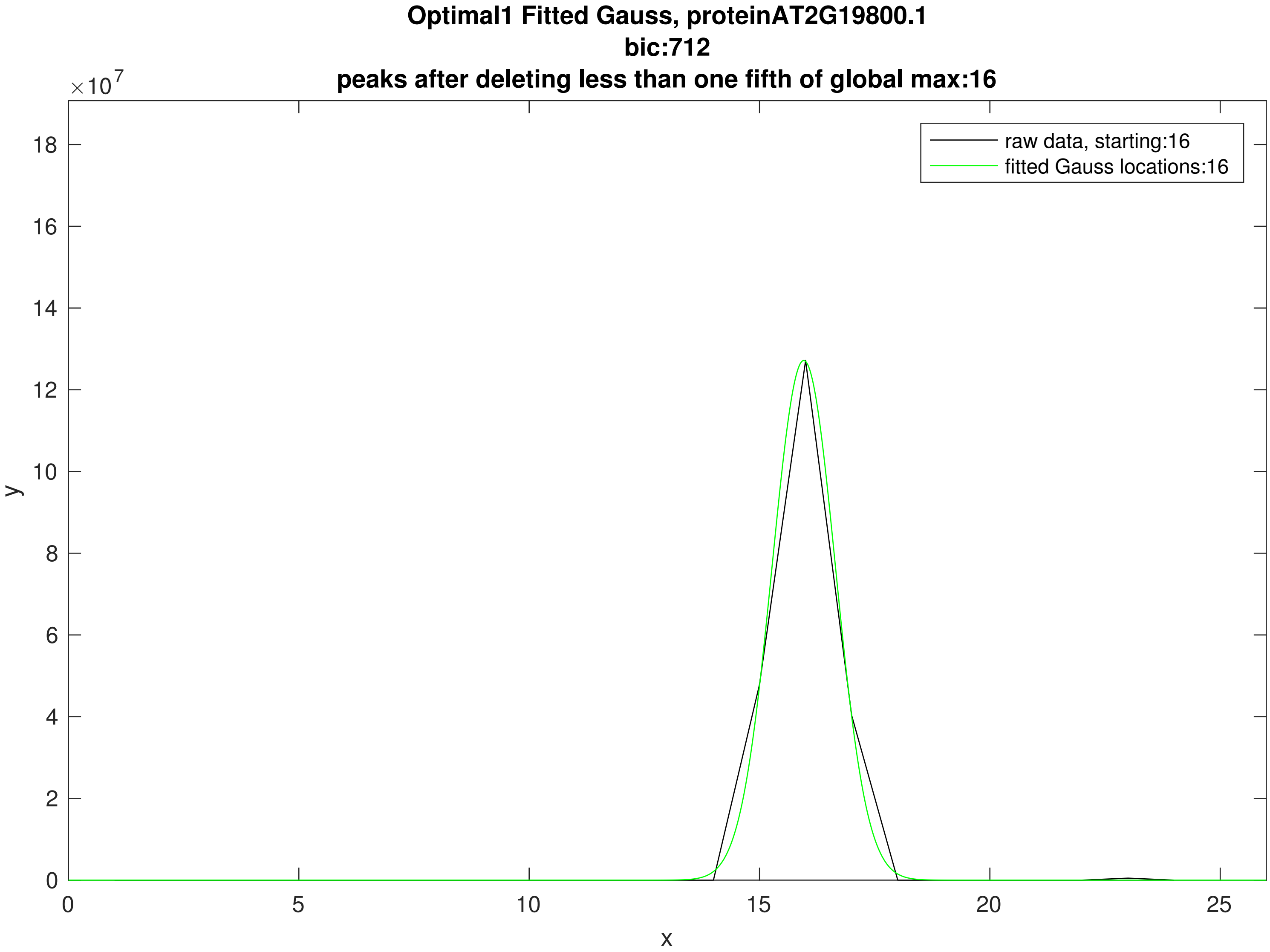
Optimal1 Fitted Gauss, proteinAT2G19800.1
bic:712
peaks after deleting less than one fifth of global max:16
×10^7
raw data, starting:16
fitted Gauss locations:16
y
x
0
2
4
6
8
10
12
14
16
18
0
5
10
15
20
25

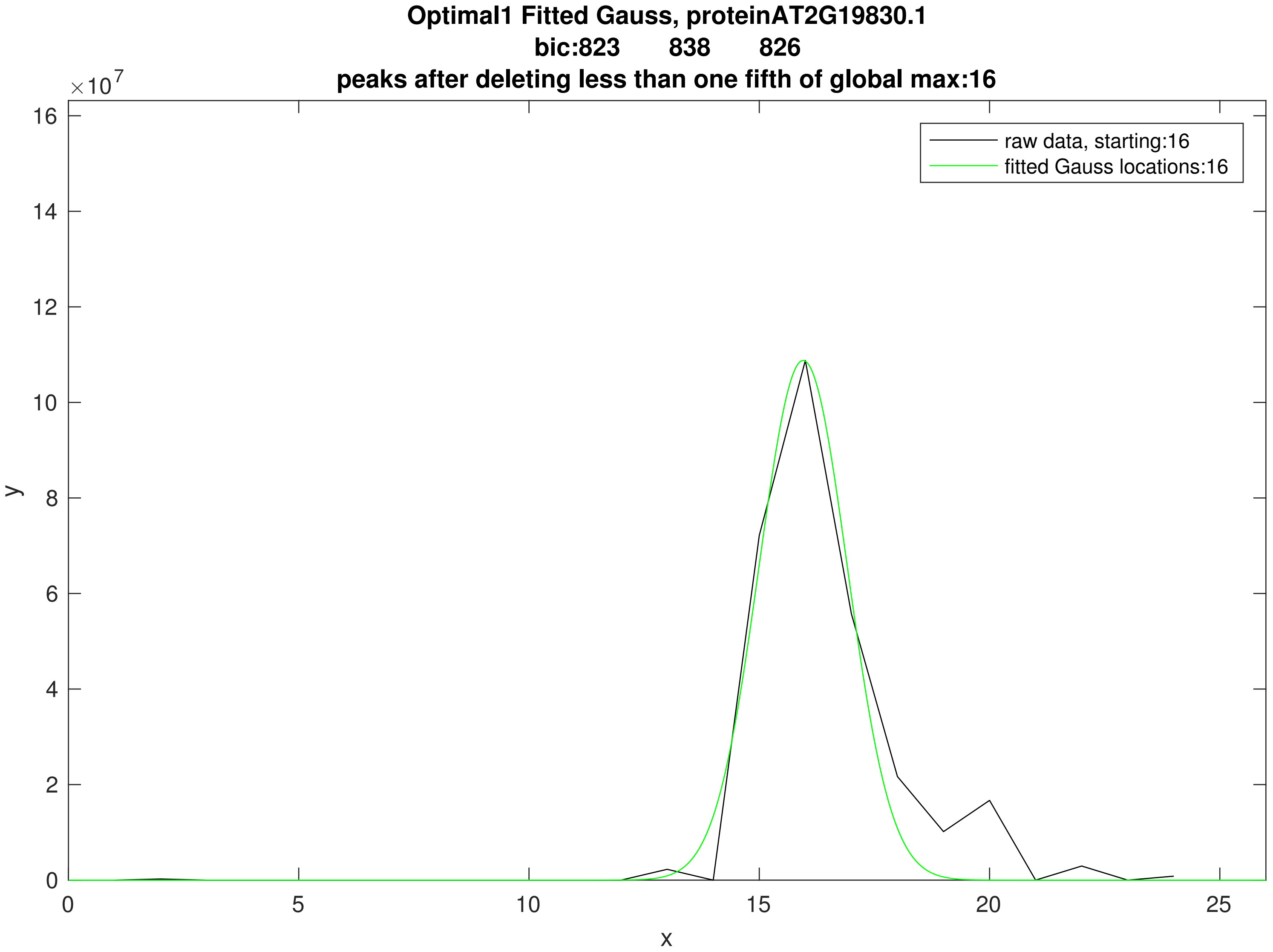

Optimal1 Fitted Gauss, proteinAT2G19830.1
bic:823 838 826
peaks after deleting less than one fifth of global max:16
×10^7
16
14
12
10
8
6
4
2
0
y
0
5
10
15
20
25
x
raw data, starting:16
fitted Gauss locations:16

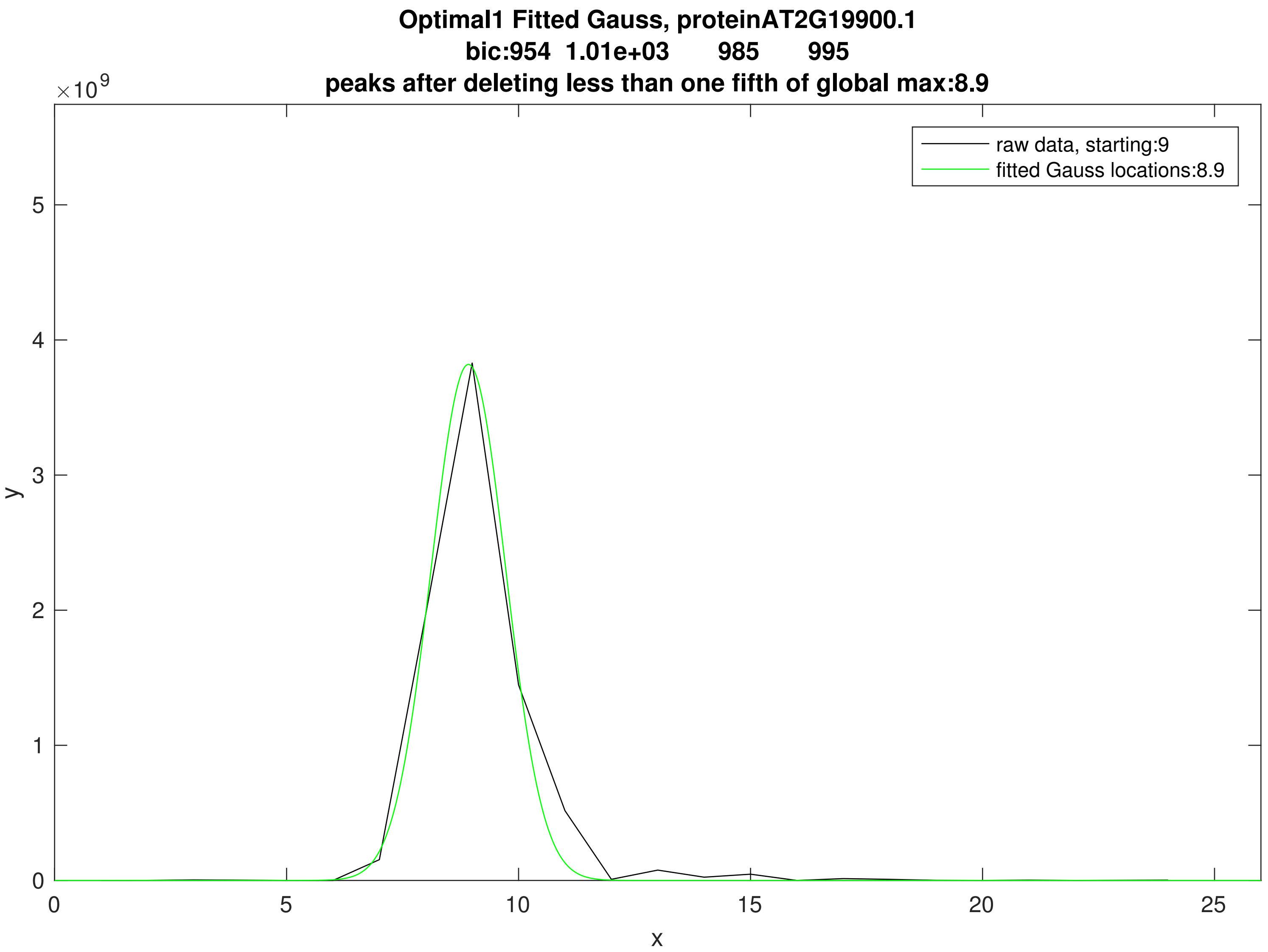
Optimal1 Fitted Gauss, proteinAT2G19900.1
bic:954 1.01e+03 985 995
peaks after deleting less than one fifth of global max:8.9
×10^9
raw data, starting:9
fitted Gauss locations:8.9
y
x
0
1
2
3
4
5
10
15
20
25

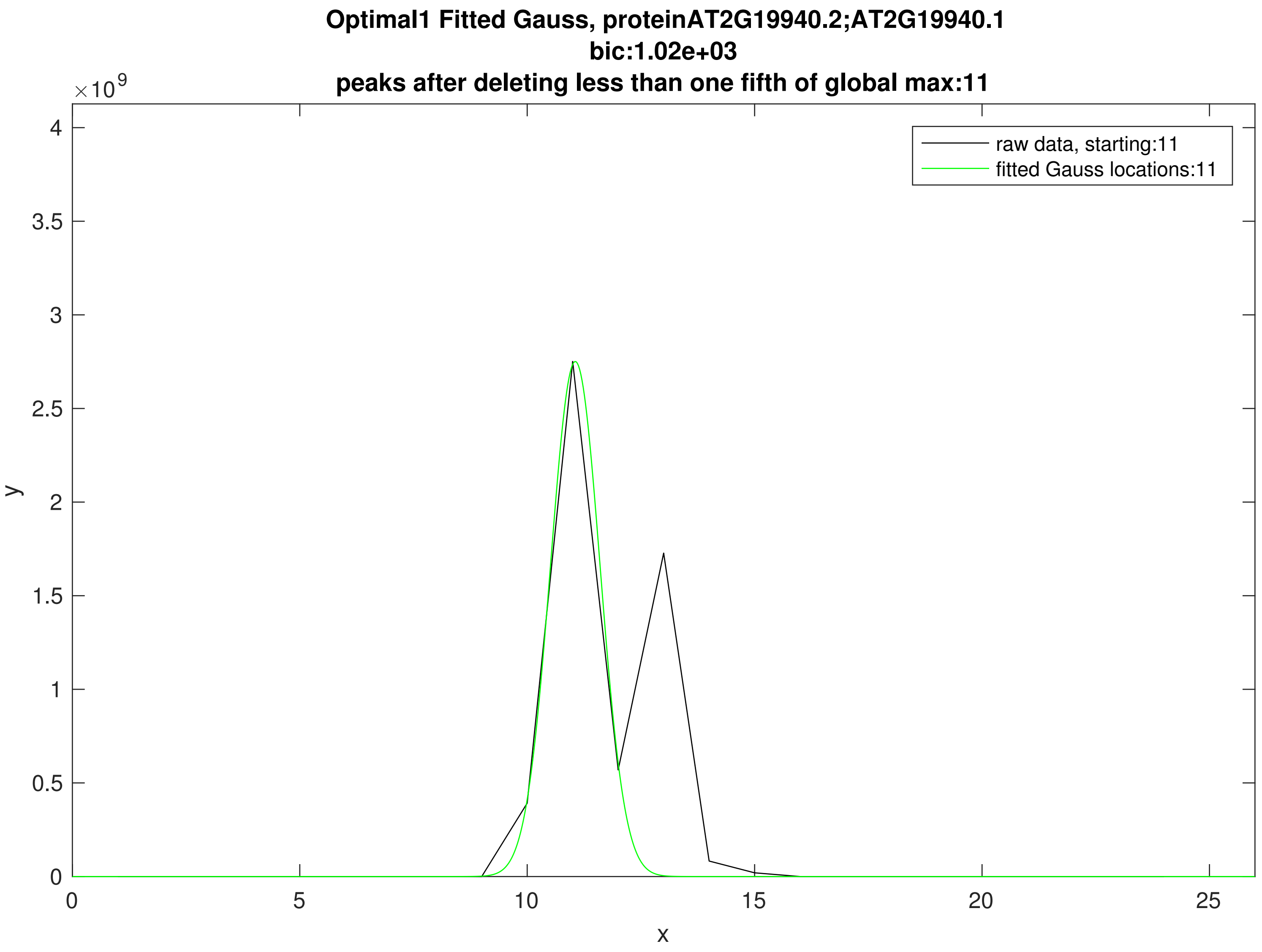
Optimal1 Fitted Gauss, proteinAT2G19940.2;AT2G19940.1
bic:1.02e+03
peaks after deleting less than one fifth of global max:11
×10^9
raw data, starting:11
fitted Gauss locations:11
y
x
0
0.5
1
1.5
2
2.5
3
3.5
4
5
10
15
20
25

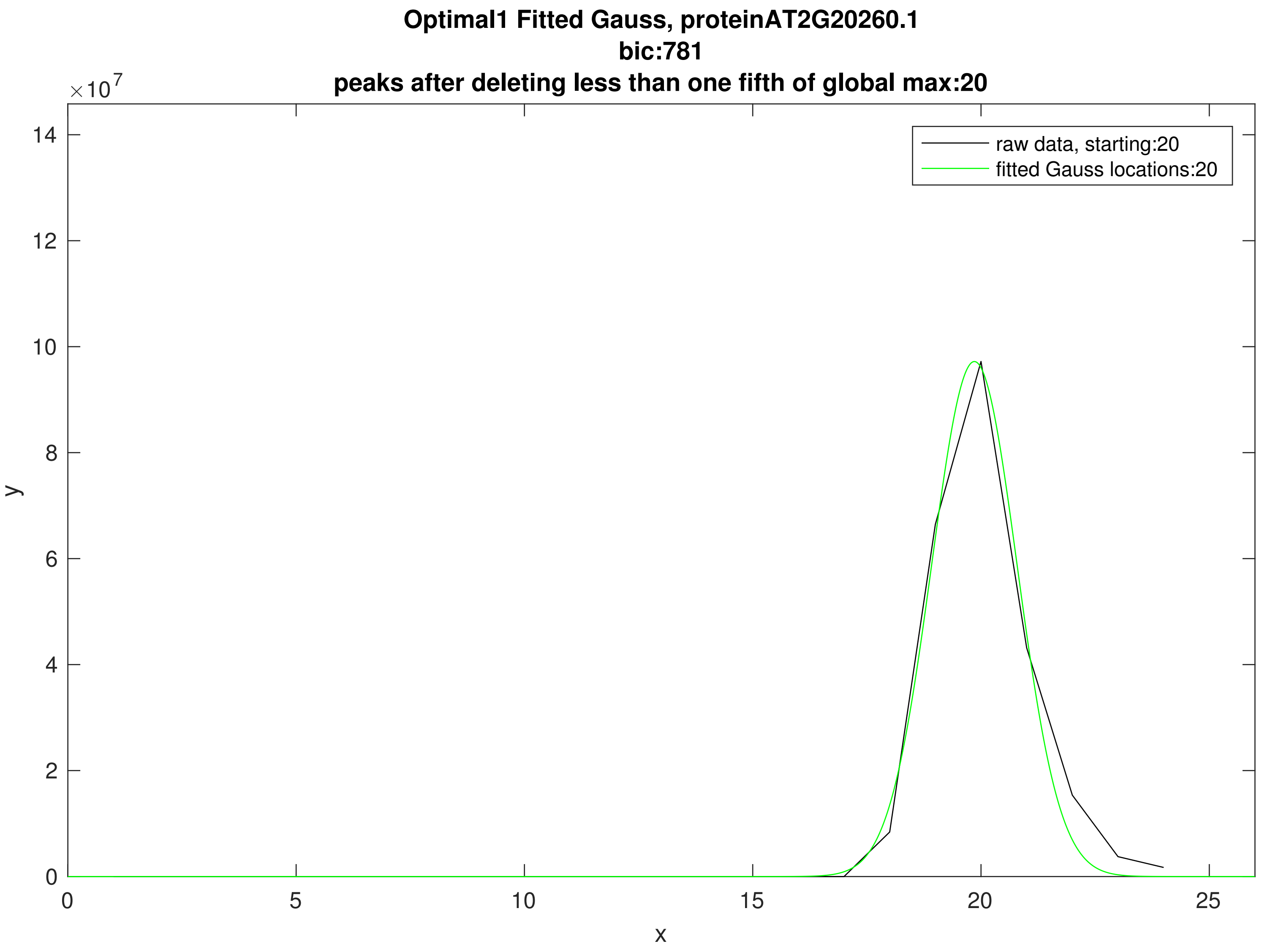
Optimal1 Fitted Gauss, proteinAT2G20260.1
bic:781
peaks after deleting less than one fifth of global max:20
×10^7
raw data, starting:20
fitted Gauss locations:20
y
x
0
2
4
6
8
10
12
14
0
5
10
15
20
25

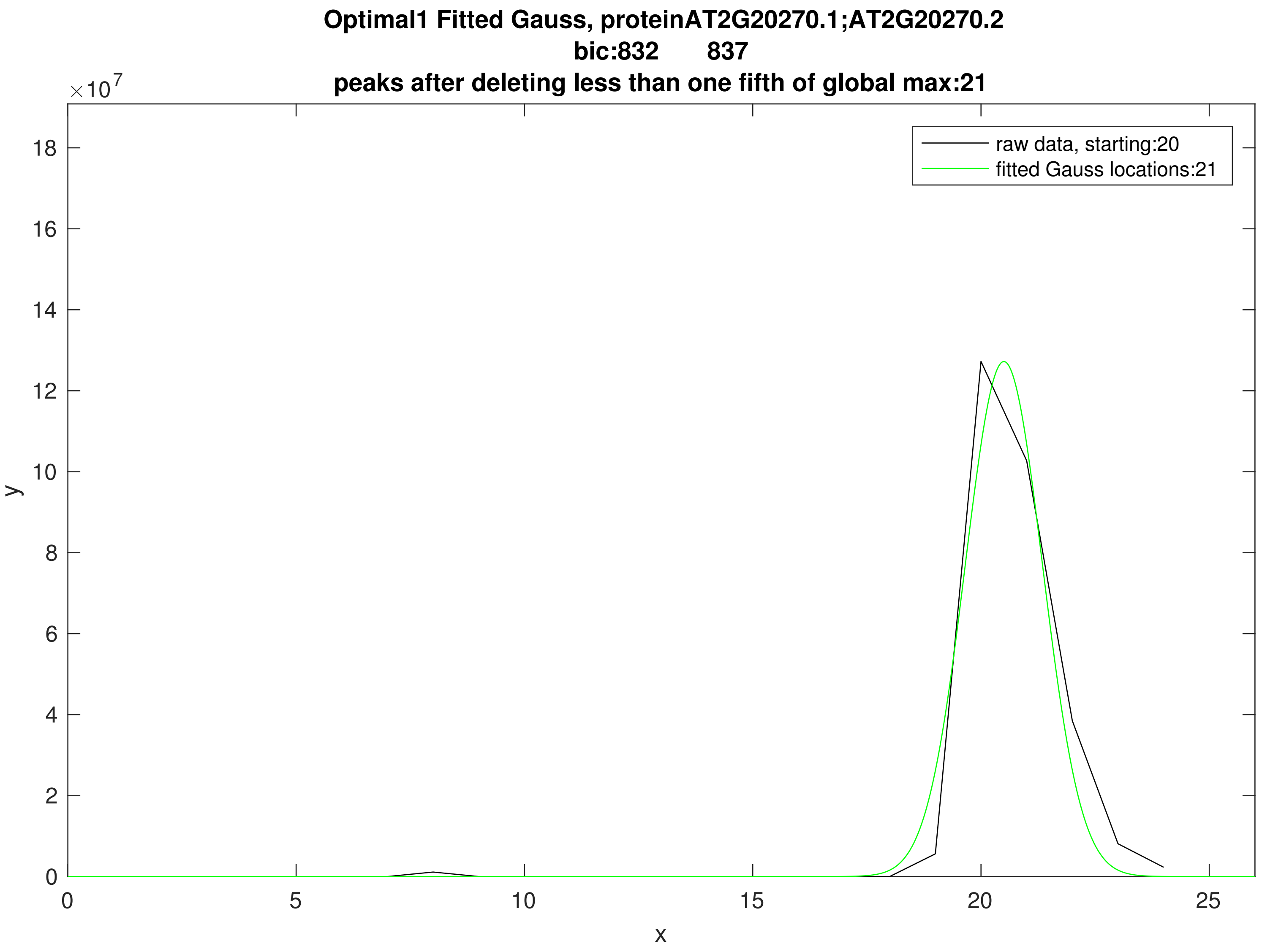

Optimal1 Fitted Gauss, proteinAT2G20270.1;AT2G20270.2
bic:832 837
peaks after deleting less than one fifth of global max:21
×10^7
raw data, starting:20
fitted Gauss locations:21
y
x
0
2
4
6
8
10
12
14
16
18
0
5
10
15
20
25

**Optimal1 Fitted Gauss, proteinAT2G20410.1**
**bic:816 819**
**peaks after deleting less than one fifth of global max:19**

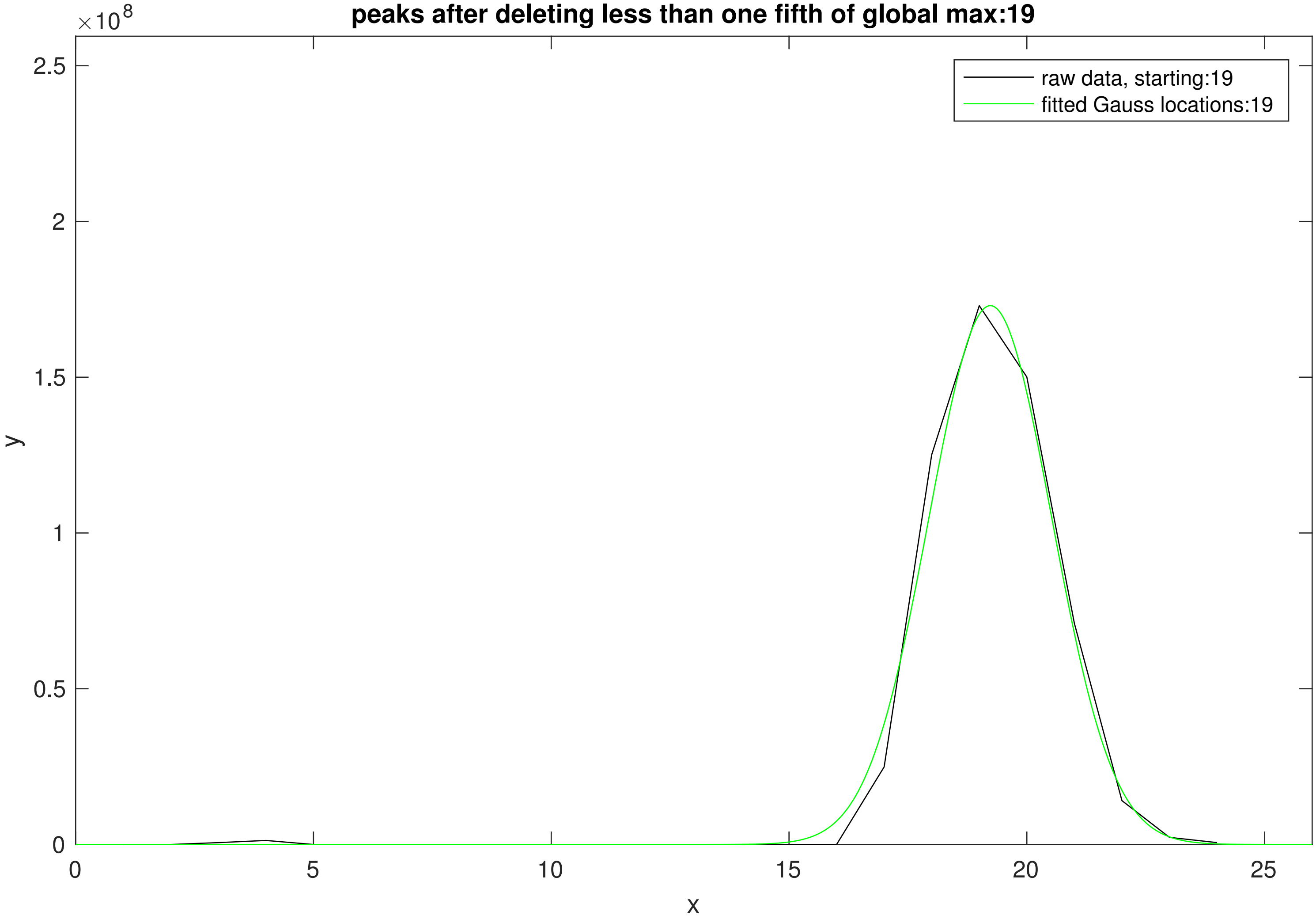

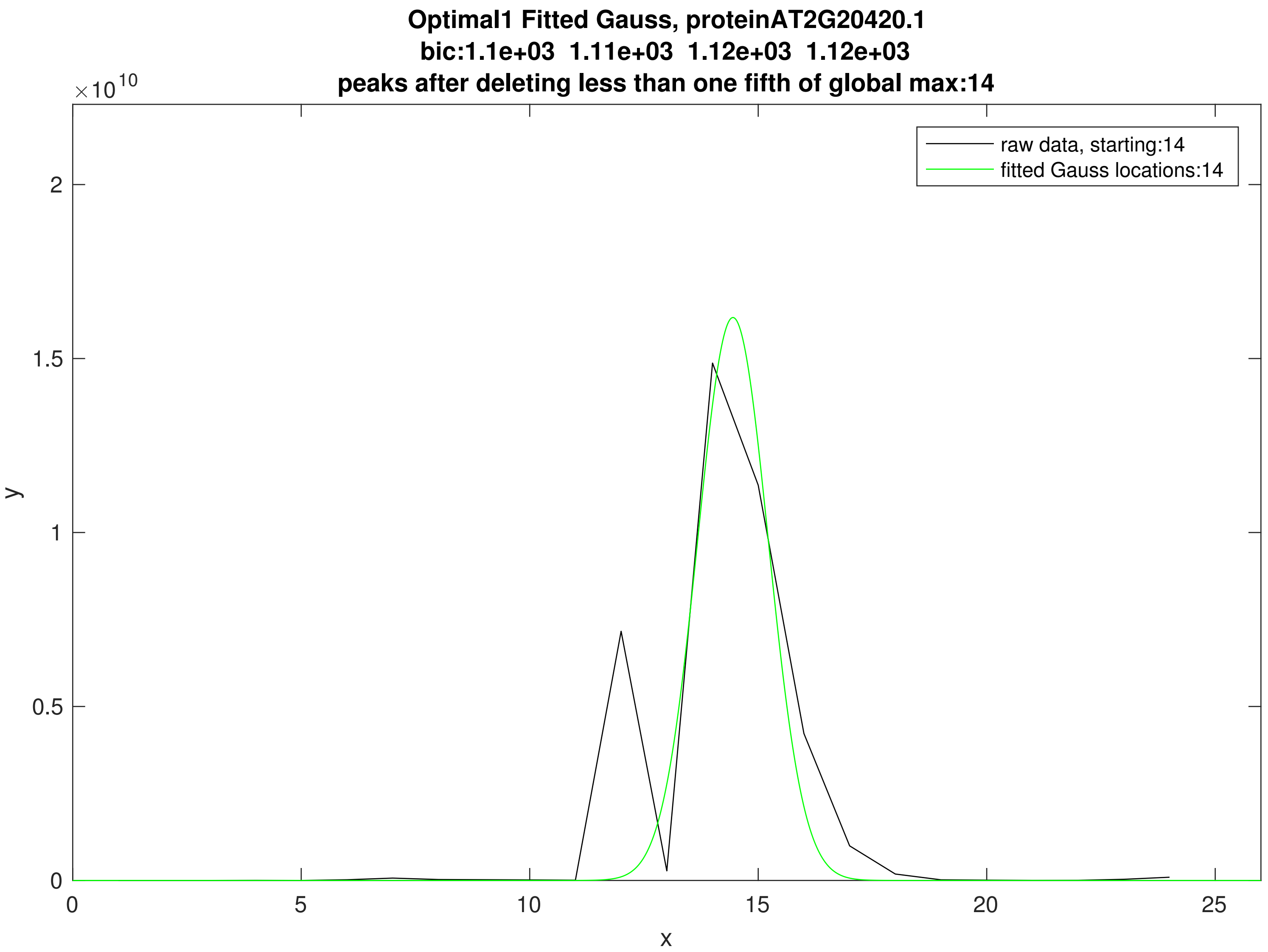
Optimal1 Fitted Gauss, proteinAT2G20420.1
bic:1.1e+03  1.11e+03  1.12e+03  1.12e+03
peaks after deleting less than one fifth of global max:14
×10^10
raw data, starting:14
fitted Gauss locations:14
2
1.5
1
0.5
0
y
0
5
10
15
20
25
x

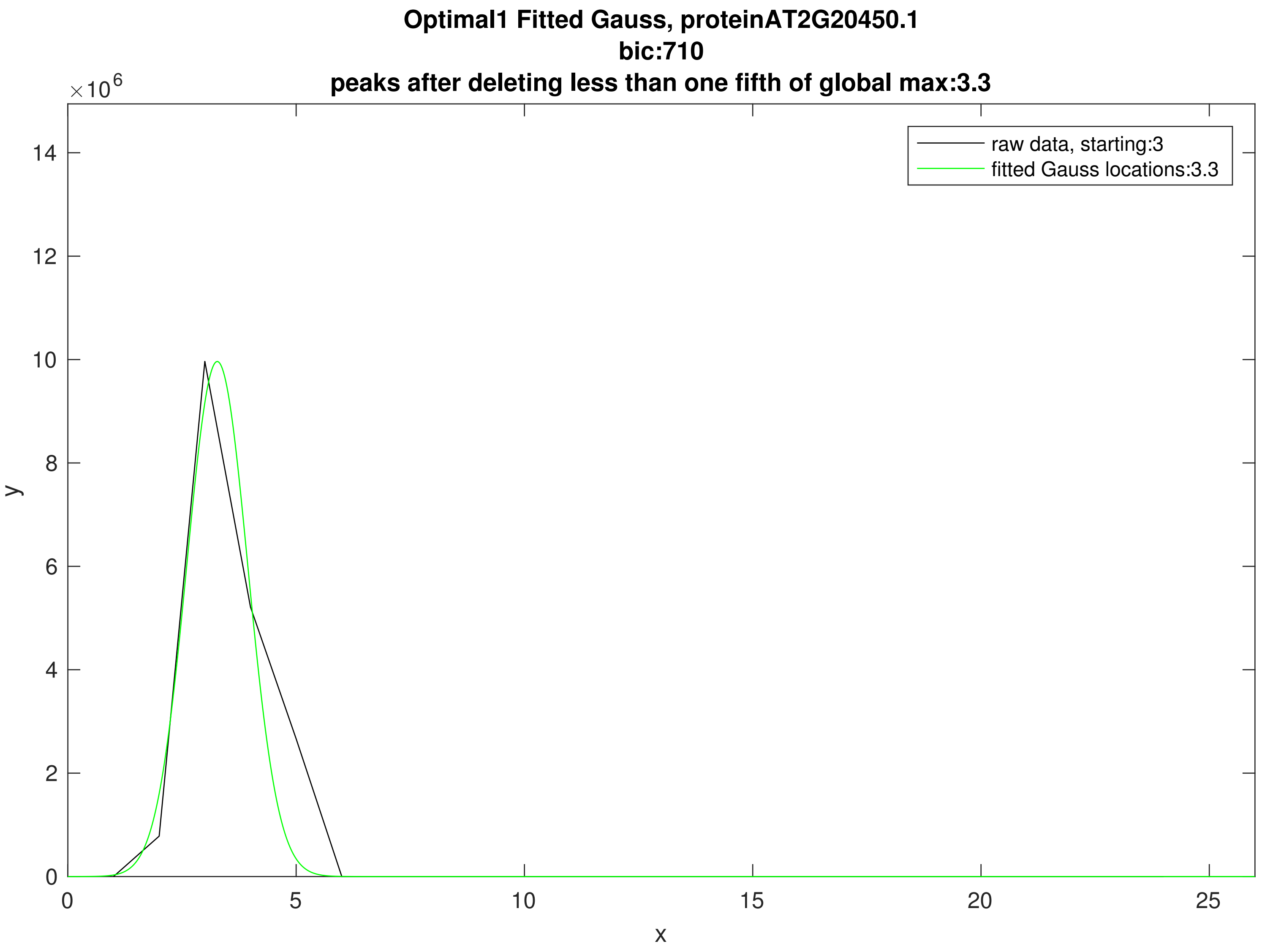
Optimal1 Fitted Gauss, proteinAT2G20450.1
bic:710
peaks after deleting less than one fifth of global max:3.3
×10^6
raw data, starting:3
fitted Gauss locations:3.3
y
x
0
2
4
6
8
10
12
14
0
5
10
15
20
25

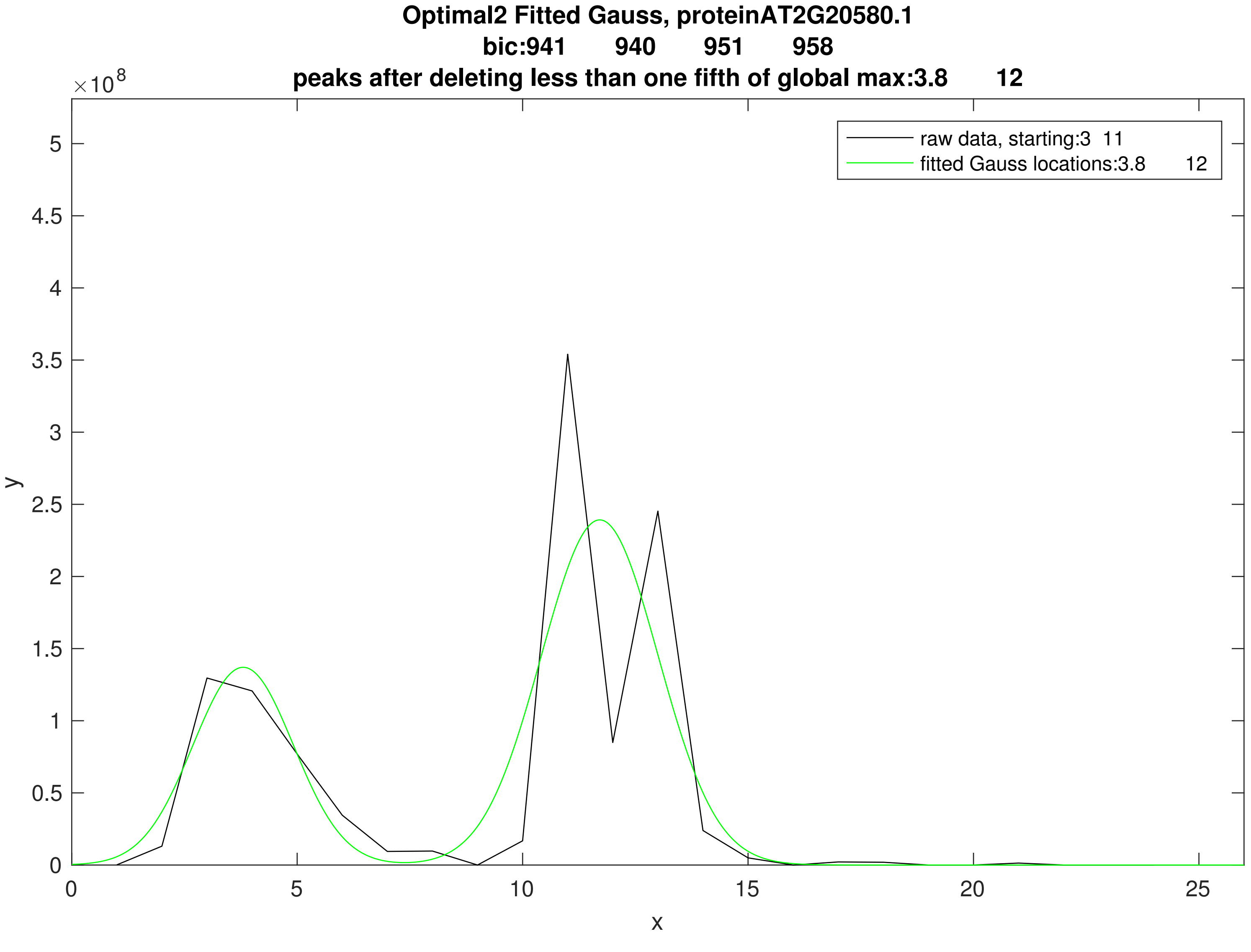
Optimal2 Fitted Gauss, proteinAT2G20580.1
bic:941 940 951 958
peaks after deleting less than one fifth of global max:3.8 12
×10^8
raw data, starting:3 11
fitted Gauss locations:3.8 12
0
0.5
1
1.5
2
2.5
3
3.5
4
4.5
5
y
0
5
10
15
20
25
x

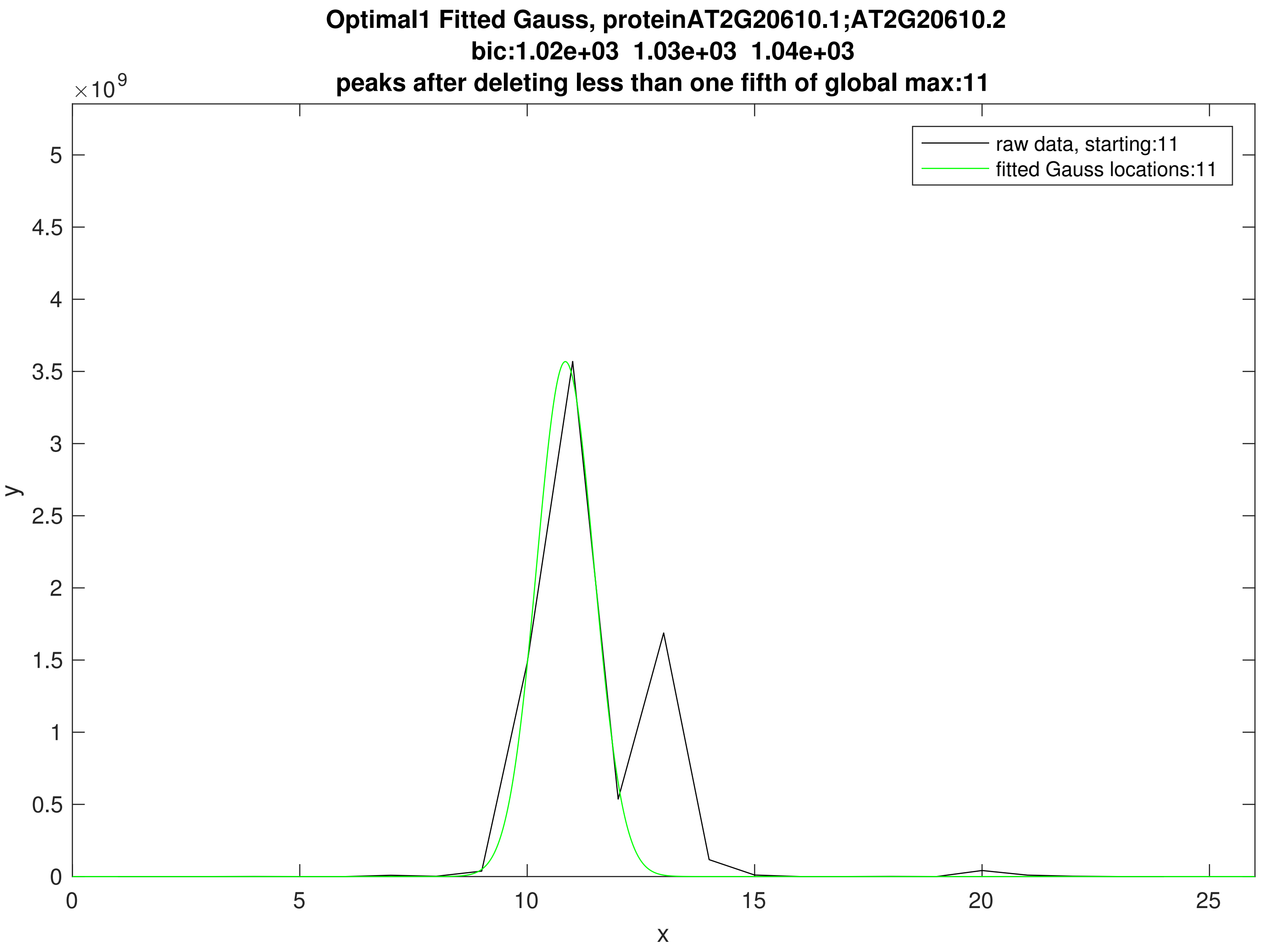
Optimal1 Fitted Gauss, proteinAT2G20610.1;AT2G20610.2
bic:1.02e+03 1.03e+03 1.04e+03
peaks after deleting less than one fifth of global max:11
×10^9
raw data, starting:11
fitted Gauss locations:11
5
4.5
4
3.5
3
2.5
2
1.5
1
0.5
0
y
0
5
10
15
20
25
x

**Optimal1 Fitted Gauss, proteinAT2G20630.1;AT2G20630.2**
**bic:819 829**
**peaks after deleting less than one fifth of global max:18**

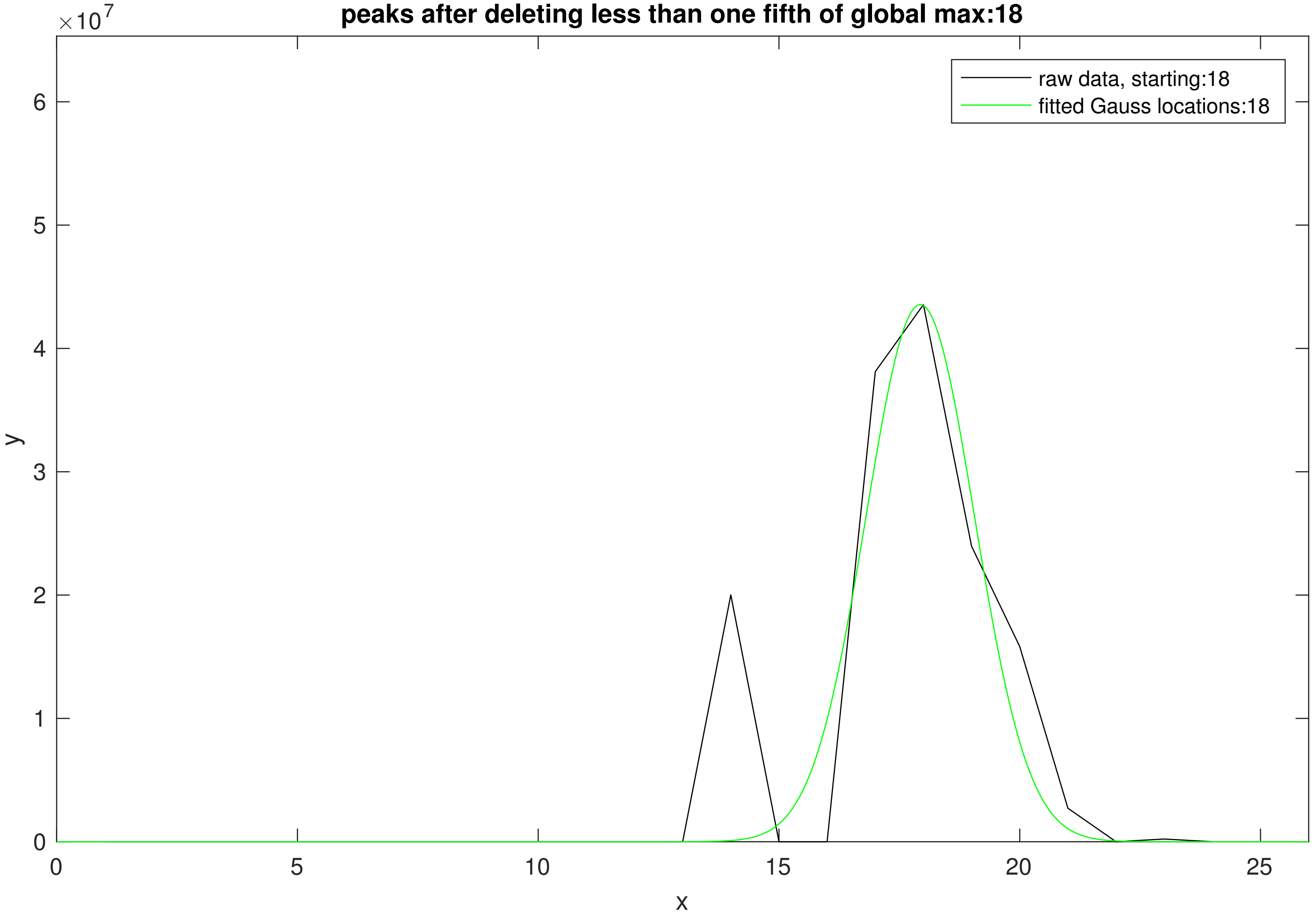

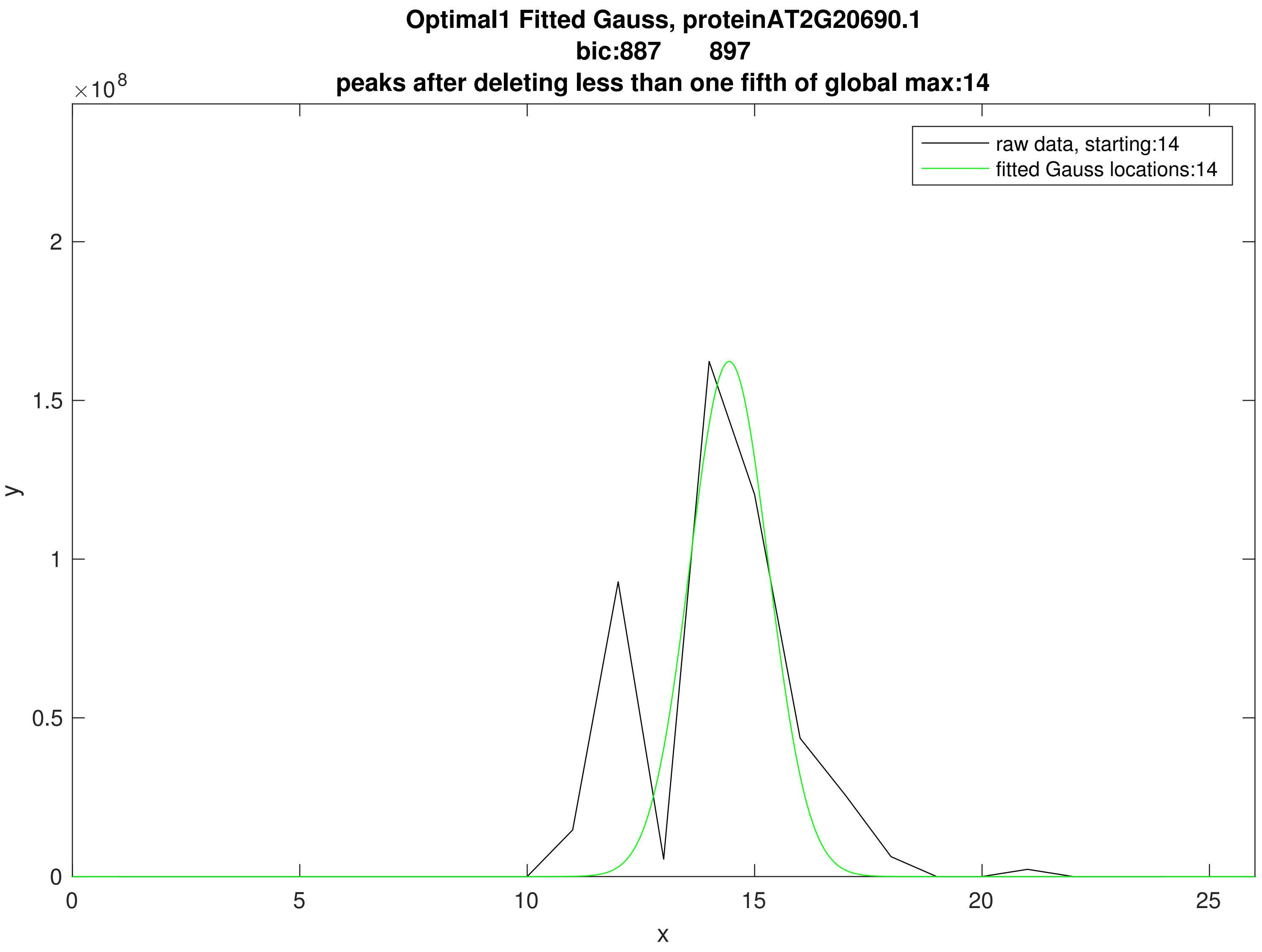

Optimal1 Fitted Gauss, proteinAT2G20690.1
bic:887 897
peaks after deleting less than one fifth of global max:14
×10^8
raw data, starting:14
fitted Gauss locations:14
y
x
0
0.5
1
1.5
2
0
5
10
15
20
25

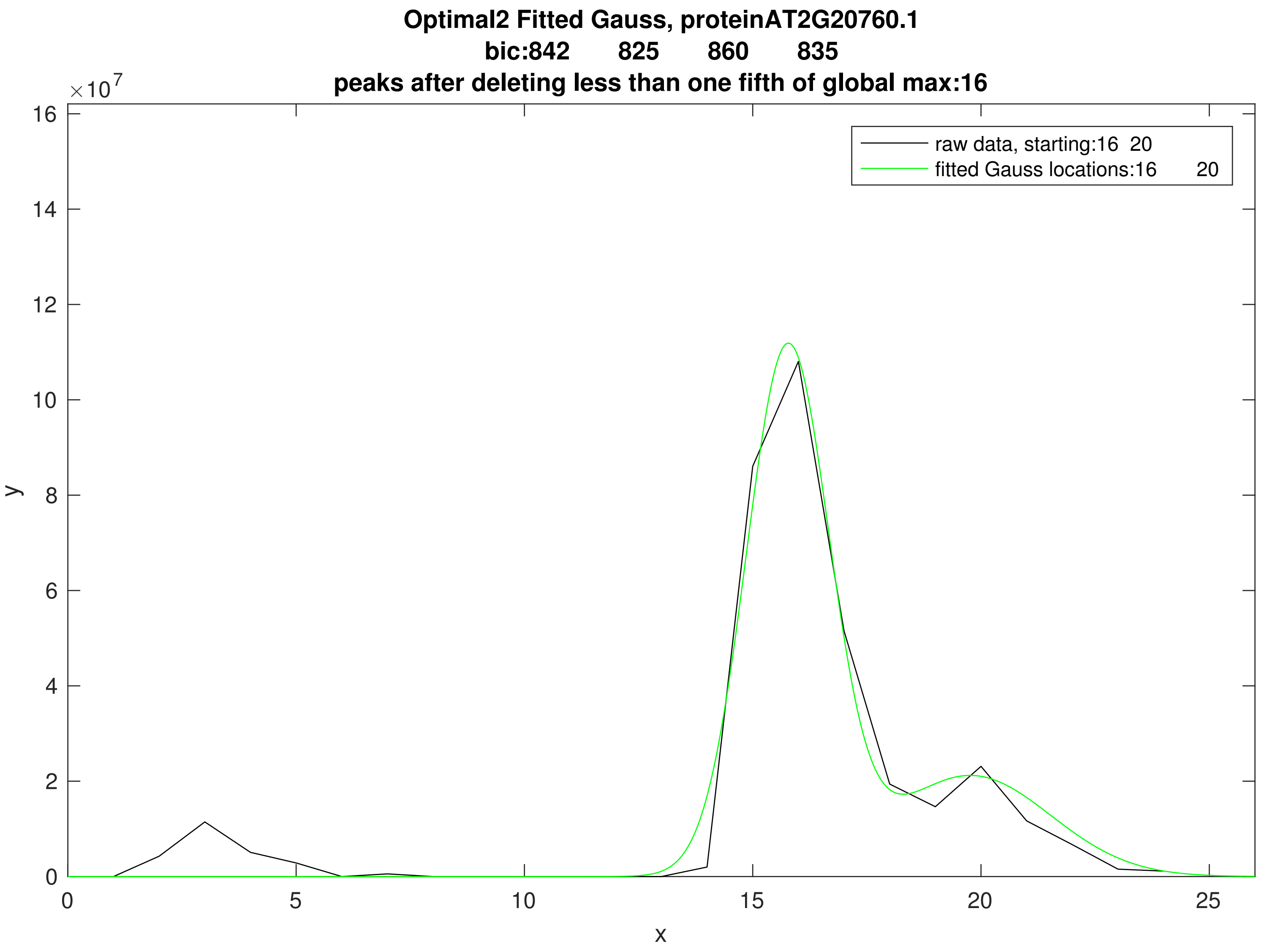

Optimal2 Fitted Gauss, proteinAT2G20760.1
bic:842 825 860 835
peaks after deleting less than one fifth of global max:16
×10^7
raw data, starting:16 20
fitted Gauss locations:16 20
y
x
0
2
4
6
8
10
12
14
16
0
5
10
15
20
25

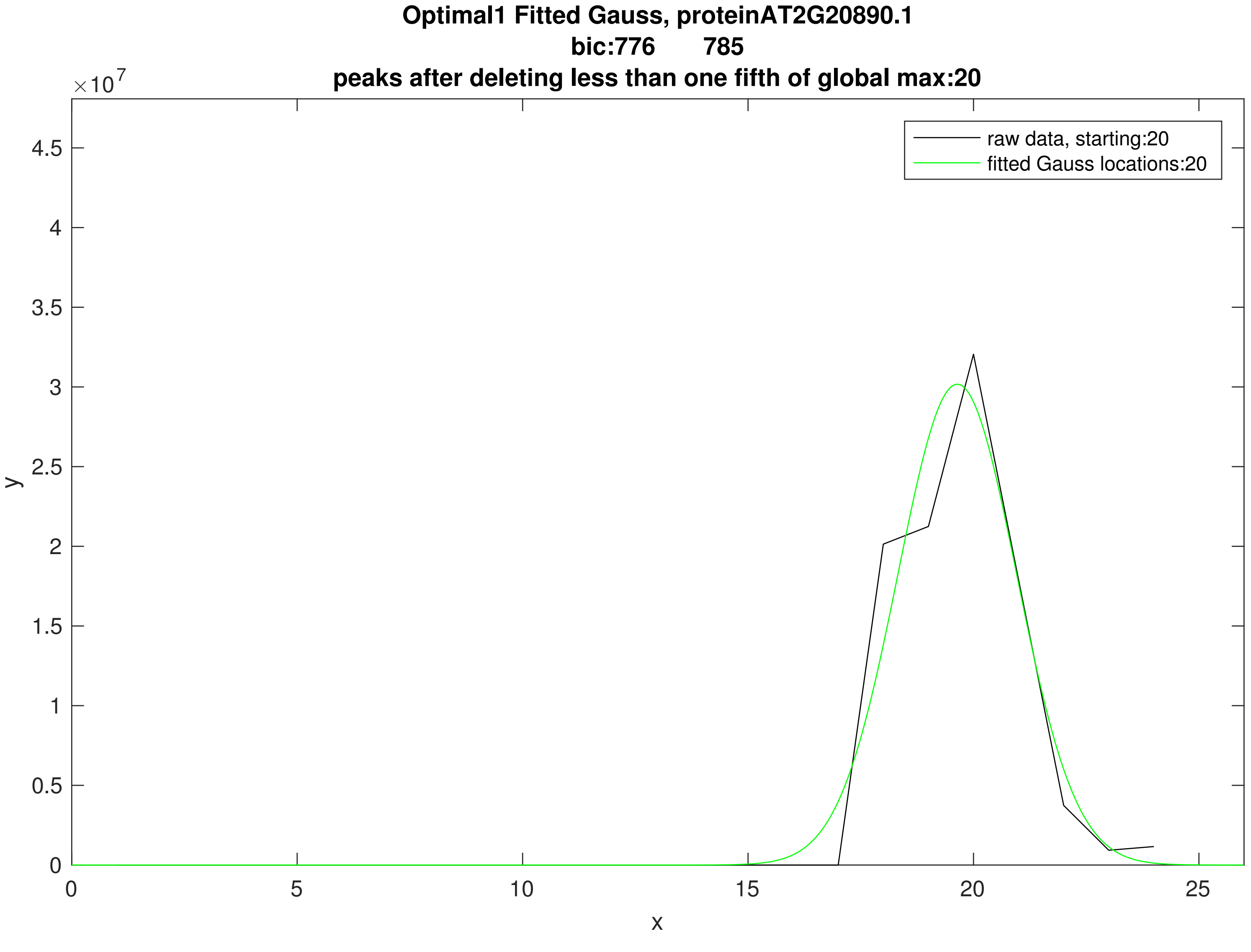
Optimal1 Fitted Gauss, proteinAT2G20890.1
bic:776 785
peaks after deleting less than one fifth of global max:20
×10^7
raw data, starting:20
fitted Gauss locations:20
y
x
0
0.5
1
1.5
2
2.5
3
3.5
4
4.5
5
10
15
20
25

**Optimal1 Fitted Gauss, proteinAT2G21045.1**
**bic:756**
**peaks after deleting less than one fifth of global max:20**

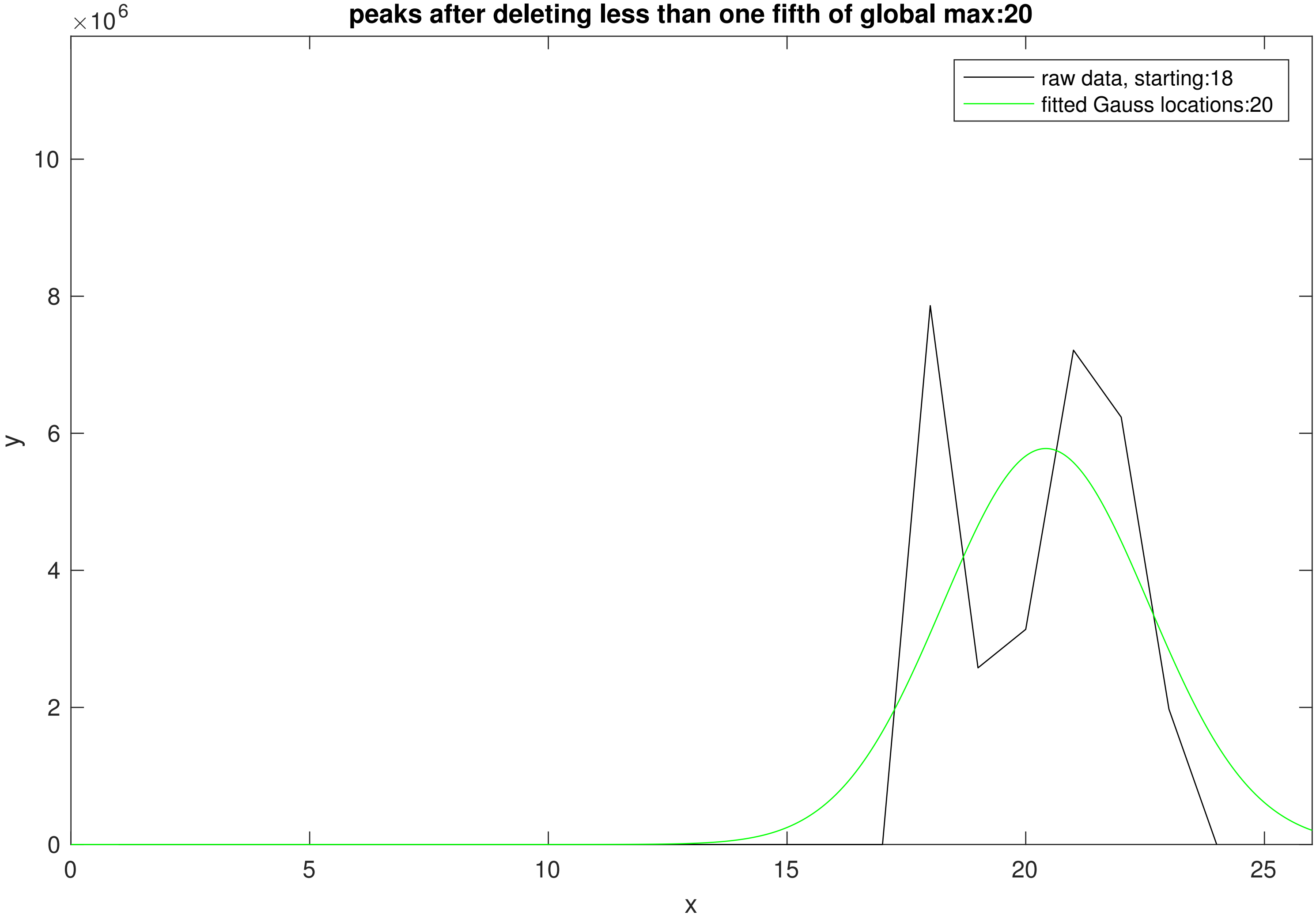

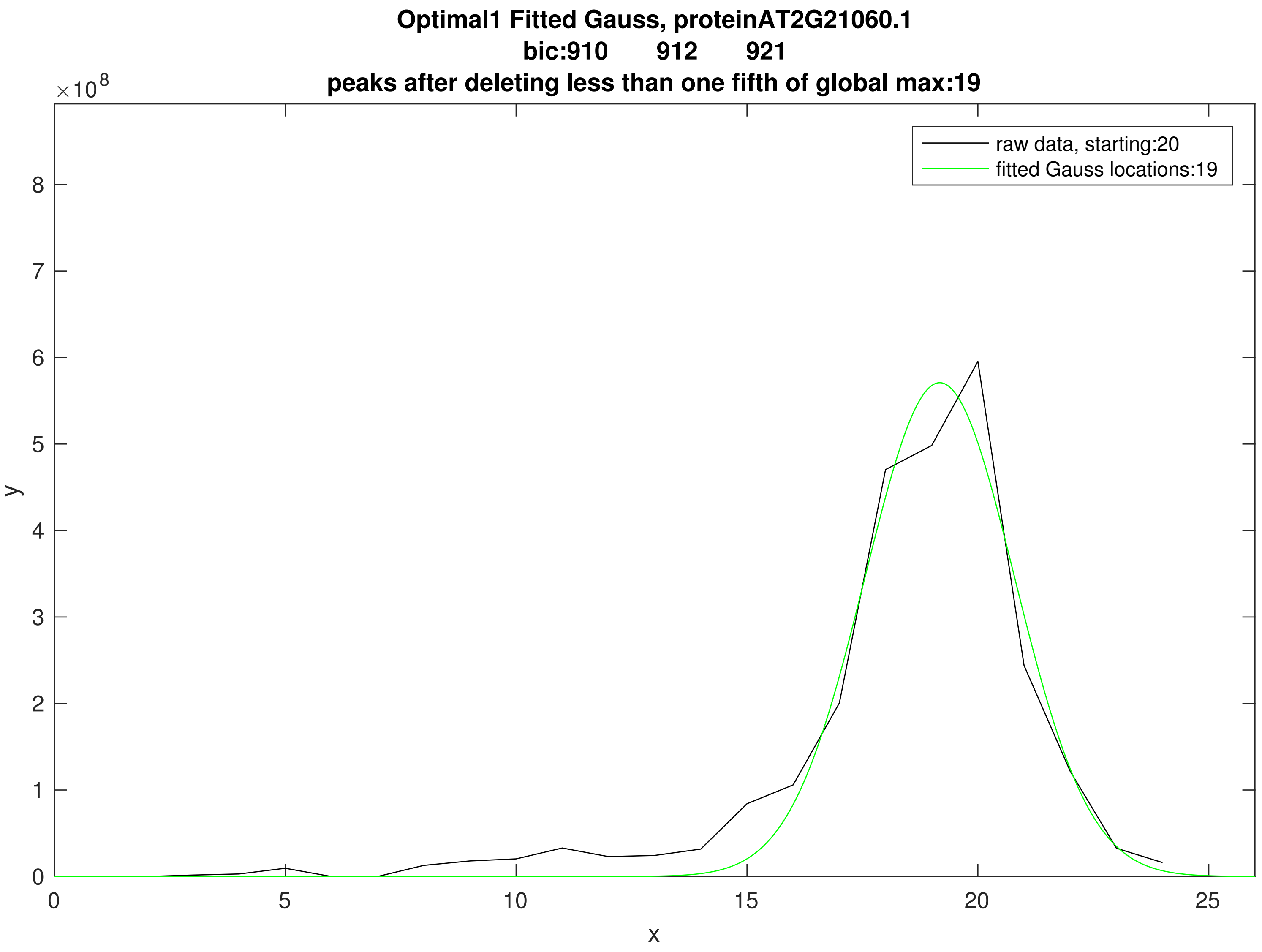
Optimal1 Fitted Gauss, proteinAT2G21060.1
bic:910 912 921
peaks after deleting less than one fifth of global max:19
×10^8
raw data, starting:20
fitted Gauss locations:19
y
x
0
1
2
3
4
5
6
7
8
0
5
10
15
20
25

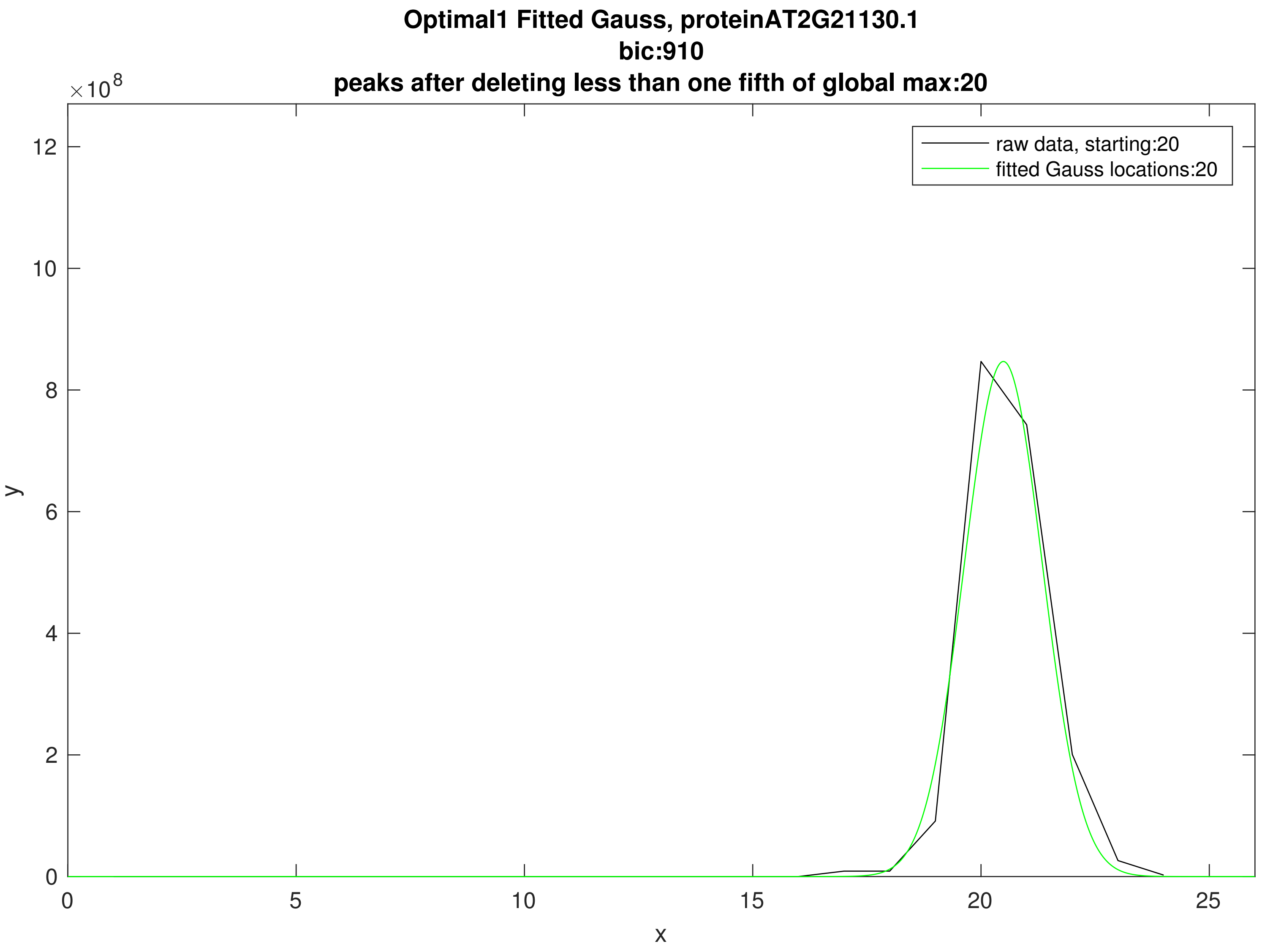

Optimal1 Fitted Gauss, proteinAT2G21130.1
bic:910
peaks after deleting less than one fifth of global max:20
×10^8
12
10
8
6
4
2
0
y
0
5
10
15
20
25
x
raw data, starting:20
fitted Gauss locations:20

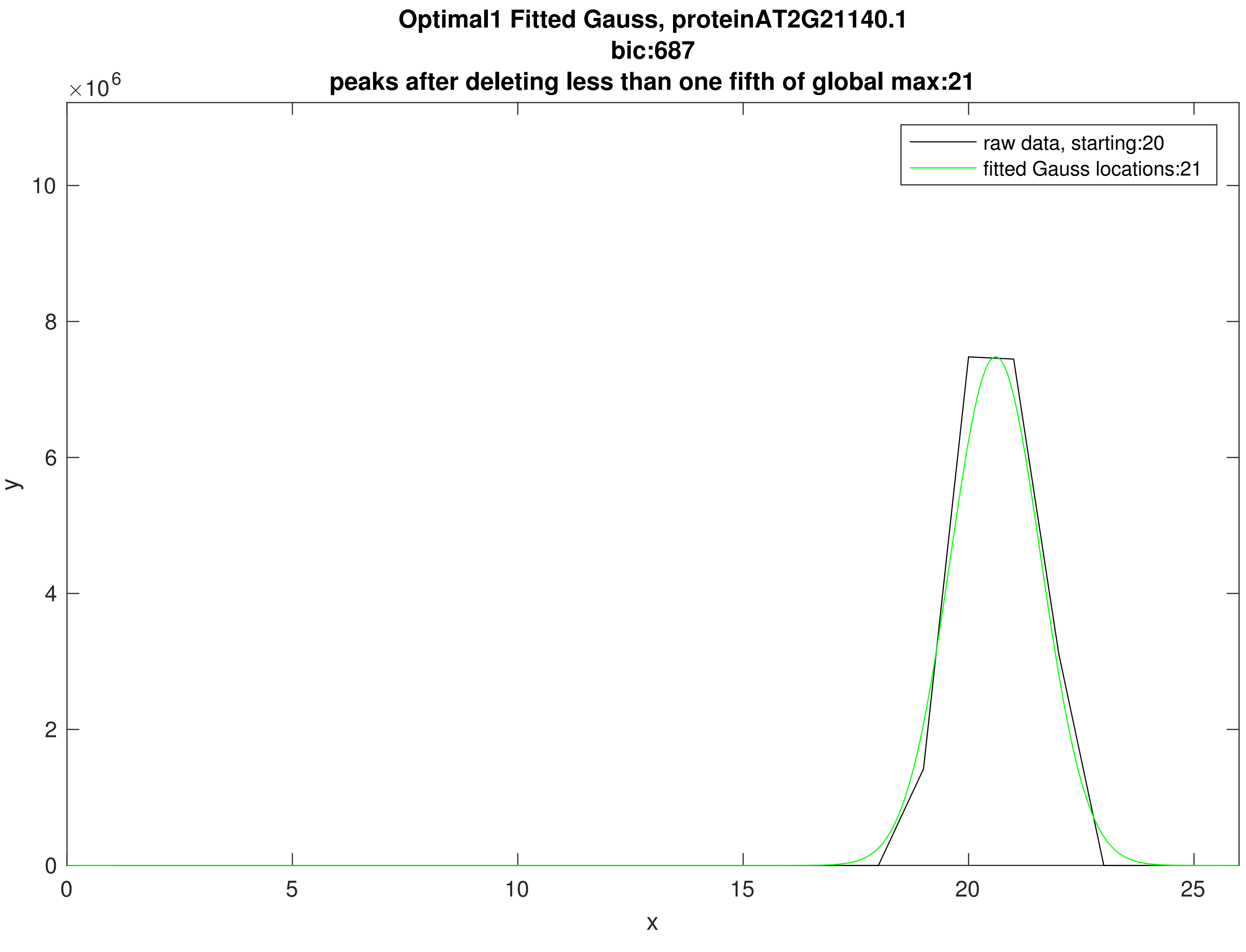

Optimal1 Fitted Gauss, proteinAT2G21140.1
bic:687
peaks after deleting less than one fifth of global max:21
$\times 10^6$
raw data, starting:20
fitted Gauss locations:21
10
8
6
4
2
0
y
0
5
10
15
20
25
x

**Optimal1 Fitted Gauss, proteinAT2G21170.1;AT2G21170.2**
**bic:1.01e+03**
**peaks after deleting less than one fifth of global max:15**

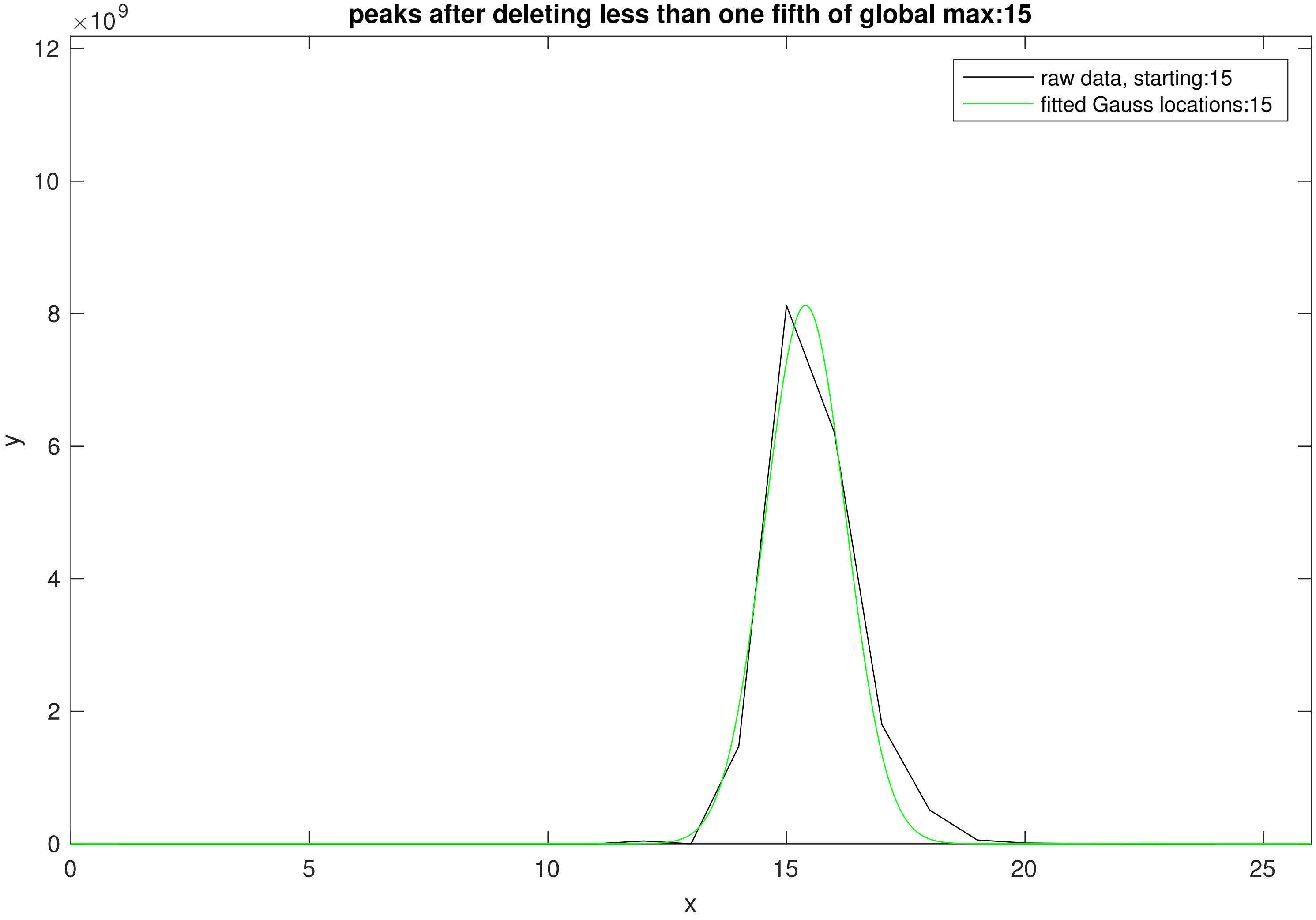

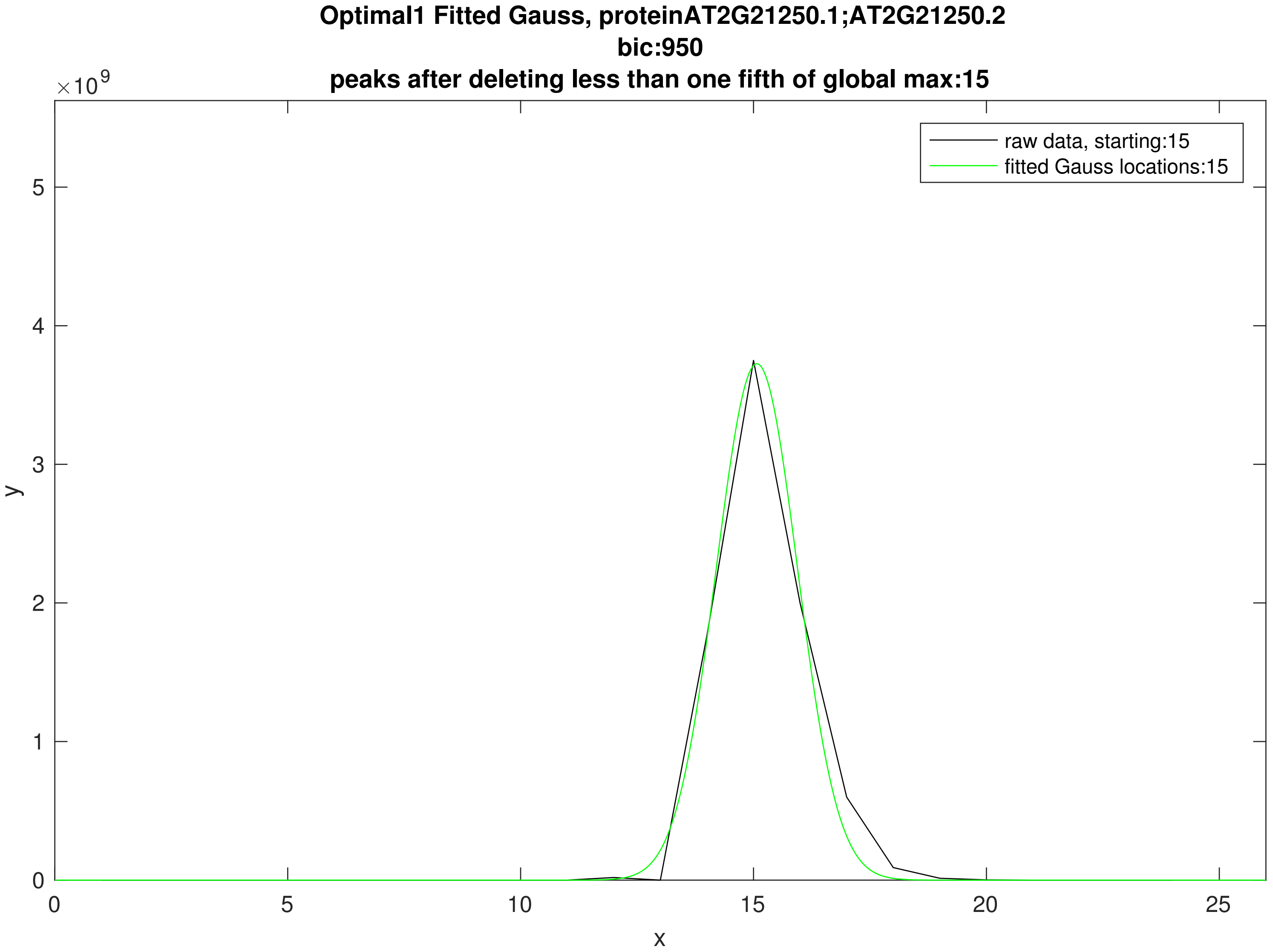

Optimal1 Fitted Gauss, proteinAT2G21250.1;AT2G21250.2
bic:950
peaks after deleting less than one fifth of global max:15
×10^9
raw data, starting:15
fitted Gauss locations:15
y
x
0
1
2
3
4
5
0
5
10
15
20
25

**Optimal1 Fitted Gauss, proteinAT2G21270.2;AT2G21270.1;AT2G21270.3**
**bic:817 833**
**peaks after deleting less than one fifth of global max:17**

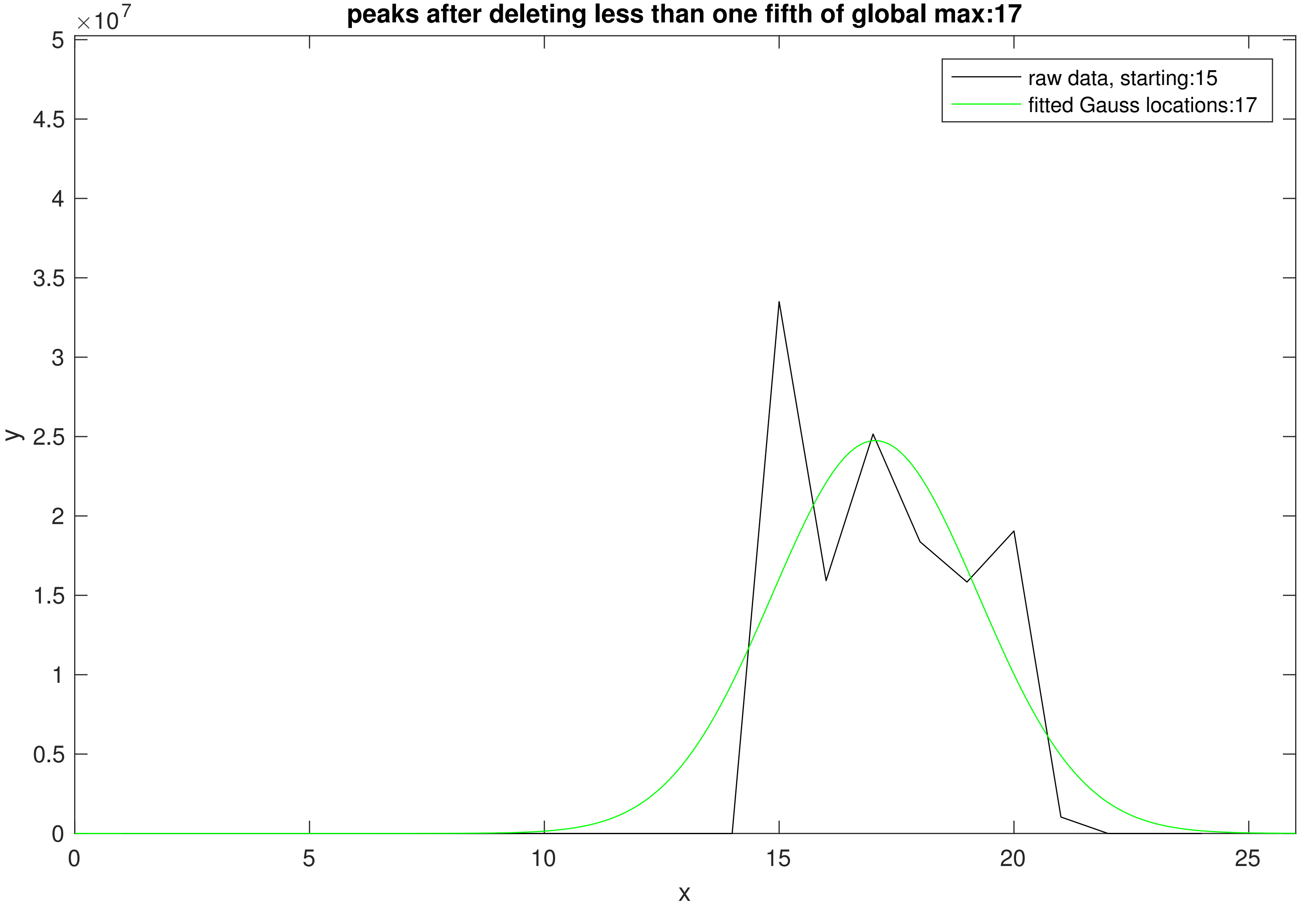

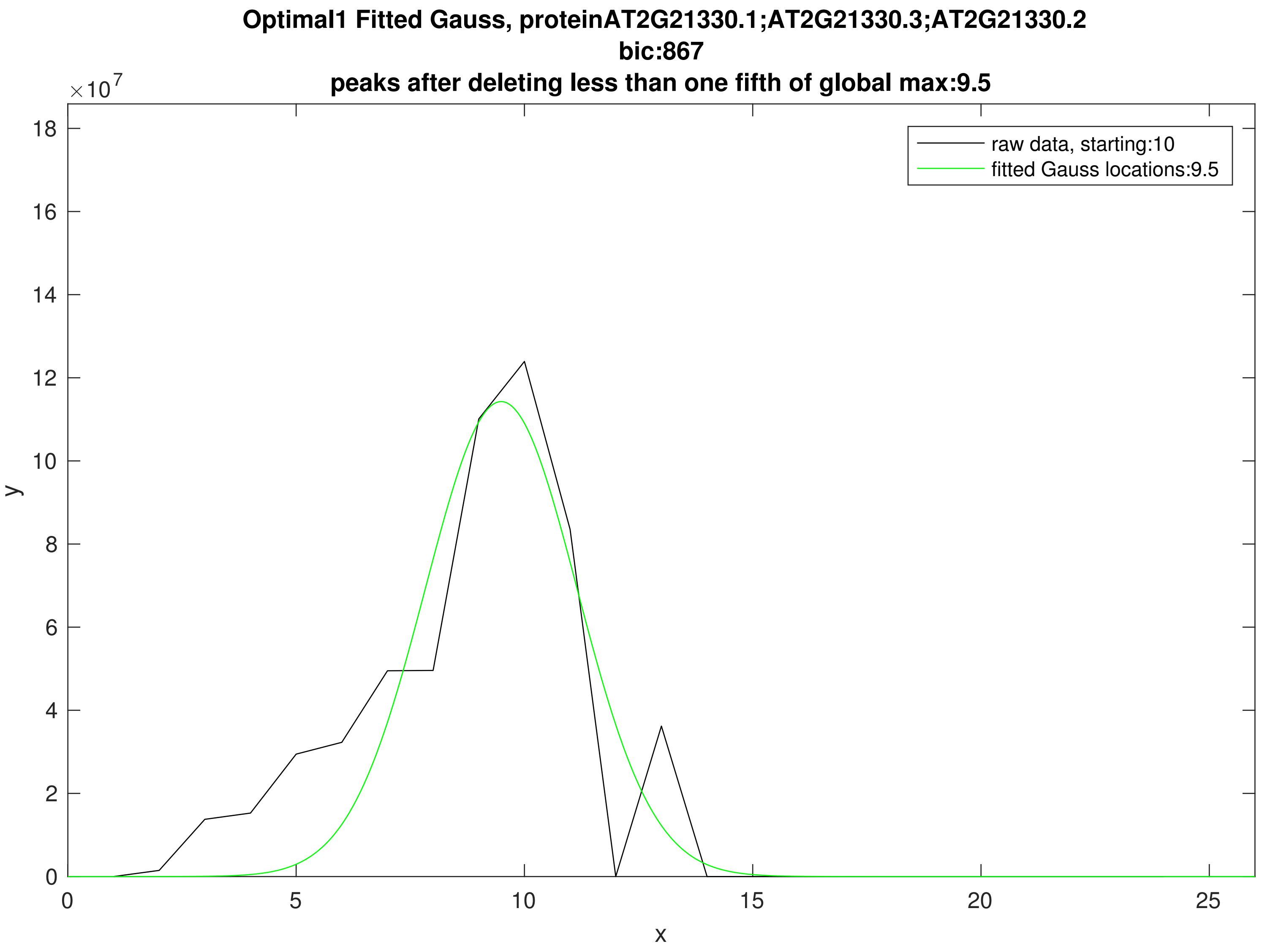

Optimal1 Fitted Gauss, proteinAT2G21330.1;AT2G21330.3;AT2G21330.2
bic:867
peaks after deleting less than one fifth of global max:9.5
×10^7
raw data, starting:10
fitted Gauss locations:9.5
y
x
0
2
4
6
8
10
12
14
16
18
5
15
20
25

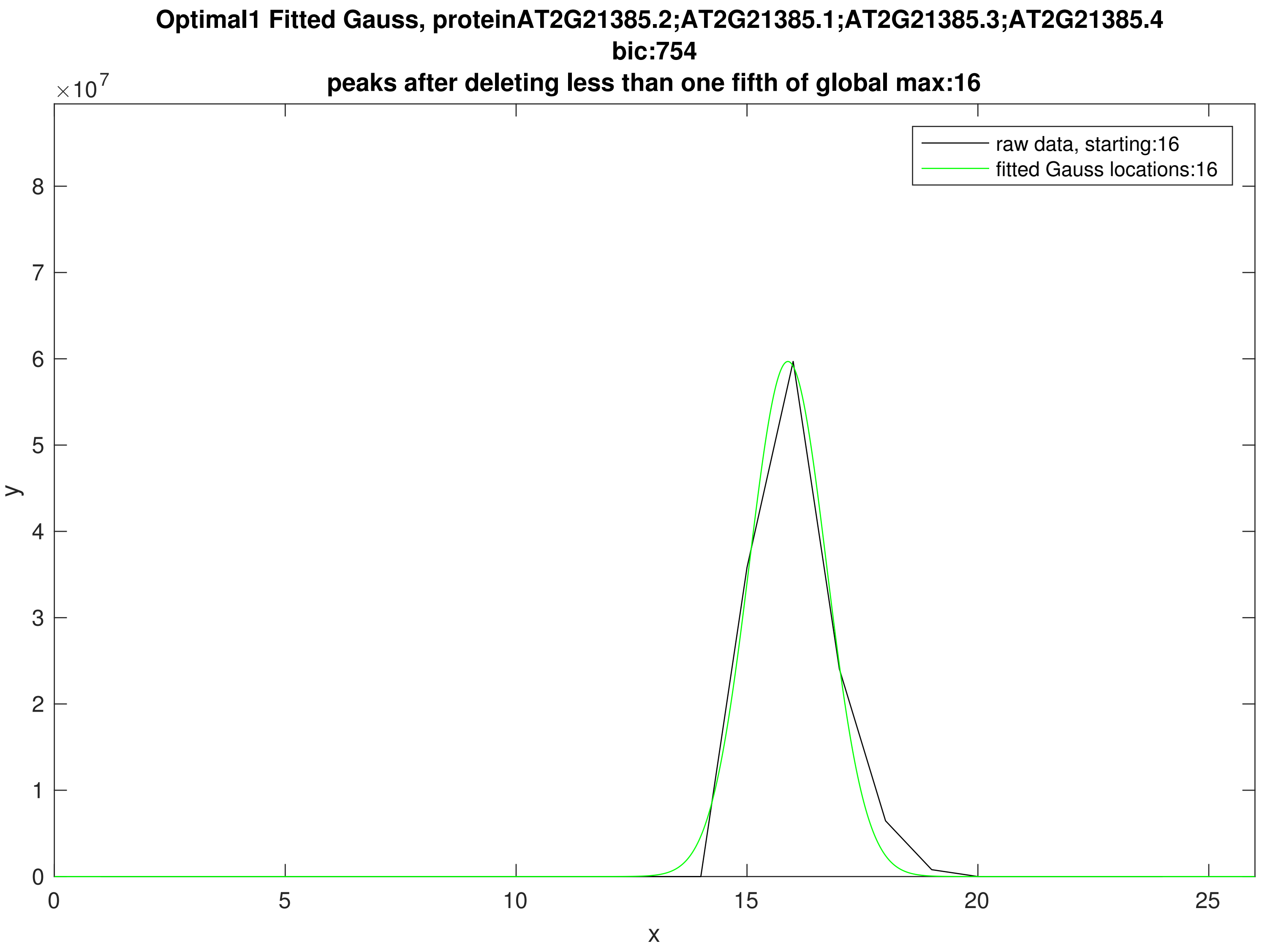

Optimal1 Fitted Gauss, proteinAT2G21385.2;AT2G21385.1;AT2G21385.3;AT2G21385.4
bic:754
peaks after deleting less than one fifth of global max:16
×10^7
raw data, starting:16
fitted Gauss locations:16
y
x
0
1
2
3
4
5
6
7
8
0
5
10
15
20
25

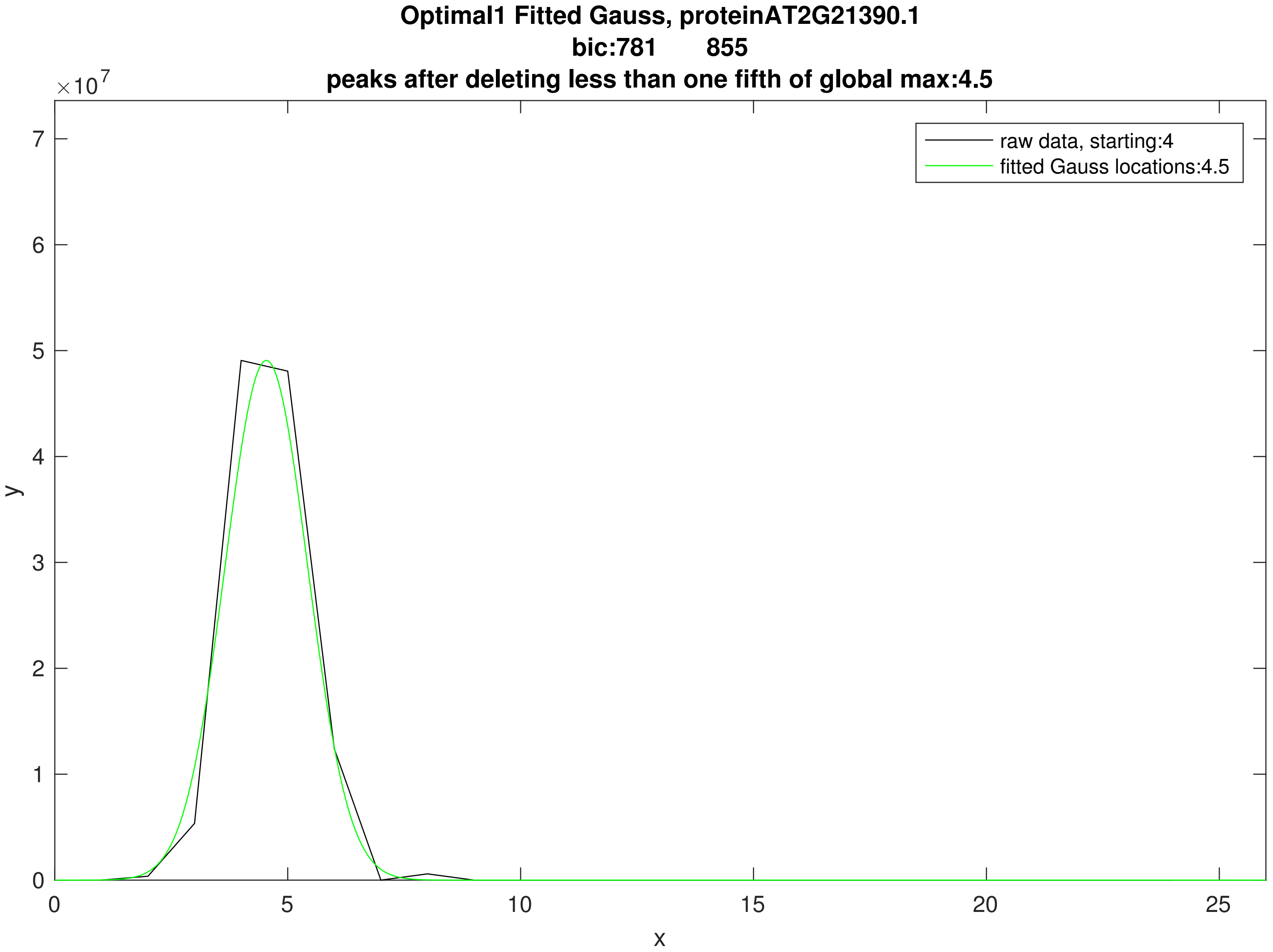

Optimal1 Fitted Gauss, proteinAT2G21390.1
bic:781 855
peaks after deleting less than one fifth of global max:4.5
×10^7
raw data, starting:4
fitted Gauss locations:4.5
y
x
0
1
2
3
4
5
6
7
0
5
10
15
20
25

Optimal1 Fitted Gauss, proteinAT2G21470.1;AT2G21470.3;AT2G21470.2
bic:814
peaks after deleting less than one fifth of global max:11

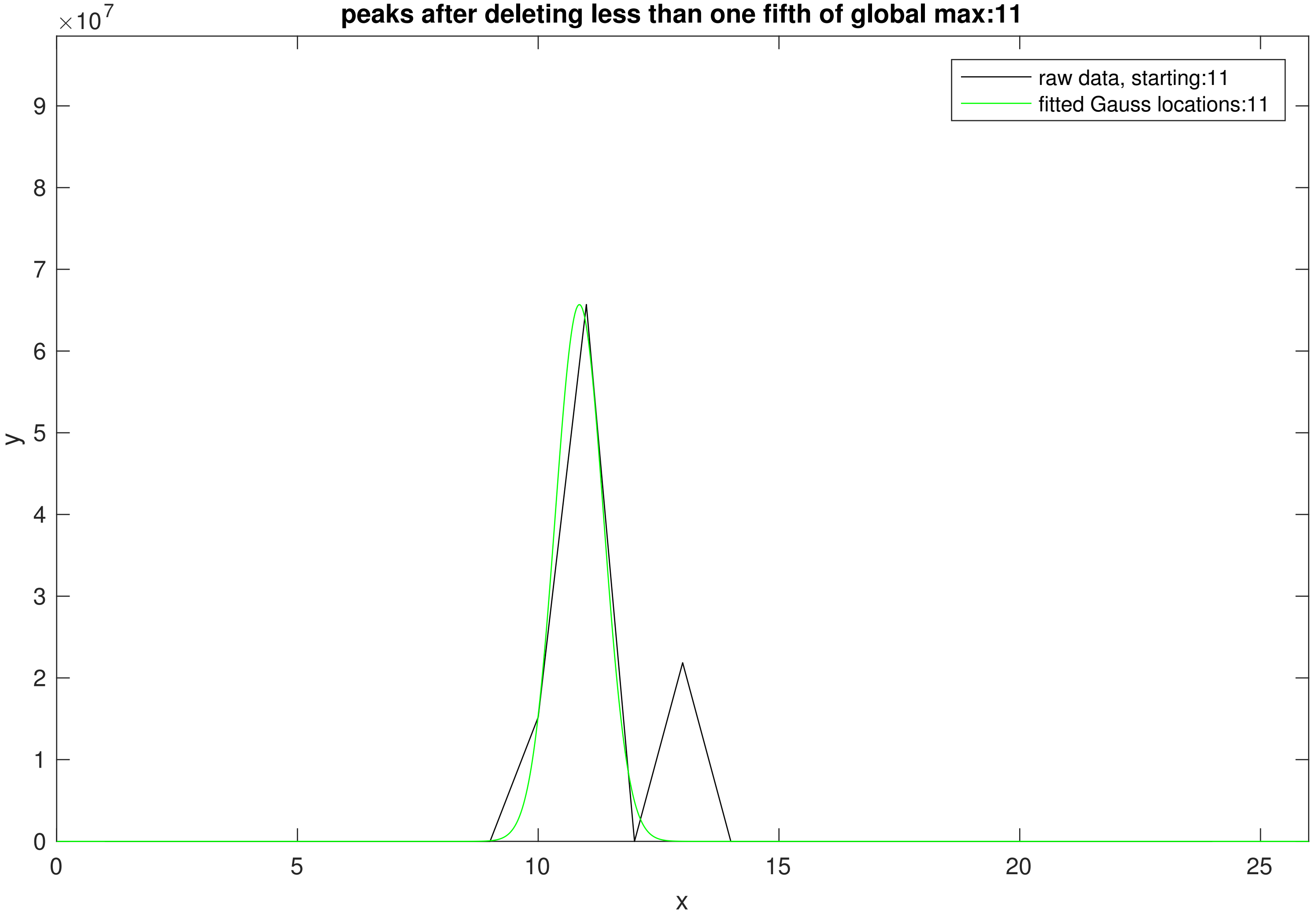

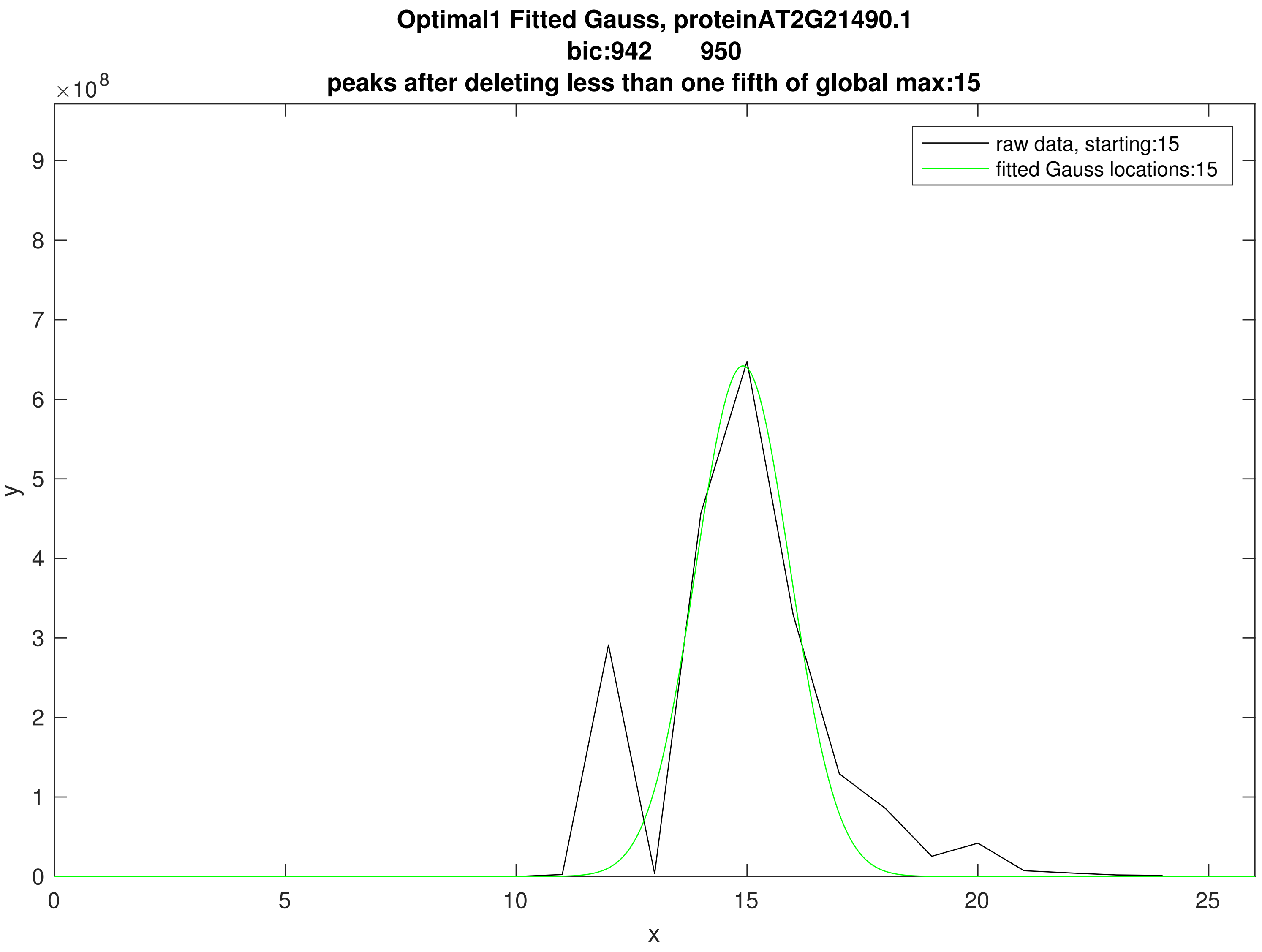

Optimal1 Fitted Gauss, proteinAT2G21490.1
bic:942 950
peaks after deleting less than one fifth of global max:15
×10^8
raw data, starting:15
fitted Gauss locations:15
y
x
0
1
2
3
4
5
6
7
8
9
0
5
10
15
20
25

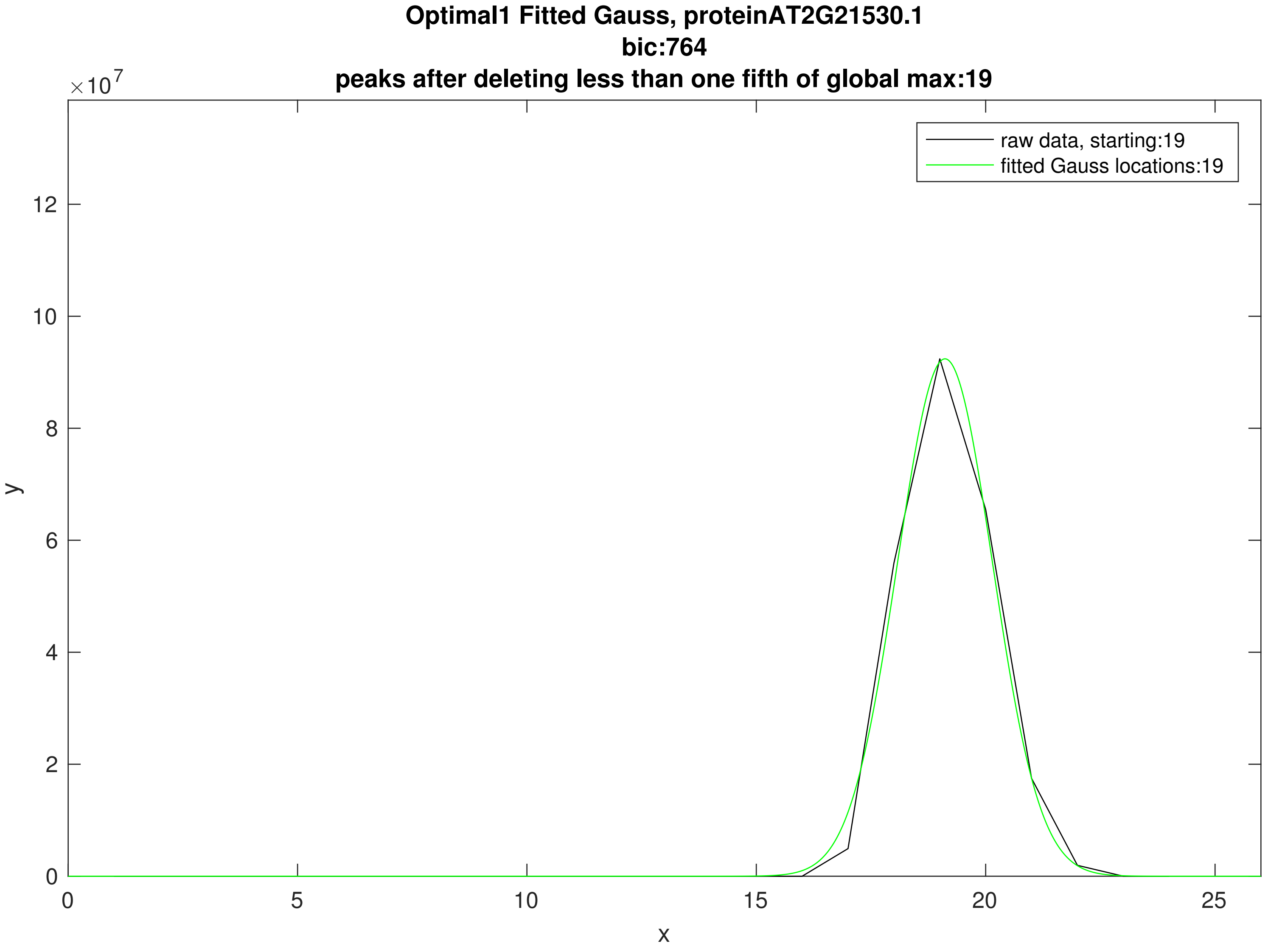
Optimal1 Fitted Gauss, proteinAT2G21530.1
bic:764
peaks after deleting less than one fifth of global max:19
$\times 10^7$
raw data, starting:19
fitted Gauss locations:19
y
x
0
2
4
6
8
10
12
0
5
10
15
20
25

#### Optimal1 Fitted Gauss, proteinAT2G21580.2;AT2G21580.1
#### bic:791
#### peaks after deleting less than one fifth of global max:3.6

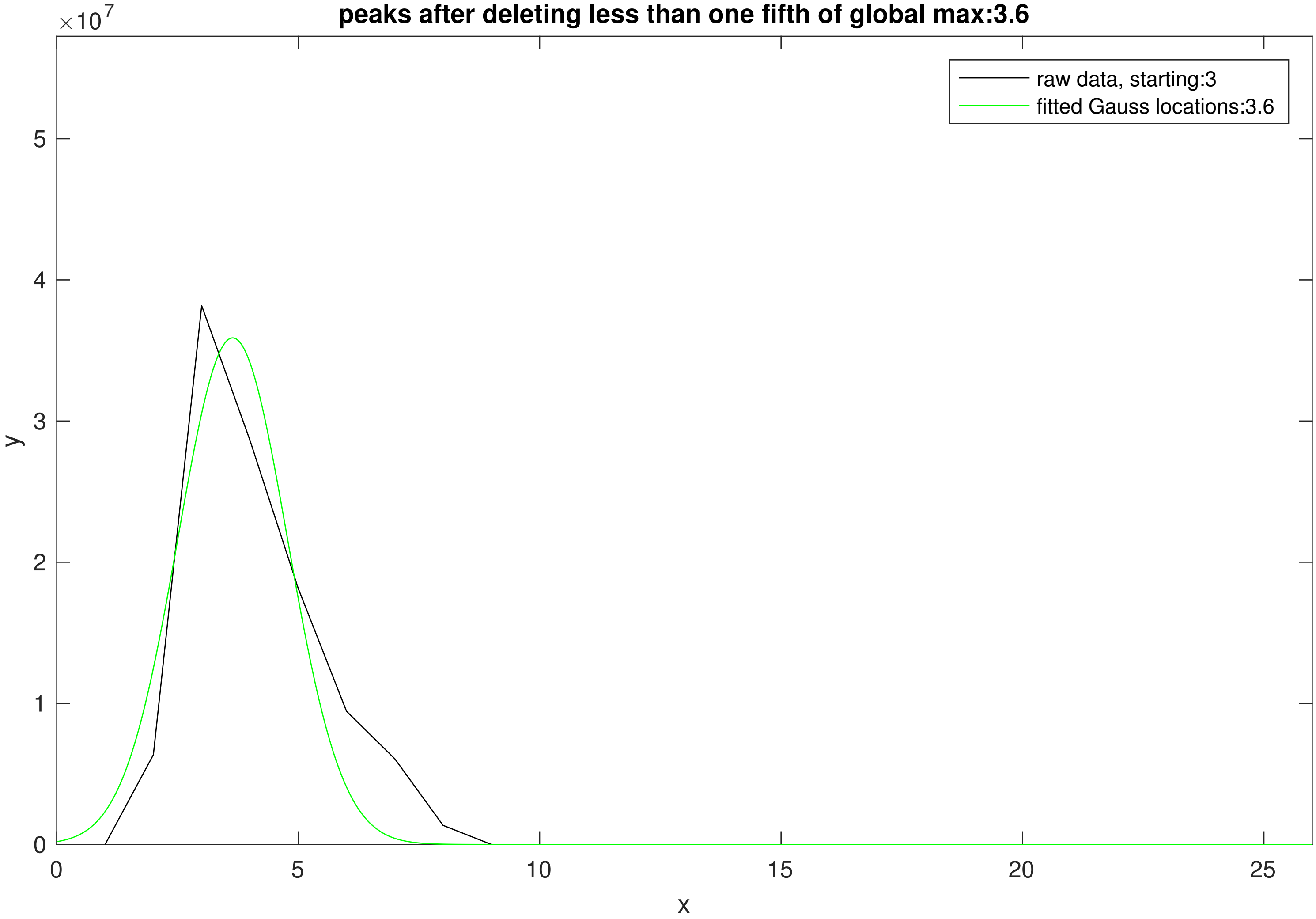

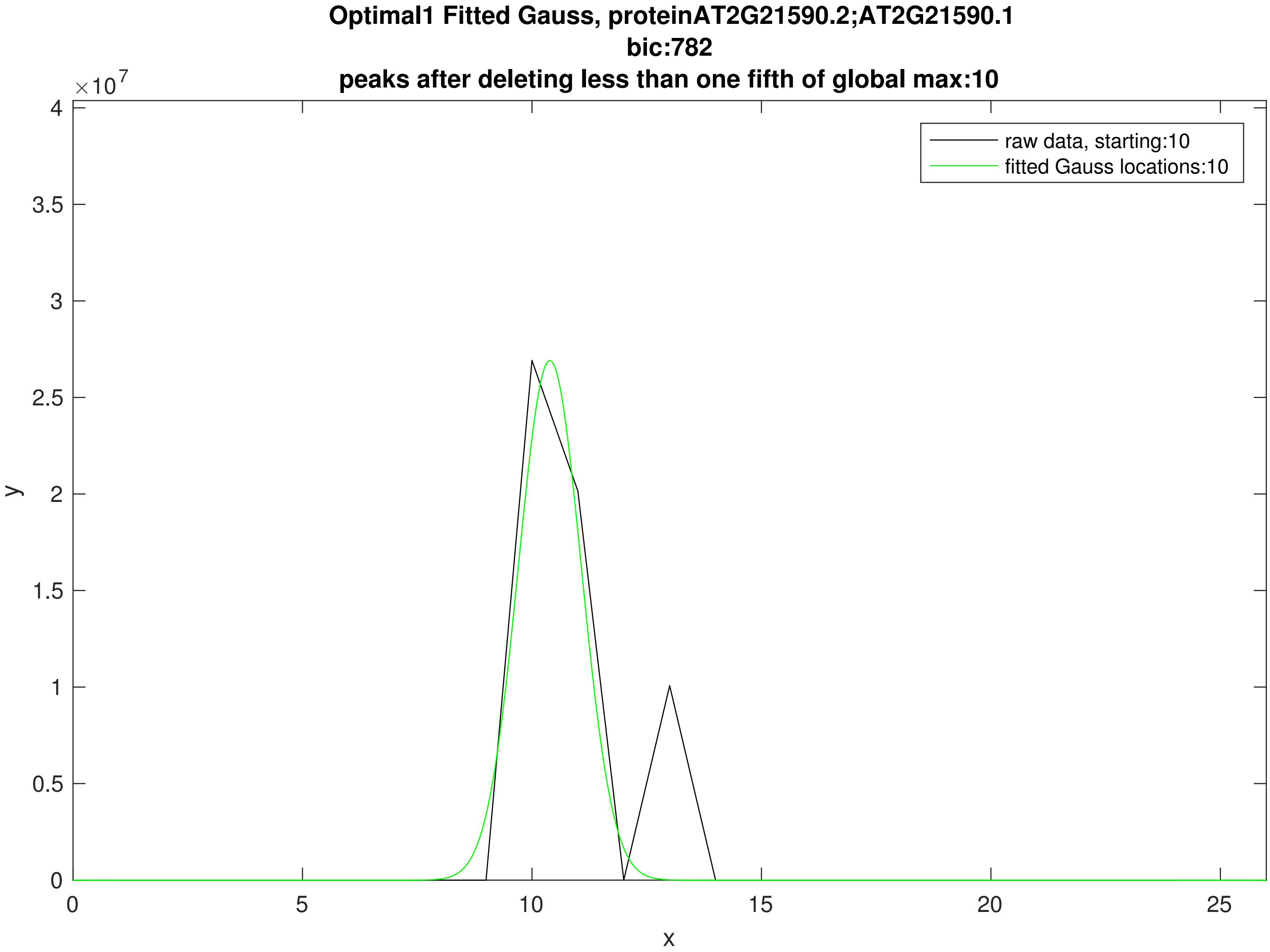

Optimal1 Fitted Gauss, proteinAT2G21590.2;AT2G21590.1
bic:782
peaks after deleting less than one fifth of global max:10
×10^7
4
3.5
3
2.5
2
1.5
1
0.5
0
y
0
5
10
15
20
25
x
raw data, starting:10
fitted Gauss locations:10

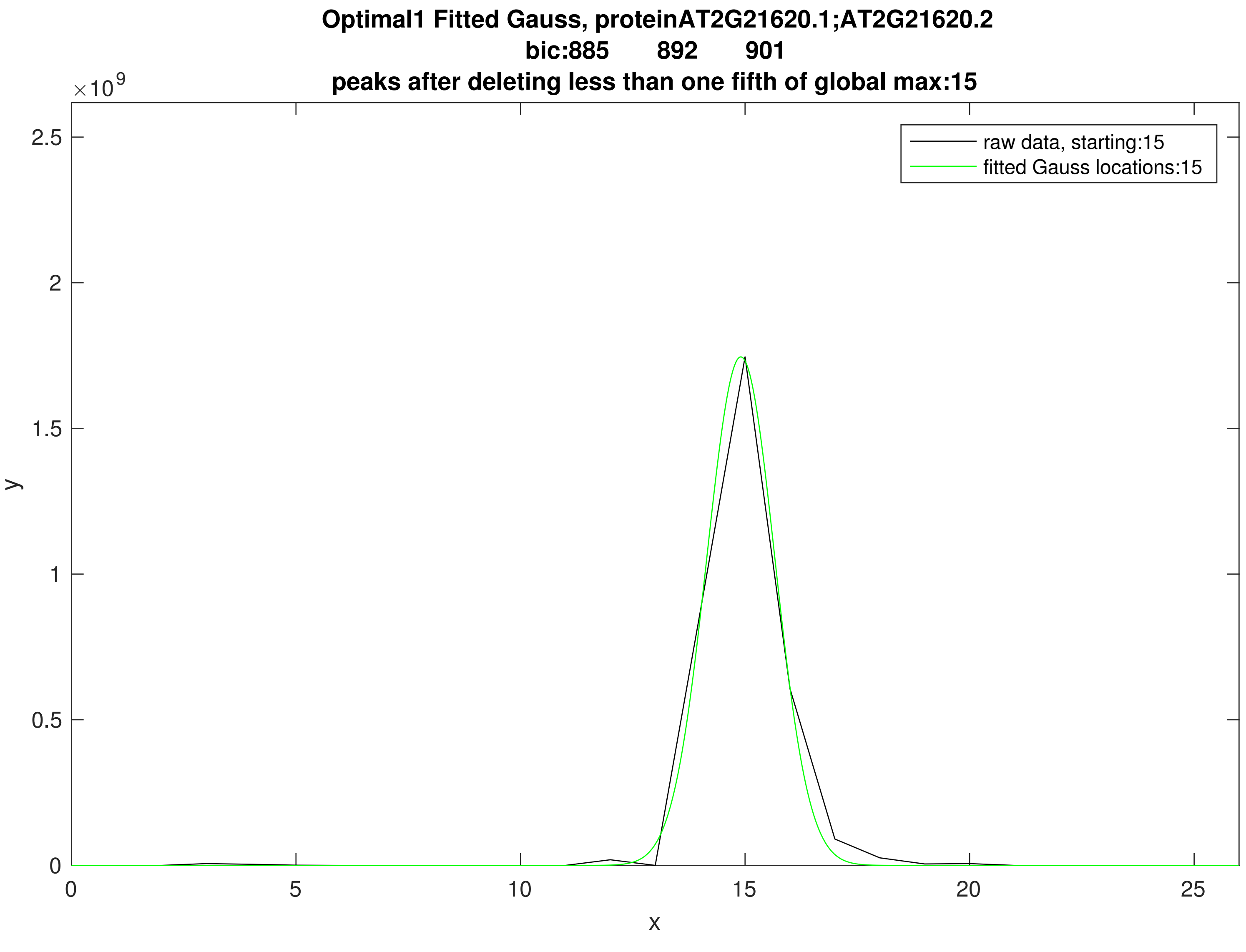

Optimal1 Fitted Gauss, proteinAT2G21620.1;AT2G21620.2
bic:885 892 901
peaks after deleting less than one fifth of global max:15
×10^9
2.5
2
1.5
1
0.5
0
y
0
5
10
15
20
25
x
raw data, starting:15
fitted Gauss locations:15

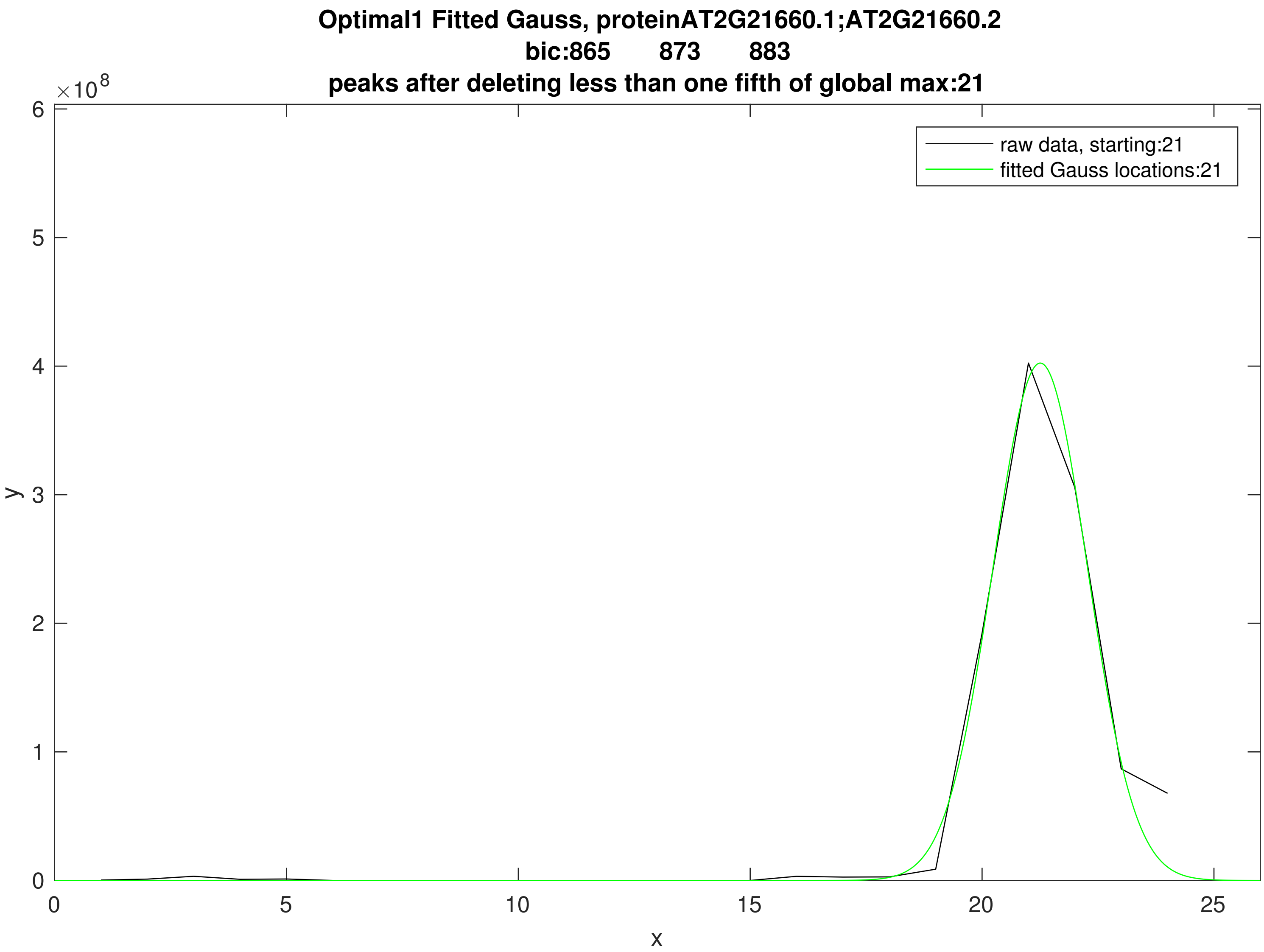

Optimal1 Fitted Gauss, proteinAT2G21660.1;AT2G21660.2
bic:865 873 883
peaks after deleting less than one fifth of global max:21
×10^8
raw data, starting:21
fitted Gauss locations:21
y
x
0
1
2
3
4
5
6
0
5
10
15
20
25

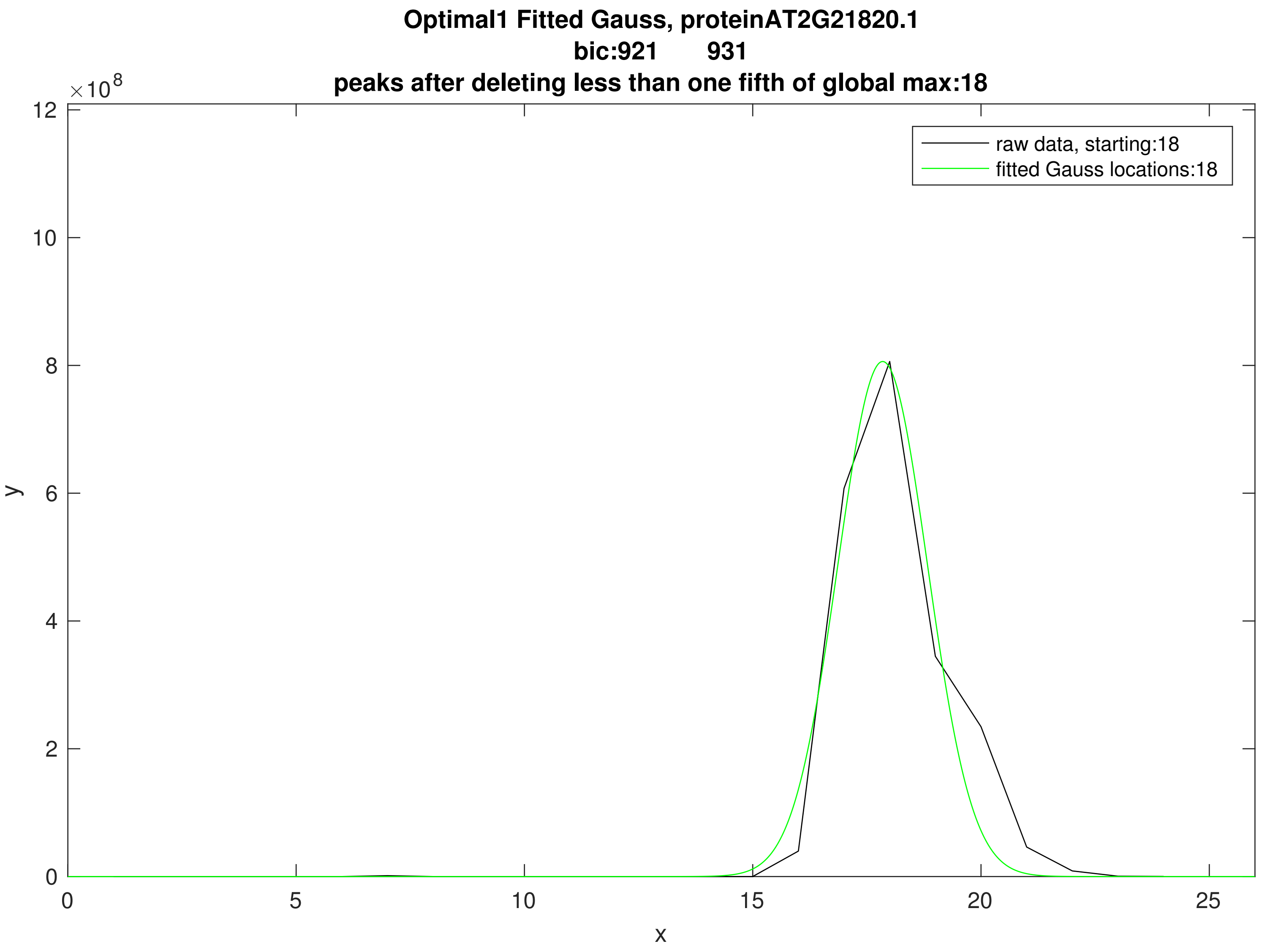

Optimal1 Fitted Gauss, proteinAT2G21820.1
bic:921 931
peaks after deleting less than one fifth of global max:18
×10^8
12
10
8
6
4
2
0
y
0
5
10
15
20
25
x
raw data, starting:18
fitted Gauss locations:18

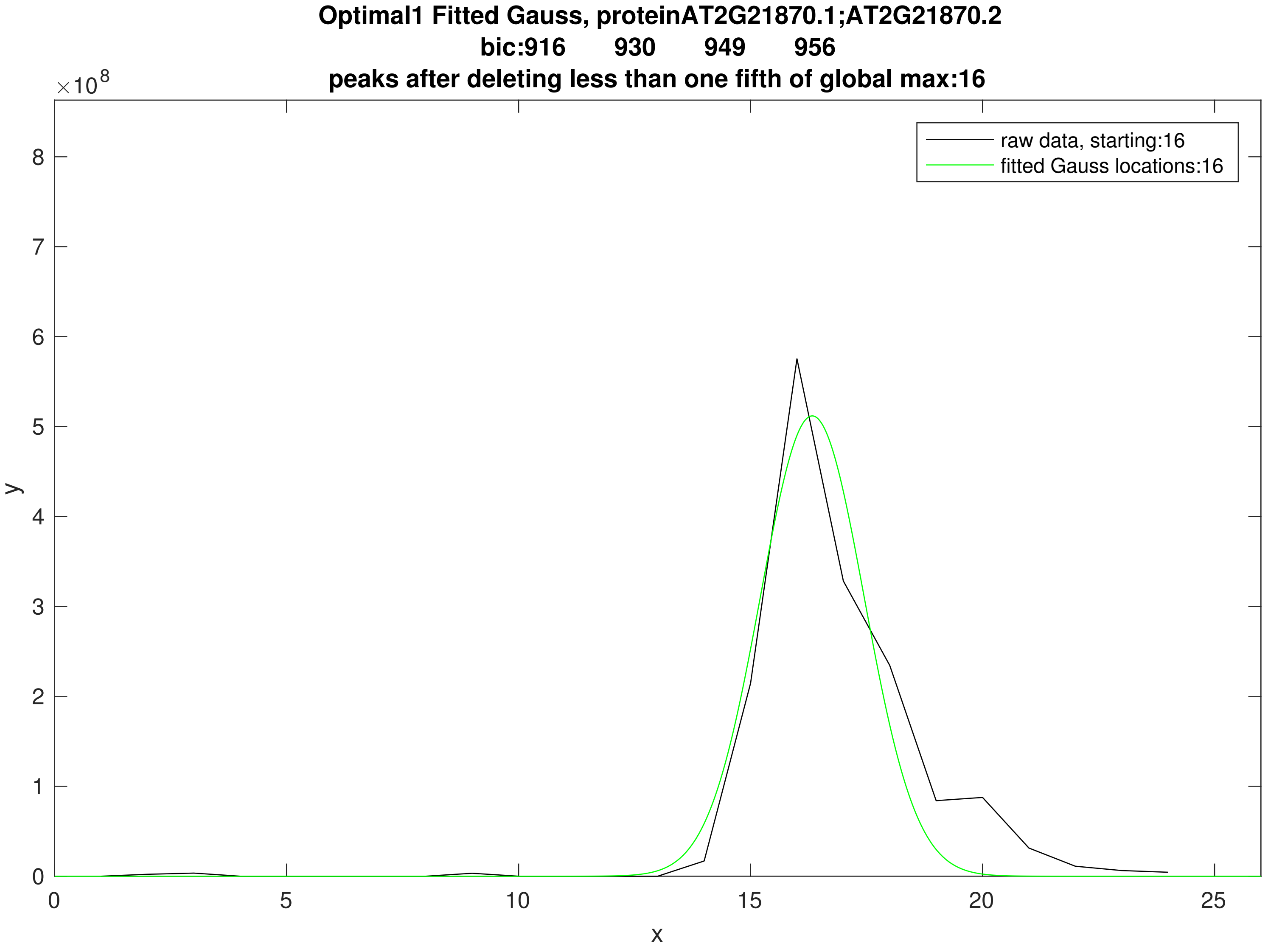

Optimal1 Fitted Gauss, proteinAT2G21870.1;AT2G21870.2
bic:916 930 949 956
peaks after deleting less than one fifth of global max:16
×10^8
raw data, starting:16
fitted Gauss locations:16
y
x
0
1
2
3
4
5
6
7
8
0
5
10
15
20
25

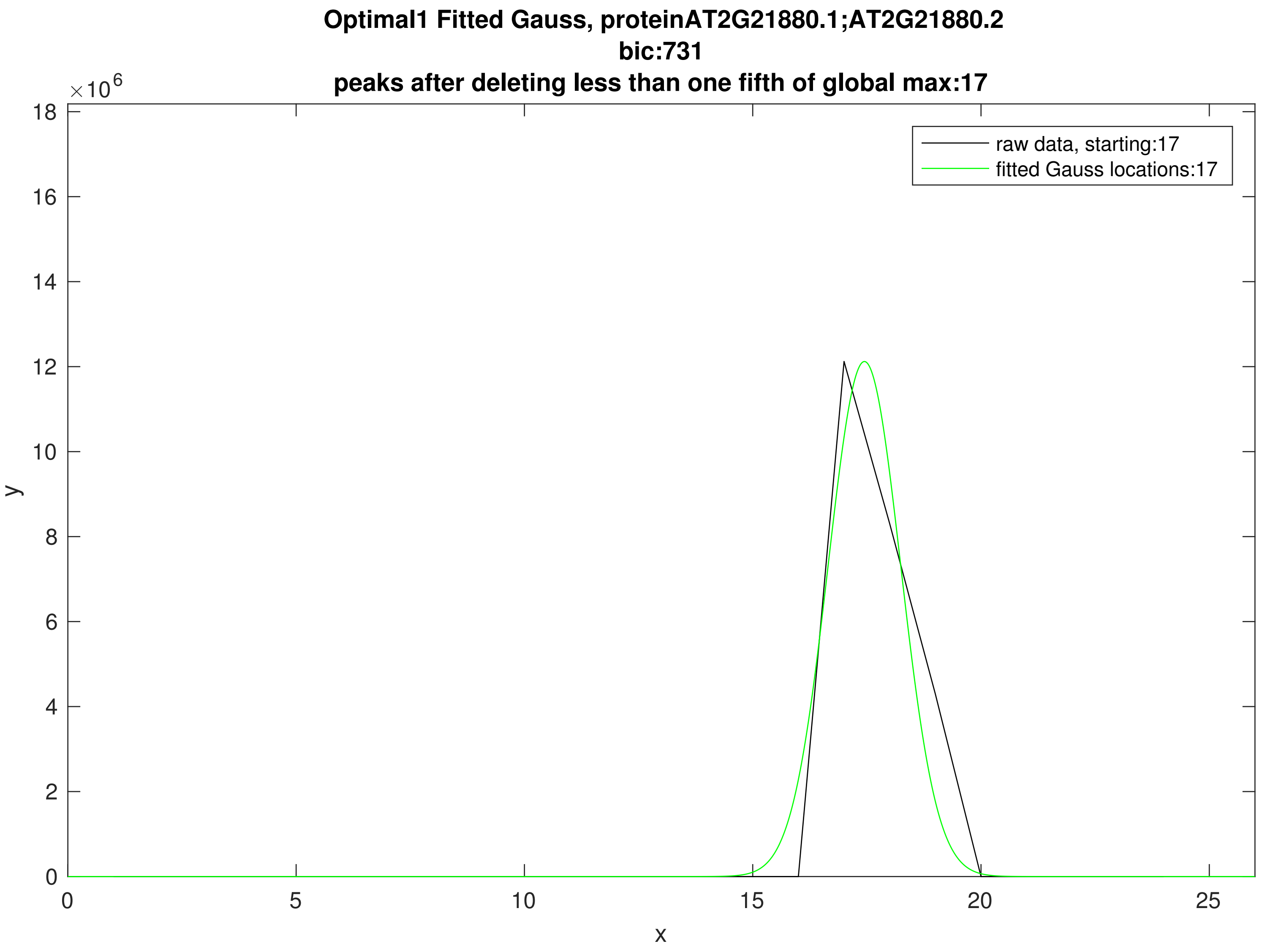

Optimal1 Fitted Gauss, proteinAT2G21880.1;AT2G21880.2
bic:731
peaks after deleting less than one fifth of global max:17
×10^6
raw data, starting:17
fitted Gauss locations:17
y
x
0
2
4
6
8
10
12
14
16
18
0
5
10
15
20
25

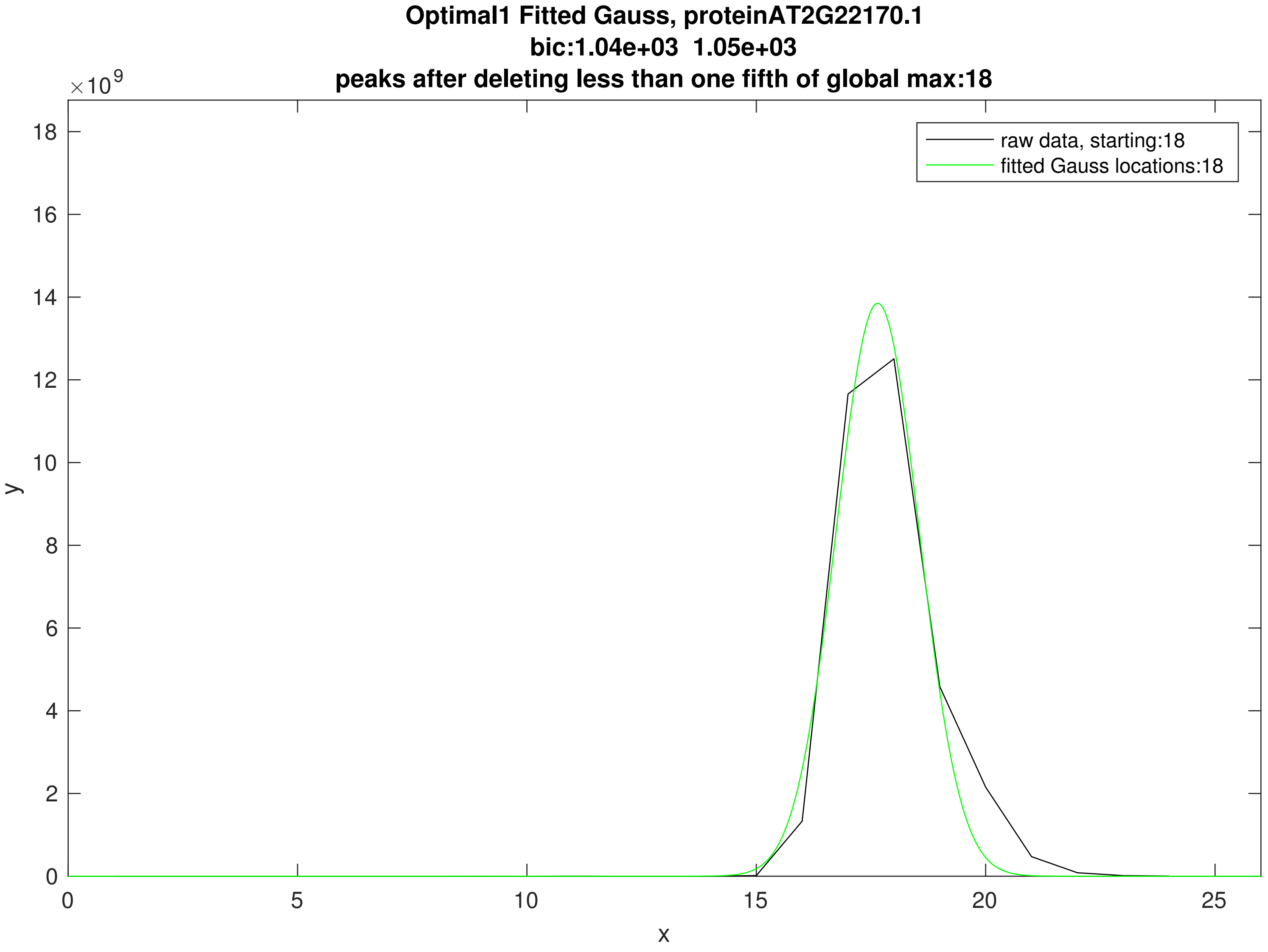

Optimal1 Fitted Gauss, proteinAT2G22170.1
bic:1.04e+03 1.05e+03
peaks after deleting less than one fifth of global max:18
×10^9
raw data, starting:18
fitted Gauss locations:18
y
x
0
2
4
6
8
10
12
14
16
18
0
5
10
15
20
25

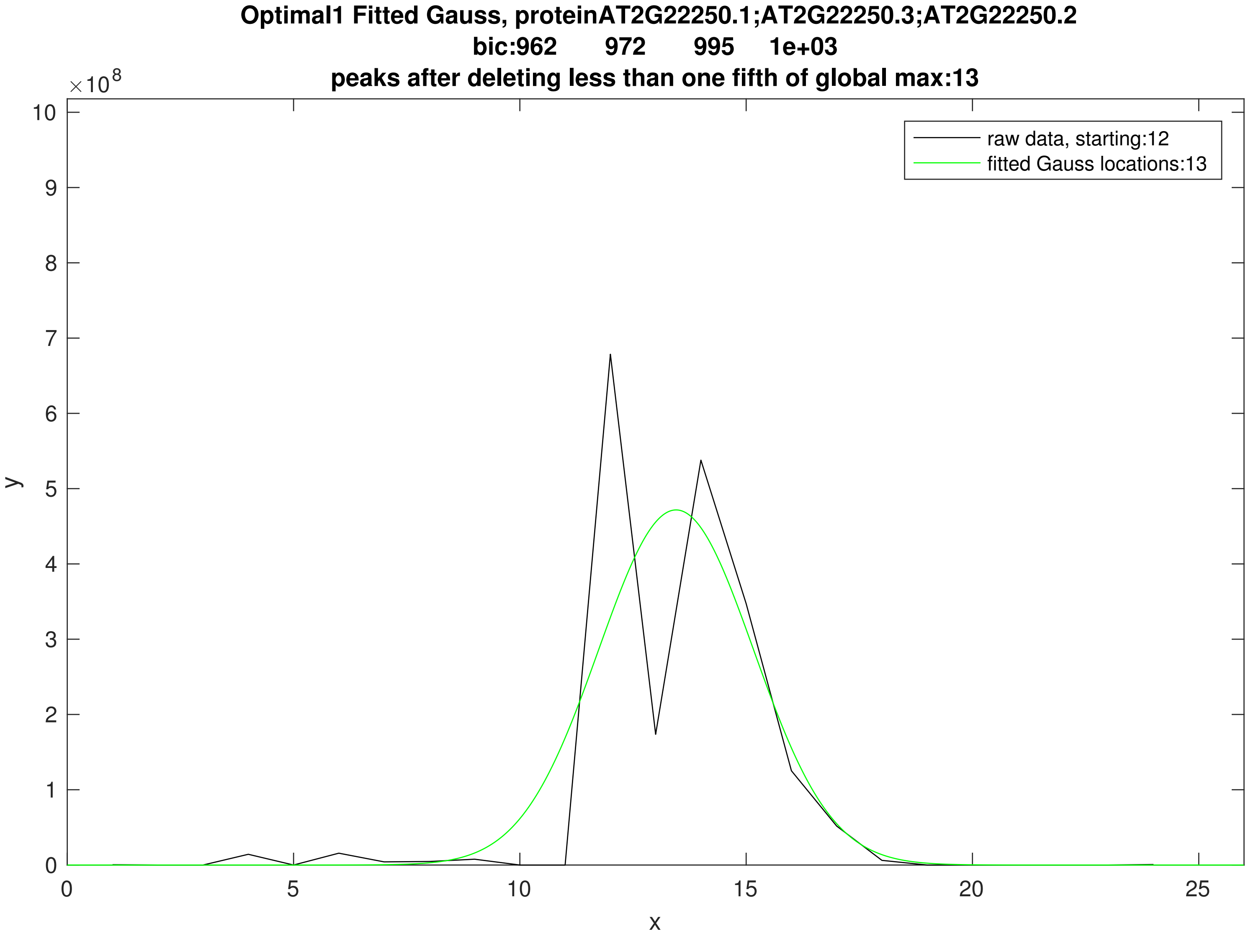
Optimal1 Fitted Gauss, proteinAT2G22250.1;AT2G22250.3;AT2G22250.2
bic:962 972 995 1e+03
peaks after deleting less than one fifth of global max:13
×10^8
10
9
8
7
6
5
4
3
2
1
0
y
0
5
10
15
20
25
x
raw data, starting:12
fitted Gauss locations:13

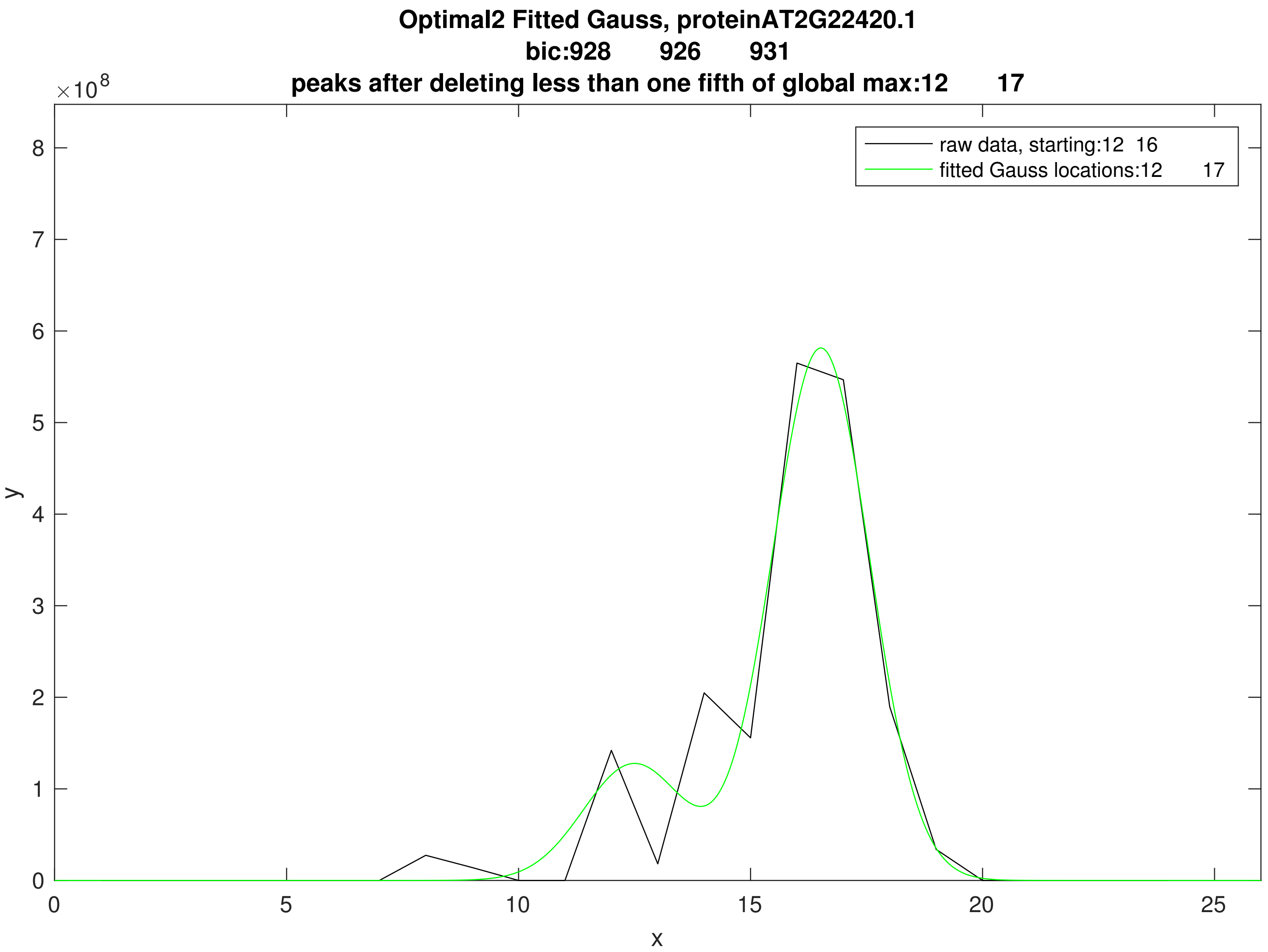
Optimal2 Fitted Gauss, proteinAT2G22420.1
bic:928 926 931
peaks after deleting less than one fifth of global max:12 17
×10^8
raw data, starting:12 16
fitted Gauss locations:12 17
y
x
0
1
2
3
4
5
6
7
8
0
5
10
15
20
25

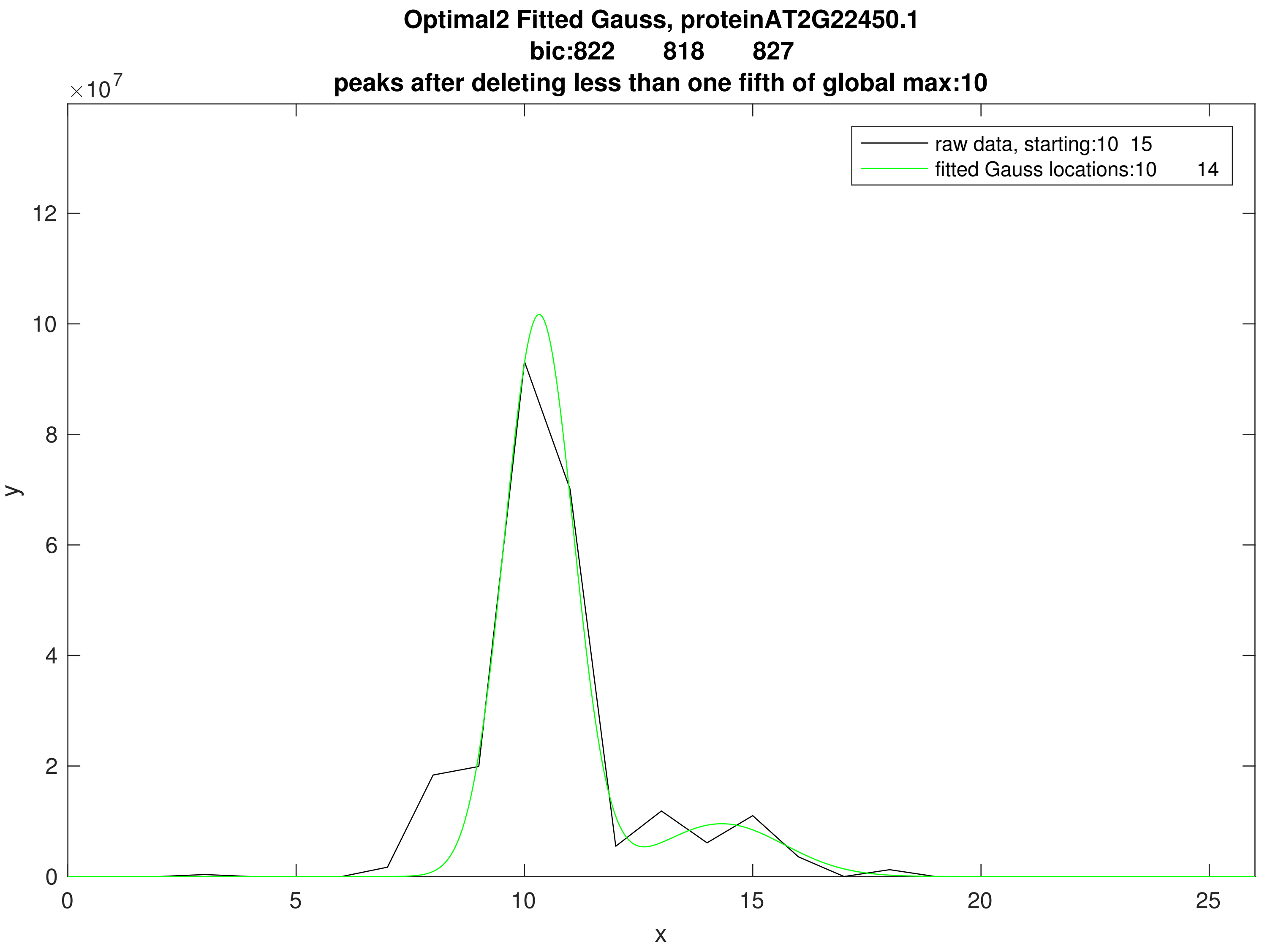

Optimal2 Fitted Gauss, proteinAT2G22450.1
bic:822 818 827
peaks after deleting less than one fifth of global max:10
×10^7
raw data, starting:10 15
fitted Gauss locations:10 14
y
x
0
2
4
6
8
10
12
0
5
10
15
20
25

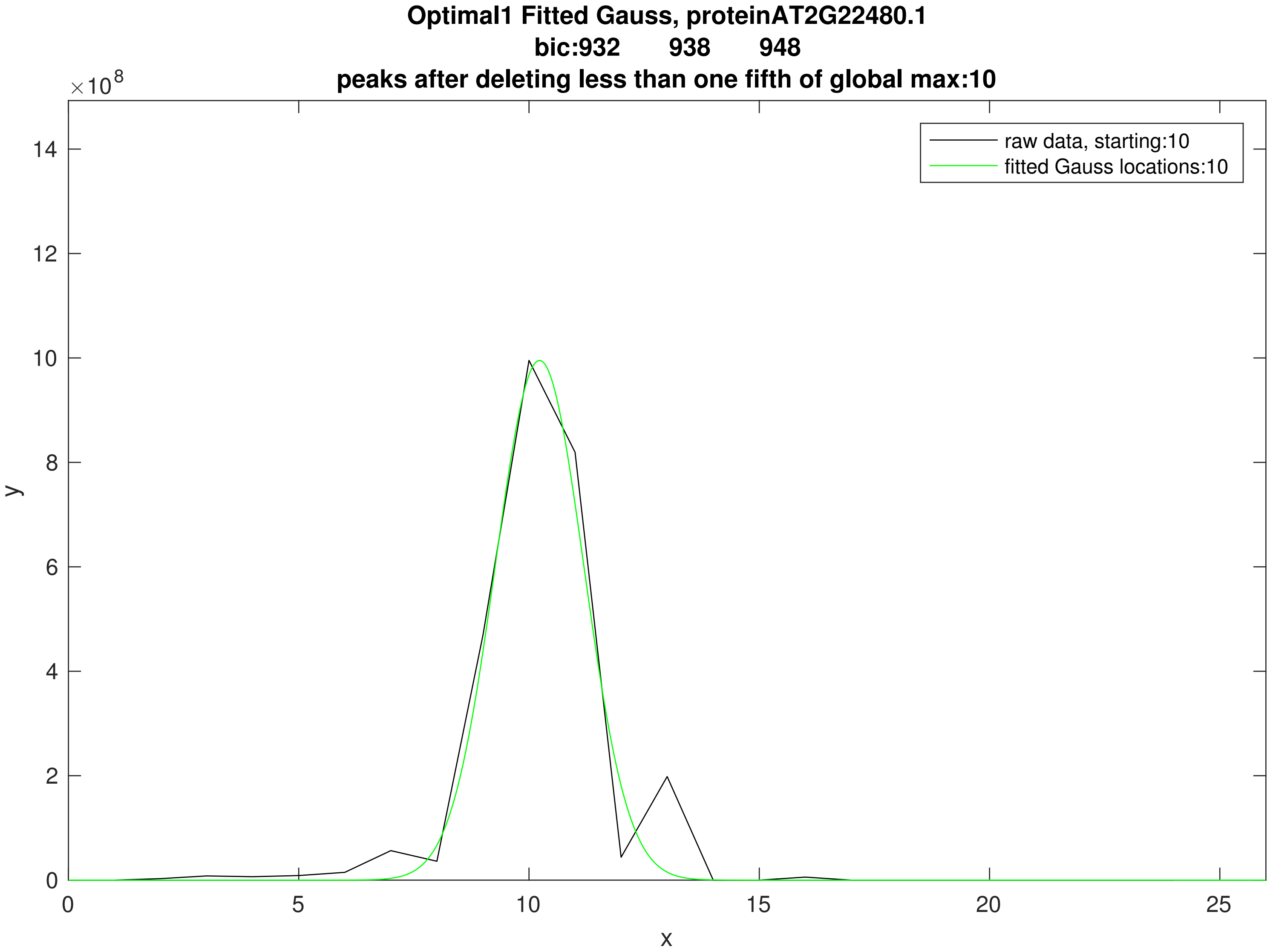

Optimal1 Fitted Gauss, proteinAT2G22480.1
bic:932 938 948
peaks after deleting less than one fifth of global max:10
×10^8
raw data, starting:10
fitted Gauss locations:10
y
x
0
2
4
6
8
10
12
14
0
5
10
15
20
25

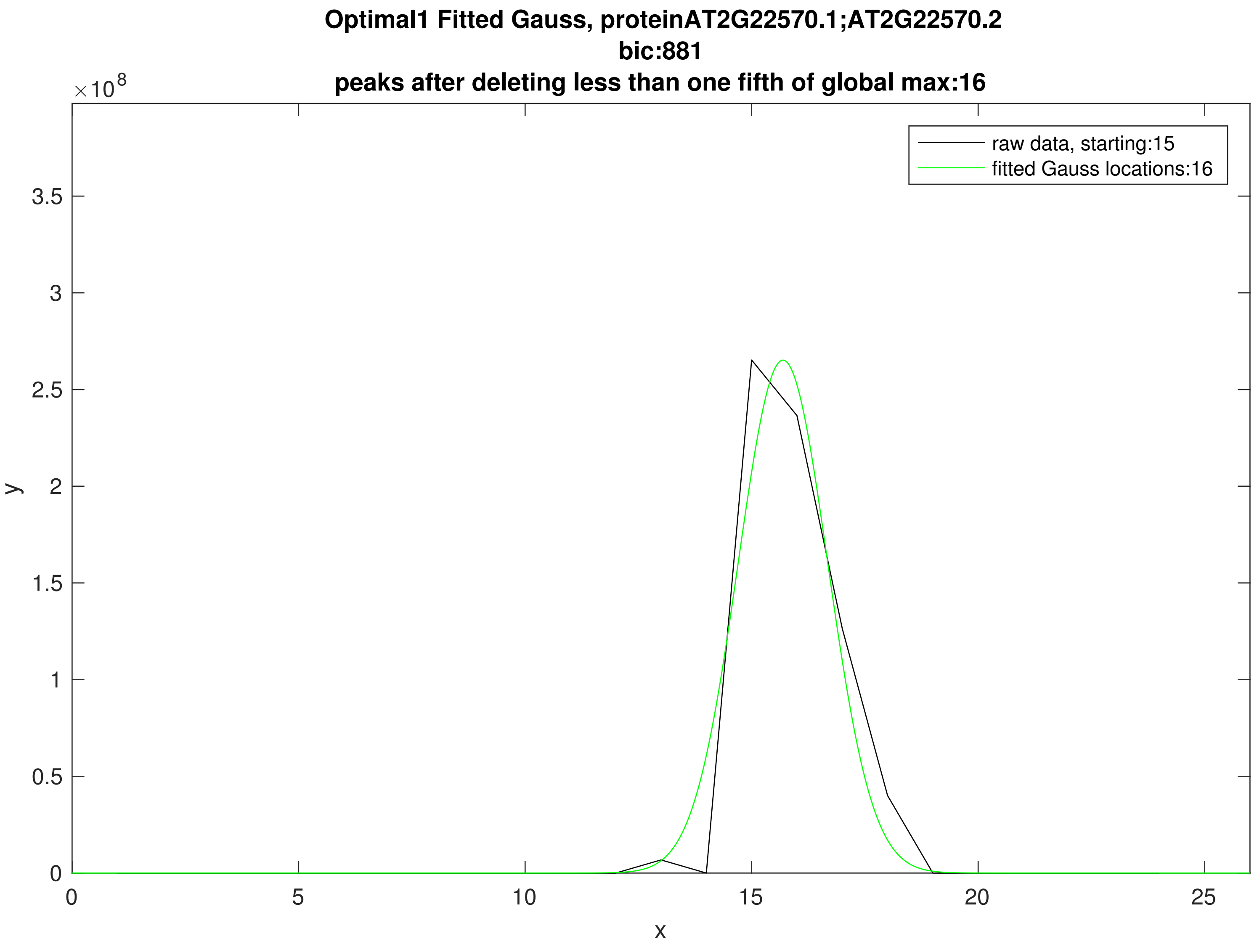
Optimal1 Fitted Gauss, proteinAT2G22570.1;AT2G22570.2
bic:881
peaks after deleting less than one fifth of global max:16
×10^8
3.5
3
2.5
2
1.5
1
0.5
0
y
0
5
10
15
20
25
x
raw data, starting:15
fitted Gauss locations:16

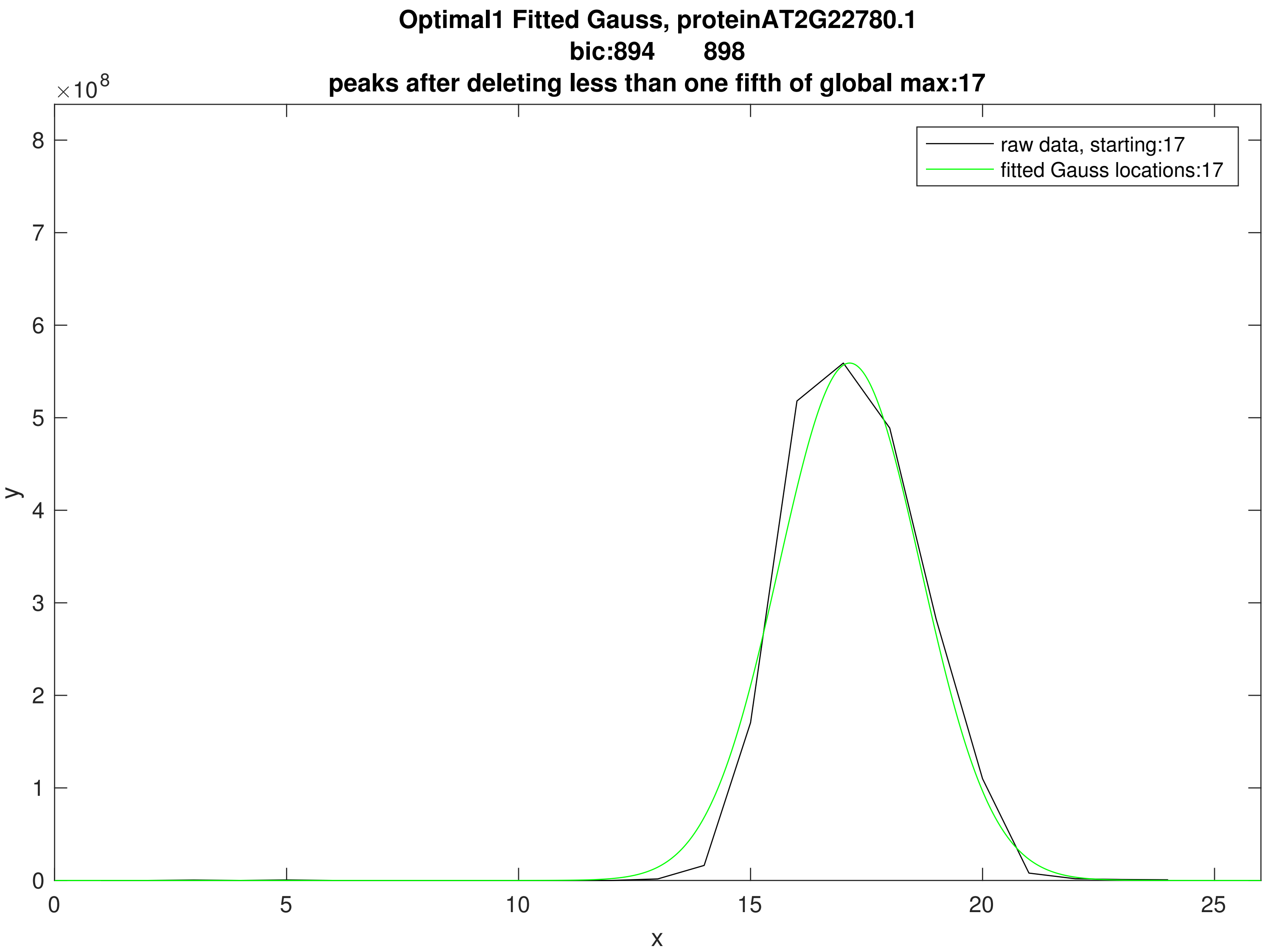

Optimal1 Fitted Gauss, proteinAT2G22780.1
bic:894 898
peaks after deleting less than one fifth of global max:17
×10^8
raw data, starting:17
fitted Gauss locations:17
y
x
0
1
2
3
4
5
6
7
8
0
5
10
15
20
25

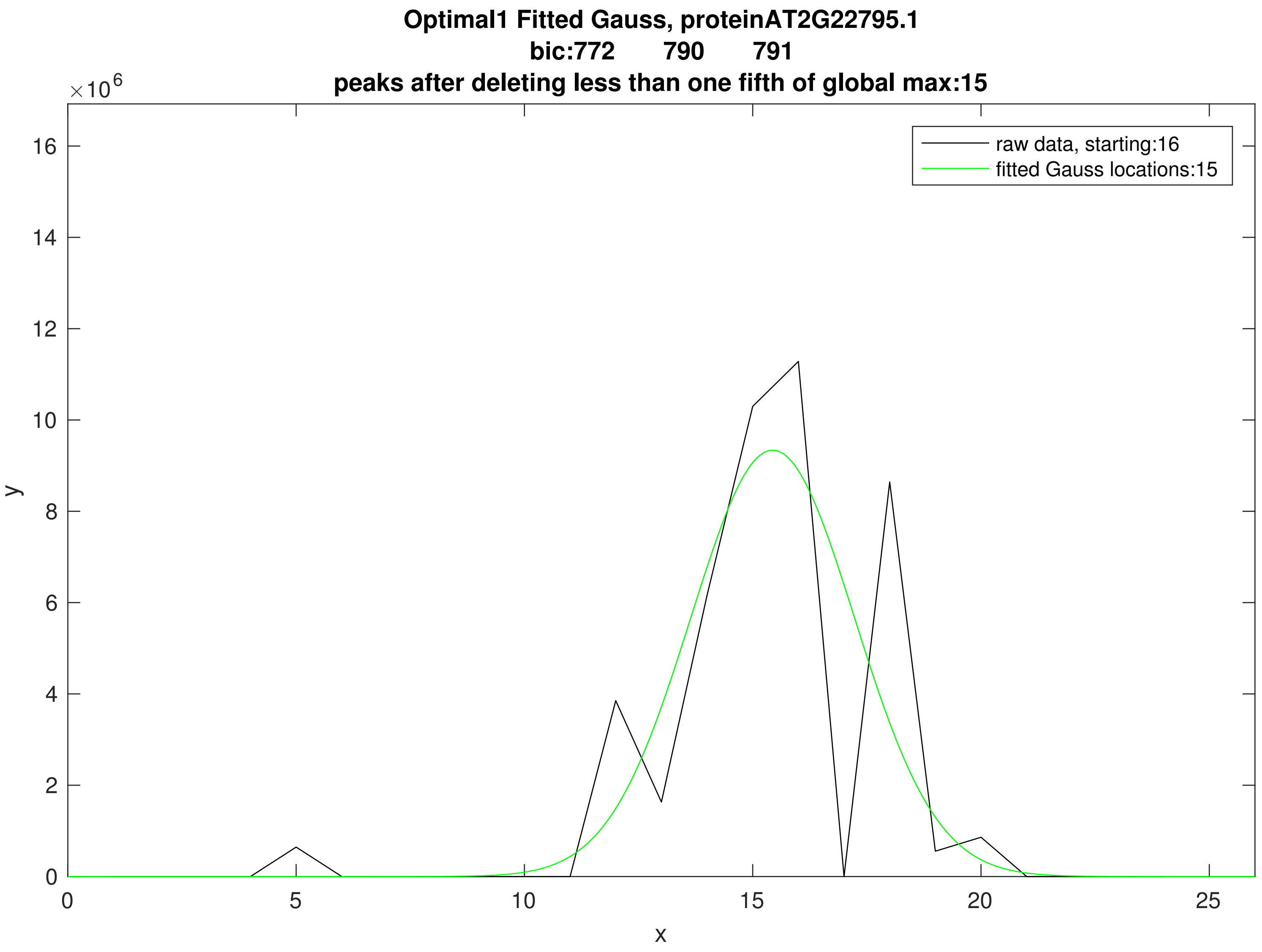

Optimal1 Fitted Gauss, proteinAT2G22795.1
bic:772 790 791
peaks after deleting less than one fifth of global max:15
×10^6
raw data, starting:16
fitted Gauss locations:15
y
x
0
2
4
6
8
10
12
14
16
0
5
10
15
20
25

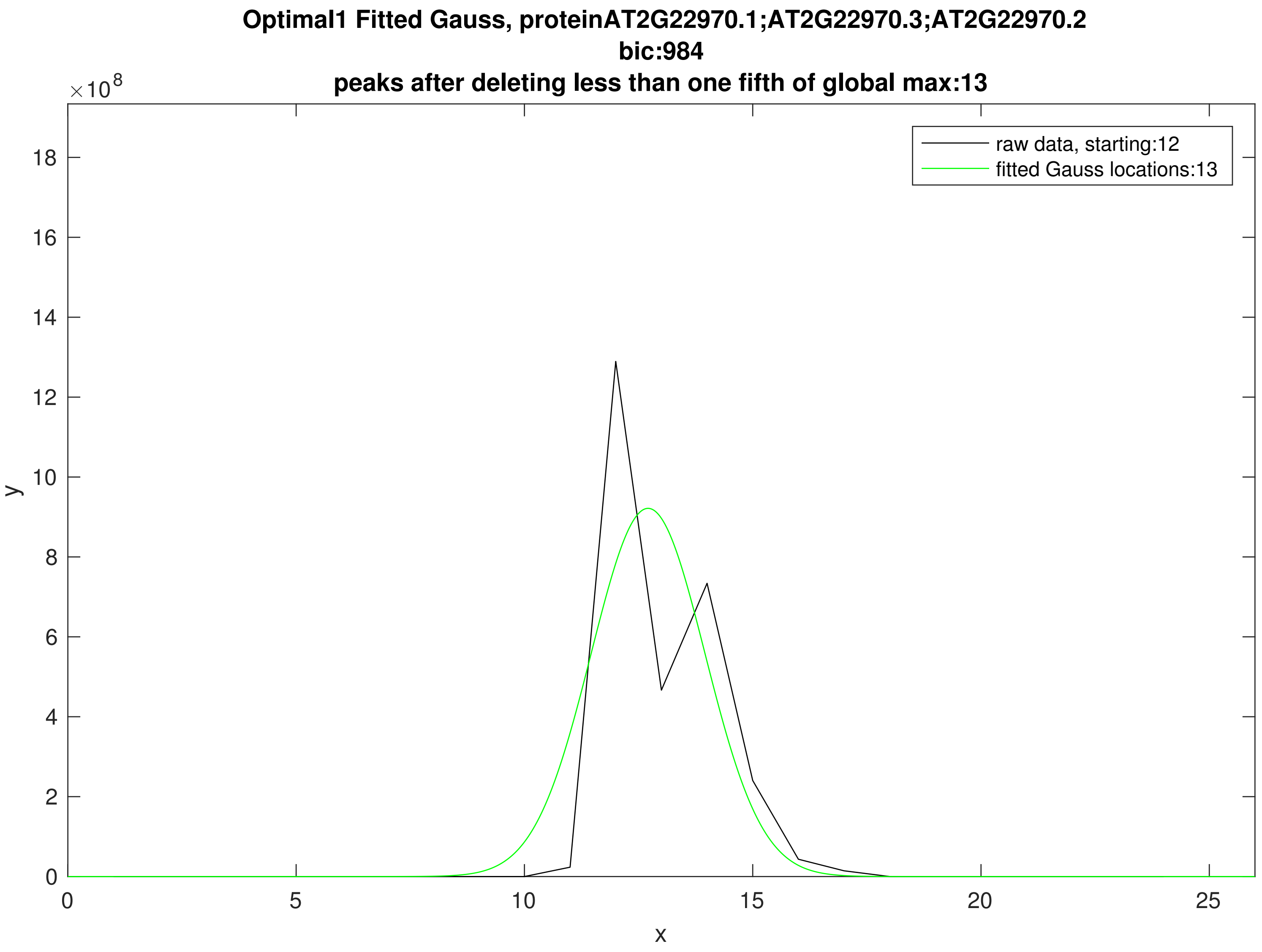
Optimal1 Fitted Gauss, proteinAT2G22970.1;AT2G22970.3;AT2G22970.2
bic:984
peaks after deleting less than one fifth of global max:13
×10^8
raw data, starting:12
fitted Gauss locations:13
y
x
0
2
4
6
8
10
12
14
16
18
0
5
10
15
20
25

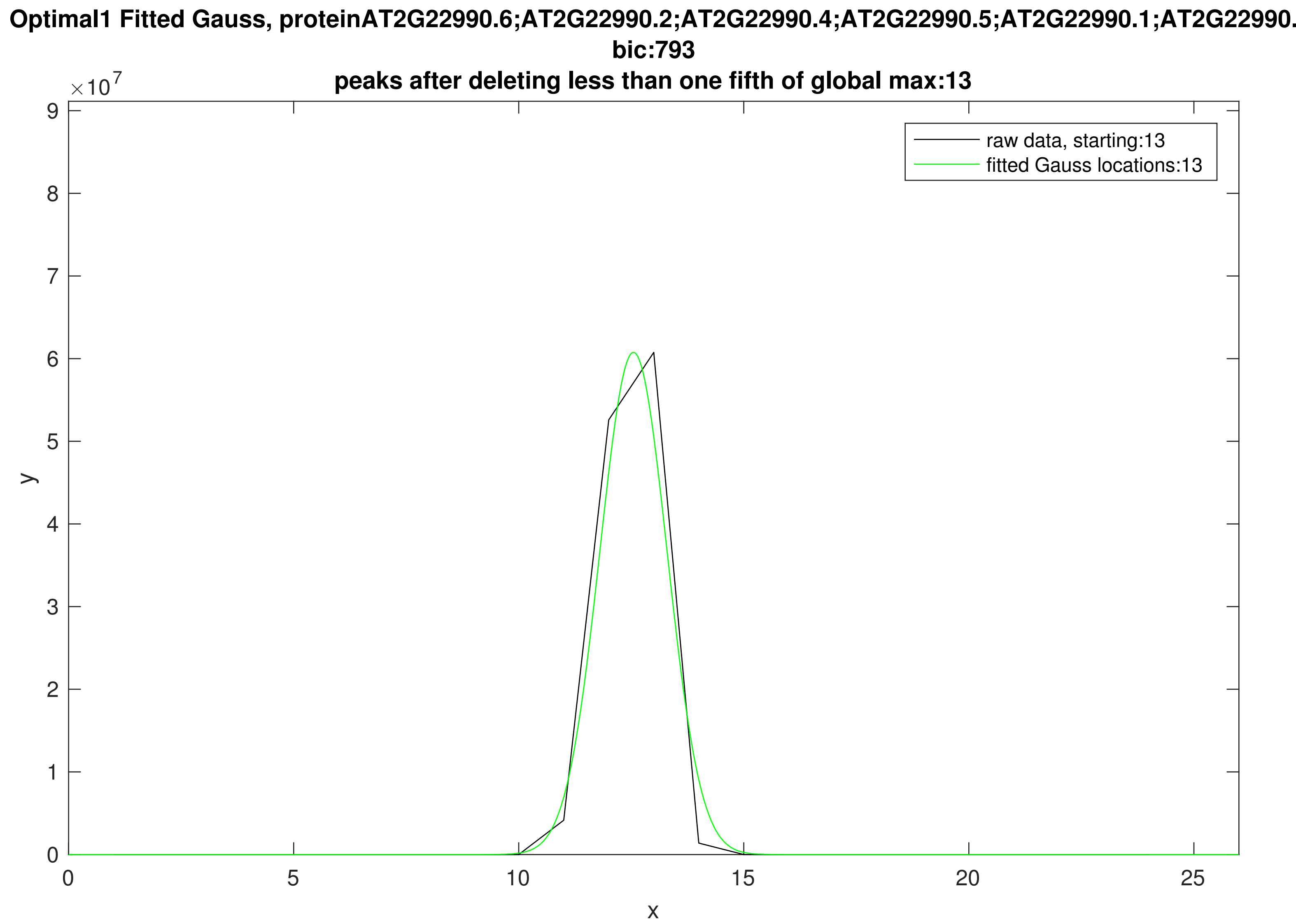
Optimal1 Fitted Gauss, proteinAT2G22990.6;AT2G22990.2;AT2G22990.4;AT2G22990.5;AT2G22990.1;AT2G22990.
bic:793
peaks after deleting less than one fifth of global max:13
×10^7
raw data, starting:13
fitted Gauss locations:13
y
x
0
1
2
3
4
5
6
7
8
9
0
5
10
15
20
25

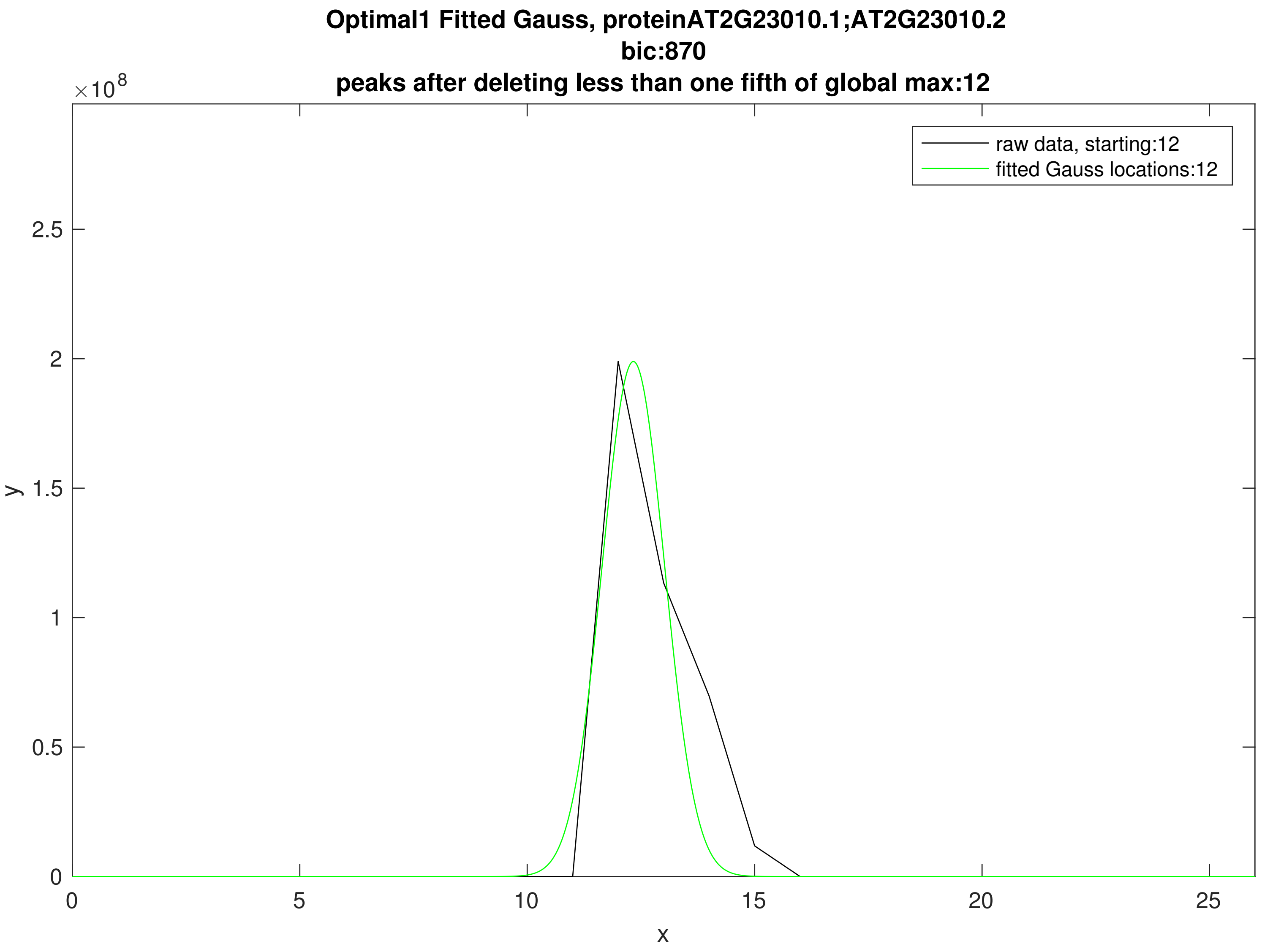
Optimal1 Fitted Gauss, proteinAT2G23010.1;AT2G23010.2
bic:870
peaks after deleting less than one fifth of global max:12
×10^8
raw data, starting:12
fitted Gauss locations:12
2.5
2
y 1.5
1
0.5
0
0
5
10
15
20
25
x

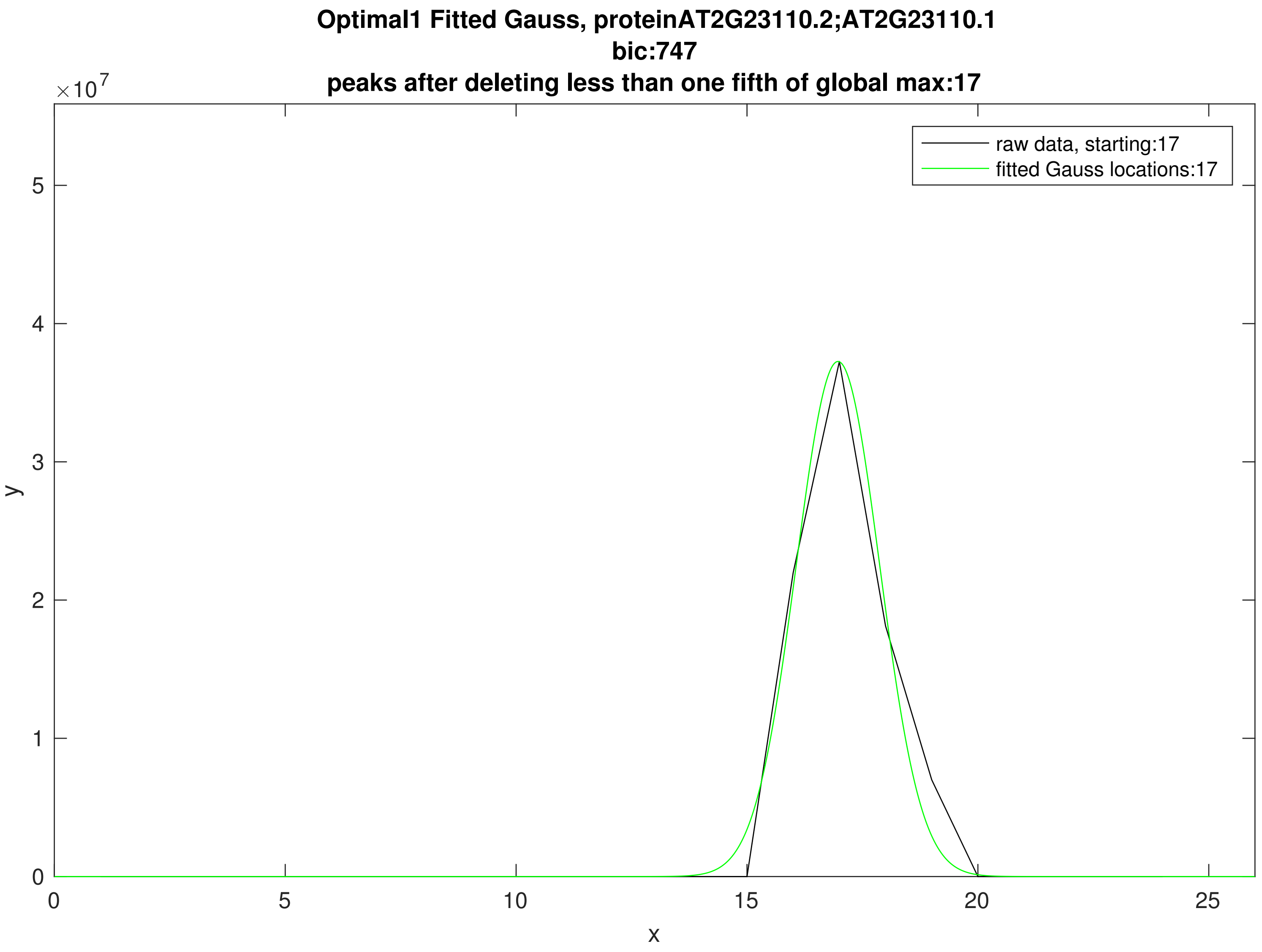
Optimal1 Fitted Gauss, proteinAT2G23110.2;AT2G23110.1
bic:747
peaks after deleting less than one fifth of global max:17
×10^7
raw data, starting:17
fitted Gauss locations:17
y
x
0
1
2
3
4
5
0
5
10
15
20
25

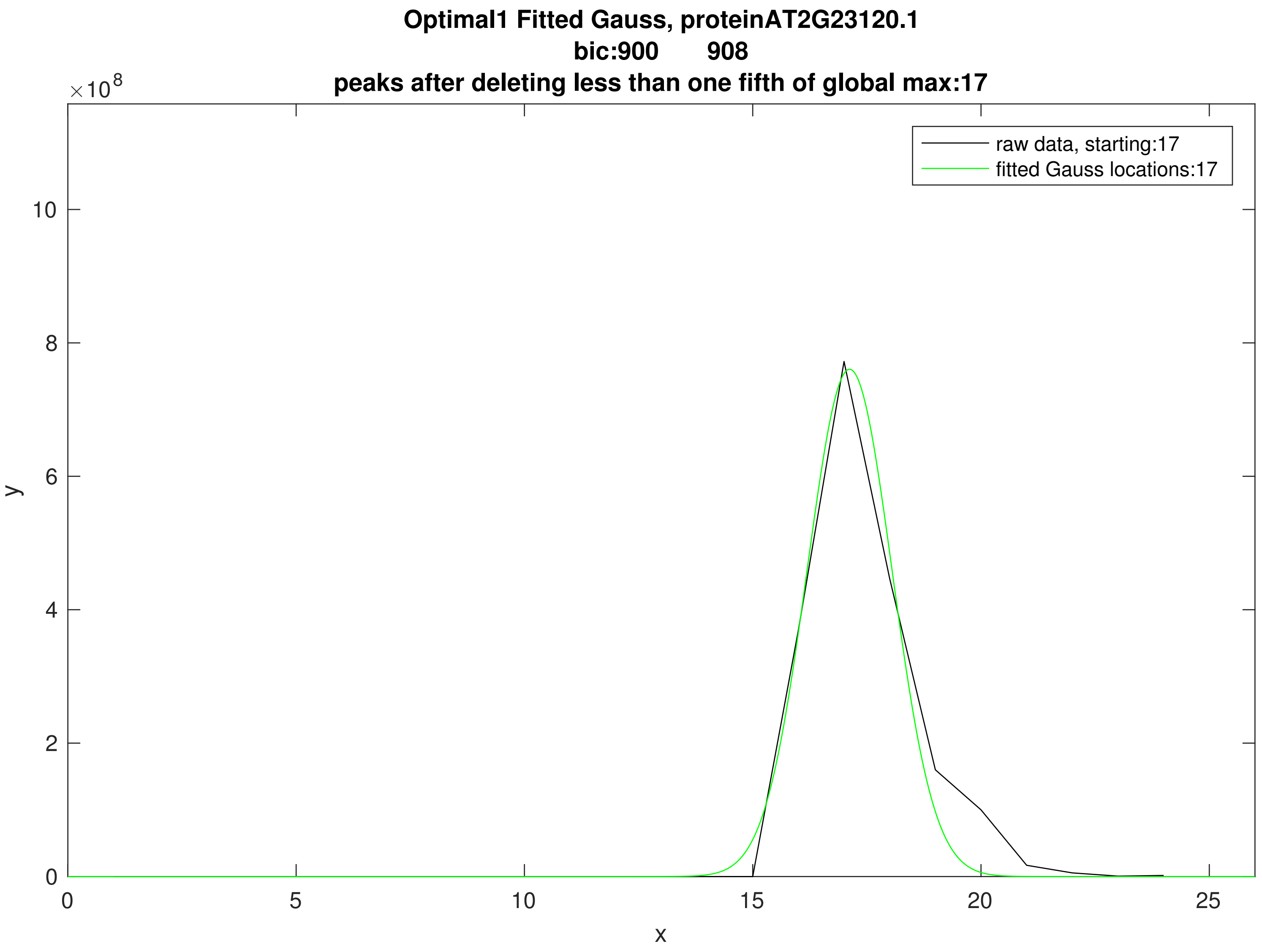

Optimal1 Fitted Gauss, proteinAT2G23120.1
bic:900 908
peaks after deleting less than one fifth of global max:17
×10^8
raw data, starting:17
fitted Gauss locations:17
y
x
0
2
4
6
8
10
0
5
10
15
20
25

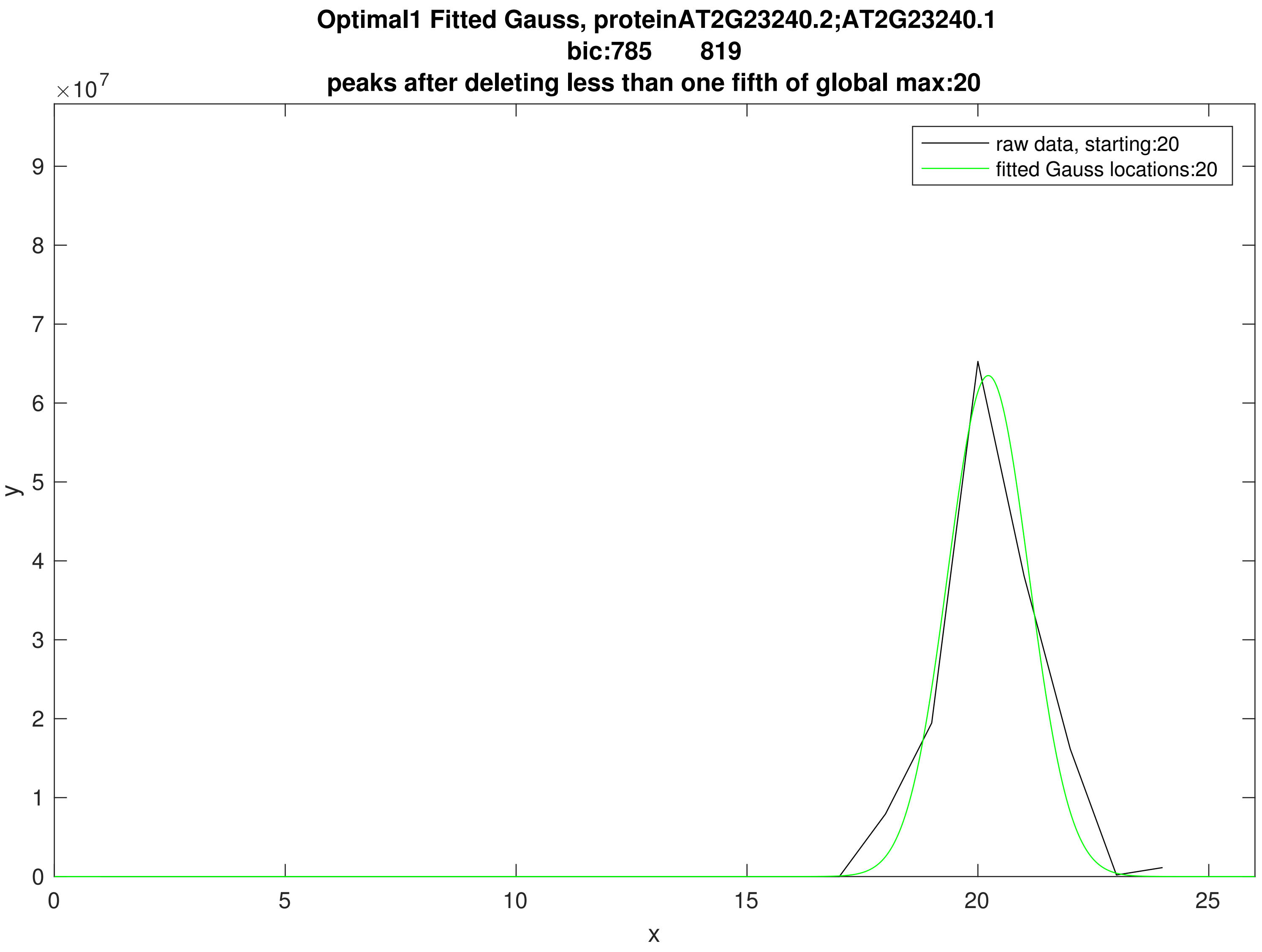
Optimal1 Fitted Gauss, proteinAT2G23240.2;AT2G23240.1
bic:785 819
peaks after deleting less than one fifth of global max:20
×10^7
raw data, starting:20
fitted Gauss locations:20
y
x
0
1
2
3
4
5
6
7
8
9
0
5
10
15
20
25

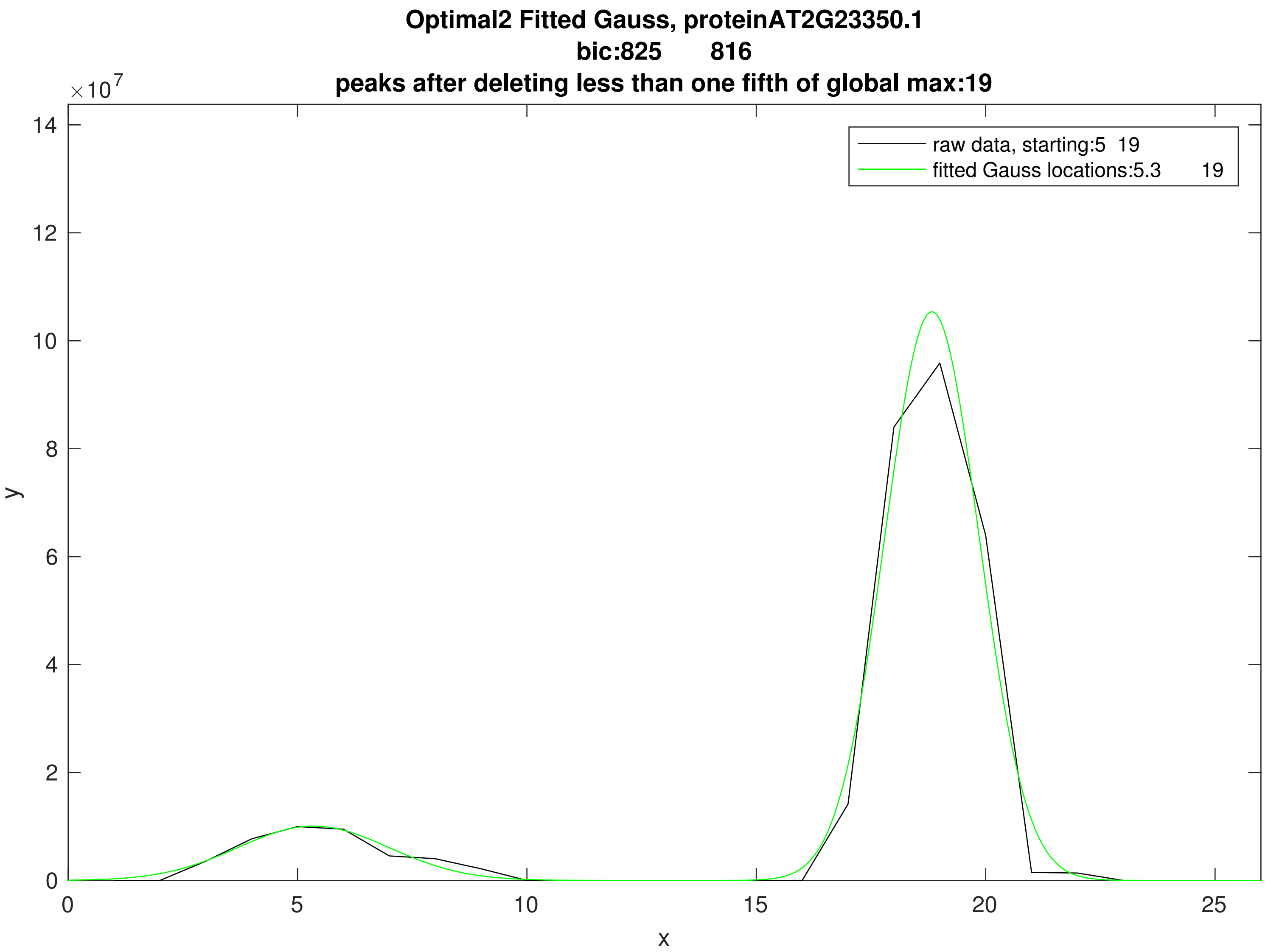
Optimal2 Fitted Gauss, proteinAT2G23350.1
bic:825 816
peaks after deleting less than one fifth of global max:19
×10^7
raw data, starting:5 19
fitted Gauss locations:5.3 19
y
x
0
2
4
6
8
10
12
14
5
15
20
25

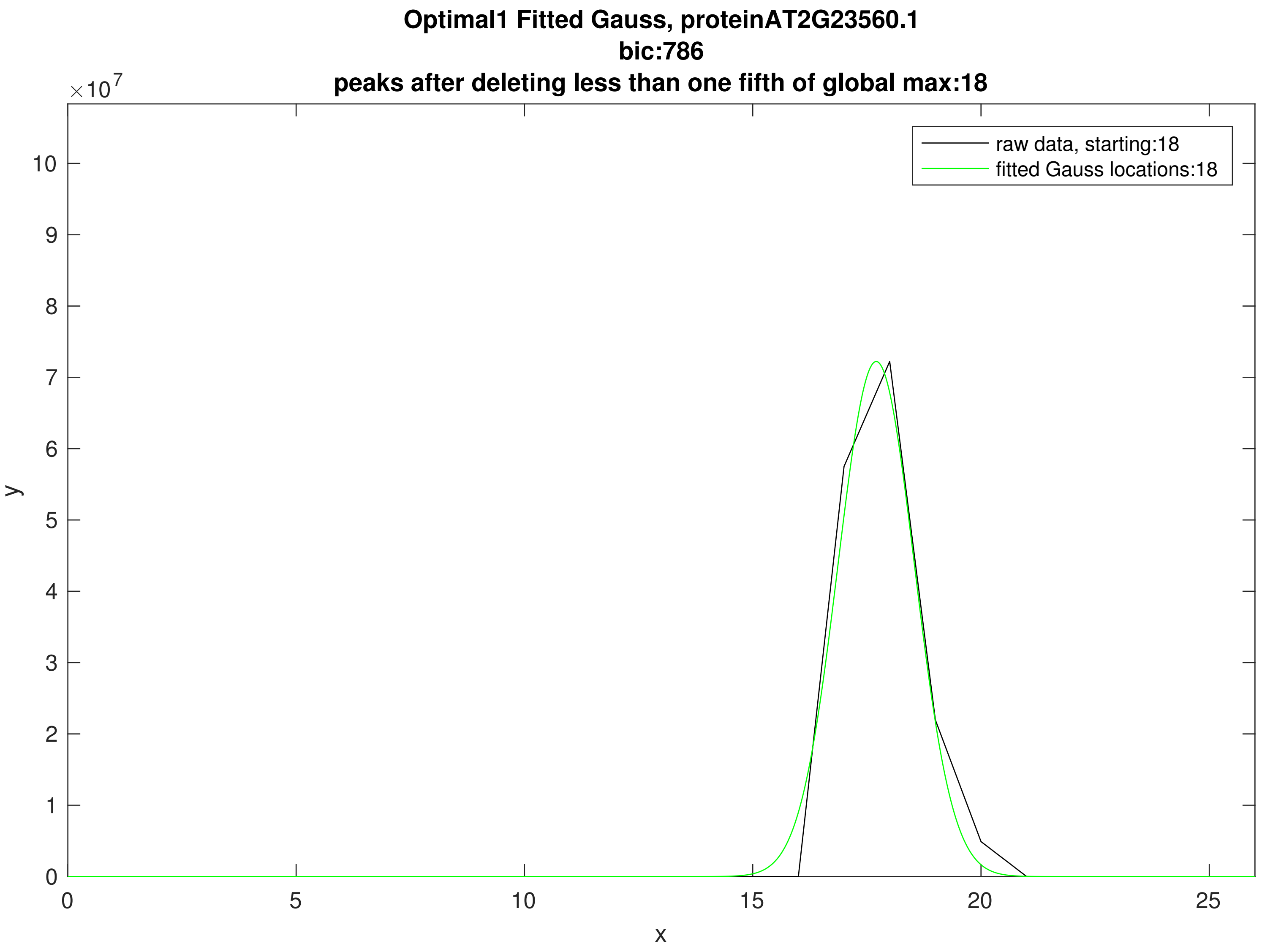

Optimal1 Fitted Gauss, proteinAT2G23560.1
bic:786
peaks after deleting less than one fifth of global max:18
×10^7
raw data, starting:18
fitted Gauss locations:18
y
x
0
1
2
3
4
5
6
7
8
9
10
0
5
10
15
20
25

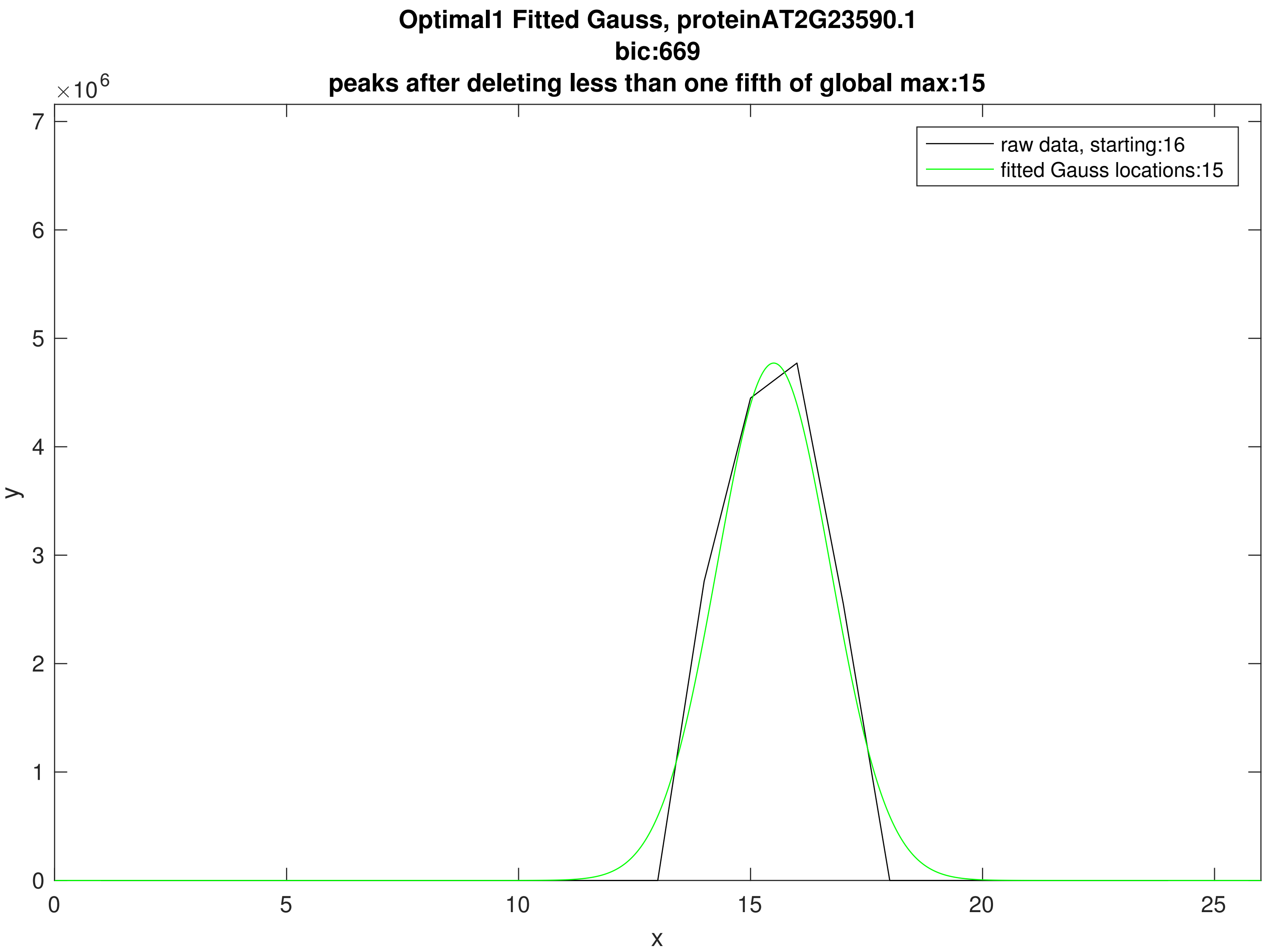
Optimal1 Fitted Gauss, proteinAT2G23590.1
bic:669
peaks after deleting less than one fifth of global max:15
×10^6
raw data, starting:16
fitted Gauss locations:15
y
x
0
1
2
3
4
5
6
7
0
5
10
15
20
25

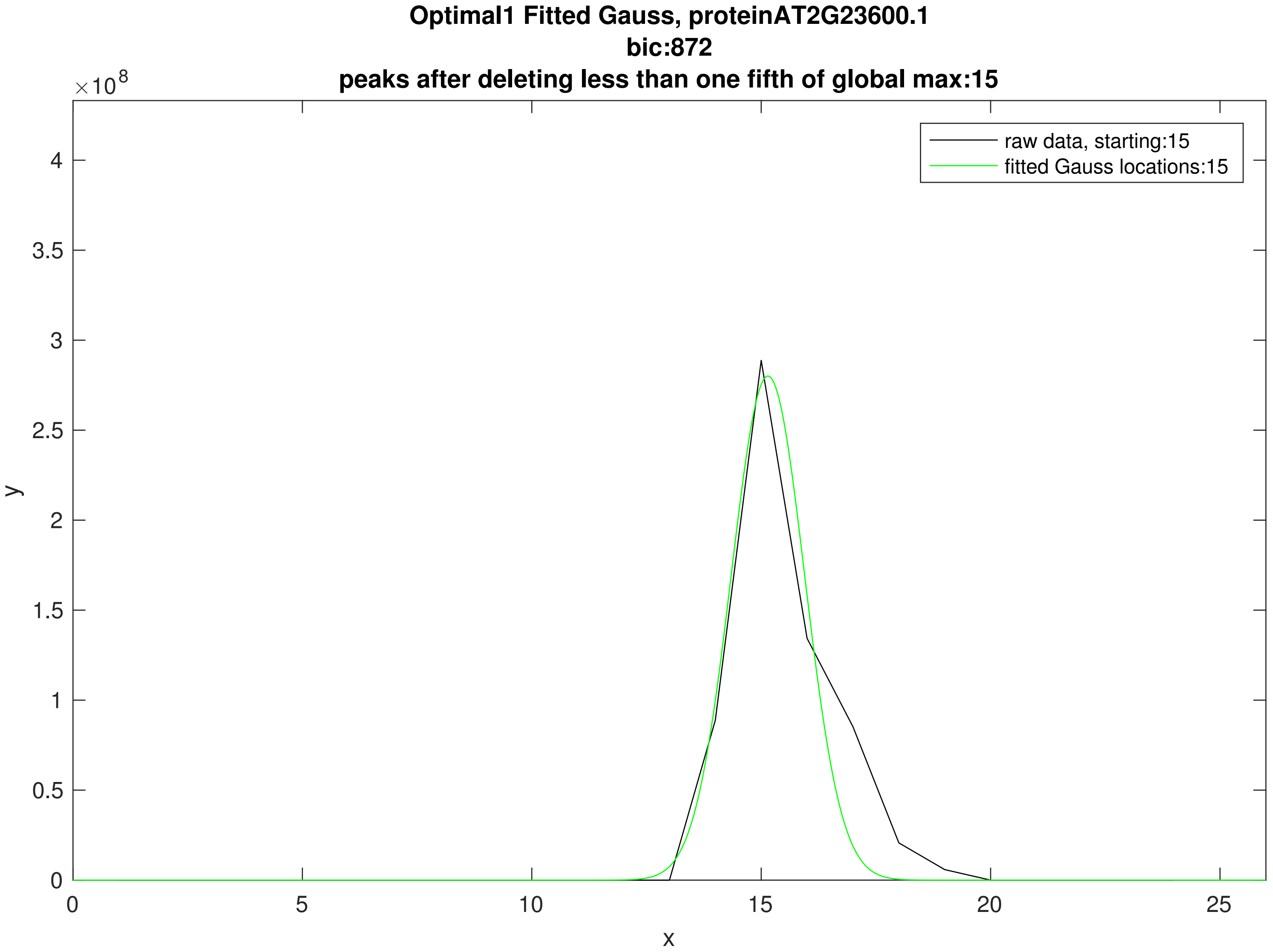
Optimal1 Fitted Gauss, proteinAT2G23600.1
bic:872
peaks after deleting less than one fifth of global max:15
×10^8
raw data, starting:15
fitted Gauss locations:15
4
3.5
3
2.5
2
1.5
1
0.5
0
y
0
5
10
15
20
25
x

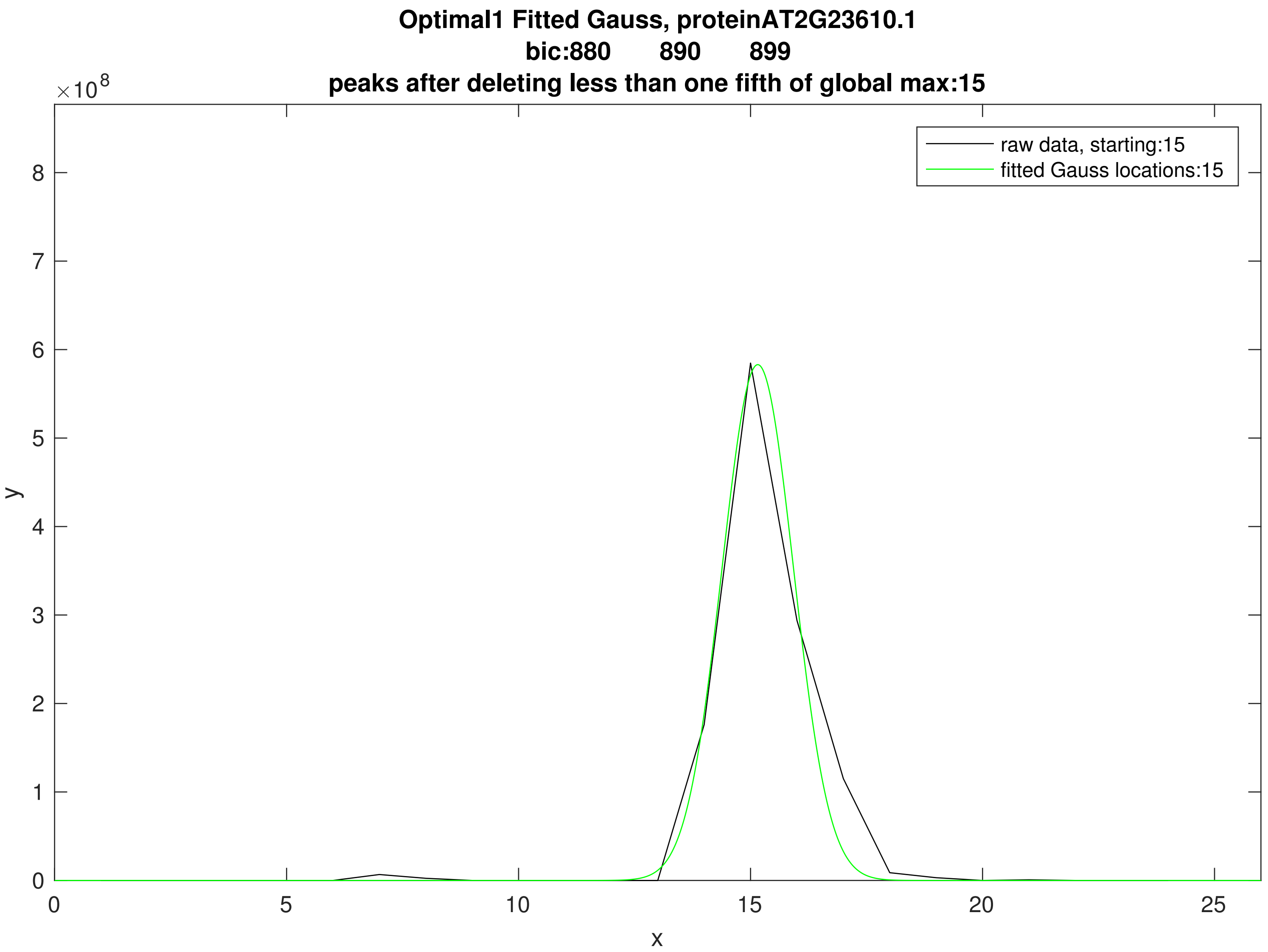

Optimal1 Fitted Gauss, proteinAT2G23610.1
bic:880 890 899
peaks after deleting less than one fifth of global max:15
×10^8
raw data, starting:15
fitted Gauss locations:15
y
x
0
1
2
3
4
5
6
7
8
0
5
10
15
20
25

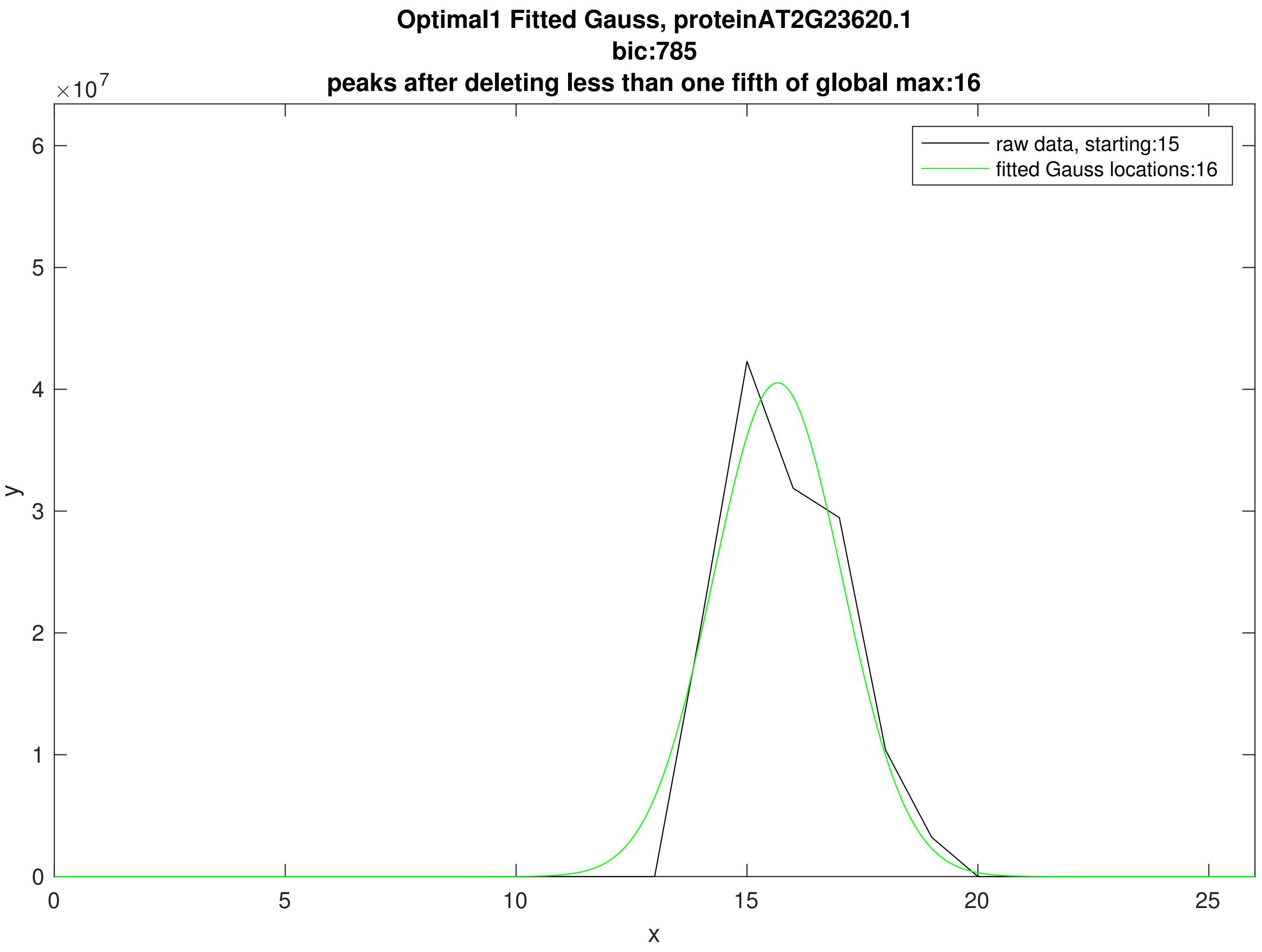

Optimal1 Fitted Gauss, proteinAT2G23620.1
bic:785
peaks after deleting less than one fifth of global max:16
×10^7
raw data, starting:15
fitted Gauss locations:16
y
x
0
1
2
3
4
5
6
0
5
10
15
20
25

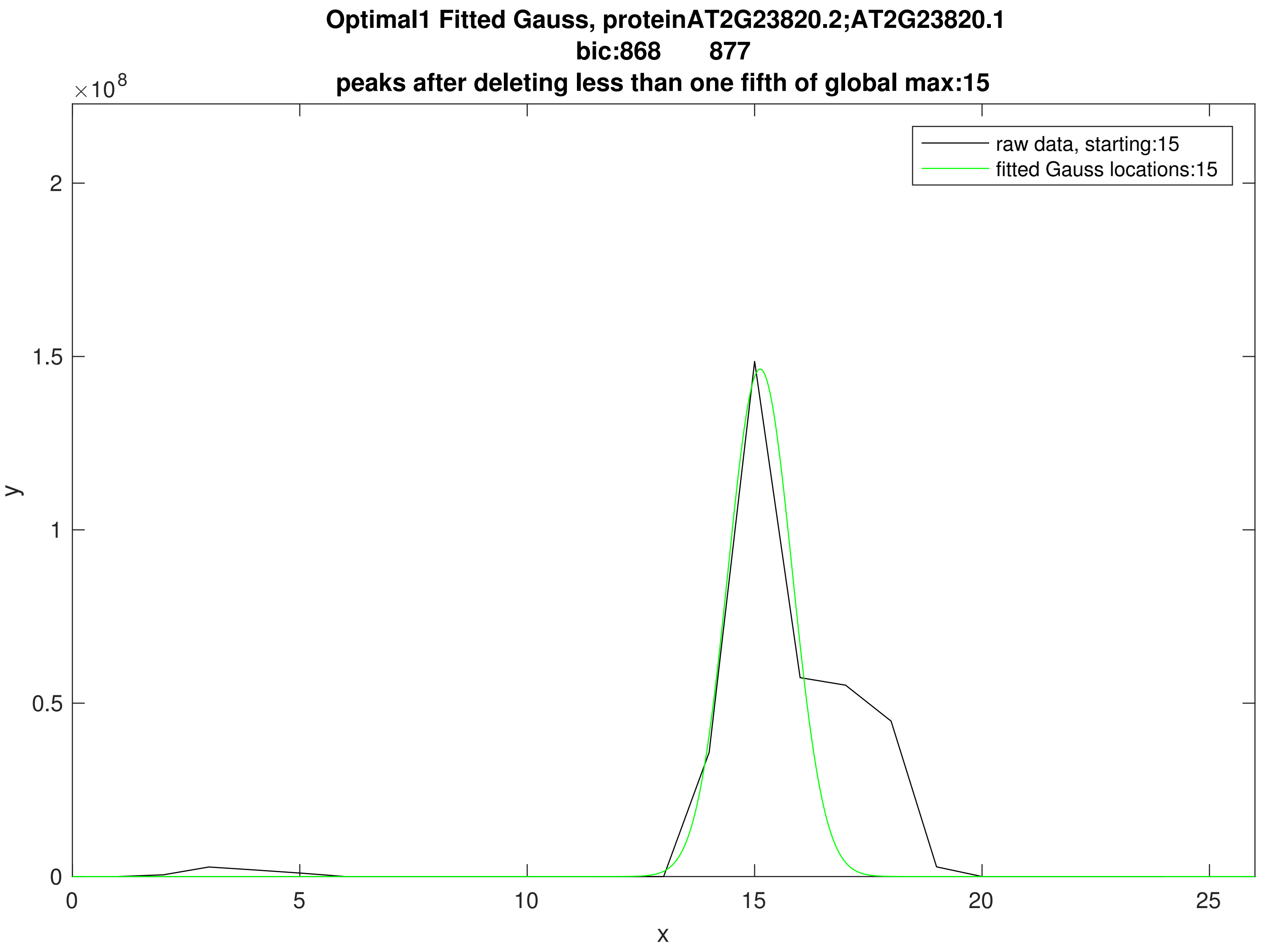

Optimal1 Fitted Gauss, proteinAT2G23820.2;AT2G23820.1
bic:868 877
peaks after deleting less than one fifth of global max:15
×10^8
raw data, starting:15
fitted Gauss locations:15
y
x
0
0.5
1
1.5
2
5
10
15
20
25

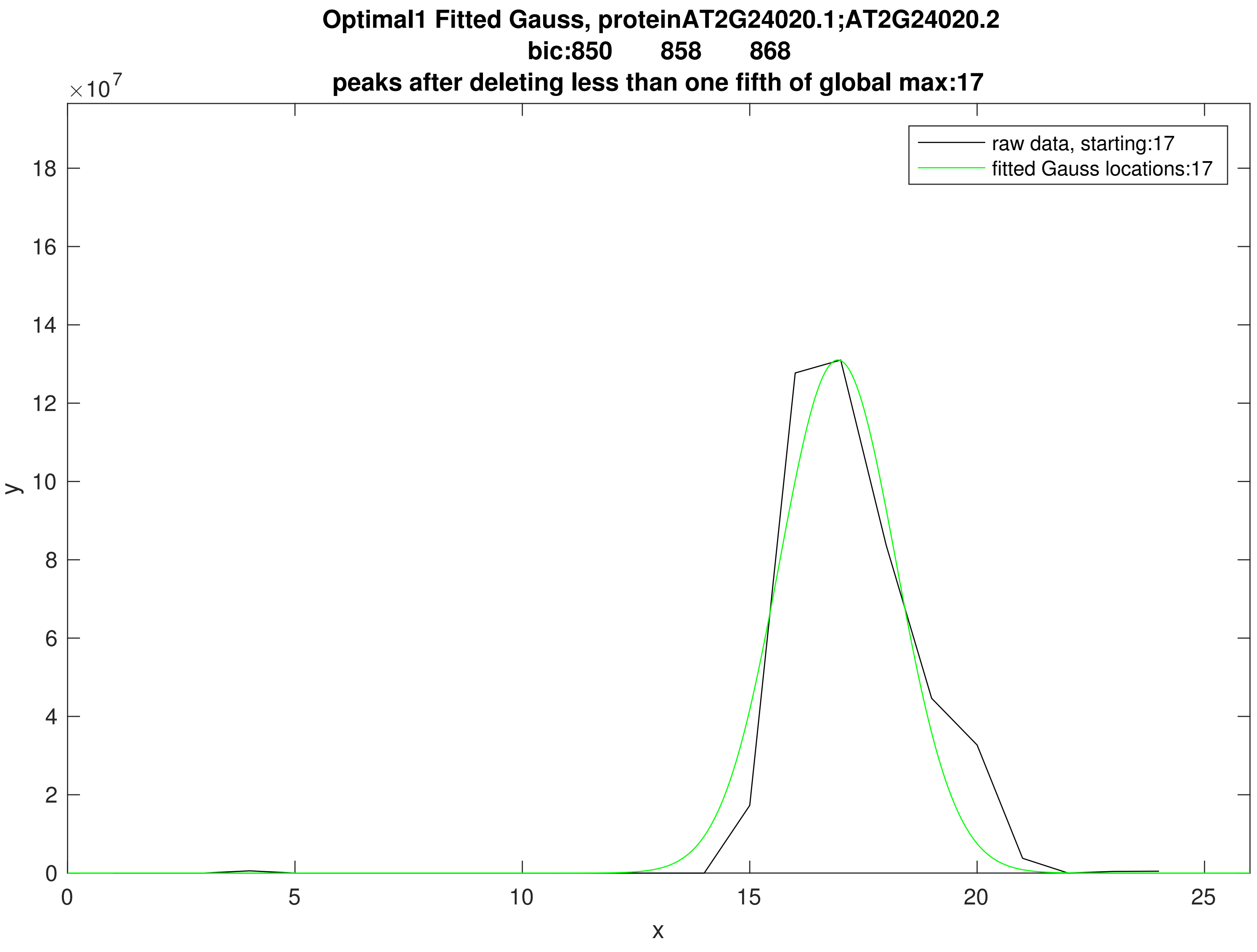

Optimal1 Fitted Gauss, proteinAT2G24020.1;AT2G24020.2
bic:850 858 868
peaks after deleting less than one fifth of global max:17
×10^7
raw data, starting:17
fitted Gauss locations:17
y
x
0
2
4
6
8
10
12
14
16
18
0
5
10
15
20
25

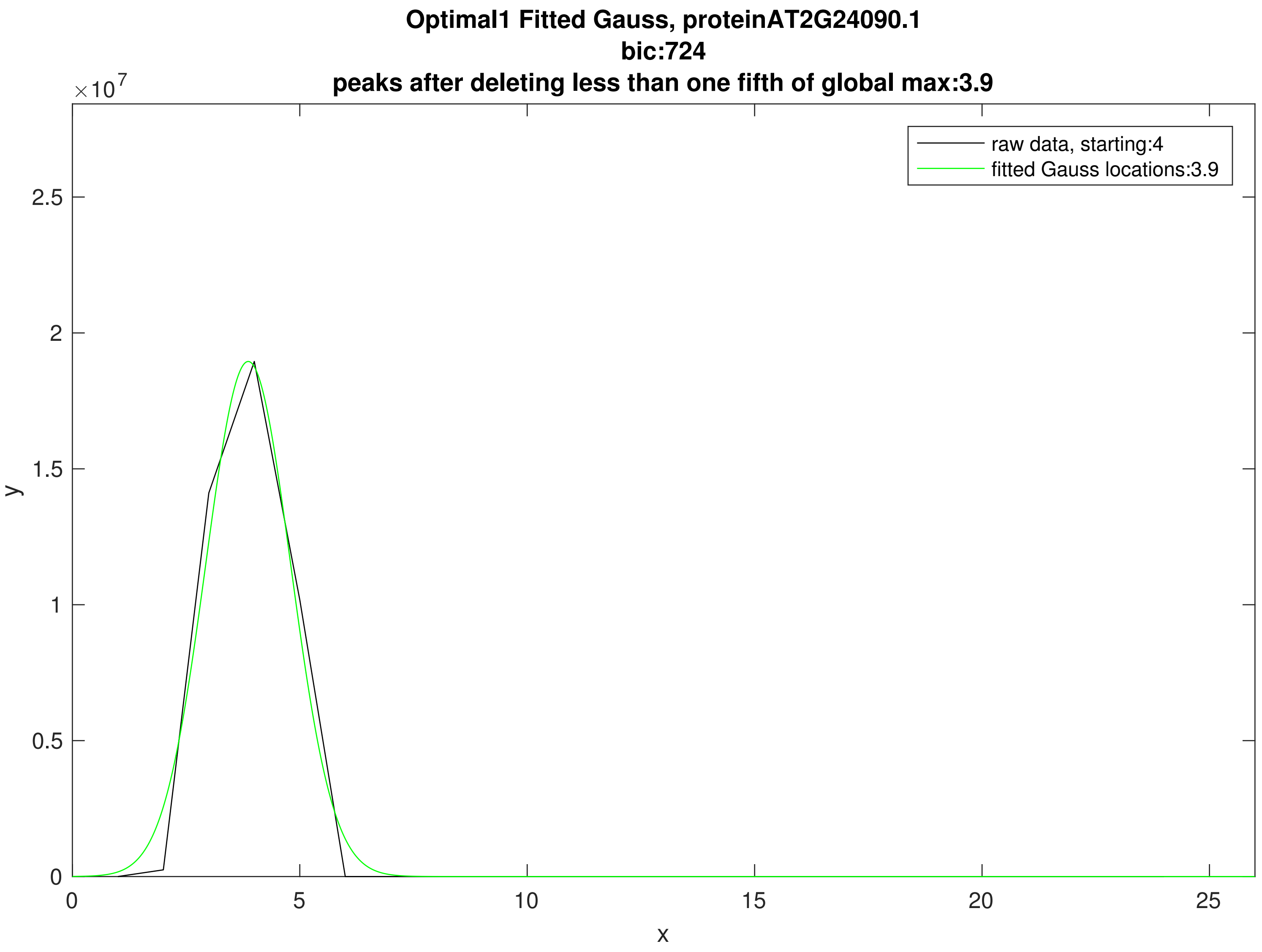
Optimal1 Fitted Gauss, proteinAT2G24090.1
bic:724
peaks after deleting less than one fifth of global max:3.9
×10^7
raw data, starting:4
fitted Gauss locations:3.9
y
x
0
0.5
1
1.5
2
2.5
0
5
10
15
20
25

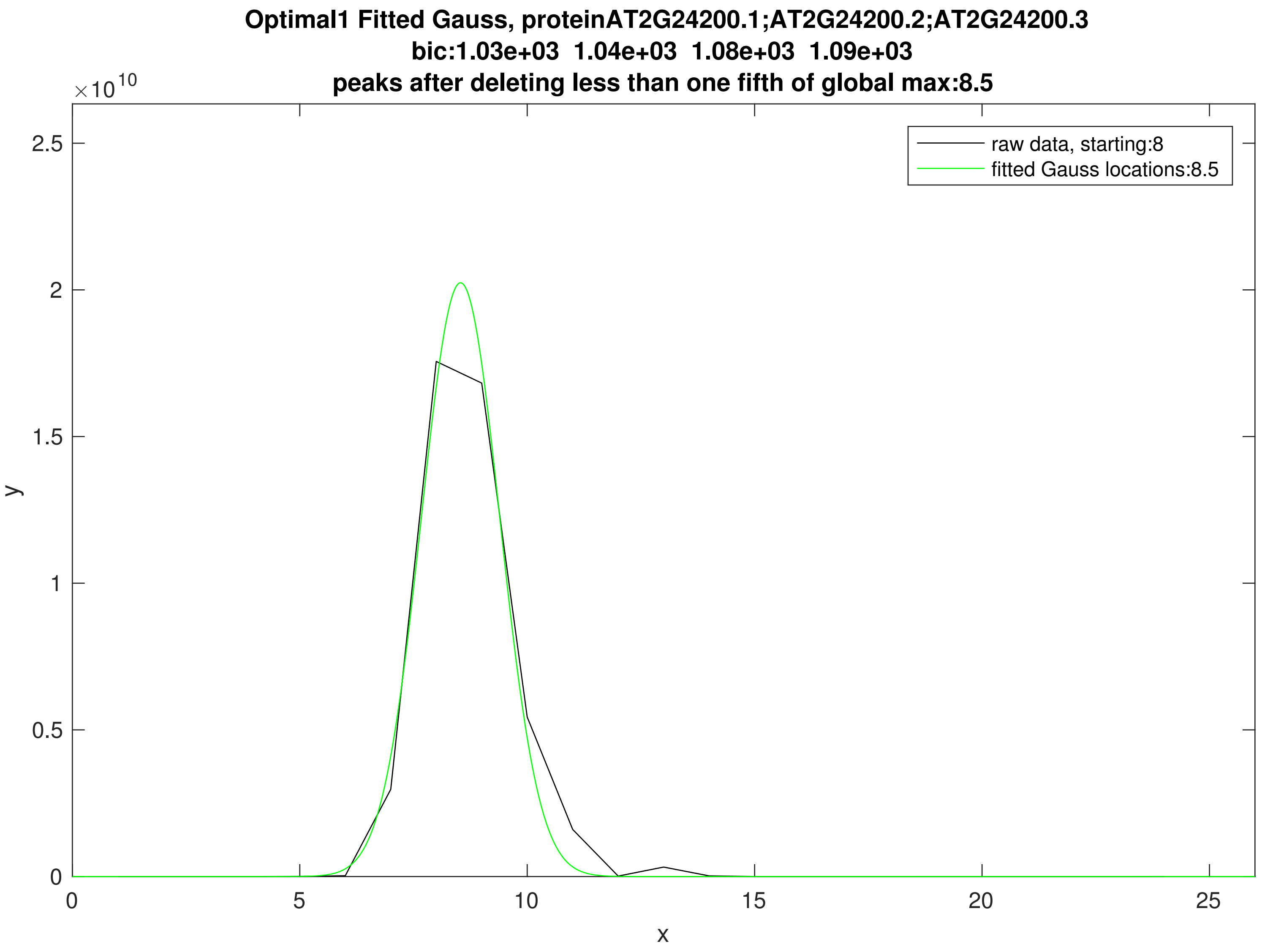

Optimal1 Fitted Gauss, proteinAT2G24200.1;AT2G24200.2;AT2G24200.3
bic:1.03e+03  1.04e+03  1.08e+03  1.09e+03
peaks after deleting less than one fifth of global max:8.5
×10^10
2.5
2
1.5
1
0.5
0
y
0
5
10
15
20
25
x
raw data, starting:8
fitted Gauss locations:8.5

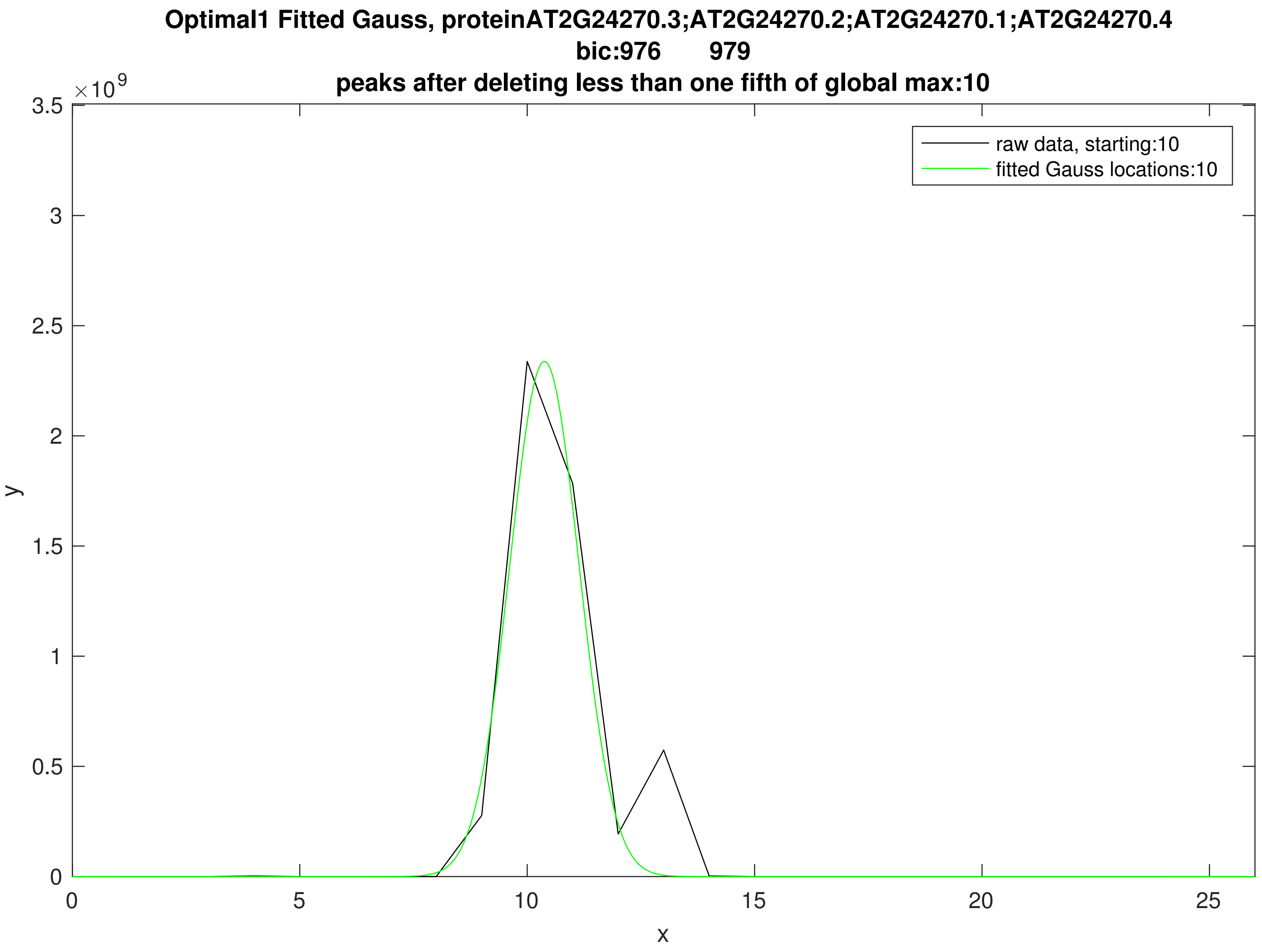
Optimal1 Fitted Gauss, proteinAT2G24270.3;AT2G24270.2;AT2G24270.1;AT2G24270.4
bic:976 979
peaks after deleting less than one fifth of global max:10
raw data, starting:10
fitted Gauss locations:10
×10^9
y
x

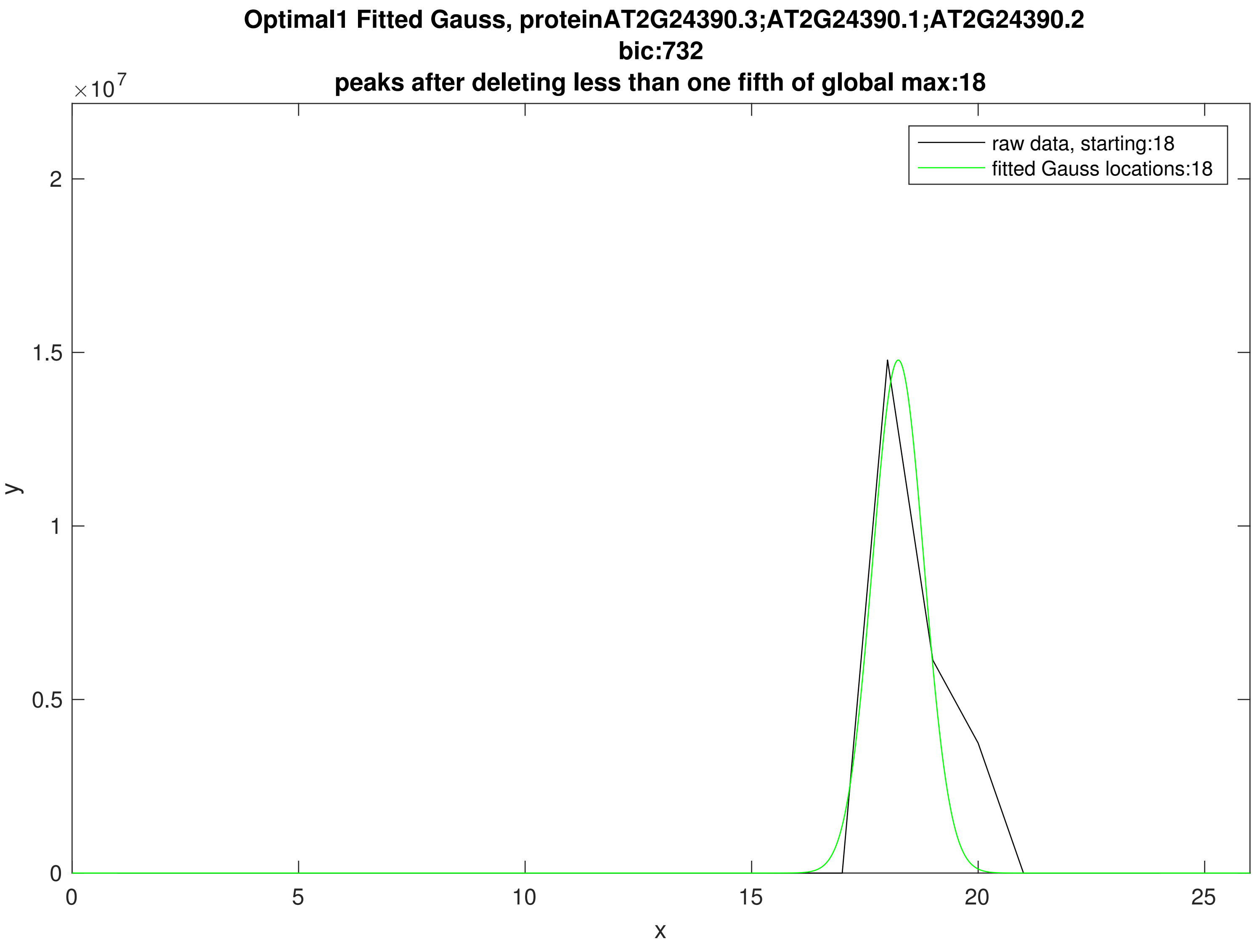

Optimal1 Fitted Gauss, proteinAT2G24390.3;AT2G24390.1;AT2G24390.2
bic:732
peaks after deleting less than one fifth of global max:18
×10^7
raw data, starting:18
fitted Gauss locations:18
y
x
0
0.5
1
1.5
2
0
5
10
15
20
25

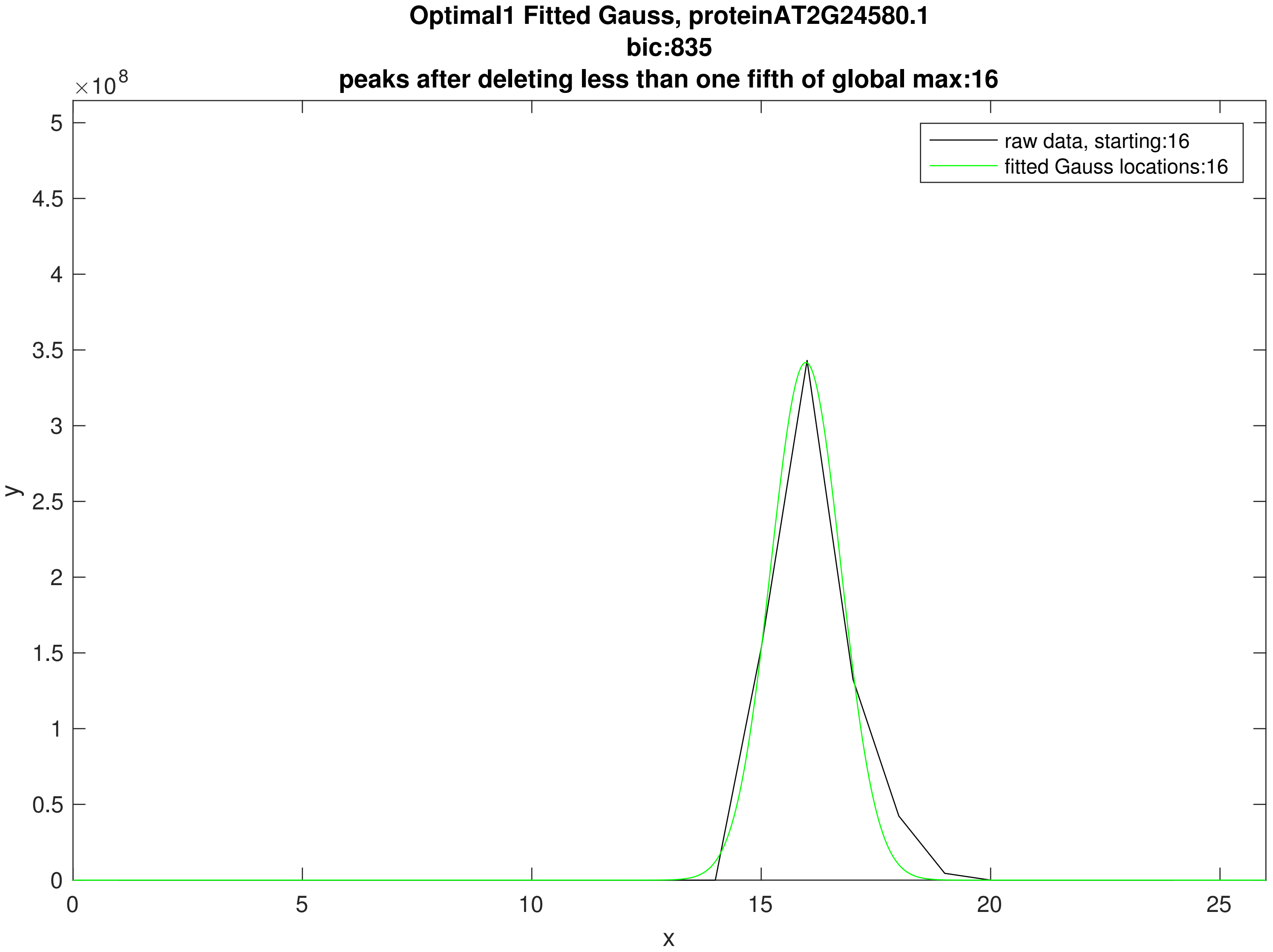
Optimal1 Fitted Gauss, proteinAT2G24580.1
bic:835
peaks after deleting less than one fifth of global max:16
×10^8
raw data, starting:16
fitted Gauss locations:16
y
x
0
0.5
1
1.5
2
2.5
3
3.5
4
4.5
5
0
5
10
15
20
25

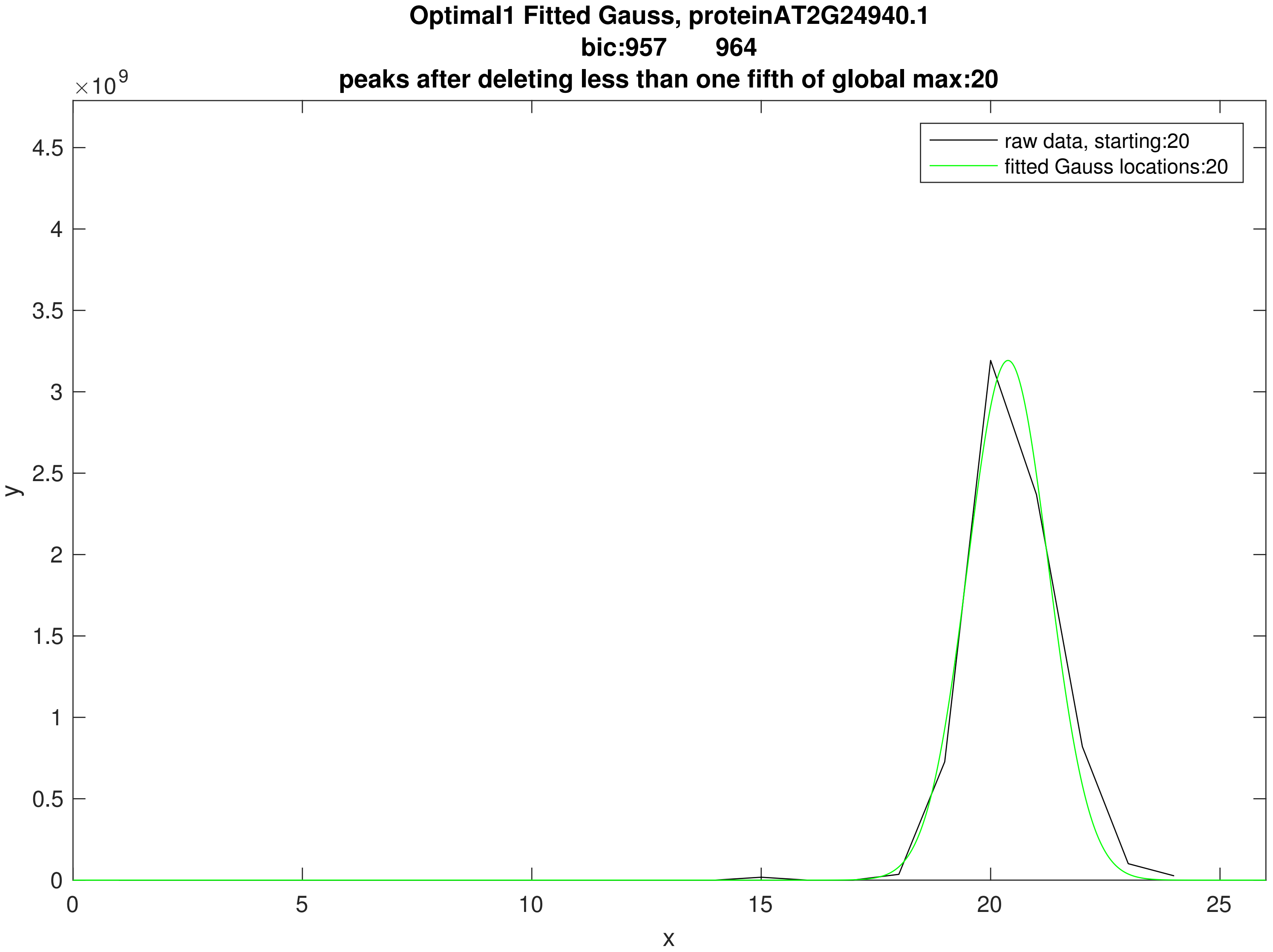

Optimal1 Fitted Gauss, proteinAT2G24940.1
bic:957 964
peaks after deleting less than one fifth of global max:20
×10^9
4.5
4
3.5
3
2.5
2
1.5
1
0.5
0
y
0
5
10
15
20
25
x
raw data, starting:20
fitted Gauss locations:20

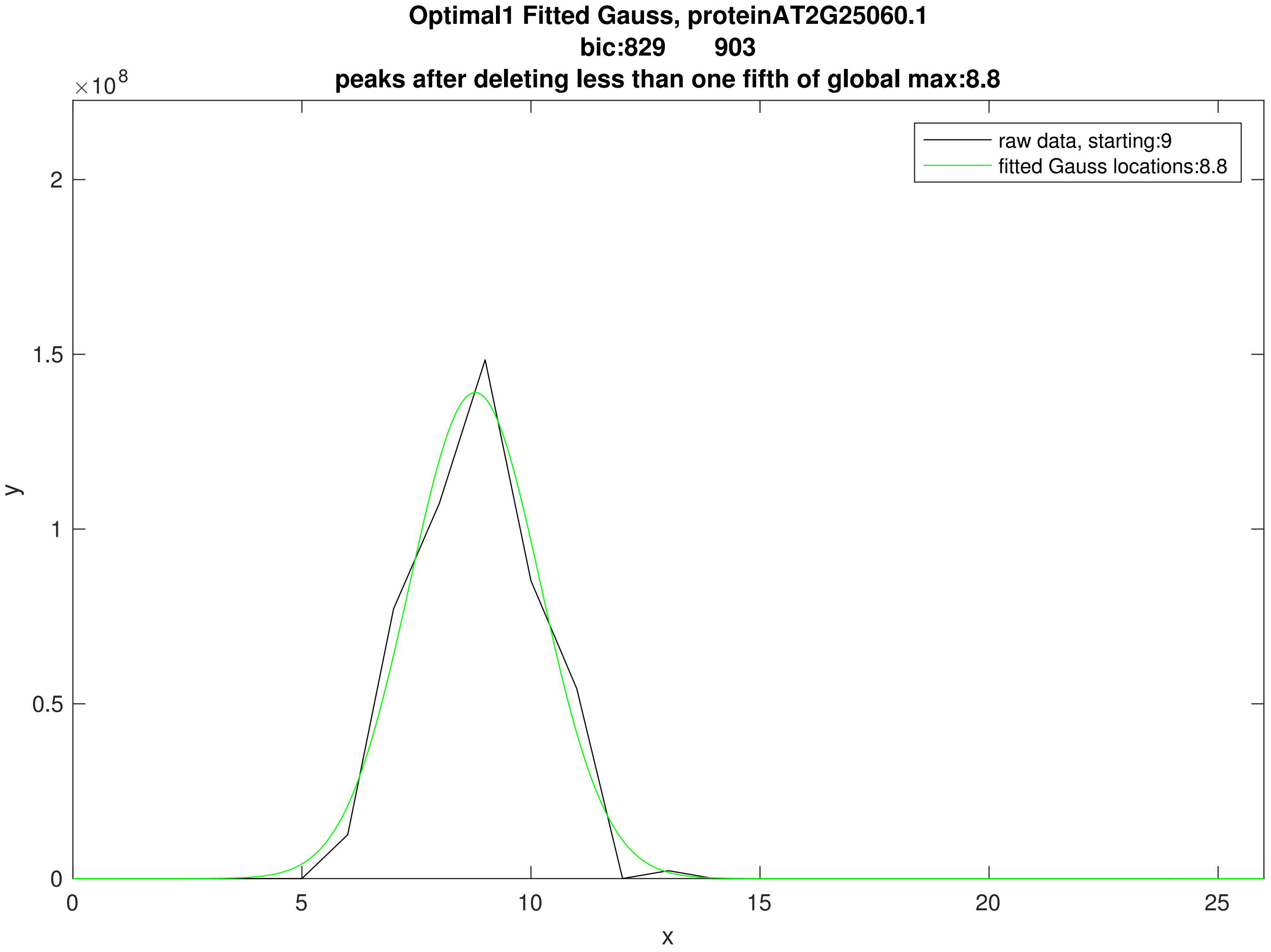
Optimal1 Fitted Gauss, proteinAT2G25060.1
bic:829 903
peaks after deleting less than one fifth of global max:8.8
×10^8
raw data, starting:9
fitted Gauss locations:8.8
y
x
0
0.5
1
1.5
2
0
5
10
15
20
25

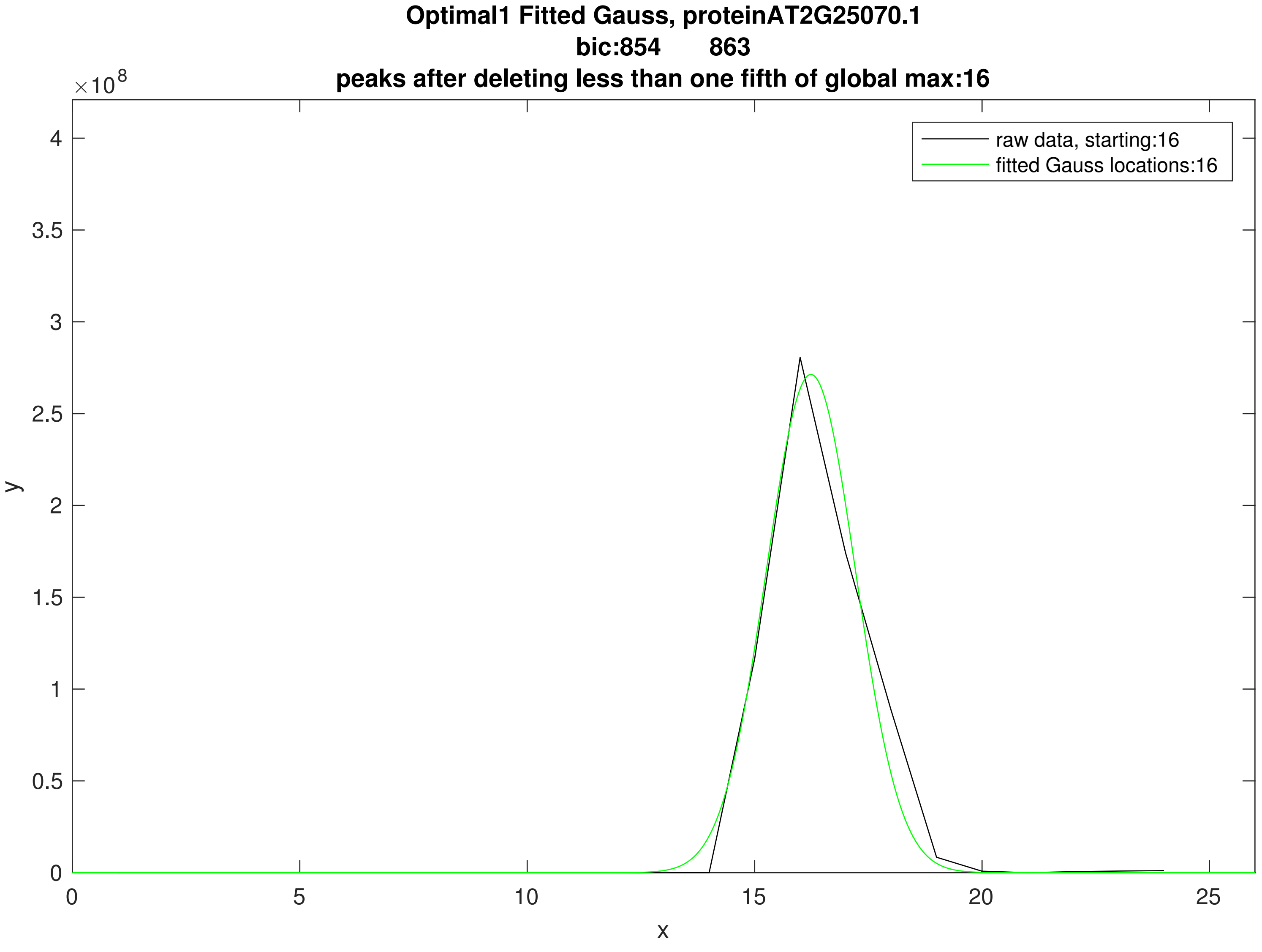

Optimal1 Fitted Gauss, proteinAT2G25070.1
bic:854 863
peaks after deleting less than one fifth of global max:16
×10^8
raw data, starting:16
fitted Gauss locations:16
y
x
0
0.5
1
1.5
2
2.5
3
3.5
4
0
5
10
15
20
25

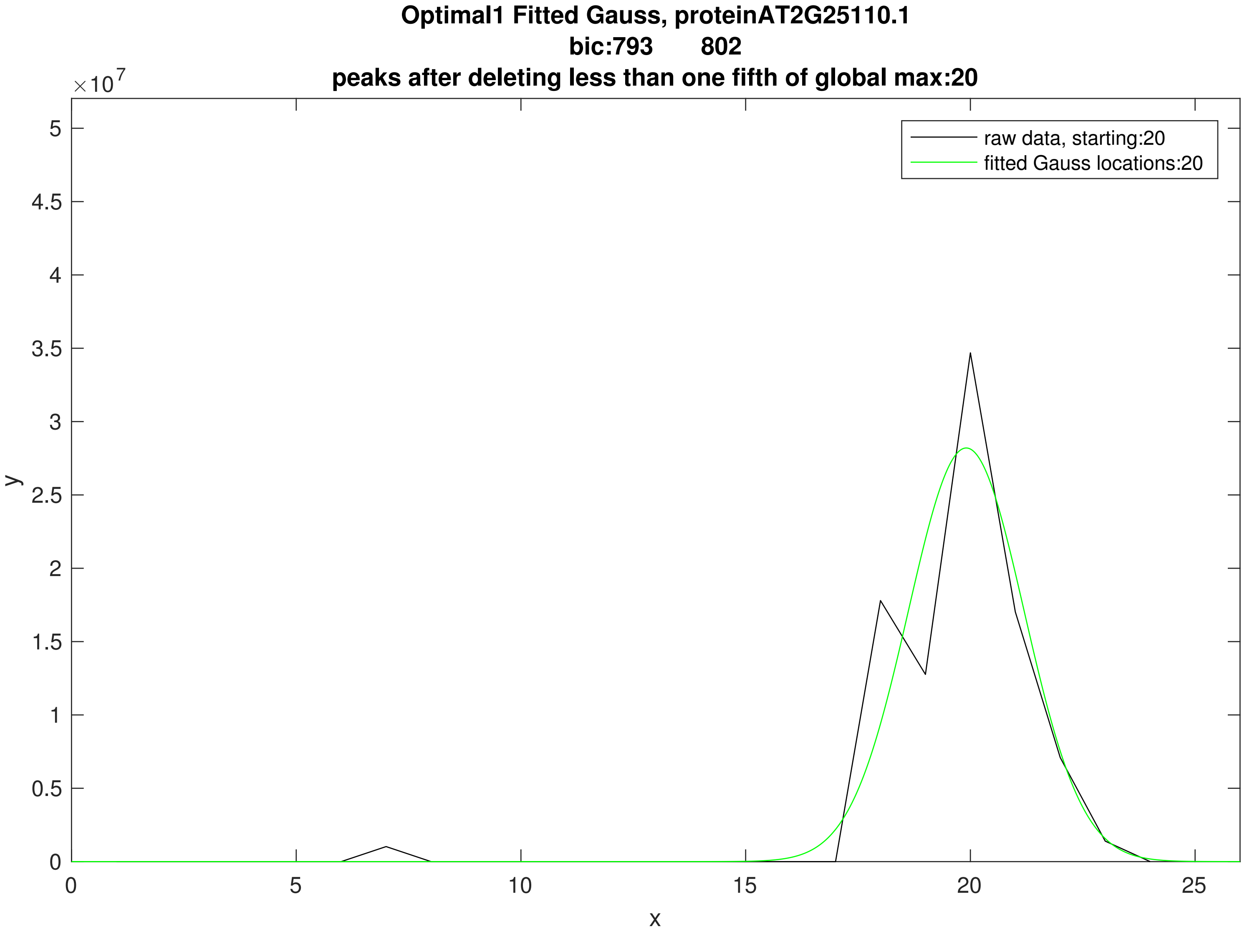
Optimal1 Fitted Gauss, proteinAT2G25110.1
bic:793 802
peaks after deleting less than one fifth of global max:20
×10^7
5
4.5
4
3.5
3
2.5
2
1.5
1
0.5
0
y
0
5
10
15
20
25
x
raw data, starting:20
fitted Gauss locations:20

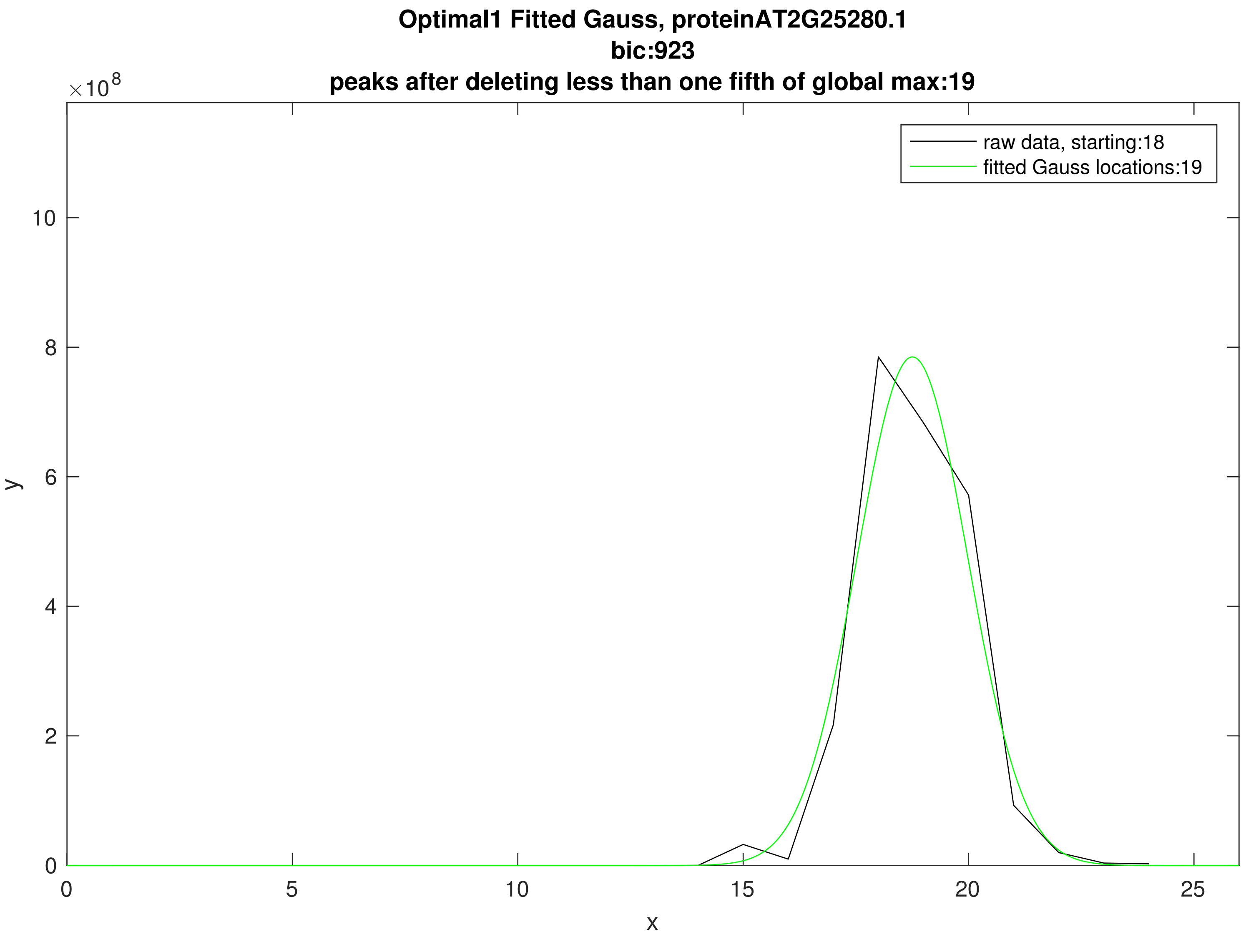

Optimal1 Fitted Gauss, proteinAT2G25280.1
bic:923
peaks after deleting less than one fifth of global max:19
×10^8
raw data, starting:18
fitted Gauss locations:19
10
8
6
4
2
0
y
0
5
10
15
20
25
x

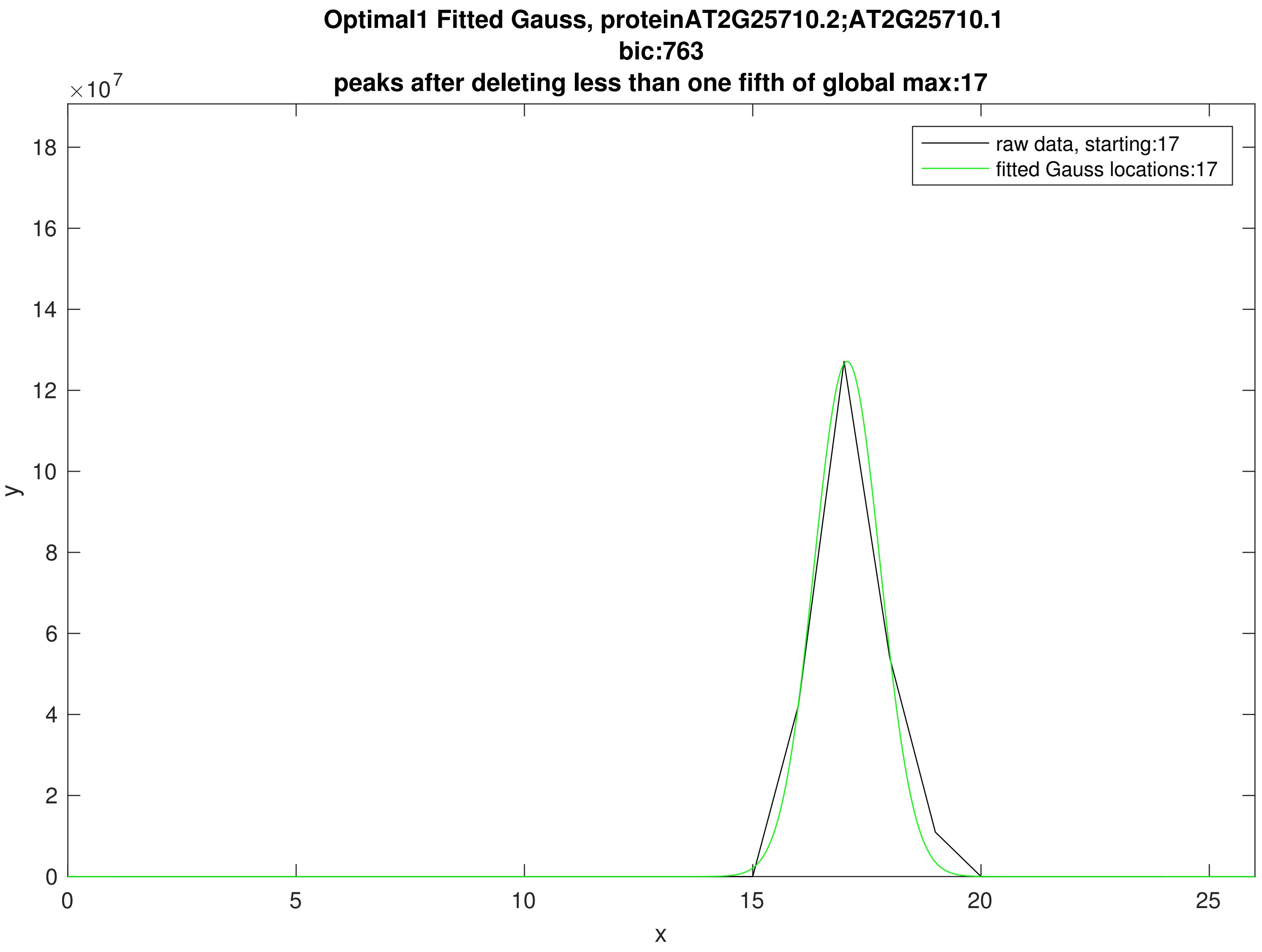
Optimal1 Fitted Gauss, proteinAT2G25710.2;AT2G25710.1
bic:763
peaks after deleting less than one fifth of global max:17
×10^7
raw data, starting:17
fitted Gauss locations:17
y
x
0
2
4
6
8
10
12
14
16
18
0
5
10
15
20
25

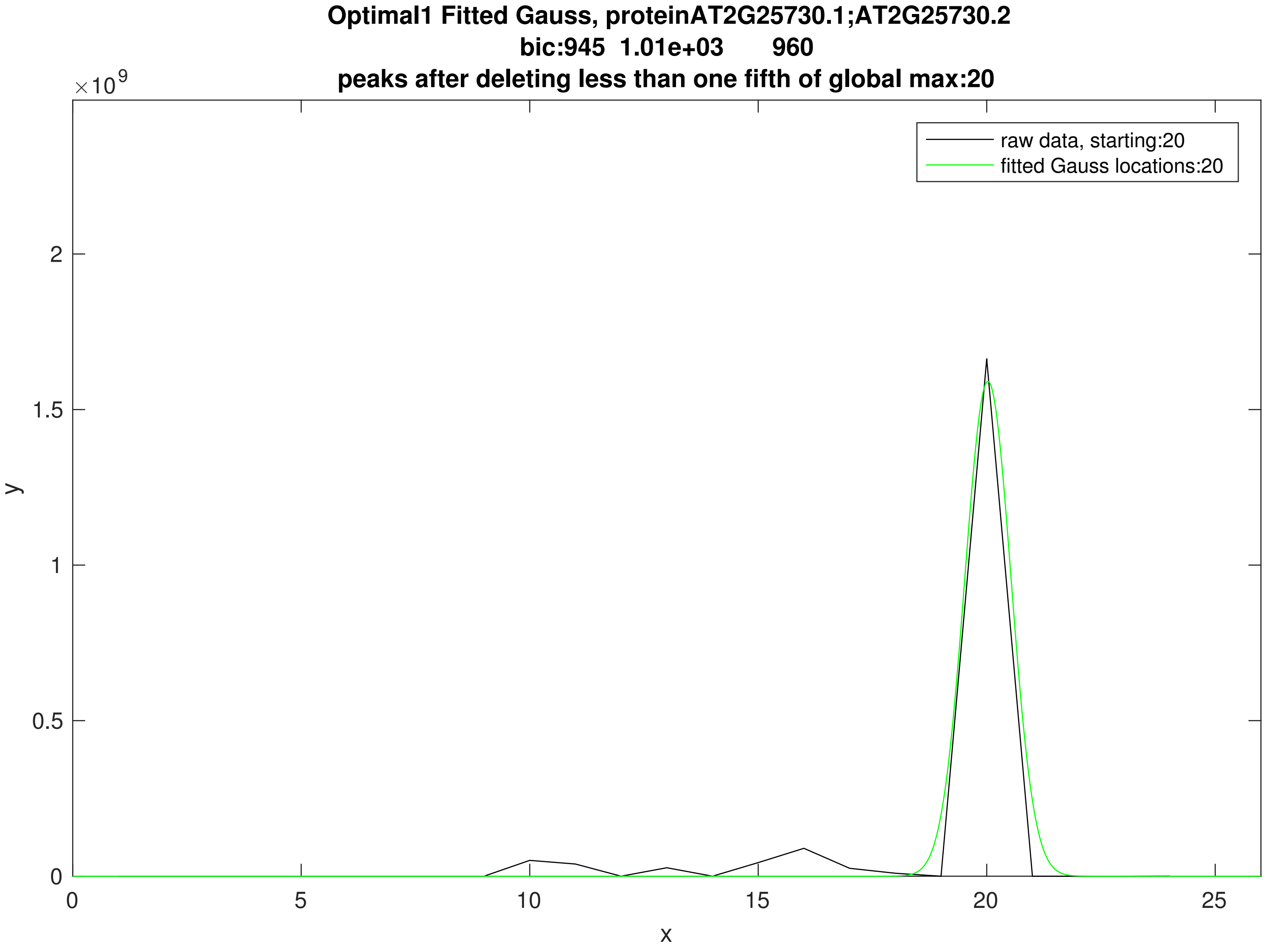
Optimal1 Fitted Gauss, proteinAT2G25730.1;AT2G25730.2
bic:945  1.01e+03       960
peaks after deleting less than one fifth of global max:20
×10^9
raw data, starting:20
fitted Gauss locations:20
2
1.5
1
0.5
0
y
0
5
10
15
20
25
x

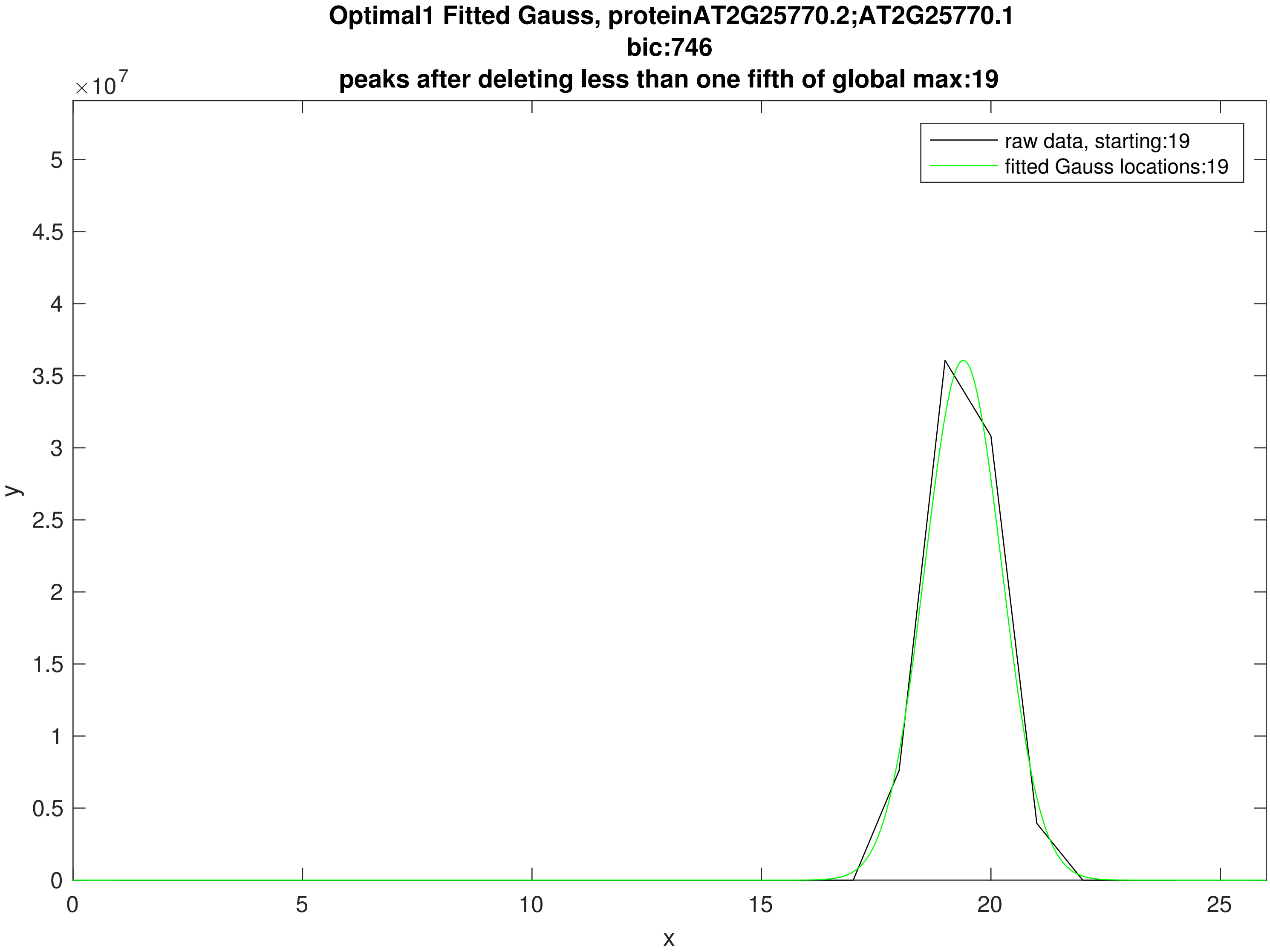

Optimal1 Fitted Gauss, proteinAT2G25770.2;AT2G25770.1
bic:746
peaks after deleting less than one fifth of global max:19
×10^7
raw data, starting:19
fitted Gauss locations:19
y
x
0
0.5
1
1.5
2
2.5
3
3.5
4
4.5
5
0
5
10
15
20
25

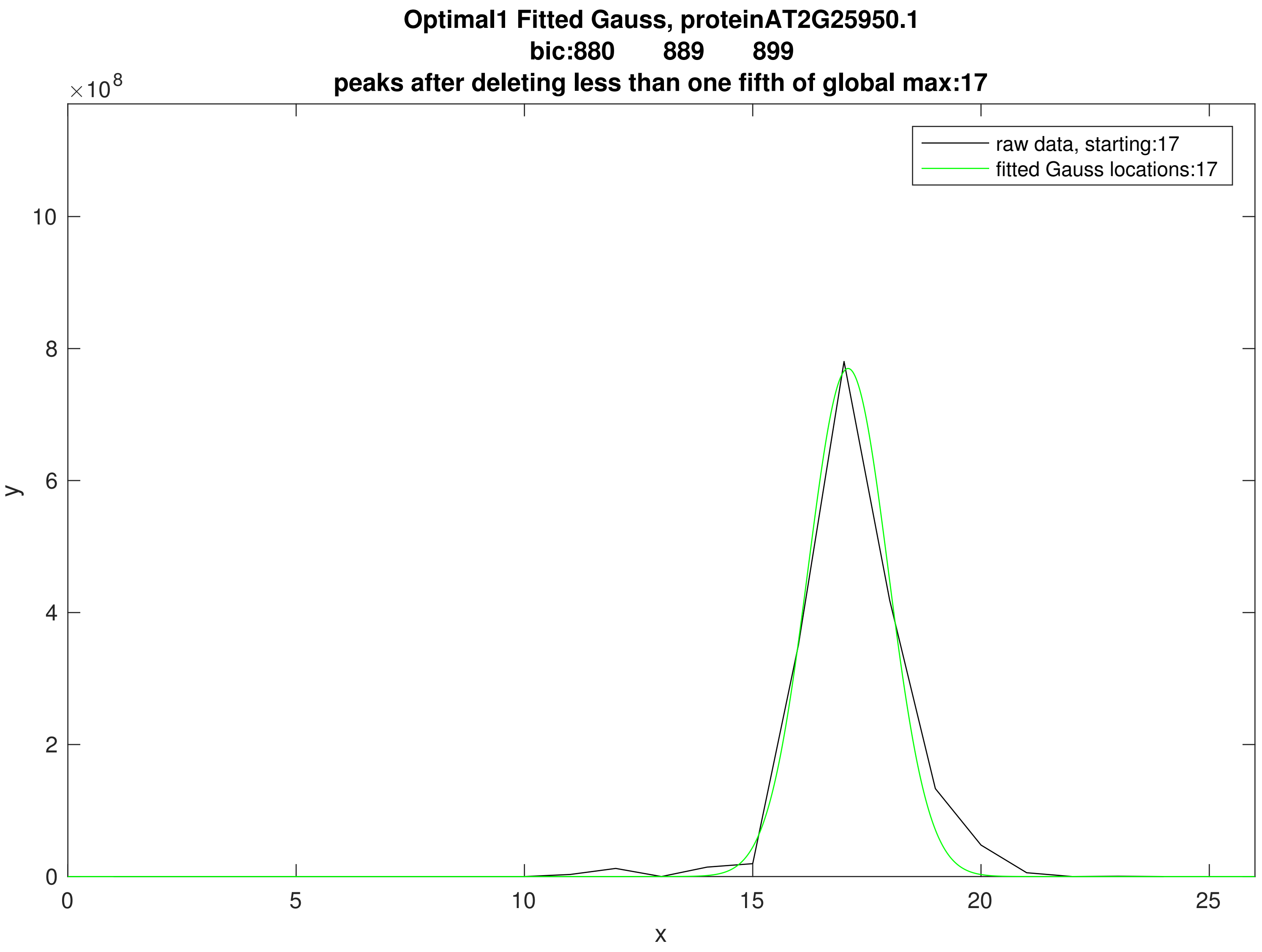
Optimal1 Fitted Gauss, proteinAT2G25950.1
bic:880 889 899
peaks after deleting less than one fifth of global max:17
×10^8
raw data, starting:17
fitted Gauss locations:17
y
x
0
2
4
6
8
10
0
5
10
15
20
25

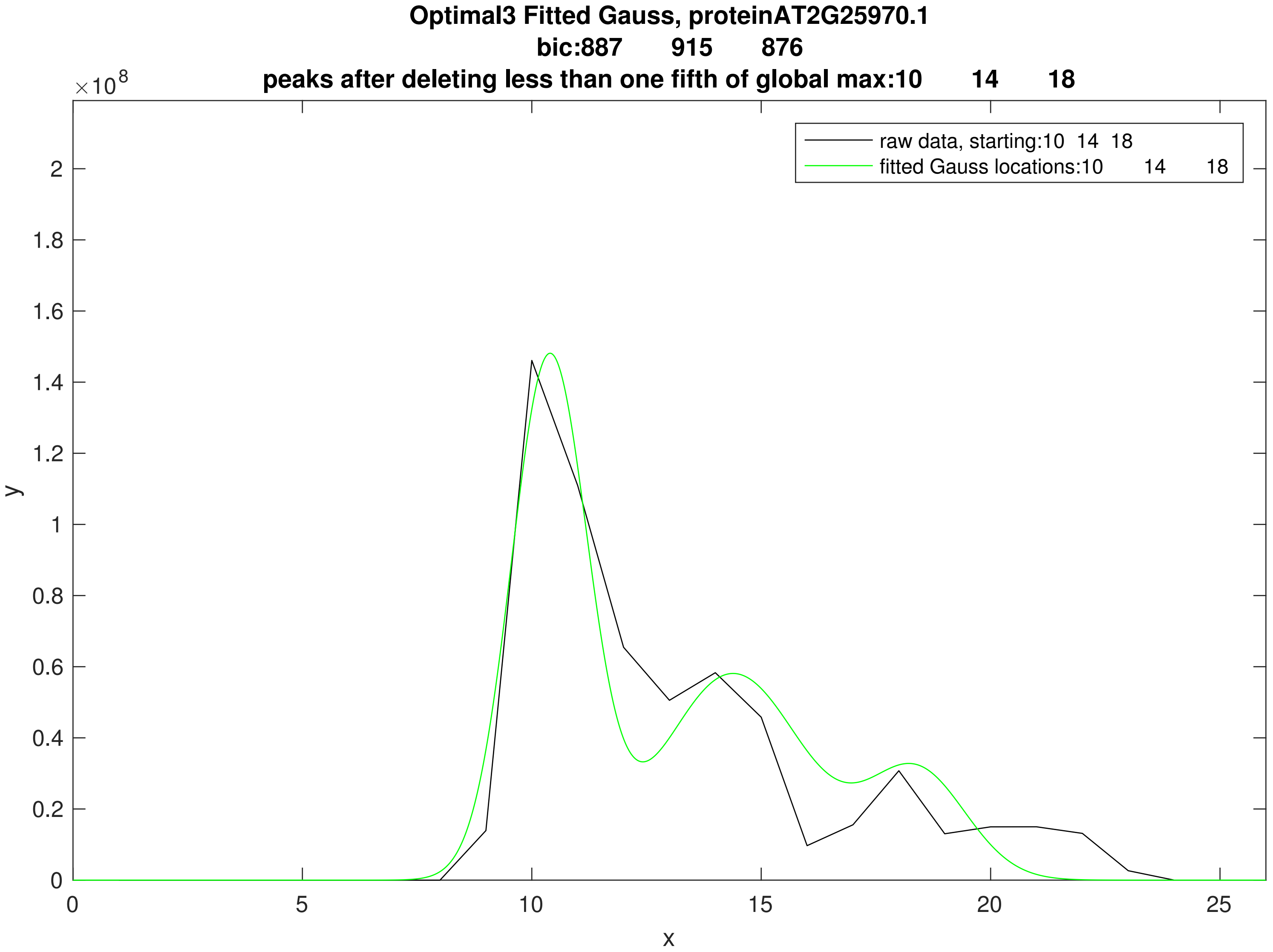
Optimal3 Fitted Gauss, proteinAT2G25970.1
bic:887 915 876
peaks after deleting less than one fifth of global max:10 14 18
×10^8
raw data, starting:10 14 18
fitted Gauss locations:10 14 18
y
x
0
0.2
0.4
0.6
0.8
1
1.2
1.4
1.6
1.8
2
0
5
10
15
20
25

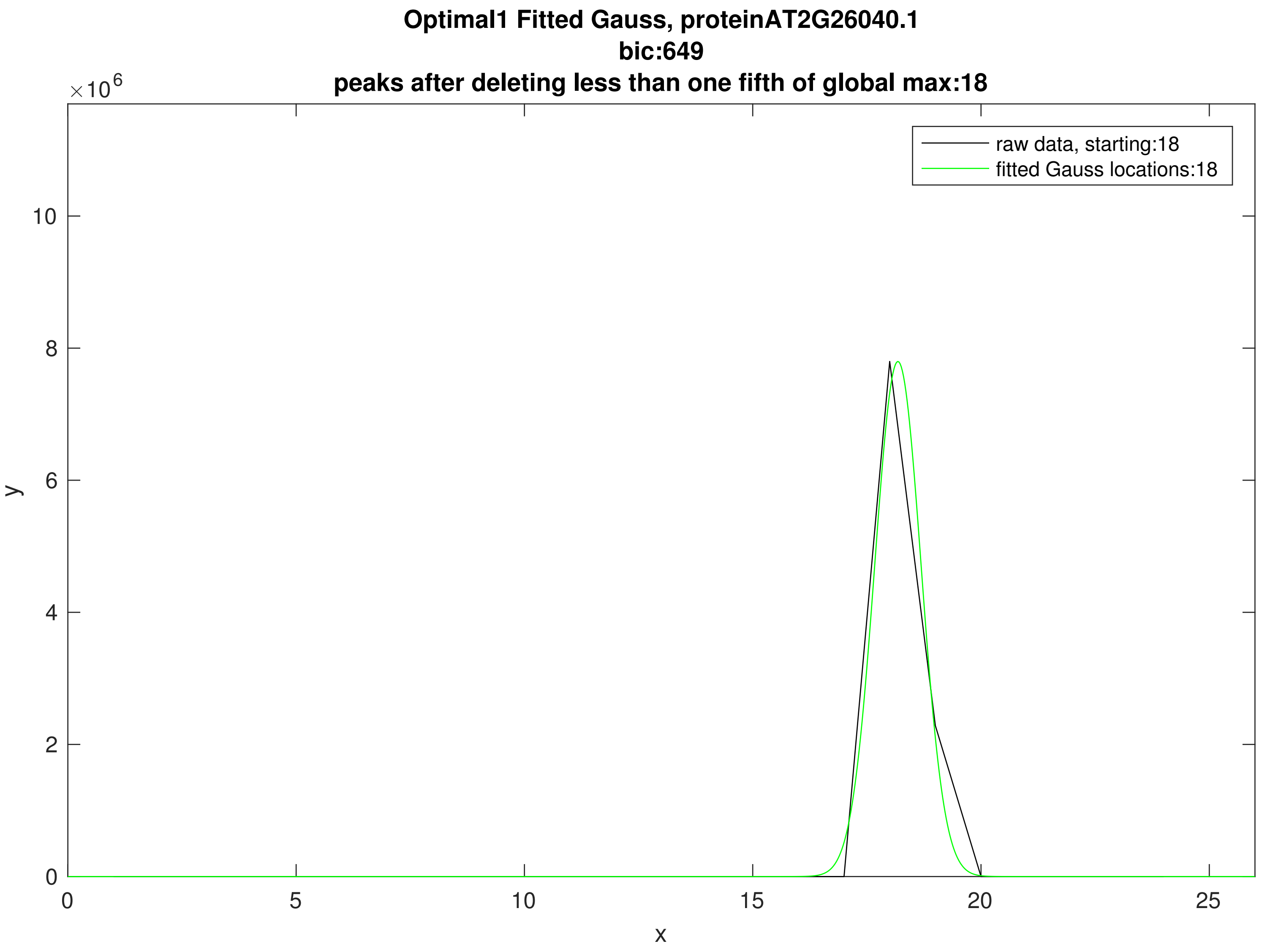
Optimal1 Fitted Gauss, proteinAT2G26040.1
bic:649
peaks after deleting less than one fifth of global max:18
×10^6
raw data, starting:18
fitted Gauss locations:18
10
8
6
4
2
0
y
0
5
10
15
20
25
x

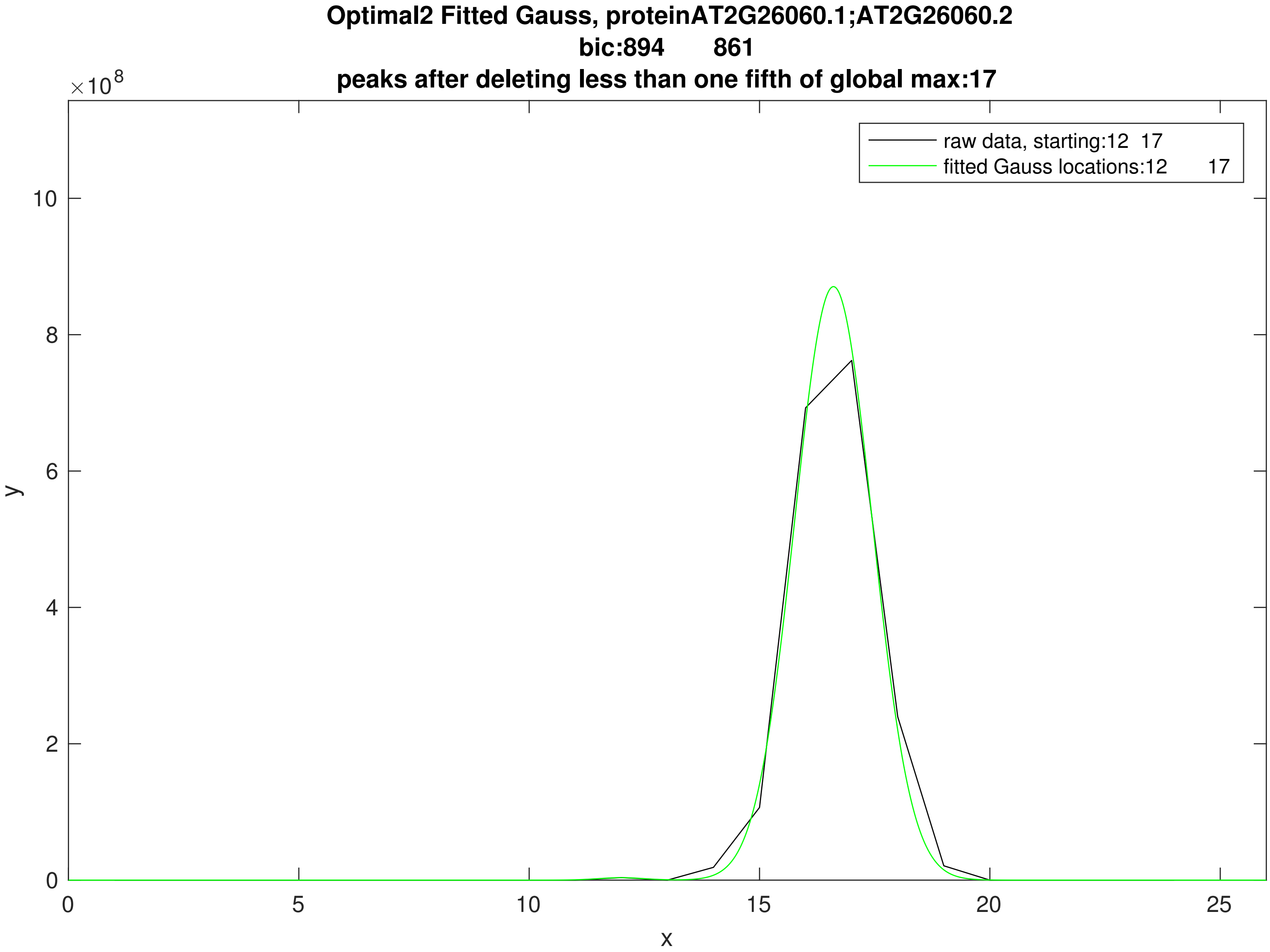

Optimal2 Fitted Gauss, proteinAT2G26060.1;AT2G26060.2
bic:894 861
peaks after deleting less than one fifth of global max:17
×10^8
raw data, starting:12 17
fitted Gauss locations:12 17
10
8
6
4
2
0
y
0
5
10
15
20
25
x

Optimal1 Fitted Gauss, proteinAT2G26080.1
bic:1.06e+03  1.07e+03  1.07e+03
peaks after deleting less than one fifth of global max:11

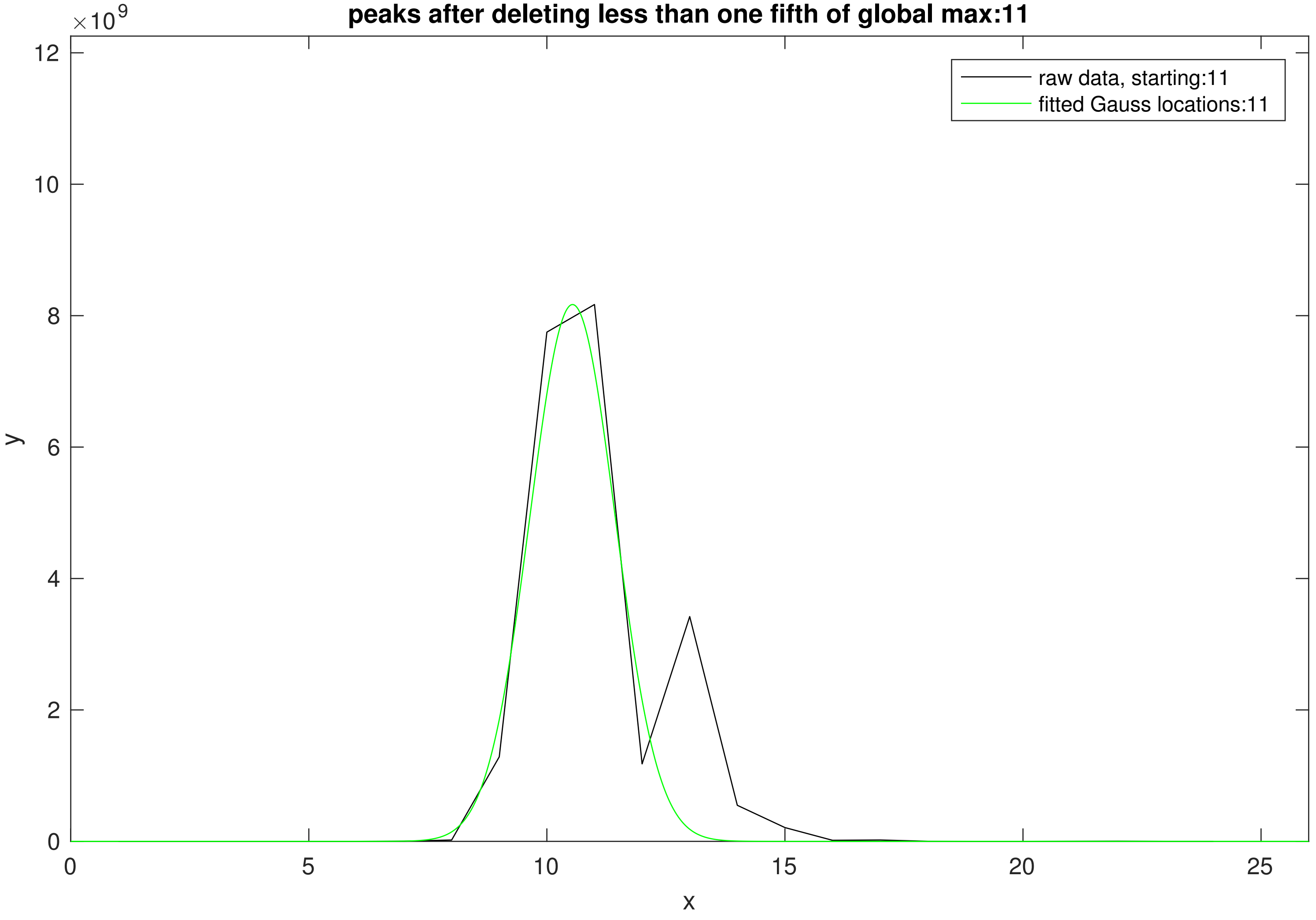

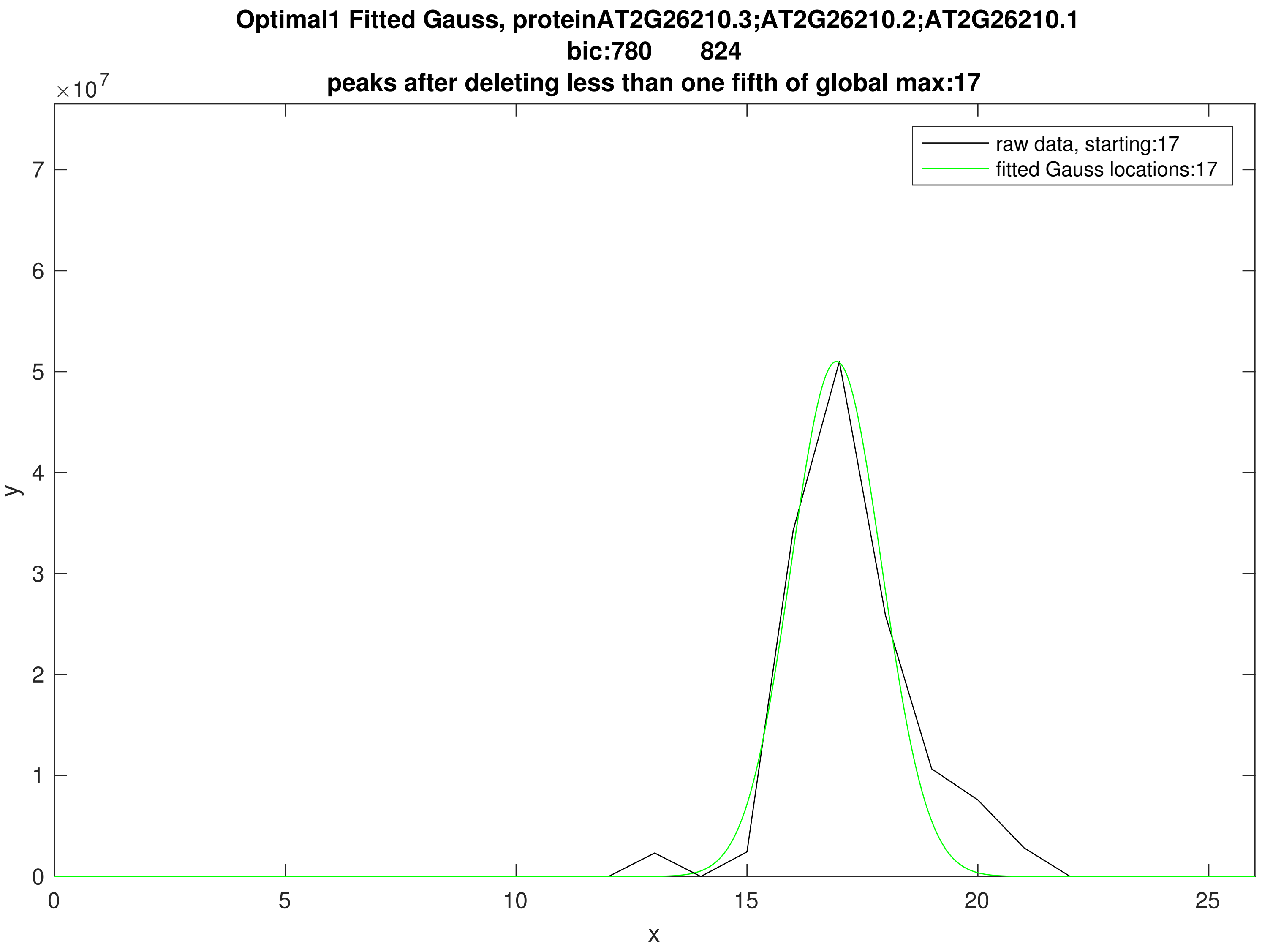

Optimal1 Fitted Gauss, proteinAT2G26210.3;AT2G26210.2;AT2G26210.1
bic:780 824
peaks after deleting less than one fifth of global max:17
×10^7
raw data, starting:17
fitted Gauss locations:17
y
x
0
1
2
3
4
5
6
7
0
5
10
15
20
25

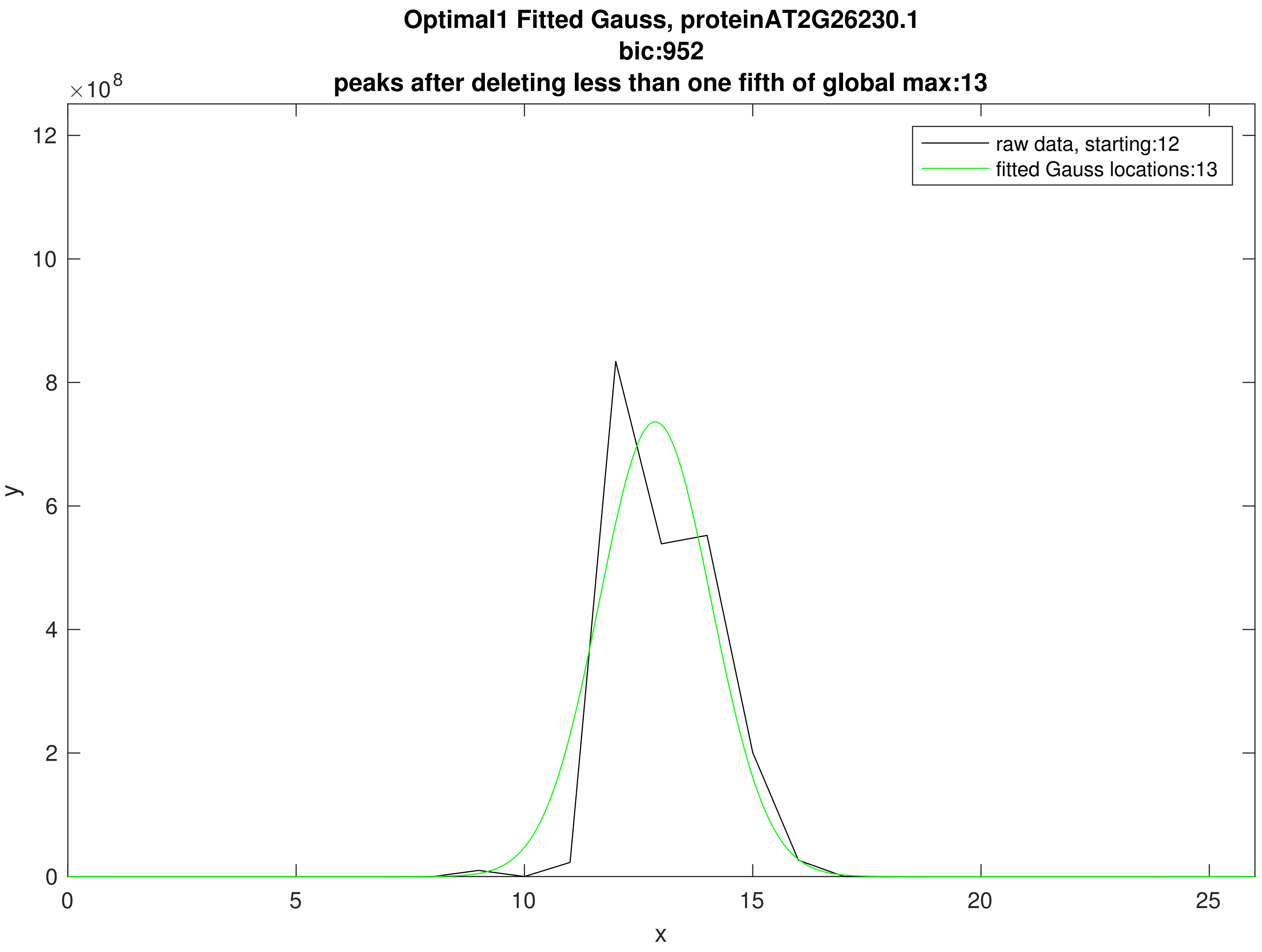
Optimal1 Fitted Gauss, proteinAT2G26230.1
bic:952
peaks after deleting less than one fifth of global max:13
×10^8
12
10
8
6
4
2
0
y
0
5
10
15
20
25
x
raw data, starting:12
fitted Gauss locations:13

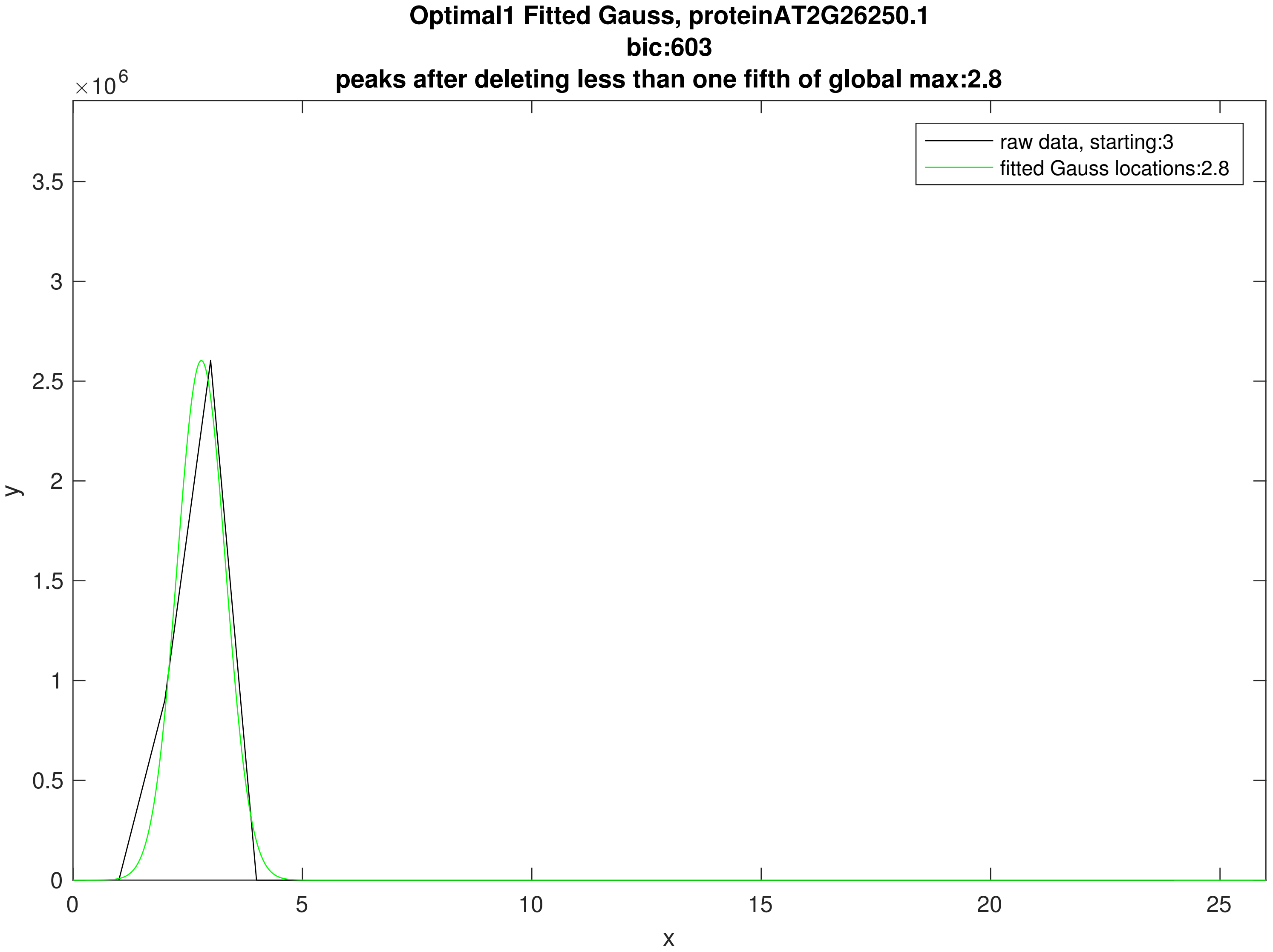

Optimal1 Fitted Gauss, proteinAT2G26250.1
bic:603
peaks after deleting less than one fifth of global max:2.8
×10^6
raw data, starting:3
fitted Gauss locations:2.8
3.5
3
2.5
2
1.5
1
0.5
0
y
0
5
10
15
20
25
x

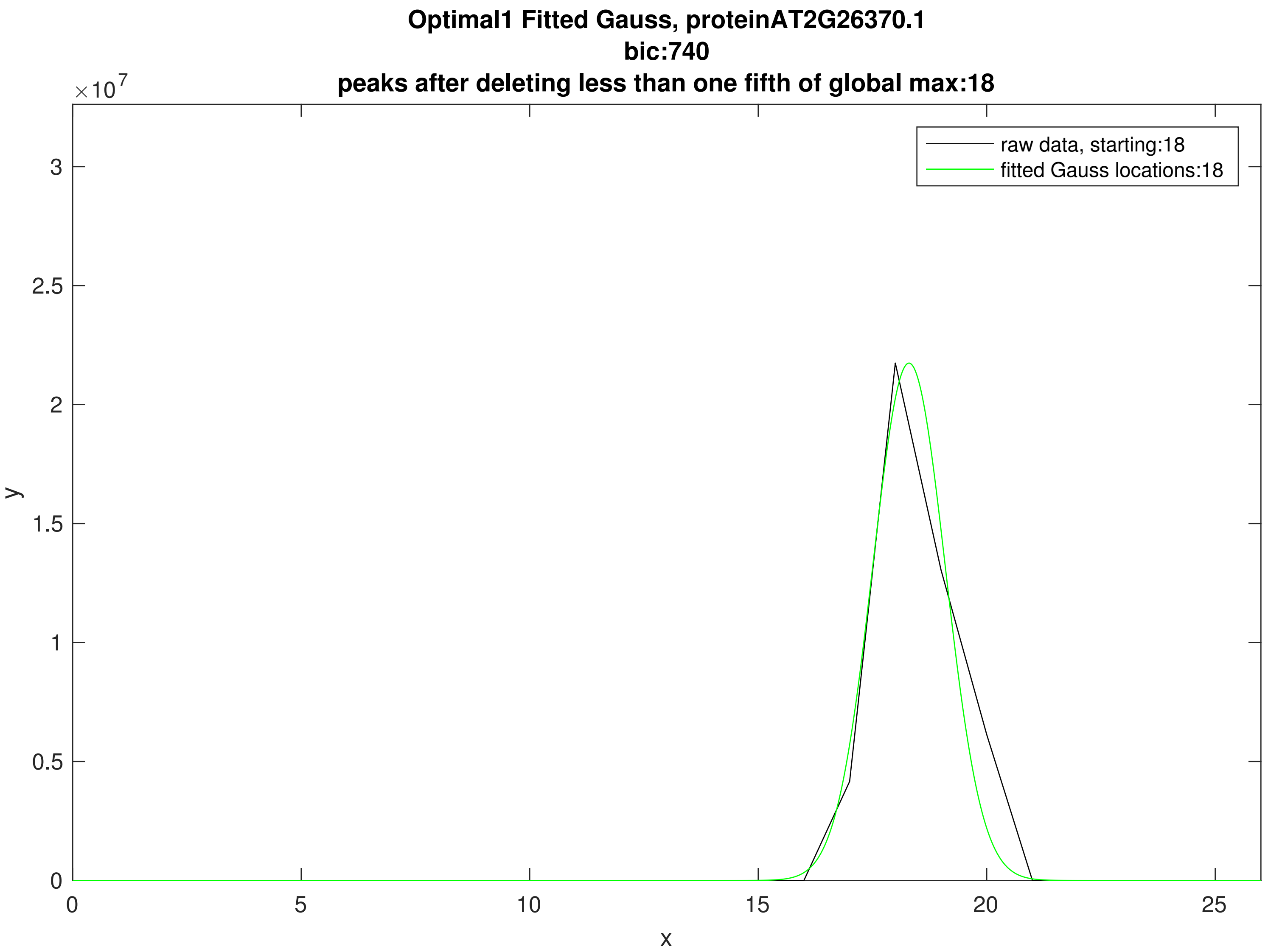
Optimal1 Fitted Gauss, proteinAT2G26370.1
bic:740
peaks after deleting less than one fifth of global max:18
×10^7
3
2.5
2
1.5
1
0.5
0
y
0
5
10
15
20
25
x
raw data, starting:18
fitted Gauss locations:18

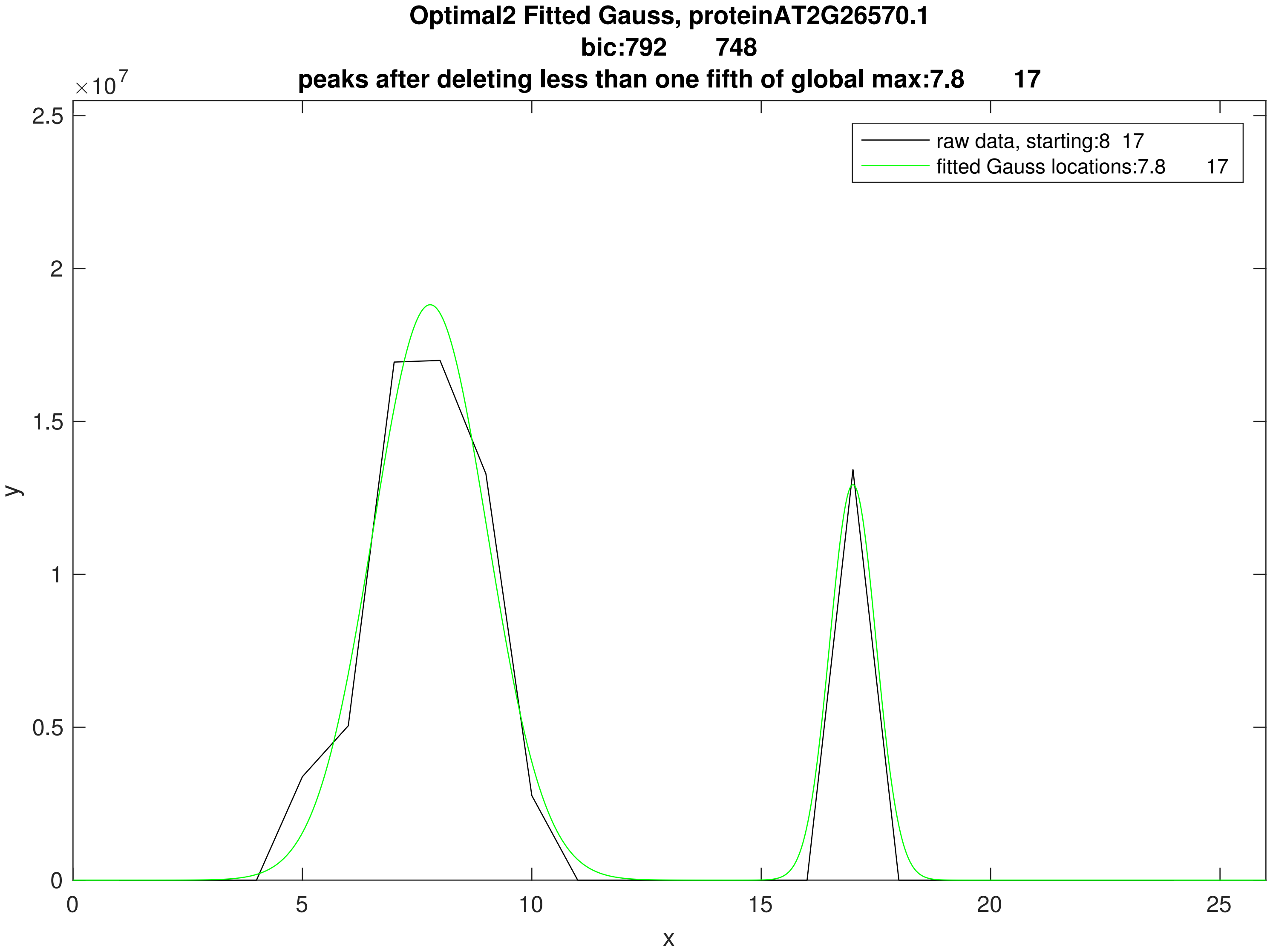

Optimal2 Fitted Gauss, proteinAT2G26570.1
bic:792 748
peaks after deleting less than one fifth of global max:7.8 17
×10^7
2.5
2
1.5
1
0.5
0
y
0
5
10
15
20
25
x
raw data, starting:8 17
fitted Gauss locations:7.8 17

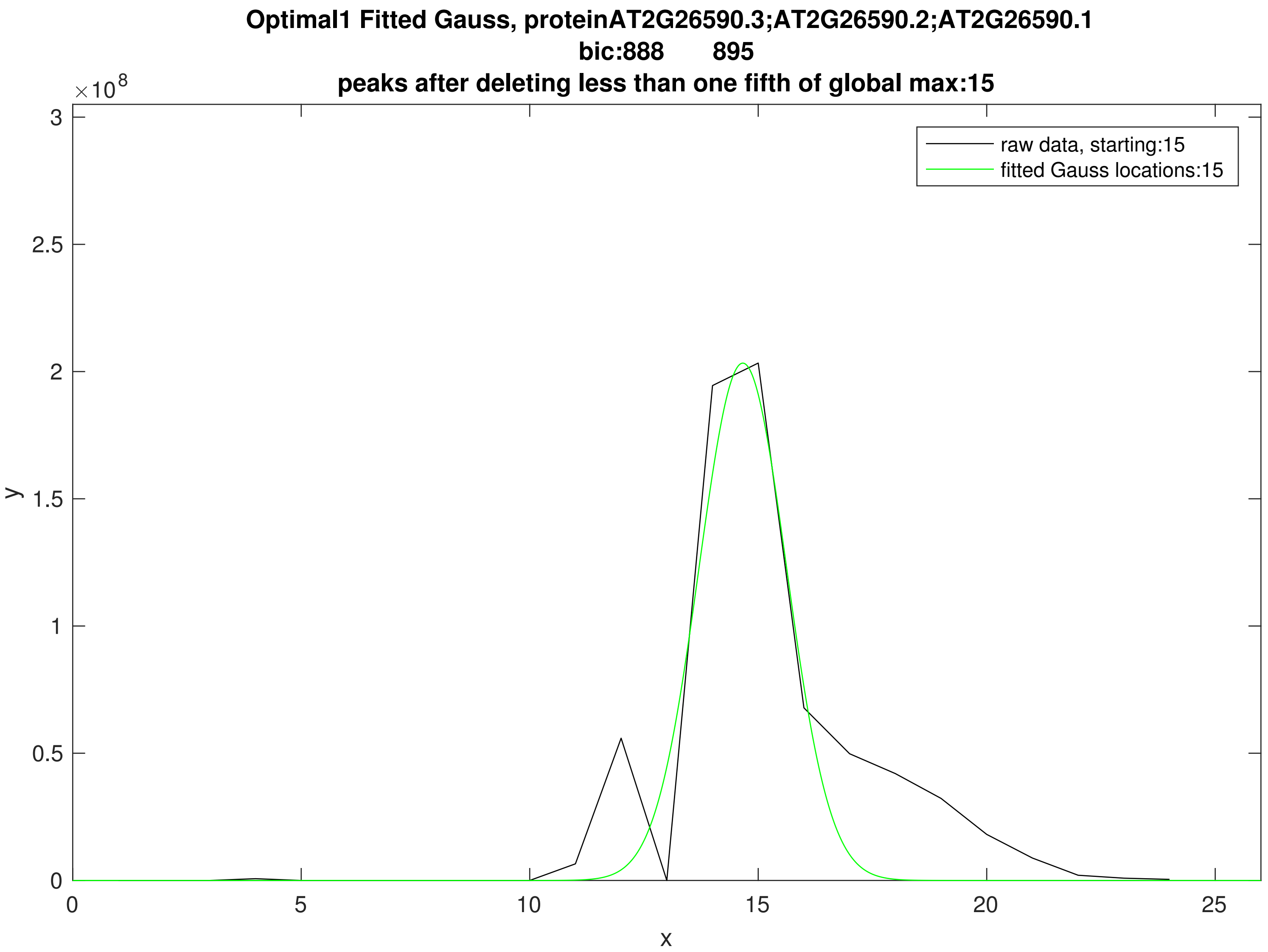

Optimal1 Fitted Gauss, proteinAT2G26590.3;AT2G26590.2;AT2G26590.1
bic:888 895
peaks after deleting less than one fifth of global max:15
×10^8
3
2.5
2
1.5
1
0.5
0
y
0
5
10
15
20
25
x
raw data, starting:15
fitted Gauss locations:15

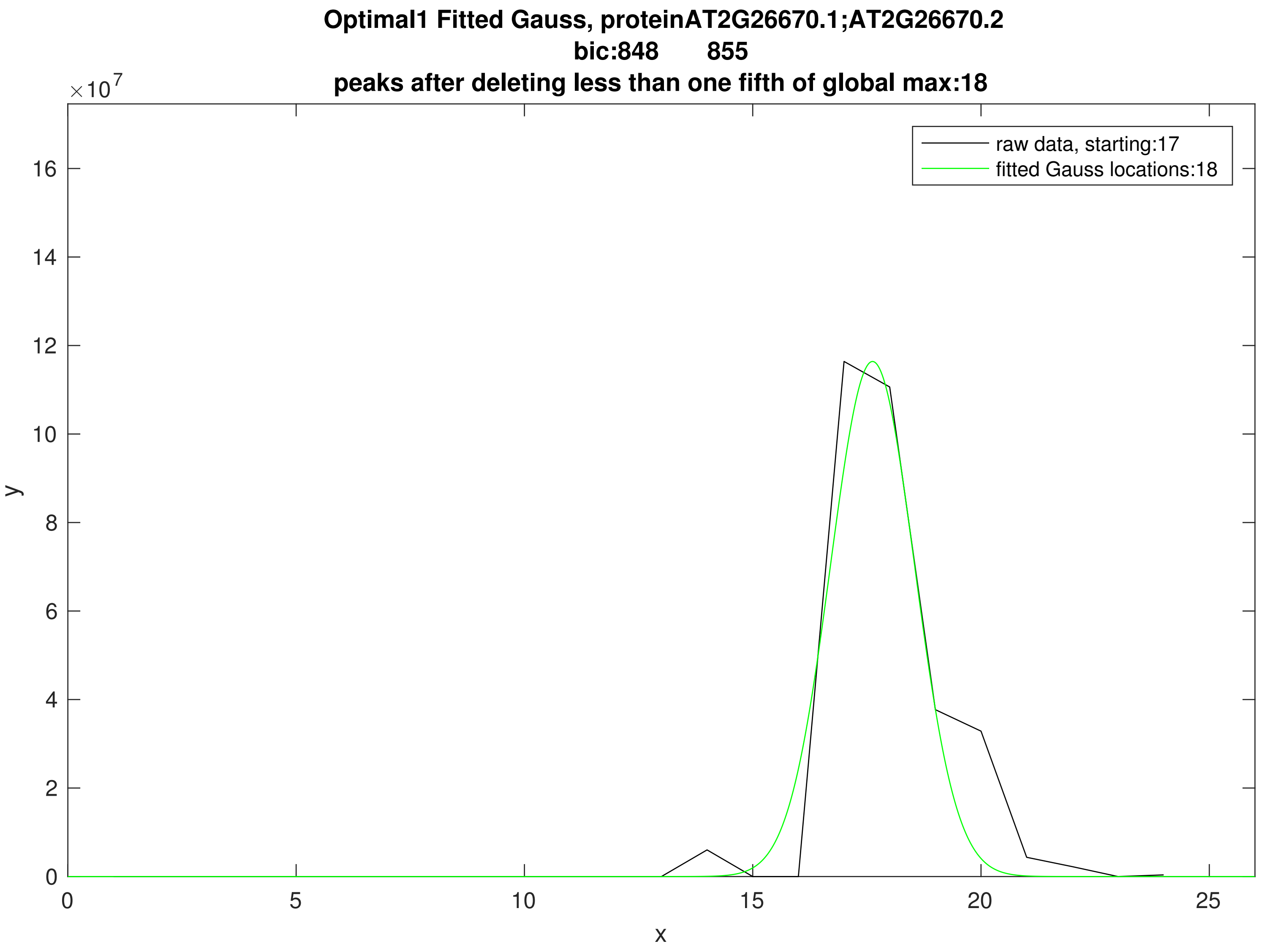
Optimal1 Fitted Gauss, proteinAT2G26670.1;AT2G26670.2
bic:848 855
peaks after deleting less than one fifth of global max:18
×10^7
raw data, starting:17
fitted Gauss locations:18
16
14
12
10
8
6
4
2
0
y
0
5
10
15
20
25
x

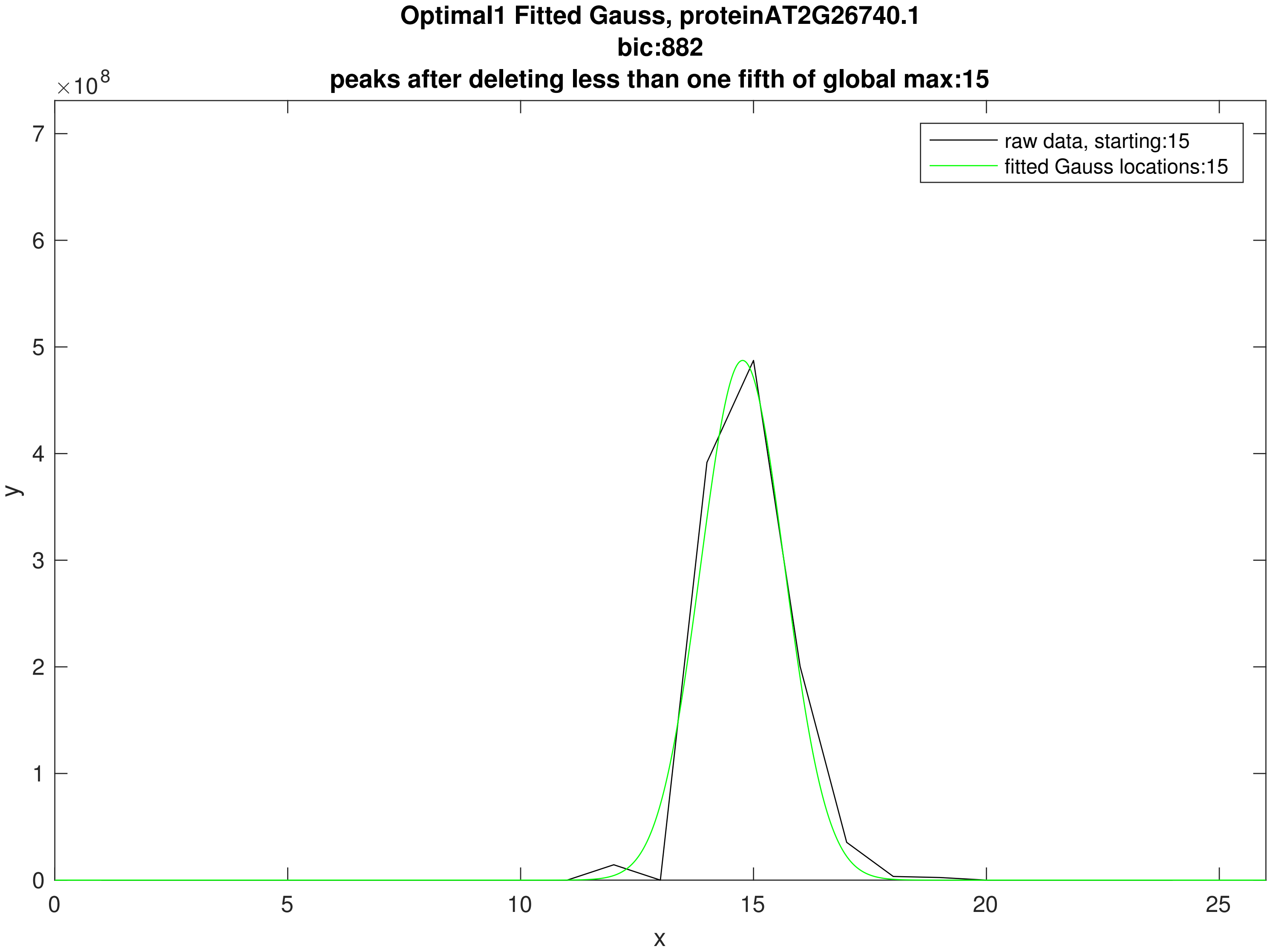
Optimal1 Fitted Gauss, proteinAT2G26740.1
bic:882
peaks after deleting less than one fifth of global max:15
×10^8
raw data, starting:15
fitted Gauss locations:15
y
x
0
1
2
3
4
5
6
7
0
5
10
15
20
25

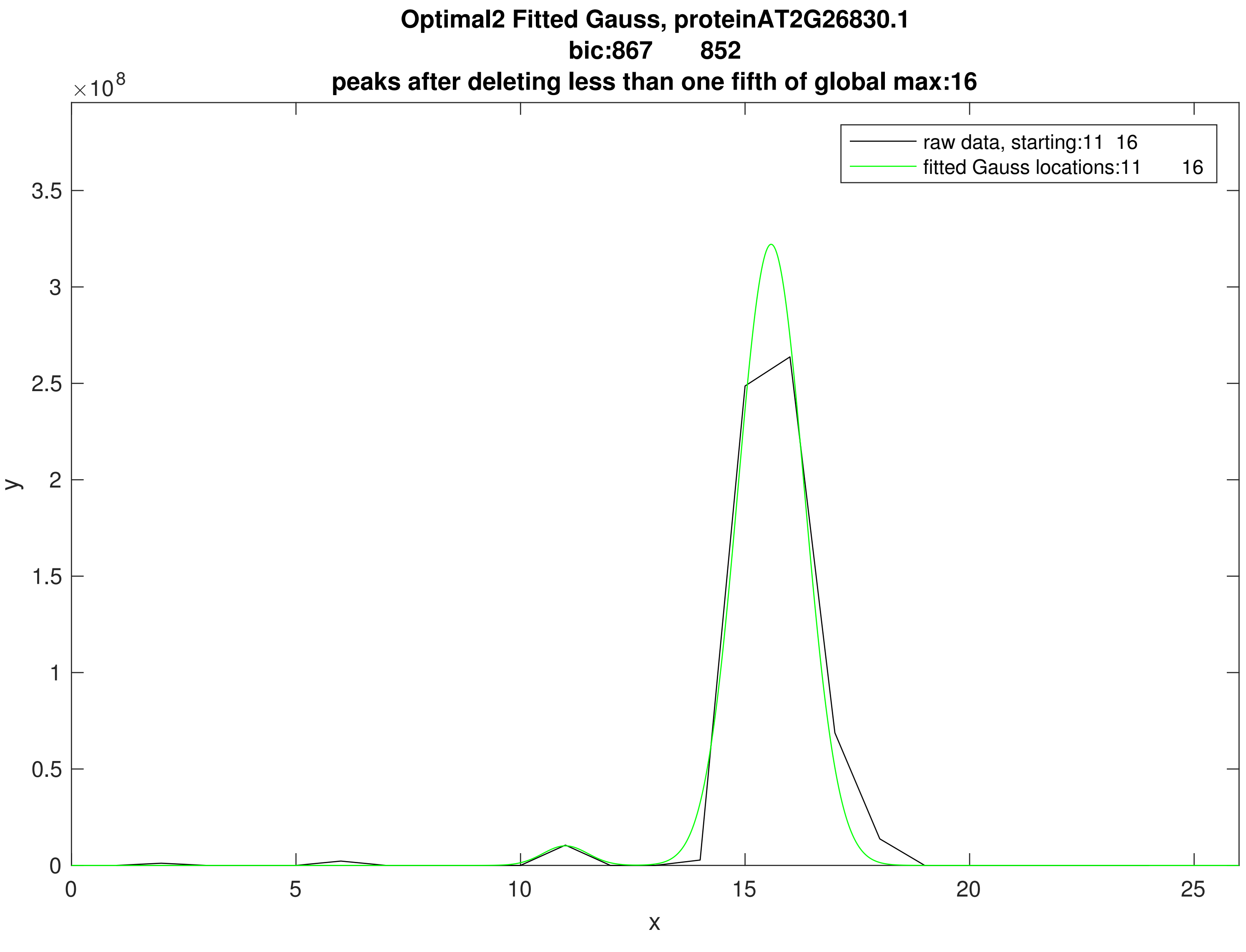
Optimal2 Fitted Gauss, proteinAT2G26830.1
bic:867 852
peaks after deleting less than one fifth of global max:16
raw data, starting:11 16
fitted Gauss locations:11 16
×10^8
3.5
3
2.5
2
1.5
1
0.5
0
y
0
5
10
15
20
25
x

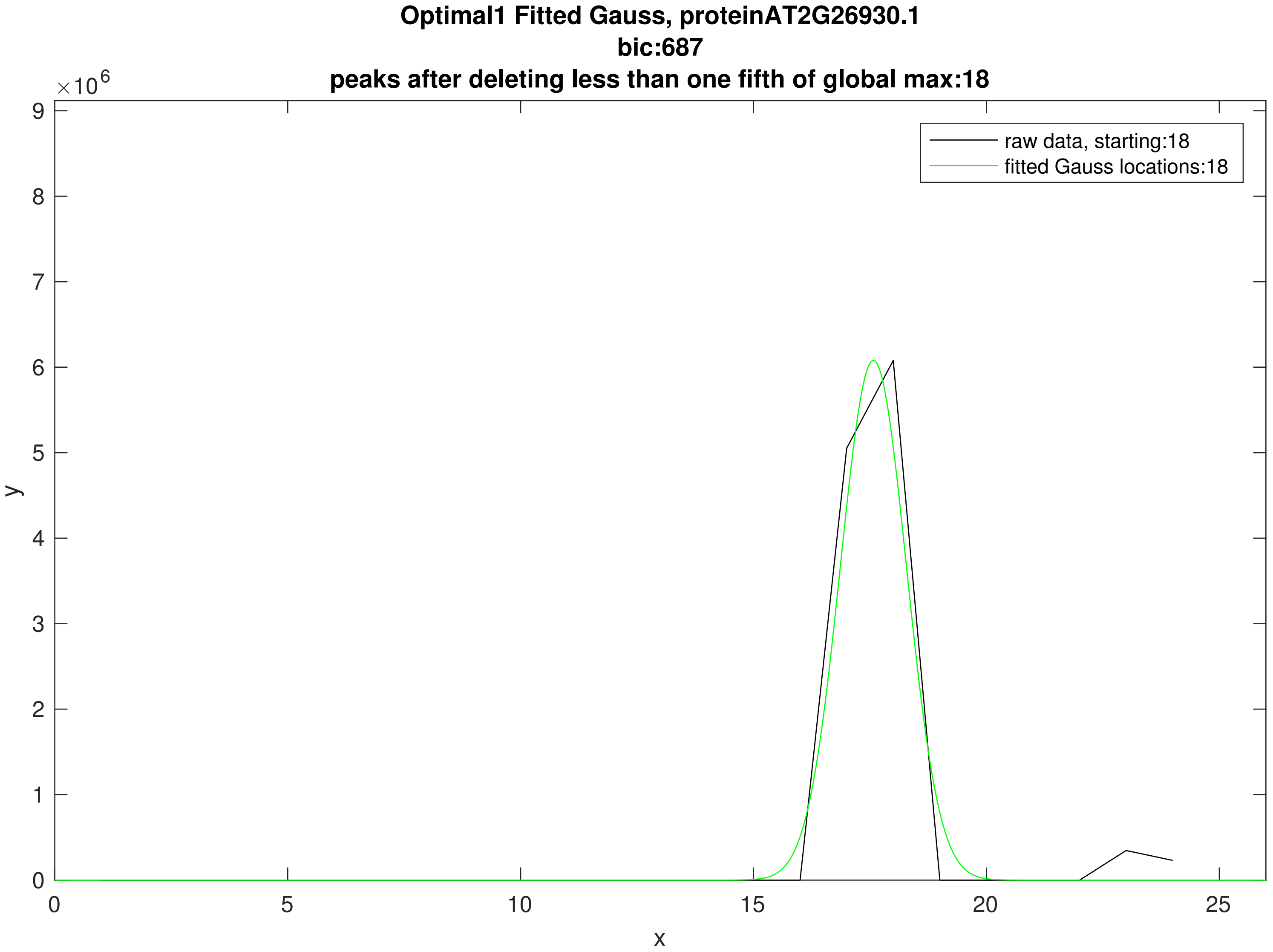

Optimal1 Fitted Gauss, proteinAT2G26930.1
bic:687
peaks after deleting less than one fifth of global max:18
×10^6
raw data, starting:18
fitted Gauss locations:18
y
x
0
1
2
3
4
5
6
7
8
9
0
5
10
15
20
25

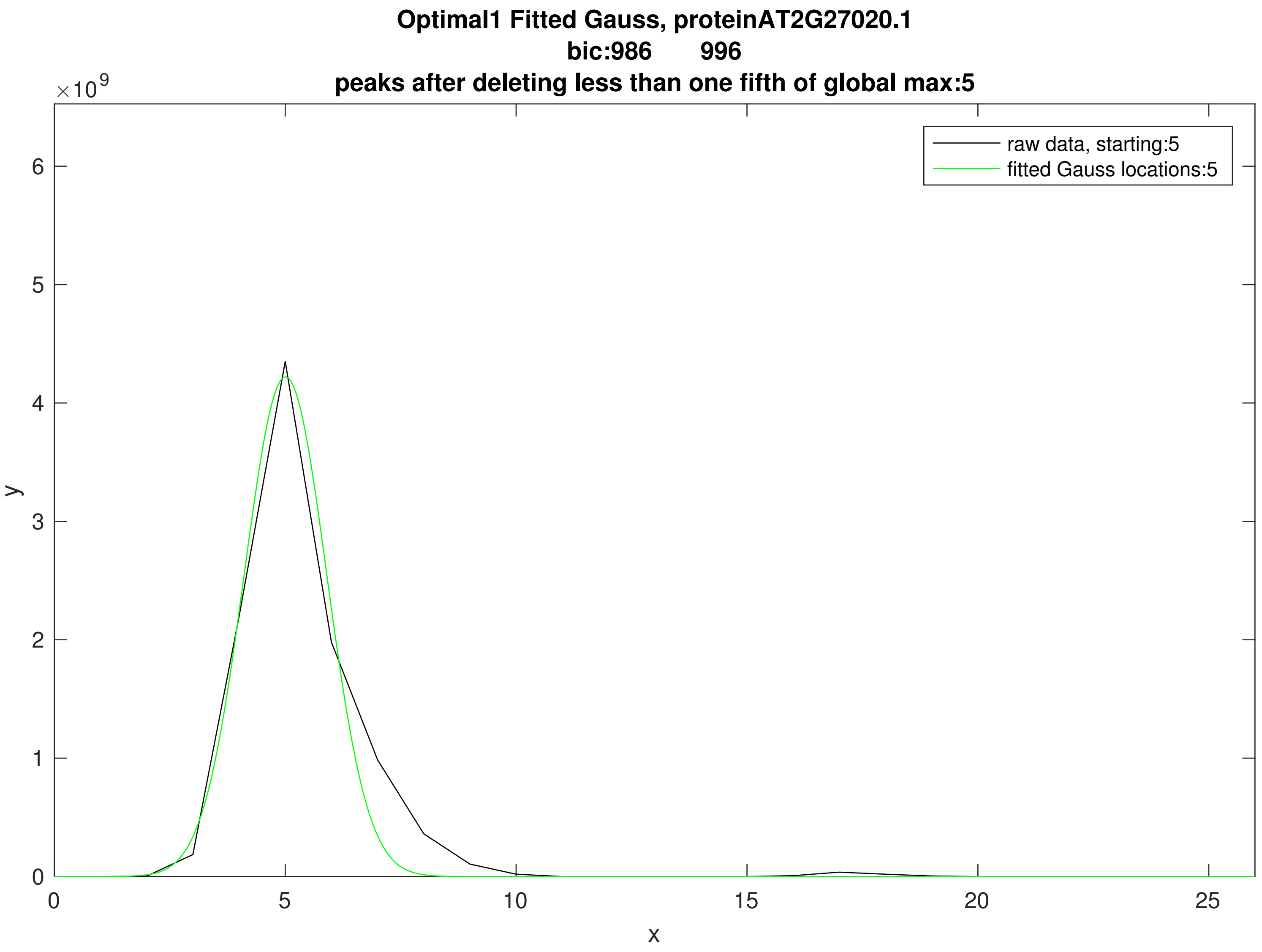

Optimal1 Fitted Gauss, proteinAT2G27020.1
bic:986 996
peaks after deleting less than one fifth of global max:5
×10^9
raw data, starting:5
fitted Gauss locations:5
y
x
0
1
2
3
4
5
6
0
5
10
15
20
25

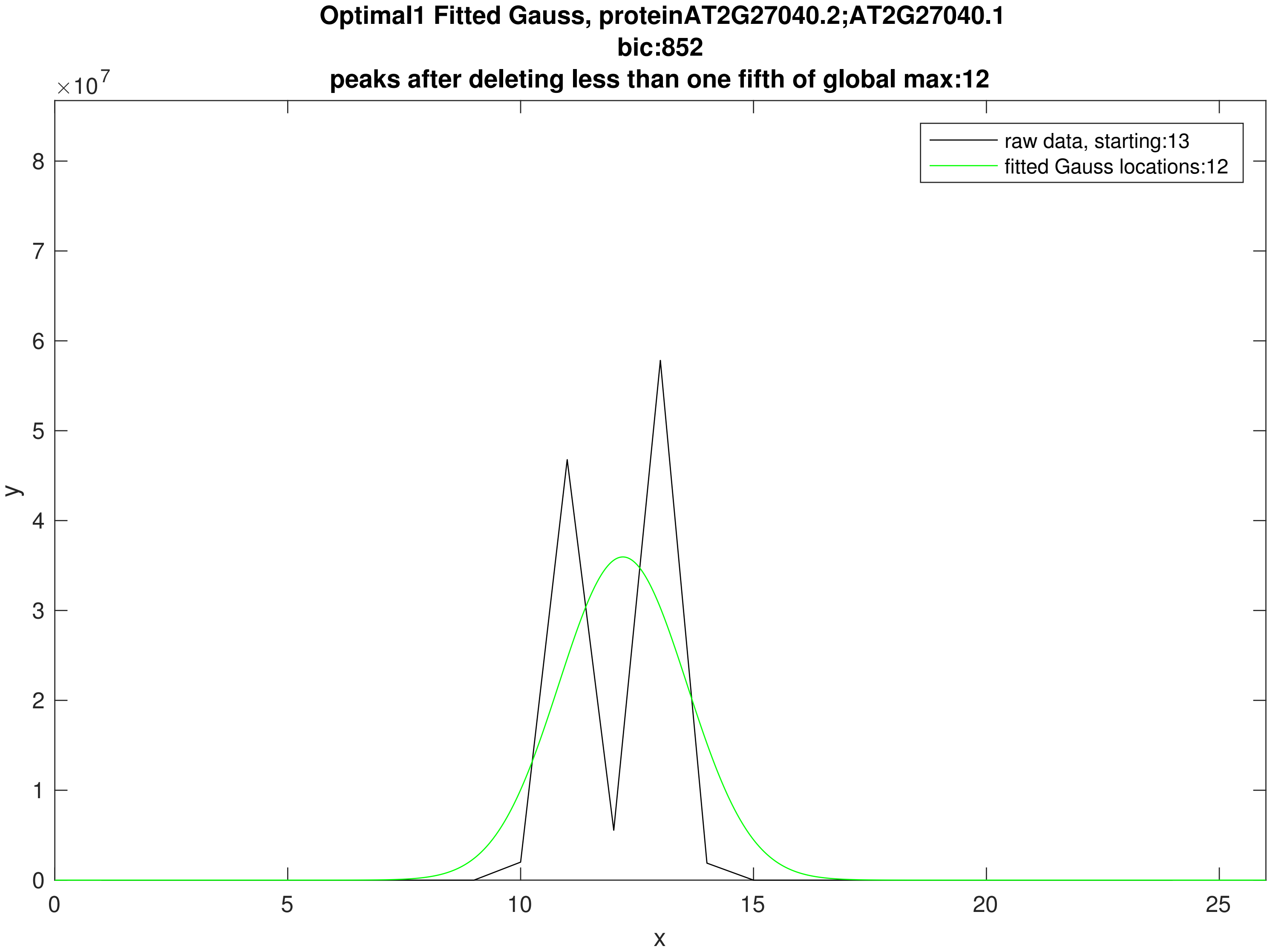

Optimal1 Fitted Gauss, proteinAT2G27040.2;AT2G27040.1
bic:852
peaks after deleting less than one fifth of global max:12
×10^7
raw data, starting:13
fitted Gauss locations:12
y
x
0
1
2
3
4
5
6
7
8
0
5
10
15
20
25

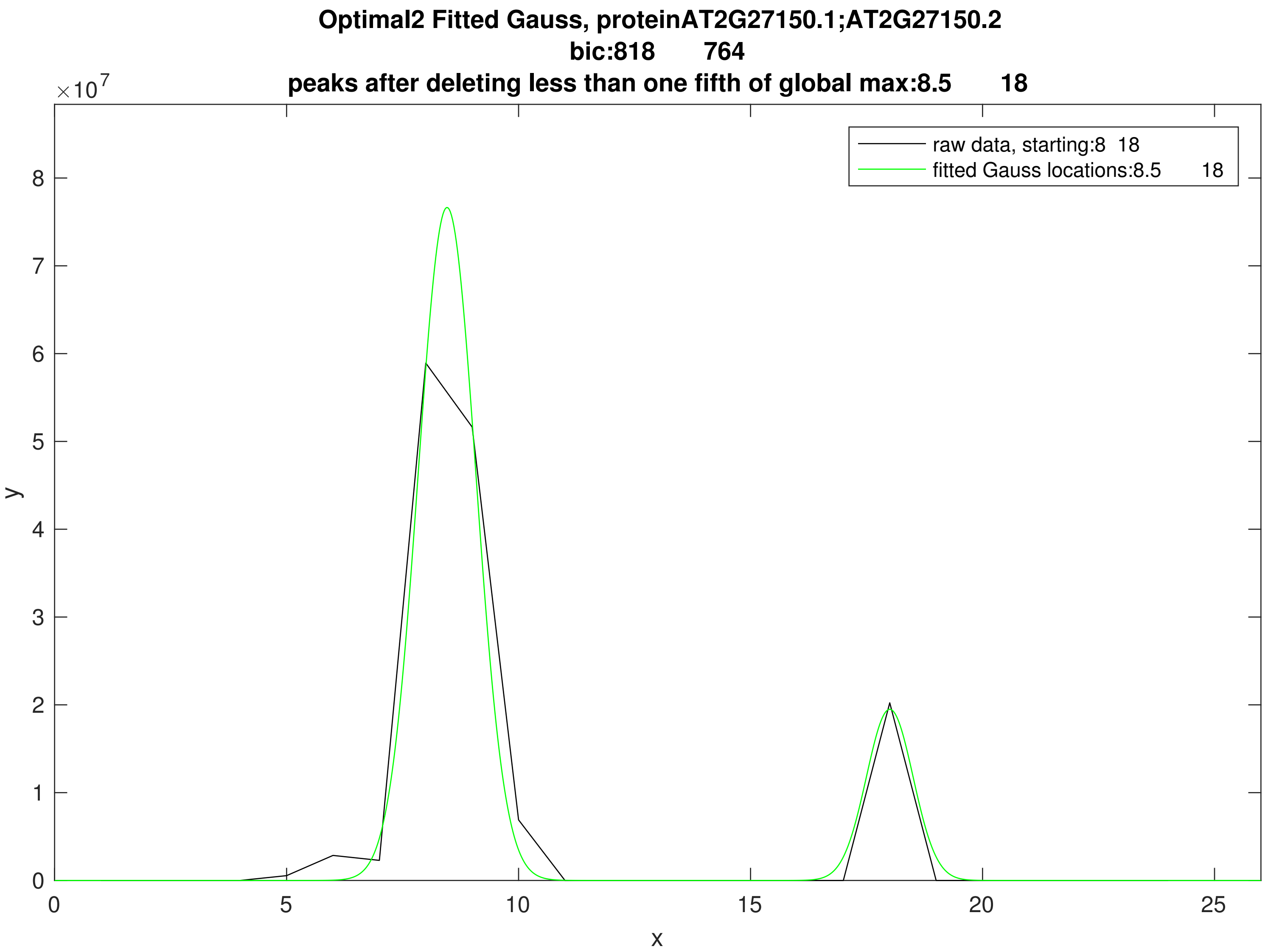

Optimal2 Fitted Gauss, proteinAT2G27150.1;AT2G27150.2
bic:818 764
peaks after deleting less than one fifth of global max:8.5 18
×10^7
raw data, starting:8 18
fitted Gauss locations:8.5 18
y
x
0
1
2
3
4
5
6
7
8
0
5
10
15
20
25

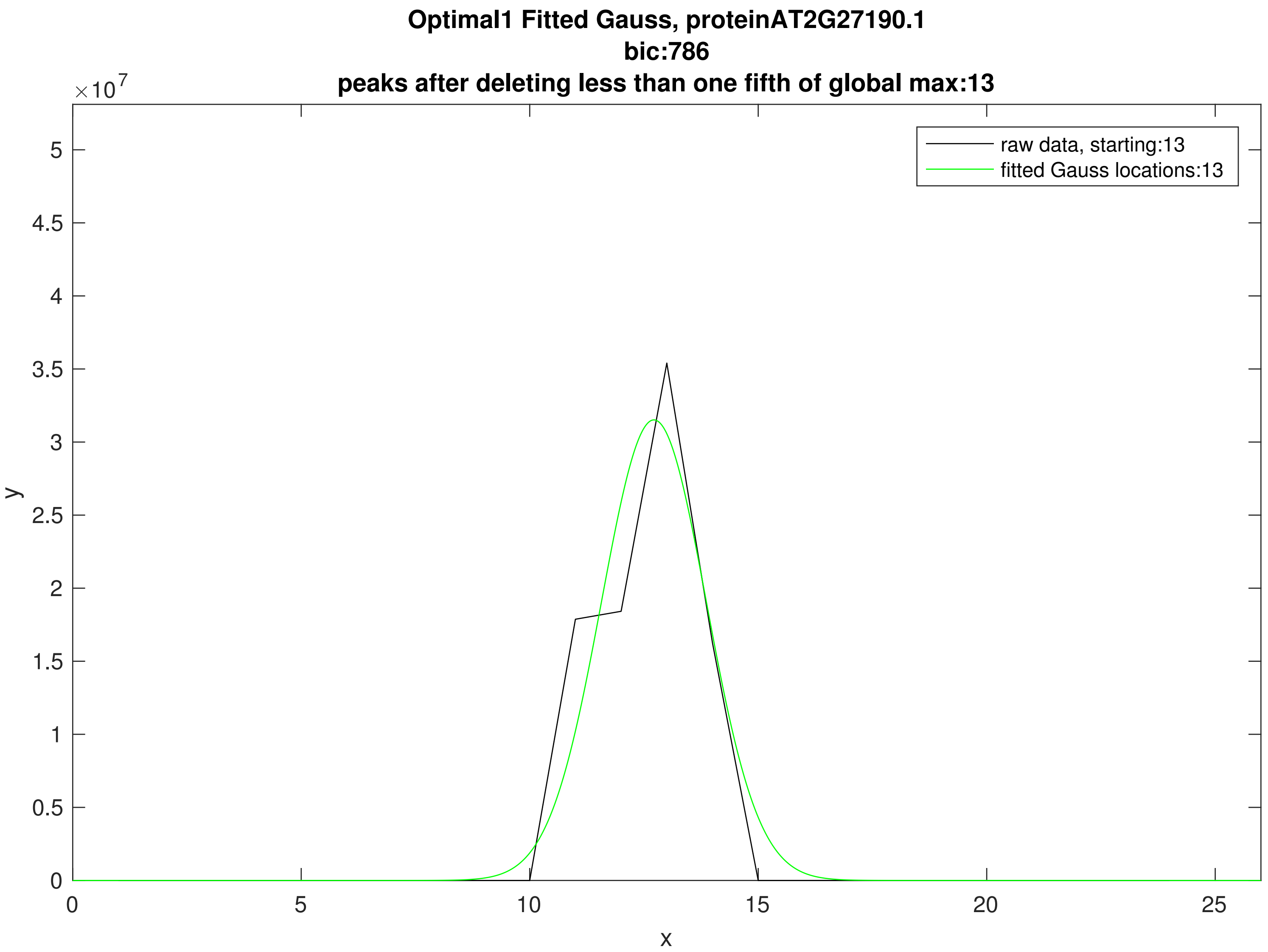
Optimal1 Fitted Gauss, proteinAT2G27190.1
bic:786
peaks after deleting less than one fifth of global max:13
×10^7
raw data, starting:13
fitted Gauss locations:13
y
x
0
0.5
1
1.5
2
2.5
3
3.5
4
4.5
5
0
5
10
15
20
25

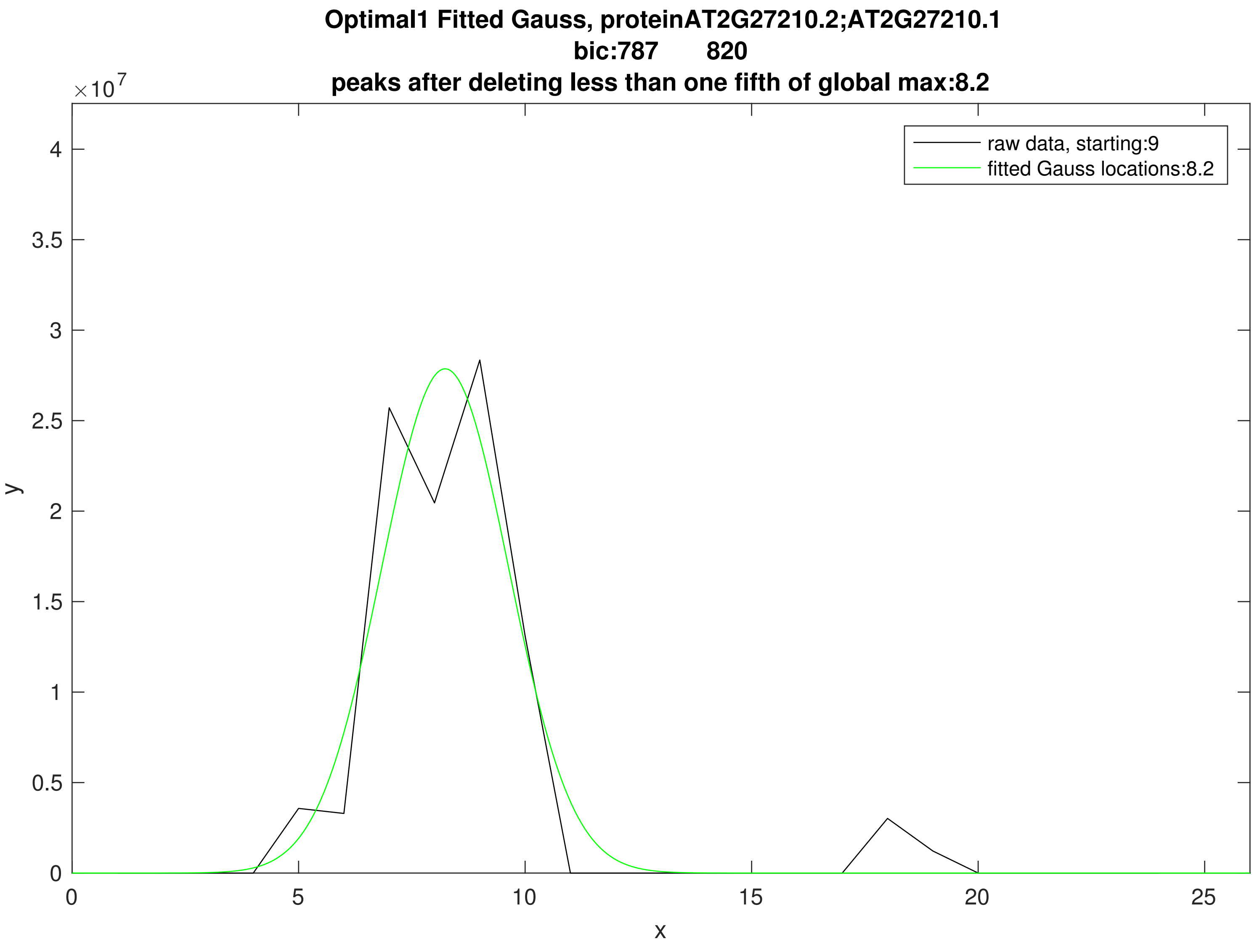
Optimal1 Fitted Gauss, proteinAT2G27210.2;AT2G27210.1
bic:787 820
peaks after deleting less than one fifth of global max:8.2
×10^7
raw data, starting:9
fitted Gauss locations:8.2
y
x
0
0.5
1
1.5
2
2.5
3
3.5
4
0
5
10
15
20
25

**Optimal1 Fitted Gauss, proteinAT2G27360.1**
**bic:850**
**peaks after deleting less than one fifth of global max:16**

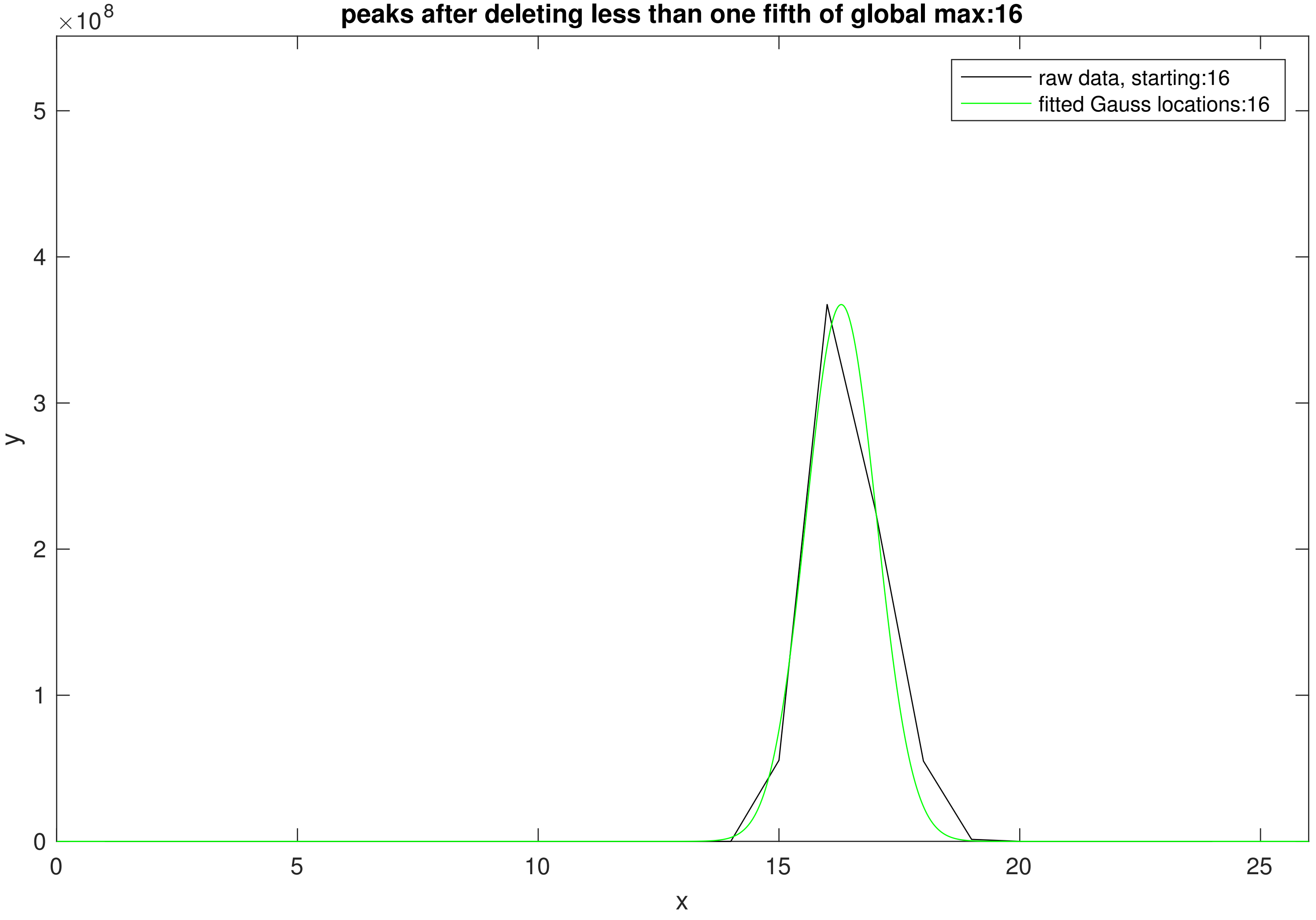

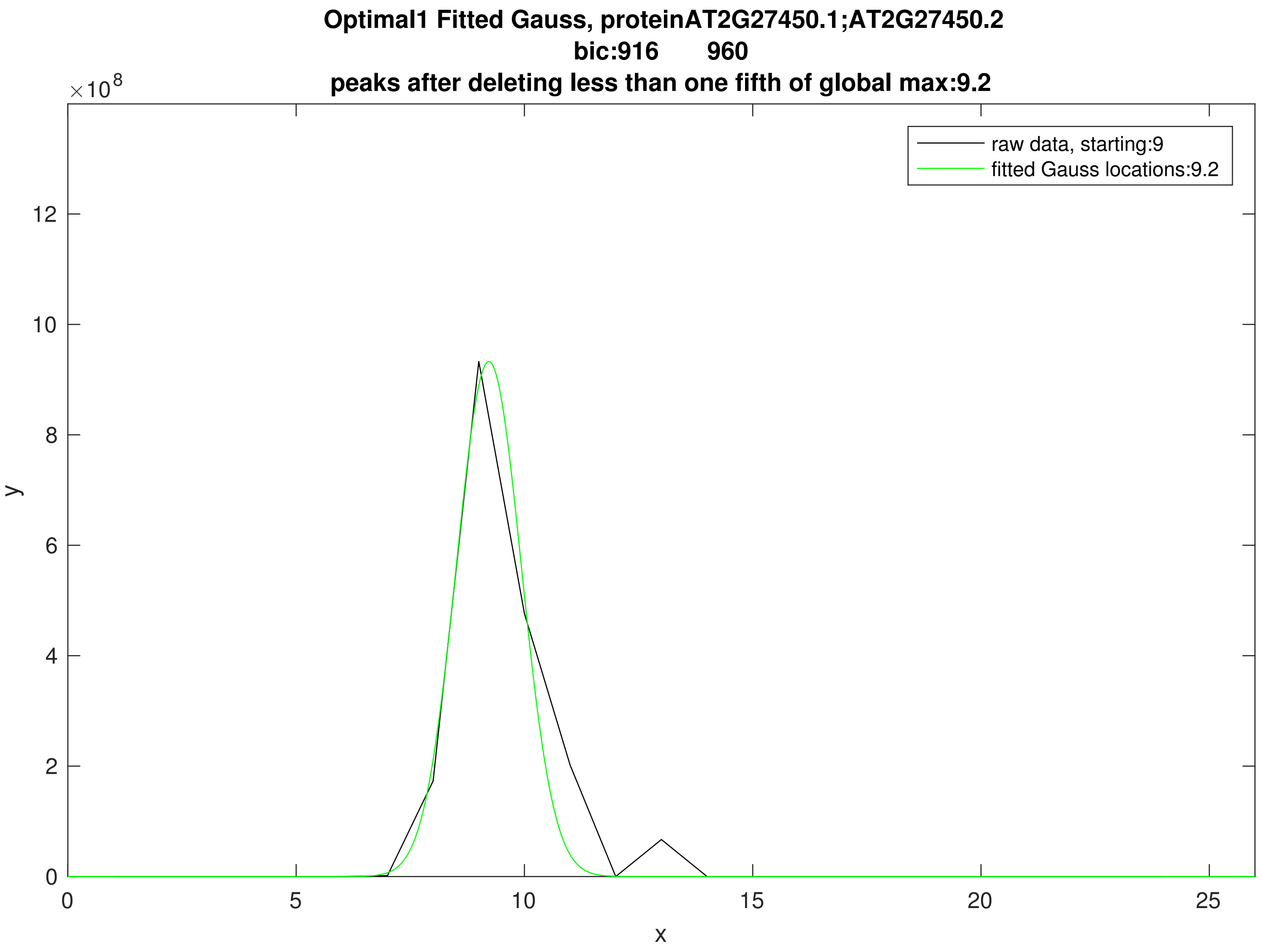

Optimal1 Fitted Gauss, proteinAT2G27450.1;AT2G27450.2
bic:916 960
peaks after deleting less than one fifth of global max:9.2
×10^8
raw data, starting:9
fitted Gauss locations:9.2
y
x
0
2
4
6
8
10
12
0
5
10
15
20
25

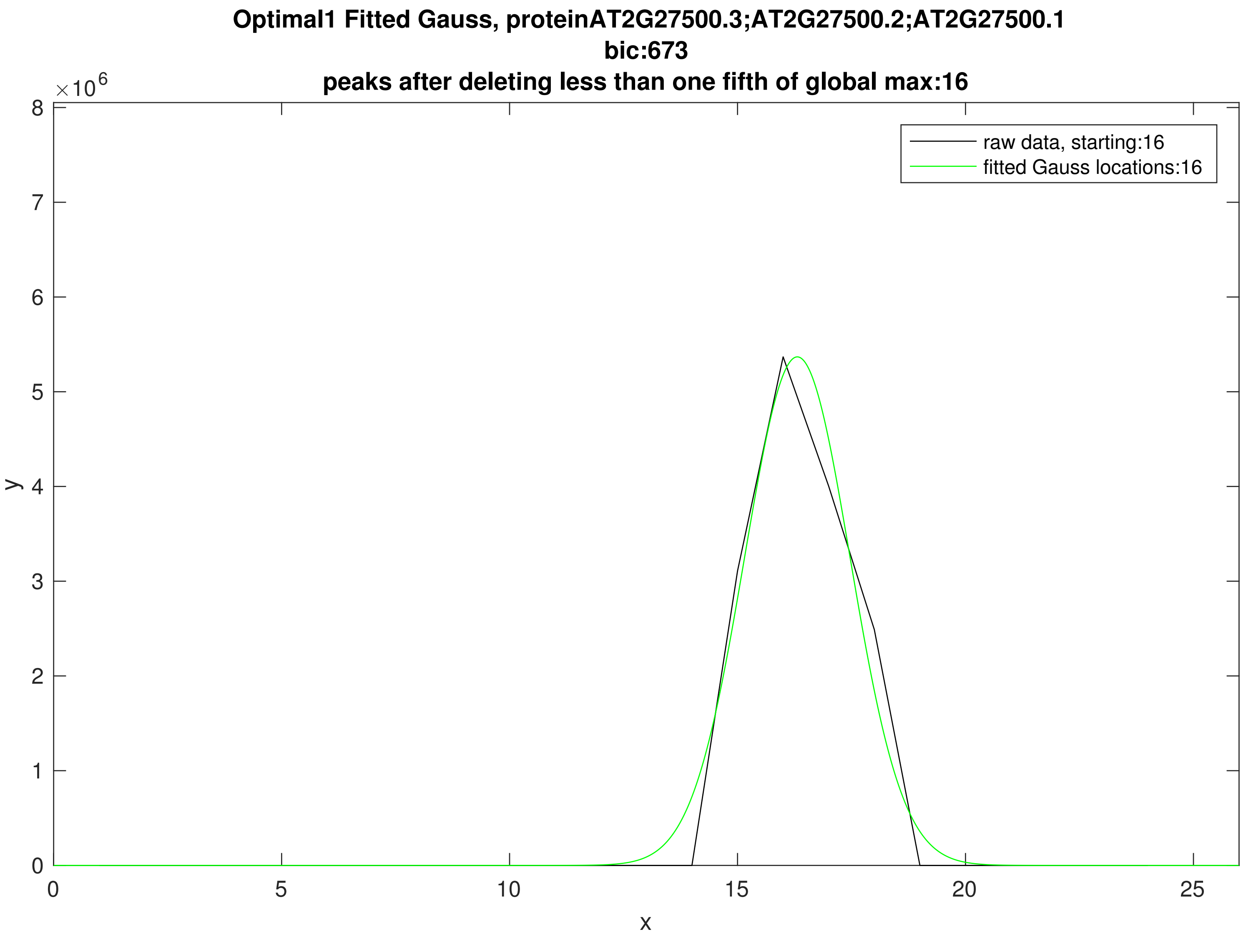
Optimal1 Fitted Gauss, proteinAT2G27500.3;AT2G27500.2;AT2G27500.1
bic:673
peaks after deleting less than one fifth of global max:16
×10^6
raw data, starting:16
fitted Gauss locations:16
y
x
0
1
2
3
4
5
6
7
8
0
5
10
15
20
25

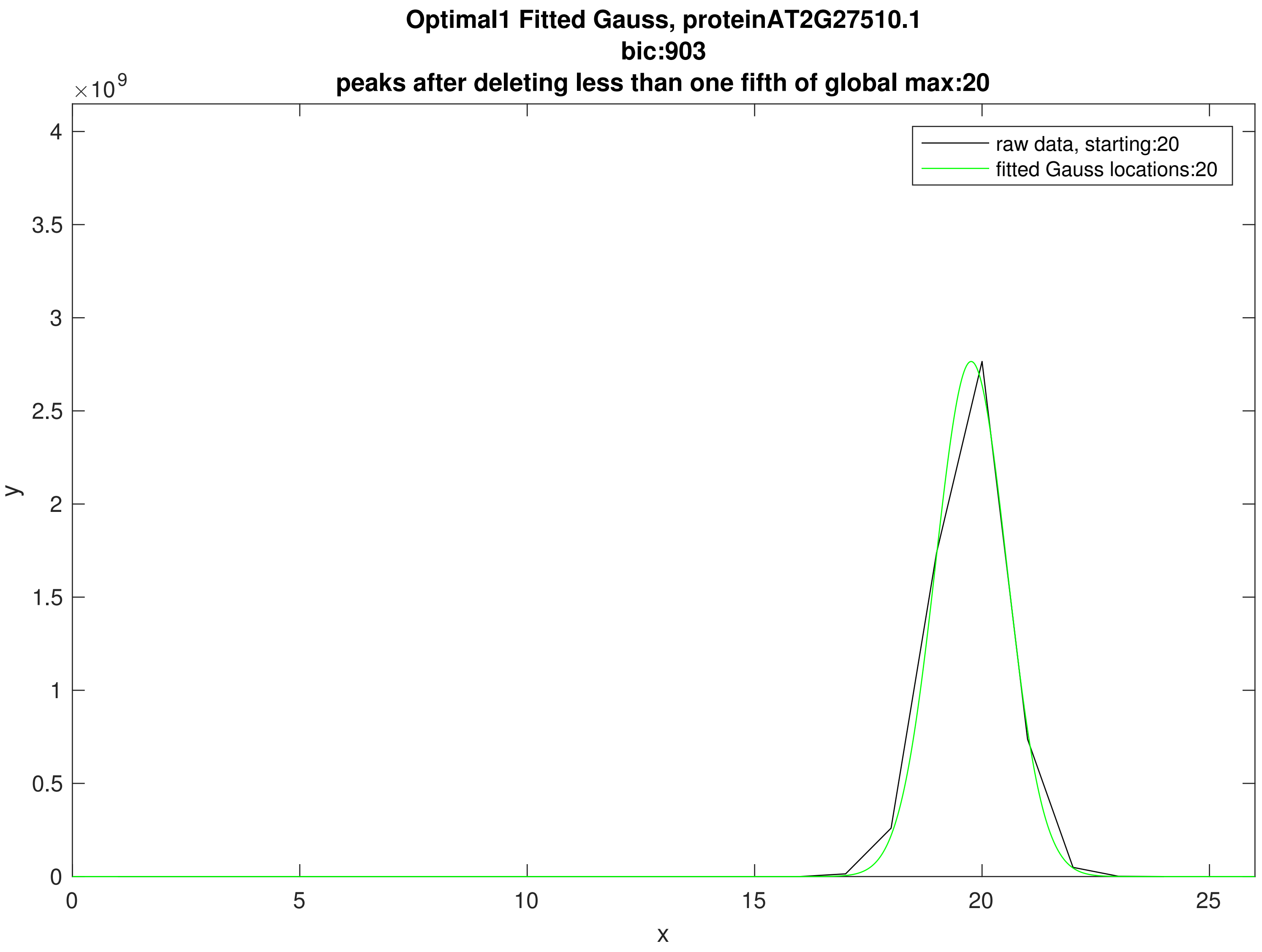

Optimal1 Fitted Gauss, proteinAT2G27510.1
bic:903
peaks after deleting less than one fifth of global max:20
×10^9
4
3.5
3
2.5
2
1.5
1
0.5
0
y
0
5
10
15
20
25
x
raw data, starting:20
fitted Gauss locations:20

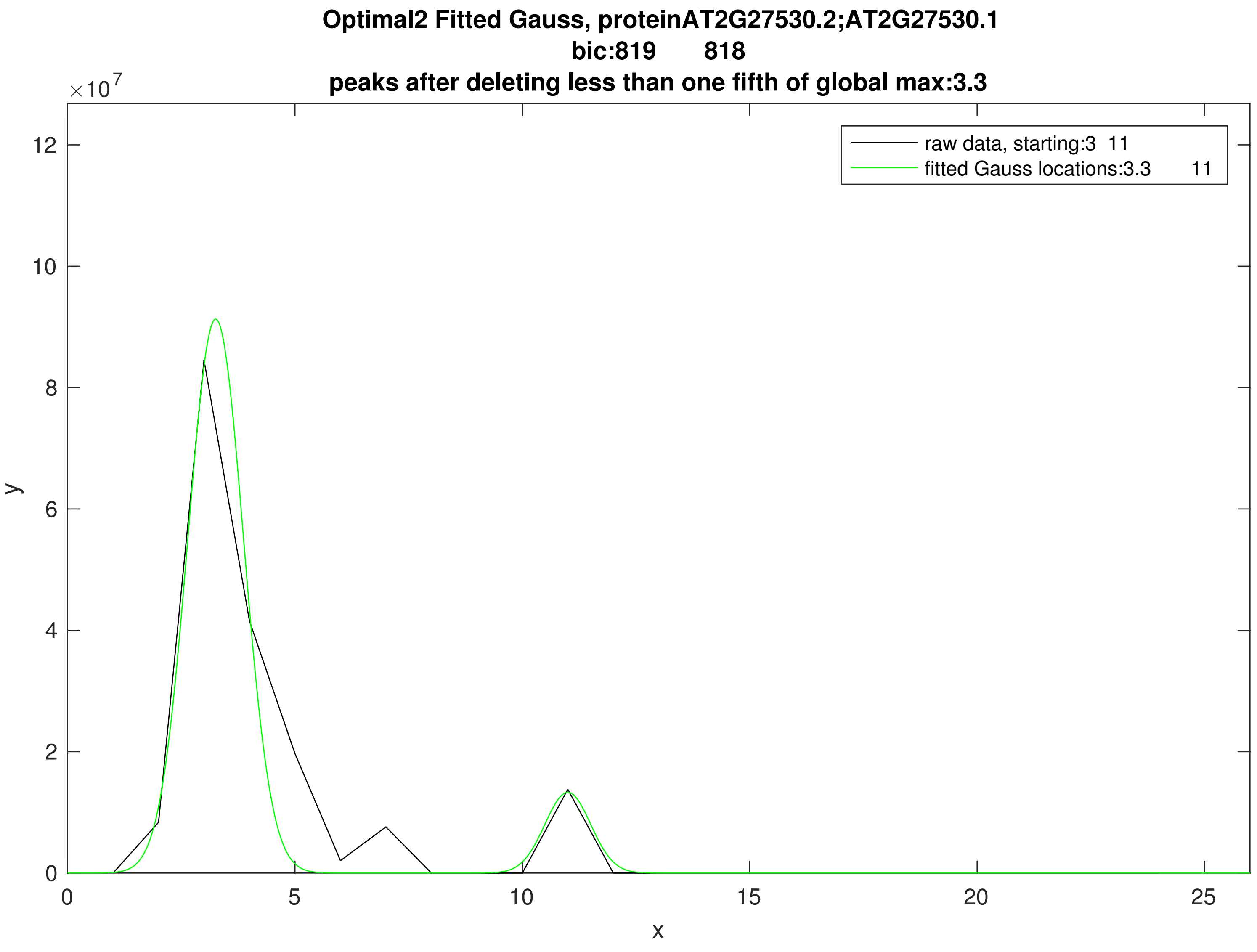

Optimal2 Fitted Gauss, proteinAT2G27530.2;AT2G27530.1
bic:819 818
peaks after deleting less than one fifth of global max:3.3
×10^7
raw data, starting:3 11
fitted Gauss locations:3.3 11
y
x
0
2
4
6
8
10
12
0
5
10
15
20
25

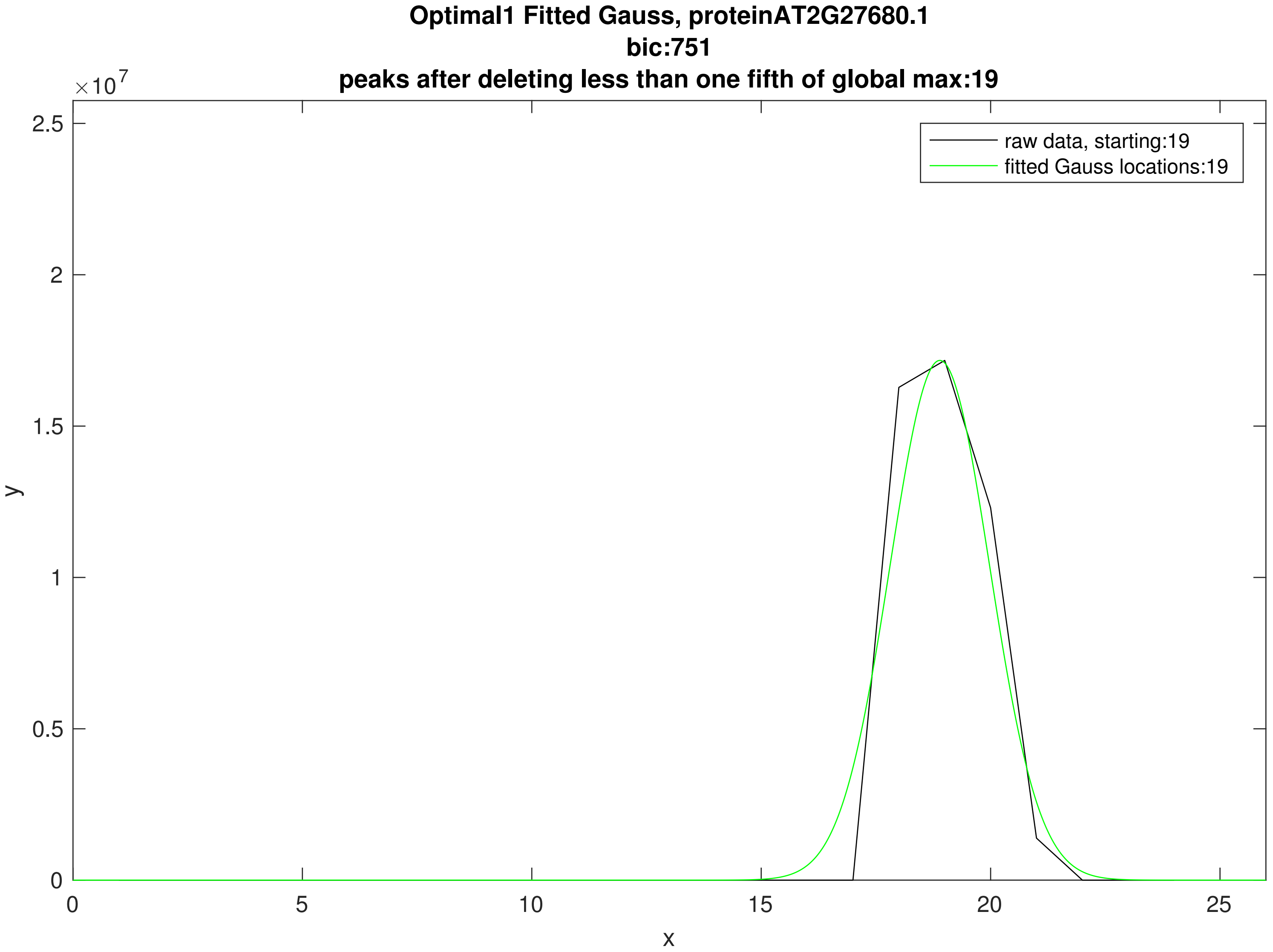

Optimal1 Fitted Gauss, proteinAT2G27680.1
bic:751
peaks after deleting less than one fifth of global max:19
×10^7
2.5
2
1.5
1
0.5
0
y
0
5
10
15
20
25
x
raw data, starting:19
fitted Gauss locations:19

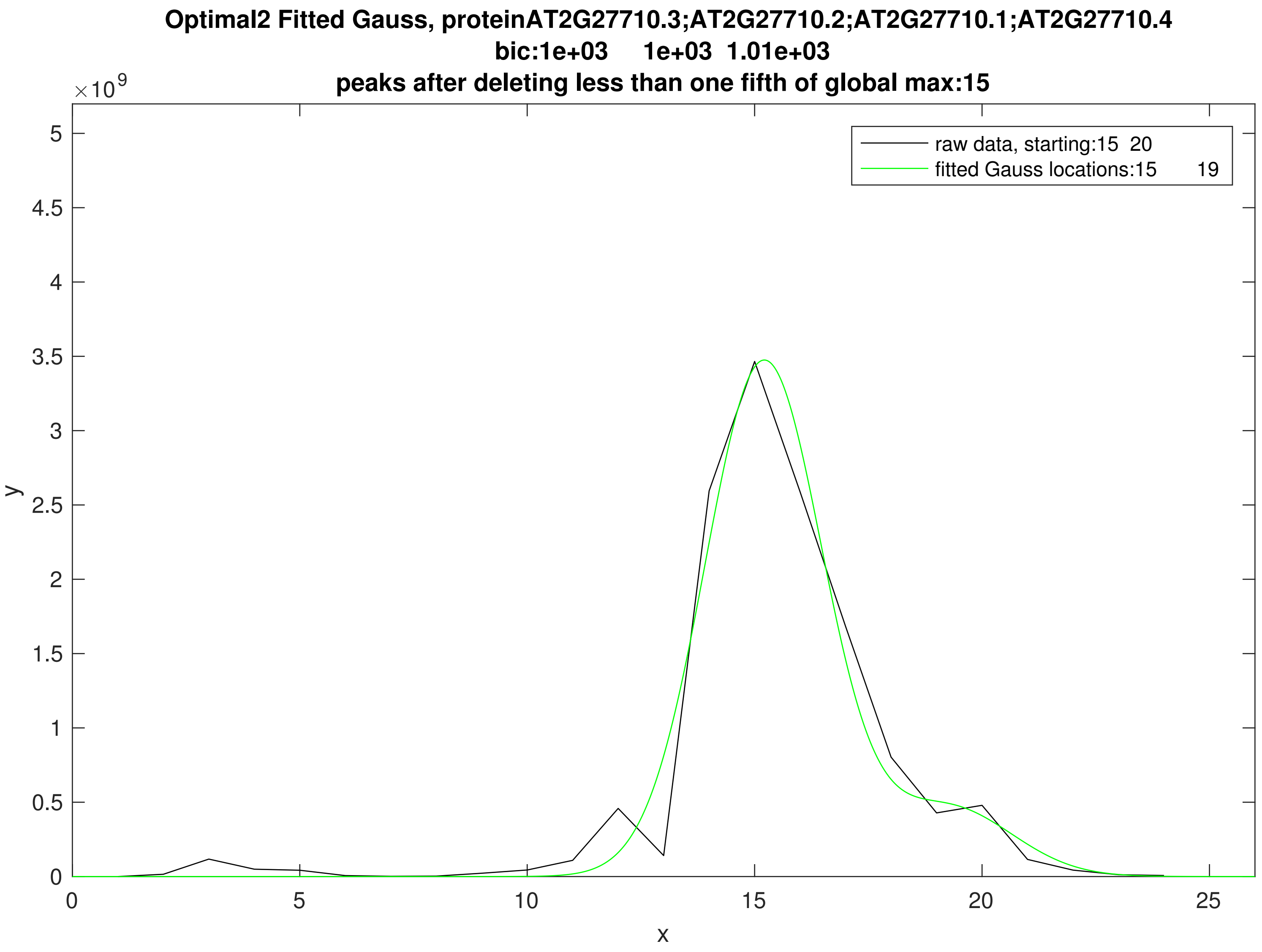

Optimal2 Fitted Gauss, proteinAT2G27710.3;AT2G27710.2;AT2G27710.1;AT2G27710.4
bic:1e+03 1e+03 1.01e+03
peaks after deleting less than one fifth of global max:15
×10^9
raw data, starting:15 20
fitted Gauss locations:15 19
y
x
0
0.5
1
1.5
2
2.5
3
3.5
4
4.5
5
0
5
10
15
20
25

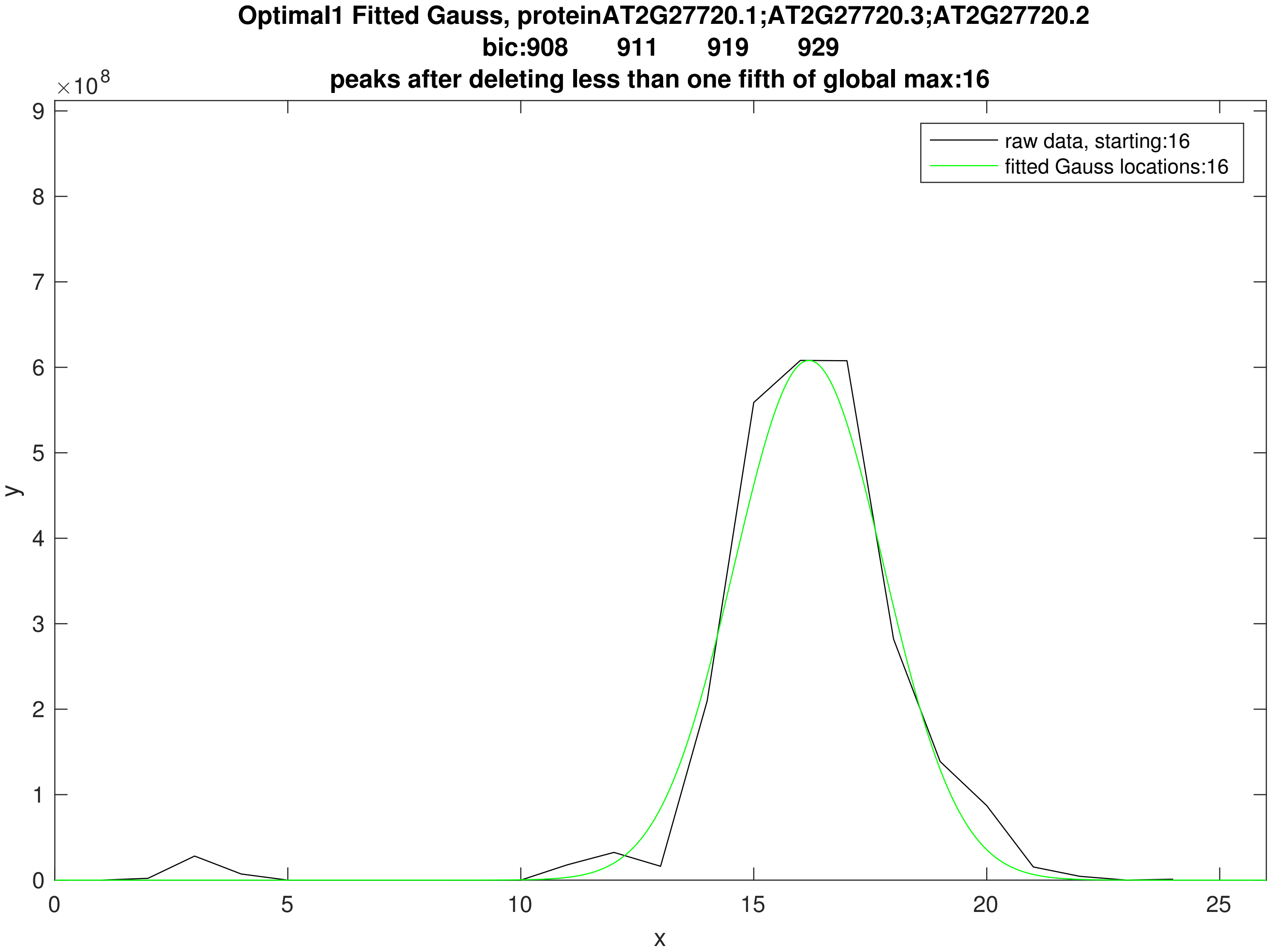

Optimal1 Fitted Gauss, proteinAT2G27720.1;AT2G27720.3;AT2G27720.2
bic:908 911 919 929
peaks after deleting less than one fifth of global max:16
×10^8
raw data, starting:16
fitted Gauss locations:16
y
x
0
1
2
3
4
5
6
7
8
9
10
15
20
25

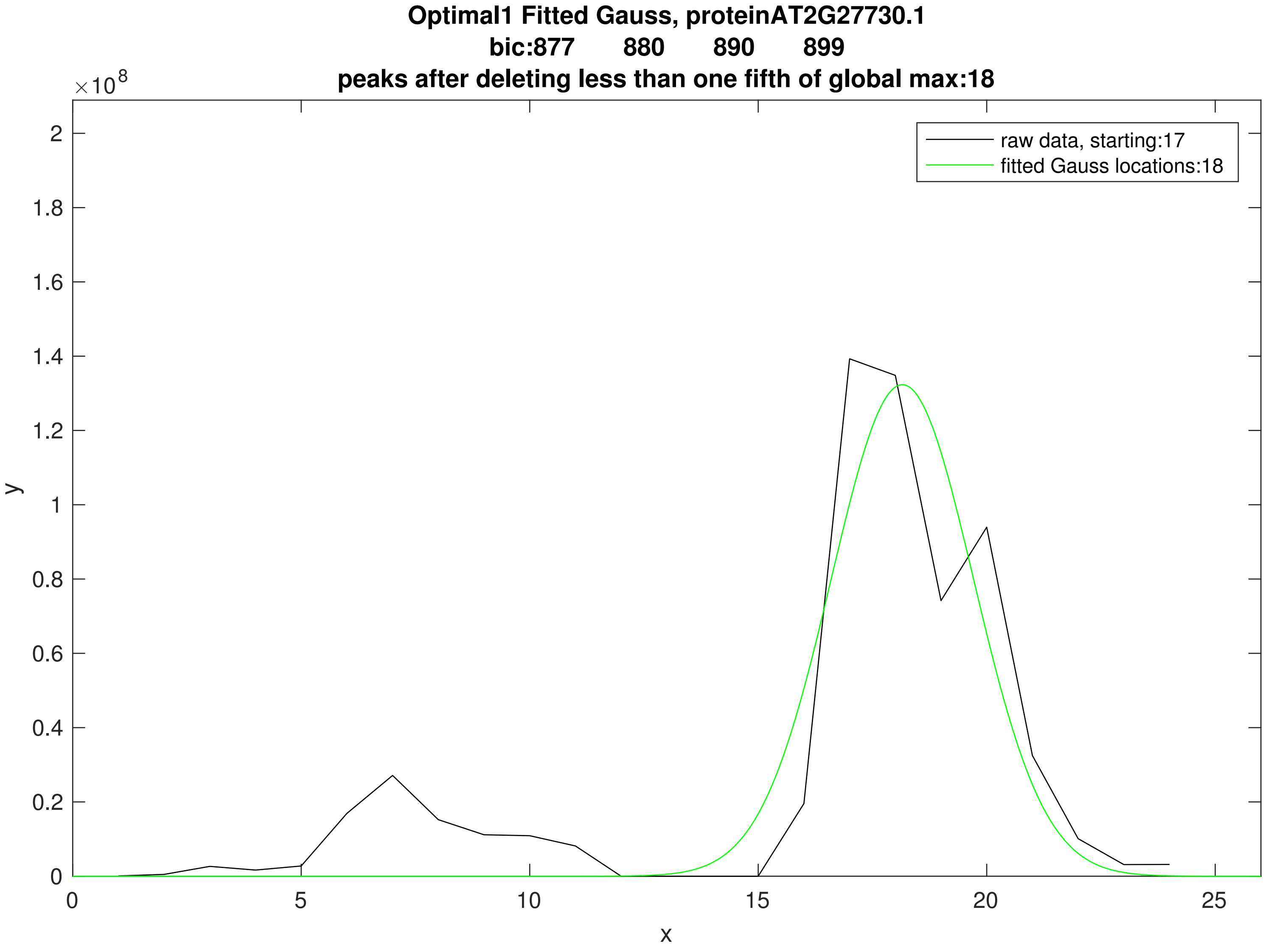
Optimal1 Fitted Gauss, proteinAT2G27730.1
bic:877 880 890 899
peaks after deleting less than one fifth of global max:18
×10^8
raw data, starting:17
fitted Gauss locations:18
y
x
0
0.2
0.4
0.6
0.8
1
1.2
1.4
1.6
1.8
2
0
5
10
15
20
25

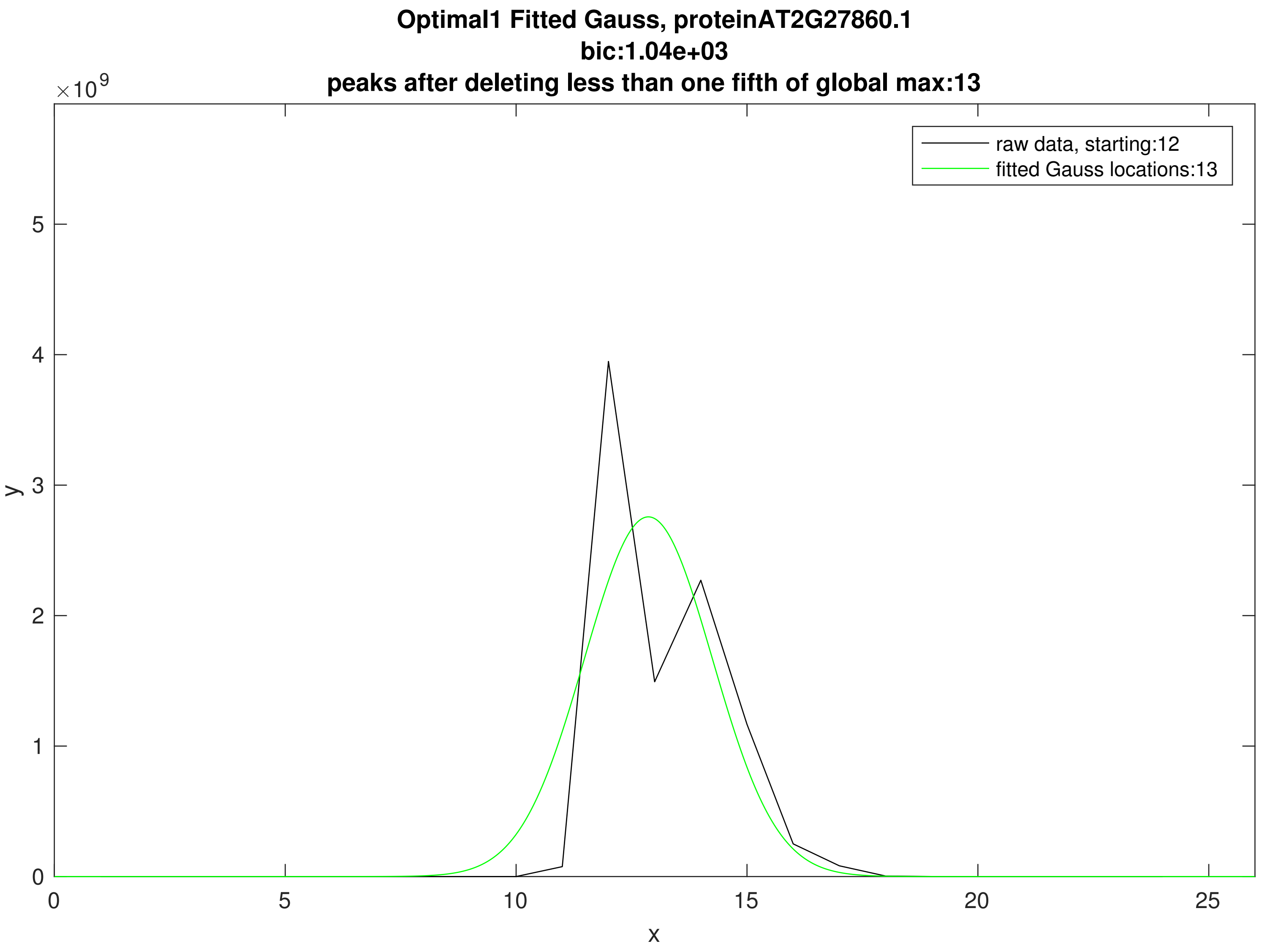
Optimal1 Fitted Gauss, proteinAT2G27860.1
bic:1.04e+03
peaks after deleting less than one fifth of global max:13
×10^9
raw data, starting:12
fitted Gauss locations:13
y
x
0
1
2
3
4
5
0
5
10
15
20
25

**Optimal1 Fitted Gauss, proteinAT2G27920.1;AT2G27920.2;AT2G27920.3**
**bic:790**
**peaks after deleting less than one fifth of global max:12**

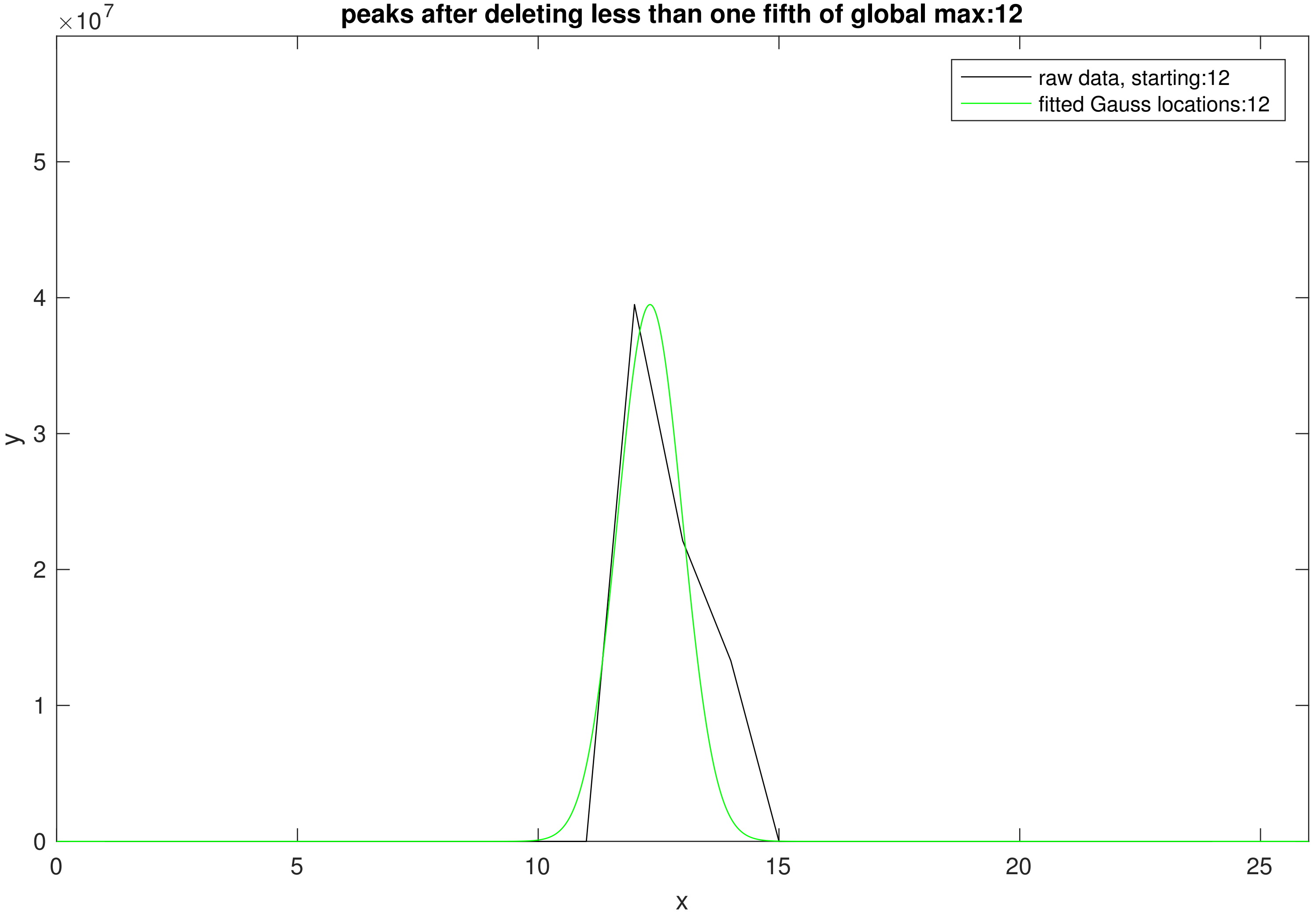

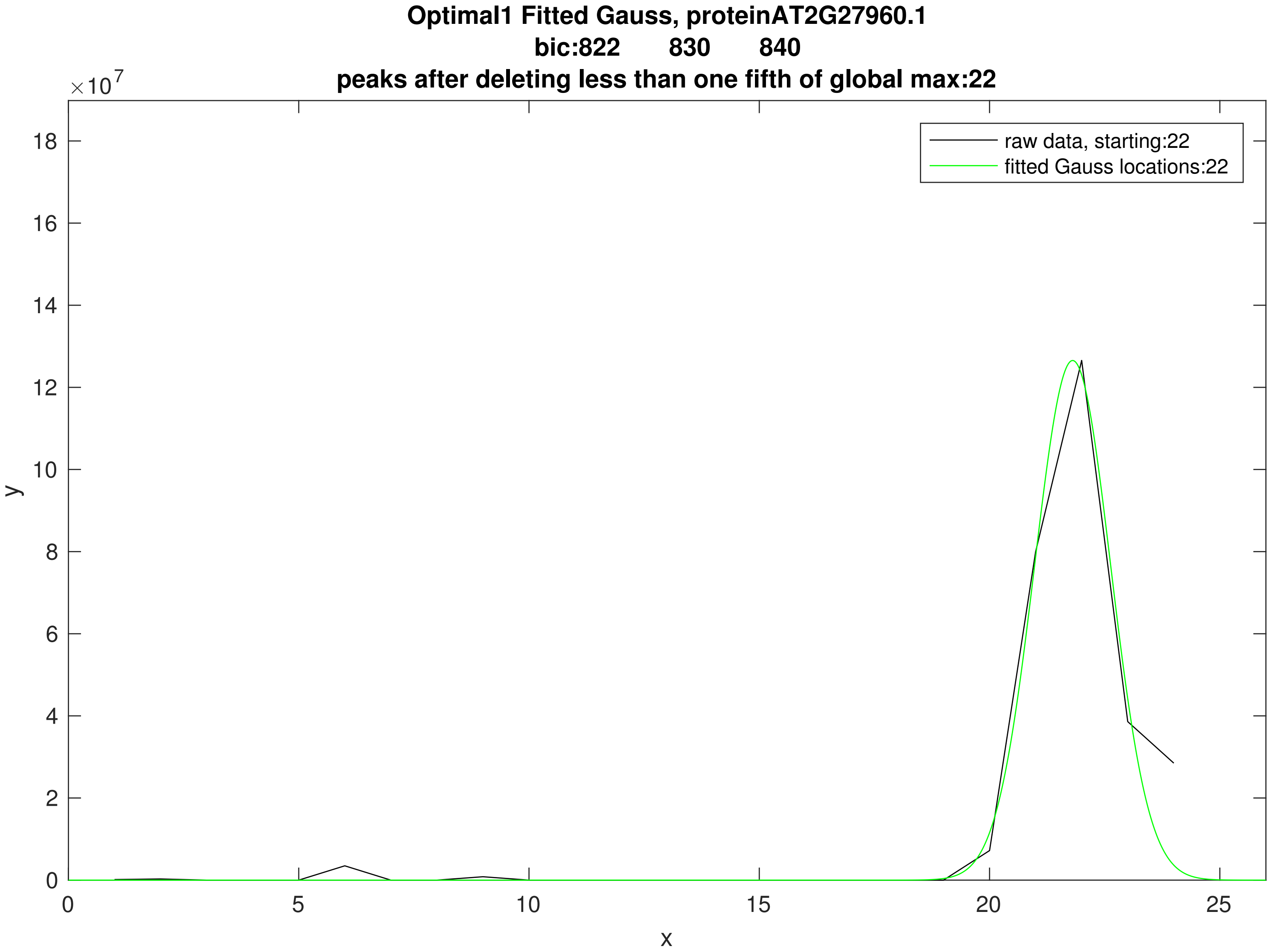
Optimal1 Fitted Gauss, proteinAT2G27960.1
bic:822 830 840
peaks after deleting less than one fifth of global max:22
×10^7
18
16
14
12
10
8
6
4
2
0
y
0
5
10
15
20
25
x
raw data, starting:22
fitted Gauss locations:22

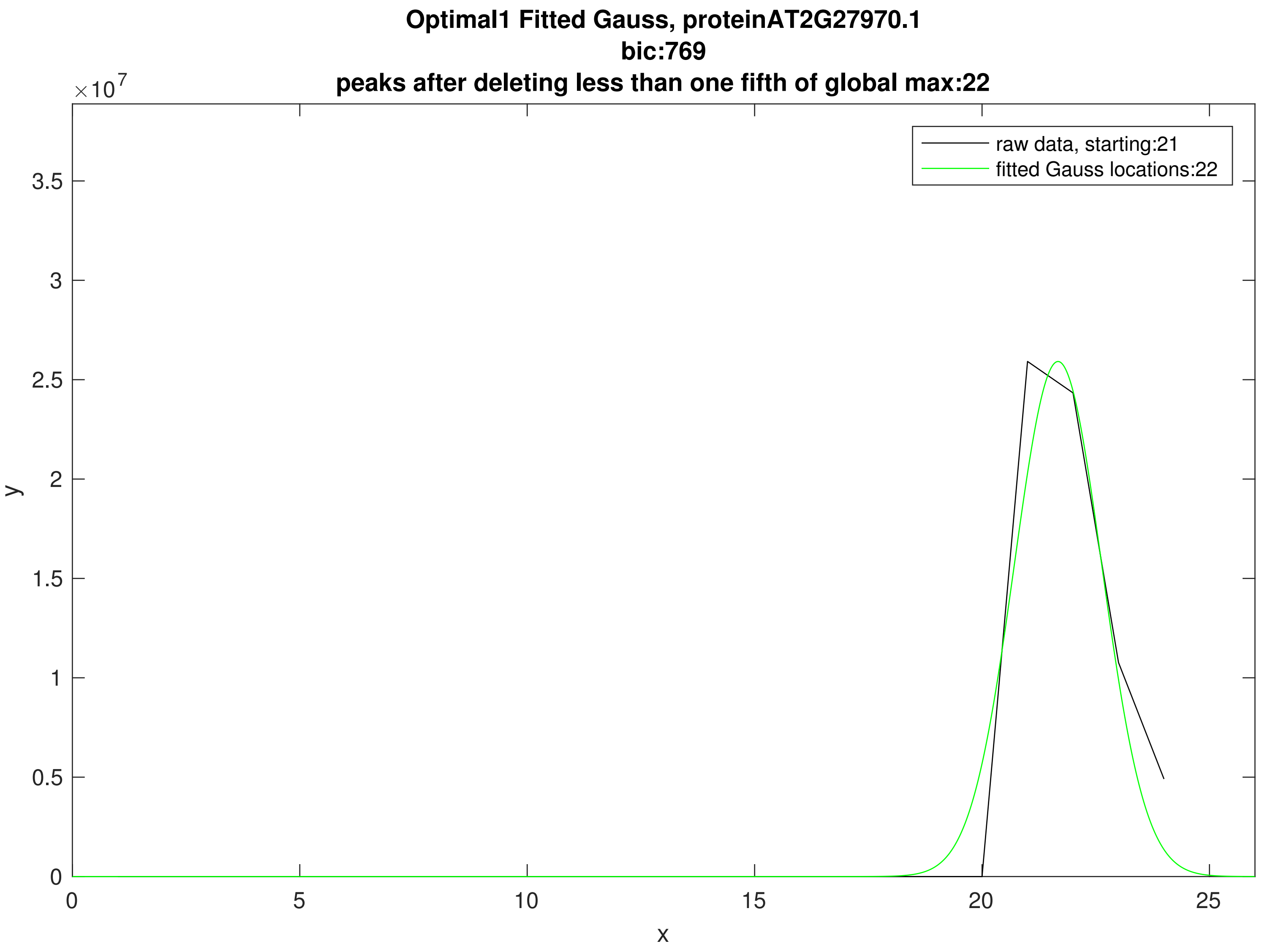

Optimal1 Fitted Gauss, proteinAT2G27970.1
bic:769
peaks after deleting less than one fifth of global max:22
×10^7
raw data, starting:21
fitted Gauss locations:22
y
x
0
0.5
1
1.5
2
2.5
3
3.5
0
5
10
15
20
25

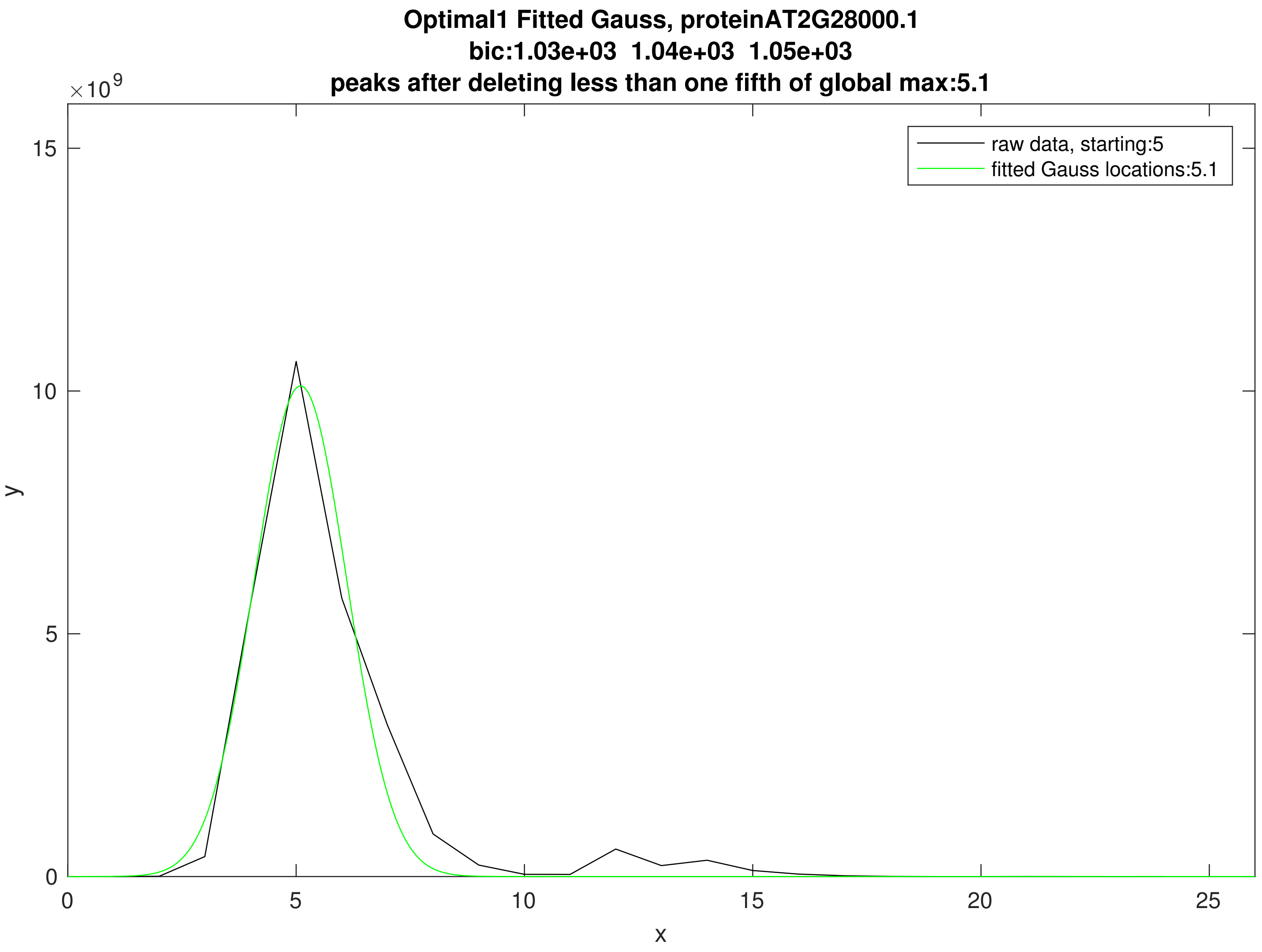

Optimal1 Fitted Gauss, proteinAT2G28000.1
bic:1.03e+03 1.04e+03 1.05e+03
peaks after deleting less than one fifth of global max:5.1
×10^9
raw data, starting:5
fitted Gauss locations:5.1
15
10
5
0
y
0
5
10
15
20
25
x

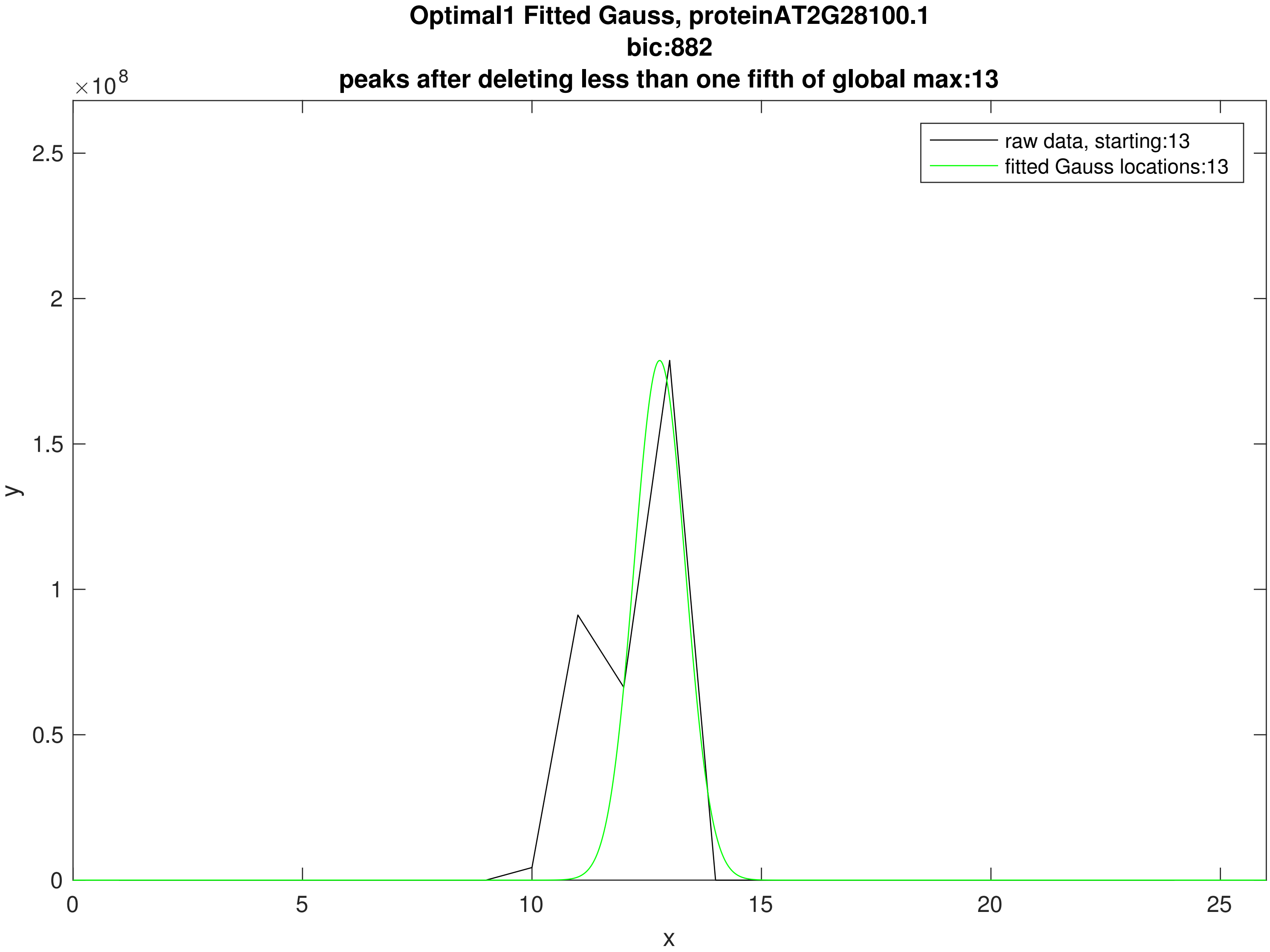

Optimal1 Fitted Gauss, proteinAT2G28100.1
bic:882
peaks after deleting less than one fifth of global max:13
×10^8
2.5
2
1.5
1
0.5
0
y
0
5
10
15
20
25
x
raw data, starting:13
fitted Gauss locations:13

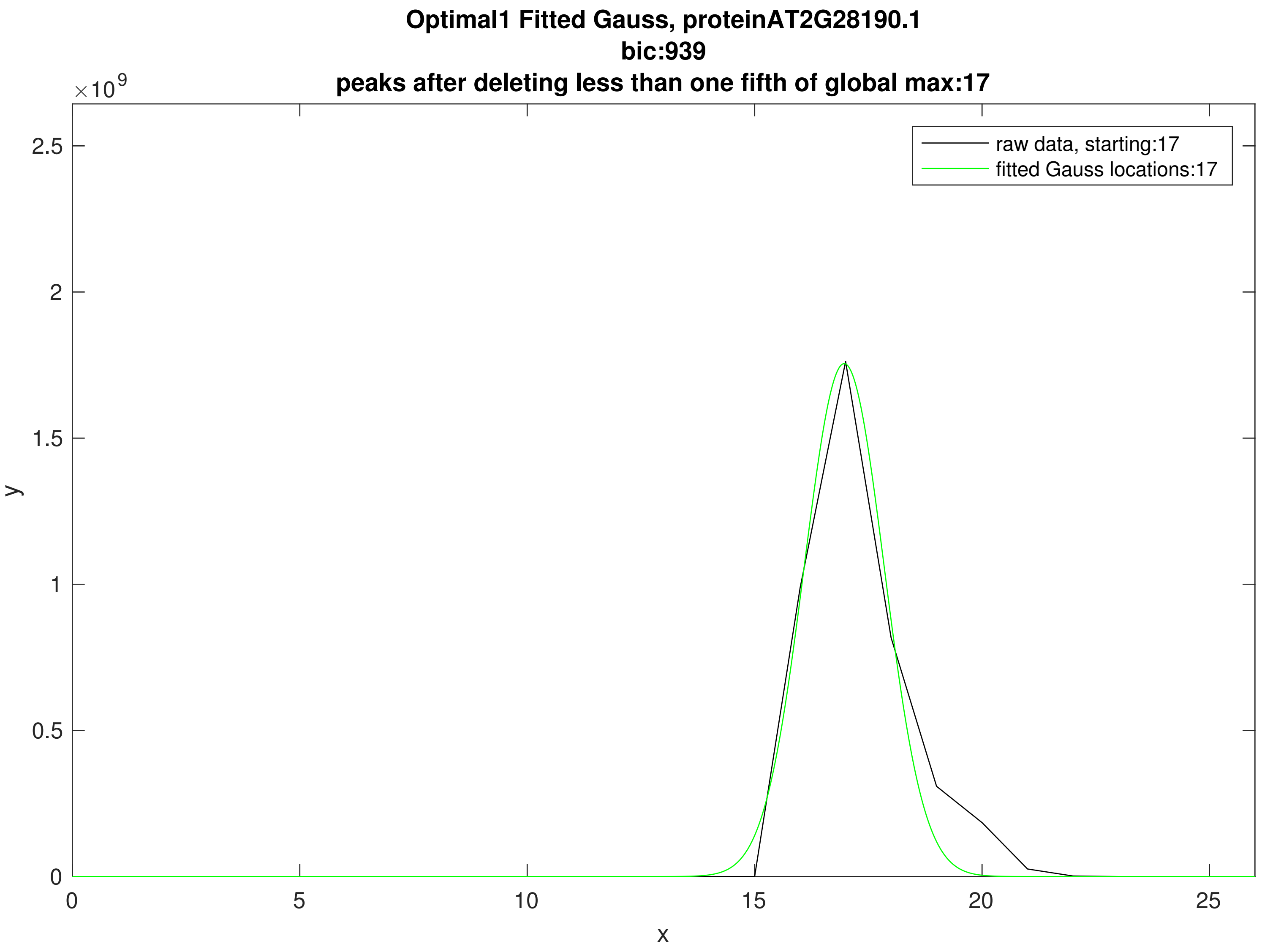

Optimal1 Fitted Gauss, proteinAT2G28190.1
bic:939
peaks after deleting less than one fifth of global max:17
×10^9
2.5
2
1.5
1
0.5
0
y
0
5
10
15
20
25
x
raw data, starting:17
fitted Gauss locations:17

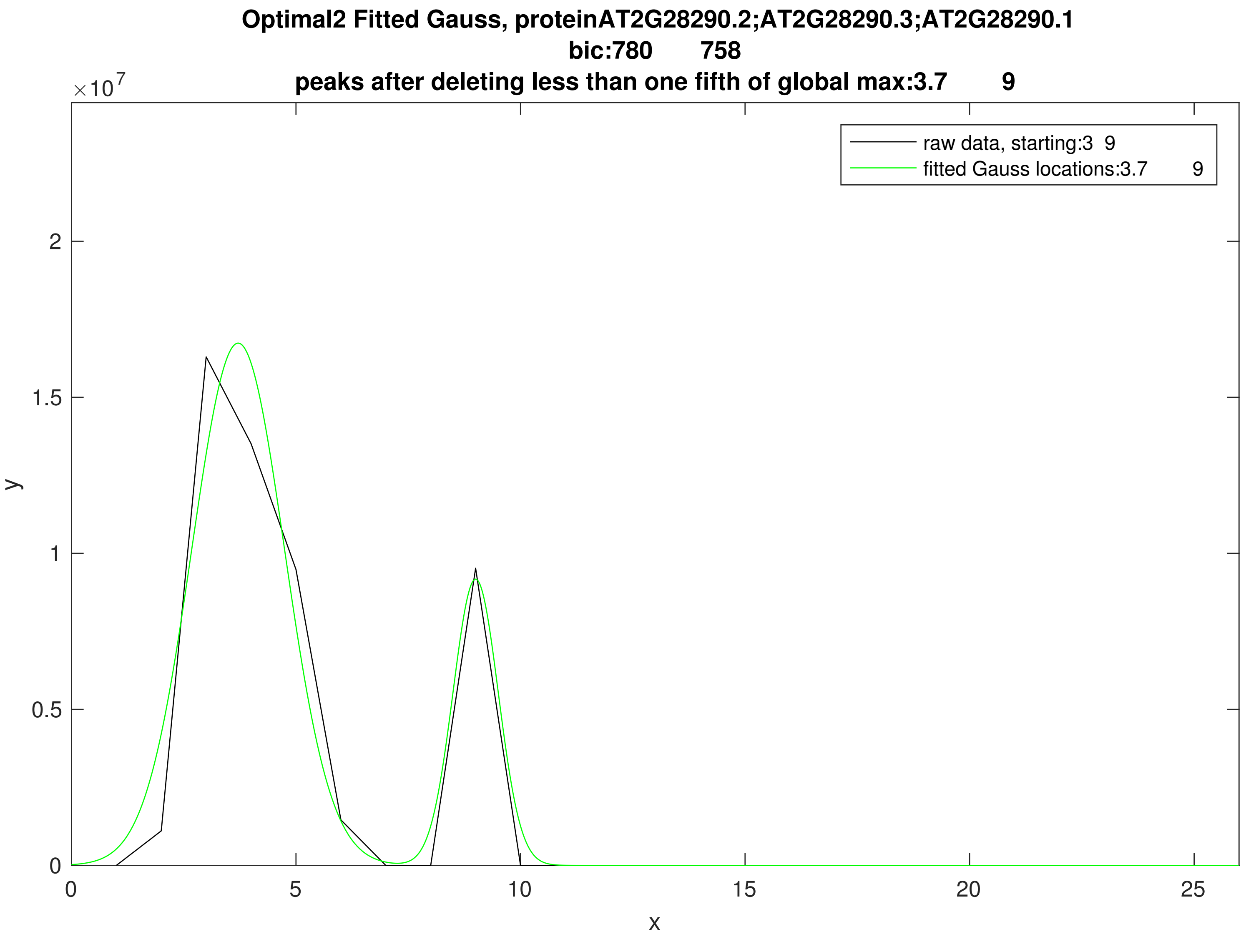

Optimal2 Fitted Gauss, proteinAT2G28290.2;AT2G28290.3;AT2G28290.1
bic:780 758
peaks after deleting less than one fifth of global max:3.7 9
×10^7
raw data, starting:3 9
fitted Gauss locations:3.7 9
y
x
0
0.5
1
1.5
2
5
10
15
20
25

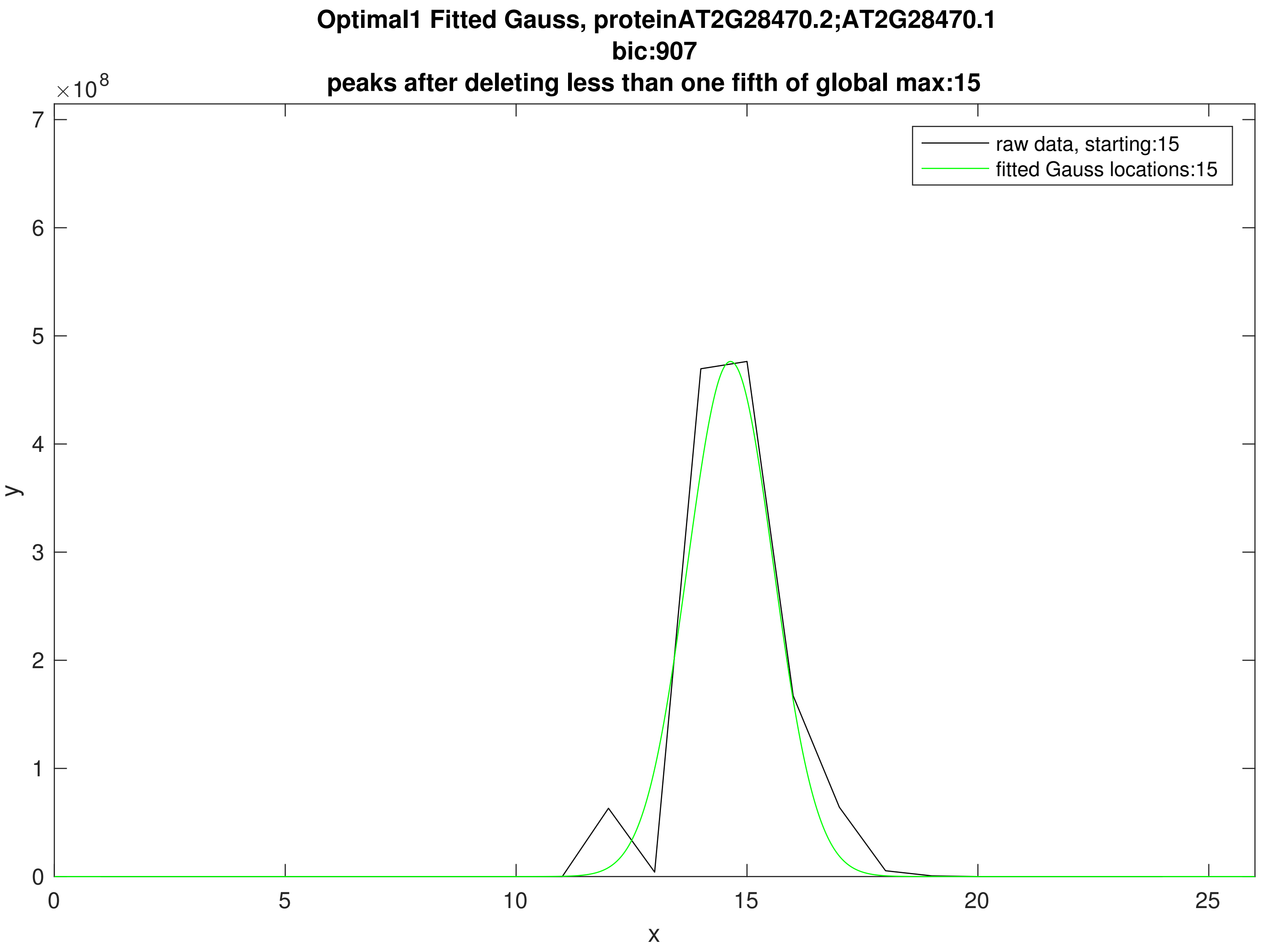
Optimal1 Fitted Gauss, proteinAT2G28470.2;AT2G28470.1
bic:907
peaks after deleting less than one fifth of global max:15
×10^8
raw data, starting:15
fitted Gauss locations:15
y
x
0
1
2
3
4
5
6
7
0
5
10
15
20
25

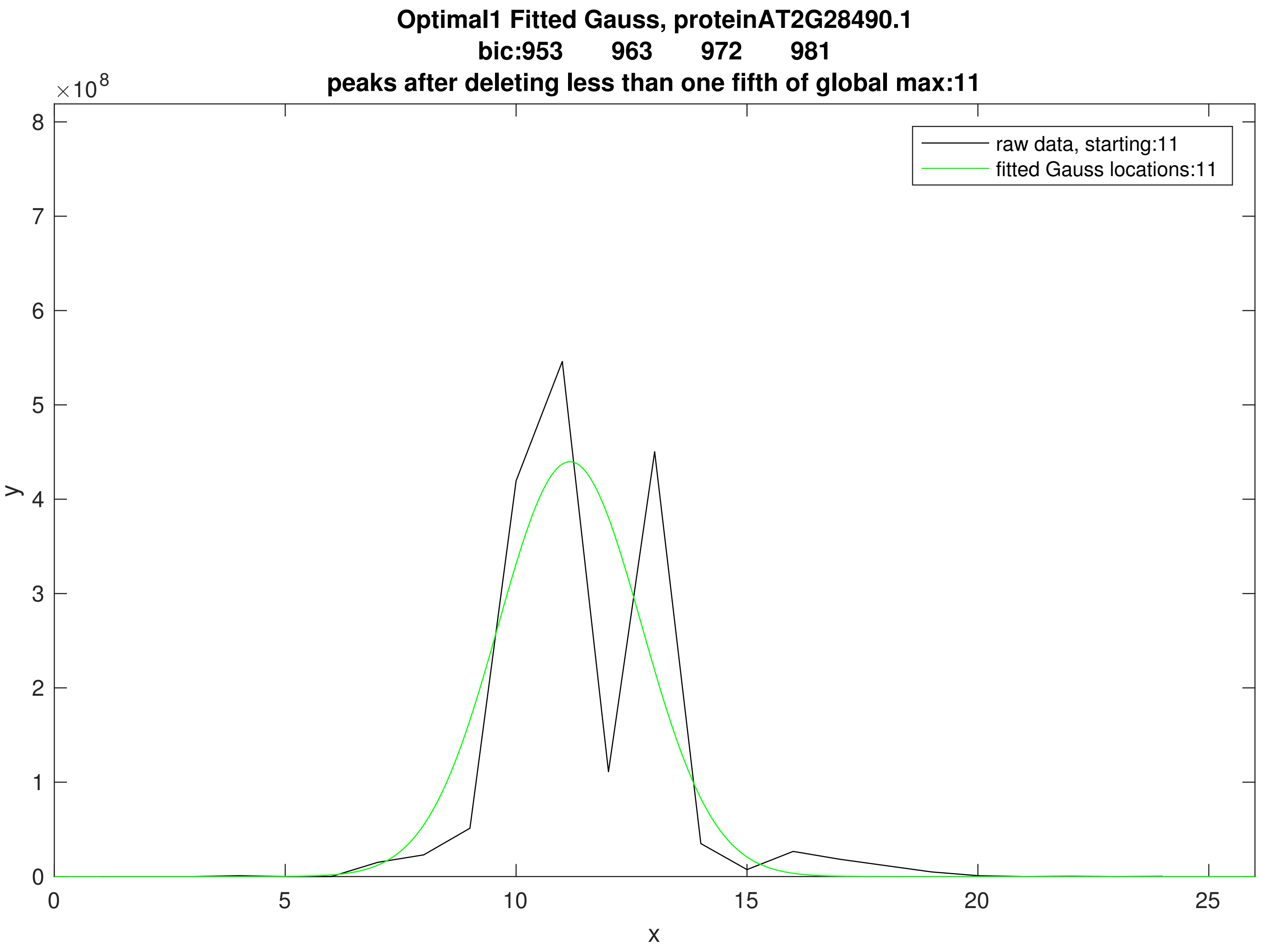
Optimal1 Fitted Gauss, proteinAT2G28490.1
bic:953 963 972 981
peaks after deleting less than one fifth of global max:11
×10^8
raw data, starting:11
fitted Gauss locations:11
y
x

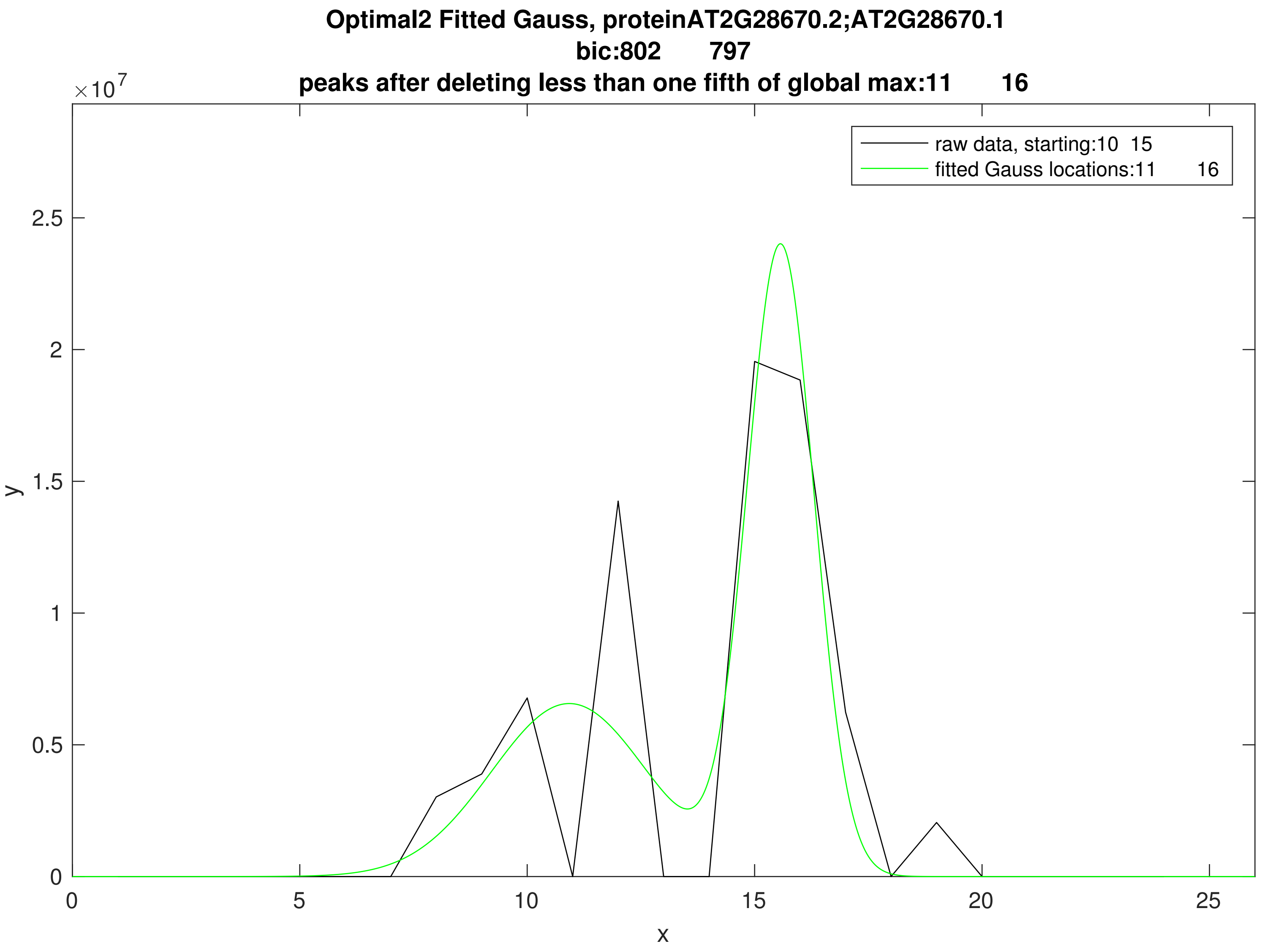
Optimal2 Fitted Gauss, proteinAT2G28670.2;AT2G28670.1
bic:802 797
peaks after deleting less than one fifth of global max:11 16
×10^7
raw data, starting:10 15
fitted Gauss locations:11 16
y
x
0
0.5
1
1.5
2
2.5
5
10
15
20
25

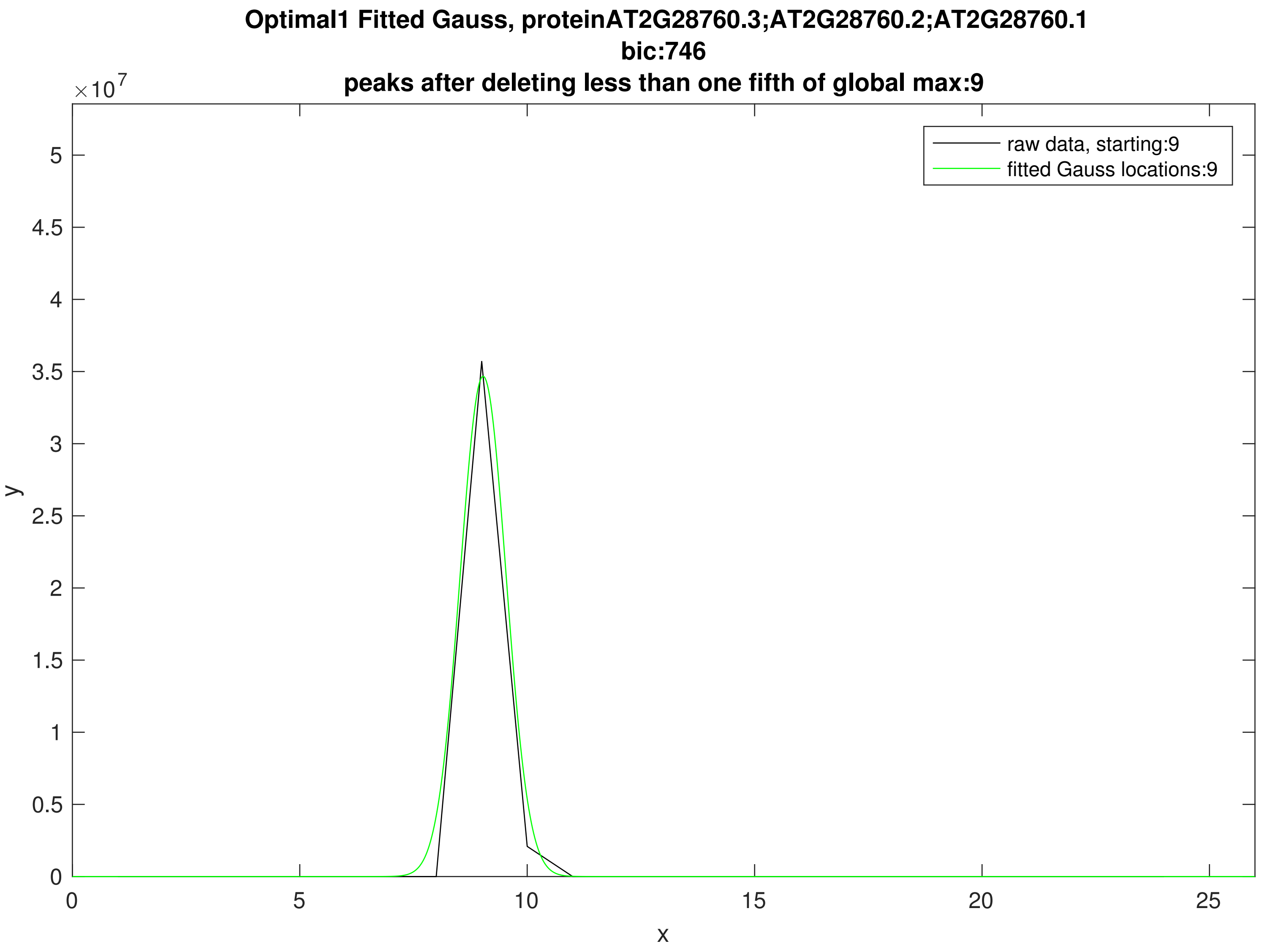

Optimal1 Fitted Gauss, proteinAT2G28760.3;AT2G28760.2;AT2G28760.1
bic:746
peaks after deleting less than one fifth of global max:9
×10^7
5
4.5
4
3.5
3
2.5
2
1.5
1
0.5
0
y
0
5
10
15
20
25
x
raw data, starting:9
fitted Gauss locations:9

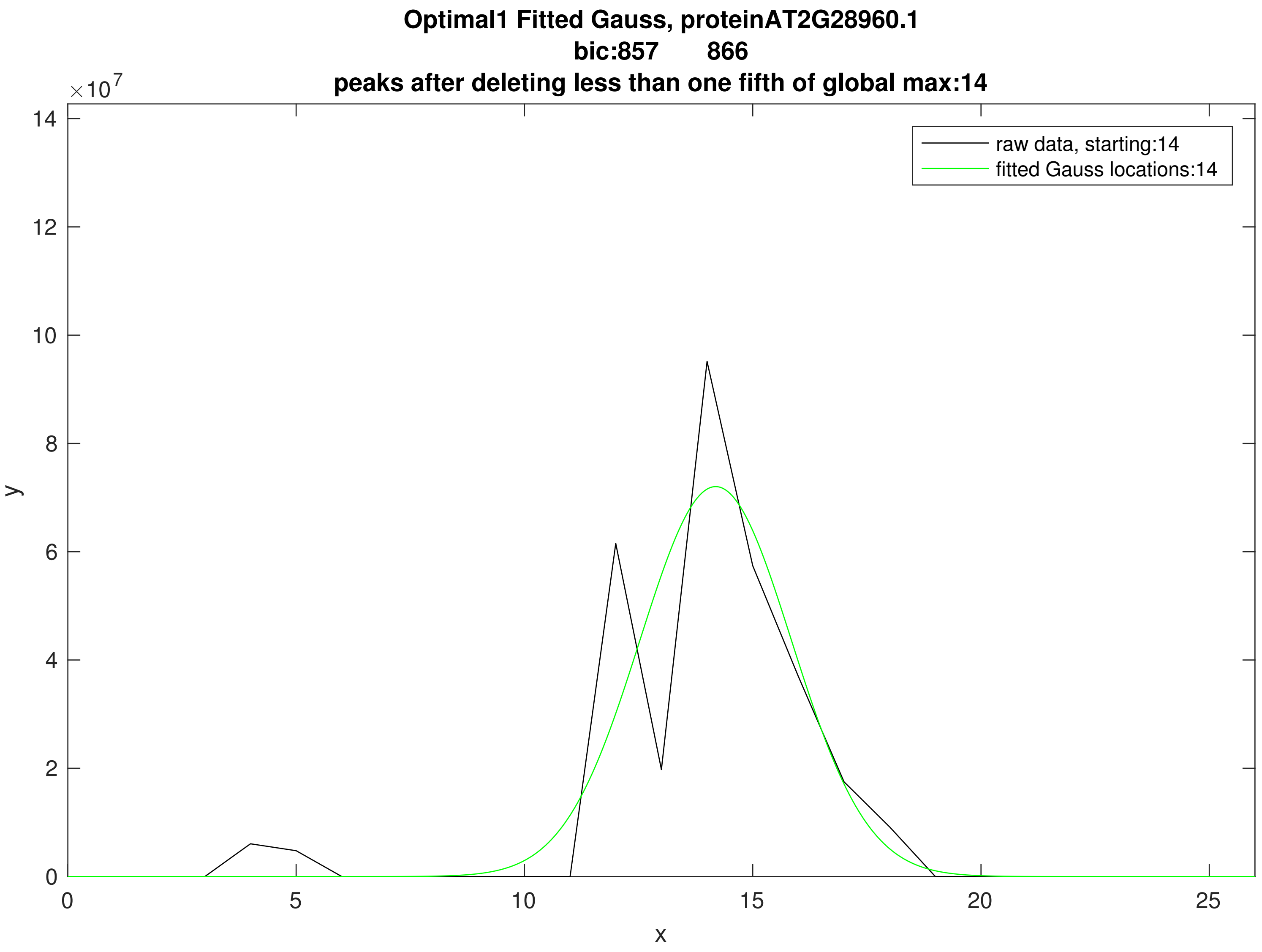

Optimal1 Fitted Gauss, proteinAT2G28960.1
bic:857 866
peaks after deleting less than one fifth of global max:14
×10^7
raw data, starting:14
fitted Gauss locations:14
y
x
0
2
4
6
8
10
12
14
0
5
10
15
20
25

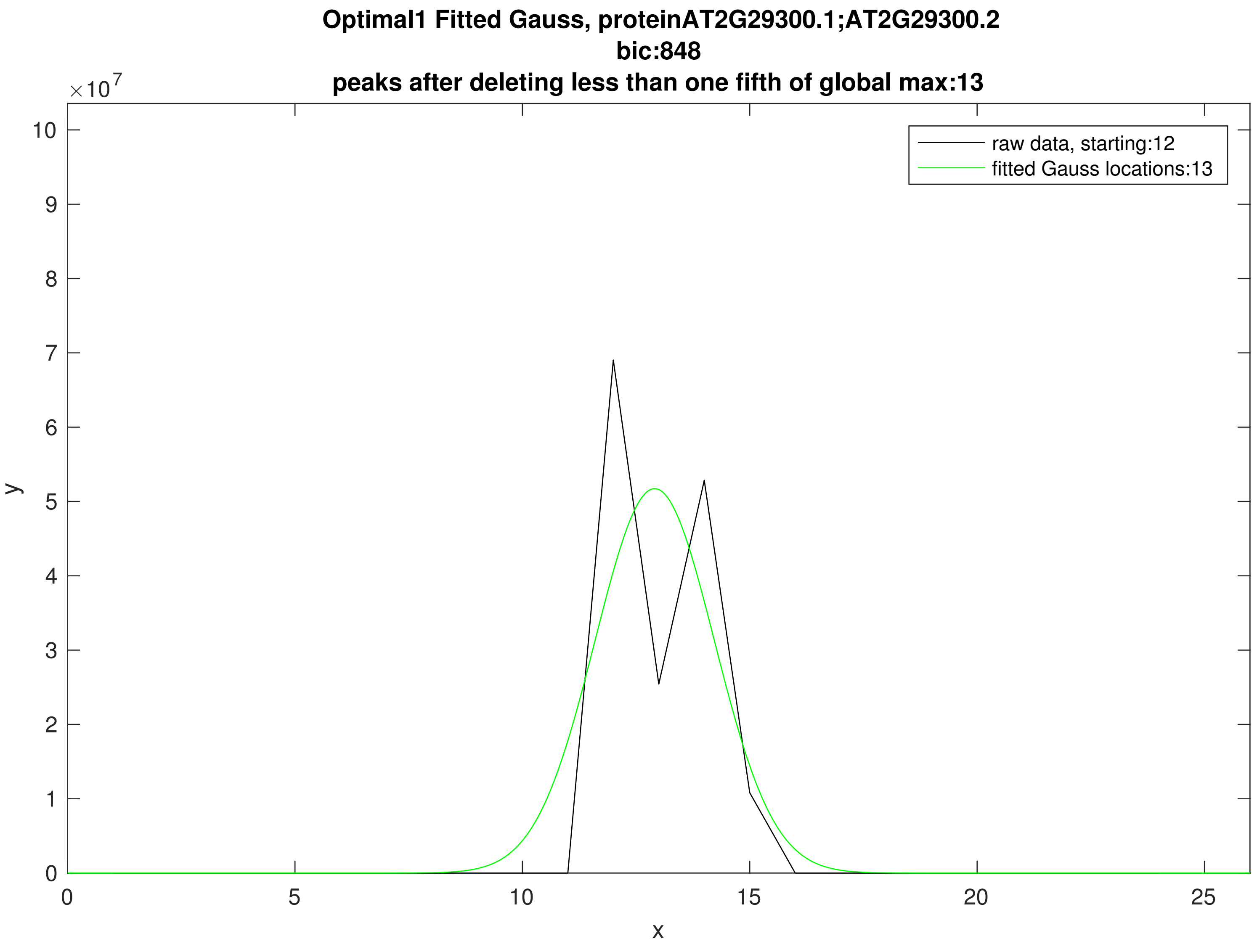

Optimal1 Fitted Gauss, proteinAT2G29300.1;AT2G29300.2
bic:848
peaks after deleting less than one fifth of global max:13
×10^7
10
9
8
7
6
5
4
3
2
1
0
y
0
5
10
15
20
25
x
raw data, starting:12
fitted Gauss locations:13

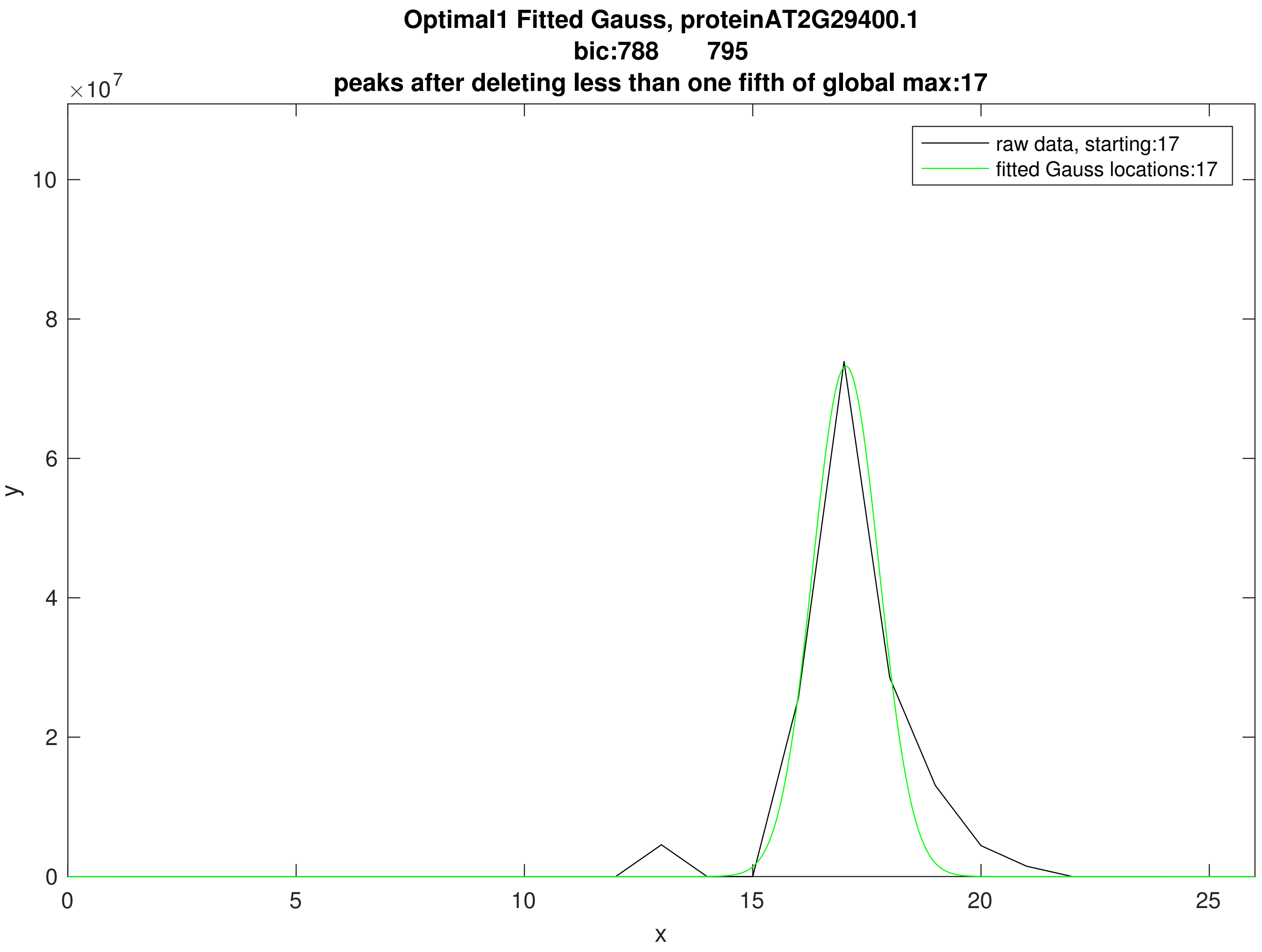
Optimal1 Fitted Gauss, proteinAT2G29400.1
bic:788 795
peaks after deleting less than one fifth of global max:17
×10^7
raw data, starting:17
fitted Gauss locations:17
y
x
0
2
4
6
8
10
0
5
10
15
20
25

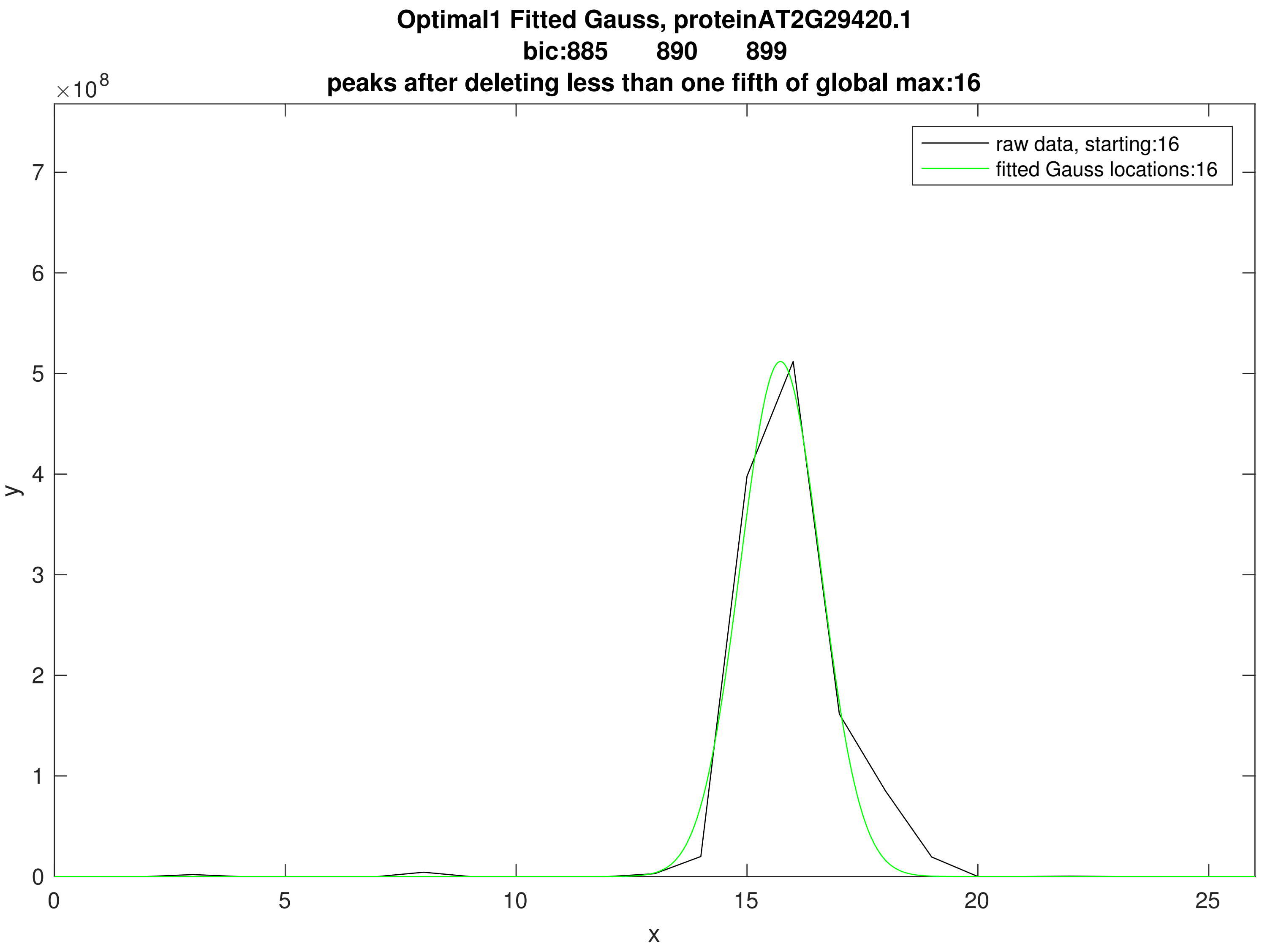

Optimal1 Fitted Gauss, proteinAT2G29420.1
bic:885 890 899
peaks after deleting less than one fifth of global max:16
×10^8
raw data, starting:16
fitted Gauss locations:16
y
x
0
1
2
3
4
5
6
7
0
5
10
15
20
25

**Optimal1 Fitted Gauss, proteinAT2G29440.1**
**bic:798**
**peaks after deleting less than one fifth of global max:16**

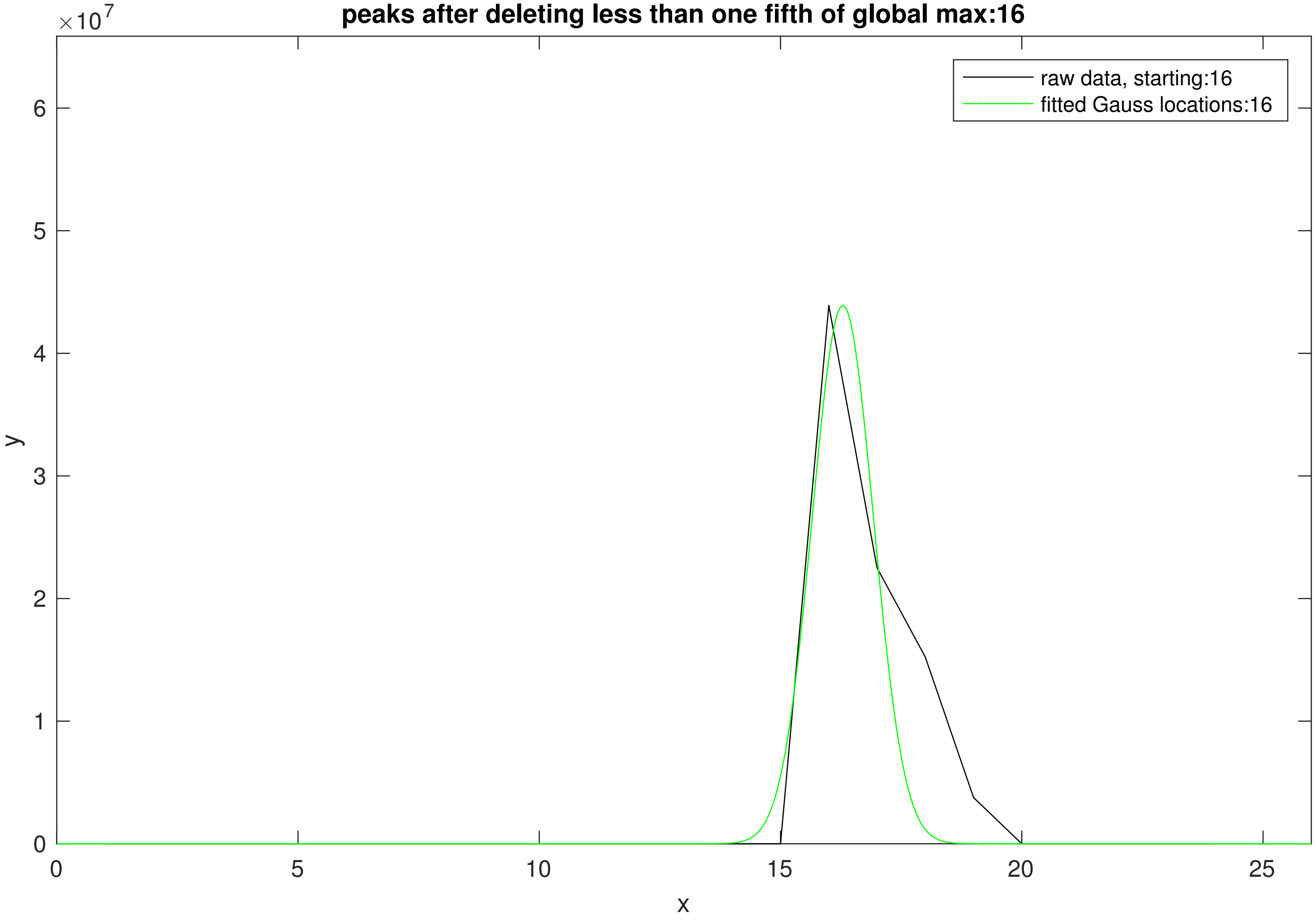

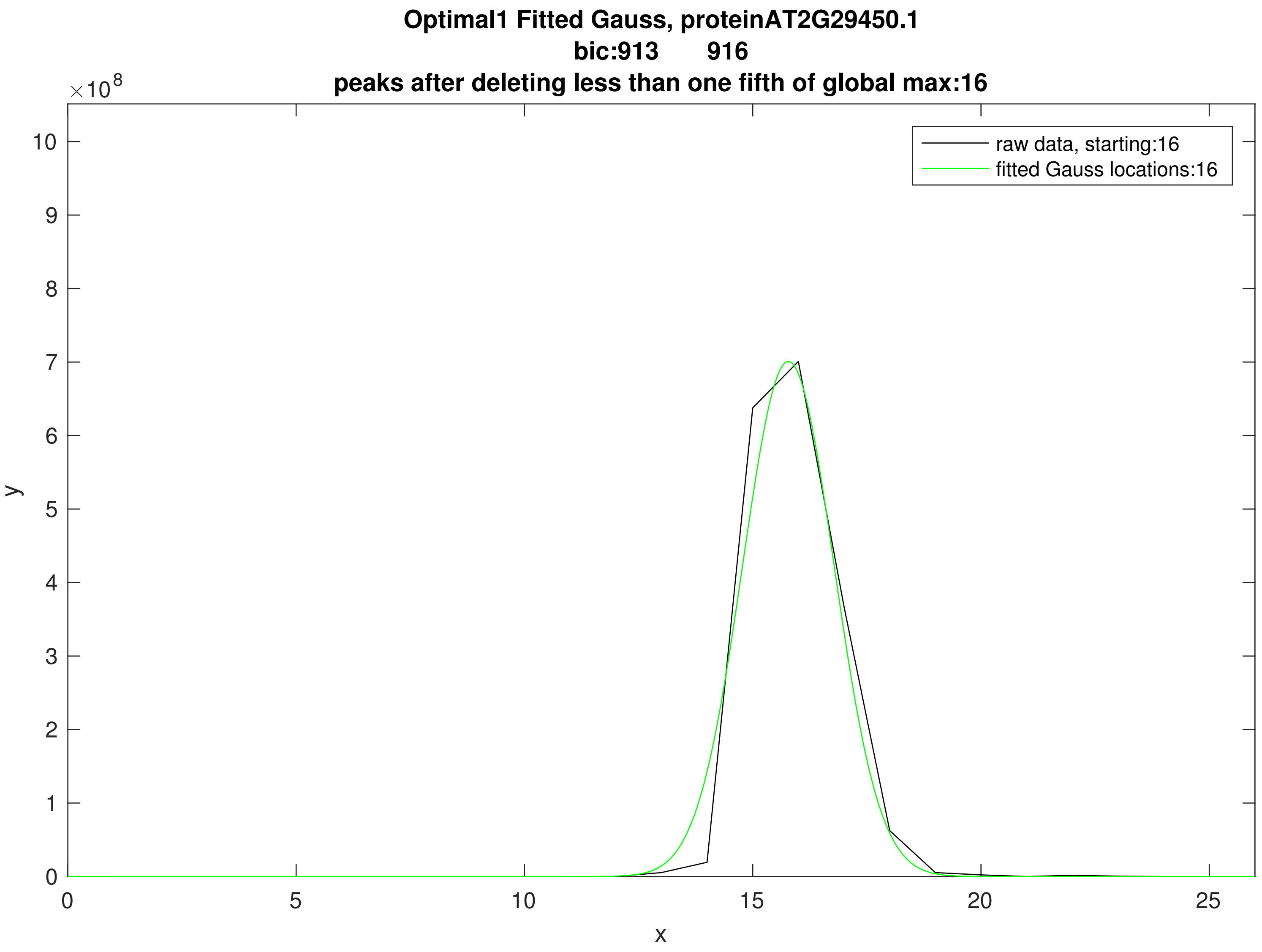

Optimal1 Fitted Gauss, proteinAT2G29450.1
bic:913 916
peaks after deleting less than one fifth of global max:16
×10^8
raw data, starting:16
fitted Gauss locations:16
y
x
0
1
2
3
4
5
6
7
8
9
10
0
5
10
15
20
25

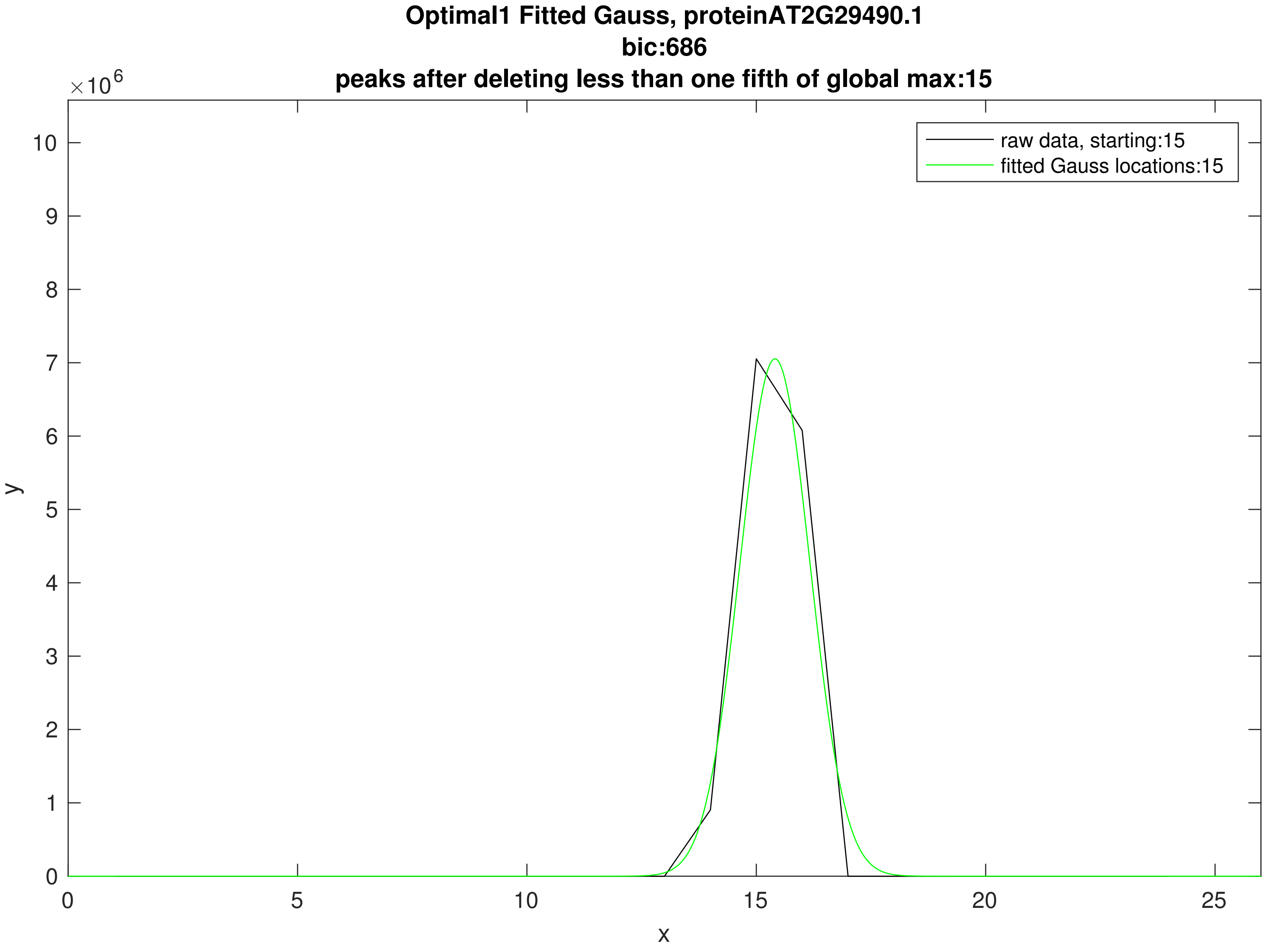

Optimal1 Fitted Gauss, proteinAT2G29490.1
bic:686
peaks after deleting less than one fifth of global max:15
×10^6
raw data, starting:15
fitted Gauss locations:15
y
x
0
1
2
3
4
5
6
7
8
9
10
0
5
10
15
20
25

**Optimal1 Fitted Gauss, proteinAT2G29530.1;AT2G29530.2;AT2G29530.3**
**bic:830**
**peaks after deleting less than one fifth of global max:20**

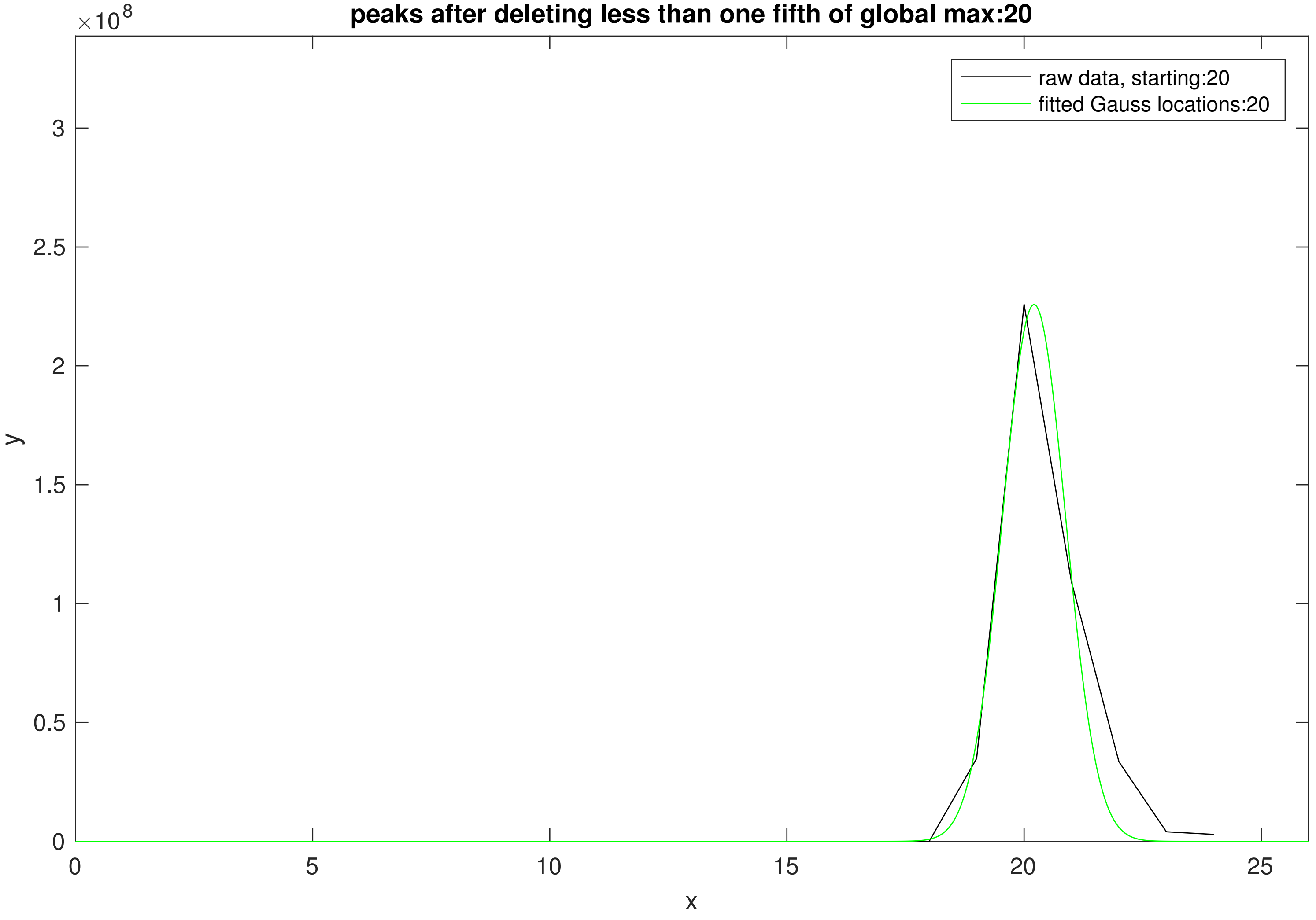

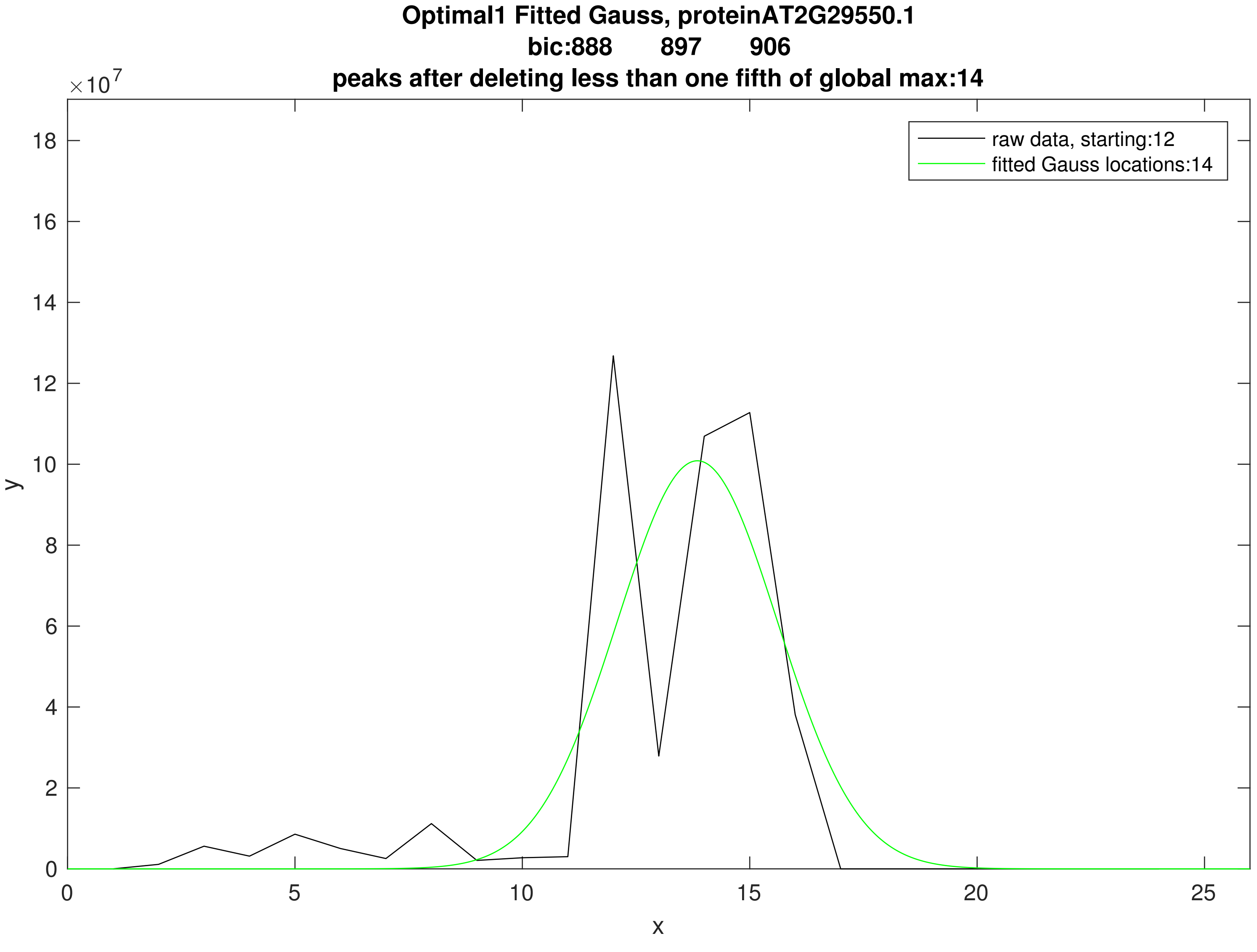
Optimal1 Fitted Gauss, proteinAT2G29550.1
bic:888 897 906
peaks after deleting less than one fifth of global max:14
×10^7
raw data, starting:12
fitted Gauss locations:14
y
x
0
2
4
6
8
10
12
14
16
18
0
5
10
15
20
25

**Optimal1 Fitted Gauss, proteinAT2G29560.1**
**bic:1e+03  1.01e+03**
**peaks after deleting less than one fifth of global max:13**

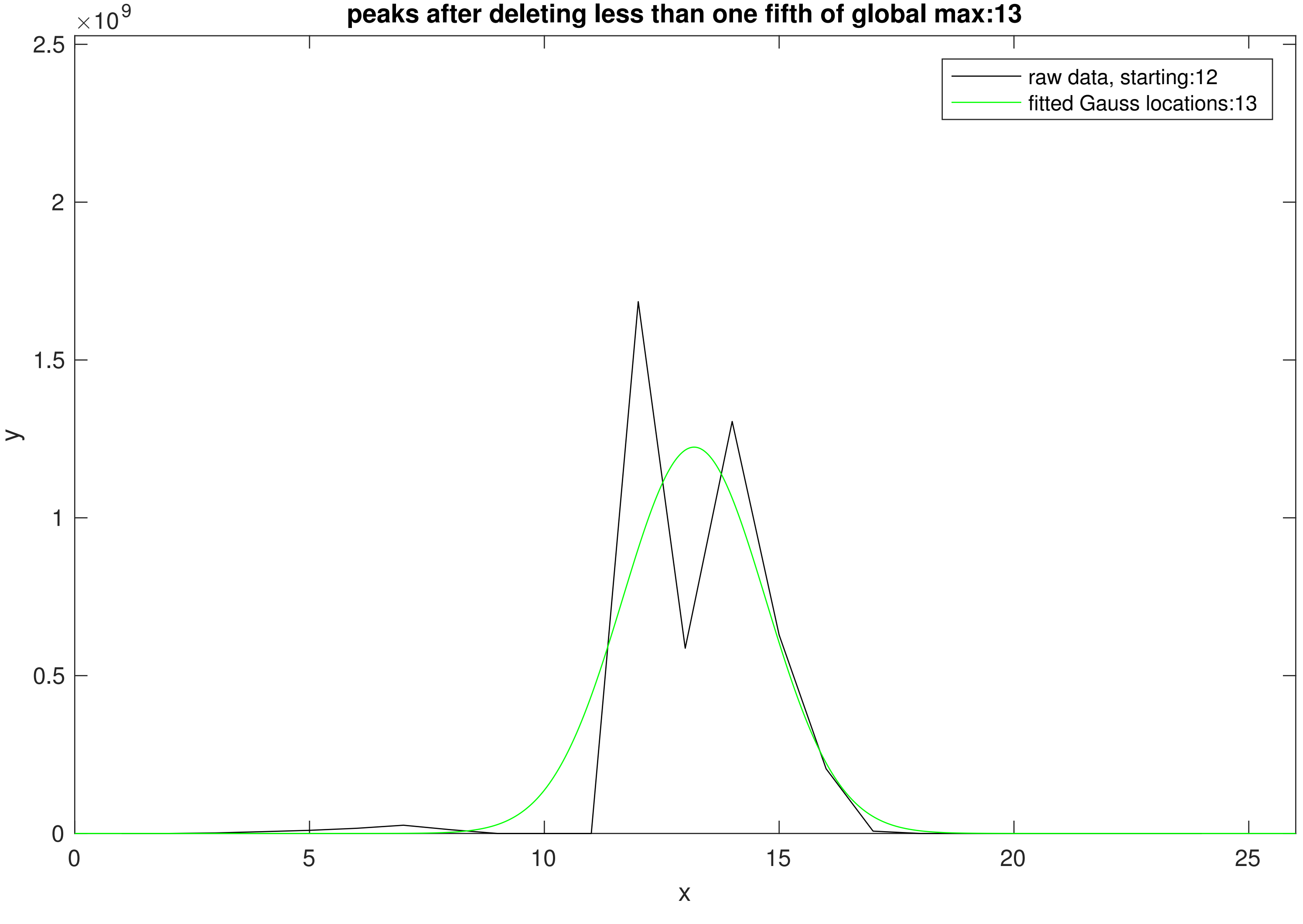

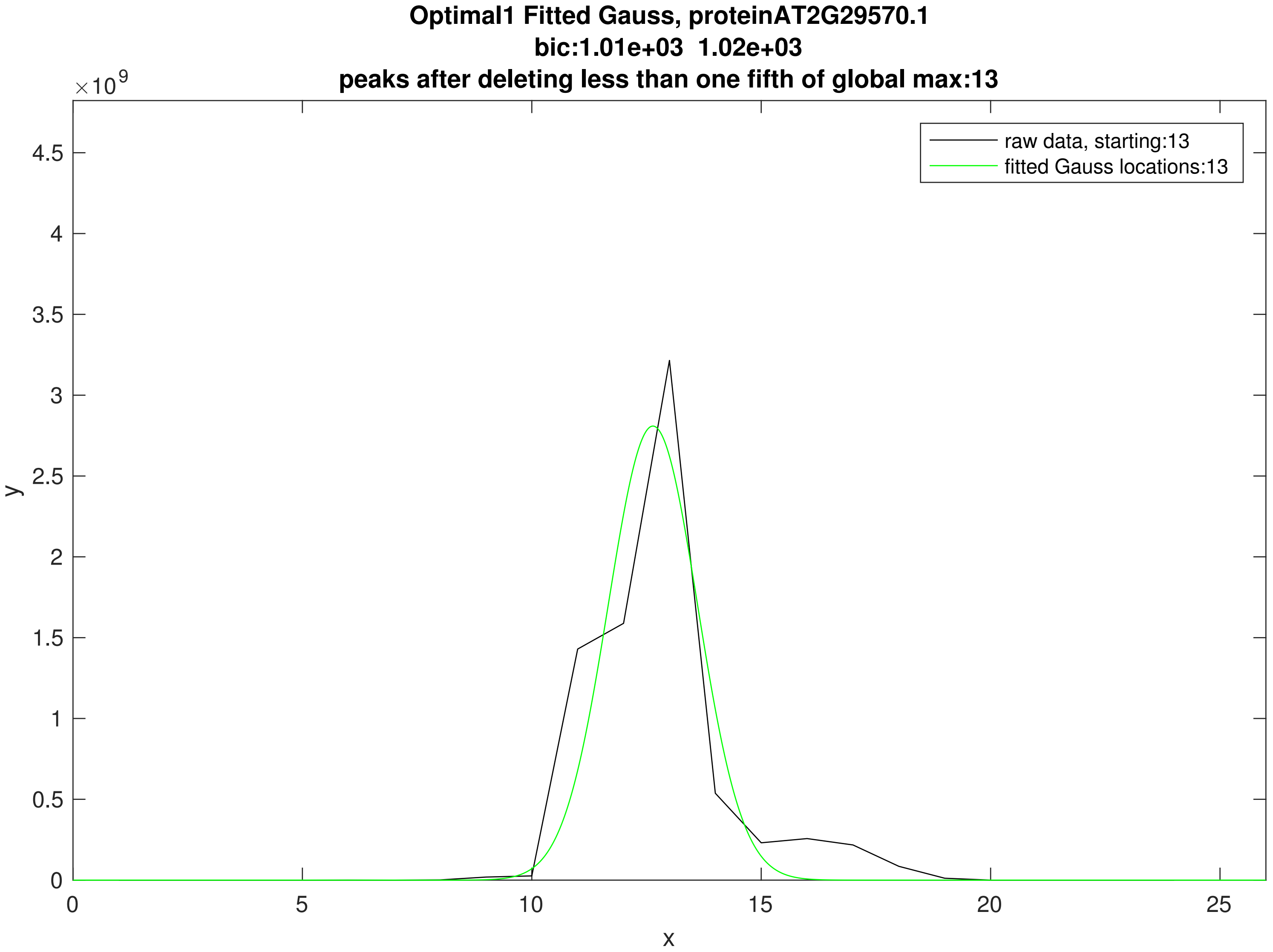

Optimal1 Fitted Gauss, proteinAT2G29570.1
bic:1.01e+03  1.02e+03
peaks after deleting less than one fifth of global max:13
×10^9
raw data, starting:13
fitted Gauss locations:13
y
x
0
0.5
1
1.5
2
2.5
3
3.5
4
4.5
5
10
15
20
25

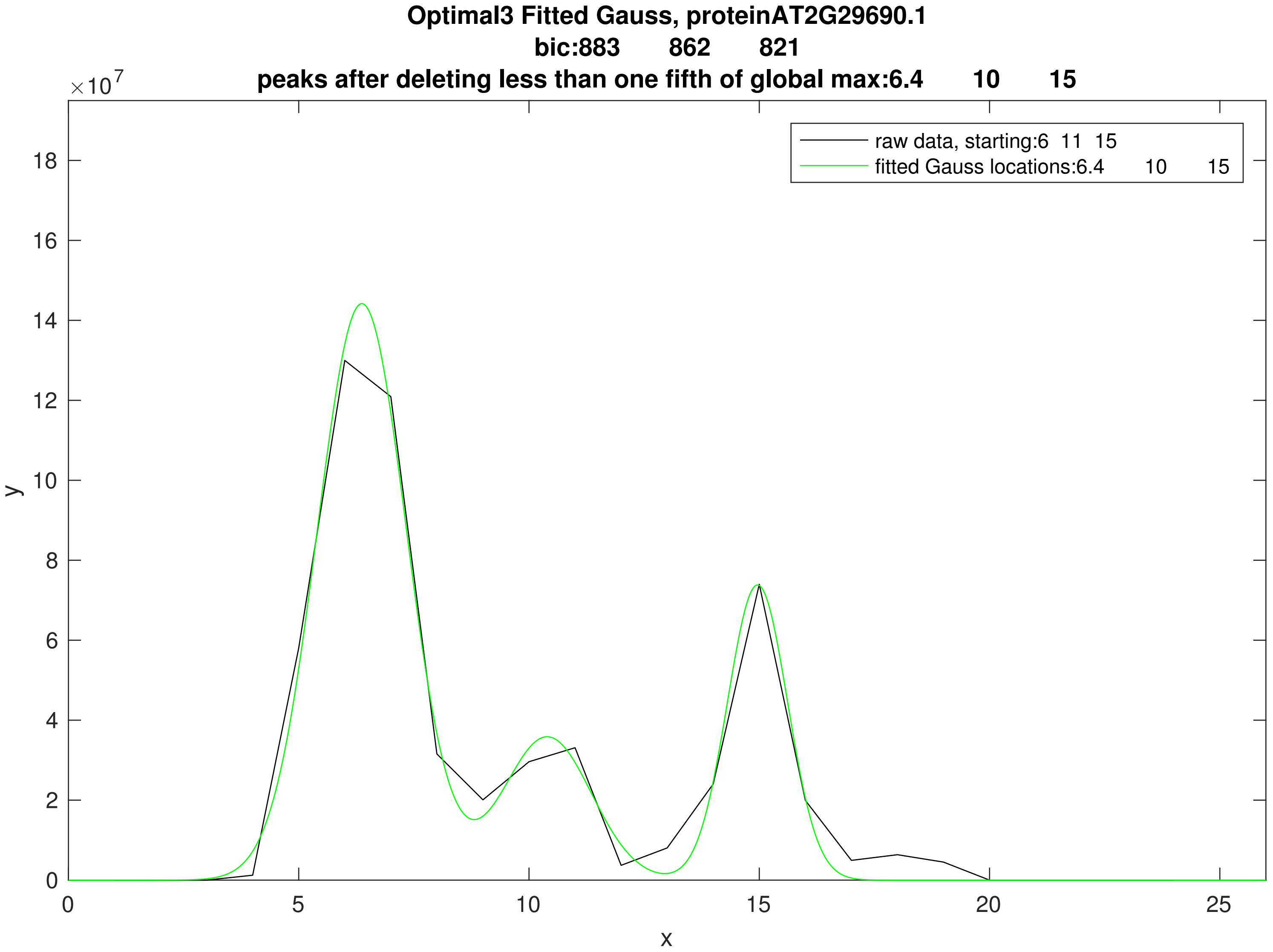

Optimal3 Fitted Gauss, proteinAT2G29690.1
bic:883 862 821
peaks after deleting less than one fifth of global max:6.4 10 15
×10^7
raw data, starting:6 11 15
fitted Gauss locations:6.4 10 15
y
x
0
2
4
6
8
10
12
14
16
18
0
5
10
15
20
25

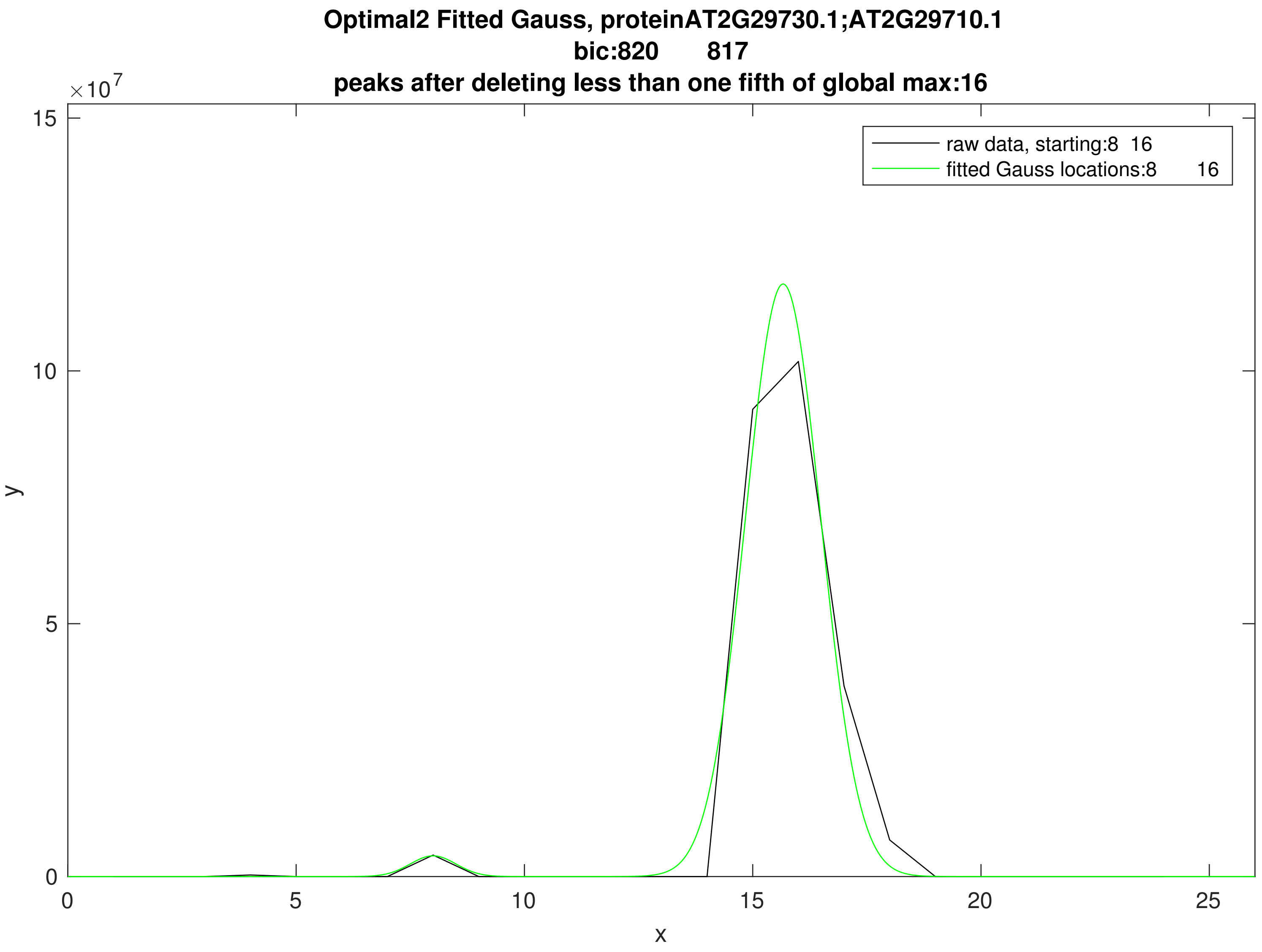

Optimal2 Fitted Gauss, proteinAT2G29730.1;AT2G29710.1
bic:820 817
peaks after deleting less than one fifth of global max:16
×10^7
15
10
5
0
y
0
5
10
15
20
25
x
raw data, starting:8 16
fitted Gauss locations:8 16

**Optimal1 Fitted Gauss, proteinAT2G29750.1**
**bic:894**
**peaks after deleting less than one fifth of global max:16**

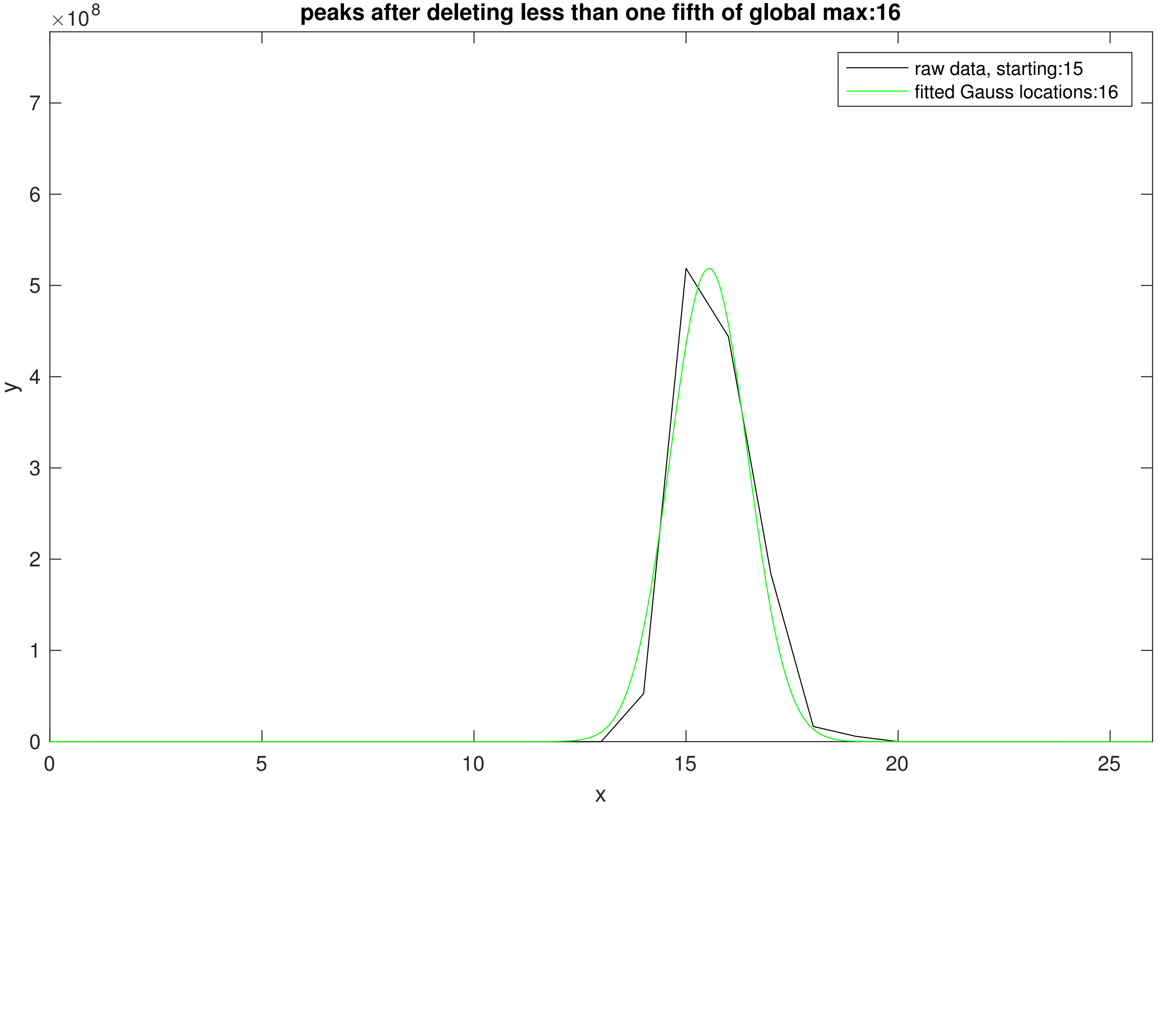

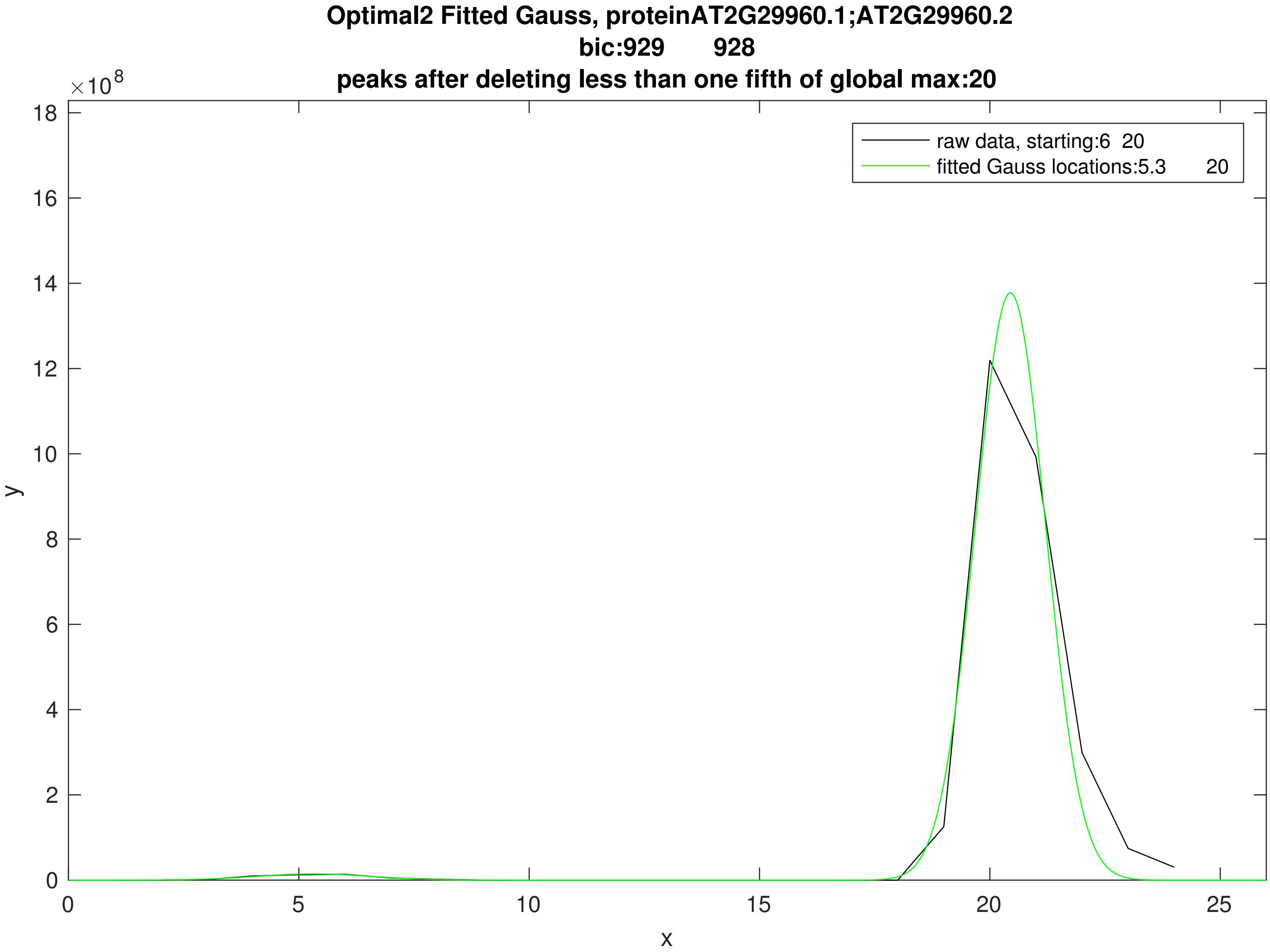

Optimal2 Fitted Gauss, proteinAT2G29960.1;AT2G29960.2
bic:929 928
peaks after deleting less than one fifth of global max:20
×10^8
18
16
14
12
10
8
6
4
2
0
y
0
5
10
15
20
25
x
raw data, starting:6 20
fitted Gauss locations:5.3 20

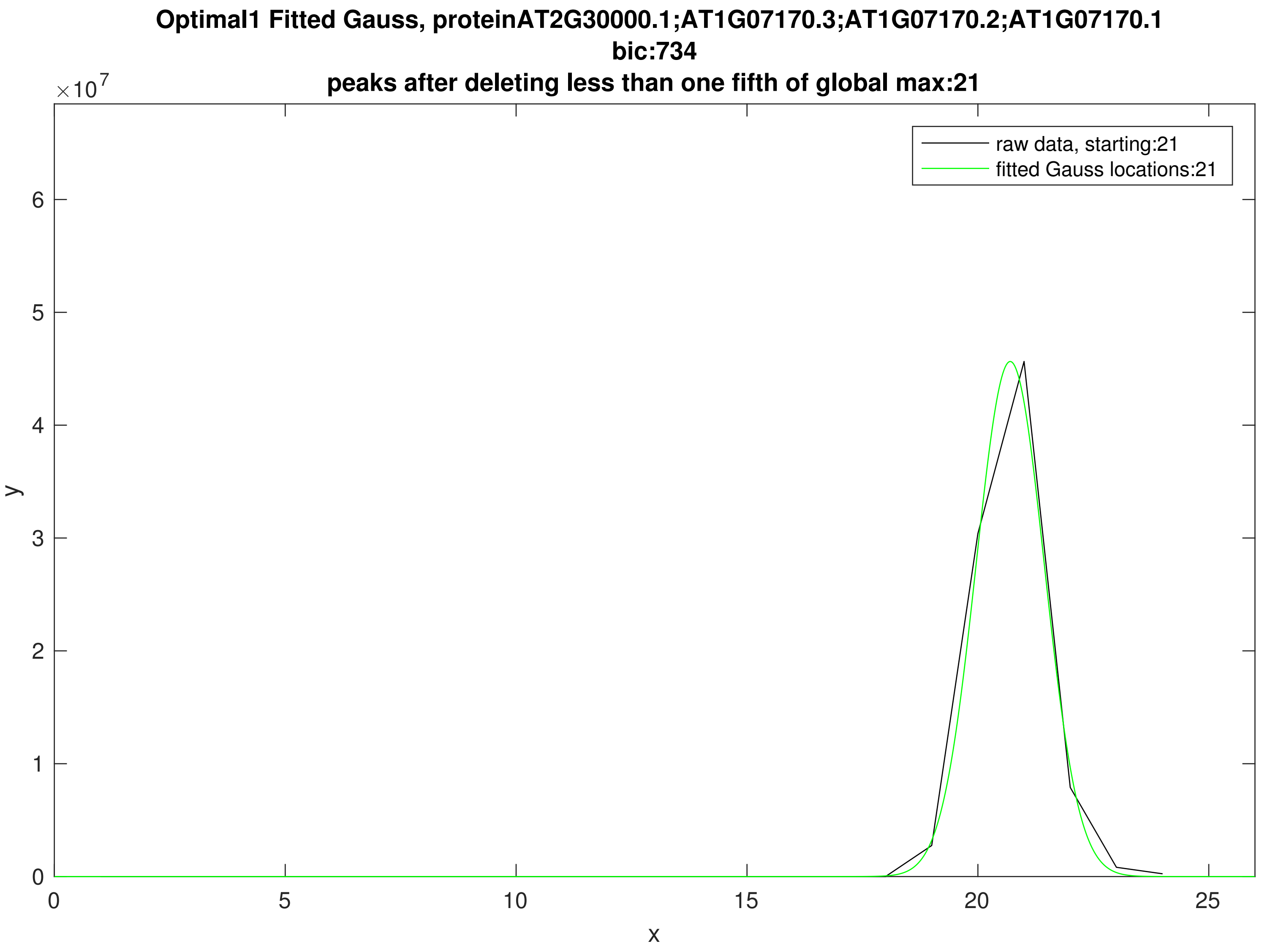

Optimal1 Fitted Gauss, proteinAT2G30000.1;AT1G07170.3;AT1G07170.2;AT1G07170.1
bic:734
peaks after deleting less than one fifth of global max:21
×10^7
raw data, starting:21
fitted Gauss locations:21
y
x
0
1
2
3
4
5
6
0
5
10
15
20
25

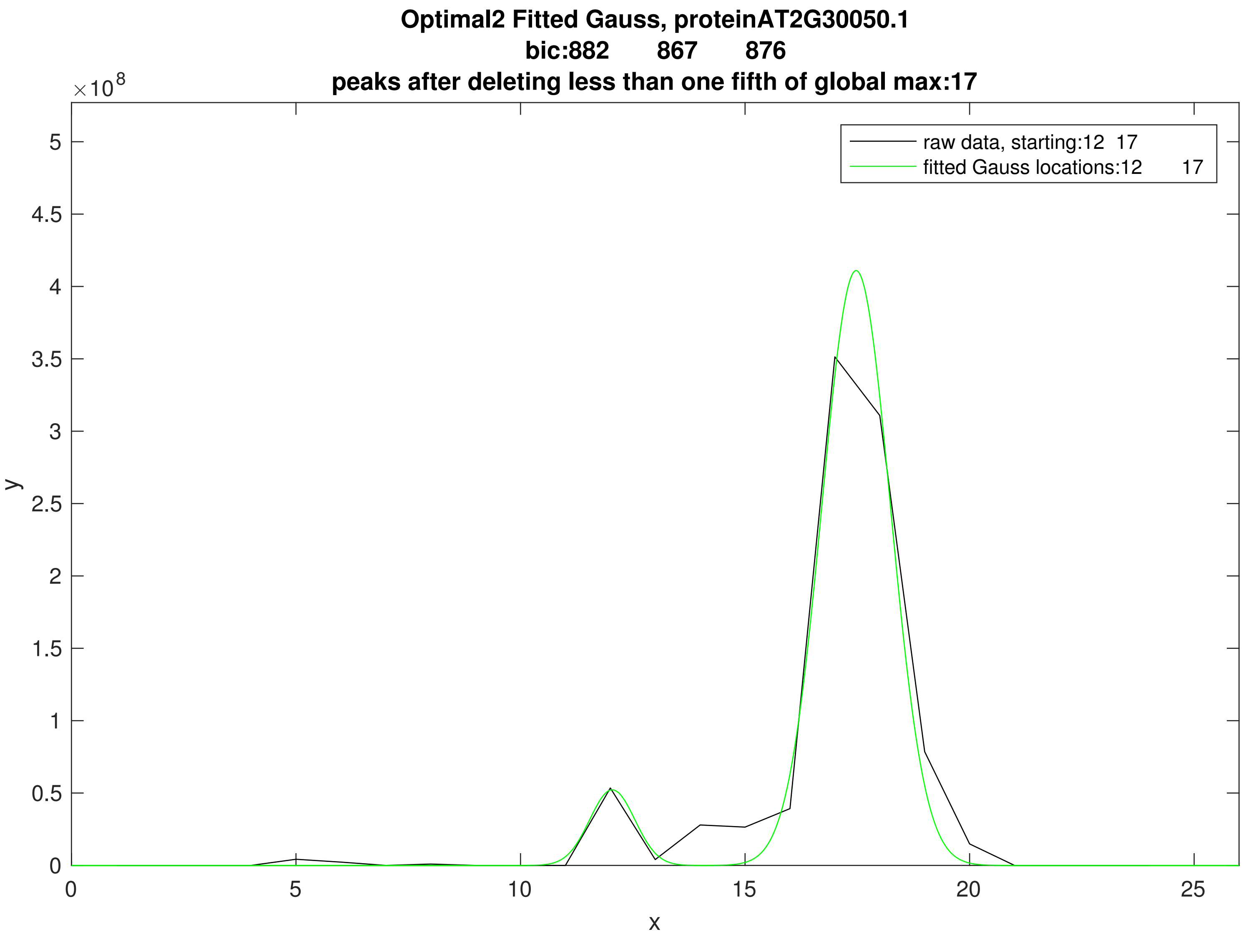
Optimal2 Fitted Gauss, proteinAT2G30050.1
bic:882 867 876
peaks after deleting less than one fifth of global max:17
×10^8
raw data, starting:12 17
fitted Gauss locations:12 17
y
x
0
0.5
1
1.5
2
2.5
3
3.5
4
4.5
5
0
5
10
15
20
25

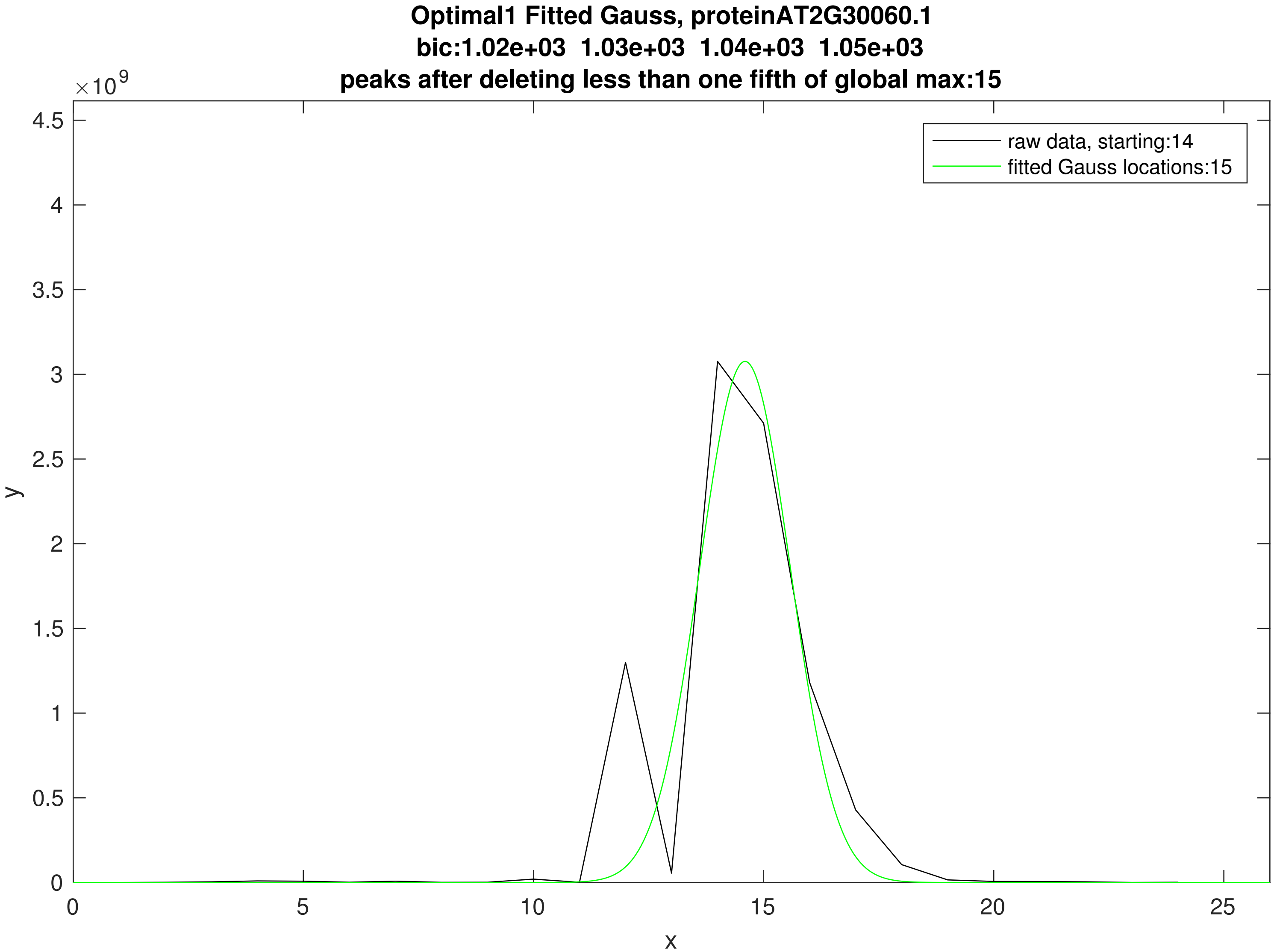
Optimal1 Fitted Gauss, proteinAT2G30060.1
bic:1.02e+03  1.03e+03  1.04e+03  1.05e+03
peaks after deleting less than one fifth of global max:15
×10^9
raw data, starting:14
fitted Gauss locations:15
y
x
0
0.5
1
1.5
2
2.5
3
3.5
4
4.5
5
10
15
20
25

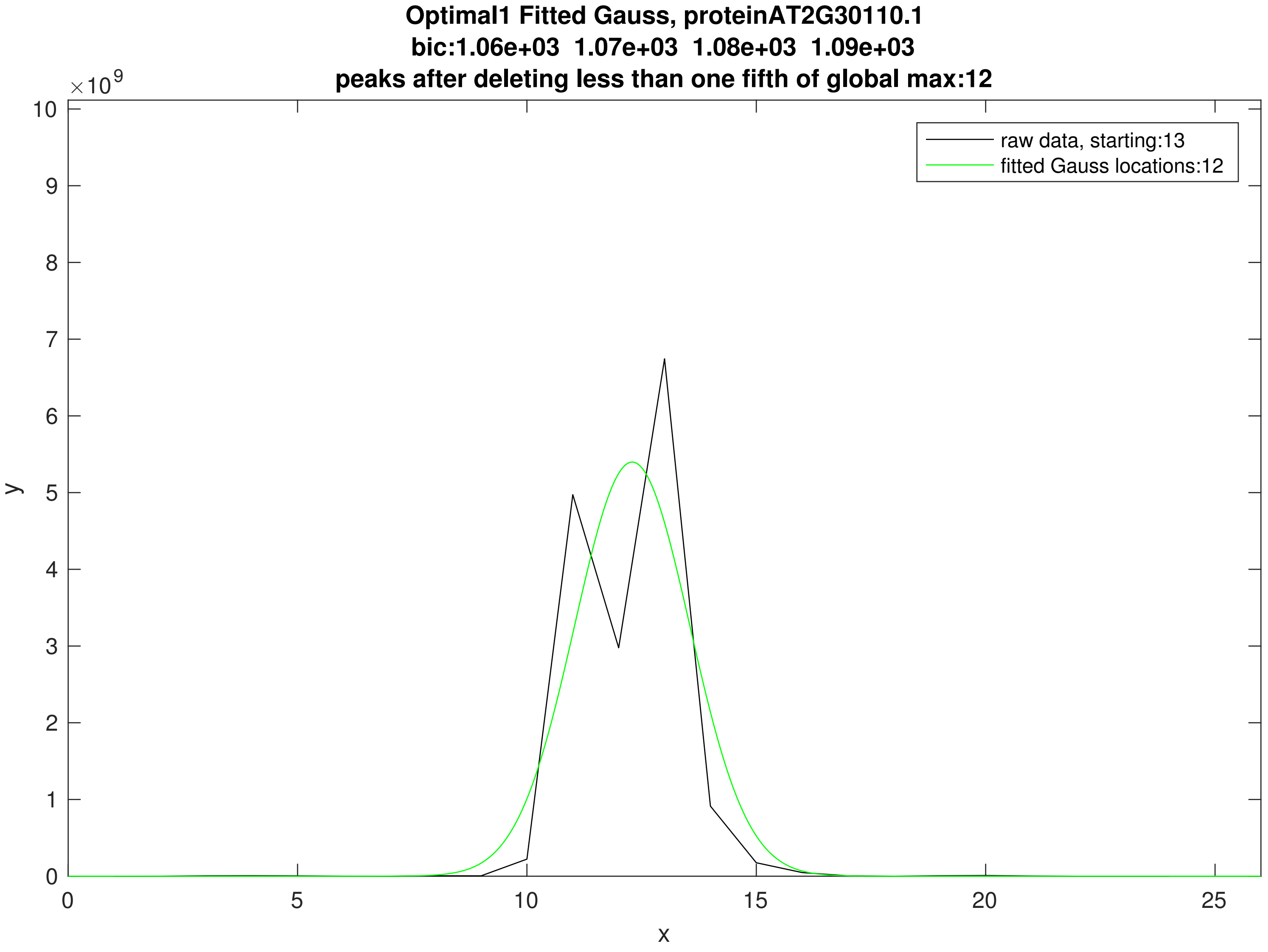

Optimal1 Fitted Gauss, proteinAT2G30110.1
bic:1.06e+03  1.07e+03  1.08e+03  1.09e+03
peaks after deleting less than one fifth of global max:12
×10^9
raw data, starting:13
fitted Gauss locations:12
y
x
0
1
2
3
4
5
6
7
8
9
10
0
5
10
15
20
25

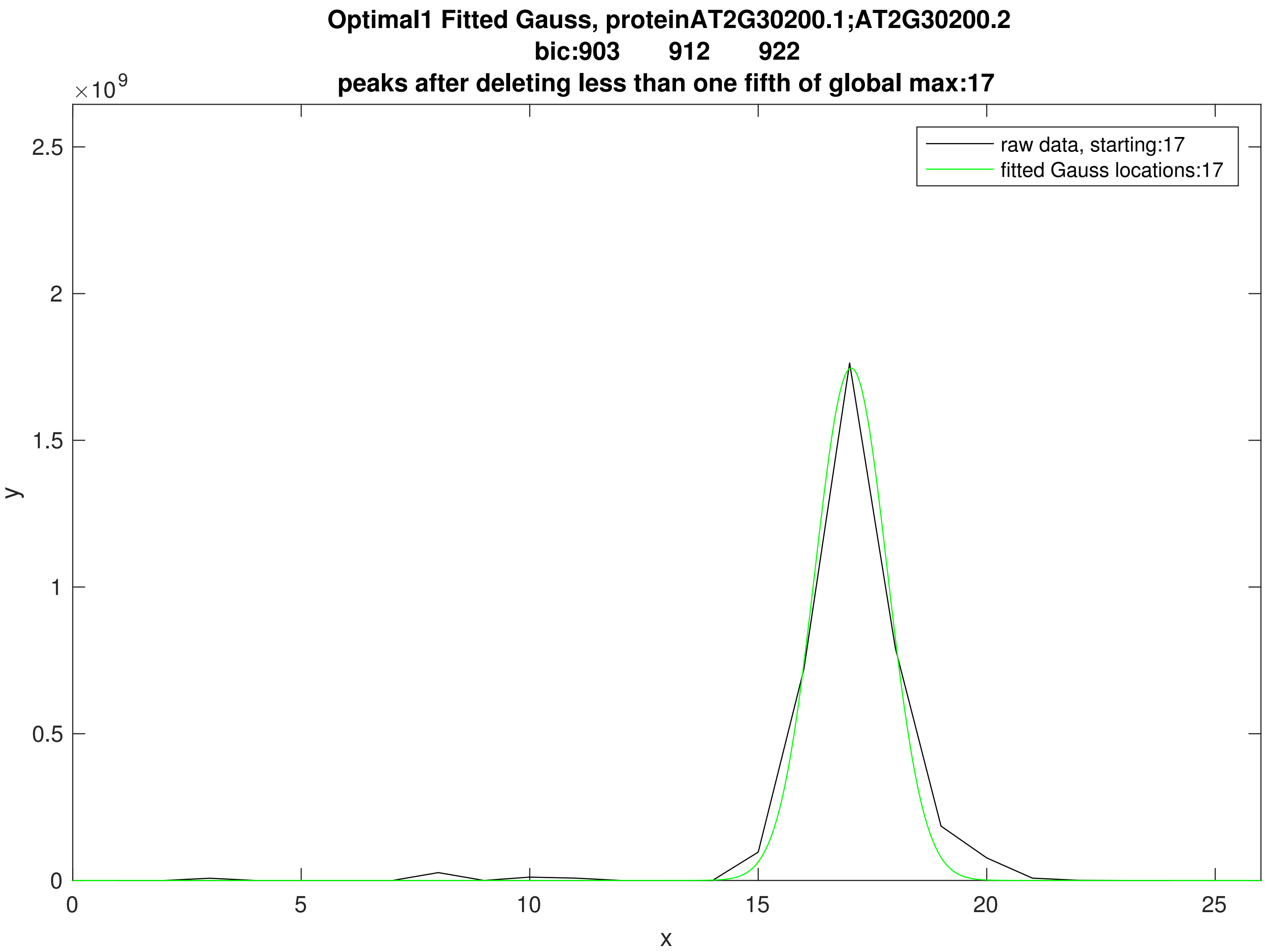

Optimal1 Fitted Gauss, proteinAT2G30200.1;AT2G30200.2
bic:903 912 922
peaks after deleting less than one fifth of global max:17
×10^9
2.5
2
1.5
1
0.5
0
y
0
5
10
15
20
25
x
raw data, starting:17
fitted Gauss locations:17

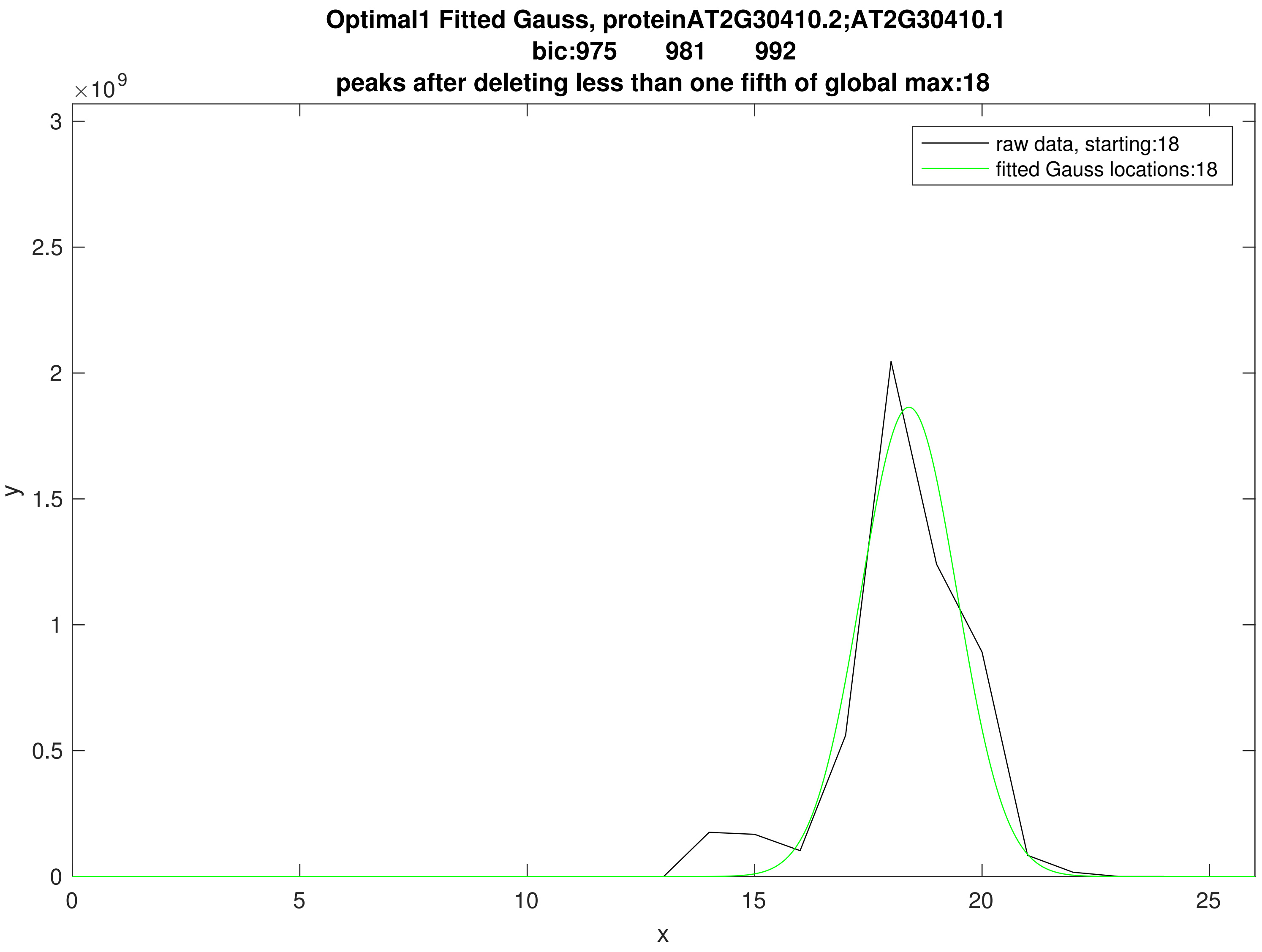
Optimal1 Fitted Gauss, proteinAT2G30410.2;AT2G30410.1
bic:975 981 992
peaks after deleting less than one fifth of global max:18
×10^9
raw data, starting:18
fitted Gauss locations:18
y
x
0
0.5
1
1.5
2
2.5
3
0
5
10
15
20
25

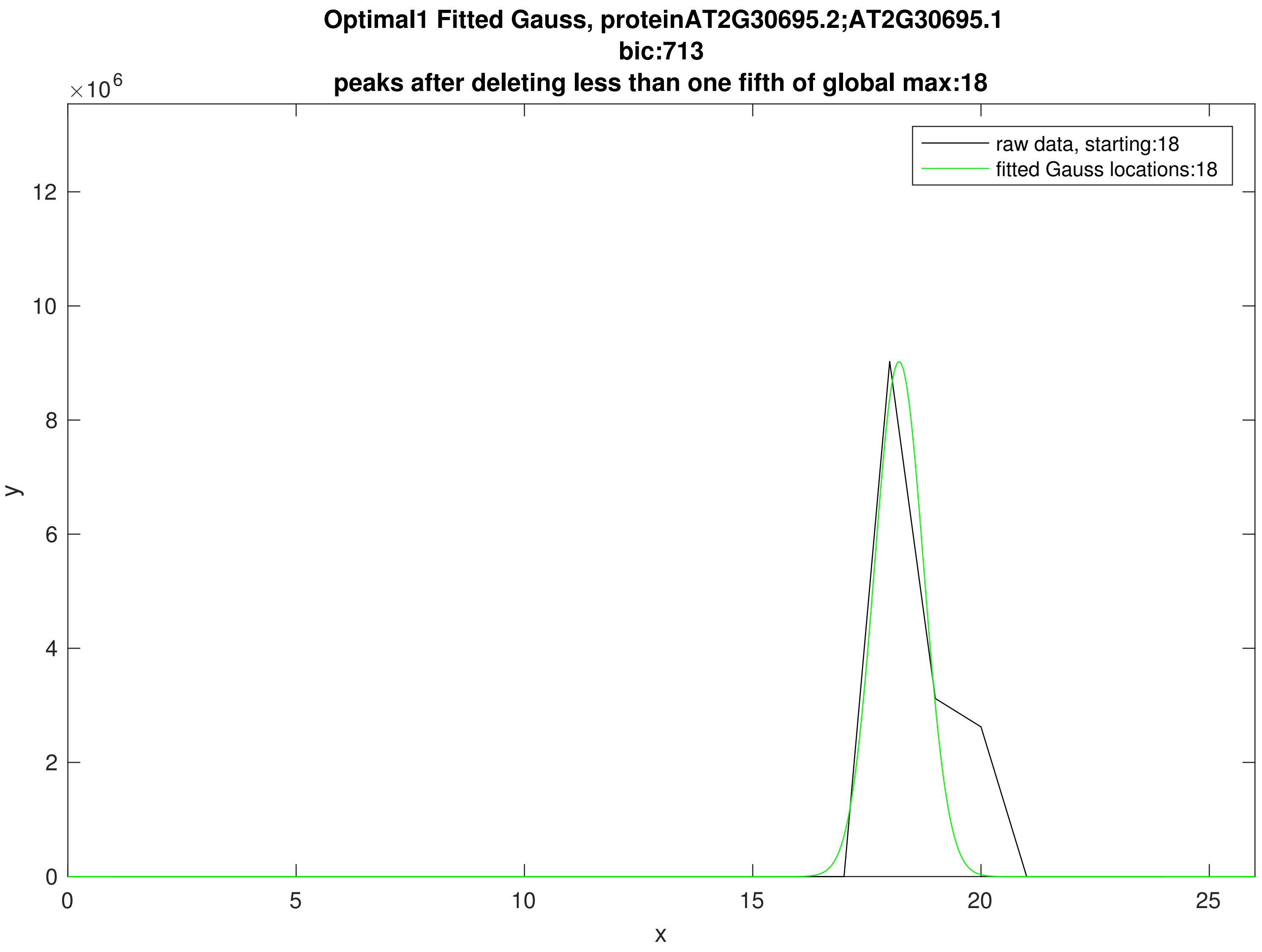

Optimal1 Fitted Gauss, proteinAT2G30695.2;AT2G30695.1
bic:713
peaks after deleting less than one fifth of global max:18
×10^6
raw data, starting:18
fitted Gauss locations:18
12
10
8
6
4
2
0
y
0
5
10
15
20
25
x

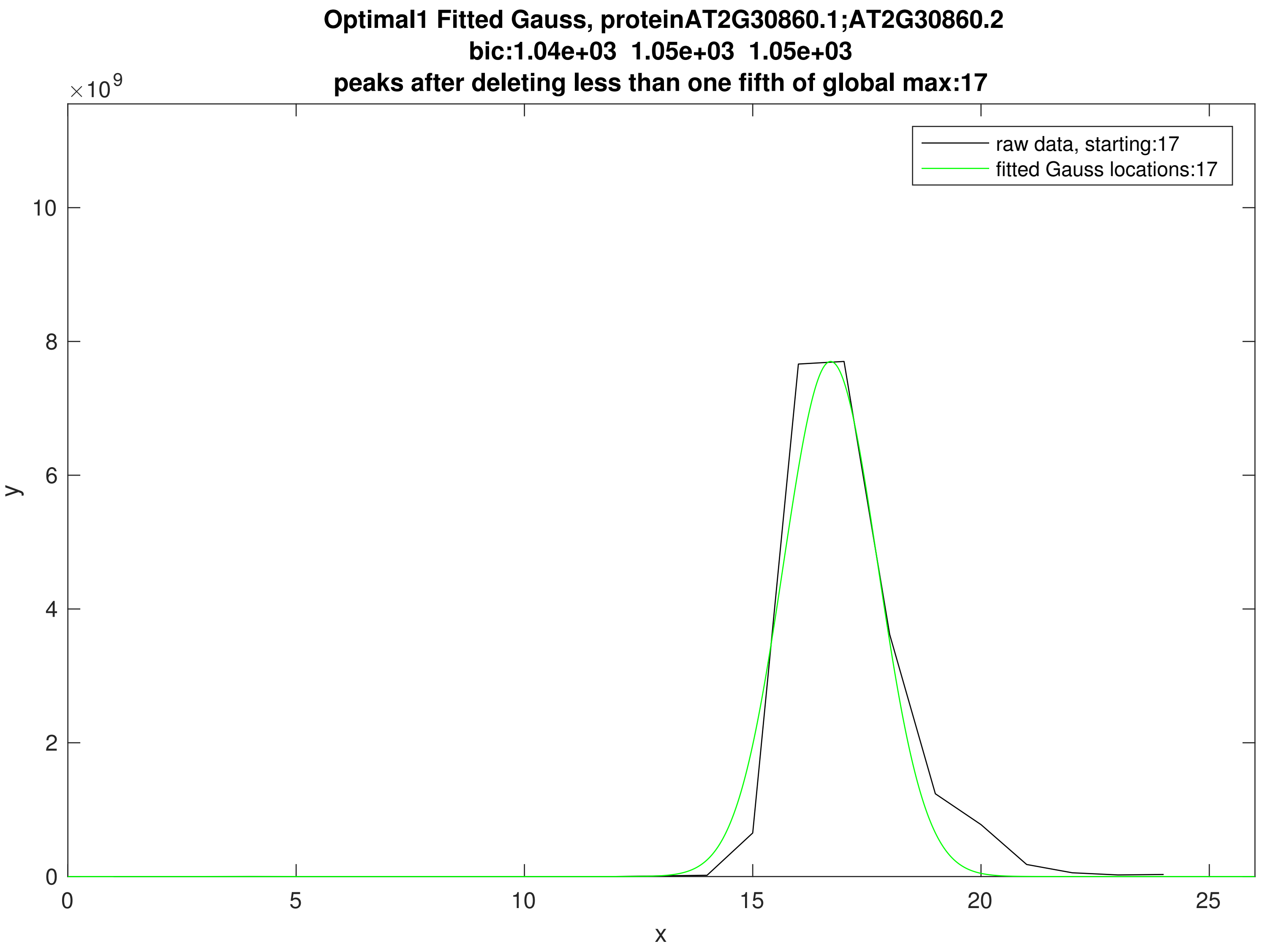

Optimal1 Fitted Gauss, proteinAT2G30860.1;AT2G30860.2
bic:1.04e+03  1.05e+03  1.05e+03
peaks after deleting less than one fifth of global max:17
×10^9
raw data, starting:17
fitted Gauss locations:17
10
8
6
4
2
0
y
0
5
10
15
20
25
x

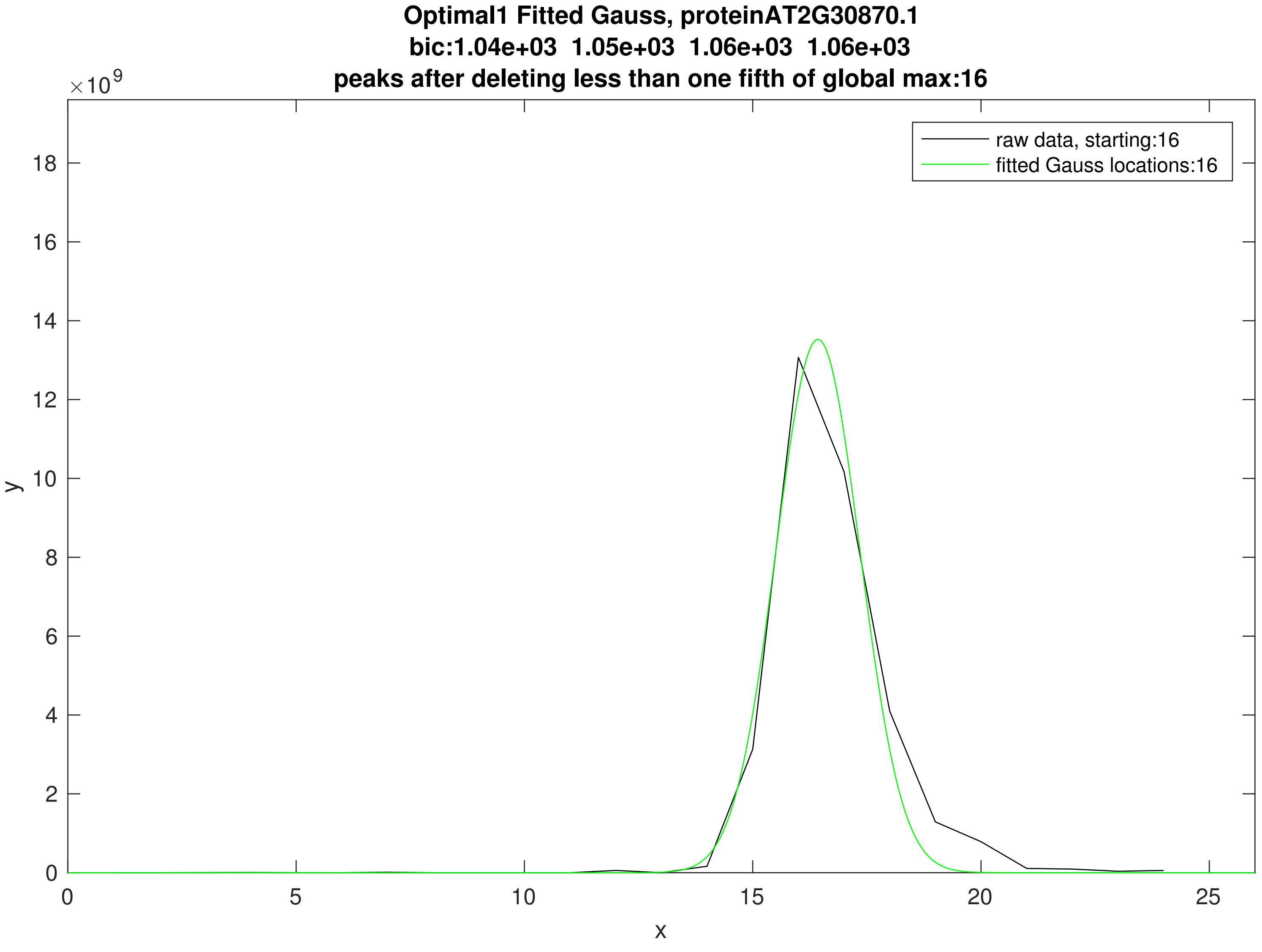

Optimal1 Fitted Gauss, proteinAT2G30870.1
bic:1.04e+03 1.05e+03 1.06e+03 1.06e+03
peaks after deleting less than one fifth of global max:16
×10^9
y
x
raw data, starting:16
fitted Gauss locations:16
0
2
4
6
8
10
12
14
16
18
0
5
10
15
20
25

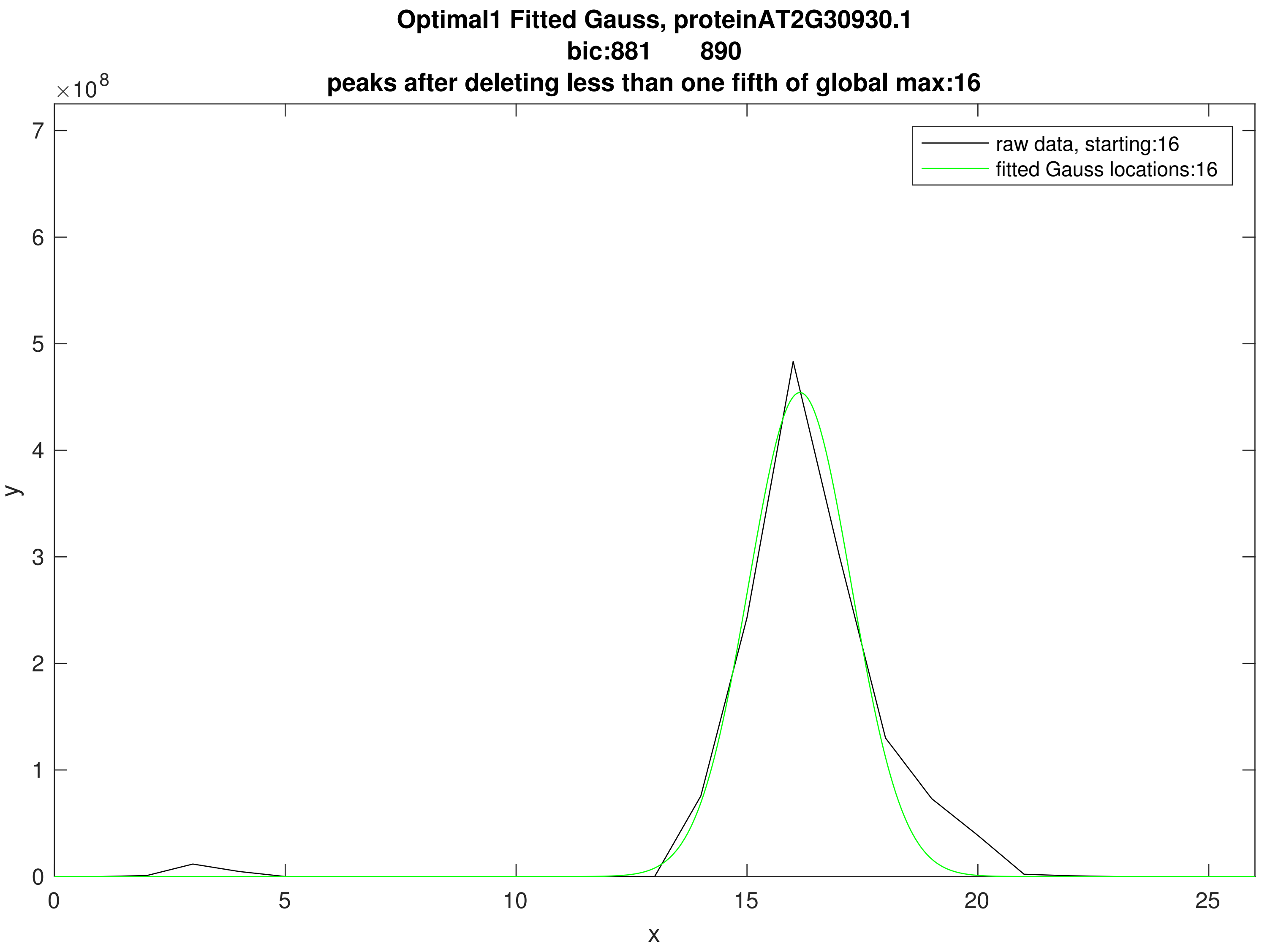
Optimal1 Fitted Gauss, proteinAT2G30930.1
bic:881 890
peaks after deleting less than one fifth of global max:16
×10^8
raw data, starting:16
fitted Gauss locations:16
y
x
0
1
2
3
4
5
6
7
0
5
10
15
20
25

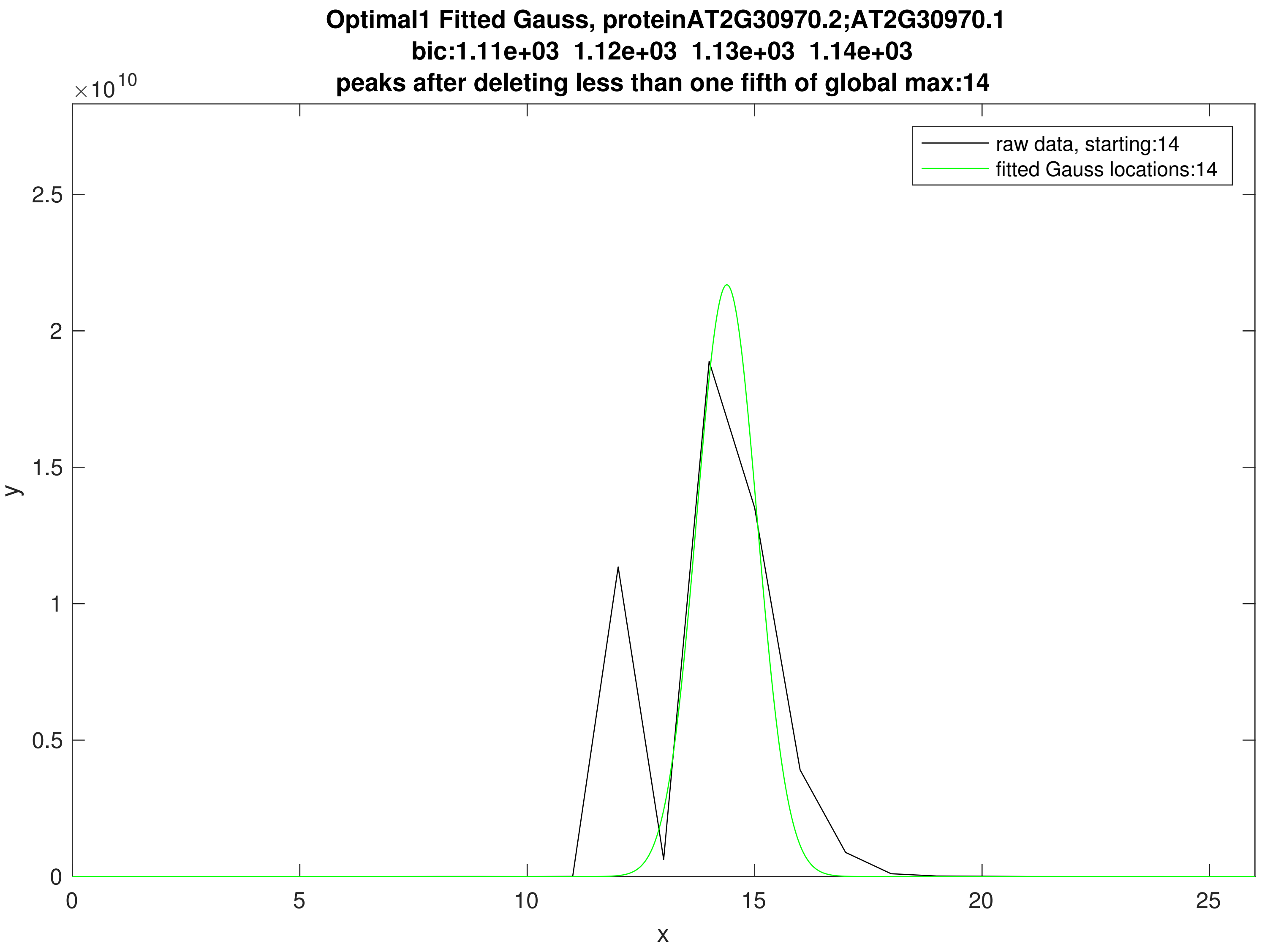

Optimal1 Fitted Gauss, proteinAT2G30970.2;AT2G30970.1
bic:1.11e+03 1.12e+03 1.13e+03 1.14e+03
peaks after deleting less than one fifth of global max:14
×10^10
2.5
2
1.5
1
0.5
0
y
0
5
10
15
20
25
x
raw data, starting:14
fitted Gauss locations:14

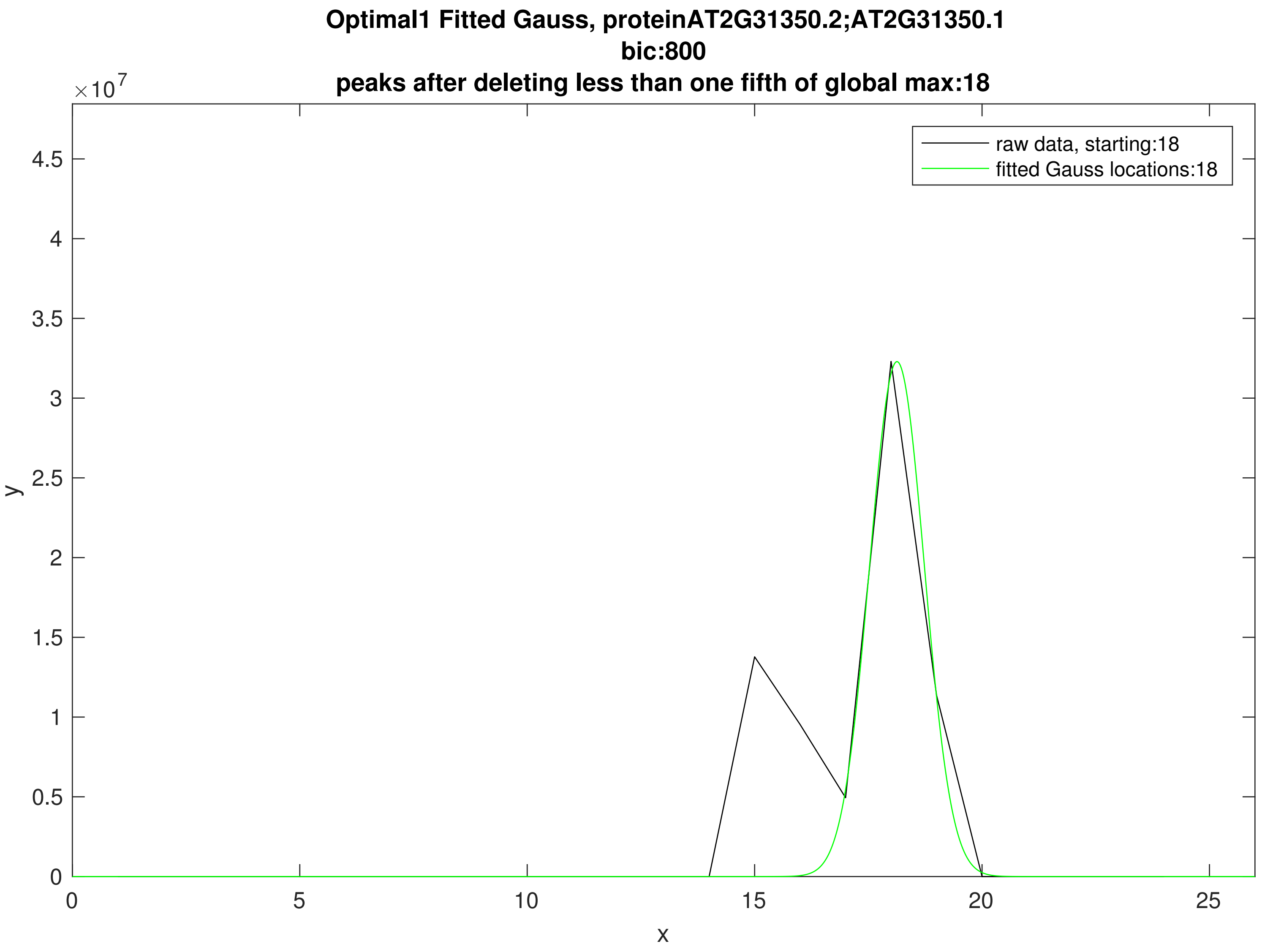
Optimal1 Fitted Gauss, proteinAT2G31350.2;AT2G31350.1
bic:800
peaks after deleting less than one fifth of global max:18
×10^7
raw data, starting:18
fitted Gauss locations:18
y
x
0
0.5
1
1.5
2
2.5
3
3.5
4
4.5
5
10
15
20
25

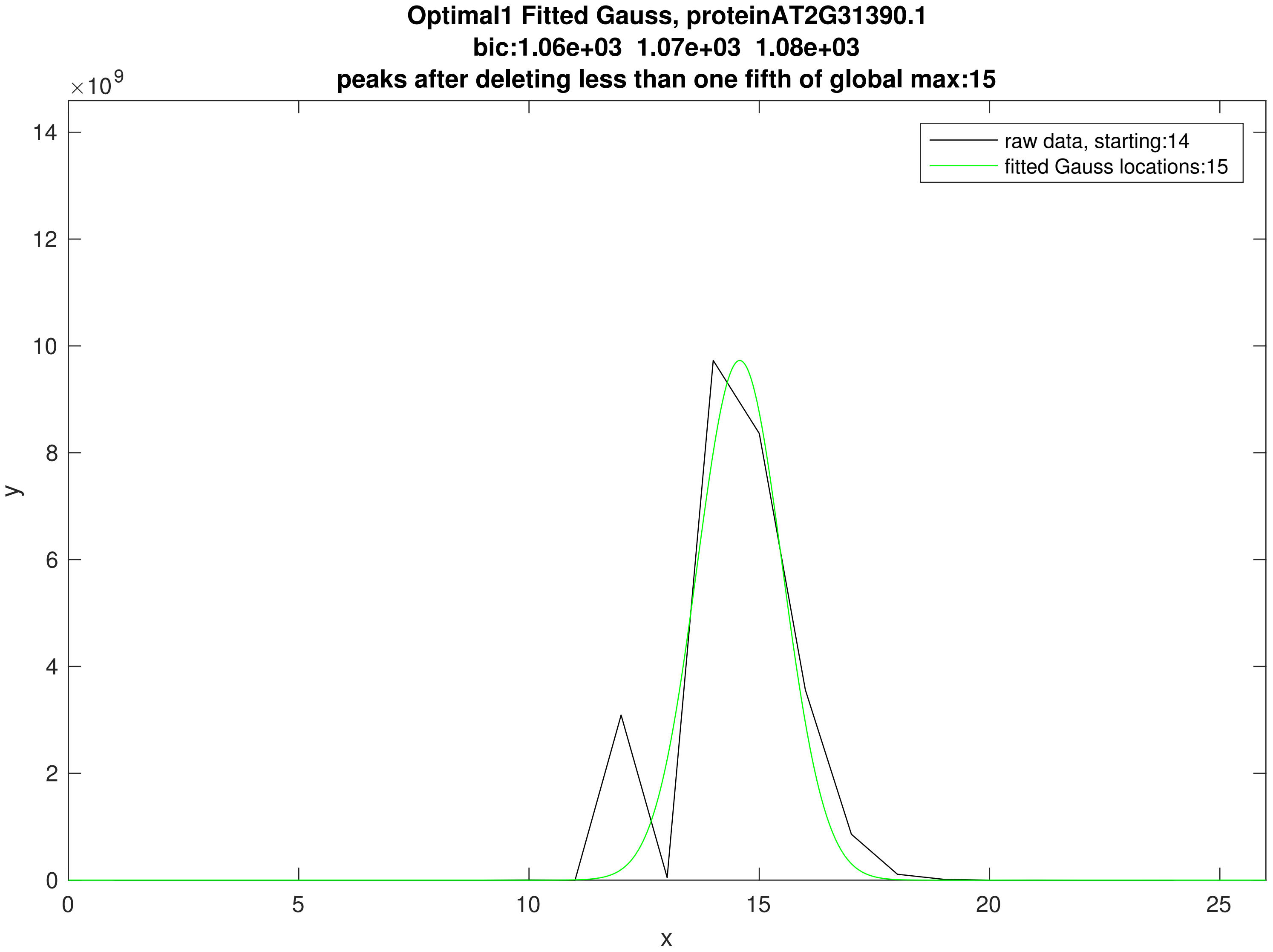

Optimal1 Fitted Gauss, proteinAT2G31390.1
bic:1.06e+03 1.07e+03 1.08e+03
peaks after deleting less than one fifth of global max:15
×10^9
14
12
10
8
6
4
2
0
y
0
5
10
15
20
25
x
raw data, starting:14
fitted Gauss locations:15

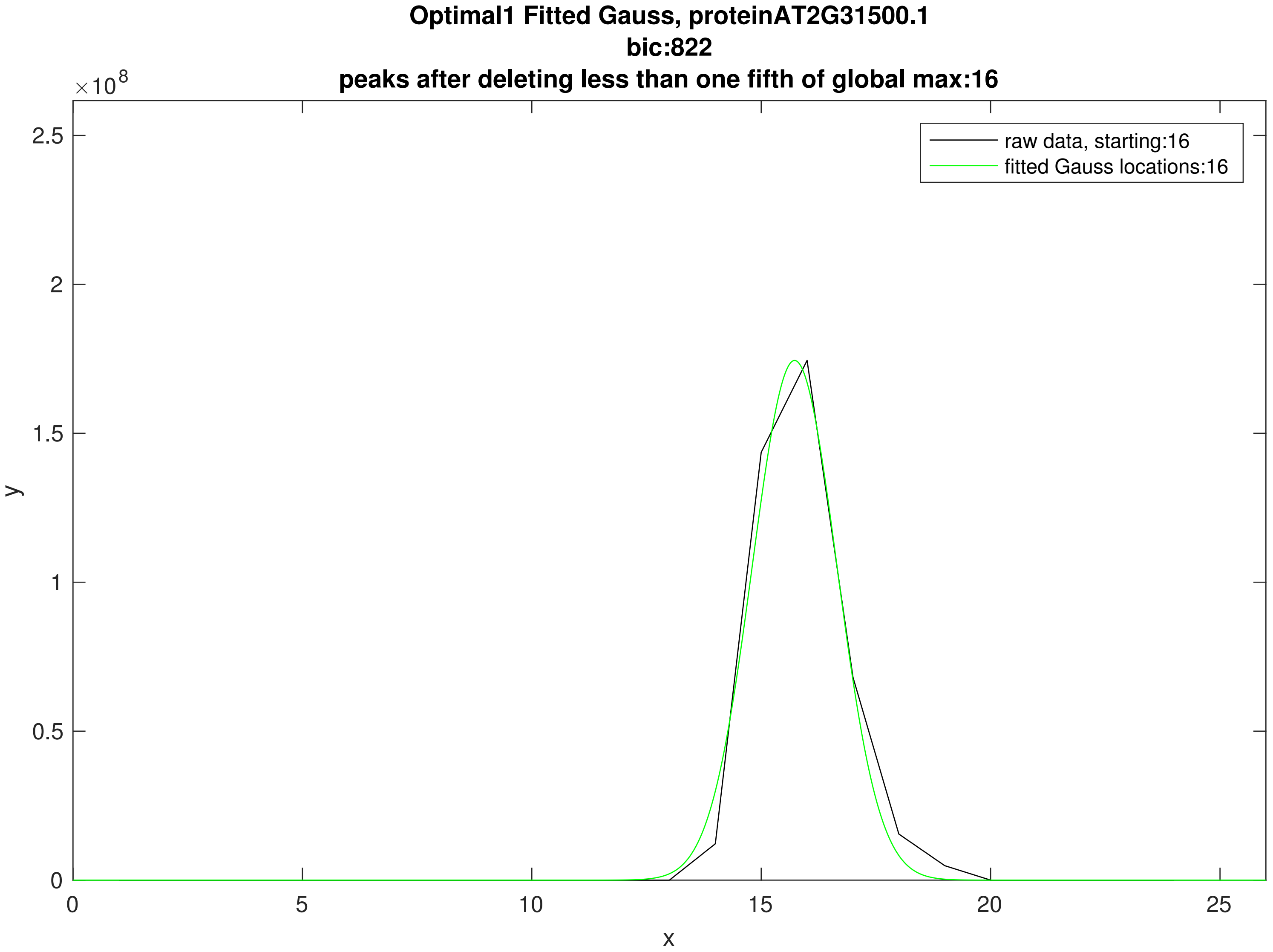

Optimal1 Fitted Gauss, proteinAT2G31500.1
bic:822
peaks after deleting less than one fifth of global max:16
×10^8
2.5
2
1.5
1
0.5
0
y
0
5
10
15
20
25
x
raw data, starting:16
fitted Gauss locations:16

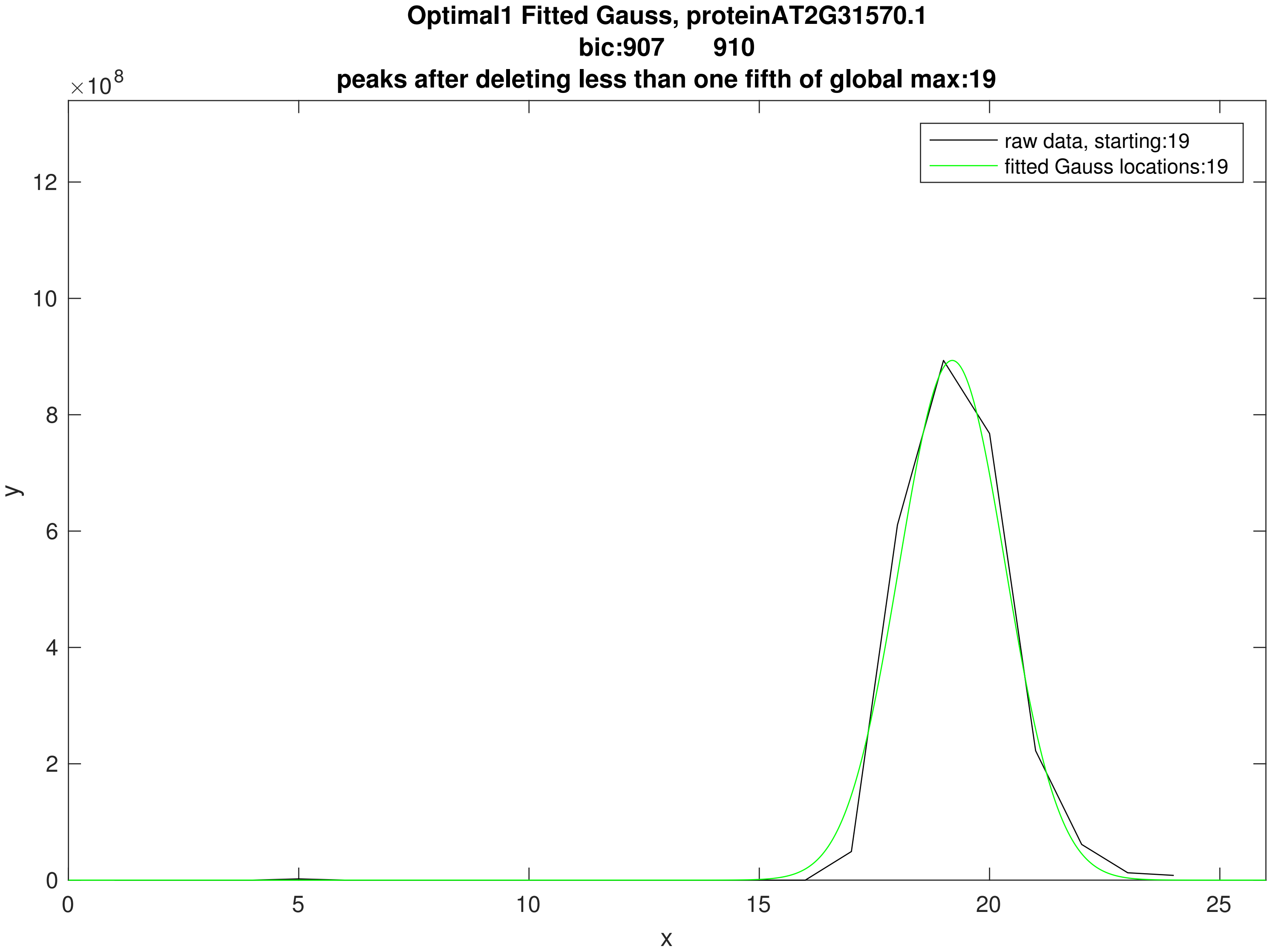
Optimal1 Fitted Gauss, proteinAT2G31570.1
bic:907 910
peaks after deleting less than one fifth of global max:19
×10^8
raw data, starting:19
fitted Gauss locations:19
y
x
0
2
4
6
8
10
12
0
5
10
15
20
25

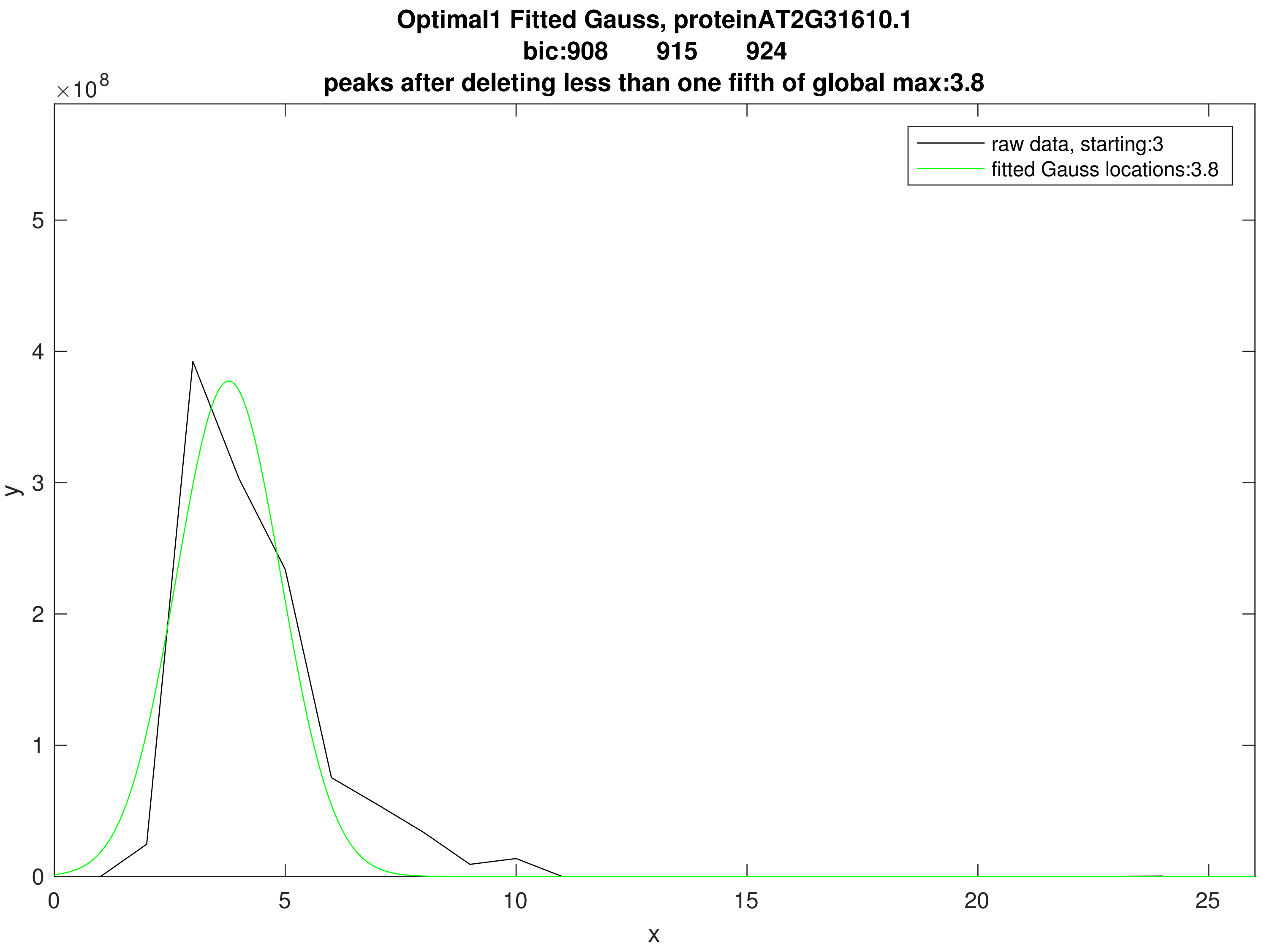

Optimal1 Fitted Gauss, proteinAT2G31610.1
bic:908 915 924
peaks after deleting less than one fifth of global max:3.8
×10^8
raw data, starting:3
fitted Gauss locations:3.8
y
x
0
1
2
3
4
5
0
5
10
15
20
25

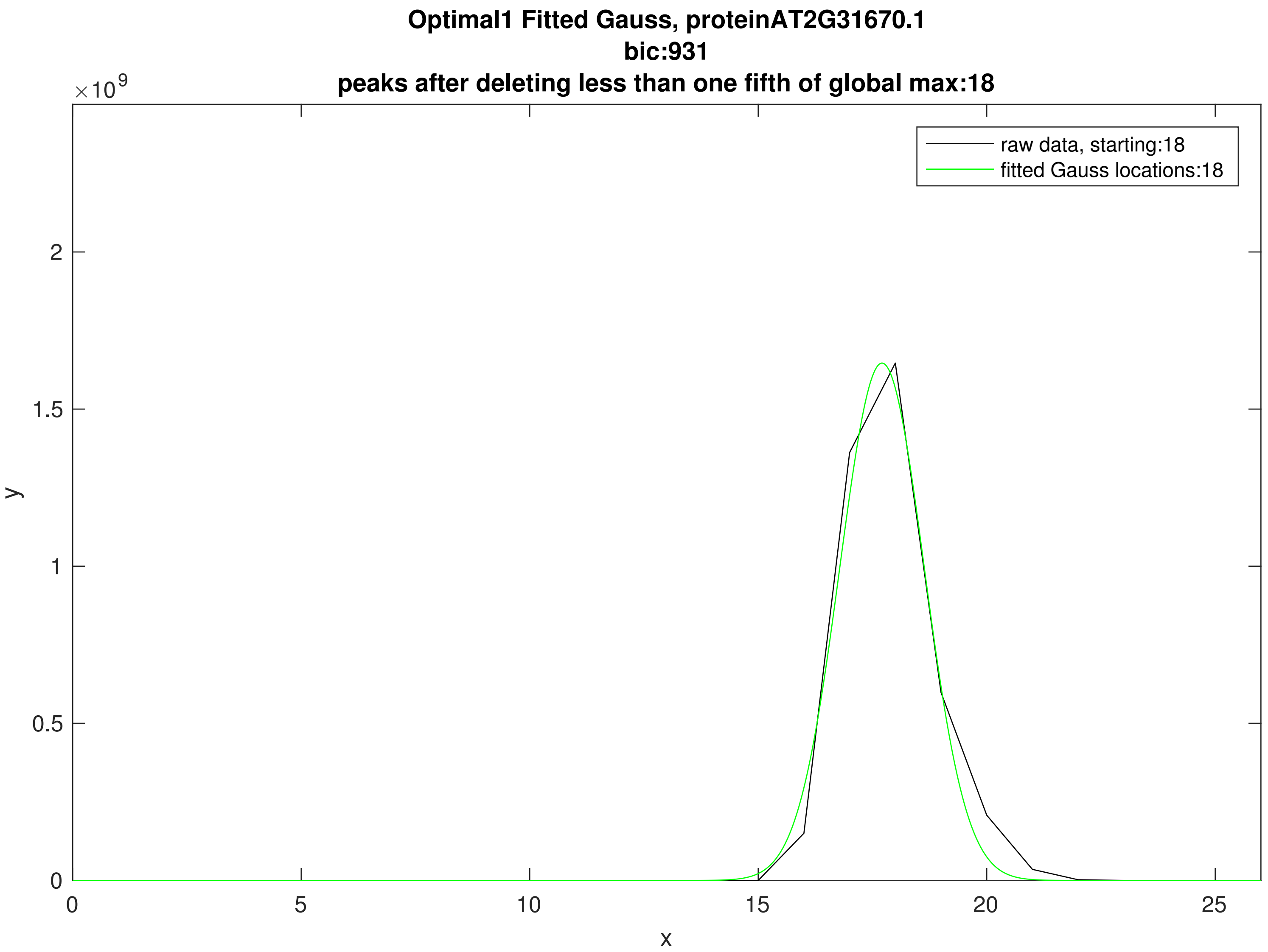

Optimal1 Fitted Gauss, proteinAT2G31670.1
bic:931
peaks after deleting less than one fifth of global max:18
×10^9
raw data, starting:18
fitted Gauss locations:18
2
1.5
1
0.5
0
y
0
5
10
15
20
25
x

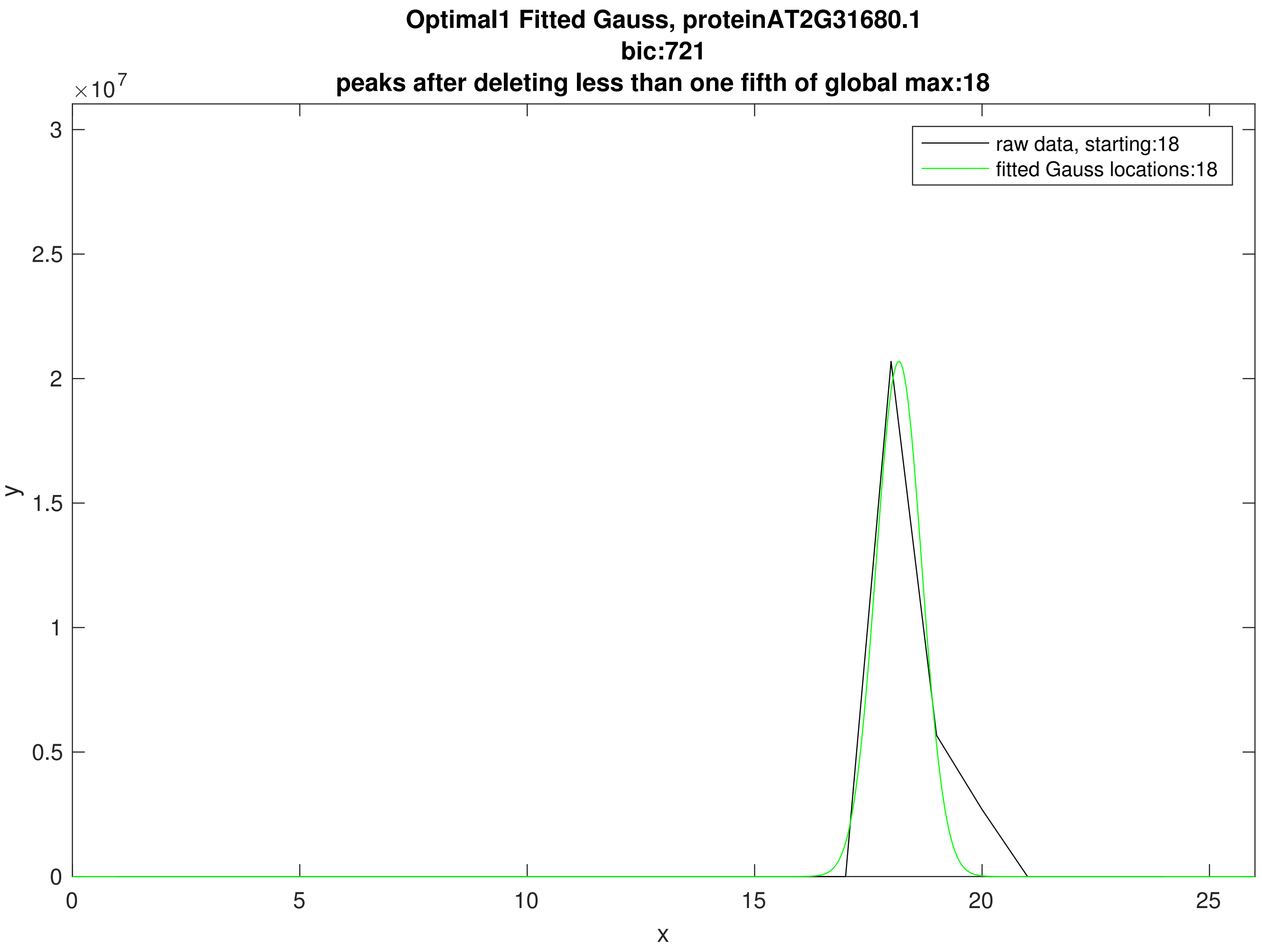
Optimal1 Fitted Gauss, proteinAT2G31680.1
bic:721
peaks after deleting less than one fifth of global max:18
×10^7
0
0.5
1
1.5
2
2.5
3
y
0
5
10
15
20
25
x
raw data, starting:18
fitted Gauss locations:18

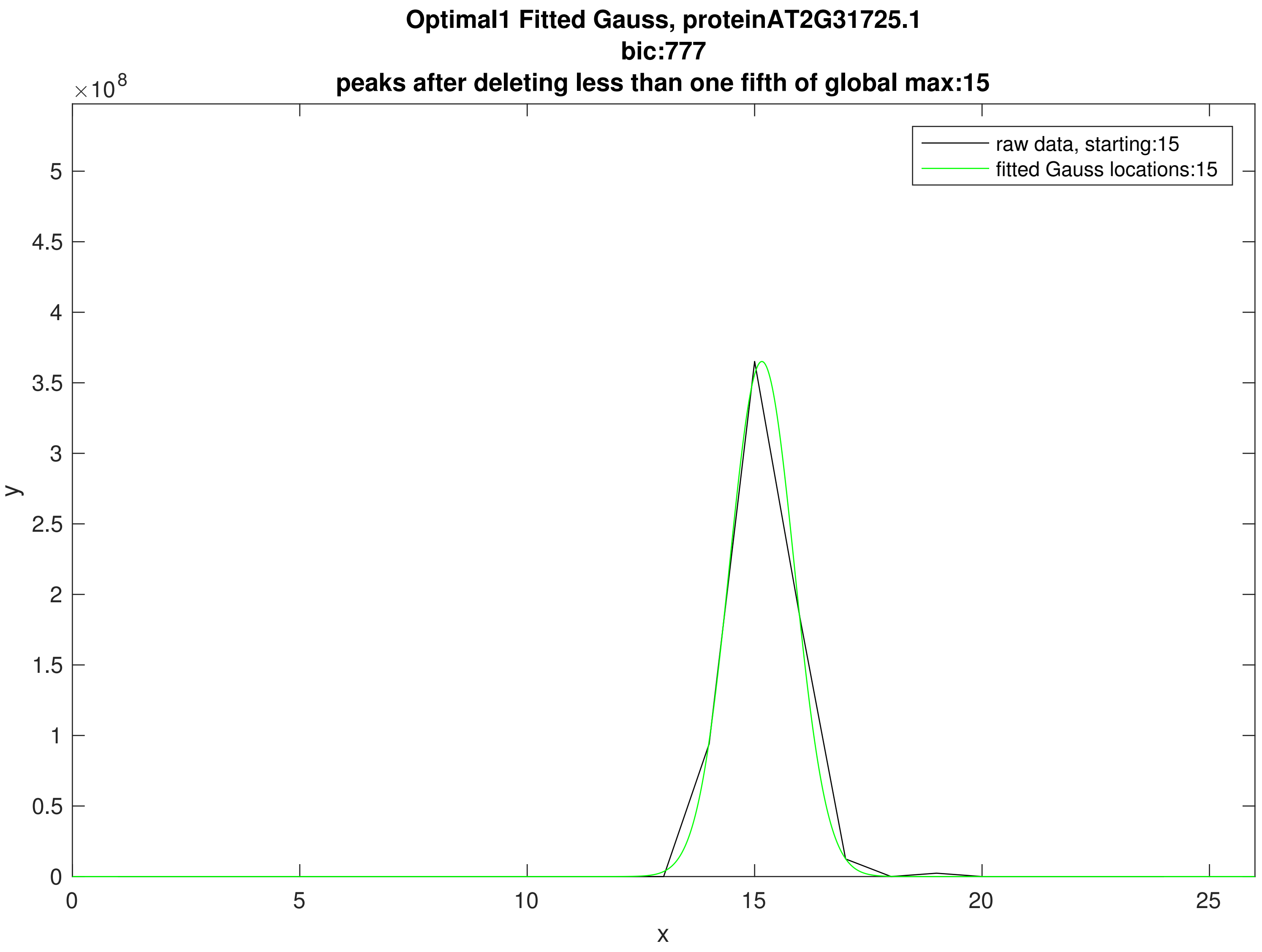

Optimal1 Fitted Gauss, proteinAT2G31725.1
bic:777
peaks after deleting less than one fifth of global max:15
×10^8
raw data, starting:15
fitted Gauss locations:15
5
4.5
4
3.5
3
2.5
2
1.5
1
0.5
0
y
0
5
10
15
20
25
x

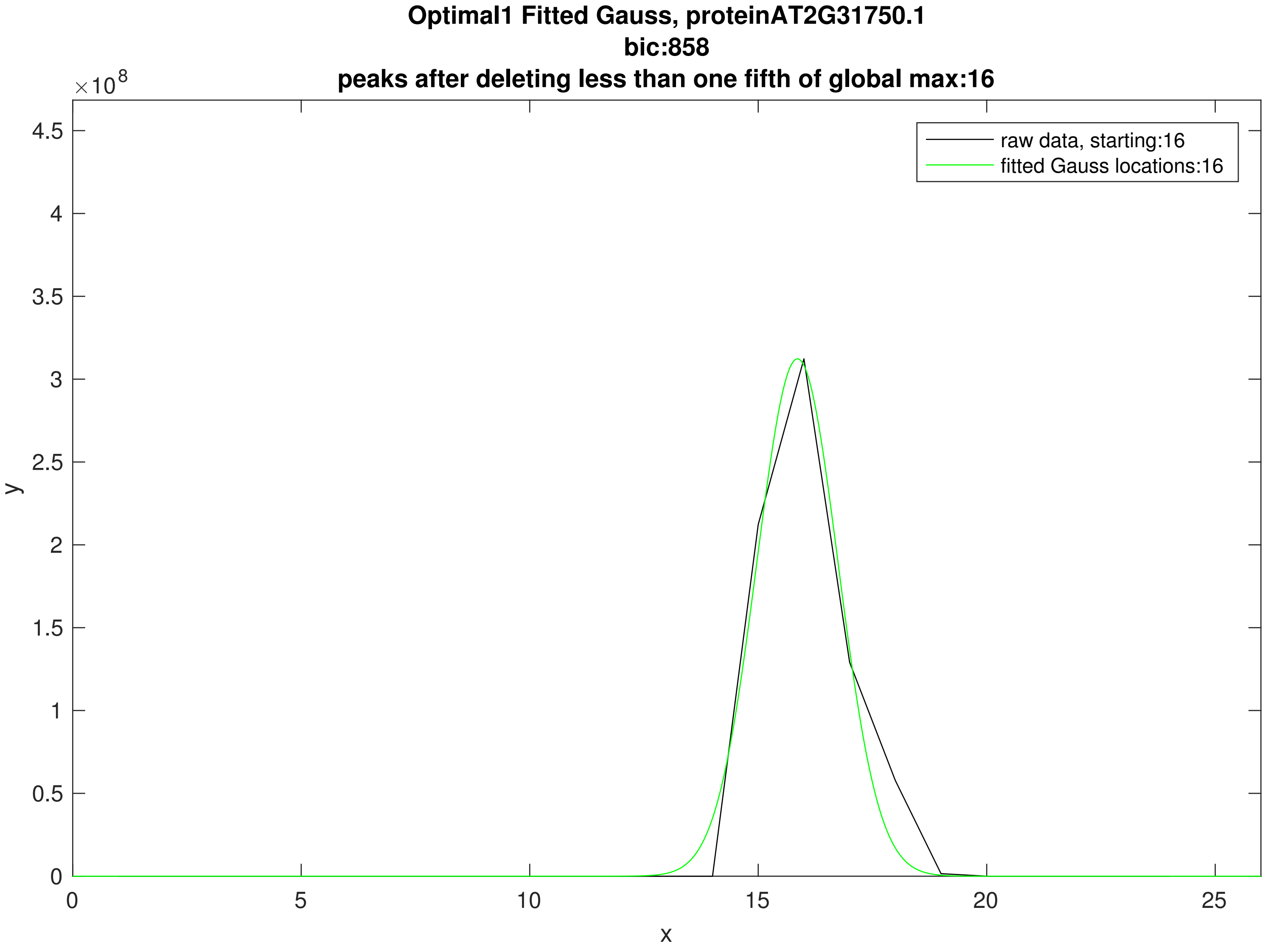
Optimal1 Fitted Gauss, proteinAT2G31750.1
bic:858
peaks after deleting less than one fifth of global max:16
×10^8
4.5
4
3.5
3
2.5
2
1.5
1
0.5
0
y
0
5
10
15
20
25
x
raw data, starting:16
fitted Gauss locations:16

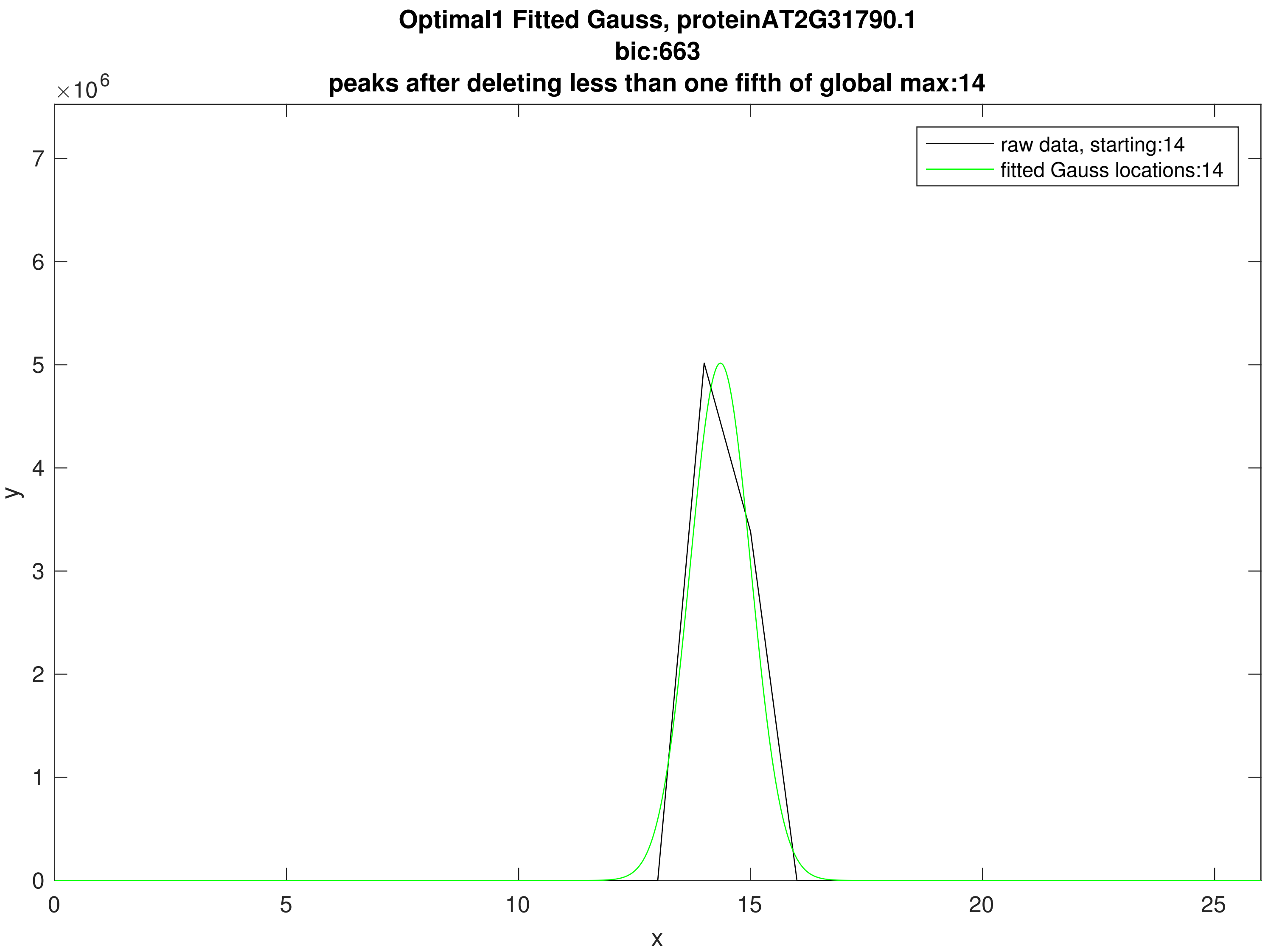
Optimal1 Fitted Gauss, proteinAT2G31790.1
bic:663
peaks after deleting less than one fifth of global max:14
×10^6
raw data, starting:14
fitted Gauss locations:14
y
x
0
1
2
3
4
5
6
7
0
5
10
15
20
25

**Optimal1 Fitted Gauss, proteinAT2G31810.2;AT2G31810.1;AT2G31810.3**
**bic:695**
**peaks after deleting less than one fifth of global max:5.3**

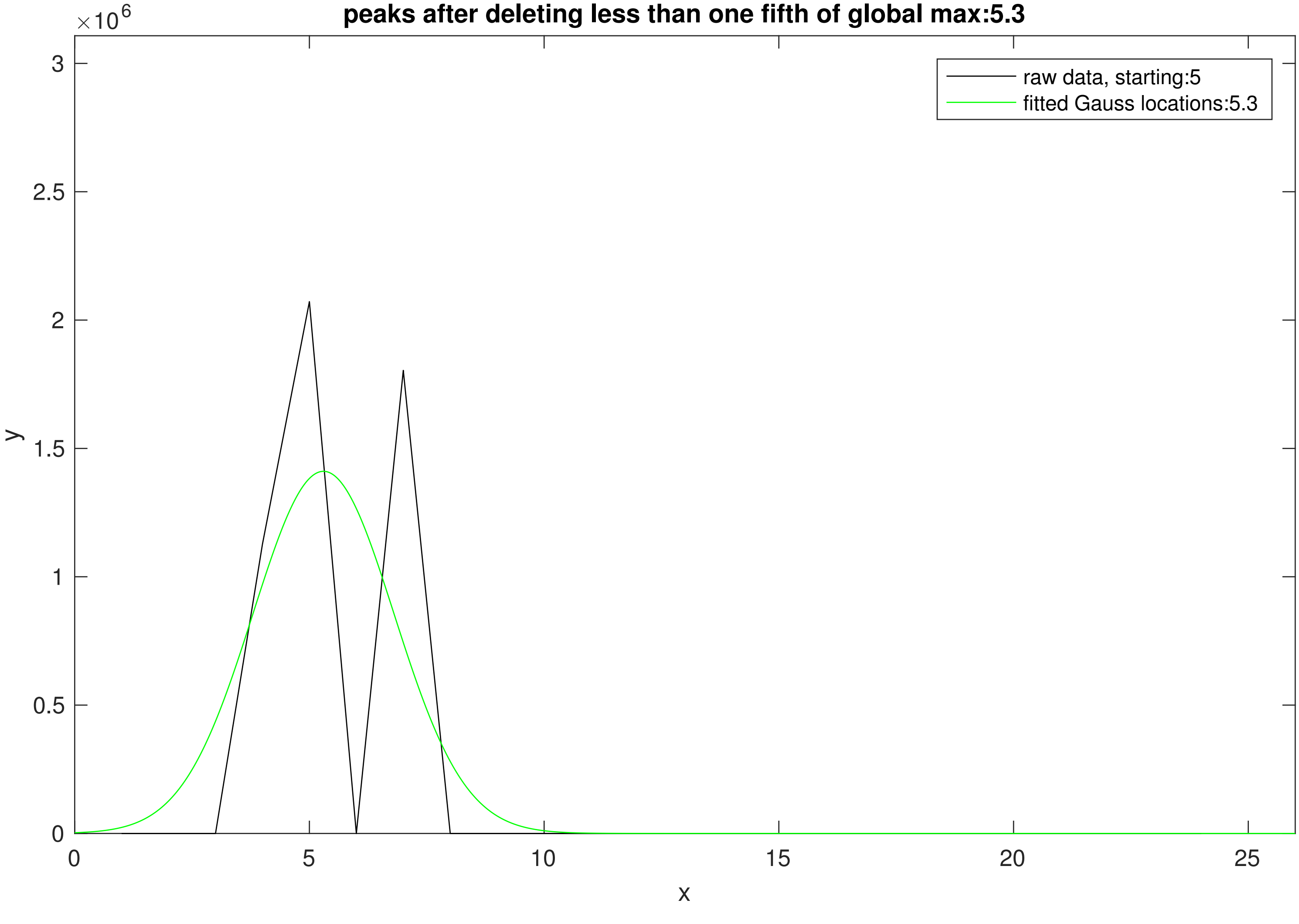

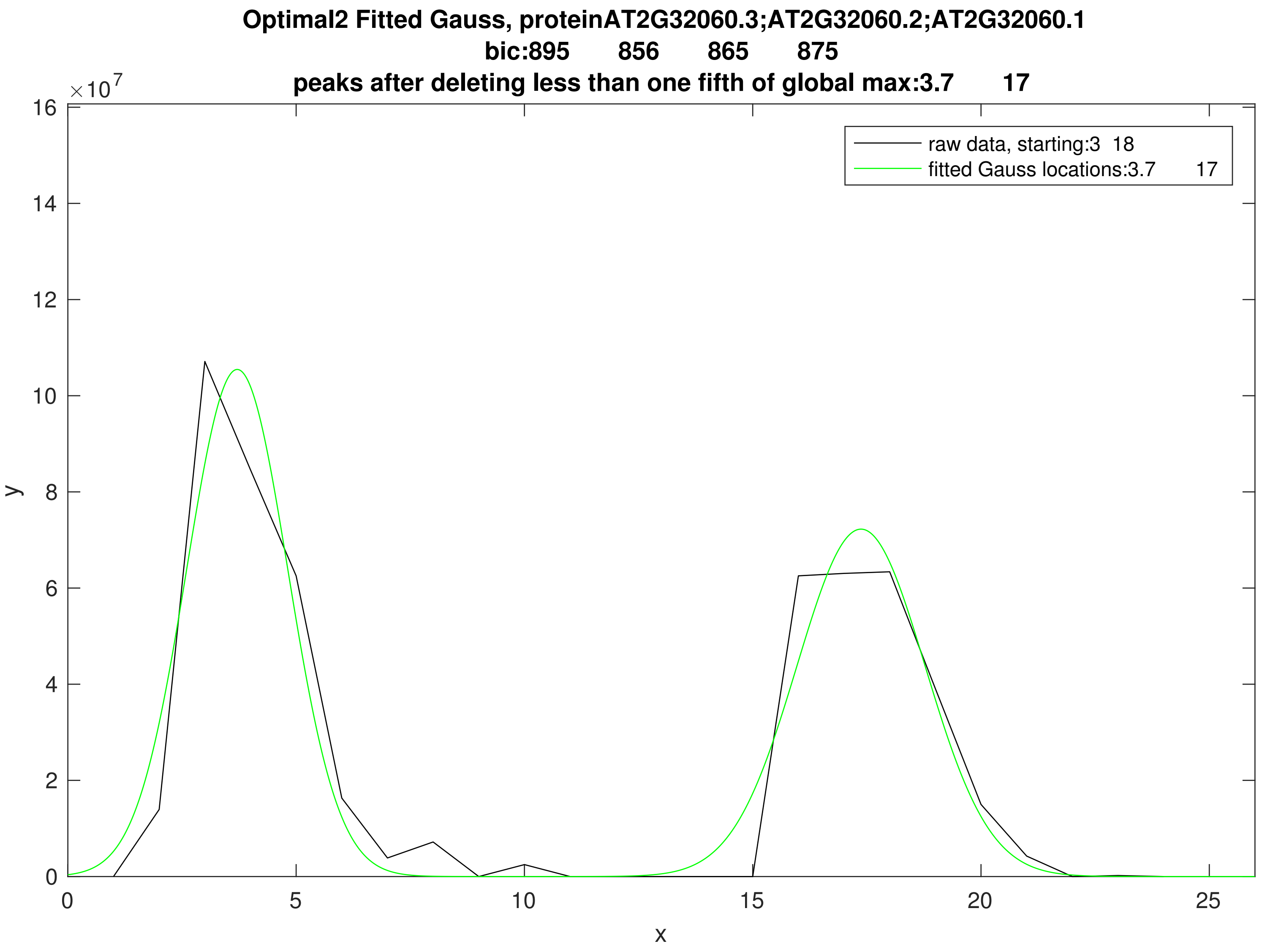

Optimal2 Fitted Gauss, proteinAT2G32060.3;AT2G32060.2;AT2G32060.1
bic:895 856 865 875
peaks after deleting less than one fifth of global max:3.7 17
×10^7
16
14
12
10
8
6
4
2
0
y
0
5
10
15
20
25
x
raw data, starting:3 18
fitted Gauss locations:3.7 17

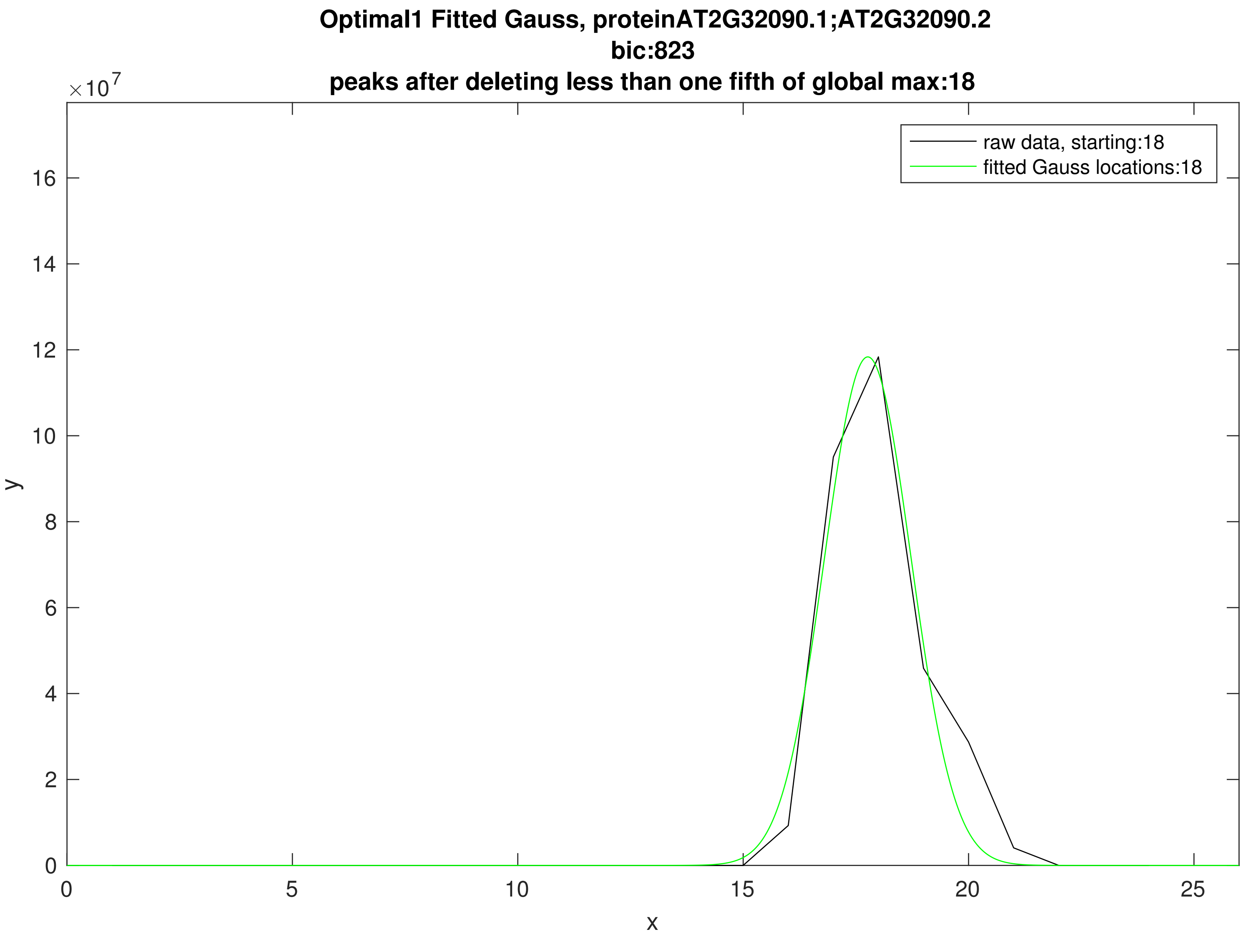
Optimal1 Fitted Gauss, proteinAT2G32090.1;AT2G32090.2
bic:823
peaks after deleting less than one fifth of global max:18
×10^7
raw data, starting:18
fitted Gauss locations:18
y
x
0
2
4
6
8
10
12
14
16
0
5
10
15
20
25

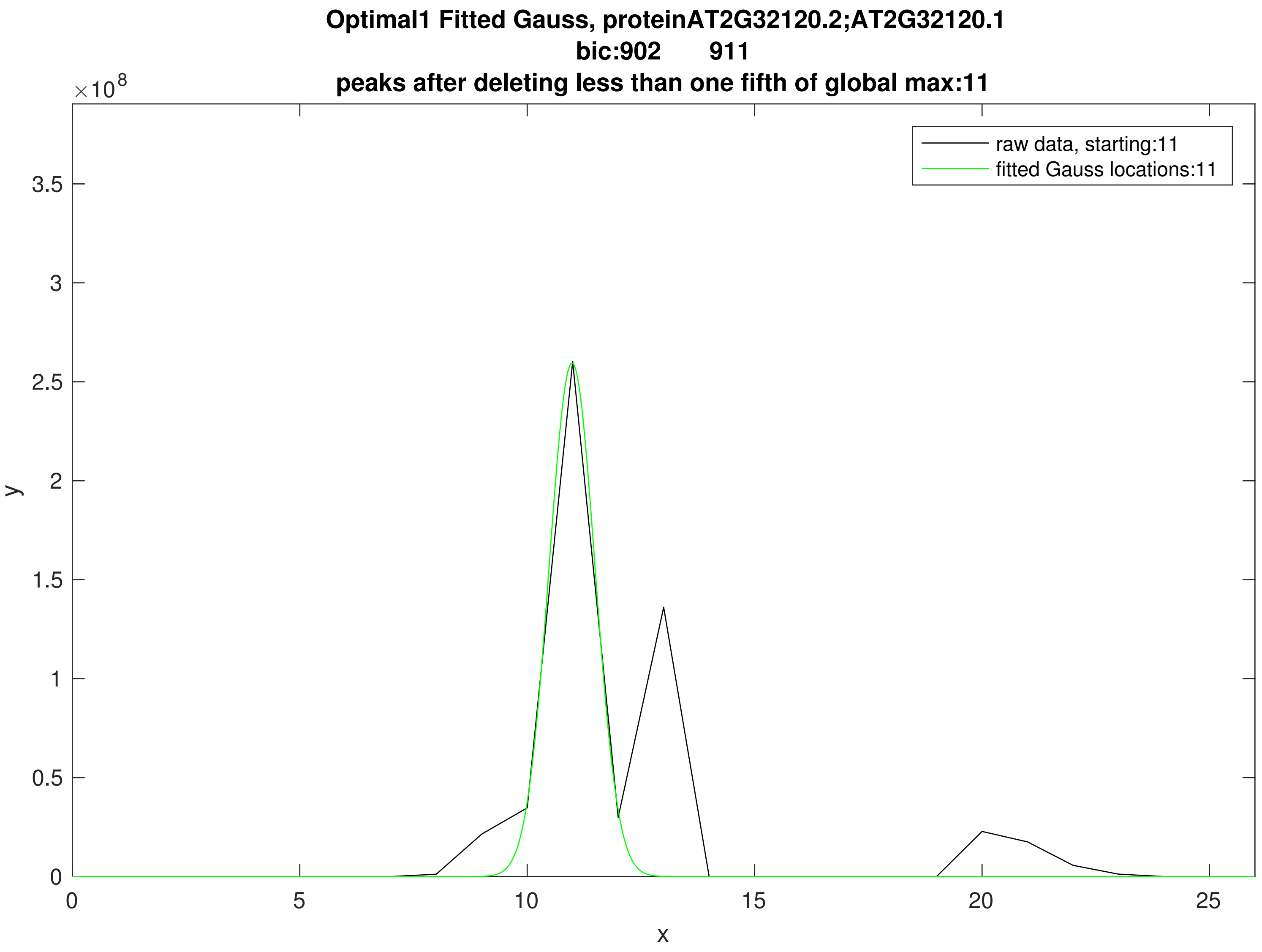

Optimal1 Fitted Gauss, proteinAT2G32120.2;AT2G32120.1
bic:902 911
peaks after deleting less than one fifth of global max:11
×10^8
raw data, starting:11
fitted Gauss locations:11
y
x
0
0.5
1
1.5
2
2.5
3
3.5
0
5
10
15
20
25

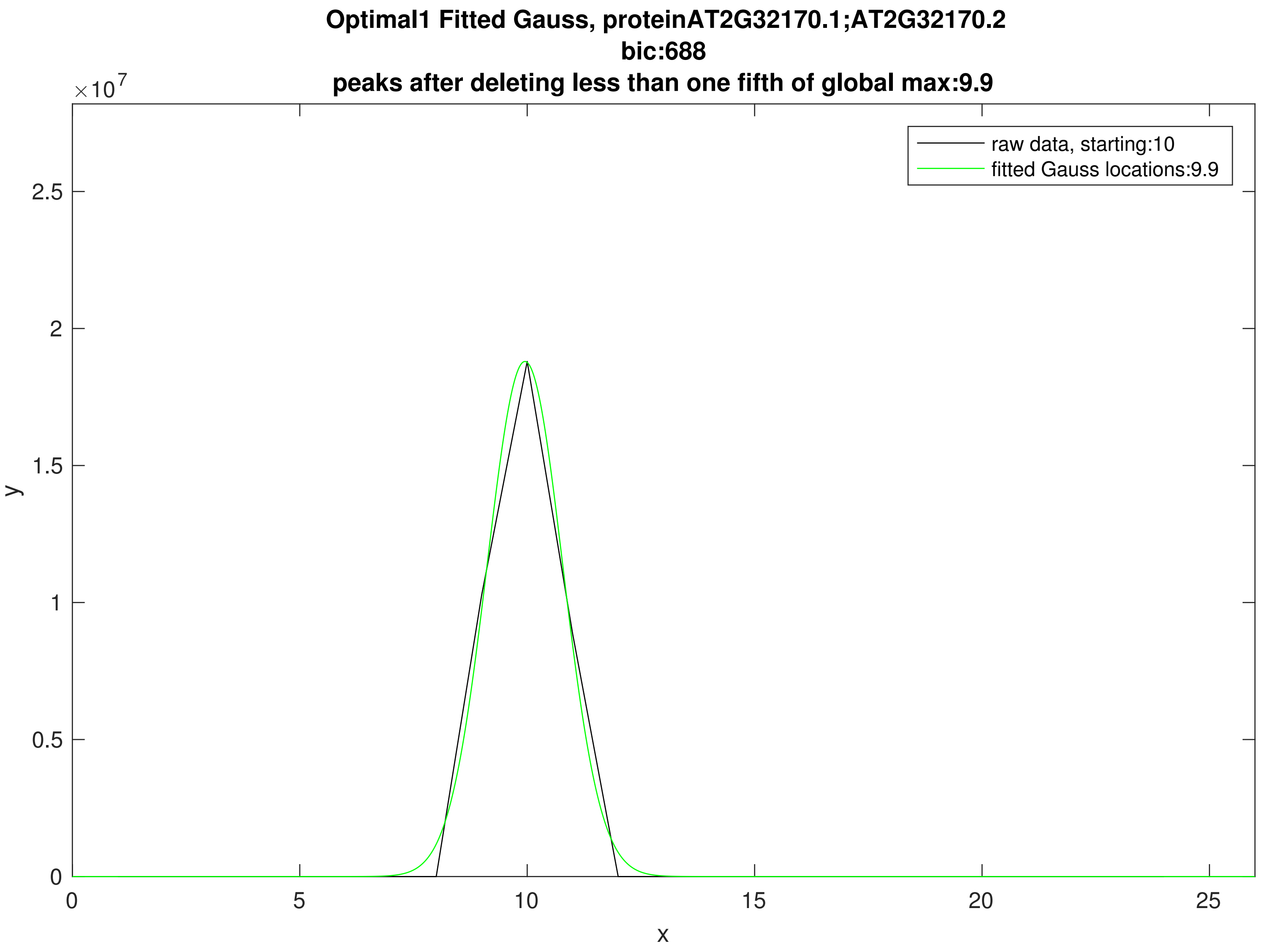

Optimal1 Fitted Gauss, proteinAT2G32170.1;AT2G32170.2
bic:688
peaks after deleting less than one fifth of global max:9.9
×10^7
raw data, starting:10
fitted Gauss locations:9.9
y
x
0
0.5
1
1.5
2
2.5
0
5
10
15
20
25

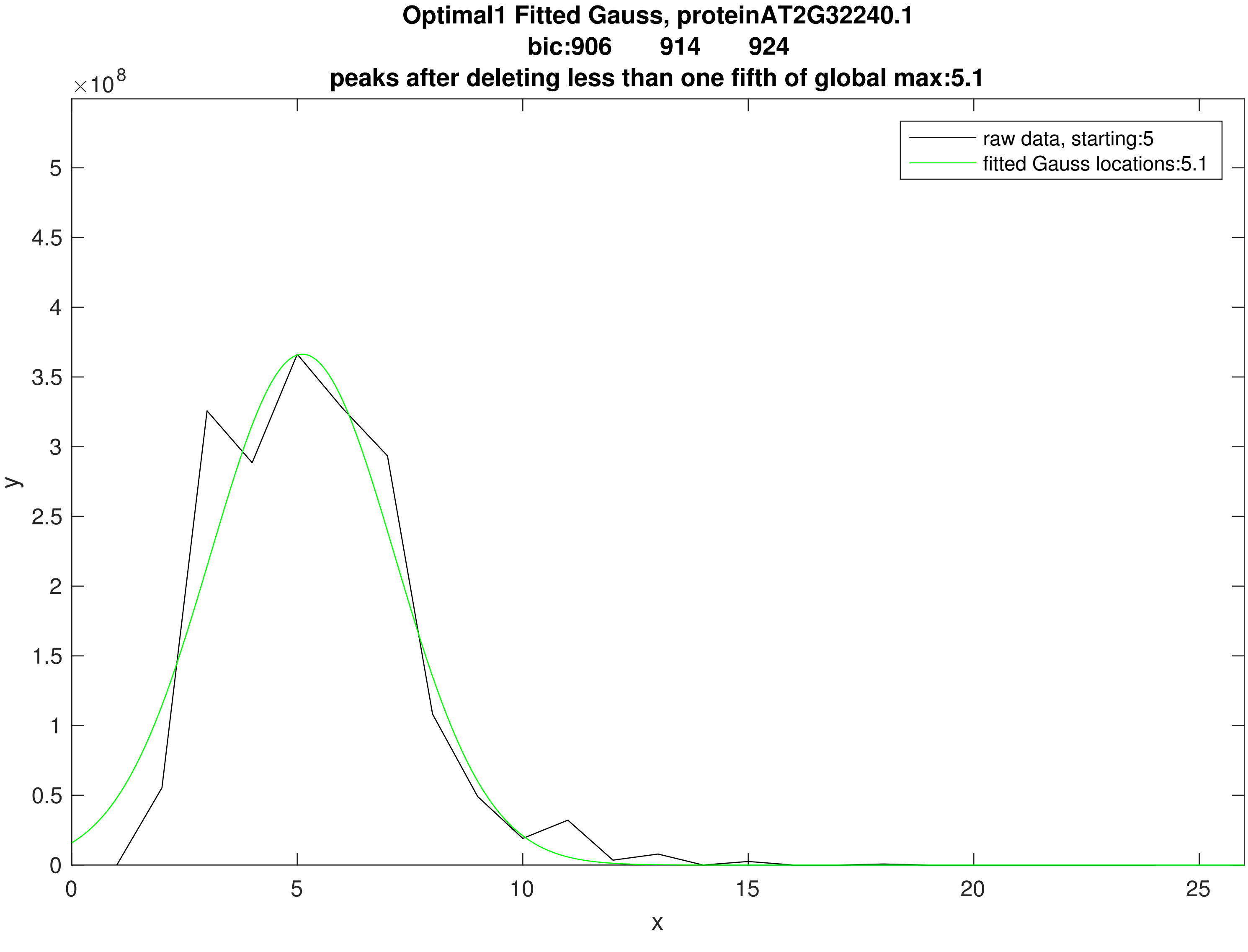
Optimal1 Fitted Gauss, proteinAT2G32240.1
bic:906 914 924
peaks after deleting less than one fifth of global max:5.1
×10^8
raw data, starting:5
fitted Gauss locations:5.1
y
x
0
0.5
1
1.5
2
2.5
3
3.5
4
4.5
5
0
5
10
15
20
25

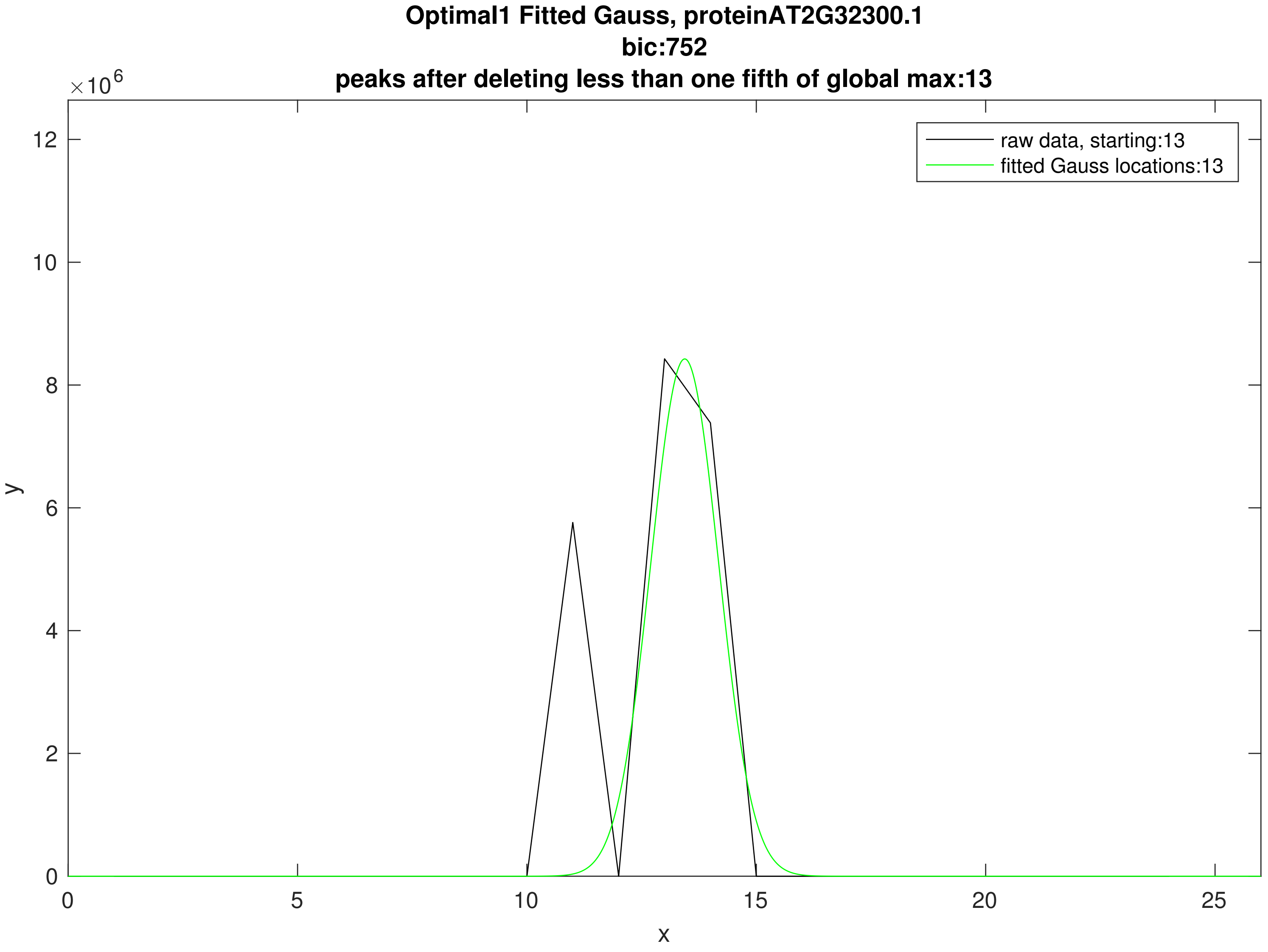

Optimal1 Fitted Gauss, proteinAT2G32300.1
bic:752
peaks after deleting less than one fifth of global max:13
×10^6
12
10
8
6
4
2
0
y
0
5
10
15
20
25
x
raw data, starting:13
fitted Gauss locations:13

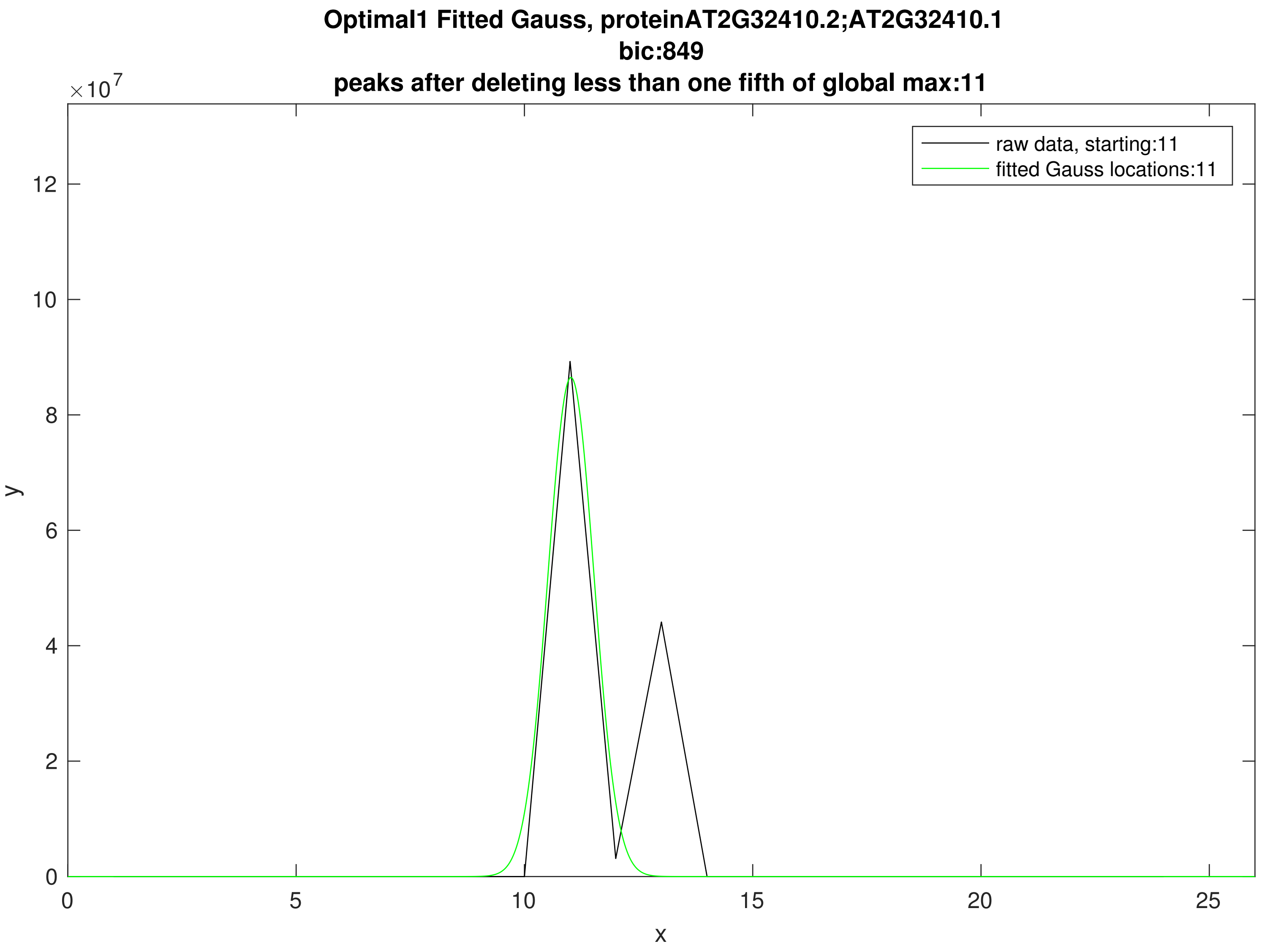

Optimal1 Fitted Gauss, proteinAT2G32410.2;AT2G32410.1
bic:849
peaks after deleting less than one fifth of global max:11
$\times 10^7$
raw data, starting:11
fitted Gauss locations:11
y
x
0
2
4
6
8
10
12
5
15
20
25

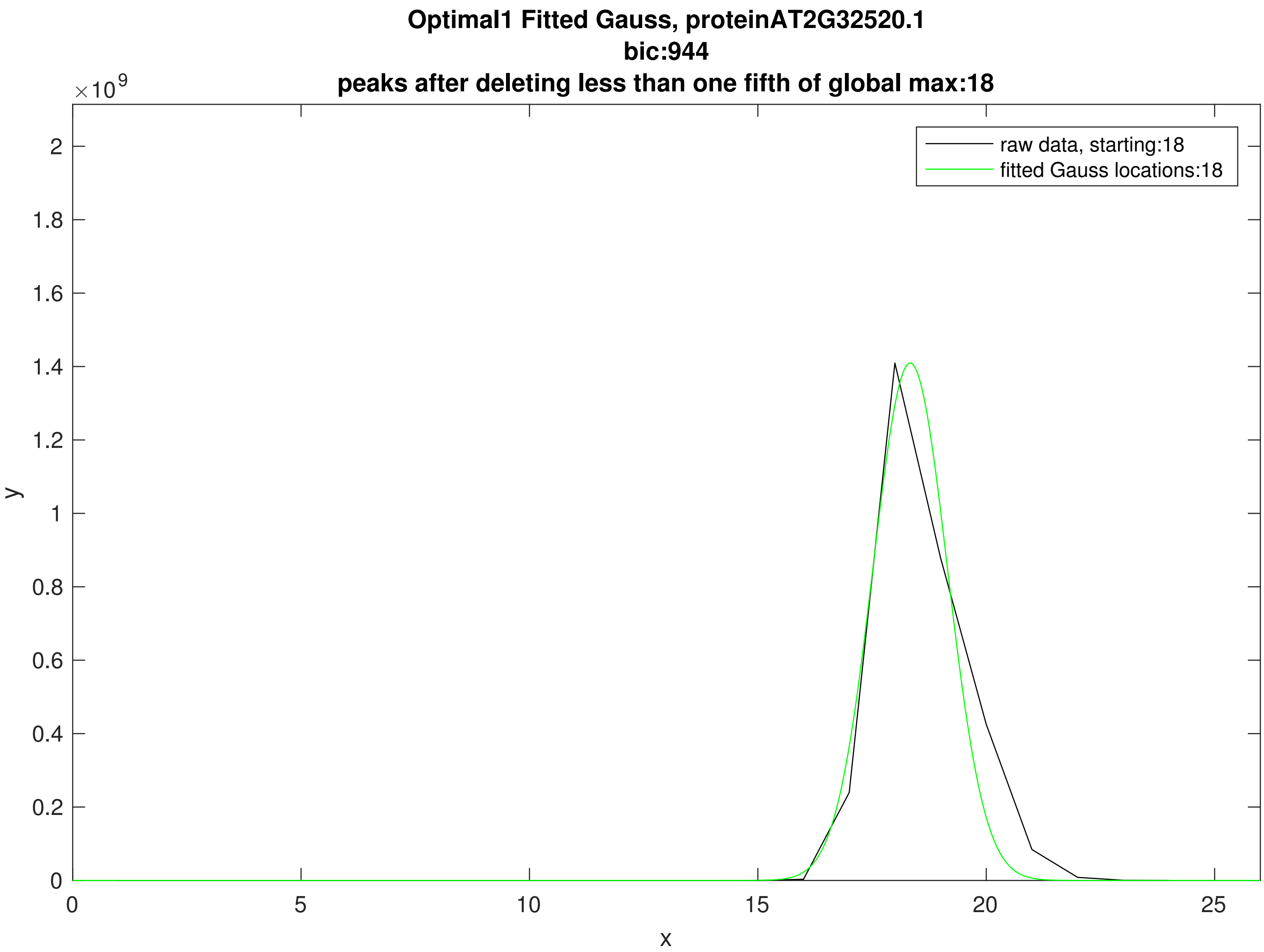

Optimal1 Fitted Gauss, proteinAT2G32520.1
bic:944
peaks after deleting less than one fifth of global max:18
×10^9
2
1.8
1.6
1.4
1.2
1
0.8
0.6
0.4
0.2
0
y
0
5
10
15
20
25
x
raw data, starting:18
fitted Gauss locations:18

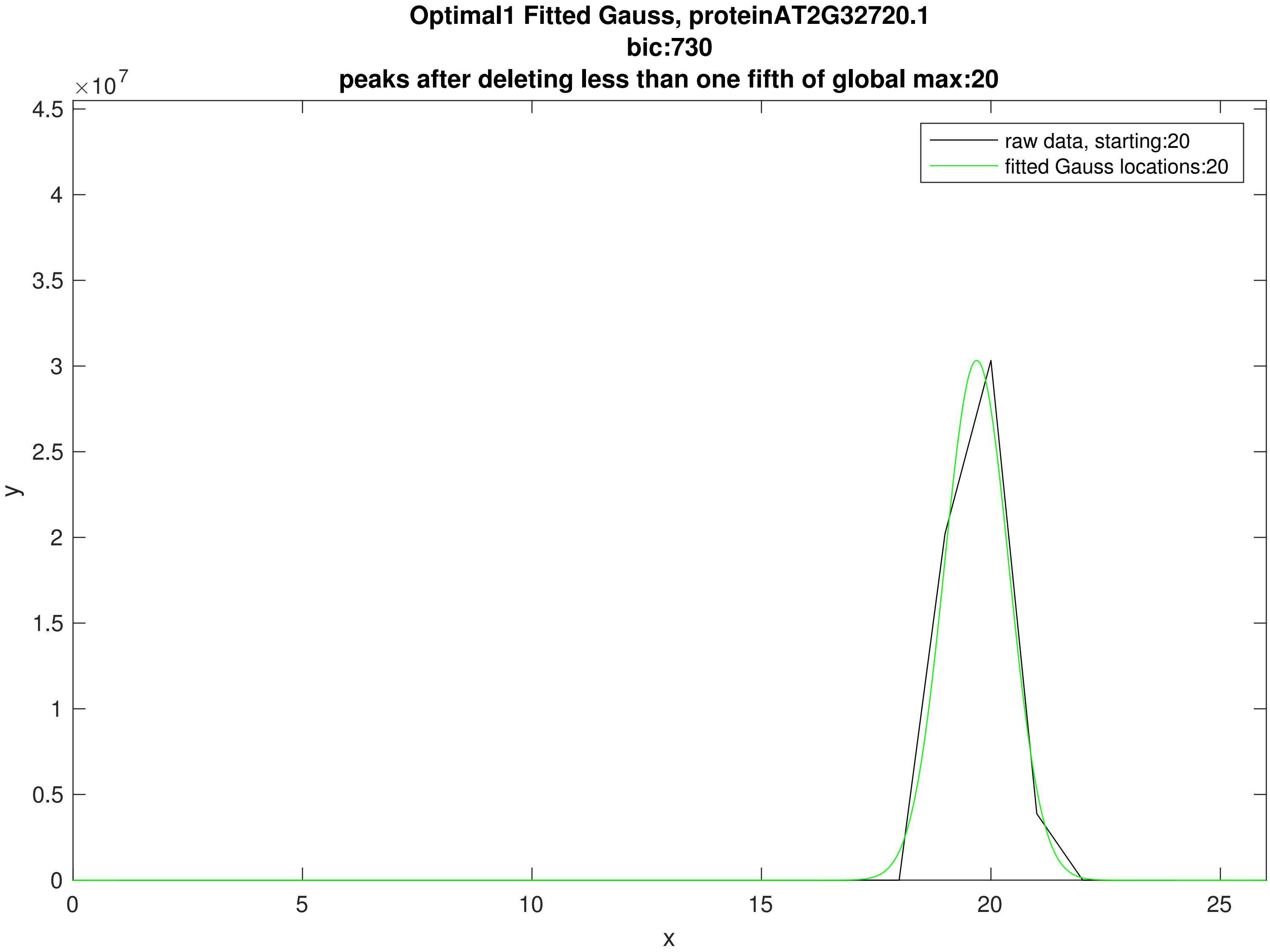

Optimal1 Fitted Gauss, proteinAT2G32720.1
bic:730
peaks after deleting less than one fifth of global max:20
×10^7
4.5
4
3.5
3
2.5
2
1.5
1
0.5
0
y
0
5
10
15
20
25
x
raw data, starting:20
fitted Gauss locations:20

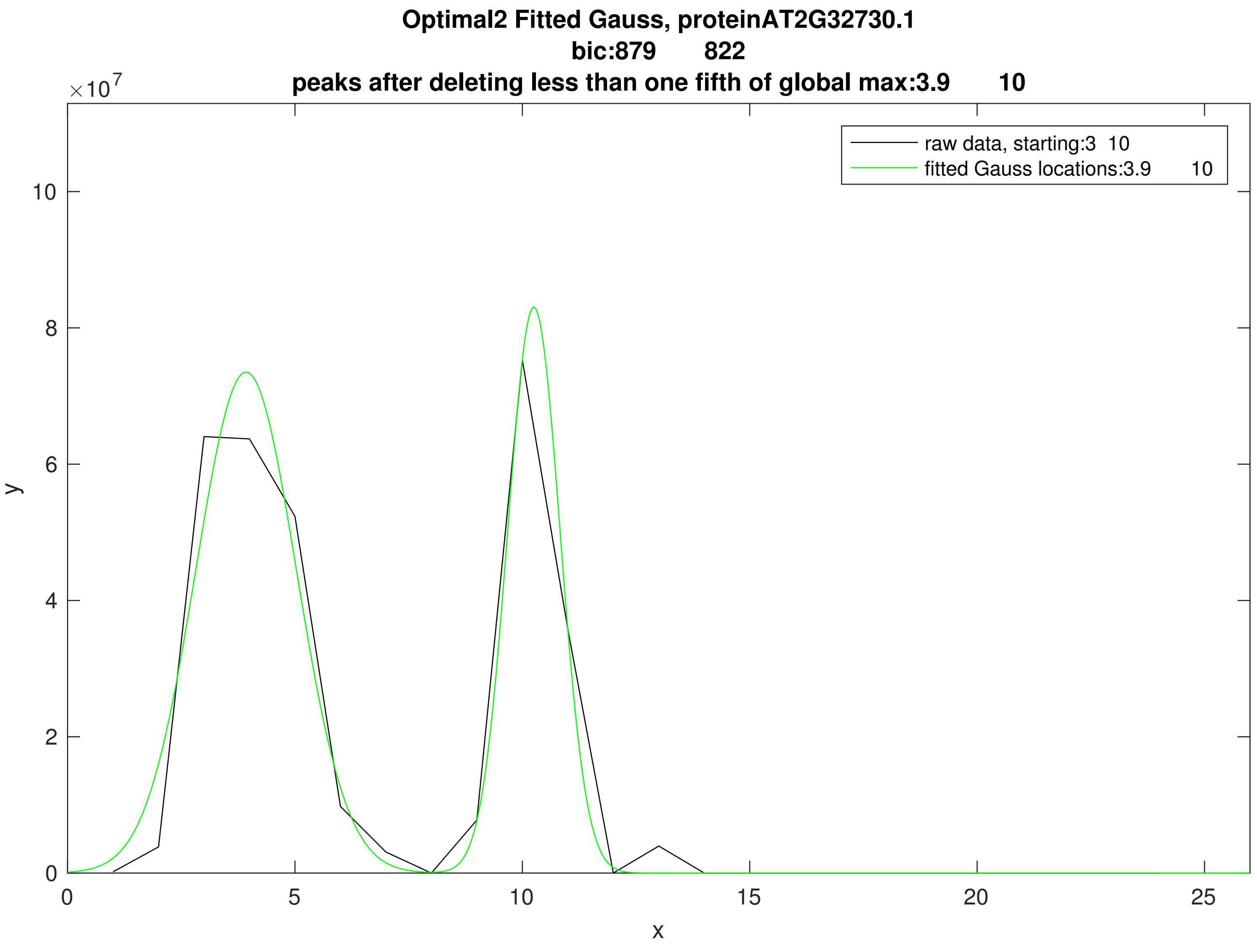

Optimal2 Fitted Gauss, proteinAT2G32730.1
bic:879 822
peaks after deleting less than one fifth of global max:3.9 10
×10^7
raw data, starting:3 10
fitted Gauss locations:3.9 10
10
8
6
4
2
0
y
0
5
10
15
20
25
x

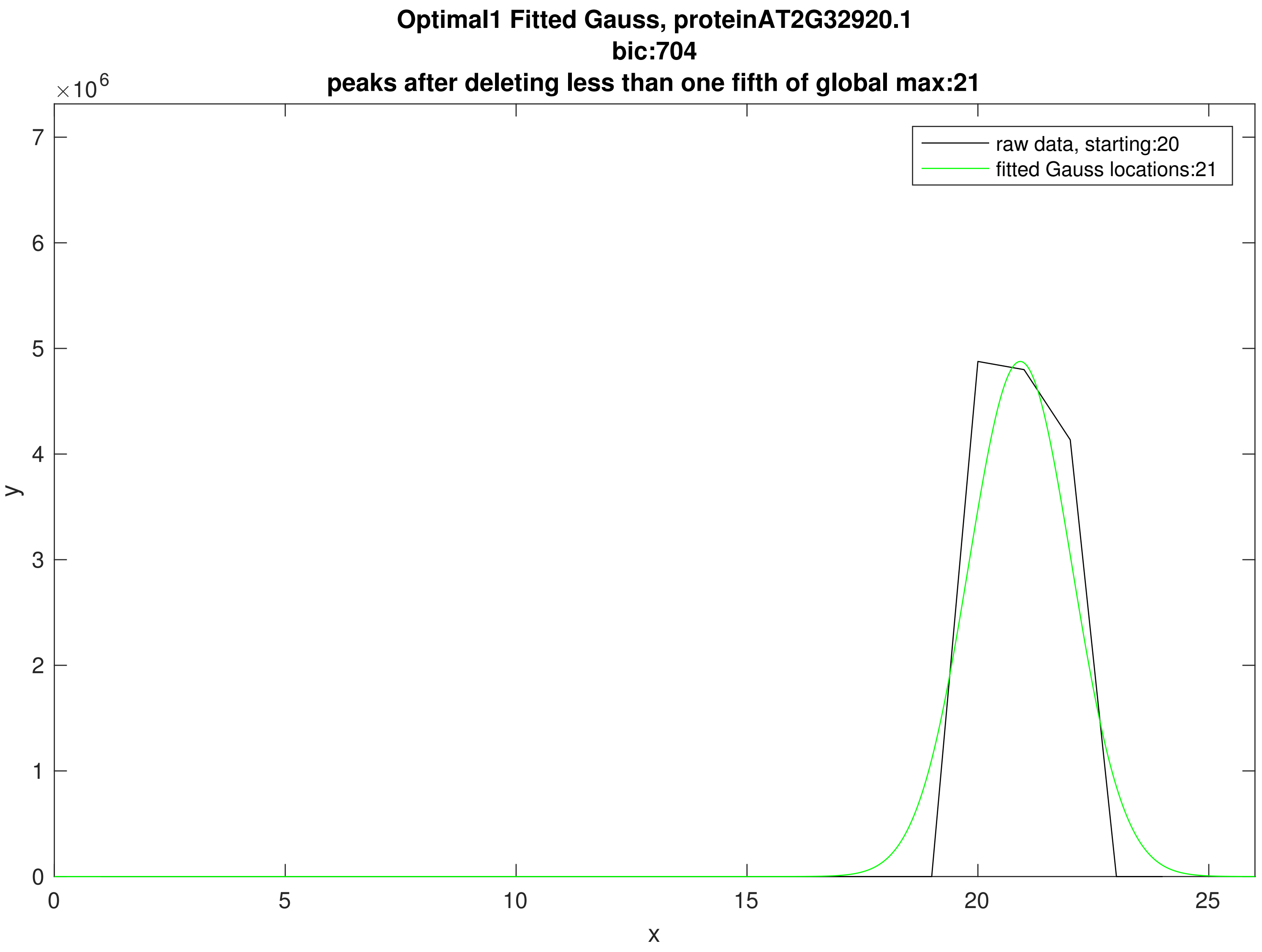
Optimal1 Fitted Gauss, proteinAT2G32920.1
bic:704
peaks after deleting less than one fifth of global max:21
×10^6
raw data, starting:20
fitted Gauss locations:21
y
x
0
1
2
3
4
5
6
7
0
5
10
15
20
25

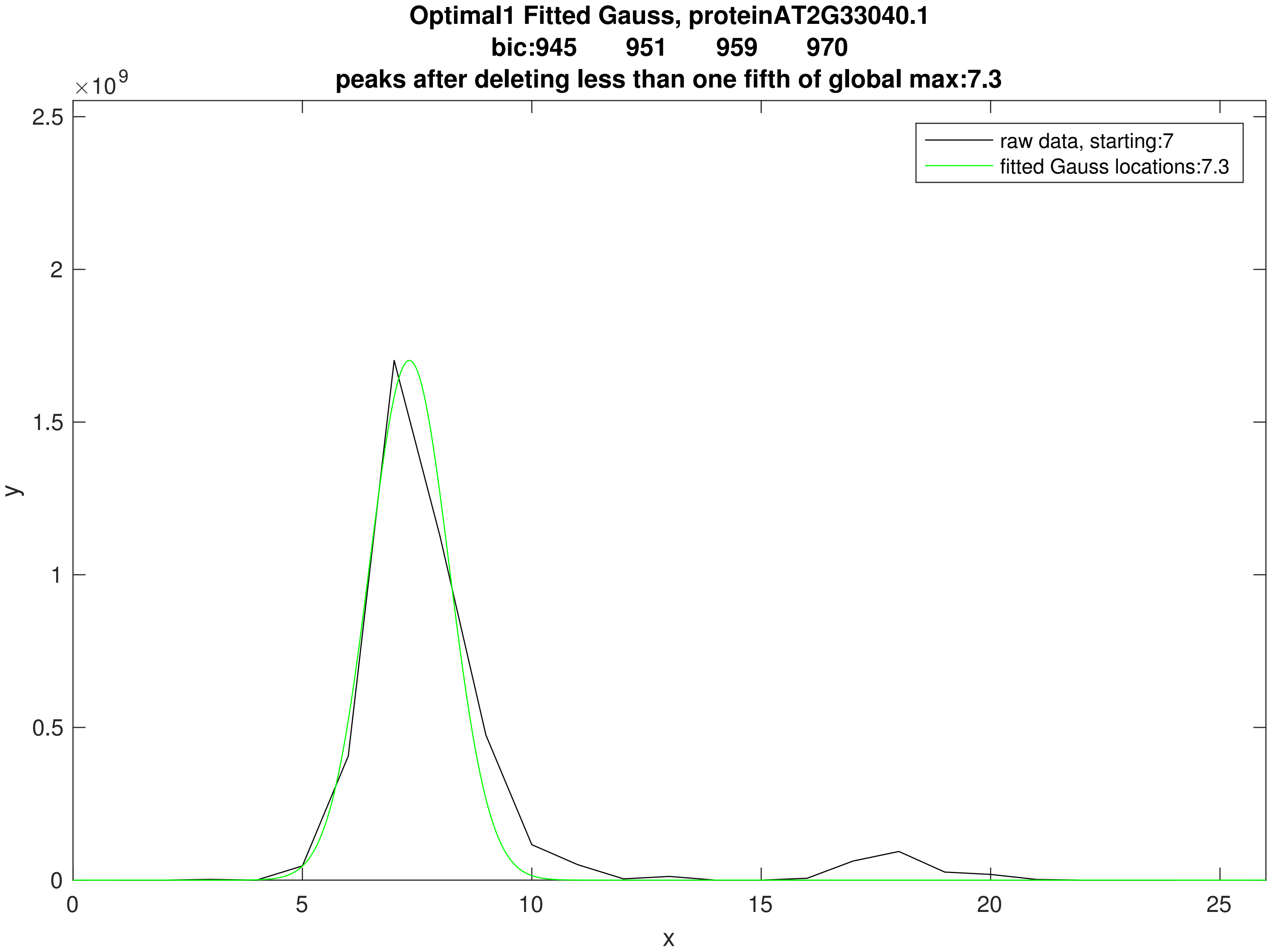
Optimal1 Fitted Gauss, proteinAT2G33040.1
bic:945 951 959 970
peaks after deleting less than one fifth of global max:7.3
×10^9
2.5
2
1.5
1
0.5
0
y
0
5
10
15
20
25
x
raw data, starting:7
fitted Gauss locations:7.3

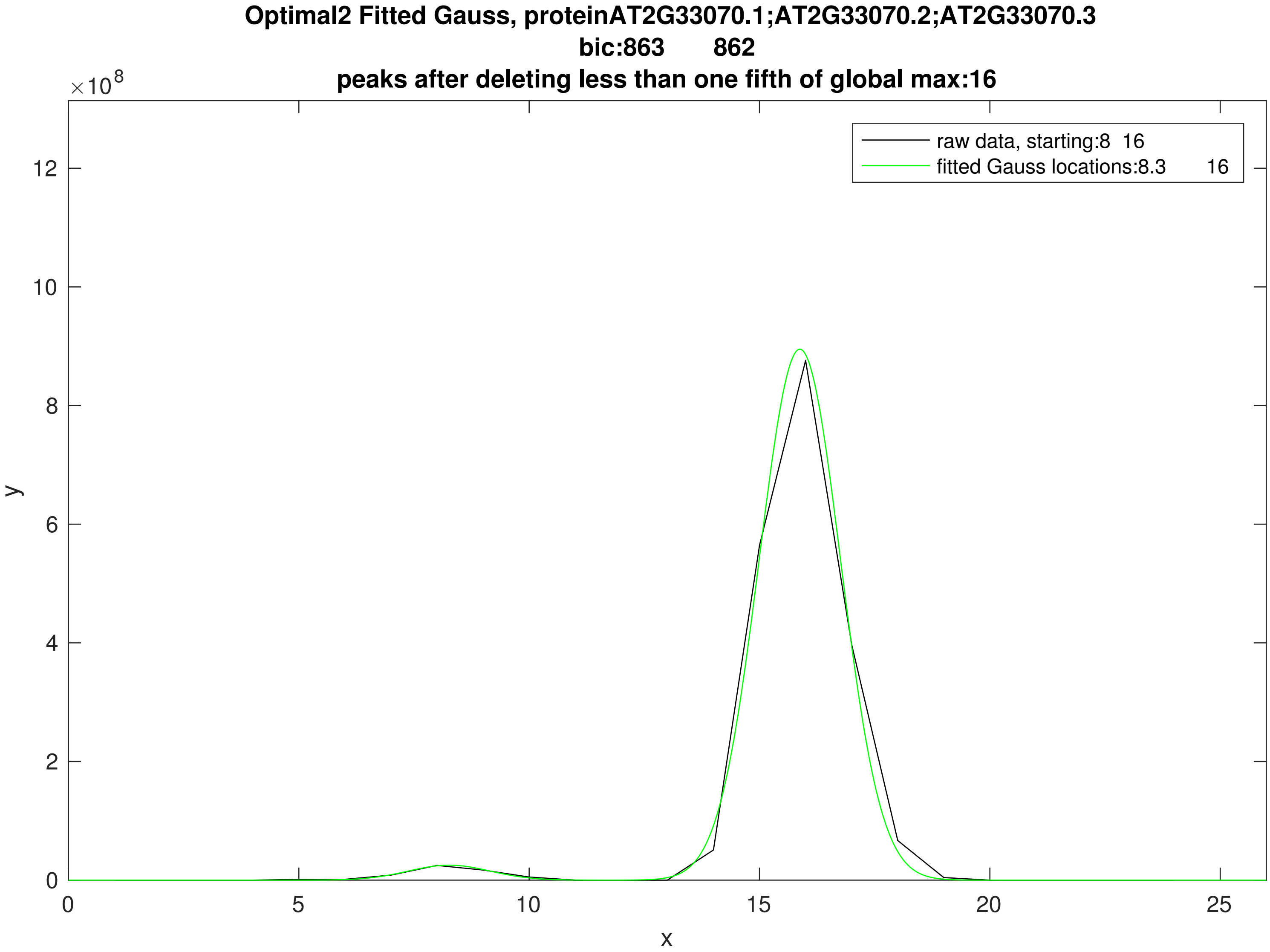
Optimal2 Fitted Gauss, proteinAT2G33070.1;AT2G33070.2;AT2G33070.3
bic:863 862
peaks after deleting less than one fifth of global max:16
×10^8
raw data, starting:8 16
fitted Gauss locations:8.3 16
y
x
0
2
4
6
8
10
12
0
5
10
15
20
25

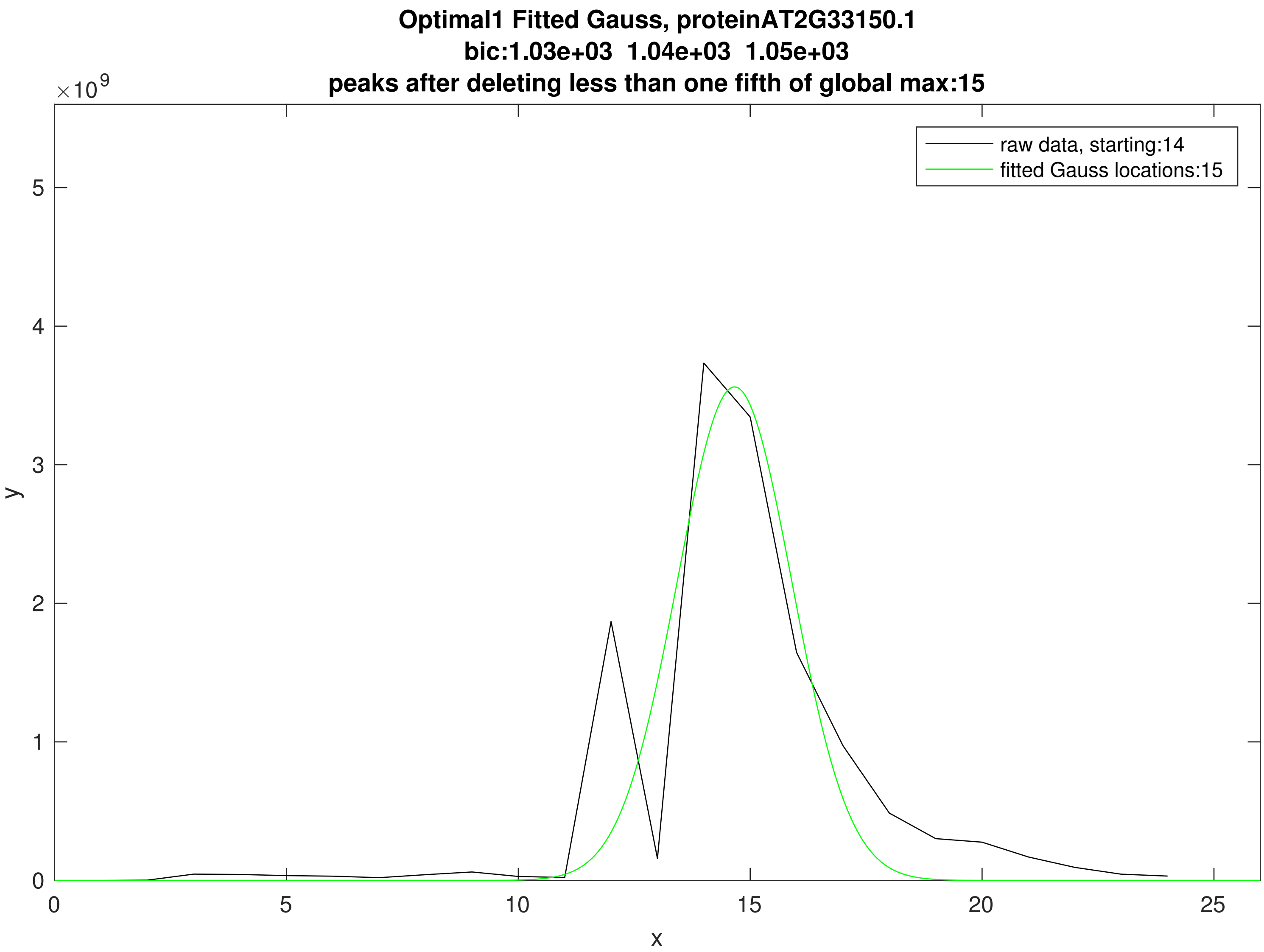

Optimal1 Fitted Gauss, proteinAT2G33150.1
bic:1.03e+03  1.04e+03  1.05e+03
peaks after deleting less than one fifth of global max:15
×10^9
raw data, starting:14
fitted Gauss locations:15
y
x
0
1
2
3
4
5
0
5
10
15
20
25

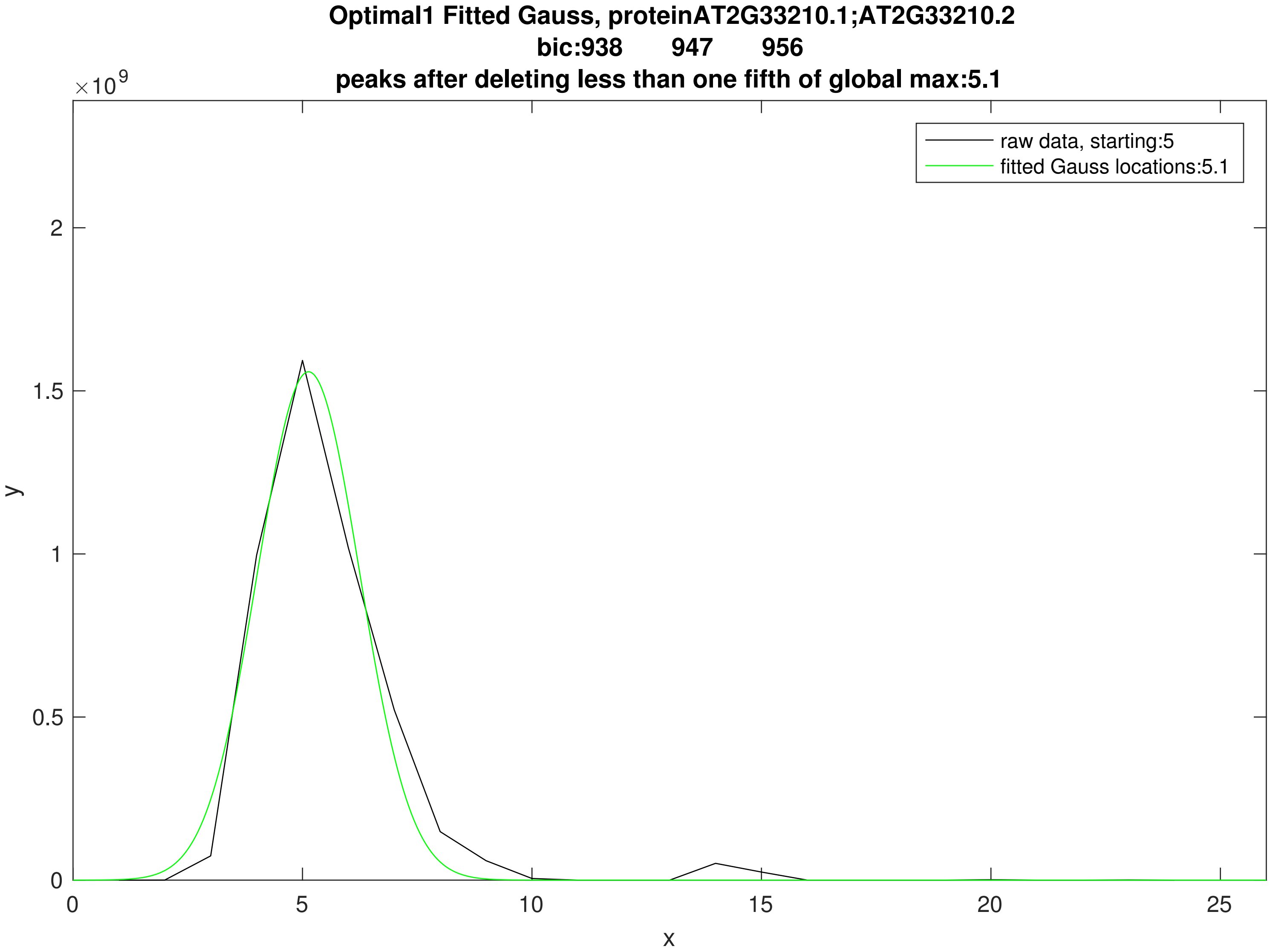

Optimal1 Fitted Gauss, proteinAT2G33210.1;AT2G33210.2
bic:938 947 956
peaks after deleting less than one fifth of global max:5.1
×10^9
raw data, starting:5
fitted Gauss locations:5.1
y
x
0
0.5
1
1.5
2
5
10
15
20
25

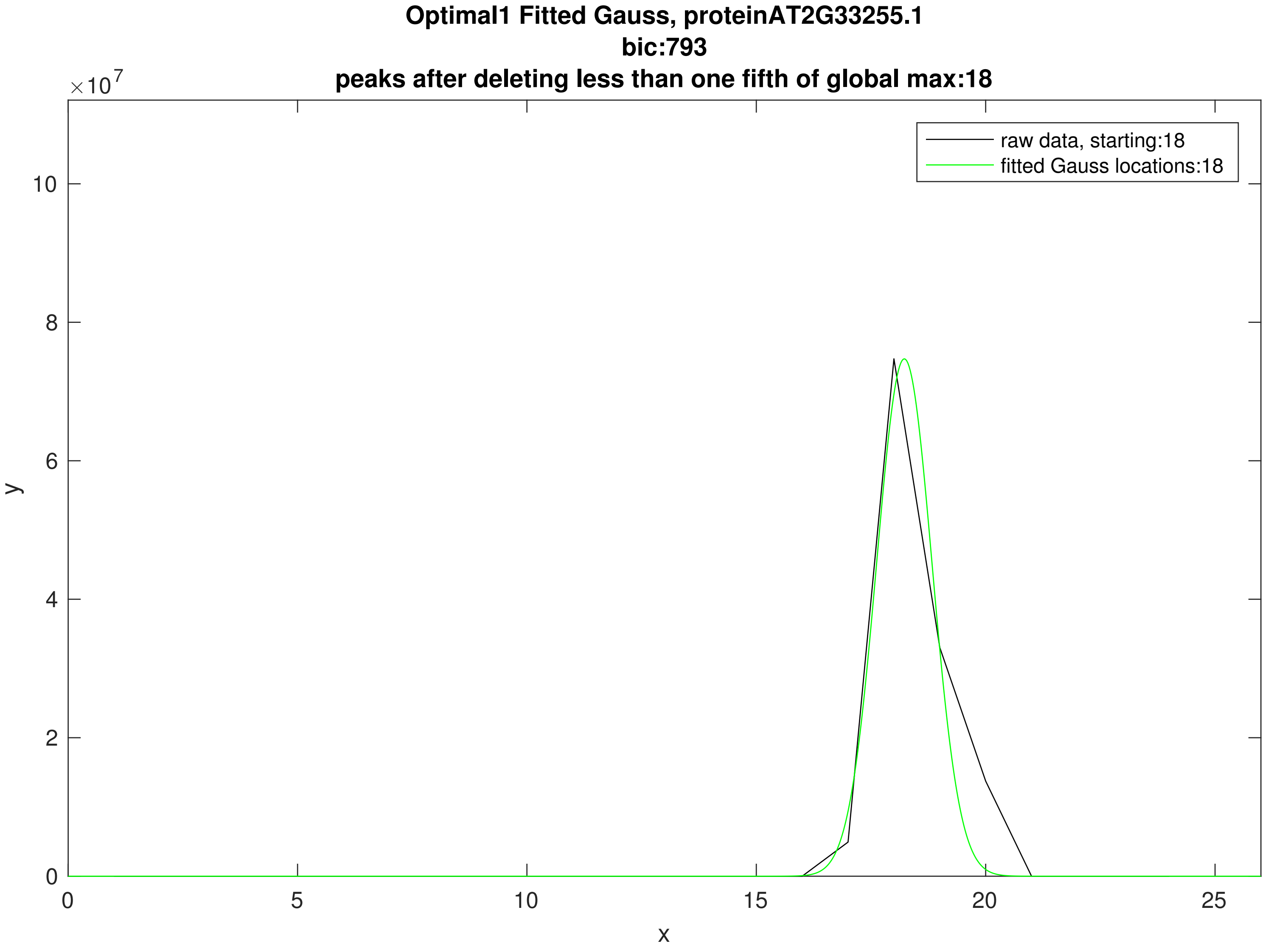

Optimal1 Fitted Gauss, proteinAT2G33255.1
bic:793
peaks after deleting less than one fifth of global max:18
×10^7
raw data, starting:18
fitted Gauss locations:18
10
8
6
4
2
0
y
0
5
10
15
20
25
x

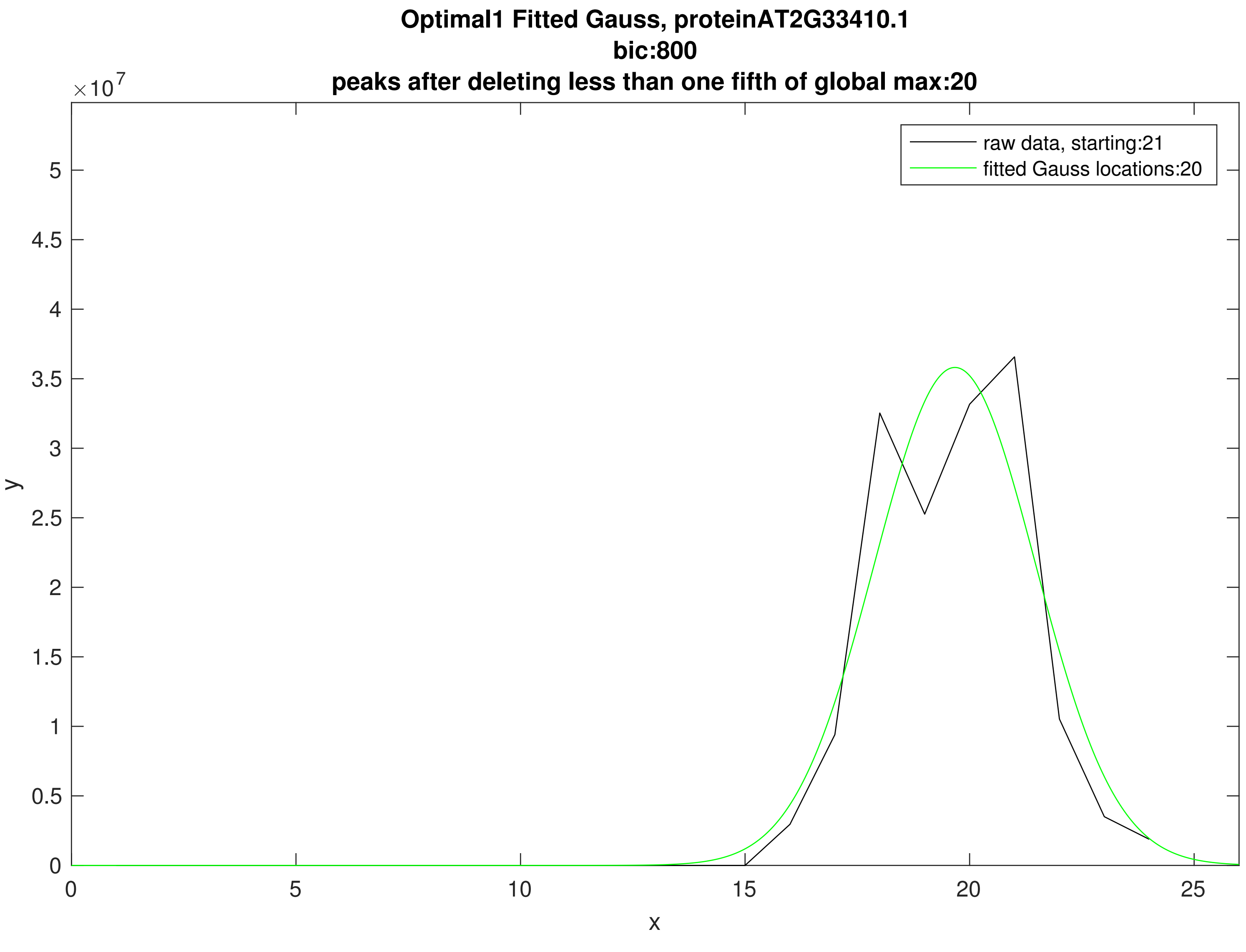

Optimal1 Fitted Gauss, proteinAT2G33410.1
bic:800
peaks after deleting less than one fifth of global max:20
×10^7
raw data, starting:21
fitted Gauss locations:20
5
4.5
4
3.5
3
2.5
2
1.5
1
0.5
0
y
0
5
10
15
20
25
x

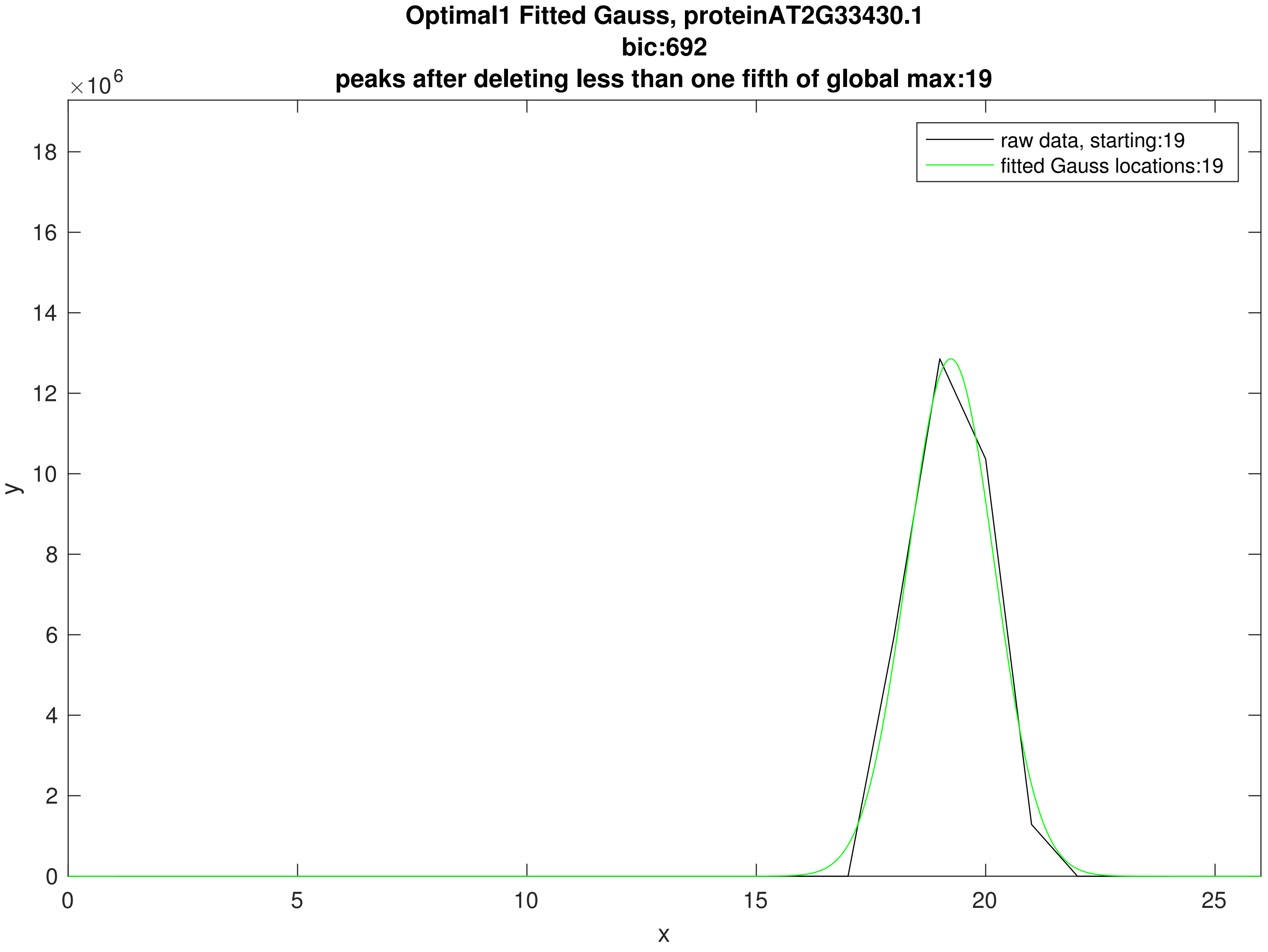
Optimal1 Fitted Gauss, proteinAT2G33430.1
bic:692
peaks after deleting less than one fifth of global max:19
×10^6
raw data, starting:19
fitted Gauss locations:19
y
x
0
2
4
6
8
10
12
14
16
18
0
5
10
15
20
25

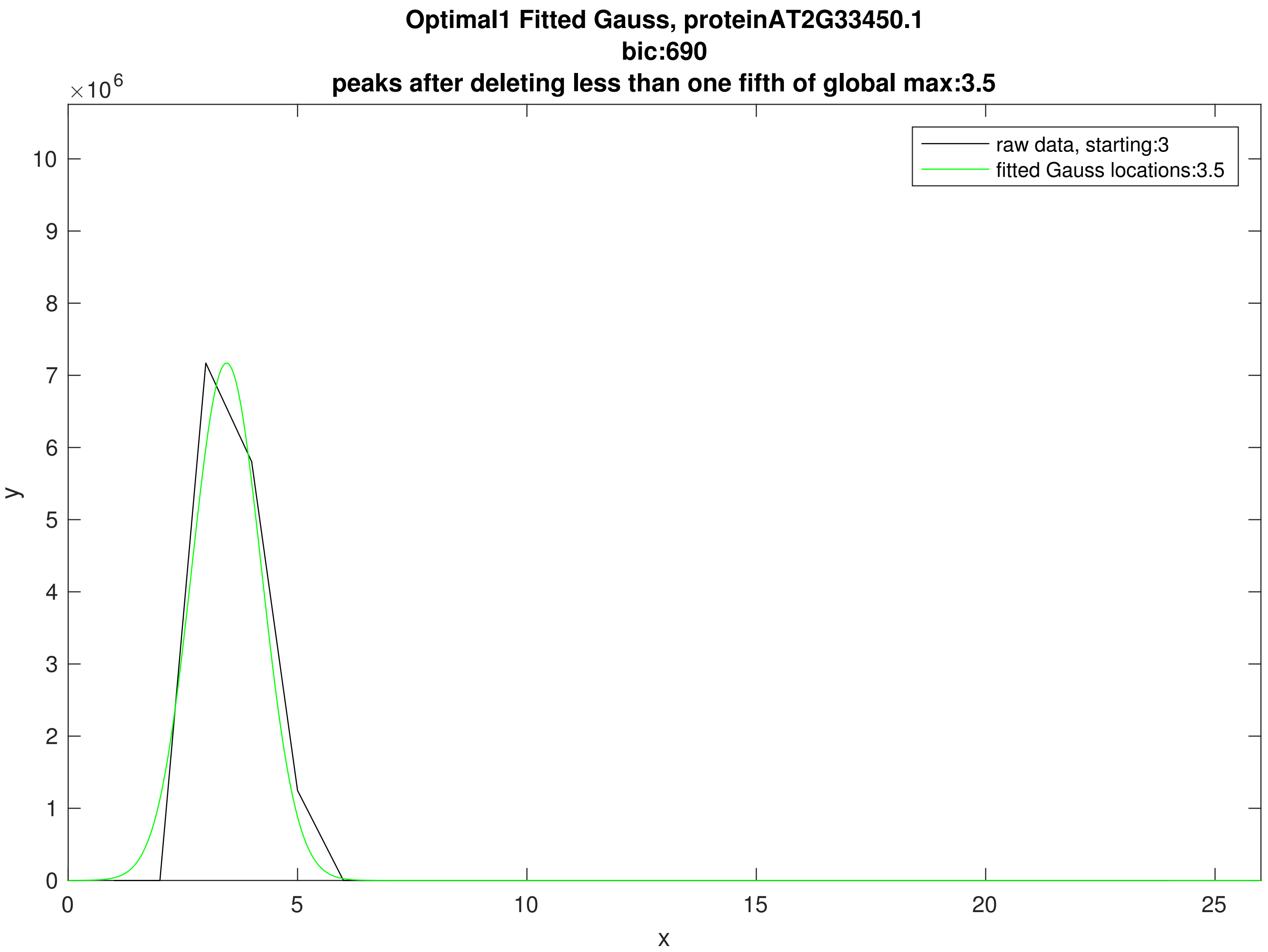
Optimal1 Fitted Gauss, proteinAT2G33450.1
bic:690
peaks after deleting less than one fifth of global max:3.5
×10^6
raw data, starting:3
fitted Gauss locations:3.5
y
x
0
1
2
3
4
5
6
7
8
9
10
0
5
10
15
20
25

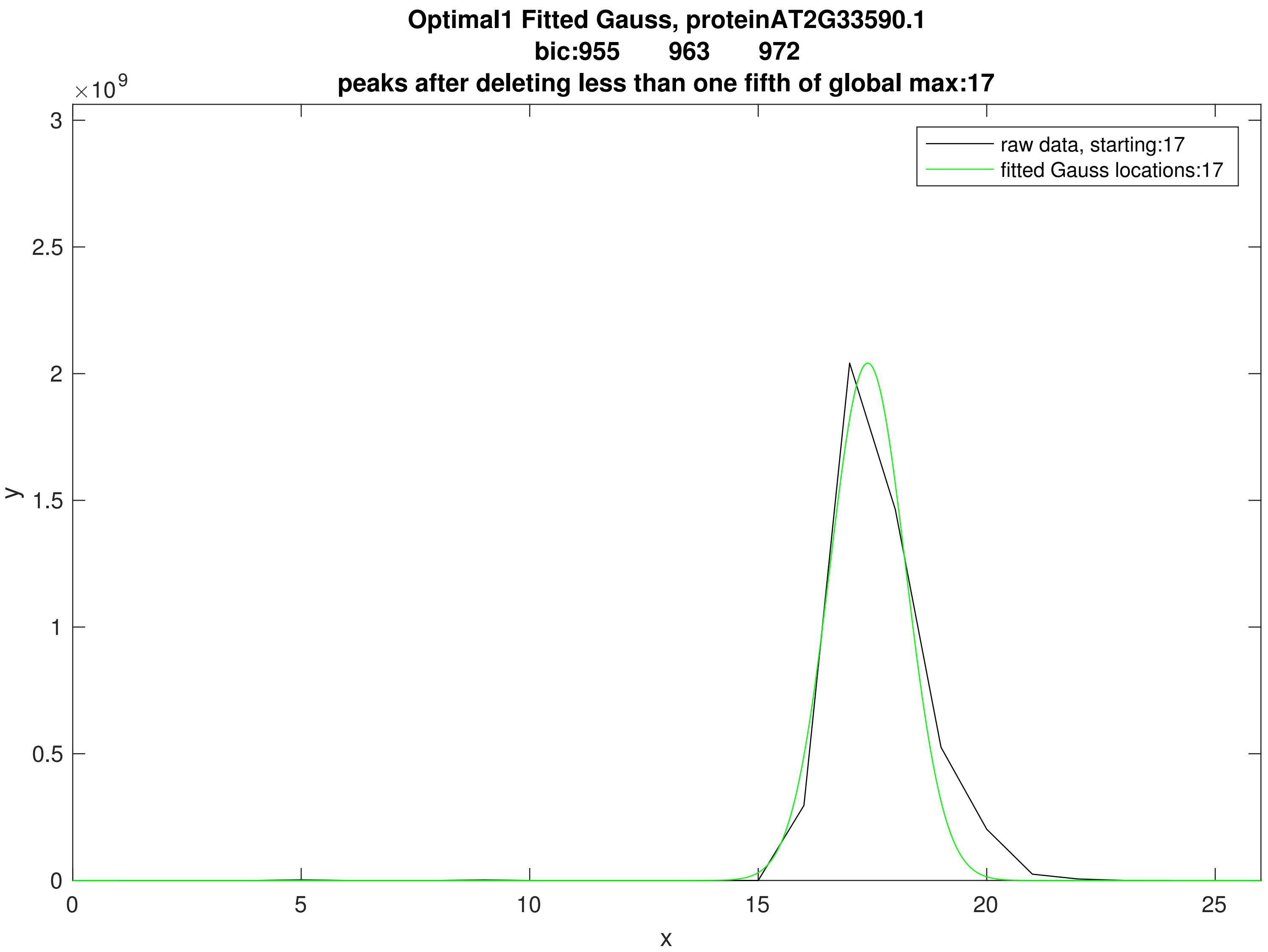
Optimal1 Fitted Gauss, proteinAT2G33590.1
bic:955 963 972
peaks after deleting less than one fifth of global max:17
×10^9
3
2.5
2
1.5
1
0.5
0
y
0
5
10
15
20
25
x
raw data, starting:17
fitted Gauss locations:17

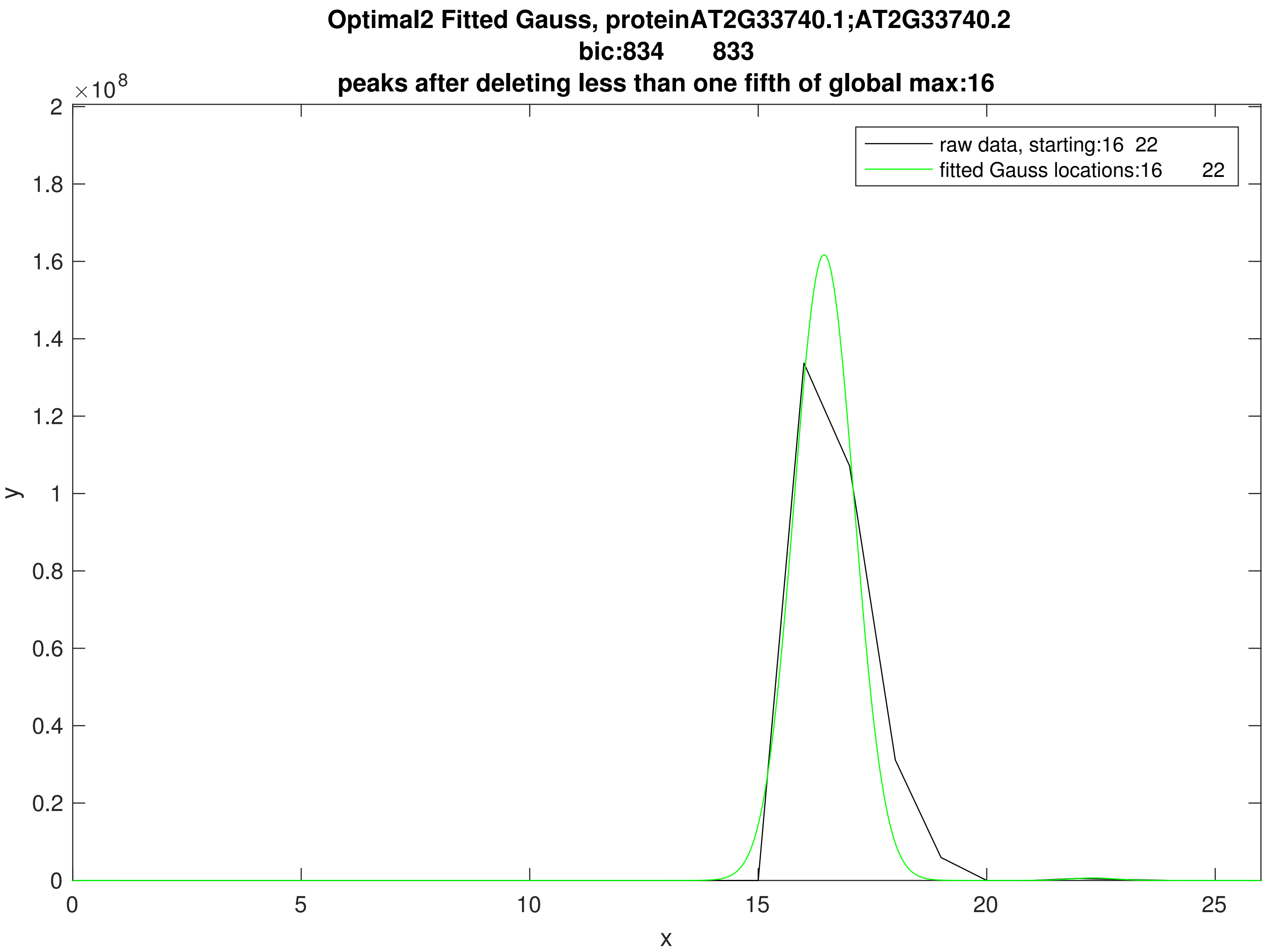
Optimal2 Fitted Gauss, proteinAT2G33740.1;AT2G33740.2
bic:834 833
peaks after deleting less than one fifth of global max:16
raw data, starting:16 22
fitted Gauss locations:16 22
×10^8
y
x

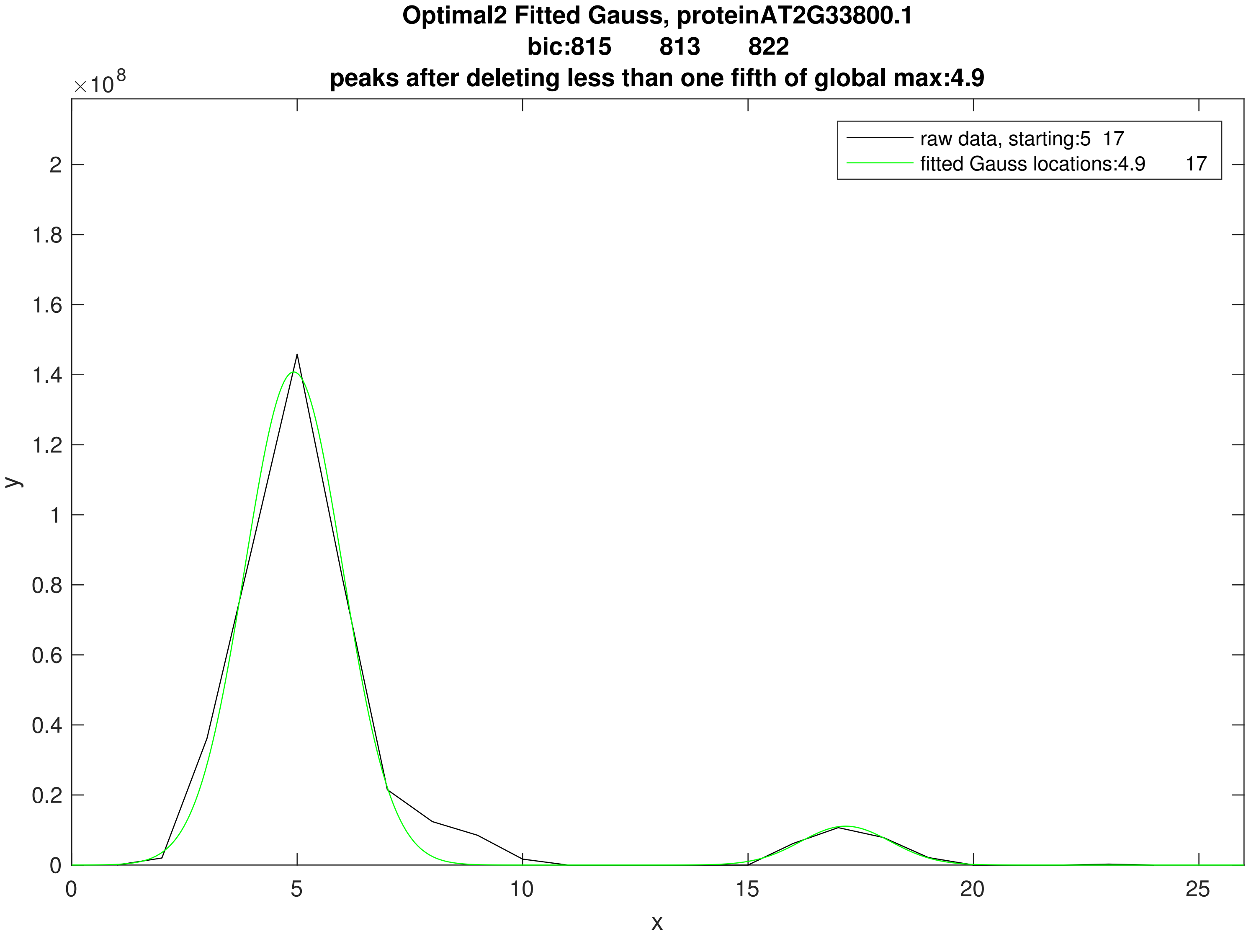
Optimal2 Fitted Gauss, proteinAT2G33800.1
bic:815 813 822
peaks after deleting less than one fifth of global max:4.9
×10^8
raw data, starting:5 17
fitted Gauss locations:4.9 17
y
x
0
0.2
0.4
0.6
0.8
1
1.2
1.4
1.6
1.8
2
0
5
10
15
20
25

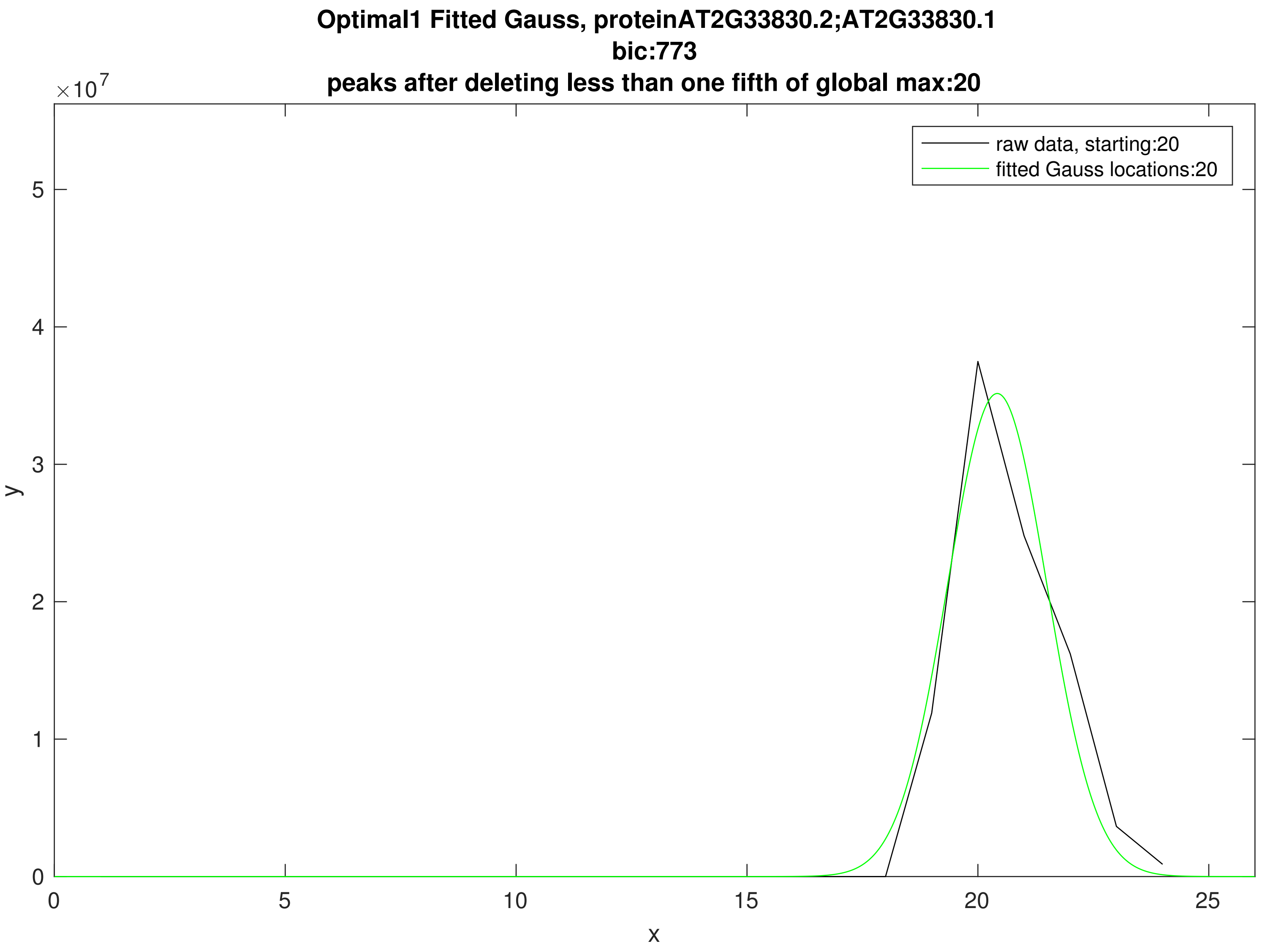
Optimal1 Fitted Gauss, proteinAT2G33830.2;AT2G33830.1
bic:773
peaks after deleting less than one fifth of global max:20
×10^7
raw data, starting:20
fitted Gauss locations:20
y
x
0
1
2
3
4
5
0
5
10
15
20
25

**Optimal1 Fitted Gauss, proteinAT2G33840.1**
**bic:824**
**peaks after deleting less than one fifth of global max:11**

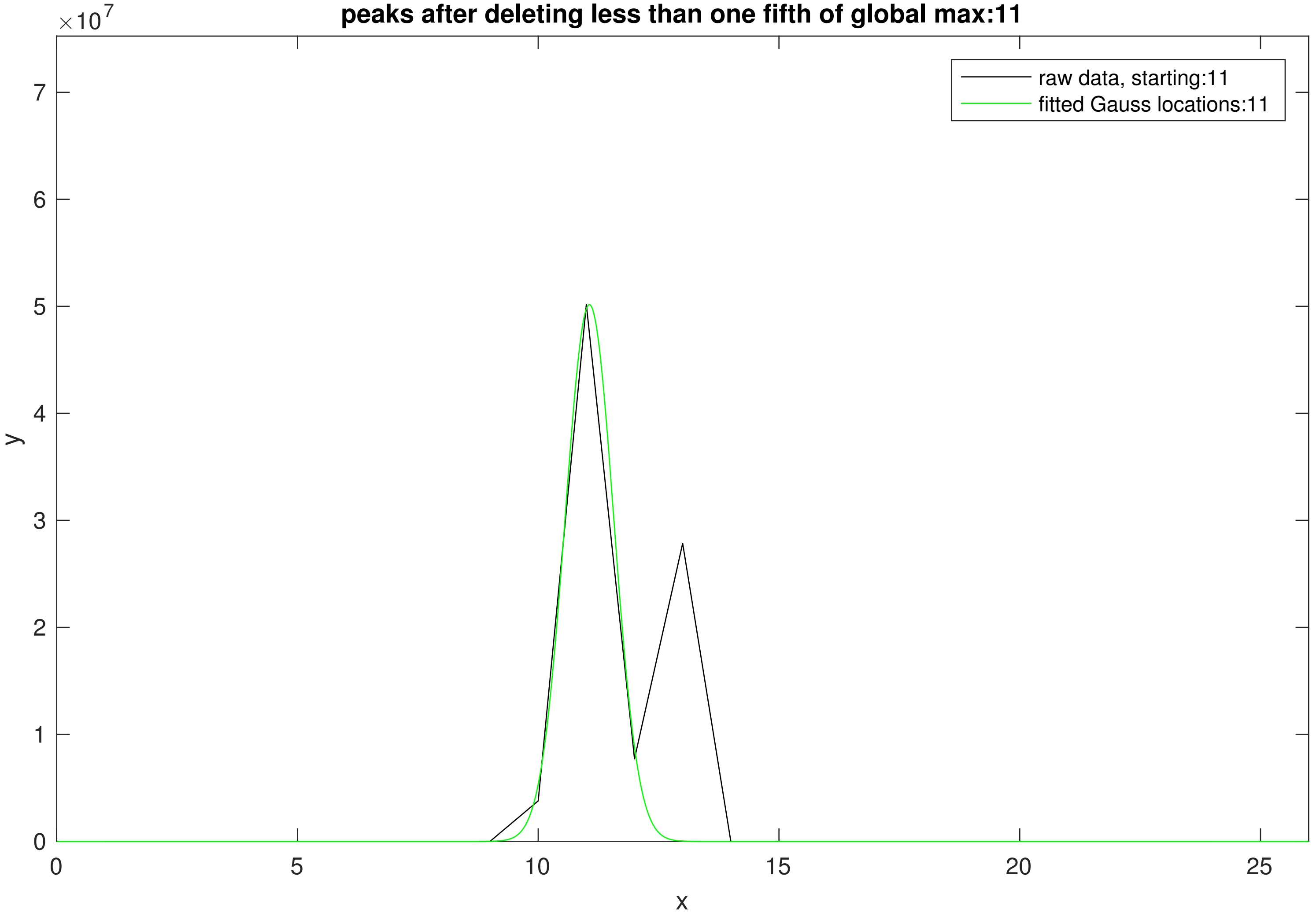

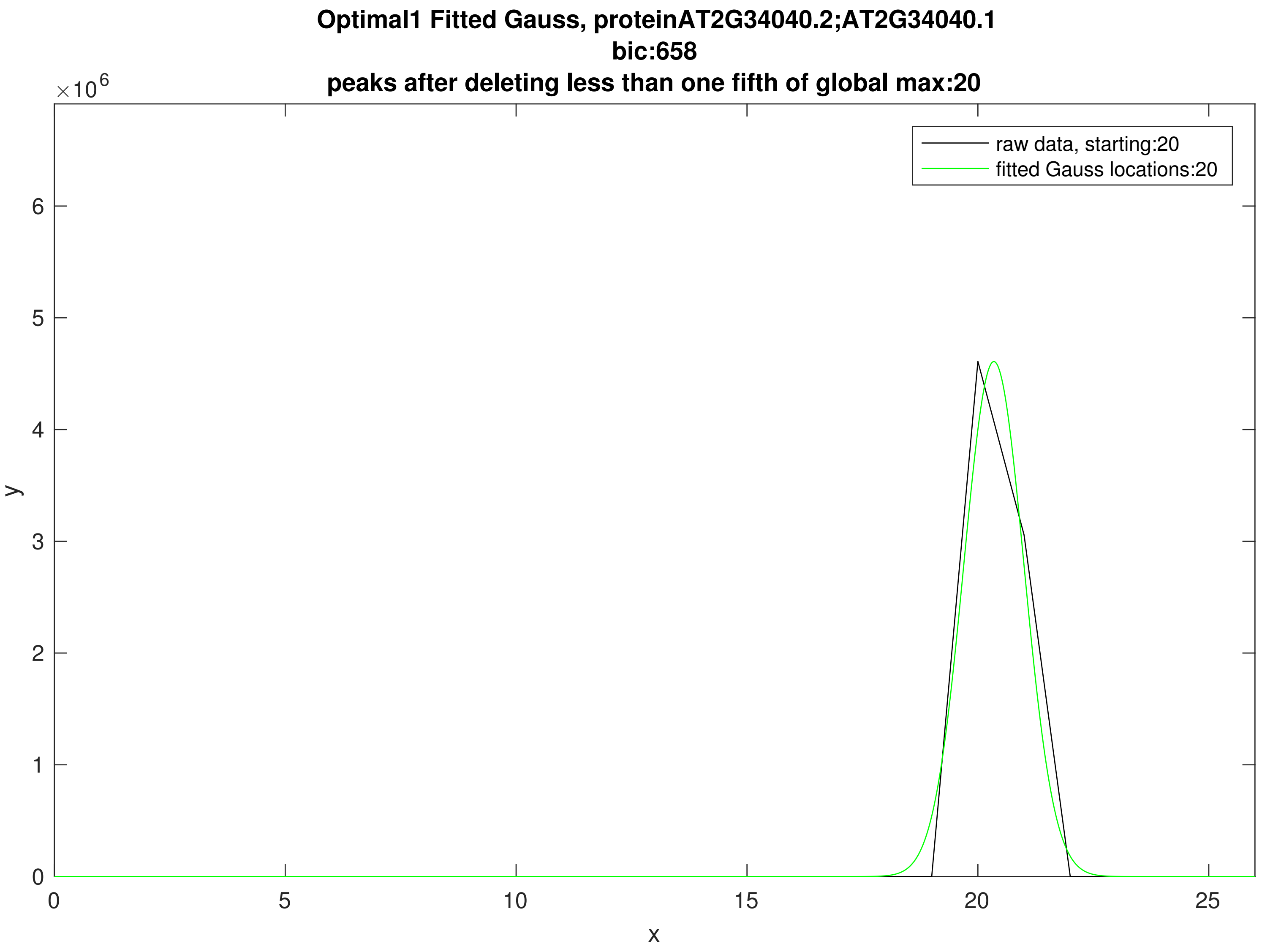
Optimal1 Fitted Gauss, proteinAT2G34040.2;AT2G34040.1
bic:658
peaks after deleting less than one fifth of global max:20
×10^6
raw data, starting:20
fitted Gauss locations:20
y
x
0
1
2
3
4
5
6
0
5
10
15
20
25

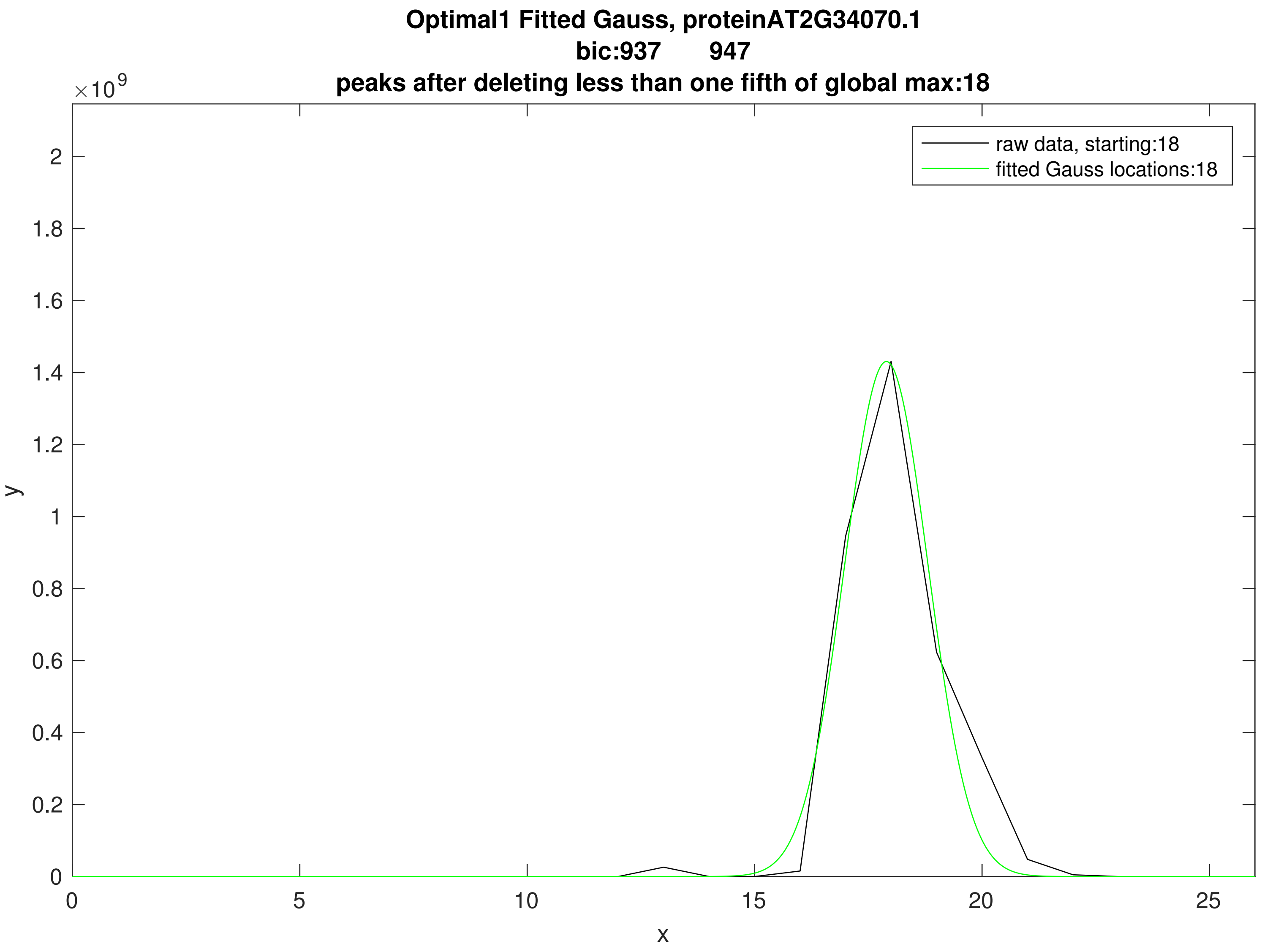
Optimal1 Fitted Gauss, proteinAT2G34070.1
bic:937 947
peaks after deleting less than one fifth of global max:18
×10^9
raw data, starting:18
fitted Gauss locations:18
y
x
0
0.2
0.4
0.6
0.8
1
1.2
1.4
1.6
1.8
2
5
10
15
20
25

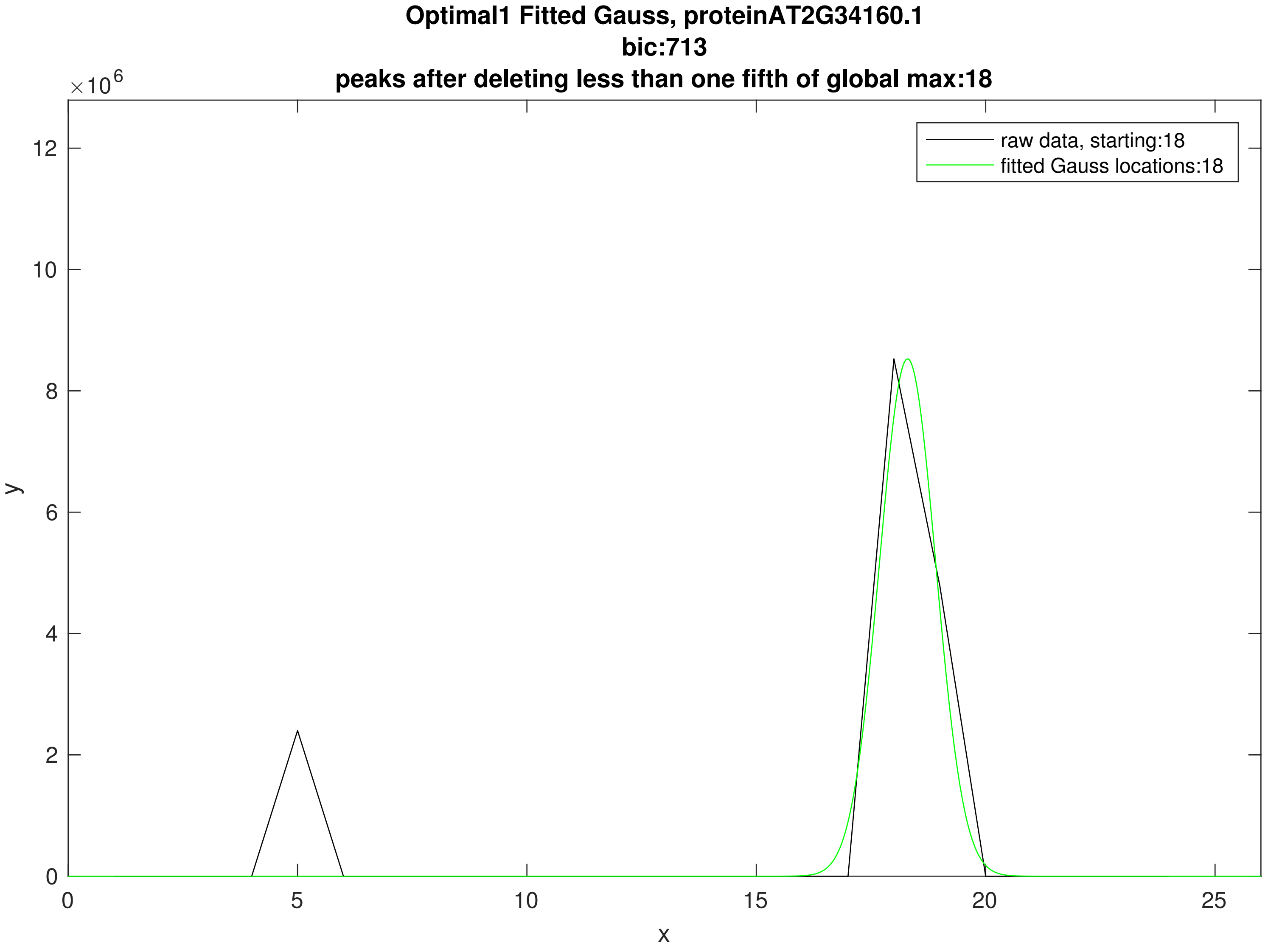
Optimal1 Fitted Gauss, proteinAT2G34160.1
bic:713
peaks after deleting less than one fifth of global max:18
×10^6
raw data, starting:18
fitted Gauss locations:18
y
x
0
2
4
6
8
10
12
0
5
10
15
20
25

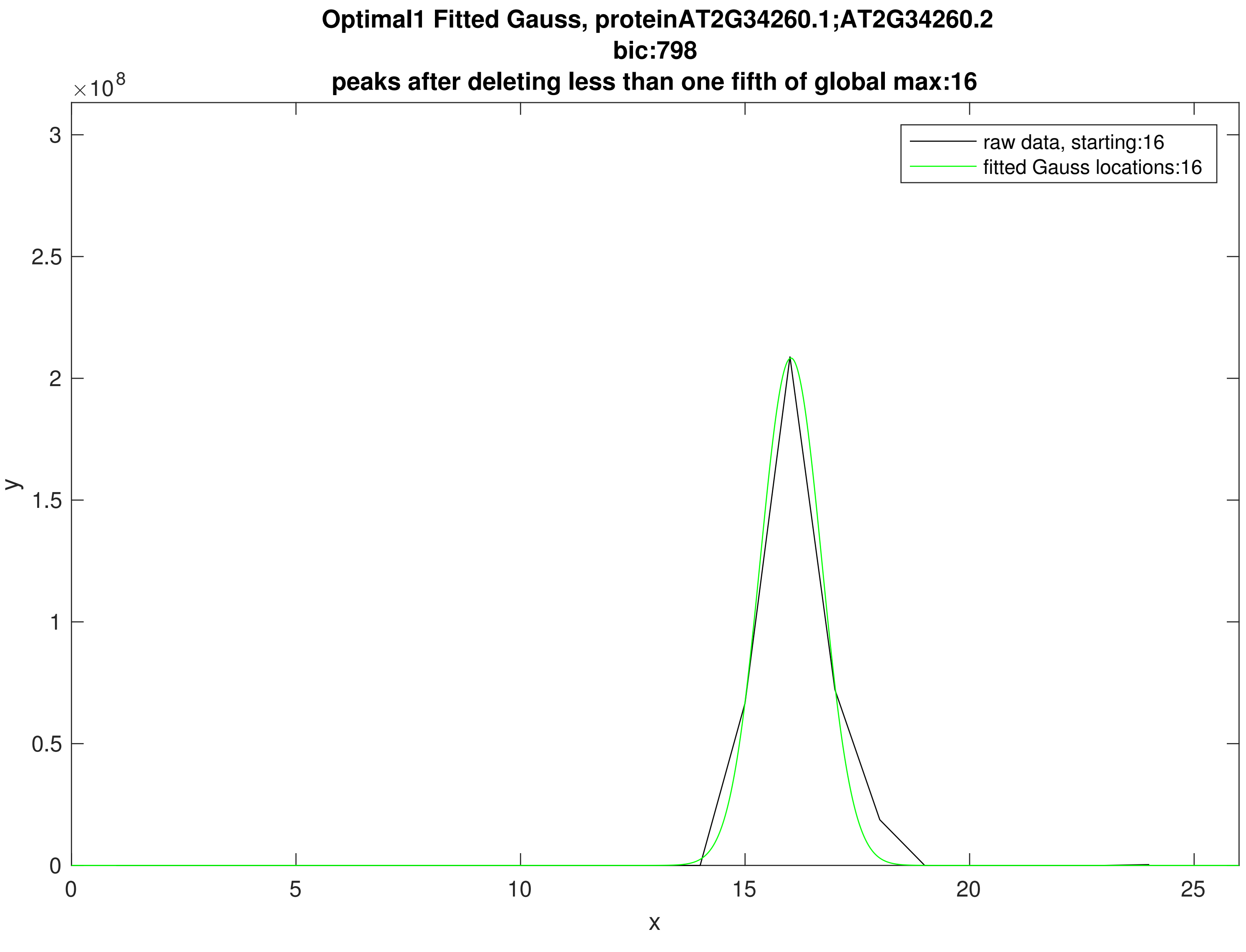

Optimal1 Fitted Gauss, proteinAT2G34260.1;AT2G34260.2
bic:798
peaks after deleting less than one fifth of global max:16
×10^8
3
2.5
2
1.5
1
0.5
0
y
0
5
10
15
20
25
x
raw data, starting:16
fitted Gauss locations:16

**Optimal1 Fitted Gauss, proteinAT2G34470.1;AT2G34470.2**
**bic:707**
**peaks after deleting less than one fifth of global max:17**

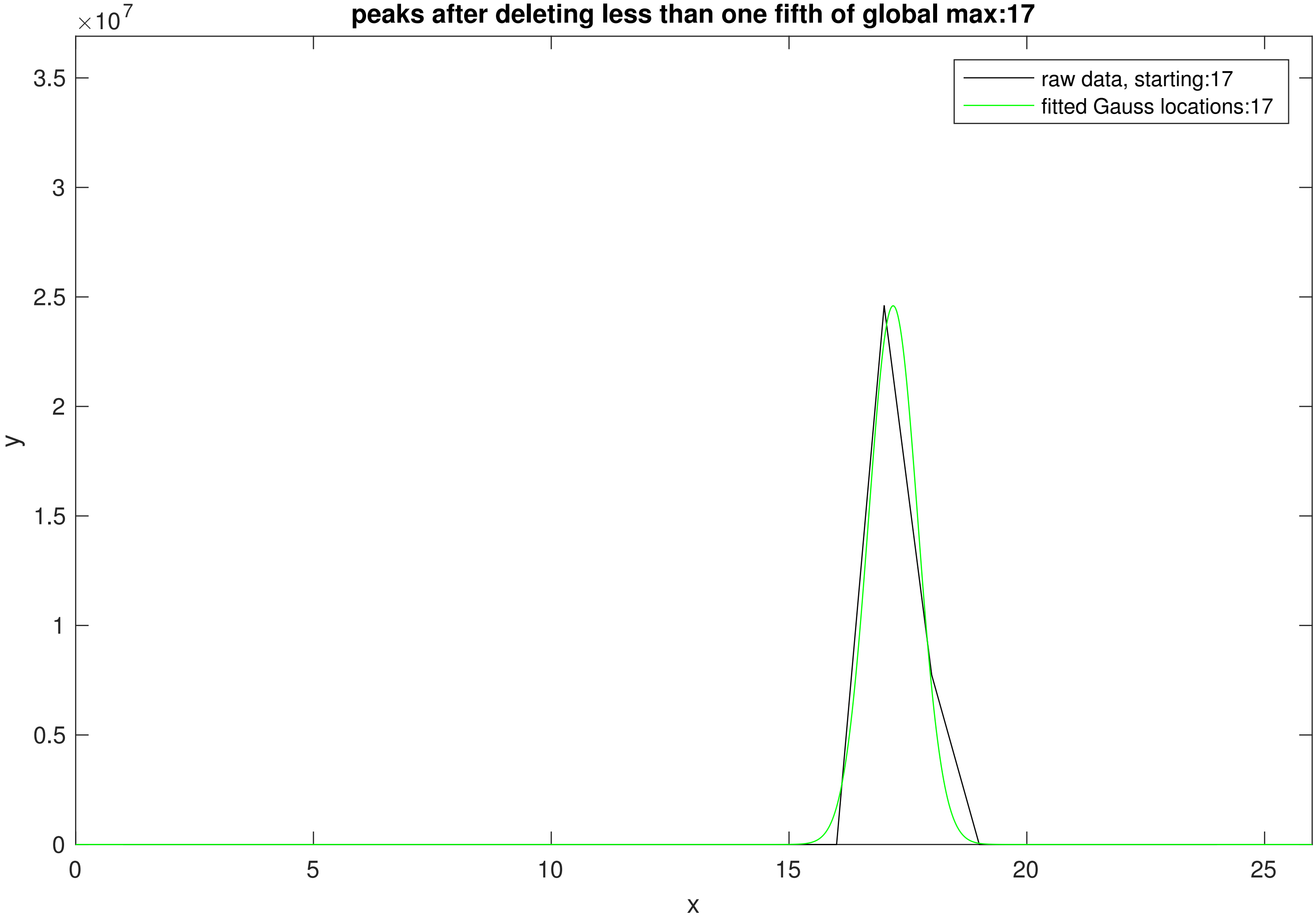

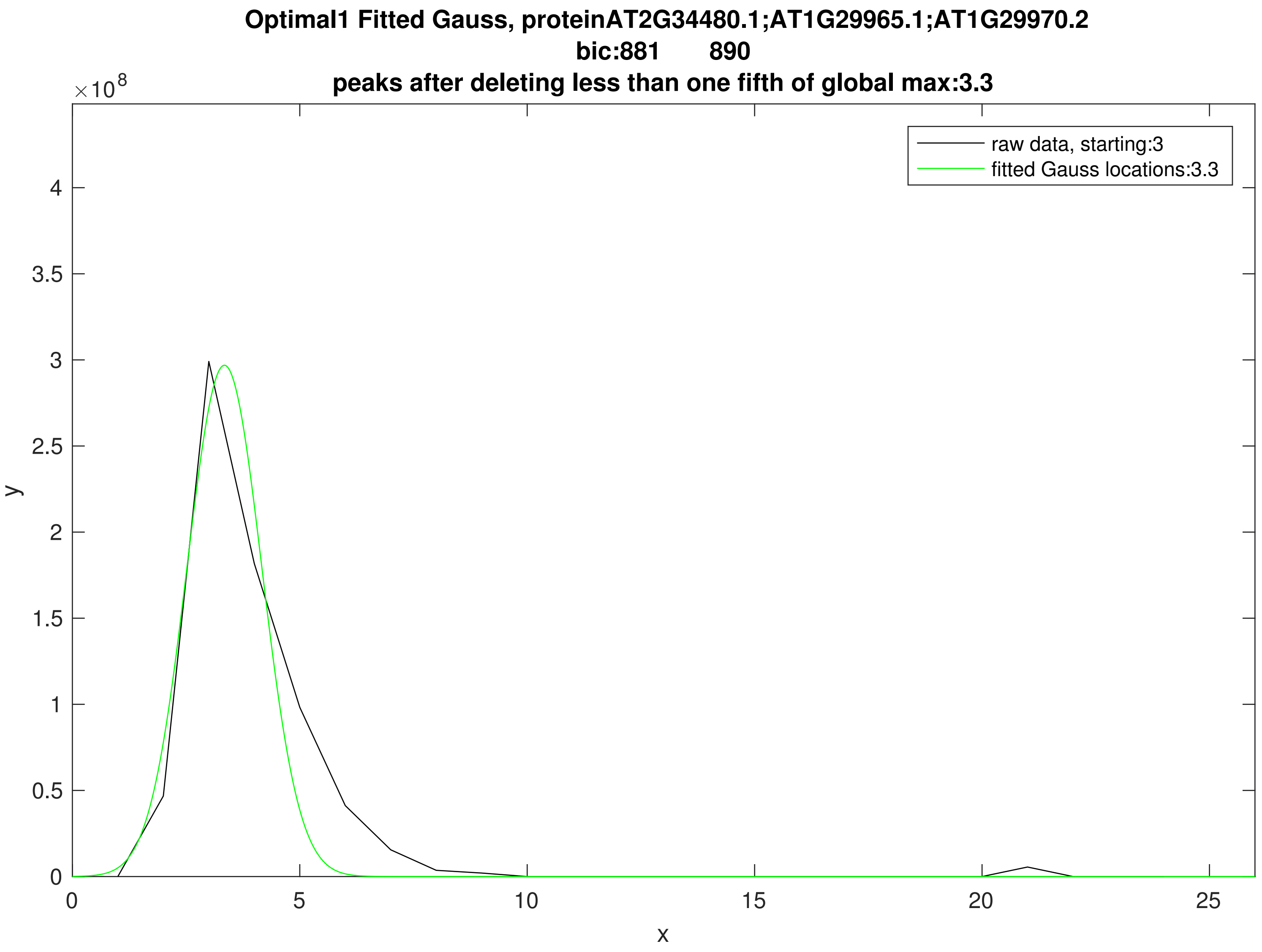

Optimal1 Fitted Gauss, proteinAT2G34480.1;AT1G29965.1;AT1G29970.2
bic:881 890
peaks after deleting less than one fifth of global max:3.3
×10^8
raw data, starting:3
fitted Gauss locations:3.3
y
x
0
0.5
1
1.5
2
2.5
3
3.5
4
5
10
15
20
25

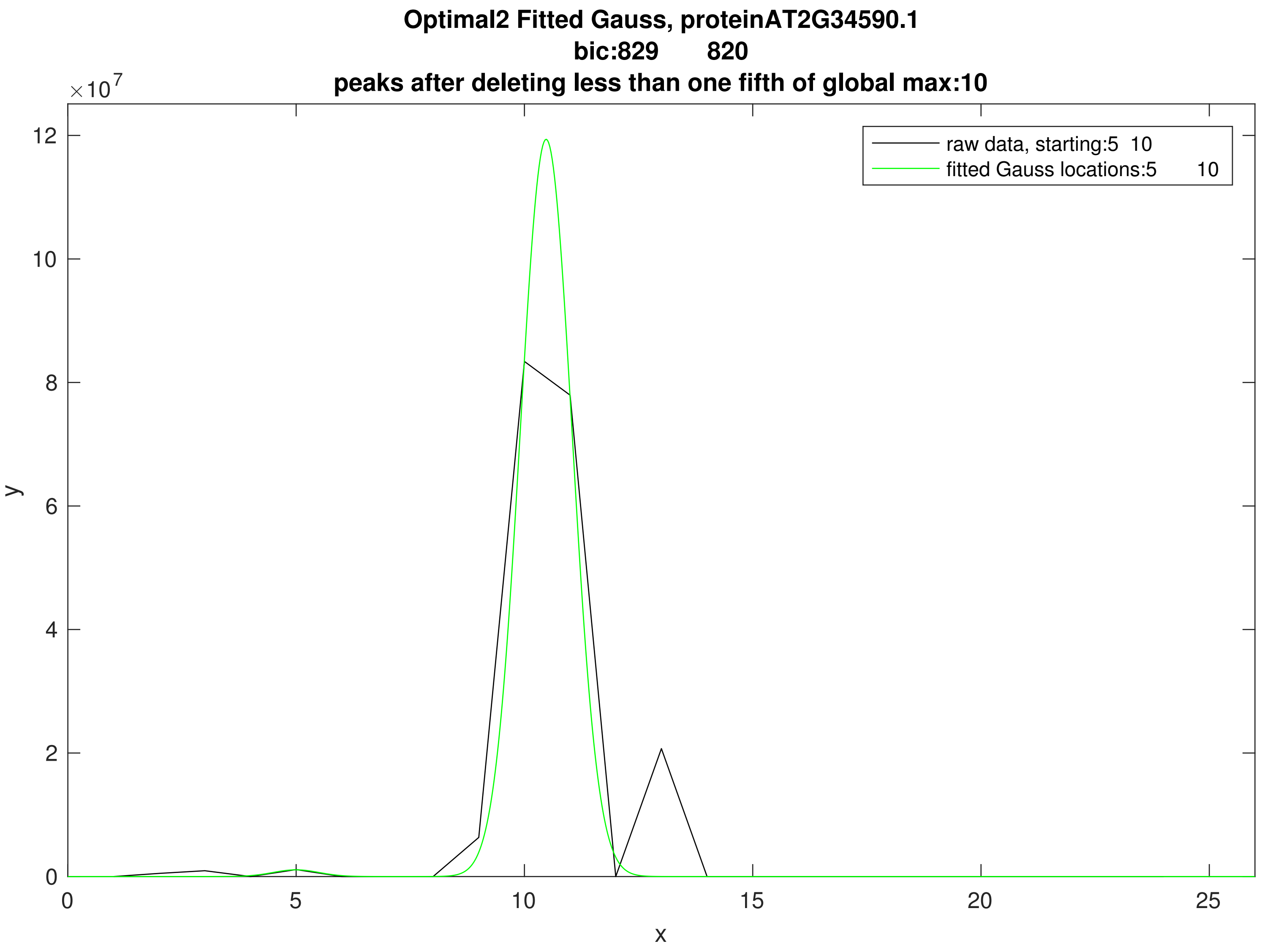
Optimal2 Fitted Gauss, proteinAT2G34590.1
bic:829 820
peaks after deleting less than one fifth of global max:10
×10^7
raw data, starting:5 10
fitted Gauss locations:5 10
y
x
0
2
4
6
8
10
12
0
5
10
15
20
25

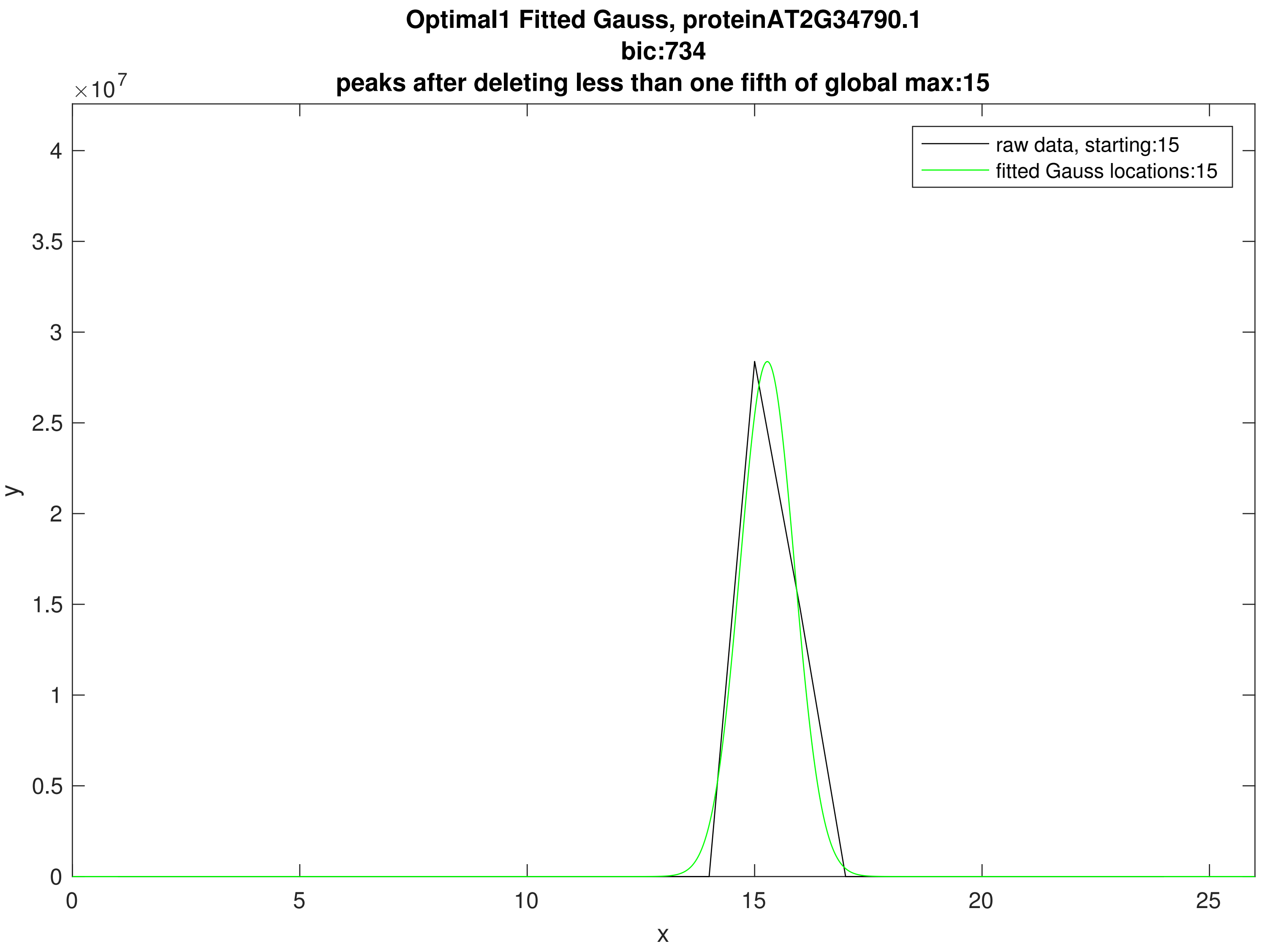
Optimal1 Fitted Gauss, proteinAT2G34790.1
bic:734
peaks after deleting less than one fifth of global max:15
×10^7
raw data, starting:15
fitted Gauss locations:15
y
x
0
0.5
1
1.5
2
2.5
3
3.5
4
0
5
10
15
20
25

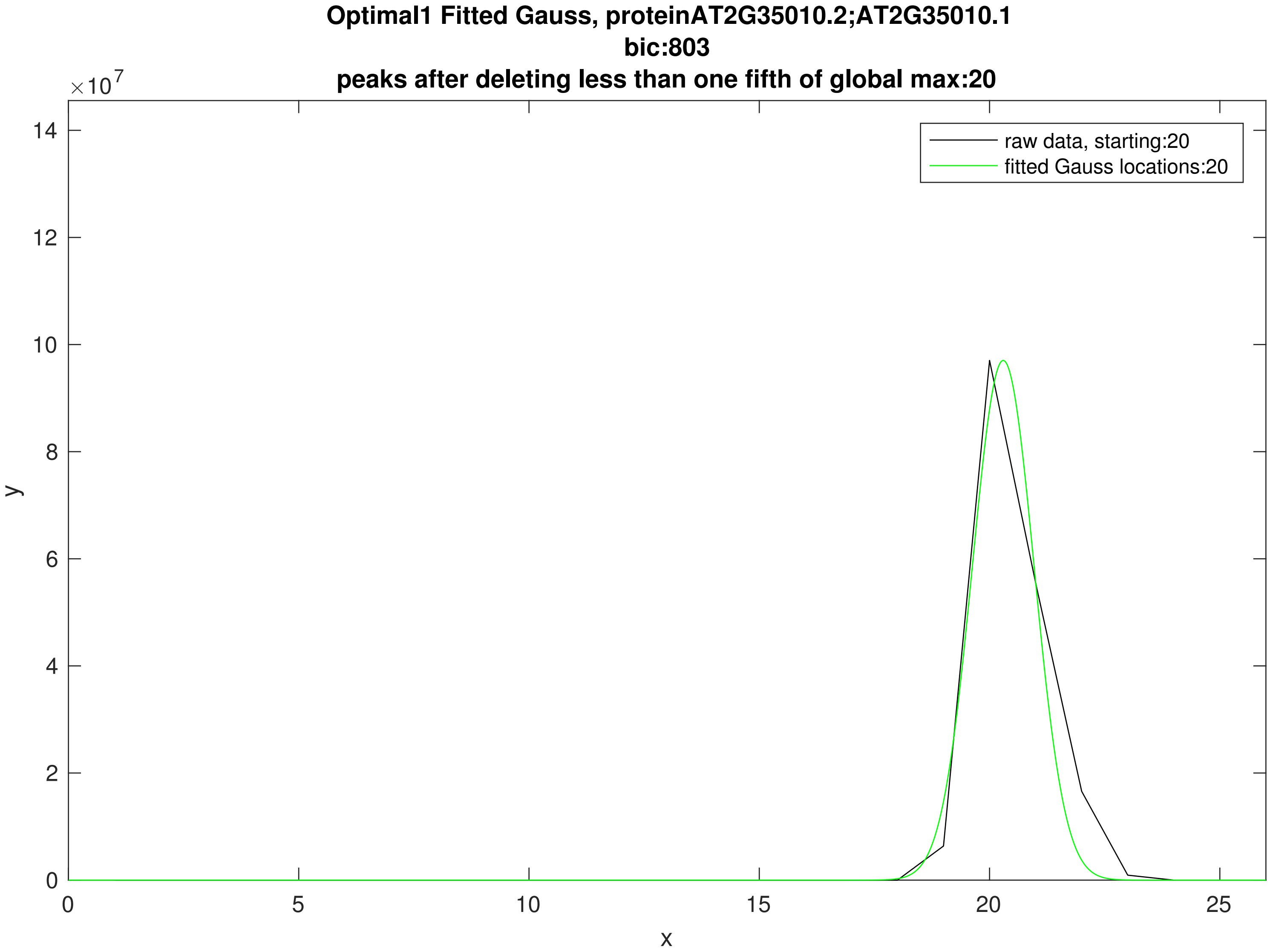
Optimal1 Fitted Gauss, proteinAT2G35010.2;AT2G35010.1
bic:803
peaks after deleting less than one fifth of global max:20
×10^7
raw data, starting:20
fitted Gauss locations:20
y
x
0
2
4
6
8
10
12
14
0
5
10
15
20
25

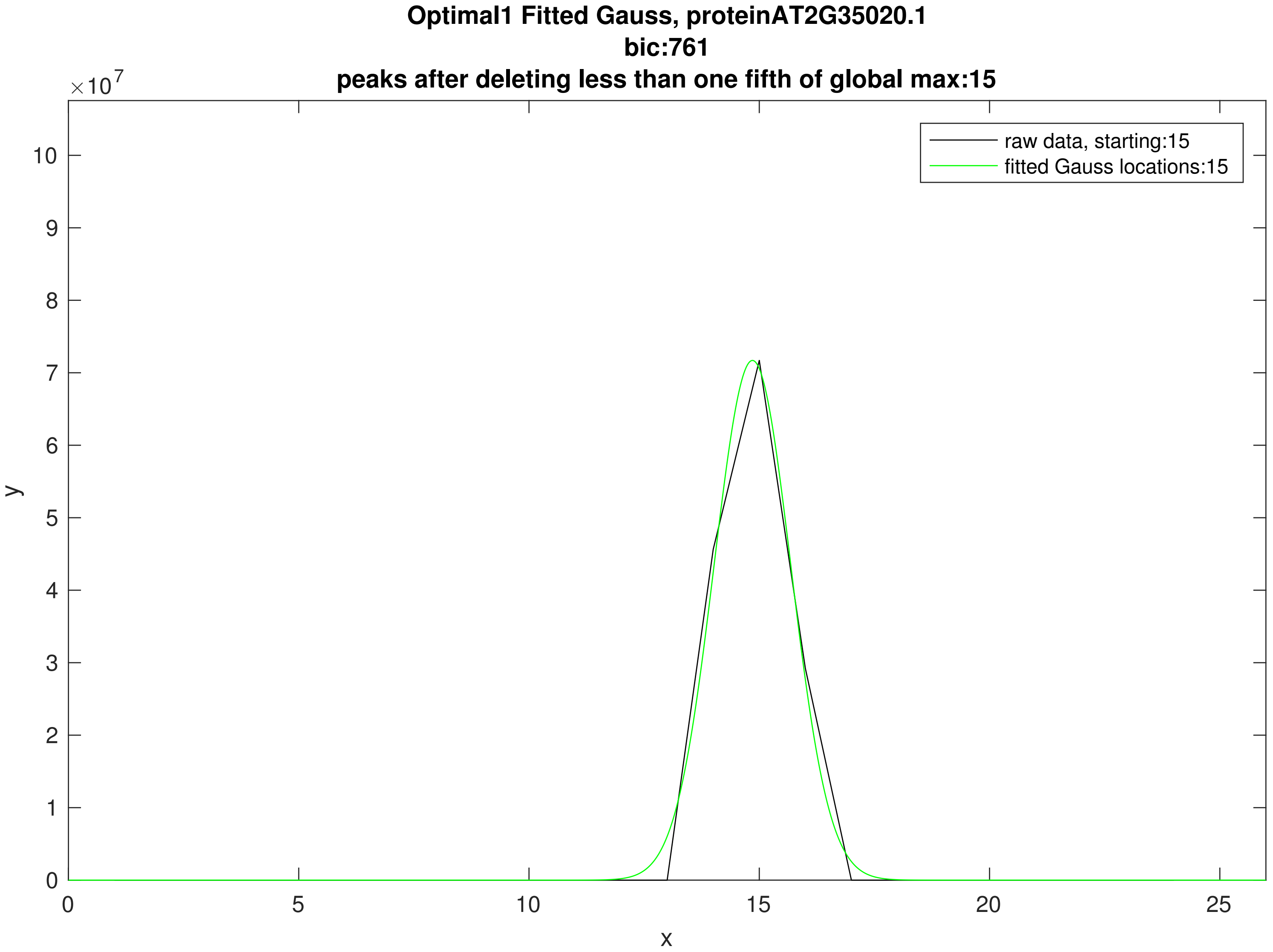
Optimal1 Fitted Gauss, proteinAT2G35020.1
bic:761
peaks after deleting less than one fifth of global max:15
×10^7
raw data, starting:15
fitted Gauss locations:15
y
x
0
1
2
3
4
5
6
7
8
9
10
0
5
10
15
20
25

Optimal1 Fitted Gauss, proteinAT2G35040.2;AT2G35040.1
bic:890
peaks after deleting less than one fifth of global max:13

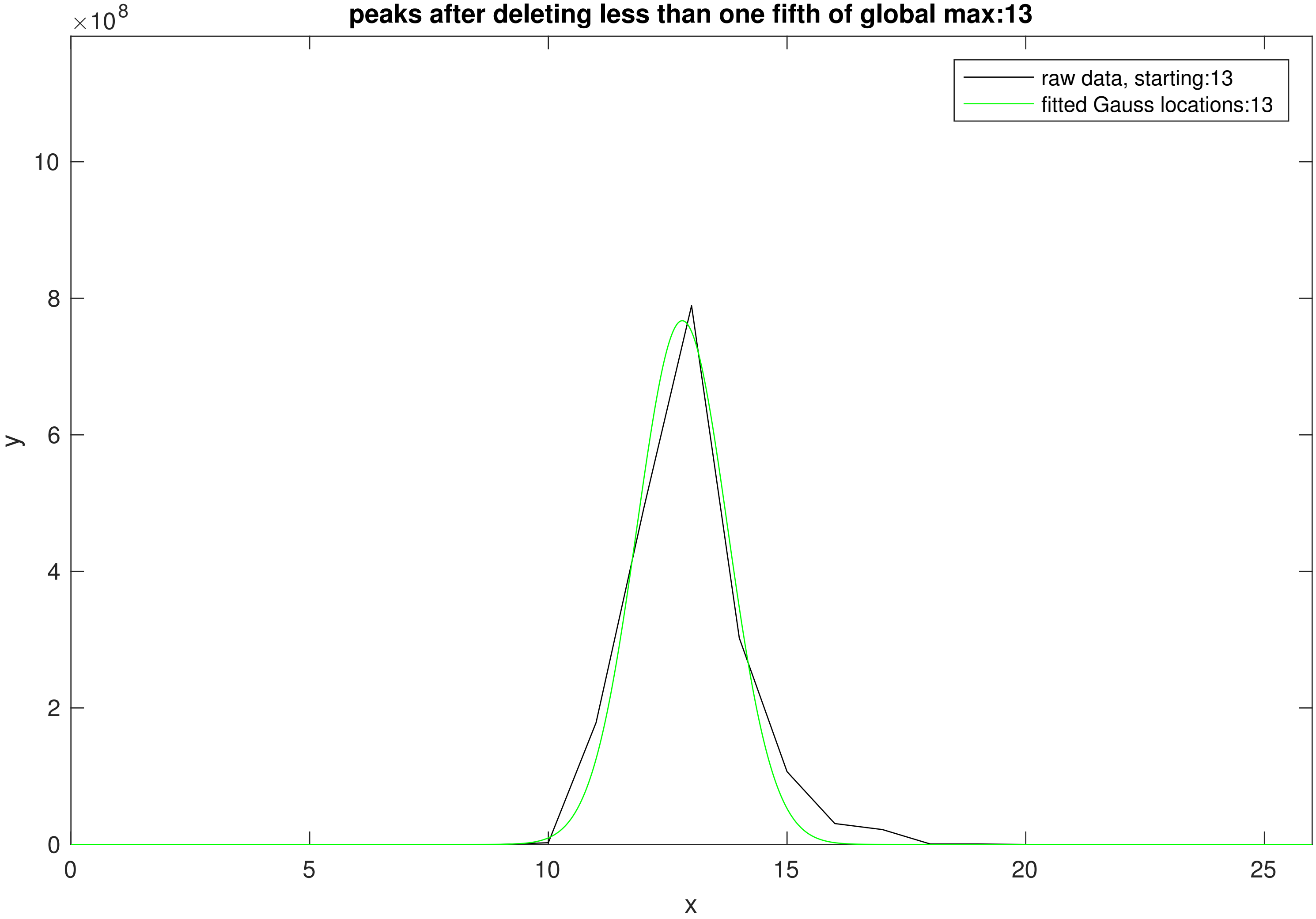

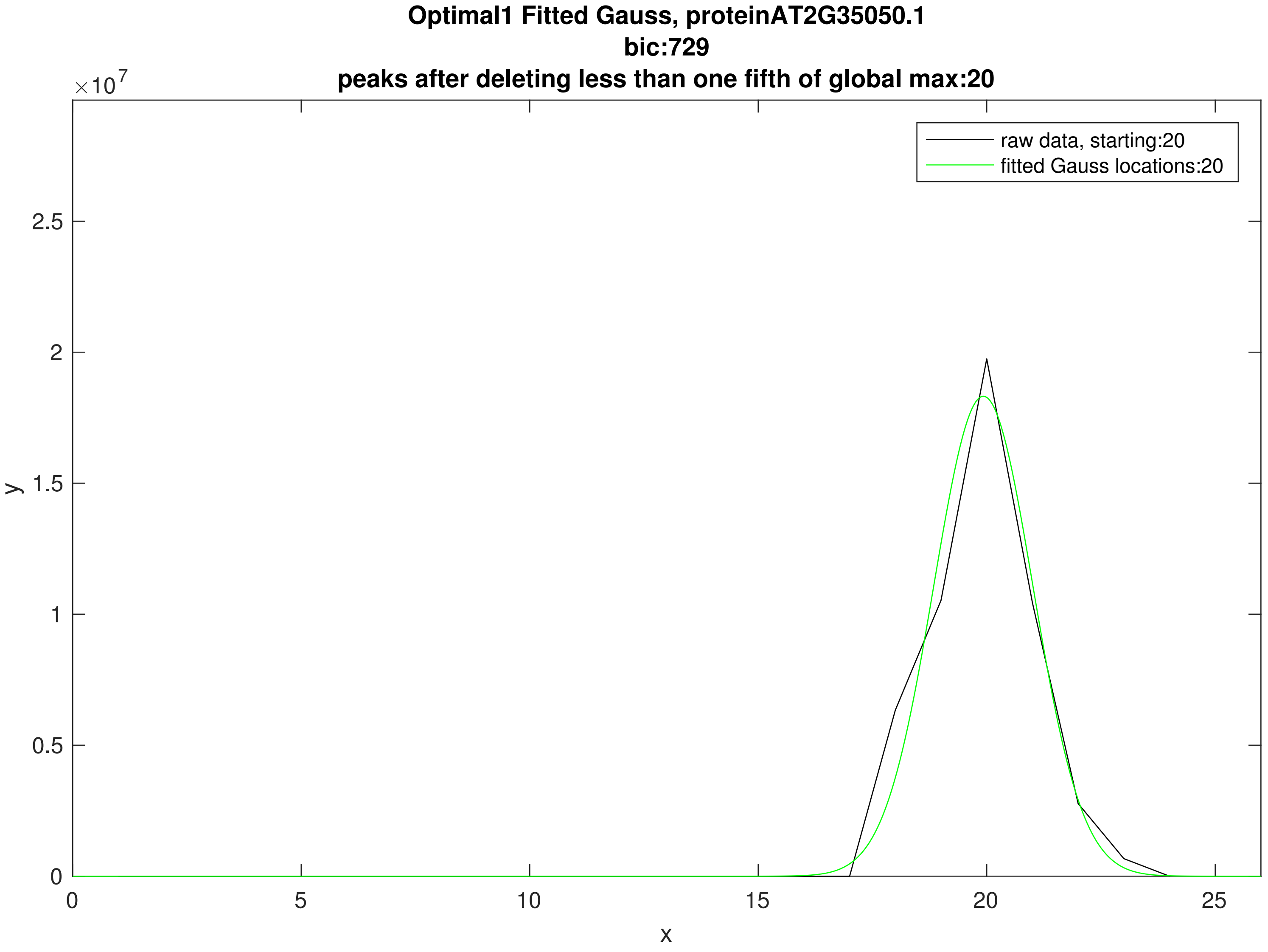

Optimal1 Fitted Gauss, proteinAT2G35050.1
bic:729
peaks after deleting less than one fifth of global max:20
×10^7
raw data, starting:20
fitted Gauss locations:20
2.5
2
1.5
1
0.5
0
y
0
5
10
15
20
25
x

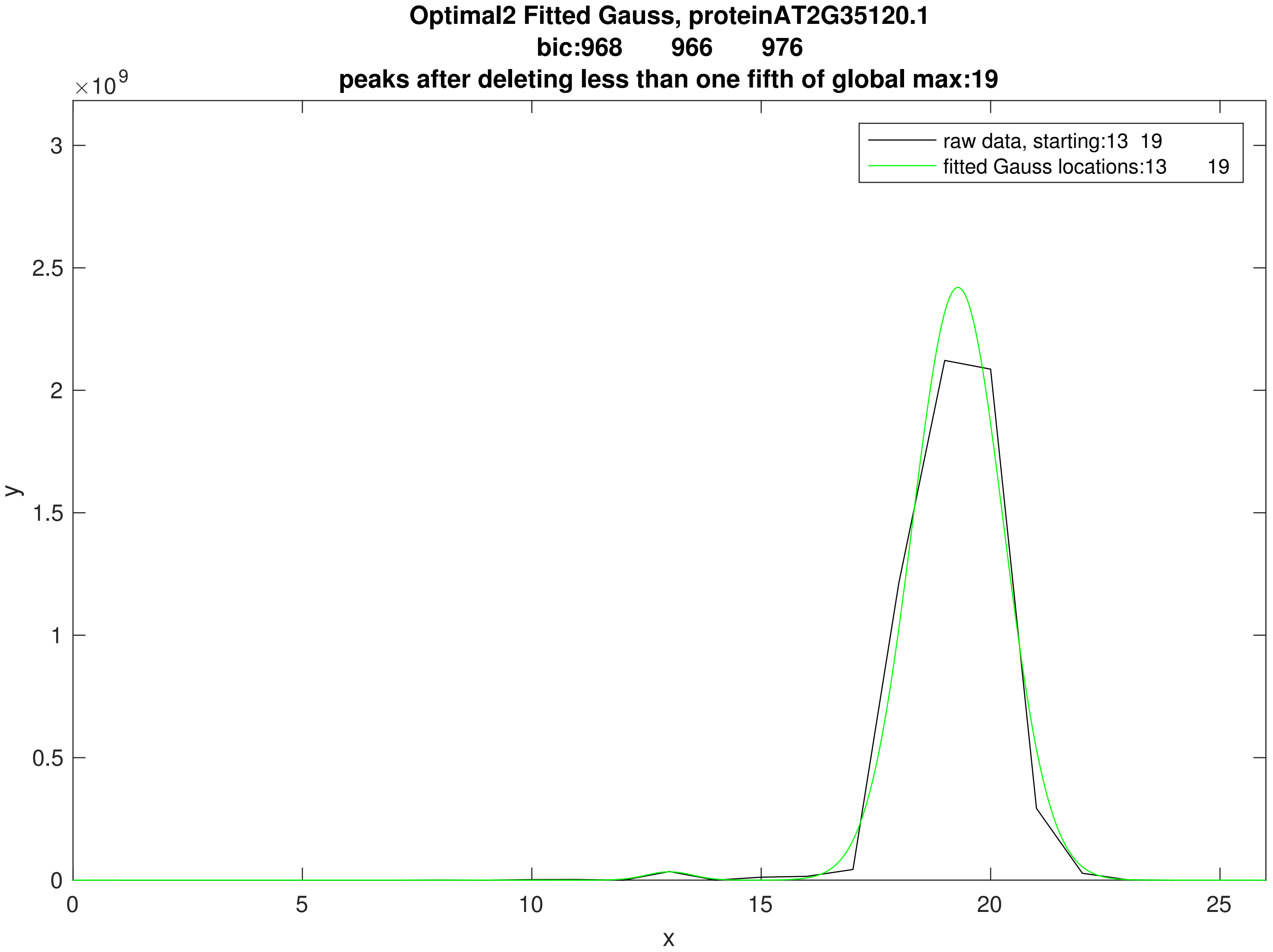

Optimal2 Fitted Gauss, proteinAT2G35120.1
bic:968 966 976
peaks after deleting less than one fifth of global max:19
×10^9
raw data, starting:13 19
fitted Gauss locations:13 19
y
x
0
0.5
1
1.5
2
2.5
3
0
5
10
15
20
25

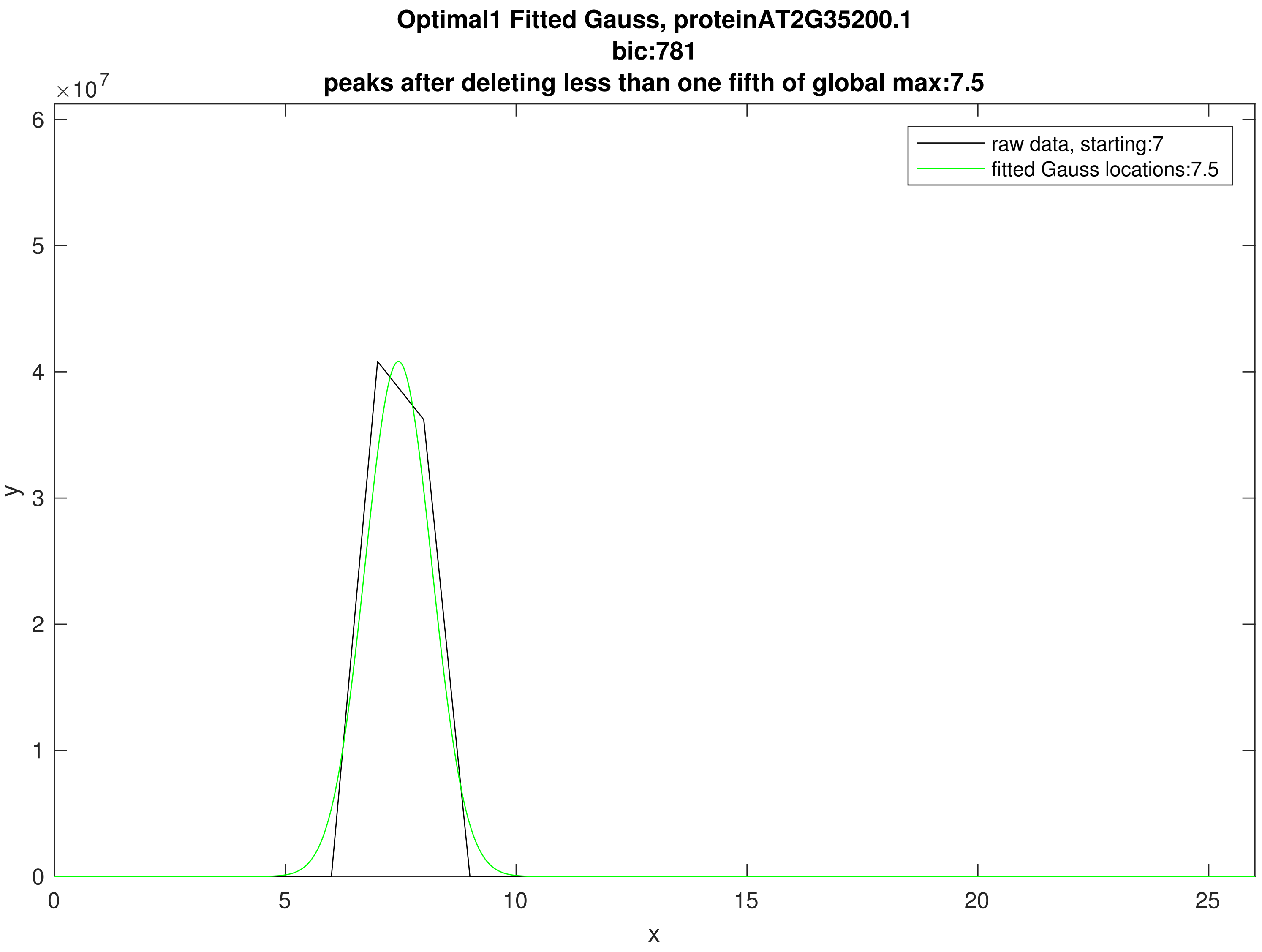

Optimal1 Fitted Gauss, proteinAT2G35200.1
bic:781
peaks after deleting less than one fifth of global max:7.5
×10^7
raw data, starting:7
fitted Gauss locations:7.5
y
x
0
1
2
3
4
5
6
0
5
10
15
20
25

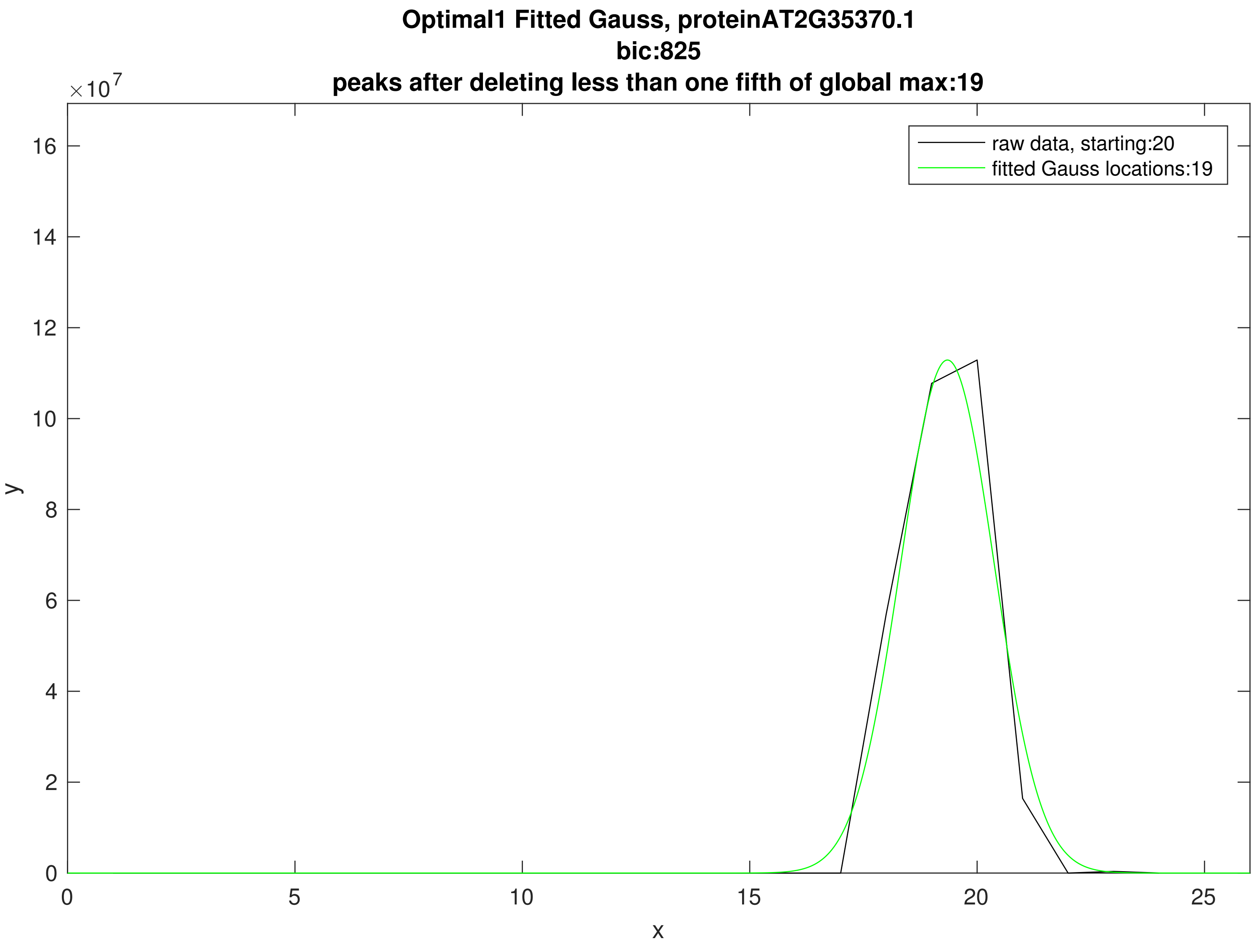
Optimal1 Fitted Gauss, proteinAT2G35370.1
bic:825
peaks after deleting less than one fifth of global max:19
×10^7
raw data, starting:20
fitted Gauss locations:19
y
x
0
2
4
6
8
10
12
14
16
0
5
10
15
20
25

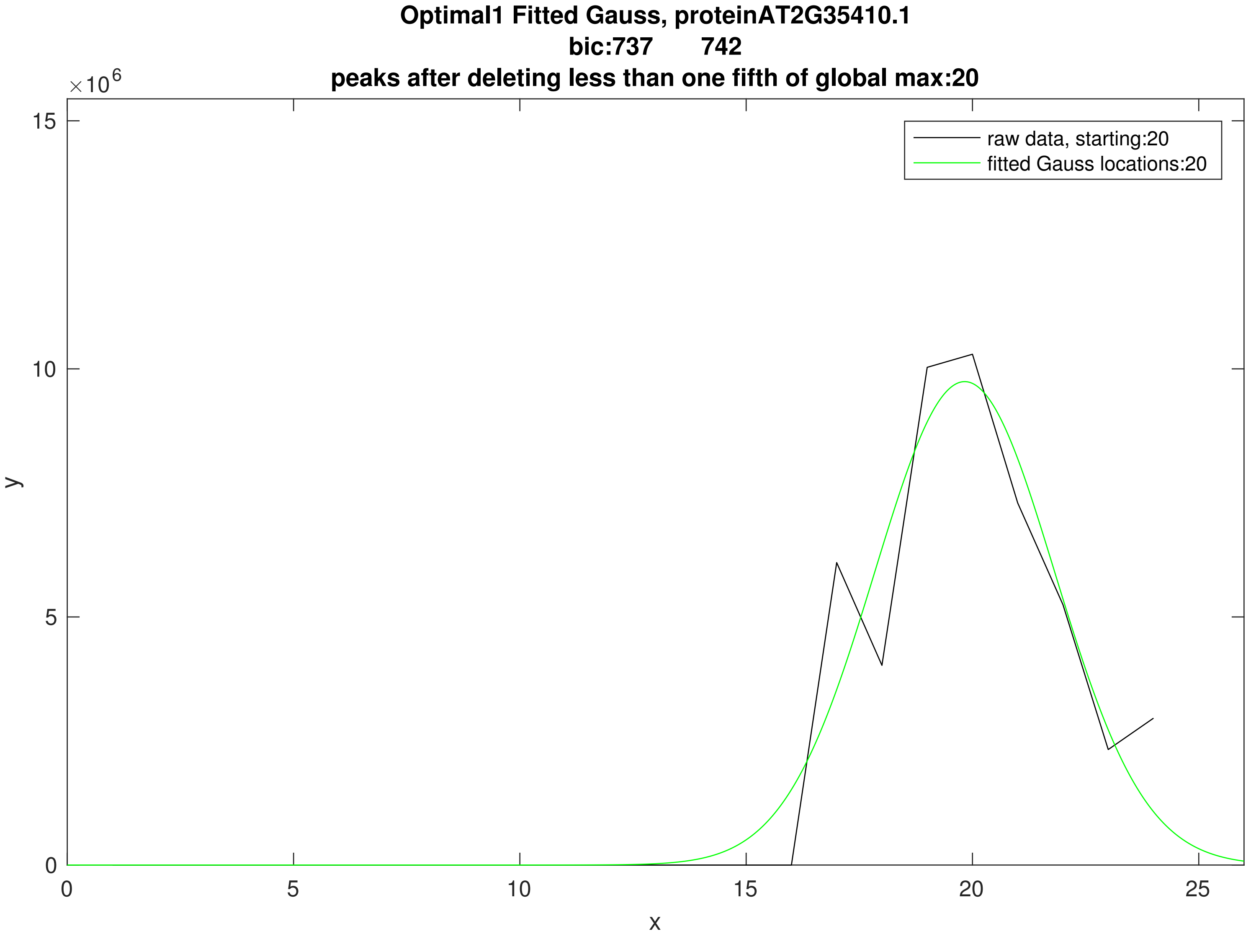
Optimal1 Fitted Gauss, proteinAT2G35410.1
bic:737 742
peaks after deleting less than one fifth of global max:20
×10^6
15
10
5
0
y
0
5
10
15
20
25
x
raw data, starting:20
fitted Gauss locations:20

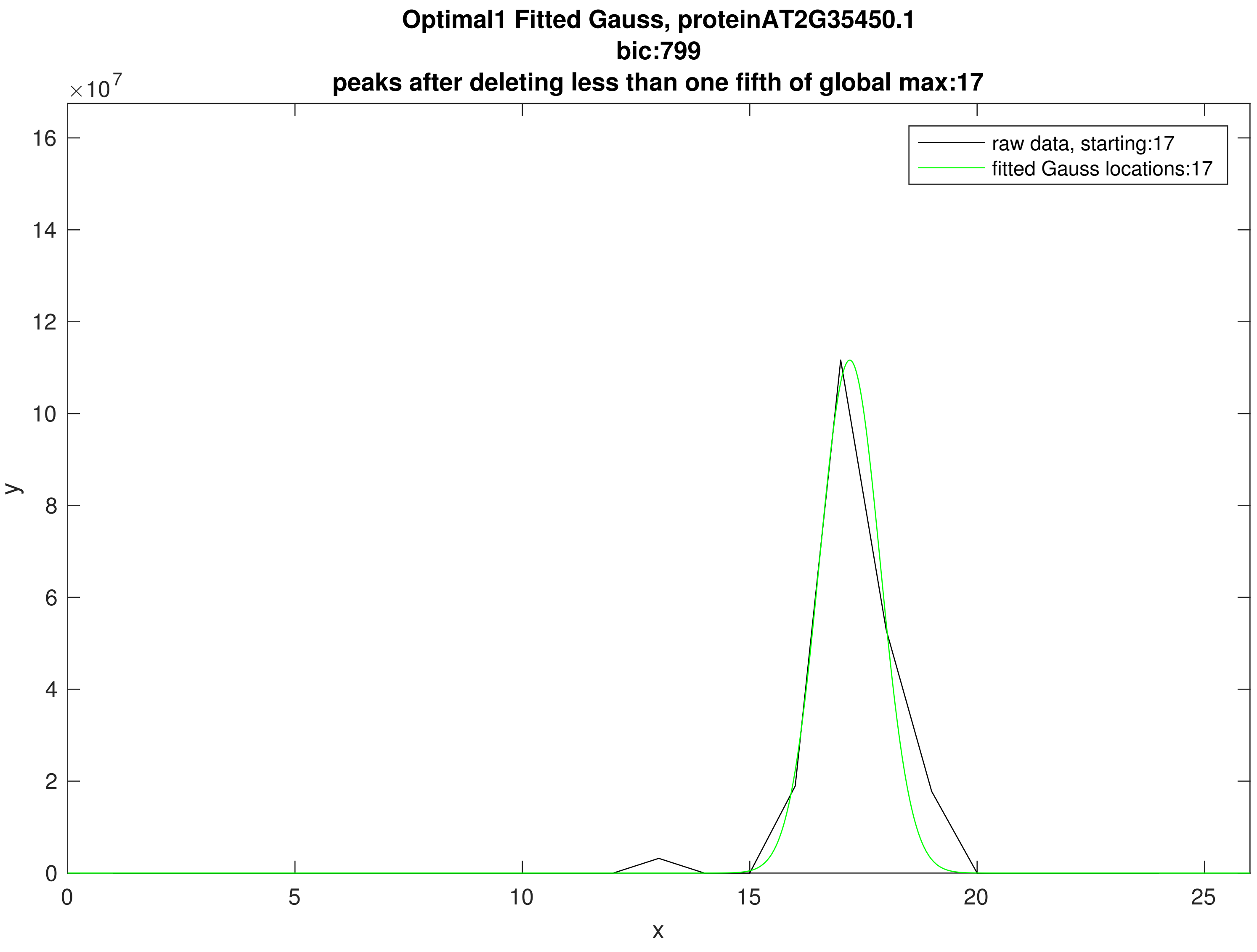
Optimal1 Fitted Gauss, proteinAT2G35450.1
bic:799
peaks after deleting less than one fifth of global max:17
×10^7
raw data, starting:17
fitted Gauss locations:17
y
x
0
2
4
6
8
10
12
14
16
0
5
10
15
20
25

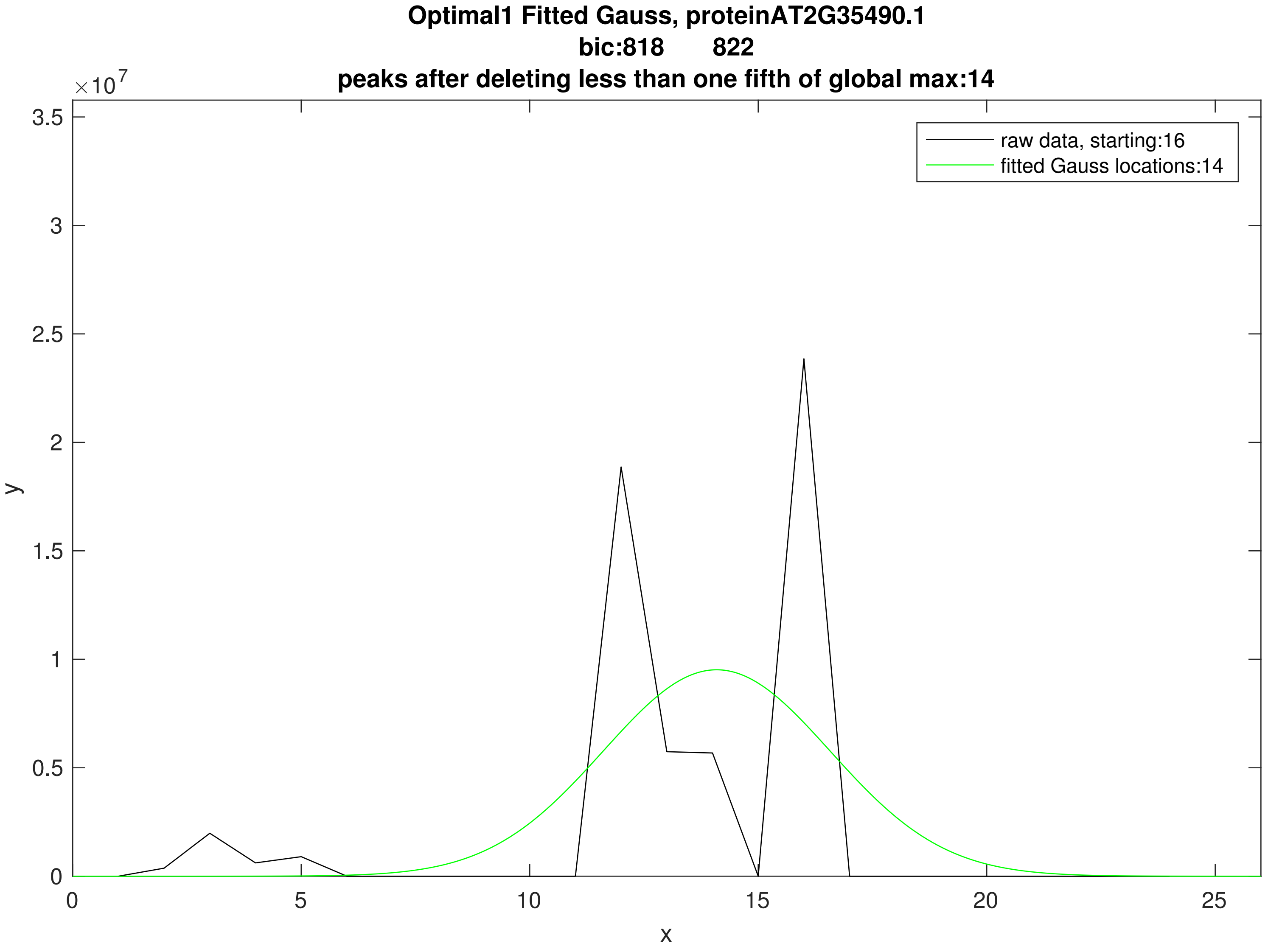

Optimal1 Fitted Gauss, proteinAT2G35490.1
bic:818 822
peaks after deleting less than one fifth of global max:14
×10^7
3.5
3
2.5
2
1.5
1
0.5
0
y
0
5
10
15
20
25
x
raw data, starting:16
fitted Gauss locations:14

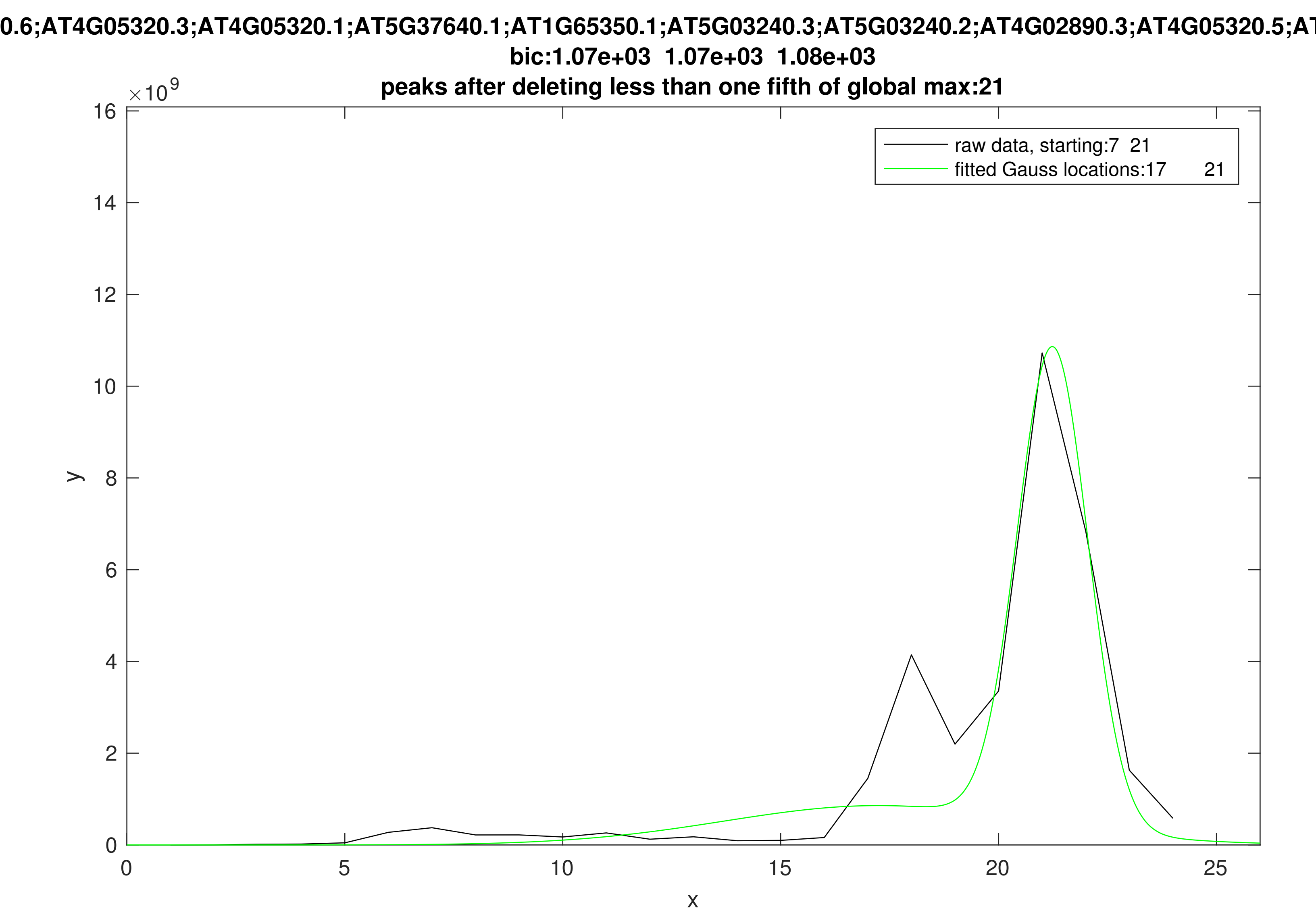
0.6;AT4G05320.3;AT4G05320.1;AT5G37640.1;AT1G65350.1;AT5G03240.3;AT5G03240.2;AT4G02890.3;AT4G05320.5;AT
bic:1.07e+03  1.07e+03  1.08e+03
peaks after deleting less than one fifth of global max:21
raw data, starting:7  21
fitted Gauss locations:17  21
×10^9
y
x

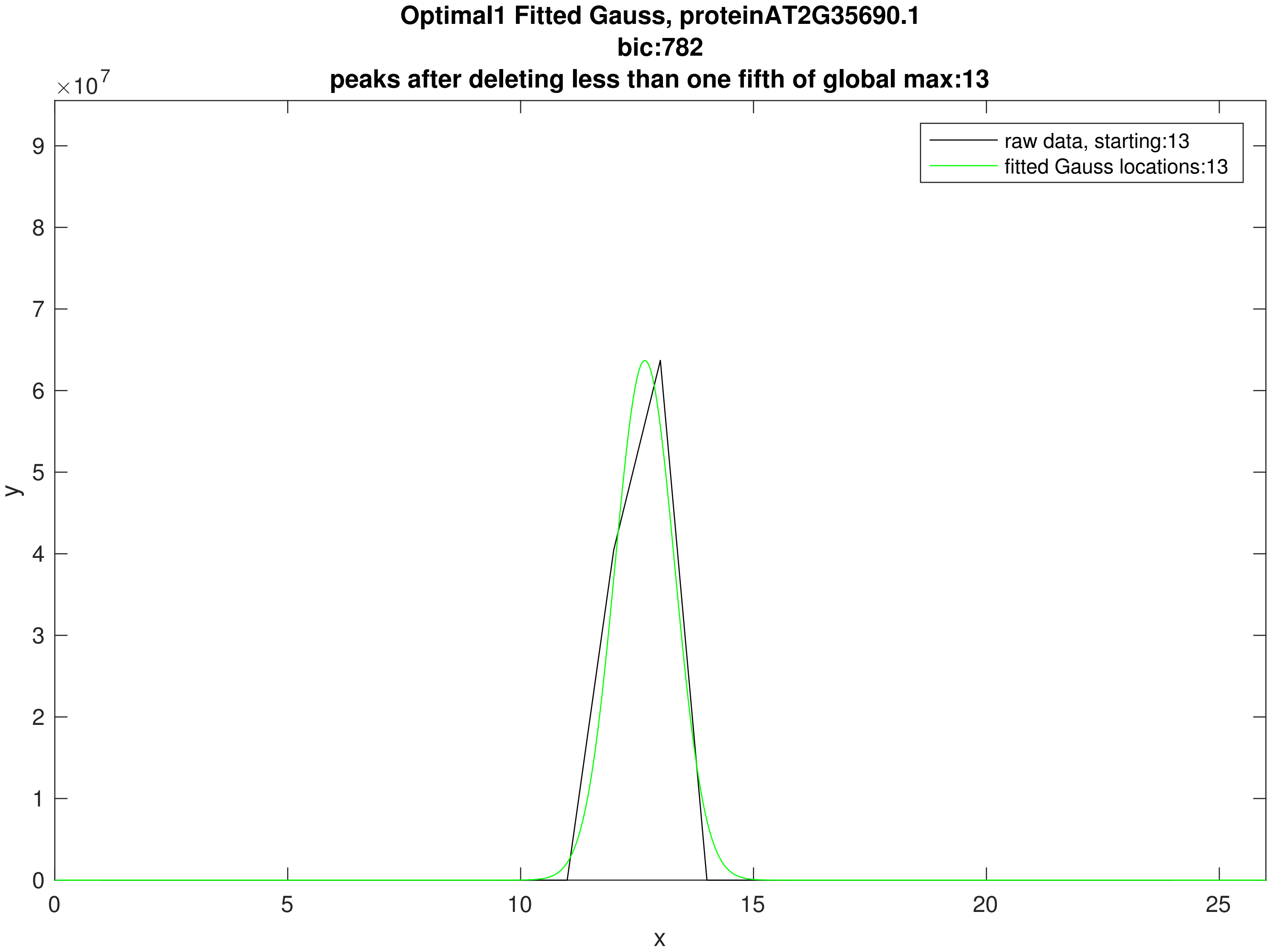

Optimal1 Fitted Gauss, proteinAT2G35690.1
bic:782
peaks after deleting less than one fifth of global max:13
×10^7
raw data, starting:13
fitted Gauss locations:13
y
x
0
1
2
3
4
5
6
7
8
9
0
5
10
15
20
25

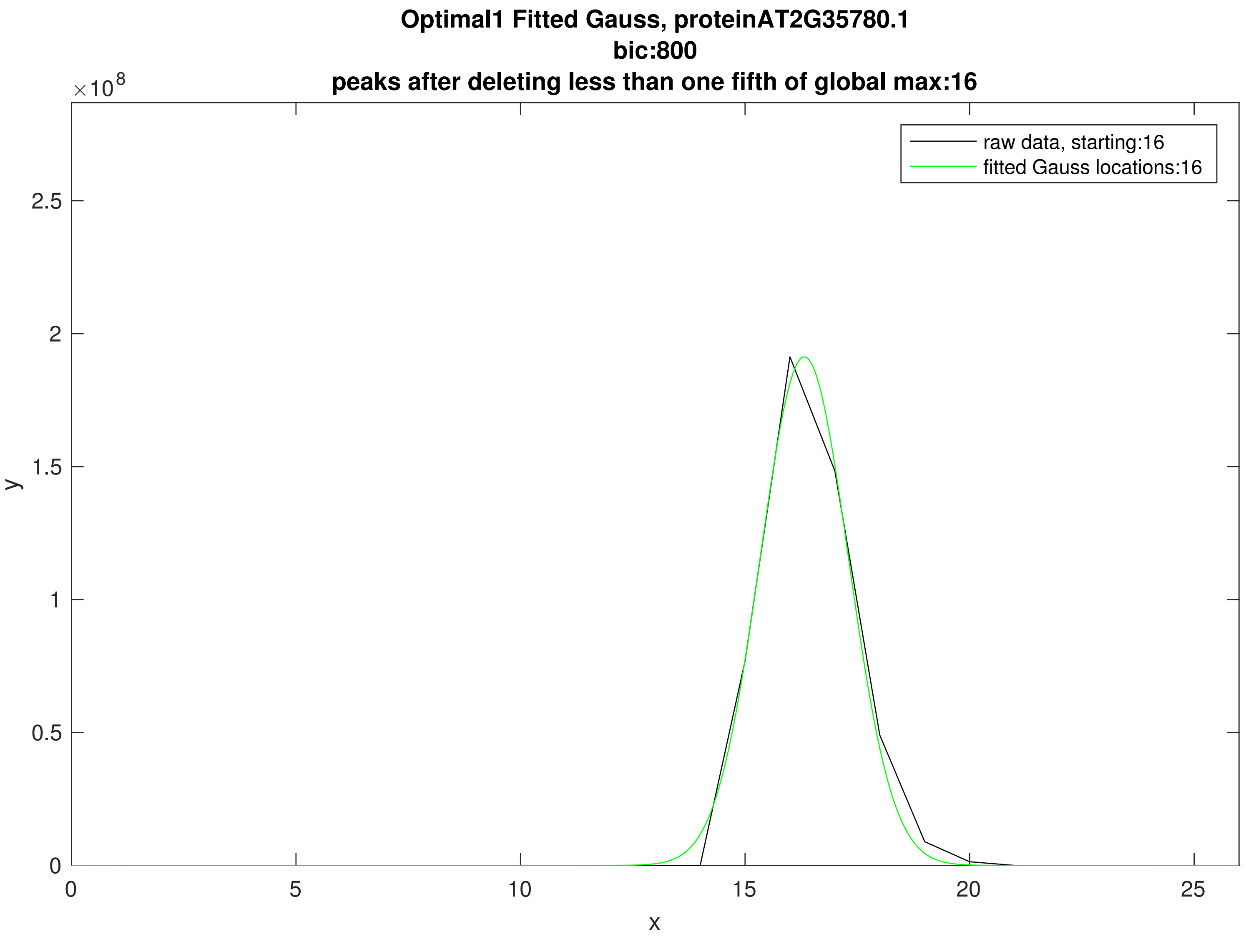
Optimal1 Fitted Gauss, proteinAT2G35780.1
bic:800
peaks after deleting less than one fifth of global max:16
×10^8
raw data, starting:16
fitted Gauss locations:16
y
x
0
0.5
1
1.5
2
2.5
5
10
15
20
25

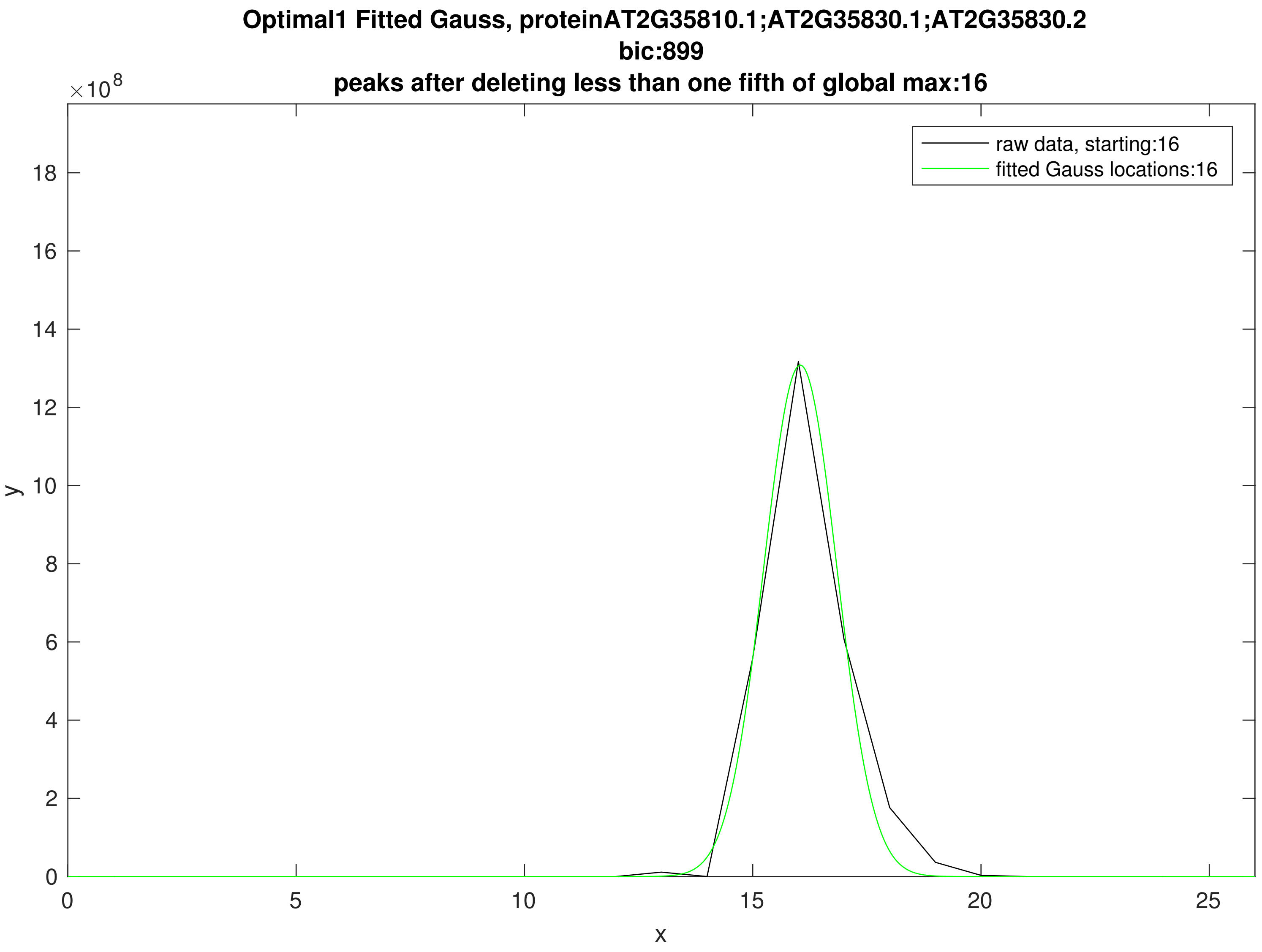

Optimal1 Fitted Gauss, proteinAT2G35810.1;AT2G35830.1;AT2G35830.2
bic:899
peaks after deleting less than one fifth of global max:16
×10^8
raw data, starting:16
fitted Gauss locations:16
y
x
0
2
4
6
8
10
12
14
16
18
0
5
10
15
20
25

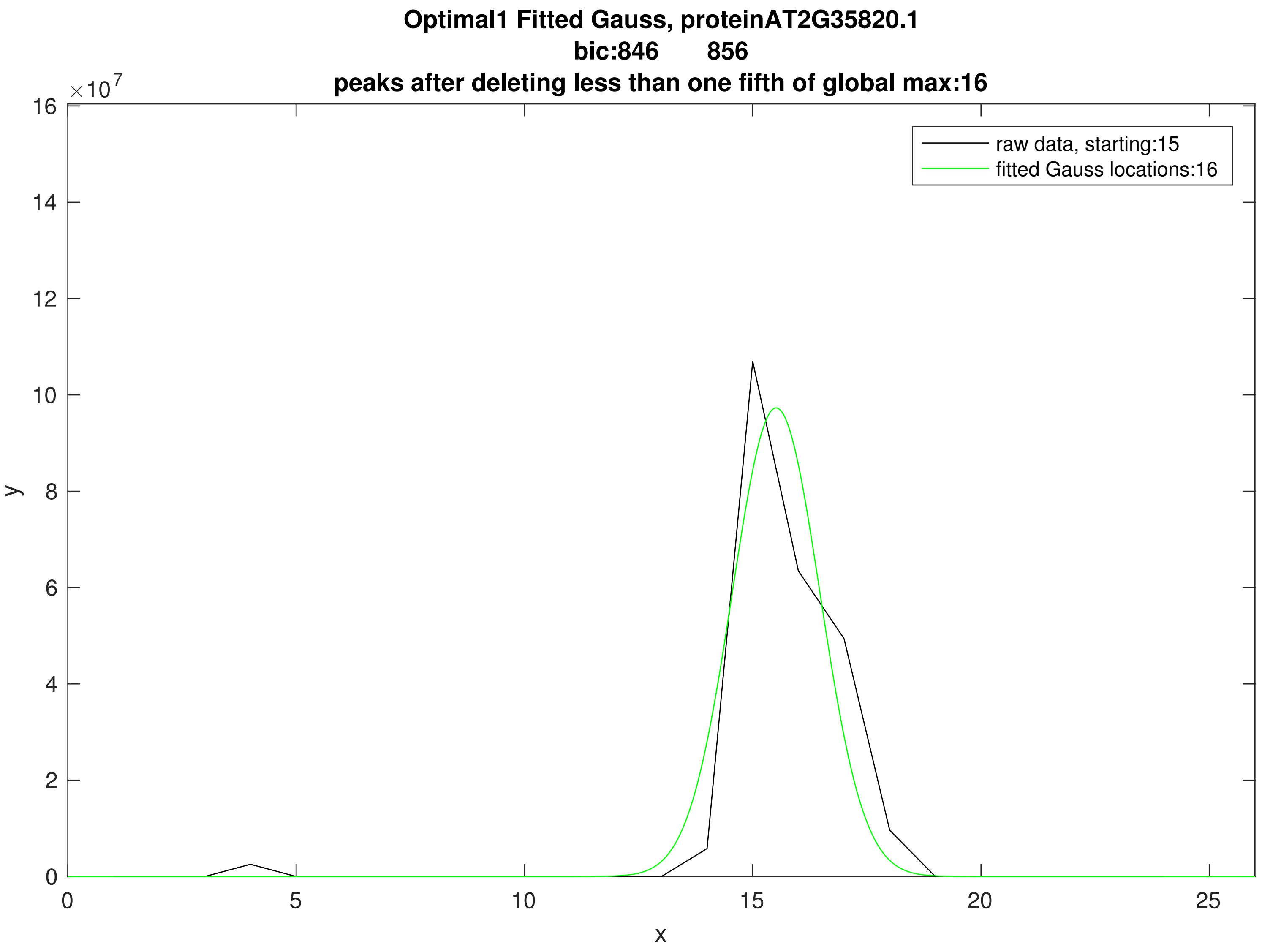
Optimal1 Fitted Gauss, proteinAT2G35820.1
bic:846 856
peaks after deleting less than one fifth of global max:16
×10^7
16
14
12
10
8
6
4
2
0
y
0
5
10
15
20
25
x
raw data, starting:15
fitted Gauss locations:16

#### Optimal1 Fitted Gauss, proteinAT2G35840.3;AT2G35840.2;AT2G35840.1
#### bic:960
#### peaks after deleting less than one fifth of global max:17

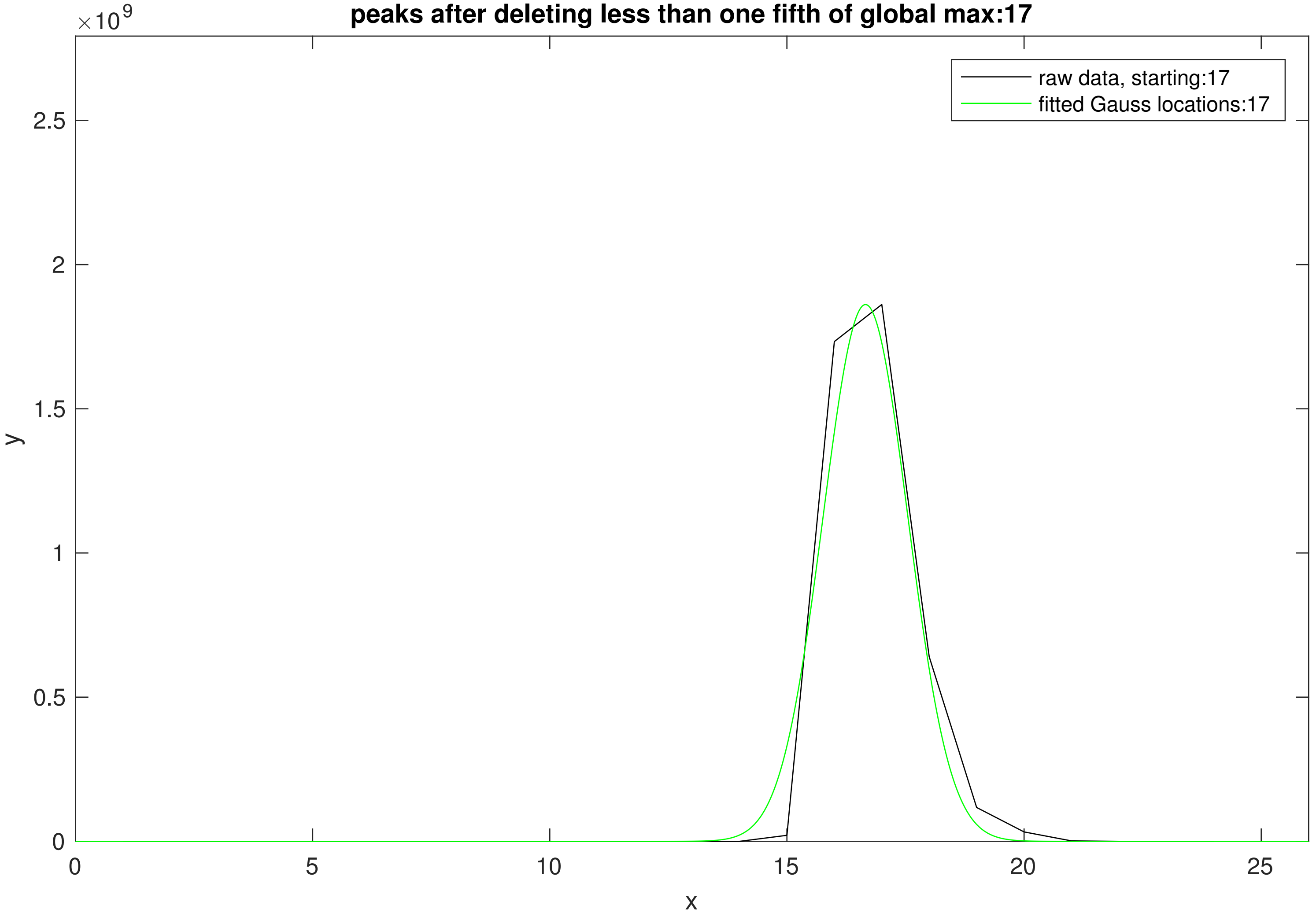

**Optimal1 Fitted Gauss, proteinAT2G35860.1**
**bic:910 973 939**
**peaks after deleting less than one fifth of global max:5.2**

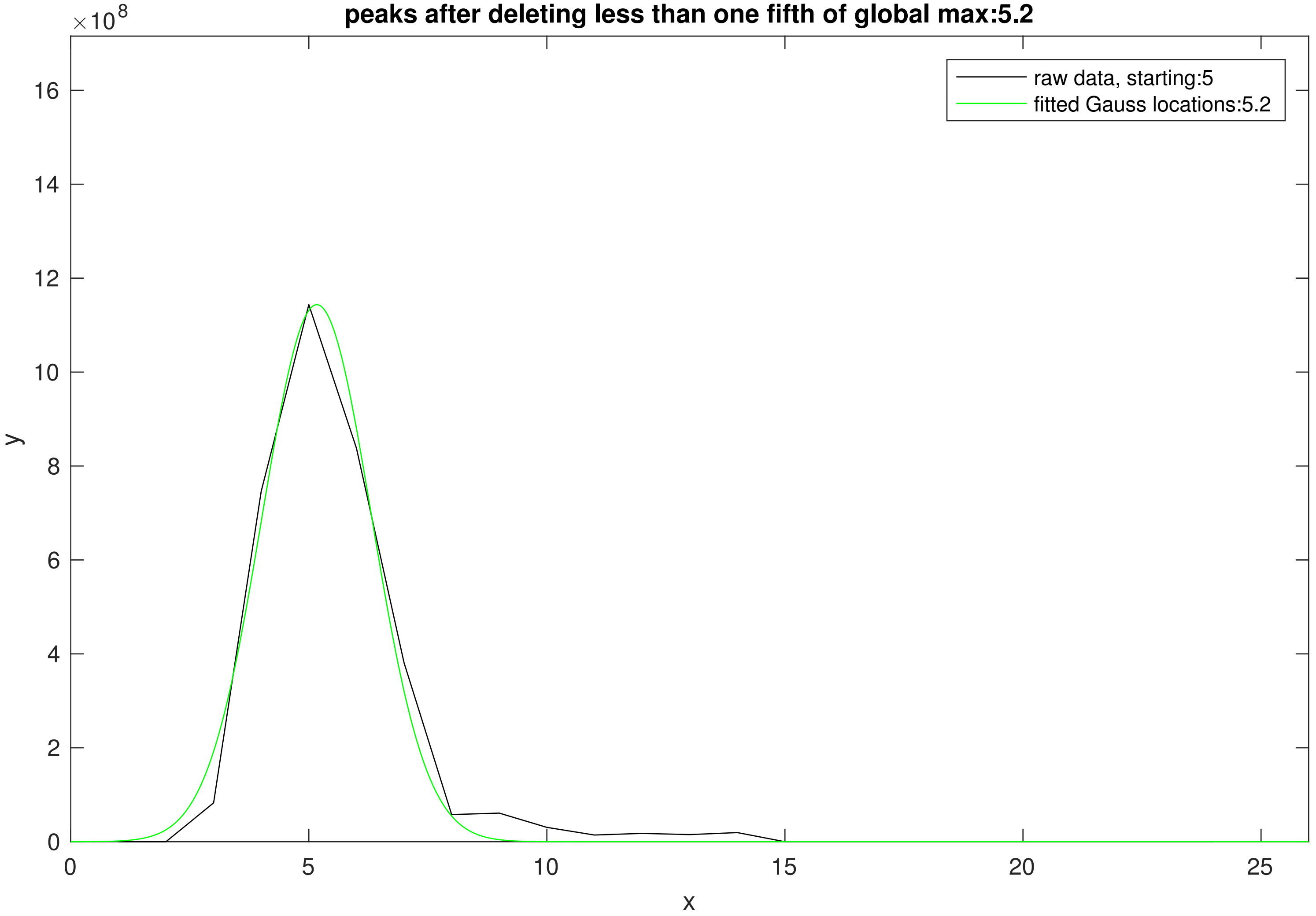

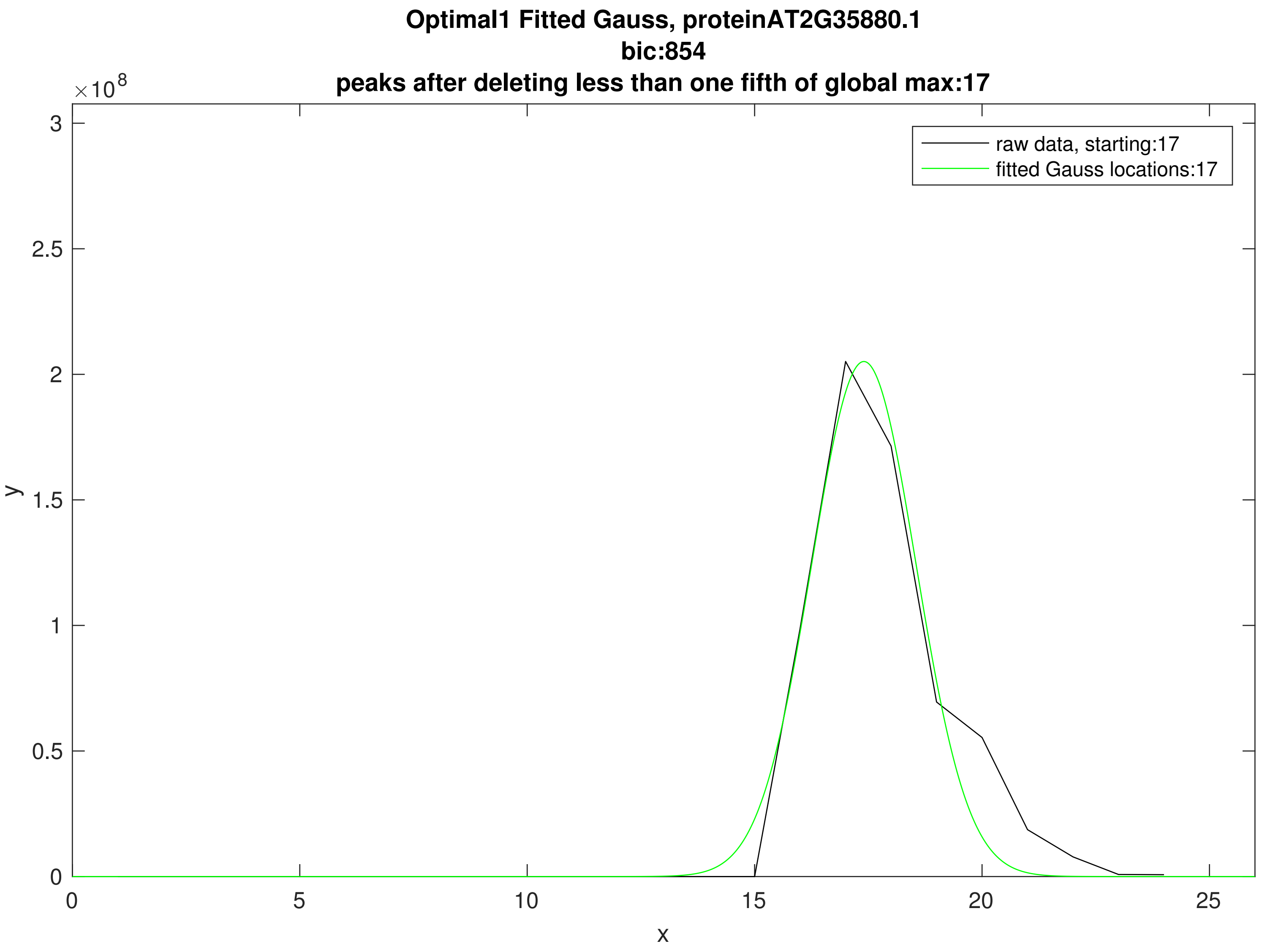

Optimal1 Fitted Gauss, proteinAT2G35880.1
bic:854
peaks after deleting less than one fifth of global max:17
×10^8
raw data, starting:17
fitted Gauss locations:17
y
x
0
0.5
1
1.5
2
2.5
3
0
5
10
15
20
25

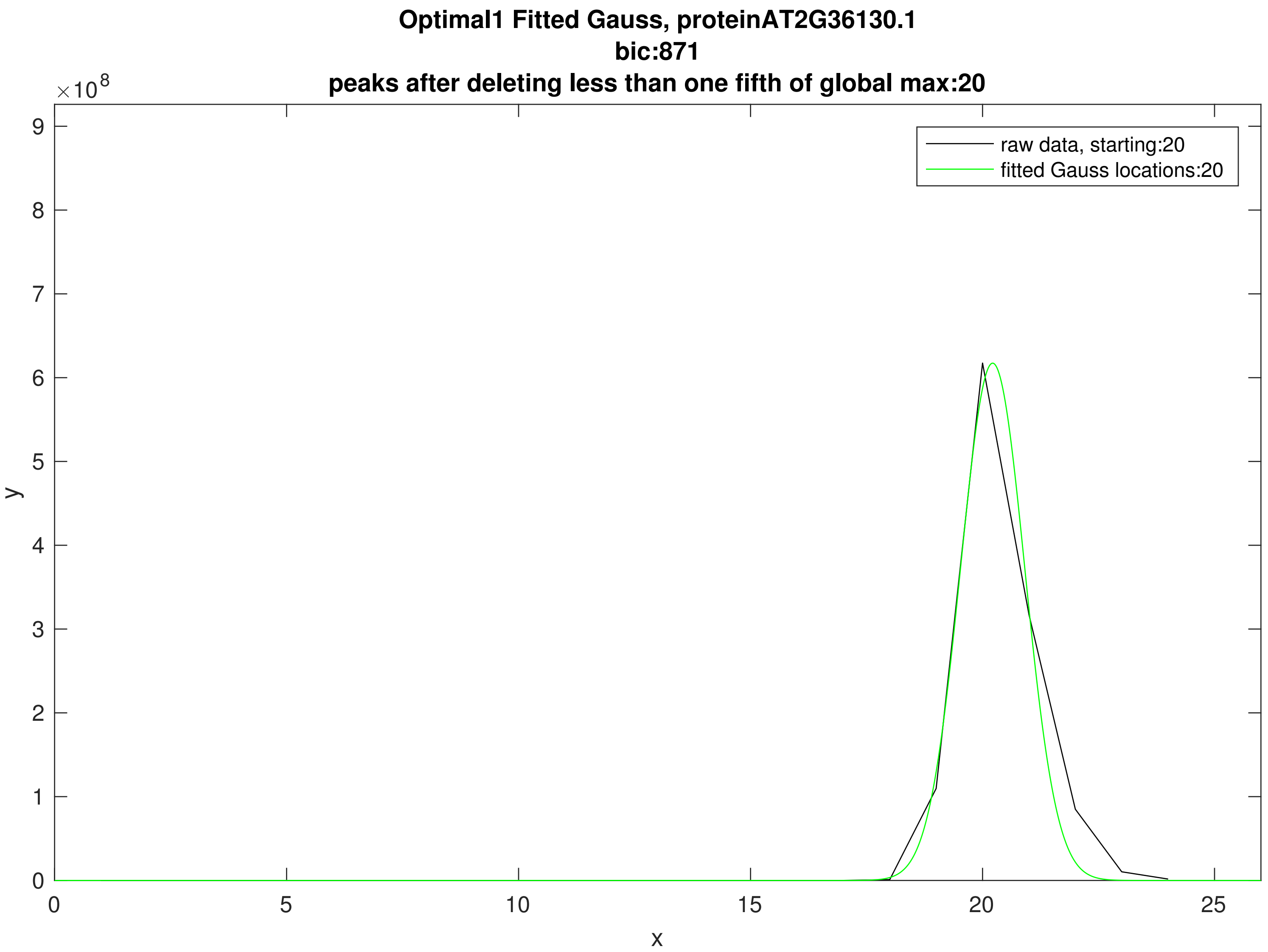
Optimal1 Fitted Gauss, proteinAT2G36130.1
bic:871
peaks after deleting less than one fifth of global max:20
×10^8
raw data, starting:20
fitted Gauss locations:20
y
x
0
1
2
3
4
5
6
7
8
9
0
5
10
15
20
25

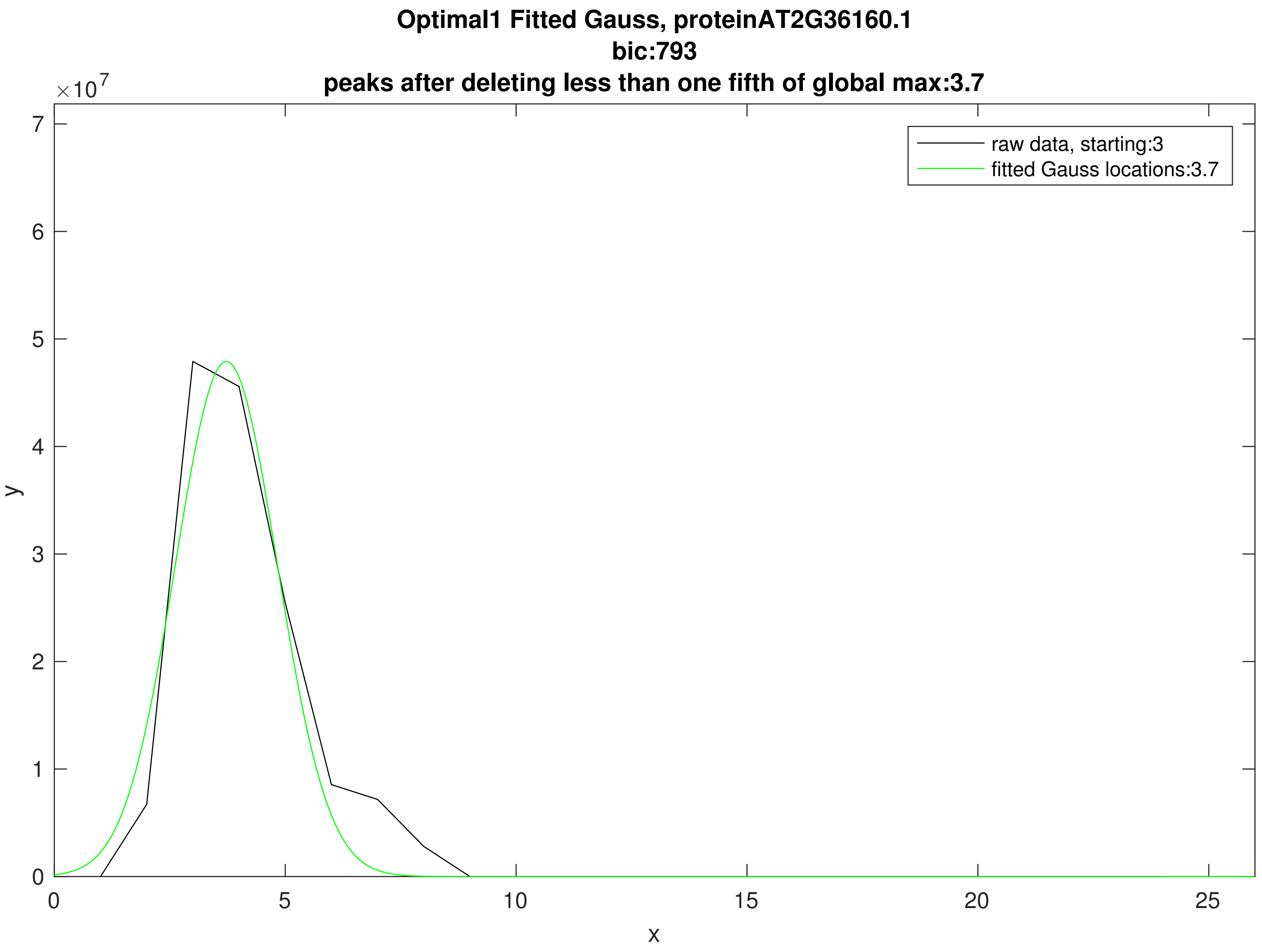
Optimal1 Fitted Gauss, proteinAT2G36160.1
bic:793
peaks after deleting less than one fifth of global max:3.7
×10^7
raw data, starting:3
fitted Gauss locations:3.7
y
x
0
1
2
3
4
5
6
7
0
5
10
15
20
25

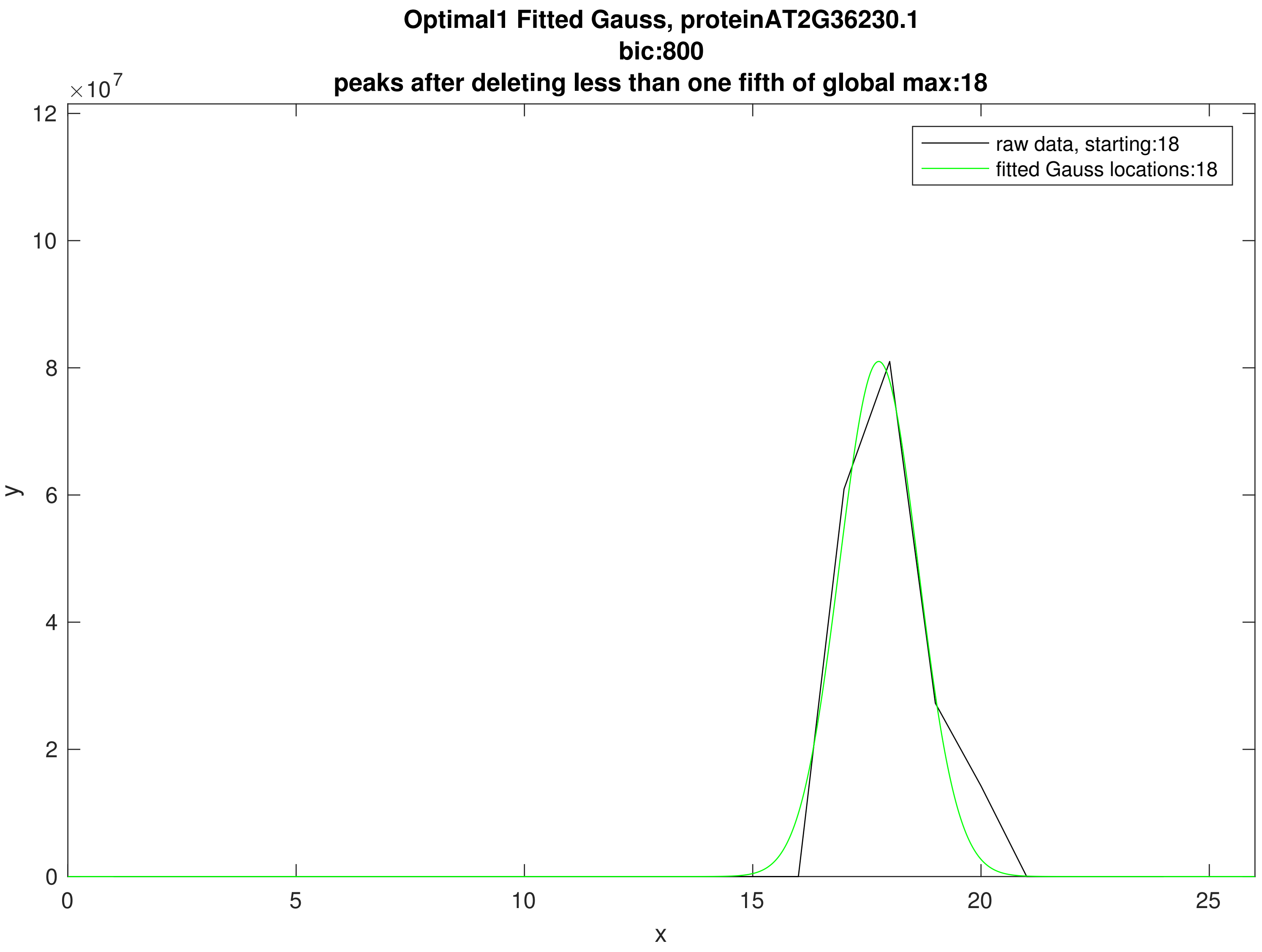

Optimal1 Fitted Gauss, proteinAT2G36230.1
bic:800
peaks after deleting less than one fifth of global max:18
×10^7
12
10
8
6
4
2
0
y
0
5
10
15
20
25
x
raw data, starting:18
fitted Gauss locations:18

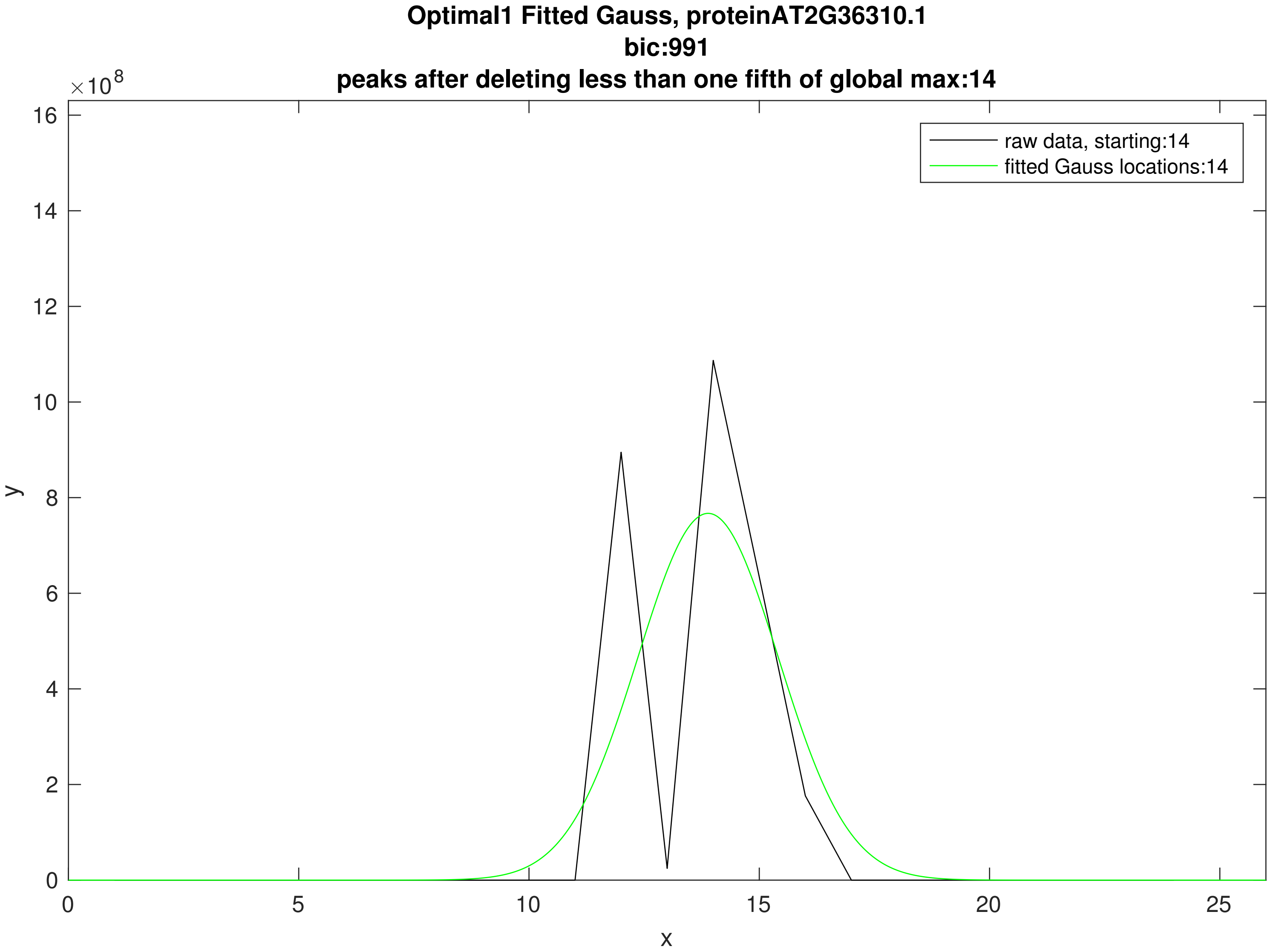
Optimal1 Fitted Gauss, proteinAT2G36310.1
bic:991
peaks after deleting less than one fifth of global max:14
×10^8
16
14
12
10
8
6
4
2
0
y
0
5
10
15
20
25
x
raw data, starting:14
fitted Gauss locations:14

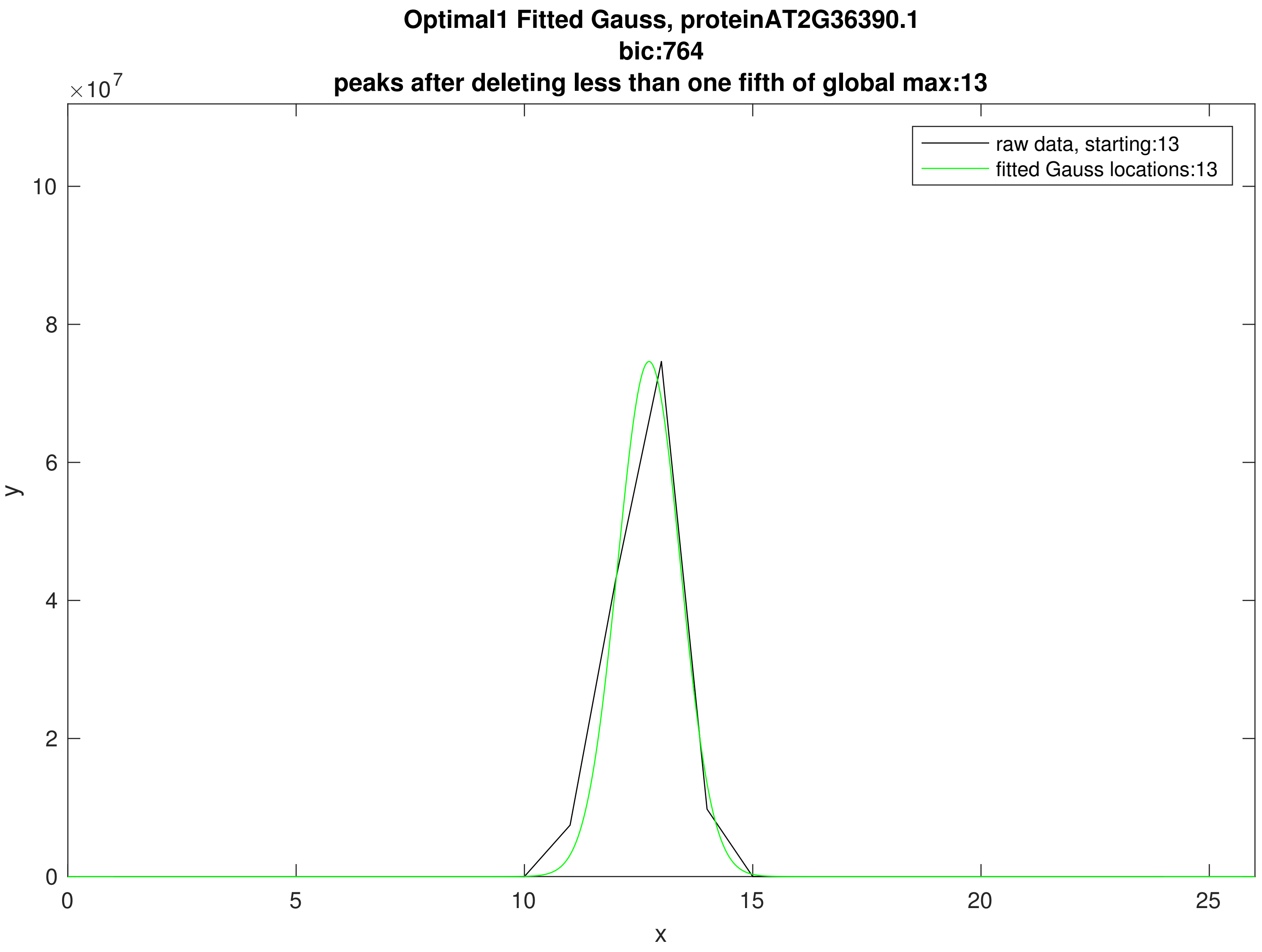

Optimal1 Fitted Gauss, proteinAT2G36390.1
bic:764
peaks after deleting less than one fifth of global max:13
×10^7
raw data, starting:13
fitted Gauss locations:13
10
8
6
4
2
0
y
0
5
10
15
20
25
x

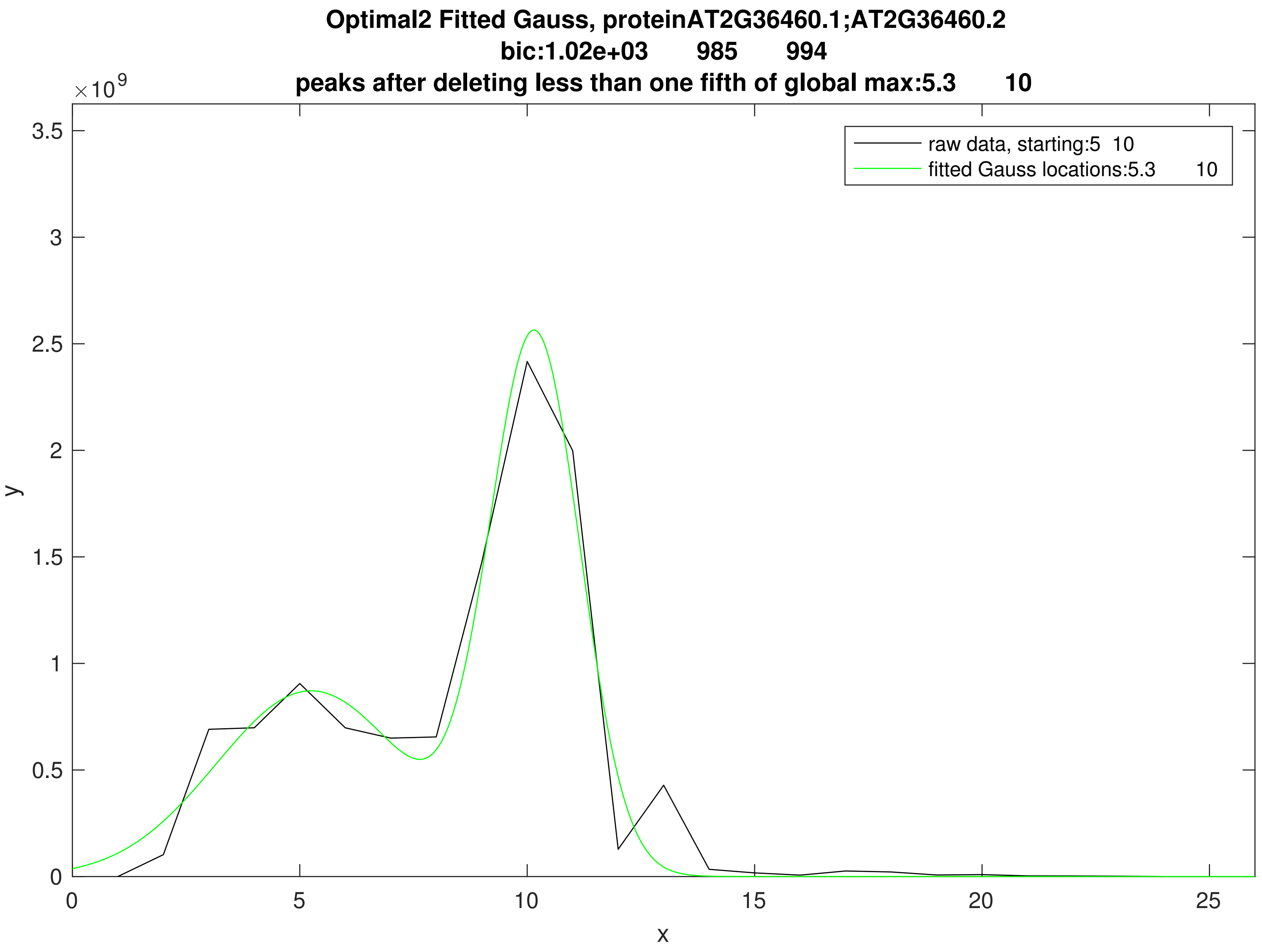
Optimal2 Fitted Gauss, proteinAT2G36460.1;AT2G36460.2
bic:1.02e+03 985 994
peaks after deleting less than one fifth of global max:5.3 10
×10^9
raw data, starting:5 10
fitted Gauss locations:5.3 10
3.5
3
2.5
2
1.5
1
0.5
0
y
0
5
10
15
20
25
x

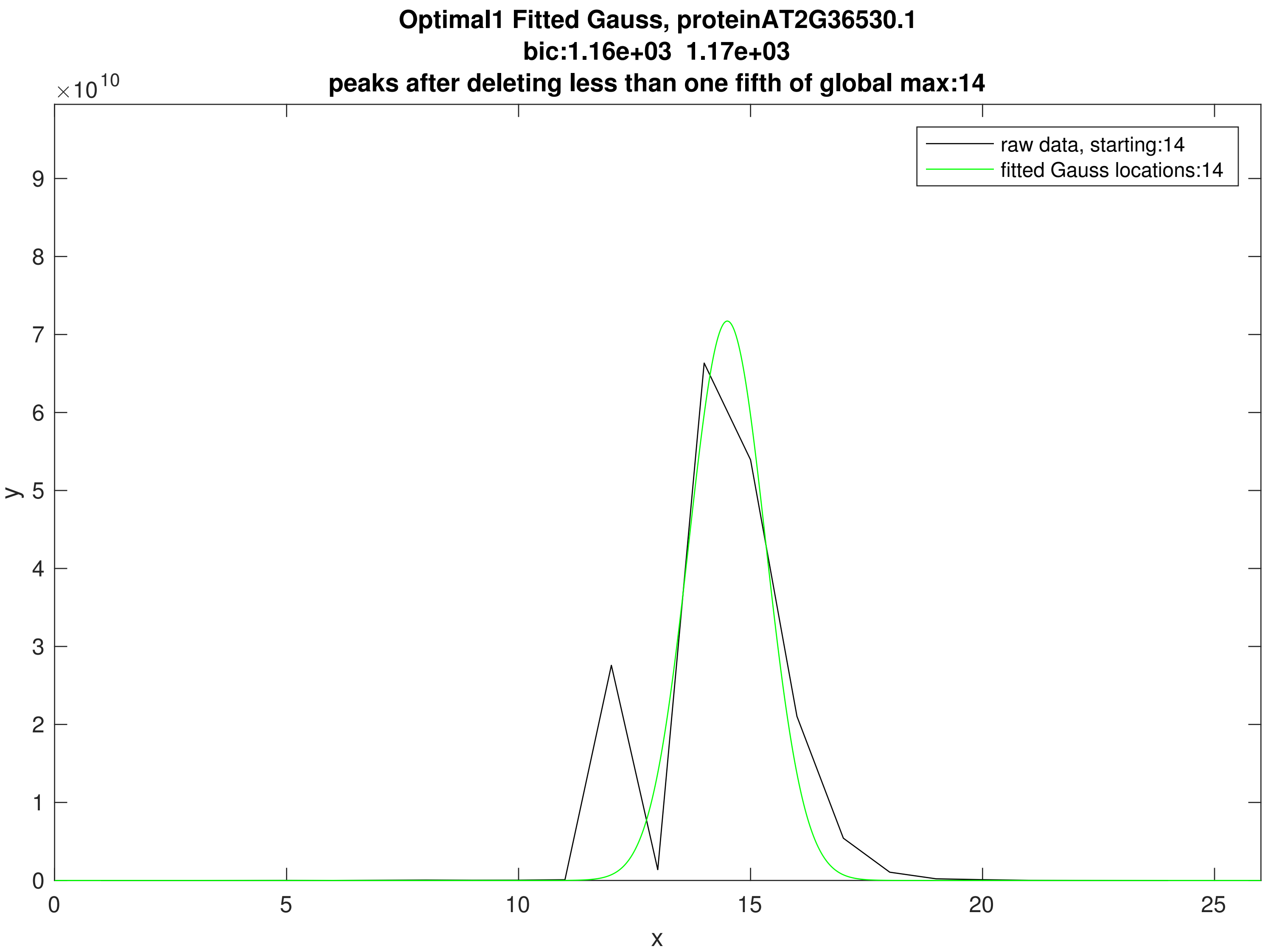
Optimal1 Fitted Gauss, proteinAT2G36530.1
bic:1.16e+03  1.17e+03
peaks after deleting less than one fifth of global max:14
×10^10
raw data, starting:14
fitted Gauss locations:14
y
x
0
1
2
3
4
5
6
7
8
9
0
5
10
15
20
25

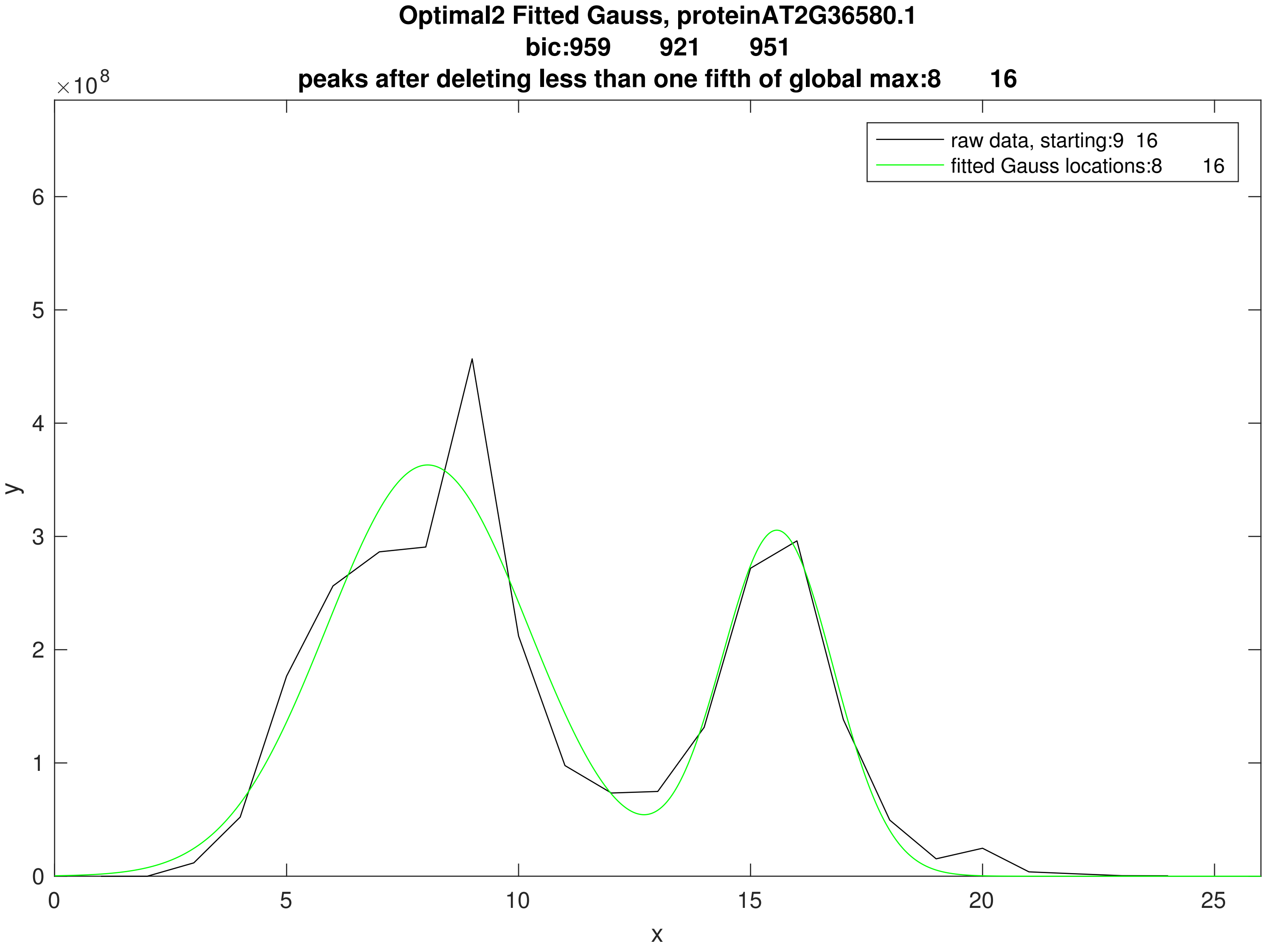
Optimal2 Fitted Gauss, proteinAT2G36580.1
bic:959 921 951
peaks after deleting less than one fifth of global max:8 16
×10^8
raw data, starting:9 16
fitted Gauss locations:8 16
y
x
0
1
2
3
4
5
6
0
5
10
15
20
25

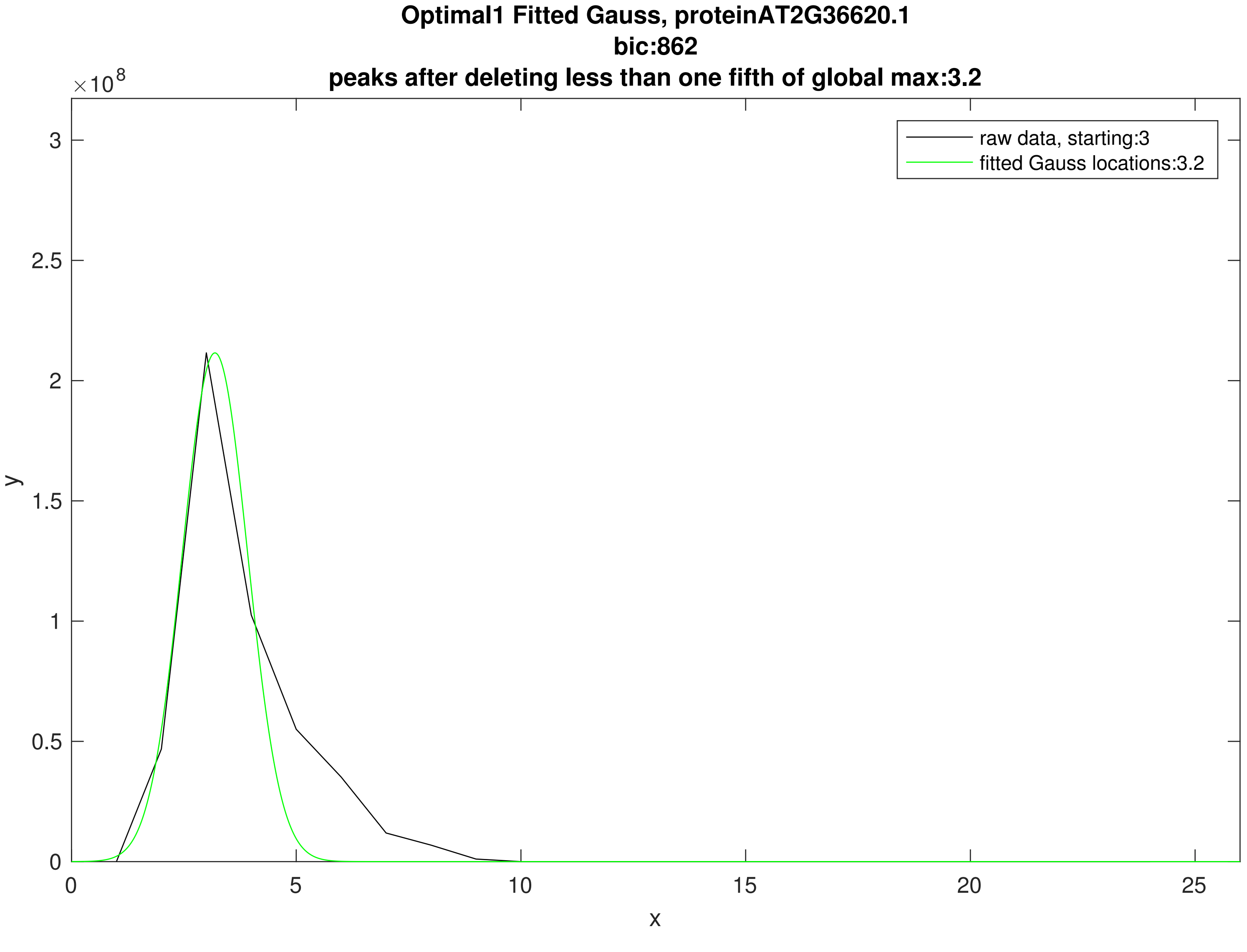
Optimal1 Fitted Gauss, proteinAT2G36620.1
bic:862
peaks after deleting less than one fifth of global max:3.2
×10^8
raw data, starting:3
fitted Gauss locations:3.2
y
x
0
0.5
1
1.5
2
2.5
3
5
10
15
20
25

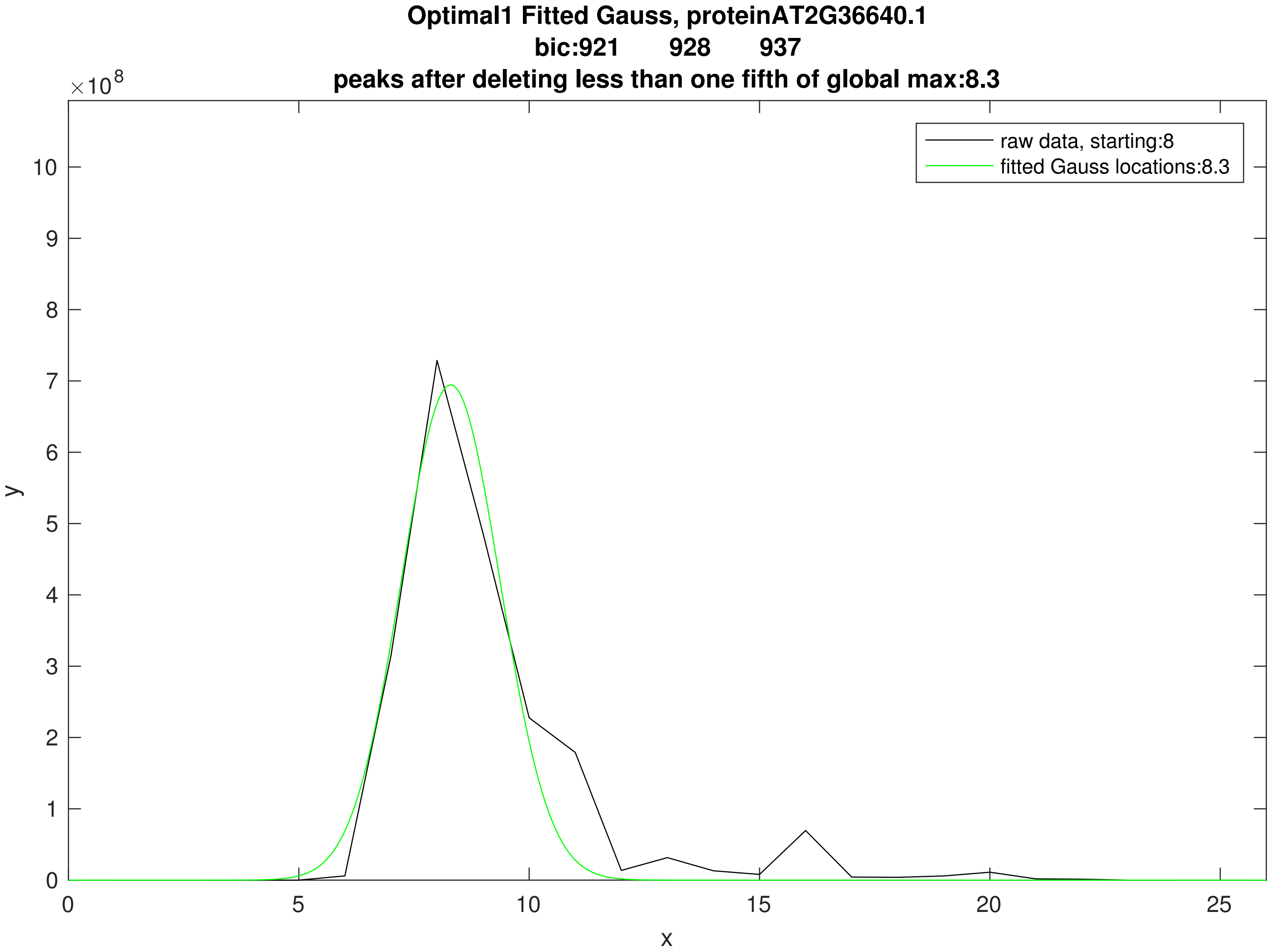
Optimal1 Fitted Gauss, proteinAT2G36640.1
bic:921 928 937
peaks after deleting less than one fifth of global max:8.3
×10^8
raw data, starting:8
fitted Gauss locations:8.3
y
x
0
1
2
3
4
5
6
7
8
9
10
0
5
10
15
20
25

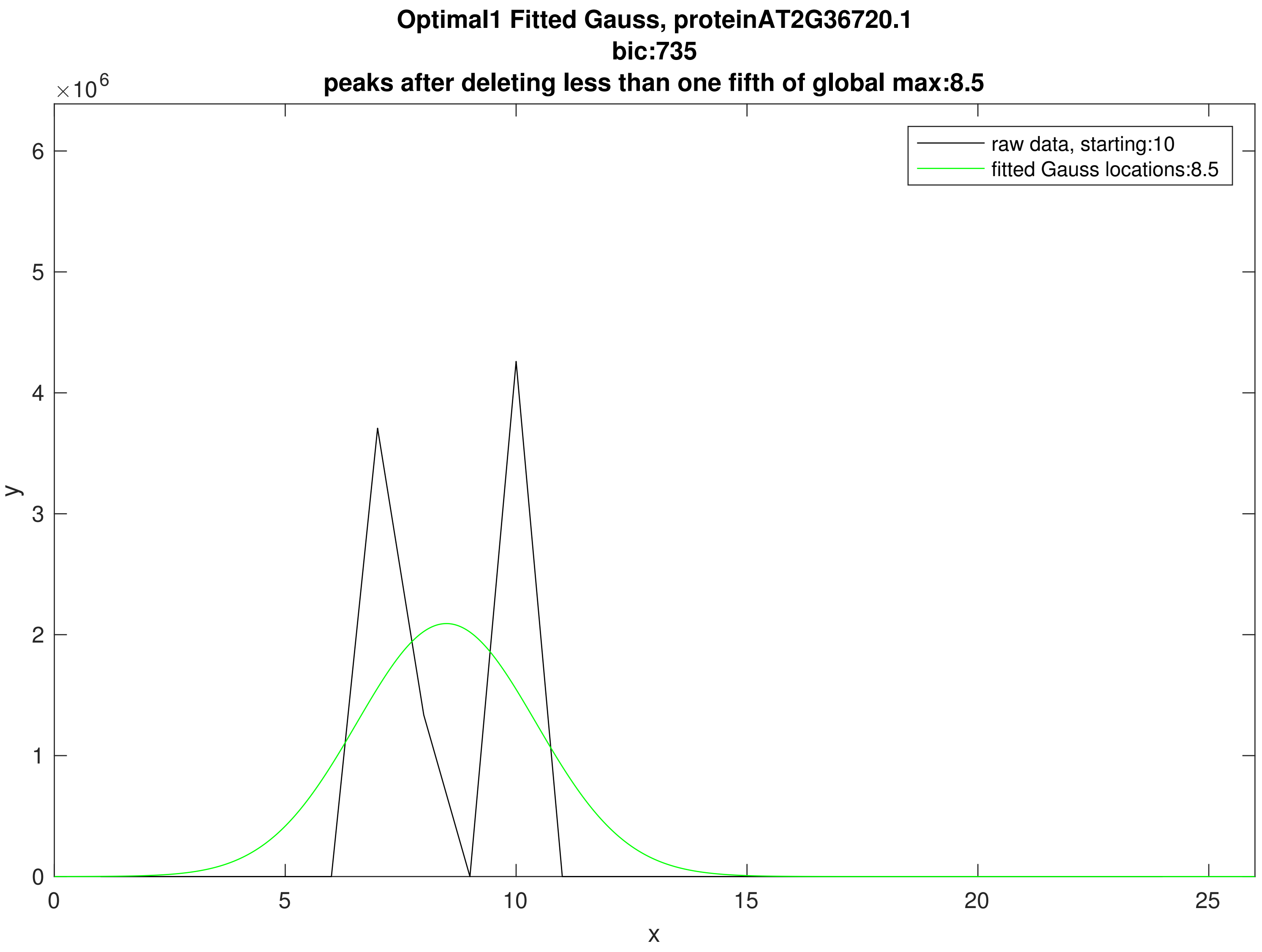

Optimal1 Fitted Gauss, proteinAT2G36720.1
bic:735
peaks after deleting less than one fifth of global max:8.5
×10^6
raw data, starting:10
fitted Gauss locations:8.5
y
x
0
1
2
3
4
5
6
0
5
10
15
20
25

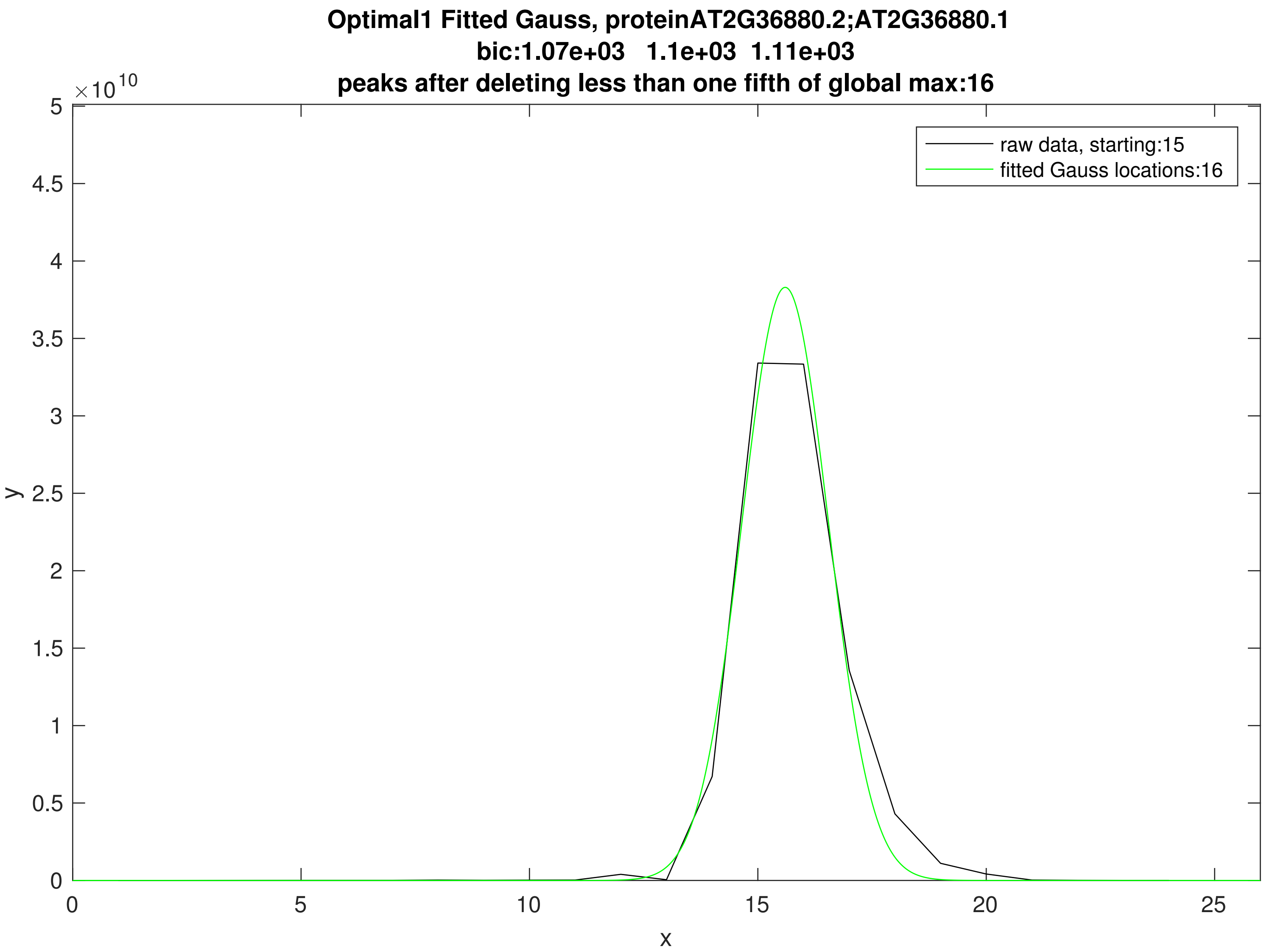
Optimal1 Fitted Gauss, proteinAT2G36880.2;AT2G36880.1
bic:1.07e+03 1.1e+03 1.11e+03
peaks after deleting less than one fifth of global max:16
×10^10
5
4.5
4
3.5
3
2.5
2
1.5
1
0.5
0
y
0
5
10
15
20
25
x
raw data, starting:15
fitted Gauss locations:16

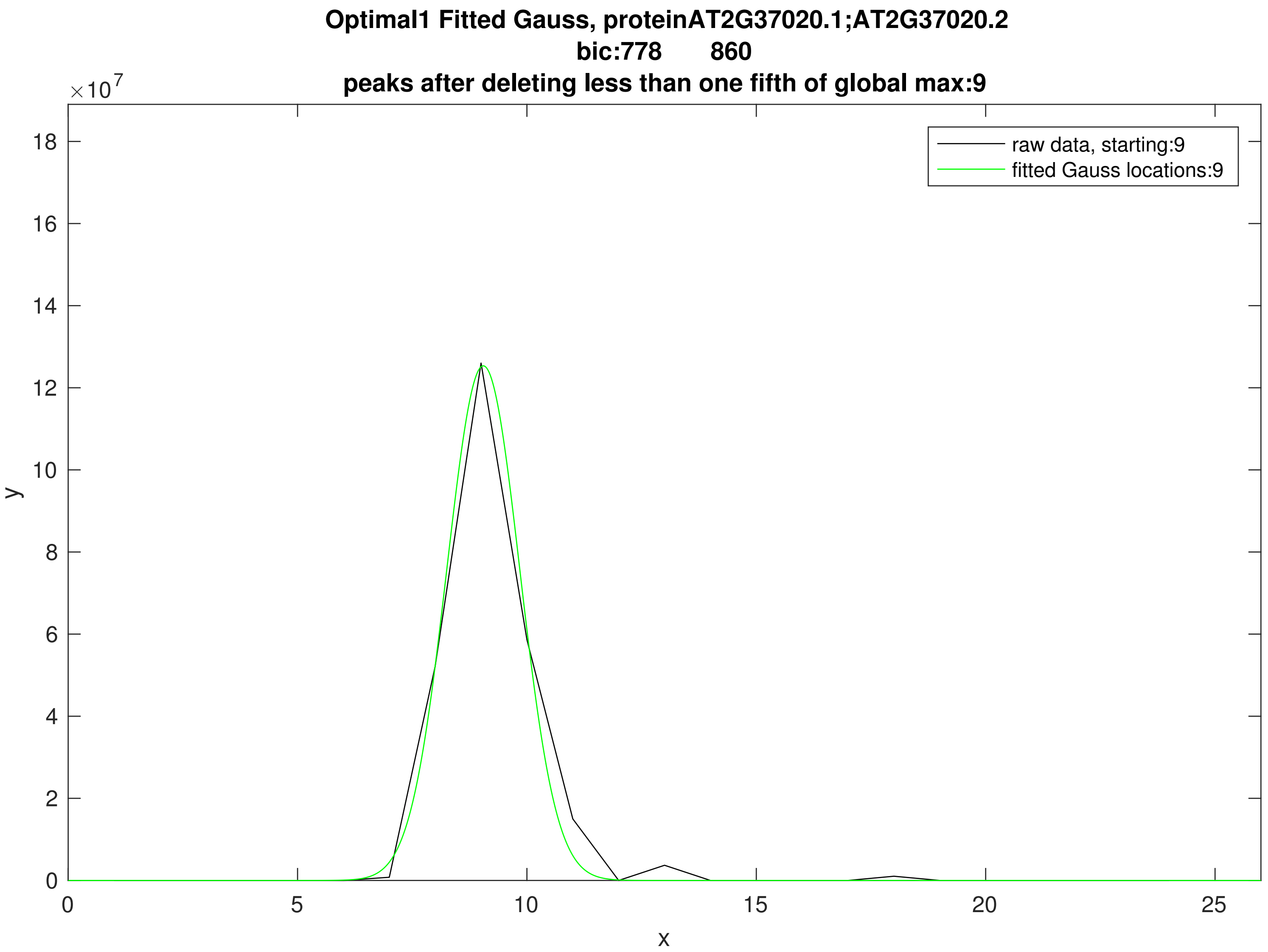

Optimal1 Fitted Gauss, proteinAT2G37020.1;AT2G37020.2
bic:778 860
peaks after deleting less than one fifth of global max:9
×10^7
raw data, starting:9
fitted Gauss locations:9
18
16
14
12
10
8
6
4
2
0
y
0
5
10
15
20
25
x

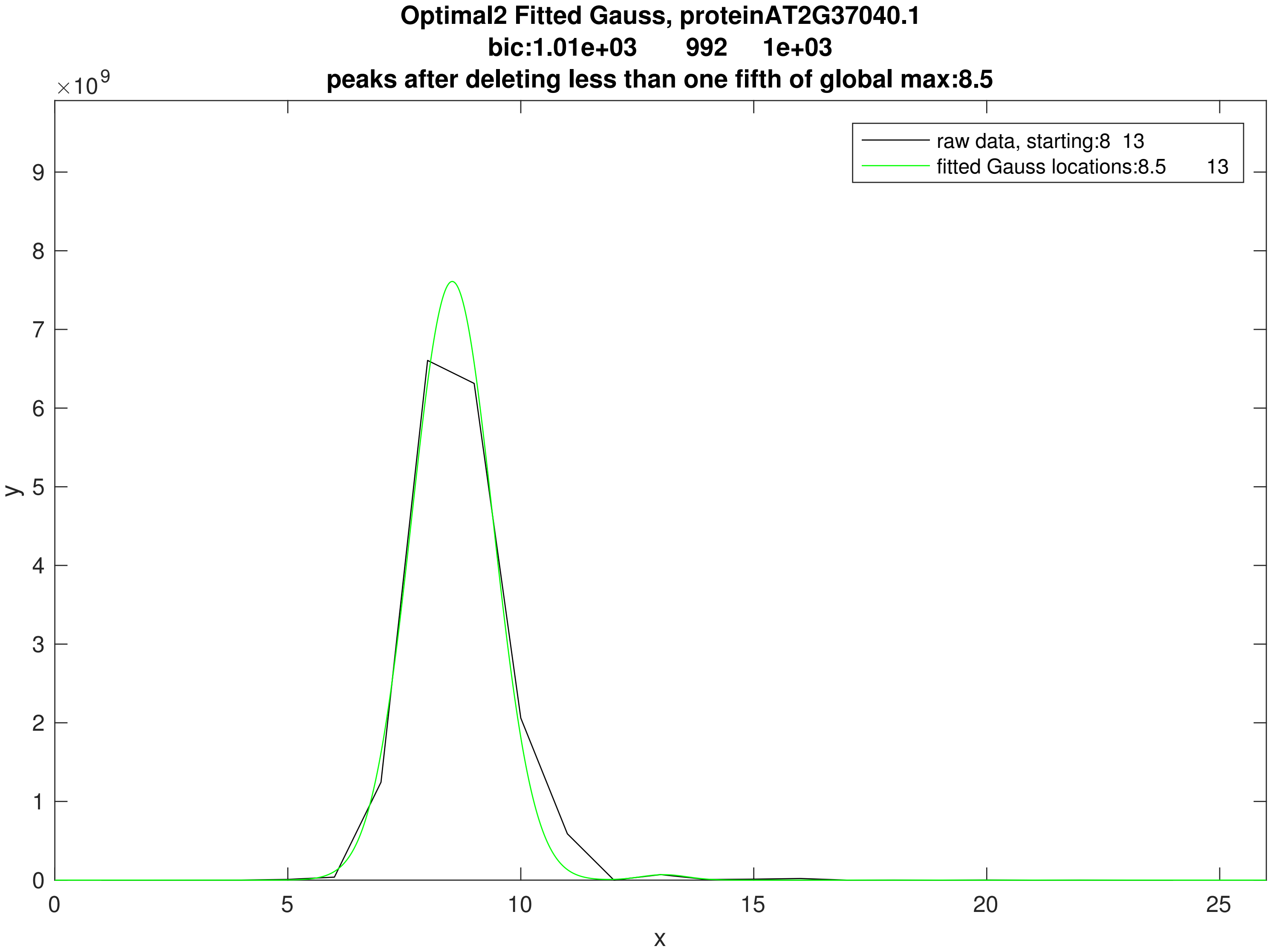
Optimal2 Fitted Gauss, proteinAT2G37040.1
bic:1.01e+03 992 1e+03
peaks after deleting less than one fifth of global max:8.5
×10^9
raw data, starting:8 13
fitted Gauss locations:8.5 13
y
x
0
1
2
3
4
5
6
7
8
9
0
5
10
15
20
25

Optimal1 Fitted Gauss, proteinAT2G37130.2;AT2G37130.1
bic:1.03e+03  1.04e+03  1.05e+03  1.06e+03
peaks after deleting less than one fifth of global max:18

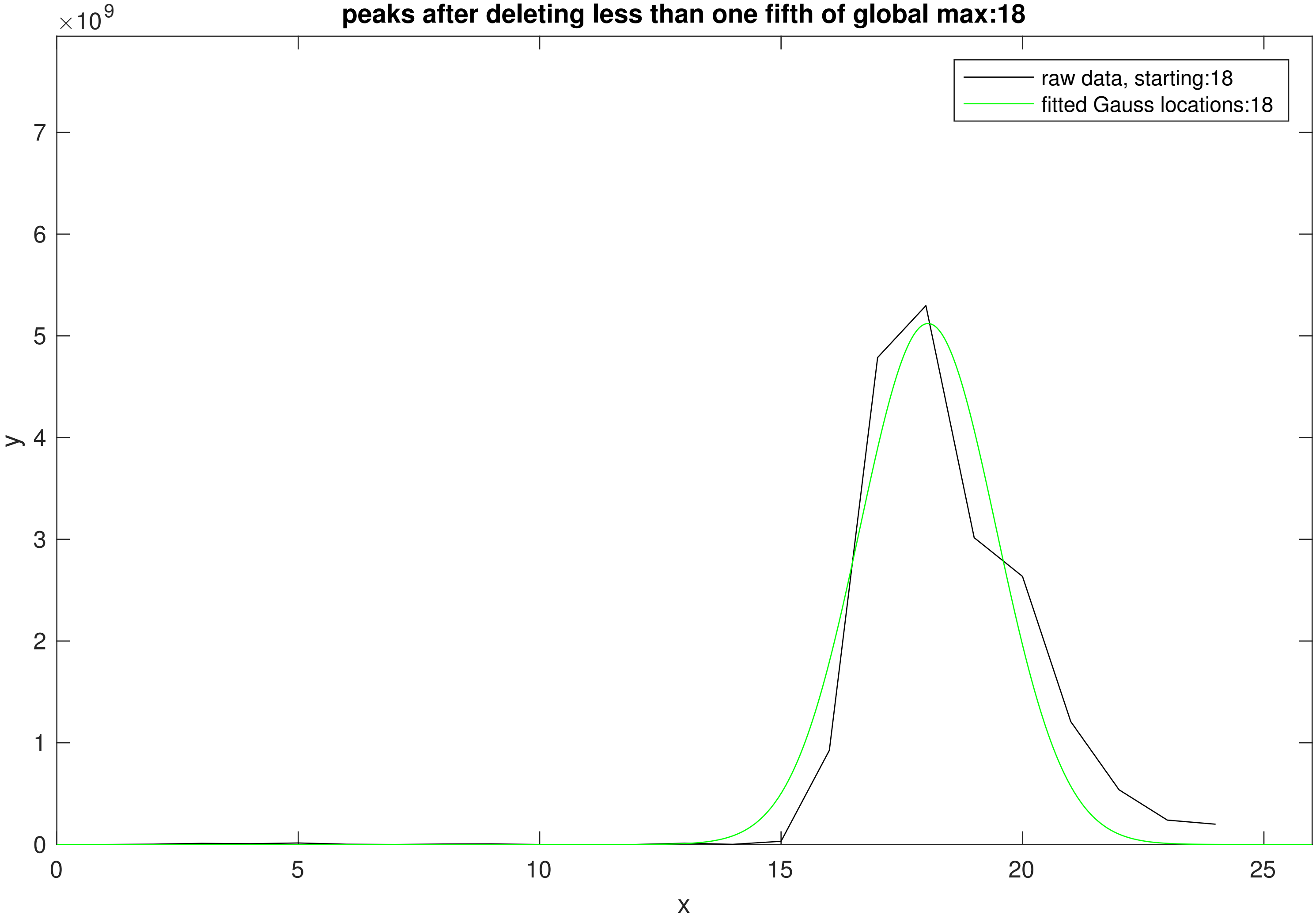

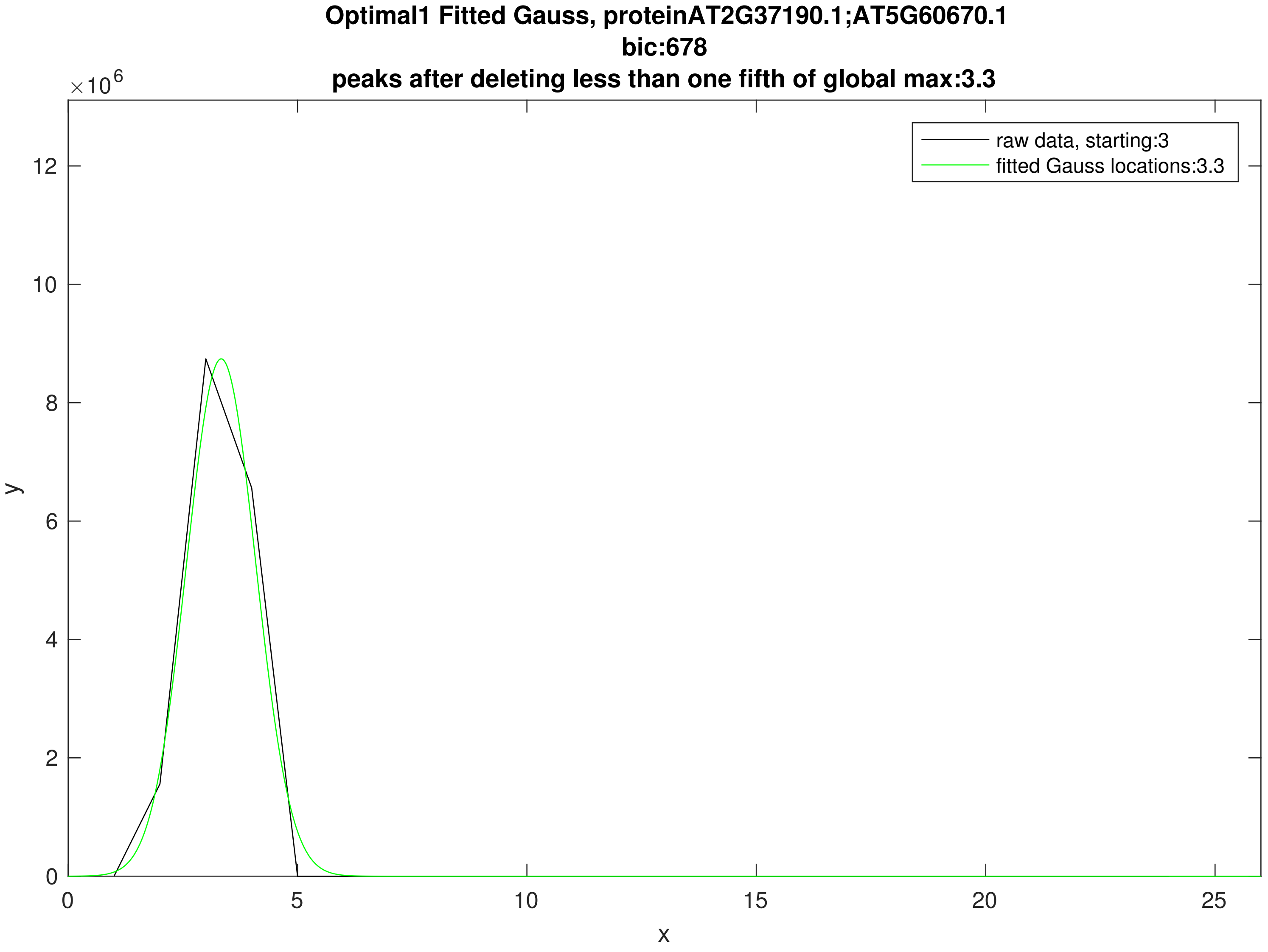
Optimal1 Fitted Gauss, proteinAT2G37190.1;AT5G60670.1
bic:678
peaks after deleting less than one fifth of global max:3.3
×10^6
raw data, starting:3
fitted Gauss locations:3.3
y
x
0
2
4
6
8
10
12
0
5
10
15
20
25

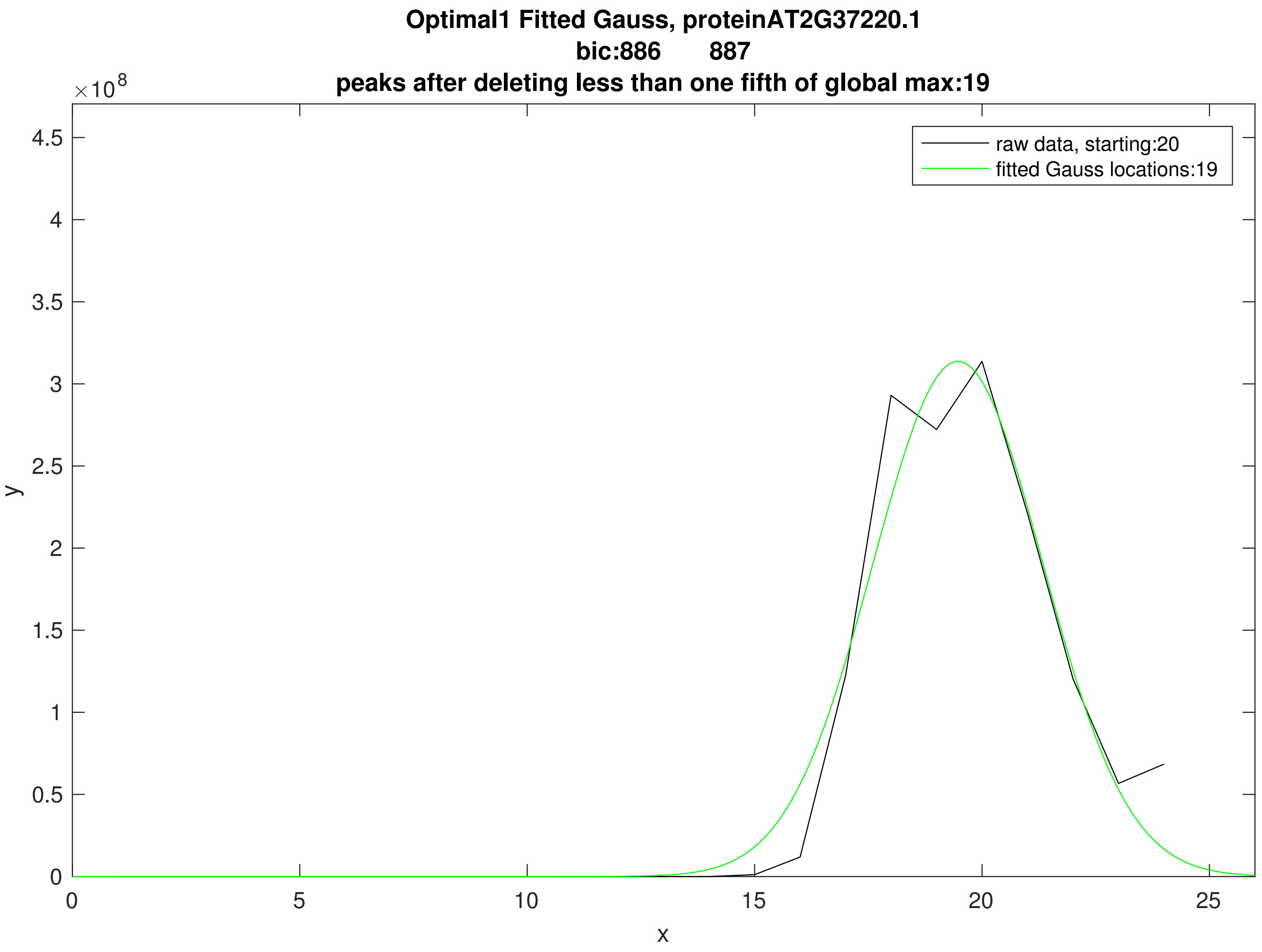
Optimal1 Fitted Gauss, proteinAT2G37220.1
bic:886 887
peaks after deleting less than one fifth of global max:19
×10^8
raw data, starting:20
fitted Gauss locations:19
y
x
0
0.5
1
1.5
2
2.5
3
3.5
4
4.5
5
10
15
20
25

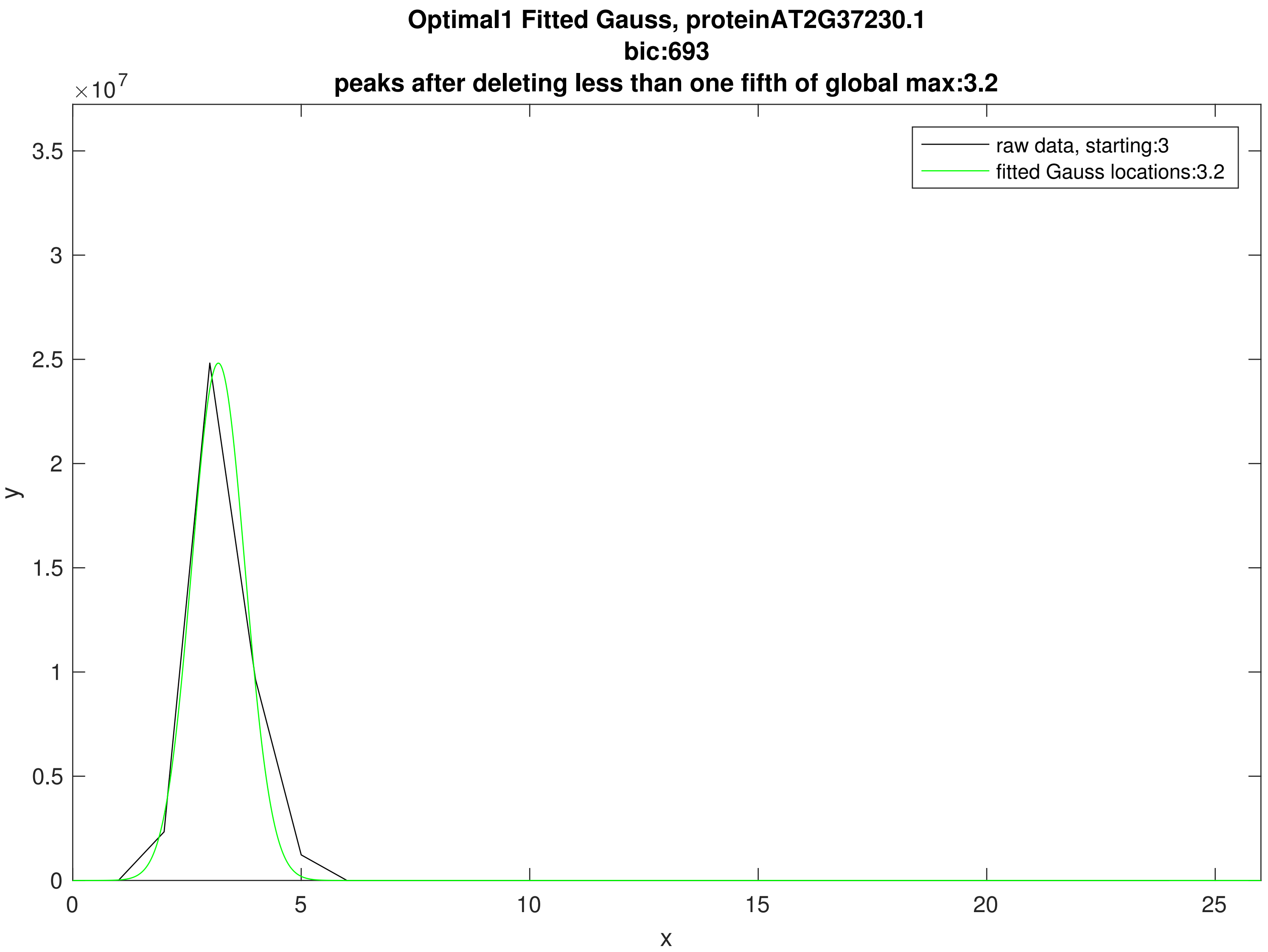
Optimal1 Fitted Gauss, proteinAT2G37230.1
bic:693
peaks after deleting less than one fifth of global max:3.2
×10^7
raw data, starting:3
fitted Gauss locations:3.2
3.5
3
2.5
2
1.5
1
0.5
0
y
0
5
10
15
20
25
x

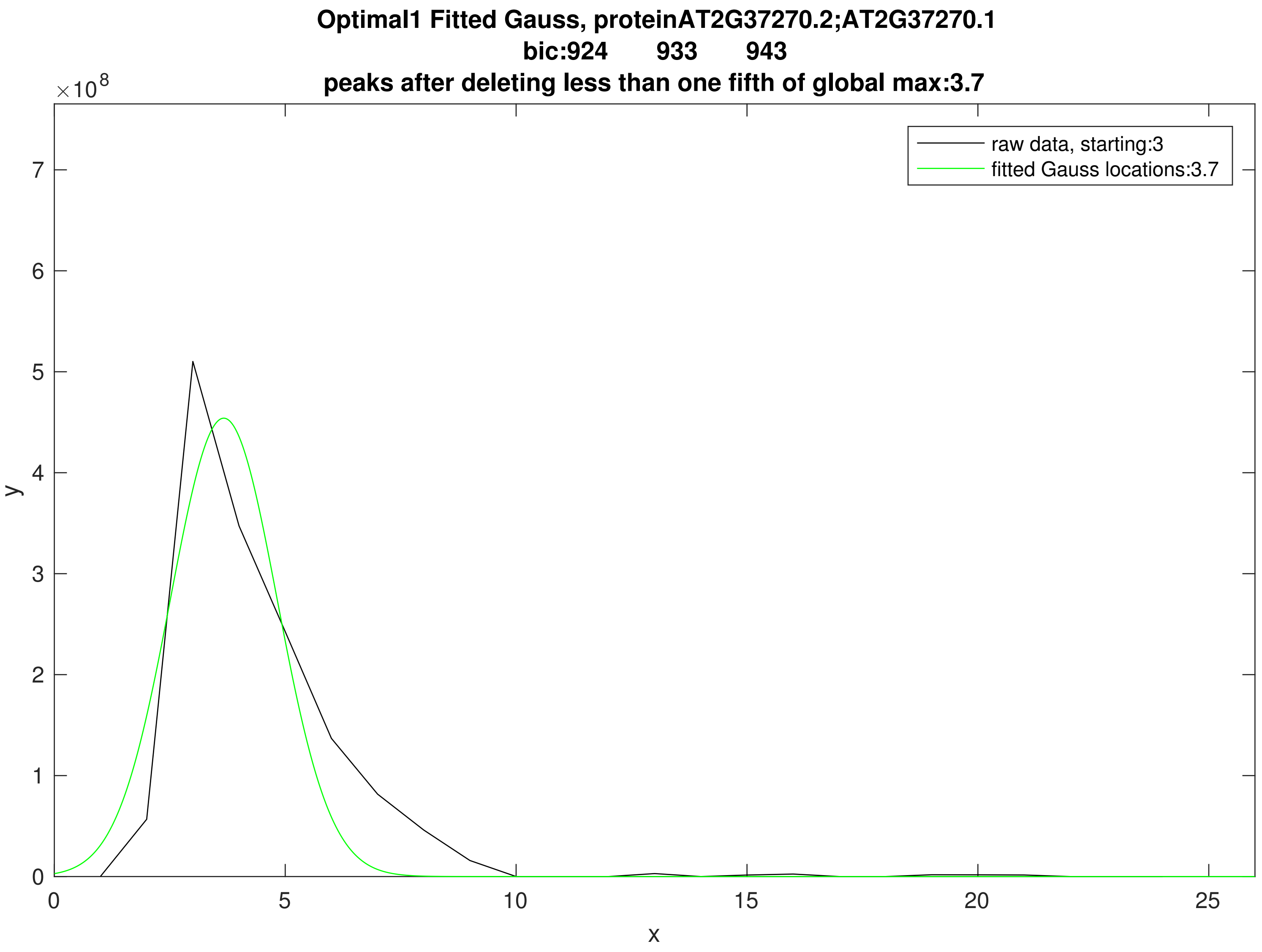
Optimal1 Fitted Gauss, proteinAT2G37270.2;AT2G37270.1
bic:924 933 943
peaks after deleting less than one fifth of global max:3.7
×10^8
raw data, starting:3
fitted Gauss locations:3.7
y
x
0
1
2
3
4
5
6
7
0
5
10
15
20
25

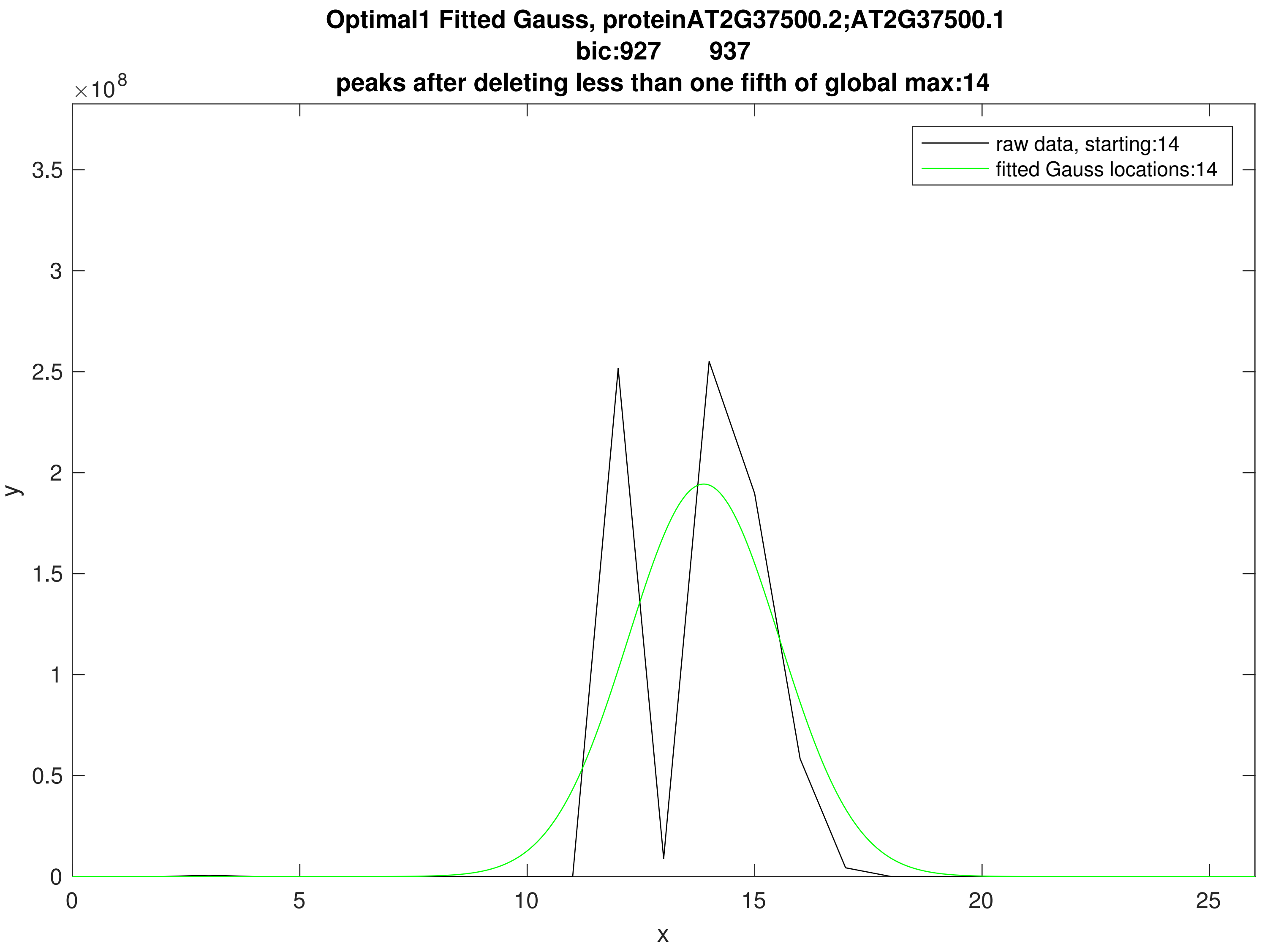
Optimal1 Fitted Gauss, proteinAT2G37500.2;AT2G37500.1
bic:927 937
peaks after deleting less than one fifth of global max:14
×10^8
raw data, starting:14
fitted Gauss locations:14
3.5
3
2.5
2
1.5
1
0.5
0
y
0
5
10
15
20
25
x

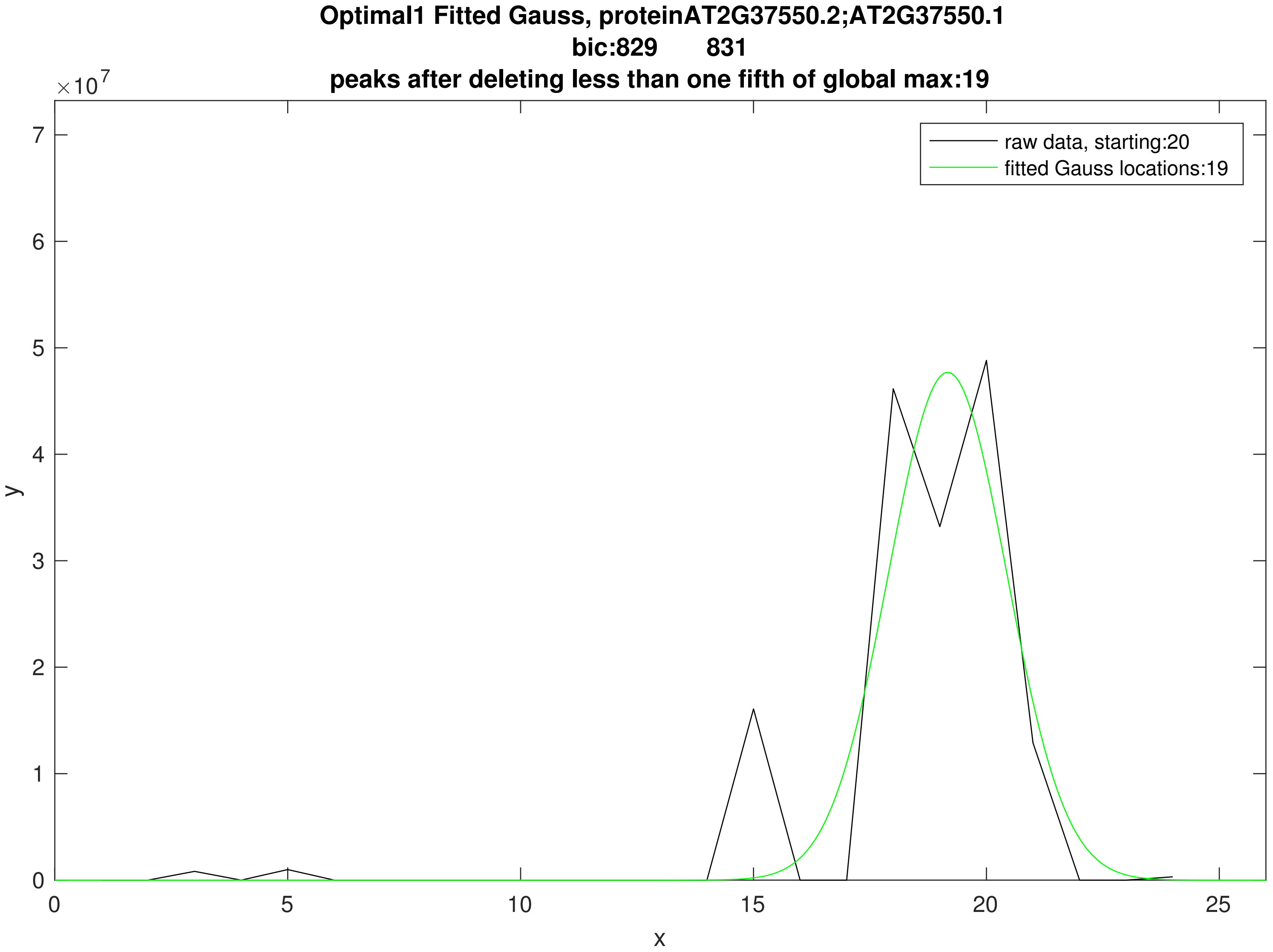

Optimal1 Fitted Gauss, proteinAT2G37550.2;AT2G37550.1
bic:829 831
peaks after deleting less than one fifth of global max:19
×10^7
raw data, starting:20
fitted Gauss locations:19
y
x
0
1
2
3
4
5
6
7
0
5
10
15
20
25

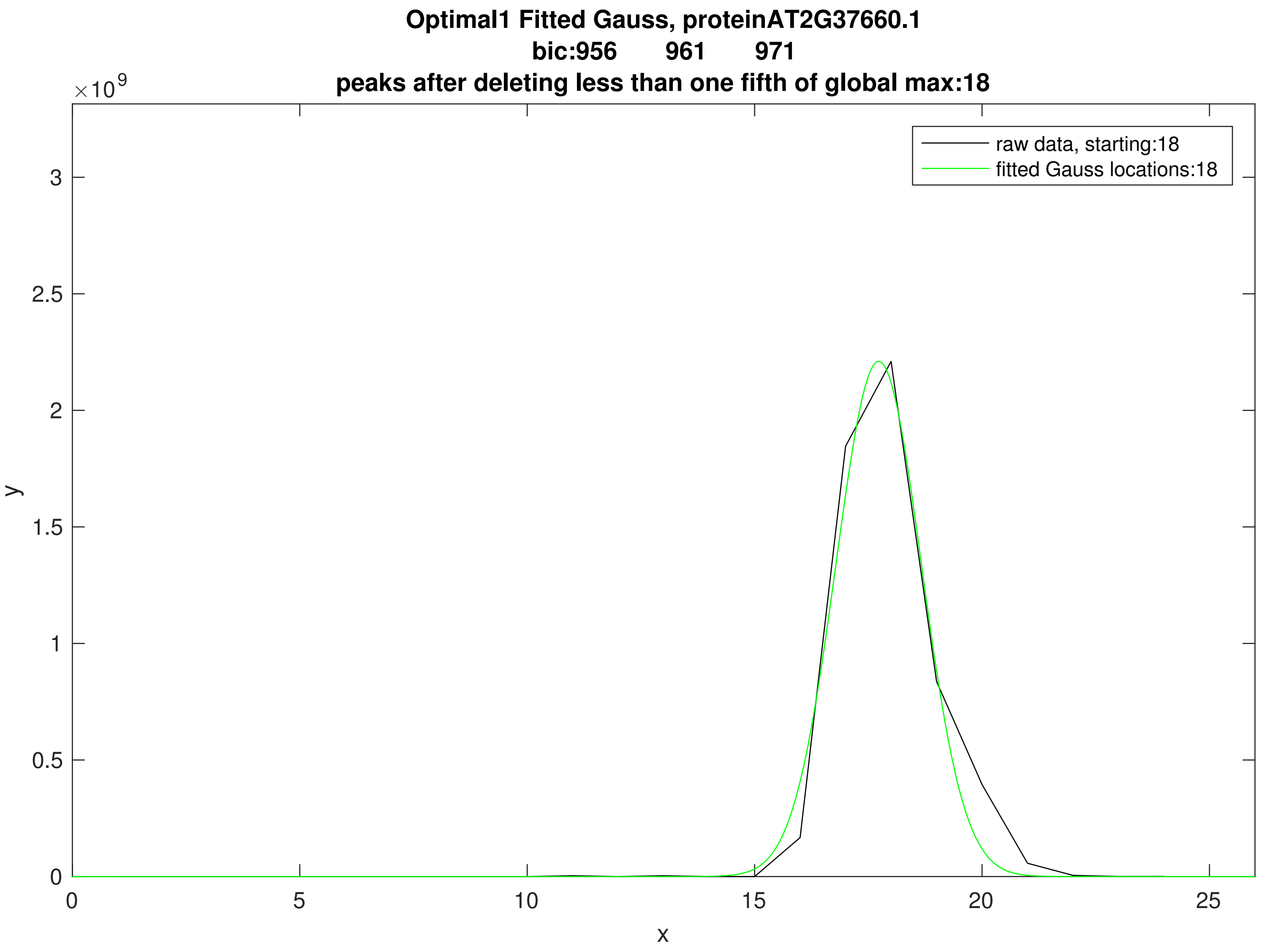
Optimal1 Fitted Gauss, proteinAT2G37660.1
bic:956 961 971
peaks after deleting less than one fifth of global max:18
×10^9
3
2.5
2
1.5
1
0.5
0
y
0
5
10
15
20
25
x
raw data, starting:18
fitted Gauss locations:18

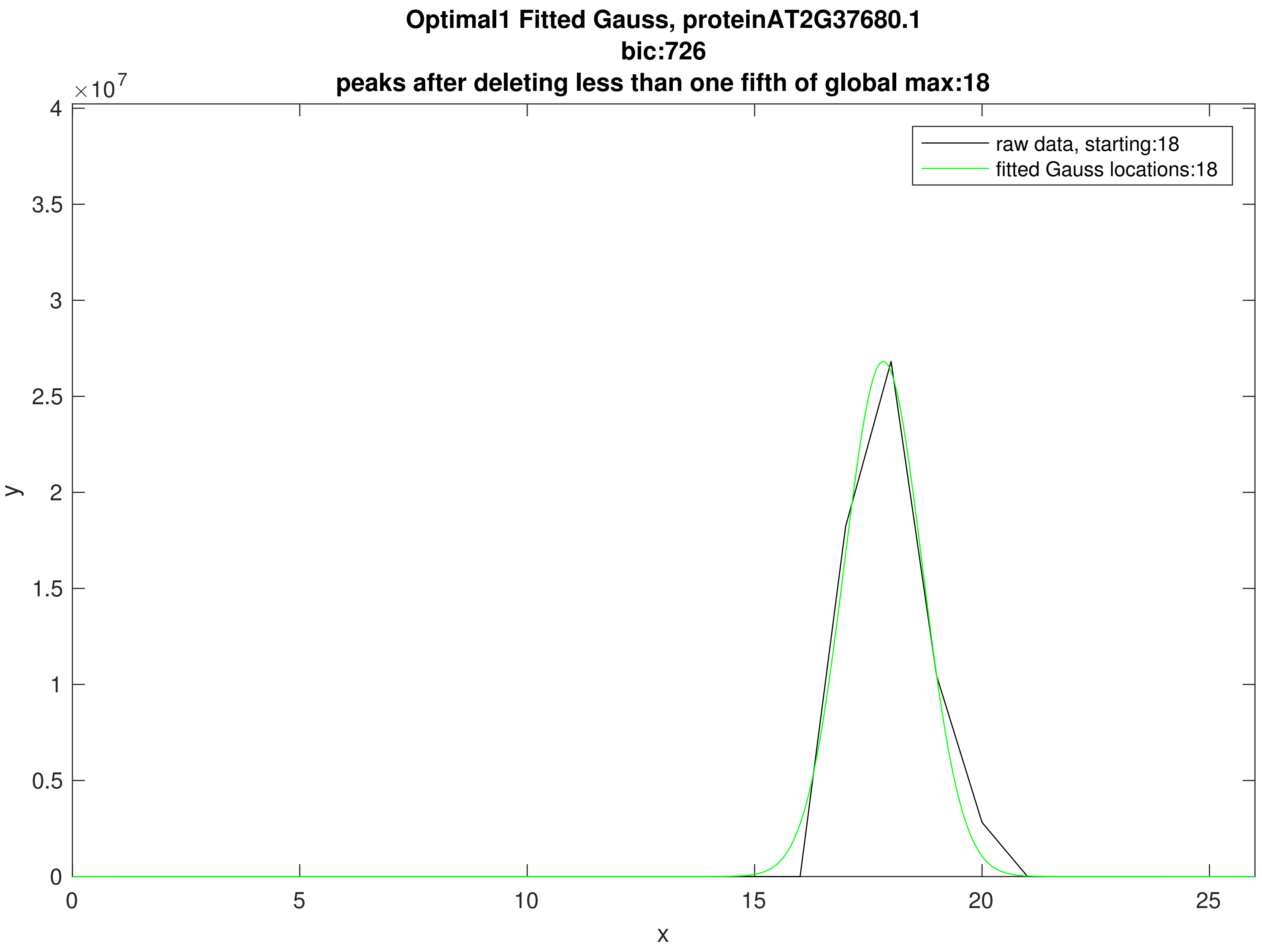
Optimal1 Fitted Gauss, proteinAT2G37680.1
bic:726
peaks after deleting less than one fifth of global max:18
×10^7
raw data, starting:18
fitted Gauss locations:18
y
x
0
0.5
1
1.5
2
2.5
3
3.5
4
0
5
10
15
20
25

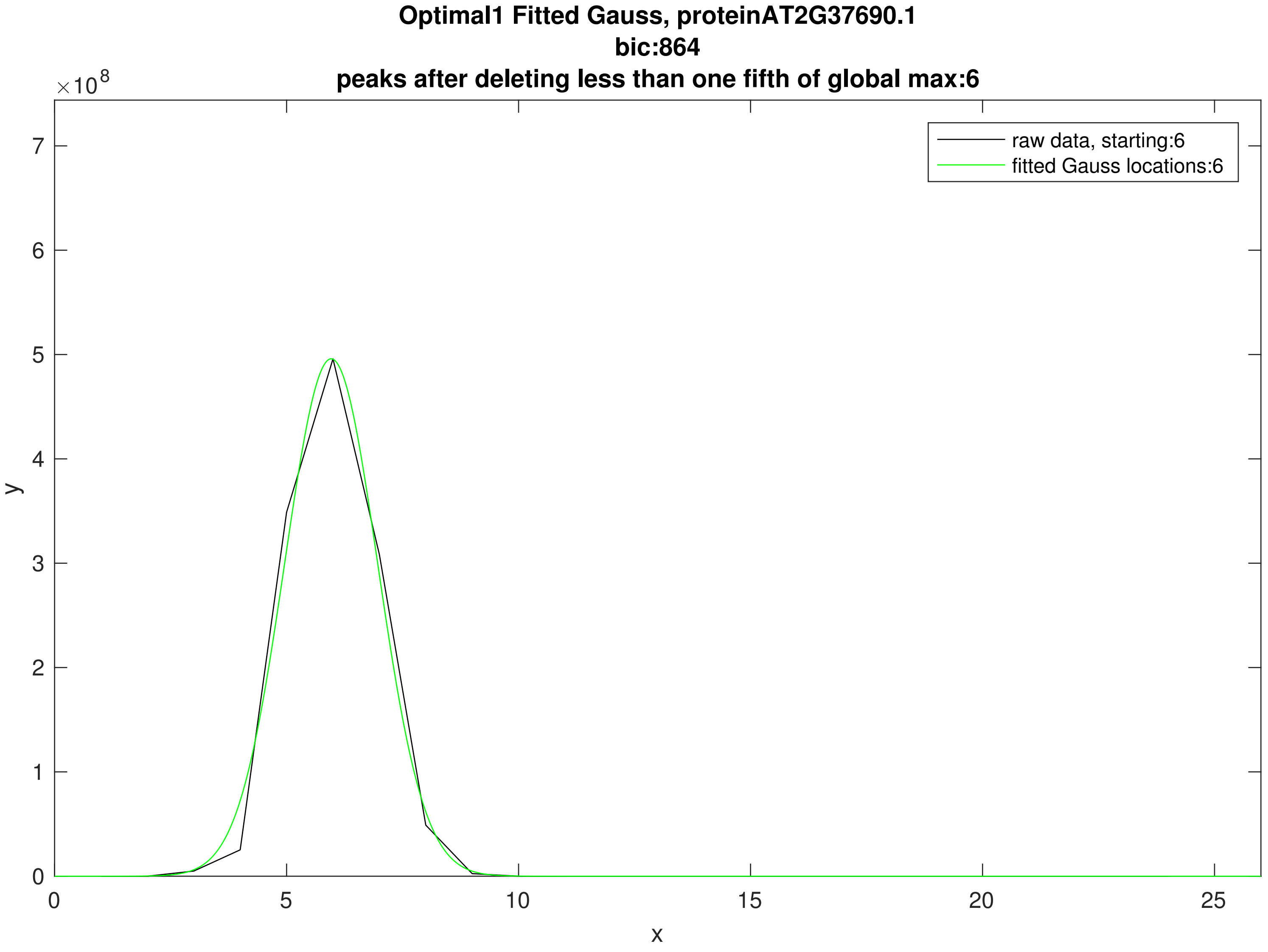
Optimal1 Fitted Gauss, proteinAT2G37690.1
bic:864
peaks after deleting less than one fifth of global max:6
×10^8
raw data, starting:6
fitted Gauss locations:6
y
x
0
1
2
3
4
5
6
7
0
5
10
15
20
25

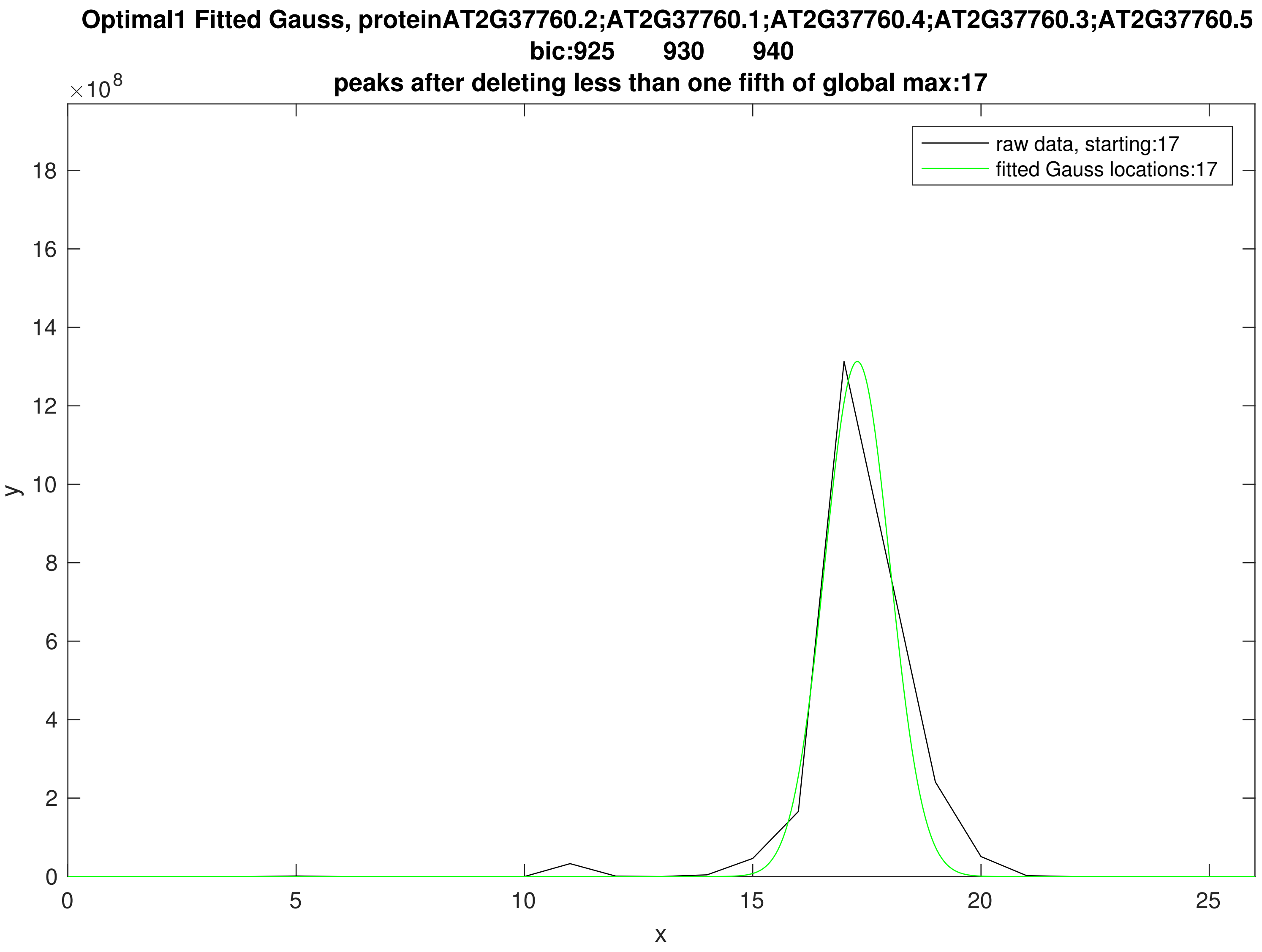
Optimal1 Fitted Gauss, proteinAT2G37760.2;AT2G37760.1;AT2G37760.4;AT2G37760.3;AT2G37760.5
bic:925 930 940
peaks after deleting less than one fifth of global max:17
×10^8
raw data, starting:17
fitted Gauss locations:17
y
x
0
2
4
6
8
10
12
14
16
18
0
5
10
15
20
25

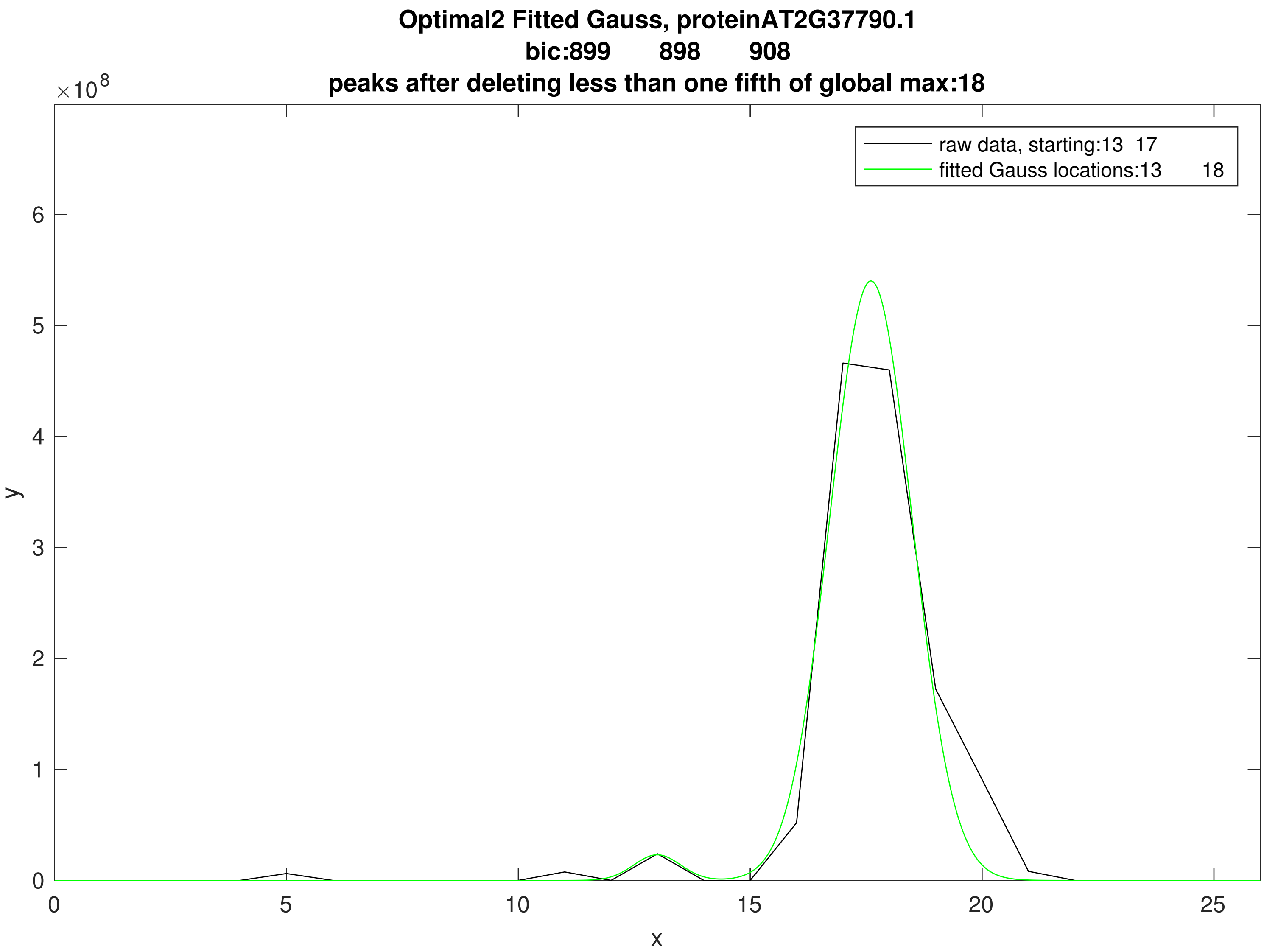
Optimal2 Fitted Gauss, proteinAT2G37790.1
bic:899 898 908
peaks after deleting less than one fifth of global max:18
×10^8
raw data, starting:13 17
fitted Gauss locations:13 18
y
x
0
1
2
3
4
5
6
0
5
10
15
20
25

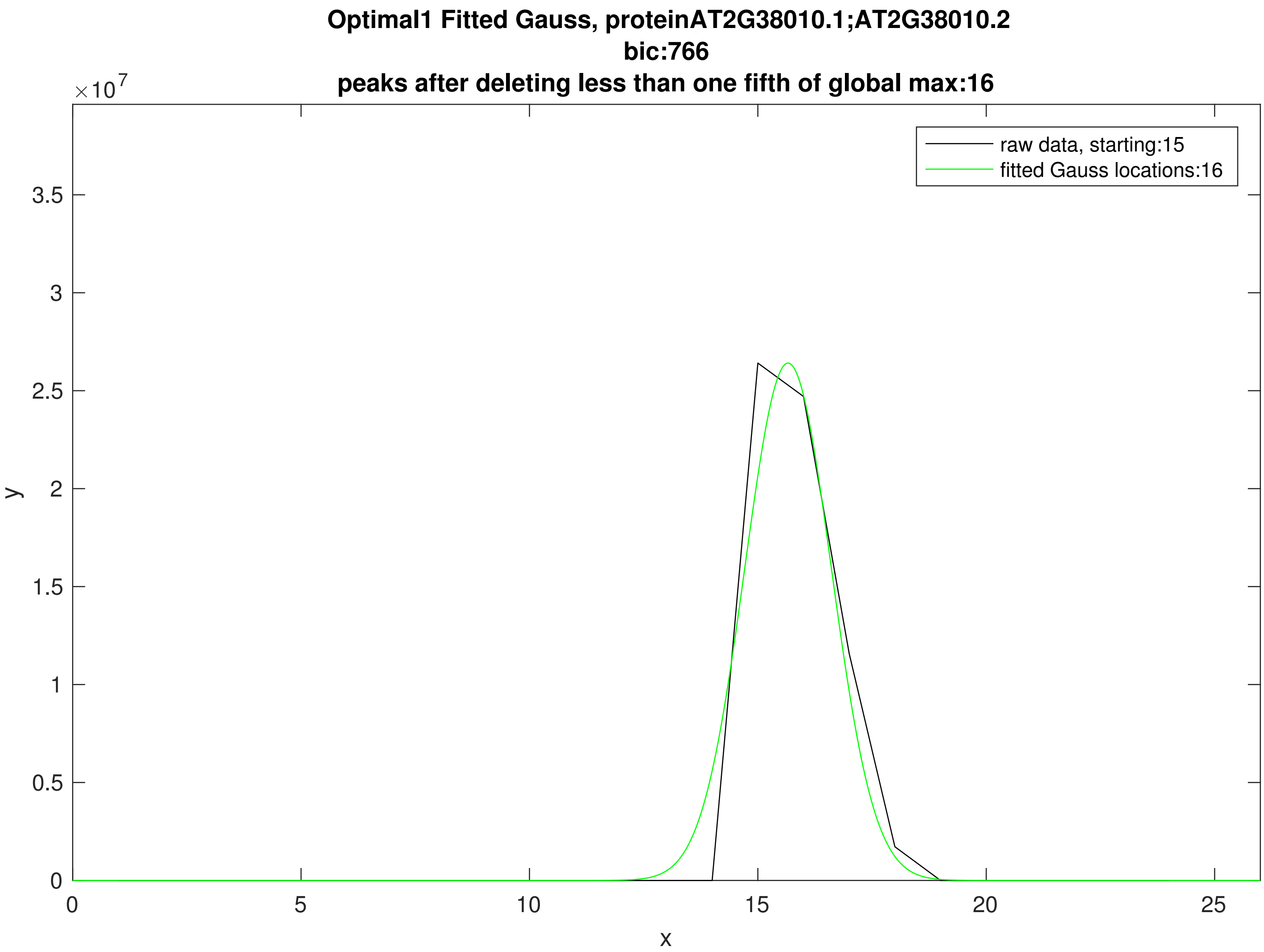
Optimal1 Fitted Gauss, proteinAT2G38010.1;AT2G38010.2
bic:766
peaks after deleting less than one fifth of global max:16
×10^7
raw data, starting:15
fitted Gauss locations:16
3.5
3
2.5
2
1.5
1
0.5
0
y
0
5
10
15
20
25
x

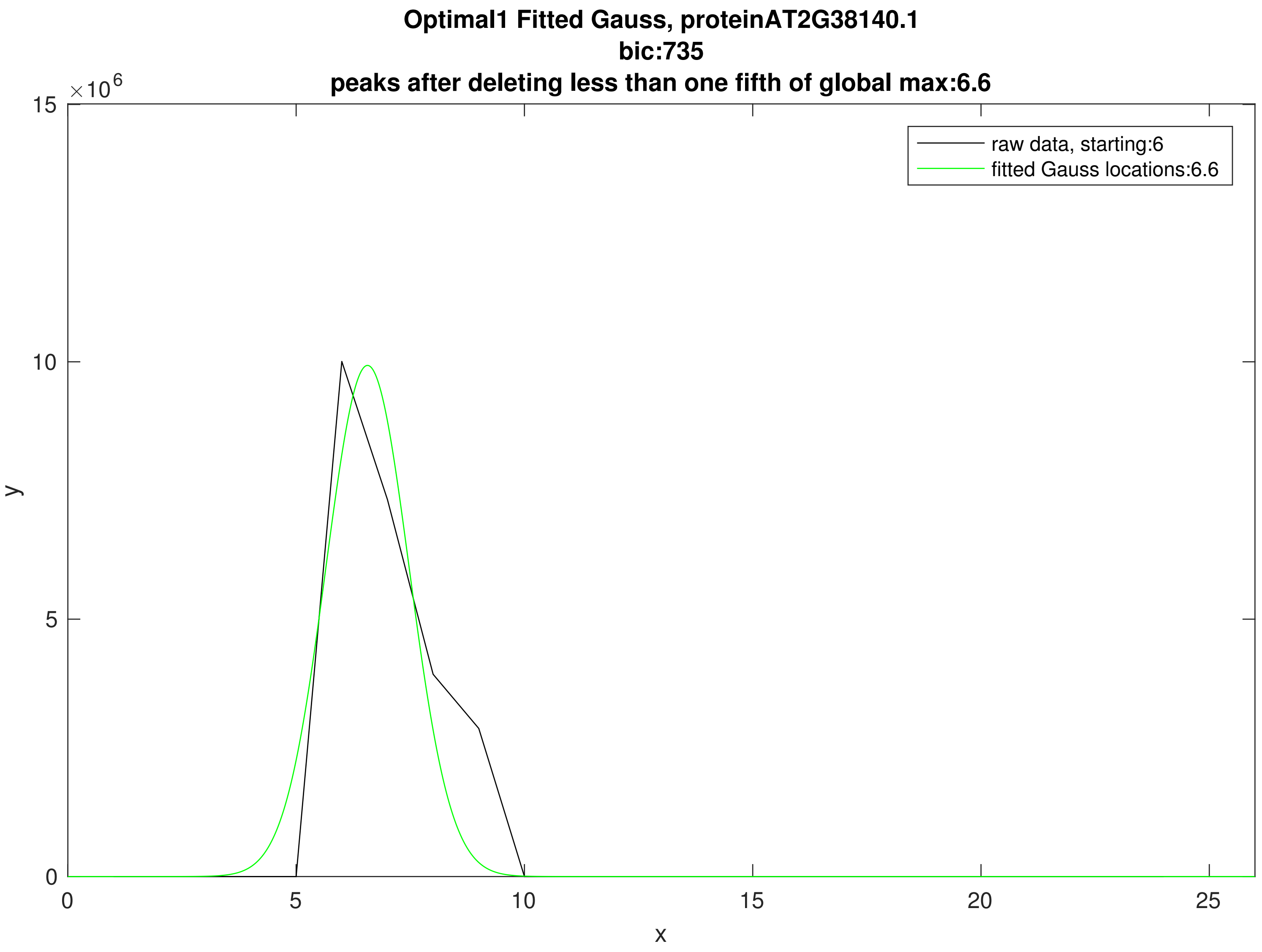

Optimal1 Fitted Gauss, proteinAT2G38140.1
bic:735
peaks after deleting less than one fifth of global max:6.6
×10^6
raw data, starting:6
fitted Gauss locations:6.6
y
x
0
5
10
15
20
25

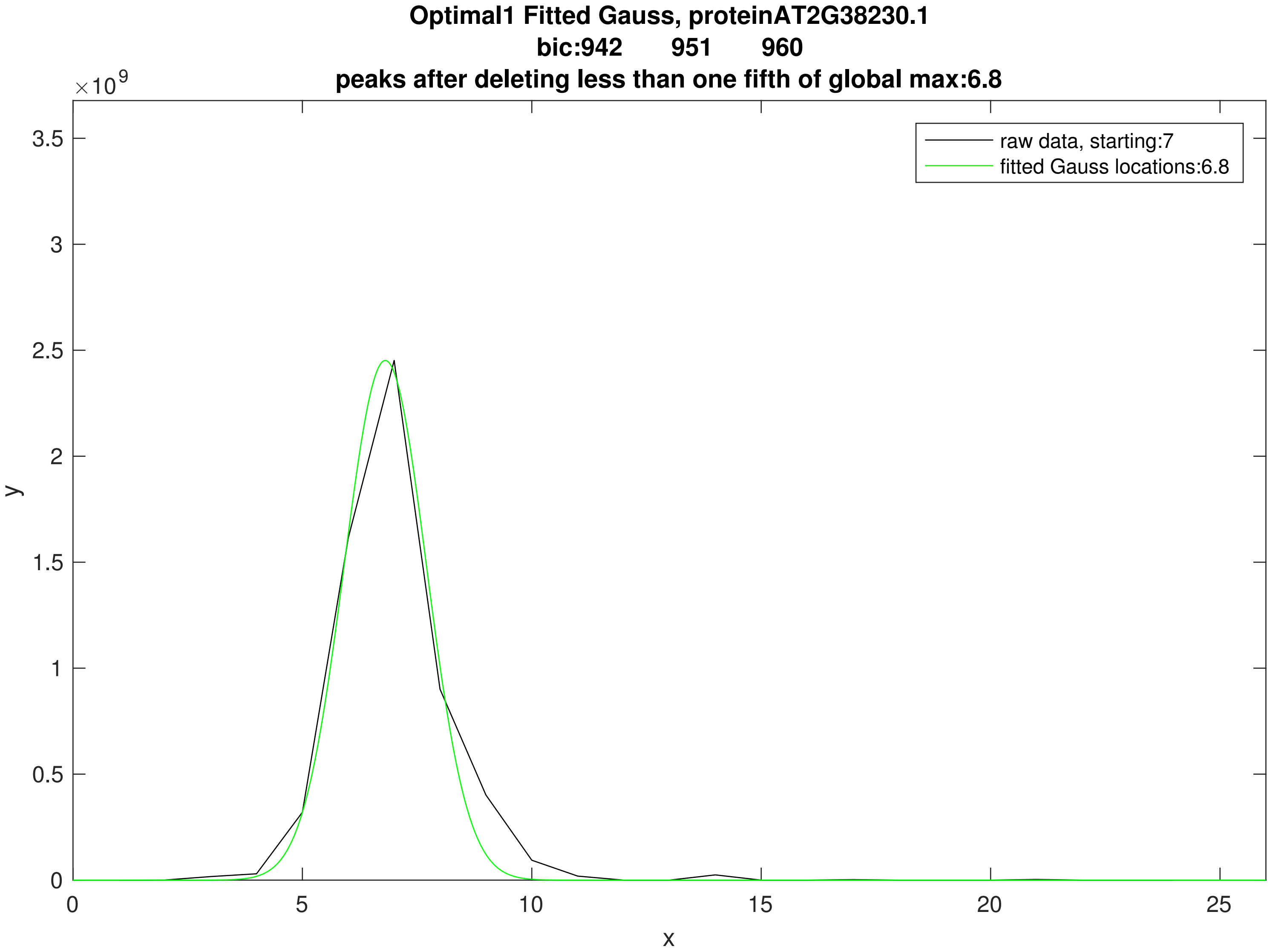
Optimal1 Fitted Gauss, proteinAT2G38230.1
bic:942 951 960
peaks after deleting less than one fifth of global max:6.8
×10^9
3.5
3
2.5
2
1.5
1
0.5
0
y
0
5
10
15
20
25
x
raw data, starting:7
fitted Gauss locations:6.8

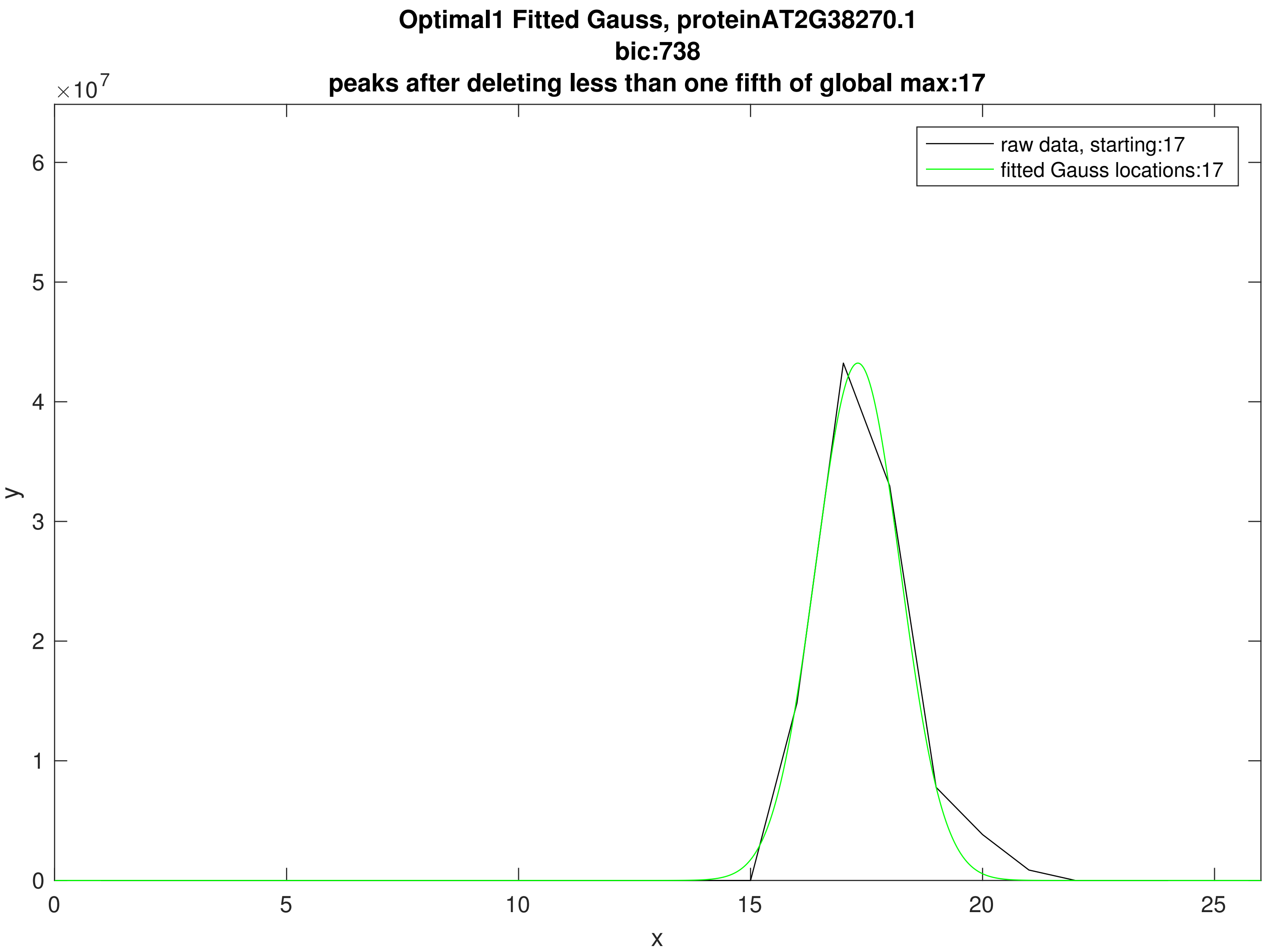
Optimal1 Fitted Gauss, proteinAT2G38270.1
bic:738
peaks after deleting less than one fifth of global max:17
×10^7
raw data, starting:17
fitted Gauss locations:17
y
x
0
1
2
3
4
5
6
0
5
10
15
20
25

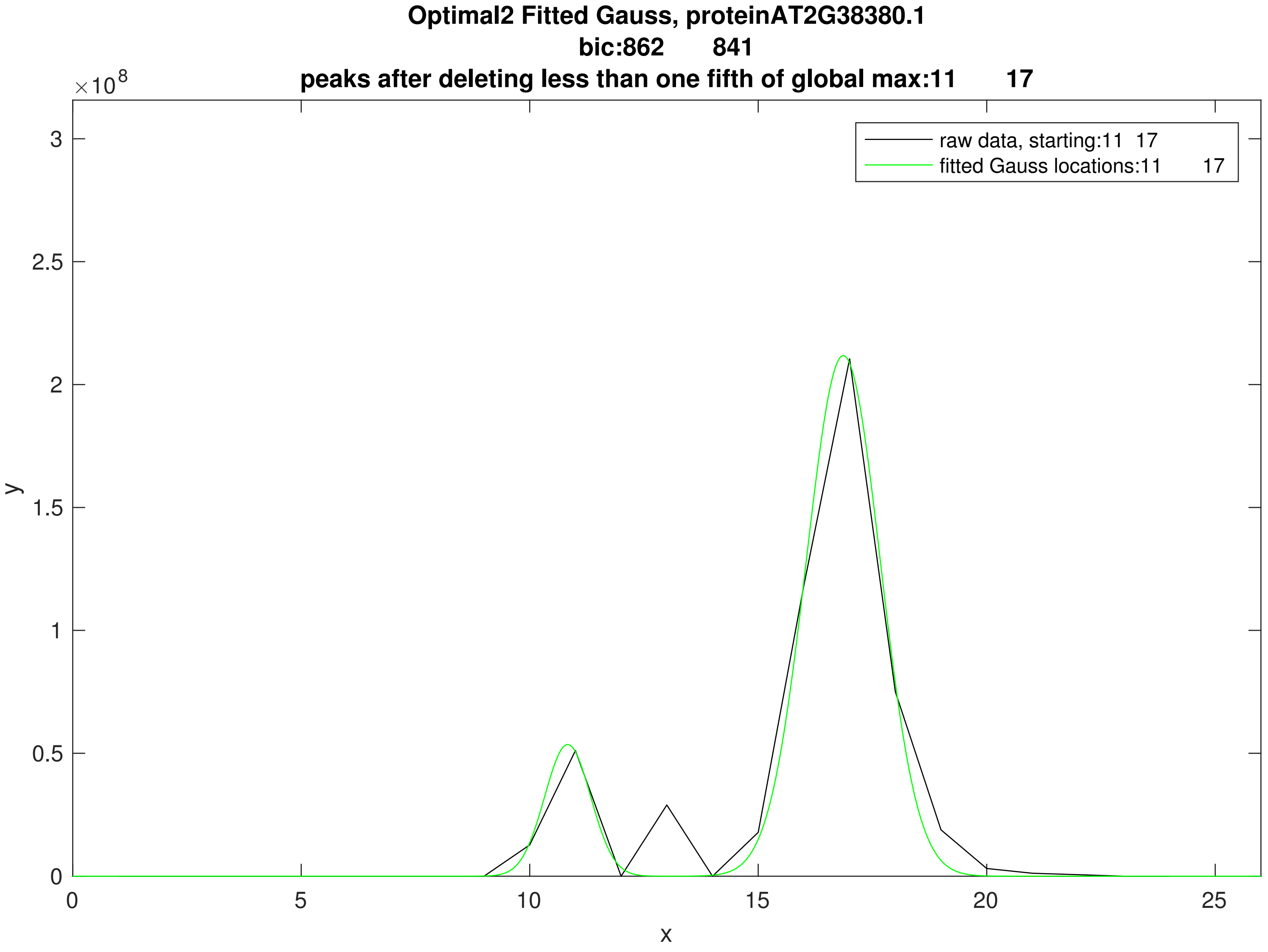

Optimal2 Fitted Gauss, proteinAT2G38380.1
bic:862 841
peaks after deleting less than one fifth of global max:11 17
×10^8
raw data, starting:11 17
fitted Gauss locations:11 17
y
x
0
0.5
1
1.5
2
2.5
3
0
5
10
15
20
25

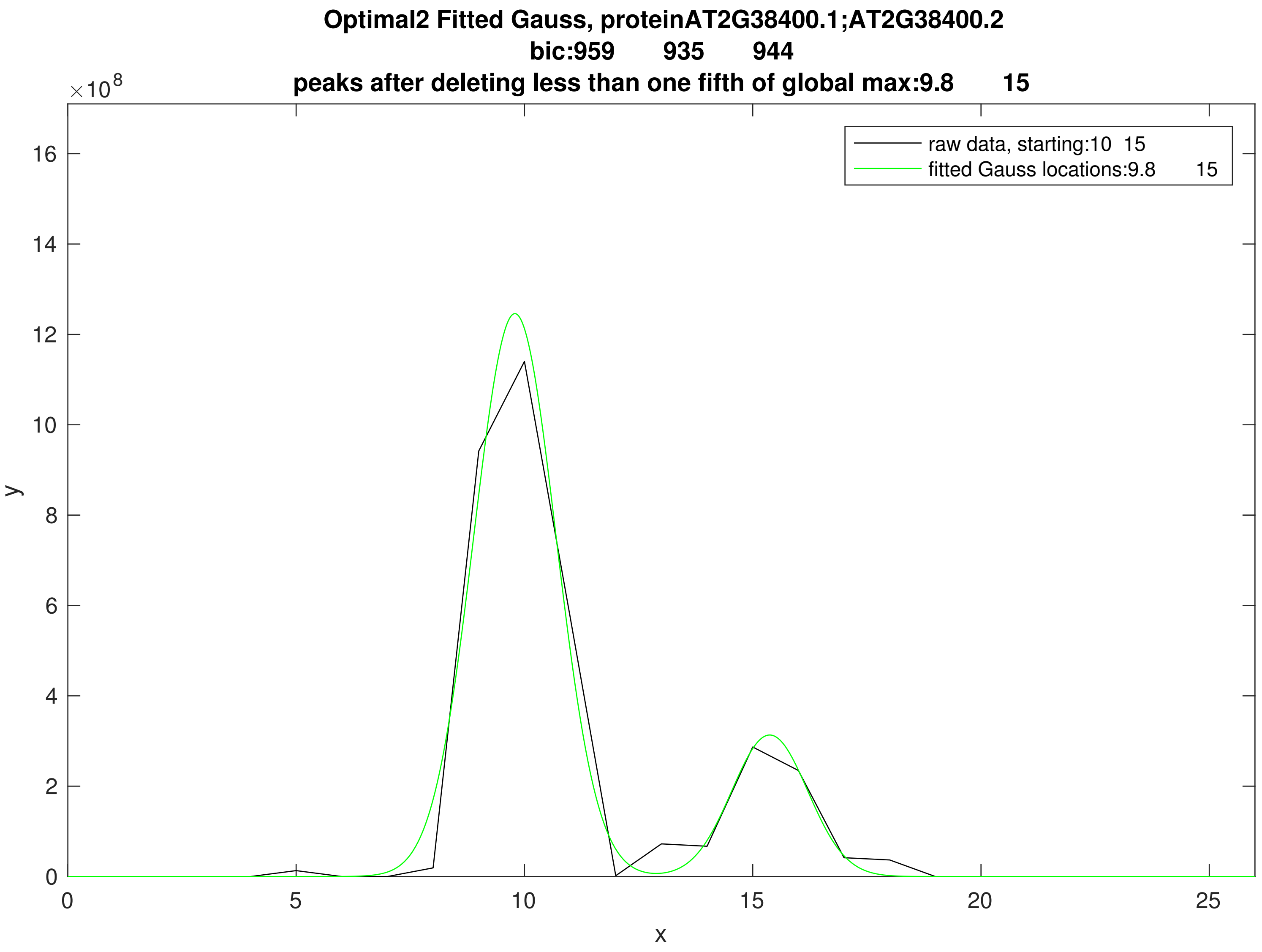

Optimal2 Fitted Gauss, proteinAT2G38400.1;AT2G38400.2
bic:959 935 944
peaks after deleting less than one fifth of global max:9.8 15
×10^8
raw data, starting:10 15
fitted Gauss locations:9.8 15
y
x
0
2
4
6
8
10
12
14
16
0
5
10
15
20
25

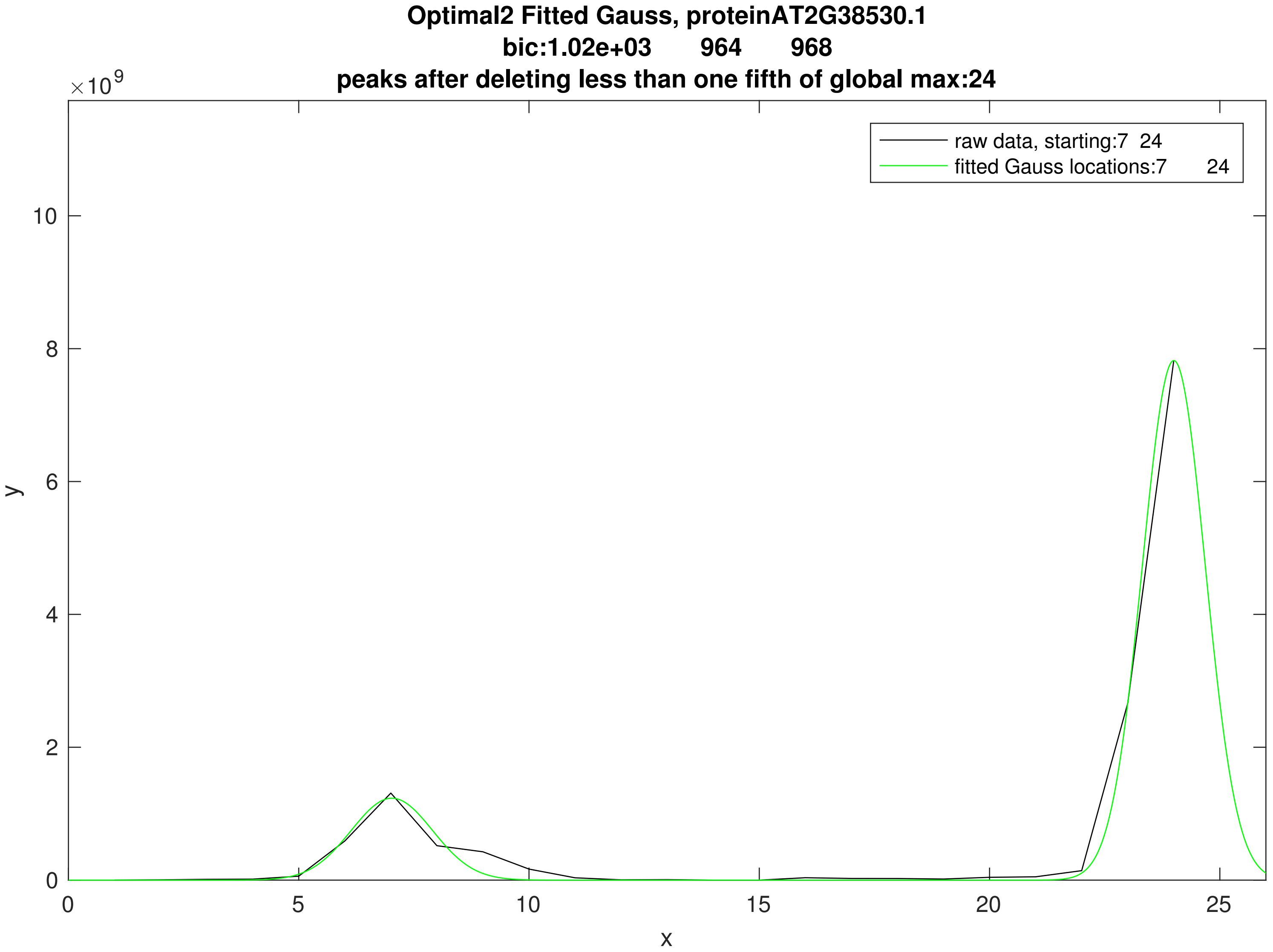
Optimal2 Fitted Gauss, proteinAT2G38530.1
bic:1.02e+03 964 968
peaks after deleting less than one fifth of global max:24
×10^9
raw data, starting:7 24
fitted Gauss locations:7 24
y
x
0
2
4
6
8
10
0
5
10
15
20
25

Optimal1 Fitted Gauss, proteinAT2G38540.1
bic:806 812
peaks after deleting less than one fifth of global max:6.9

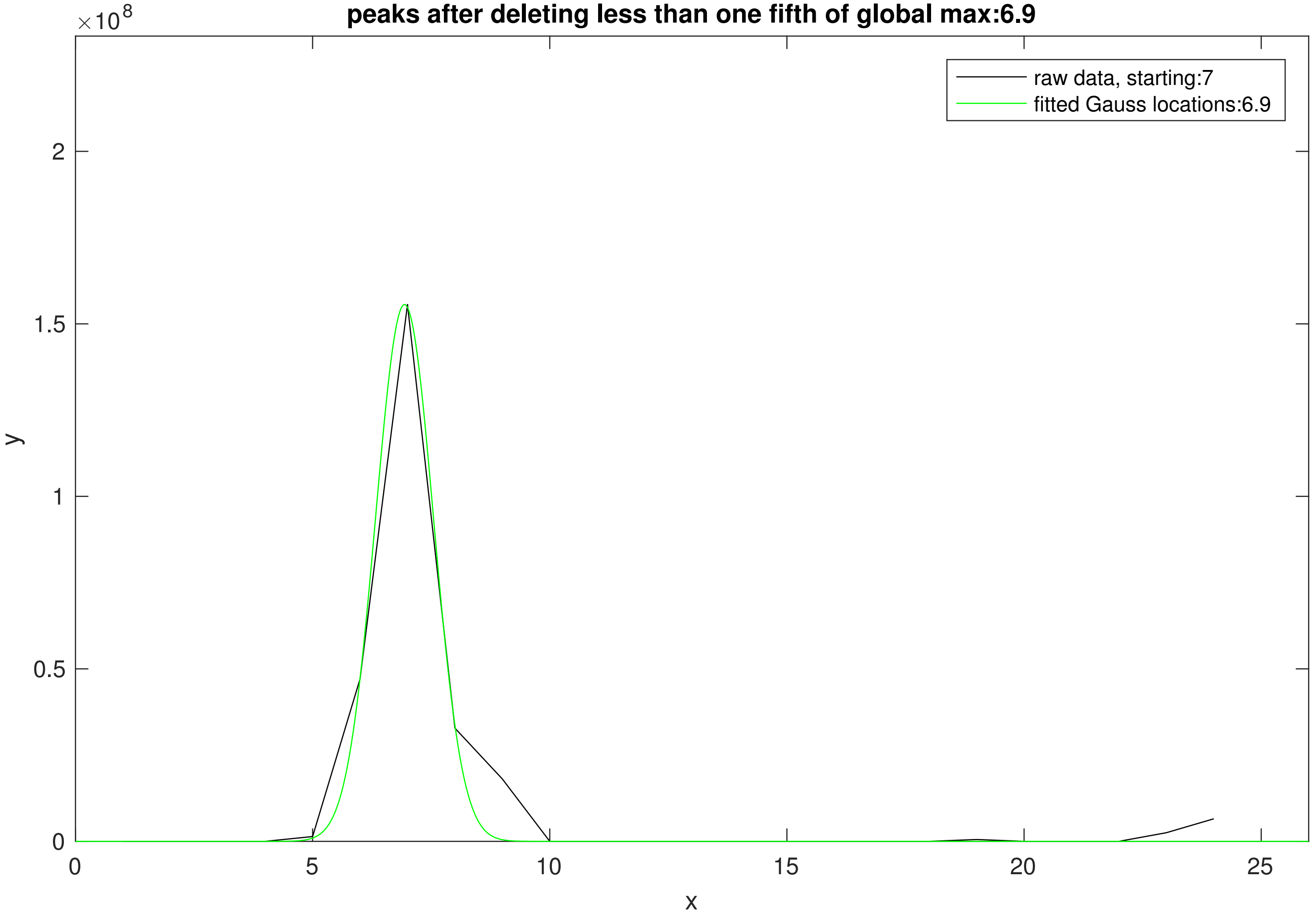

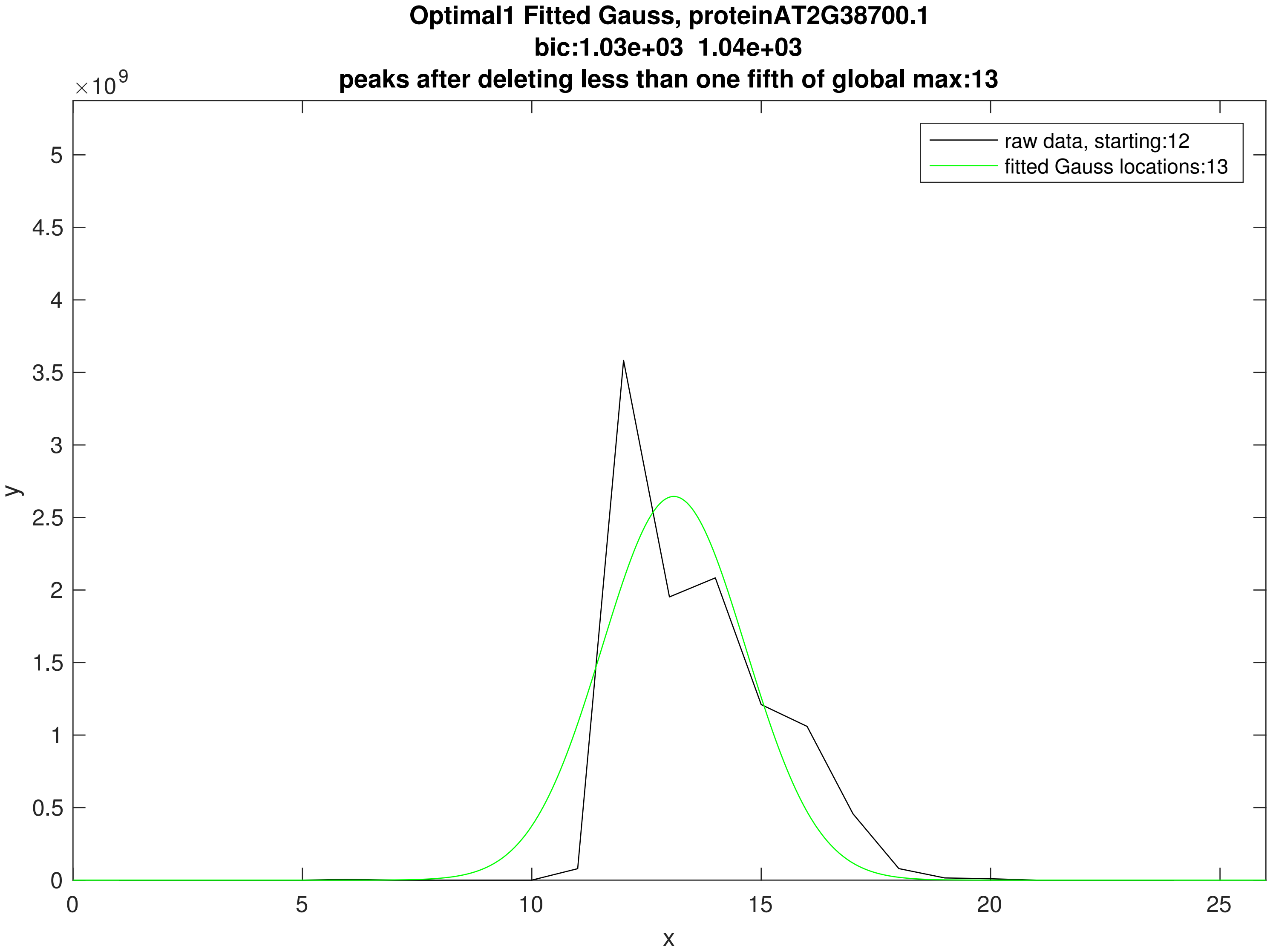
Optimal1 Fitted Gauss, proteinAT2G38700.1
bic:1.03e+03 1.04e+03
peaks after deleting less than one fifth of global max:13
×10^9
raw data, starting:12
fitted Gauss locations:13
y
x
0
0.5
1
1.5
2
2.5
3
3.5
4
4.5
5
0
5
10
15
20
25

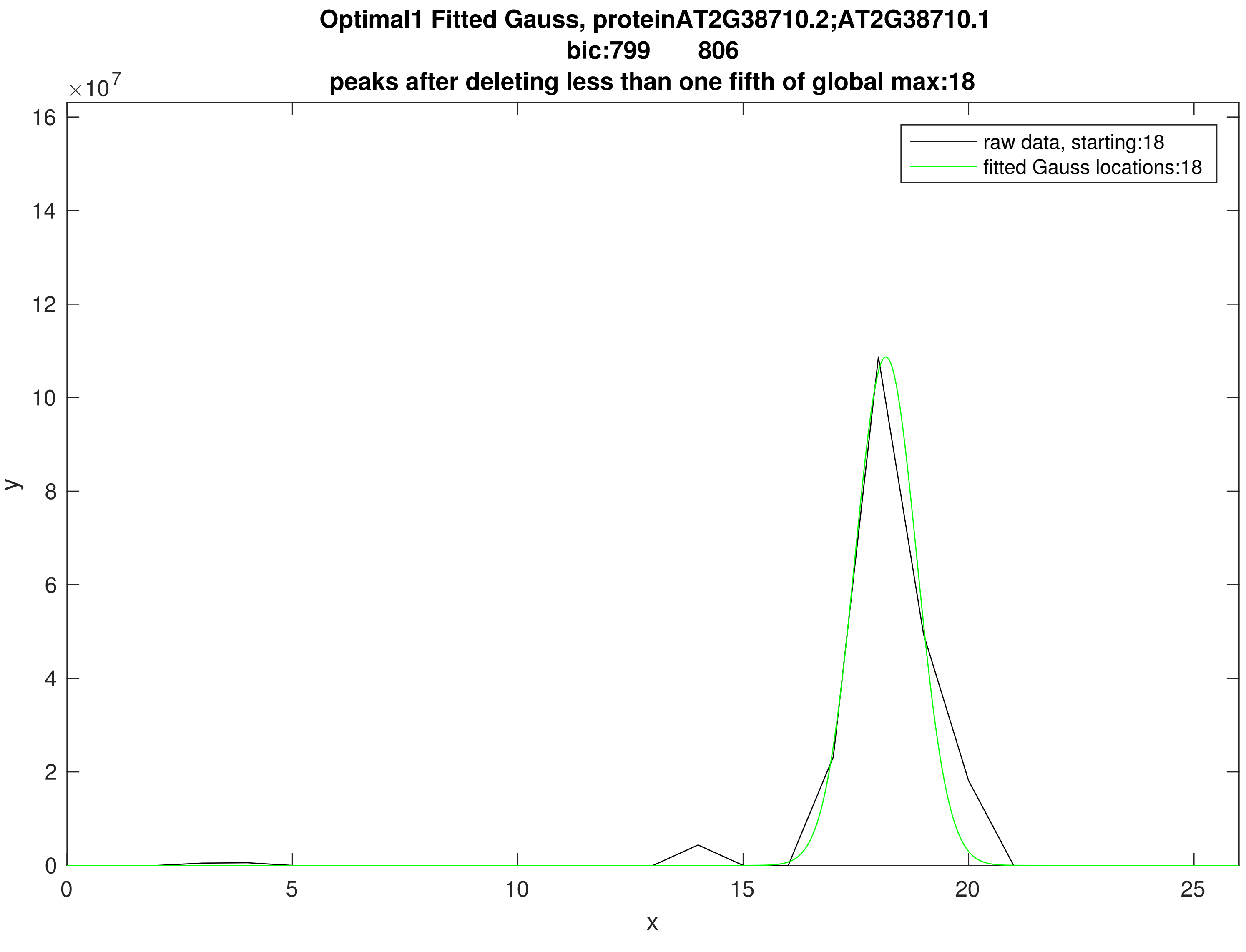
Optimal1 Fitted Gauss, proteinAT2G38710.2;AT2G38710.1
bic:799 806
peaks after deleting less than one fifth of global max:18
×10^7
16
14
12
10
8
6
4
2
0
y
0
5
10
15
20
25
x
raw data, starting:18
fitted Gauss locations:18

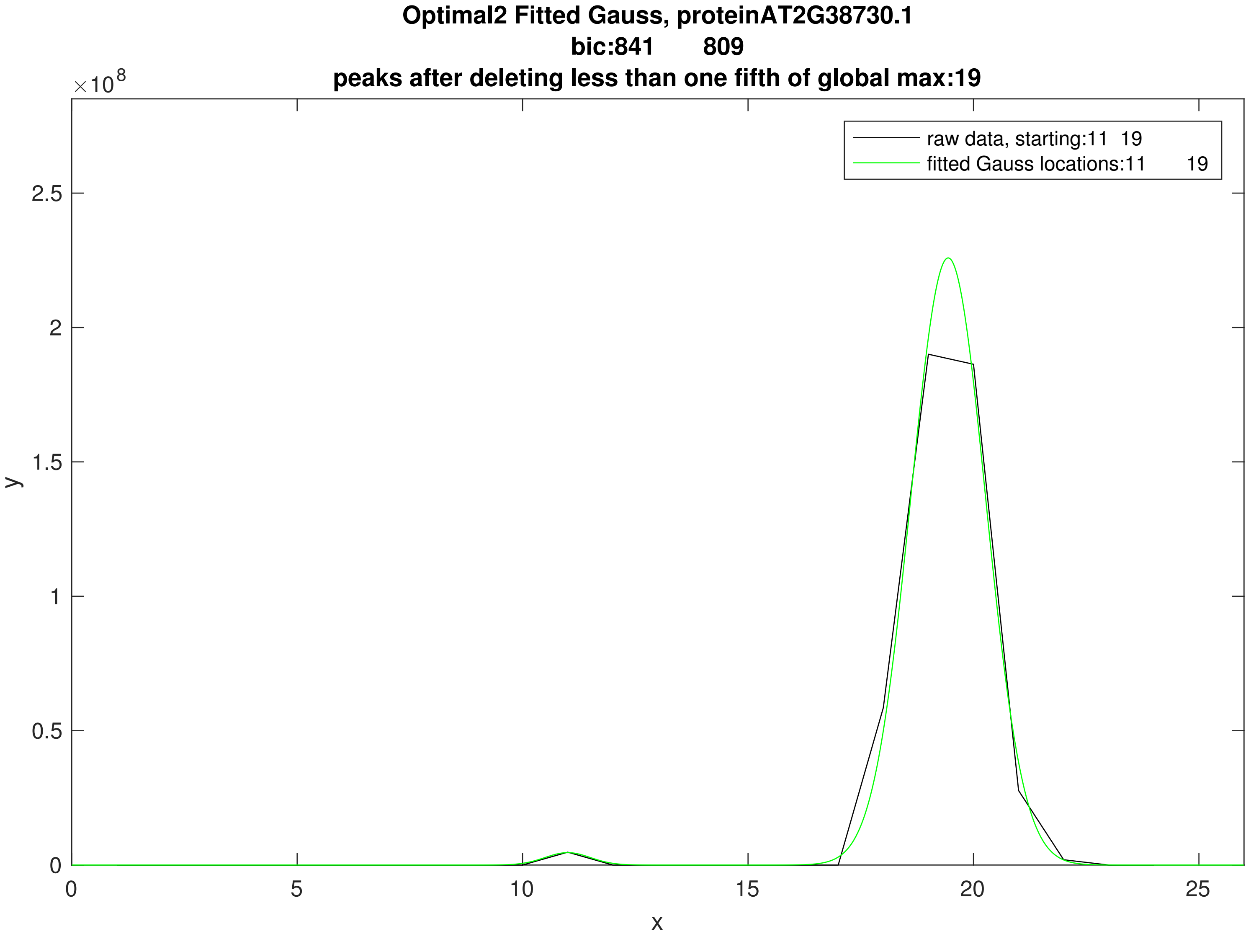

Optimal2 Fitted Gauss, proteinAT2G38730.1
bic:841 809
peaks after deleting less than one fifth of global max:19
×10^8
raw data, starting:11 19
fitted Gauss locations:11 19
2.5
2
1.5
1
0.5
0
y
0
5
10
15
20
25
x

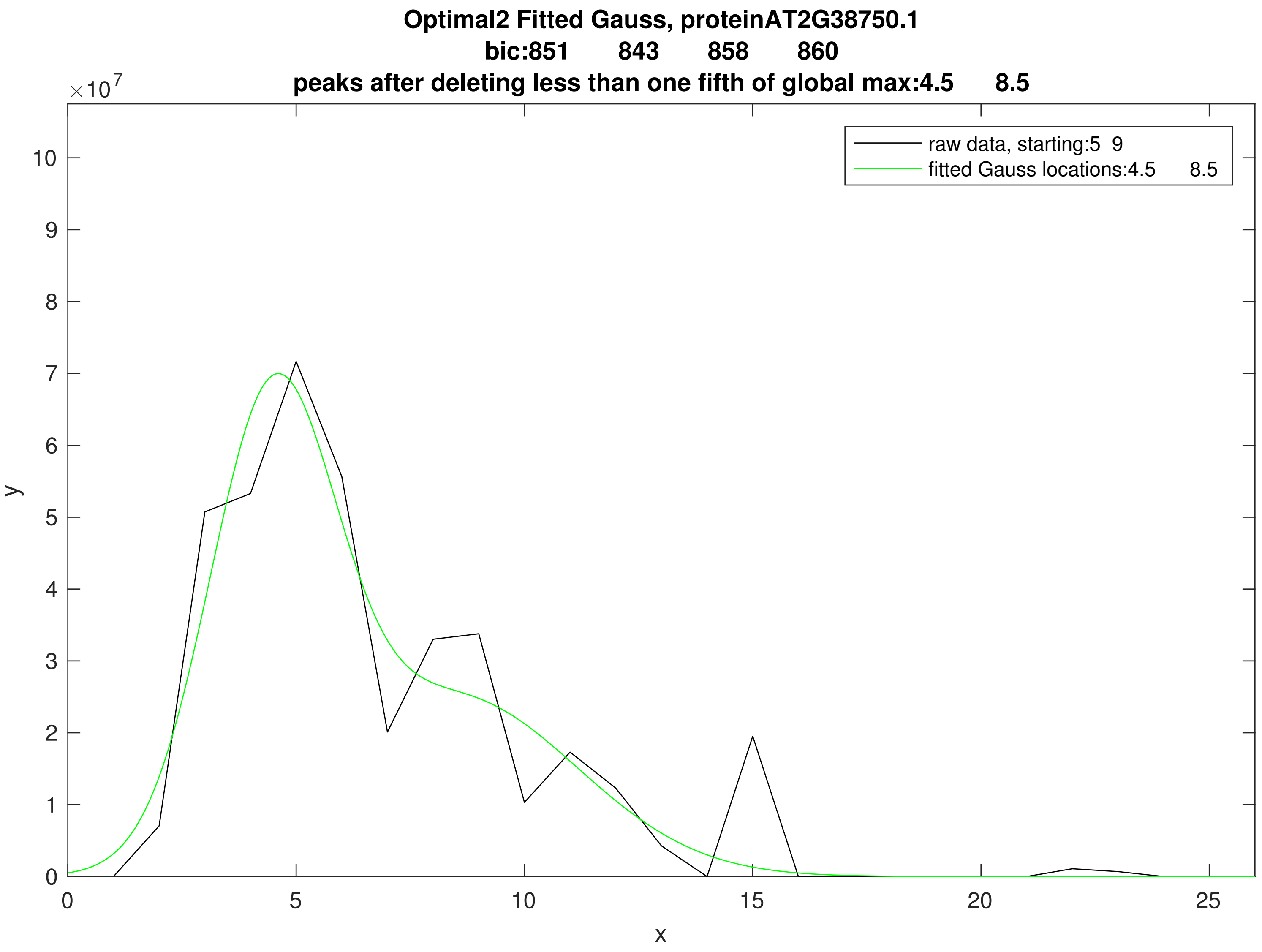
Optimal2 Fitted Gauss, proteinAT2G38750.1
bic:851 843 858 860
peaks after deleting less than one fifth of global max:4.5 8.5
×10^7
raw data, starting:5 9
fitted Gauss locations:4.5 8.5
y
x
0
1
2
3
4
5
6
7
8
9
10
15
20
25

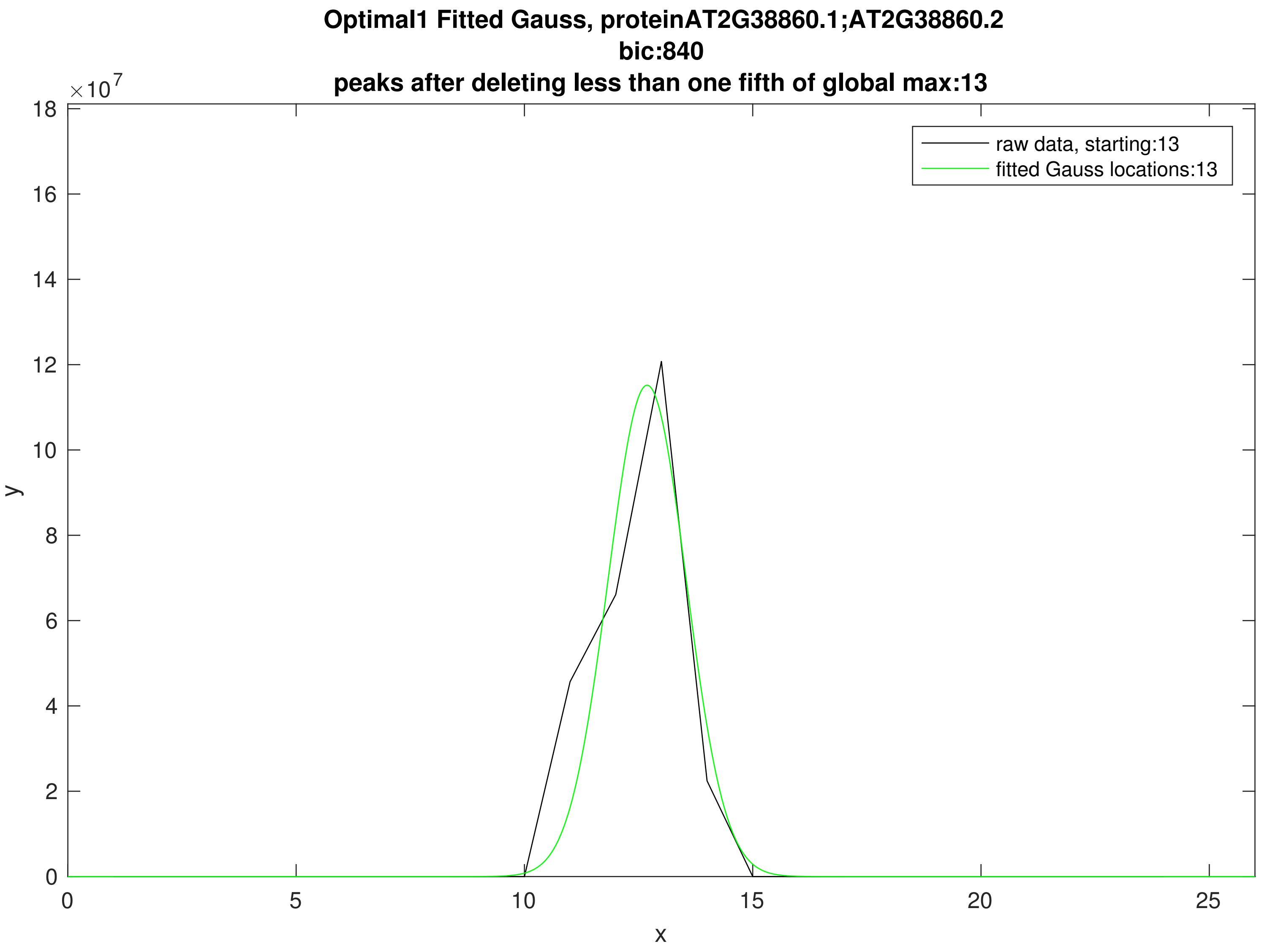
Optimal1 Fitted Gauss, proteinAT2G38860.1;AT2G38860.2
bic:840
peaks after deleting less than one fifth of global max:13
×10^7
raw data, starting:13
fitted Gauss locations:13
y
x
0
2
4
6
8
10
12
14
16
18
0
5
10
15
20
25

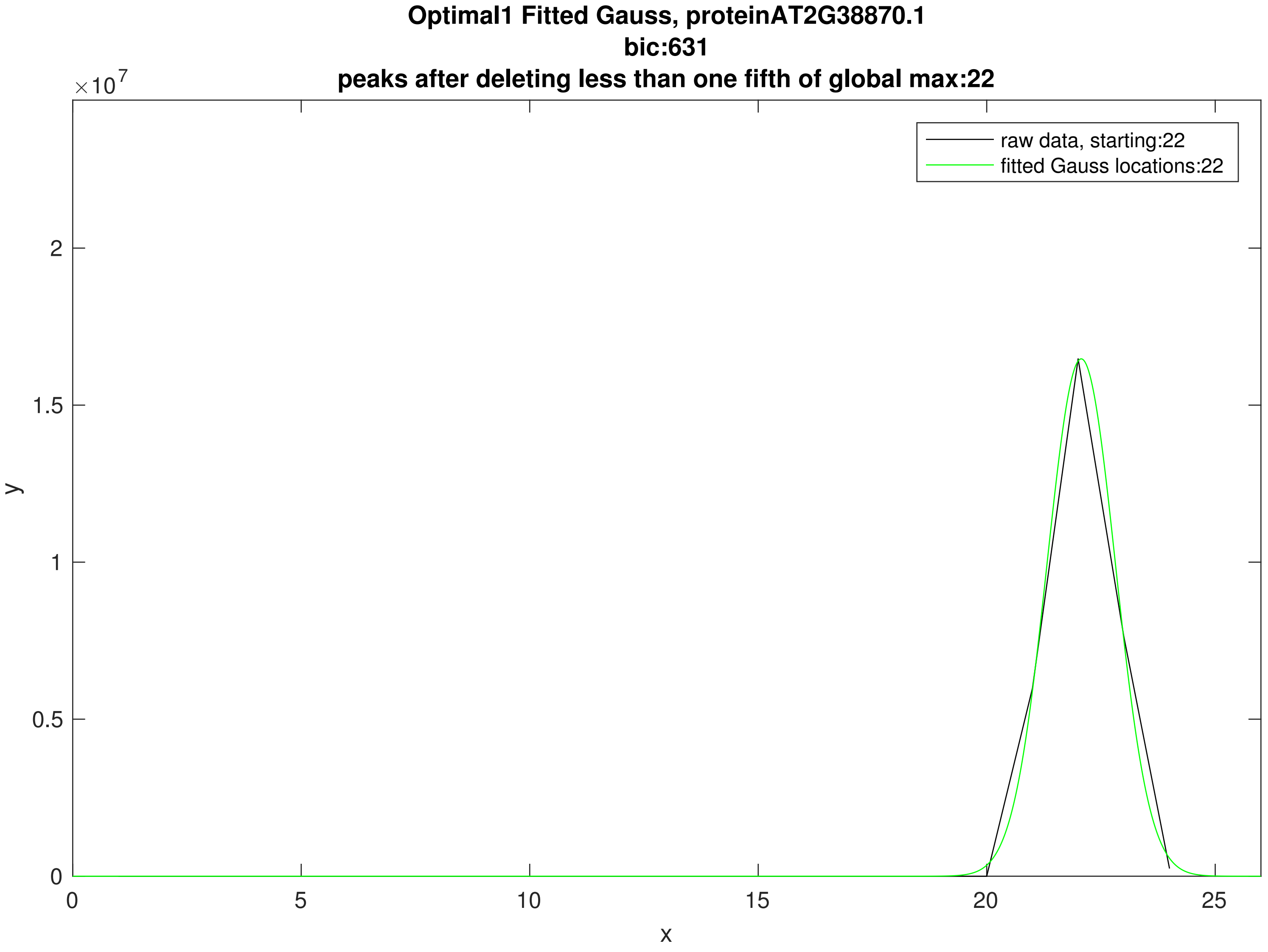
Optimal1 Fitted Gauss, proteinAT2G38870.1
bic:631
peaks after deleting less than one fifth of global max:22
×10^7
raw data, starting:22
fitted Gauss locations:22
y
x
0
0.5
1
1.5
2
0
5
10
15
20
25

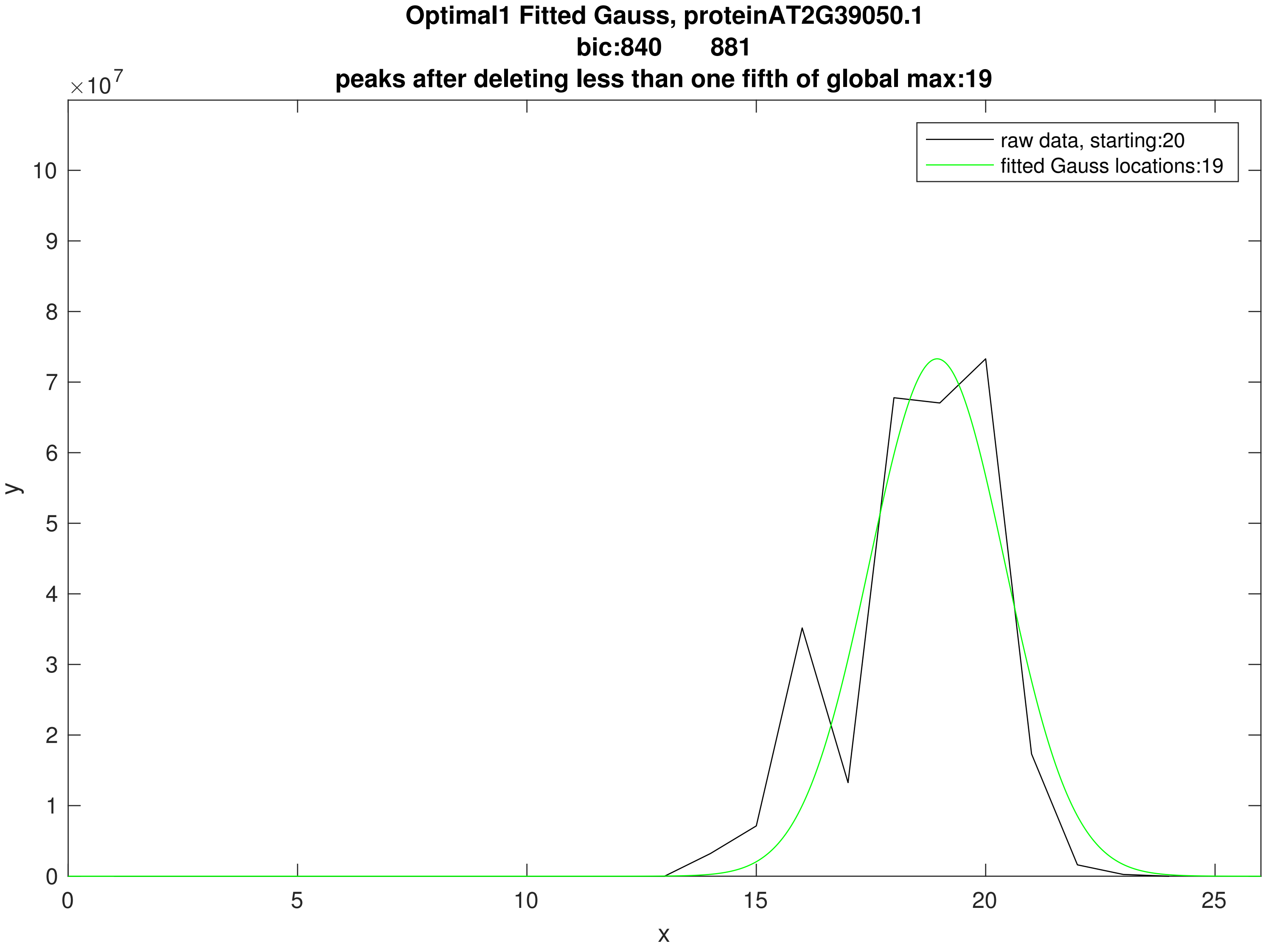
Optimal1 Fitted Gauss, proteinAT2G39050.1
bic:840 881
peaks after deleting less than one fifth of global max:19
×10^7
raw data, starting:20
fitted Gauss locations:19
y
x
0
1
2
3
4
5
6
7
8
9
10
0
5
10
15
20
25

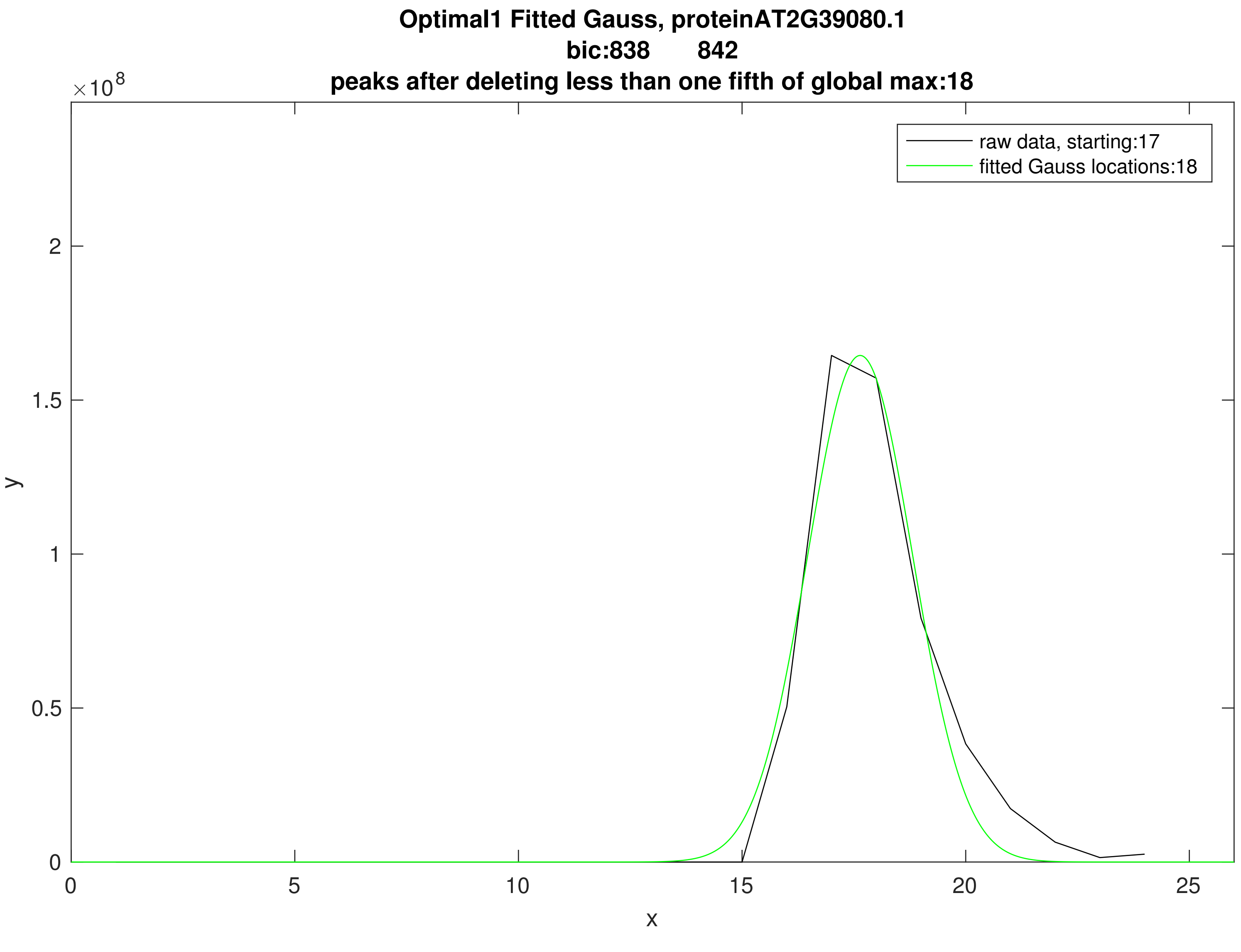
Optimal1 Fitted Gauss, proteinAT2G39080.1
bic:838 842
peaks after deleting less than one fifth of global max:18
×10^8
raw data, starting:17
fitted Gauss locations:18
y
x
0
0.5
1
1.5
2
5
10
15
20
25

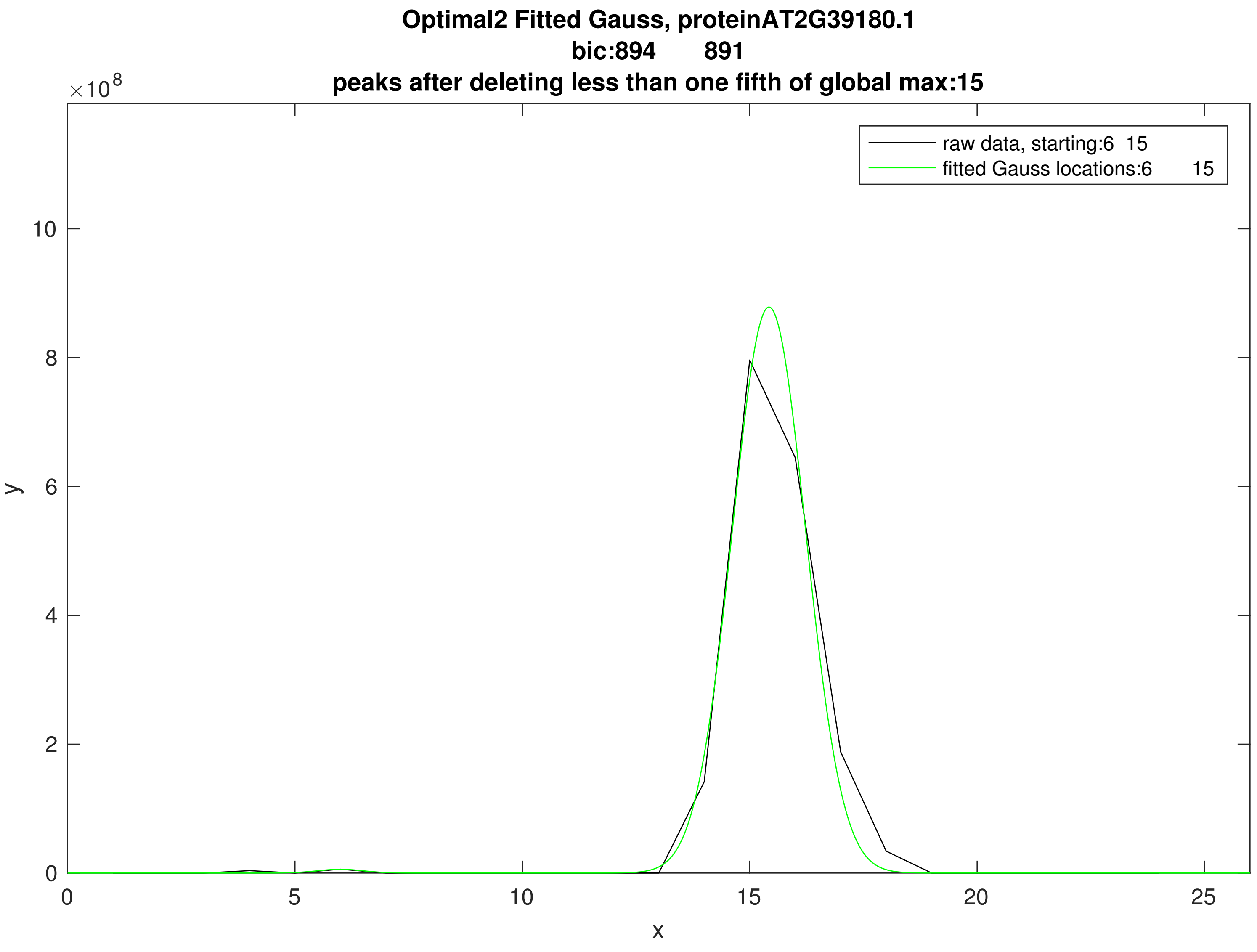

Optimal2 Fitted Gauss, proteinAT2G39180.1
bic:894 891
peaks after deleting less than one fifth of global max:15
×10^8
raw data, starting:6 15
fitted Gauss locations:6 15
10
8
6
4
2
0
y
0
5
10
15
20
25
x

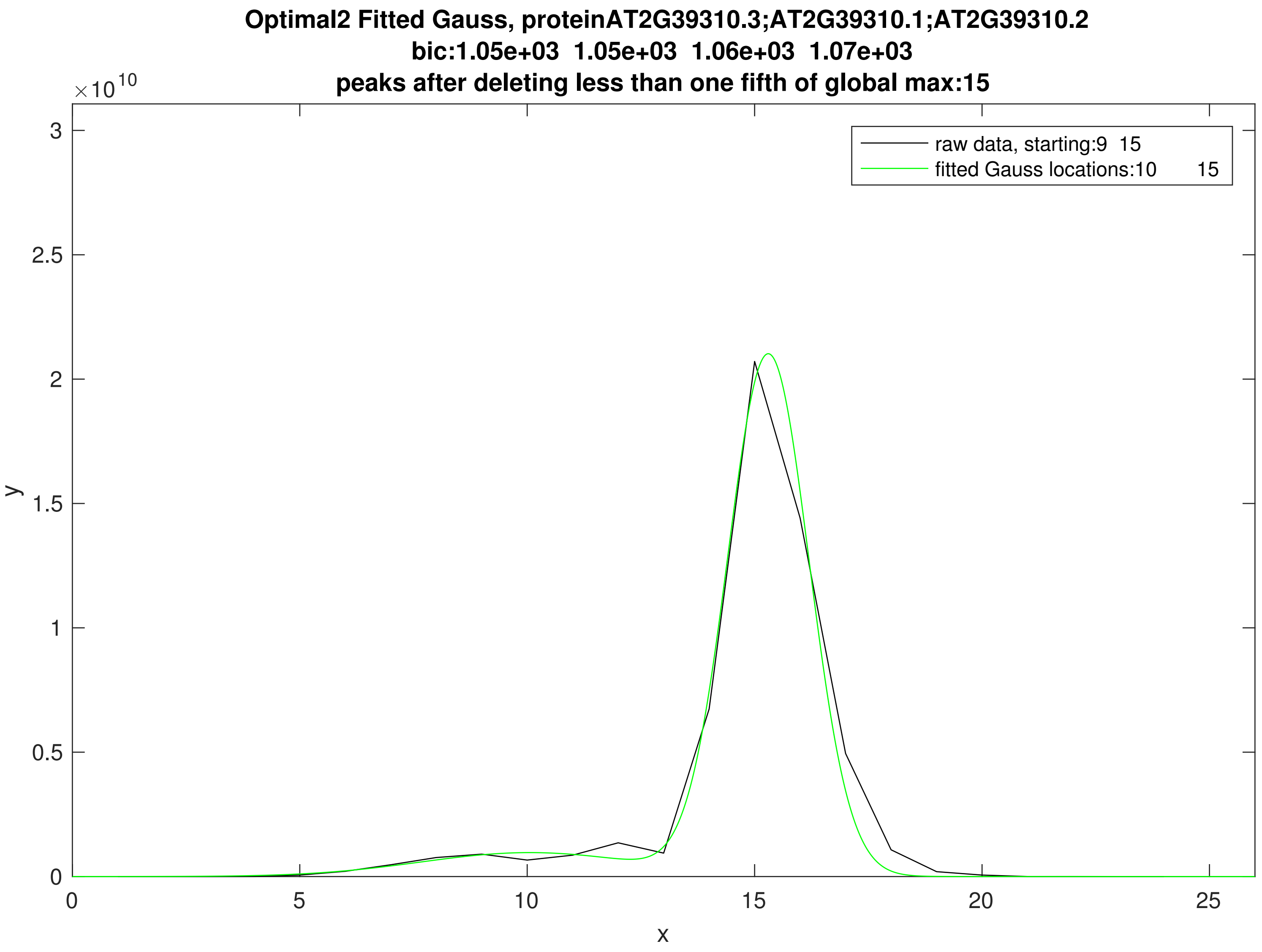

Optimal2 Fitted Gauss, proteinAT2G39310.3;AT2G39310.1;AT2G39310.2
bic:1.05e+03 1.05e+03 1.06e+03 1.07e+03
peaks after deleting less than one fifth of global max:15
×10^10
raw data, starting:9 15
fitted Gauss locations:10 15
y
x
0
0.5
1
1.5
2
2.5
3
5
10
15
20
25

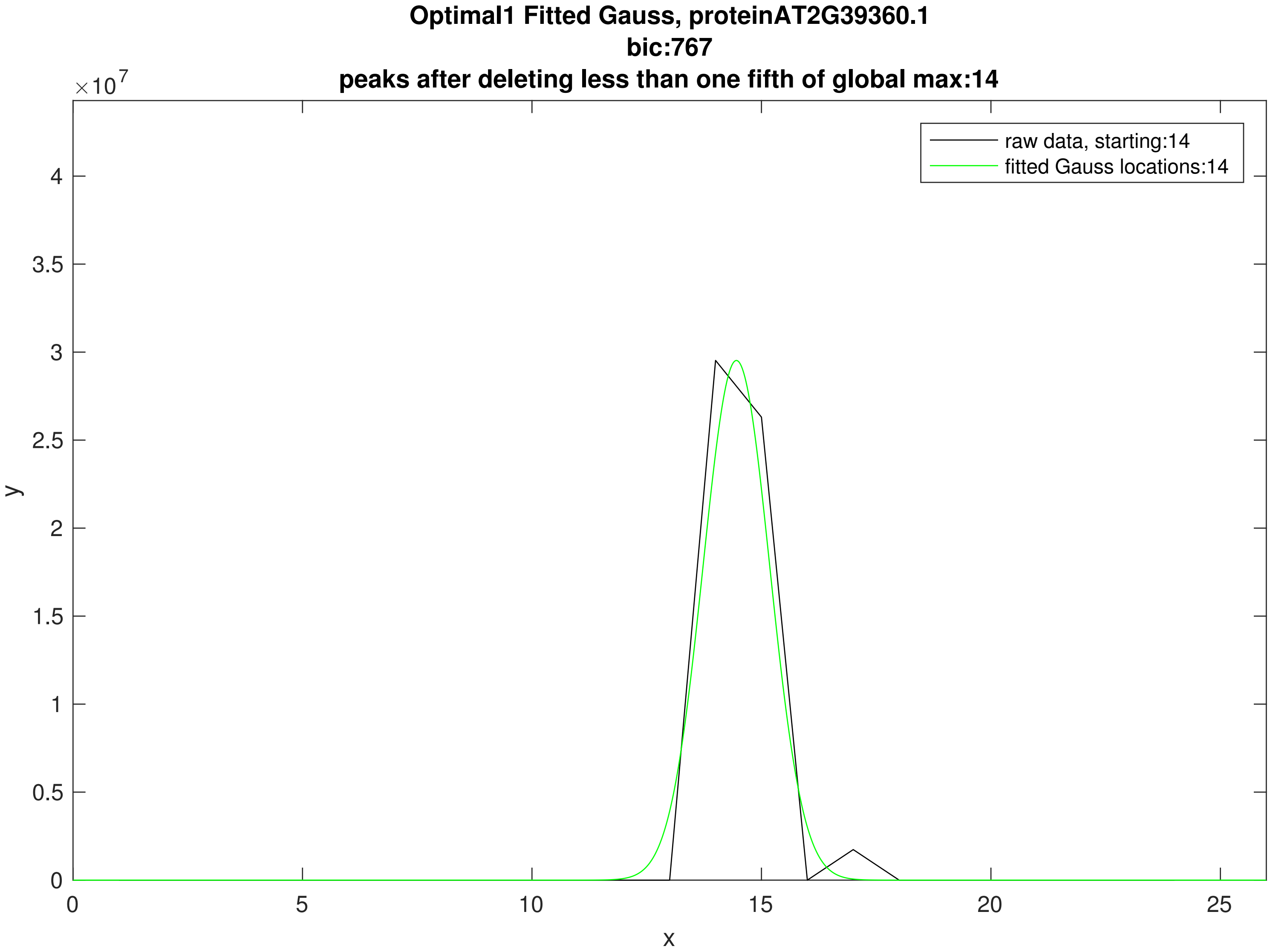

Optimal1 Fitted Gauss, proteinAT2G39360.1
bic:767
peaks after deleting less than one fifth of global max:14
×10^7
raw data, starting:14
fitted Gauss locations:14
y
x
0
0.5
1
1.5
2
2.5
3
3.5
4
5
10
15
20
25

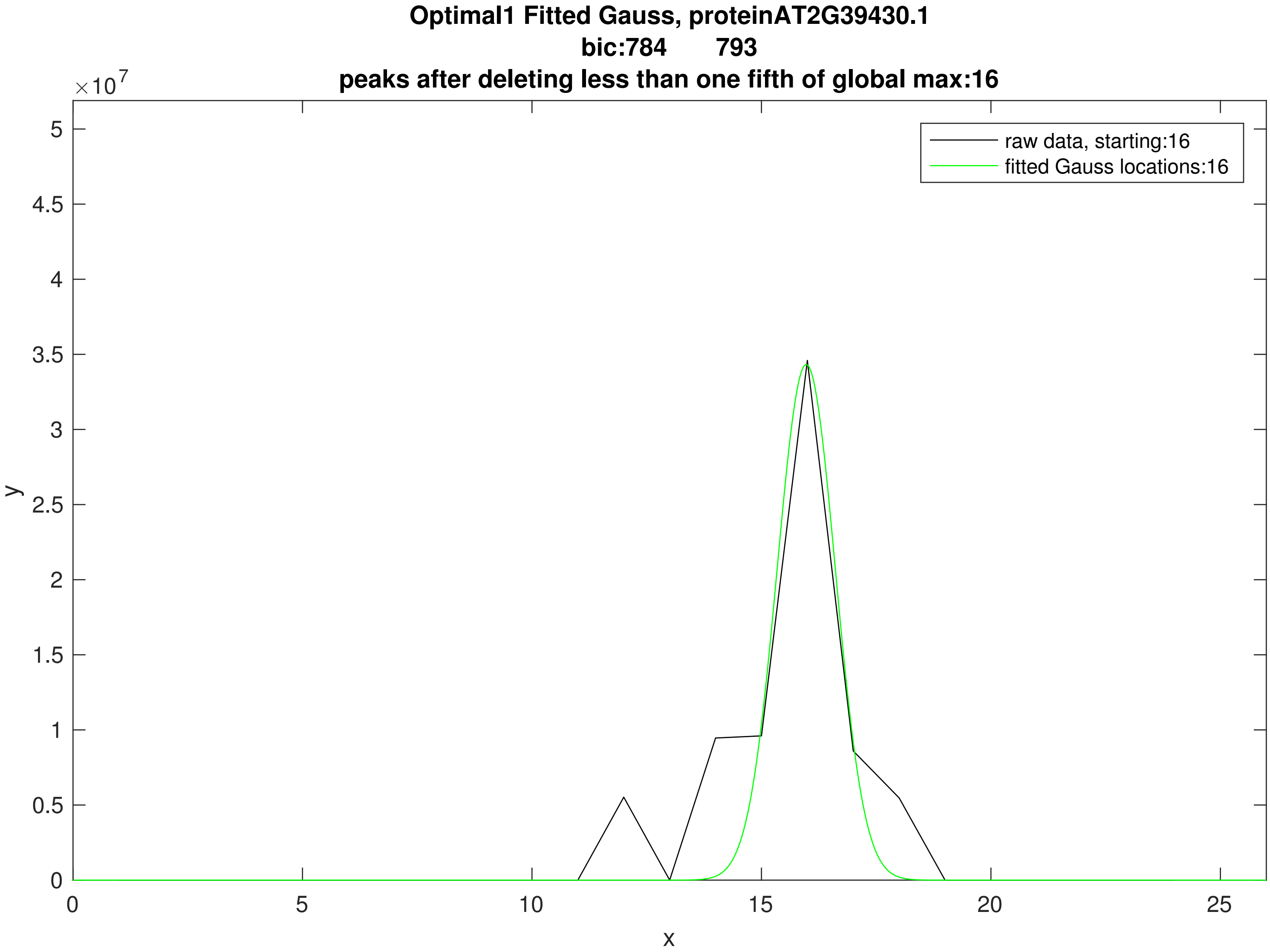
Optimal1 Fitted Gauss, proteinAT2G39430.1
bic:784 793
peaks after deleting less than one fifth of global max:16
×10^7
5
4.5
4
3.5
3
2.5
2
1.5
1
0.5
0
y
0
5
10
15
20
25
x
raw data, starting:16
fitted Gauss locations:16

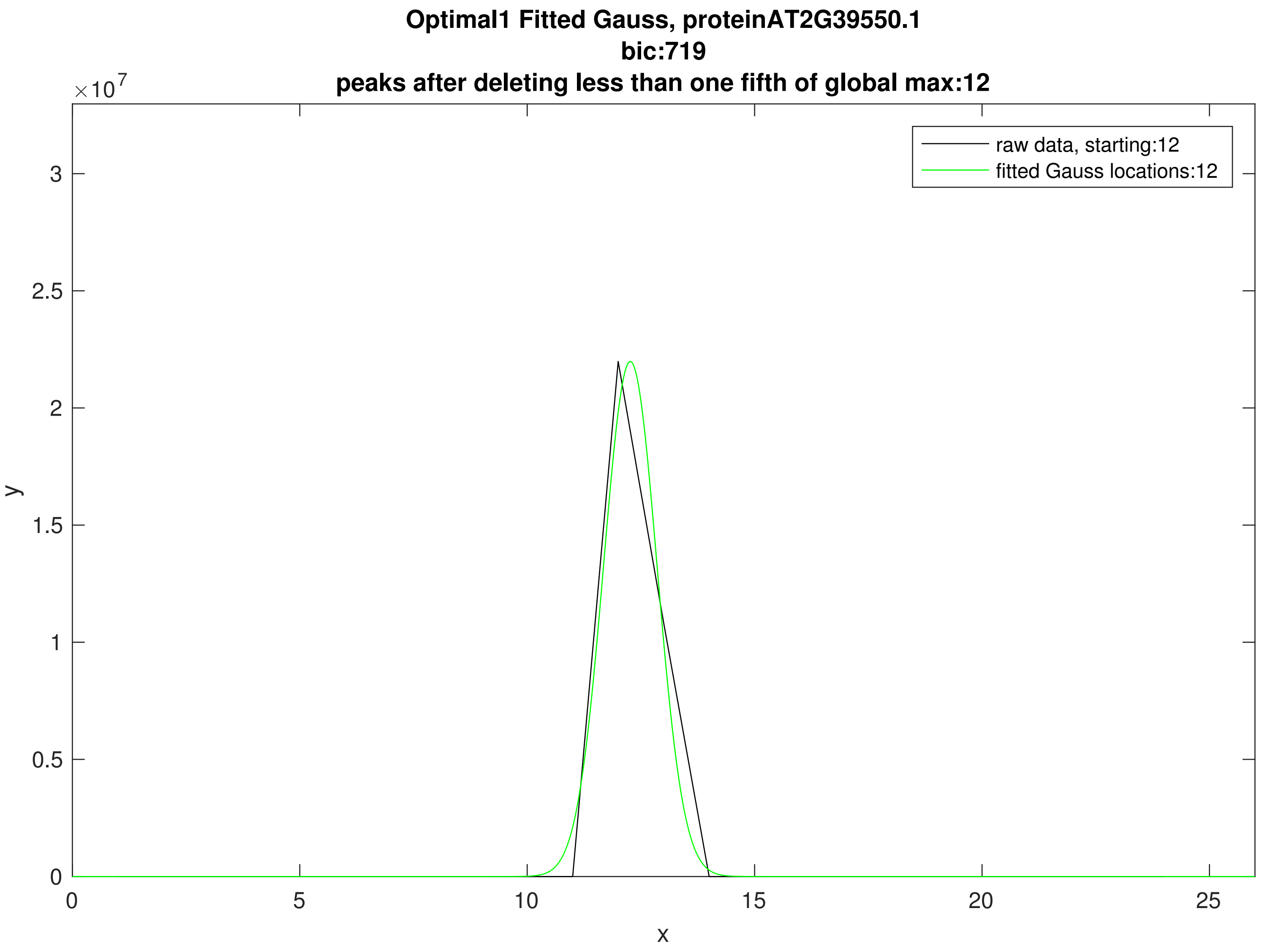
Optimal1 Fitted Gauss, proteinAT2G39550.1
bic:719
peaks after deleting less than one fifth of global max:12
×10^7
3
2.5
2
1.5
1
0.5
0
y
0
5
10
15
20
25
x
raw data, starting:12
fitted Gauss locations:12

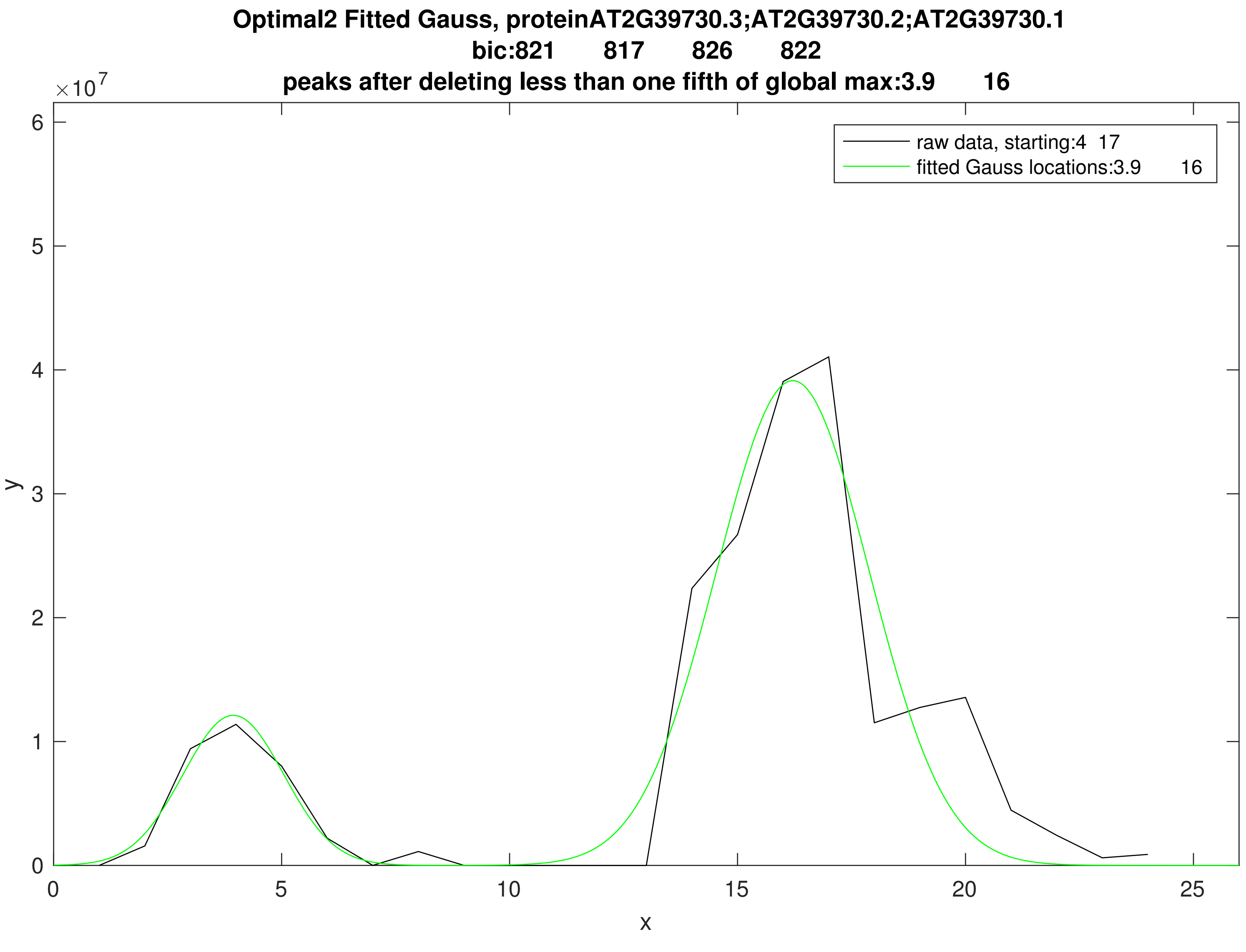
Optimal2 Fitted Gauss, proteinAT2G39730.3;AT2G39730.2;AT2G39730.1
bic:821 817 826 822
peaks after deleting less than one fifth of global max:3.9 16
×10^7
raw data, starting:4 17
fitted Gauss locations:3.9 16
y
x
0
1
2
3
4
5
6
0
5
10
15
20
25

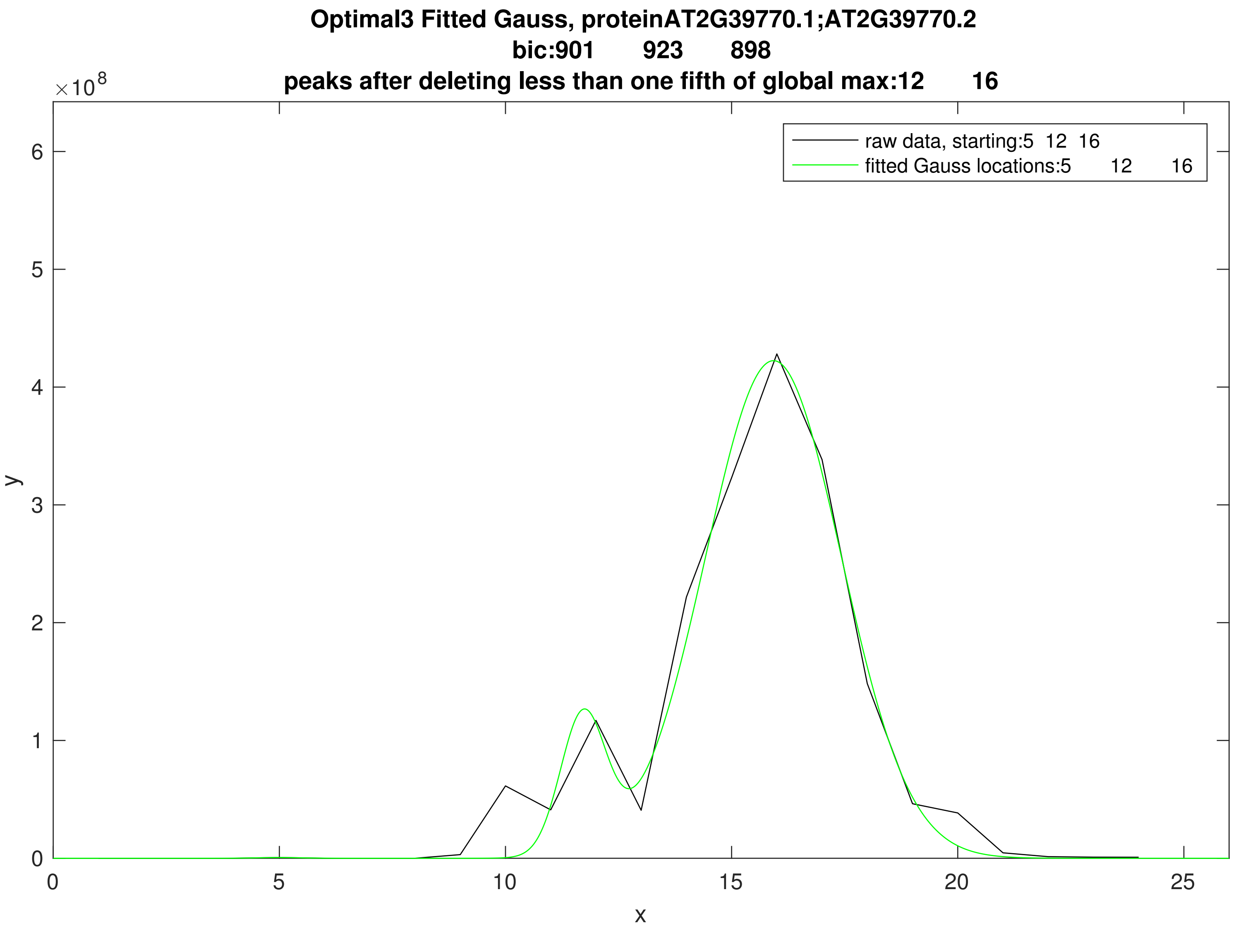
Optimal3 Fitted Gauss, proteinAT2G39770.1;AT2G39770.2
bic:901 923 898
peaks after deleting less than one fifth of global max:12 16
×10^8
raw data, starting:5 12 16
fitted Gauss locations:5 12 16
y
x
0
1
2
3
4
5
6
0
5
10
15
20
25

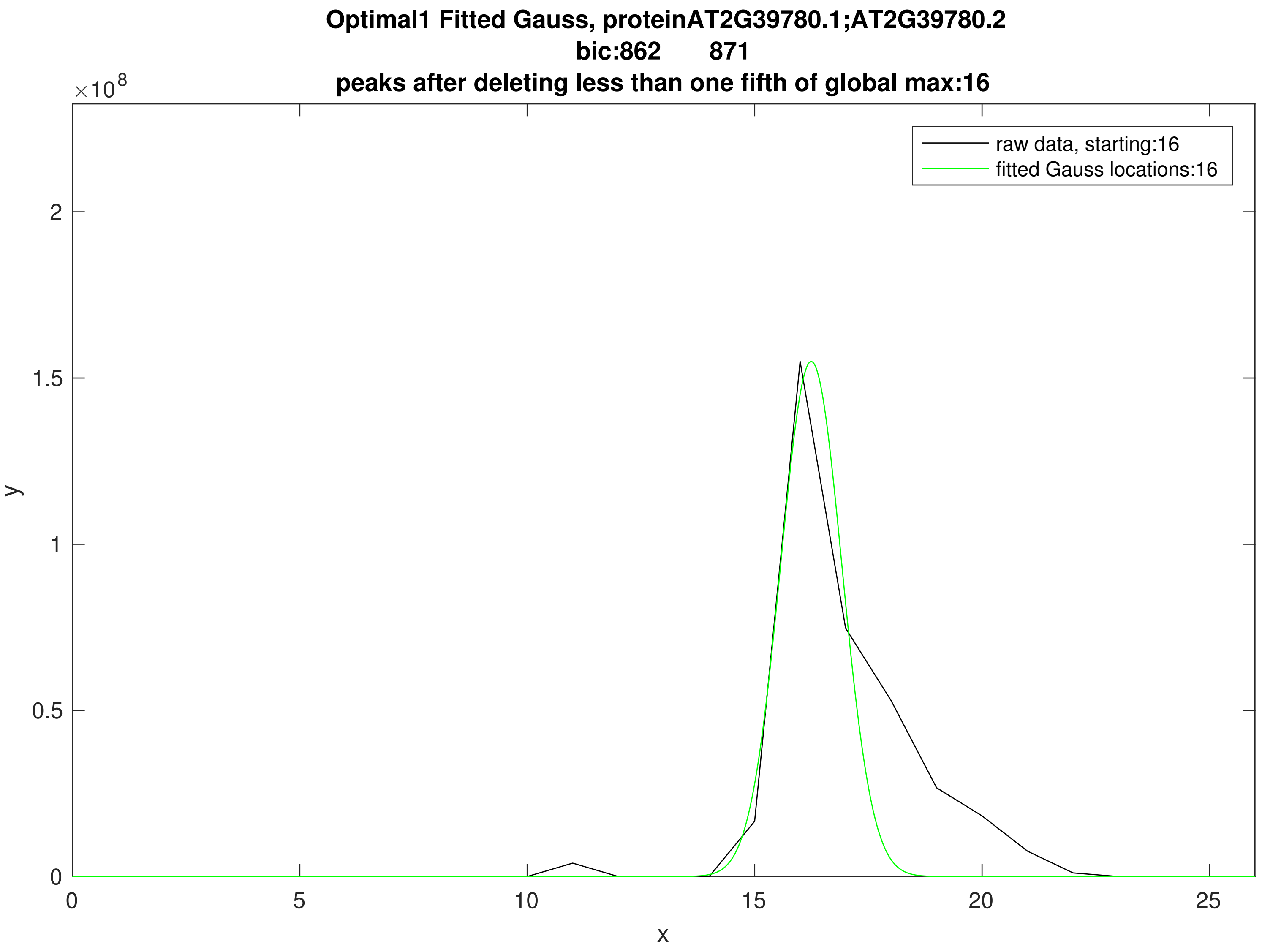

Optimal1 Fitted Gauss, proteinAT2G39780.1;AT2G39780.2
bic:862 871
peaks after deleting less than one fifth of global max:16
×10^8
raw data, starting:16
fitted Gauss locations:16
y
x
0
0.5
1
1.5
2
0
5
10
15
20
25

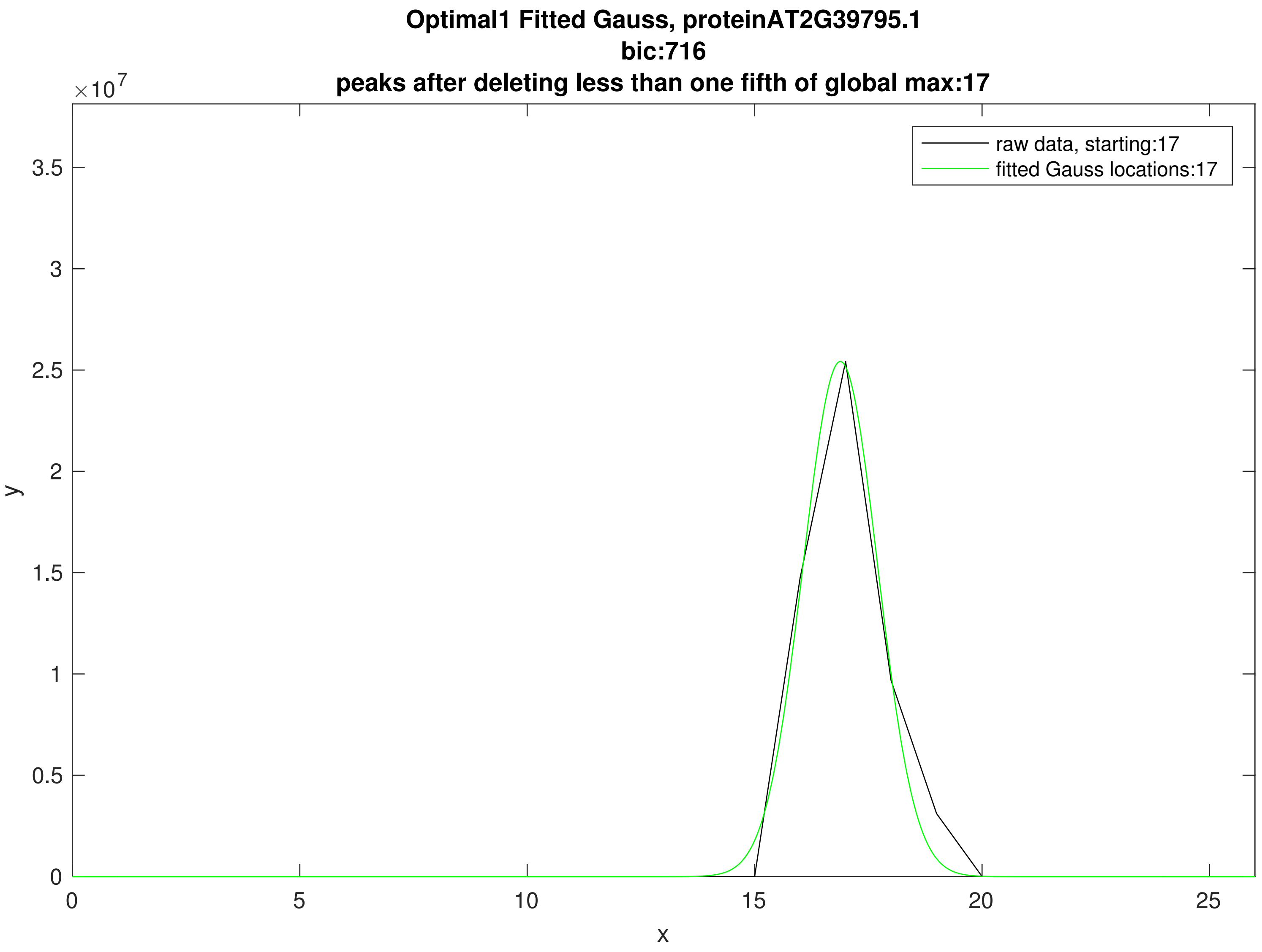
Optimal1 Fitted Gauss, proteinAT2G39795.1
bic:716
peaks after deleting less than one fifth of global max:17
×10^7
3.5
3
2.5
2
1.5
1
0.5
0
y
0
5
10
15
20
25
x
raw data, starting:17
fitted Gauss locations:17

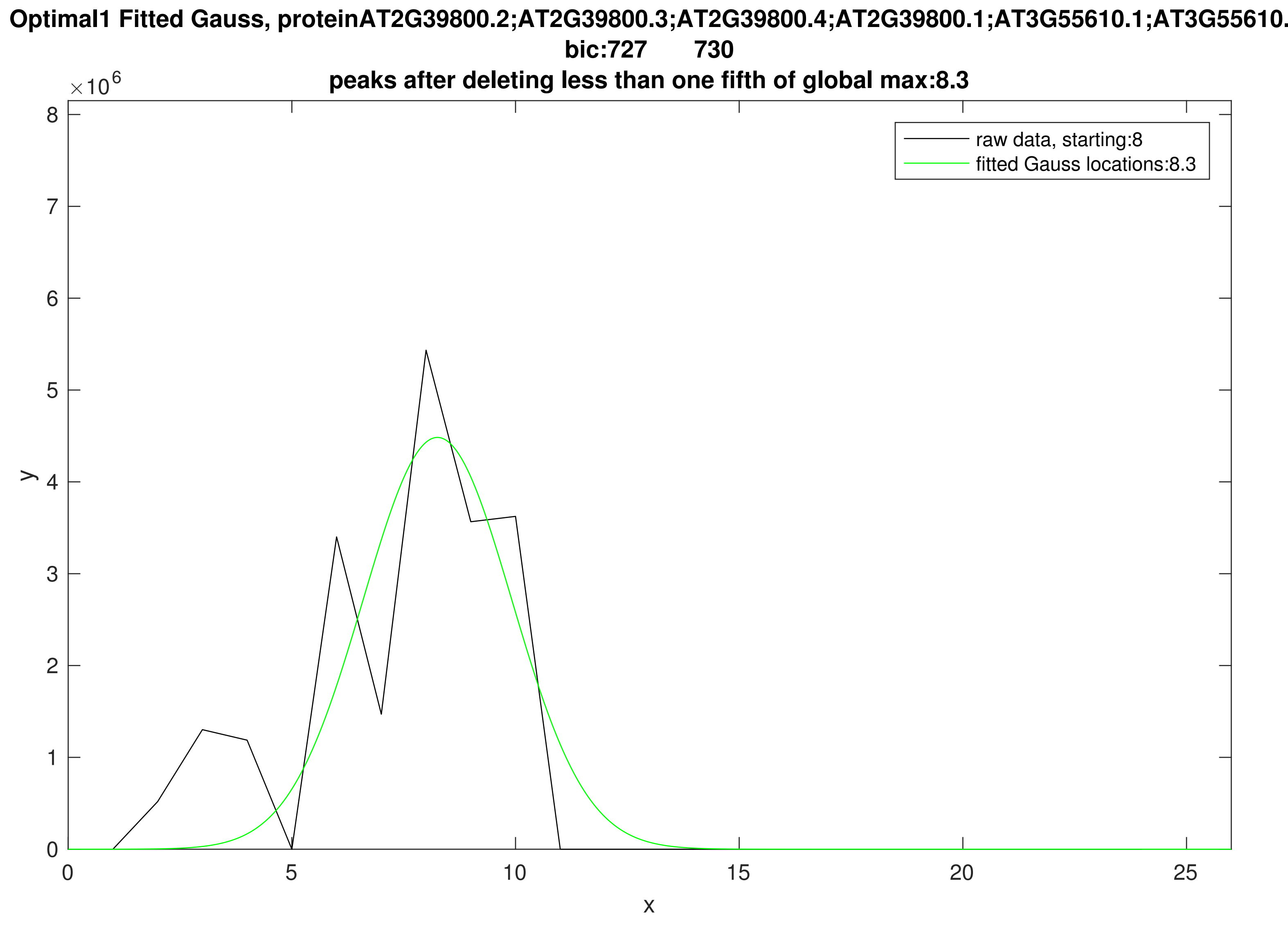

Optimal1 Fitted Gauss, proteinAT2G39800.2;AT2G39800.3;AT2G39800.4;AT2G39800.1;AT3G55610.1;AT3G55610.
bic:727 730
peaks after deleting less than one fifth of global max:8.3
×10^6
raw data, starting:8
fitted Gauss locations:8.3
y
x
0
1
2
3
4
5
6
7
8
0
5
10
15
20
25

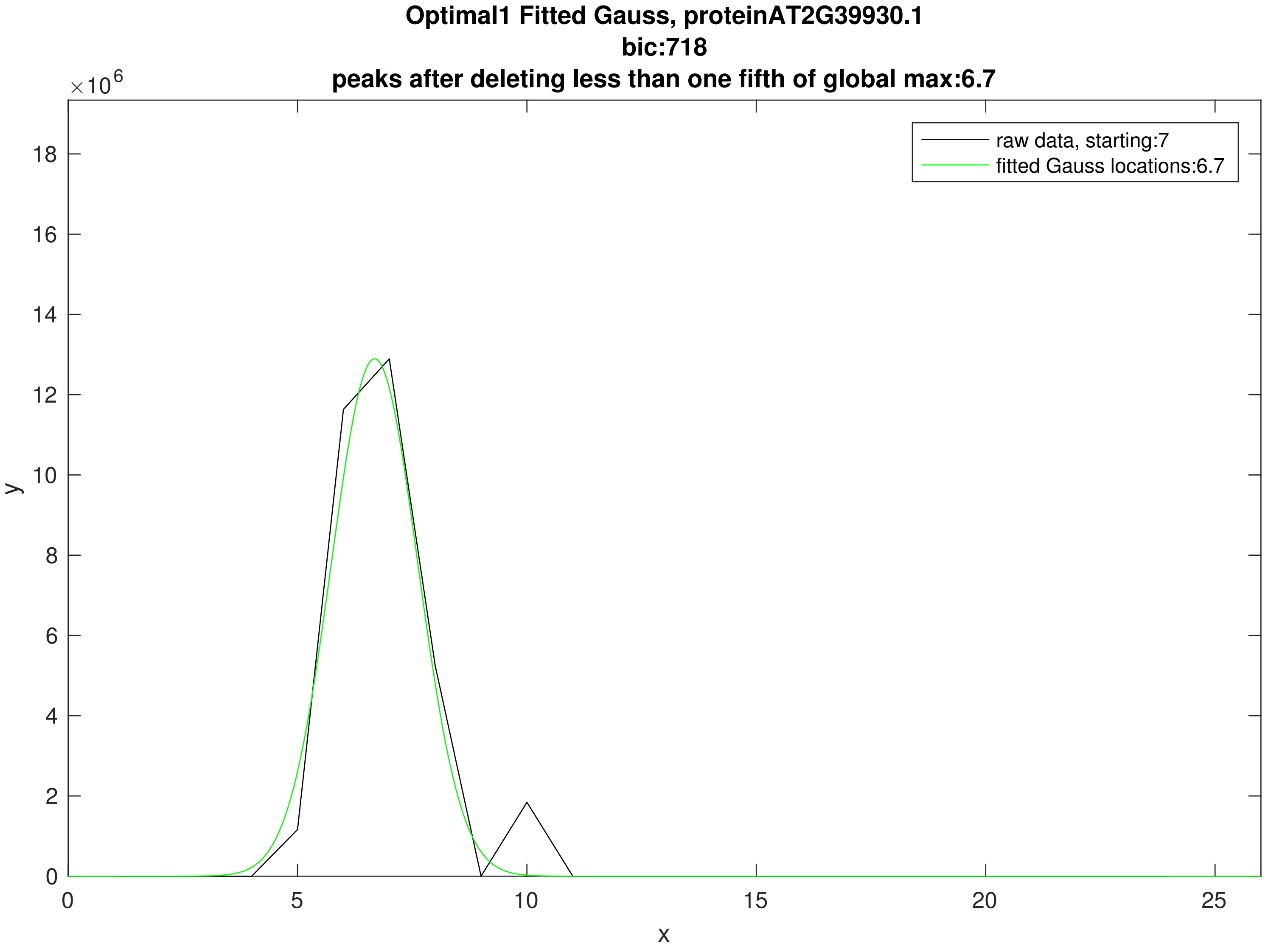

Optimal1 Fitted Gauss, proteinAT2G39930.1
bic:718
peaks after deleting less than one fifth of global max:6.7
×10^6
raw data, starting:7
fitted Gauss locations:6.7
18
16
14
12
10
8
6
4
2
0
y
0
5
10
15
20
25
x

#### Optimal2 Fitted Gauss, proteinAT2G39940.1
#### bic:749 702
#### peaks after deleting less than one fifth of global max:13

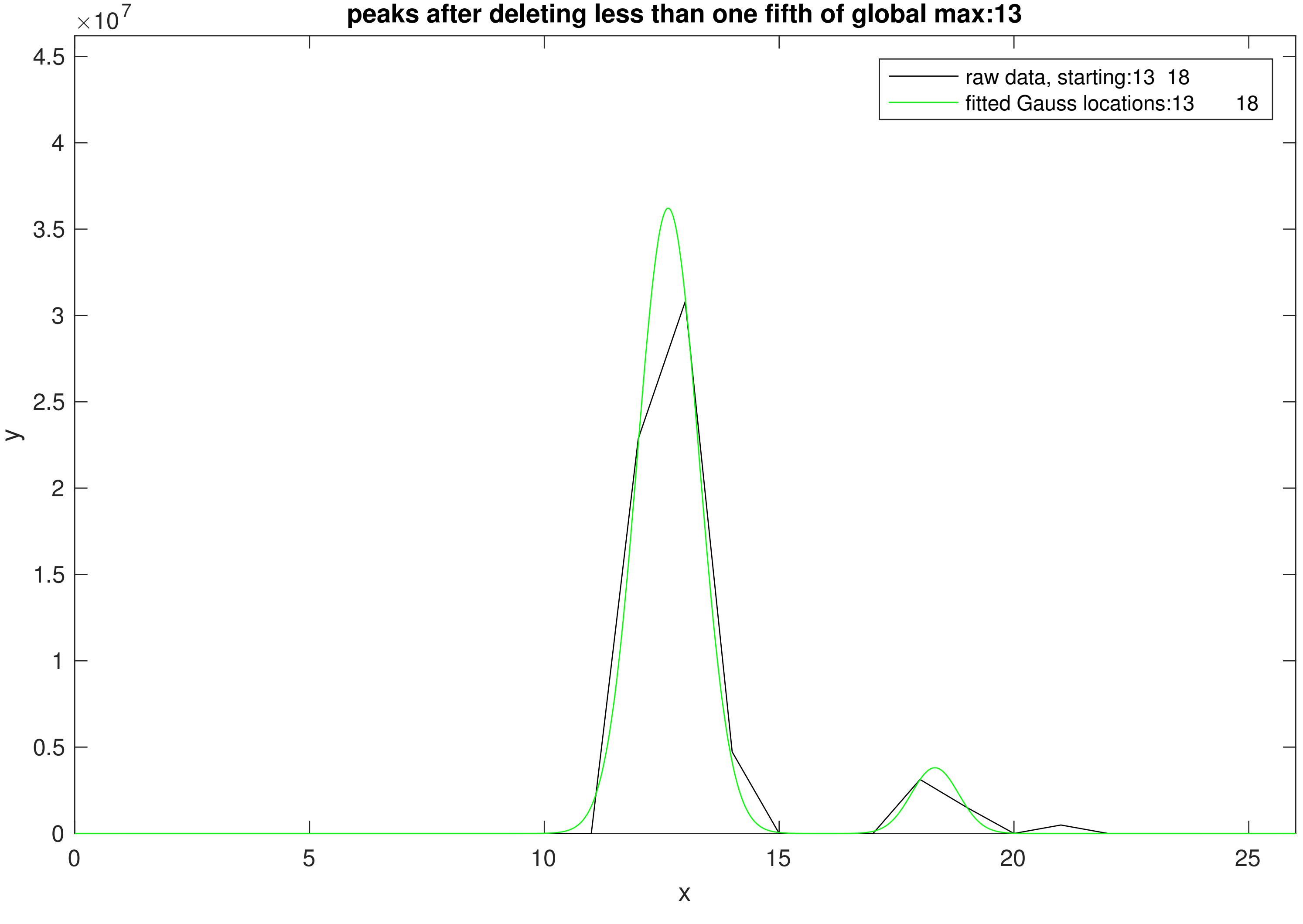

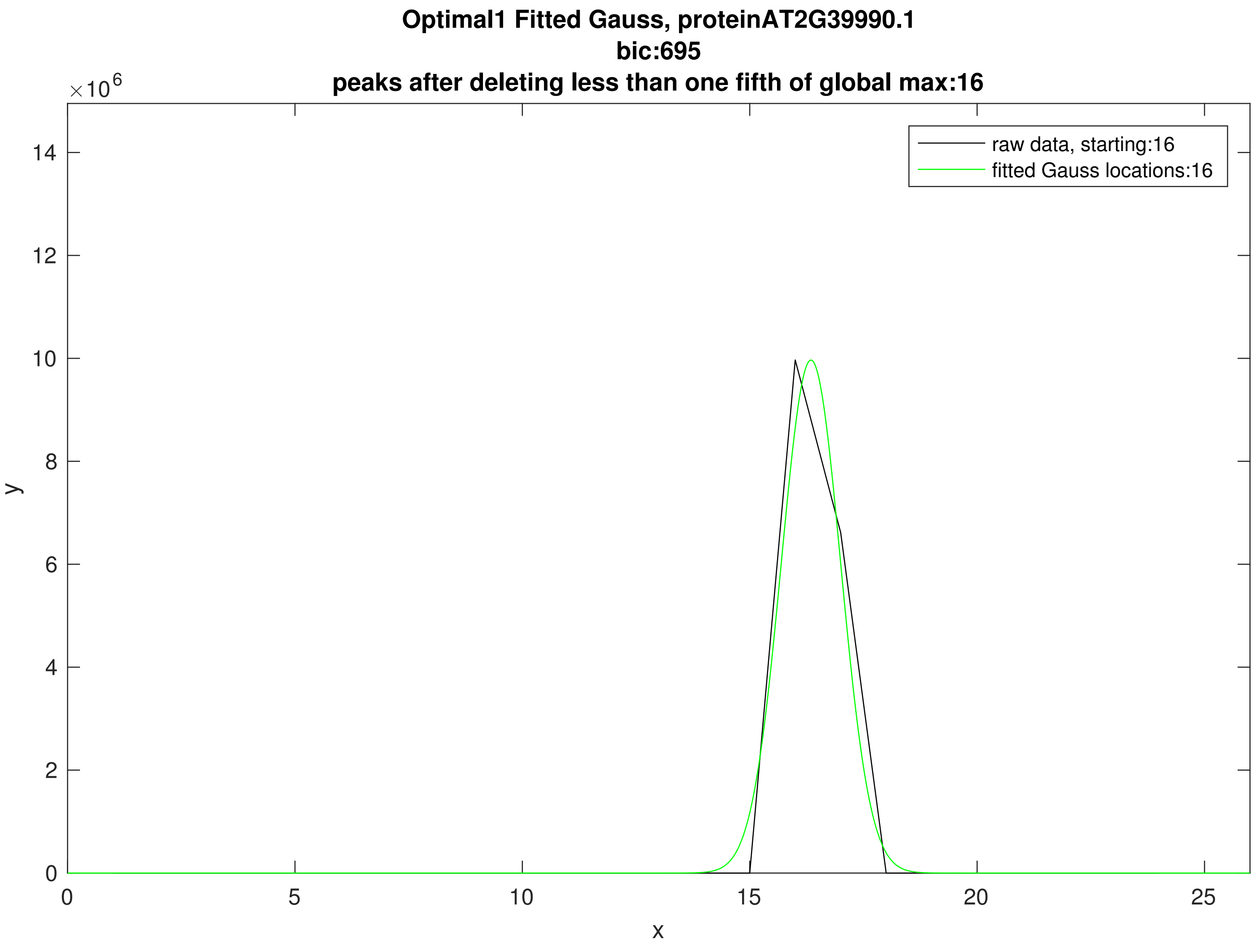

Optimal1 Fitted Gauss, proteinAT2G39990.1
bic:695
peaks after deleting less than one fifth of global max:16
×10^6
raw data, starting:16
fitted Gauss locations:16
14
12
10
8
6
4
2
0
y
0
5
10
15
20
25
x

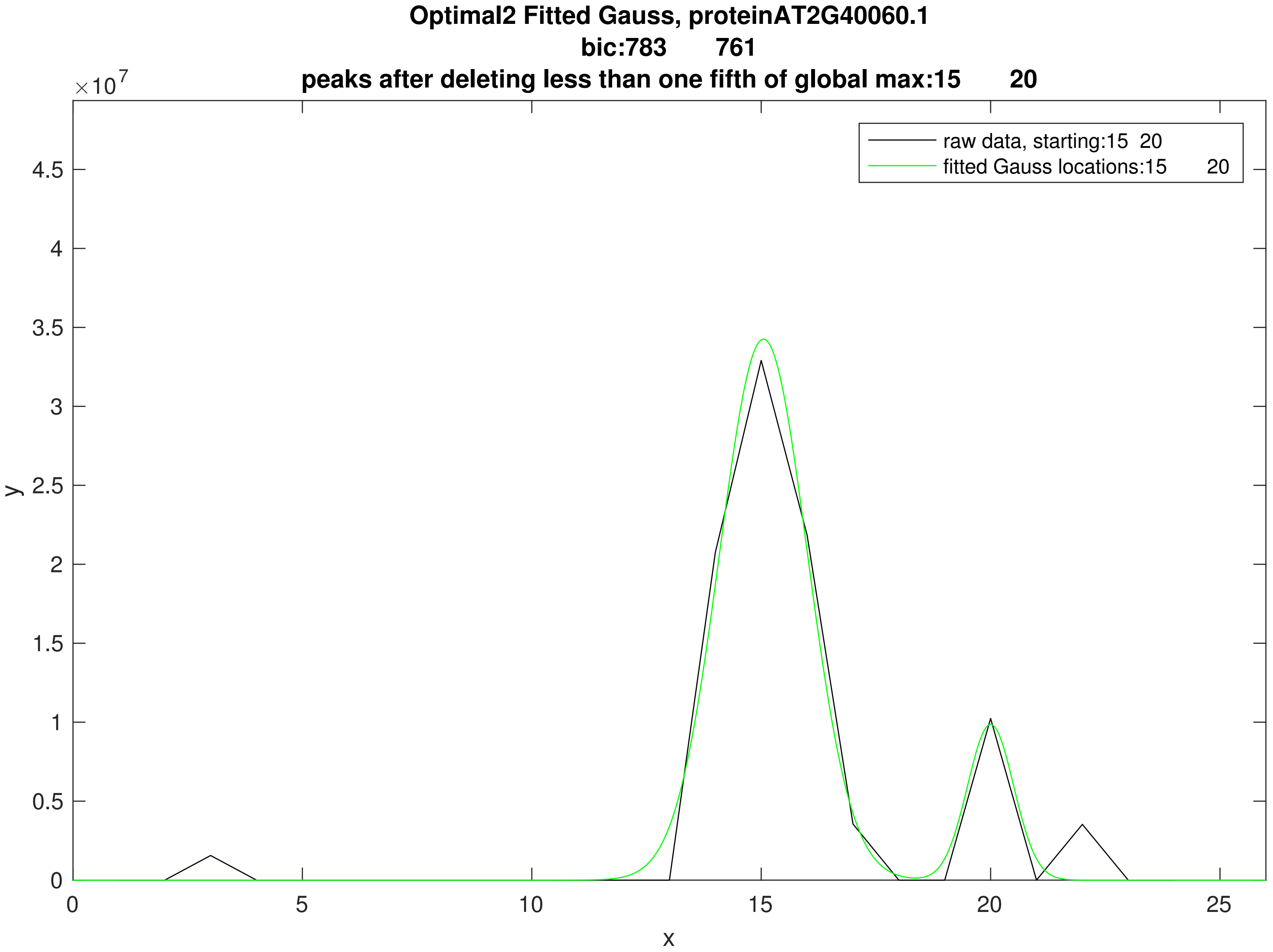
Optimal2 Fitted Gauss, proteinAT2G40060.1
bic:783 761
peaks after deleting less than one fifth of global max:15 20
raw data, starting:15 20
fitted Gauss locations:15 20
×10^7
y
x
0
0.5
1
1.5
2
2.5
3
3.5
4
4.5
5
10
15
20
25

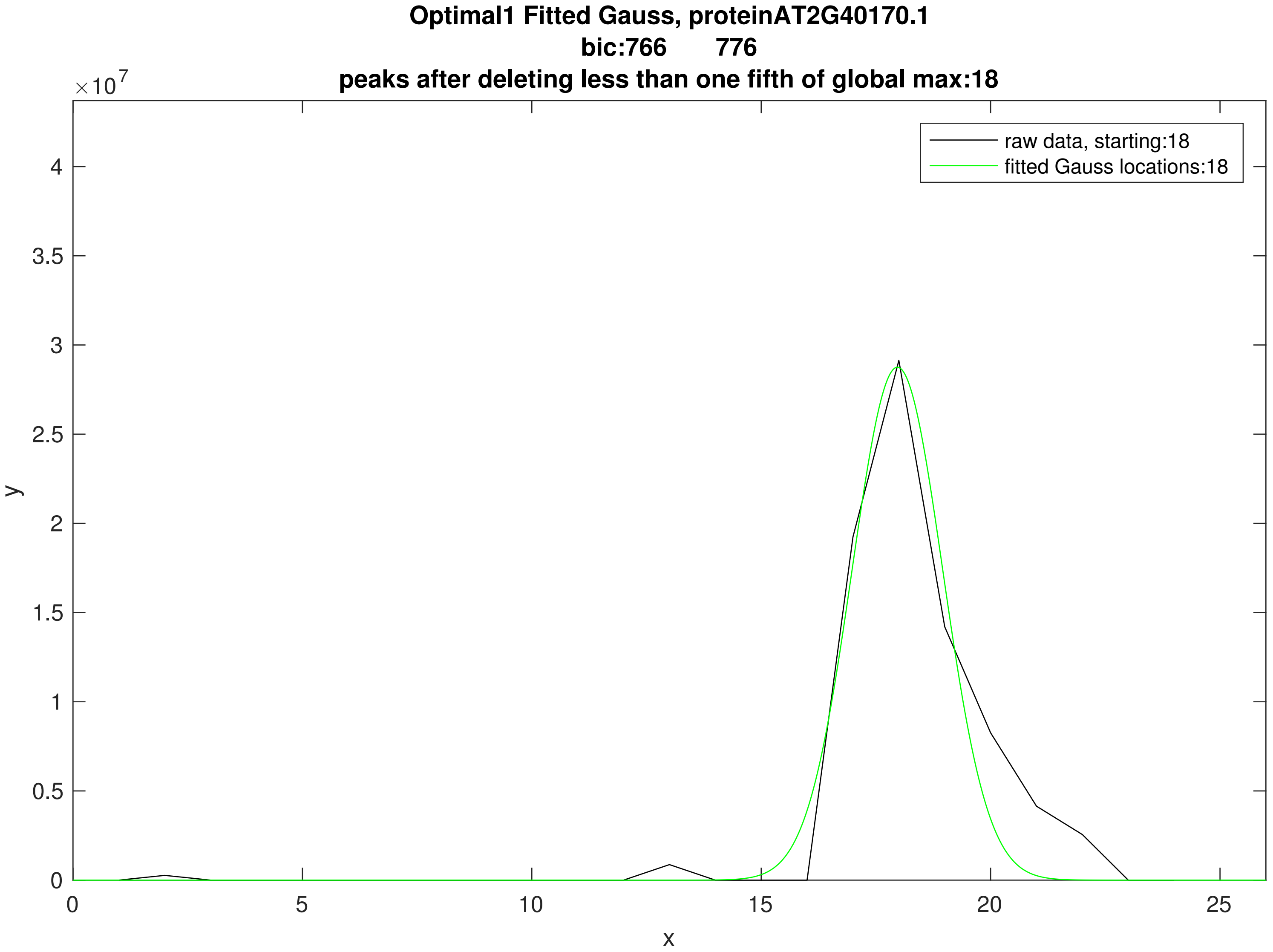
Optimal1 Fitted Gauss, proteinAT2G40170.1
bic:766 776
peaks after deleting less than one fifth of global max:18
×10^7
raw data, starting:18
fitted Gauss locations:18
y
x
0
0.5
1
1.5
2
2.5
3
3.5
4
0
5
10
15
20
25

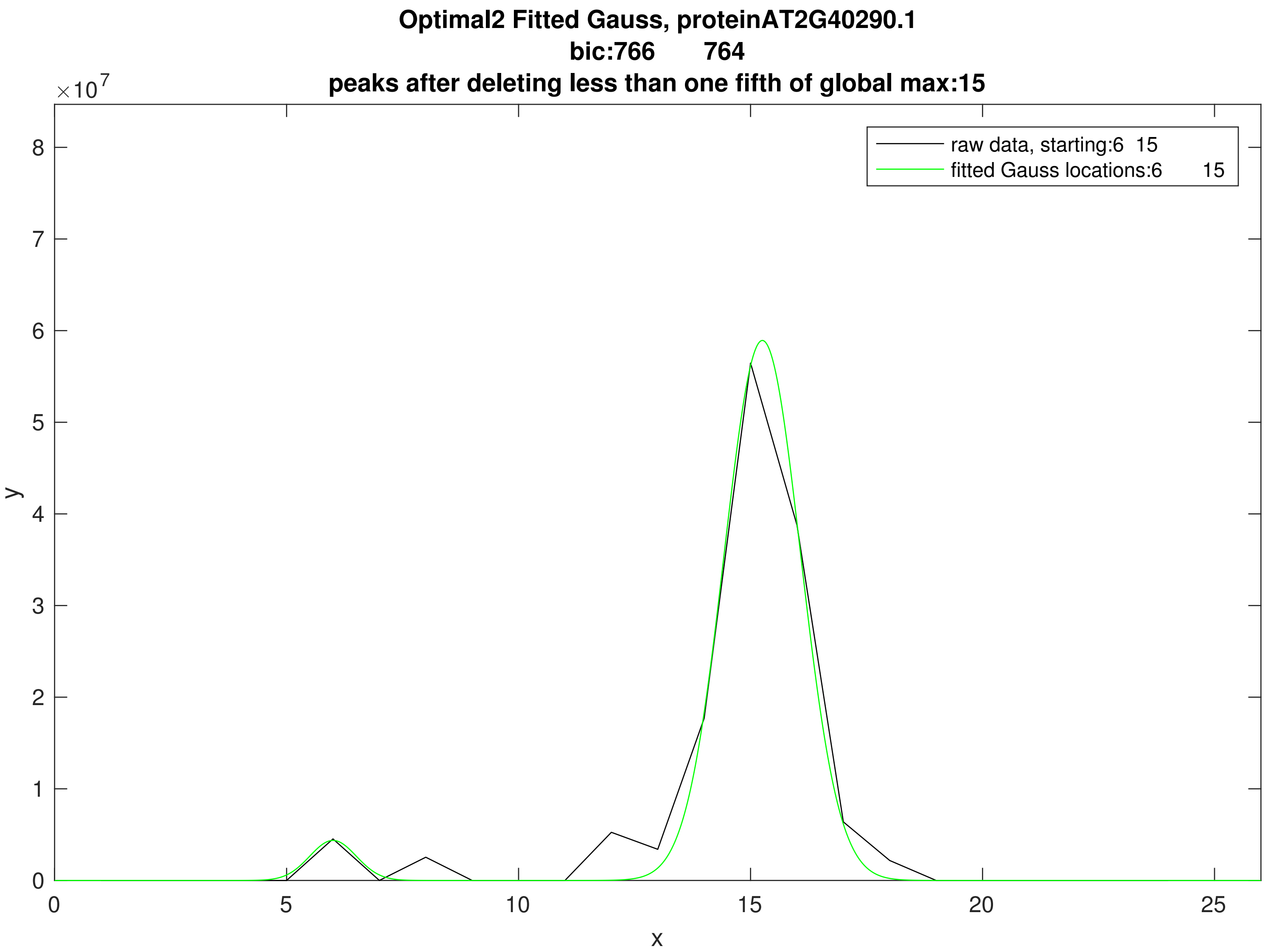
Optimal2 Fitted Gauss, proteinAT2G40290.1
bic:766 764
peaks after deleting less than one fifth of global max:15
×10^7
raw data, starting:6 15
fitted Gauss locations:6 15
y
x
0
1
2
3
4
5
6
7
8
0
5
10
15
20
25

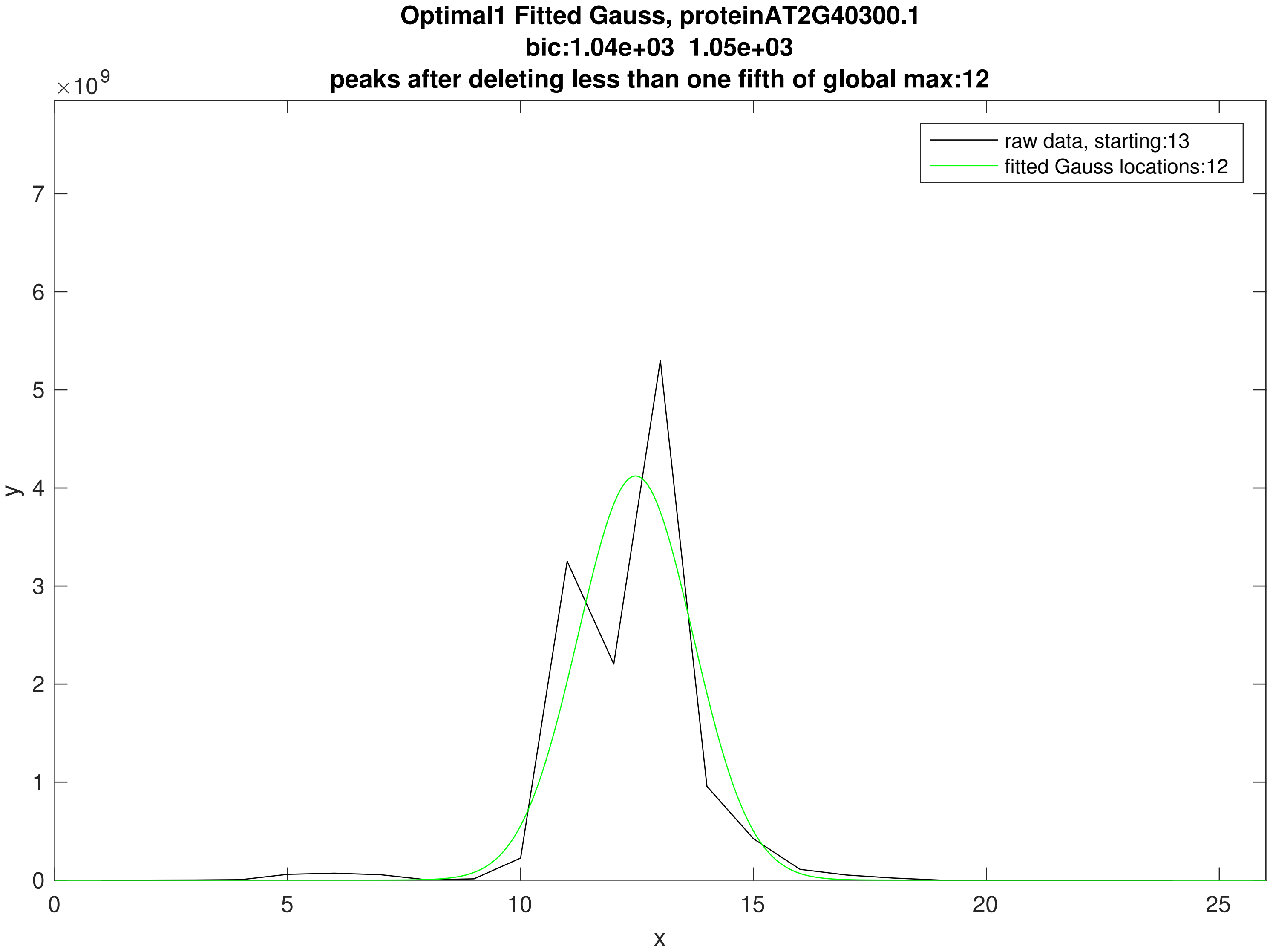

Optimal1 Fitted Gauss, proteinAT2G40300.1
bic:1.04e+03 1.05e+03
peaks after deleting less than one fifth of global max:12
×10^9
raw data, starting:13
fitted Gauss locations:12
y
x
0
1
2
3
4
5
6
7
0
5
10
15
20
25

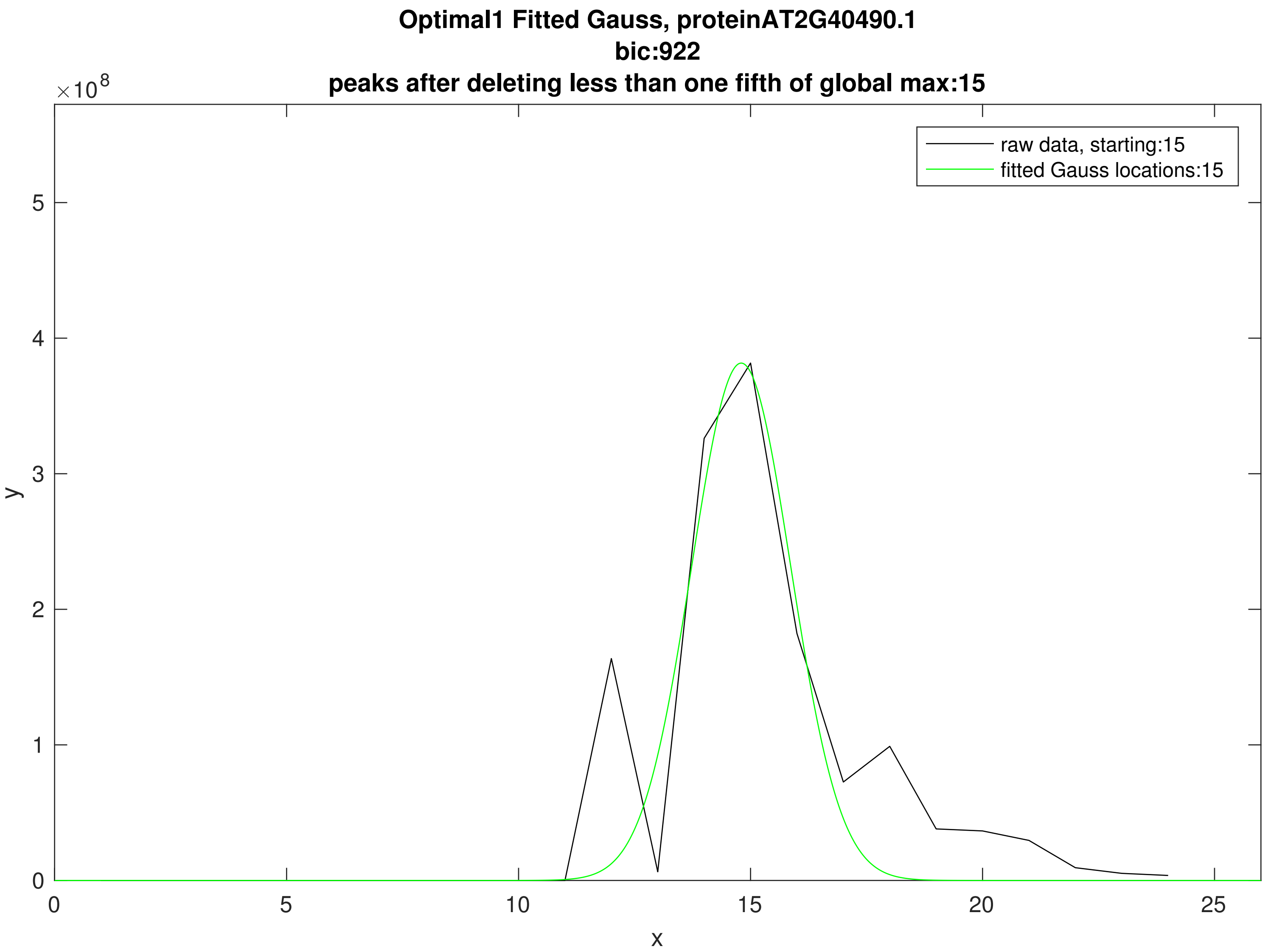

Optimal1 Fitted Gauss, proteinAT2G40490.1
bic:922
peaks after deleting less than one fifth of global max:15
×10^8
raw data, starting:15
fitted Gauss locations:15
y
x
0
1
2
3
4
5
0
5
10
15
20
25

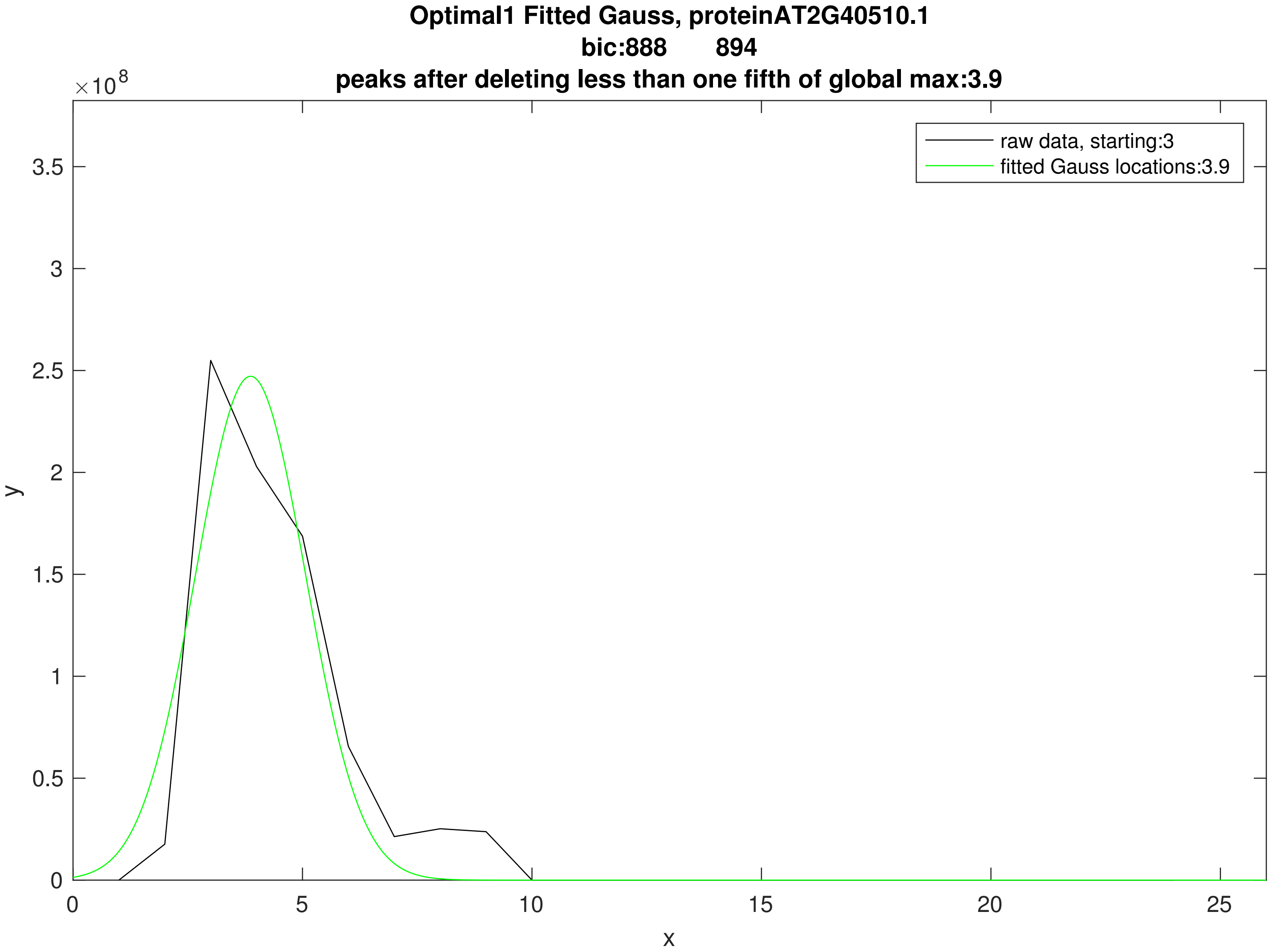

Optimal1 Fitted Gauss, proteinAT2G40510.1
bic:888 894
peaks after deleting less than one fifth of global max:3.9
×10^8
3.5
3
2.5
2
1.5
1
0.5
0
y
0
5
10
15
20
25
x
raw data, starting:3
fitted Gauss locations:3.9

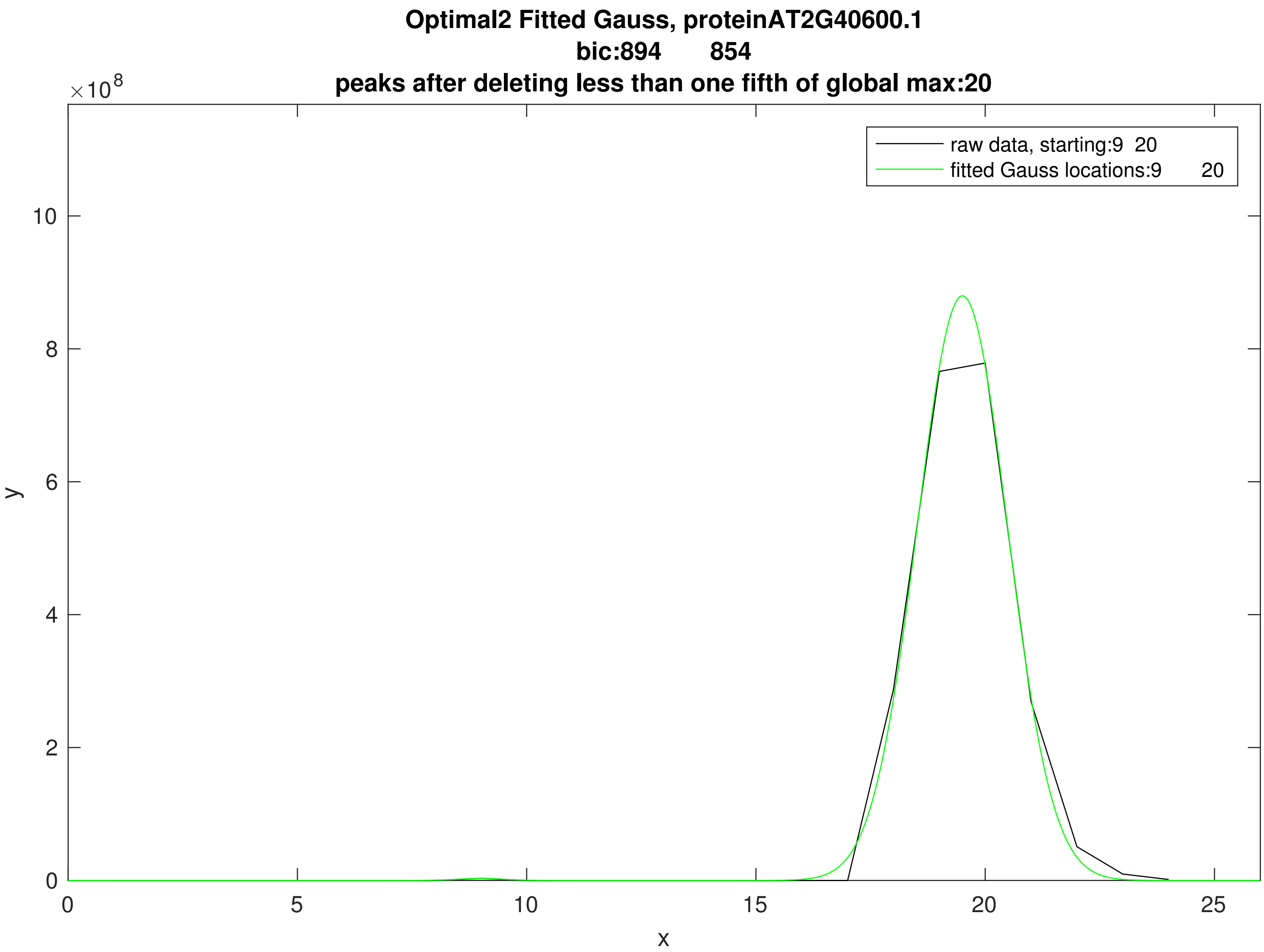

Optimal2 Fitted Gauss, proteinAT2G40600.1
bic:894 854
peaks after deleting less than one fifth of global max:20
×10^8
raw data, starting:9 20
fitted Gauss locations:9 20
10
8
6
4
2
0
y
0
5
10
15
20
25
x

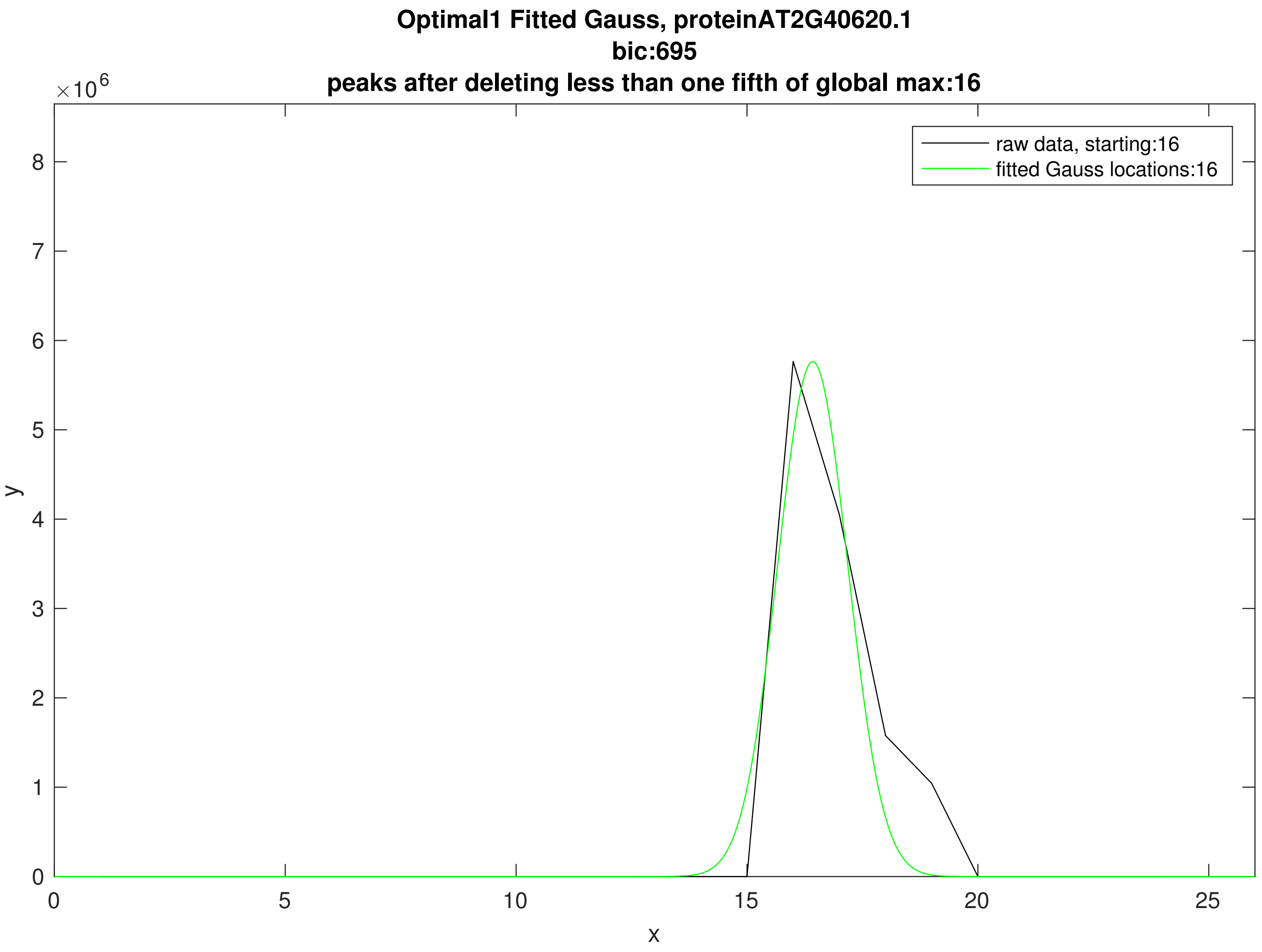

Optimal1 Fitted Gauss, proteinAT2G40620.1
bic:695
peaks after deleting less than one fifth of global max:16
×10^6
raw data, starting:16
fitted Gauss locations:16
y
x
0
1
2
3
4
5
6
7
8
0
5
10
15
20
25

**Optimal1 Fitted Gauss, proteinAT2G40660.1**
**bic:742**
**peaks after deleting less than one fifth of global max:4**

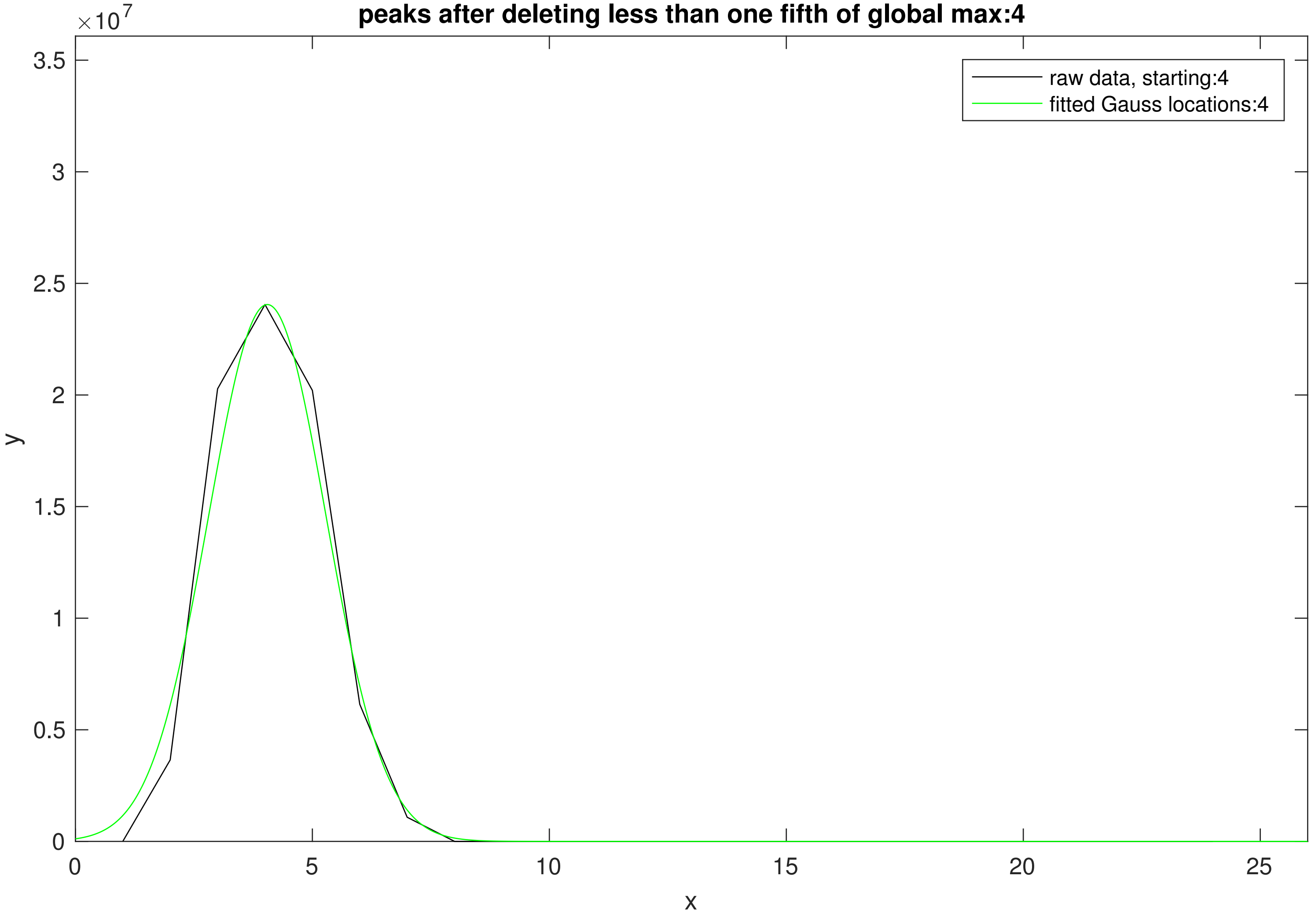

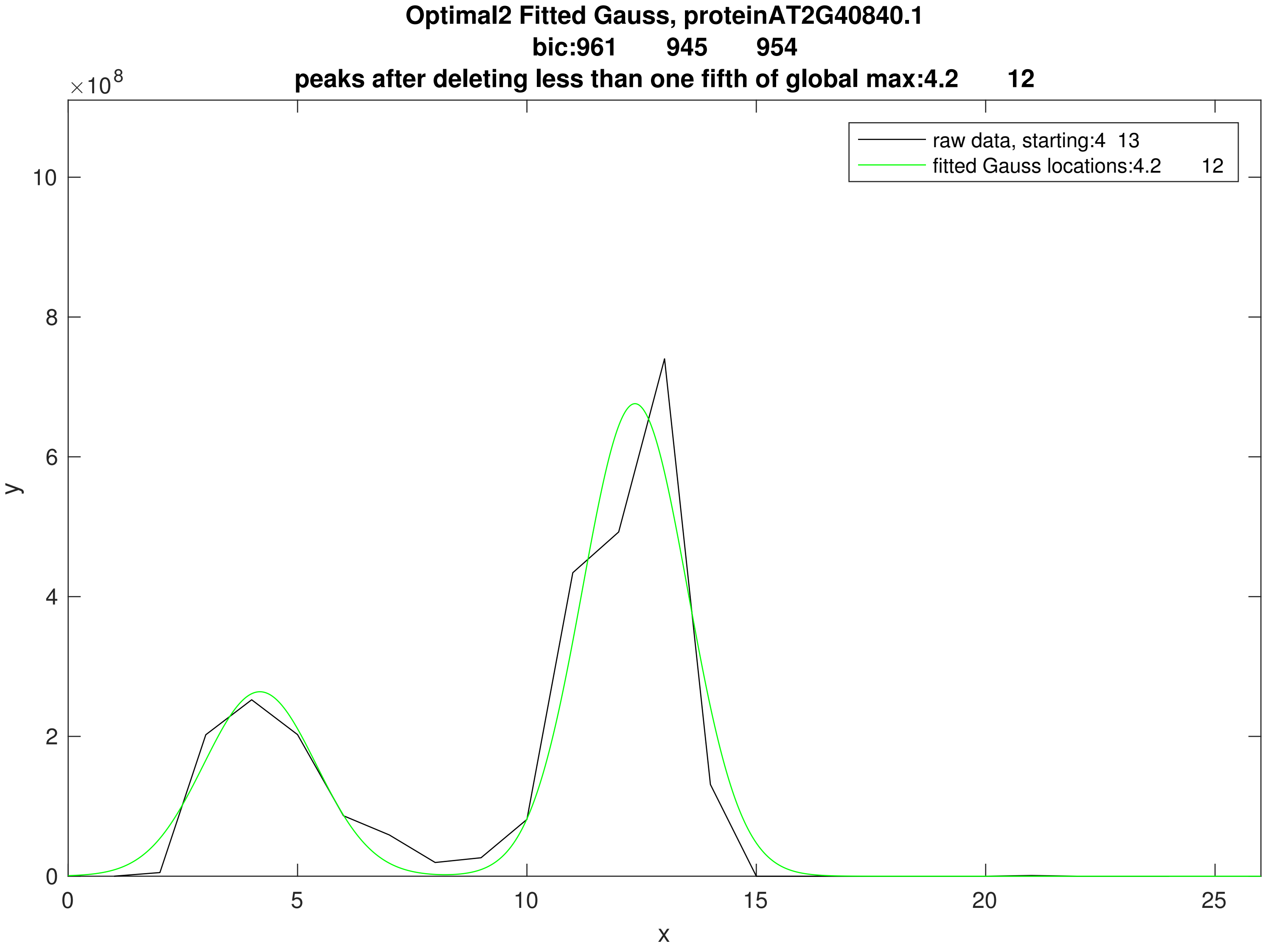
Optimal2 Fitted Gauss, proteinAT2G40840.1
bic:961 945 954
peaks after deleting less than one fifth of global max:4.2 12
×10^8
raw data, starting:4 13
fitted Gauss locations:4.2 12
y
x
0
2
4
6
8
10
0
5
10
15
20
25

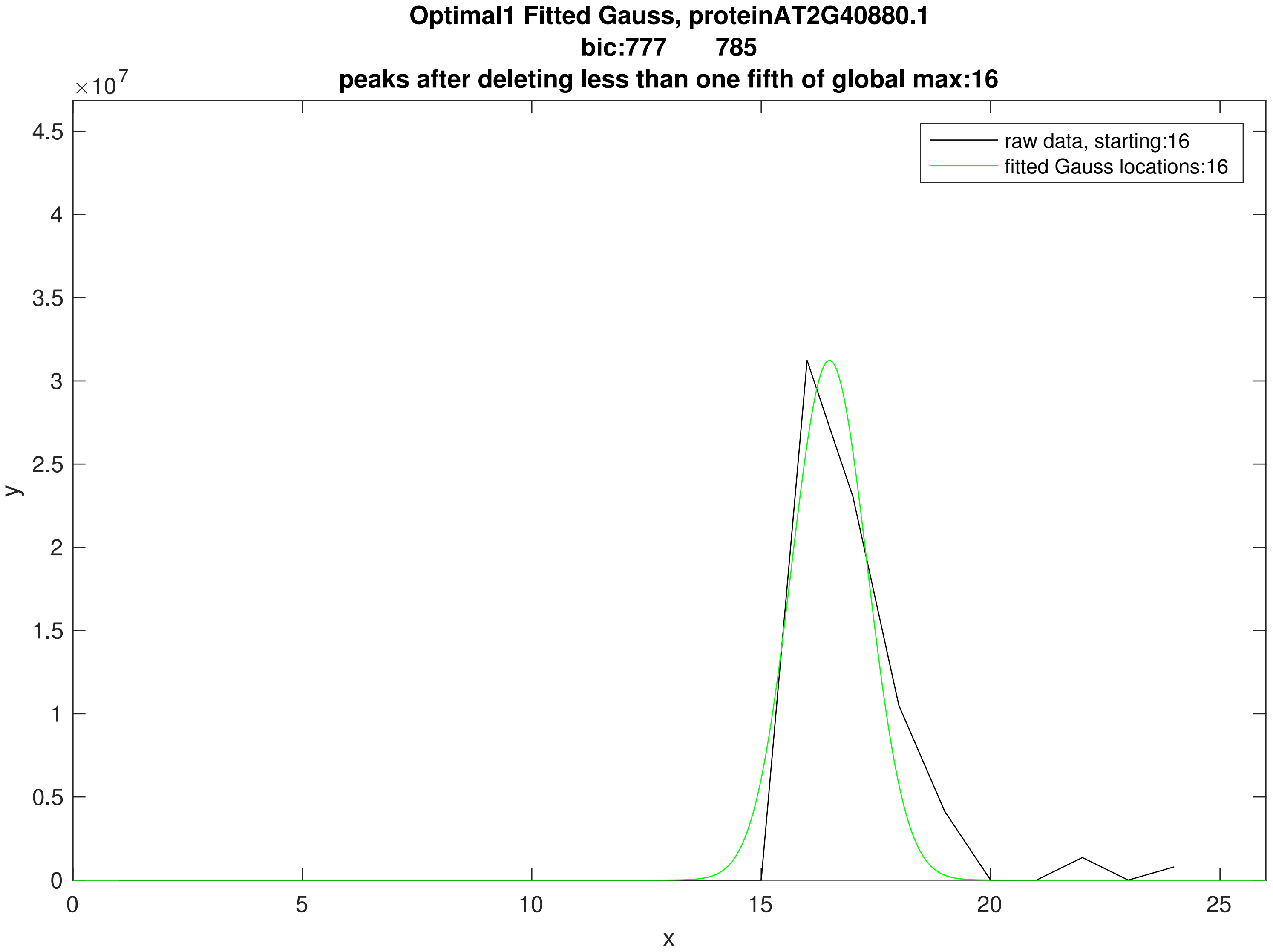

Optimal1 Fitted Gauss, proteinAT2G40880.1
bic:777 785
peaks after deleting less than one fifth of global max:16
×10^7
4.5
4
3.5
3
2.5
2
1.5
1
0.5
0
y
0
5
10
15
20
25
x
raw data, starting:16
fitted Gauss locations:16

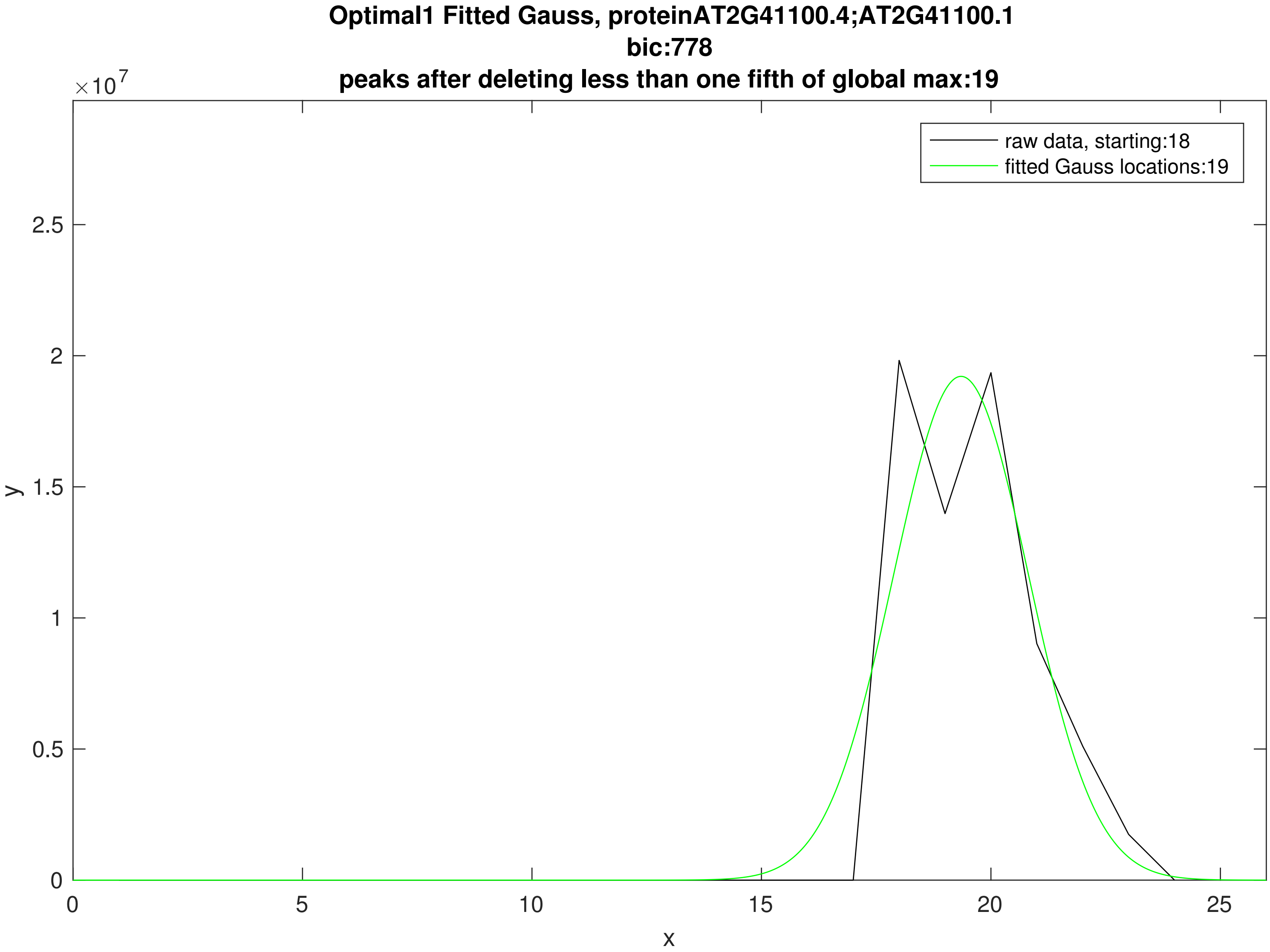
Optimal1 Fitted Gauss, proteinAT2G41100.4;AT2G41100.1
bic:778
peaks after deleting less than one fifth of global max:19
×10^7
raw data, starting:18
fitted Gauss locations:19
2.5
2
1.5
1
0.5
0
y
0
5
10
15
20
25
x

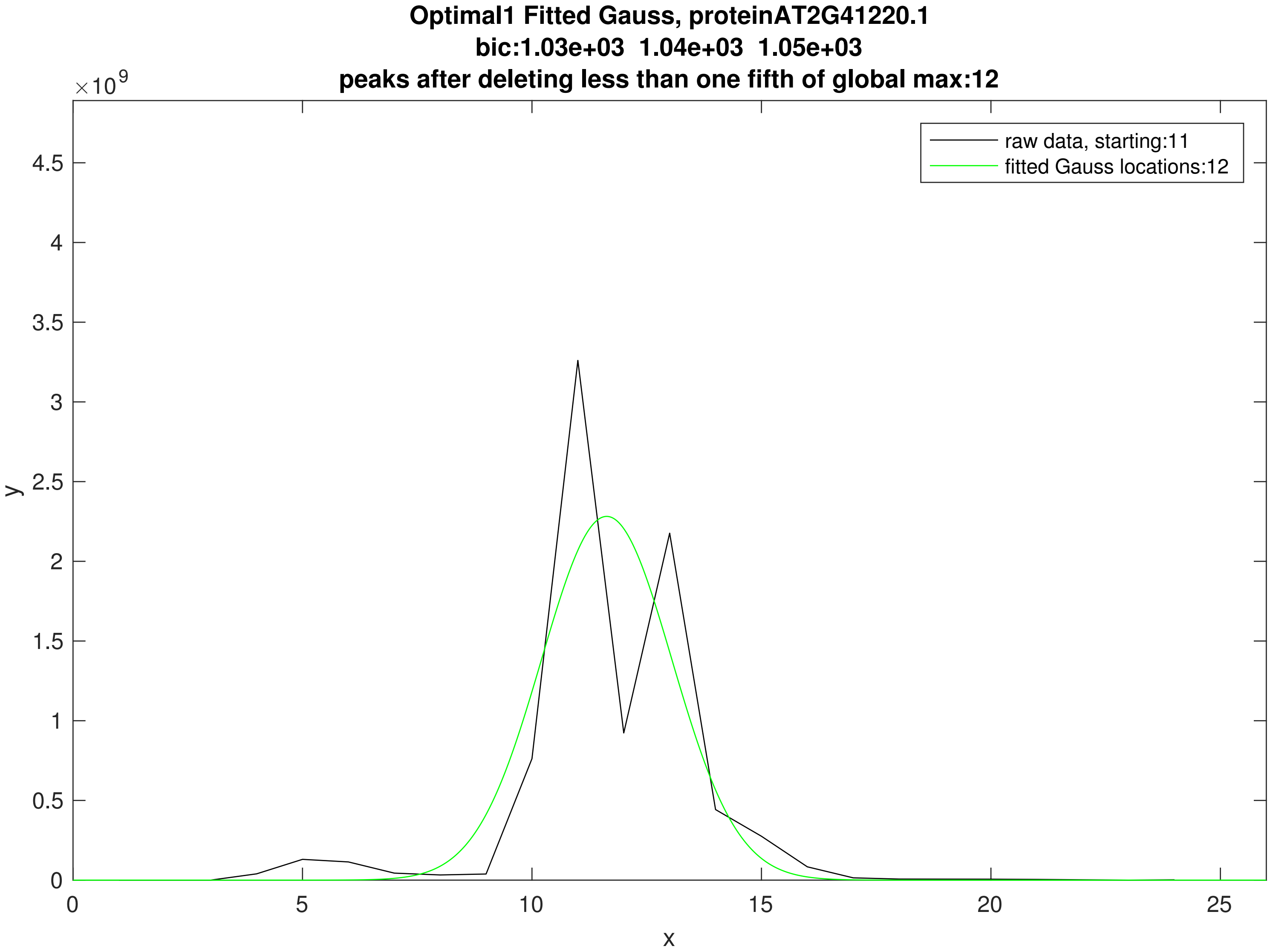

Optimal1 Fitted Gauss, proteinAT2G41220.1
bic:1.03e+03 1.04e+03 1.05e+03
peaks after deleting less than one fifth of global max:12
×10^9
4.5
4
3.5
3
2.5
2
1.5
1
0.5
0
y
0
5
10
15
20
25
x
raw data, starting:11
fitted Gauss locations:12

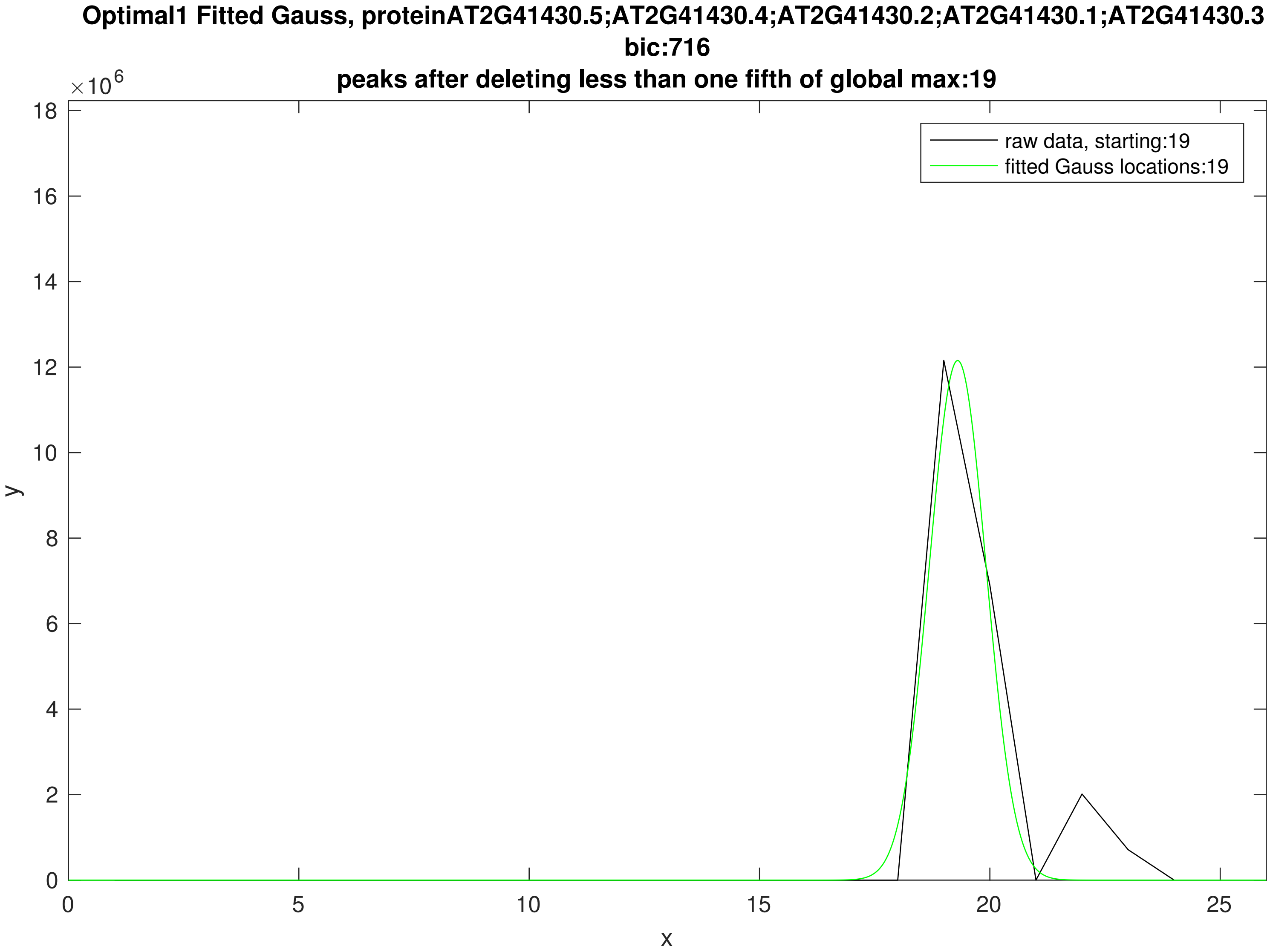
Optimal1 Fitted Gauss, proteinAT2G41430.5;AT2G41430.4;AT2G41430.2;AT2G41430.1;AT2G41430.3
bic:716
peaks after deleting less than one fifth of global max:19
×10^6
raw data, starting:19
fitted Gauss locations:19
y
x
0
2
4
6
8
10
12
14
16
18
5
15
20
25

**Optimal1 Fitted Gauss, proteinAT2G41530.1**
**bic:981 990**
**peaks after deleting less than one fifth of global max:15**

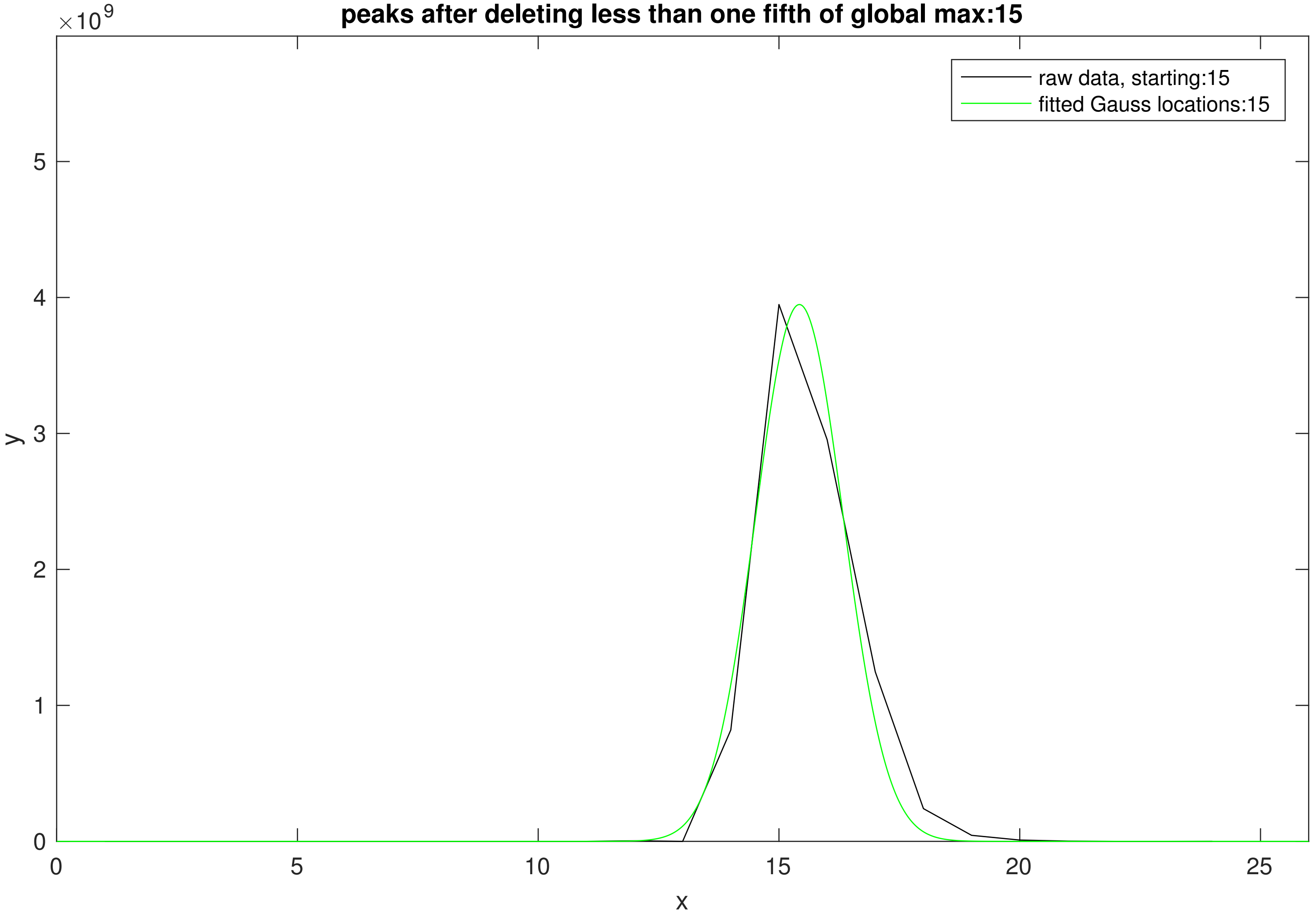

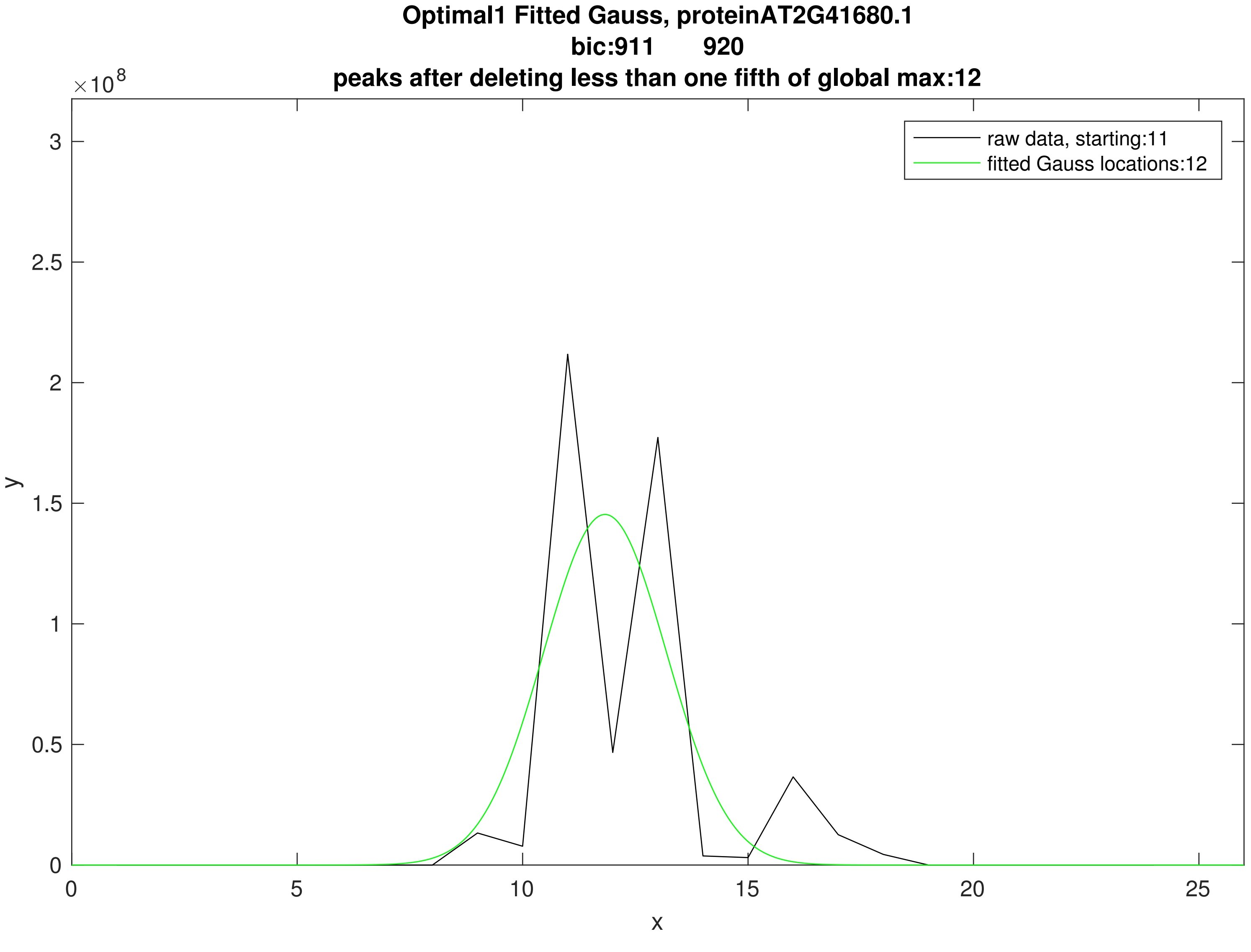
Optimal1 Fitted Gauss, proteinAT2G41680.1
bic:911 920
peaks after deleting less than one fifth of global max:12
×10^8
raw data, starting:11
fitted Gauss locations:12
y
x
0
0.5
1
1.5
2
2.5
3
0
5
10
15
20
25

#### Optimal2 Fitted Gauss, proteinAT2G41740.1
#### bic:870 870
#### peaks after deleting less than one fifth of global max:16

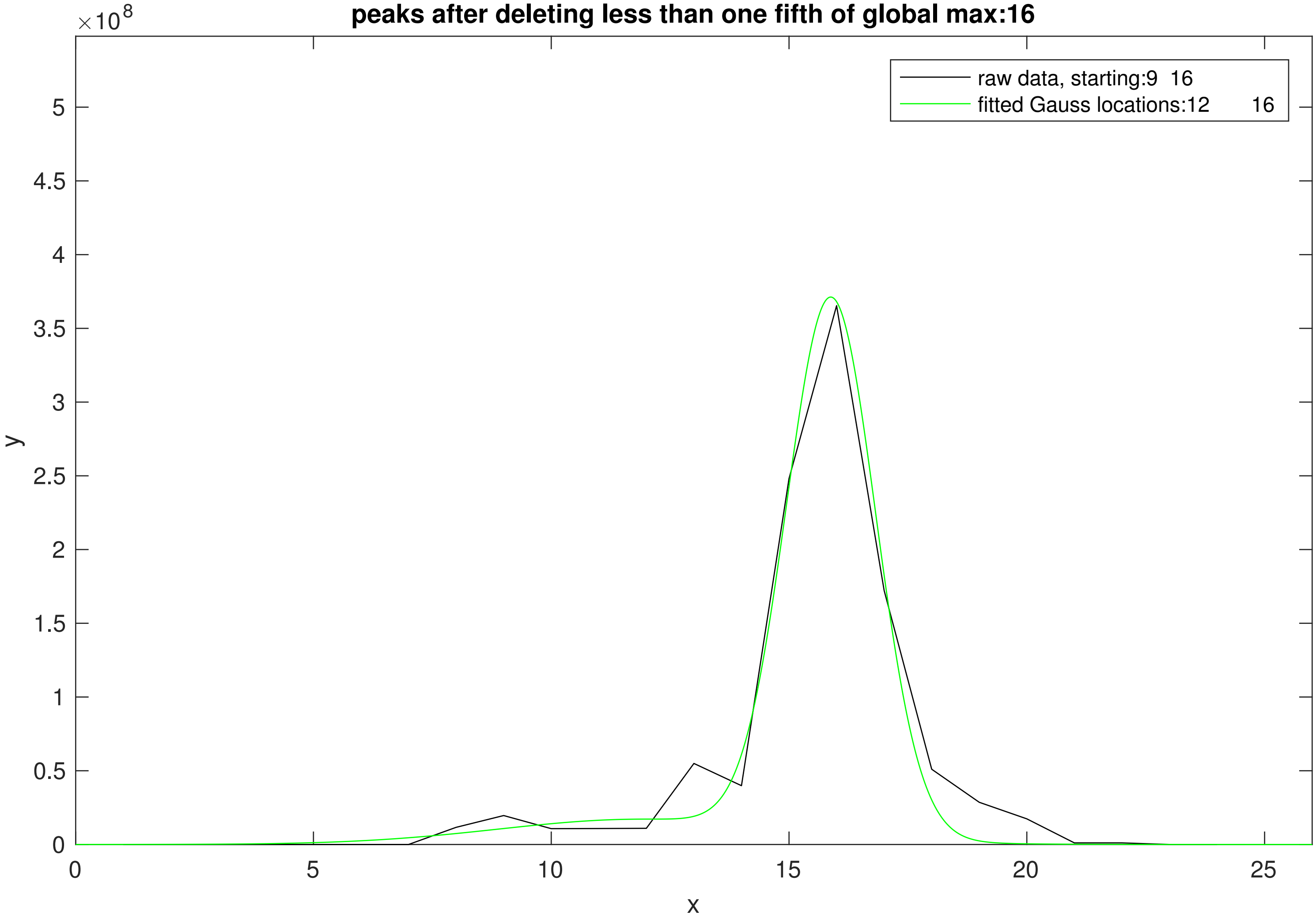

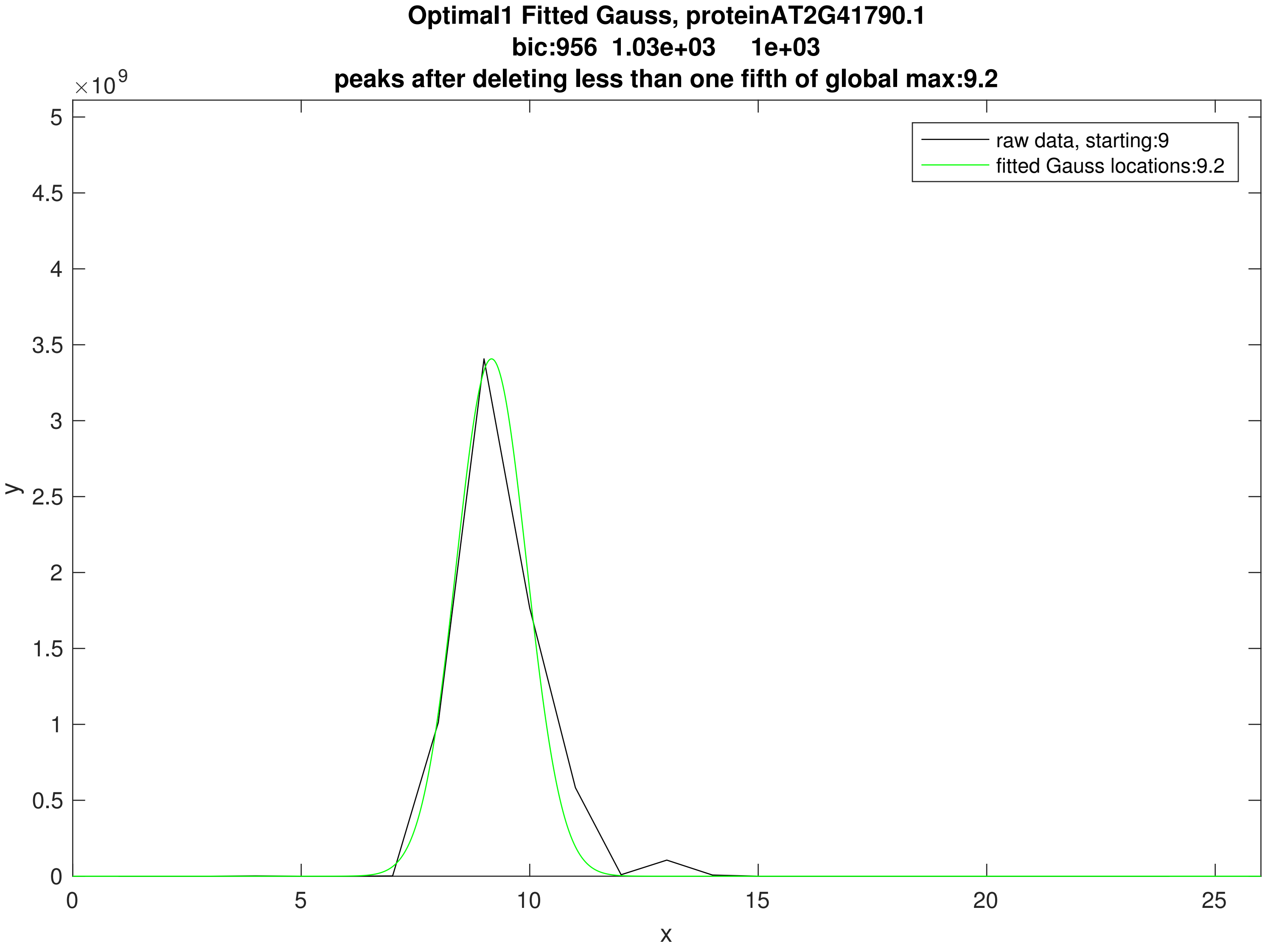
Optimal1 Fitted Gauss, proteinAT2G41790.1
bic:956 1.03e+03 1e+03
peaks after deleting less than one fifth of global max:9.2
×10^9
y
x
raw data, starting:9
fitted Gauss locations:9.2

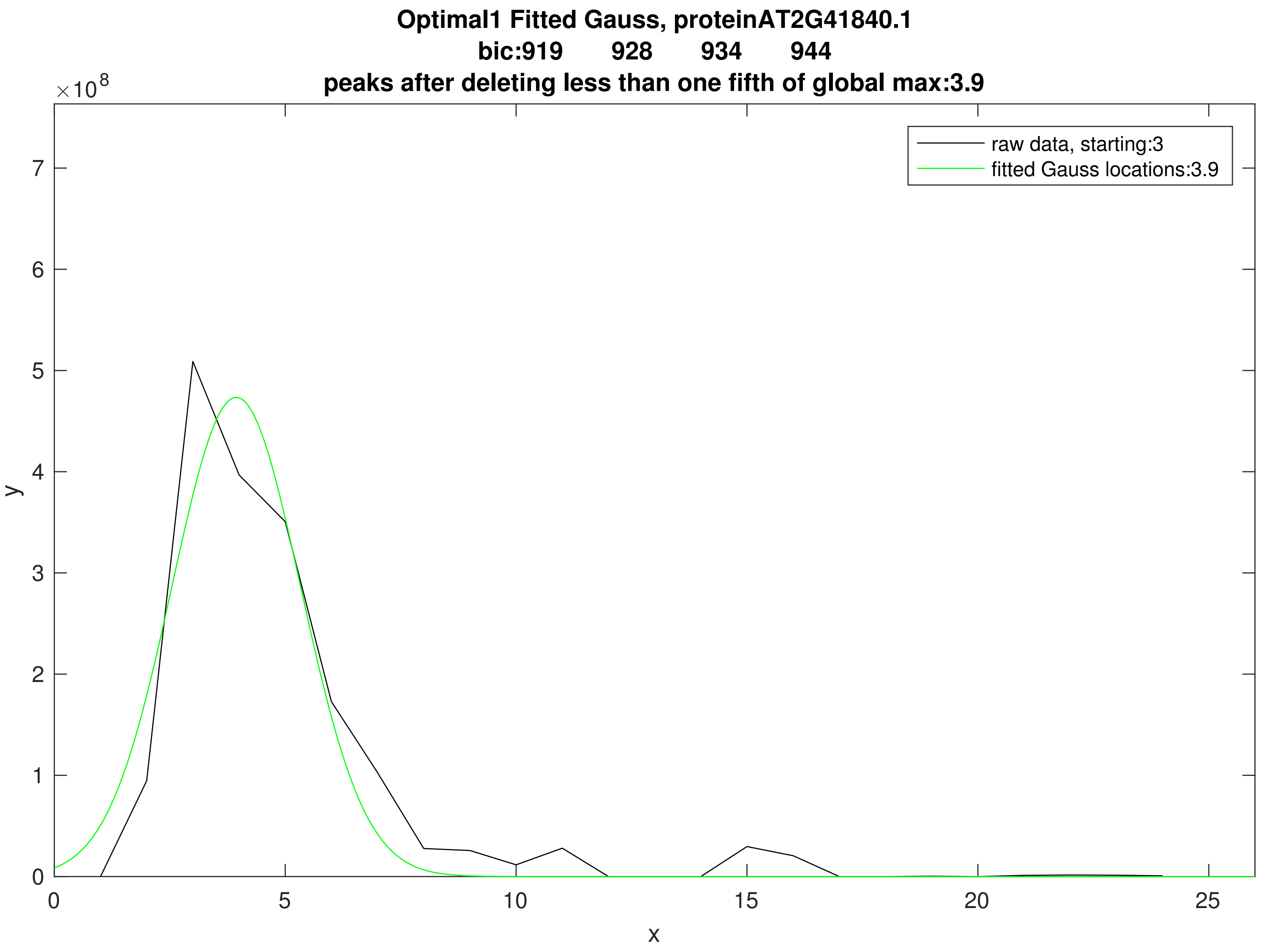
Optimal1 Fitted Gauss, proteinAT2G41840.1
bic:919 928 934 944
peaks after deleting less than one fifth of global max:3.9
×10^8
raw data, starting:3
fitted Gauss locations:3.9
y
x
0
1
2
3
4
5
6
7
0
5
10
15
20
25

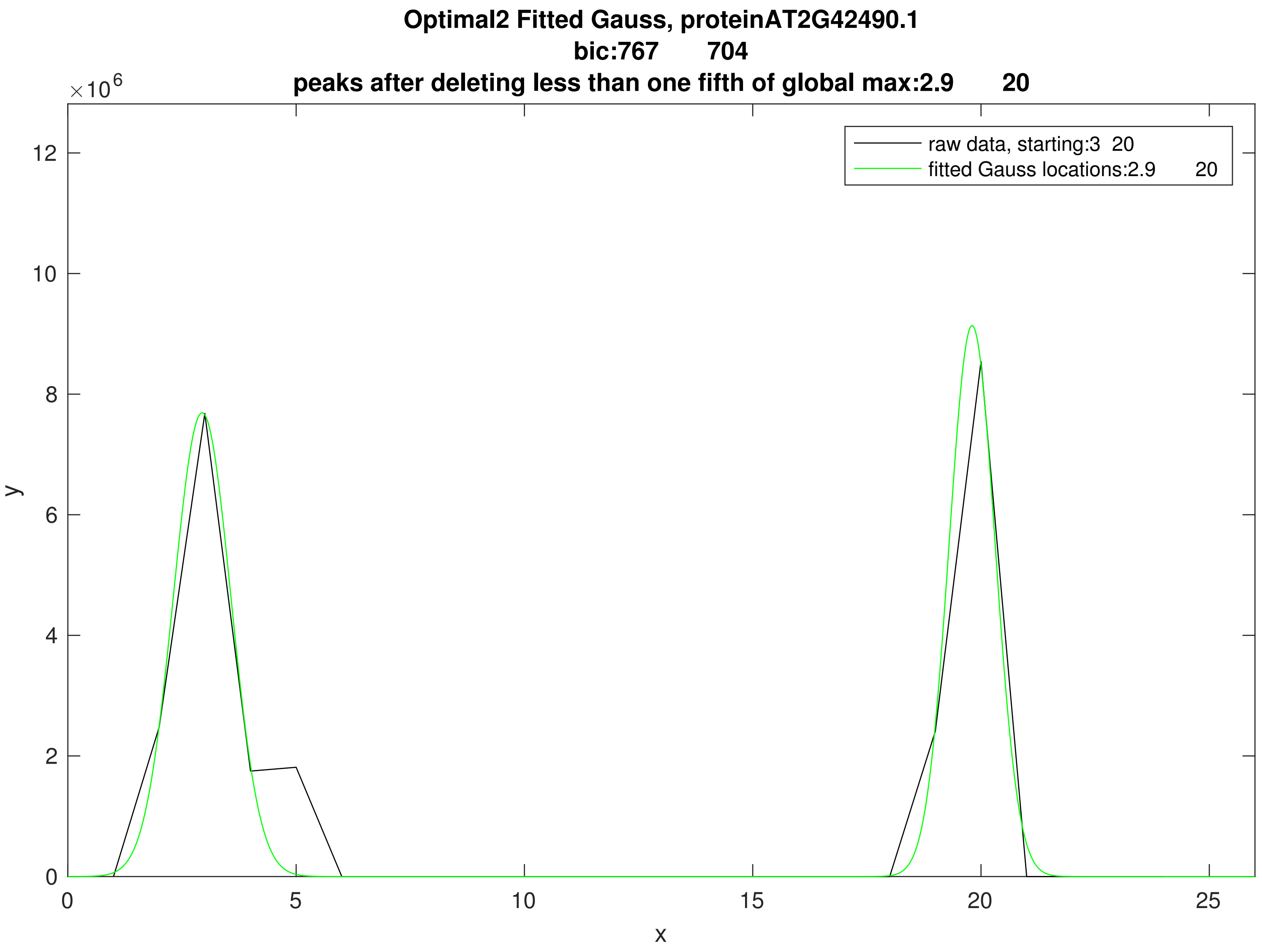

Optimal2 Fitted Gauss, proteinAT2G42490.1
bic:767 704
peaks after deleting less than one fifth of global max:2.9 20
×10^6
raw data, starting:3 20
fitted Gauss locations:2.9 20
12
10
8
6
4
2
0
y
0
5
10
15
20
25
x

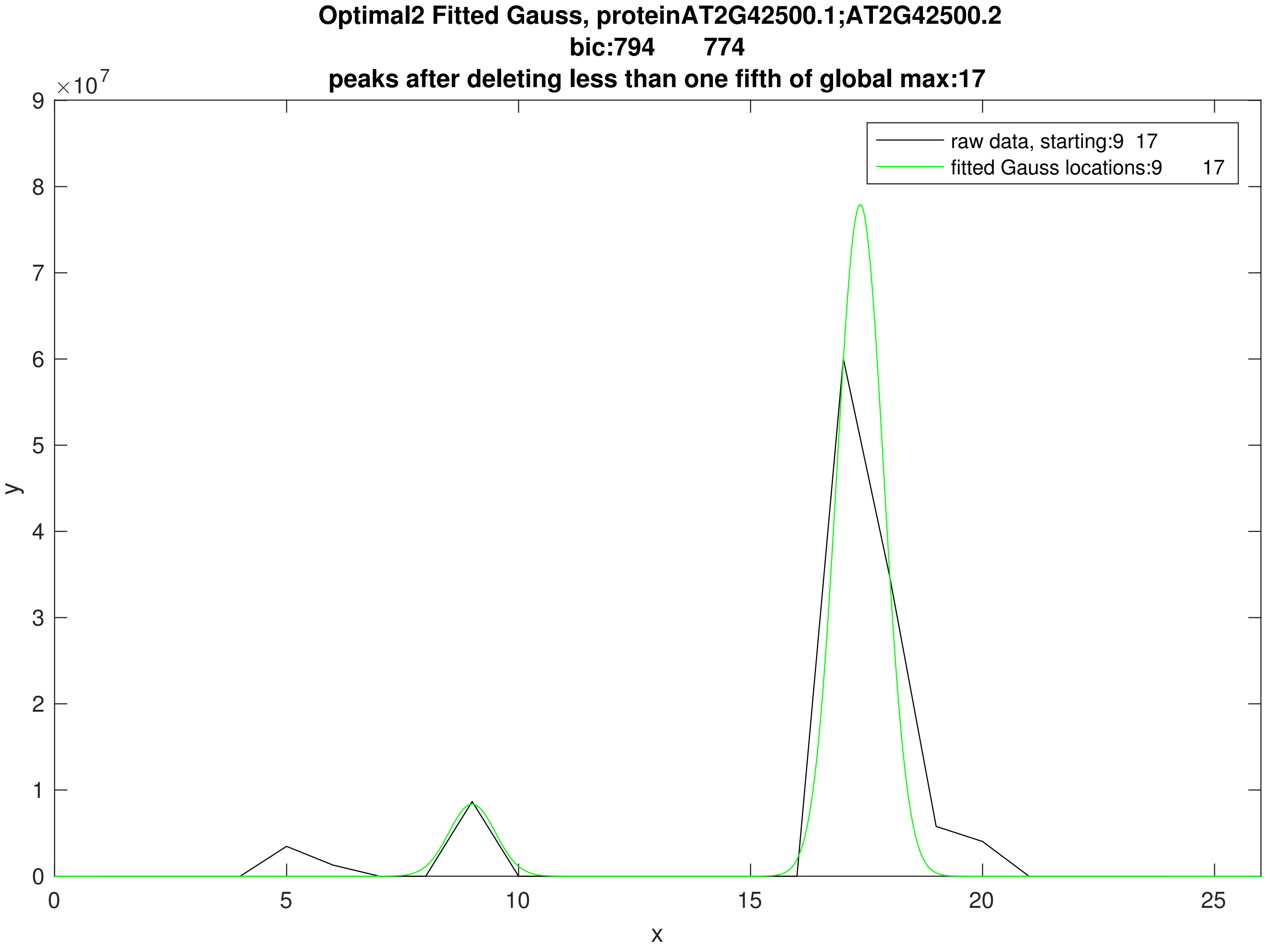

Optimal2 Fitted Gauss, proteinAT2G42500.1;AT2G42500.2
bic:794 774
peaks after deleting less than one fifth of global max:17
×10^7
raw data, starting:9 17
fitted Gauss locations:9 17
y
x
0
1
2
3
4
5
6
7
8
9
0
5
10
15
20
25

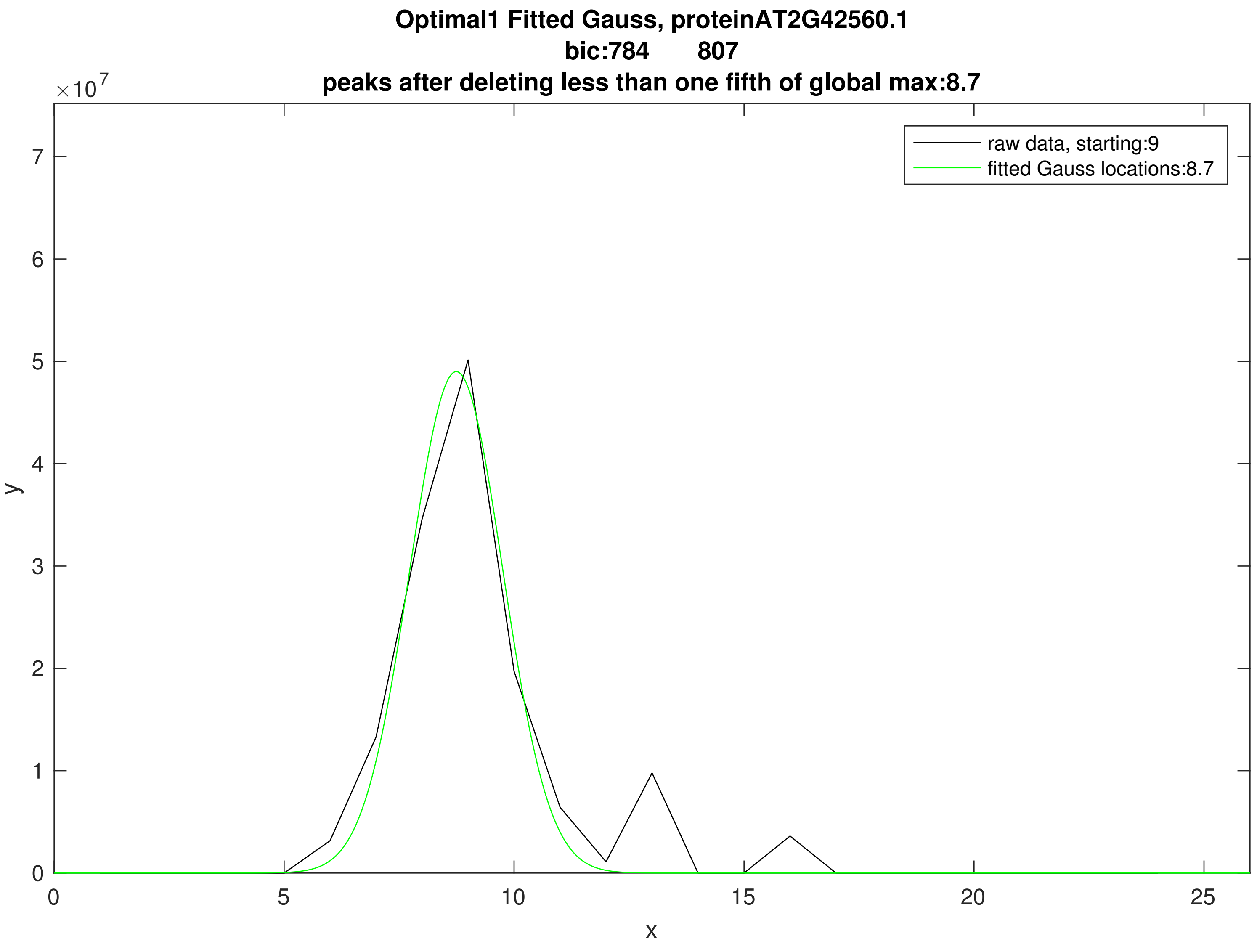
Optimal1 Fitted Gauss, proteinAT2G42560.1
bic:784 807
peaks after deleting less than one fifth of global max:8.7
×10^7
raw data, starting:9
fitted Gauss locations:8.7
y
x
0
1
2
3
4
5
6
7
0
5
10
15
20
25

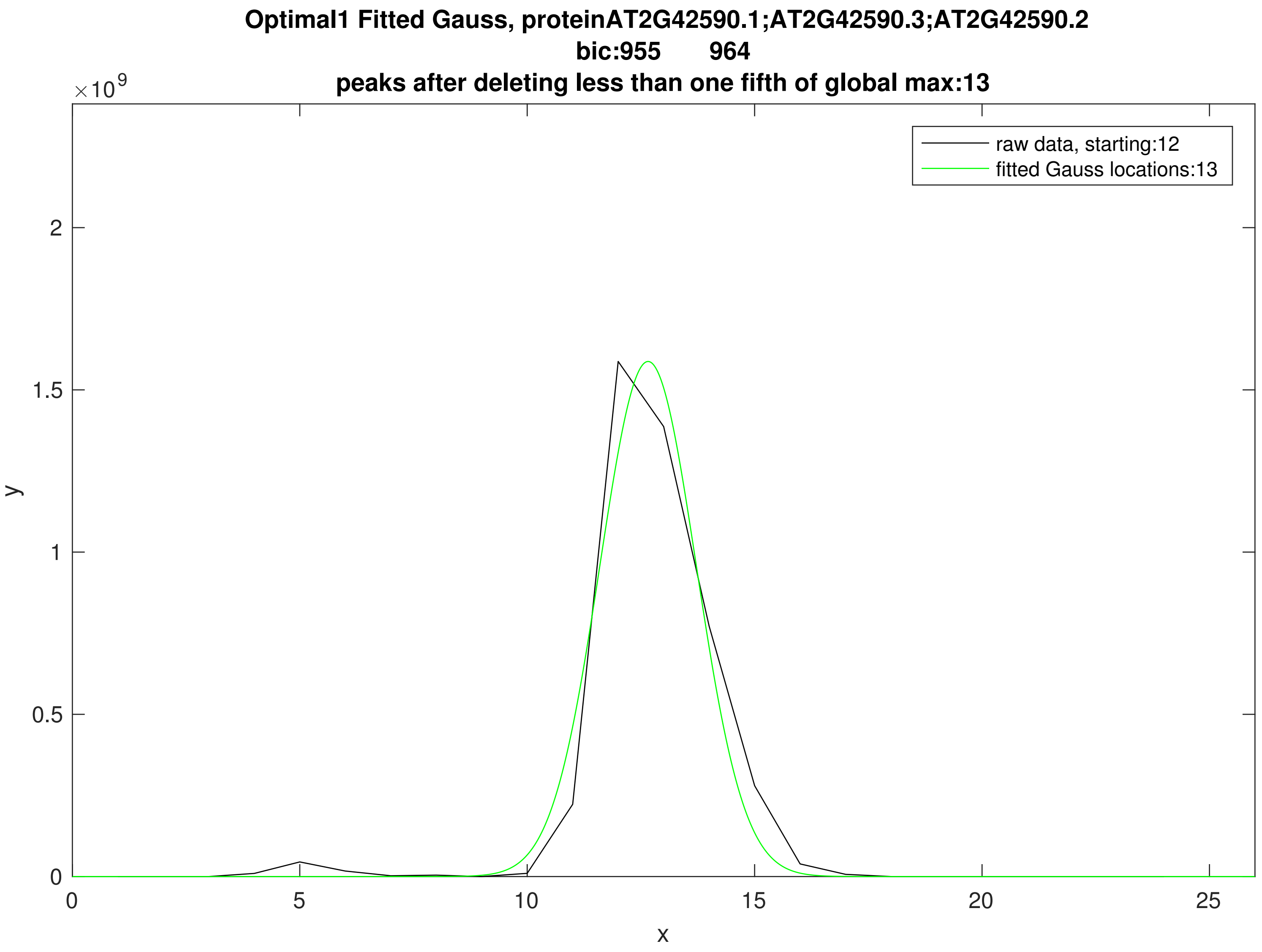

Optimal1 Fitted Gauss, proteinAT2G42590.1;AT2G42590.3;AT2G42590.2
bic:955 964
peaks after deleting less than one fifth of global max:13
×10^9
raw data, starting:12
fitted Gauss locations:13
y
x
0
0.5
1
1.5
2
0
5
10
15
20
25

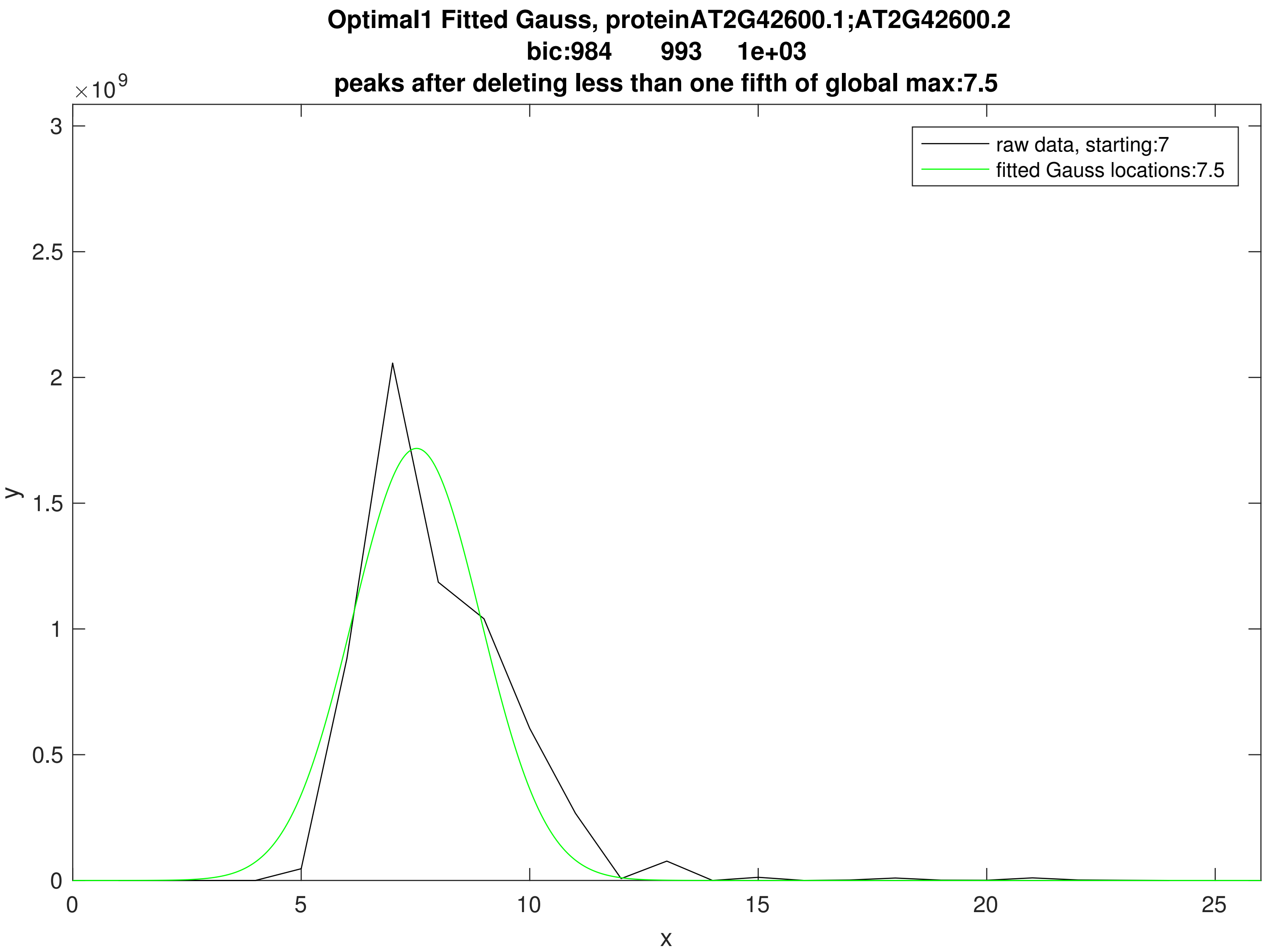
Optimal1 Fitted Gauss, proteinAT2G42600.1;AT2G42600.2
bic:984 993 1e+03
peaks after deleting less than one fifth of global max:7.5
×10^9
3
2.5
2
1.5
1
0.5
0
y
0
5
10
15
20
25
x
raw data, starting:7
fitted Gauss locations:7.5

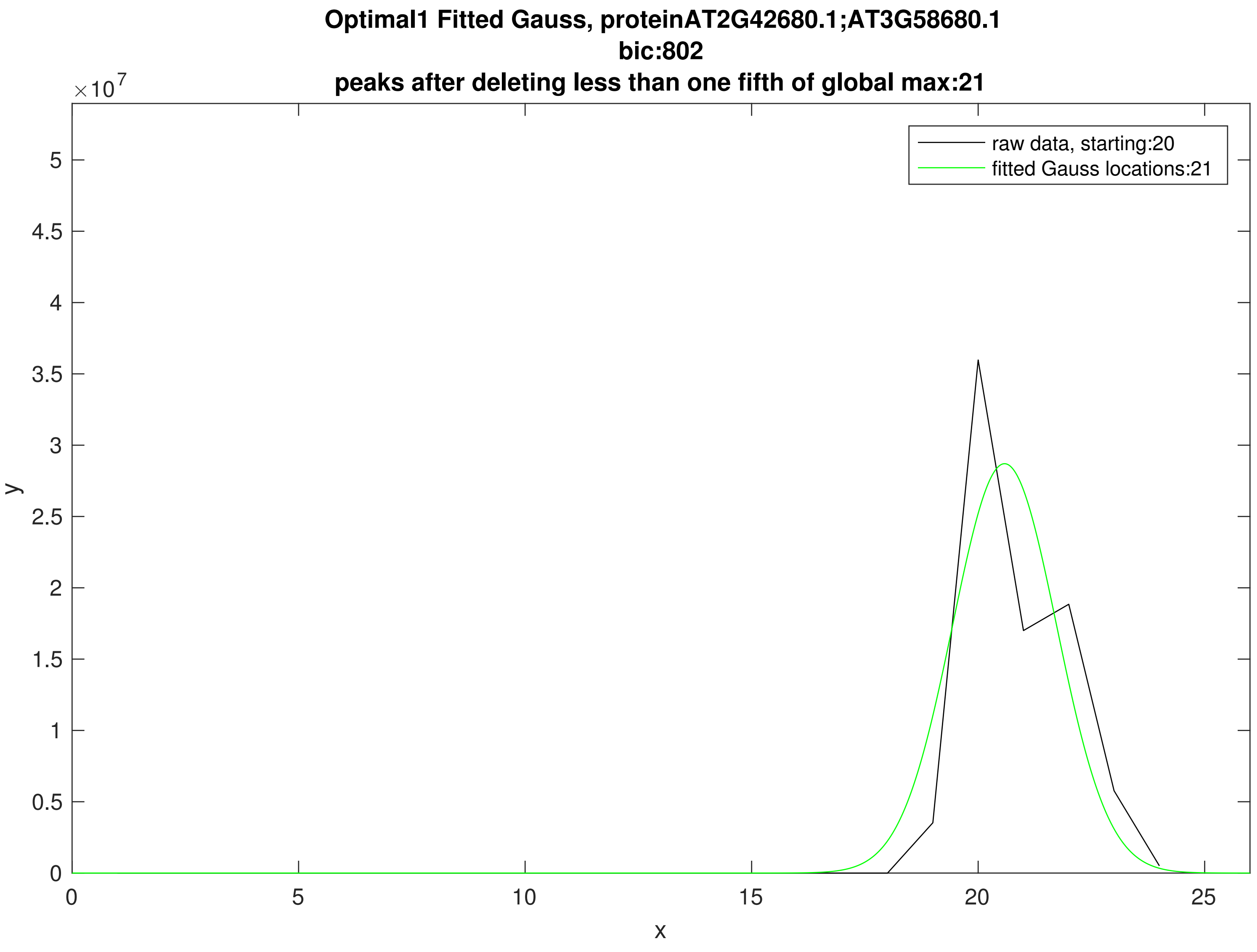
Optimal1 Fitted Gauss, proteinAT2G42680.1;AT3G58680.1
bic:802
peaks after deleting less than one fifth of global max:21
×10^7
raw data, starting:20
fitted Gauss locations:21
y
x
0
0.5
1
1.5
2
2.5
3
3.5
4
4.5
5
0
5
10
15
20
25

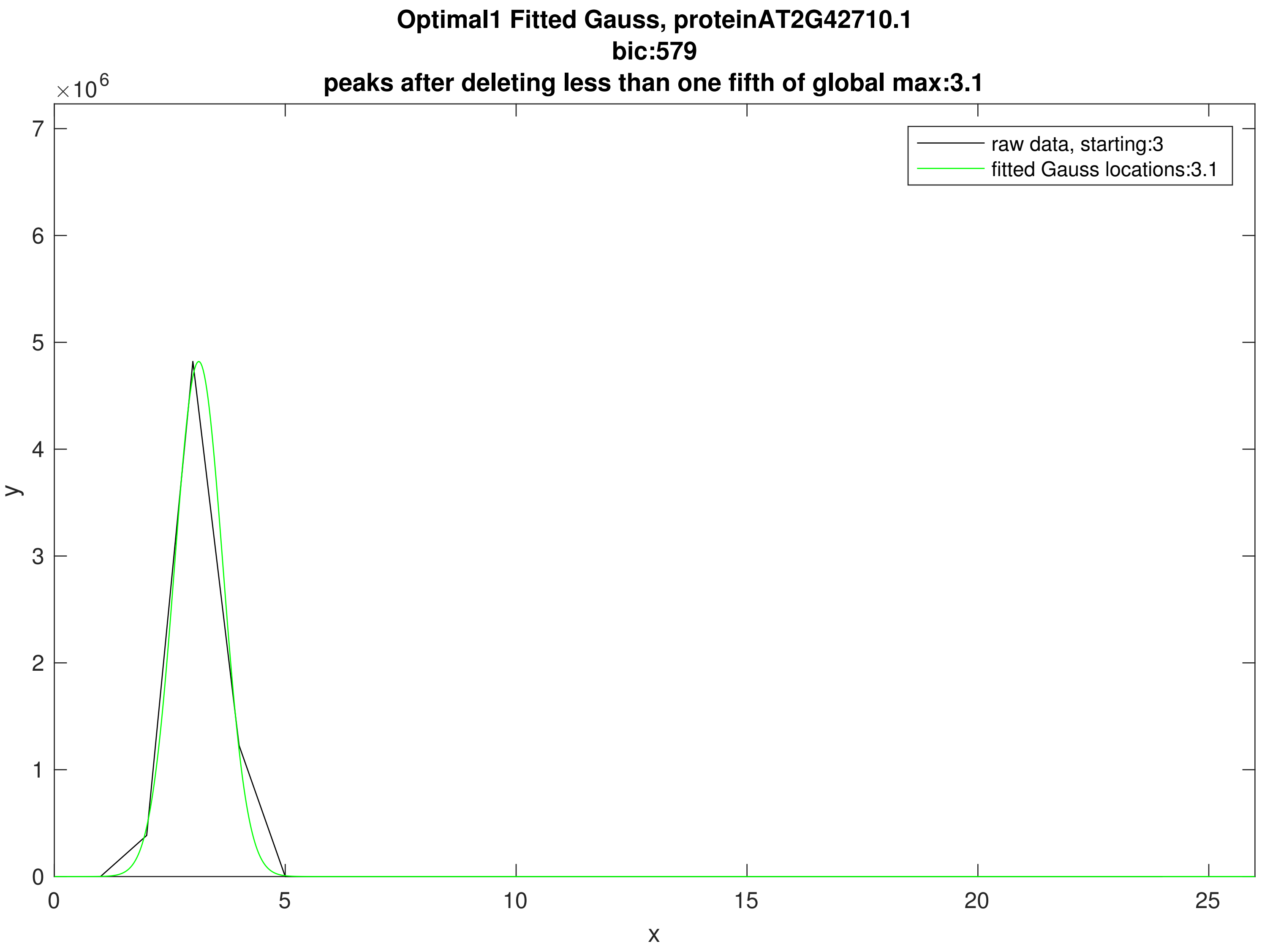

Optimal1 Fitted Gauss, proteinAT2G42710.1
bic:579
peaks after deleting less than one fifth of global max:3.1
×10^6
raw data, starting:3
fitted Gauss locations:3.1
y
x

**Optimal1 Fitted Gauss, proteinAT2G42790.1**
**bic:1.04e+03  1.05e+03  1.06e+03**
**peaks after deleting less than one fifth of global max:13**

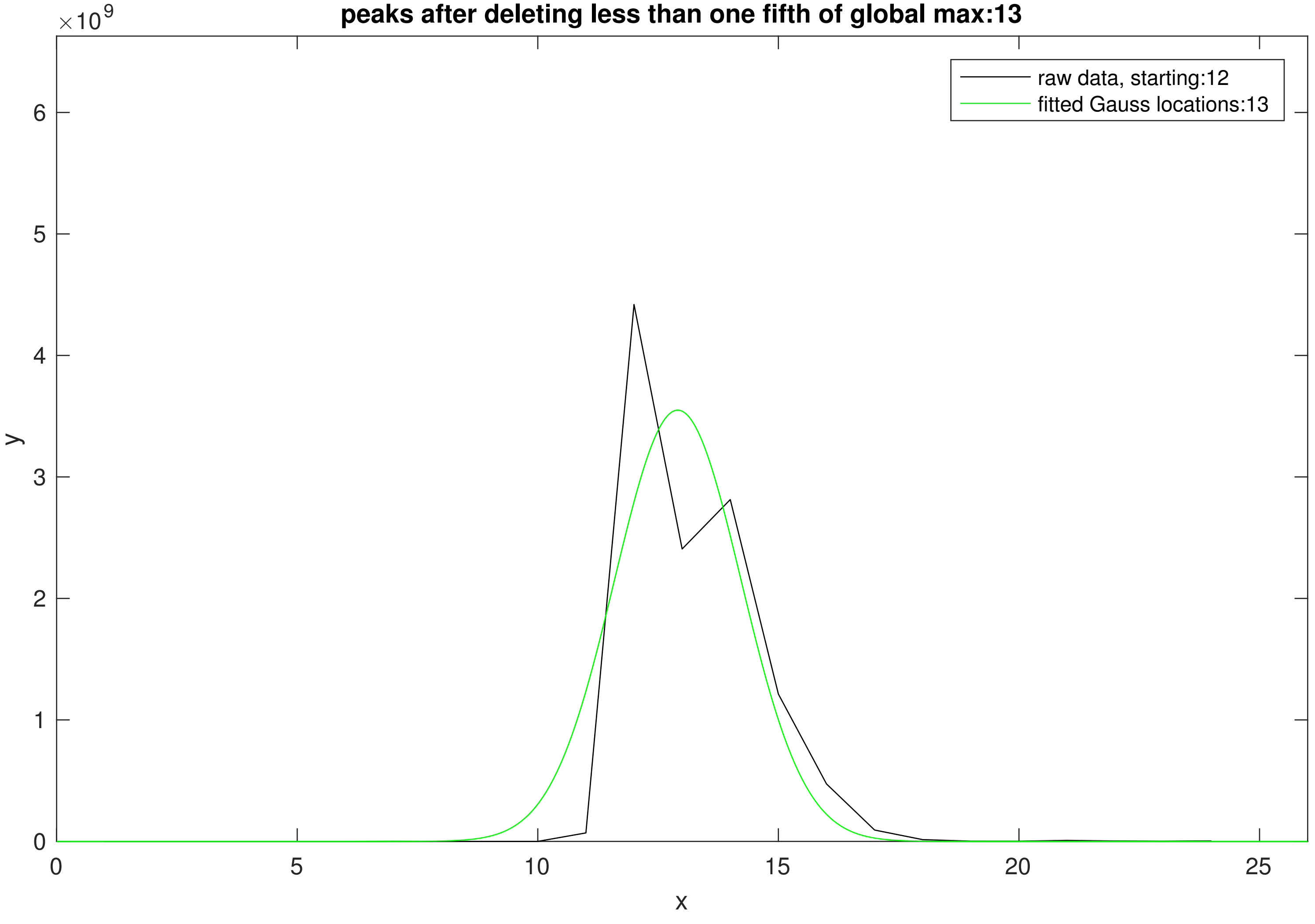

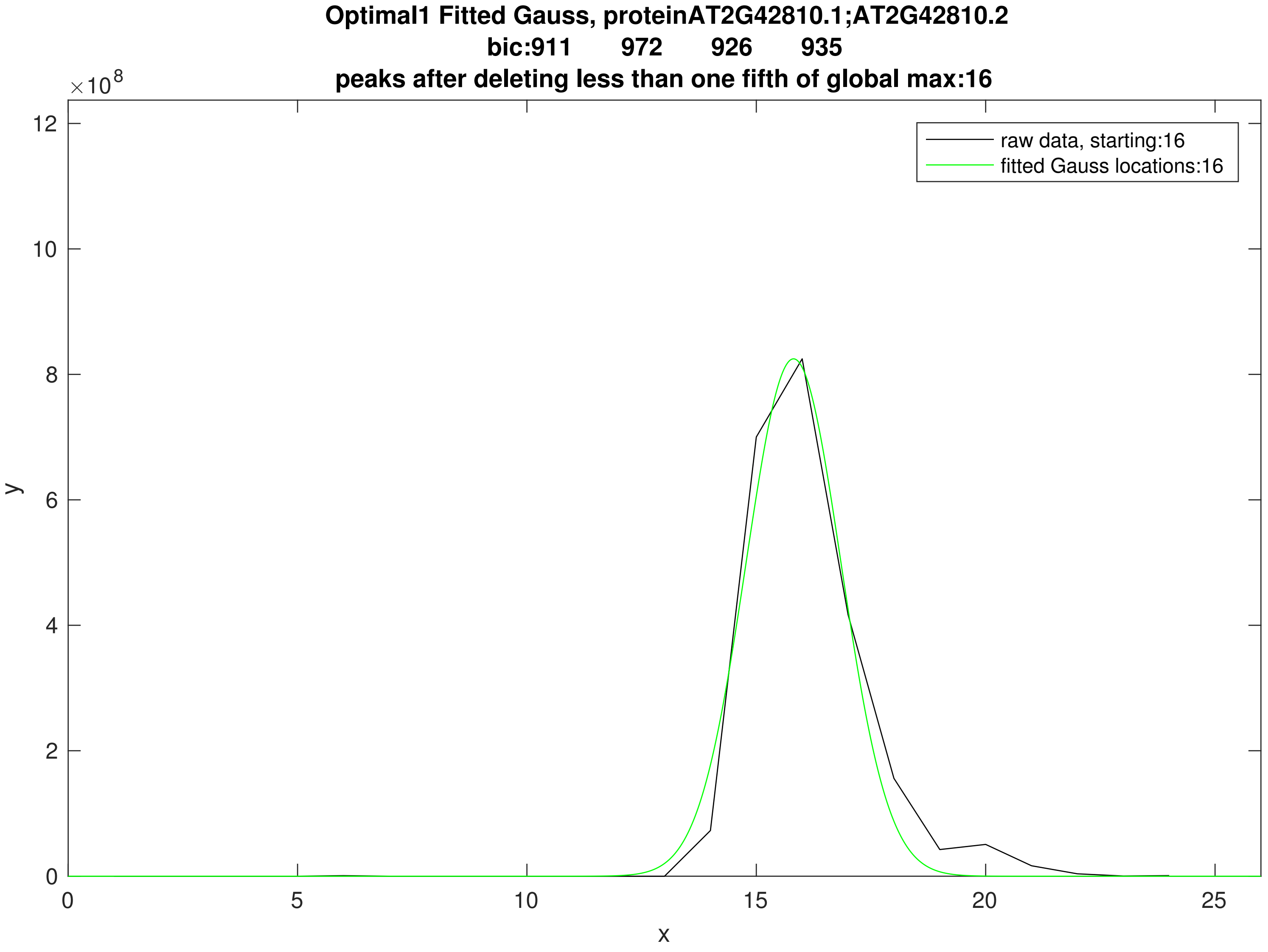

Optimal1 Fitted Gauss, proteinAT2G42810.1;AT2G42810.2
bic:911 972 926 935
peaks after deleting less than one fifth of global max:16
×10^8
raw data, starting:16
fitted Gauss locations:16
y
x
0
2
4
6
8
10
12
0
5
10
15
20
25

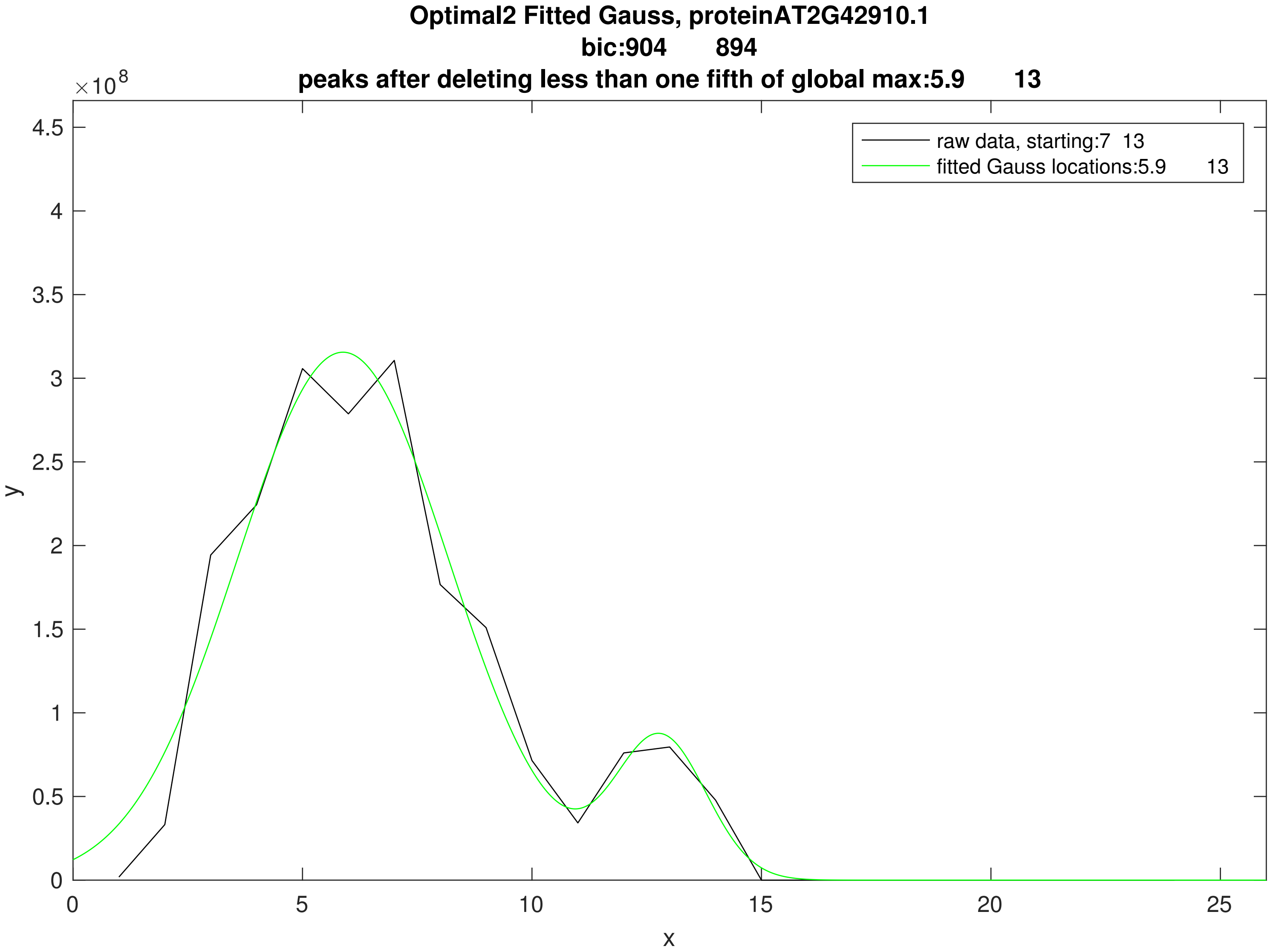

Optimal2 Fitted Gauss, proteinAT2G42910.1
bic:904 894
peaks after deleting less than one fifth of global max:5.9 13
×10^8
4.5
4
3.5
3
2.5
2
1.5
1
0.5
0
y
raw data, starting:7 13
fitted Gauss locations:5.9 13
0
5
10
15
20
25
x

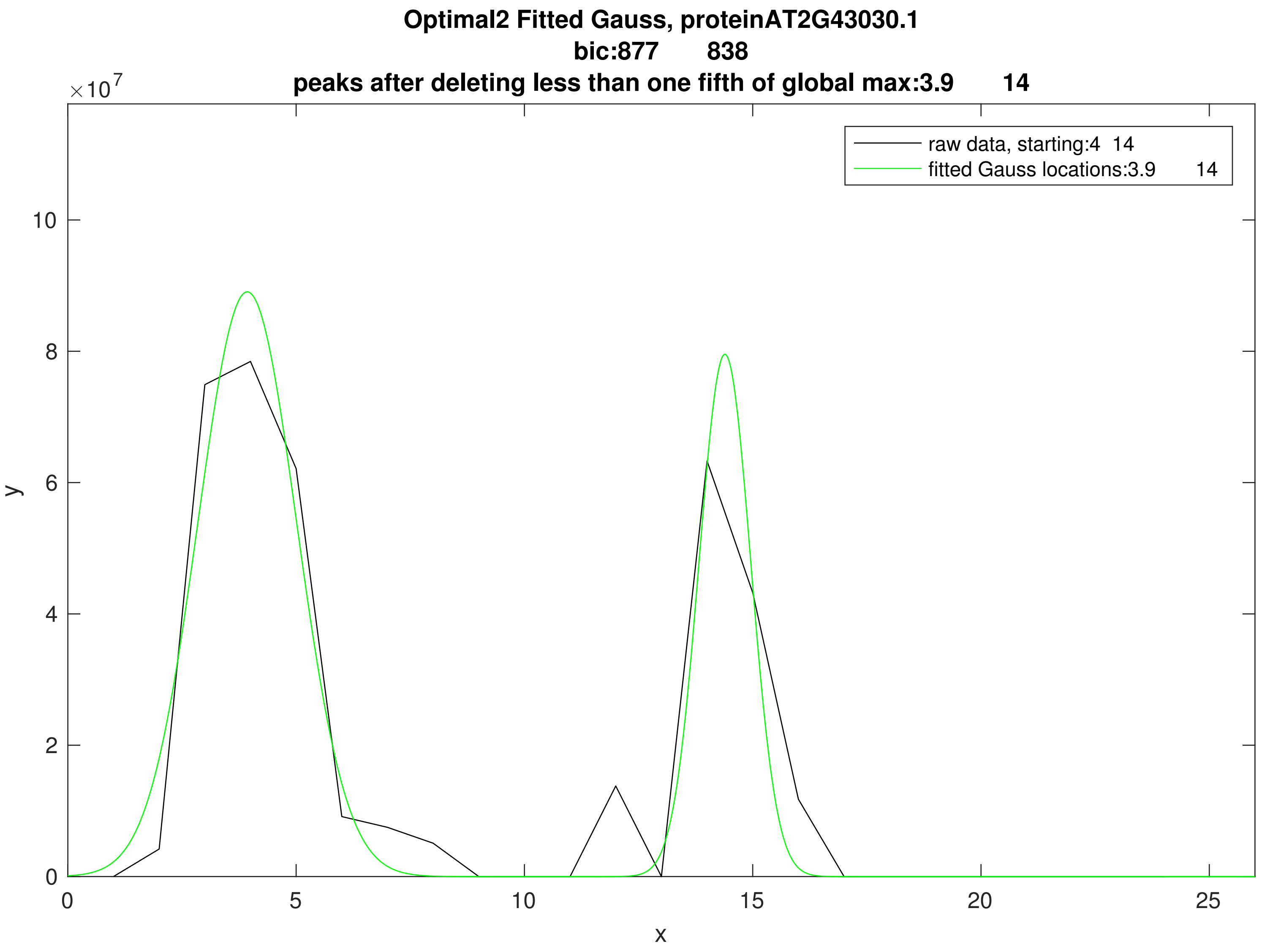
Optimal2 Fitted Gauss, proteinAT2G43030.1
bic:877 838
peaks after deleting less than one fifth of global max:3.9 14
×10^7
raw data, starting:4 14
fitted Gauss locations:3.9 14
y
x
0
2
4
6
8
10
0
5
10
15
20
25

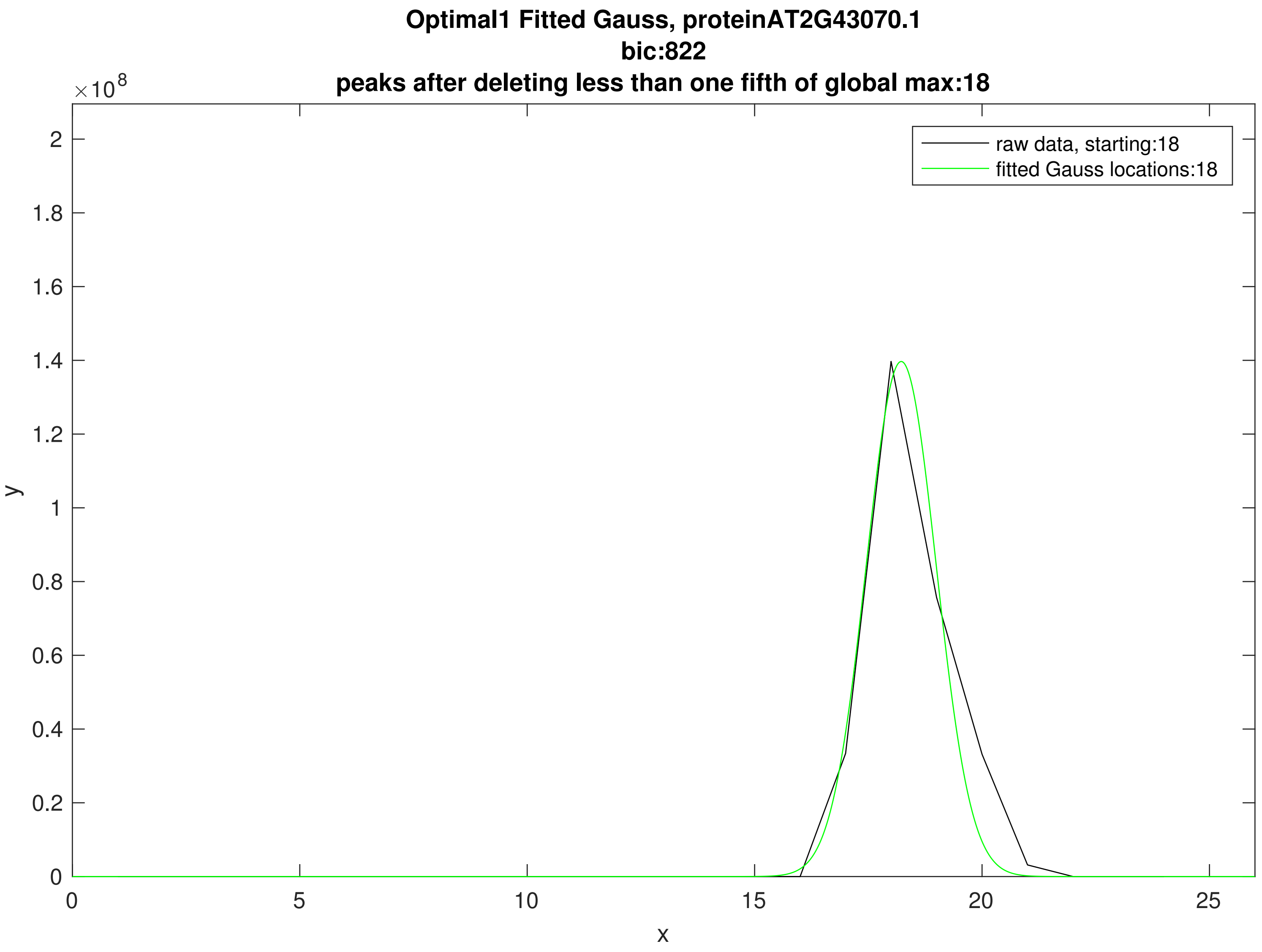

Optimal1 Fitted Gauss, proteinAT2G43070.1
bic:822
peaks after deleting less than one fifth of global max:18
×10^8
2
1.8
1.6
1.4
1.2
1
0.8
0.6
0.4
0.2
0
y
0
5
10
15
20
25
x
raw data, starting:18
fitted Gauss locations:18

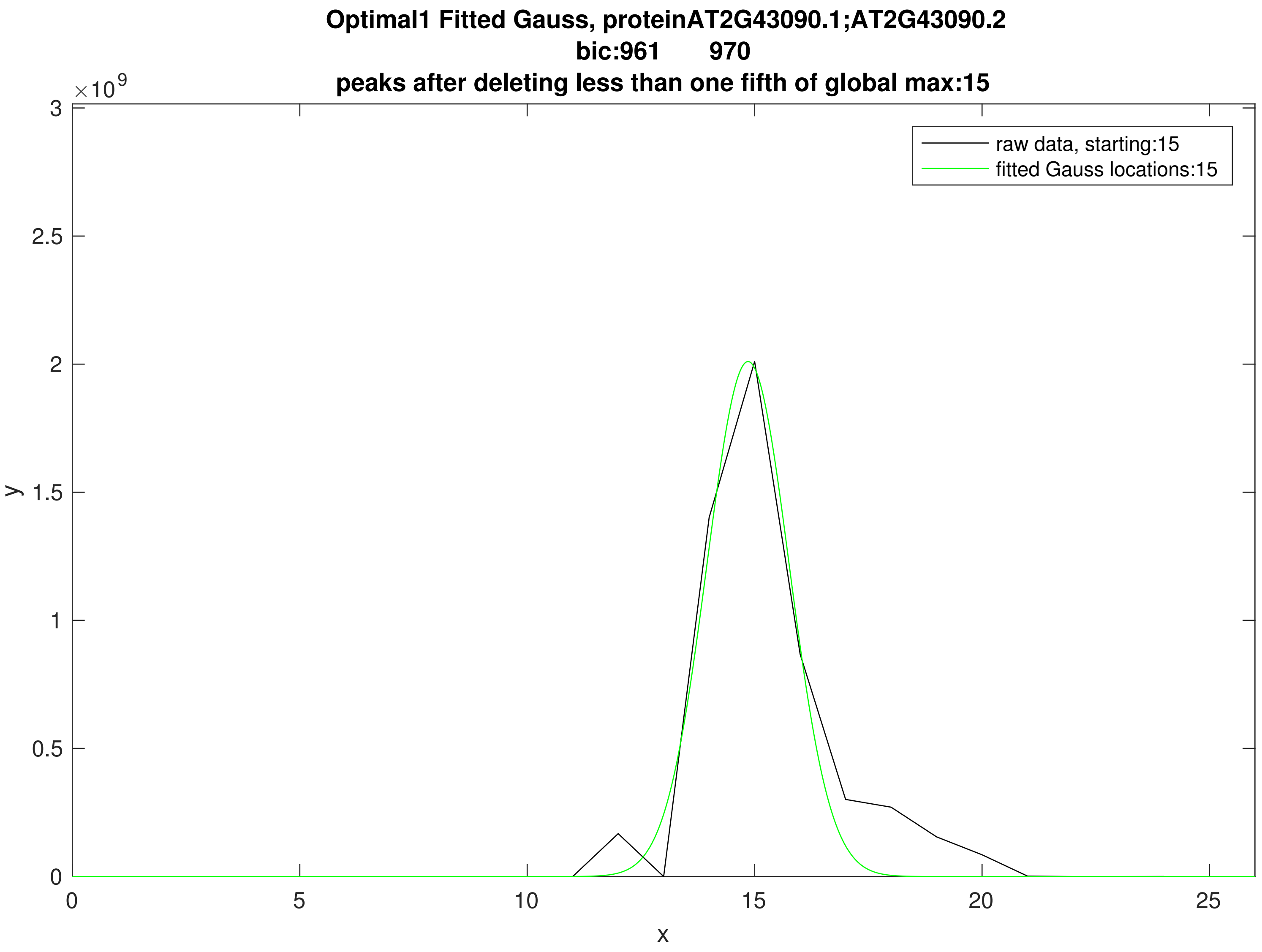
Optimal1 Fitted Gauss, proteinAT2G43090.1;AT2G43090.2
bic:961 970
peaks after deleting less than one fifth of global max:15
raw data, starting:15
fitted Gauss locations:15
×10^9
y
x
0
0.5
1
1.5
2
2.5
3
5
10
15
20
25

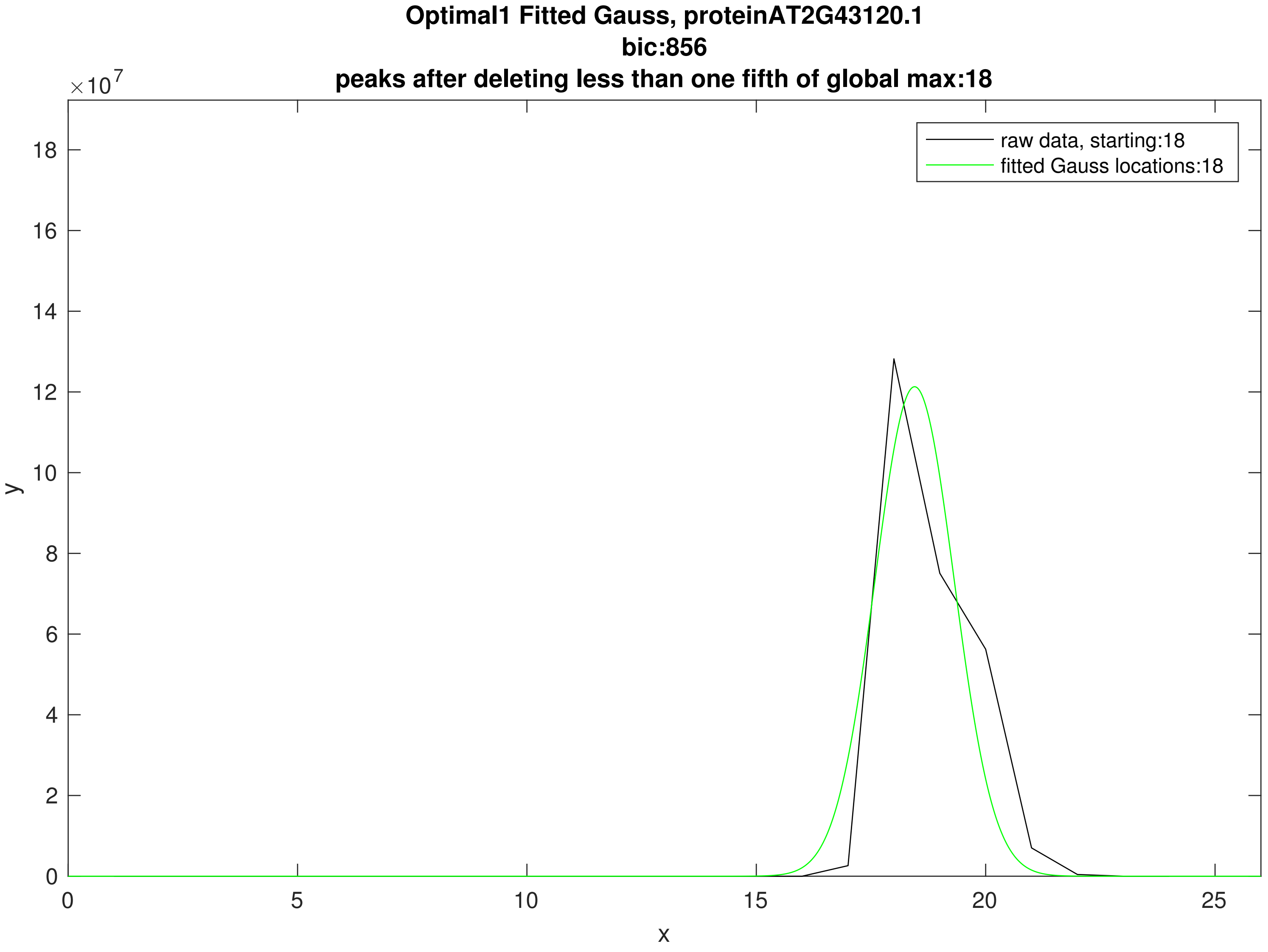
Optimal1 Fitted Gauss, proteinAT2G43120.1
bic:856
peaks after deleting less than one fifth of global max:18
×10^7
raw data, starting:18
fitted Gauss locations:18
y
x
0
2
4
6
8
10
12
14
16
18
0
5
10
15
20
25

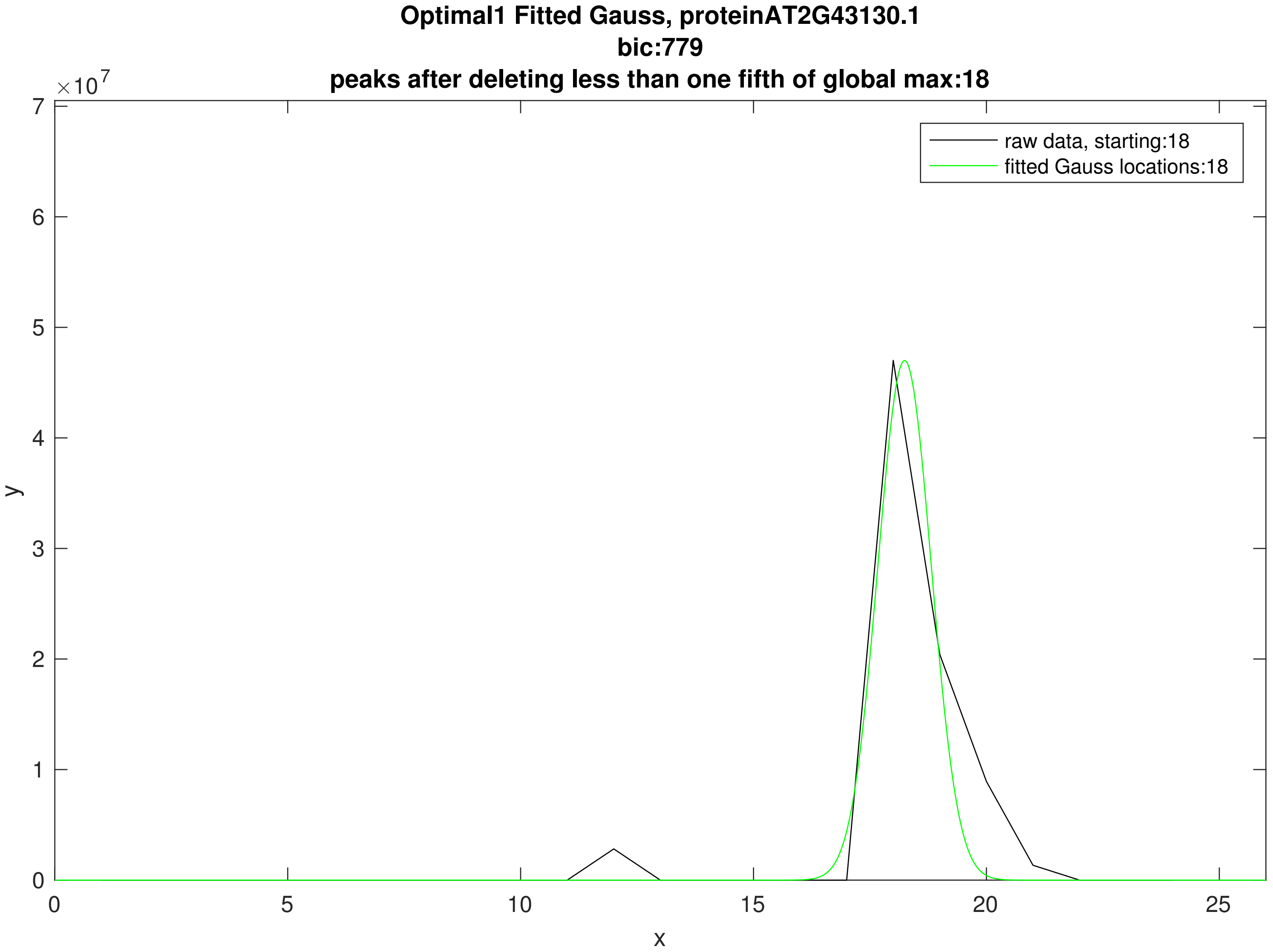

Optimal1 Fitted Gauss, proteinAT2G43130.1
bic:779
peaks after deleting less than one fifth of global max:18
×10^7
raw data, starting:18
fitted Gauss locations:18
y
x
0
1
2
3
4
5
6
7
0
5
10
15
20
25

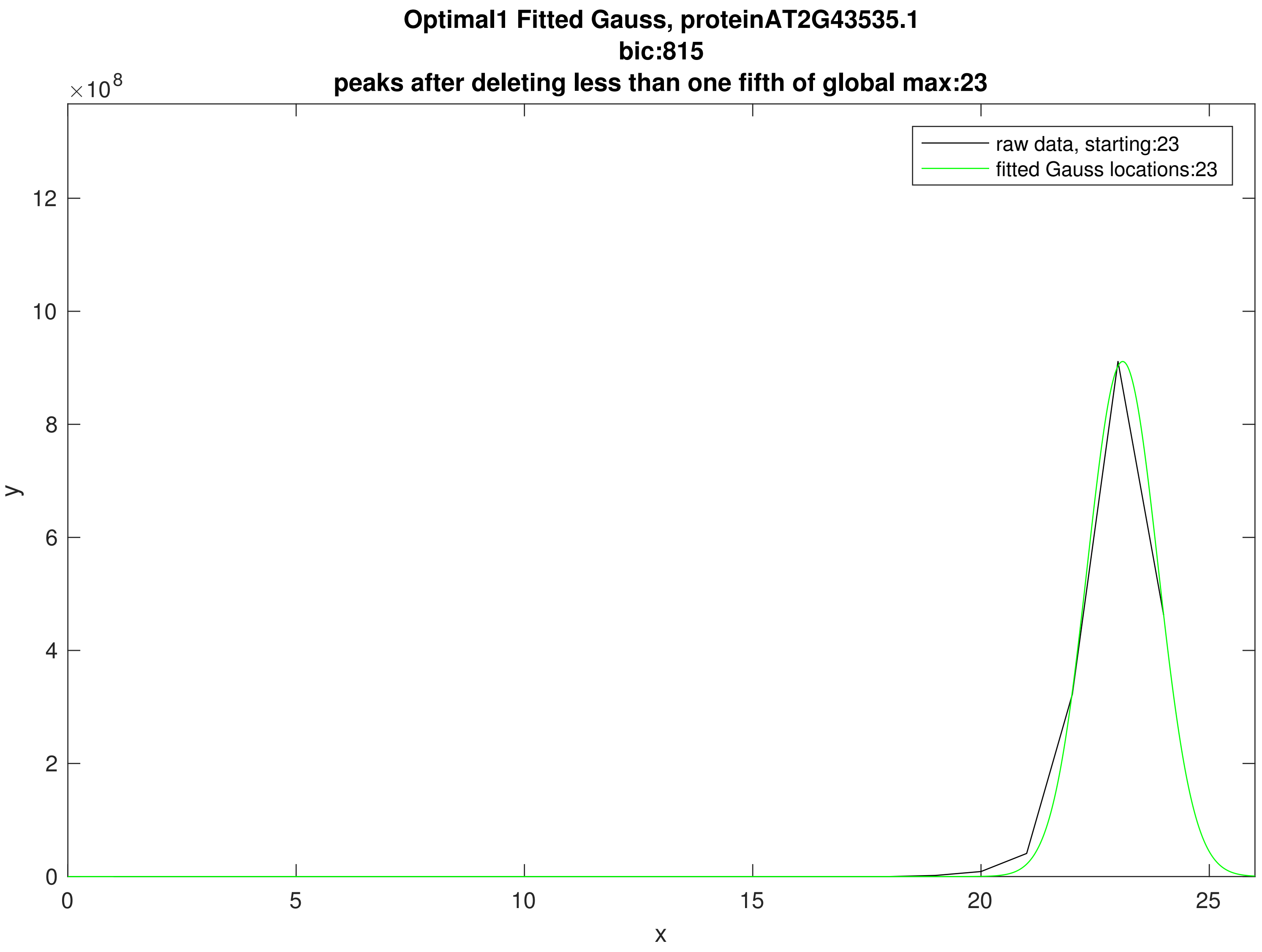

Optimal1 Fitted Gauss, proteinAT2G43535.1
bic:815
peaks after deleting less than one fifth of global max:23
×10^8
raw data, starting:23
fitted Gauss locations:23
y
x
0
2
4
6
8
10
12
0
5
10
15
20
25

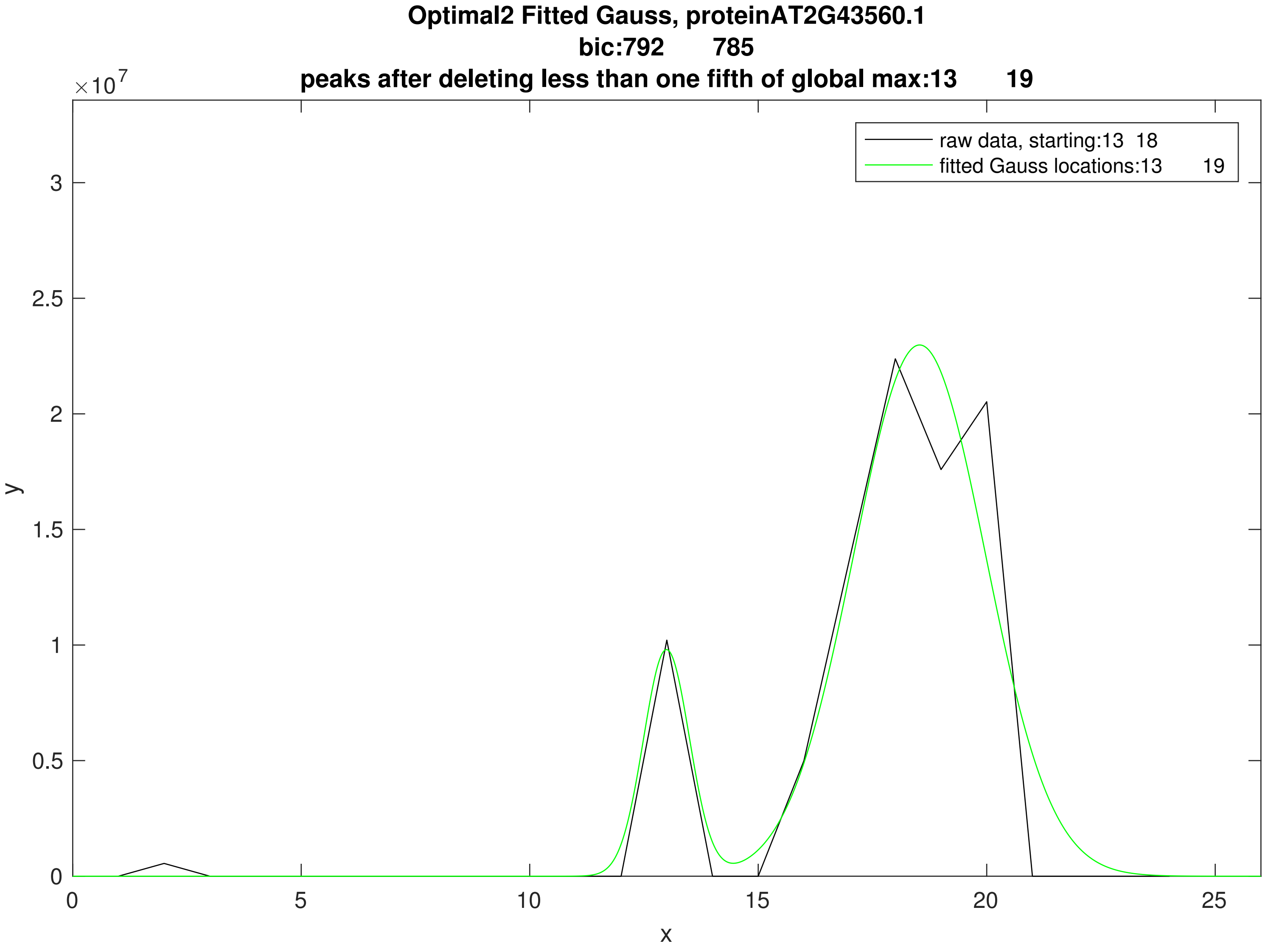
Optimal2 Fitted Gauss, proteinAT2G43560.1
bic:792 785
peaks after deleting less than one fifth of global max:13 19
×10^7
raw data, starting:13 18
fitted Gauss locations:13 19
y
x
0
0.5
1
1.5
2
2.5
3
5
10
15
20
25

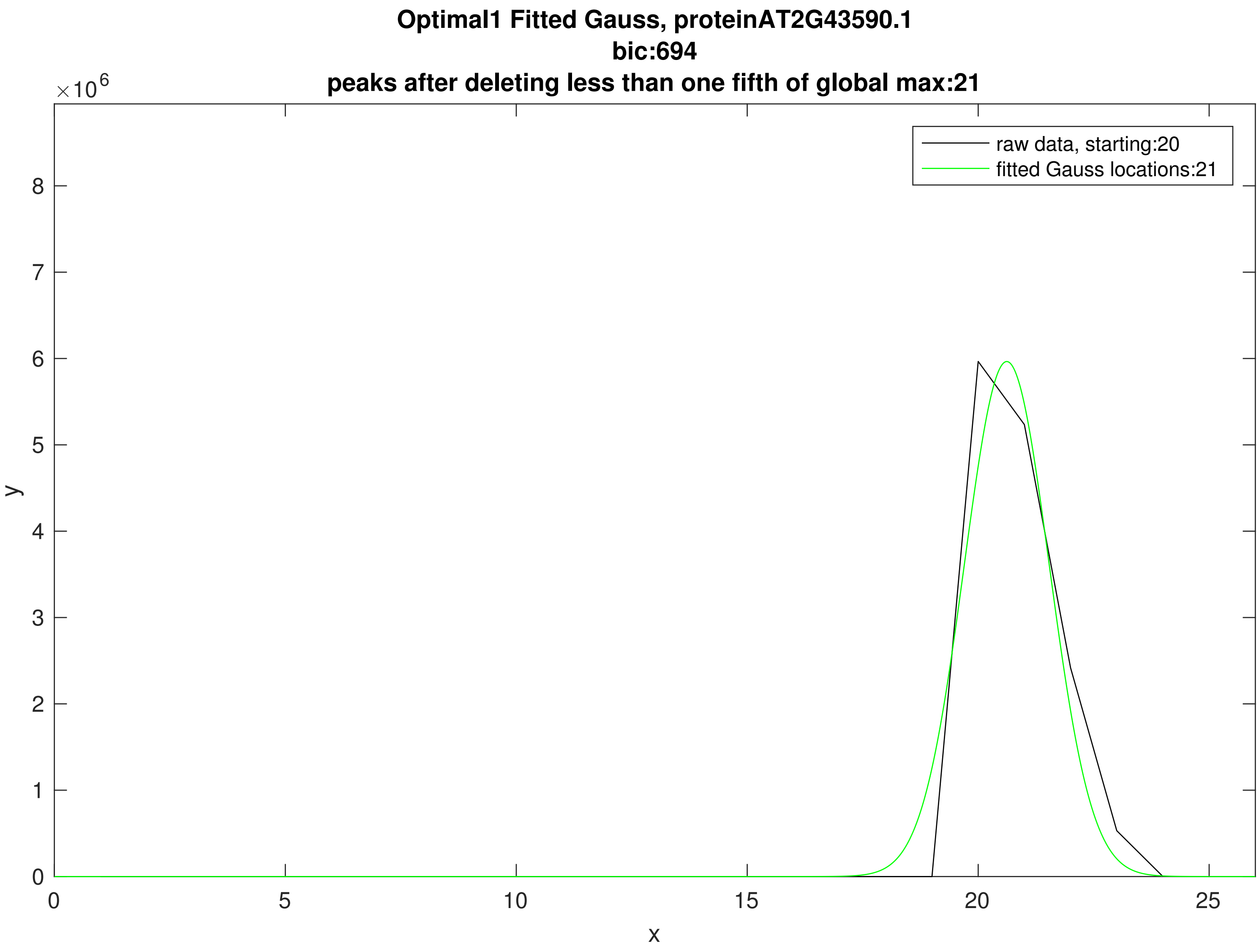
Optimal1 Fitted Gauss, proteinAT2G43590.1
bic:694
peaks after deleting less than one fifth of global max:21
×10^6
raw data, starting:20
fitted Gauss locations:21
0
1
2
3
4
5
6
7
8
y
0
5
10
15
20
25
x

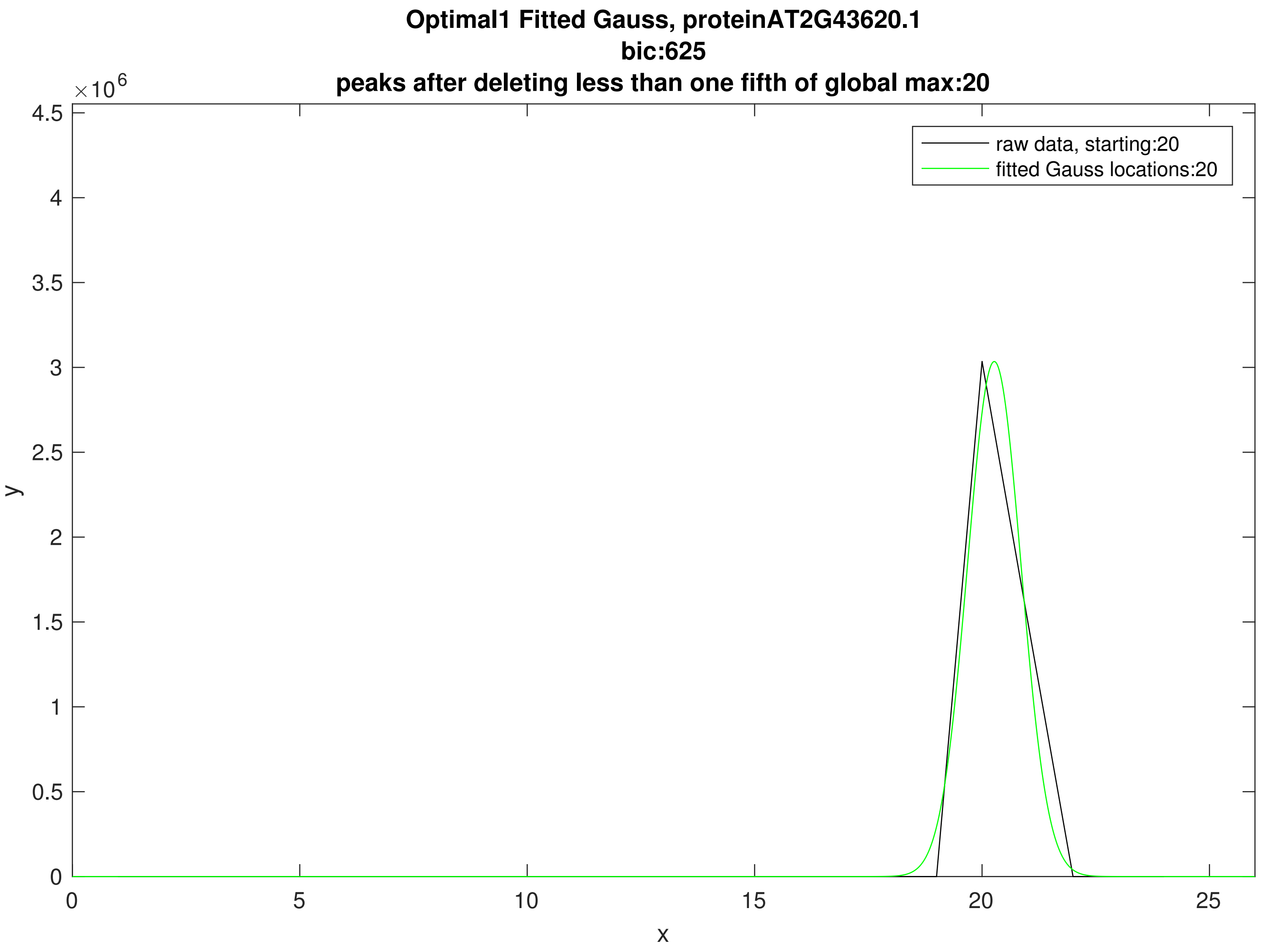

Optimal1 Fitted Gauss, proteinAT2G43620.1
bic:625
peaks after deleting less than one fifth of global max:20
×10^6
4.5
4
3.5
3
2.5
2
1.5
1
0.5
0
y
0
5
10
15
20
25
x
raw data, starting:20
fitted Gauss locations:20

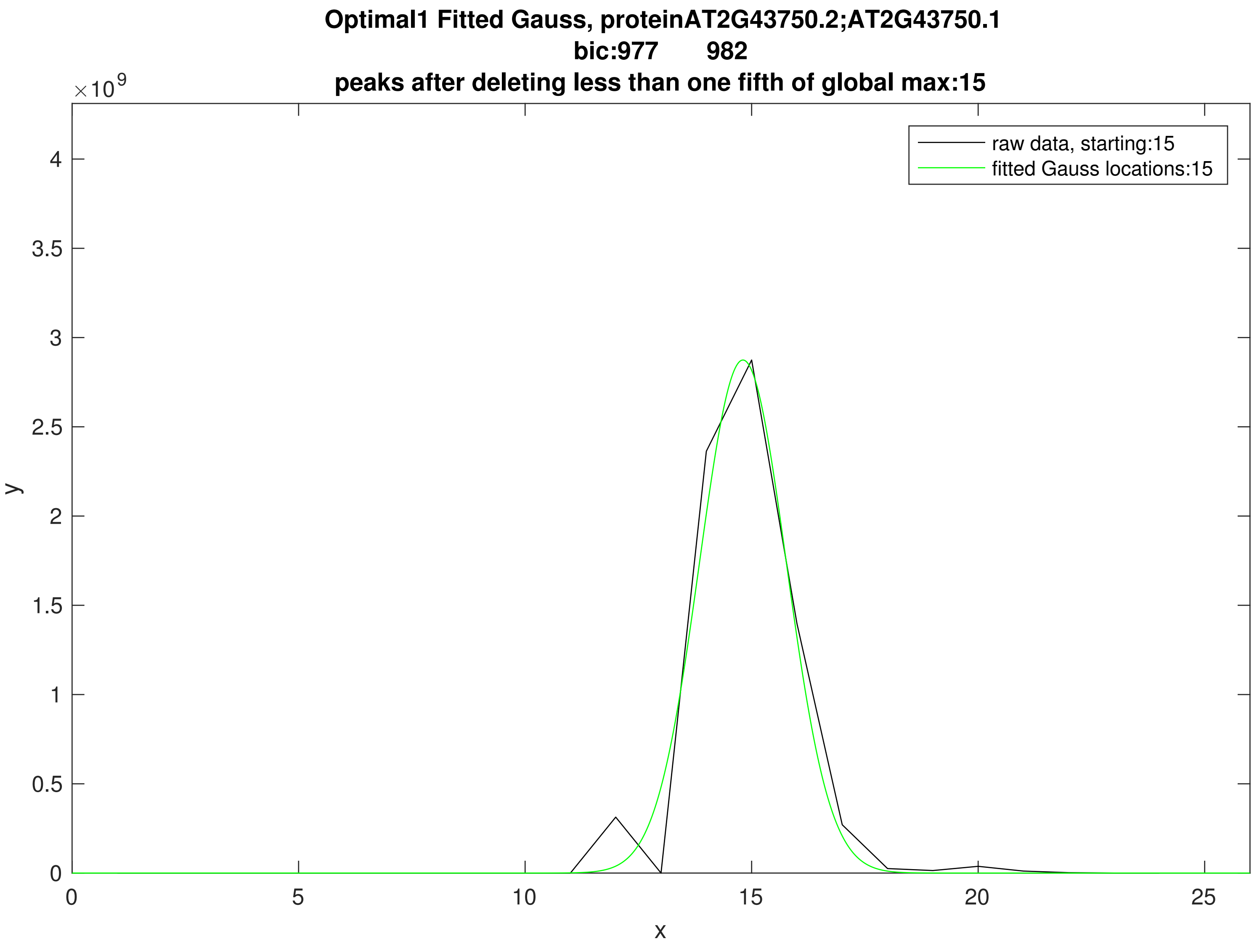
Optimal1 Fitted Gauss, proteinAT2G43750.2;AT2G43750.1
bic:977 982
peaks after deleting less than one fifth of global max:15
×10^9
raw data, starting:15
fitted Gauss locations:15
y
x
4
3.5
3
2.5
2
1.5
1
0.5
0
0
5
10
15
20
25

**Optimal1 Fitted Gauss, proteinAT2G43760.3;AT2G43760.2;AT2G43760.1**
**bic:826**
**peaks after deleting less than one fifth of global max:15**

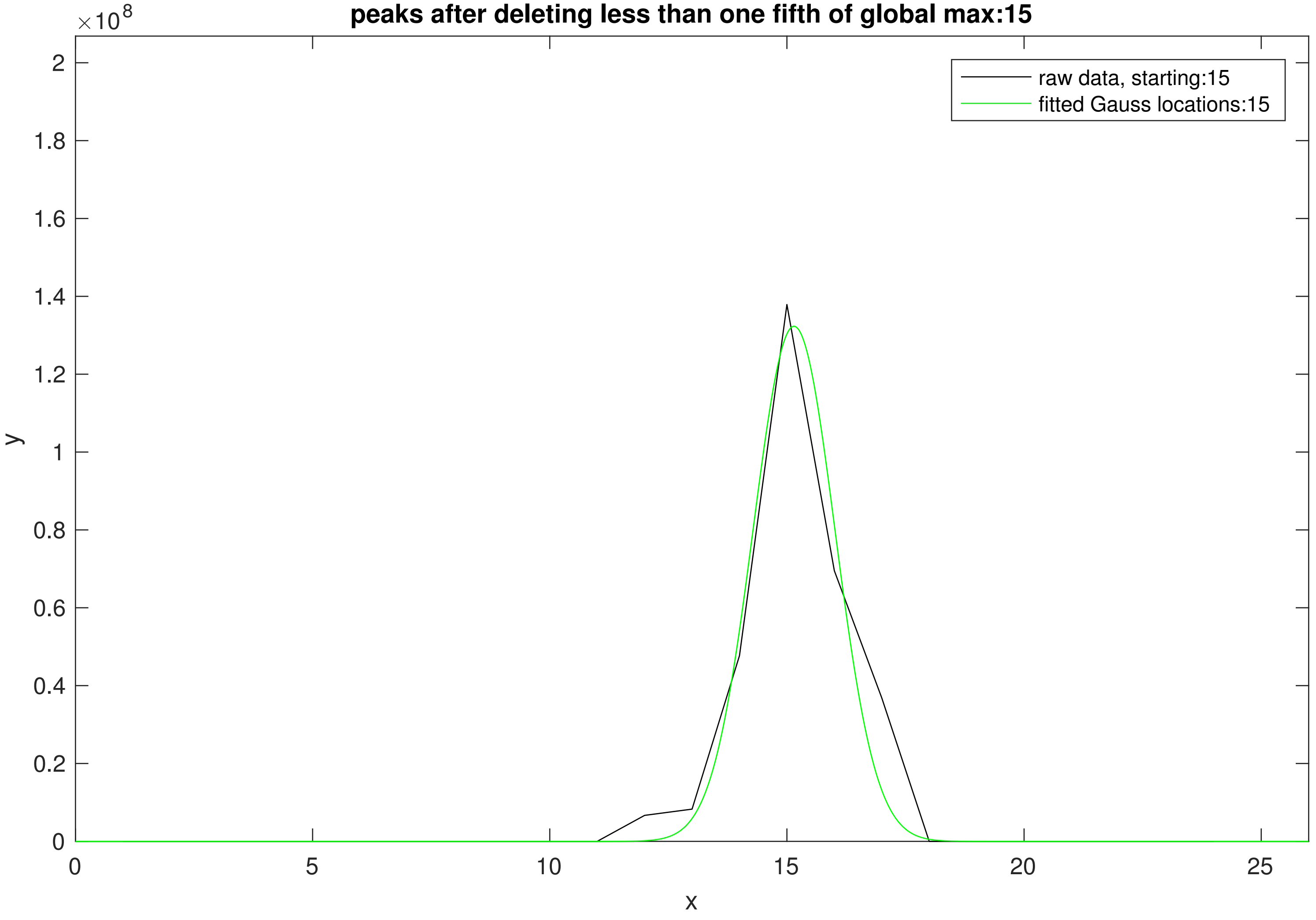

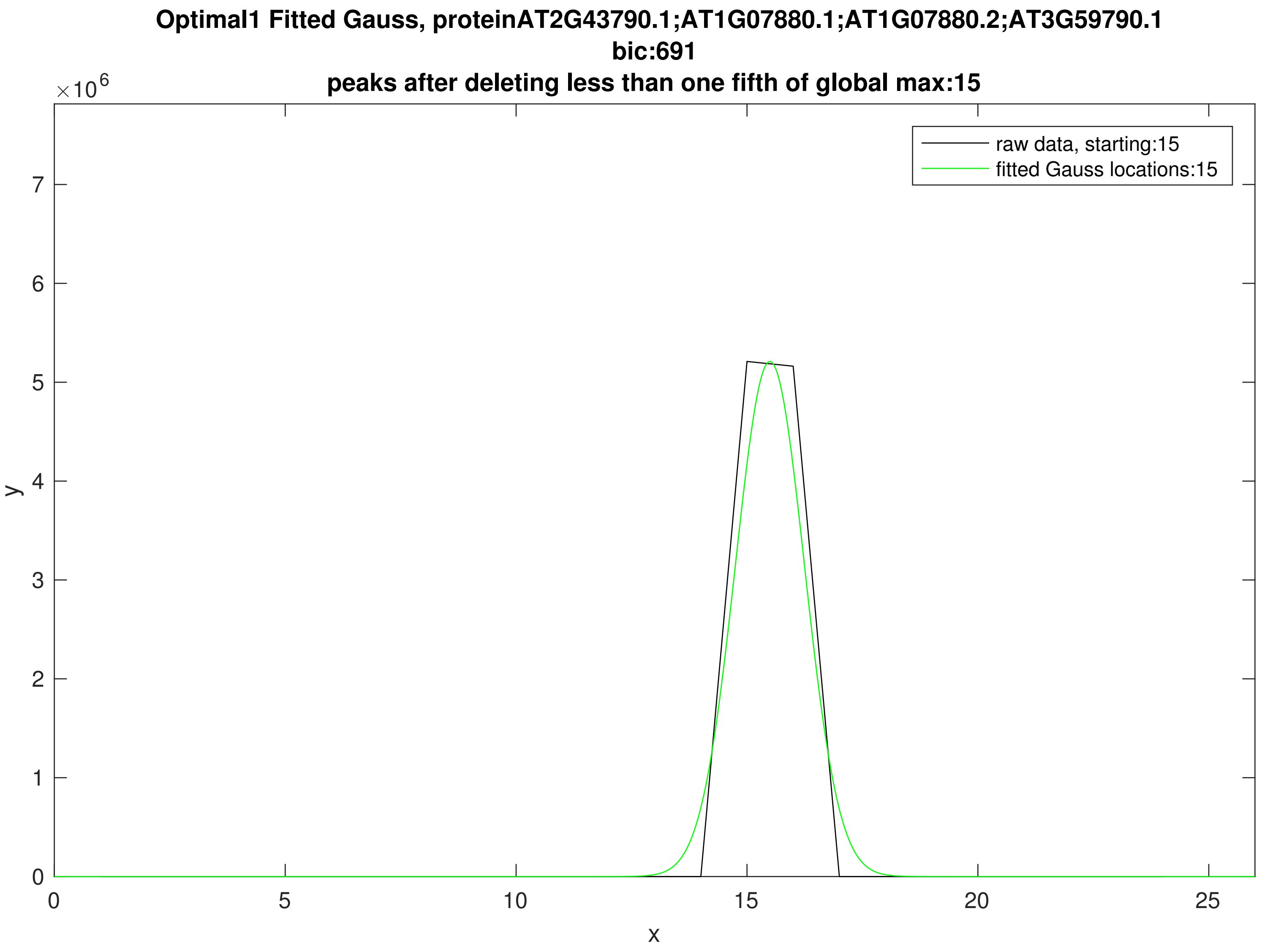
Optimal1 Fitted Gauss, proteinAT2G43790.1;AT1G07880.1;AT1G07880.2;AT3G59790.1
bic:691
peaks after deleting less than one fifth of global max:15
×10^6
raw data, starting:15
fitted Gauss locations:15
y
x
0
1
2
3
4
5
6
7
0
5
10
15
20
25

Optimal1 Fitted Gauss, proteinAT2G43810.2;AT2G43810.1
bic:827 836
peaks after deleting less than one fifth of global max:14

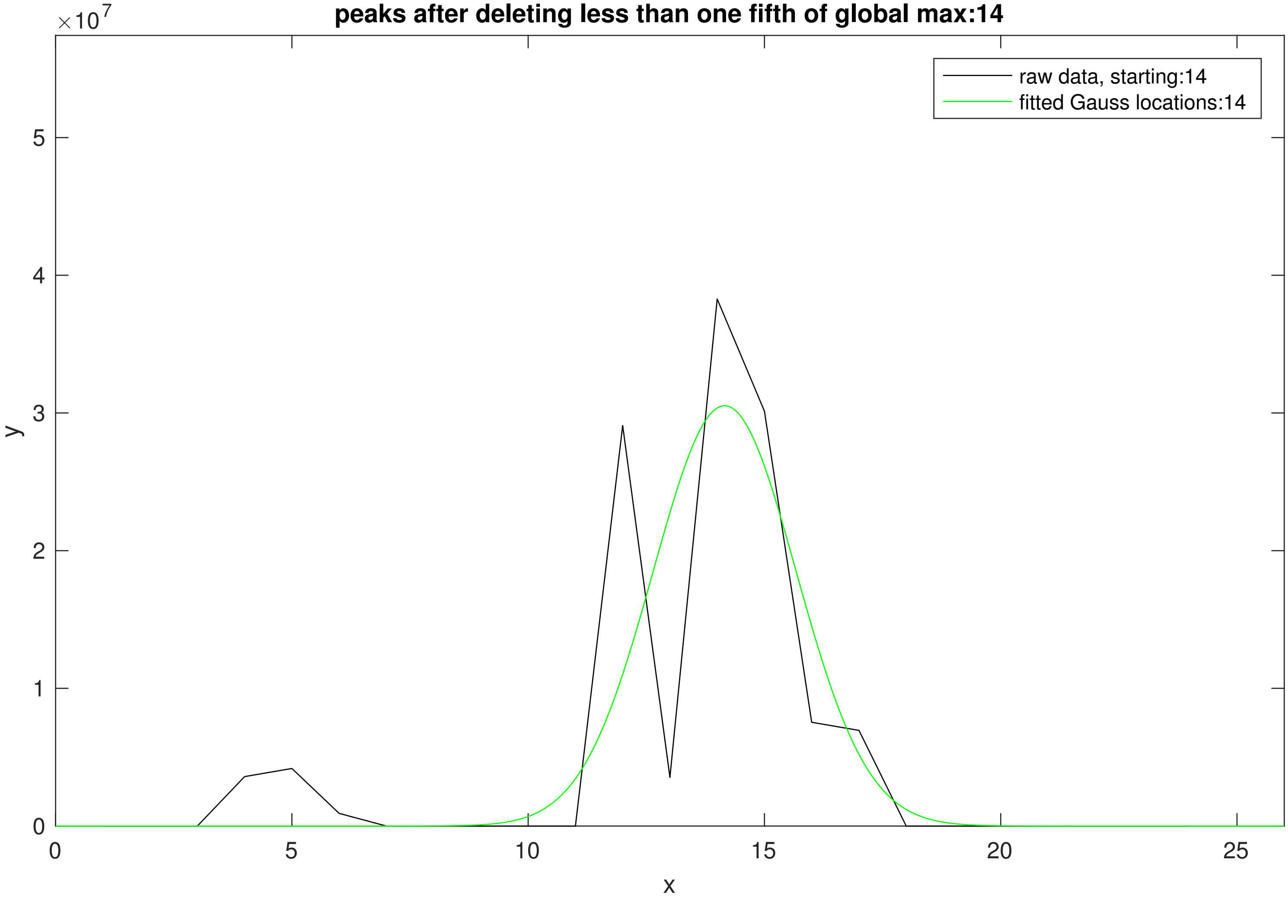

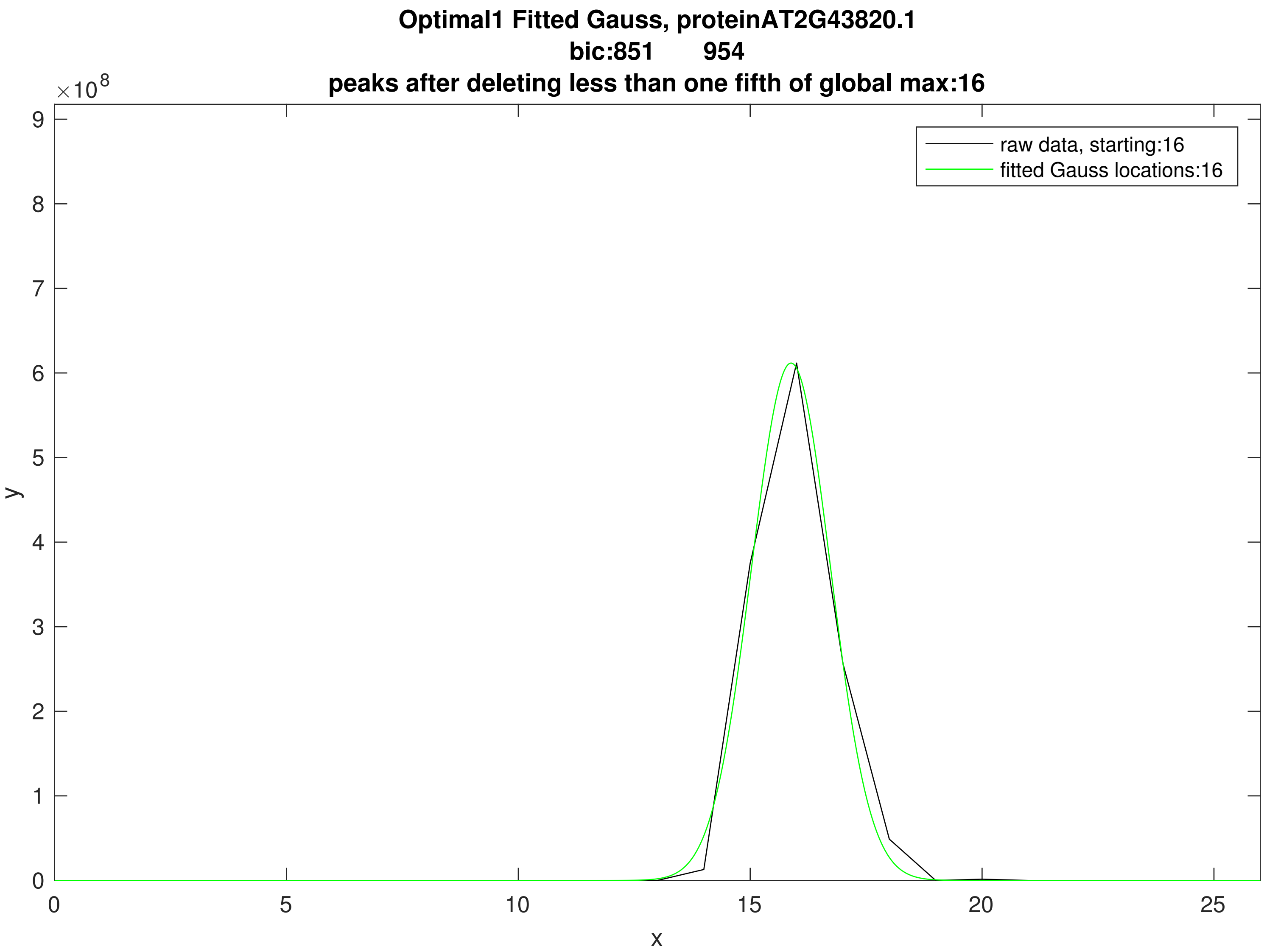
Optimal1 Fitted Gauss, proteinAT2G43820.1
bic:851 954
peaks after deleting less than one fifth of global max:16
×10^8
raw data, starting:16
fitted Gauss locations:16
y
x
0
1
2
3
4
5
6
7
8
9
0
5
10
15
20
25

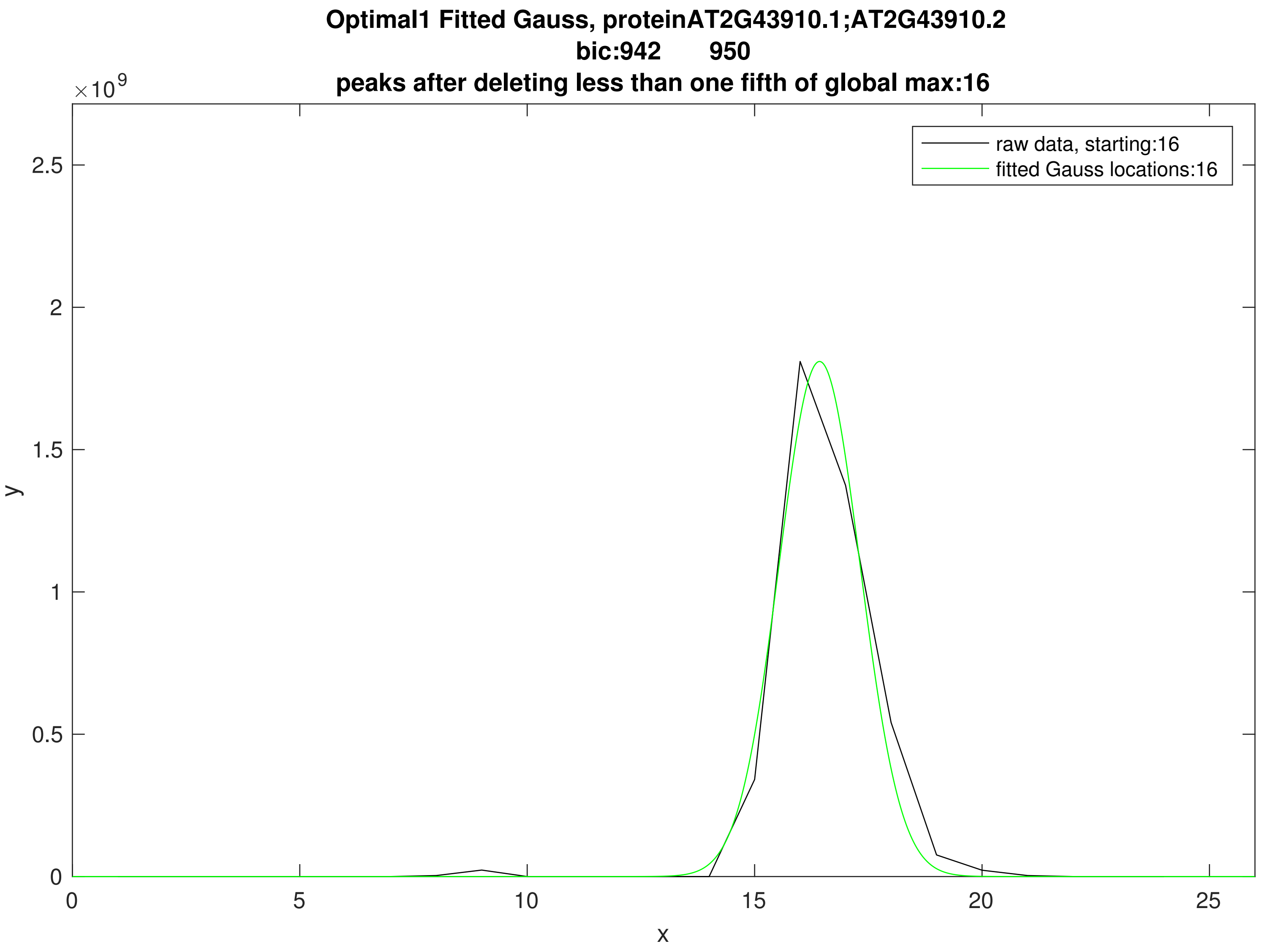
Optimal1 Fitted Gauss, proteinAT2G43910.1;AT2G43910.2
bic:942 950
peaks after deleting less than one fifth of global max:16
×10^9
2.5
2
1.5
1
0.5
0
y
0
5
10
15
20
25
x
raw data, starting:16
fitted Gauss locations:16

Optimal1 Fitted Gauss, proteinAT2G43940.1
bic:688
peaks after deleting less than one fifth of global max:18

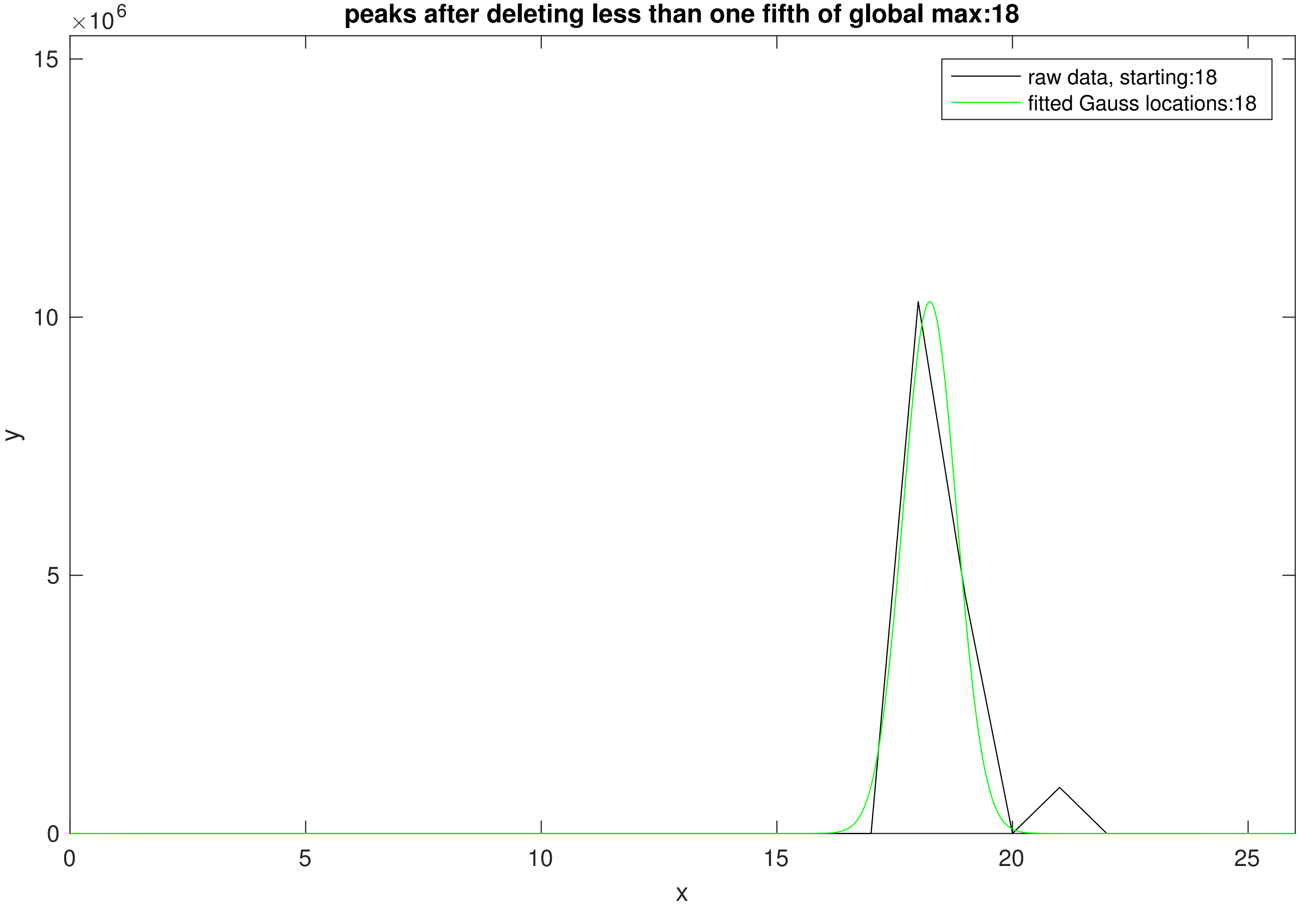

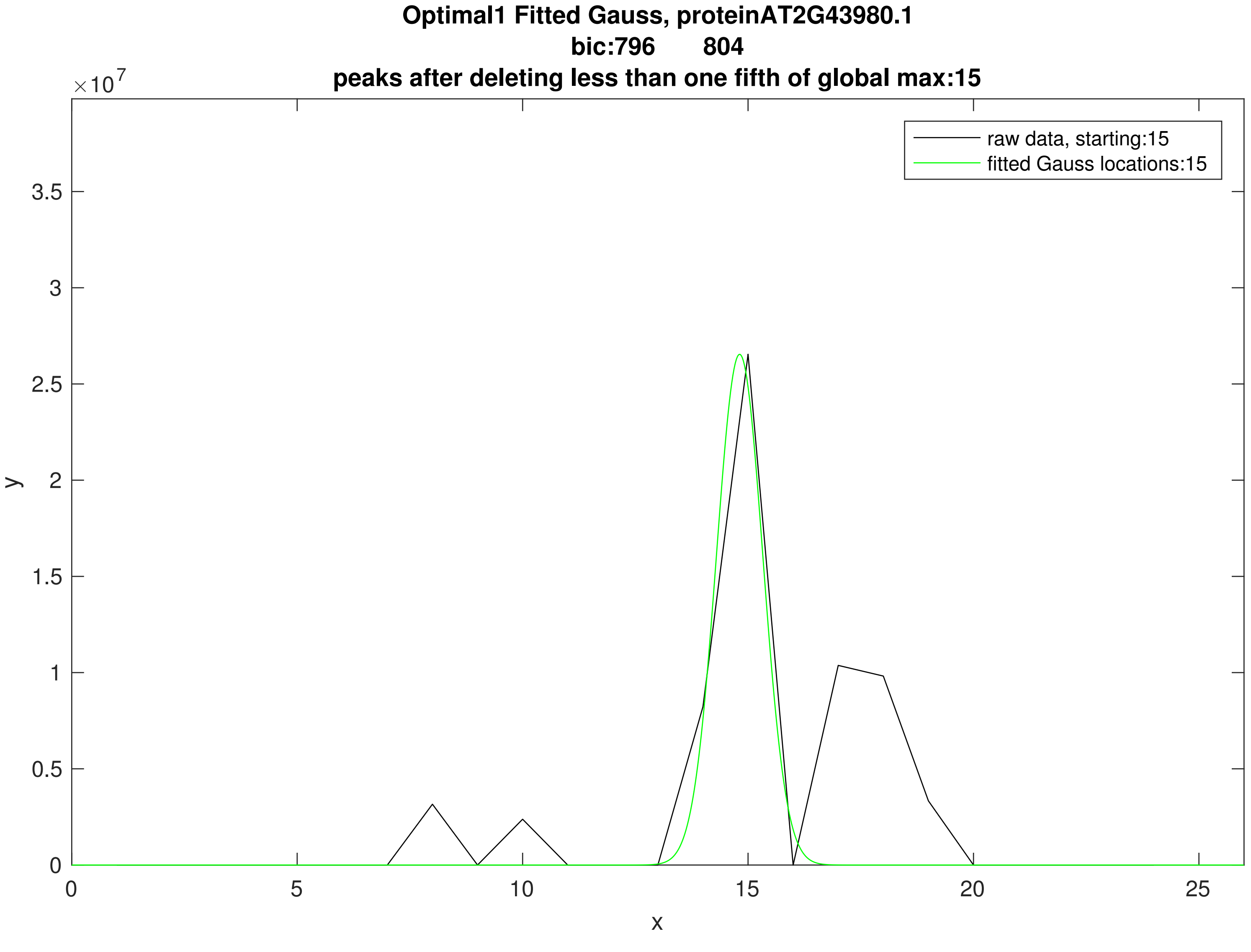

Optimal1 Fitted Gauss, proteinAT2G43980.1
bic:796 804
peaks after deleting less than one fifth of global max:15
×10^7
3.5
3
2.5
2
1.5
1
0.5
0
y
0
5
10
15
20
25
x
raw data, starting:15
fitted Gauss locations:15

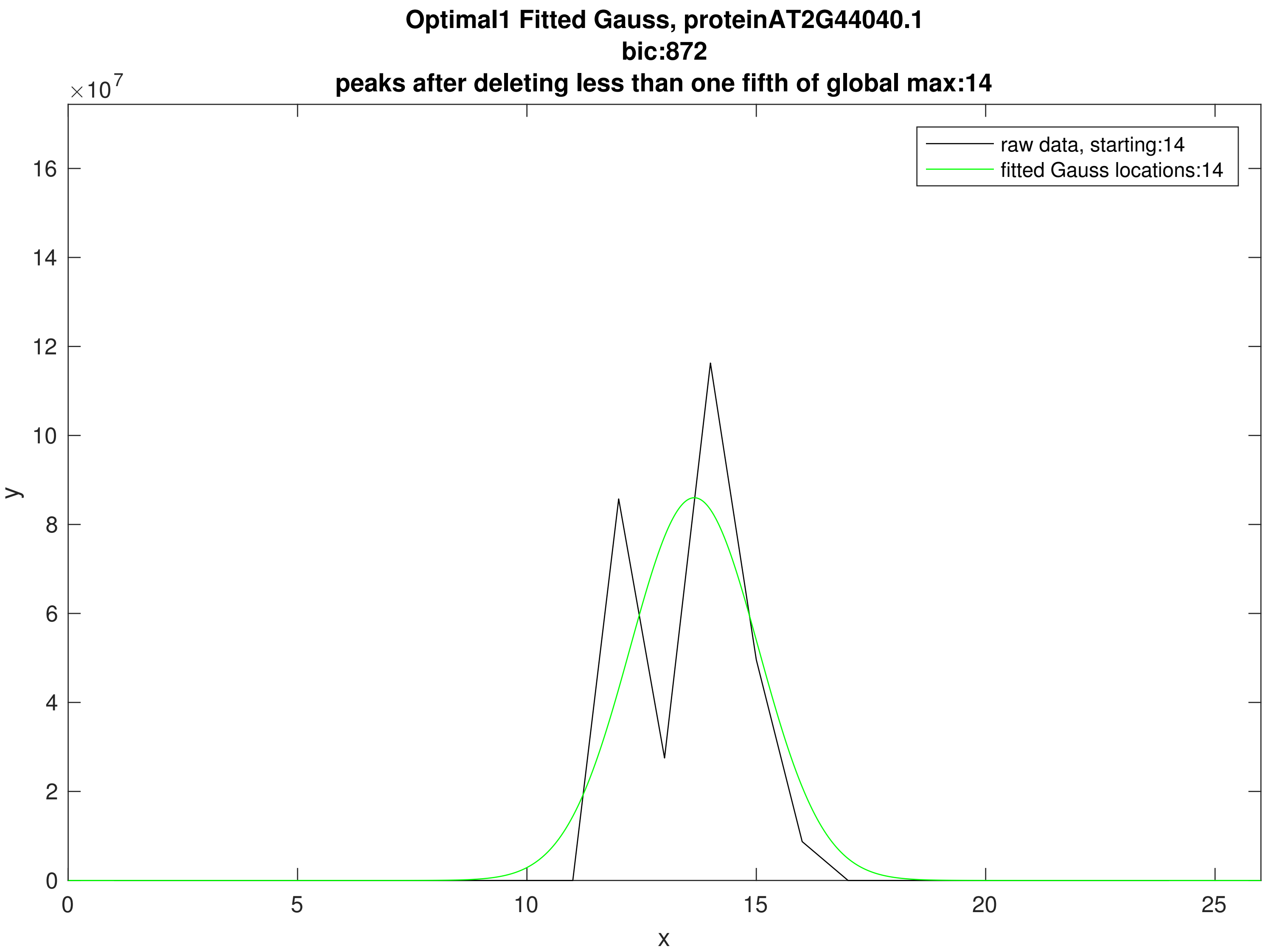
Optimal1 Fitted Gauss, proteinAT2G44040.1
bic:872
peaks after deleting less than one fifth of global max:14
×10^7
raw data, starting:14
fitted Gauss locations:14
y
x
0
2
4
6
8
10
12
14
16
0
5
10
15
20
25

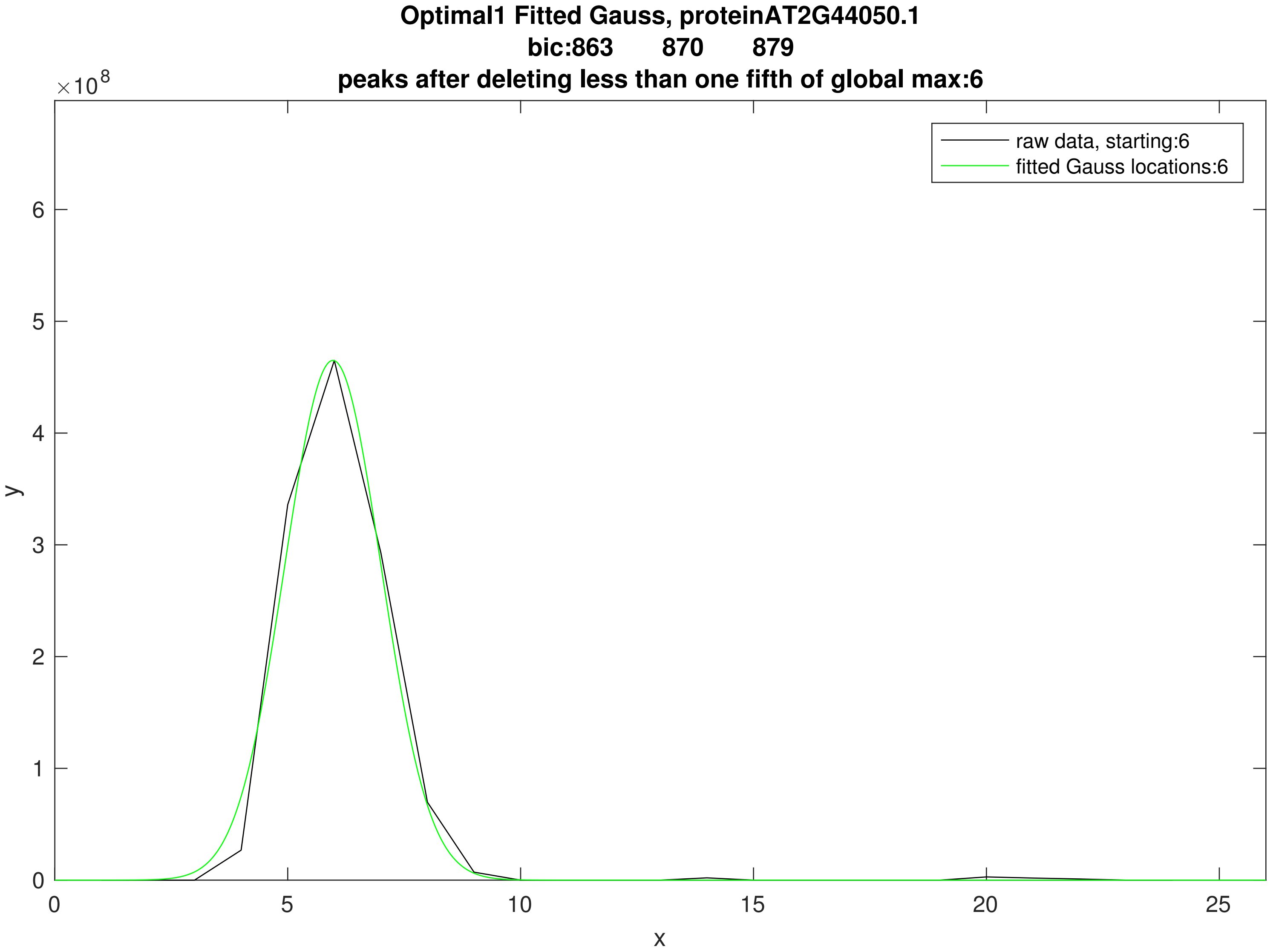
Optimal1 Fitted Gauss, proteinAT2G44050.1
bic:863 870 879
peaks after deleting less than one fifth of global max:6
×10^8
raw data, starting:6
fitted Gauss locations:6
y
x
0
1
2
3
4
5
6
0
5
10
15
20
25

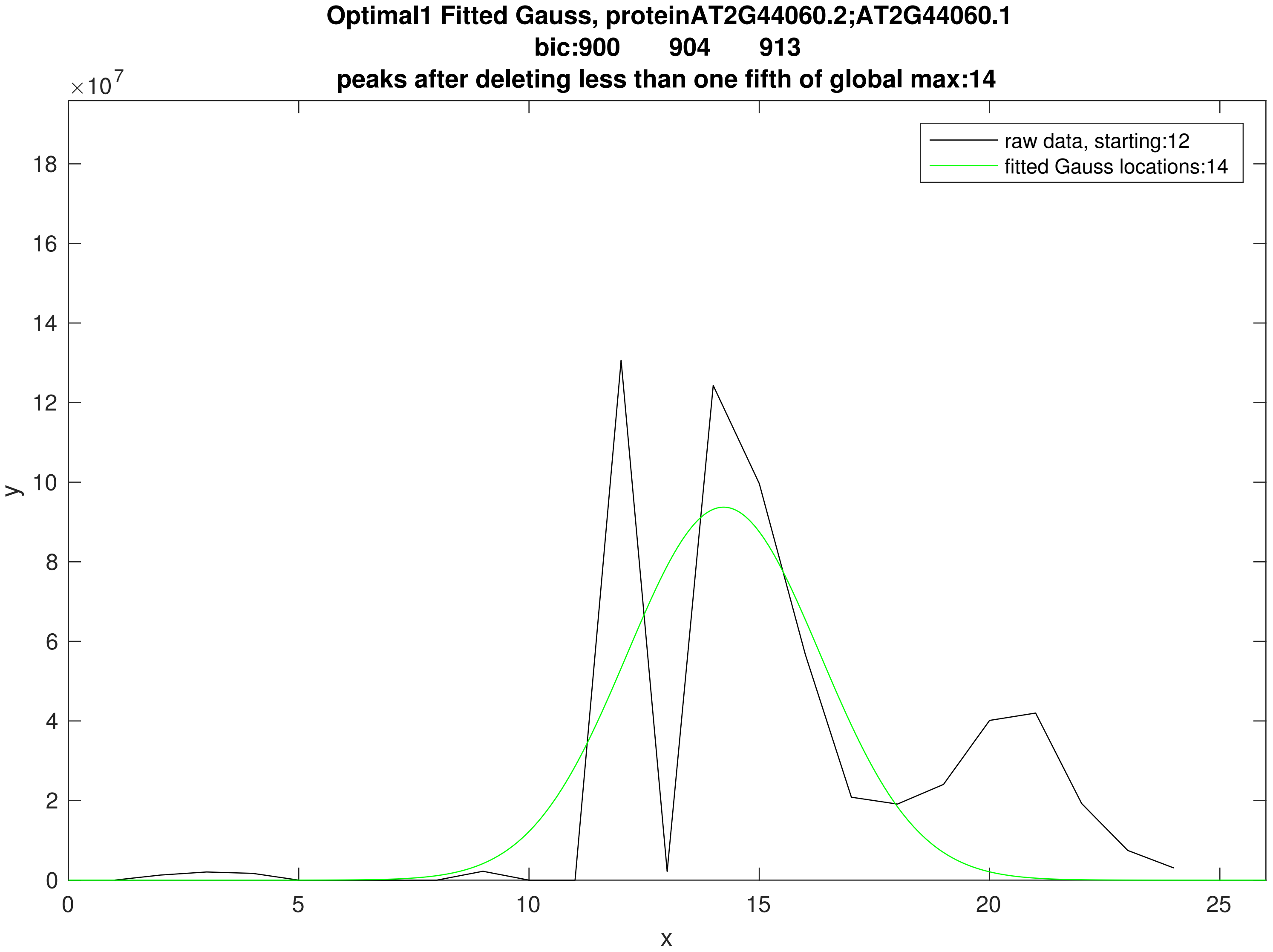
Optimal1 Fitted Gauss, proteinAT2G44060.2;AT2G44060.1
bic:900 904 913
peaks after deleting less than one fifth of global max:14
×10^7
raw data, starting:12
fitted Gauss locations:14
y
x
0
2
4
6
8
10
12
14
16
18
0
5
10
15
20
25

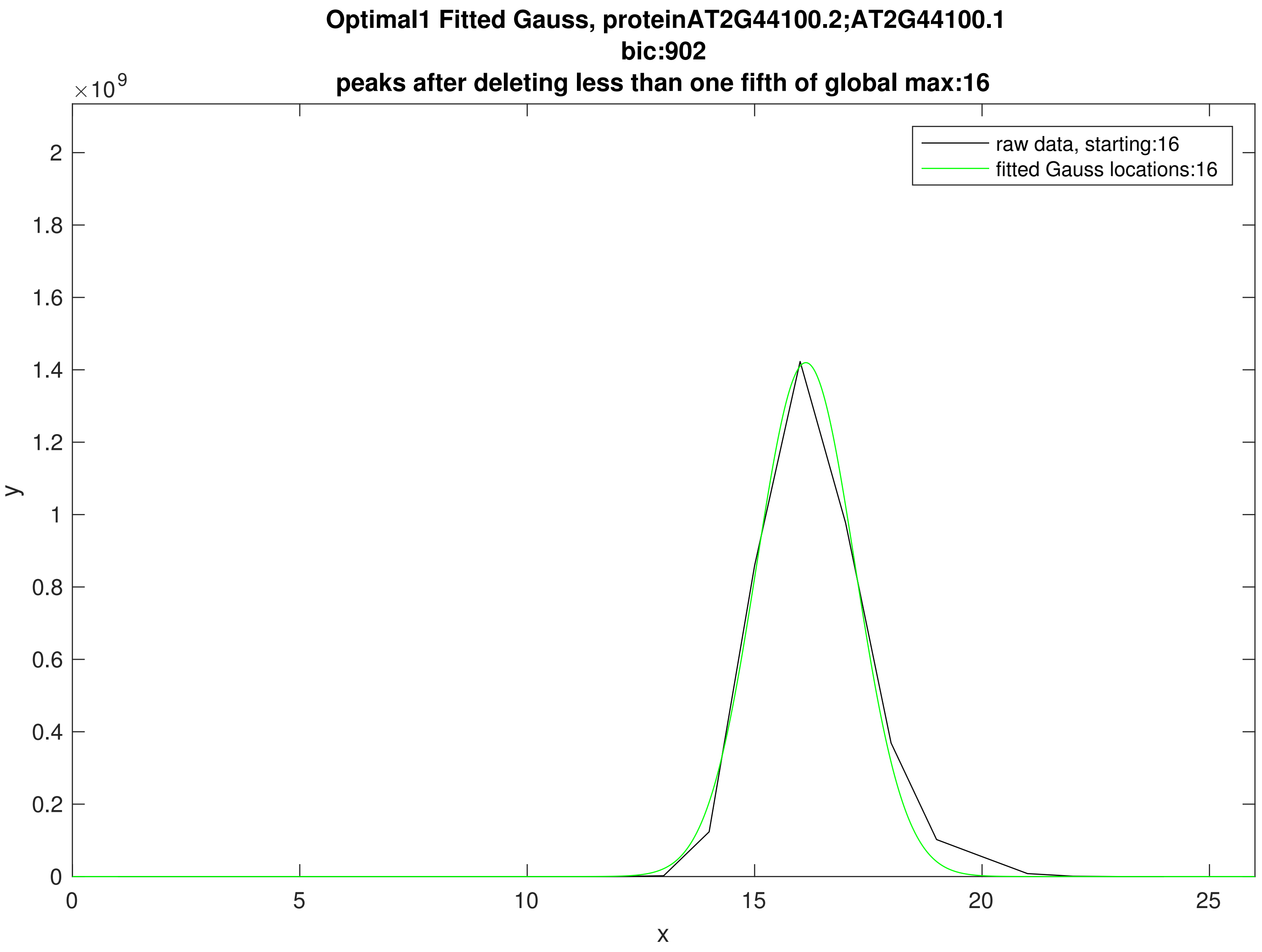

Optimal1 Fitted Gauss, proteinAT2G44100.2;AT2G44100.1
bic:902
peaks after deleting less than one fifth of global max:16
×10^9
y
x
raw data, starting:16
fitted Gauss locations:16

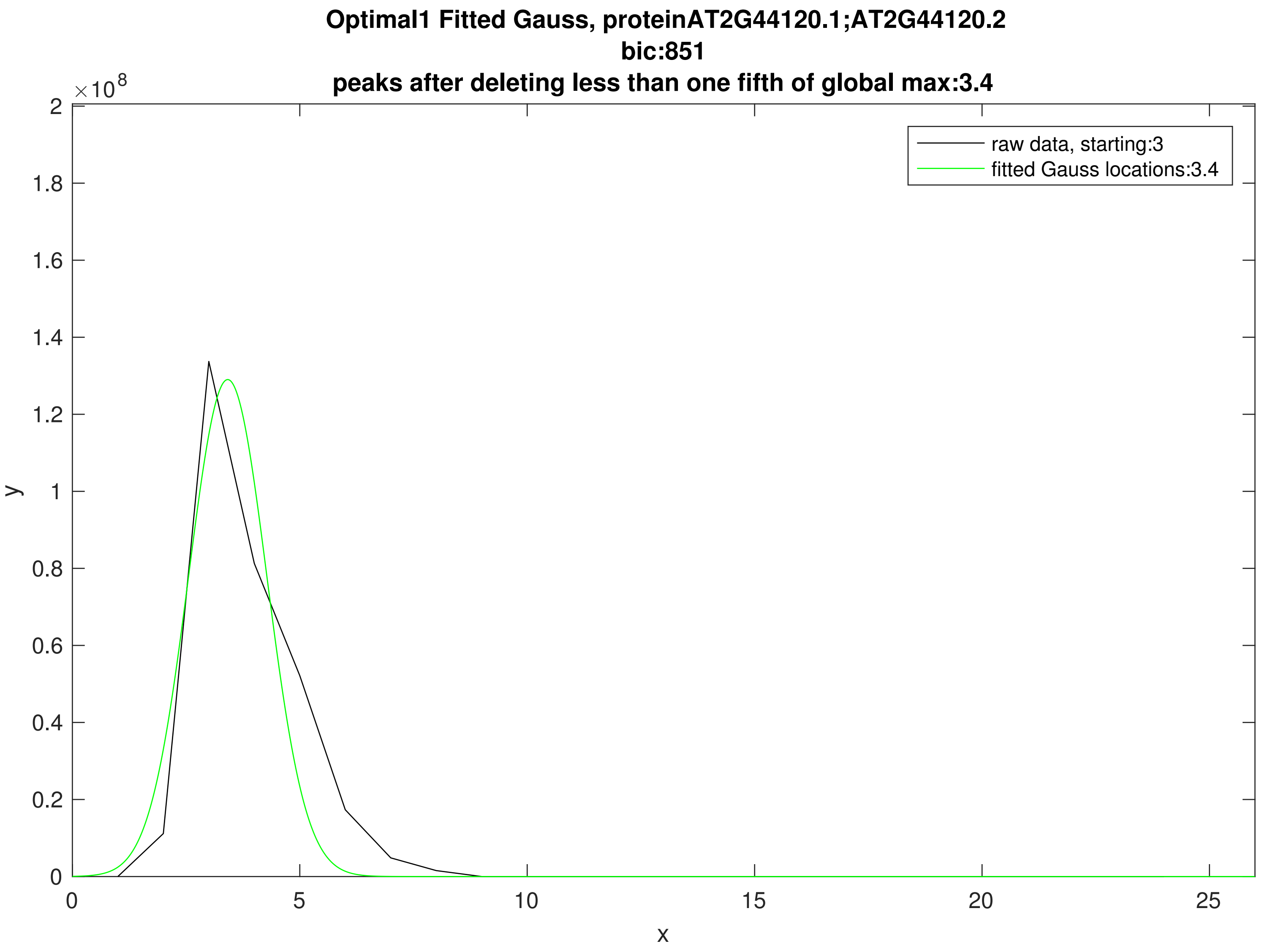
Optimal1 Fitted Gauss, proteinAT2G44120.1;AT2G44120.2
bic:851
peaks after deleting less than one fifth of global max:3.4
×10^8
raw data, starting:3
fitted Gauss locations:3.4
y
x
0
0.2
0.4
0.6
0.8
1
1.2
1.4
1.6
1.8
2
5
10
15
20
25

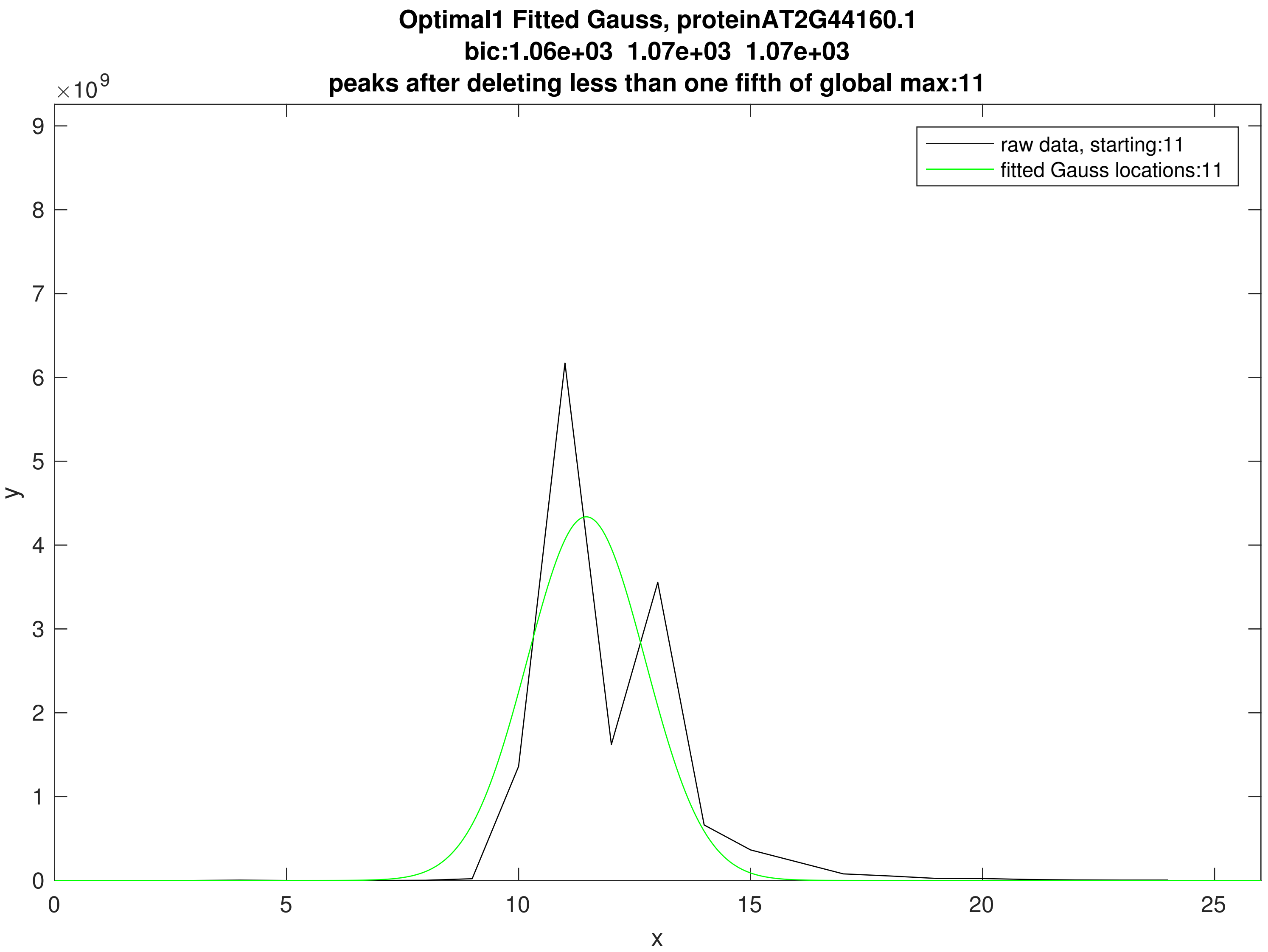

Optimal1 Fitted Gauss, proteinAT2G44160.1
bic:1.06e+03 1.07e+03 1.07e+03
peaks after deleting less than one fifth of global max:11
×10^9
raw data, starting:11
fitted Gauss locations:11
y
x
0
1
2
3
4
5
6
7
8
9
0
5
10
15
20
25

**Optimal1 Fitted Gauss, proteinAT2G44310.1**
**bic:915 925 934**
**peaks after deleting less than one fifth of global max:18**

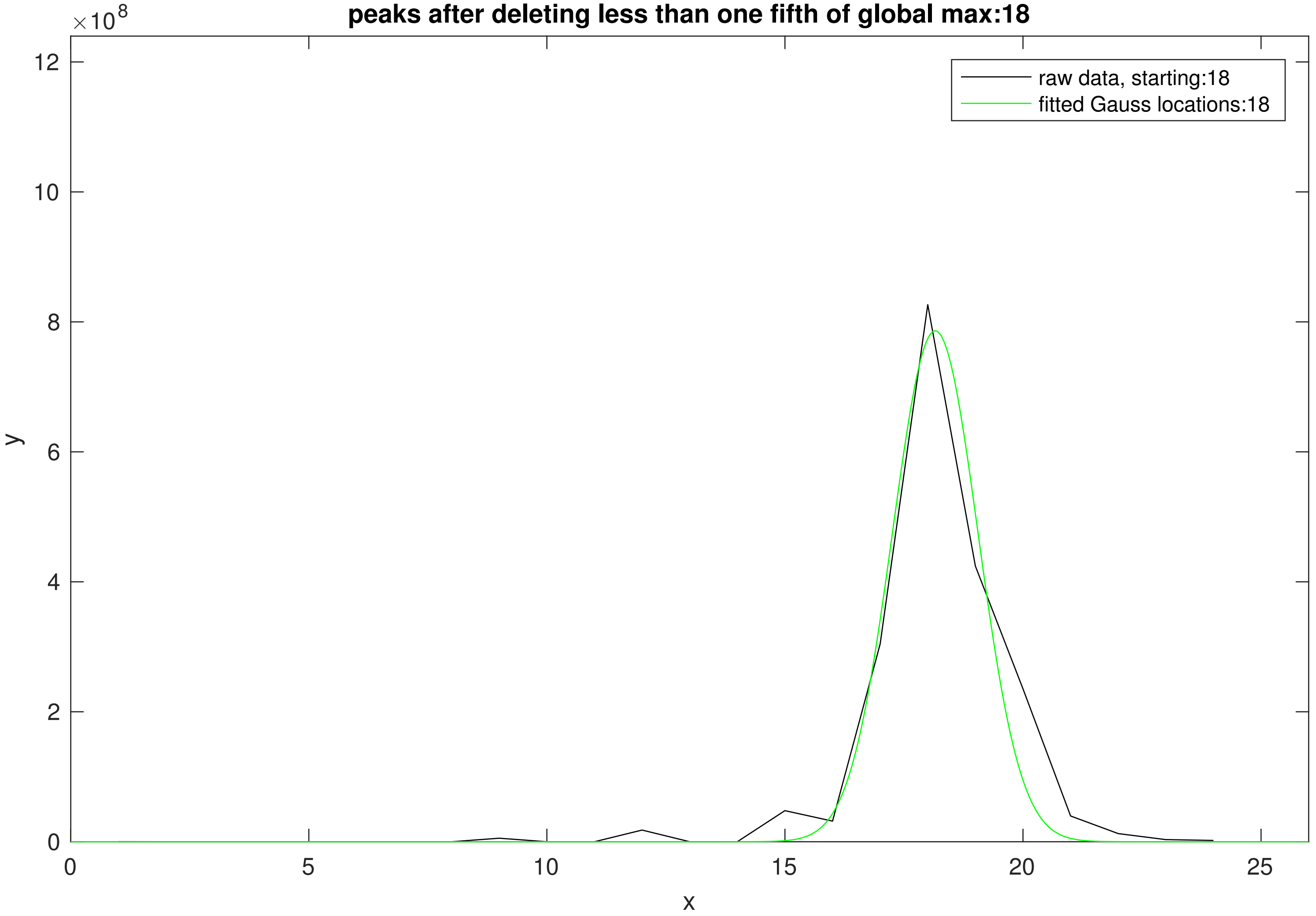

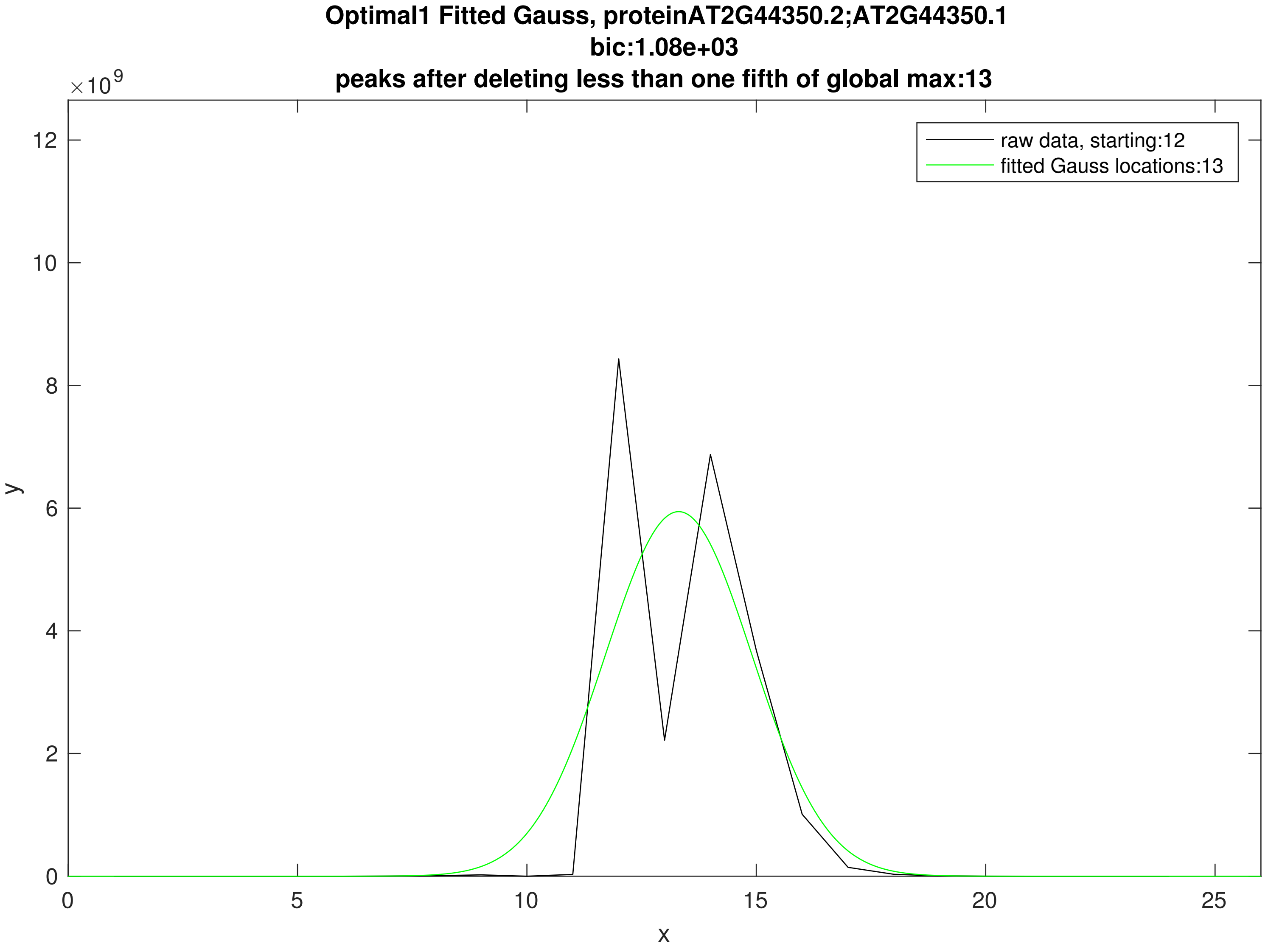

Optimal1 Fitted Gauss, proteinAT2G44350.2;AT2G44350.1
bic:1.08e+03
peaks after deleting less than one fifth of global max:13
×10^9
12
10
8
6
4
2
0
y
0
5
10
15
20
25
x
raw data, starting:12
fitted Gauss locations:13

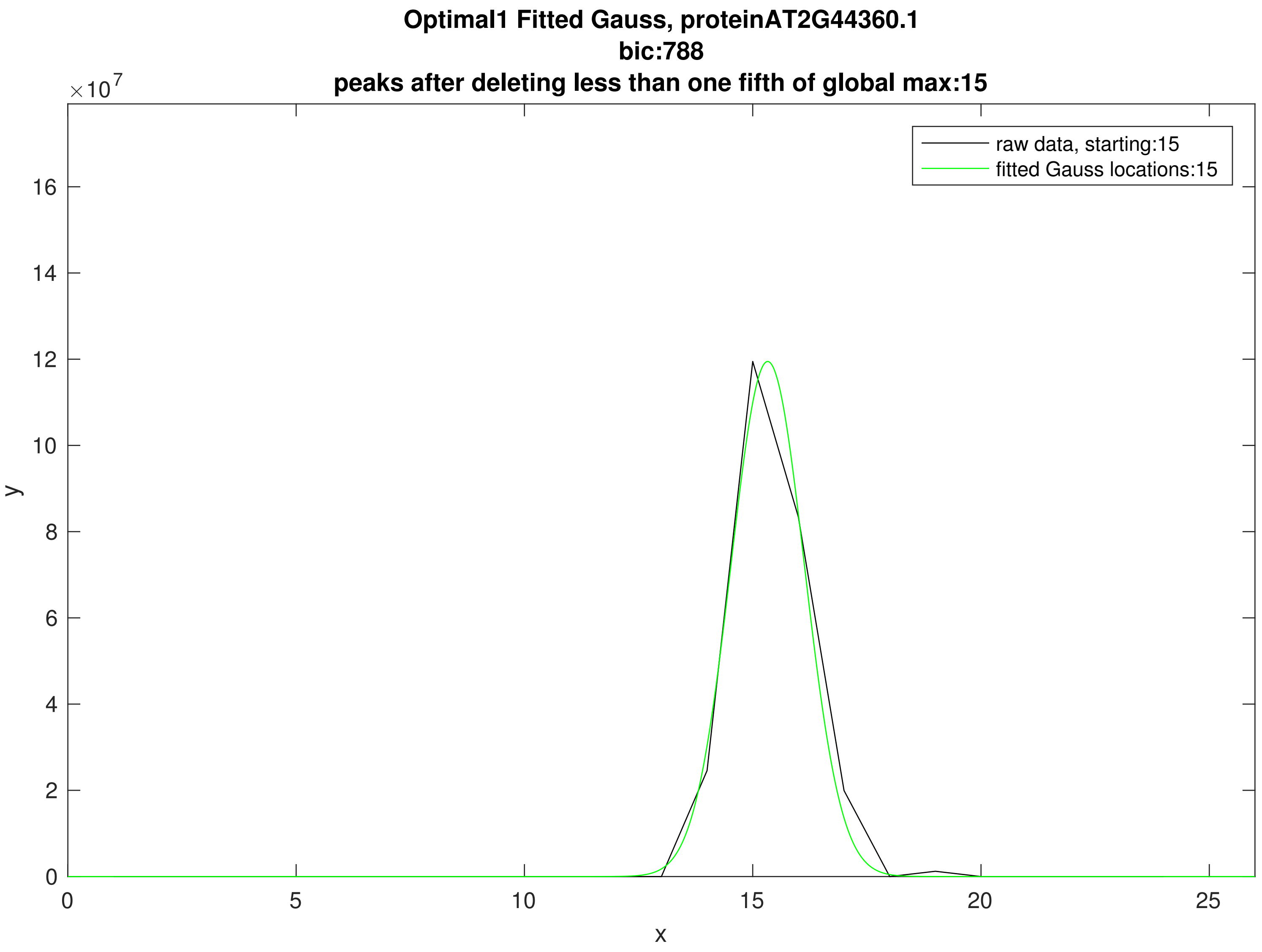

Optimal1 Fitted Gauss, proteinAT2G44360.1
bic:788
peaks after deleting less than one fifth of global max:15
×10^7
raw data, starting:15
fitted Gauss locations:15
y
x

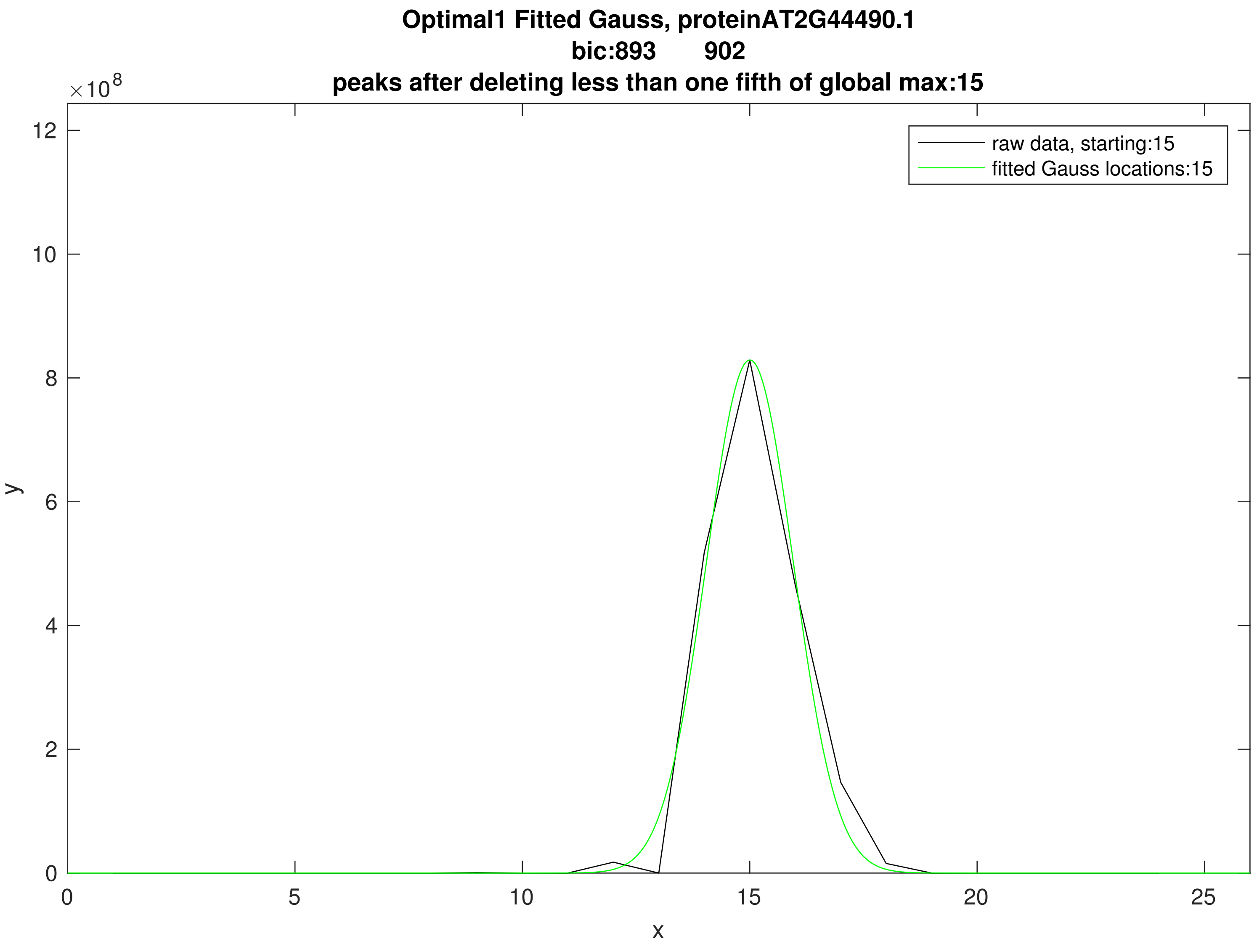

Optimal1 Fitted Gauss, proteinAT2G44490.1
bic:893 902
peaks after deleting less than one fifth of global max:15
×10^8
12
10
8
6
4
2
0
y
0
5
10
15
20
25
x
raw data, starting:15
fitted Gauss locations:15

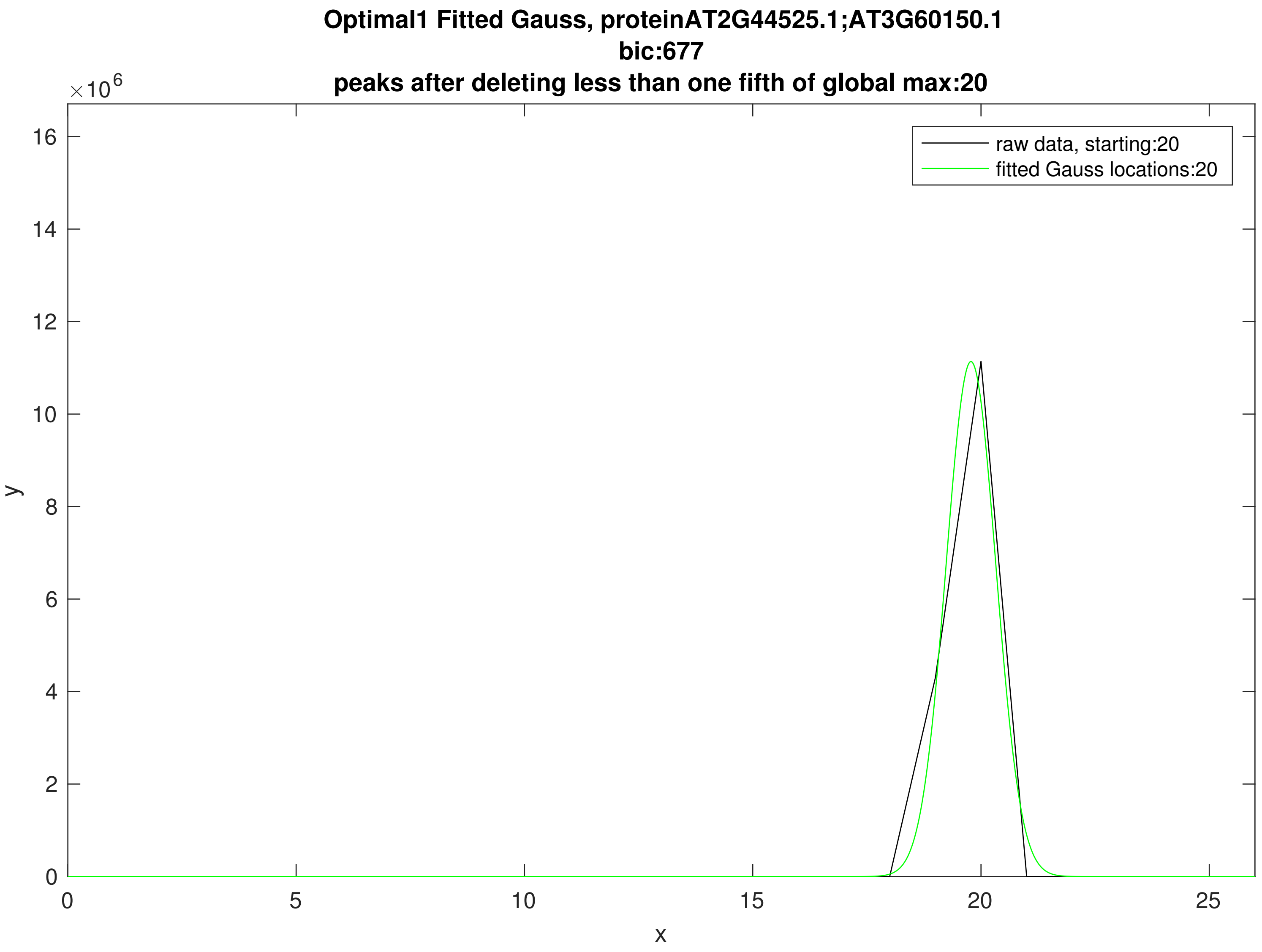
Optimal1 Fitted Gauss, proteinAT2G44525.1;AT3G60150.1
bic:677
peaks after deleting less than one fifth of global max:20
×10^6
raw data, starting:20
fitted Gauss locations:20
y
x
0
2
4
6
8
10
12
14
16
0
5
10
15
20
25

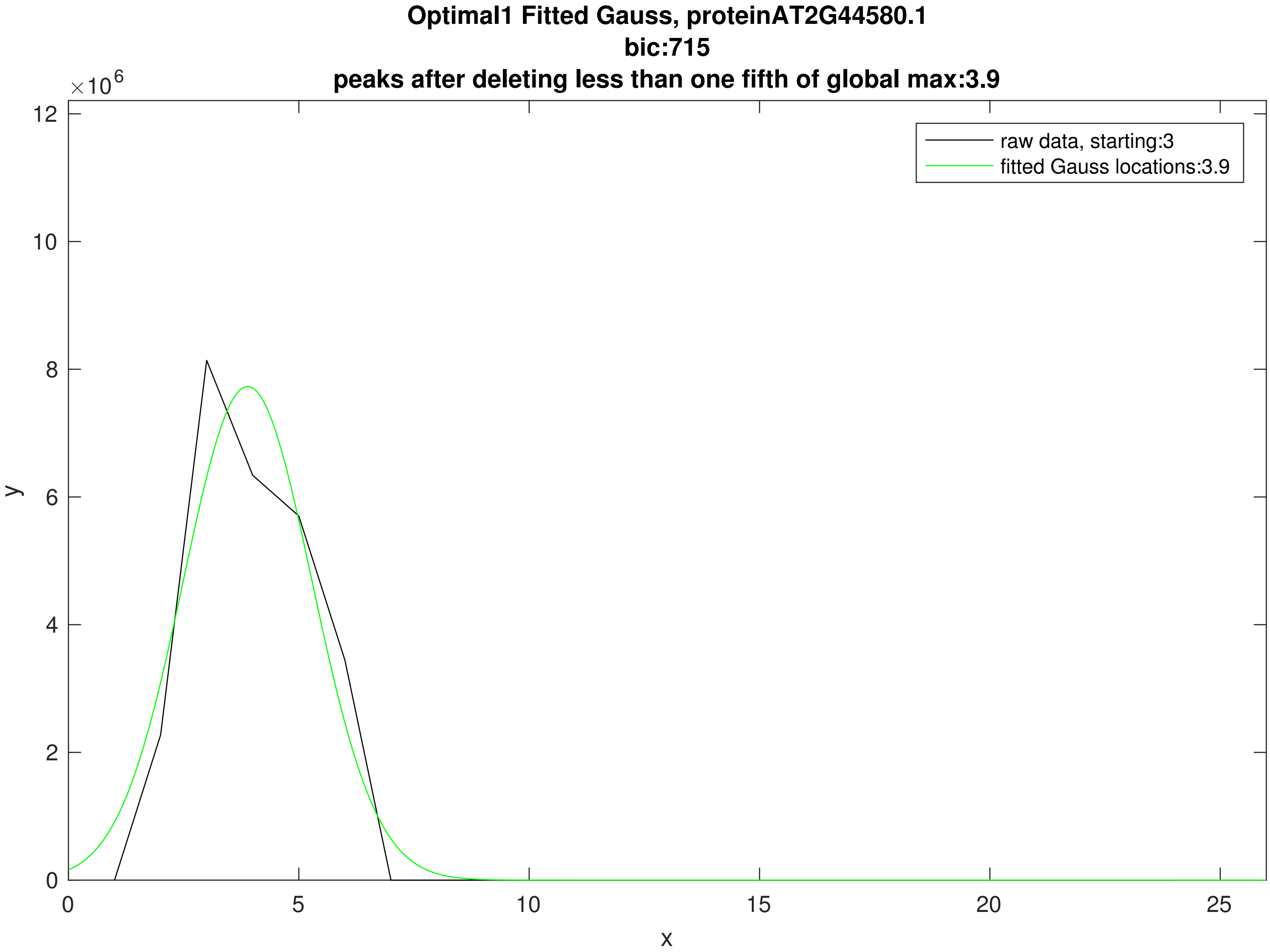

Optimal1 Fitted Gauss, proteinAT2G44580.1
bic:715
peaks after deleting less than one fifth of global max:3.9
×10^6
12
10
8
6
4
2
0
y
0
5
10
15
20
25
x
raw data, starting:3
fitted Gauss locations:3.9

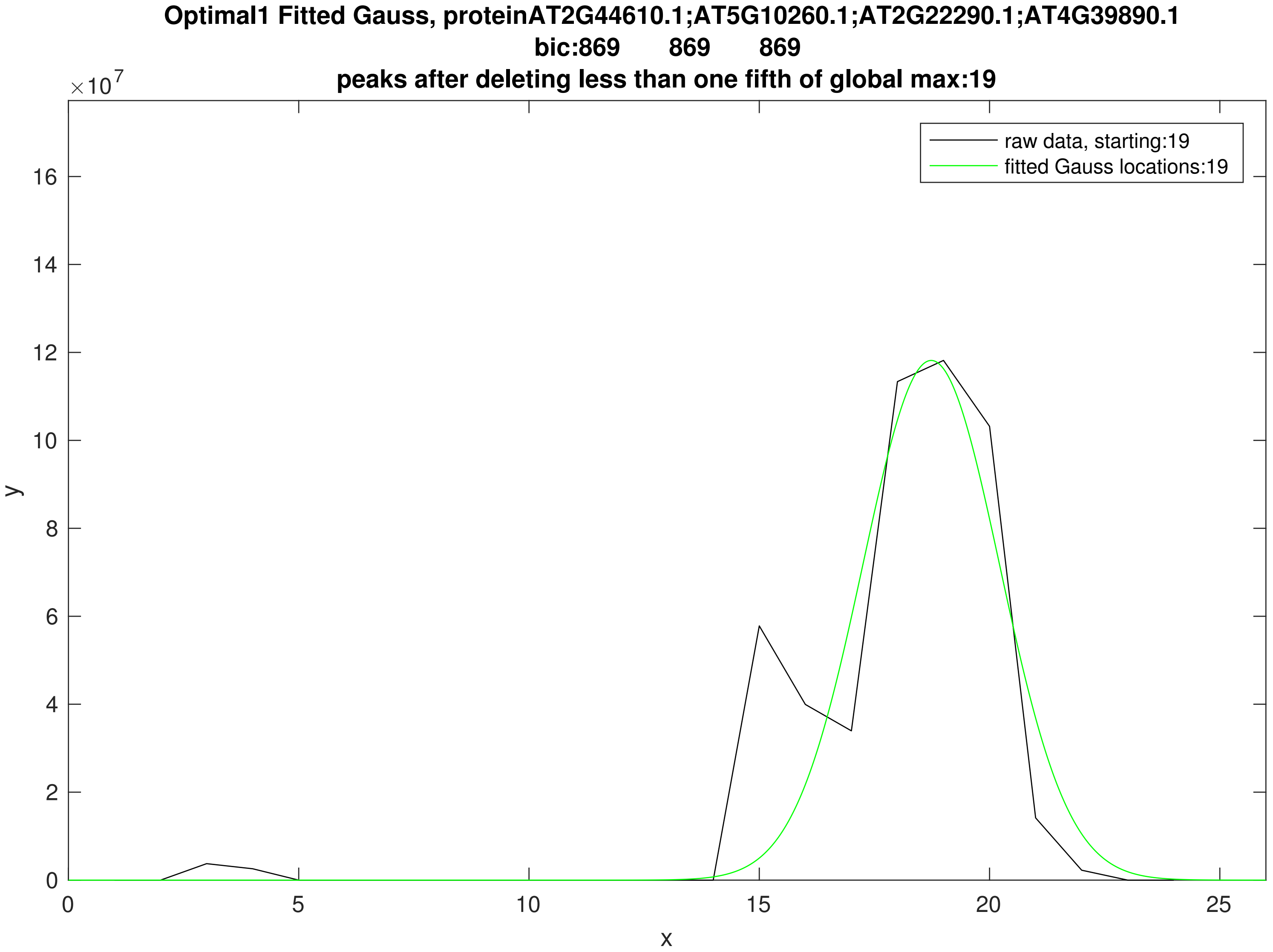

Optimal1 Fitted Gauss, proteinAT2G44610.1;AT5G10260.1;AT2G22290.1;AT4G39890.1
bic:869 869 869
peaks after deleting less than one fifth of global max:19
×10^7
raw data, starting:19
fitted Gauss locations:19
y
x
0
2
4
6
8
10
12
14
16
0
5
10
15
20
25

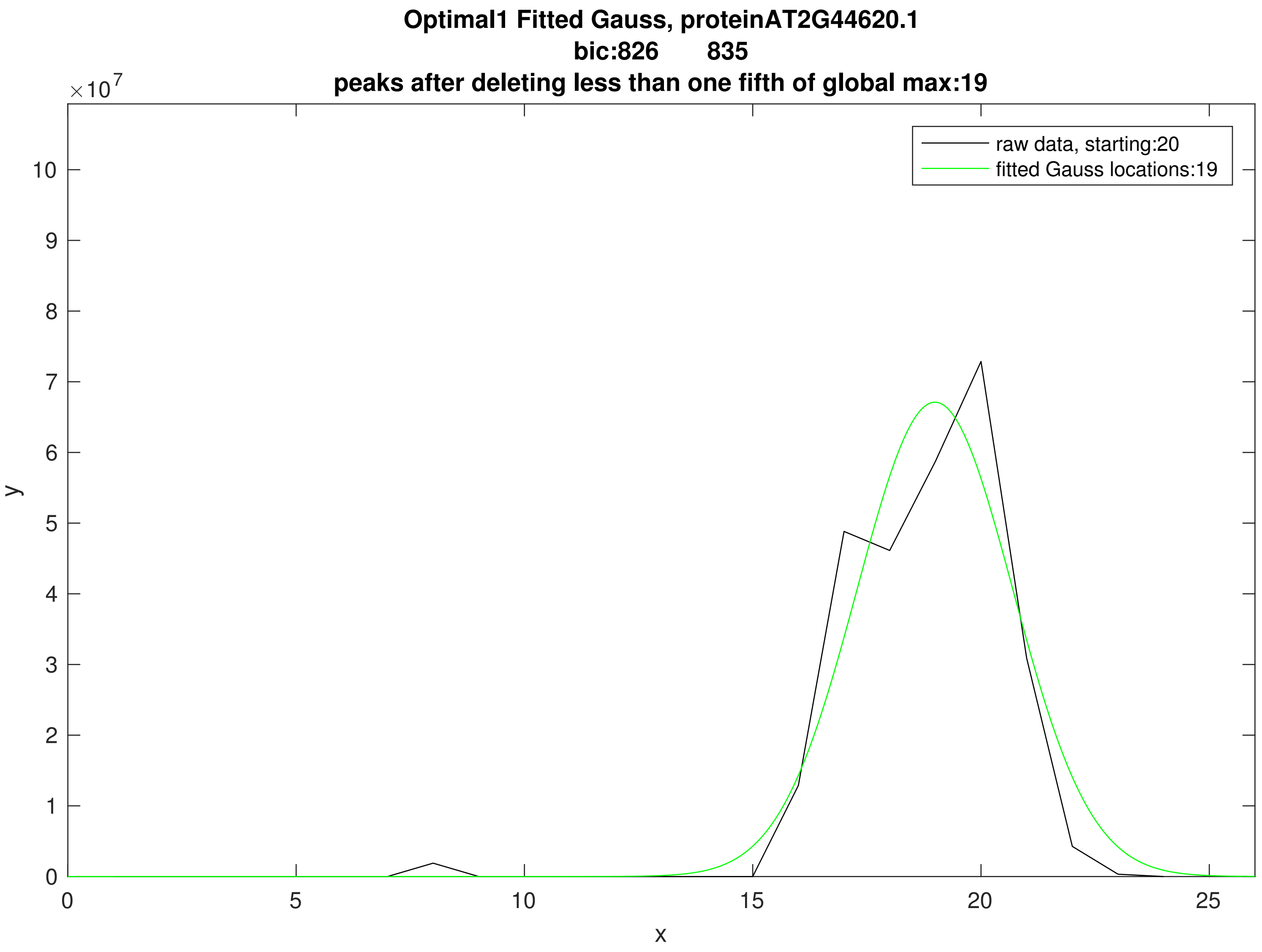

Optimal1 Fitted Gauss, proteinAT2G44620.1
bic:826 835
peaks after deleting less than one fifth of global max:19
×10^7
raw data, starting:20
fitted Gauss locations:19
y
x
0
1
2
3
4
5
6
7
8
9
10
0
5
10
15
20
25

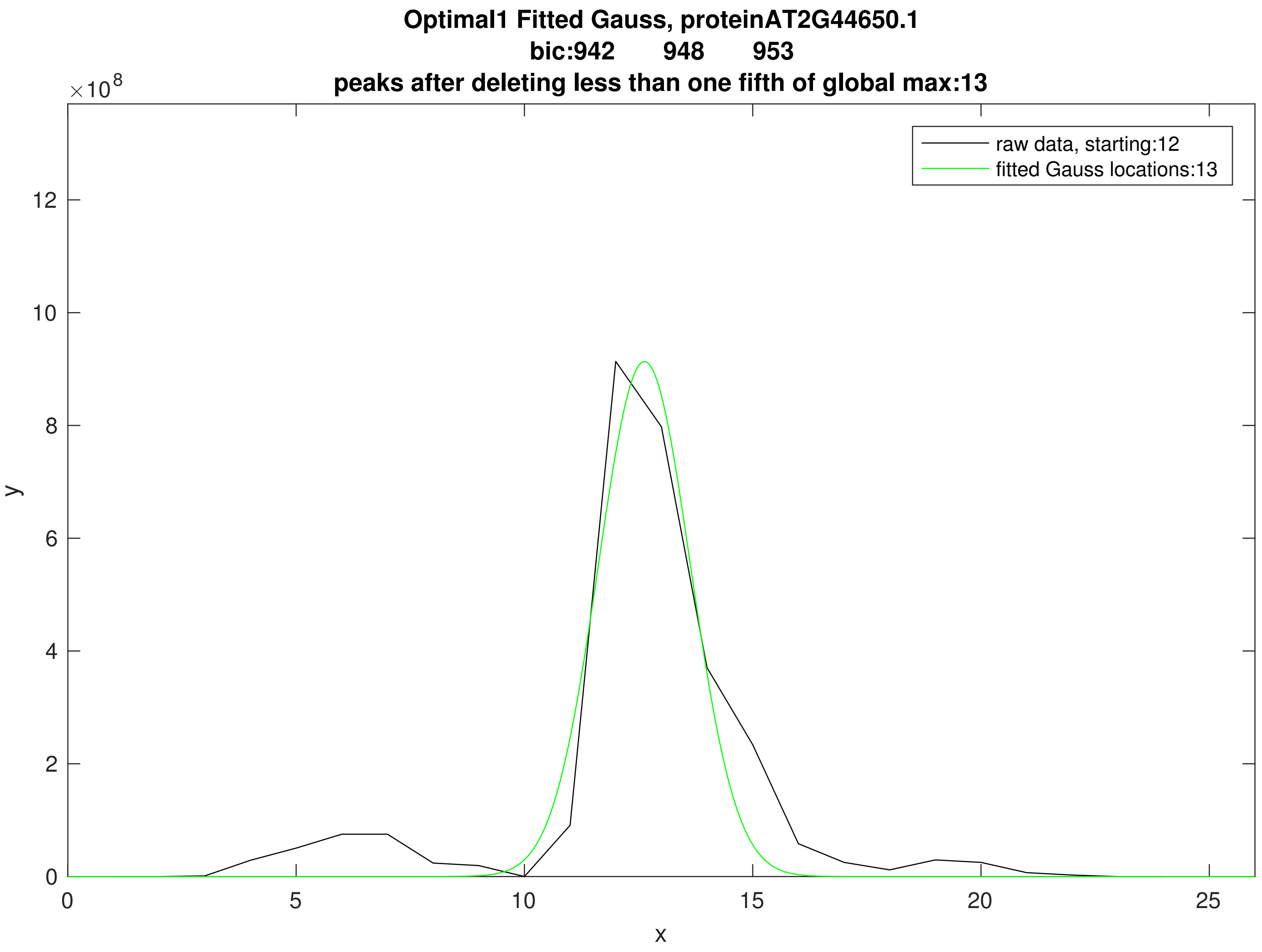
Optimal1 Fitted Gauss, proteinAT2G44650.1
bic:942 948 953
peaks after deleting less than one fifth of global max:13
×10^8
raw data, starting:12
fitted Gauss locations:13
y
x
0
2
4
6
8
10
12
0
5
10
15
20
25

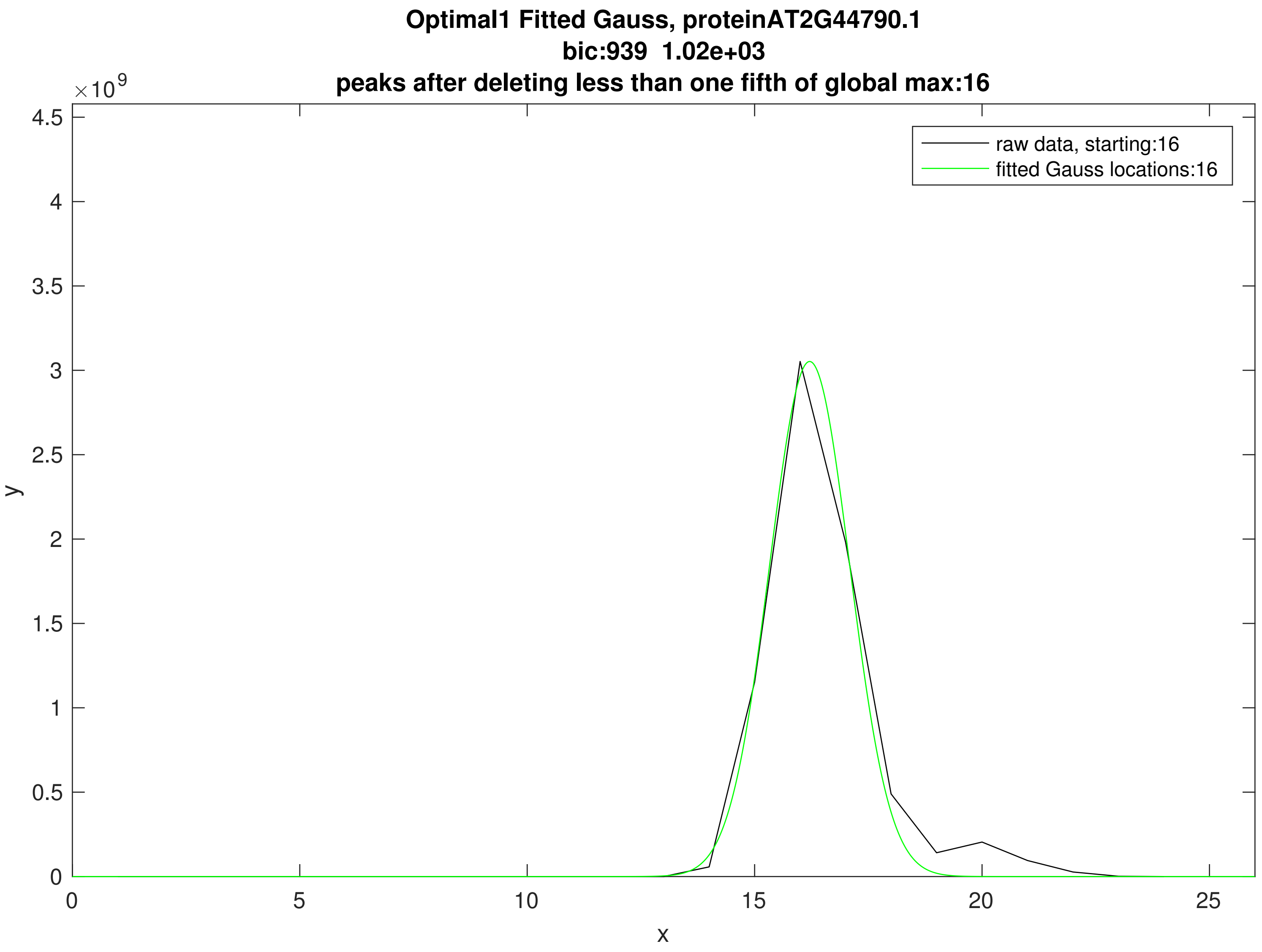

Optimal1 Fitted Gauss, proteinAT2G44790.1
bic:939 1.02e+03
peaks after deleting less than one fifth of global max:16
×10^9
4.5
4
3.5
3
2.5
2
1.5
1
0.5
0
y
0
5
10
15
20
25
x
raw data, starting:16
fitted Gauss locations:16

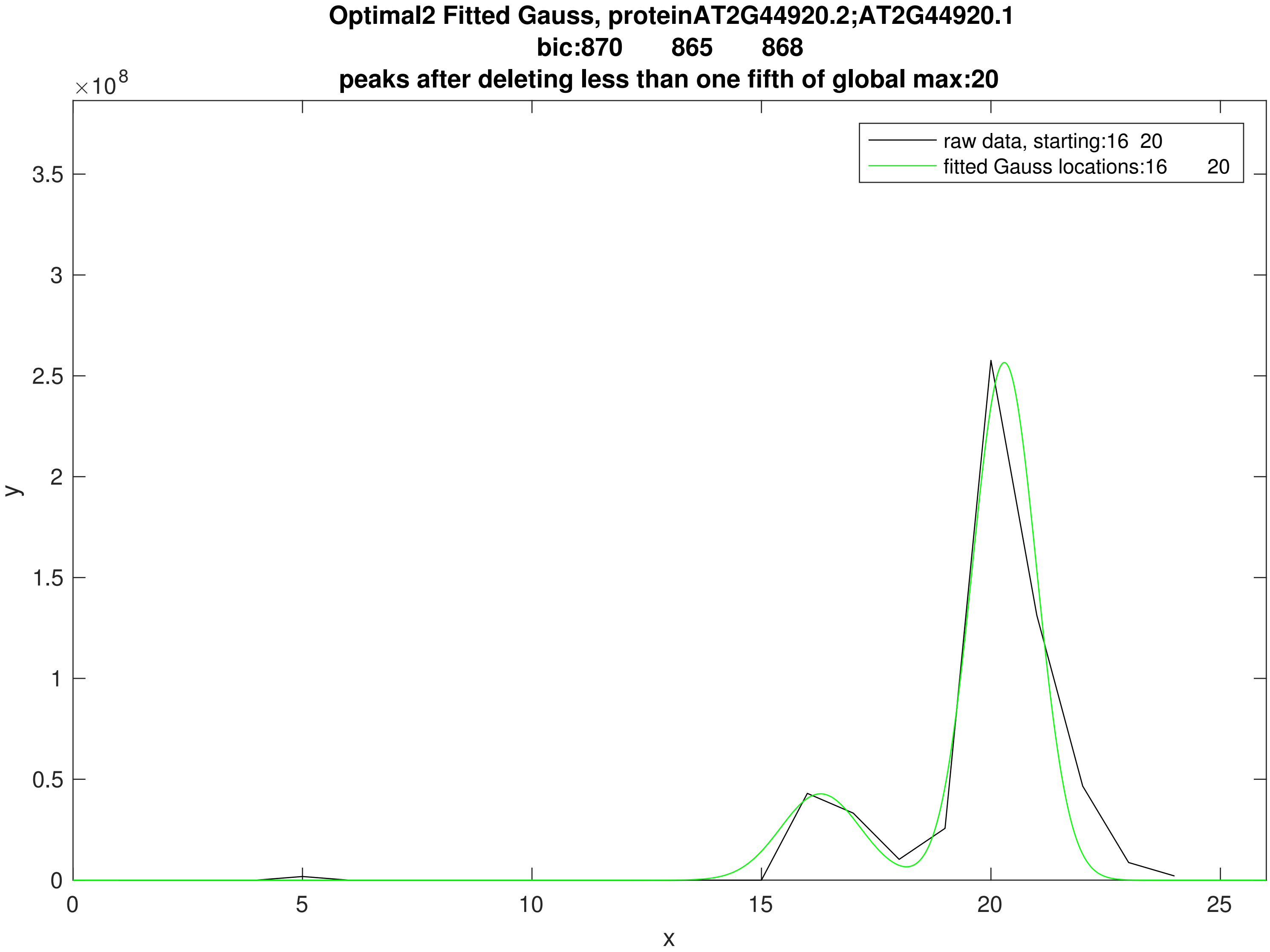
Optimal2 Fitted Gauss, proteinAT2G44920.2;AT2G44920.1
bic:870 865 868
peaks after deleting less than one fifth of global max:20
×10^8
raw data, starting:16 20
fitted Gauss locations:16 20
3.5
3
2.5
2
1.5
1
0.5
0
y
0
5
10
15
20
25
x

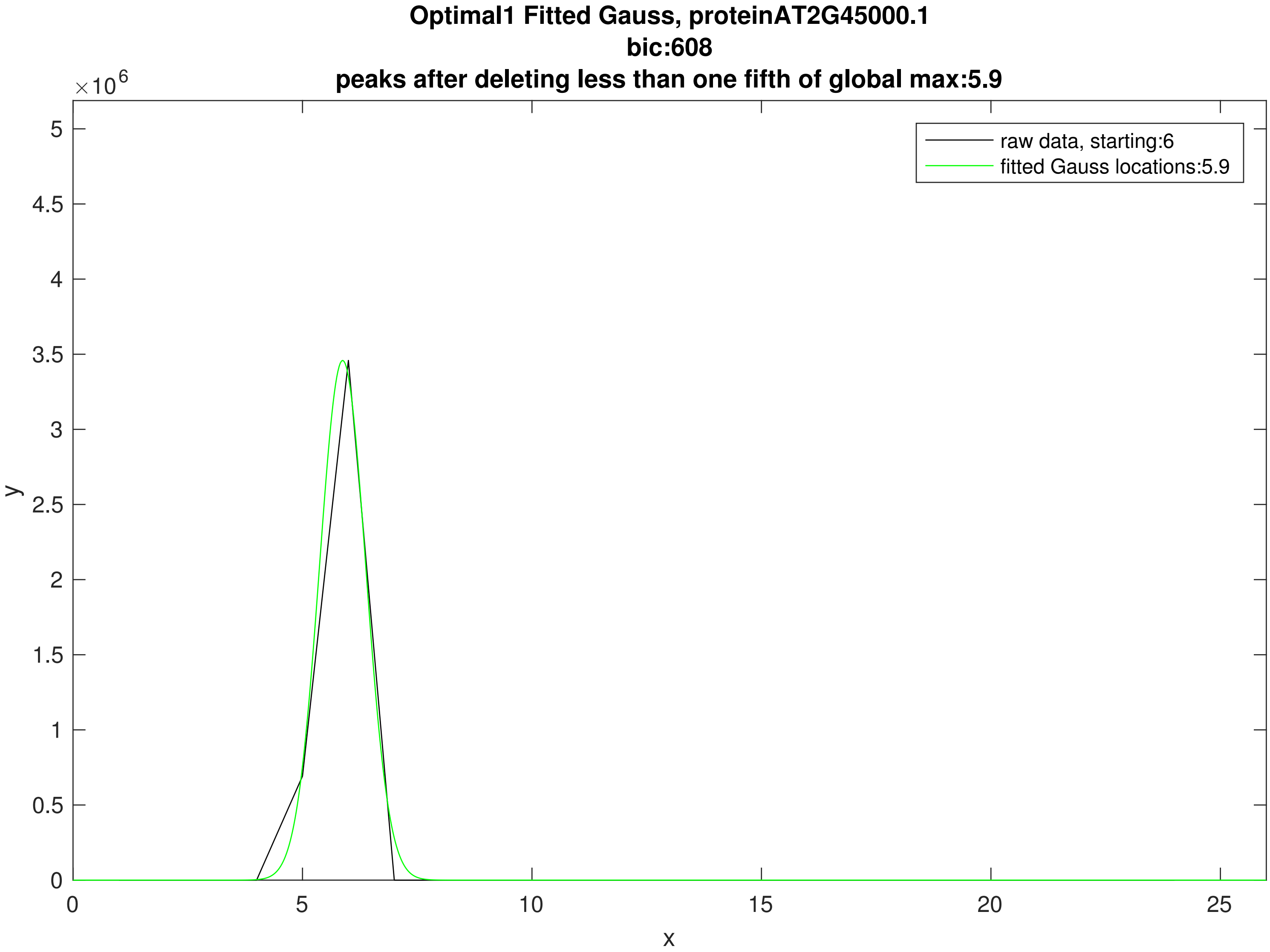

Optimal1 Fitted Gauss, proteinAT2G45000.1
bic:608
peaks after deleting less than one fifth of global max:5.9
×10^6
raw data, starting:6
fitted Gauss locations:5.9
y
x

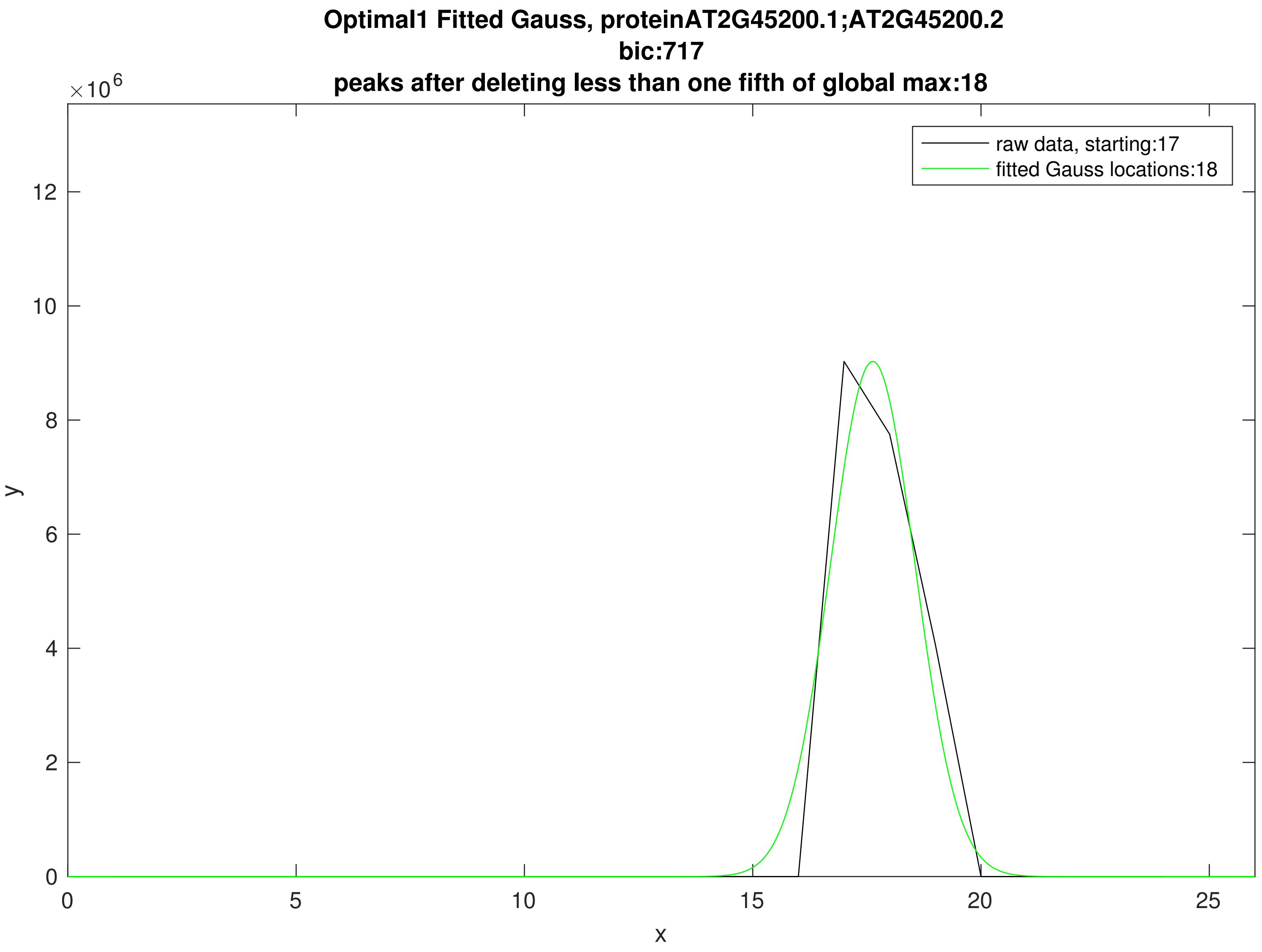
Optimal1 Fitted Gauss, proteinAT2G45200.1;AT2G45200.2
bic:717
peaks after deleting less than one fifth of global max:18
×10^6
raw data, starting:17
fitted Gauss locations:18
y
x
0
2
4
6
8
10
12
0
5
10
15
20
25

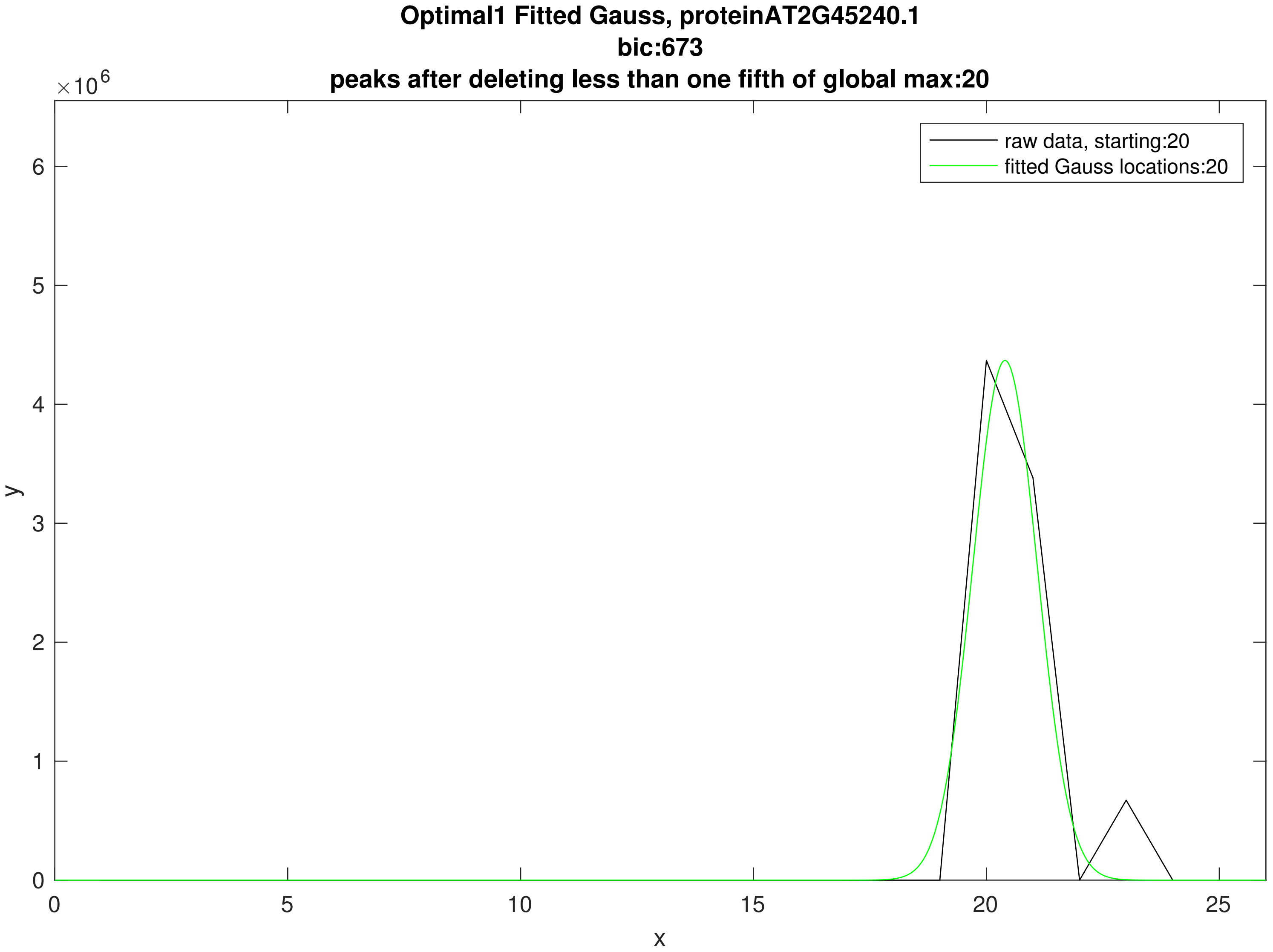
Optimal1 Fitted Gauss, proteinAT2G45240.1
bic:673
peaks after deleting less than one fifth of global max:20
×10^6
raw data, starting:20
fitted Gauss locations:20
y
x
0
1
2
3
4
5
6
0
5
10
15
20
25

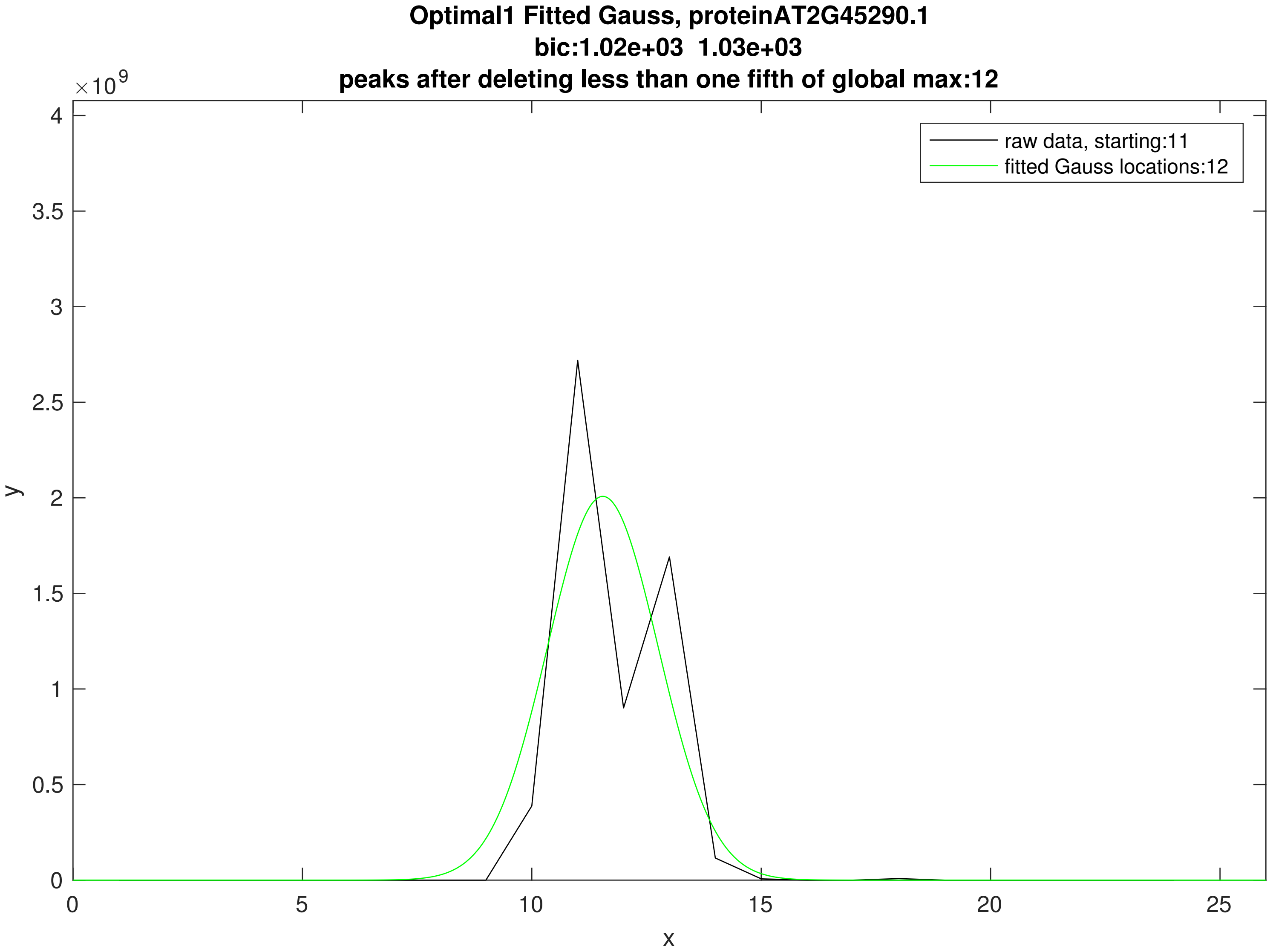
Optimal1 Fitted Gauss, proteinAT2G45290.1
bic:1.02e+03  1.03e+03
peaks after deleting less than one fifth of global max:12
×10^9
raw data, starting:11
fitted Gauss locations:12
y
x
0
0.5
1
1.5
2
2.5
3
3.5
4
0
5
10
15
20
25

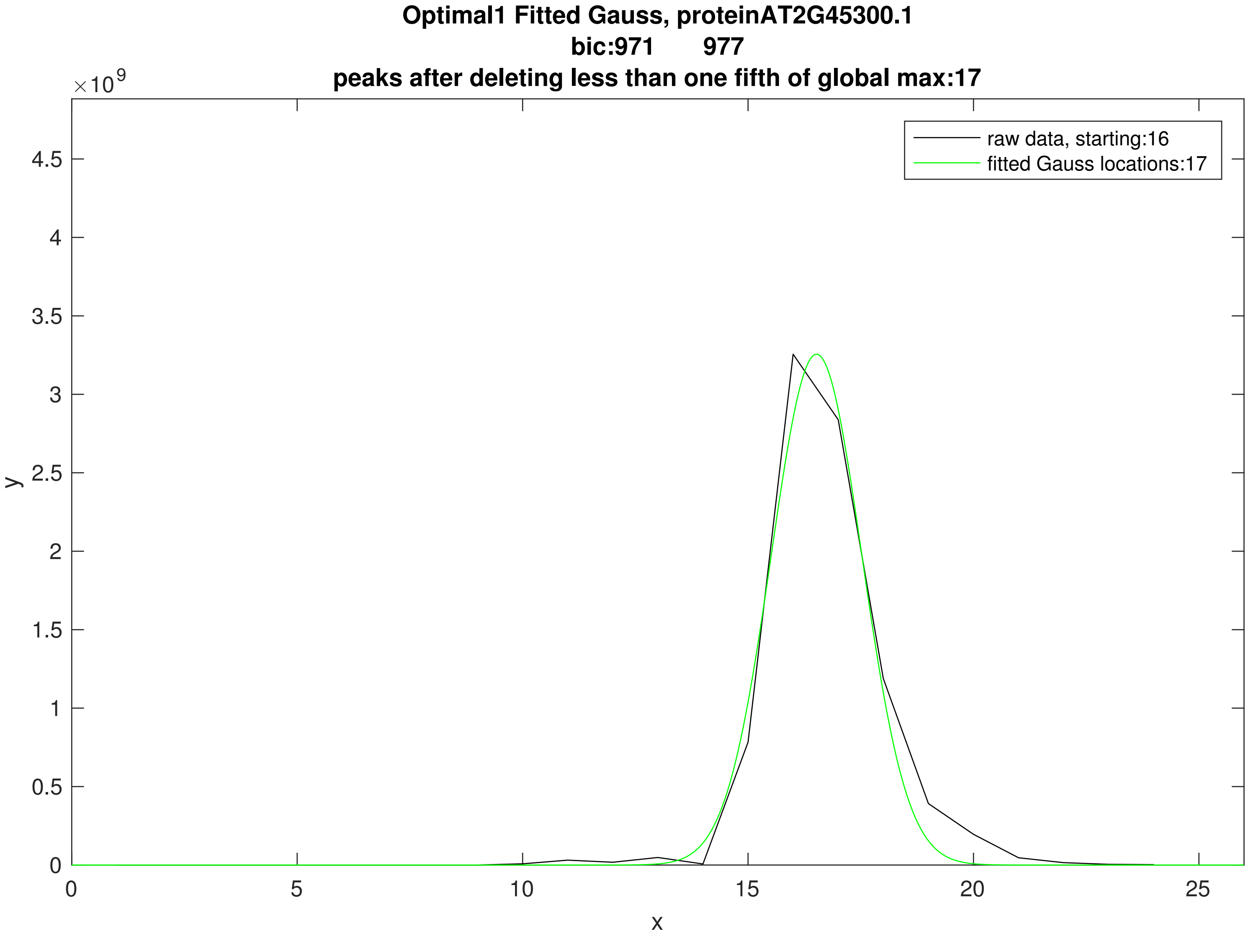
Optimal1 Fitted Gauss, proteinAT2G45300.1
bic:971 977
peaks after deleting less than one fifth of global max:17
×10^9
raw data, starting:16
fitted Gauss locations:17
y
x
0
0.5
1
1.5
2
2.5
3
3.5
4
4.5
0
5
10
15
20
25

**Optimal1 Fitted Gauss, proteinAT2G45440.1**
**bic:842**
**peaks after deleting less than one fifth of global max:13**

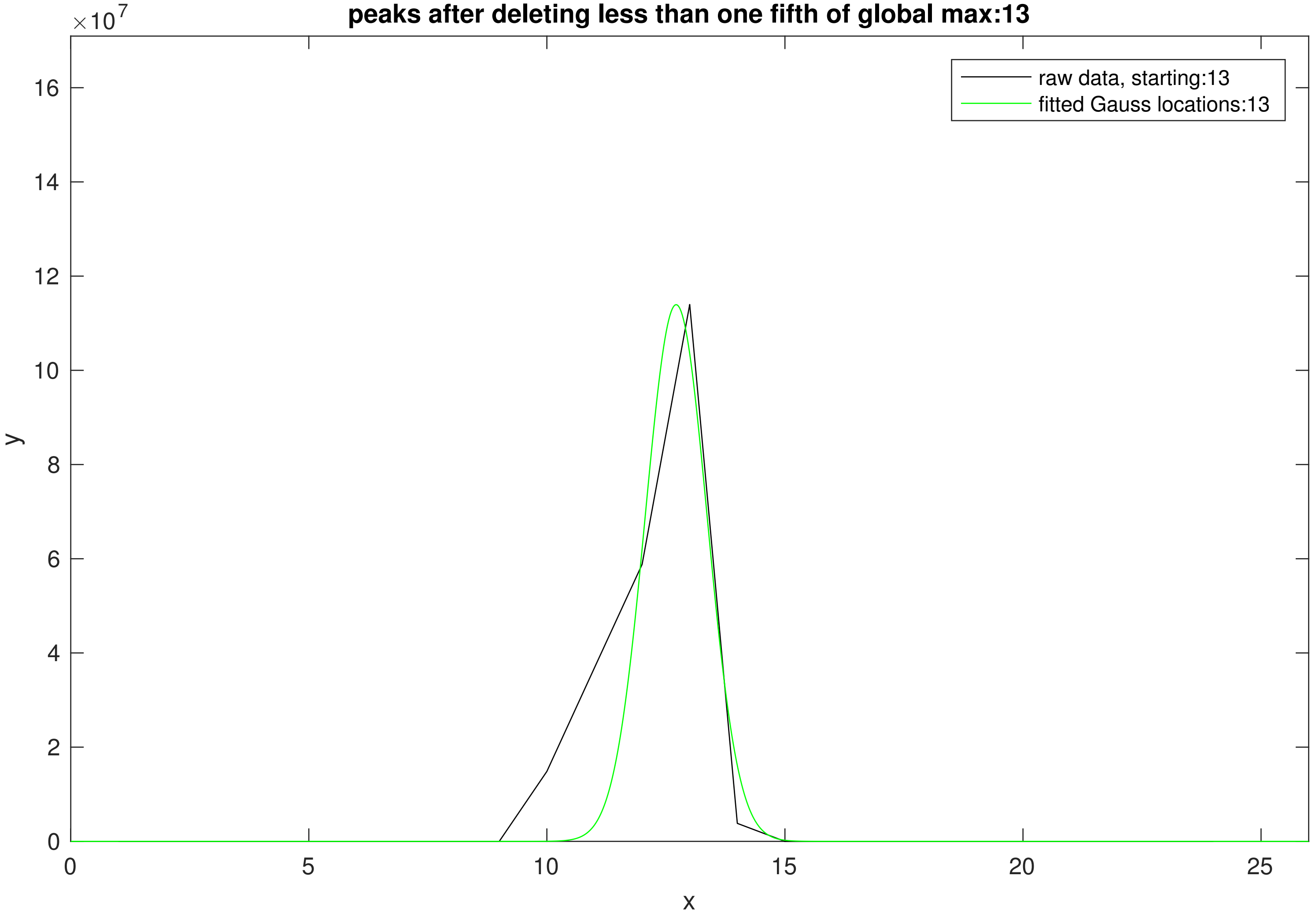

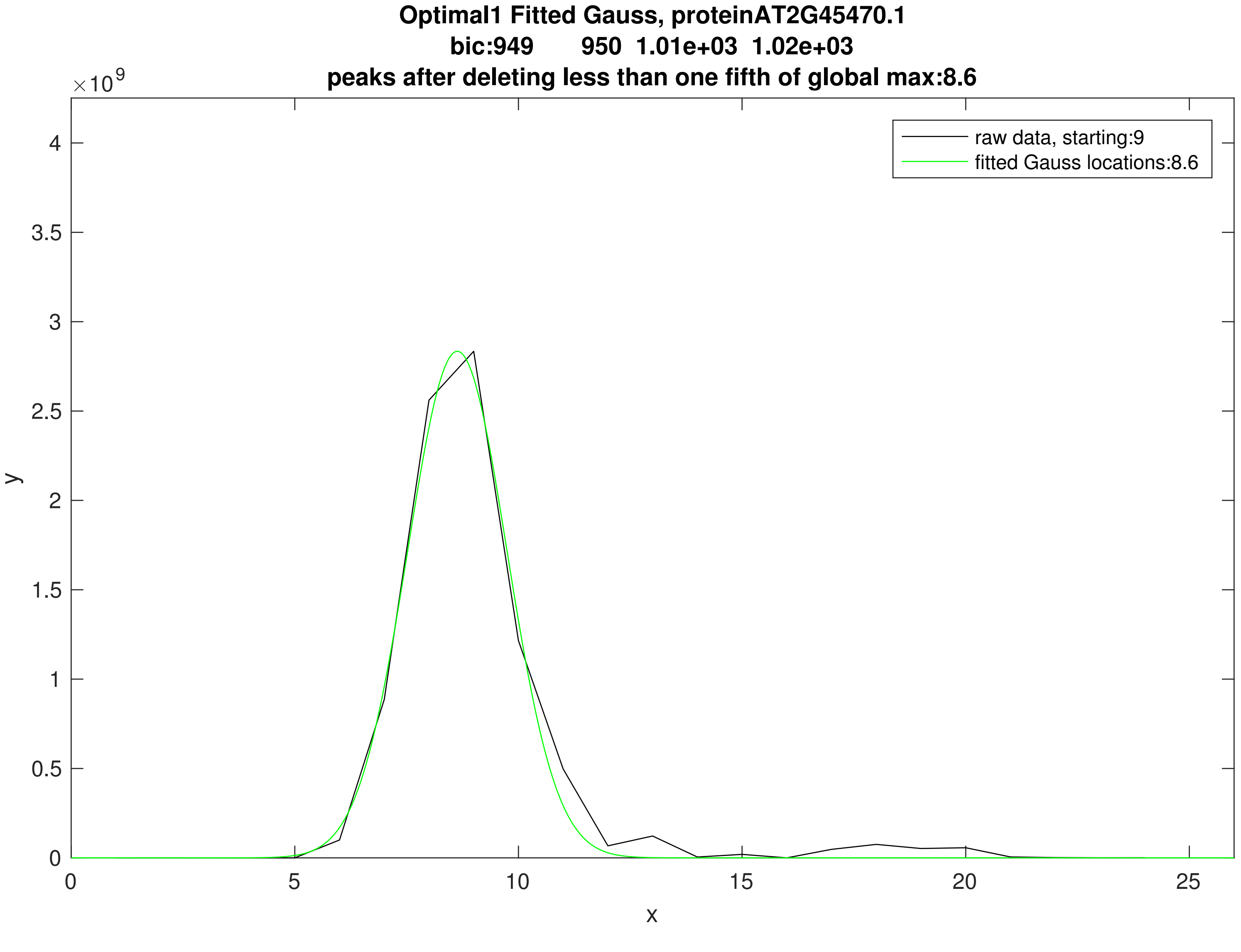
Optimal1 Fitted Gauss, proteinAT2G45470.1
bic:949 950 1.01e+03 1.02e+03
peaks after deleting less than one fifth of global max:8.6
×10^9
raw data, starting:9
fitted Gauss locations:8.6
y
x
0
0.5
1
1.5
2
2.5
3
3.5
4
5
10
15
20
25

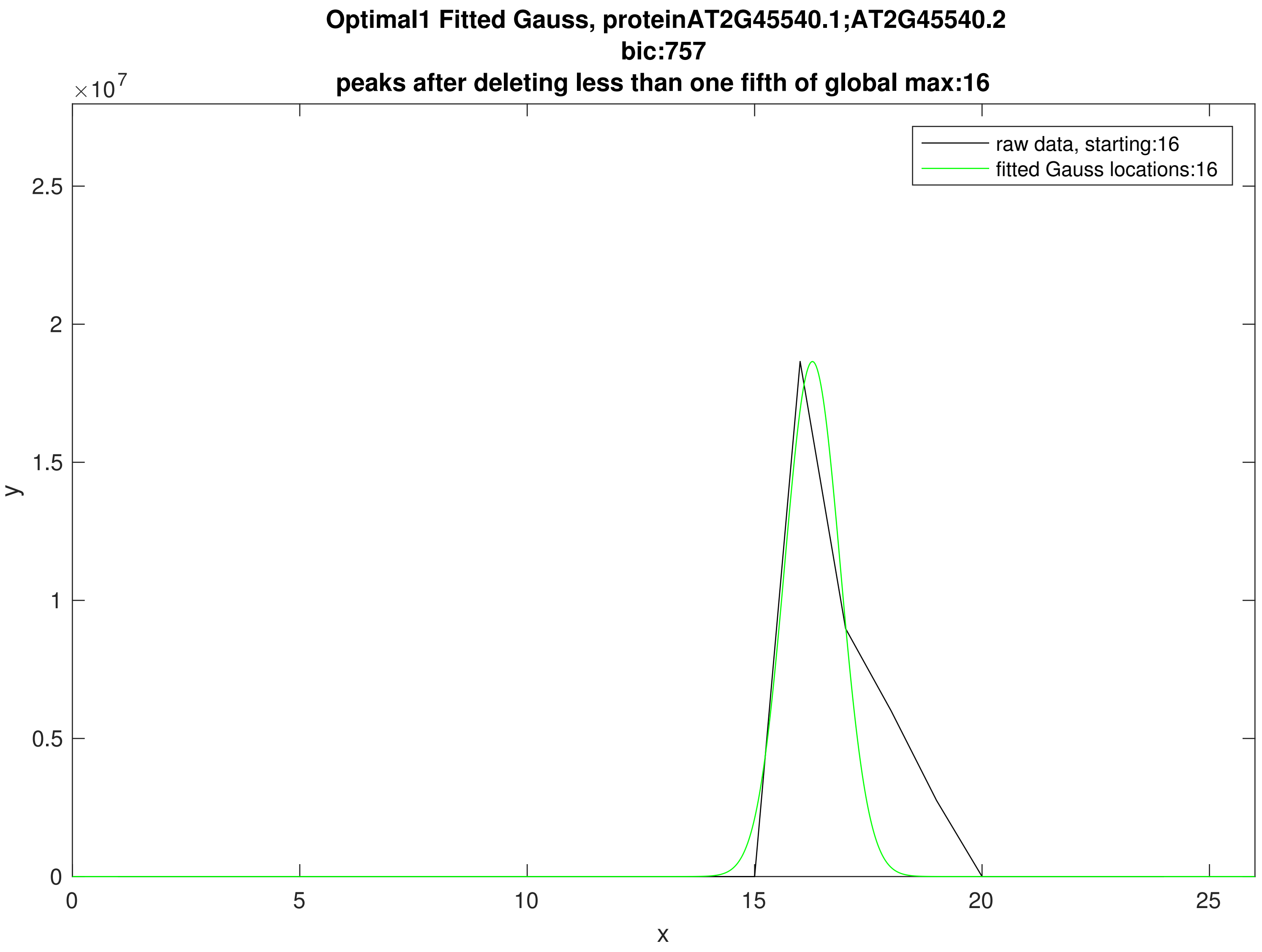
Optimal1 Fitted Gauss, proteinAT2G45540.1;AT2G45540.2
bic:757
peaks after deleting less than one fifth of global max:16
×10^7
raw data, starting:16
fitted Gauss locations:16
y
x
0
0.5
1
1.5
2
2.5
0
5
10
15
20
25

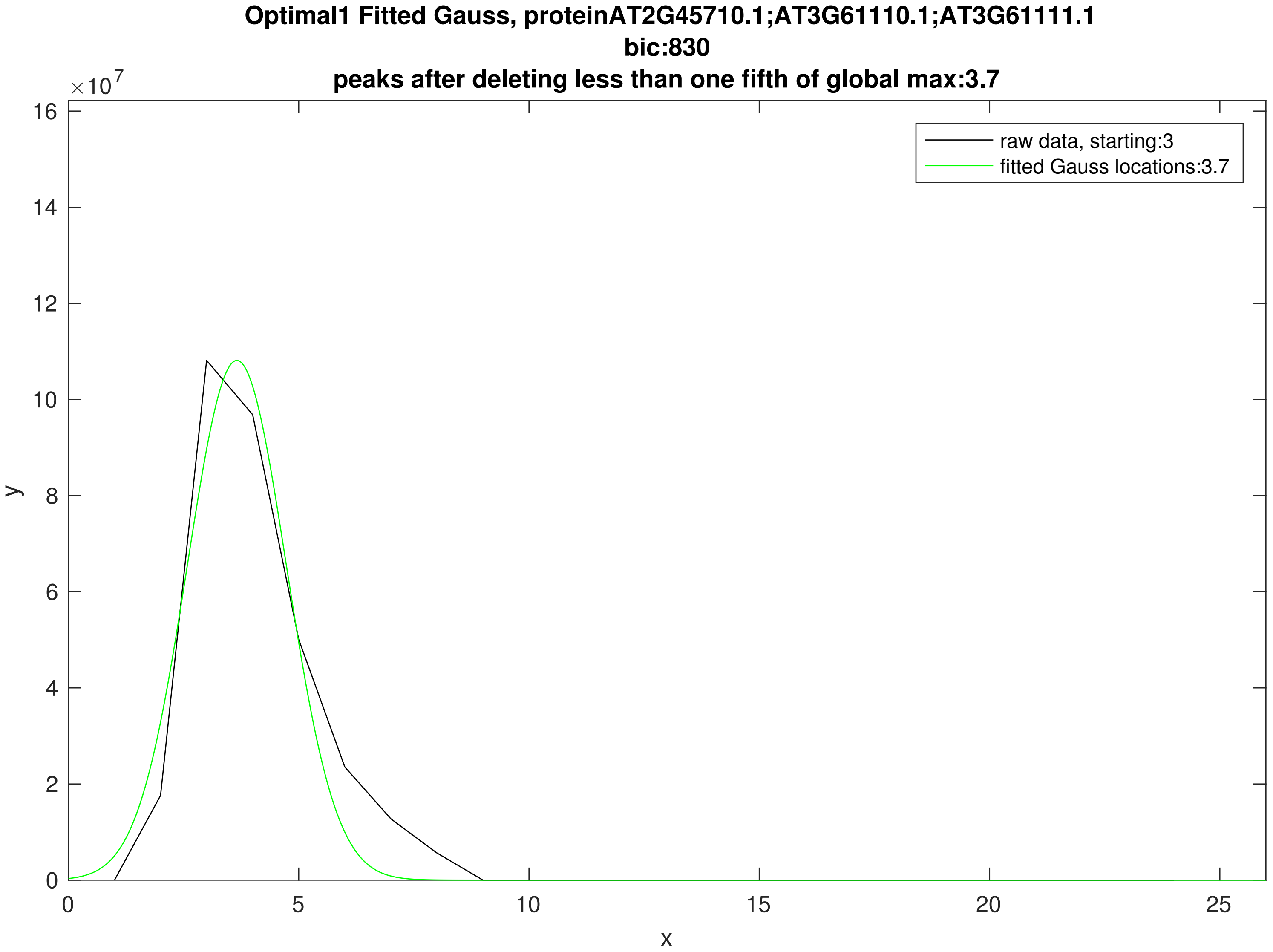

Optimal1 Fitted Gauss, proteinAT2G45710.1;AT3G61110.1;AT3G61111.1
bic:830
peaks after deleting less than one fifth of global max:3.7
×10^7
16
14
12
10
8
6
4
2
0
y
0
5
10
15
20
25
x
raw data, starting:3
fitted Gauss locations:3.7

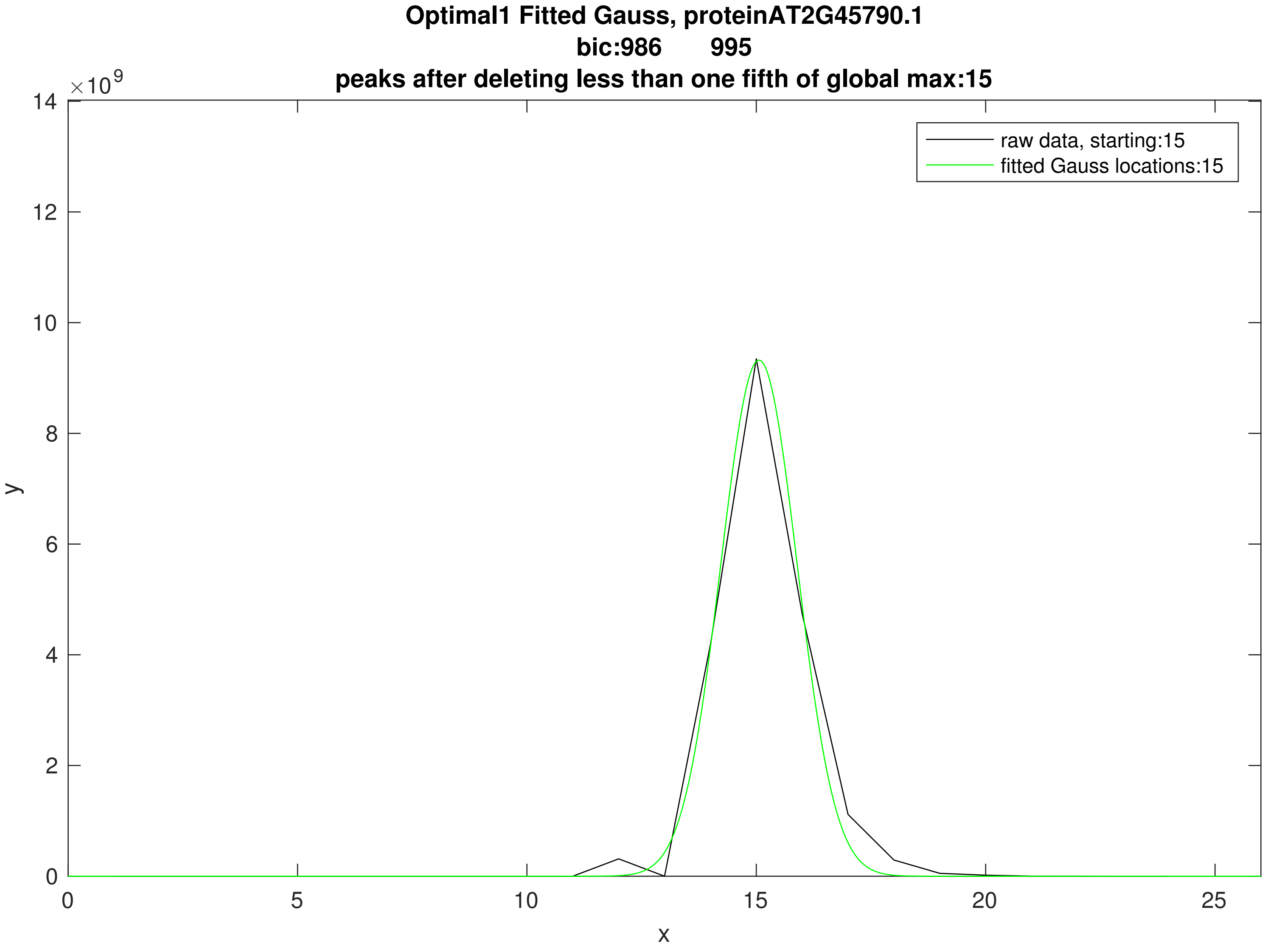

Optimal1 Fitted Gauss, proteinAT2G45790.1
bic:986 995
peaks after deleting less than one fifth of global max:15
×10^9
14
12
10
8
6
4
2
0
y
0
5
10
15
20
25
x
raw data, starting:15
fitted Gauss locations:15

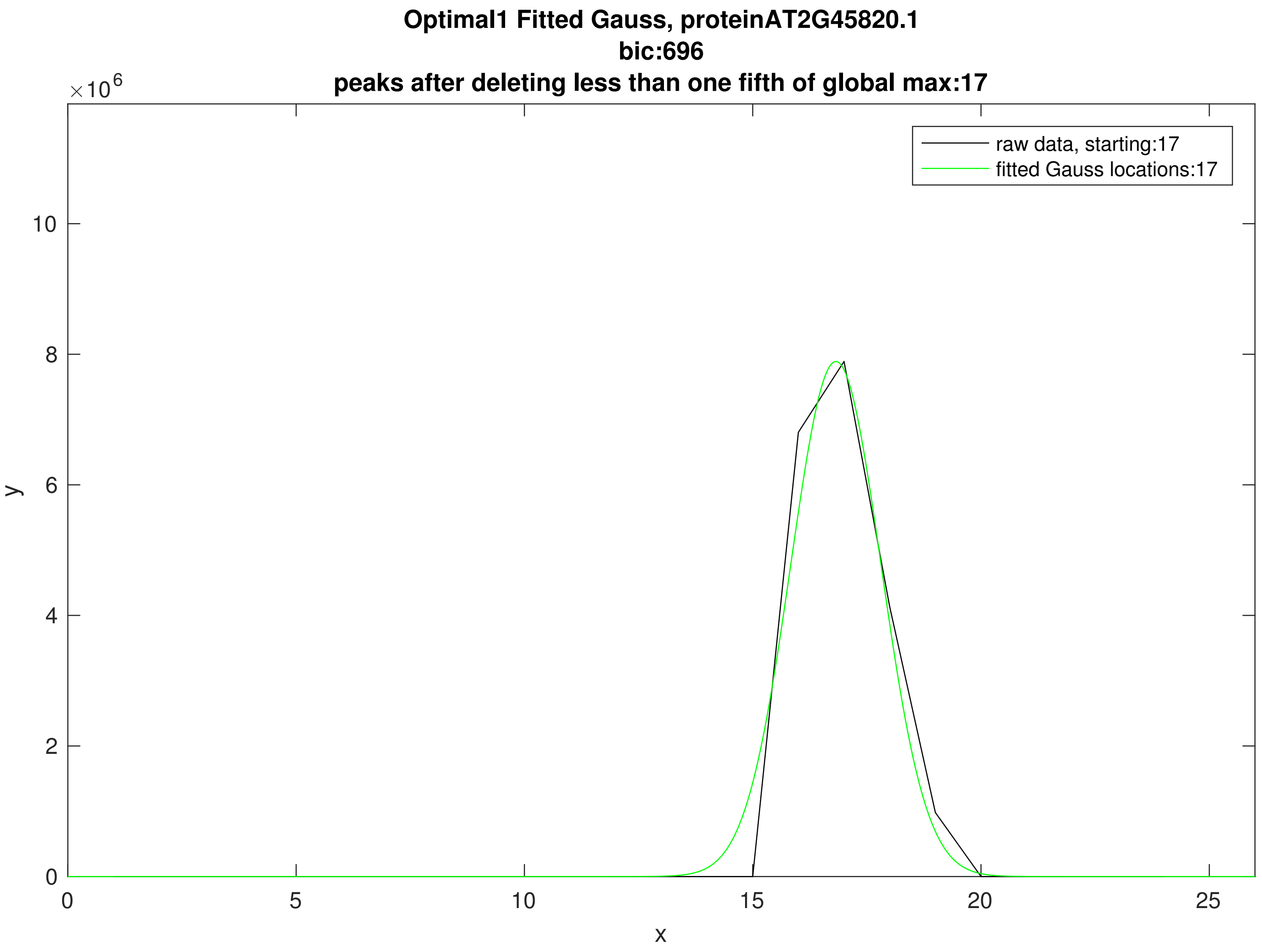
Optimal1 Fitted Gauss, proteinAT2G45820.1
bic:696
peaks after deleting less than one fifth of global max:17
×10^6
raw data, starting:17
fitted Gauss locations:17
y
x
0
2
4
6
8
10
0
5
10
15
20
25

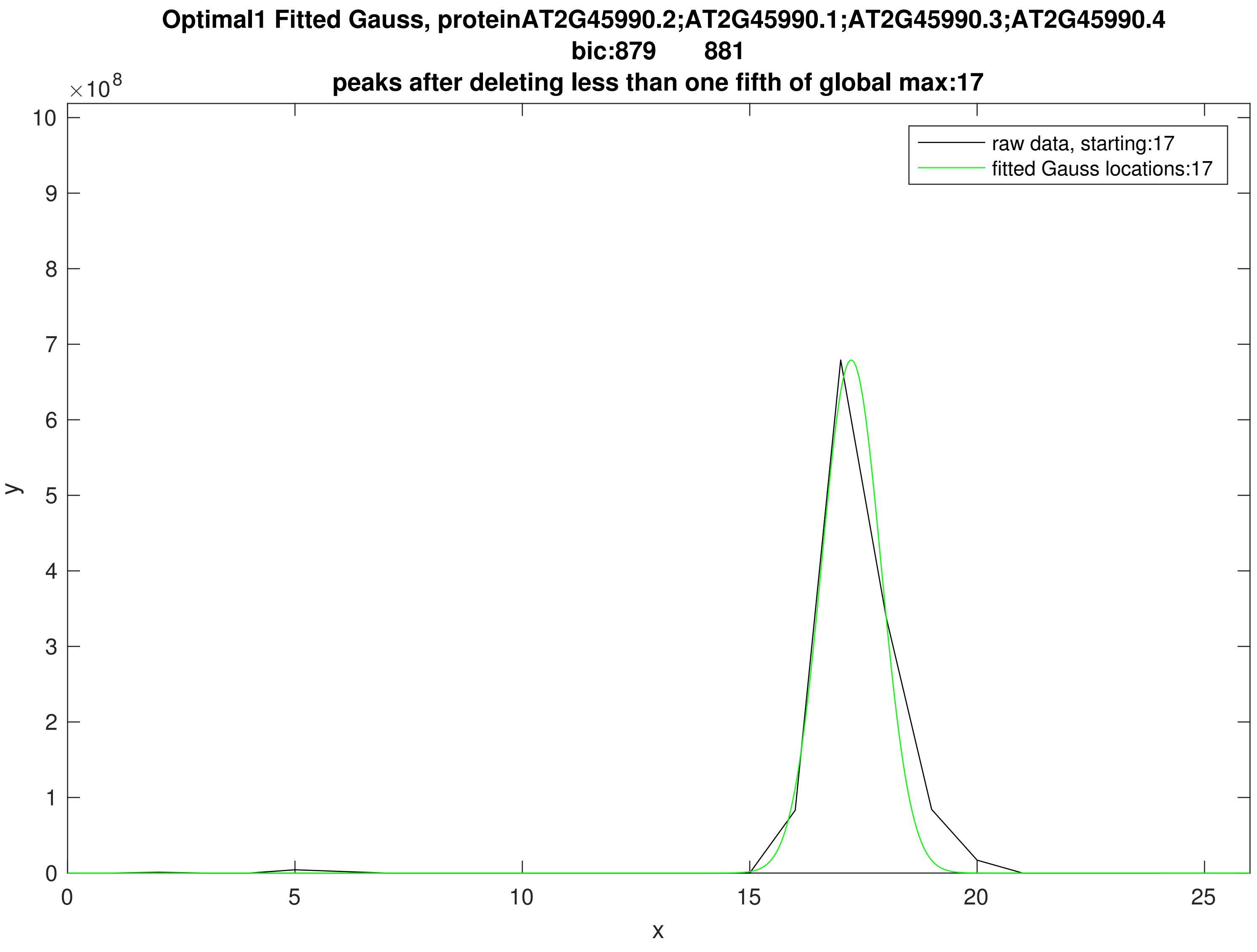
Optimal1 Fitted Gauss, proteinAT2G45990.2;AT2G45990.1;AT2G45990.3;AT2G45990.4
bic:879 881
peaks after deleting less than one fifth of global max:17
×10^8
y
x
raw data, starting:17
fitted Gauss locations:17

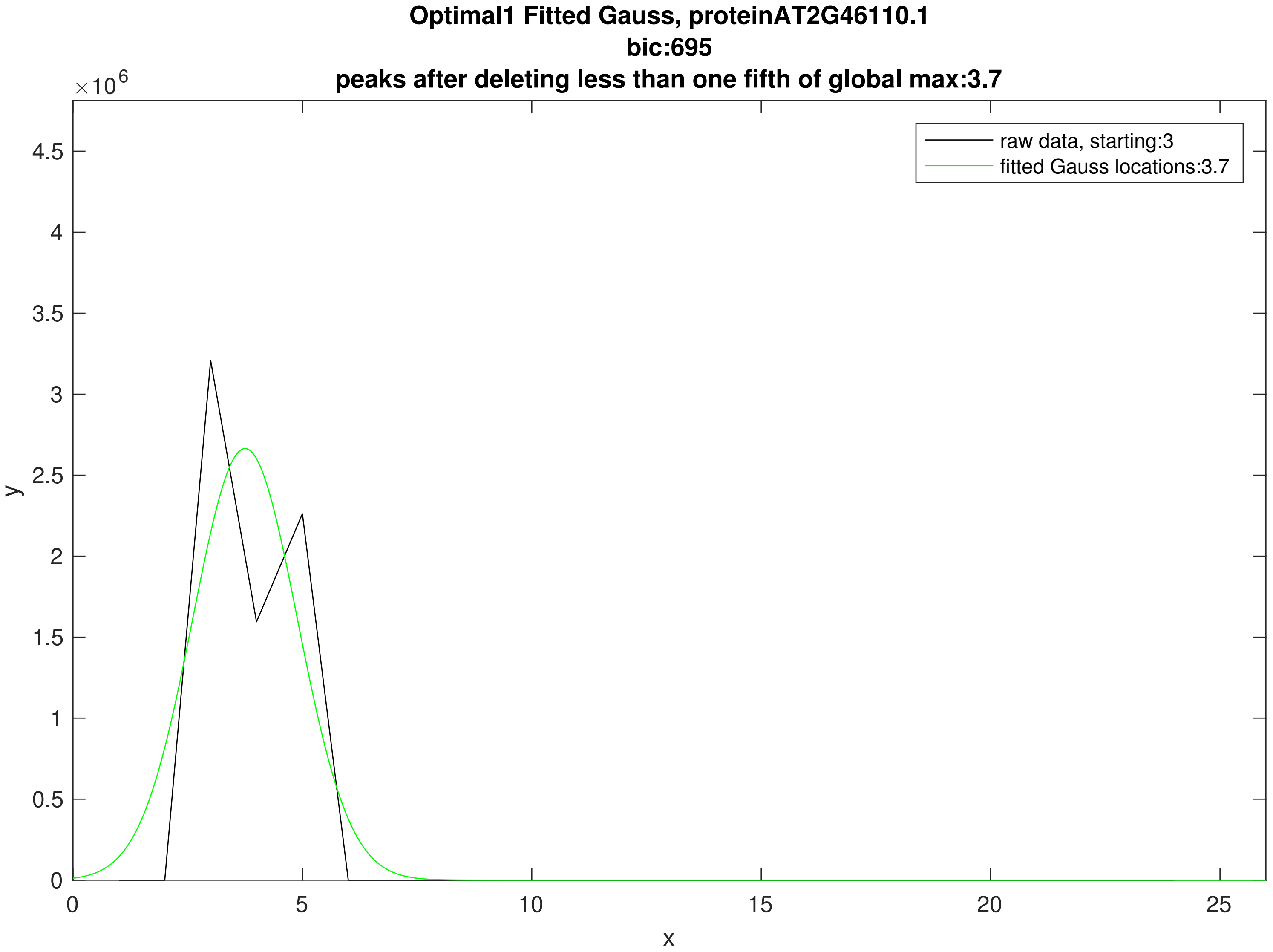

Optimal1 Fitted Gauss, proteinAT2G46110.1
bic:695
peaks after deleting less than one fifth of global max:3.7
×10^6
raw data, starting:3
fitted Gauss locations:3.7
4.5
4
3.5
3
2.5
2
1.5
1
0.5
0
y
0
5
10
15
20
25
x

**Optimal1 Fitted Gauss, proteinAT2G46140.1**
**bic:821**
**peaks after deleting less than one fifth of global max:19**

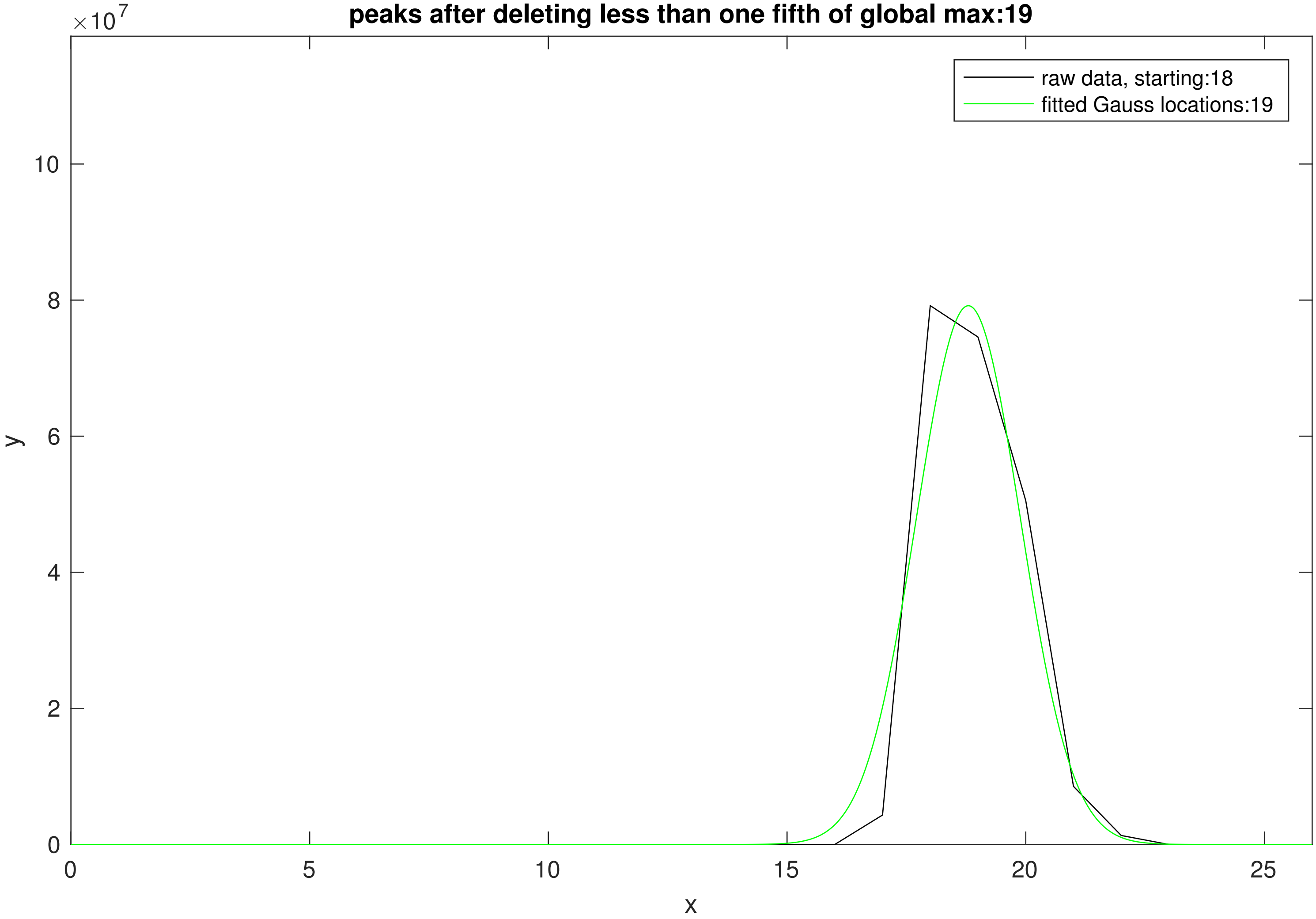

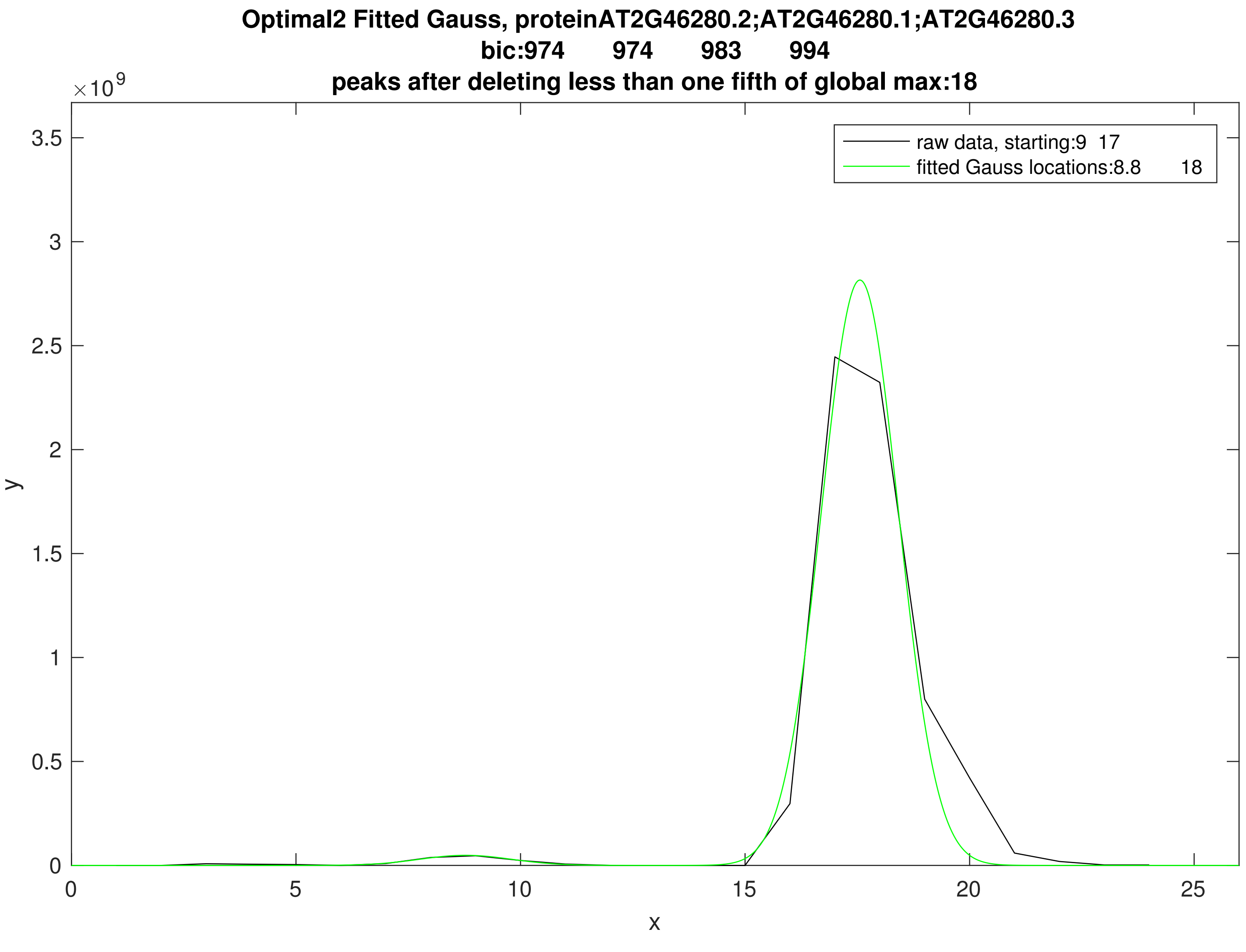

Optimal2 Fitted Gauss, proteinAT2G46280.2;AT2G46280.1;AT2G46280.3
bic:974 974 983 994
peaks after deleting less than one fifth of global max:18
×10^9
raw data, starting:9 17
fitted Gauss locations:8.8 18
3.5
3
2.5
2
1.5
1
0.5
0
y
0
5
10
15
20
25
x

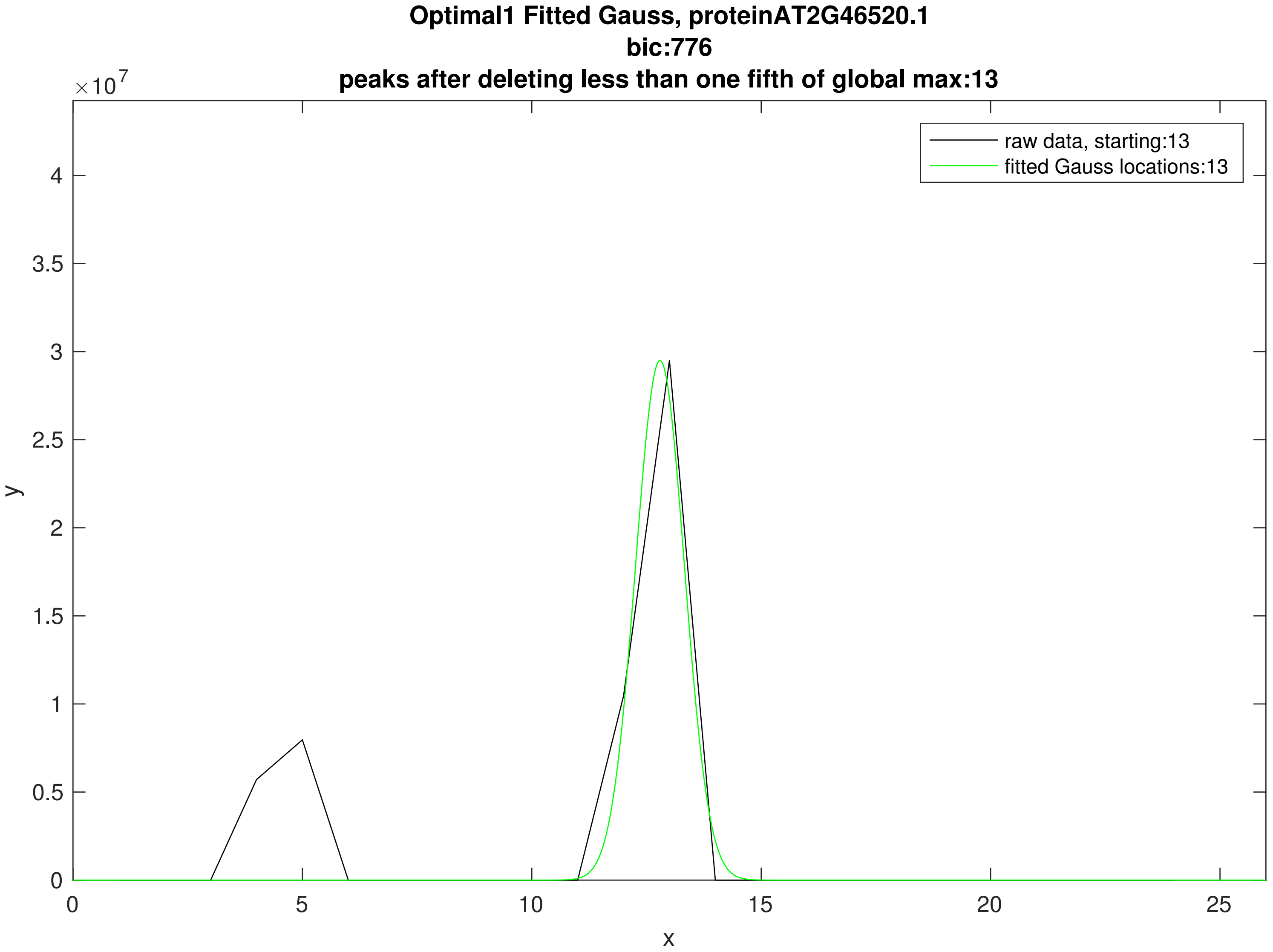

Optimal1 Fitted Gauss, proteinAT2G46520.1
bic:776
peaks after deleting less than one fifth of global max:13
×10^7
raw data, starting:13
fitted Gauss locations:13
y
x
0
0.5
1
1.5
2
2.5
3
3.5
4
0
5
10
15
20
25

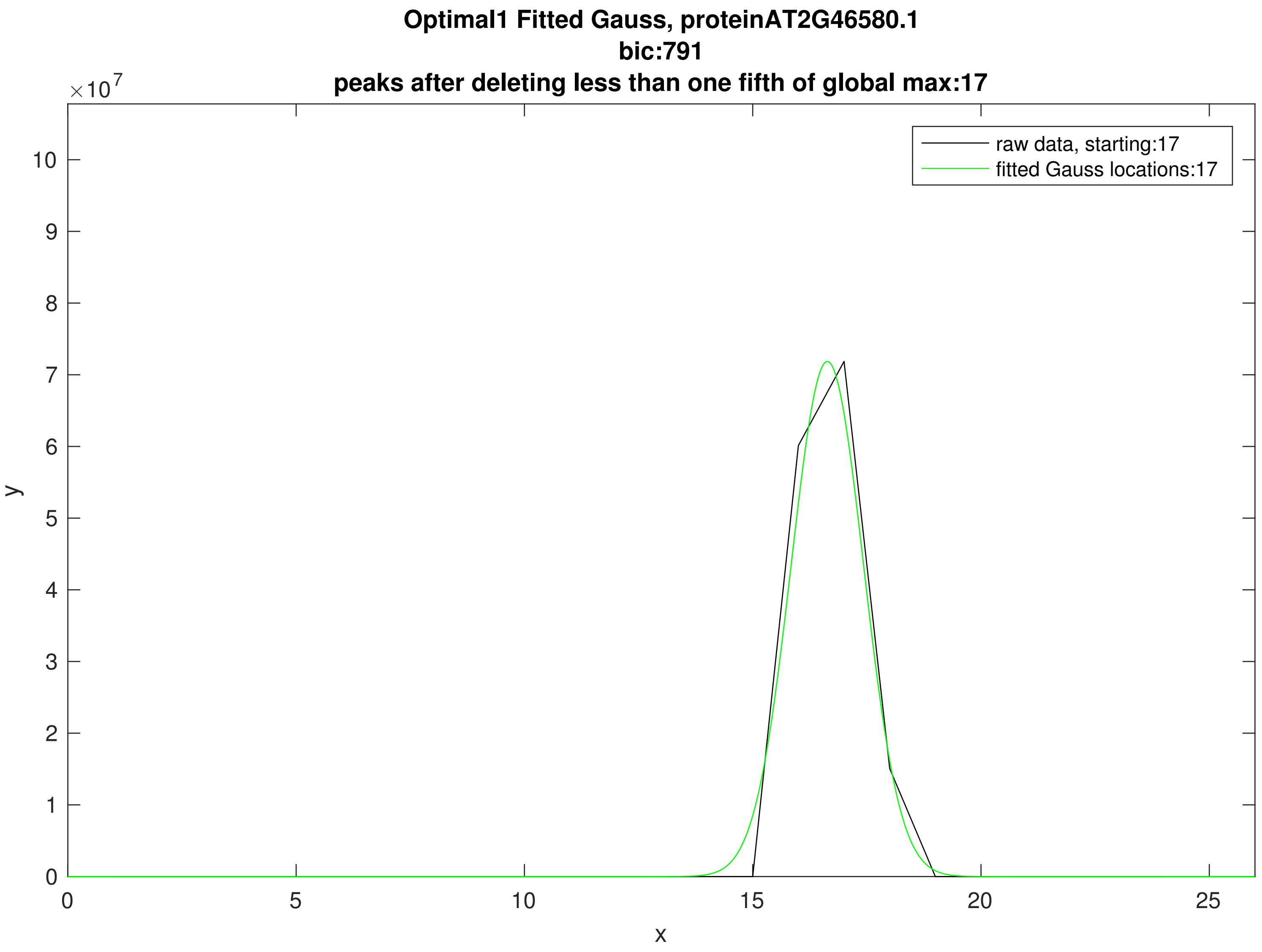
Optimal1 Fitted Gauss, proteinAT2G46580.1
bic:791
peaks after deleting less than one fifth of global max:17
×10^7
raw data, starting:17
fitted Gauss locations:17
y
x
0
1
2
3
4
5
6
7
8
9
10
15
20
25

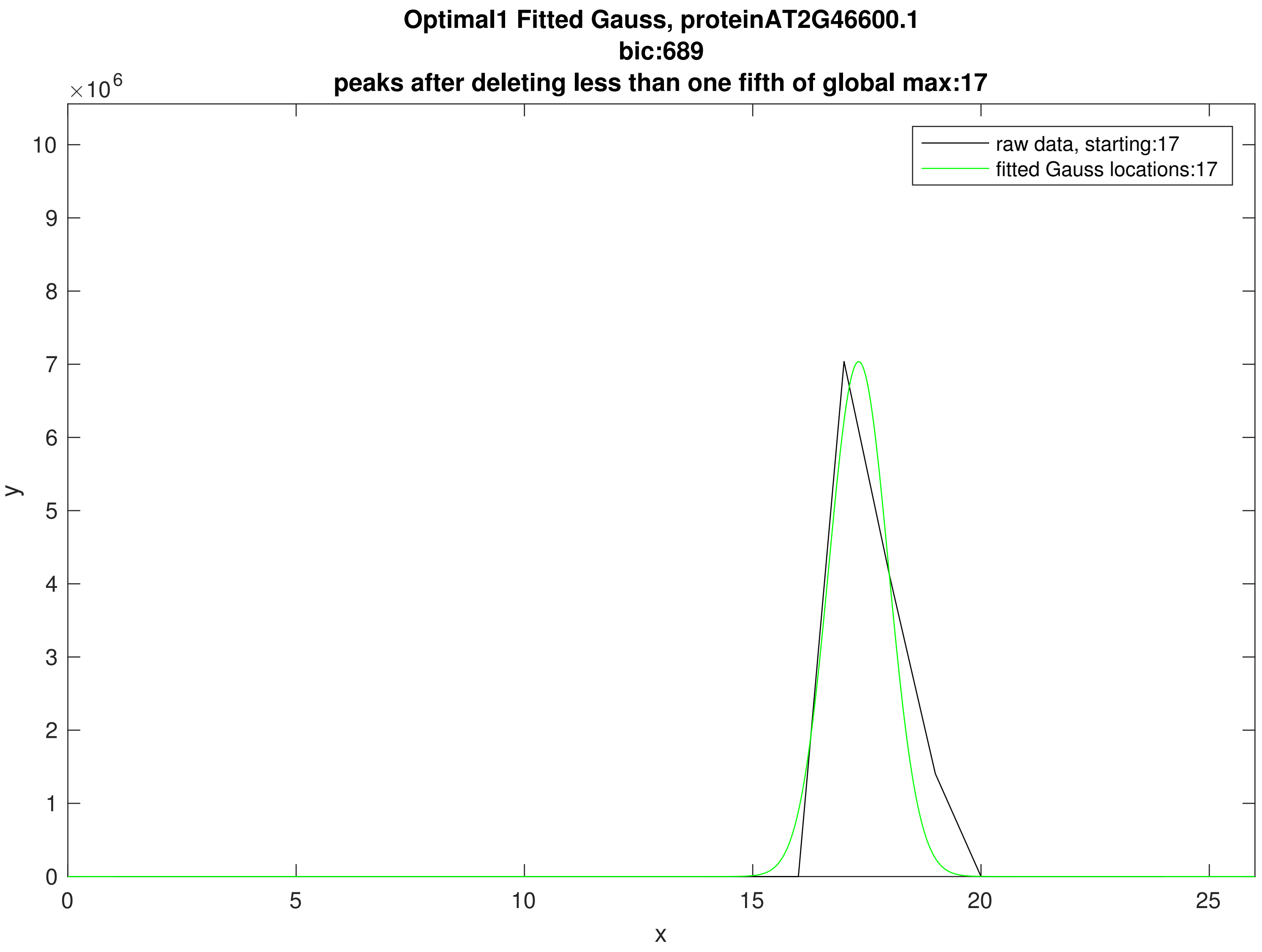
Optimal1 Fitted Gauss, proteinAT2G46600.1
bic:689
peaks after deleting less than one fifth of global max:17
×10^6
raw data, starting:17
fitted Gauss locations:17
y
x
0
1
2
3
4
5
6
7
8
9
10
0
5
10
15
20
25

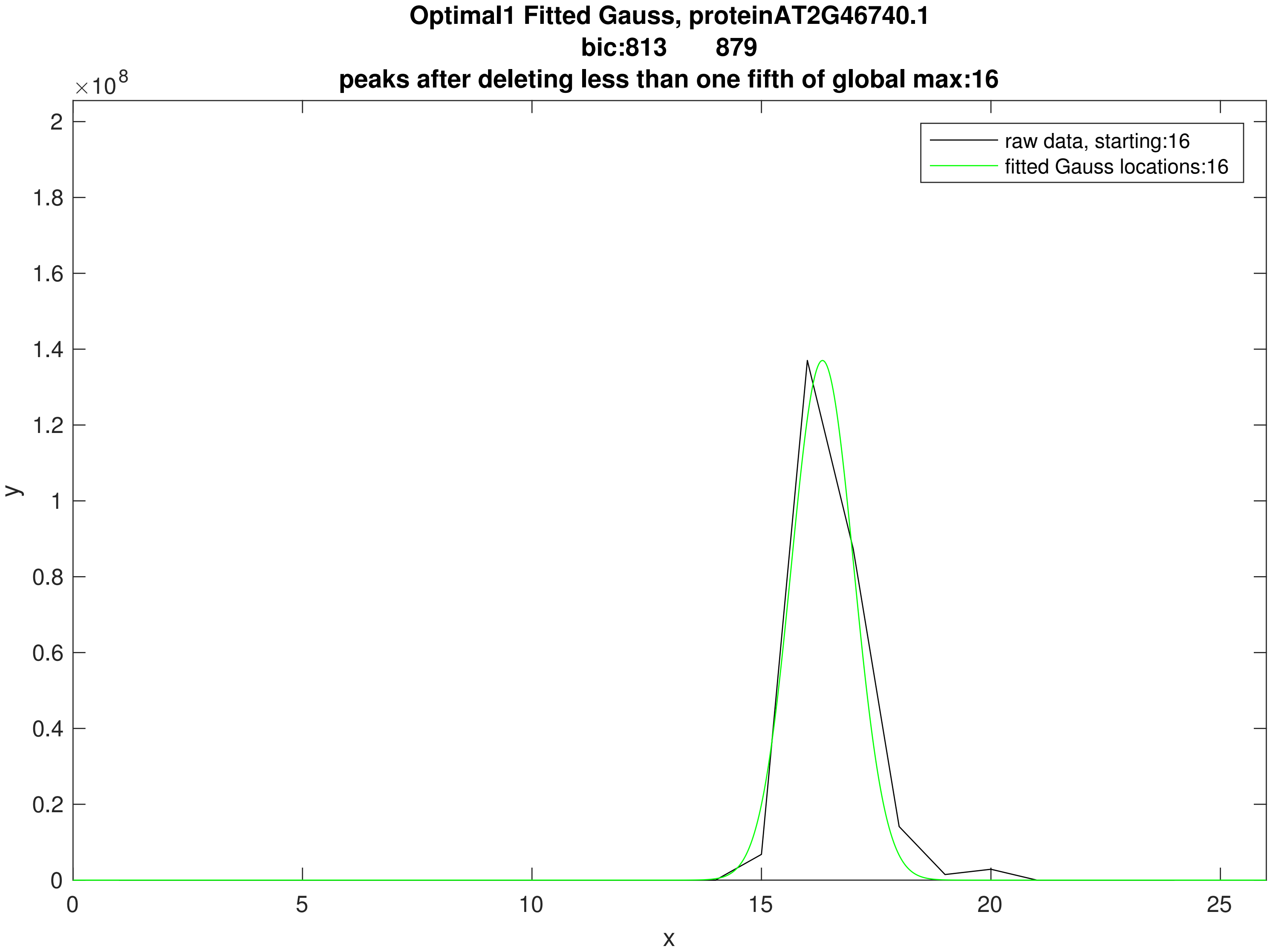
Optimal1 Fitted Gauss, proteinAT2G46740.1
bic:813 879
peaks after deleting less than one fifth of global max:16
×10^8
y
x
raw data, starting:16
fitted Gauss locations:16
0
0.2
0.4
0.6
0.8
1
1.2
1.4
1.6
1.8
2
5
10
15
20
25

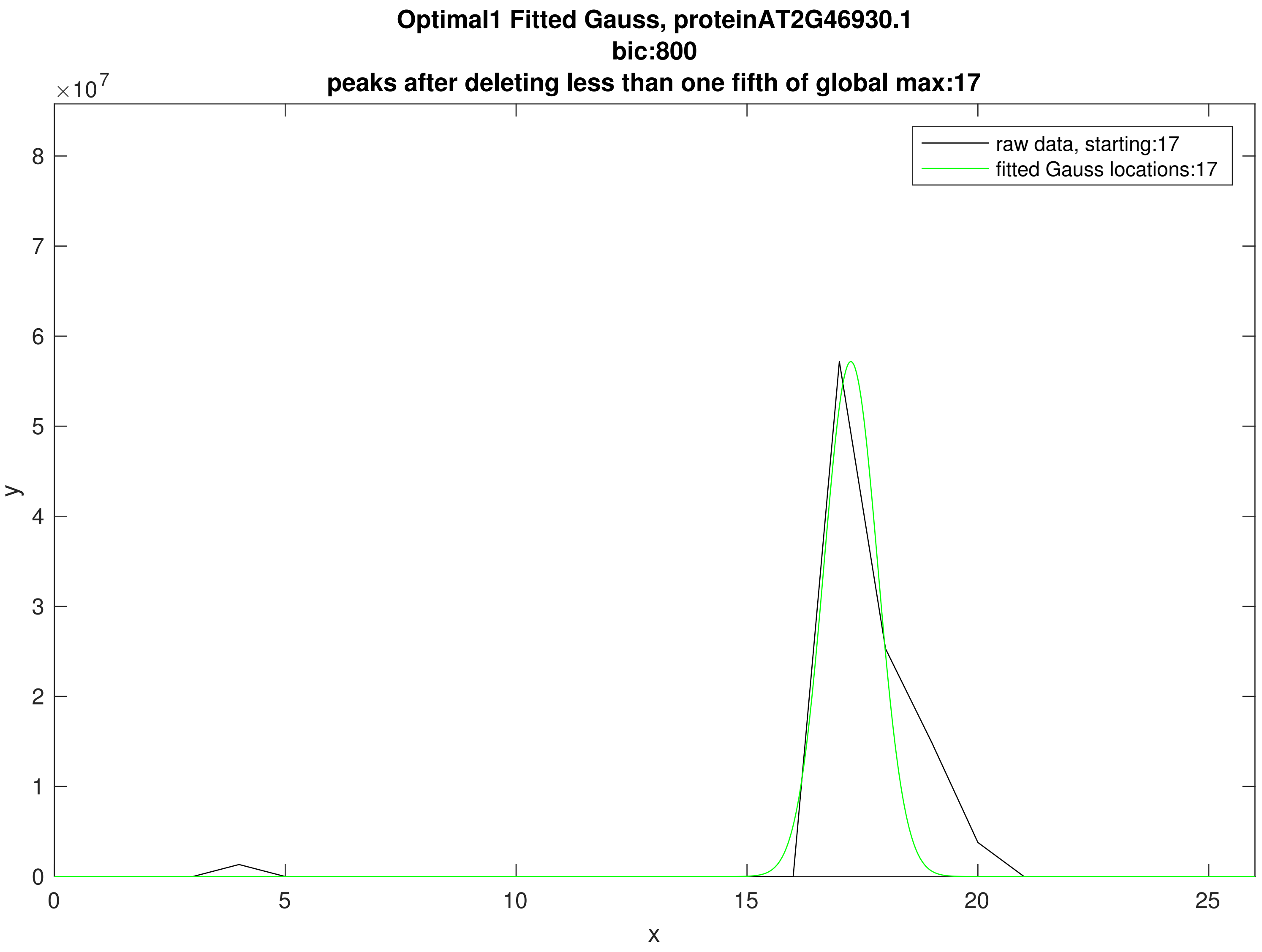

Optimal1 Fitted Gauss, proteinAT2G46930.1
bic:800
peaks after deleting less than one fifth of global max:17
×10^7
raw data, starting:17
fitted Gauss locations:17
y
x
0
1
2
3
4
5
6
7
8
0
5
10
15
20
25

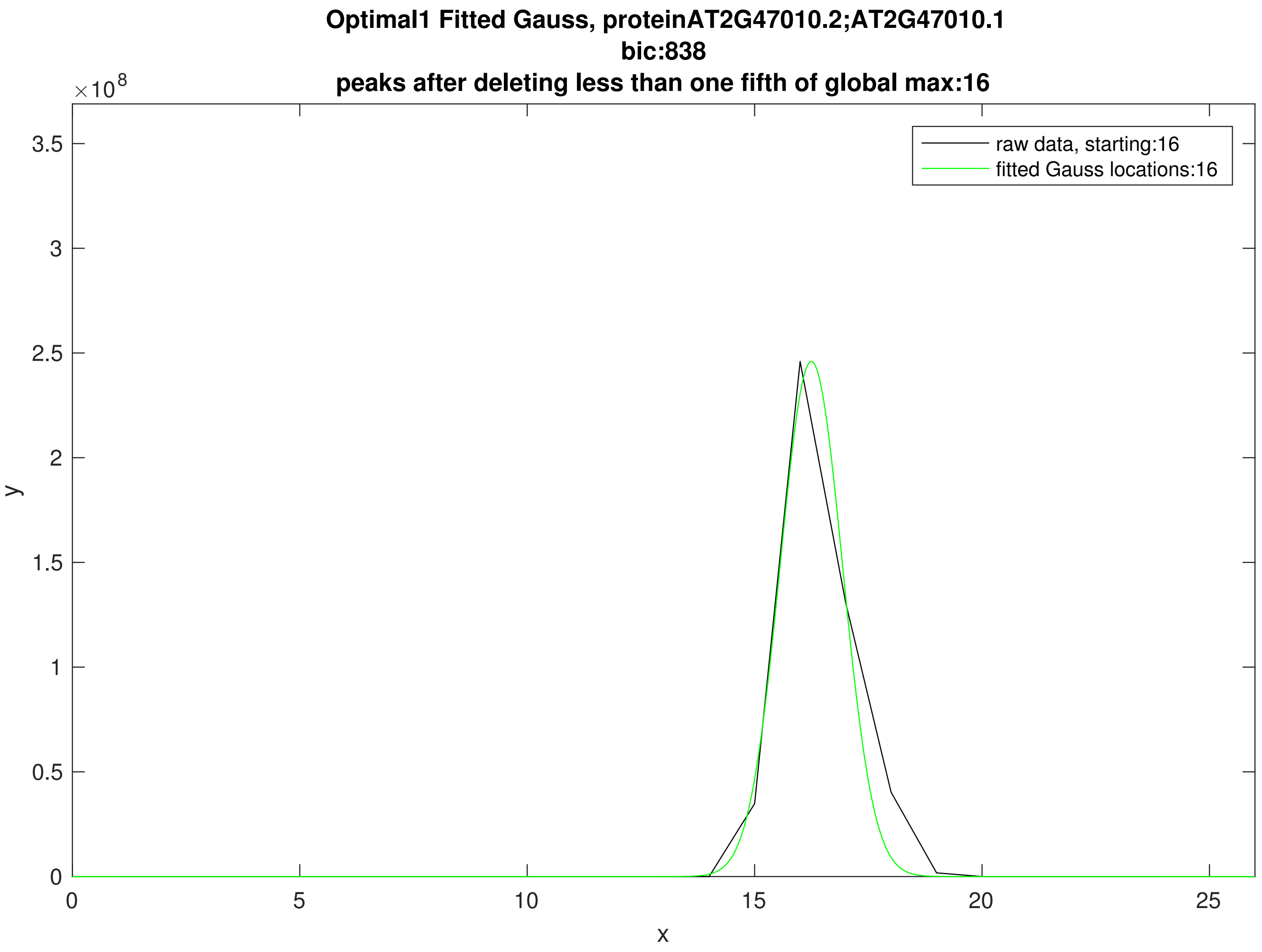

Optimal1 Fitted Gauss, proteinAT2G47010.2;AT2G47010.1
bic:838
peaks after deleting less than one fifth of global max:16
×10^8
3.5
3
2.5
2
1.5
1
0.5
0
y
0
5
10
15
20
25
x
raw data, starting:16
fitted Gauss locations:16

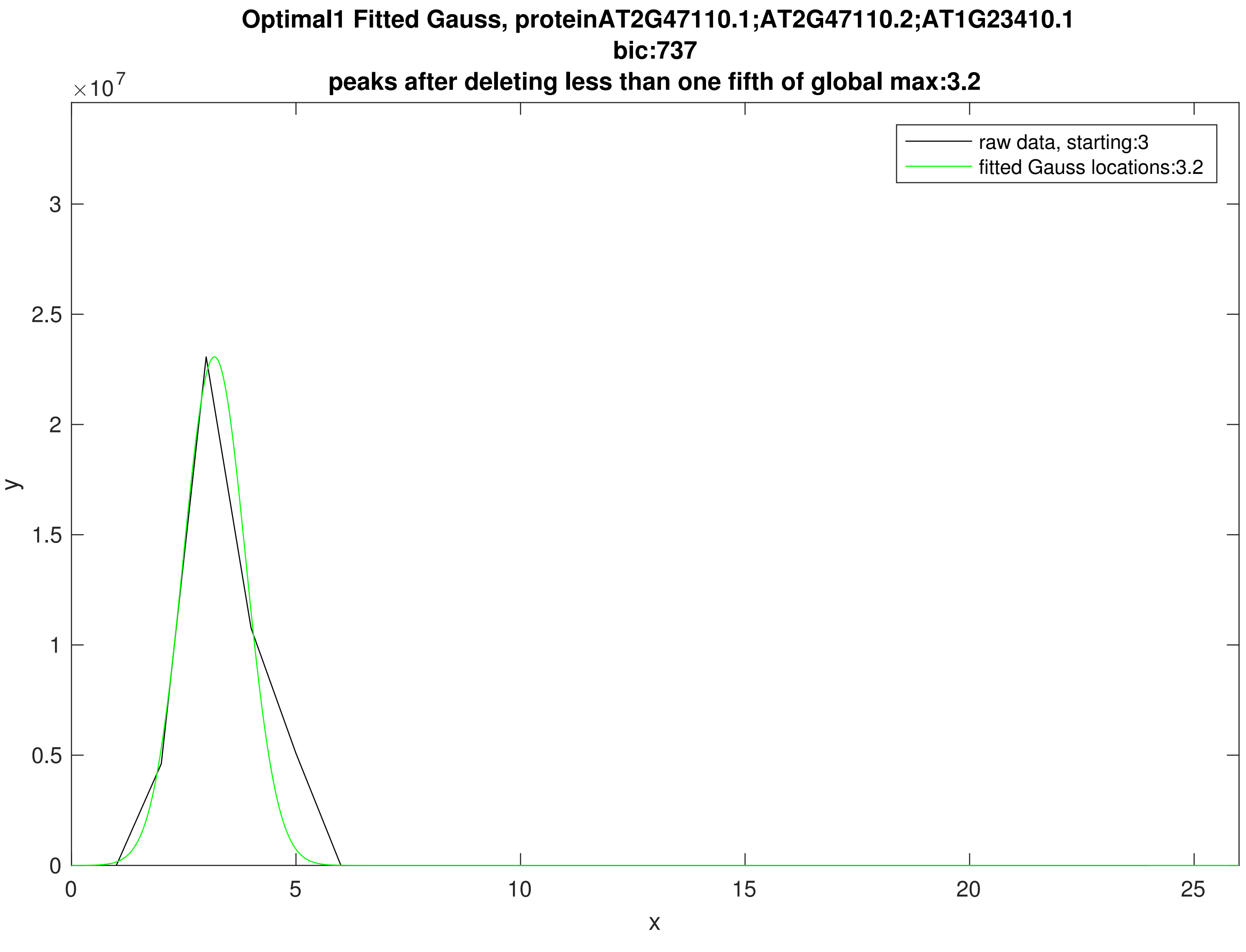
Optimal1 Fitted Gauss, proteinAT2G47110.1;AT2G47110.2;AT1G23410.1
bic:737
peaks after deleting less than one fifth of global max:3.2
×10^7
raw data, starting:3
fitted Gauss locations:3.2
y
x
0
0.5
1
1.5
2
2.5
3
0
5
10
15
20
25

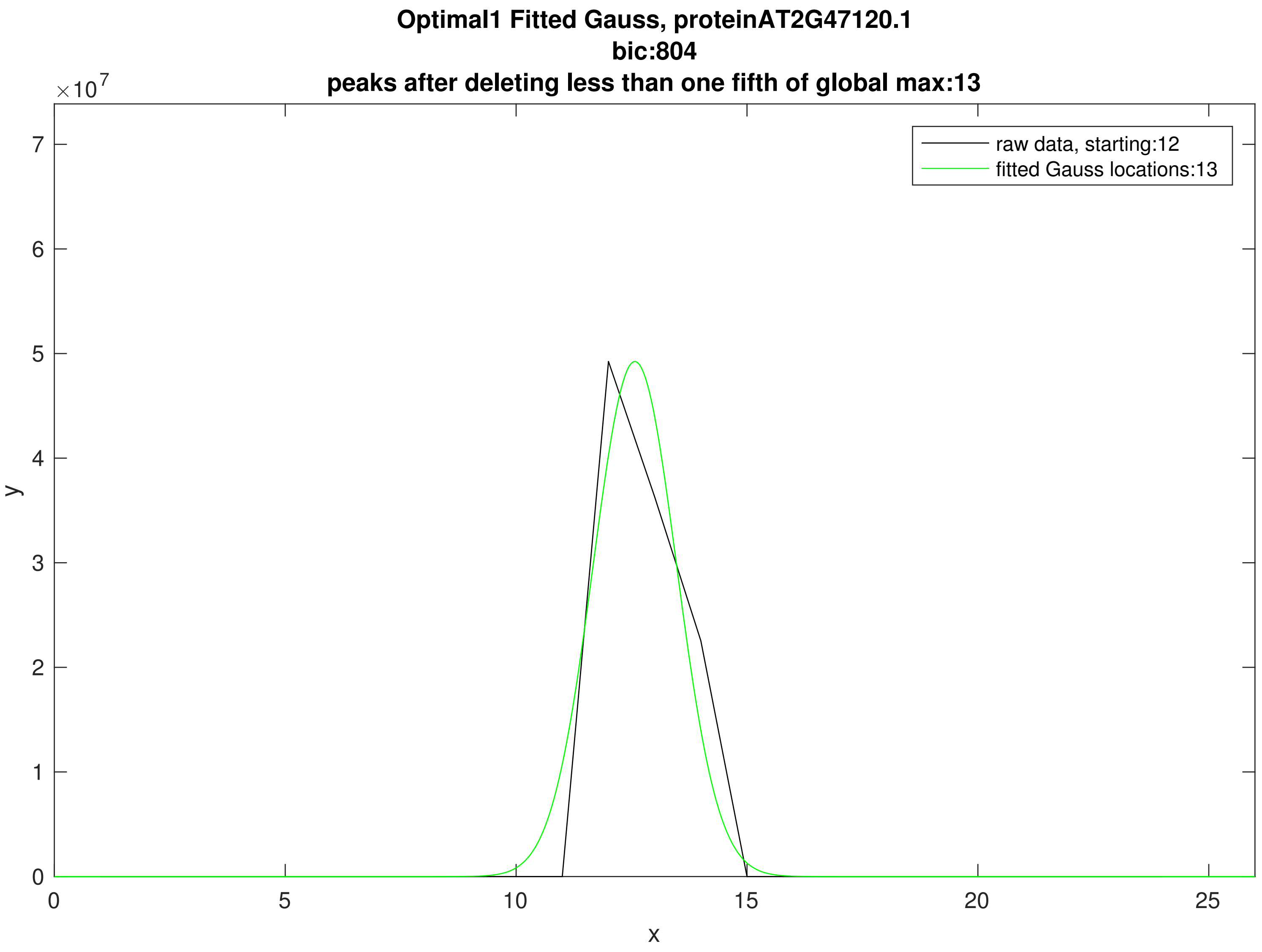
Optimal1 Fitted Gauss, proteinAT2G47120.1
bic:804
peaks after deleting less than one fifth of global max:13
×10^7
raw data, starting:12
fitted Gauss locations:13
y
x
0
1
2
3
4
5
6
7
0
5
10
15
20
25

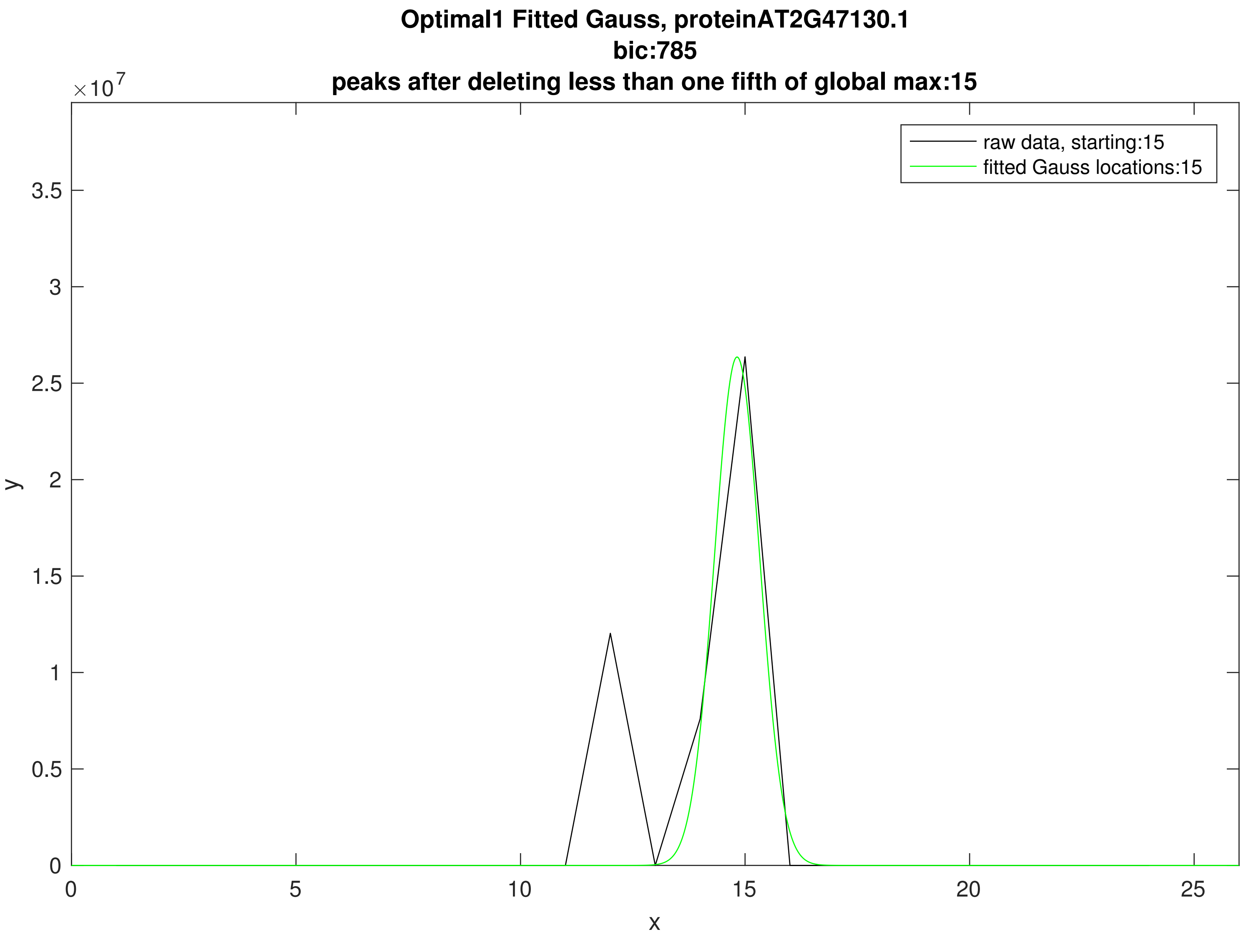
Optimal1 Fitted Gauss, proteinAT2G47130.1
bic:785
peaks after deleting less than one fifth of global max:15
×10^7
raw data, starting:15
fitted Gauss locations:15
3.5
3
2.5
2
1.5
1
0.5
0
y
0
5
10
15
20
25
x

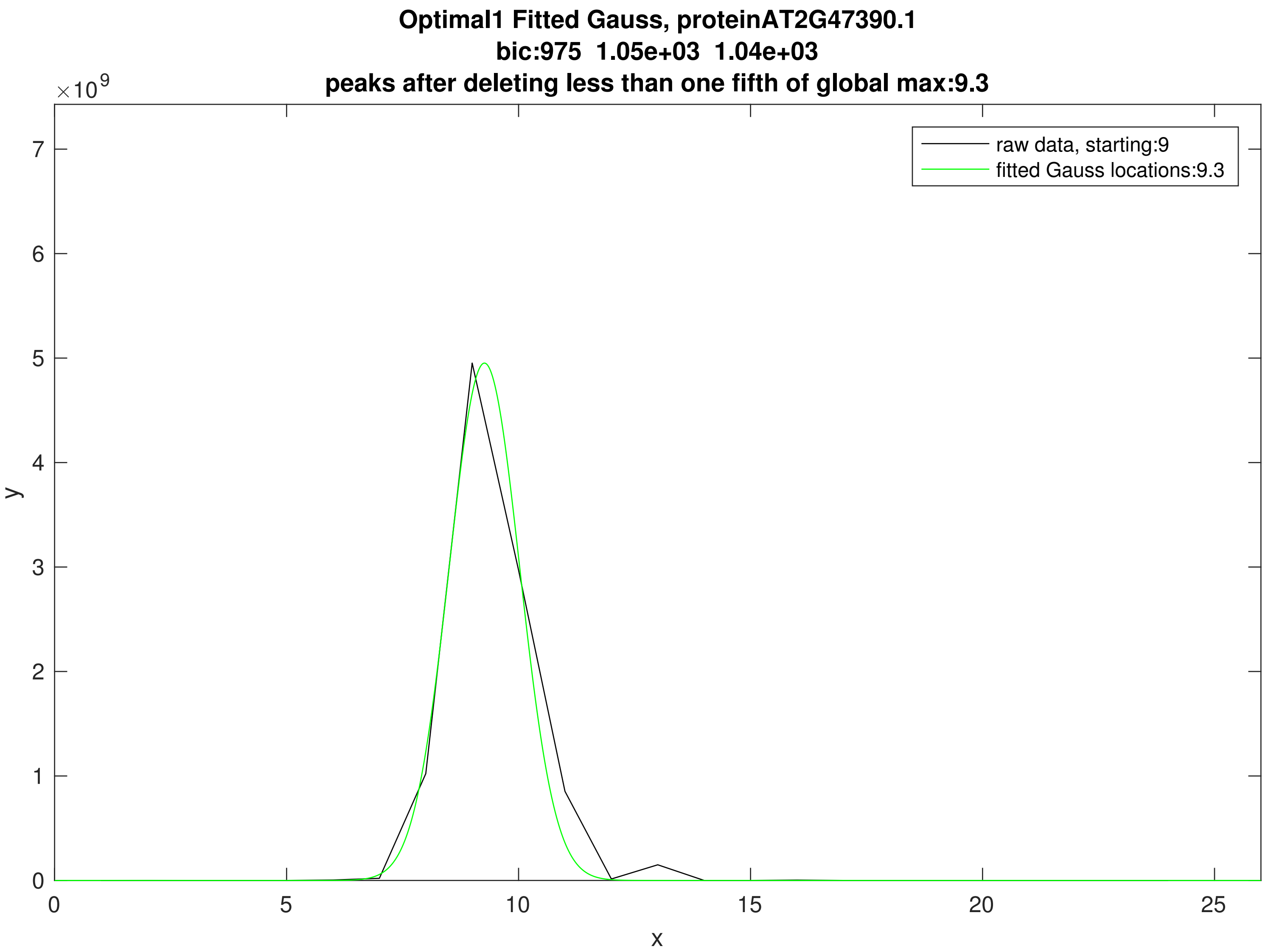
Optimal1 Fitted Gauss, proteinAT2G47390.1
bic:975 1.05e+03 1.04e+03
peaks after deleting less than one fifth of global max:9.3
×10^9
raw data, starting:9
fitted Gauss locations:9.3
y
x
0
1
2
3
4
5
6
7
0
5
10
15
20
25

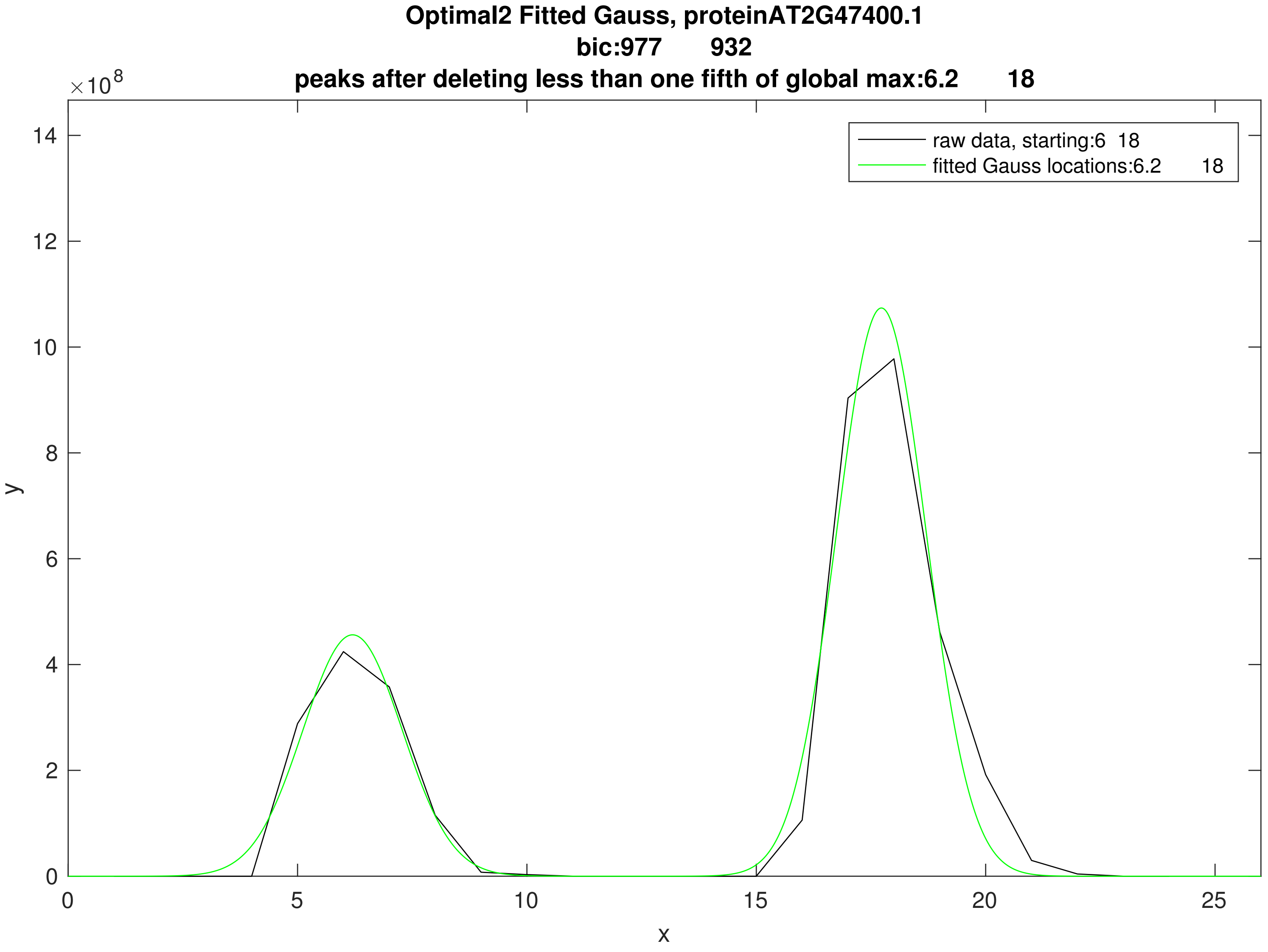

Optimal2 Fitted Gauss, proteinAT2G47400.1
bic:977 932
peaks after deleting less than one fifth of global max:6.2 18
×10^8
raw data, starting:6 18
fitted Gauss locations:6.2 18
14
12
10
8
6
4
2
0
y
0
5
10
15
20
25
x

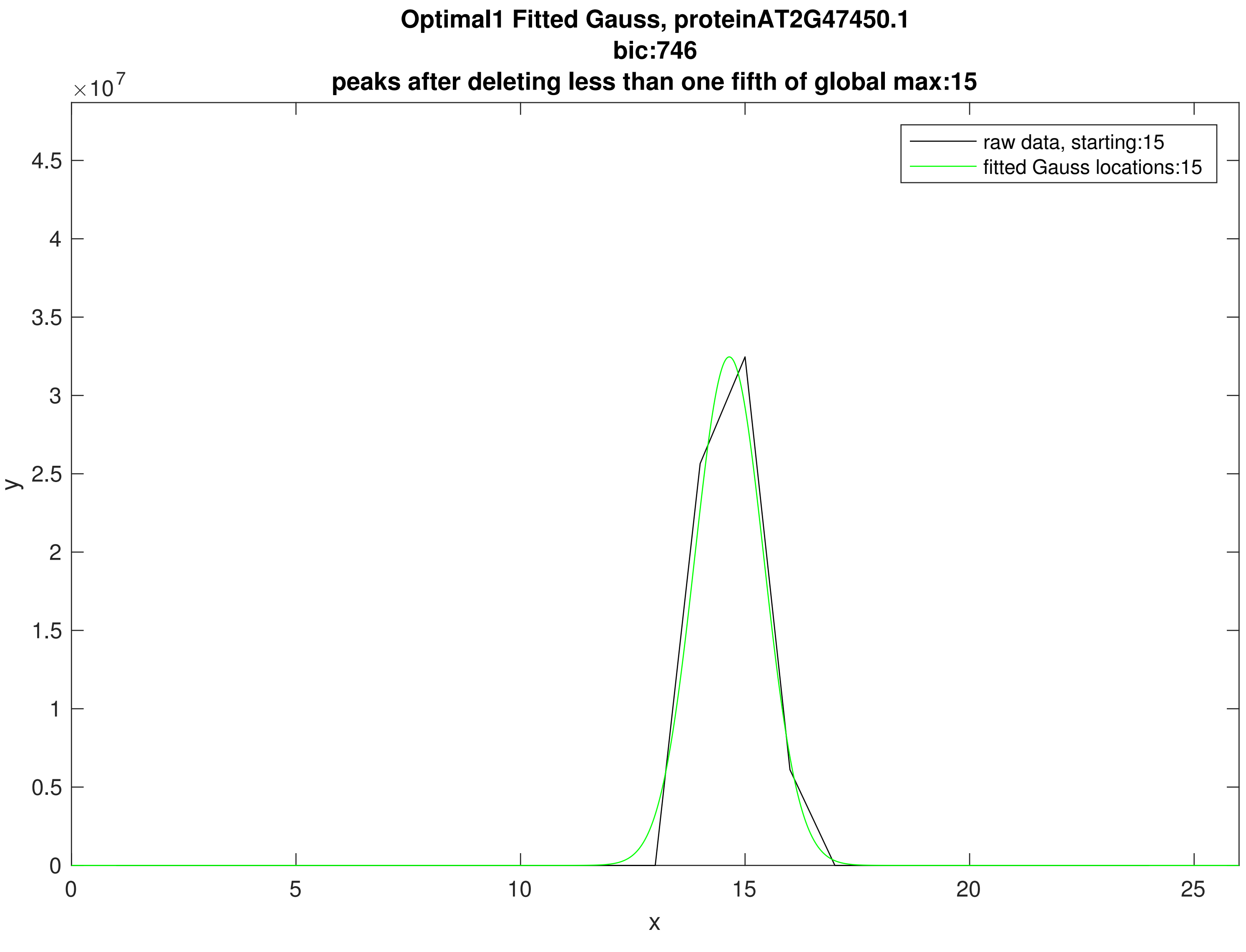

Optimal1 Fitted Gauss, proteinAT2G47450.1
bic:746
peaks after deleting less than one fifth of global max:15
×10^7
4.5
4
3.5
3
2.5
2
1.5
1
0.5
0
y
0
5
10
15
20
25
x
raw data, starting:15
fitted Gauss locations:15

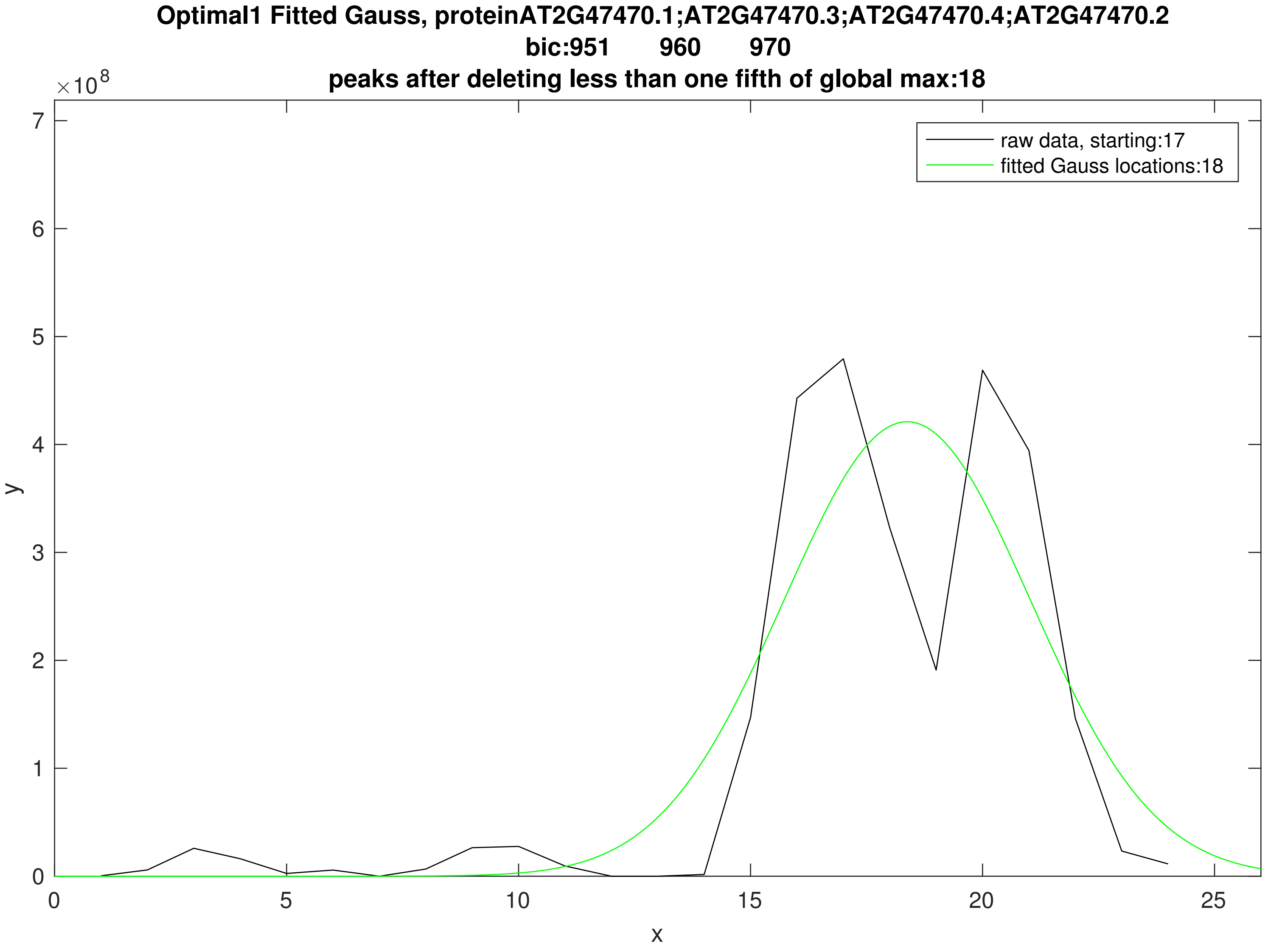

Optimal1 Fitted Gauss, proteinAT2G47470.1;AT2G47470.3;AT2G47470.4;AT2G47470.2
bic:951 960 970
peaks after deleting less than one fifth of global max:18
×10^8
raw data, starting:17
fitted Gauss locations:18
y
x
0
1
2
3
4
5
6
7
10
15
20
25

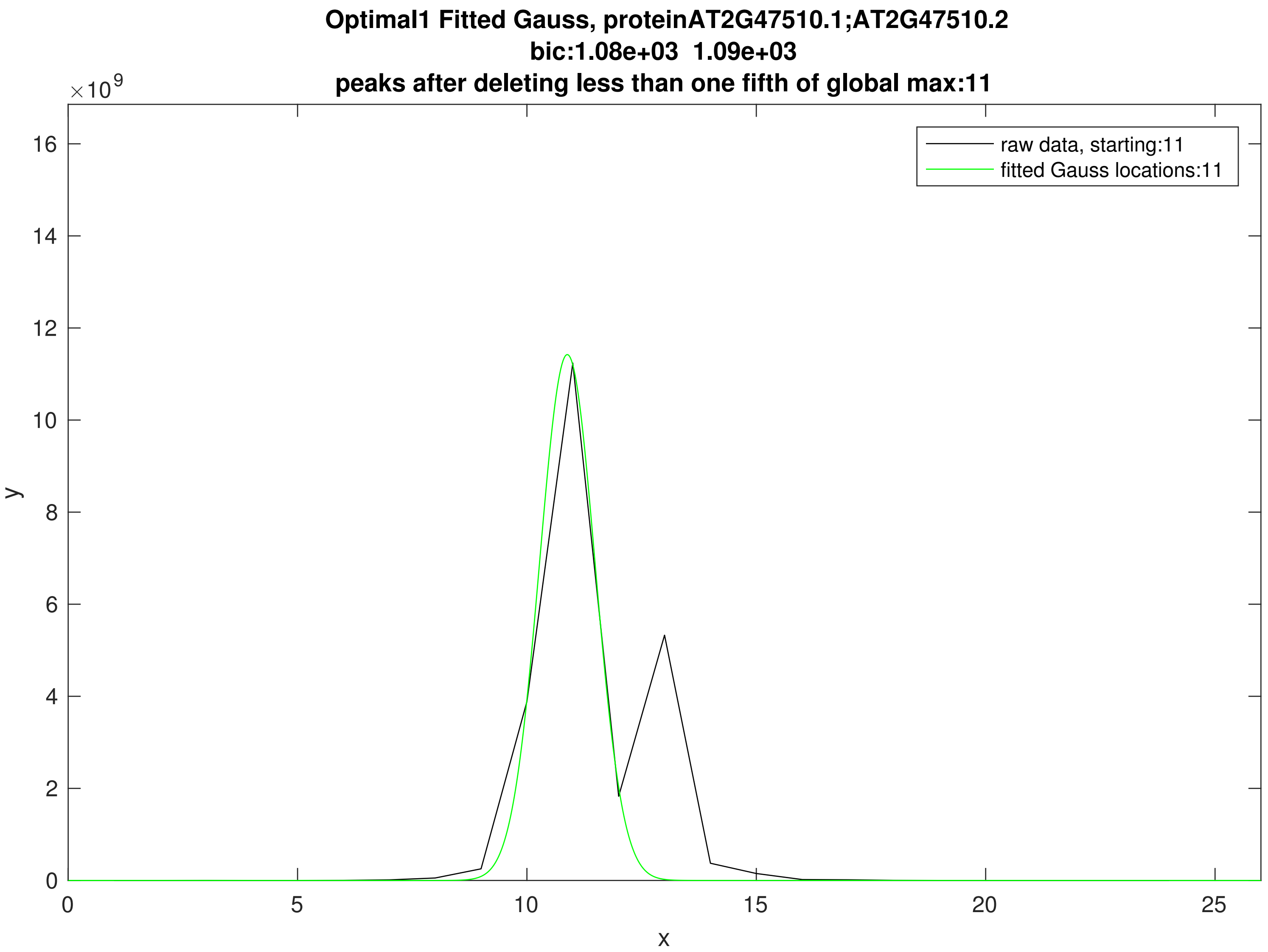

Optimal1 Fitted Gauss, proteinAT2G47510.1;AT2G47510.2
bic:1.08e+03  1.09e+03
peaks after deleting less than one fifth of global max:11
×10^9
raw data, starting:11
fitted Gauss locations:11
y
x

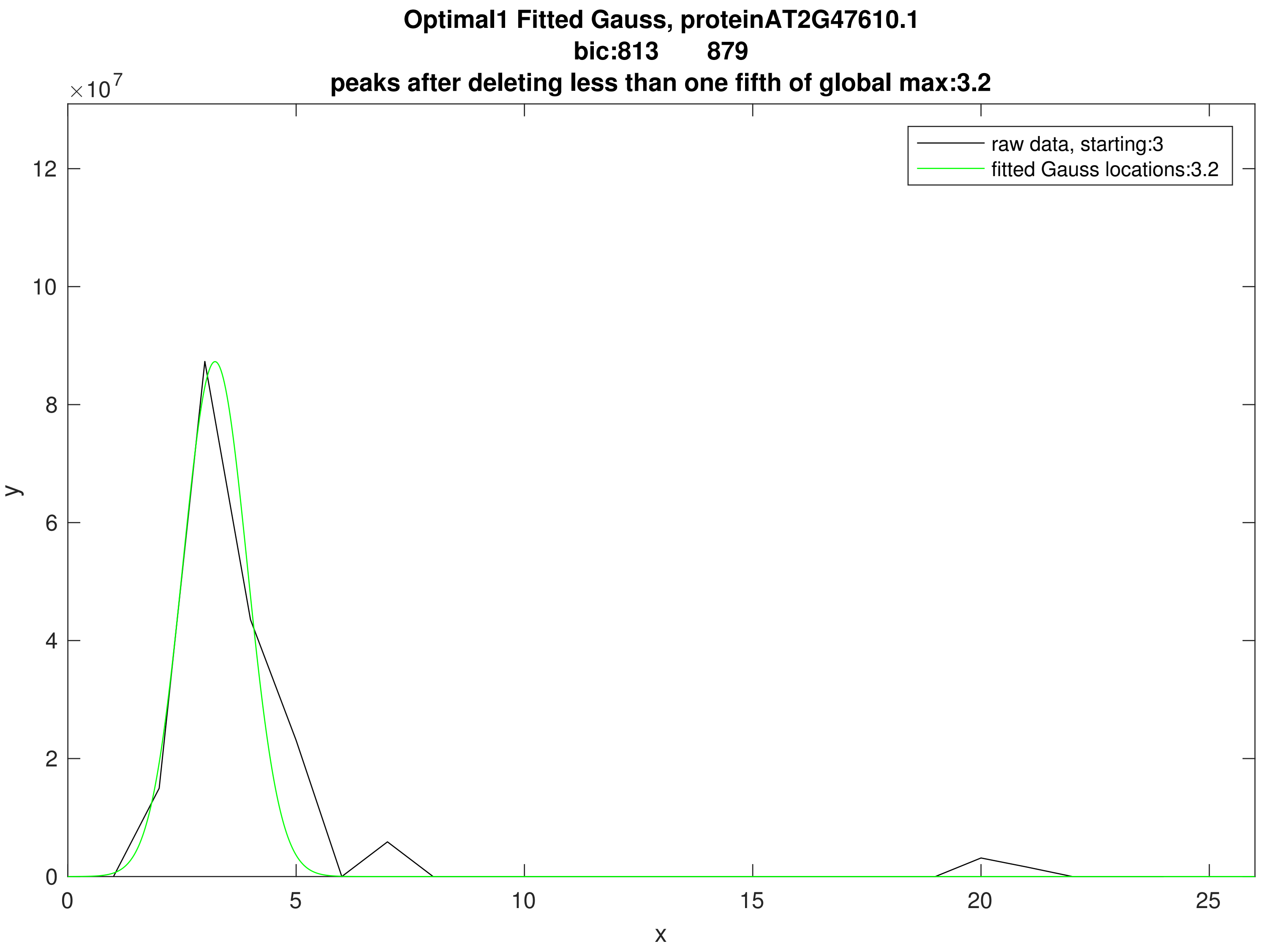

Optimal1 Fitted Gauss, proteinAT2G47610.1
bic:813 879
peaks after deleting less than one fifth of global max:3.2
×10^7
12
10
8
6
4
2
0
y
0
5
10
15
20
25
x
raw data, starting:3
fitted Gauss locations:3.2

**Optimal1 Fitted Gauss, proteinAT2G47730.1**
**bic:963  1.01e+03  1.02e+03  1.03e+03**
**peaks after deleting less than one fifth of global max:17**

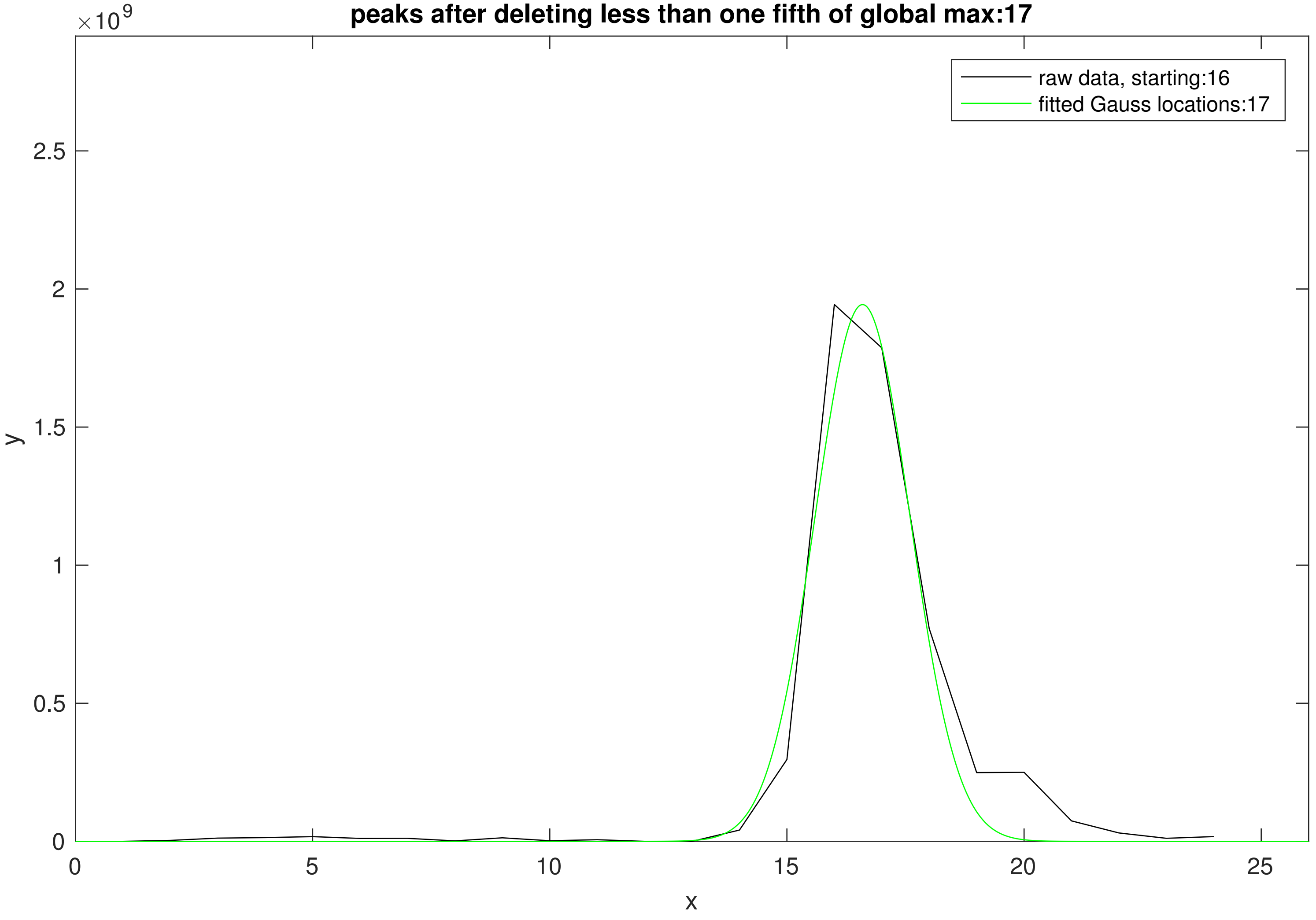

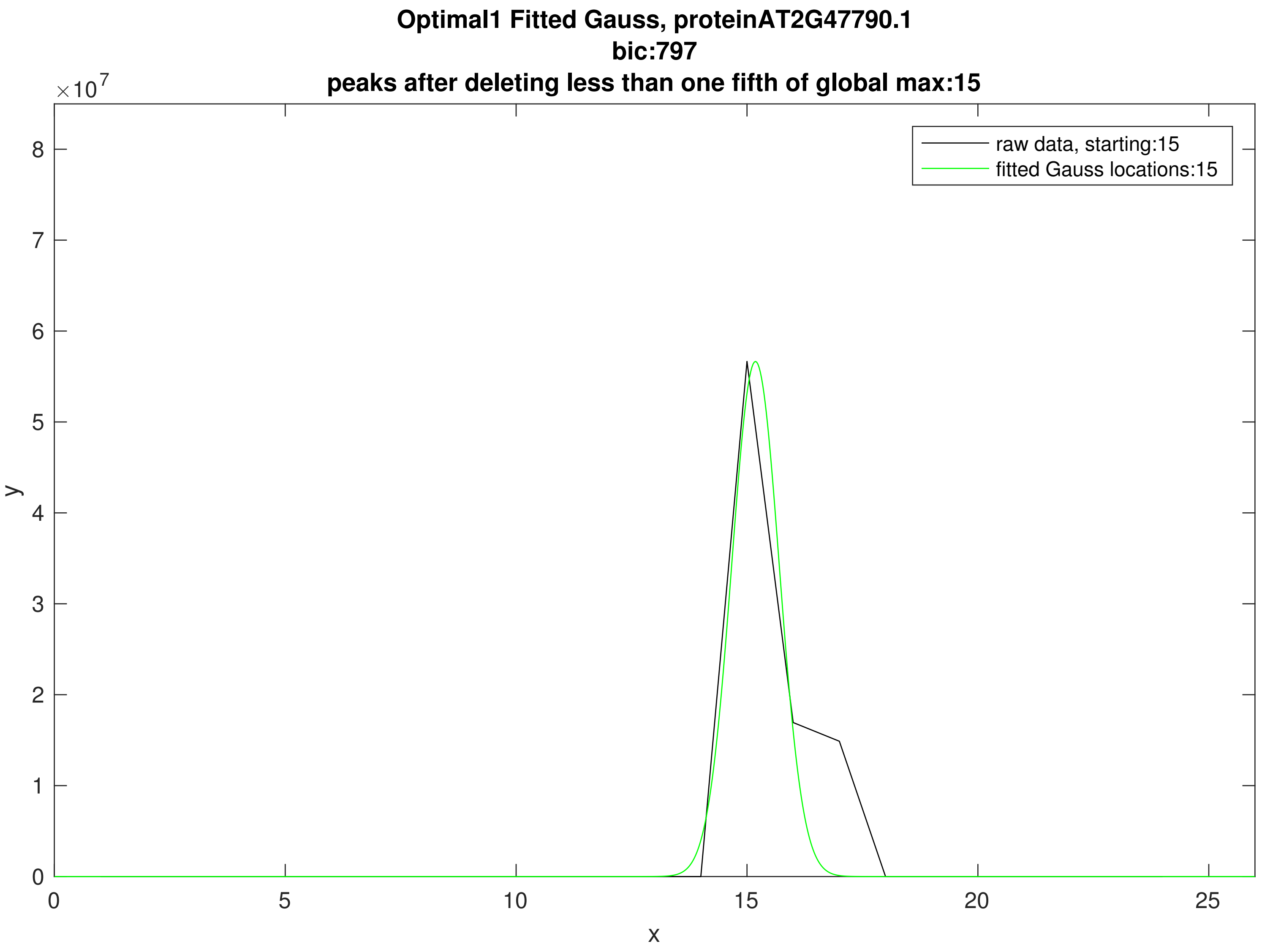
Optimal1 Fitted Gauss, proteinAT2G47790.1
bic:797
peaks after deleting less than one fifth of global max:15
×10^7
raw data, starting:15
fitted Gauss locations:15
y
x
0
1
2
3
4
5
6
7
8
0
5
10
15
20
25

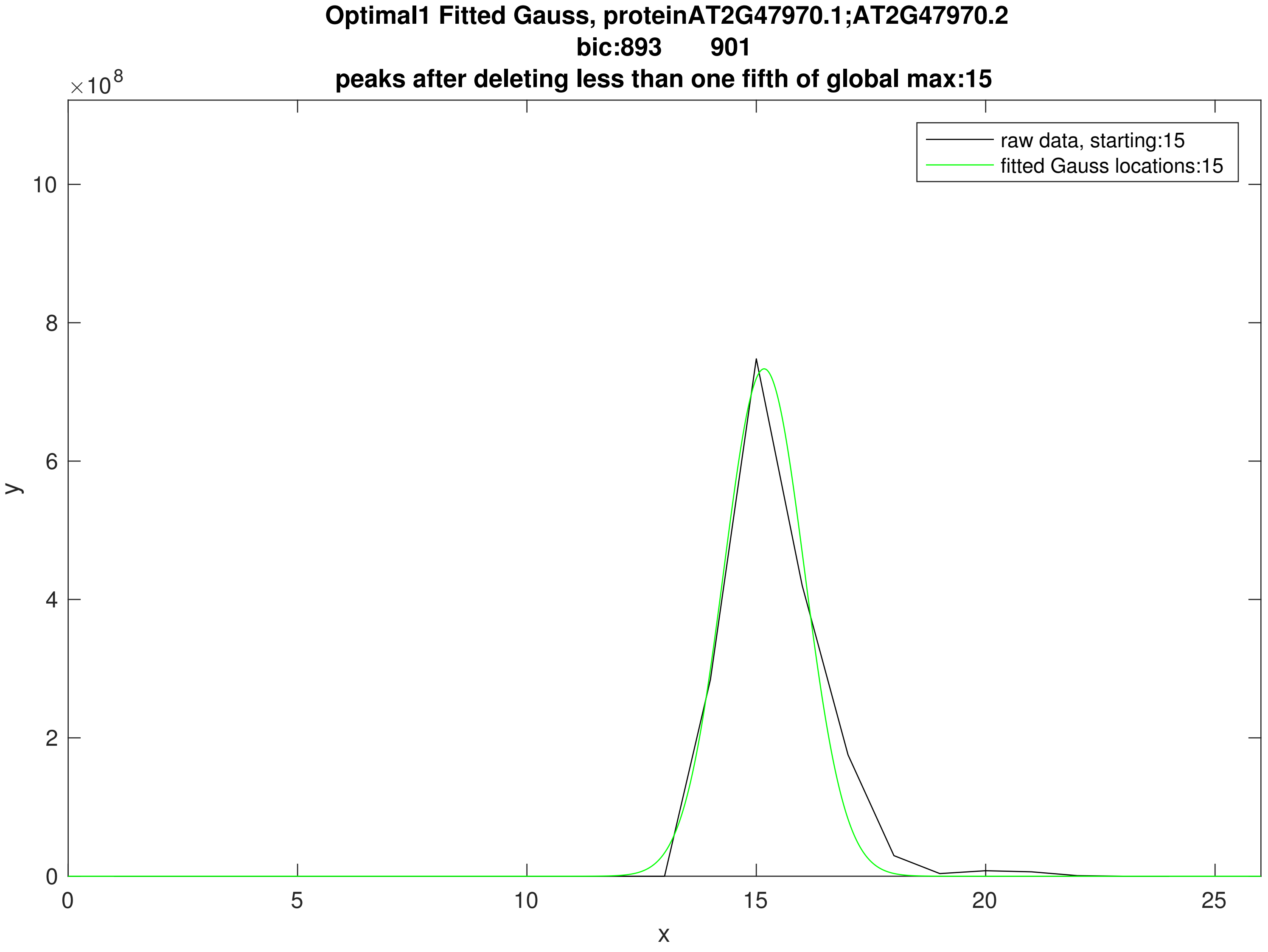
Optimal1 Fitted Gauss, proteinAT2G47970.1;AT2G47970.2
bic:893 901
peaks after deleting less than one fifth of global max:15
×10^8
raw data, starting:15
fitted Gauss locations:15
y
x
0
2
4
6
8
10
0
5
10
15
20
25

Optimal1 Fitted Gauss, proteinAT2G48120.2;AT2G48120.1
bic:578
peaks after deleting less than one fifth of global max:15

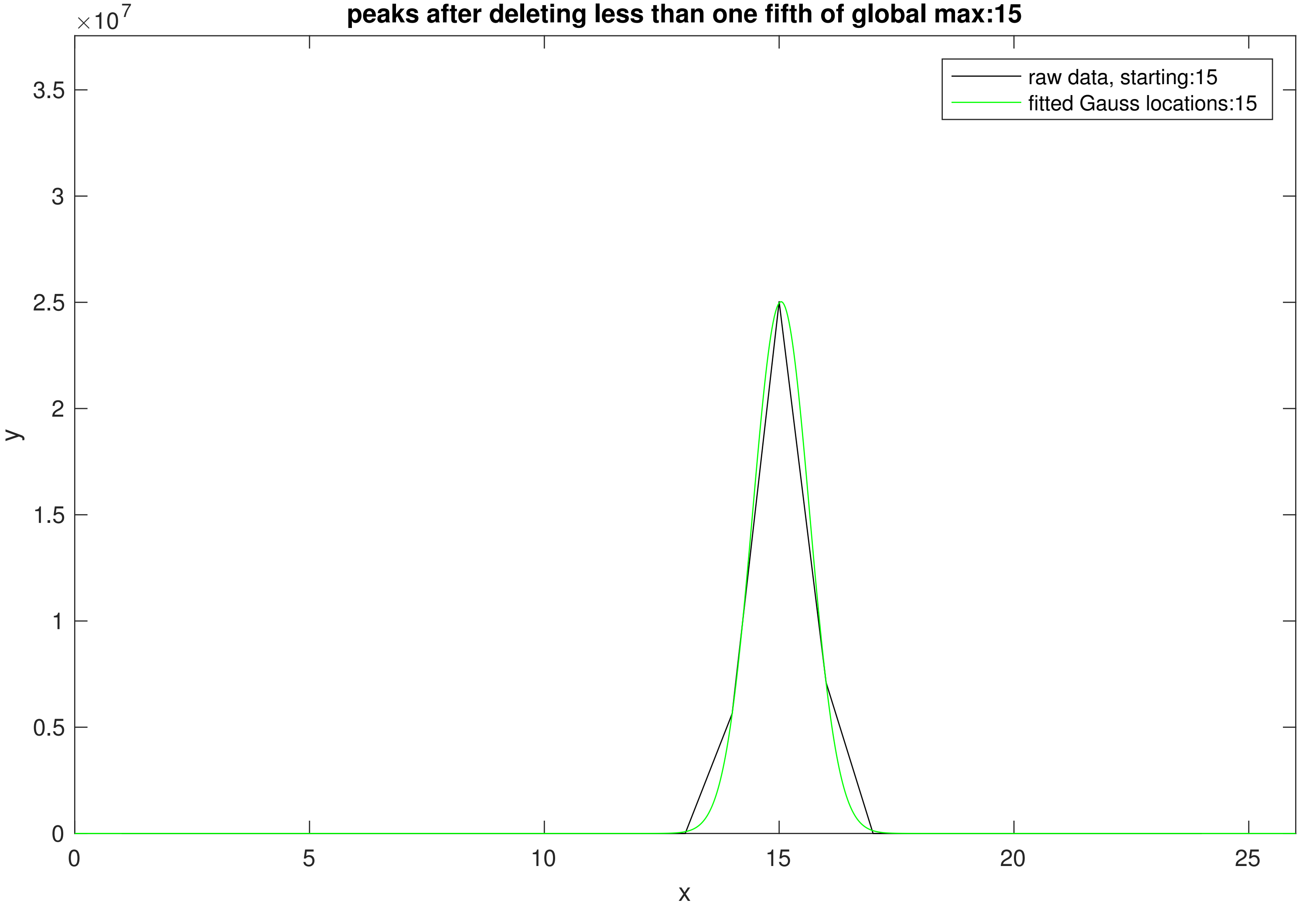

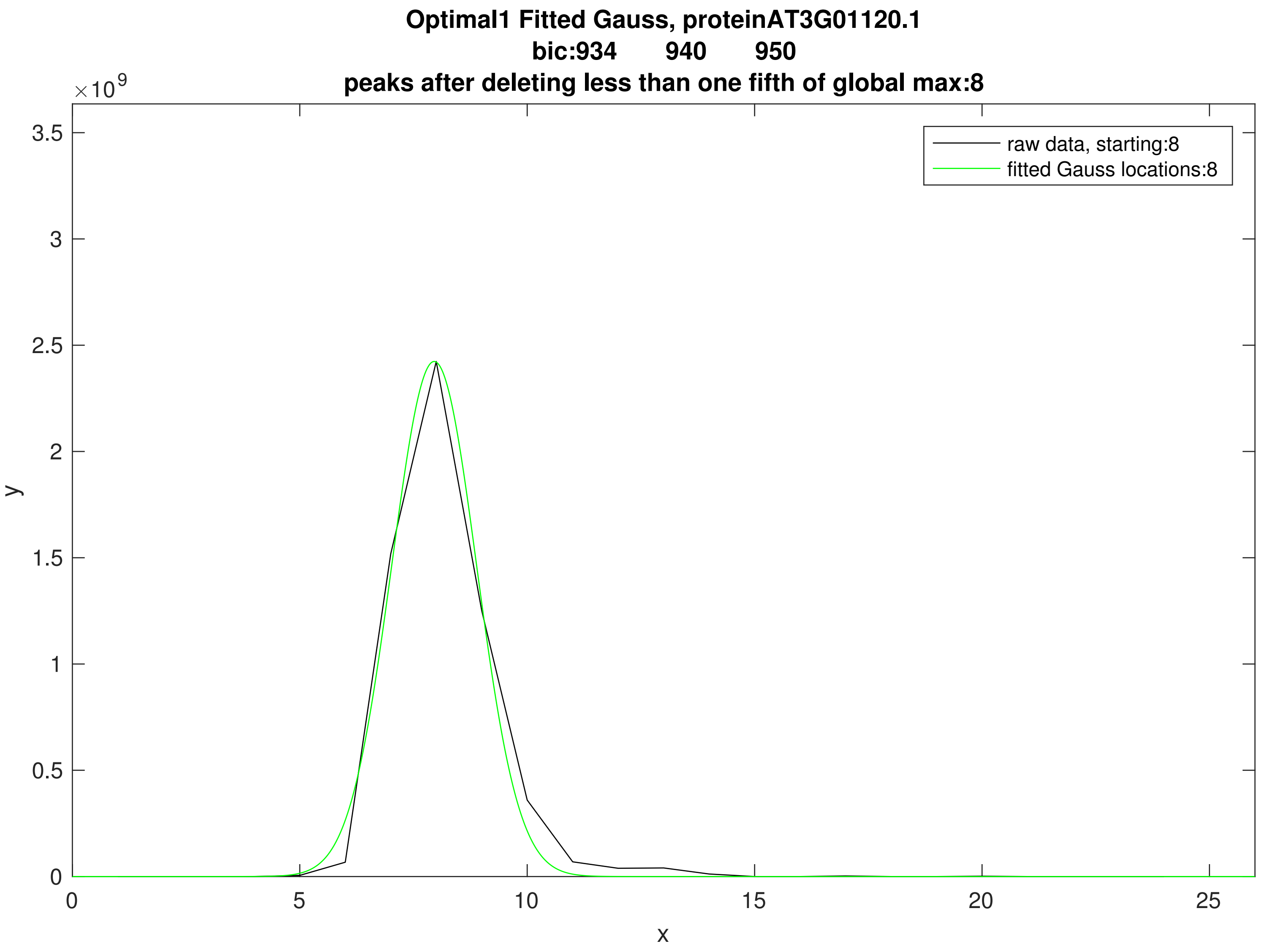
Optimal1 Fitted Gauss, proteinAT3G01120.1
bic:934 940 950
peaks after deleting less than one fifth of global max:8
×10^9
3.5
3
2.5
2
1.5
1
0.5
0
y
0
5
10
15
20
25
x
raw data, starting:8
fitted Gauss locations:8

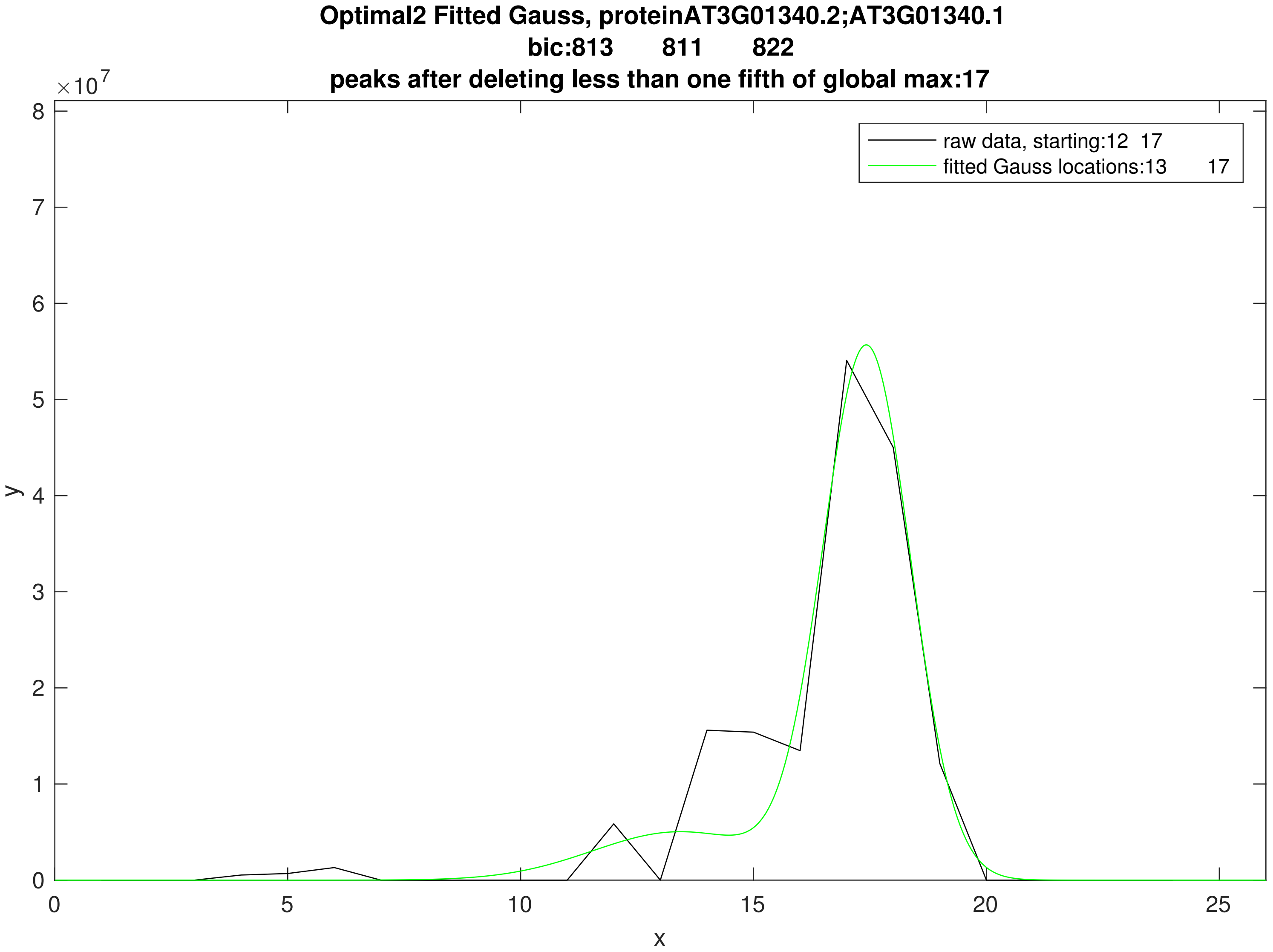
Optimal2 Fitted Gauss, proteinAT3G01340.2;AT3G01340.1
bic:813 811 822
peaks after deleting less than one fifth of global max:17
×10^7
raw data, starting:12 17
fitted Gauss locations:13 17
y
x
0
1
2
3
4
5
6
7
8
0
5
10
15
20
25

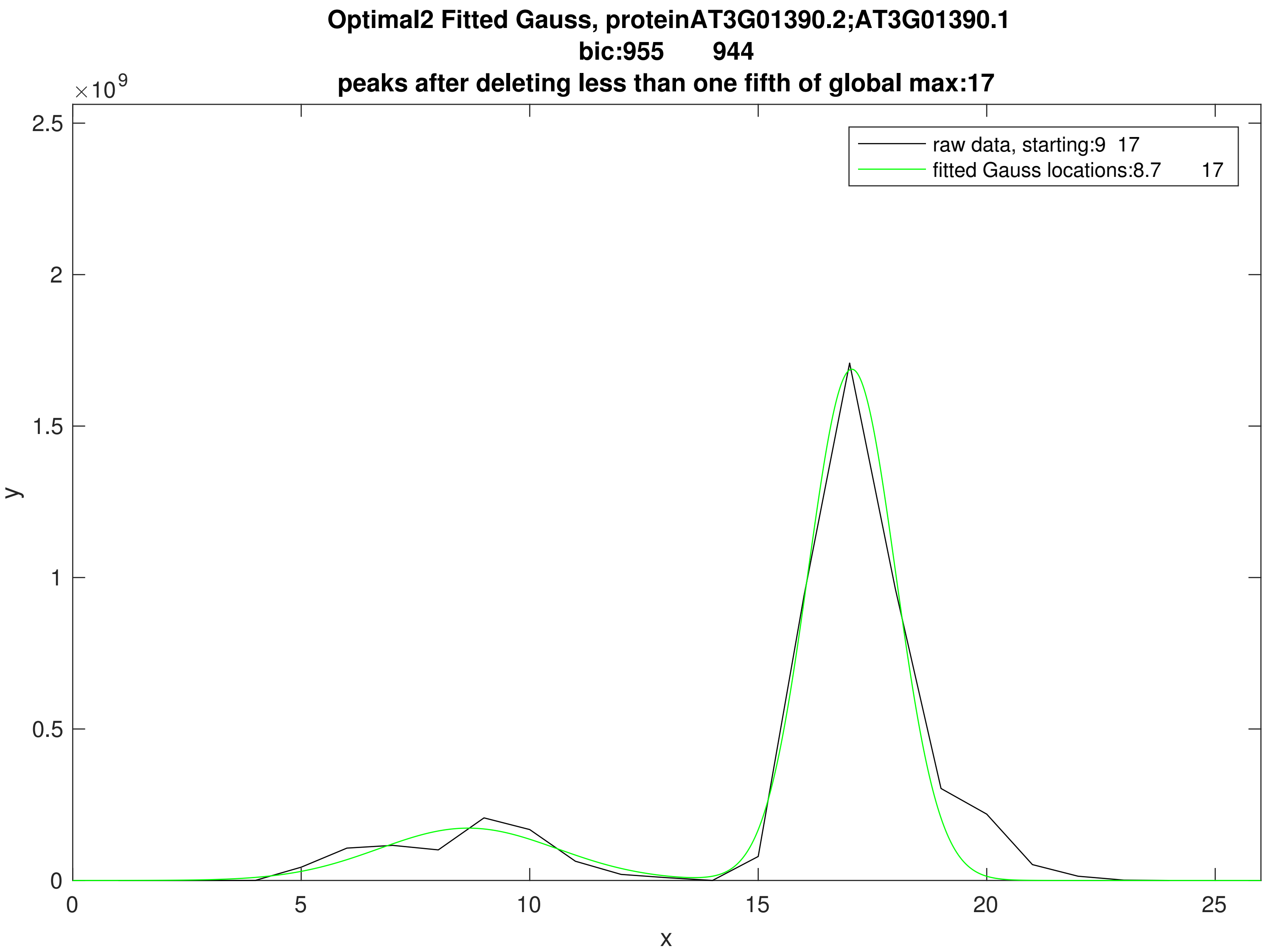
Optimal2 Fitted Gauss, proteinAT3G01390.2;AT3G01390.1
bic:955 944
peaks after deleting less than one fifth of global max:17
×10^9
2.5
2
1.5
1
0.5
0
y
0
5
10
15
20
25
x
raw data, starting:9 17
fitted Gauss locations:8.7 17

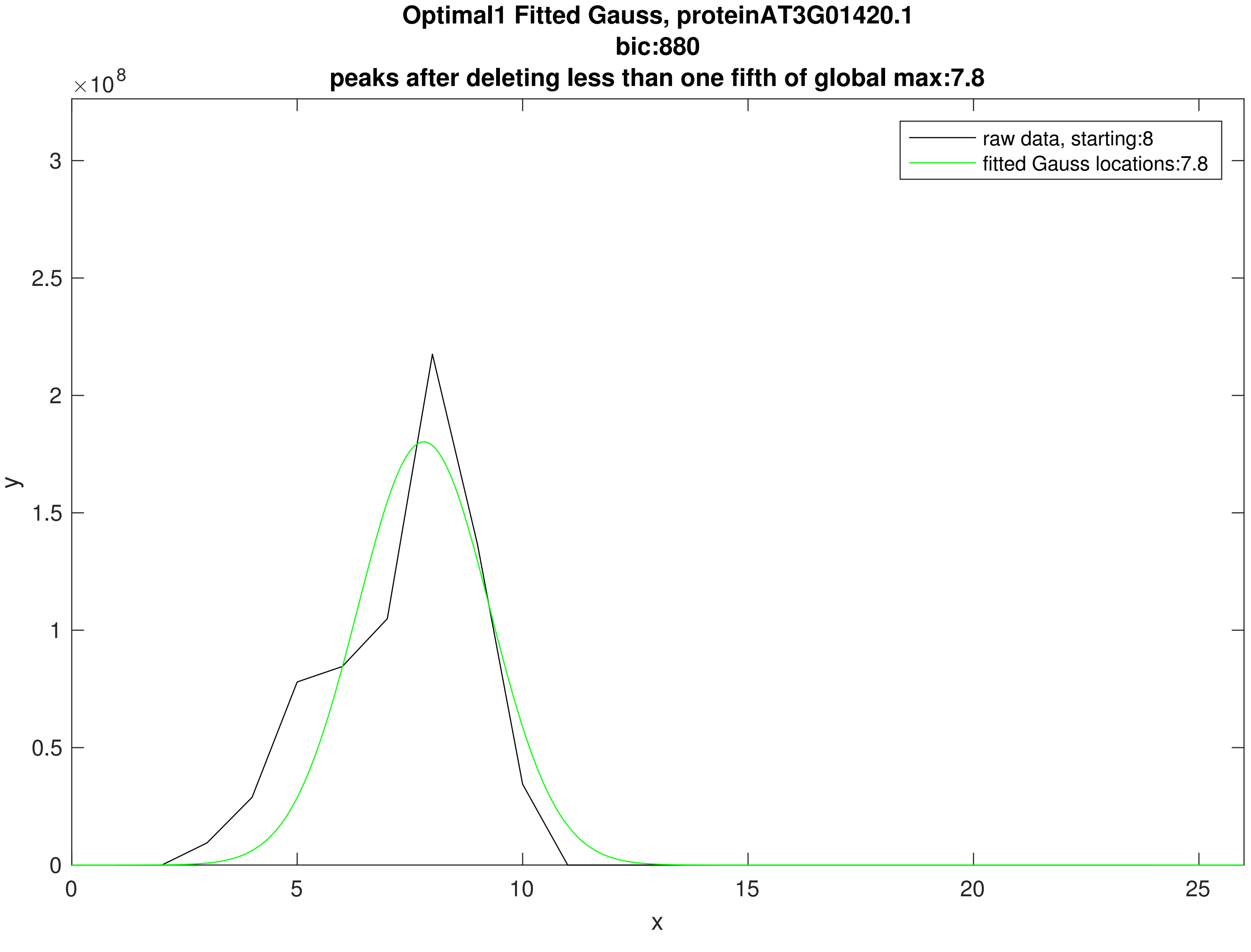
Optimal1 Fitted Gauss, proteinAT3G01420.1
bic:880
peaks after deleting less than one fifth of global max:7.8
×10^8
raw data, starting:8
fitted Gauss locations:7.8
y
x
0
0.5
1
1.5
2
2.5
3
0
5
10
15
20
25

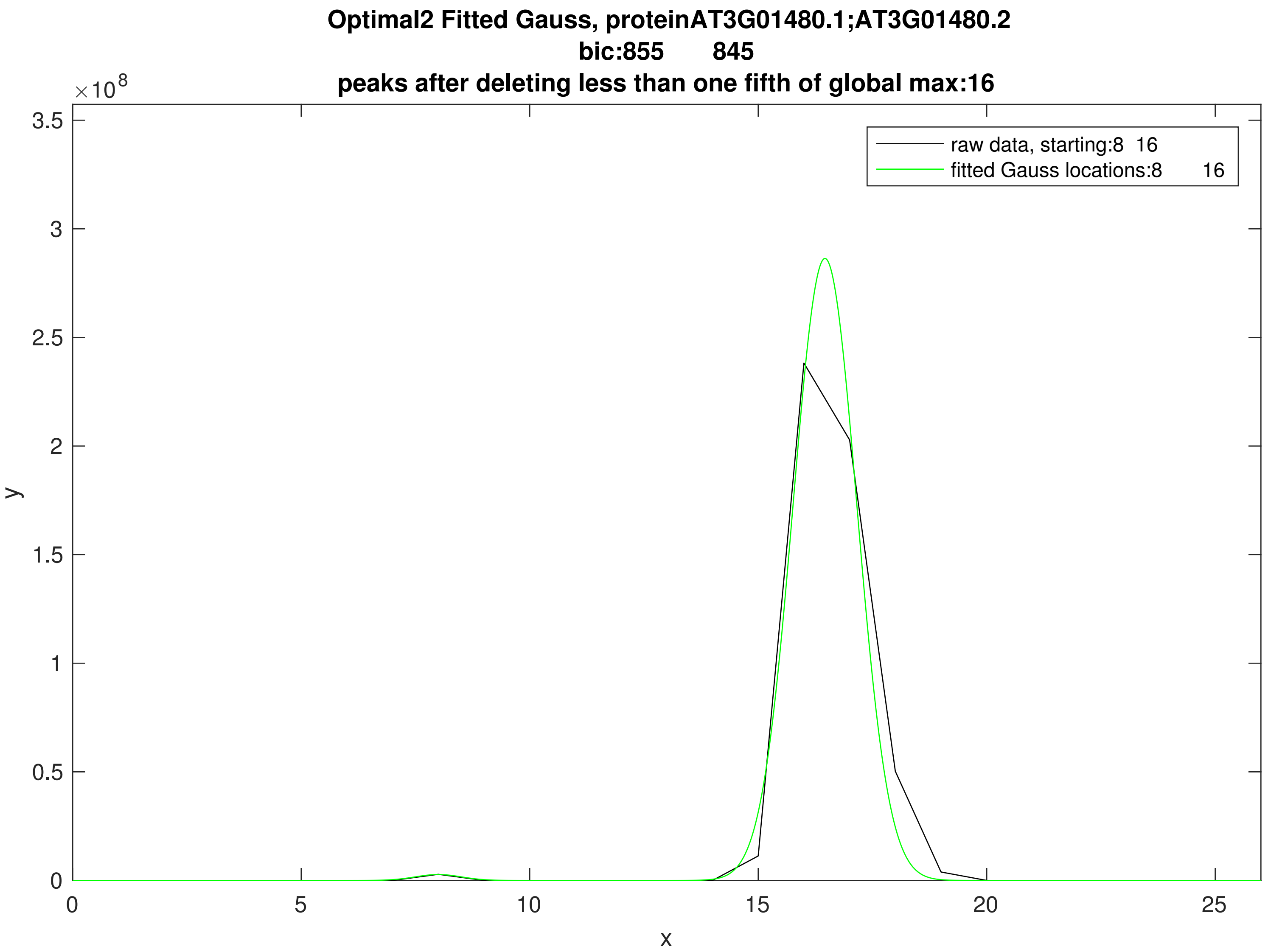

Optimal2 Fitted Gauss, proteinAT3G01480.1;AT3G01480.2
bic:855 845
peaks after deleting less than one fifth of global max:16
×10^8
raw data, starting:8 16
fitted Gauss locations:8 16
3.5
3
2.5
2
1.5
1
0.5
0
y
0
5
10
15
20
25
x

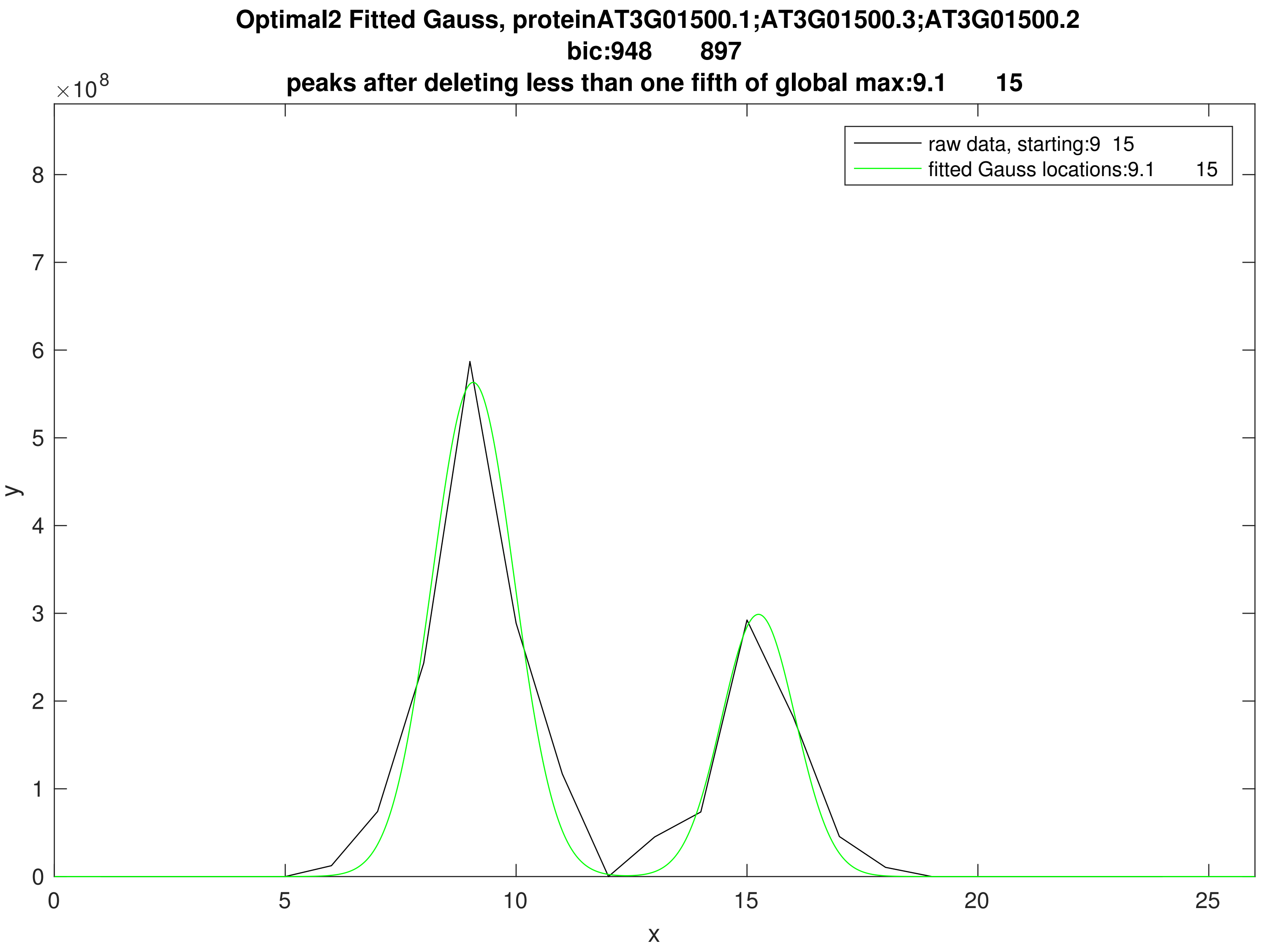
Optimal2 Fitted Gauss, proteinAT3G01500.1;AT3G01500.3;AT3G01500.2
bic:948 897
peaks after deleting less than one fifth of global max:9.1 15
raw data, starting:9 15
fitted Gauss locations:9.1 15
×10^8
y
x
0
1
2
3
4
5
6
7
8
0
5
10
15
20
25

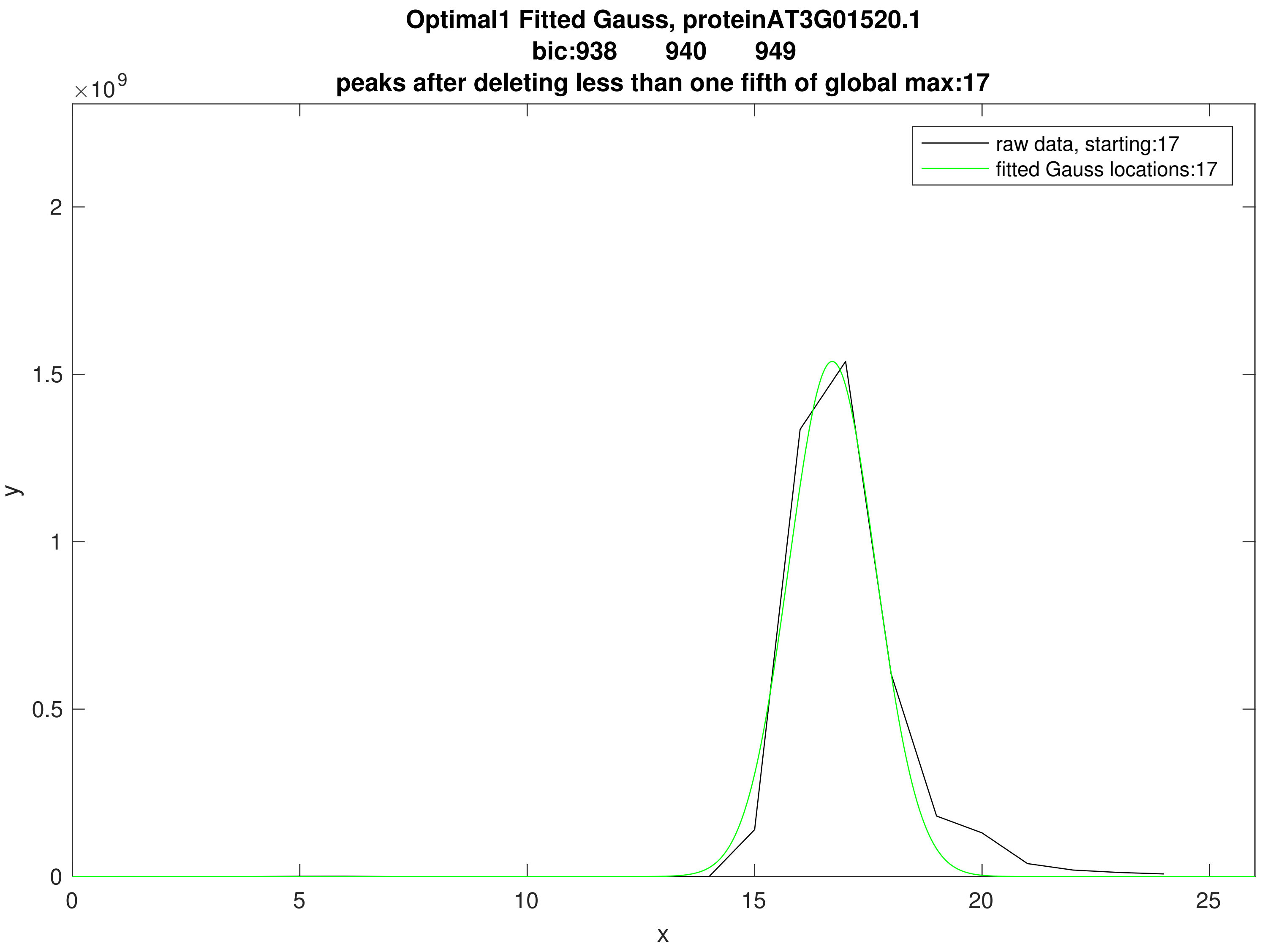

Optimal1 Fitted Gauss, proteinAT3G01520.1
bic:938 940 949
peaks after deleting less than one fifth of global max:17
×10^9
raw data, starting:17
fitted Gauss locations:17
y
x
0
0.5
1
1.5
2
0
5
10
15
20
25

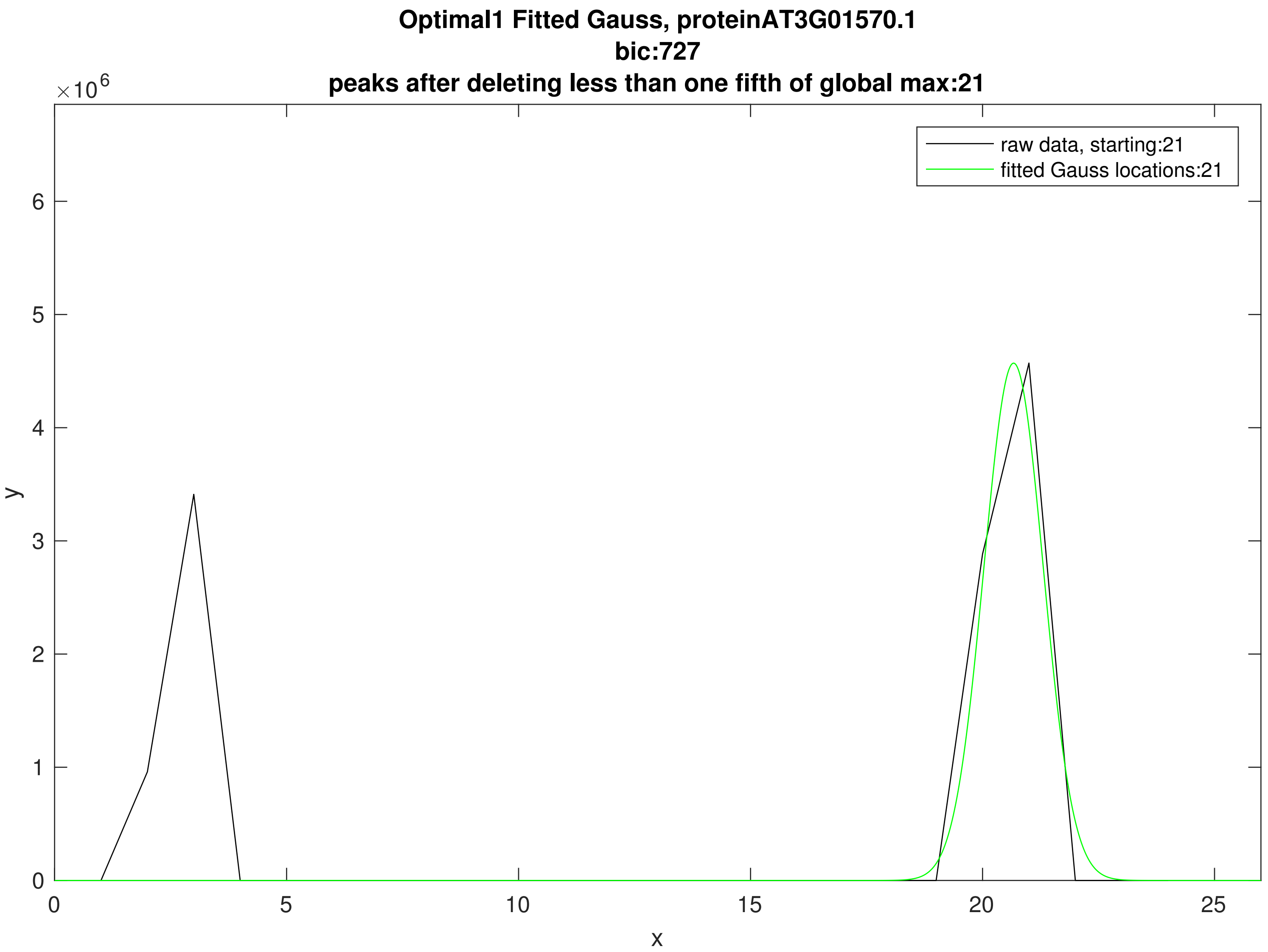

Optimal1 Fitted Gauss, proteinAT3G01570.1
bic:727
peaks after deleting less than one fifth of global max:21
×10^6
raw data, starting:21
fitted Gauss locations:21
y
x
0
1
2
3
4
5
6
0
5
10
15
20
25

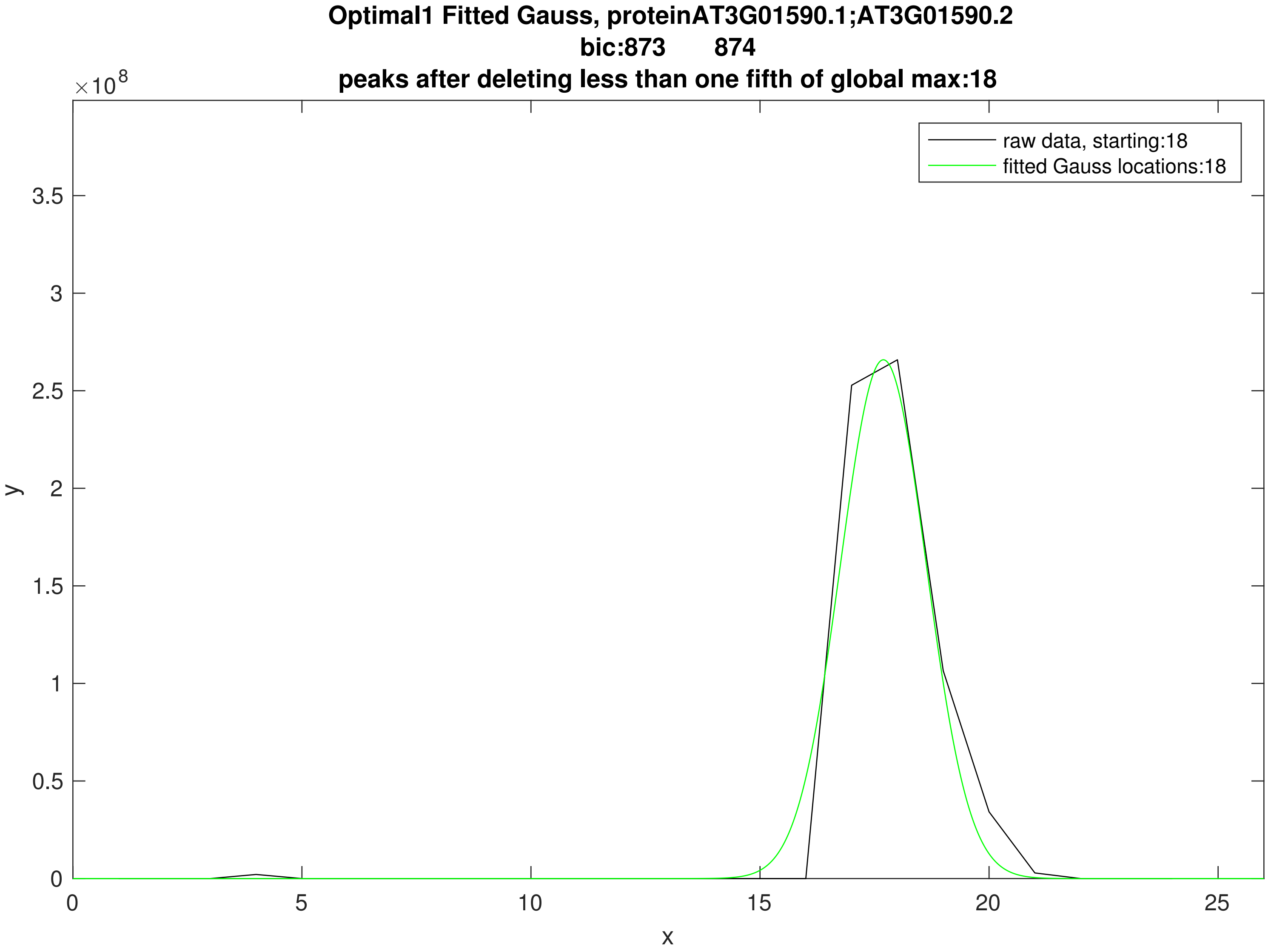

Optimal1 Fitted Gauss, proteinAT3G01590.1;AT3G01590.2
bic:873 874
peaks after deleting less than one fifth of global max:18
×10^8
raw data, starting:18
fitted Gauss locations:18
y
x
0
0.5
1
1.5
2
2.5
3
3.5
5
10
15
20
25

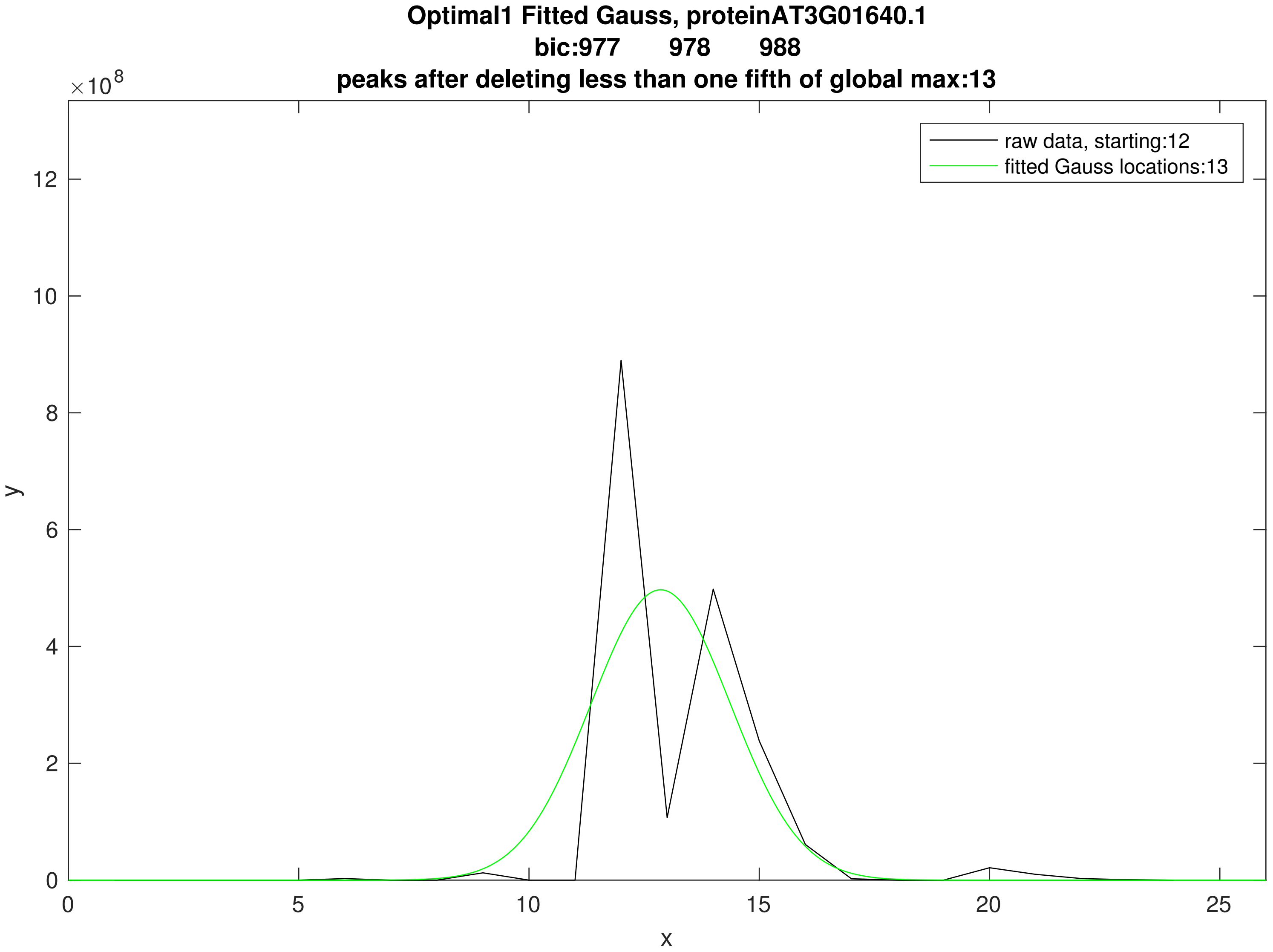
Optimal1 Fitted Gauss, proteinAT3G01640.1
bic:977 978 988
peaks after deleting less than one fifth of global max:13
×10^8
raw data, starting:12
fitted Gauss locations:13
y
x
0
2
4
6
8
10
12
0
5
10
15
20
25

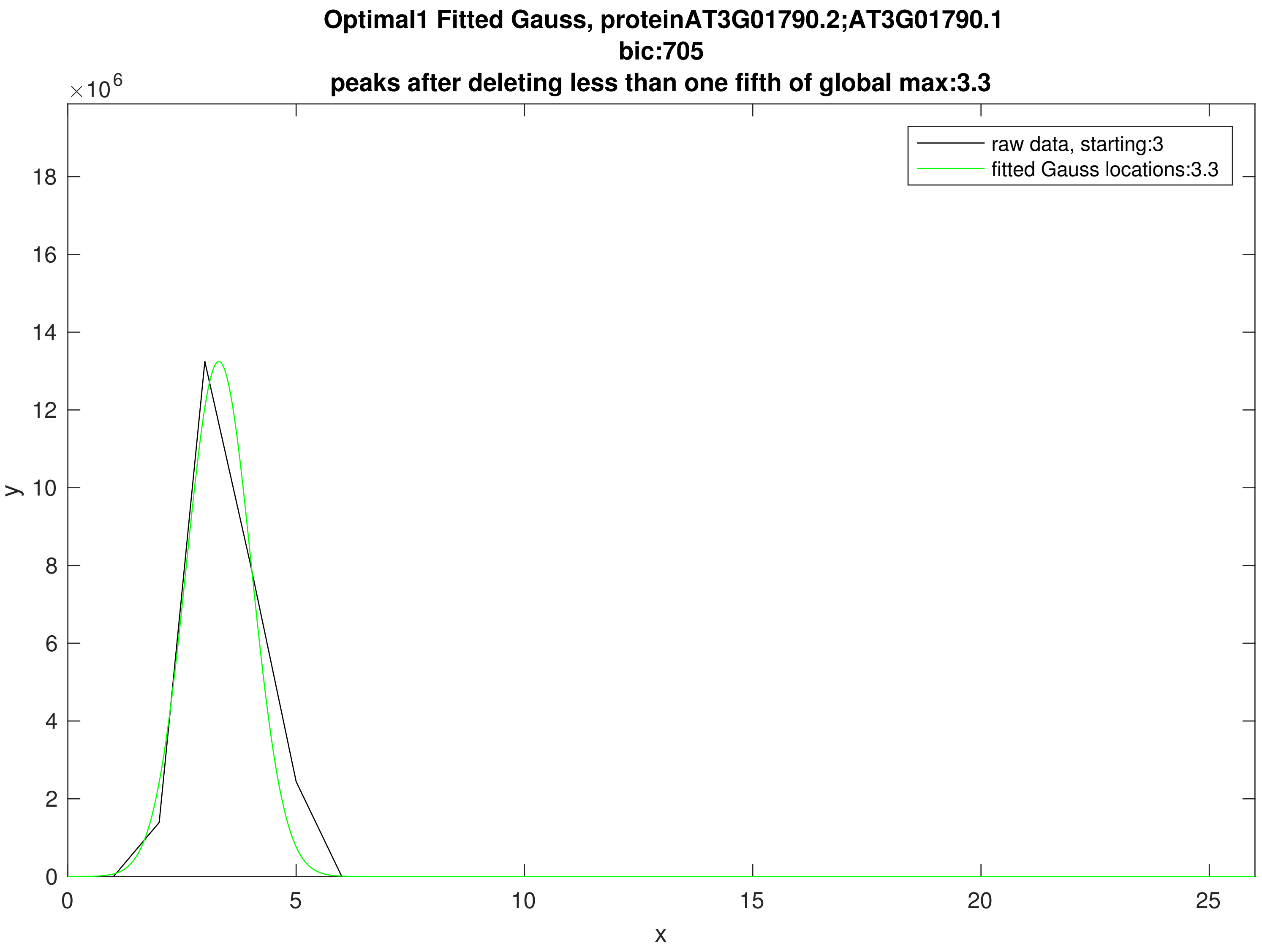
Optimal1 Fitted Gauss, proteinAT3G01790.2;AT3G01790.1
bic:705
peaks after deleting less than one fifth of global max:3.3
×10^6
raw data, starting:3
fitted Gauss locations:3.3
y
x
0
2
4
6
8
10
12
14
16
18
0
5
10
15
20
25

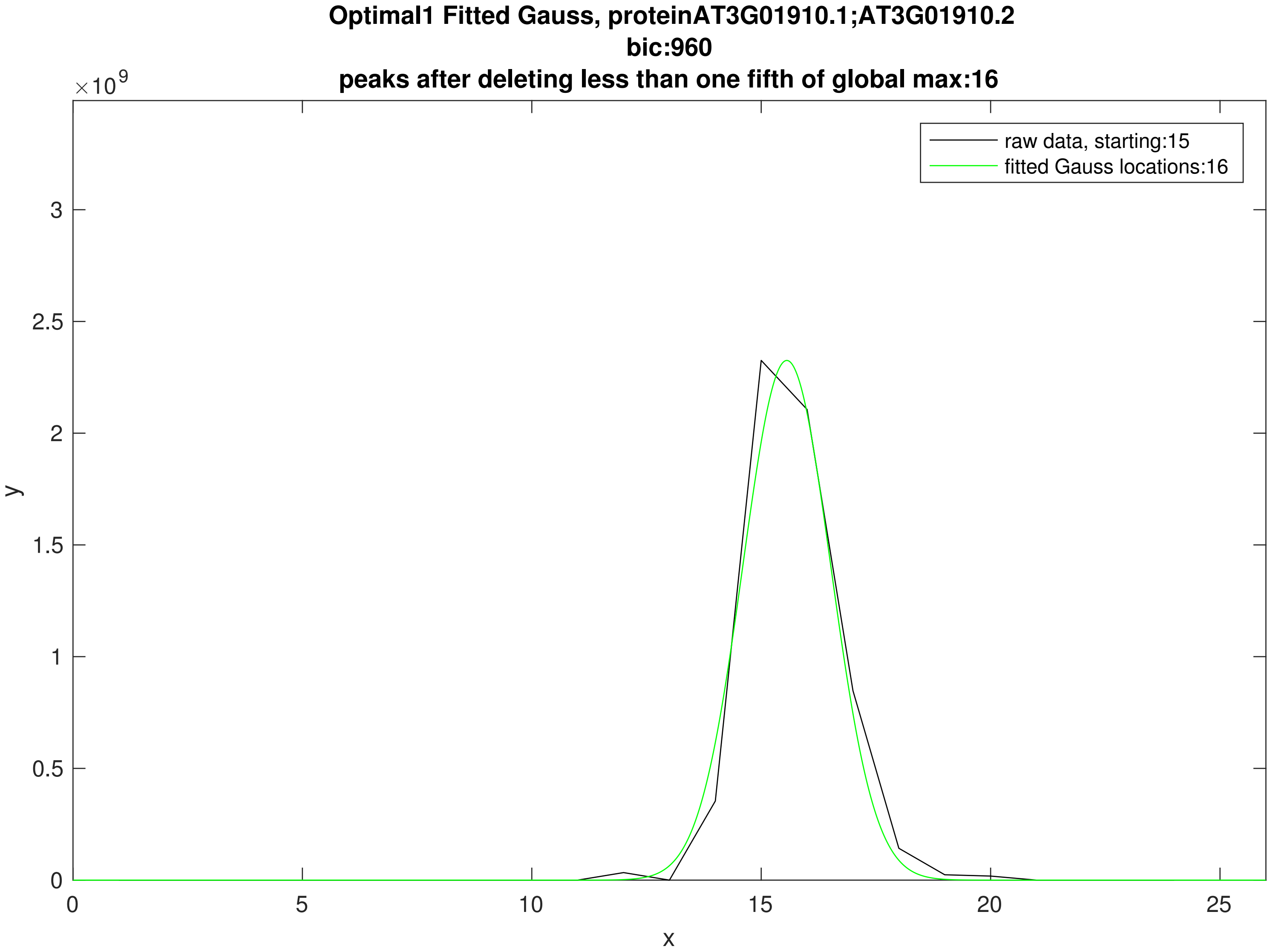

Optimal1 Fitted Gauss, proteinAT3G01910.1;AT3G01910.2
bic:960
peaks after deleting less than one fifth of global max:16
×10^9
raw data, starting:15
fitted Gauss locations:16
y
x
0
0.5
1
1.5
2
2.5
3
0
5
10
15
20
25

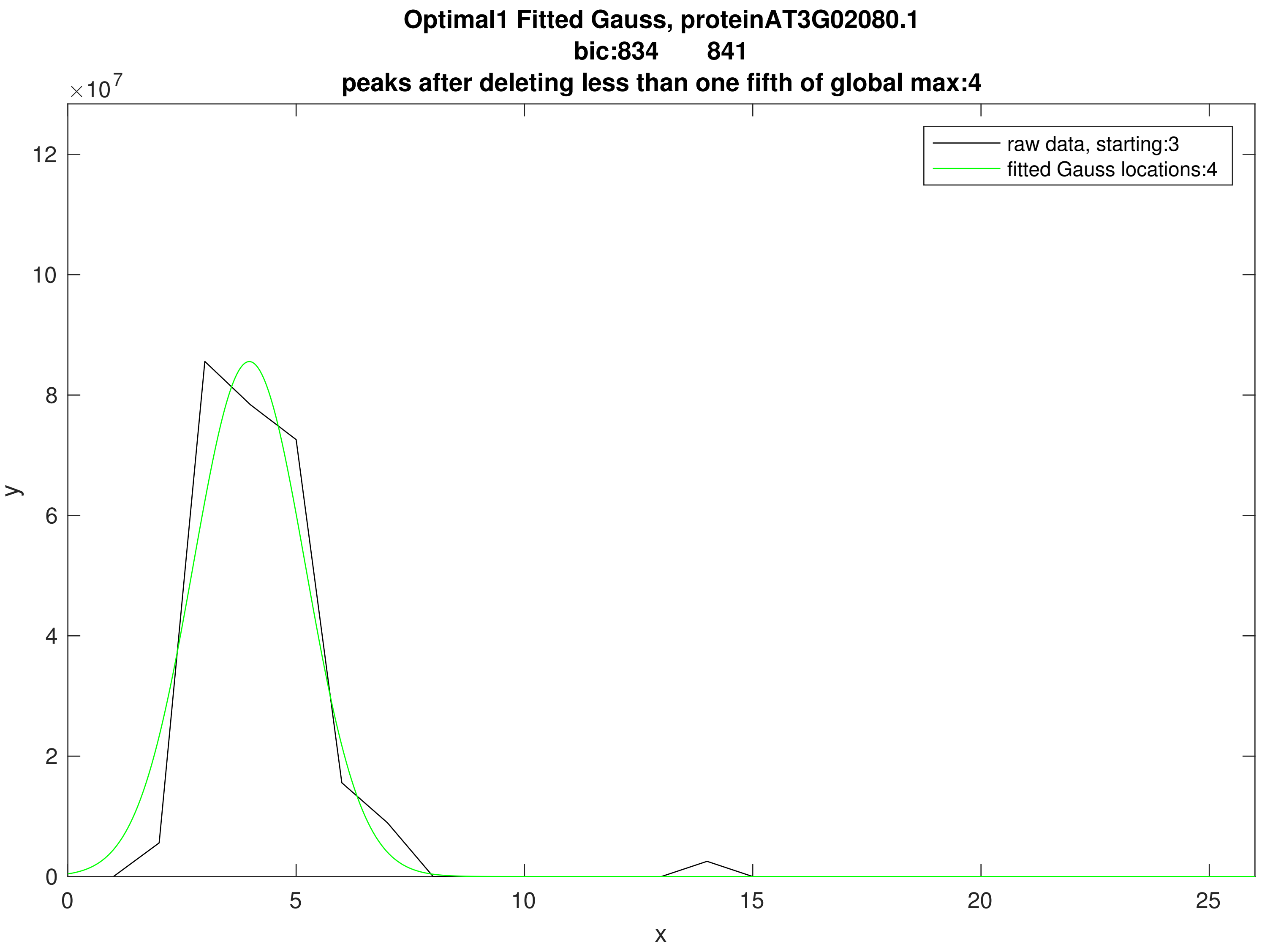
Optimal1 Fitted Gauss, proteinAT3G02080.1
bic:834 841
peaks after deleting less than one fifth of global max:4
×10^7
raw data, starting:3
fitted Gauss locations:4
y
x
0
2
4
6
8
10
12
0
5
10
15
20
25

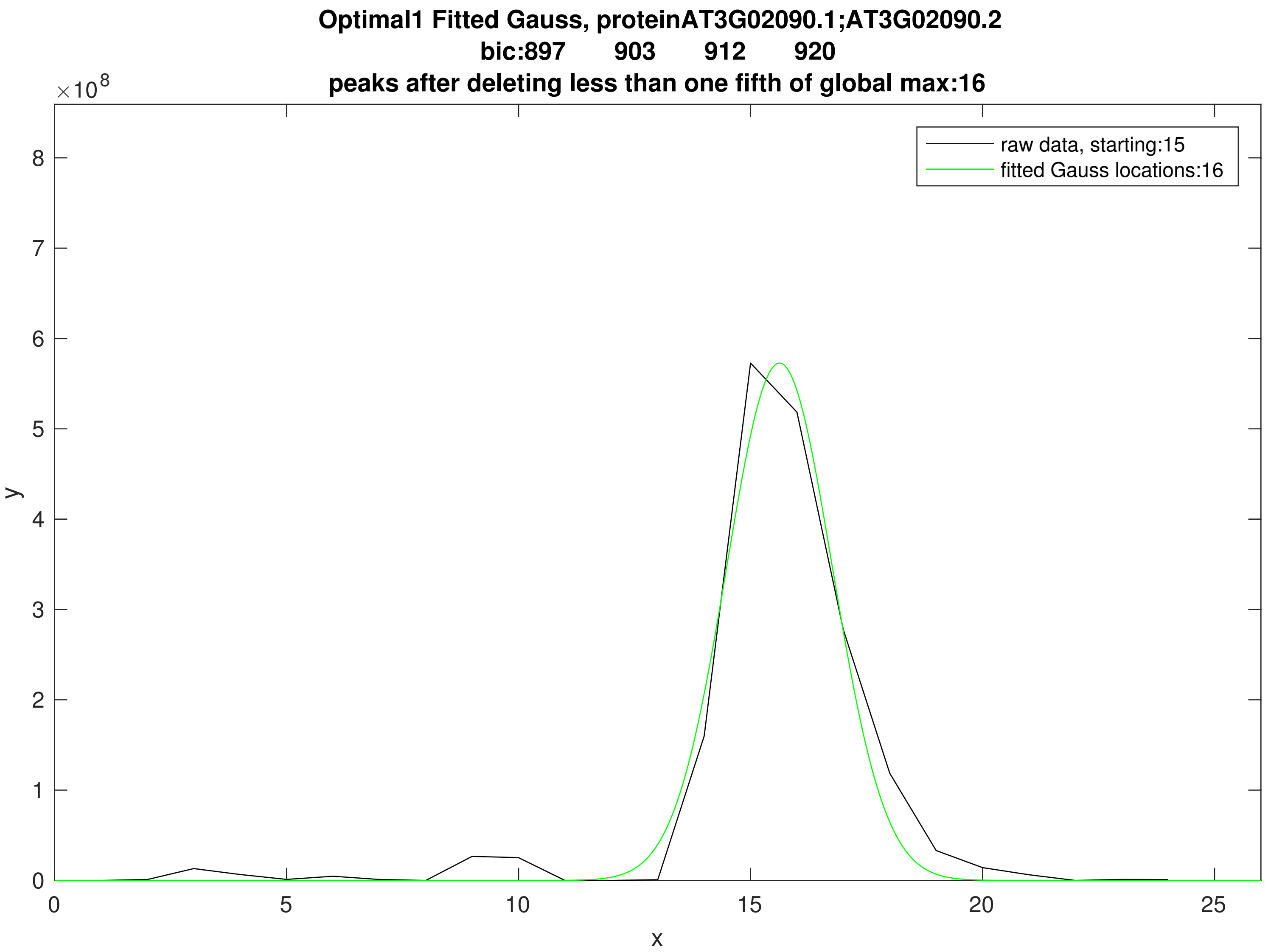
Optimal1 Fitted Gauss, proteinAT3G02090.1;AT3G02090.2
bic:897 903 912 920
peaks after deleting less than one fifth of global max:16
×10^8
raw data, starting:15
fitted Gauss locations:16
y
x
0
1
2
3
4
5
6
7
8
0
5
10
15
20
25

Optimal1 Fitted Gauss, proteinAT3G02180.1;AT3G02180.2;AT3G02180.3
bic:747
peaks after deleting less than one fifth of global max:19

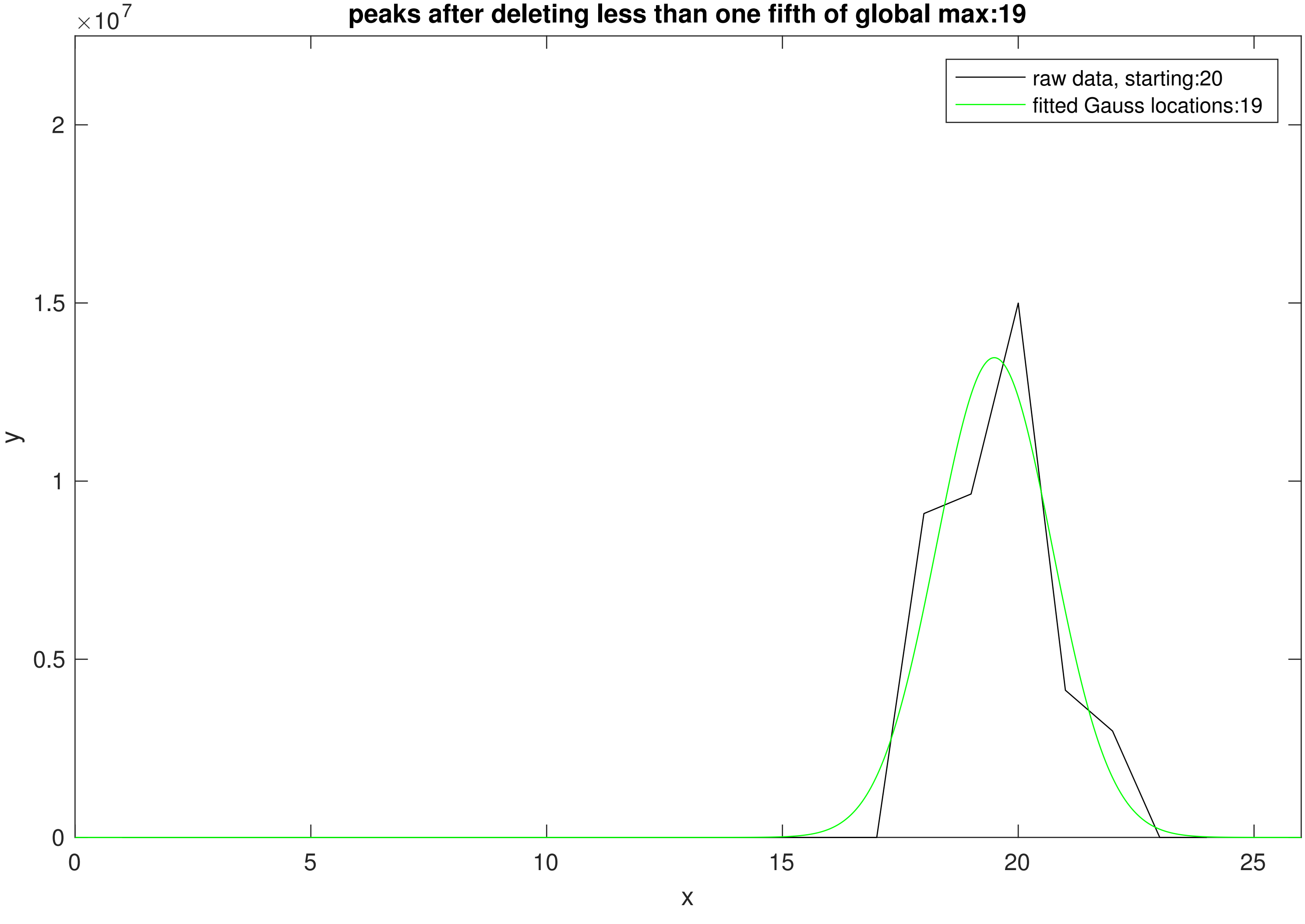

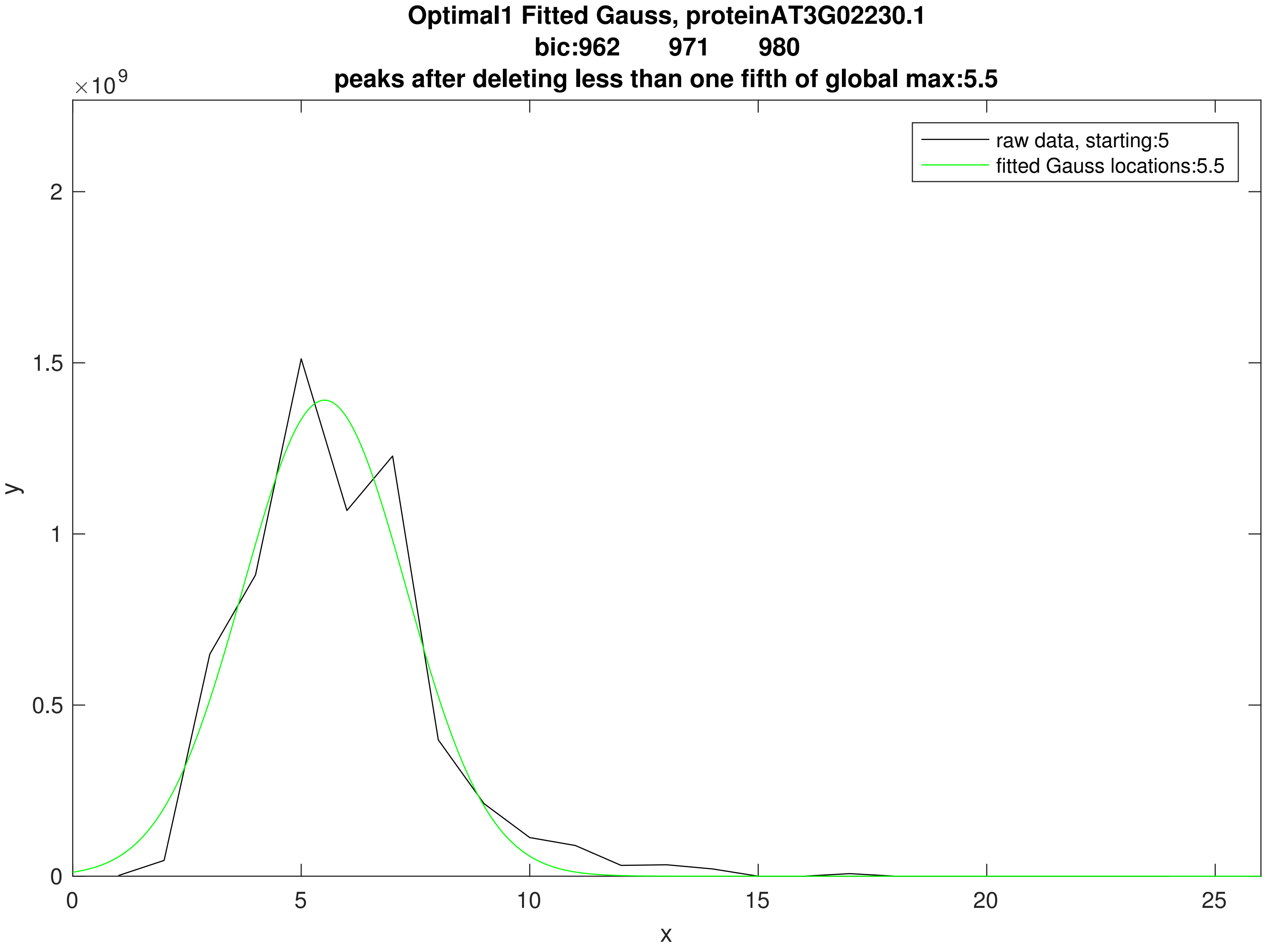
Optimal1 Fitted Gauss, proteinAT3G02230.1
bic:962 971 980
peaks after deleting less than one fifth of global max:5.5
×10^9
raw data, starting:5
fitted Gauss locations:5.5
y
x
0
0.5
1
1.5
2
0
5
10
15
20
25

**Optimal1 Fitted Gauss, proteinAT3G02360.2;AT3G02360.1**
**bic:1.08e+03  1.11e+03  1.12e+03**
**peaks after deleting less than one fifth of global max:13**

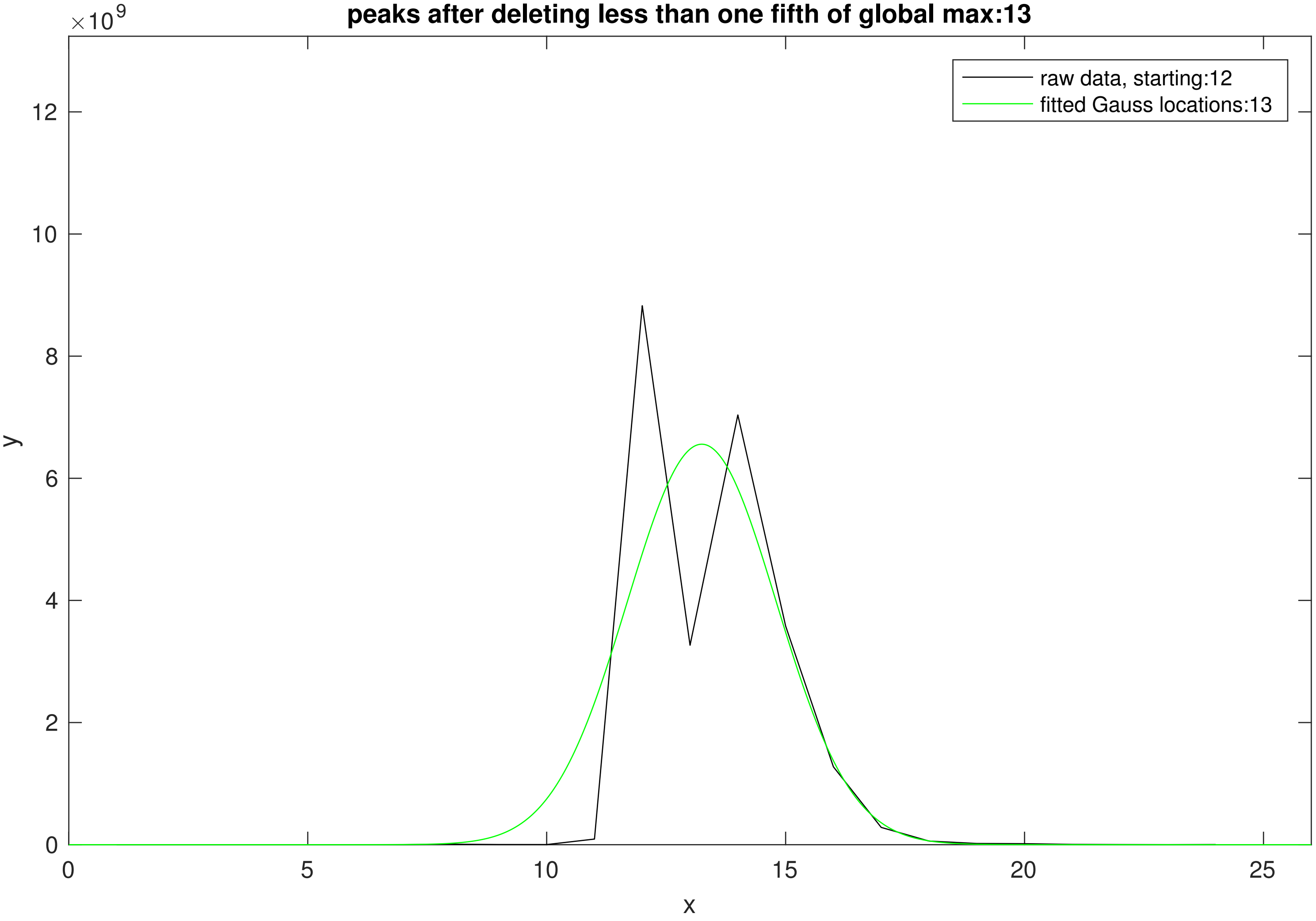

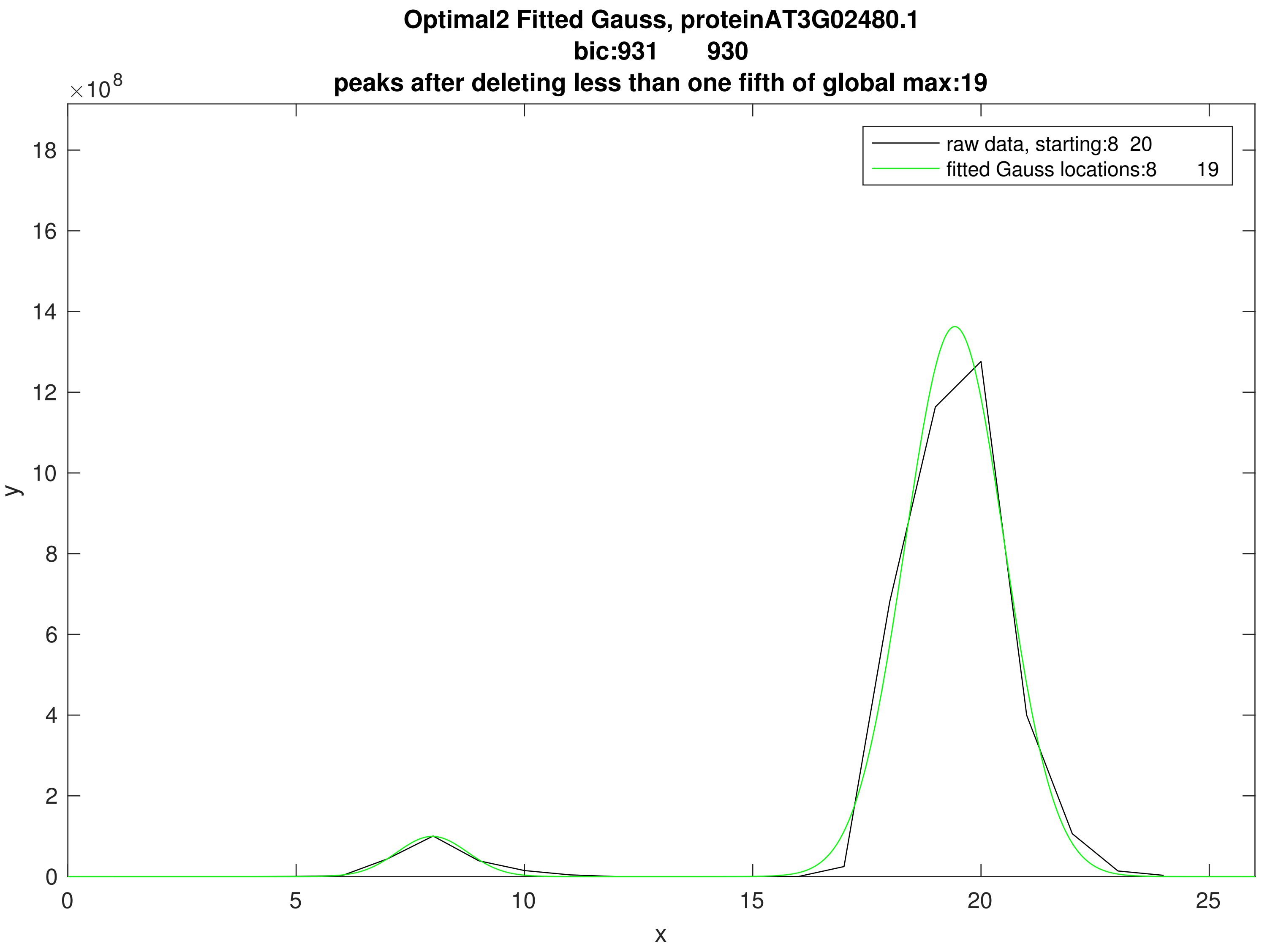

Optimal2 Fitted Gauss, proteinAT3G02480.1
bic:931 930
peaks after deleting less than one fifth of global max:19
×10^8
raw data, starting:8 20
fitted Gauss locations:8 19
y
x
0
2
4
6
8
10
12
14
16
18
0
5
10
15
20
25

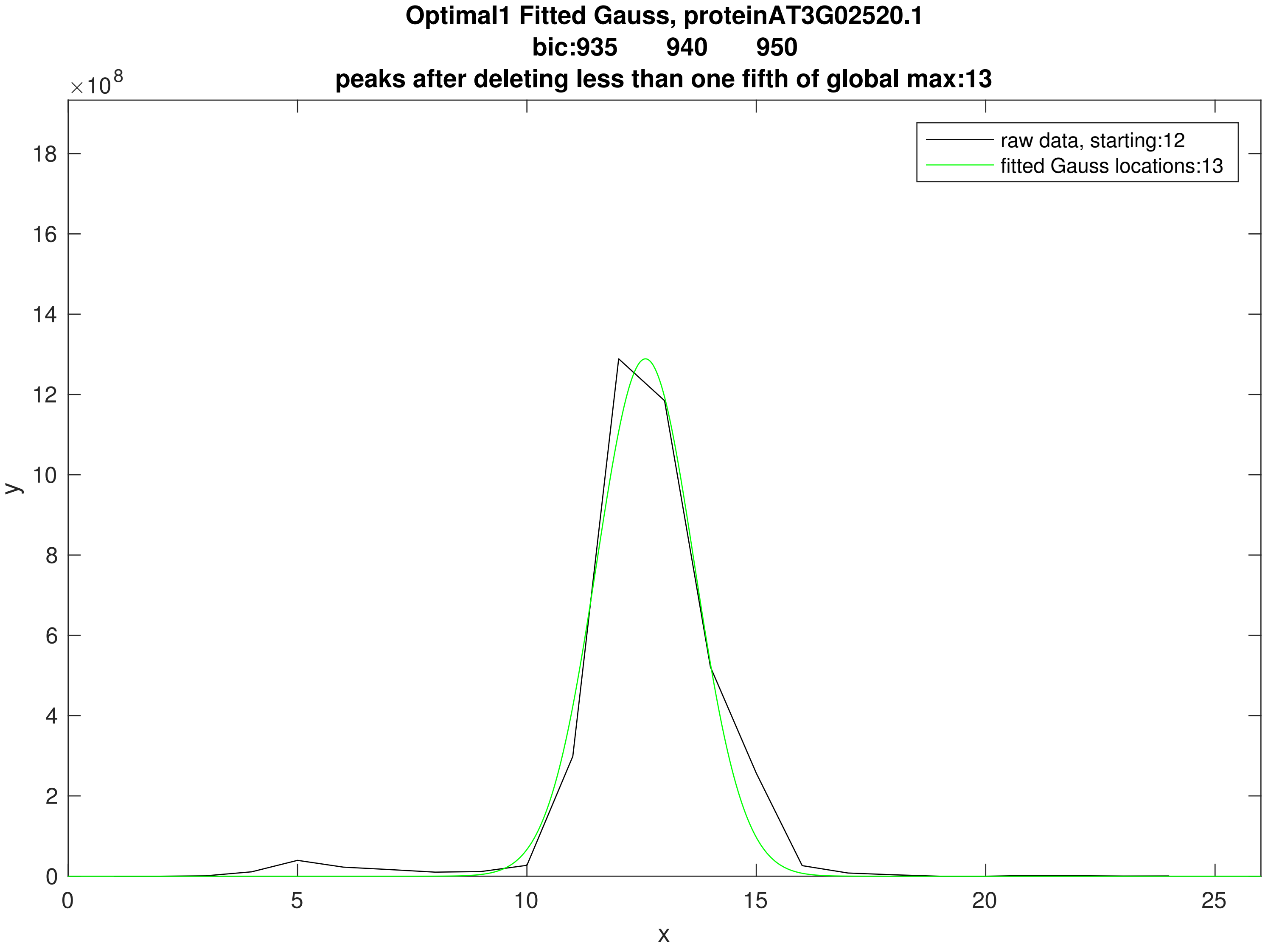

Optimal1 Fitted Gauss, proteinAT3G02520.1
bic:935 940 950
peaks after deleting less than one fifth of global max:13
×10^8
raw data, starting:12
fitted Gauss locations:13
y
x
0
2
4
6
8
10
12
14
16
18
0
5
10
15
20
25

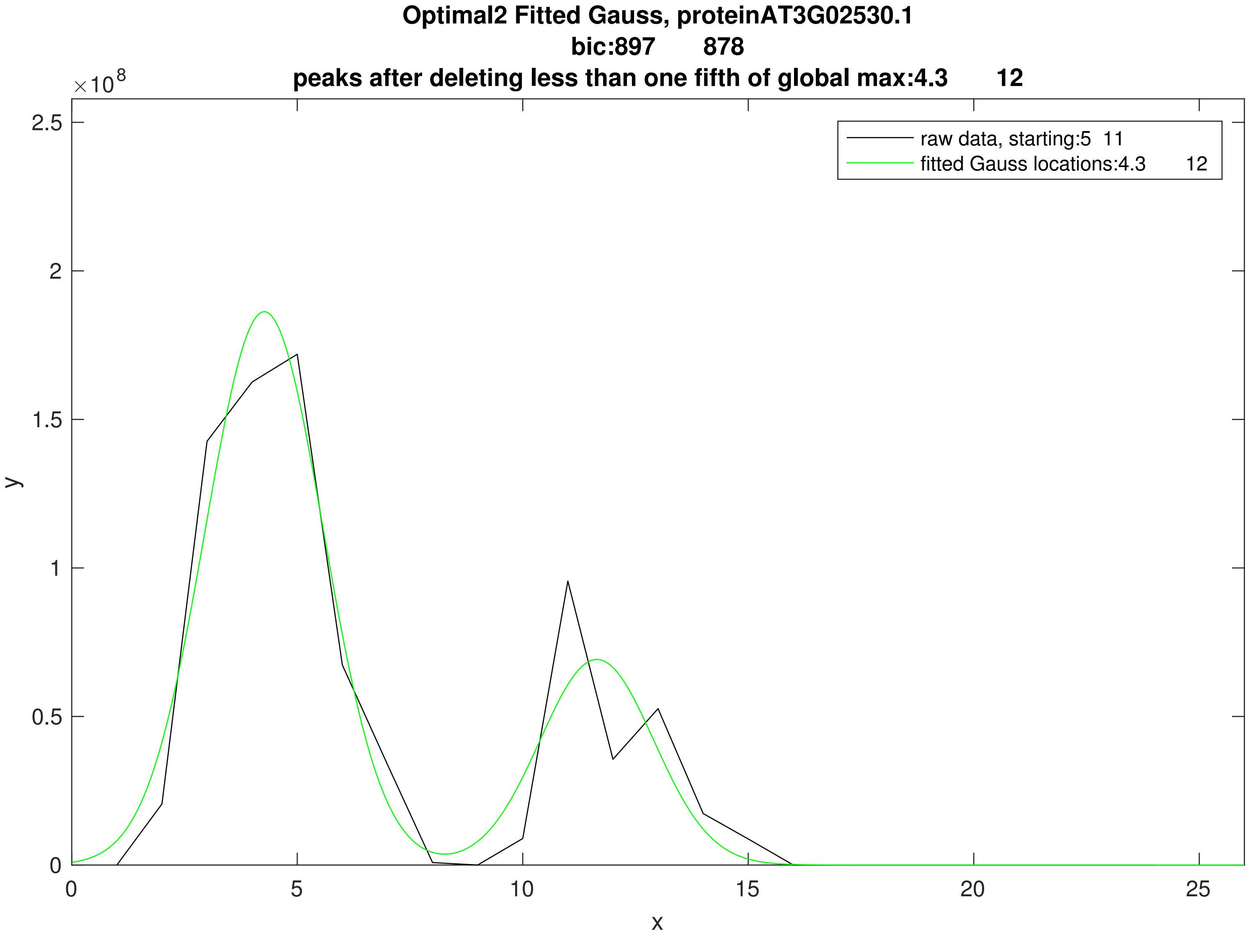
Optimal2 Fitted Gauss, proteinAT3G02530.1
bic:897 878
peaks after deleting less than one fifth of global max:4.3 12
×10^8
2.5
2
1.5
1
0.5
0
y
0
5
10
15
20
25
x
raw data, starting:5 11
fitted Gauss locations:4.3 12

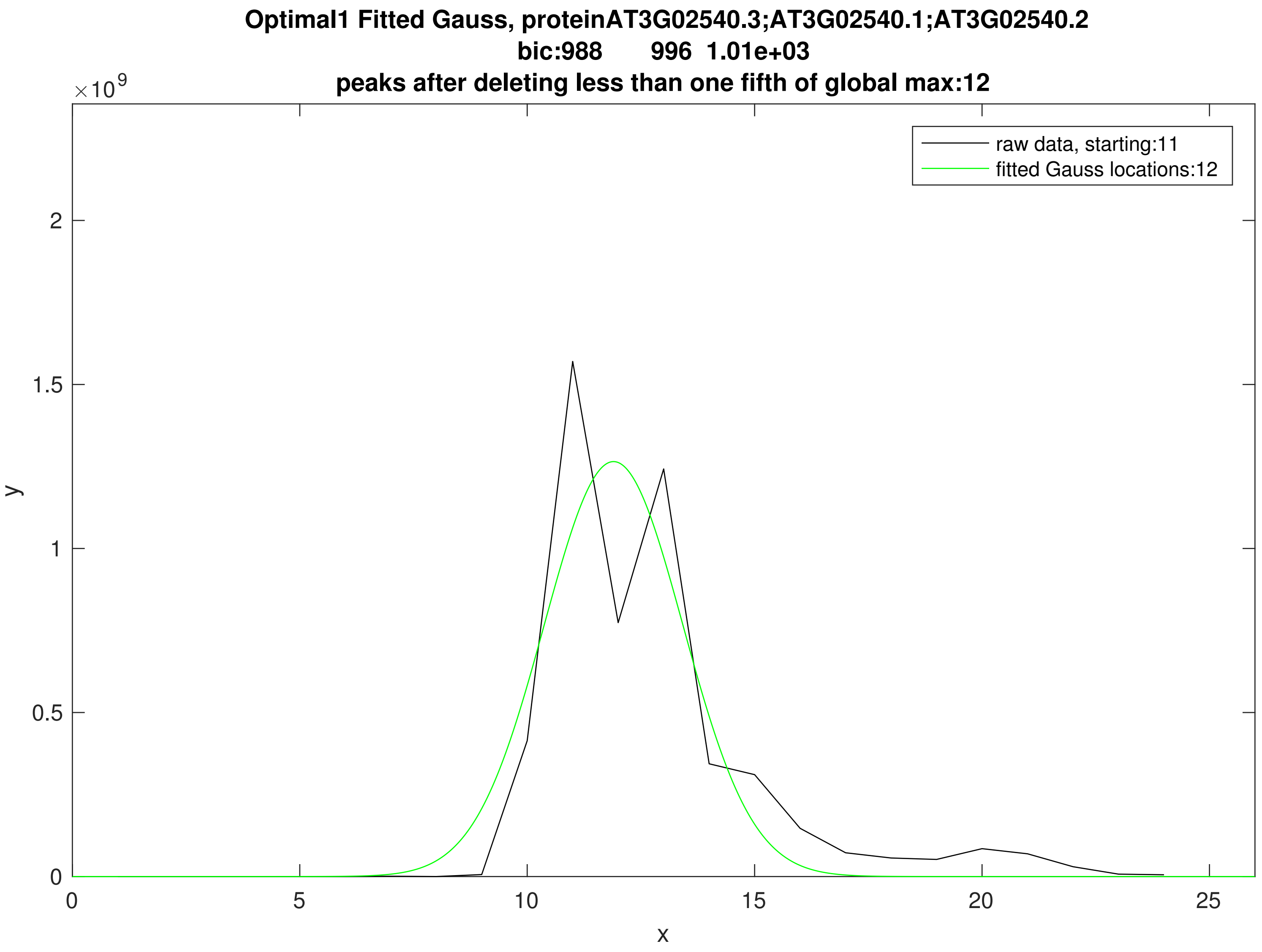

Optimal1 Fitted Gauss, proteinAT3G02540.3;AT3G02540.1;AT3G02540.2
bic:988 996 1.01e+03
peaks after deleting less than one fifth of global max:12
×10^9
raw data, starting:11
fitted Gauss locations:12
y
x
0
0.5
1
1.5
2
0
5
10
15
20
25

**Optimal1 Fitted Gauss, proteinAT3G02560.2;AT3G02560.1**
**bic:813 816**
**peaks after deleting less than one fifth of global max:3.8**

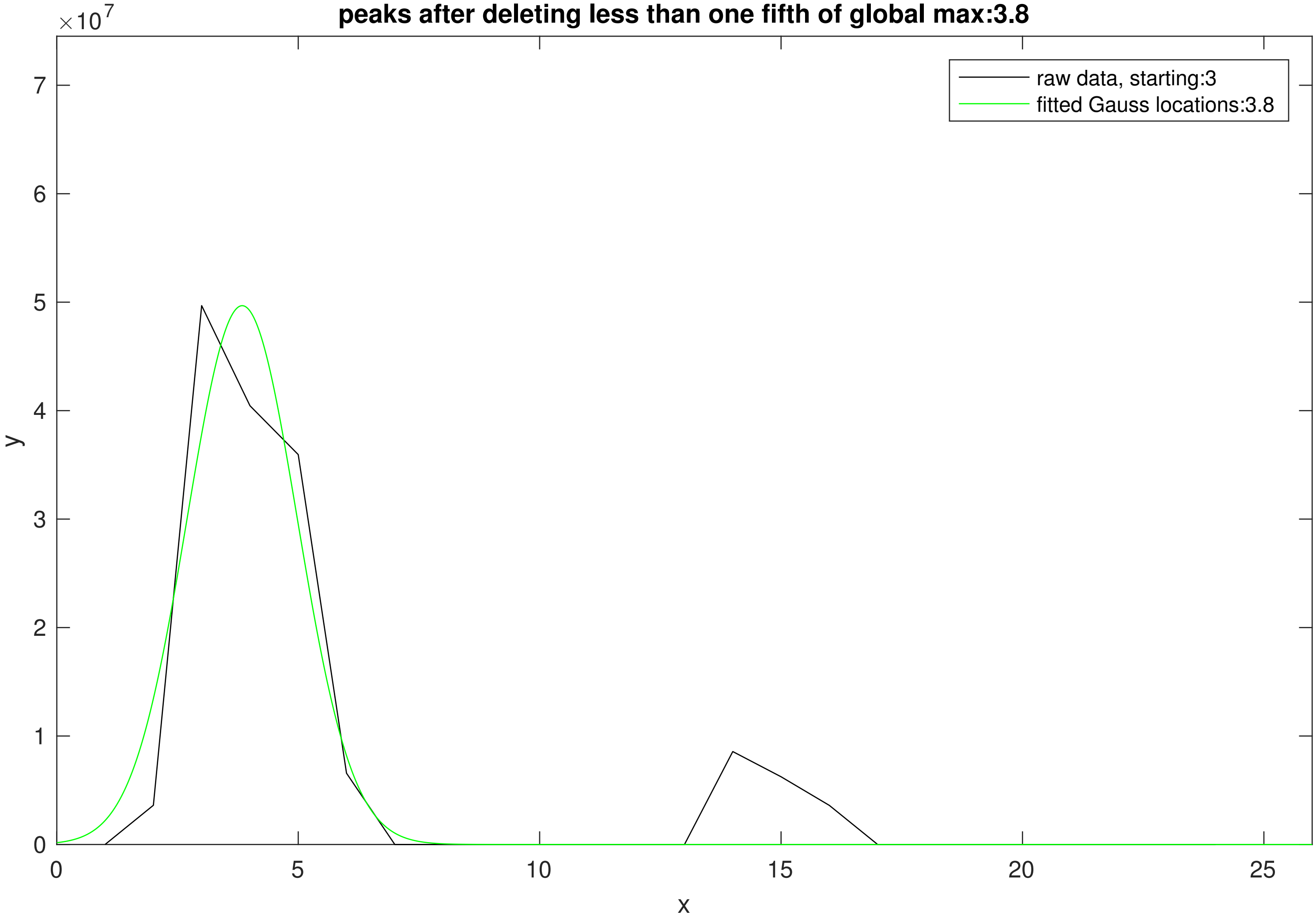

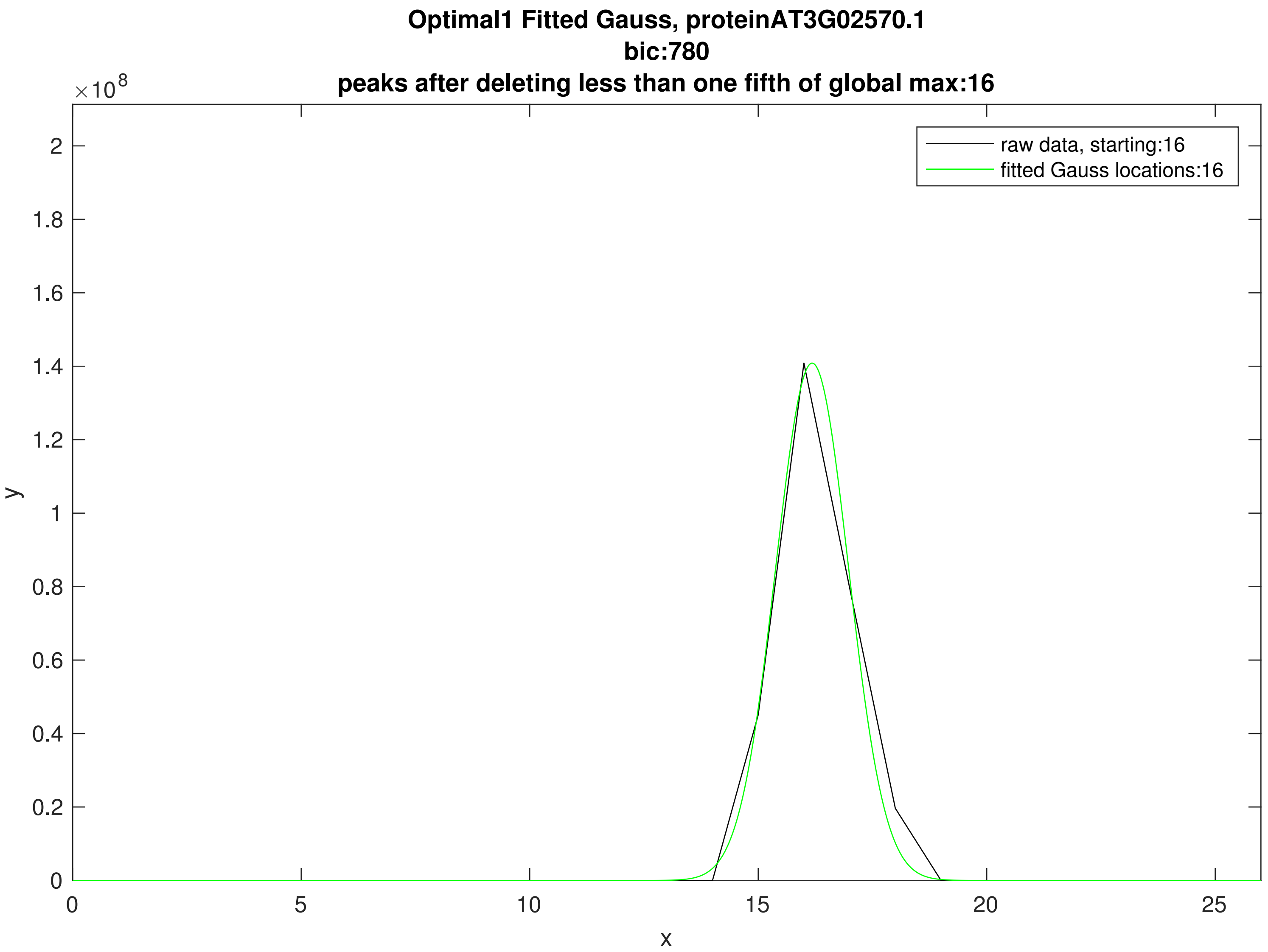
Optimal1 Fitted Gauss, proteinAT3G02570.1
bic:780
peaks after deleting less than one fifth of global max:16
×10^8
2
1.8
1.6
1.4
1.2
1
0.8
0.6
0.4
0.2
0
y
0
5
10
15
20
25
x
raw data, starting:16
fitted Gauss locations:16

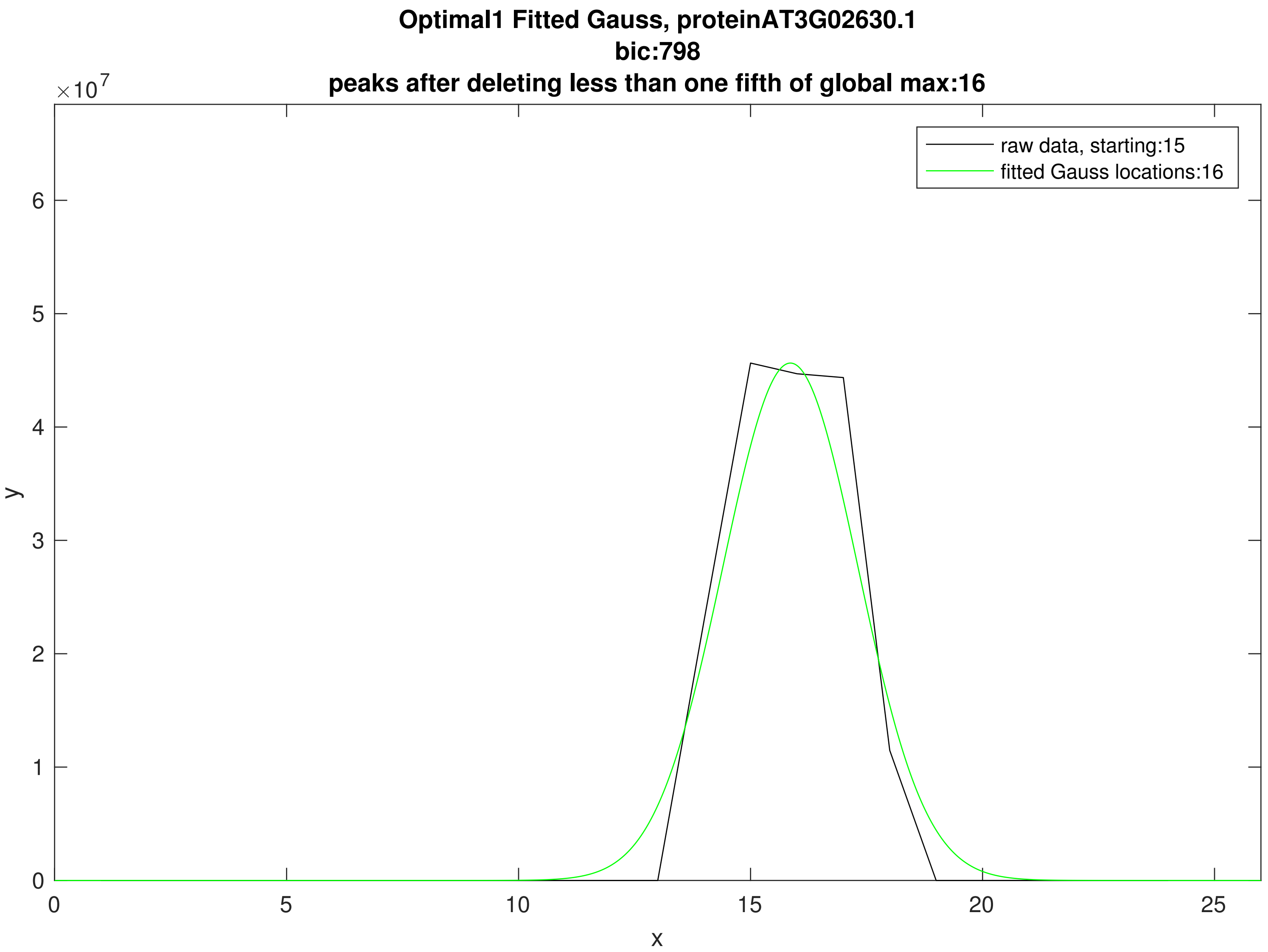

Optimal1 Fitted Gauss, proteinAT3G02630.1
bic:798
peaks after deleting less than one fifth of global max:16
×10^7
raw data, starting:15
fitted Gauss locations:16
y
x
0
1
2
3
4
5
6
0
5
10
15
20
25

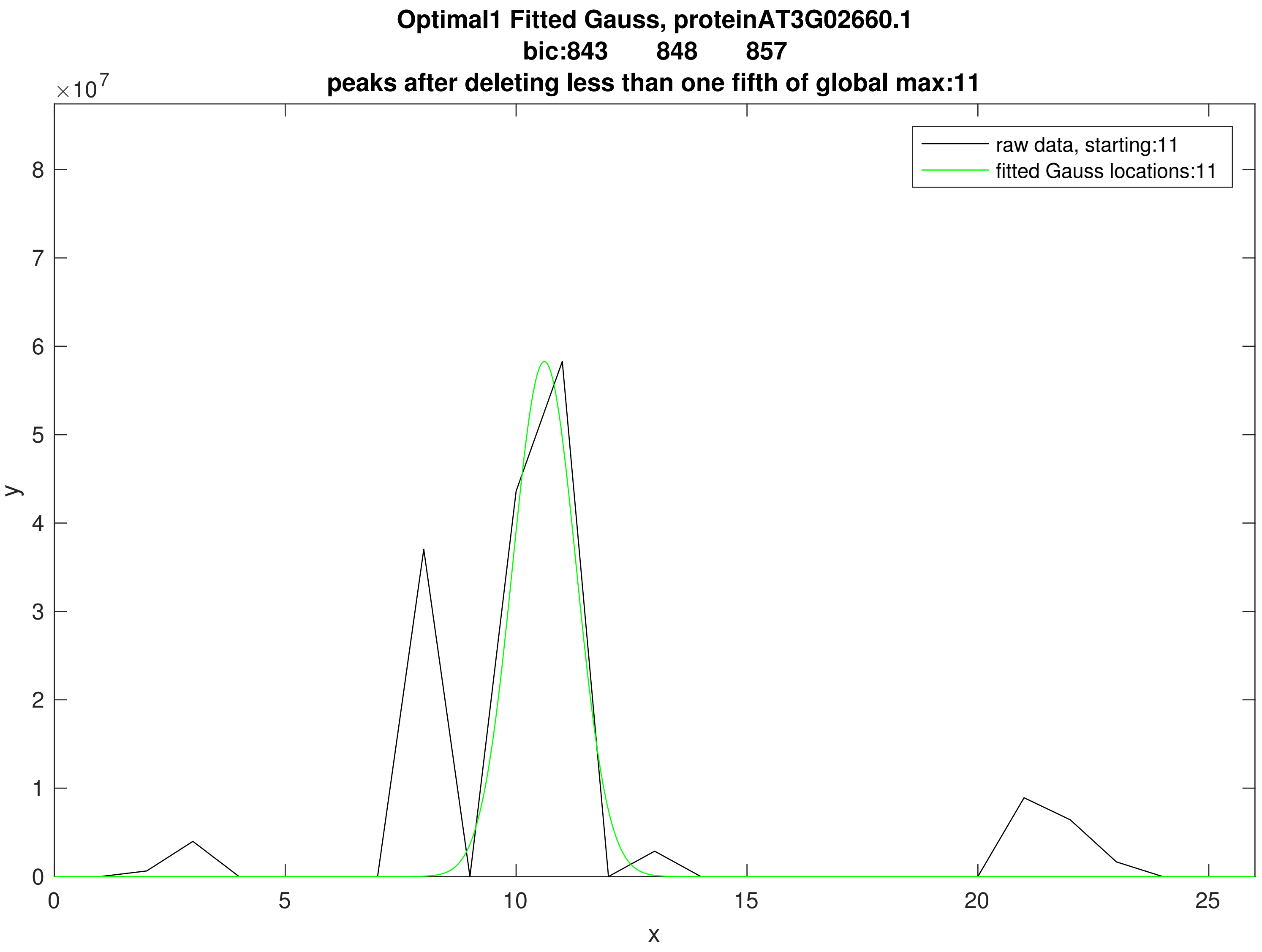
Optimal1 Fitted Gauss, proteinAT3G02660.1
bic:843 848 857
peaks after deleting less than one fifth of global max:11
×10^7
raw data, starting:11
fitted Gauss locations:11
y
x

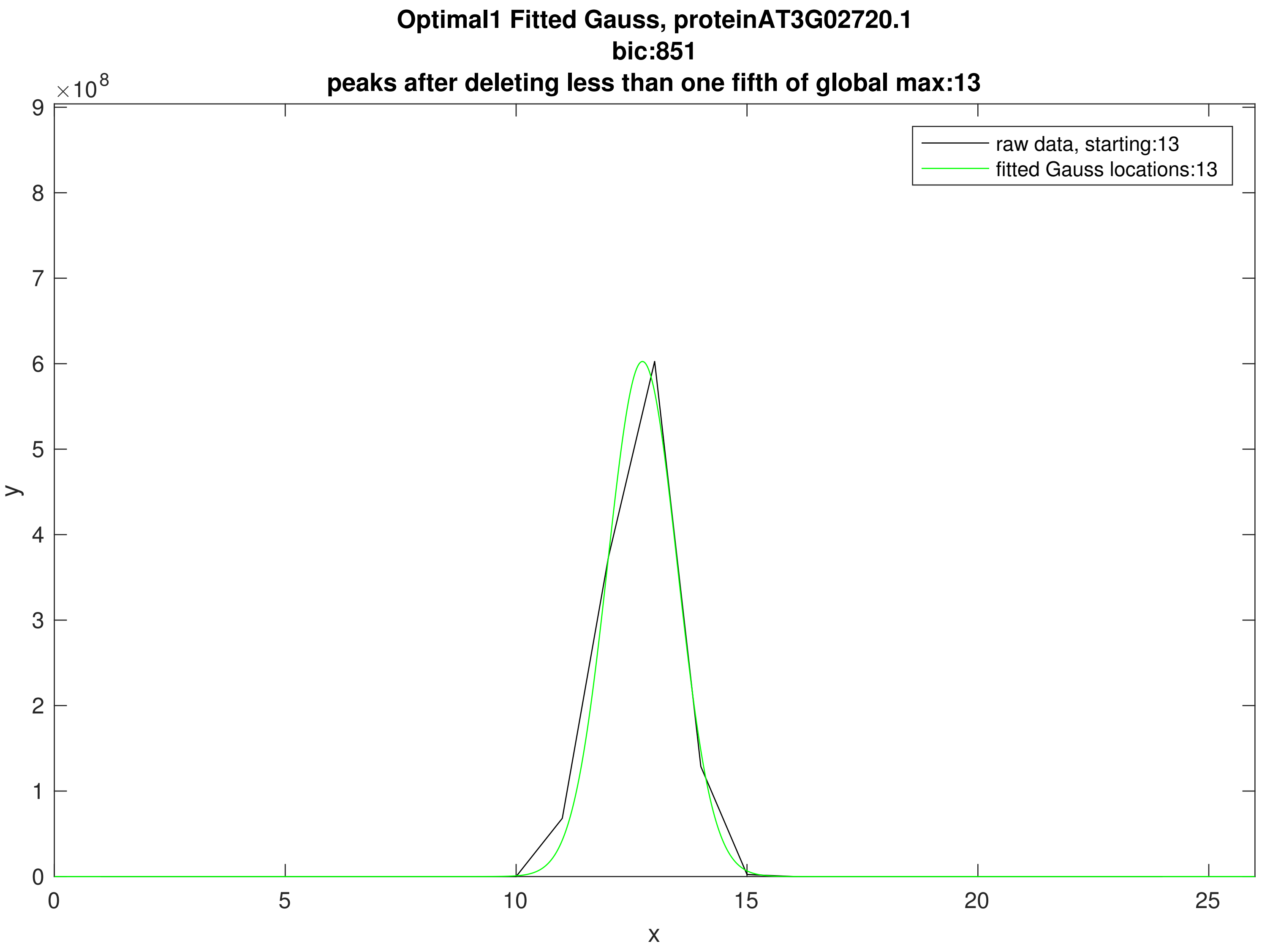
Optimal1 Fitted Gauss, proteinAT3G02720.1
bic:851
peaks after deleting less than one fifth of global max:13
×10^8
9
8
7
6
5
4
3
2
1
0
y
0
5
10
15
20
25
x
raw data, starting:13
fitted Gauss locations:13

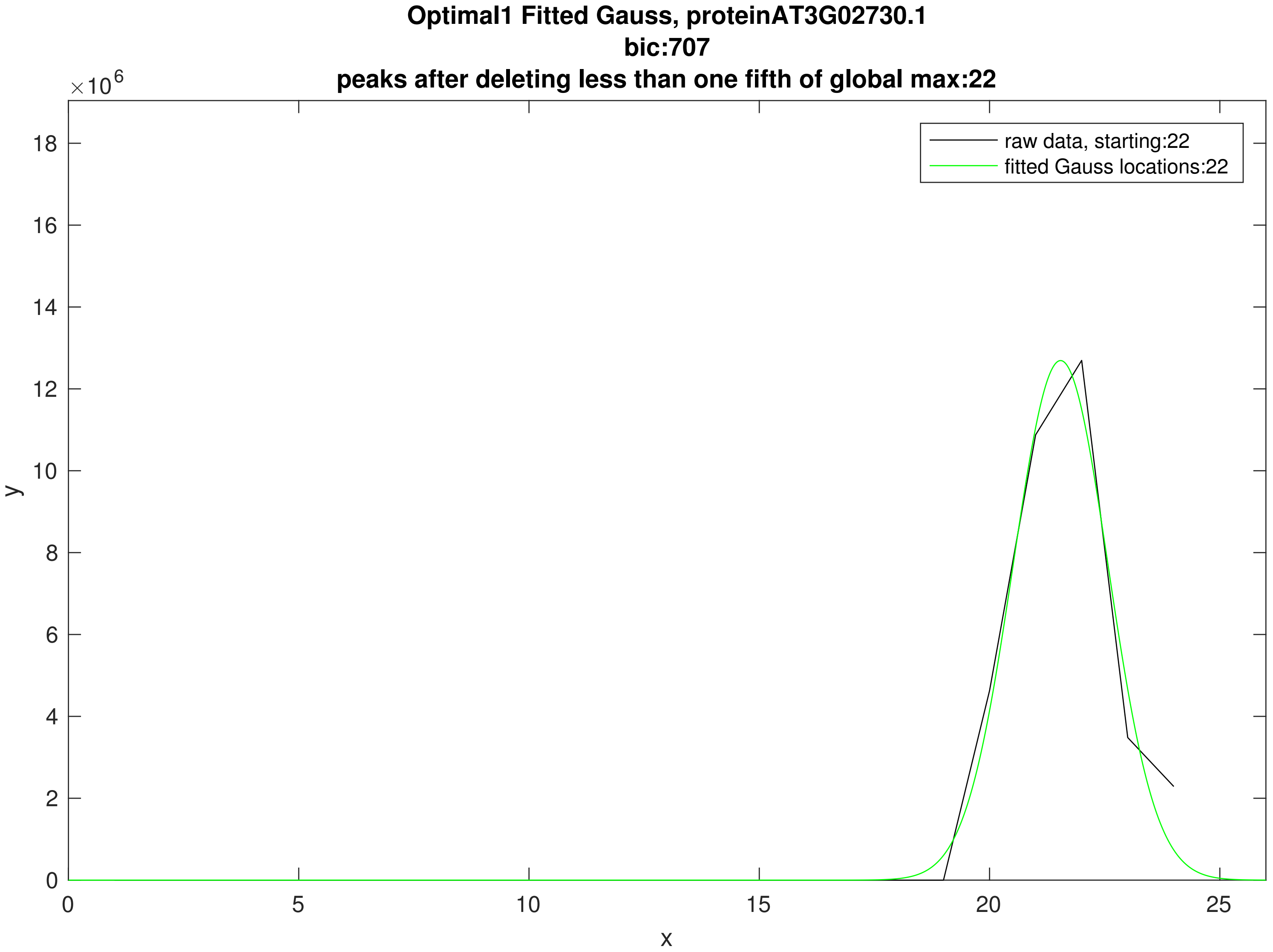

Optimal1 Fitted Gauss, proteinAT3G02730.1
bic:707
peaks after deleting less than one fifth of global max:22
×10^6
raw data, starting:22
fitted Gauss locations:22
y
x
0
2
4
6
8
10
12
14
16
18
0
5
10
15
20
25

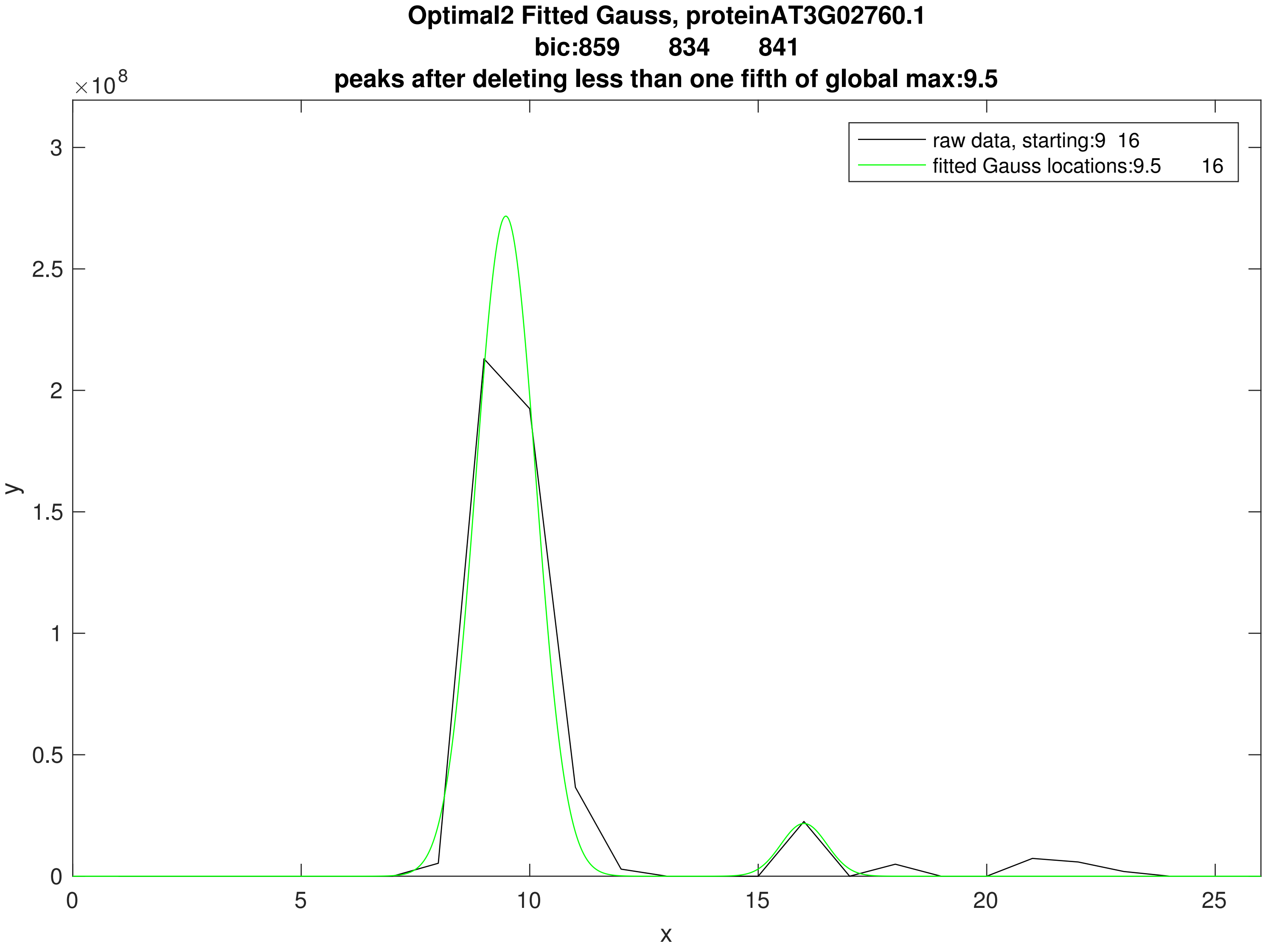
Optimal2 Fitted Gauss, proteinAT3G02760.1
bic:859 834 841
peaks after deleting less than one fifth of global max:9.5
×10^8
3
2.5
2
1.5
1
0.5
0
y
0
5
10
15
20
25
x
raw data, starting:9 16
fitted Gauss locations:9.5 16

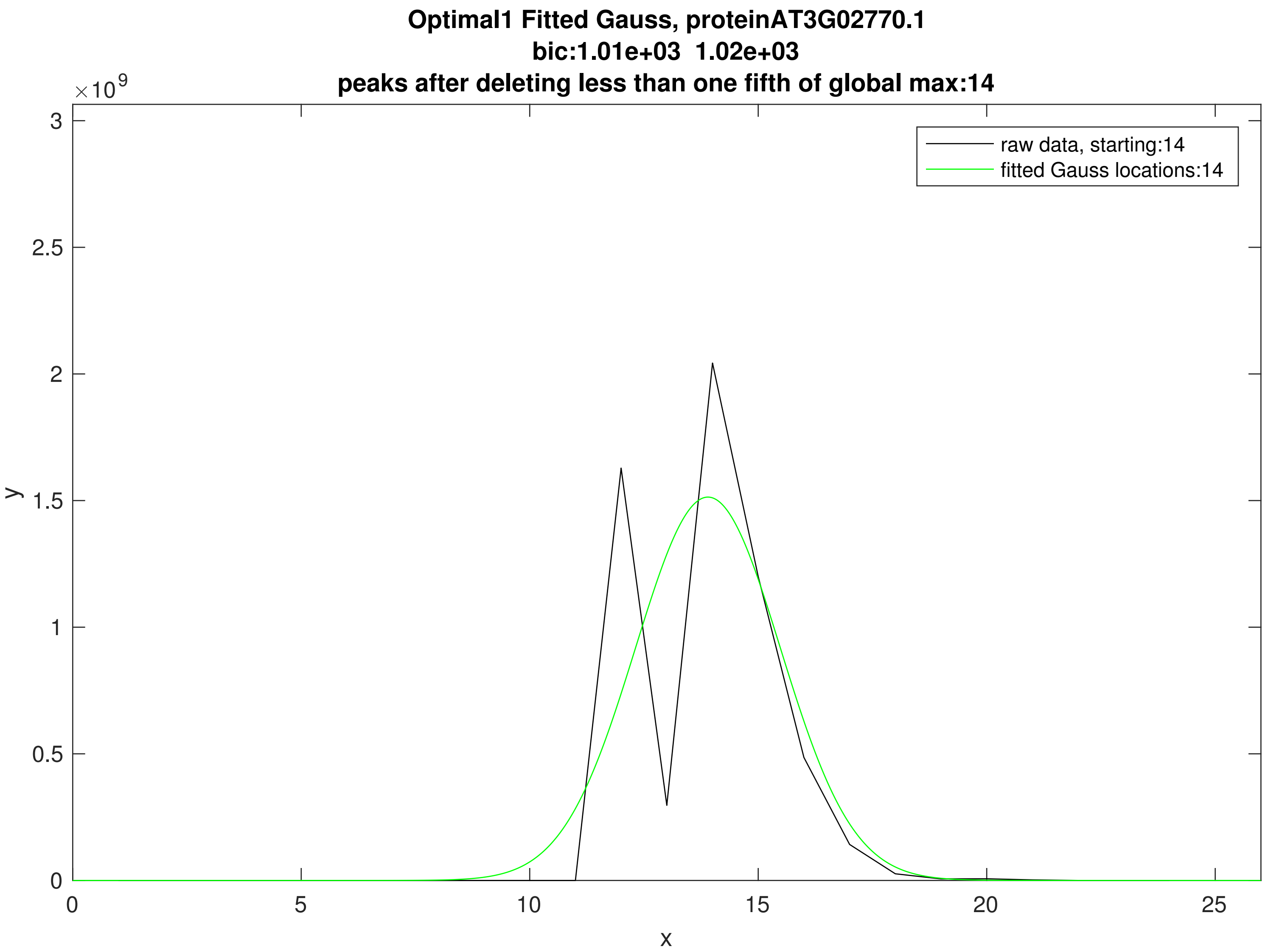
Optimal1 Fitted Gauss, proteinAT3G02770.1
bic:1.01e+03  1.02e+03
peaks after deleting less than one fifth of global max:14
×10^9
3
2.5
2
1.5
1
0.5
0
y
0
5
10
15
20
25
x
raw data, starting:14
fitted Gauss locations:14

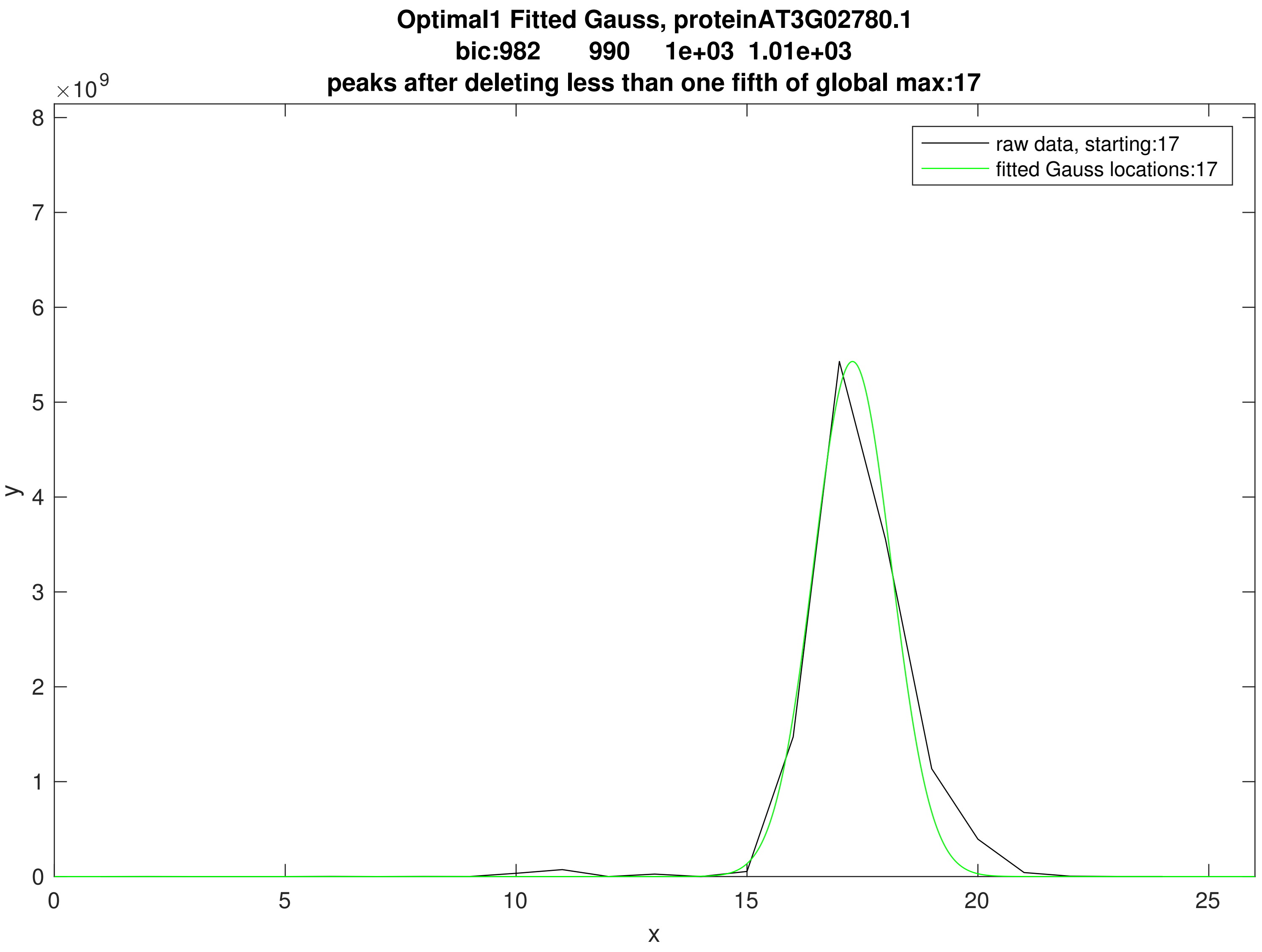
Optimal1 Fitted Gauss, proteinAT3G02780.1
bic:982 990 1e+03 1.01e+03
peaks after deleting less than one fifth of global max:17
×10^9
raw data, starting:17
fitted Gauss locations:17
y
x
0
1
2
3
4
5
6
7
8
0
5
10
15
20
25

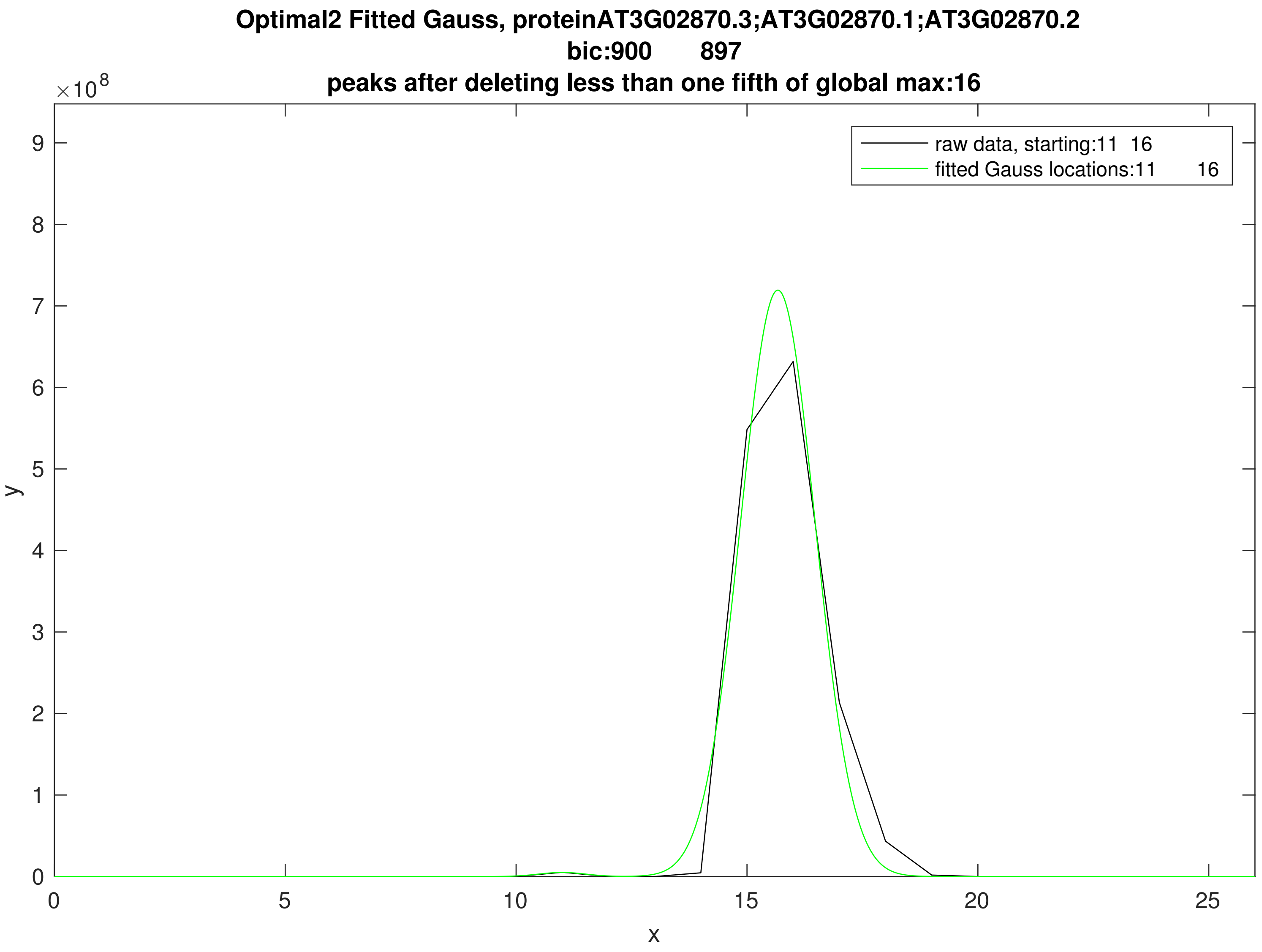
Optimal2 Fitted Gauss, proteinAT3G02870.3;AT3G02870.1;AT3G02870.2
bic:900 897
peaks after deleting less than one fifth of global max:16
raw data, starting:11 16
fitted Gauss locations:11 16
×10^8
y
x

Optimal1 Fitted Gauss, proteinAT3G02875.1
bic:800 920
peaks after deleting less than one fifth of global max:9.1

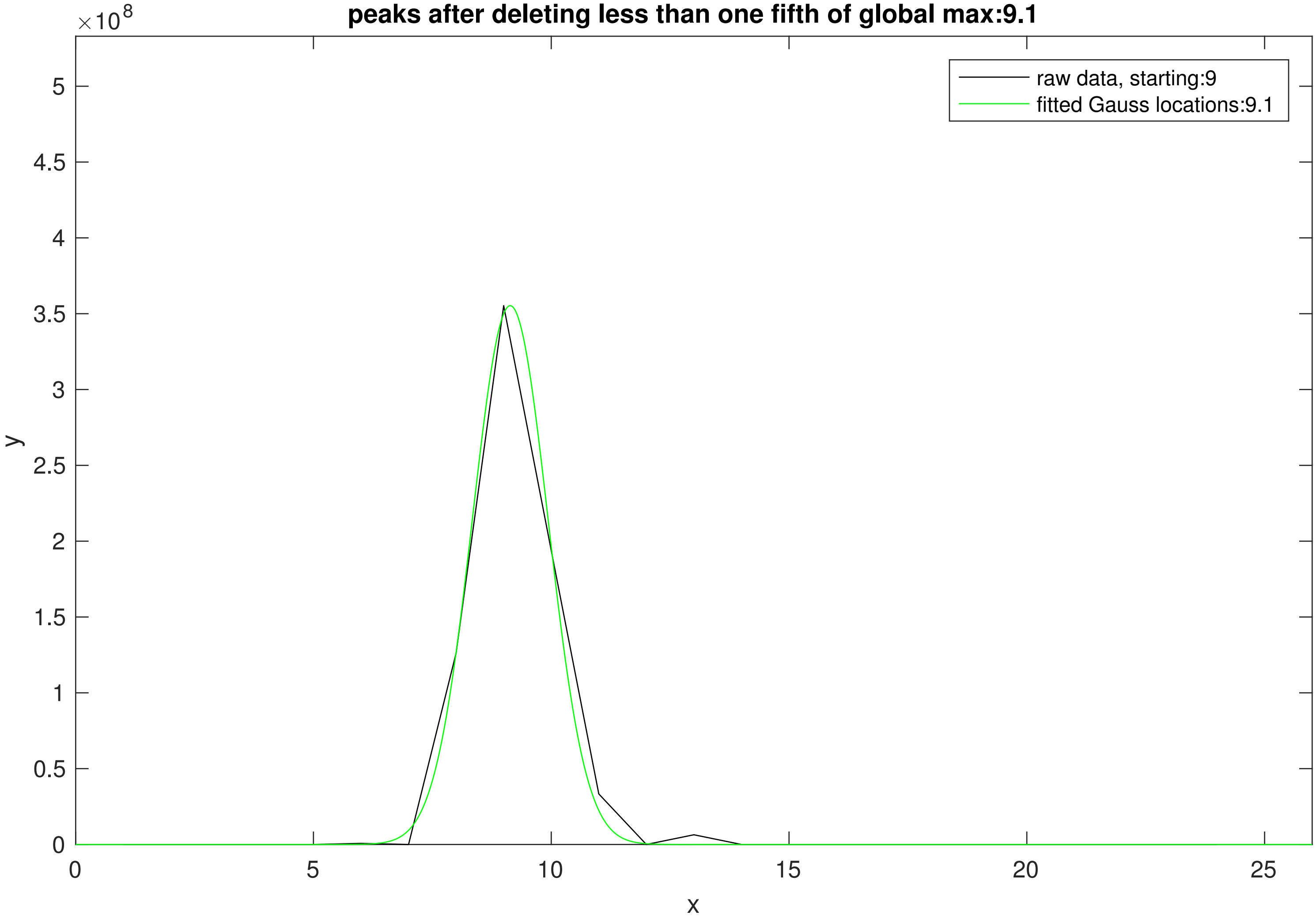

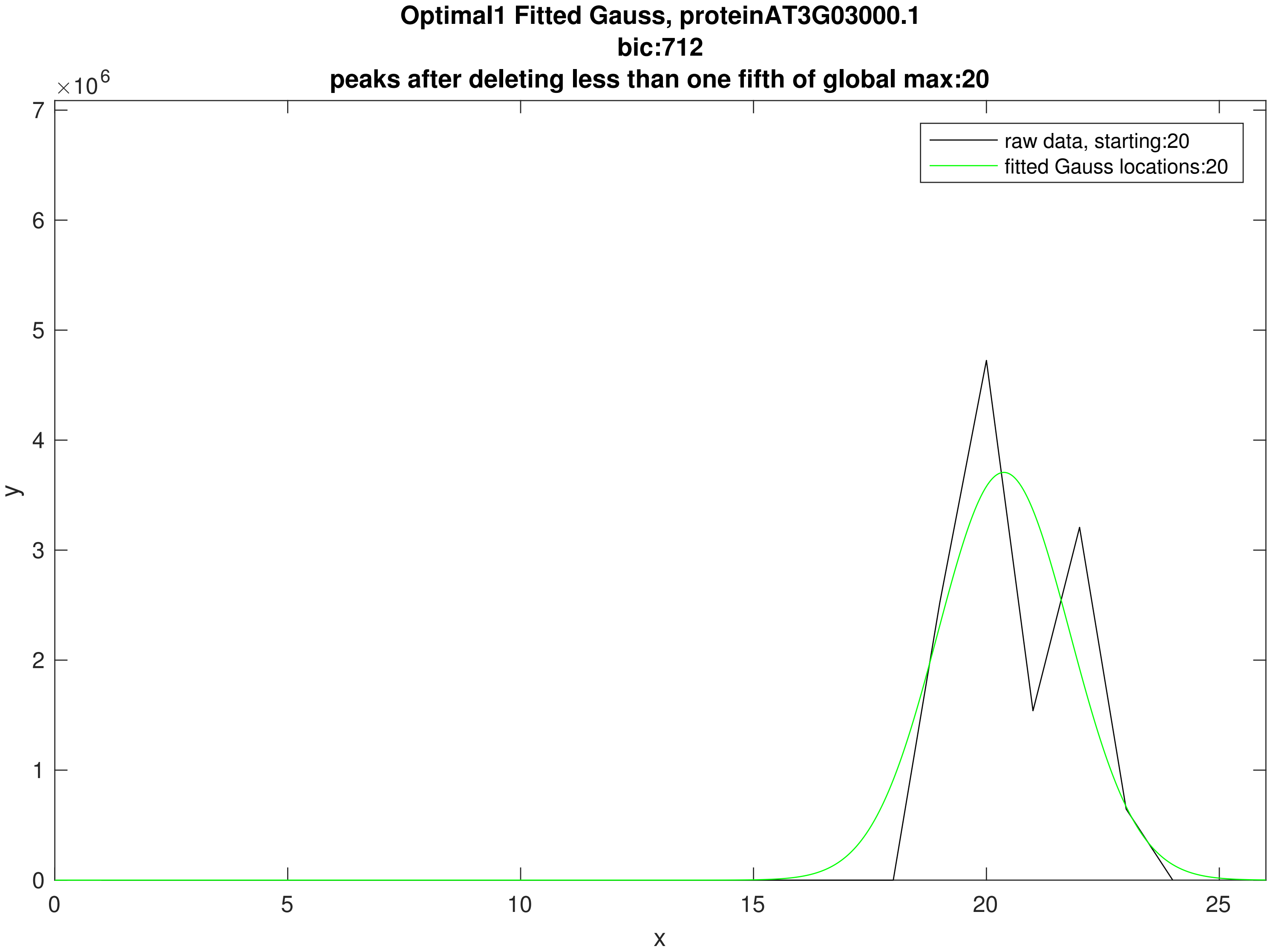

Optimal1 Fitted Gauss, proteinAT3G03000.1
bic:712
peaks after deleting less than one fifth of global max:20
×10^6
raw data, starting:20
fitted Gauss locations:20
y
x
0
1
2
3
4
5
6
7
0
5
10
15
20
25

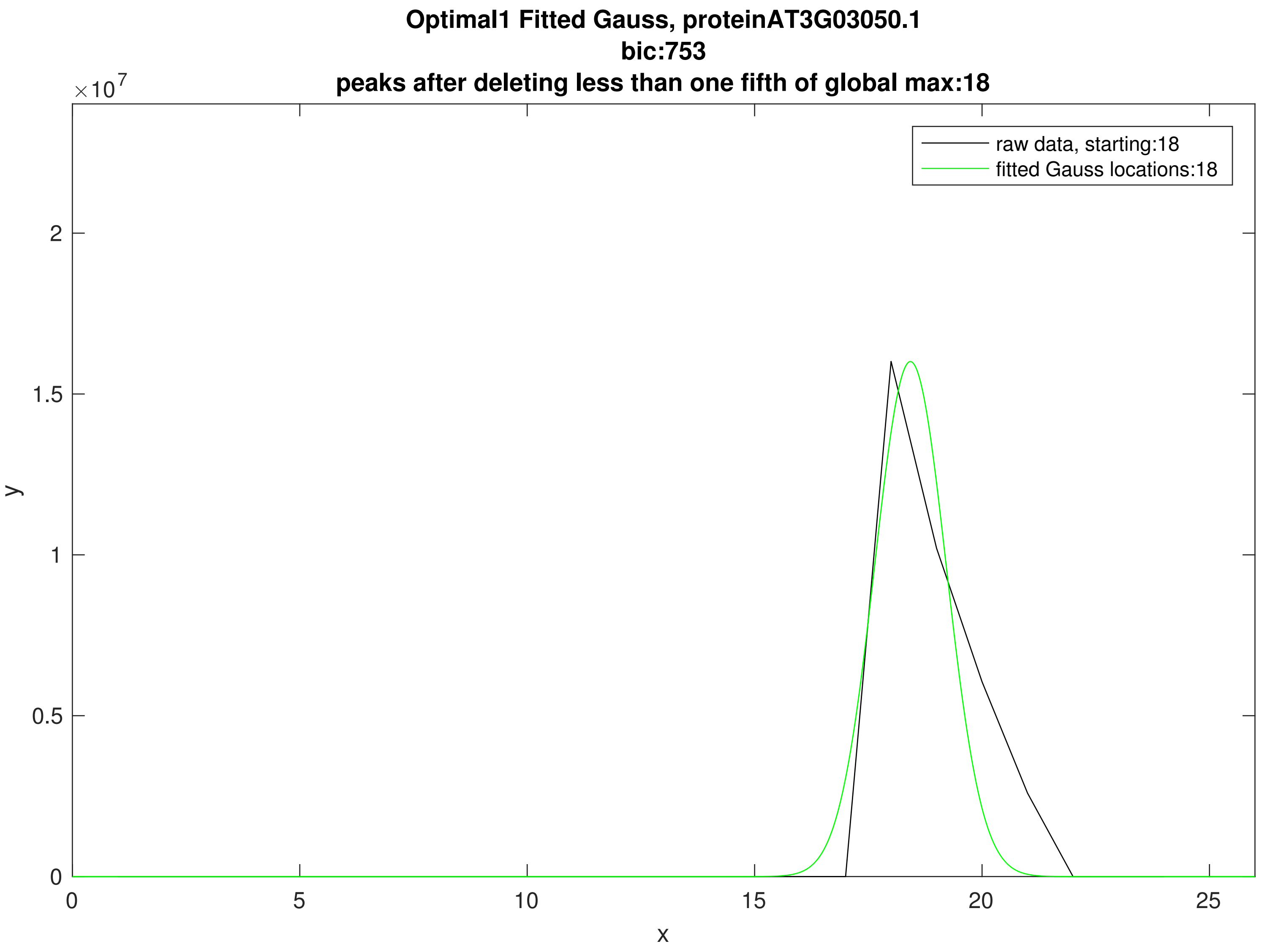

Optimal1 Fitted Gauss, proteinAT3G03050.1
bic:753
peaks after deleting less than one fifth of global max:18
×10^7
raw data, starting:18
fitted Gauss locations:18
y
x
0
0.5
1
1.5
2
0
5
10
15
20
25

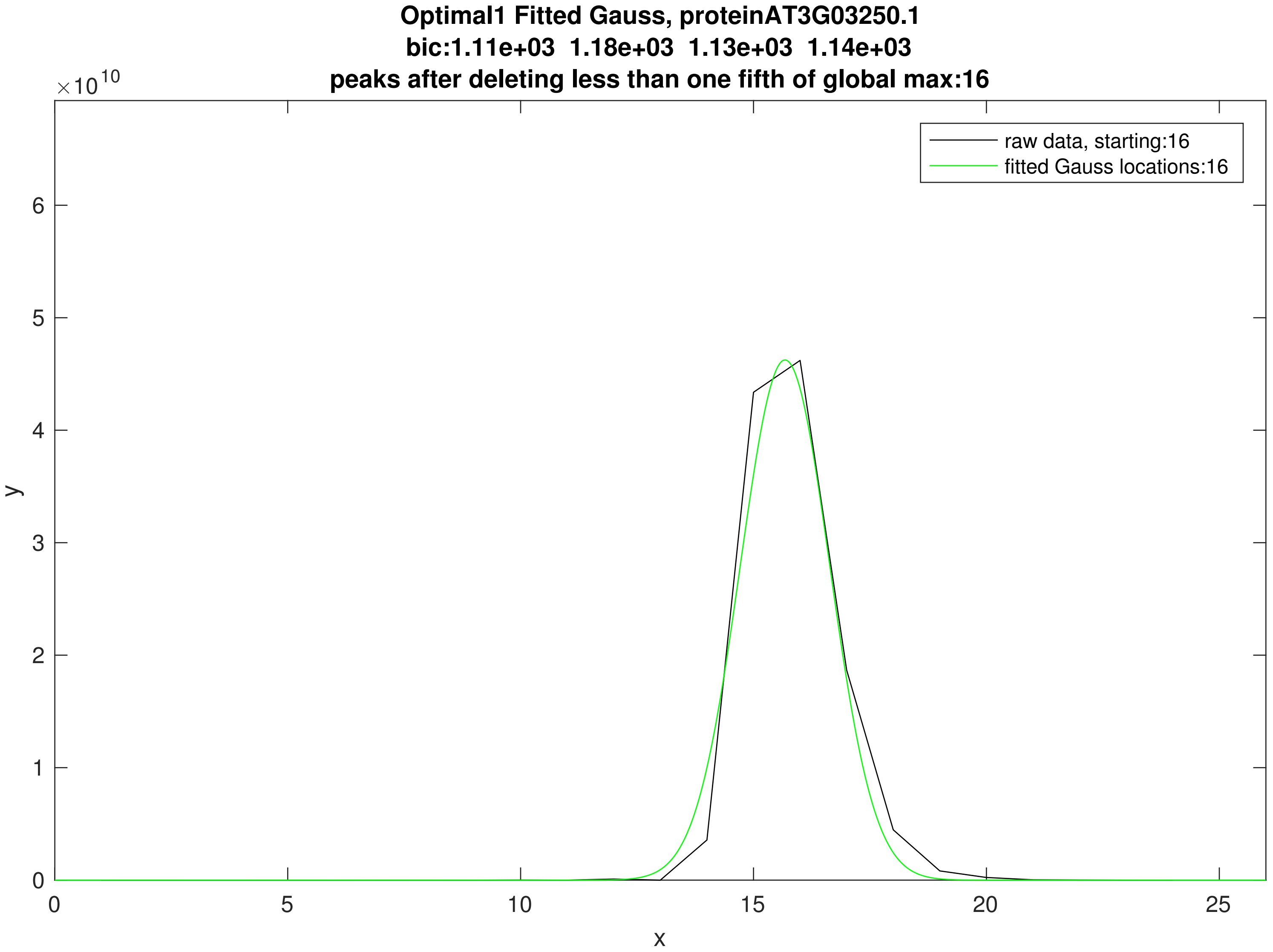

Optimal1 Fitted Gauss, proteinAT3G03250.1
bic:1.11e+03  1.18e+03  1.13e+03  1.14e+03
peaks after deleting less than one fifth of global max:16
×10^10
raw data, starting:16
fitted Gauss locations:16
y
x
0
1
2
3
4
5
6
0
5
10
15
20
25

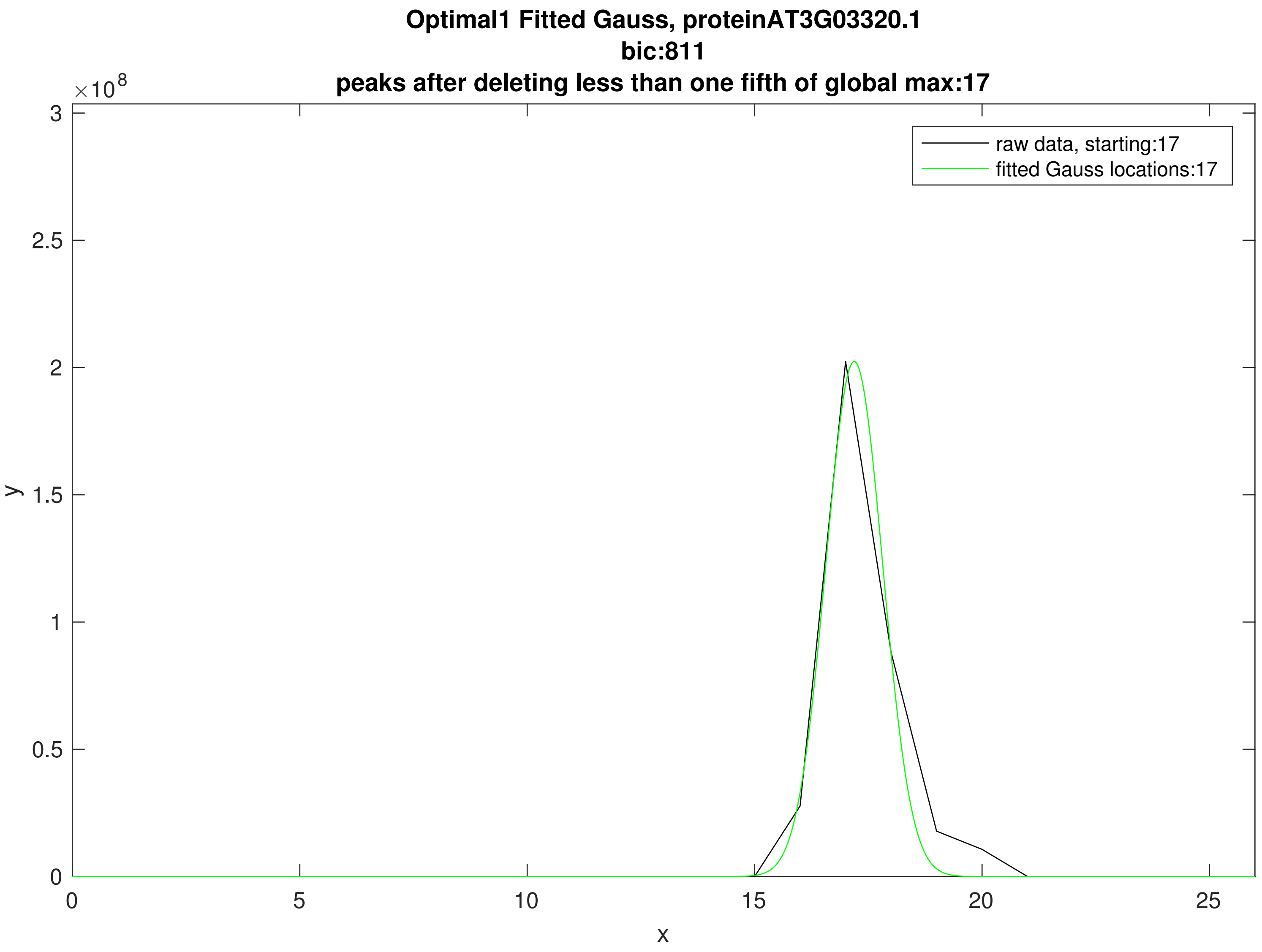
Optimal1 Fitted Gauss, proteinAT3G03320.1
bic:811
peaks after deleting less than one fifth of global max:17
×10^8
3
2.5
2
1.5
1
0.5
0
y
0
5
10
15
20
25
x
raw data, starting:17
fitted Gauss locations:17

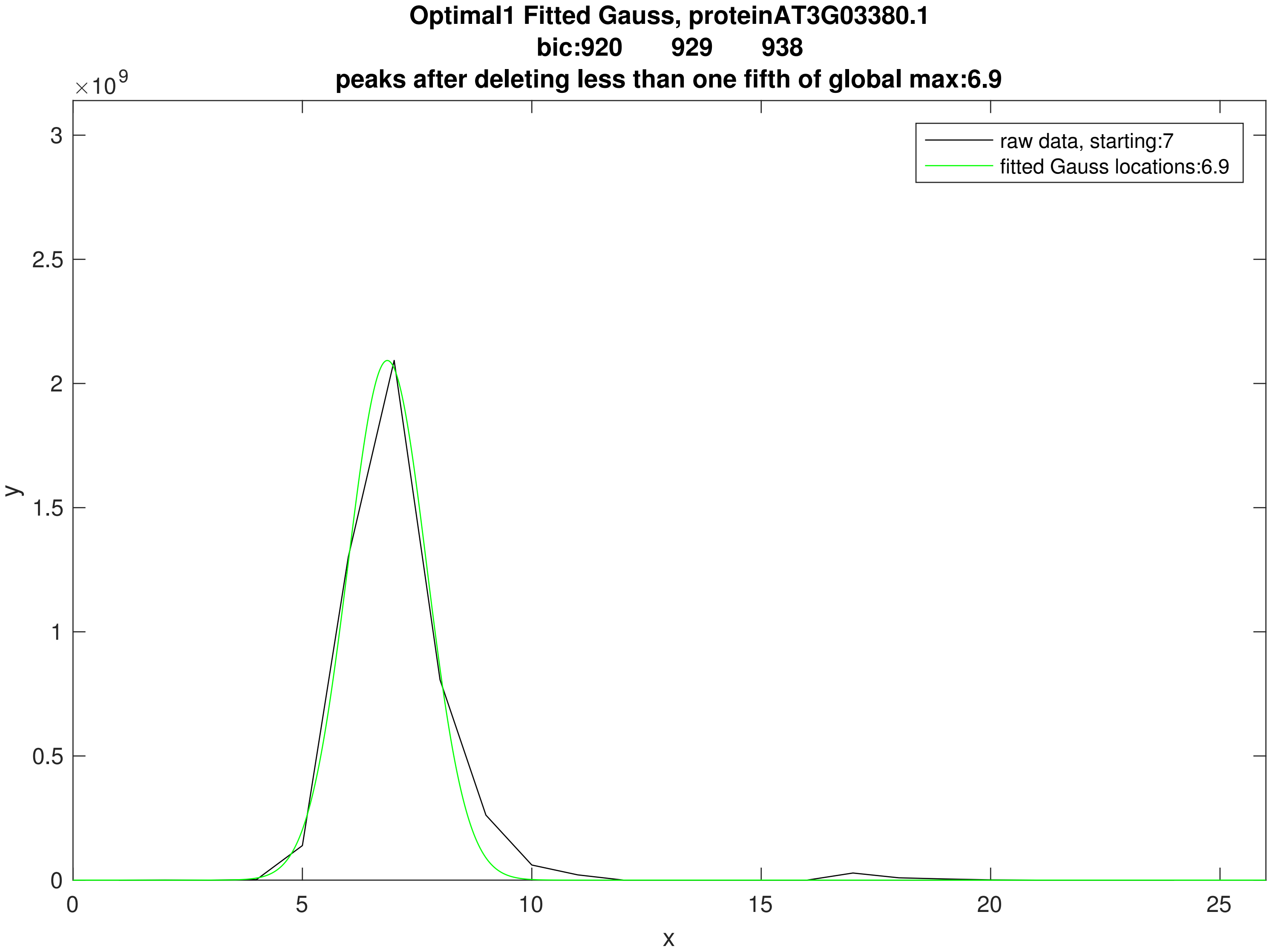
Optimal1 Fitted Gauss, proteinAT3G03380.1
bic:920 929 938
peaks after deleting less than one fifth of global max:6.9
×10^9
raw data, starting:7
fitted Gauss locations:6.9
y
x
0
0.5
1
1.5
2
2.5
3
0
5
10
15
20
25

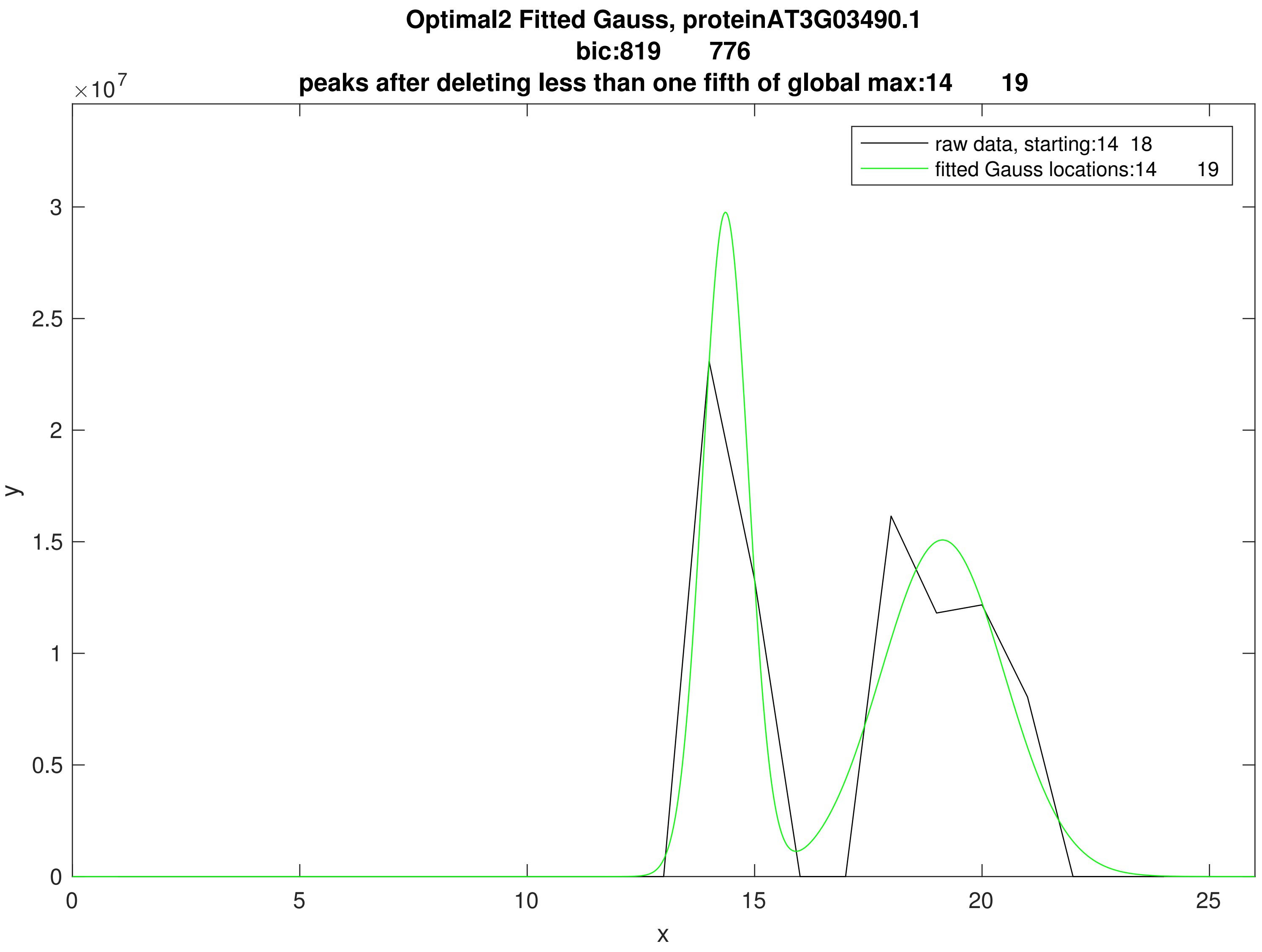
Optimal2 Fitted Gauss, proteinAT3G03490.1
bic:819 776
peaks after deleting less than one fifth of global max:14 19
×10^7
raw data, starting:14 18
fitted Gauss locations:14 19
y
x
0
0.5
1
1.5
2
2.5
3
0
5
10
15
20
25

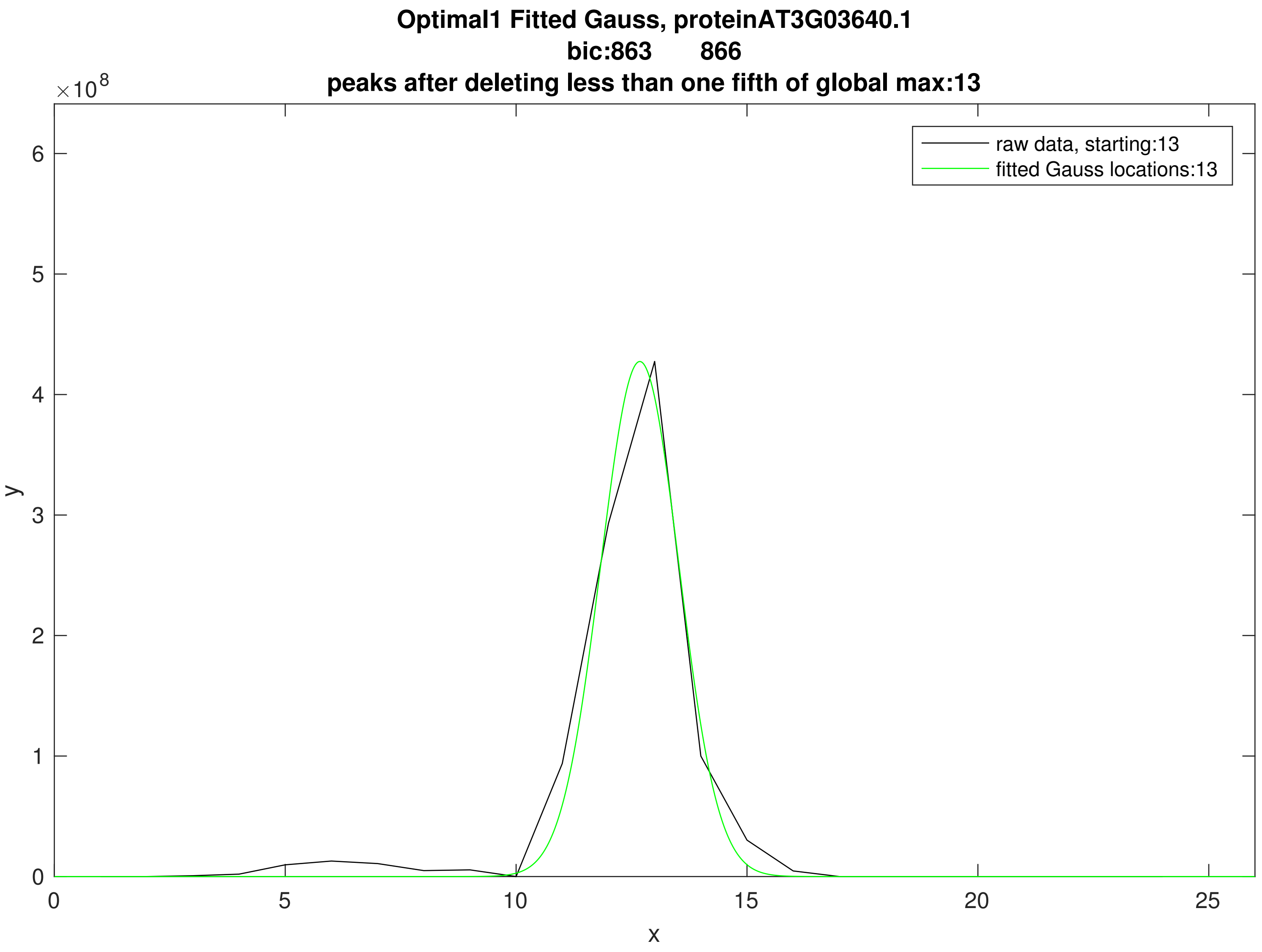
Optimal1 Fitted Gauss, proteinAT3G03640.1
bic:863 866
peaks after deleting less than one fifth of global max:13
$\times 10^8$
raw data, starting:13
fitted Gauss locations:13
y
x
0
1
2
3
4
5
6
0
5
10
15
20
25

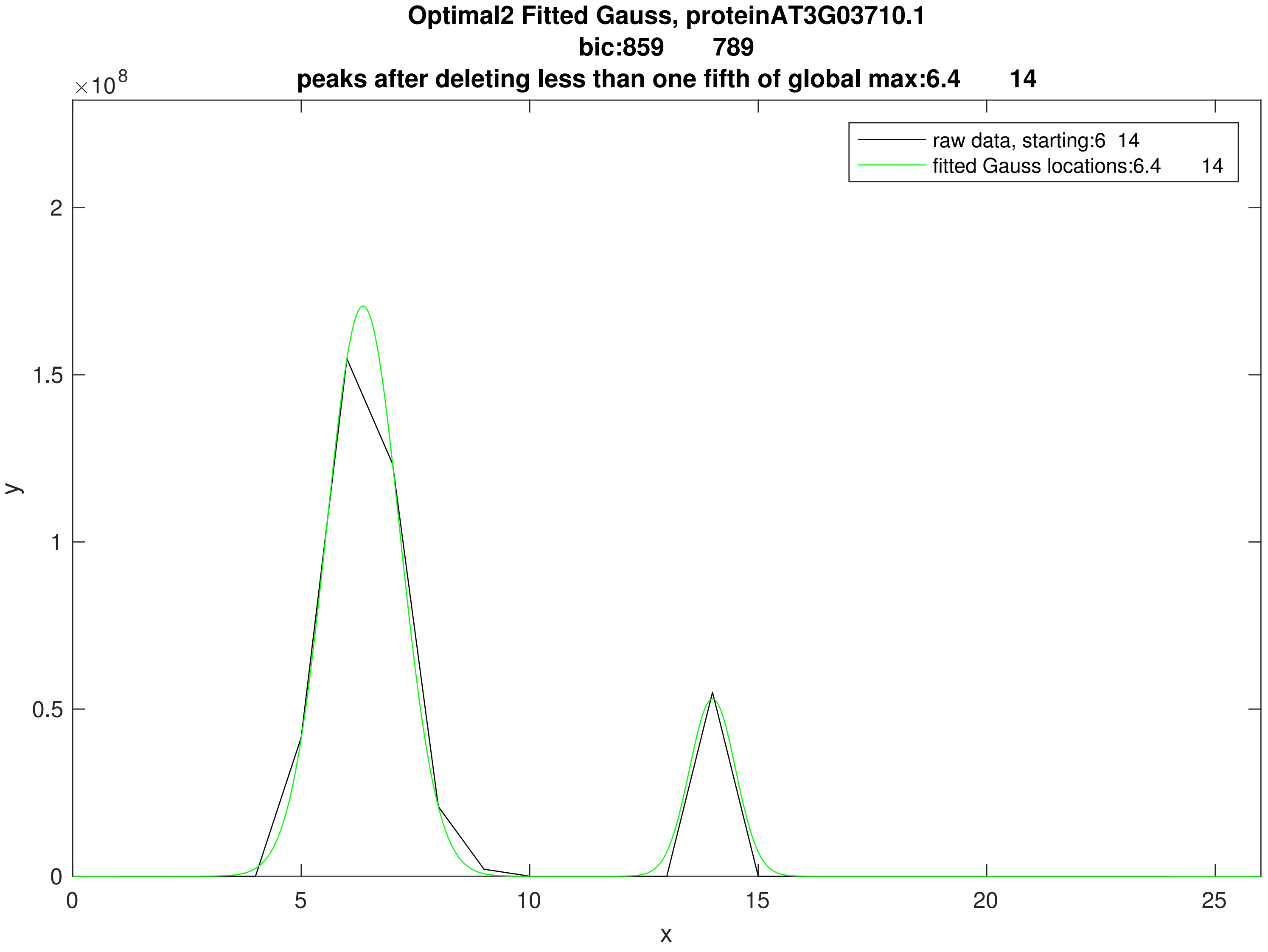
Optimal2 Fitted Gauss, proteinAT3G03710.1
bic:859 789
peaks after deleting less than one fifth of global max:6.4 14
×10^8
raw data, starting:6 14
fitted Gauss locations:6.4 14
y
x
0
0.5
1
1.5
2
0
5
10
15
20
25

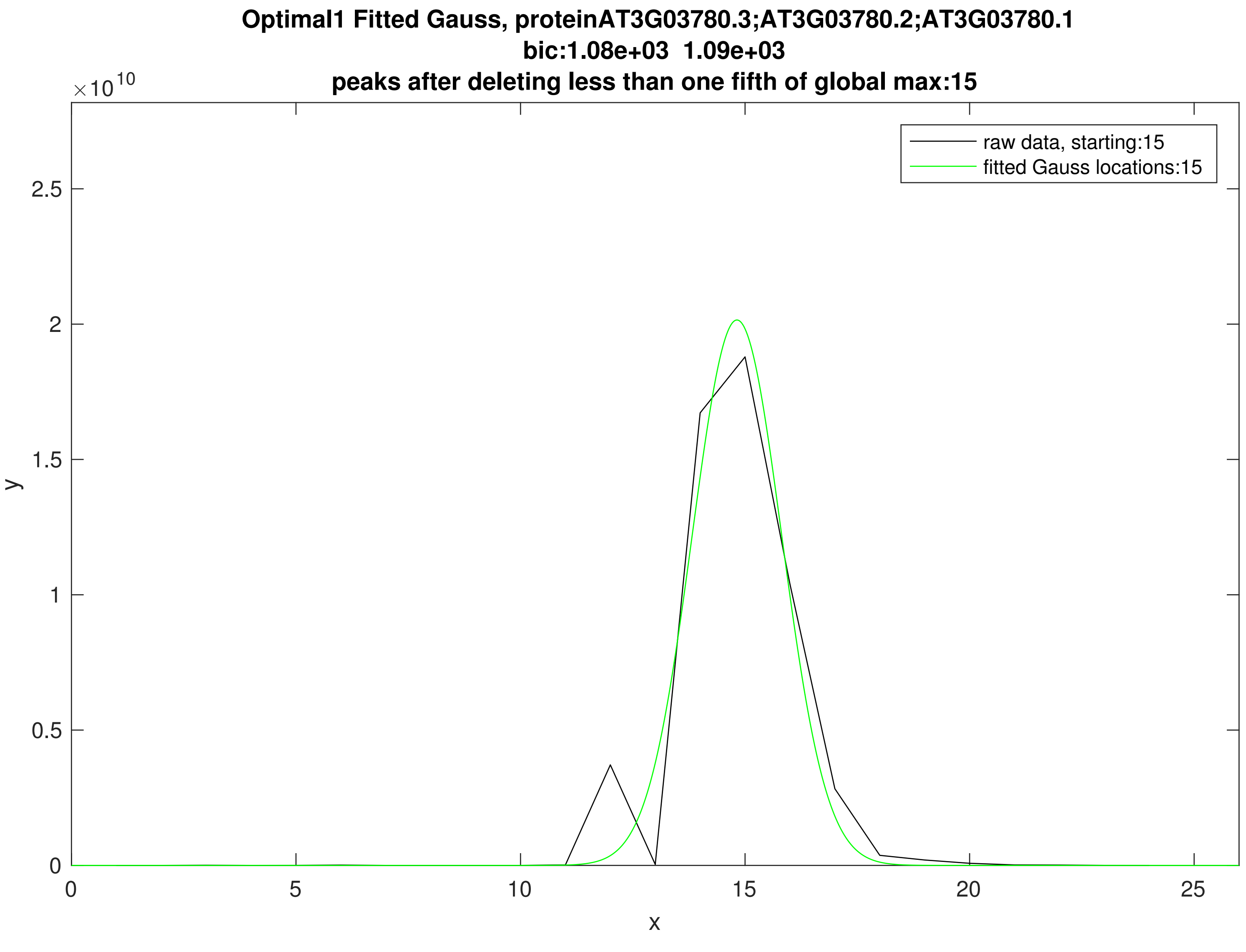

Optimal1 Fitted Gauss, proteinAT3G03780.3;AT3G03780.2;AT3G03780.1
bic:1.08e+03 1.09e+03
peaks after deleting less than one fifth of global max:15
×10^10
raw data, starting:15
fitted Gauss locations:15
y
x
0
0.5
1
1.5
2
2.5
0
5
10
15
20
25

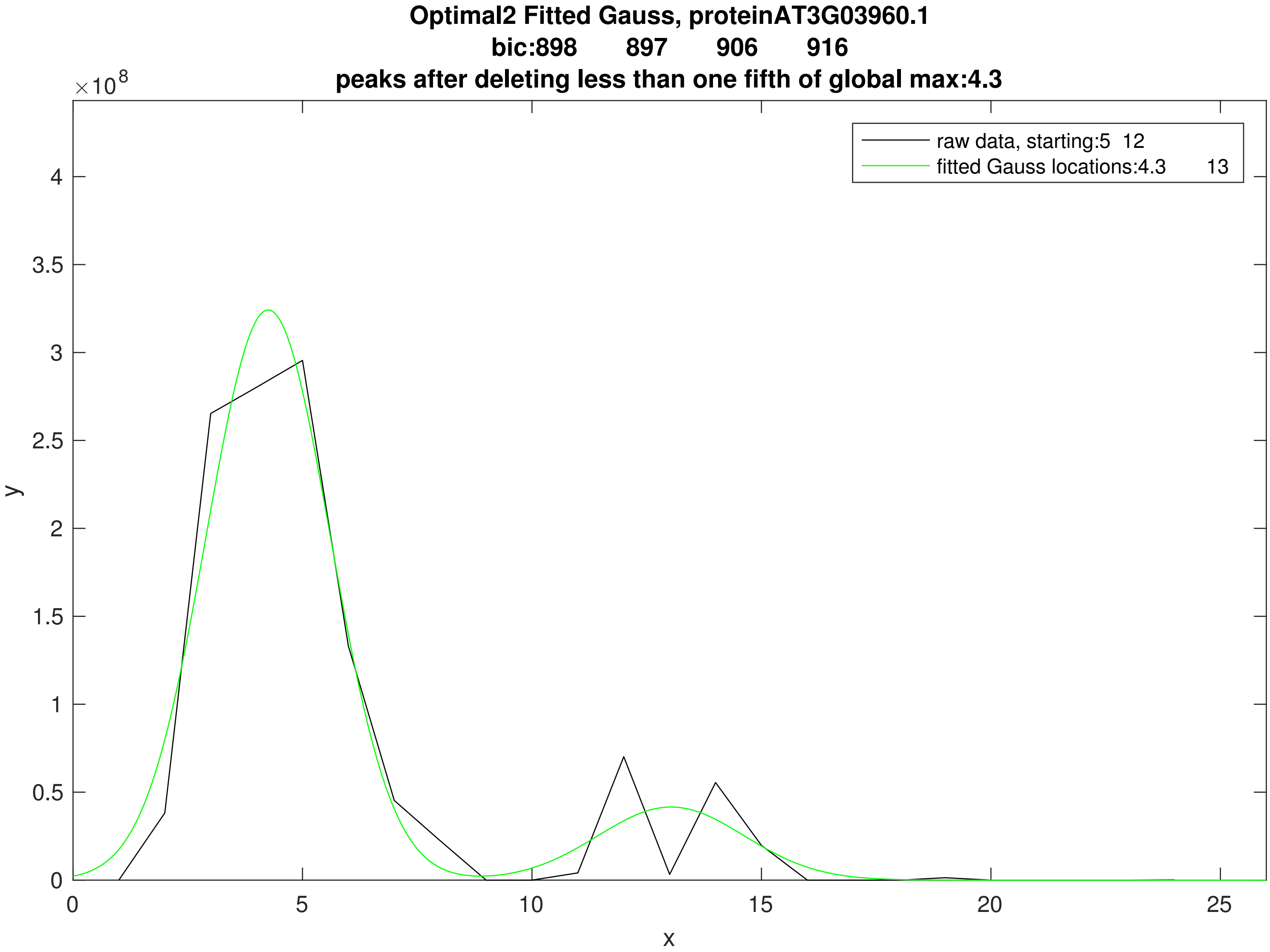

Optimal2 Fitted Gauss, proteinAT3G03960.1
bic:898 897 906 916
peaks after deleting less than one fifth of global max:4.3
×10^8
raw data, starting:5 12
fitted Gauss locations:4.3 13
y
x
0
0.5
1
1.5
2
2.5
3
3.5
4
5
10
15
20
25

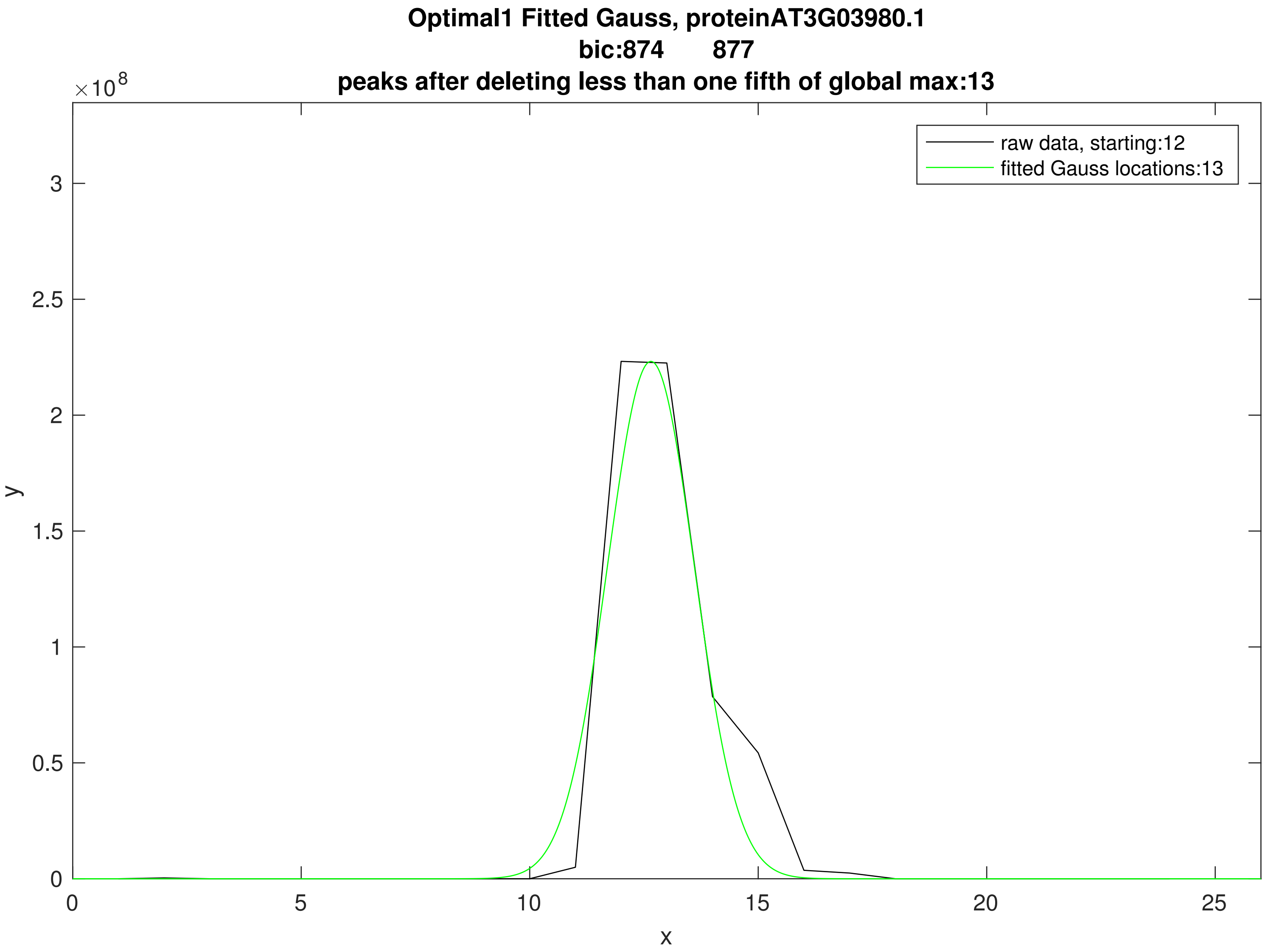
Optimal1 Fitted Gauss, proteinAT3G03980.1
bic:874 877
peaks after deleting less than one fifth of global max:13
×10^8
raw data, starting:12
fitted Gauss locations:13
y
x
0
0.5
1
1.5
2
2.5
3
0
5
10
15
20
25

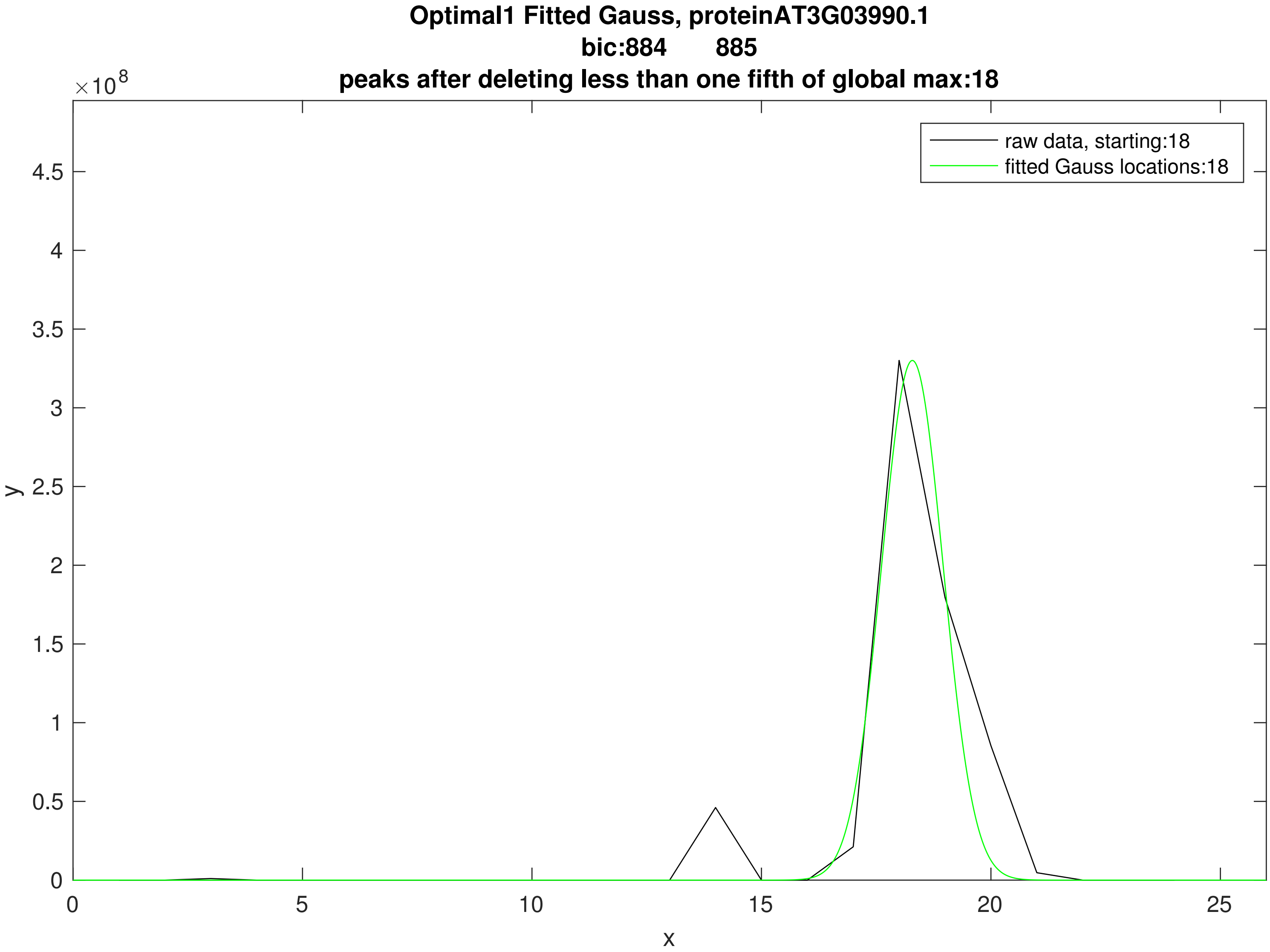
Optimal1 Fitted Gauss, proteinAT3G03990.1
bic:884 885
peaks after deleting less than one fifth of global max:18
×10^8
4.5
4
3.5
3
2.5
2
1.5
1
0.5
0
y
0
5
10
15
20
25
x
raw data, starting:18
fitted Gauss locations:18

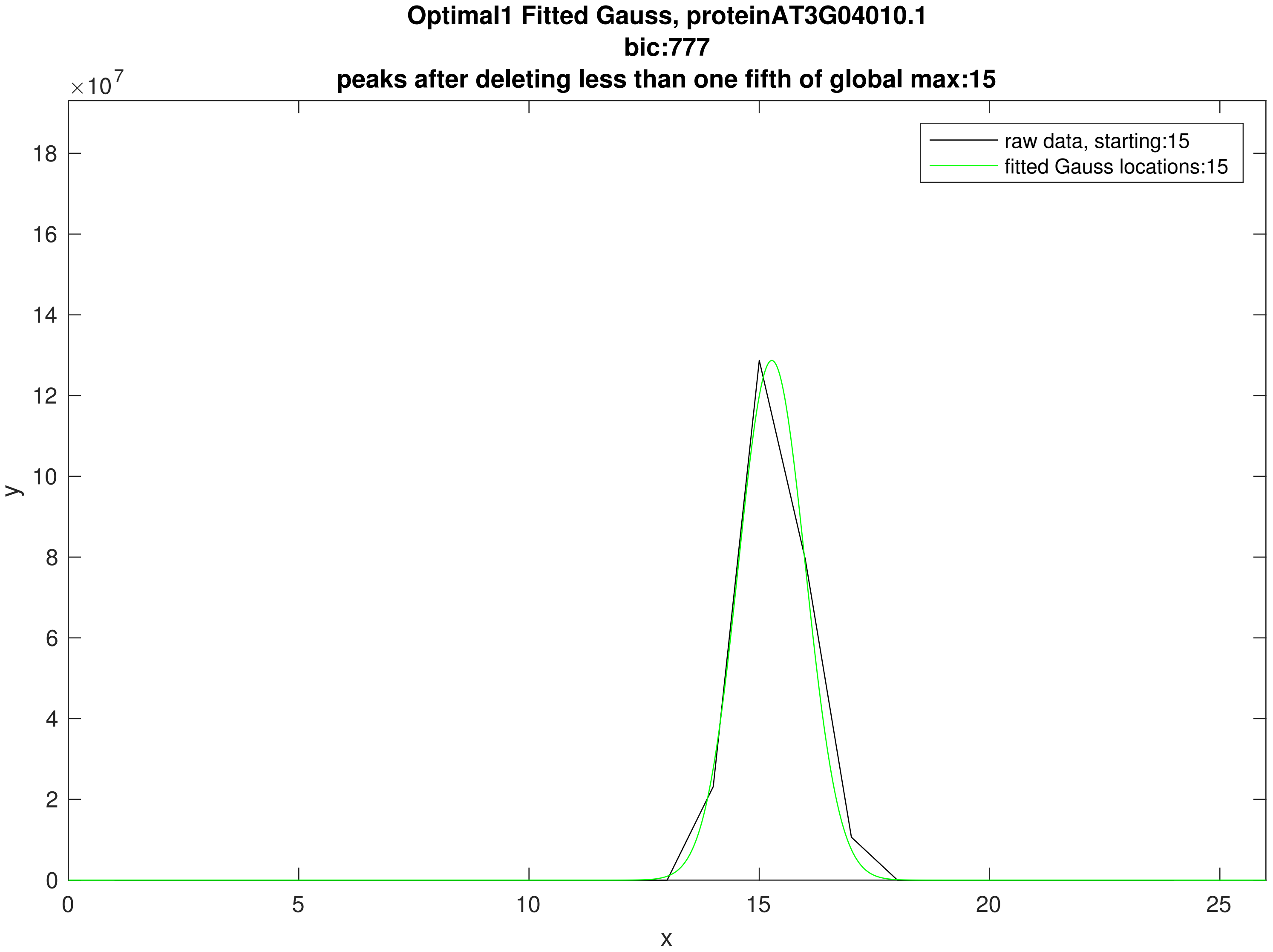

Optimal1 Fitted Gauss, proteinAT3G04010.1
bic:777
peaks after deleting less than one fifth of global max:15
×10^7
raw data, starting:15
fitted Gauss locations:15
y
x
0
2
4
6
8
10
12
14
16
18
0
5
10
15
20
25

**Optimal1 Fitted Gauss, proteinAT3G04120.1**
**bic:1.2e+03  1.21e+03  1.22e+03**
**peaks after deleting less than one fifth of global max:12**

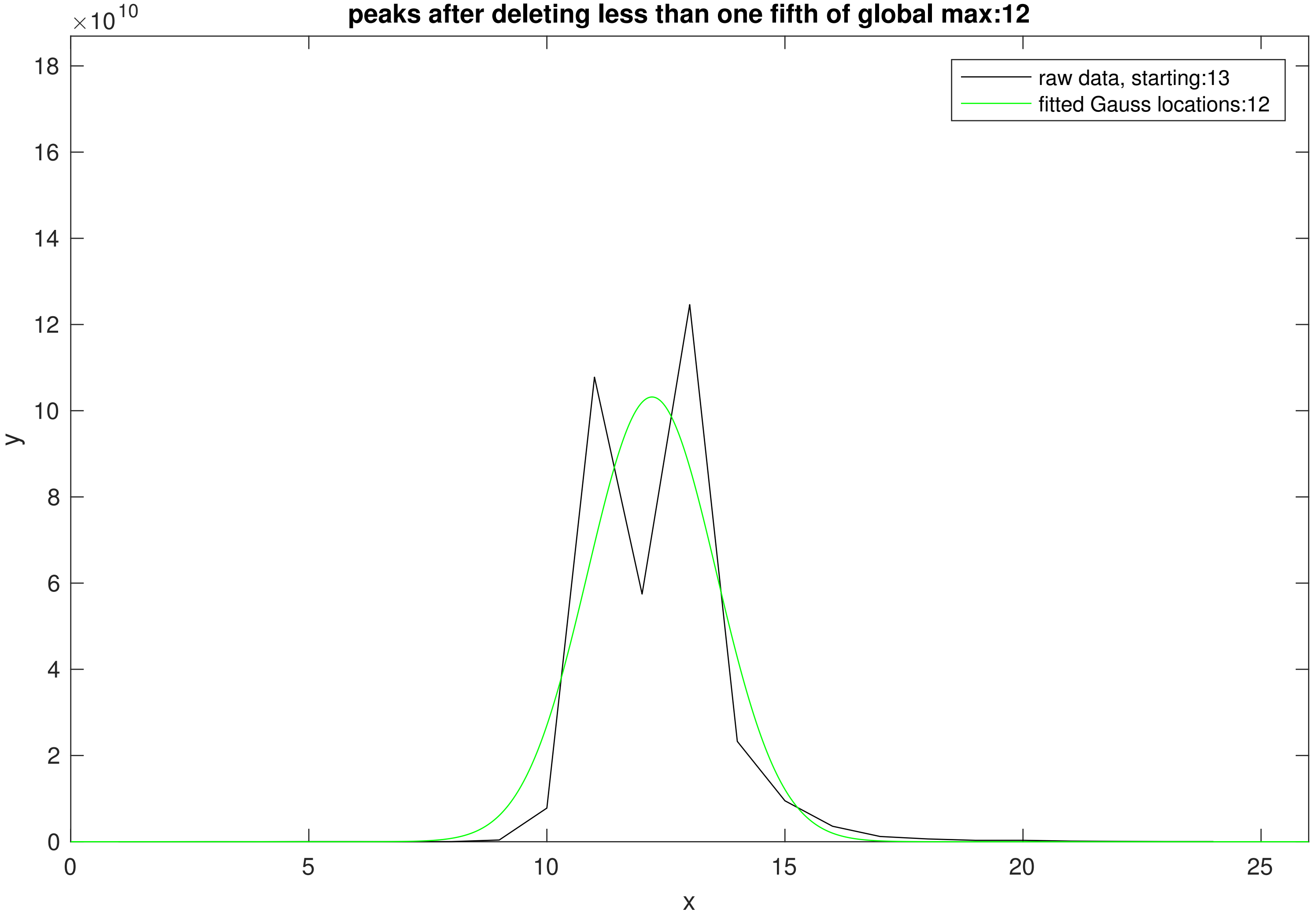

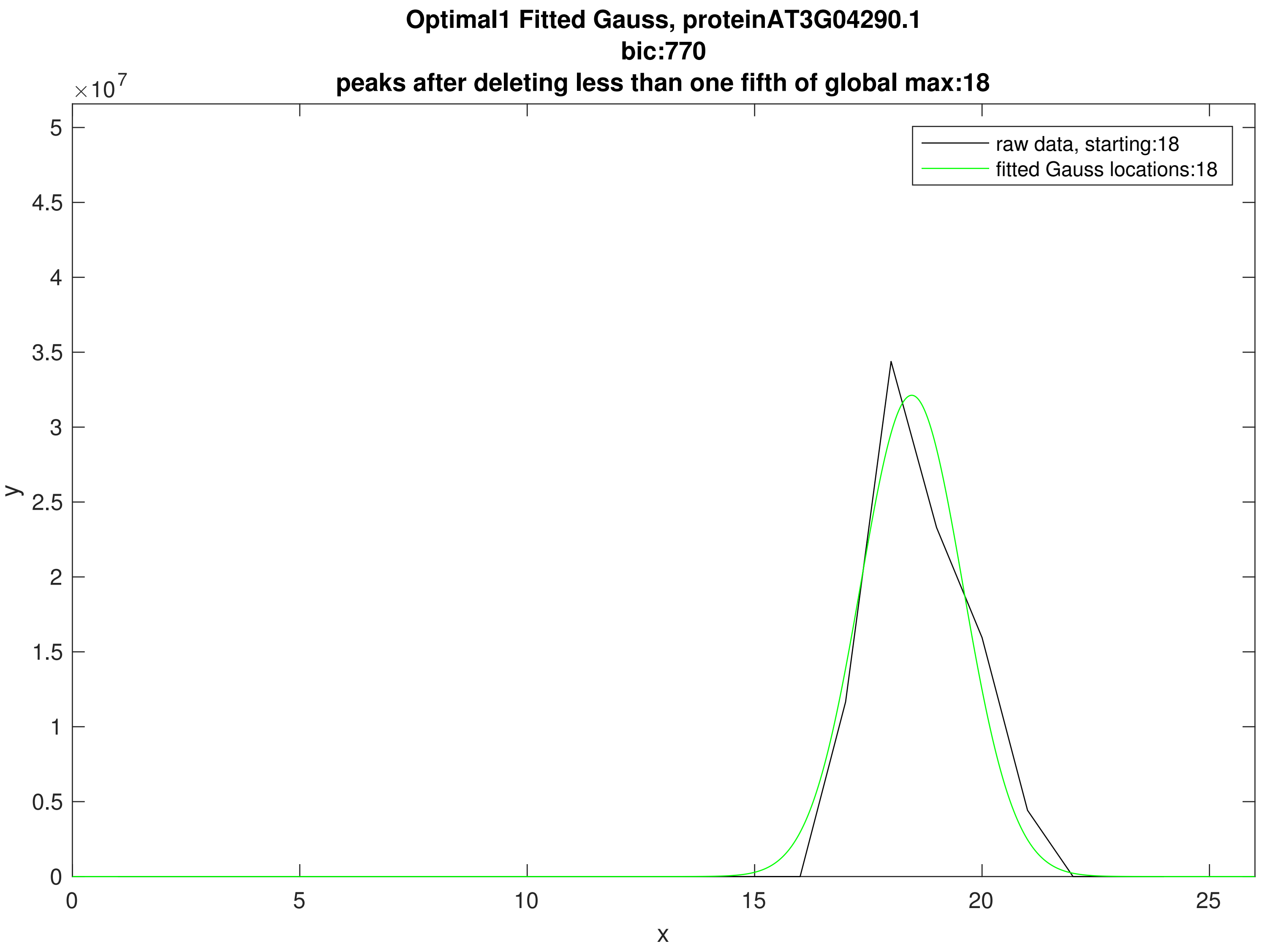
Optimal1 Fitted Gauss, proteinAT3G04290.1
bic:770
peaks after deleting less than one fifth of global max:18
×10^7
raw data, starting:18
fitted Gauss locations:18
y
x
0
0.5
1
1.5
2
2.5
3
3.5
4
4.5
5
0
5
10
15
20
25

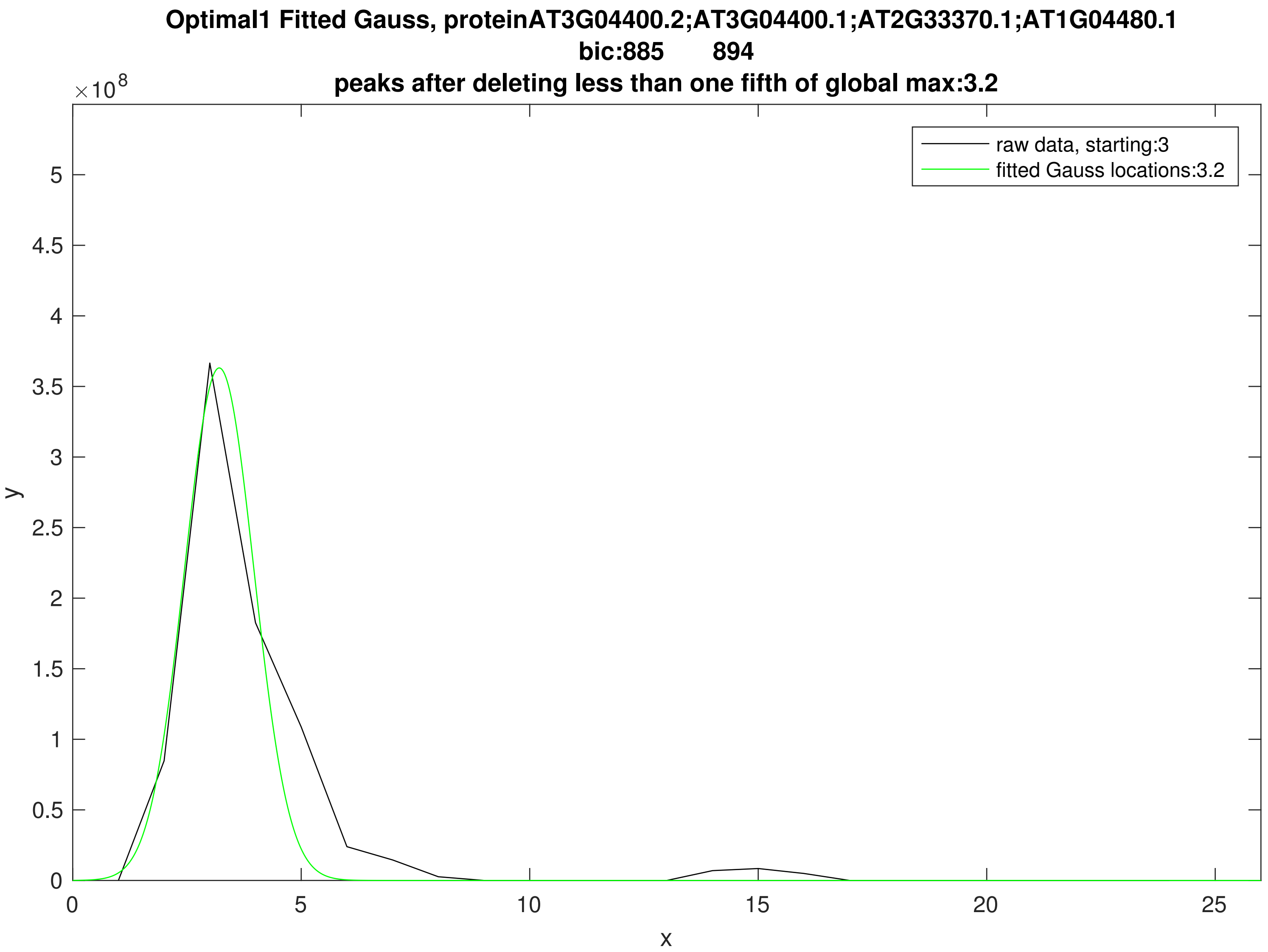

Optimal1 Fitted Gauss, proteinAT3G04400.2;AT3G04400.1;AT2G33370.1;AT1G04480.1
bic:885 894
peaks after deleting less than one fifth of global max:3.2
×10^8
raw data, starting:3
fitted Gauss locations:3.2
y
x
0
0.5
1
1.5
2
2.5
3
3.5
4
4.5
5
0
5
10
15
20
25

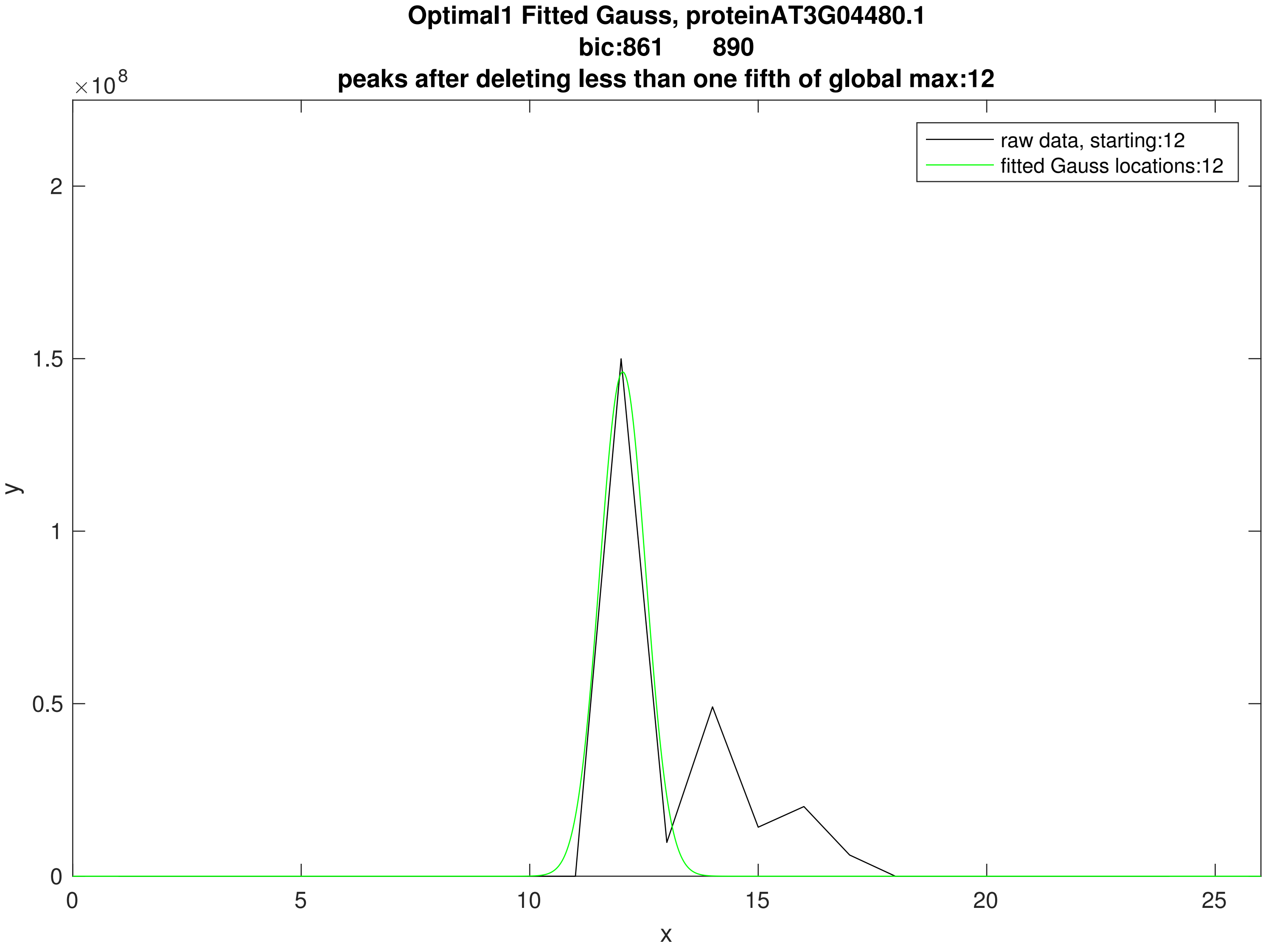
Optimal1 Fitted Gauss, proteinAT3G04480.1
bic:861 890
peaks after deleting less than one fifth of global max:12
×10^8
raw data, starting:12
fitted Gauss locations:12
y
x
0
0.5
1
1.5
2
5
10
15
20
25

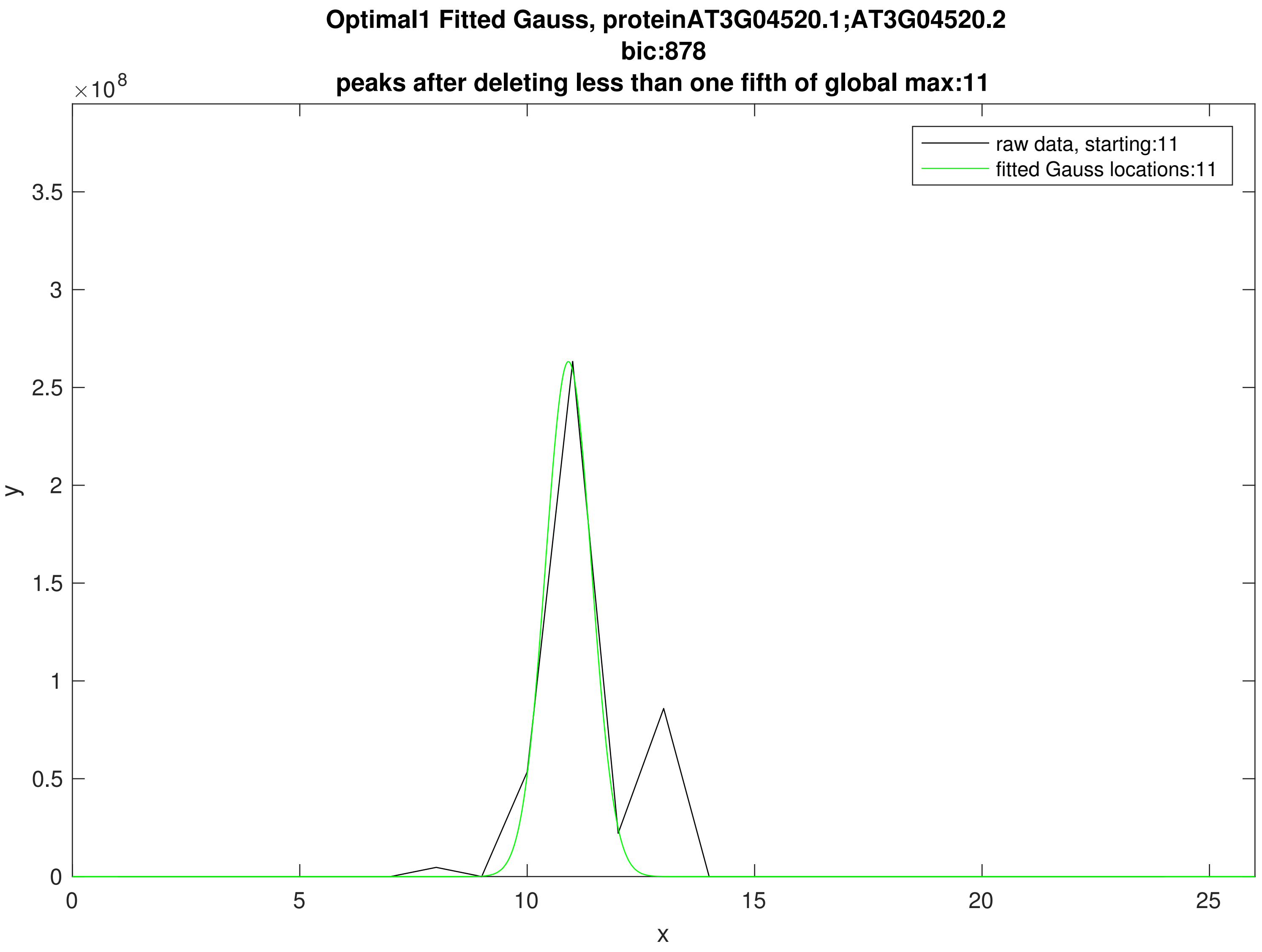
Optimal1 Fitted Gauss, proteinAT3G04520.1;AT3G04520.2
bic:878
peaks after deleting less than one fifth of global max:11
×10^8
raw data, starting:11
fitted Gauss locations:11
3.5
3
2.5
2
1.5
1
0.5
0
y
0
5
10
15
20
25
x

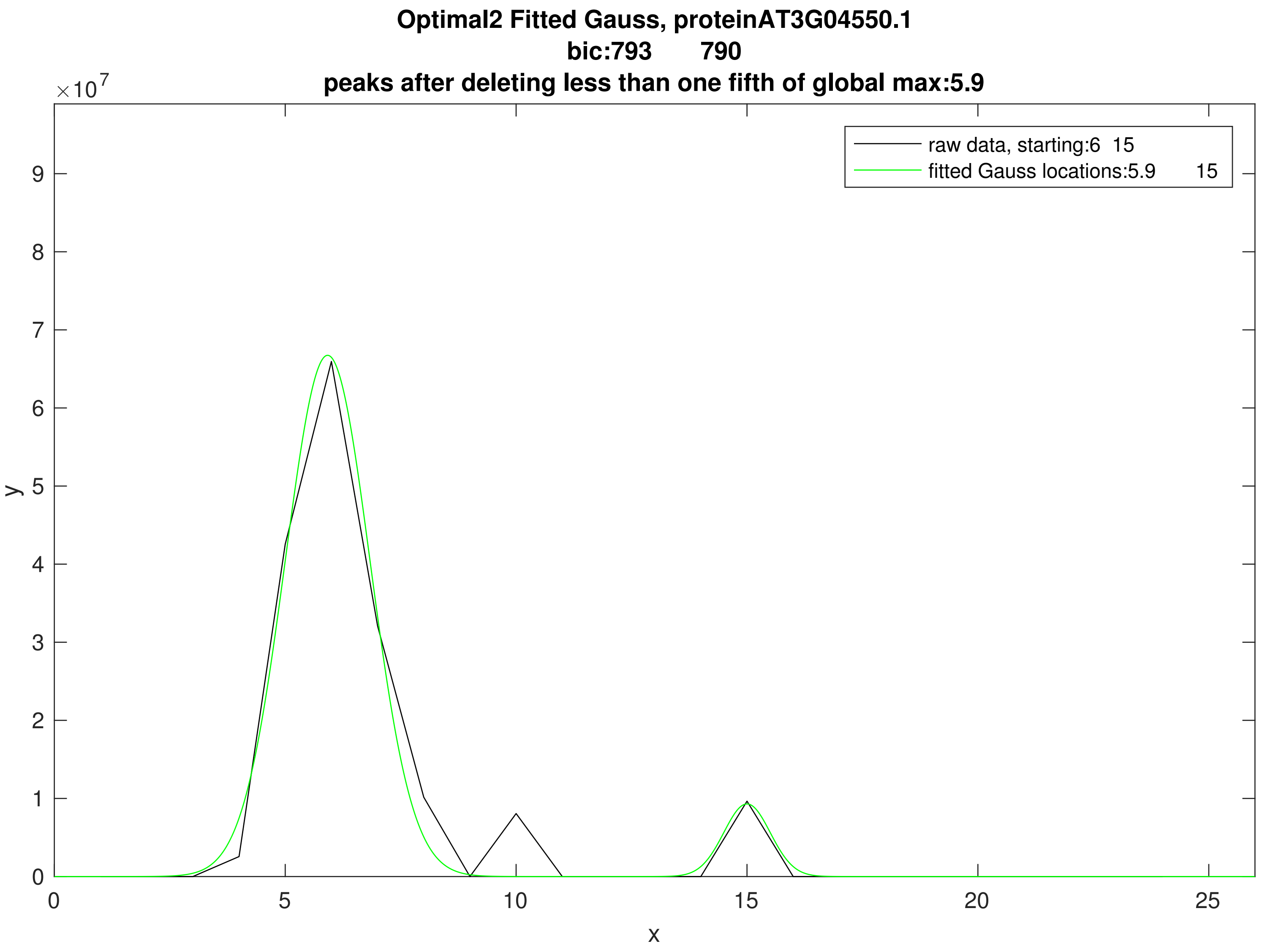

Optimal2 Fitted Gauss, proteinAT3G04550.1
bic:793 790
peaks after deleting less than one fifth of global max:5.9
×10^7
raw data, starting:6 15
fitted Gauss locations:5.9 15
y
x
0
1
2
3
4
5
6
7
8
9
0
5
10
15
20
25

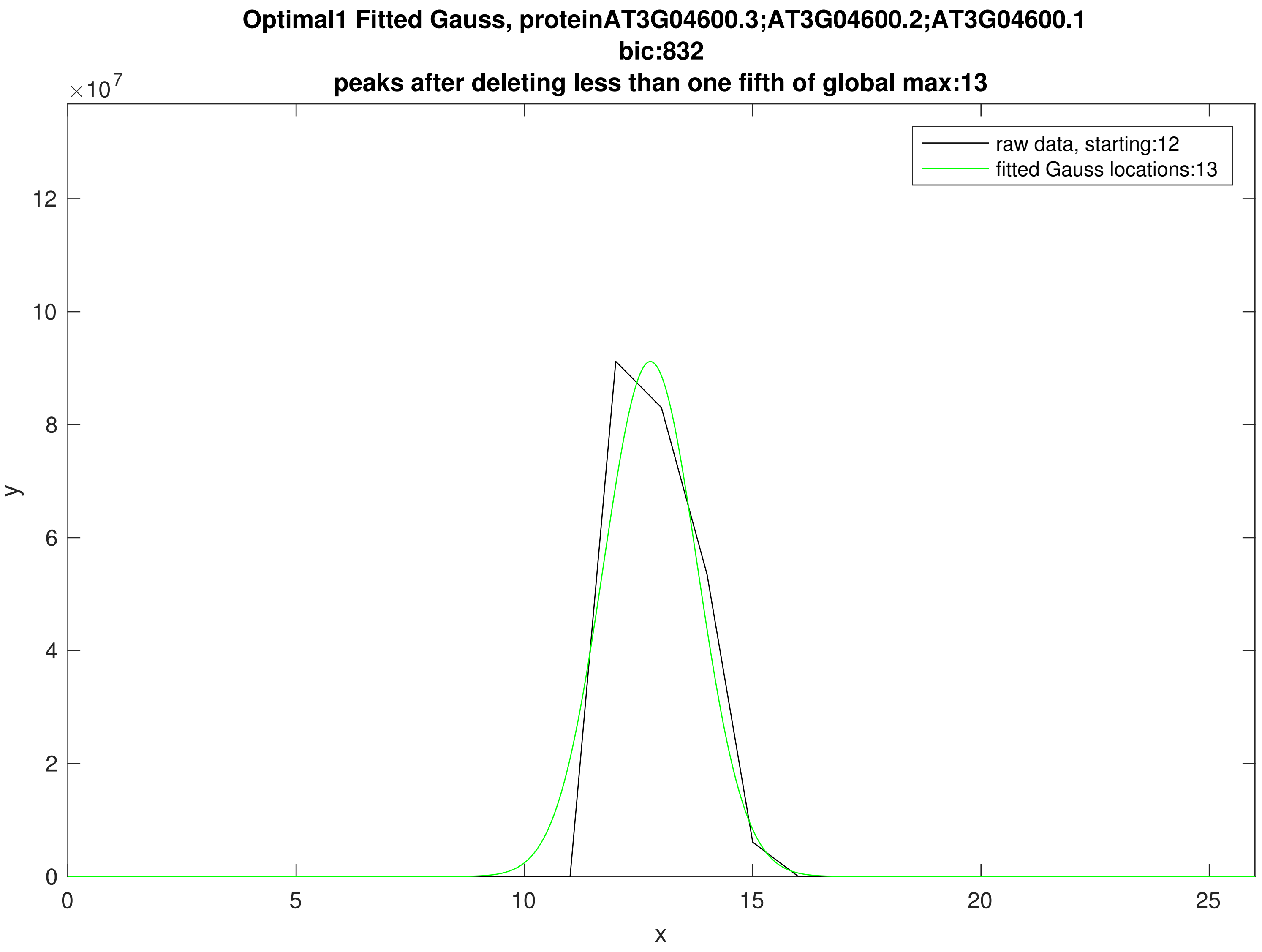

Optimal1 Fitted Gauss, proteinAT3G04600.3;AT3G04600.2;AT3G04600.1
bic:832
peaks after deleting less than one fifth of global max:13
×10^7
raw data, starting:12
fitted Gauss locations:13
y
x
0
2
4
6
8
10
12
0
5
10
15
20
25

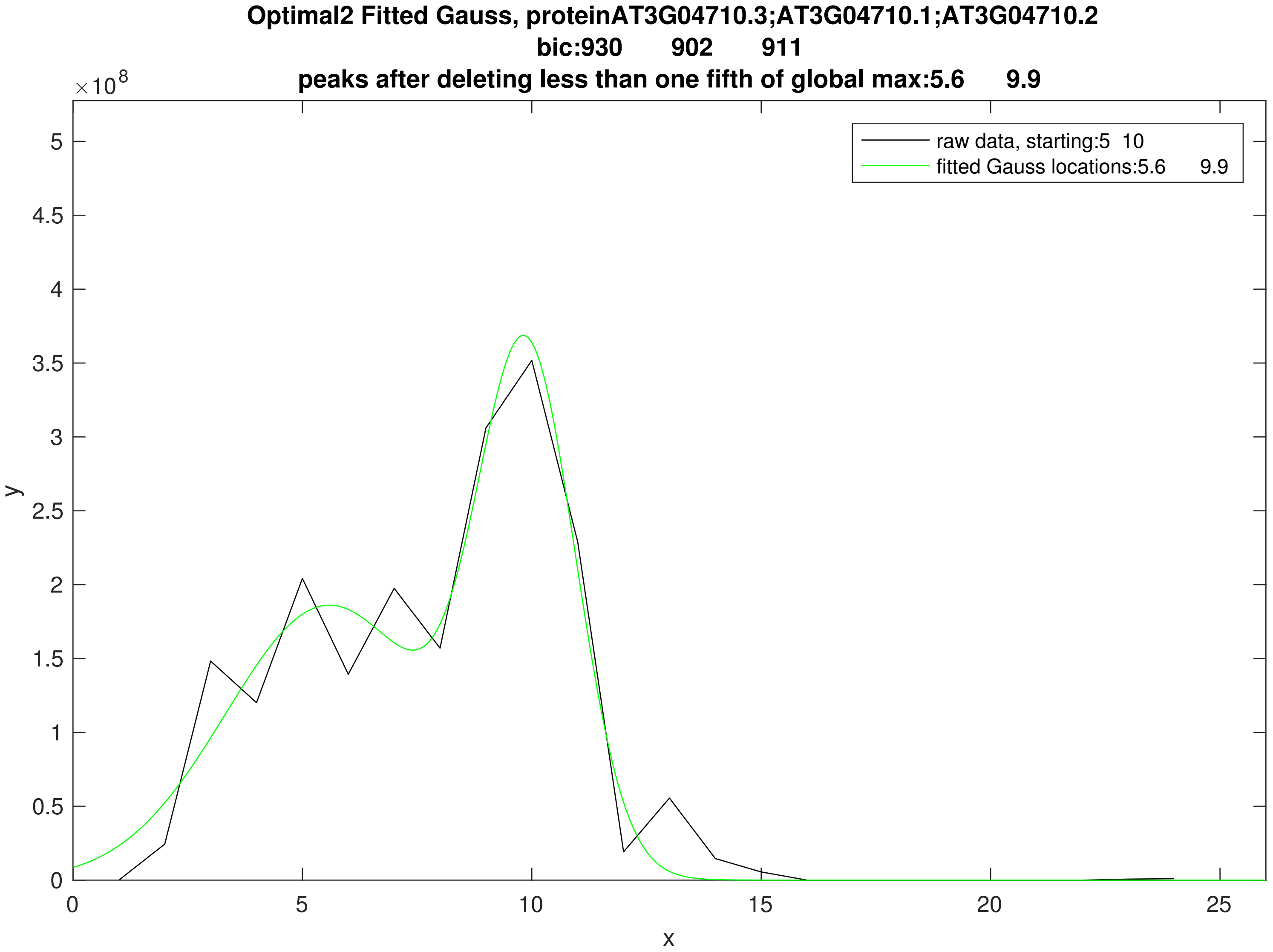
Optimal2 Fitted Gauss, proteinAT3G04710.3;AT3G04710.1;AT3G04710.2
bic:930 902 911
peaks after deleting less than one fifth of global max:5.6 9.9
×10^8
raw data, starting:5 10
fitted Gauss locations:5.6 9.9
y
x
0
0.5
1
1.5
2
2.5
3
3.5
4
4.5
5
0
5
10
15
20
25

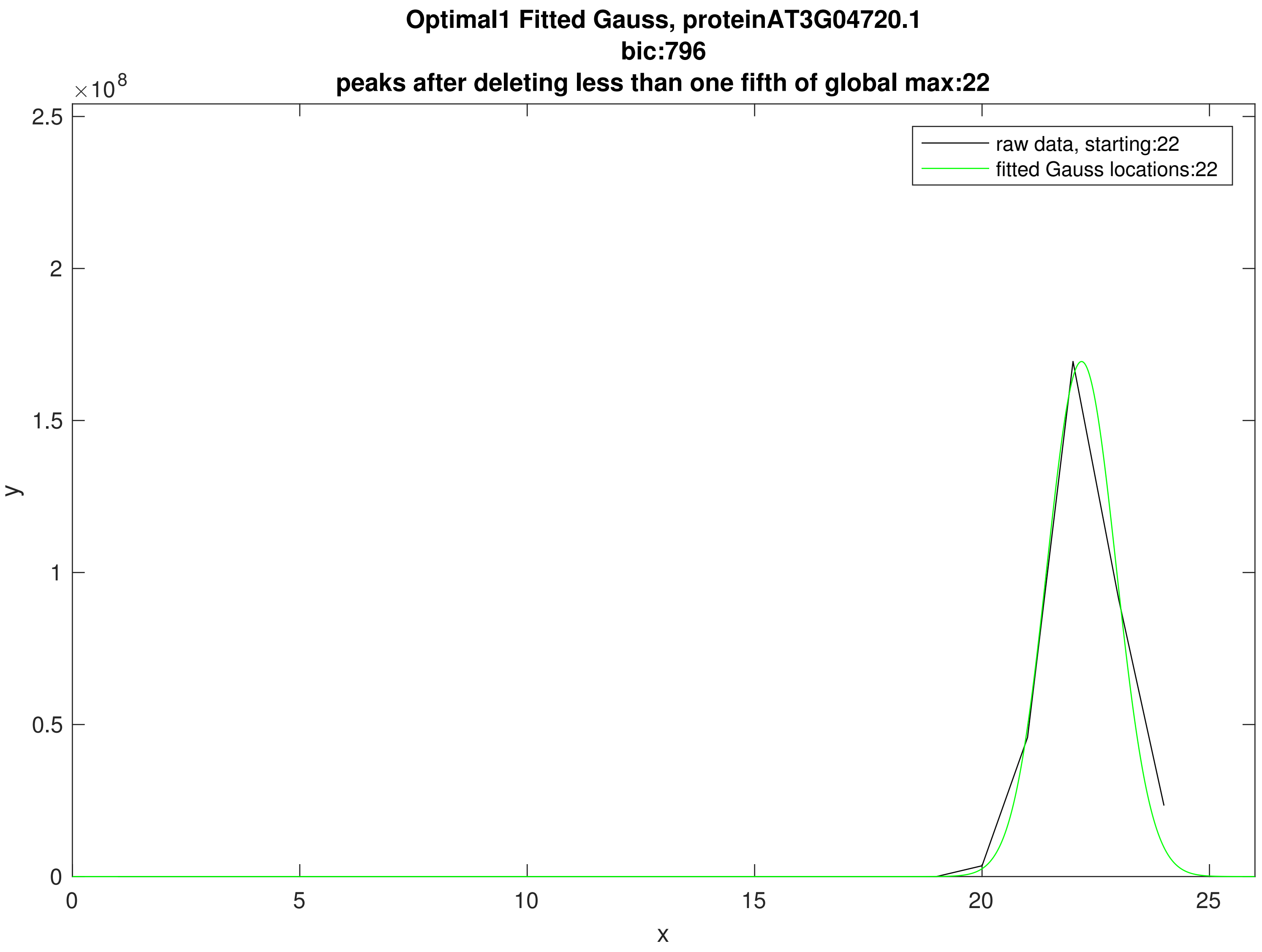
Optimal1 Fitted Gauss, proteinAT3G04720.1
bic:796
peaks after deleting less than one fifth of global max:22
$\times 10^8$
2.5
2
1.5
1
0.5
0
y
0
5
10
15
20
25
x
raw data, starting:22
fitted Gauss locations:22

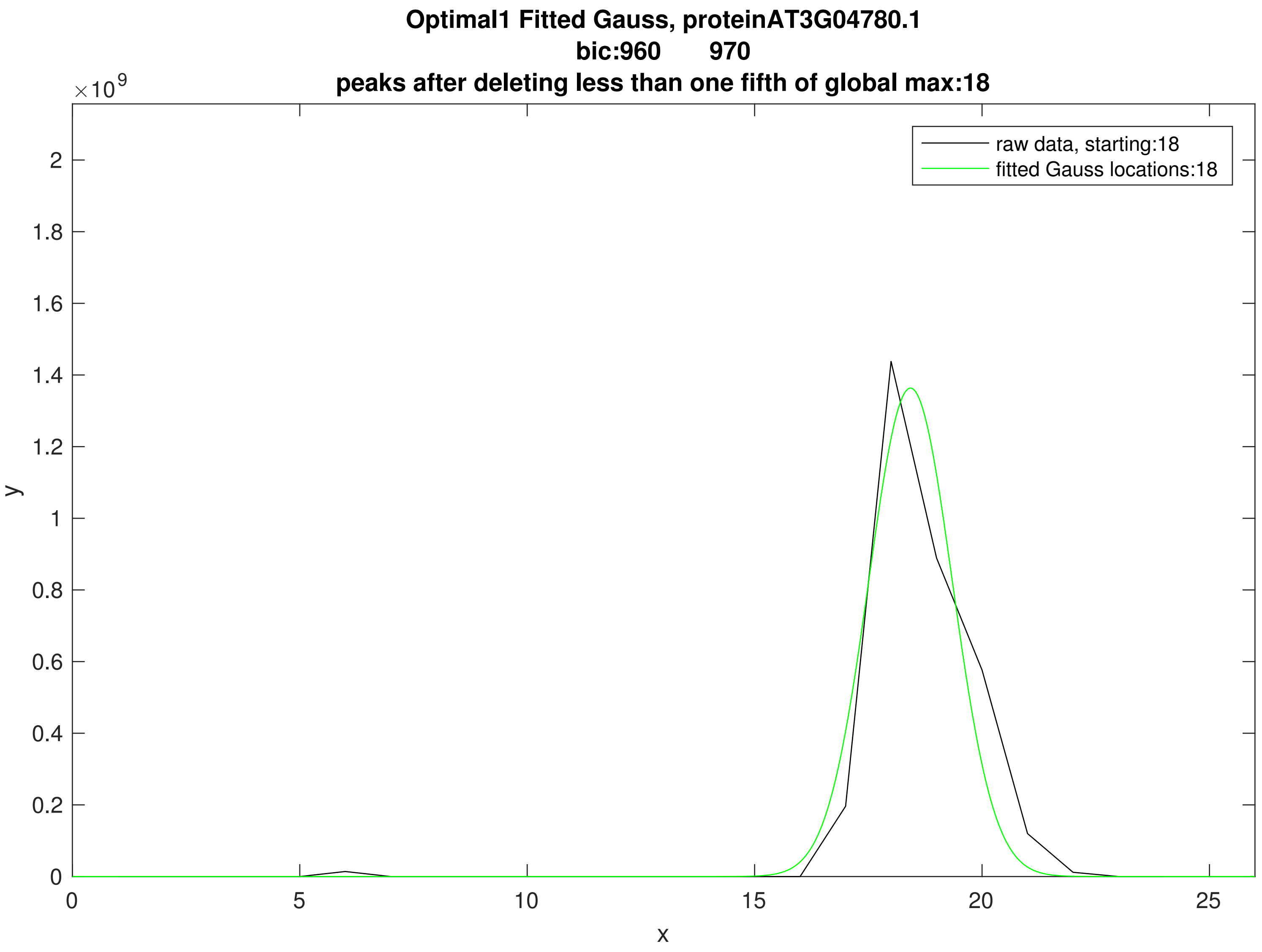

Optimal1 Fitted Gauss, proteinAT3G04780.1
bic:960 970
peaks after deleting less than one fifth of global max:18
×10^9
2
1.8
1.6
1.4
1.2
1
0.8
0.6
0.4
0.2
0
y
0
5
10
15
20
25
x
raw data, starting:18
fitted Gauss locations:18

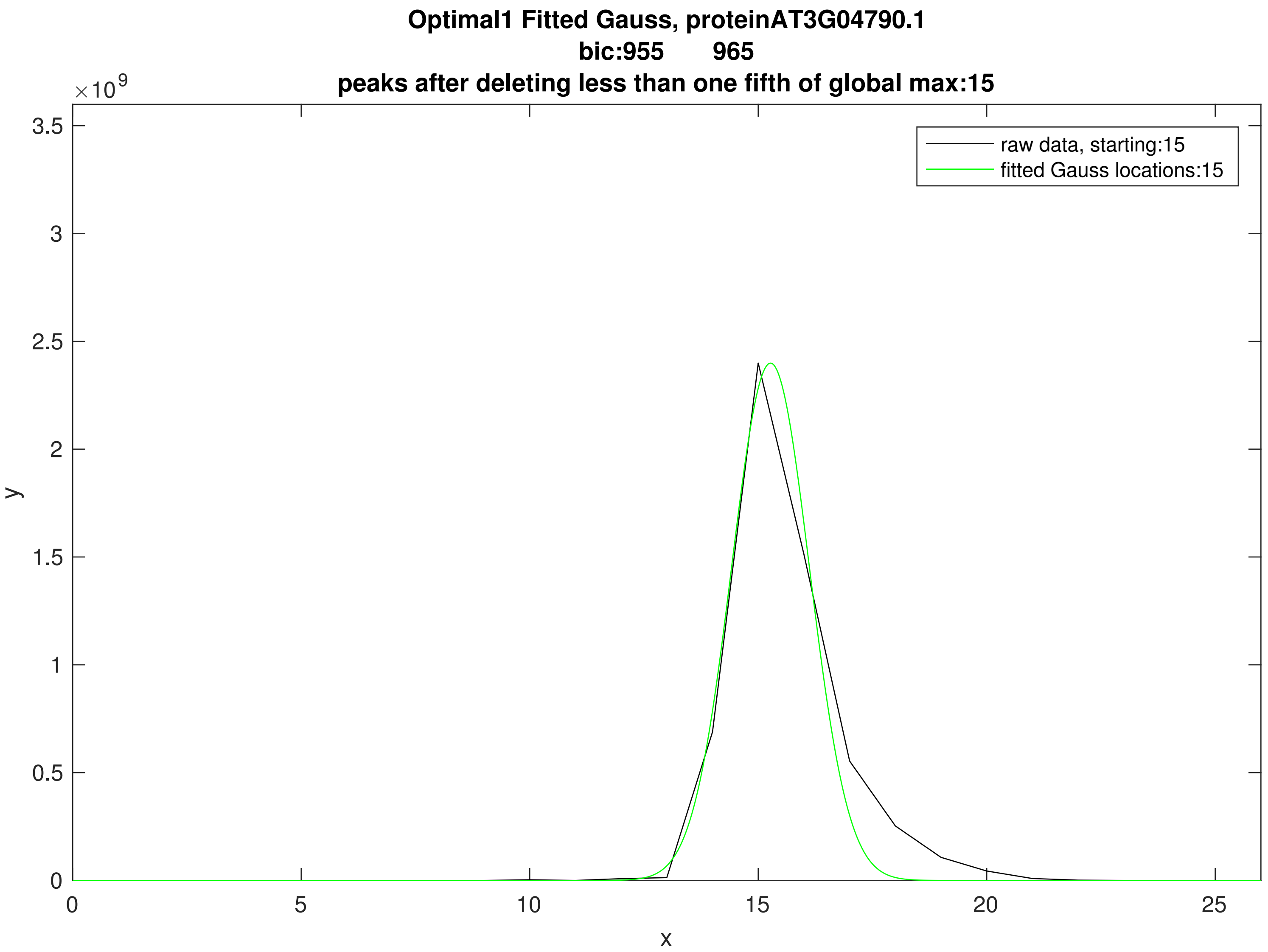

Optimal1 Fitted Gauss, proteinAT3G04790.1
bic:955 965
peaks after deleting less than one fifth of global max:15
×10^9
3.5
3
2.5
2
1.5
1
0.5
0
y
0
5
10
15
20
25
x
raw data, starting:15
fitted Gauss locations:15

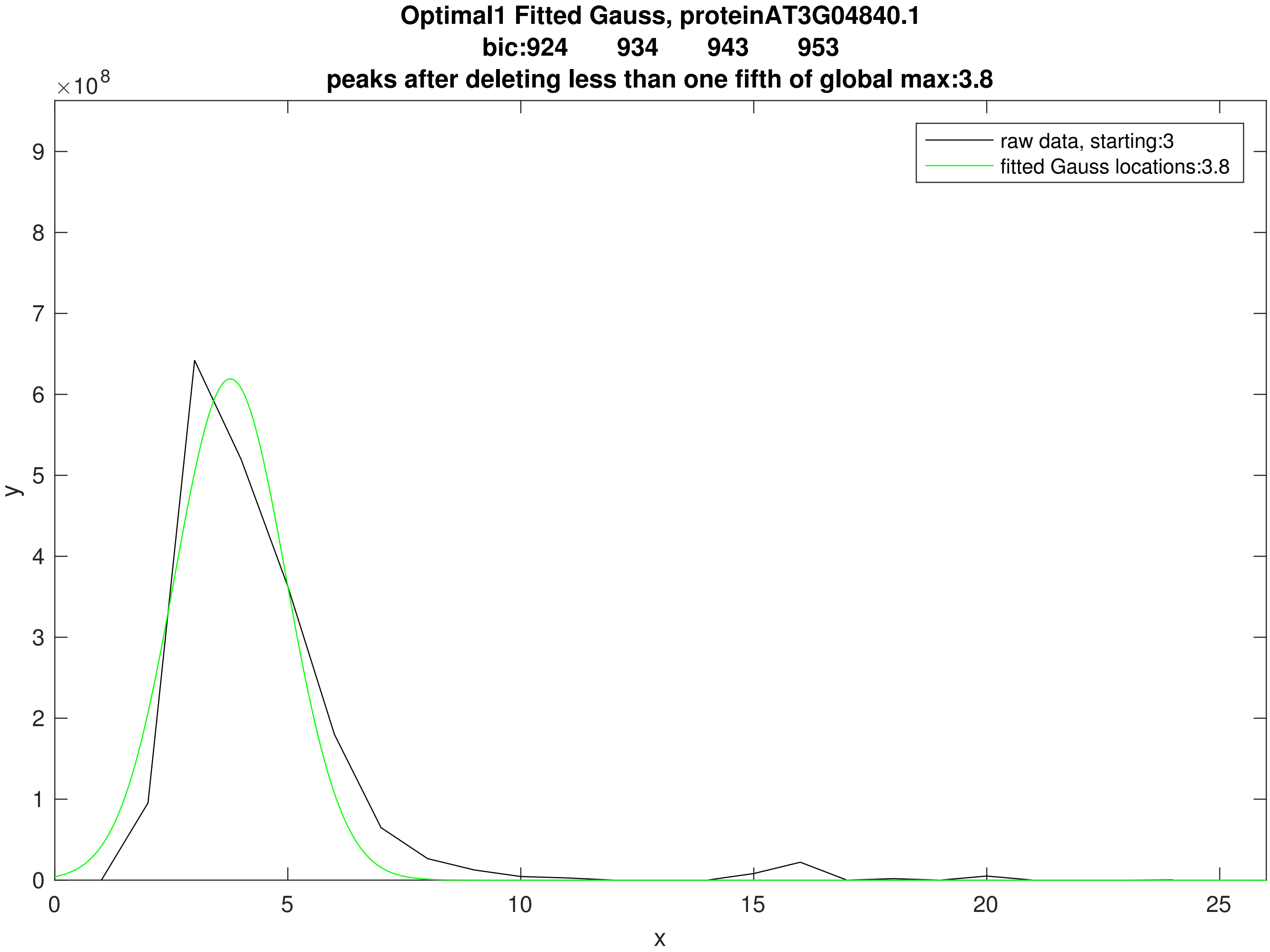

Optimal1 Fitted Gauss, proteinAT3G04840.1
bic:924 934 943 953
peaks after deleting less than one fifth of global max:3.8
×10^8
raw data, starting:3
fitted Gauss locations:3.8
y
x
0
1
2
3
4
5
6
7
8
9
0
5
10
15
20
25

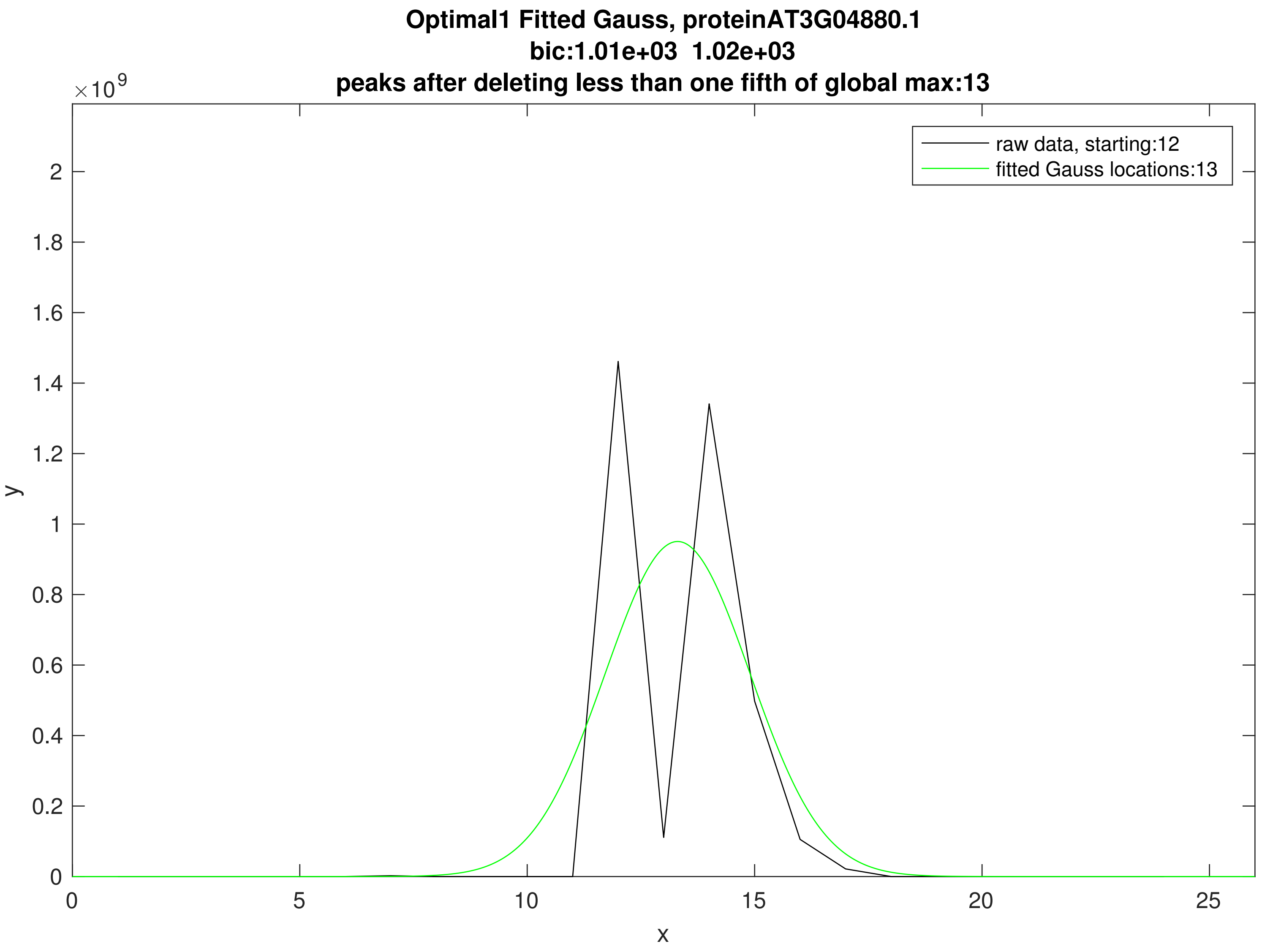
Optimal1 Fitted Gauss, proteinAT3G04880.1
bic:1.01e+03  1.02e+03
peaks after deleting less than one fifth of global max:13
×10^9
raw data, starting:12
fitted Gauss locations:13
y
x
0
0.2
0.4
0.6
0.8
1
1.2
1.4
1.6
1.8
2
5
10
15
20
25

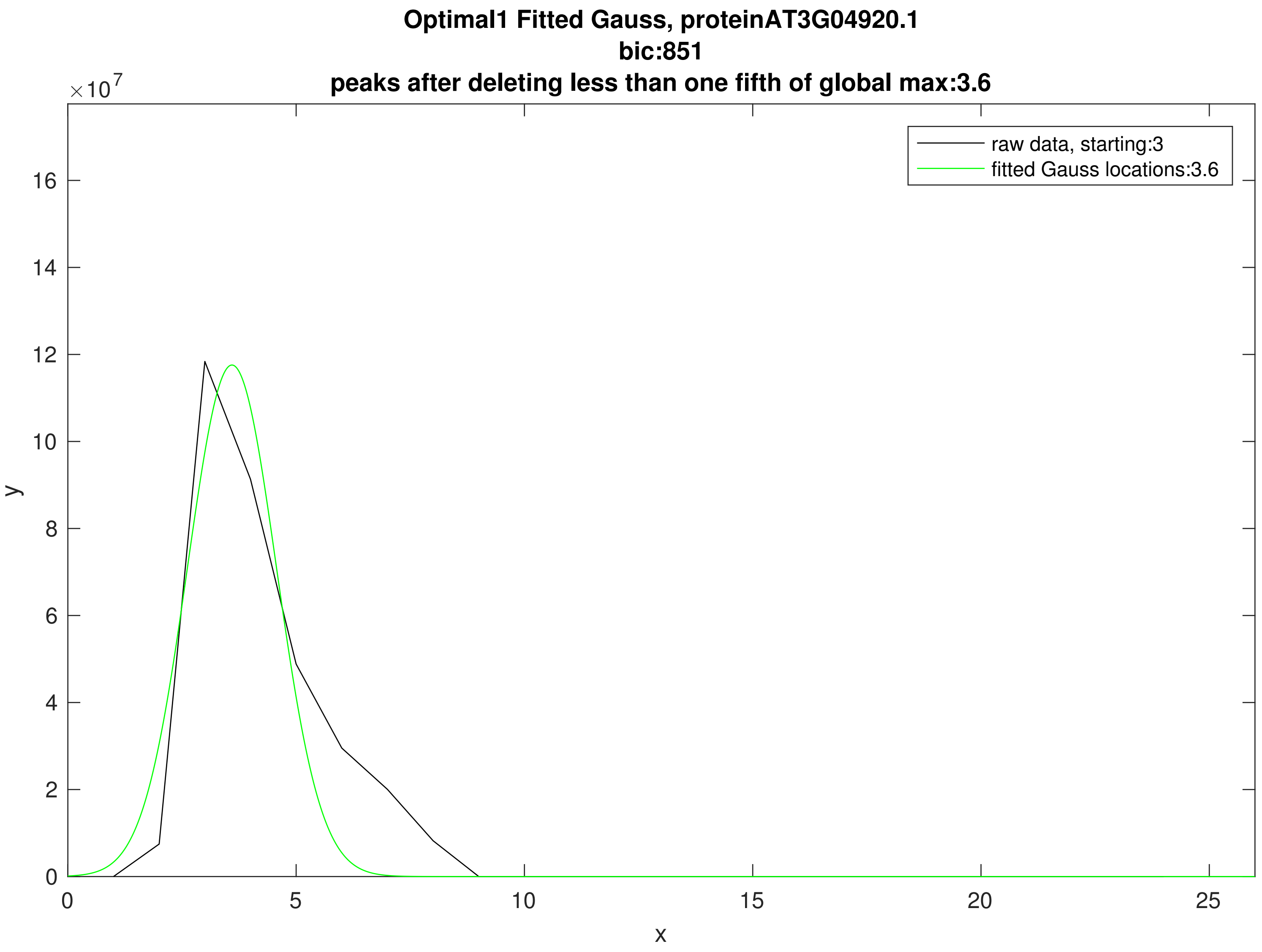
Optimal1 Fitted Gauss, proteinAT3G04920.1
bic:851
peaks after deleting less than one fifth of global max:3.6
×10^7
raw data, starting:3
fitted Gauss locations:3.6
y
x
0
2
4
6
8
10
12
14
16
0
5
10
15
20
25

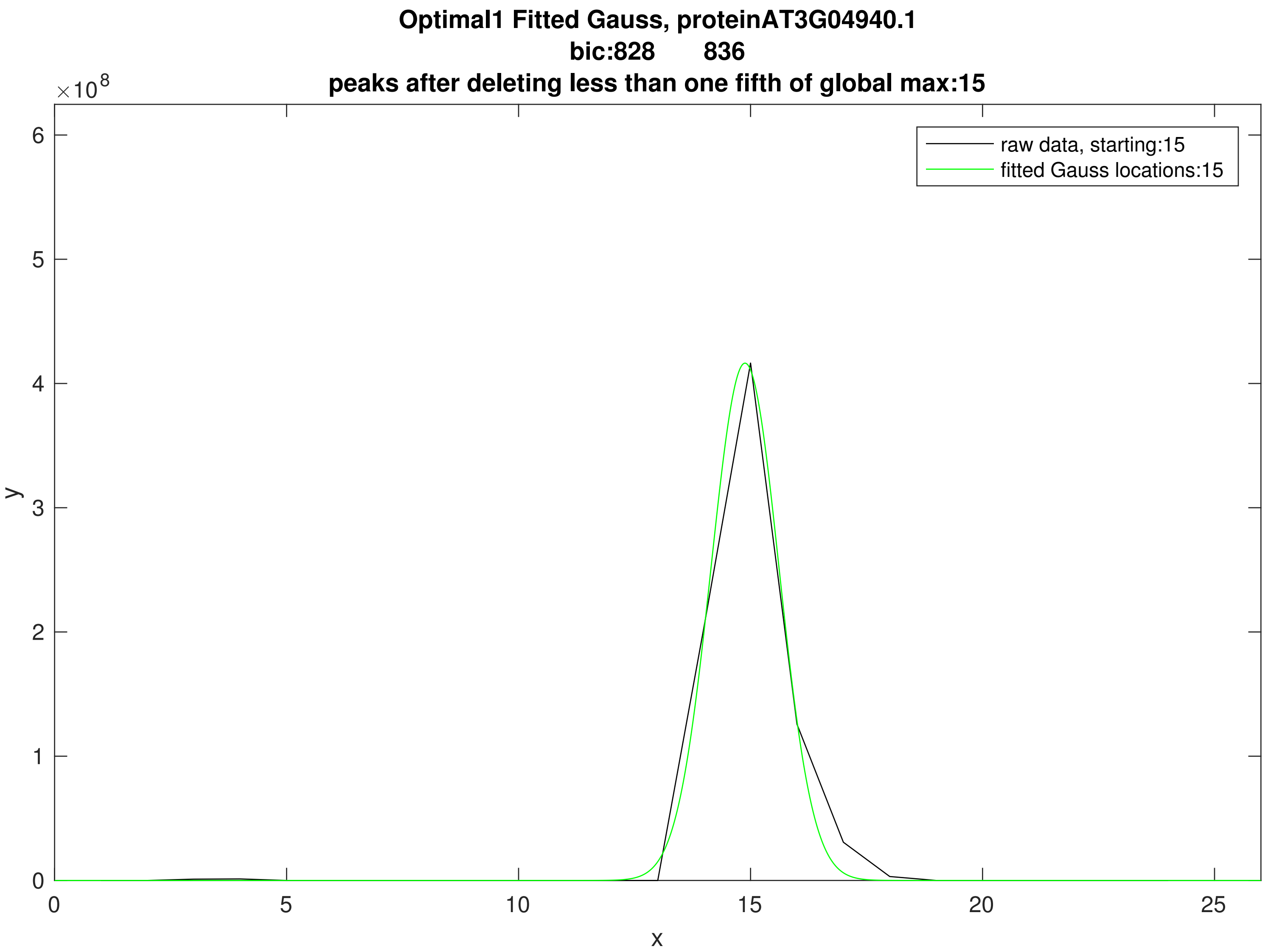

Optimal1 Fitted Gauss, proteinAT3G04940.1
bic:828 836
peaks after deleting less than one fifth of global max:15
×10^8
raw data, starting:15
fitted Gauss locations:15
y
x
0
1
2
3
4
5
6
0
5
10
15
20
25

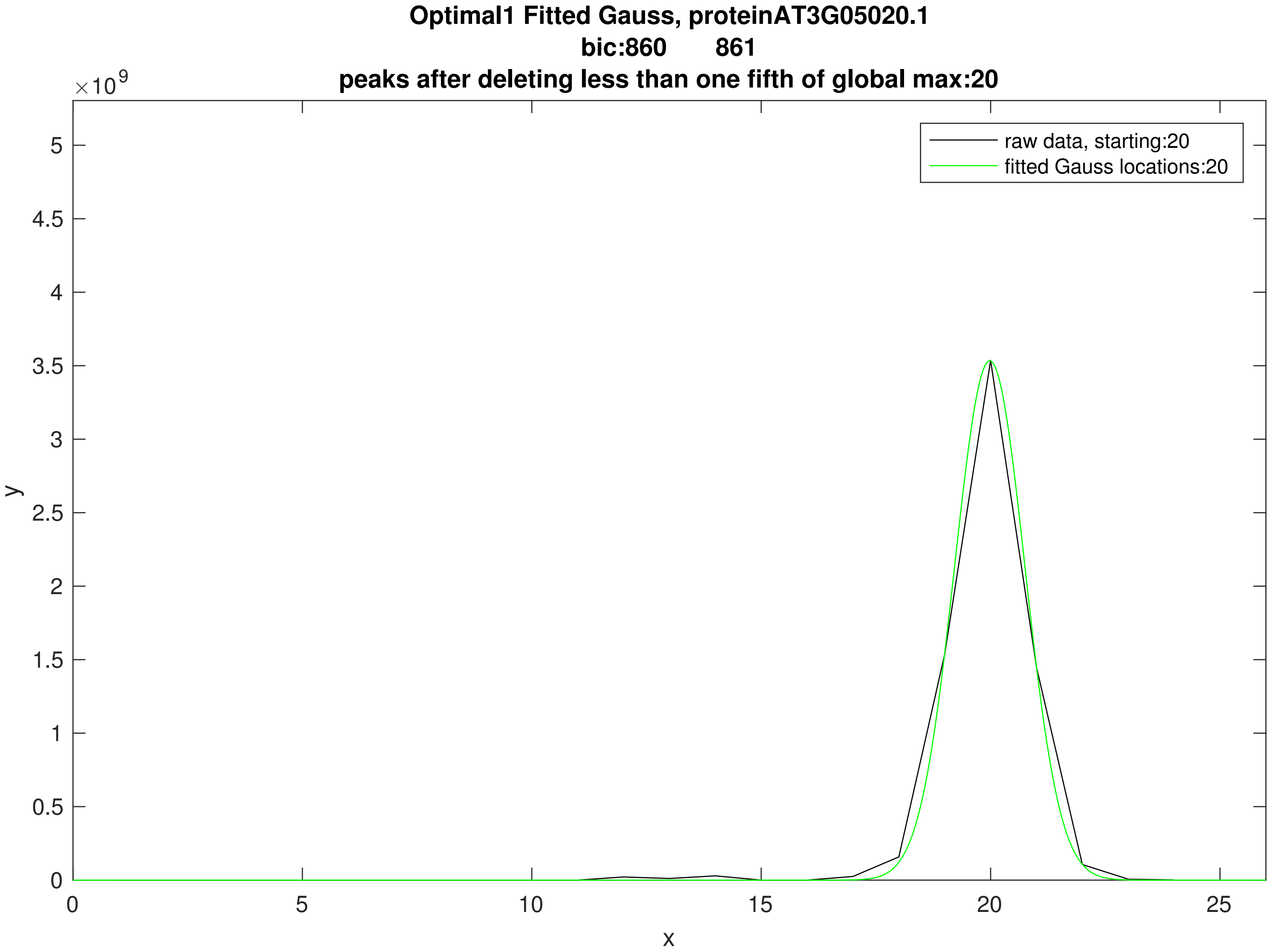
Optimal1 Fitted Gauss, proteinAT3G05020.1
bic:860 861
peaks after deleting less than one fifth of global max:20
×10^9
raw data, starting:20
fitted Gauss locations:20
y
x
0
0.5
1
1.5
2
2.5
3
3.5
4
4.5
5
0
5
10
15
20
25

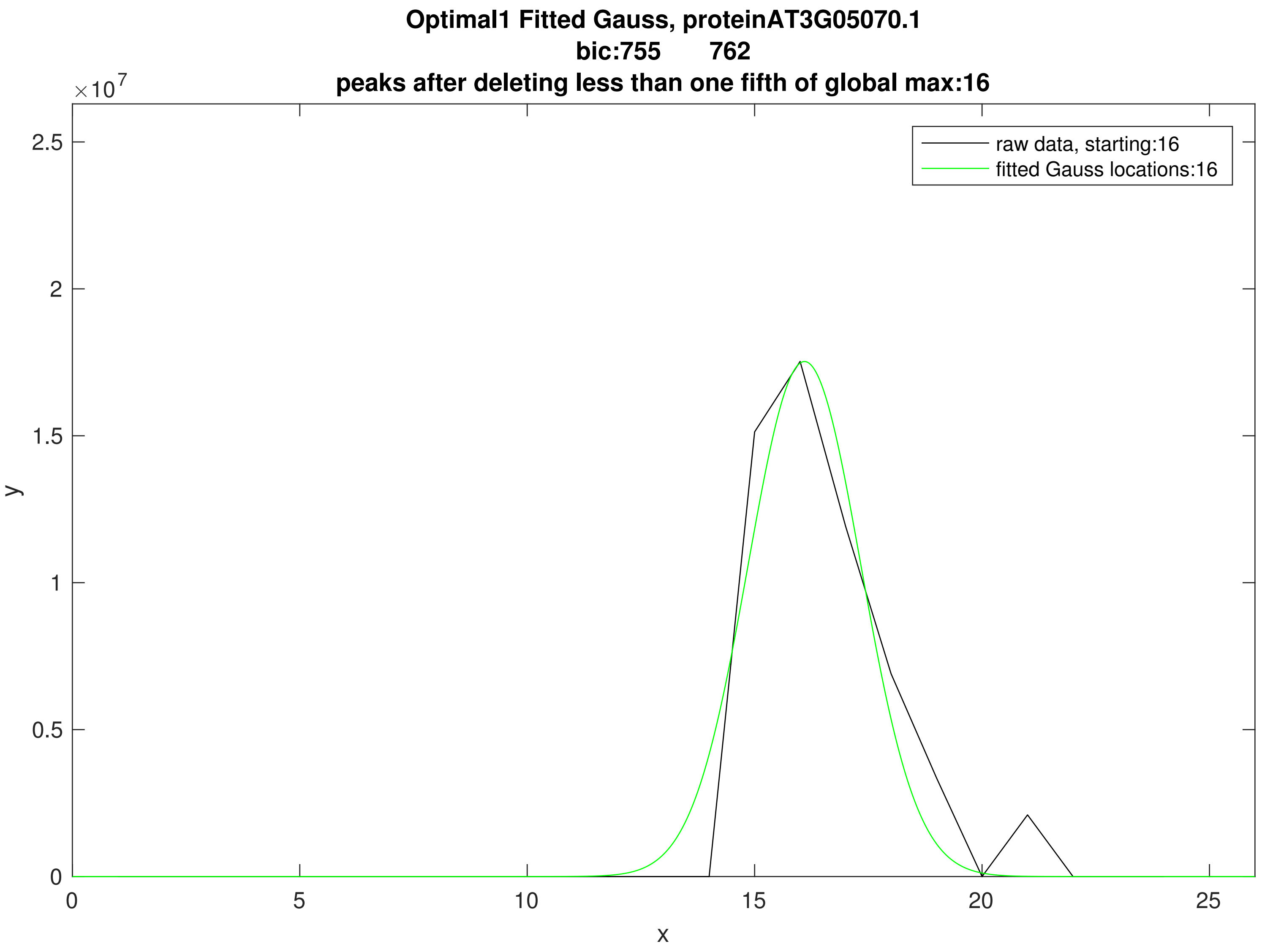
Optimal1 Fitted Gauss, proteinAT3G05070.1
bic:755 762
peaks after deleting less than one fifth of global max:16
×10^7
2.5
2
1.5
1
0.5
0
y
0
5
10
15
20
25
x
raw data, starting:16
fitted Gauss locations:16

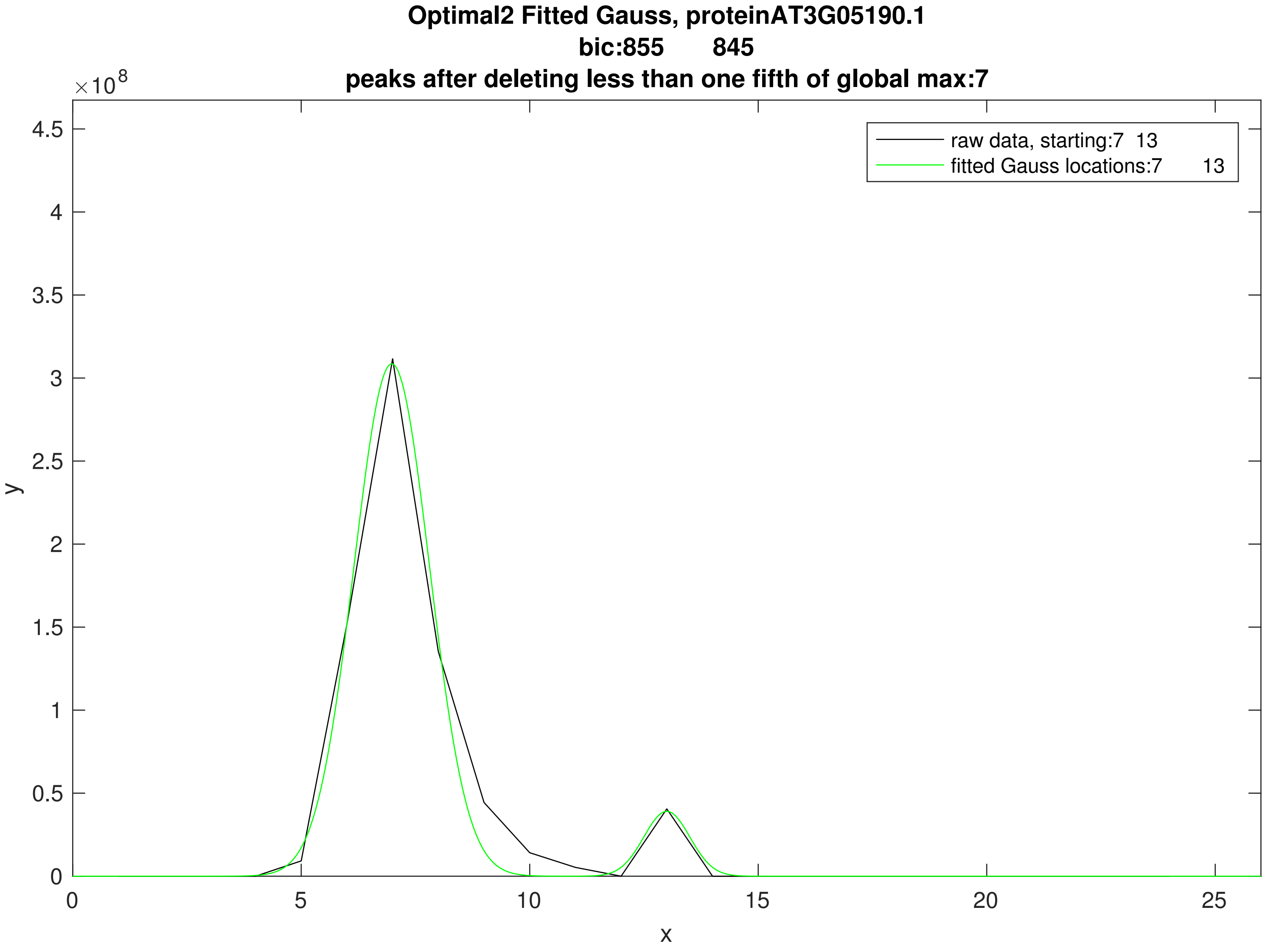

Optimal2 Fitted Gauss, proteinAT3G05190.1
bic:855 845
peaks after deleting less than one fifth of global max:7
×10^8
4.5
4
3.5
3
2.5
2
1.5
1
0.5
0
y
0
5
10
15
20
25
x
raw data, starting:7 13
fitted Gauss locations:7 13

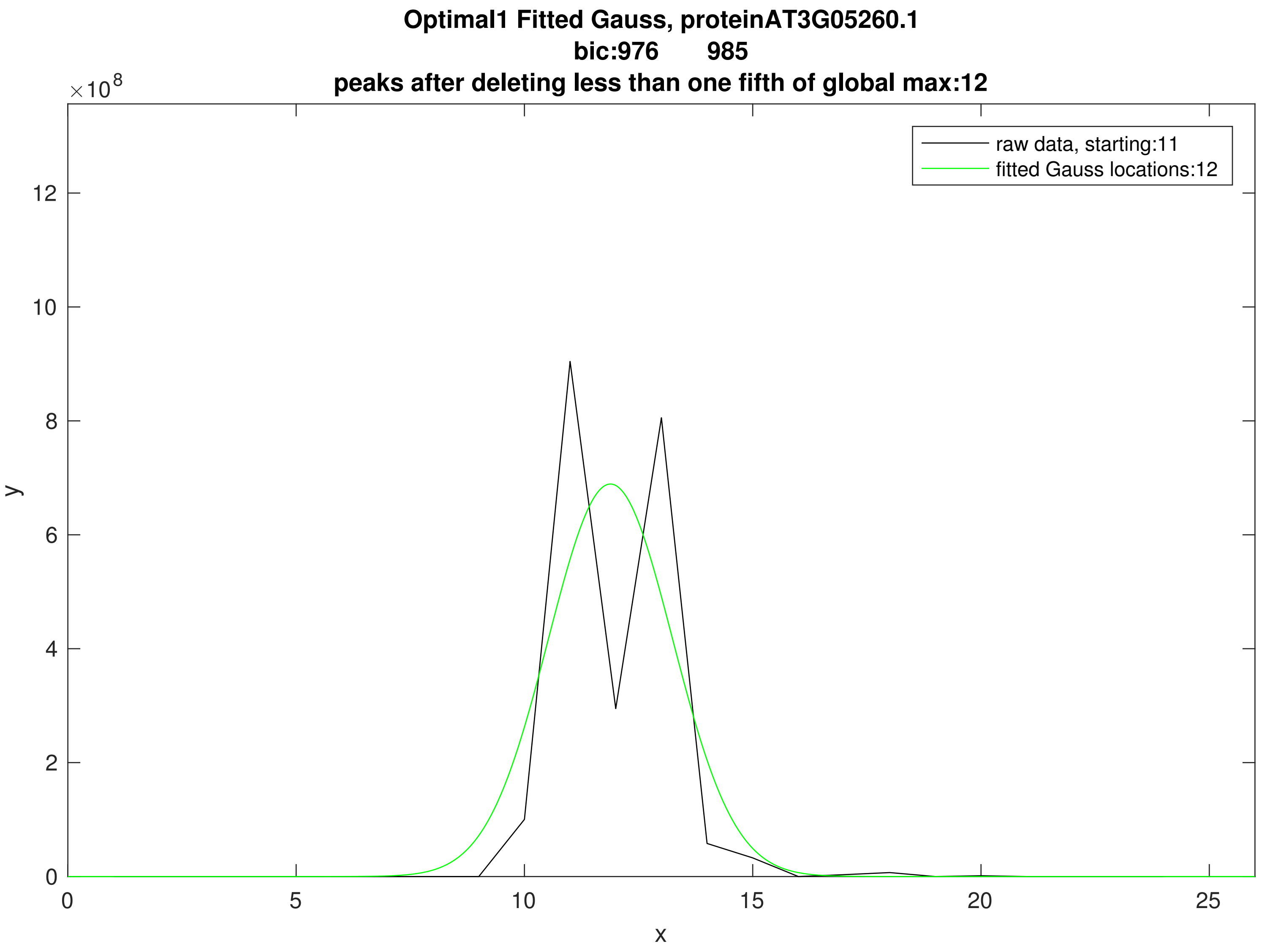
Optimal1 Fitted Gauss, proteinAT3G05260.1
bic:976 985
peaks after deleting less than one fifth of global max:12
×10^8
raw data, starting:11
fitted Gauss locations:12
12
10
8
6
4
2
0
y
0
5
10
15
20
25
x

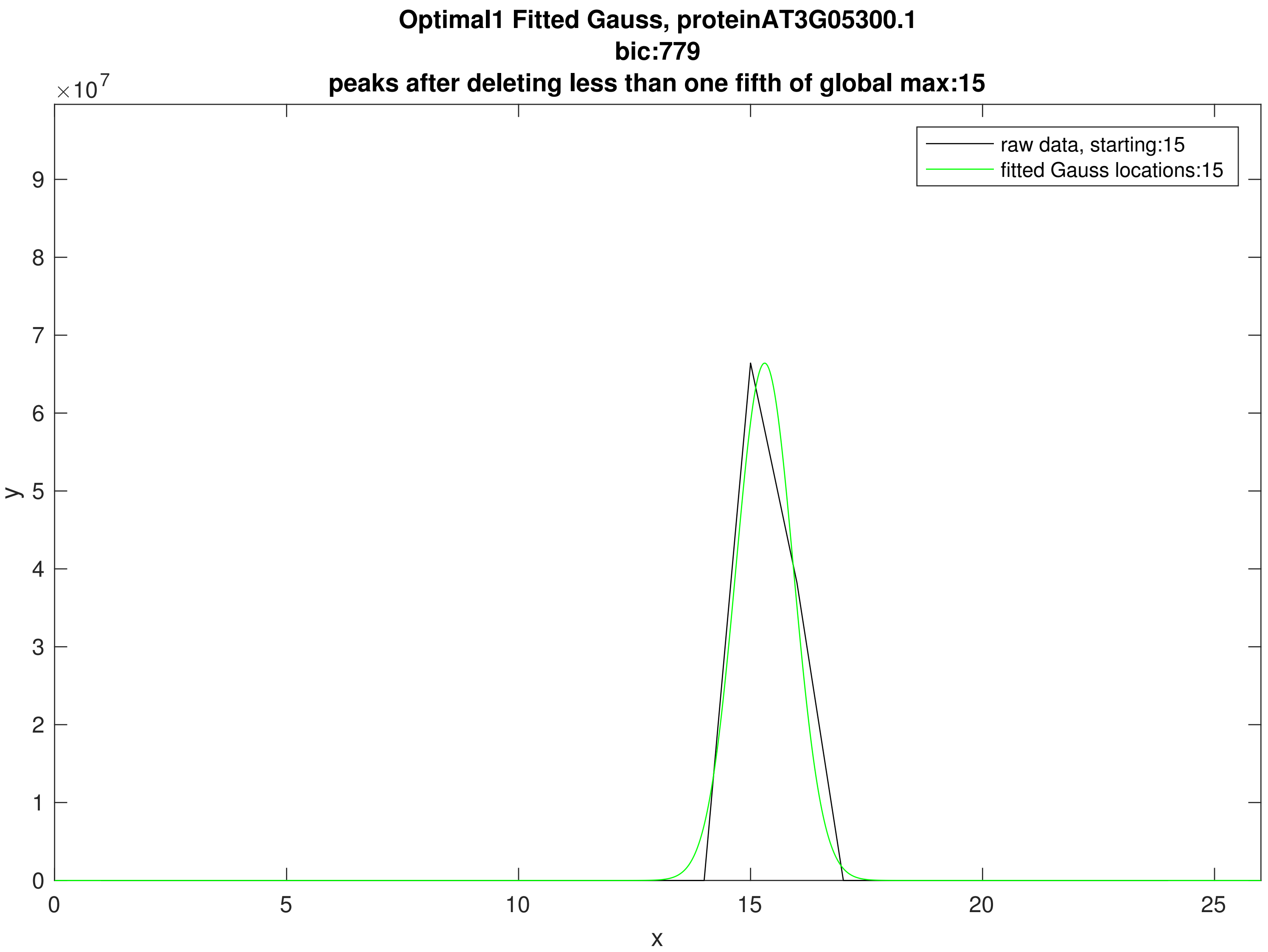

Optimal1 Fitted Gauss, proteinAT3G05300.1
bic:779
peaks after deleting less than one fifth of global max:15
×10^7
raw data, starting:15
fitted Gauss locations:15
y
x
0
1
2
3
4
5
6
7
8
9
0
5
10
15
20
25

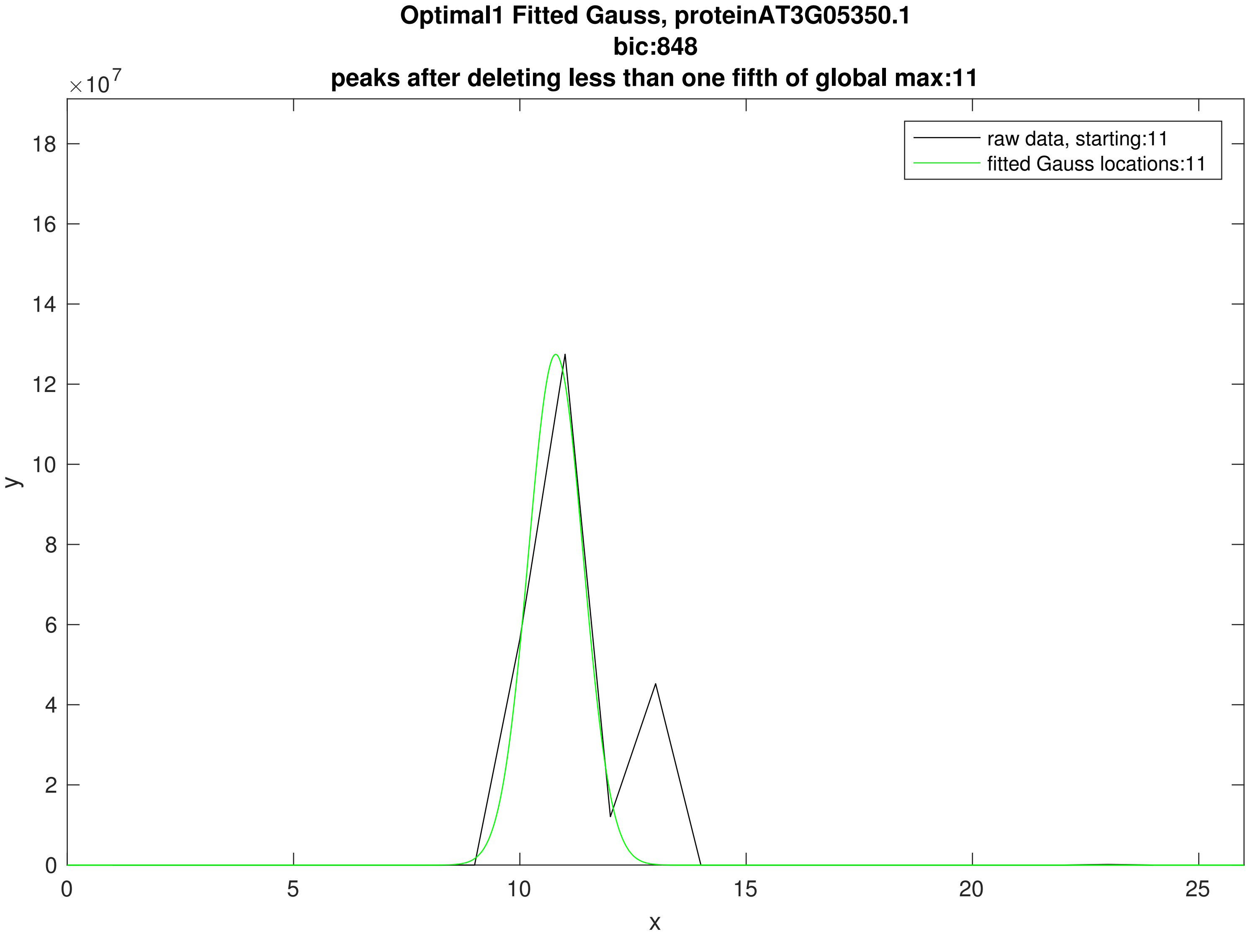
Optimal1 Fitted Gauss, proteinAT3G05350.1
bic:848
peaks after deleting less than one fifth of global max:11
×10^7
raw data, starting:11
fitted Gauss locations:11
18
16
14
12
10
8
6
4
2
0
y
0
5
10
15
20
25
x

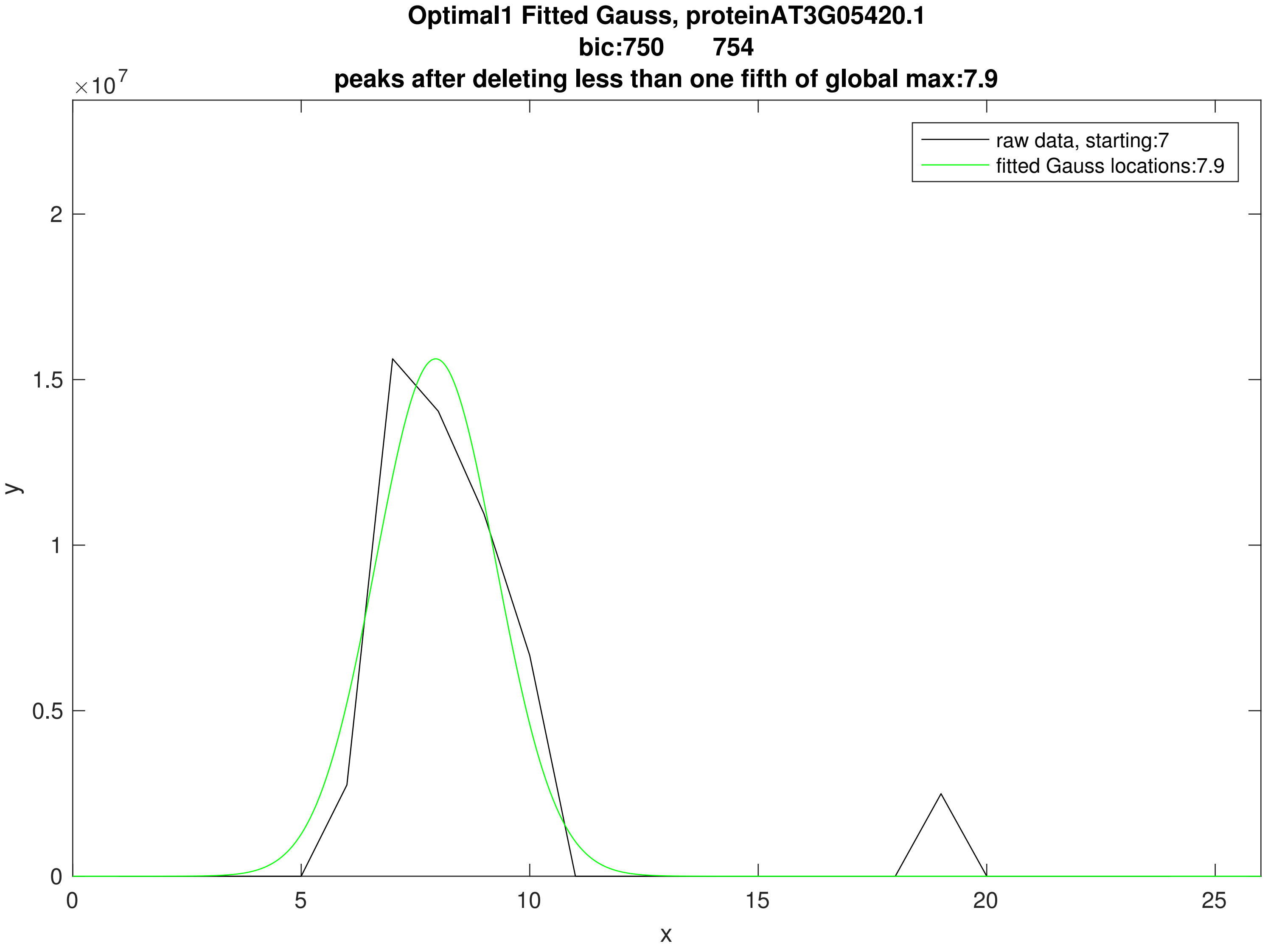
Optimal1 Fitted Gauss, proteinAT3G05420.1
bic:750 754
peaks after deleting less than one fifth of global max:7.9
×10^7
raw data, starting:7
fitted Gauss locations:7.9
y
x
0
0.5
1
1.5
2
0
5
10
15
20
25

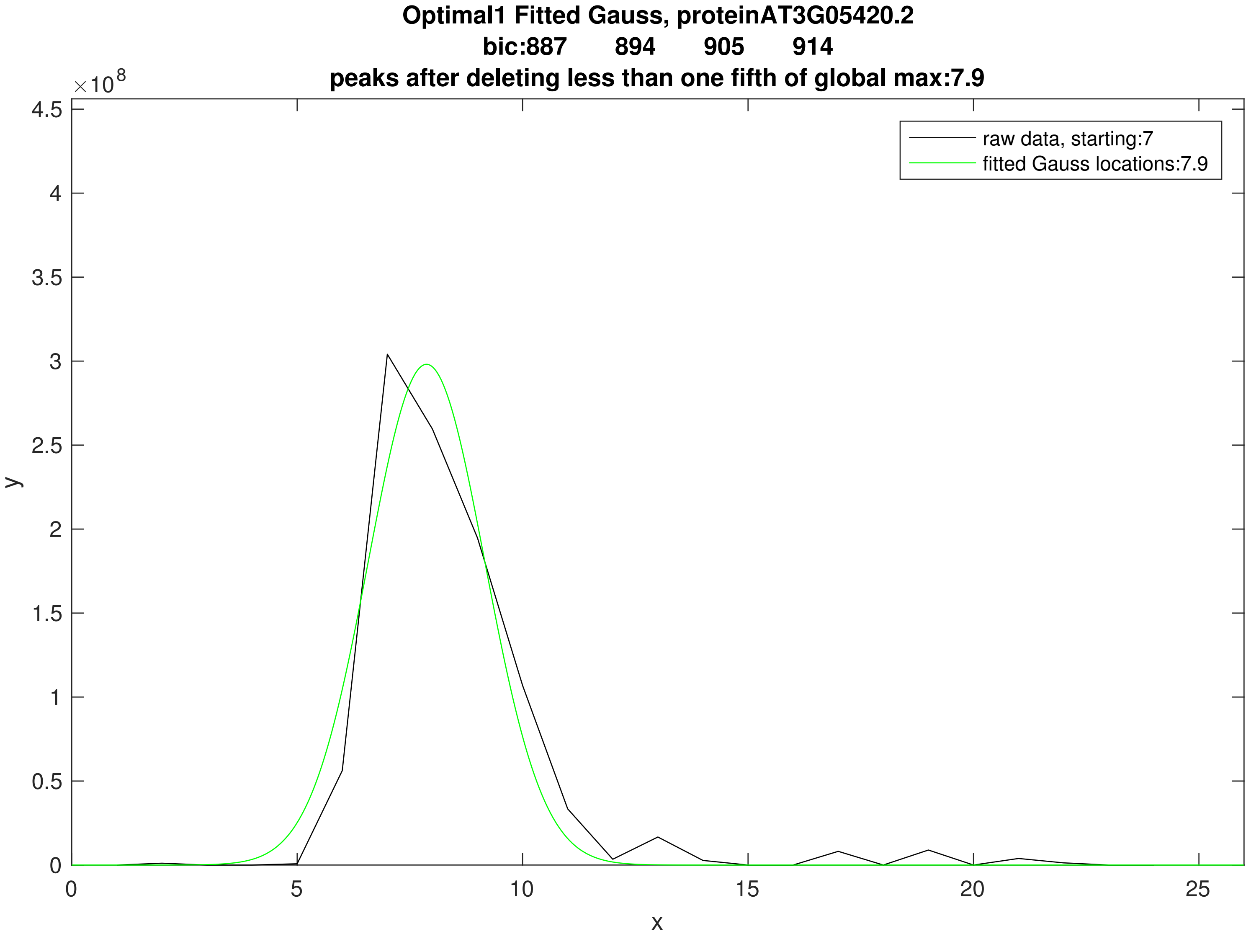
Optimal1 Fitted Gauss, proteinAT3G05420.2
bic:887 894 905 914
peaks after deleting less than one fifth of global max:7.9
raw data, starting:7
fitted Gauss locations:7.9
y
x

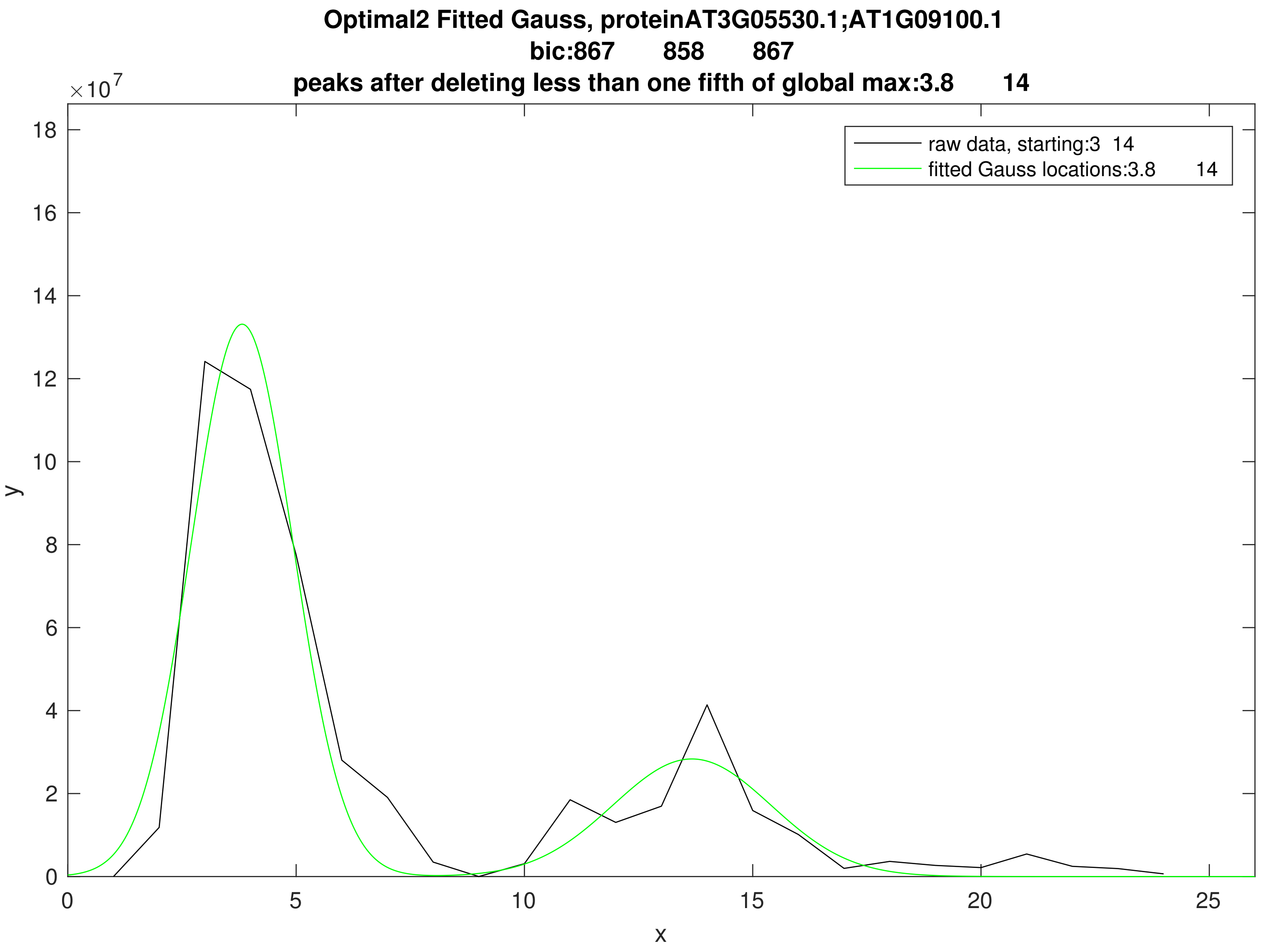

Optimal2 Fitted Gauss, proteinAT3G05530.1;AT1G09100.1
bic:867 858 867
peaks after deleting less than one fifth of global max:3.8 14
×10^7
raw data, starting:3 14
fitted Gauss locations:3.8 14
y
x
0
2
4
6
8
10
12
14
16
18
5
15
20
25

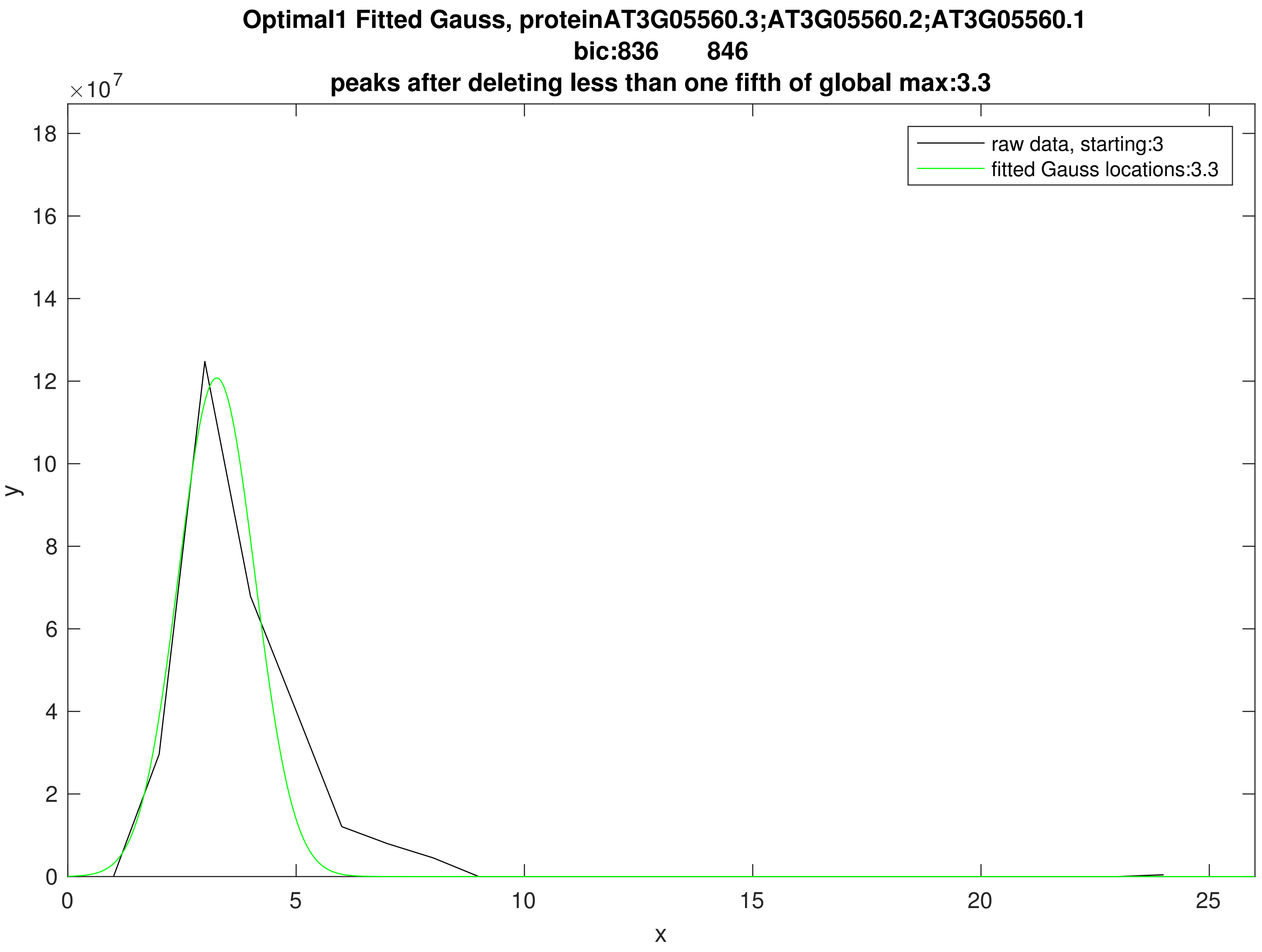
Optimal1 Fitted Gauss, proteinAT3G05560.3;AT3G05560.2;AT3G05560.1
bic:836 846
peaks after deleting less than one fifth of global max:3.3
×10^7
raw data, starting:3
fitted Gauss locations:3.3
y
x
0
2
4
6
8
10
12
14
16
18
0
5
10
15
20
25

**Optimal1 Fitted Gauss, proteinAT3G05580.1**
**bic:679**
**peaks after deleting less than one fifth of global max:17**

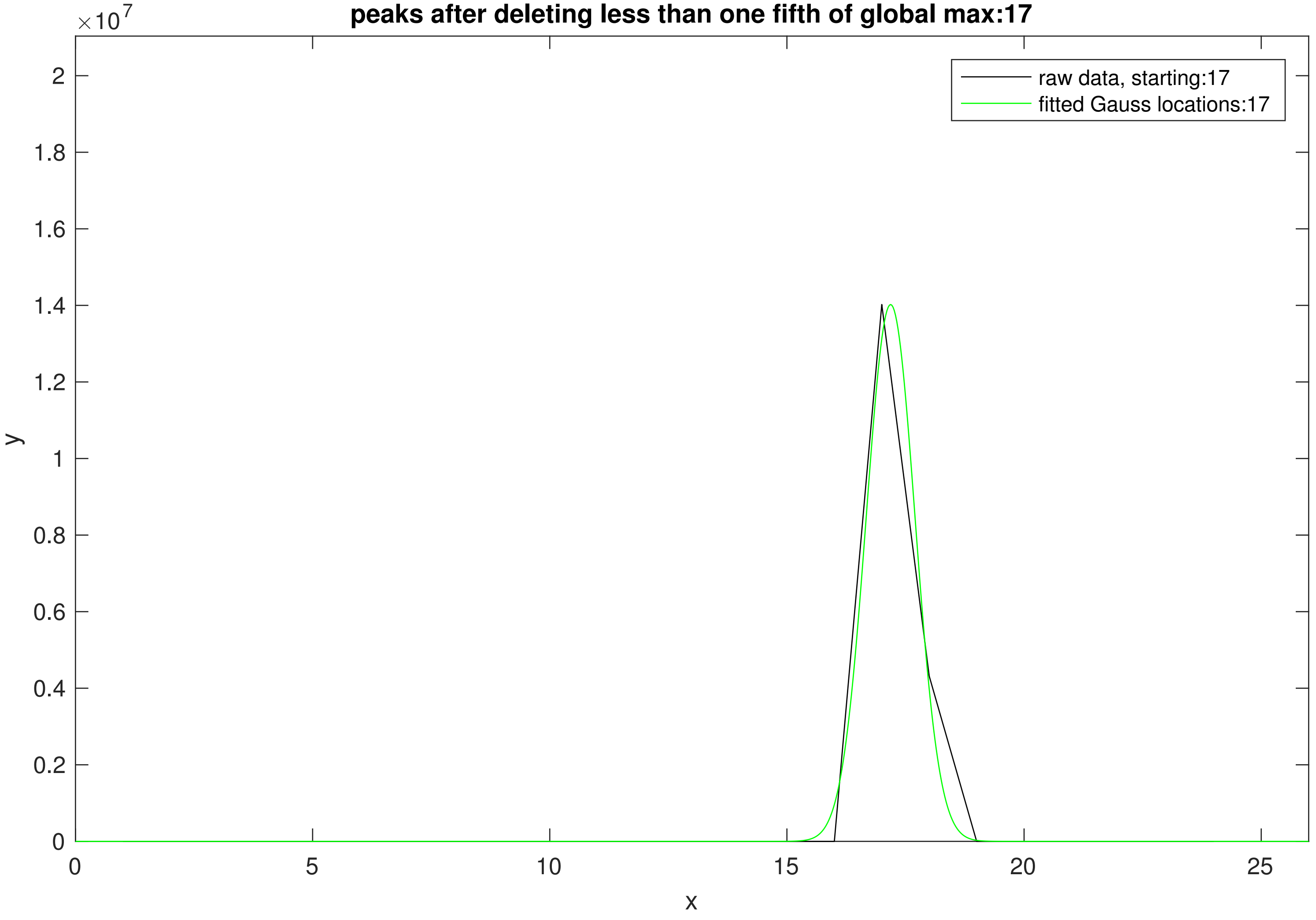

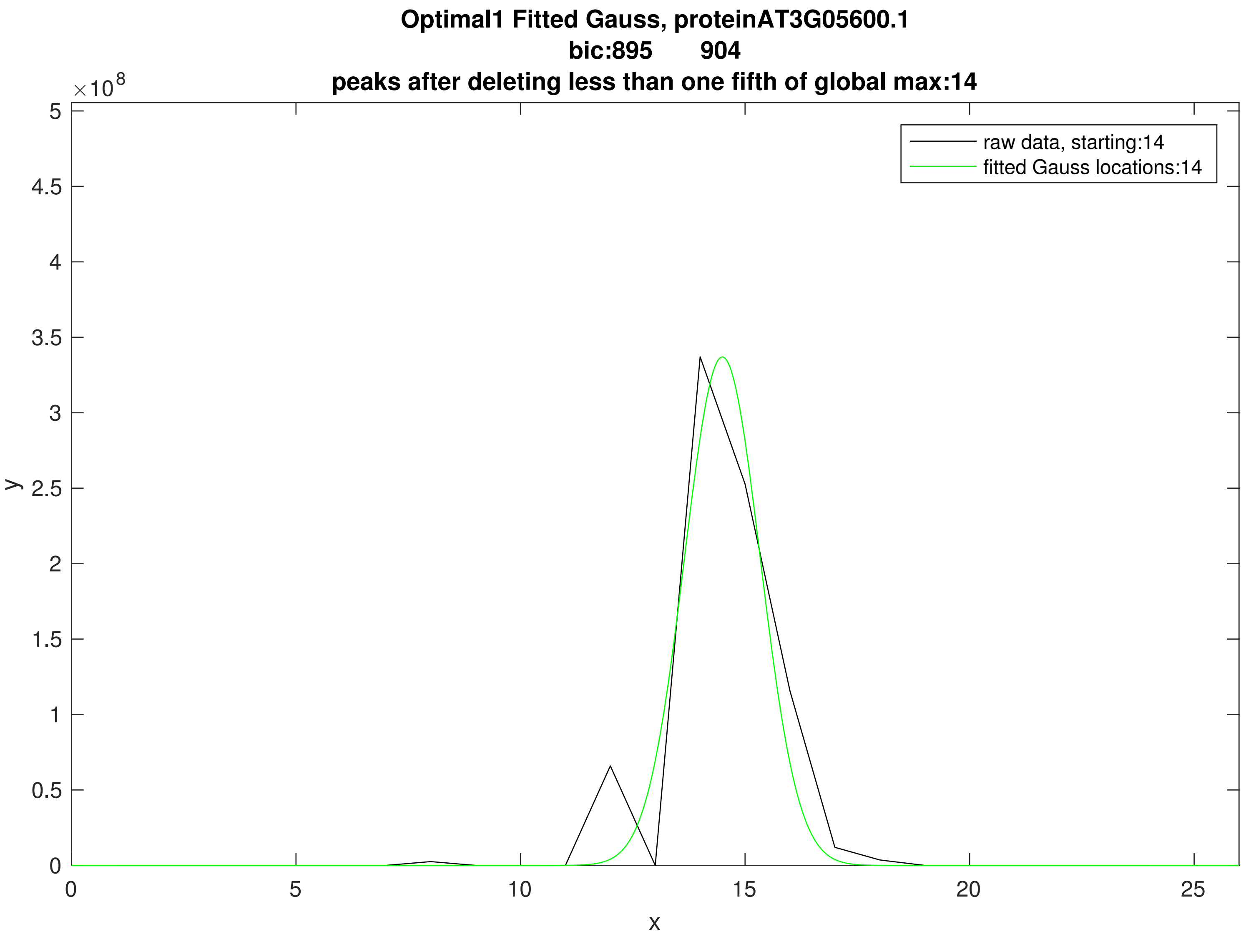
Optimal1 Fitted Gauss, proteinAT3G05600.1
bic:895 904
peaks after deleting less than one fifth of global max:14
×10^8
5
4.5
4
3.5
3
2.5
2
1.5
1
0.5
0
y
0
5
10
15
20
25
x
raw data, starting:14
fitted Gauss locations:14

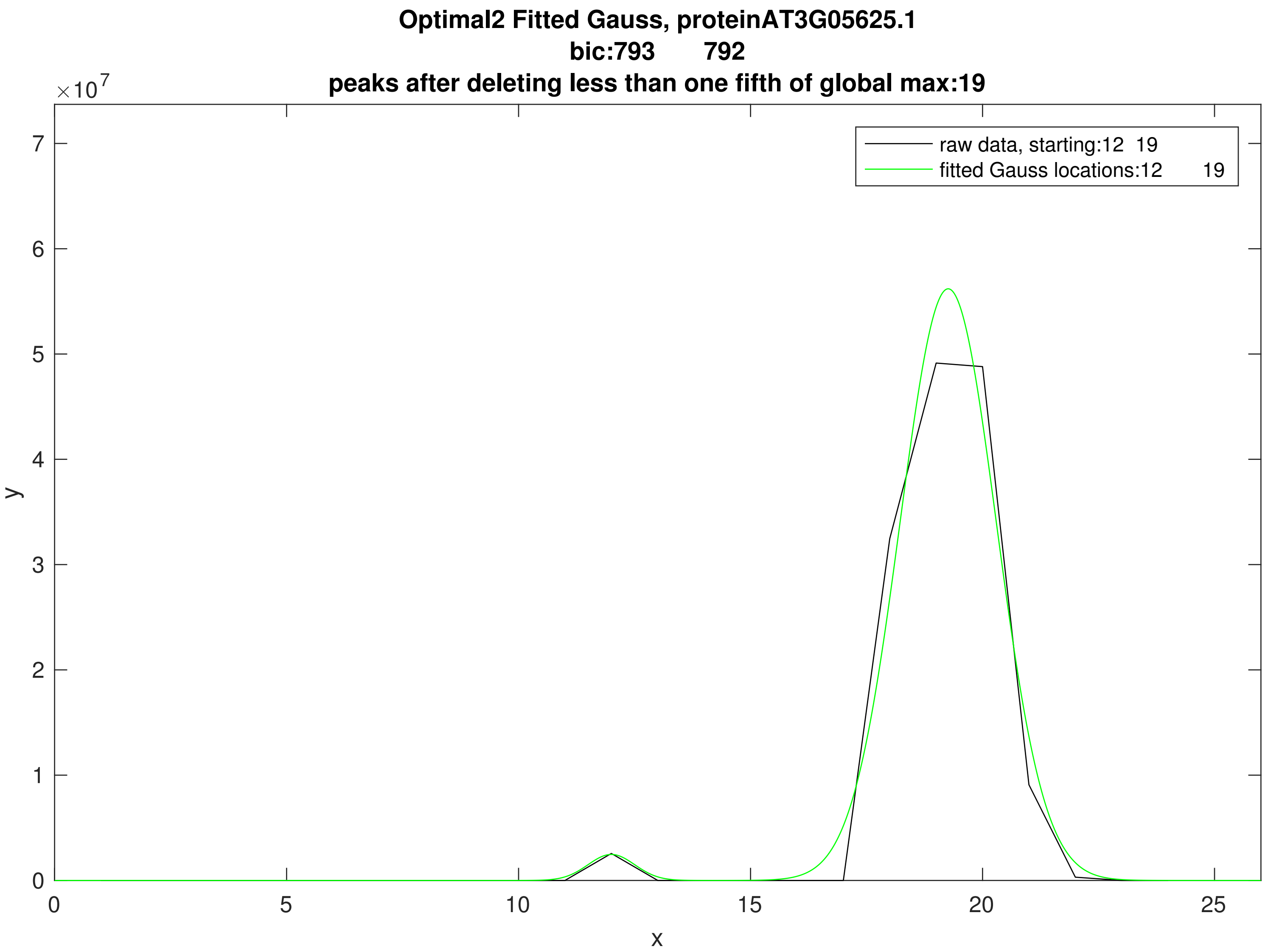
Optimal2 Fitted Gauss, proteinAT3G05625.1
bic:793 792
peaks after deleting less than one fifth of global max:19
×10^7
raw data, starting:12 19
fitted Gauss locations:12 19
y
x
0
1
2
3
4
5
6
7
0
5
10
15
20
25

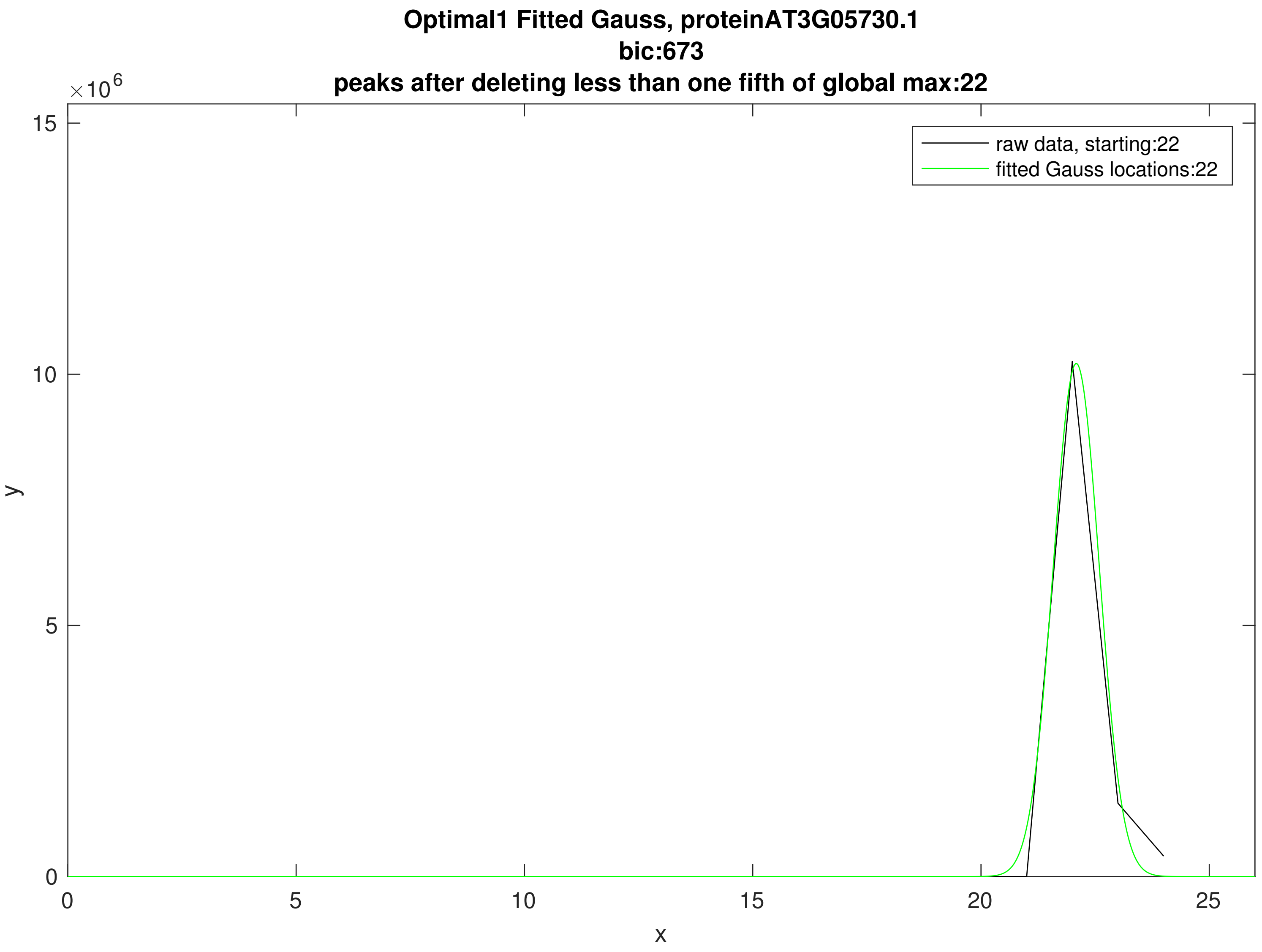
Optimal1 Fitted Gauss, proteinAT3G05730.1
bic:673
peaks after deleting less than one fifth of global max:22
×10^6
raw data, starting:22
fitted Gauss locations:22
y
x
0
5
10
15
20
25
0
5
10
15

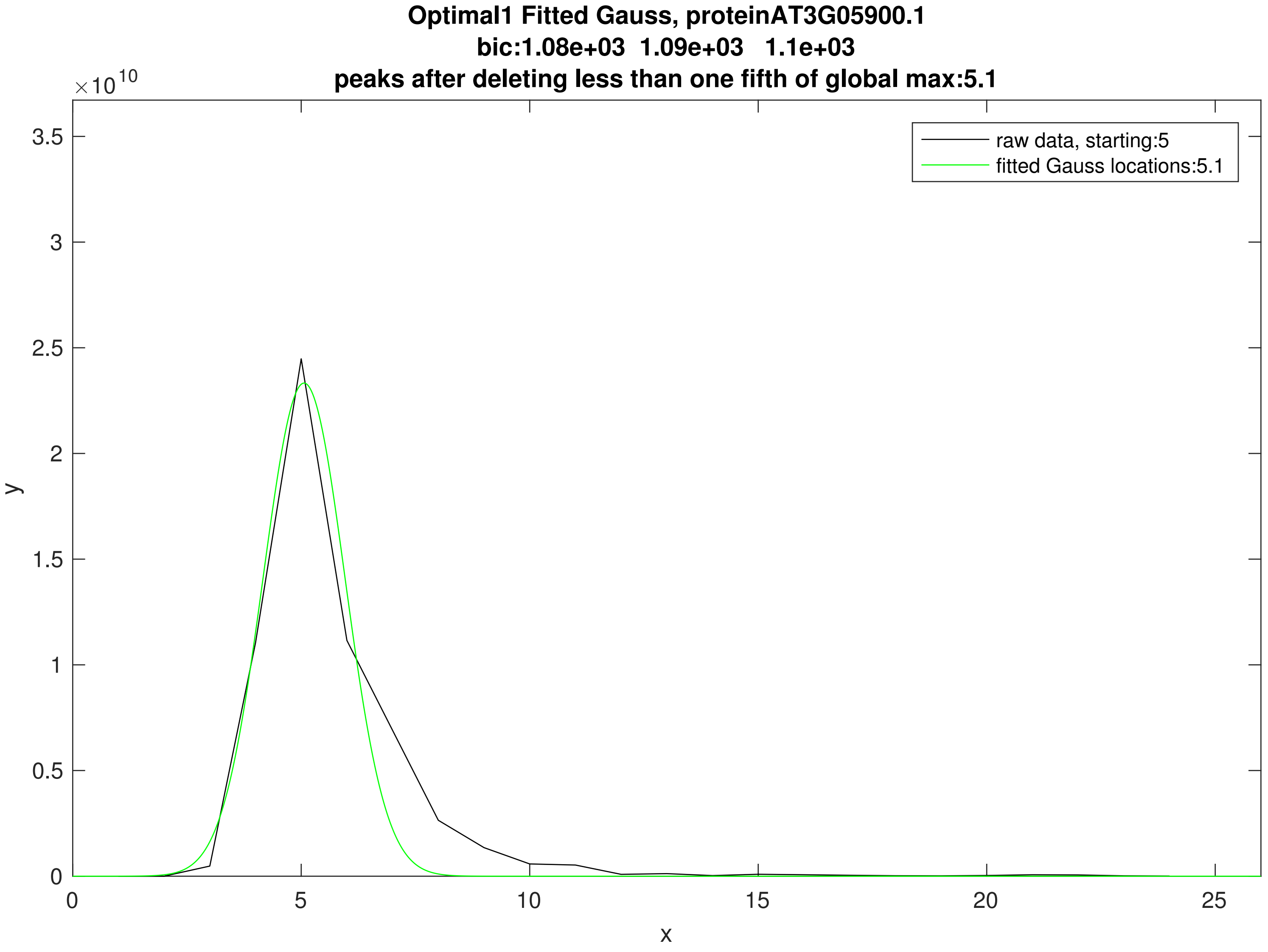

Optimal1 Fitted Gauss, proteinAT3G05900.1
bic:1.08e+03 1.09e+03 1.1e+03
peaks after deleting less than one fifth of global max:5.1
×10^10
3.5
3
2.5
2
1.5
1
0.5
0
y
0
5
10
15
20
25
x
raw data, starting:5
fitted Gauss locations:5.1

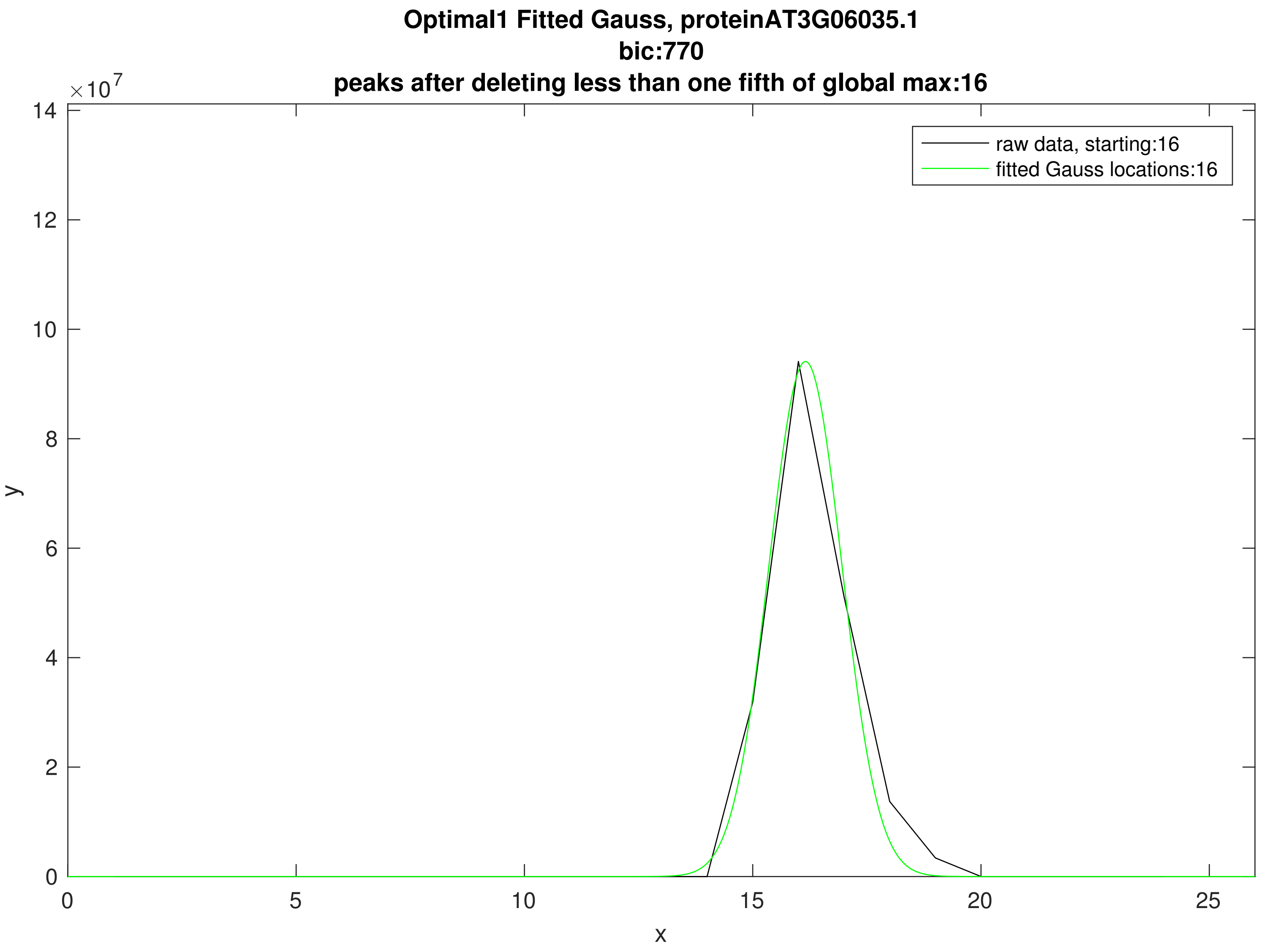

Optimal1 Fitted Gauss, proteinAT3G06035.1
bic:770
peaks after deleting less than one fifth of global max:16
×10^7
14
12
10
8
6
4
2
0
y
0
5
10
15
20
25
x
raw data, starting:16
fitted Gauss locations:16

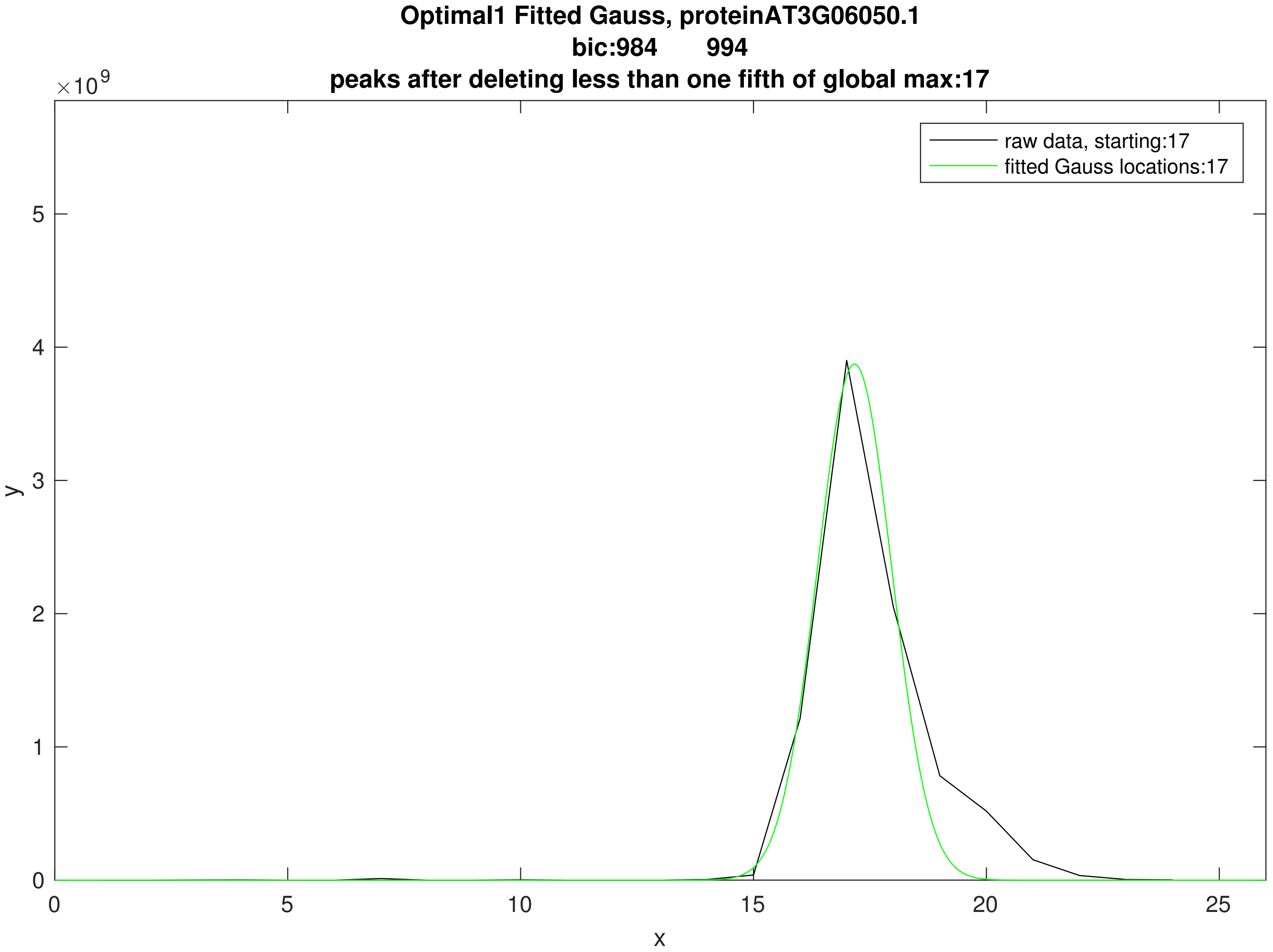

Optimal1 Fitted Gauss, proteinAT3G06050.1
bic:984 994
peaks after deleting less than one fifth of global max:17
×10^9
raw data, starting:17
fitted Gauss locations:17
y
x
0
1
2
3
4
5
0
5
10
15
20
25

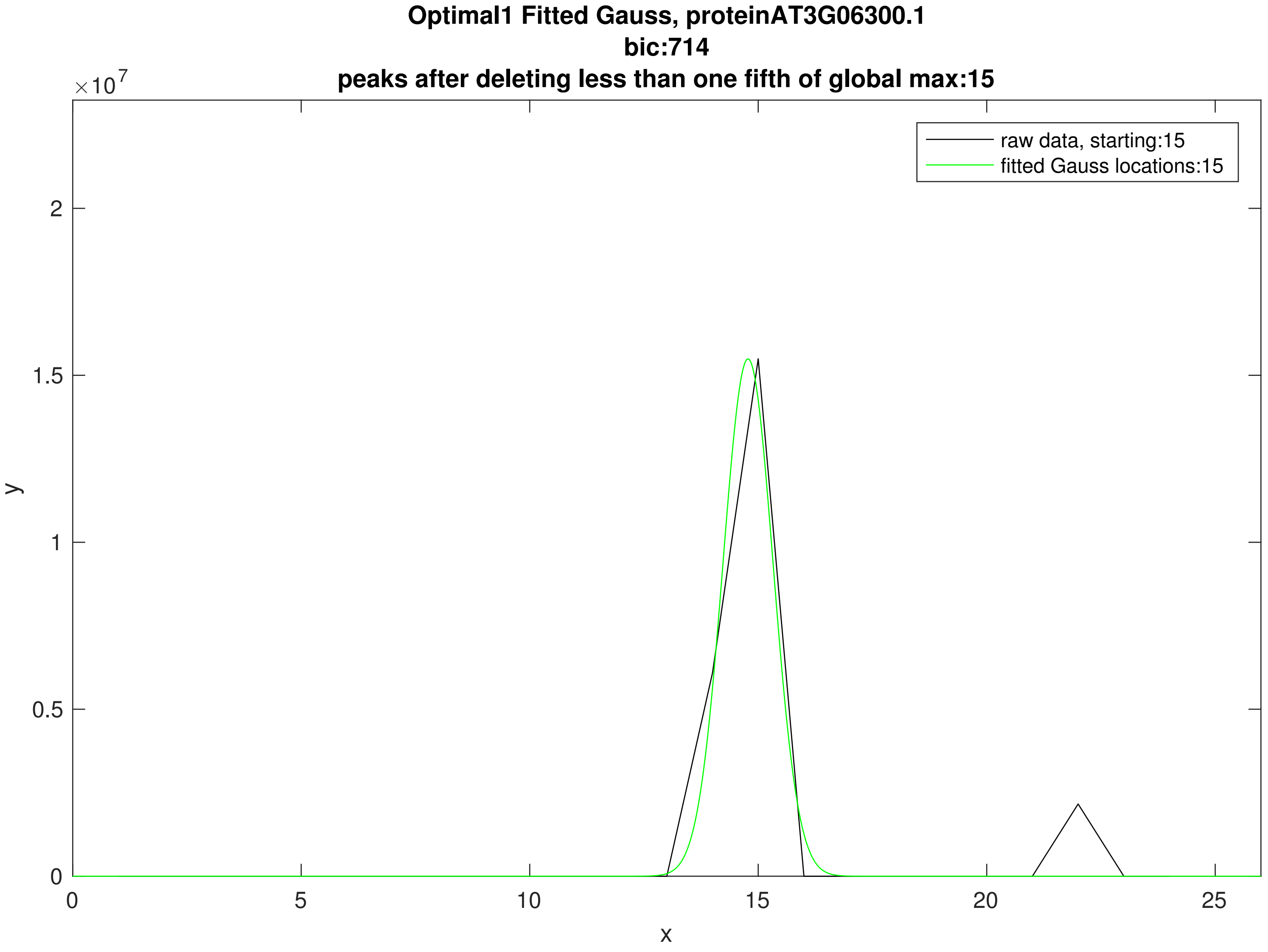
Optimal1 Fitted Gauss, proteinAT3G06300.1
bic:714
peaks after deleting less than one fifth of global max:15
×10^7
raw data, starting:15
fitted Gauss locations:15
2
1.5
1
0.5
0
y
0
5
10
15
20
25
x

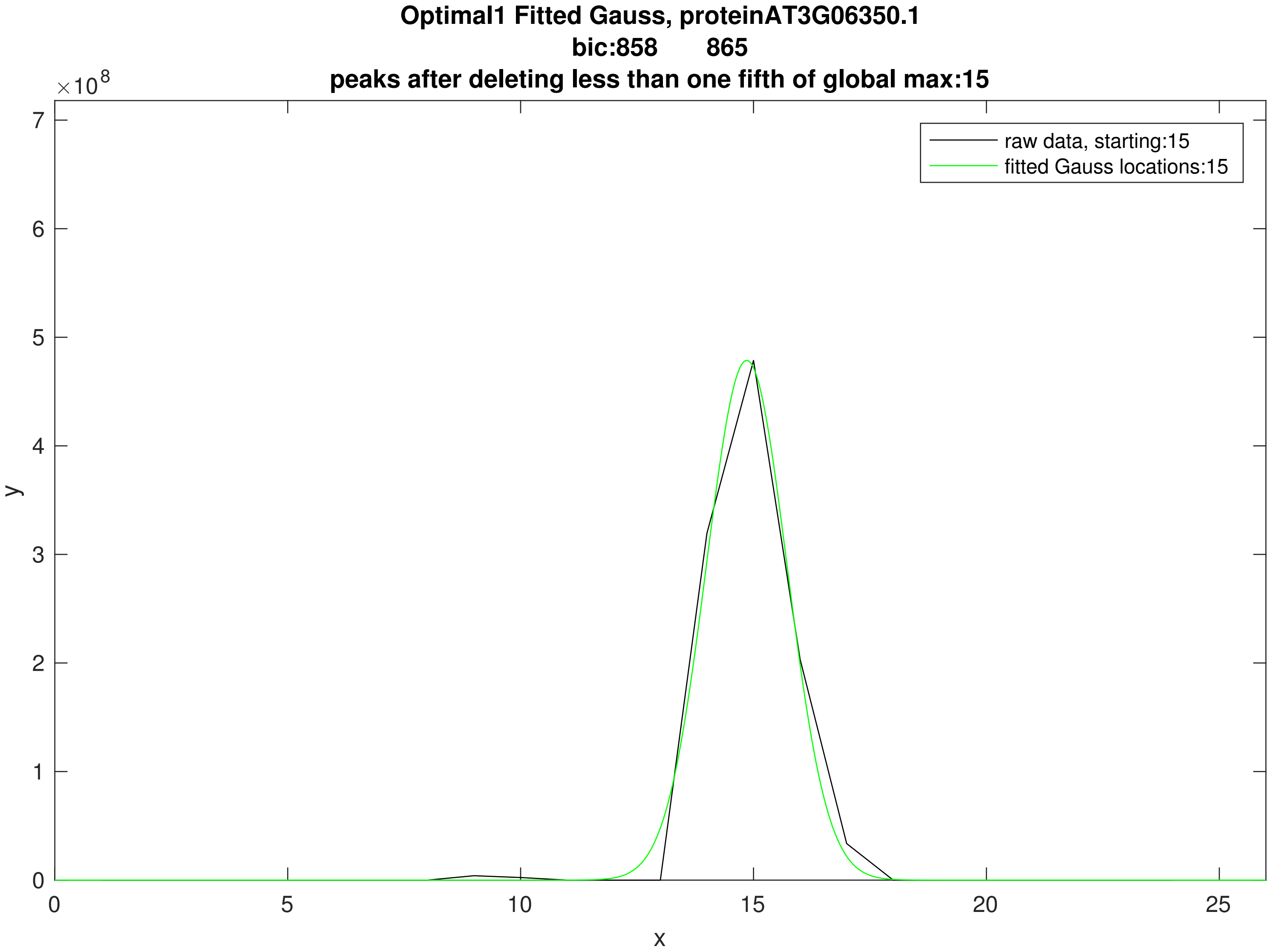
Optimal1 Fitted Gauss, proteinAT3G06350.1
bic:858 865
peaks after deleting less than one fifth of global max:15
×10^8
raw data, starting:15
fitted Gauss locations:15
y
x
0
1
2
3
4
5
6
7
0
5
10
15
20
25

**Optimal1 Fitted Gauss, proteinAT3G06540.1**
**bic:769**
**peaks after deleting less than one fifth of global max:13**

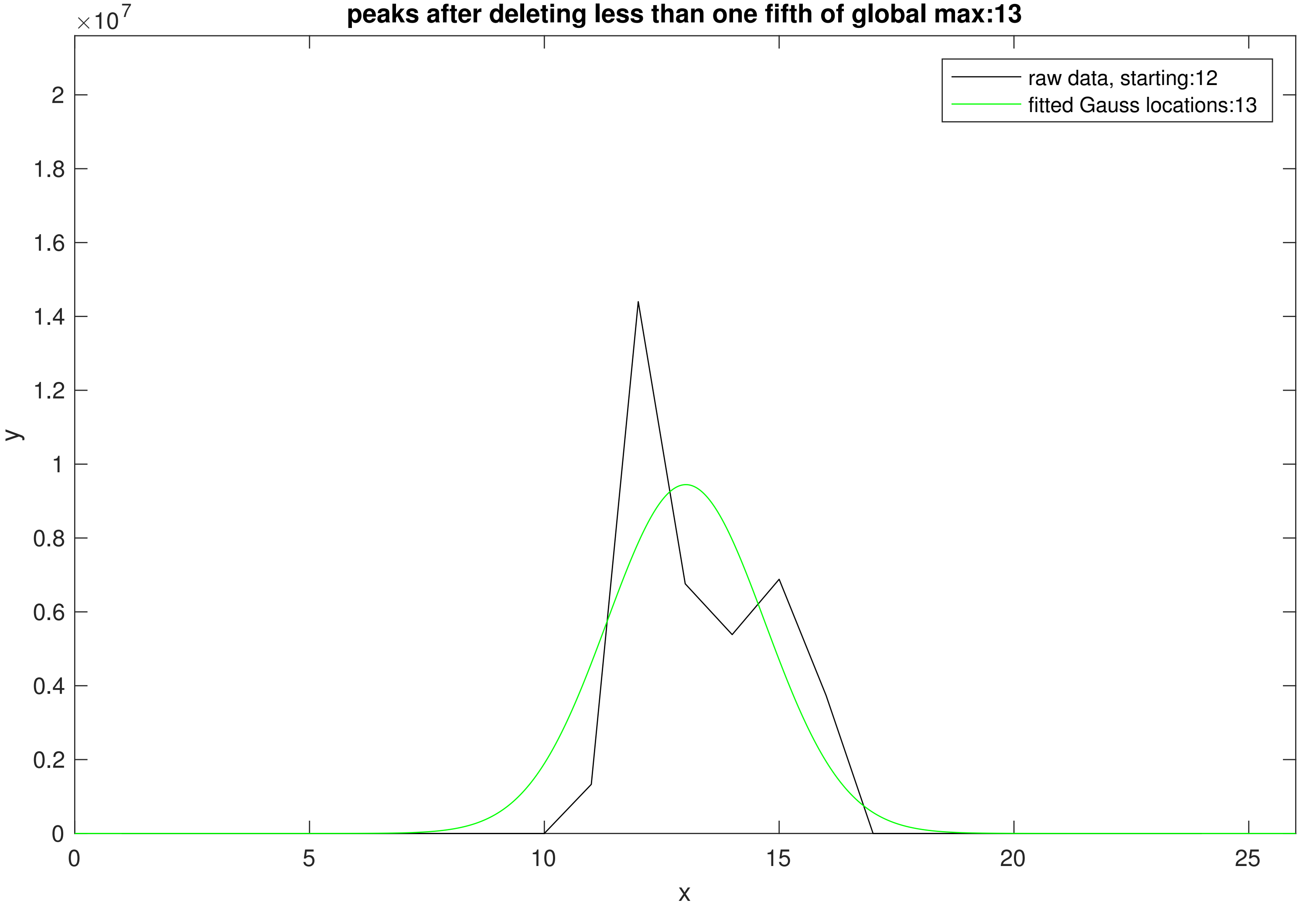

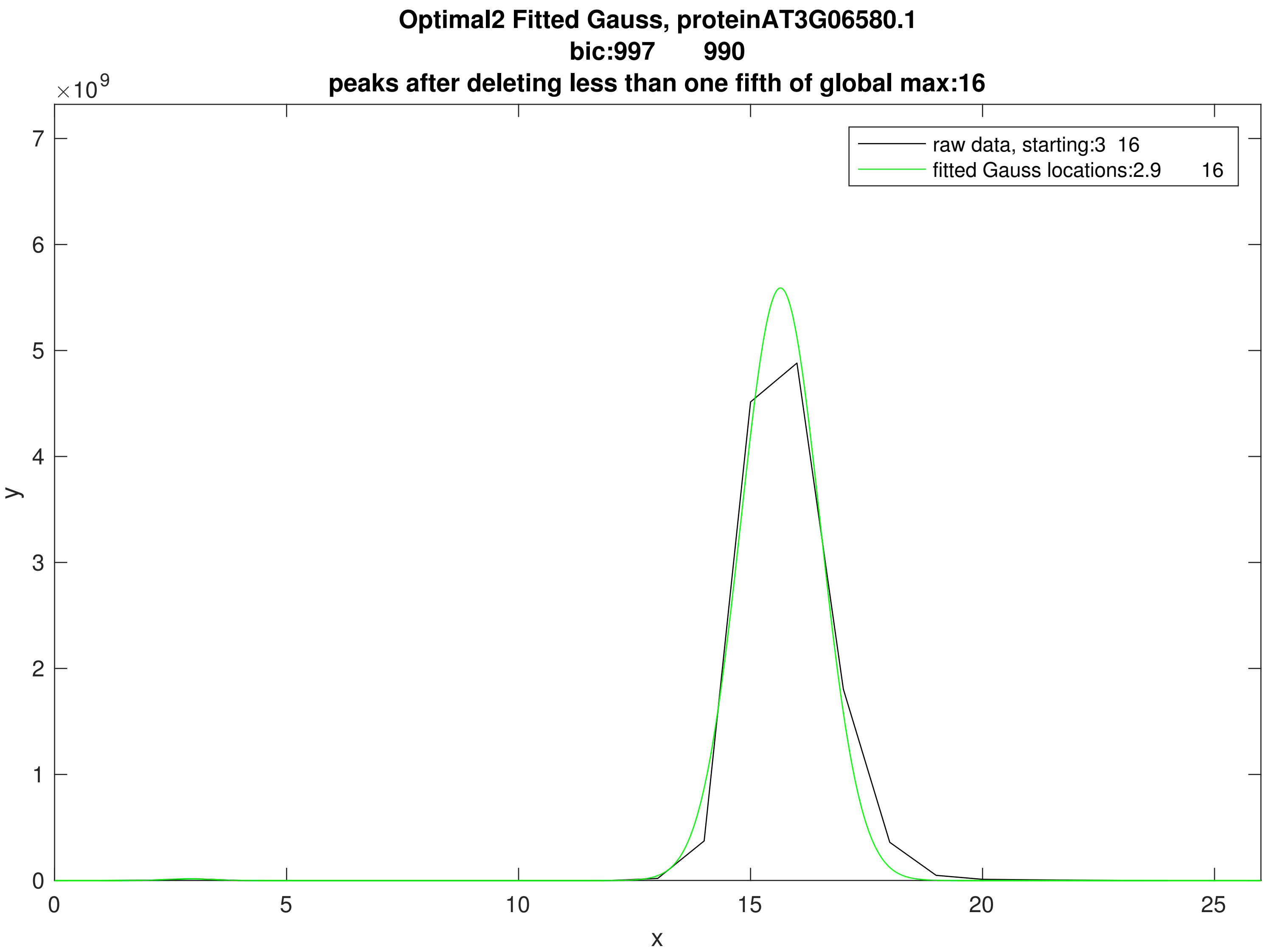

Optimal2 Fitted Gauss, proteinAT3G06580.1
bic:997 990
peaks after deleting less than one fifth of global max:16
×10^9
raw data, starting:3 16
fitted Gauss locations:2.9 16
y
x
0
1
2
3
4
5
6
7
0
5
10
15
20
25

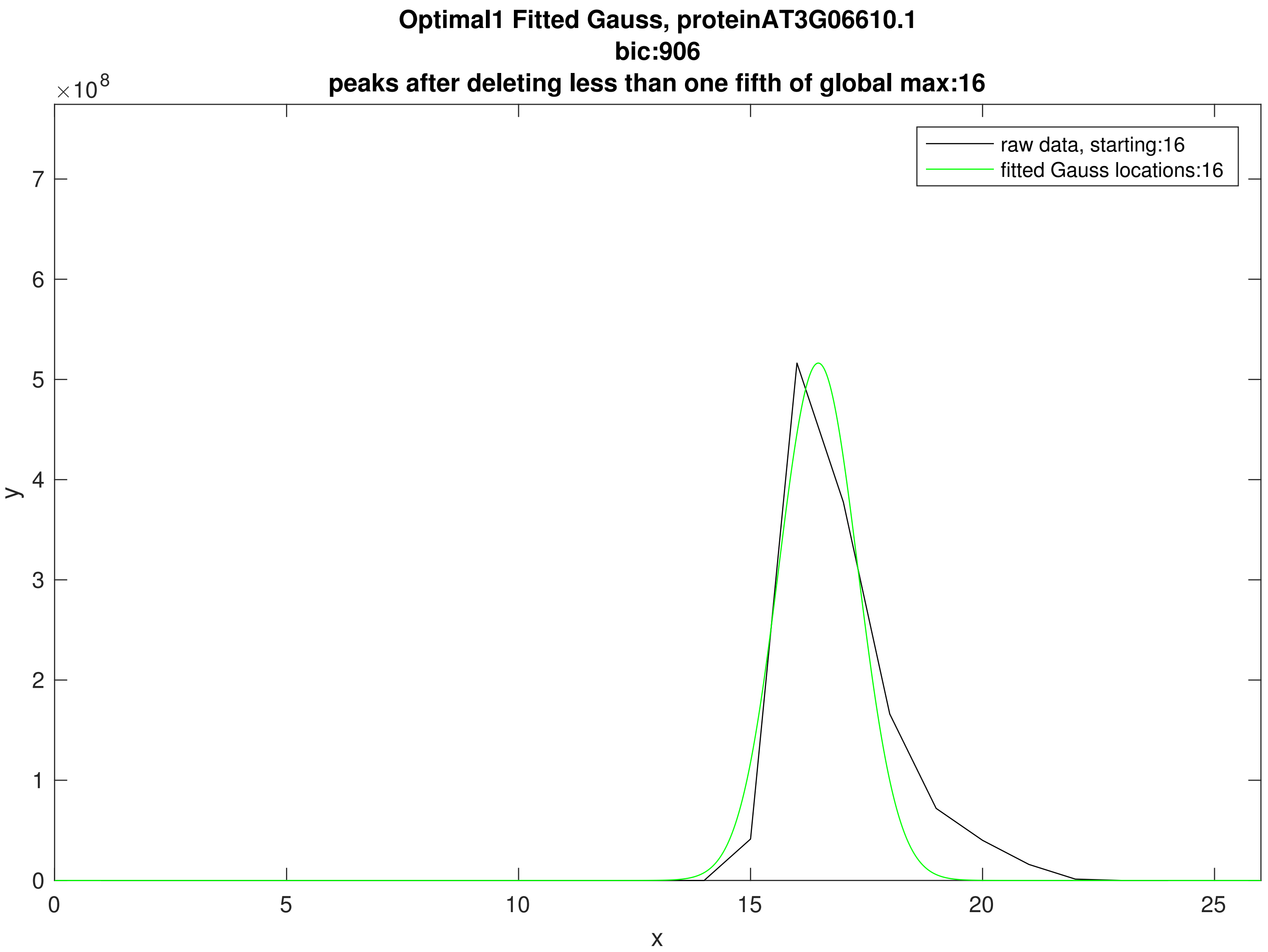

Optimal1 Fitted Gauss, proteinAT3G06610.1
bic:906
peaks after deleting less than one fifth of global max:16
×10^8
raw data, starting:16
fitted Gauss locations:16
y
x
0
1
2
3
4
5
6
7
0
5
10
15
20
25

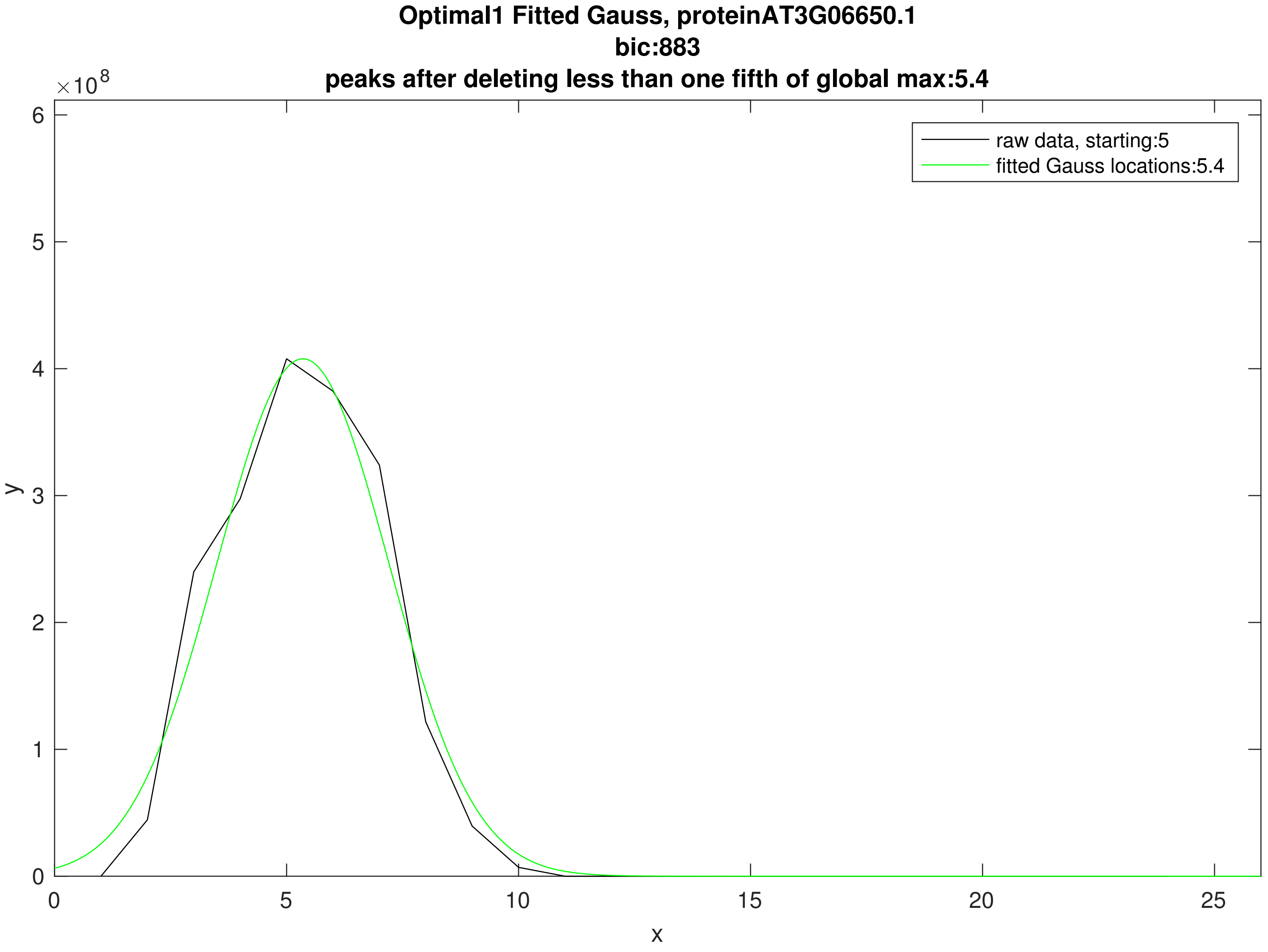

Optimal1 Fitted Gauss, proteinAT3G06650.1
bic:883
peaks after deleting less than one fifth of global max:5.4
×10^8
6
5
4
3
2
1
0
y
0
5
10
15
20
25
x
raw data, starting:5
fitted Gauss locations:5.4

#### Optimal1 Fitted Gauss, proteinAT3G06770.2;AT3G06770.3;AT3G06770.1
#### bic:914
#### peaks after deleting less than one fifth of global max:16

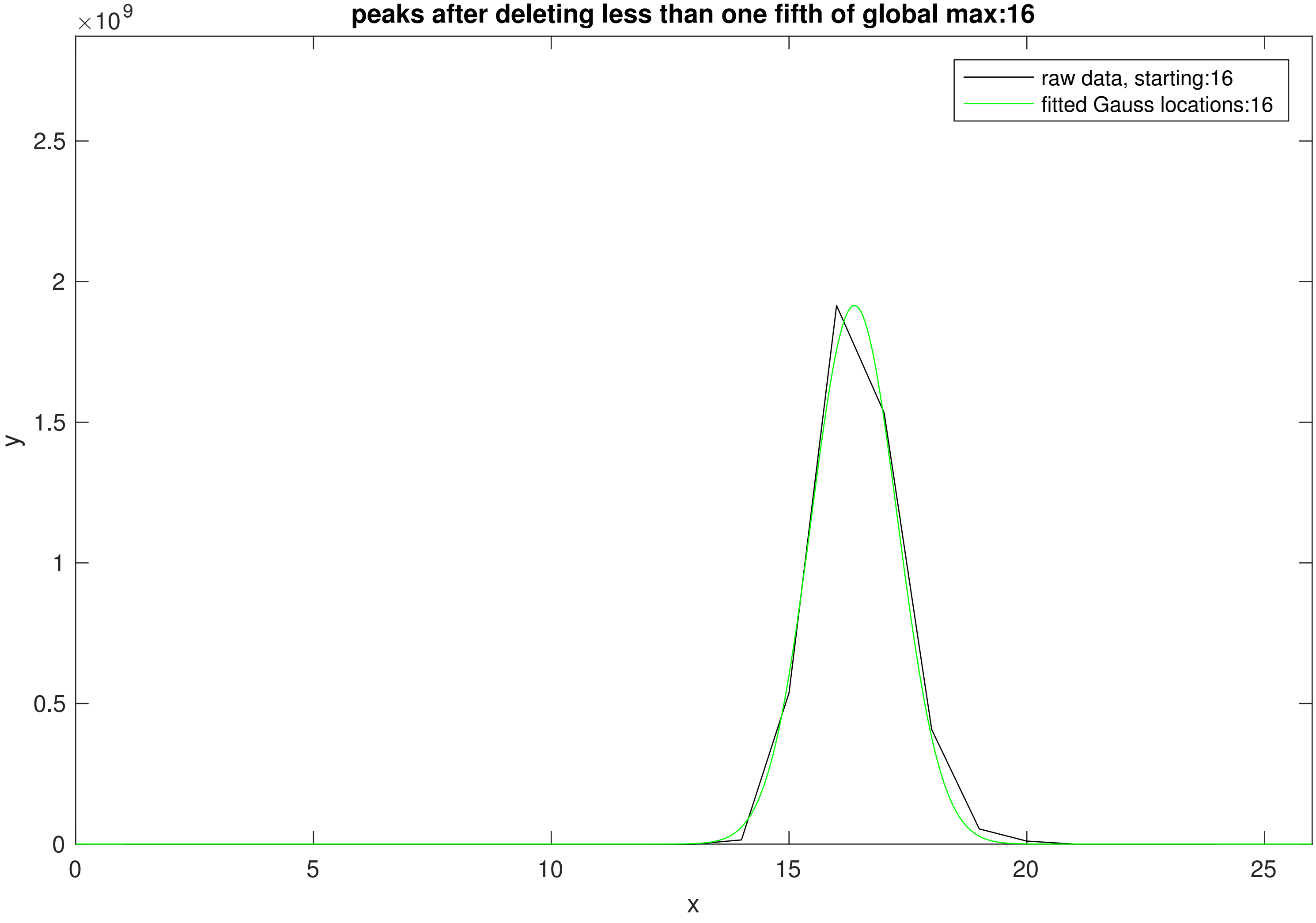

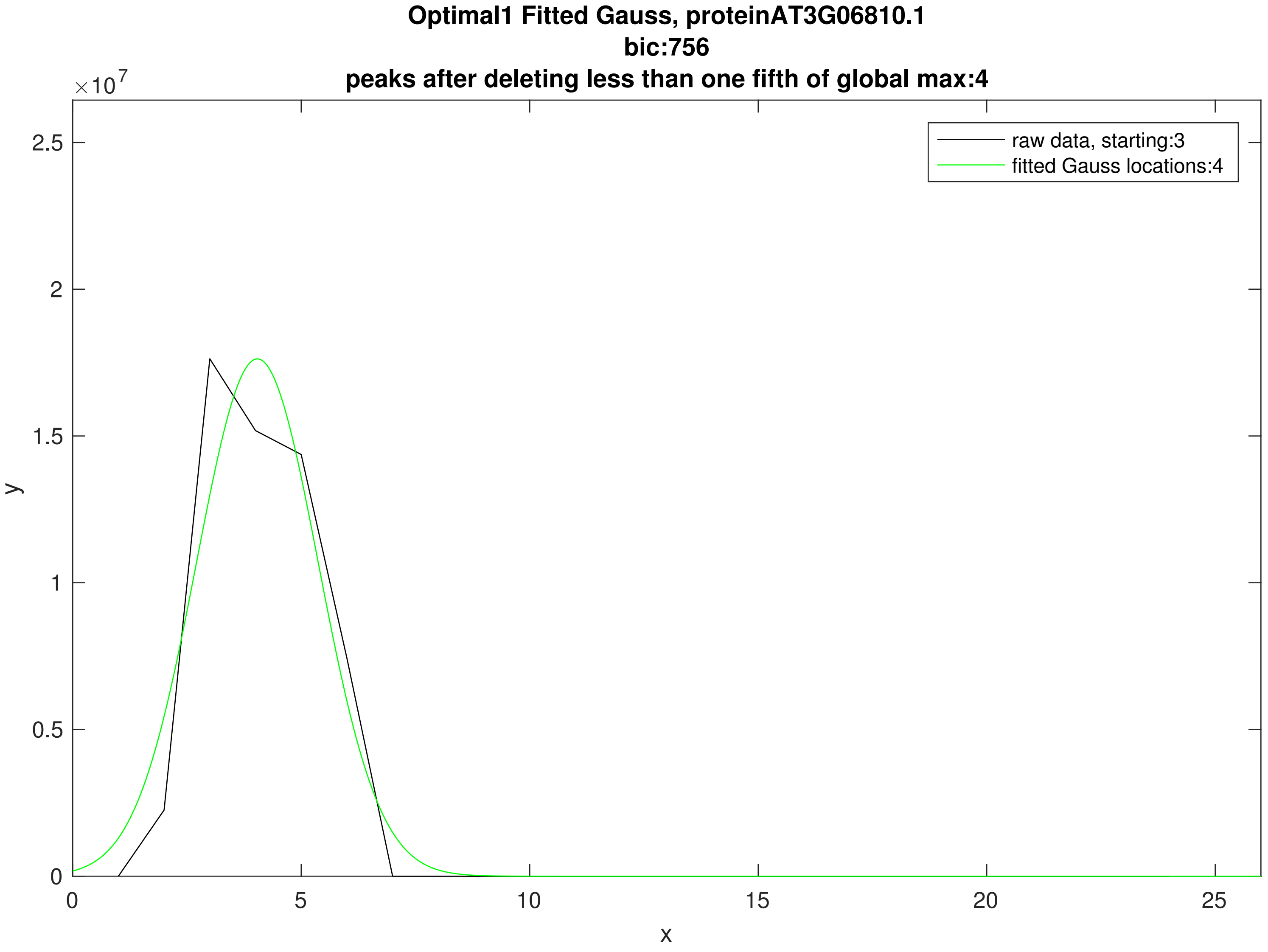

Optimal1 Fitted Gauss, proteinAT3G06810.1
bic:756
peaks after deleting less than one fifth of global max:4
×10^7
2.5
2
1.5
1
0.5
0
y
0
5
10
15
20
25
x
raw data, starting:3
fitted Gauss locations:4

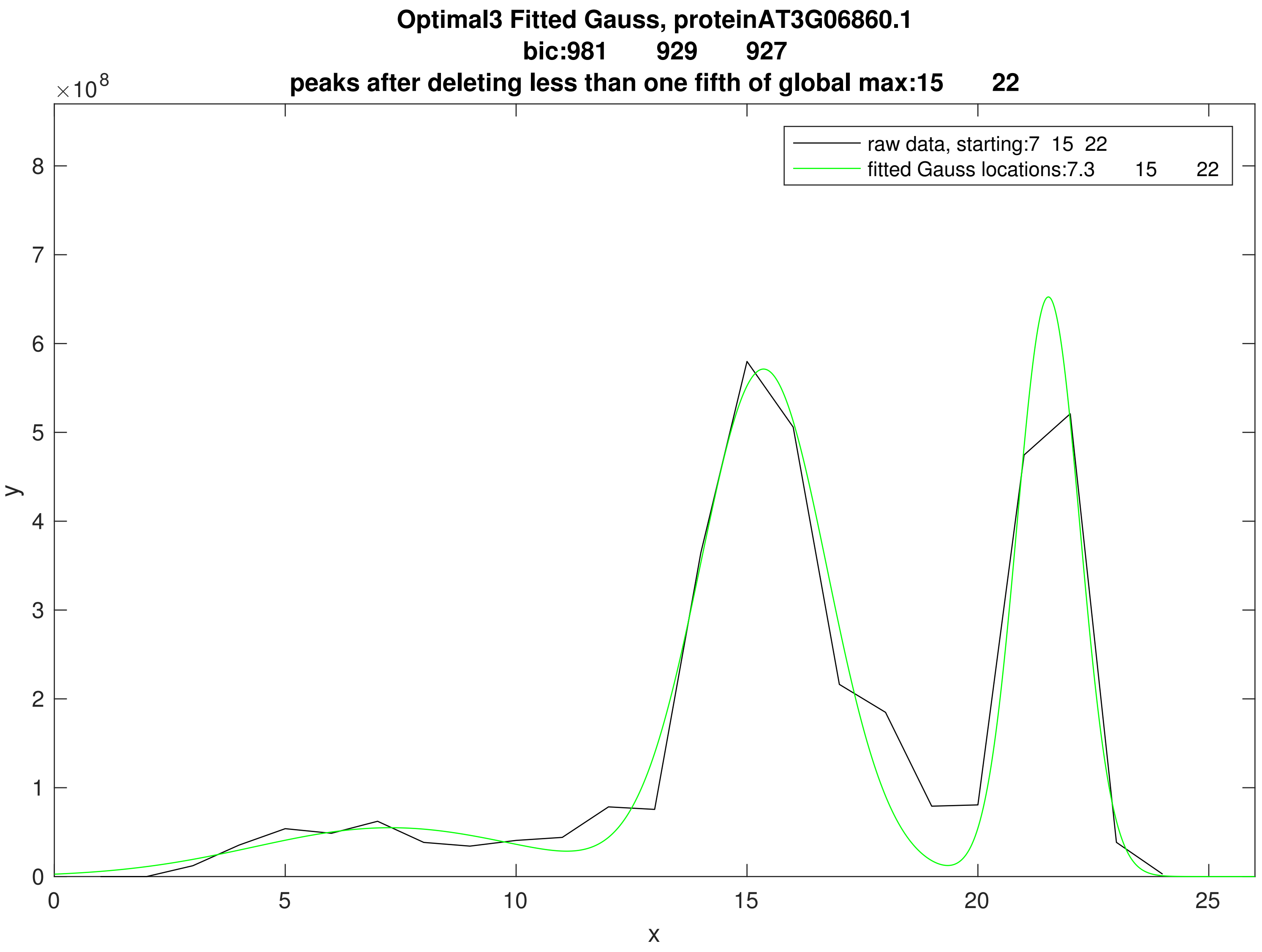

Optimal3 Fitted Gauss, proteinAT3G06860.1
bic:981 929 927
peaks after deleting less than one fifth of global max:15 22
×10^8
raw data, starting:7 15 22
fitted Gauss locations:7.3 15 22
y
x
0
1
2
3
4
5
6
7
8
0
5
10
15
20
25

Optimal2 Fitted Gauss, proteinAT3G06930.1;AT3G06930.2
bic:814 806
peaks after deleting less than one fifth of global max:6.4

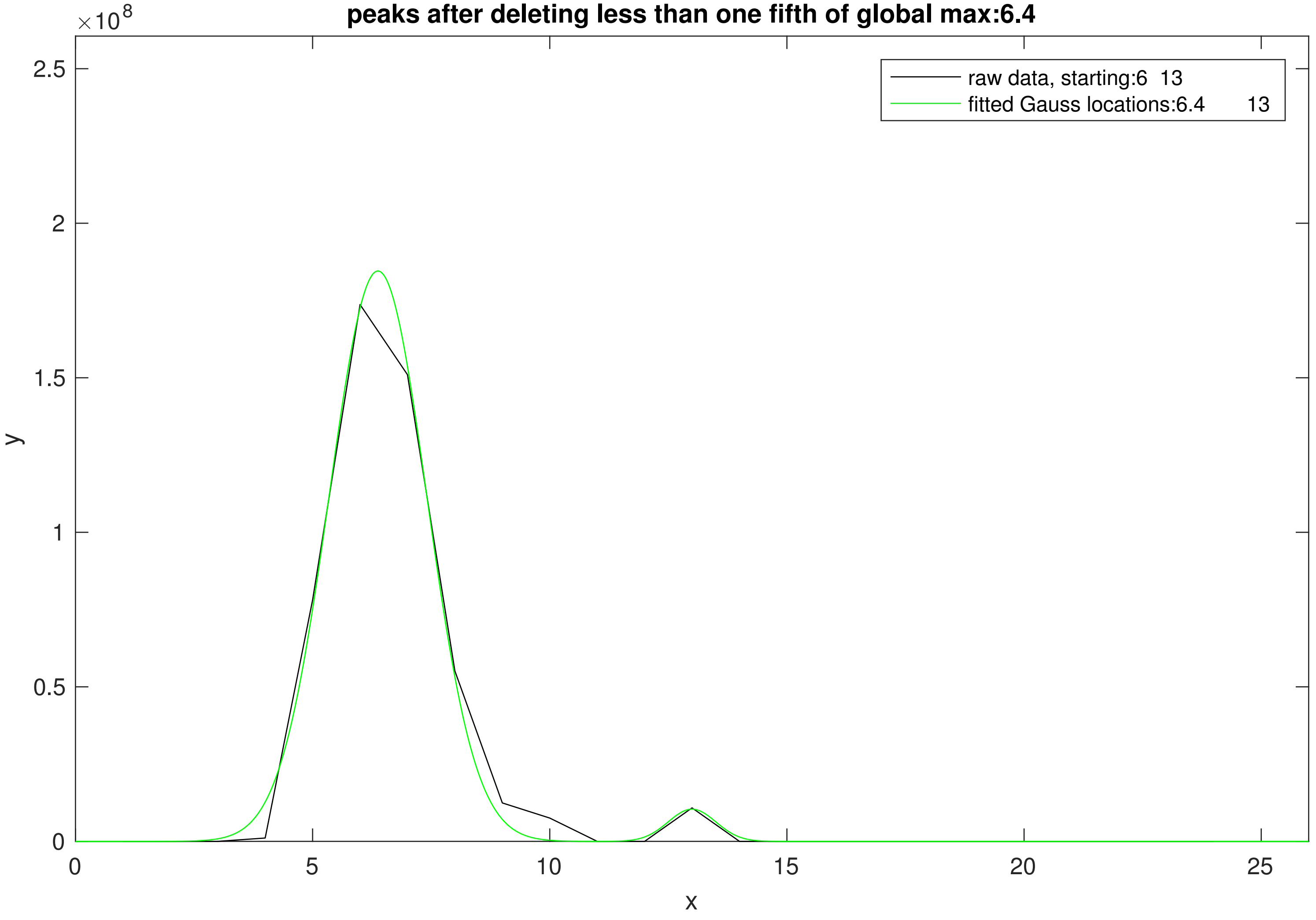

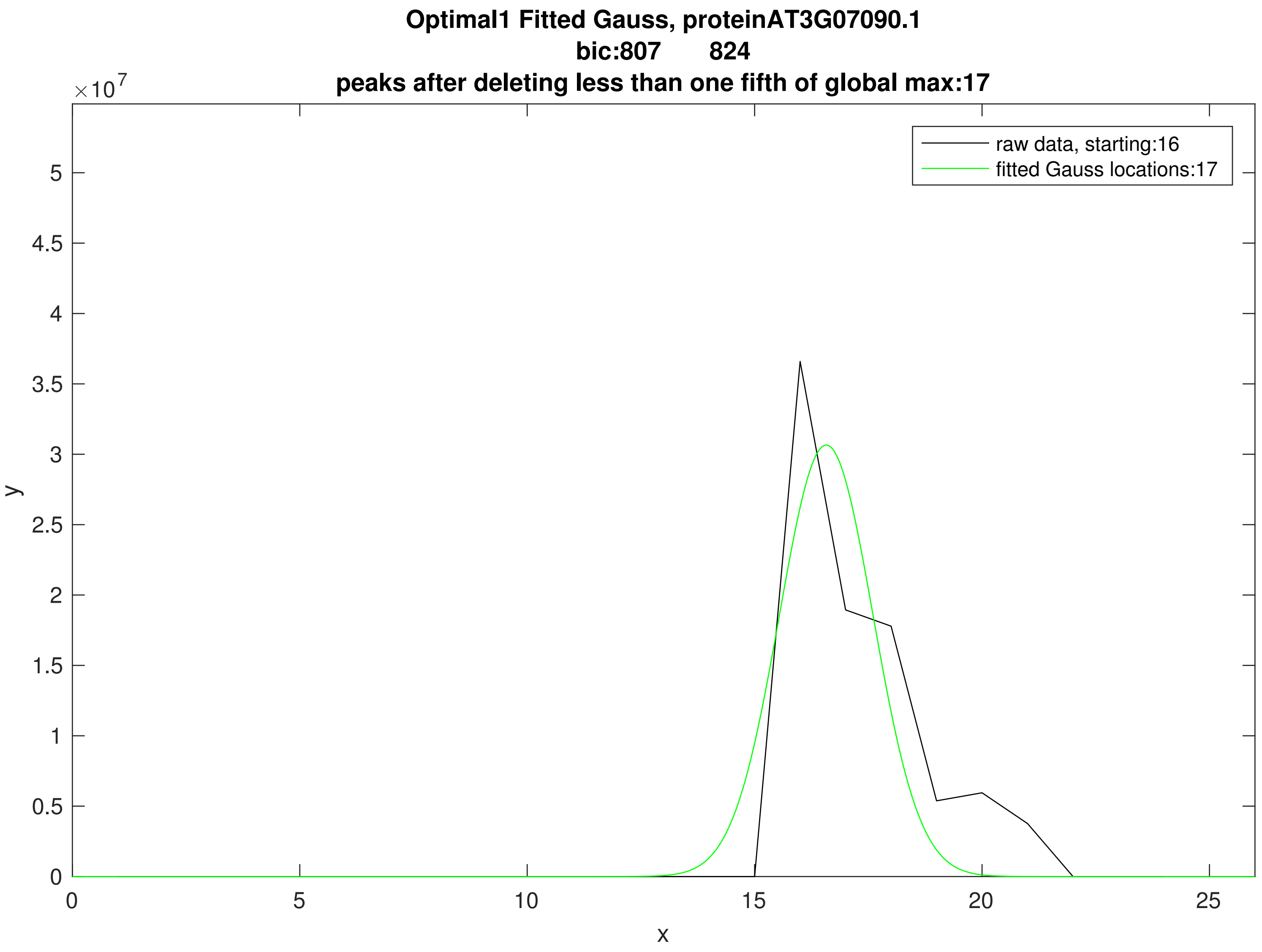

Optimal1 Fitted Gauss, proteinAT3G07090.1
bic:807 824
peaks after deleting less than one fifth of global max:17
×10^7
raw data, starting:16
fitted Gauss locations:17
y
x
0
0.5
1
1.5
2
2.5
3
3.5
4
4.5
5
0
5
10
15
20
25

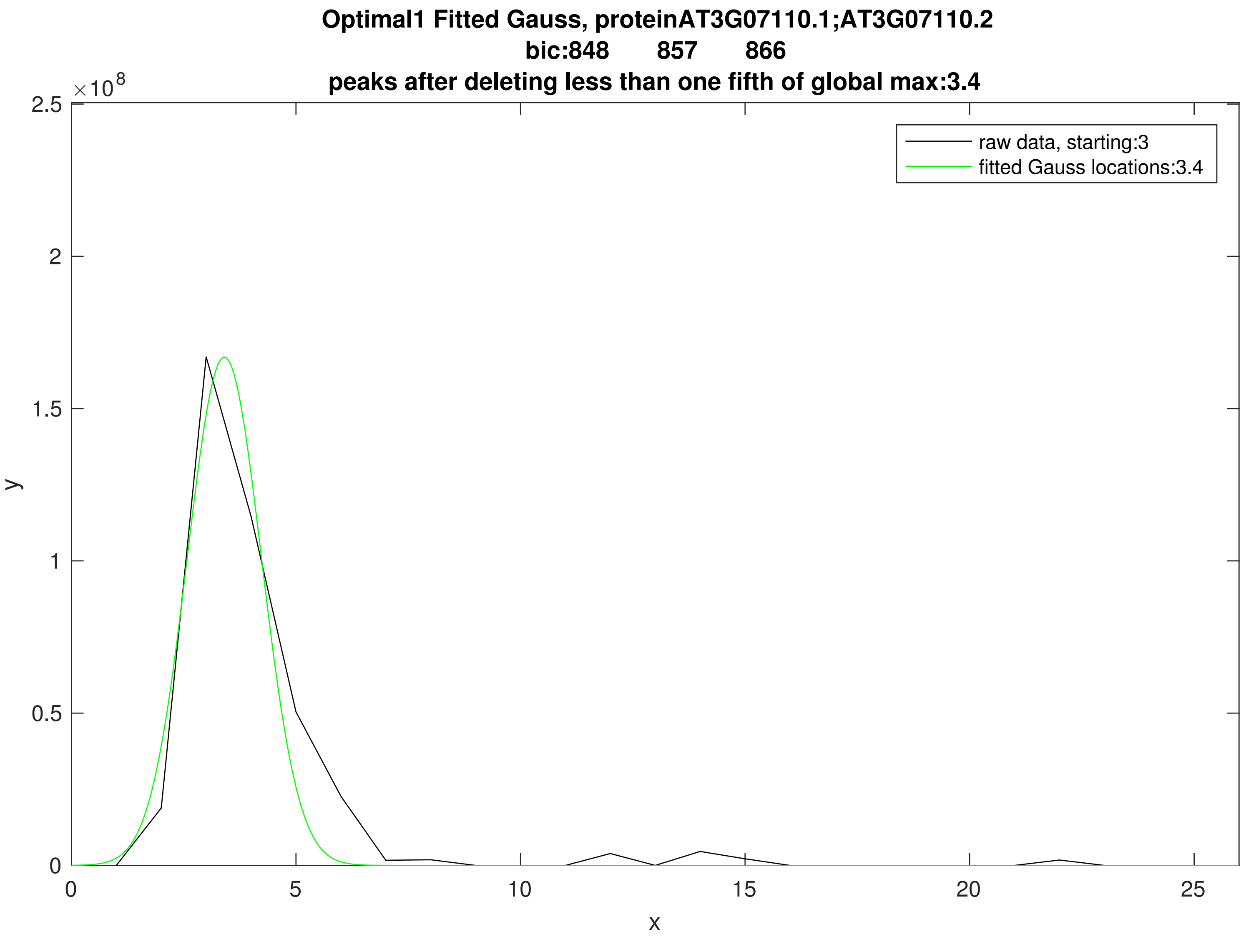
Optimal1 Fitted Gauss, proteinAT3G07110.1;AT3G07110.2
bic:848 857 866
peaks after deleting less than one fifth of global max:3.4
×10^8
2.5
2
1.5
1
0.5
0
y
0
5
10
15
20
25
x
raw data, starting:3
fitted Gauss locations:3.4

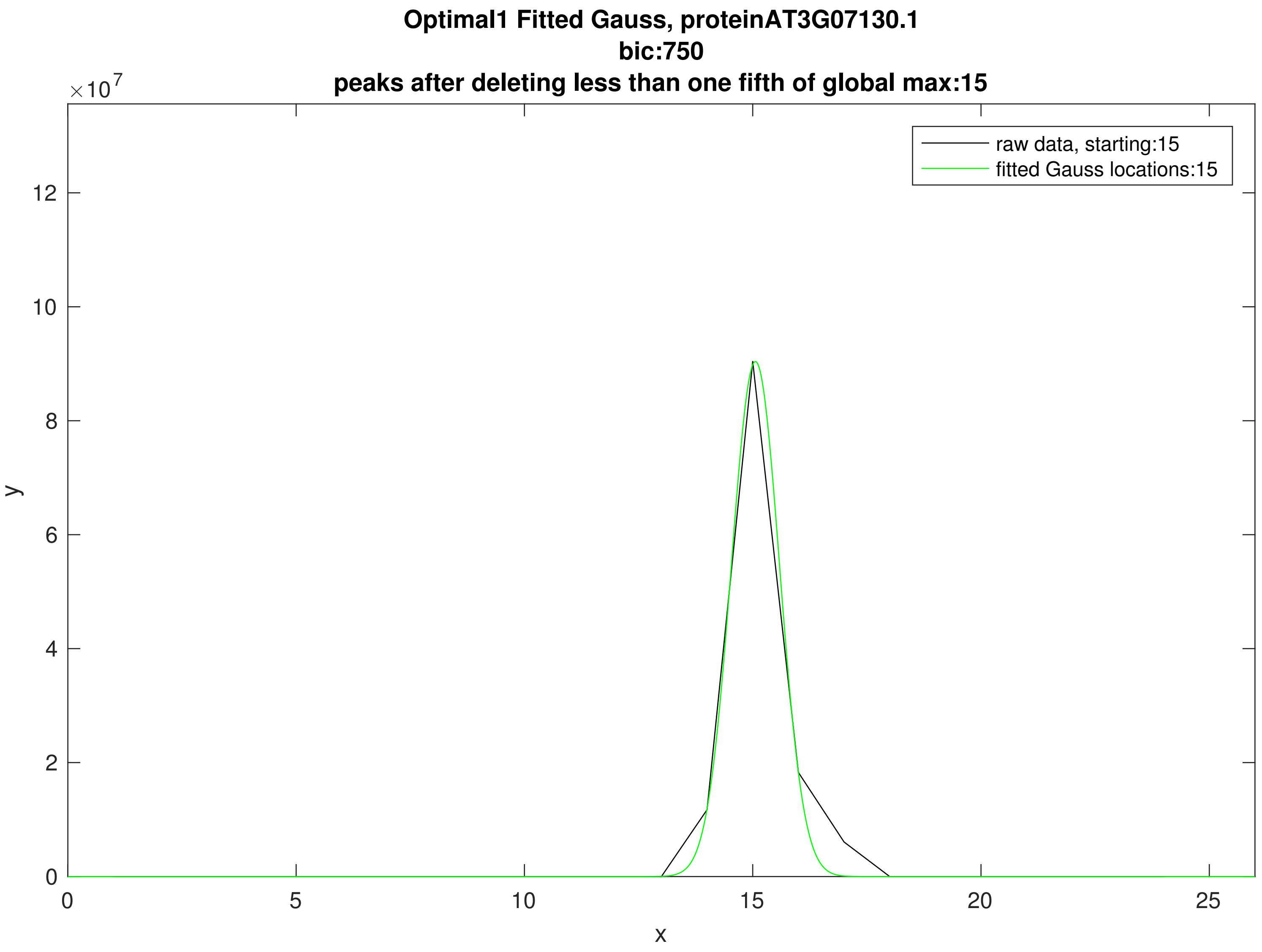
Optimal1 Fitted Gauss, proteinAT3G07130.1
bic:750
peaks after deleting less than one fifth of global max:15
×10^7
raw data, starting:15
fitted Gauss locations:15
12
10
8
6
4
2
0
y
0
5
10
15
20
25
x

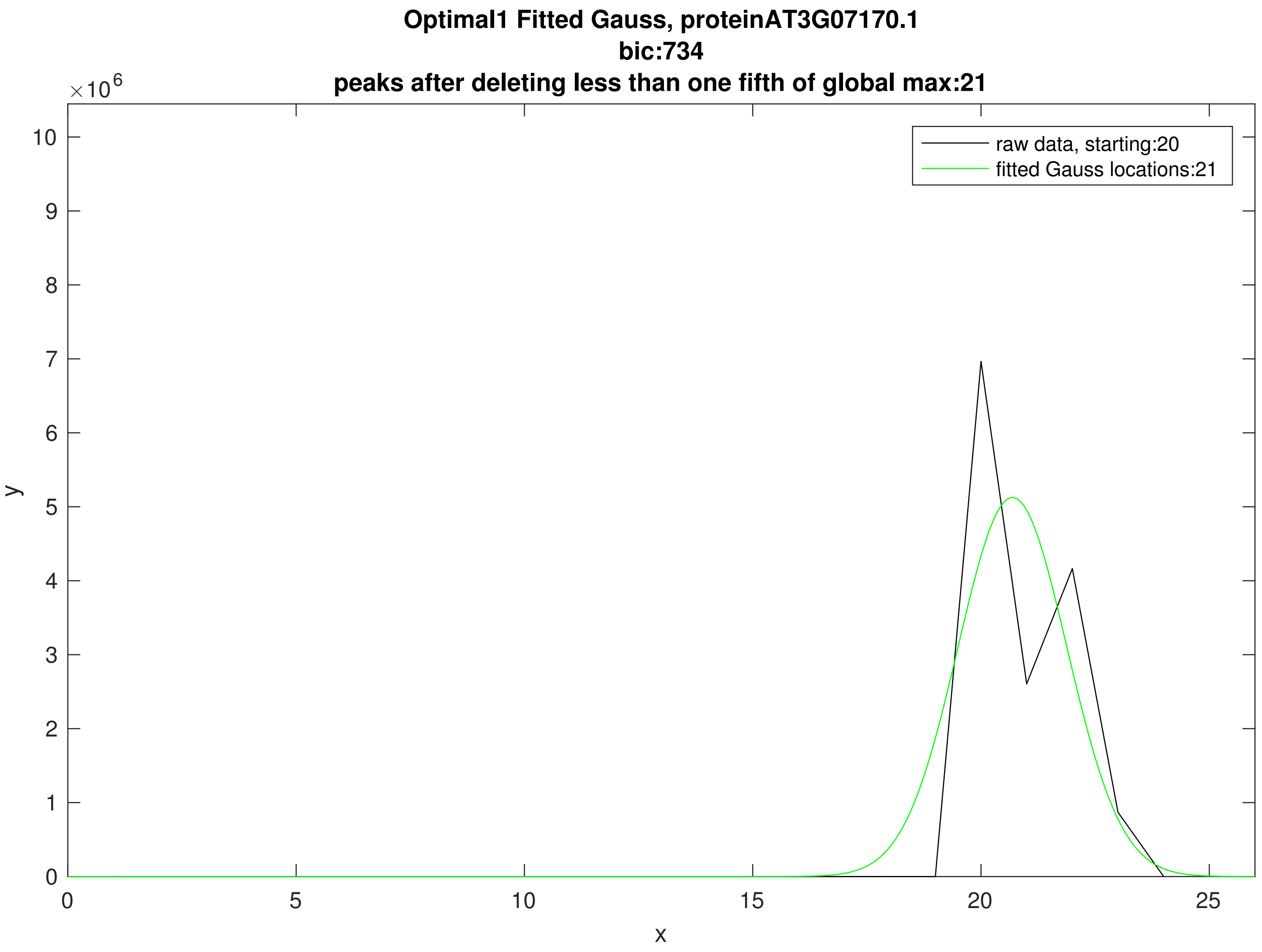
Optimal1 Fitted Gauss, proteinAT3G07170.1
bic:734
peaks after deleting less than one fifth of global max:21
×10^6
raw data, starting:20
fitted Gauss locations:21
y
x
0
1
2
3
4
5
6
7
8
9
10
0
5
10
15
20
25

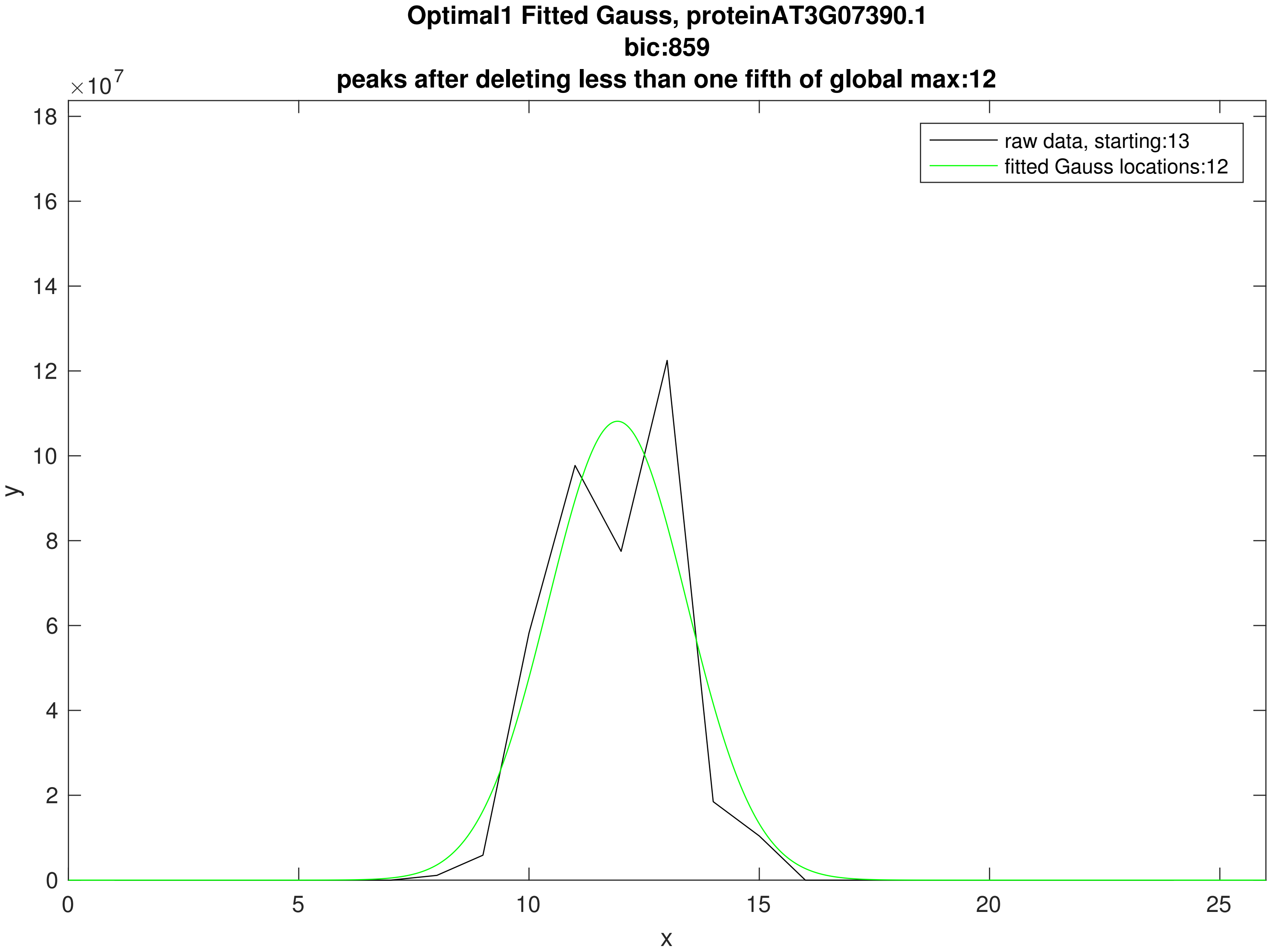
Optimal1 Fitted Gauss, proteinAT3G07390.1
bic:859
peaks after deleting less than one fifth of global max:12
×10^7
raw data, starting:13
fitted Gauss locations:12
y
x
0
2
4
6
8
10
12
14
16
18
0
5
10
15
20
25

**Optimal1 Fitted Gauss, proteinAT3G07460.1;AT3G07460.2**
**bic:858 863 872**
**peaks after deleting less than one fifth of global max:20**

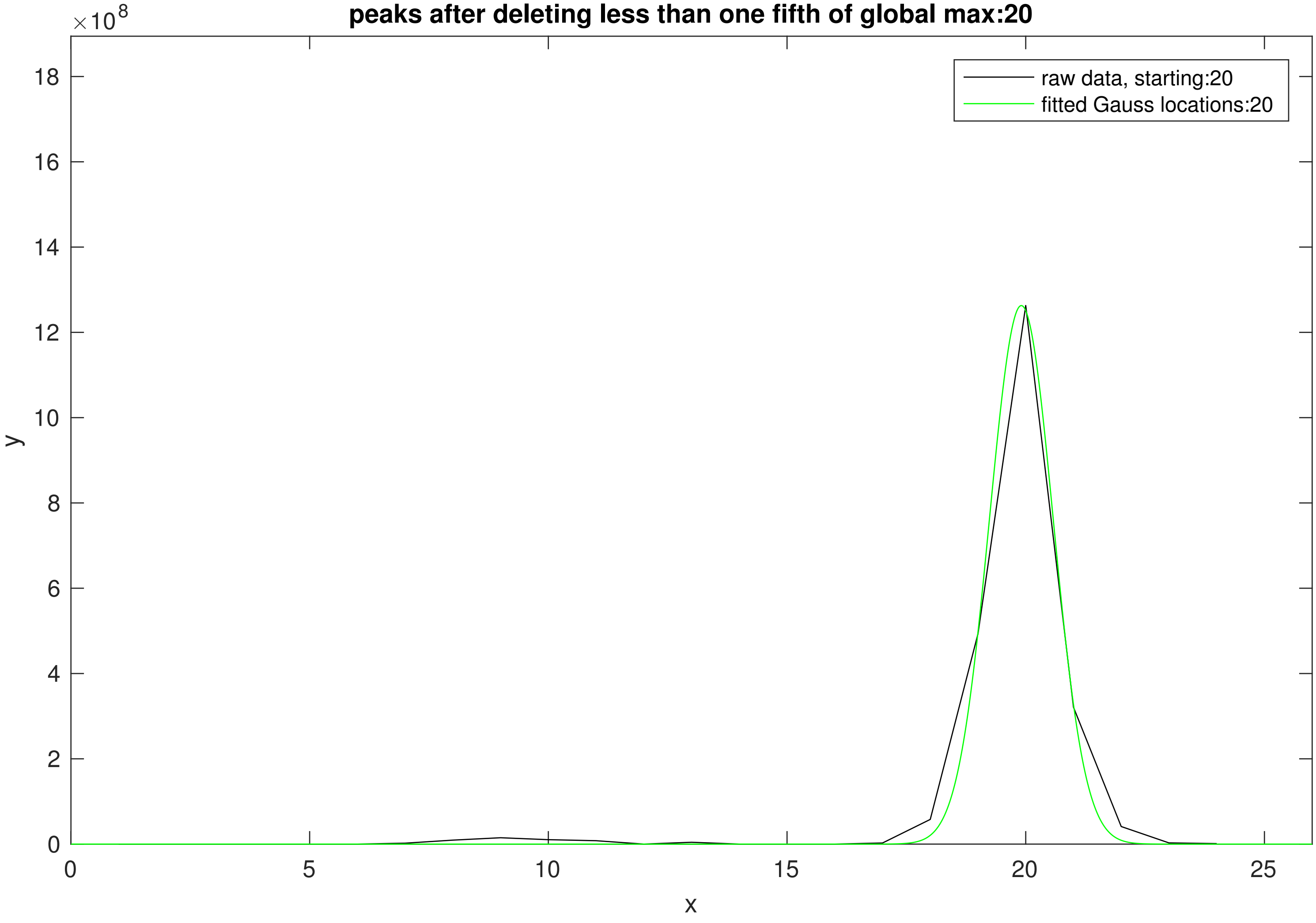

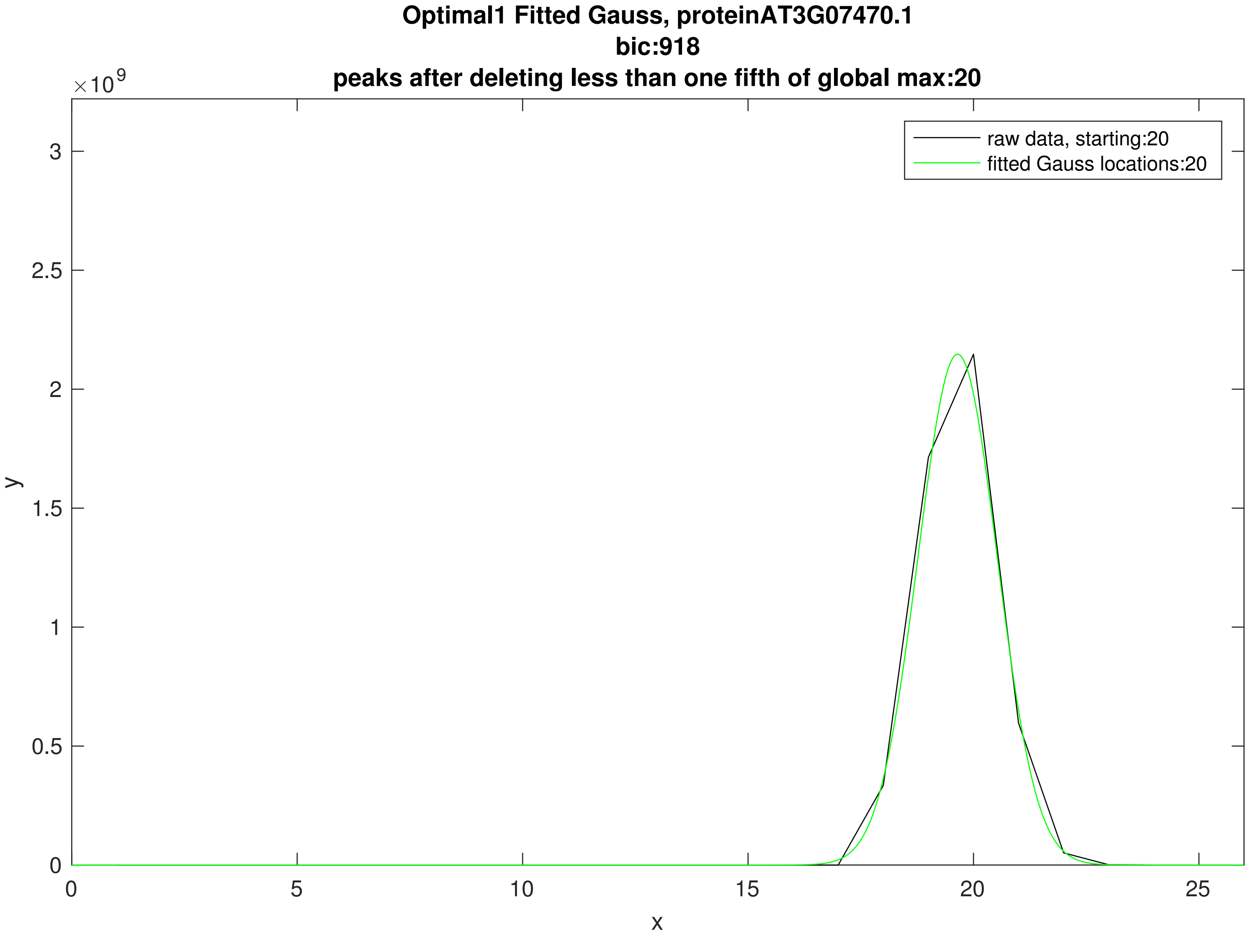
Optimal1 Fitted Gauss, proteinAT3G07470.1
bic:918
peaks after deleting less than one fifth of global max:20
×10^9
raw data, starting:20
fitted Gauss locations:20
y
x
0
0.5
1
1.5
2
2.5
3
0
5
10
15
20
25

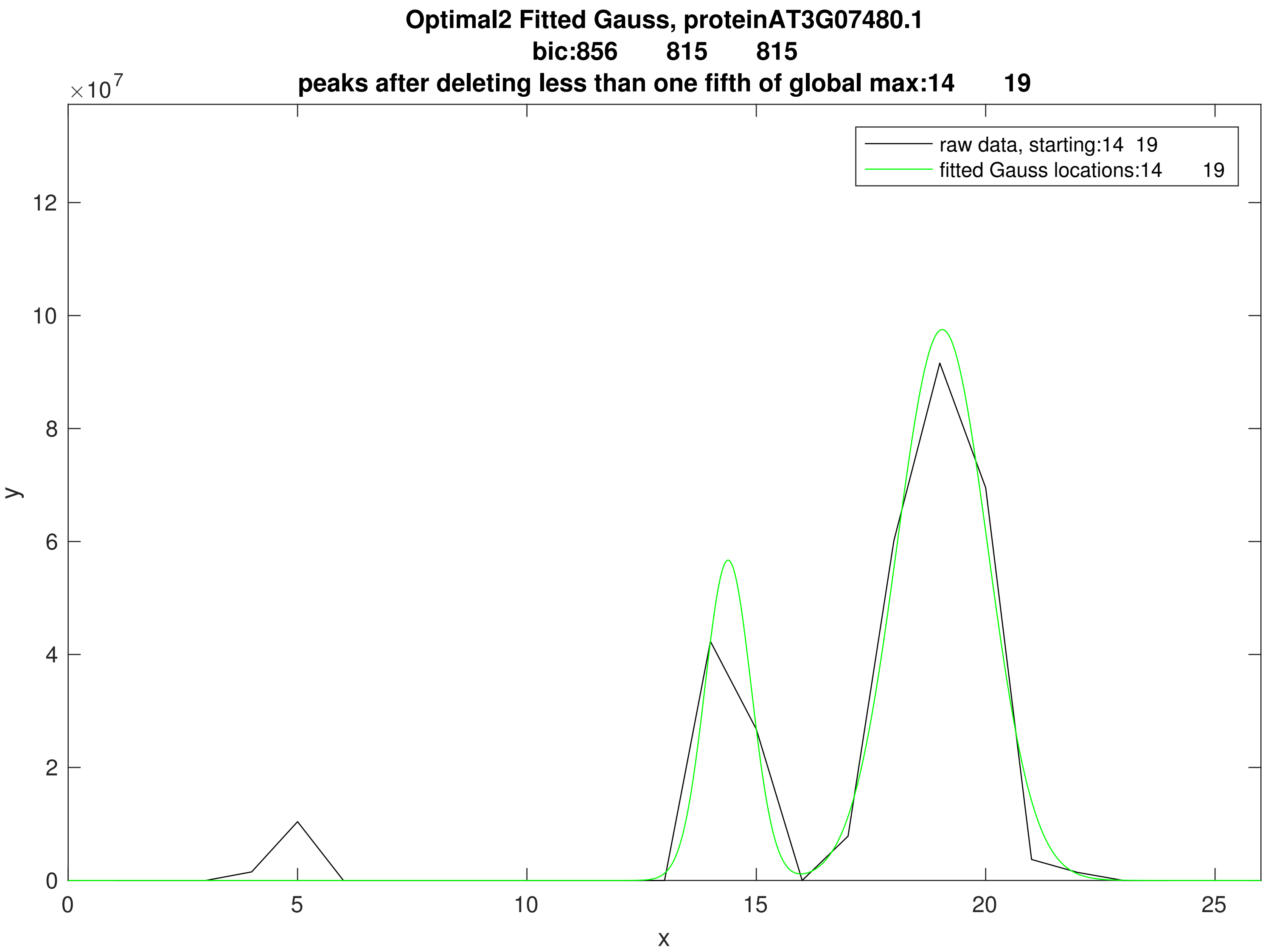
Optimal2 Fitted Gauss, proteinAT3G07480.1
bic:856 815 815
peaks after deleting less than one fifth of global max:14 19
×10^7
raw data, starting:14 19
fitted Gauss locations:14 19
y
x
0
2
4
6
8
10
12
0
5
10
15
20
25

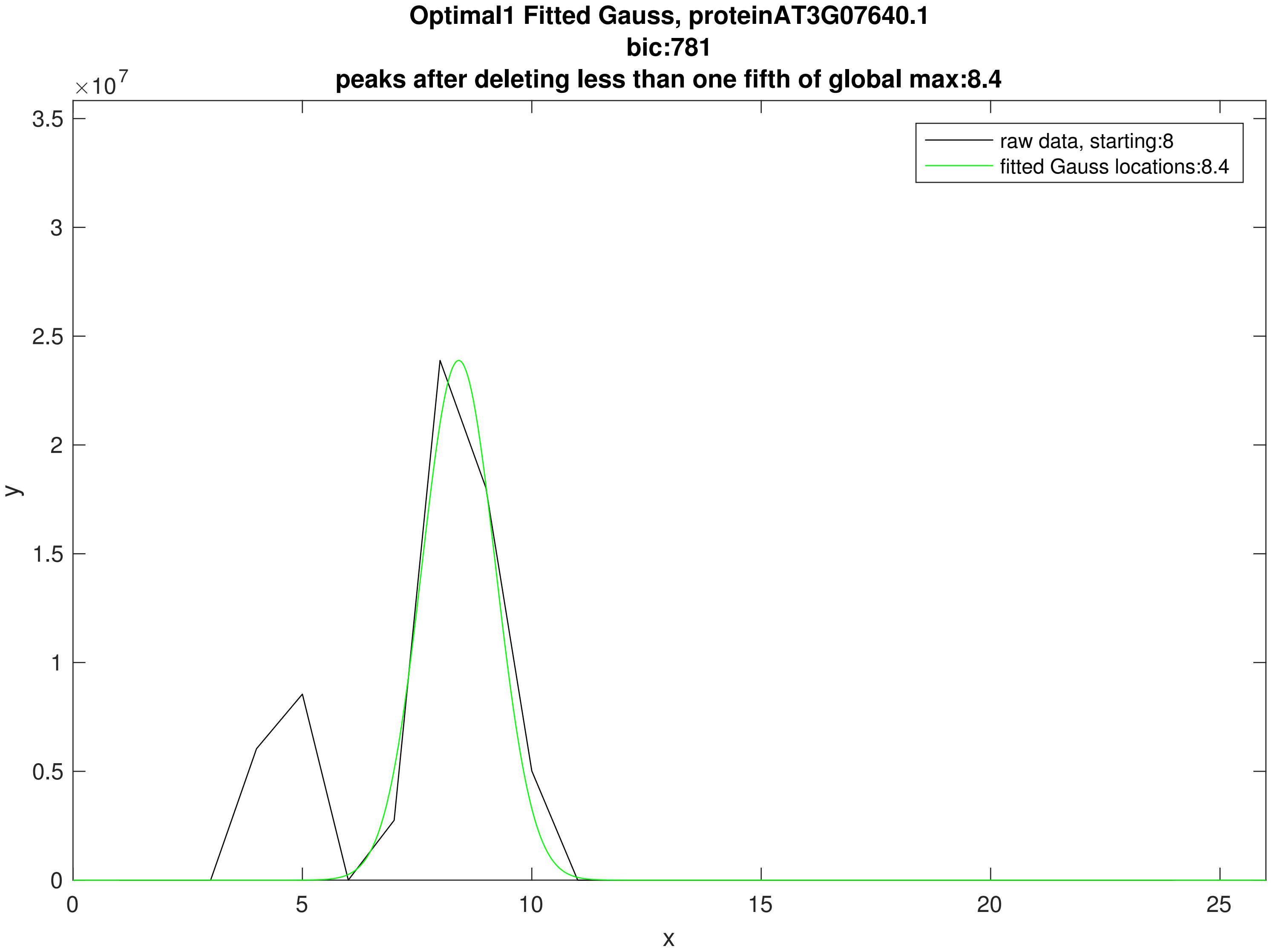
Optimal1 Fitted Gauss, proteinAT3G07640.1
bic:781
peaks after deleting less than one fifth of global max:8.4
×10^7
3.5
3
2.5
2
1.5
1
0.5
0
y
0
5
10
15
20
25
x
raw data, starting:8
fitted Gauss locations:8.4

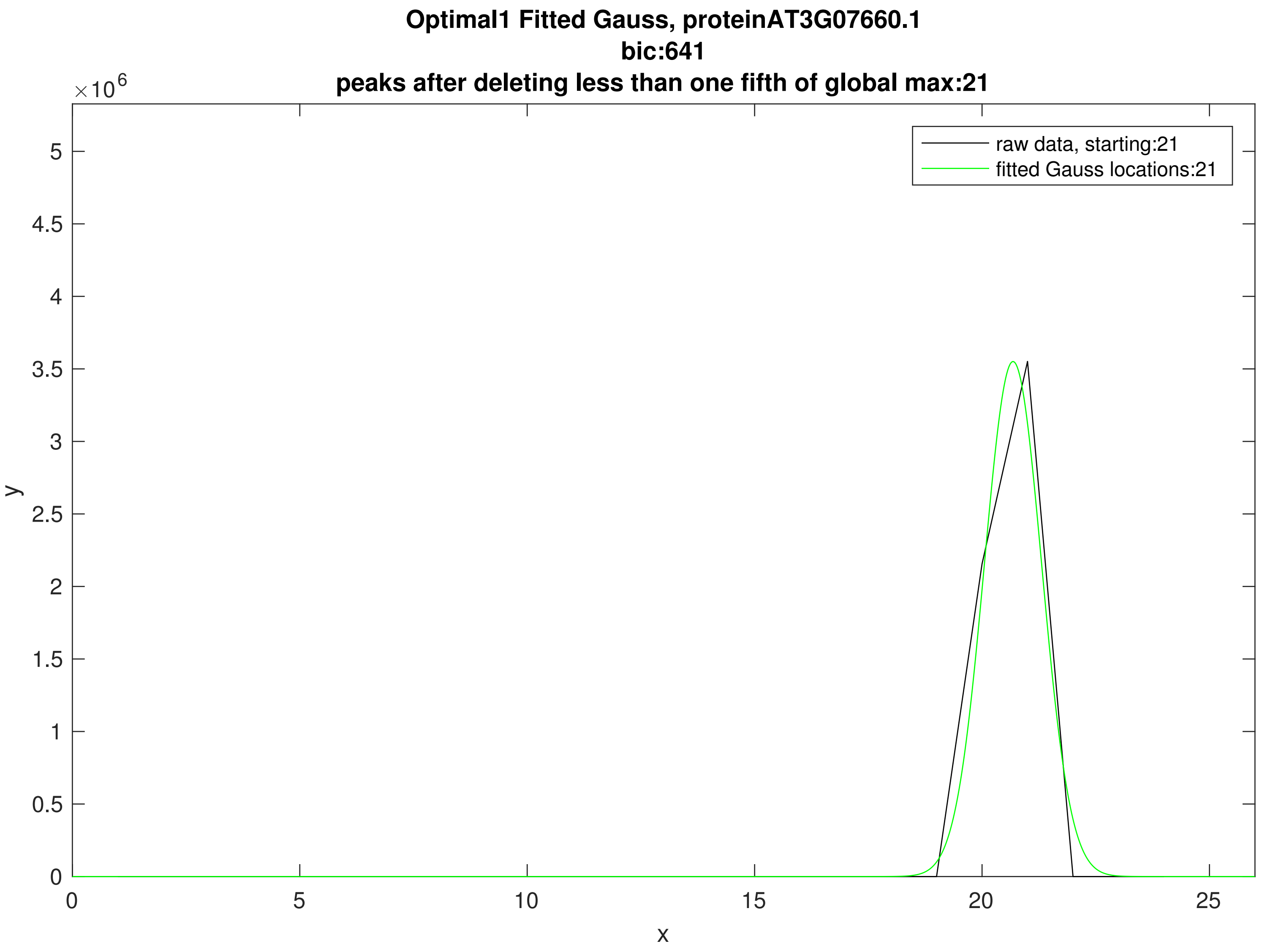
Optimal1 Fitted Gauss, proteinAT3G07660.1
bic:641
peaks after deleting less than one fifth of global max:21
×10^6
raw data, starting:21
fitted Gauss locations:21
y
x
0
0.5
1
1.5
2
2.5
3
3.5
4
4.5
5
0
5
10
15
20
25

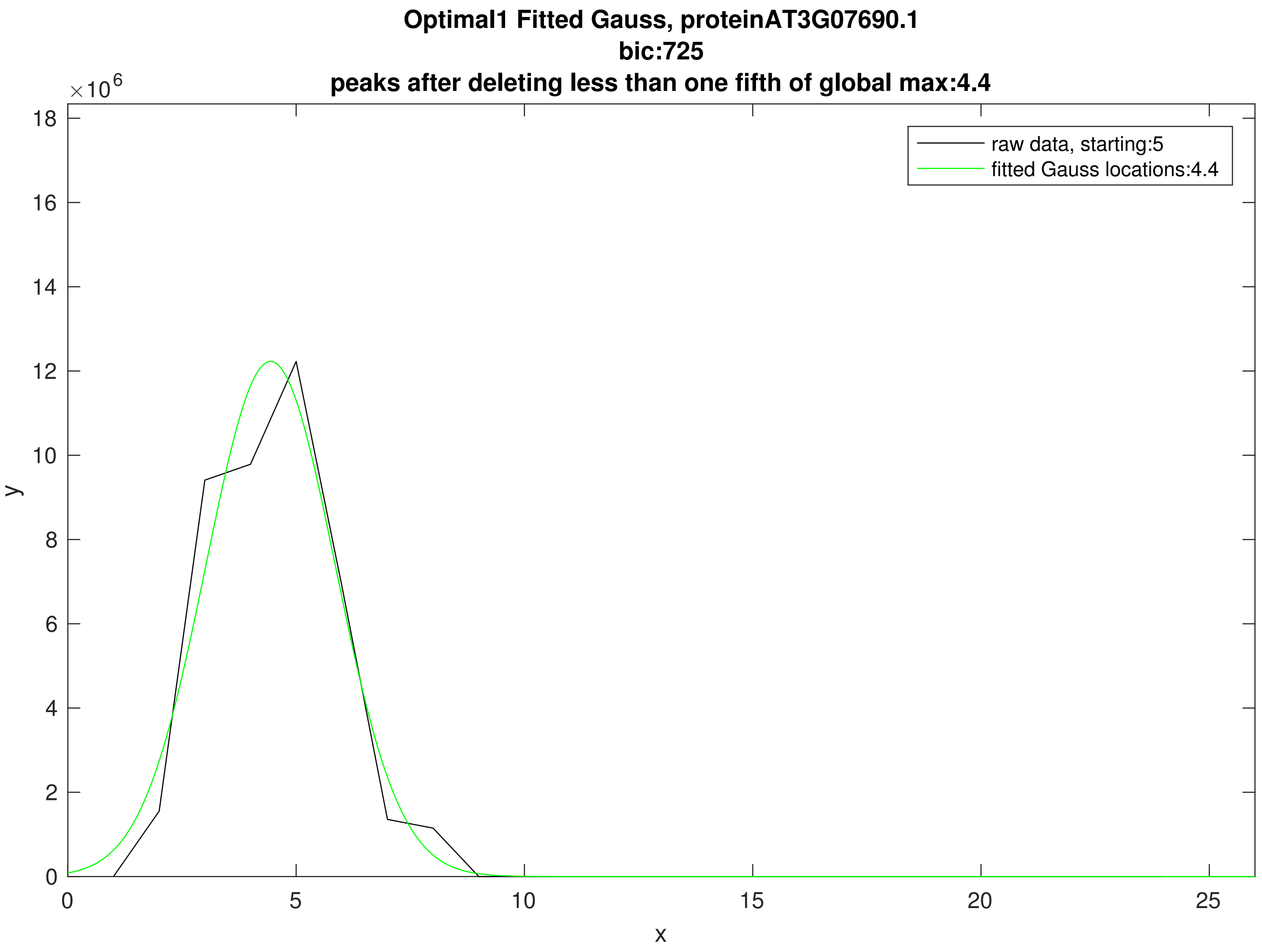
Optimal1 Fitted Gauss, proteinAT3G07690.1
bic:725
peaks after deleting less than one fifth of global max:4.4
×10^6
18
16
14
12
10
8
6
4
2
0
y
0
5
10
15
20
25
x
raw data, starting:5
fitted Gauss locations:4.4

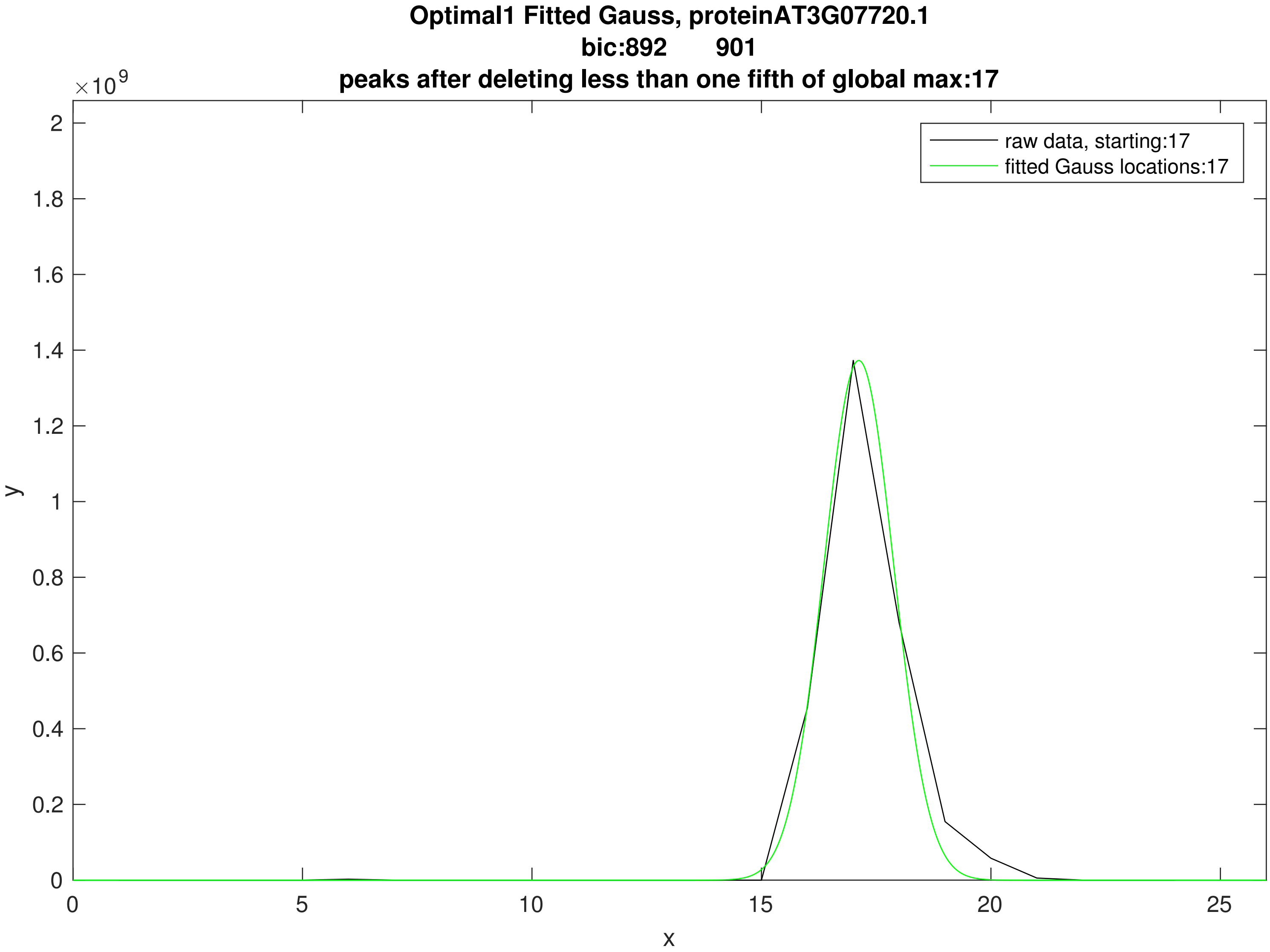
Optimal1 Fitted Gauss, proteinAT3G07720.1
bic:892 901
peaks after deleting less than one fifth of global max:17
×10^9
raw data, starting:17
fitted Gauss locations:17
y
x
0
0.2
0.4
0.6
0.8
1
1.2
1.4
1.6
1.8
2
5
10
15
20
25

**Optimal1 Fitted Gauss, proteinAT3G07770.1**
**bic:817 824**
**peaks after deleting less than one fifth of global max:11**

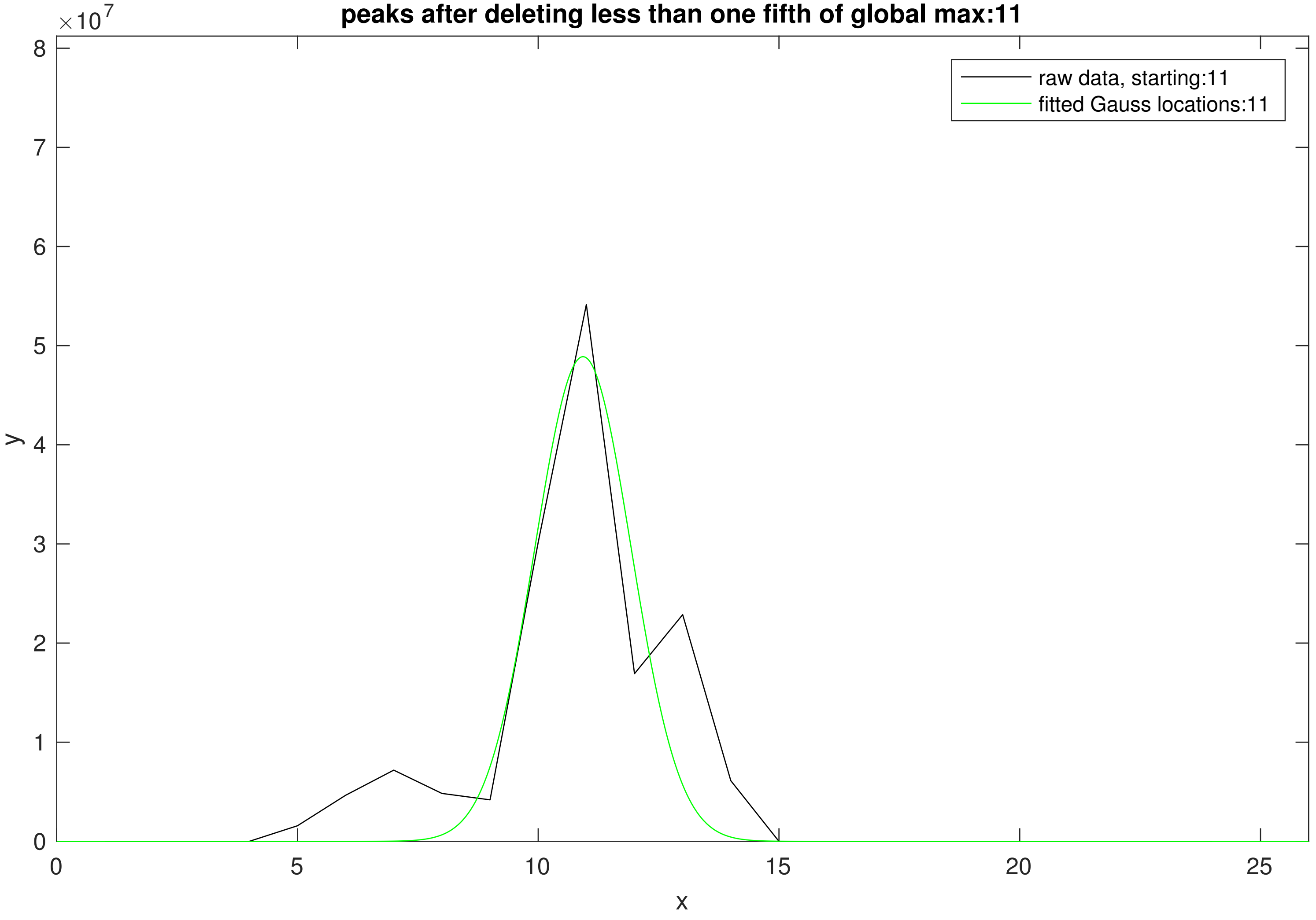

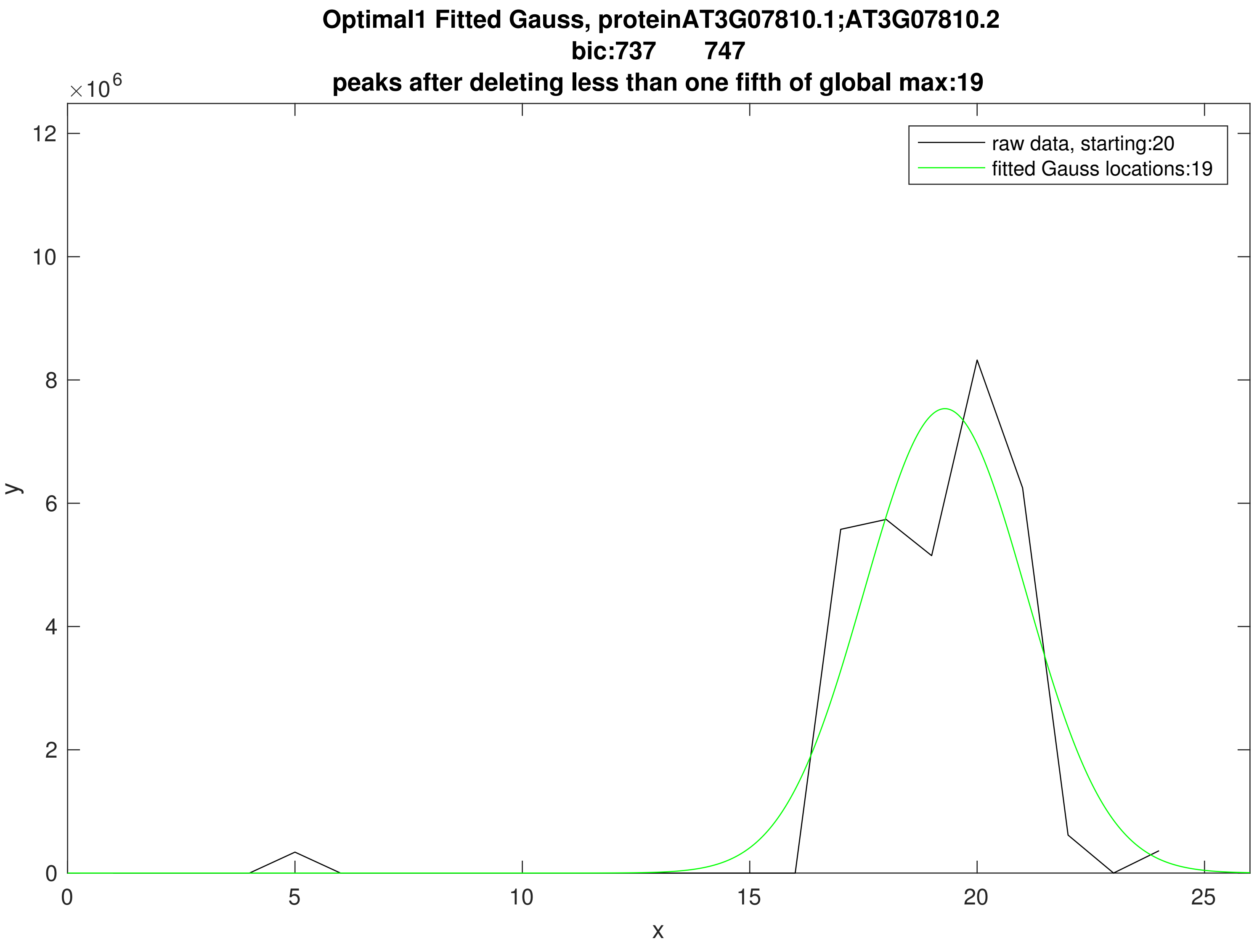

Optimal1 Fitted Gauss, proteinAT3G07810.1;AT3G07810.2
bic:737 747
peaks after deleting less than one fifth of global max:19
×10^6
raw data, starting:20
fitted Gauss locations:19
y
x
0
2
4
6
8
10
12
0
5
10
15
20
25

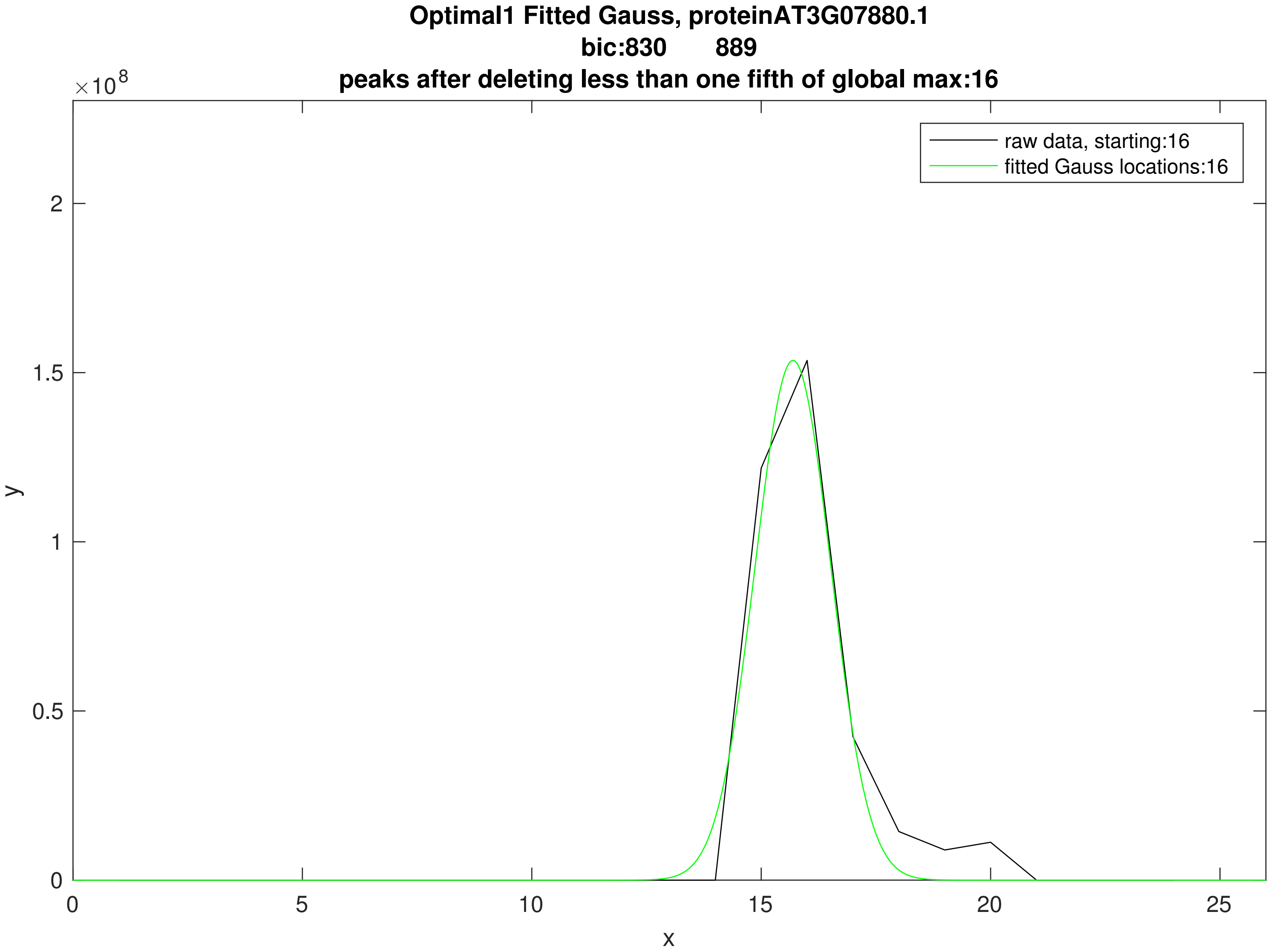

Optimal1 Fitted Gauss, proteinAT3G07880.1
bic:830 889
peaks after deleting less than one fifth of global max:16
×10^8
raw data, starting:16
fitted Gauss locations:16
y
x
0
0.5
1
1.5
2
0
5
10
15
20
25

Optimal1 Fitted Gauss, proteinAT3G08030.1;AT3G08030.2
bic:1.05e+03 1.06e+03 1.07e+03
peaks after deleting less than one fifth of global max:17

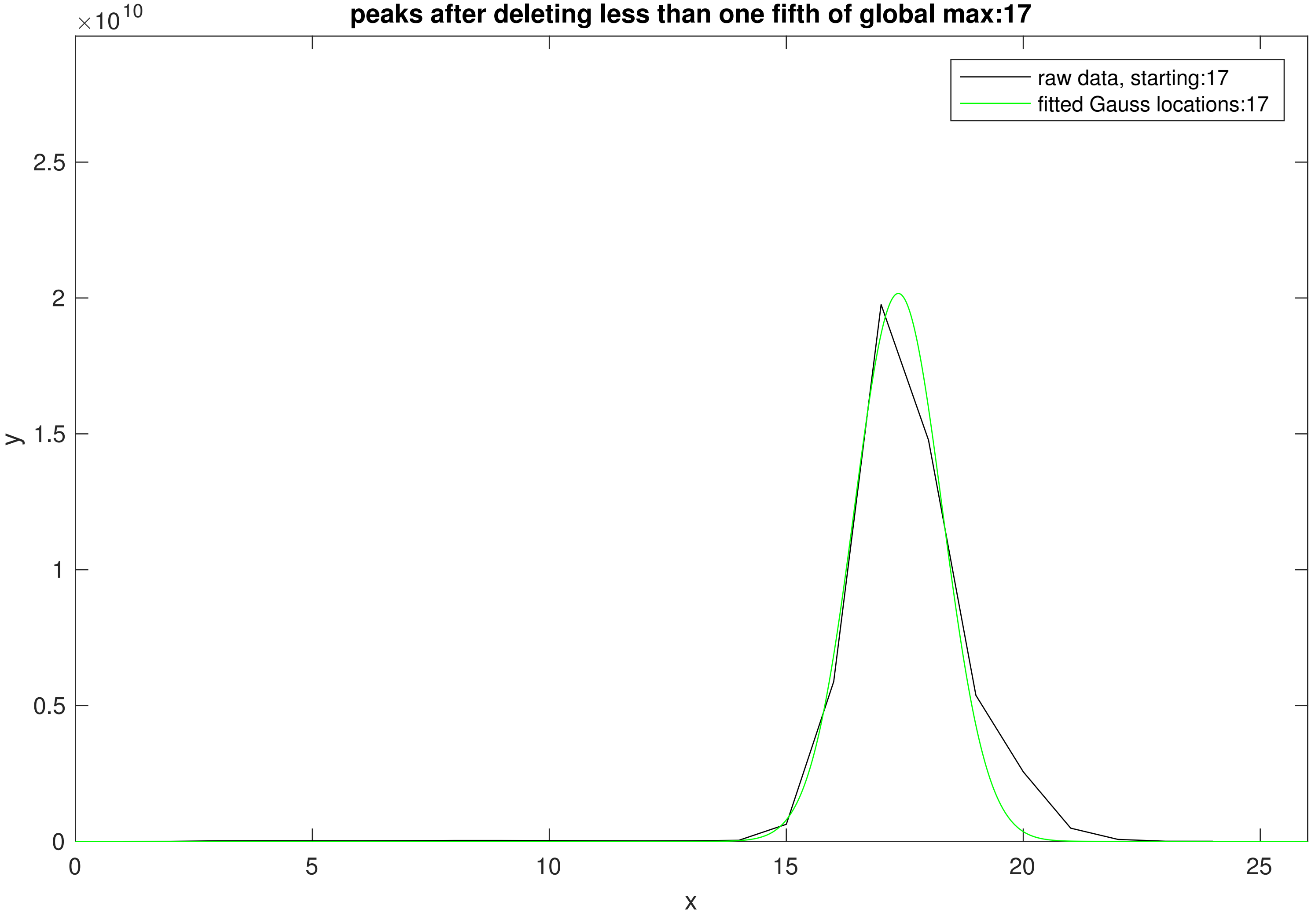

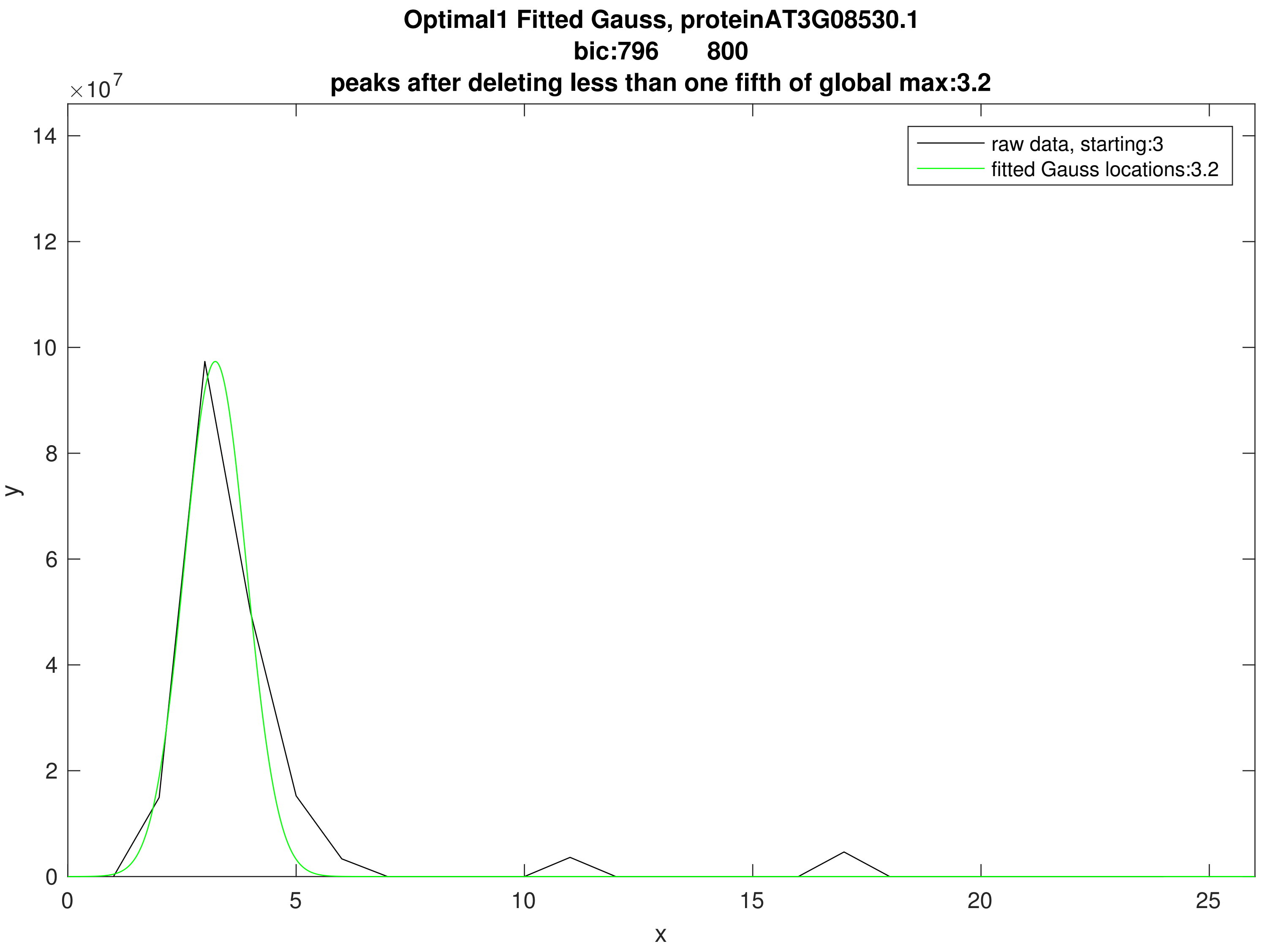
Optimal1 Fitted Gauss, proteinAT3G08530.1
bic:796 800
peaks after deleting less than one fifth of global max:3.2
×10^7
raw data, starting:3
fitted Gauss locations:3.2
y
x
0
2
4
6
8
10
12
14
0
5
10
15
20
25

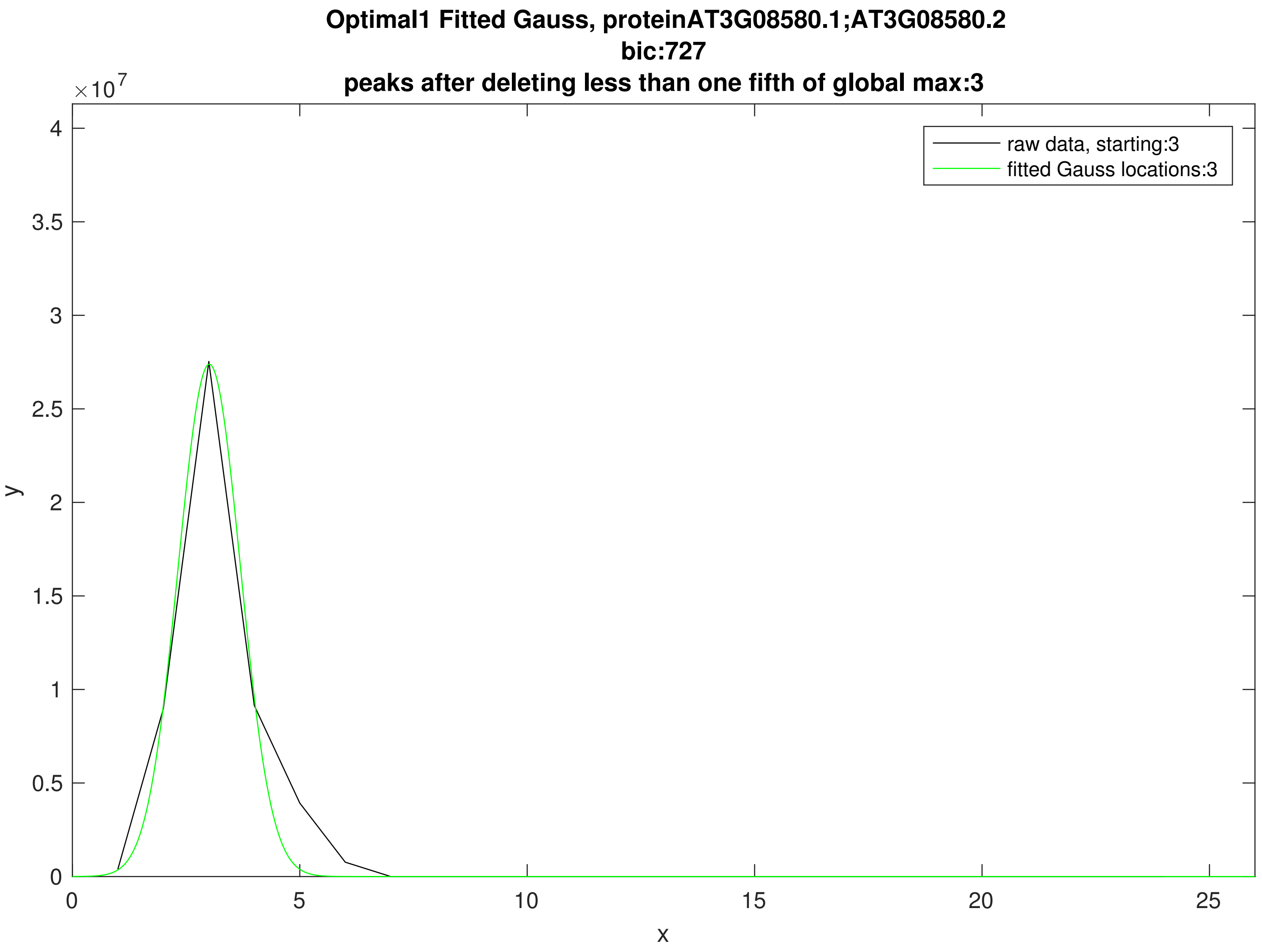

Optimal1 Fitted Gauss, proteinAT3G08580.1;AT3G08580.2
bic:727
peaks after deleting less than one fifth of global max:3
$\times 10^7$
raw data, starting:3
fitted Gauss locations:3
y
x
0
0.5
1
1.5
2
2.5
3
3.5
4
0
5
10
15
20
25

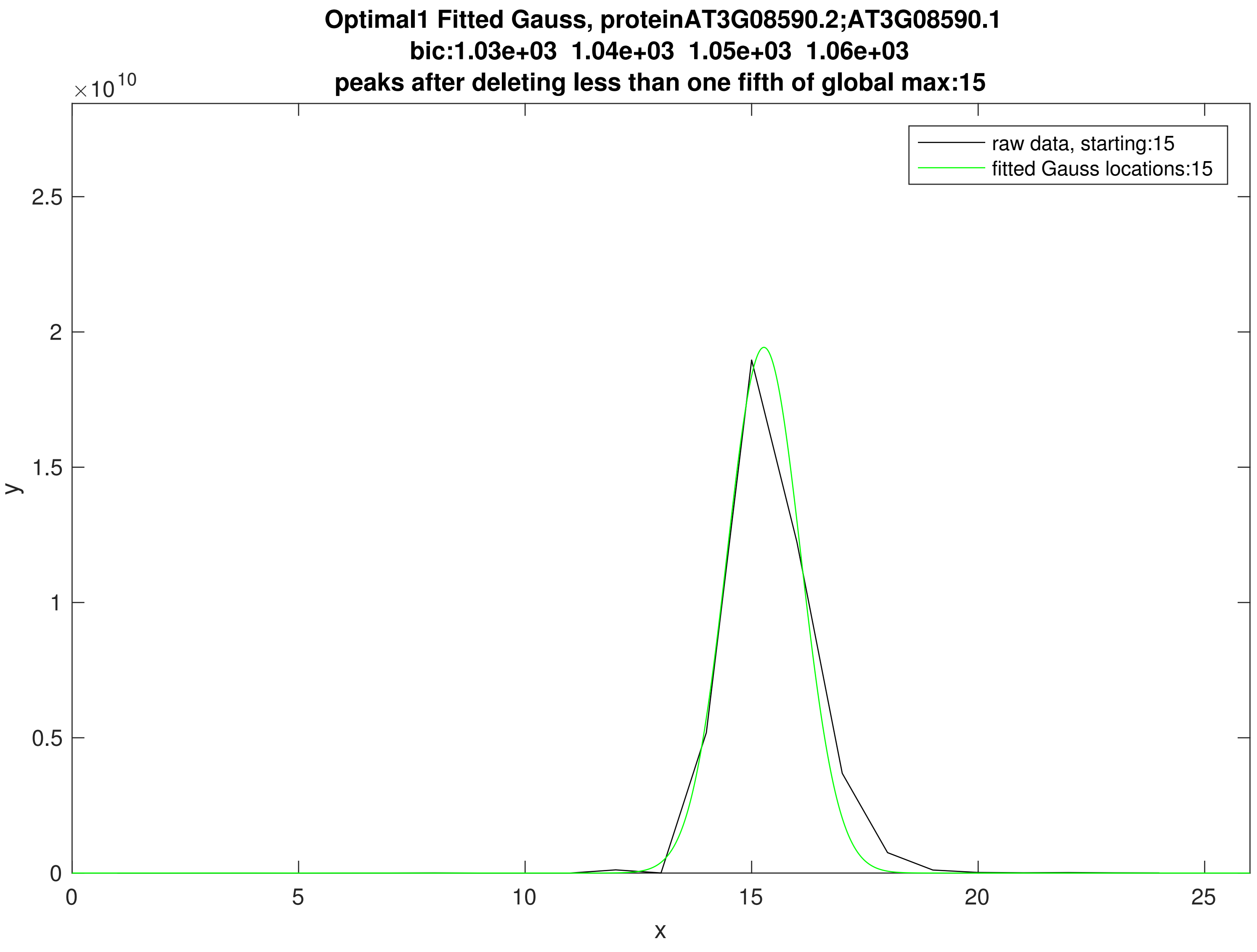
Optimal1 Fitted Gauss, proteinAT3G08590.2;AT3G08590.1
bic:1.03e+03  1.04e+03  1.05e+03  1.06e+03
peaks after deleting less than one fifth of global max:15
×10^10
raw data, starting:15
fitted Gauss locations:15
y
x
0
0.5
1
1.5
2
2.5
0
5
10
15
20
25

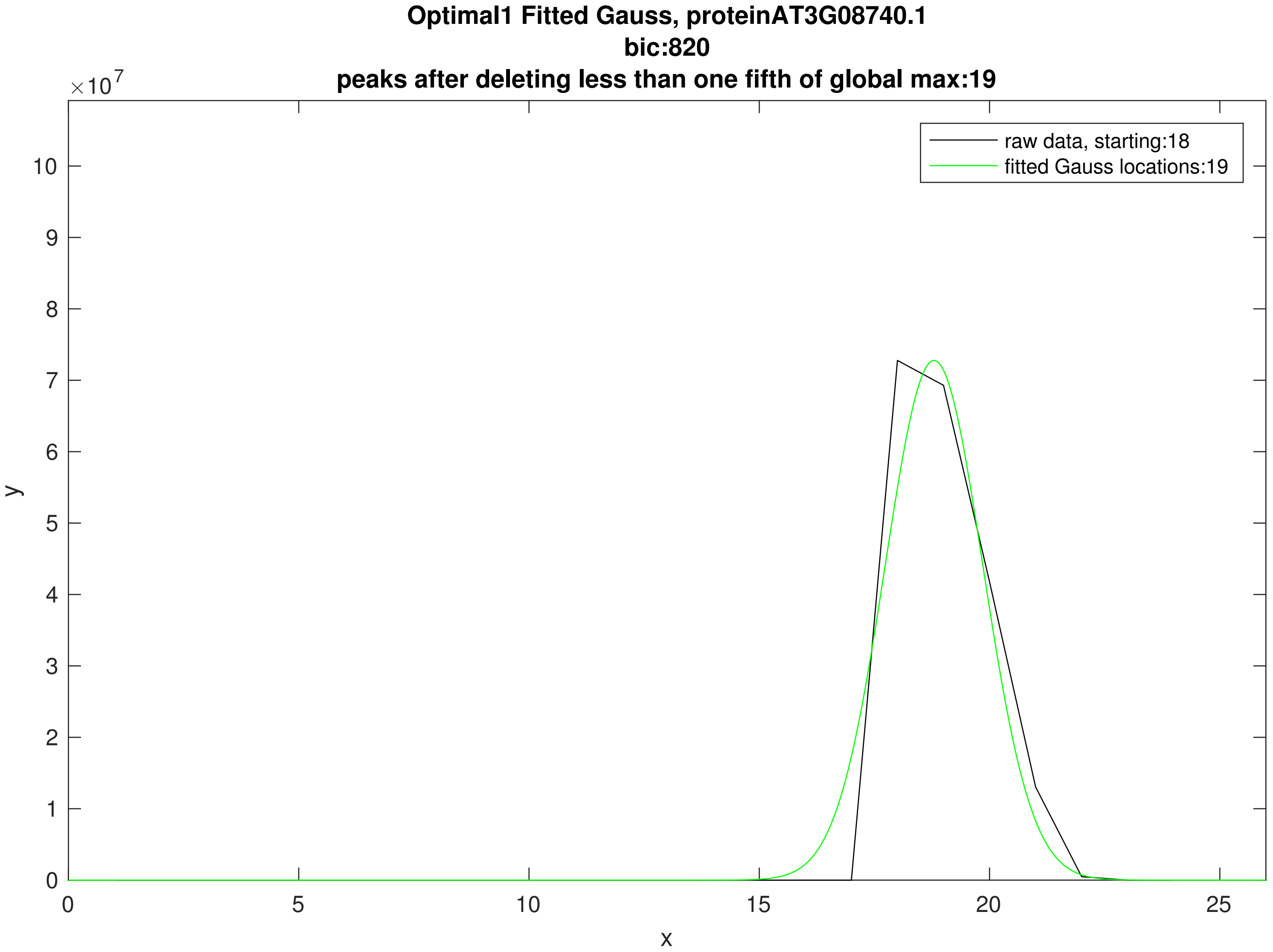
Optimal1 Fitted Gauss, proteinAT3G08740.1
bic:820
peaks after deleting less than one fifth of global max:19
×10^7
raw data, starting:18
fitted Gauss locations:19
y
x
0
1
2
3
4
5
6
7
8
9
10
0
5
10
15
20
25

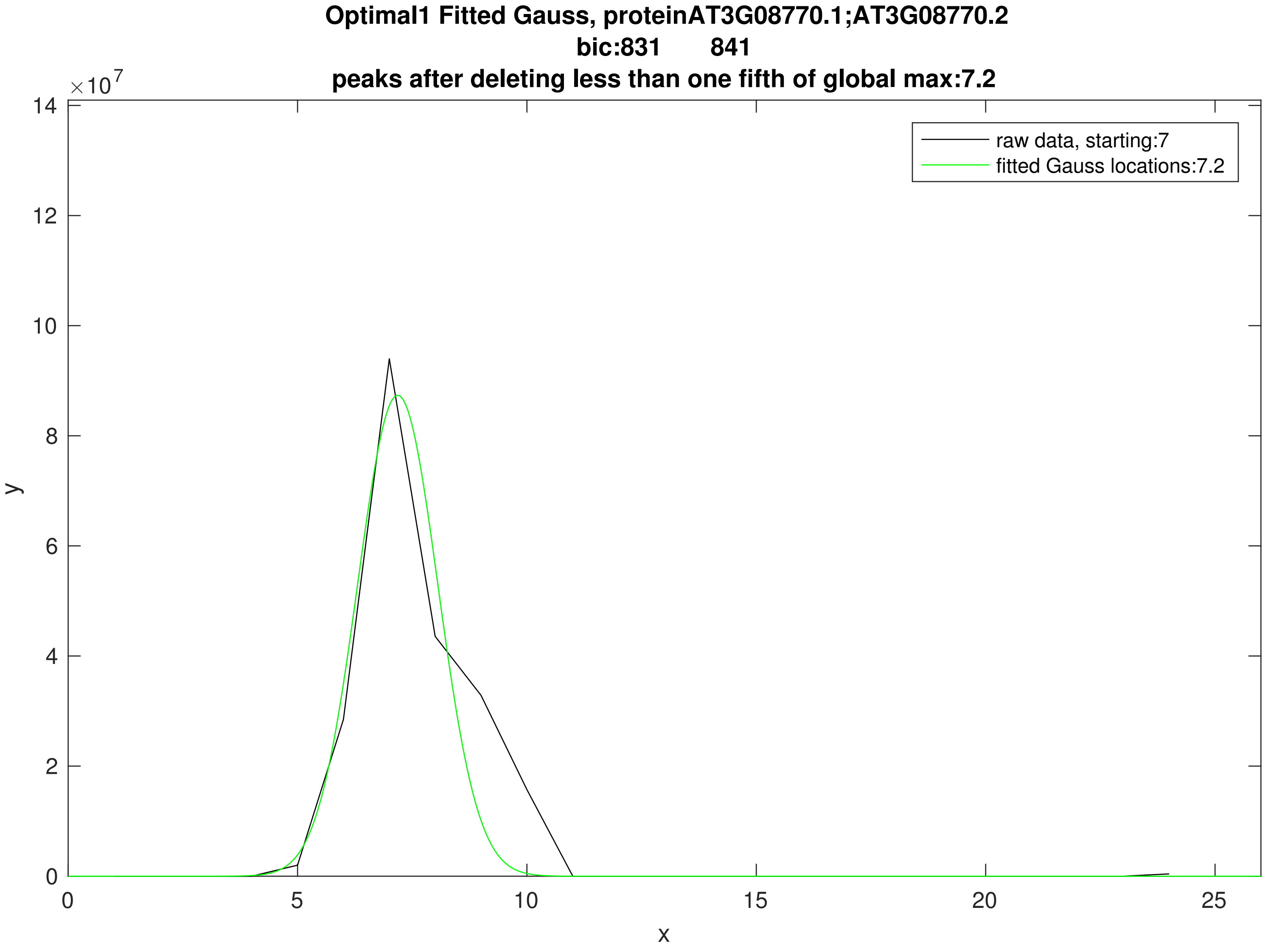
Optimal1 Fitted Gauss, proteinAT3G08770.1;AT3G08770.2
bic:831 841
peaks after deleting less than one fifth of global max:7.2
×10^7
14
12
10
8
6
4
2
0
y
0
5
10
15
20
25
x
raw data, starting:7
fitted Gauss locations:7.2

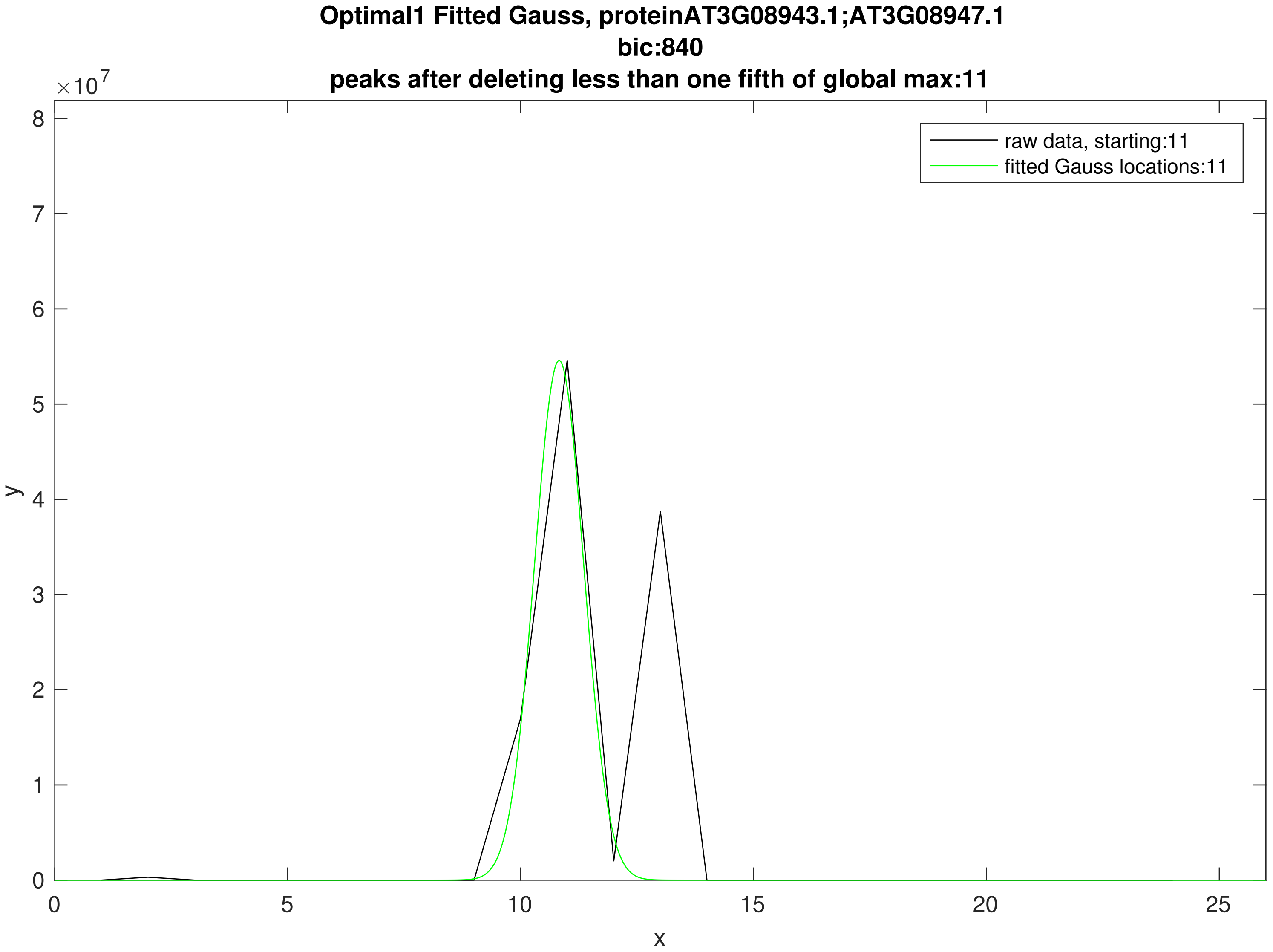
Optimal1 Fitted Gauss, proteinAT3G08943.1;AT3G08947.1
bic:840
peaks after deleting less than one fifth of global max:11
×10^7
raw data, starting:11
fitted Gauss locations:11
y
x
0
1
2
3
4
5
6
7
8
0
5
10
15
20
25

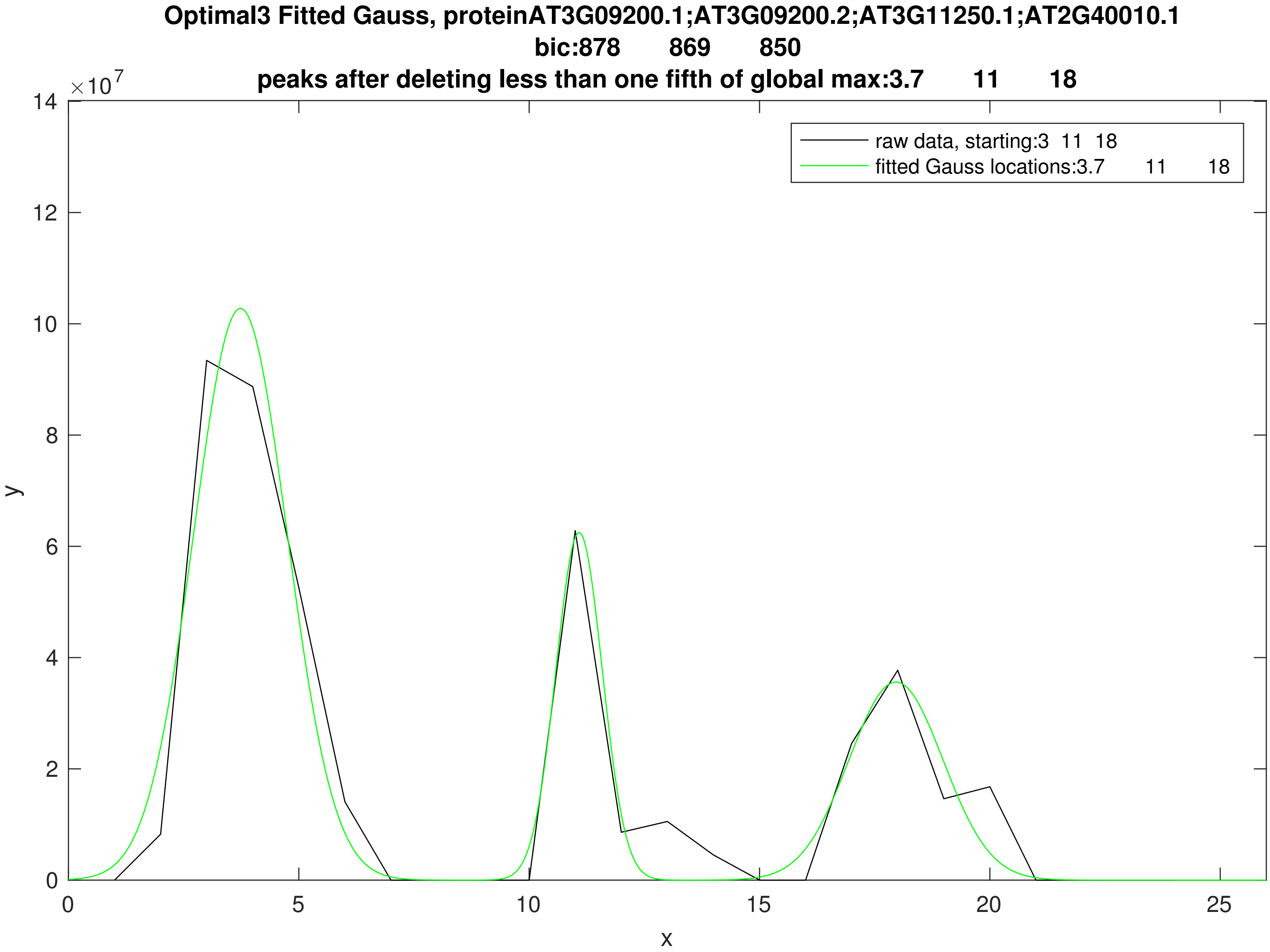
Optimal3 Fitted Gauss, proteinAT3G09200.1;AT3G09200.2;AT3G11250.1;AT2G40010.1
bic:878 869 850
peaks after deleting less than one fifth of global max:3.7 11 18
×10^7
14
12
10
8
6
4
2
0
y
0
5
10
15
20
25
x
raw data, starting:3 11 18
fitted Gauss locations:3.7 11 18

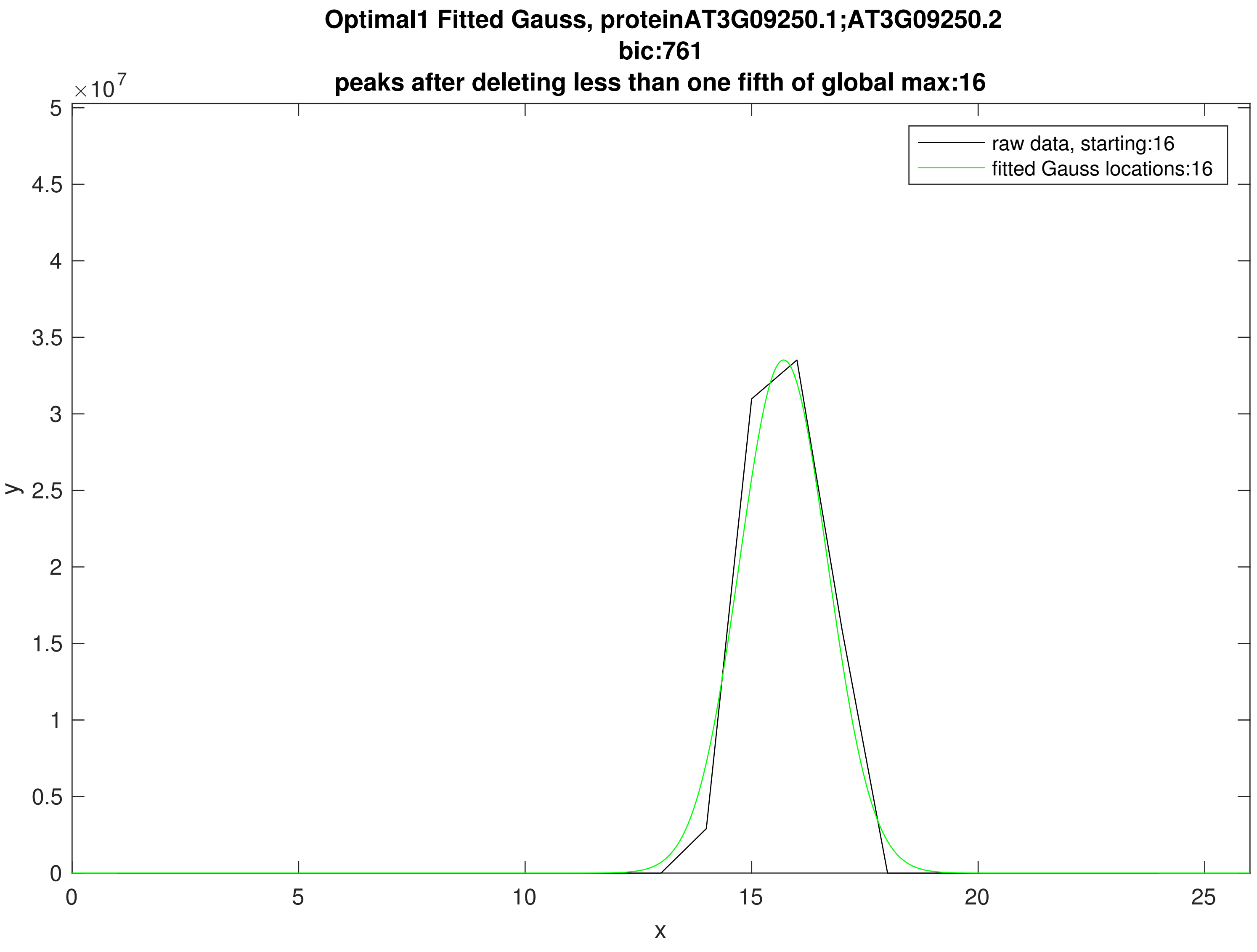
Optimal1 Fitted Gauss, proteinAT3G09250.1;AT3G09250.2
bic:761
peaks after deleting less than one fifth of global max:16
×10^7
raw data, starting:16
fitted Gauss locations:16
y
x
0
0.5
1
1.5
2
2.5
3
3.5
4
4.5
5
0
5
10
15
20
25

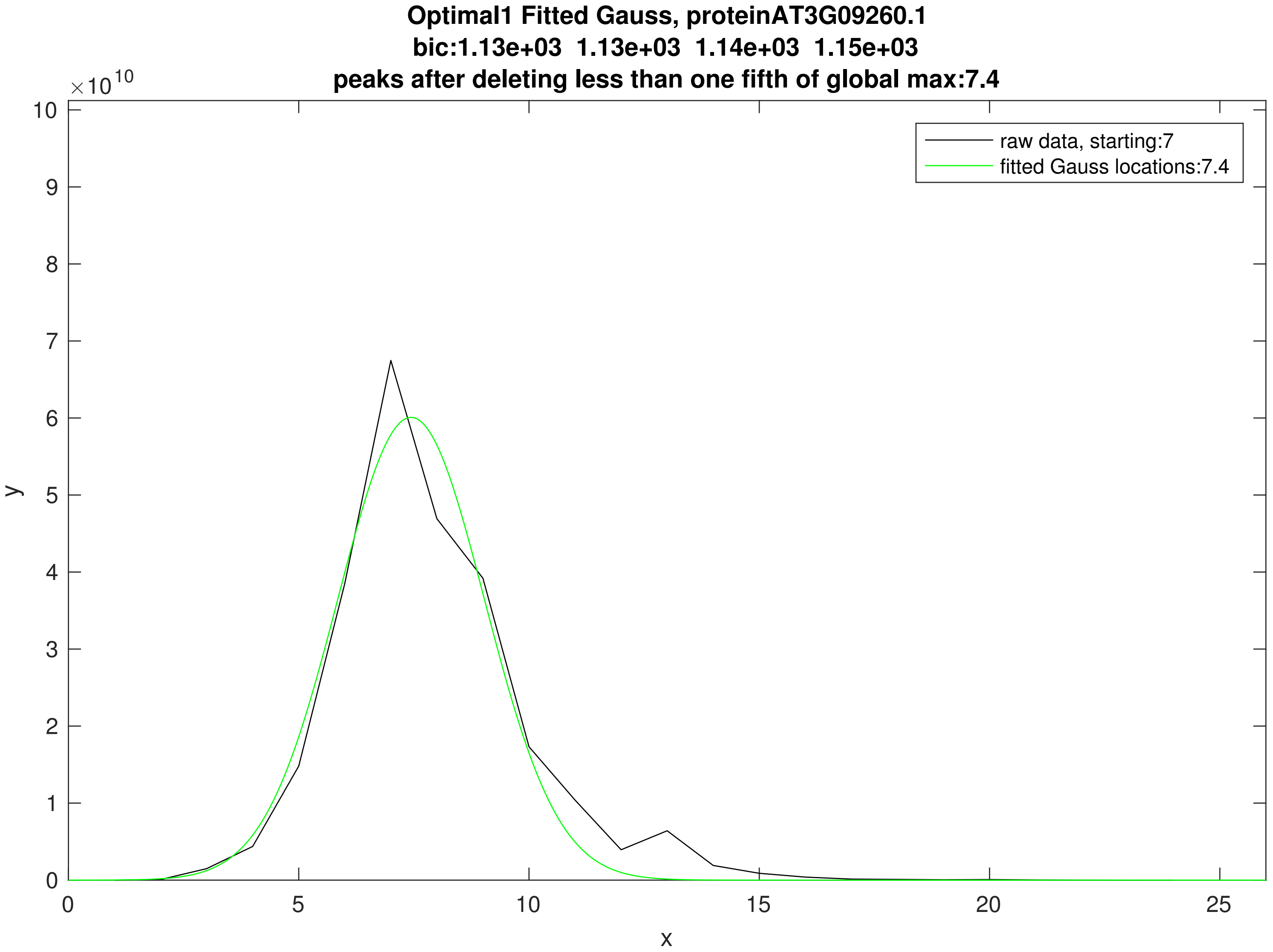
Optimal1 Fitted Gauss, proteinAT3G09260.1
bic:1.13e+03 1.13e+03 1.14e+03 1.15e+03
peaks after deleting less than one fifth of global max:7.4
×10^10
y
x
raw data, starting:7
fitted Gauss locations:7.4
0
1
2
3
4
5
6
7
8
9
10
0
5
10
15
20
25

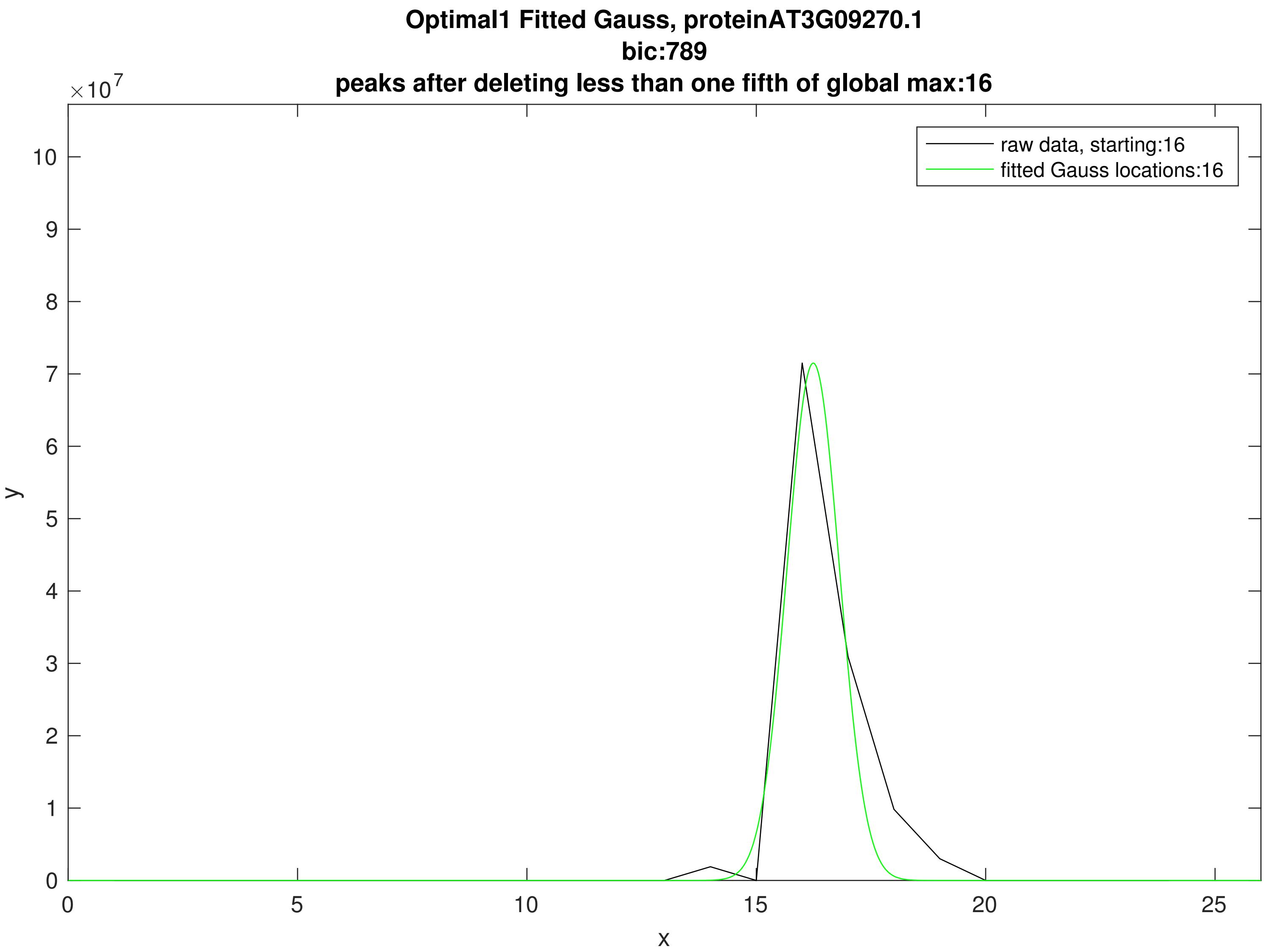
Optimal1 Fitted Gauss, proteinAT3G09270.1
bic:789
peaks after deleting less than one fifth of global max:16
×10^7
raw data, starting:16
fitted Gauss locations:16
y
x
0
1
2
3
4
5
6
7
8
9
10
0
5
10
15
20
25

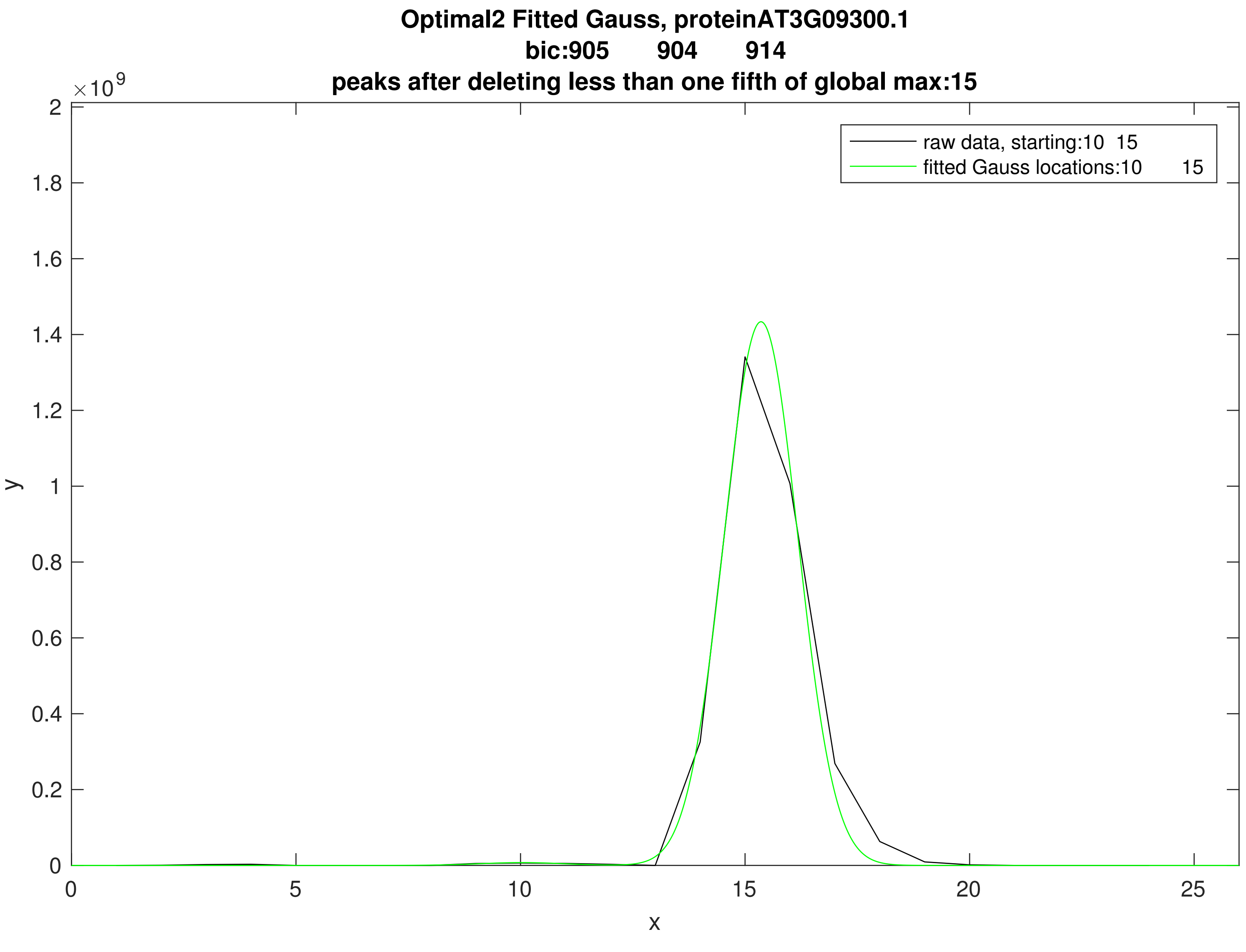
Optimal2 Fitted Gauss, proteinAT3G09300.1
bic:905 904 914
peaks after deleting less than one fifth of global max:15
×10^9
raw data, starting:10 15
fitted Gauss locations:10 15
y
x
0
0.2
0.4
0.6
0.8
1
1.2
1.4
1.6
1.8
2
0
5
10
15
20
25

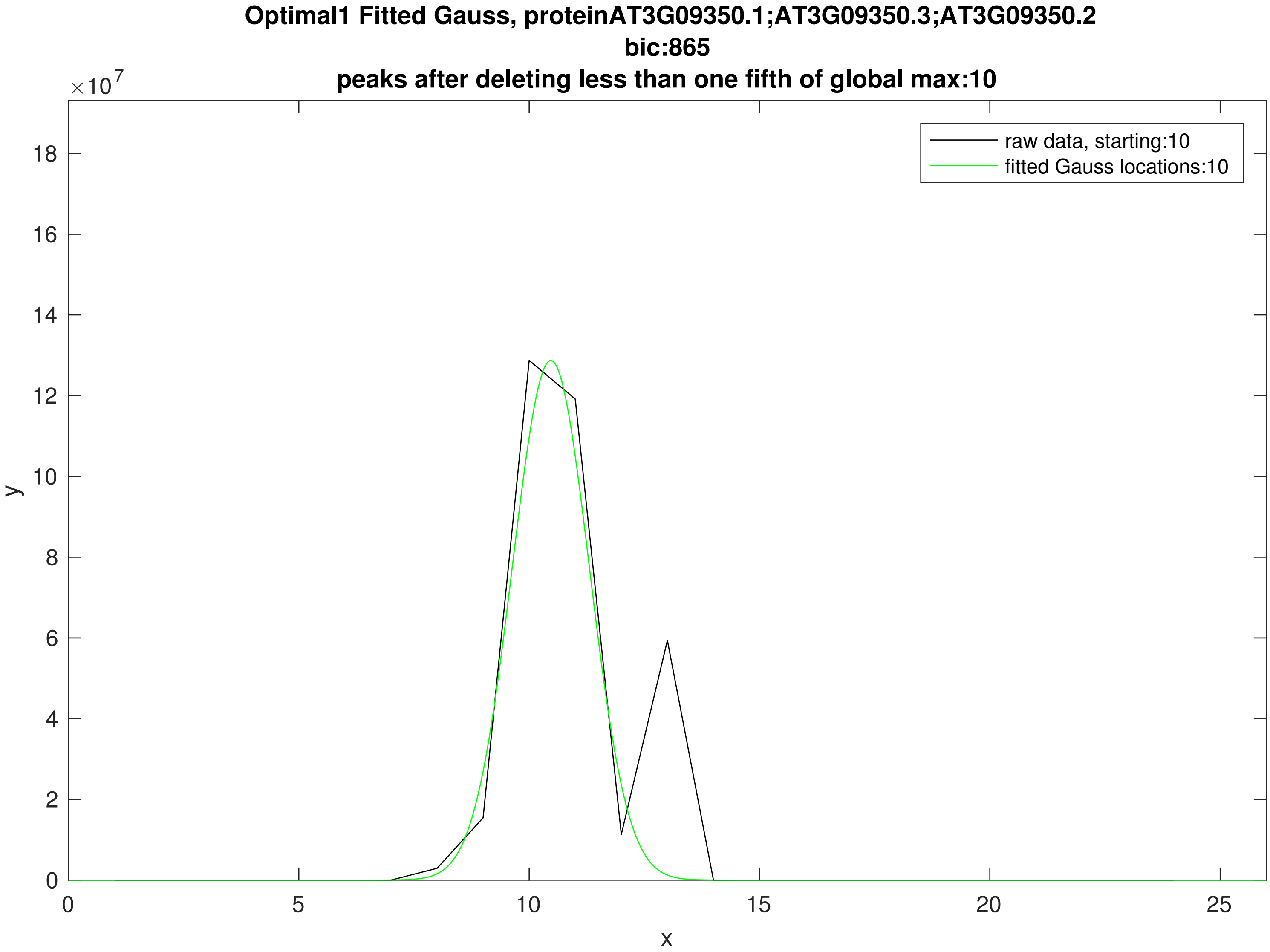

Optimal1 Fitted Gauss, proteinAT3G09350.1;AT3G09350.3;AT3G09350.2
bic:865
peaks after deleting less than one fifth of global max:10
×10^7
18
16
14
12
10
8
6
4
2
0
y
0
5
10
15
20
25
x
raw data, starting:10
fitted Gauss locations:10

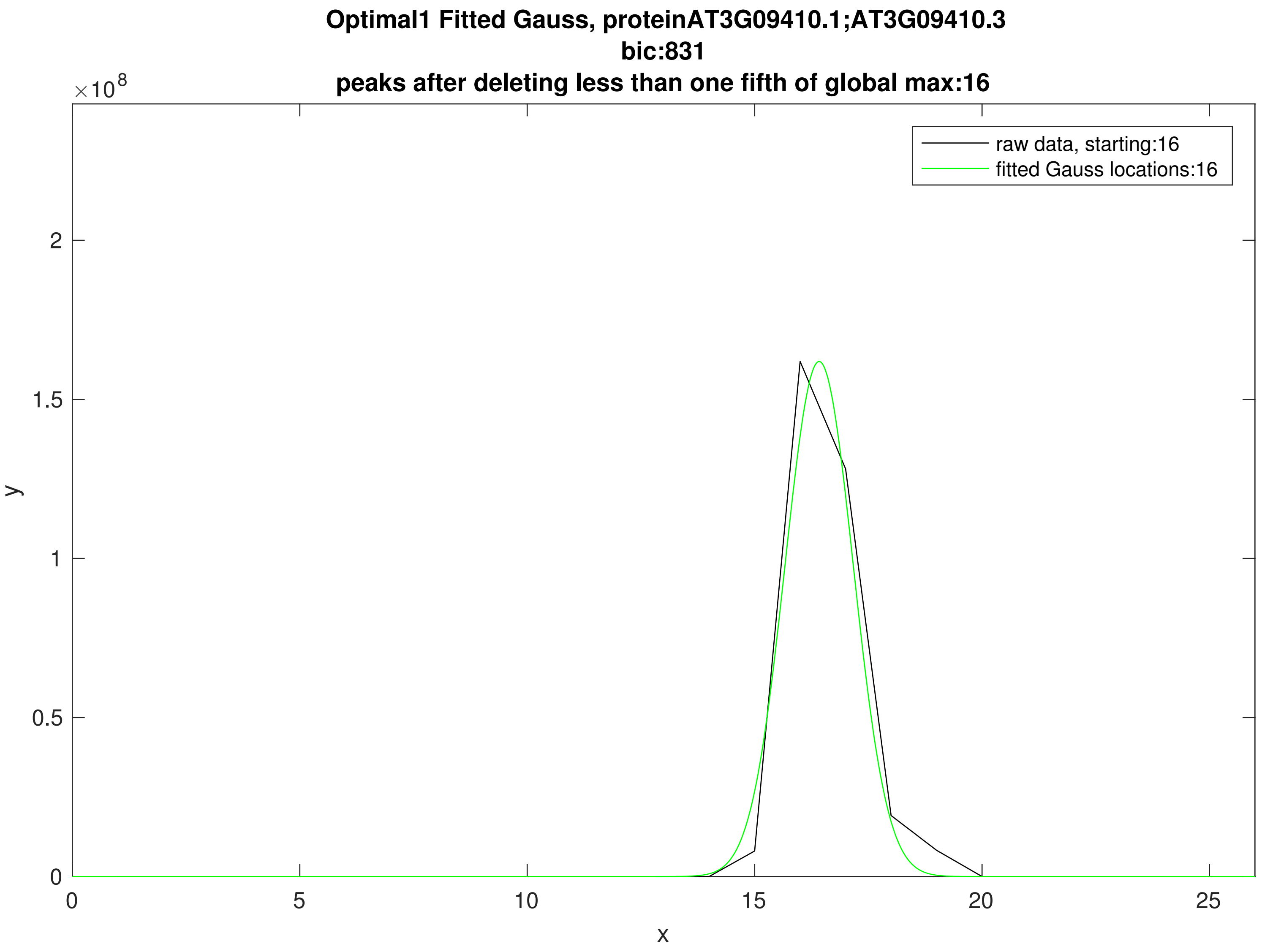
Optimal1 Fitted Gauss, proteinAT3G09410.1;AT3G09410.3
bic:831
peaks after deleting less than one fifth of global max:16
×10^8
raw data, starting:16
fitted Gauss locations:16
2
1.5
1
0.5
0
y
0
5
10
15
20
25
x

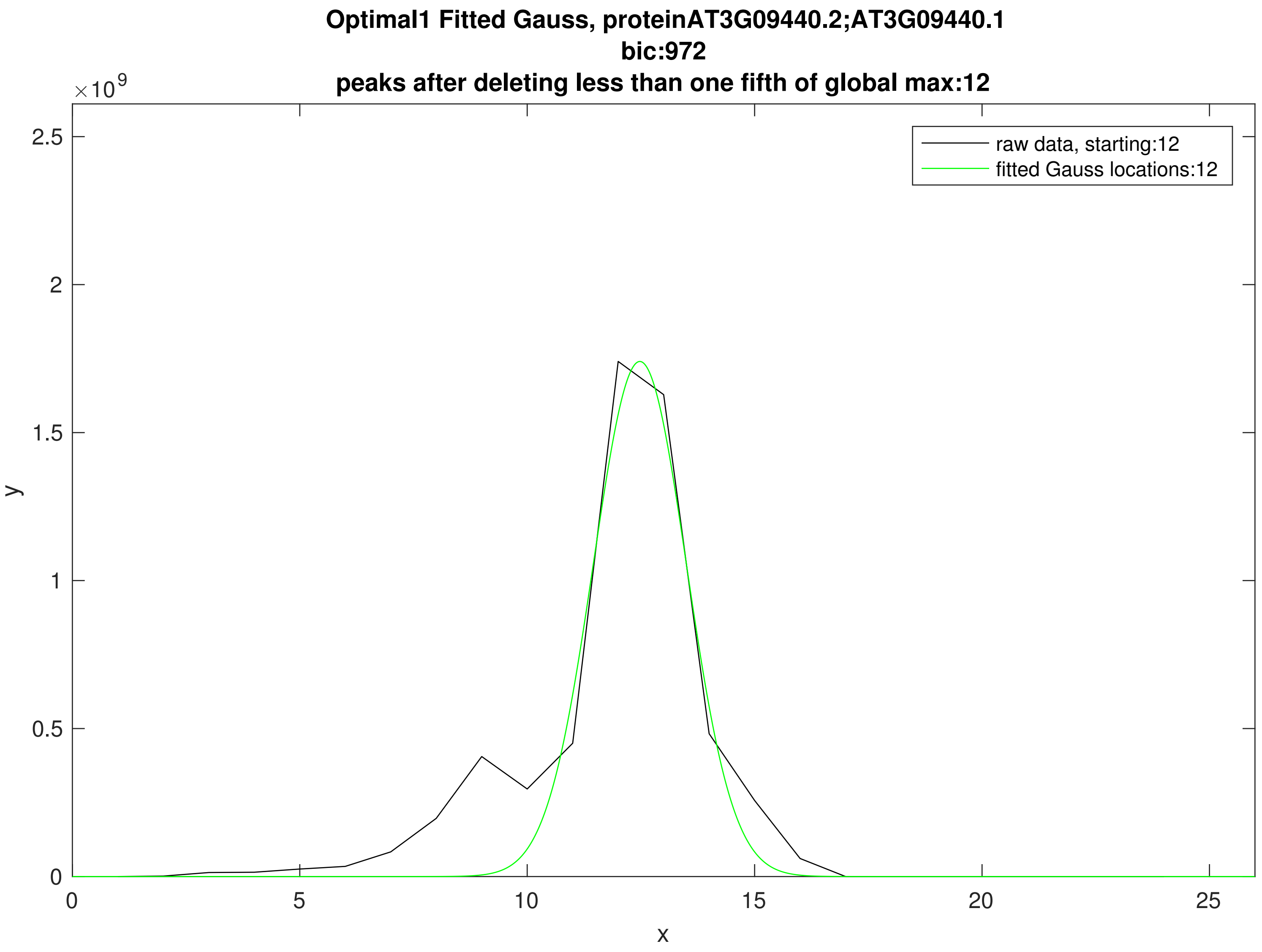
Optimal1 Fitted Gauss, proteinAT3G09440.2;AT3G09440.1
bic:972
peaks after deleting less than one fifth of global max:12
×10^9
2.5
2
1.5
1
0.5
0
y
0
5
10
15
20
25
x
raw data, starting:12
fitted Gauss locations:12

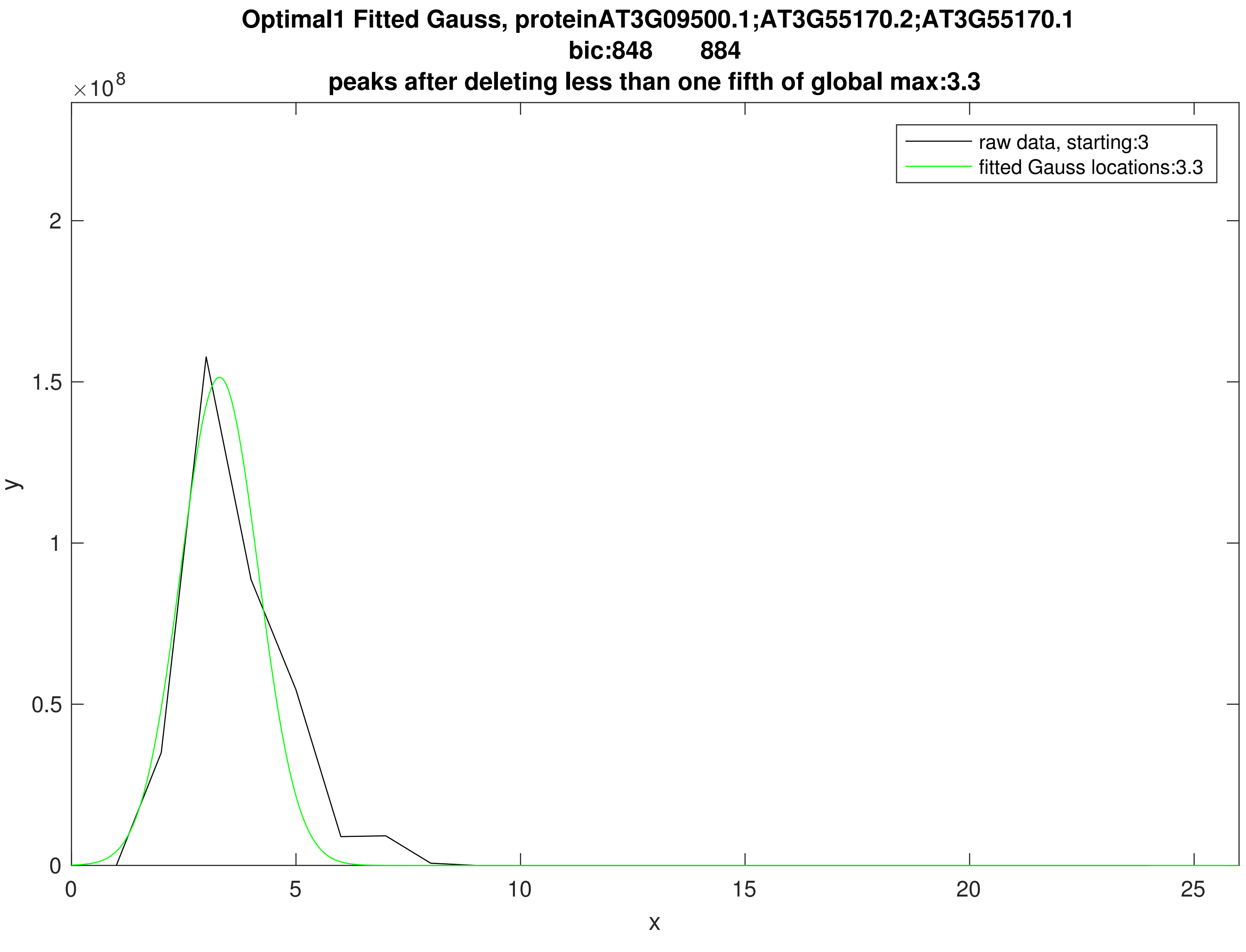

Optimal1 Fitted Gauss, proteinAT3G09500.1;AT3G55170.2;AT3G55170.1
bic:848 884
peaks after deleting less than one fifth of global max:3.3
×10^8
raw data, starting:3
fitted Gauss locations:3.3
y
x
0
0.5
1
1.5
2
0
5
10
15
20
25

**Optimal1 Fitted Gauss, proteinAT3G09630.1;AT3G09630.2**
**bic:927 937 946**
**peaks after deleting less than one fifth of global max:3.5**

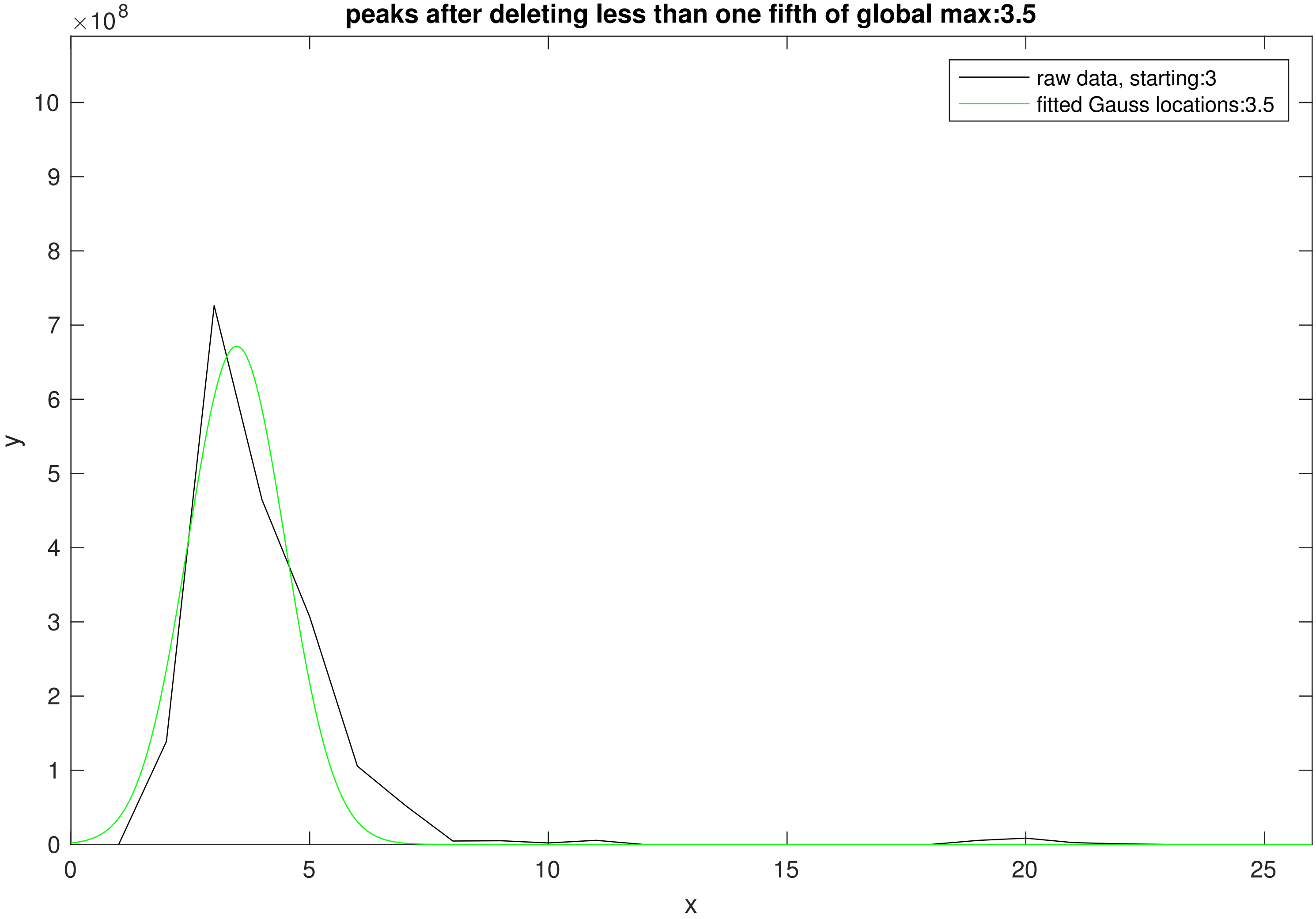

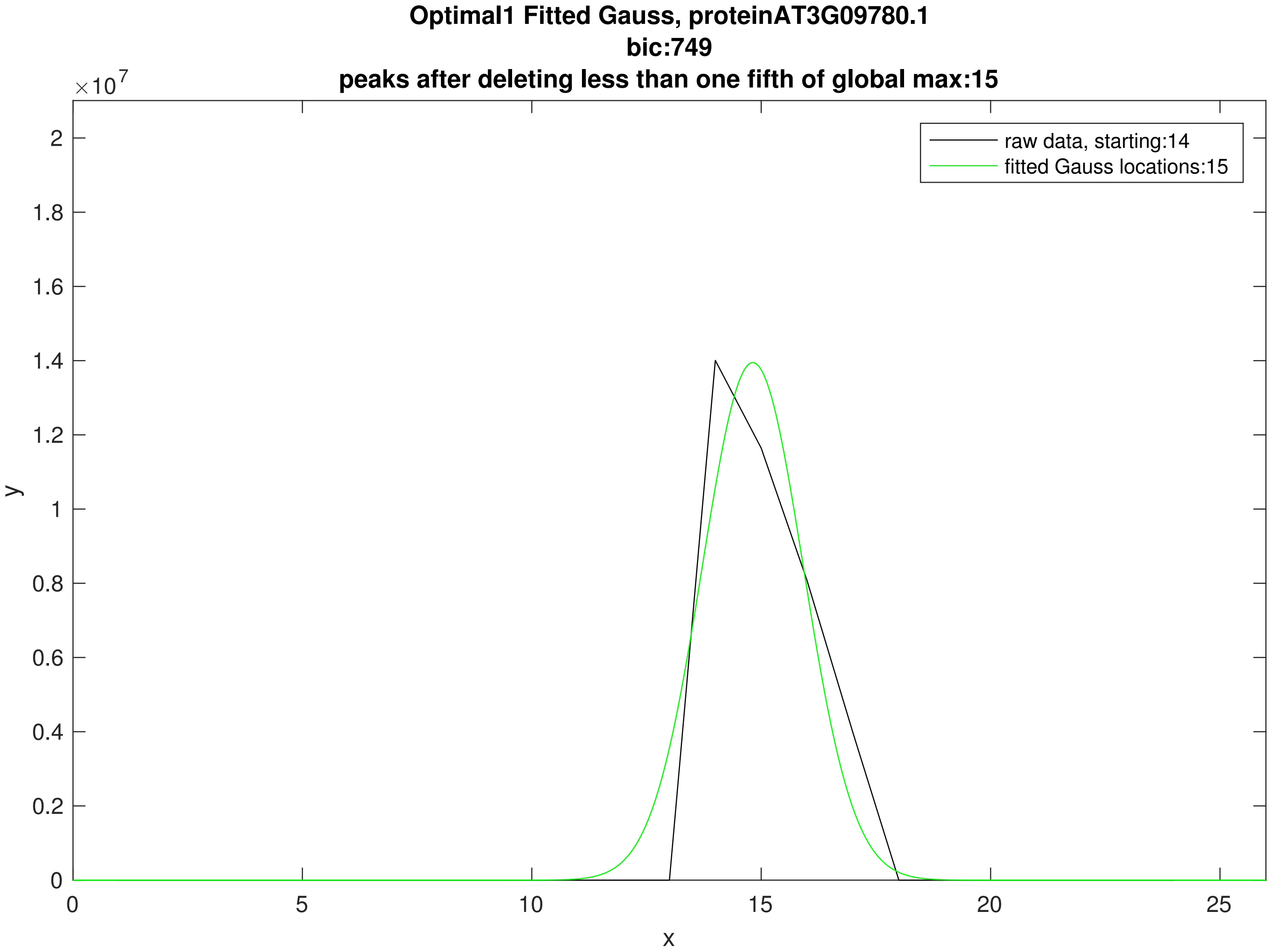

Optimal1 Fitted Gauss, proteinAT3G09780.1
bic:749
peaks after deleting less than one fifth of global max:15
×10^7
raw data, starting:14
fitted Gauss locations:15
y
x
0
0.2
0.4
0.6
0.8
1
1.2
1.4
1.6
1.8
2
0
5
10
15
20
25

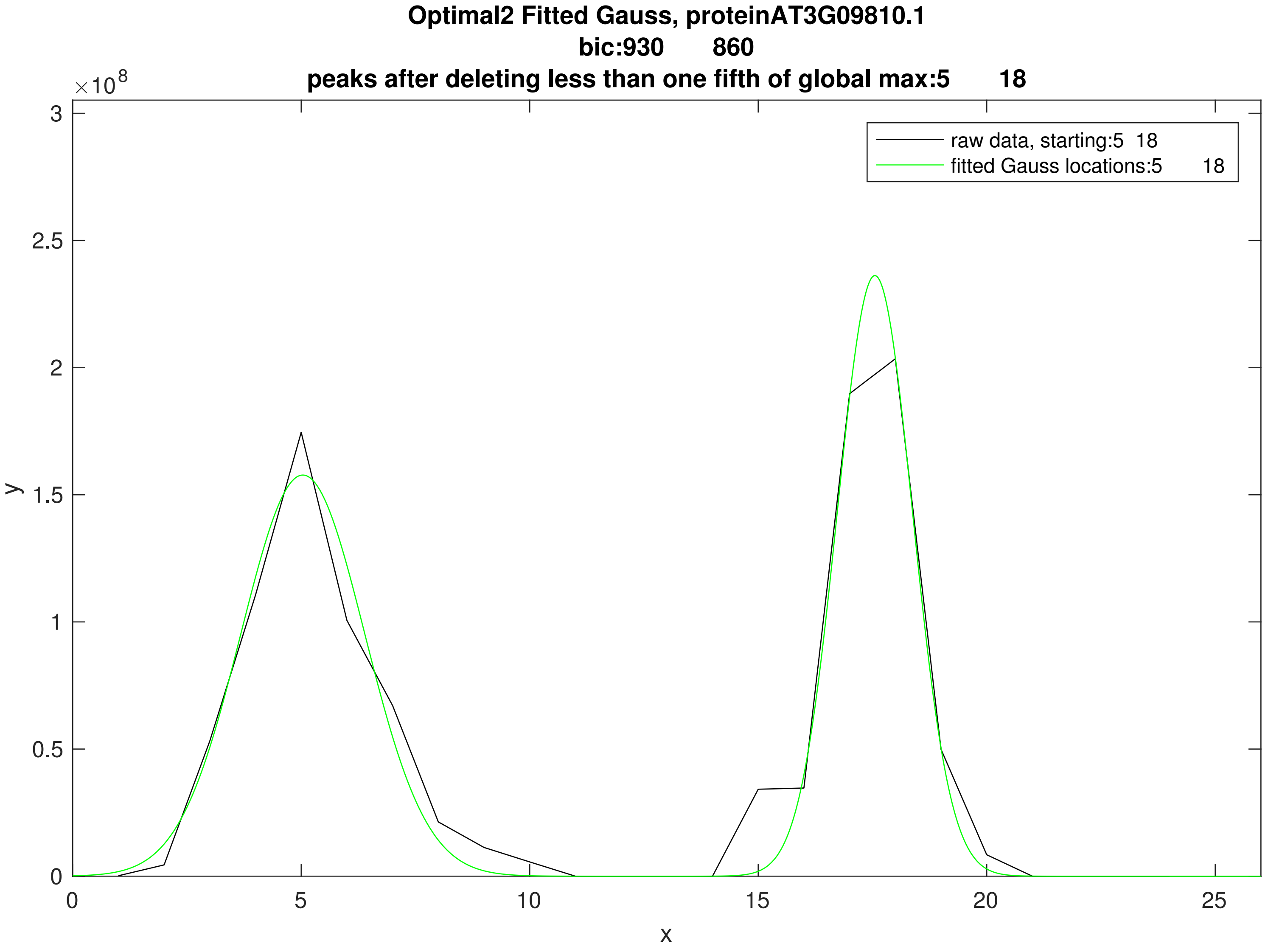

Optimal2 Fitted Gauss, proteinAT3G09810.1
bic:930 860
peaks after deleting less than one fifth of global max:5 18
raw data, starting:5 18
fitted Gauss locations:5 18
×10^8
y
x

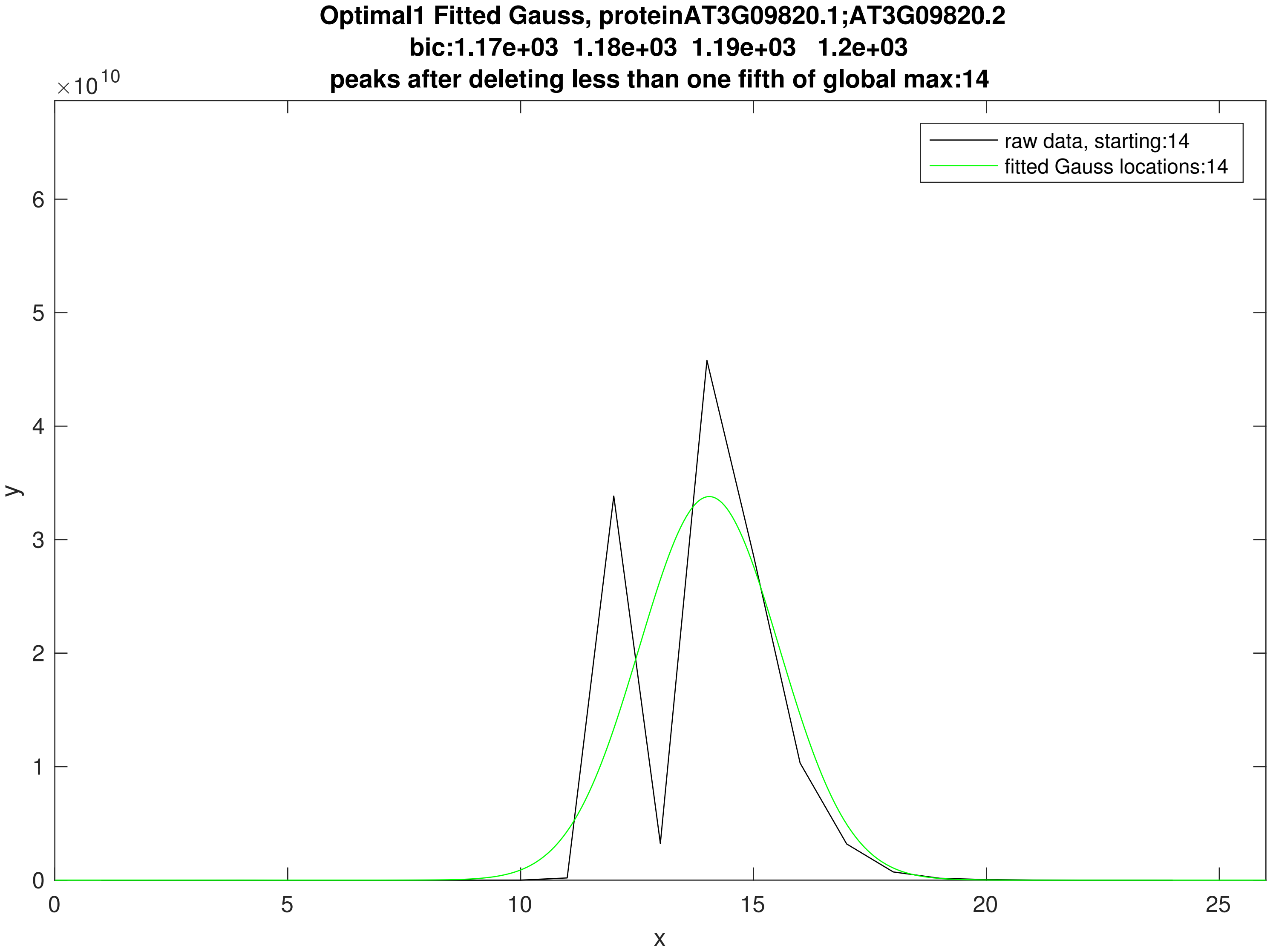

Optimal1 Fitted Gauss, proteinAT3G09820.1;AT3G09820.2
bic:1.17e+03 1.18e+03 1.19e+03 1.2e+03
peaks after deleting less than one fifth of global max:14
×10^10
raw data, starting:14
fitted Gauss locations:14
y
x
0
1
2
3
4
5
6
0
5
10
15
20
25

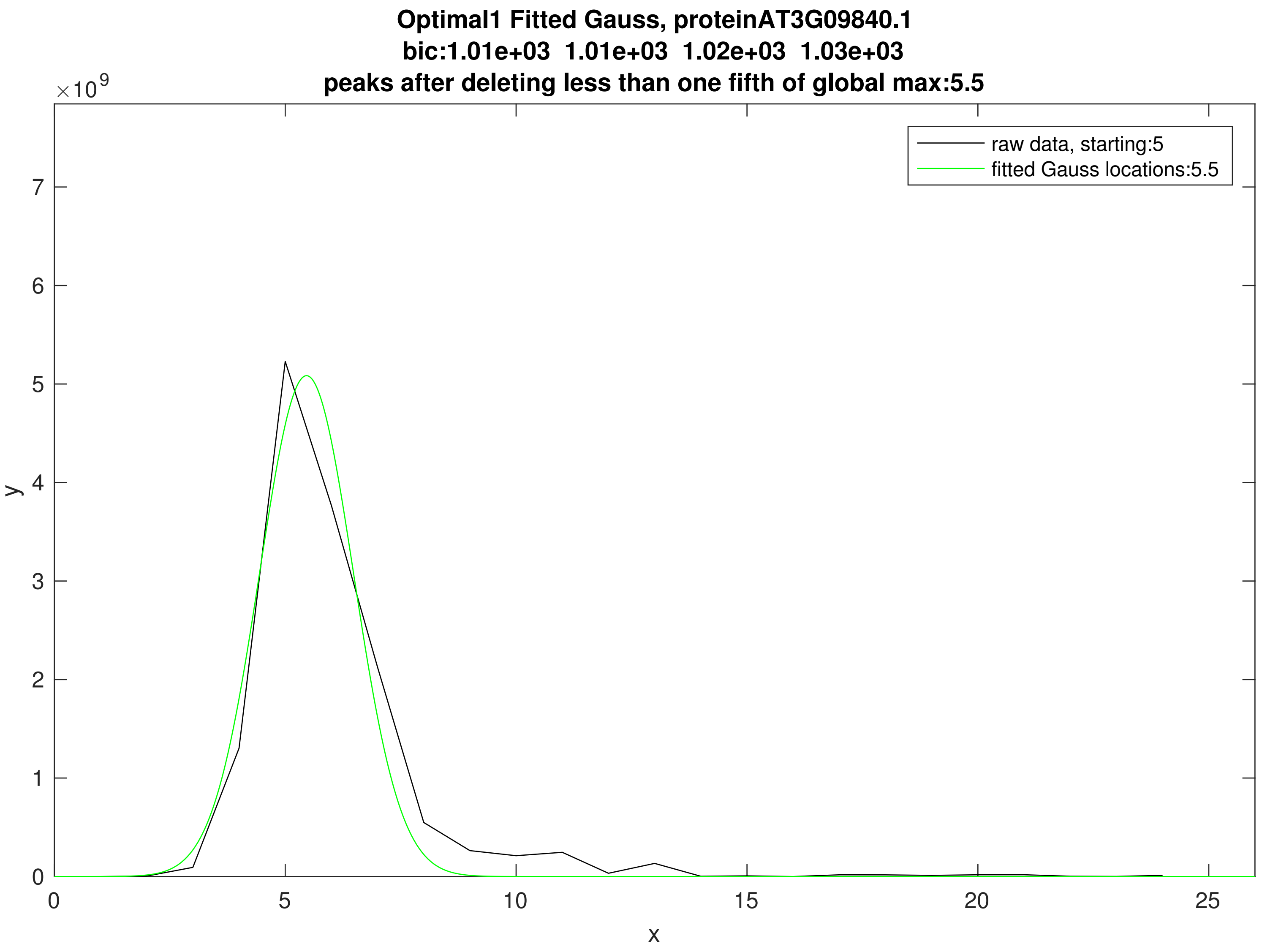
Optimal1 Fitted Gauss, proteinAT3G09840.1
bic:1.01e+03 1.01e+03 1.02e+03 1.03e+03
peaks after deleting less than one fifth of global max:5.5
×10^9
raw data, starting:5
fitted Gauss locations:5.5
y
x
0
1
2
3
4
5
6
7
10
15
20
25

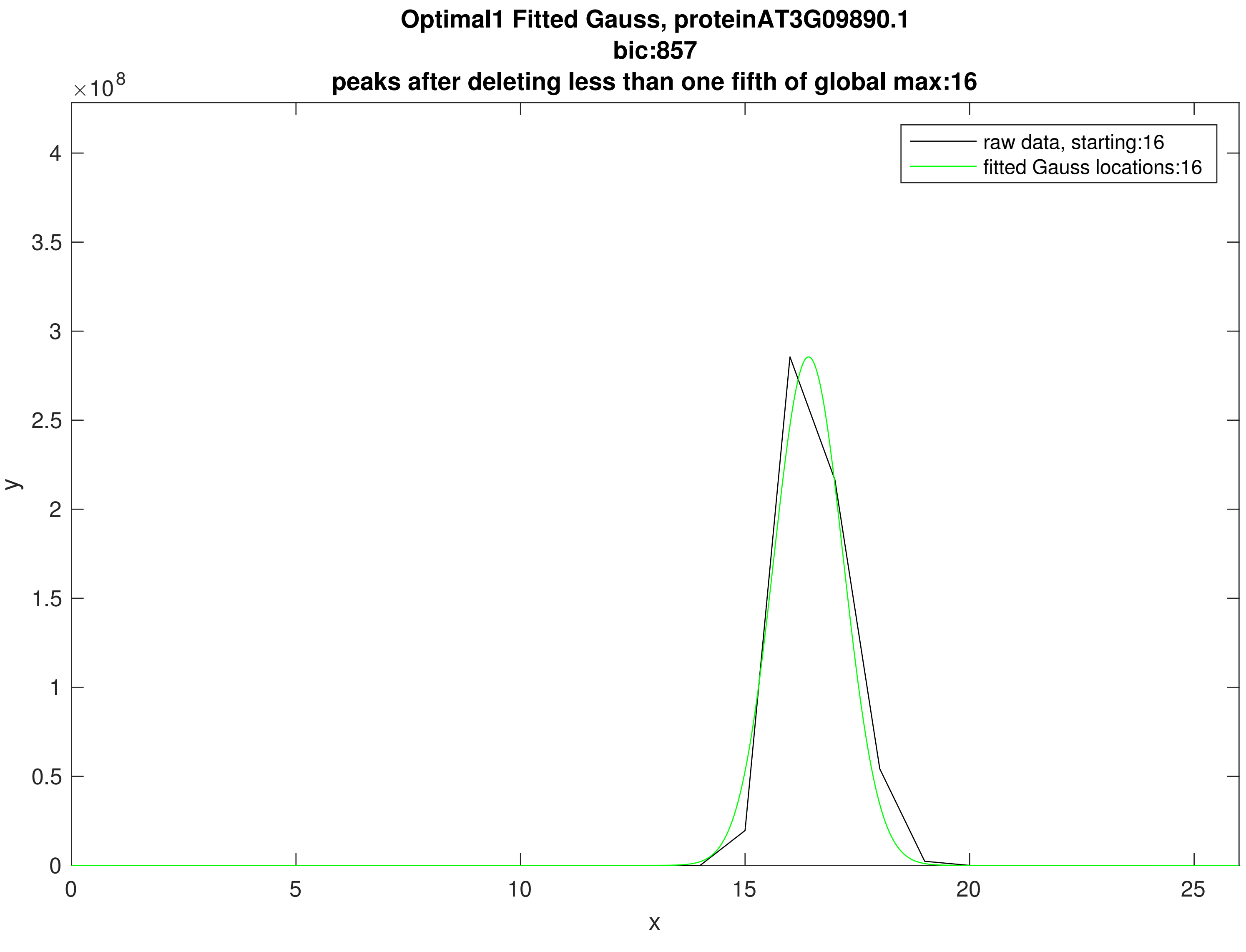

Optimal1 Fitted Gauss, proteinAT3G09890.1
bic:857
peaks after deleting less than one fifth of global max:16
×10^8
4
3.5
3
2.5
2
1.5
1
0.5
0
y
0
5
10
15
20
25
x
raw data, starting:16
fitted Gauss locations:16

Optimal1 Fitted Gauss, proteinAT3G09940.1;AT3G09940.2
bic:946 953 963
peaks after deleting less than one fifth of global max:16

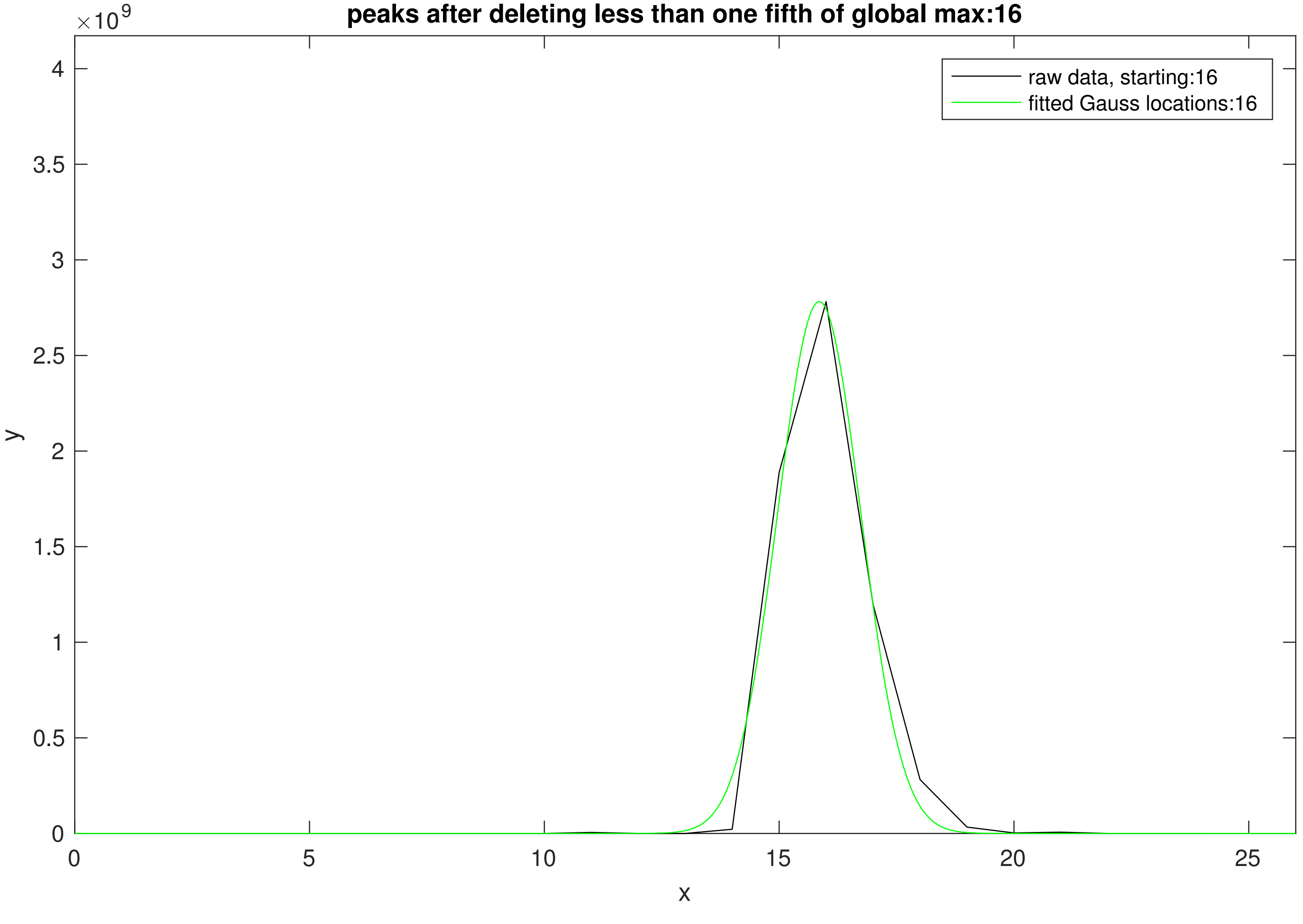

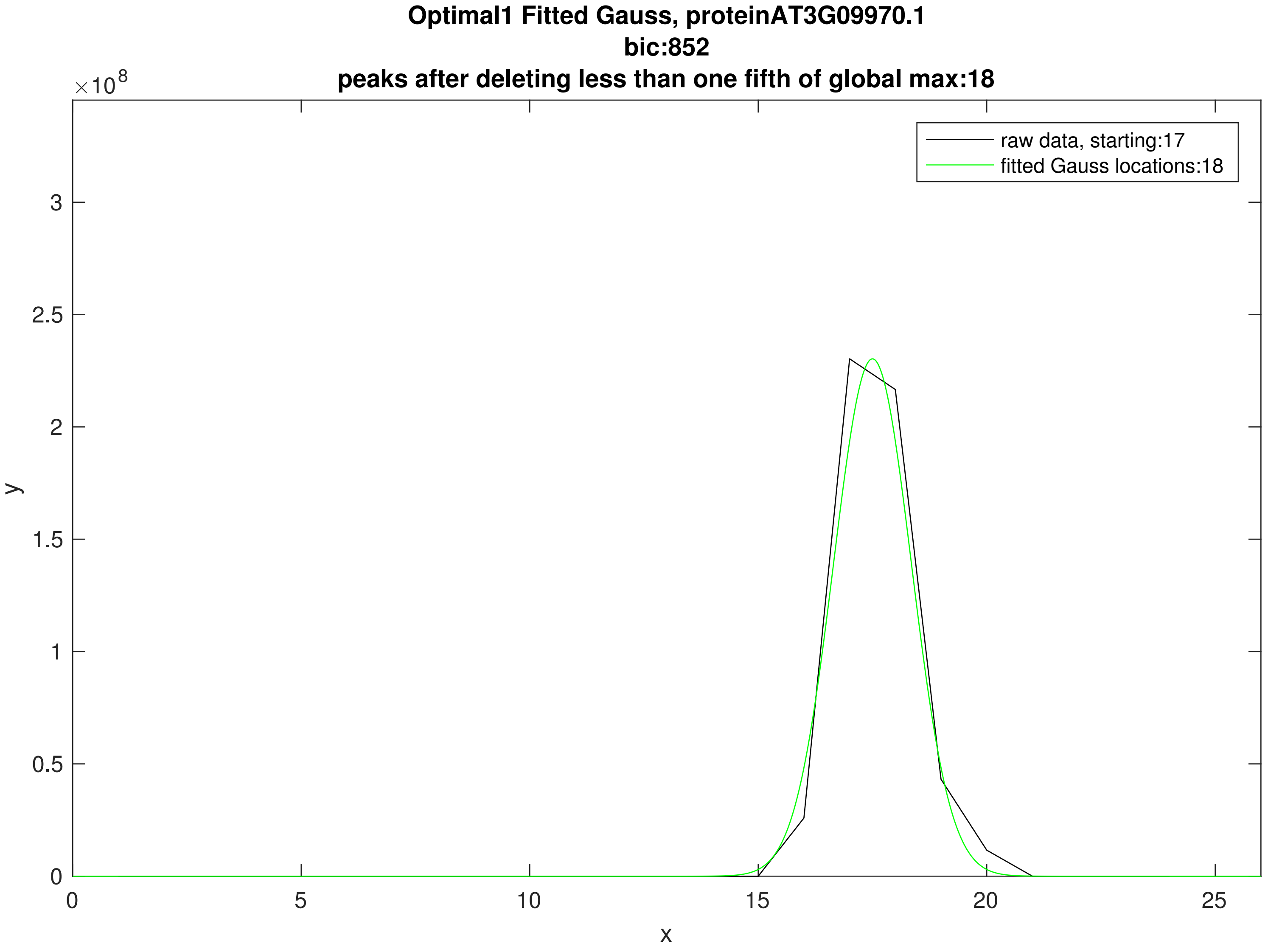

Optimal1 Fitted Gauss, proteinAT3G09970.1
bic:852
peaks after deleting less than one fifth of global max:18
×10^8
raw data, starting:17
fitted Gauss locations:18
y
x
0
0.5
1
1.5
2
2.5
3
0
5
10
15
20
25

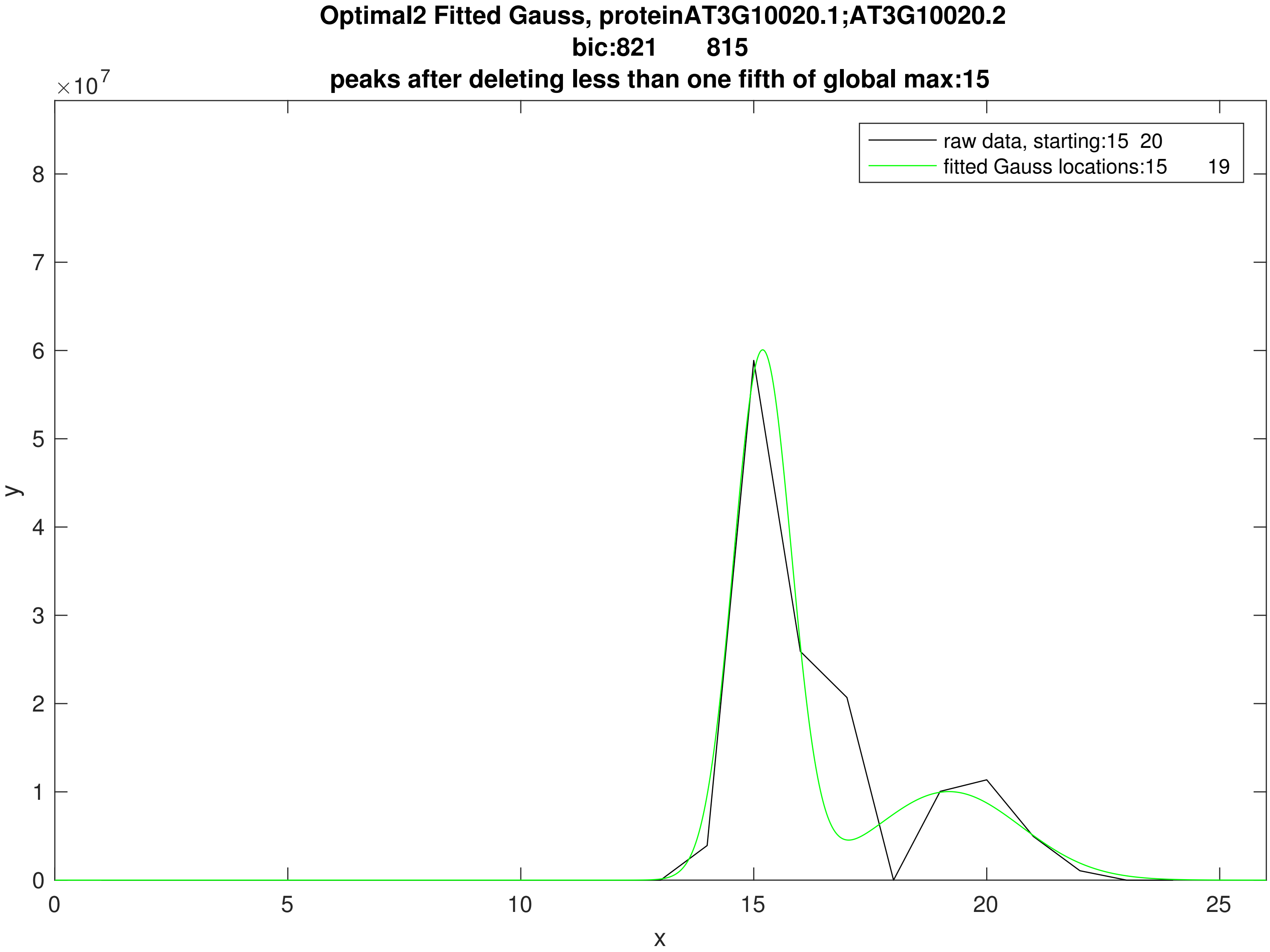
Optimal2 Fitted Gauss, proteinAT3G10020.1;AT3G10020.2
bic:821 815
peaks after deleting less than one fifth of global max:15
×10^7
raw data, starting:15 20
fitted Gauss locations:15 19
y
x
0
1
2
3
4
5
6
7
8
0
5
10
15
20
25

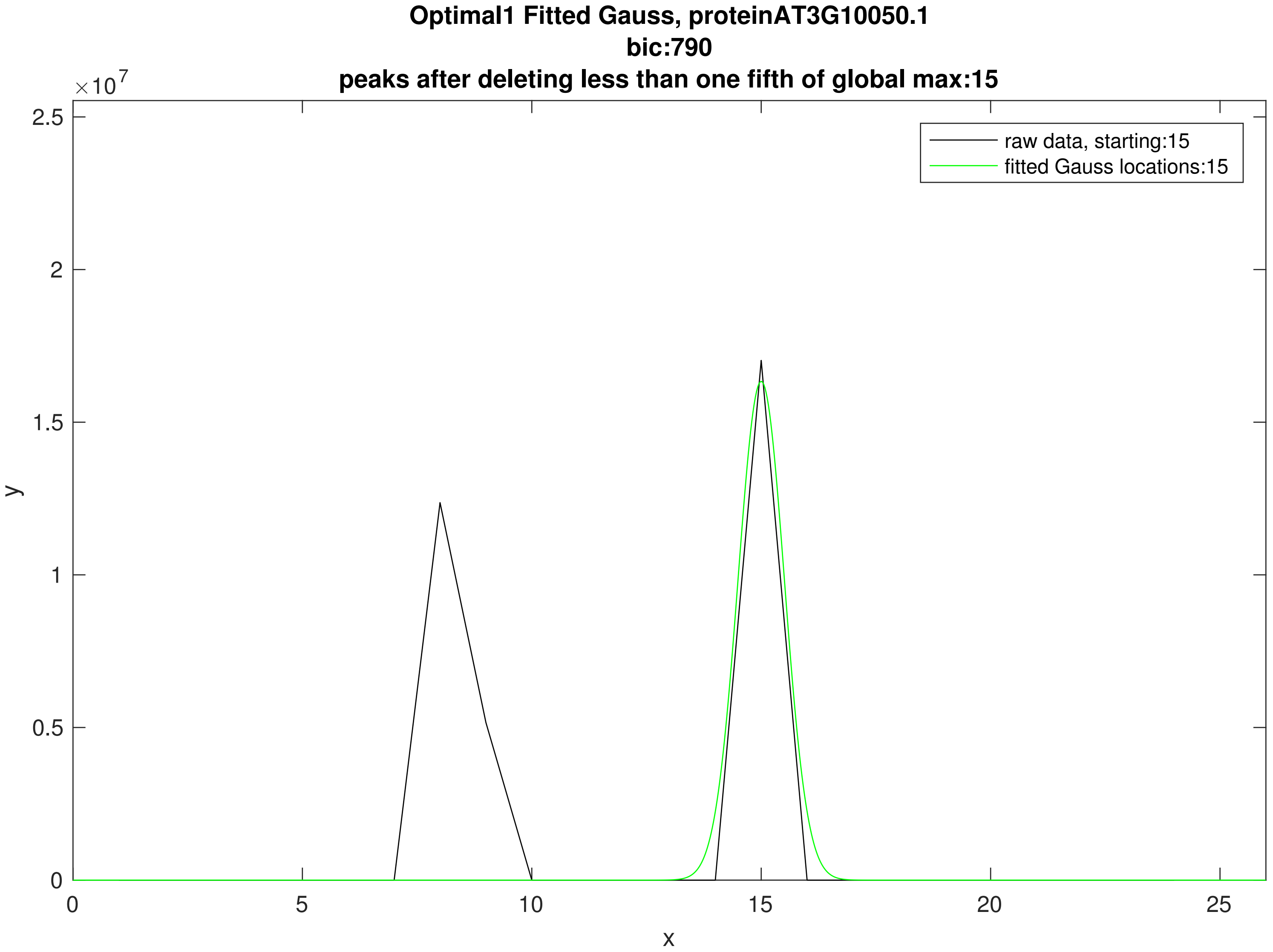
Optimal1 Fitted Gauss, proteinAT3G10050.1
bic:790
peaks after deleting less than one fifth of global max:15
×10^7
raw data, starting:15
fitted Gauss locations:15
y
x
0
0.5
1
1.5
2
2.5
0
5
10
15
20
25

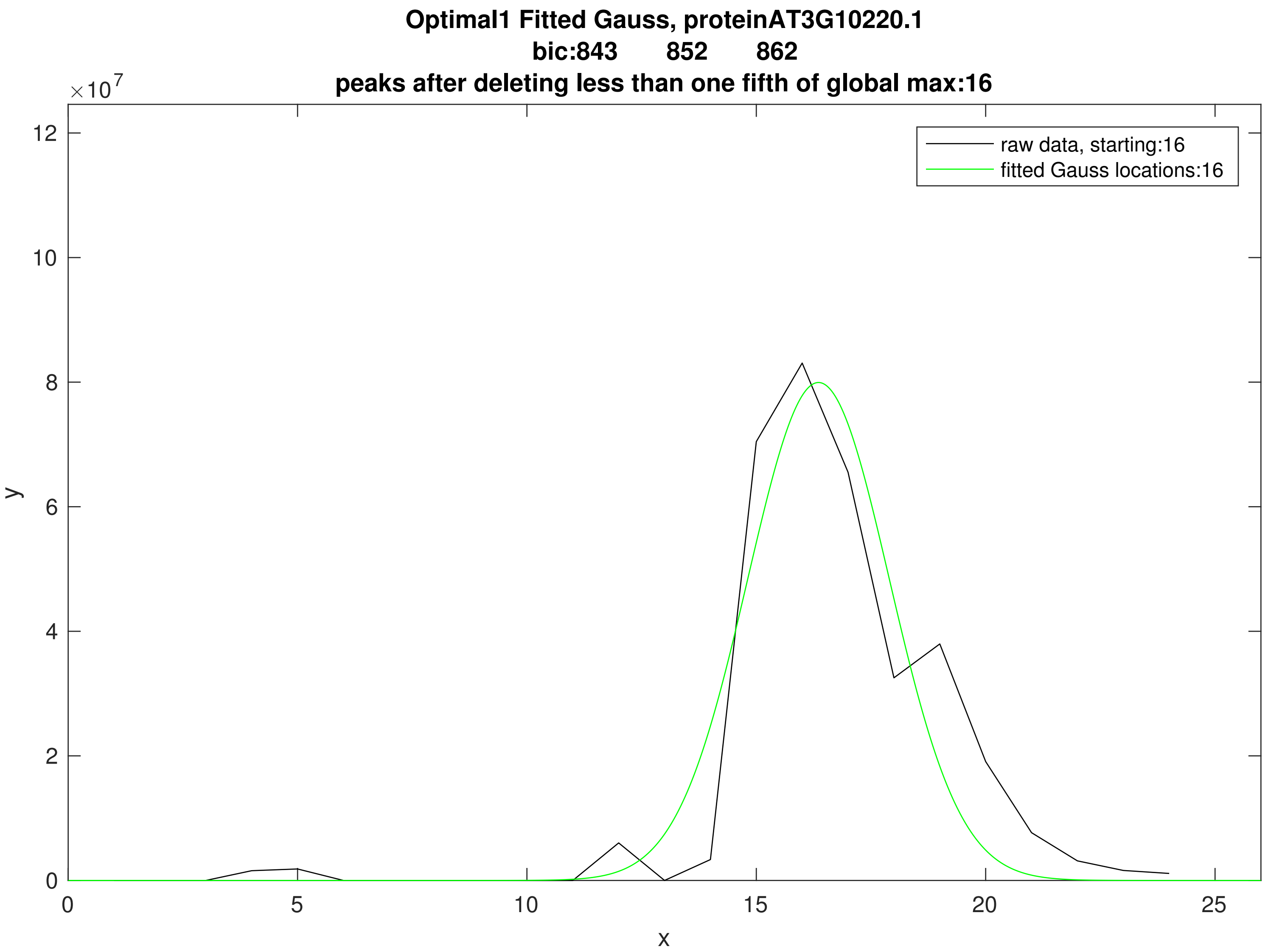

Optimal1 Fitted Gauss, proteinAT3G10220.1
bic:843 852 862
peaks after deleting less than one fifth of global max:16
×10^7
raw data, starting:16
fitted Gauss locations:16
y
x
0
2
4
6
8
10
12
0
5
10
15
20
25

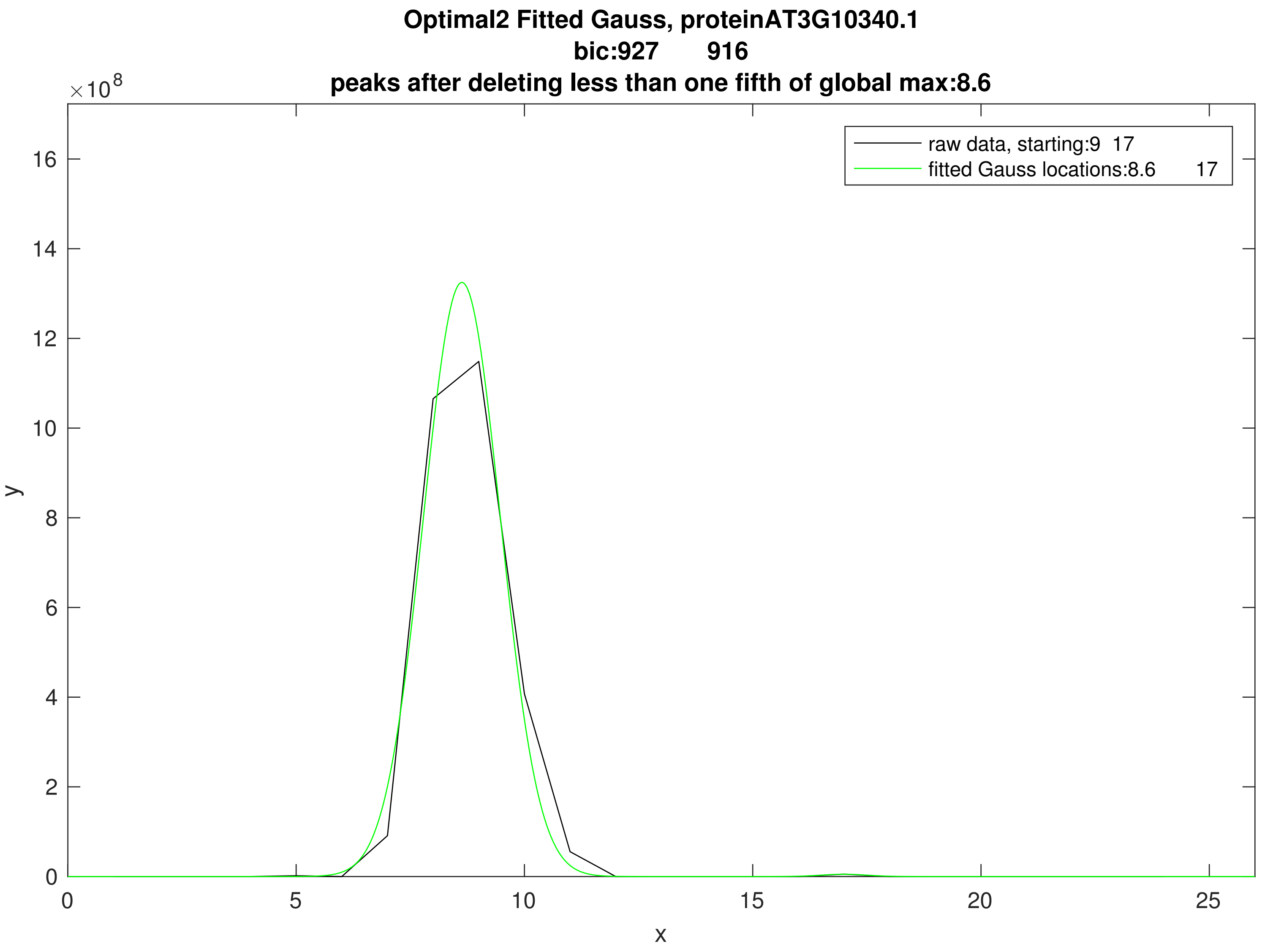
Optimal2 Fitted Gauss, proteinAT3G10340.1
bic:927 916
peaks after deleting less than one fifth of global max:8.6
×10^8
raw data, starting:9 17
fitted Gauss locations:8.6 17
y
x
0
2
4
6
8
10
12
14
16
0
5
10
15
20
25

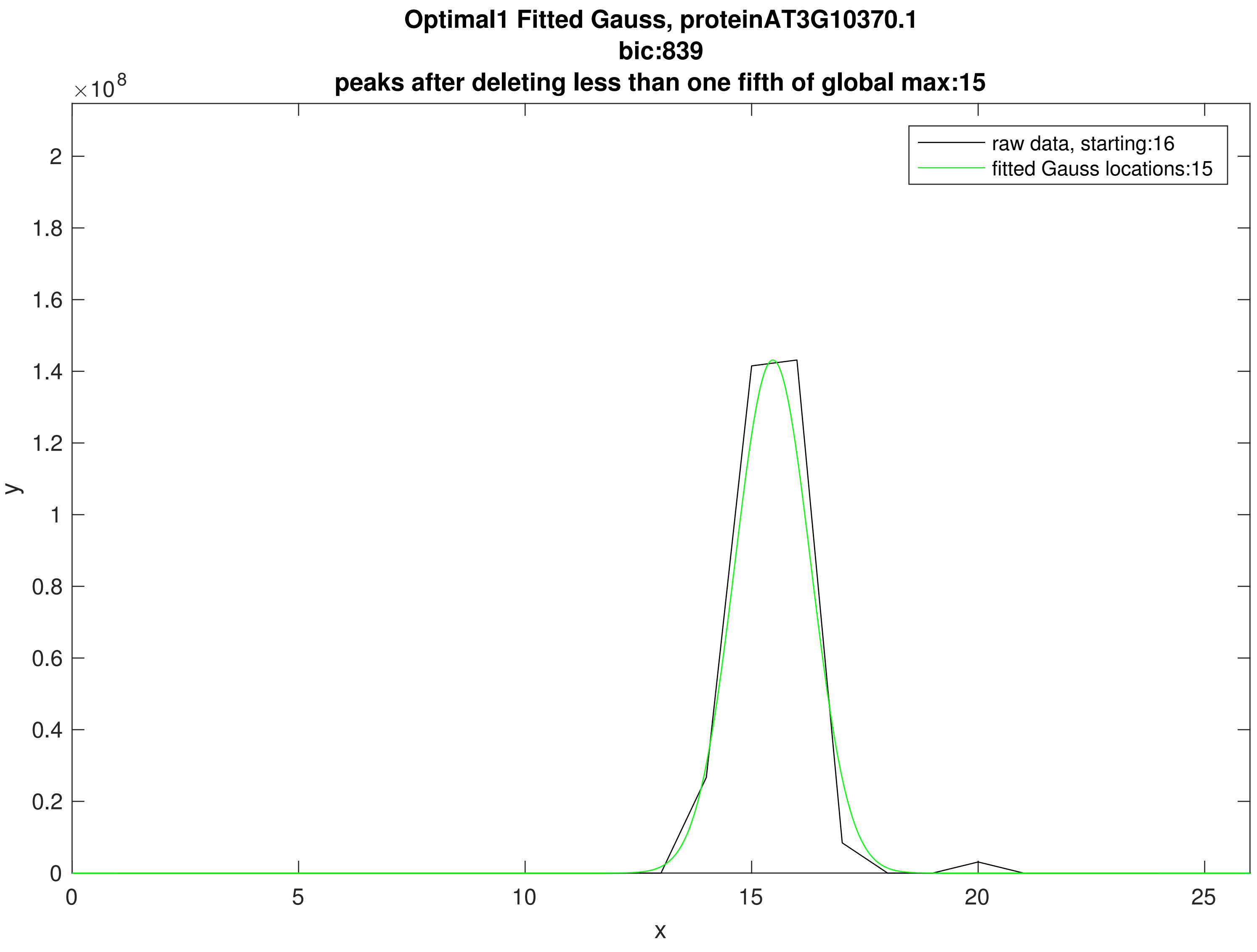

Optimal1 Fitted Gauss, proteinAT3G10370.1
bic:839
peaks after deleting less than one fifth of global max:15
×10^8
raw data, starting:16
fitted Gauss locations:15
y
x
0
0.2
0.4
0.6
0.8
1
1.2
1.4
1.6
1.8
2
5
10
15
20
25

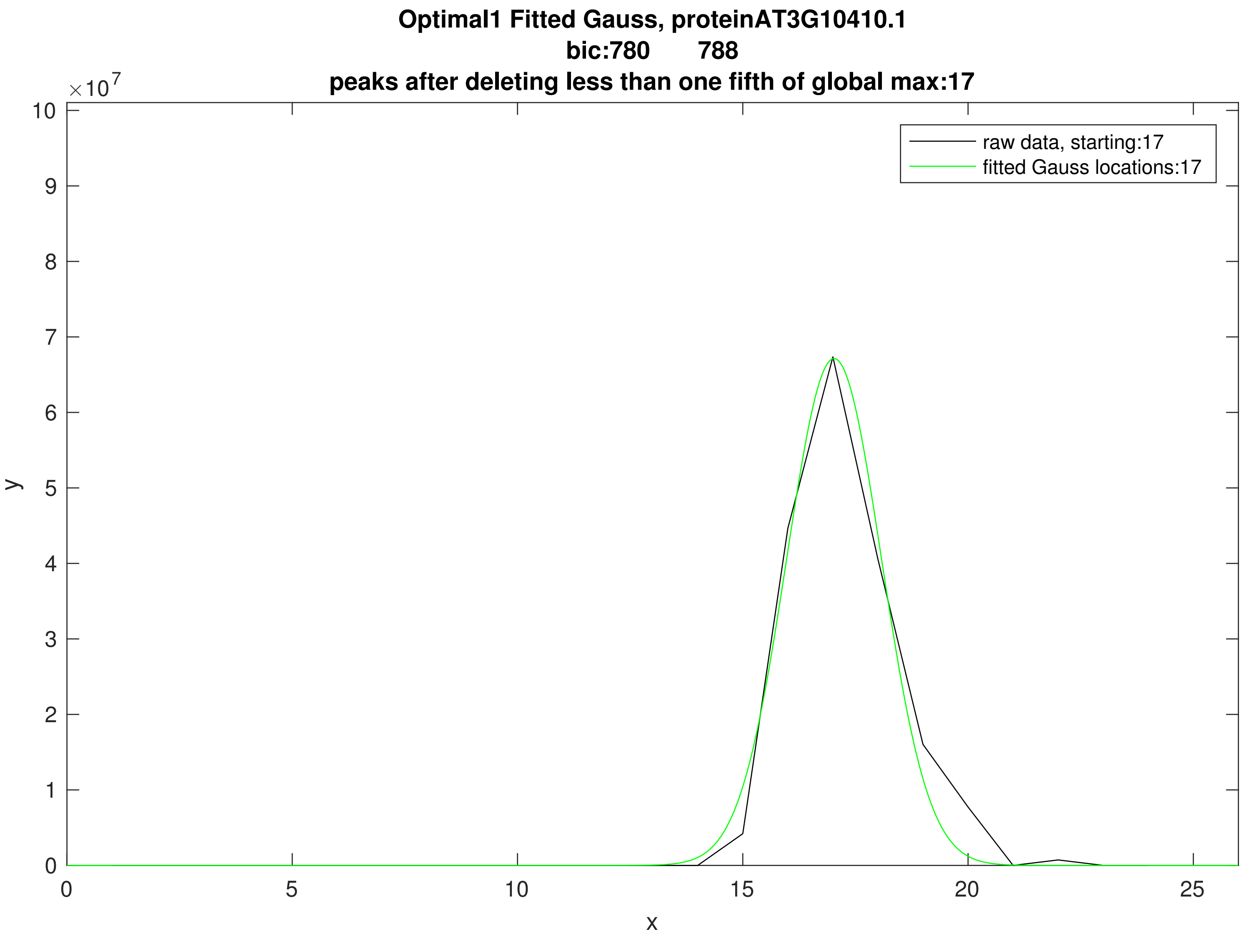

Optimal1 Fitted Gauss, proteinAT3G10410.1
bic:780 788
peaks after deleting less than one fifth of global max:17
×10^7
raw data, starting:17
fitted Gauss locations:17
y
x
0
1
2
3
4
5
6
7
8
9
10
0
5
10
15
20
25

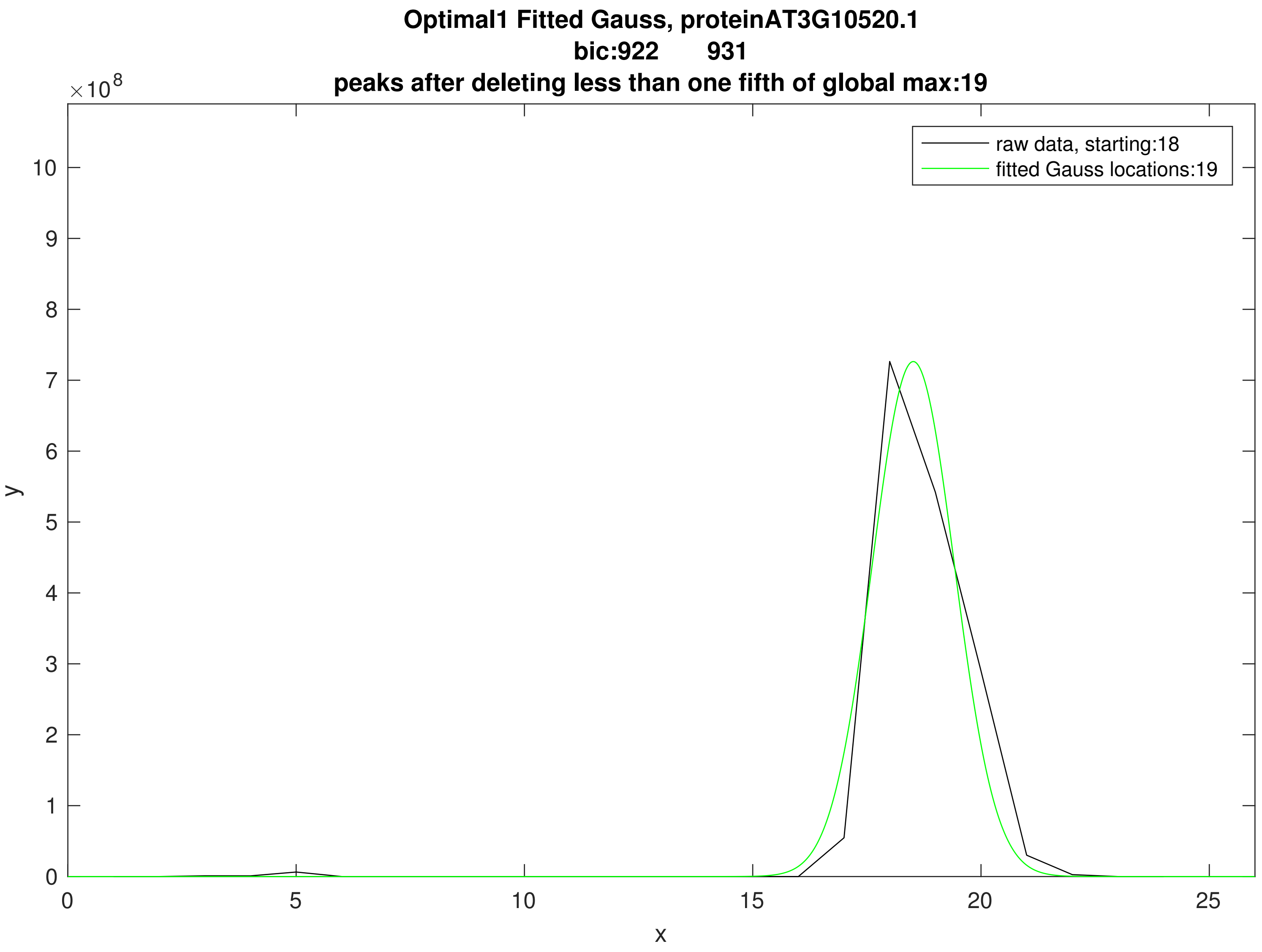
Optimal1 Fitted Gauss, proteinAT3G10520.1
bic:922 931
peaks after deleting less than one fifth of global max:19
×10^8
raw data, starting:18
fitted Gauss locations:19
y
x
0
1
2
3
4
5
6
7
8
9
10
0
5
10
15
20
25

**1 Fitted Gauss, proteinAT3G10610.1;AT2G05220.2;AT2G05220.1;AT5G04800.4;AT5G04800.3;AT5G04800.2;AT5G0480**

**bic:871 876**

**peaks after deleting less than one fifth of global max:3.7**

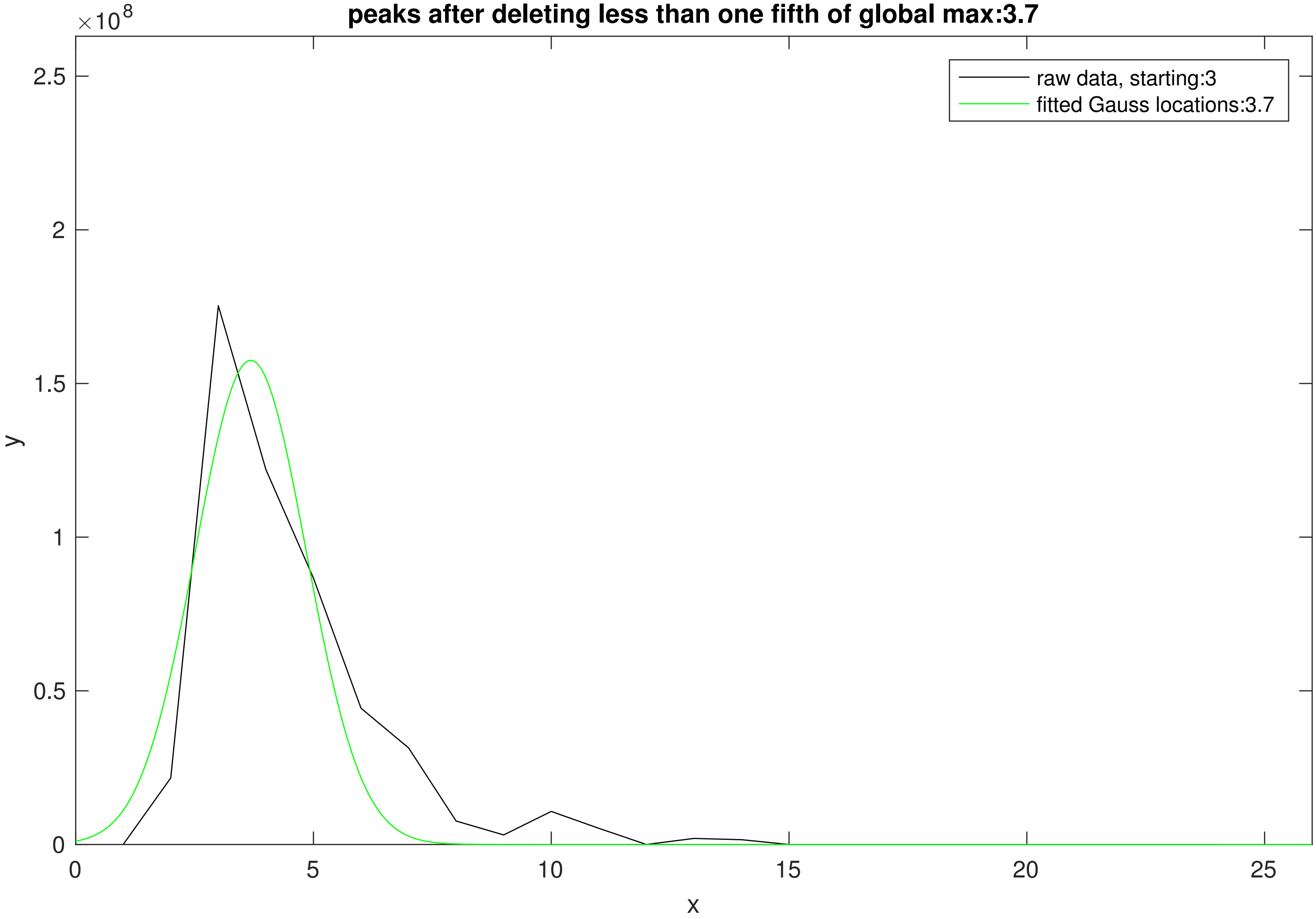

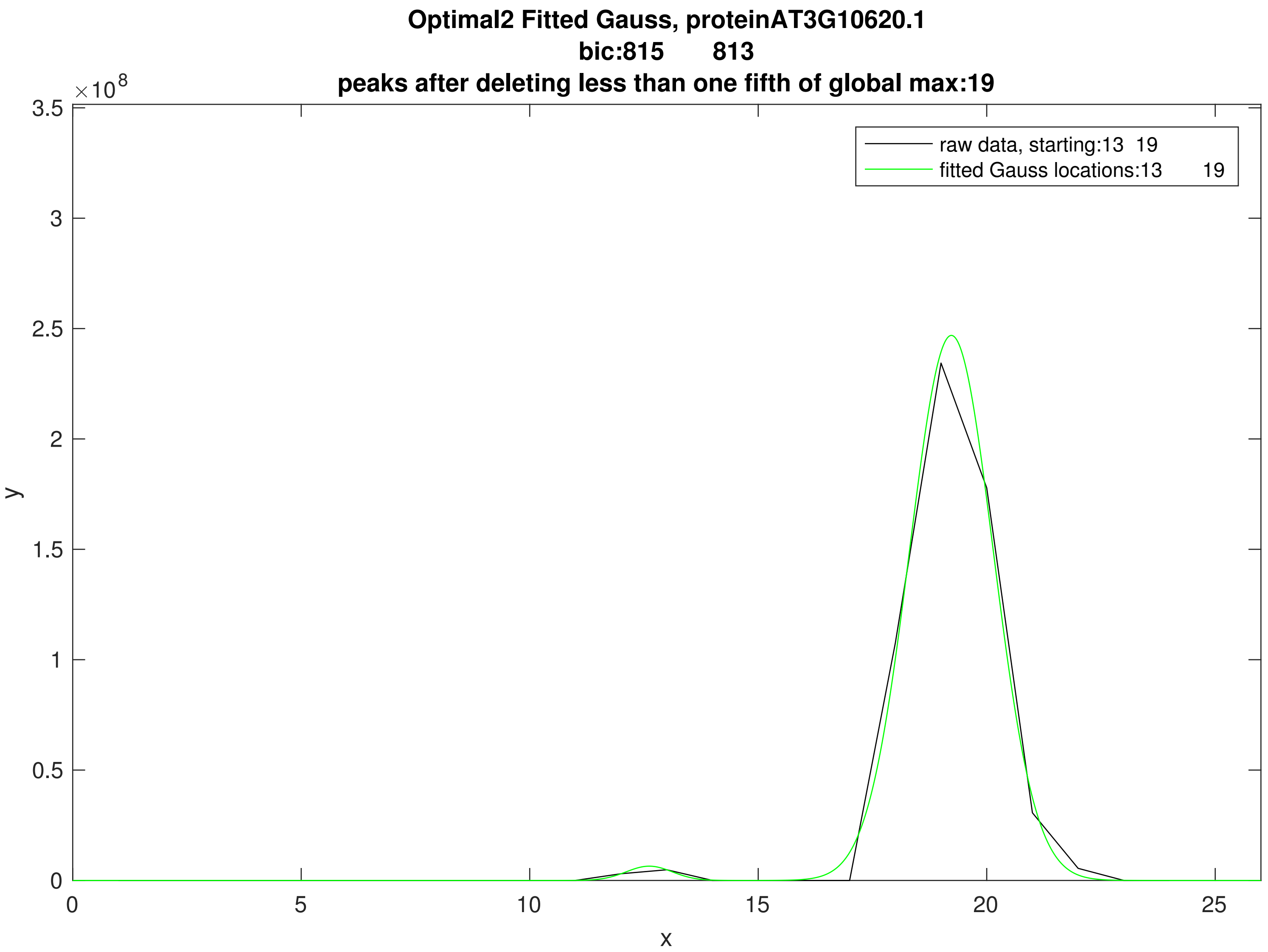

Optimal2 Fitted Gauss, proteinAT3G10620.1
bic:815 813
peaks after deleting less than one fifth of global max:19
×10^8
3.5
3
2.5
2
1.5
1
0.5
0
y
0
5
10
15
20
25
x
raw data, starting:13 19
fitted Gauss locations:13 19

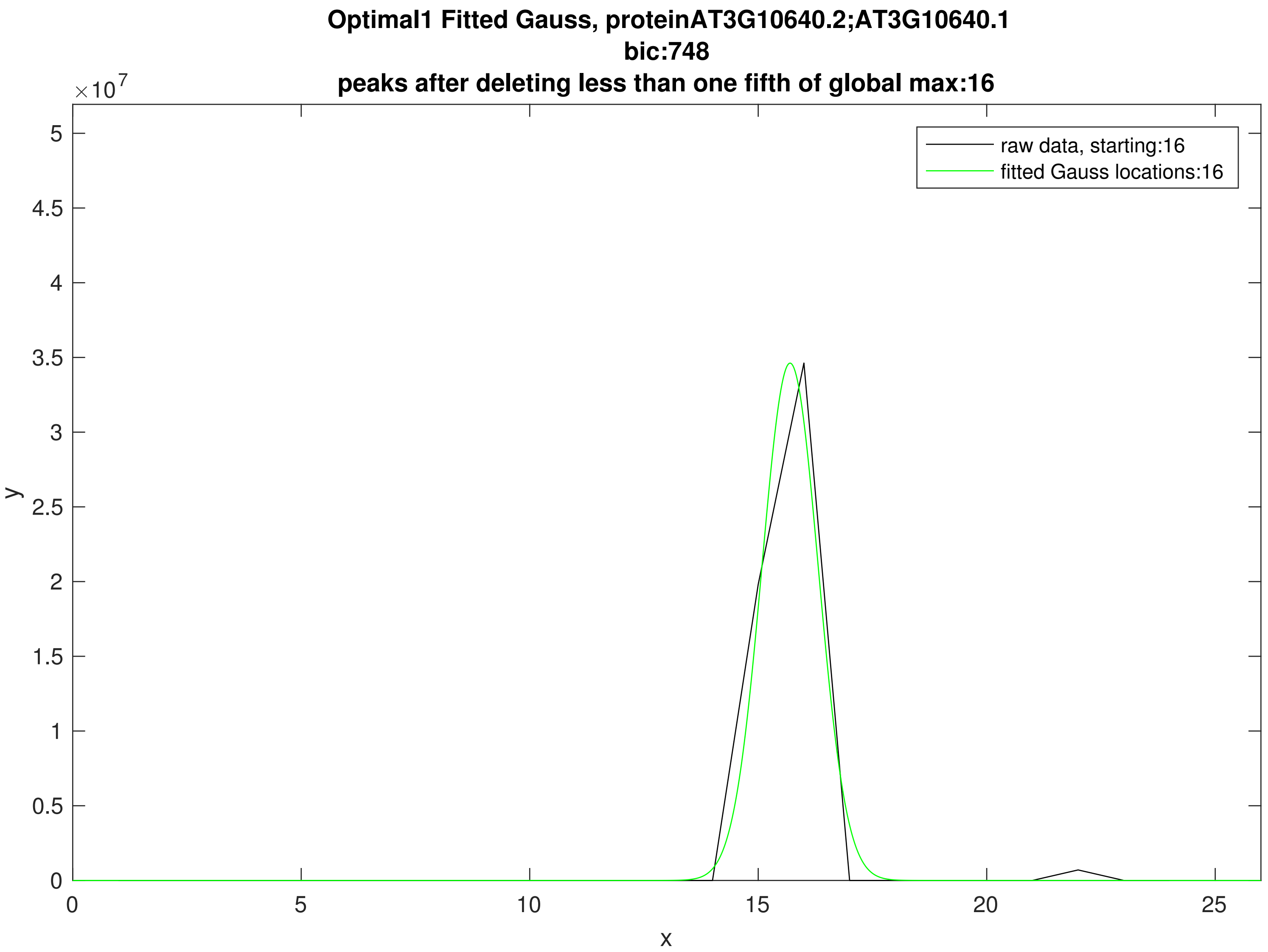
Optimal1 Fitted Gauss, proteinAT3G10640.2;AT3G10640.1
bic:748
peaks after deleting less than one fifth of global max:16
×10^7
raw data, starting:16
fitted Gauss locations:16
y
x
0
0.5
1
1.5
2
2.5
3
3.5
4
4.5
5
0
5
10
15
20
25

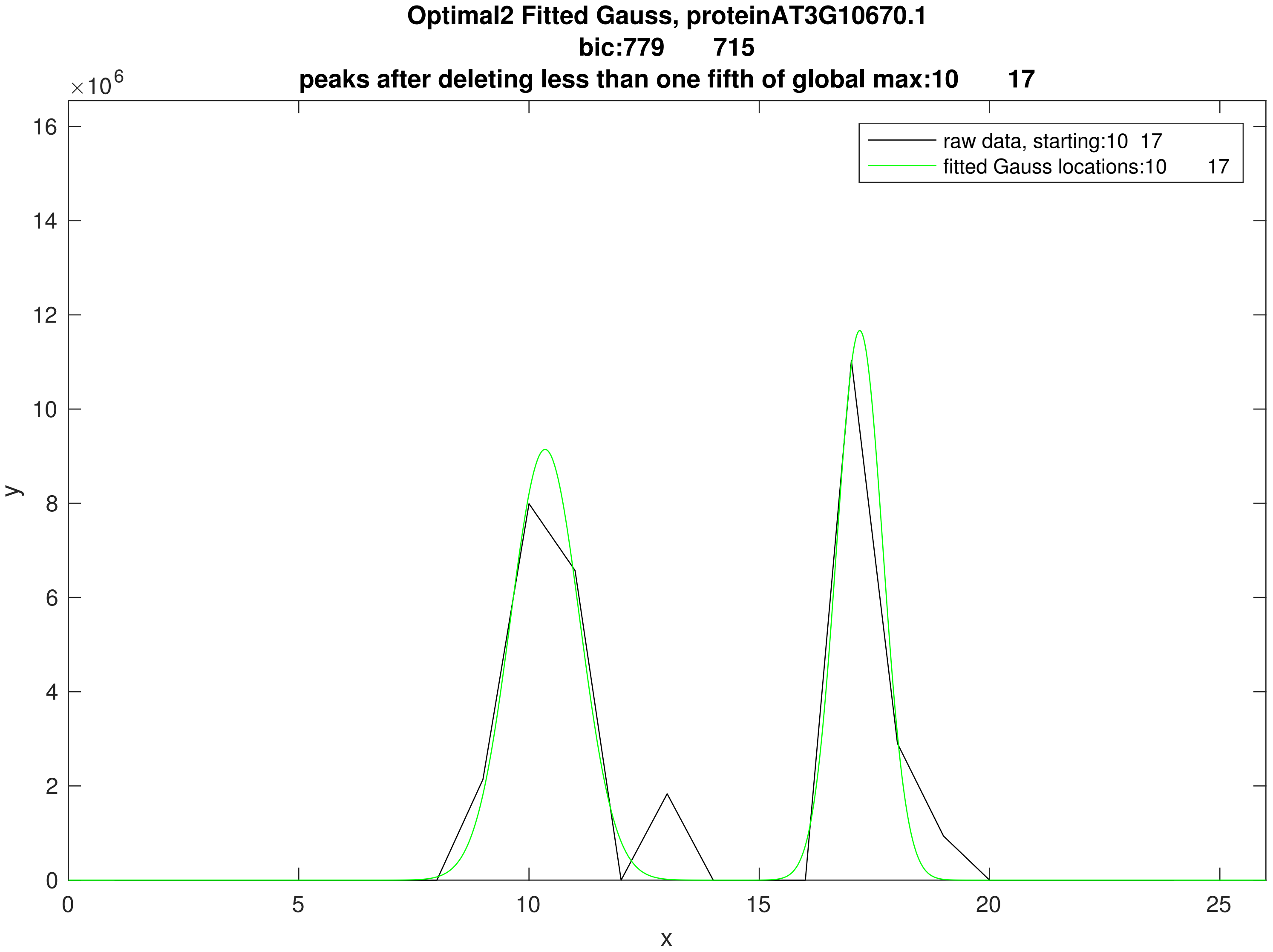

Optimal2 Fitted Gauss, proteinAT3G10670.1
bic:779 715
peaks after deleting less than one fifth of global max:10 17
×10^6
raw data, starting:10 17
fitted Gauss locations:10 17
y
x
0
2
4
6
8
10
12
14
16
0
5
10
15
20
25

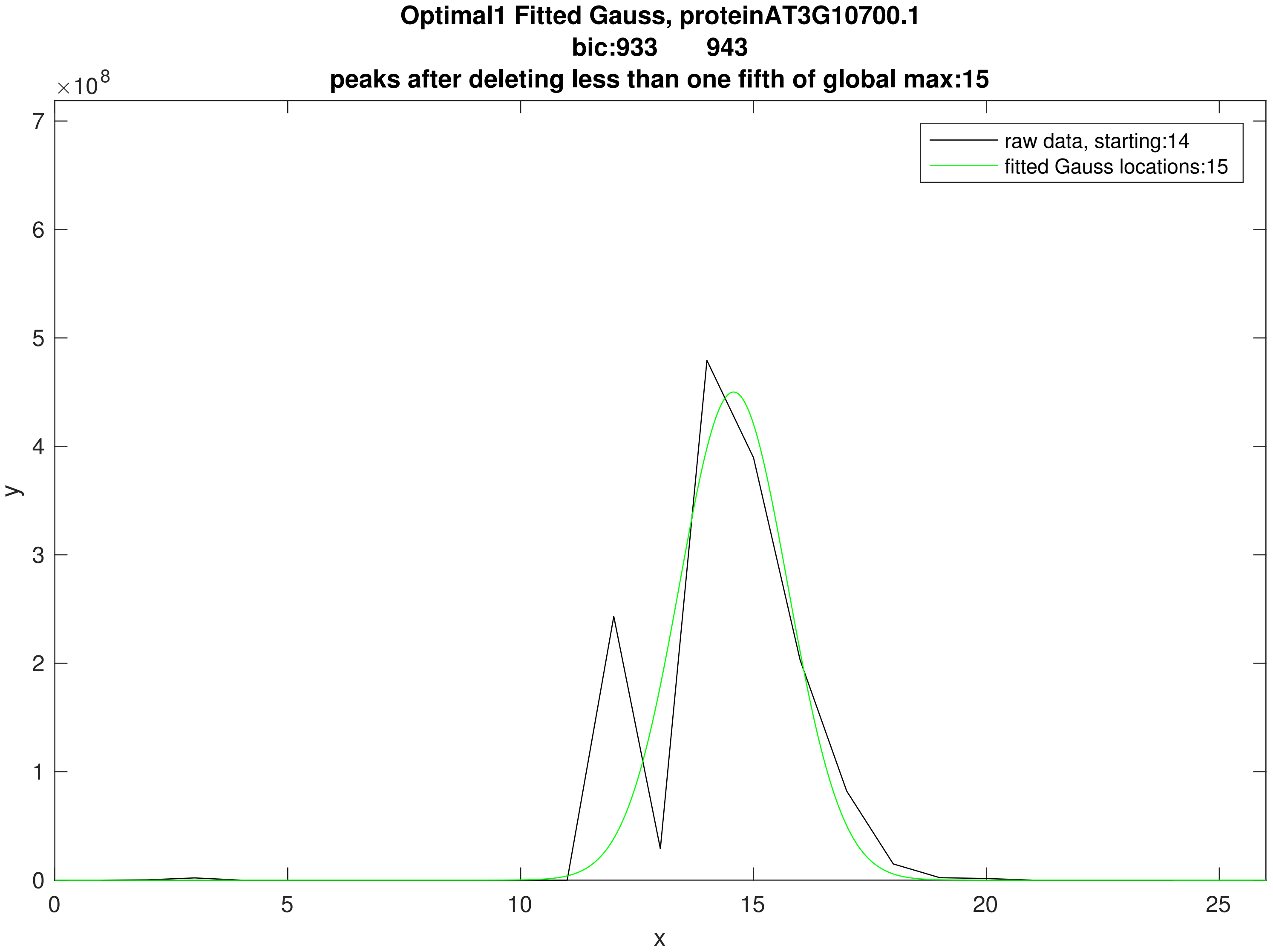
Optimal1 Fitted Gauss, proteinAT3G10700.1
bic:933 943
peaks after deleting less than one fifth of global max:15
×10^8
raw data, starting:14
fitted Gauss locations:15
y
x
0
1
2
3
4
5
6
7
0
5
10
15
20
25

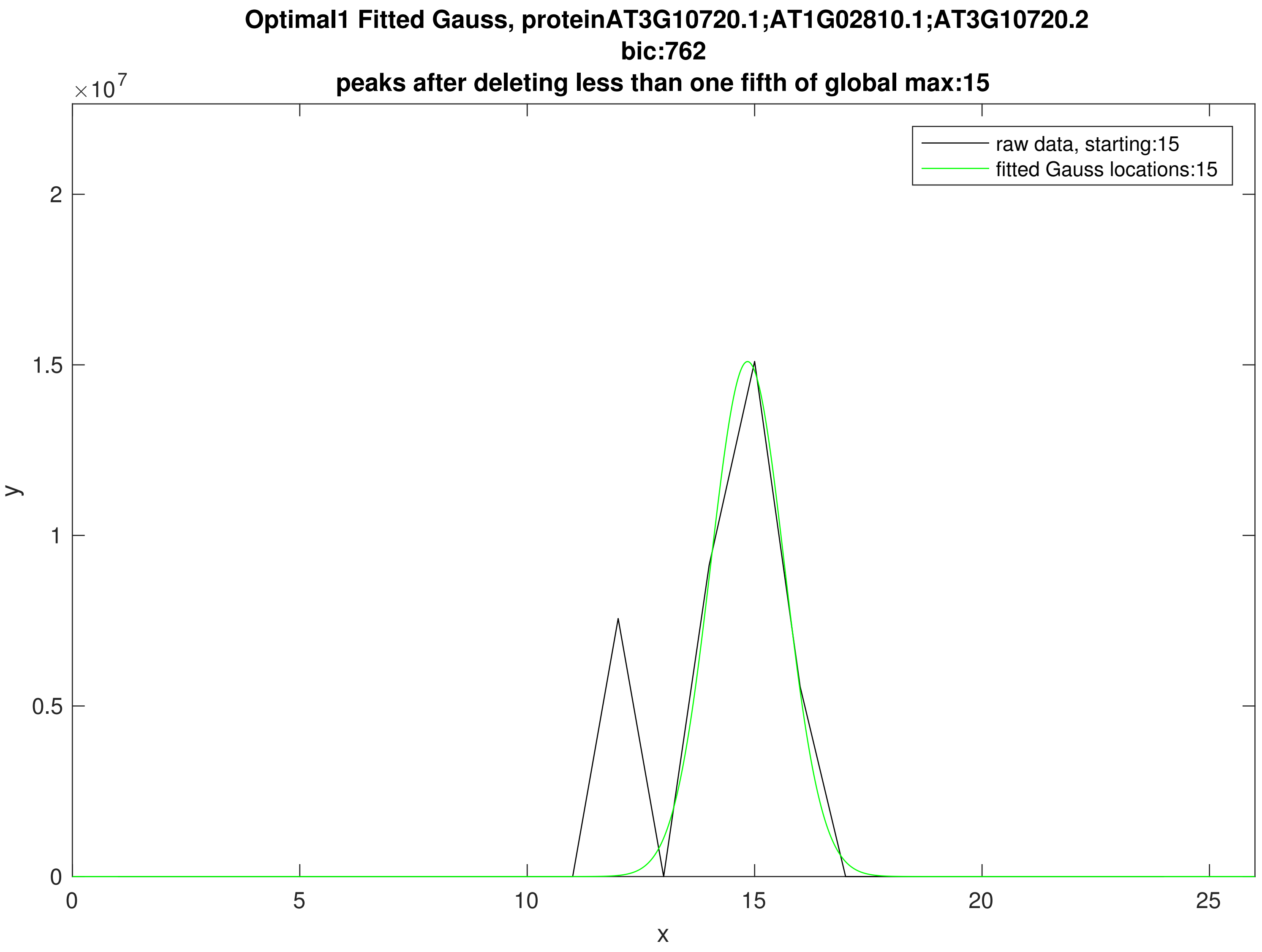

Optimal1 Fitted Gauss, proteinAT3G10720.1;AT1G02810.1;AT3G10720.2
bic:762
peaks after deleting less than one fifth of global max:15
×10^7
raw data, starting:15
fitted Gauss locations:15
2
1.5
1
0.5
0
y
0
5
10
15
20
25
x

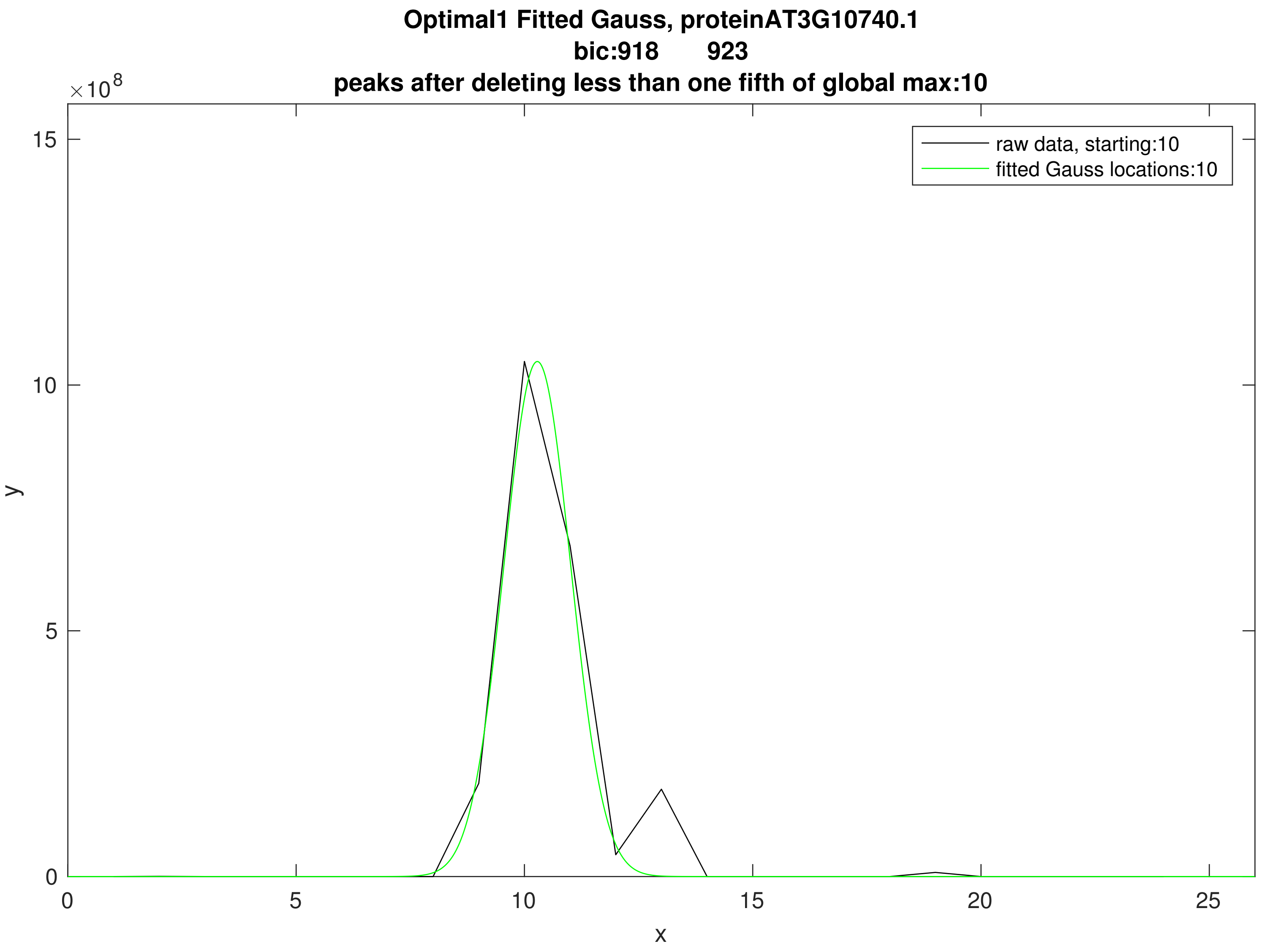
Optimal1 Fitted Gauss, proteinAT3G10740.1
bic:918 923
peaks after deleting less than one fifth of global max:10
×10^8
15
10
5
0
y
0
5
10
15
20
25
x
raw data, starting:10
fitted Gauss locations:10

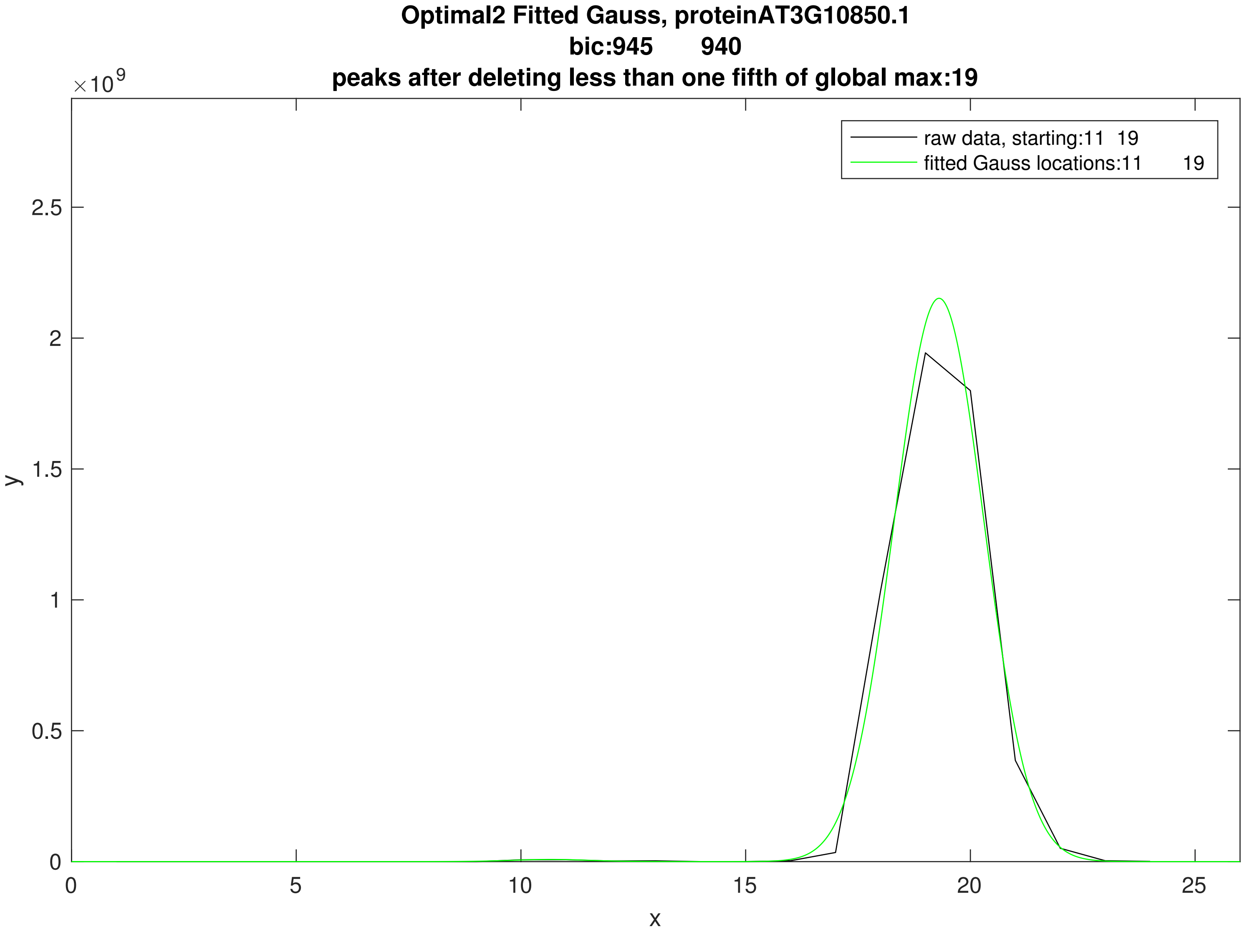

Optimal2 Fitted Gauss, proteinAT3G10850.1
bic:945 940
peaks after deleting less than one fifth of global max:19
×10^9
raw data, starting:11 19
fitted Gauss locations:11 19
2.5
2
1.5
1
0.5
0
y
0
5
10
15
20
25
x

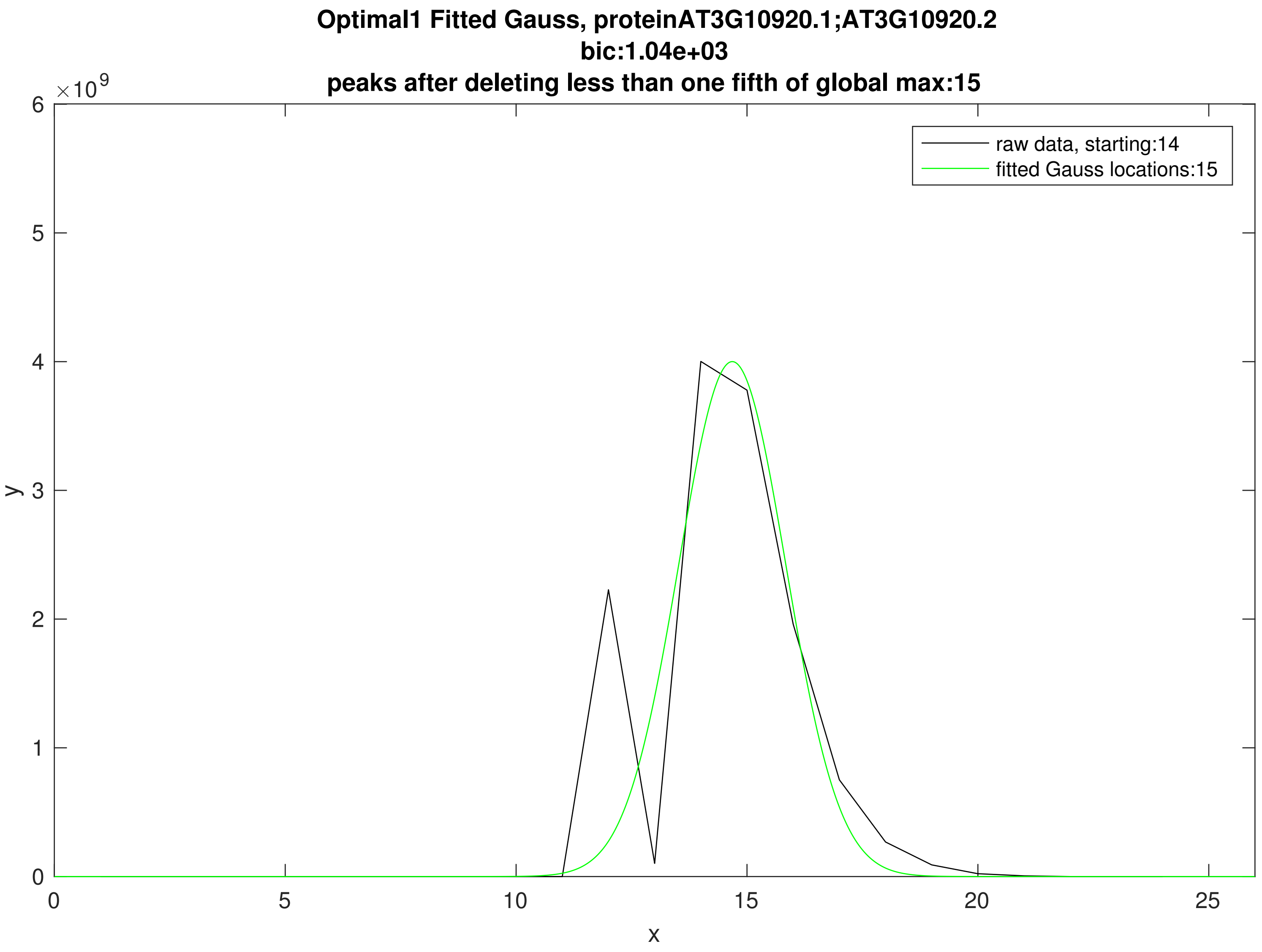
Optimal1 Fitted Gauss, proteinAT3G10920.1;AT3G10920.2
bic:1.04e+03
peaks after deleting less than one fifth of global max:15
×10^9
raw data, starting:14
fitted Gauss locations:15
y
x
0
1
2
3
4
5
6
0
5
10
15
20
25

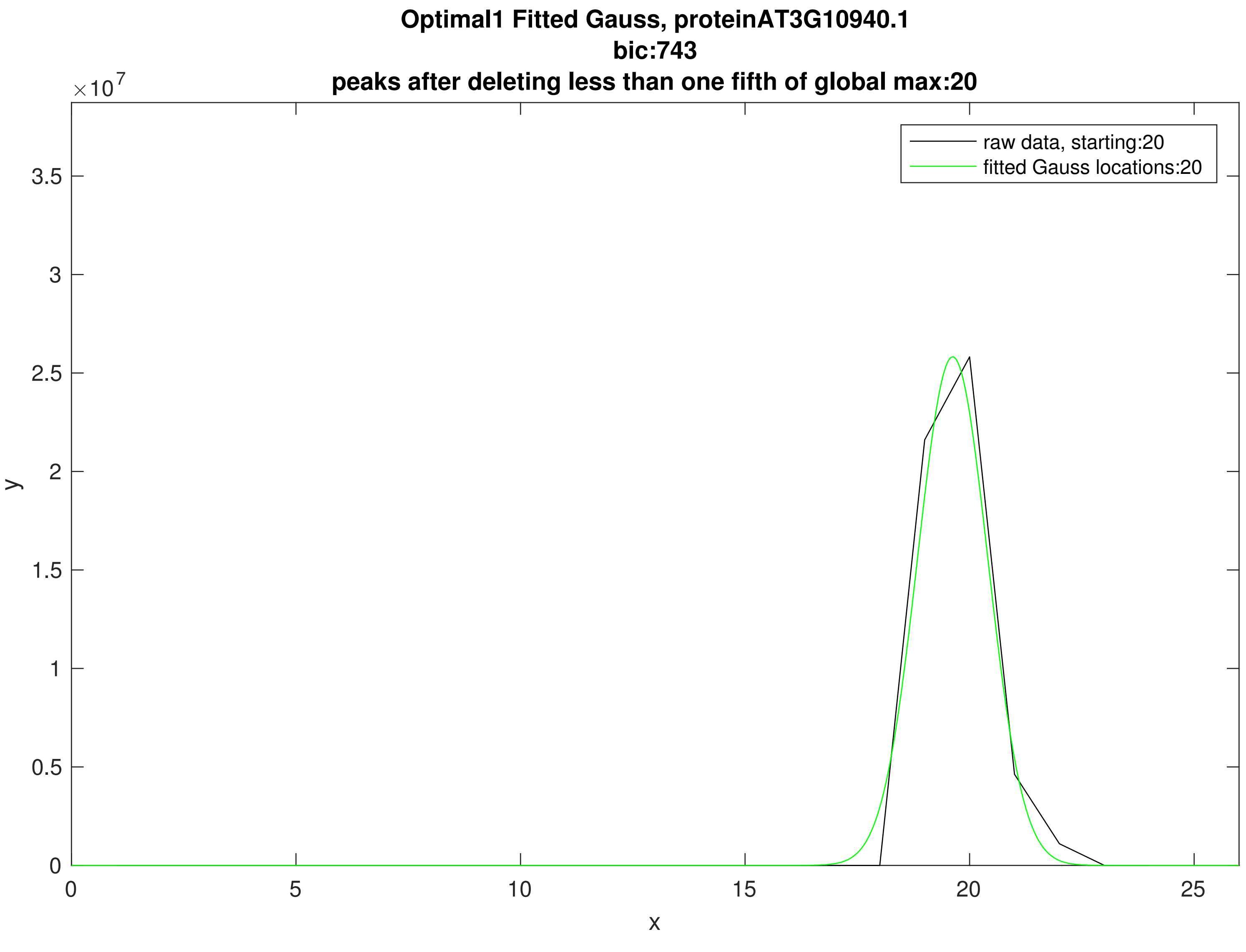
Optimal1 Fitted Gauss, proteinAT3G10940.1
bic:743
peaks after deleting less than one fifth of global max:20
×10^7
raw data, starting:20
fitted Gauss locations:20
3.5
3
2.5
2
1.5
1
0.5
0
y
0
5
10
15
20
25
x

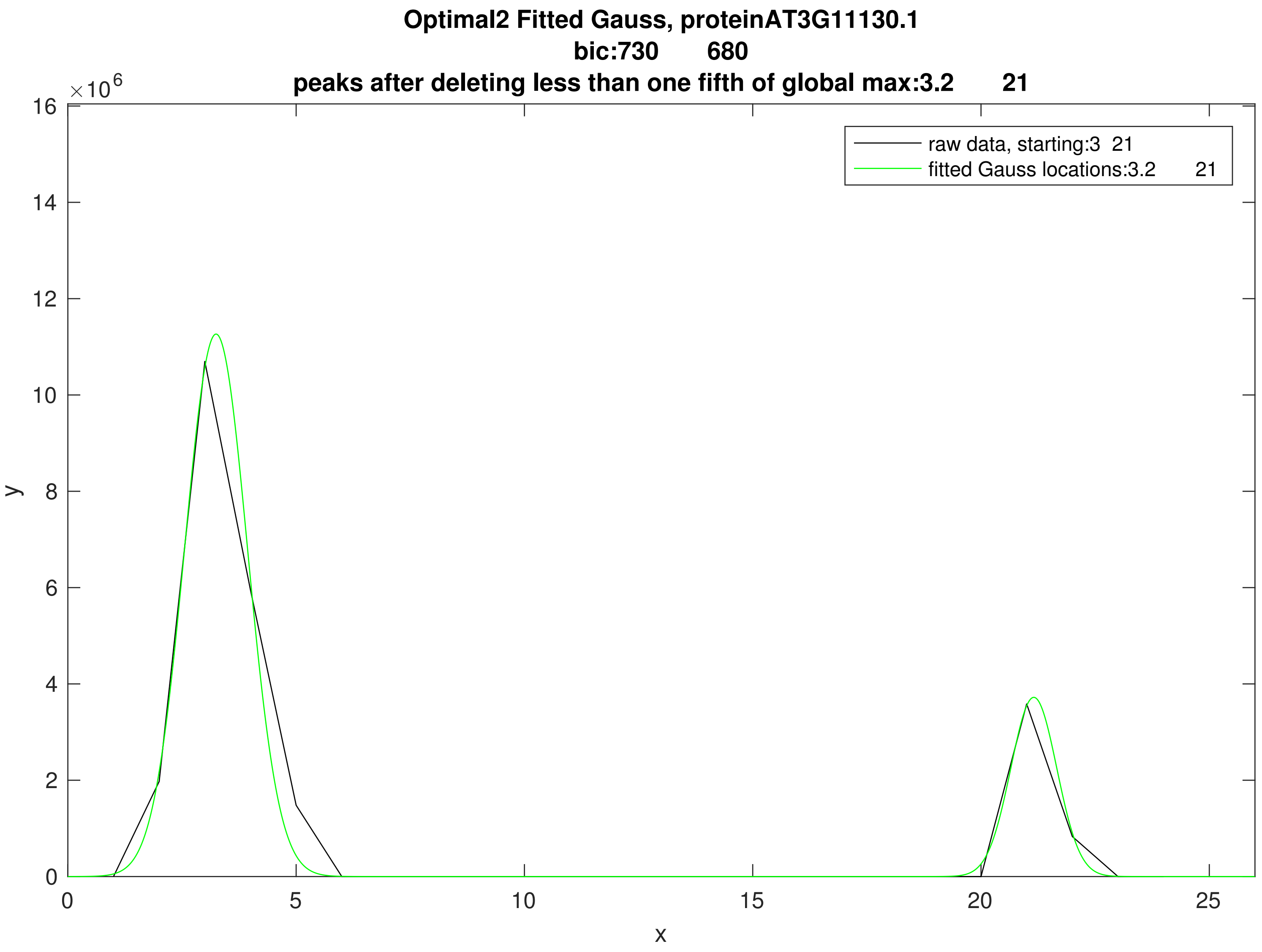
Optimal2 Fitted Gauss, proteinAT3G11130.1
bic:730 680
peaks after deleting less than one fifth of global max:3.2 21
raw data, starting:3 21
fitted Gauss locations:3.2 21
×10^6
16
14
12
10
8
6
4
2
0
y
0
5
10
15
20
25
x

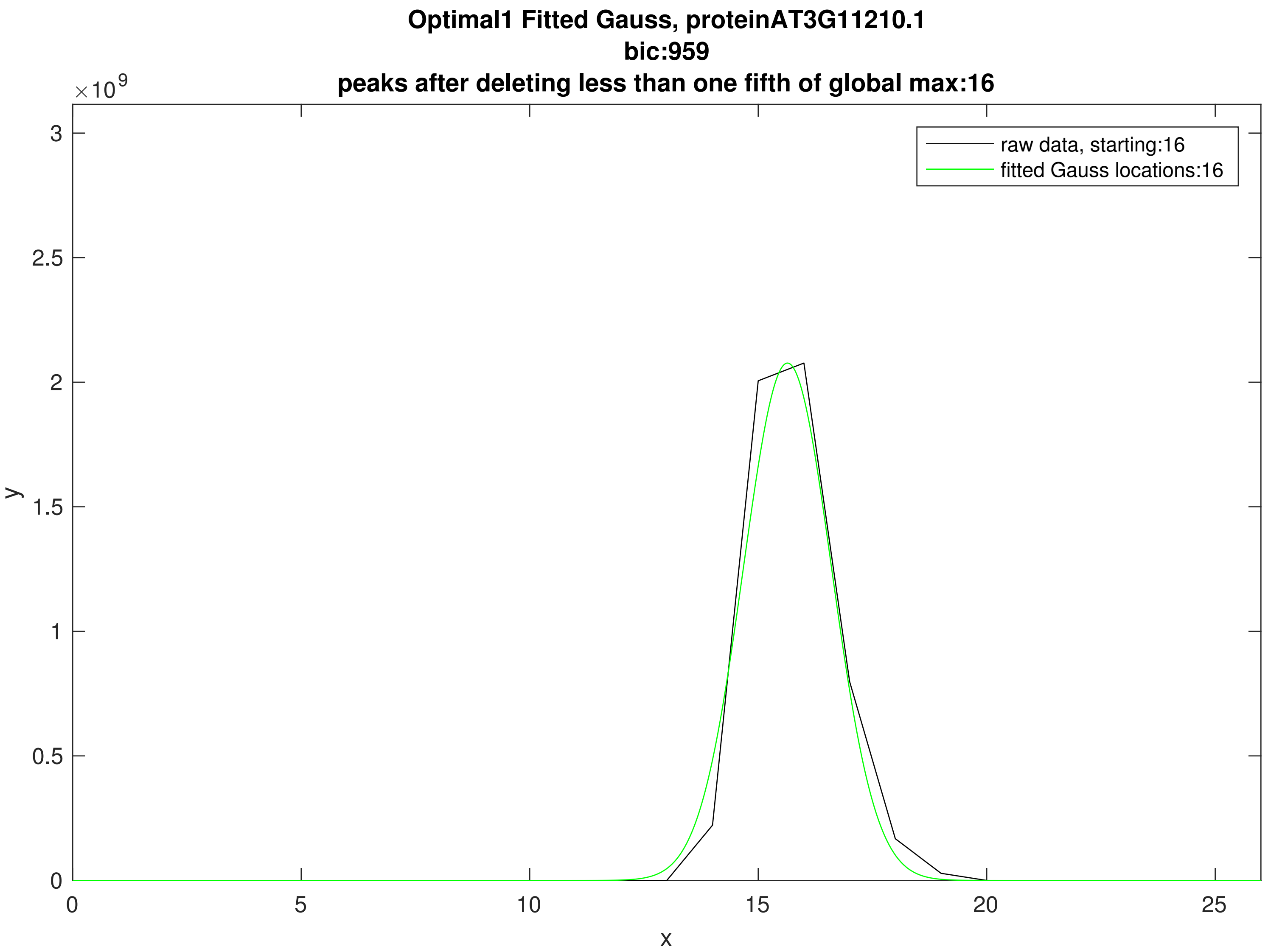
Optimal1 Fitted Gauss, proteinAT3G11210.1
bic:959
peaks after deleting less than one fifth of global max:16
×10^9
raw data, starting:16
fitted Gauss locations:16
y
x
0
0.5
1
1.5
2
2.5
3
0
5
10
15
20
25

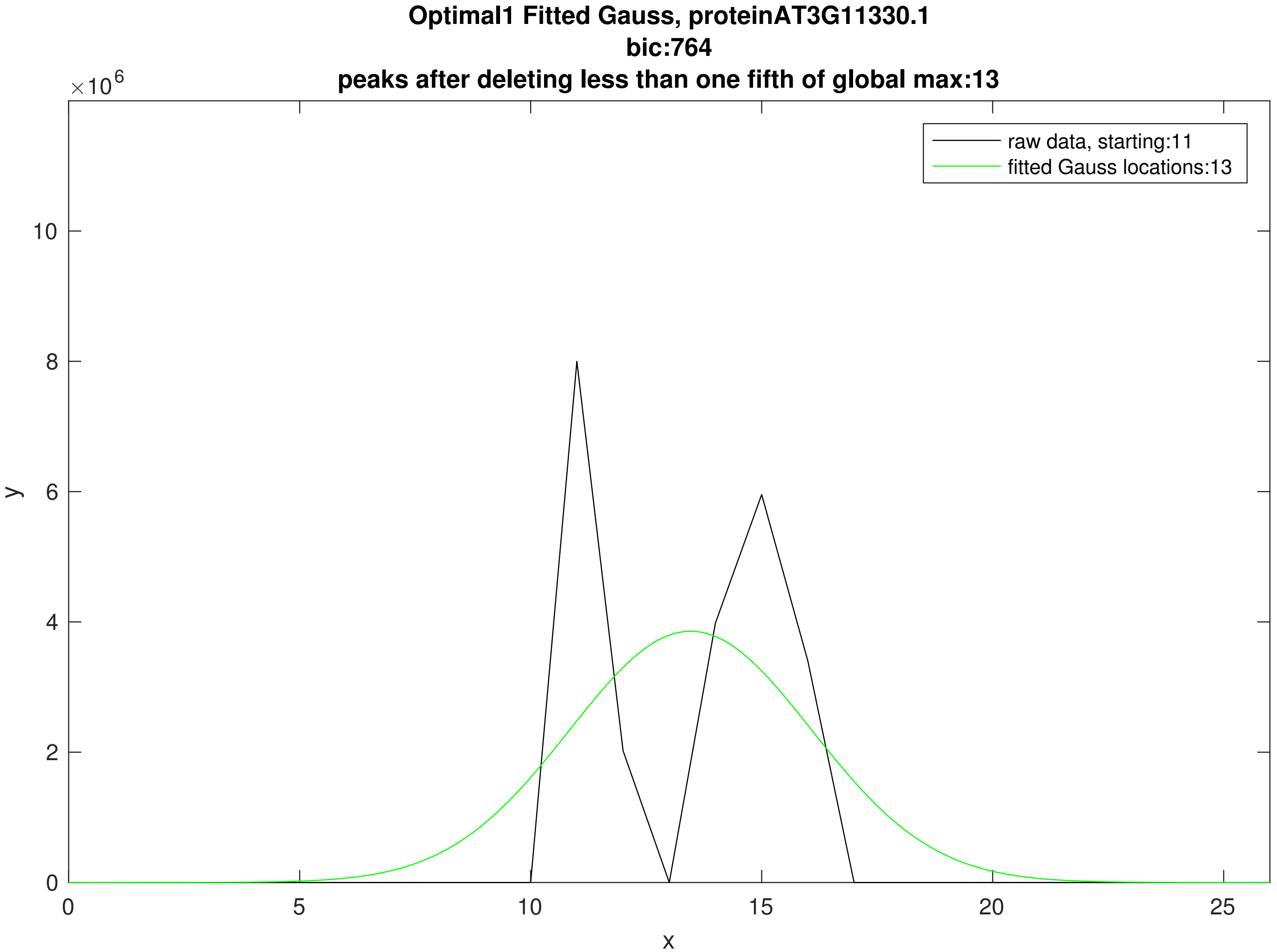
Optimal1 Fitted Gauss, proteinAT3G11330.1
bic:764
peaks after deleting less than one fifth of global max:13
×10^6
raw data, starting:11
fitted Gauss locations:13
y
x
0
2
4
6
8
10
0
5
10
15
20
25

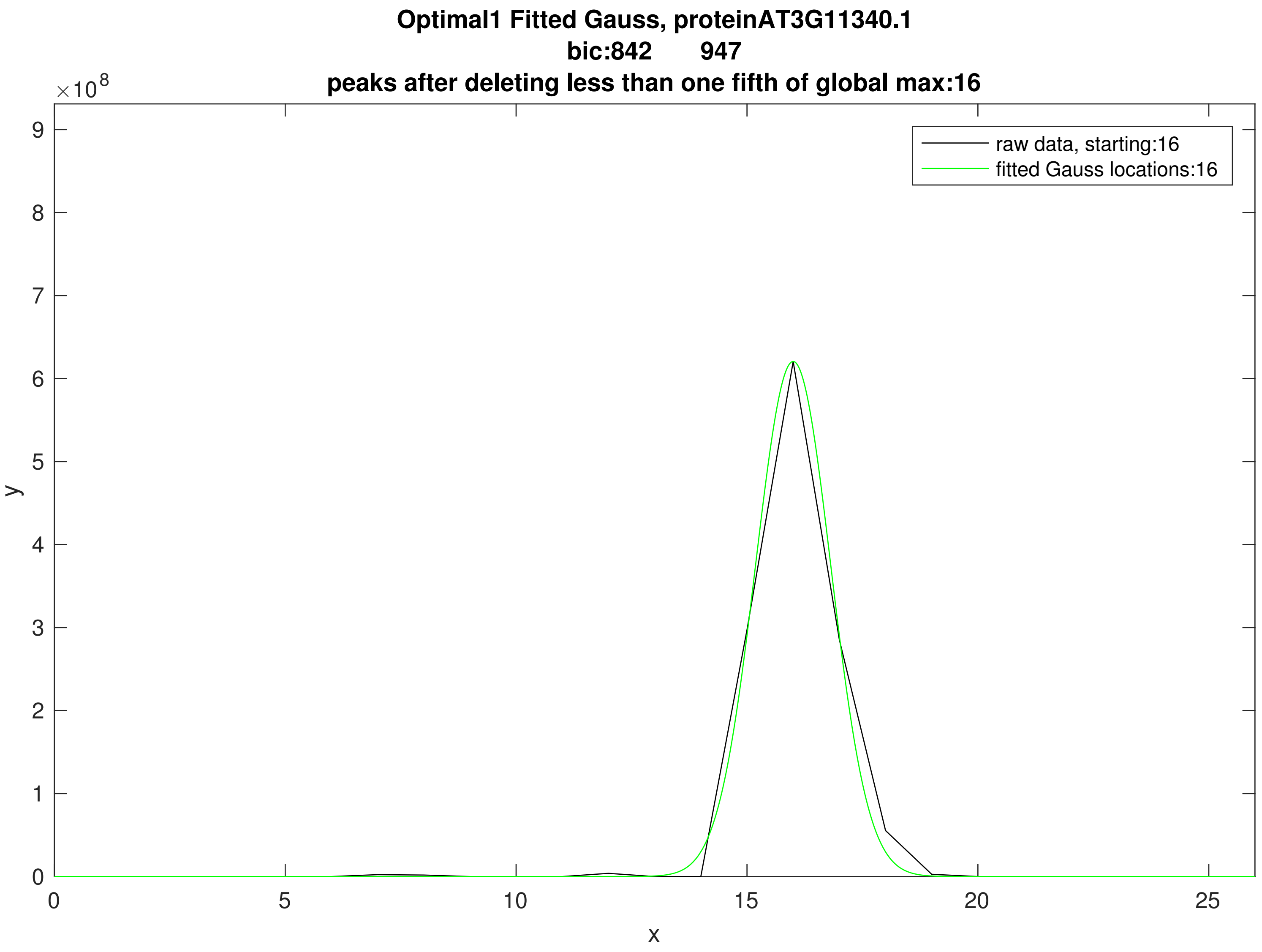

Optimal1 Fitted Gauss, proteinAT3G11340.1
bic:842 947
peaks after deleting less than one fifth of global max:16
×10^8
raw data, starting:16
fitted Gauss locations:16
y
x
0
1
2
3
4
5
6
7
8
9
0
5
10
15
20
25

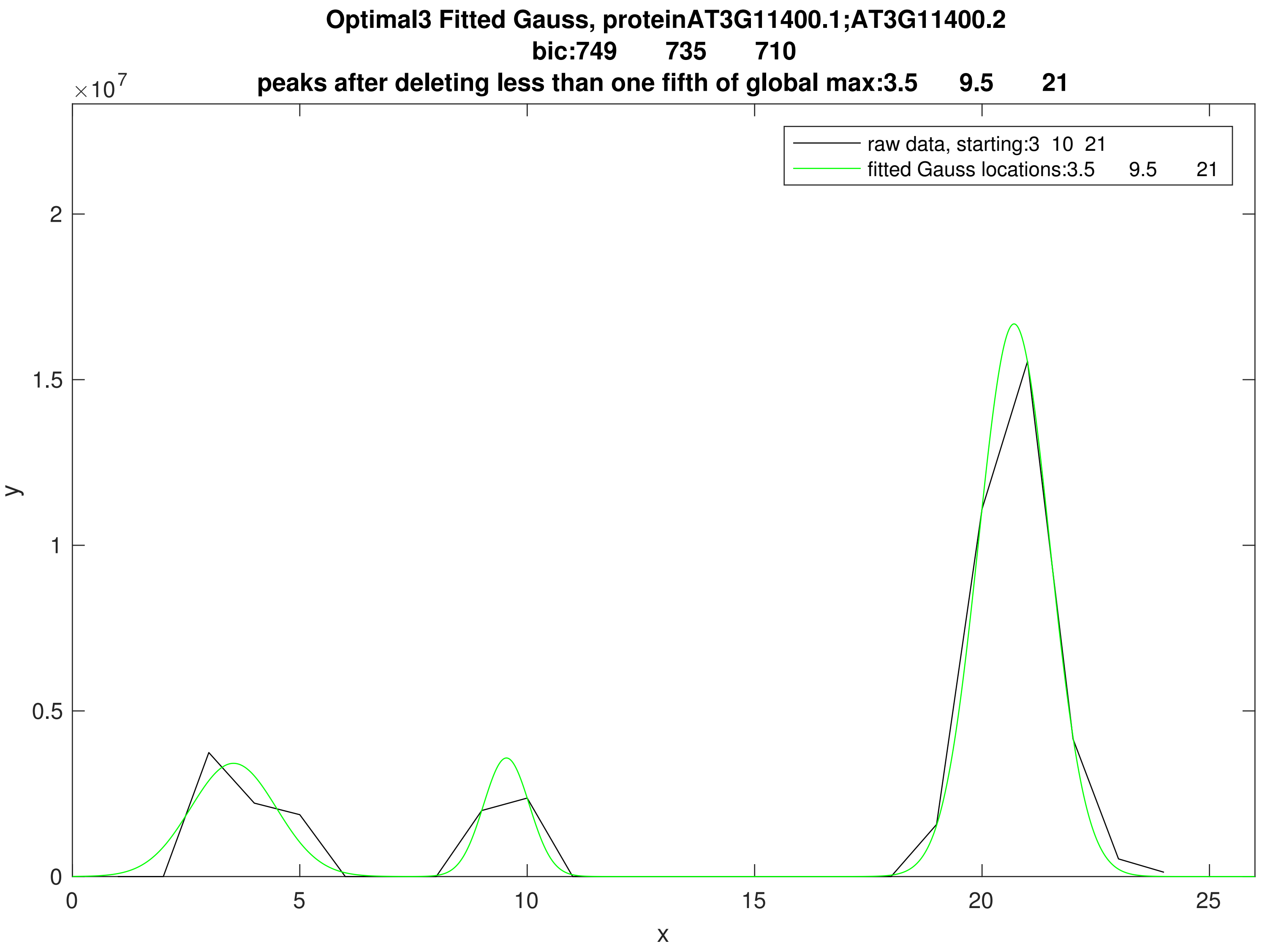
Optimal3 Fitted Gauss, proteinAT3G11400.1;AT3G11400.2
bic:749 735 710
peaks after deleting less than one fifth of global max:3.5 9.5 21
×10^7
raw data, starting:3 10 21
fitted Gauss locations:3.5 9.5 21
y
x
0
0.5
1
1.5
2
5
10
15
20
25

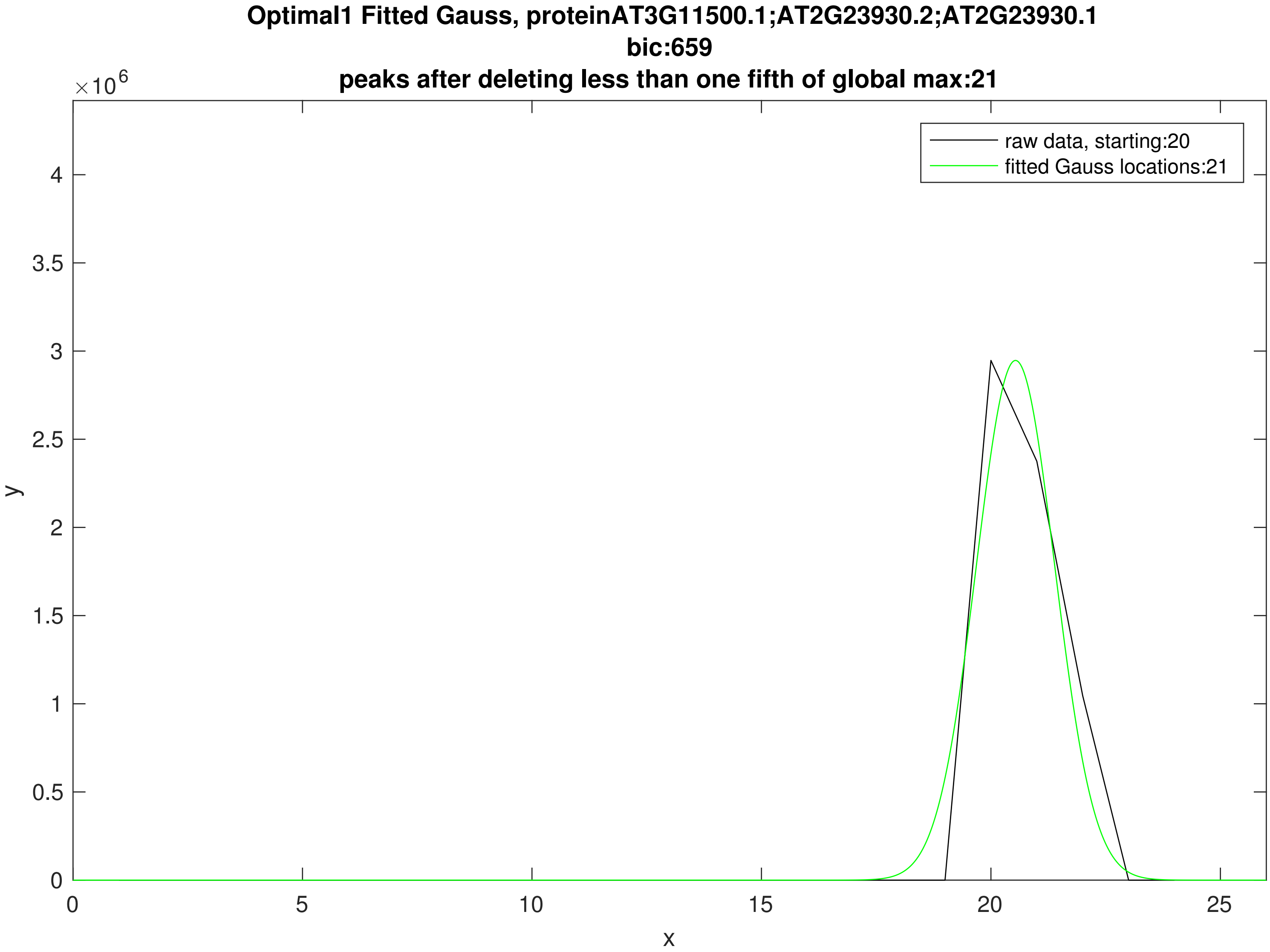

Optimal1 Fitted Gauss, proteinAT3G11500.1;AT2G23930.2;AT2G23930.1
bic:659
peaks after deleting less than one fifth of global max:21
×10^6
raw data, starting:20
fitted Gauss locations:21
y
x
0
0.5
1
1.5
2
2.5
3
3.5
4
0
5
10
15
20
25

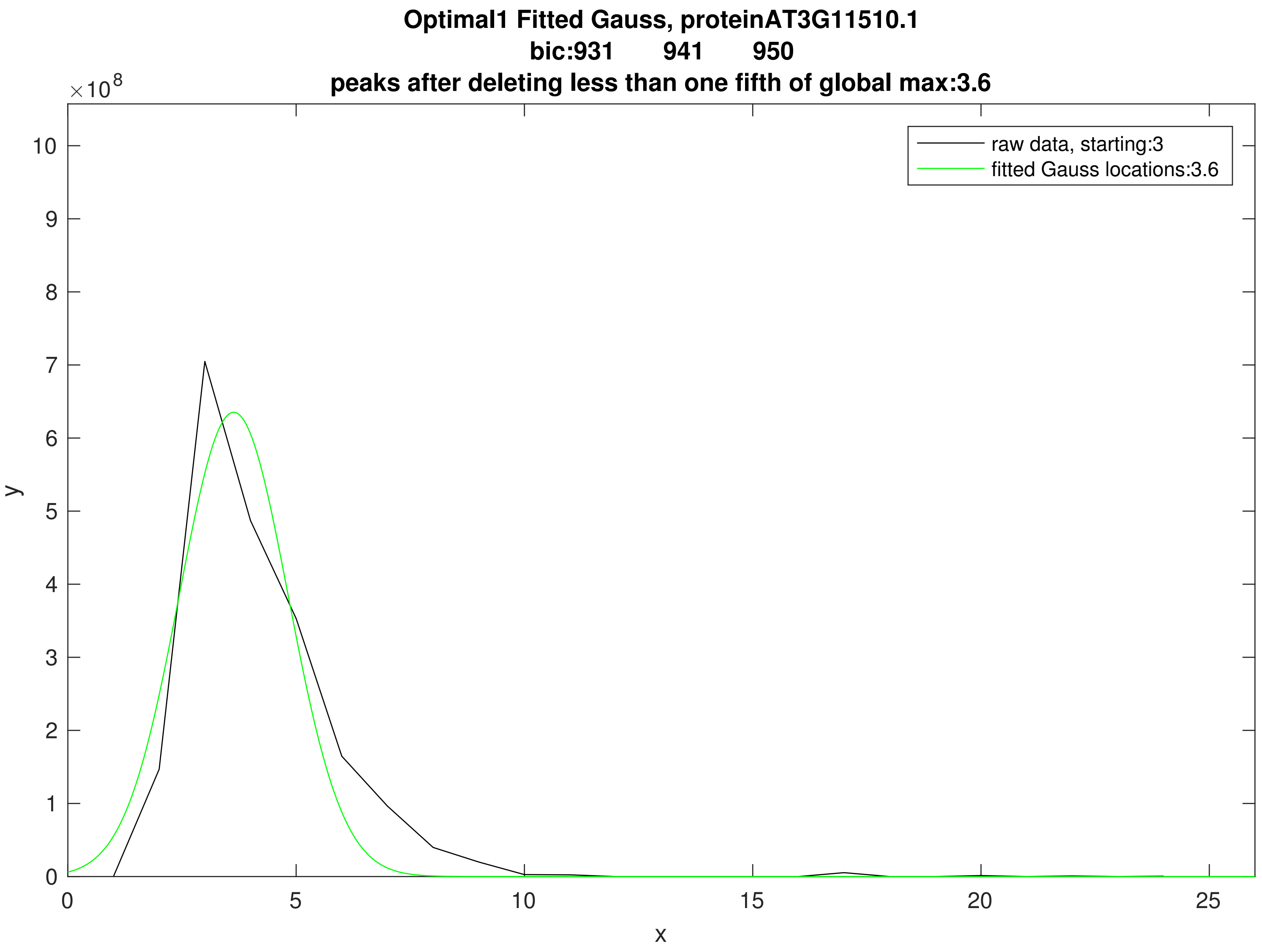

Optimal1 Fitted Gauss, proteinAT3G11510.1
bic:931 941 950
peaks after deleting less than one fifth of global max:3.6
×10^8
raw data, starting:3
fitted Gauss locations:3.6
y
x
0
1
2
3
4
5
6
7
8
9
10
0
5
10
15
20
25

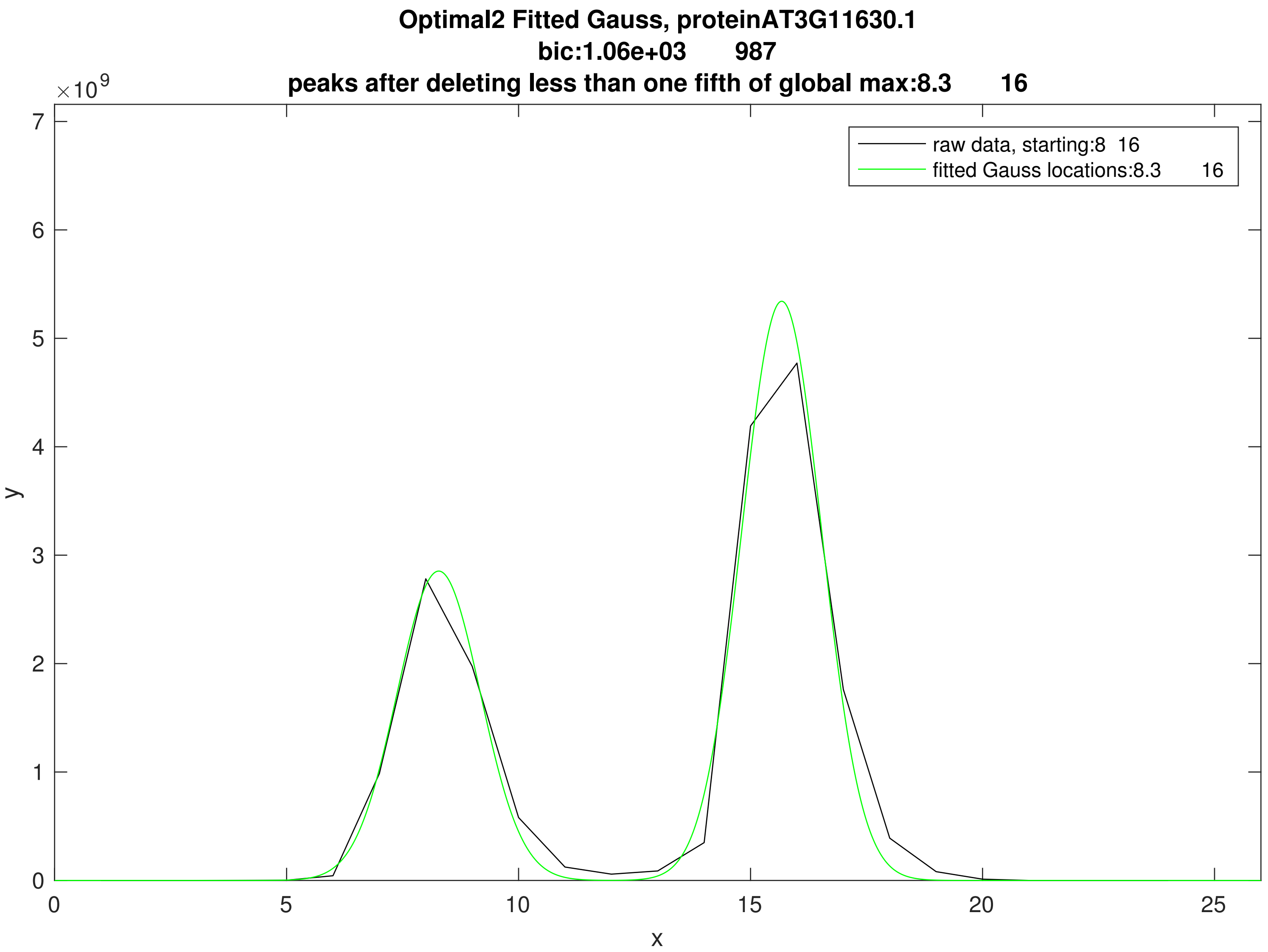

Optimal2 Fitted Gauss, proteinAT3G11630.1
bic:1.06e+03 987
peaks after deleting less than one fifth of global max:8.3 16
×10^9
7
6
5
4
3
2
1
0
y
0
5
10
15
20
25
x
raw data, starting:8 16
fitted Gauss locations:8.3 16

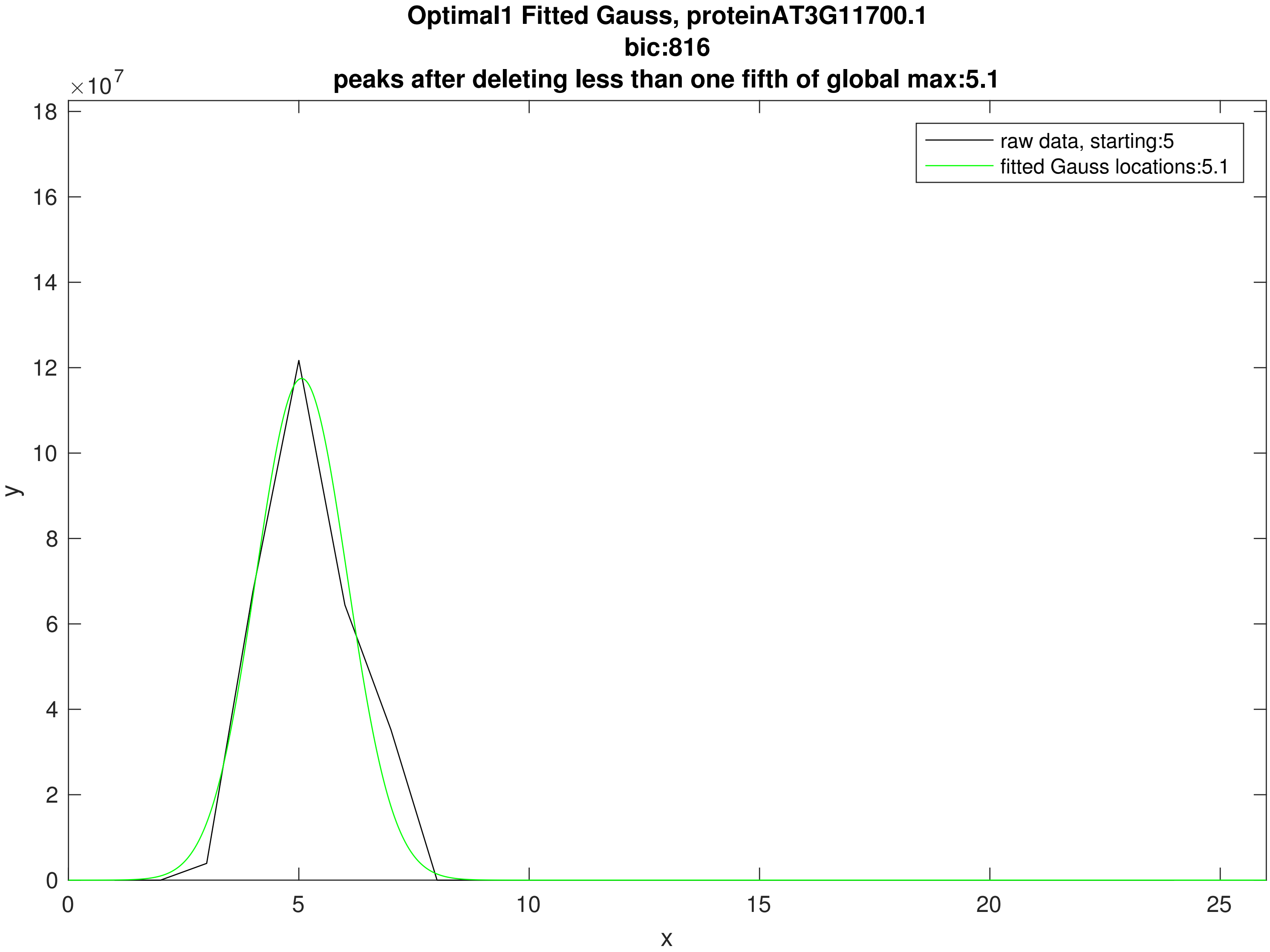
Optimal1 Fitted Gauss, proteinAT3G11700.1
bic:816
peaks after deleting less than one fifth of global max:5.1
raw data, starting:5
fitted Gauss locations:5.1
×10^7
y
x
0
2
4
6
8
10
12
14
16
18
0
5
10
15
20
25

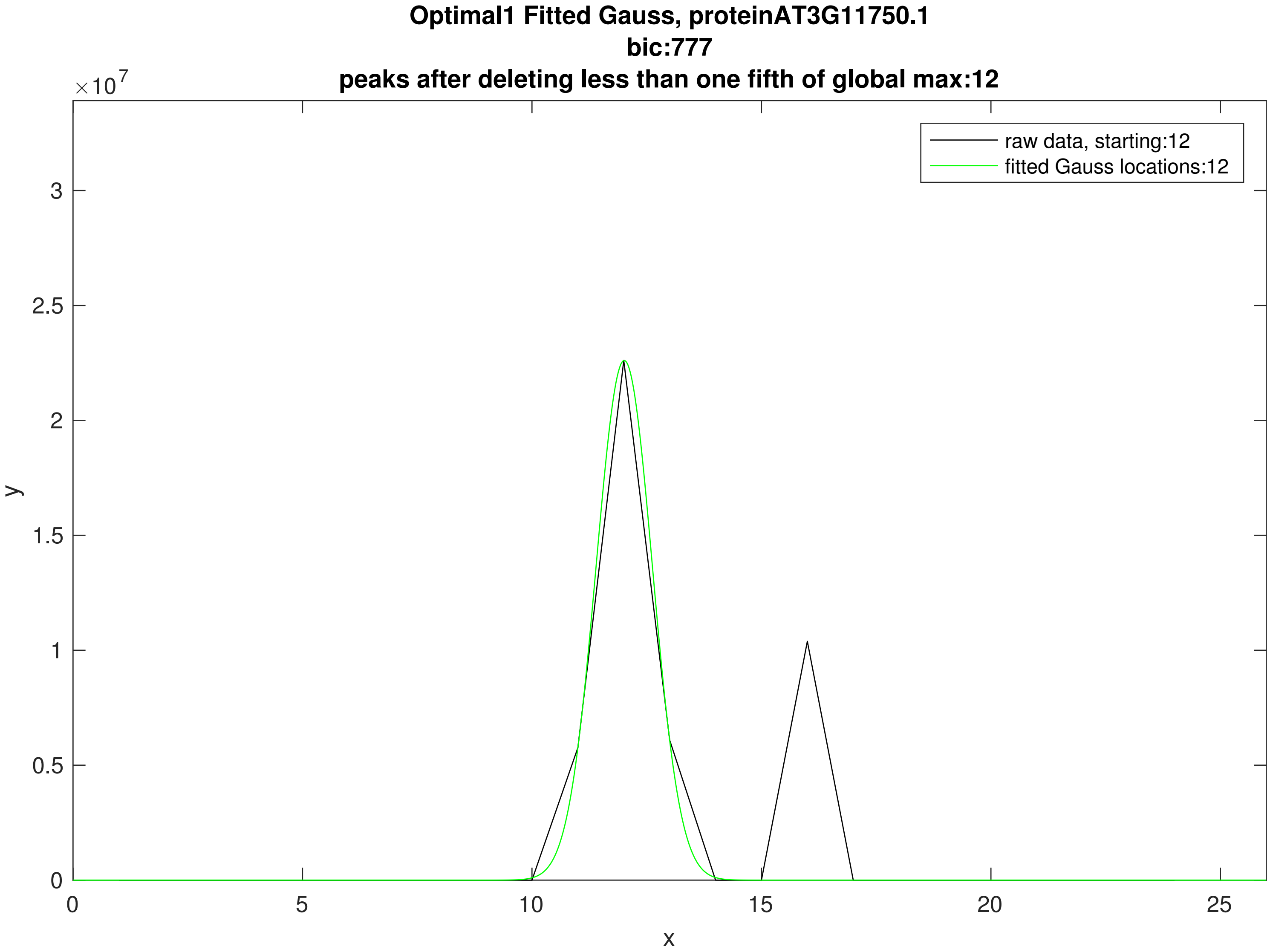

Optimal1 Fitted Gauss, proteinAT3G11750.1
bic:777
peaks after deleting less than one fifth of global max:12
×10^7
raw data, starting:12
fitted Gauss locations:12
y
x
0
0.5
1
1.5
2
2.5
3
0
5
10
15
20
25

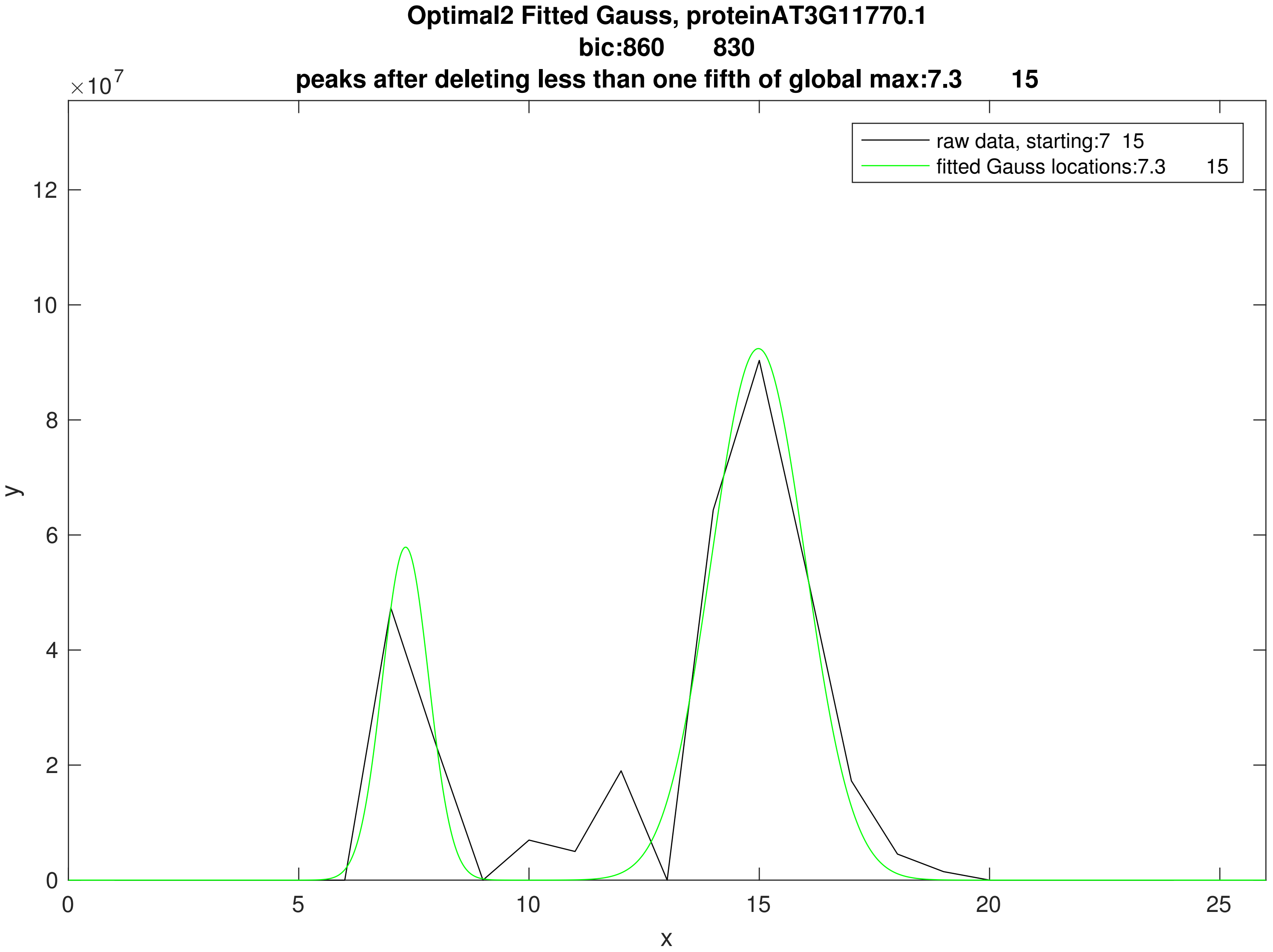
Optimal2 Fitted Gauss, proteinAT3G11770.1
bic:860 830
peaks after deleting less than one fifth of global max:7.3 15
×10^7
raw data, starting:7 15
fitted Gauss locations:7.3 15
y
x
0
2
4
6
8
10
12
0
5
10
15
20
25

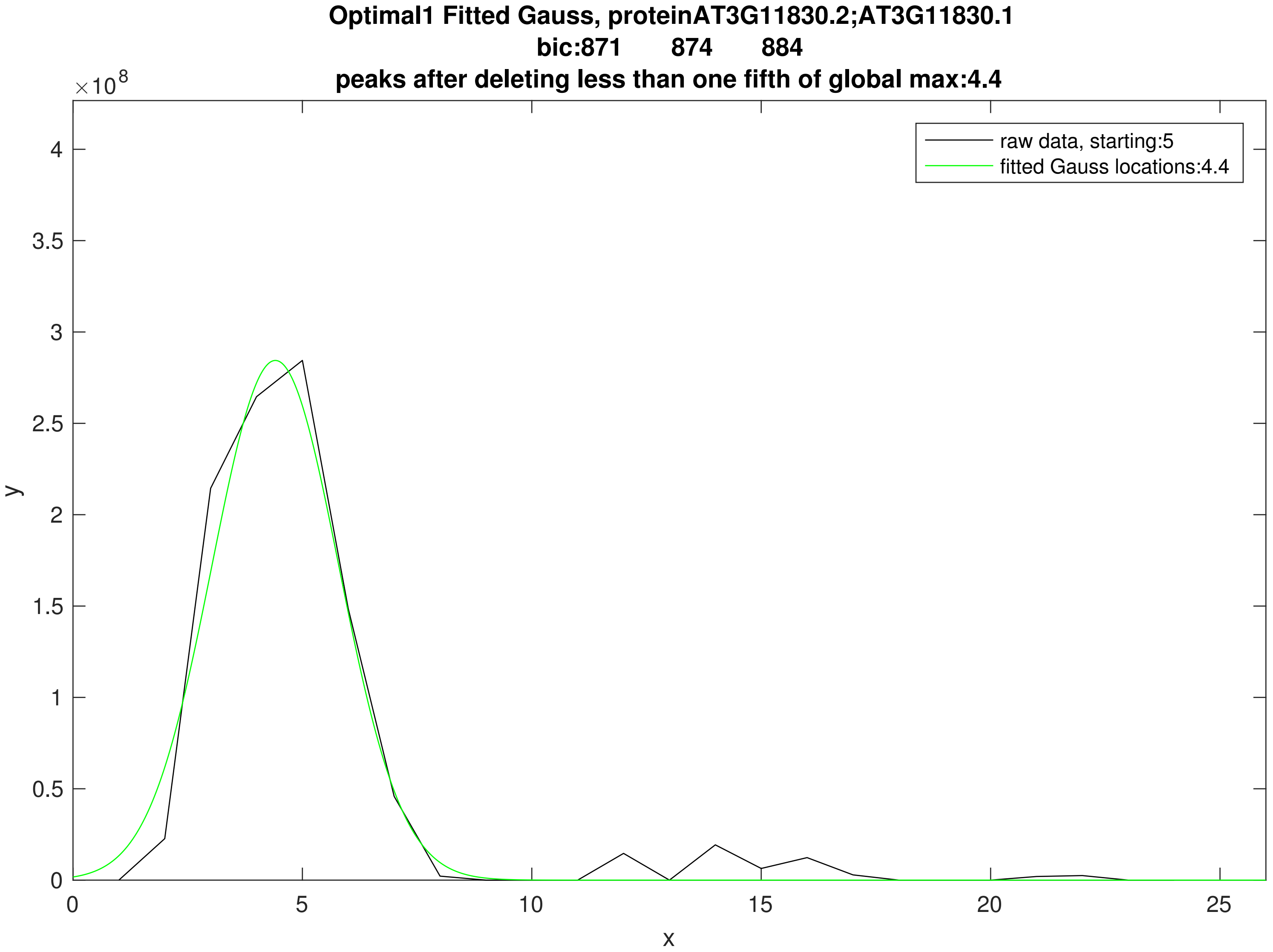

Optimal1 Fitted Gauss, proteinAT3G11830.2;AT3G11830.1
bic:871 874 884
peaks after deleting less than one fifth of global max:4.4
×10^8
raw data, starting:5
fitted Gauss locations:4.4
y
x
0
0.5
1
1.5
2
2.5
3
3.5
4
0
5
10
15
20
25

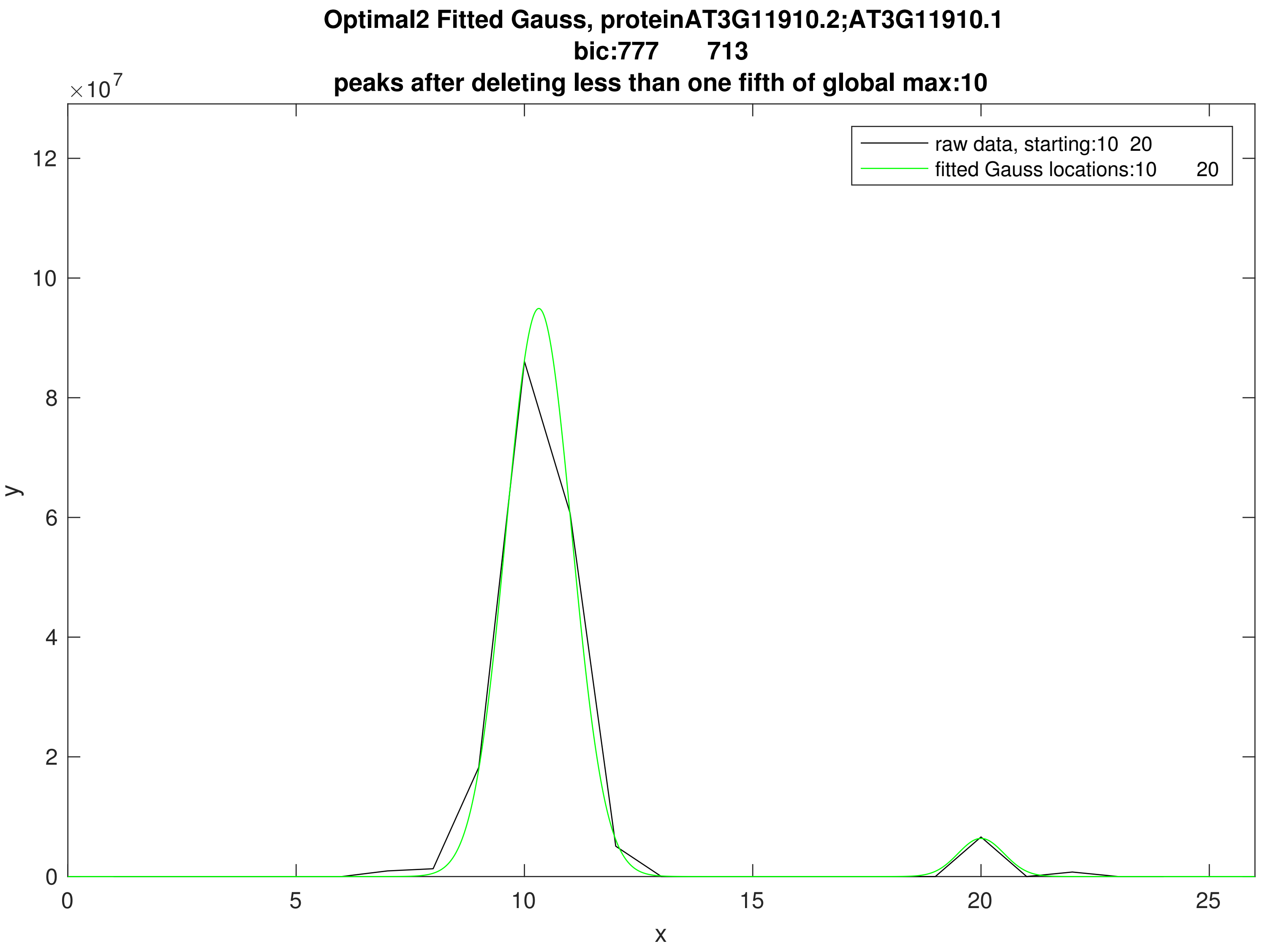

Optimal2 Fitted Gauss, proteinAT3G11910.2;AT3G11910.1
bic:777 713
peaks after deleting less than one fifth of global max:10
×10^7
raw data, starting:10 20
fitted Gauss locations:10 20
y
x
0
2
4
6
8
10
12
0
5
10
15
20
25

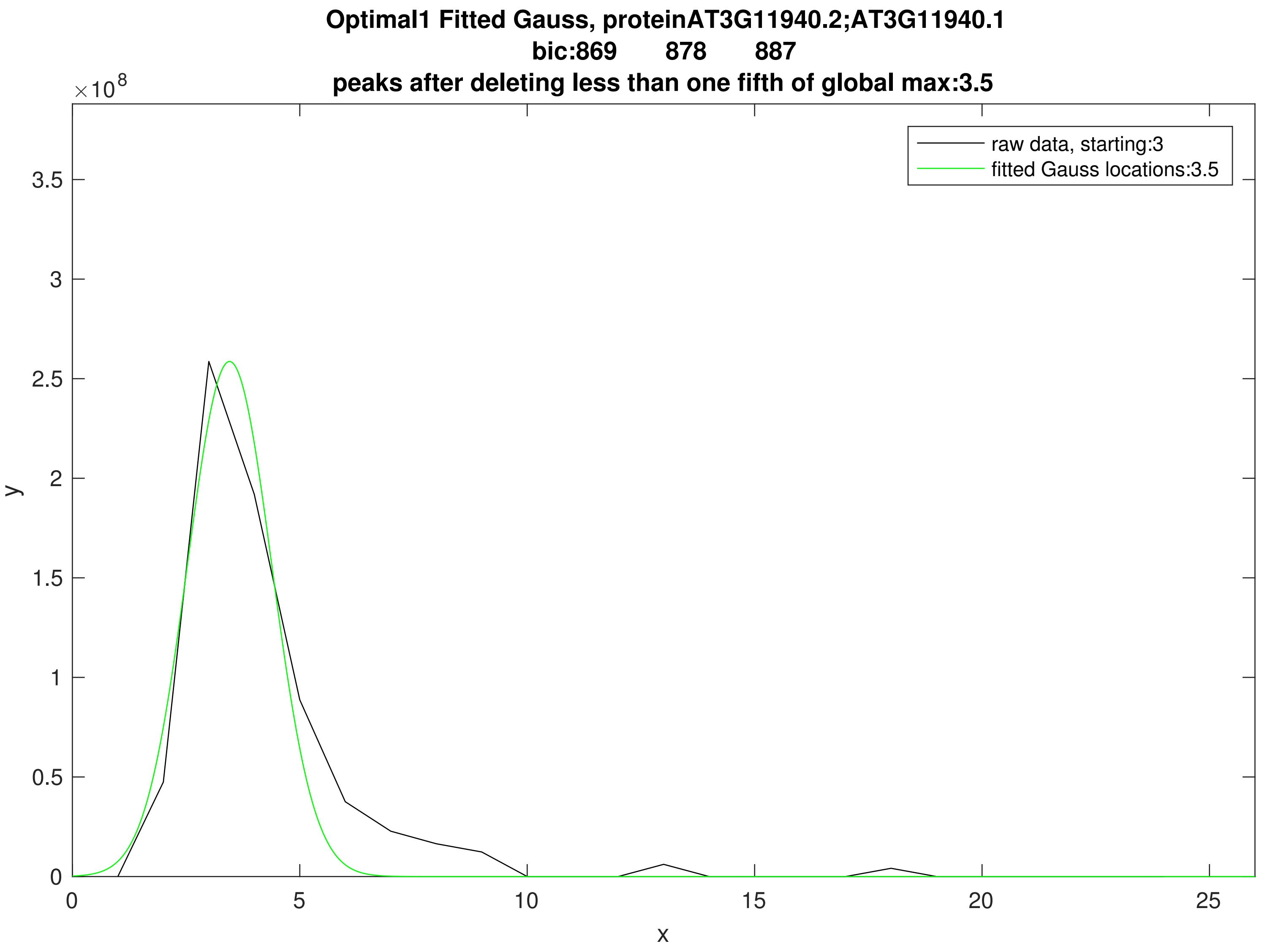

Optimal1 Fitted Gauss, proteinAT3G11940.2;AT3G11940.1
bic:869 878 887
peaks after deleting less than one fifth of global max:3.5
×10^8
raw data, starting:3
fitted Gauss locations:3.5
y
x
0
0.5
1
1.5
2
2.5
3
3.5
0
5
10
15
20
25

**Optimal1 Fitted Gauss, proteinAT3G12110.1**
**bic:733**
**peaks after deleting less than one fifth of global max:4.6**

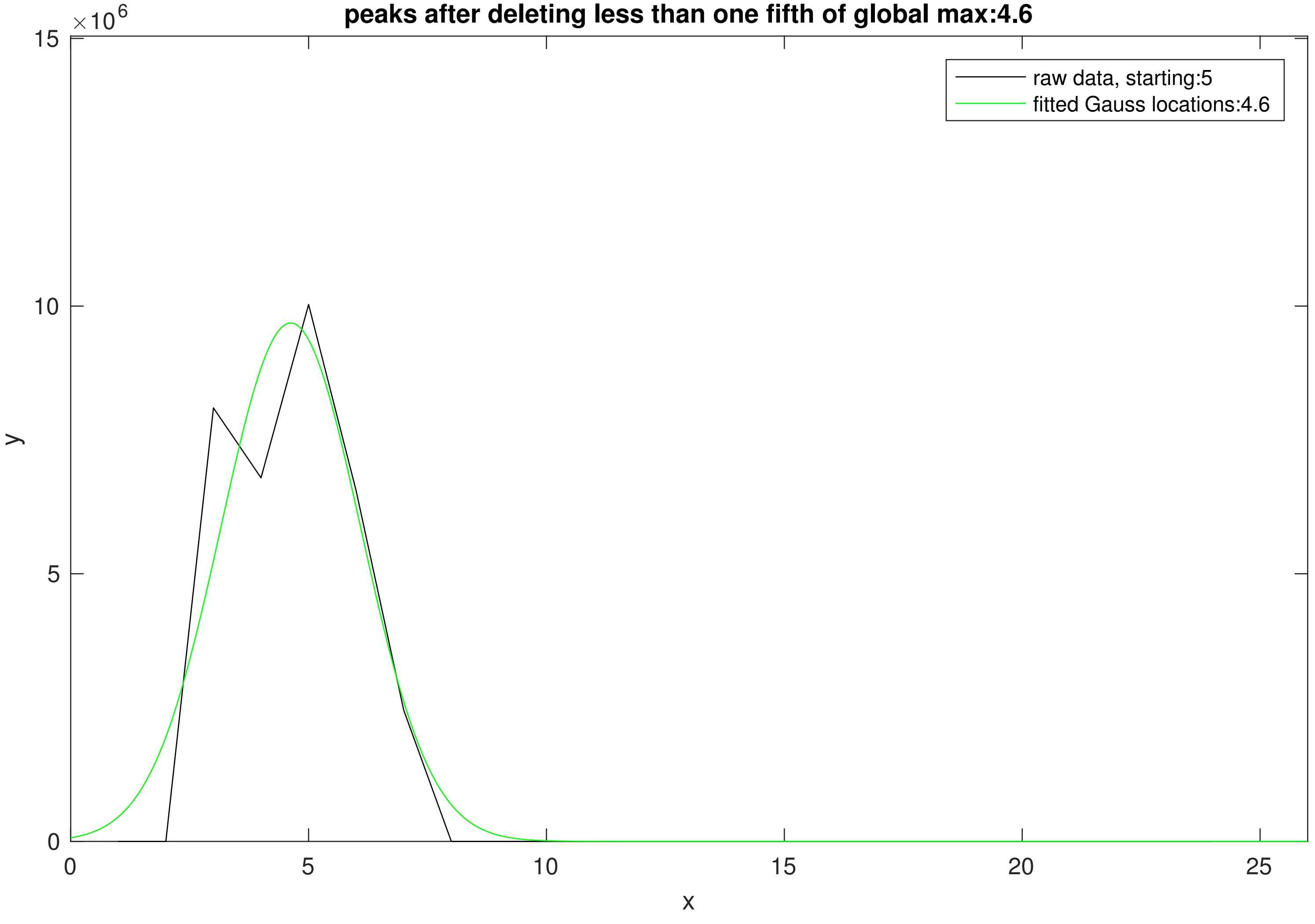

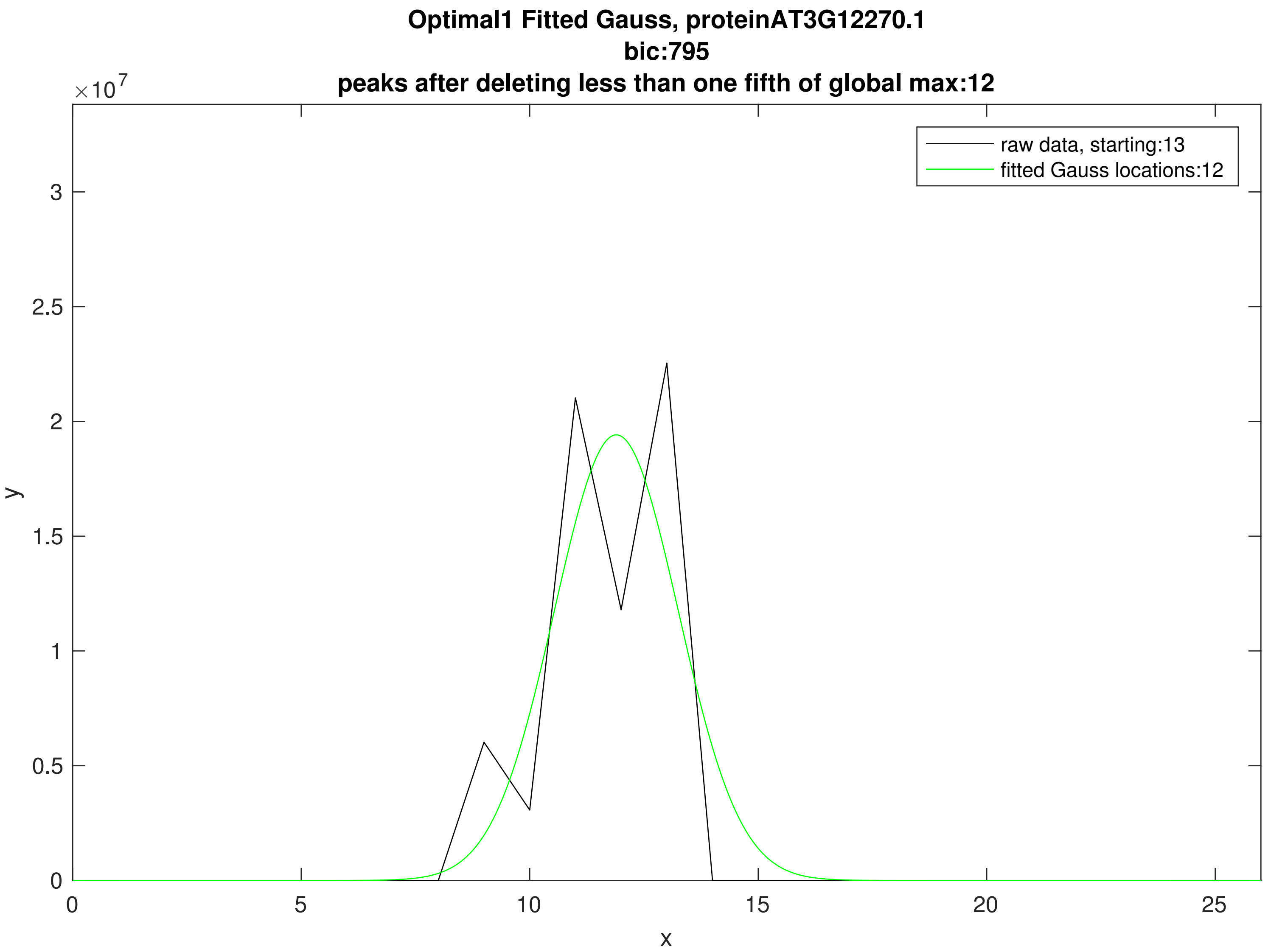

Optimal1 Fitted Gauss, proteinAT3G12270.1
bic:795
peaks after deleting less than one fifth of global max:12
×10^7
raw data, starting:13
fitted Gauss locations:12
y
x
0
0.5
1
1.5
2
2.5
3
0
5
10
15
20
25

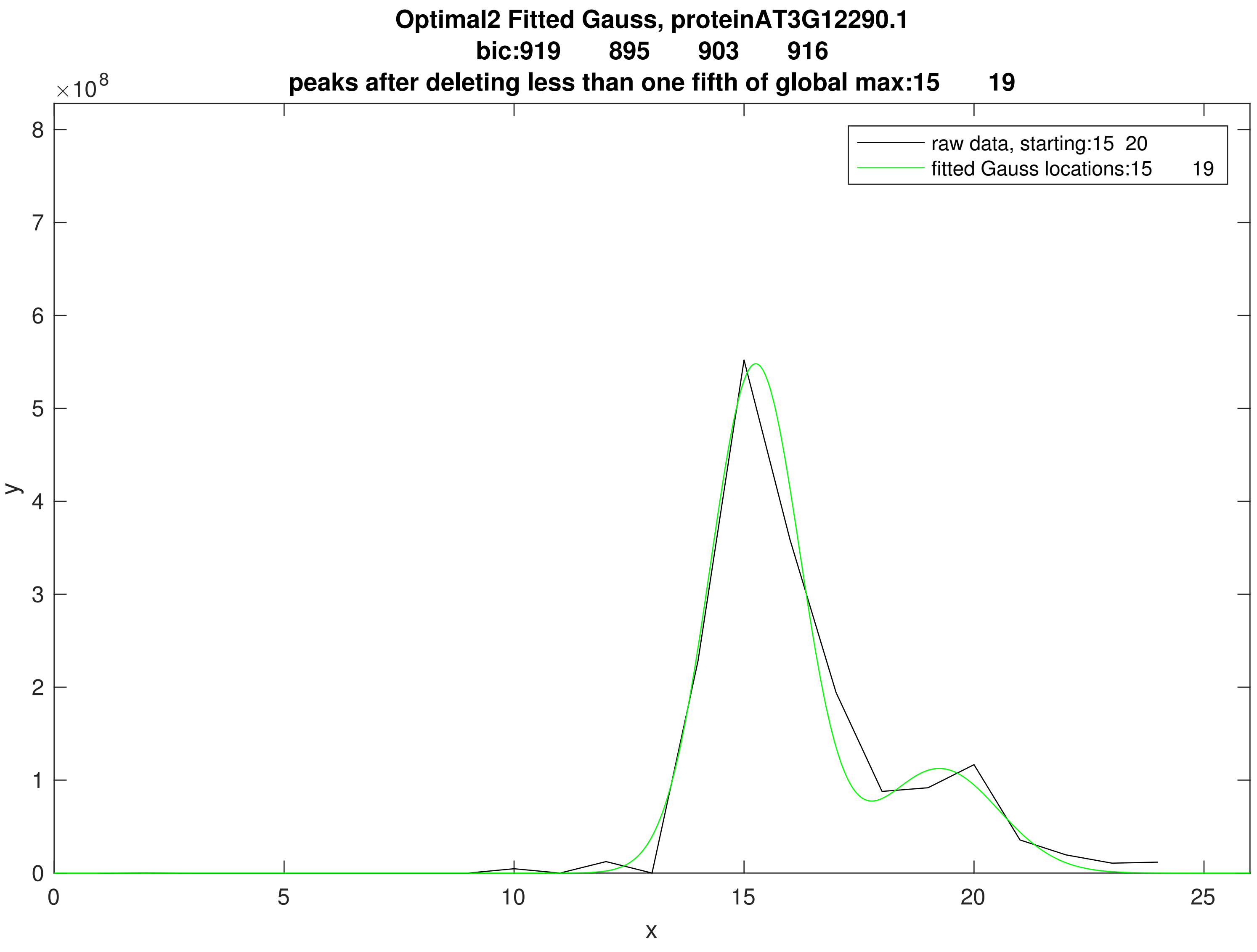
Optimal2 Fitted Gauss, proteinAT3G12290.1
bic:919 895 903 916
peaks after deleting less than one fifth of global max:15 19
×10^8
raw data, starting:15 20
fitted Gauss locations:15 19
y
x
0
1
2
3
4
5
6
7
8
0
5
10
15
20
25

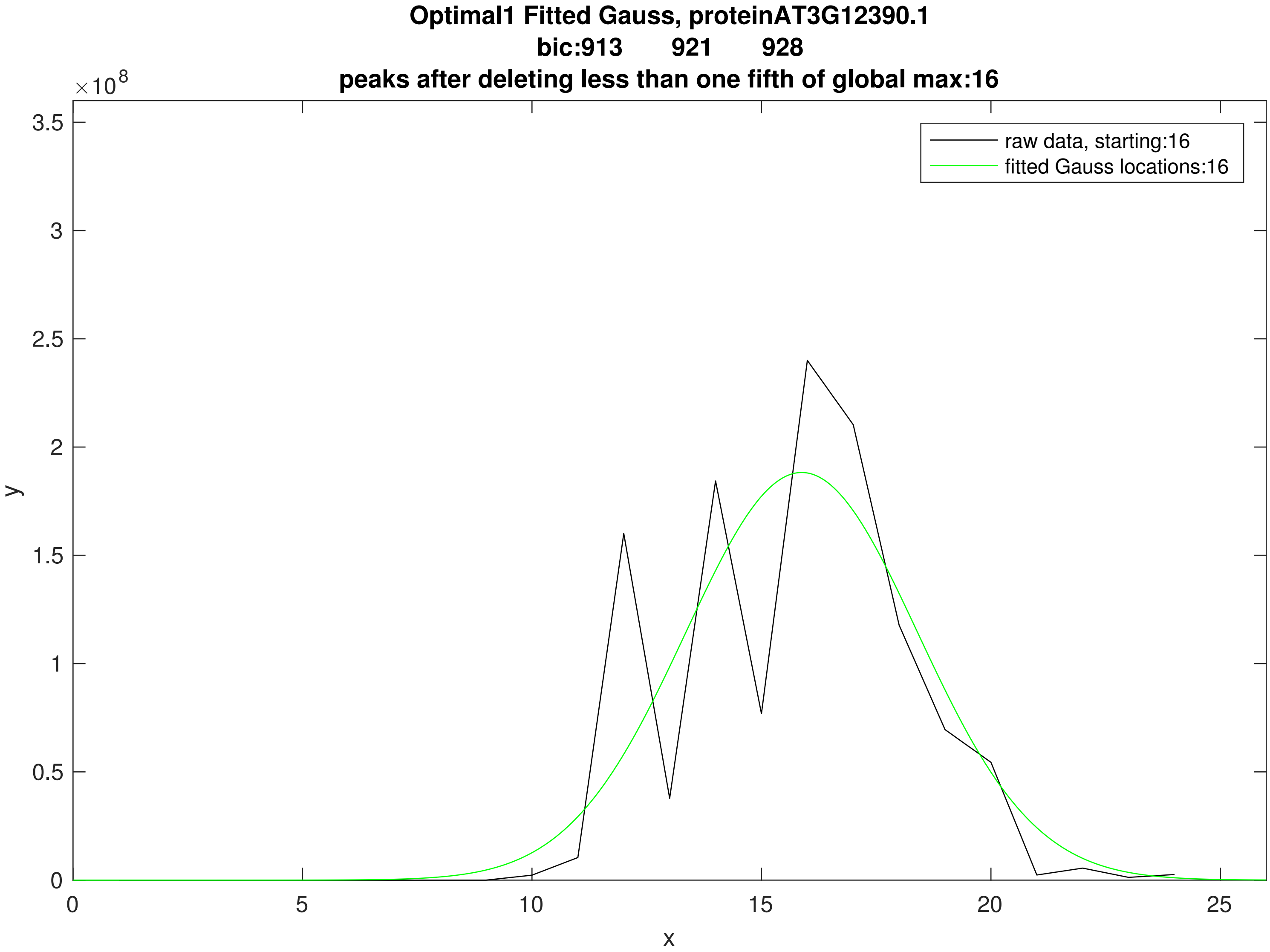

Optimal1 Fitted Gauss, proteinAT3G12390.1
bic:913 921 928
peaks after deleting less than one fifth of global max:16
×10^8
3.5
3
2.5
2
1.5
1
0.5
0
y
0
5
10
15
20
25
x
raw data, starting:16
fitted Gauss locations:16

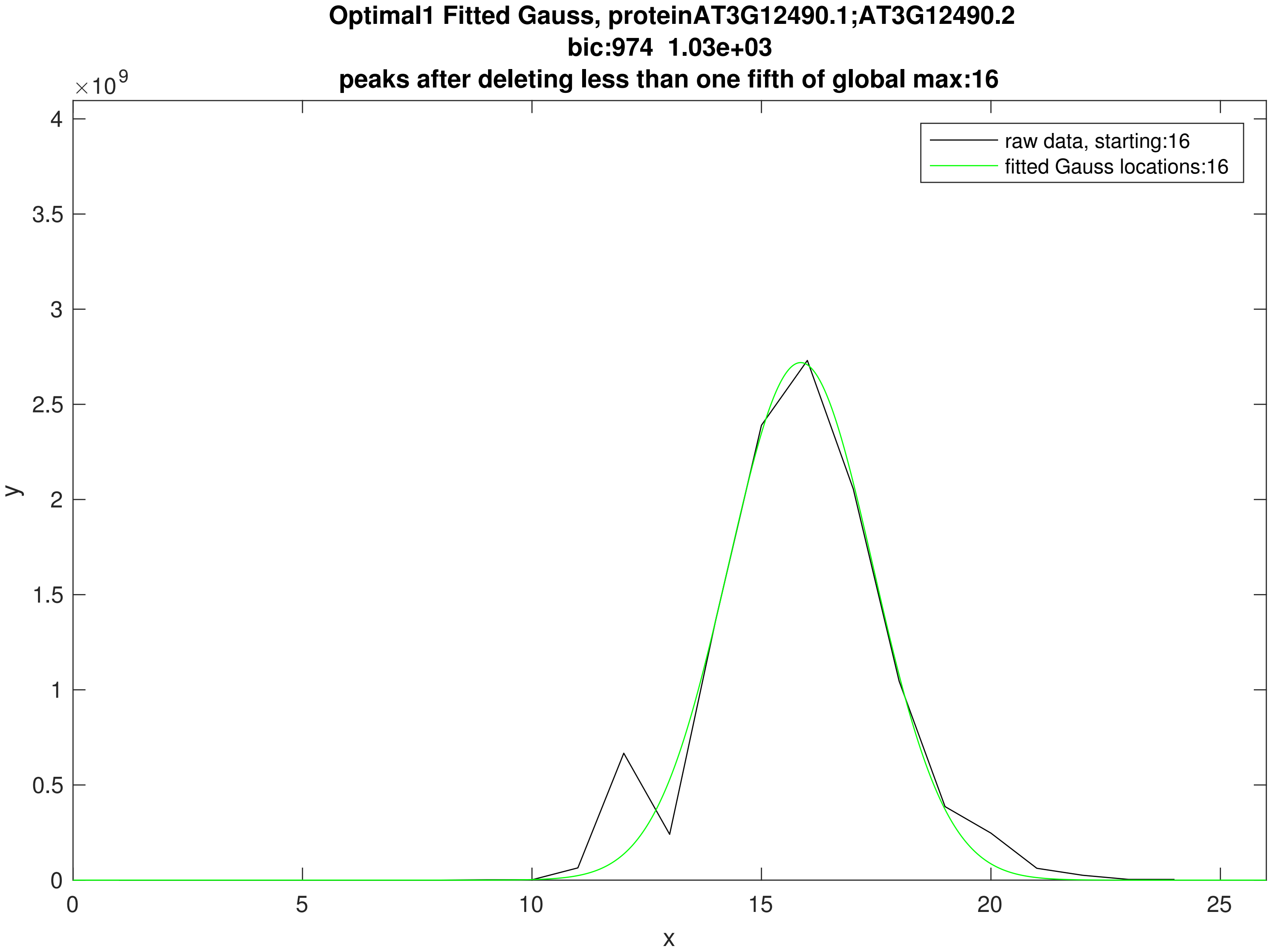
Optimal1 Fitted Gauss, proteinAT3G12490.1;AT3G12490.2
bic:974 1.03e+03
peaks after deleting less than one fifth of global max:16
×10^9
raw data, starting:16
fitted Gauss locations:16
y
x
0
0.5
1
1.5
2
2.5
3
3.5
4
0
5
10
15
20
25

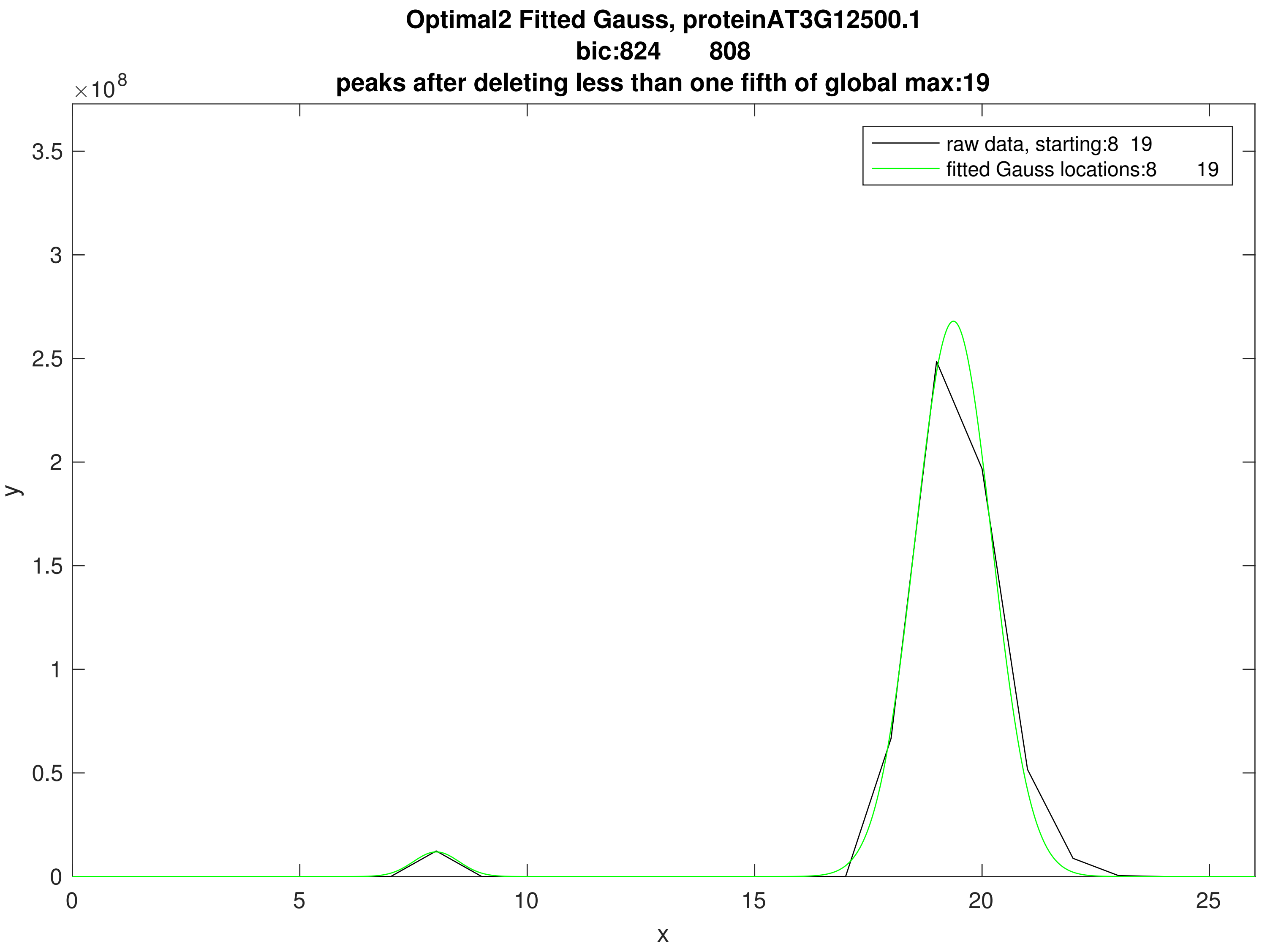

Optimal2 Fitted Gauss, proteinAT3G12500.1
bic:824 808
peaks after deleting less than one fifth of global max:19
×10^8
3.5
3
2.5
2
1.5
1
0.5
0
y
0
5
10
15
20
25
x
raw data, starting:8 19
fitted Gauss locations:8 19

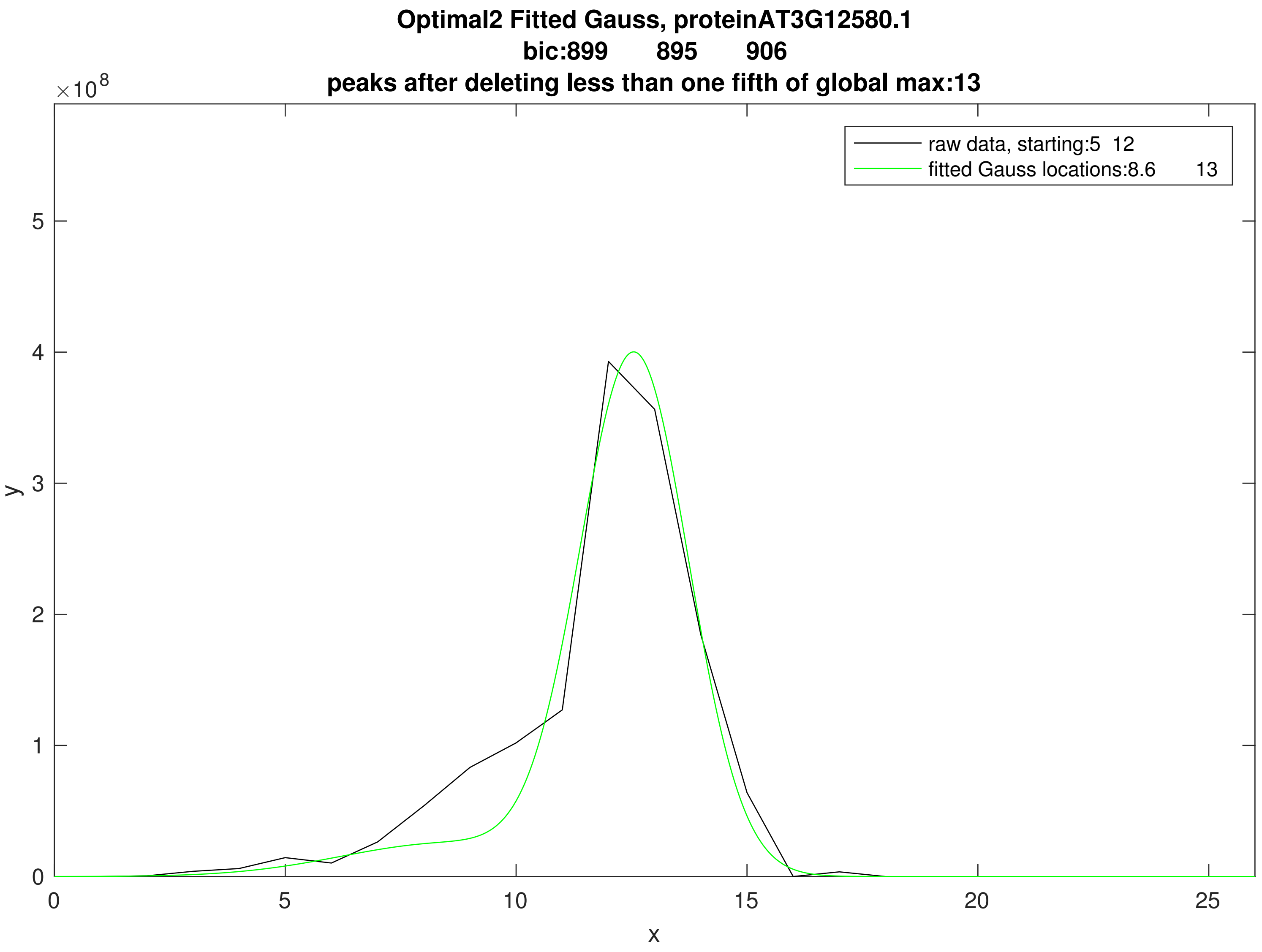
Optimal2 Fitted Gauss, proteinAT3G12580.1
bic:899 895 906
peaks after deleting less than one fifth of global max:13
×10^8
raw data, starting:5 12
fitted Gauss locations:8.6 13
y
x
0
1
2
3
4
5
0
5
10
15
20
25

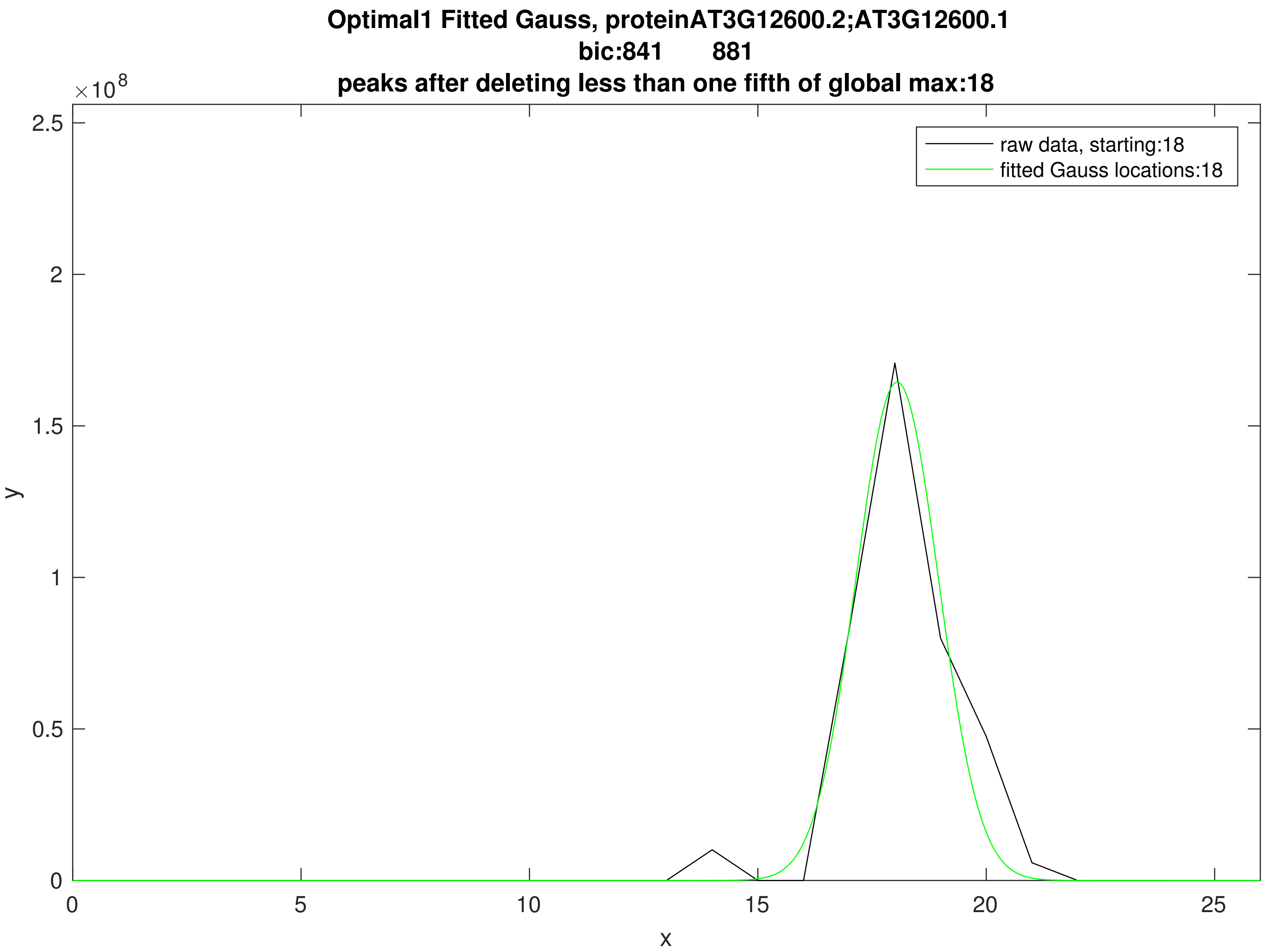

Optimal1 Fitted Gauss, proteinAT3G12600.2;AT3G12600.1
bic:841 881
peaks after deleting less than one fifth of global max:18
×10^8
2.5
2
1.5
1
0.5
0
y
0
5
10
15
20
25
x
raw data, starting:18
fitted Gauss locations:18

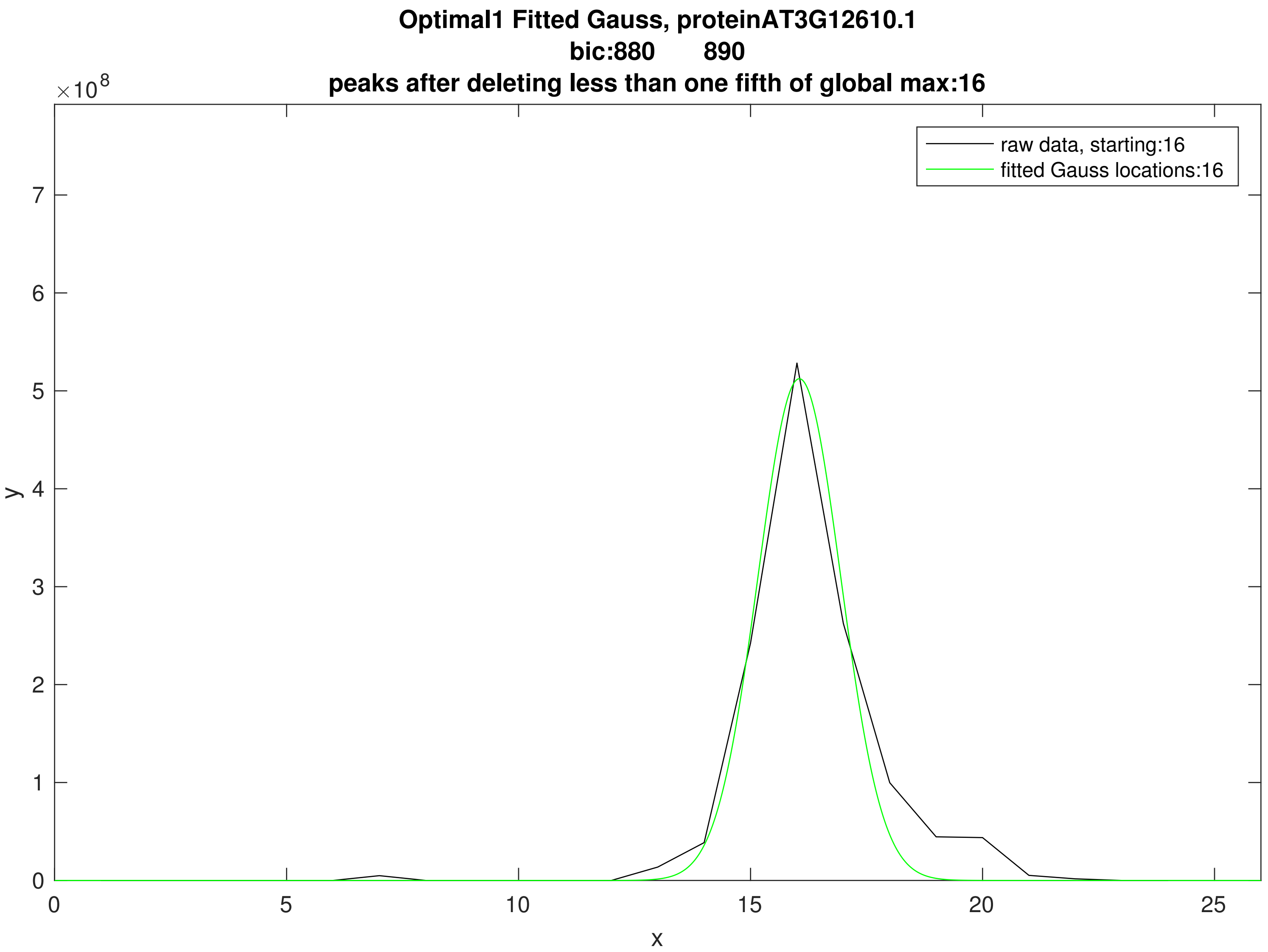

Optimal1 Fitted Gauss, proteinAT3G12610.1
bic:880 890
peaks after deleting less than one fifth of global max:16
×10^8
raw data, starting:16
fitted Gauss locations:16
y
x
0
1
2
3
4
5
6
7
0
5
10
15
20
25

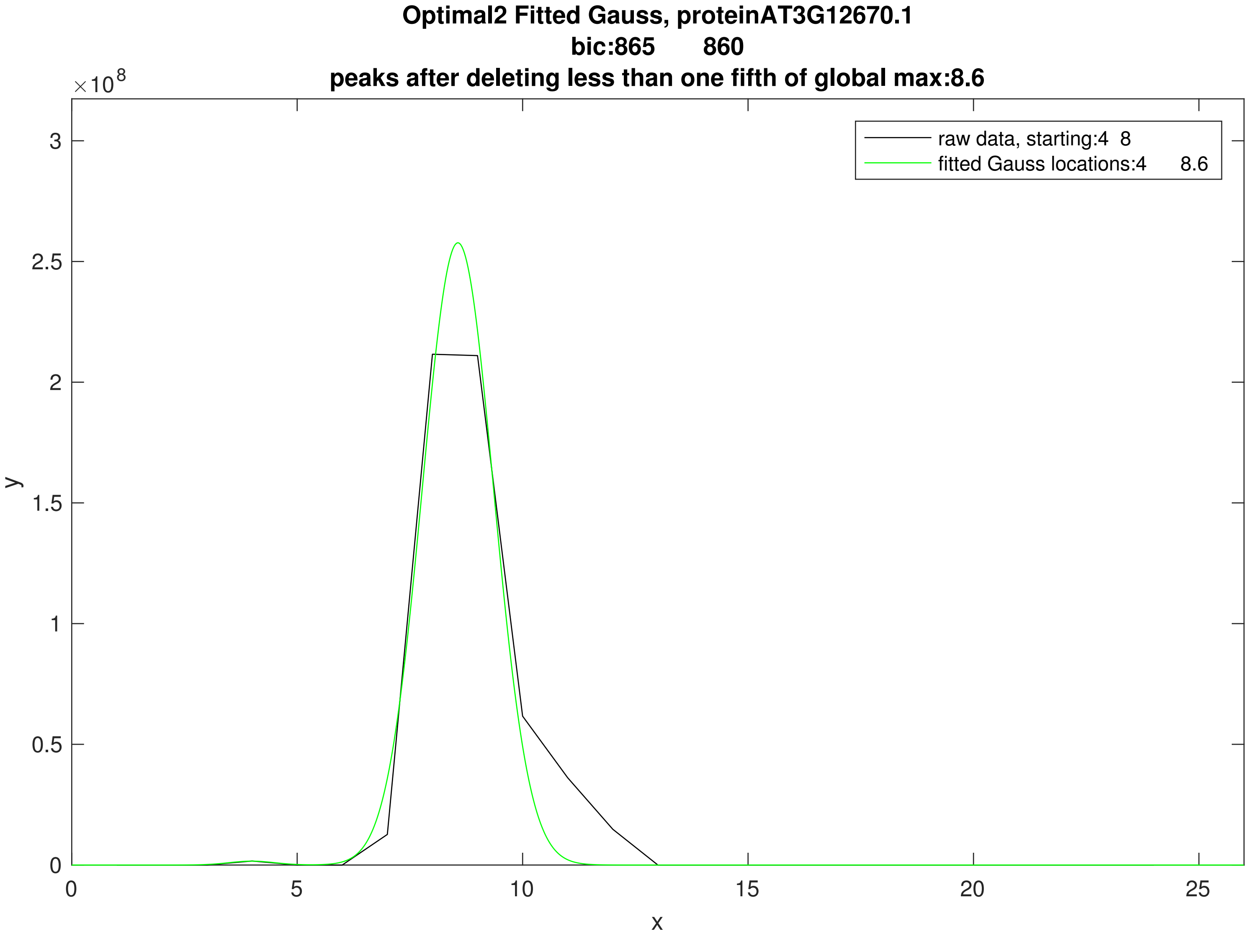
Optimal2 Fitted Gauss, proteinAT3G12670.1
bic:865 860
peaks after deleting less than one fifth of global max:8.6
×10^8
raw data, starting:4 8
fitted Gauss locations:4 8.6
y
x
0
0.5
1
1.5
2
2.5
3
0
5
10
15
20
25

Optimal1 Fitted Gauss, proteinAT3G12780.1
bic:1.02e+03  1.03e+03  1.05e+03
peaks after deleting less than one fifth of global max:16

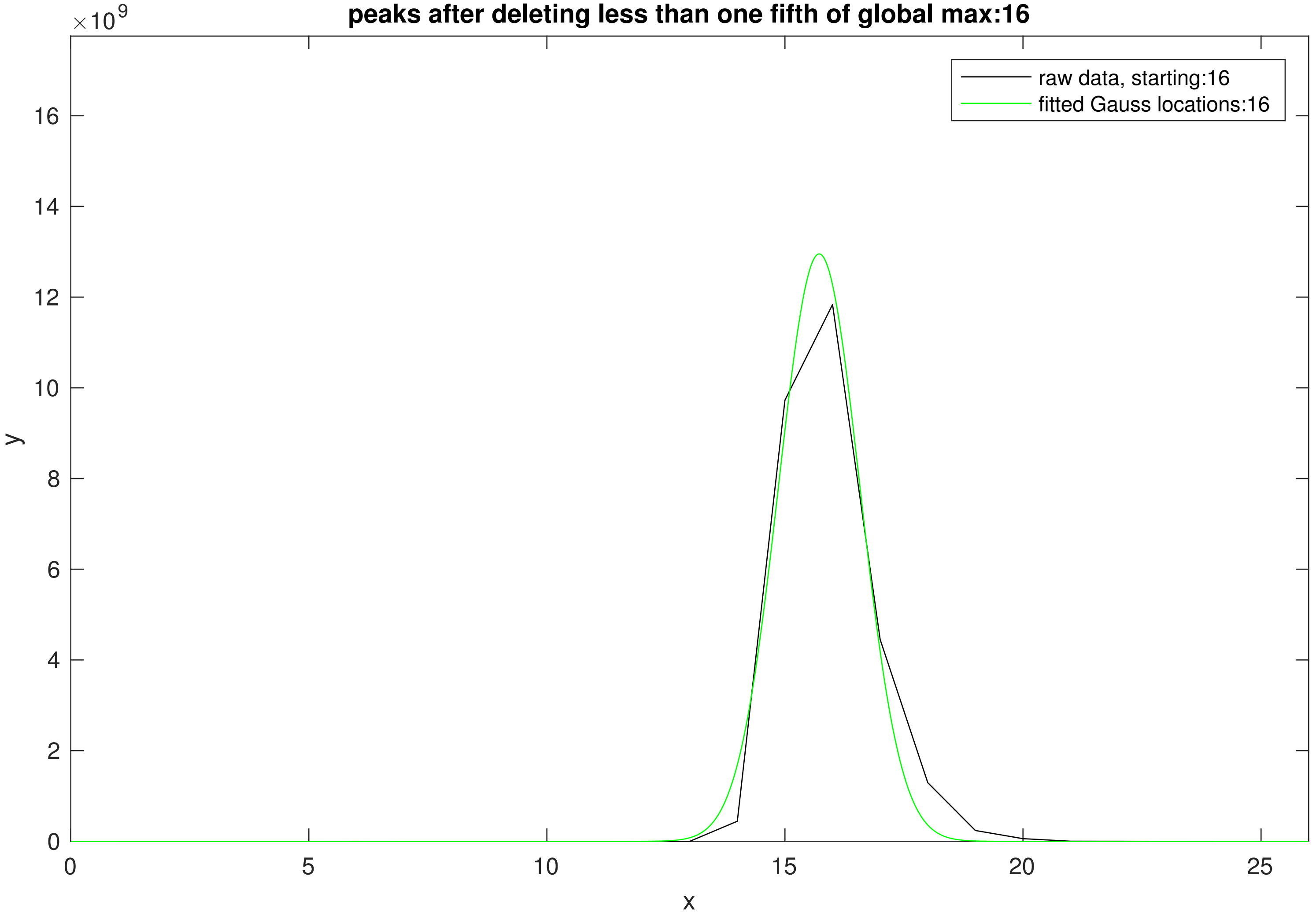

**Optimal2 Fitted Gauss, proteinAT3G12800.1**
**bic:799 753 787**
**peaks after deleting less than one fifth of global max:4.7 13**

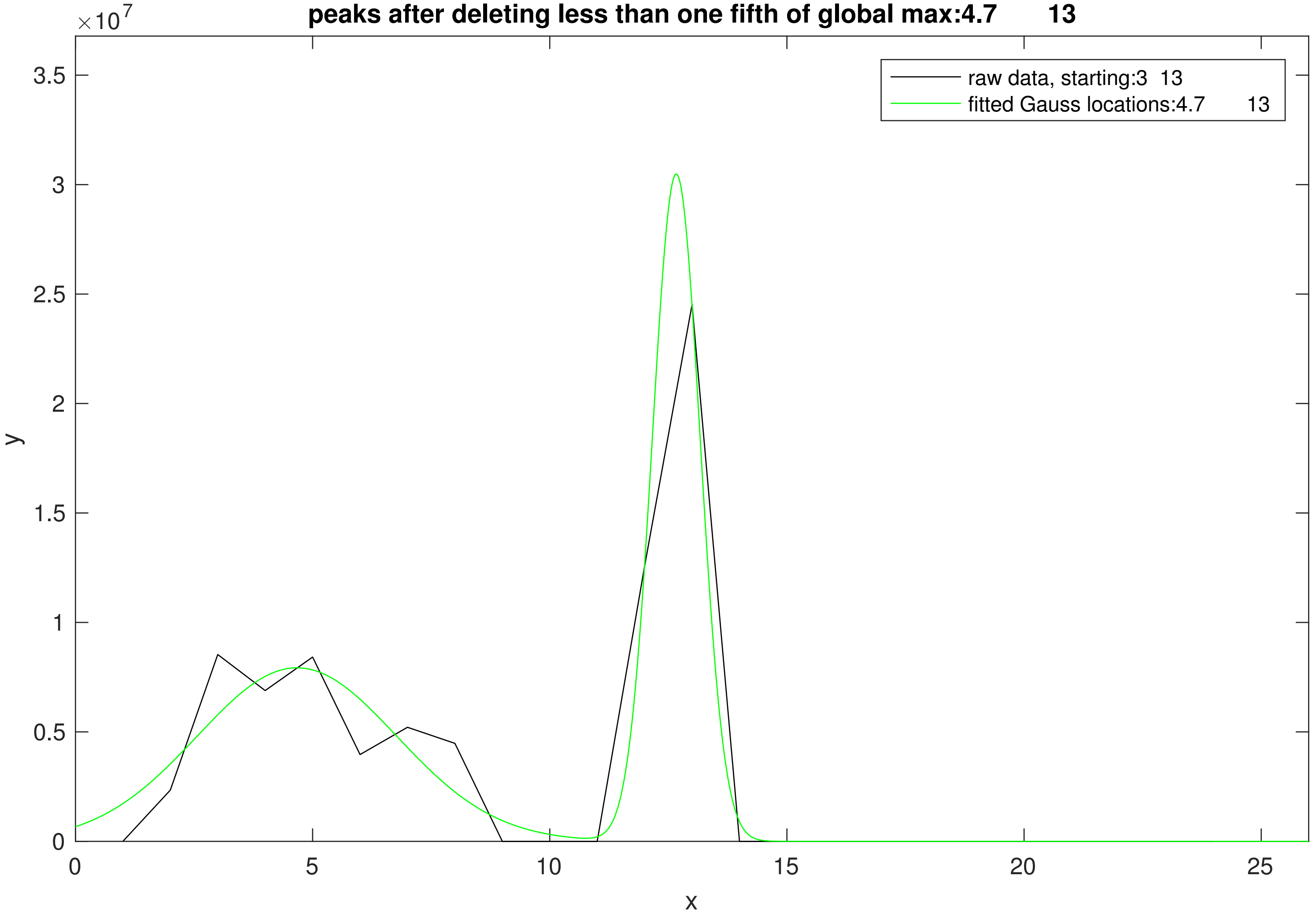

Optimal1 Fitted Gauss, proteinAT3G12915.1;AT3G12915.2
bic:827
peaks after deleting less than one fifth of global max:13

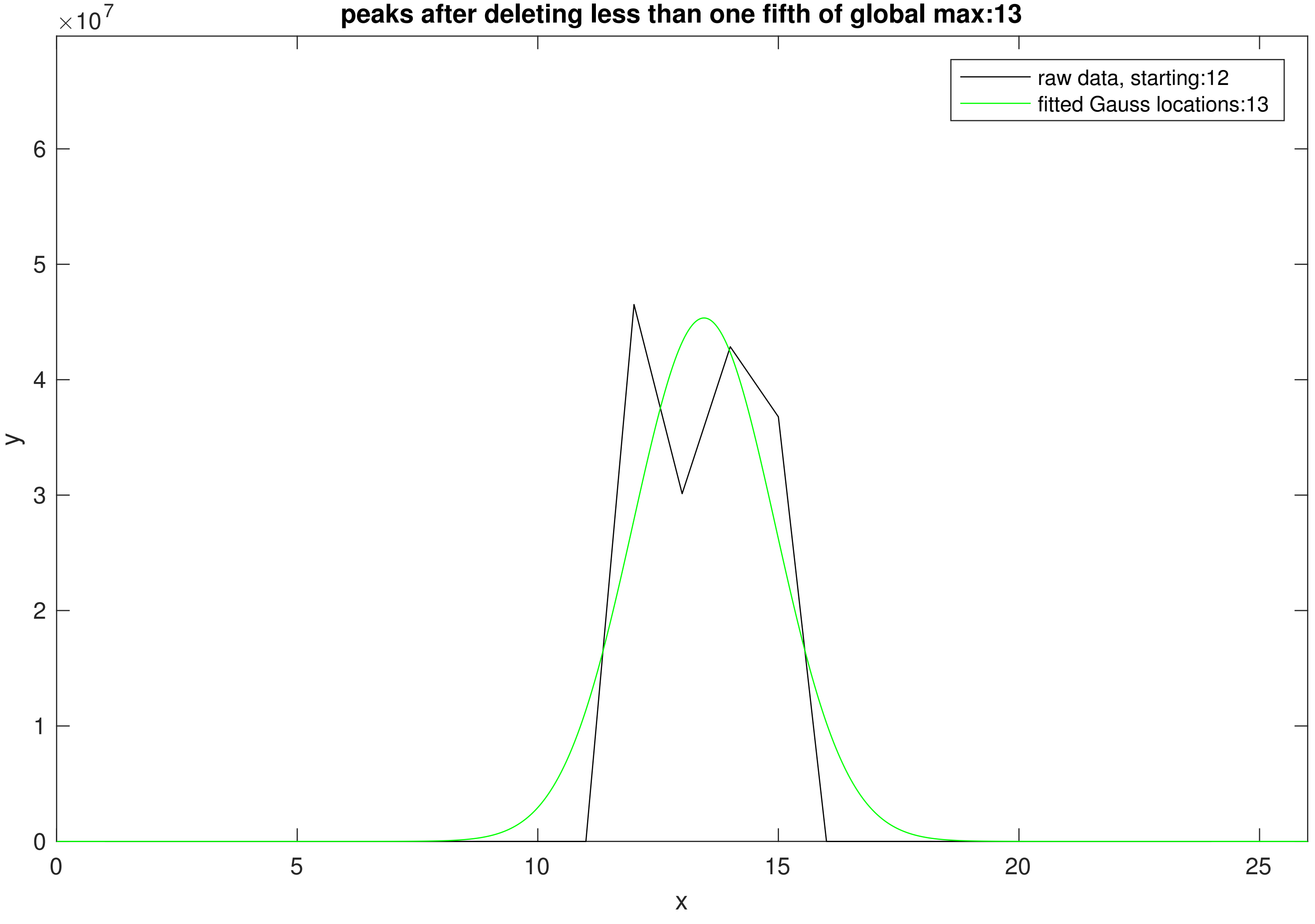

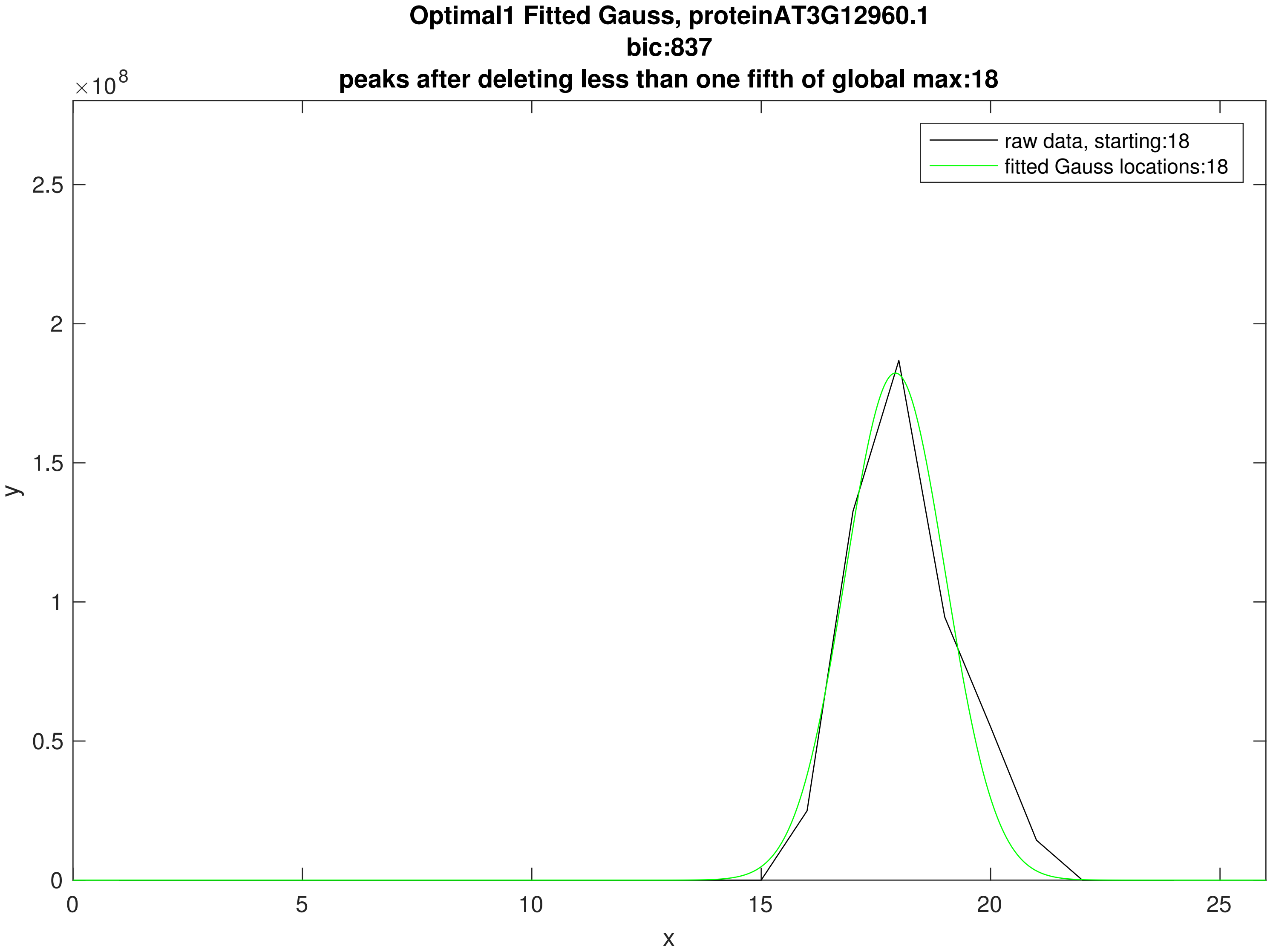
Optimal1 Fitted Gauss, proteinAT3G12960.1
bic:837
peaks after deleting less than one fifth of global max:18
×10^8
raw data, starting:18
fitted Gauss locations:18
y
x
0
0.5
1
1.5
2
2.5
0
5
10
15
20
25

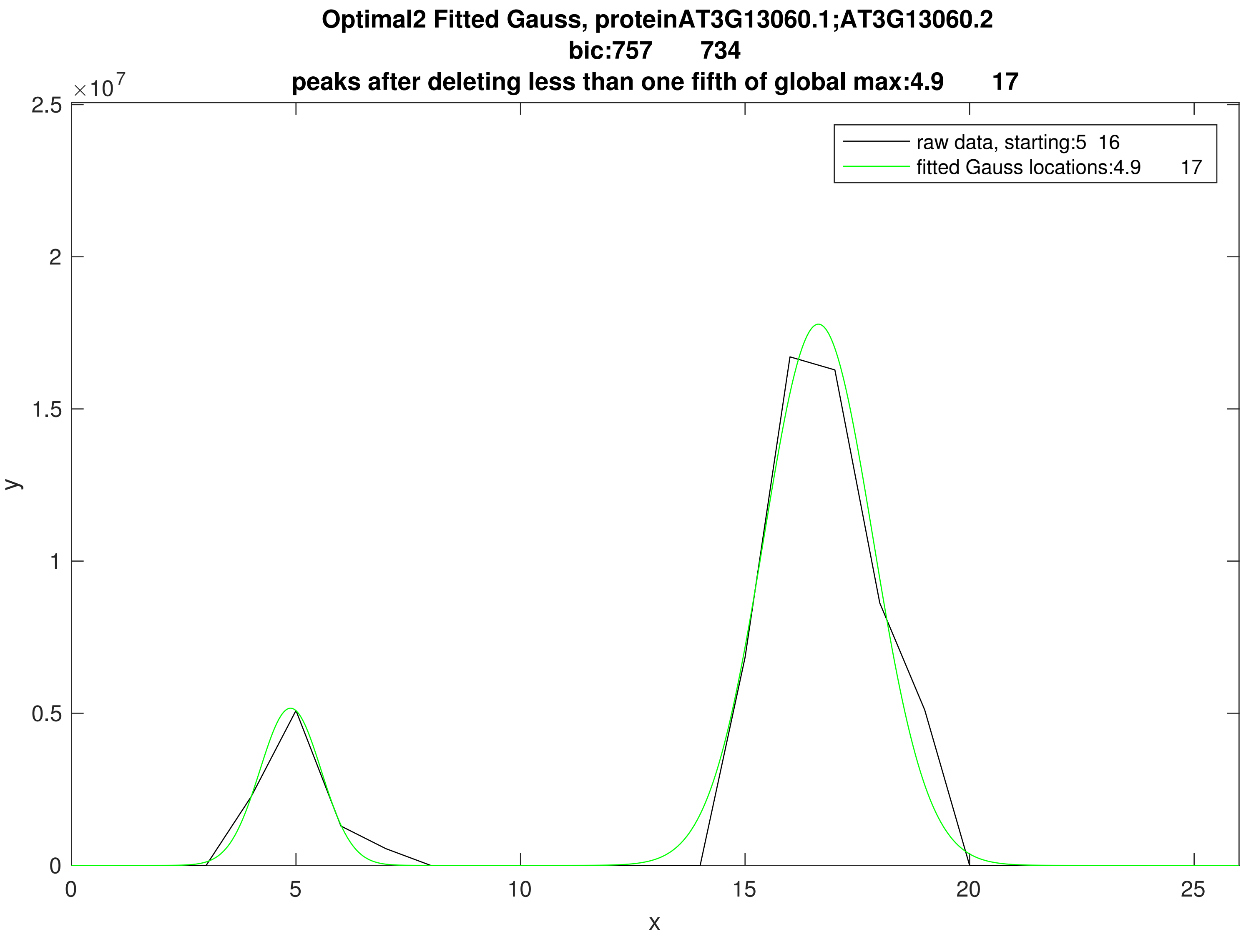

Optimal2 Fitted Gauss, proteinAT3G13060.1;AT3G13060.2
bic:757 734
peaks after deleting less than one fifth of global max:4.9 17
×10^7
2.5
2
1.5
1
0.5
0
y
0
5
10
15
20
25
x
raw data, starting:5 16
fitted Gauss locations:4.9 17

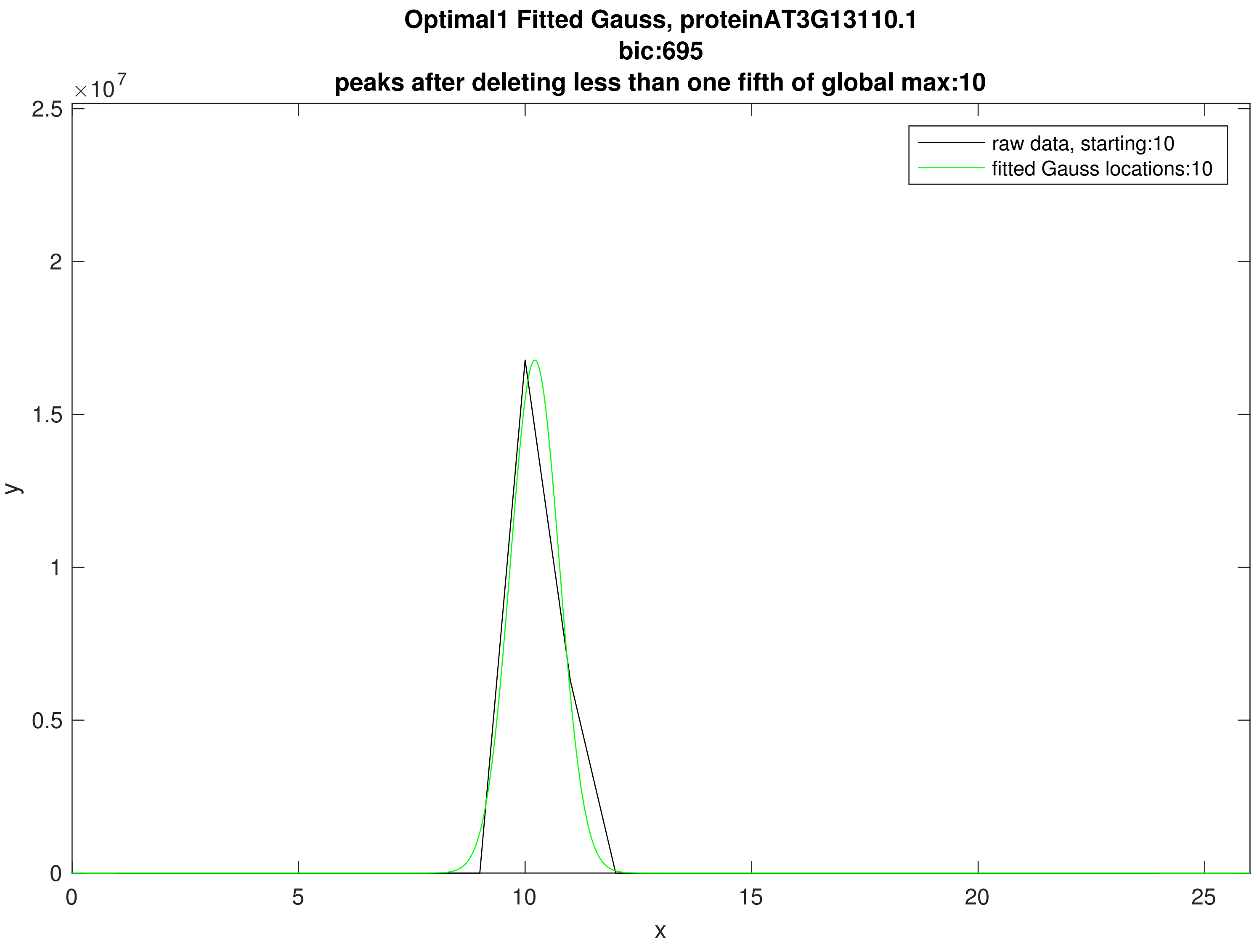

Optimal1 Fitted Gauss, proteinAT3G13110.1
bic:695
peaks after deleting less than one fifth of global max:10
×10^7
2.5
2
1.5
1
0.5
0
y
0
5
10
15
20
25
x
raw data, starting:10
fitted Gauss locations:10

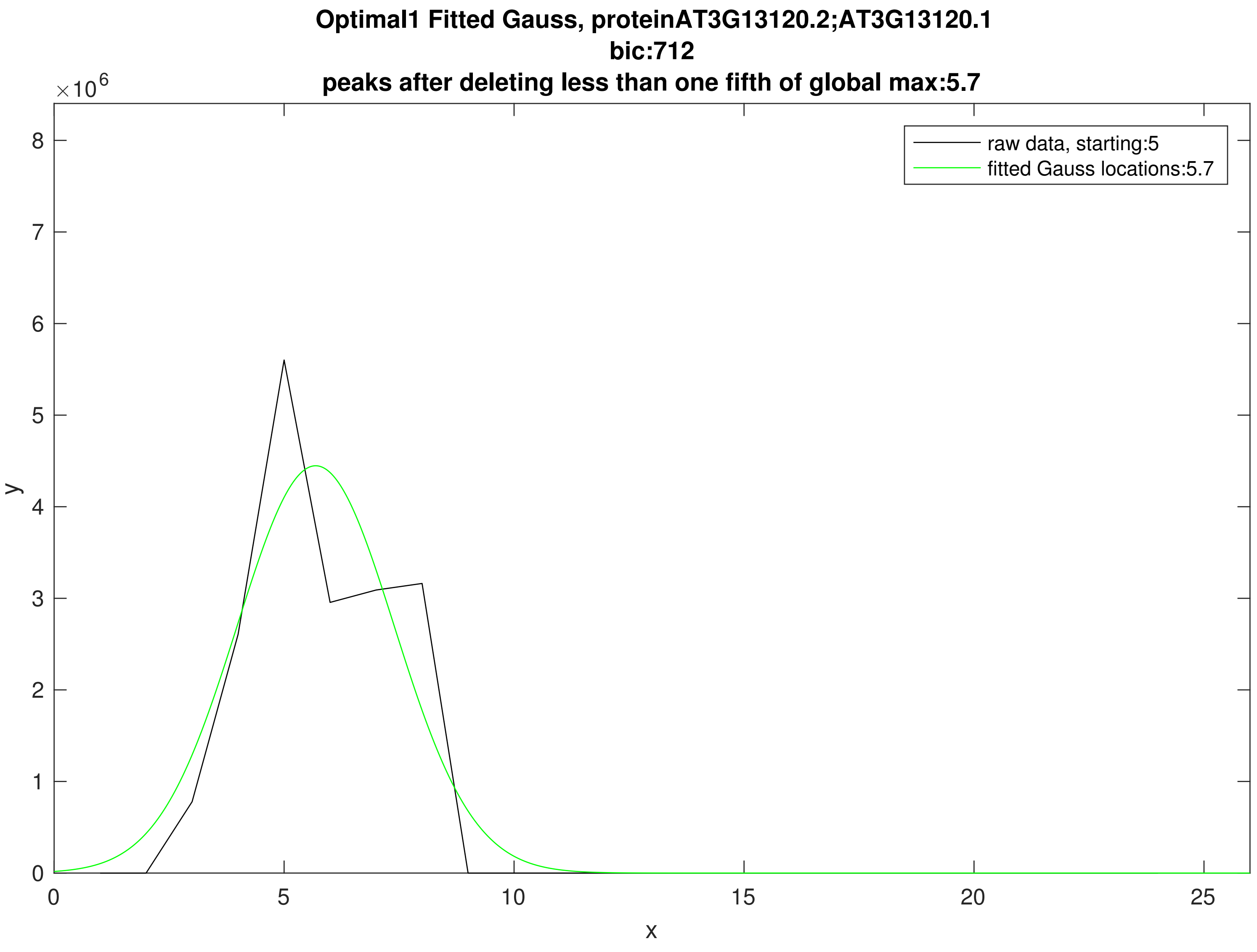
Optimal1 Fitted Gauss, proteinAT3G13120.2;AT3G13120.1
bic:712
peaks after deleting less than one fifth of global max:5.7
×10^6
raw data, starting:5
fitted Gauss locations:5.7
y
x
0
1
2
3
4
5
6
7
8
0
5
10
15
20
25

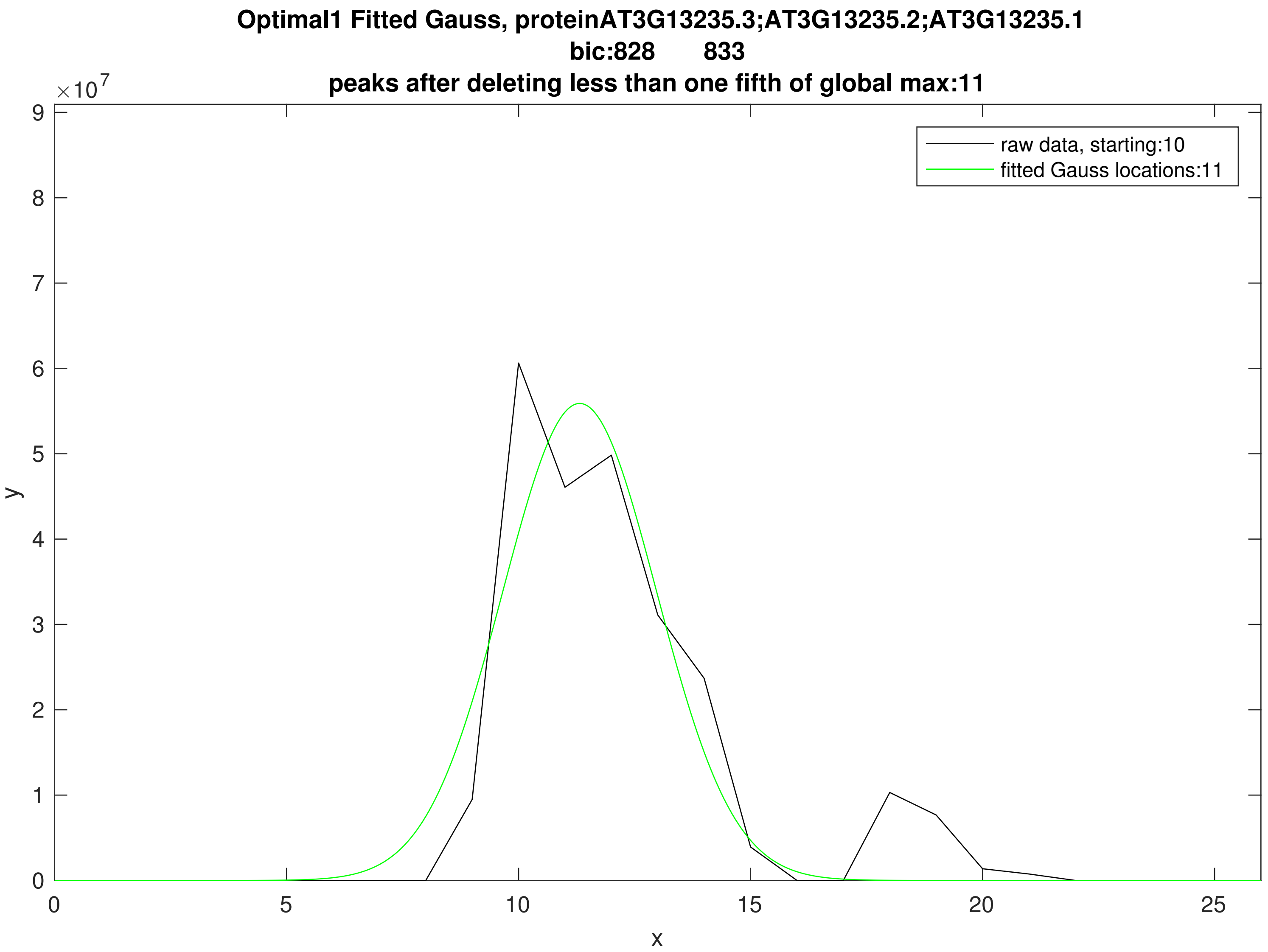

Optimal1 Fitted Gauss, proteinAT3G13235.3;AT3G13235.2;AT3G13235.1
bic:828 833
peaks after deleting less than one fifth of global max:11
×10^7
raw data, starting:10
fitted Gauss locations:11
y
x
0
1
2
3
4
5
6
7
8
9
0
5
10
15
20
25

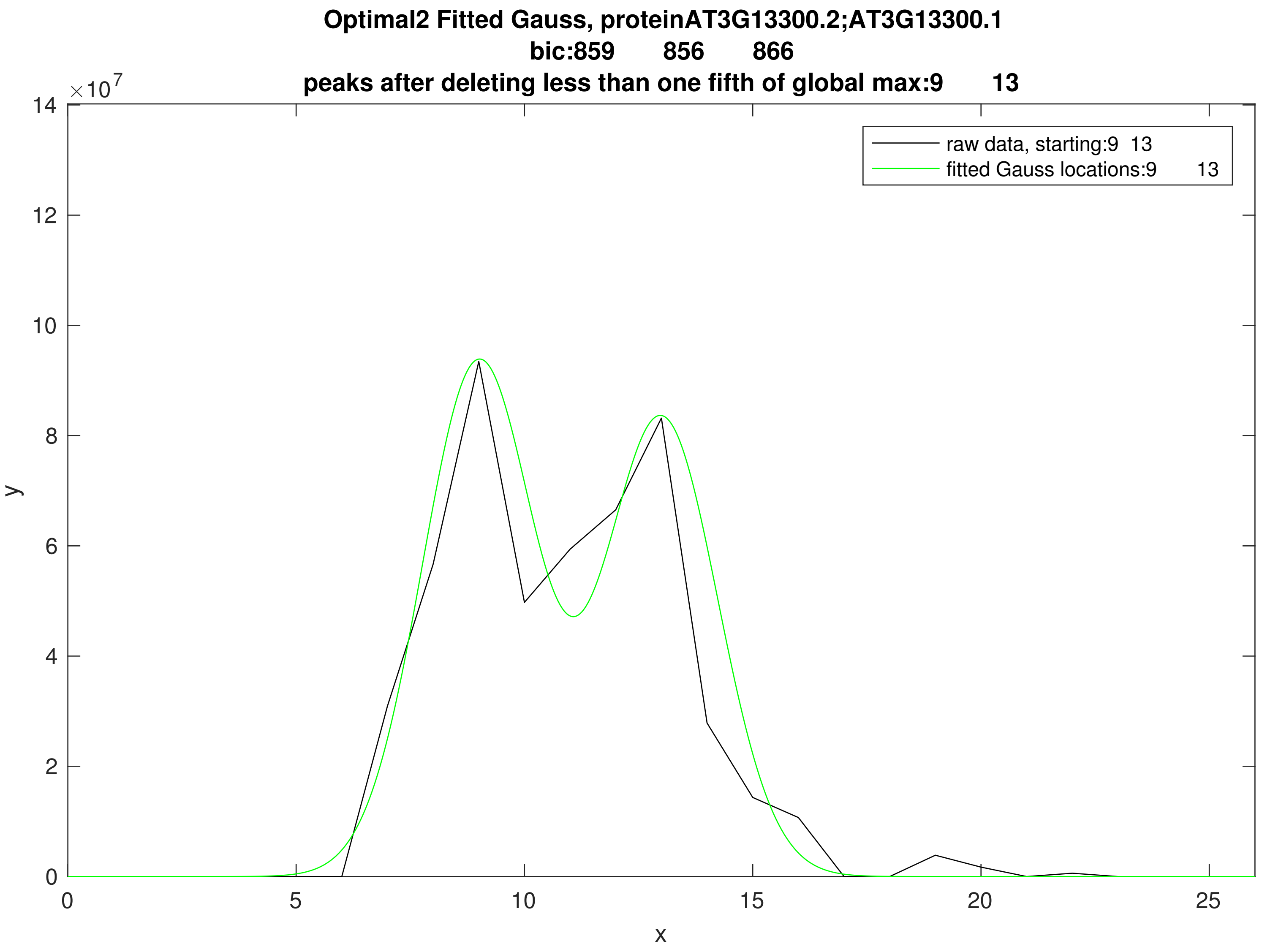

Optimal2 Fitted Gauss, proteinAT3G13300.2;AT3G13300.1
bic:859 856 866
peaks after deleting less than one fifth of global max:9 13
×10^7
raw data, starting:9 13
fitted Gauss locations:9 13
y
x
0
2
4
6
8
10
12
14
5
15
20
25

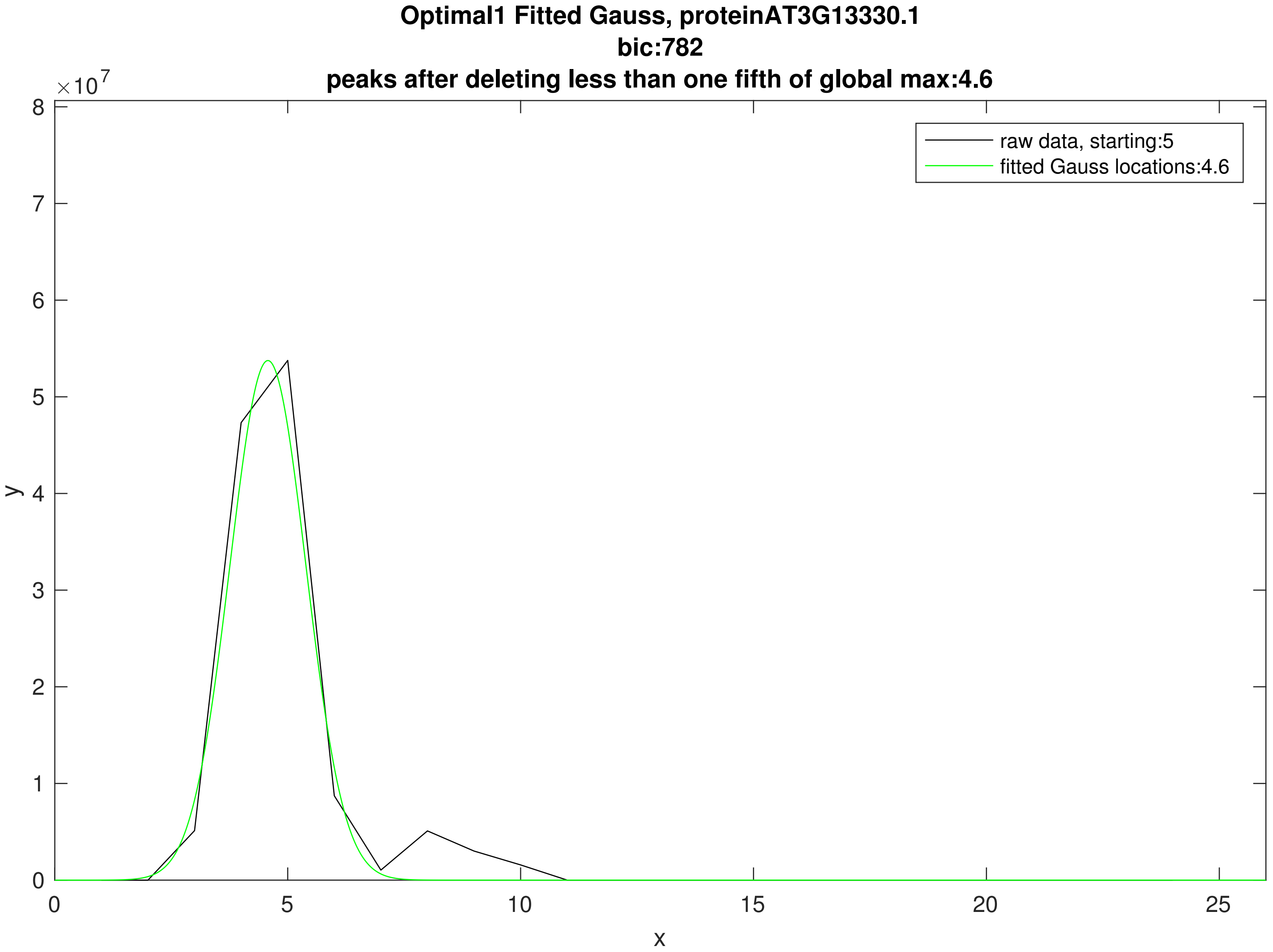
Optimal1 Fitted Gauss, proteinAT3G13330.1
bic:782
peaks after deleting less than one fifth of global max:4.6
×10^7
y
x
raw data, starting:5
fitted Gauss locations:4.6
0
1
2
3
4
5
6
7
8
0
5
10
15
20
25

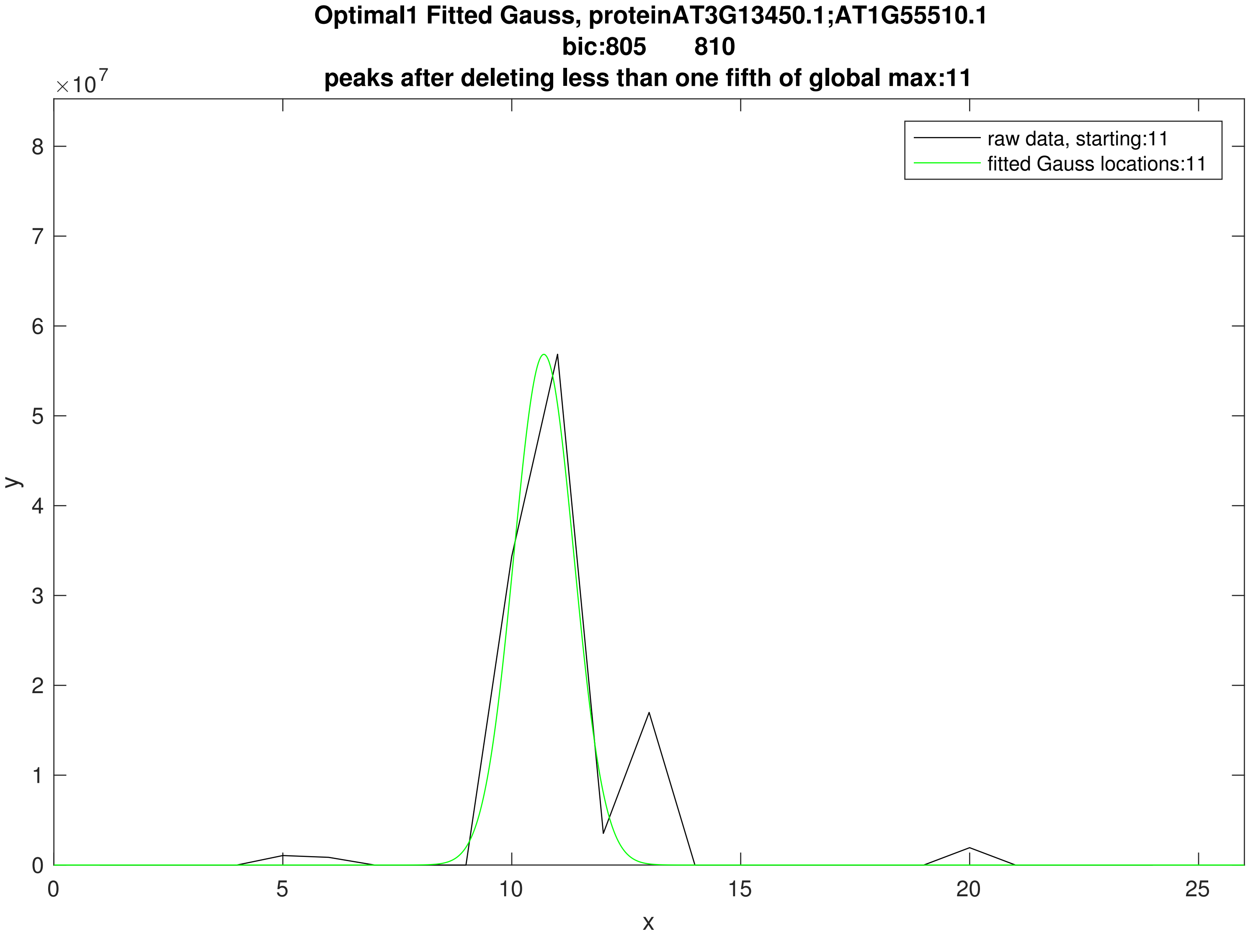
Optimal1 Fitted Gauss, proteinAT3G13450.1;AT1G55510.1
bic:805 810
peaks after deleting less than one fifth of global max:11
×10^7
raw data, starting:11
fitted Gauss locations:11
y
x
0
1
2
3
4
5
6
7
8
0
5
10
15
20
25

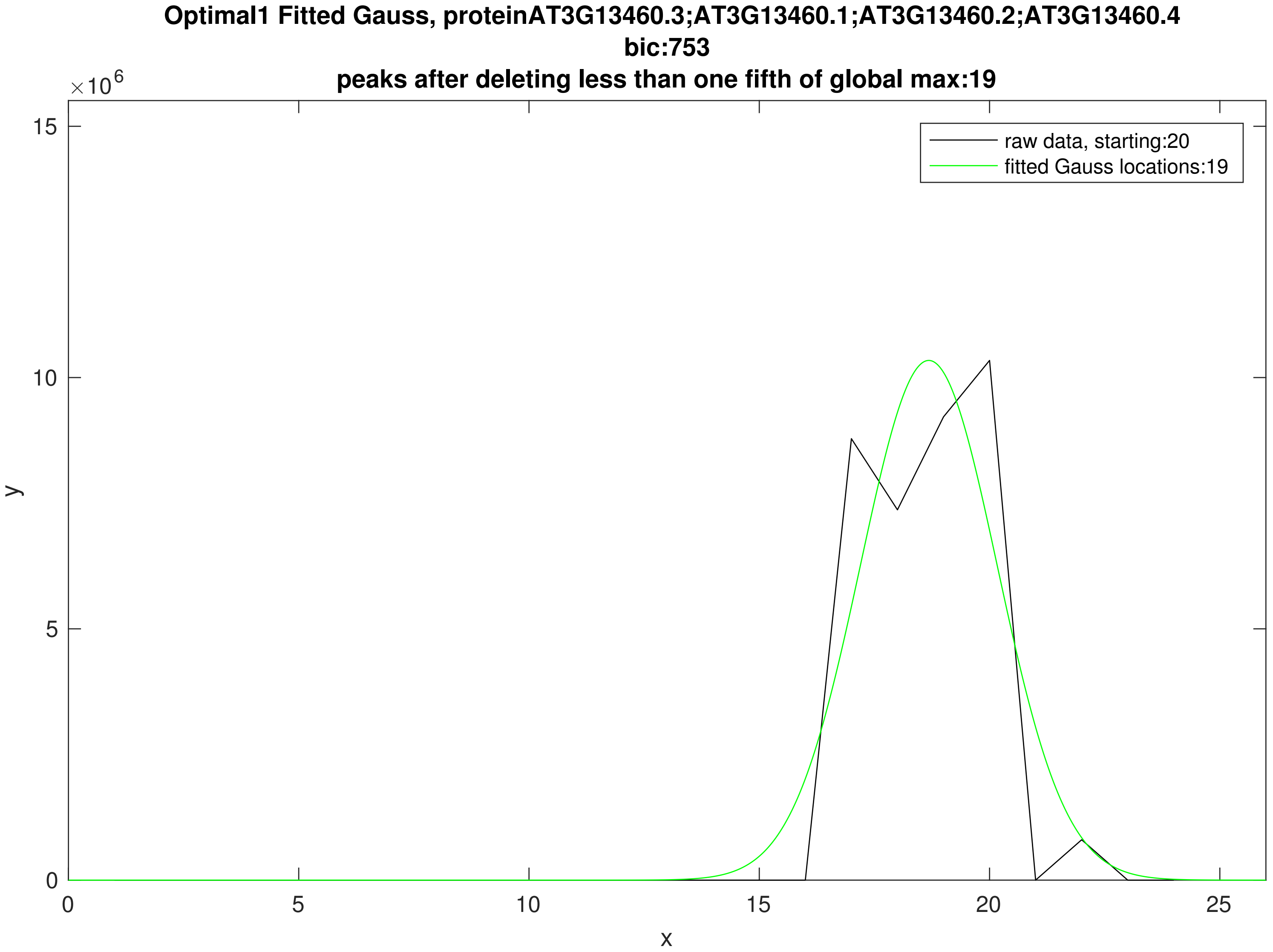

Optimal1 Fitted Gauss, proteinAT3G13460.3;AT3G13460.1;AT3G13460.2;AT3G13460.4
bic:753
peaks after deleting less than one fifth of global max:19
×10^6
raw data, starting:20
fitted Gauss locations:19
y
x
0
5
10
15
20
25

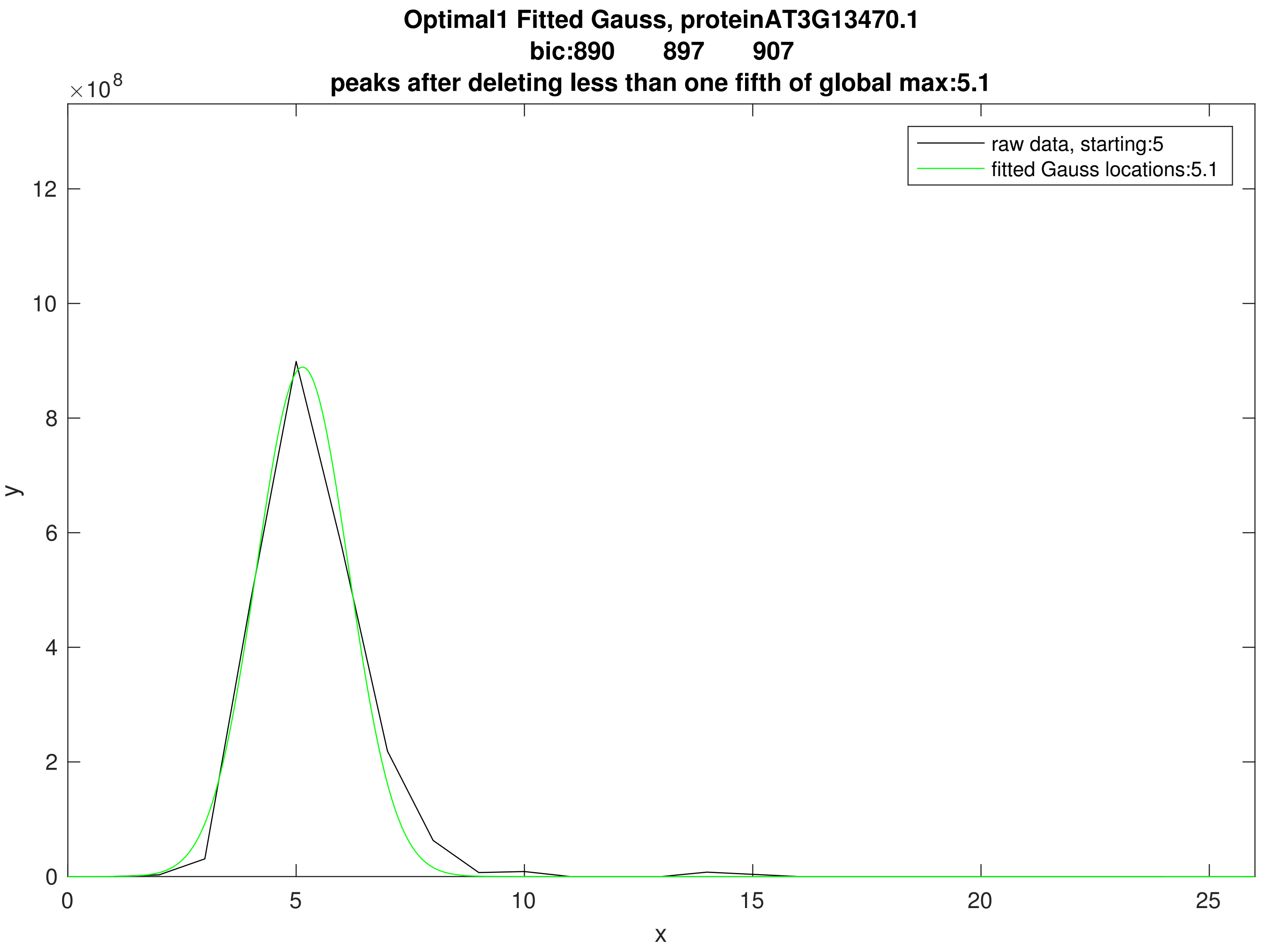

Optimal1 Fitted Gauss, proteinAT3G13470.1
bic:890 897 907
peaks after deleting less than one fifth of global max:5.1
×10^8
raw data, starting:5
fitted Gauss locations:5.1
y
x
0
2
4
6
8
10
12
0
5
10
15
20
25

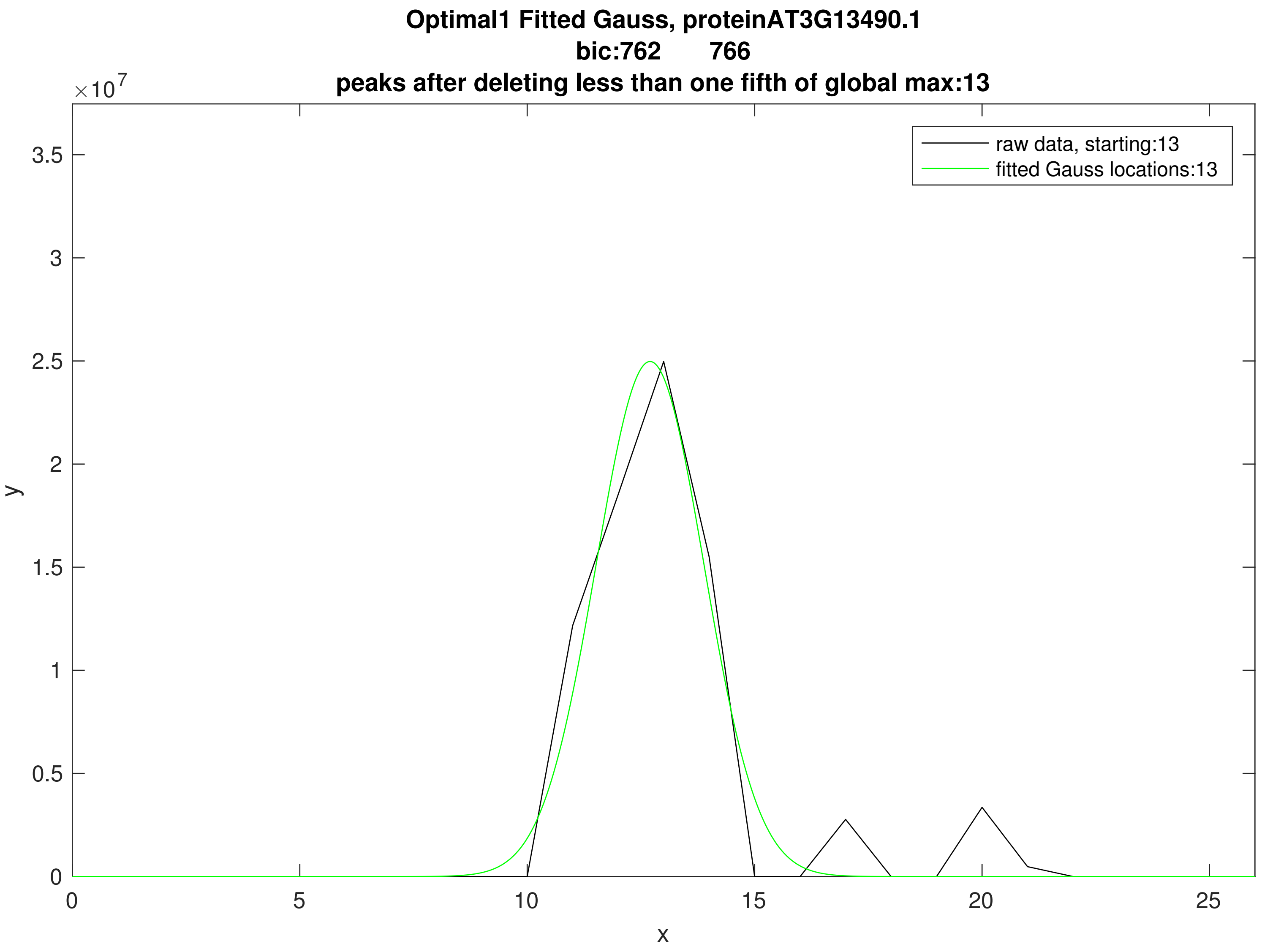
Optimal1 Fitted Gauss, proteinAT3G13490.1
bic:762 766
peaks after deleting less than one fifth of global max:13
×10^7
3.5
3
2.5
2
1.5
1
0.5
0
y
0
5
10
15
20
25
x
raw data, starting:13
fitted Gauss locations:13

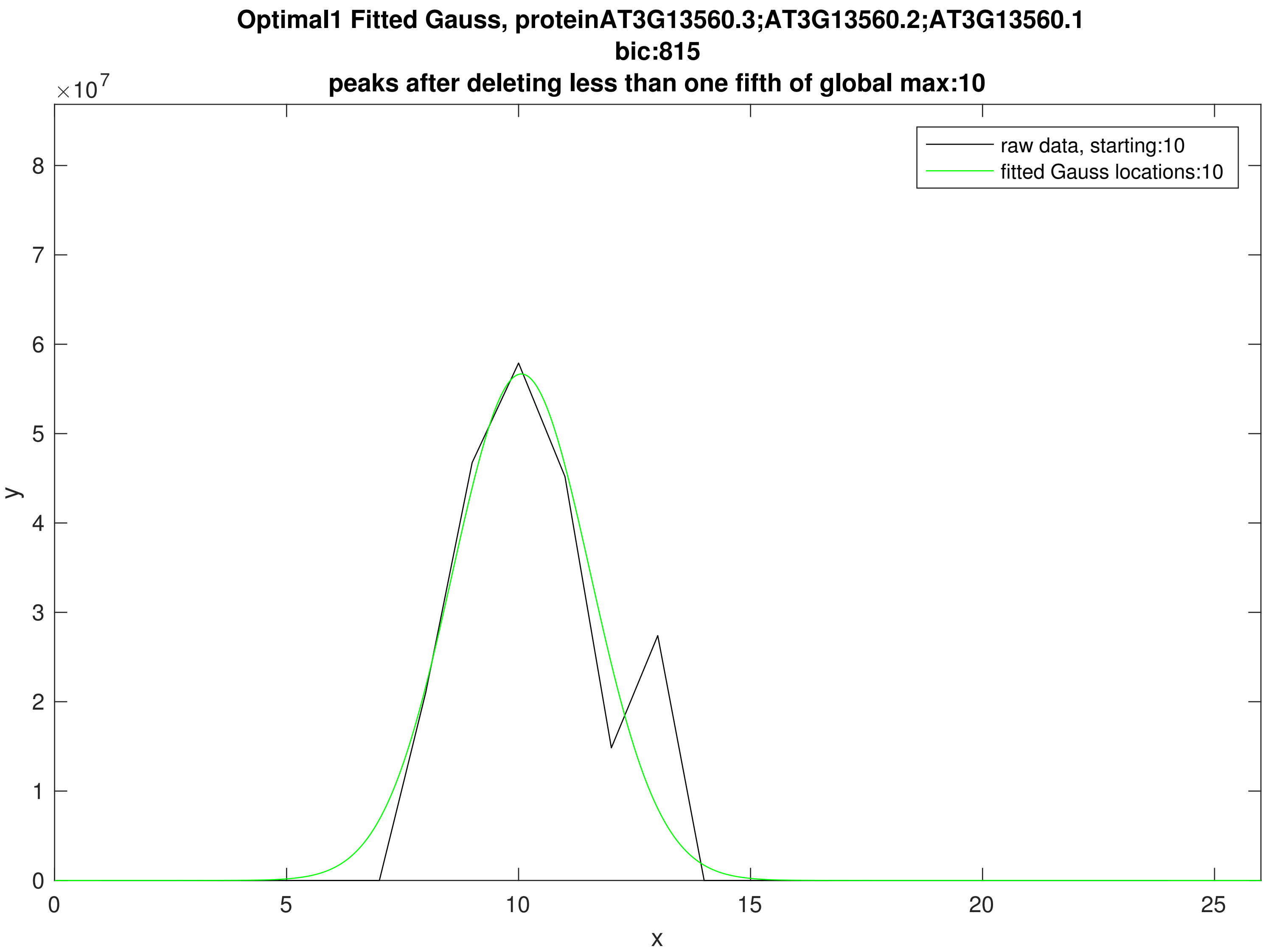
Optimal1 Fitted Gauss, proteinAT3G13560.3;AT3G13560.2;AT3G13560.1
bic:815
peaks after deleting less than one fifth of global max:10
×10^7
raw data, starting:10
fitted Gauss locations:10
y
x
0
1
2
3
4
5
6
7
8
0
5
10
15
20
25

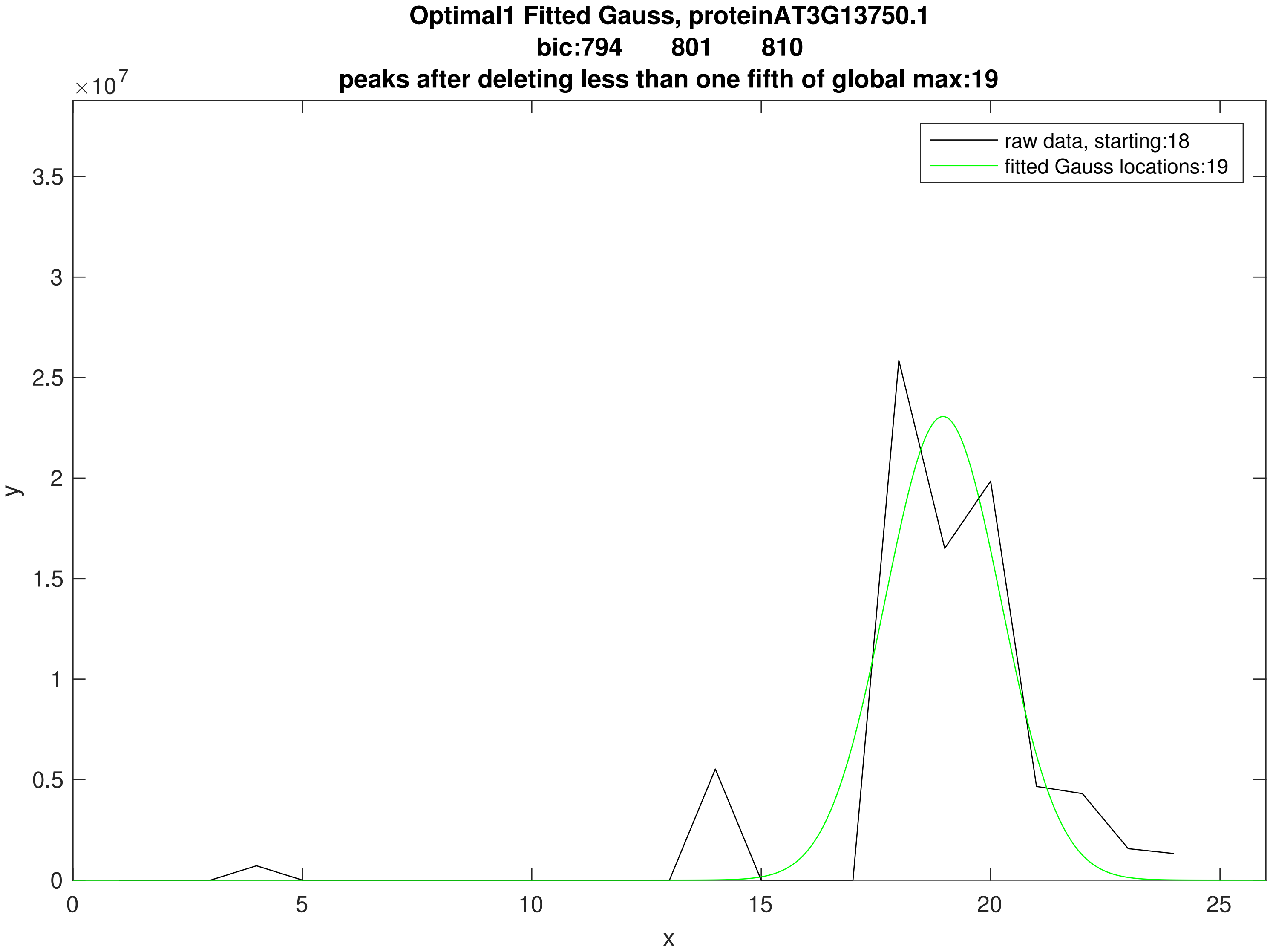
Optimal1 Fitted Gauss, proteinAT3G13750.1
bic:794 801 810
peaks after deleting less than one fifth of global max:19
×10^7
raw data, starting:18
fitted Gauss locations:19
y
x
0
0.5
1
1.5
2
2.5
3
3.5
5
10
15
20
25

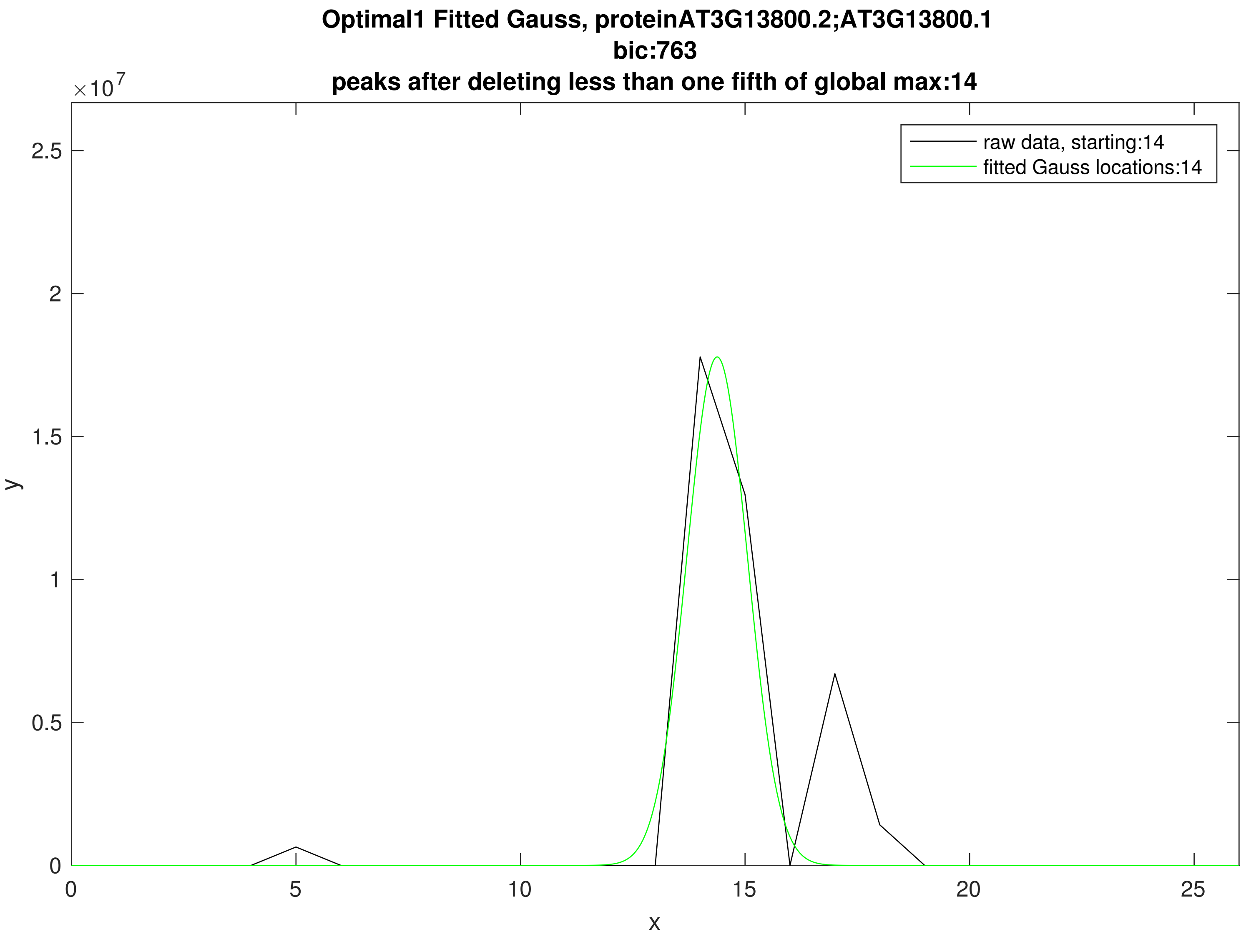
Optimal1 Fitted Gauss, proteinAT3G13800.2;AT3G13800.1
bic:763
peaks after deleting less than one fifth of global max:14
×10^7
2.5
2
1.5
1
0.5
0
y
0
5
10
15
20
25
x
raw data, starting:14
fitted Gauss locations:14

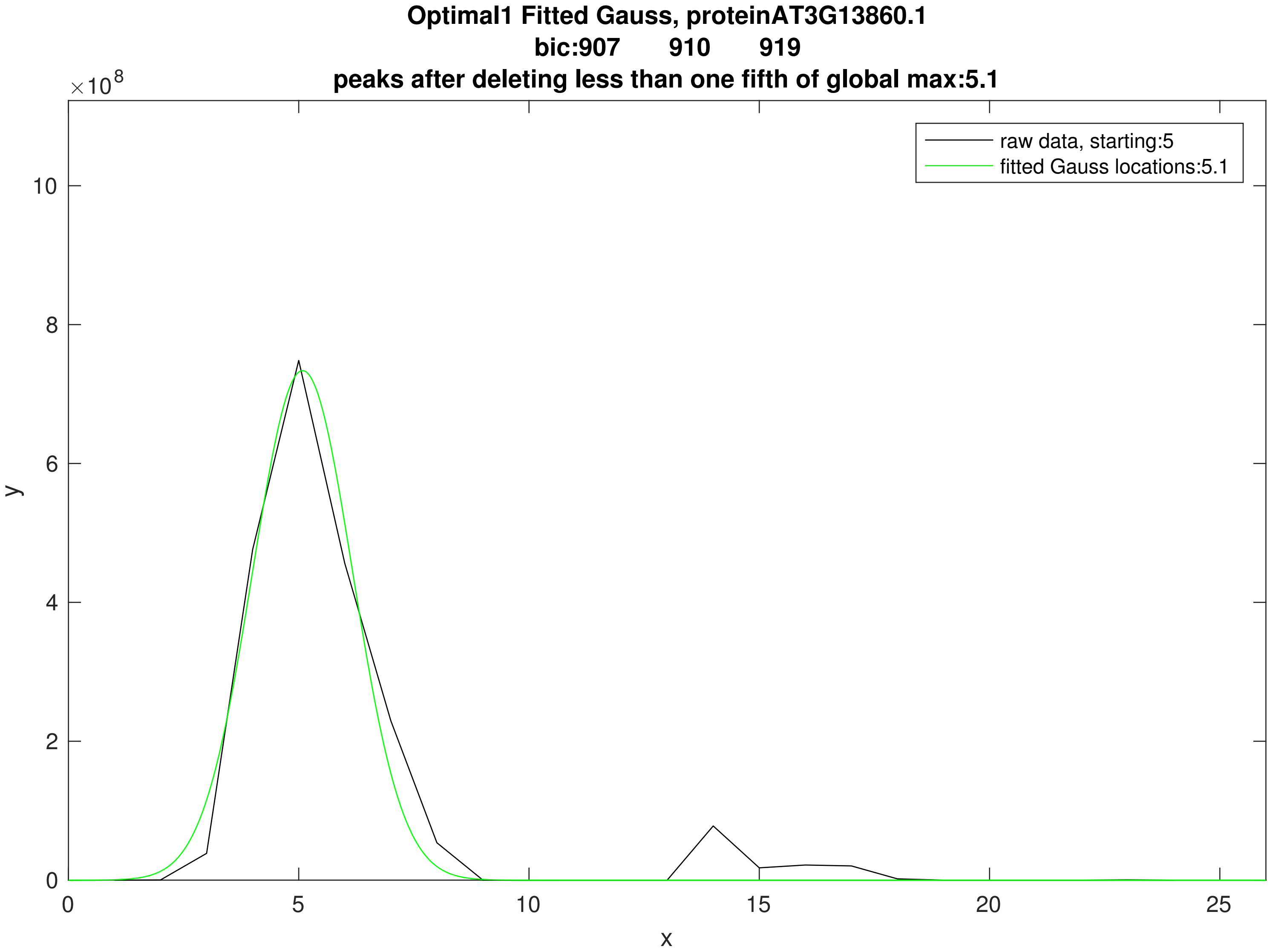

Optimal1 Fitted Gauss, proteinAT3G13860.1
bic:907 910 919
peaks after deleting less than one fifth of global max:5.1
×10^8
y
x
raw data, starting:5
fitted Gauss locations:5.1
0
2
4
6
8
10
0
5
10
15
20
25

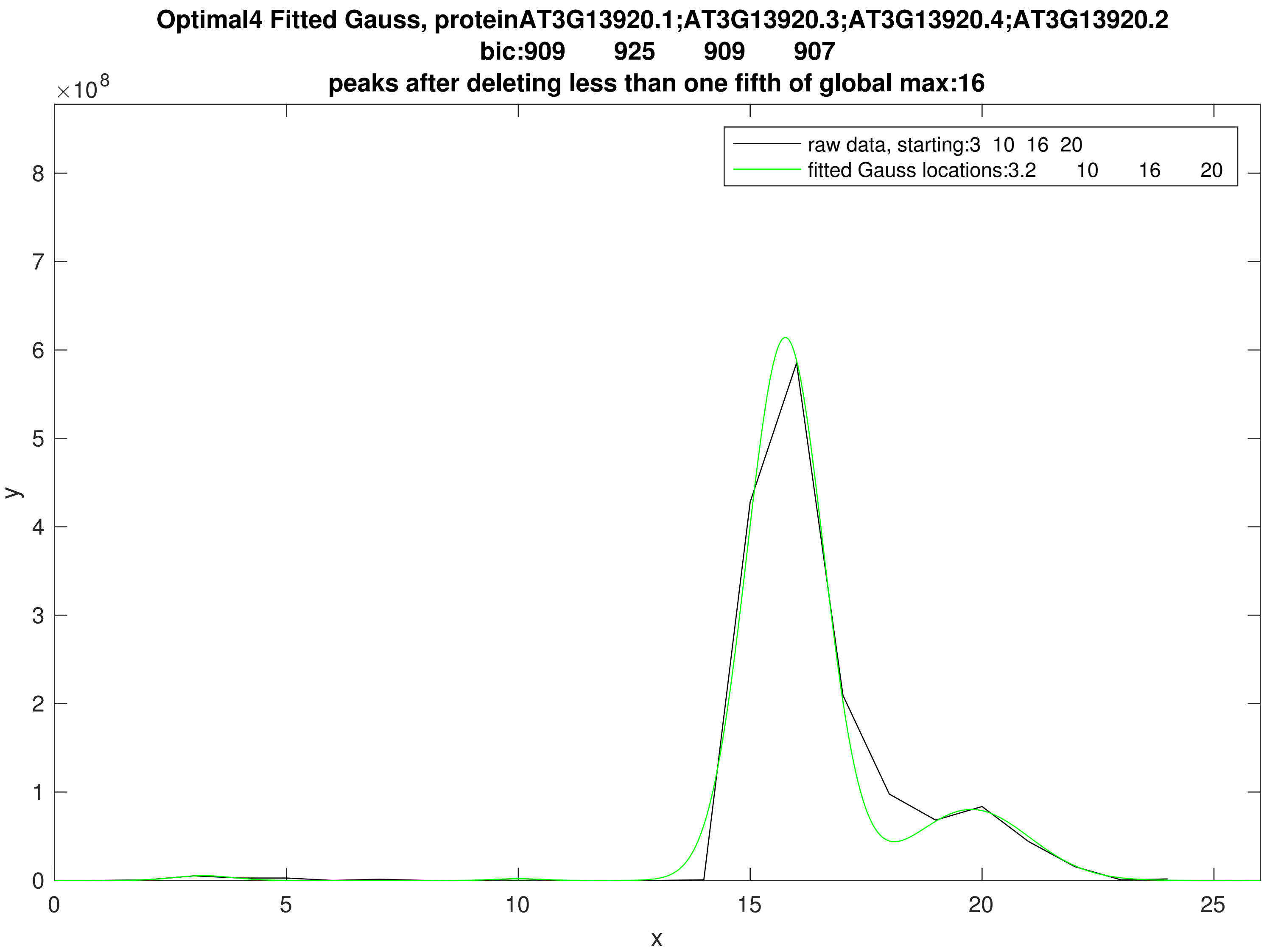

Optimal4 Fitted Gauss, proteinAT3G13920.1;AT3G13920.3;AT3G13920.4;AT3G13920.2
bic:909 925 909 907
peaks after deleting less than one fifth of global max:16
×10^8
raw data, starting:3 10 16 20
fitted Gauss locations:3.2 10 16 20
y
x
0
1
2
3
4
5
6
7
8
10
15
20
25

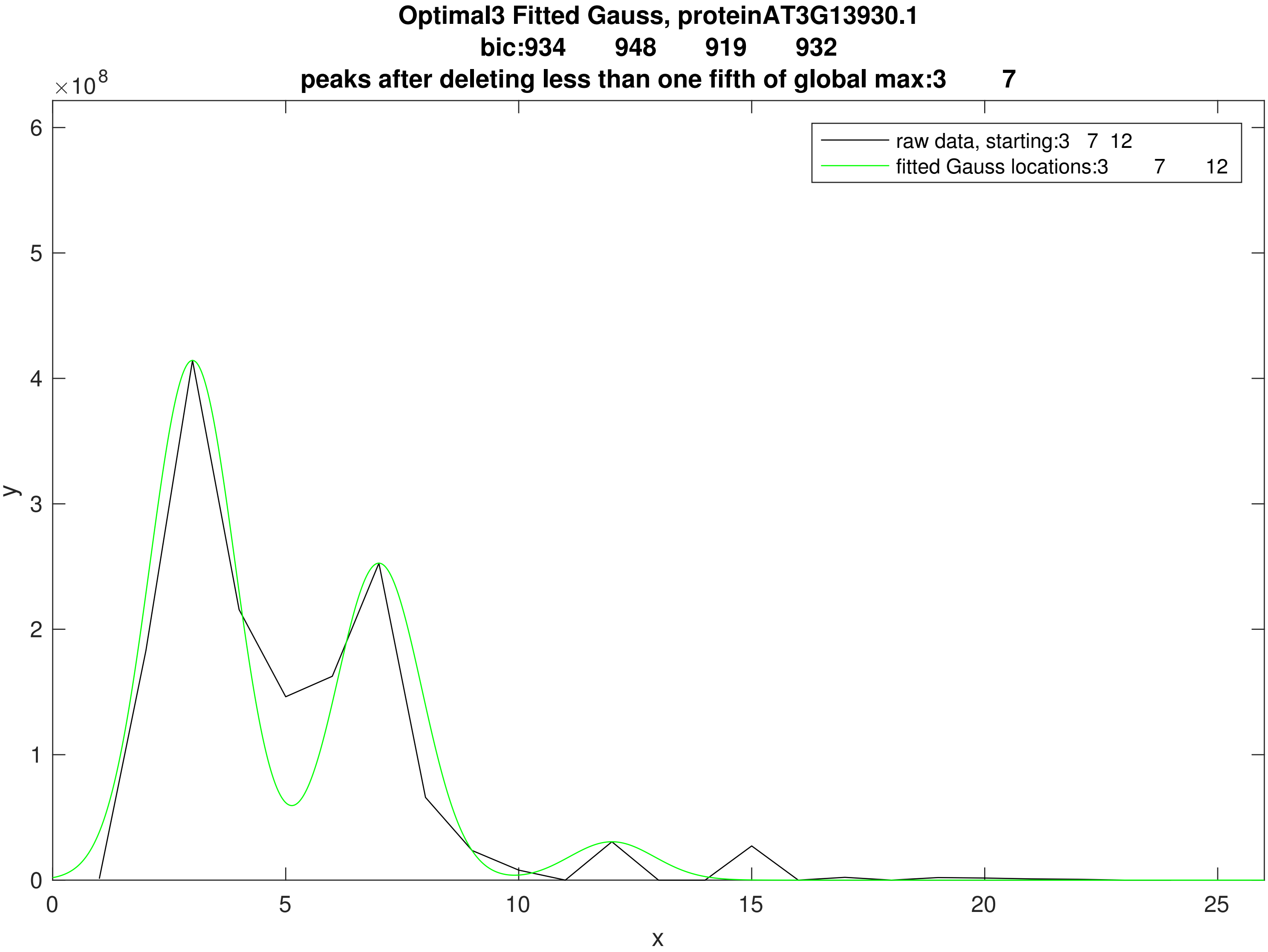

Optimal3 Fitted Gauss, proteinAT3G13930.1
bic:934 948 919 932
peaks after deleting less than one fifth of global max:3 7
×10^8
raw data, starting:3 7 12
fitted Gauss locations:3 7 12
y
x
0
1
2
3
4
5
6
0
5
10
15
20
25

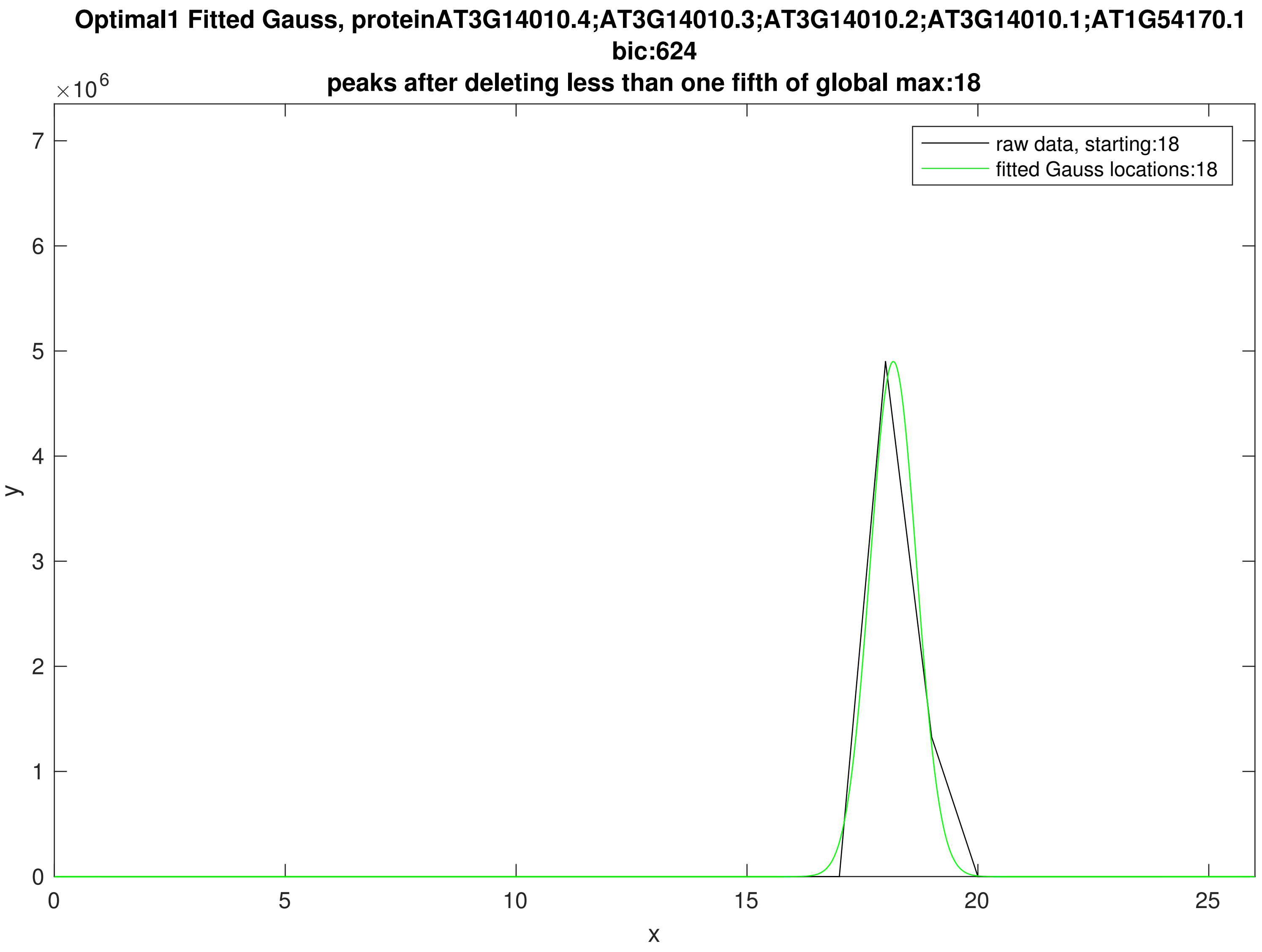

Optimal1 Fitted Gauss, proteinAT3G14010.4;AT3G14010.3;AT3G14010.2;AT3G14010.1;AT1G54170.1
bic:624
peaks after deleting less than one fifth of global max:18
×10^6
7
6
5
4
3
2
1
0
y
0
5
10
15
20
25
x
raw data, starting:18
fitted Gauss locations:18

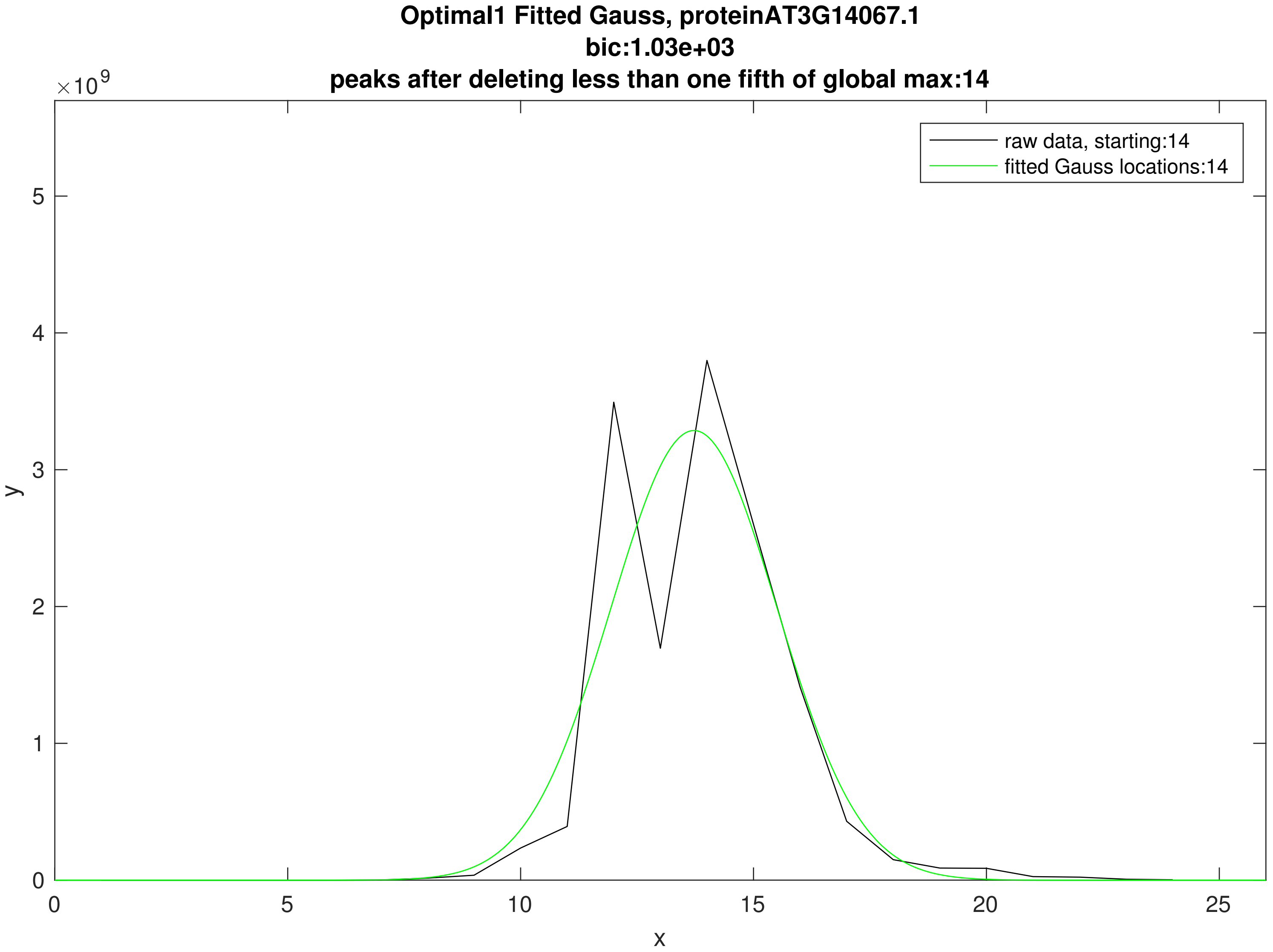
Optimal1 Fitted Gauss, proteinAT3G14067.1
bic:1.03e+03
peaks after deleting less than one fifth of global max:14
×10^9
raw data, starting:14
fitted Gauss locations:14
y
x
0
1
2
3
4
5
0
5
10
15
20
25

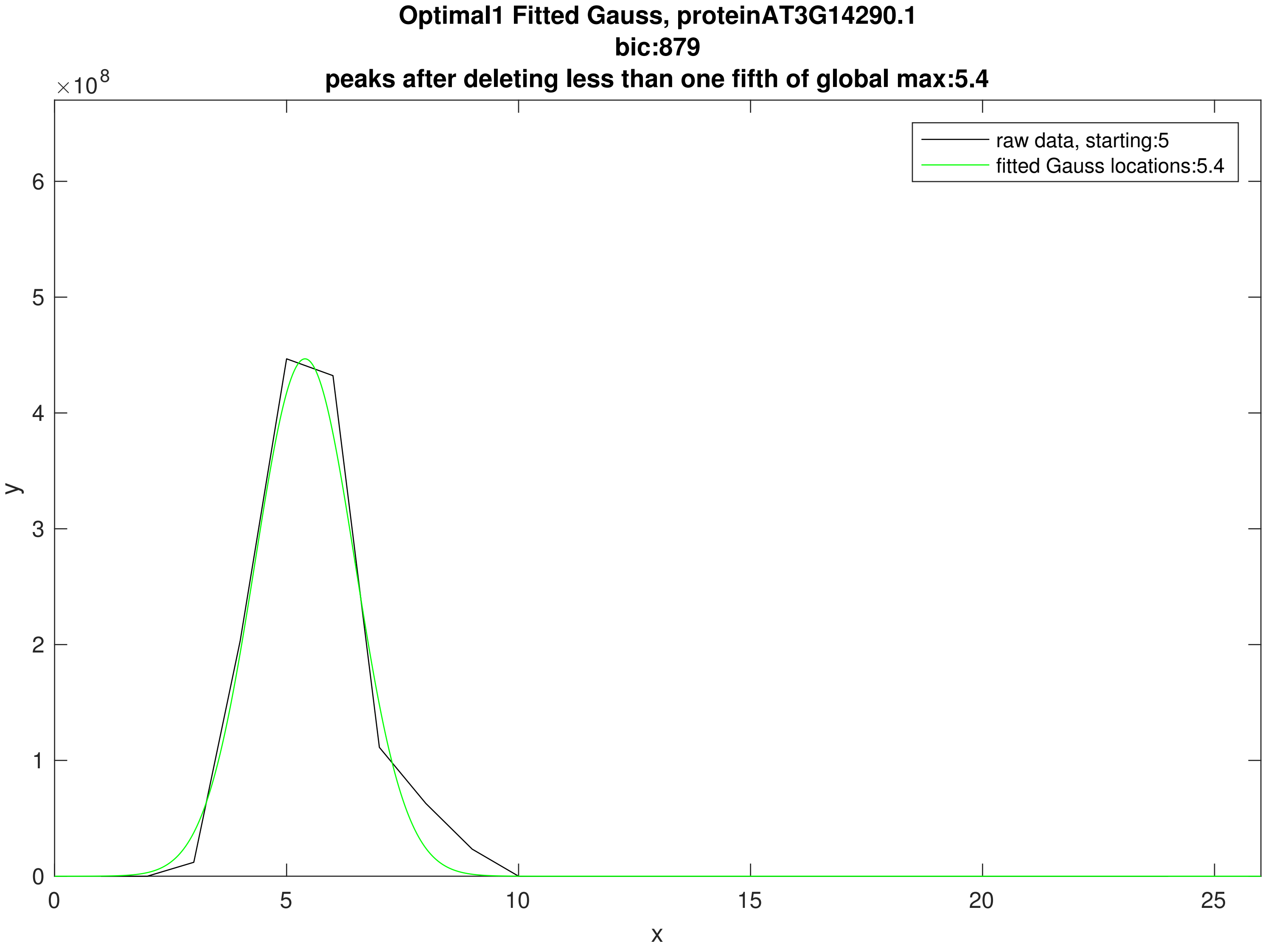
Optimal1 Fitted Gauss, proteinAT3G14290.1
bic:879
peaks after deleting less than one fifth of global max:5.4
×10^8
raw data, starting:5
fitted Gauss locations:5.4
y
x
0
1
2
3
4
5
6
10
15
20
25

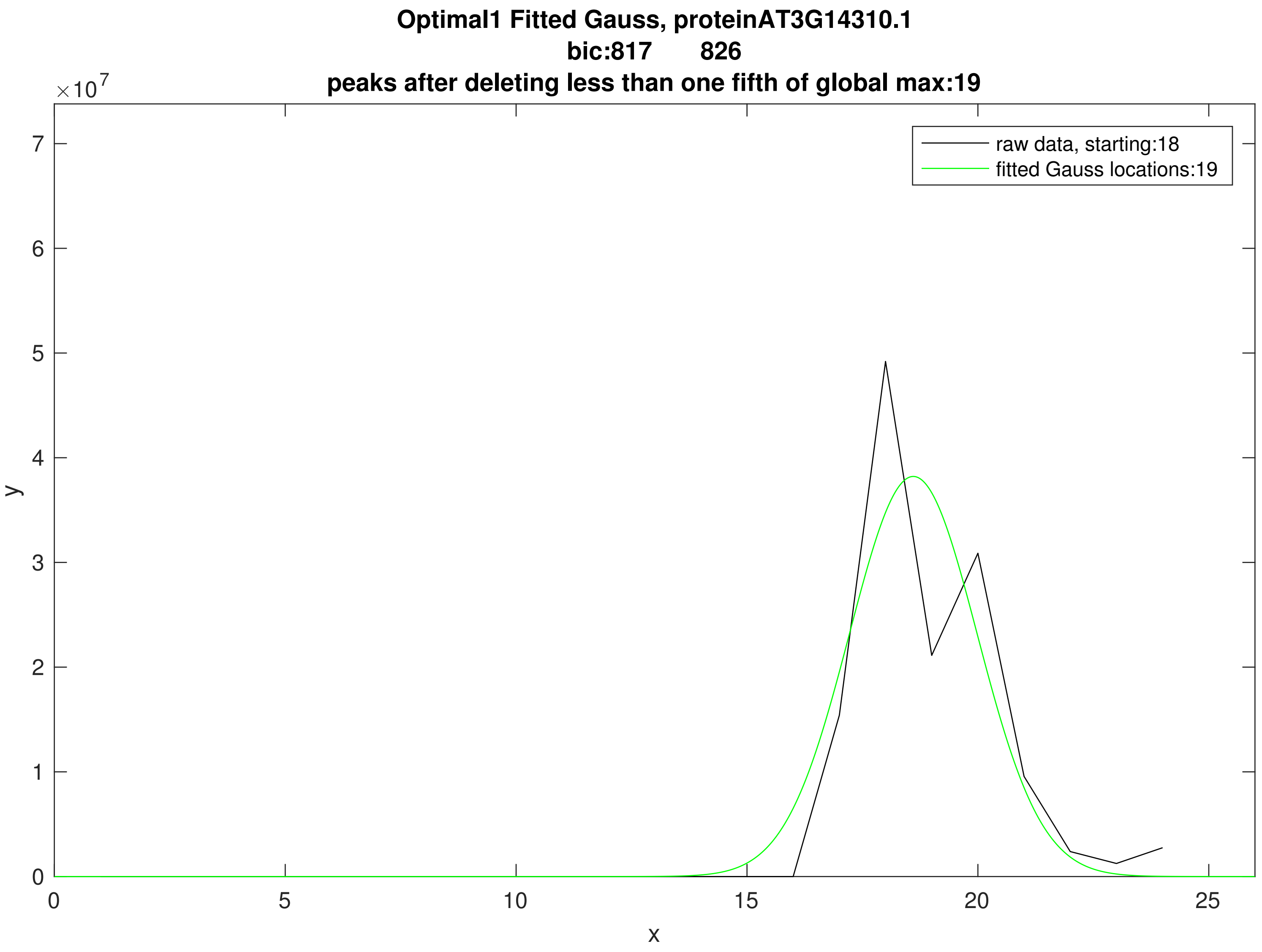
Optimal1 Fitted Gauss, proteinAT3G14310.1
bic:817 826
peaks after deleting less than one fifth of global max:19
×10^7
raw data, starting:18
fitted Gauss locations:19
y
x
0
1
2
3
4
5
6
7
0
5
10
15
20
25

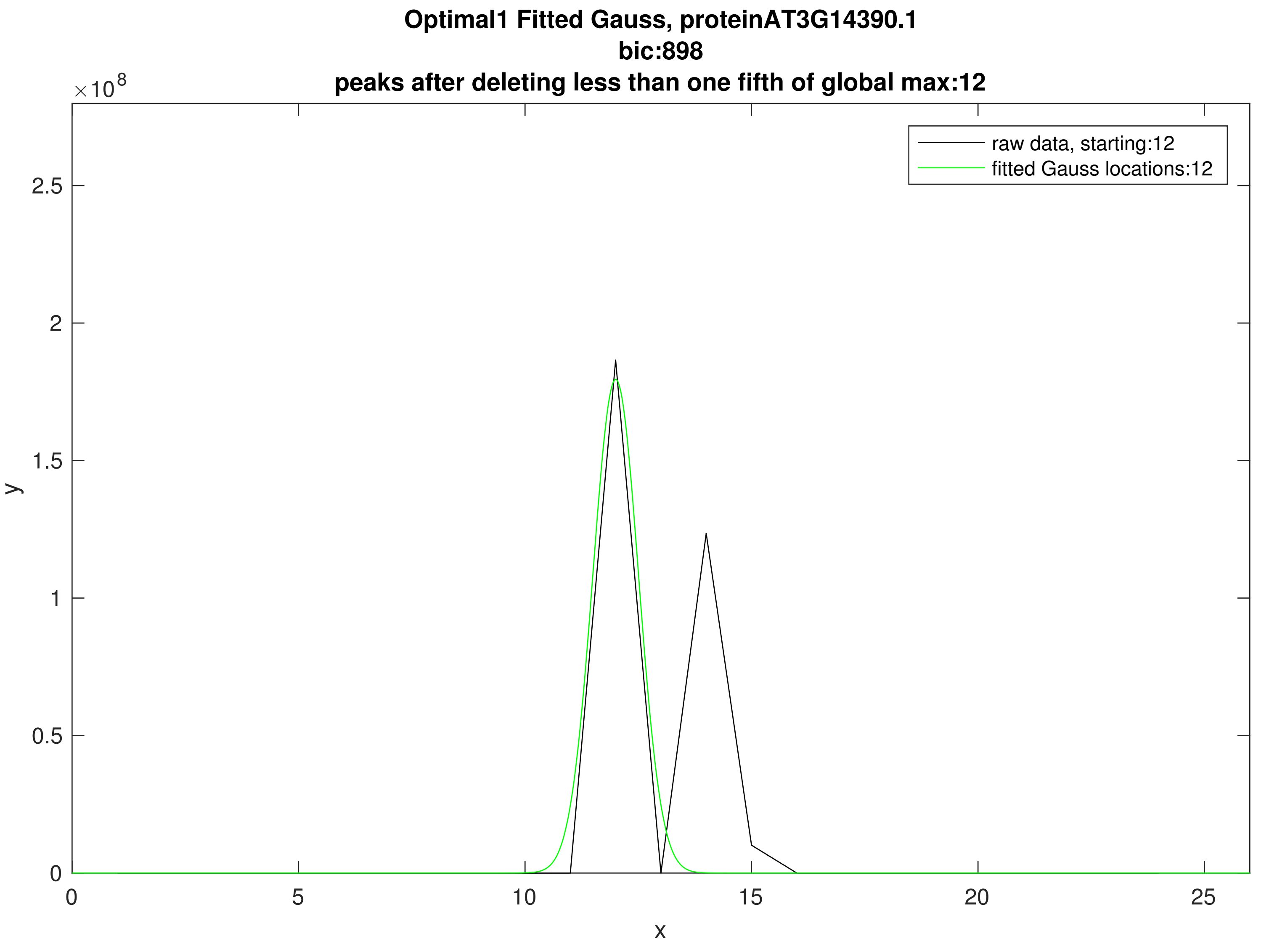

Optimal1 Fitted Gauss, proteinAT3G14390.1
bic:898
peaks after deleting less than one fifth of global max:12
×10^8
raw data, starting:12
fitted Gauss locations:12
y
x
0
0.5
1
1.5
2
2.5
0
5
10
15
20
25

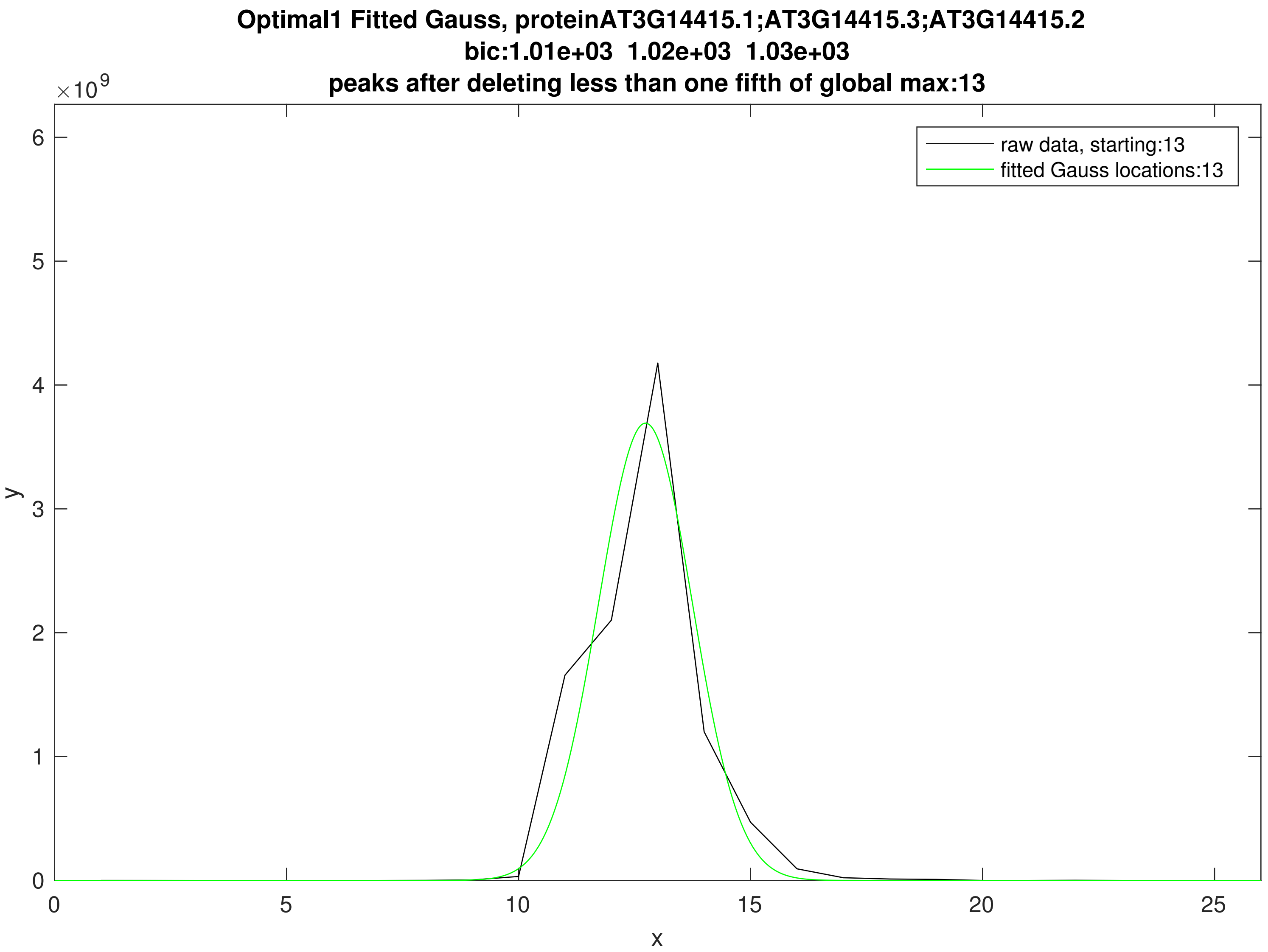

Optimal1 Fitted Gauss, proteinAT3G14415.1;AT3G14415.3;AT3G14415.2
bic:1.01e+03 1.02e+03 1.03e+03
peaks after deleting less than one fifth of global max:13
×10^9
raw data, starting:13
fitted Gauss locations:13
y
x

**Optimal1 Fitted Gauss, proteinAT3G14420.2;AT3G14420.1;AT3G14420.4;AT3G14420.3;AT3G14420.5;AT3G14420.**
**bic:894 951**
**peaks after deleting less than one fifth of global max:12**

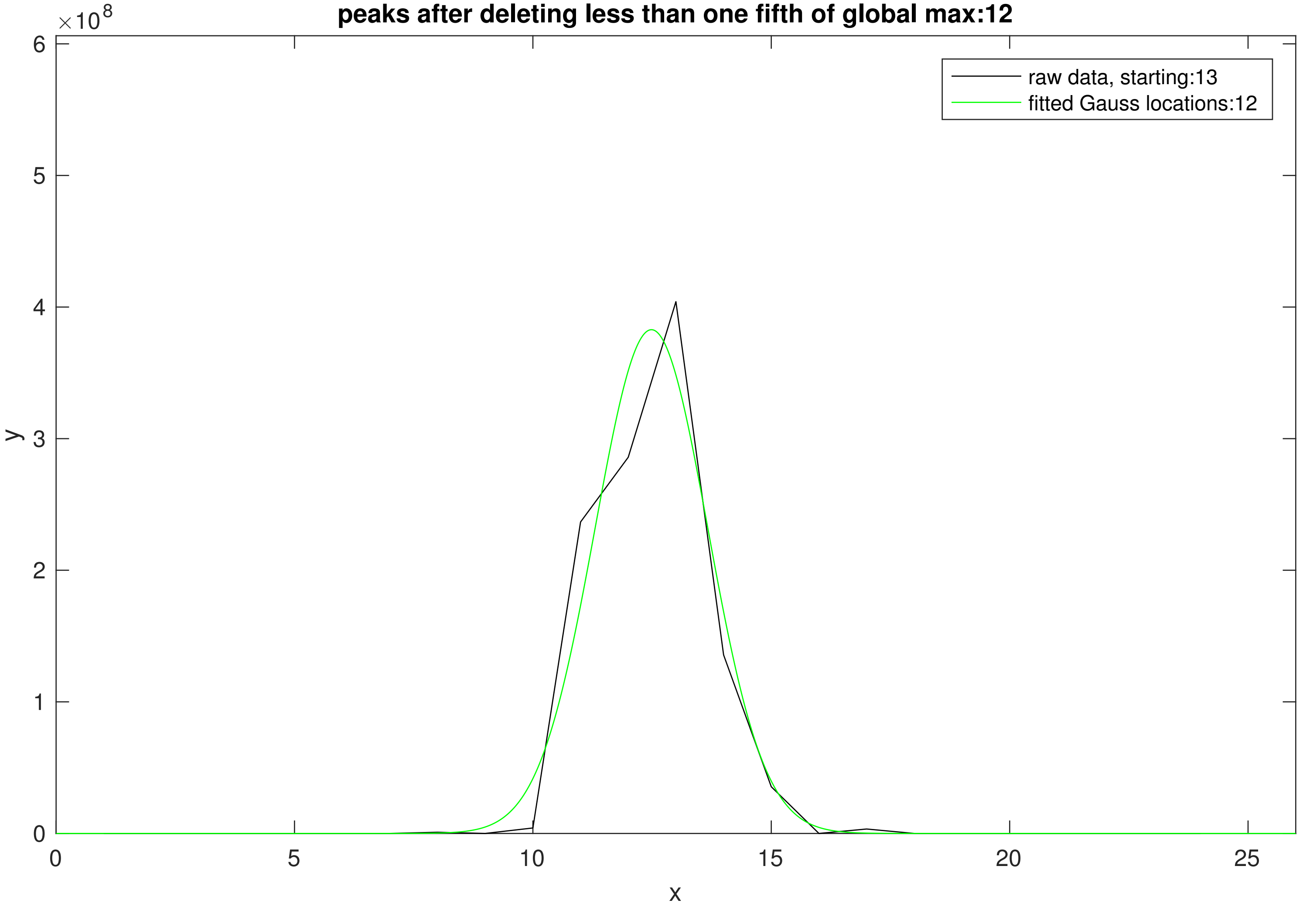

**Optimal1 Fitted Gauss, proteinAT3G14600.1**
**bic:652**
**peaks after deleting less than one fifth of global max:3**

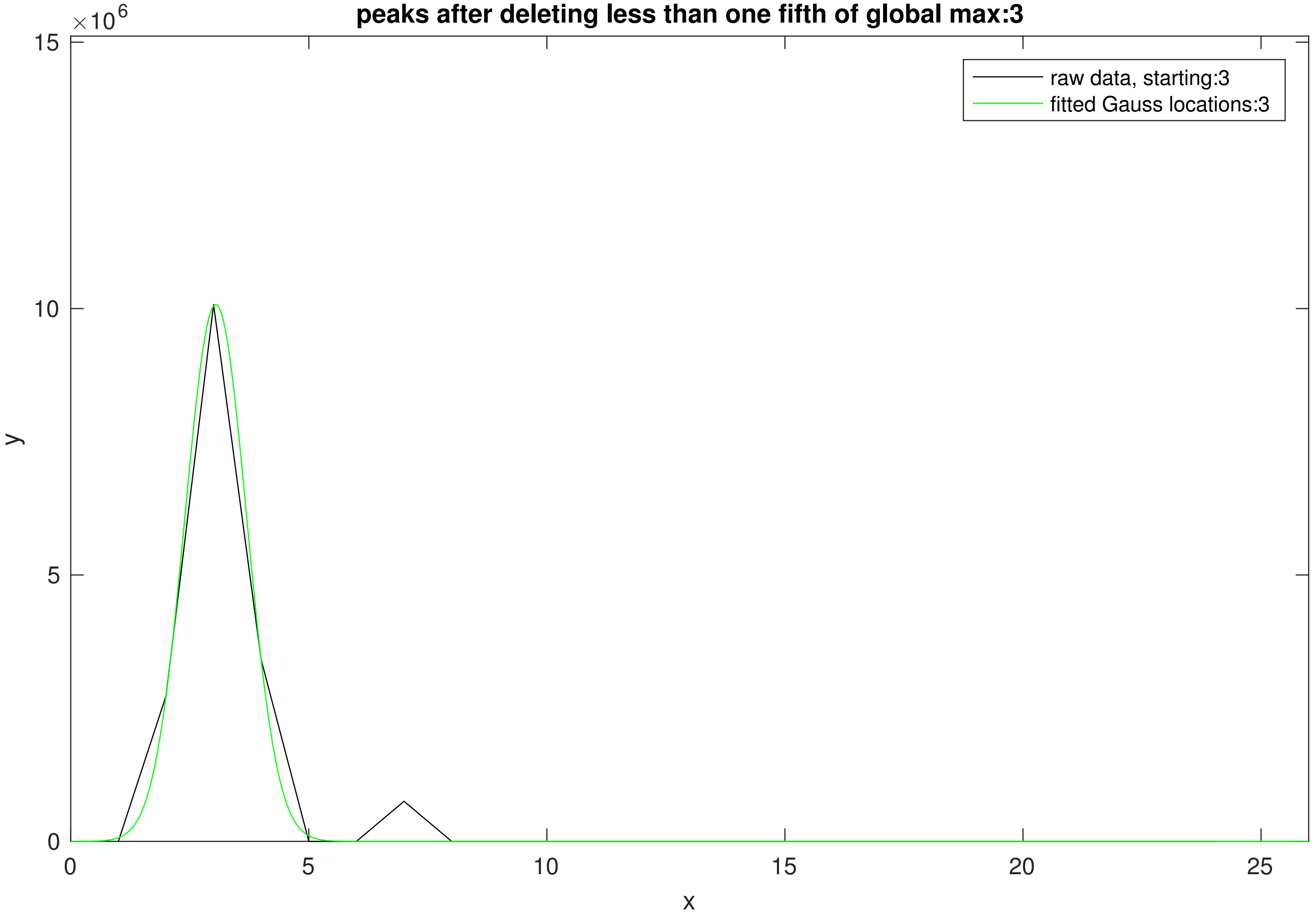

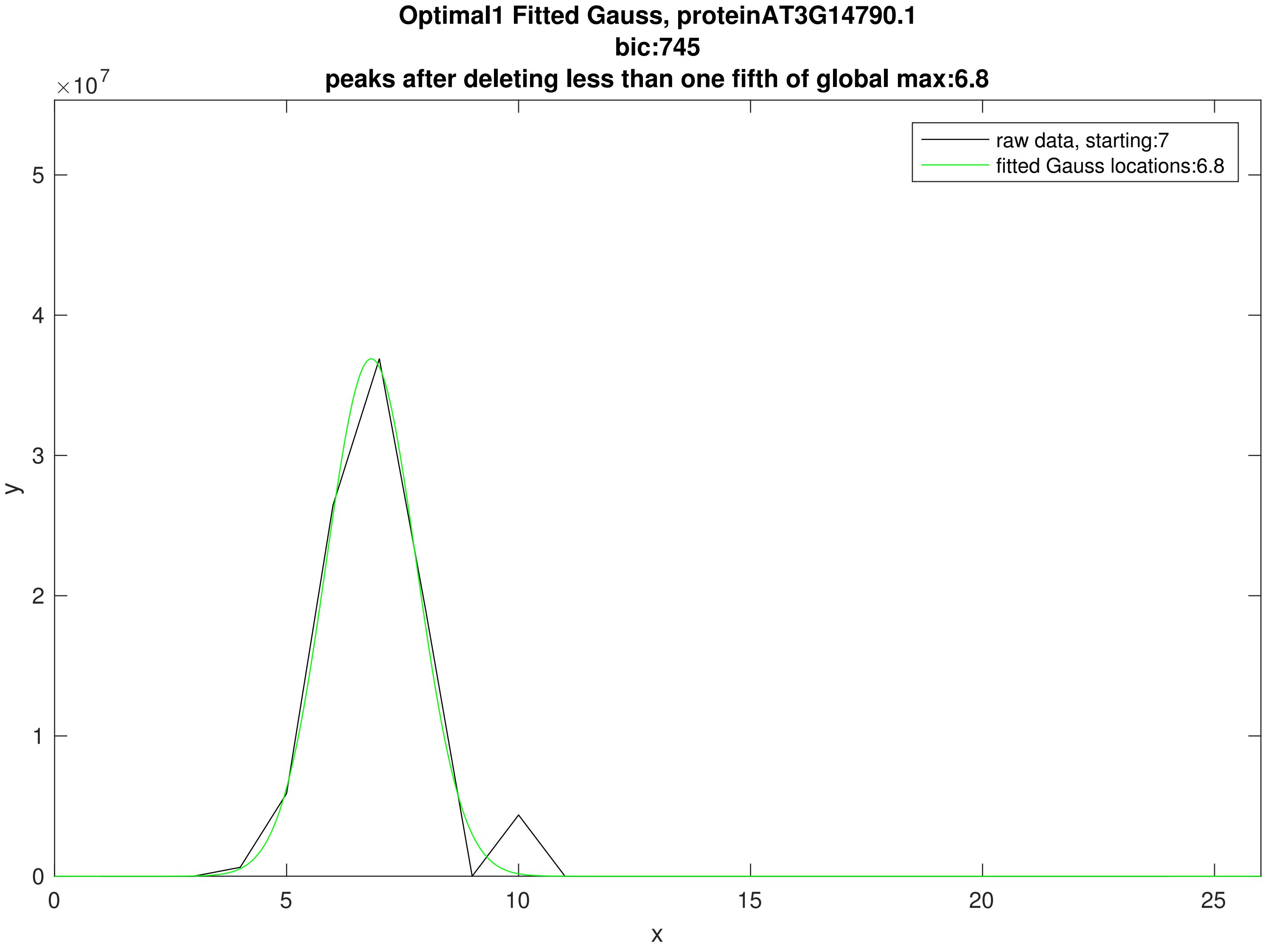
Optimal1 Fitted Gauss, proteinAT3G14790.1
bic:745
peaks after deleting less than one fifth of global max:6.8
×10^7
raw data, starting:7
fitted Gauss locations:6.8
y
x
0
1
2
3
4
5
0
5
10
15
20
25

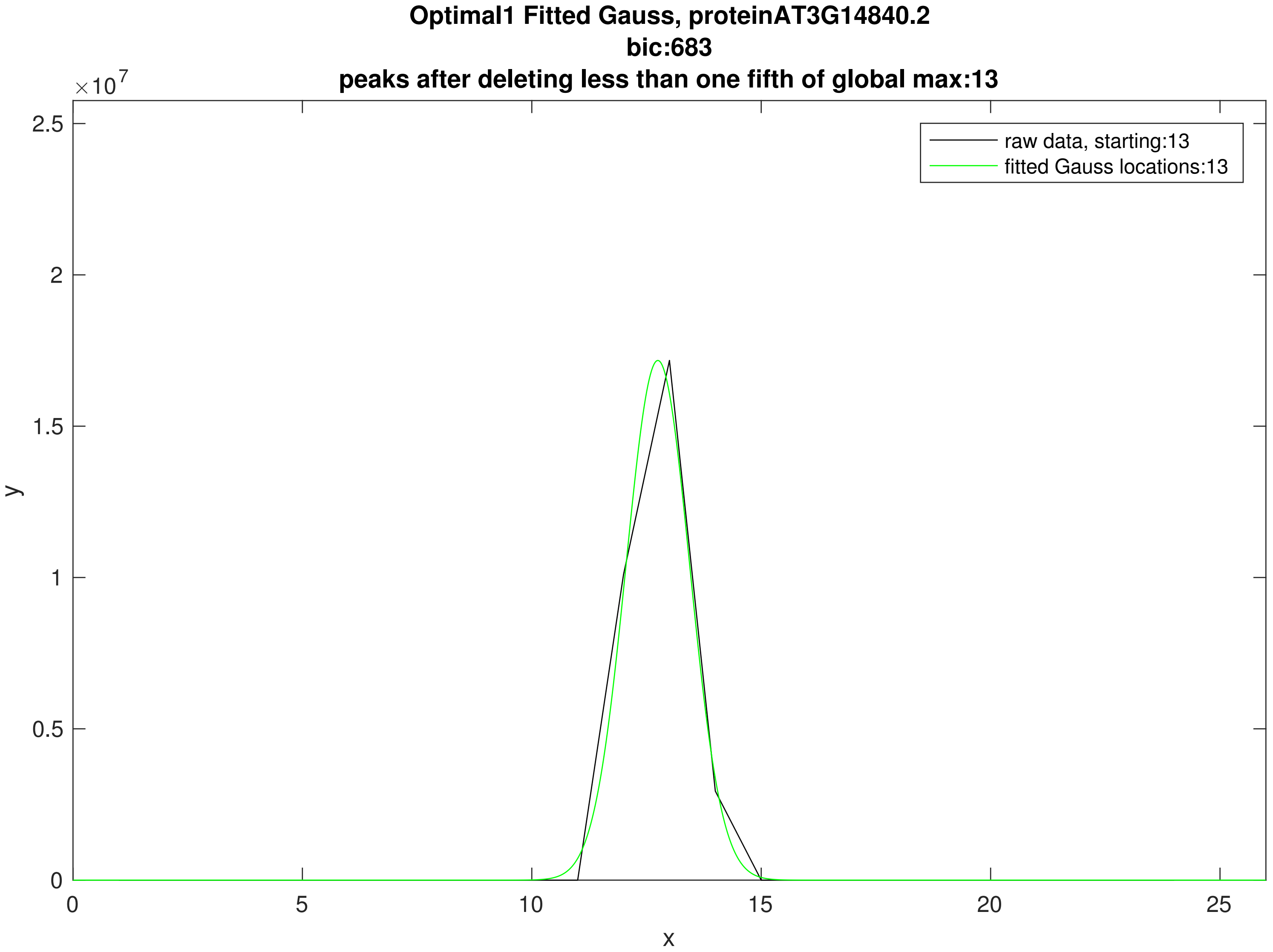

Optimal1 Fitted Gauss, proteinAT3G14840.2
bic:683
peaks after deleting less than one fifth of global max:13
×10^7
2.5
2
1.5
1
0.5
0
y
0
5
10
15
20
25
x
raw data, starting:13
fitted Gauss locations:13

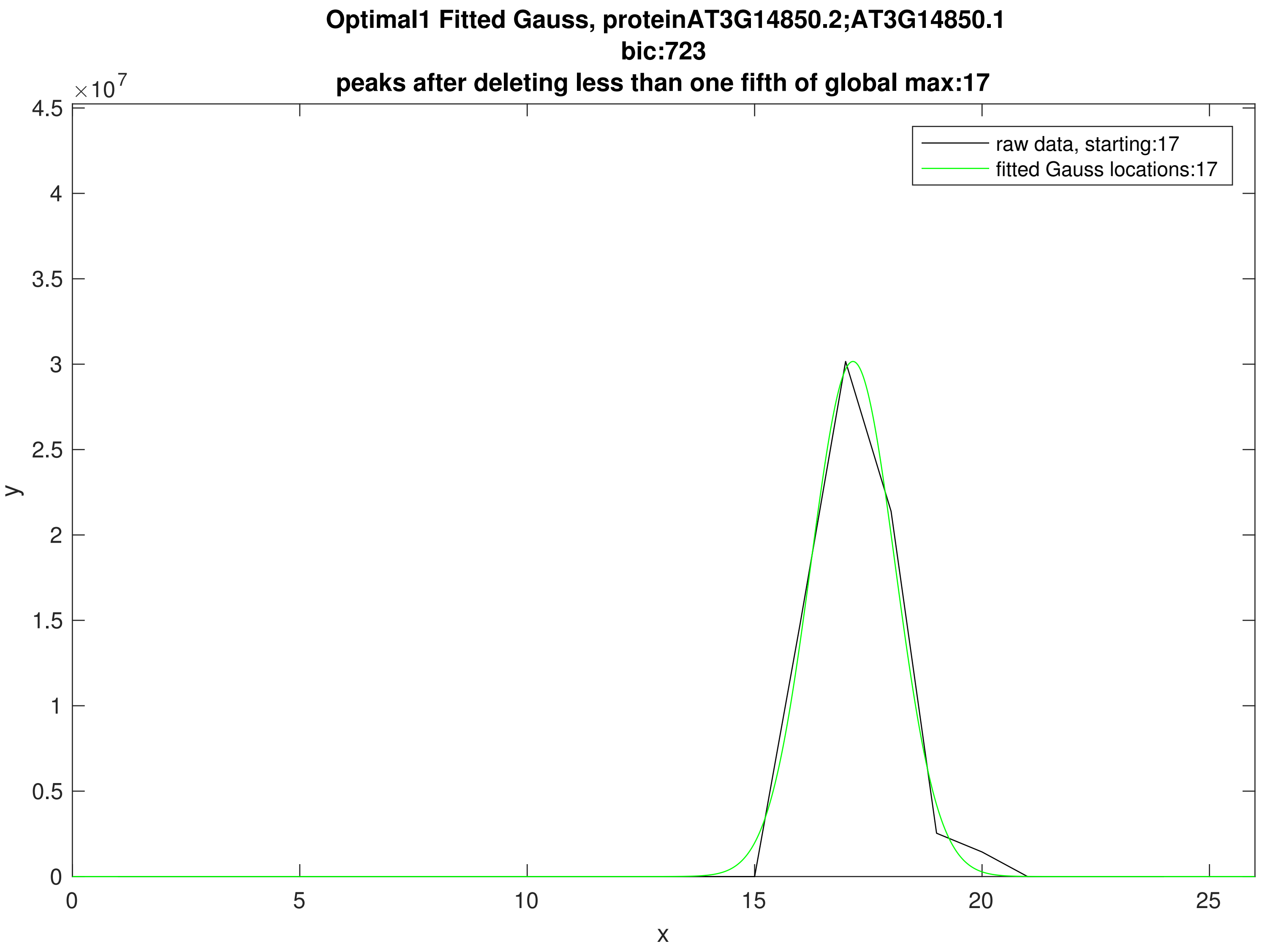
Optimal1 Fitted Gauss, proteinAT3G14850.2;AT3G14850.1
bic:723
peaks after deleting less than one fifth of global max:17
×10^7
4.5
4
3.5
3
2.5
2
1.5
1
0.5
0
y
0
5
10
15
20
25
x
raw data, starting:17
fitted Gauss locations:17

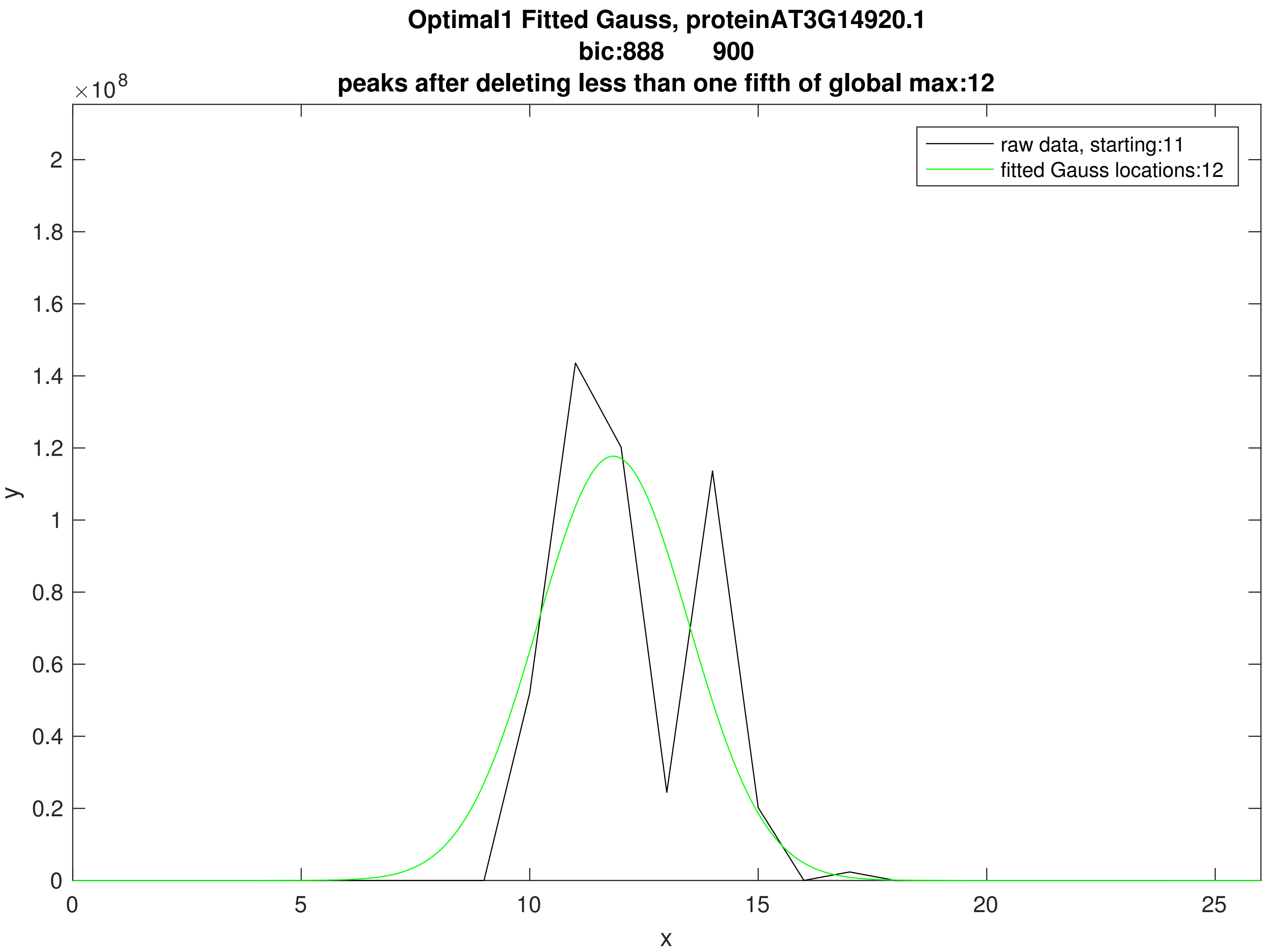
Optimal1 Fitted Gauss, proteinAT3G14920.1
bic:888 900
peaks after deleting less than one fifth of global max:12
×10^8
raw data, starting:11
fitted Gauss locations:12
y
x
0
0.2
0.4
0.6
0.8
1
1.2
1.4
1.6
1.8
2
5
10
15
20
25

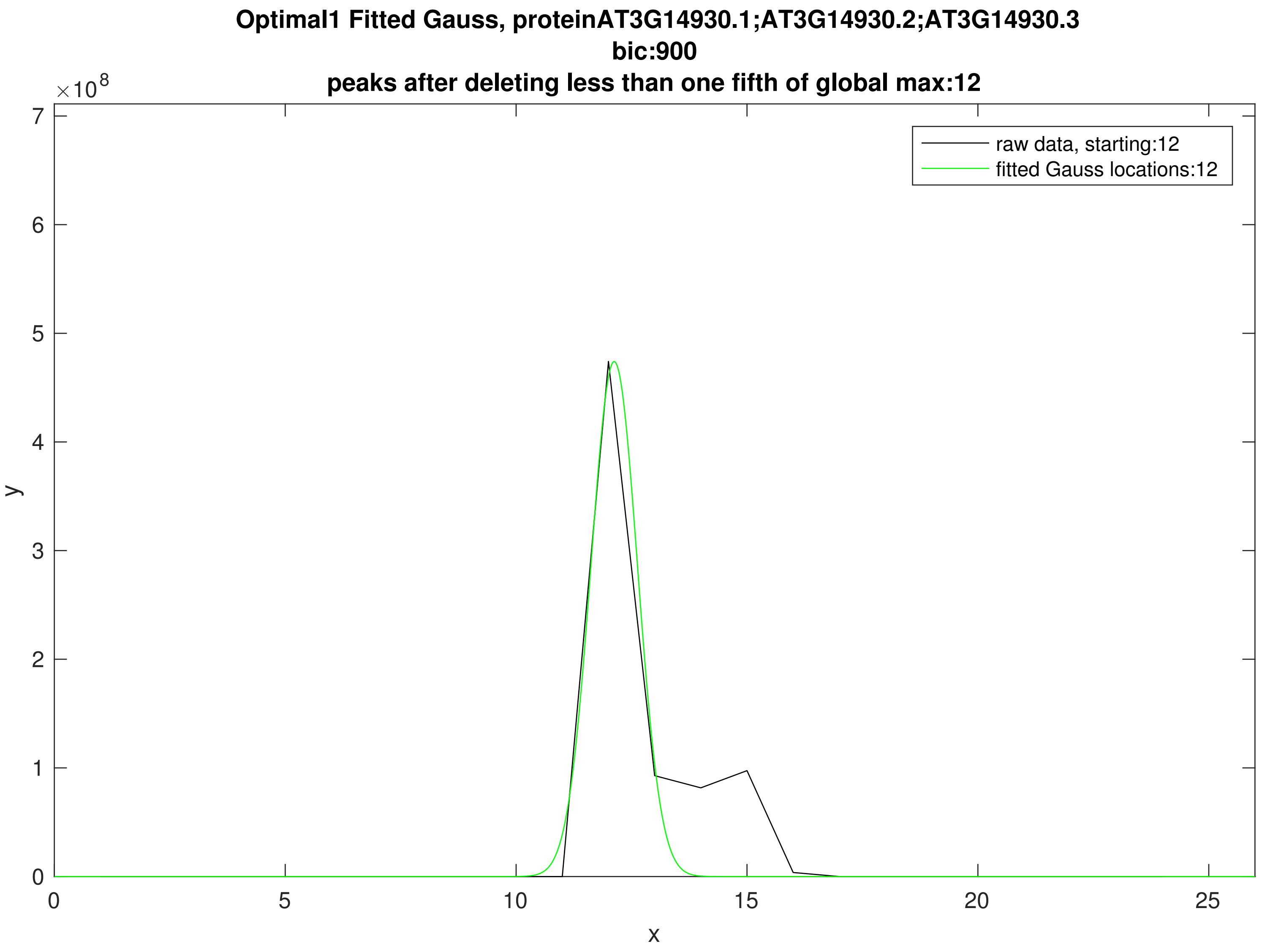

Optimal1 Fitted Gauss, proteinAT3G14930.1;AT3G14930.2;AT3G14930.3
bic:900
peaks after deleting less than one fifth of global max:12
×10^8
7
6
5
4
3
2
1
0
y
0
5
10
15
20
25
x
raw data, starting:12
fitted Gauss locations:12

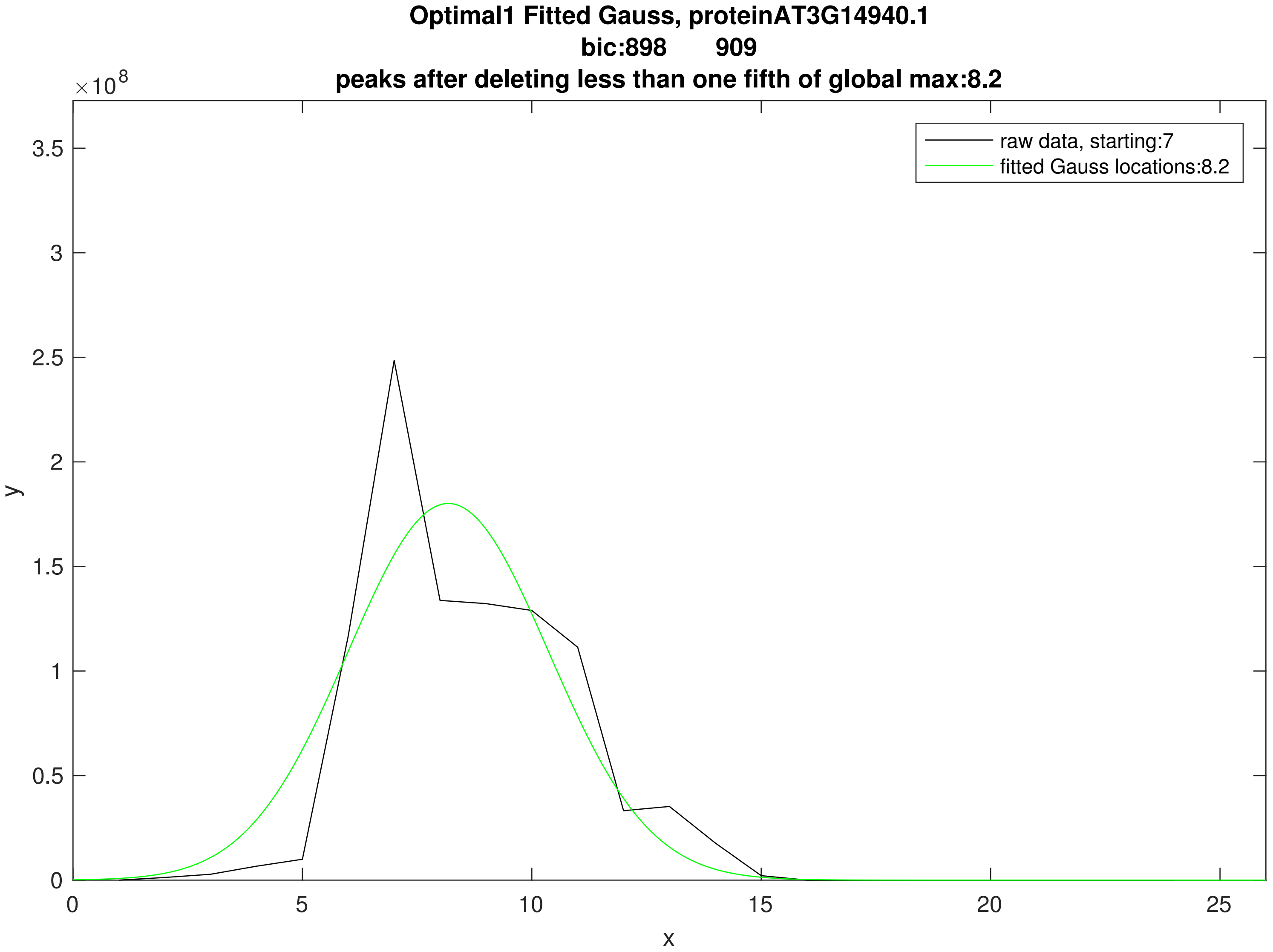
Optimal1 Fitted Gauss, proteinAT3G14940.1
bic:898 909
peaks after deleting less than one fifth of global max:8.2
×10^8
3.5
3
2.5
2
1.5
1
0.5
0
y
0
5
10
15
20
25
x
raw data, starting:7
fitted Gauss locations:8.2

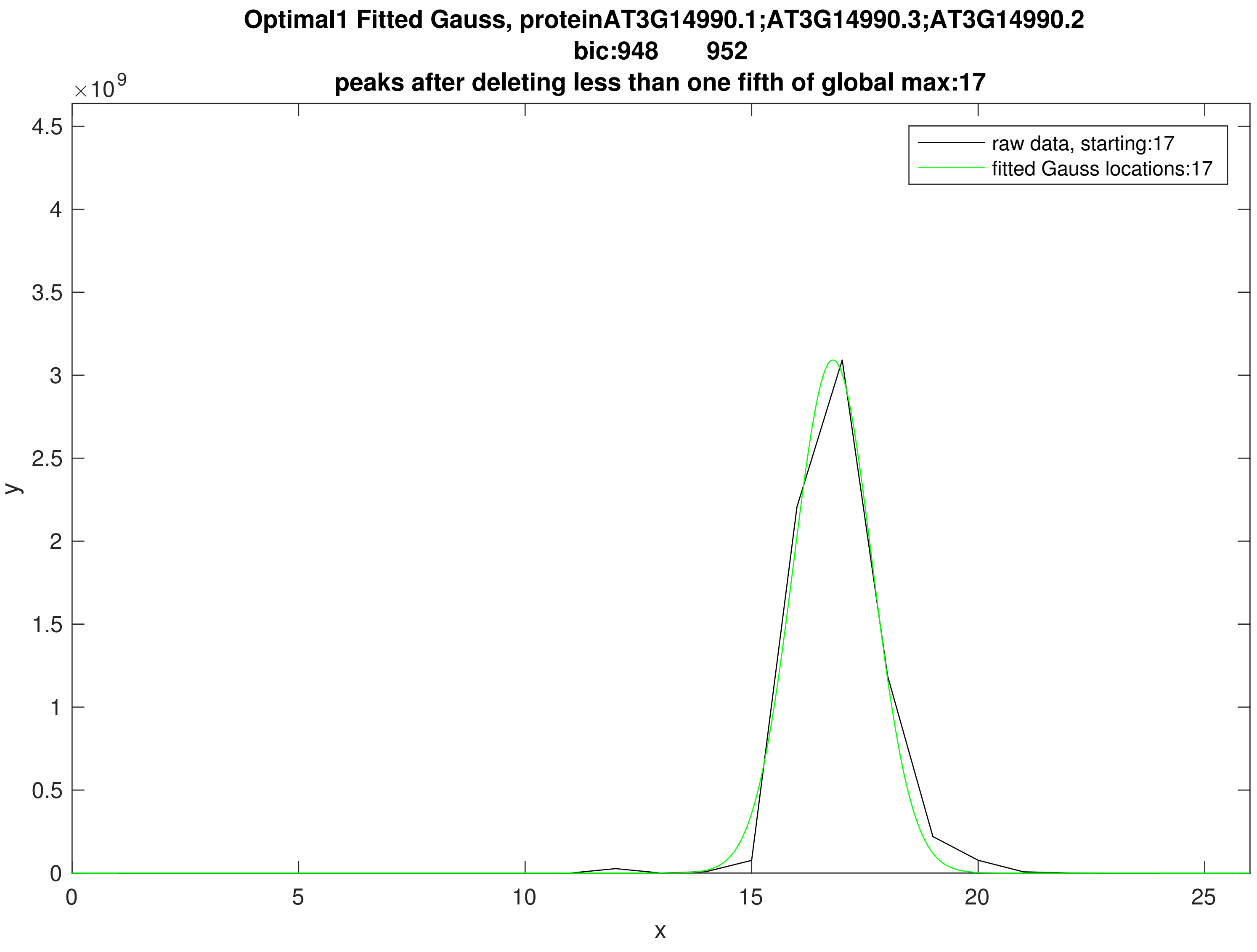

Optimal1 Fitted Gauss, proteinAT3G14990.1;AT3G14990.3;AT3G14990.2
bic:948 952
peaks after deleting less than one fifth of global max:17
×10^9
raw data, starting:17
fitted Gauss locations:17
y
x
0
0.5
1
1.5
2
2.5
3
3.5
4
4.5
5
10
15
20
25

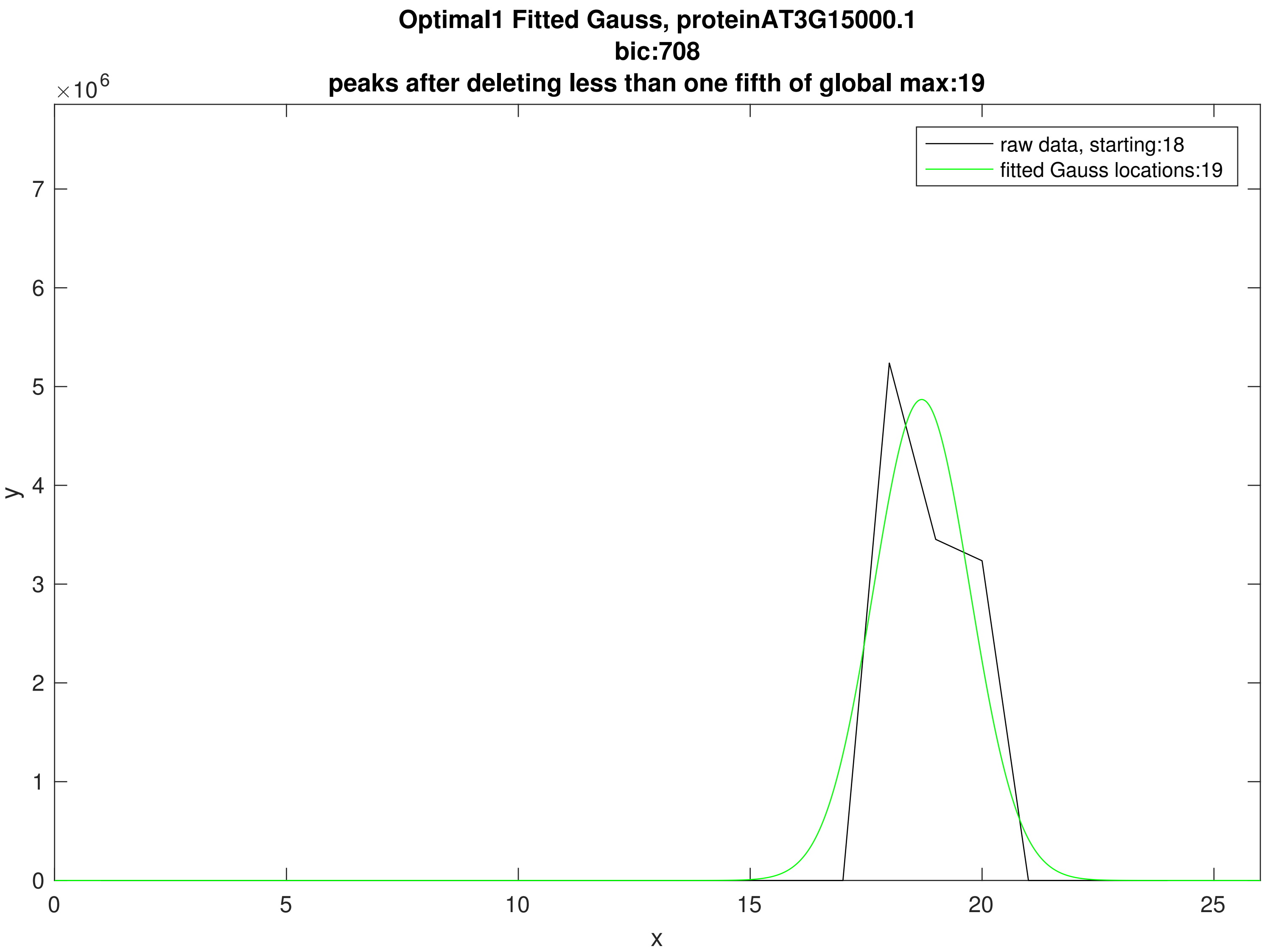
Optimal1 Fitted Gauss, proteinAT3G15000.1
bic:708
peaks after deleting less than one fifth of global max:19
×10^6
raw data, starting:18
fitted Gauss locations:19
y
x
0
1
2
3
4
5
6
7
0
5
10
15
20
25

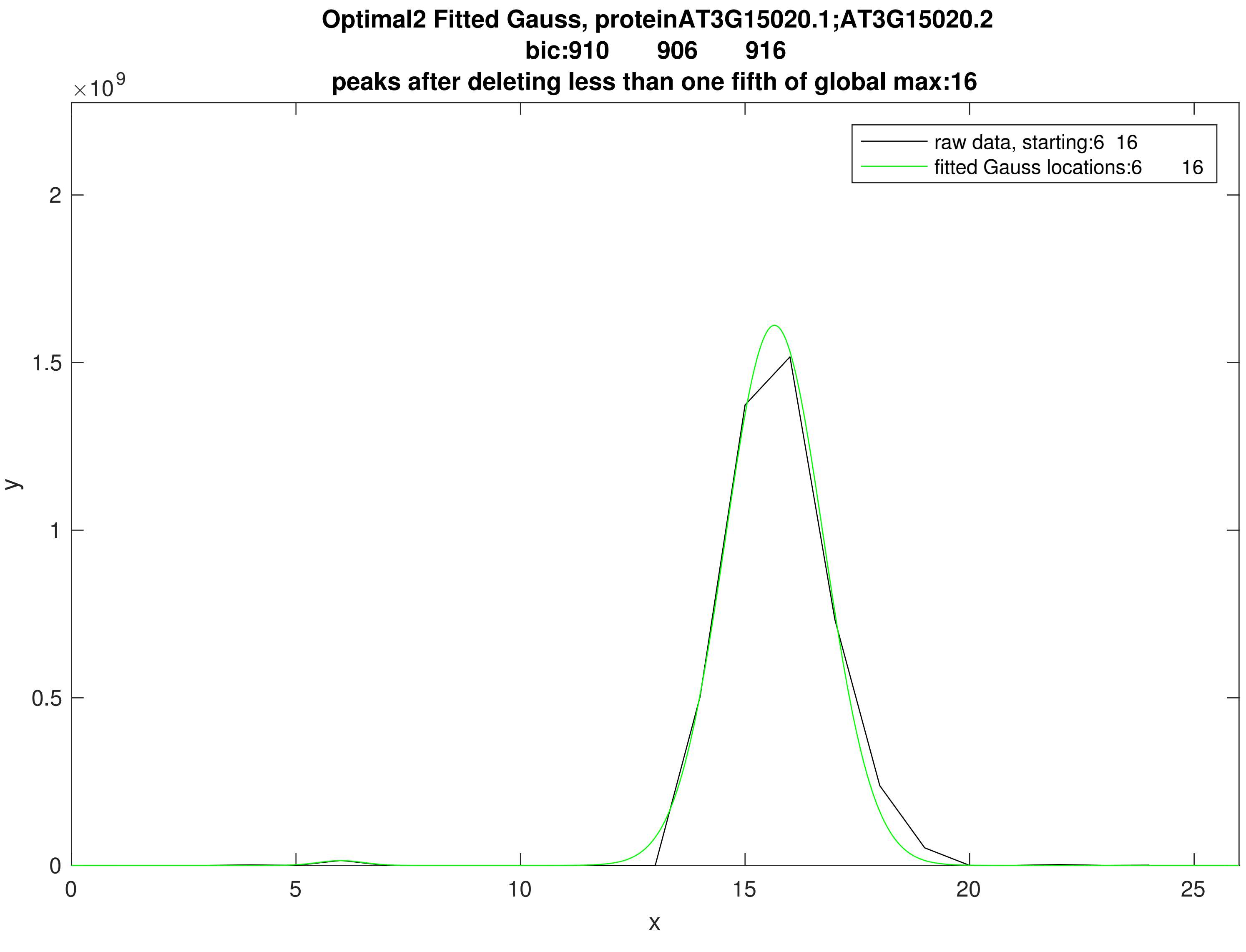

Optimal2 Fitted Gauss, proteinAT3G15020.1;AT3G15020.2
bic:910 906 916
peaks after deleting less than one fifth of global max:16
×10^9
raw data, starting:6 16
fitted Gauss locations:6 16
y
x
0
0.5
1
1.5
2
0
5
10
15
20
25

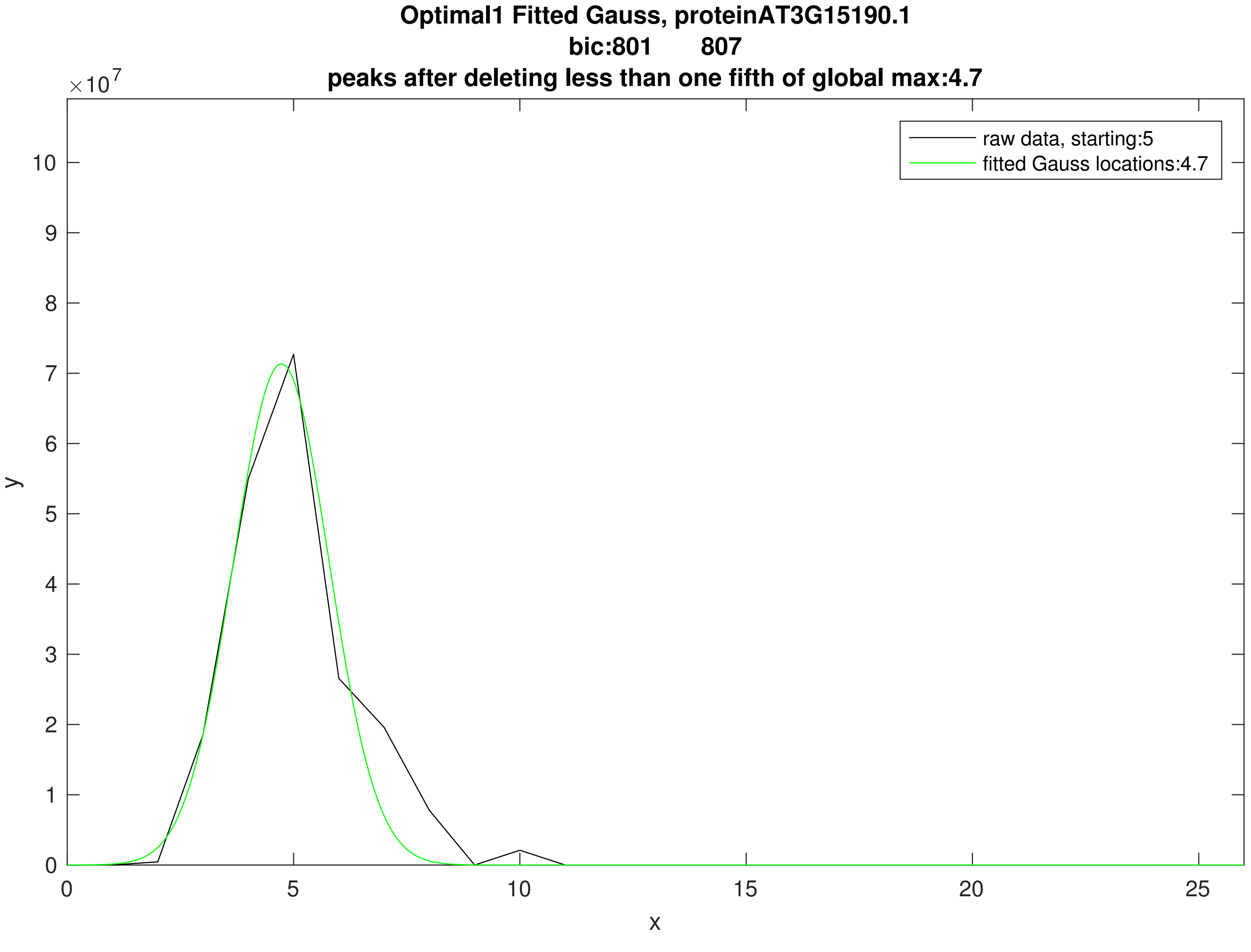
Optimal1 Fitted Gauss, proteinAT3G15190.1
bic:801 807
peaks after deleting less than one fifth of global max:4.7
×10^7
raw data, starting:5
fitted Gauss locations:4.7
y
x
0
1
2
3
4
5
6
7
8
9
10
0
5
10
15
20
25

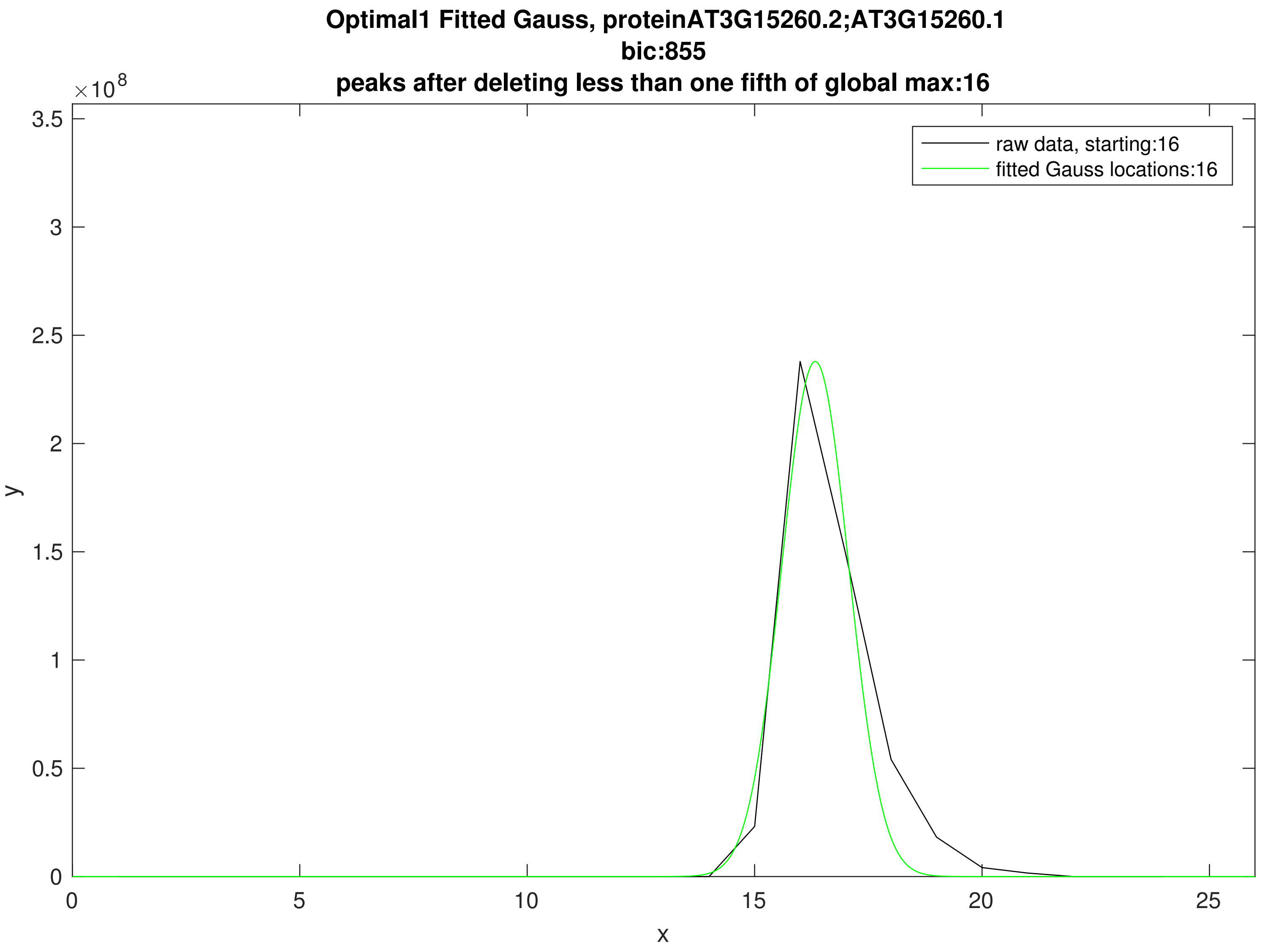
Optimal1 Fitted Gauss, proteinAT3G15260.2;AT3G15260.1
bic:855
peaks after deleting less than one fifth of global max:16
×10^8
3.5
3
2.5
2
1.5
1
0.5
0
y
0
5
10
15
20
25
x
raw data, starting:16
fitted Gauss locations:16

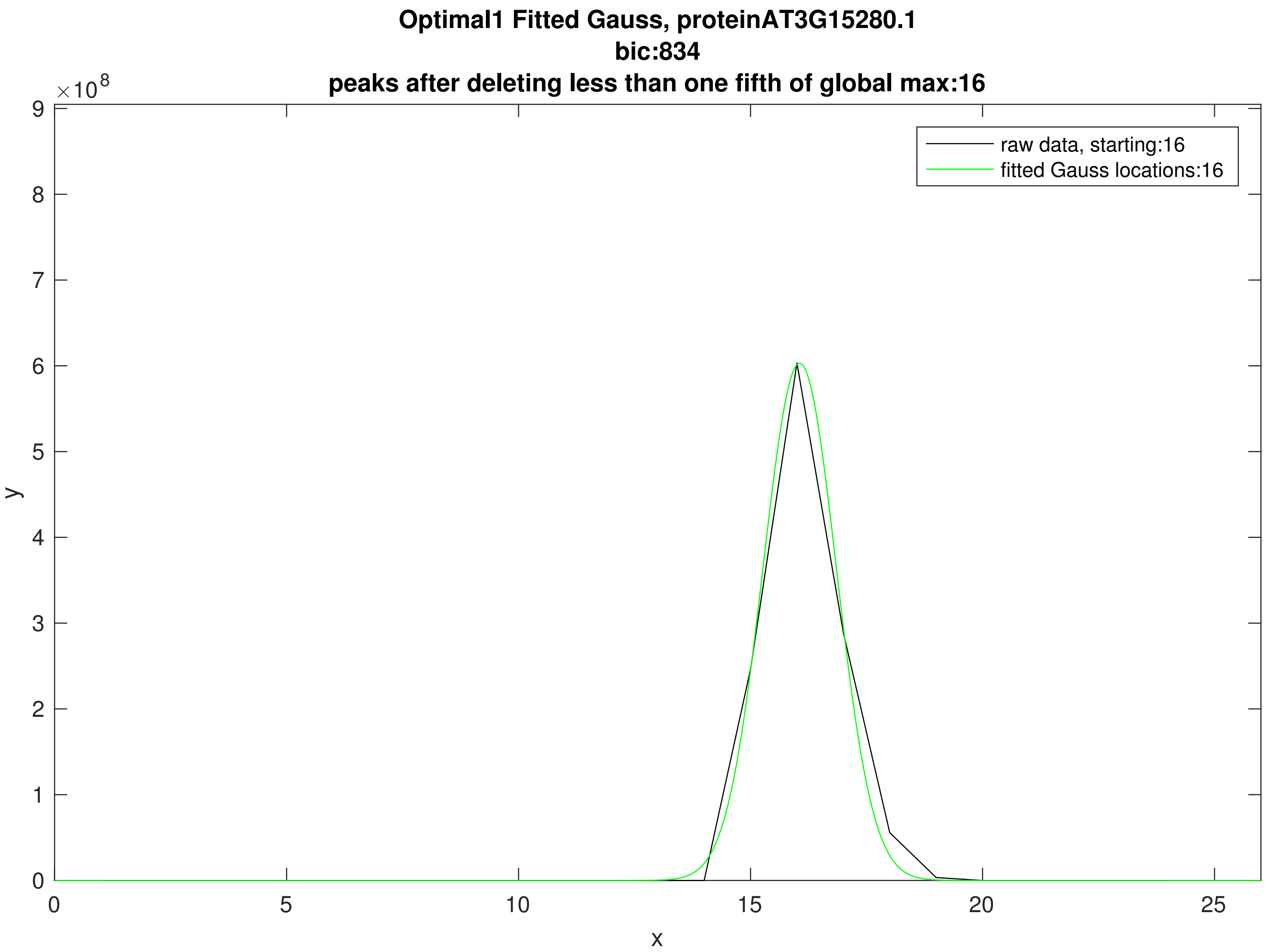
Optimal1 Fitted Gauss, proteinAT3G15280.1
bic:834
peaks after deleting less than one fifth of global max:16
×10^8
raw data, starting:16
fitted Gauss locations:16
y
x
0
1
2
3
4
5
6
7
8
9
0
5
10
15
20
25

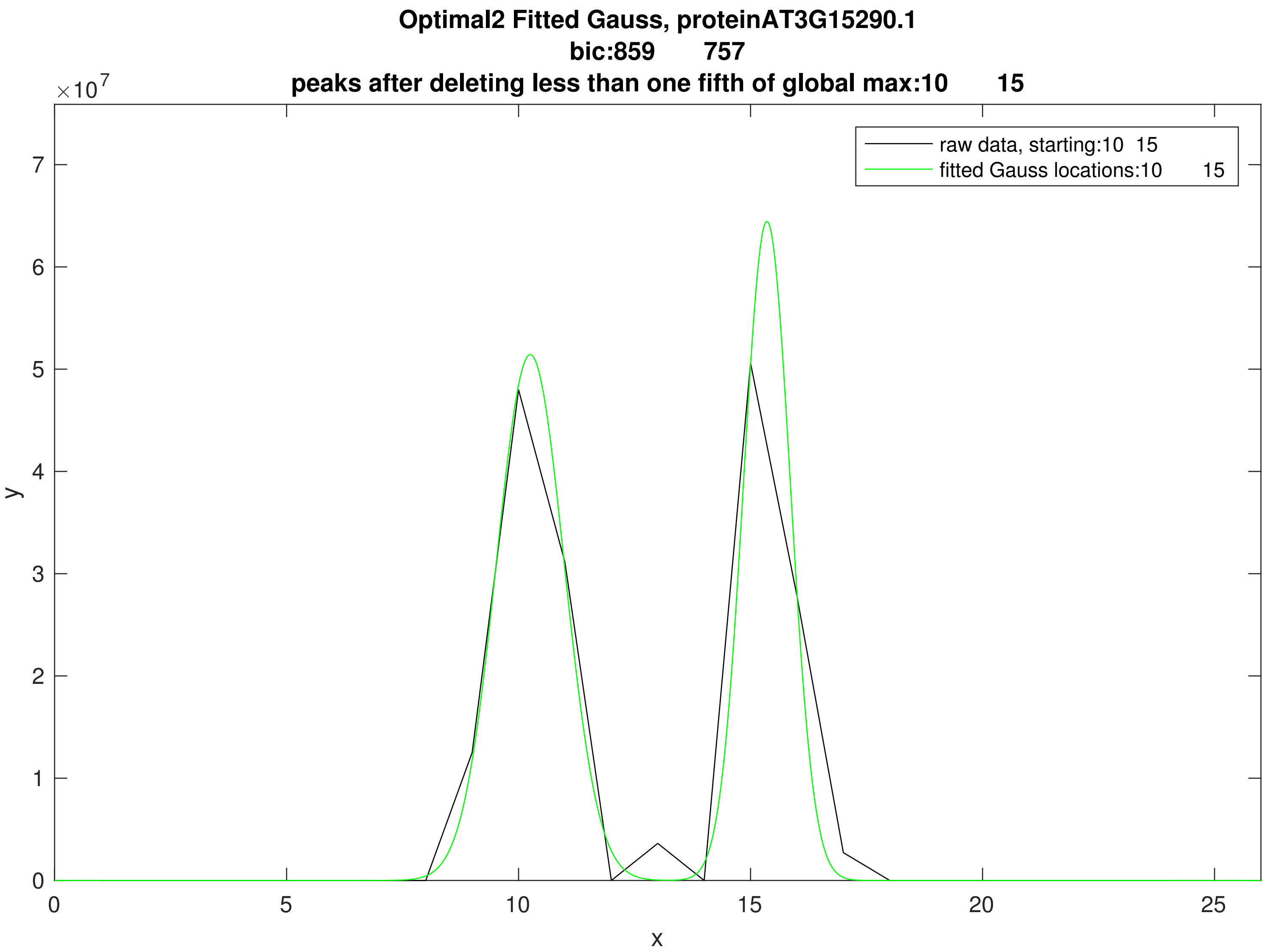
Optimal2 Fitted Gauss, proteinAT3G15290.1
bic:859 757
peaks after deleting less than one fifth of global max:10 15
×10^7
raw data, starting:10 15
fitted Gauss locations:10 15
y
x
0
1
2
3
4
5
6
7
0
5
10
15
20
25

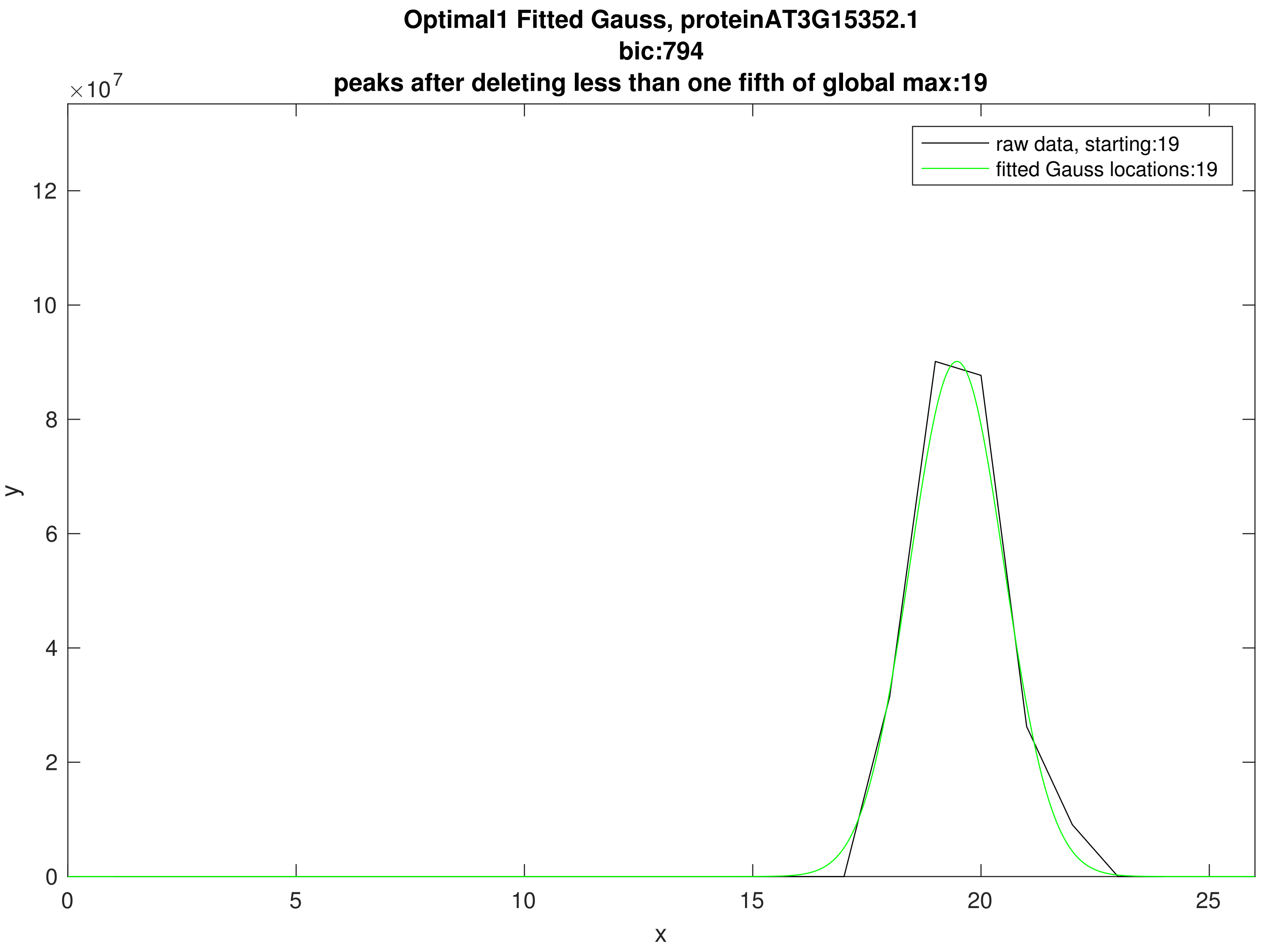

Optimal1 Fitted Gauss, proteinAT3G15352.1
bic:794
peaks after deleting less than one fifth of global max:19
$\times 10^7$
raw data, starting:19
fitted Gauss locations:19
12
10
8
6
4
2
0
y
0
5
10
15
20
25
x

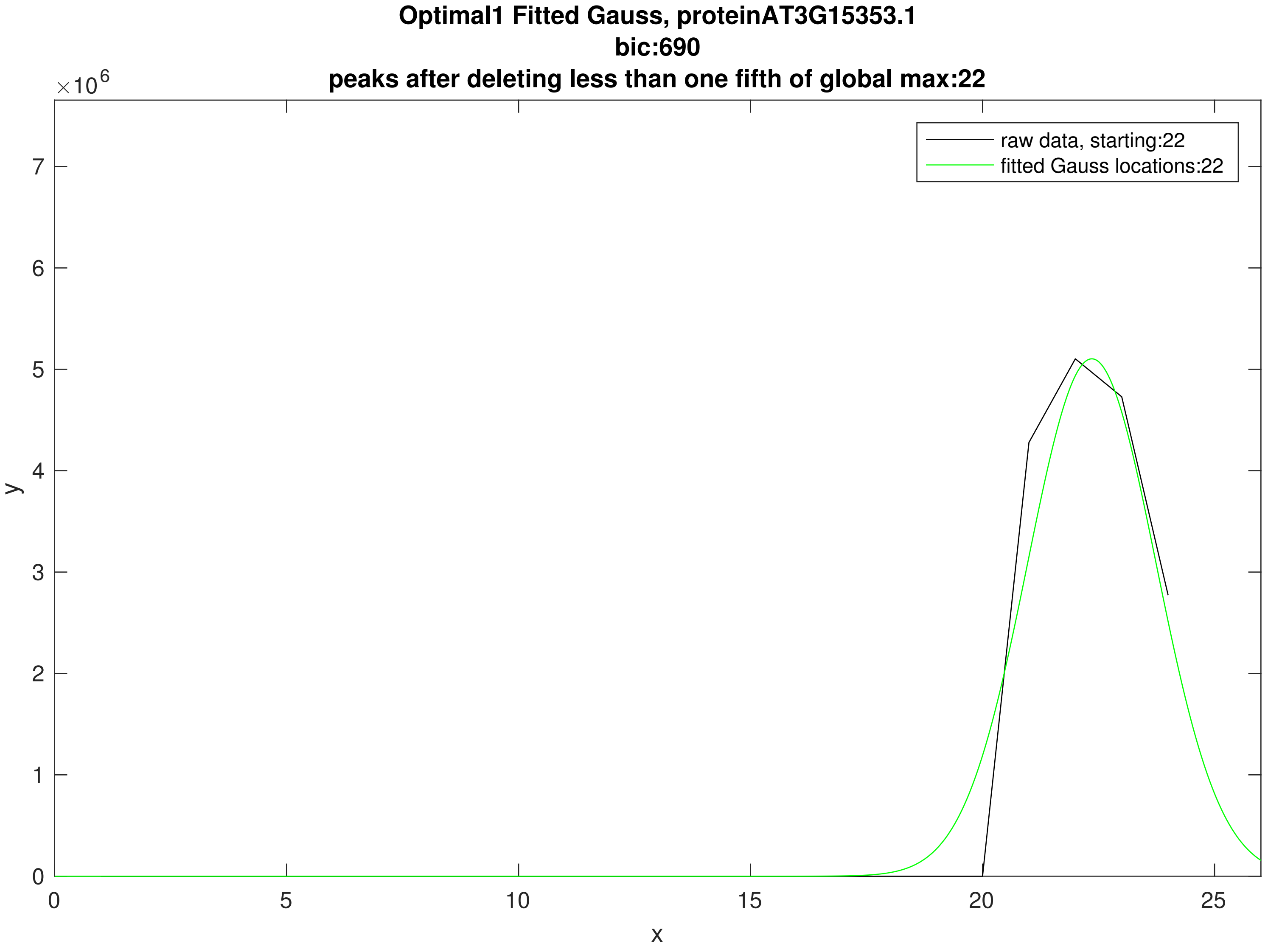

Optimal1 Fitted Gauss, proteinAT3G15353.1
bic:690
peaks after deleting less than one fifth of global max:22
×10^6
raw data, starting:22
fitted Gauss locations:22
y
x
0
1
2
3
4
5
6
7
0
5
10
15
20
25

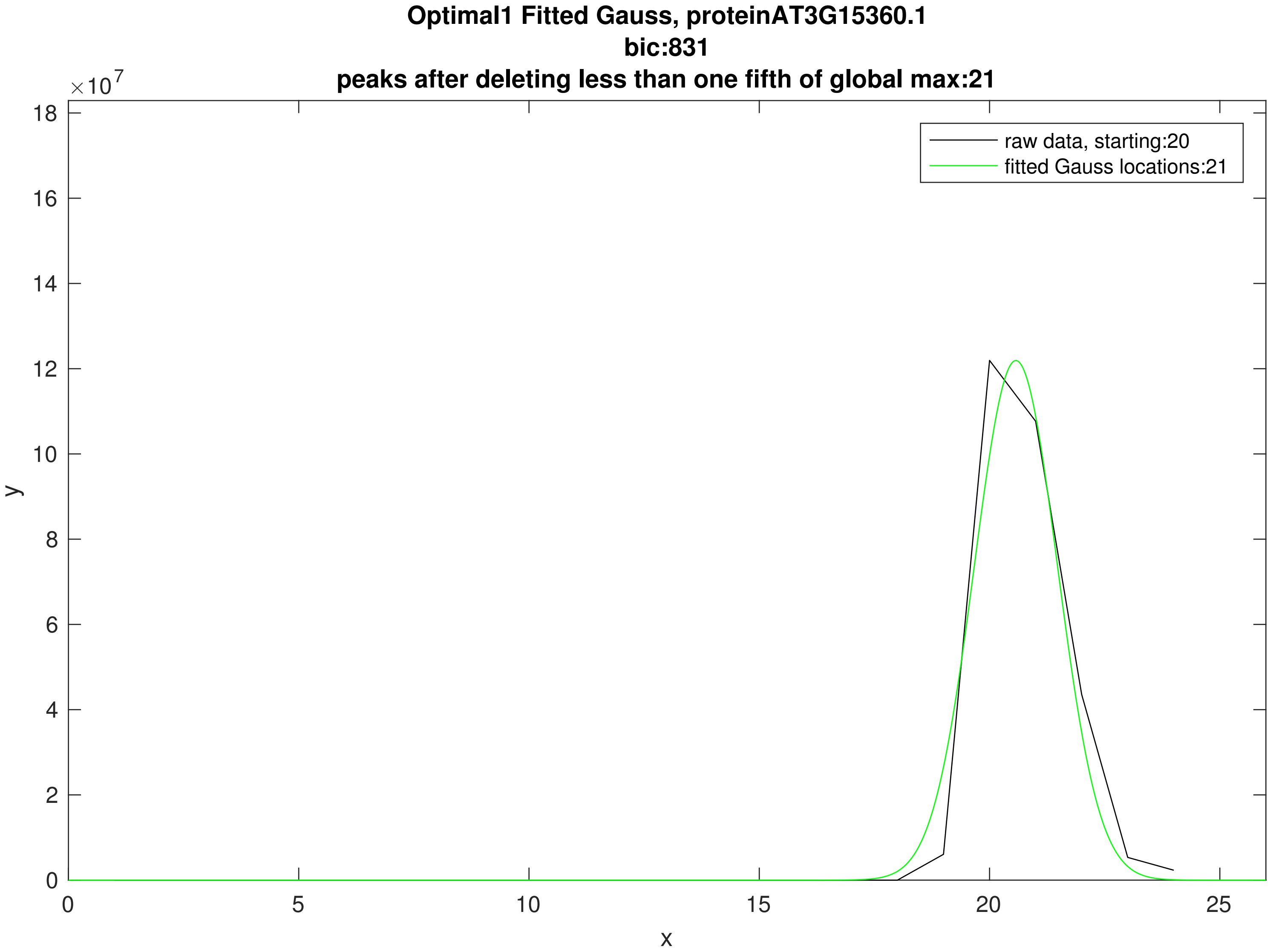
Optimal1 Fitted Gauss, proteinAT3G15360.1
bic:831
peaks after deleting less than one fifth of global max:21
×10^7
18
16
14
12
10
8
6
4
2
0
y
0
5
10
15
20
25
x
raw data, starting:20
fitted Gauss locations:21

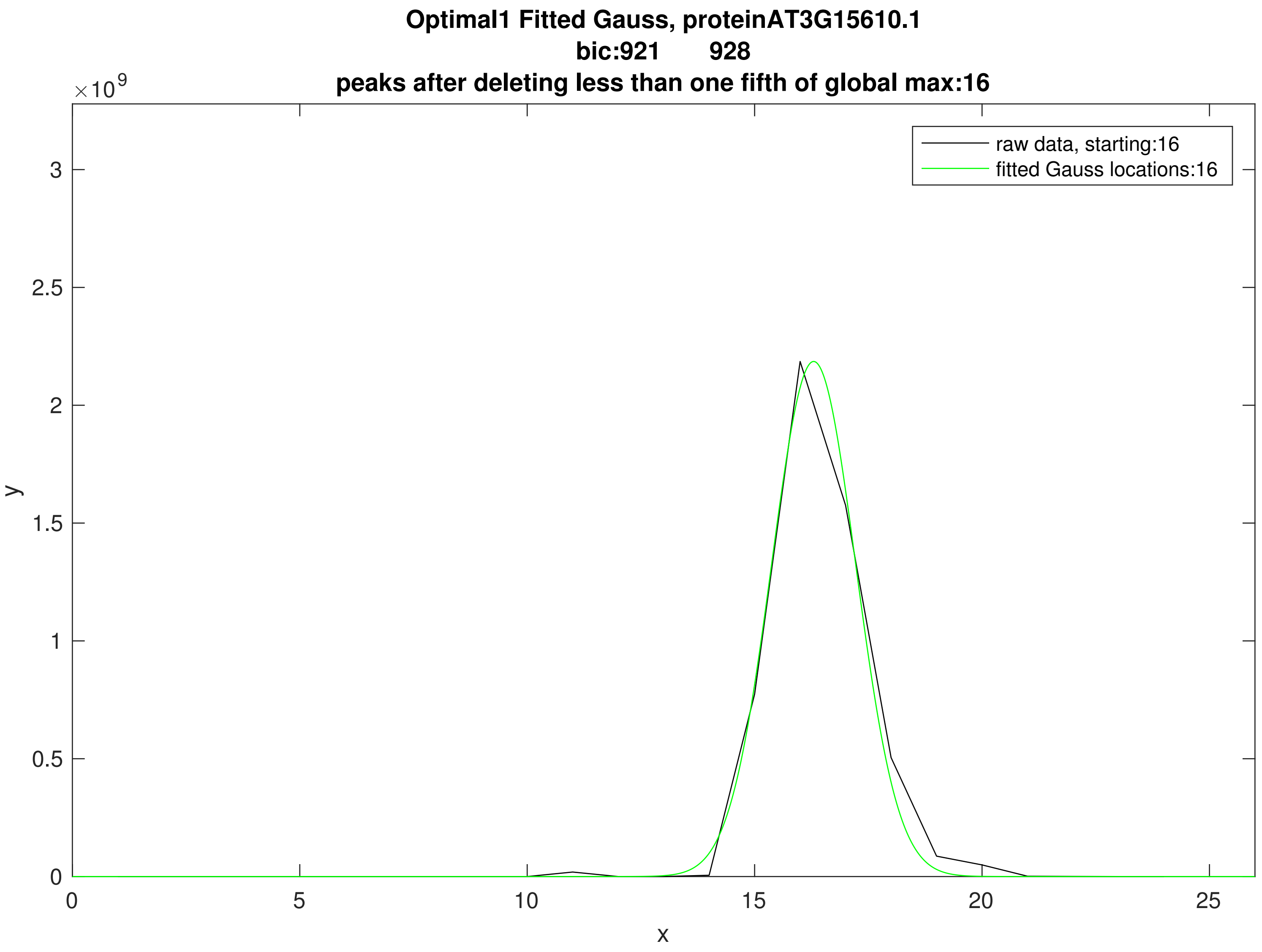
Optimal1 Fitted Gauss, proteinAT3G15610.1
bic:921 928
peaks after deleting less than one fifth of global max:16
×10^9
raw data, starting:16
fitted Gauss locations:16
y
x
0
0.5
1
1.5
2
2.5
3
0
5
10
15
20
25

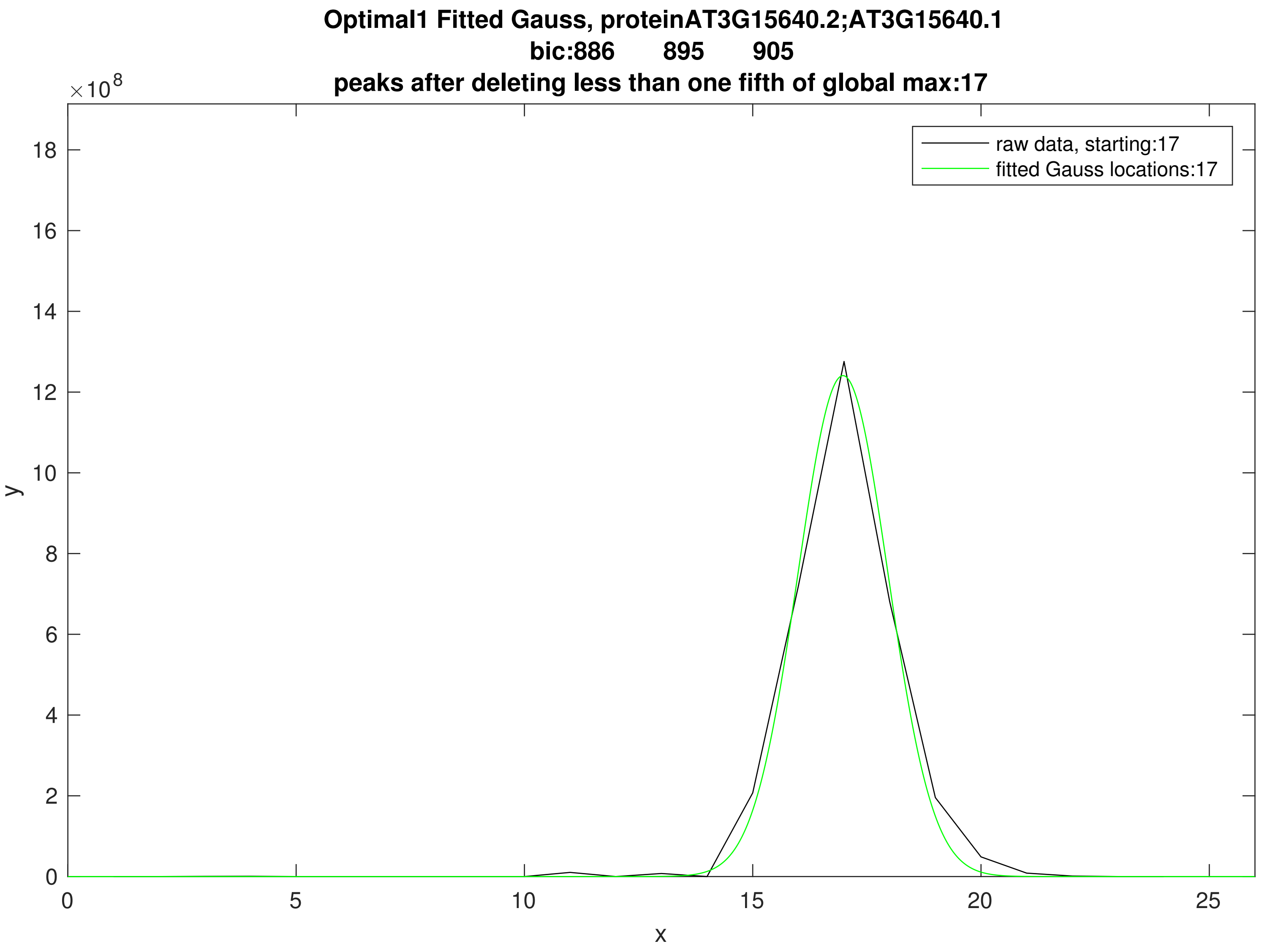
Optimal1 Fitted Gauss, proteinAT3G15640.2;AT3G15640.1
bic:886 895 905
peaks after deleting less than one fifth of global max:17
×10^8
raw data, starting:17
fitted Gauss locations:17
y
x
0
2
4
6
8
10
12
14
16
18
0
5
10
15
20
25

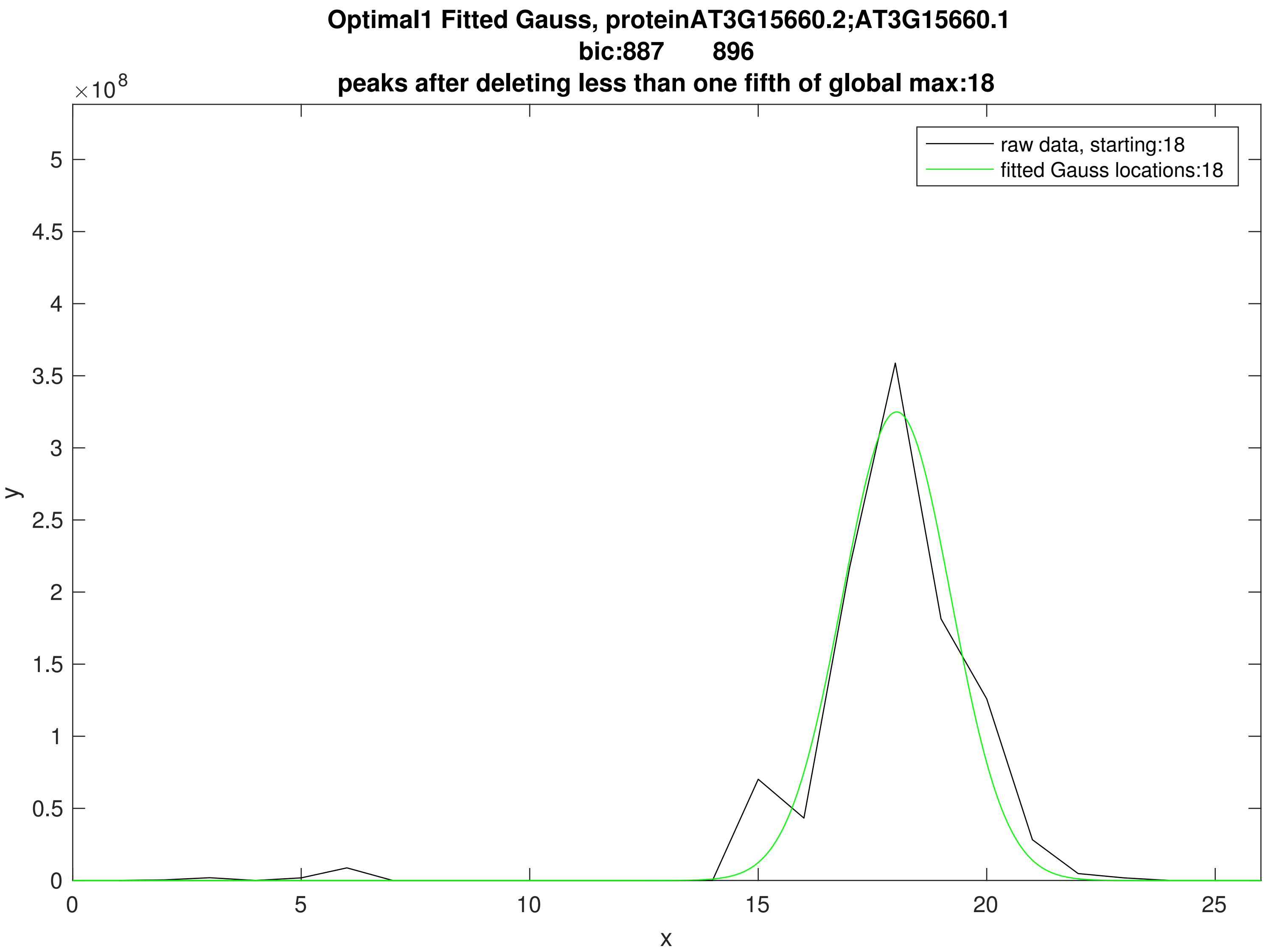

Optimal1 Fitted Gauss, proteinAT3G15660.2;AT3G15660.1
bic:887 896
peaks after deleting less than one fifth of global max:18
×10^8
5
4.5
4
3.5
3
2.5
2
1.5
1
0.5
0
y
0
5
10
15
20
25
x
raw data, starting:18
fitted Gauss locations:18

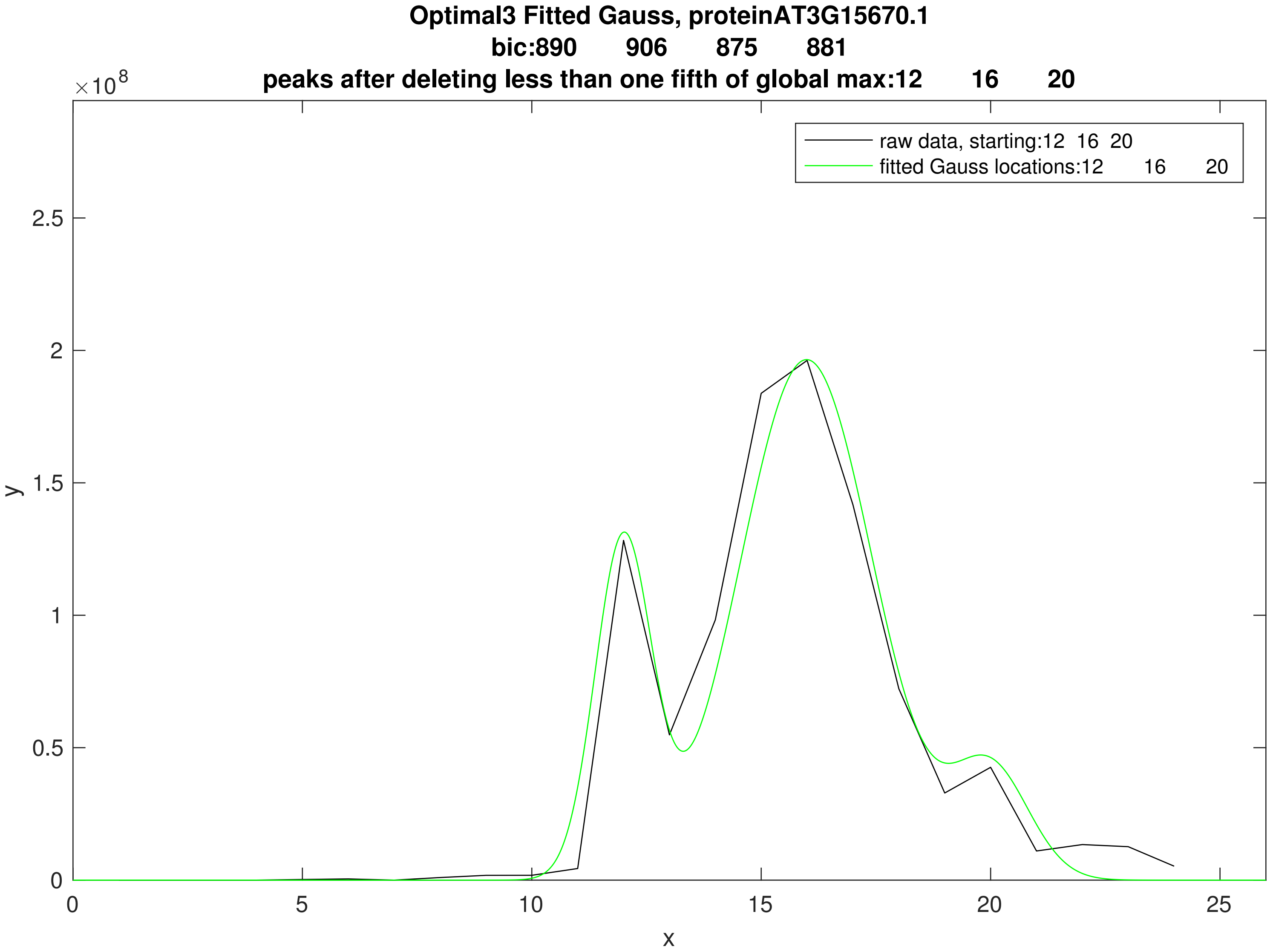

Optimal3 Fitted Gauss, proteinAT3G15670.1
bic:890 906 875 881
peaks after deleting less than one fifth of global max:12 16 20
×10^8
raw data, starting:12 16 20
fitted Gauss locations:12 16 20
2.5
2
1.5
1
0.5
0
y
0
5
10
15
20
25
x

**Optimal1 Fitted Gauss, proteinAT3G15730.1**
**bic:1.07e+03  1.07e+03  1.09e+03**
**peaks after deleting less than one fifth of global max:9.9**

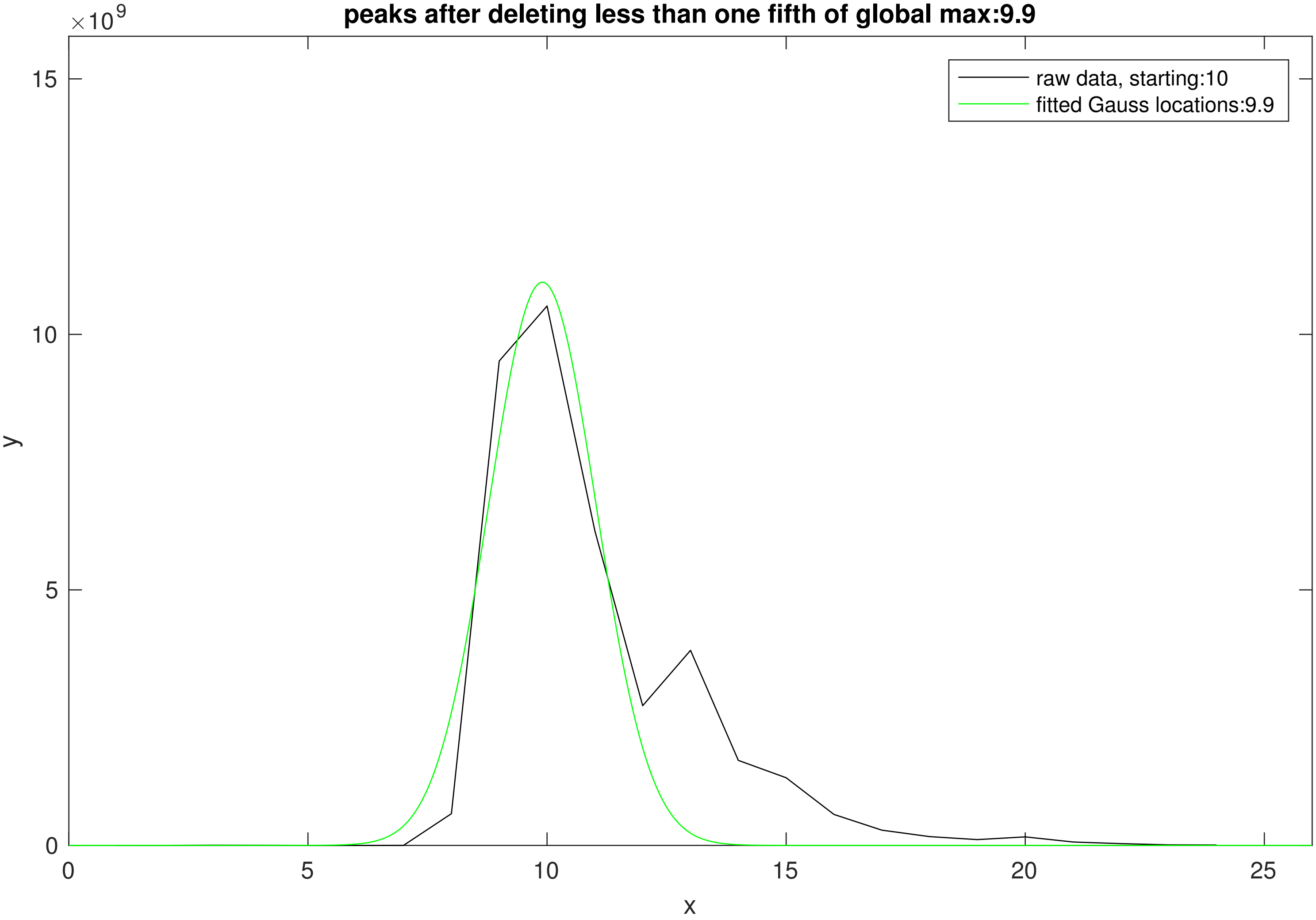

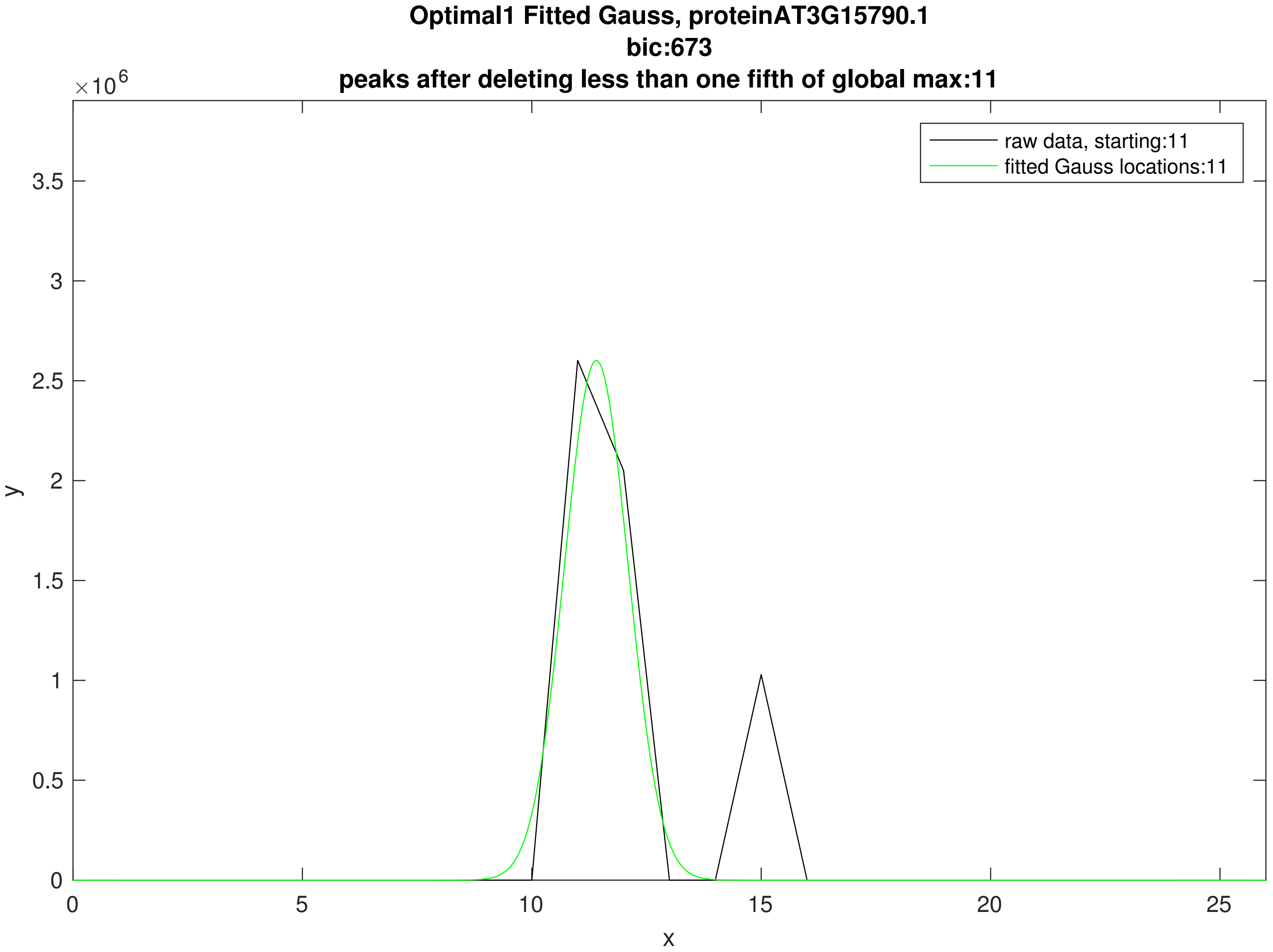
Optimal1 Fitted Gauss, proteinAT3G15790.1
bic:673
peaks after deleting less than one fifth of global max:11
×10^6
3.5
3
2.5
2
1.5
1
0.5
0
y
0
5
10
15
20
25
x
raw data, starting:11
fitted Gauss locations:11

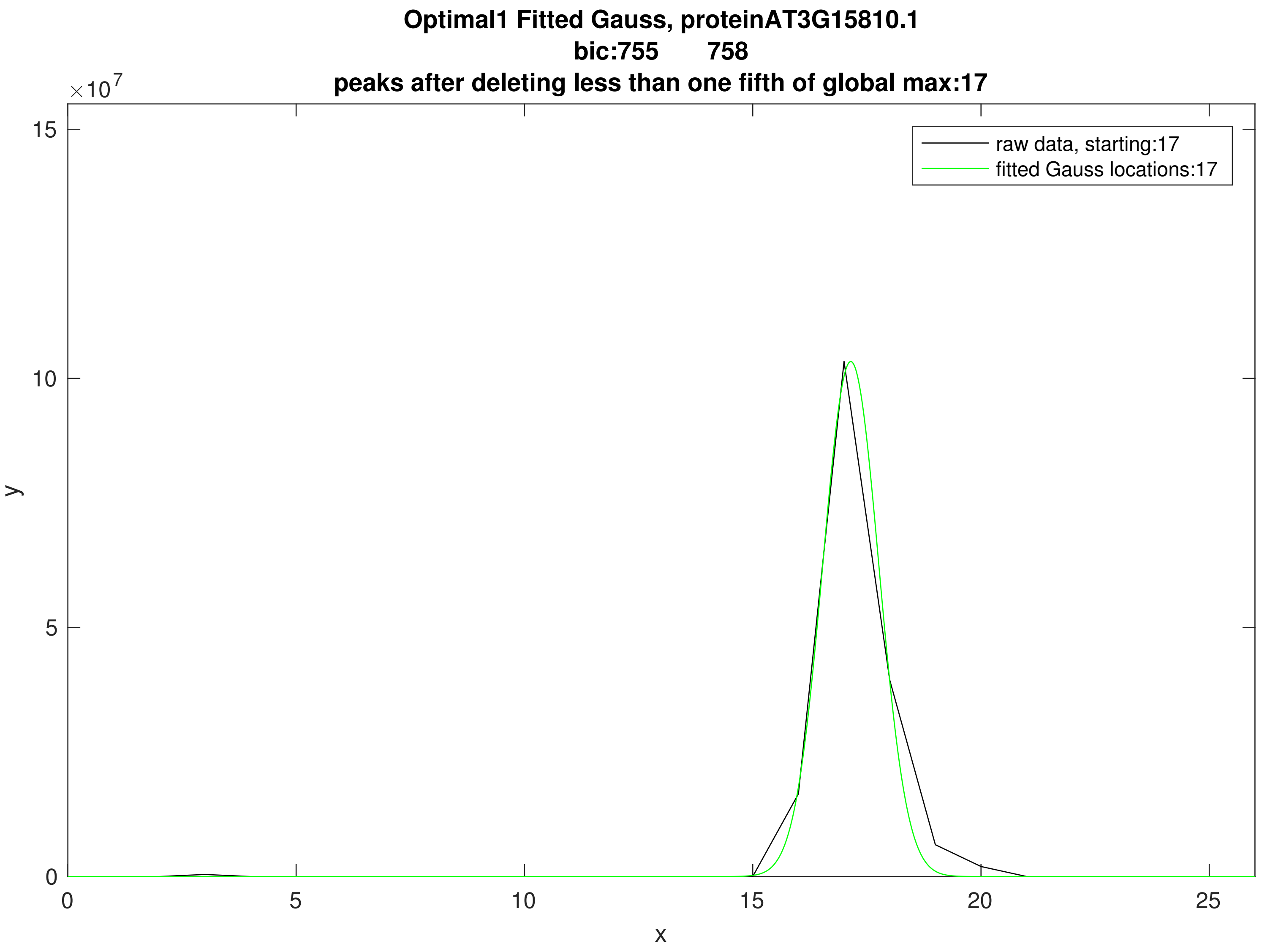

Optimal1 Fitted Gauss, proteinAT3G15810.1
bic:755 758
peaks after deleting less than one fifth of global max:17
×10^7
15
10
5
0
y
0
5
10
15
20
25
x
raw data, starting:17
fitted Gauss locations:17

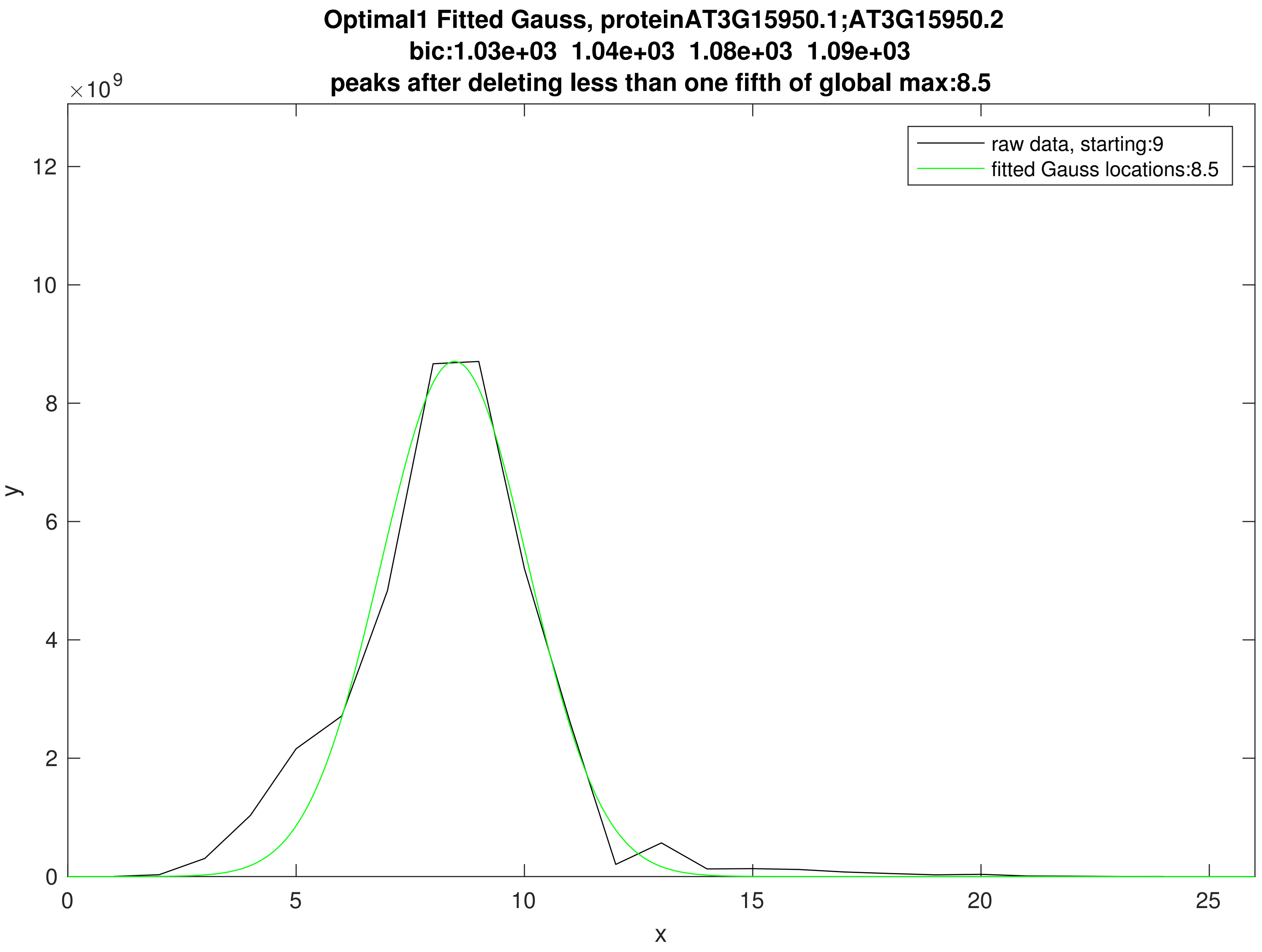
Optimal1 Fitted Gauss, proteinAT3G15950.1;AT3G15950.2
bic:1.03e+03  1.04e+03  1.08e+03  1.09e+03
peaks after deleting less than one fifth of global max:8.5
×10^9
raw data, starting:9
fitted Gauss locations:8.5
y
x
0
2
4
6
8
10
12
5
15
20
25

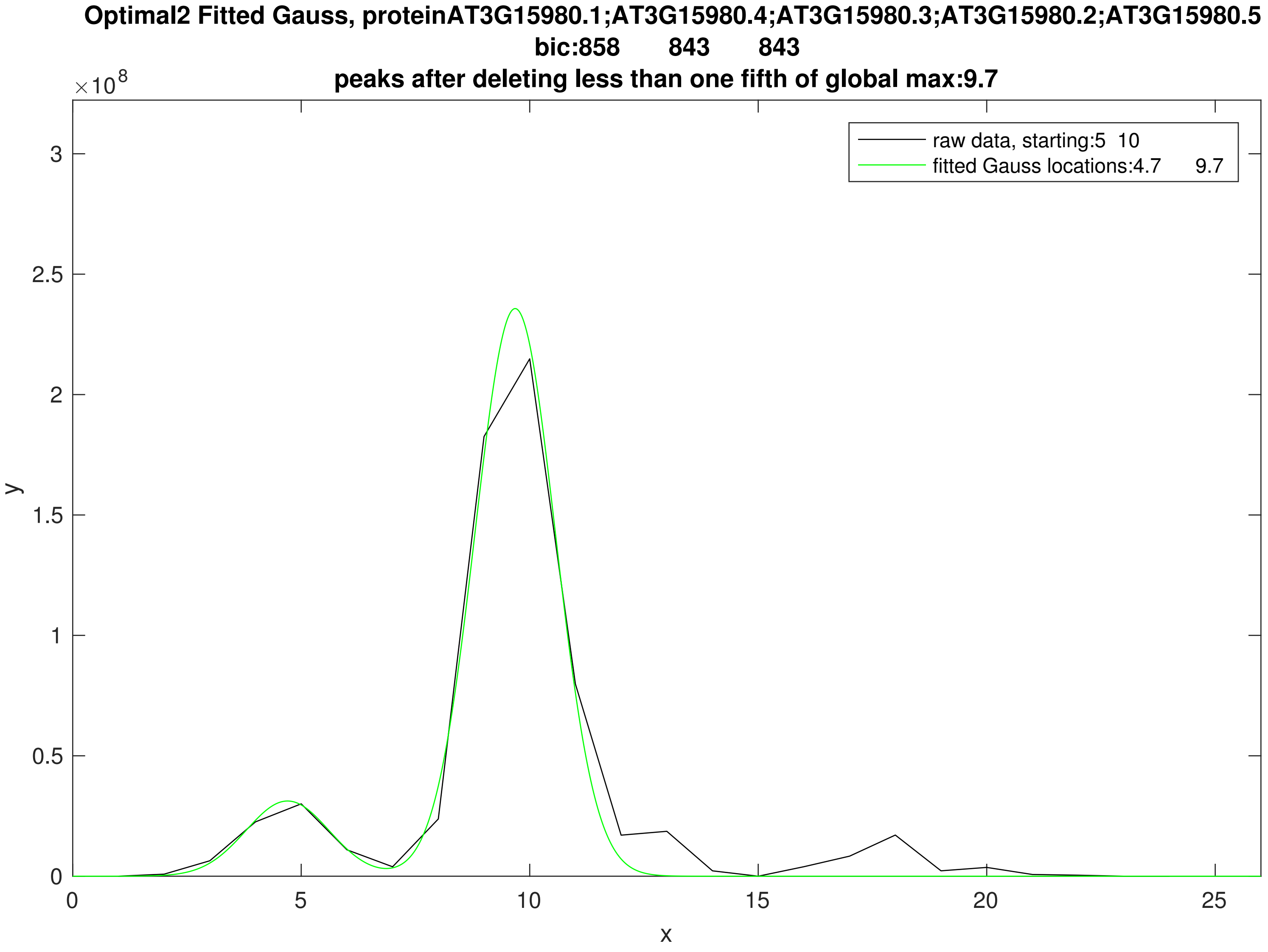

Optimal2 Fitted Gauss, proteinAT3G15980.1;AT3G15980.4;AT3G15980.3;AT3G15980.2;AT3G15980.5
bic:858 843 843
peaks after deleting less than one fifth of global max:9.7
×10^8
raw data, starting:5 10
fitted Gauss locations:4.7 9.7
y
x
0
0.5
1
1.5
2
2.5
3
5
10
15
20
25

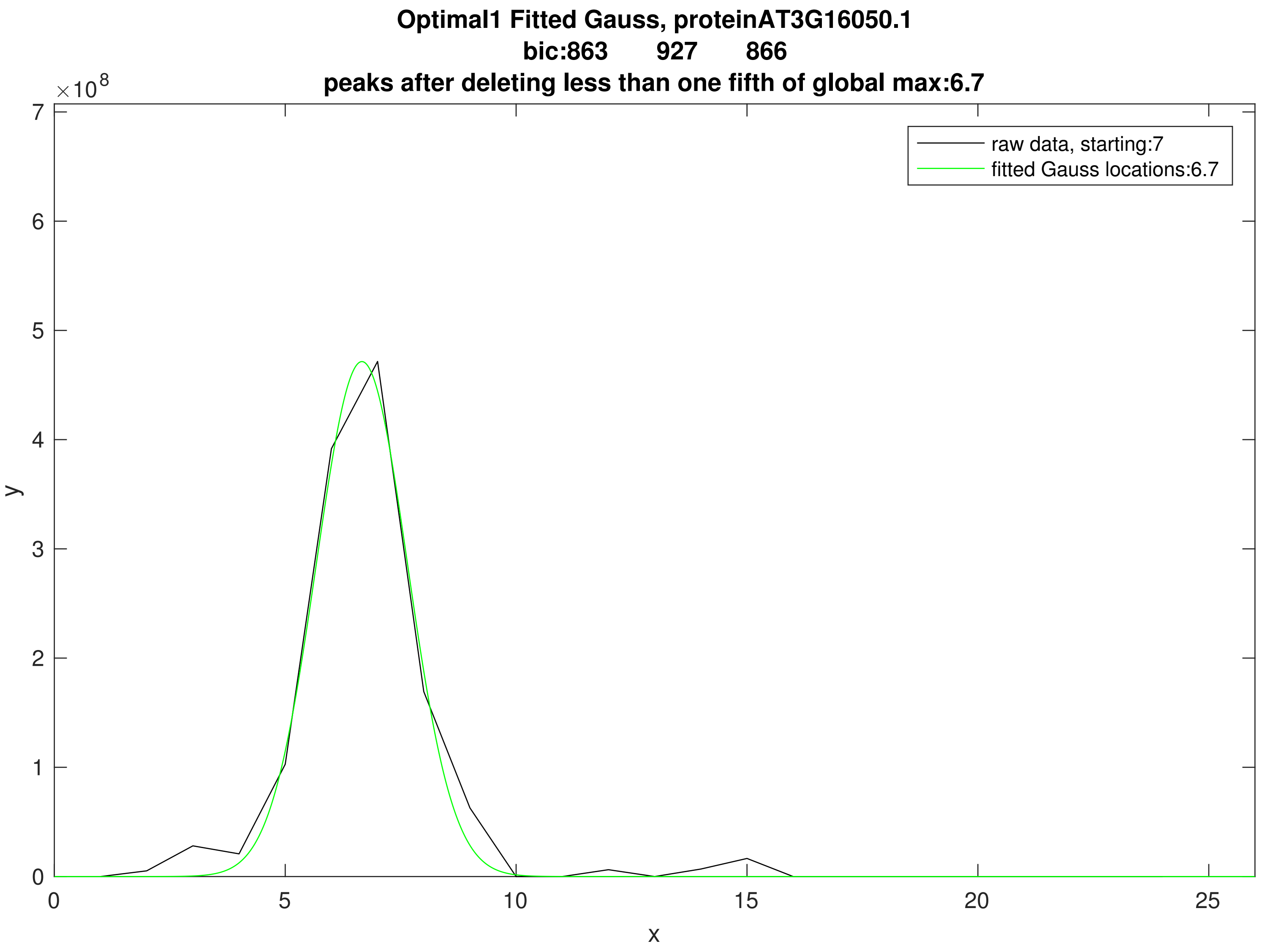
Optimal1 Fitted Gauss, proteinAT3G16050.1
bic:863 927 866
peaks after deleting less than one fifth of global max:6.7
×10^8
y
x
raw data, starting:7
fitted Gauss locations:6.7
0
1
2
3
4
5
6
7
0
5
10
15
20
25

Optimal1 Fitted Gauss, proteinAT3G16110.1;AT1G52260.1
bic:757
peaks after deleting less than one fifth of global max:13

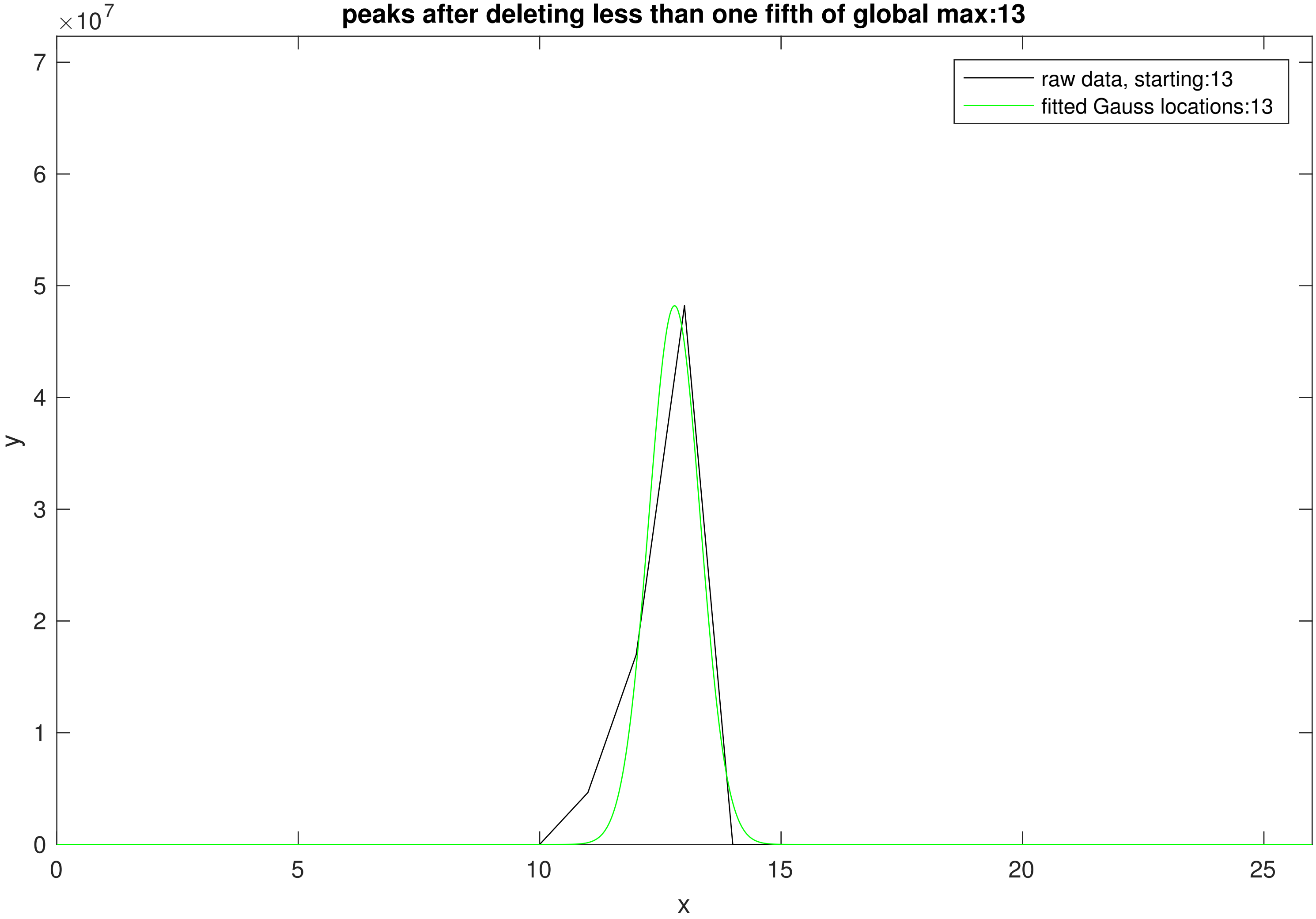

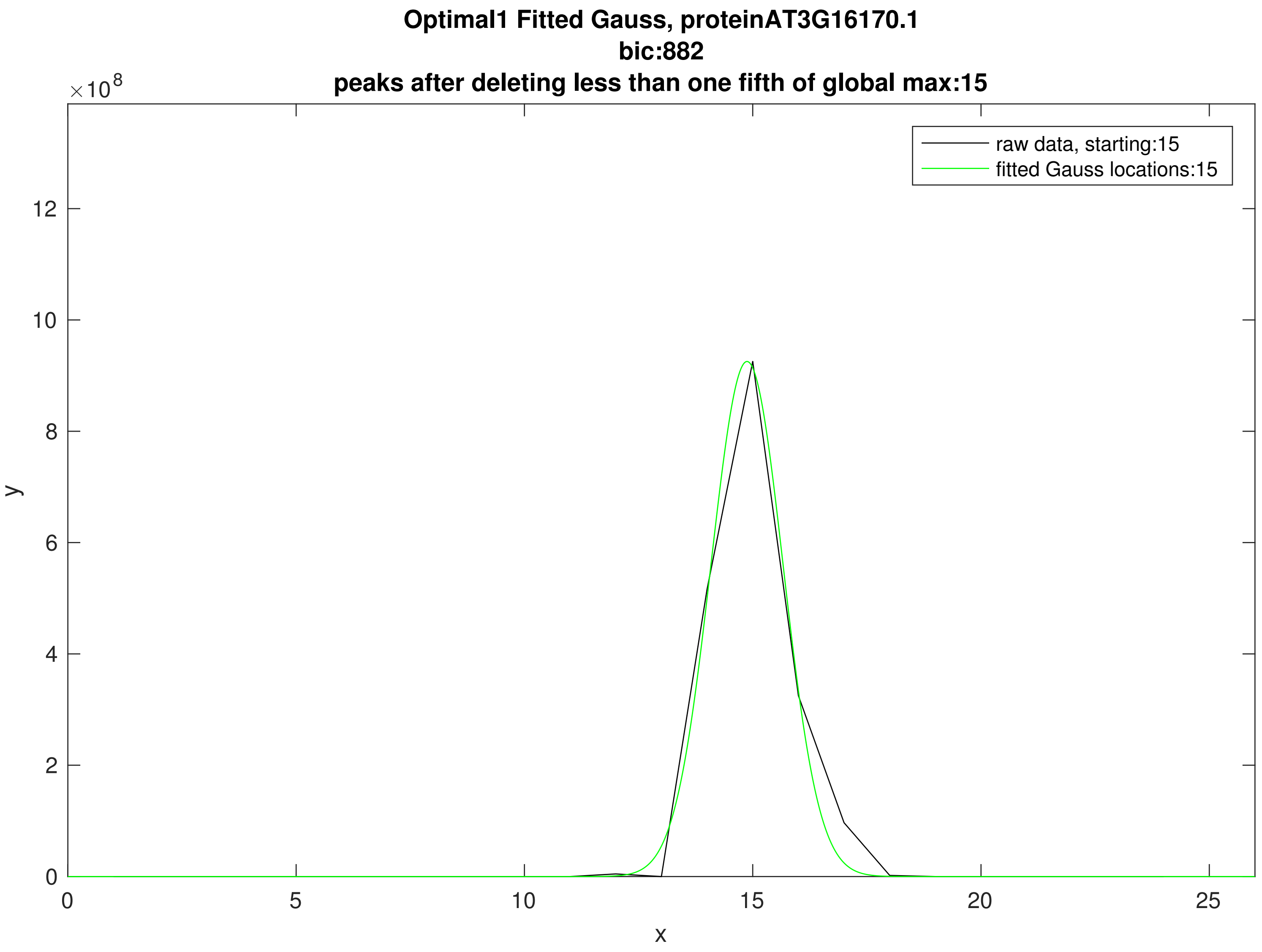

Optimal1 Fitted Gauss, proteinAT3G16170.1
bic:882
peaks after deleting less than one fifth of global max:15
×10^8
raw data, starting:15
fitted Gauss locations:15
12
10
8
6
4
2
0
y
0
5
10
15
20
25
x

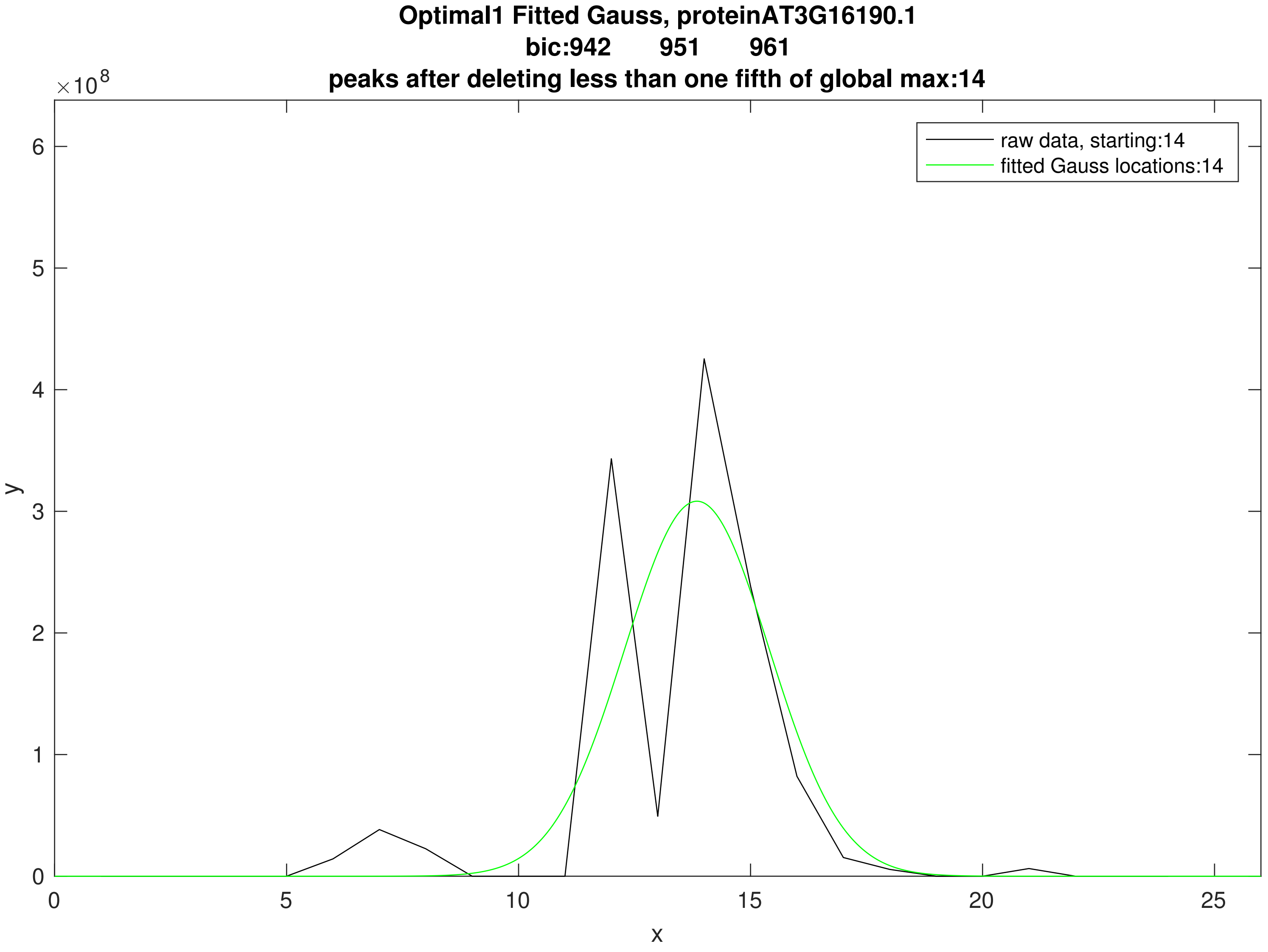

Optimal1 Fitted Gauss, proteinAT3G16190.1
bic:942 951 961
peaks after deleting less than one fifth of global max:14
×10^8
raw data, starting:14
fitted Gauss locations:14
y
x
0
1
2
3
4
5
6
0
5
10
15
20
25

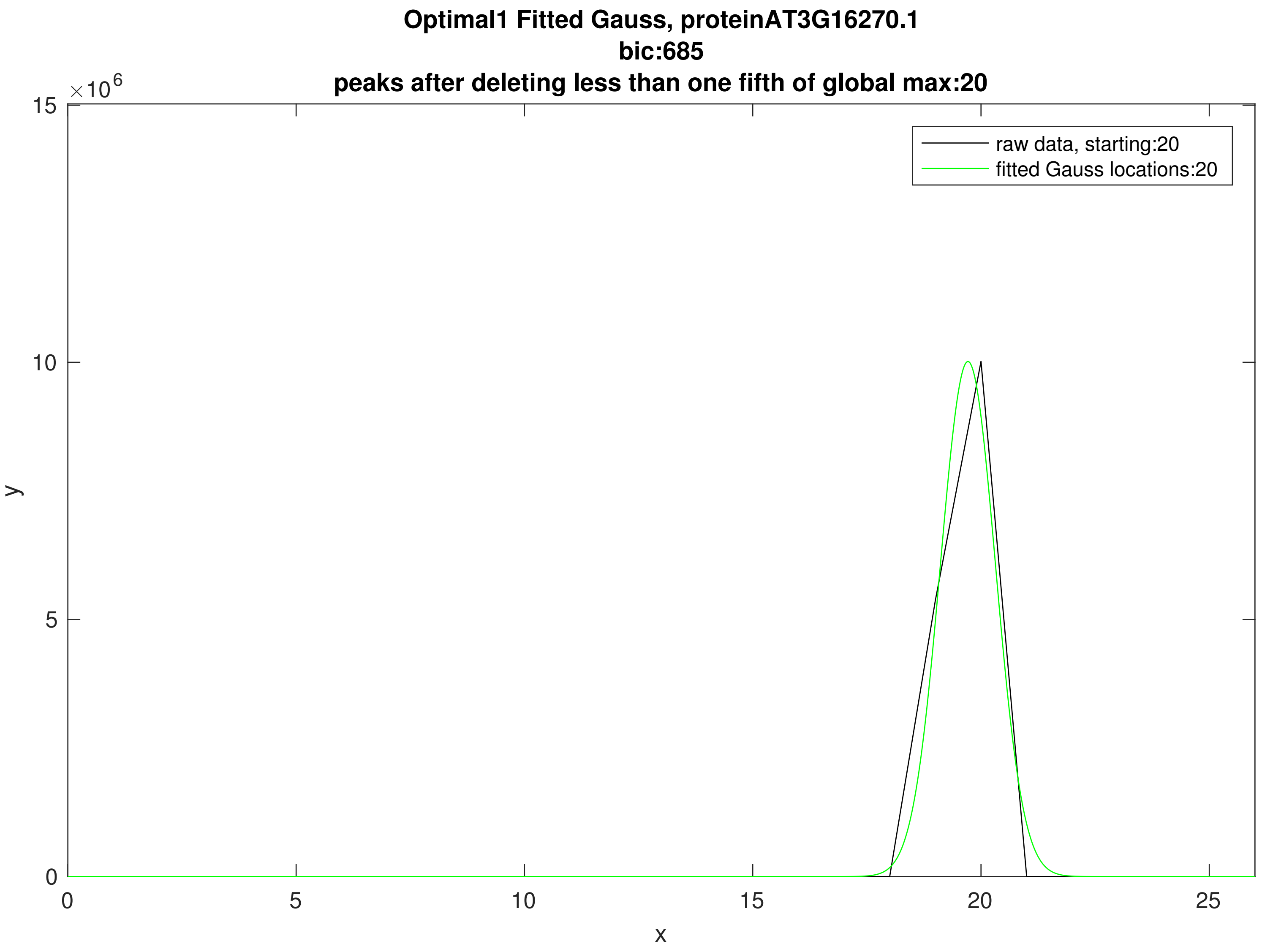

Optimal1 Fitted Gauss, proteinAT3G16270.1
bic:685
peaks after deleting less than one fifth of global max:20
×10^6
15
10
5
0
y
0
5
10
15
20
25
x
raw data, starting:20
fitted Gauss locations:20

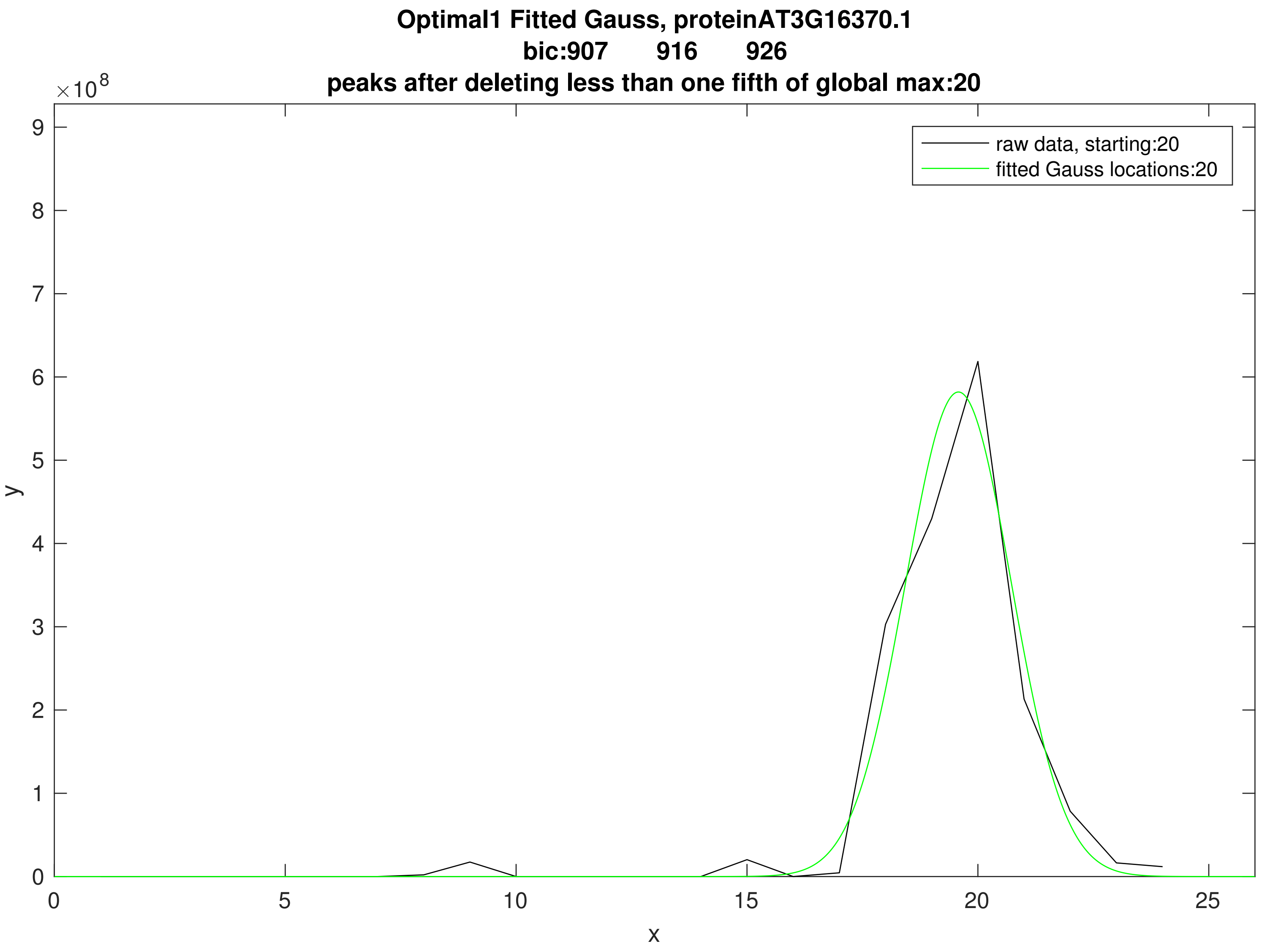

Optimal1 Fitted Gauss, proteinAT3G16370.1
bic:907 916 926
peaks after deleting less than one fifth of global max:20
×10^8
raw data, starting:20
fitted Gauss locations:20
y
x
0
1
2
3
4
5
6
7
8
9
0
5
10
15
20
25

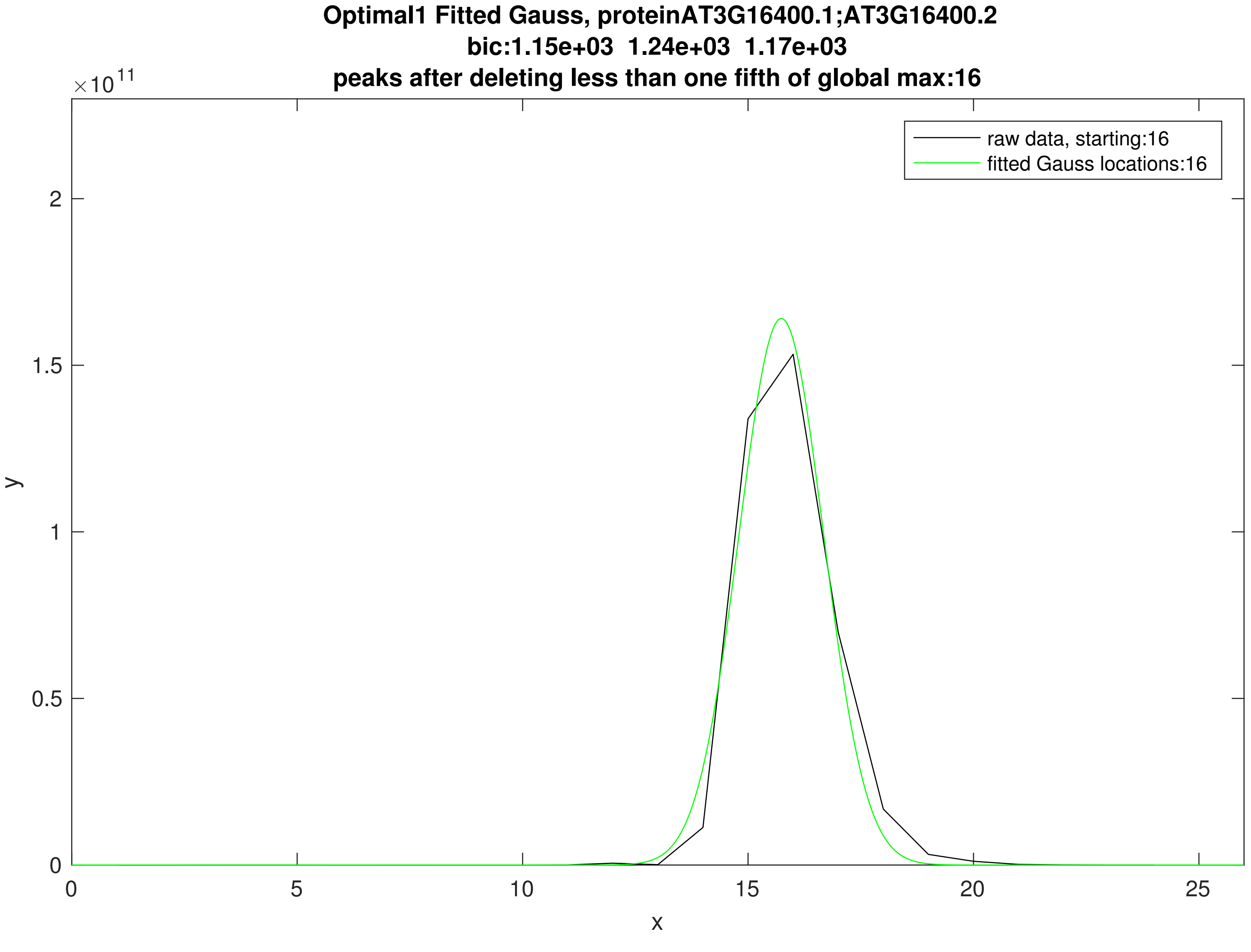
Optimal1 Fitted Gauss, proteinAT3G16400.1;AT3G16400.2
bic:1.15e+03  1.24e+03  1.17e+03
peaks after deleting less than one fifth of global max:16
×10^11
raw data, starting:16
fitted Gauss locations:16
y
x
0
0.5
1
1.5
2
5
10
15
20
25

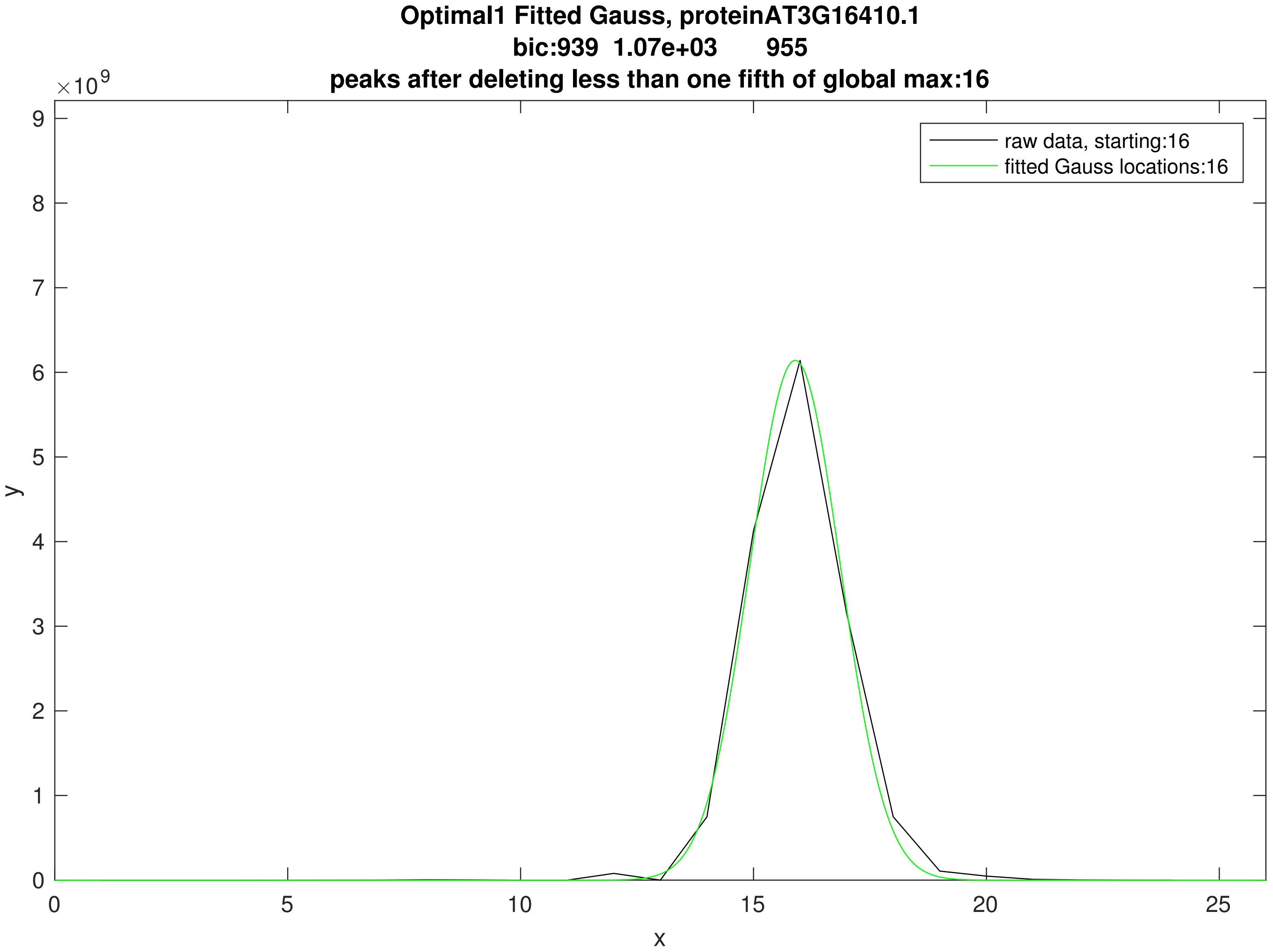
Optimal1 Fitted Gauss, proteinAT3G16410.1
bic:939 1.07e+03 955
peaks after deleting less than one fifth of global max:16
×10^9
raw data, starting:16
fitted Gauss locations:16
y
x

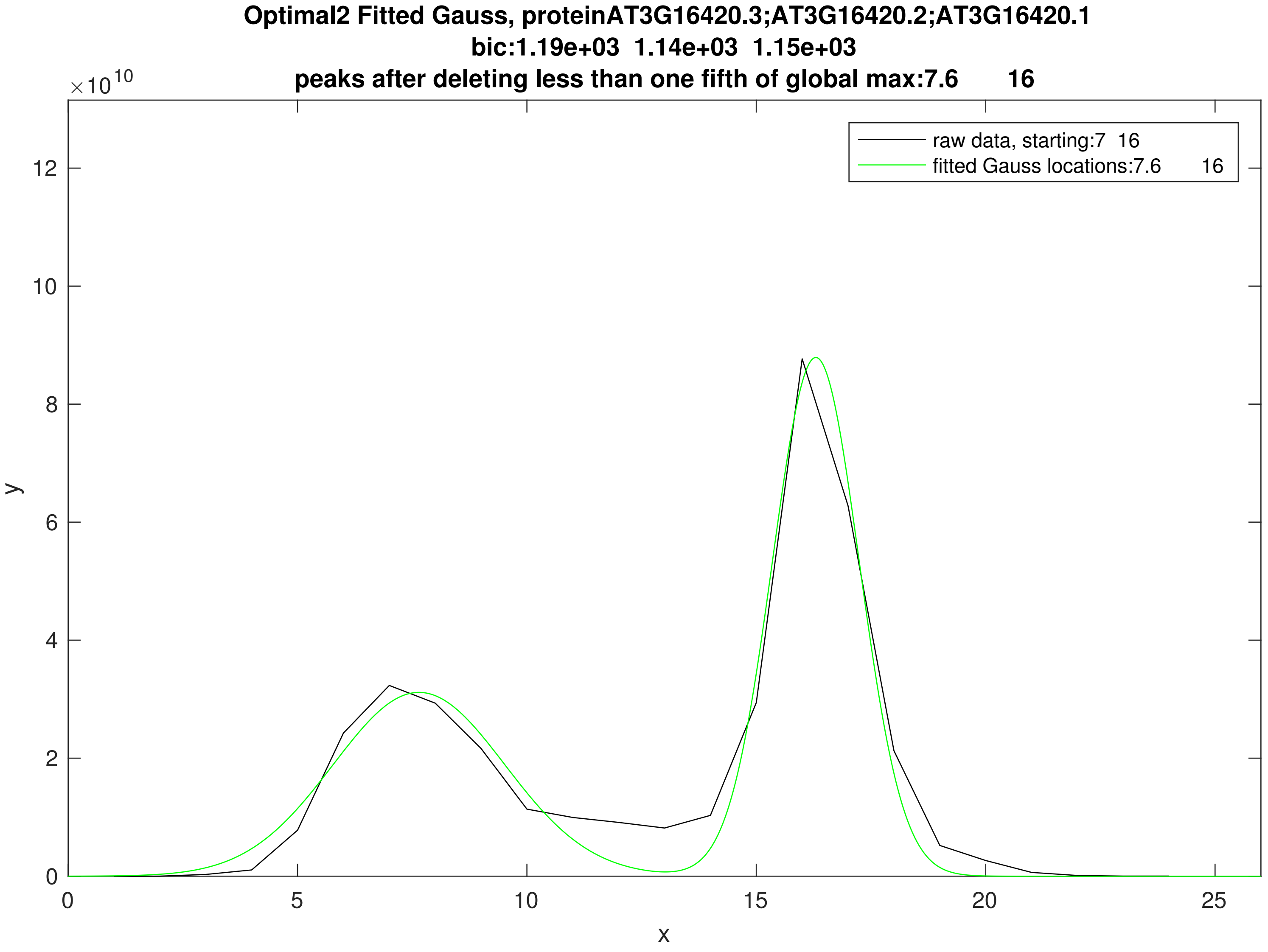
Optimal2 Fitted Gauss, proteinAT3G16420.3;AT3G16420.2;AT3G16420.1
bic:1.19e+03  1.14e+03  1.15e+03
peaks after deleting less than one fifth of global max:7.6   16
×10^10
raw data, starting:7  16
fitted Gauss locations:7.6   16
y
x
0
2
4
6
8
10
12
0
5
10
15
20
25

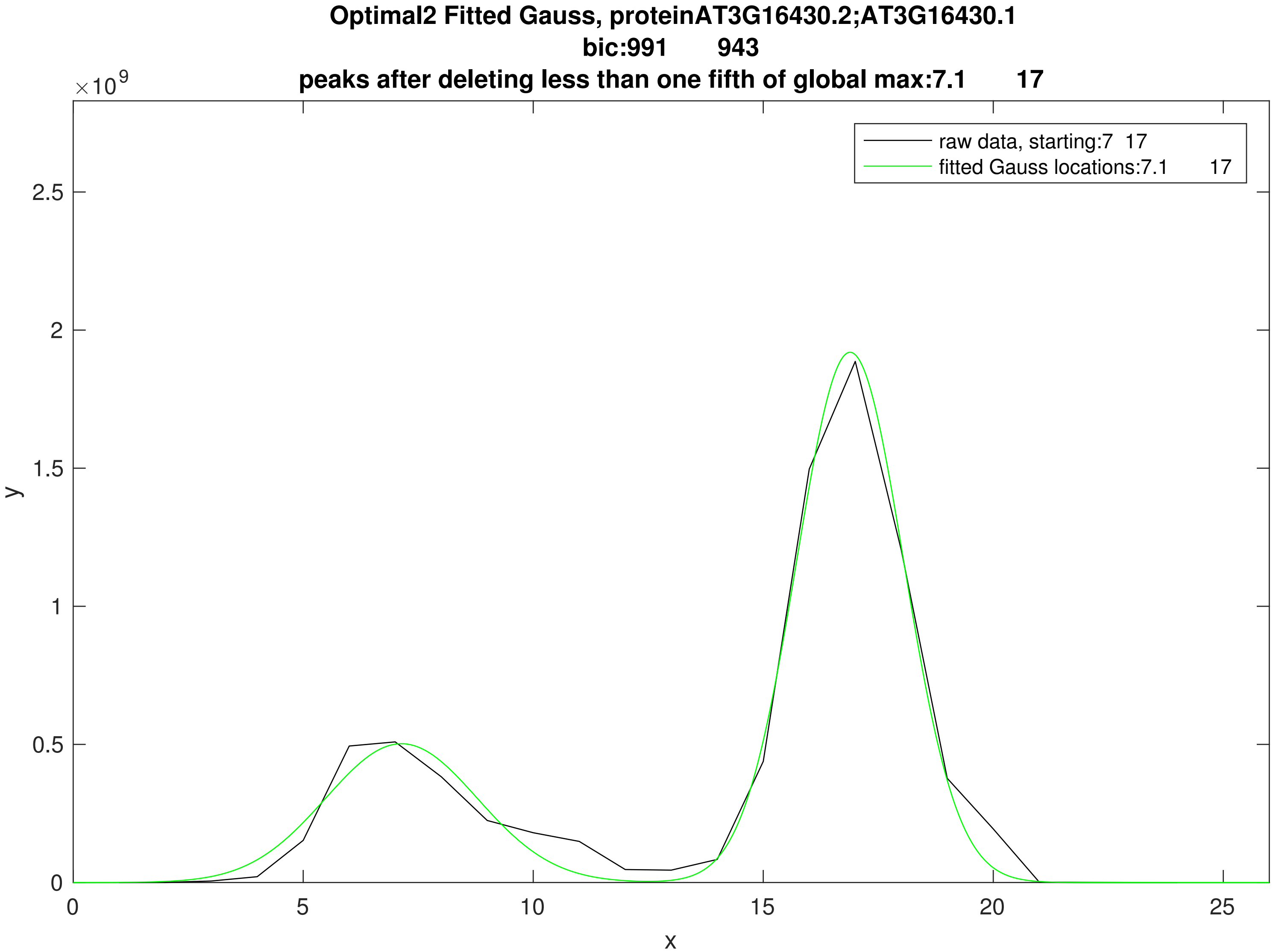

Optimal2 Fitted Gauss, proteinAT3G16430.2;AT3G16430.1
bic:991 943
peaks after deleting less than one fifth of global max:7.1 17
×10^9
raw data, starting:7 17
fitted Gauss locations:7.1 17
y
x
0
0.5
1
1.5
2
2.5
5
10
15
20
25

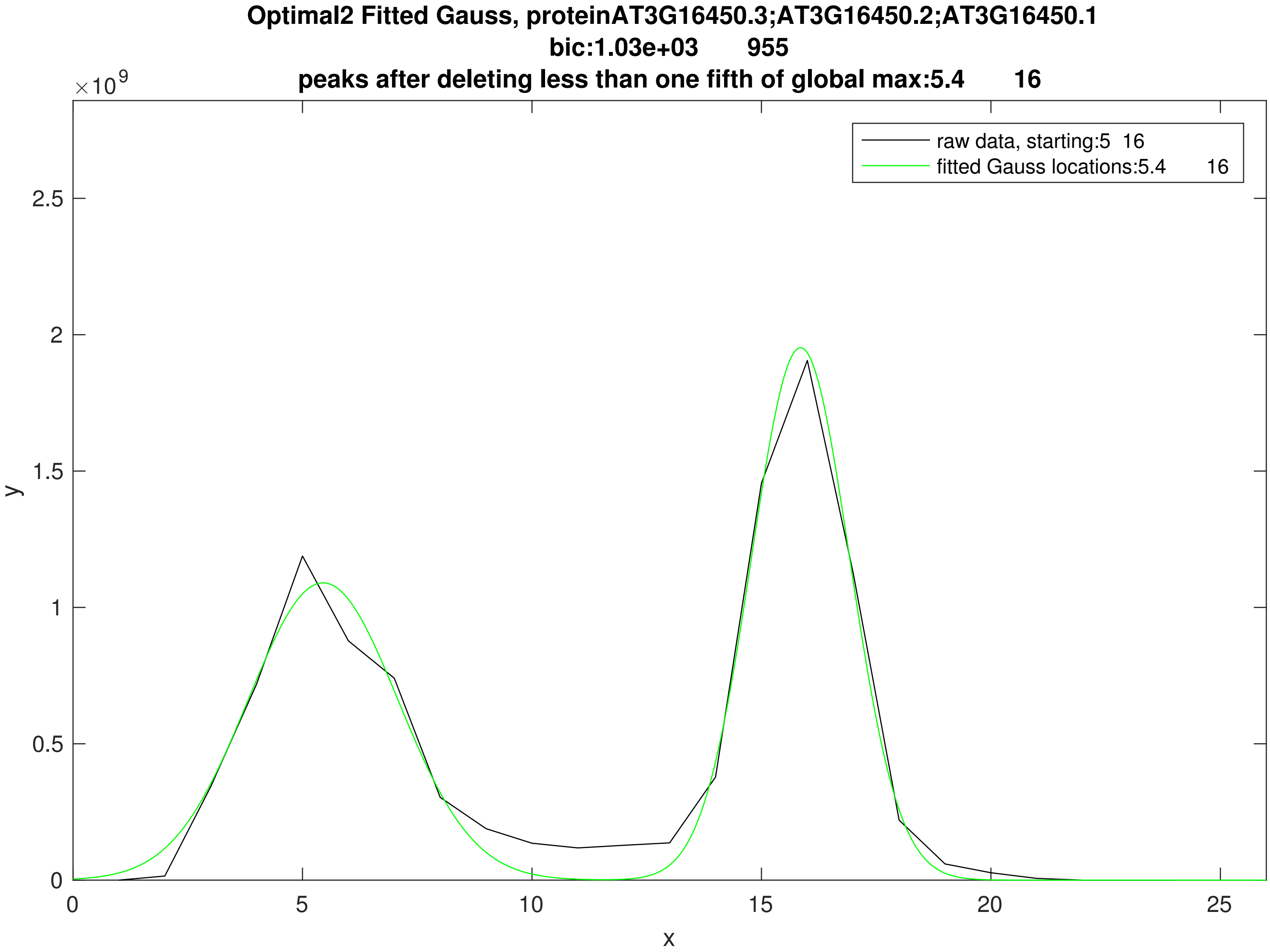
Optimal2 Fitted Gauss, proteinAT3G16450.3;AT3G16450.2;AT3G16450.1
bic:1.03e+03 955
peaks after deleting less than one fifth of global max:5.4 16
×10^9
raw data, starting:5 16
fitted Gauss locations:5.4 16
y
x
0
0.5
1
1.5
2
2.5
5
10
15
20
25

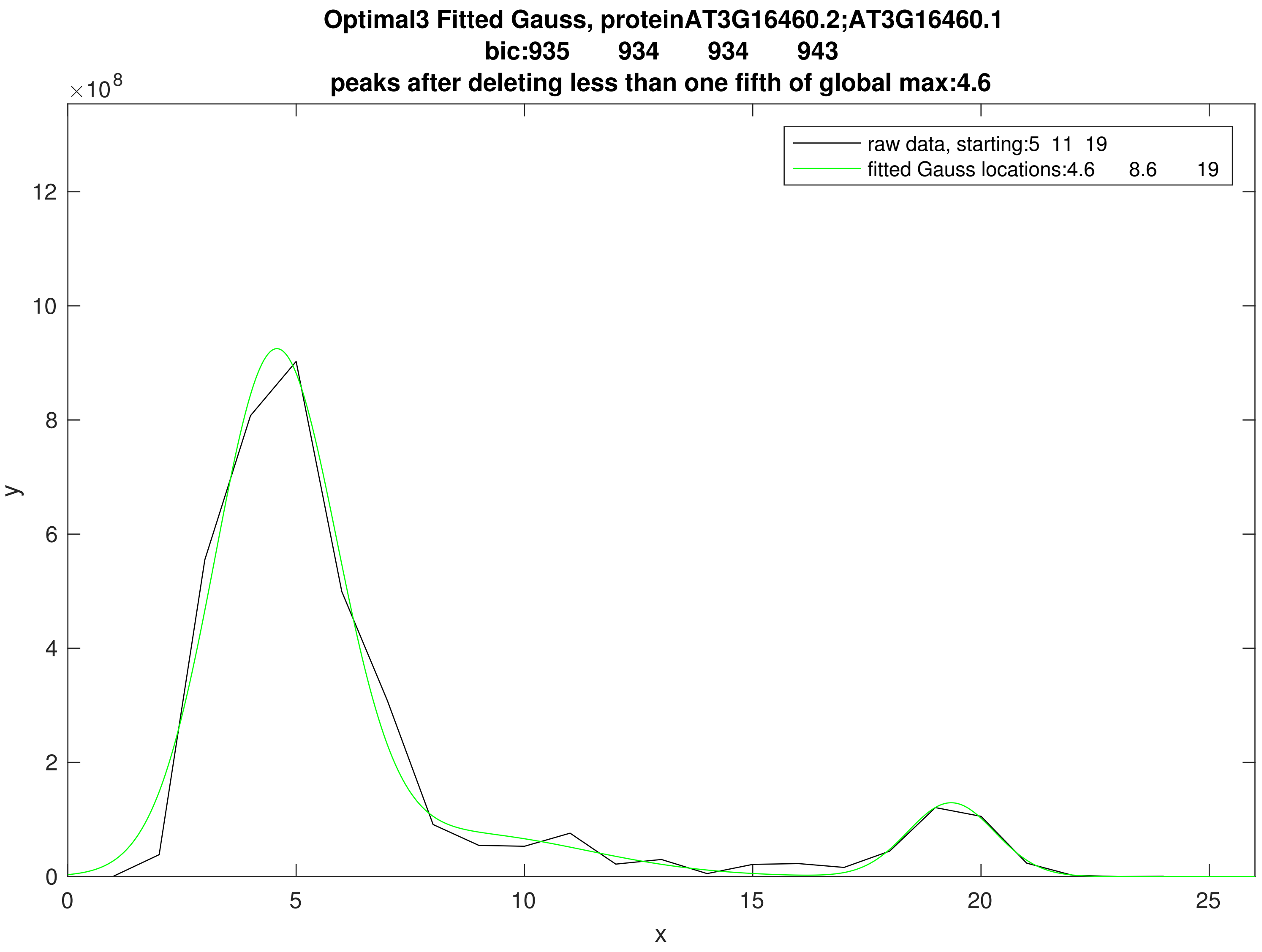

Optimal3 Fitted Gauss, proteinAT3G16460.2;AT3G16460.1
bic:935 934 934 943
peaks after deleting less than one fifth of global max:4.6
×10^8
raw data, starting:5 11 19
fitted Gauss locations:4.6 8.6 19
y
x
0
2
4
6
8
10
12
0
5
10
15
20
25

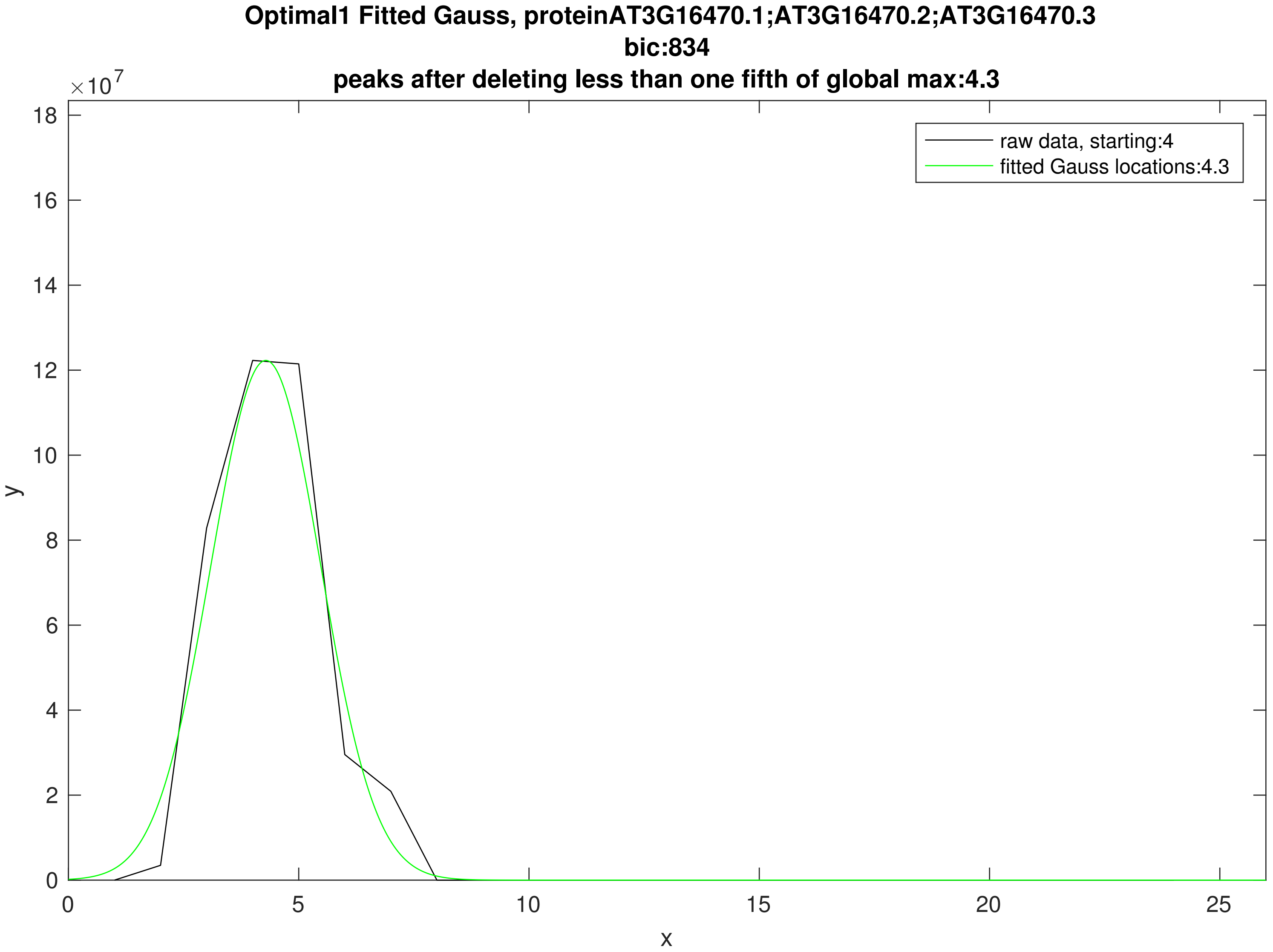
Optimal1 Fitted Gauss, proteinAT3G16470.1;AT3G16470.2;AT3G16470.3
bic:834
peaks after deleting less than one fifth of global max:4.3
×10^7
raw data, starting:4
fitted Gauss locations:4.3
y
x
0
2
4
6
8
10
12
14
16
18
0
5
10
15
20
25

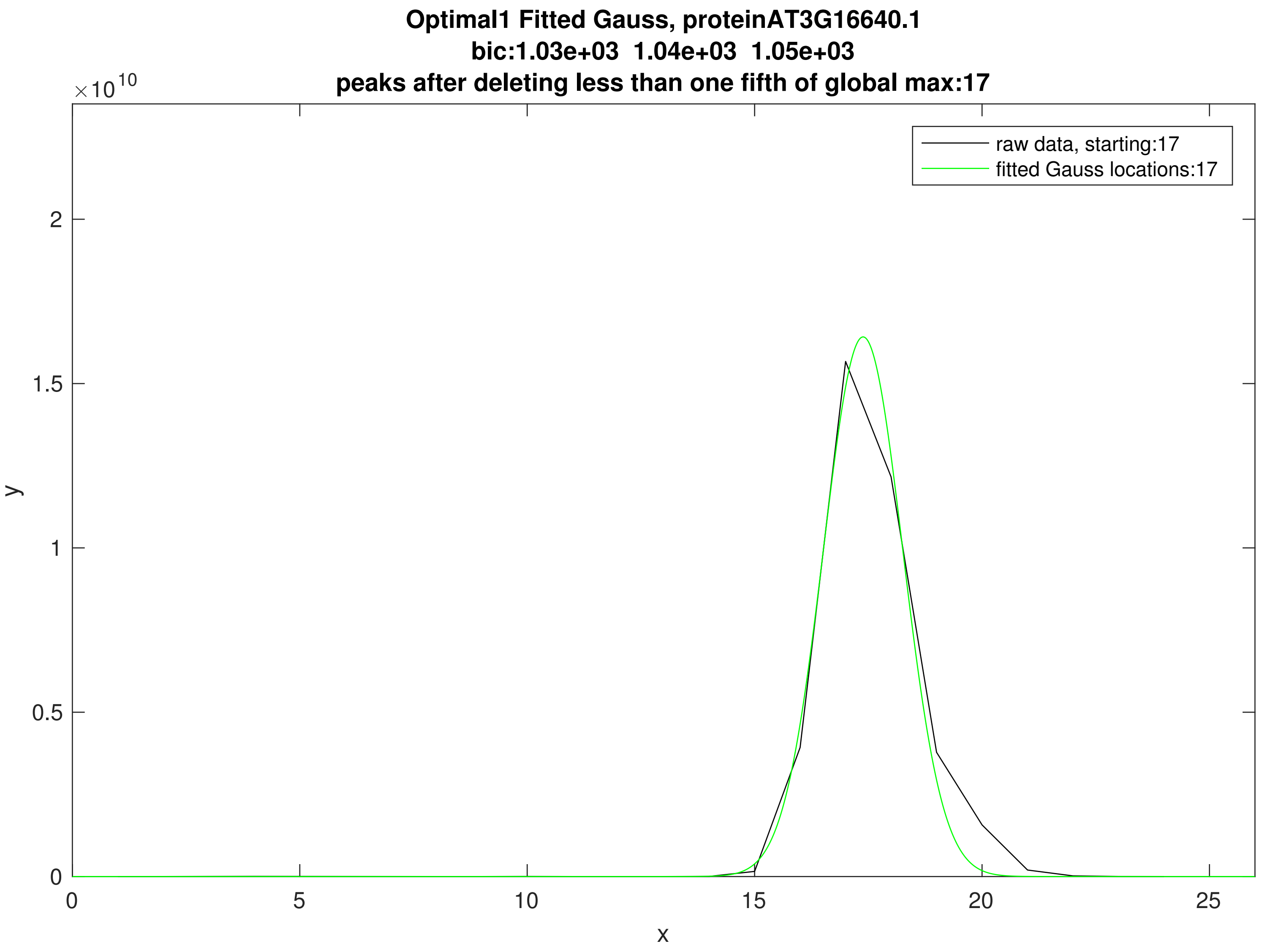
Optimal1 Fitted Gauss, proteinAT3G16640.1
bic:1.03e+03 1.04e+03 1.05e+03
peaks after deleting less than one fifth of global max:17
×10^10
raw data, starting:17
fitted Gauss locations:17
y
x
0
0.5
1
1.5
2
5
10
15
20
25

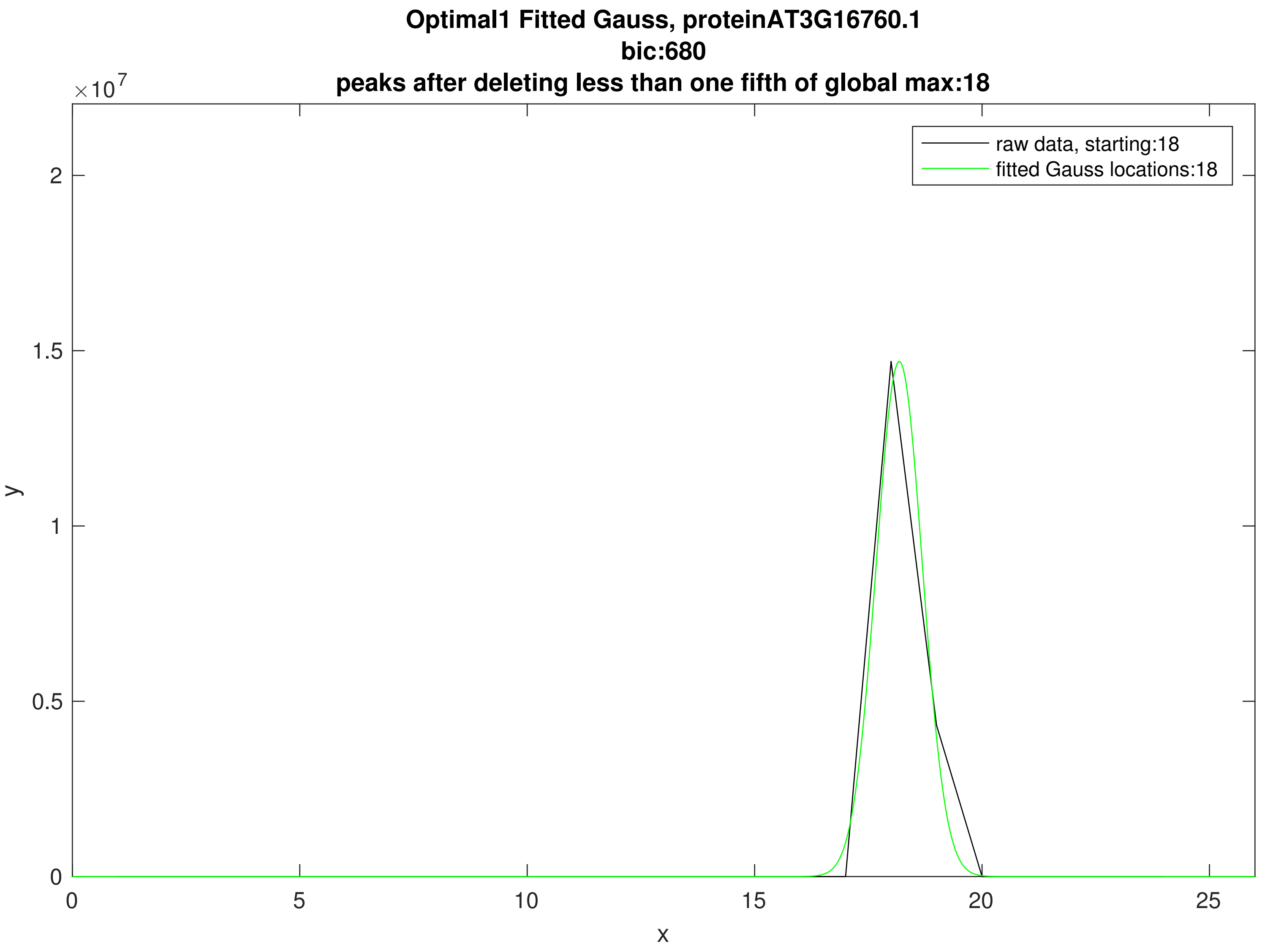
Optimal1 Fitted Gauss, proteinAT3G16760.1
bic:680
peaks after deleting less than one fifth of global max:18
×10^7
2
1.5
1
0.5
0
y
0
5
10
15
20
25
x
raw data, starting:18
fitted Gauss locations:18

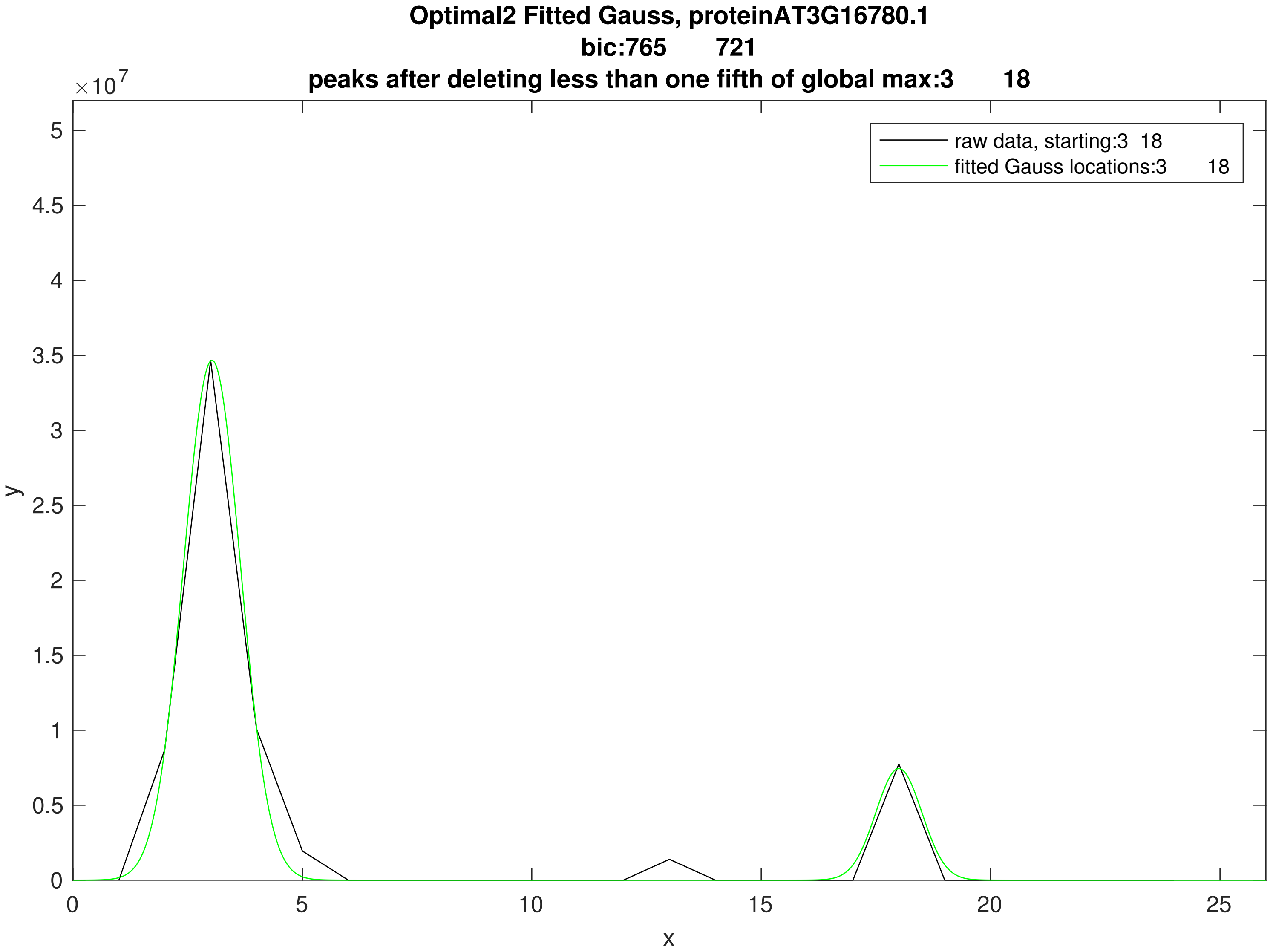
Optimal2 Fitted Gauss, proteinAT3G16780.1
bic:765 721
peaks after deleting less than one fifth of global max:3 18
×10^7
raw data, starting:3 18
fitted Gauss locations:3 18
y
x
0
0.5
1
1.5
2
2.5
3
3.5
4
4.5
5
0
5
10
15
20
25

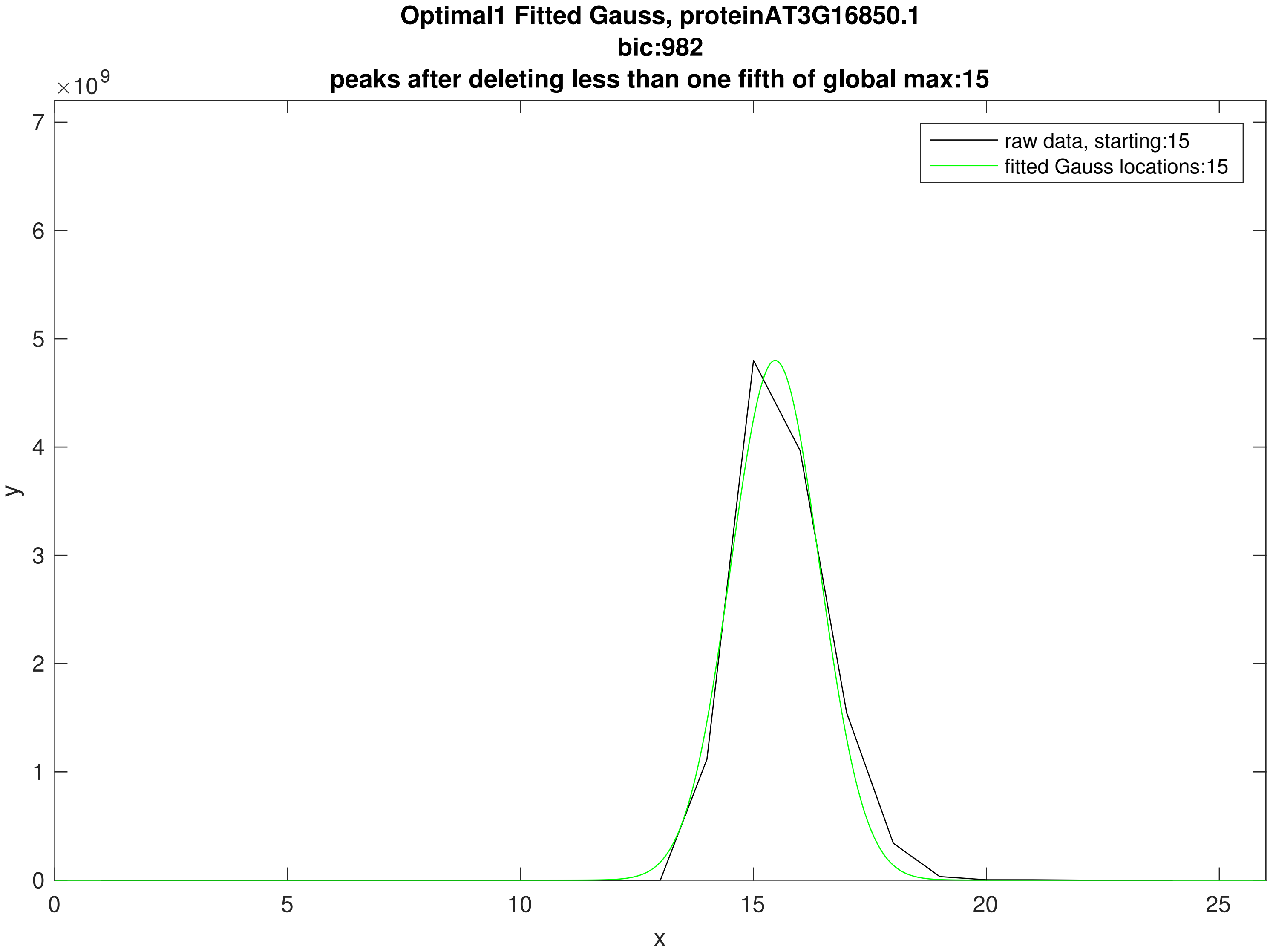
Optimal1 Fitted Gauss, proteinAT3G16850.1
bic:982
peaks after deleting less than one fifth of global max:15
×10^9
raw data, starting:15
fitted Gauss locations:15
y
x
0
1
2
3
4
5
6
7
0
5
10
15
20
25

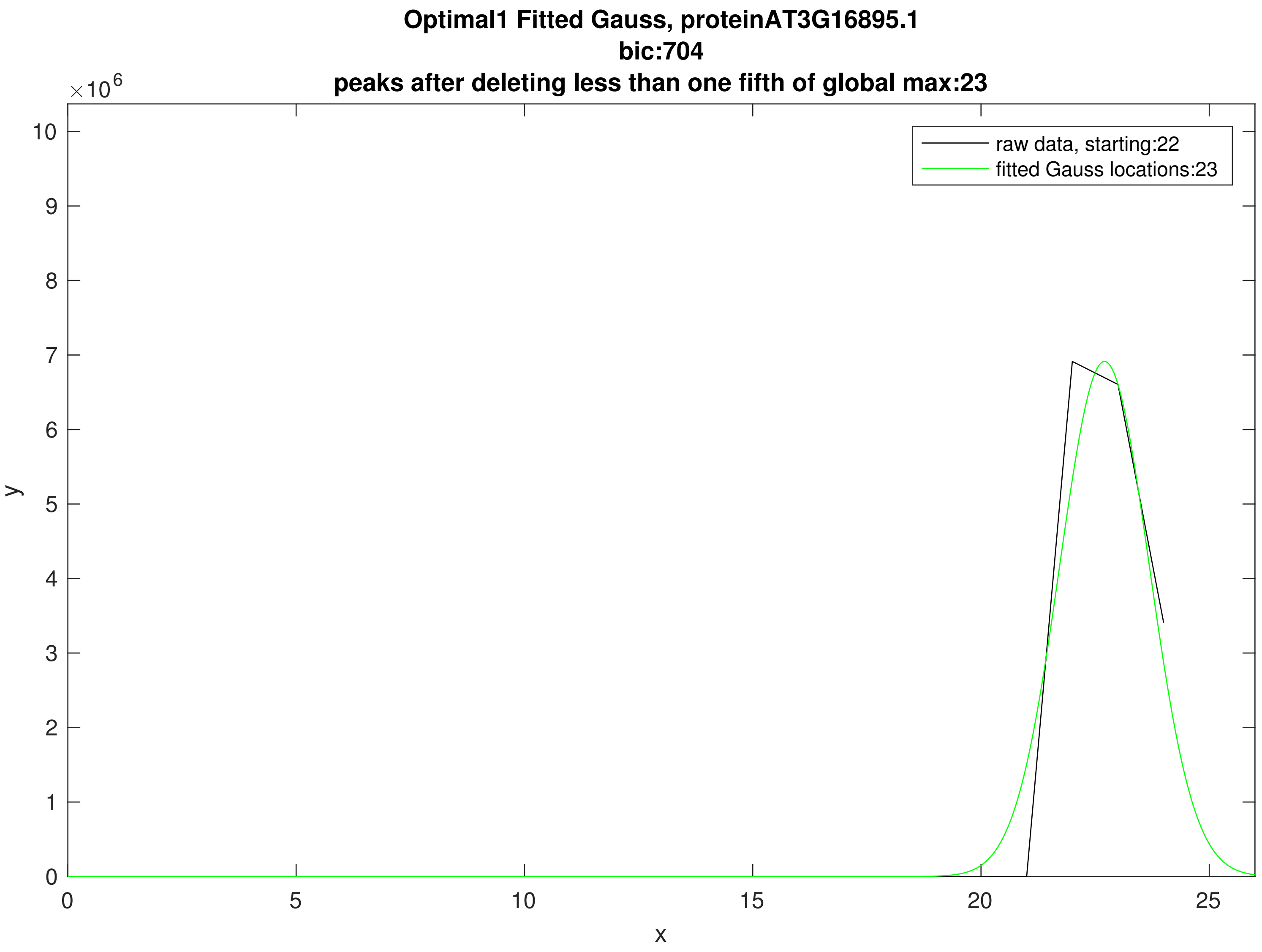

Optimal1 Fitted Gauss, proteinAT3G16895.1
bic:704
peaks after deleting less than one fifth of global max:23
$\times 10^6$
raw data, starting:22
fitted Gauss locations:23
y
x
0
1
2
3
4
5
6
7
8
9
10
0
5
10
15
20
25

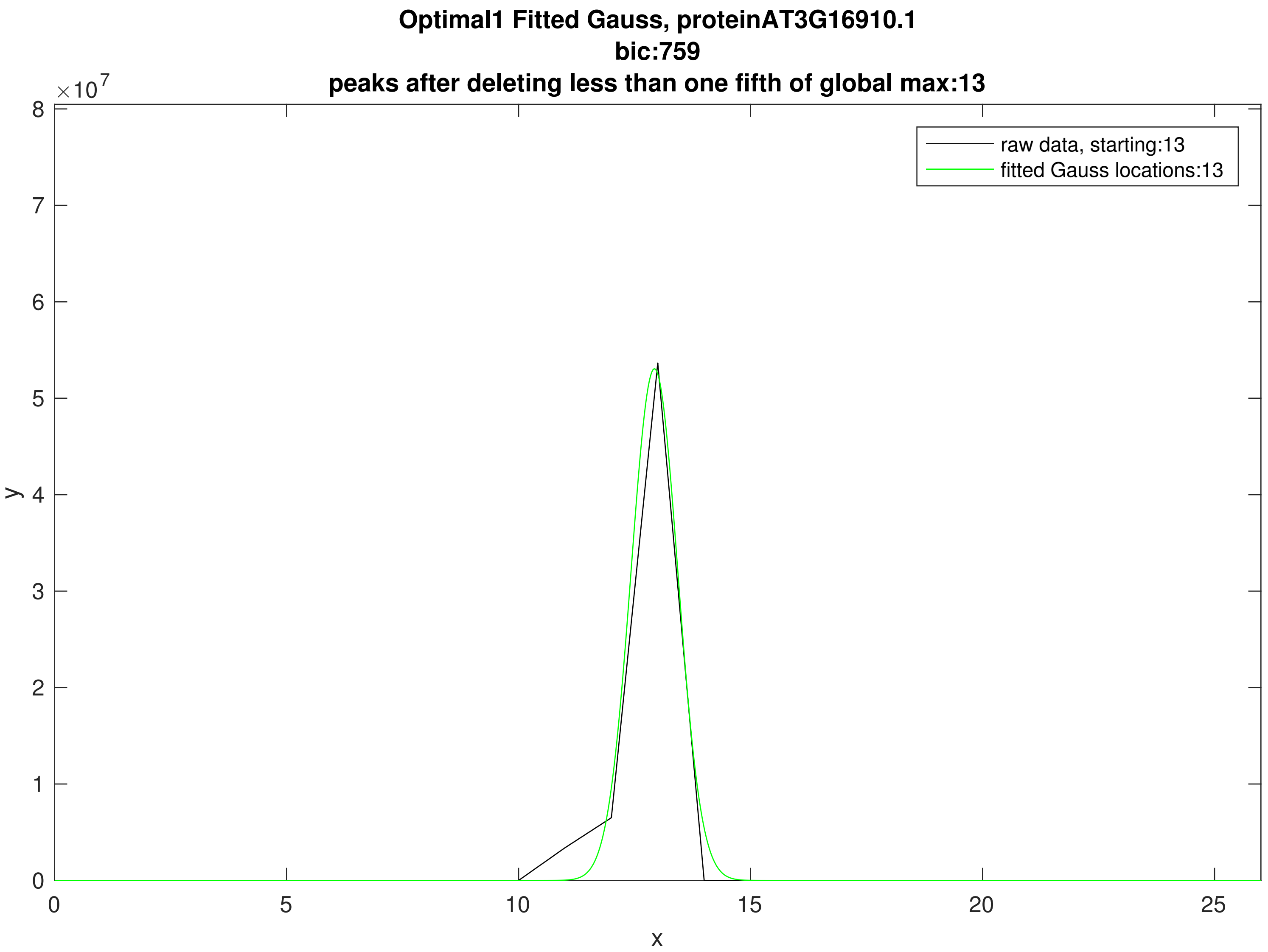
Optimal1 Fitted Gauss, proteinAT3G16910.1
bic:759
peaks after deleting less than one fifth of global max:13
×10^7
8
7
6
5
4
3
2
1
0
y
0
5
10
15
20
25
x
raw data, starting:13
fitted Gauss locations:13

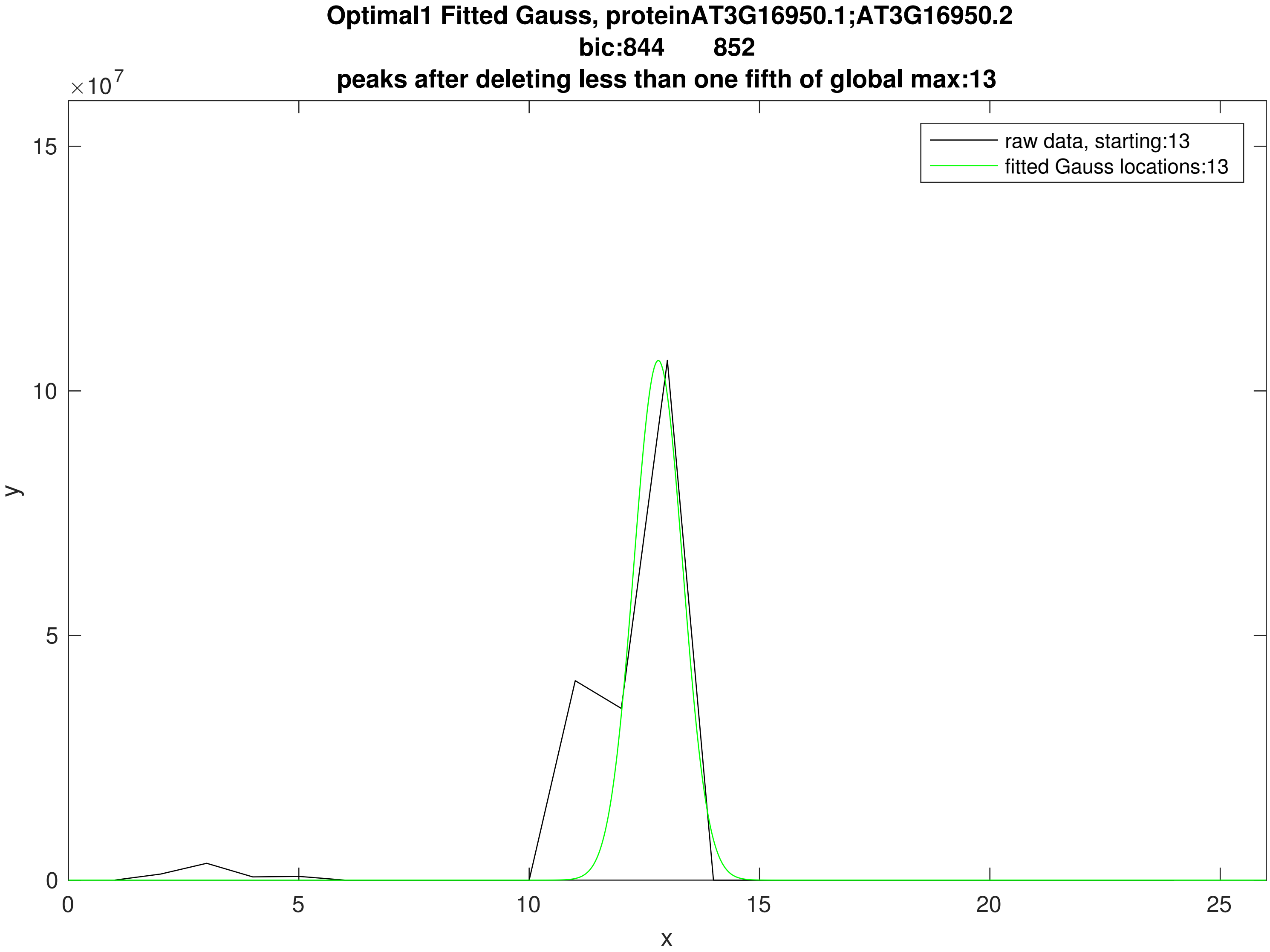
Optimal1 Fitted Gauss, proteinAT3G16950.1;AT3G16950.2
bic:844 852
peaks after deleting less than one fifth of global max:13
×10^7
raw data, starting:13
fitted Gauss locations:13
y
x
0
5
10
15
20
25

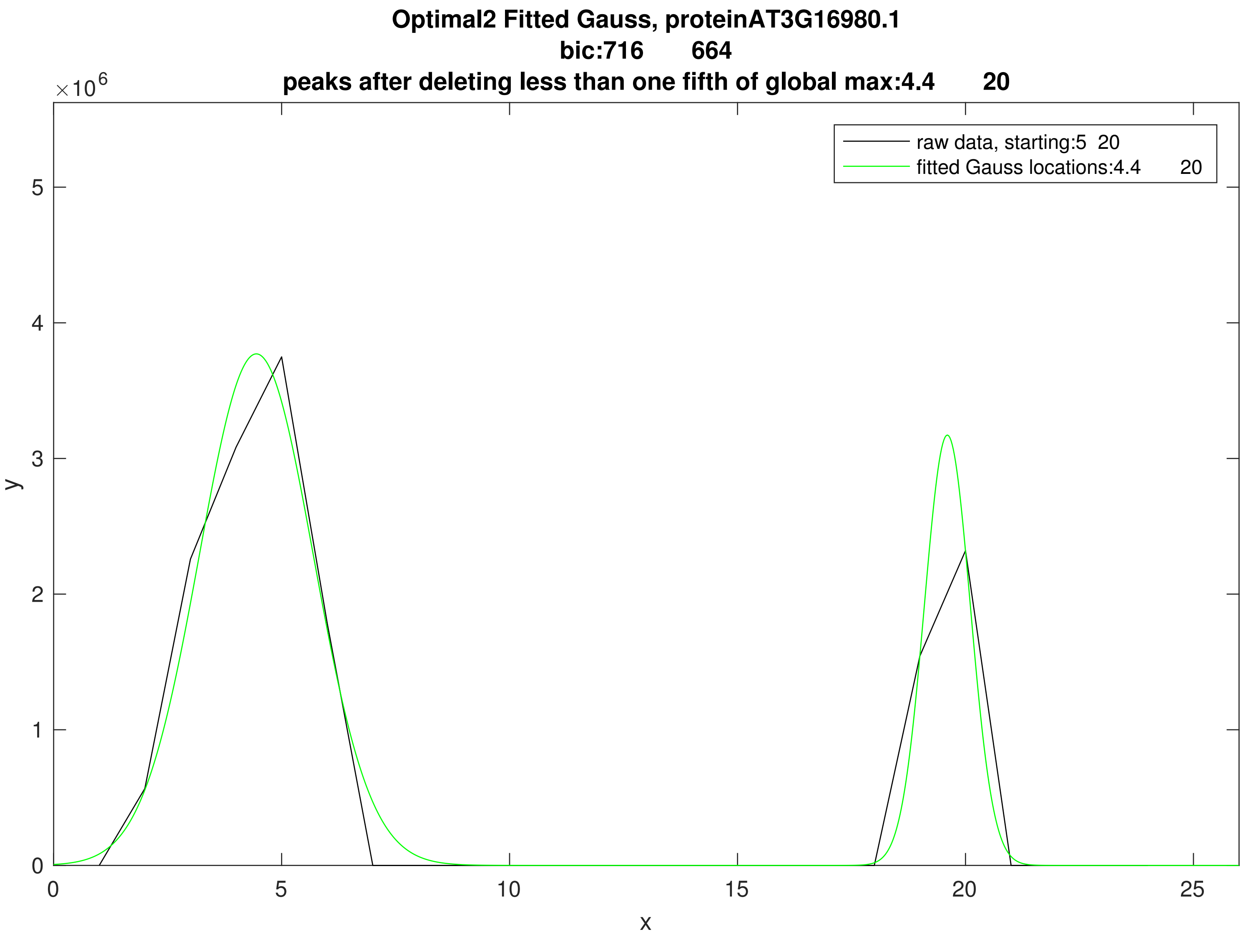
Optimal2 Fitted Gauss, proteinAT3G16980.1
bic:716 664
peaks after deleting less than one fifth of global max:4.4 20
×10^6
raw data, starting:5 20
fitted Gauss locations:4.4 20
y
x
0
1
2
3
4
5
0
5
10
15
20
25

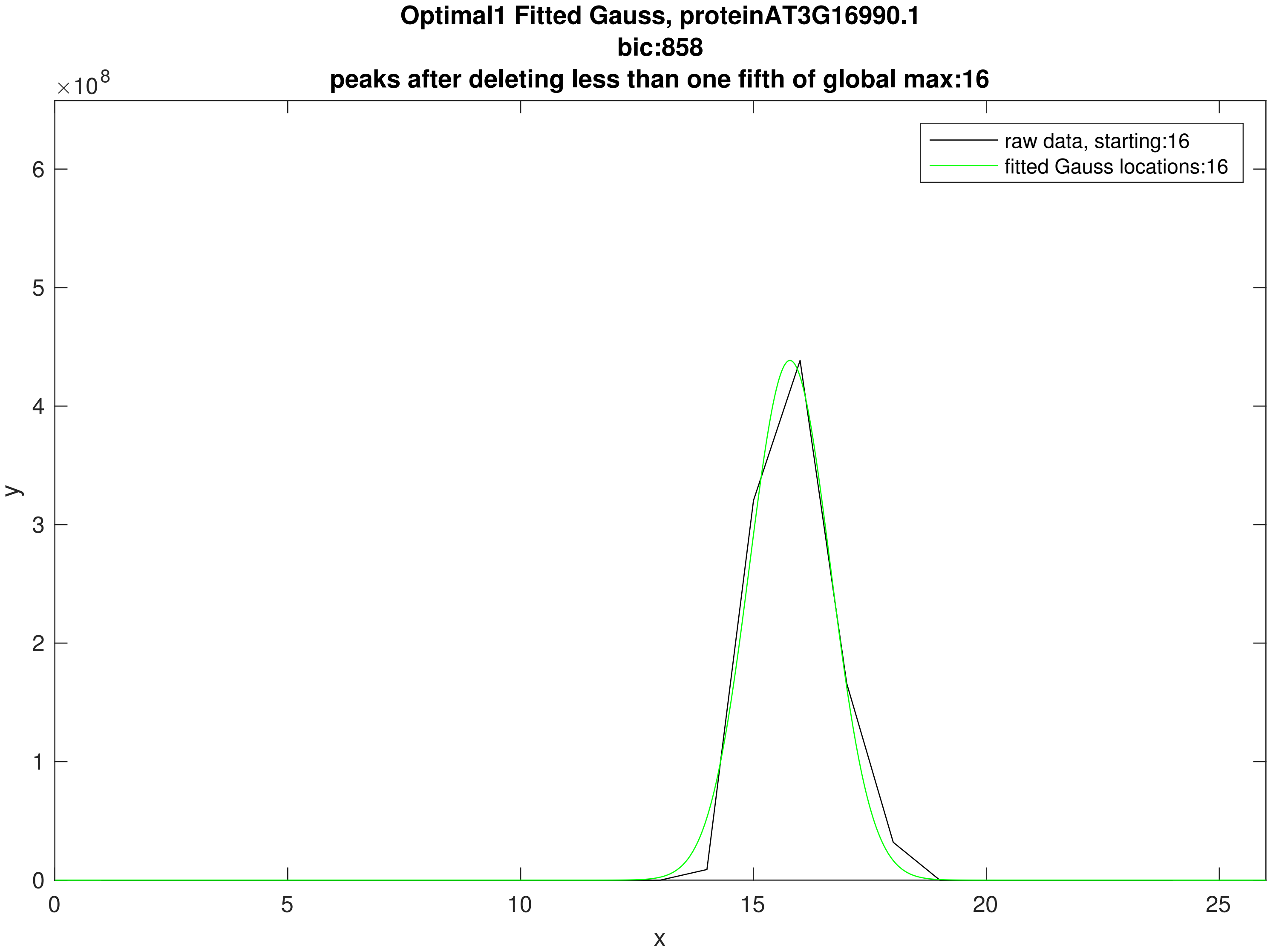
Optimal1 Fitted Gauss, proteinAT3G16990.1
bic:858
peaks after deleting less than one fifth of global max:16
×10^8
raw data, starting:16
fitted Gauss locations:16
y
x
0
1
2
3
4
5
6
0
5
10
15
20
25

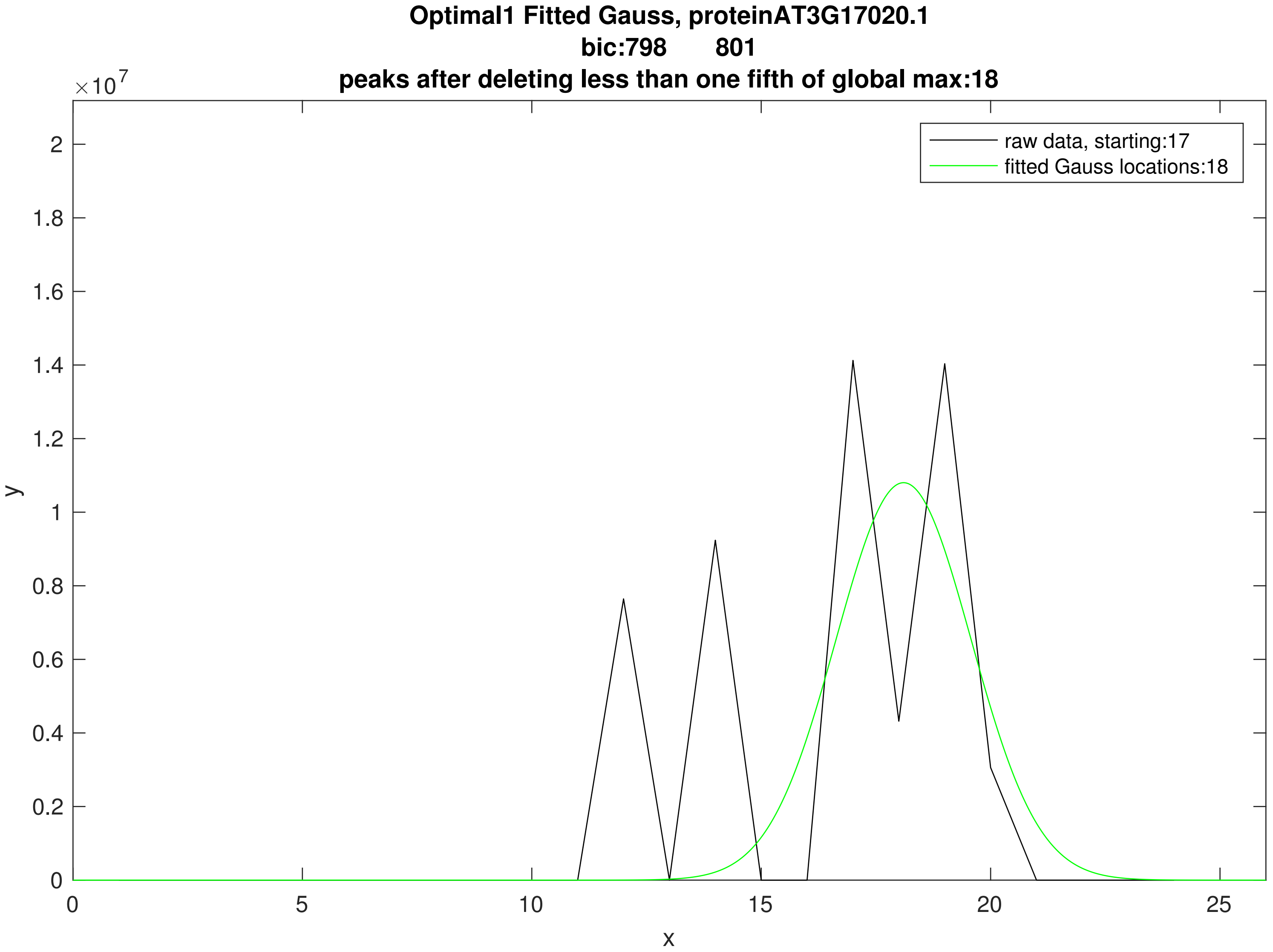
Optimal1 Fitted Gauss, proteinAT3G17020.1
bic:798 801
peaks after deleting less than one fifth of global max:18
×10^7
raw data, starting:17
fitted Gauss locations:18
y
x
0
0.2
0.4
0.6
0.8
1
1.2
1.4
1.6
1.8
2
0
5
10
15
20
25

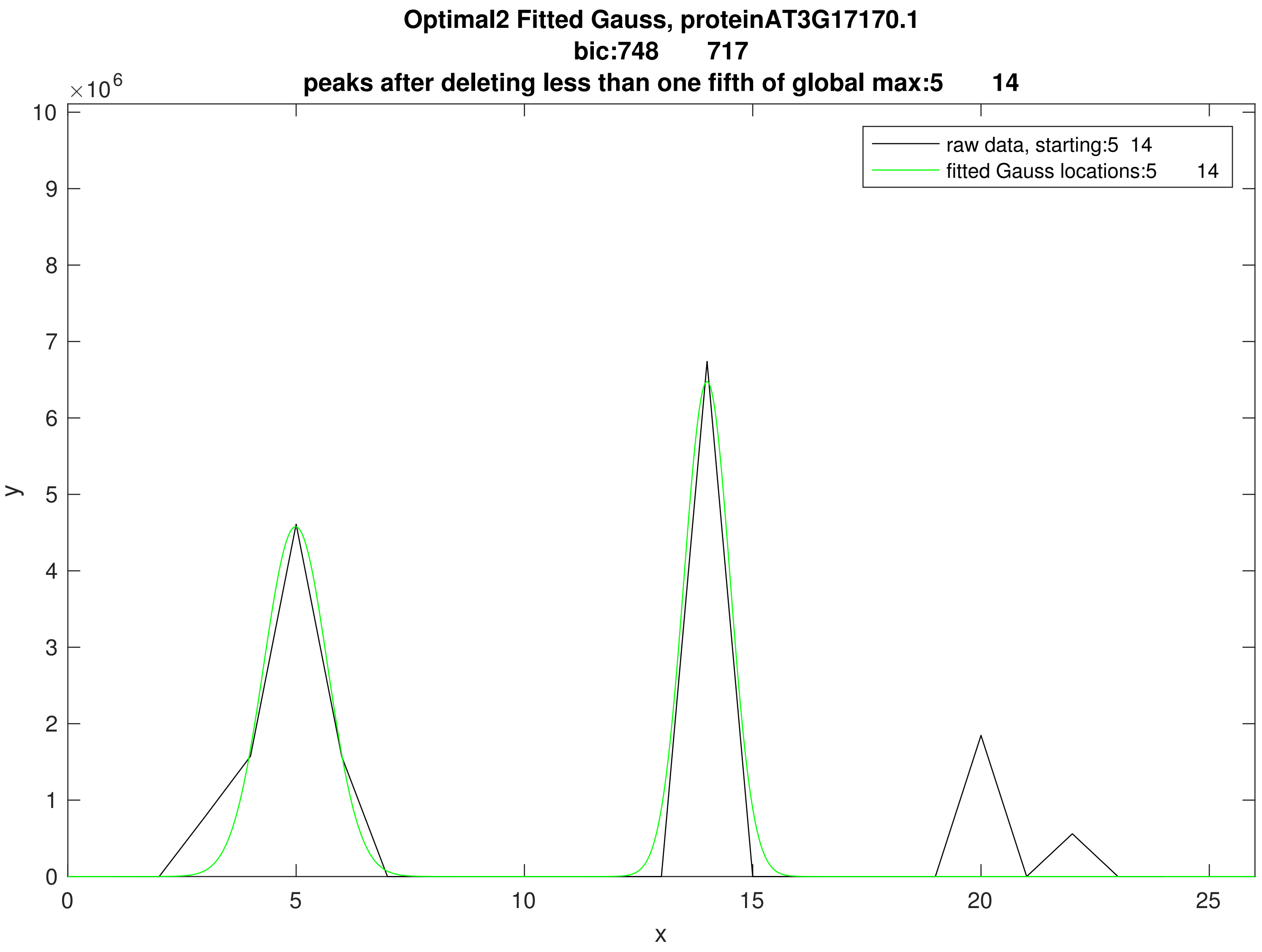

Optimal2 Fitted Gauss, proteinAT3G17170.1
bic:748 717
peaks after deleting less than one fifth of global max:5 14
×10^6
raw data, starting:5 14
fitted Gauss locations:5 14
y
x
0
1
2
3
4
5
6
7
8
9
10
0
5
10
15
20
25

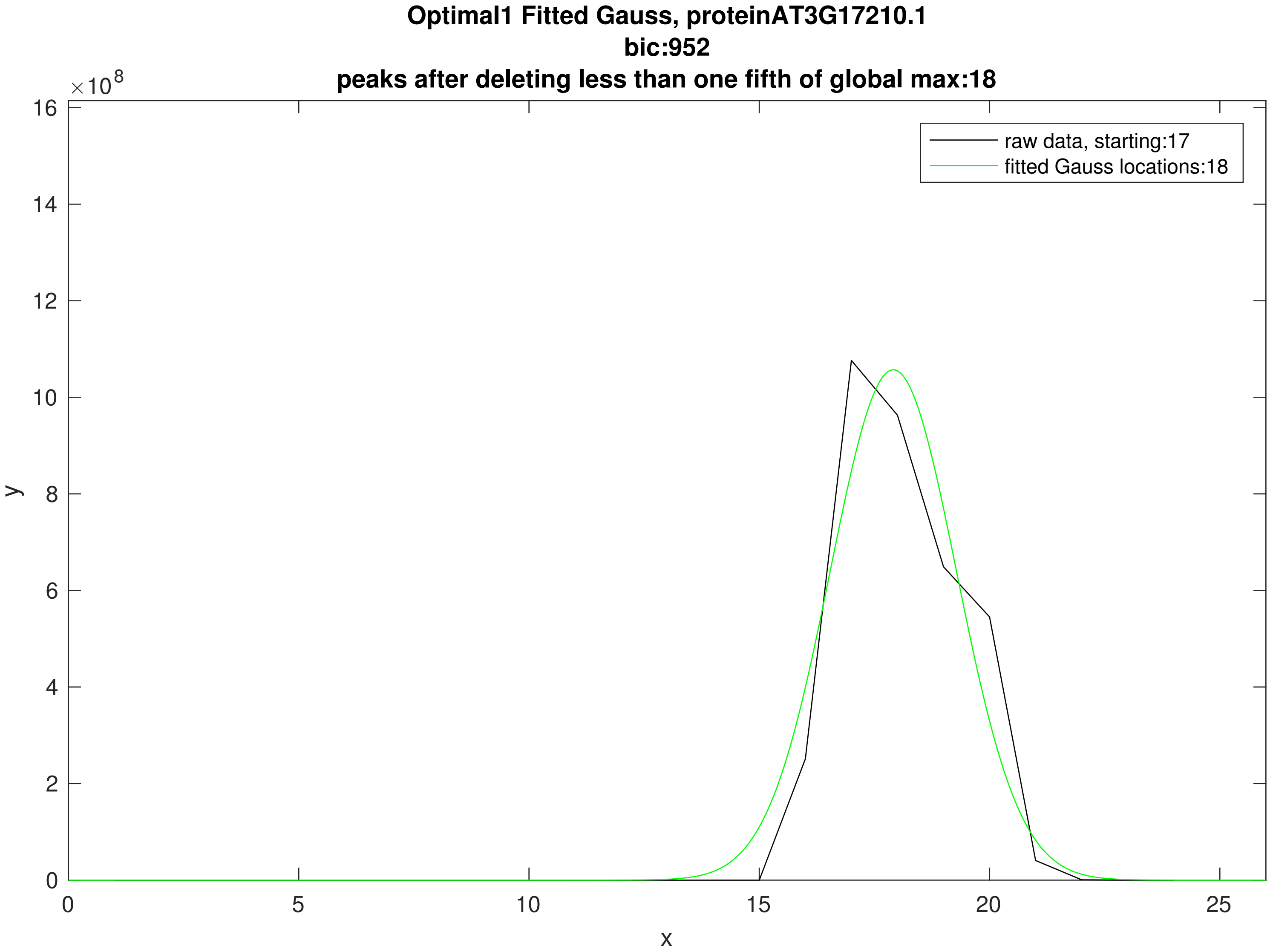
Optimal1 Fitted Gauss, proteinAT3G17210.1
bic:952
peaks after deleting less than one fifth of global max:18
×10^8
16
14
12
10
8
6
4
2
0
y
0
5
10
15
20
25
x
raw data, starting:17
fitted Gauss locations:18

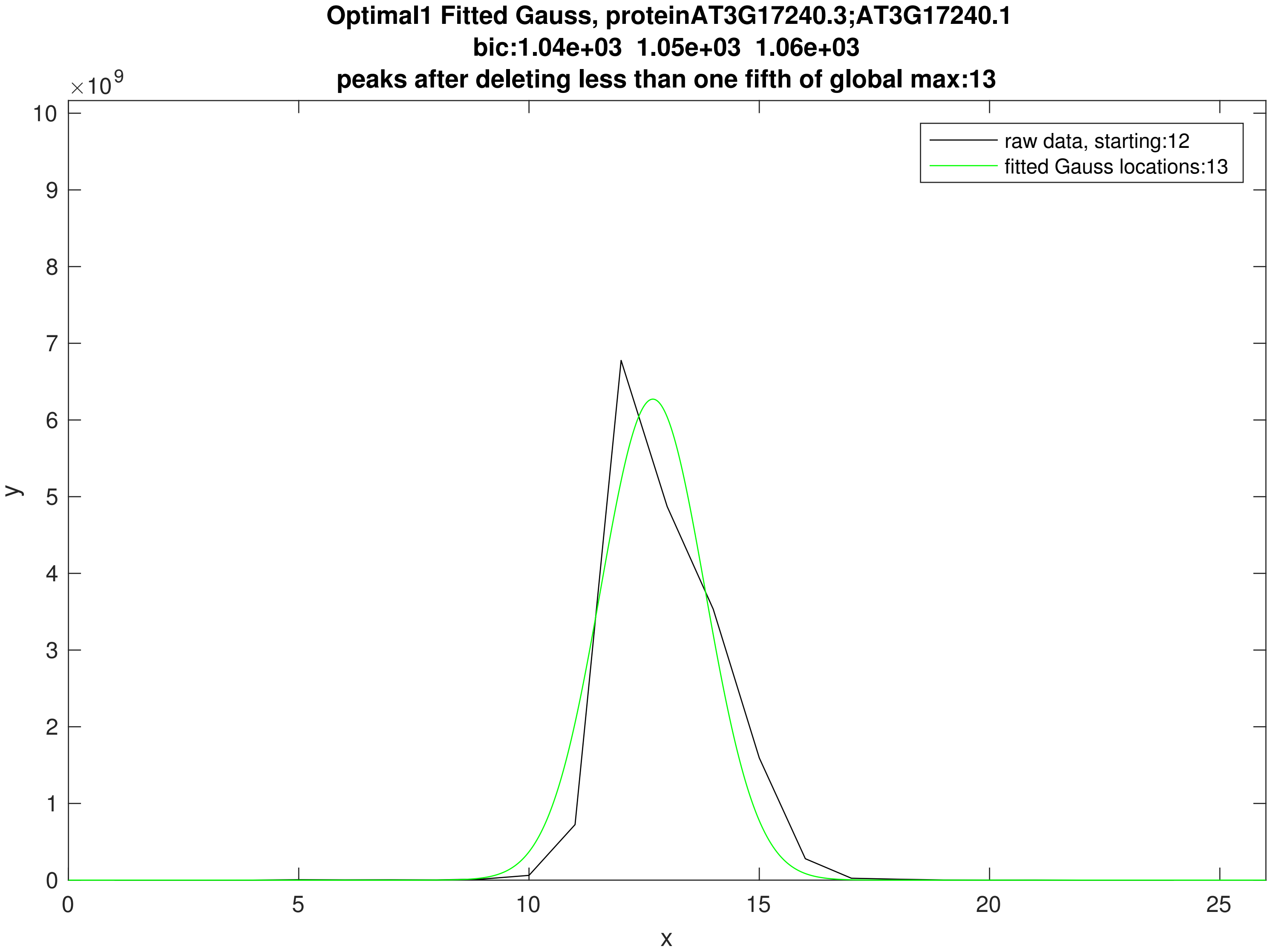

Optimal1 Fitted Gauss, proteinAT3G17240.3;AT3G17240.1
bic:1.04e+03  1.05e+03  1.06e+03
peaks after deleting less than one fifth of global max:13
×10^9
y
x
raw data, starting:12
fitted Gauss locations:13

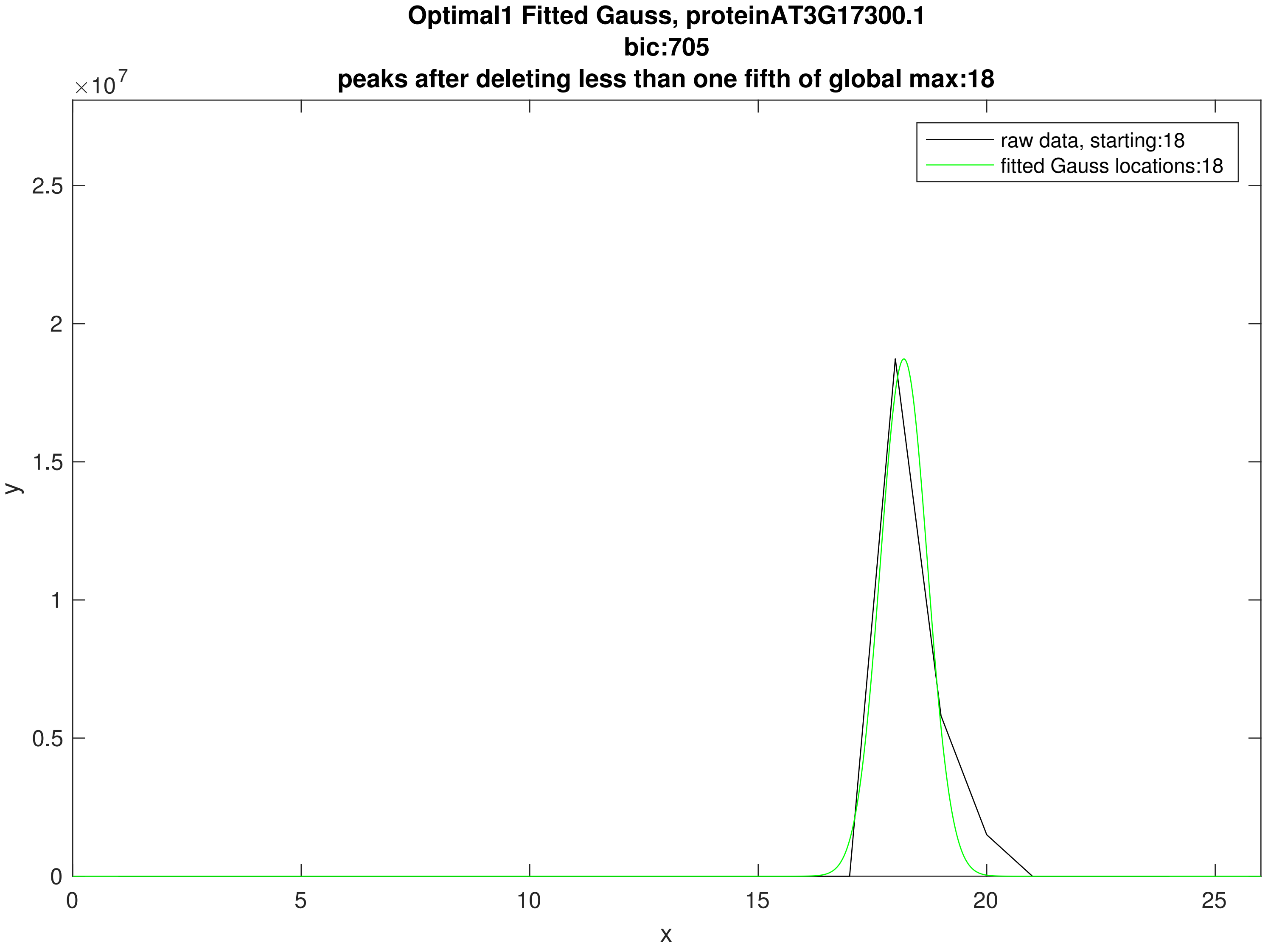
Optimal1 Fitted Gauss, proteinAT3G17300.1
bic:705
peaks after deleting less than one fifth of global max:18
×10^7
raw data, starting:18
fitted Gauss locations:18
2.5
2
1.5
1
0.5
0
y
0
5
10
15
20
25
x

Optimal1 Fitted Gauss, proteinAT3G17390.1
bic:1.13e+03  1.15e+03  1.16e+03
peaks after deleting less than one fifth of global max:16

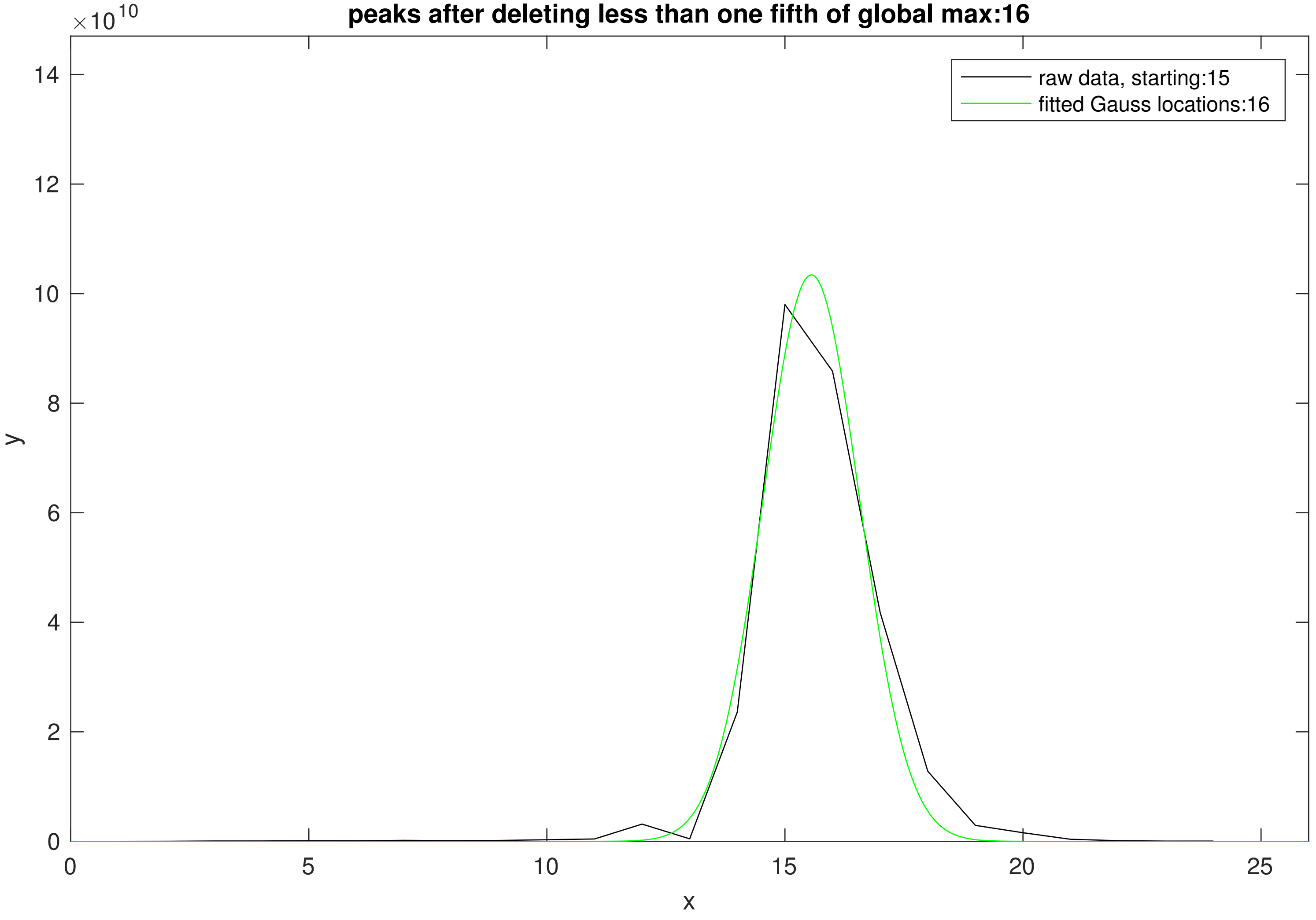

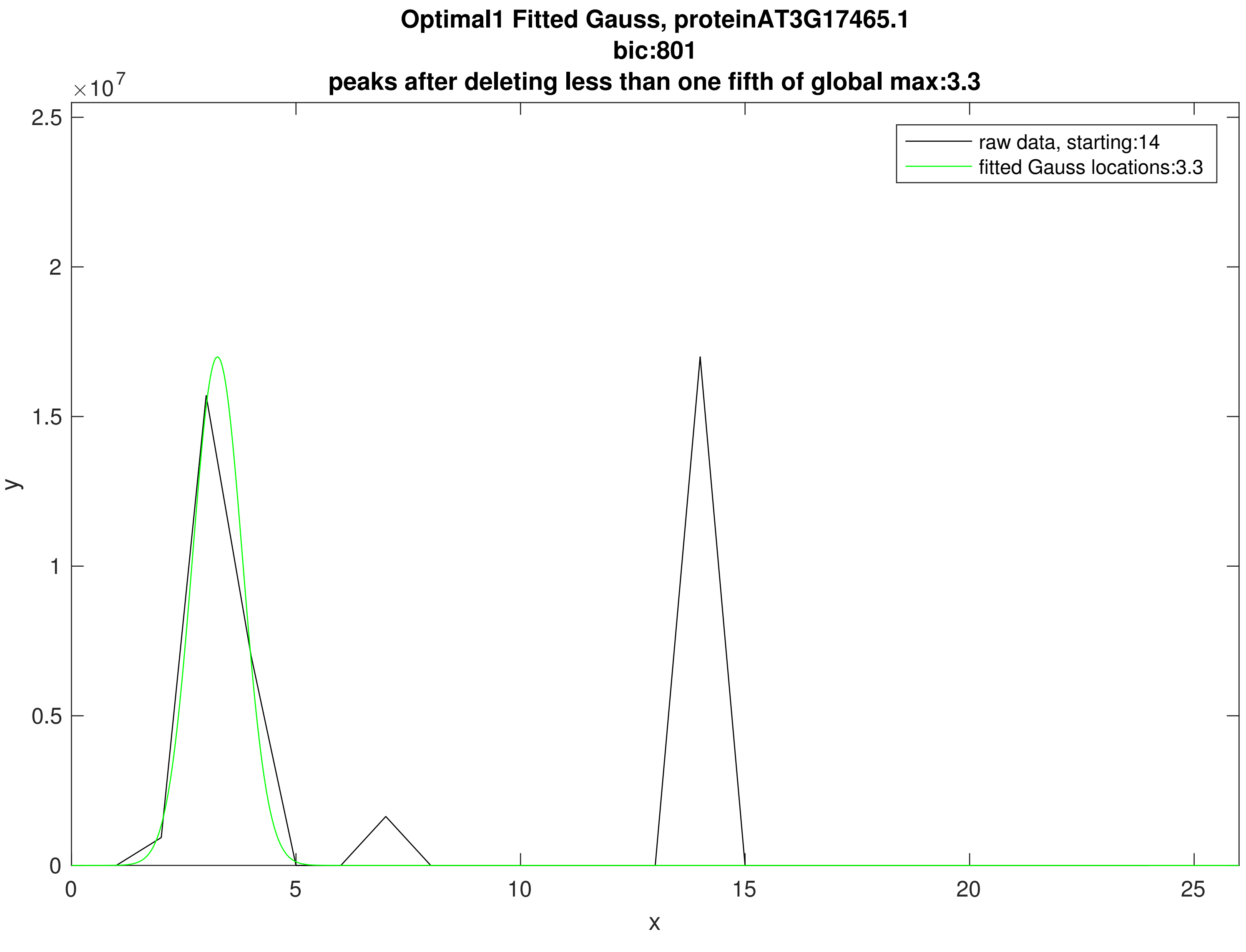
Optimal1 Fitted Gauss, proteinAT3G17465.1
bic:801
peaks after deleting less than one fifth of global max:3.3
×10^7
2.5
2
1.5
1
0.5
0
y
0
5
10
15
20
25
x
raw data, starting:14
fitted Gauss locations:3.3

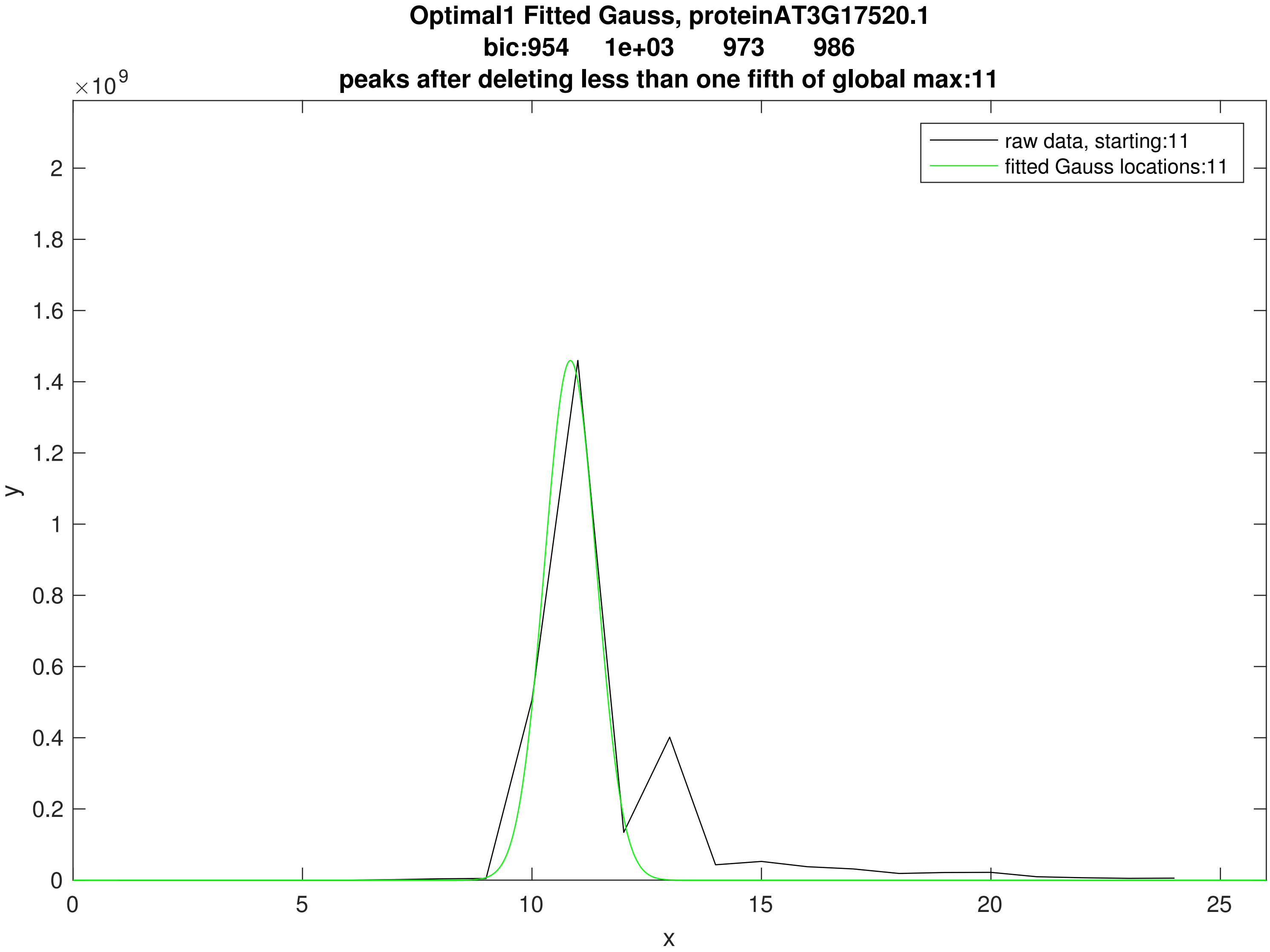
Optimal1 Fitted Gauss, proteinAT3G17520.1
bic:954 1e+03 973 986
peaks after deleting less than one fifth of global max:11
×10^9
raw data, starting:11
fitted Gauss locations:11
y
x
0
0.2
0.4
0.6
0.8
1
1.2
1.4
1.6
1.8
2
0
5
10
15
20
25

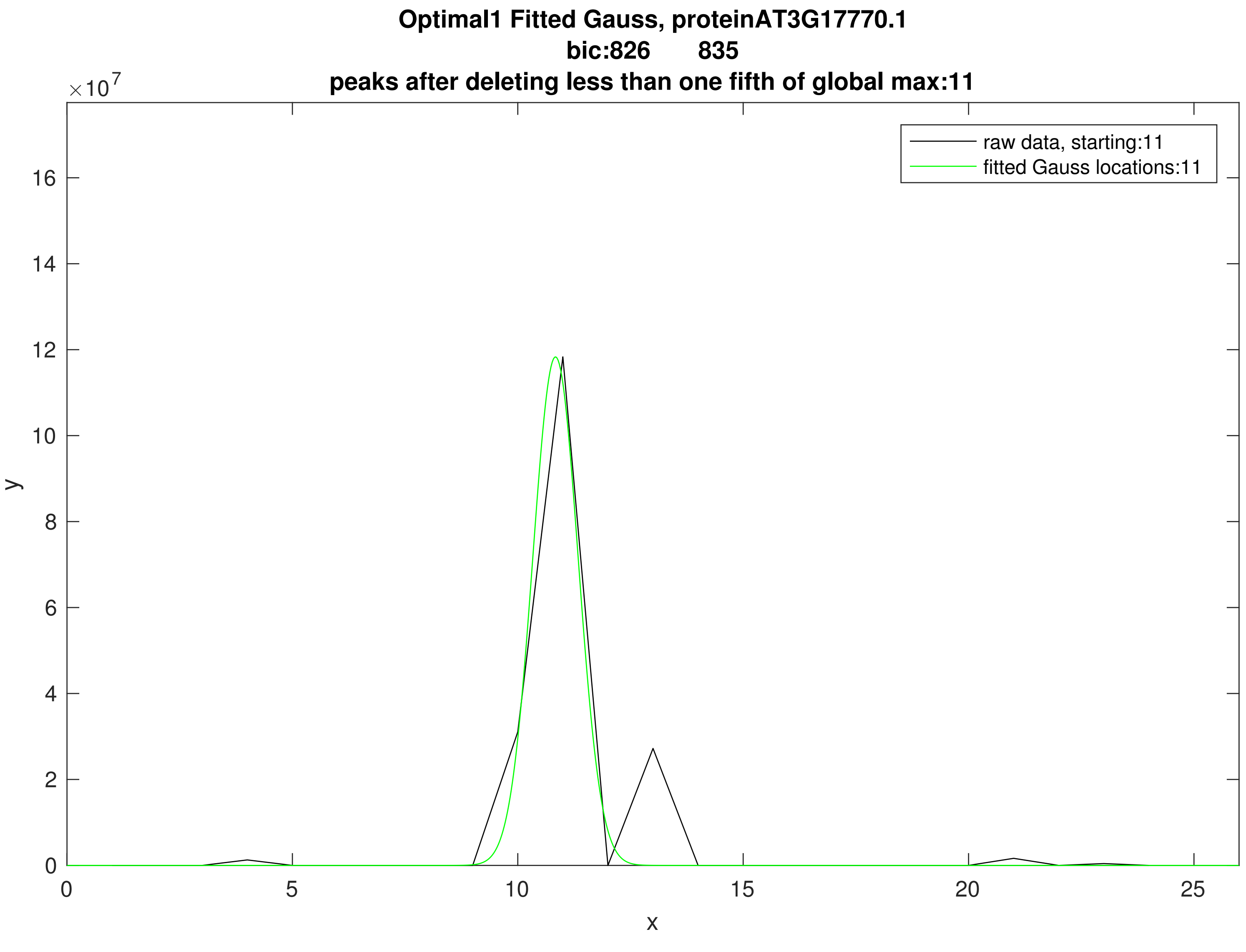

Optimal1 Fitted Gauss, proteinAT3G17770.1
bic:826 835
peaks after deleting less than one fifth of global max:11
×10^7
16
14
12
10
8
6
4
2
0
y
0
5
10
15
20
25
x
raw data, starting:11
fitted Gauss locations:11

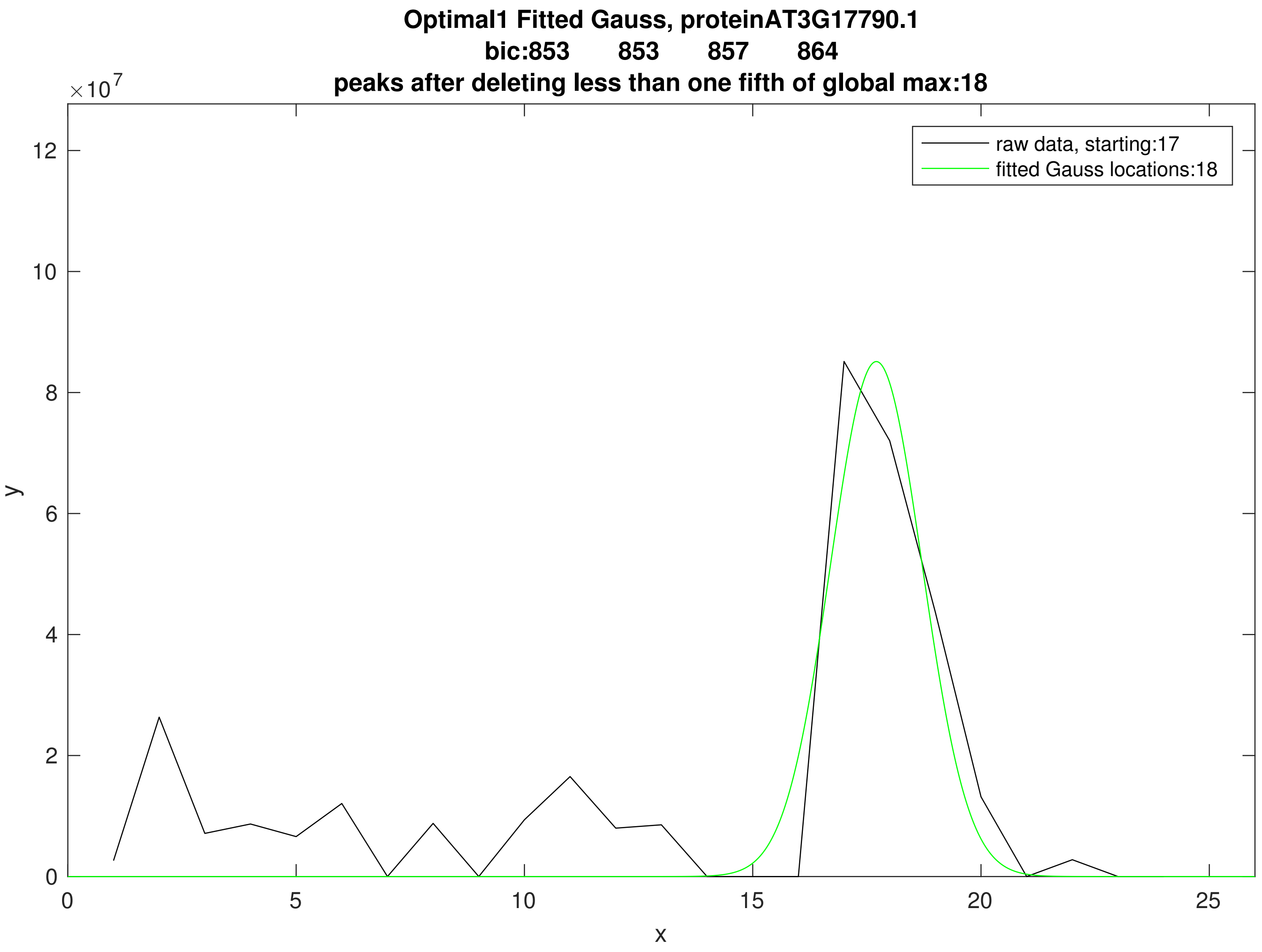

Optimal1 Fitted Gauss, proteinAT3G17790.1
bic:853 853 857 864
peaks after deleting less than one fifth of global max:18
×10^7
raw data, starting:17
fitted Gauss locations:18
y
x
0
2
4
6
8
10
12
0
5
10
15
20
25

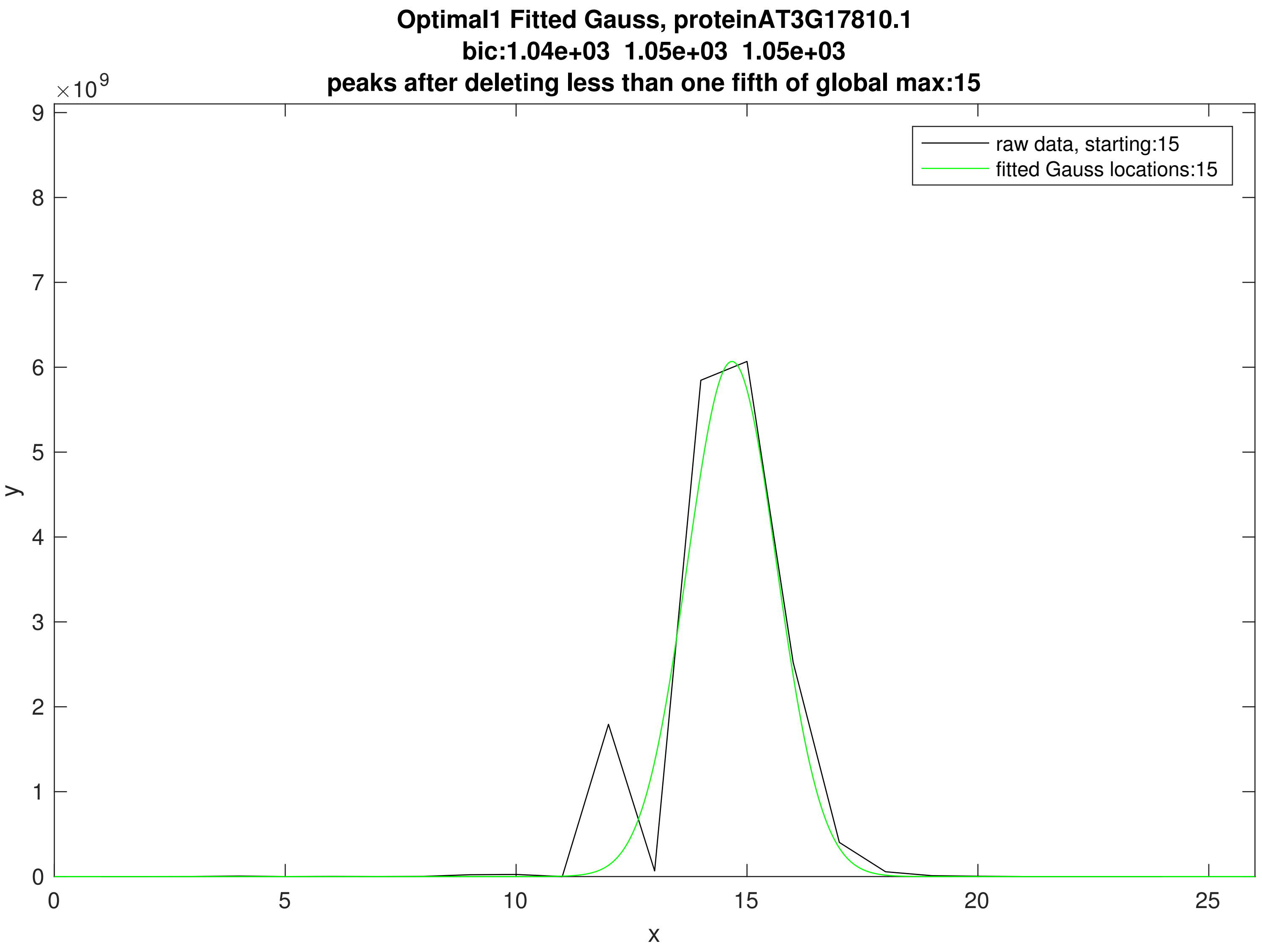

Optimal1 Fitted Gauss, proteinAT3G17810.1
bic:1.04e+03 1.05e+03 1.05e+03
peaks after deleting less than one fifth of global max:15
×10^9
raw data, starting:15
fitted Gauss locations:15
y
x
0
1
2
3
4
5
6
7
8
9
0
5
10
15
20
25

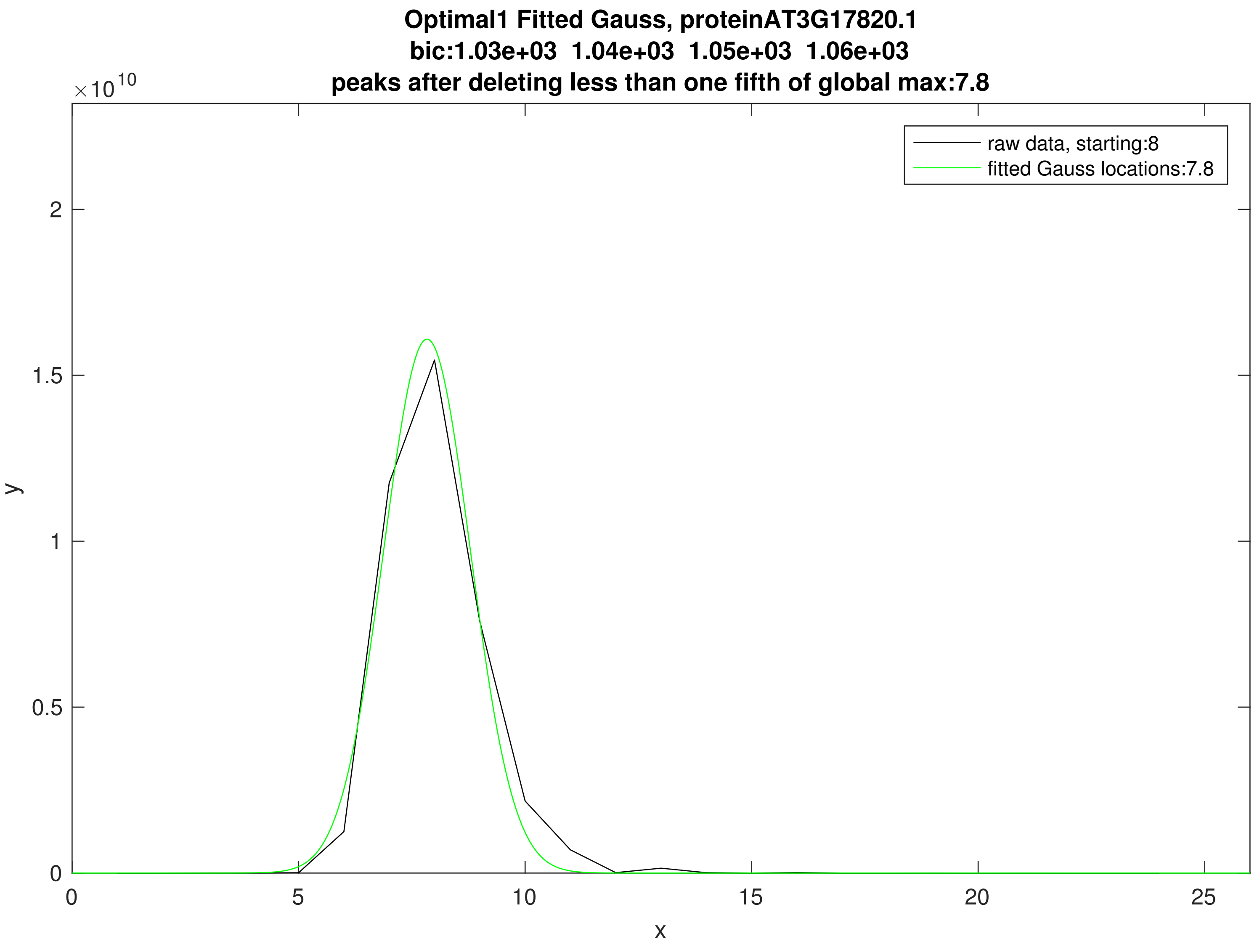

Optimal1 Fitted Gauss, proteinAT3G17820.1
bic:1.03e+03 1.04e+03 1.05e+03 1.06e+03
peaks after deleting less than one fifth of global max:7.8
×10^10
raw data, starting:8
fitted Gauss locations:7.8
y
x
0
0.5
1
1.5
2
5
10
15
20
25

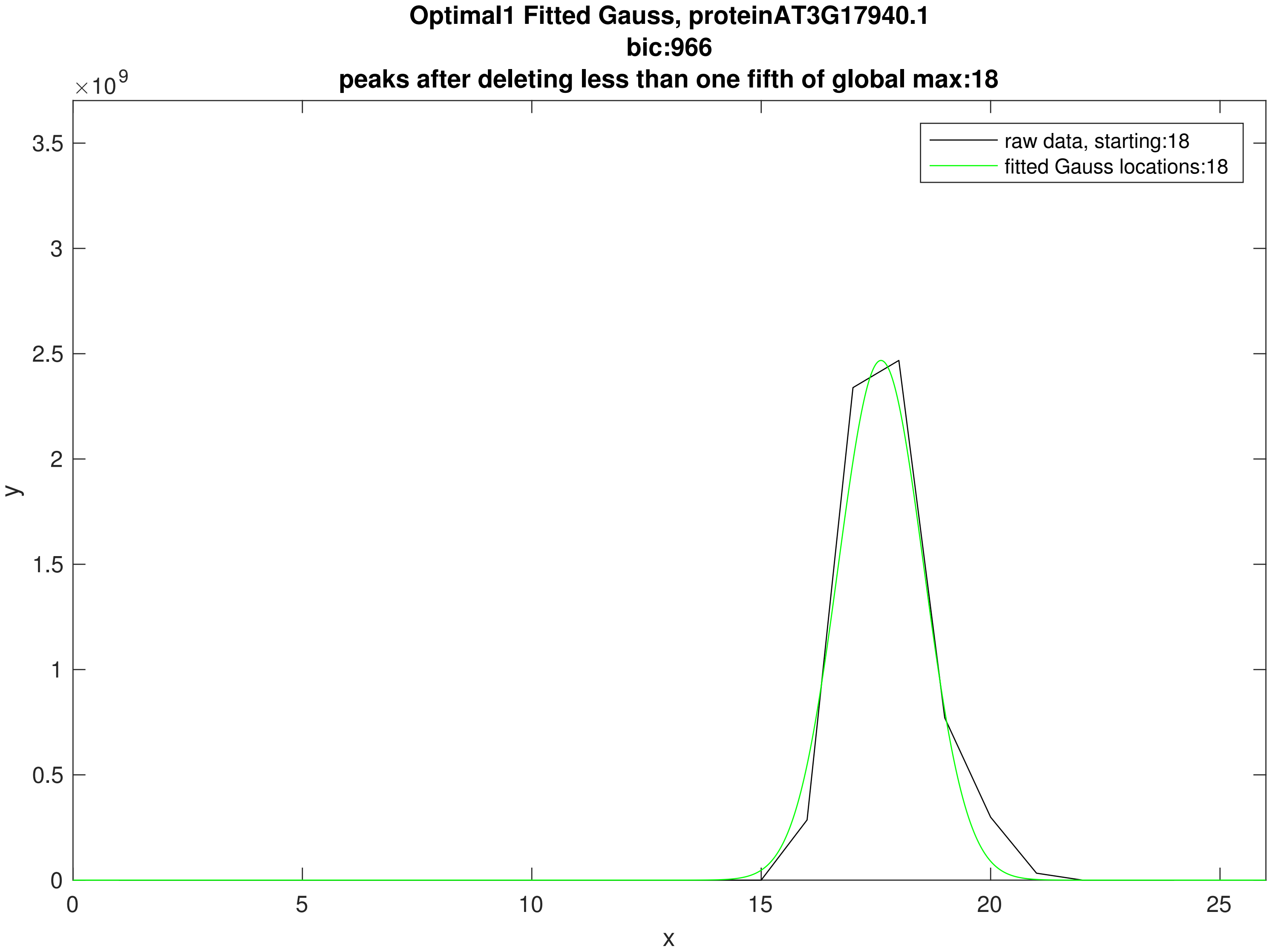
Optimal1 Fitted Gauss, proteinAT3G17940.1
bic:966
peaks after deleting less than one fifth of global max:18
×10^9
raw data, starting:18
fitted Gauss locations:18
y
x
0
0.5
1
1.5
2
2.5
3
3.5
0
5
10
15
20
25

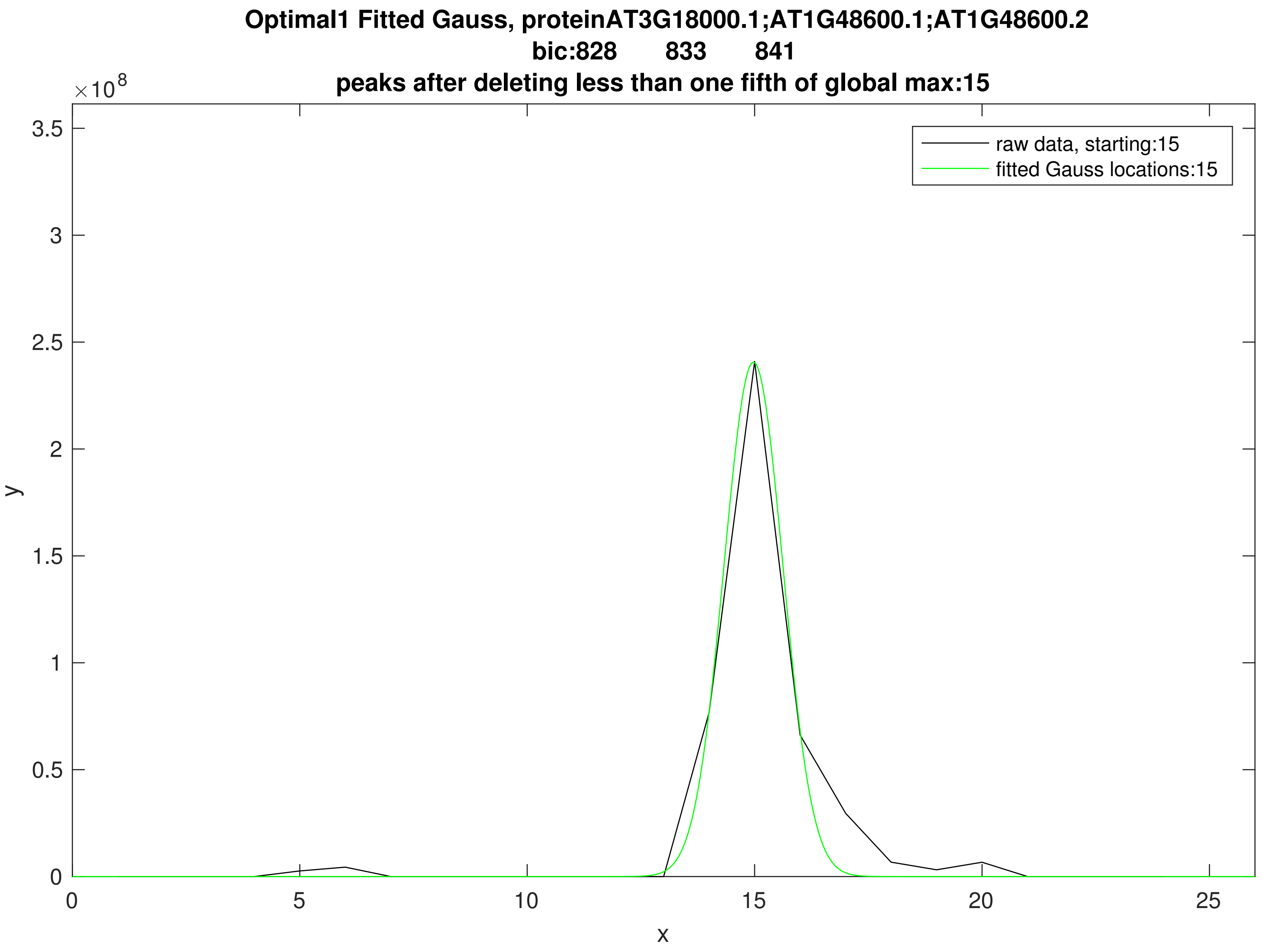

Optimal1 Fitted Gauss, proteinAT3G18000.1;AT1G48600.1;AT1G48600.2
bic:828 833 841
peaks after deleting less than one fifth of global max:15
×10^8
3.5
3
2.5
2
1.5
1
0.5
0
y
0
5
10
15
20
25
x
raw data, starting:15
fitted Gauss locations:15

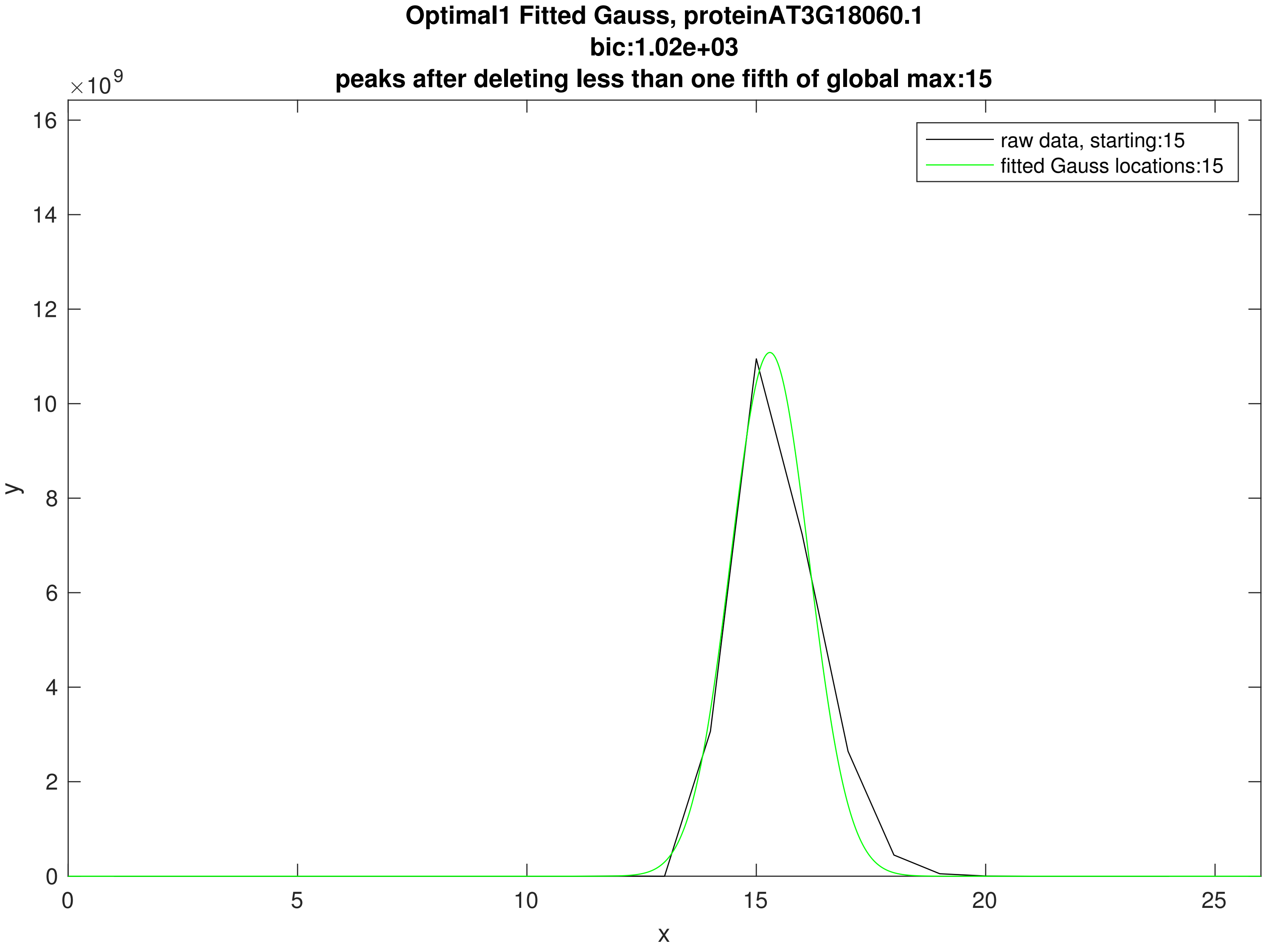
Optimal1 Fitted Gauss, proteinAT3G18060.1
bic:1.02e+03
peaks after deleting less than one fifth of global max:15
×10^9
raw data, starting:15
fitted Gauss locations:15
y
x
0
2
4
6
8
10
12
14
16
0
5
10
15
20
25

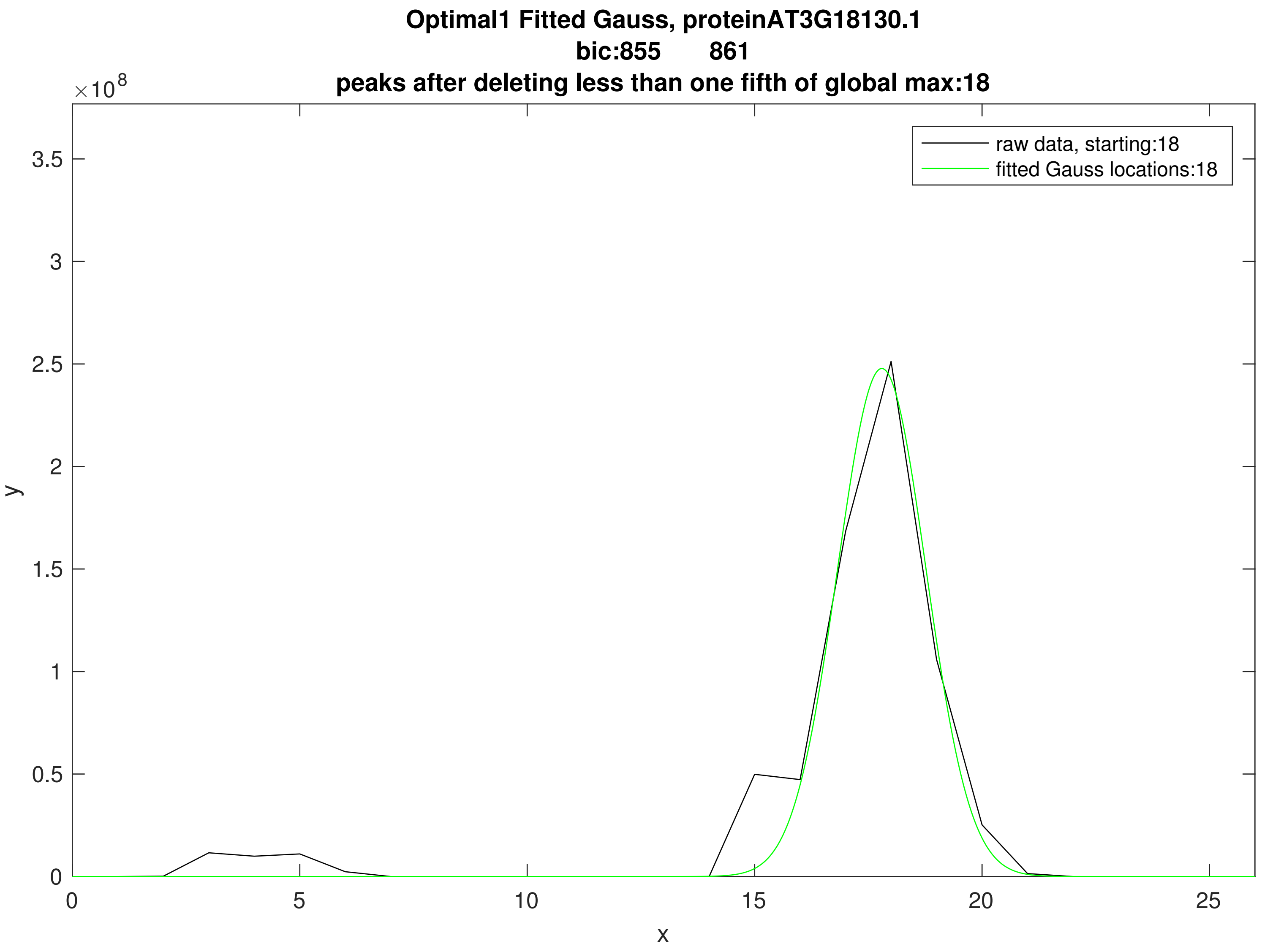

Optimal1 Fitted Gauss, proteinAT3G18130.1
bic:855 861
peaks after deleting less than one fifth of global max:18
×10^8
3.5
3
2.5
2
1.5
1
0.5
0
y
0
5
10
15
20
25
x
raw data, starting:18
fitted Gauss locations:18

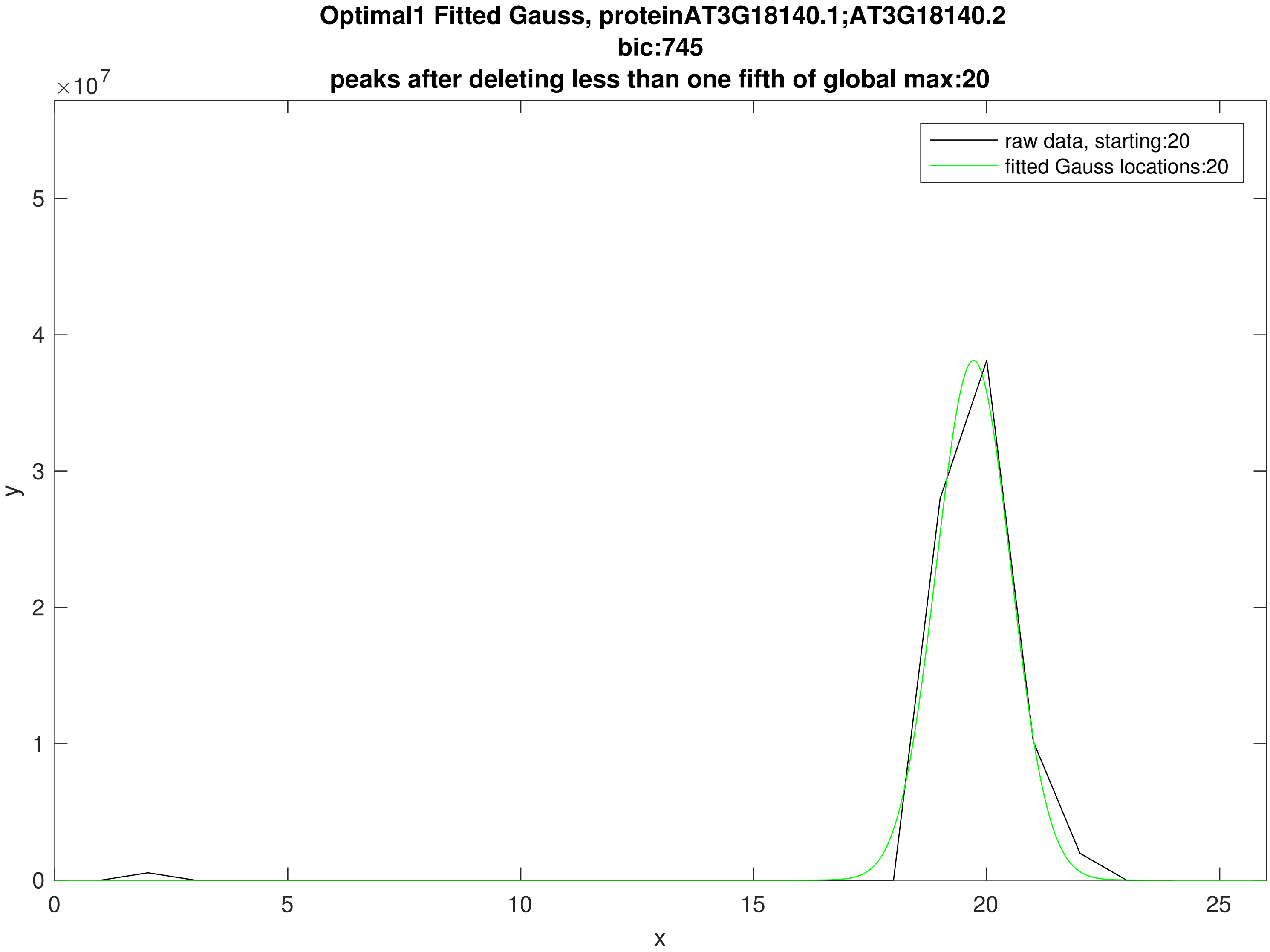

Optimal1 Fitted Gauss, proteinAT3G18140.1;AT3G18140.2
bic:745
peaks after deleting less than one fifth of global max:20
×10^7
raw data, starting:20
fitted Gauss locations:20
y
x
0
1
2
3
4
5
0
5
10
15
20
25

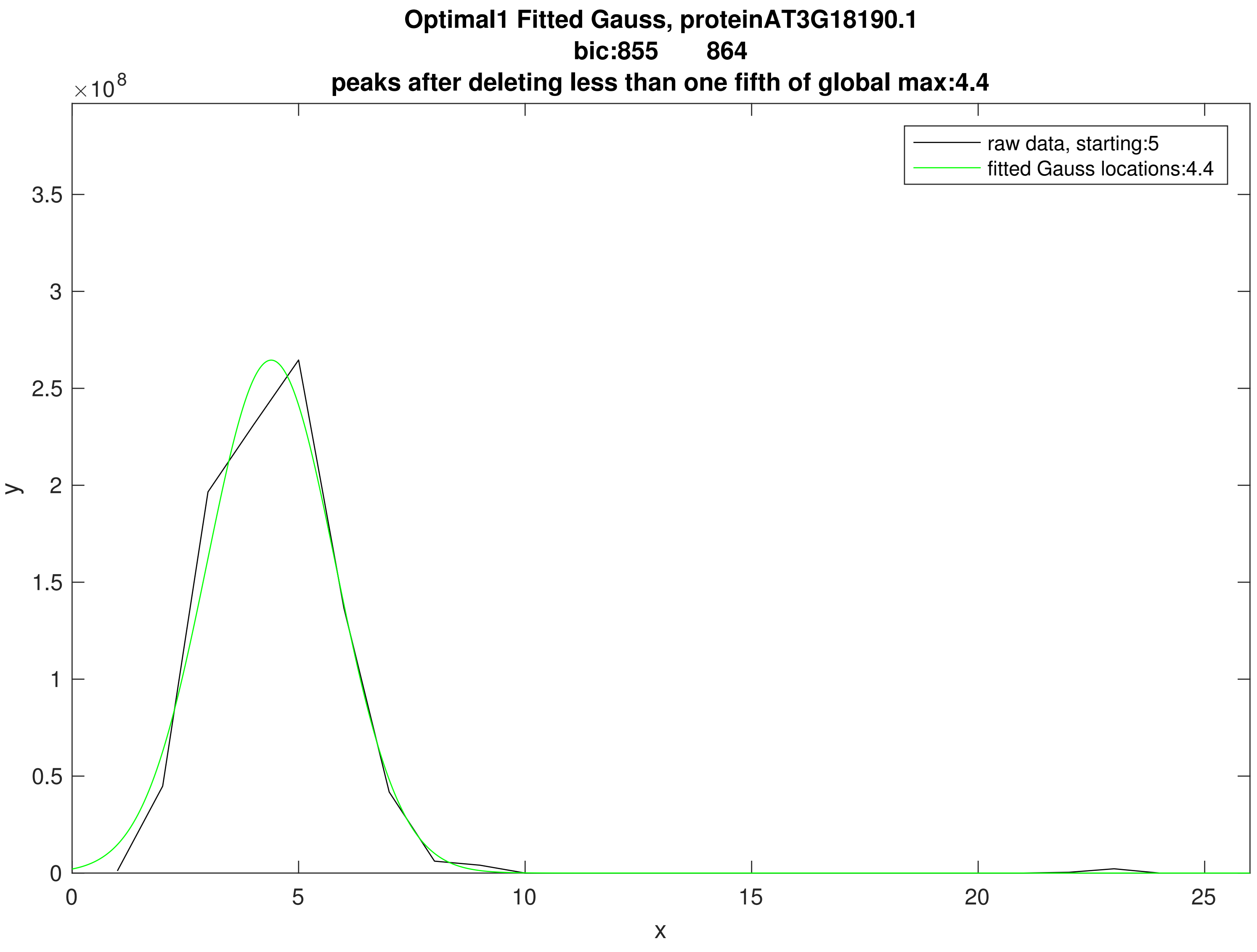

Optimal1 Fitted Gauss, proteinAT3G18190.1
bic:855 864
peaks after deleting less than one fifth of global max:4.4
×10^8
raw data, starting:5
fitted Gauss locations:4.4
3.5
3
2.5
2
1.5
1
0.5
0
y
0
5
10
15
20
25
x

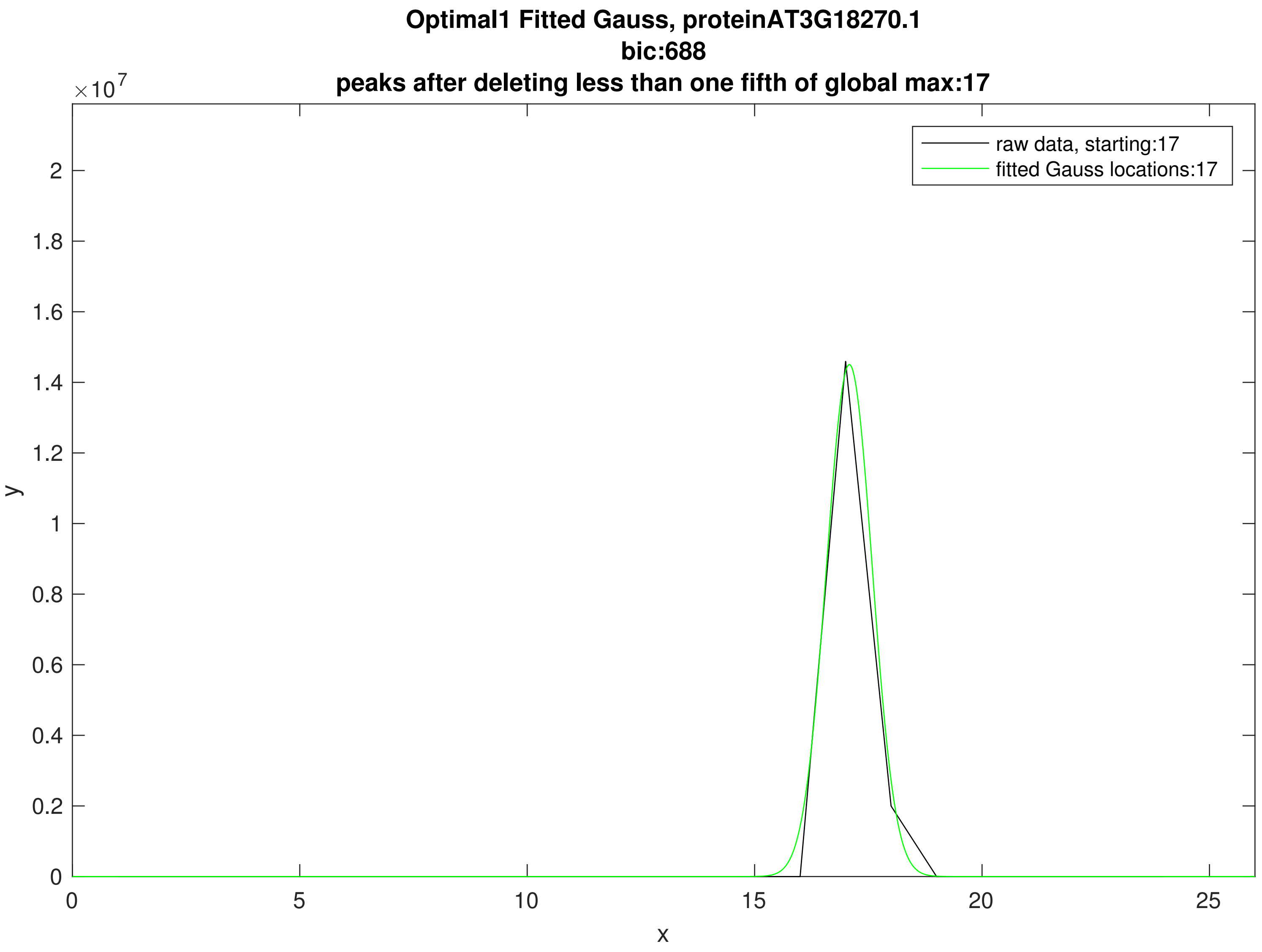
Optimal1 Fitted Gauss, proteinAT3G18270.1
bic:688
peaks after deleting less than one fifth of global max:17
×10^7
raw data, starting:17
fitted Gauss locations:17
y
x
0
0.2
0.4
0.6
0.8
1
1.2
1.4
1.6
1.8
2
0
5
10
15
20
25

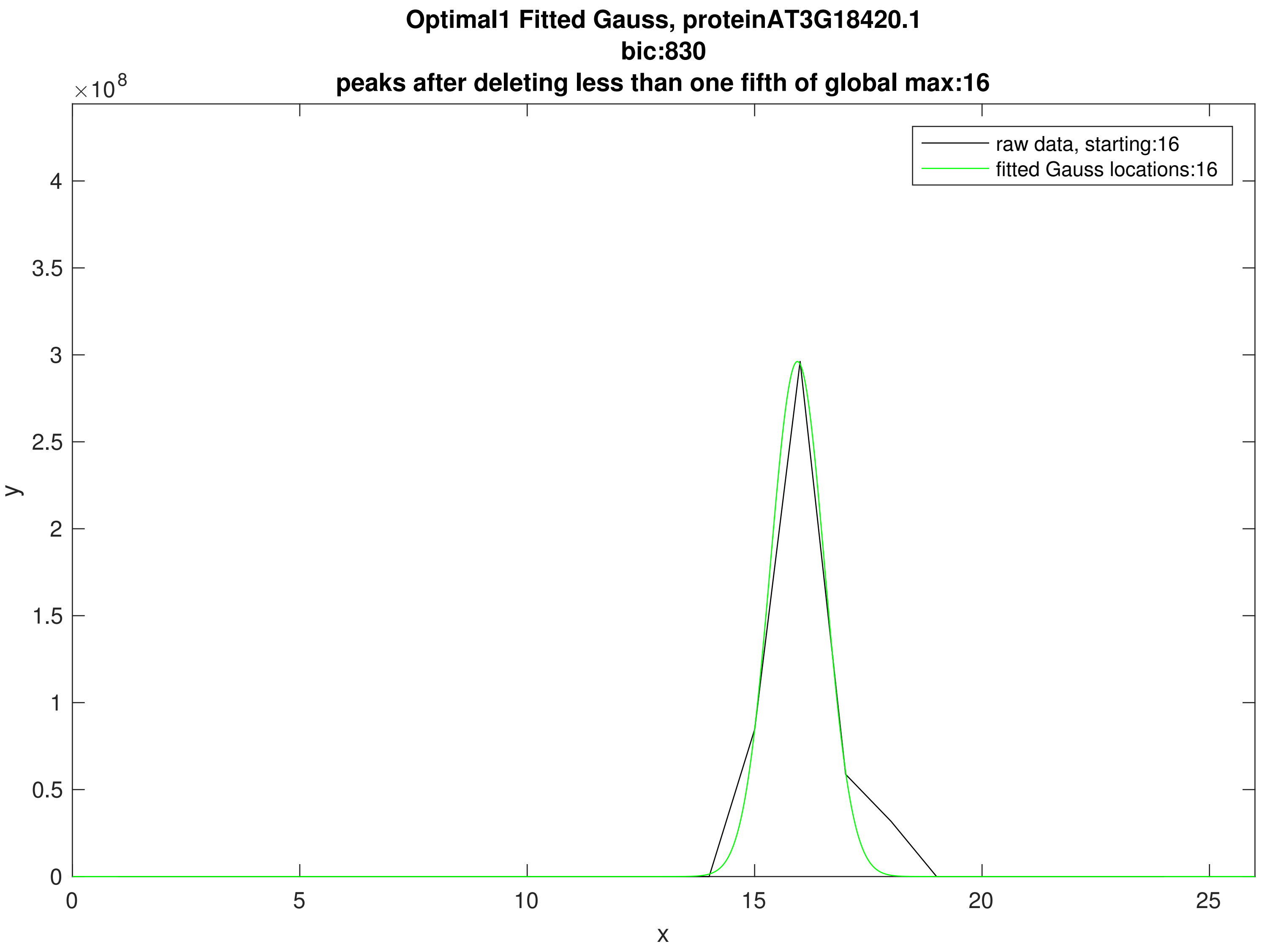

Optimal1 Fitted Gauss, proteinAT3G18420.1
bic:830
peaks after deleting less than one fifth of global max:16
×10^8
raw data, starting:16
fitted Gauss locations:16
0
0.5
1
1.5
2
2.5
3
3.5
4
y
0
5
10
15
20
25
x

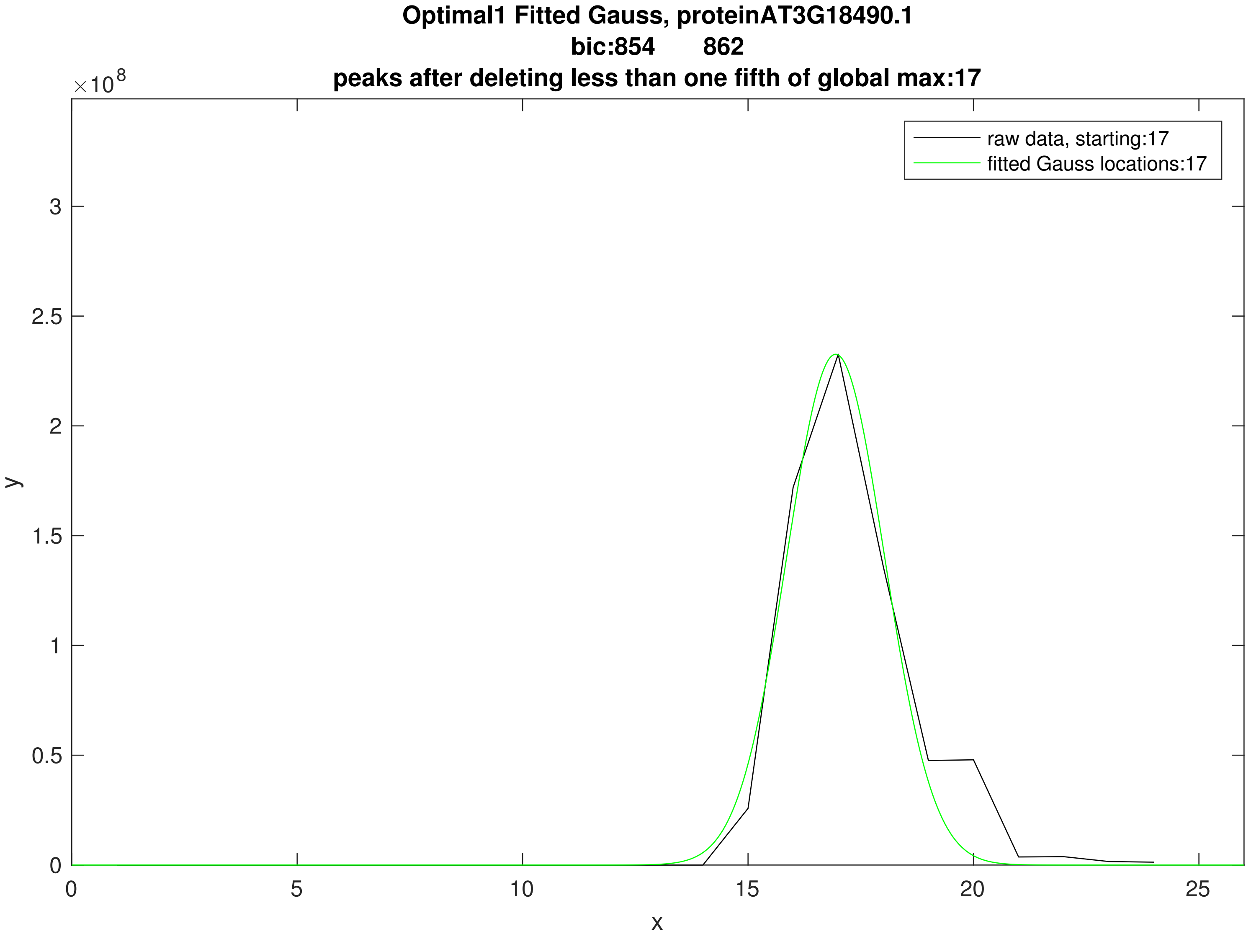
Optimal1 Fitted Gauss, proteinAT3G18490.1
bic:854 862
peaks after deleting less than one fifth of global max:17
$\times 10^8$
raw data, starting:17
fitted Gauss locations:17
y
x
0
0.5
1
1.5
2
2.5
3
0
5
10
15
20
25

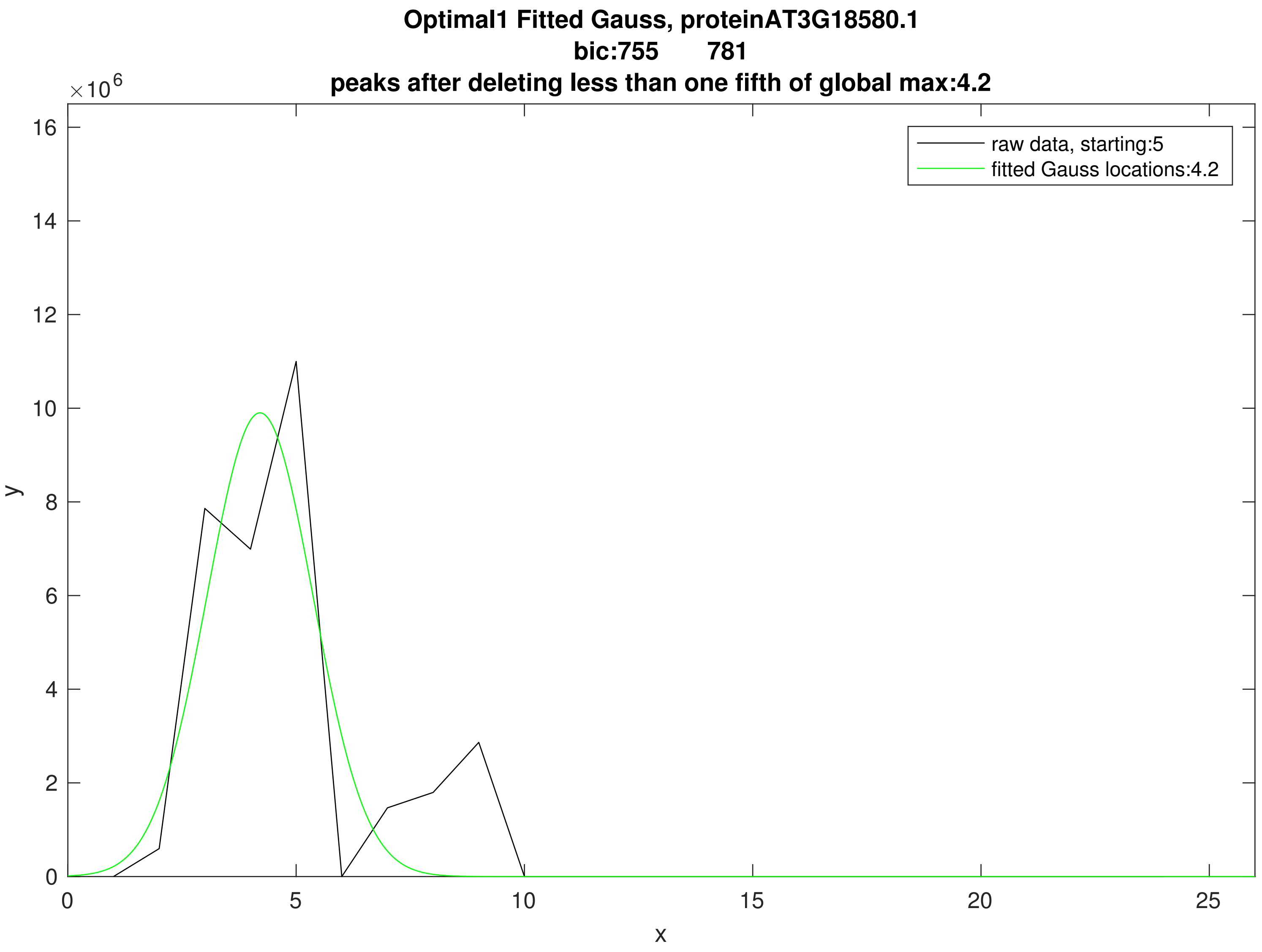
Optimal1 Fitted Gauss, proteinAT3G18580.1
bic:755 781
peaks after deleting less than one fifth of global max:4.2
×10^6
raw data, starting:5
fitted Gauss locations:4.2
y
x
0
2
4
6
8
10
12
14
16
0
5
10
15
20
25

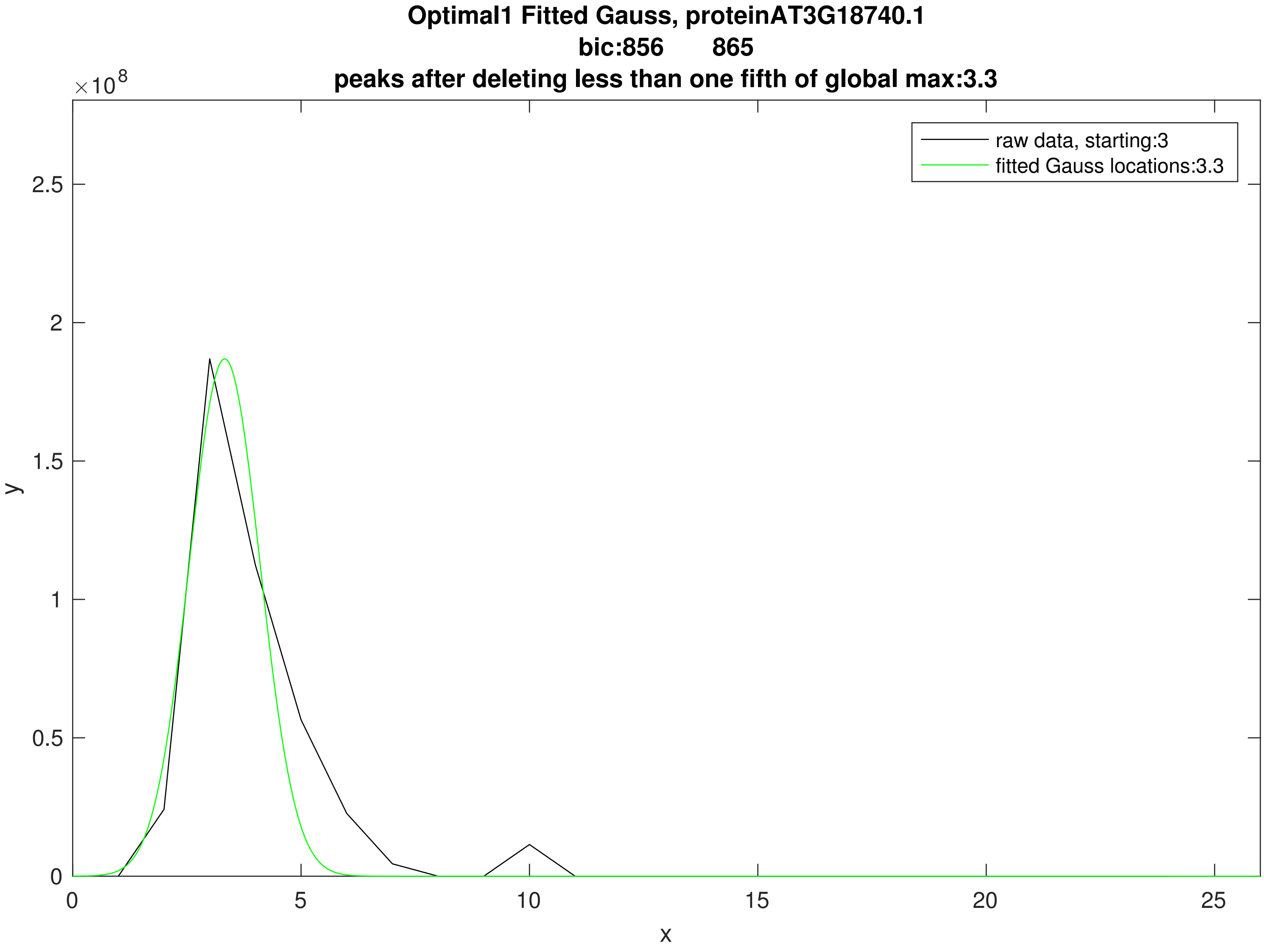

Optimal1 Fitted Gauss, proteinAT3G18740.1
bic:856 865
peaks after deleting less than one fifth of global max:3.3
×10^8
2.5
2
1.5
1
0.5
0
y
0
5
10
15
20
25
x
raw data, starting:3
fitted Gauss locations:3.3

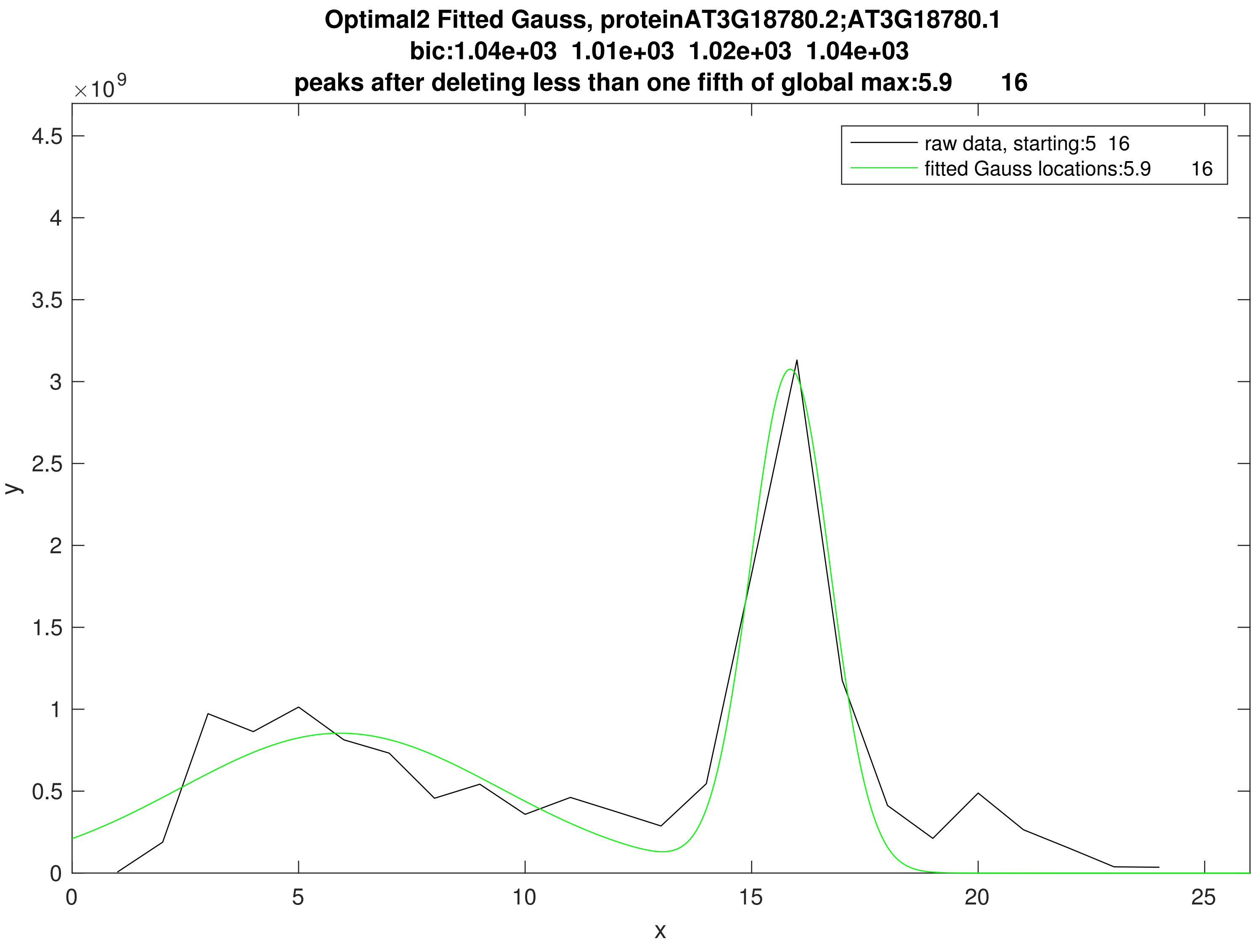
Optimal2 Fitted Gauss, proteinAT3G18780.2;AT3G18780.1
bic:1.04e+03 1.01e+03 1.02e+03 1.04e+03
peaks after deleting less than one fifth of global max:5.9 16
×10^9
raw data, starting:5 16
fitted Gauss locations:5.9 16
y
x
0
0.5
1
1.5
2
2.5
3
3.5
4
4.5
5
10
15
20
25

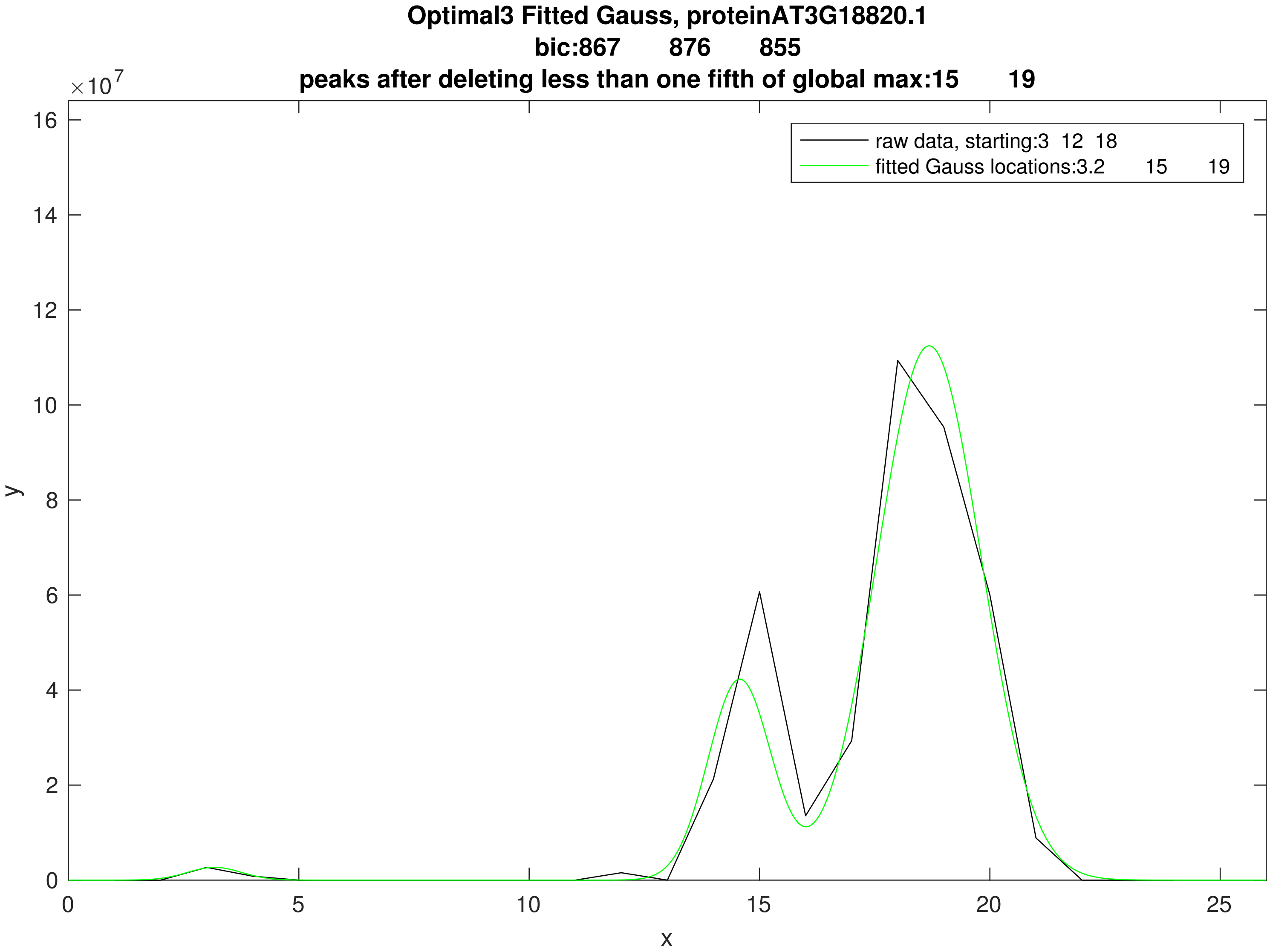
Optimal3 Fitted Gauss, proteinAT3G18820.1
bic:867 876 855
peaks after deleting less than one fifth of global max:15 19
×10^7
raw data, starting:3 12 18
fitted Gauss locations:3.2 15 19
y
x
0
2
4
6
8
10
12
14
16
0
5
10
15
20
25

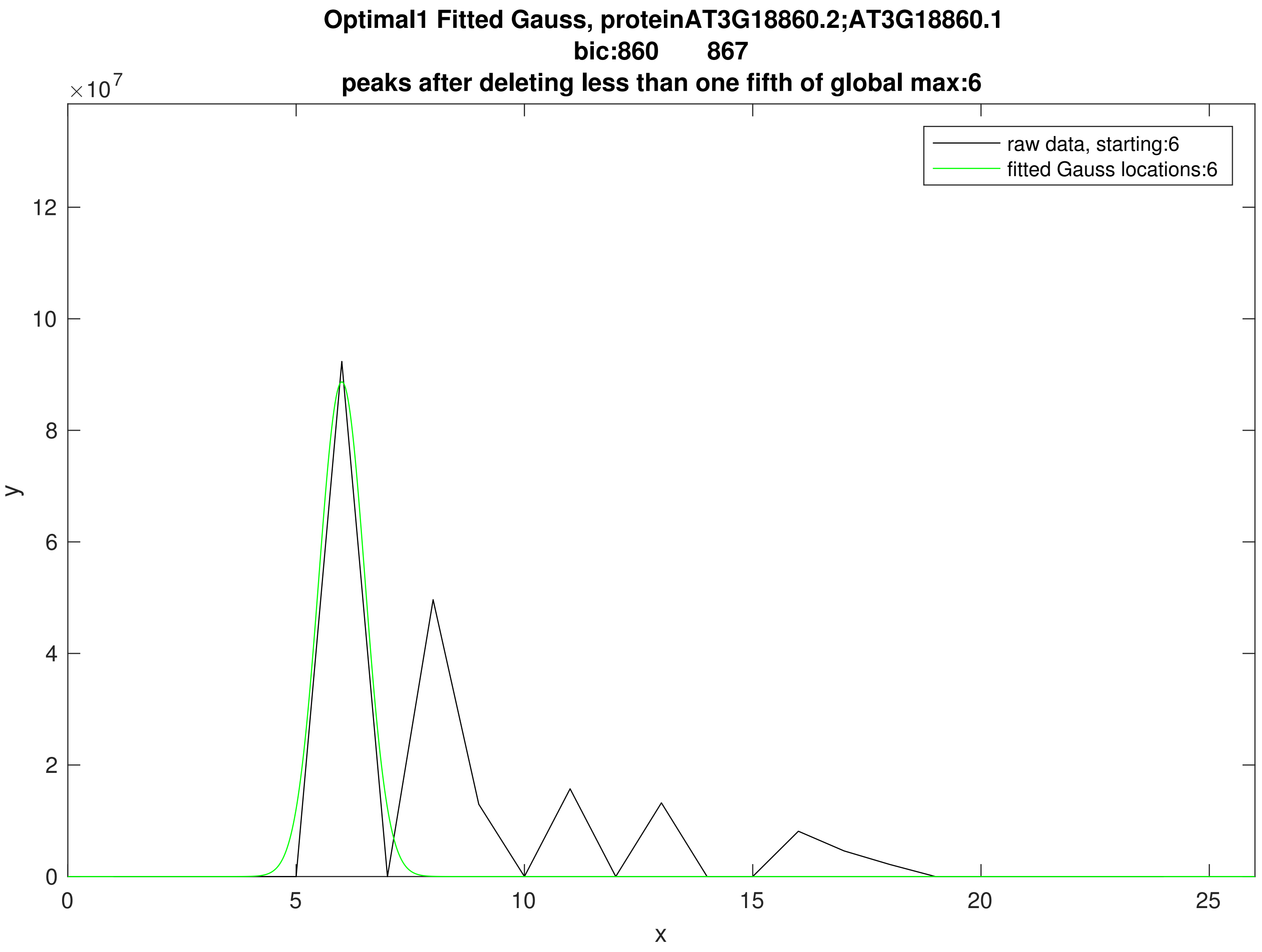
Optimal1 Fitted Gauss, proteinAT3G18860.2;AT3G18860.1
bic:860 867
peaks after deleting less than one fifth of global max:6
×10^7
raw data, starting:6
fitted Gauss locations:6
y
x
0
2
4
6
8
10
12
0
5
10
15
20
25

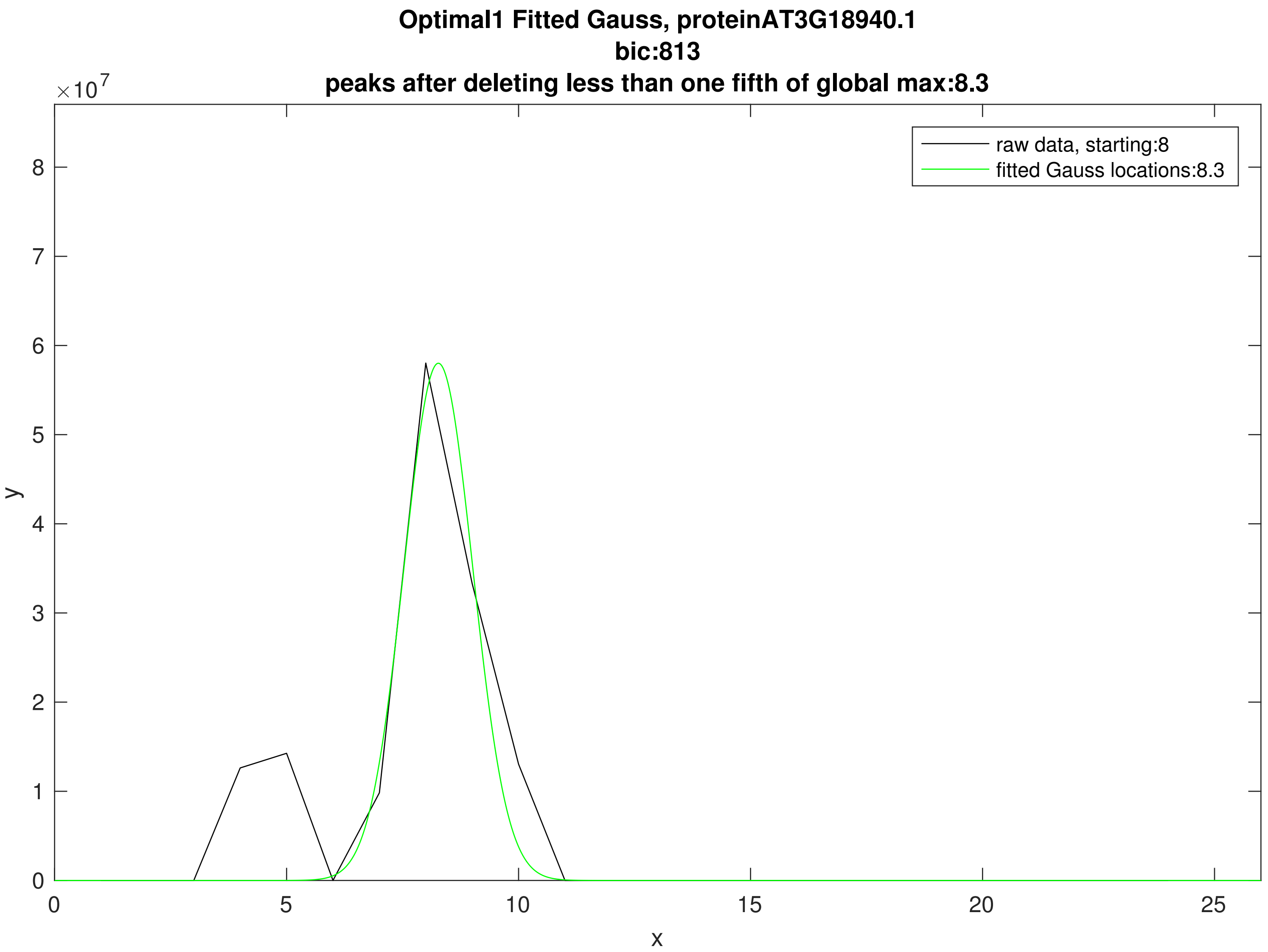
Optimal1 Fitted Gauss, proteinAT3G18940.1
bic:813
peaks after deleting less than one fifth of global max:8.3
×10^7
raw data, starting:8
fitted Gauss locations:8.3
y
x
0
1
2
3
4
5
6
7
8
0
5
10
15
20
25

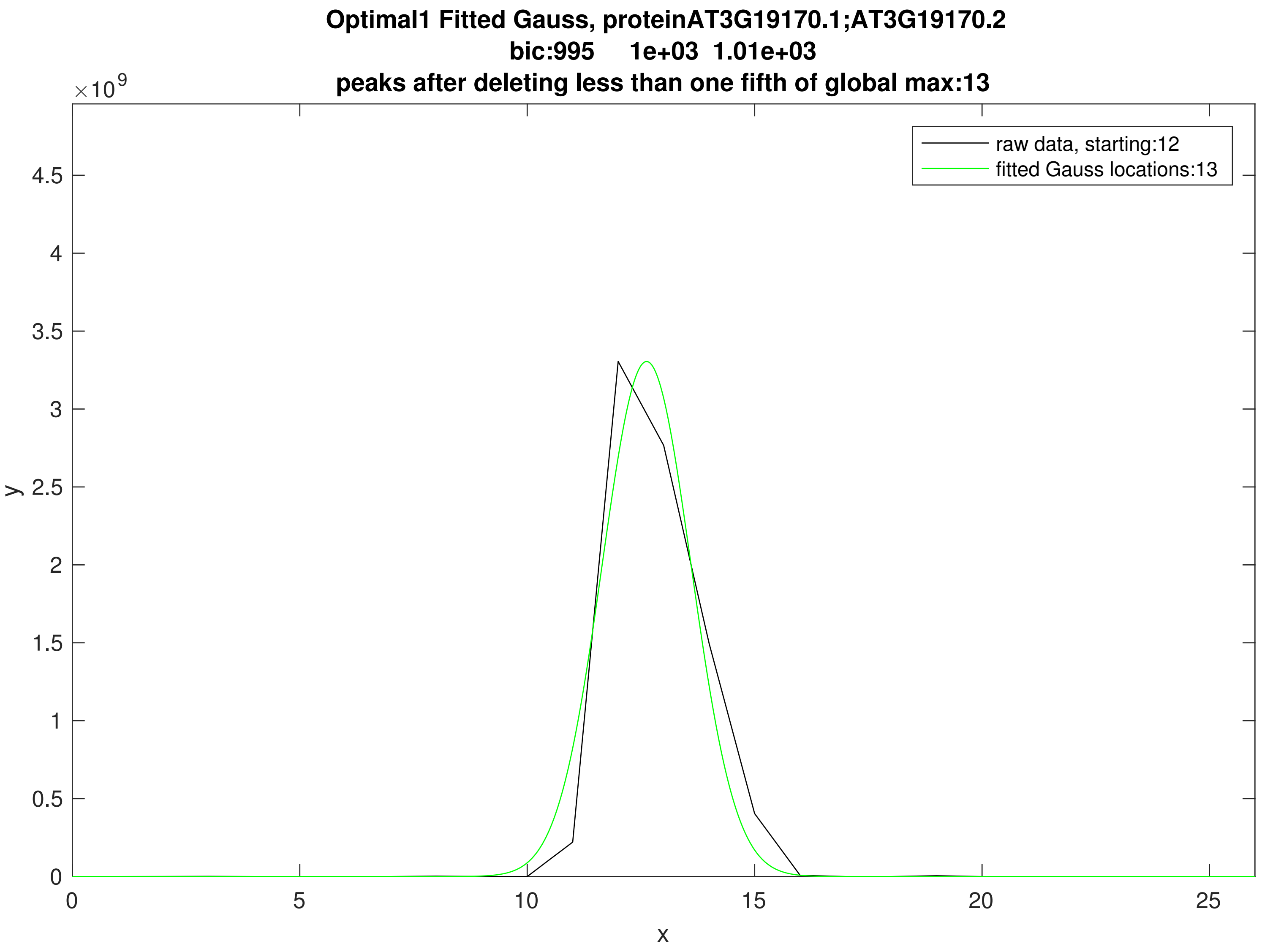

Optimal1 Fitted Gauss, proteinAT3G19170.1;AT3G19170.2
bic:995 1e+03 1.01e+03
peaks after deleting less than one fifth of global max:13
×10^9
raw data, starting:12
fitted Gauss locations:13
y
x
0
0.5
1
1.5
2
2.5
3
3.5
4
4.5
5
10
15
20
25

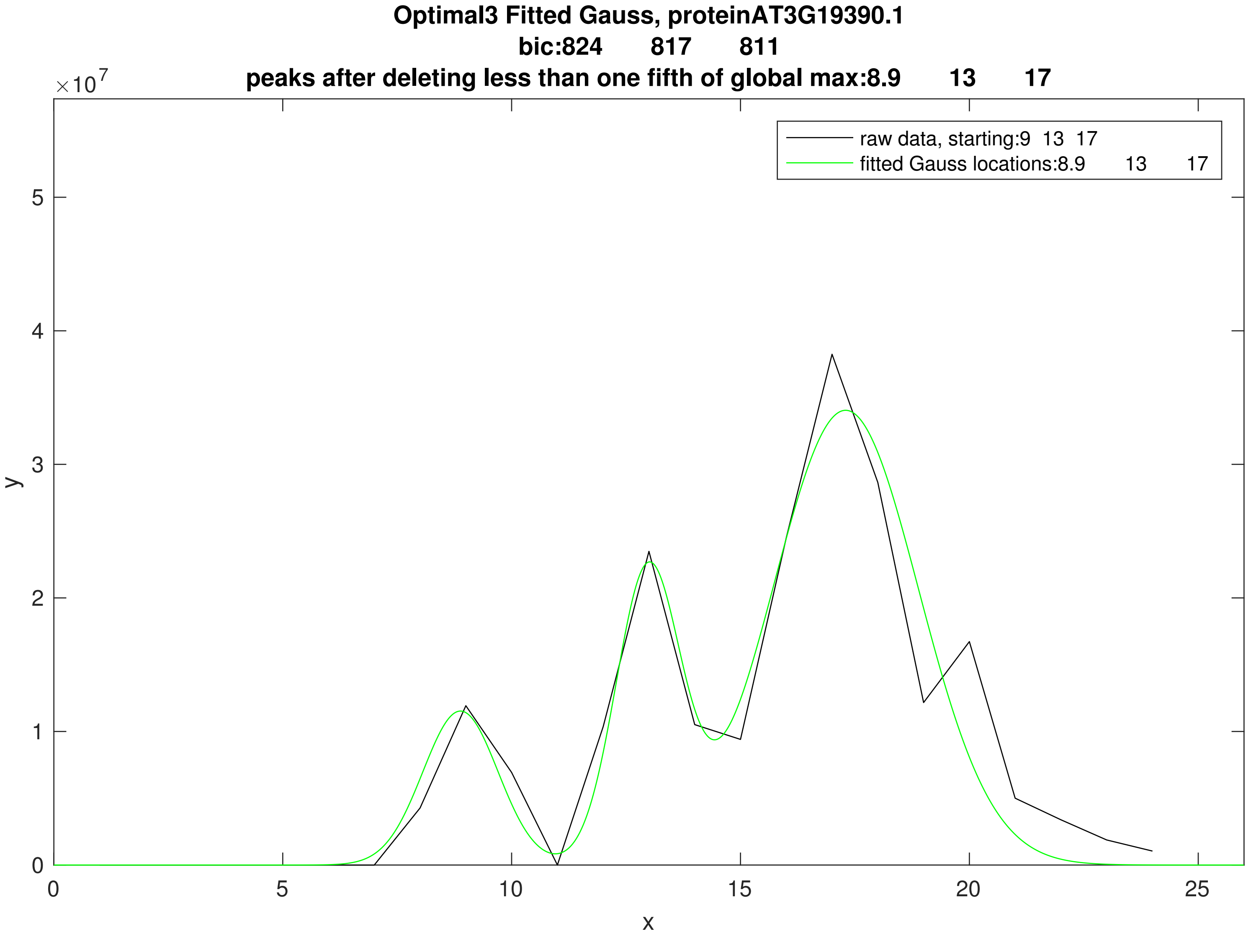
Optimal3 Fitted Gauss, proteinAT3G19390.1
bic:824 817 811
peaks after deleting less than one fifth of global max:8.9 13 17
×10^7
raw data, starting:9 13 17
fitted Gauss locations:8.9 13 17
y
x
0
1
2
3
4
5
0
5
10
15
20
25

Optimal1 Fitted Gauss, proteinAT3G19400.1;AT3G19400.2
bic:854 858
peaks after deleting less than one fifth of global max:17

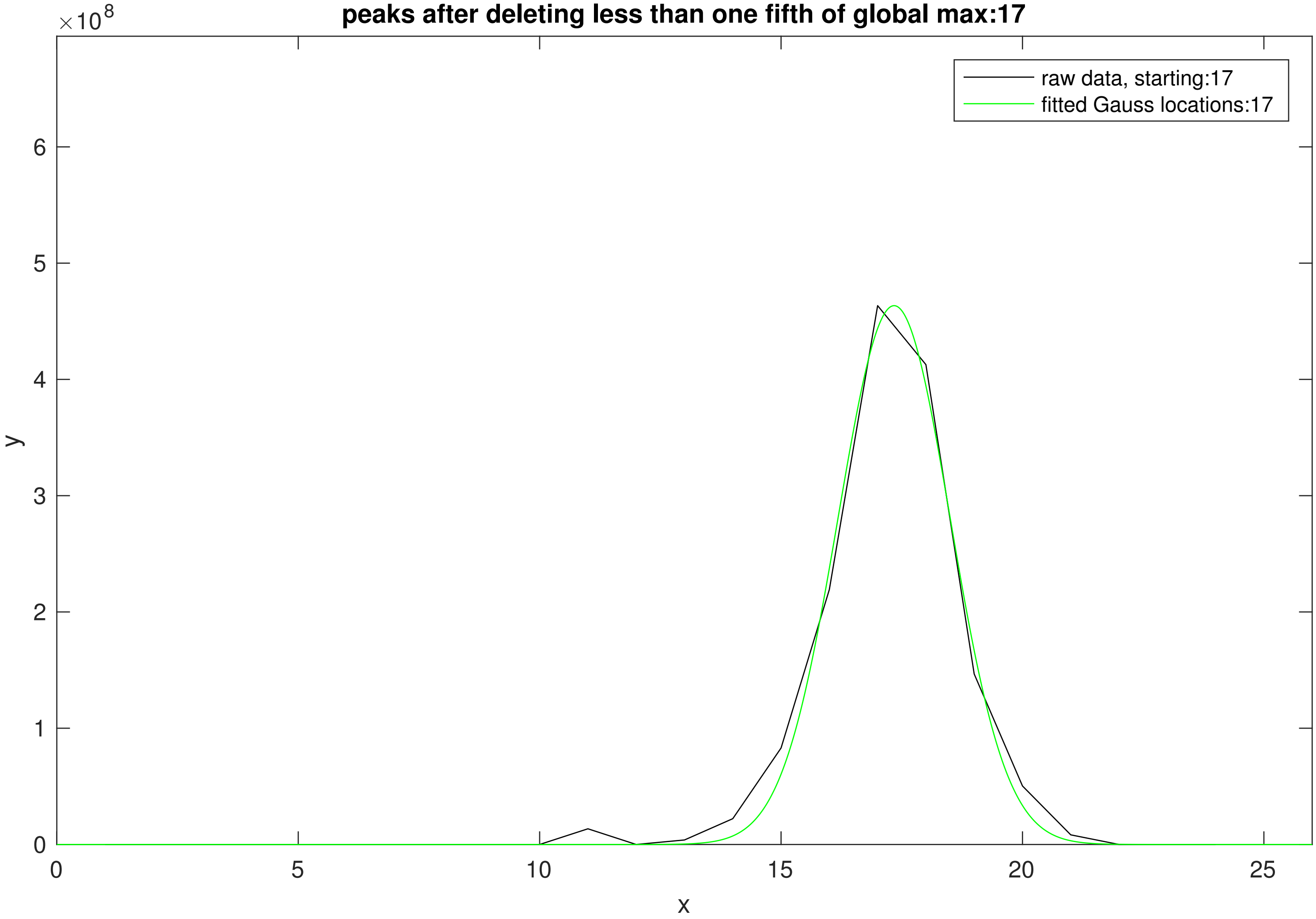

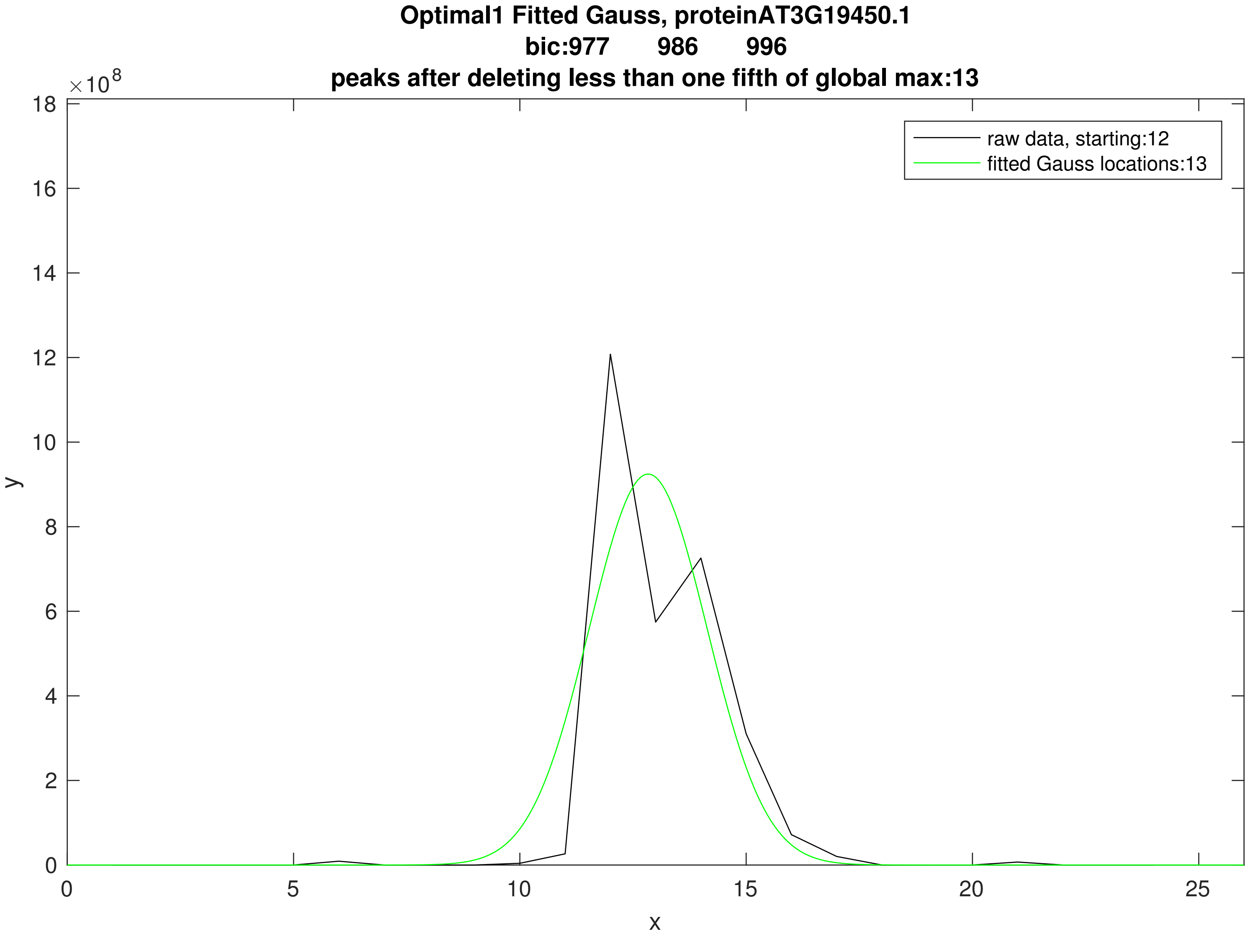

Optimal1 Fitted Gauss, proteinAT3G19450.1
bic:977 986 996
peaks after deleting less than one fifth of global max:13
×10^8
18
16
14
12
10
8
6
4
2
0
y
raw data, starting:12
fitted Gauss locations:13
0
5
10
15
20
25
x

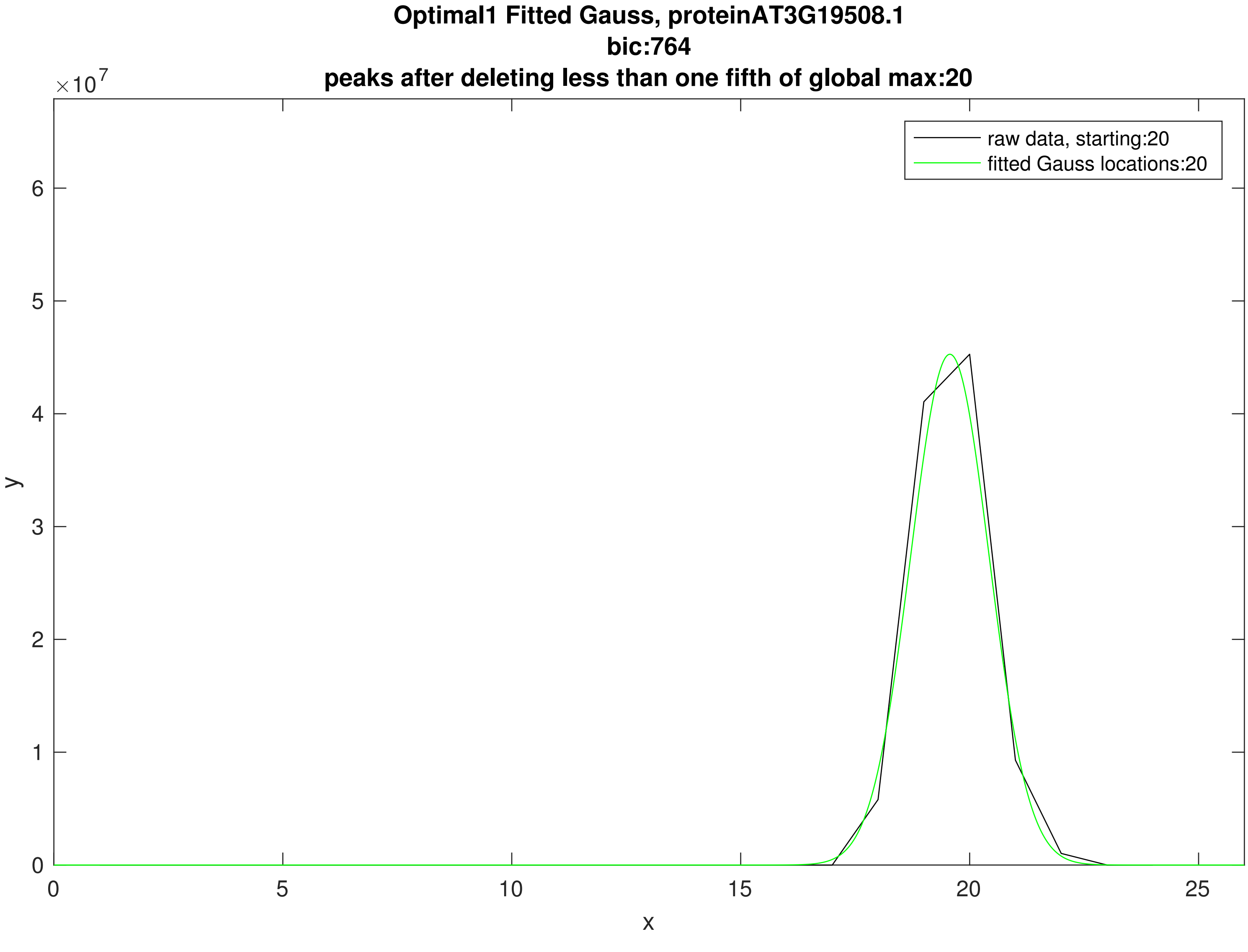
Optimal1 Fitted Gauss, proteinAT3G19508.1
bic:764
peaks after deleting less than one fifth of global max:20
×10^7
raw data, starting:20
fitted Gauss locations:20
y
x
0
1
2
3
4
5
6
0
5
10
15
20
25

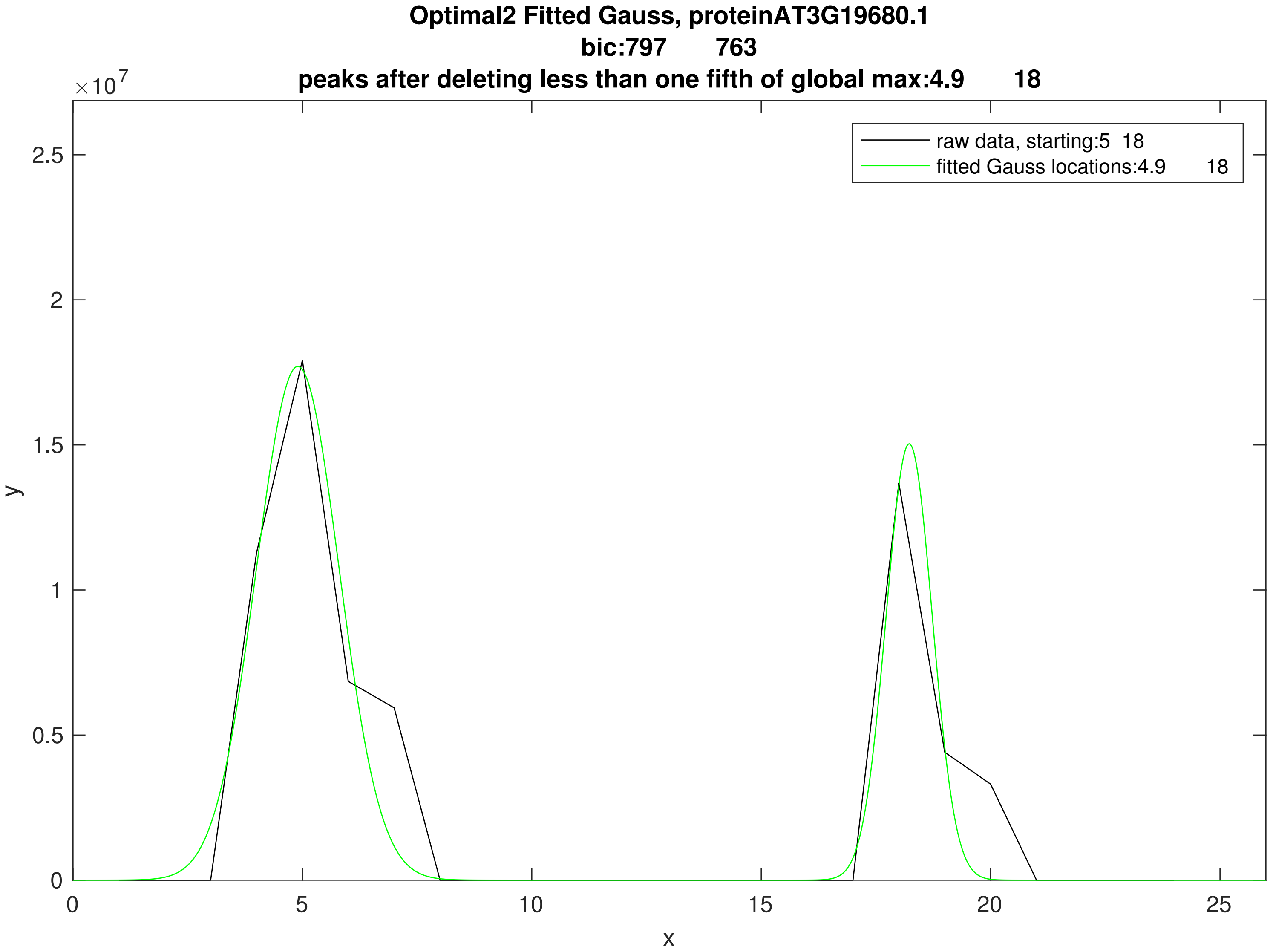

Optimal2 Fitted Gauss, proteinAT3G19680.1
bic:797 763
peaks after deleting less than one fifth of global max:4.9 18
×10^7
2.5
2
1.5
1
0.5
0
y
0
5
10
15
20
25
x
raw data, starting:5 18
fitted Gauss locations:4.9 18

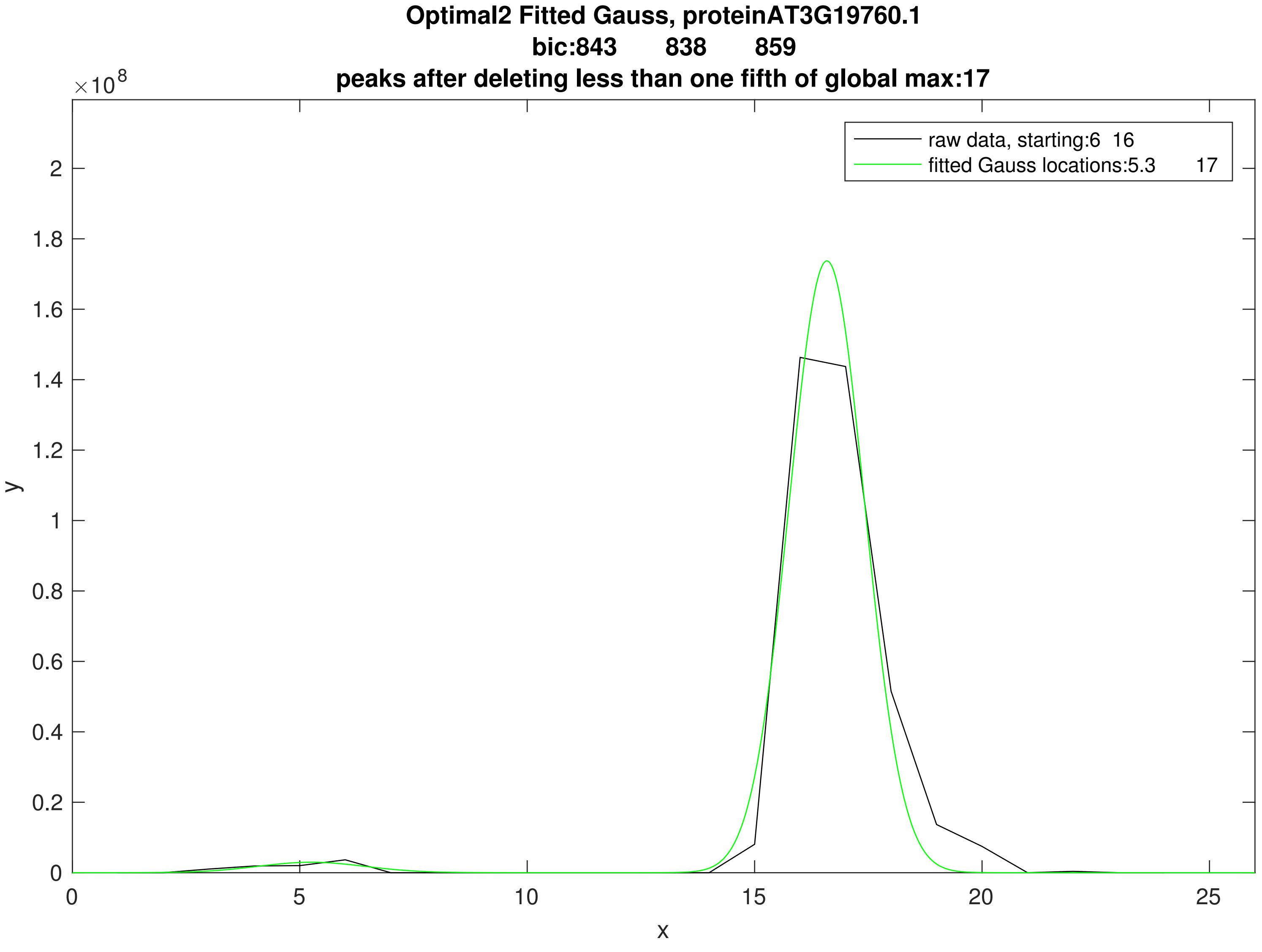

Optimal2 Fitted Gauss, proteinAT3G19760.1
bic:843 838 859
peaks after deleting less than one fifth of global max:17
×10^8
raw data, starting:6 16
fitted Gauss locations:5.3 17
y
x
0
0.2
0.4
0.6
0.8
1
1.2
1.4
1.6
1.8
2
0
5
10
15
20
25

**Optimal1 Fitted Gauss, proteinAT3G19820.3;AT3G19820.2;AT3G19820.1**
**bic:651**
**peaks after deleting less than one fifth of global max:3.4**

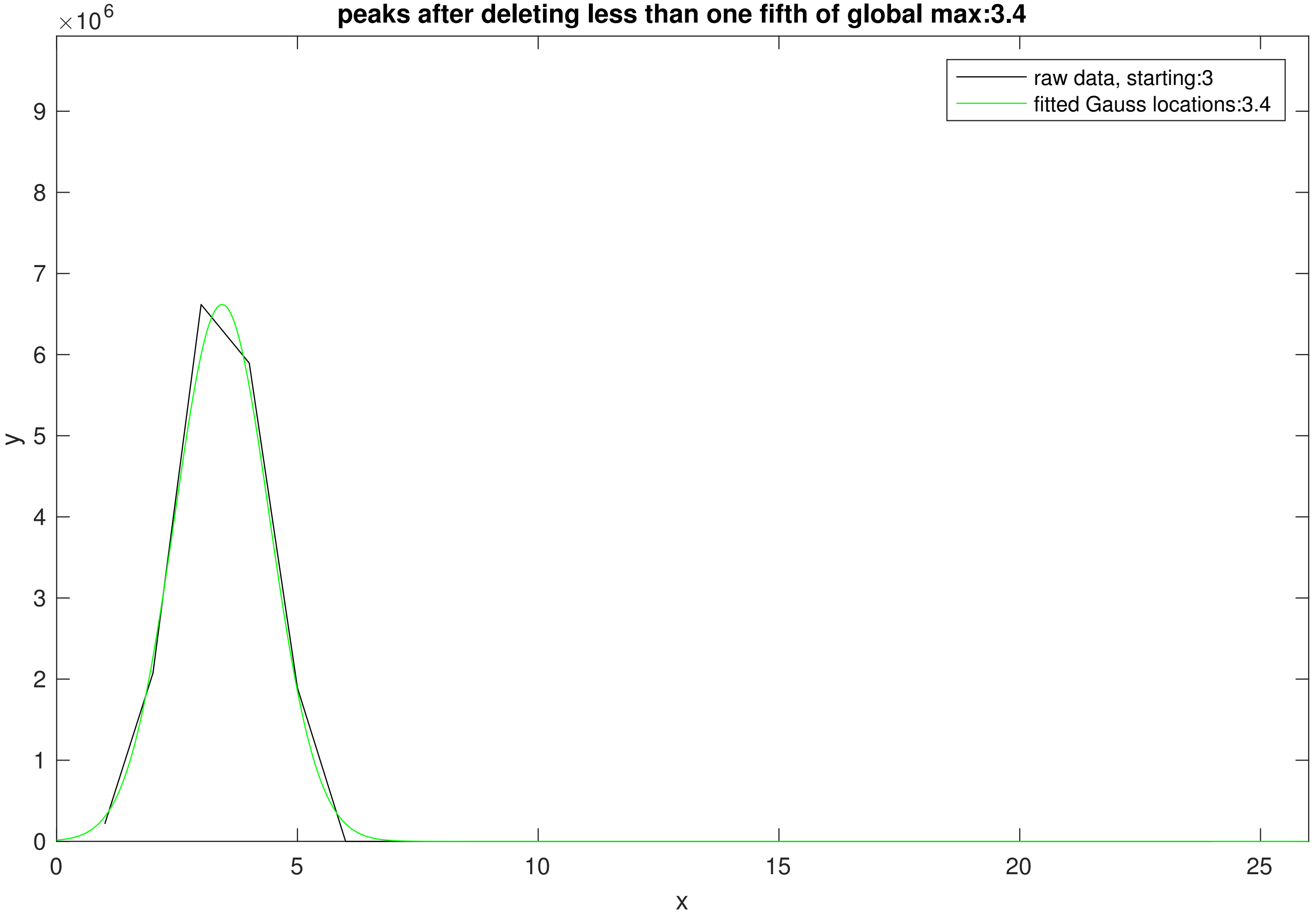

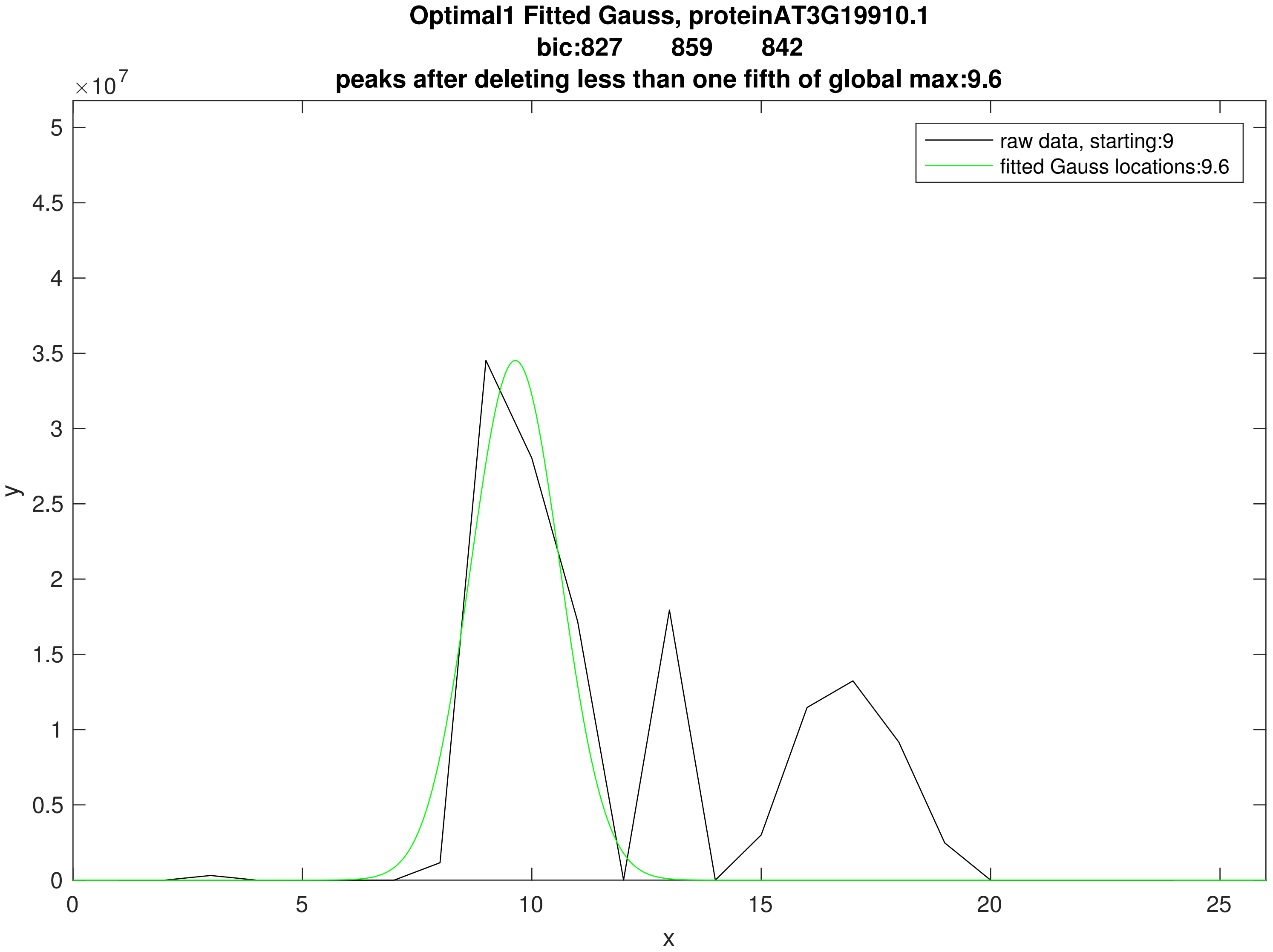

Optimal1 Fitted Gauss, proteinAT3G19910.1
bic:827 859 842
peaks after deleting less than one fifth of global max:9.6
×10^7
raw data, starting:9
fitted Gauss locations:9.6
y
x
0
0.5
1
1.5
2
2.5
3
3.5
4
4.5
5
0
5
10
15
20
25

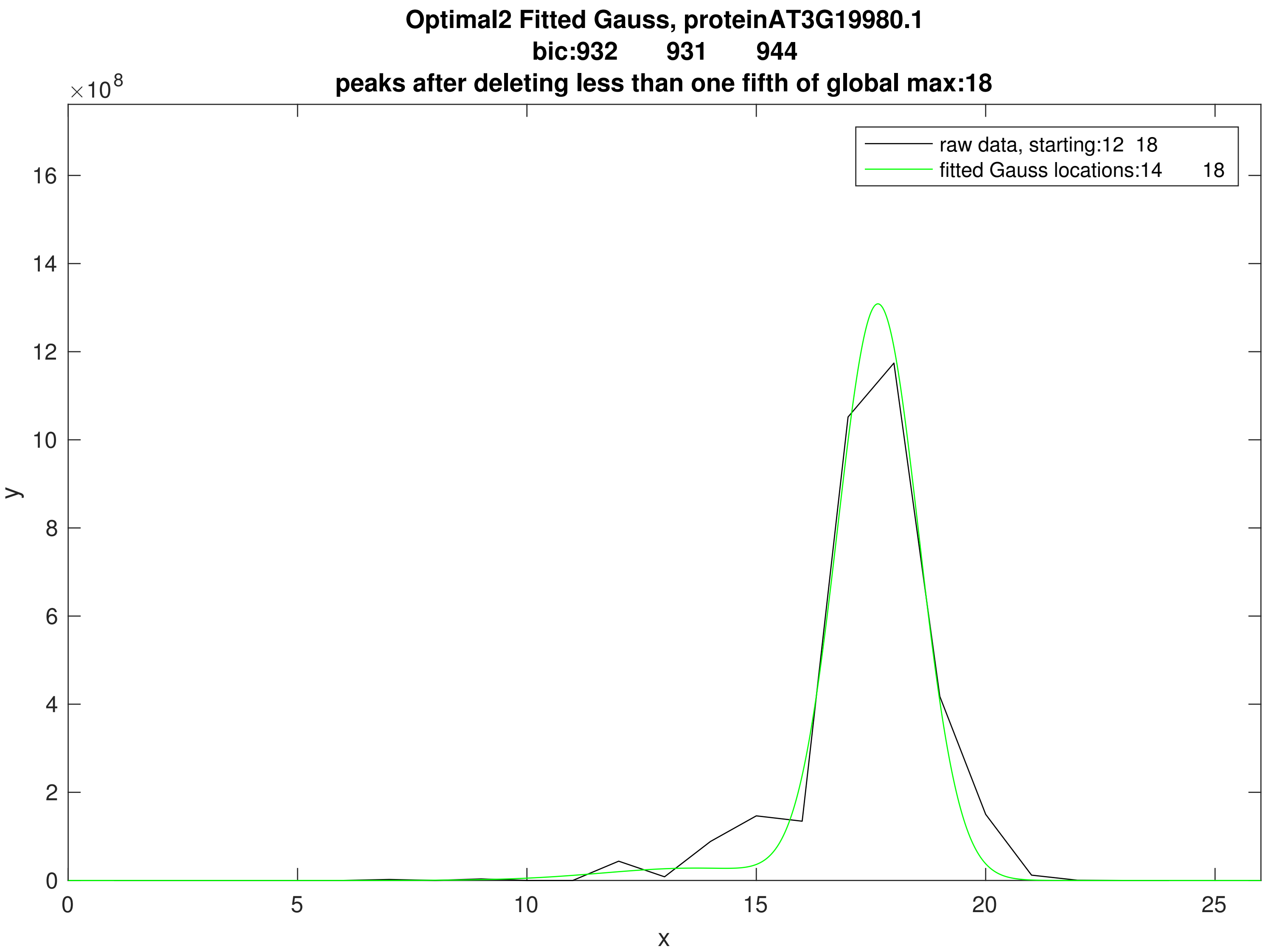

Optimal2 Fitted Gauss, proteinAT3G19980.1
bic:932 931 944
peaks after deleting less than one fifth of global max:18
×10^8
raw data, starting:12 18
fitted Gauss locations:14 18
y
x
0
2
4
6
8
10
12
14
16
0
5
10
15
20
25

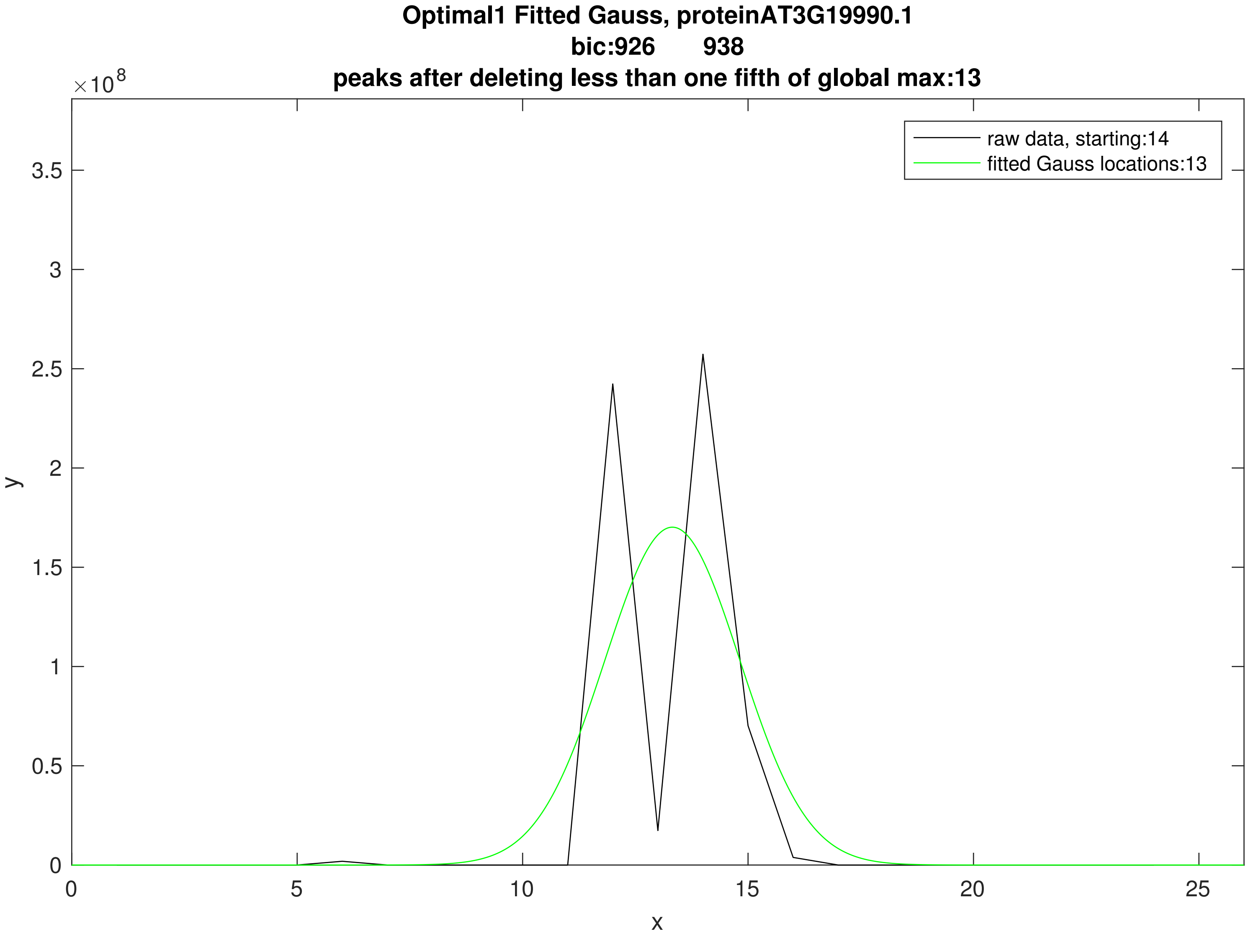
Optimal1 Fitted Gauss, proteinAT3G19990.1
bic:926 938
peaks after deleting less than one fifth of global max:13
×10^8
raw data, starting:14
fitted Gauss locations:13
y
x
0
0.5
1
1.5
2
2.5
3
3.5
0
5
10
15
20
25

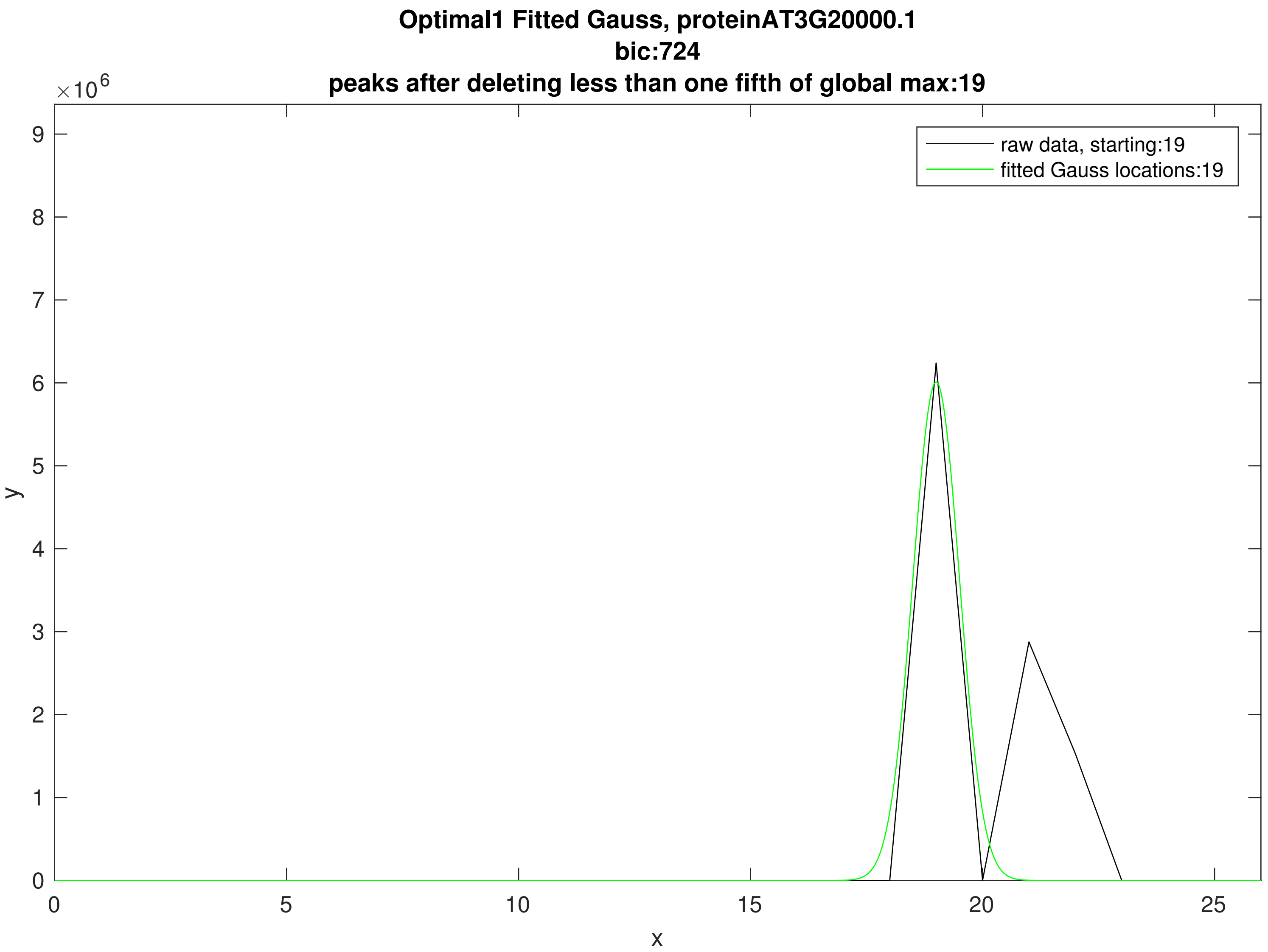
Optimal1 Fitted Gauss, proteinAT3G20000.1
bic:724
peaks after deleting less than one fifth of global max:19
×10^6
9
8
7
6
5
4
3
2
1
0
y
0
5
10
15
20
25
x
raw data, starting:19
fitted Gauss locations:19

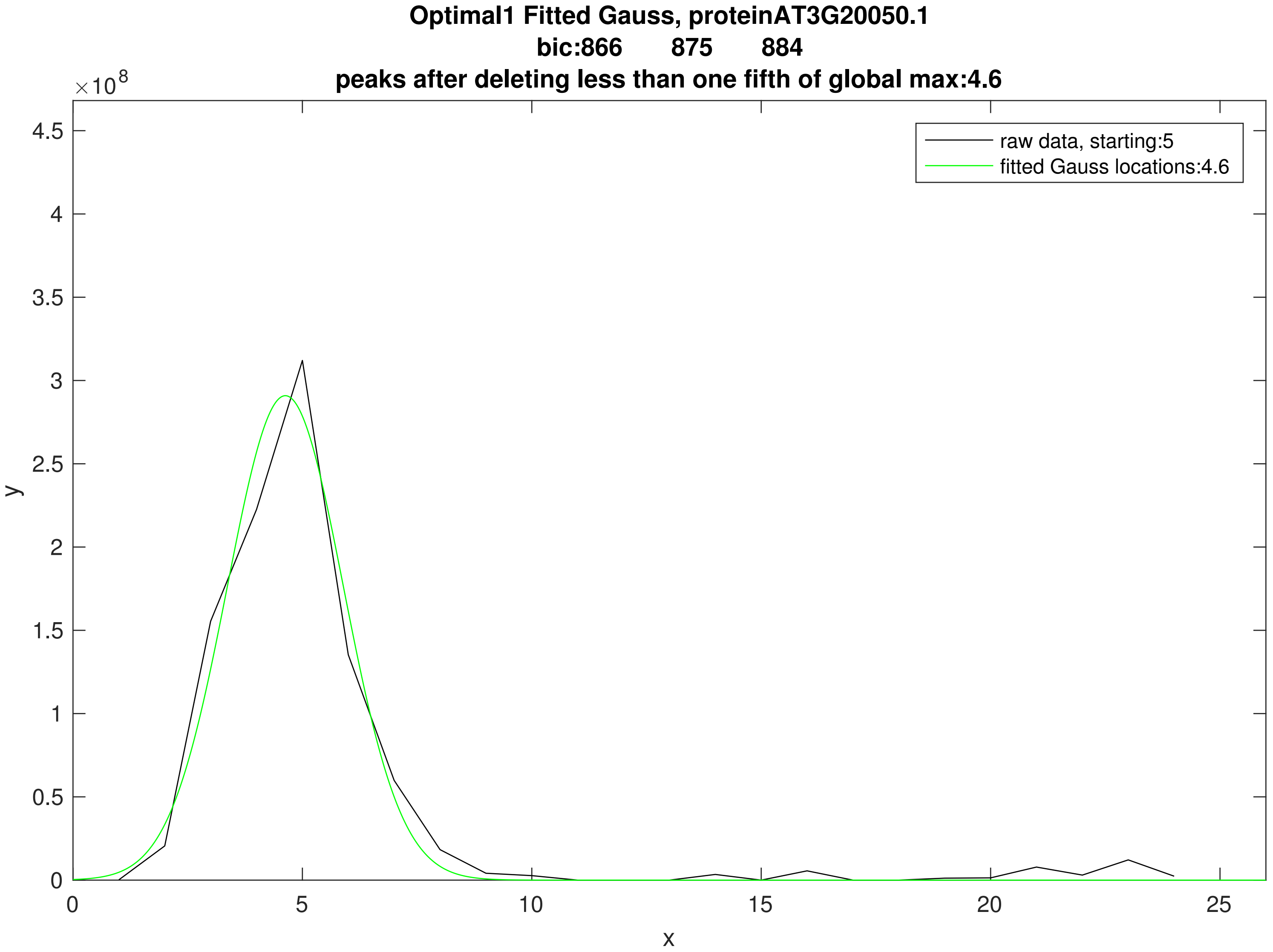

Optimal1 Fitted Gauss, proteinAT3G20050.1
bic:866 875 884
peaks after deleting less than one fifth of global max:4.6
×10^8
4.5
4
3.5
3
2.5
2
1.5
1
0.5
0
y
0
5
10
15
20
25
x
raw data, starting:5
fitted Gauss locations:4.6

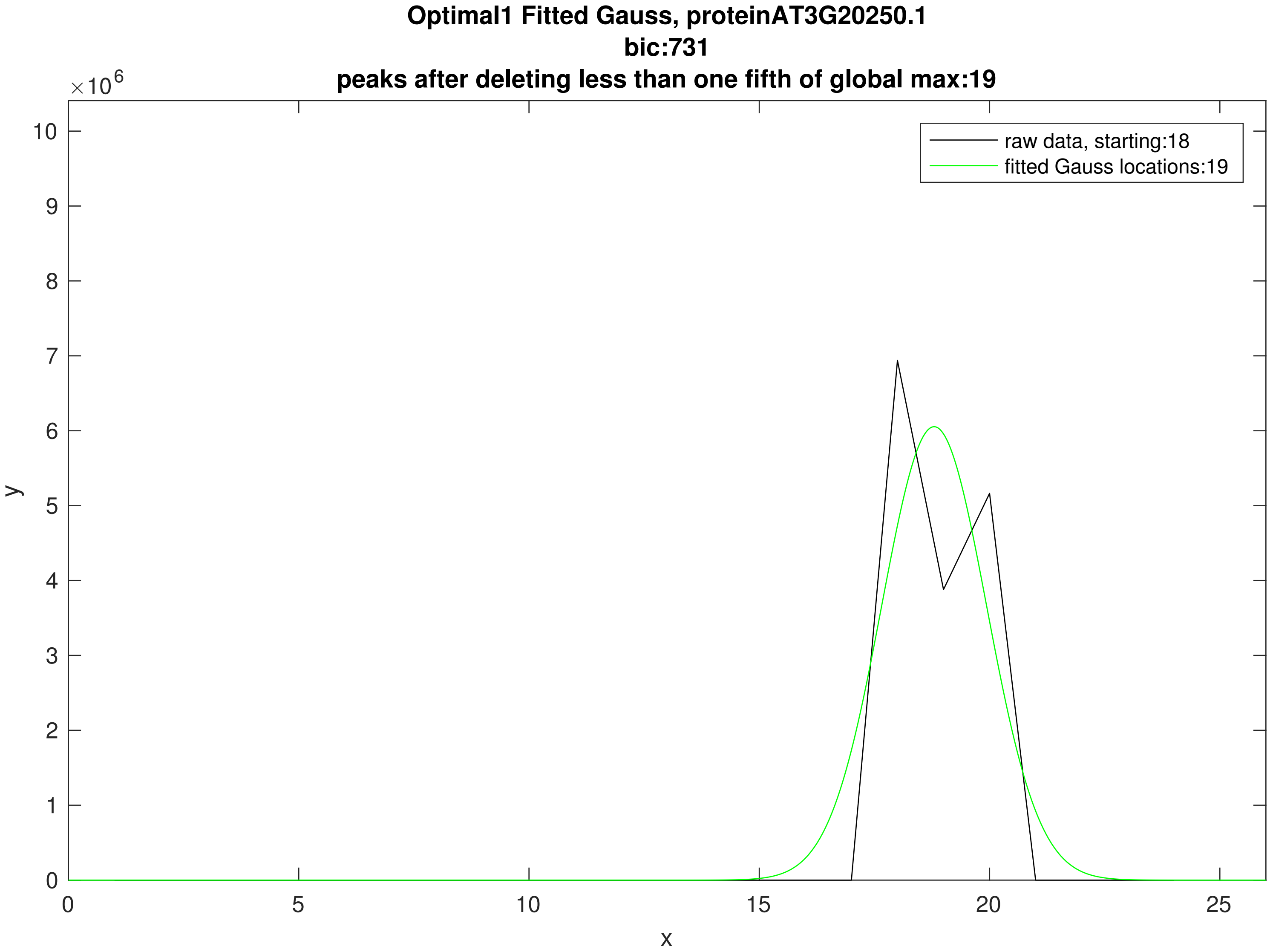
Optimal1 Fitted Gauss, proteinAT3G20250.1
bic:731
peaks after deleting less than one fifth of global max:19
×10^6
raw data, starting:18
fitted Gauss locations:19
y
x
0
1
2
3
4
5
6
7
8
9
10
0
5
10
15
20
25

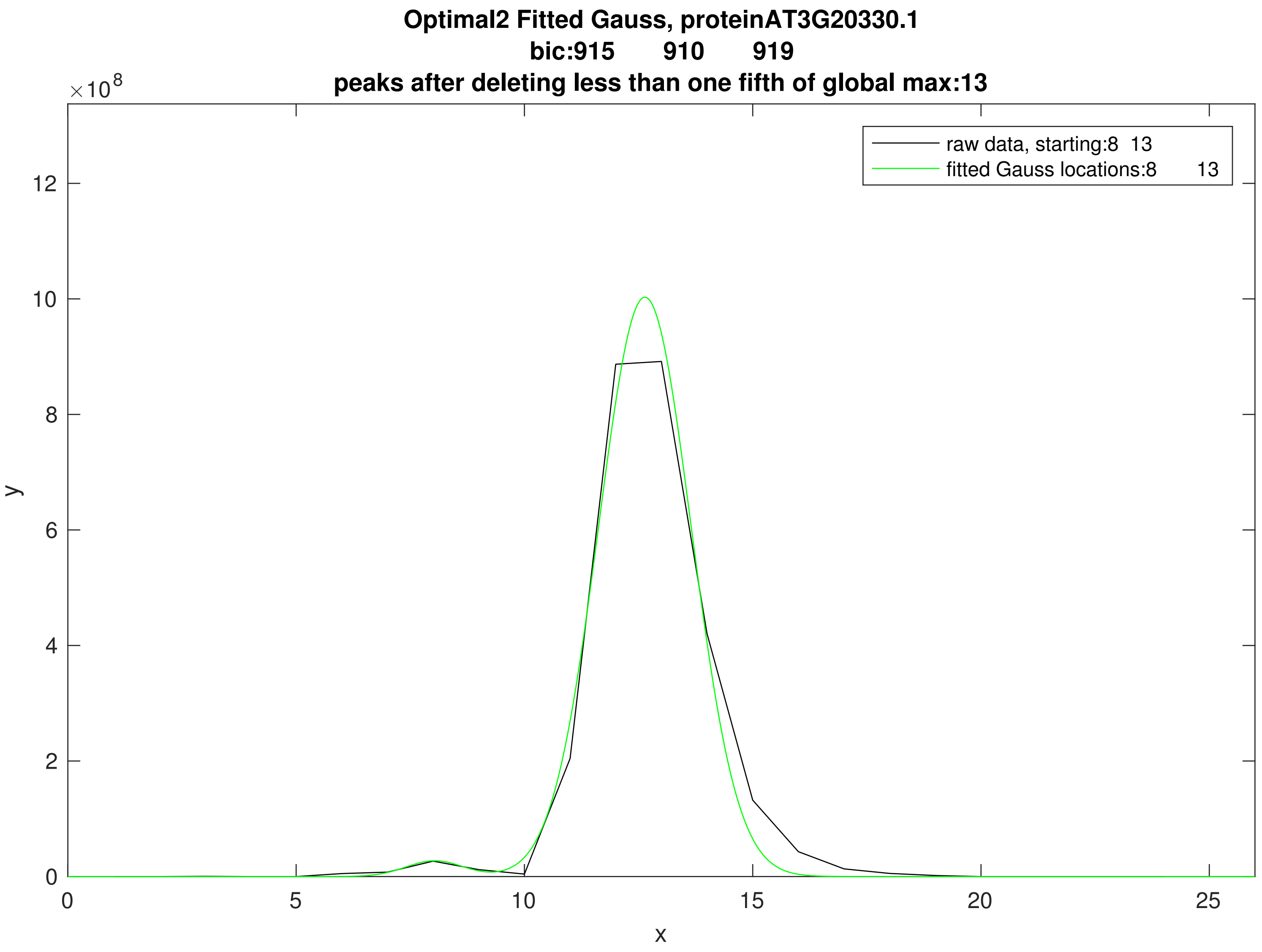
Optimal2 Fitted Gauss, proteinAT3G20330.1
bic:915 910 919
peaks after deleting less than one fifth of global max:13
×10^8
raw data, starting:8 13
fitted Gauss locations:8 13
y
x
0
2
4
6
8
10
12
0
5
10
15
20
25

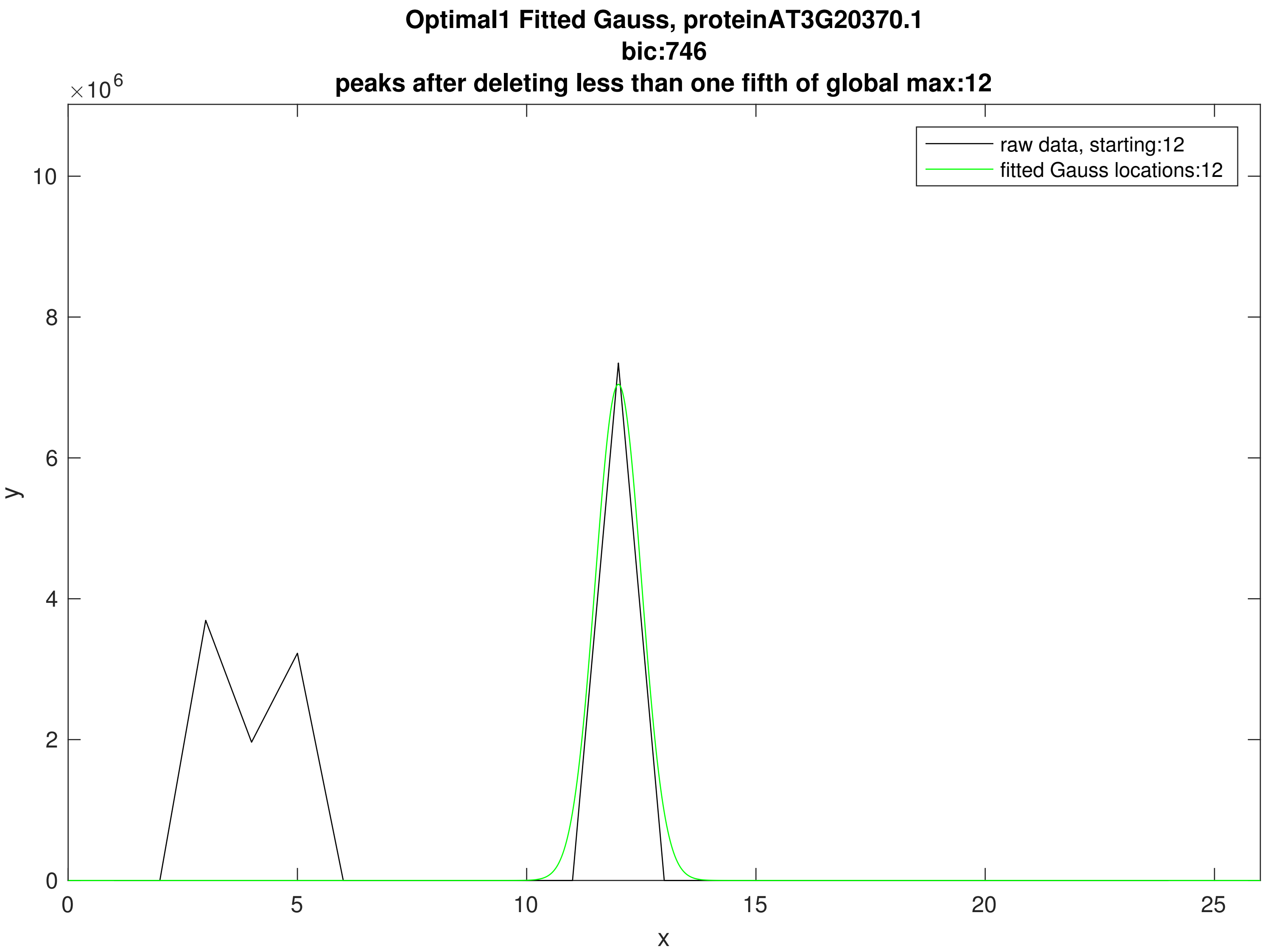

Optimal1 Fitted Gauss, proteinAT3G20370.1
bic:746
peaks after deleting less than one fifth of global max:12
×10^6
raw data, starting:12
fitted Gauss locations:12
y
x
0
2
4
6
8
10
0
5
10
15
20
25

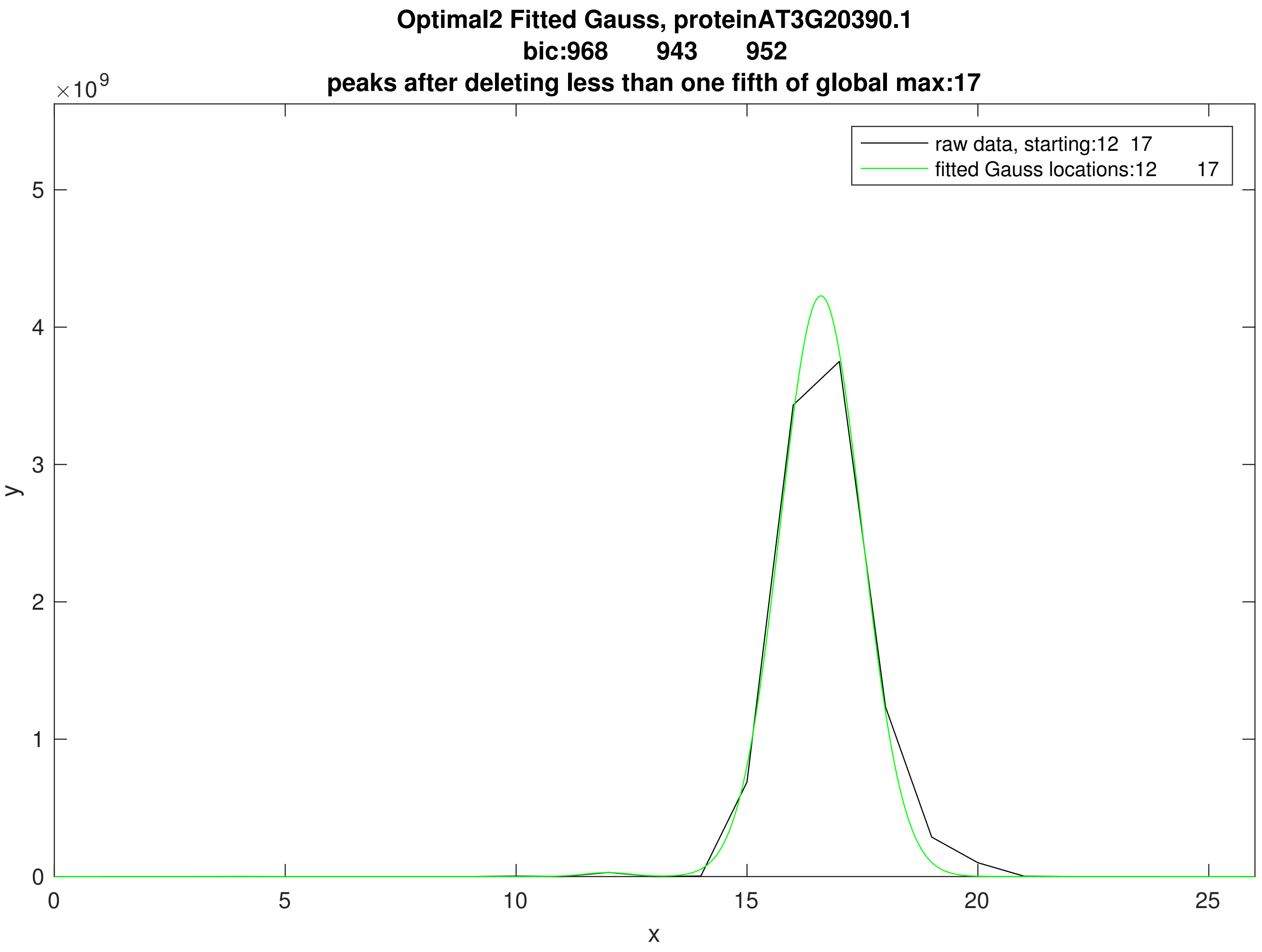

Optimal2 Fitted Gauss, proteinAT3G20390.1
bic:968 943 952
peaks after deleting less than one fifth of global max:17
×10^9
raw data, starting:12 17
fitted Gauss locations:12 17
y
x
0
1
2
3
4
5
10
15
20
25

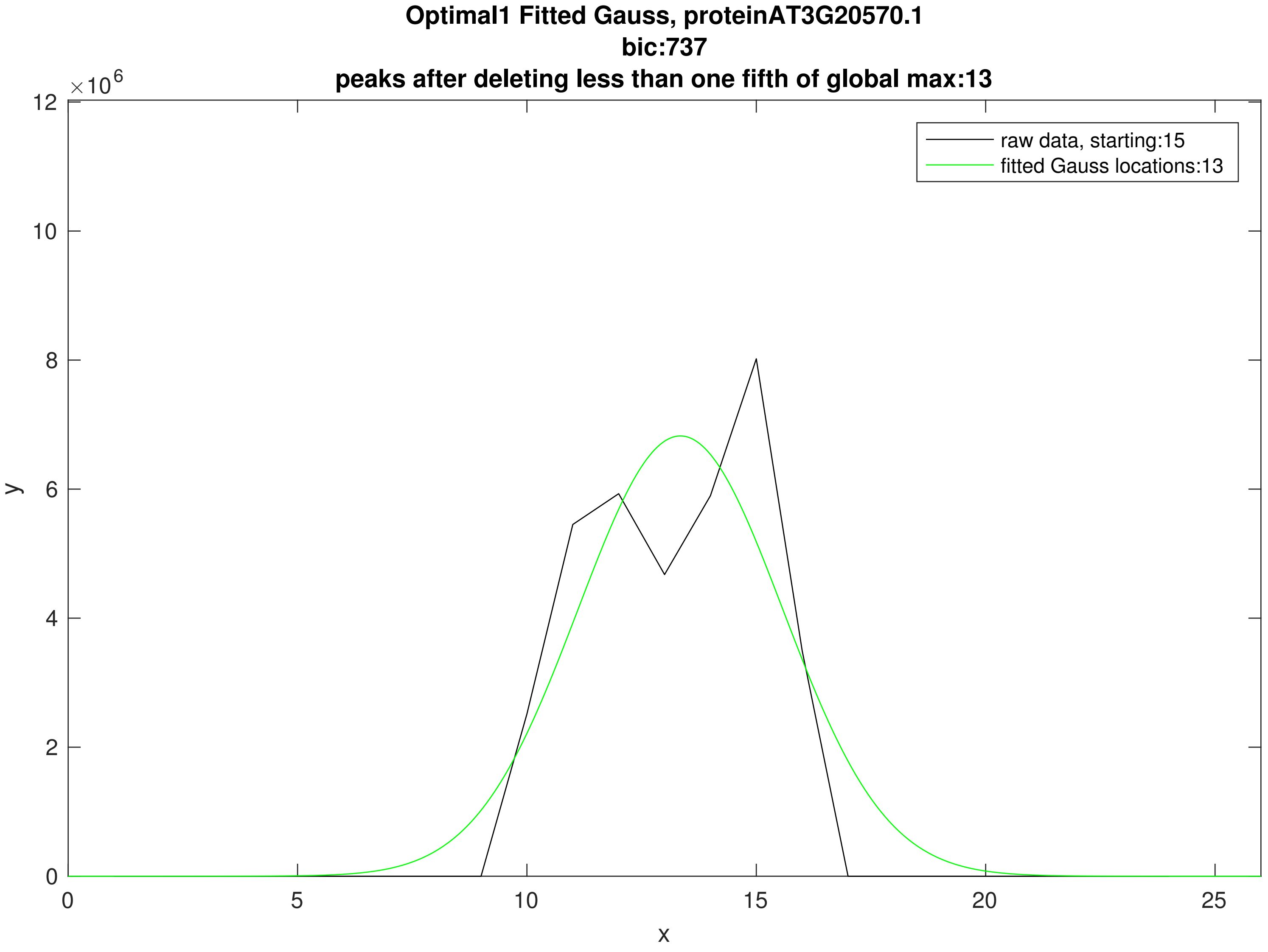
Optimal1 Fitted Gauss, proteinAT3G20570.1
bic:737
peaks after deleting less than one fifth of global max:13
$\times 10^6$
raw data, starting:15
fitted Gauss locations:13
y
x
0
2
4
6
8
10
12
0
5
10
15
20
25

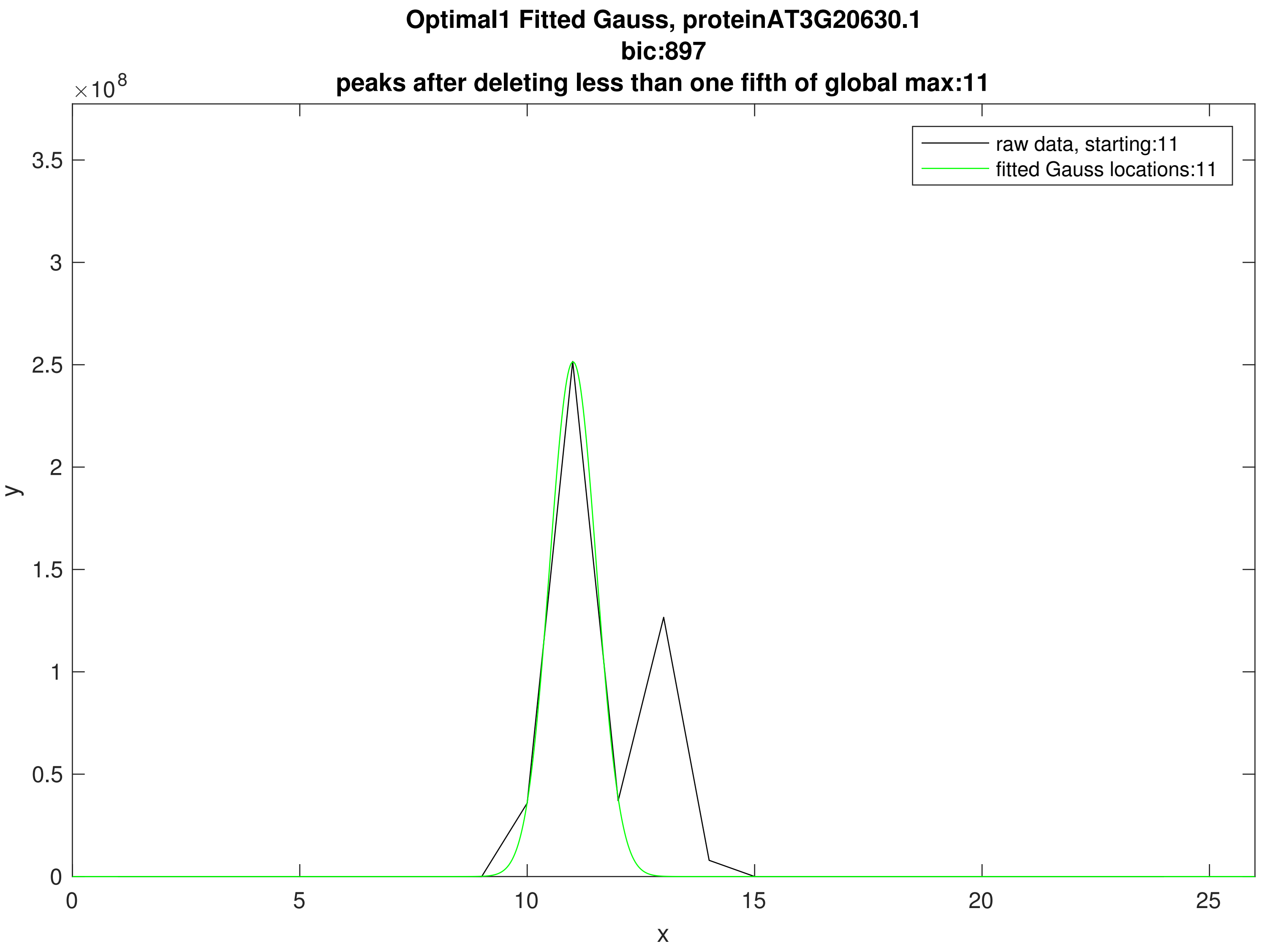

Optimal1 Fitted Gauss, proteinAT3G20630.1
bic:897
peaks after deleting less than one fifth of global max:11
×10^8
3.5
3
2.5
2
1.5
1
0.5
0
y
0
5
10
15
20
25
x
raw data, starting:11
fitted Gauss locations:11

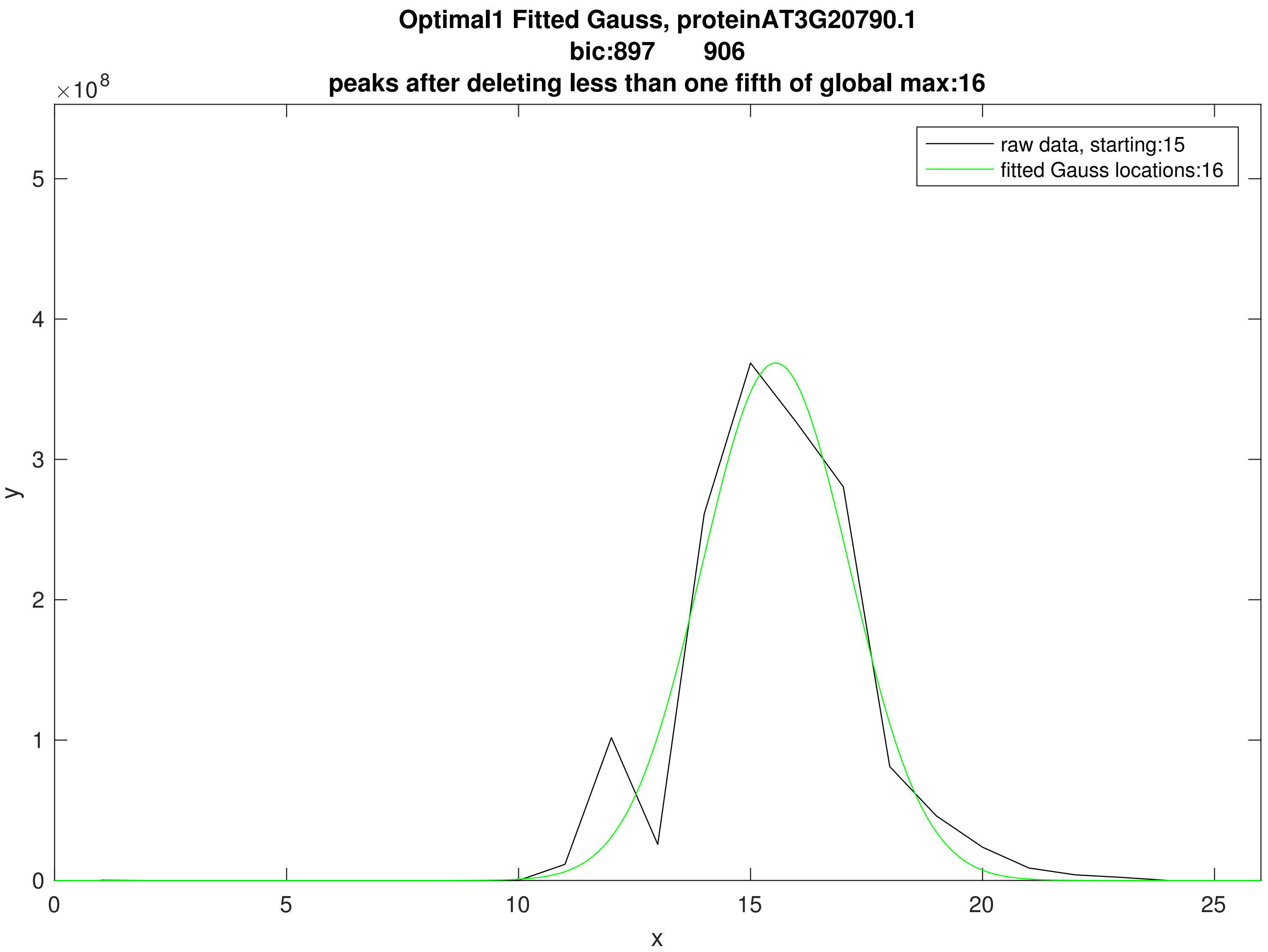
Optimal1 Fitted Gauss, proteinAT3G20790.1
bic:897 906
peaks after deleting less than one fifth of global max:16
×10^8
raw data, starting:15
fitted Gauss locations:16
y
x
0
1
2
3
4
5
0
5
10
15
20
25

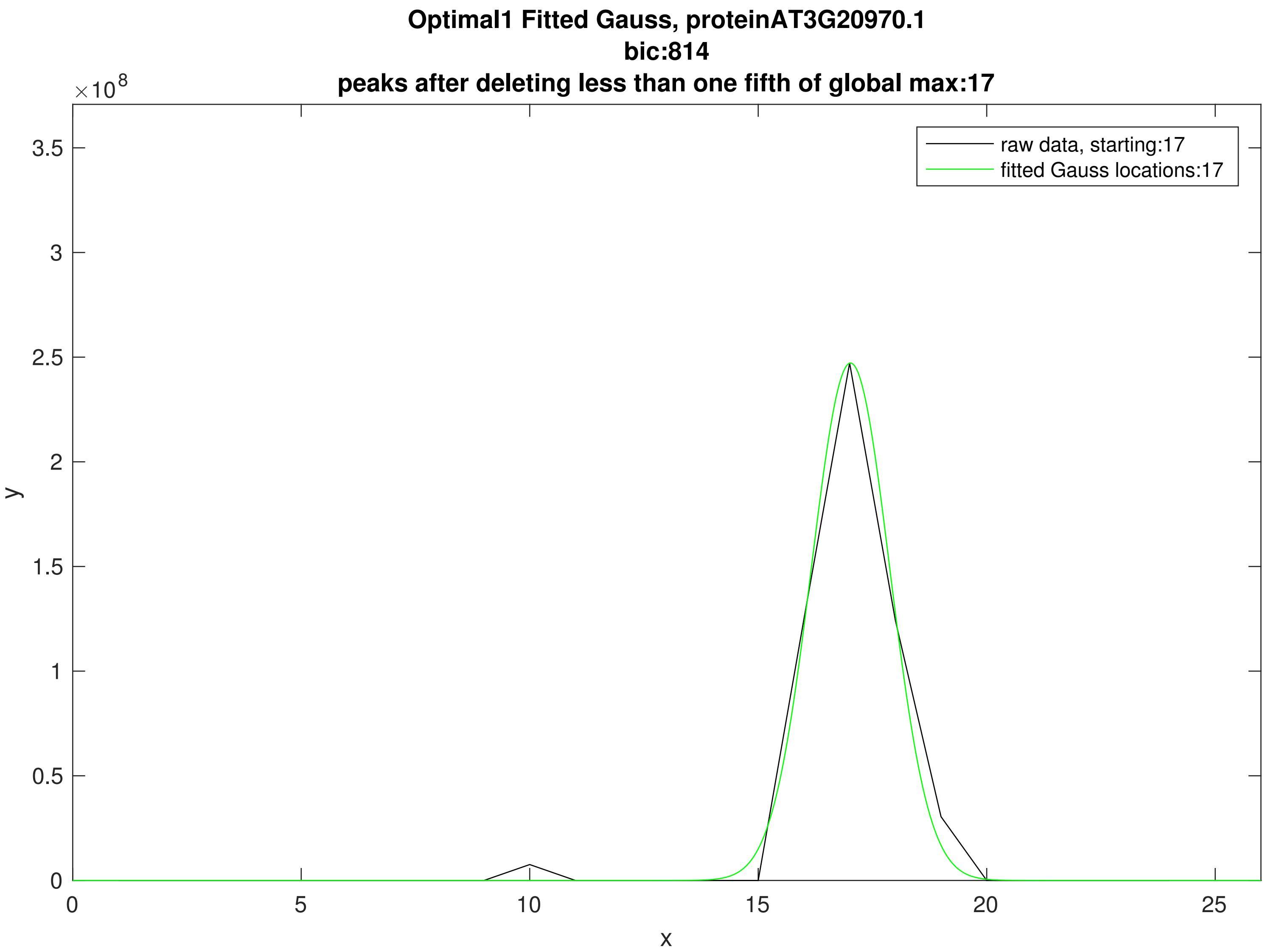
Optimal1 Fitted Gauss, proteinAT3G20970.1
bic:814
peaks after deleting less than one fifth of global max:17
×10^8
raw data, starting:17
fitted Gauss locations:17
y
x
0
0.5
1
1.5
2
2.5
3
3.5
0
5
10
15
20
25

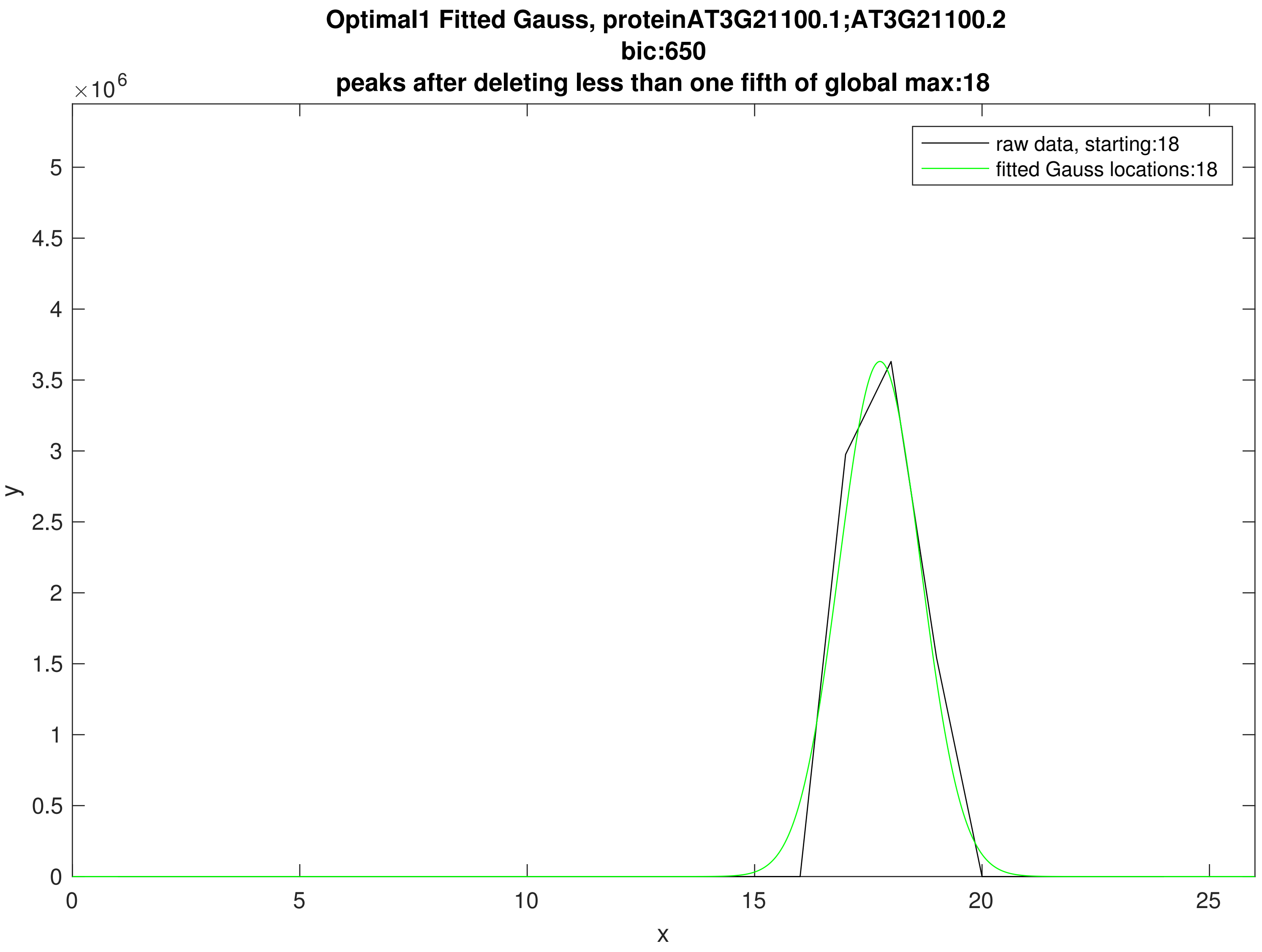
Optimal1 Fitted Gauss, proteinAT3G21100.1;AT3G21100.2
bic:650
peaks after deleting less than one fifth of global max:18
×10^6
raw data, starting:18
fitted Gauss locations:18
5
4.5
4
3.5
3
2.5
2
1.5
1
0.5
0
y
0
5
10
15
20
25
x

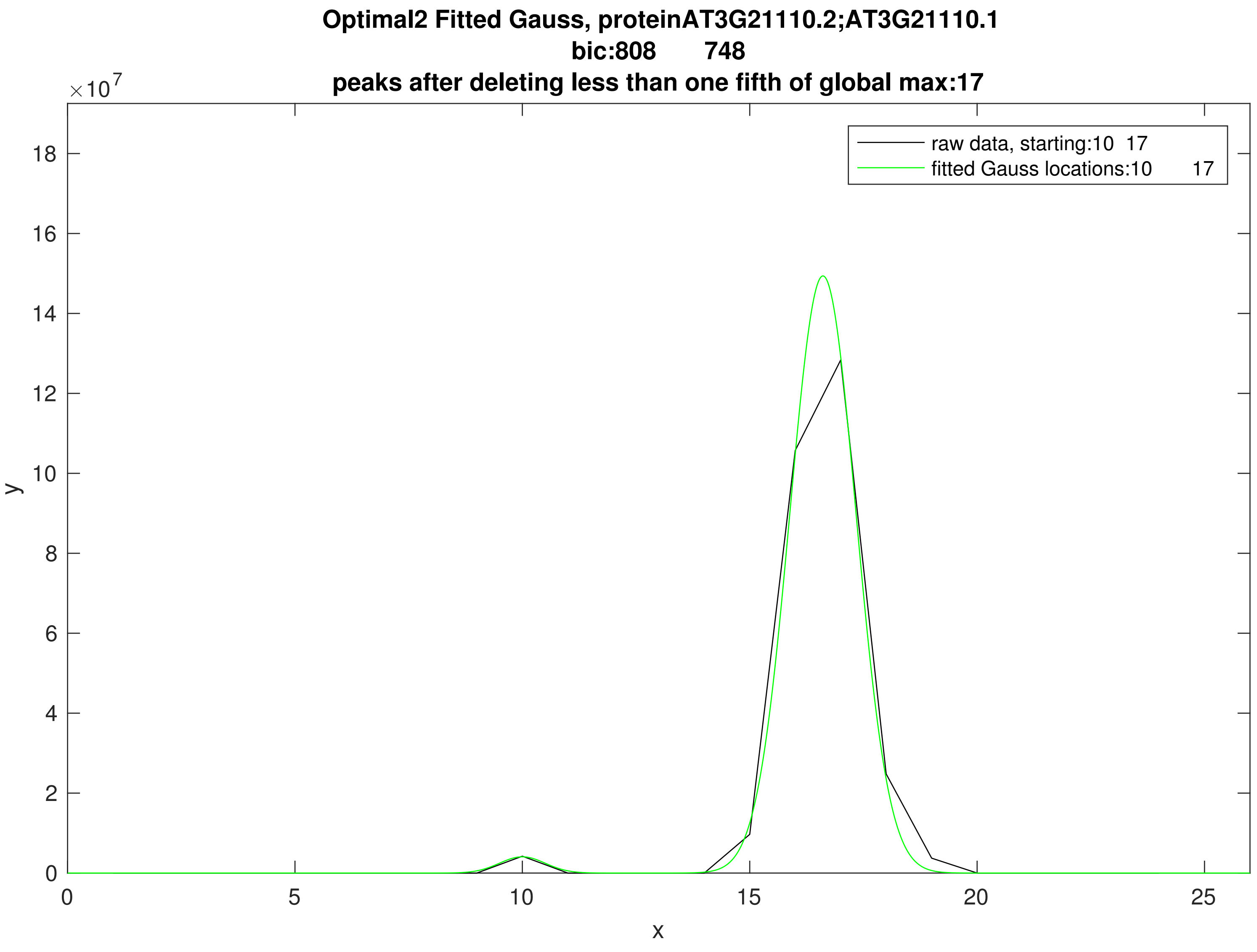

Optimal2 Fitted Gauss, proteinAT3G21110.2;AT3G21110.1
bic:808 748
peaks after deleting less than one fifth of global max:17
×10^7
raw data, starting:10 17
fitted Gauss locations:10 17
y
x
0
2
4
6
8
10
12
14
16
18
0
5
10
15
20
25

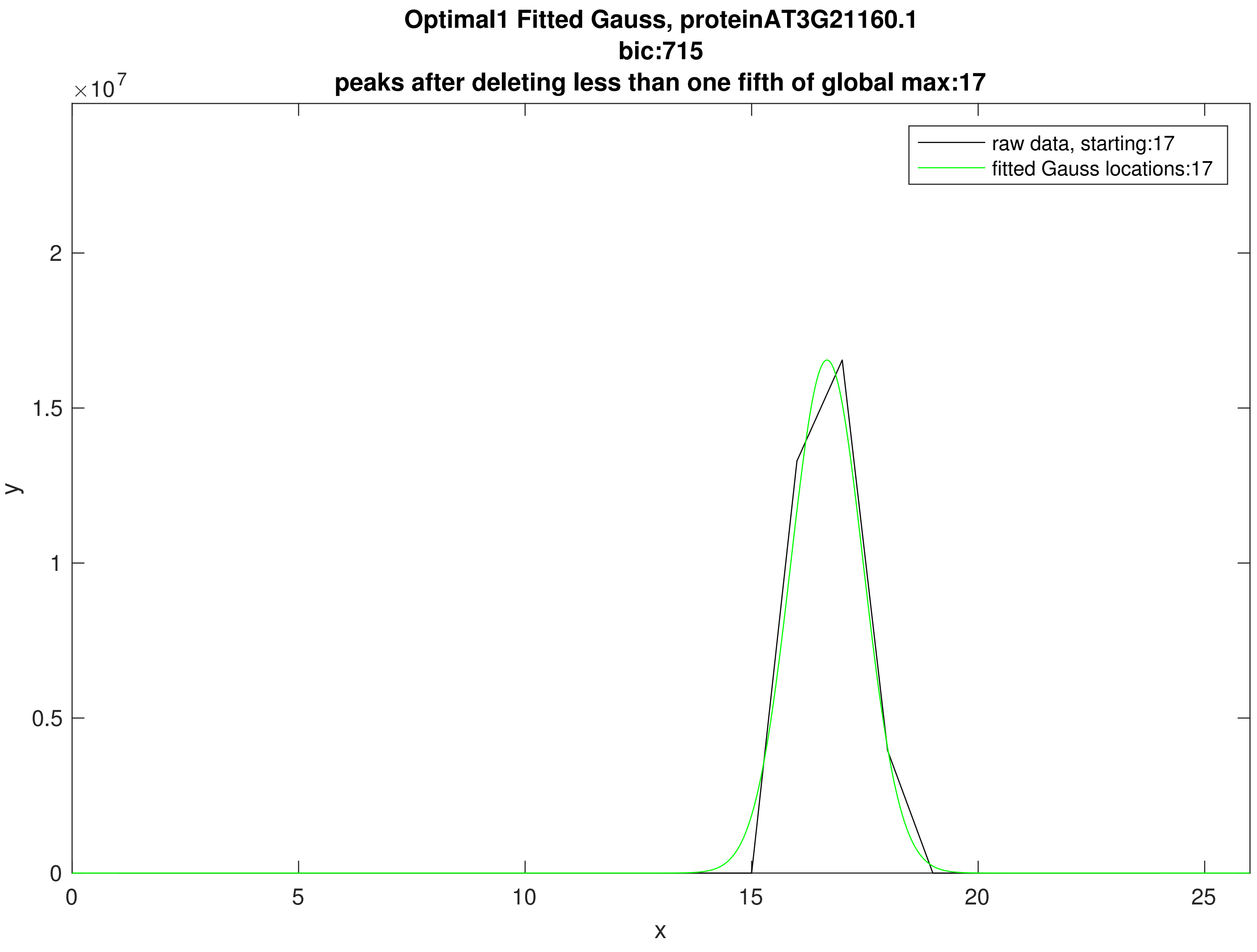

Optimal1 Fitted Gauss, proteinAT3G21160.1
bic:715
peaks after deleting less than one fifth of global max:17
×10^7
raw data, starting:17
fitted Gauss locations:17
2
1.5
1
0.5
0
y
0
5
10
15
20
25
x

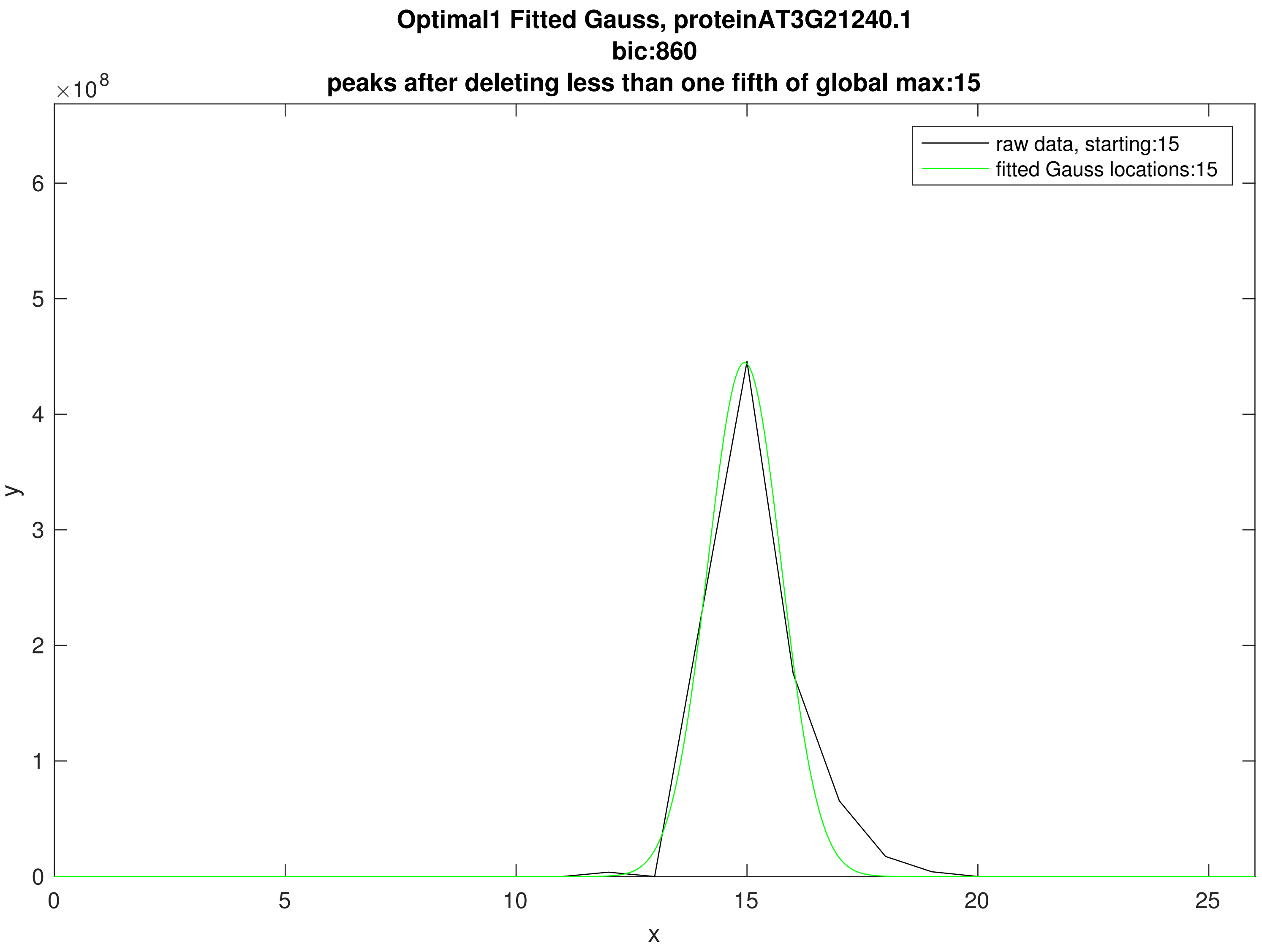
Optimal1 Fitted Gauss, proteinAT3G21240.1
bic:860
peaks after deleting less than one fifth of global max:15
×10^8
raw data, starting:15
fitted Gauss locations:15
y
x
0
1
2
3
4
5
6
0
5
10
15
20
25

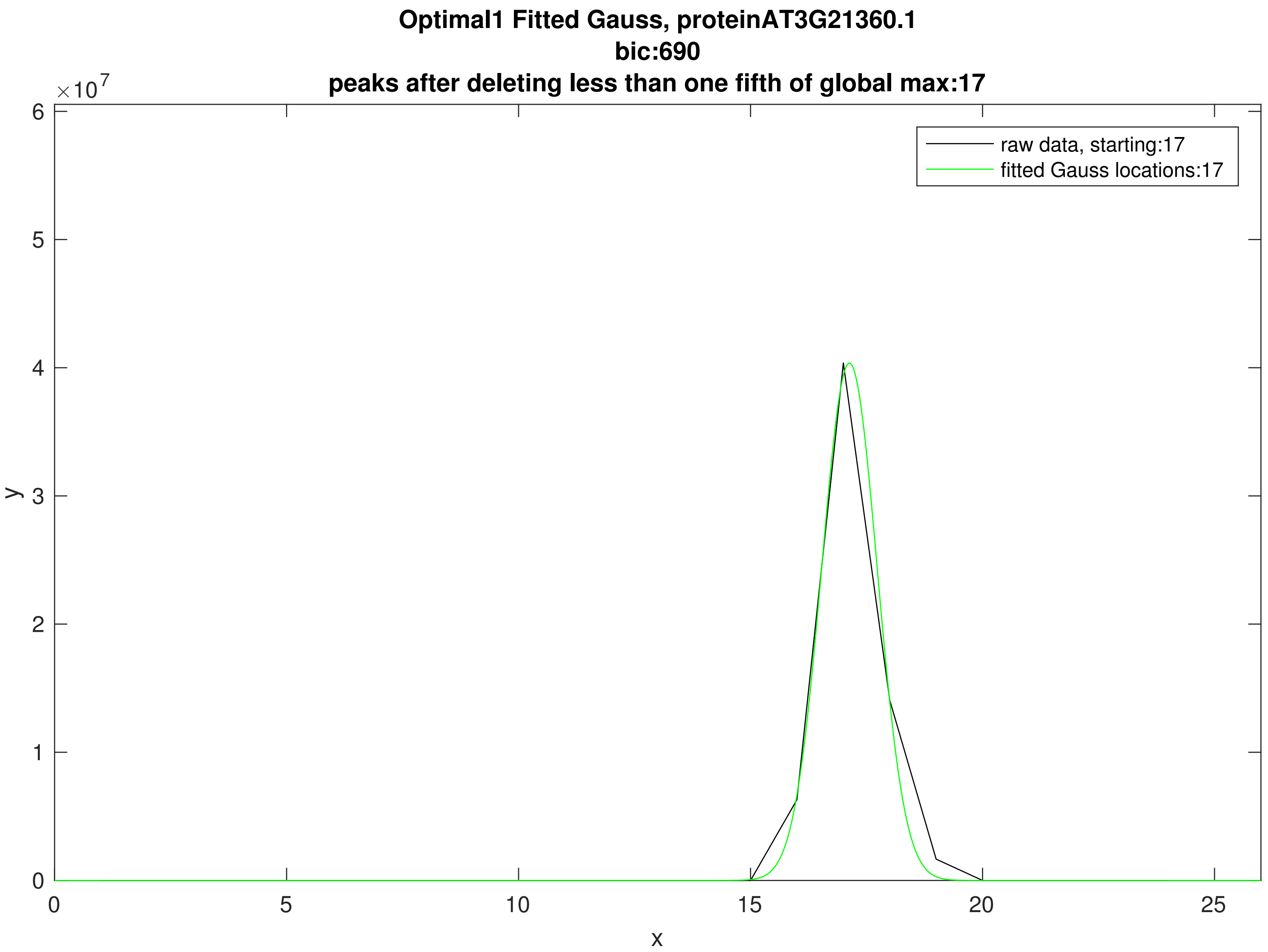

Optimal1 Fitted Gauss, proteinAT3G21360.1
bic:690
peaks after deleting less than one fifth of global max:17
×10^7
raw data, starting:17
fitted Gauss locations:17
y
x
0
1
2
3
4
5
6
0
5
10
15
20
25

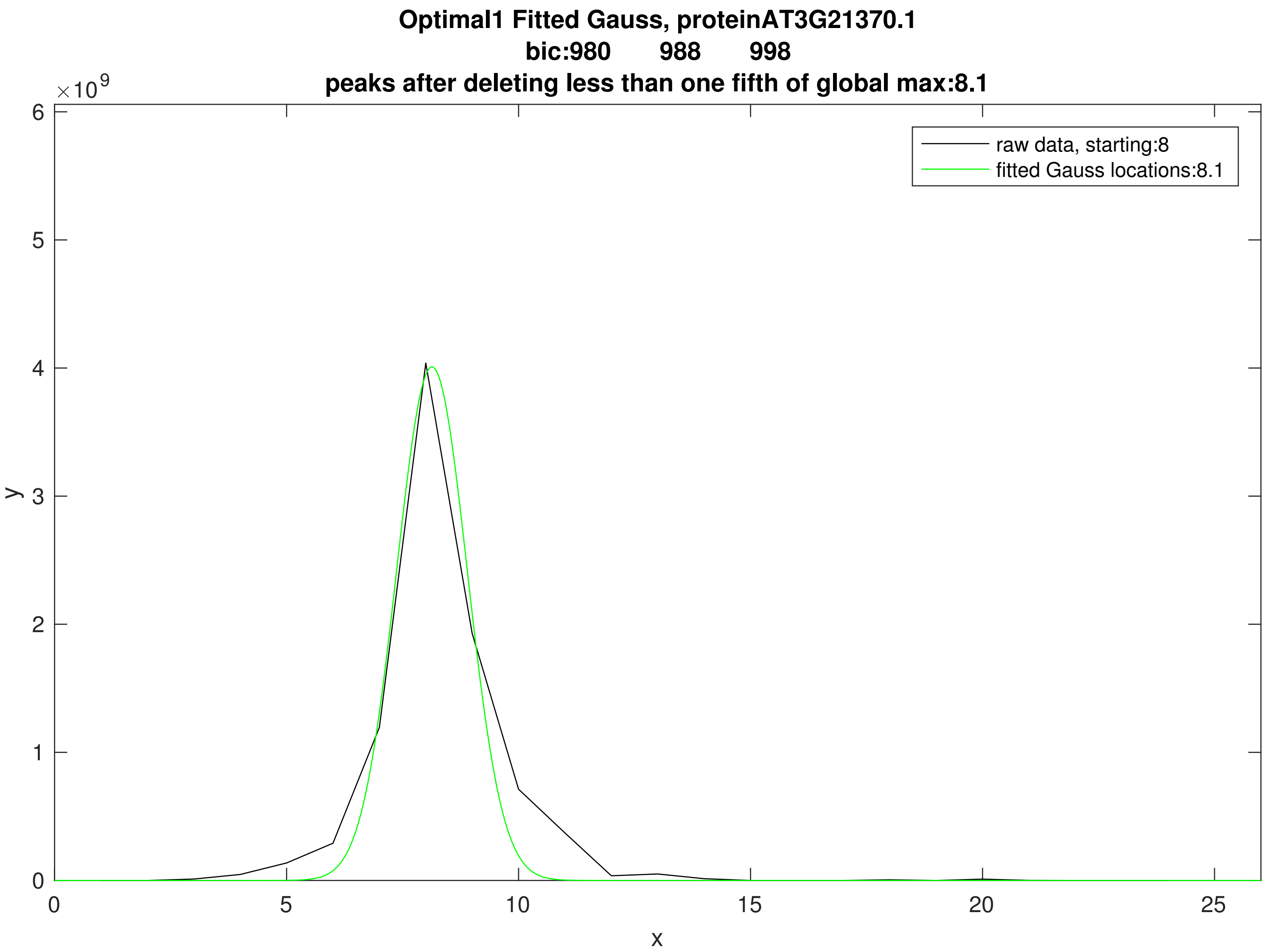
Optimal1 Fitted Gauss, proteinAT3G21370.1
bic:980 988 998
peaks after deleting less than one fifth of global max:8.1
×10^9
y
x
raw data, starting:8
fitted Gauss locations:8.1

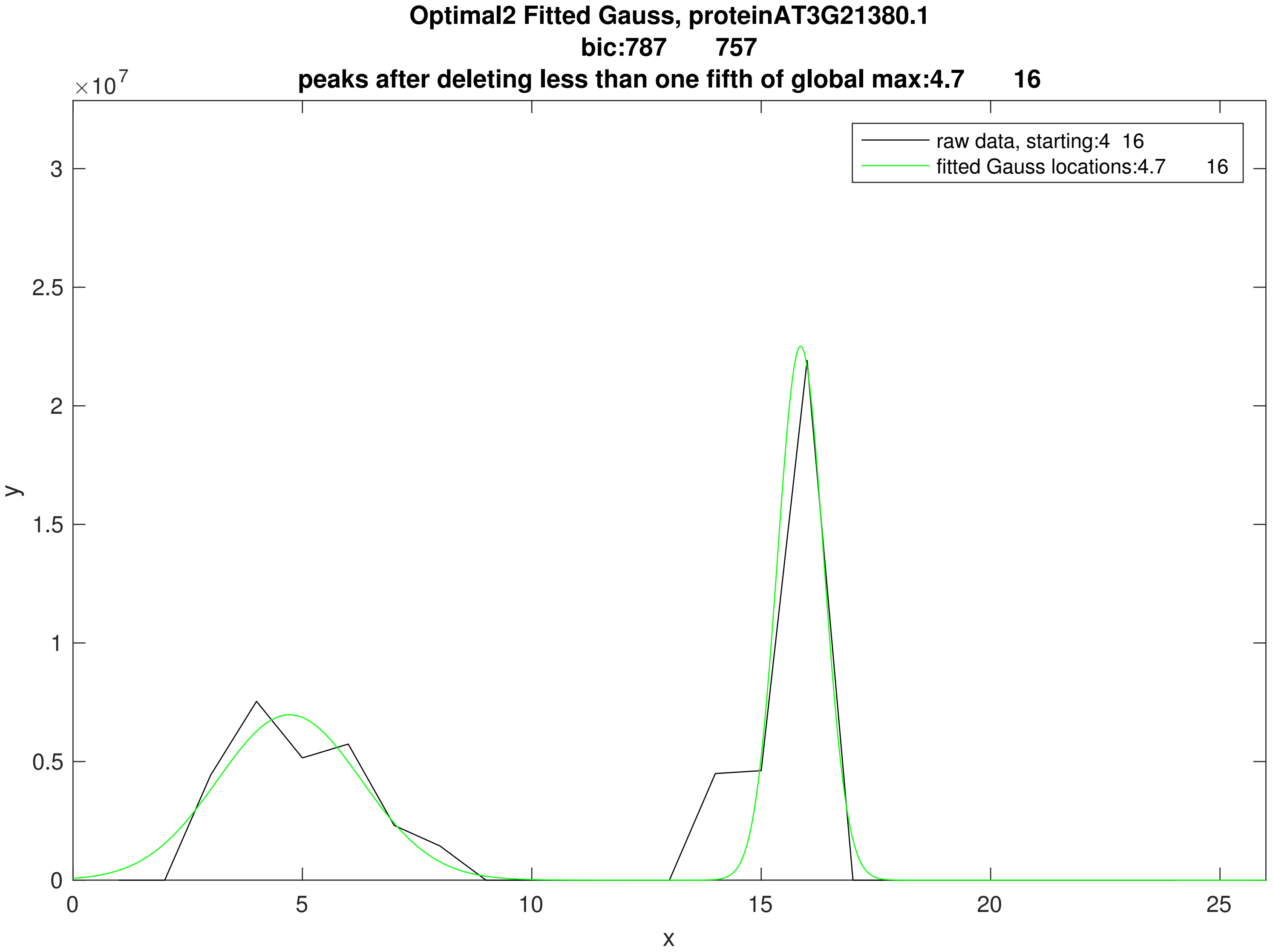
Optimal2 Fitted Gauss, proteinAT3G21380.1
bic:787 757
peaks after deleting less than one fifth of global max:4.7 16
×10^7
3
2.5
2
1.5
1
0.5
0
y
0
5
10
15
20
25
x
raw data, starting:4 16
fitted Gauss locations:4.7 16

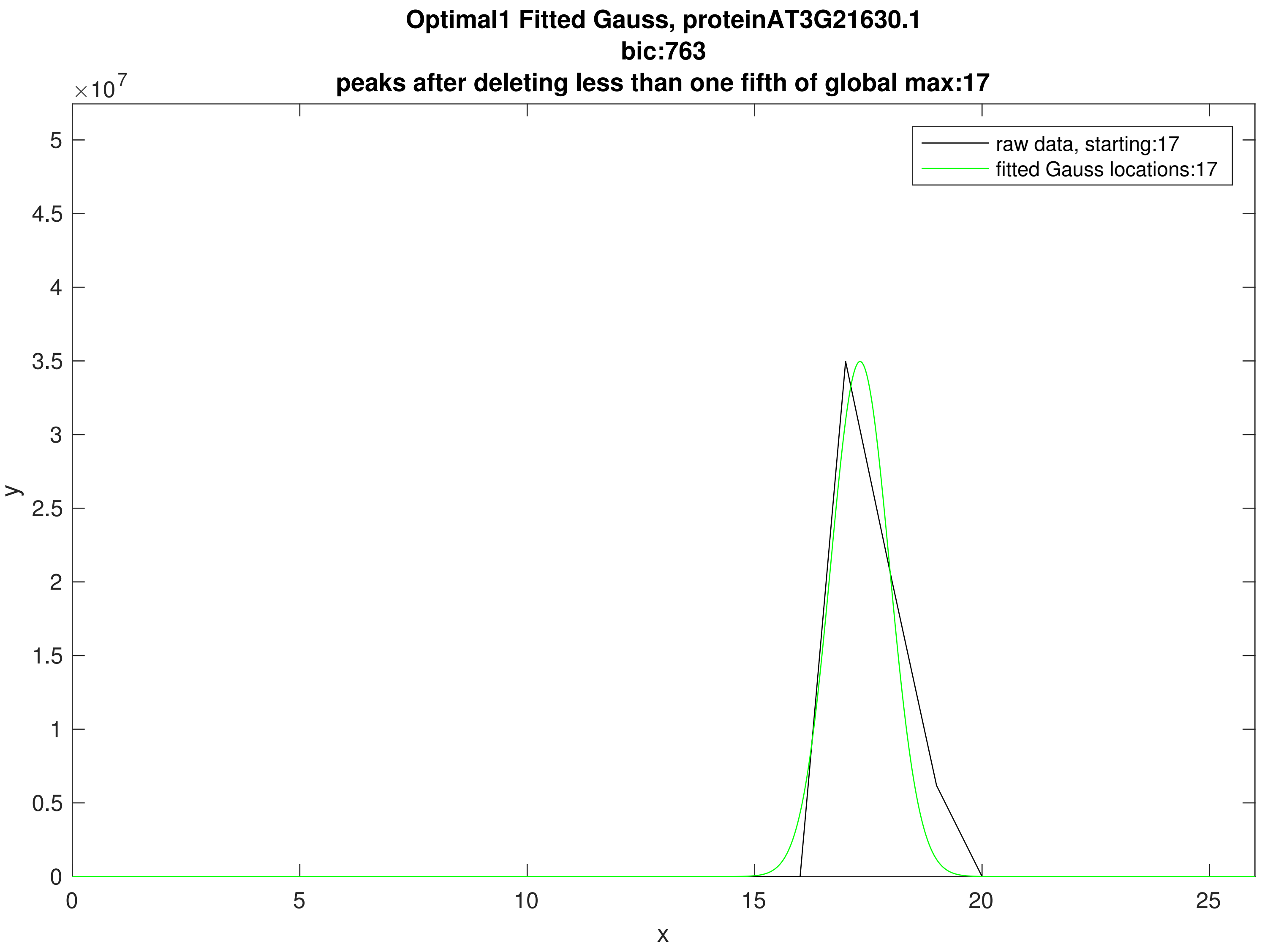

Optimal1 Fitted Gauss, proteinAT3G21630.1
bic:763
peaks after deleting less than one fifth of global max:17
×10^7
raw data, starting:17
fitted Gauss locations:17
y
x
0
0.5
1
1.5
2
2.5
3
3.5
4
4.5
5
0
5
10
15
20
25

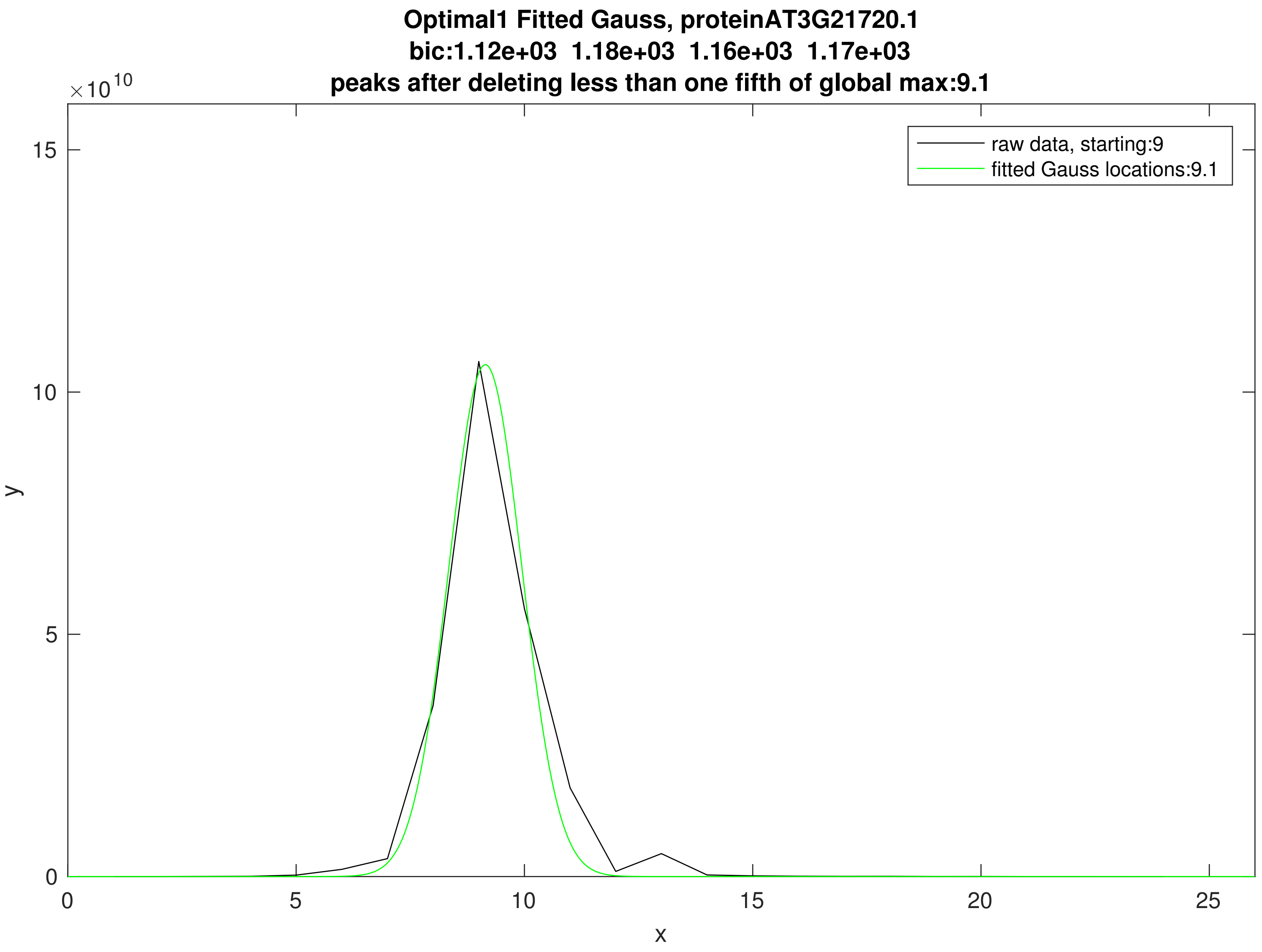
Optimal1 Fitted Gauss, proteinAT3G21720.1
bic:1.12e+03  1.18e+03  1.16e+03  1.17e+03
peaks after deleting less than one fifth of global max:9.1
×10^10
15
10
5
0
y
0
5
10
15
20
25
x
raw data, starting:9
fitted Gauss locations:9.1

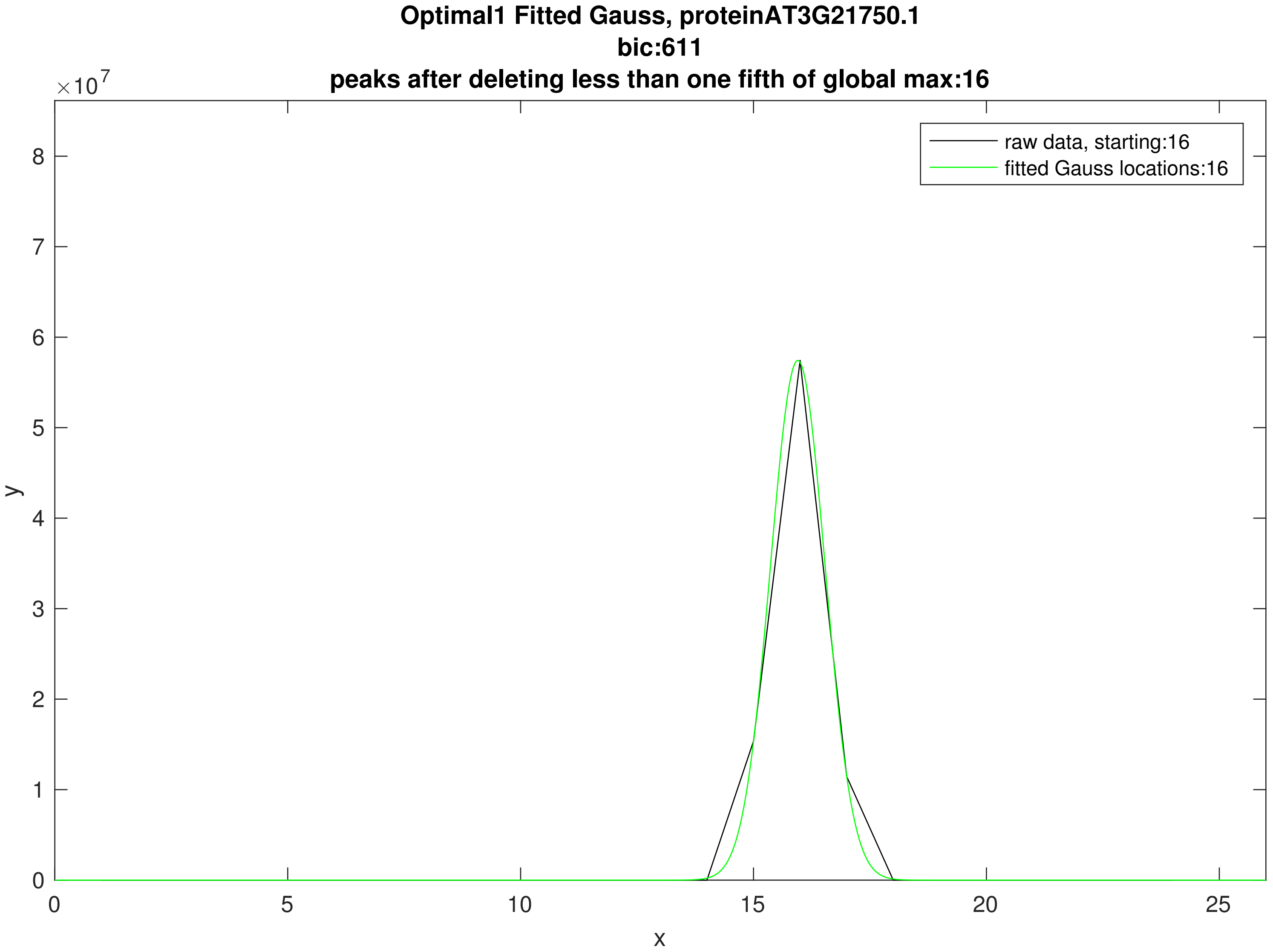
Optimal1 Fitted Gauss, proteinAT3G21750.1
bic:611
peaks after deleting less than one fifth of global max:16
×10^7
raw data, starting:16
fitted Gauss locations:16
y
x
0
1
2
3
4
5
6
7
8
0
5
10
15
20
25

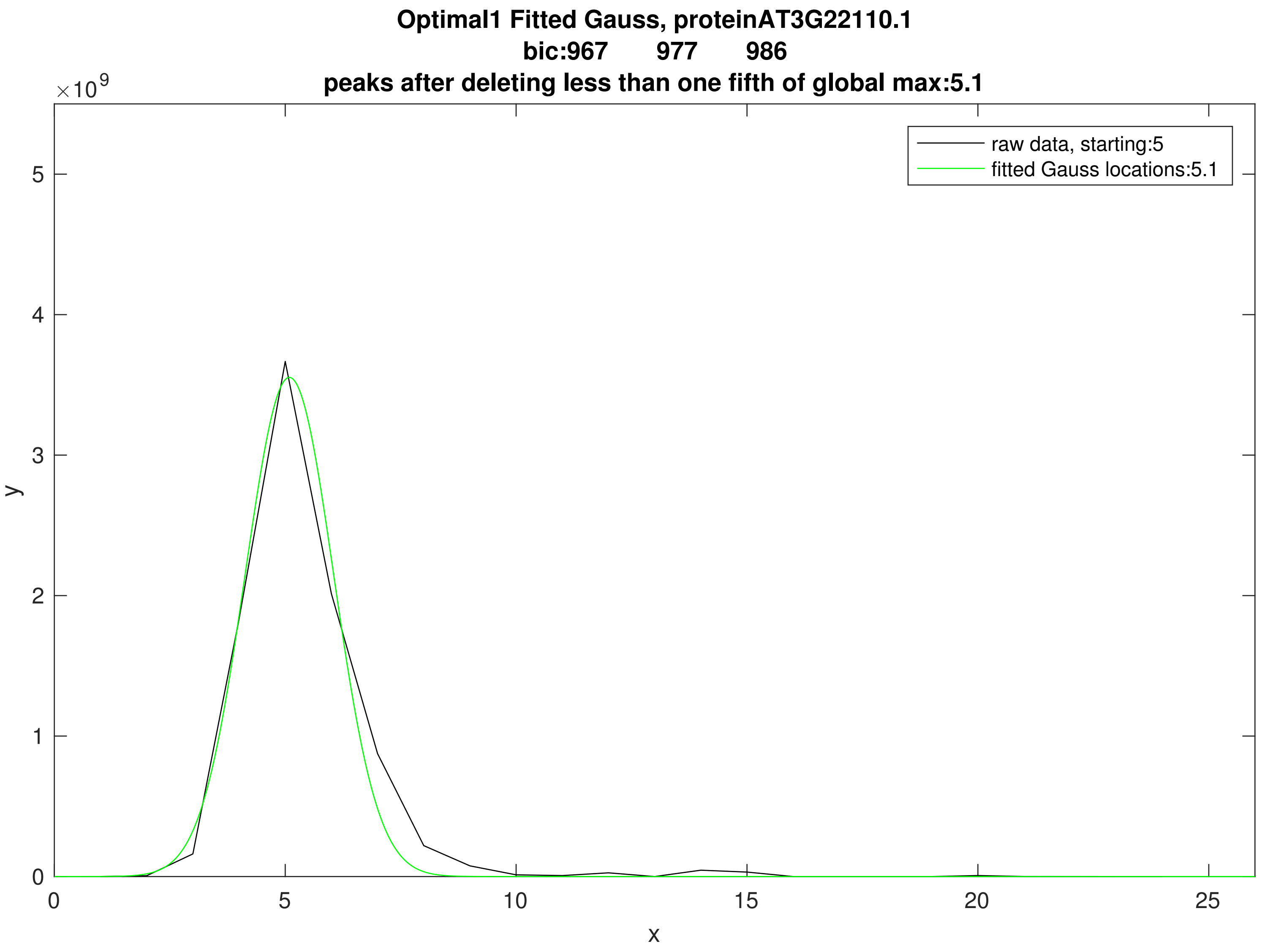

Optimal1 Fitted Gauss, proteinAT3G22110.1
bic:967 977 986
peaks after deleting less than one fifth of global max:5.1
×10^9
raw data, starting:5
fitted Gauss locations:5.1
y
x
0
1
2
3
4
5
0
5
10
15
20
25

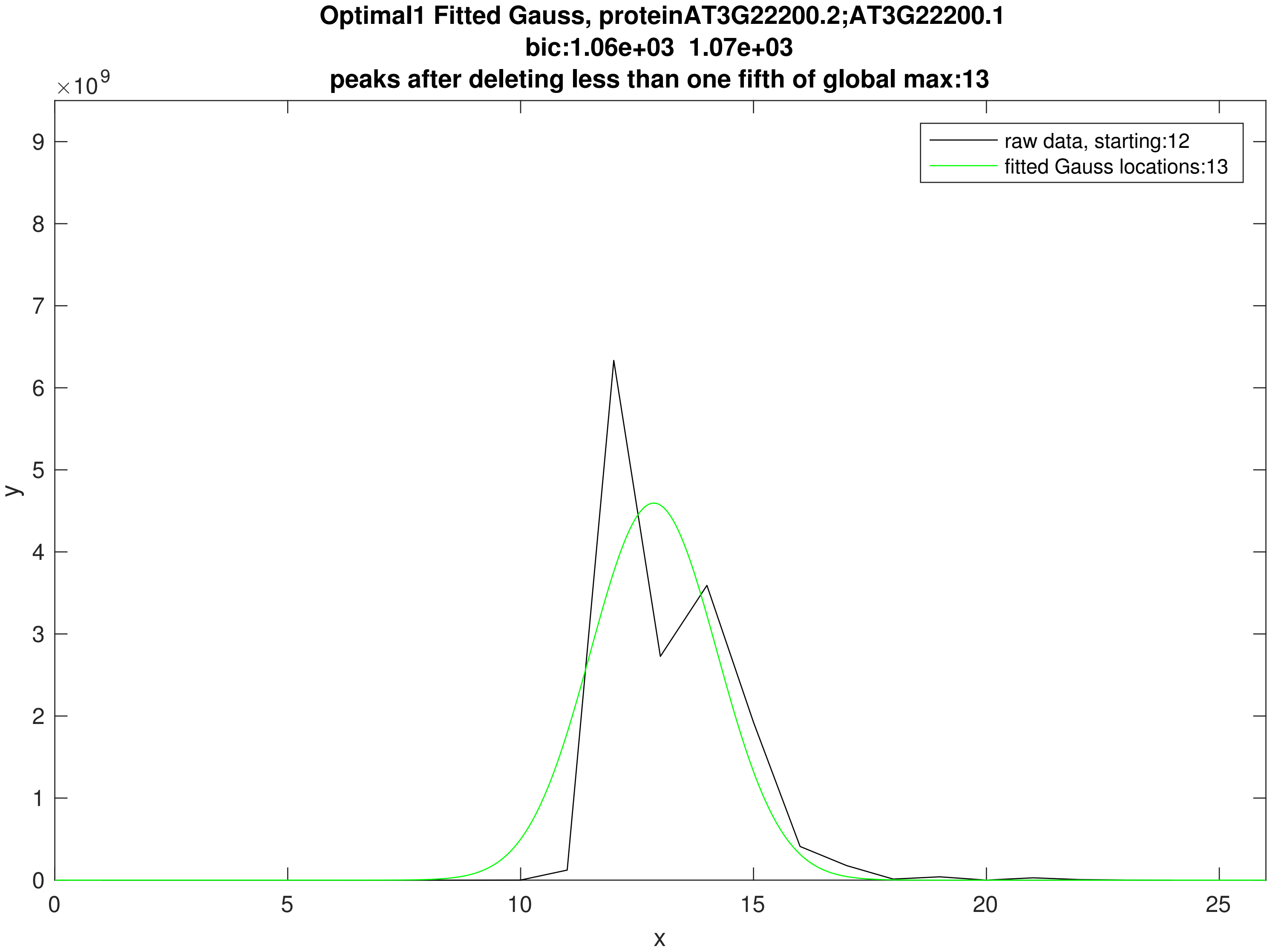
Optimal1 Fitted Gauss, proteinAT3G22200.2;AT3G22200.1
bic:1.06e+03  1.07e+03
peaks after deleting less than one fifth of global max:13
×10^9
raw data, starting:12
fitted Gauss locations:13
y
x
0
1
2
3
4
5
6
7
8
9
0
5
10
15
20
25

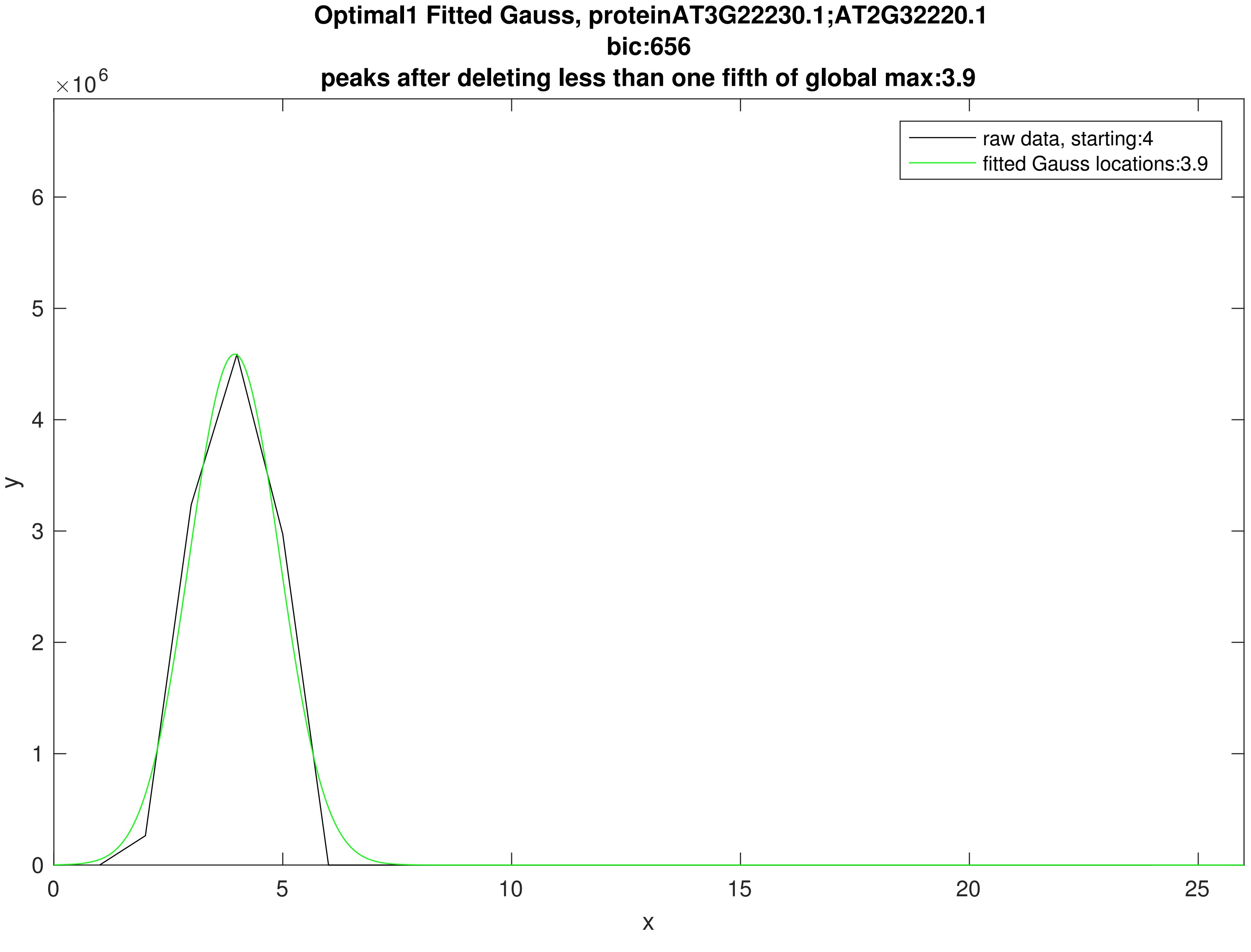
Optimal1 Fitted Gauss, proteinAT3G22230.1;AT2G32220.1
bic:656
peaks after deleting less than one fifth of global max:3.9
×10^6
raw data, starting:4
fitted Gauss locations:3.9
y
x
0
1
2
3
4
5
6
0
5
10
15
20
25

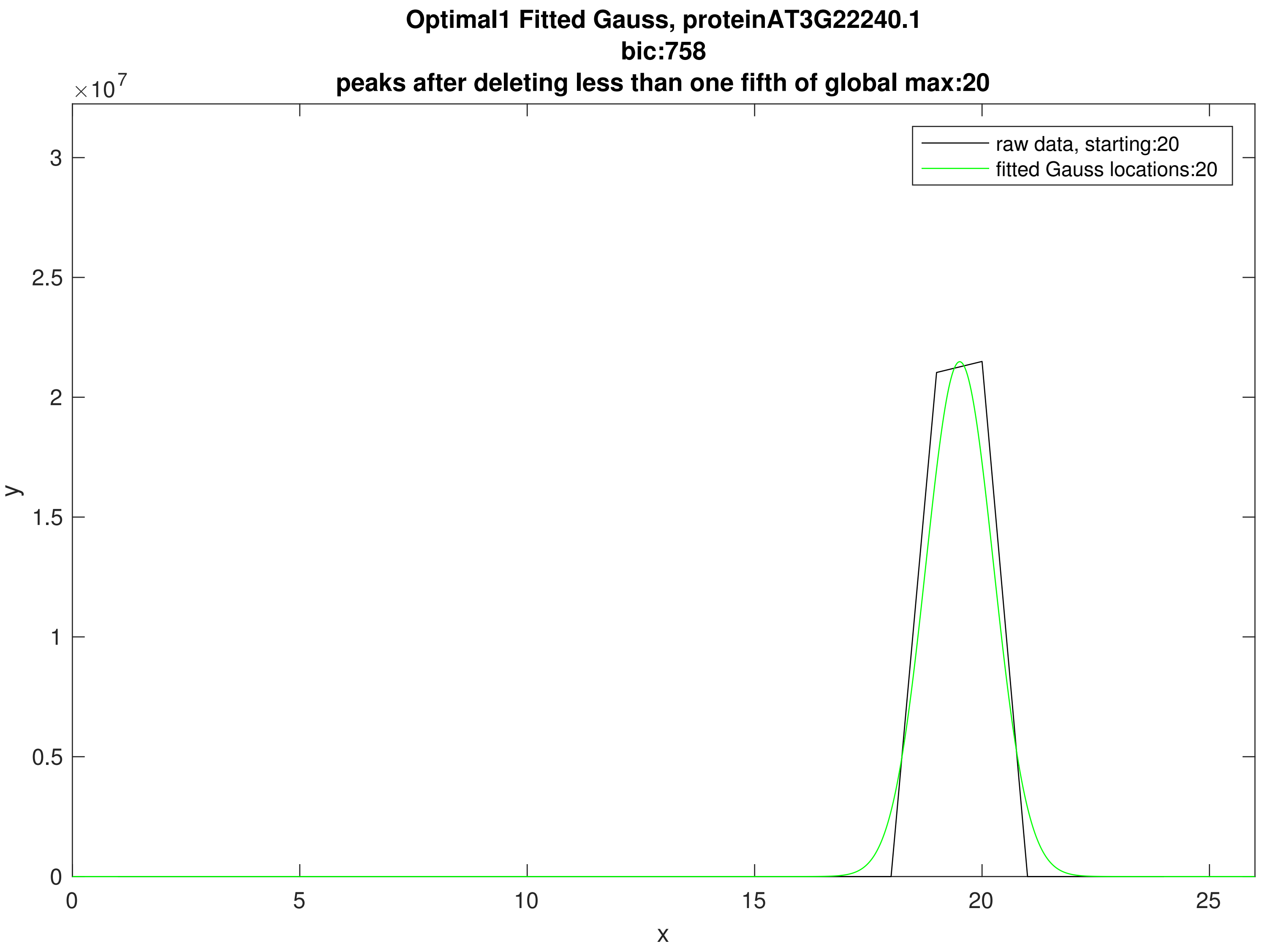
Optimal1 Fitted Gauss, proteinAT3G22240.1
bic:758
peaks after deleting less than one fifth of global max:20
×10^7
raw data, starting:20
fitted Gauss locations:20
3
2.5
2
1.5
1
0.5
0
y
0
5
10
15
20
25
x

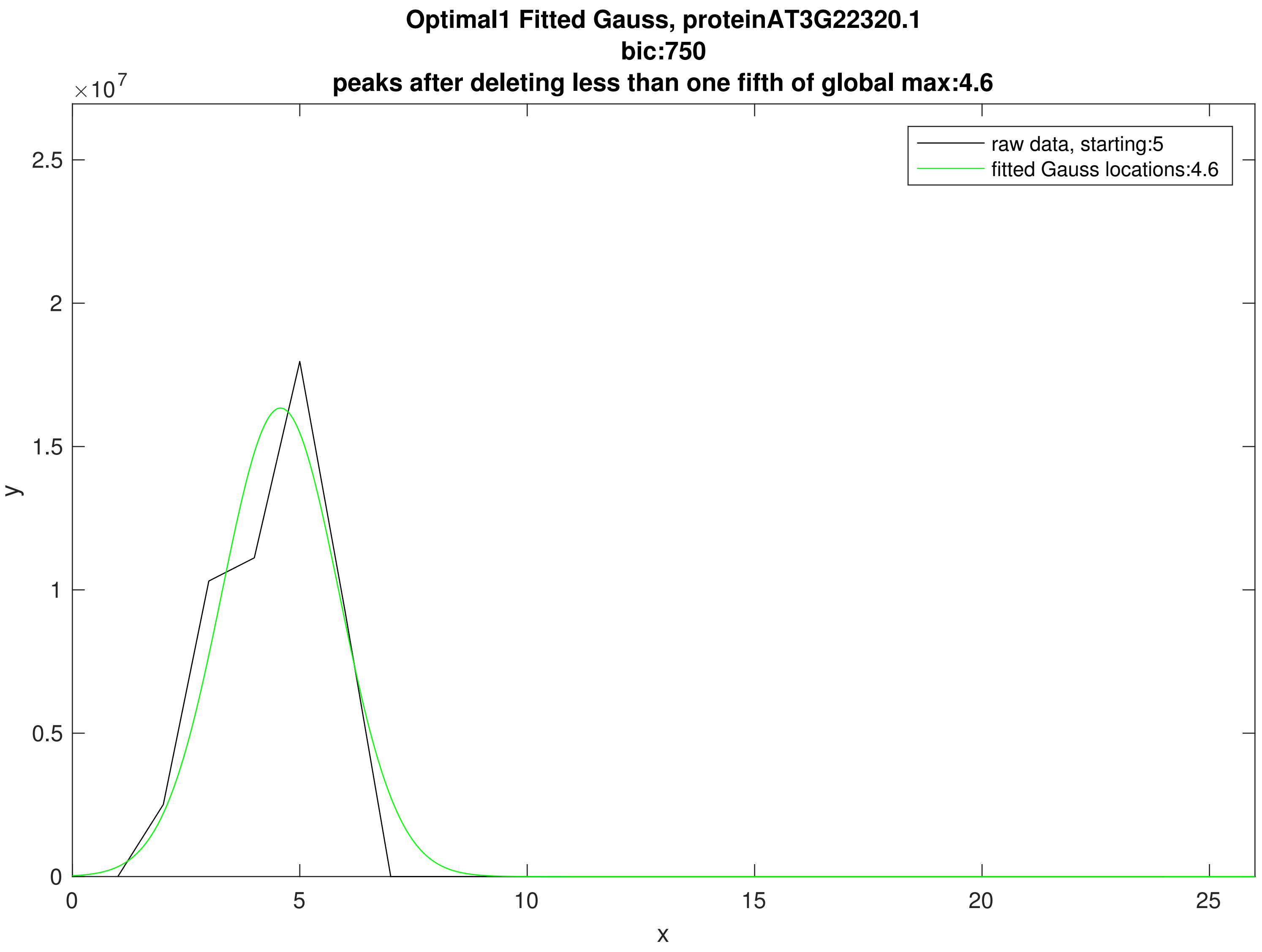
Optimal1 Fitted Gauss, proteinAT3G22320.1
bic:750
peaks after deleting less than one fifth of global max:4.6
×10^7
raw data, starting:5
fitted Gauss locations:4.6
y
x
0
0.5
1
1.5
2
2.5
5
10
15
20
25

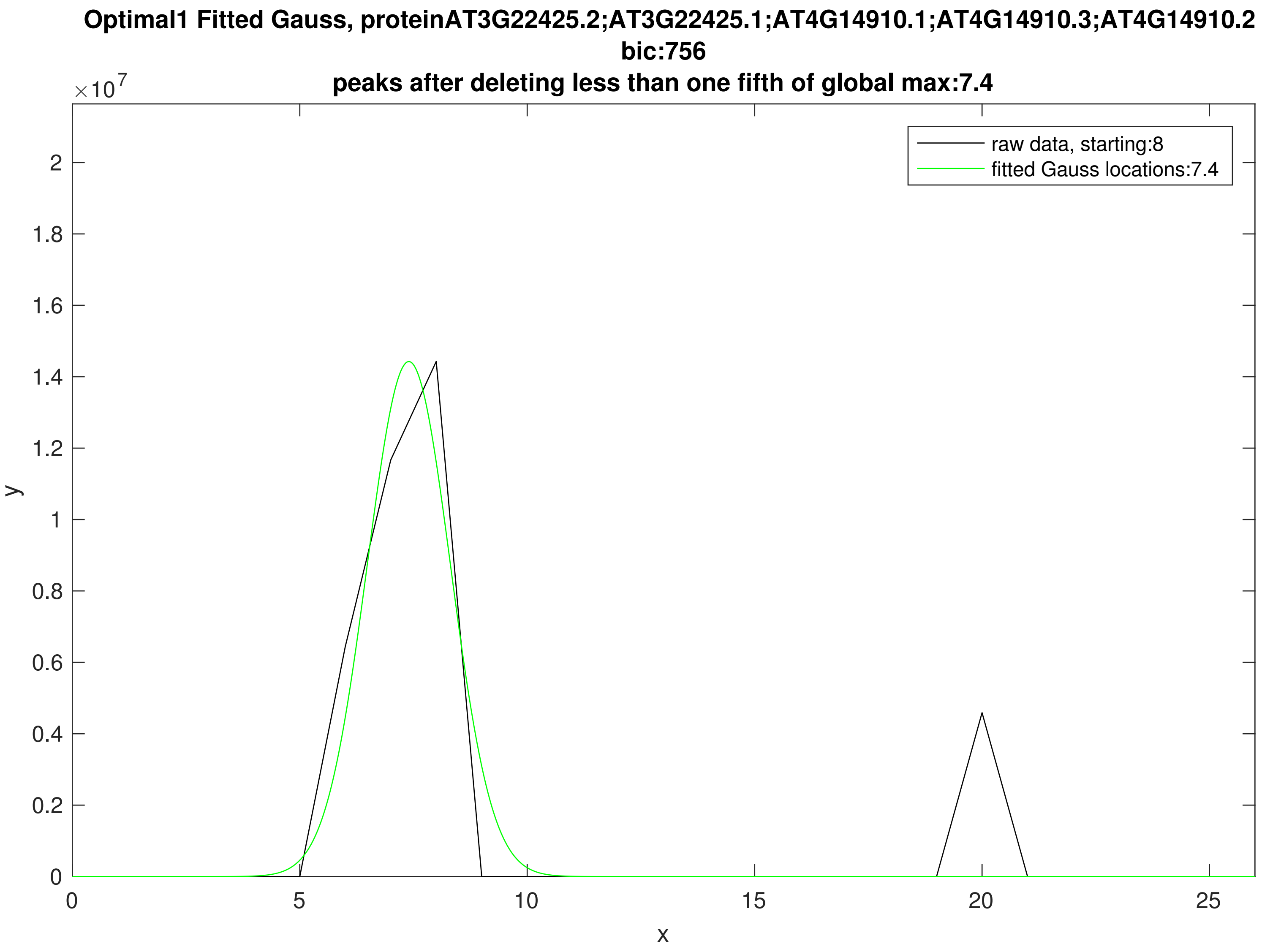
Optimal1 Fitted Gauss, proteinAT3G22425.2;AT3G22425.1;AT4G14910.1;AT4G14910.3;AT4G14910.2
bic:756
peaks after deleting less than one fifth of global max:7.4
×10^7
raw data, starting:8
fitted Gauss locations:7.4
y
x
0
0.2
0.4
0.6
0.8
1
1.2
1.4
1.6
1.8
2
5
10
15
20
25

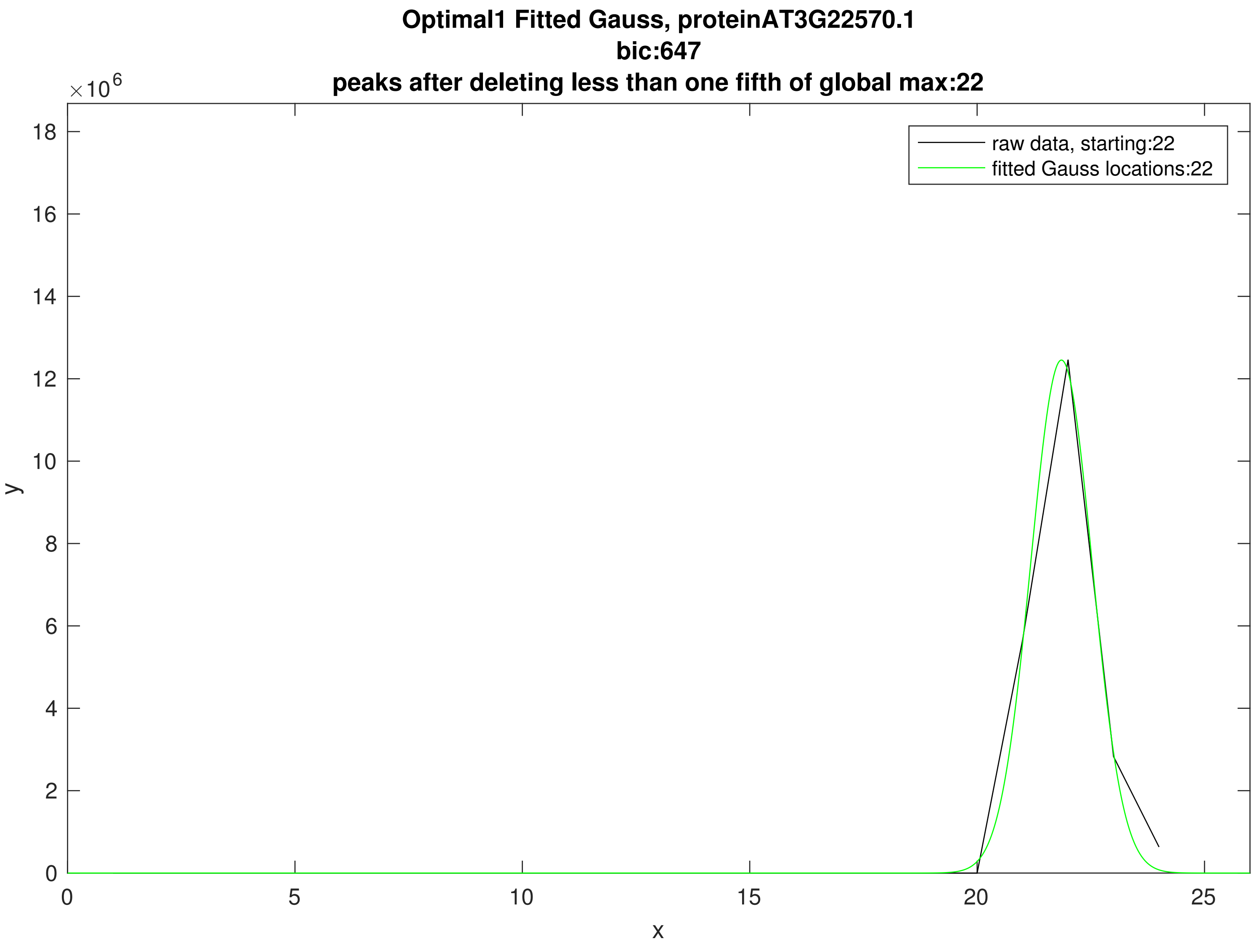

Optimal1 Fitted Gauss, proteinAT3G22570.1
bic:647
peaks after deleting less than one fifth of global max:22
×10^6
raw data, starting:22
fitted Gauss locations:22
y
x
0
2
4
6
8
10
12
14
16
18
0
5
10
15
20
25

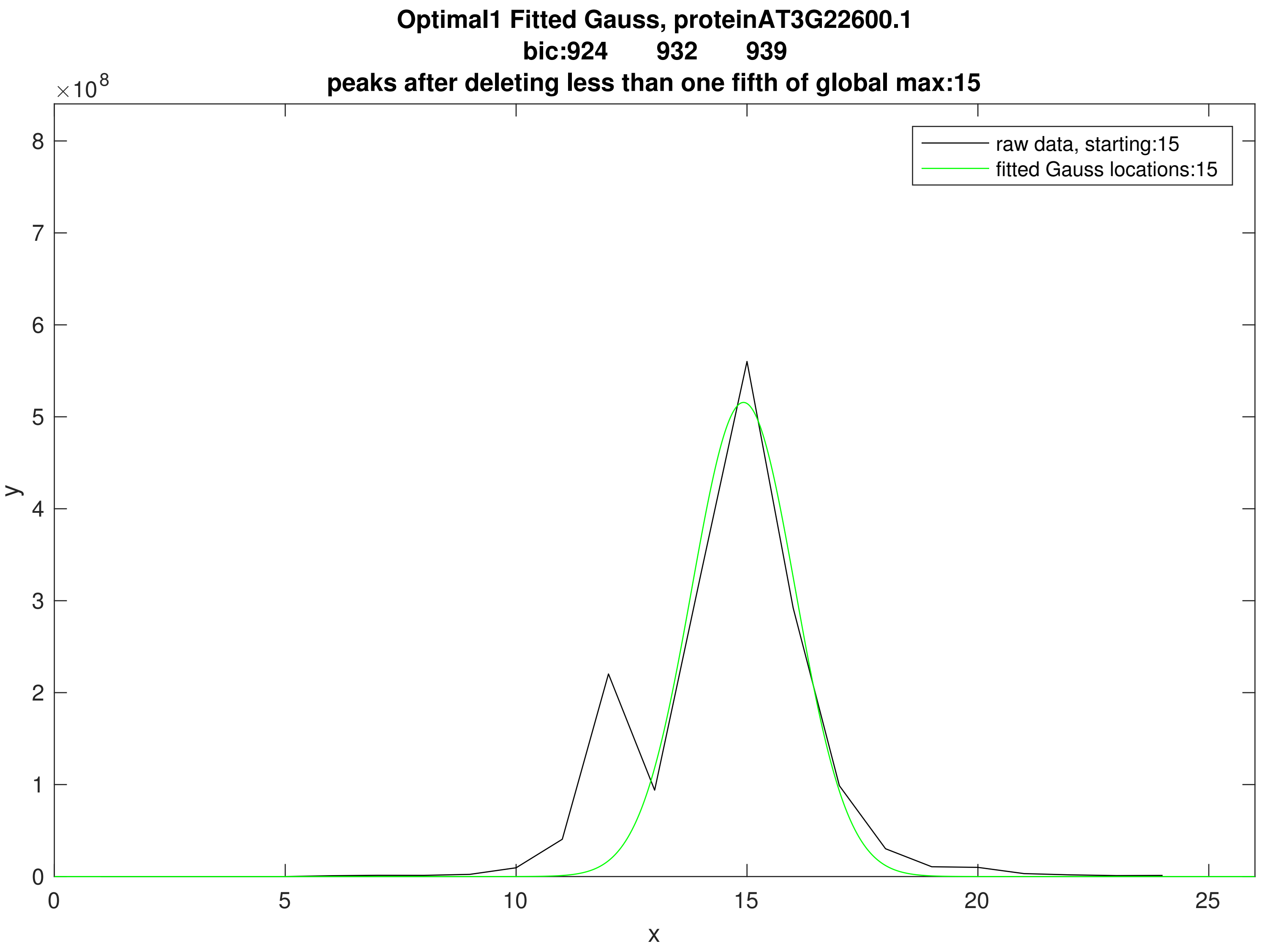
Optimal1 Fitted Gauss, proteinAT3G22600.1
bic:924 932 939
peaks after deleting less than one fifth of global max:15
×10^8
raw data, starting:15
fitted Gauss locations:15
y
x
0
1
2
3
4
5
6
7
8
0
5
10
15
20
25

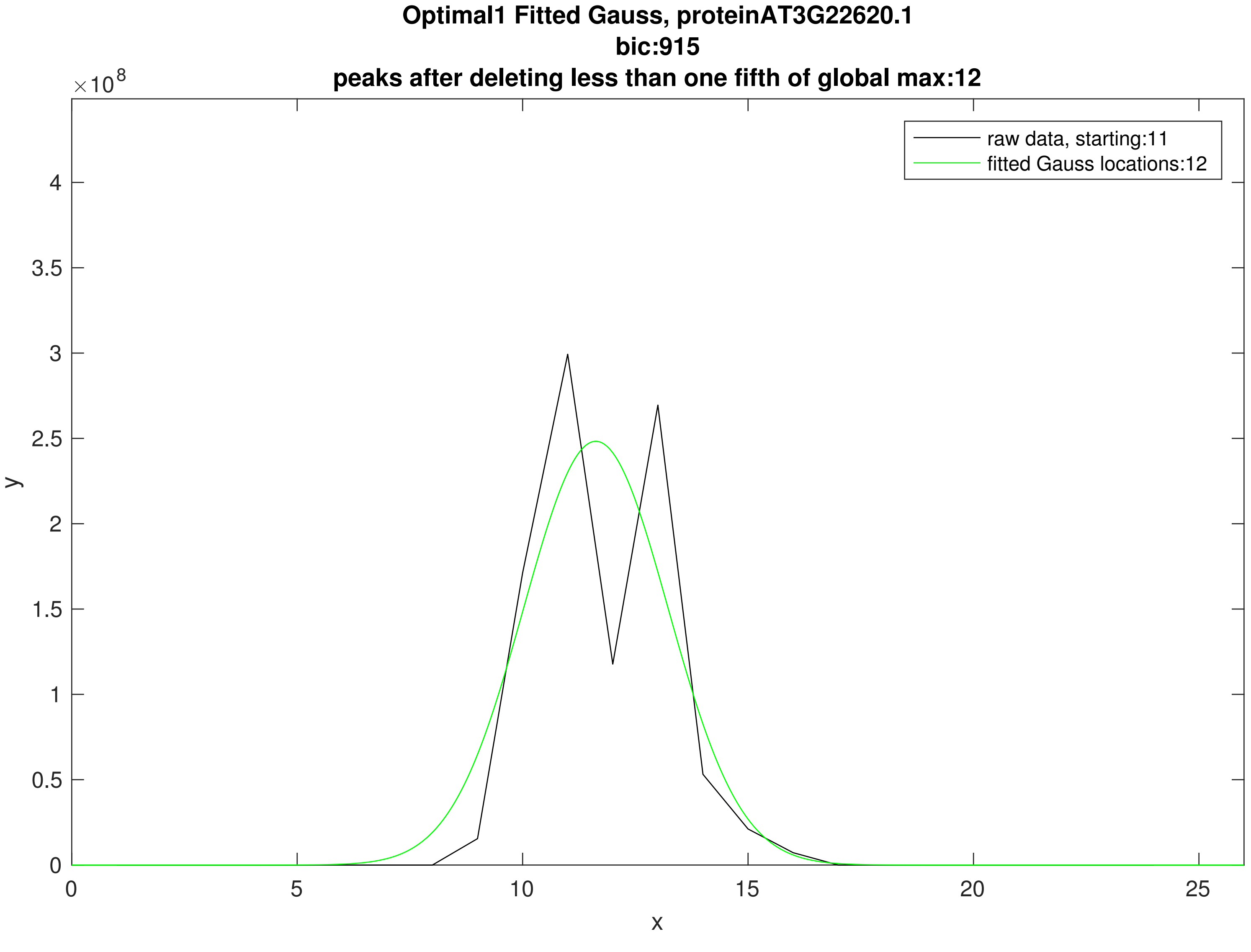

Optimal1 Fitted Gauss, proteinAT3G22620.1
bic:915
peaks after deleting less than one fifth of global max:12
×10^8
raw data, starting:11
fitted Gauss locations:12
4
3.5
3
2.5
2
1.5
1
0.5
0
y
0
5
10
15
20
25
x

**Optimal1 Fitted Gauss, proteinAT3G22630.1**
**bic:885**
**peaks after deleting less than one fifth of global max:5.1**

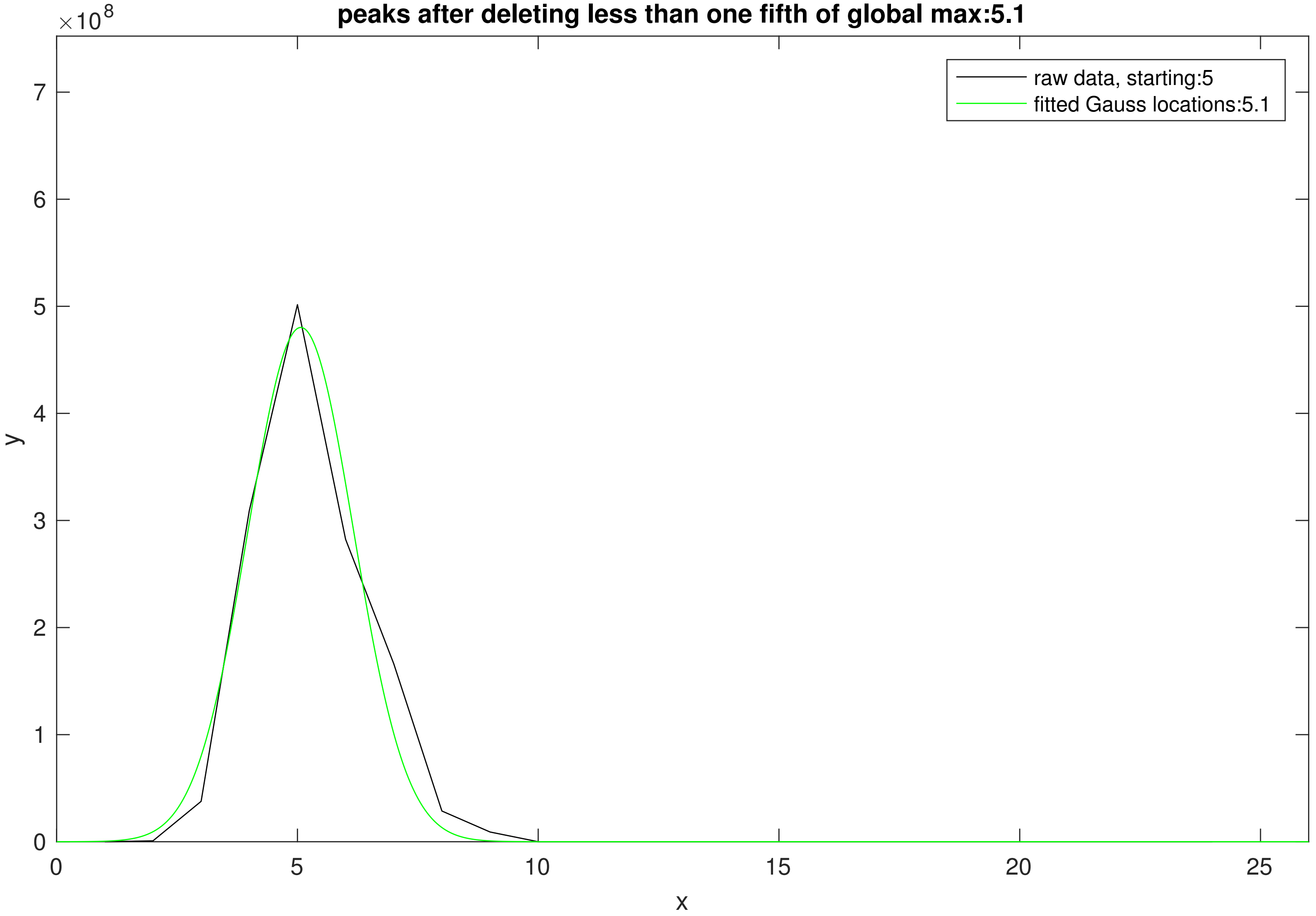

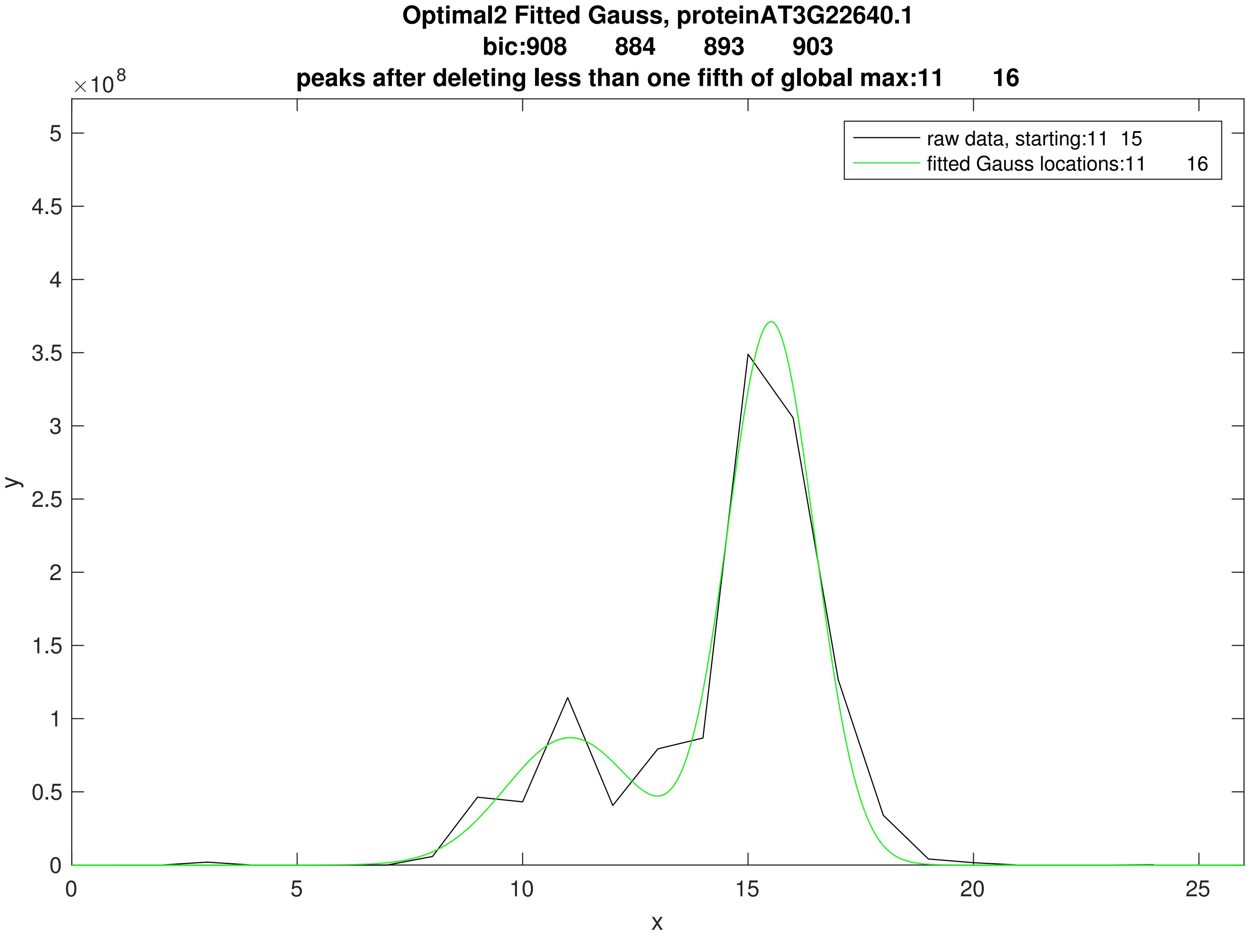

Optimal2 Fitted Gauss, proteinAT3G22640.1
bic:908 884 893 903
peaks after deleting less than one fifth of global max:11 16
×10^8
raw data, starting:11 15
fitted Gauss locations:11 16
y
x
0
0.5
1
1.5
2
2.5
3
3.5
4
4.5
5
0
5
10
15
20
25

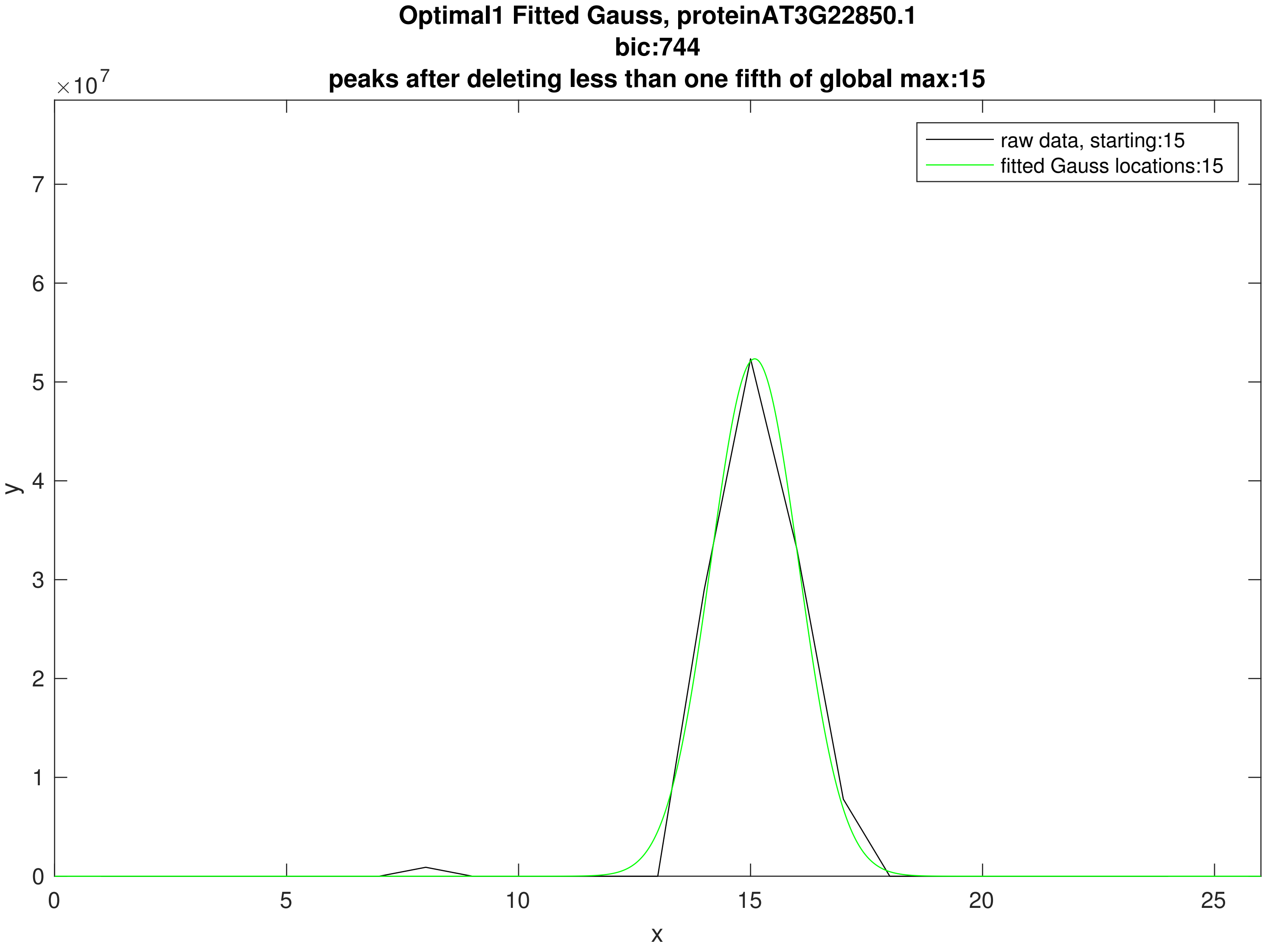
Optimal1 Fitted Gauss, proteinAT3G22850.1
bic:744
peaks after deleting less than one fifth of global max:15
×10^7
raw data, starting:15
fitted Gauss locations:15
y
x
0
1
2
3
4
5
6
7
0
5
10
15
20
25

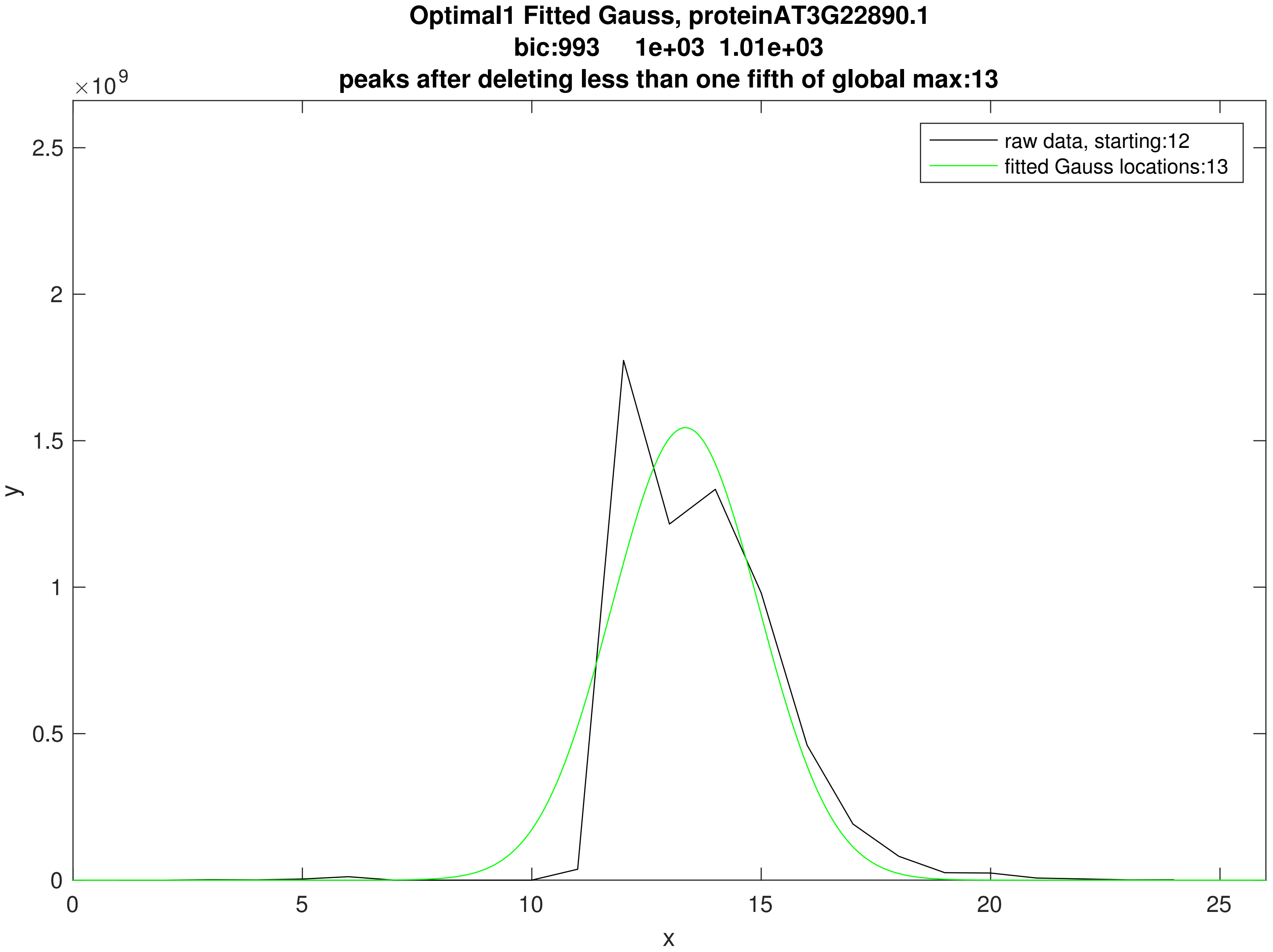

Optimal1 Fitted Gauss, proteinAT3G22890.1
bic:993 1e+03 1.01e+03
peaks after deleting less than one fifth of global max:13
×10^9
raw data, starting:12
fitted Gauss locations:13
y
x

Optimal2 Fitted Gauss, proteinAT3G22960.1
bic:915 897 907 916
peaks after deleting less than one fifth of global max:8.4

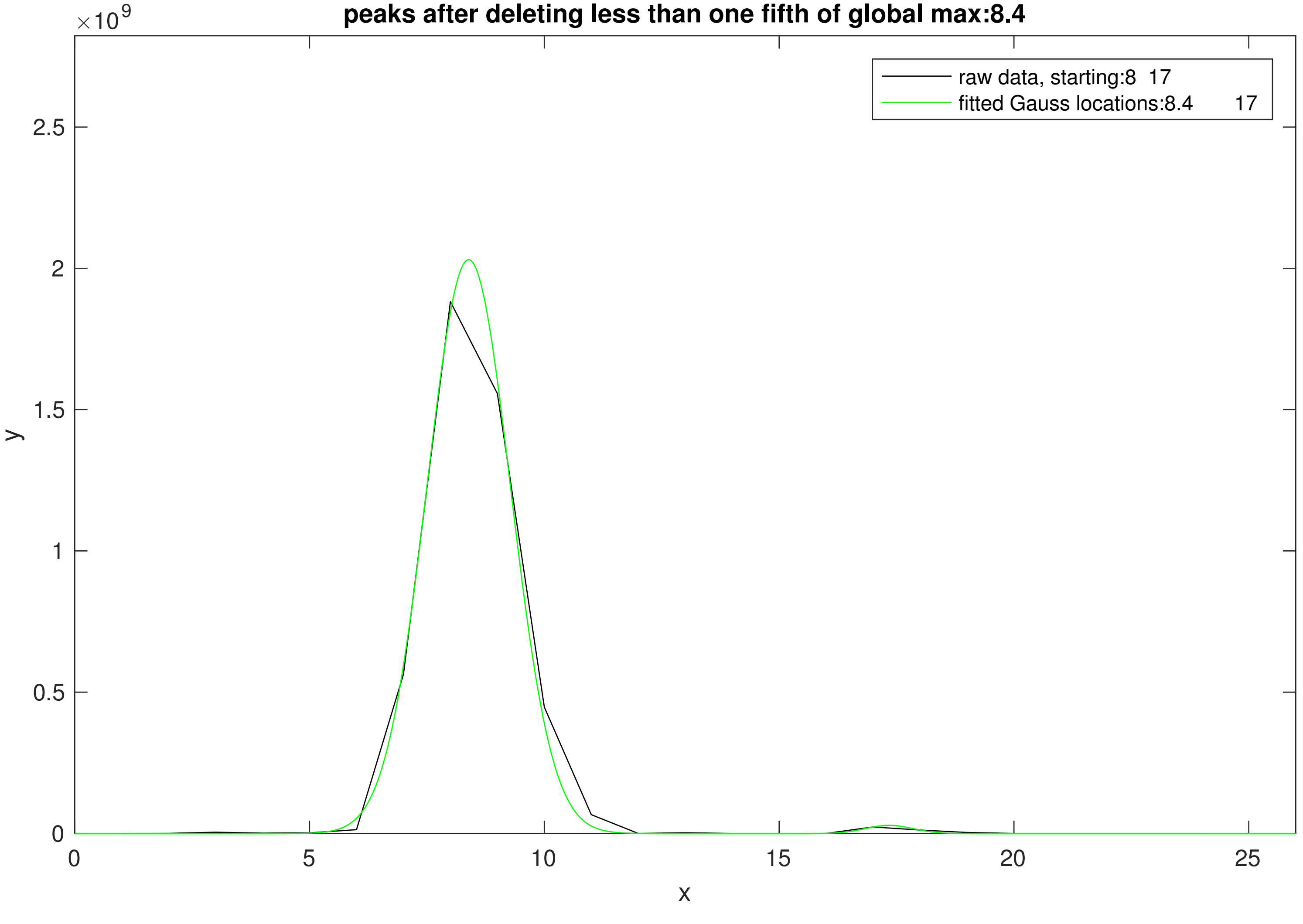

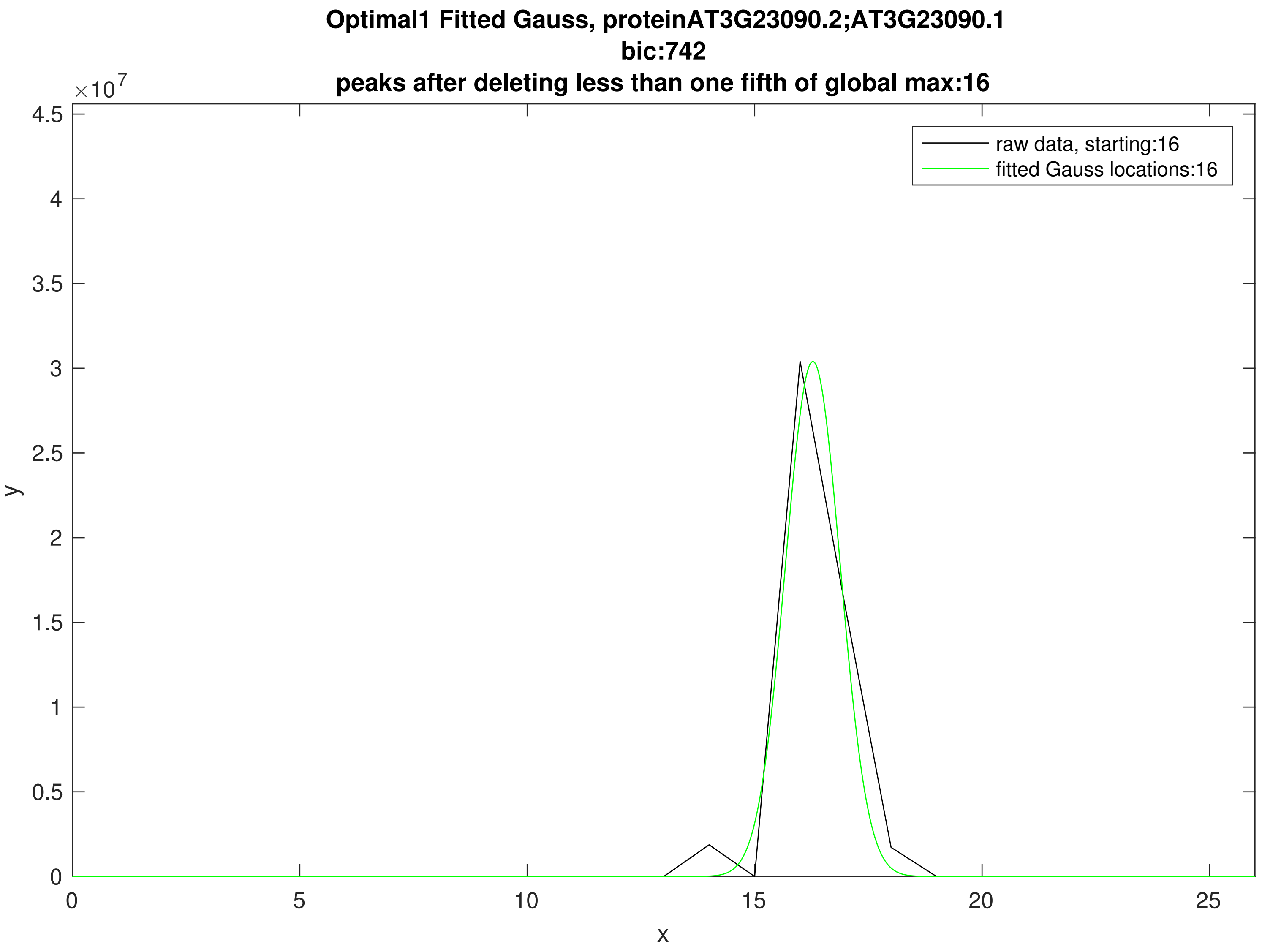
Optimal1 Fitted Gauss, proteinAT3G23090.2;AT3G23090.1
bic:742
peaks after deleting less than one fifth of global max:16
×10^7
raw data, starting:16
fitted Gauss locations:16
y
x
0
0.5
1
1.5
2
2.5
3
3.5
4
4.5
5
10
15
20
25

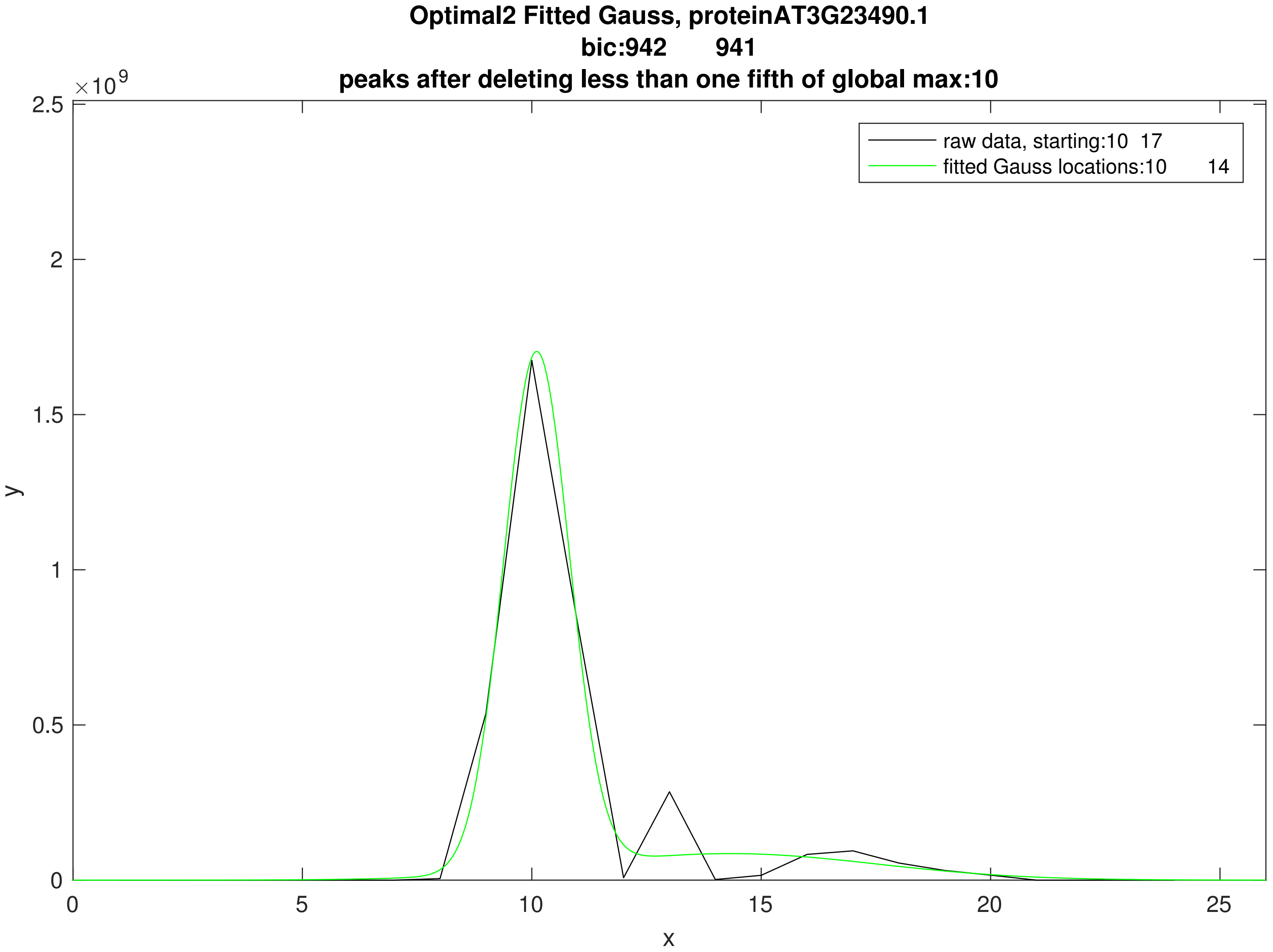
Optimal2 Fitted Gauss, proteinAT3G23490.1
bic:942 941
peaks after deleting less than one fifth of global max:10
raw data, starting:10 17
fitted Gauss locations:10 14
×10^9
2.5
2
1.5
1
0.5
0
y
0
5
10
15
20
25
x

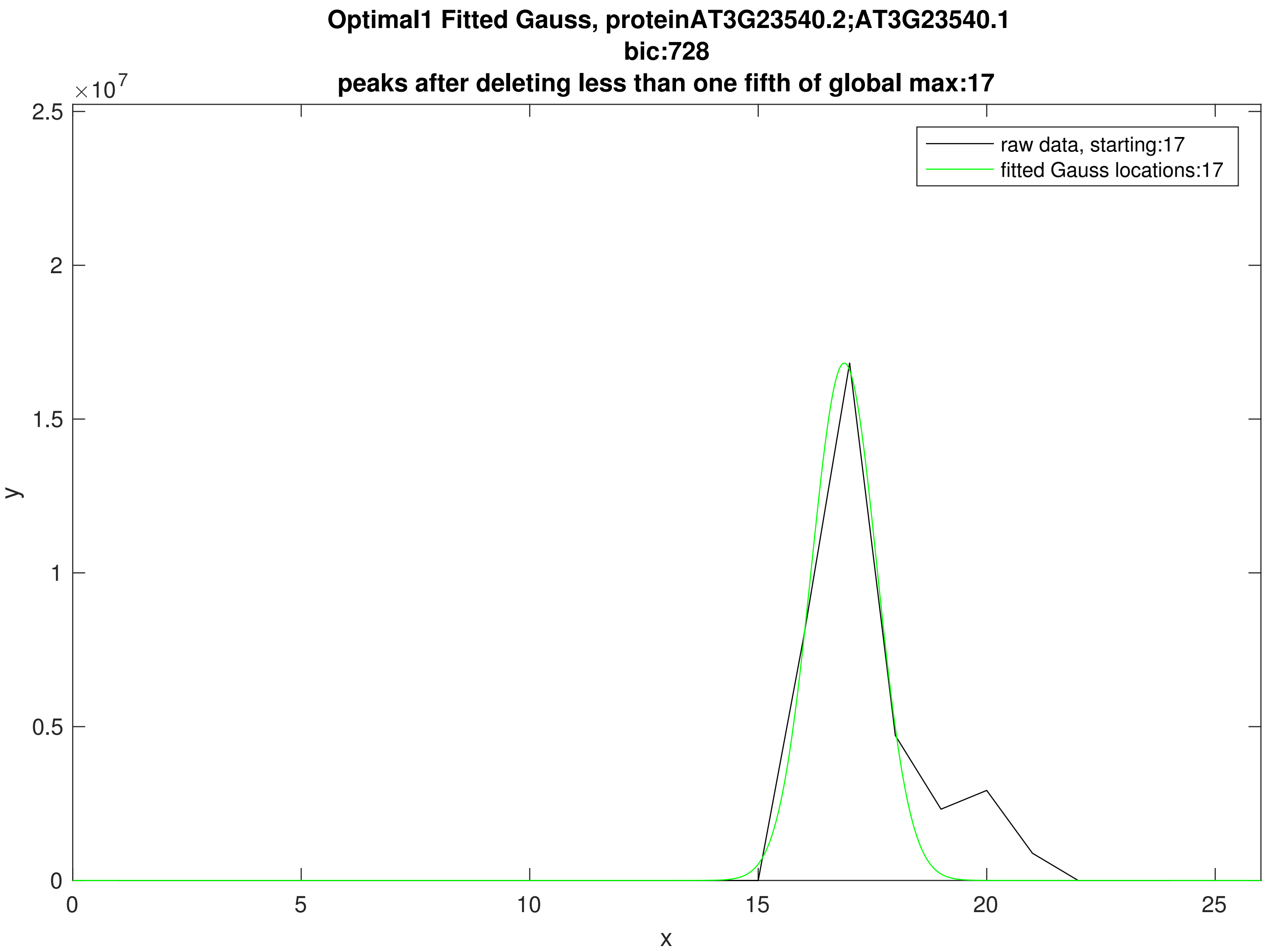

Optimal1 Fitted Gauss, proteinAT3G23540.2;AT3G23540.1
bic:728
peaks after deleting less than one fifth of global max:17
×10^7
2.5
2
1.5
1
0.5
0
y
0
5
10
15
20
25
x
raw data, starting:17
fitted Gauss locations:17

**Optimal1 Fitted Gauss, proteinAT3G23570.1**
**bic:878**
**peaks after deleting less than one fifth of global max:18**

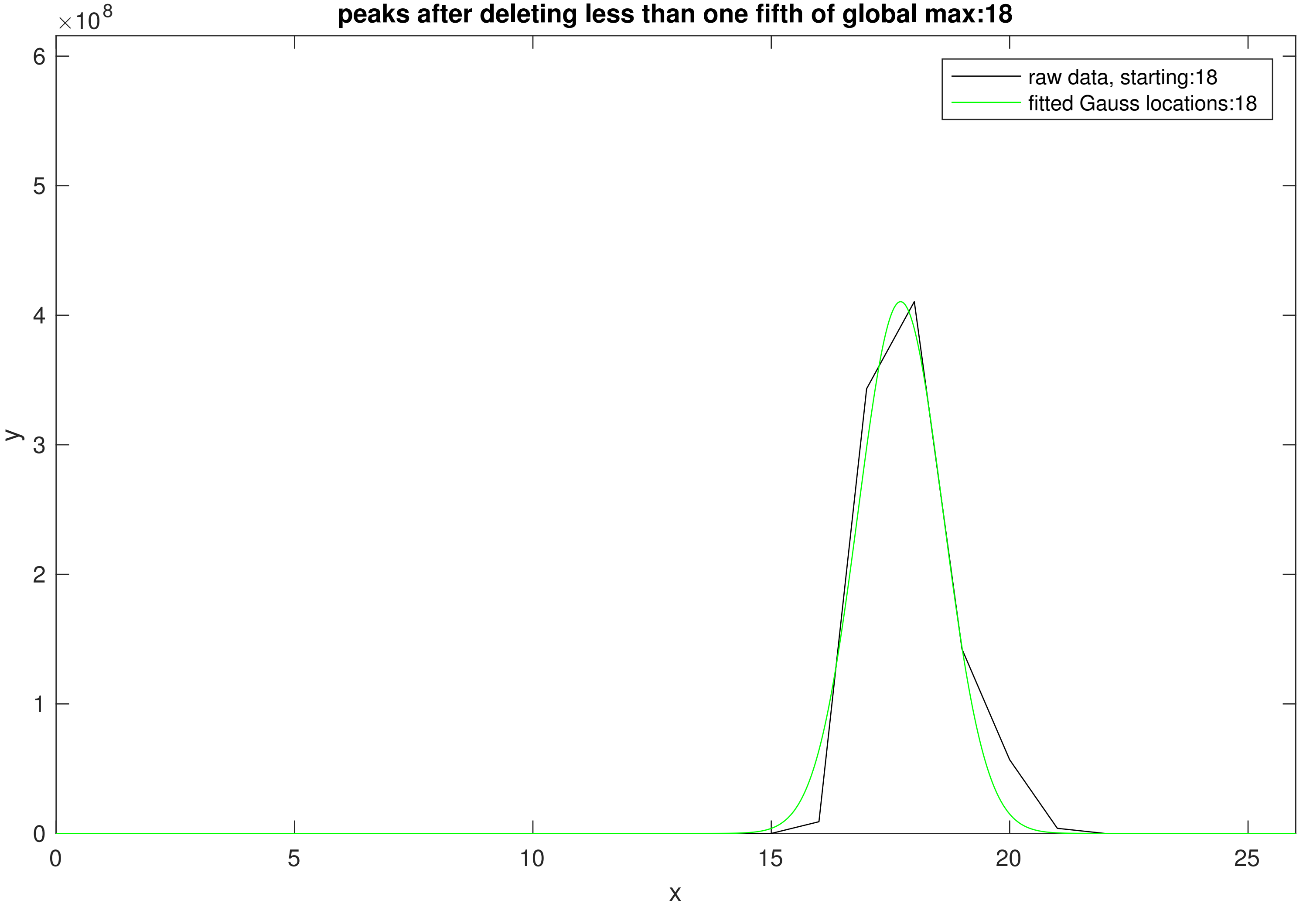

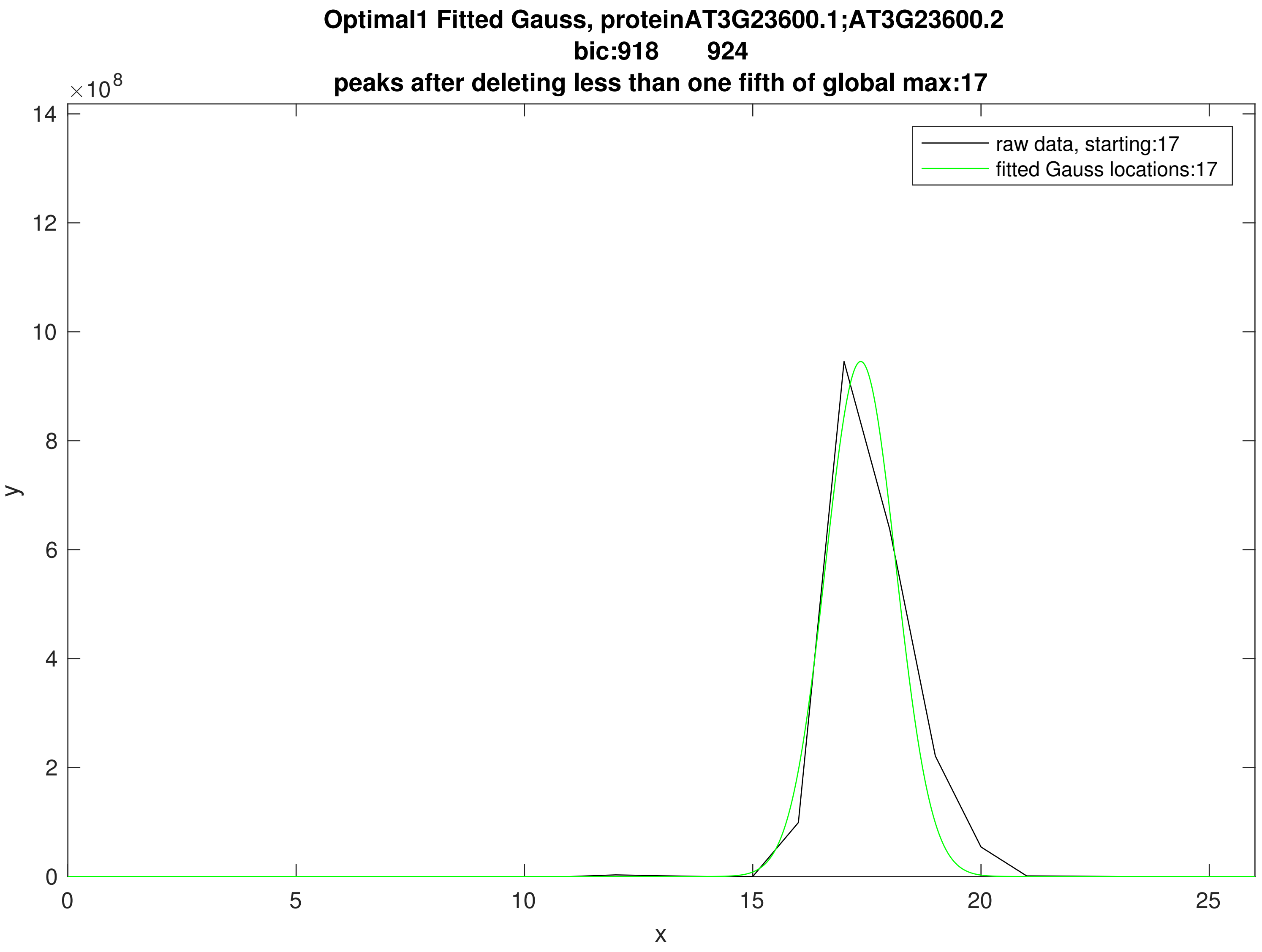

Optimal1 Fitted Gauss, proteinAT3G23600.1;AT3G23600.2
bic:918 924
peaks after deleting less than one fifth of global max:17
×10^8
14
12
10
8
6
4
2
0
y
0
5
10
15
20
25
x
raw data, starting:17
fitted Gauss locations:17

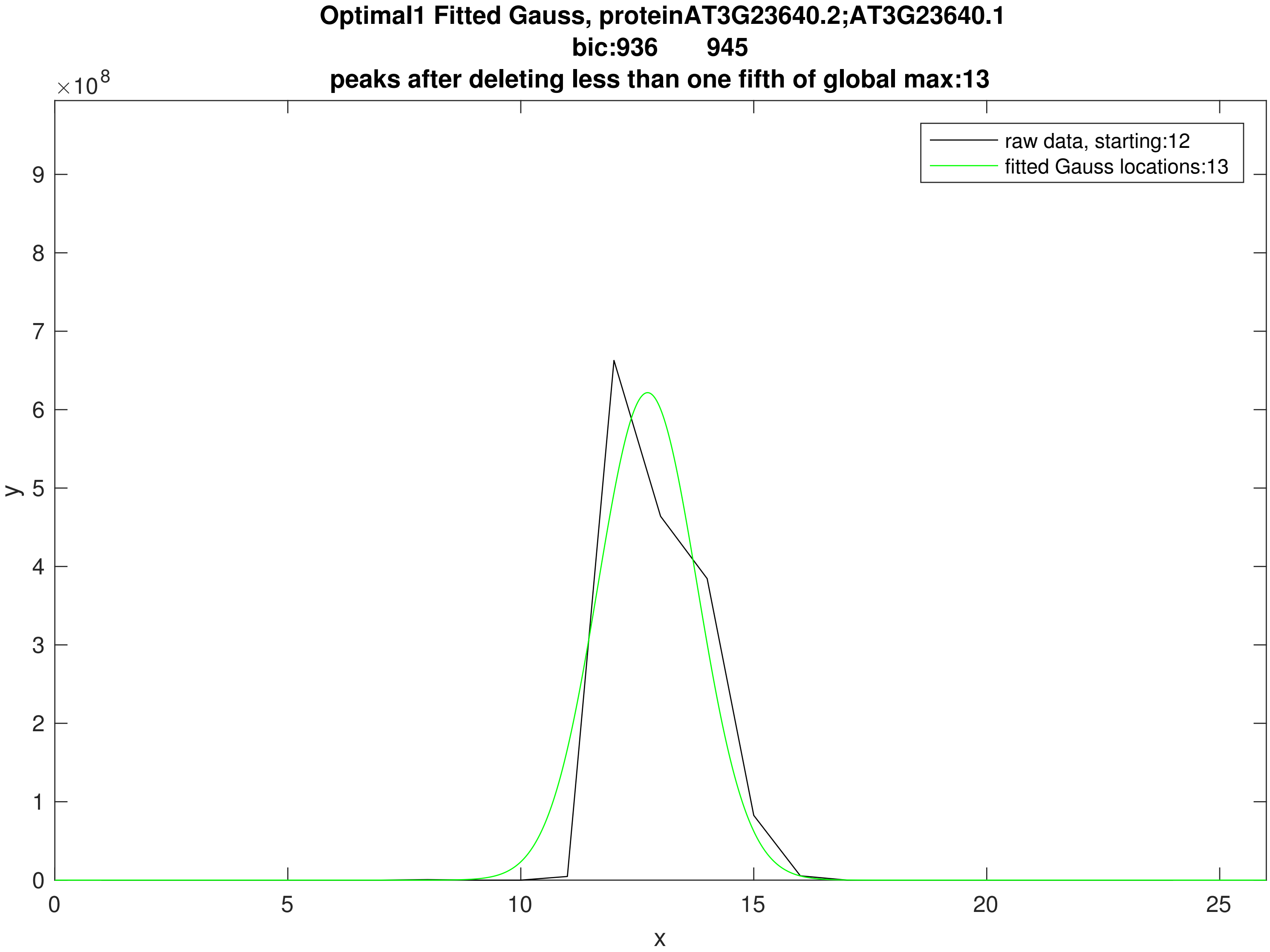
Optimal1 Fitted Gauss, proteinAT3G23640.2;AT3G23640.1
bic:936 945
peaks after deleting less than one fifth of global max:13
×10^8
raw data, starting:12
fitted Gauss locations:13
y
x
0
1
2
3
4
5
6
7
8
9
0
5
10
15
20
25

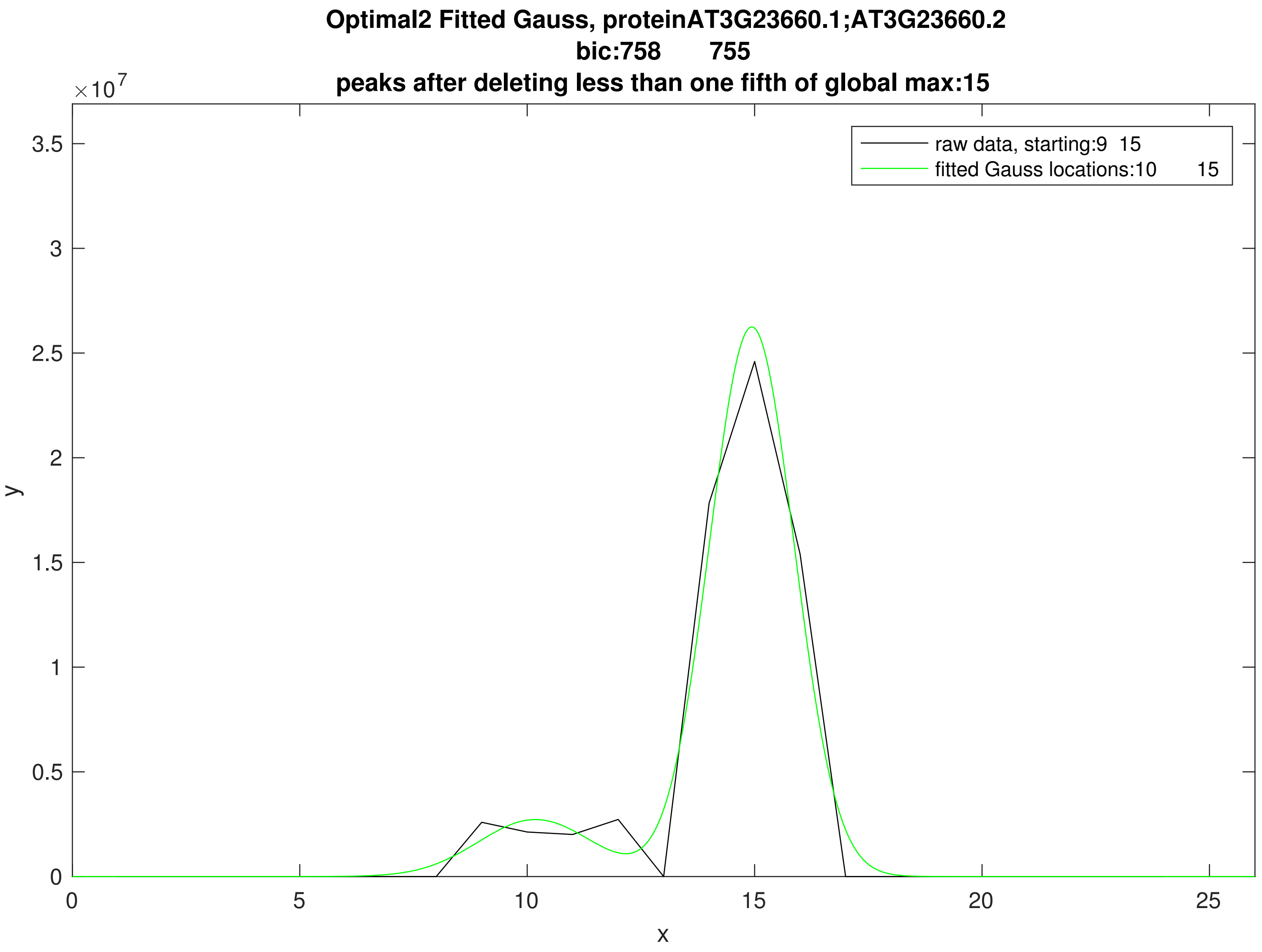
Optimal2 Fitted Gauss, proteinAT3G23660.1;AT3G23660.2
bic:758 755
peaks after deleting less than one fifth of global max:15
×10^7
3.5
3
2.5
2
1.5
1
0.5
0
y
0
5
10
15
20
25
x
raw data, starting:9 15
fitted Gauss locations:10 15

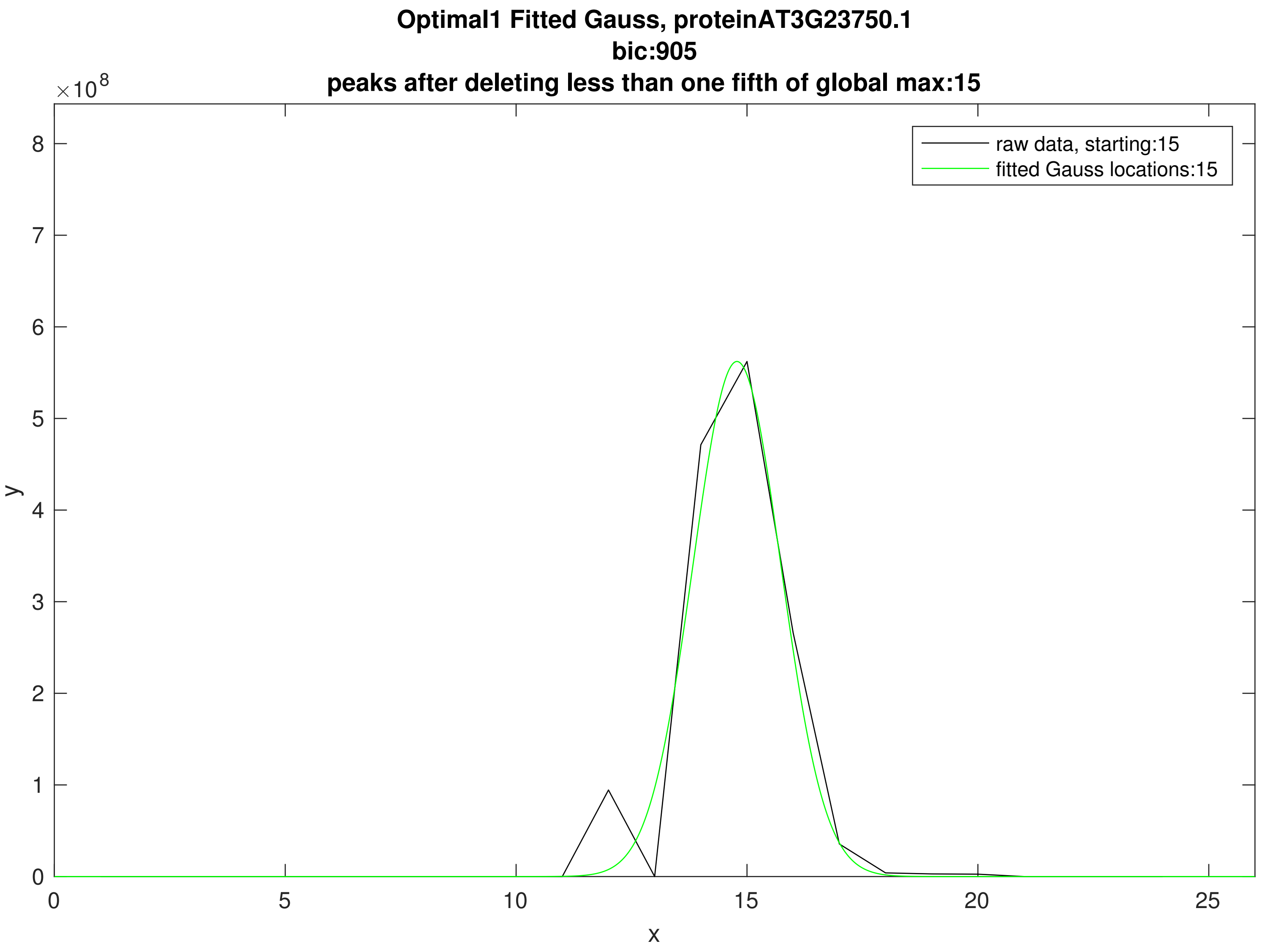
Optimal1 Fitted Gauss, proteinAT3G23750.1
bic:905
peaks after deleting less than one fifth of global max:15
×10^8
raw data, starting:15
fitted Gauss locations:15
y
x
0
1
2
3
4
5
6
7
8
0
5
10
15
20
25

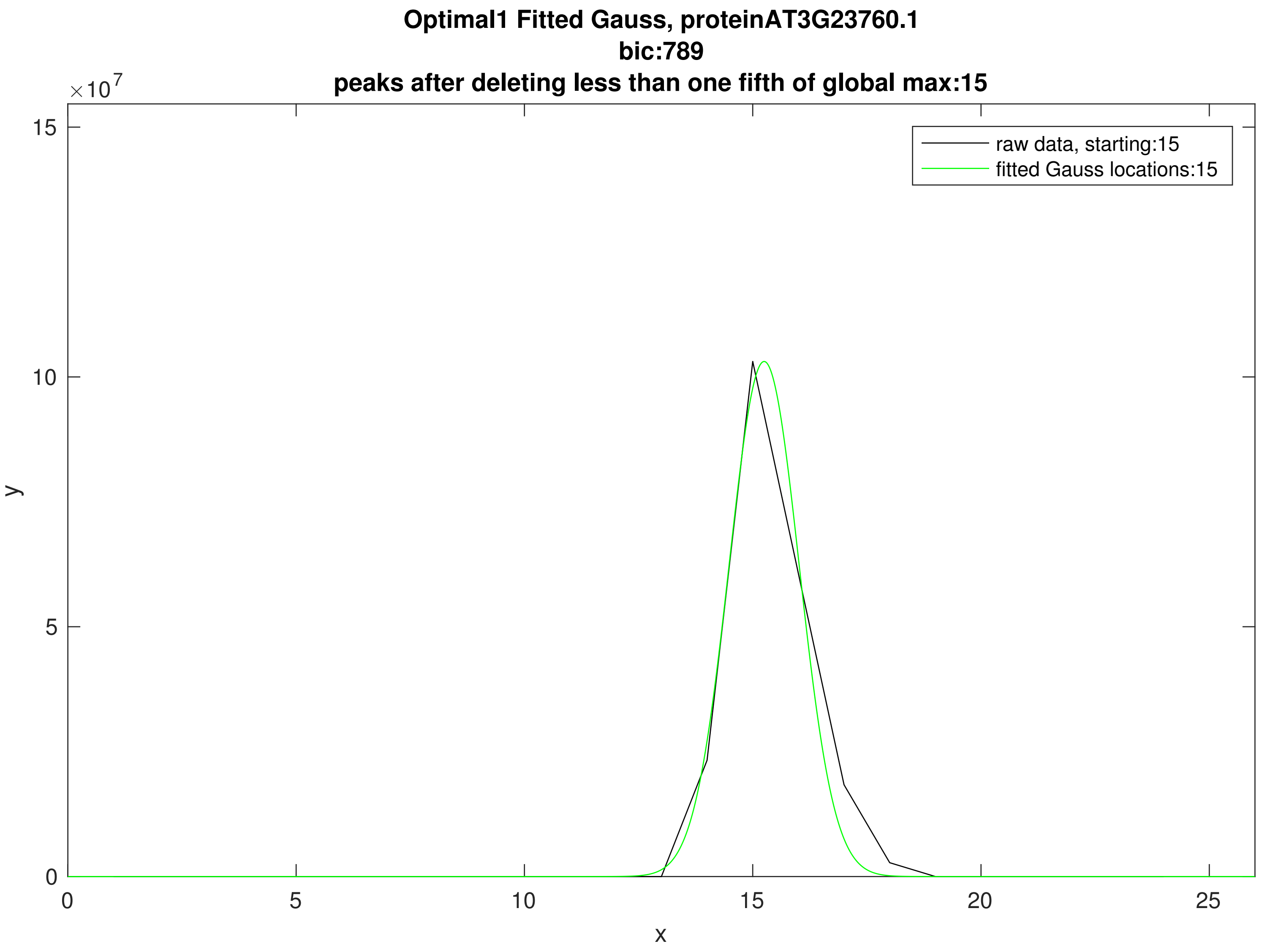
Optimal1 Fitted Gauss, proteinAT3G23760.1
bic:789
peaks after deleting less than one fifth of global max:15
$\times 10^7$
raw data, starting:15
fitted Gauss locations:15
y
x
0
5
10
15
20
25

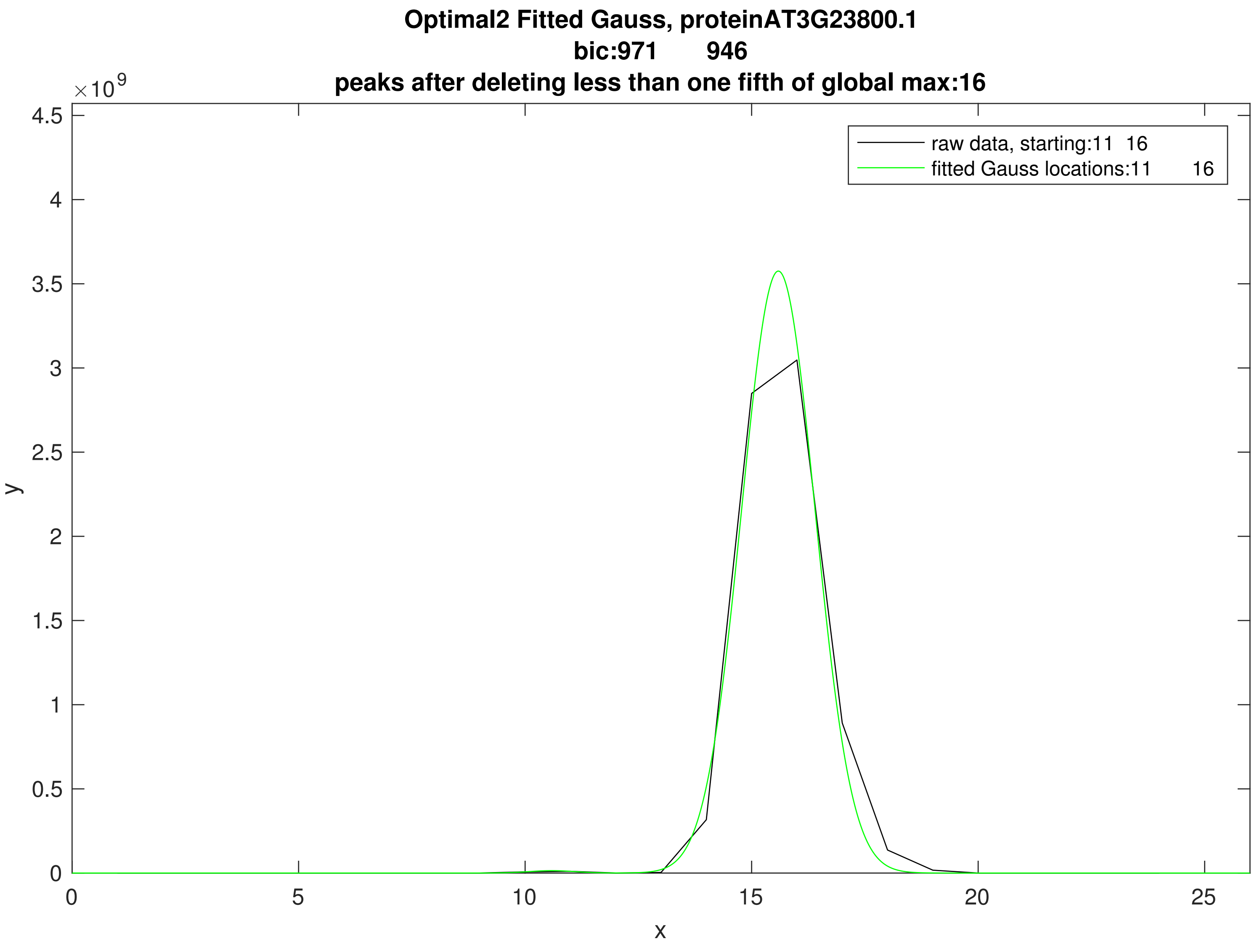
Optimal2 Fitted Gauss, proteinAT3G23800.1
bic:971 946
peaks after deleting less than one fifth of global max:16
×10^9
raw data, starting:11 16
fitted Gauss locations:11 16
y
x
0
0.5
1
1.5
2
2.5
3
3.5
4
4.5
0
5
10
15
20
25

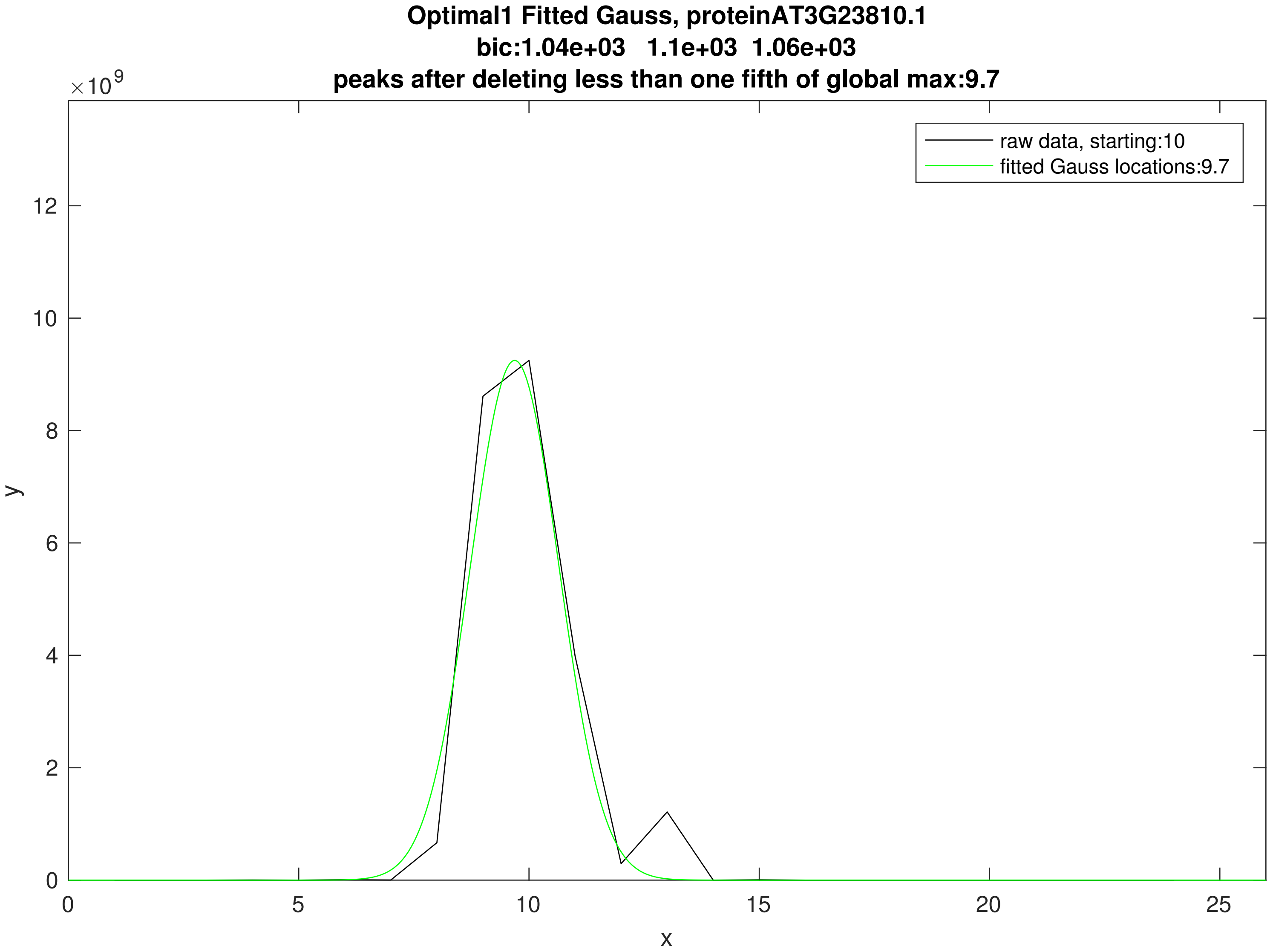
Optimal1 Fitted Gauss, proteinAT3G23810.1
bic:1.04e+03 1.1e+03 1.06e+03
peaks after deleting less than one fifth of global max:9.7
×10^9
raw data, starting:10
fitted Gauss locations:9.7
y
x
0
2
4
6
8
10
12
5
15
20
25

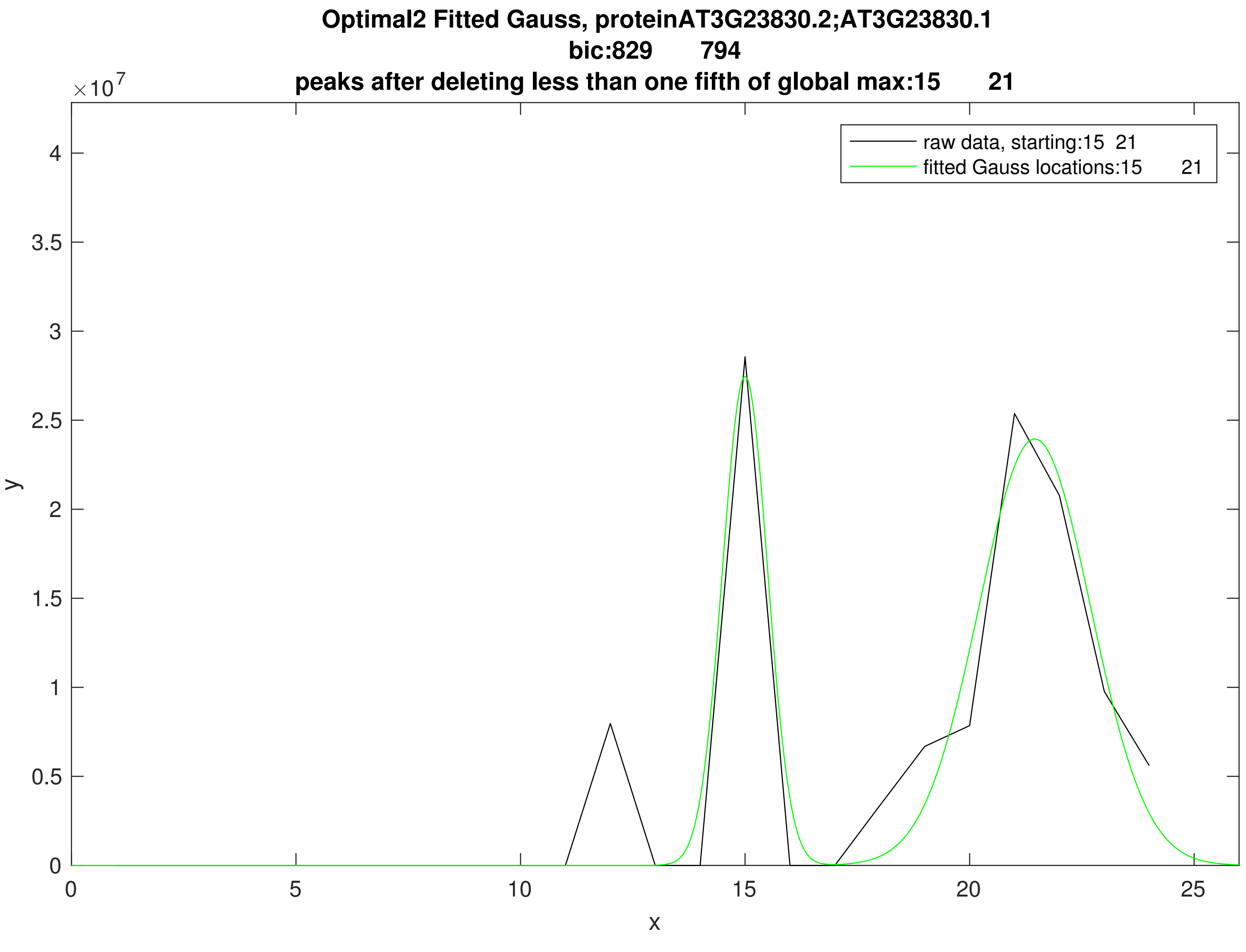
Optimal2 Fitted Gauss, proteinAT3G23830.2;AT3G23830.1
bic:829 794
peaks after deleting less than one fifth of global max:15 21
raw data, starting:15 21
fitted Gauss locations:15 21
×10^7
4
3.5
3
2.5
2
1.5
1
0.5
0
y
0
5
10
15
20
25
x

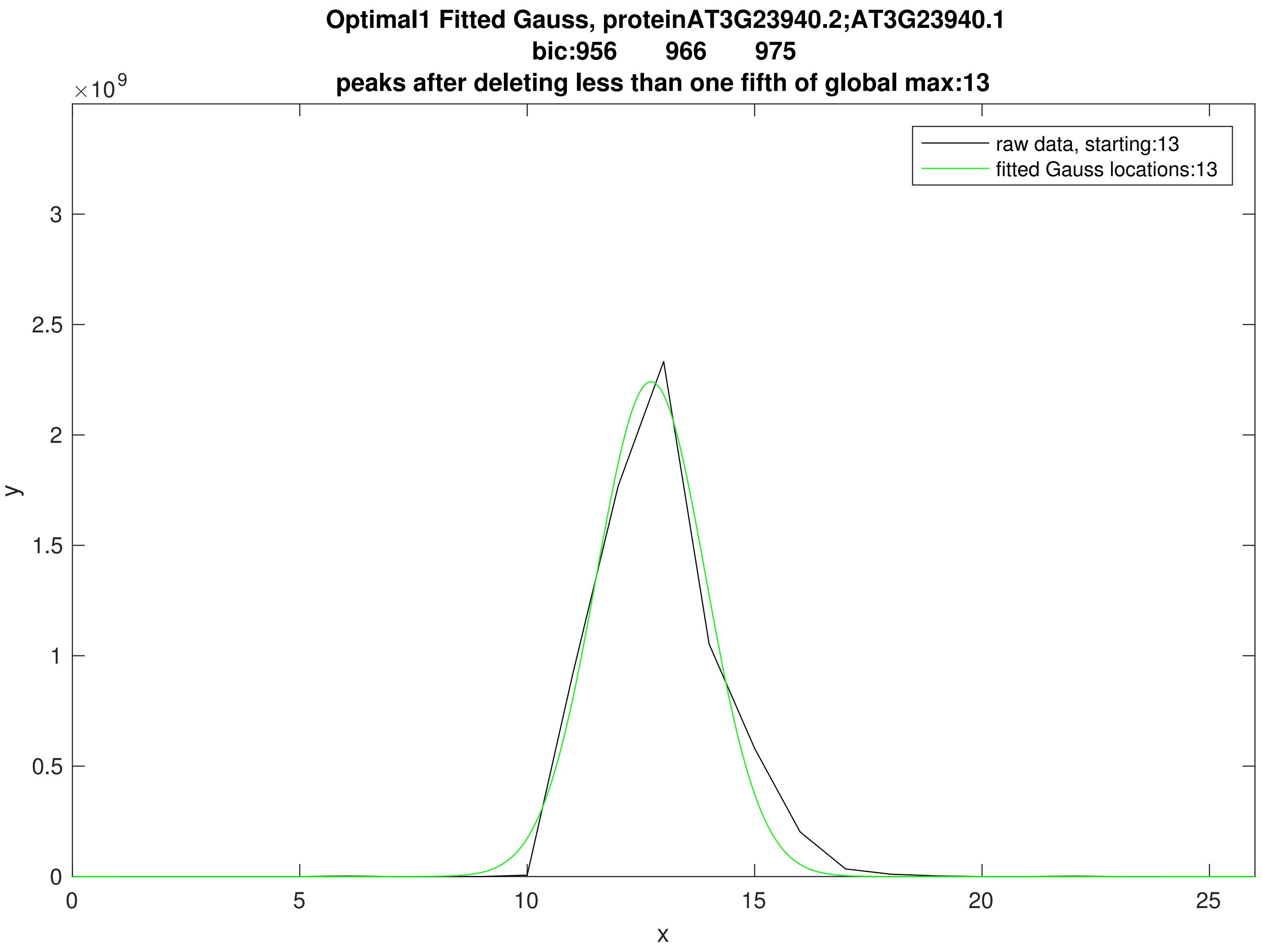
Optimal1 Fitted Gauss, proteinAT3G23940.2;AT3G23940.1
bic:956 966 975
peaks after deleting less than one fifth of global max:13
×10^9
raw data, starting:13
fitted Gauss locations:13
y
x
0
0.5
1
1.5
2
2.5
3
0
5
10
15
20
25

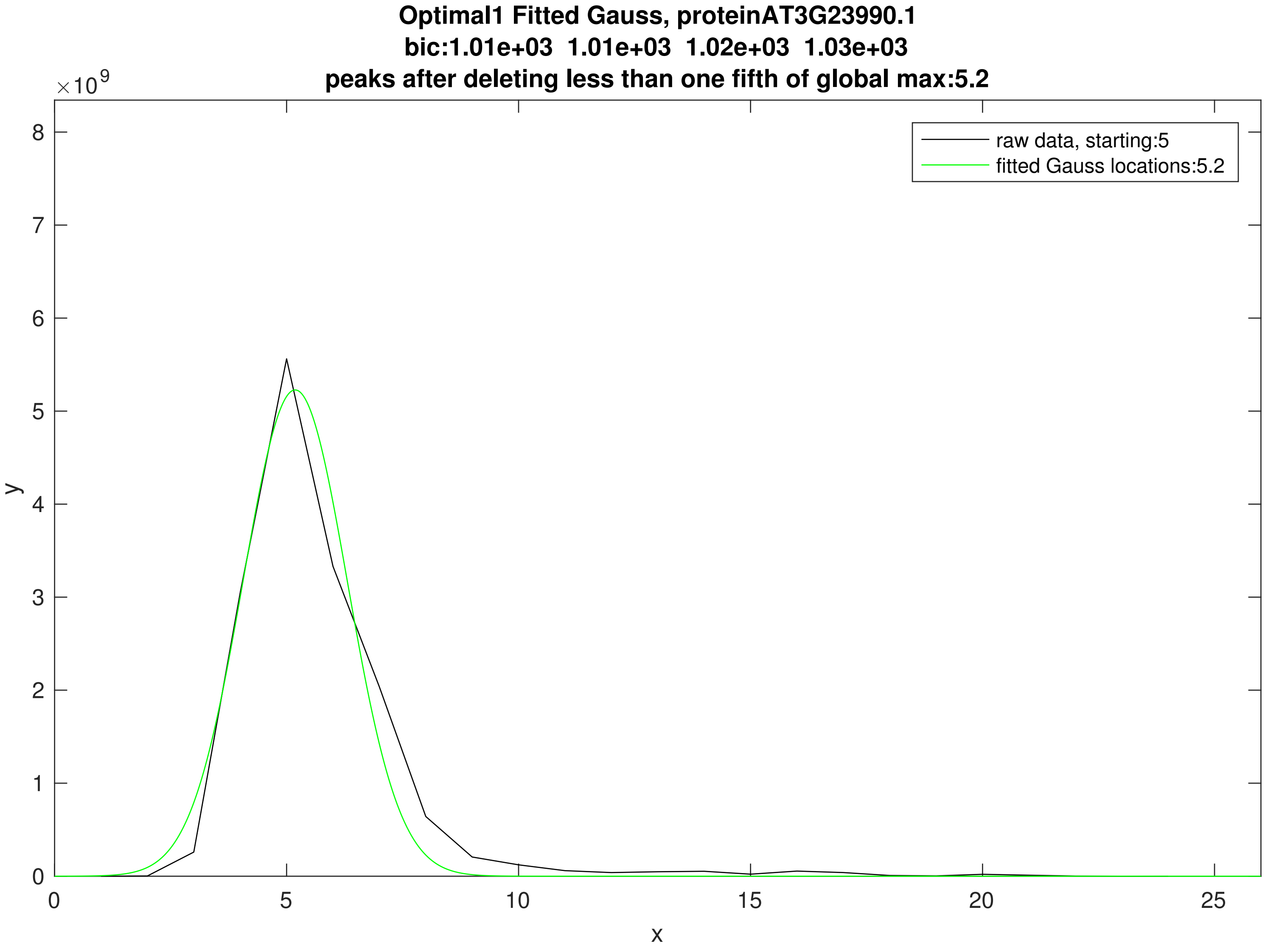
Optimal1 Fitted Gauss, proteinAT3G23990.1
bic:1.01e+03 1.01e+03 1.02e+03 1.03e+03
peaks after deleting less than one fifth of global max:5.2
×10^9
raw data, starting:5
fitted Gauss locations:5.2
y
x
0
1
2
3
4
5
6
7
8
10
15
20
25

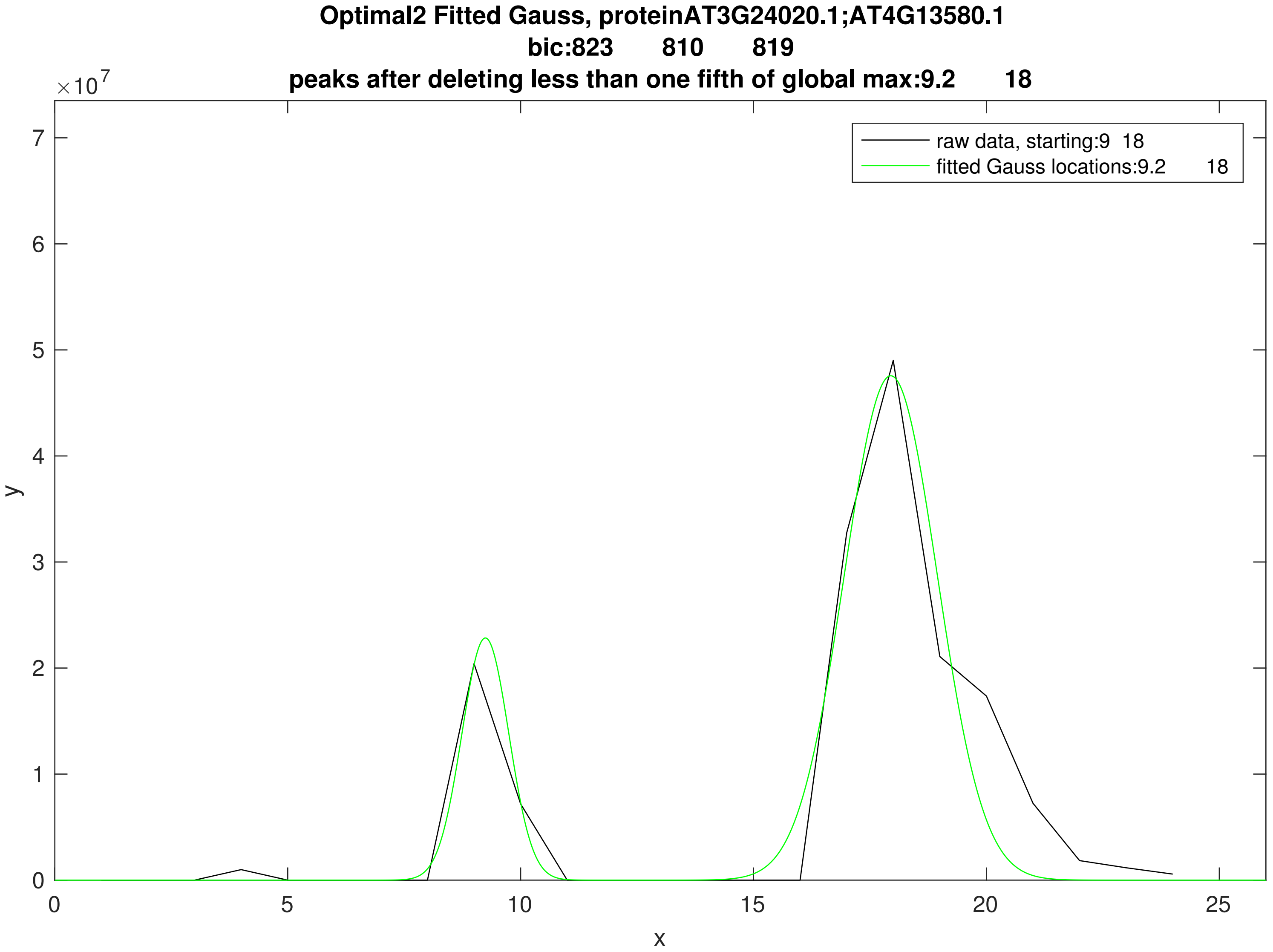
Optimal2 Fitted Gauss, proteinAT3G24020.1;AT4G13580.1
bic:823 810 819
peaks after deleting less than one fifth of global max:9.2 18
×10^7
raw data, starting:9 18
fitted Gauss locations:9.2 18
y
x

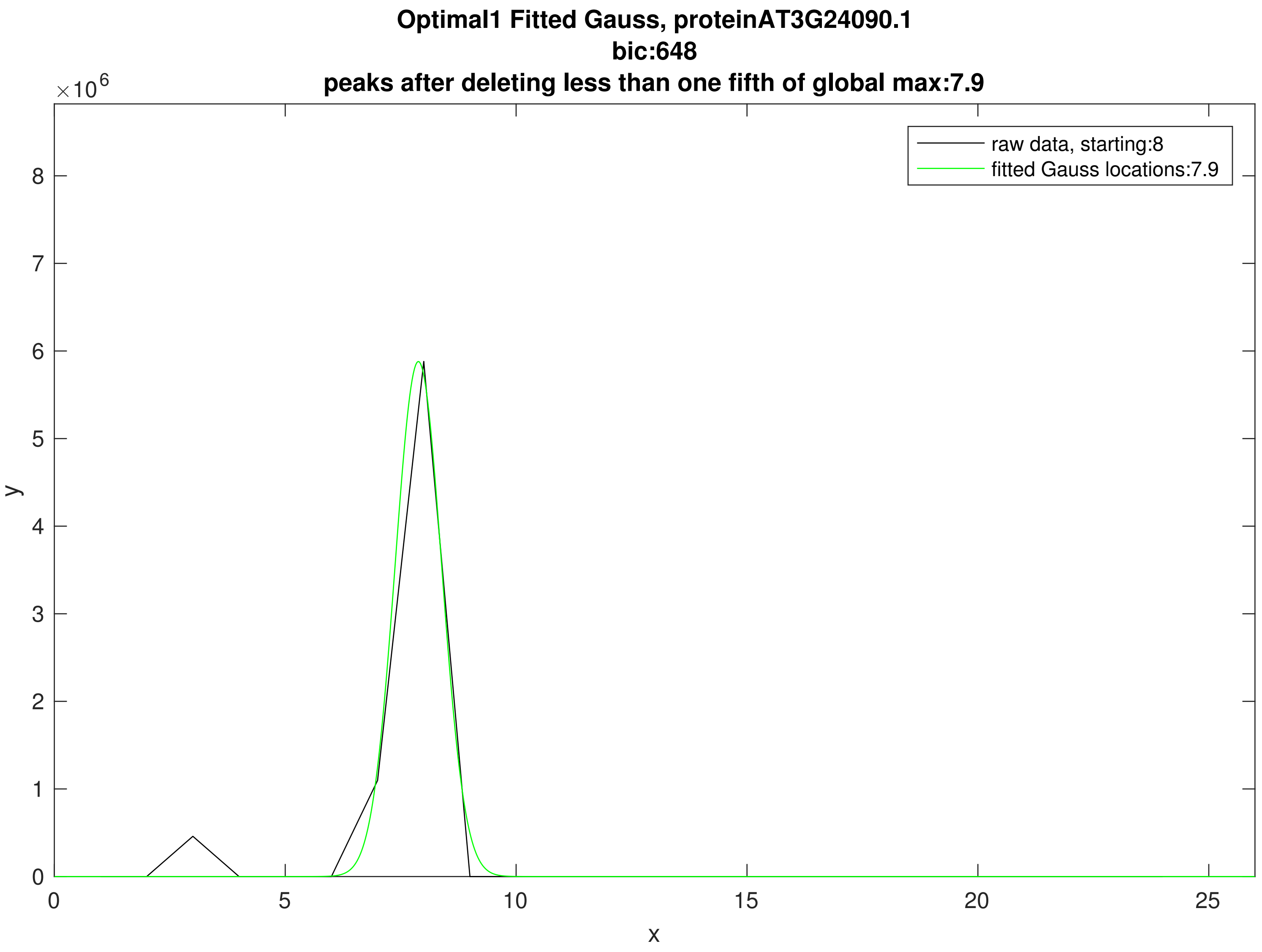
Optimal1 Fitted Gauss, proteinAT3G24090.1
bic:648
peaks after deleting less than one fifth of global max:7.9
×10^6
raw data, starting:8
fitted Gauss locations:7.9
y
x
0
1
2
3
4
5
6
7
8
0
5
10
15
20
25

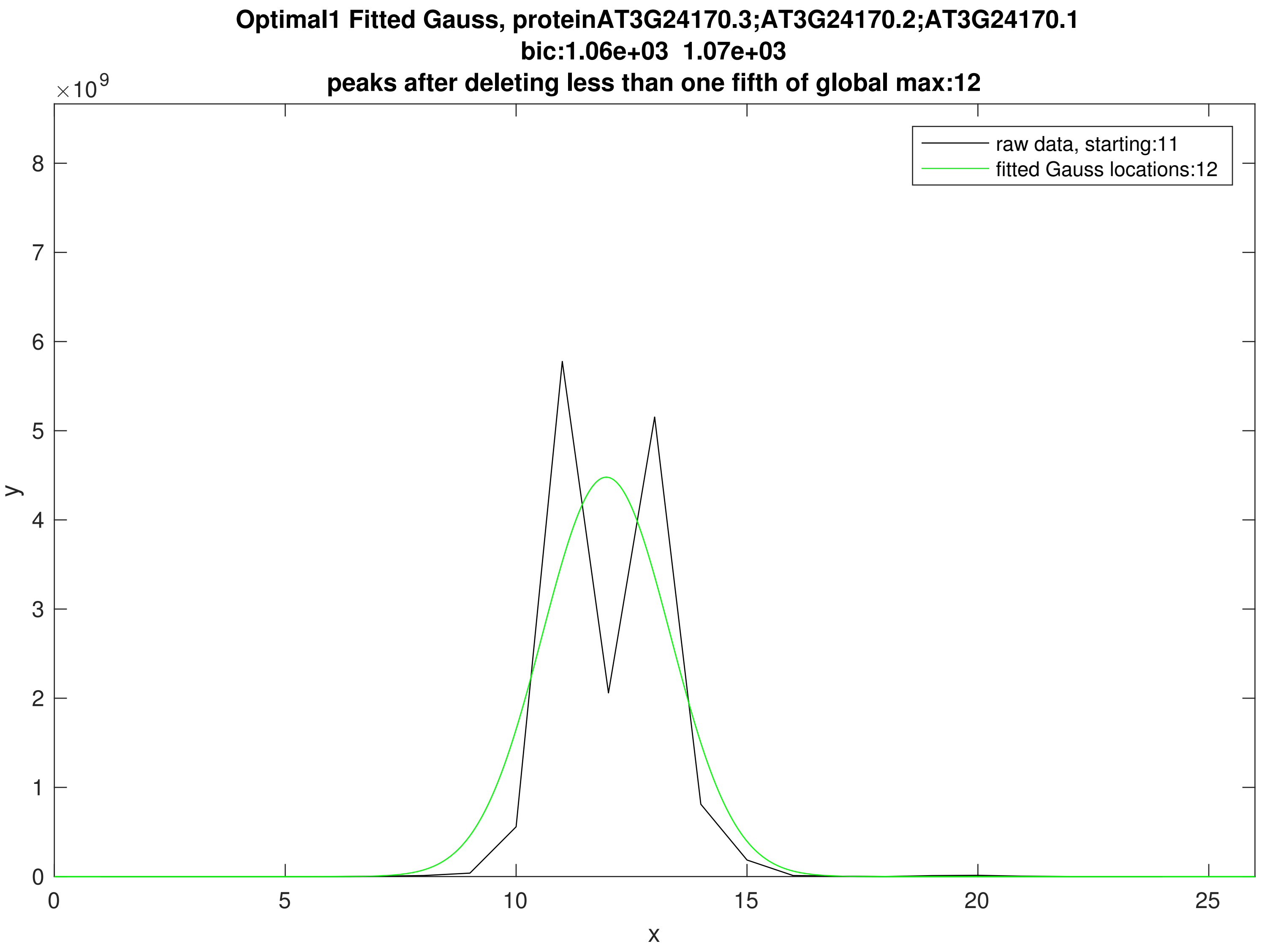
Optimal1 Fitted Gauss, proteinAT3G24170.3;AT3G24170.2;AT3G24170.1
bic:1.06e+03 1.07e+03
peaks after deleting less than one fifth of global max:12
×10^9
raw data, starting:11
fitted Gauss locations:12
y
x
0
1
2
3
4
5
6
7
8
0
5
10
15
20
25

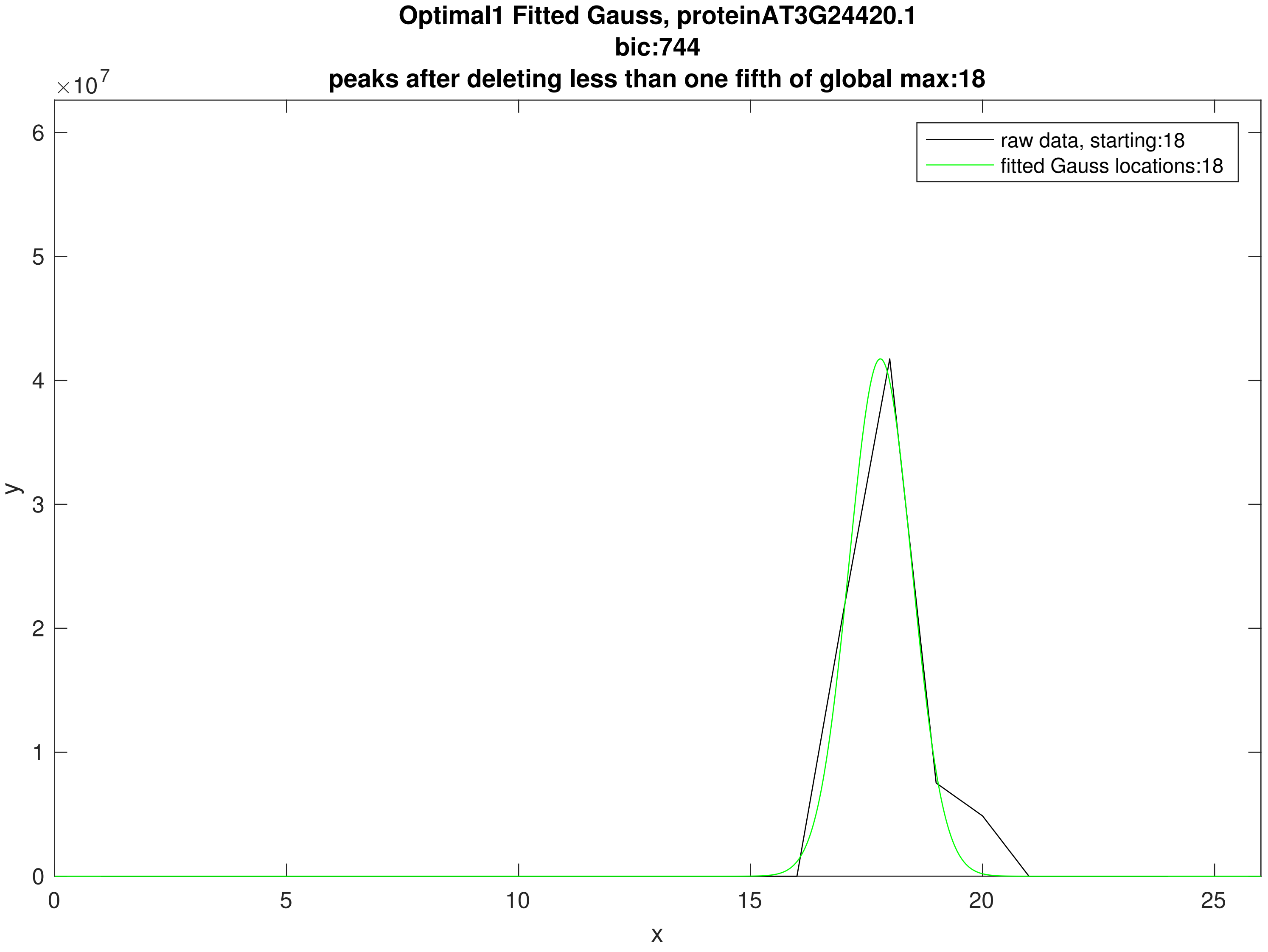
Optimal1 Fitted Gauss, proteinAT3G24420.1
bic:744
peaks after deleting less than one fifth of global max:18
×10^7
raw data, starting:18
fitted Gauss locations:18
y
x

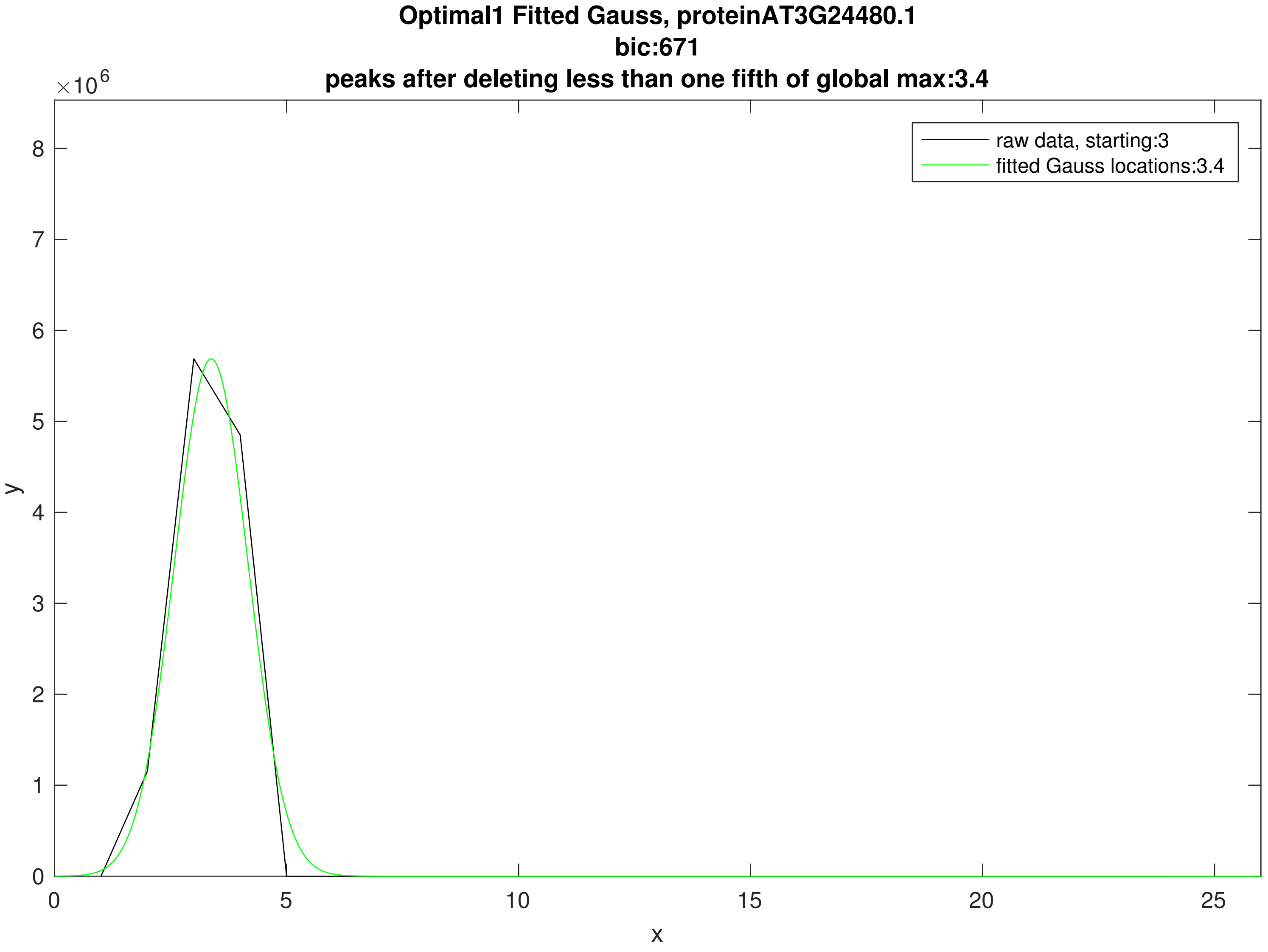
Optimal1 Fitted Gauss, proteinAT3G24480.1
bic:671
peaks after deleting less than one fifth of global max:3.4
×10^6
raw data, starting:3
fitted Gauss locations:3.4
y
x
0
1
2
3
4
5
6
7
8
0
5
10
15
20
25

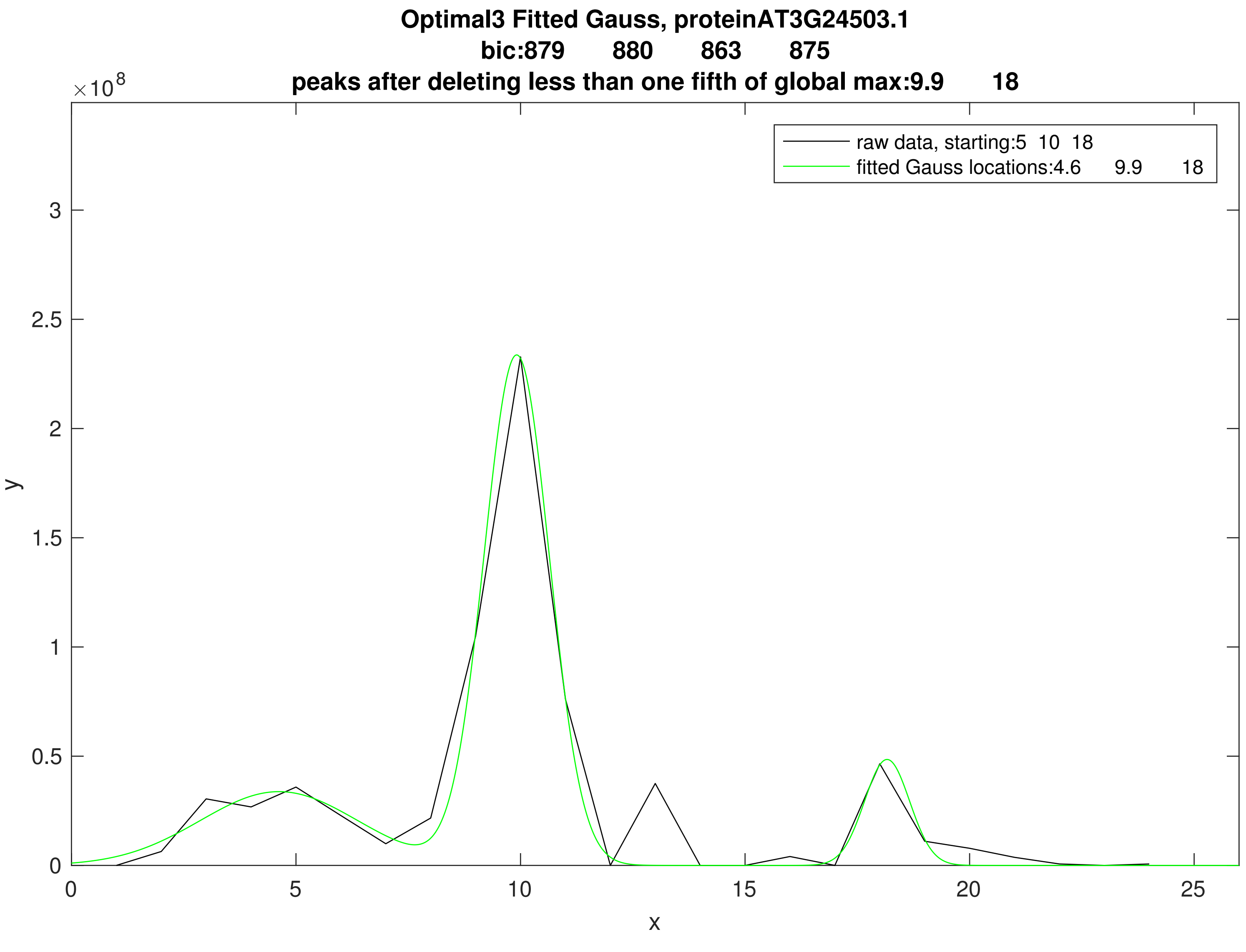

Optimal3 Fitted Gauss, proteinAT3G24503.1
bic:879 880 863 875
peaks after deleting less than one fifth of global max:9.9 18
×10^8
raw data, starting:5 10 18
fitted Gauss locations:4.6 9.9 18
0
0.5
1
1.5
2
2.5
3
y
0
5
10
15
20
25
x

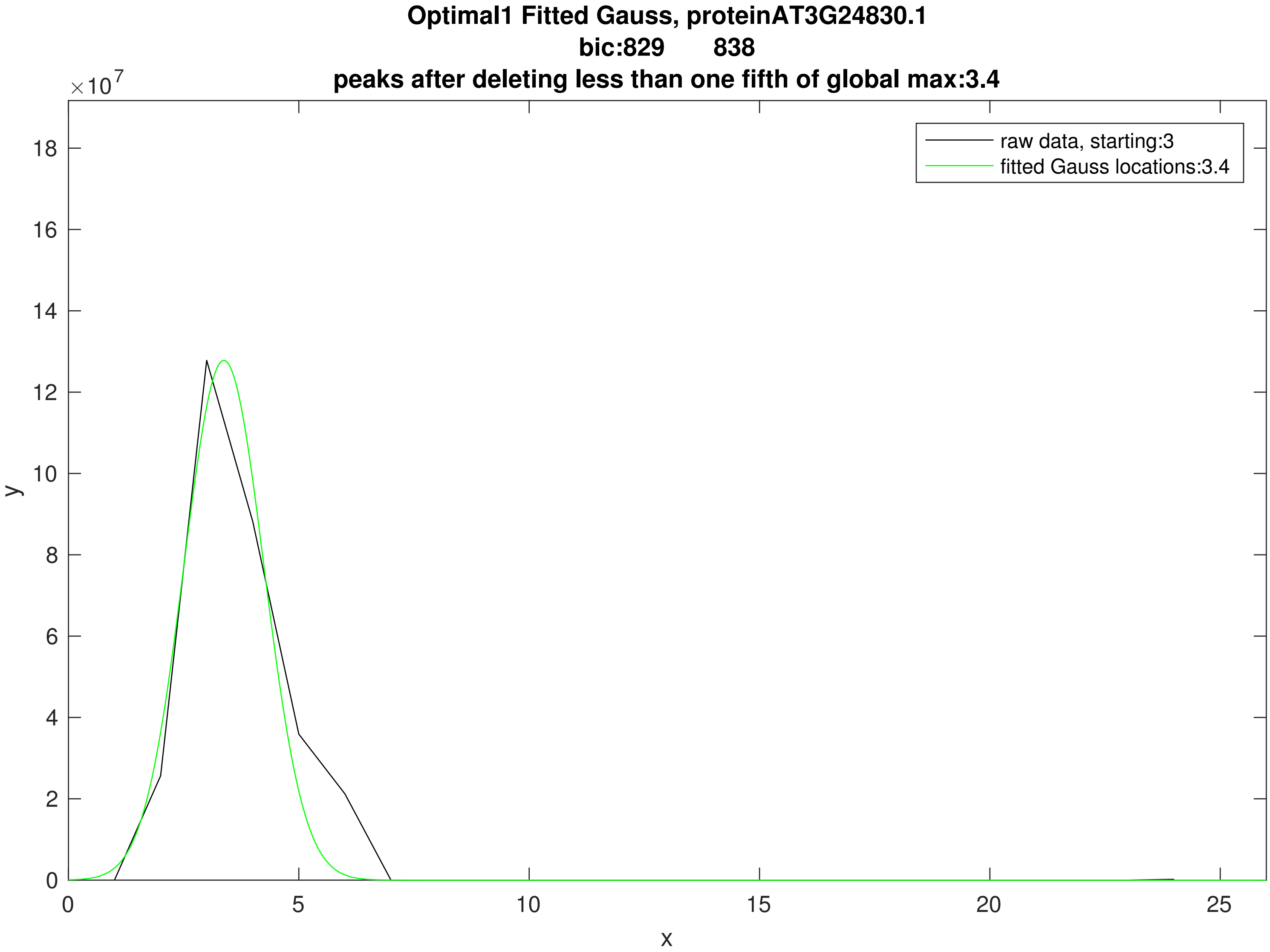

Optimal1 Fitted Gauss, proteinAT3G24830.1
bic:829 838
peaks after deleting less than one fifth of global max:3.4
×10^7
18
16
14
12
10
8
6
4
2
0
y
0
5
10
15
20
25
x
raw data, starting:3
fitted Gauss locations:3.4

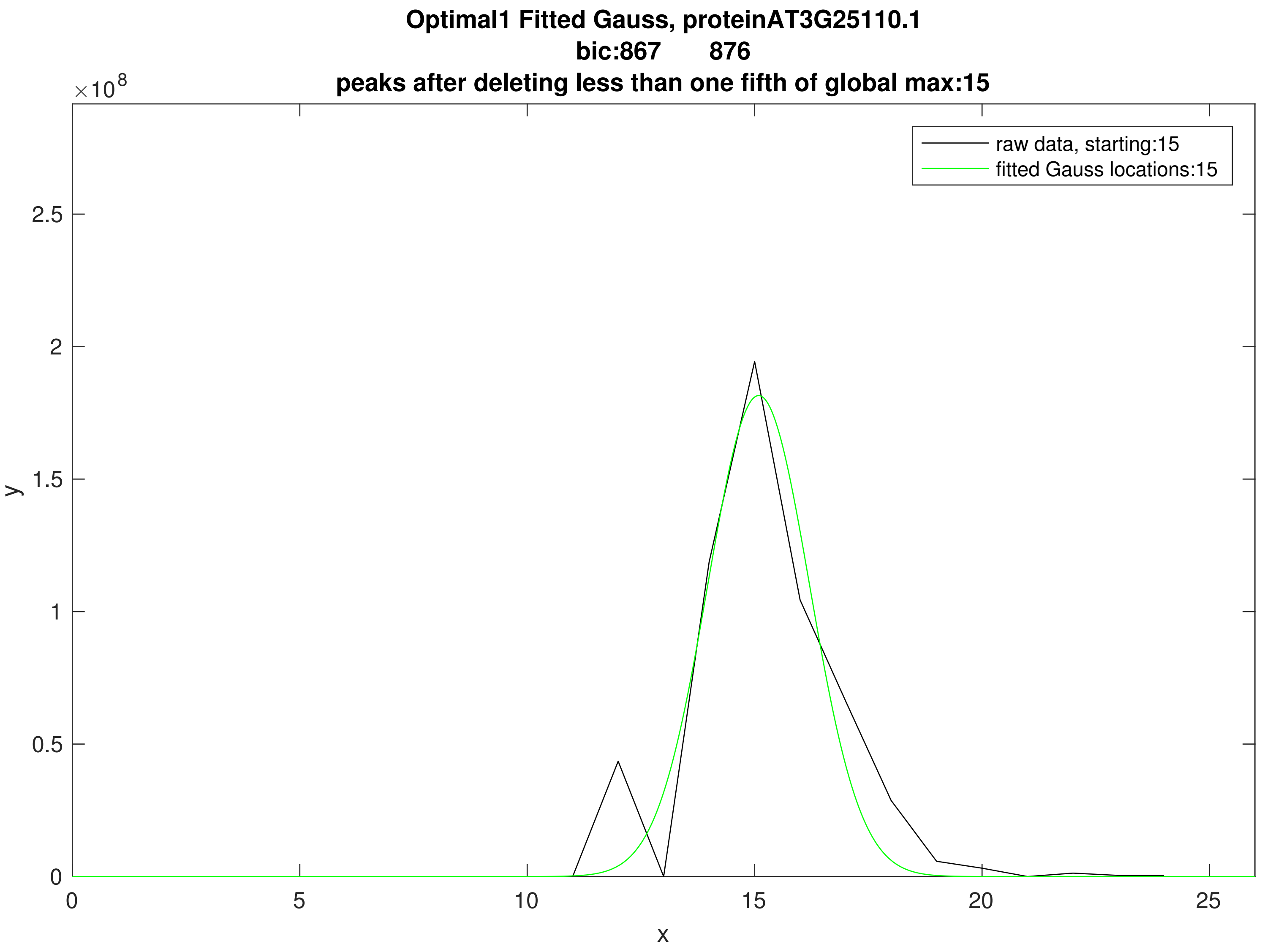
Optimal1 Fitted Gauss, proteinAT3G25110.1
bic:867 876
peaks after deleting less than one fifth of global max:15
×10^8
raw data, starting:15
fitted Gauss locations:15
2.5
2
1.5
1
0.5
0
y
0
5
10
15
20
25
x

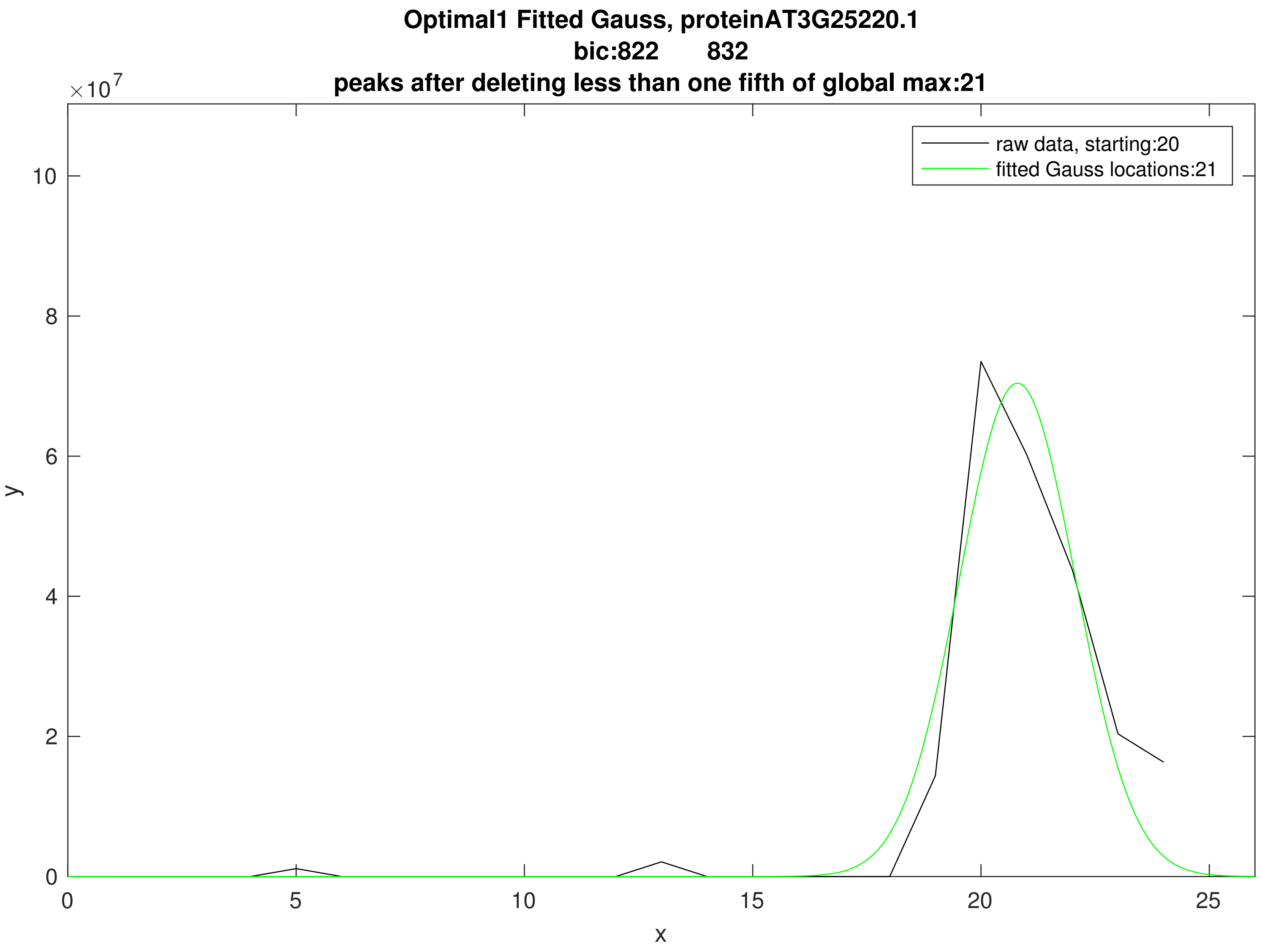
Optimal1 Fitted Gauss, proteinAT3G25220.1
bic:822 832
peaks after deleting less than one fifth of global max:21
×10^7
raw data, starting:20
fitted Gauss locations:21
y
x
0
2
4
6
8
10
5
15
20
25

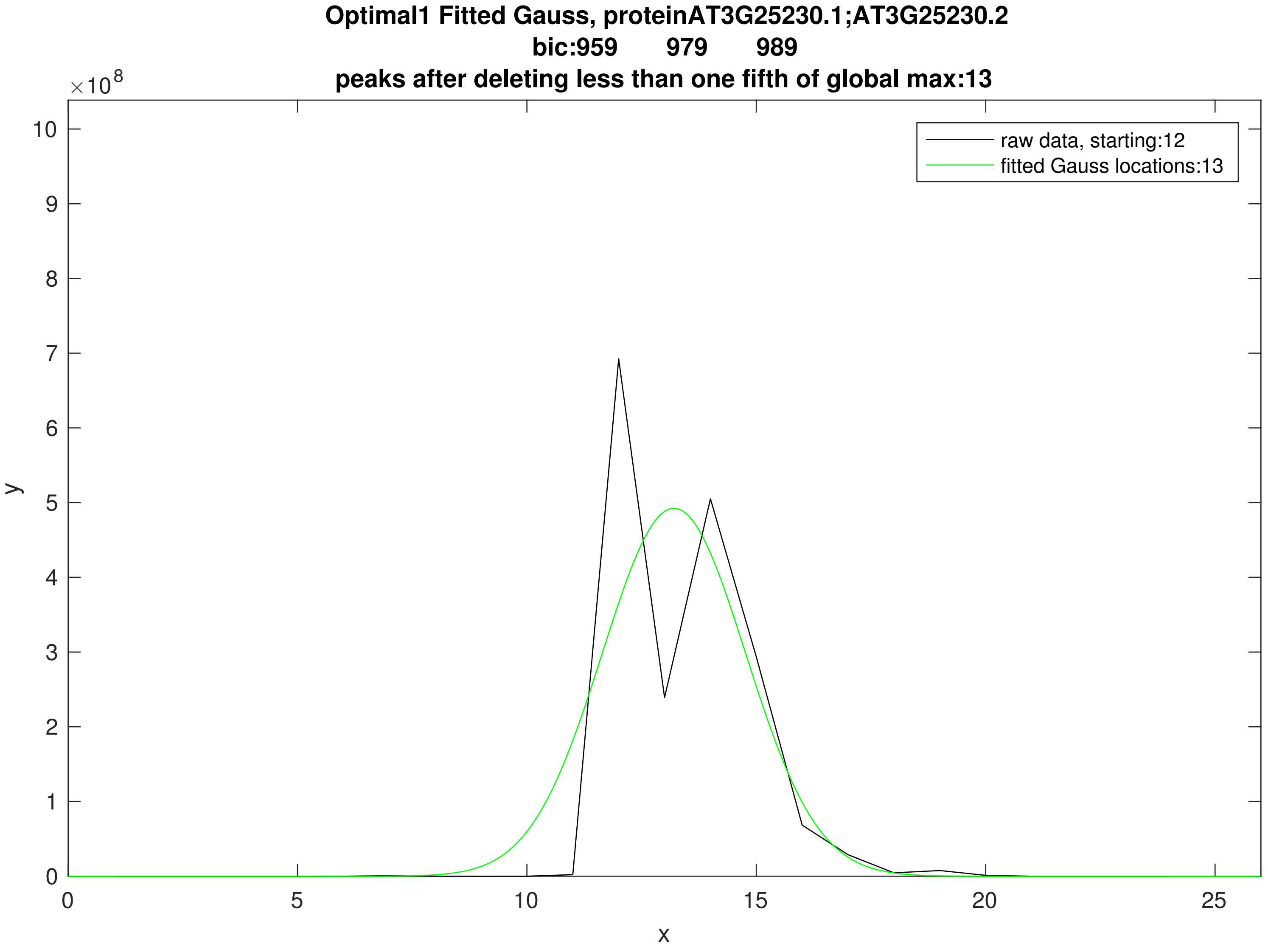
Optimal1 Fitted Gauss, proteinAT3G25230.1;AT3G25230.2
bic:959 979 989
peaks after deleting less than one fifth of global max:13
×10^8
raw data, starting:12
fitted Gauss locations:13
y
x
0
1
2
3
4
5
6
7
8
9
10
0
5
10
15
20
25

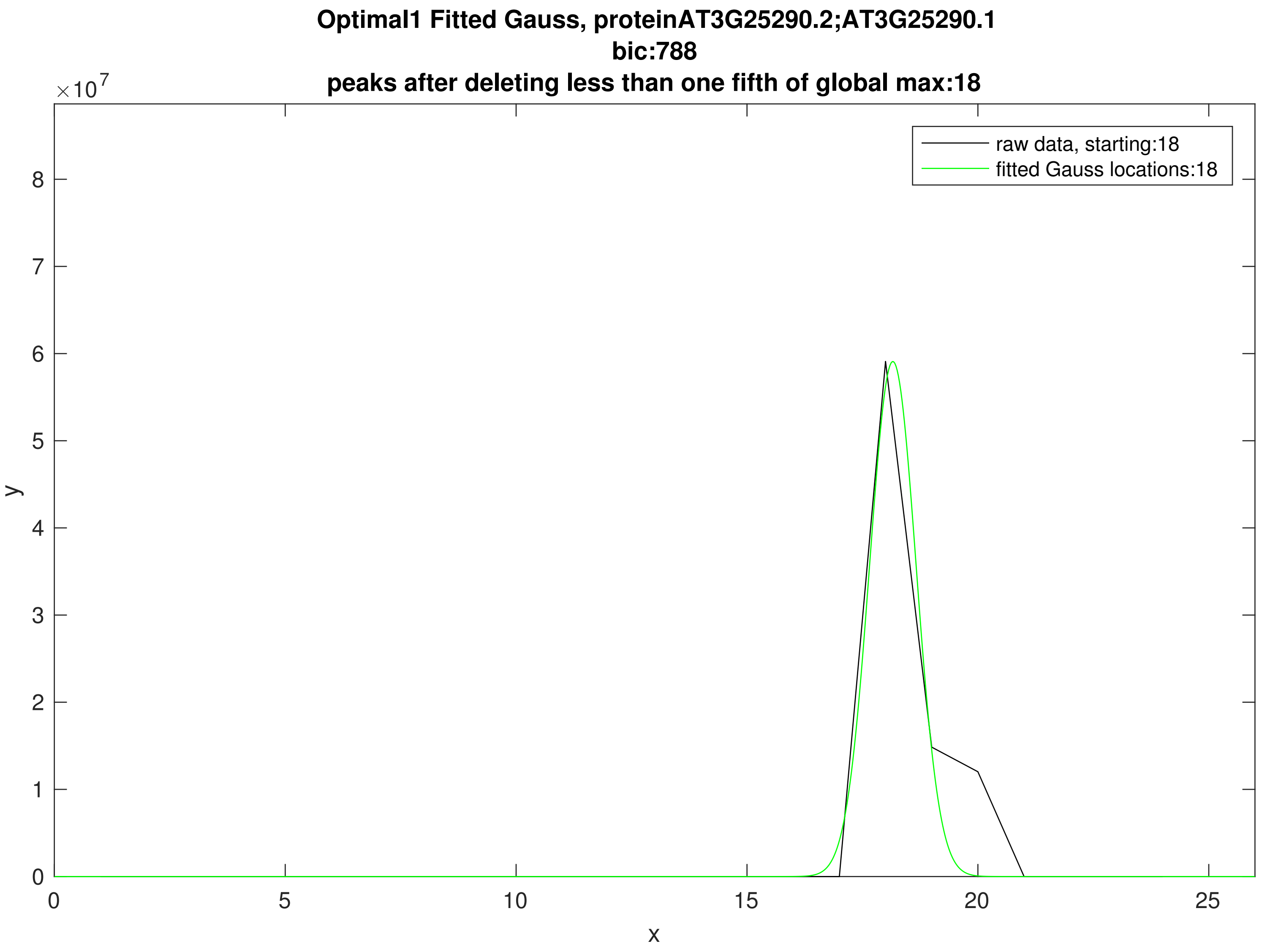

Optimal1 Fitted Gauss, proteinAT3G25290.2;AT3G25290.1
bic:788
peaks after deleting less than one fifth of global max:18
×10^7
raw data, starting:18
fitted Gauss locations:18
y
x
0
1
2
3
4
5
6
7
8
0
5
10
15
20
25

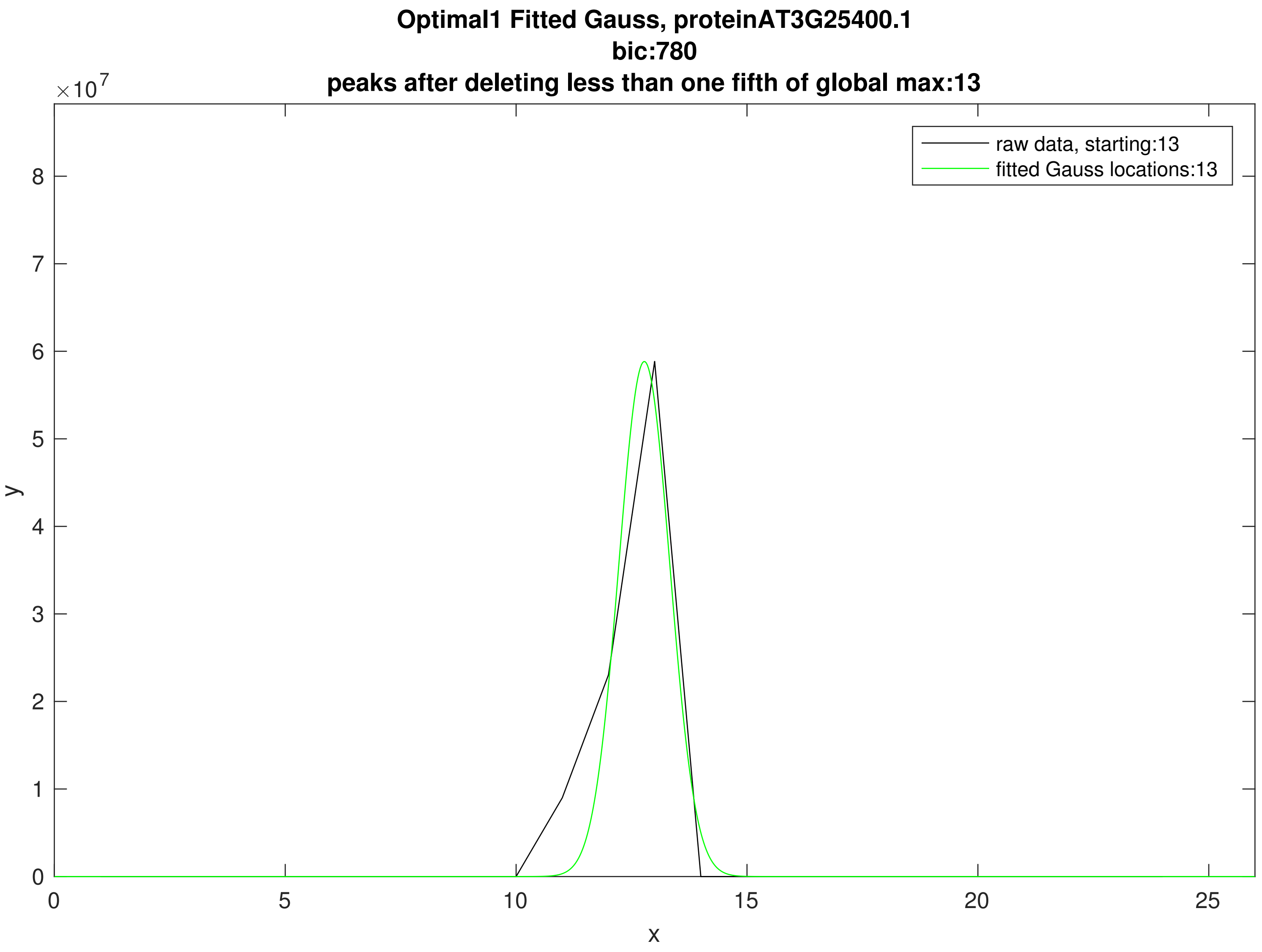
Optimal1 Fitted Gauss, proteinAT3G25400.1
bic:780
peaks after deleting less than one fifth of global max:13
×10^7
raw data, starting:13
fitted Gauss locations:13
y
x
0
1
2
3
4
5
6
7
8
0
5
10
15
20
25

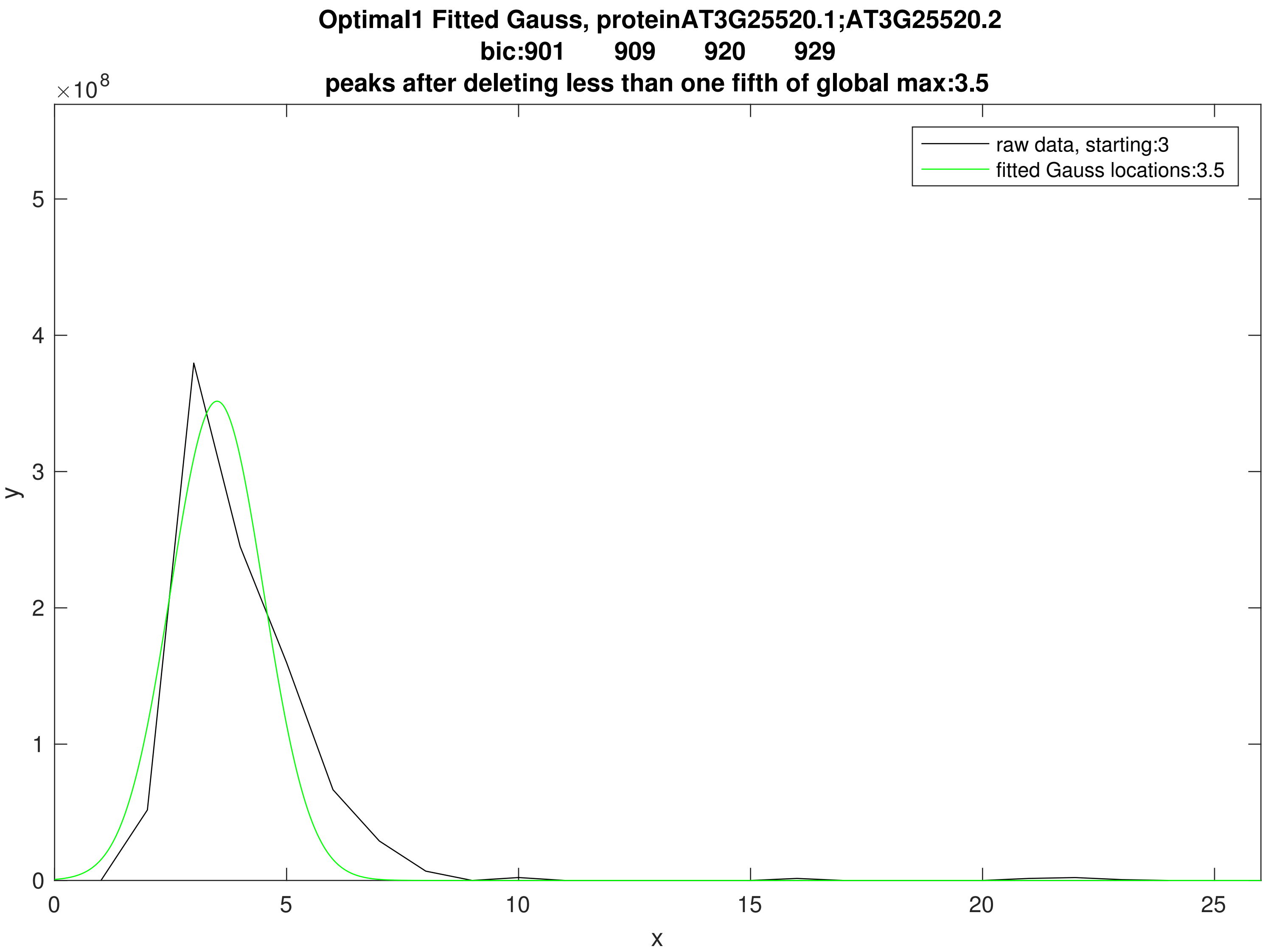

Optimal1 Fitted Gauss, proteinAT3G25520.1;AT3G25520.2
bic:901 909 920 929
peaks after deleting less than one fifth of global max:3.5
×10^8
raw data, starting:3
fitted Gauss locations:3.5
y
x
0
1
2
3
4
5
0
5
10
15
20
25

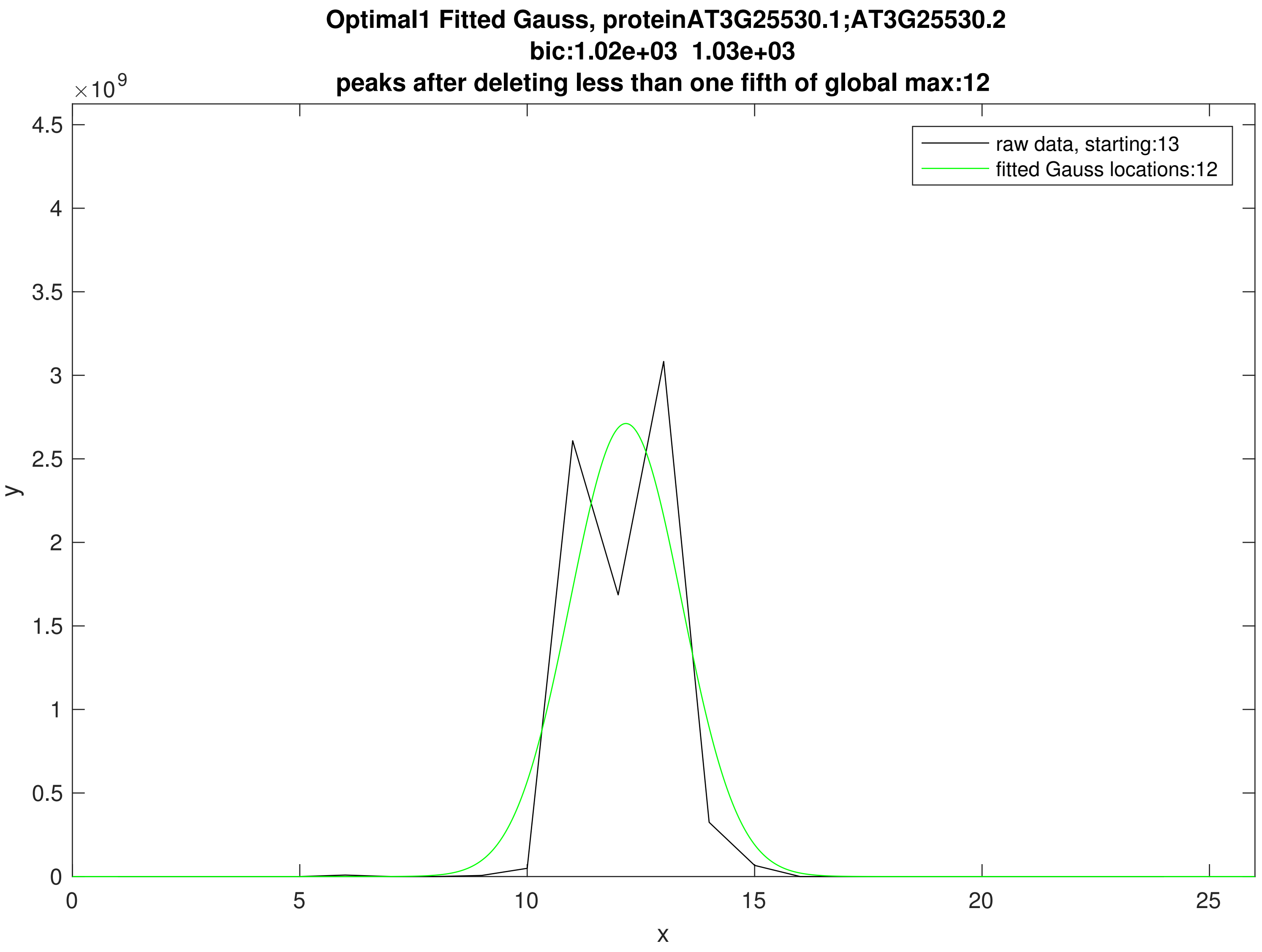

Optimal1 Fitted Gauss, proteinAT3G25530.1;AT3G25530.2
bic:1.02e+03 1.03e+03
peaks after deleting less than one fifth of global max:12
×10^9
raw data, starting:13
fitted Gauss locations:12
y
x

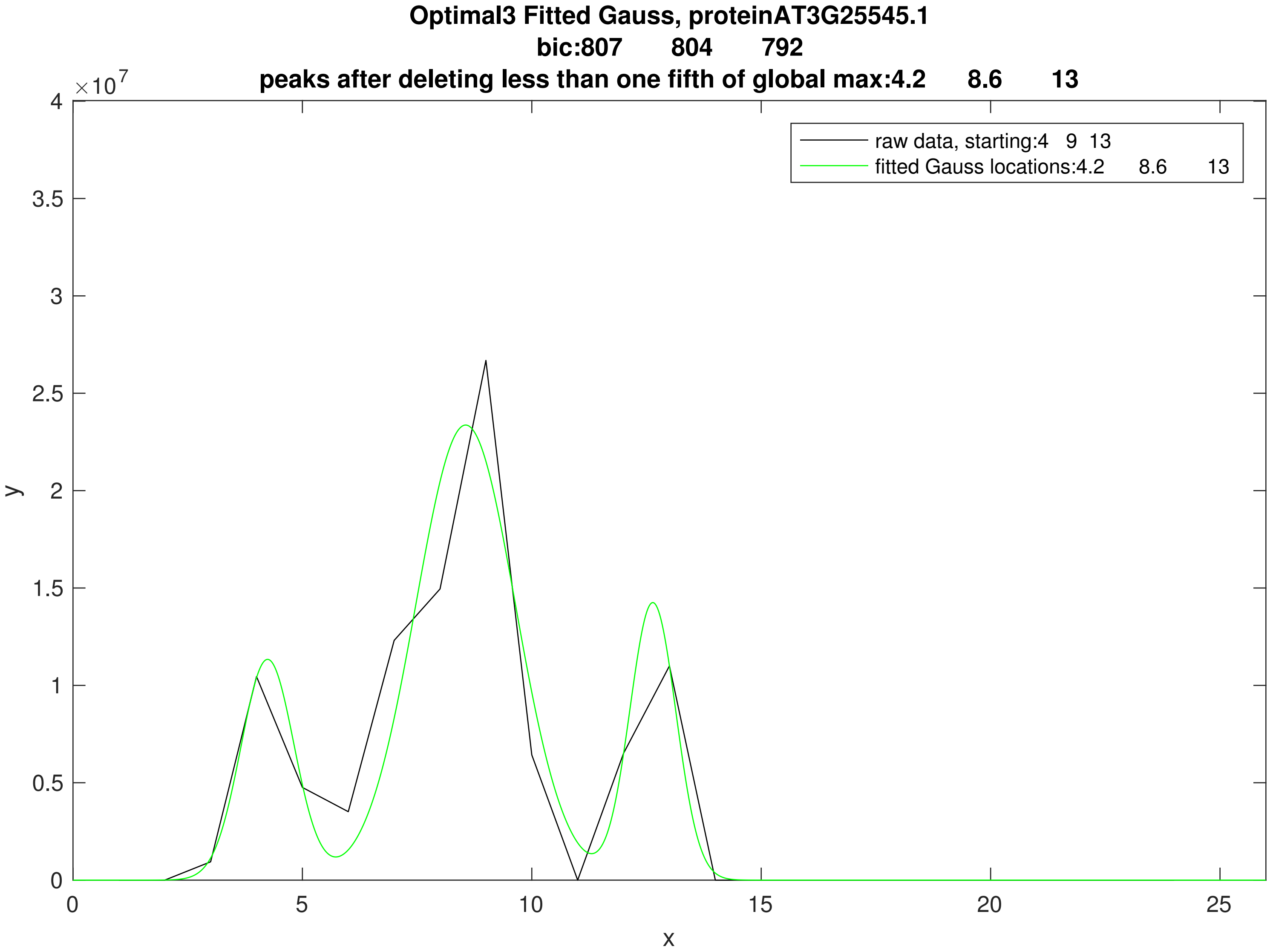

Optimal3 Fitted Gauss, proteinAT3G25545.1
bic:807 804 792
peaks after deleting less than one fifth of global max:4.2 8.6 13
raw data, starting:4 9 13
fitted Gauss locations:4.2 8.6 13
×10^7
y
x

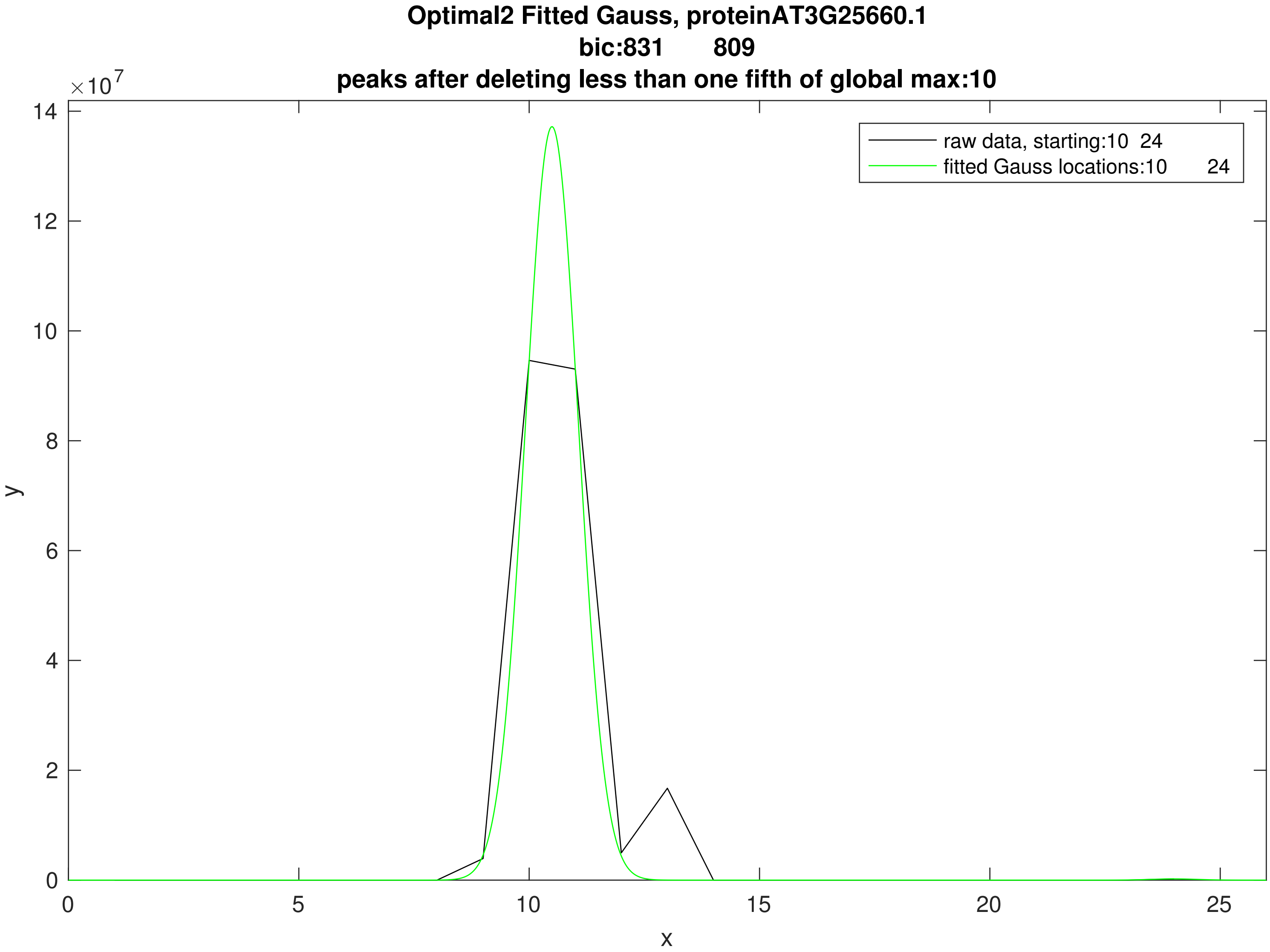

Optimal2 Fitted Gauss, proteinAT3G25660.1
bic:831 809
peaks after deleting less than one fifth of global max:10
×10^7
14
12
10
8
6
4
2
0
y
0
5
10
15
20
25
x
raw data, starting:10 24
fitted Gauss locations:10 24

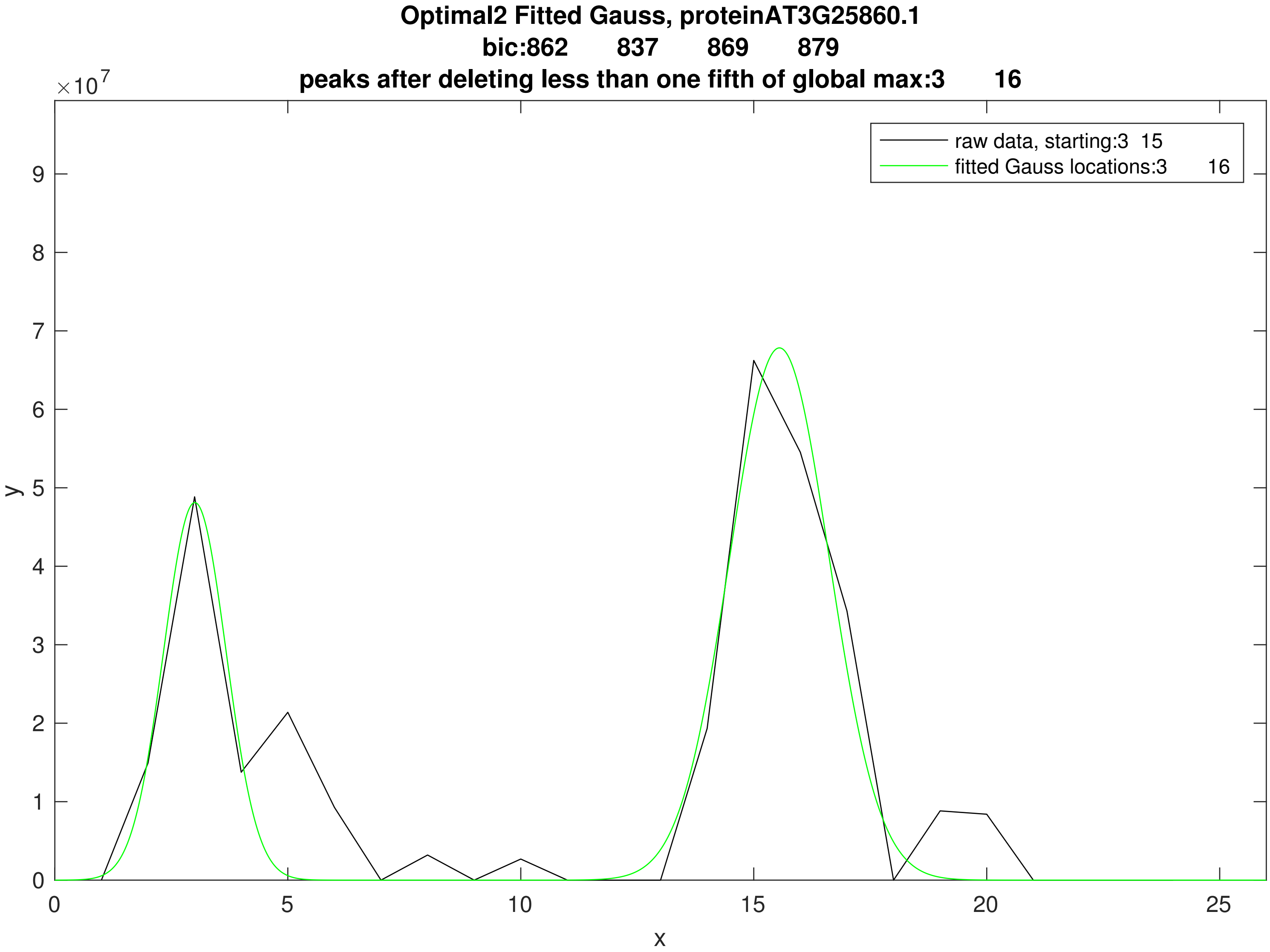

Optimal2 Fitted Gauss, proteinAT3G25860.1
bic:862 837 869 879
peaks after deleting less than one fifth of global max:3 16
×10^7
raw data, starting:3 15
fitted Gauss locations:3 16
y
x
0
1
2
3
4
5
6
7
8
9
5
10
15
20
25

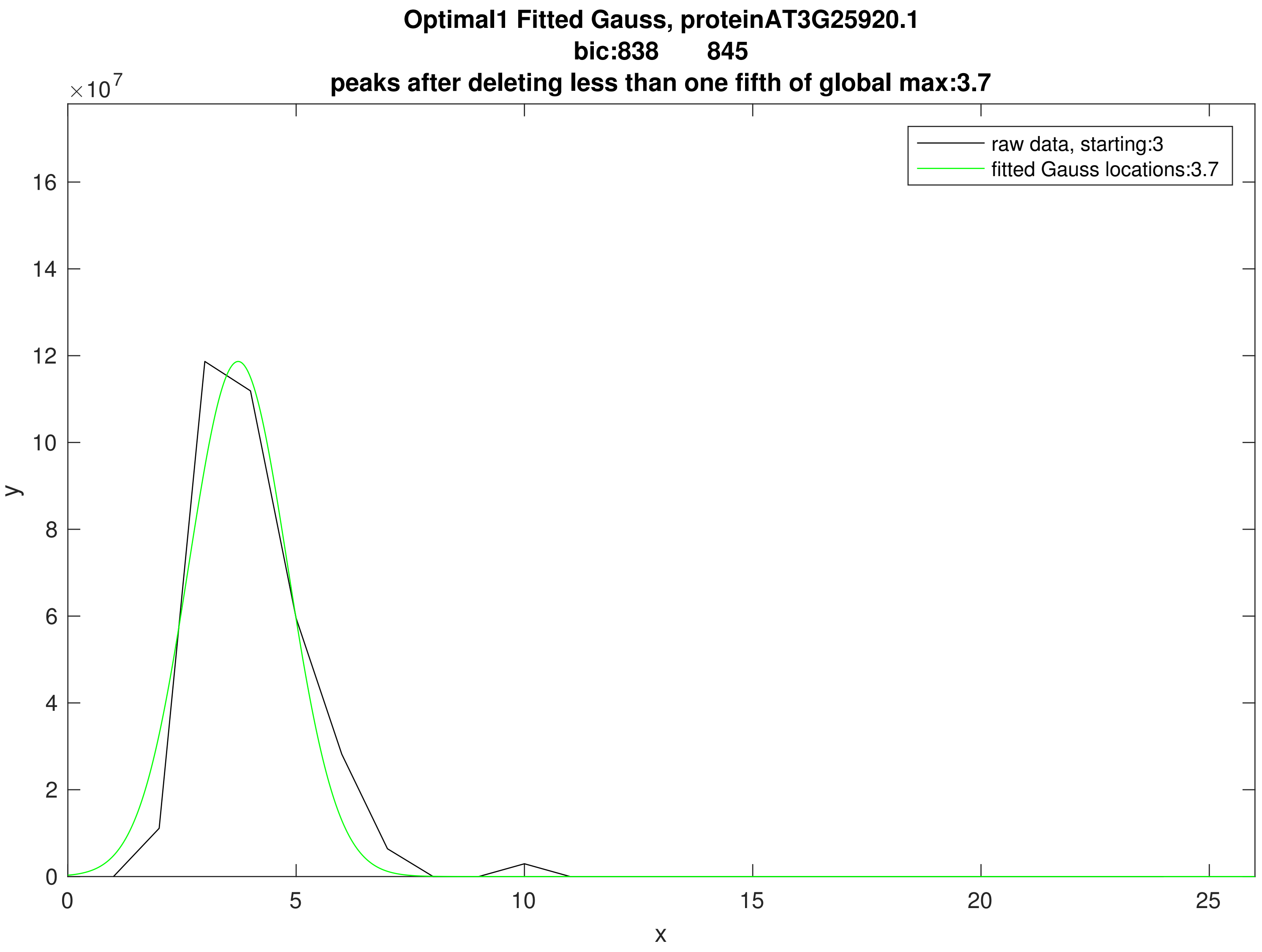
Optimal1 Fitted Gauss, proteinAT3G25920.1
bic:838 845
peaks after deleting less than one fifth of global max:3.7
×10^7
raw data, starting:3
fitted Gauss locations:3.7
y
x
0
2
4
6
8
10
12
14
16
5
15
20
25

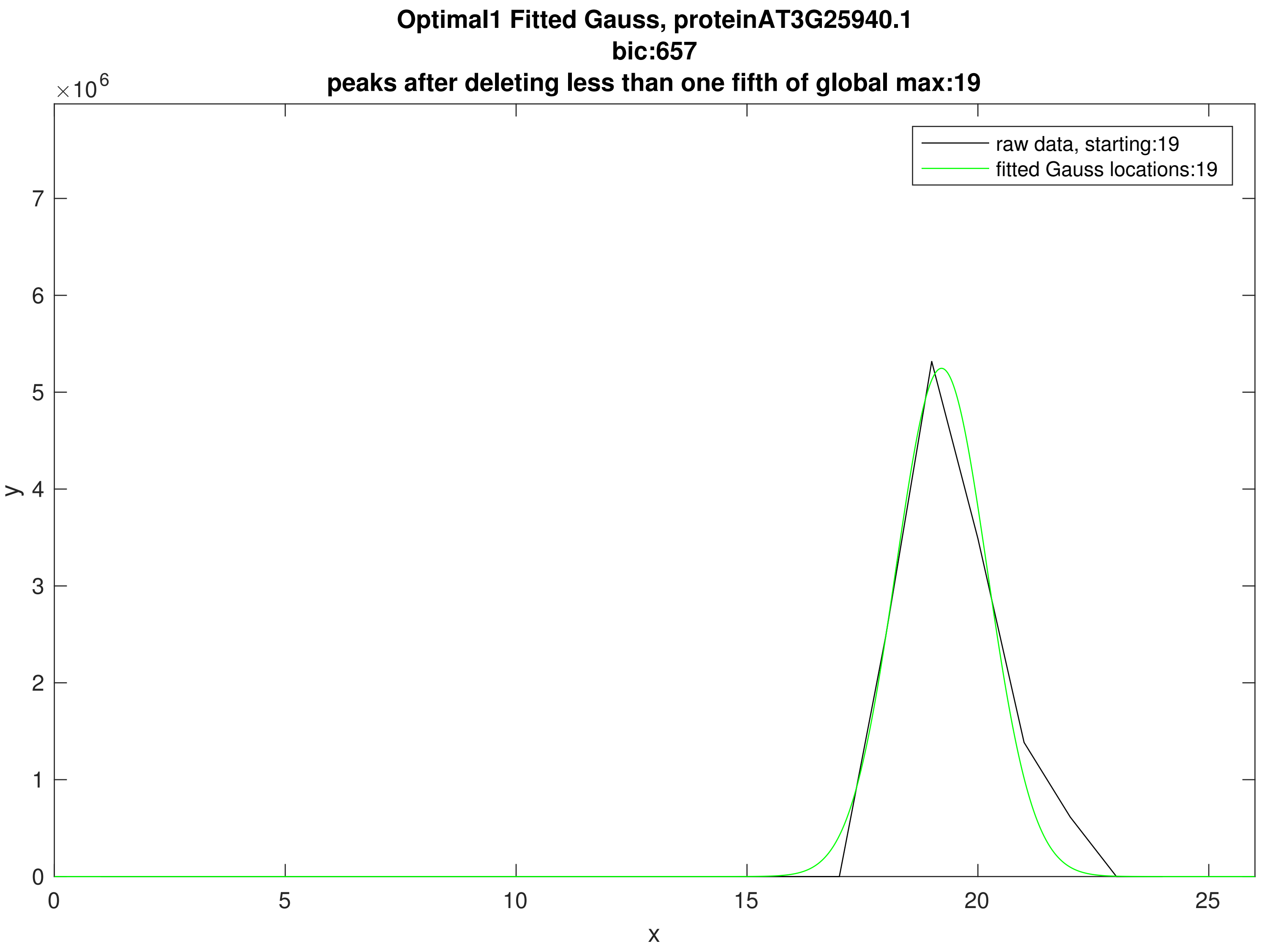
Optimal1 Fitted Gauss, proteinAT3G25940.1
bic:657
peaks after deleting less than one fifth of global max:19
×10^6
raw data, starting:19
fitted Gauss locations:19
y
x
0
1
2
3
4
5
6
7
0
5
10
15
20
25

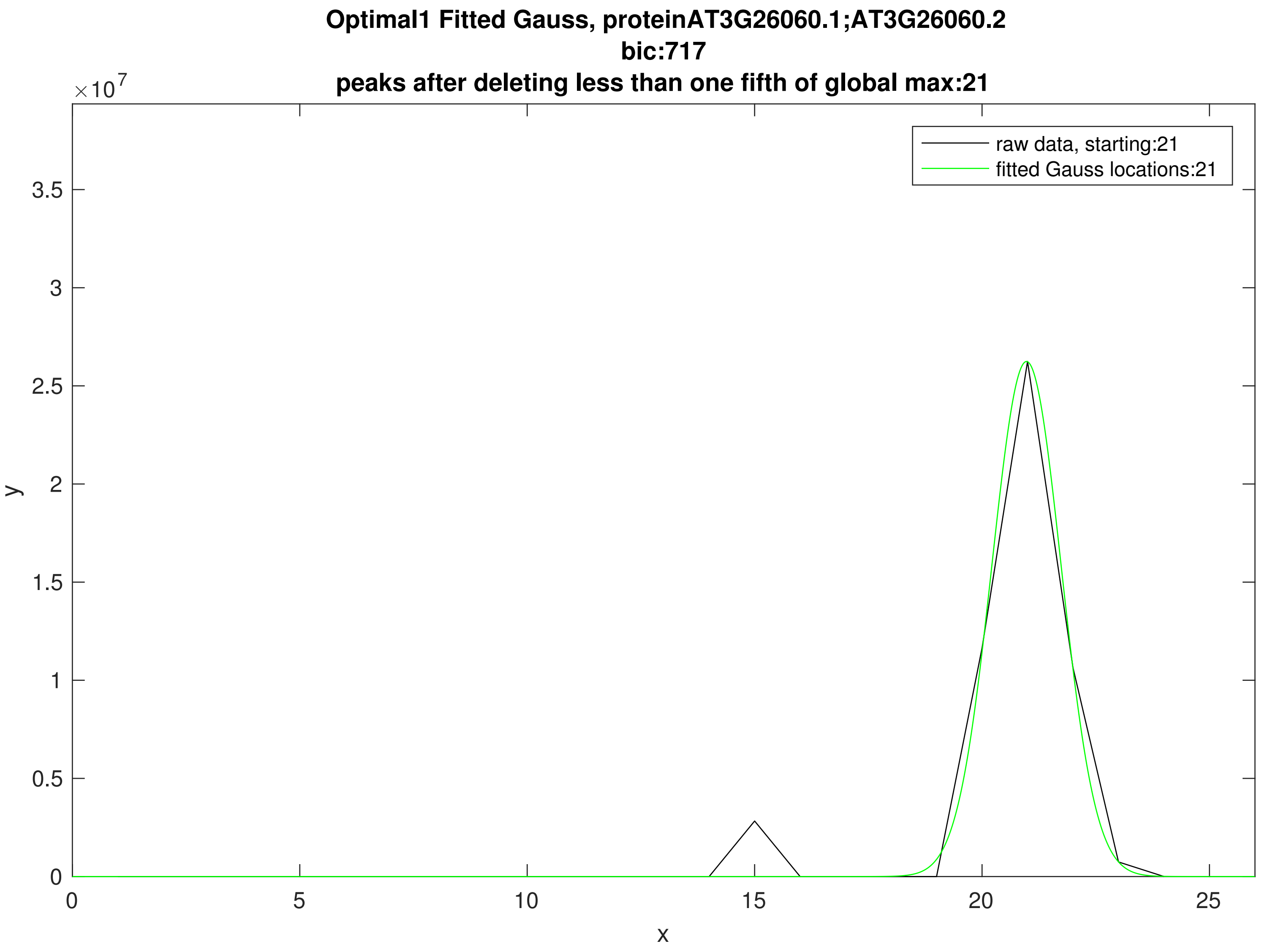

Optimal1 Fitted Gauss, proteinAT3G26060.1;AT3G26060.2
bic:717
peaks after deleting less than one fifth of global max:21
×10^7
raw data, starting:21
fitted Gauss locations:21
3.5
3
2.5
2
1.5
1
0.5
0
y
0
5
10
15
20
25
x

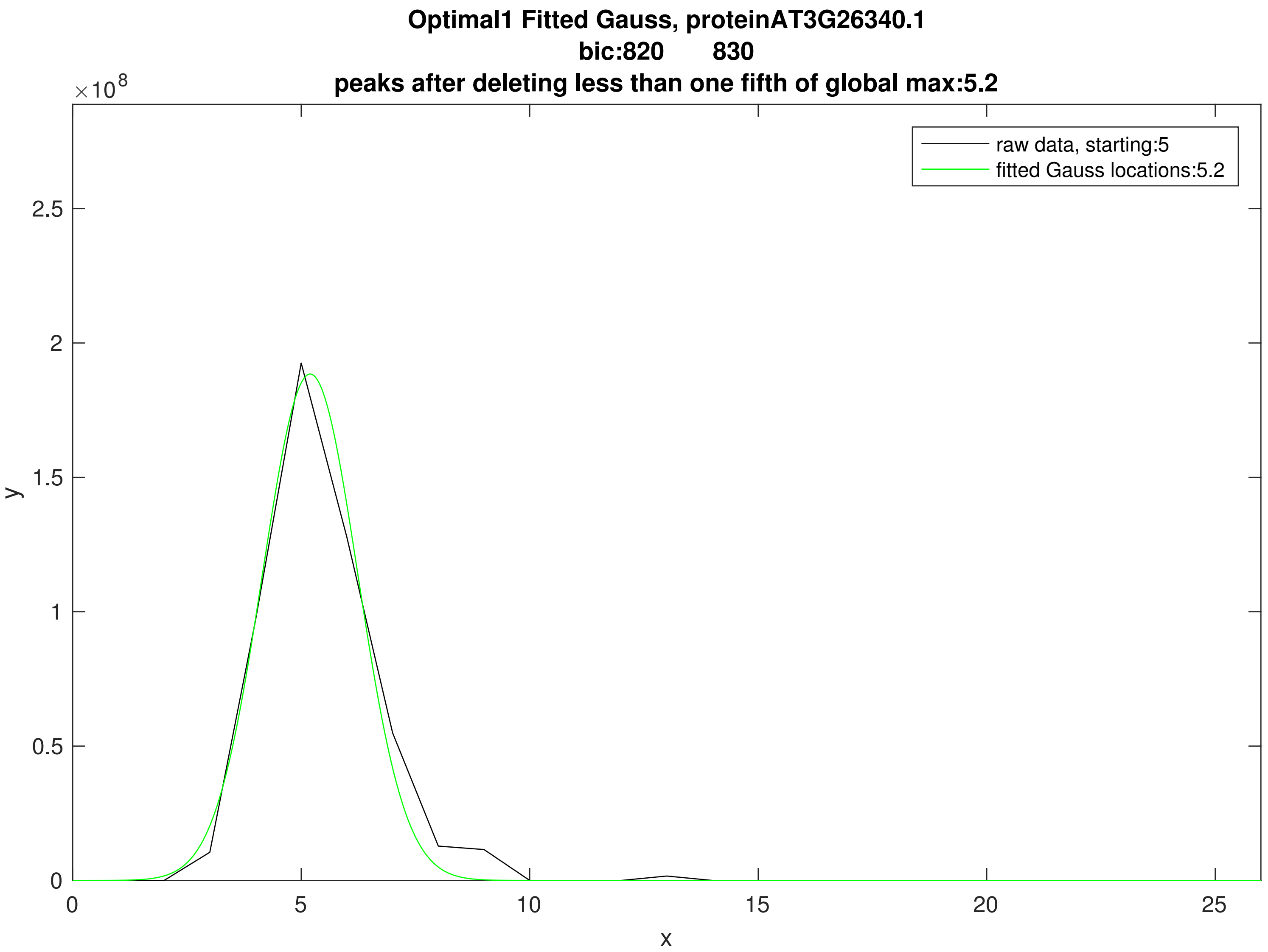

Optimal1 Fitted Gauss, proteinAT3G26340.1
bic:820 830
peaks after deleting less than one fifth of global max:5.2
raw data, starting:5
fitted Gauss locations:5.2
×10^8
0
0.5
1
1.5
2
2.5
y
0
5
10
15
20
25
x

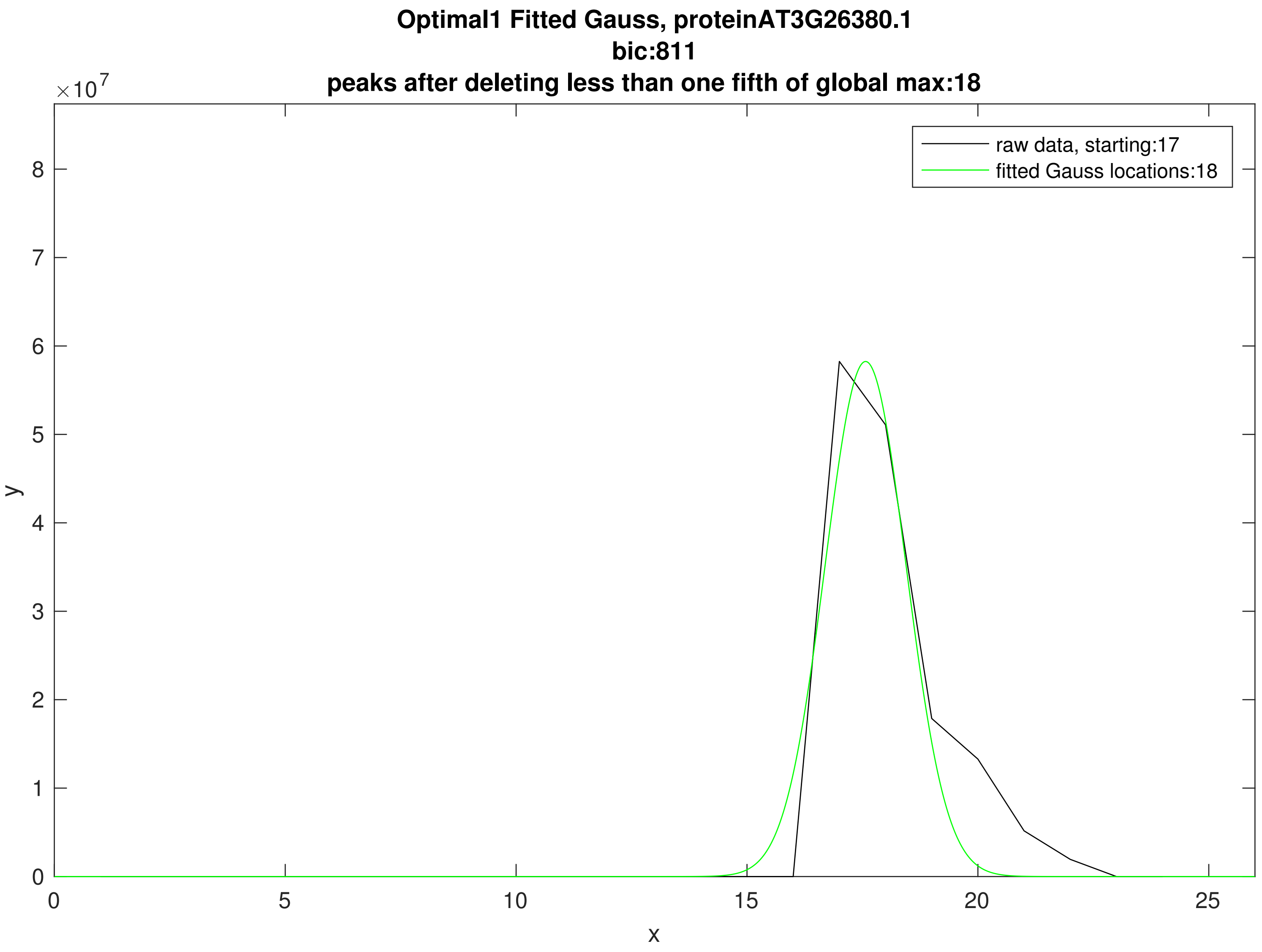
Optimal1 Fitted Gauss, proteinAT3G26380.1
bic:811
peaks after deleting less than one fifth of global max:18
×10^7
raw data, starting:17
fitted Gauss locations:18
y
x
0
1
2
3
4
5
6
7
8
0
5
10
15
20
25

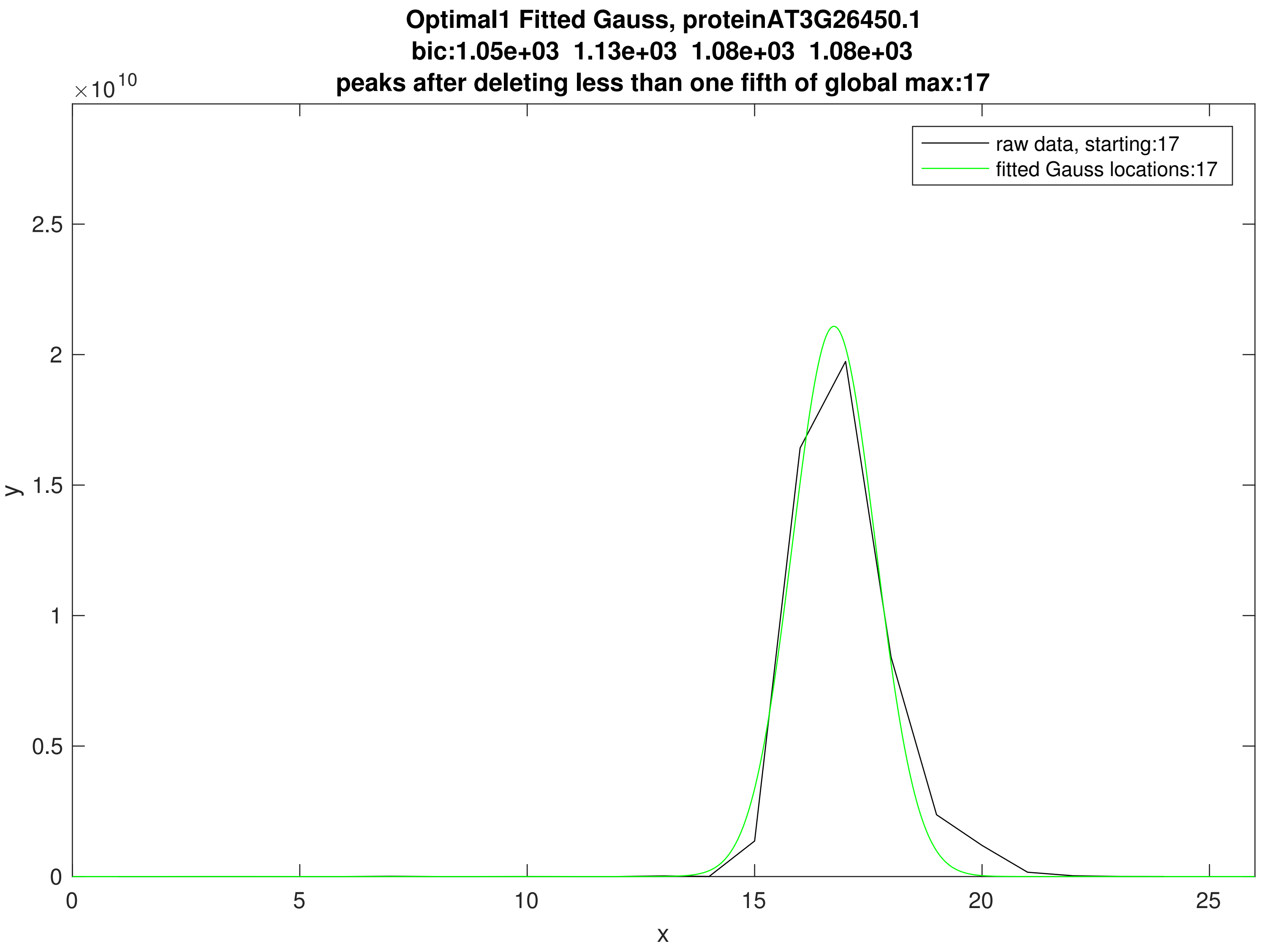

Optimal1 Fitted Gauss, proteinAT3G26450.1
bic:1.05e+03  1.13e+03  1.08e+03  1.08e+03
peaks after deleting less than one fifth of global max:17
×10^10
raw data, starting:17
fitted Gauss locations:17
y
x
0
0.5
1
1.5
2
2.5
0
5
10
15
20
25

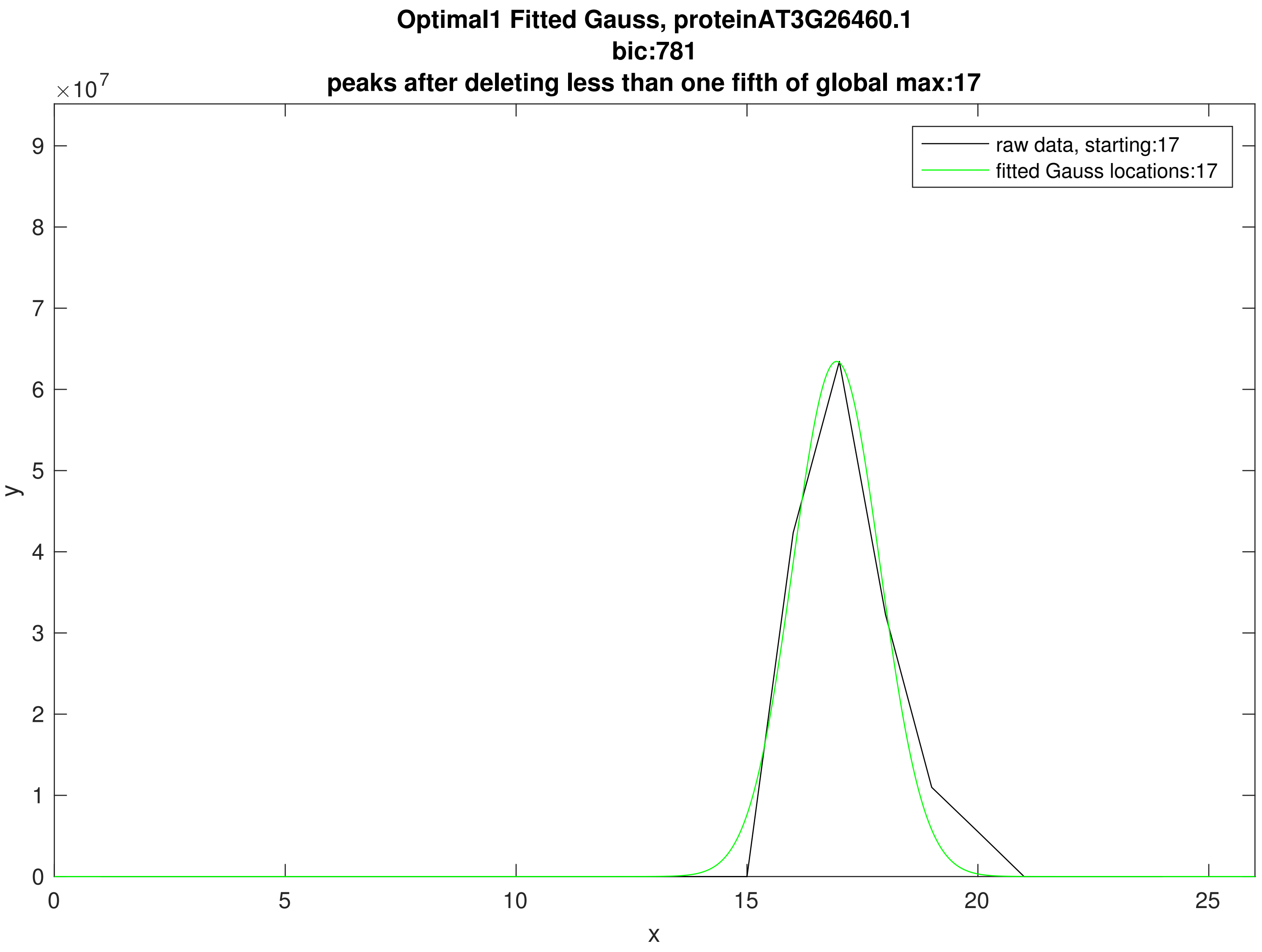
Optimal1 Fitted Gauss, proteinAT3G26460.1
bic:781
peaks after deleting less than one fifth of global max:17
×10^7
raw data, starting:17
fitted Gauss locations:17
y
x
0
1
2
3
4
5
6
7
8
9
0
5
10
15
20
25

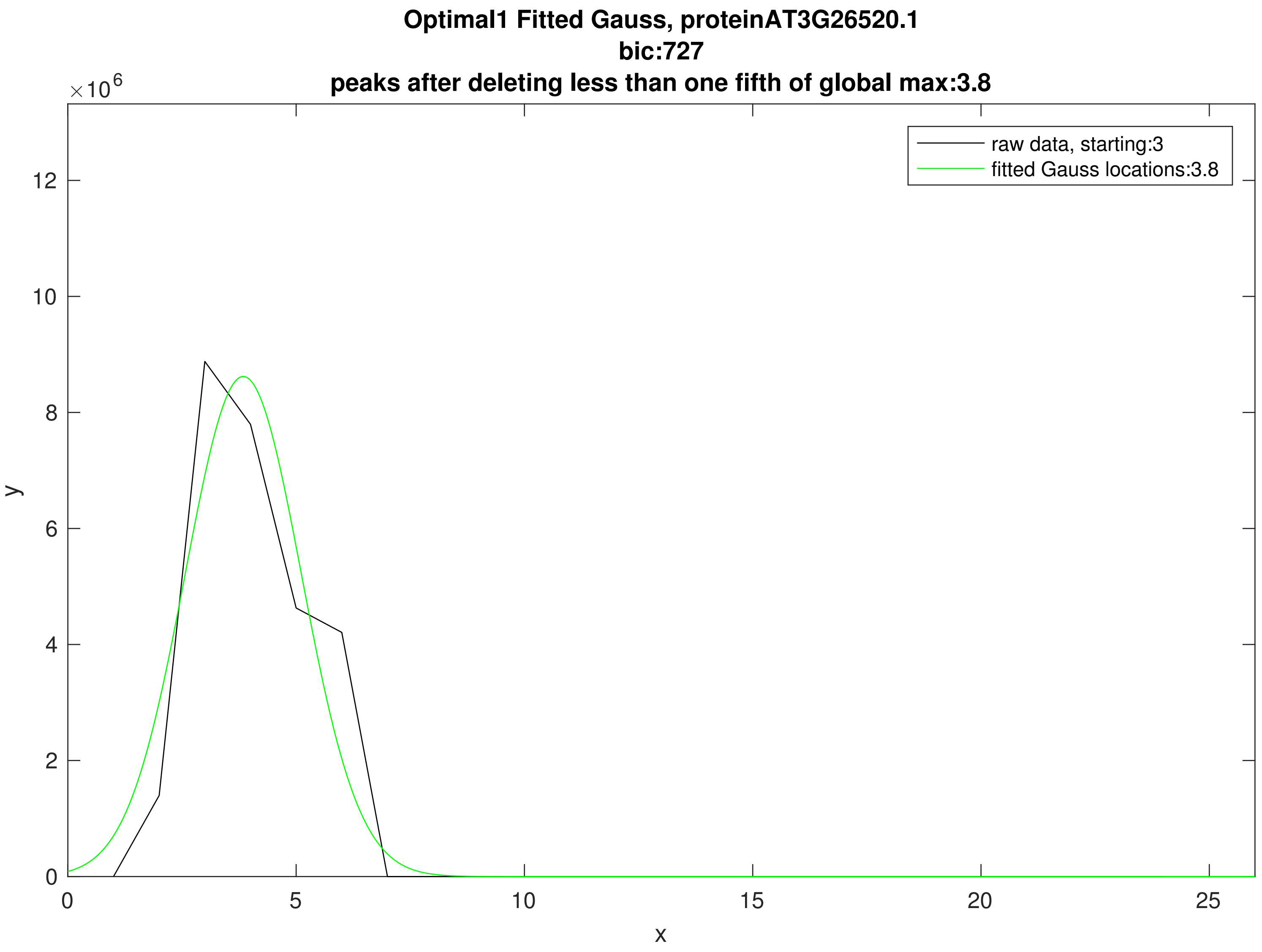
Optimal1 Fitted Gauss, proteinAT3G26520.1
bic:727
peaks after deleting less than one fifth of global max:3.8
×10^6
raw data, starting:3
fitted Gauss locations:3.8
y
x
0
2
4
6
8
10
12
0
5
10
15
20
25

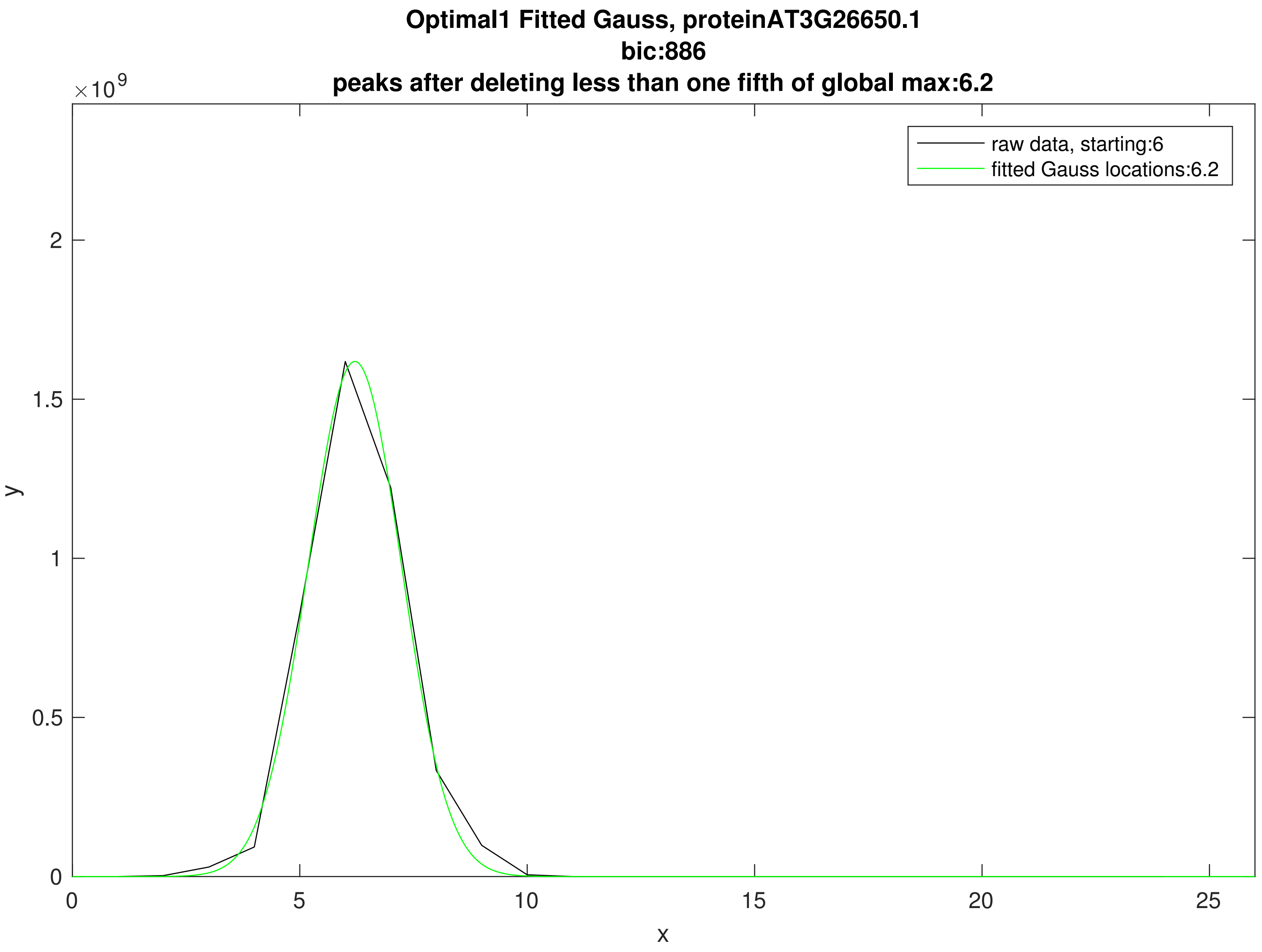

Optimal1 Fitted Gauss, proteinAT3G26650.1
bic:886
peaks after deleting less than one fifth of global max:6.2
×10^9
raw data, starting:6
fitted Gauss locations:6.2
y
x
0
0.5
1
1.5
2
5
10
15
20
25

#### Optimal1 Fitted Gauss, proteinAT3G26720.1;AT3G26720.2
#### bic:931 1e+03 948
#### peaks after deleting less than one fifth of global max:9

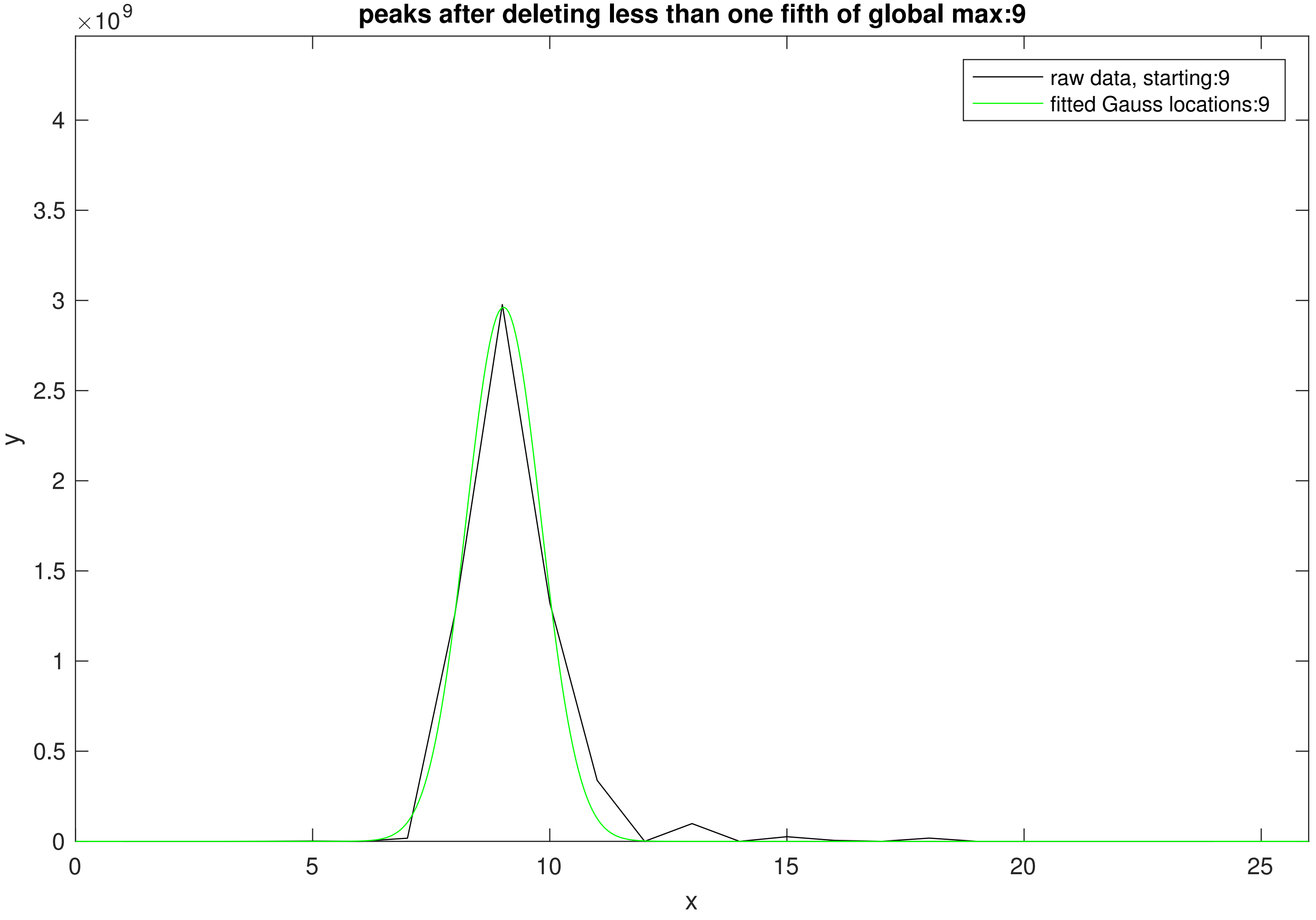

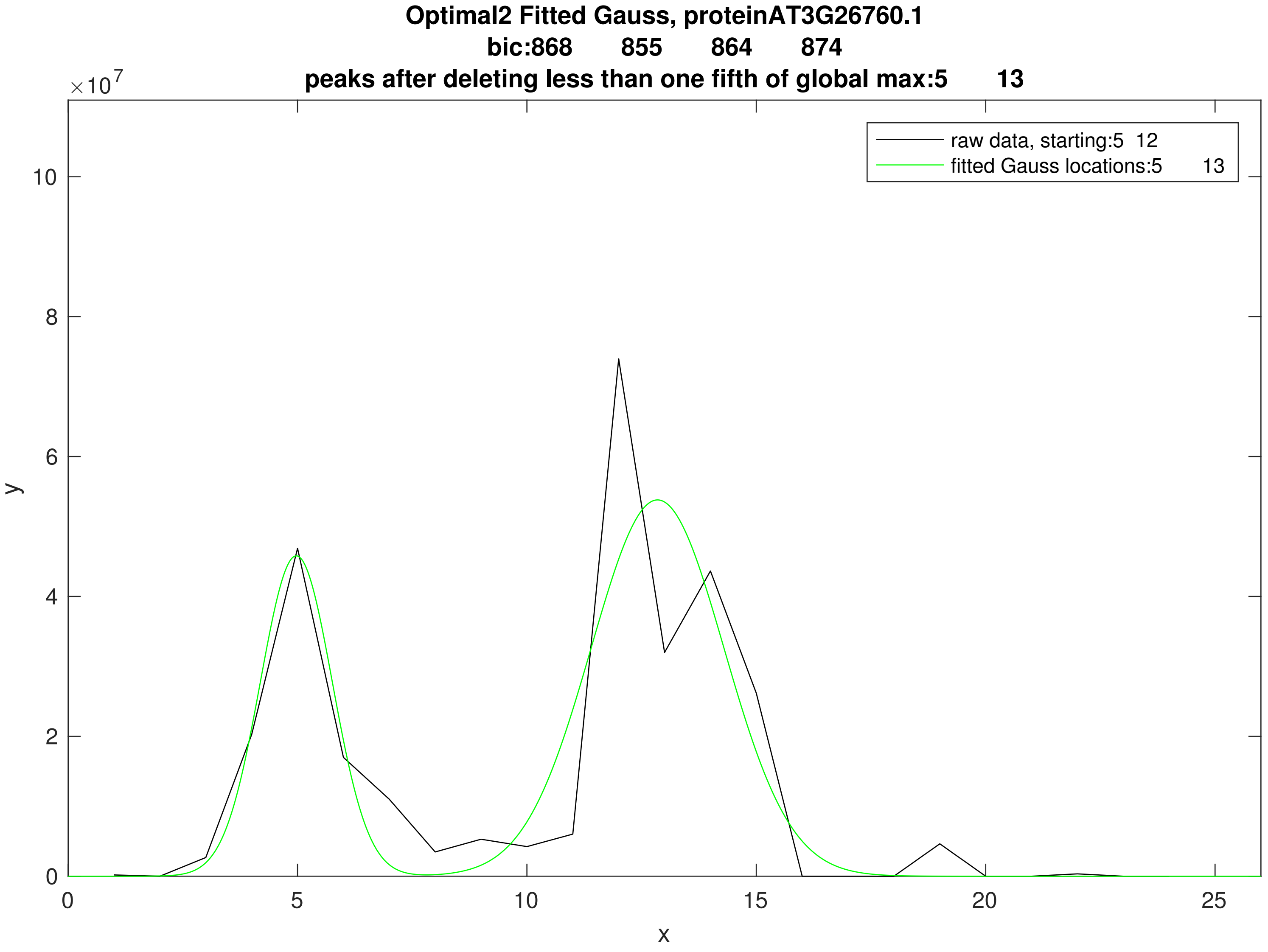
Optimal2 Fitted Gauss, proteinAT3G26760.1
bic:868 855 864 874
peaks after deleting less than one fifth of global max:5 13
×10^7
raw data, starting:5 12
fitted Gauss locations:5 13
y
x
0
2
4
6
8
10
0
5
10
15
20
25

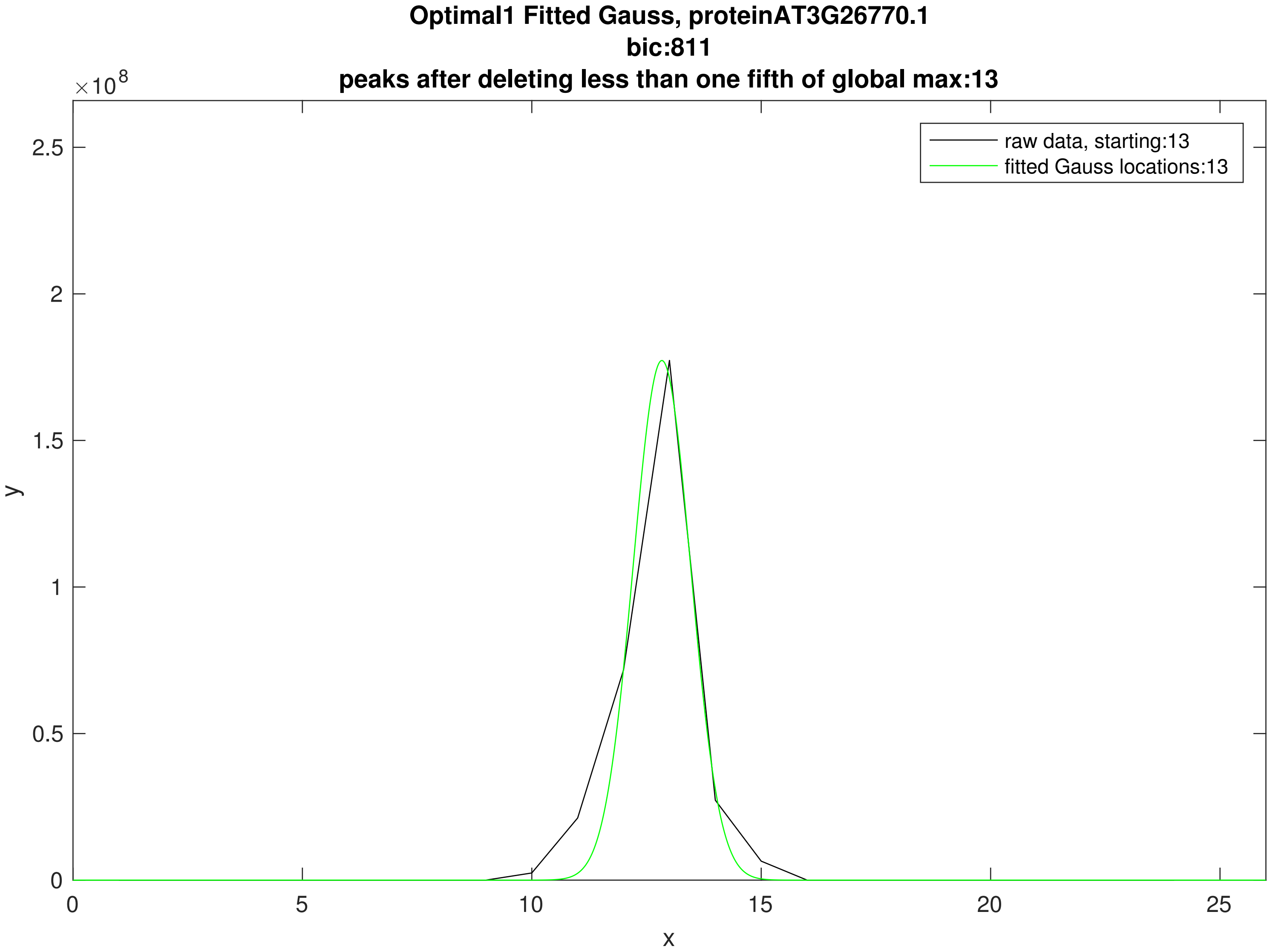
Optimal1 Fitted Gauss, proteinAT3G26770.1
bic:811
peaks after deleting less than one fifth of global max:13
×10^8
2.5
2
1.5
1
0.5
0
y
0
5
10
15
20
25
x
raw data, starting:13
fitted Gauss locations:13

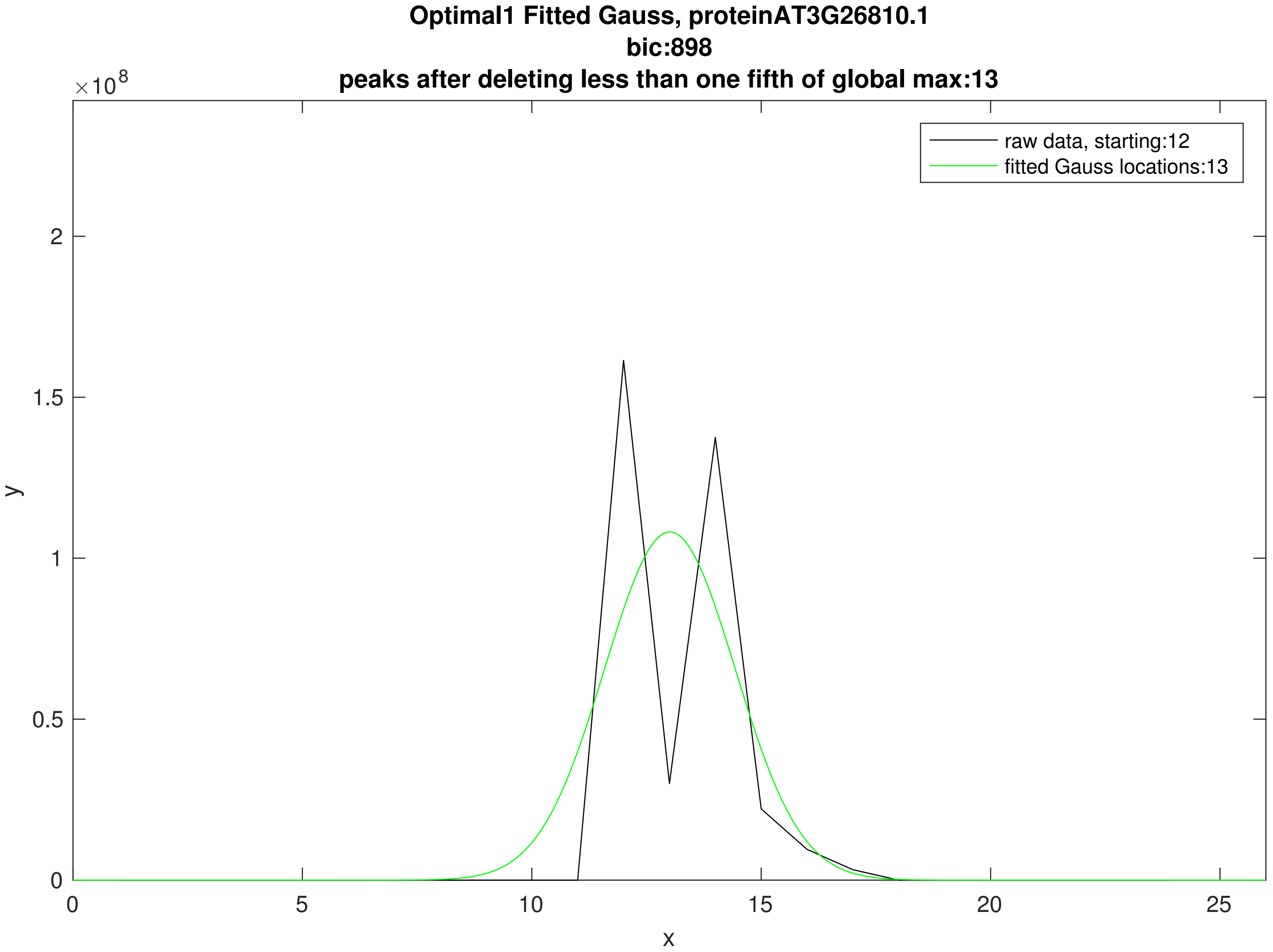

Optimal1 Fitted Gauss, proteinAT3G26810.1
bic:898
peaks after deleting less than one fifth of global max:13
×10^8
raw data, starting:12
fitted Gauss locations:13
2
1.5
1
0.5
0
y
0
5
10
15
20
25
x

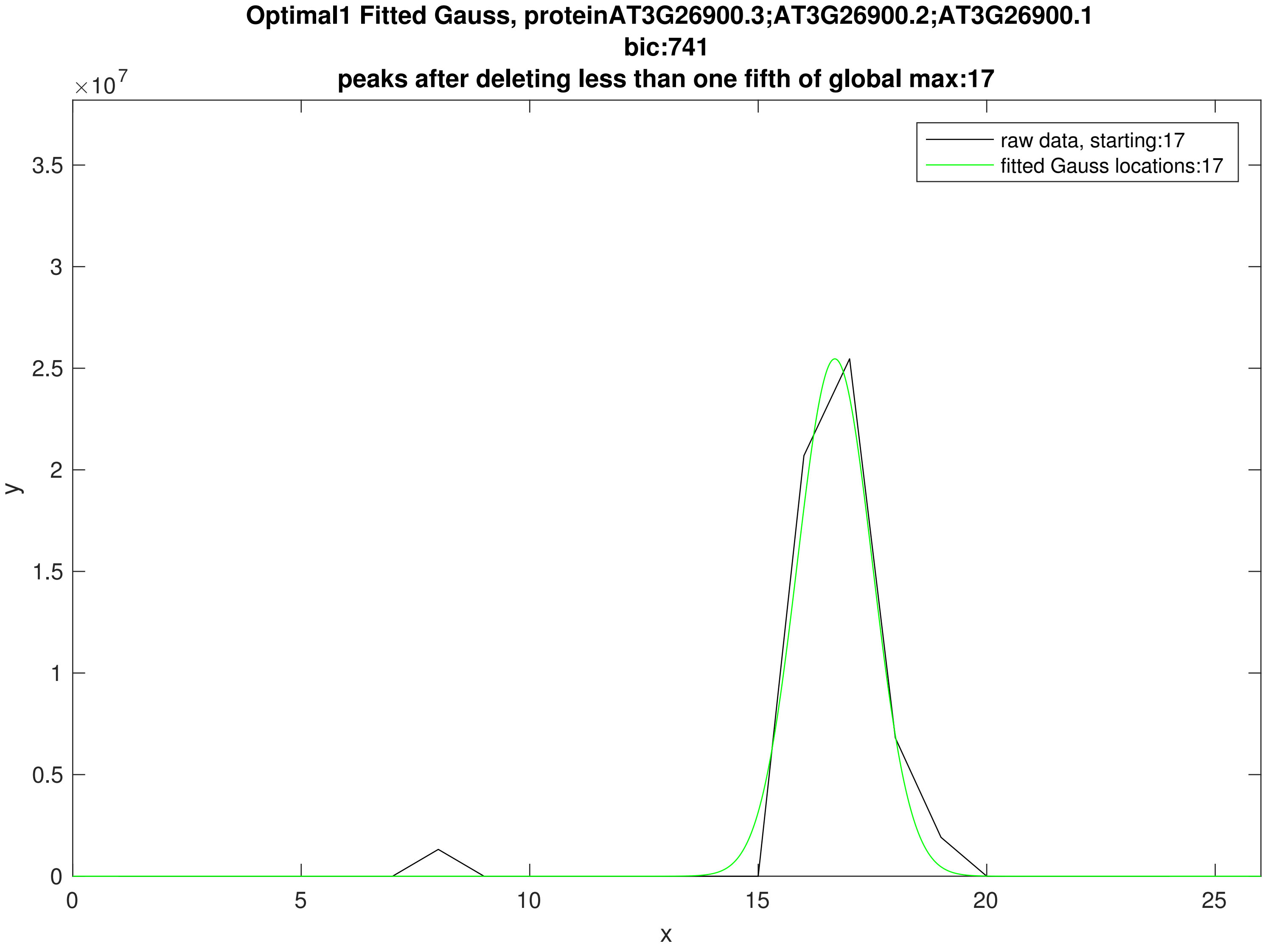
Optimal1 Fitted Gauss, proteinAT3G26900.3;AT3G26900.2;AT3G26900.1
bic:741
peaks after deleting less than one fifth of global max:17
×10^7
raw data, starting:17
fitted Gauss locations:17
y
x
0
0.5
1
1.5
2
2.5
3
3.5
0
5
10
15
20
25

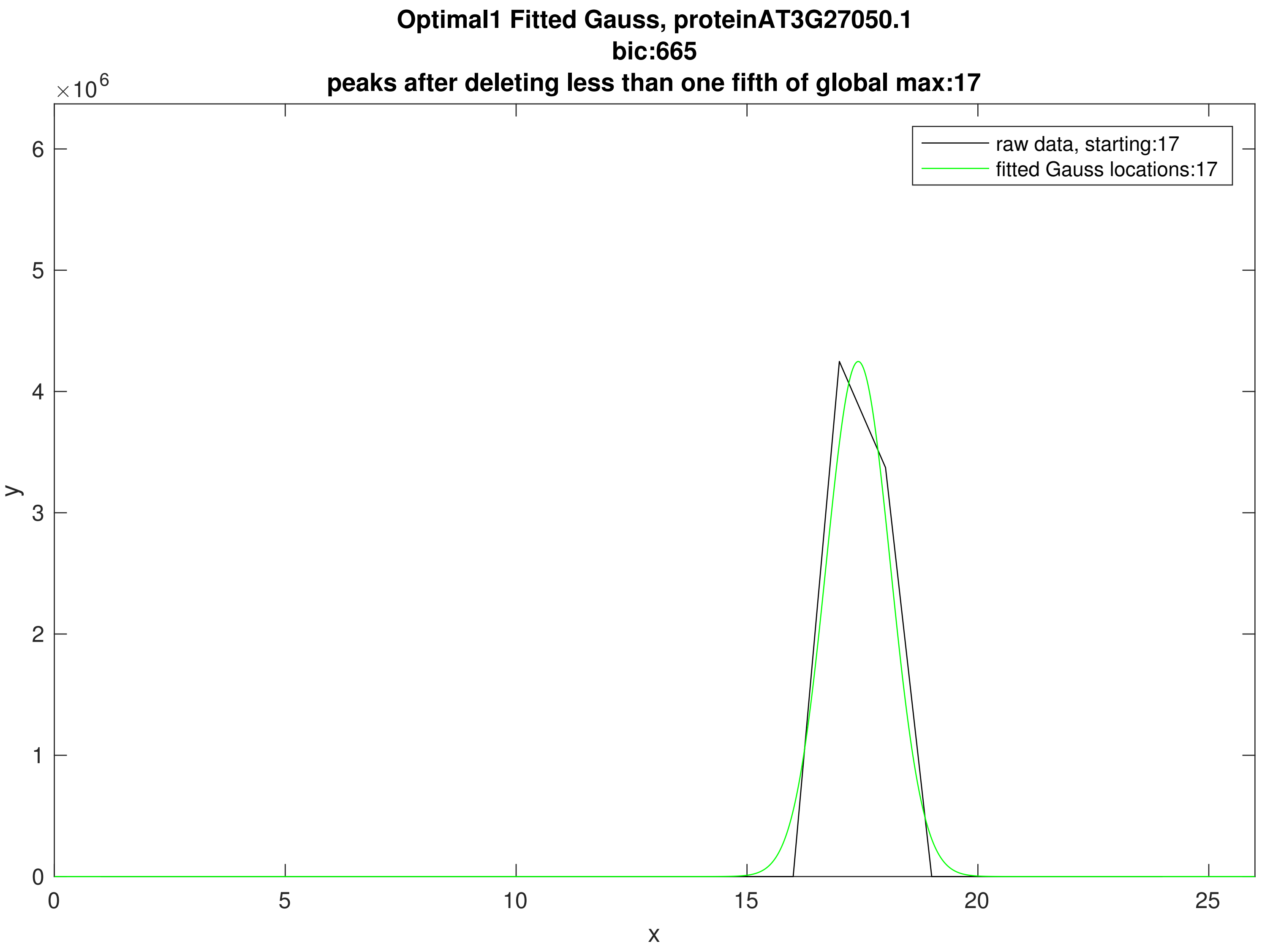

Optimal1 Fitted Gauss, proteinAT3G27050.1
bic:665
peaks after deleting less than one fifth of global max:17
×10^6
raw data, starting:17
fitted Gauss locations:17
y
x
0
1
2
3
4
5
6
0
5
10
15
20
25

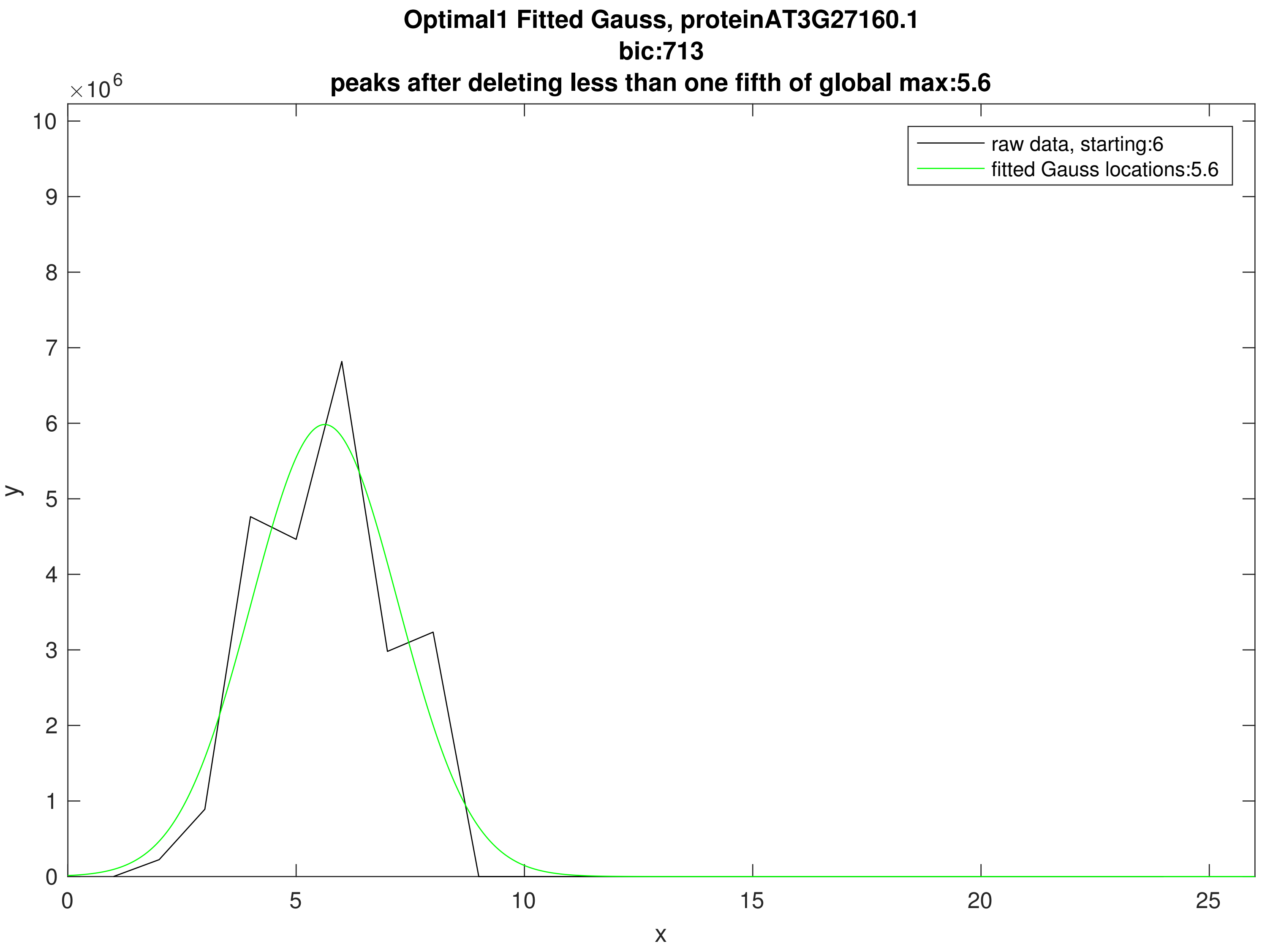

Optimal1 Fitted Gauss, proteinAT3G27160.1
bic:713
peaks after deleting less than one fifth of global max:5.6
×10^6
10
9
8
7
6
5
4
3
2
1
0
y
0
5
10
15
20
25
x
raw data, starting:6
fitted Gauss locations:5.6

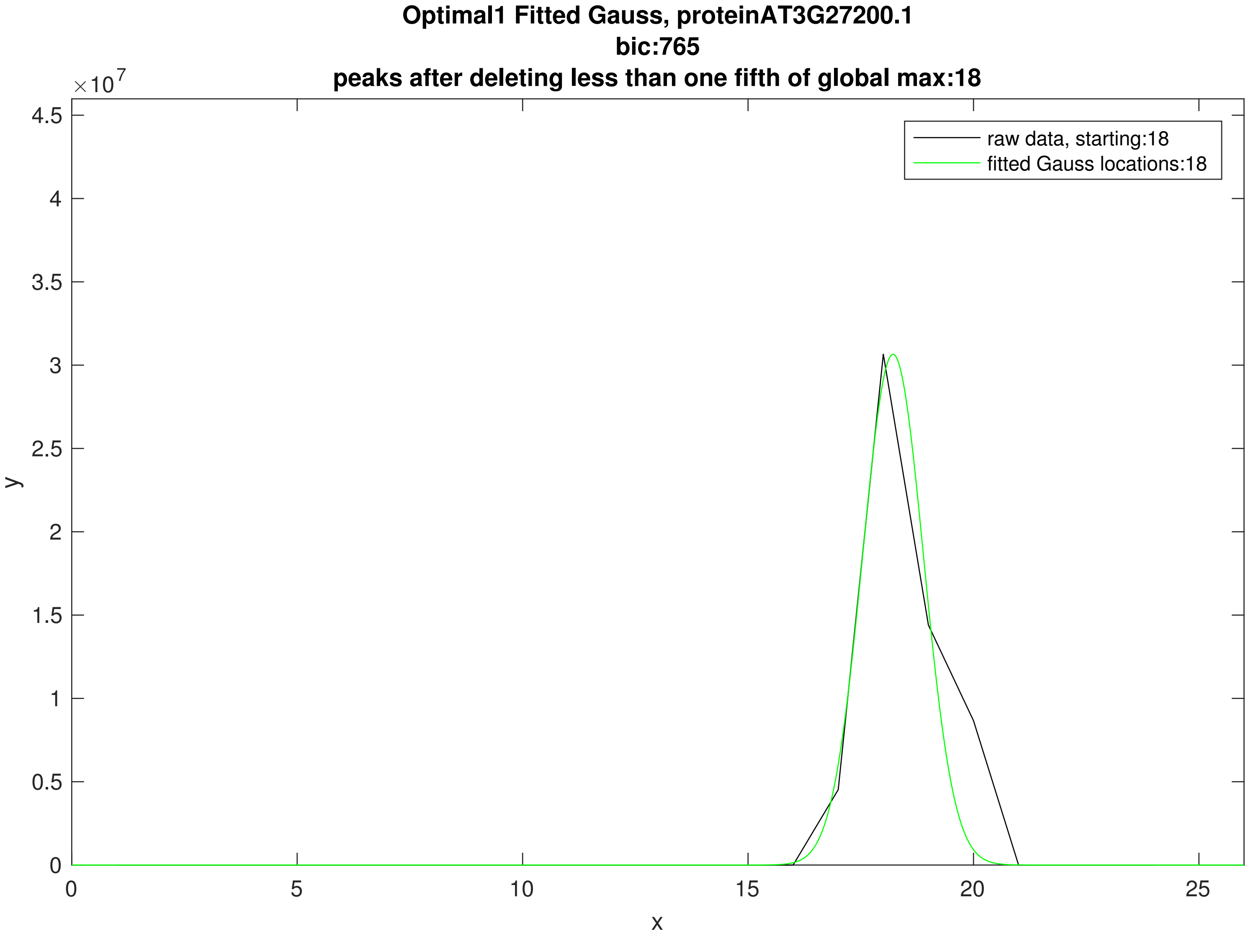

Optimal1 Fitted Gauss, proteinAT3G27200.1
bic:765
peaks after deleting less than one fifth of global max:18
×10^7
raw data, starting:18
fitted Gauss locations:18
y
x
0
0.5
1
1.5
2
2.5
3
3.5
4
4.5
0
5
10
15
20
25

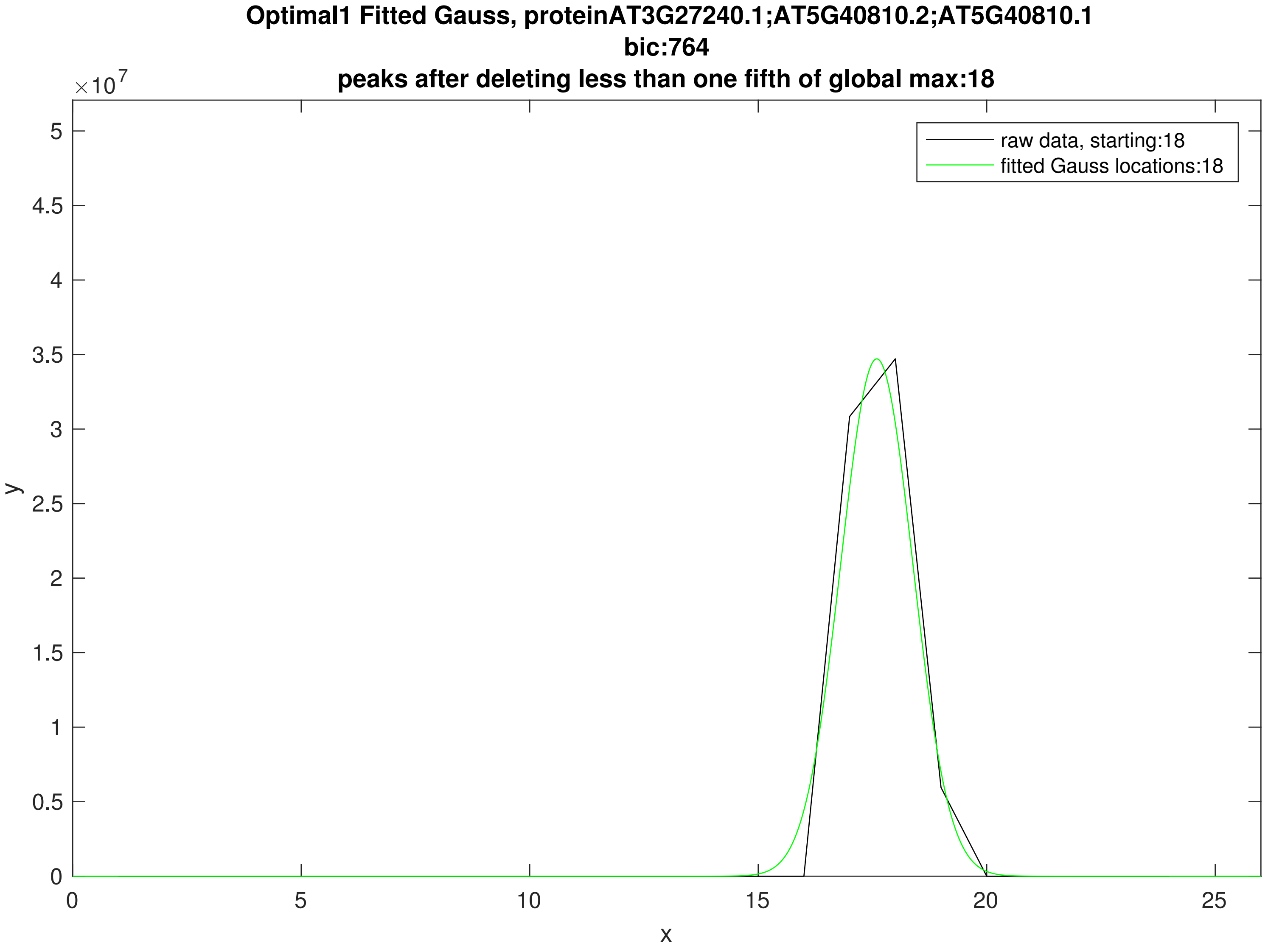
Optimal1 Fitted Gauss, proteinAT3G27240.1;AT5G40810.2;AT5G40810.1
bic:764
peaks after deleting less than one fifth of global max:18
×10^7
5
4.5
4
3.5
3
2.5
2
1.5
1
0.5
0
y
0
5
10
15
20
25
x
raw data, starting:18
fitted Gauss locations:18

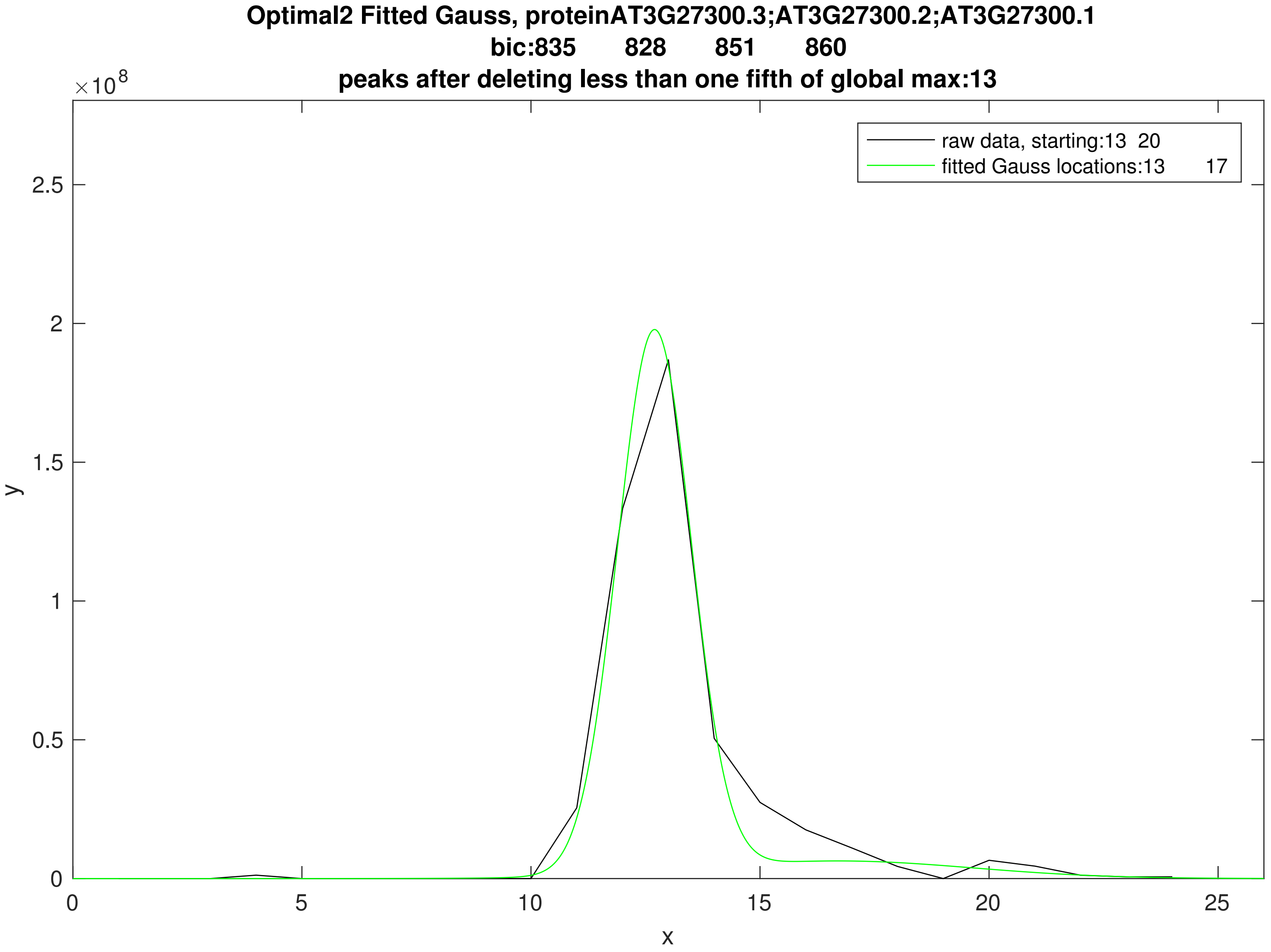
Optimal2 Fitted Gauss, proteinAT3G27300.3;AT3G27300.2;AT3G27300.1
bic:835 828 851 860
peaks after deleting less than one fifth of global max:13
×10^8
raw data, starting:13 20
fitted Gauss locations:13 17
y
x
0
0.5
1
1.5
2
2.5
0
5
10
15
20
25

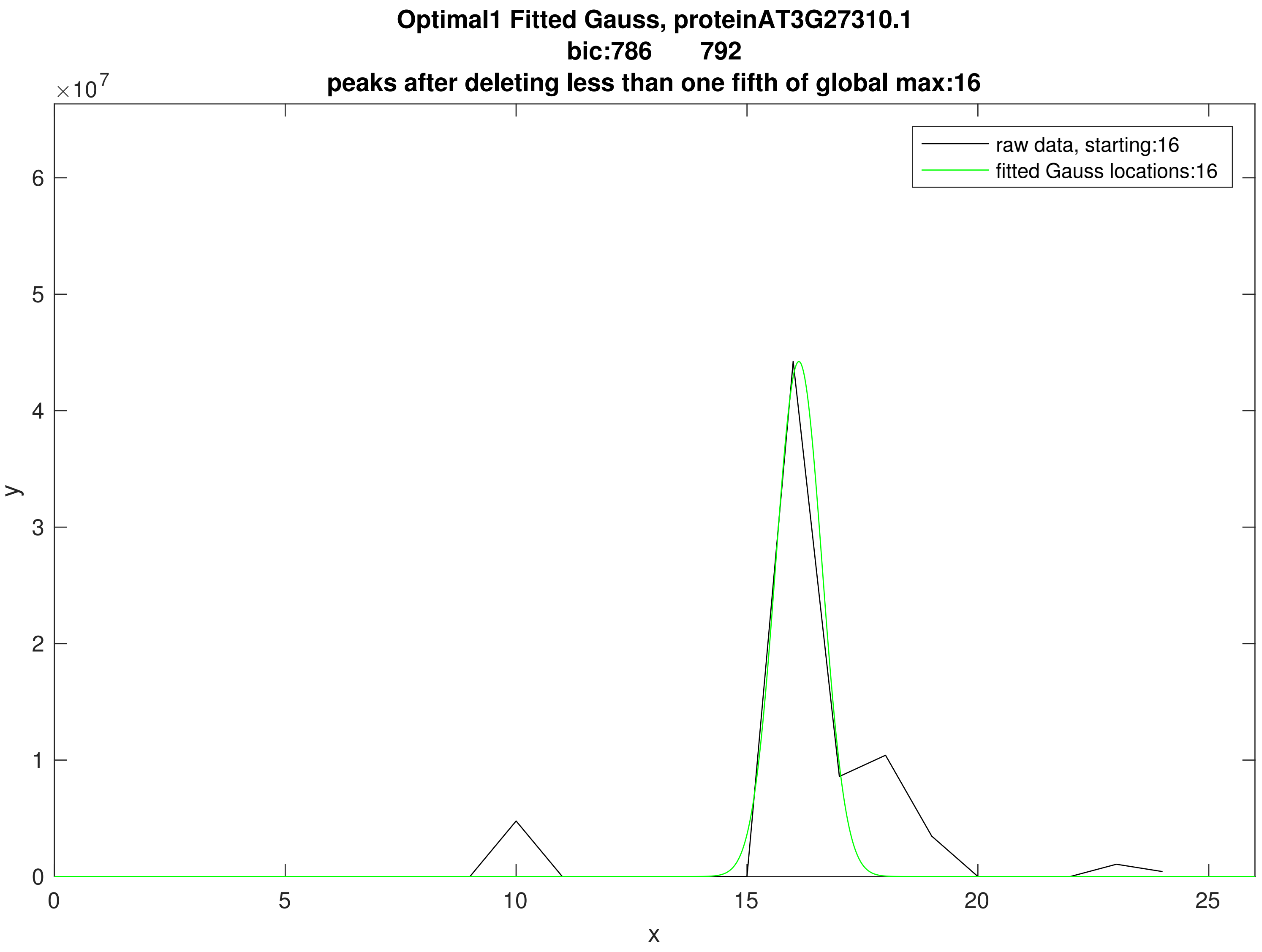
Optimal1 Fitted Gauss, proteinAT3G27310.1
bic:786 792
peaks after deleting less than one fifth of global max:16
×10^7
raw data, starting:16
fitted Gauss locations:16
y
x
0
1
2
3
4
5
6
0
5
10
15
20
25

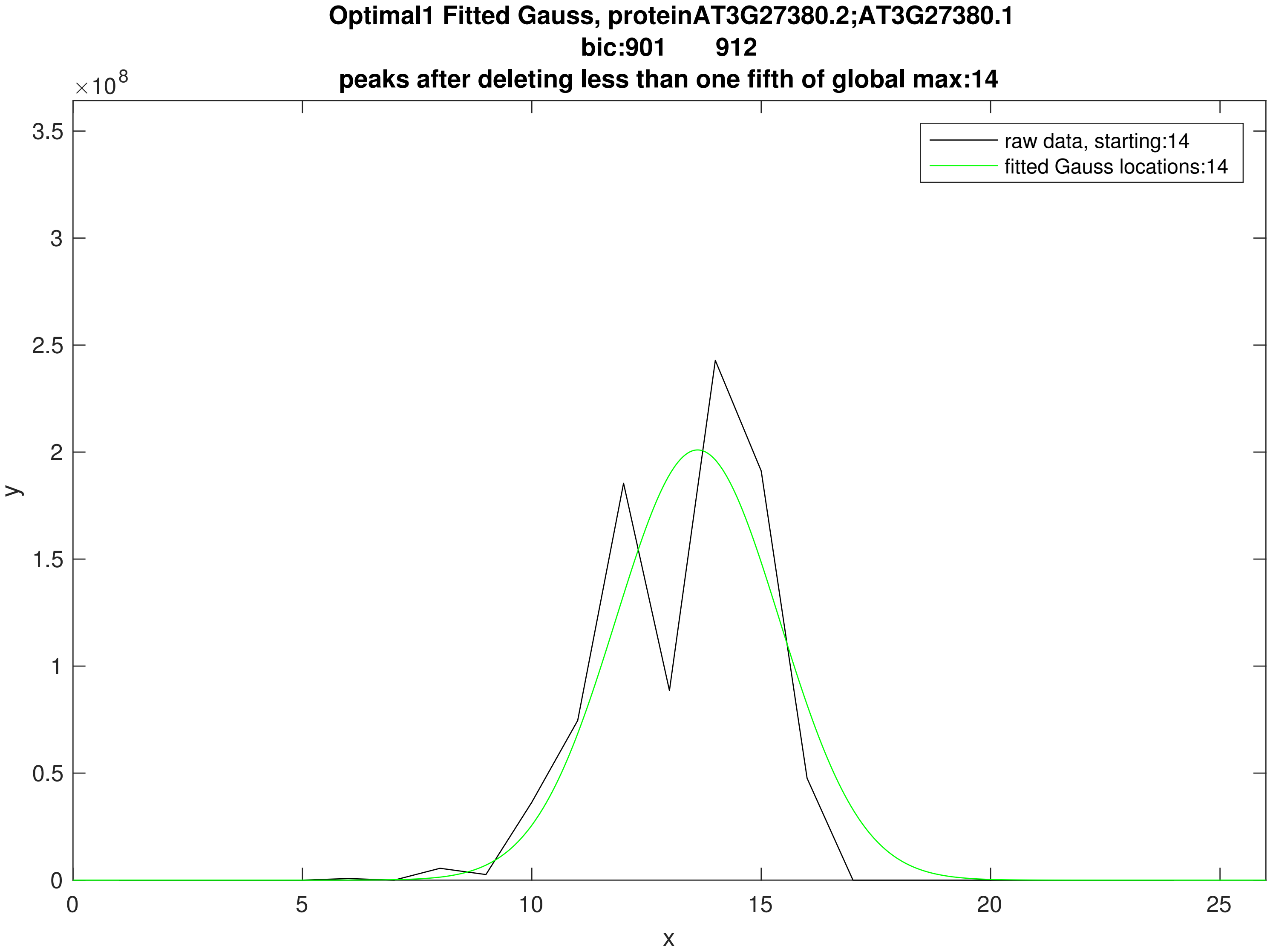
Optimal1 Fitted Gauss, proteinAT3G27380.2;AT3G27380.1
bic:901 912
peaks after deleting less than one fifth of global max:14
×10^8
3.5
3
2.5
2
1.5
1
0.5
0
y
0
5
10
15
20
25
x
raw data, starting:14
fitted Gauss locations:14

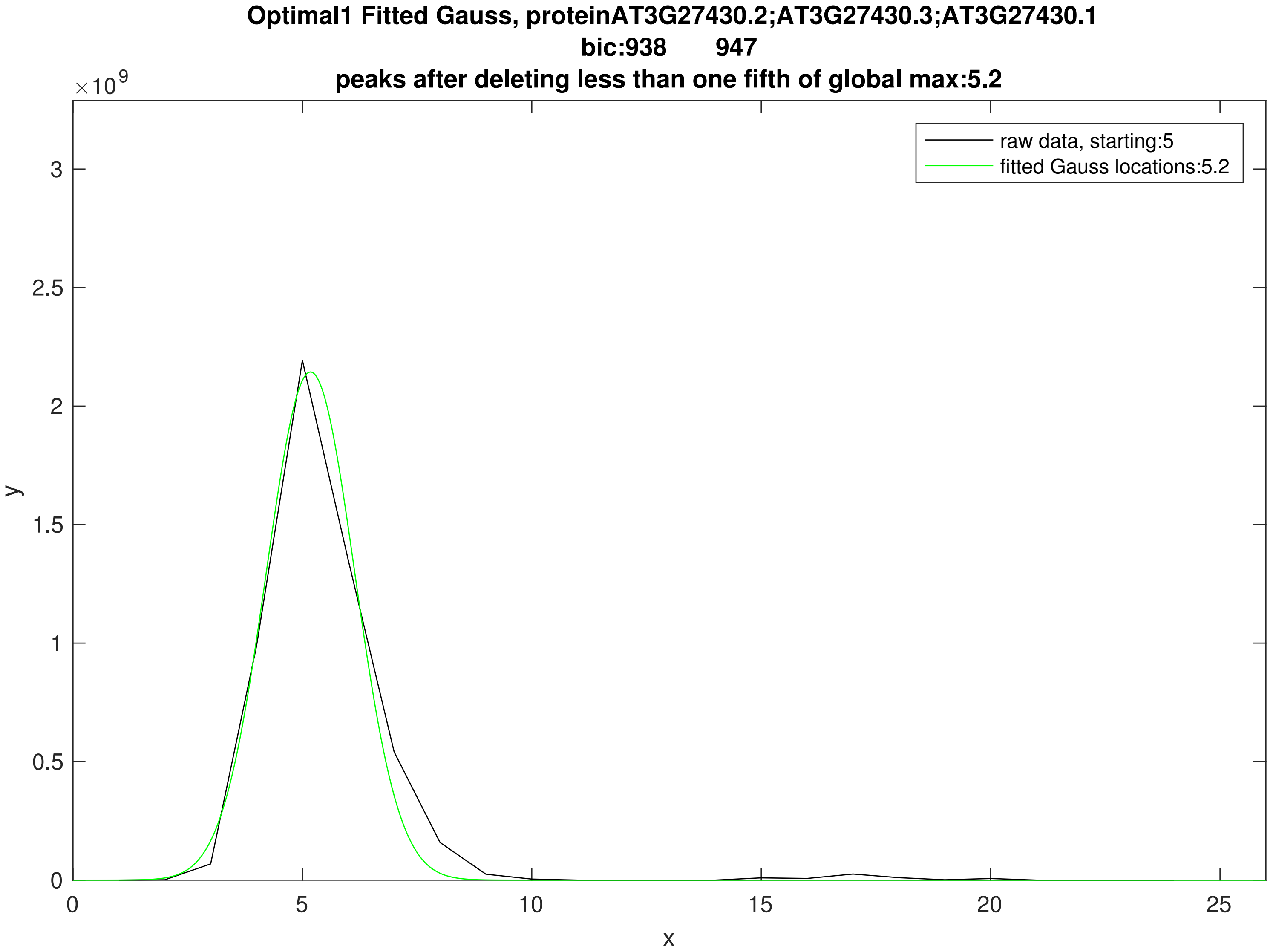

Optimal1 Fitted Gauss, proteinAT3G27430.2;AT3G27430.3;AT3G27430.1
bic:938 947
peaks after deleting less than one fifth of global max:5.2
×10^9
raw data, starting:5
fitted Gauss locations:5.2
y
x

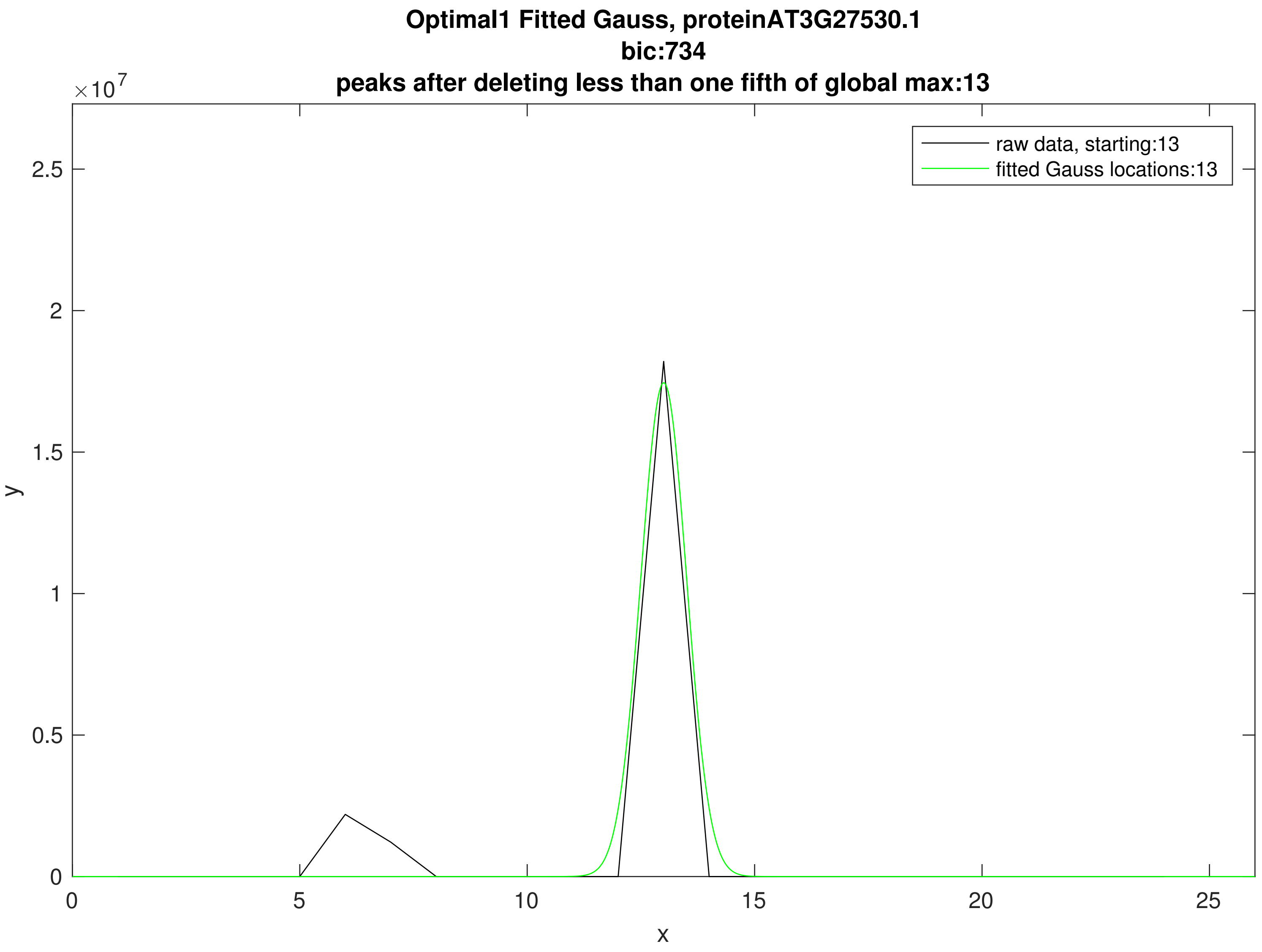
Optimal1 Fitted Gauss, proteinAT3G27530.1
bic:734
peaks after deleting less than one fifth of global max:13
×10^7
raw data, starting:13
fitted Gauss locations:13
2.5
2
1.5
1
0.5
0
y
0
5
10
15
20
25
x

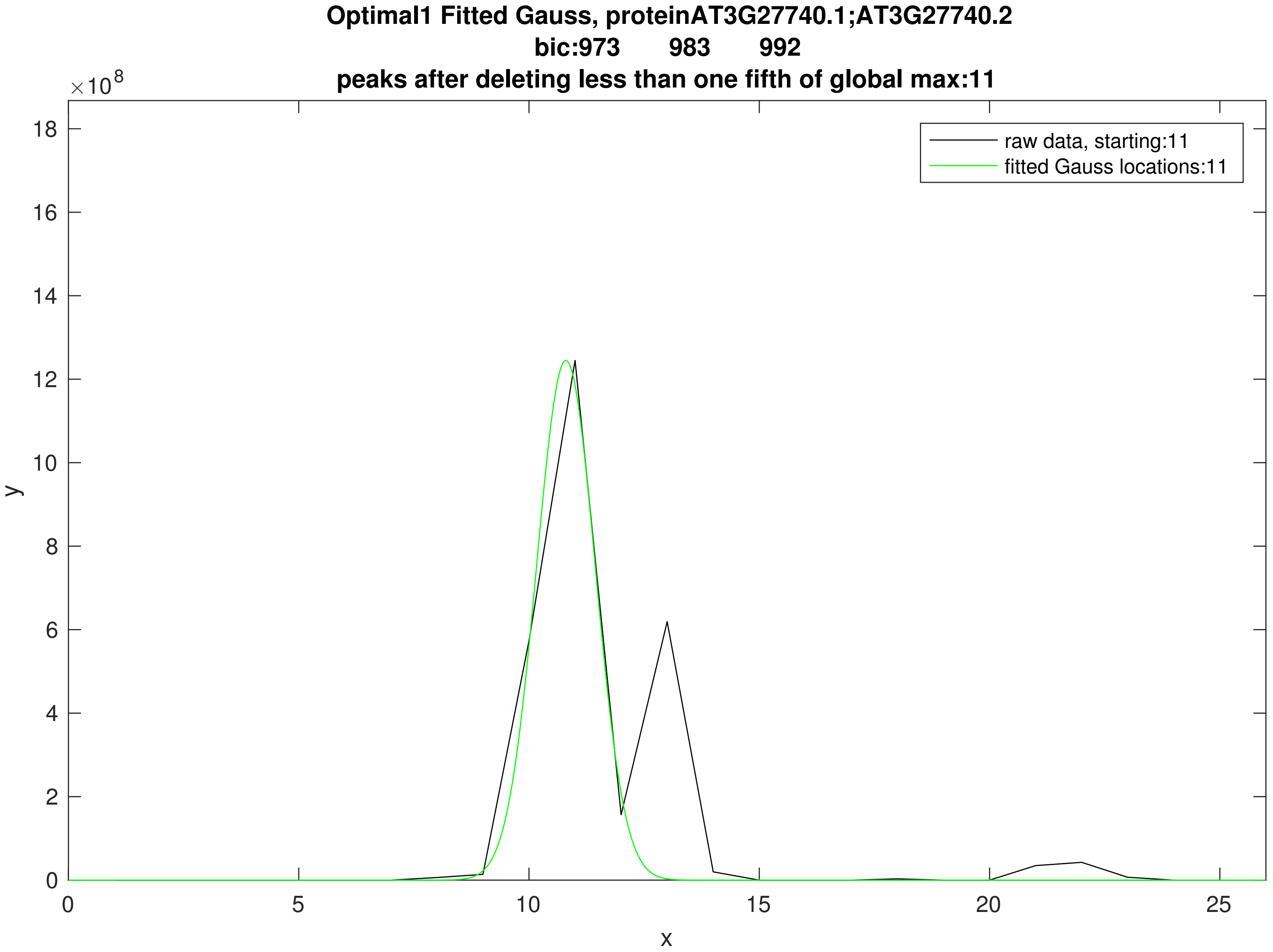
Optimal1 Fitted Gauss, proteinAT3G27740.1;AT3G27740.2
bic:973 983 992
peaks after deleting less than one fifth of global max:11
×10^8
raw data, starting:11
fitted Gauss locations:11
y
x
0
2
4
6
8
10
12
14
16
18
0
5
10
15
20
25

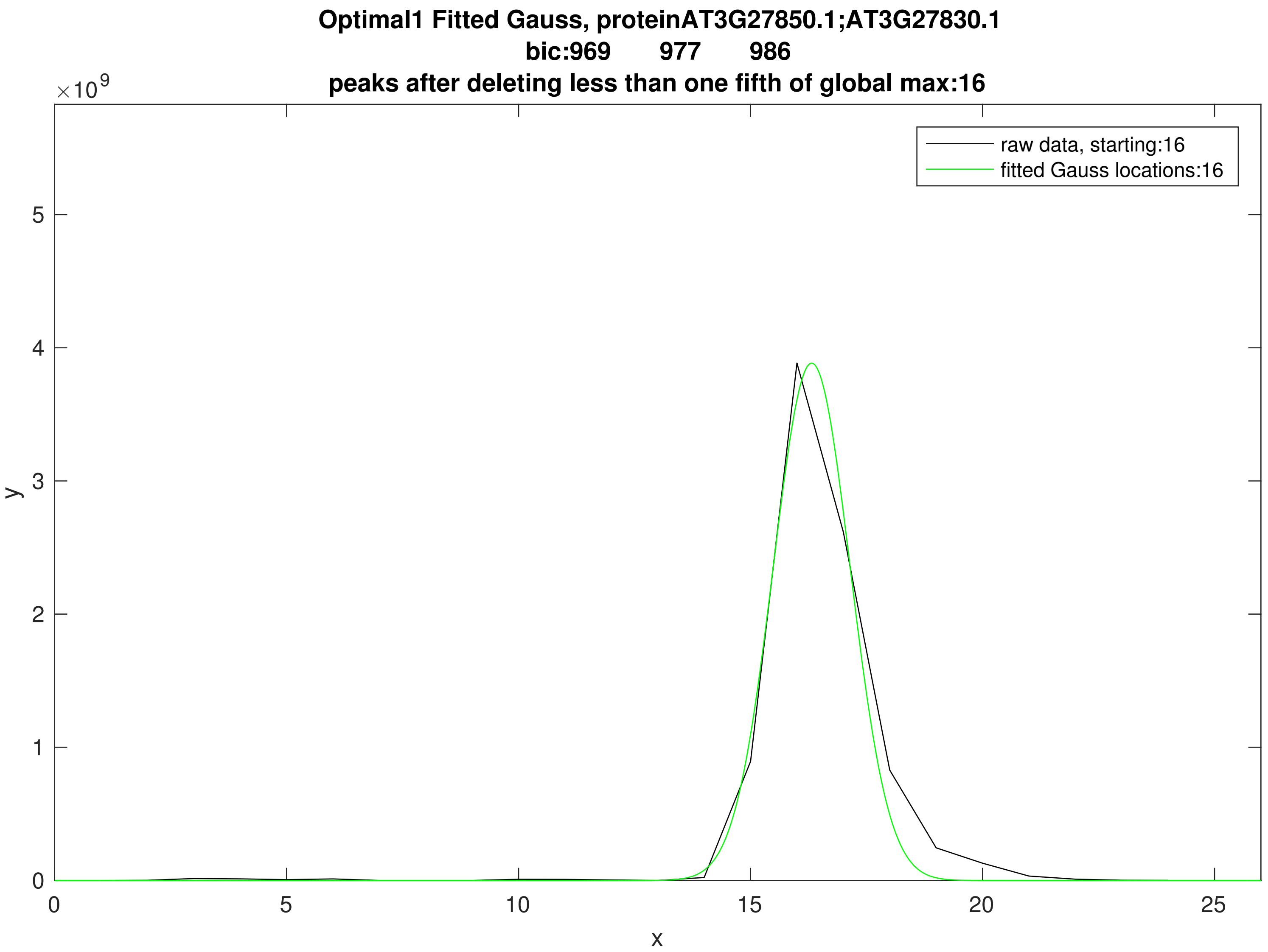

Optimal1 Fitted Gauss, proteinAT3G27850.1;AT3G27830.1
bic:969 977 986
peaks after deleting less than one fifth of global max:16
×10^9
raw data, starting:16
fitted Gauss locations:16
y
x
0
1
2
3
4
5
0
5
10
15
20
25

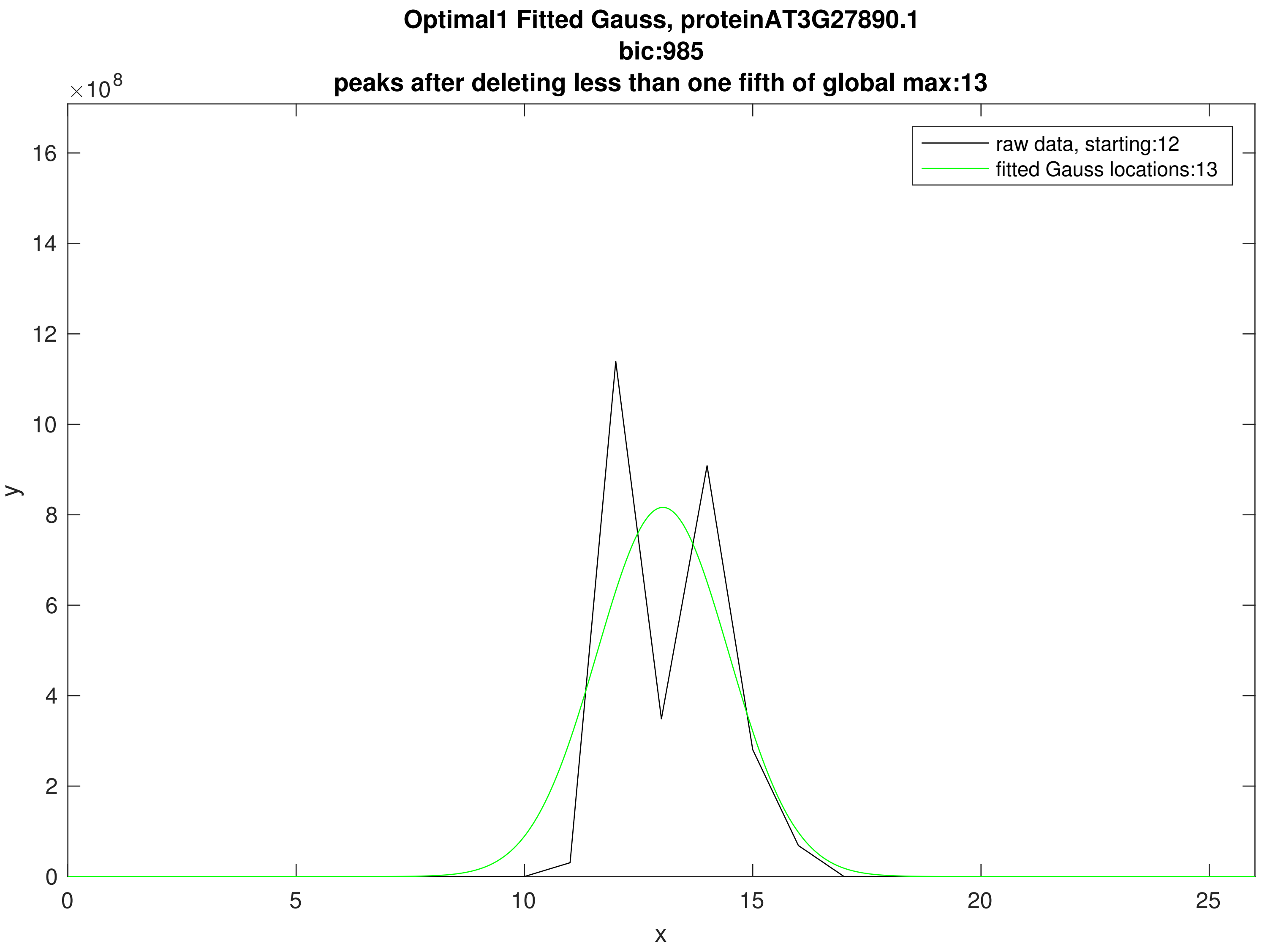

Optimal1 Fitted Gauss, proteinAT3G27890.1
bic:985
peaks after deleting less than one fifth of global max:13
×10^8
16
14
12
10
8
6
4
2
0
y
0
5
10
15
20
25
x
raw data, starting:12
fitted Gauss locations:13

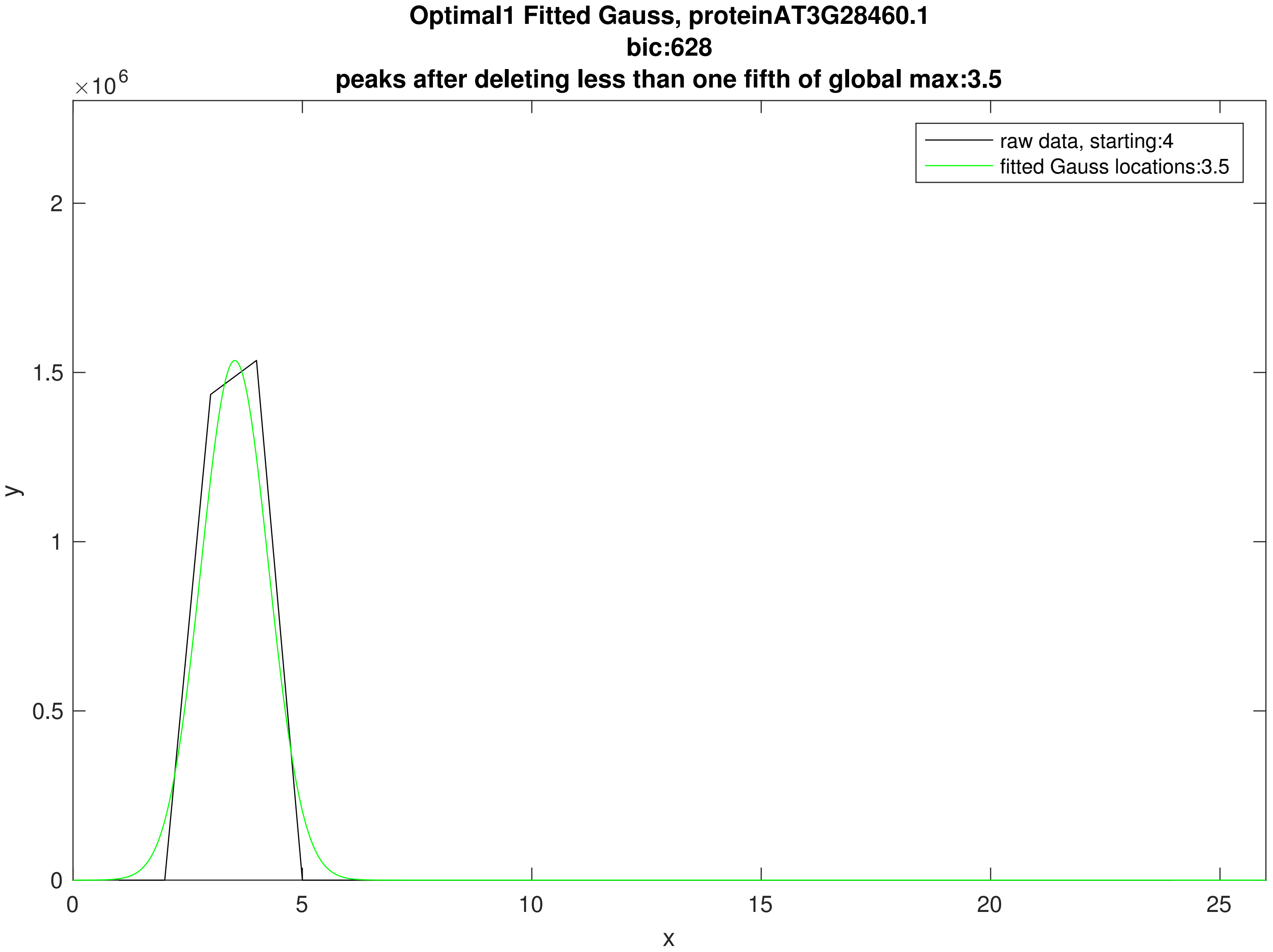

Optimal1 Fitted Gauss, proteinAT3G28460.1
bic:628
peaks after deleting less than one fifth of global max:3.5
×10^6
raw data, starting:4
fitted Gauss locations:3.5
y
x
0
0.5
1
1.5
2
0
5
10
15
20
25

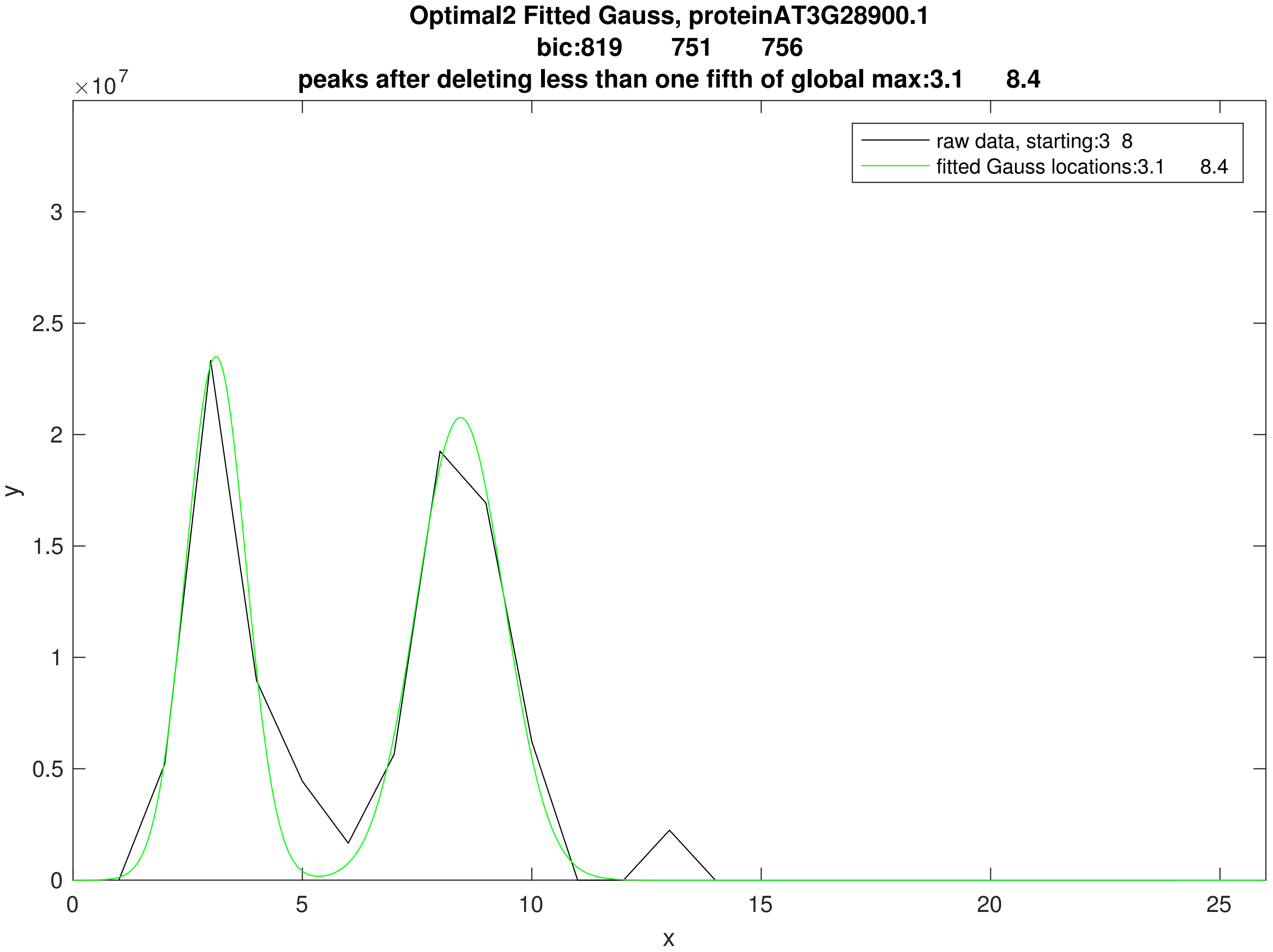
Optimal2 Fitted Gauss, proteinAT3G28900.1
bic:819 751 756
peaks after deleting less than one fifth of global max:3.1 8.4
×10^7
raw data, starting:3 8
fitted Gauss locations:3.1 8.4
3
2.5
2
1.5
1
0.5
0
y
0
5
10
15
20
25
x

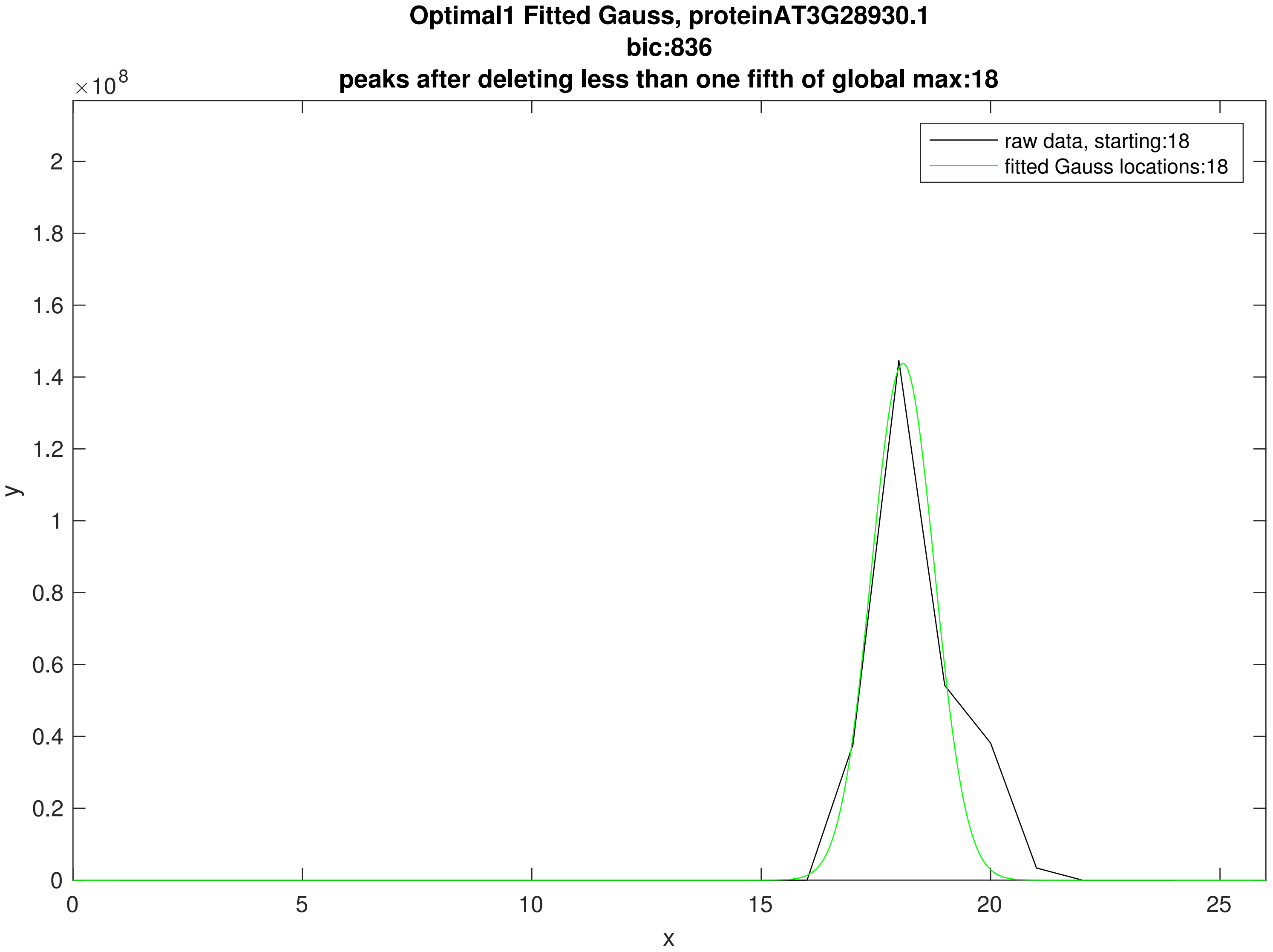
Optimal1 Fitted Gauss, proteinAT3G28930.1
bic:836
peaks after deleting less than one fifth of global max:18
×10^8
raw data, starting:18
fitted Gauss locations:18
2
1.8
1.6
1.4
1.2
1
0.8
0.6
0.4
0.2
0
y
0
5
10
15
20
25
x

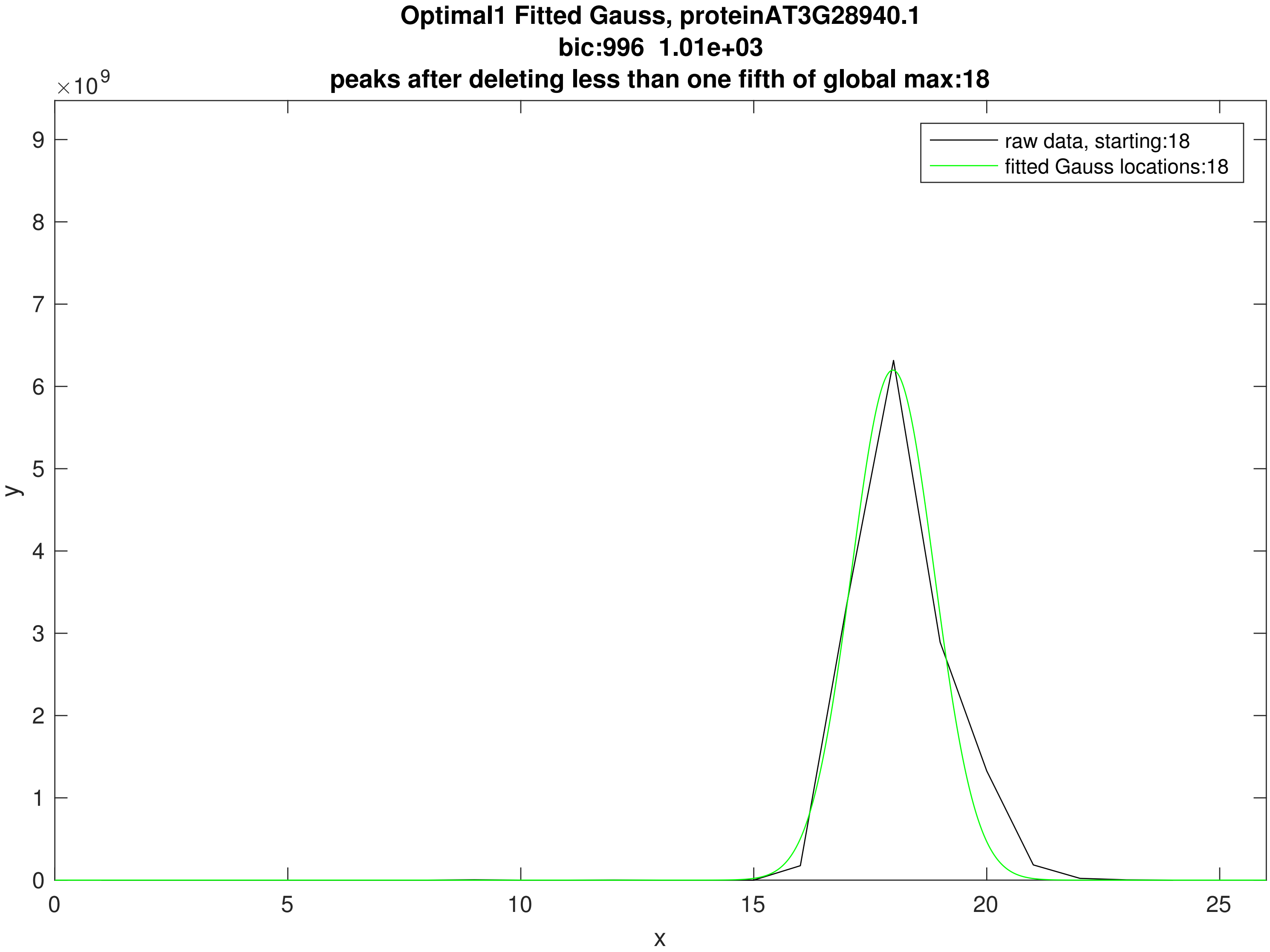
Optimal1 Fitted Gauss, proteinAT3G28940.1
bic:996  1.01e+03
peaks after deleting less than one fifth of global max:18
×10^9
9
8
7
6
5
4
3
2
1
0
y
0
5
10
15
20
25
x
raw data, starting:18
fitted Gauss locations:18

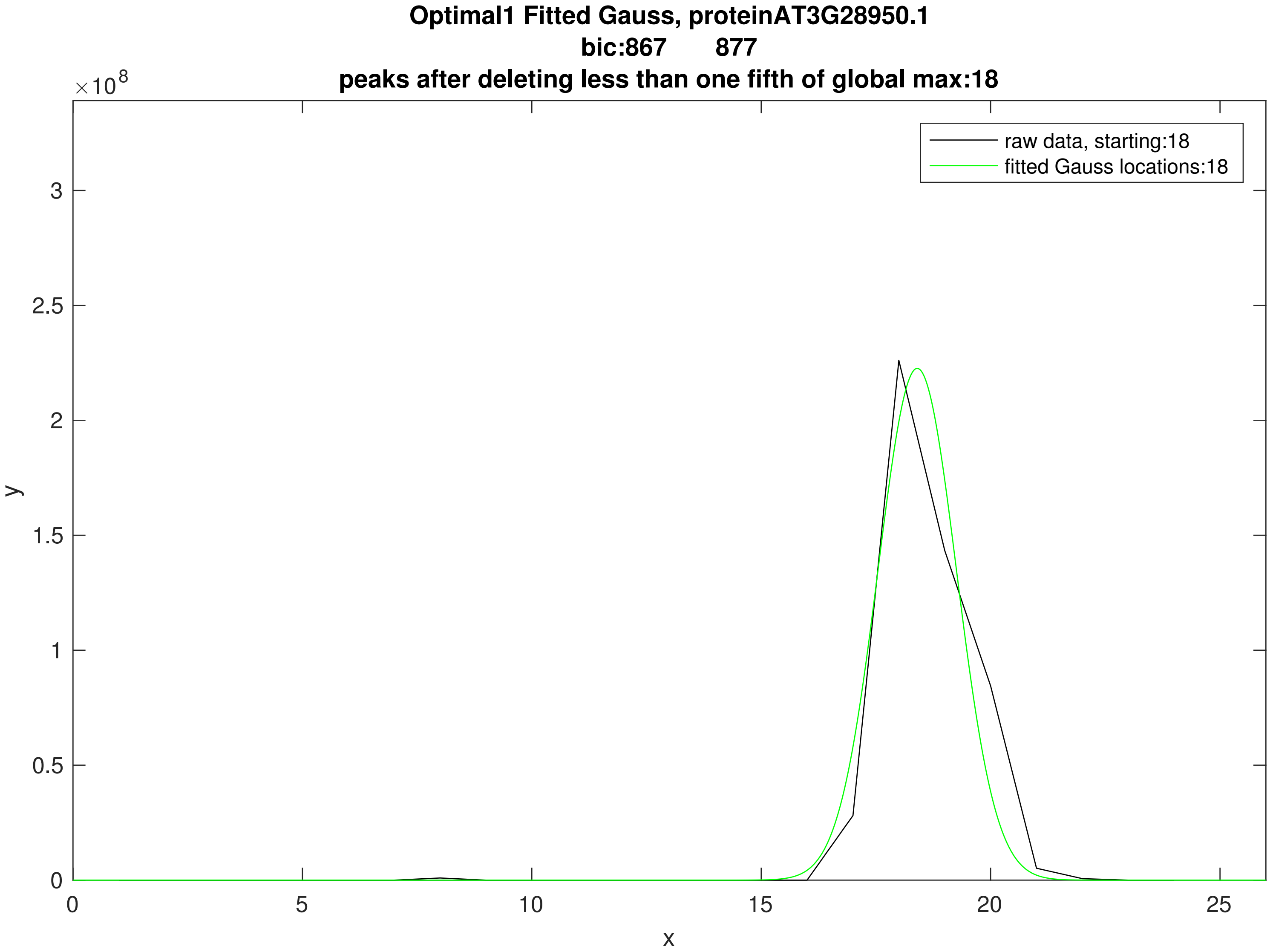

Optimal1 Fitted Gauss, proteinAT3G28950.1
bic:867 877
peaks after deleting less than one fifth of global max:18
×10^8
raw data, starting:18
fitted Gauss locations:18
y
x
0
0.5
1
1.5
2
2.5
3
0
5
10
15
20
25

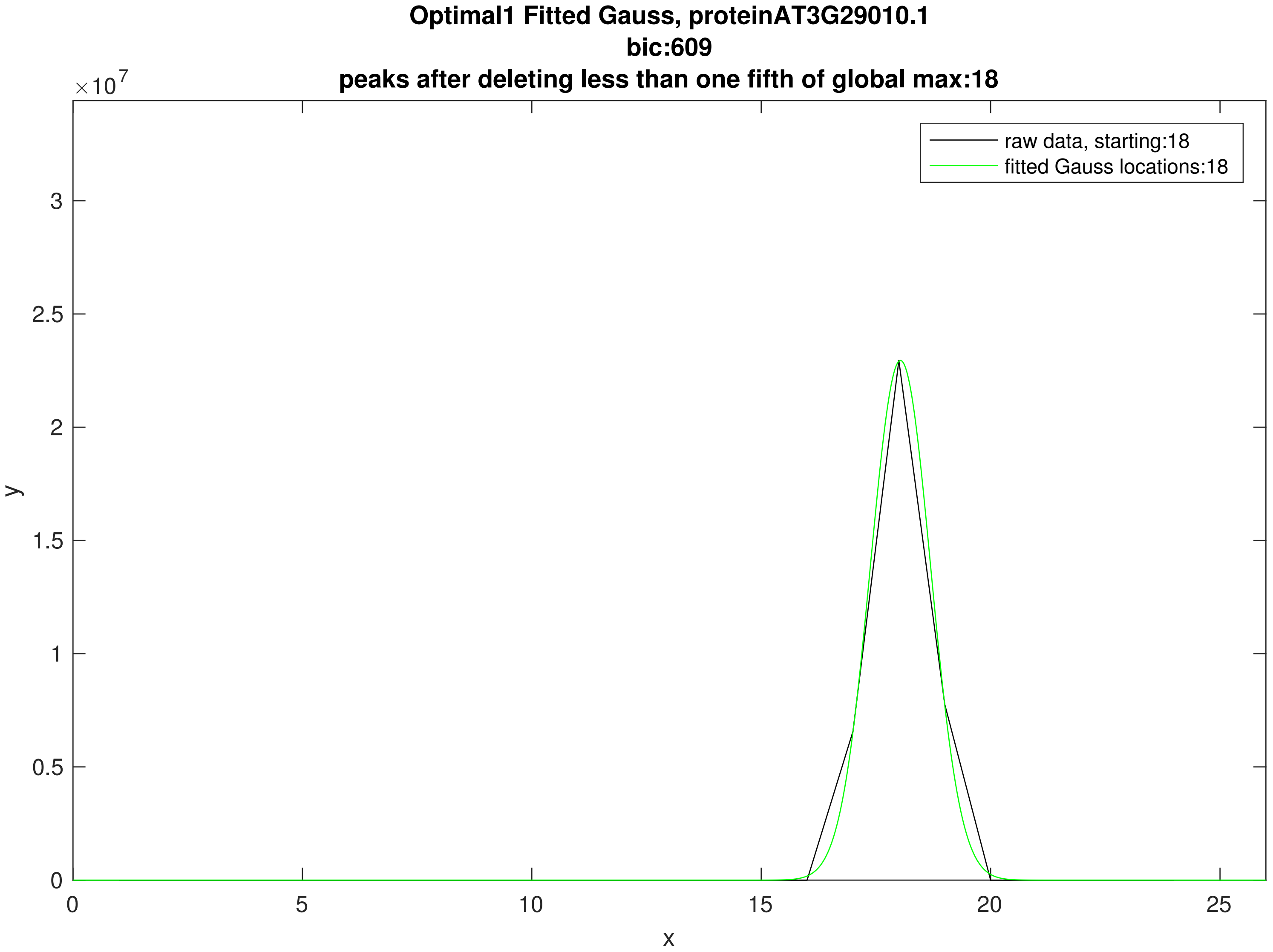

Optimal1 Fitted Gauss, proteinAT3G29010.1
bic:609
peaks after deleting less than one fifth of global max:18
×10^7
raw data, starting:18
fitted Gauss locations:18
y
x
0
0.5
1
1.5
2
2.5
3
0
5
10
15
20
25

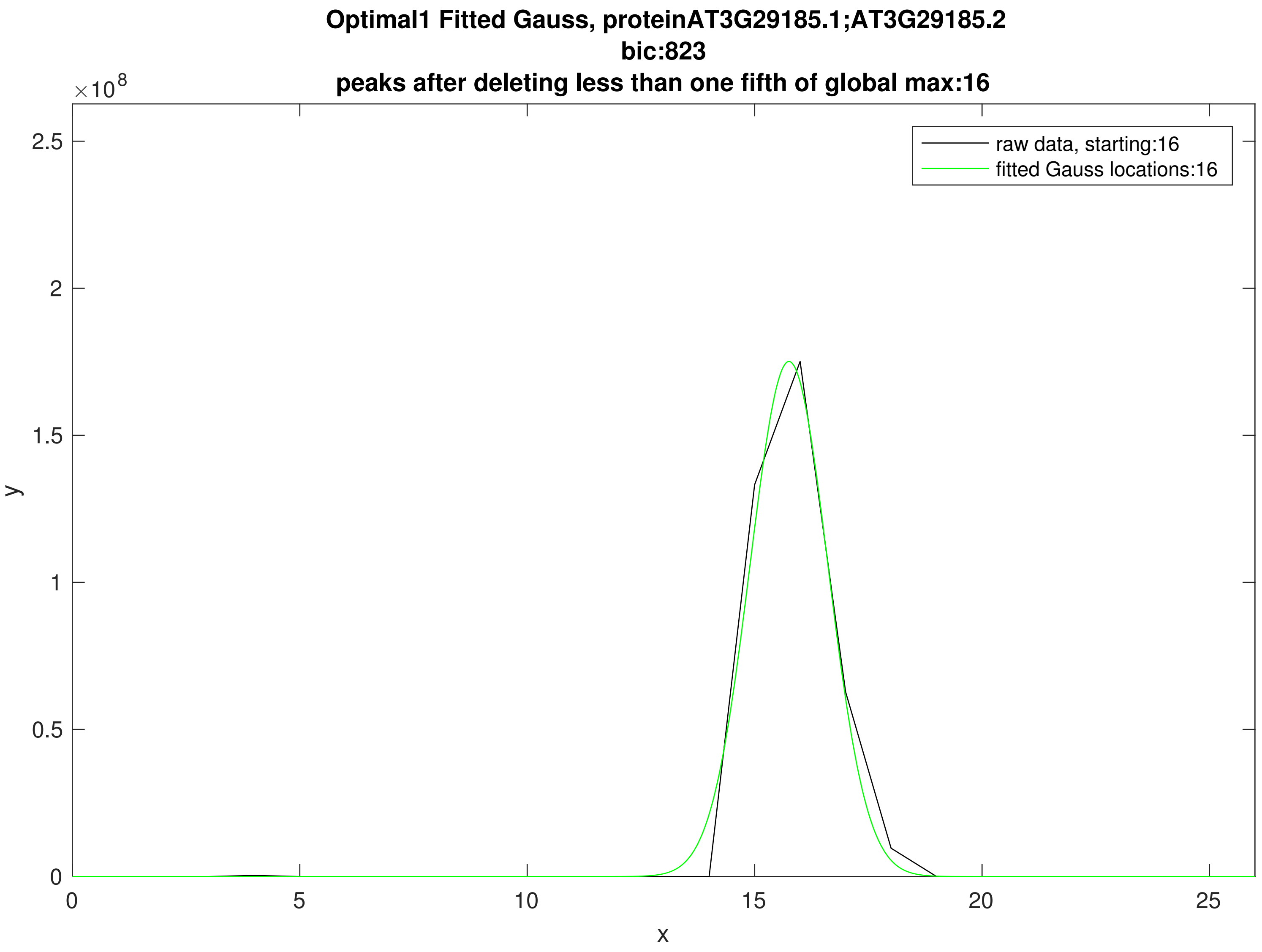

Optimal1 Fitted Gauss, proteinAT3G29185.1;AT3G29185.2
bic:823
peaks after deleting less than one fifth of global max:16
×10^8
2.5
2
1.5
1
0.5
0
y
0
5
10
15
20
25
x
raw data, starting:16
fitted Gauss locations:16

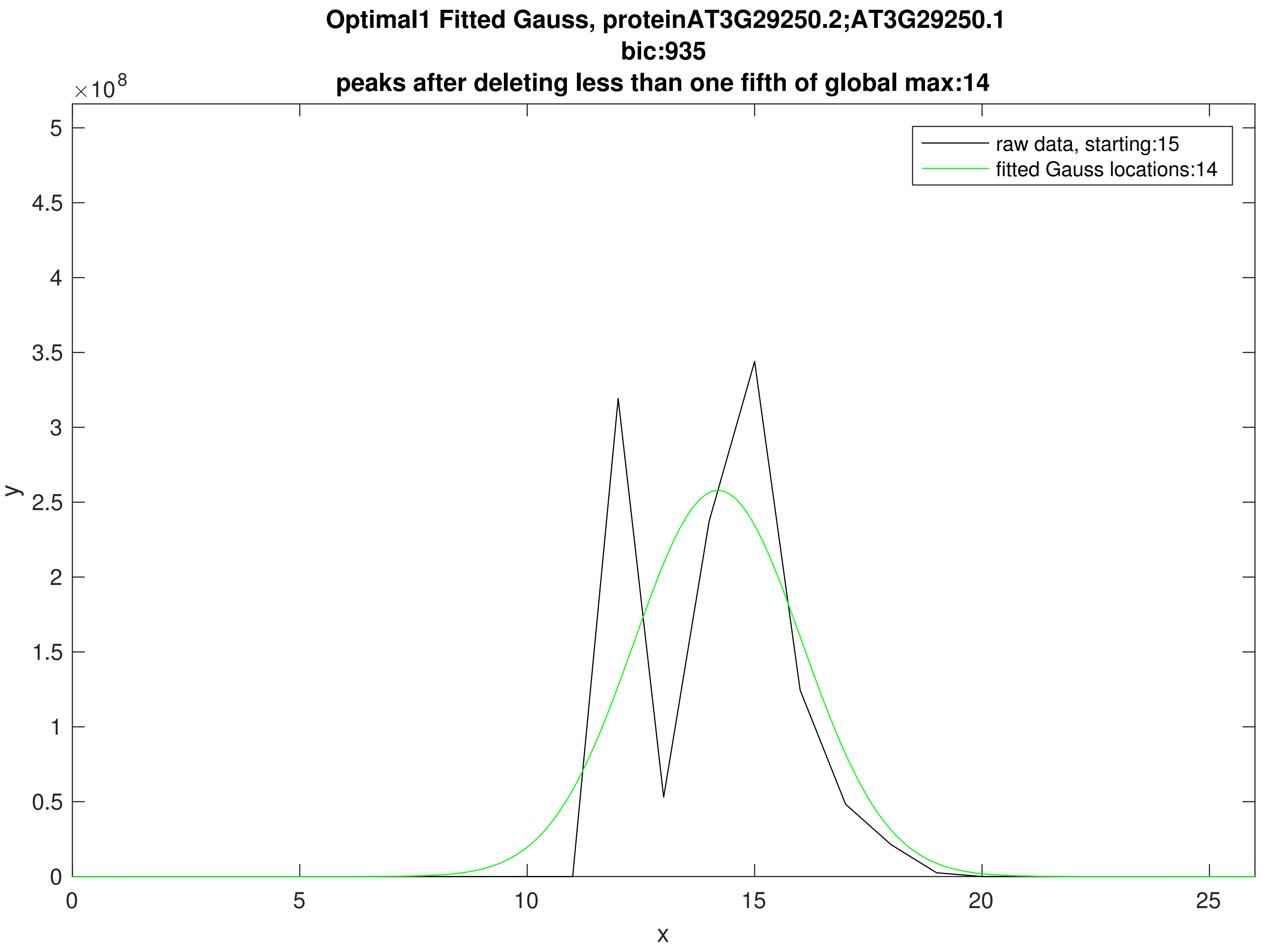
Optimal1 Fitted Gauss, proteinAT3G29250.2;AT3G29250.1
bic:935
peaks after deleting less than one fifth of global max:14
×10^8
raw data, starting:15
fitted Gauss locations:14
y
x
0
0.5
1
1.5
2
2.5
3
3.5
4
4.5
5
0
5
10
15
20
25

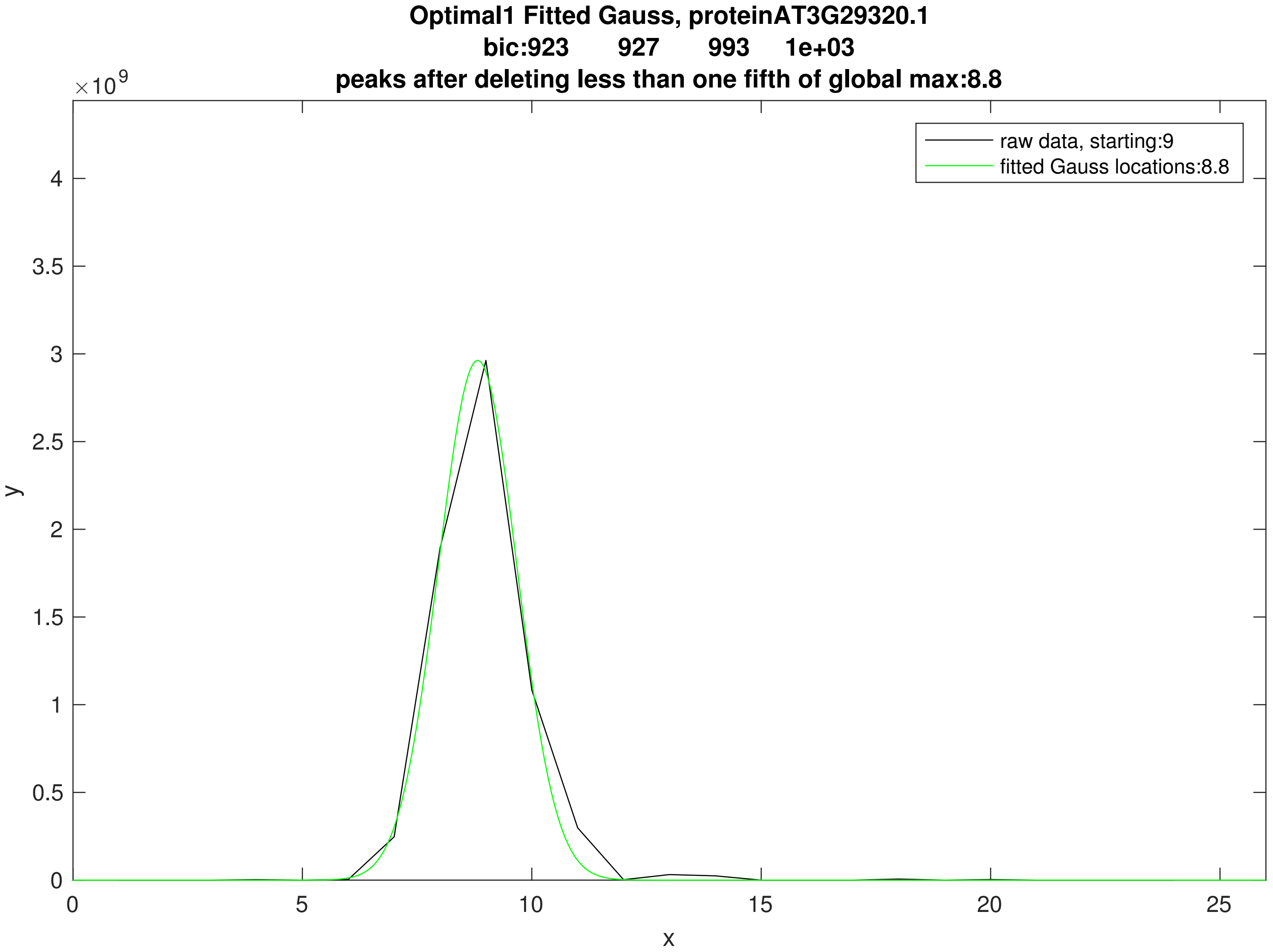

Optimal1 Fitted Gauss, proteinAT3G29320.1
bic:923 927 993 1e+03
peaks after deleting less than one fifth of global max:8.8
×10^9
raw data, starting:9
fitted Gauss locations:8.8
y
x

Optimal1 Fitted Gauss, proteinAT3G29350.1;AT3G29350.2
bic:671
peaks after deleting less than one fifth of global max:19

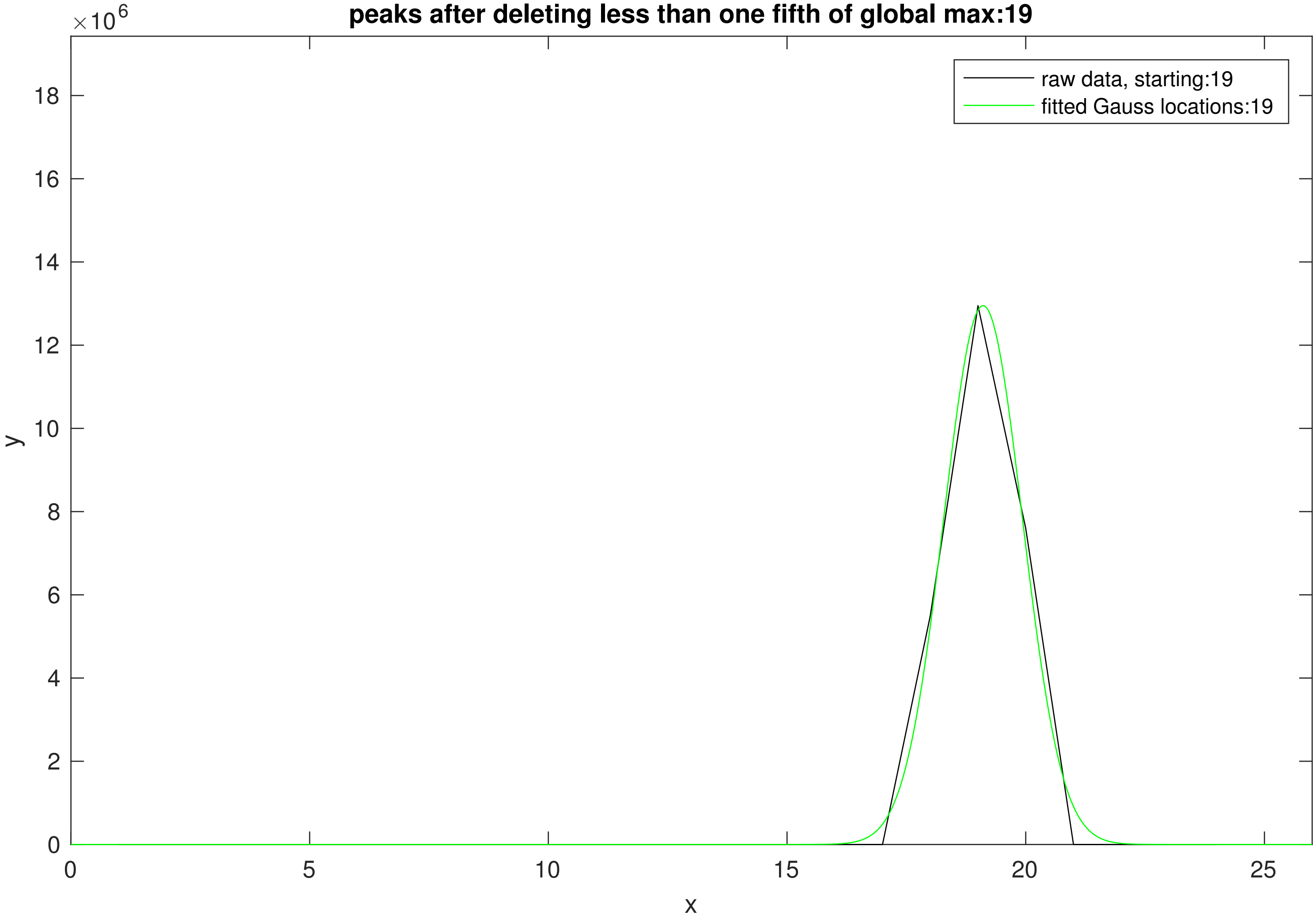

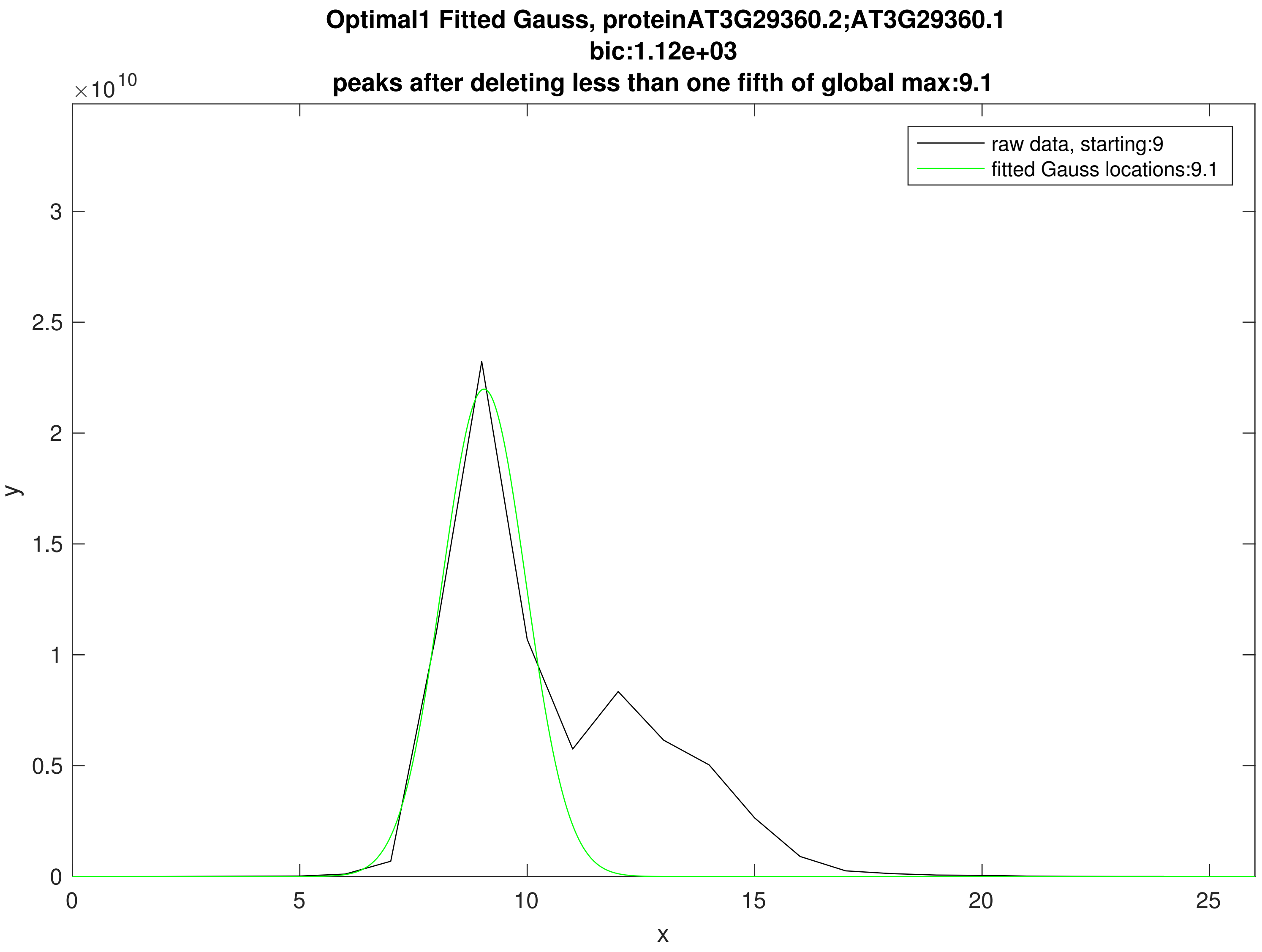
Optimal1 Fitted Gauss, proteinAT3G29360.2;AT3G29360.1
bic:1.12e+03
peaks after deleting less than one fifth of global max:9.1
×10^10
raw data, starting:9
fitted Gauss locations:9.1
y
x
0
0.5
1
1.5
2
2.5
3
5
10
15
20
25

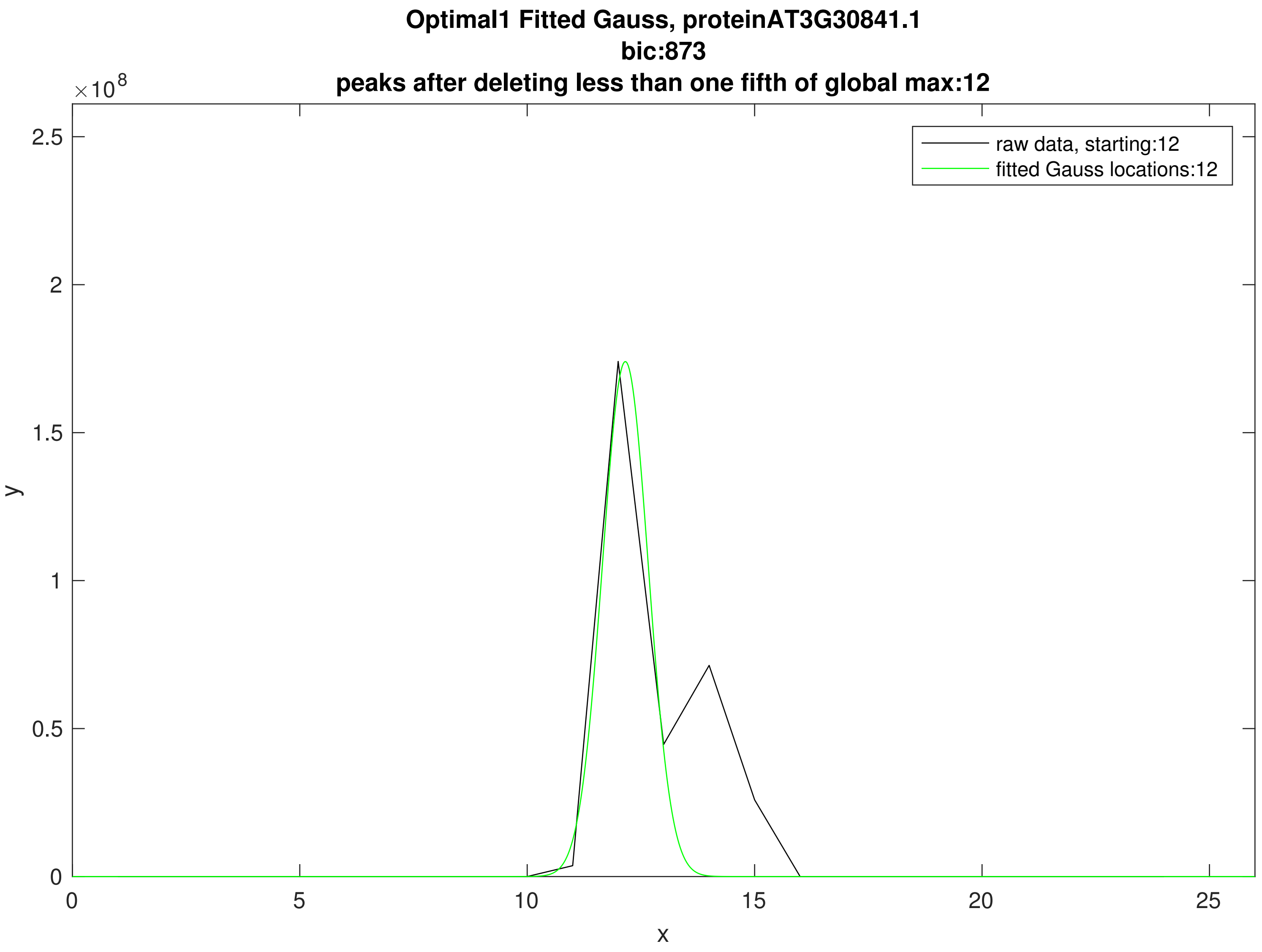
Optimal1 Fitted Gauss, proteinAT3G30841.1
bic:873
peaks after deleting less than one fifth of global max:12
×10^8
2.5
2
1.5
1
0.5
0
y
0
5
10
15
20
25
x
raw data, starting:12
fitted Gauss locations:12

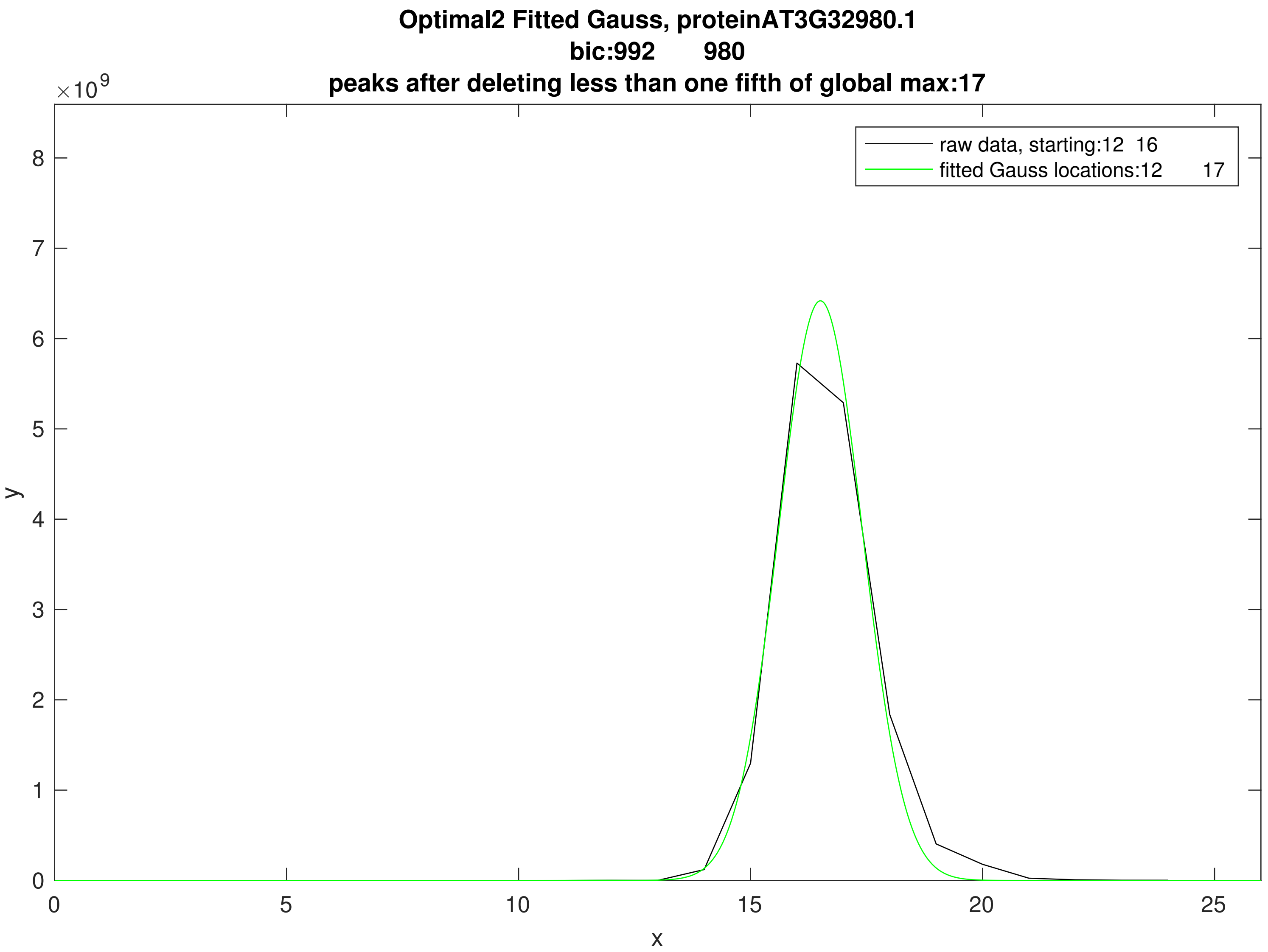
Optimal2 Fitted Gauss, proteinAT3G32980.1
bic:992 980
peaks after deleting less than one fifth of global max:17
×10^9
raw data, starting:12 16
fitted Gauss locations:12 17
y
x
0
1
2
3
4
5
6
7
8
0
5
10
15
20
25

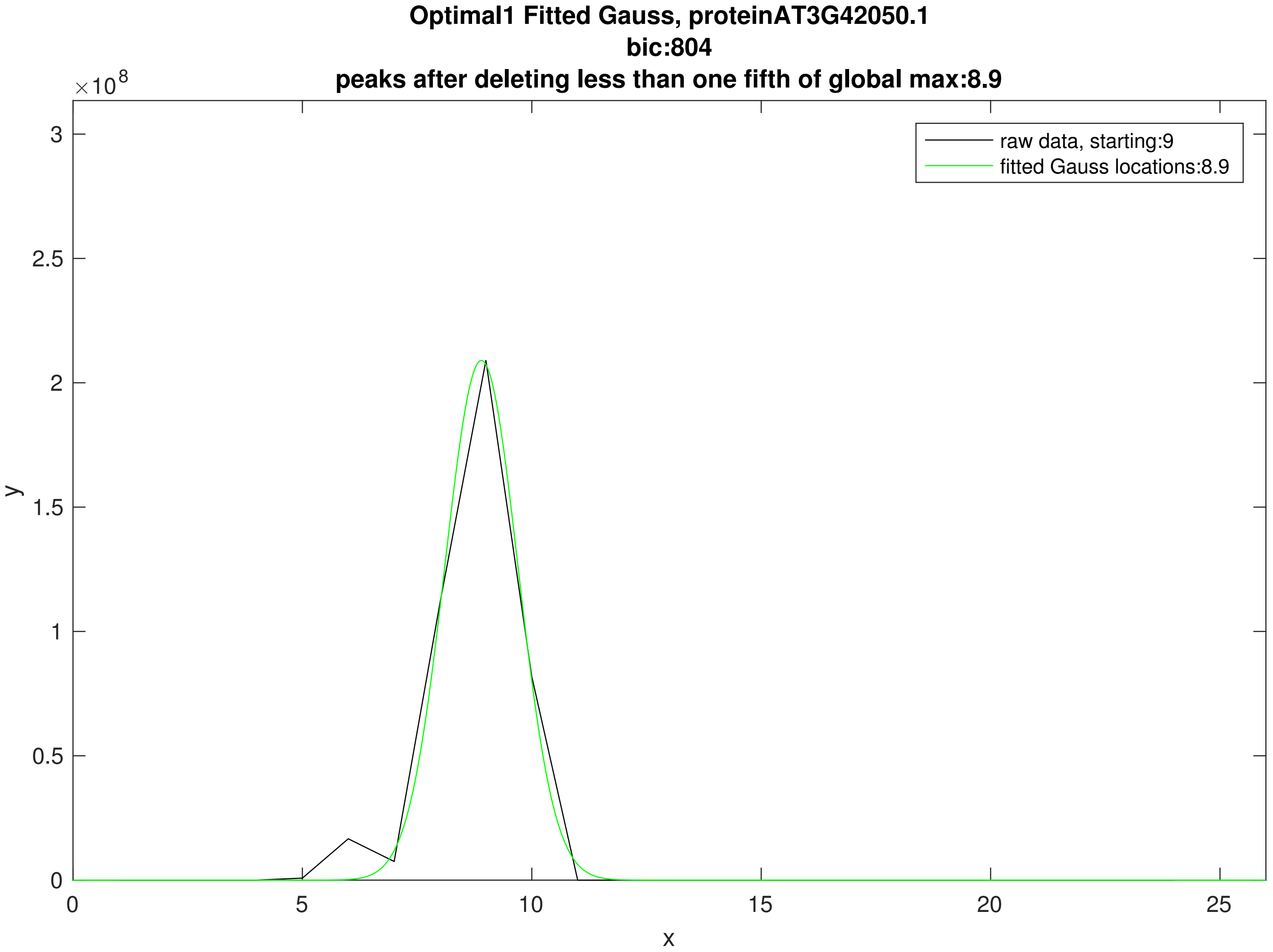

Optimal1 Fitted Gauss, proteinAT3G42050.1
bic:804
peaks after deleting less than one fifth of global max:8.9
×10^8
raw data, starting:9
fitted Gauss locations:8.9
y
x
0
0.5
1
1.5
2
2.5
3
0
5
10
15
20
25

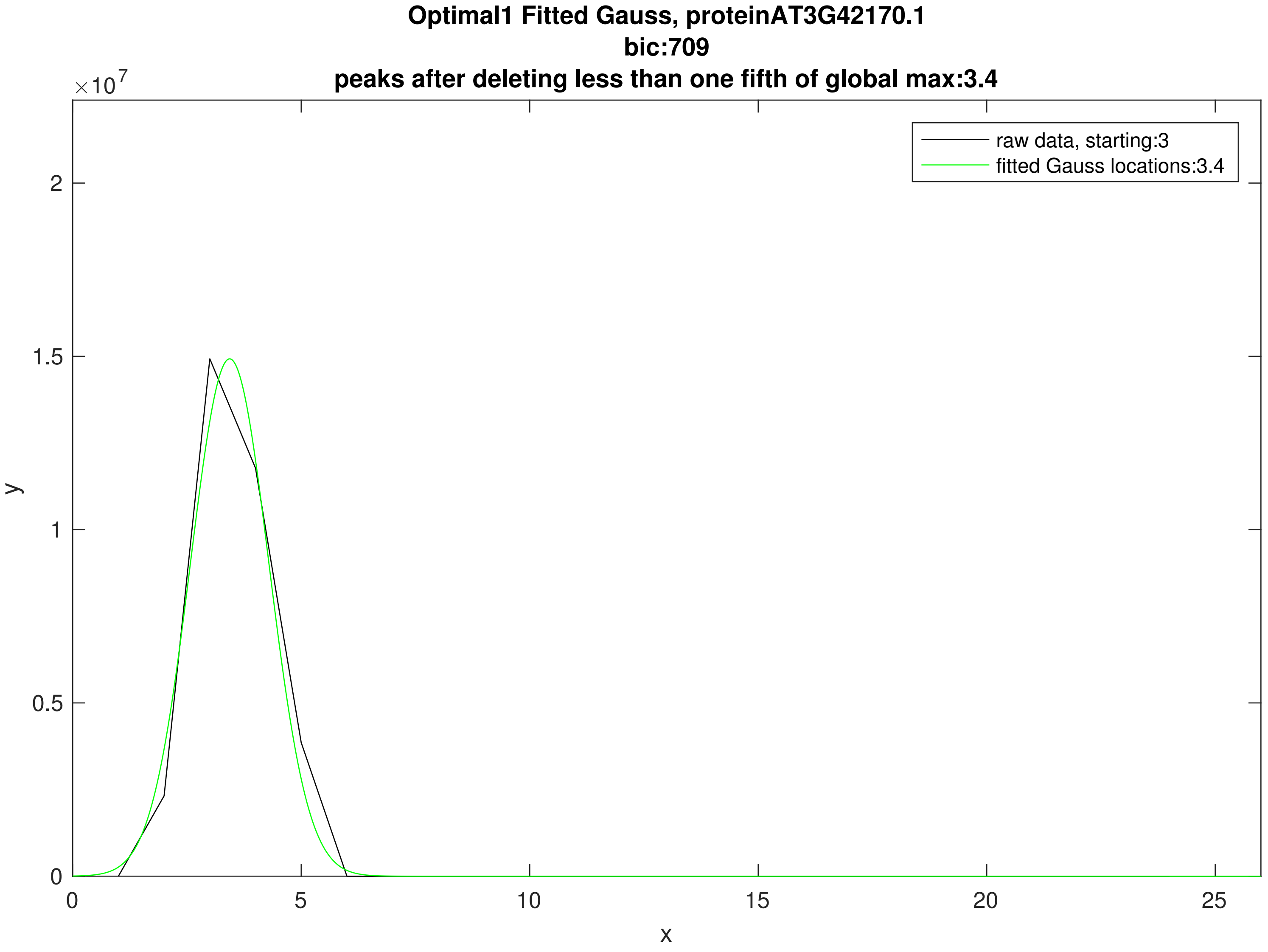
Optimal1 Fitted Gauss, proteinAT3G42170.1
bic:709
peaks after deleting less than one fifth of global max:3.4
×10^7
raw data, starting:3
fitted Gauss locations:3.4
y
x
0
0.5
1
1.5
2
0
5
10
15
20
25

Optimal1 Fitted Gauss, proteinAT3G43270.1
bic:896 922 915
peaks after deleting less than one fifth of global max:14

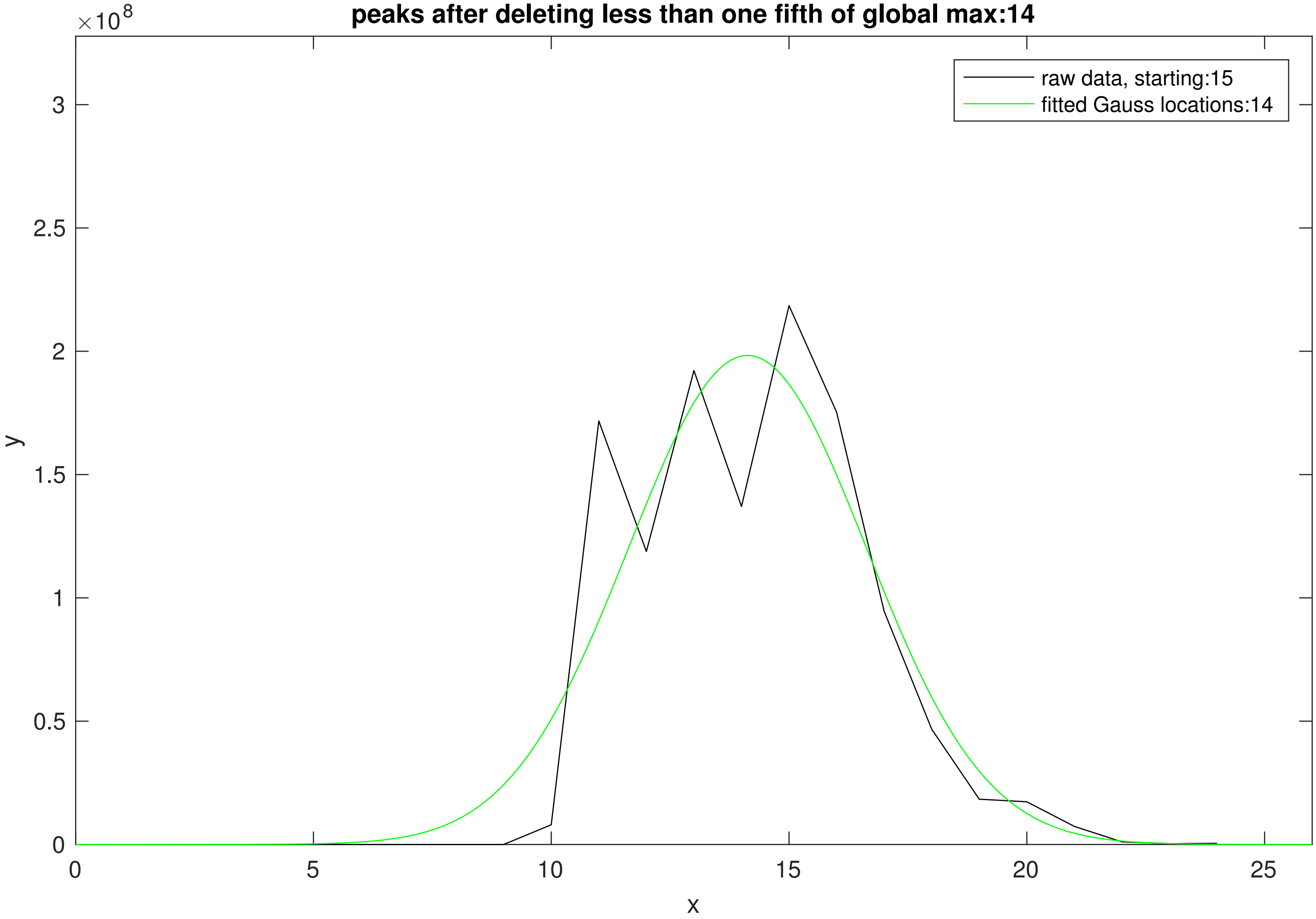

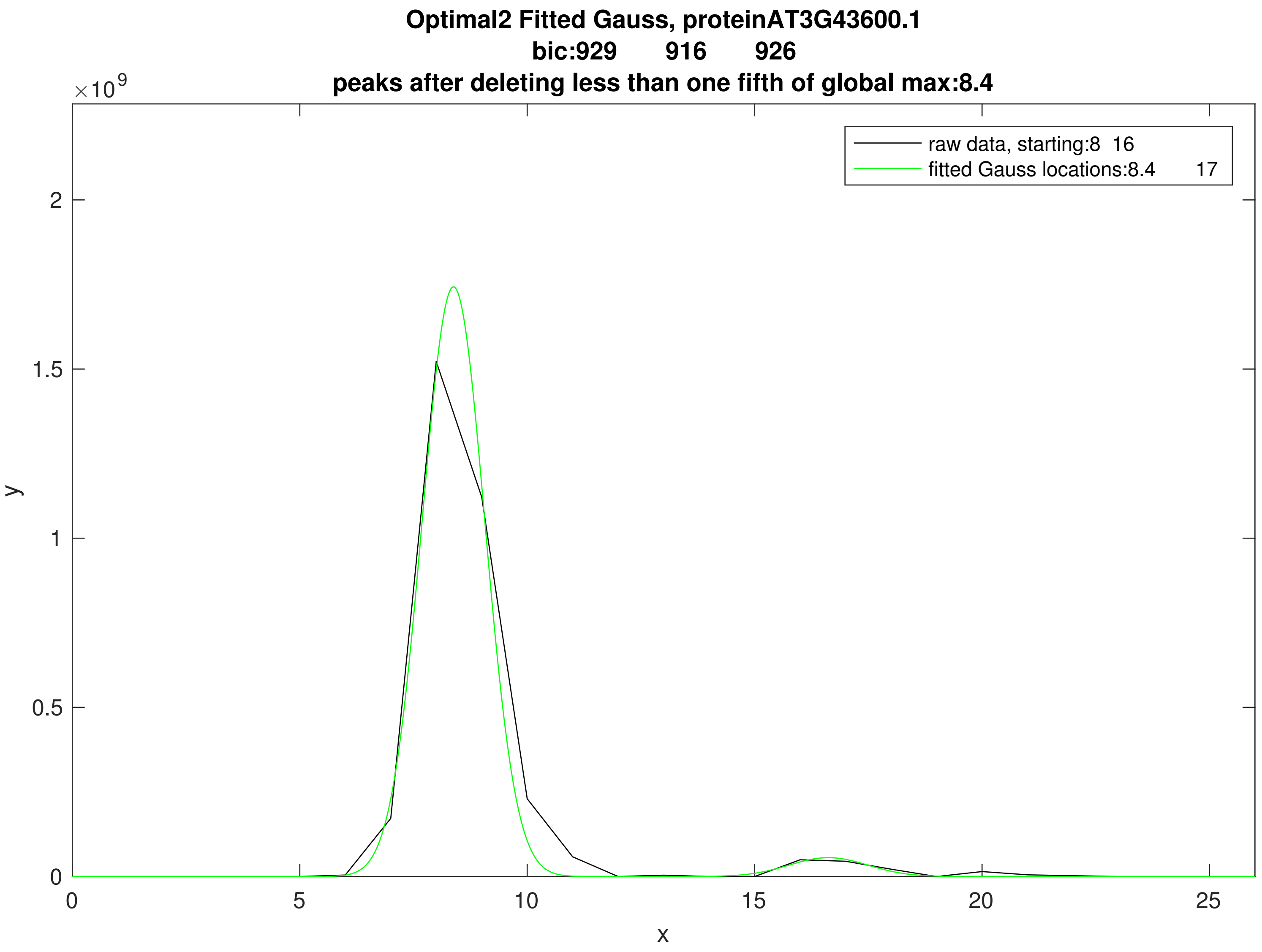
Optimal2 Fitted Gauss, proteinAT3G43600.1
bic:929 916 926
peaks after deleting less than one fifth of global max:8.4
×10^9
raw data, starting:8 16
fitted Gauss locations:8.4 17
y
x
0
0.5
1
1.5
2
5
10
15
20
25

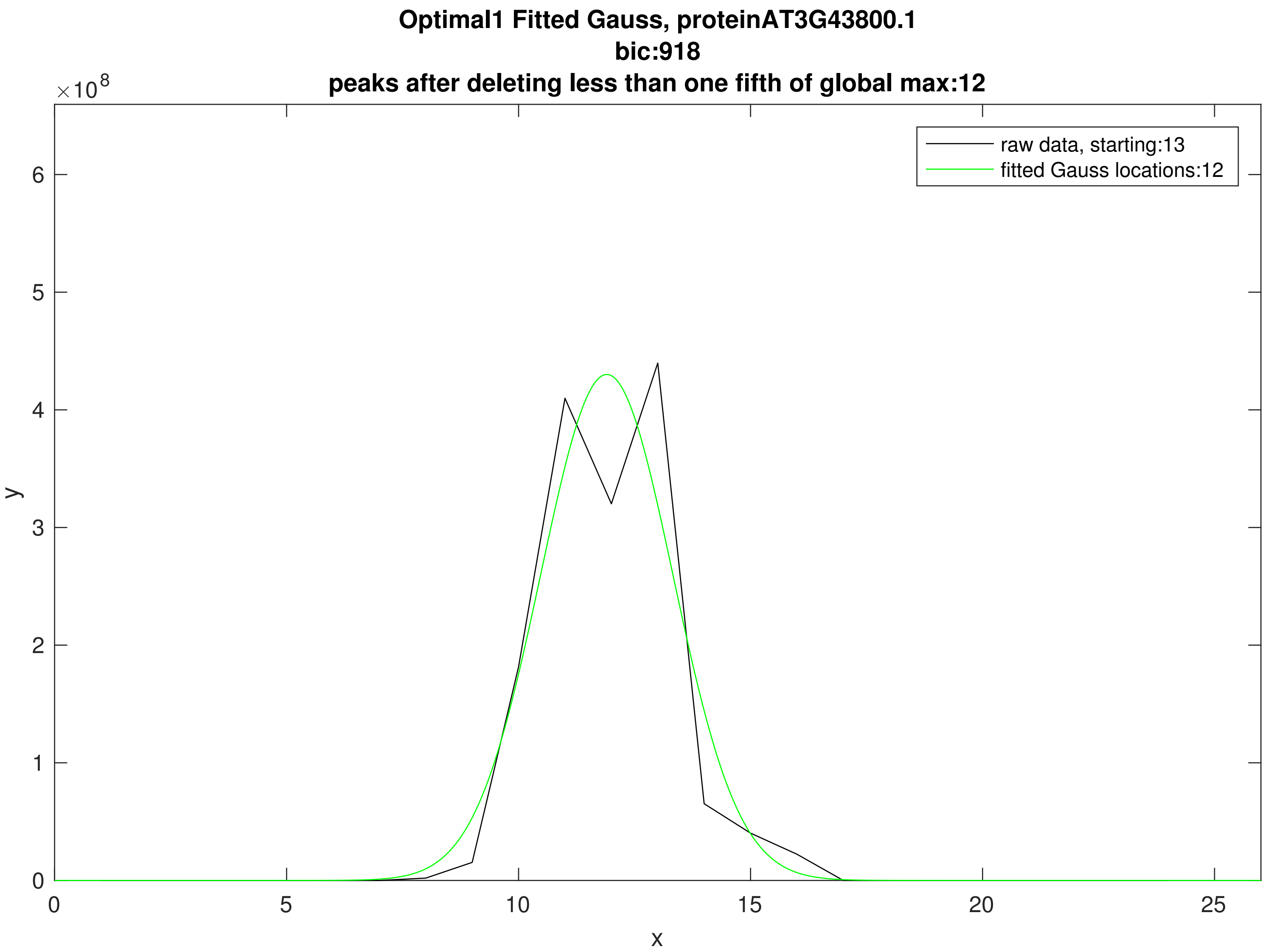

Optimal1 Fitted Gauss, proteinAT3G43800.1
bic:918
peaks after deleting less than one fifth of global max:12
×10^8
raw data, starting:13
fitted Gauss locations:12
y
x
0
1
2
3
4
5
6
0
5
10
15
20
25

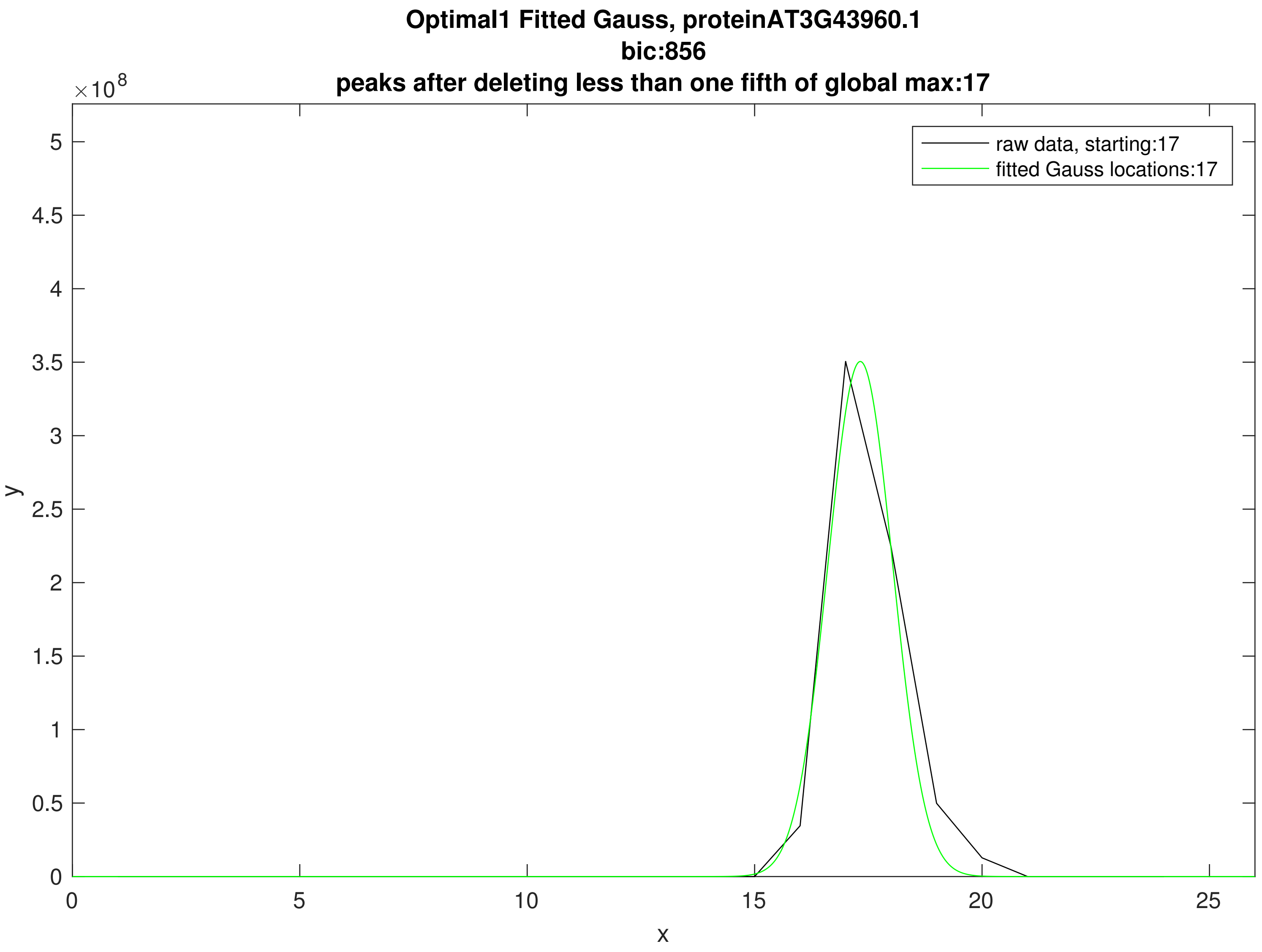
Optimal1 Fitted Gauss, proteinAT3G43960.1
bic:856
peaks after deleting less than one fifth of global max:17
×10^8
raw data, starting:17
fitted Gauss locations:17
y
x
0
0.5
1
1.5
2
2.5
3
3.5
4
4.5
5
0
5
10
15
20
25

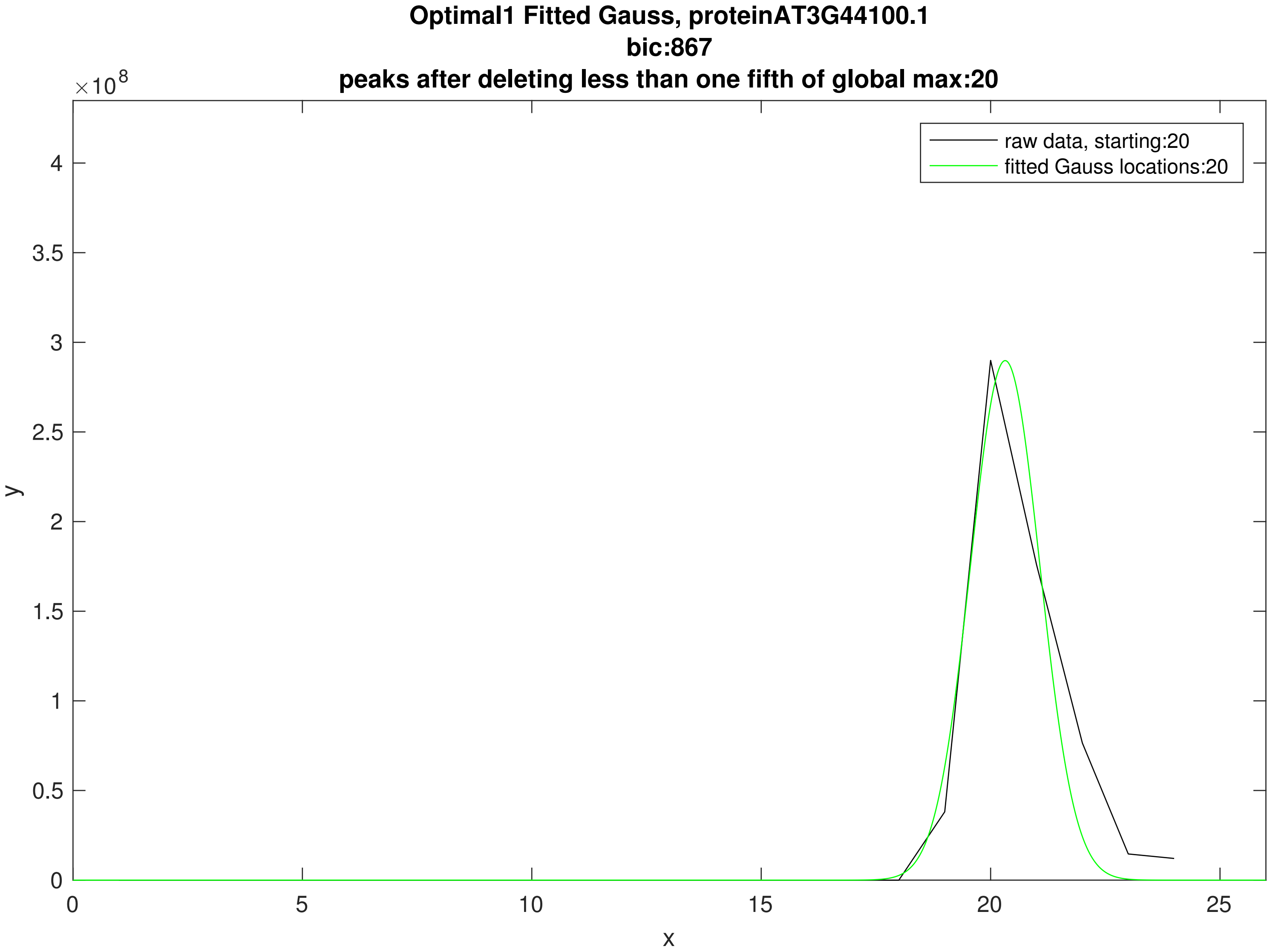
Optimal1 Fitted Gauss, proteinAT3G44100.1
bic:867
peaks after deleting less than one fifth of global max:20
×10^8
raw data, starting:20
fitted Gauss locations:20
y
x
0
0.5
1
1.5
2
2.5
3
3.5
4
0
5
10
15
20
25

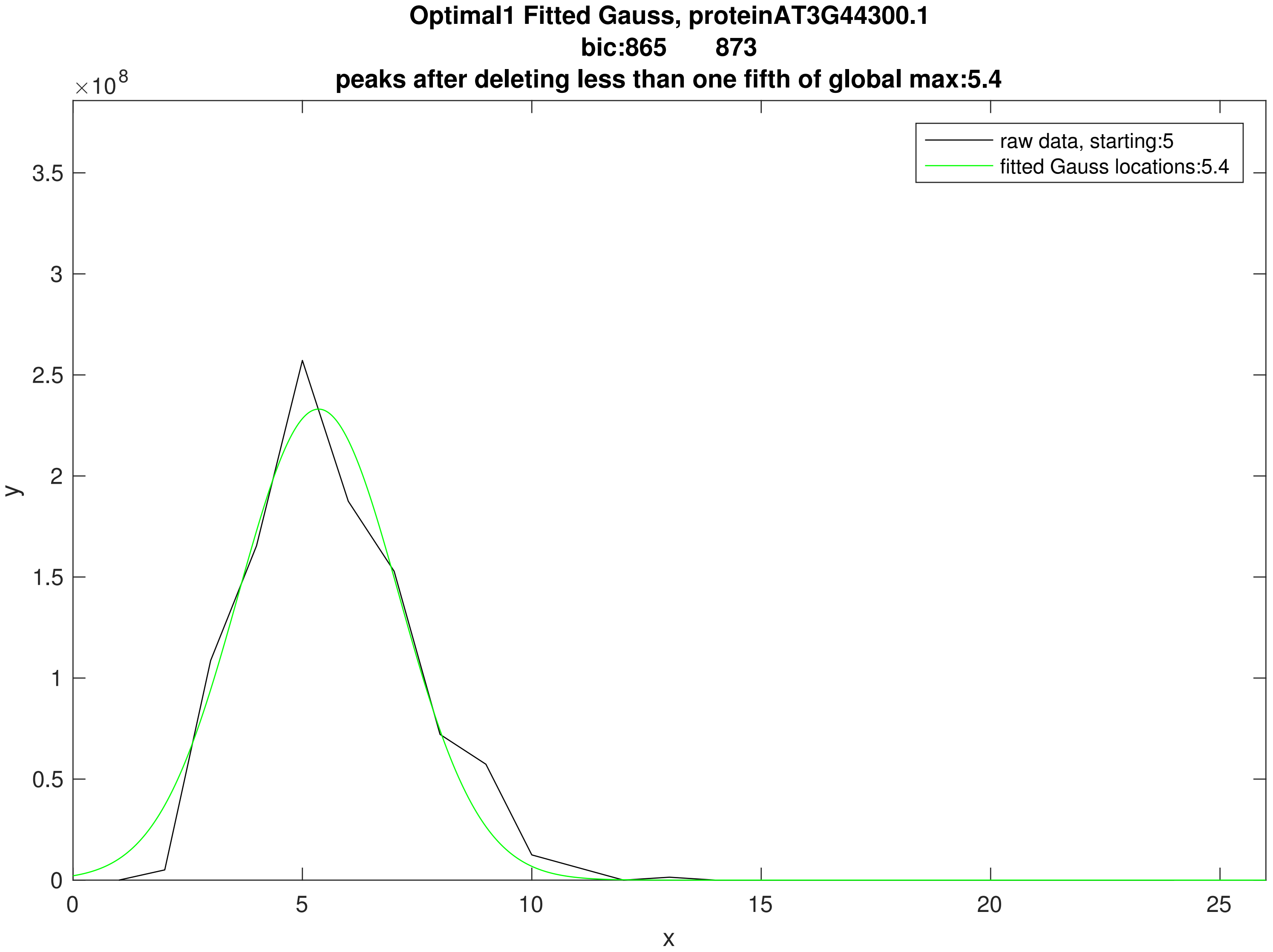
Optimal1 Fitted Gauss, proteinAT3G44300.1
bic:865 873
peaks after deleting less than one fifth of global max:5.4
×10^8
raw data, starting:5
fitted Gauss locations:5.4
y
x
0
0.5
1
1.5
2
2.5
3
3.5
0
5
10
15
20
25

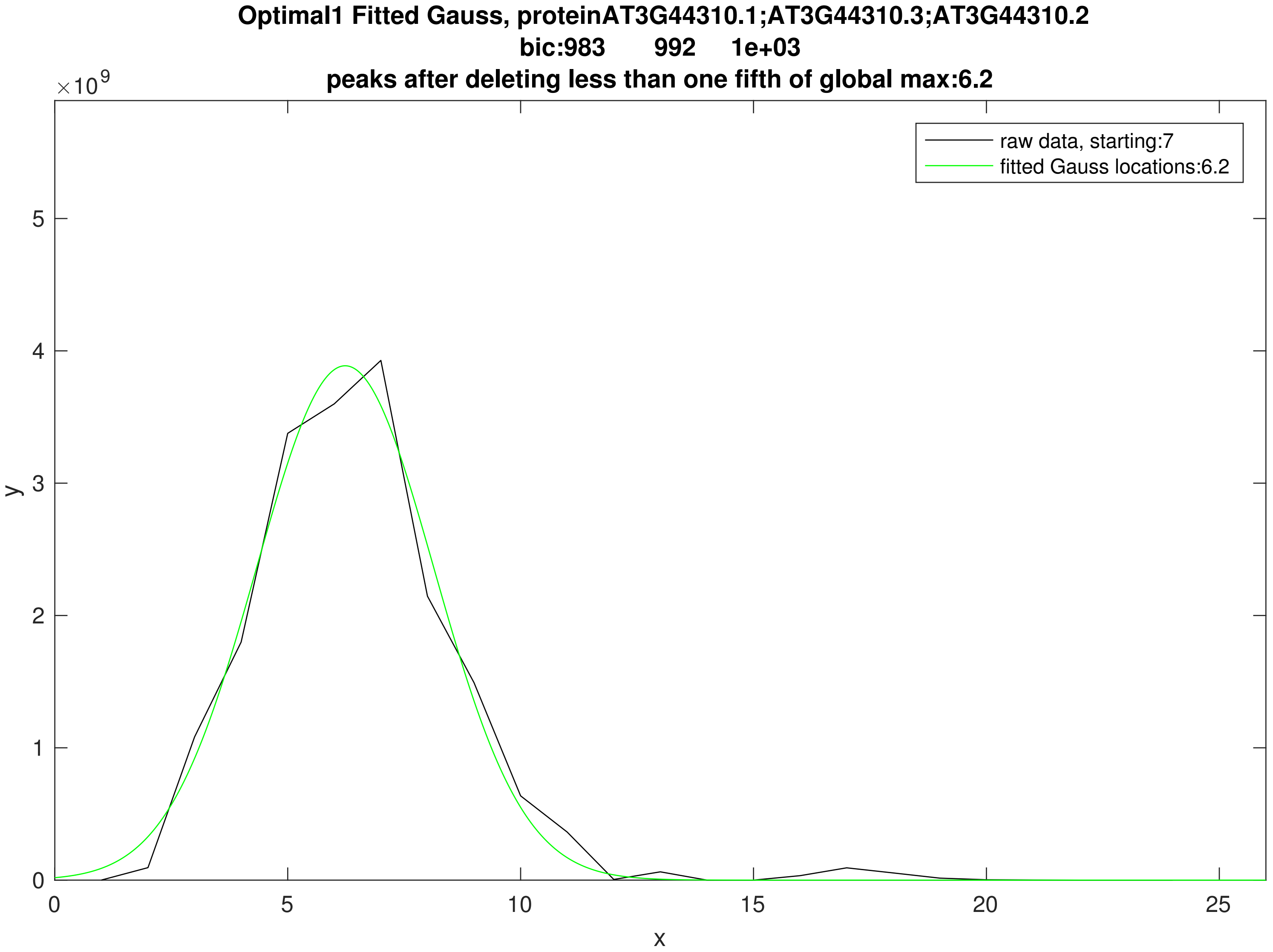

Optimal1 Fitted Gauss, proteinAT3G44310.1;AT3G44310.3;AT3G44310.2
bic:983 992 1e+03
peaks after deleting less than one fifth of global max:6.2
×10^9
raw data, starting:7
fitted Gauss locations:6.2
y
x
0
1
2
3
4
5
0
5
10
15
20
25

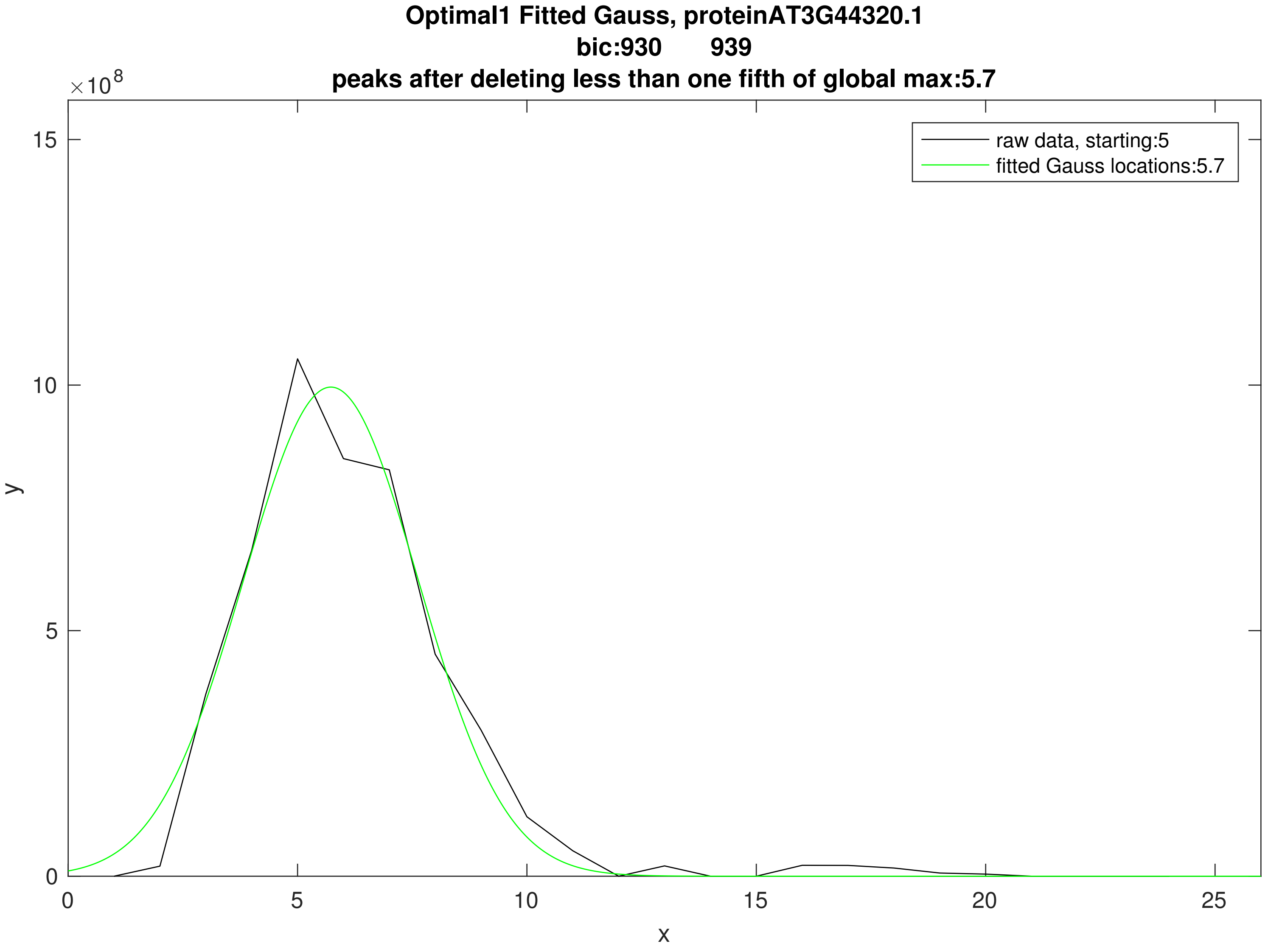
Optimal1 Fitted Gauss, proteinAT3G44320.1
bic:930 939
peaks after deleting less than one fifth of global max:5.7
×10^8
15
10
5
0
y
0
5
10
15
20
25
x
raw data, starting:5
fitted Gauss locations:5.7

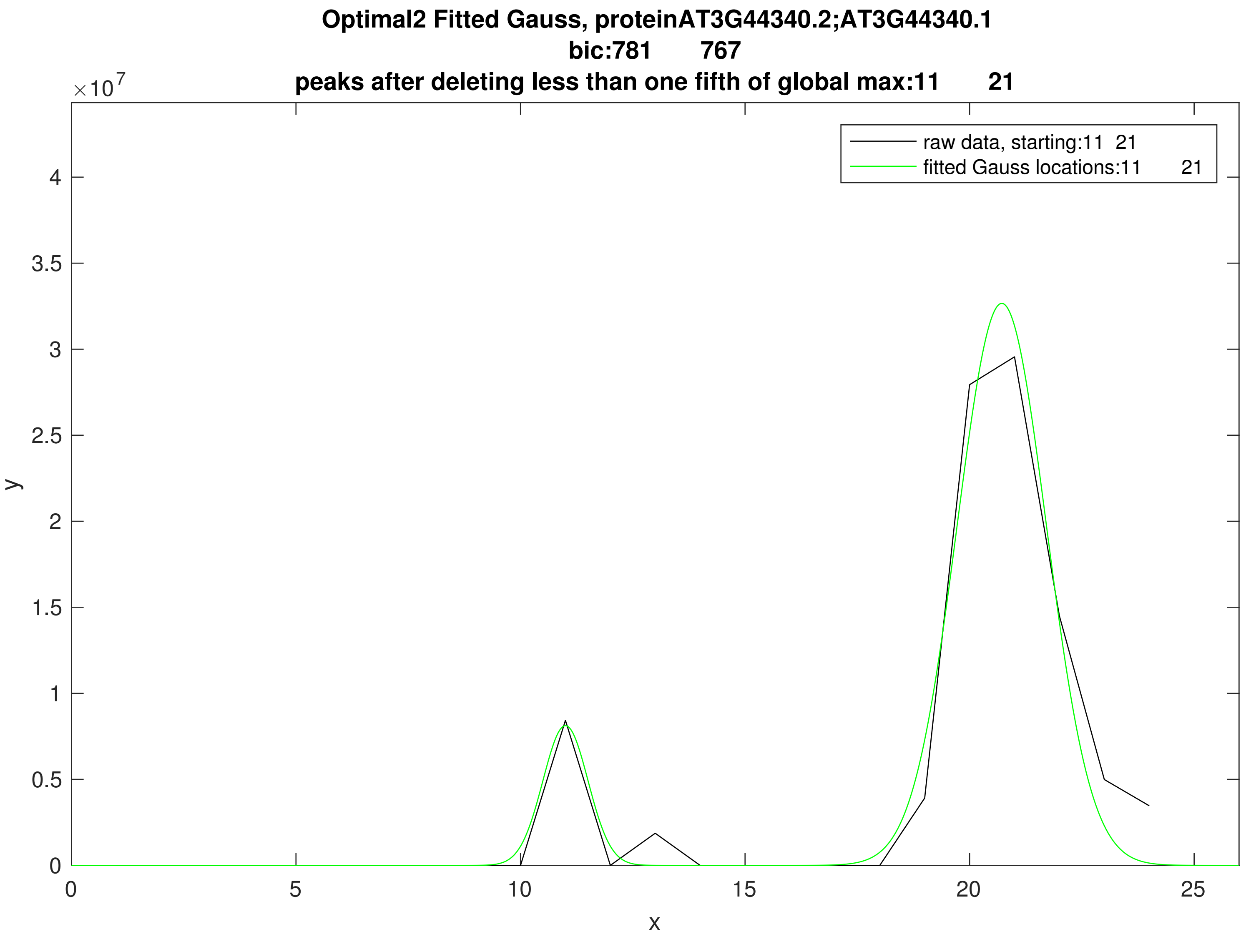
Optimal2 Fitted Gauss, proteinAT3G44340.2;AT3G44340.1
bic:781 767
peaks after deleting less than one fifth of global max:11 21
×10^7
raw data, starting:11 21
fitted Gauss locations:11 21
y
x
0
0.5
1
1.5
2
2.5
3
3.5
4
0
5
10
15
20
25

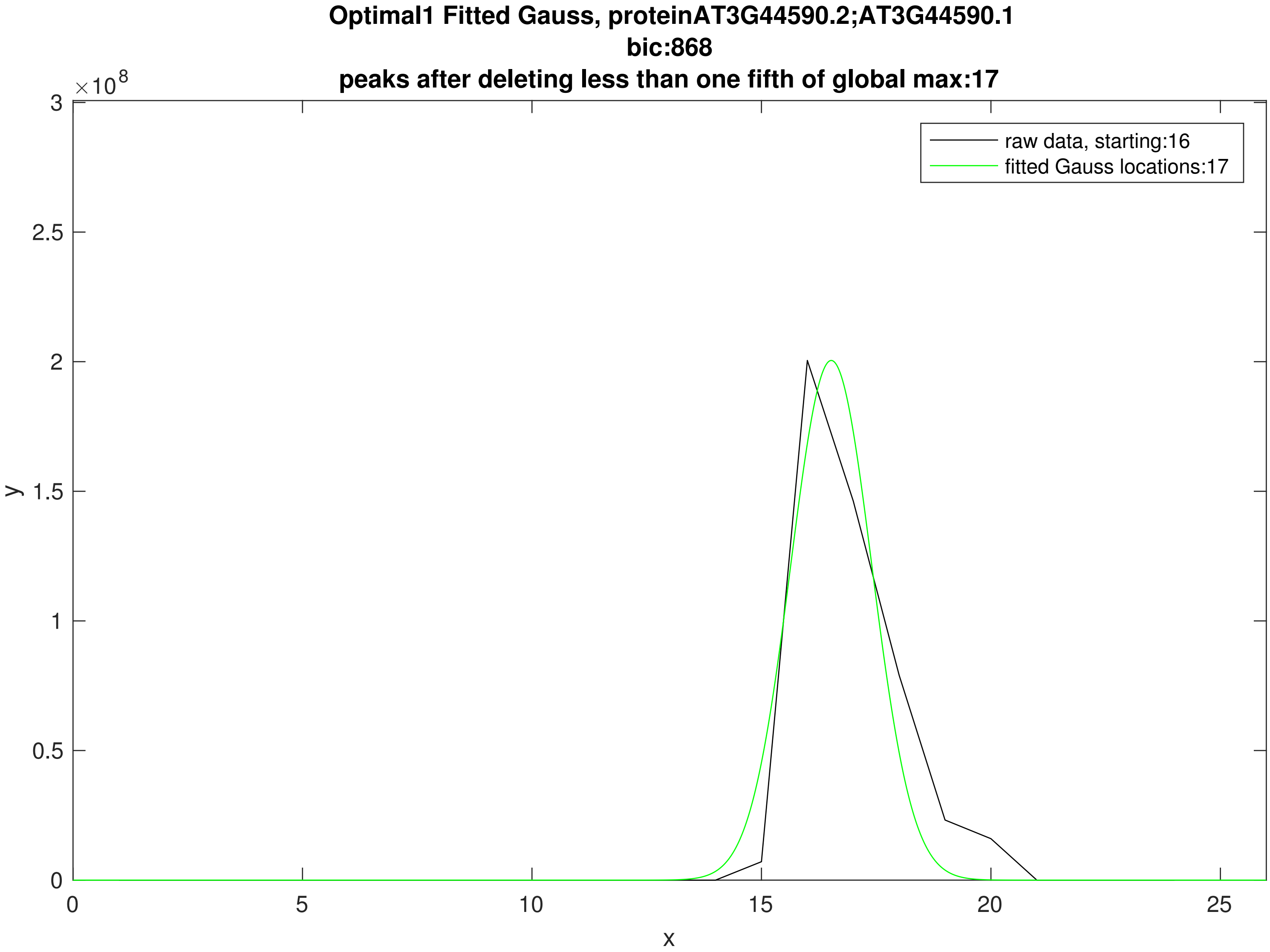
Optimal1 Fitted Gauss, proteinAT3G44590.2;AT3G44590.1
bic:868
peaks after deleting less than one fifth of global max:17
×10^8
3
2.5
2
1.5
1
0.5
0
y
0
5
10
15
20
25
x
raw data, starting:16
fitted Gauss locations:17

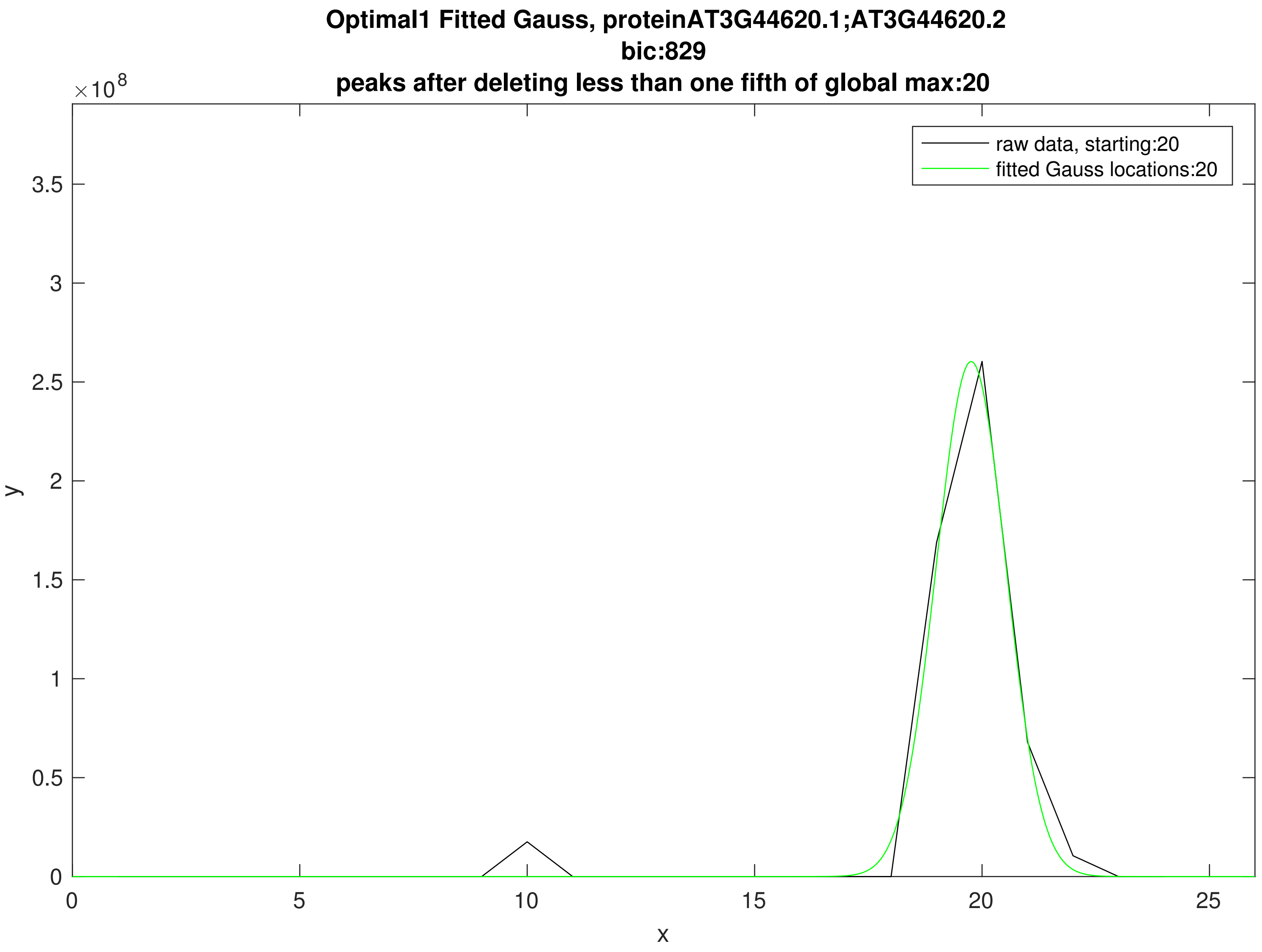

Optimal1 Fitted Gauss, proteinAT3G44620.1;AT3G44620.2
bic:829
peaks after deleting less than one fifth of global max:20
×10^8
raw data, starting:20
fitted Gauss locations:20
y
x
0
0.5
1
1.5
2
2.5
3
3.5
5
10
15
20
25

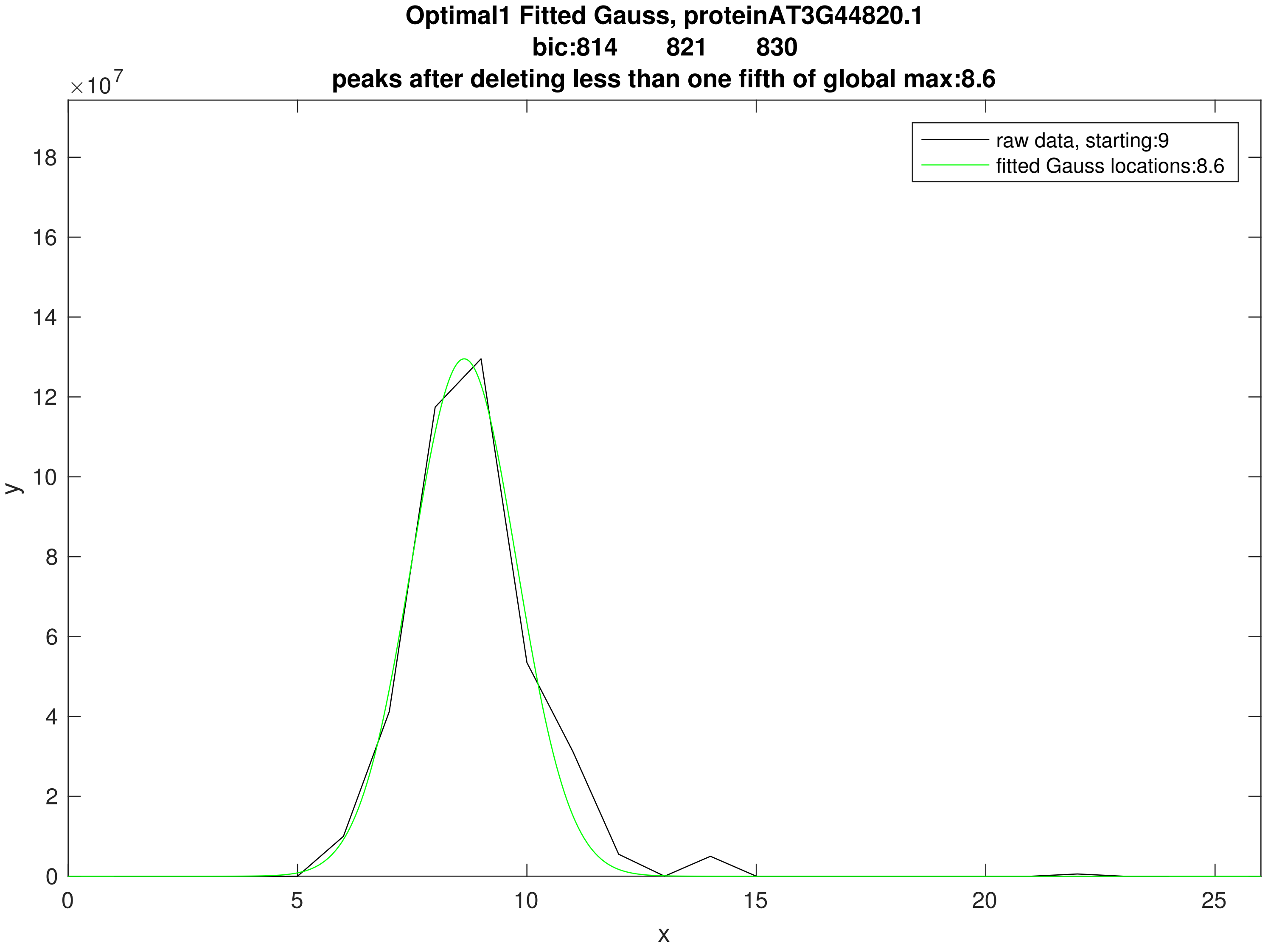

Optimal1 Fitted Gauss, proteinAT3G44820.1
bic:814 821 830
peaks after deleting less than one fifth of global max:8.6
×10^7
raw data, starting:9
fitted Gauss locations:8.6
y
x
0
2
4
6
8
10
12
14
16
18
0
5
10
15
20
25

**Optimal1 Fitted Gauss, proteinAT3G44890.1**
**bic:721**
**peaks after deleting less than one fifth of global max:3.9**

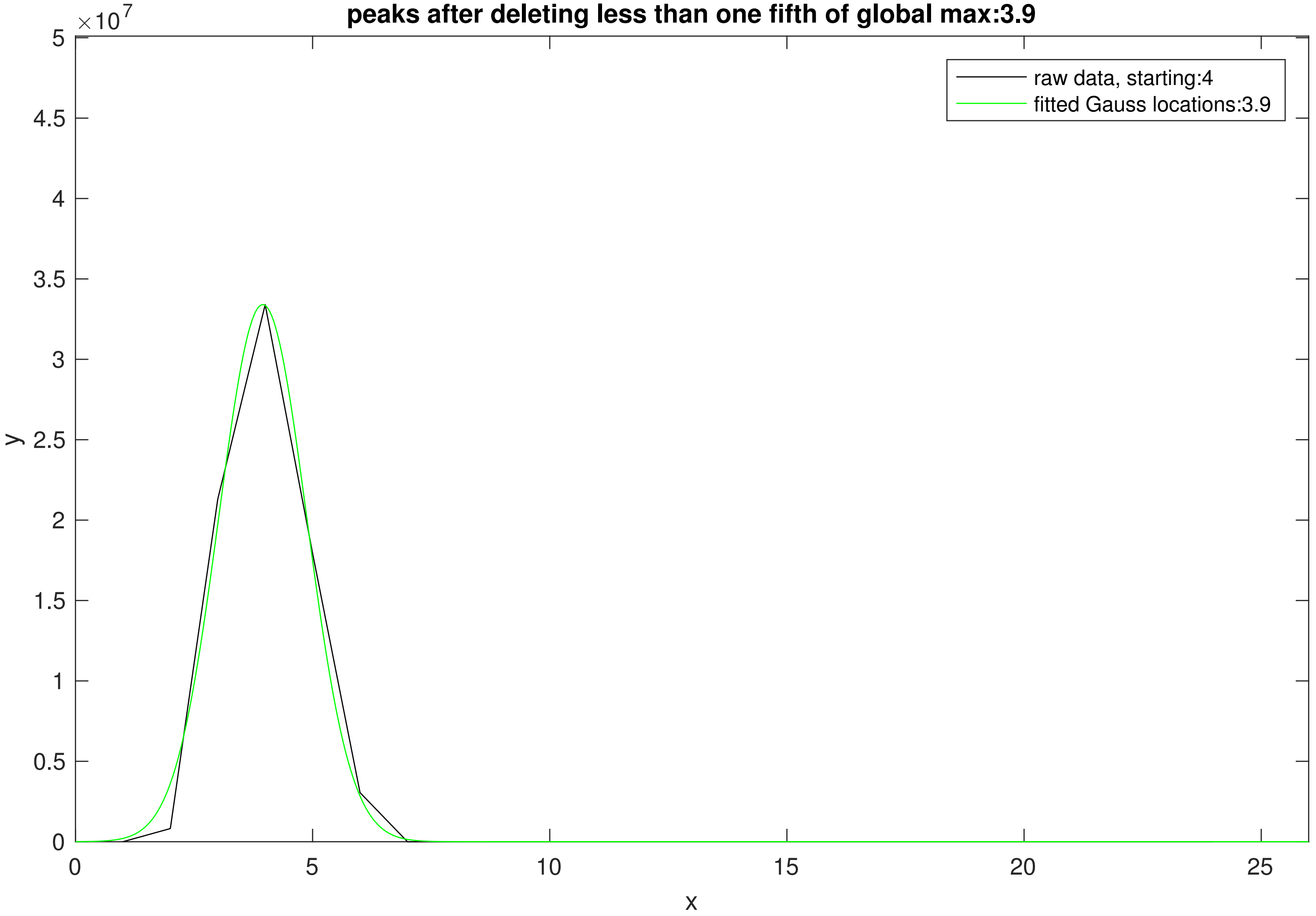

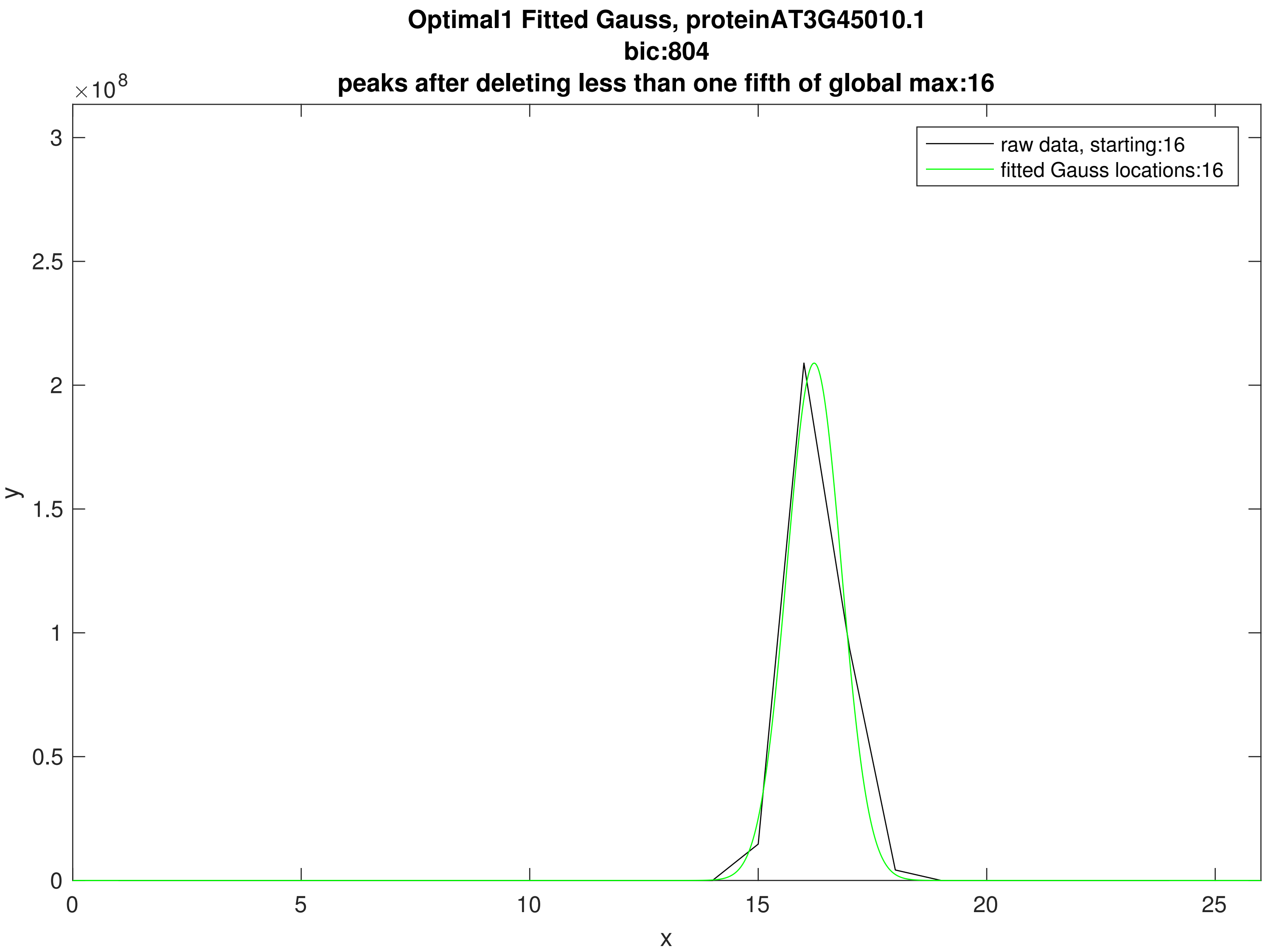

Optimal1 Fitted Gauss, proteinAT3G45010.1
bic:804
peaks after deleting less than one fifth of global max:16
×10^8
3
2.5
2
1.5
1
0.5
0
y
0
5
10
15
20
25
x
raw data, starting:16
fitted Gauss locations:16

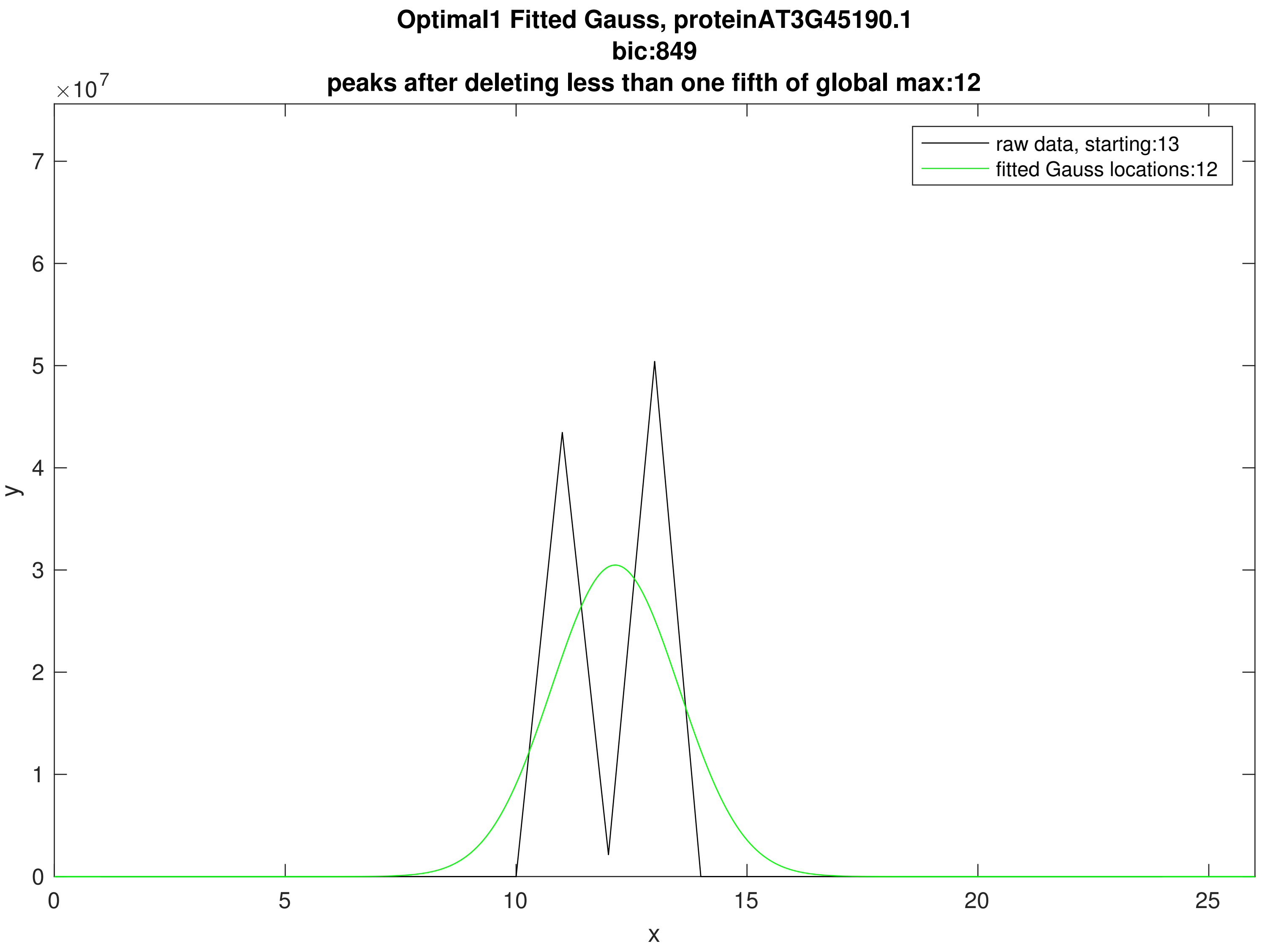
Optimal1 Fitted Gauss, proteinAT3G45190.1
bic:849
peaks after deleting less than one fifth of global max:12
×10^7
raw data, starting:13
fitted Gauss locations:12
y
x
0
1
2
3
4
5
6
7
0
5
10
15
20
25

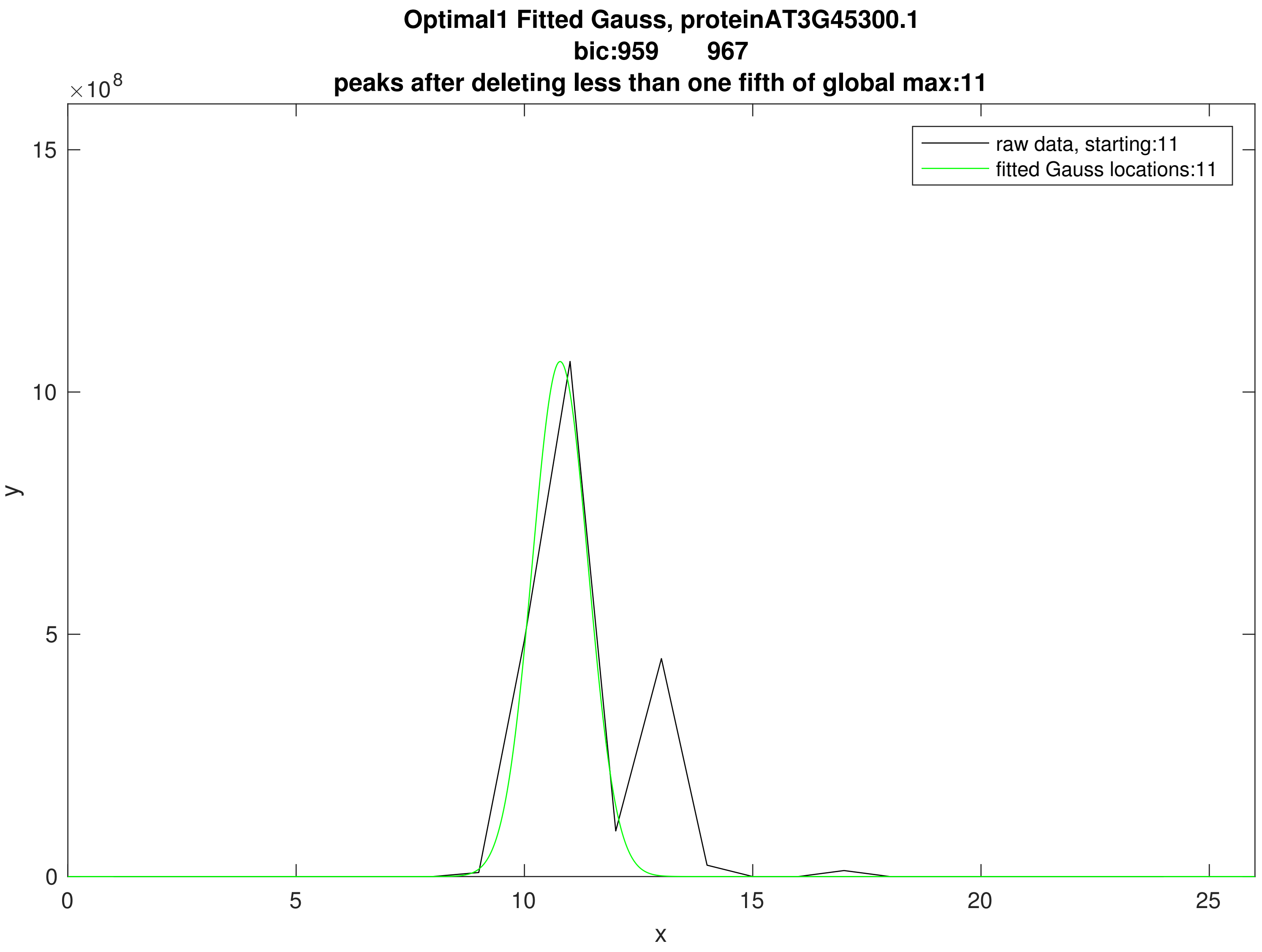
Optimal1 Fitted Gauss, proteinAT3G45300.1
bic:959 967
peaks after deleting less than one fifth of global max:11
×10^8
15
10
5
0
y
0
5
10
15
20
25
x
raw data, starting:11
fitted Gauss locations:11

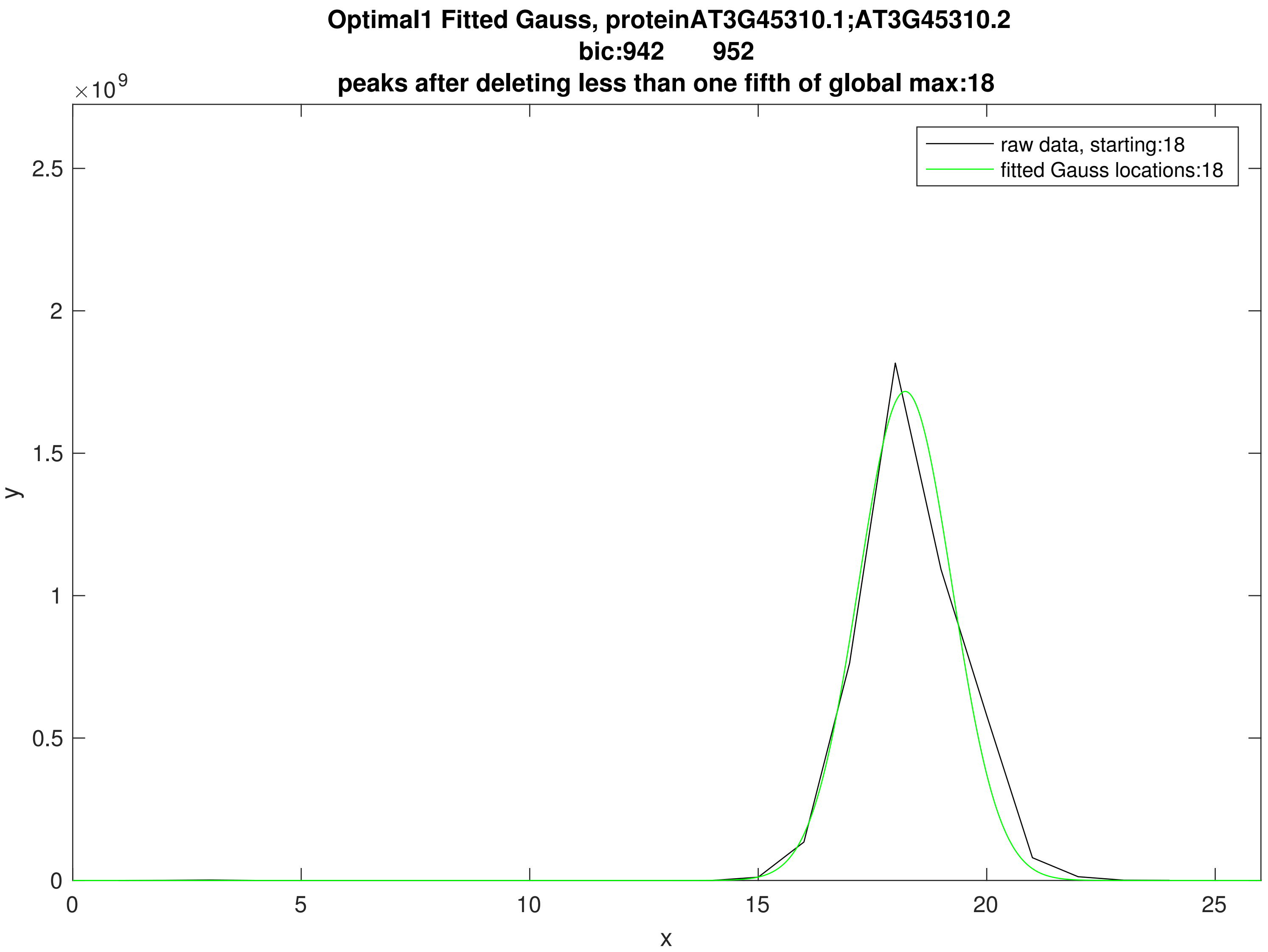
Optimal1 Fitted Gauss, proteinAT3G45310.1;AT3G45310.2
bic:942 952
peaks after deleting less than one fifth of global max:18
×10^9
2.5
2
1.5
1
0.5
0
y
0
5
10
15
20
25
x
raw data, starting:18
fitted Gauss locations:18

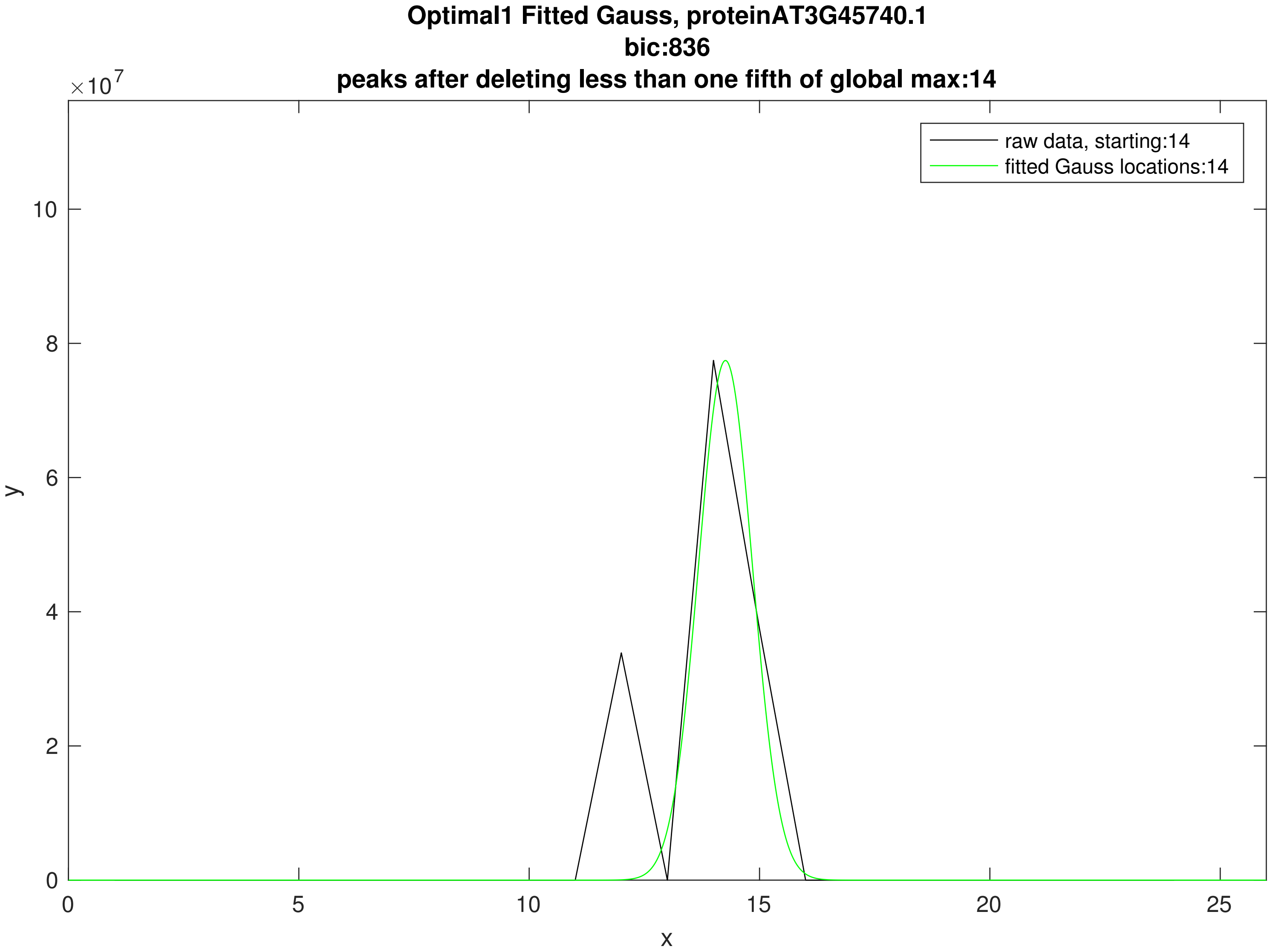

Optimal1 Fitted Gauss, proteinAT3G45740.1
bic:836
peaks after deleting less than one fifth of global max:14
×10^7
raw data, starting:14
fitted Gauss locations:14
10
8
6
4
2
0
y
0
5
10
15
20
25
x

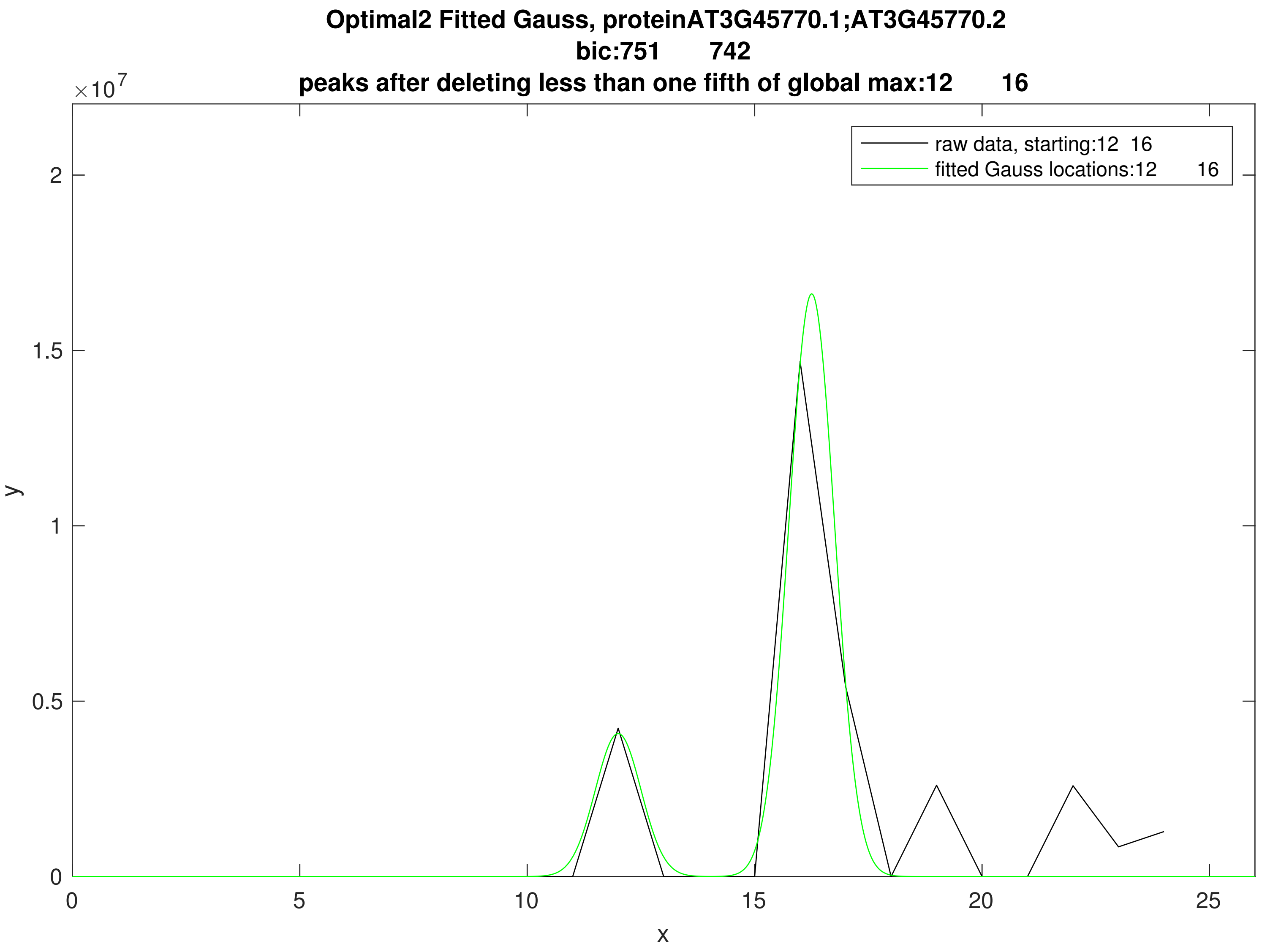
Optimal2 Fitted Gauss, proteinAT3G45770.1;AT3G45770.2
bic:751 742
peaks after deleting less than one fifth of global max:12 16
×10^7
raw data, starting:12 16
fitted Gauss locations:12 16
2
1.5
1
0.5
0
y
0
5
10
15
20
25
x

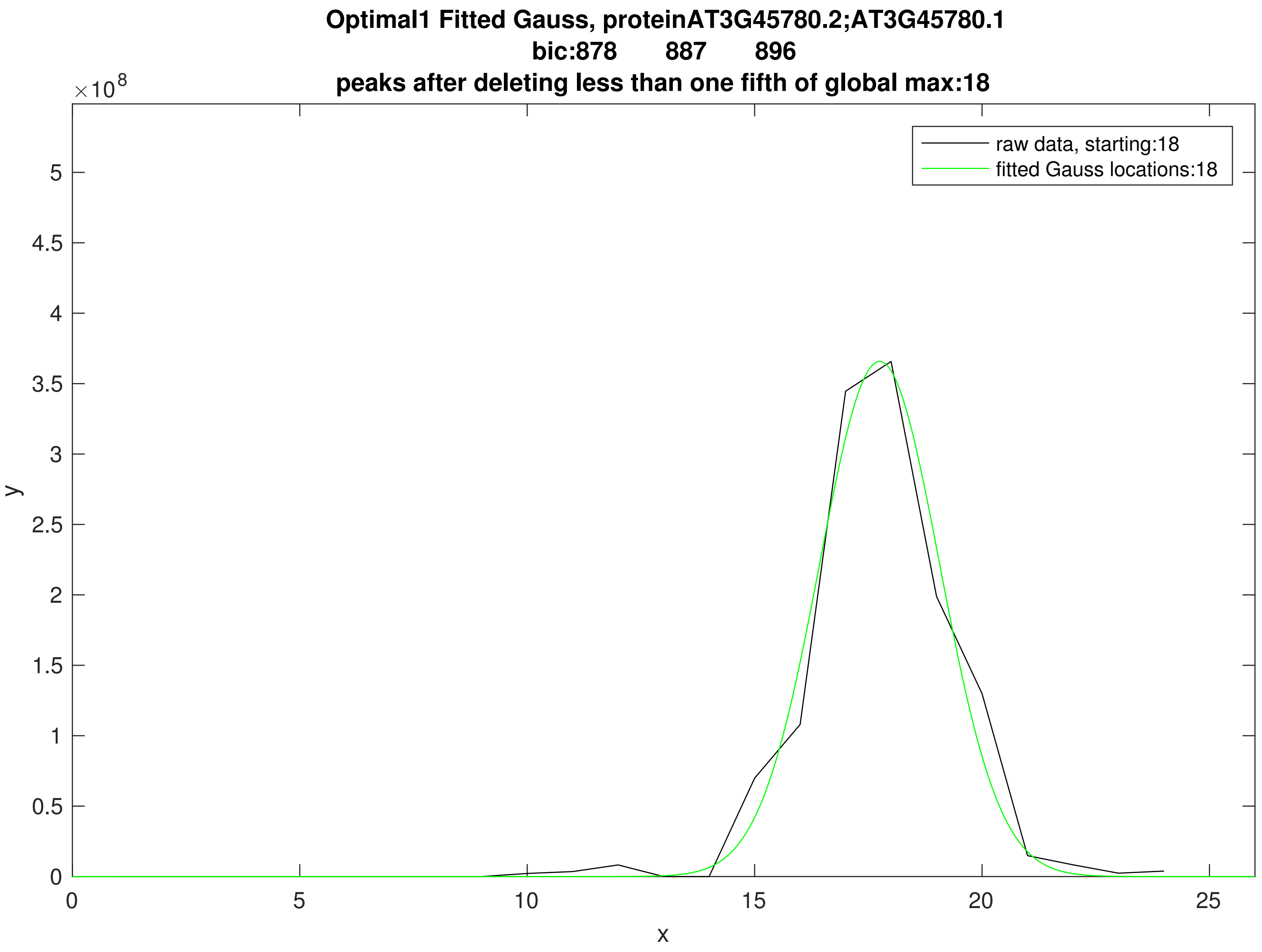

Optimal1 Fitted Gauss, proteinAT3G45780.2;AT3G45780.1
bic:878 887 896
peaks after deleting less than one fifth of global max:18
×10^8
raw data, starting:18
fitted Gauss locations:18
y
x
0
0.5
1
1.5
2
2.5
3
3.5
4
4.5
5
0
5
10
15
20
25

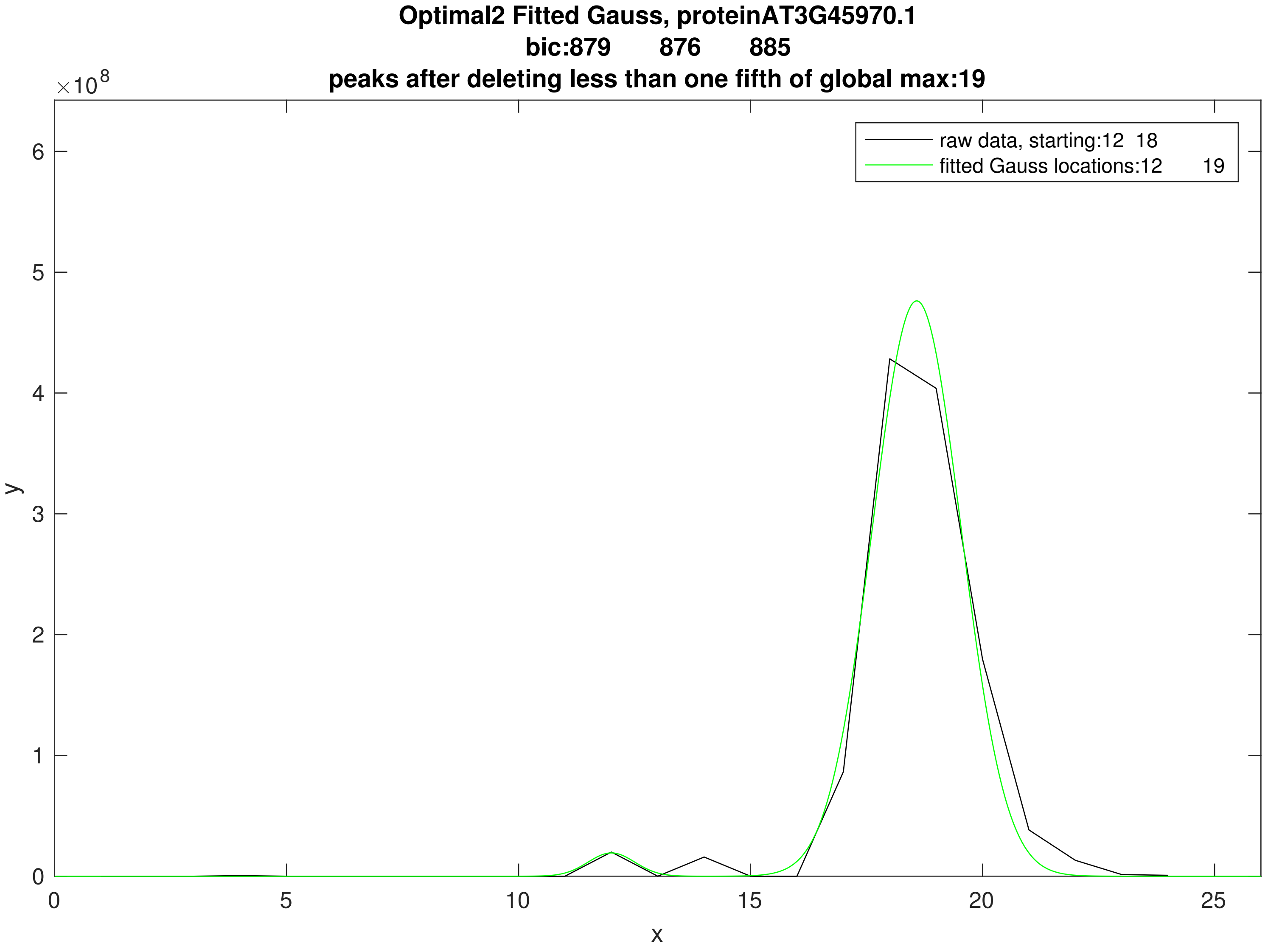
Optimal2 Fitted Gauss, proteinAT3G45970.1
bic:879 876 885
peaks after deleting less than one fifth of global max:19
×10^8
raw data, starting:12 18
fitted Gauss locations:12 19
y
x
0
1
2
3
4
5
6
0
5
10
15
20
25

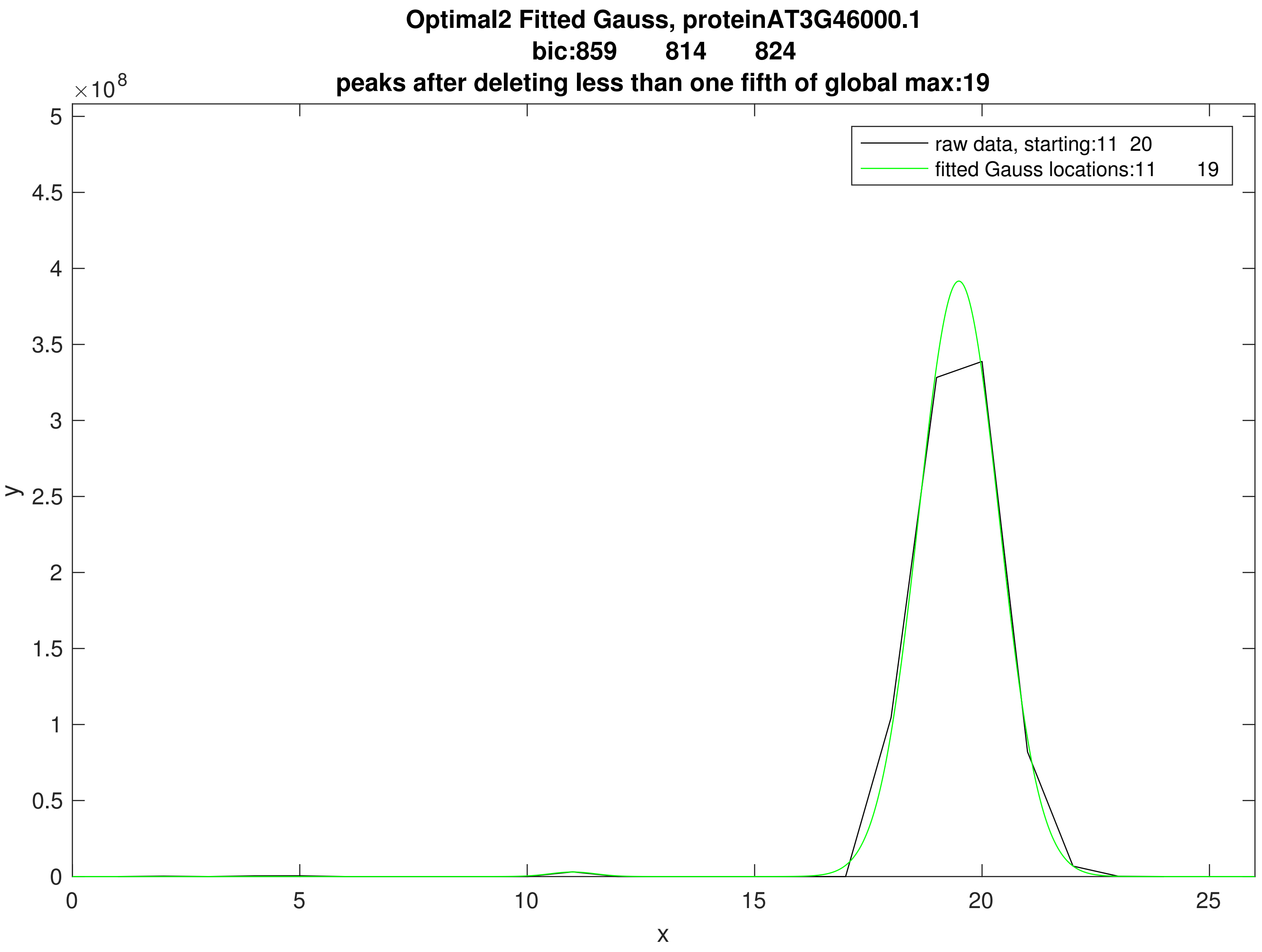

Optimal2 Fitted Gauss, proteinAT3G46000.1
bic:859 814 824
peaks after deleting less than one fifth of global max:19
×10^8
raw data, starting:11 20
fitted Gauss locations:11 19
y
x
0
0.5
1
1.5
2
2.5
3
3.5
4
4.5
5
0
5
10
15
20
25

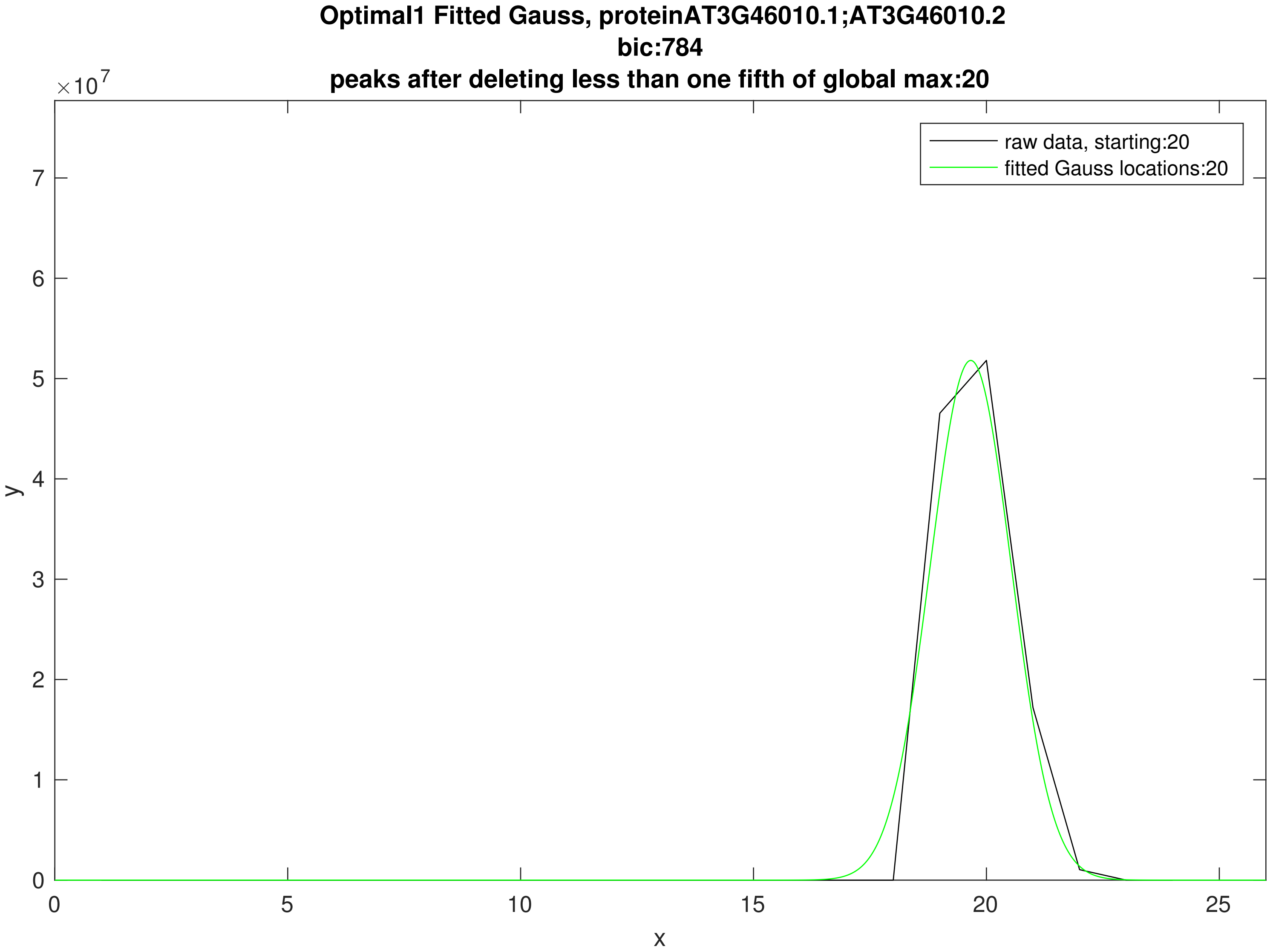
Optimal1 Fitted Gauss, proteinAT3G46010.1;AT3G46010.2
bic:784
peaks after deleting less than one fifth of global max:20
×10^7
raw data, starting:20
fitted Gauss locations:20
y
x
0
1
2
3
4
5
6
7
0
5
10
15
20
25

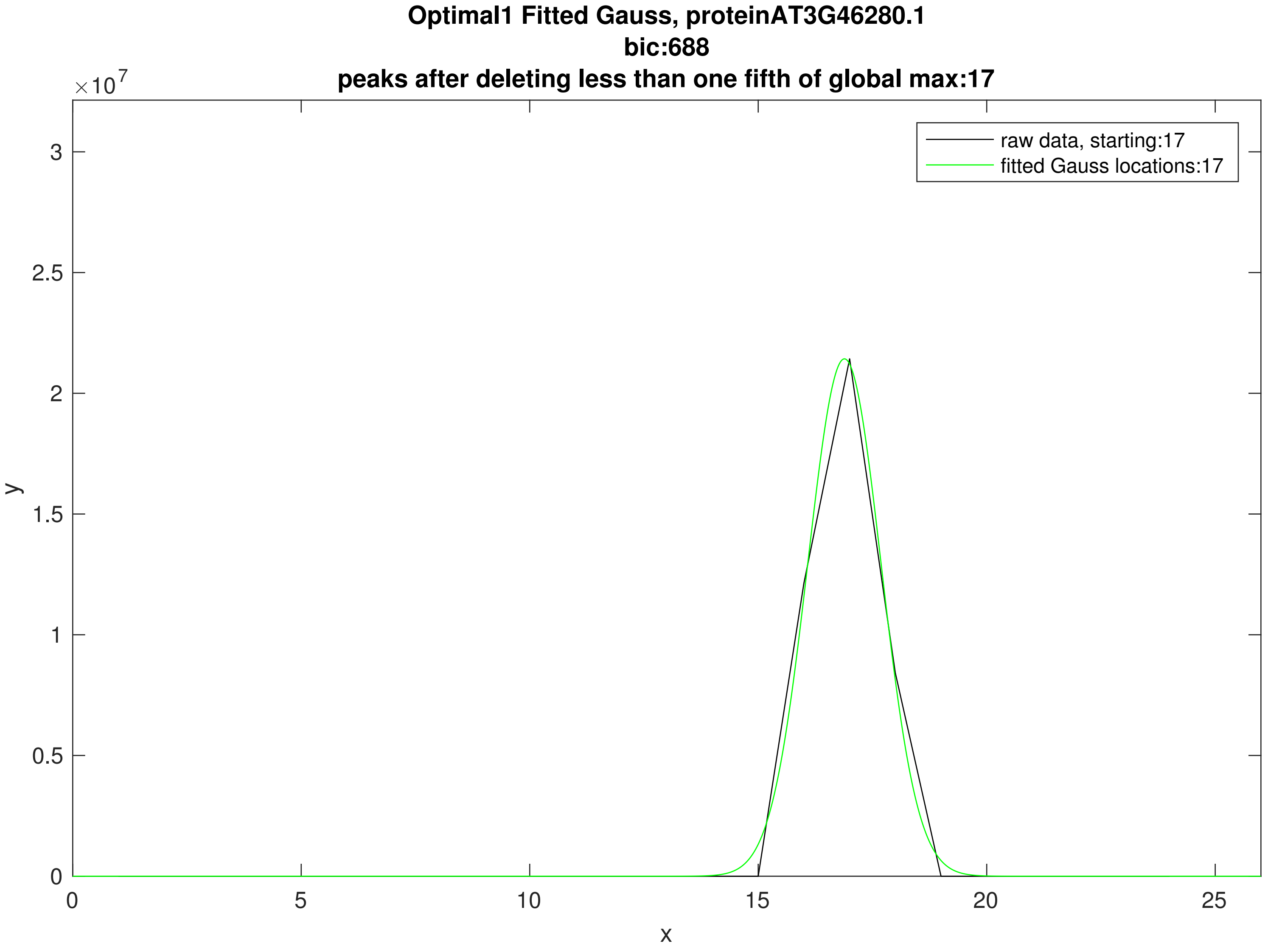

Optimal1 Fitted Gauss, proteinAT3G46280.1
bic:688
peaks after deleting less than one fifth of global max:17
×10^7
raw data, starting:17
fitted Gauss locations:17
3
2.5
2
1.5
1
0.5
0
y
0
5
10
15
20
25
x

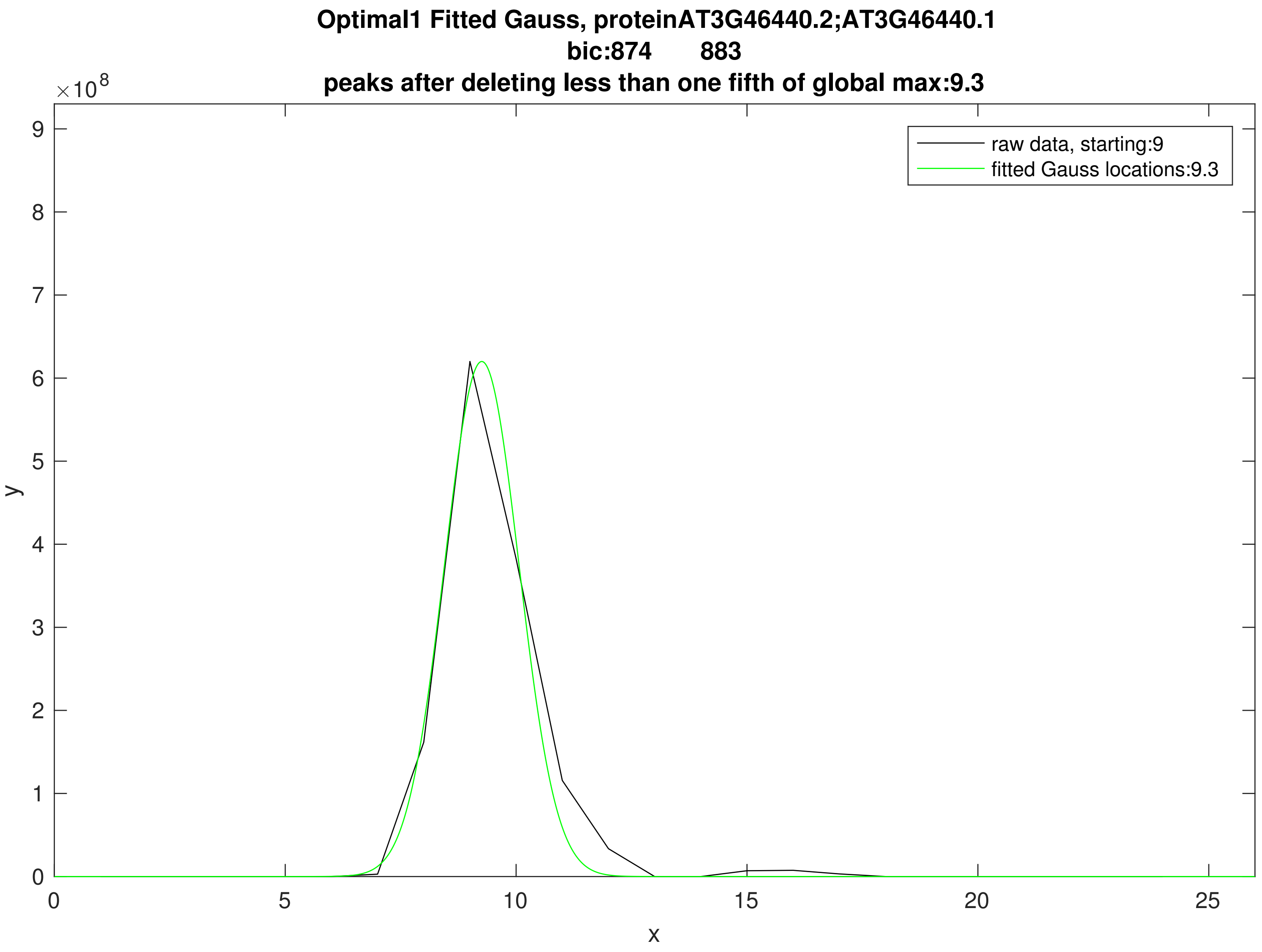
Optimal1 Fitted Gauss, proteinAT3G46440.2;AT3G46440.1
bic:874 883
peaks after deleting less than one fifth of global max:9.3
×10^8
raw data, starting:9
fitted Gauss locations:9.3
y
x
0
1
2
3
4
5
6
7
8
9
0
5
10
15
20
25

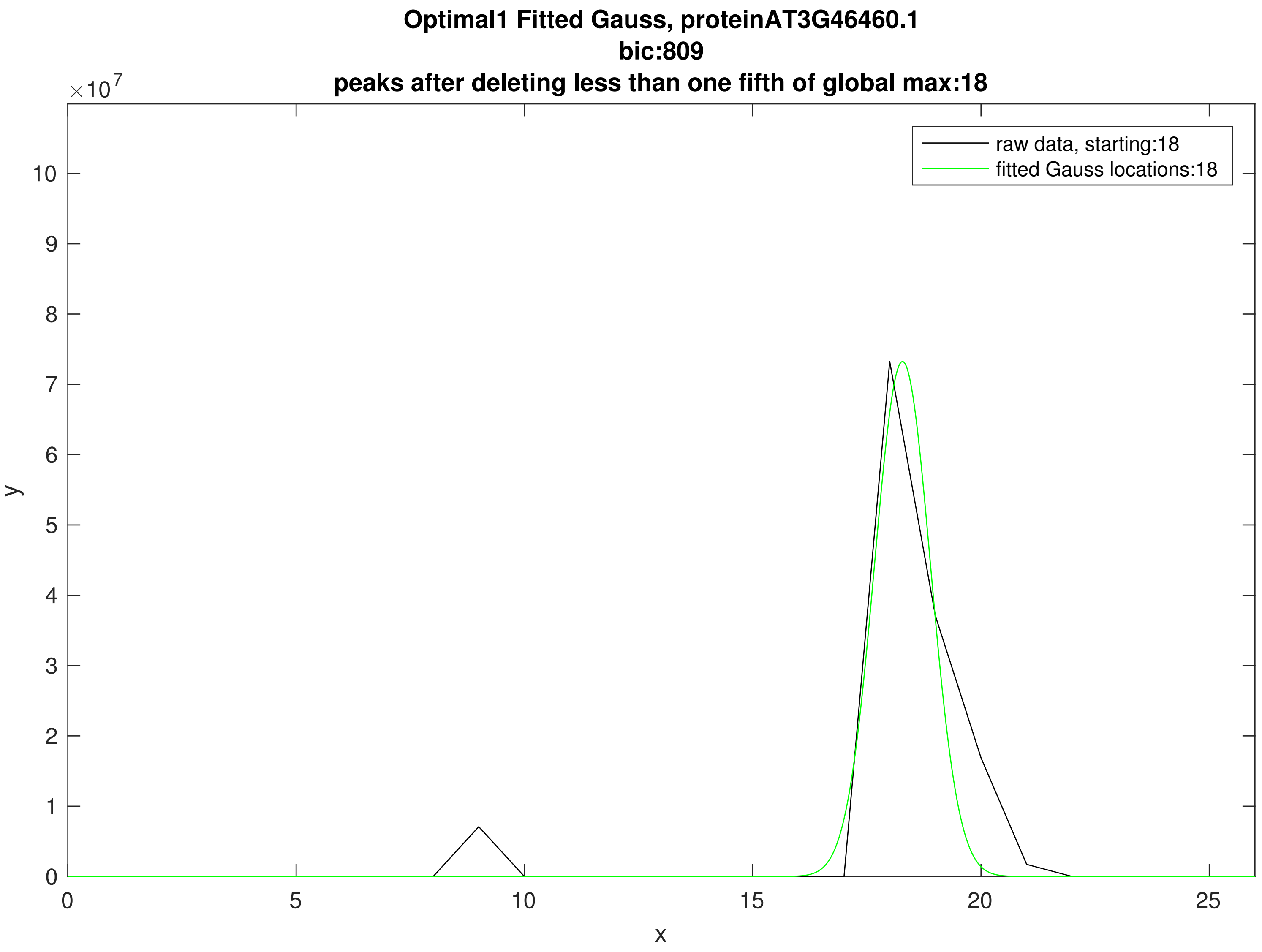
Optimal1 Fitted Gauss, proteinAT3G46460.1
bic:809
peaks after deleting less than one fifth of global max:18
×10^7
raw data, starting:18
fitted Gauss locations:18
y
x
0
1
2
3
4
5
6
7
8
9
10
0
5
10
15
20
25

#### Optimal1 Fitted Gauss, proteinAT3G46560.1
#### bic:869 870 879
#### peaks after deleting less than one fifth of global max:20

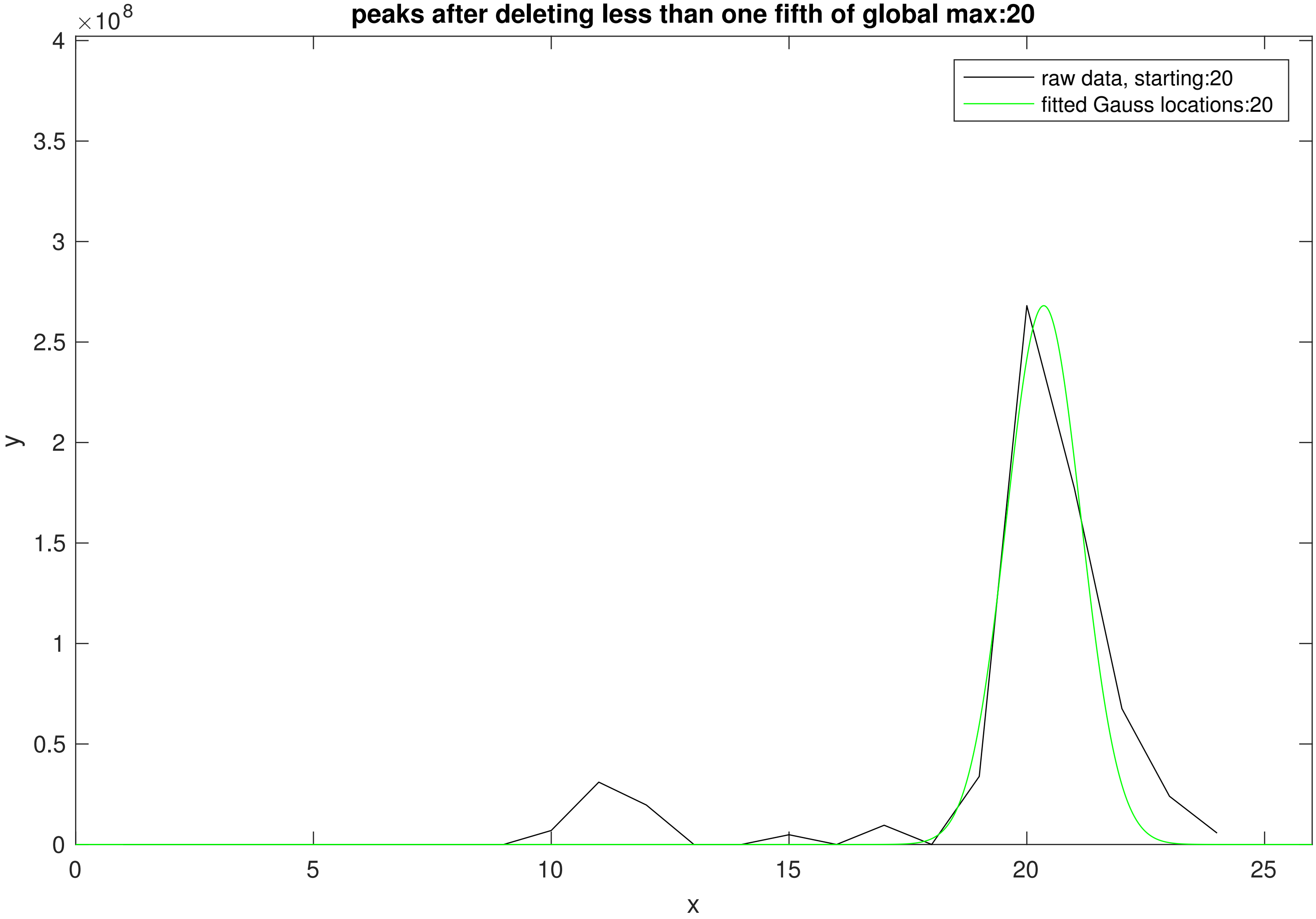

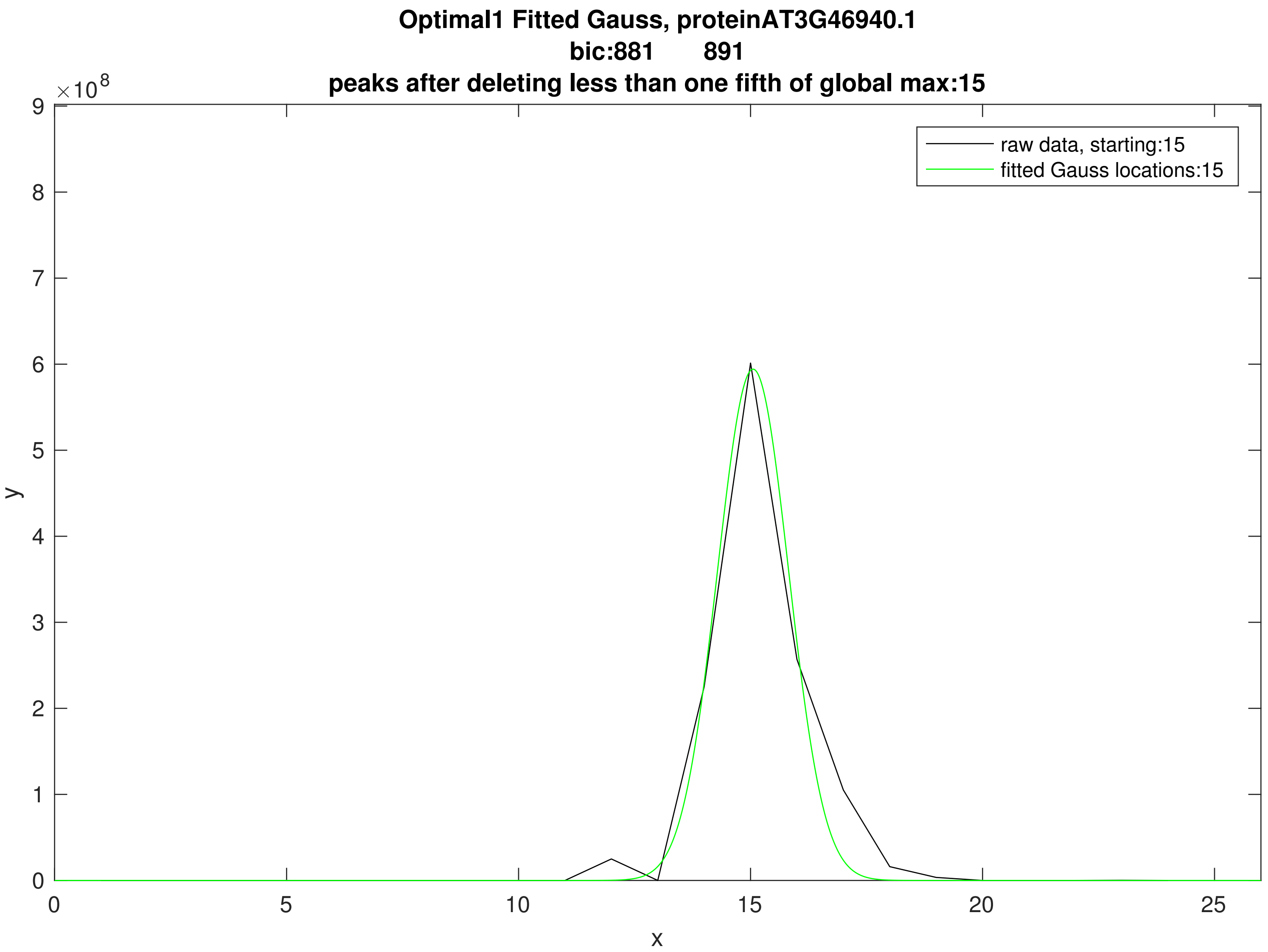
Optimal1 Fitted Gauss, proteinAT3G46940.1
bic:881 891
peaks after deleting less than one fifth of global max:15
raw data, starting:15
fitted Gauss locations:15
×10^8
y
x

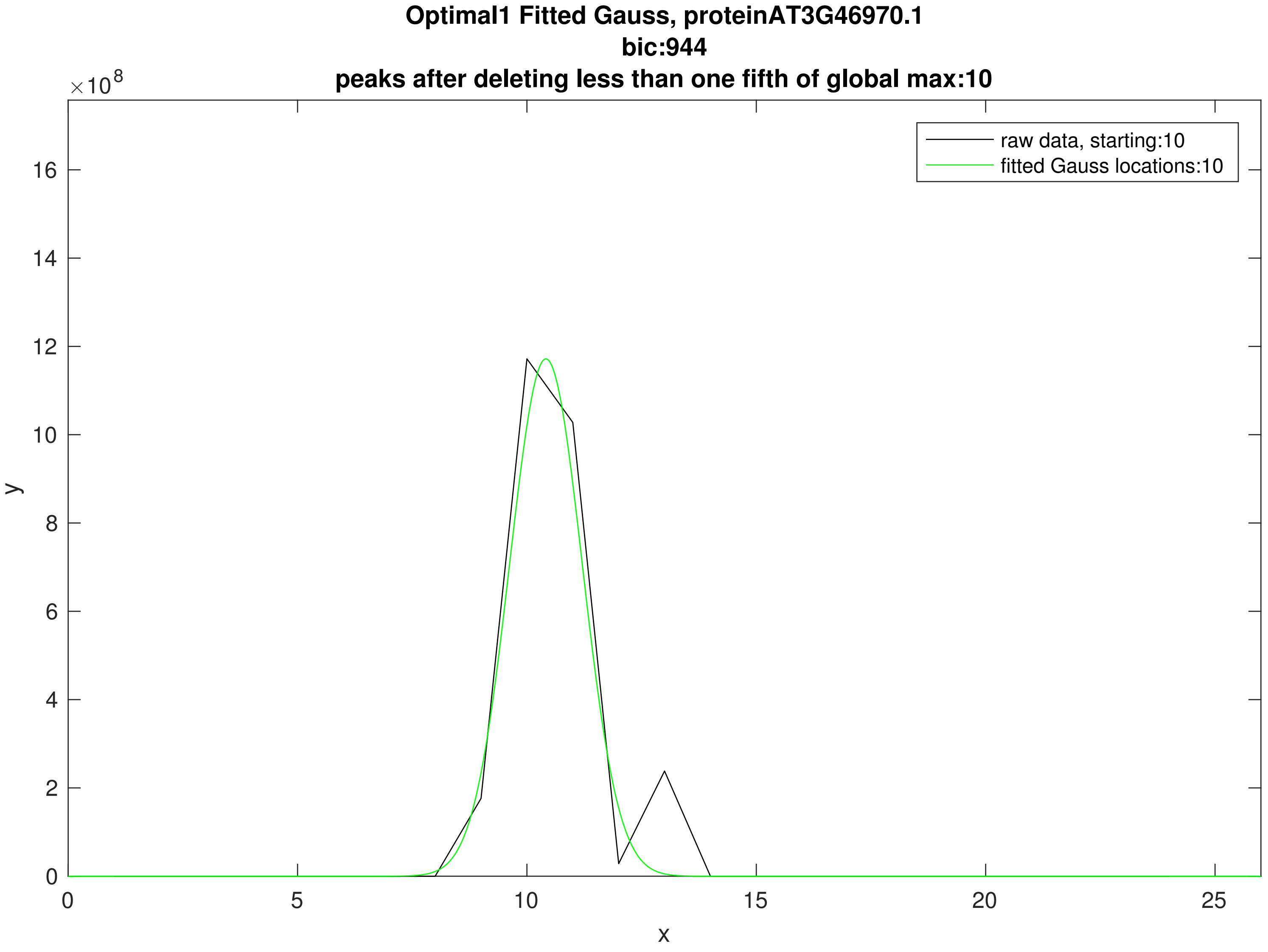

Optimal1 Fitted Gauss, proteinAT3G46970.1
bic:944
peaks after deleting less than one fifth of global max:10
×10^8
raw data, starting:10
fitted Gauss locations:10
y
x
0
2
4
6
8
10
12
14
16
0
5
10
15
20
25

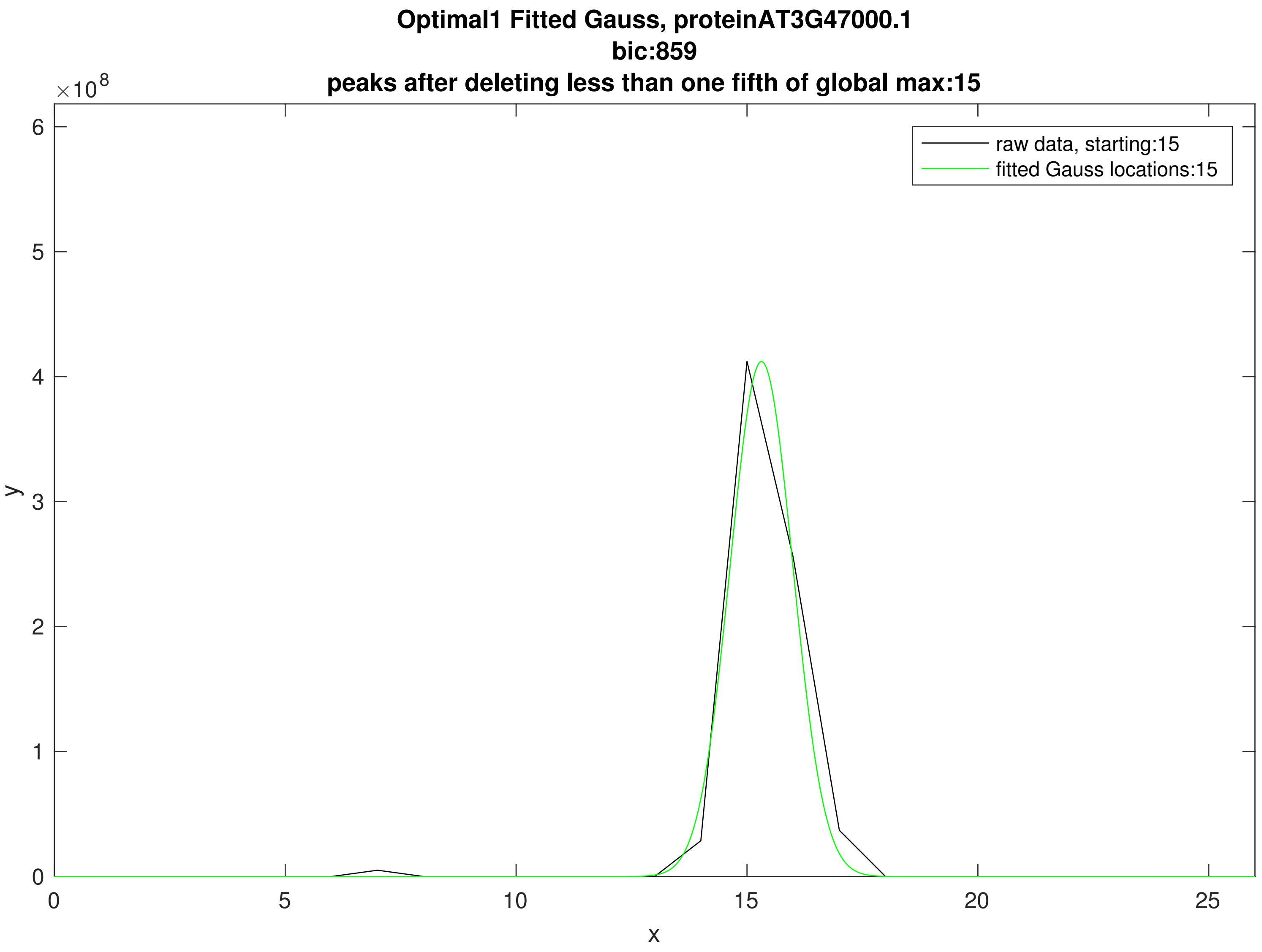
Optimal1 Fitted Gauss, proteinAT3G47000.1
bic:859
peaks after deleting less than one fifth of global max:15
×10^8
6
5
4
3
2
1
0
y
0
5
10
15
20
25
x
raw data, starting:15
fitted Gauss locations:15

#### Optimal1 Fitted Gauss, proteinAT3G47070.1
#### bic:752
#### peaks after deleting less than one fifth of global max:18

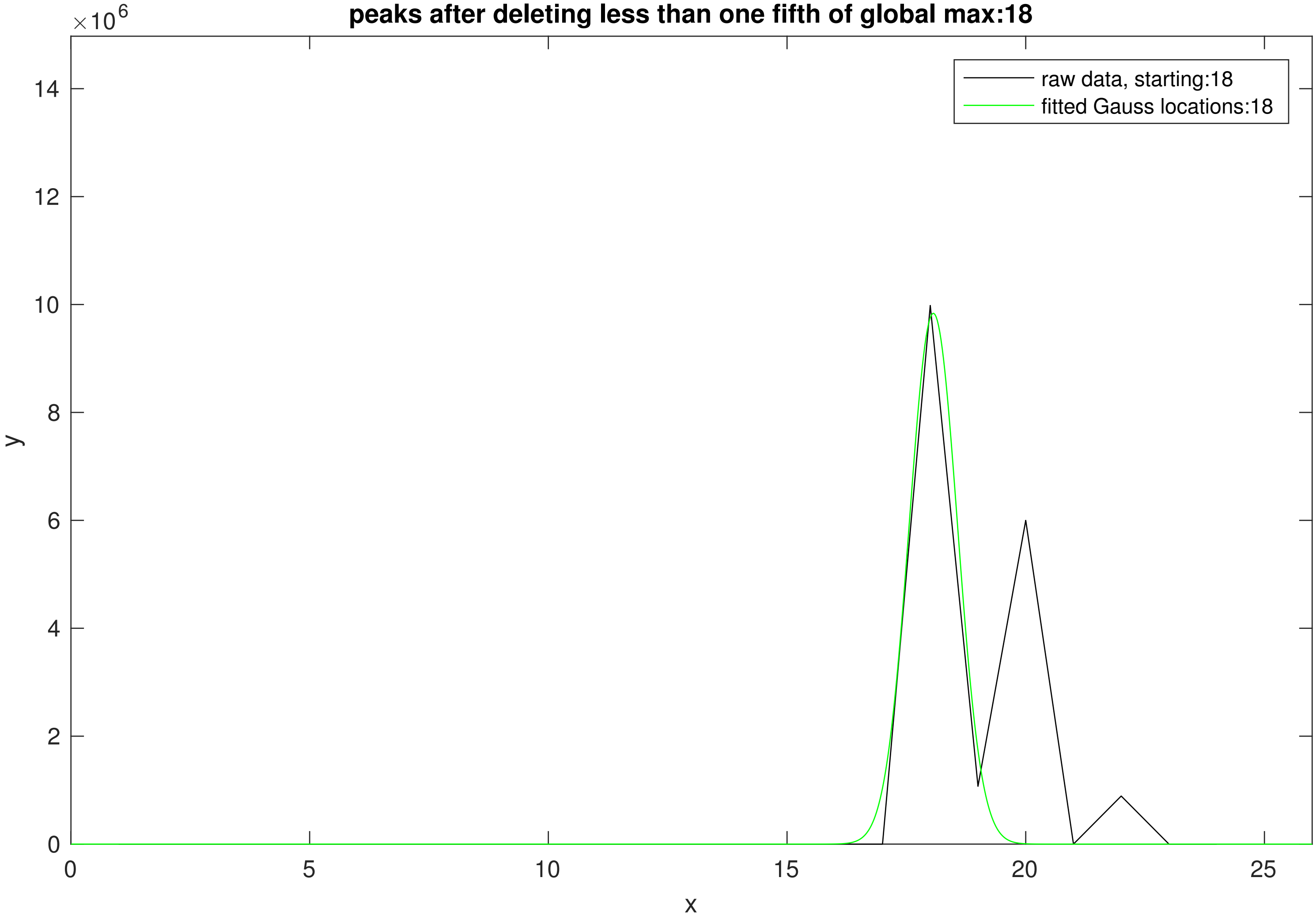

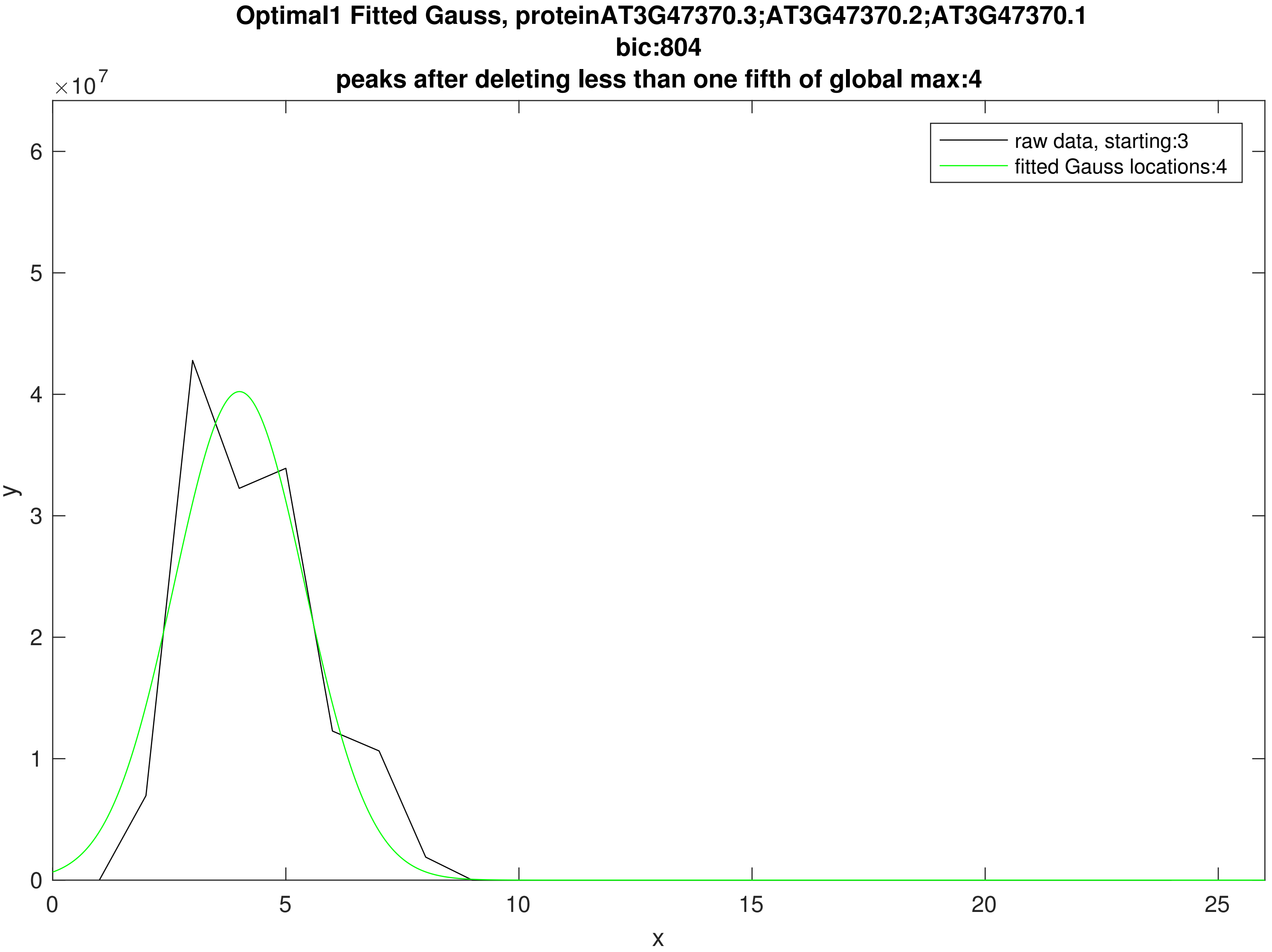
Optimal1 Fitted Gauss, proteinAT3G47370.3;AT3G47370.2;AT3G47370.1
bic:804
peaks after deleting less than one fifth of global max:4
×10^7
raw data, starting:3
fitted Gauss locations:4
y
x
0
1
2
3
4
5
6
0
5
10
15
20
25

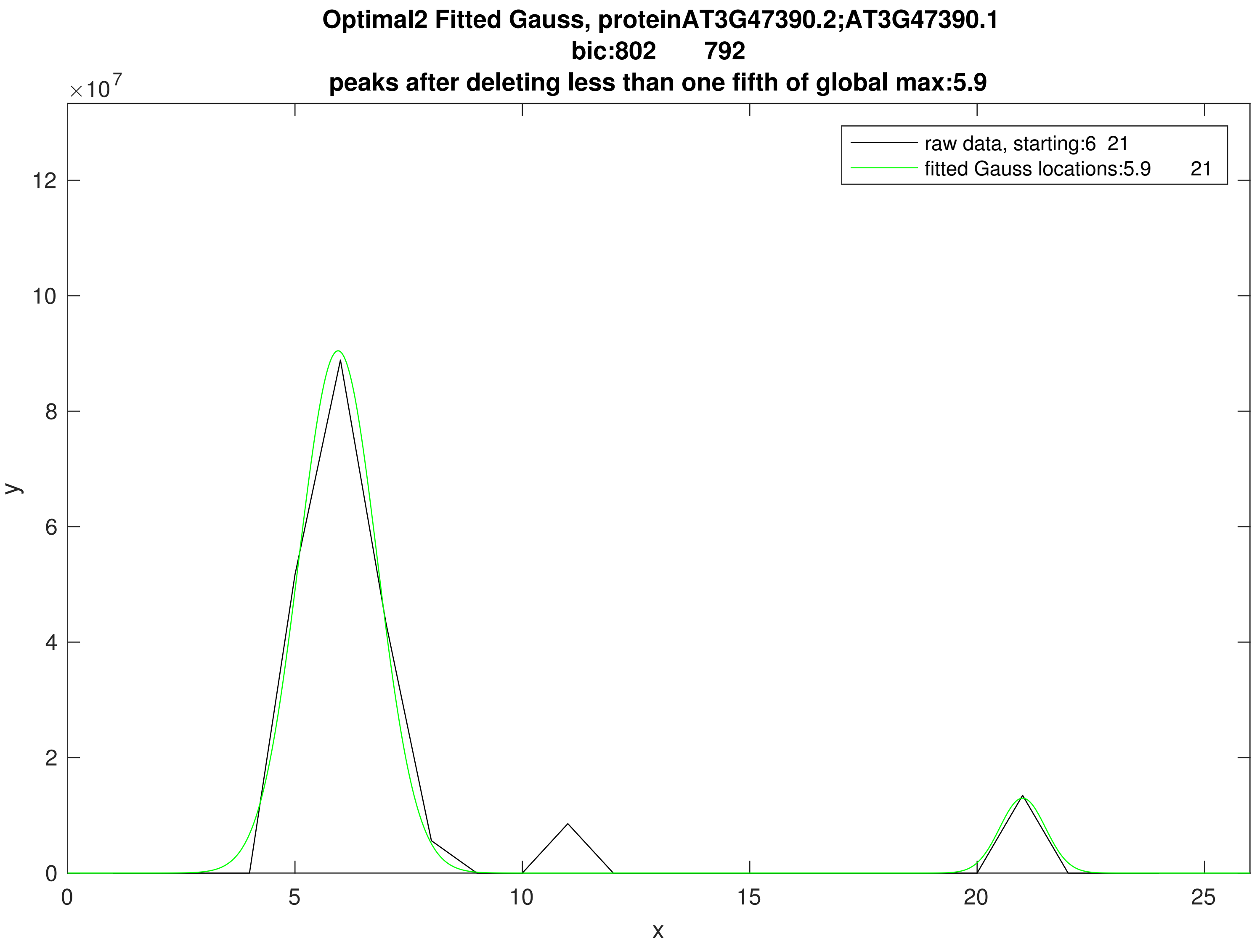
Optimal2 Fitted Gauss, proteinAT3G47390.2;AT3G47390.1
bic:802 792
peaks after deleting less than one fifth of global max:5.9
×10^7
raw data, starting:6 21
fitted Gauss locations:5.9 21
y
x
0
2
4
6
8
10
12
5
15
20
25

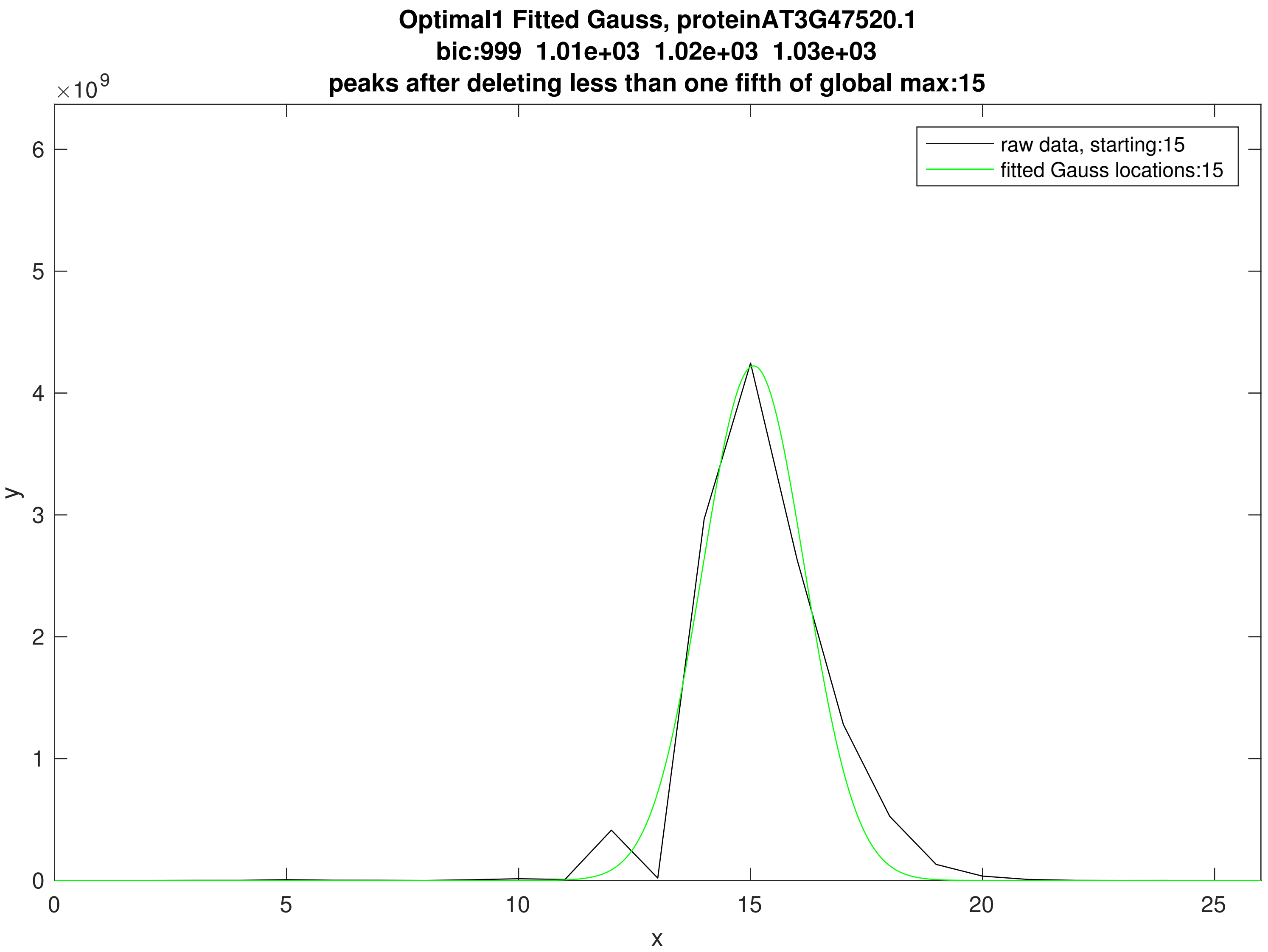
Optimal1 Fitted Gauss, proteinAT3G47520.1
bic:999 1.01e+03 1.02e+03 1.03e+03
peaks after deleting less than one fifth of global max:15
×10^9
raw data, starting:15
fitted Gauss locations:15
y
x
0
1
2
3
4
5
6
0
5
10
15
20
25

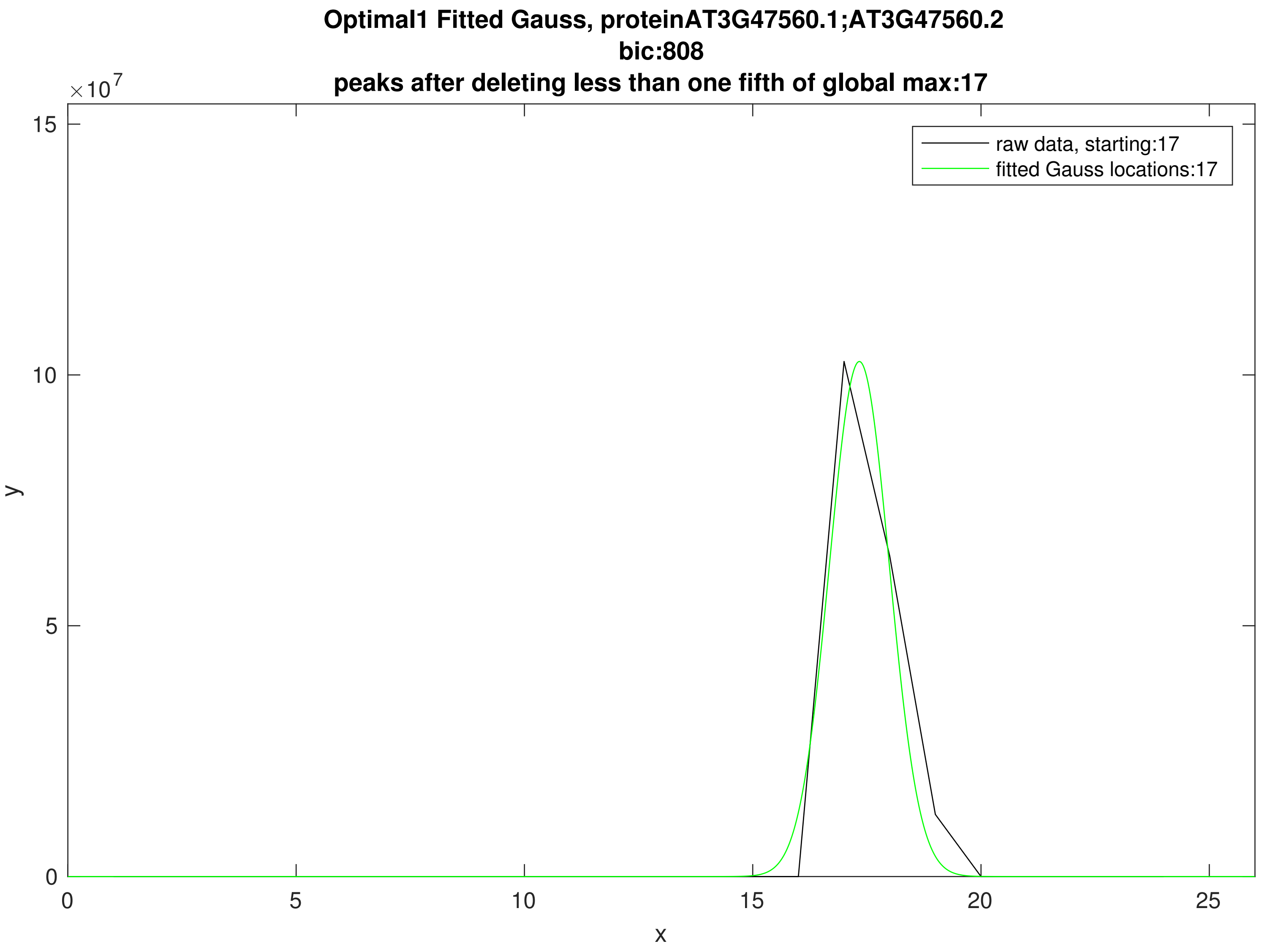
Optimal1 Fitted Gauss, proteinAT3G47560.1;AT3G47560.2
bic:808
peaks after deleting less than one fifth of global max:17
×10^7
raw data, starting:17
fitted Gauss locations:17
y
x
0
5
10
15
20
25

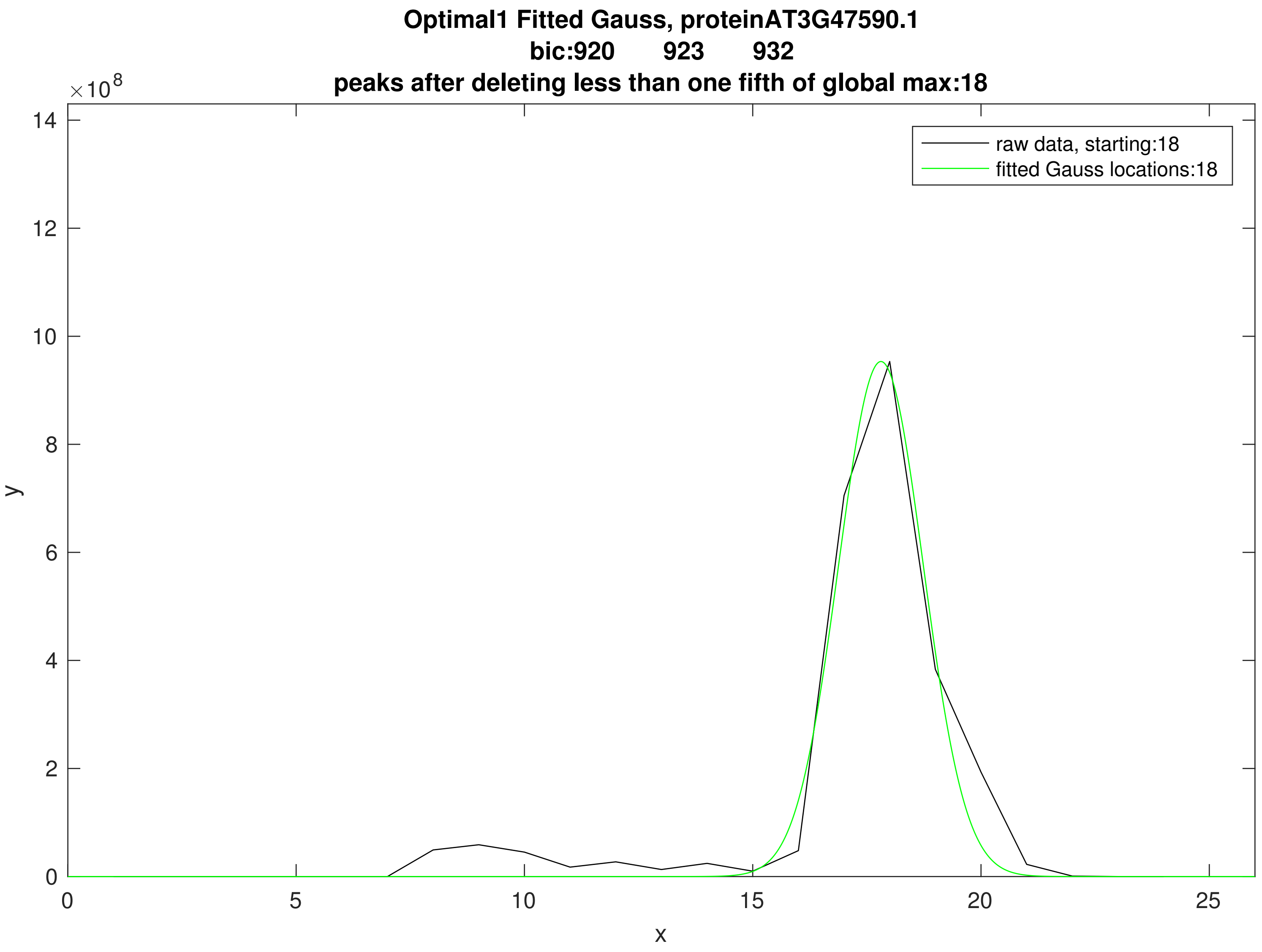
Optimal1 Fitted Gauss, proteinAT3G47590.1
bic:920 923 932
peaks after deleting less than one fifth of global max:18
×10^8
raw data, starting:18
fitted Gauss locations:18
y
x
0
2
4
6
8
10
12
14
0
5
10
15
20
25

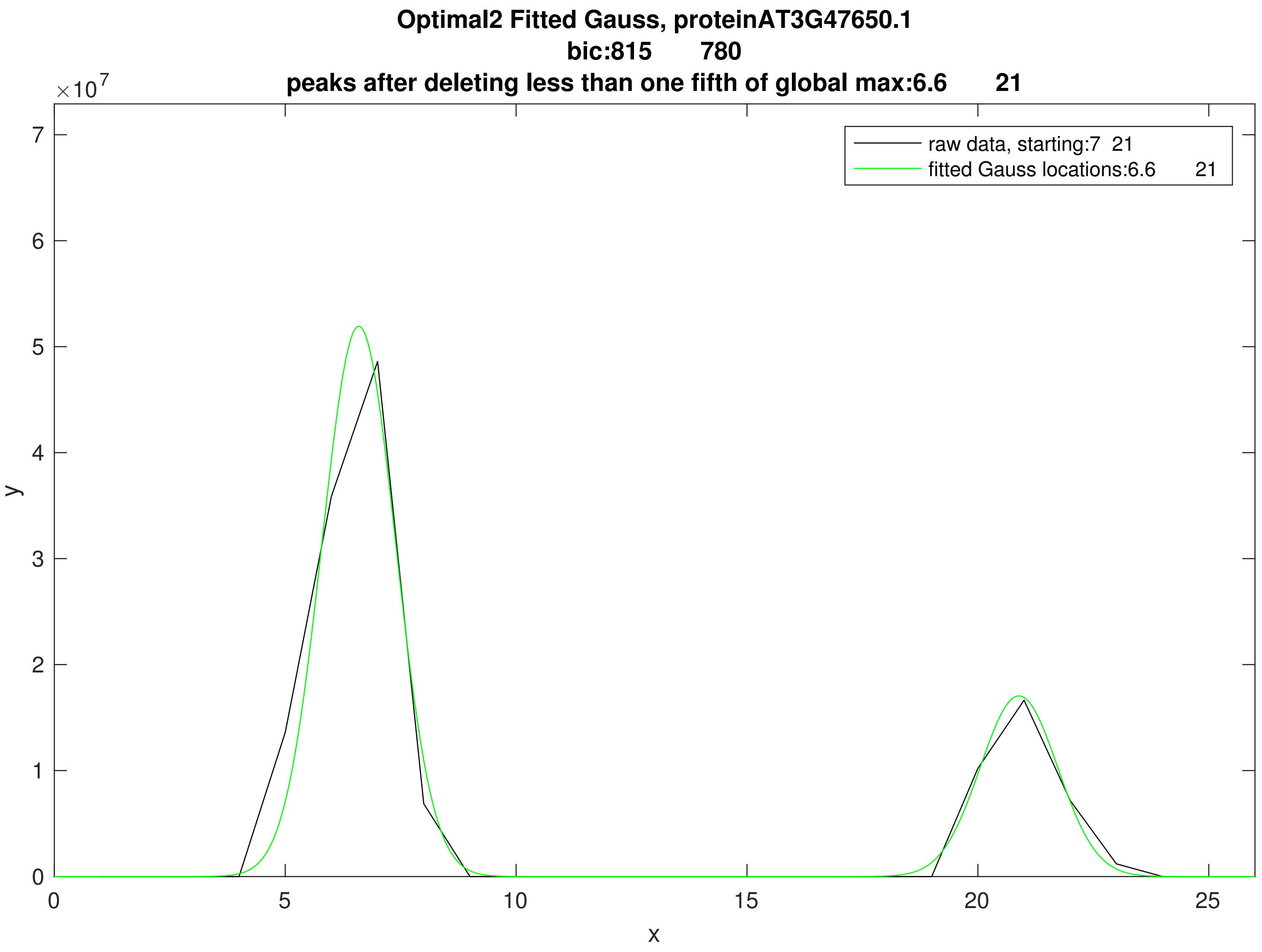
Optimal2 Fitted Gauss, proteinAT3G47650.1
bic:815 780
peaks after deleting less than one fifth of global max:6.6 21
×10^7
raw data, starting:7 21
fitted Gauss locations:6.6 21
y
x
0
1
2
3
4
5
6
7
0
5
10
15
20
25

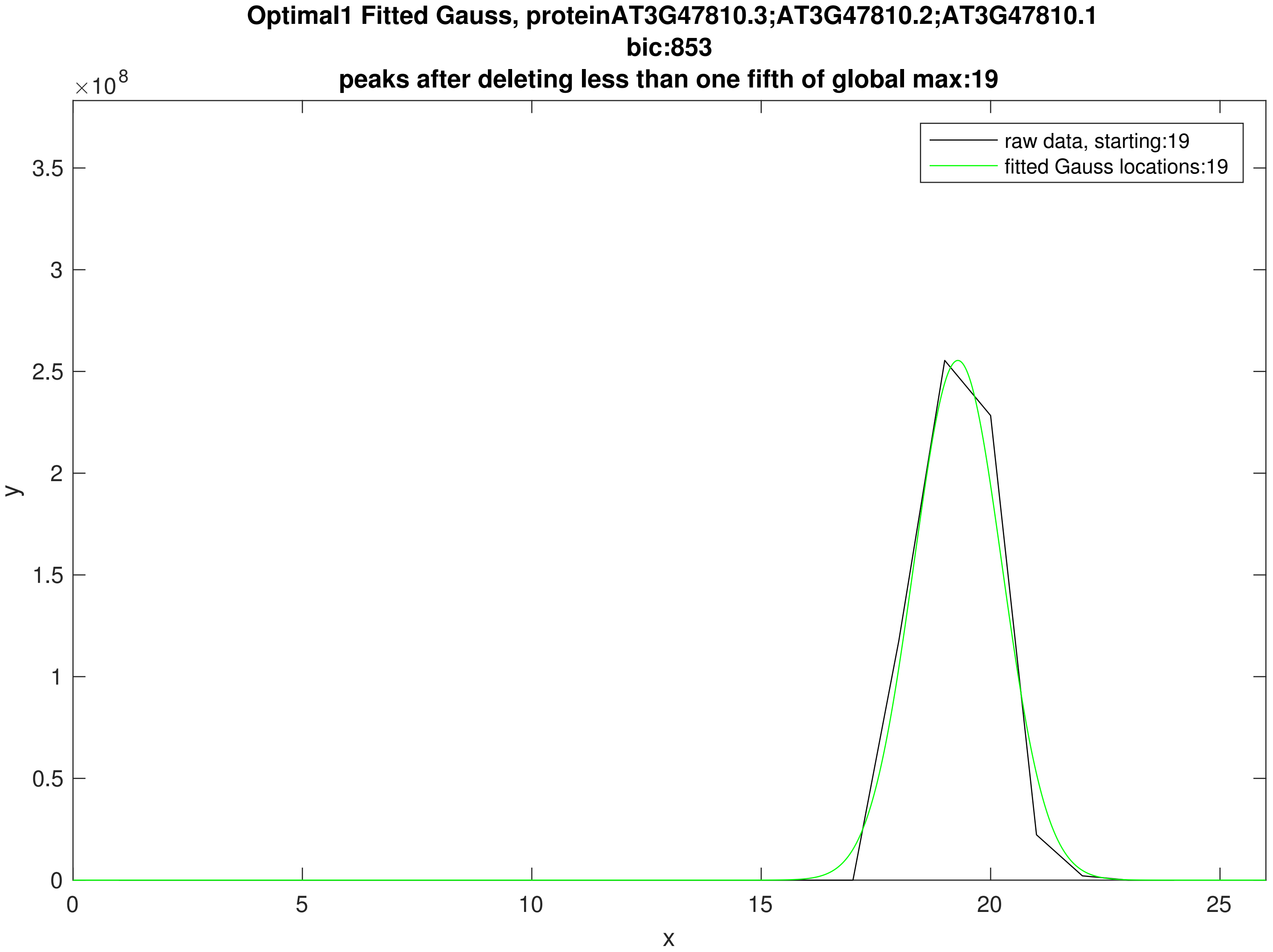

Optimal1 Fitted Gauss, proteinAT3G47810.3;AT3G47810.2;AT3G47810.1
bic:853
peaks after deleting less than one fifth of global max:19
×10^8
3.5
3
2.5
2
1.5
1
0.5
0
y
0
5
10
15
20
25
x
raw data, starting:19
fitted Gauss locations:19

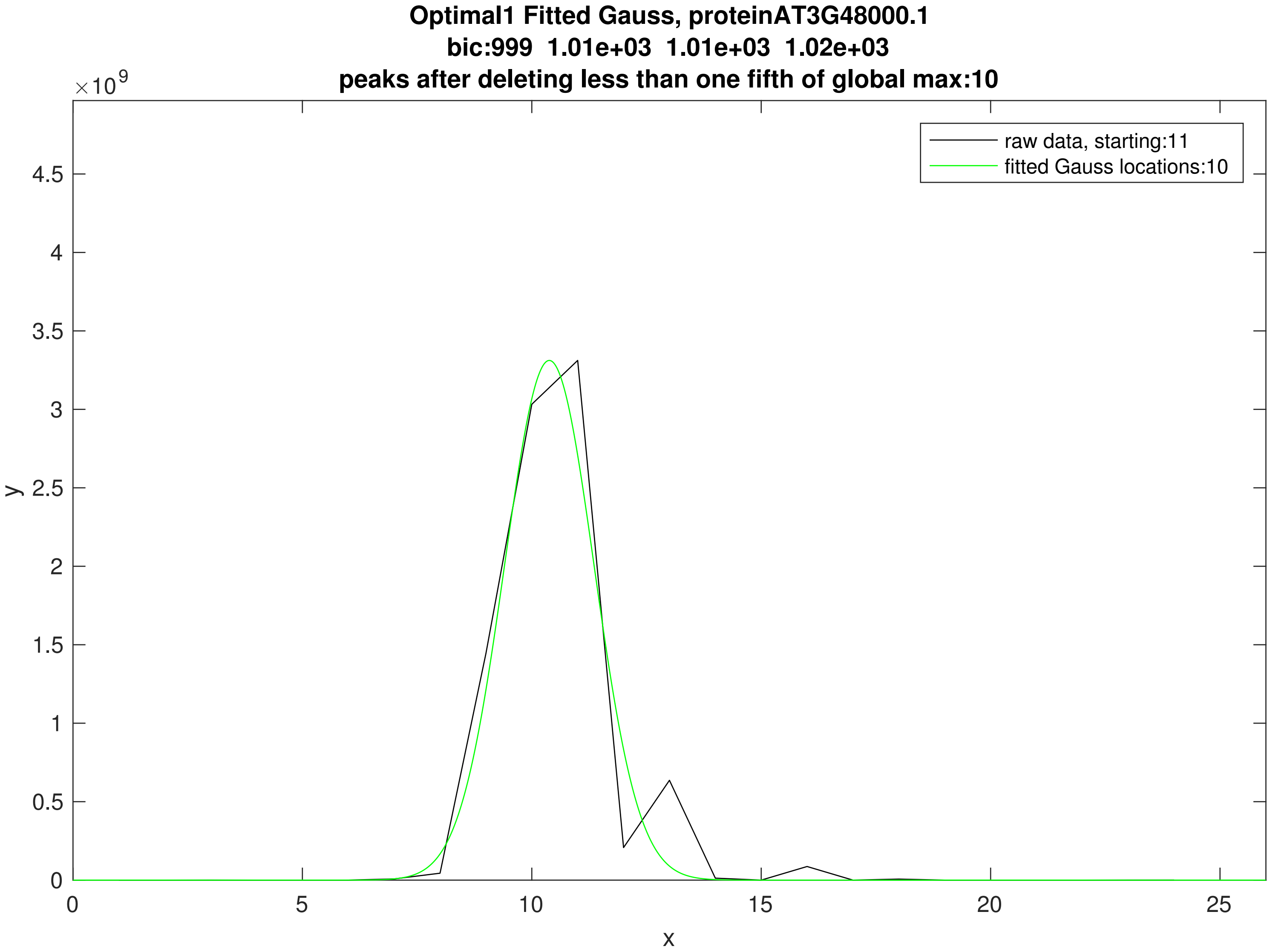
Optimal1 Fitted Gauss, proteinAT3G48000.1
bic:999 1.01e+03 1.01e+03 1.02e+03
peaks after deleting less than one fifth of global max:10
×10^9
raw data, starting:11
fitted Gauss locations:10
y
x
0
0.5
1
1.5
2
2.5
3
3.5
4
4.5
5
10
15
20
25

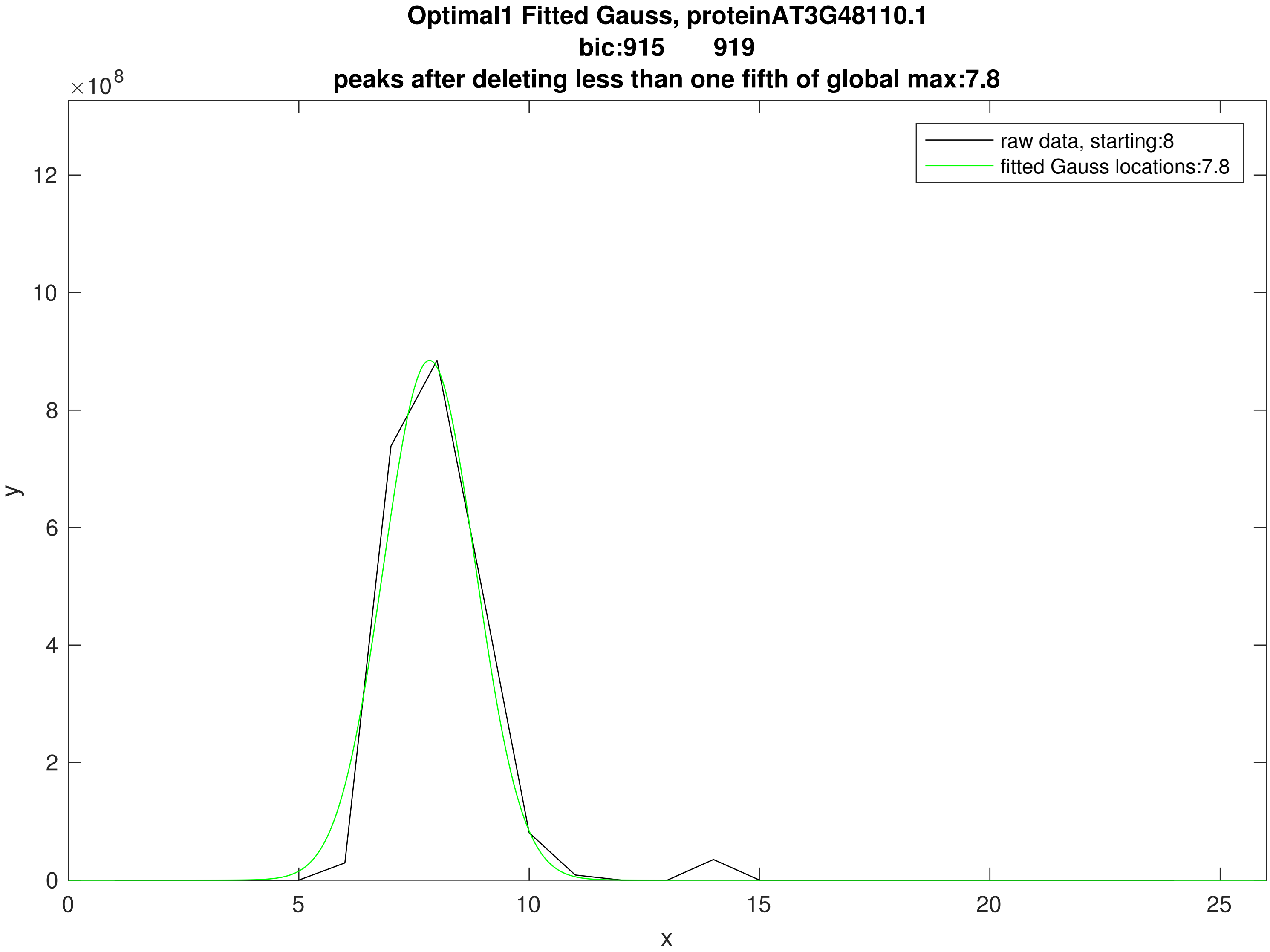

Optimal1 Fitted Gauss, proteinAT3G48110.1
bic:915 919
peaks after deleting less than one fifth of global max:7.8
×10^8
raw data, starting:8
fitted Gauss locations:7.8
12
10
8
6
4
2
0
y
0
5
10
15
20
25
x

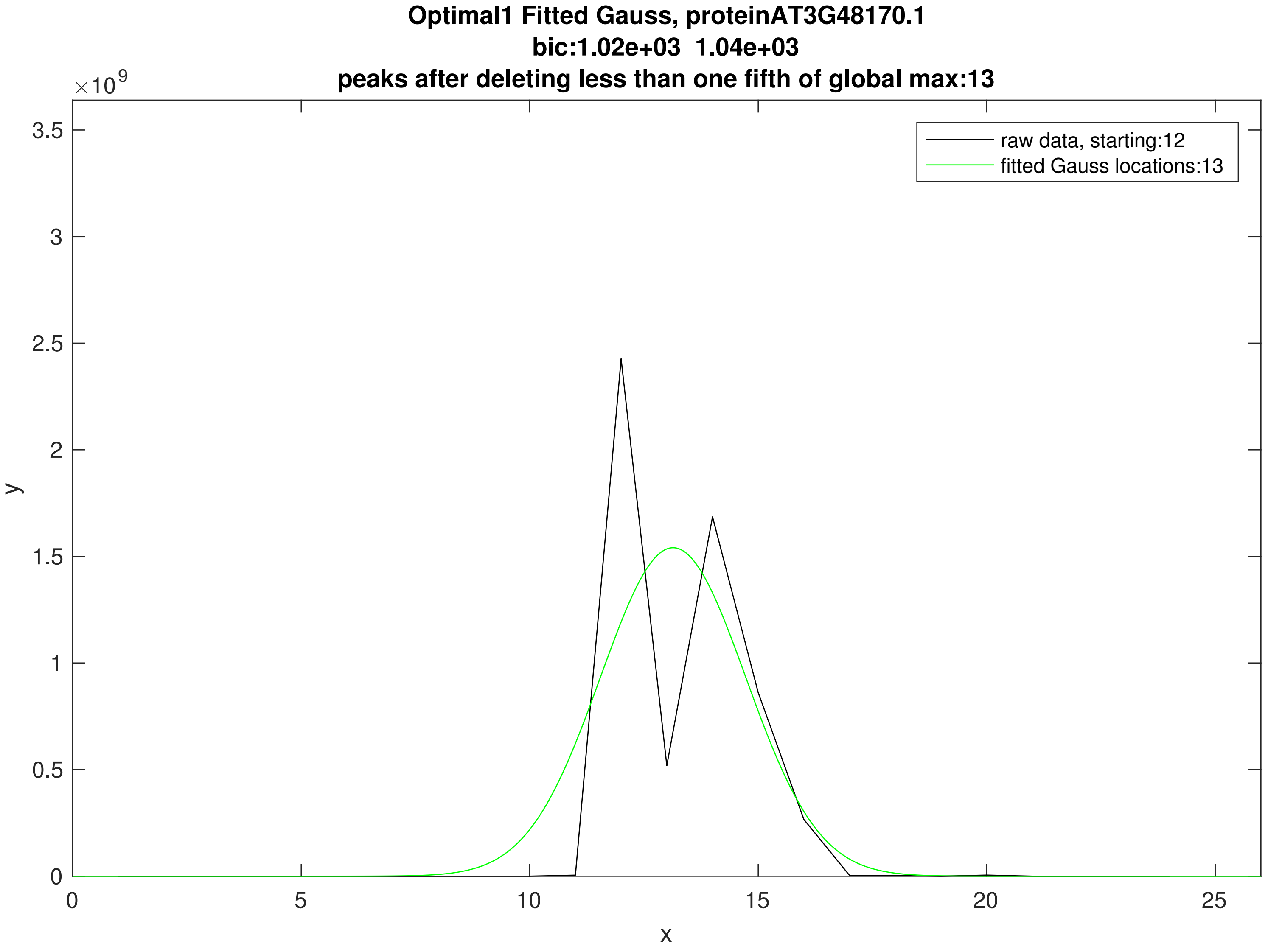

Optimal1 Fitted Gauss, proteinAT3G48170.1
bic:1.02e+03  1.04e+03
peaks after deleting less than one fifth of global max:13
$\times 10^9$
3.5
3
2.5
2
1.5
1
0.5
0
y
0
5
10
15
20
25
x
raw data, starting:12
fitted Gauss locations:13

**Optimal1 Fitted Gauss, proteinAT3G48350.1**
**bic:837 844**
**peaks after deleting less than one fifth of global max:13**

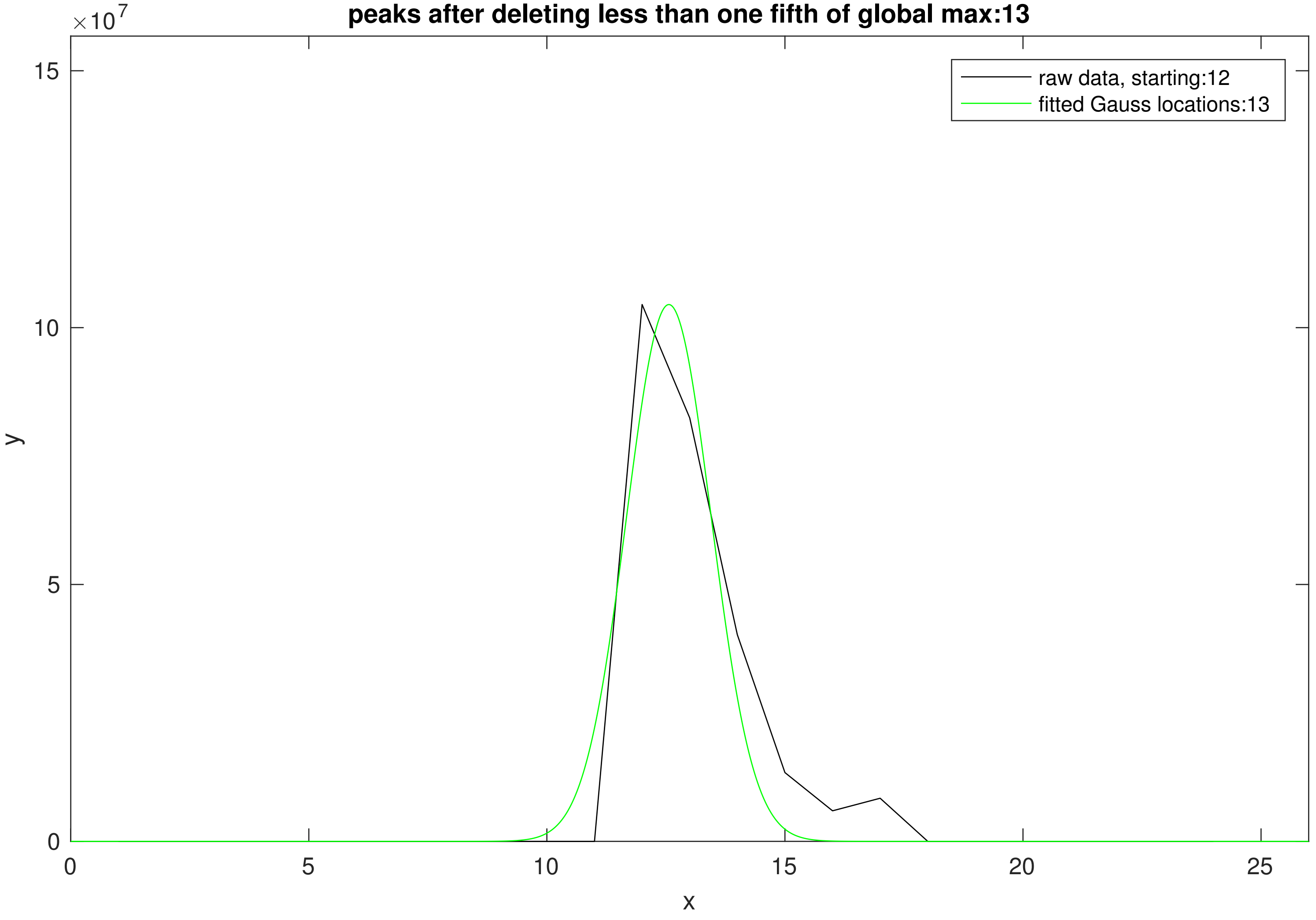

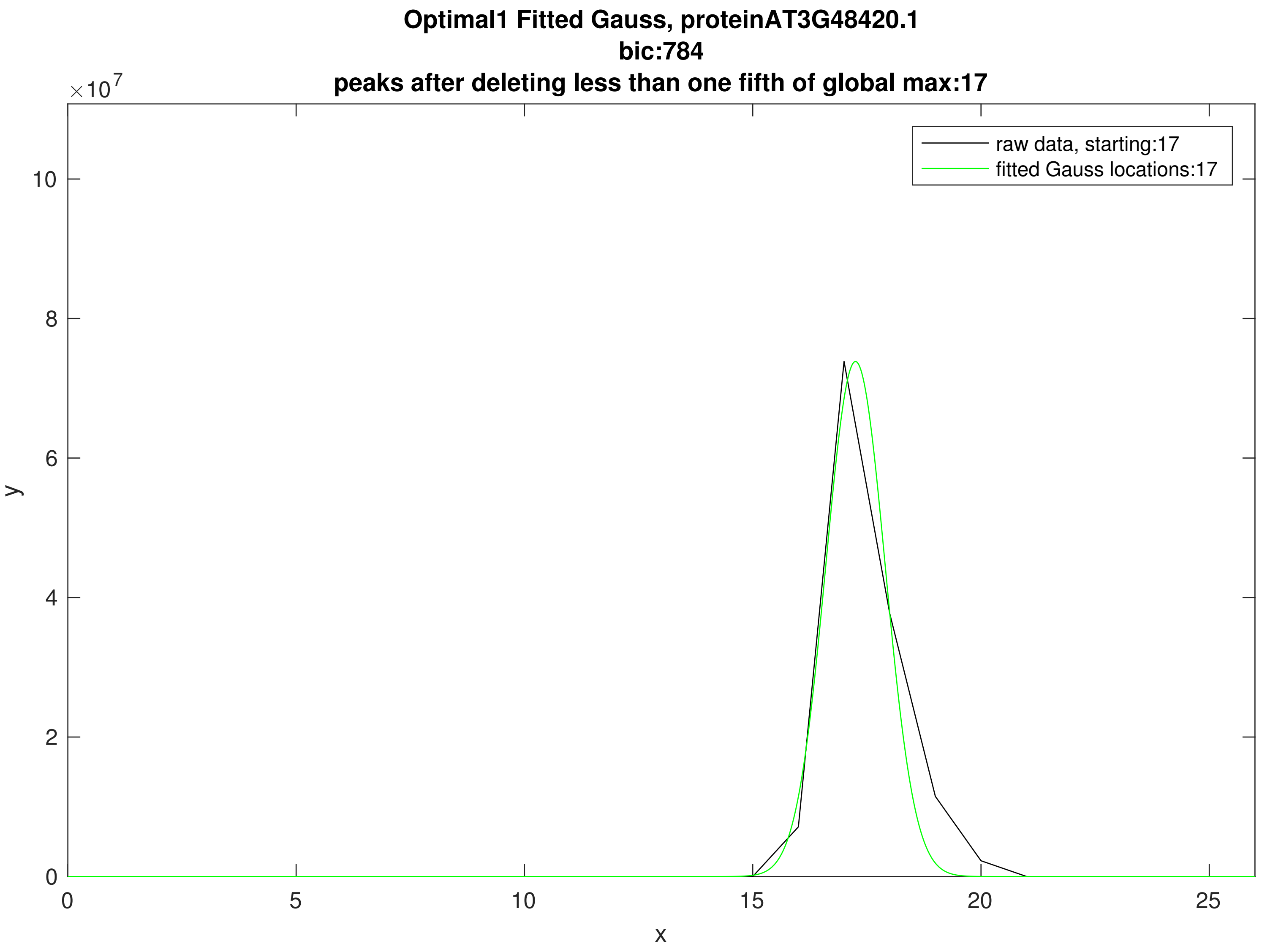

Optimal1 Fitted Gauss, proteinAT3G48420.1
bic:784
peaks after deleting less than one fifth of global max:17
×10^7
raw data, starting:17
fitted Gauss locations:17
y
x
0
2
4
6
8
10
0
5
10
15
20
25

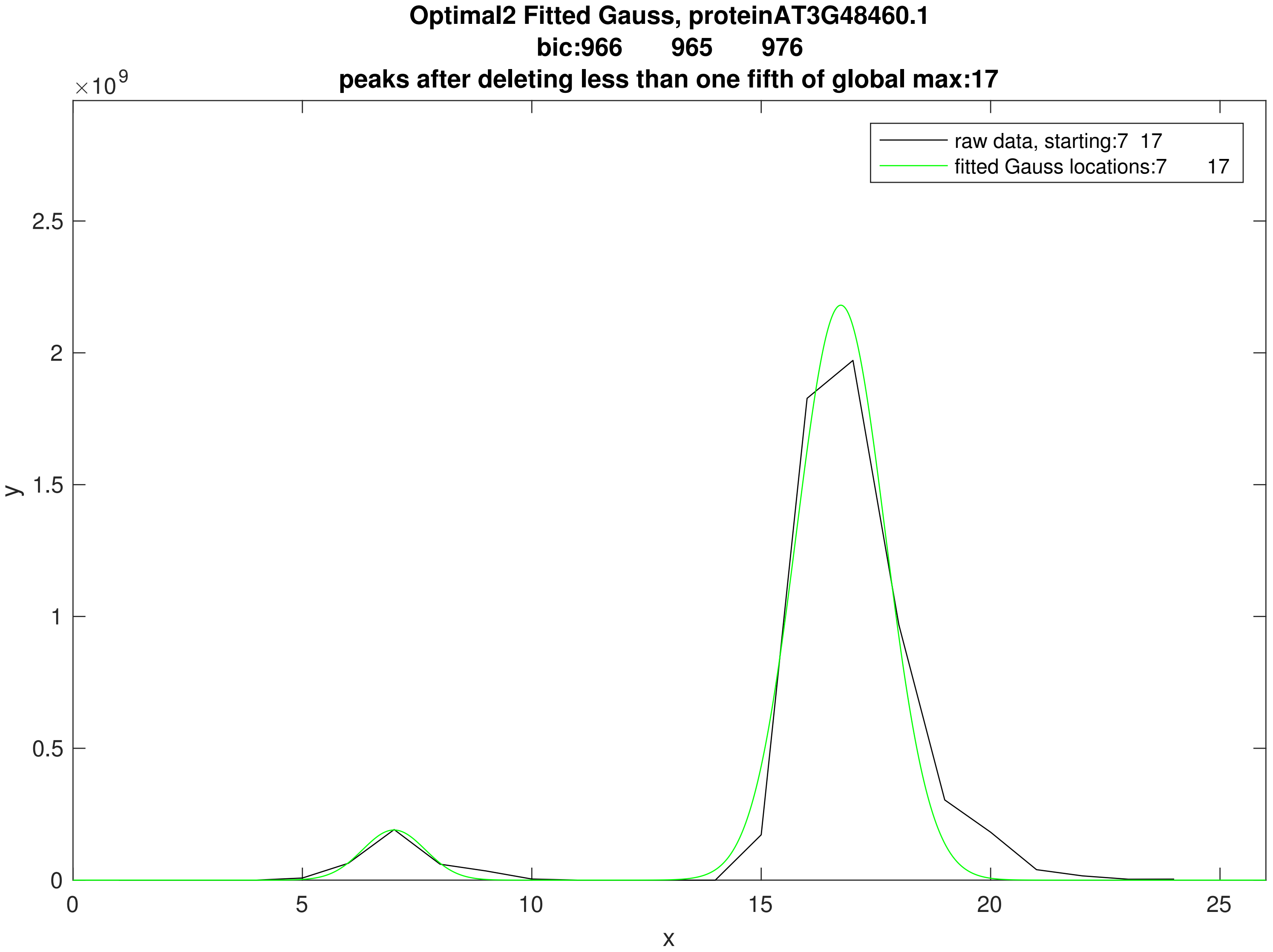

Optimal2 Fitted Gauss, proteinAT3G48460.1
bic:966 965 976
peaks after deleting less than one fifth of global max:17
×10^9
raw data, starting:7 17
fitted Gauss locations:7 17
y
x
0
0.5
1
1.5
2
2.5
5
10
15
20
25

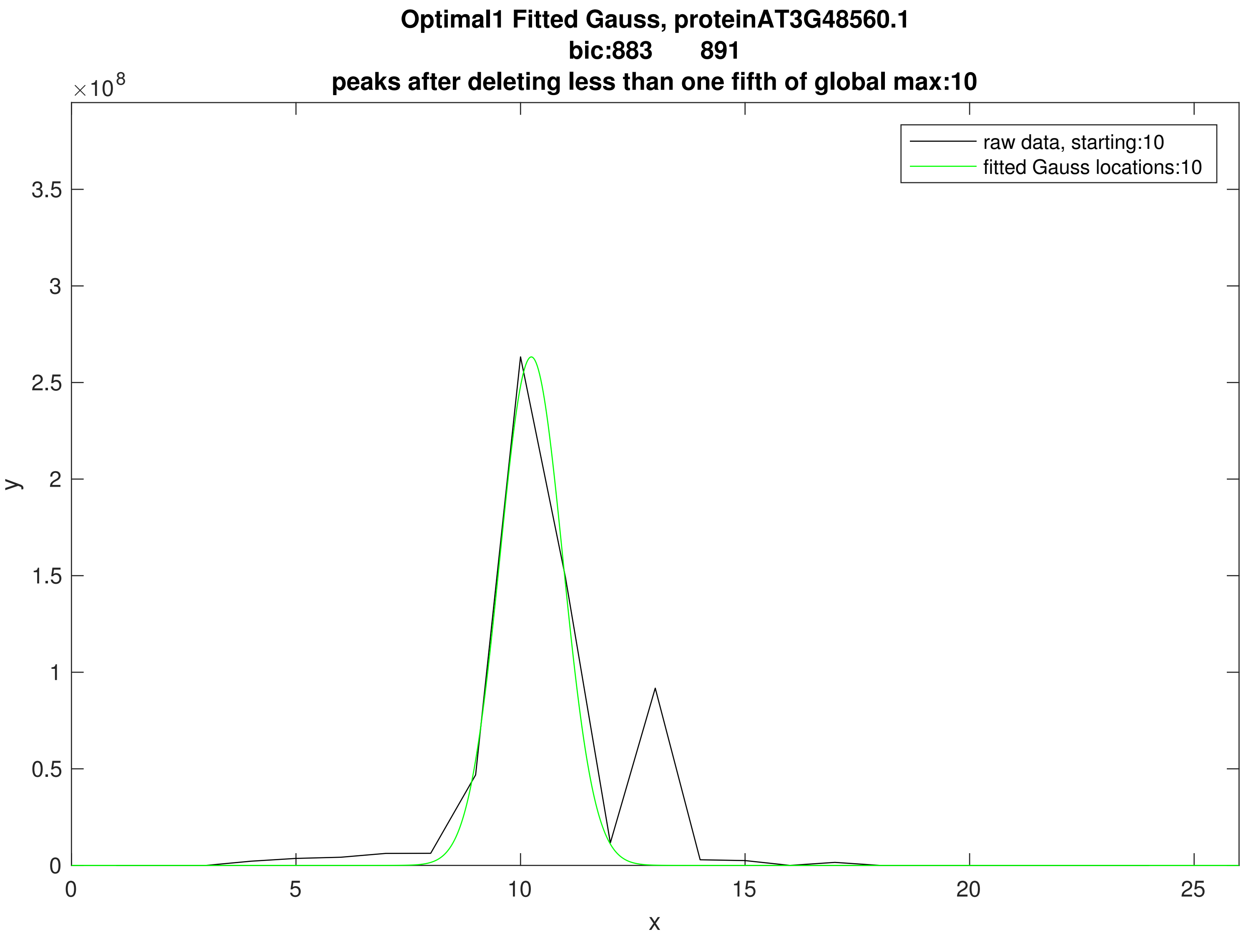

Optimal1 Fitted Gauss, proteinAT3G48560.1
bic:883 891
peaks after deleting less than one fifth of global max:10
×10^8
3.5
3
2.5
2
1.5
1
0.5
0
y
0
5
10
15
20
25
x
raw data, starting:10
fitted Gauss locations:10

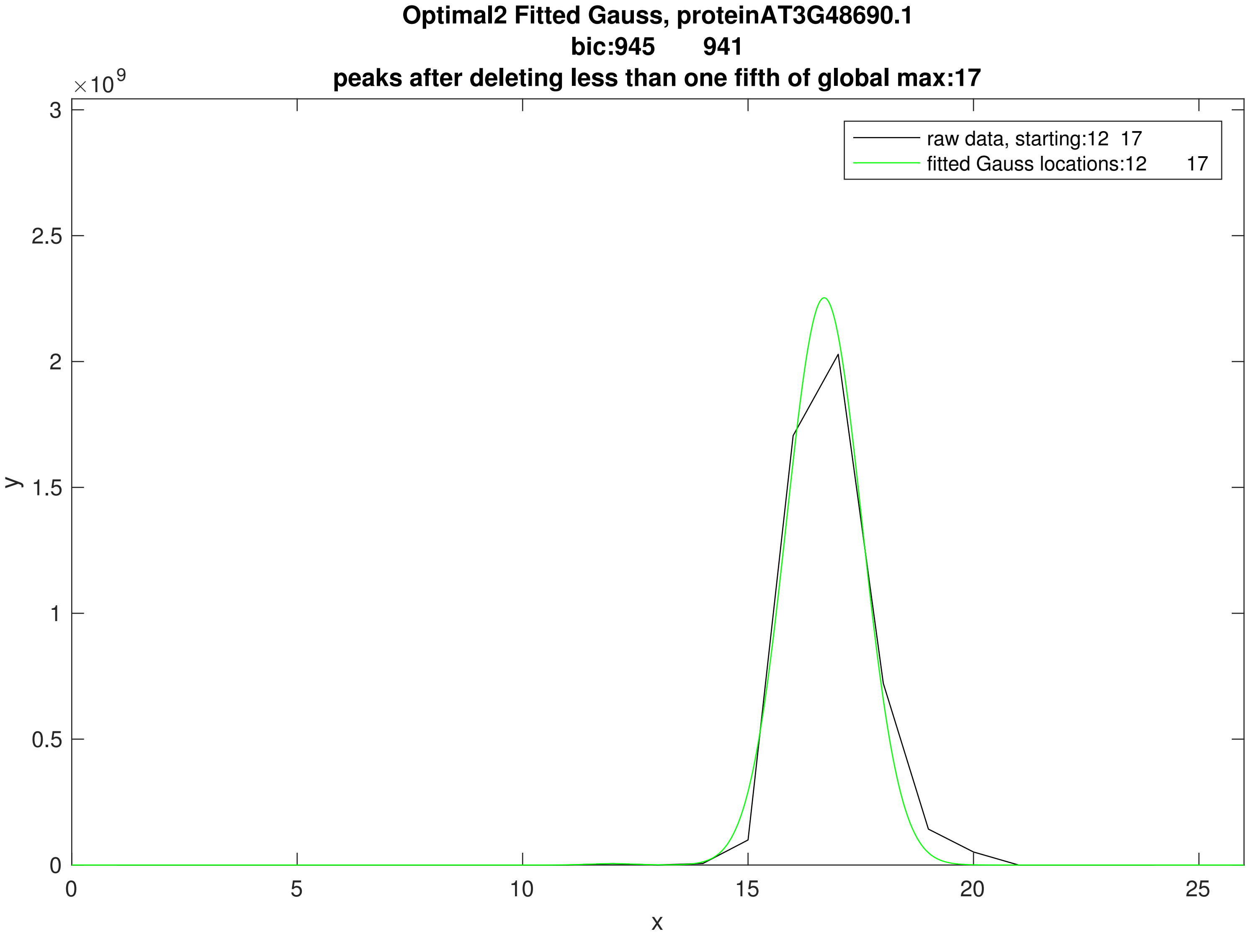
Optimal2 Fitted Gauss, proteinAT3G48690.1
bic:945 941
peaks after deleting less than one fifth of global max:17
×10^9
3
2.5
2
1.5
1
0.5
0
y
0
5
10
15
20
25
x
raw data, starting:12 17
fitted Gauss locations:12 17

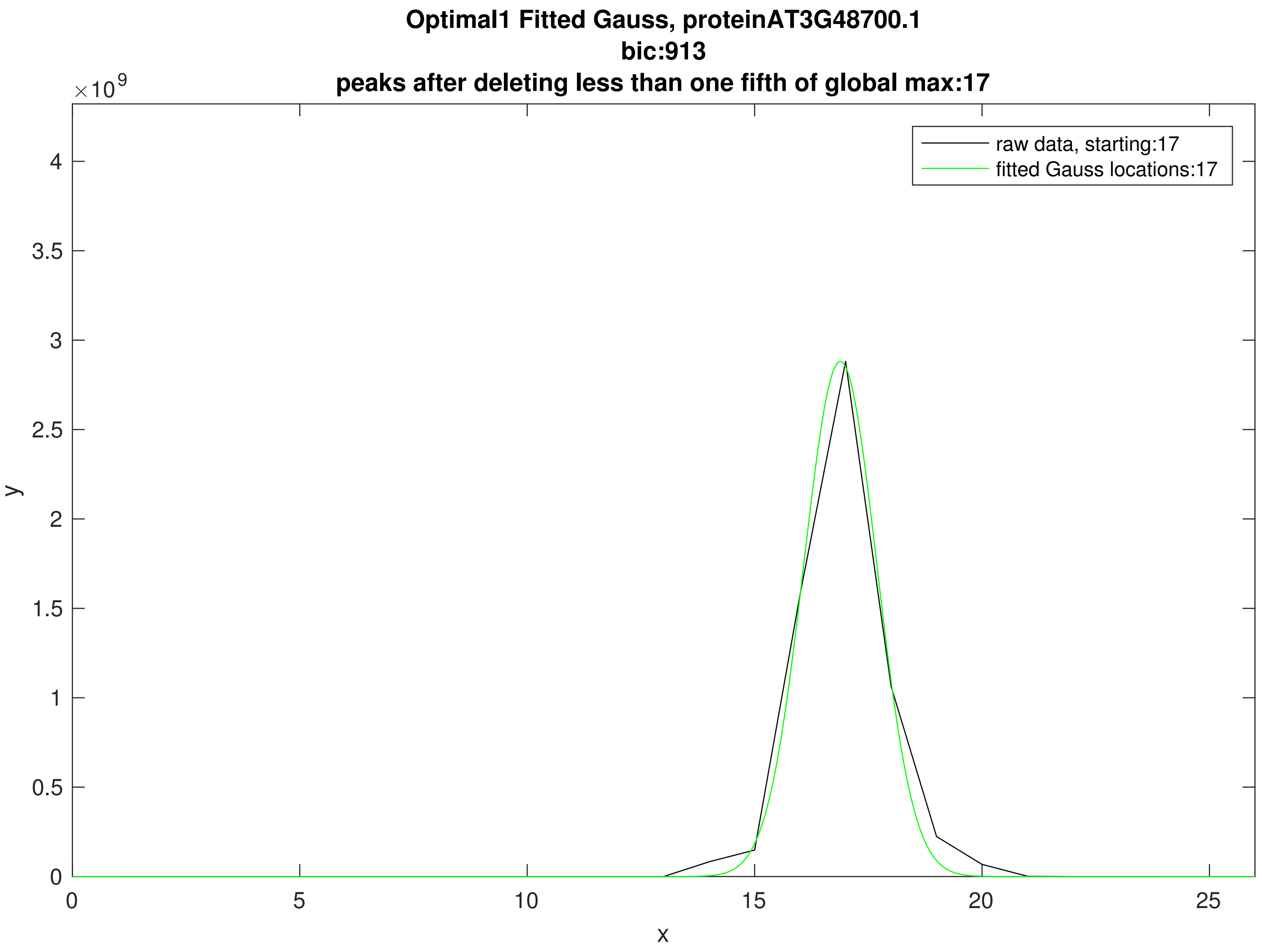
Optimal1 Fitted Gauss, proteinAT3G48700.1
bic:913
peaks after deleting less than one fifth of global max:17
×10^9
raw data, starting:17
fitted Gauss locations:17
y
x
0
0.5
1
1.5
2
2.5
3
3.5
4
5
10
15
20
25

**Optimal2 Fitted Gauss, proteinAT3G48720.1**
**bic:772 743**
**peaks after deleting less than one fifth of global max:13 17**

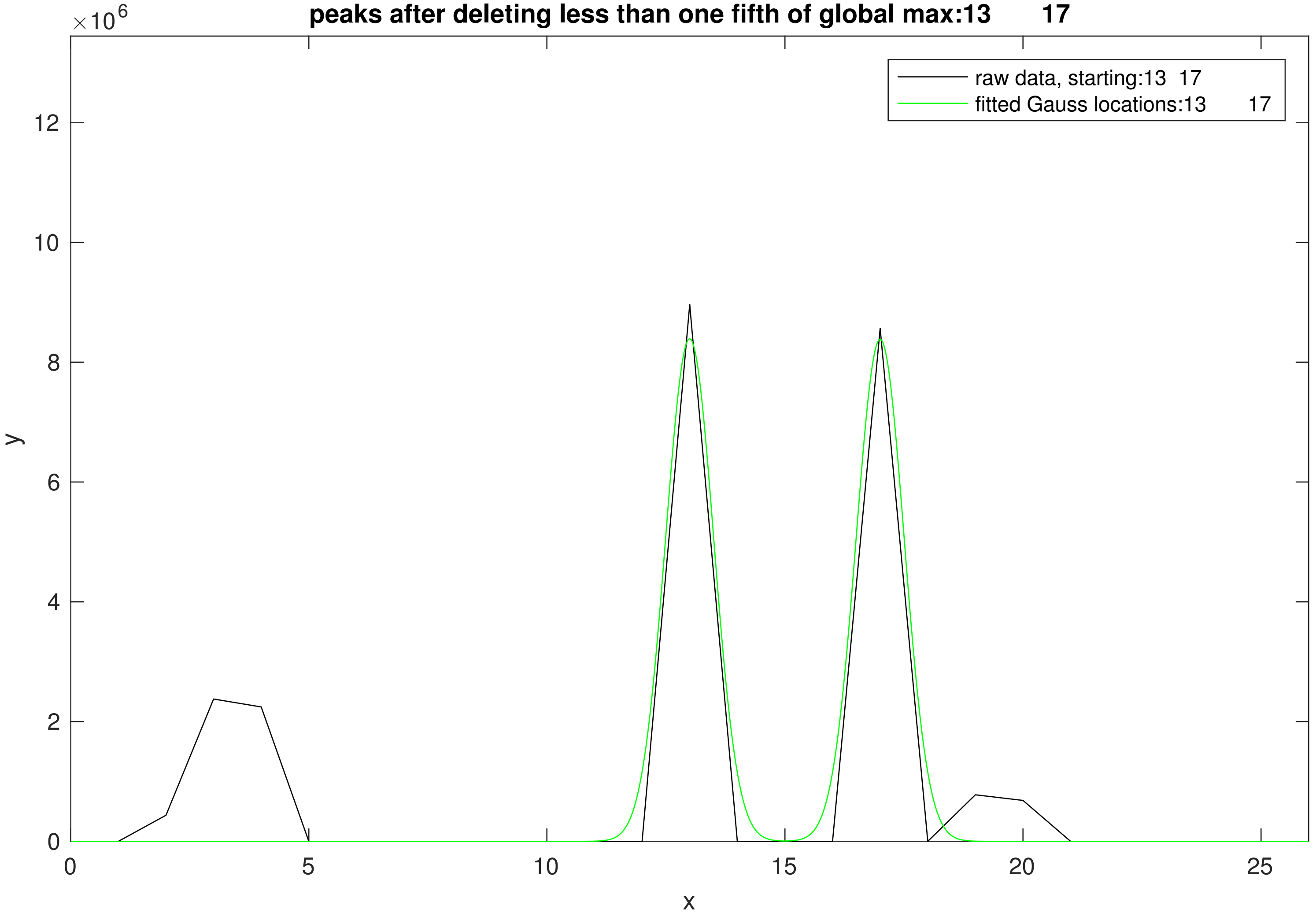

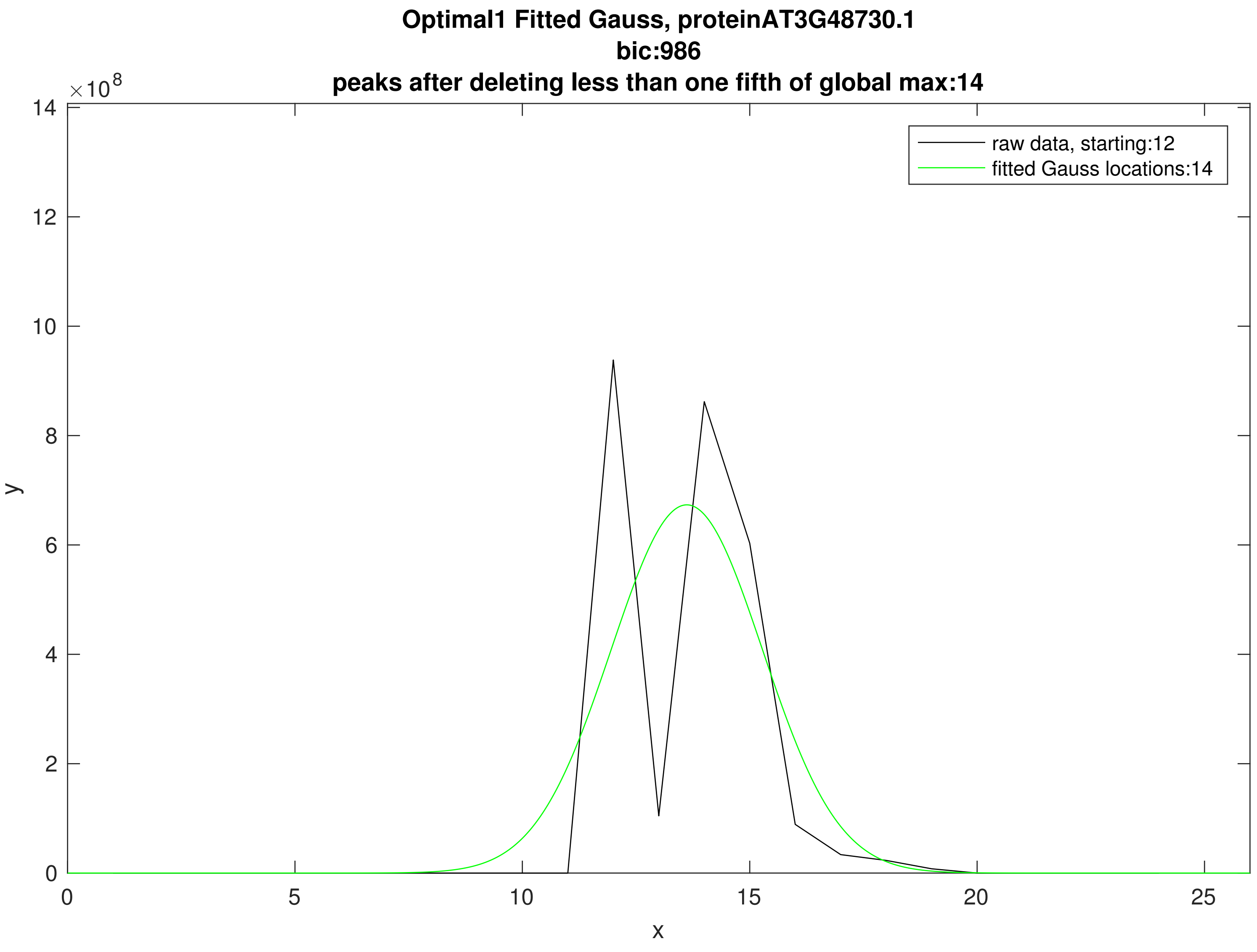
Optimal1 Fitted Gauss, proteinAT3G48730.1
bic:986
peaks after deleting less than one fifth of global max:14
×10^8
raw data, starting:12
fitted Gauss locations:14
y
x
0
2
4
6
8
10
12
14
0
5
10
15
20
25

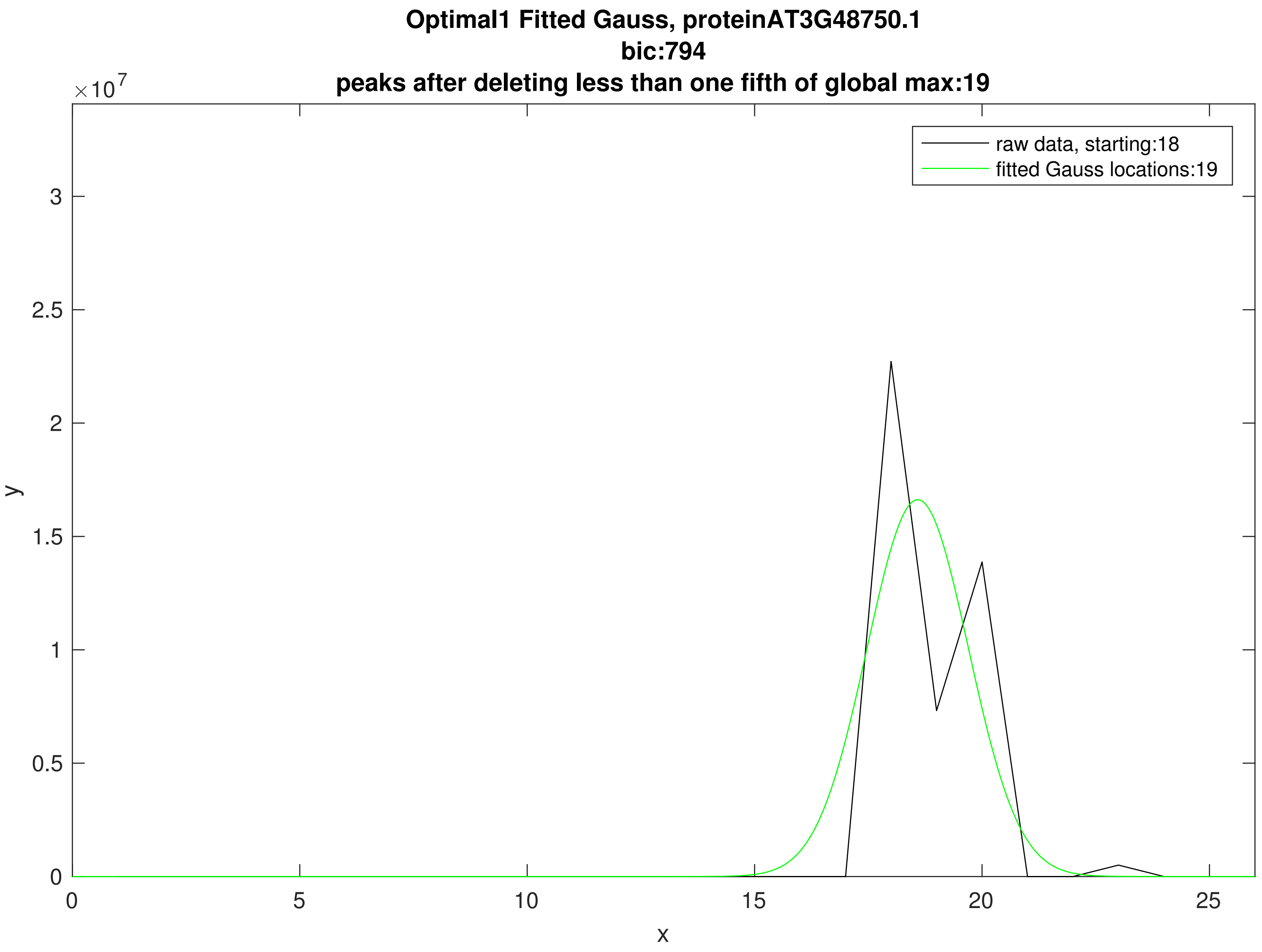

Optimal1 Fitted Gauss, proteinAT3G48750.1
bic:794
peaks after deleting less than one fifth of global max:19
×10^7
raw data, starting:18
fitted Gauss locations:19
y
x
0
0.5
1
1.5
2
2.5
3
0
5
10
15
20
25

**Optimal1 Fitted Gauss, proteinAT3G48890.1**
**bic:762**
**peaks after deleting less than one fifth of global max:16**

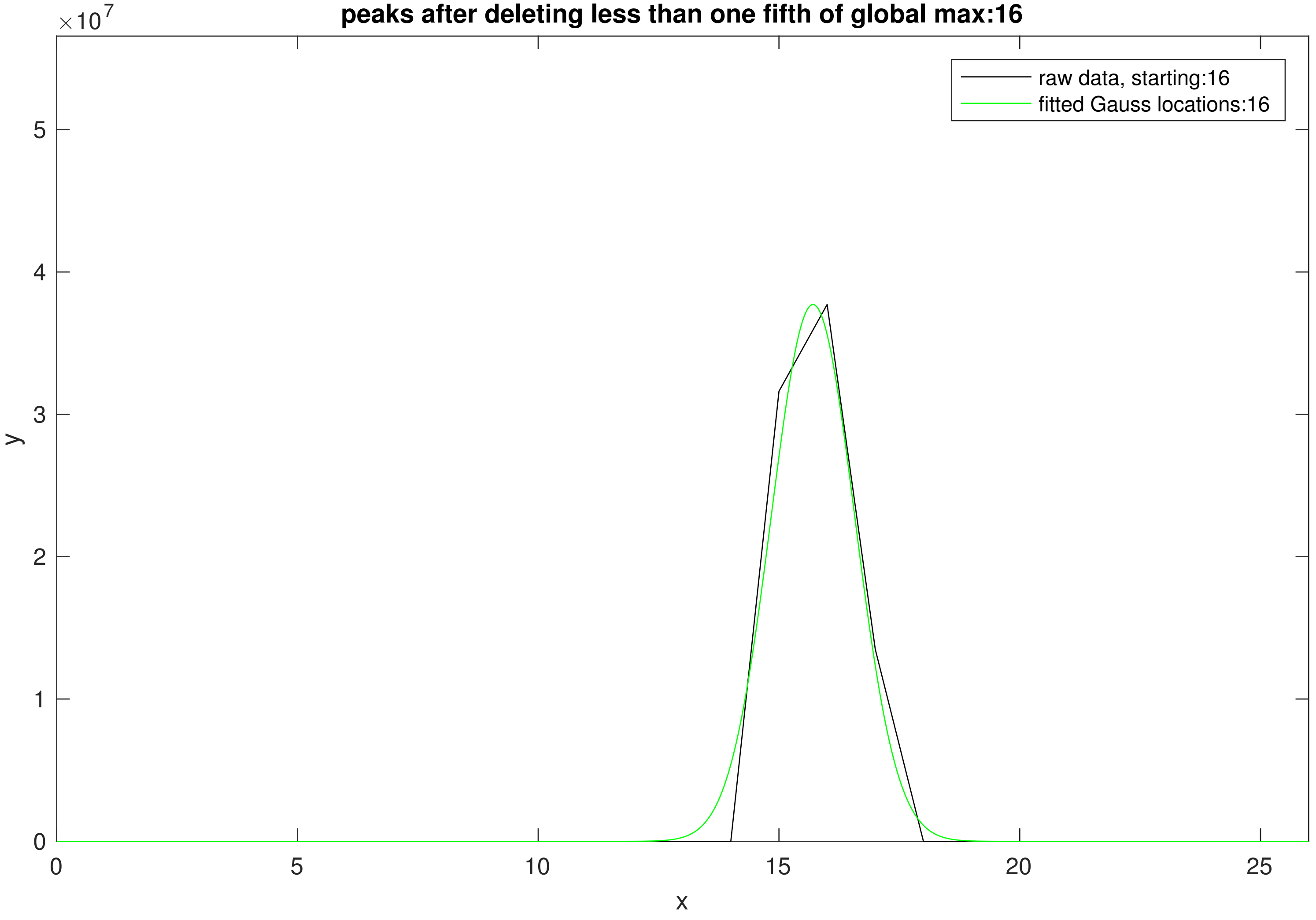

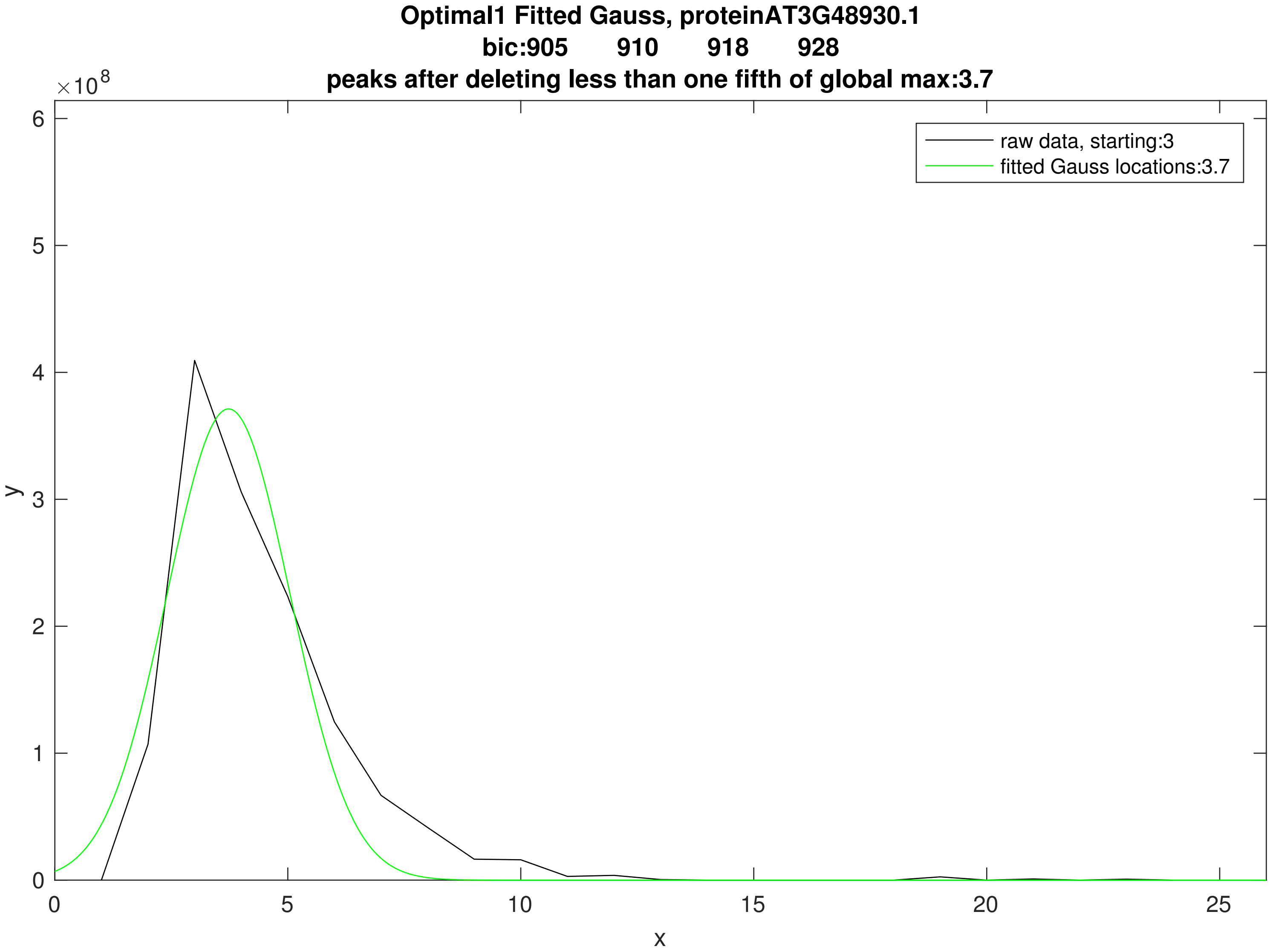
Optimal1 Fitted Gauss, proteinAT3G48930.1
bic:905 910 918 928
peaks after deleting less than one fifth of global max:3.7
×10^8
raw data, starting:3
fitted Gauss locations:3.7
y
x
0
1
2
3
4
5
6
0
5
10
15
20
25

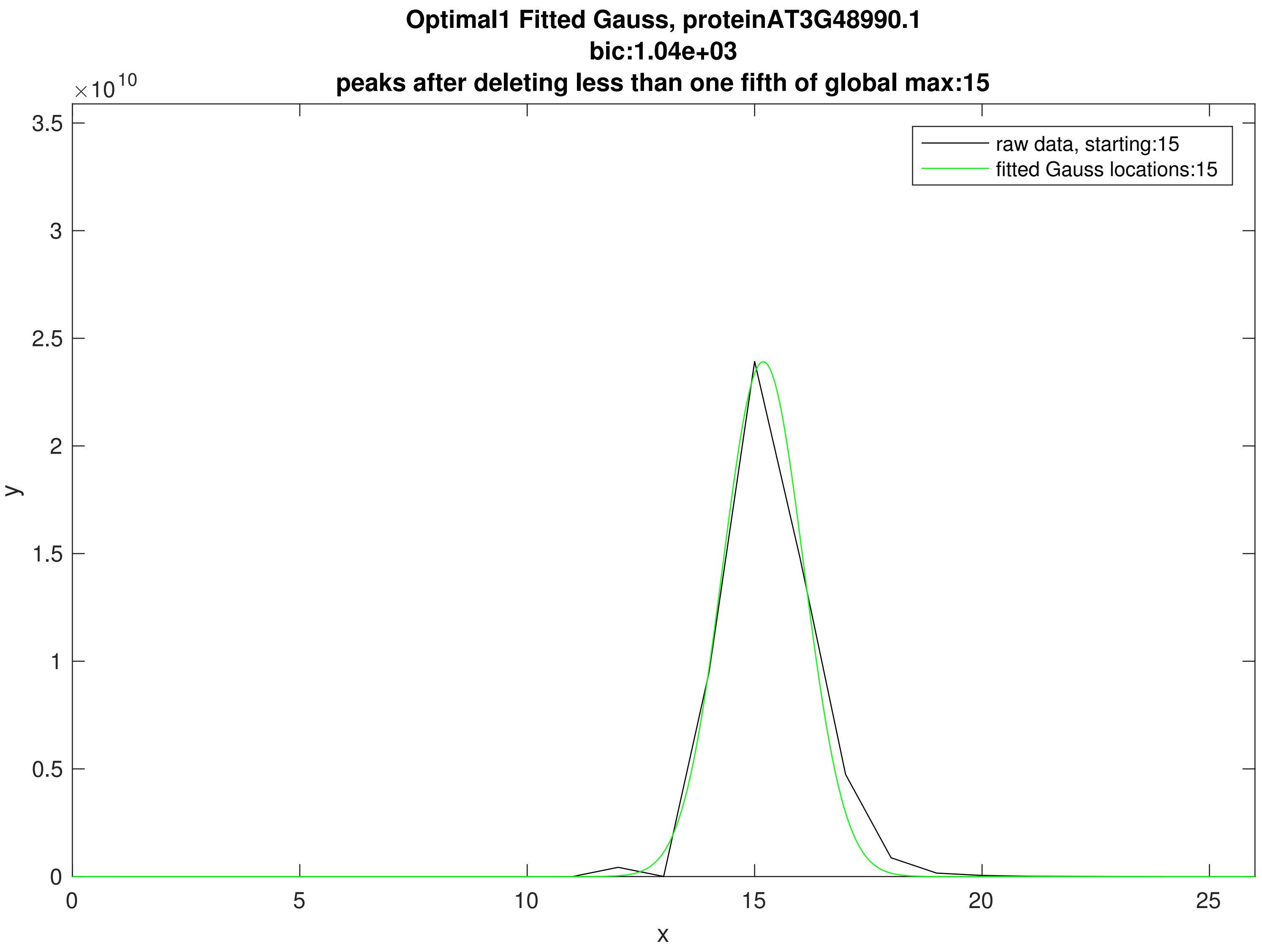

Optimal1 Fitted Gauss, proteinAT3G48990.1
bic:1.04e+03
peaks after deleting less than one fifth of global max:15
×10^10
3.5
3
2.5
2
1.5
1
0.5
0
y
0
5
10
15
20
25
x
raw data, starting:15
fitted Gauss locations:15

**Optimal1 Fitted Gauss, proteinAT3G49010.3;AT3G49010.2;AT3G49010.1;AT3G49010.4;AT3G49010.5**
**bic:913 923**
**peaks after deleting less than one fifth of global max:3.3**

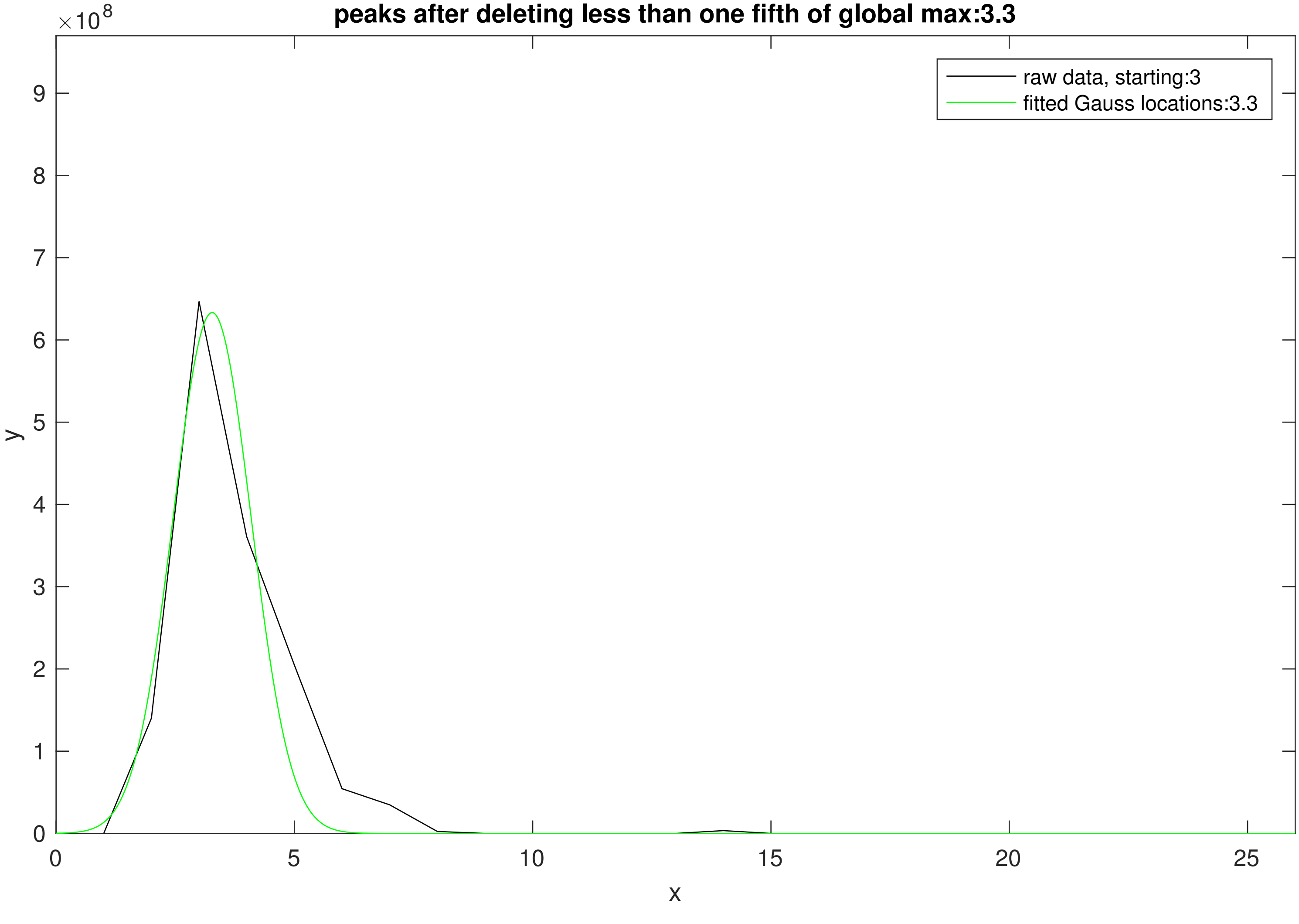

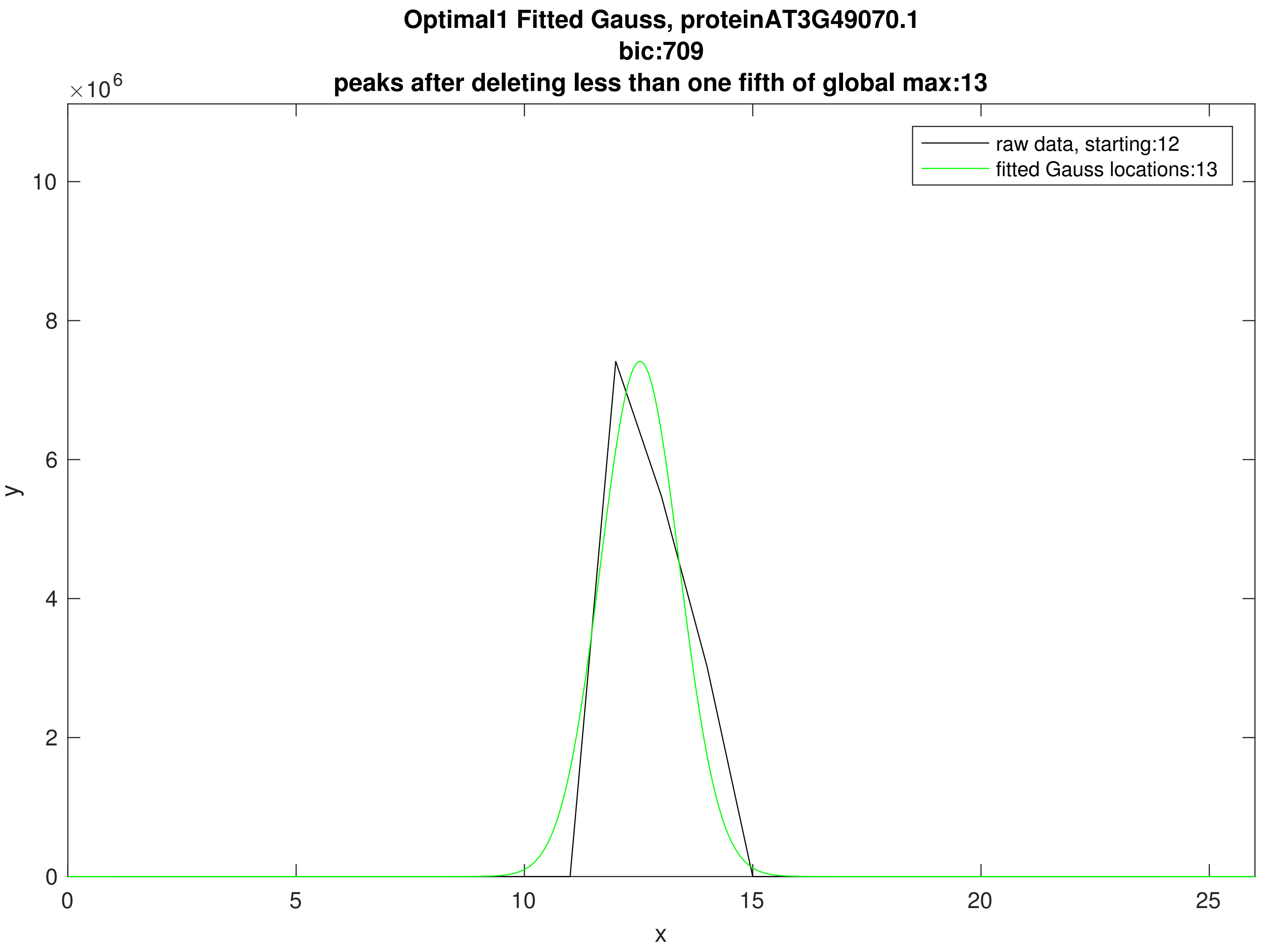
Optimal1 Fitted Gauss, proteinAT3G49070.1
bic:709
peaks after deleting less than one fifth of global max:13
×10^6
raw data, starting:12
fitted Gauss locations:13
y
x
0
2
4
6
8
10
0
5
10
15
20
25

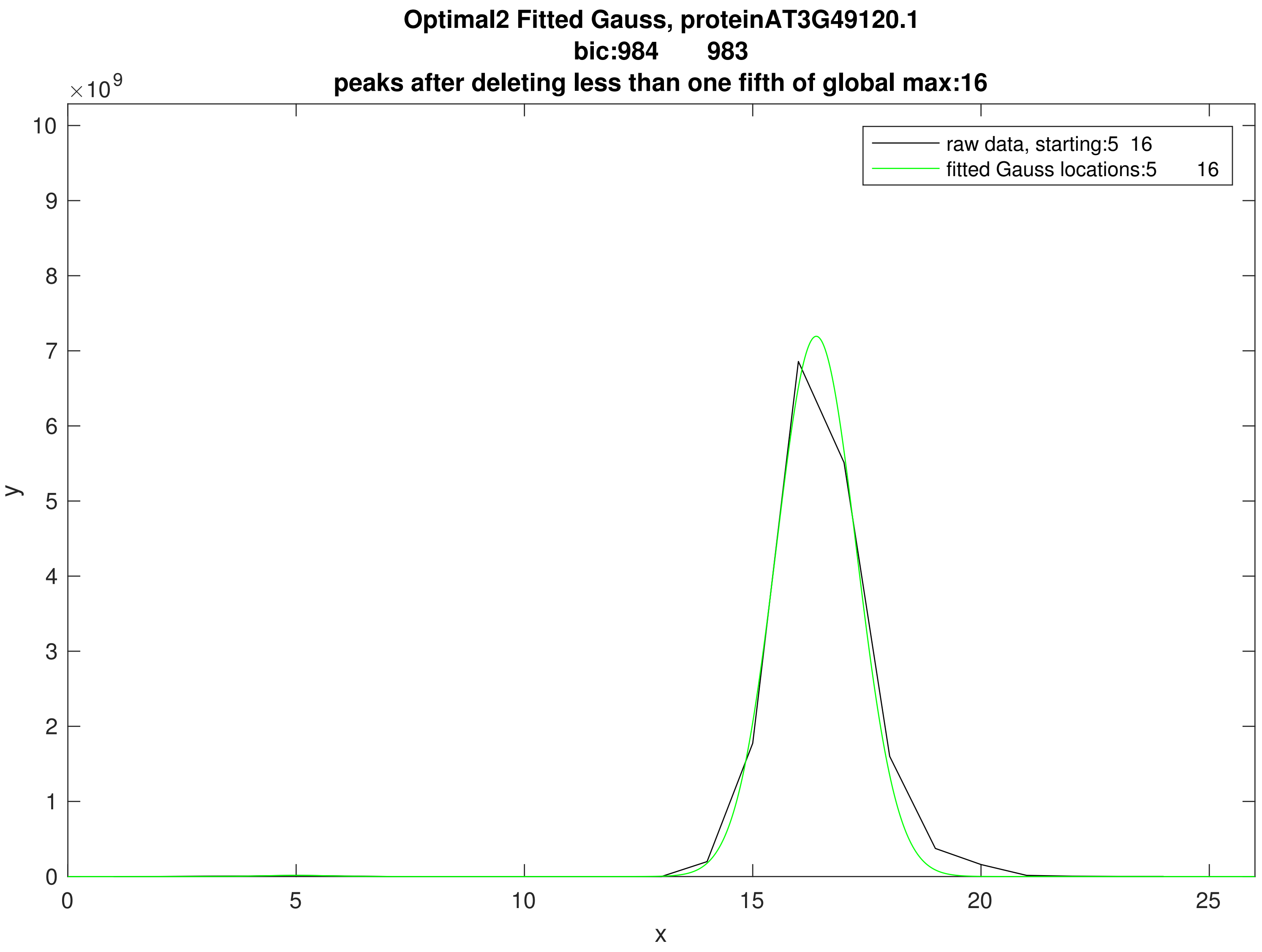

Optimal2 Fitted Gauss, proteinAT3G49120.1
bic:984 983
peaks after deleting less than one fifth of global max:16
×10^9
raw data, starting:5 16
fitted Gauss locations:5 16
y
x
0
1
2
3
4
5
6
7
8
9
10
0
5
10
15
20
25

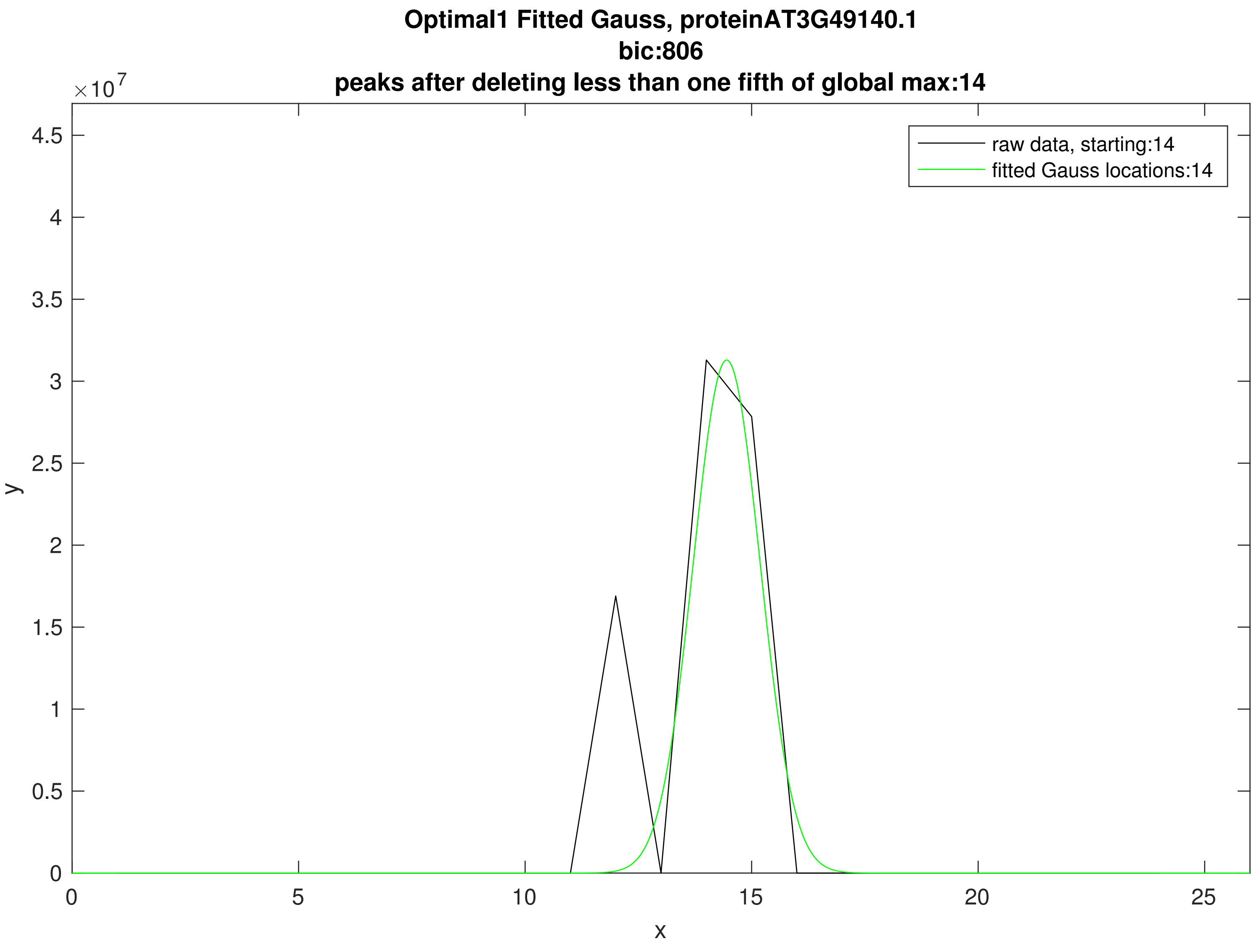
Optimal1 Fitted Gauss, proteinAT3G49140.1
bic:806
peaks after deleting less than one fifth of global max:14
×10^7
4.5
4
3.5
3
2.5
2
1.5
1
0.5
0
y
0
5
10
15
20
25
x
raw data, starting:14
fitted Gauss locations:14

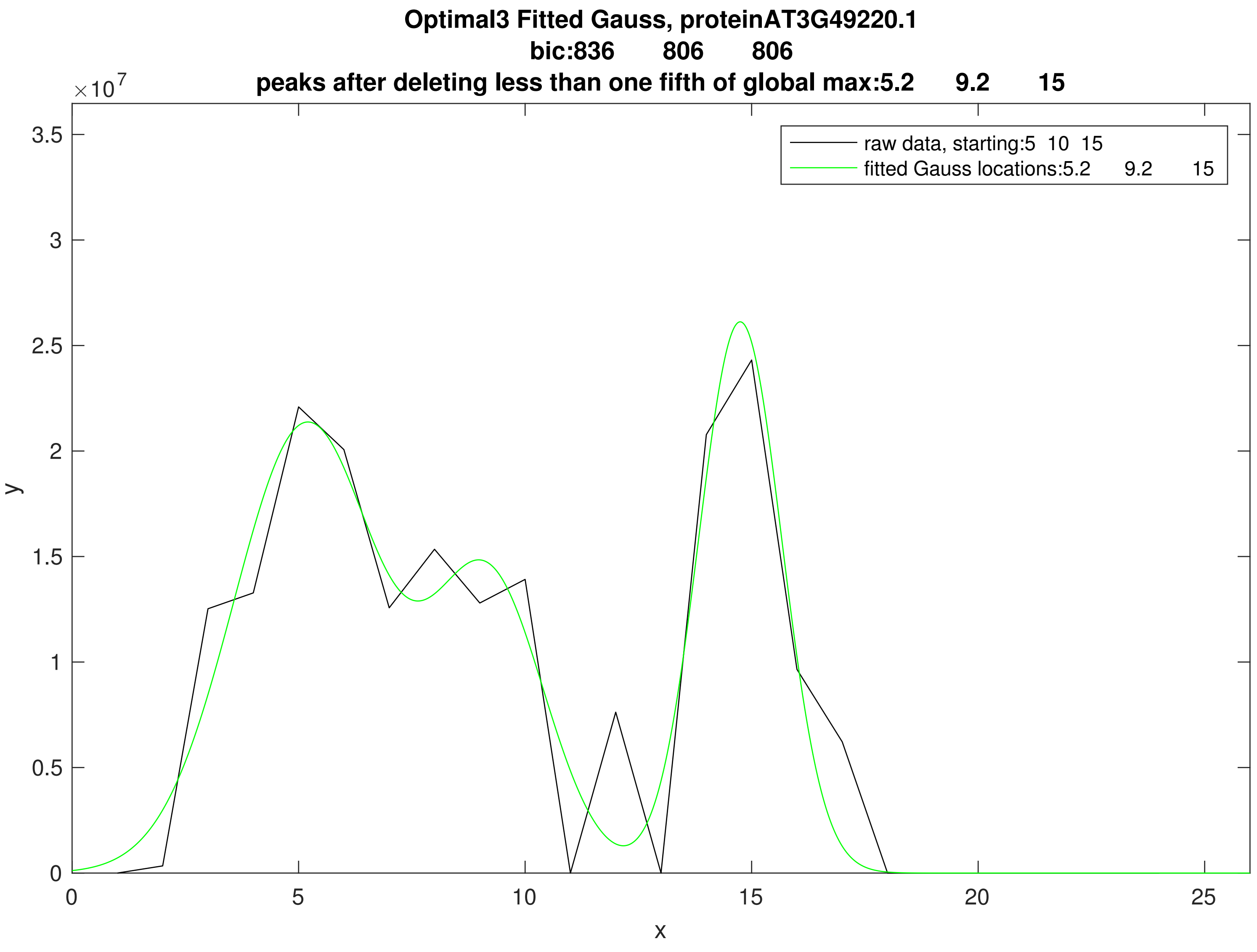
Optimal3 Fitted Gauss, proteinAT3G49220.1
bic:836 806 806
peaks after deleting less than one fifth of global max:5.2 9.2 15
×10^7
3.5
3
2.5
2
1.5
1
0.5
0
y
raw data, starting:5 10 15
fitted Gauss locations:5.2 9.2 15
0
5
10
15
20
25
x

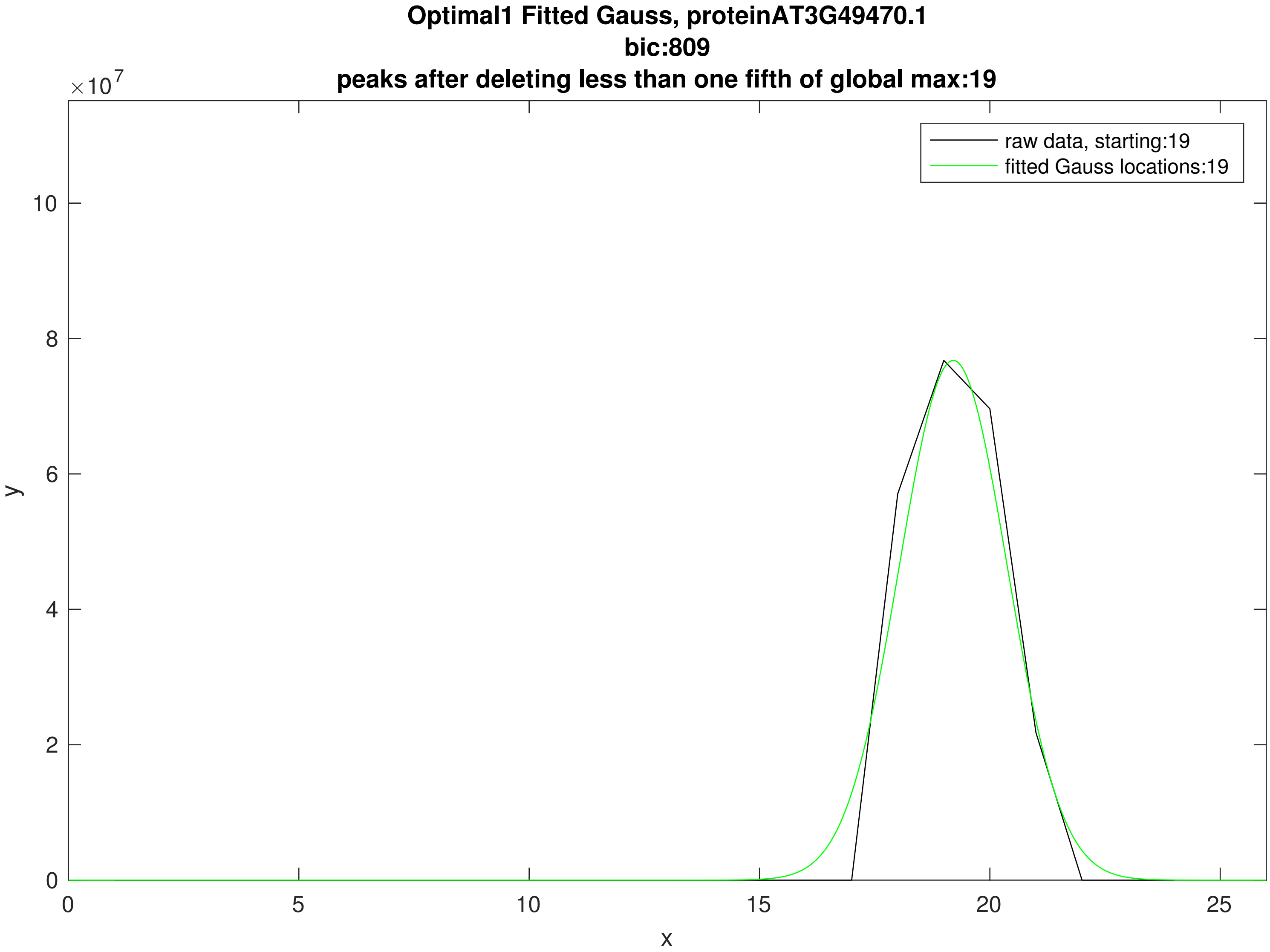
Optimal1 Fitted Gauss, proteinAT3G49470.1
bic:809
peaks after deleting less than one fifth of global max:19
×10^7
raw data, starting:19
fitted Gauss locations:19
10
8
6
4
2
0
y
0
5
10
15
20
25
x

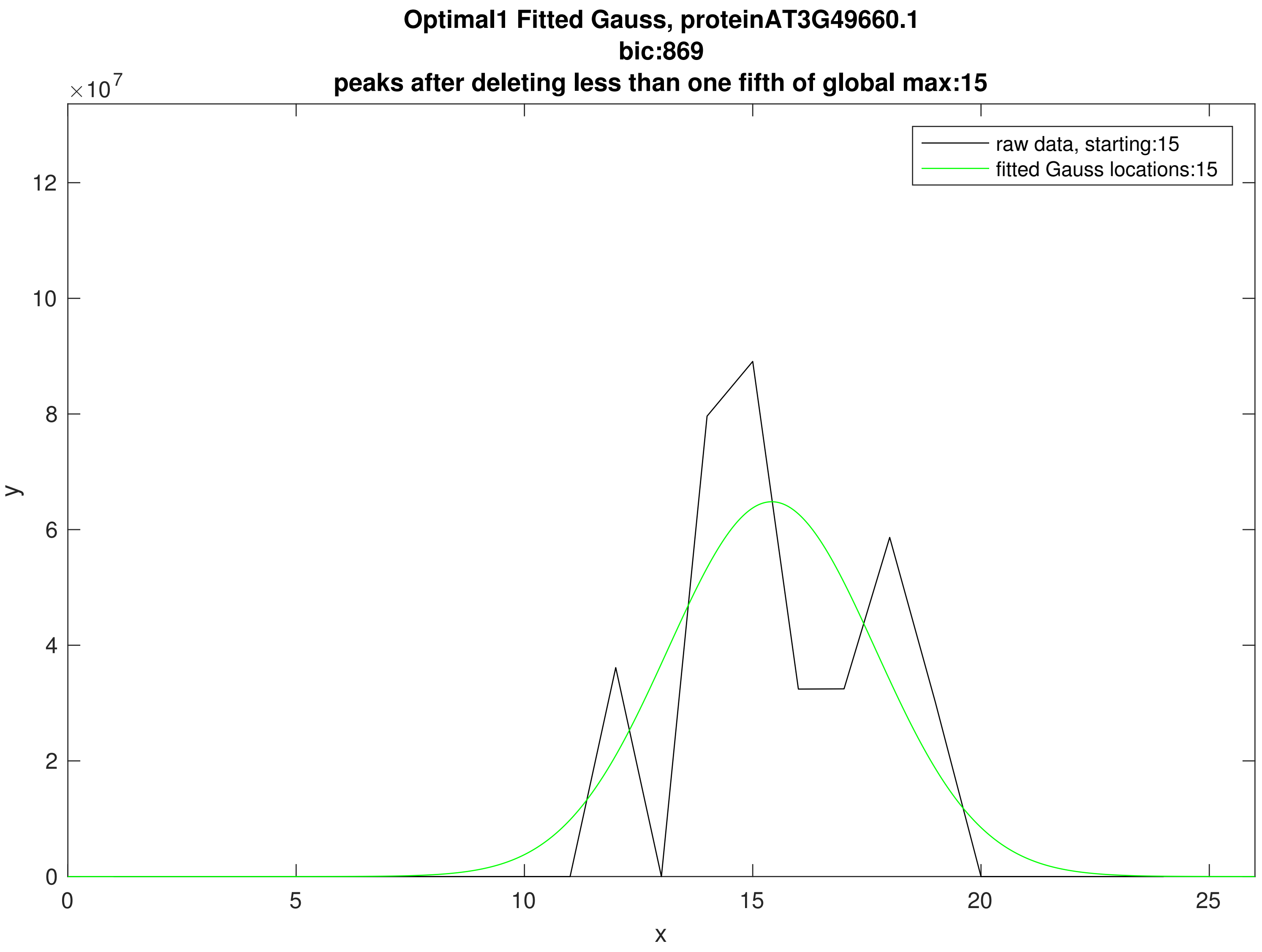
Optimal1 Fitted Gauss, proteinAT3G49660.1
bic:869
peaks after deleting less than one fifth of global max:15
×10^7
raw data, starting:15
fitted Gauss locations:15
y
x
0
2
4
6
8
10
12
0
5
10
15
20
25

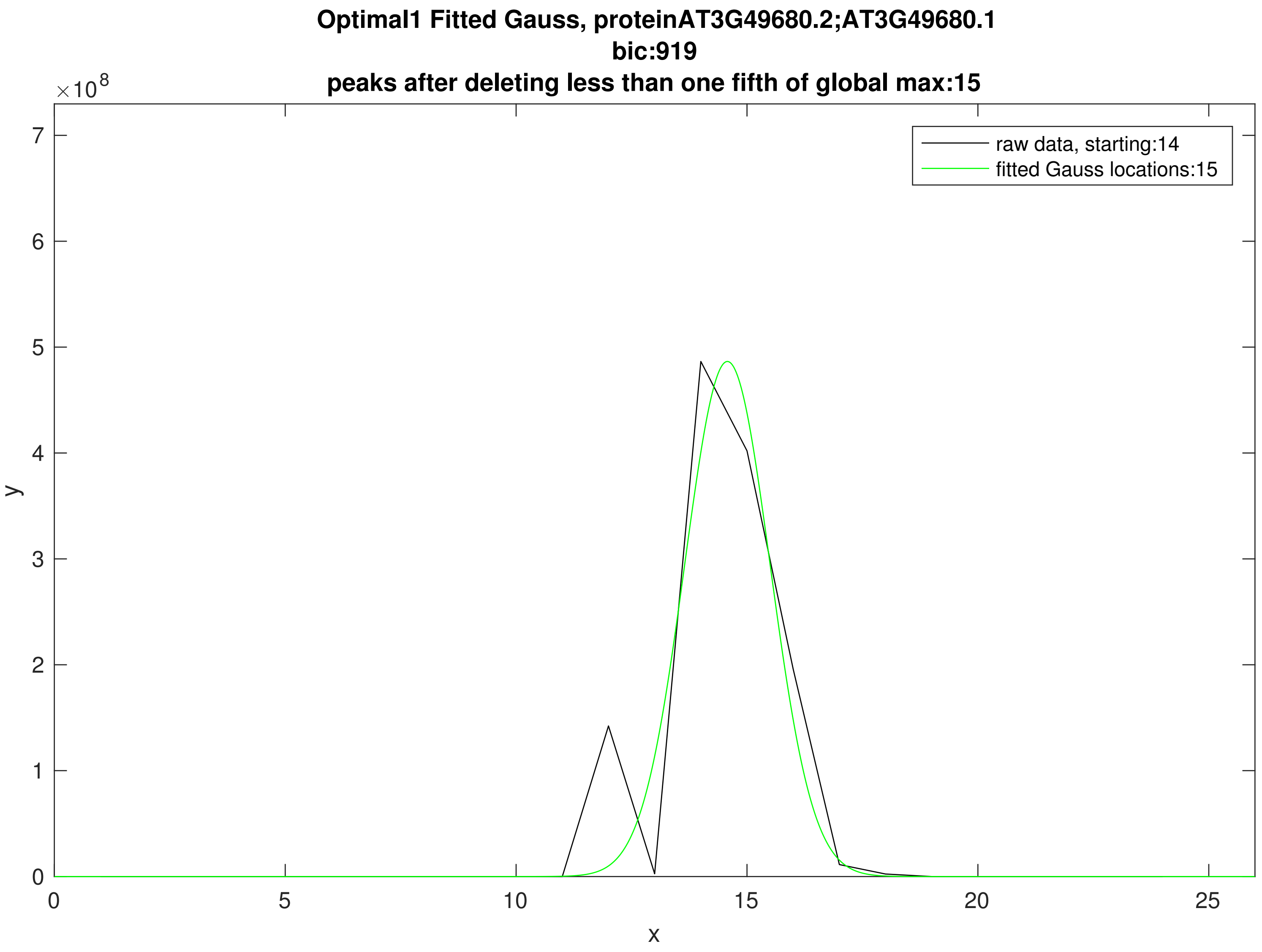
Optimal1 Fitted Gauss, proteinAT3G49680.2;AT3G49680.1
bic:919
peaks after deleting less than one fifth of global max:15
×10^8
raw data, starting:14
fitted Gauss locations:15
y
x
0
1
2
3
4
5
6
7
0
5
10
15
20
25

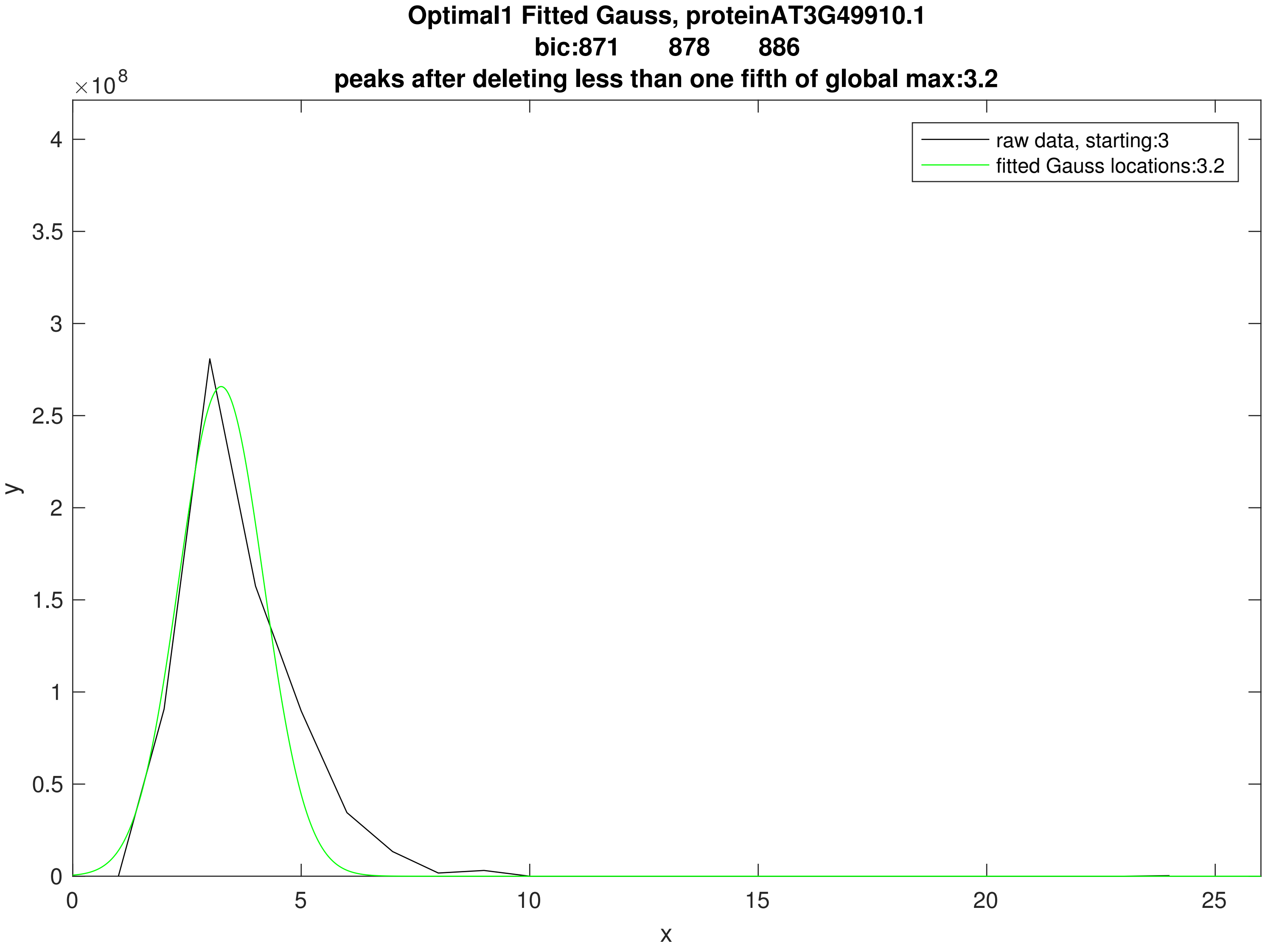
Optimal1 Fitted Gauss, proteinAT3G49910.1
bic:871 878 886
peaks after deleting less than one fifth of global max:3.2
×10^8
raw data, starting:3
fitted Gauss locations:3.2
y
x
0
0.5
1
1.5
2
2.5
3
3.5
4
5
10
15
20
25

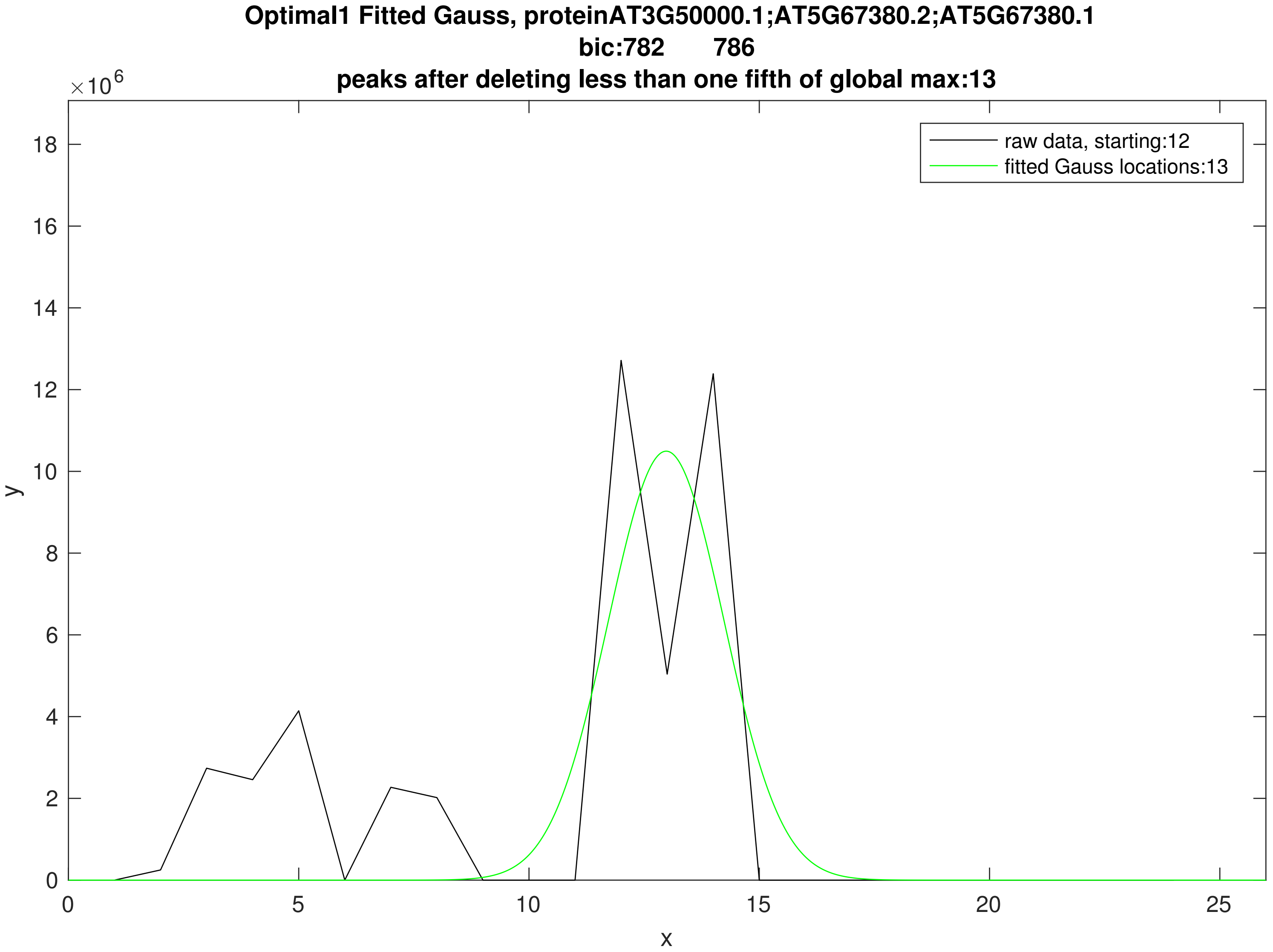

Optimal1 Fitted Gauss, proteinAT3G50000.1;AT5G67380.2;AT5G67380.1
bic:782 786
peaks after deleting less than one fifth of global max:13
$\times 10^6$
raw data, starting:12
fitted Gauss locations:13
y
x
0
2
4
6
8
10
12
14
16
18
0
5
10
15
20
25

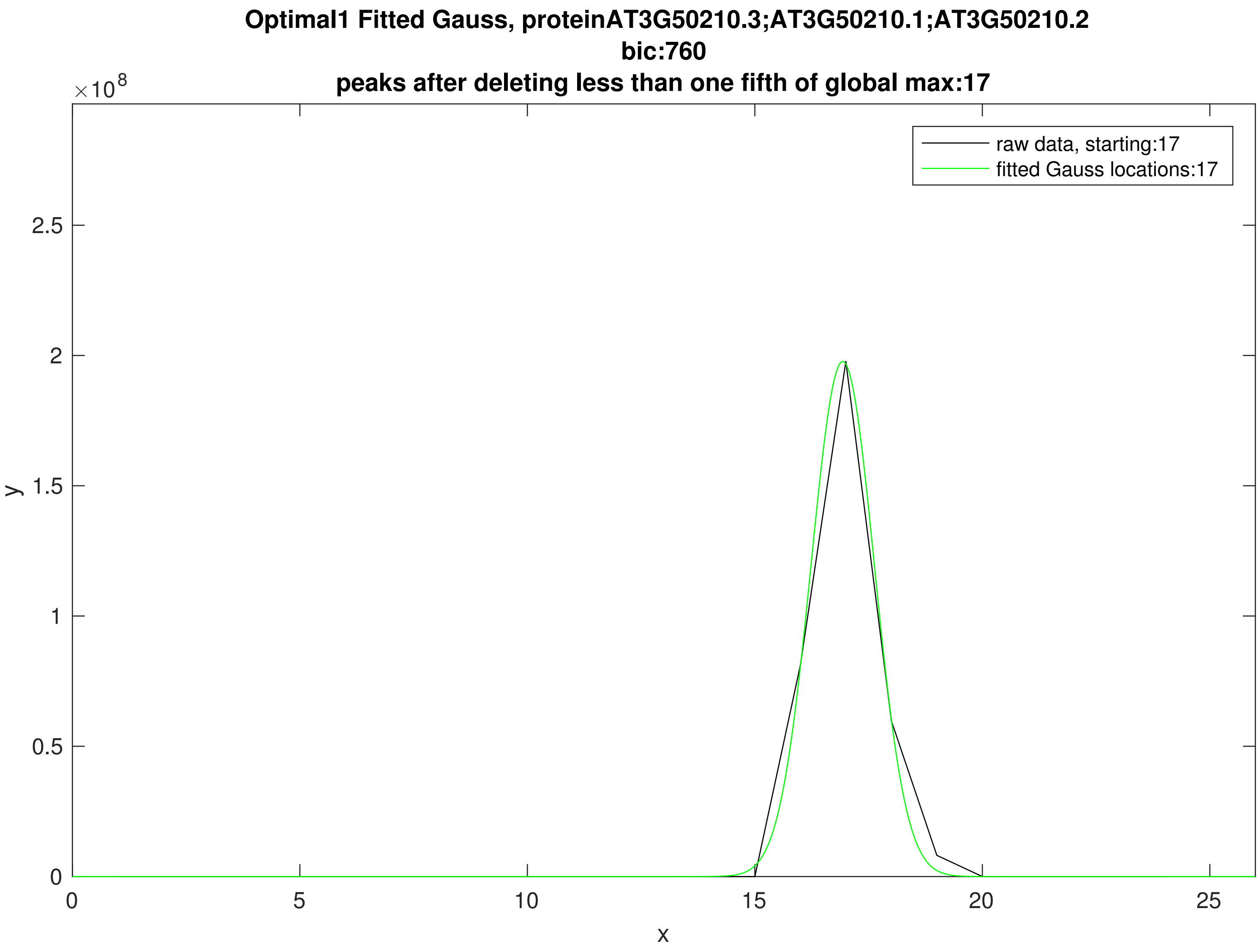
Optimal1 Fitted Gauss, proteinAT3G50210.3;AT3G50210.1;AT3G50210.2
bic:760
peaks after deleting less than one fifth of global max:17
×10^8
raw data, starting:17
fitted Gauss locations:17
y
x
0
0.5
1
1.5
2
2.5
0
5
10
15
20
25

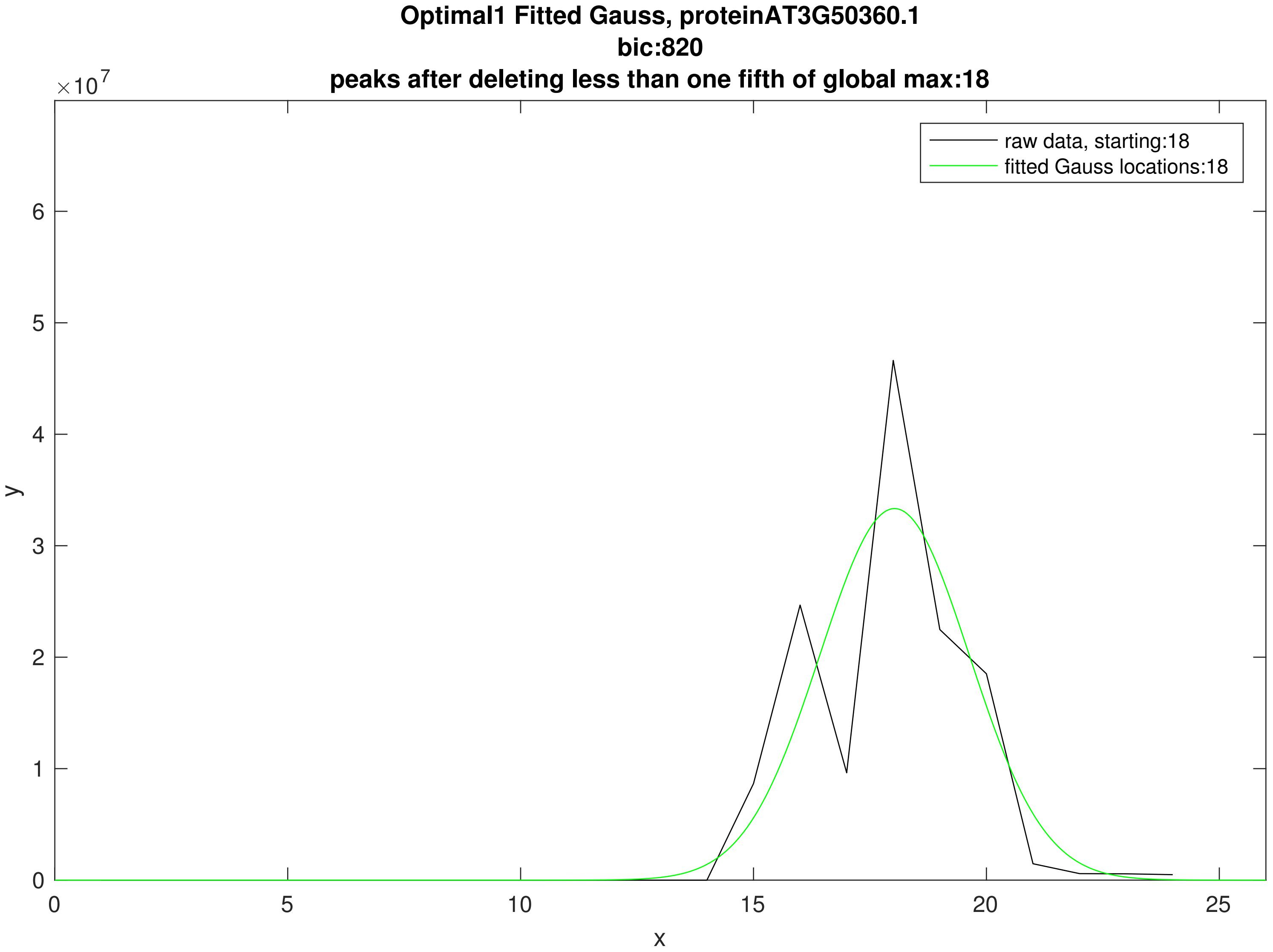
Optimal1 Fitted Gauss, proteinAT3G50360.1
bic:820
peaks after deleting less than one fifth of global max:18
×10^7
raw data, starting:18
fitted Gauss locations:18
y
0
1
2
3
4
5
6
0
5
10
15
20
25
x

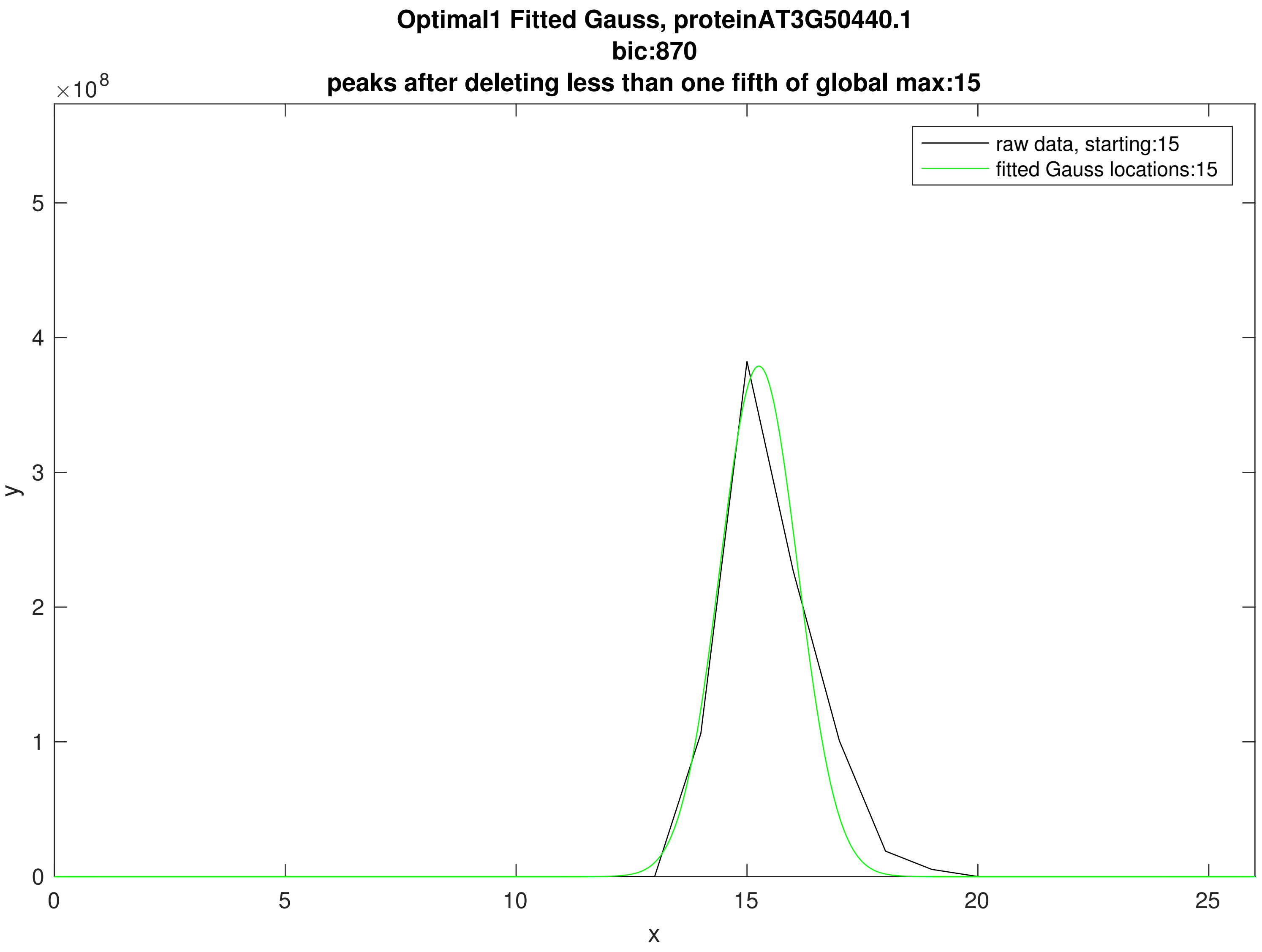
Optimal1 Fitted Gauss, proteinAT3G50440.1
bic:870
peaks after deleting less than one fifth of global max:15
×10^8
raw data, starting:15
fitted Gauss locations:15
y
x
0
1
2
3
4
5
0
5
10
15
20
25

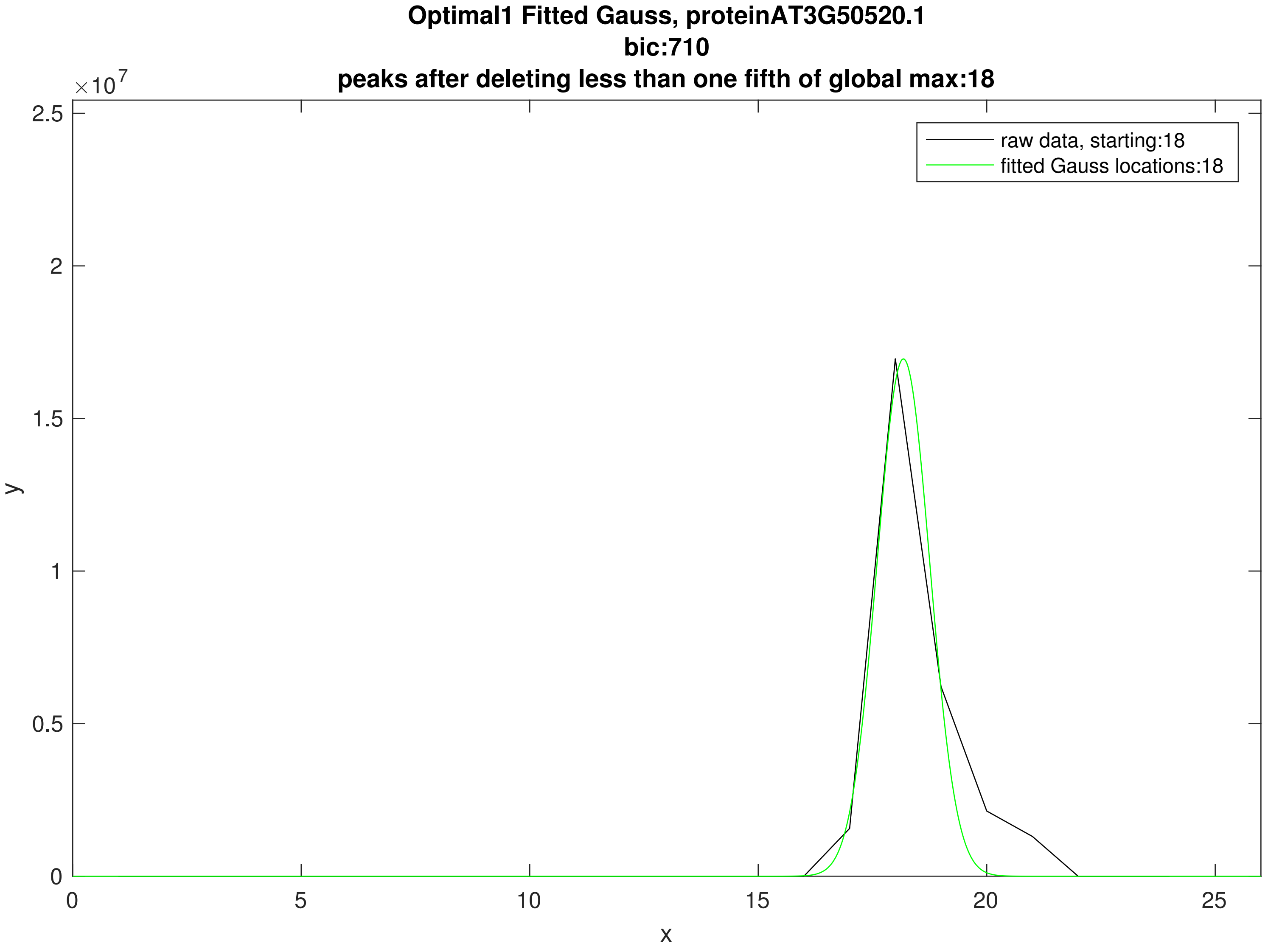
Optimal1 Fitted Gauss, proteinAT3G50520.1
bic:710
peaks after deleting less than one fifth of global max:18
×10^7
2.5
2
1.5
1
0.5
0
y
0
5
10
15
20
25
x
raw data, starting:18
fitted Gauss locations:18

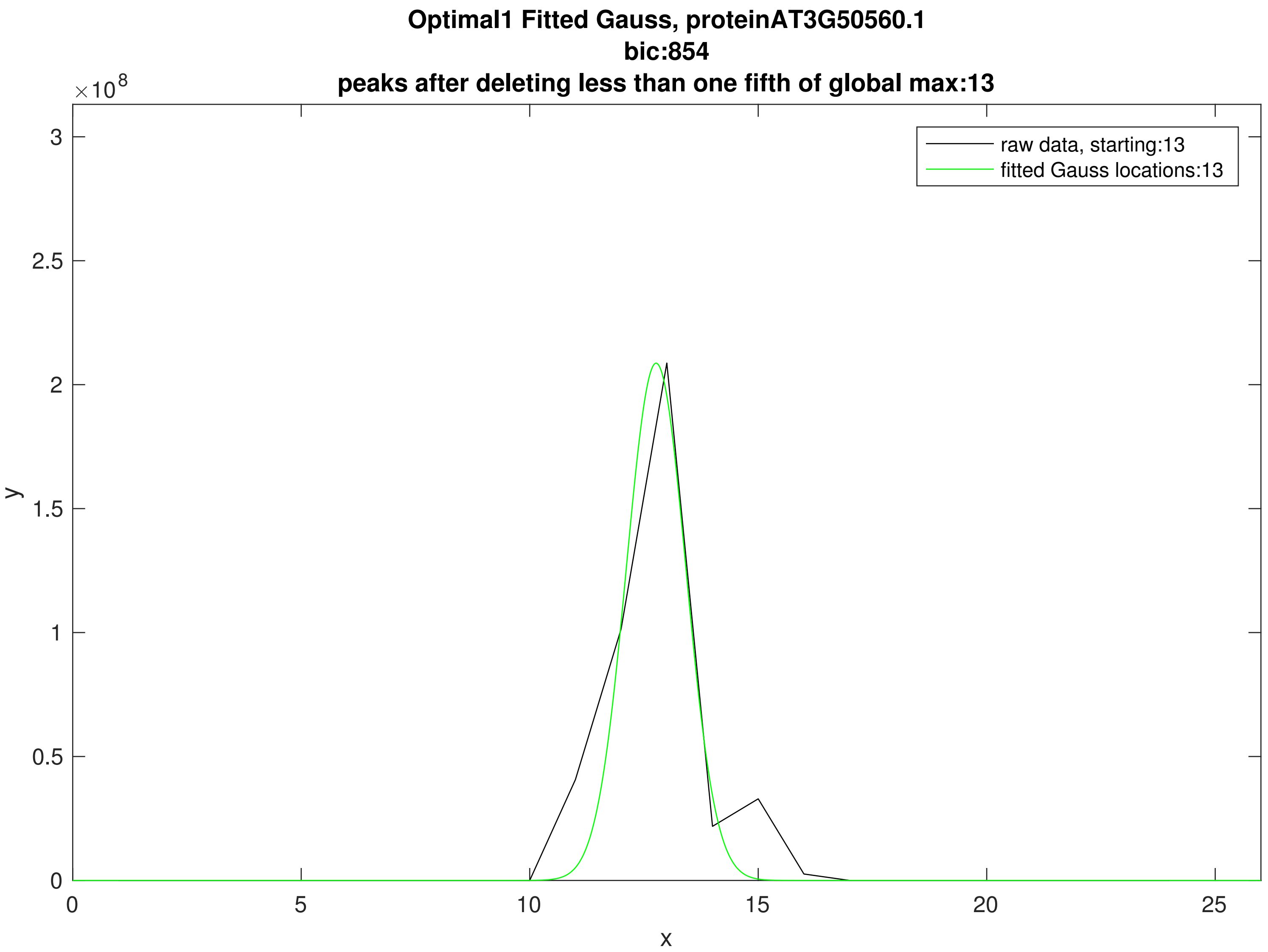

Optimal1 Fitted Gauss, proteinAT3G50560.1
bic:854
peaks after deleting less than one fifth of global max:13
×10^8
raw data, starting:13
fitted Gauss locations:13
y
x
0
0.5
1
1.5
2
2.5
3
0
5
10
15
20
25

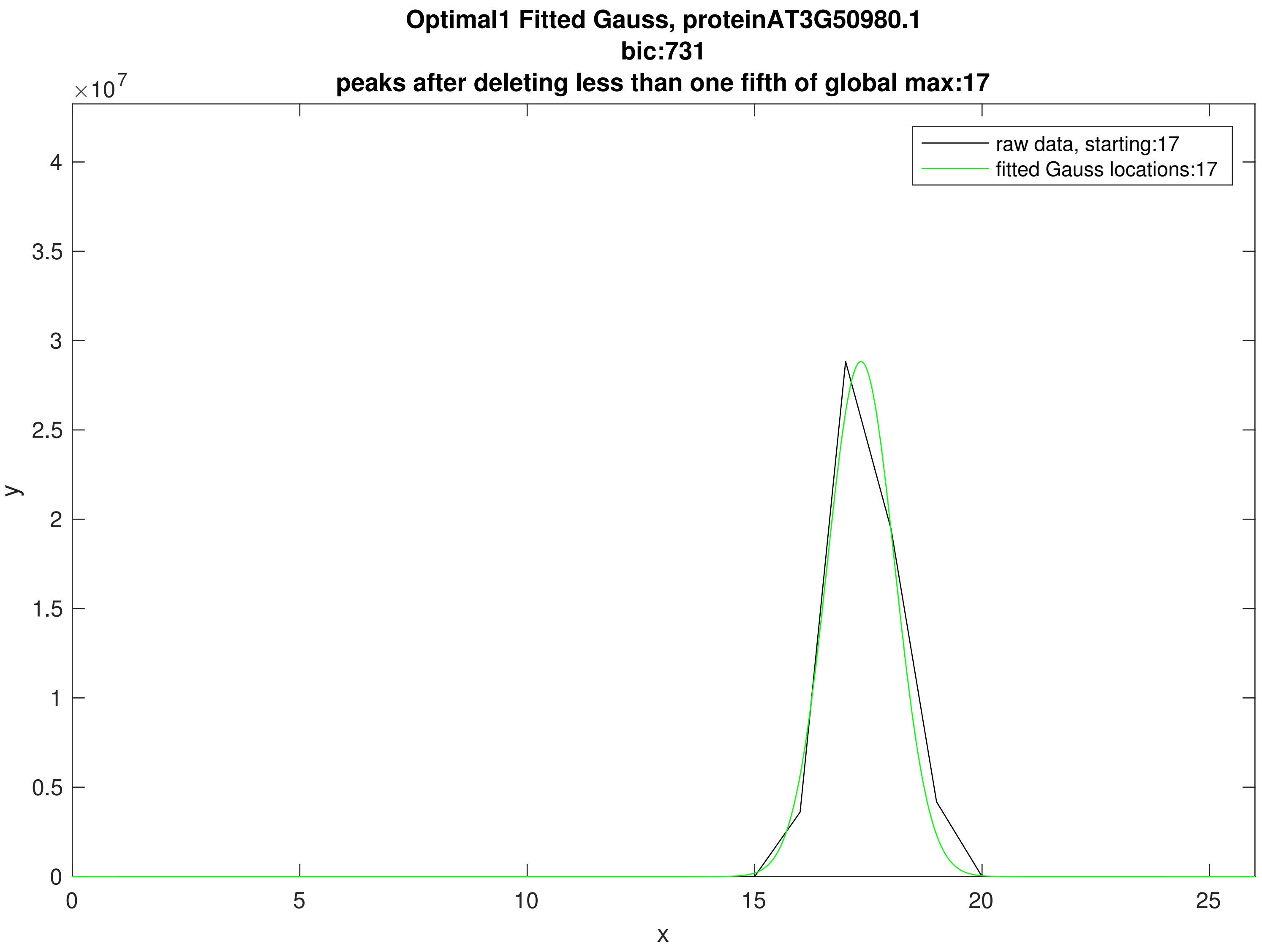
Optimal1 Fitted Gauss, proteinAT3G50980.1
bic:731
peaks after deleting less than one fifth of global max:17
×10^7
raw data, starting:17
fitted Gauss locations:17
y
x
0
0.5
1
1.5
2
2.5
3
3.5
4
0
5
10
15
20
25

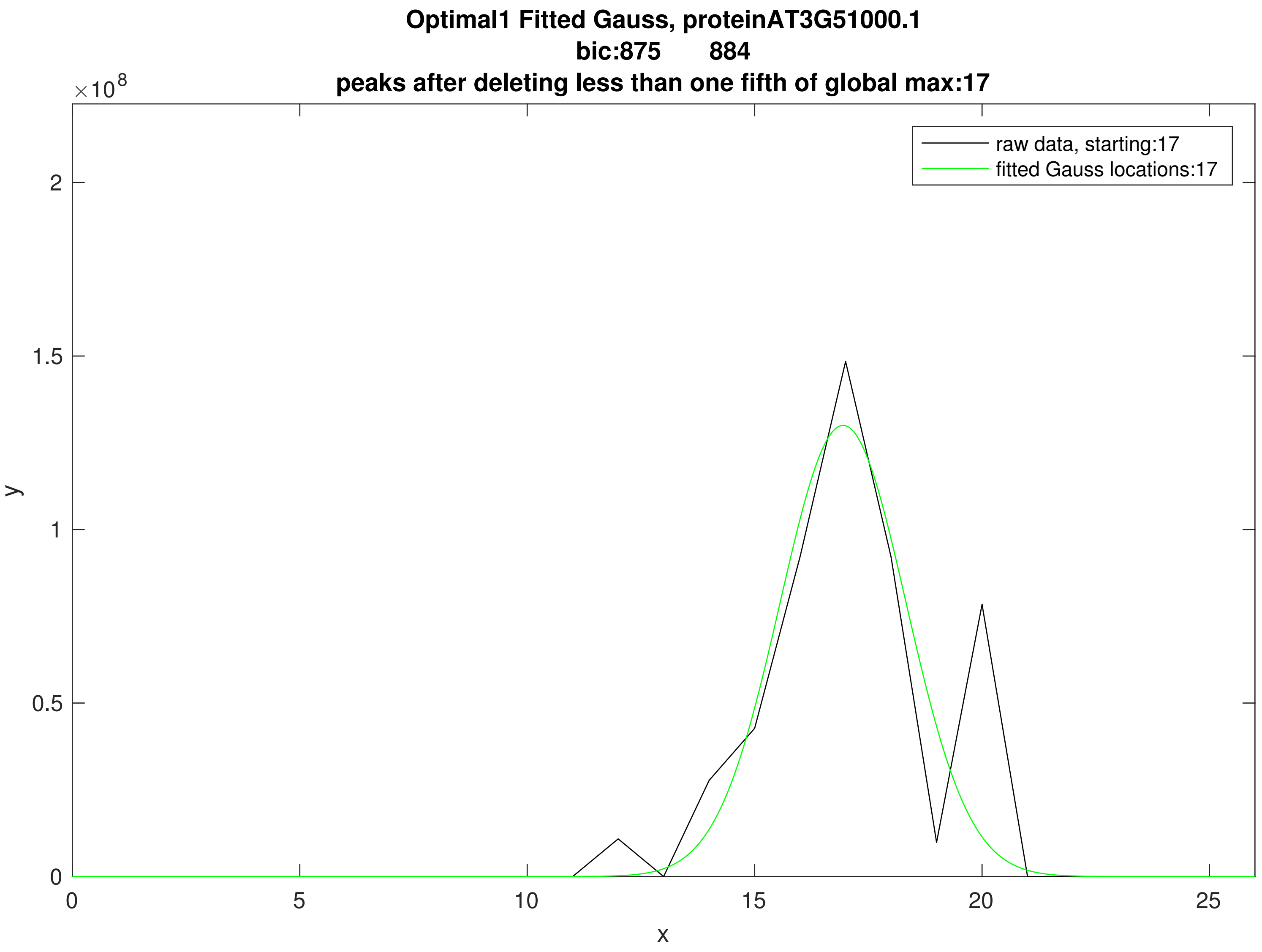
Optimal1 Fitted Gauss, proteinAT3G51000.1
bic:875 884
peaks after deleting less than one fifth of global max:17
×10^8
raw data, starting:17
fitted Gauss locations:17
y
x
0
0.5
1
1.5
2
0
5
10
15
20
25

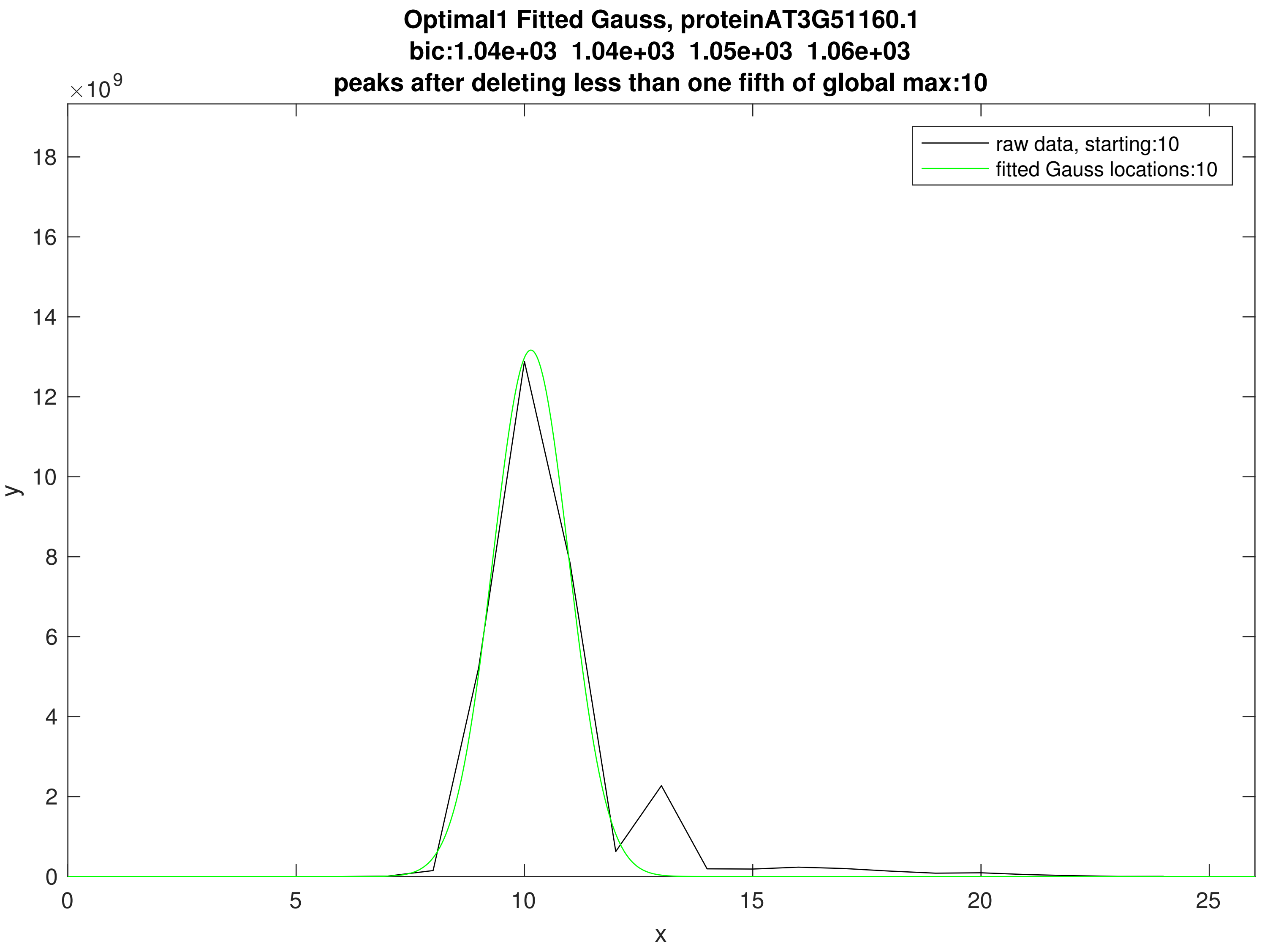
Optimal1 Fitted Gauss, proteinAT3G51160.1
bic:1.04e+03  1.04e+03  1.05e+03  1.06e+03
peaks after deleting less than one fifth of global max:10
×10^9
raw data, starting:10
fitted Gauss locations:10
y
x
0
2
4
6
8
10
12
14
16
18
0
5
10
15
20
25

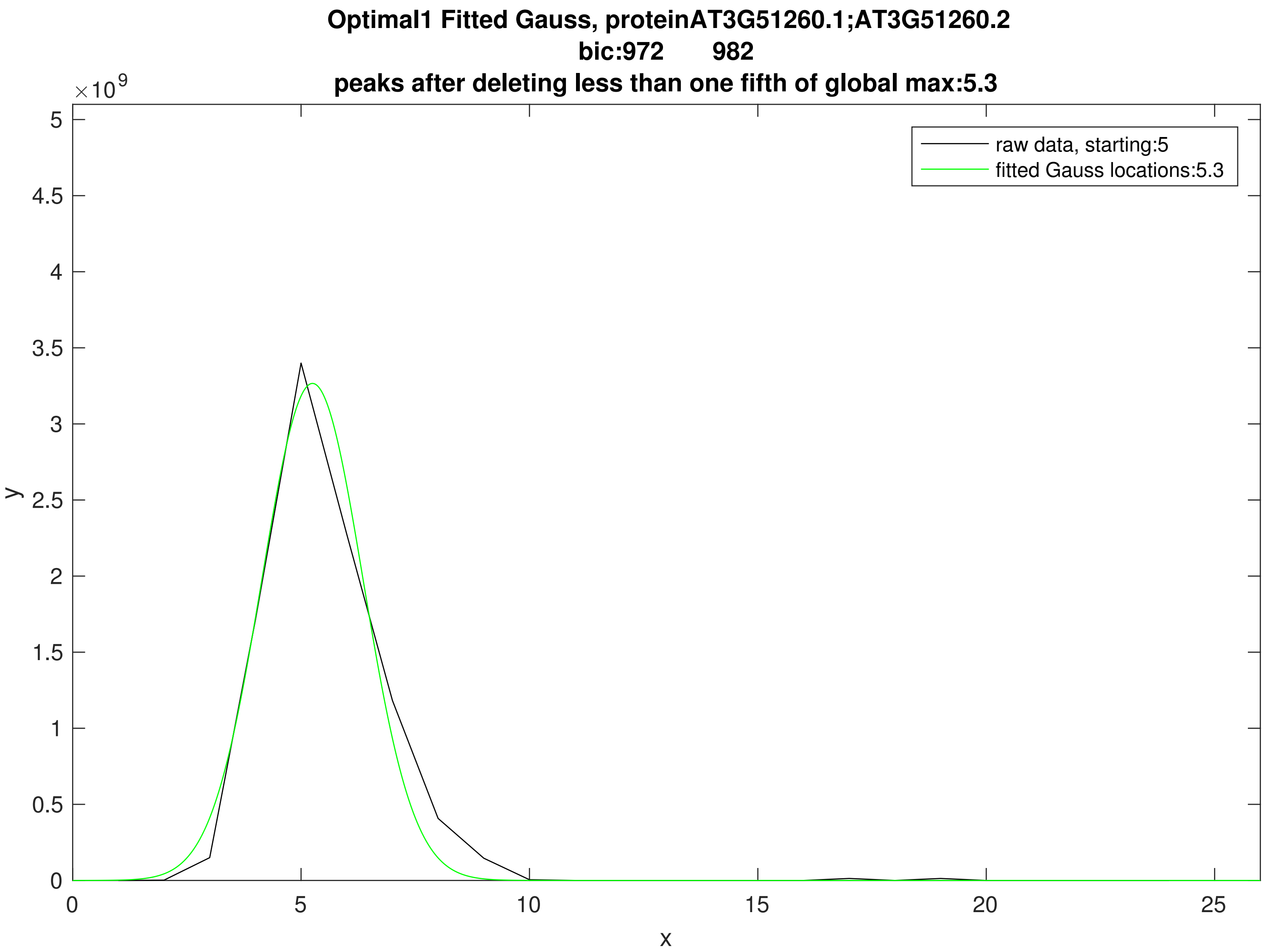
Optimal1 Fitted Gauss, proteinAT3G51260.1;AT3G51260.2
bic:972 982
peaks after deleting less than one fifth of global max:5.3
×10^9
5
4.5
4
3.5
3
2.5
2
1.5
1
0.5
0
y
0
5
10
15
20
25
x
raw data, starting:5
fitted Gauss locations:5.3

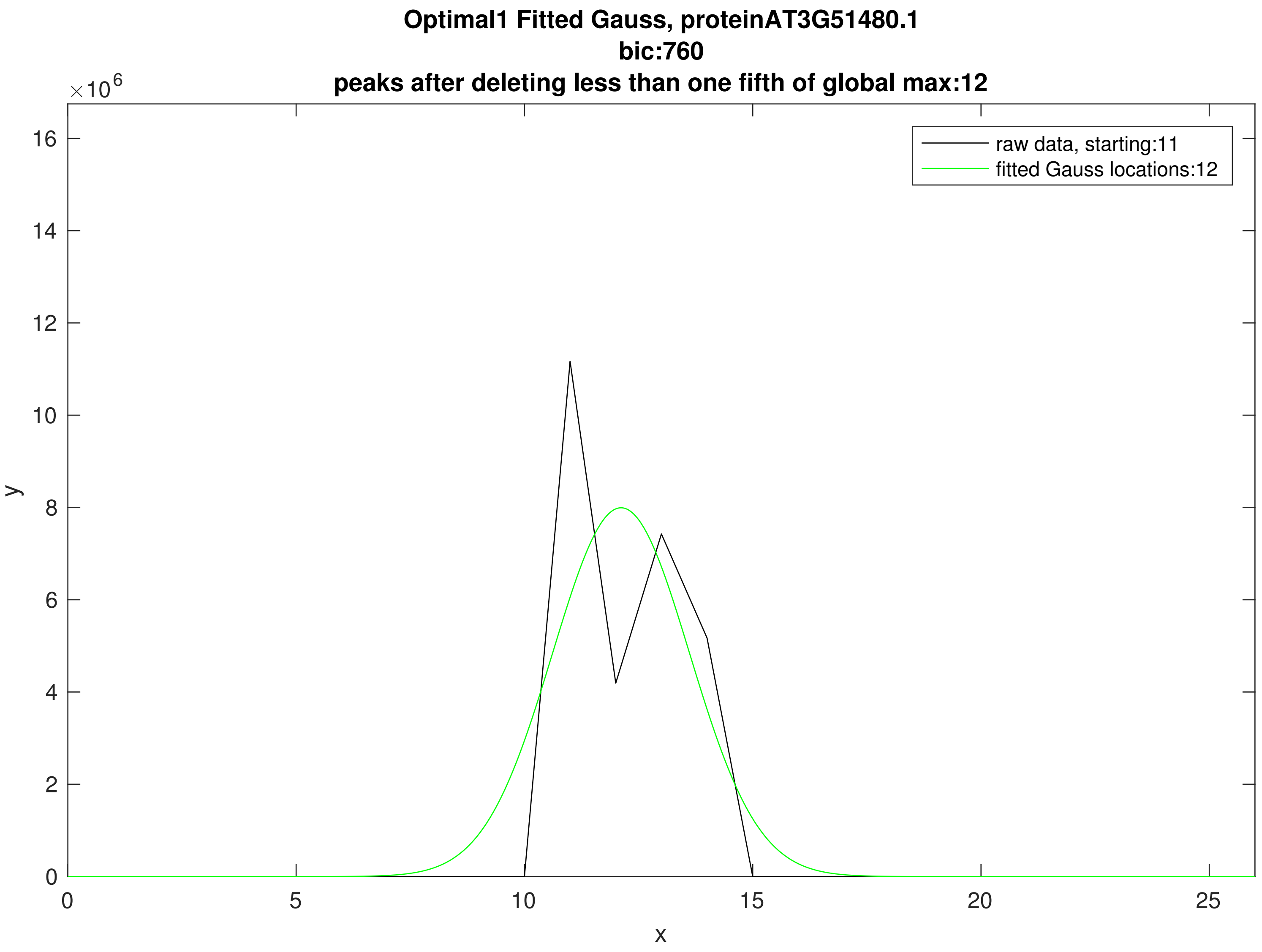
Optimal1 Fitted Gauss, proteinAT3G51480.1
bic:760
peaks after deleting less than one fifth of global max:12
×10^6
raw data, starting:11
fitted Gauss locations:12
16
14
12
10
8
6
4
2
0
y
0
5
10
15
20
25
x

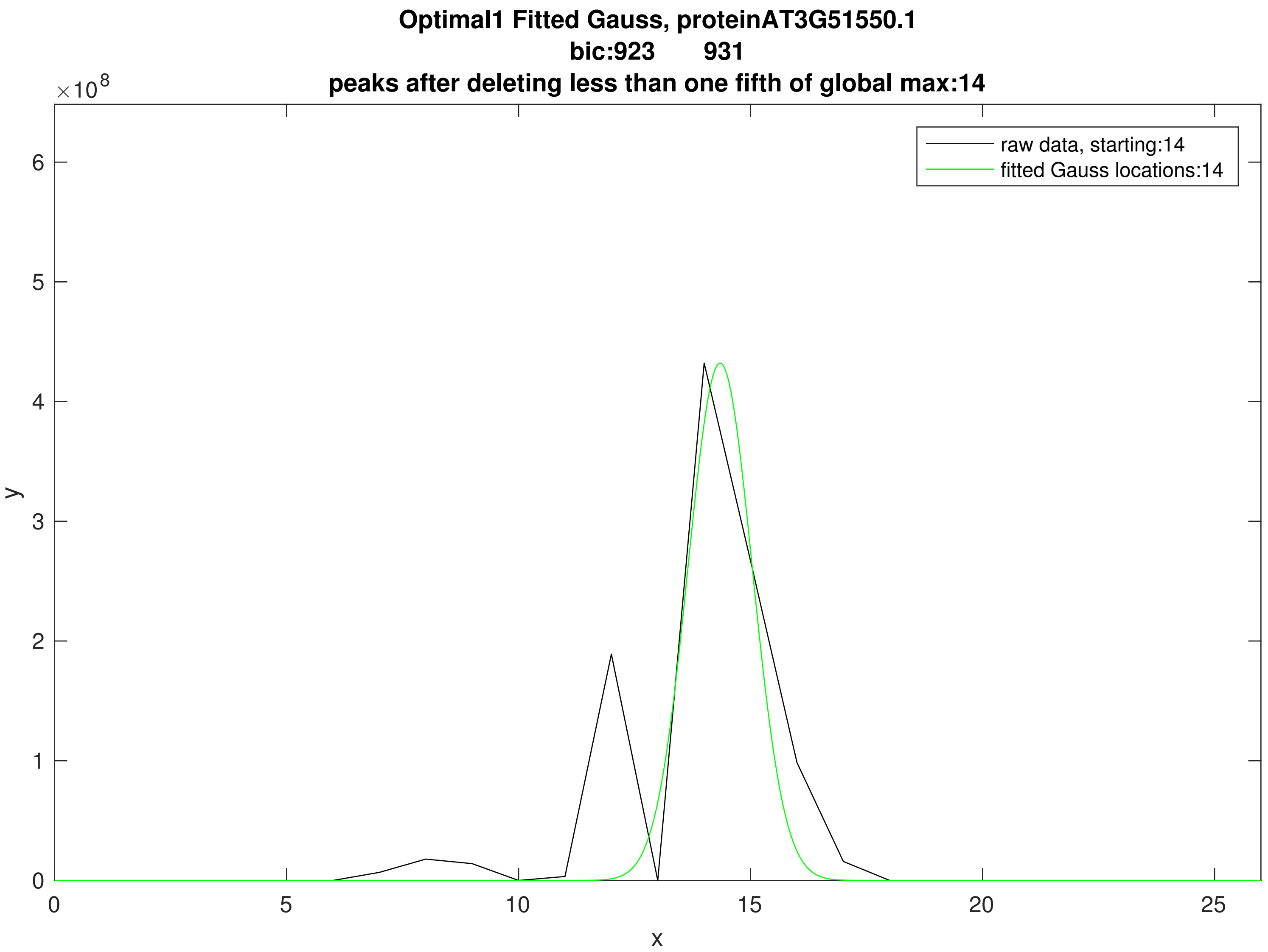

Optimal1 Fitted Gauss, proteinAT3G51550.1
bic:923 931
peaks after deleting less than one fifth of global max:14
×10^8
raw data, starting:14
fitted Gauss locations:14
y
x
0
1
2
3
4
5
6
0
5
10
15
20
25

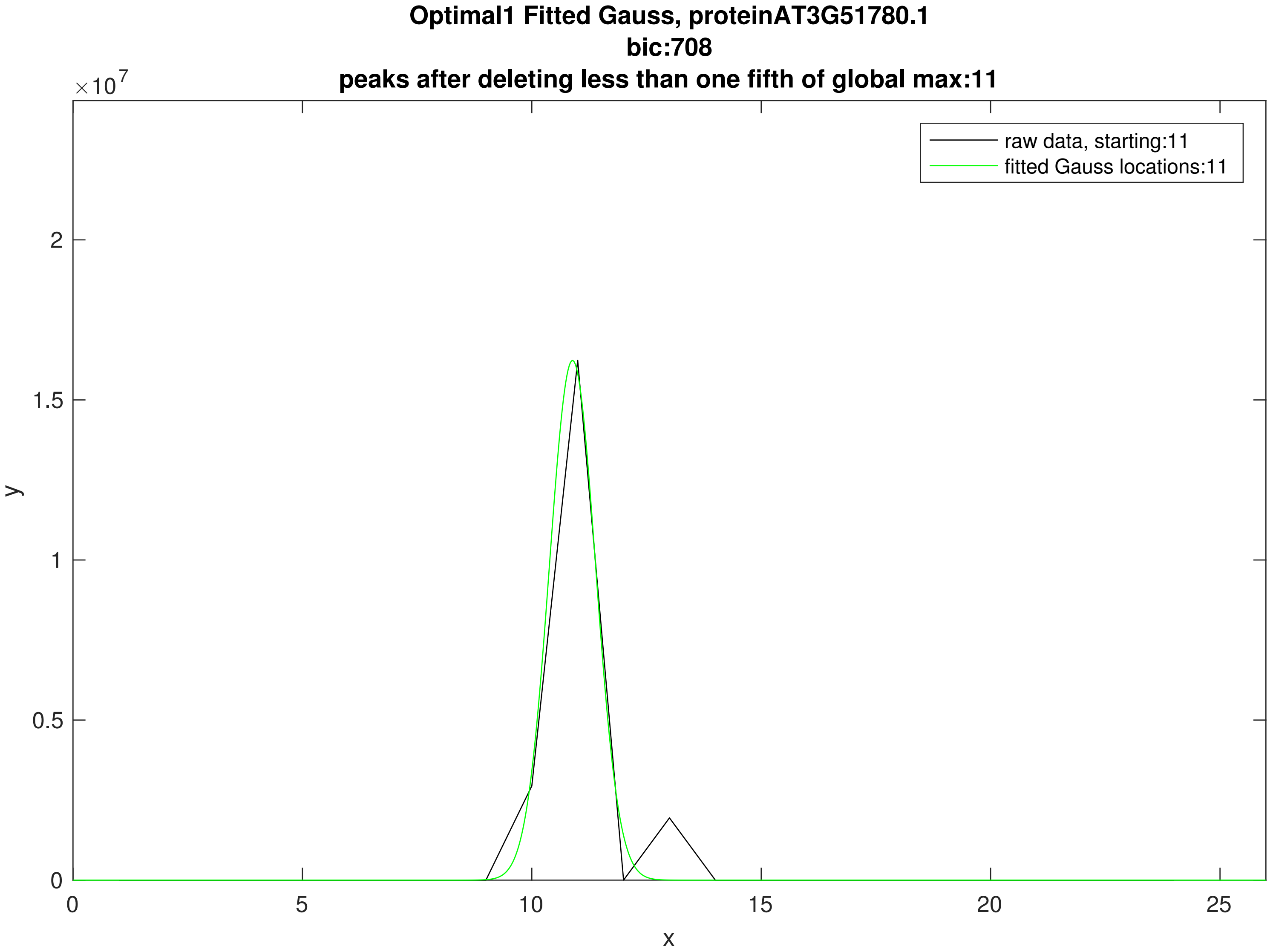
Optimal1 Fitted Gauss, proteinAT3G51780.1
bic:708
peaks after deleting less than one fifth of global max:11
×10^7
raw data, starting:11
fitted Gauss locations:11
2
1.5
1
0.5
0
y
0
5
10
15
20
25
x

**Optimal2 Fitted Gauss, proteinAT3G51800.3;AT3G51800.1;AT3G51800.2**
**bic:902 880**
**peaks after deleting less than one fifth of global max:9.8 14**

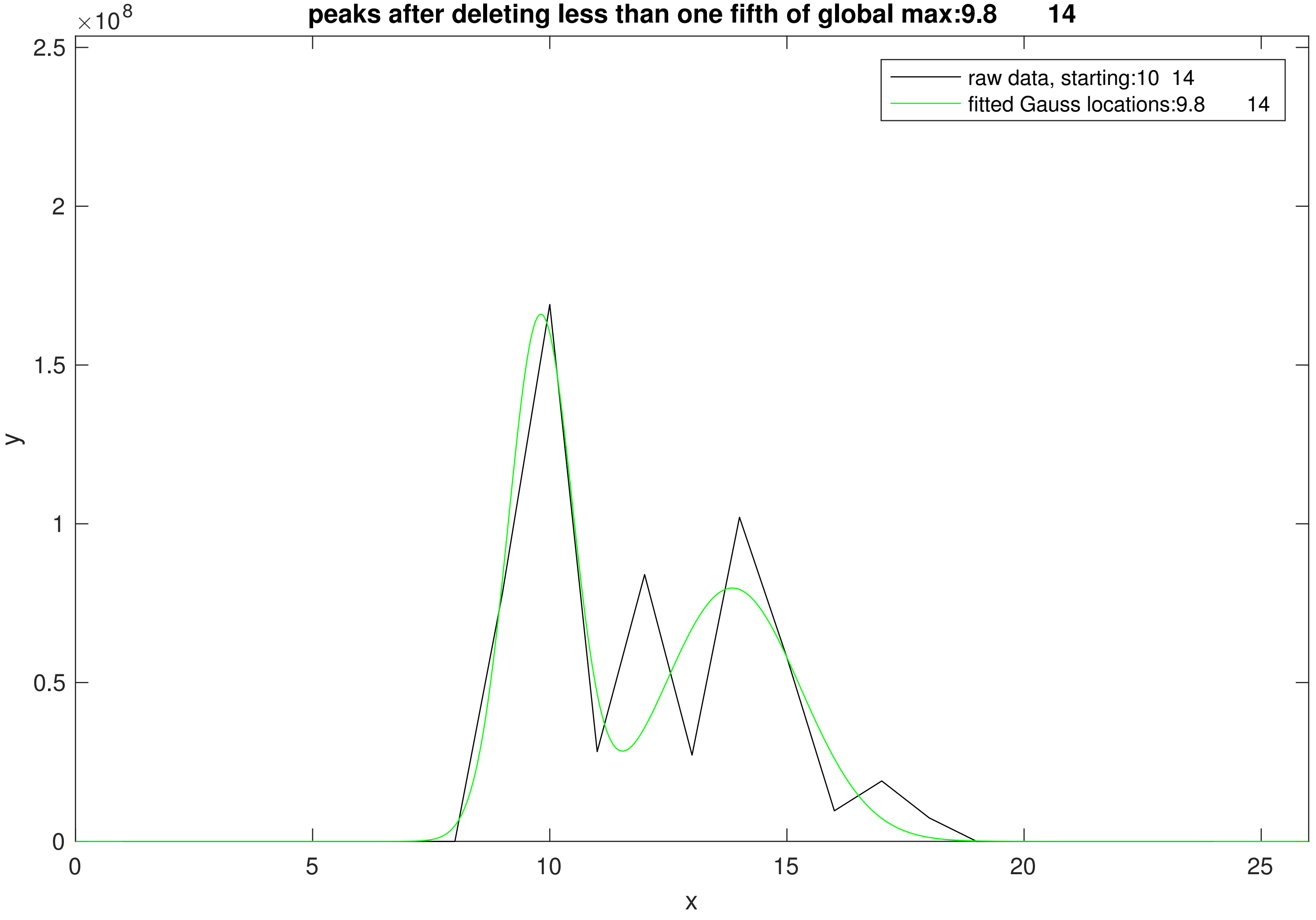

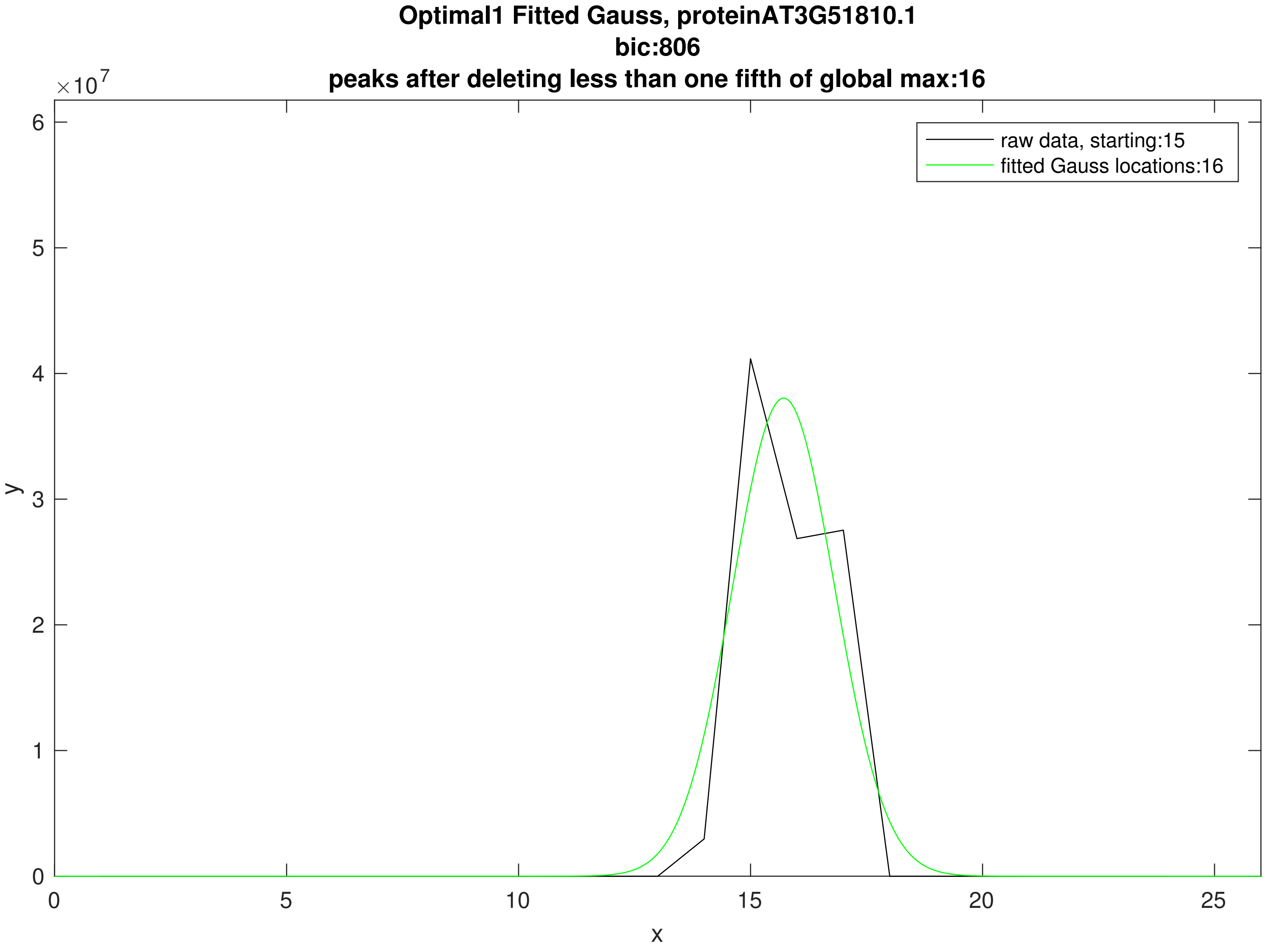

Optimal1 Fitted Gauss, proteinAT3G51810.1
bic:806
peaks after deleting less than one fifth of global max:16
×10^7
6
5
4
3
2
1
0
y
0
5
10
15
20
25
x
raw data, starting:15
fitted Gauss locations:16

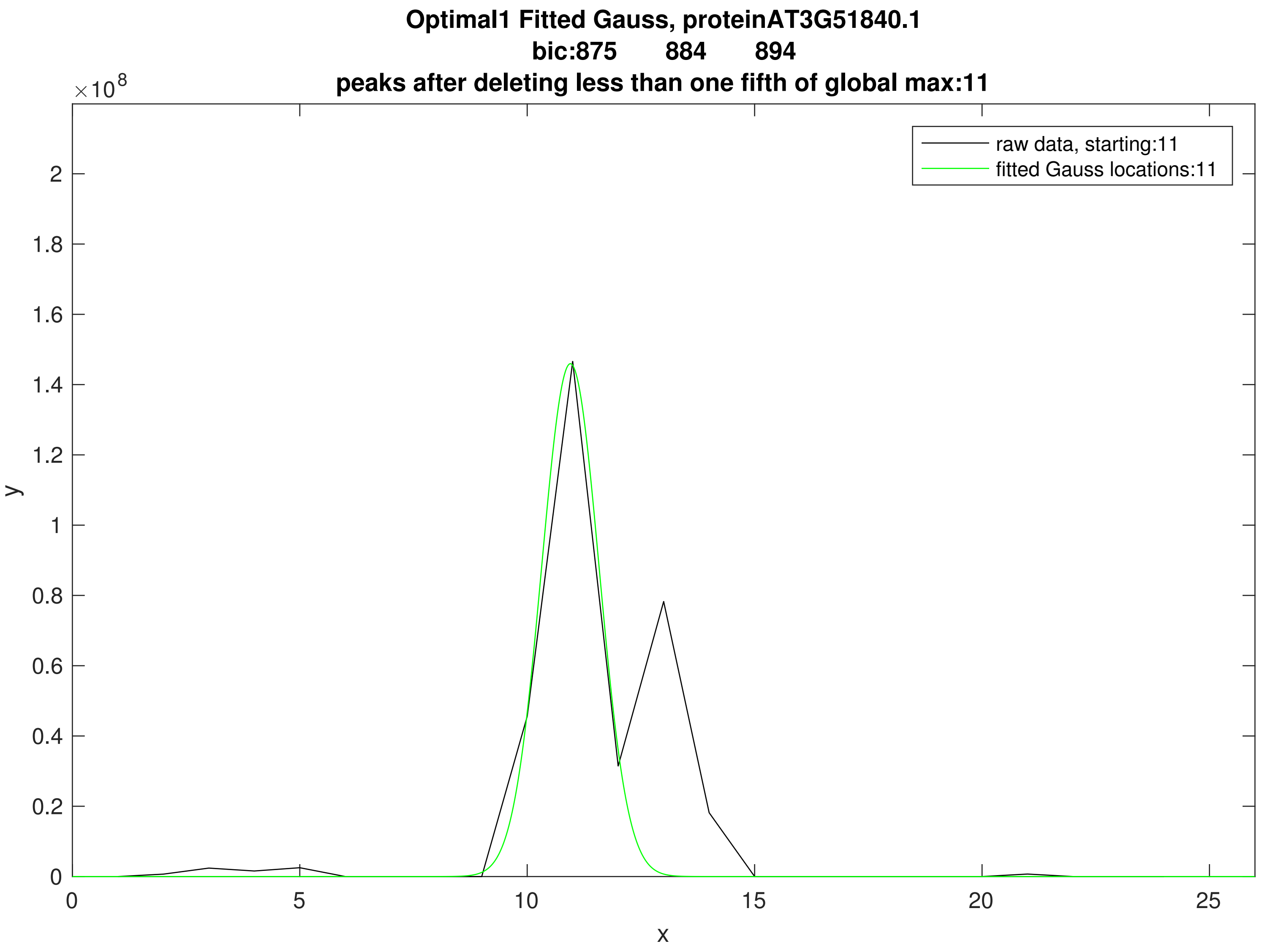

Optimal1 Fitted Gauss, proteinAT3G51840.1
bic:875 884 894
peaks after deleting less than one fifth of global max:11
×10^8
raw data, starting:11
fitted Gauss locations:11
2
1.8
1.6
1.4
1.2
1
0.8
0.6
0.4
0.2
0
y
0
5
10
15
20
25
x

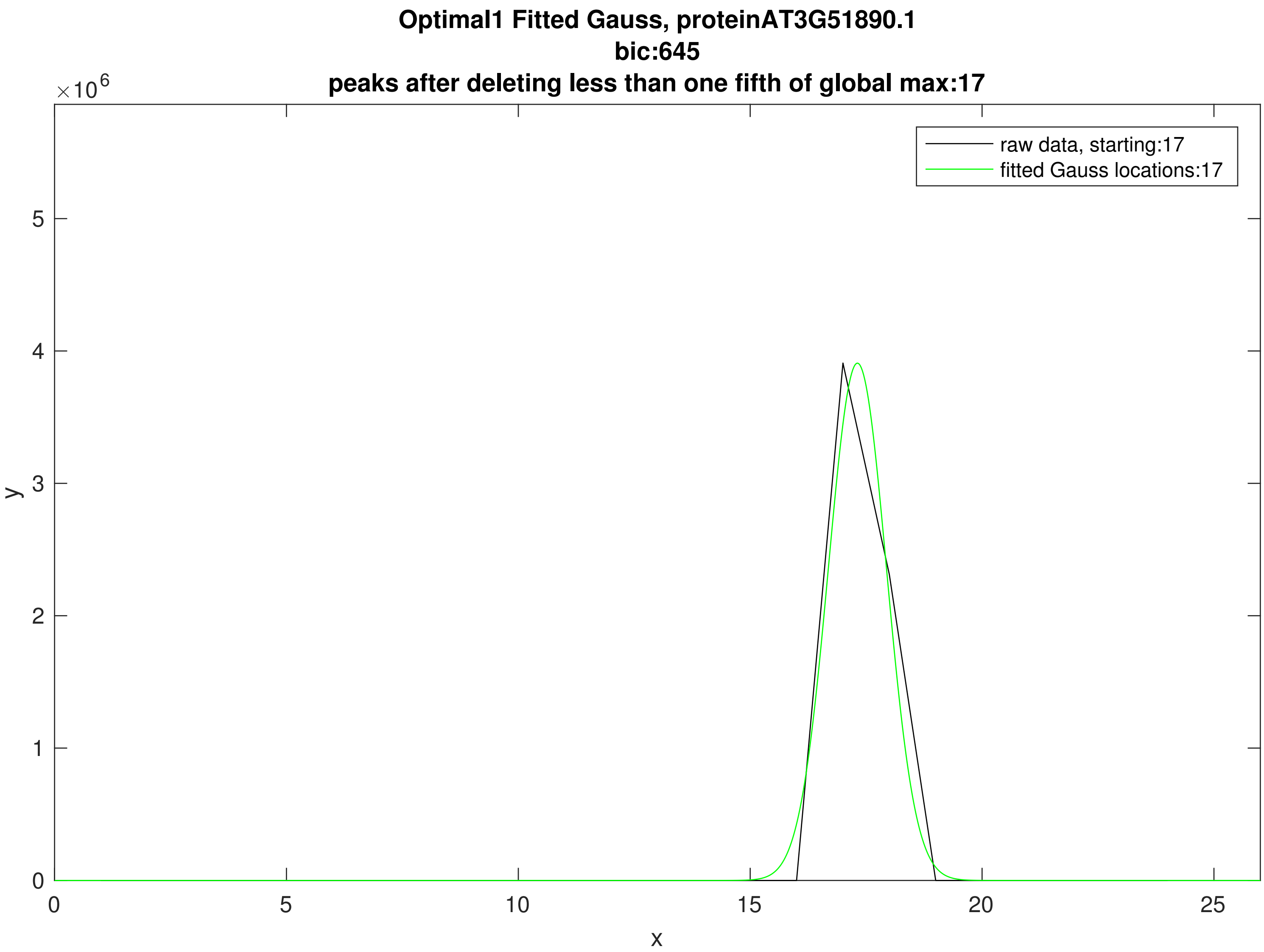

Optimal1 Fitted Gauss, proteinAT3G51890.1
bic:645
peaks after deleting less than one fifth of global max:17
×10^6
raw data, starting:17
fitted Gauss locations:17
5
4
3
2
1
0
y
0
5
10
15
20
25
x

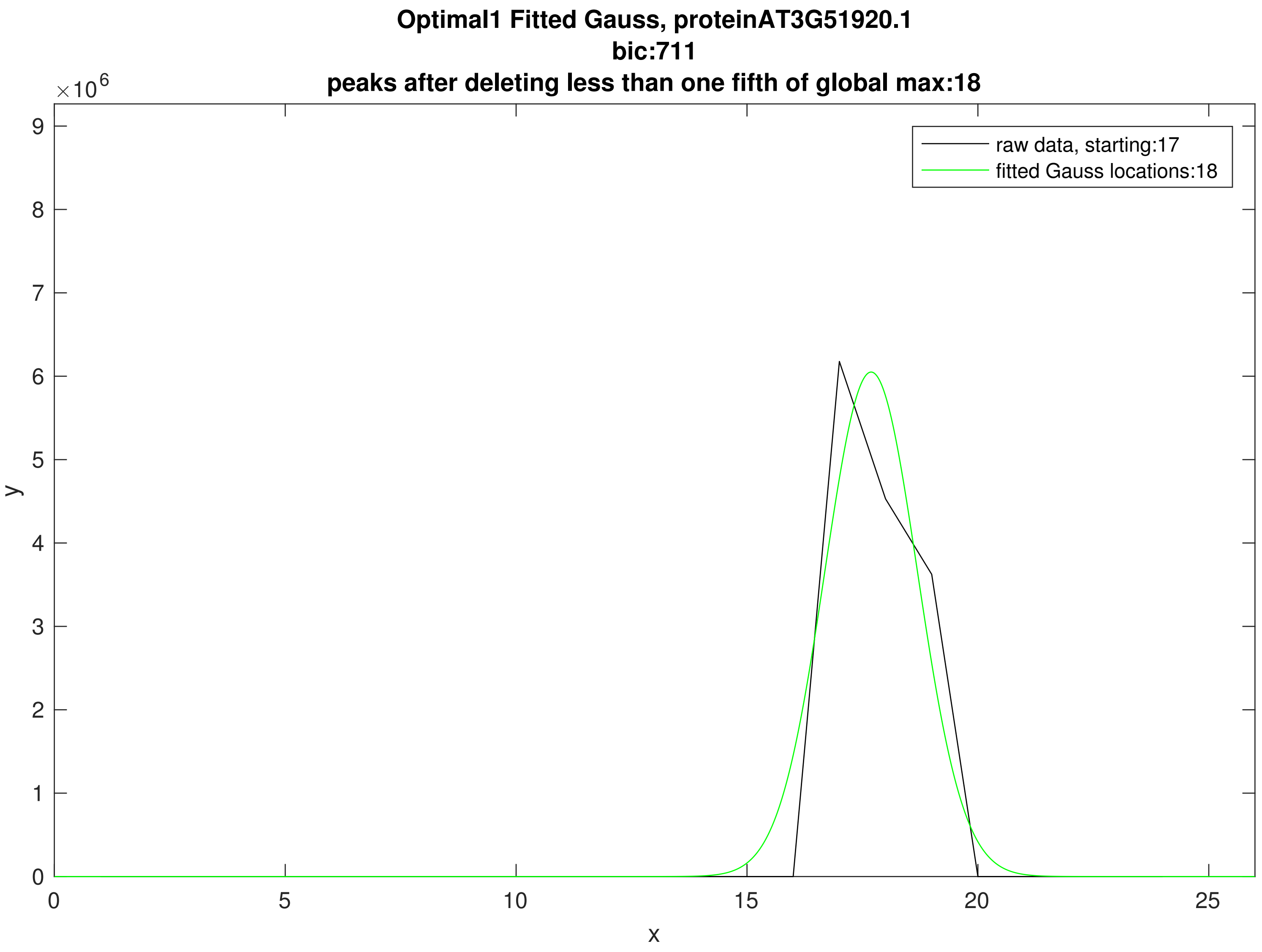

Optimal1 Fitted Gauss, proteinAT3G51920.1
bic:711
peaks after deleting less than one fifth of global max:18
×10^6
raw data, starting:17
fitted Gauss locations:18
y
x
0
1
2
3
4
5
6
7
8
9
0
5
10
15
20
25

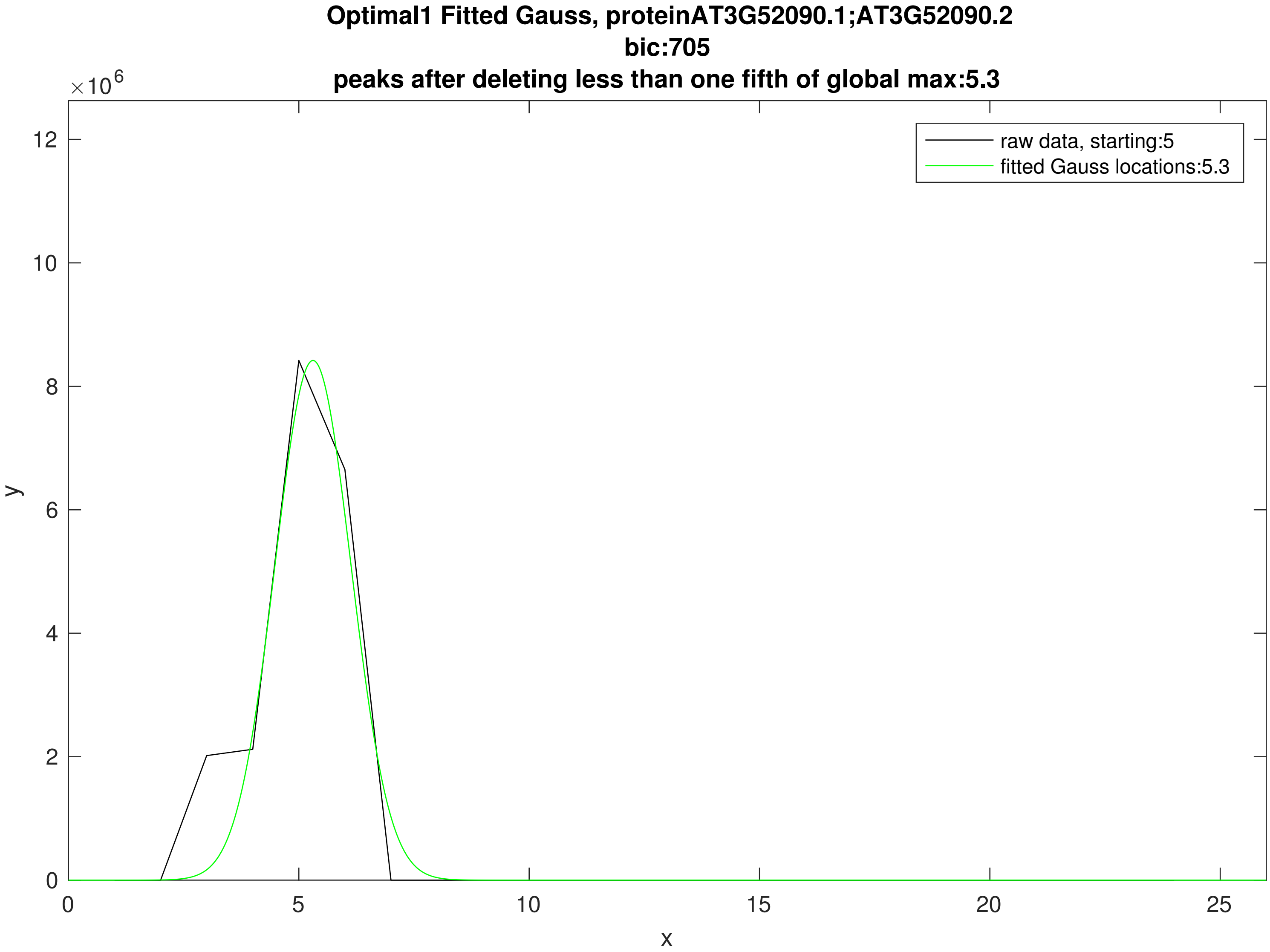
Optimal1 Fitted Gauss, proteinAT3G52090.1;AT3G52090.2
bic:705
peaks after deleting less than one fifth of global max:5.3
×10^6
raw data, starting:5
fitted Gauss locations:5.3
y
x
0
2
4
6
8
10
12
0
5
10
15
20
25

**Optimal2 Fitted Gauss, proteinAT3G52150.2;AT3G52150.1**
**bic:857 844**
**peaks after deleting less than one fifth of global max:5 19**

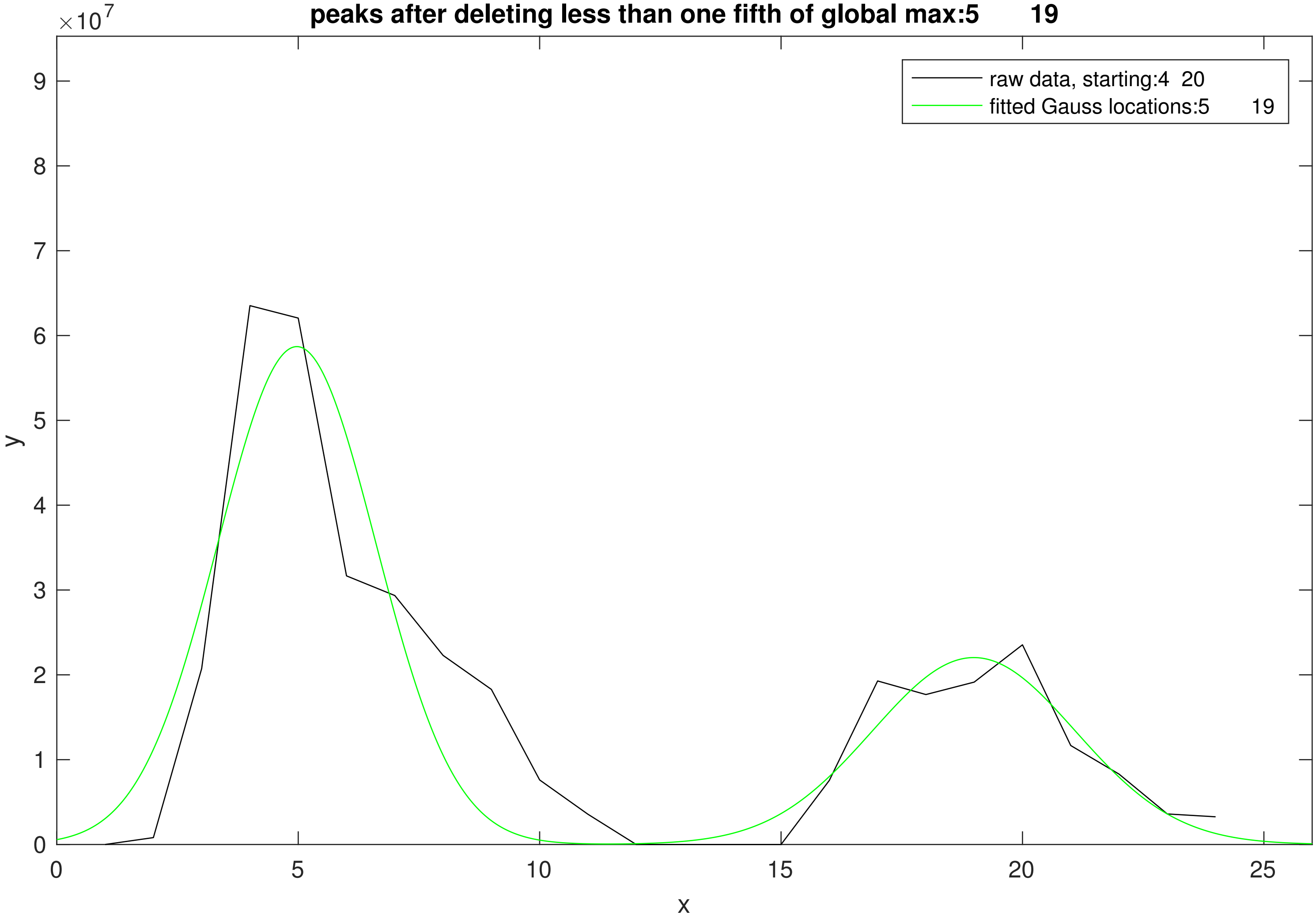

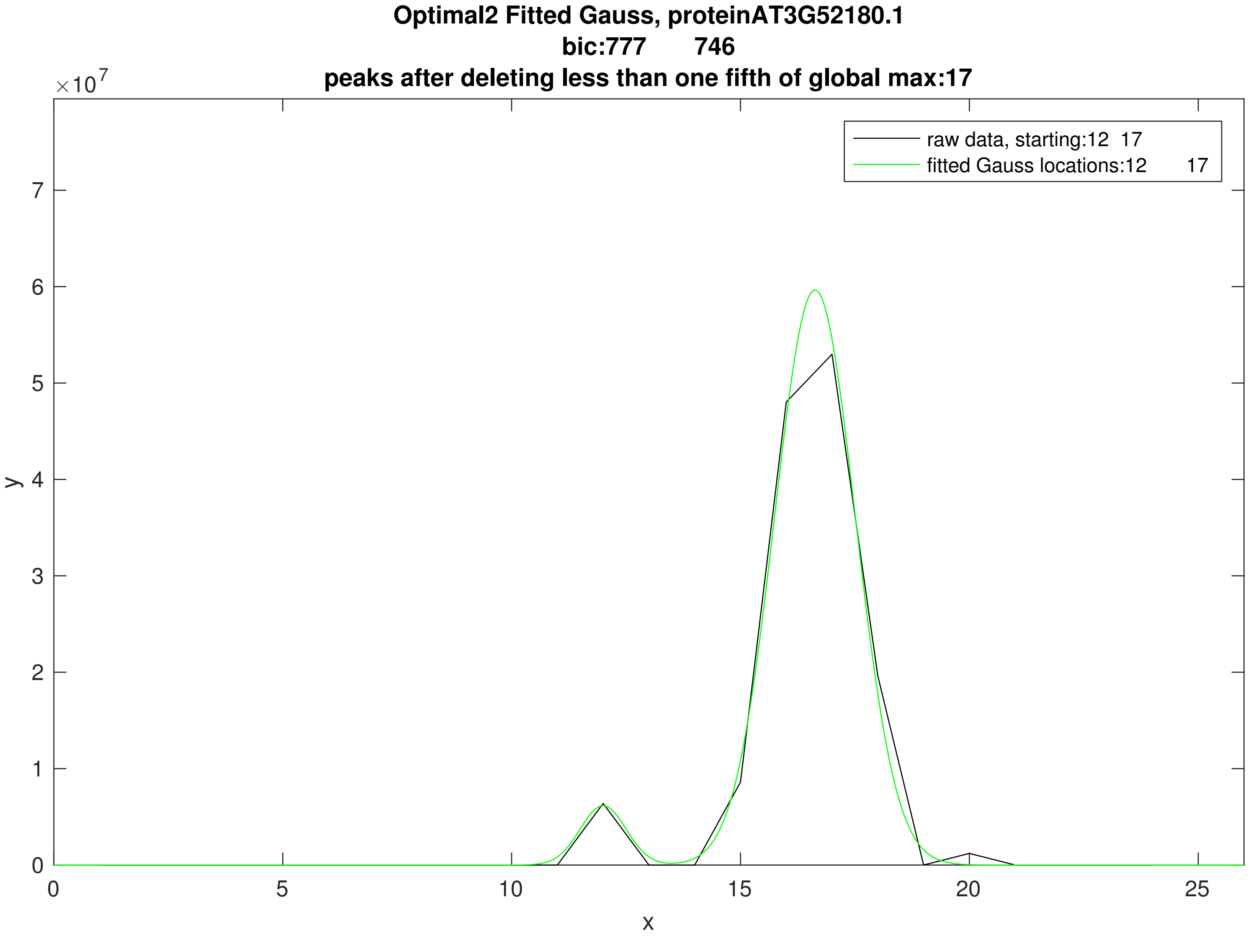

Optimal2 Fitted Gauss, proteinAT3G52180.1
bic:777 746
peaks after deleting less than one fifth of global max:17
×10^7
raw data, starting:12 17
fitted Gauss locations:12 17
y
x
0
1
2
3
4
5
6
7
0
5
10
15
20
25

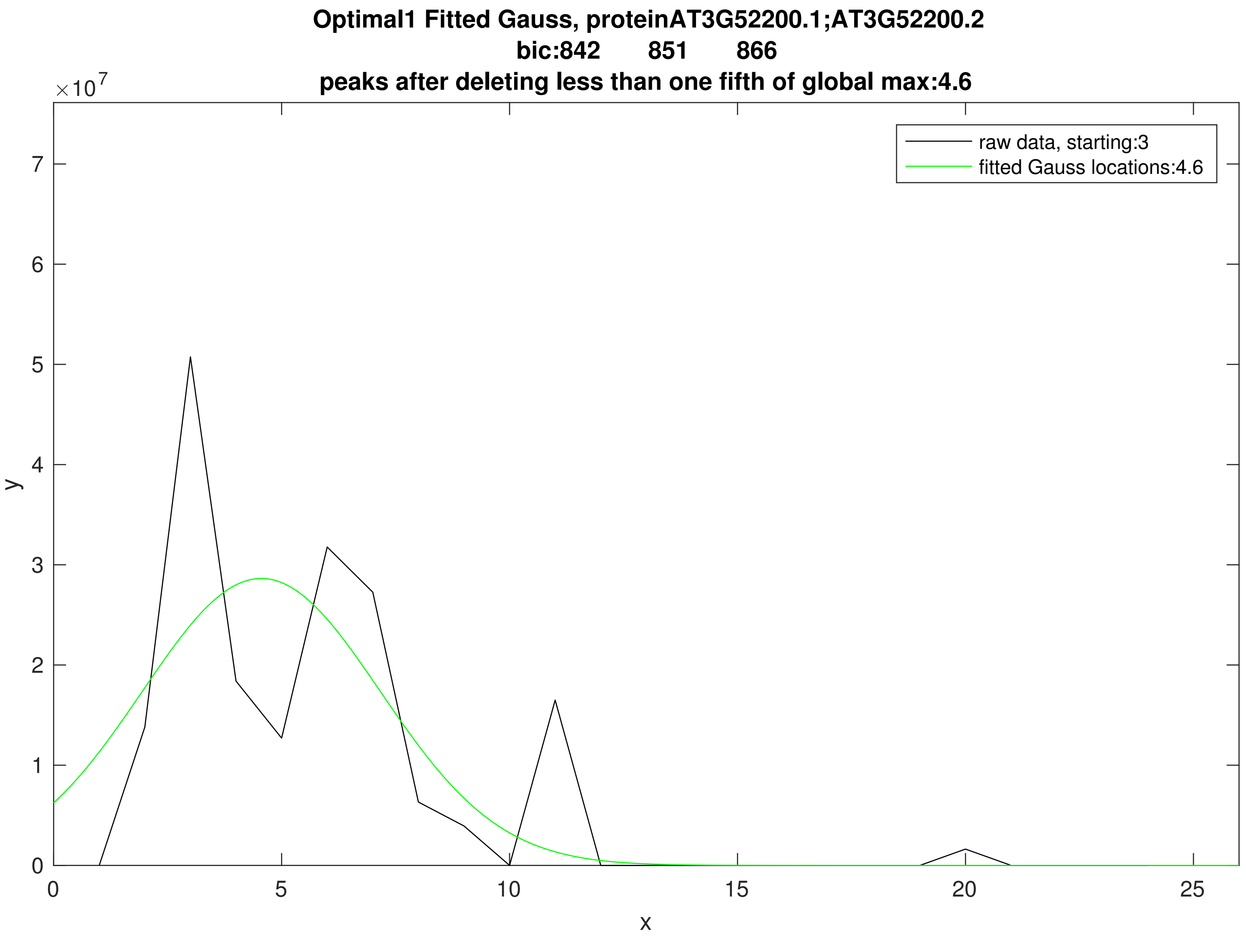

Optimal1 Fitted Gauss, proteinAT3G52200.1;AT3G52200.2
bic:842 851 866
peaks after deleting less than one fifth of global max:4.6
×10^7
raw data, starting:3
fitted Gauss locations:4.6
y
x
0
1
2
3
4
5
6
7
0
5
10
15
20
25

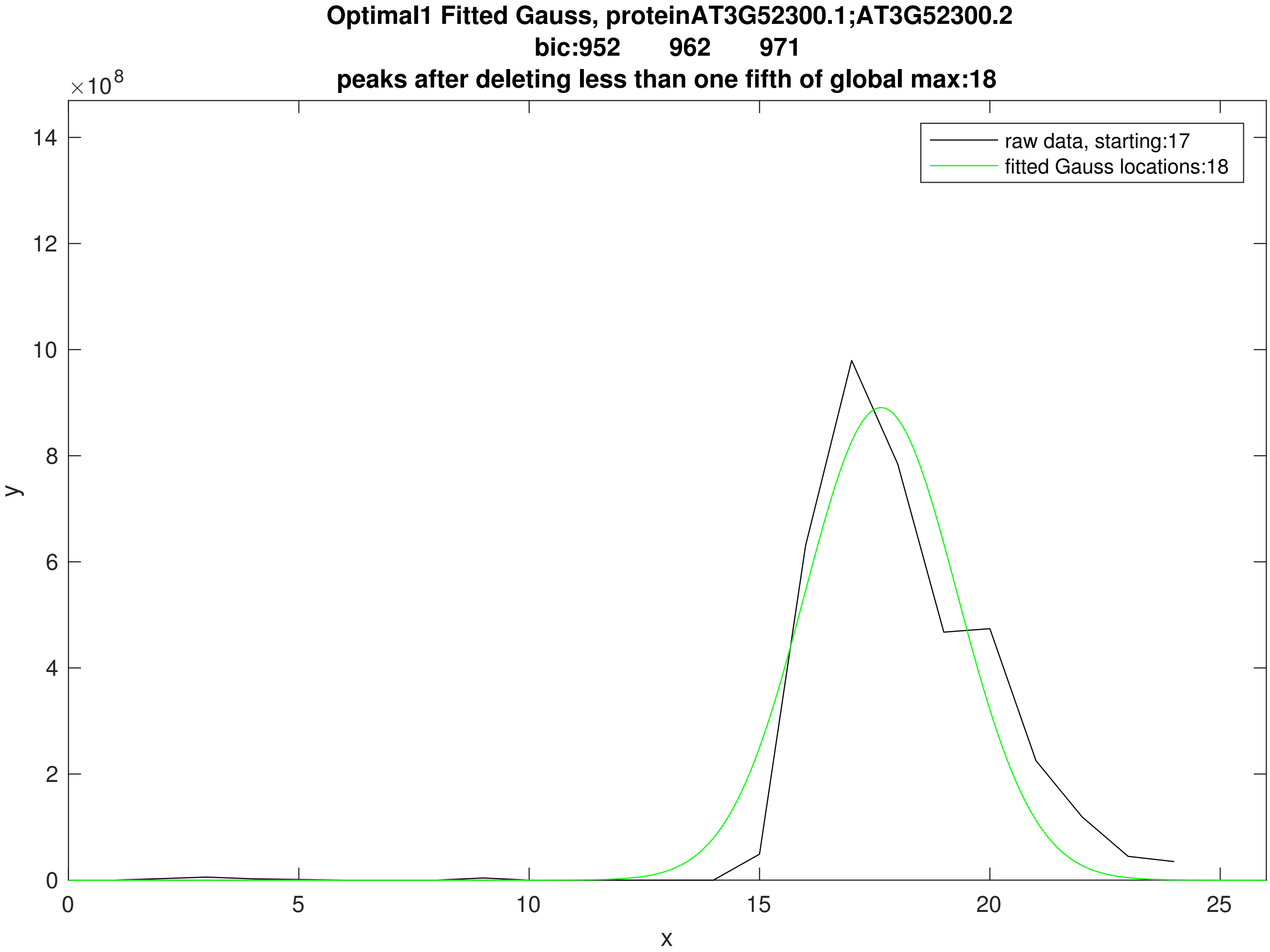

Optimal1 Fitted Gauss, proteinAT3G52300.1;AT3G52300.2
bic:952 962 971
peaks after deleting less than one fifth of global max:18
×10^8
14
12
10
8
6
4
2
0
y
0
5
10
15
20
25
x
raw data, starting:17
fitted Gauss locations:18

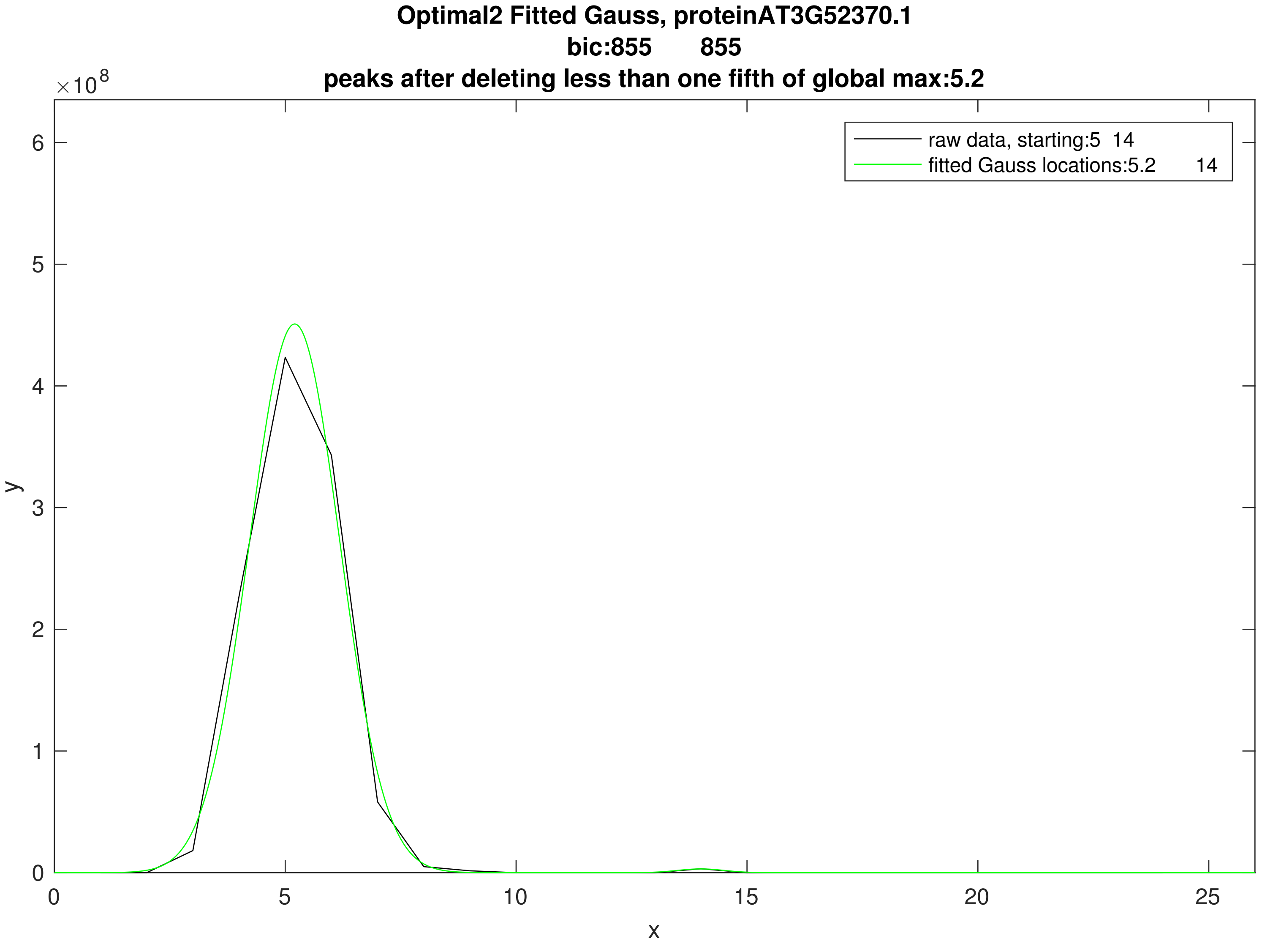

Optimal2 Fitted Gauss, proteinAT3G52370.1
bic:855 855
peaks after deleting less than one fifth of global max:5.2
×10^8
raw data, starting:5 14
fitted Gauss locations:5.2 14
y
x
0
1
2
3
4
5
6
0
5
10
15
20
25

Optimal1 Fitted Gauss, proteinAT3G52380.1
bic:930 940 951
peaks after deleting less than one fifth of global max:14

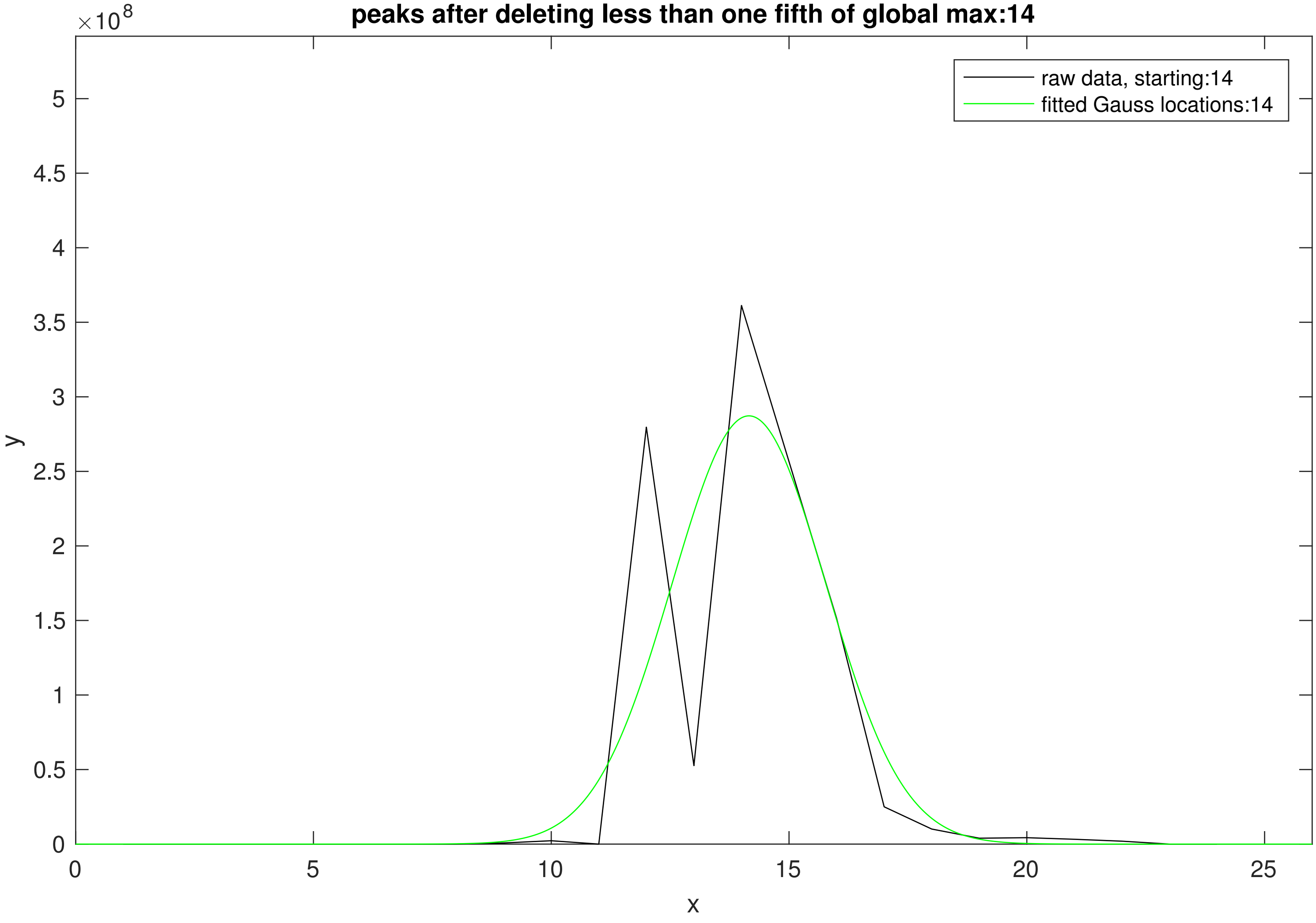

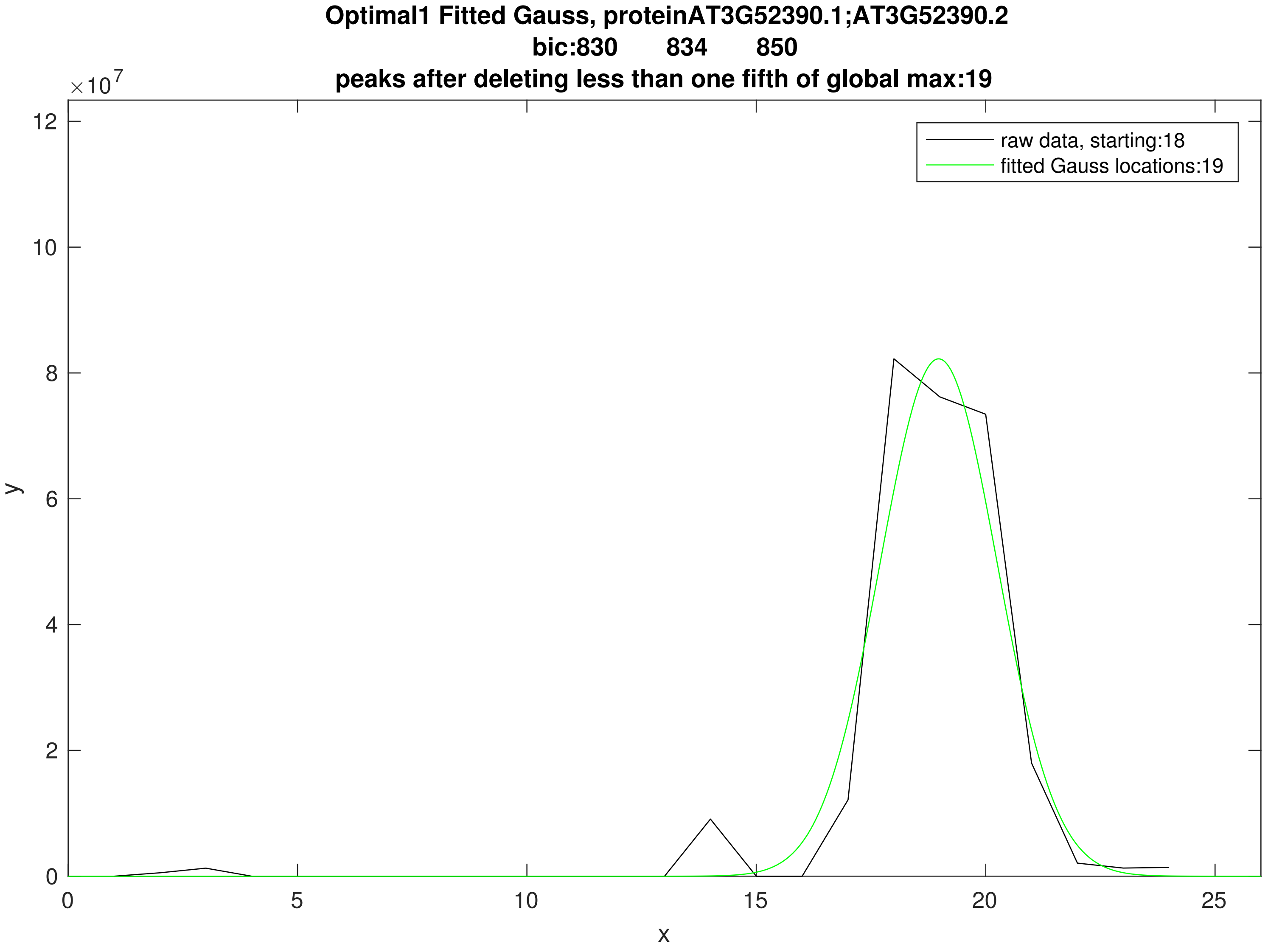
Optimal1 Fitted Gauss, proteinAT3G52390.1;AT3G52390.2
bic:830 834 850
peaks after deleting less than one fifth of global max:19
×10^7
raw data, starting:18
fitted Gauss locations:19
y
x
0
2
4
6
8
10
12
0
5
10
15
20
25

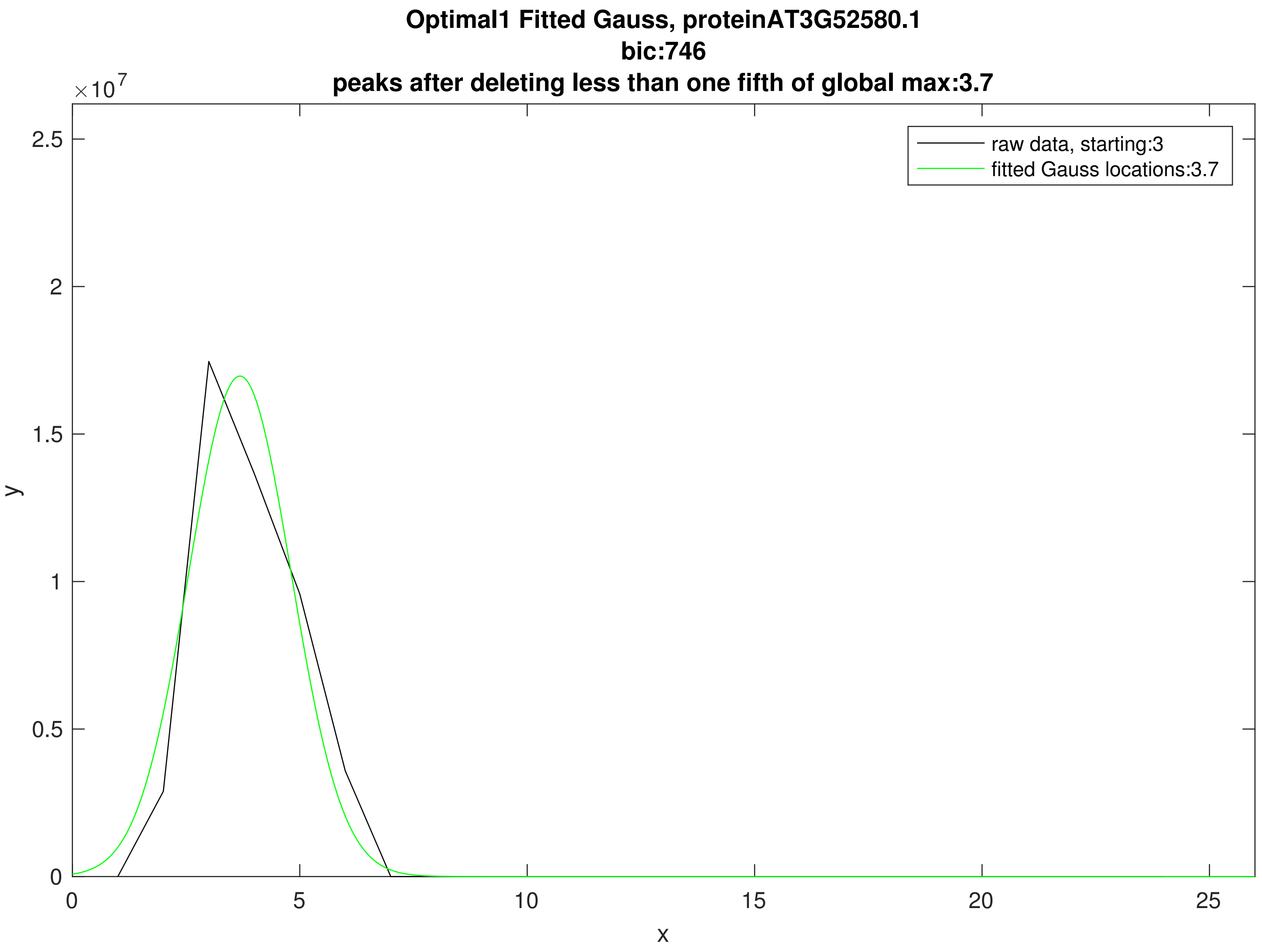
Optimal1 Fitted Gauss, proteinAT3G52580.1
bic:746
peaks after deleting less than one fifth of global max:3.7
×10^7
2.5
2
1.5
1
0.5
0
y
0
5
10
15
20
25
x
raw data, starting:3
fitted Gauss locations:3.7

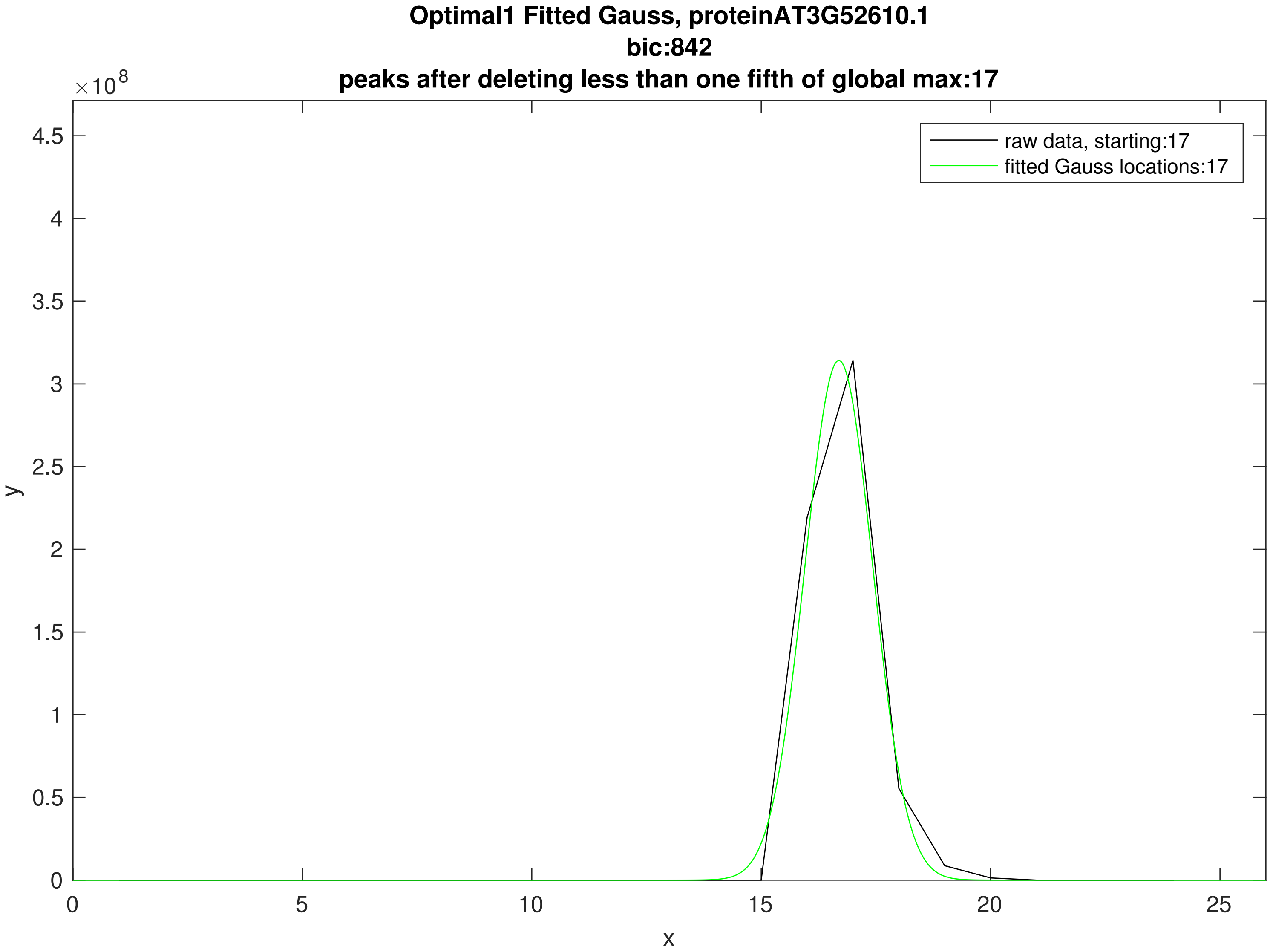
Optimal1 Fitted Gauss, proteinAT3G52610.1
bic:842
peaks after deleting less than one fifth of global max:17
×10^8
y
x
raw data, starting:17
fitted Gauss locations:17

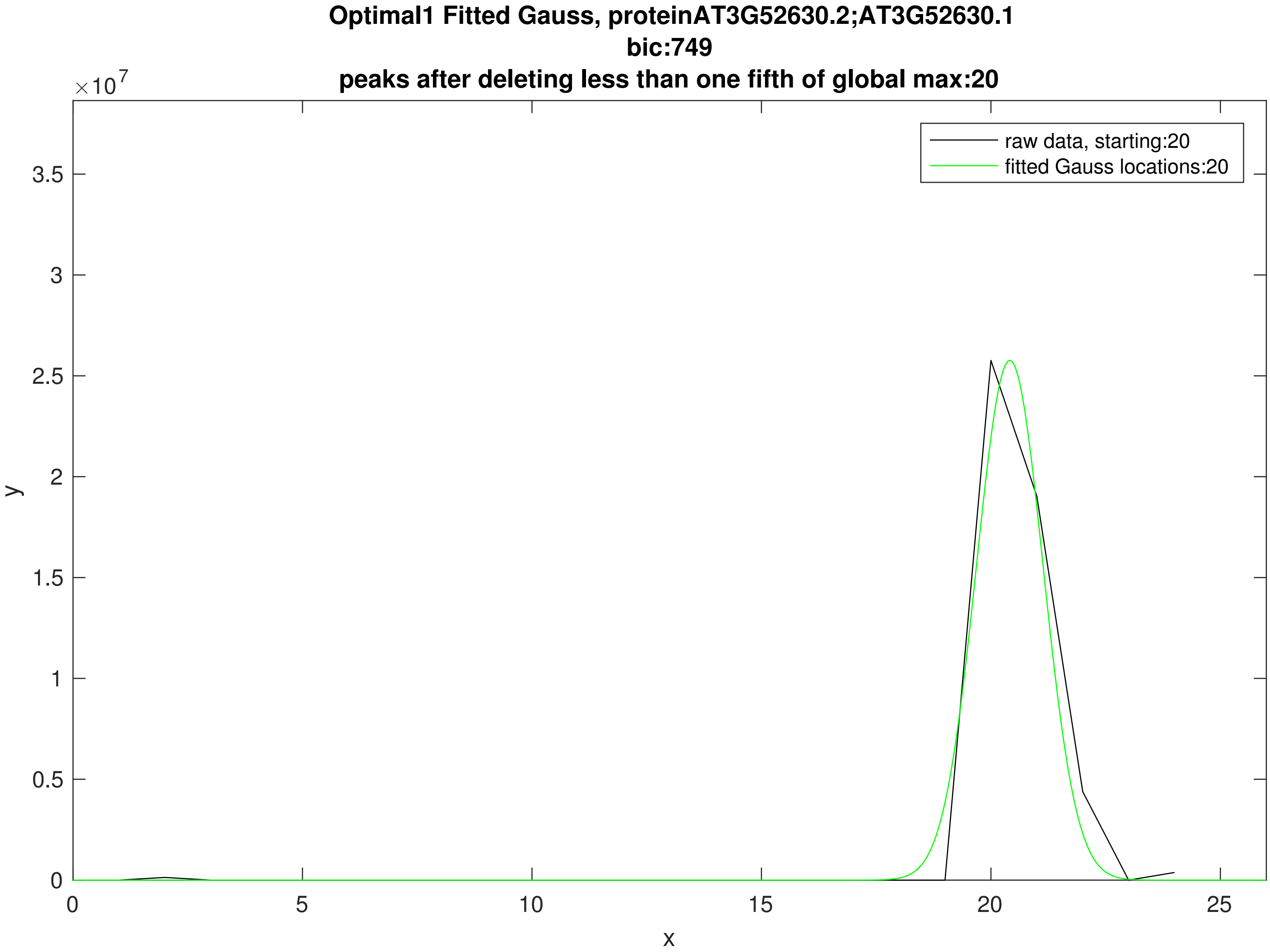
Optimal1 Fitted Gauss, proteinAT3G52630.2;AT3G52630.1
bic:749
peaks after deleting less than one fifth of global max:20
×10^7
raw data, starting:20
fitted Gauss locations:20
y
x
0
0.5
1
1.5
2
2.5
3
3.5
5
10
15
20
25

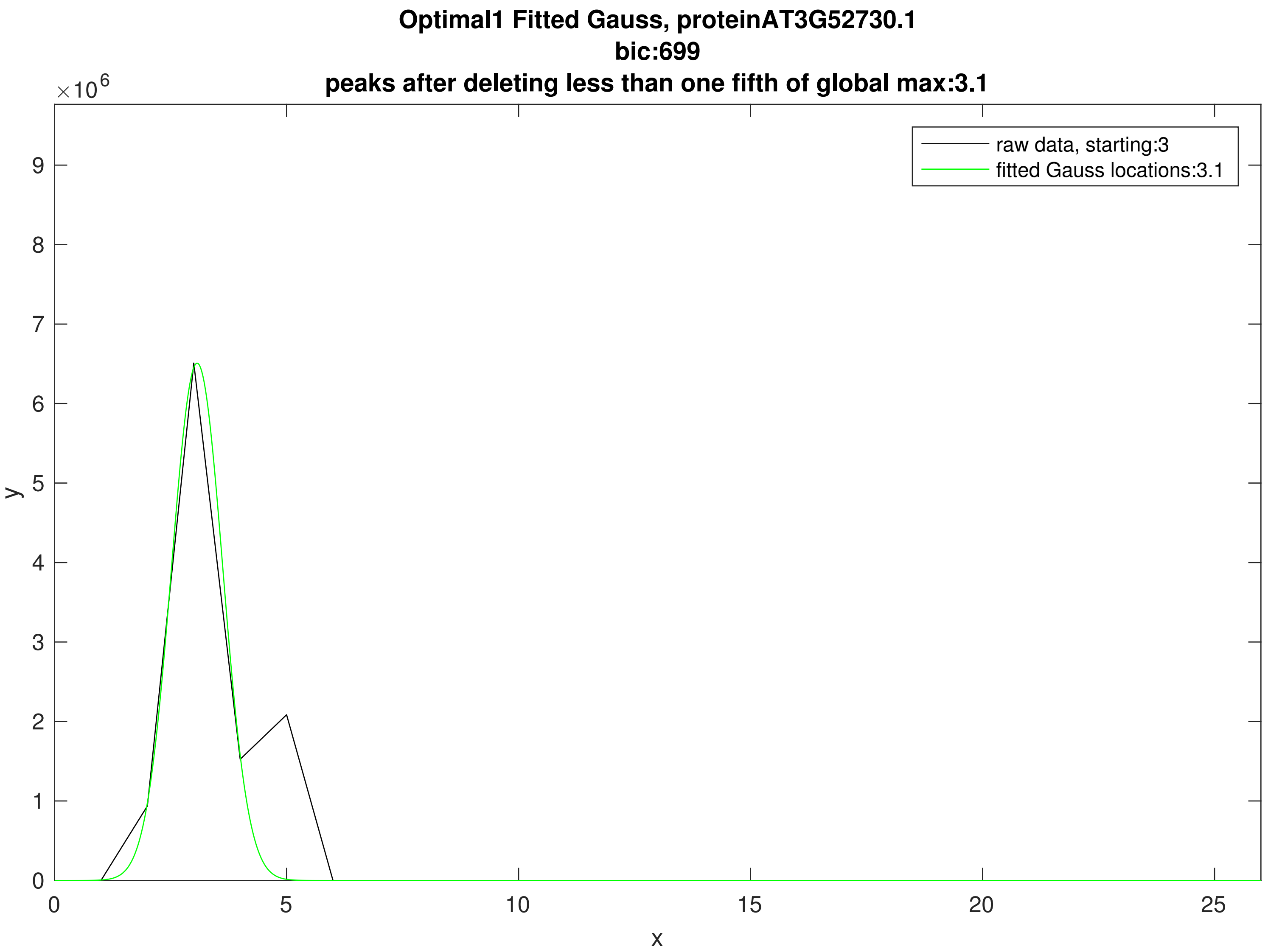

Optimal1 Fitted Gauss, proteinAT3G52730.1
bic:699
peaks after deleting less than one fifth of global max:3.1
×10^6
raw data, starting:3
fitted Gauss locations:3.1
y
x
0
1
2
3
4
5
6
7
8
9
0
5
10
15
20
25

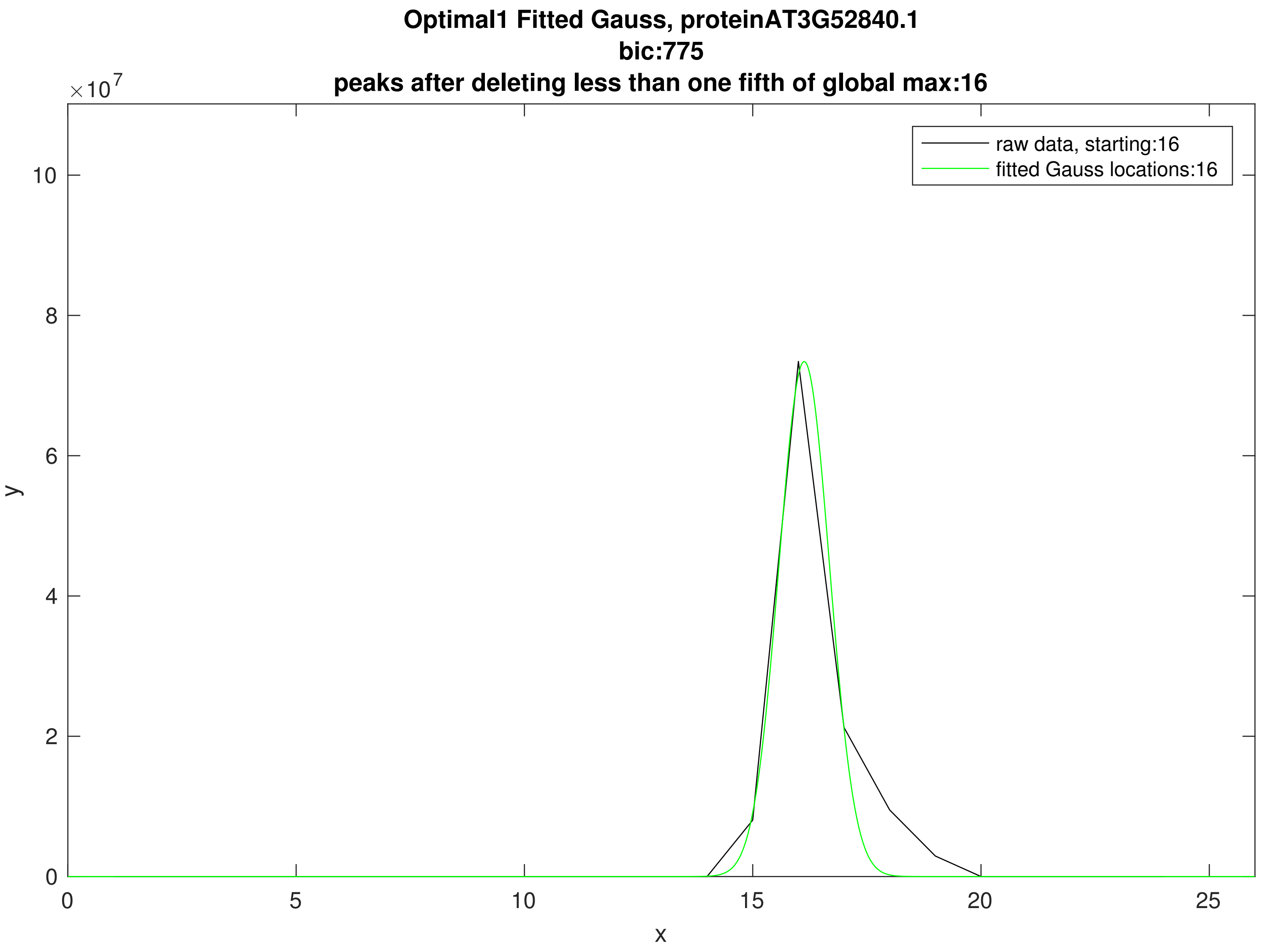

Optimal1 Fitted Gauss, proteinAT3G52840.1
bic:775
peaks after deleting less than one fifth of global max:16
×10^7
raw data, starting:16
fitted Gauss locations:16
y
x
0
2
4
6
8
10
0
5
10
15
20
25

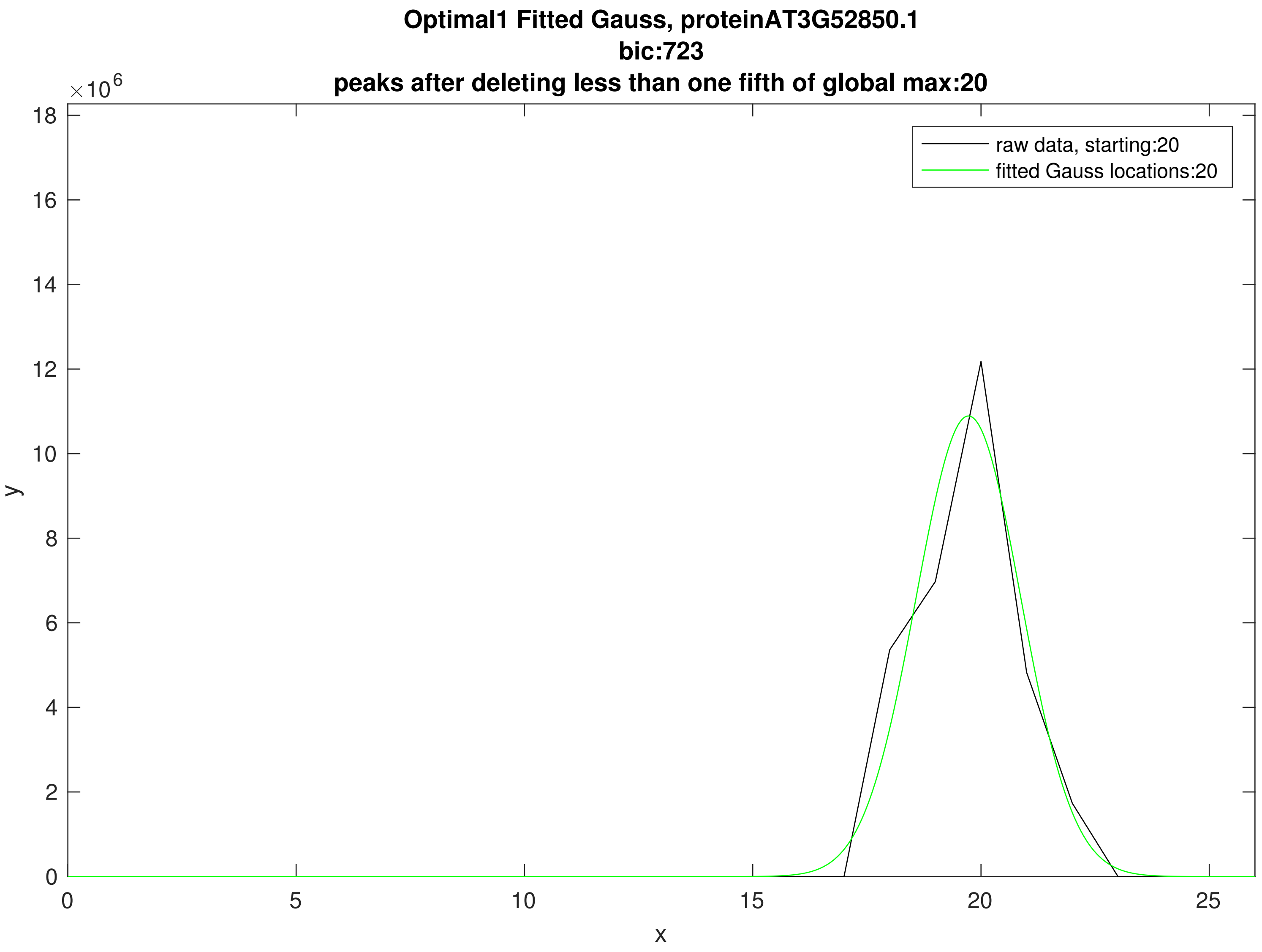
Optimal1 Fitted Gauss, proteinAT3G52850.1
bic:723
peaks after deleting less than one fifth of global max:20
raw data, starting:20
fitted Gauss locations:20
×10^6
y
x
0
2
4
6
8
10
12
14
16
18
0
5
10
15
20
25

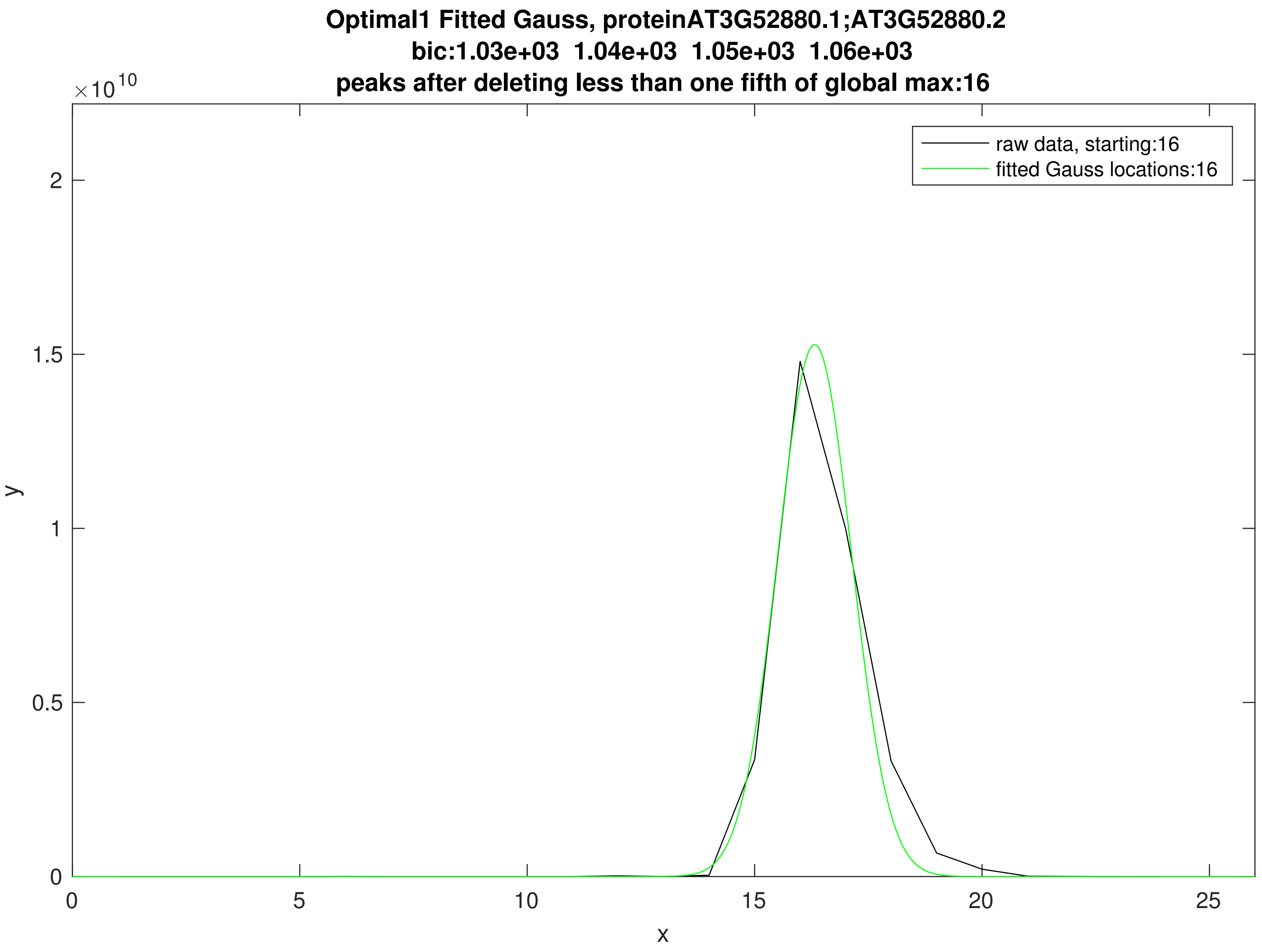
Optimal1 Fitted Gauss, proteinAT3G52880.1;AT3G52880.2
bic:1.03e+03  1.04e+03  1.05e+03  1.06e+03
peaks after deleting less than one fifth of global max:16
×10^10
raw data, starting:16
fitted Gauss locations:16
y
x
0
0.5
1
1.5
2
0
5
10
15
20
25

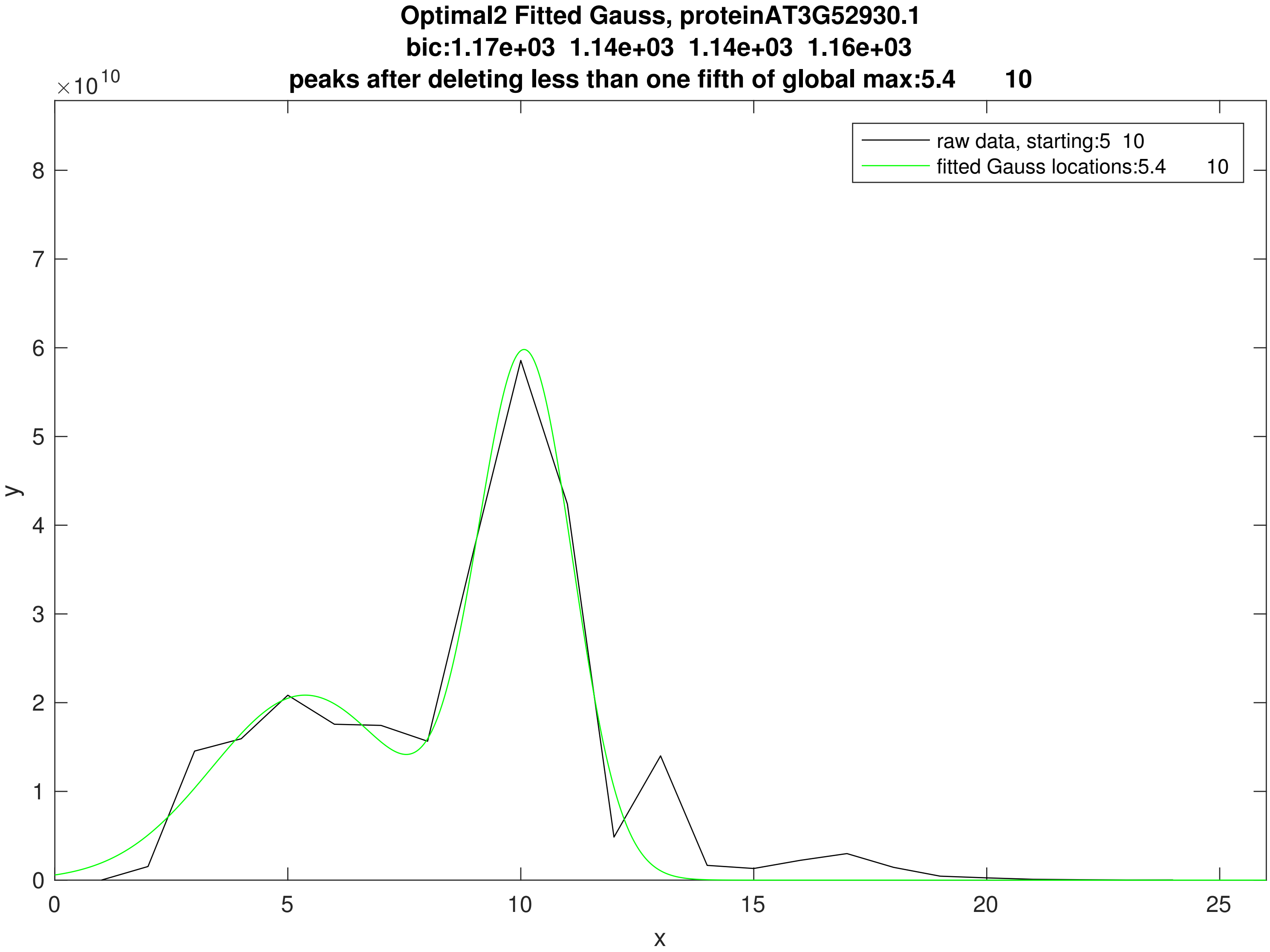

Optimal2 Fitted Gauss, proteinAT3G52930.1
bic:1.17e+03 1.14e+03 1.14e+03 1.16e+03
peaks after deleting less than one fifth of global max:5.4 10
×10^10
raw data, starting:5 10
fitted Gauss locations:5.4 10
y
x
0
1
2
3
4
5
6
7
8
0
5
10
15
20
25

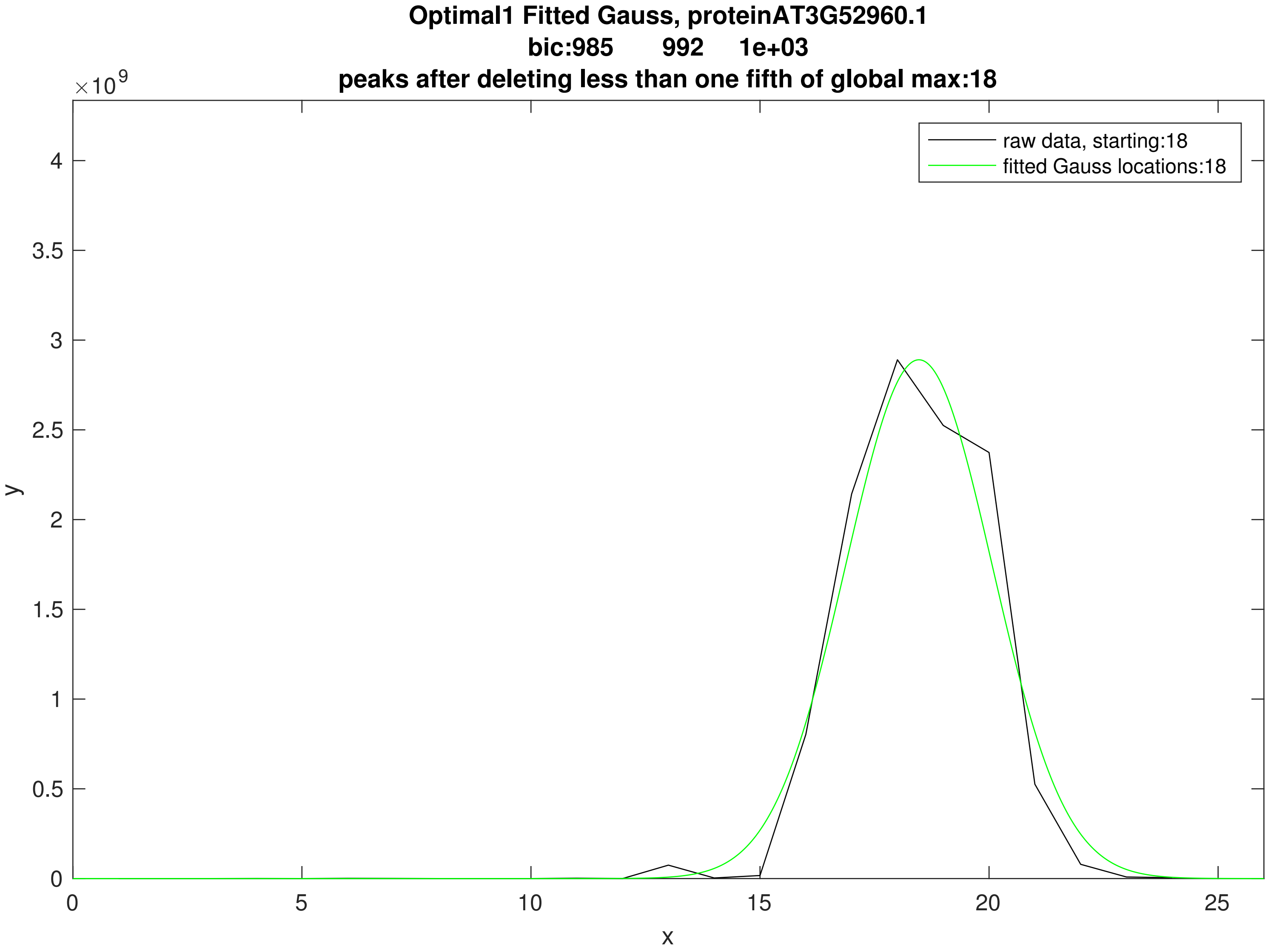

Optimal1 Fitted Gauss, proteinAT3G52960.1
bic:985 992 1e+03
peaks after deleting less than one fifth of global max:18
×10^9
raw data, starting:18
fitted Gauss locations:18
y
x

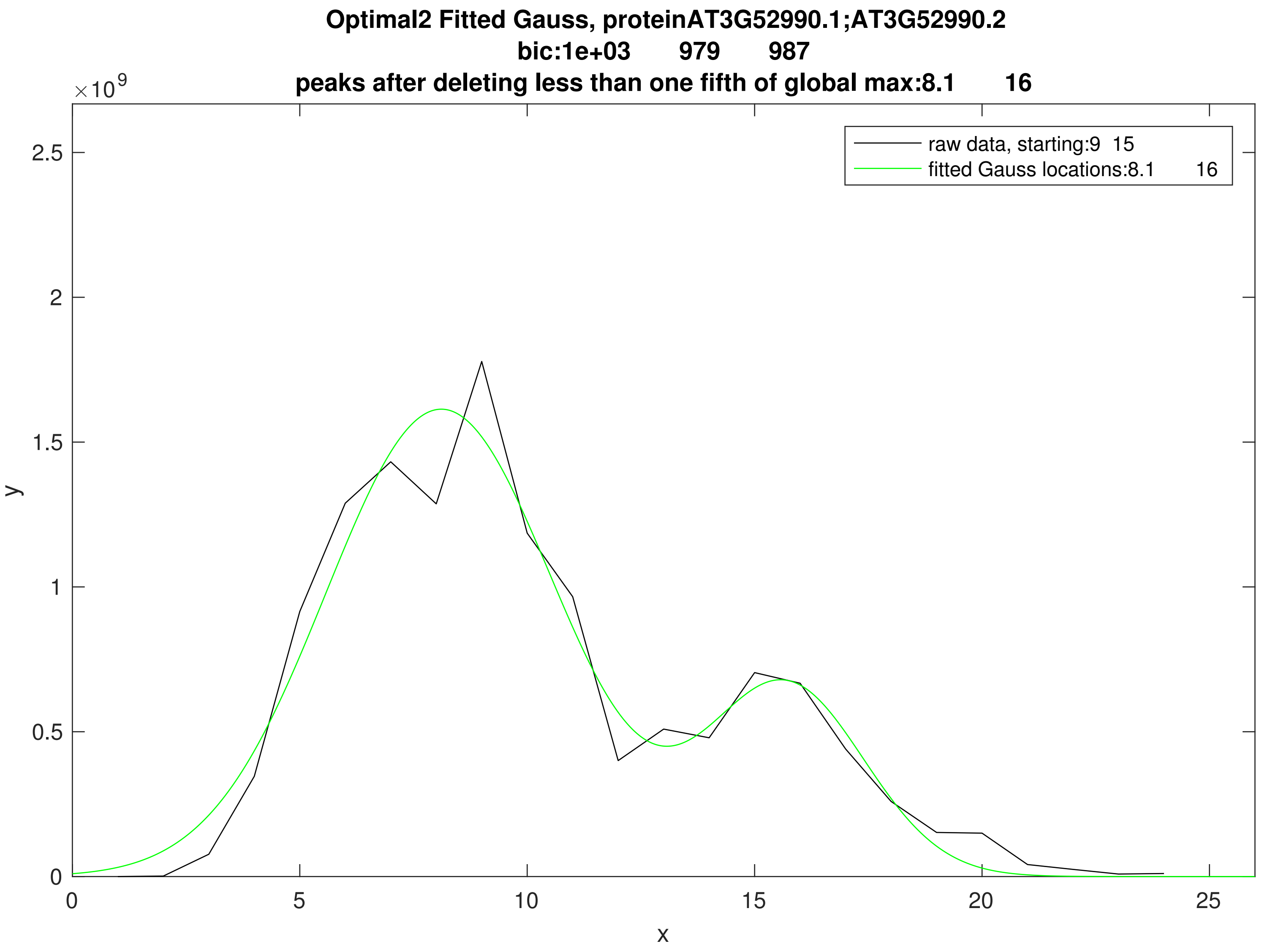

Optimal2 Fitted Gauss, proteinAT3G52990.1;AT3G52990.2
bic:1e+03 979 987
peaks after deleting less than one fifth of global max:8.1 16
×10^9
raw data, starting:9 15
fitted Gauss locations:8.1 16
y
x
0
0.5
1
1.5
2
2.5
5
10
15
20
25

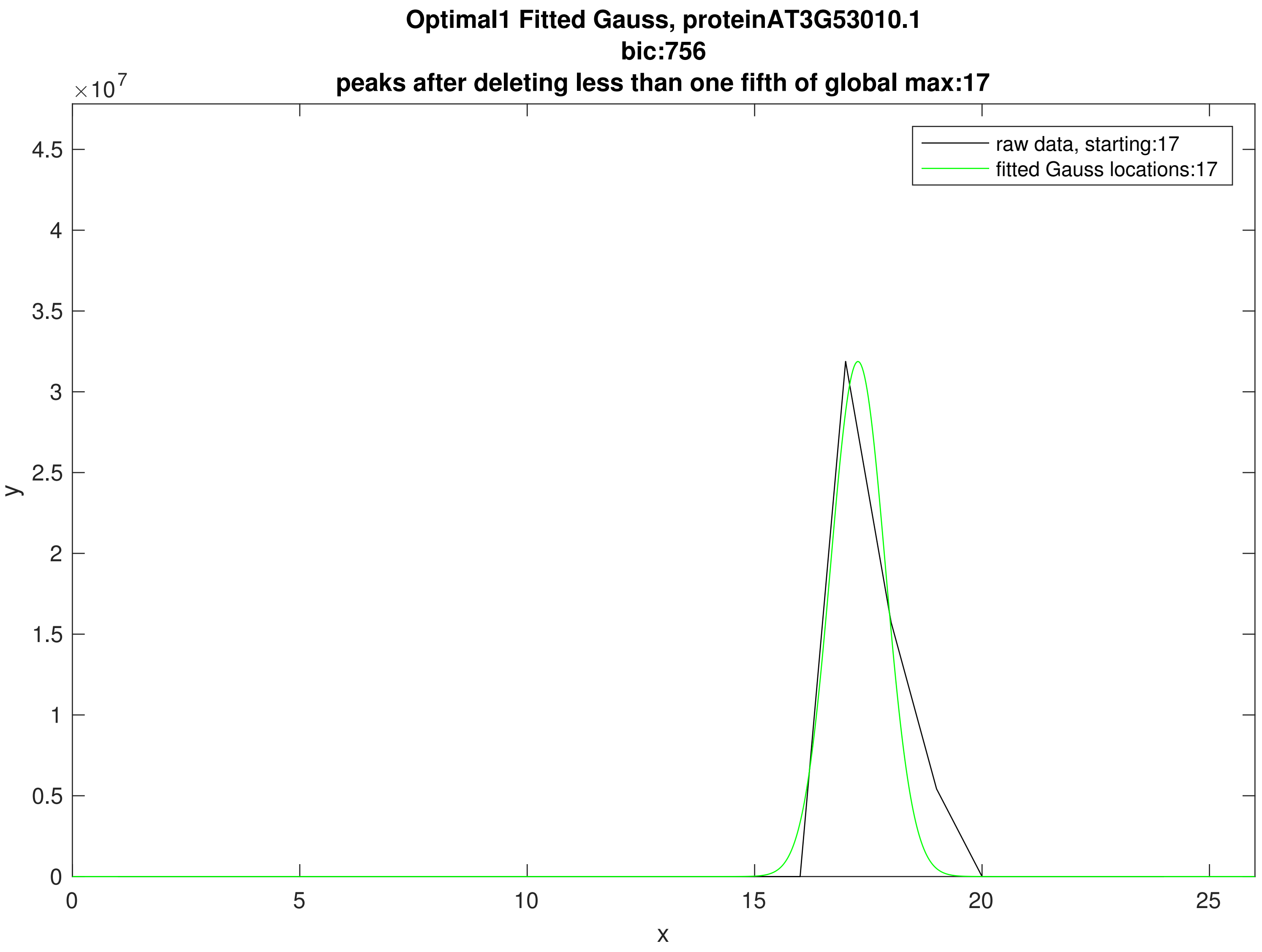

Optimal1 Fitted Gauss, proteinAT3G53010.1
bic:756
peaks after deleting less than one fifth of global max:17
×10^7
raw data, starting:17
fitted Gauss locations:17
y
x
0
0.5
1
1.5
2
2.5
3
3.5
4
4.5
5
10
15
20
25

#### Optimal1 Fitted Gauss, proteinAT3G53040.1
#### bic:955 957 972
#### peaks after deleting less than one fifth of global max:7.9

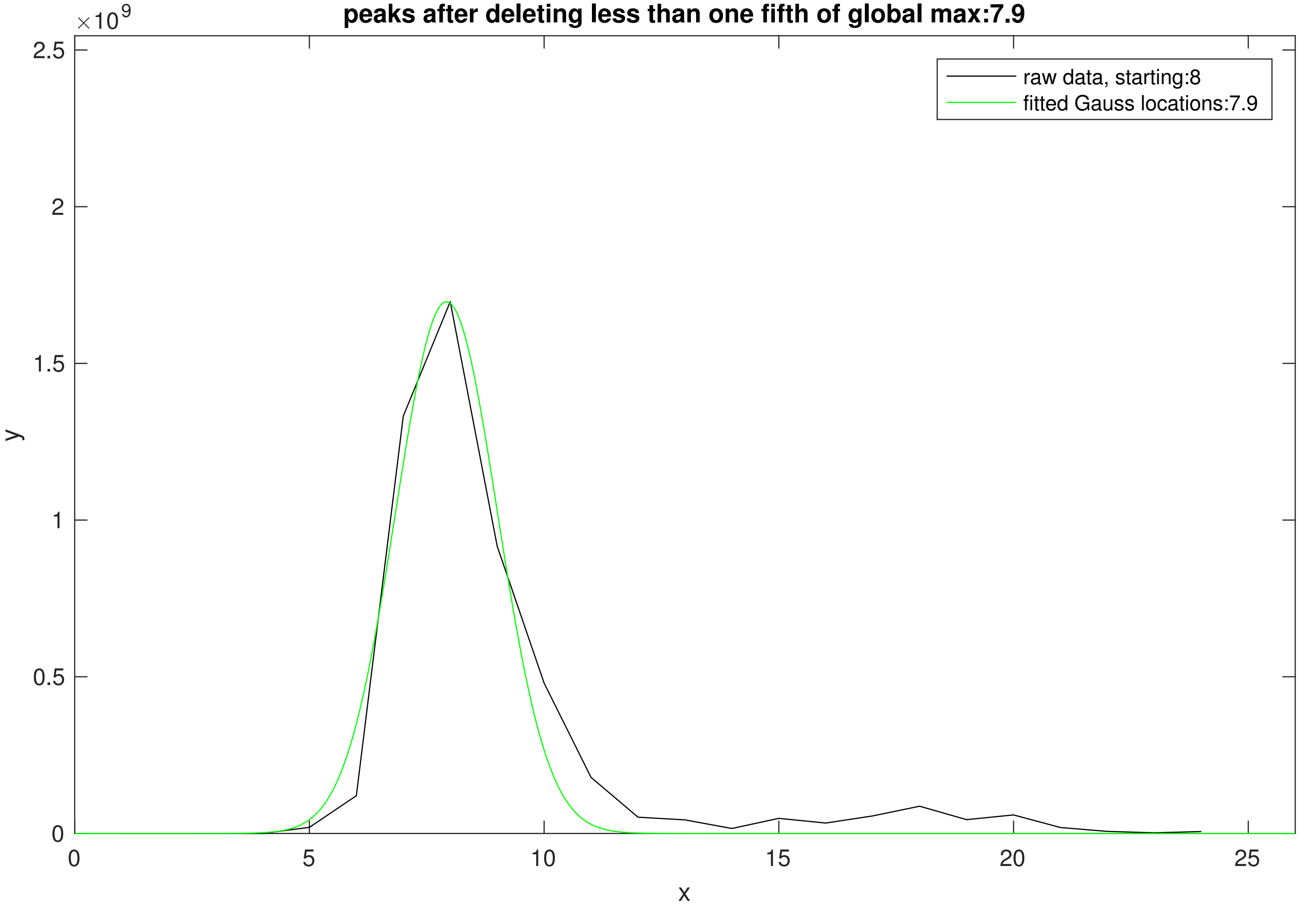

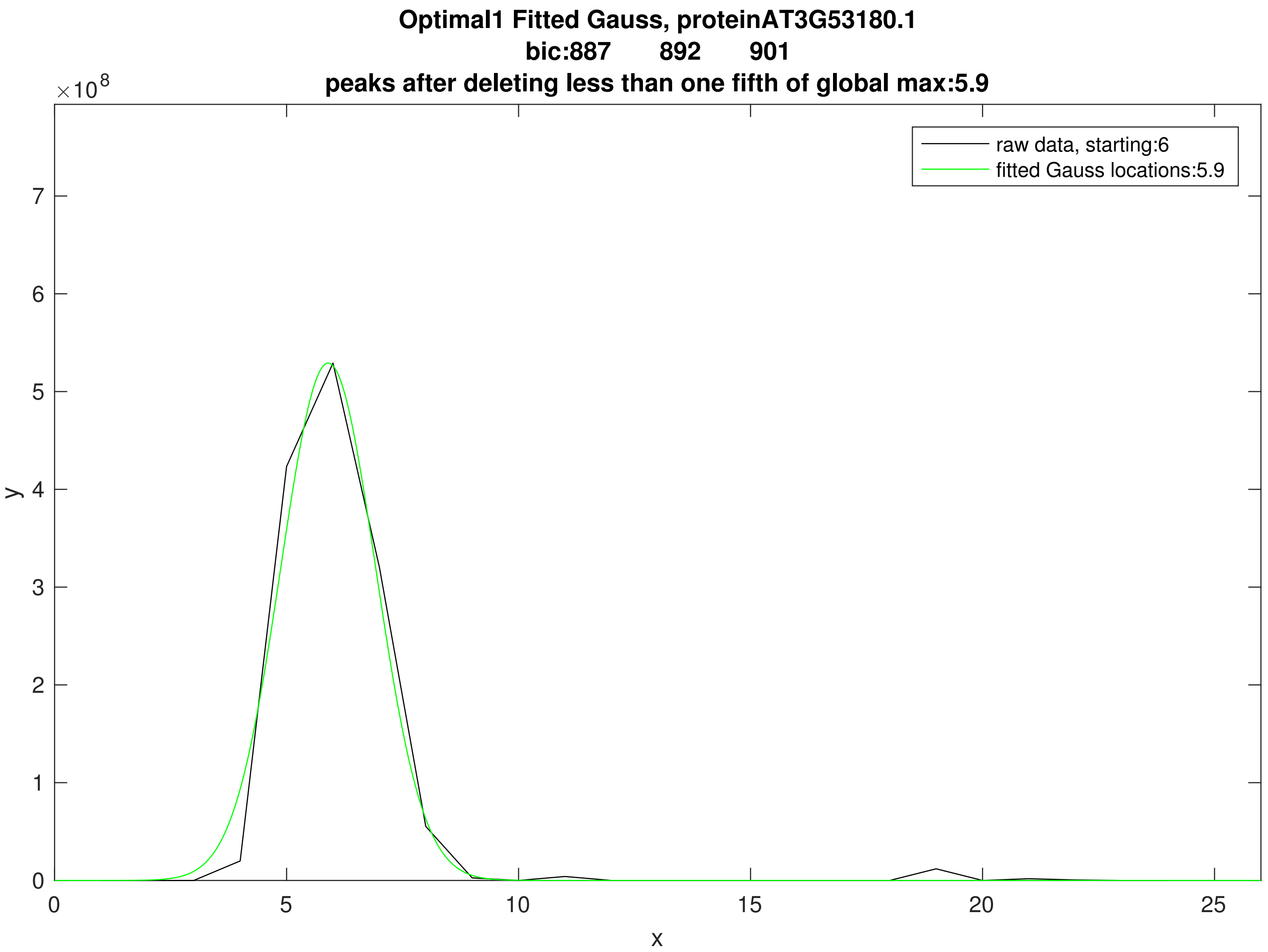
Optimal1 Fitted Gauss, proteinAT3G53180.1
bic:887 892 901
peaks after deleting less than one fifth of global max:5.9
×10^8
raw data, starting:6
fitted Gauss locations:5.9
y
x
0
1
2
3
4
5
6
7
0
5
10
15
20
25

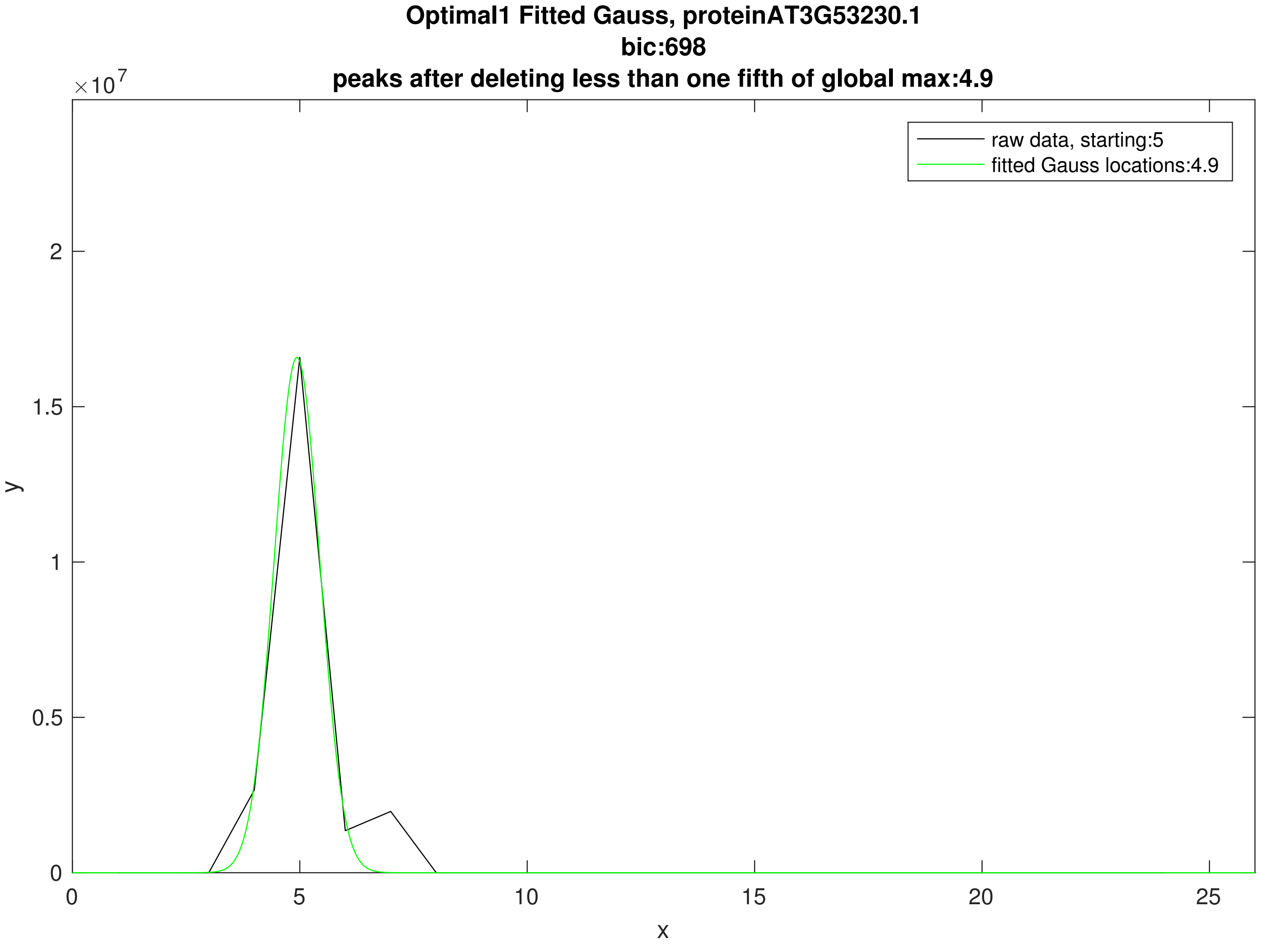

Optimal1 Fitted Gauss, proteinAT3G53230.1
bic:698
peaks after deleting less than one fifth of global max:4.9
×10^7
raw data, starting:5
fitted Gauss locations:4.9
2
1.5
1
0.5
0
y
0
5
10
15
20
25
x

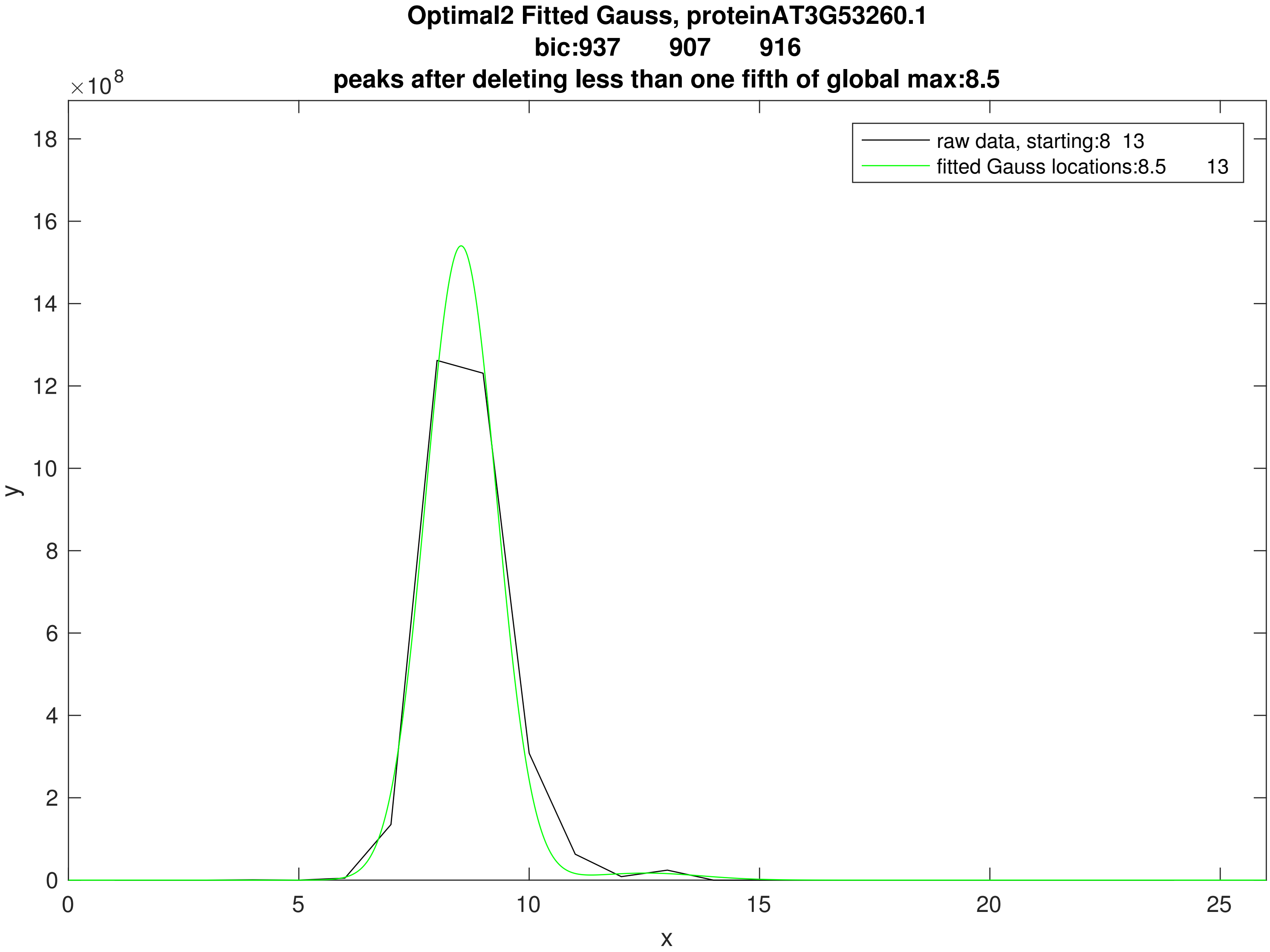

Optimal2 Fitted Gauss, proteinAT3G53260.1
bic:937 907 916
peaks after deleting less than one fifth of global max:8.5
×10^8
raw data, starting:8 13
fitted Gauss locations:8.5 13
18
16
14
12
10
8
6
4
2
0
y
0
5
10
15
20
25
x

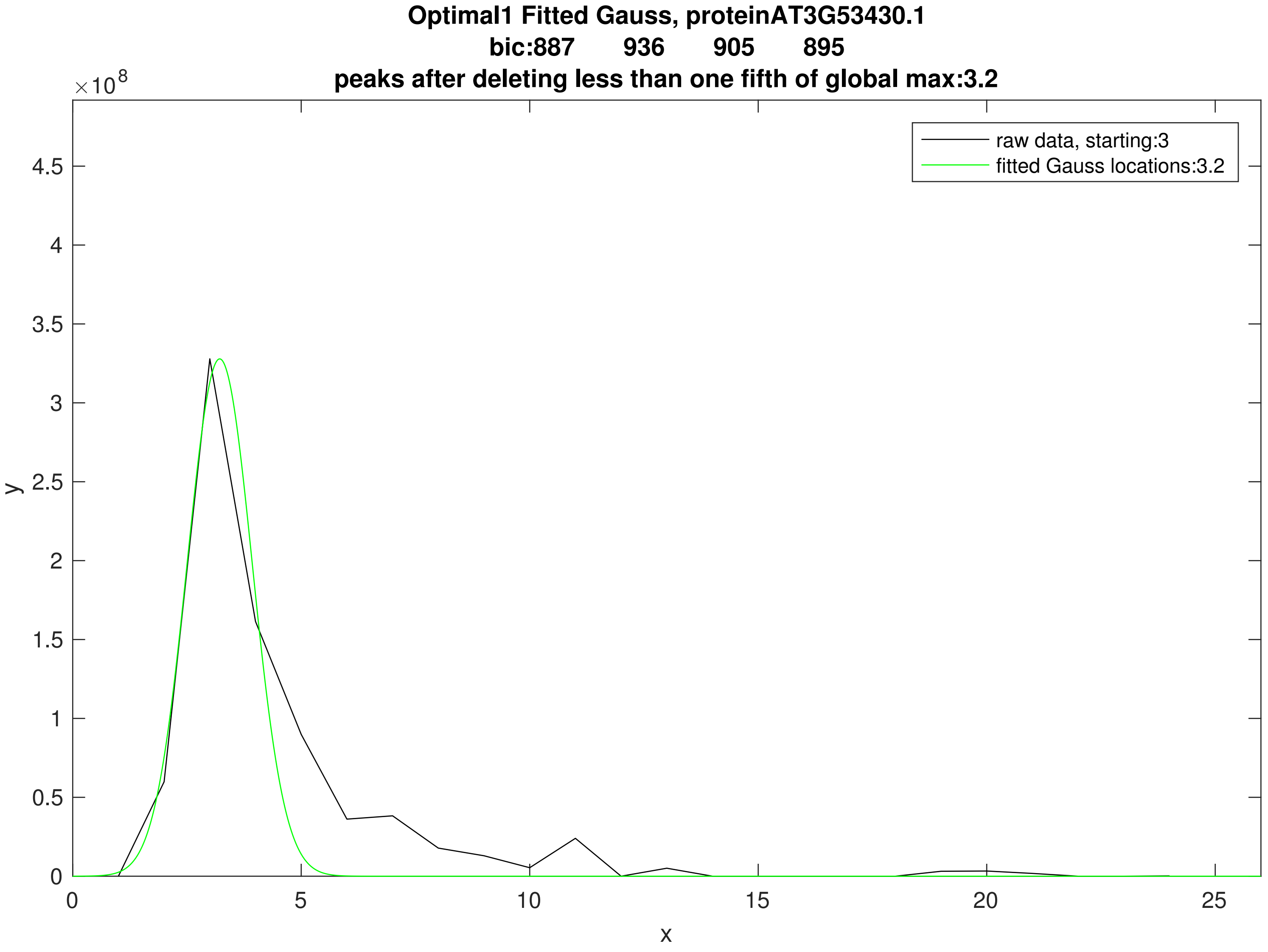

Optimal1 Fitted Gauss, proteinAT3G53430.1
bic:887 936 905 895
peaks after deleting less than one fifth of global max:3.2
×10^8
raw data, starting:3
fitted Gauss locations:3.2
4.5
4
3.5
3
2.5
2
1.5
1
0.5
0
y
0
5
10
15
20
25
x

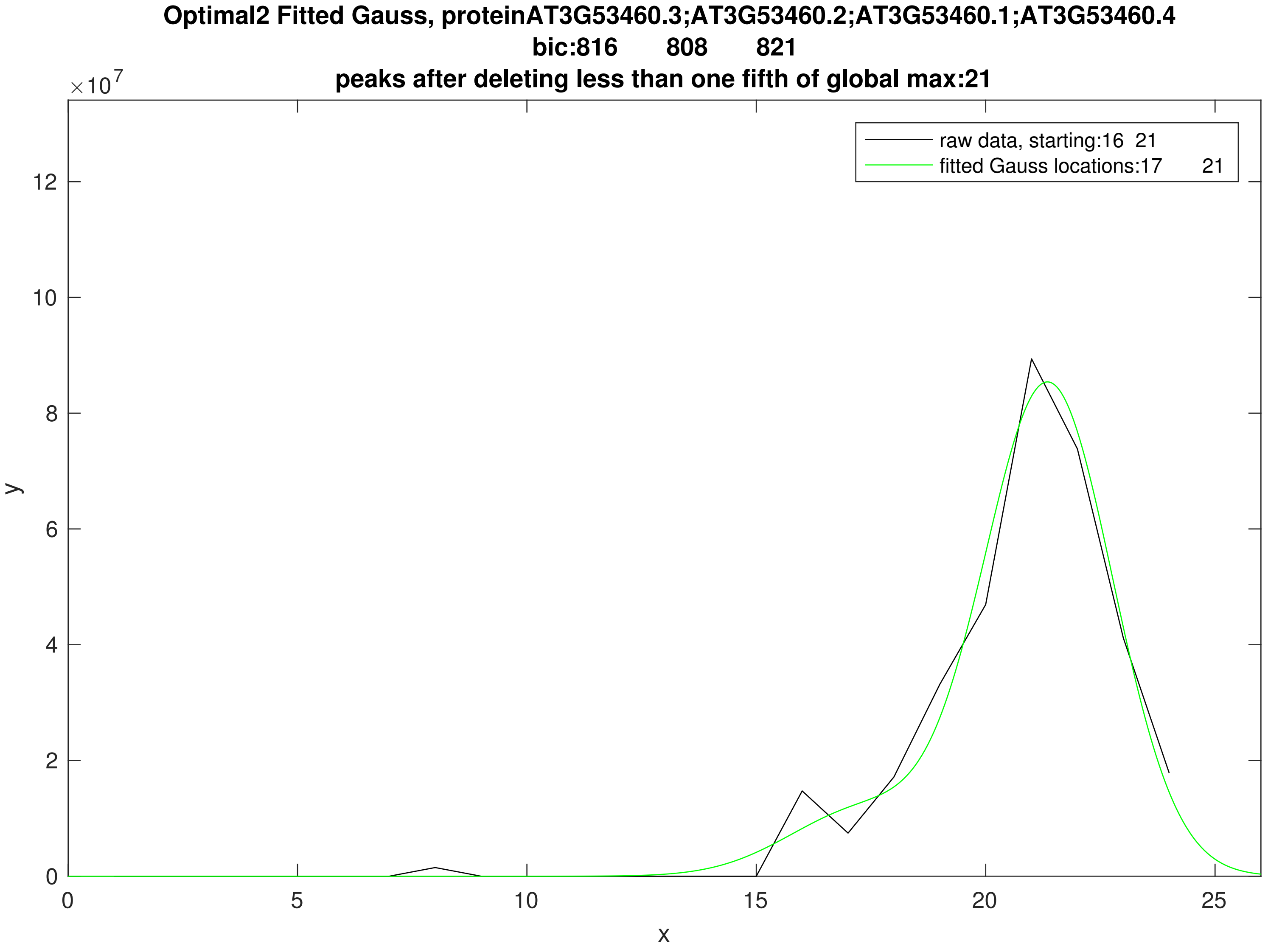

Optimal2 Fitted Gauss, proteinAT3G53460.3;AT3G53460.2;AT3G53460.1;AT3G53460.4
bic:816 808 821
peaks after deleting less than one fifth of global max:21
×10^7
raw data, starting:16 21
fitted Gauss locations:17 21
12
10
8
6
4
2
0
y
0
5
10
15
20
25
x

**Optimal1 Fitted Gauss, proteinAT3G53560.1**
**bic:890**
**peaks after deleting less than one fifth of global max:16**

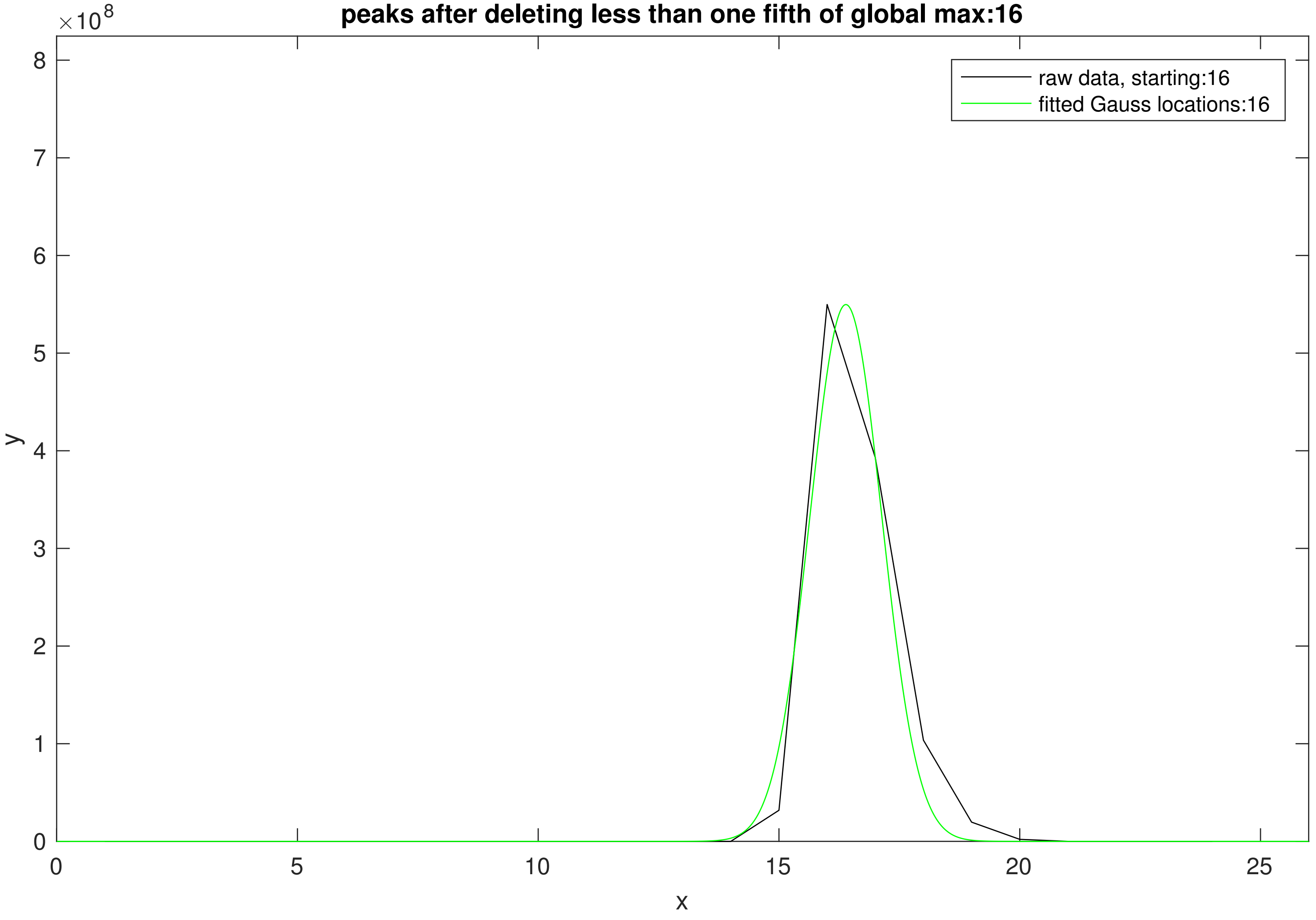

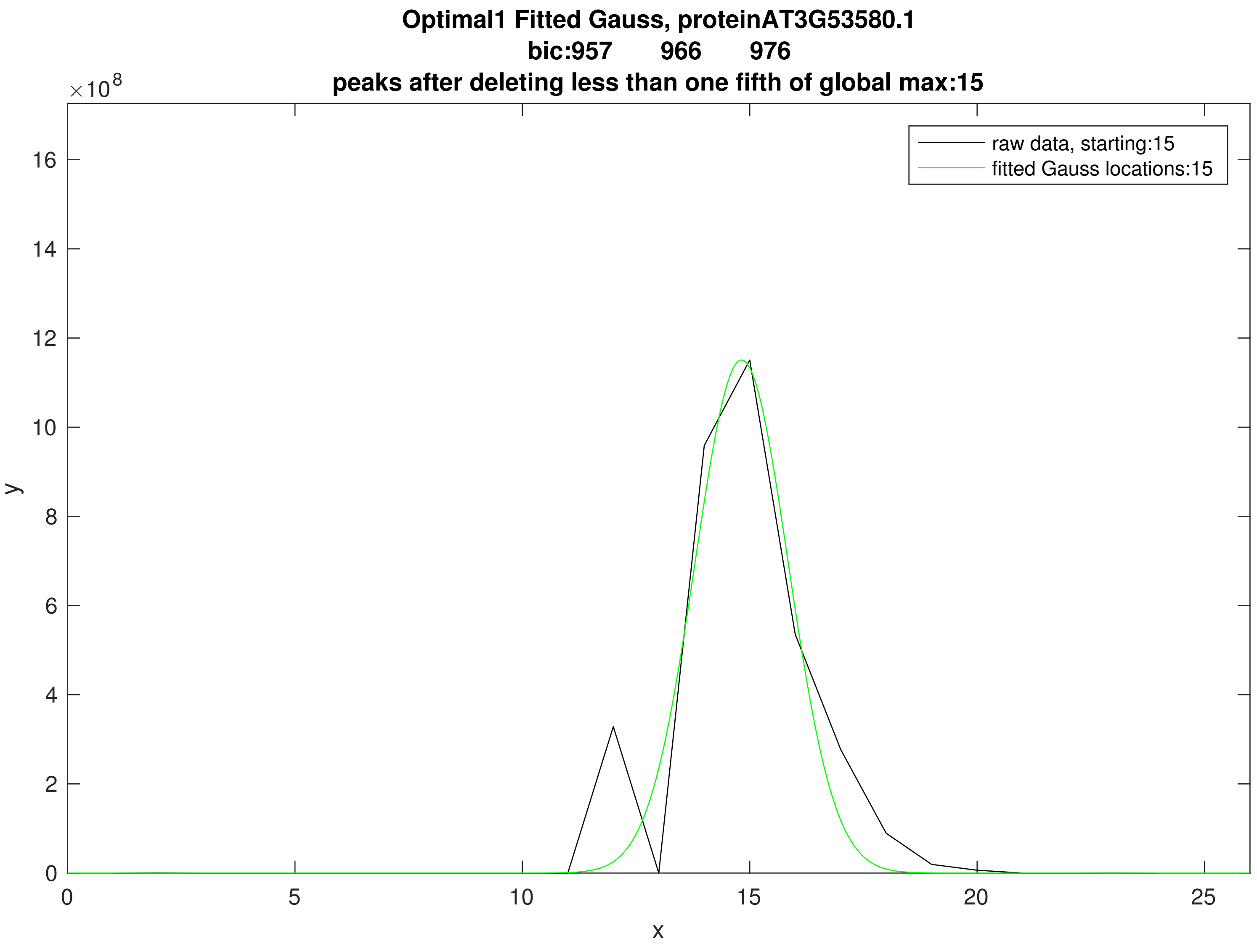
Optimal1 Fitted Gauss, proteinAT3G53580.1
bic:957 966 976
peaks after deleting less than one fifth of global max:15
×10^8
raw data, starting:15
fitted Gauss locations:15
y
x
0
2
4
6
8
10
12
14
16
0
5
10
15
20
25

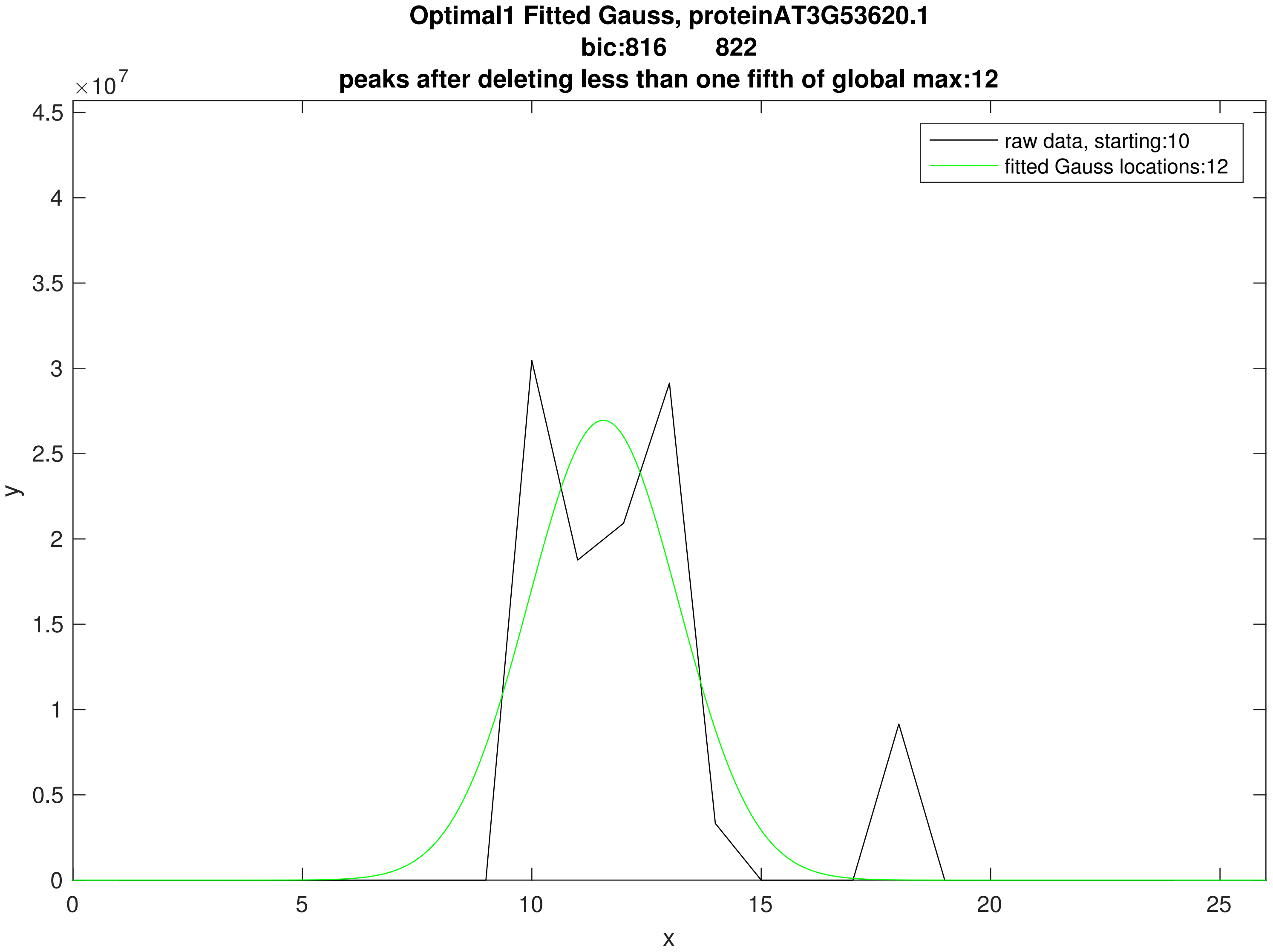

Optimal1 Fitted Gauss, proteinAT3G53620.1
bic:816 822
peaks after deleting less than one fifth of global max:12
×10^7
4.5
4
3.5
3
2.5
2
1.5
1
0.5
0
y
0
5
10
15
20
25
x
raw data, starting:10
fitted Gauss locations:12

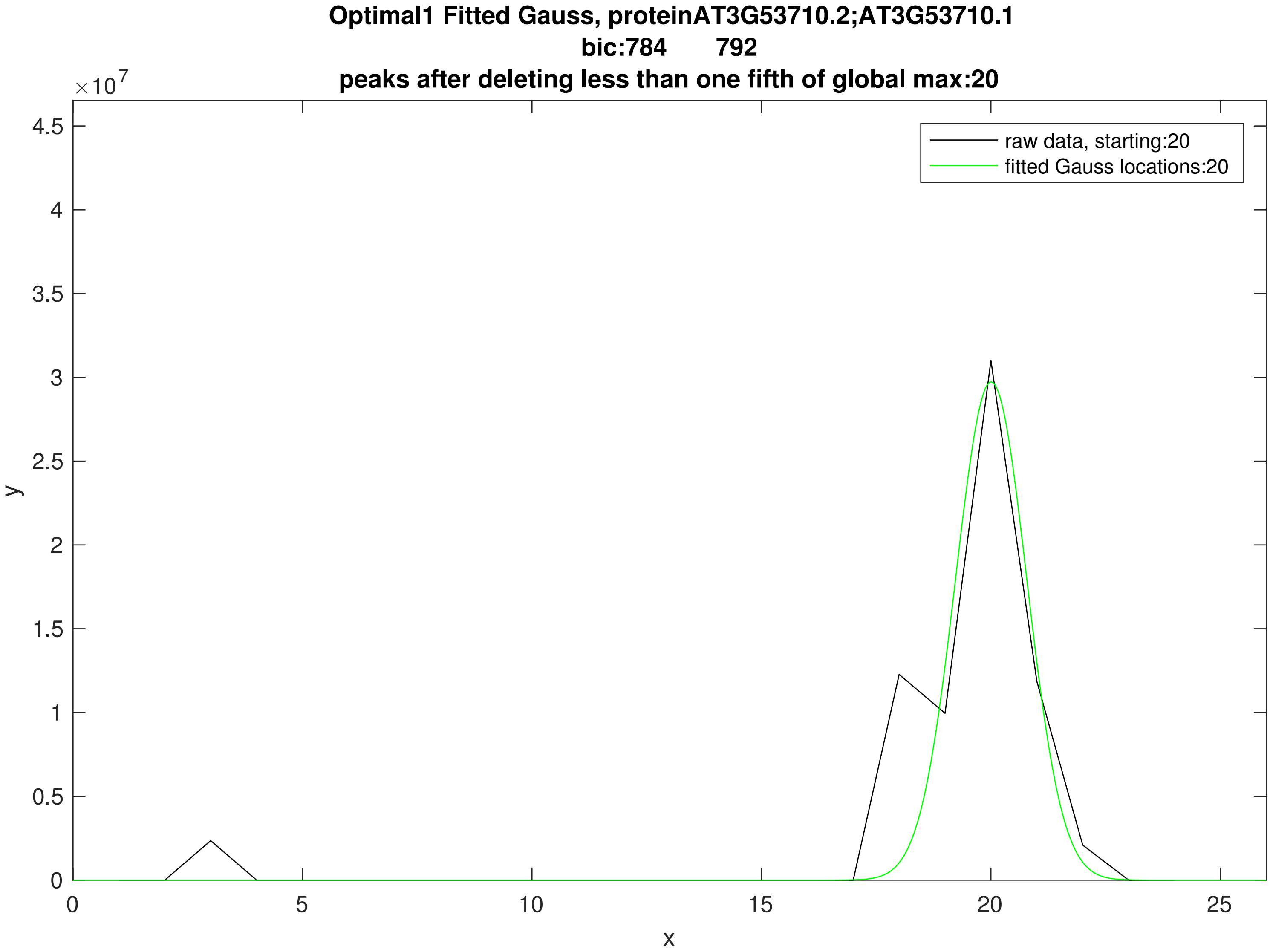

Optimal1 Fitted Gauss, proteinAT3G53710.2;AT3G53710.1
bic:784 792
peaks after deleting less than one fifth of global max:20
×10^7
4.5
4
3.5
3
2.5
2
1.5
1
0.5
0
y
0
5
10
15
20
25
x
raw data, starting:20
fitted Gauss locations:20

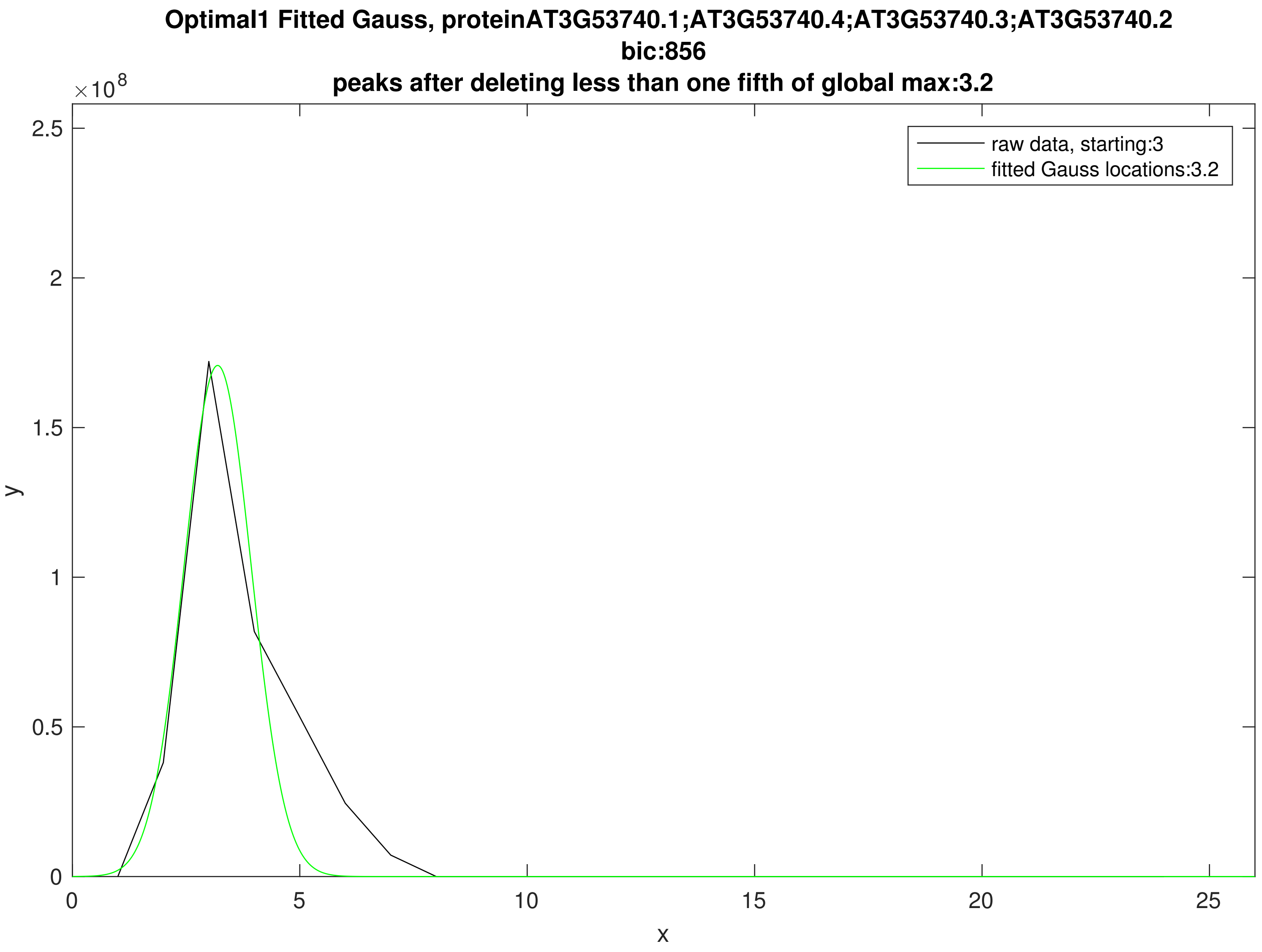

Optimal1 Fitted Gauss, proteinAT3G53740.1;AT3G53740.4;AT3G53740.3;AT3G53740.2
bic:856
peaks after deleting less than one fifth of global max:3.2
×10^8
2.5
2
1.5
1
0.5
0
y
0
5
10
15
20
25
x
raw data, starting:3
fitted Gauss locations:3.2

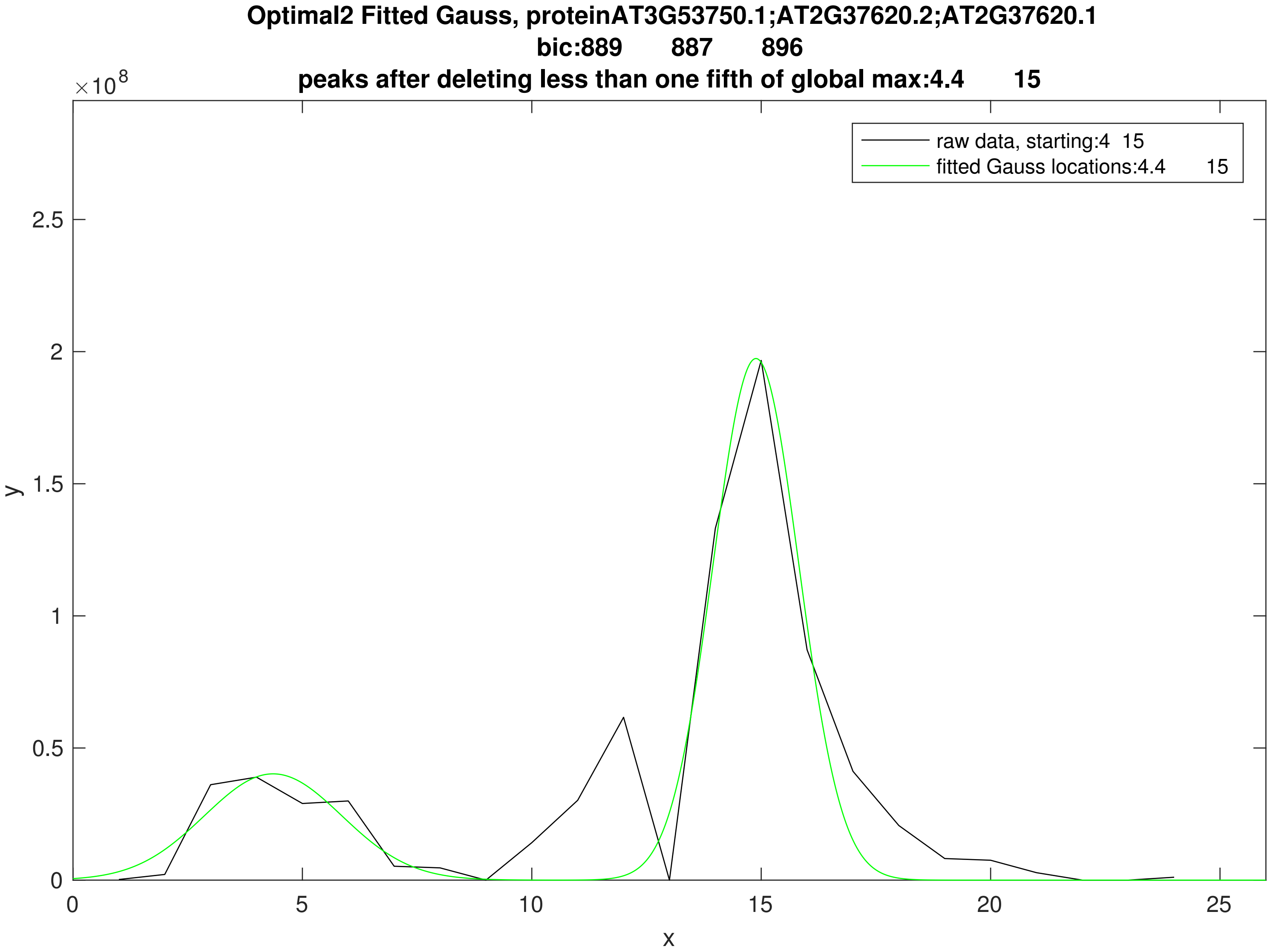
Optimal2 Fitted Gauss, proteinAT3G53750.1;AT2G37620.2;AT2G37620.1
bic:889 887 896
peaks after deleting less than one fifth of global max:4.4 15
×10^8
raw data, starting:4 15
fitted Gauss locations:4.4 15
y
x
0
0.5
1
1.5
2
2.5
5
10
15
20
25

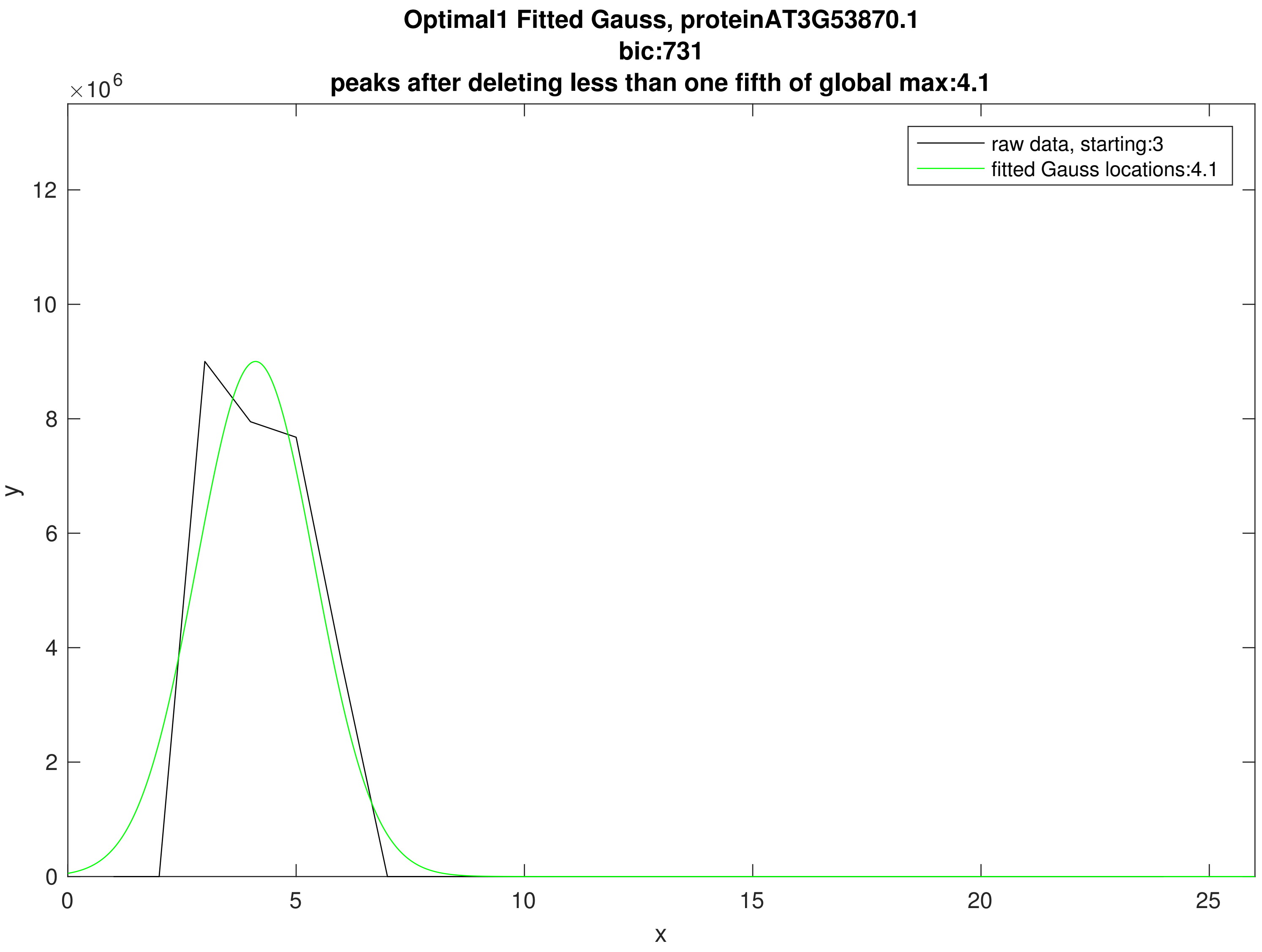
Optimal1 Fitted Gauss, proteinAT3G53870.1
bic:731
peaks after deleting less than one fifth of global max:4.1
×10^6
raw data, starting:3
fitted Gauss locations:4.1
12
10
8
6
4
2
0
y
0
5
10
15
20
25
x

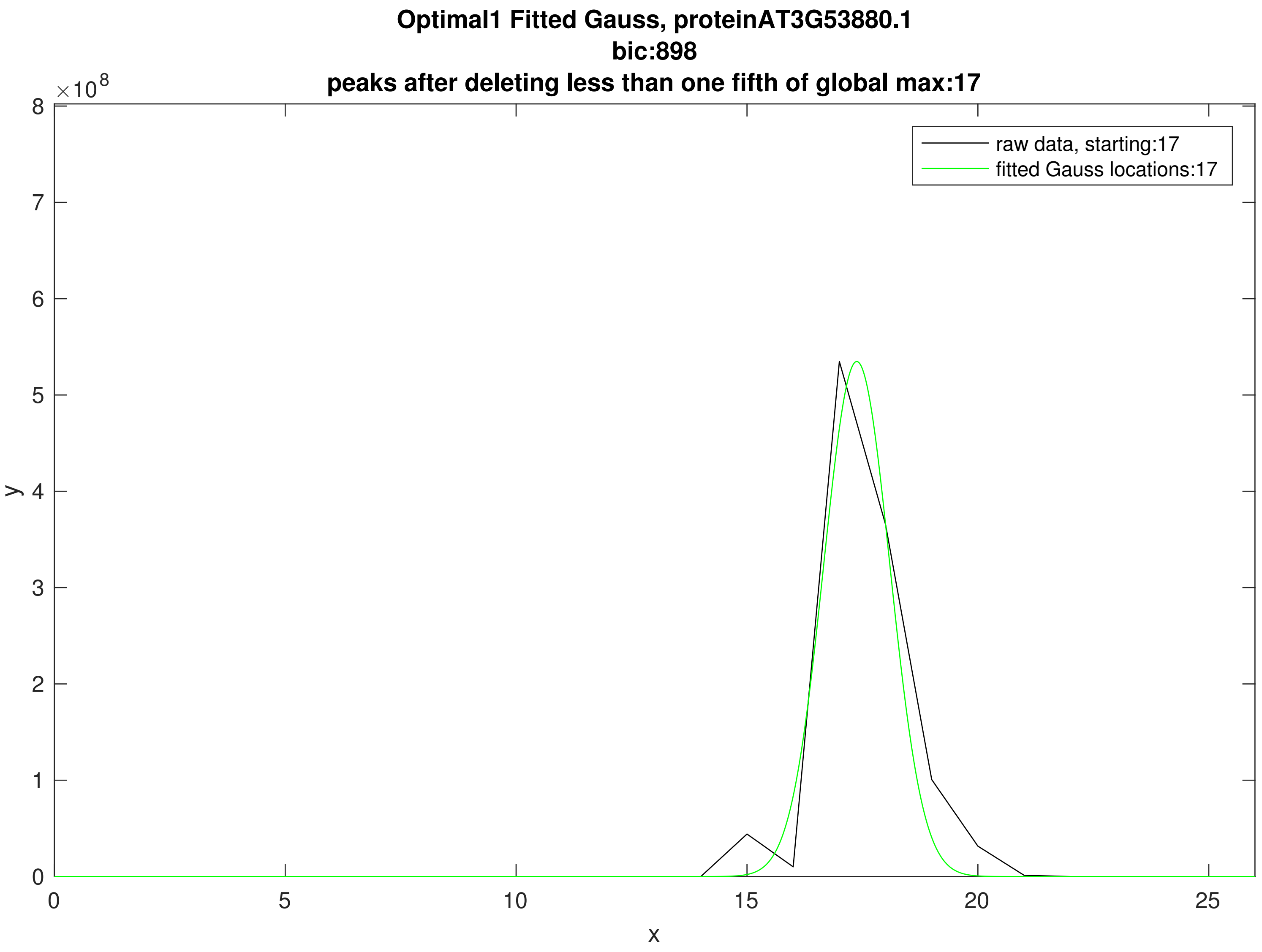
Optimal1 Fitted Gauss, proteinAT3G53880.1
bic:898
peaks after deleting less than one fifth of global max:17
×10^8
raw data, starting:17
fitted Gauss locations:17
y
x
0
1
2
3
4
5
6
7
8
0
5
10
15
20
25

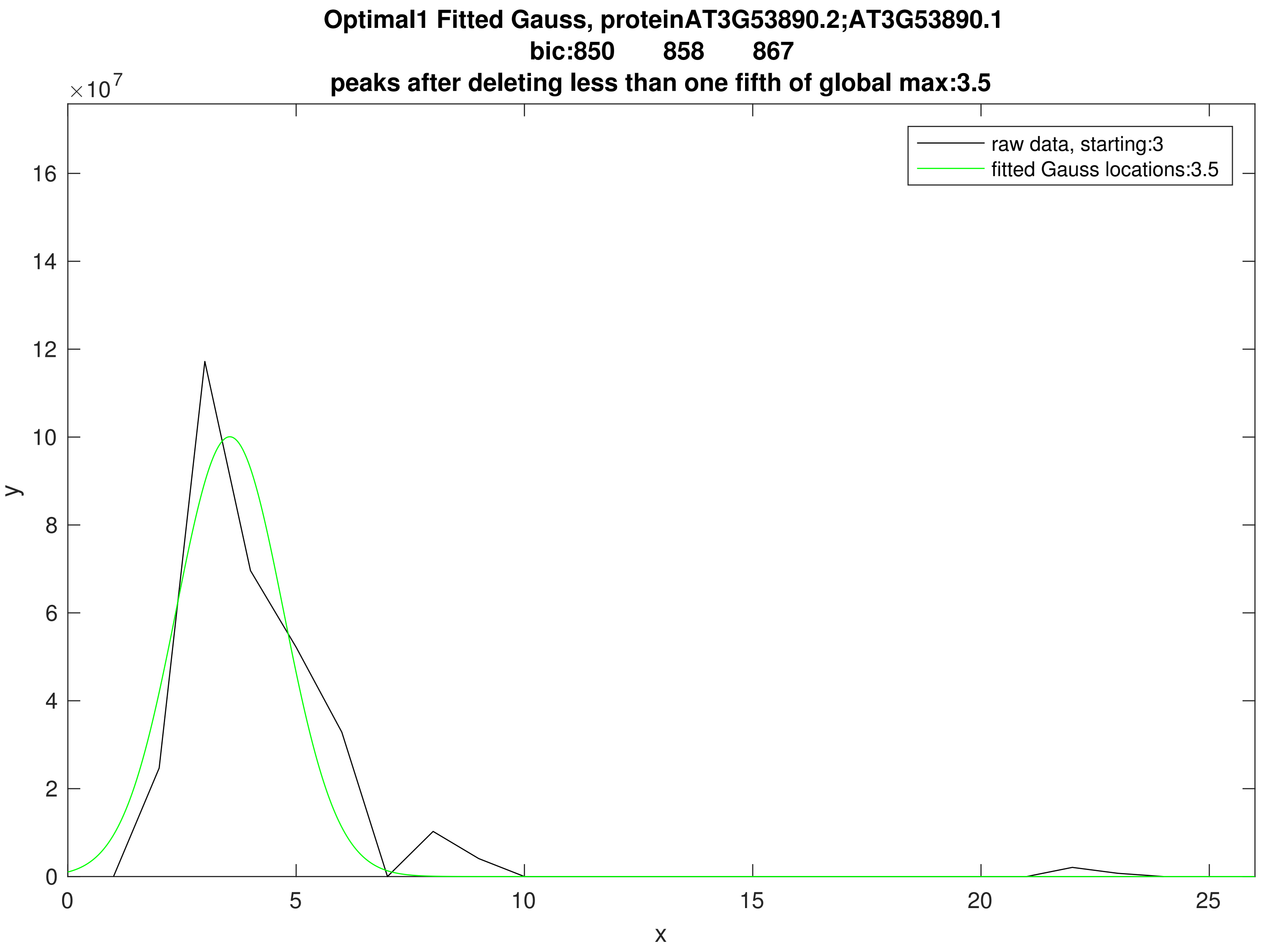

Optimal1 Fitted Gauss, proteinAT3G53890.2;AT3G53890.1
bic:850 858 867
peaks after deleting less than one fifth of global max:3.5
×10^7
raw data, starting:3
fitted Gauss locations:3.5
16
14
12
10
8
6
4
2
0
y
0
5
10
15
20
25
x

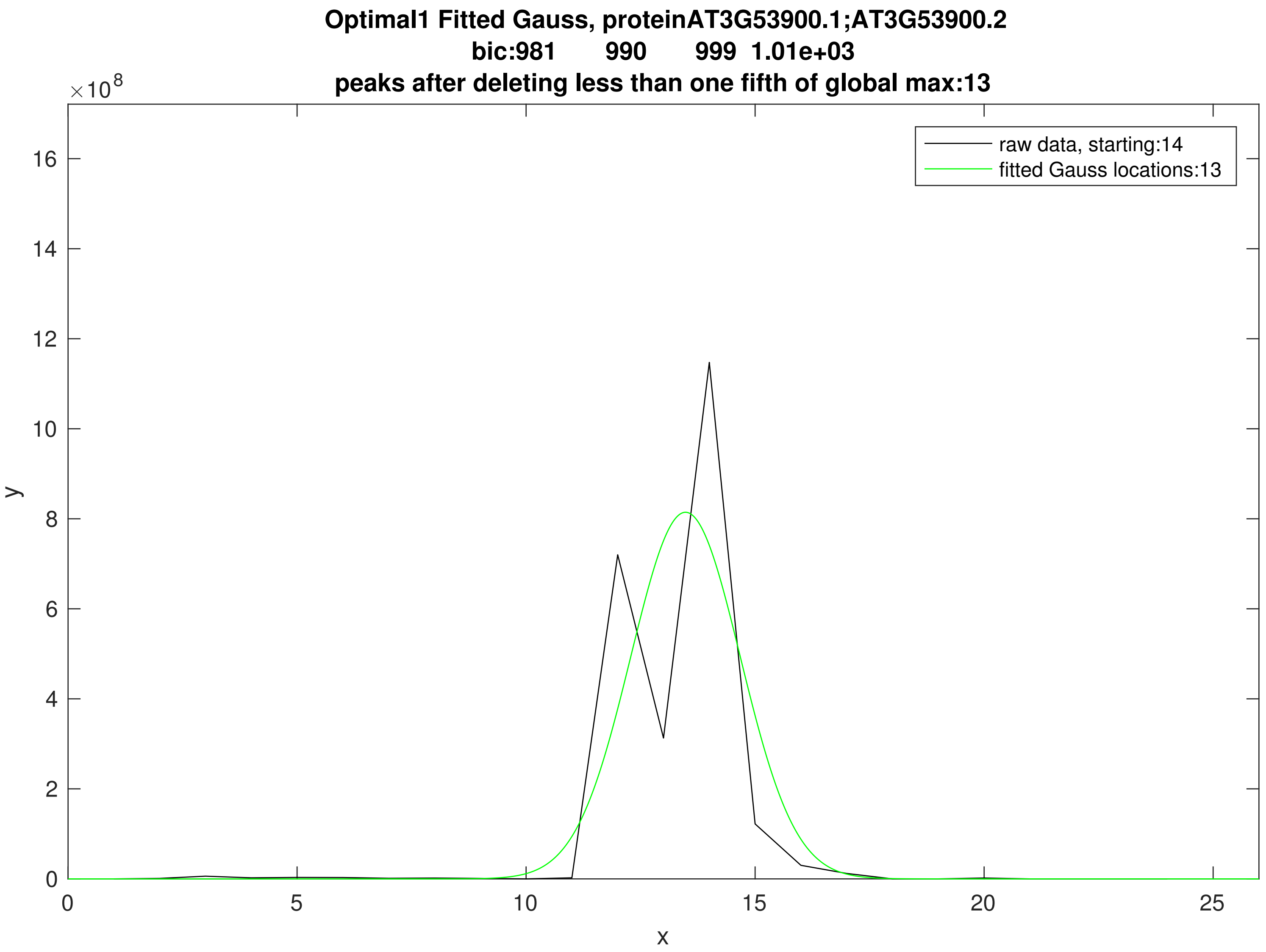
Optimal1 Fitted Gauss, proteinAT3G53900.1;AT3G53900.2
bic:981 990 999 1.01e+03
peaks after deleting less than one fifth of global max:13
×10^8
16
14
12
10
8
6
4
2
0
y
0
5
10
15
20
25
x
raw data, starting:14
fitted Gauss locations:13

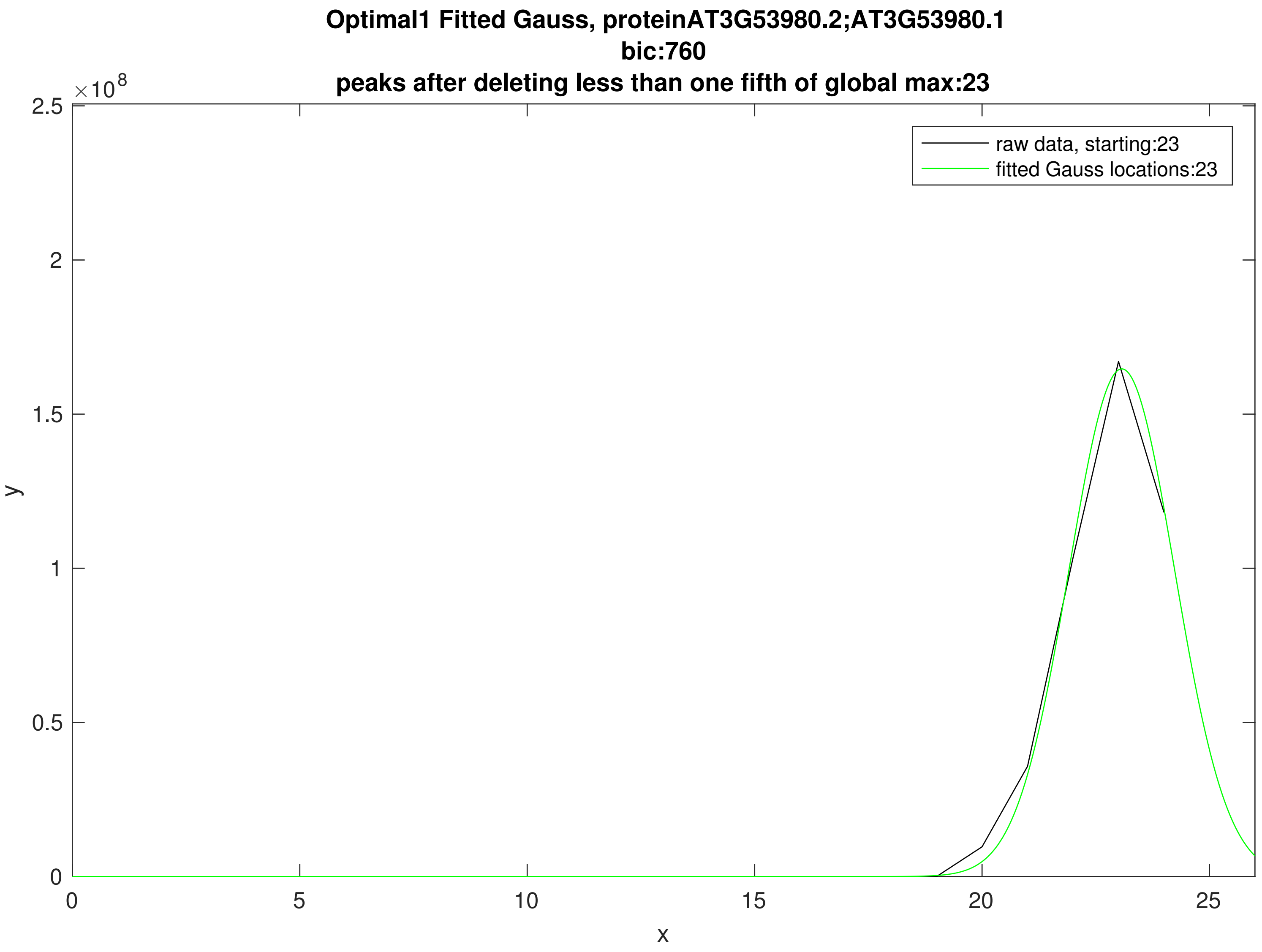
Optimal1 Fitted Gauss, proteinAT3G53980.2;AT3G53980.1
bic:760
peaks after deleting less than one fifth of global max:23
×10^8
2.5
2
1.5
1
0.5
0
y
0
5
10
15
20
25
x
raw data, starting:23
fitted Gauss locations:23

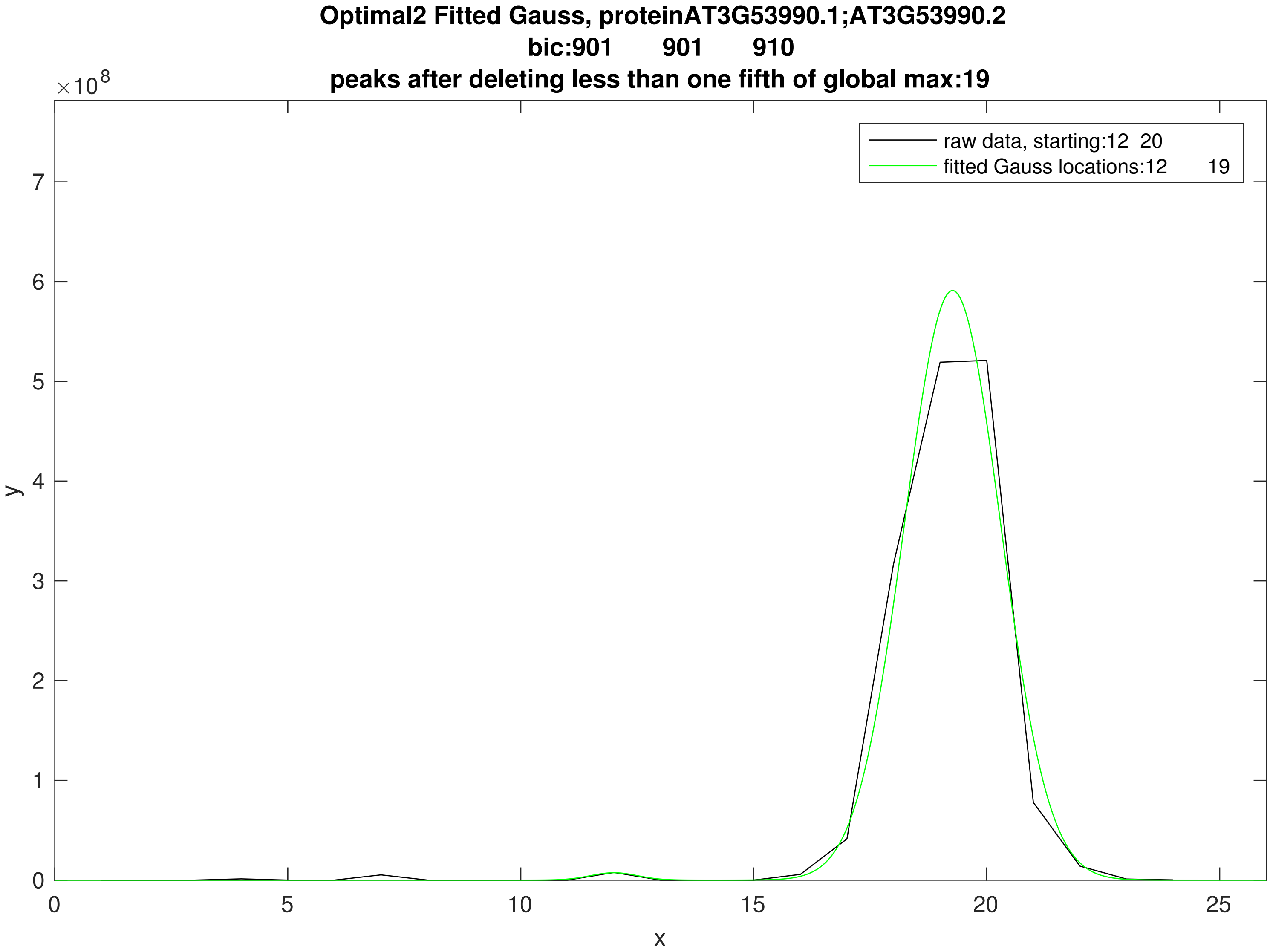
Optimal2 Fitted Gauss, proteinAT3G53990.1;AT3G53990.2
bic:901 901 910
peaks after deleting less than one fifth of global max:19
×10^8
raw data, starting:12 20
fitted Gauss locations:12 19
y
x
0
1
2
3
4
5
6
7
0
5
10
15
20
25

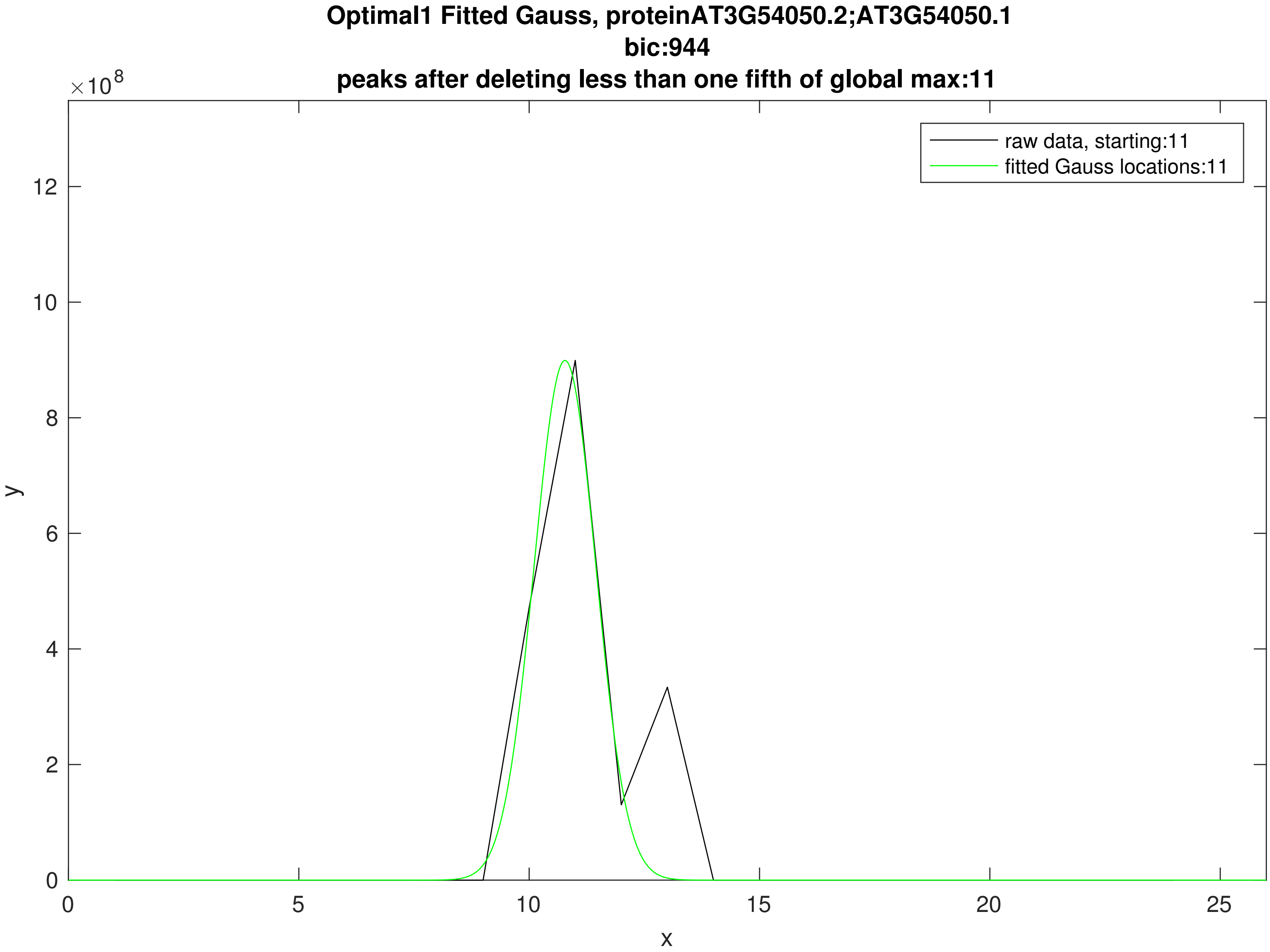

Optimal1 Fitted Gauss, proteinAT3G54050.2;AT3G54050.1
bic:944
peaks after deleting less than one fifth of global max:11
×10^8
raw data, starting:11
fitted Gauss locations:11
y
x
0
2
4
6
8
10
12
0
5
10
15
20
25

**Optimal1 Fitted Gauss, proteinAT3G54210.1**
**bic:781**
**peaks after deleting less than one fifth of global max:3.9**

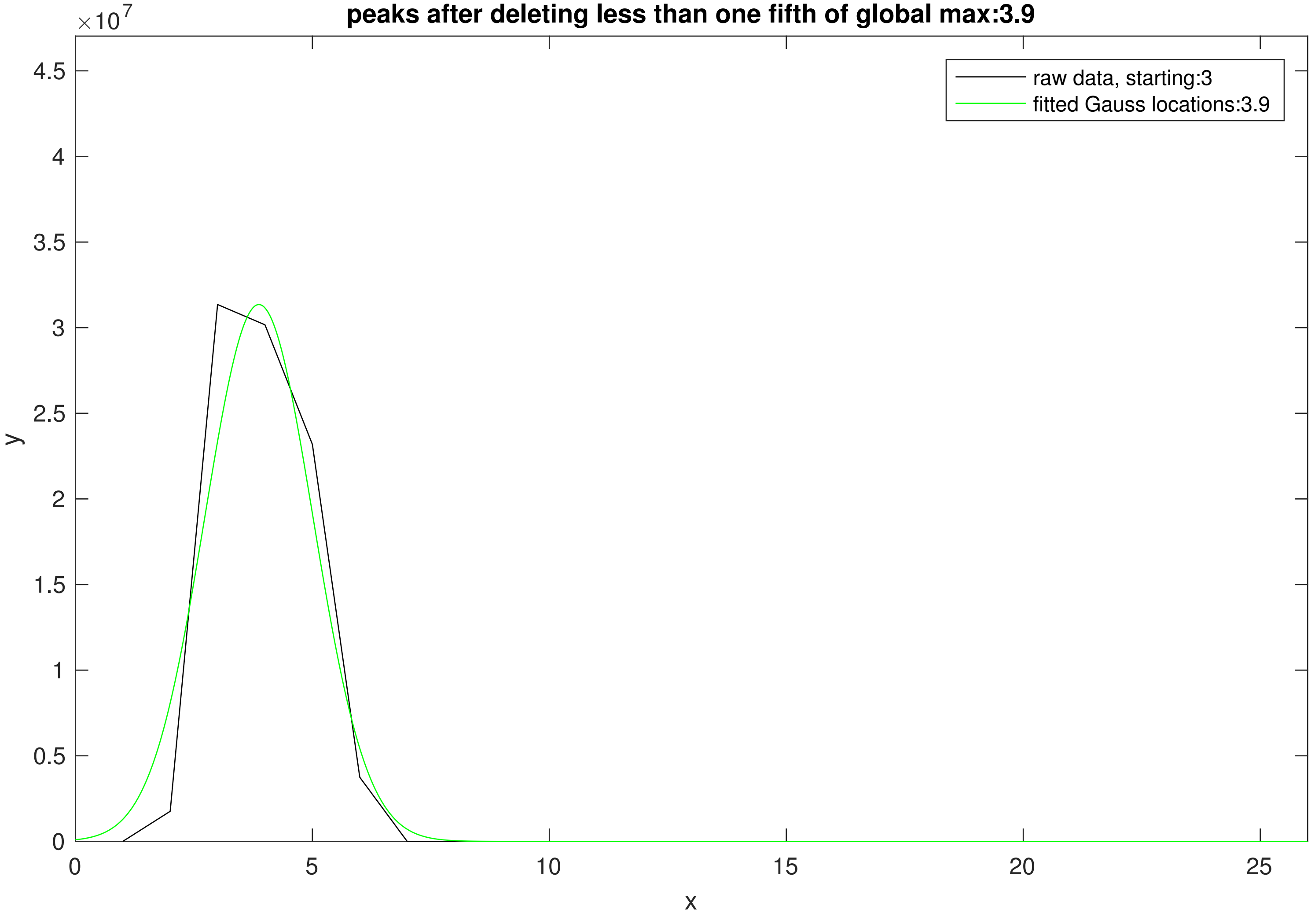

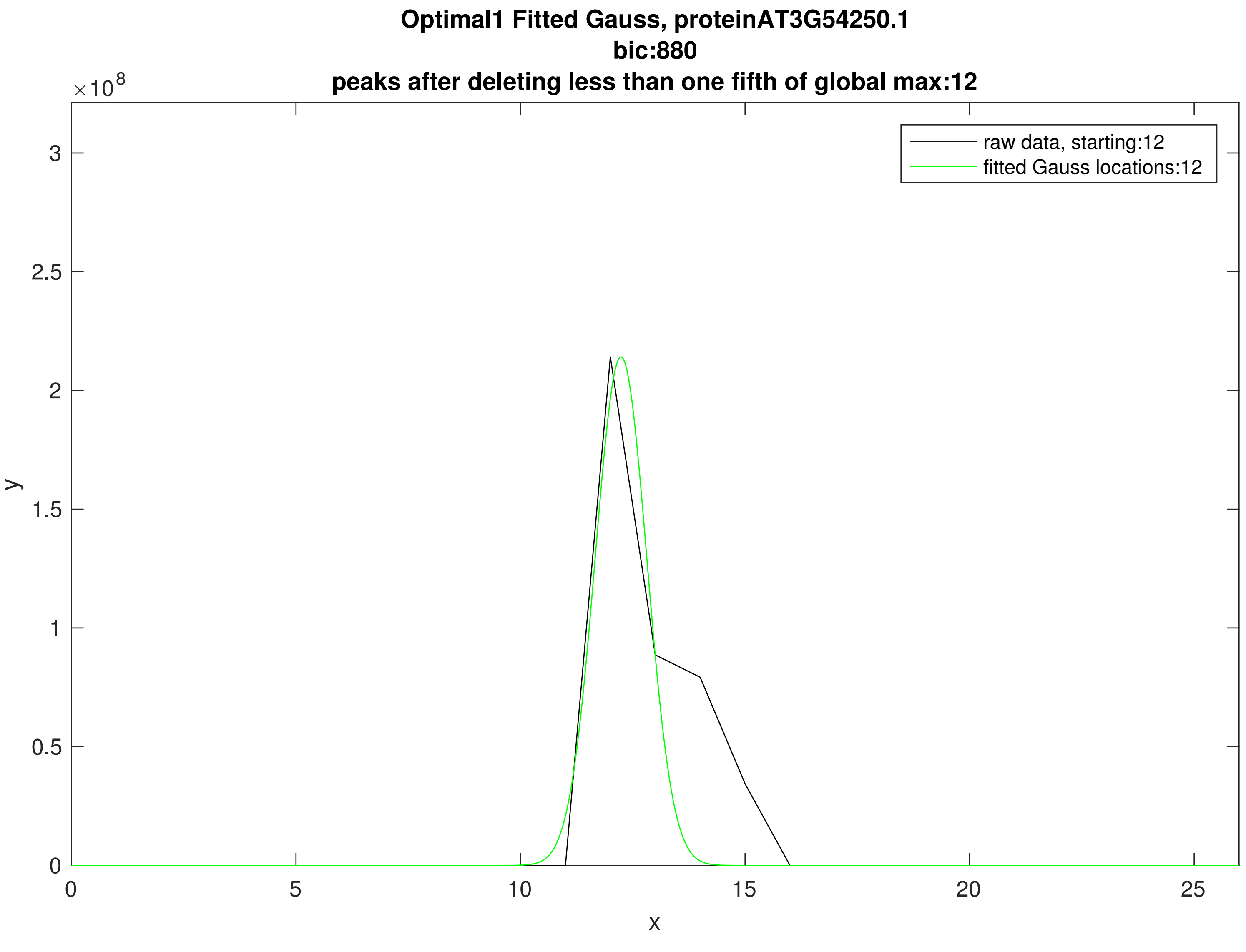
Optimal1 Fitted Gauss, proteinAT3G54250.1
bic:880
peaks after deleting less than one fifth of global max:12
×10^8
3
2.5
2
1.5
1
0.5
0
y
0
5
10
15
20
25
x
raw data, starting:12
fitted Gauss locations:12

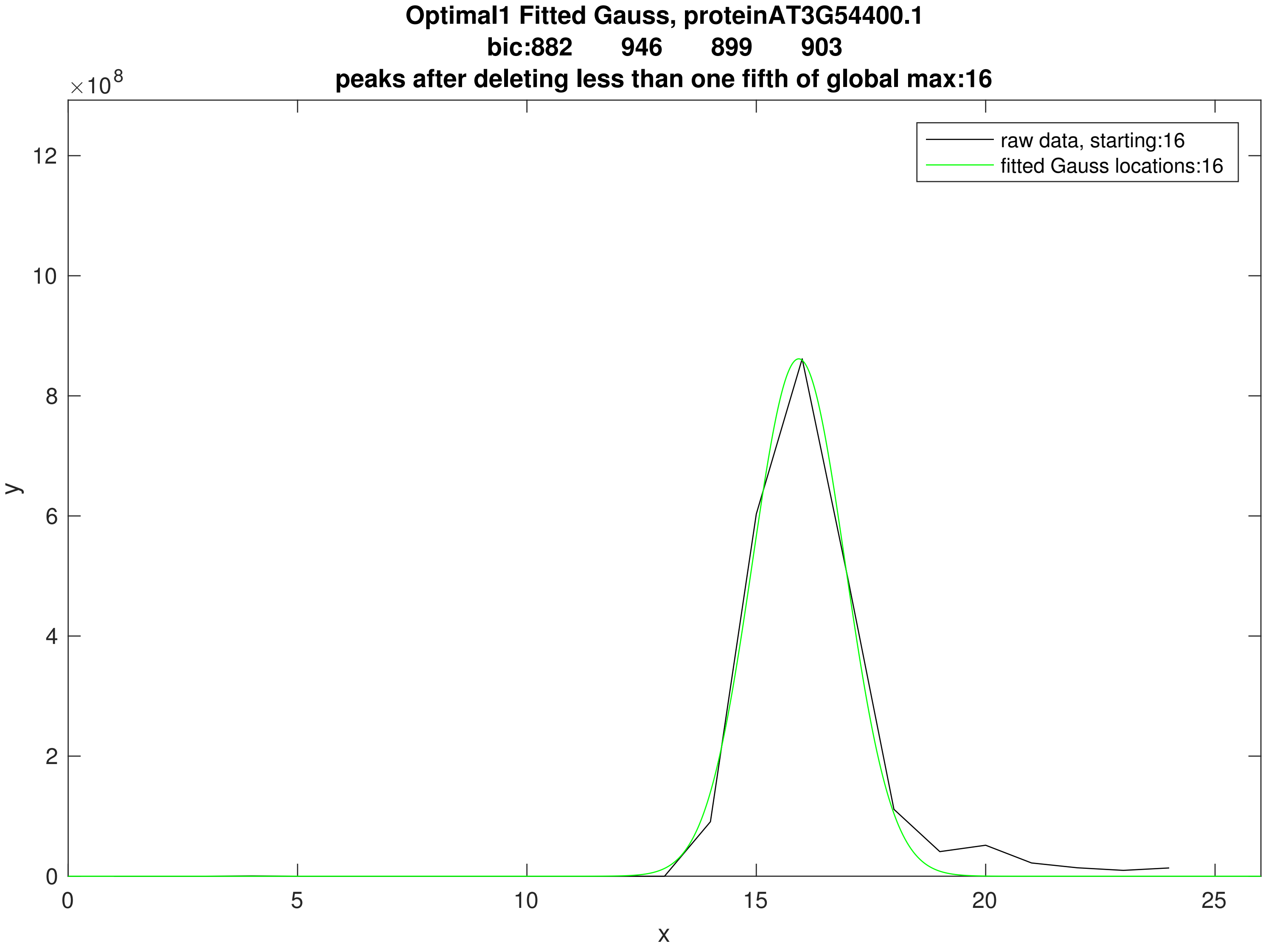

Optimal1 Fitted Gauss, proteinAT3G54400.1
bic:882 946 899 903
peaks after deleting less than one fifth of global max:16
×10^8
12
10
8
6
4
2
0
y
0
5
10
15
20
25
x
raw data, starting:16
fitted Gauss locations:16

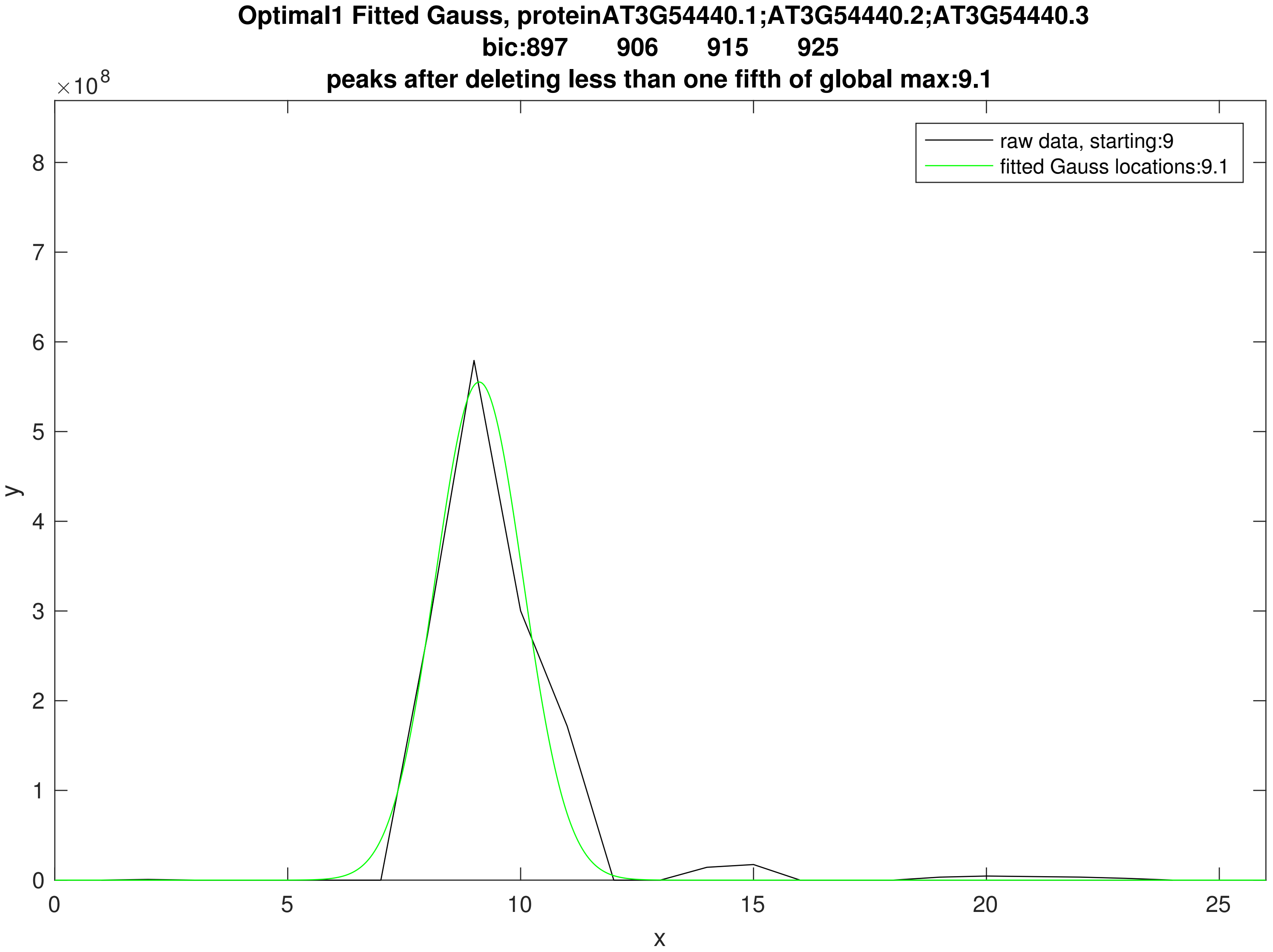

Optimal1 Fitted Gauss, proteinAT3G54440.1;AT3G54440.2;AT3G54440.3
bic:897 906 915 925
peaks after deleting less than one fifth of global max:9.1
×10^8
raw data, starting:9
fitted Gauss locations:9.1
y
x
0
1
2
3
4
5
6
7
8
0
5
10
15
20
25

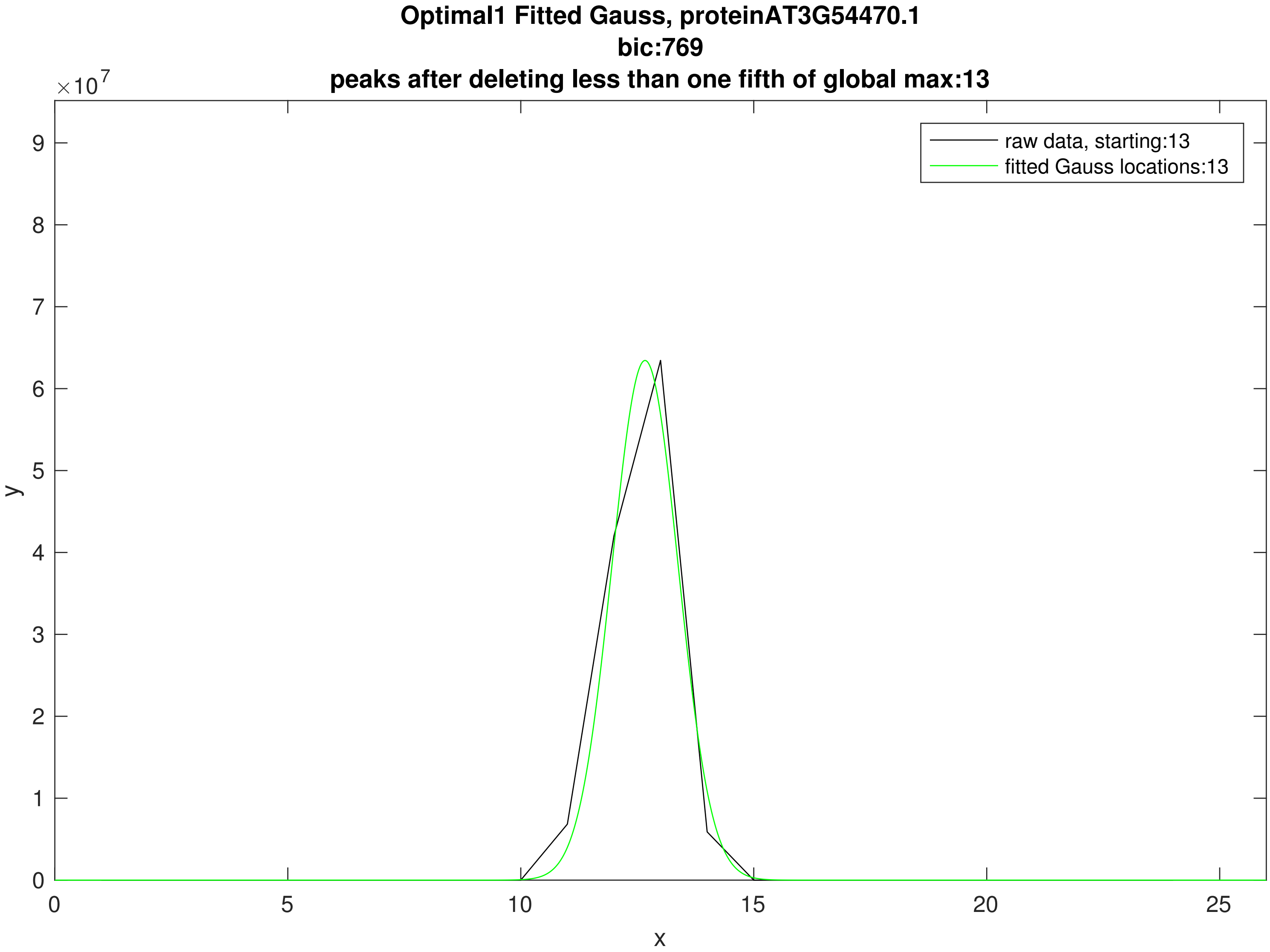

Optimal1 Fitted Gauss, proteinAT3G54470.1
bic:769
peaks after deleting less than one fifth of global max:13
×10^7
raw data, starting:13
fitted Gauss locations:13
y
x
0
1
2
3
4
5
6
7
8
9
0
5
10
15
20
25

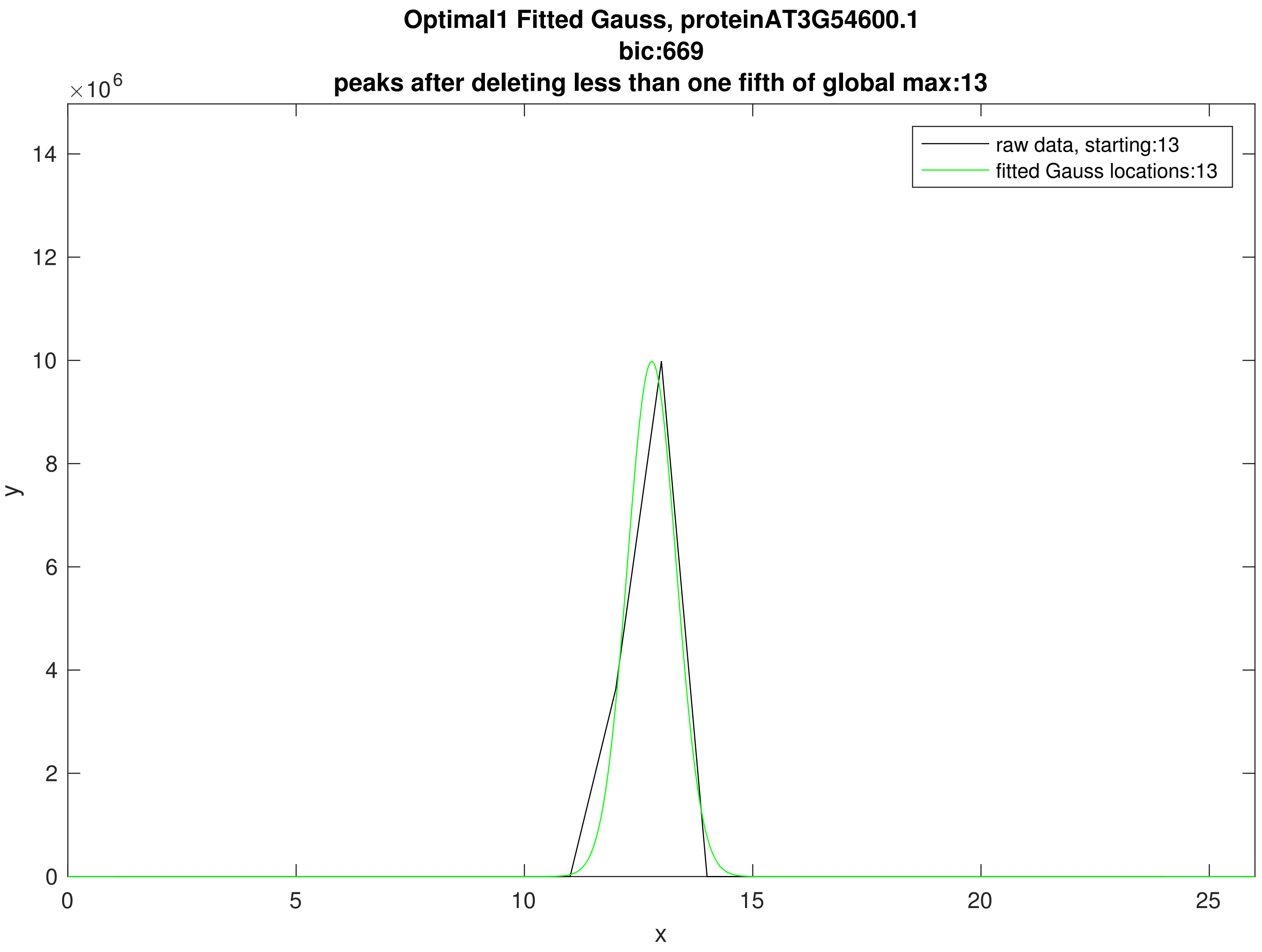
Optimal1 Fitted Gauss, proteinAT3G54600.1
bic:669
peaks after deleting less than one fifth of global max:13
×10^6
raw data, starting:13
fitted Gauss locations:13
y
x
0
2
4
6
8
10
12
14
0
5
10
15
20
25

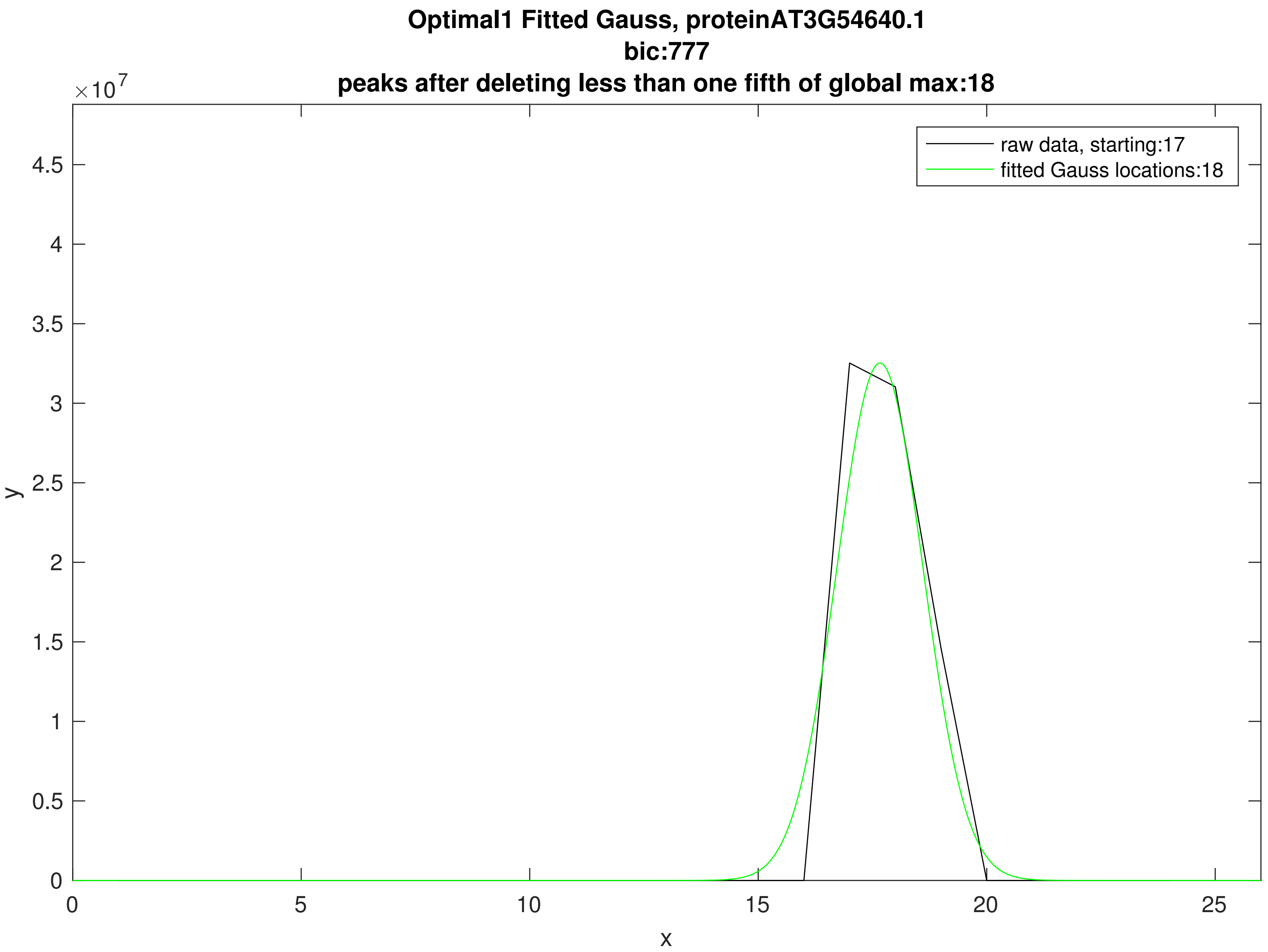

Optimal1 Fitted Gauss, proteinAT3G54640.1
bic:777
peaks after deleting less than one fifth of global max:18
×10^7
4.5
4
3.5
3
2.5
2
1.5
1
0.5
0
y
0
5
10
15
20
25
x
raw data, starting:17
fitted Gauss locations:18

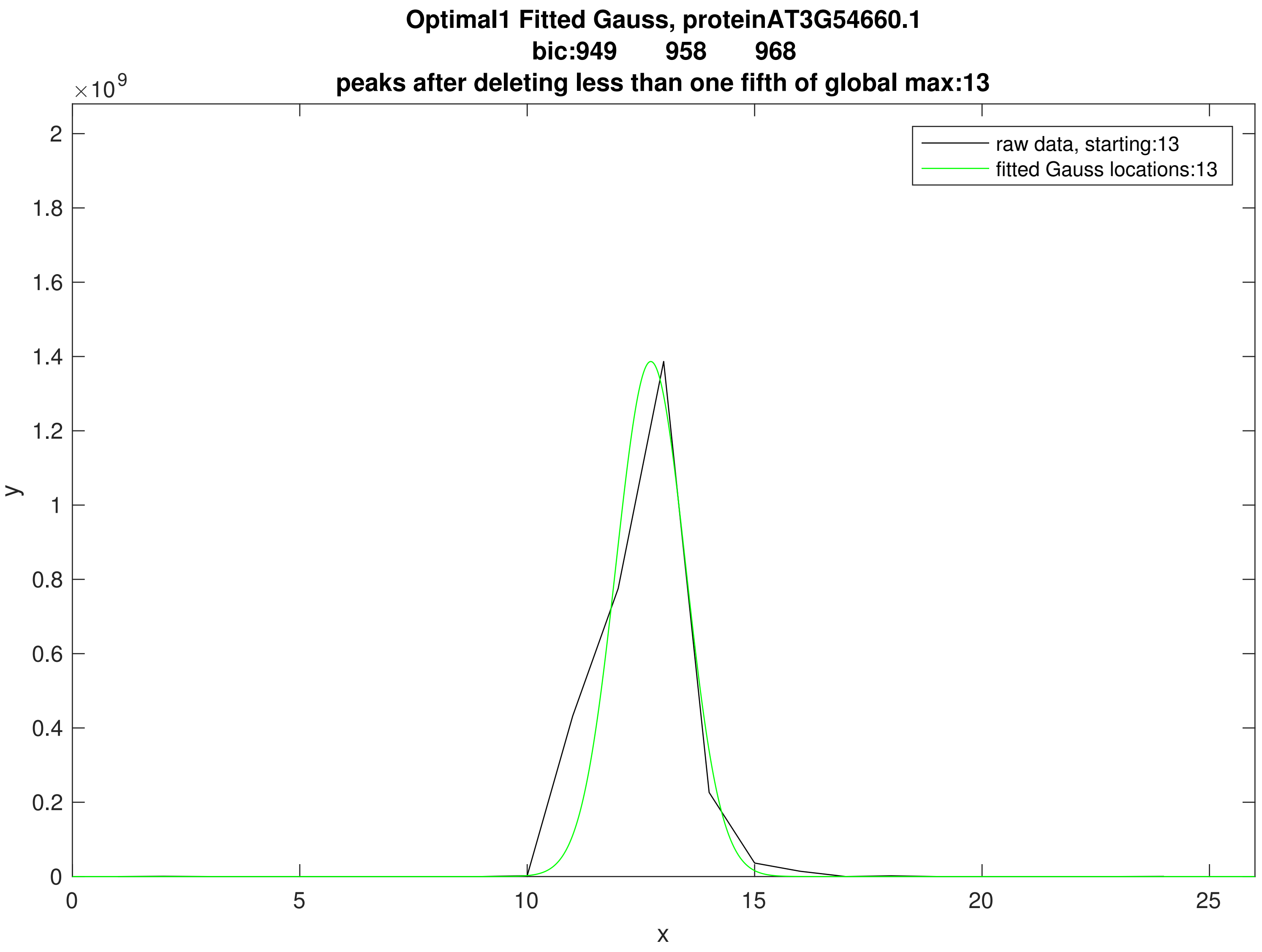
Optimal1 Fitted Gauss, proteinAT3G54660.1
bic:949 958 968
peaks after deleting less than one fifth of global max:13
×10^9
raw data, starting:13
fitted Gauss locations:13
y
x
0
0.2
0.4
0.6
0.8
1
1.2
1.4
1.6
1.8
2
0
5
10
15
20
25

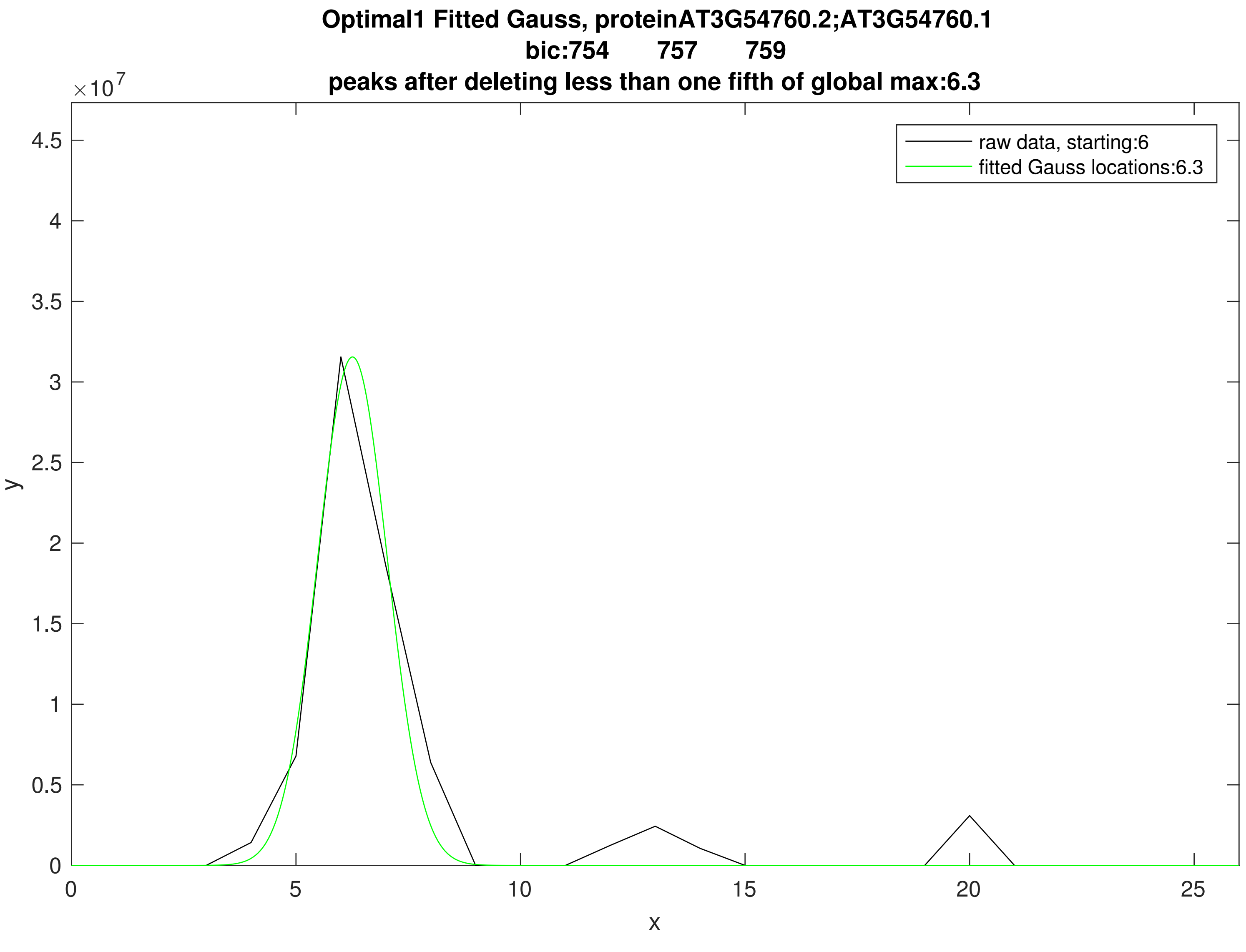
Optimal1 Fitted Gauss, proteinAT3G54760.2;AT3G54760.1
bic:754 757 759
peaks after deleting less than one fifth of global max:6.3
×10^7
raw data, starting:6
fitted Gauss locations:6.3
y
x
0
5
10
15
20
25
0.5
1
1.5
2
2.5
3
3.5
4
4.5

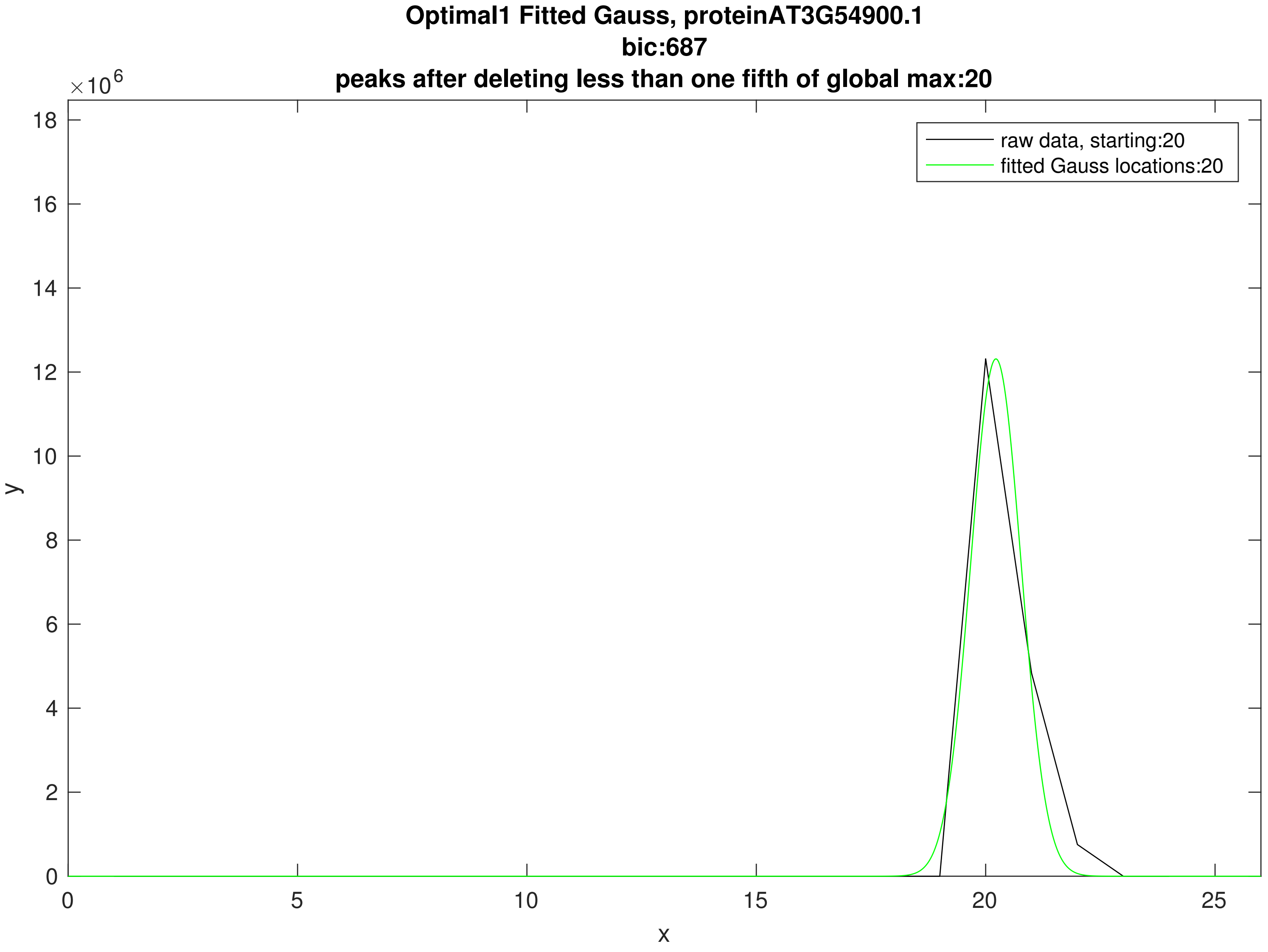

Optimal1 Fitted Gauss, proteinAT3G54900.1
bic:687
peaks after deleting less than one fifth of global max:20
×10^6
raw data, starting:20
fitted Gauss locations:20
y
x
0
2
4
6
8
10
12
14
16
18
0
5
10
15
20
25

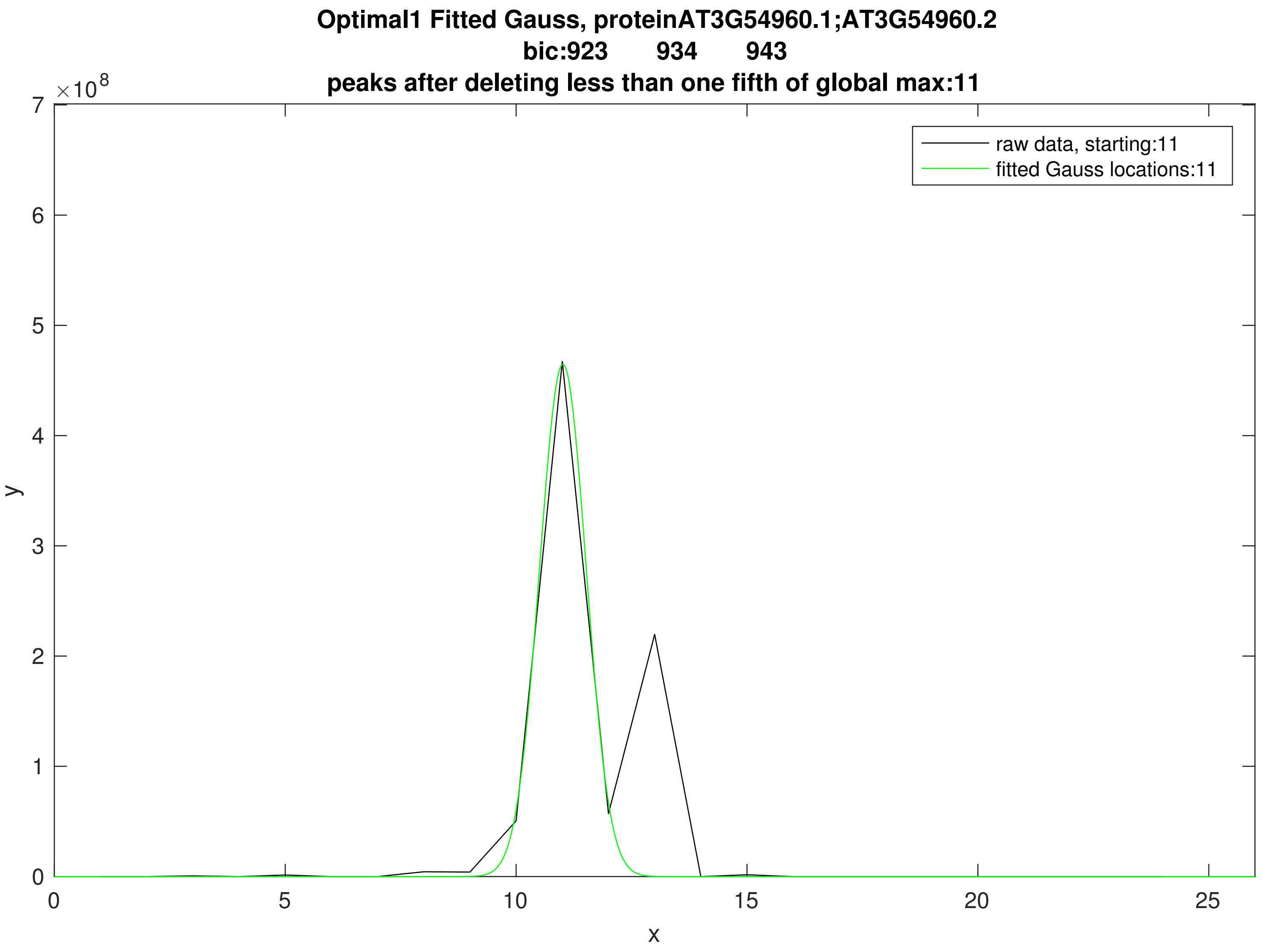
Optimal1 Fitted Gauss, proteinAT3G54960.1;AT3G54960.2
bic:923 934 943
peaks after deleting less than one fifth of global max:11
raw data, starting:11
fitted Gauss locations:11
y
x
×10^8
0
1
2
3
4
5
6
7
0
5
10
15
20
25

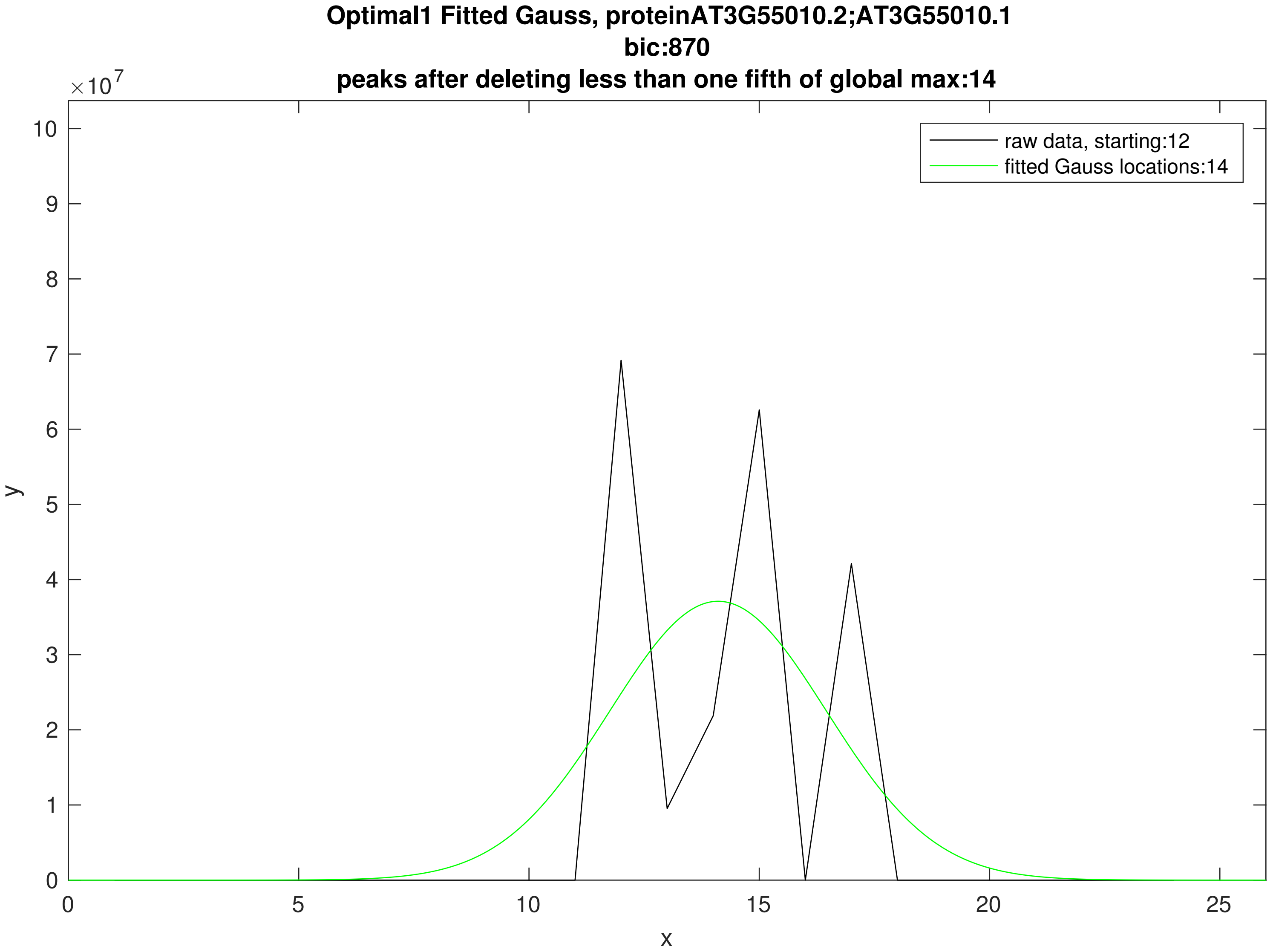

Optimal1 Fitted Gauss, proteinAT3G55010.2;AT3G55010.1
bic:870
peaks after deleting less than one fifth of global max:14
×10^7
raw data, starting:12
fitted Gauss locations:14
y
x

**Optimal1 Fitted Gauss, proteinAT3G55040.1**
**bic:753**
**peaks after deleting less than one fifth of global max:18**

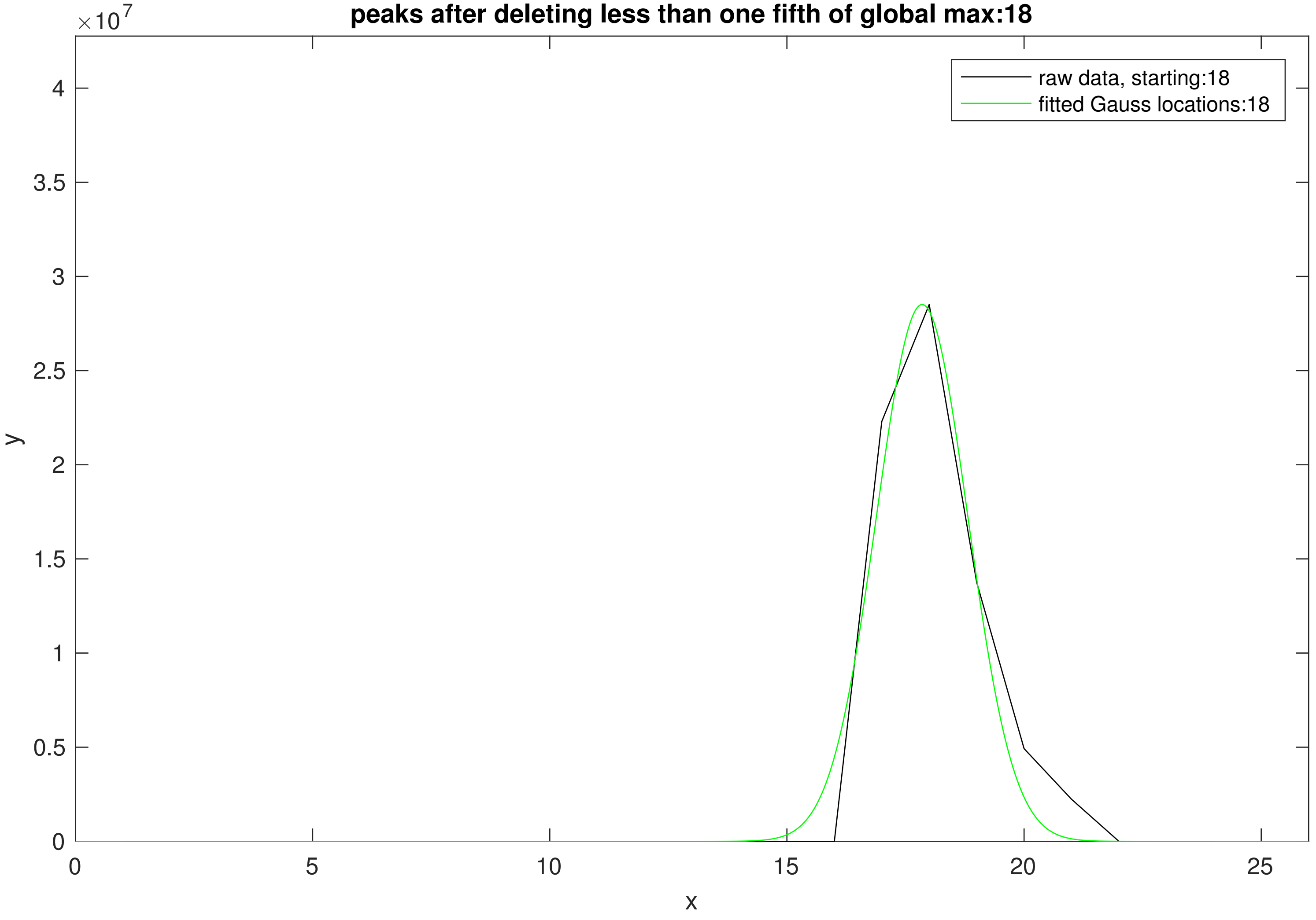

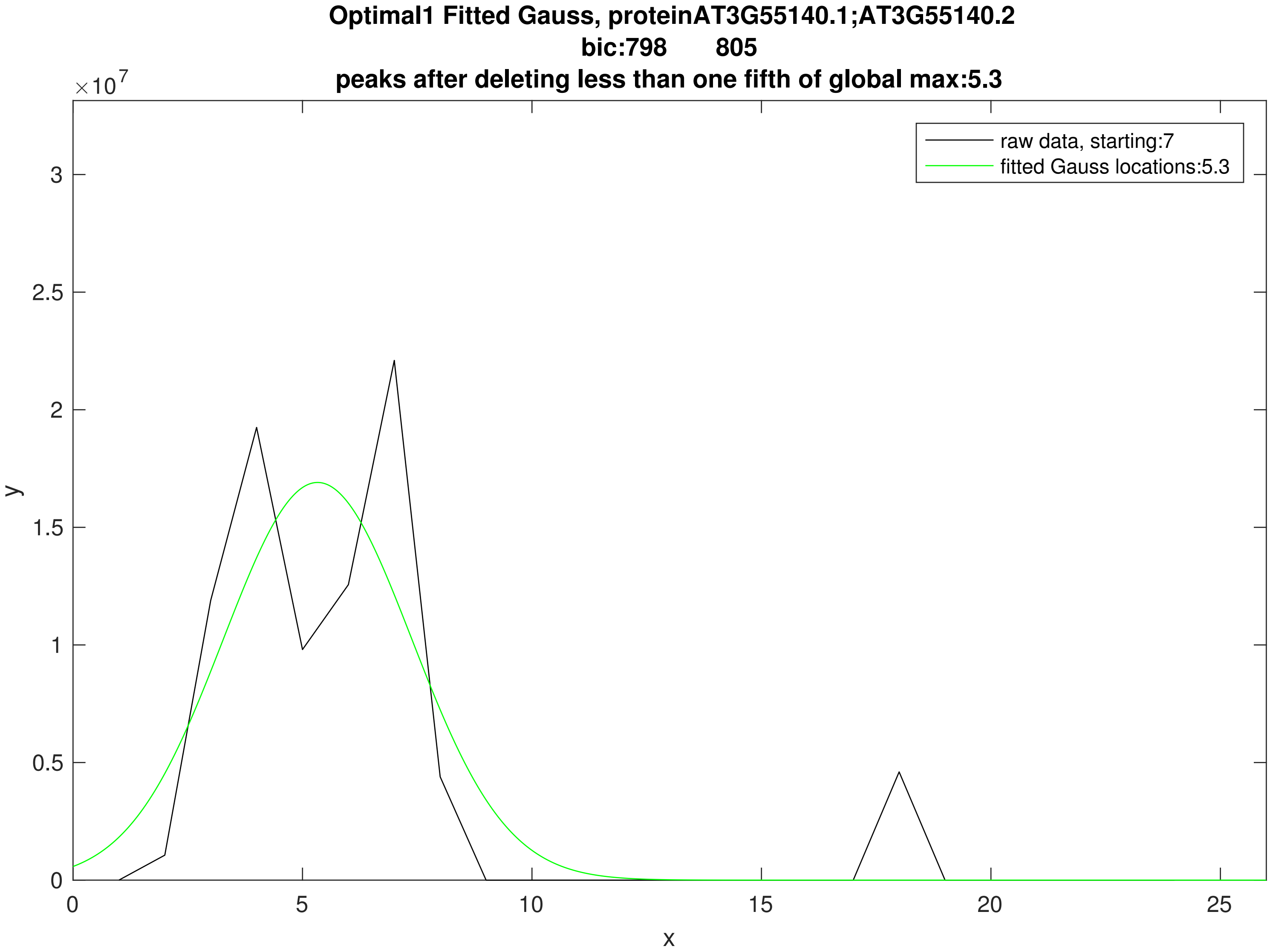

Optimal1 Fitted Gauss, proteinAT3G55140.1;AT3G55140.2
bic:798 805
peaks after deleting less than one fifth of global max:5.3
×10^7
3
2.5
2
1.5
1
0.5
0
y
0
5
10
15
20
25
x
raw data, starting:7
fitted Gauss locations:5.3

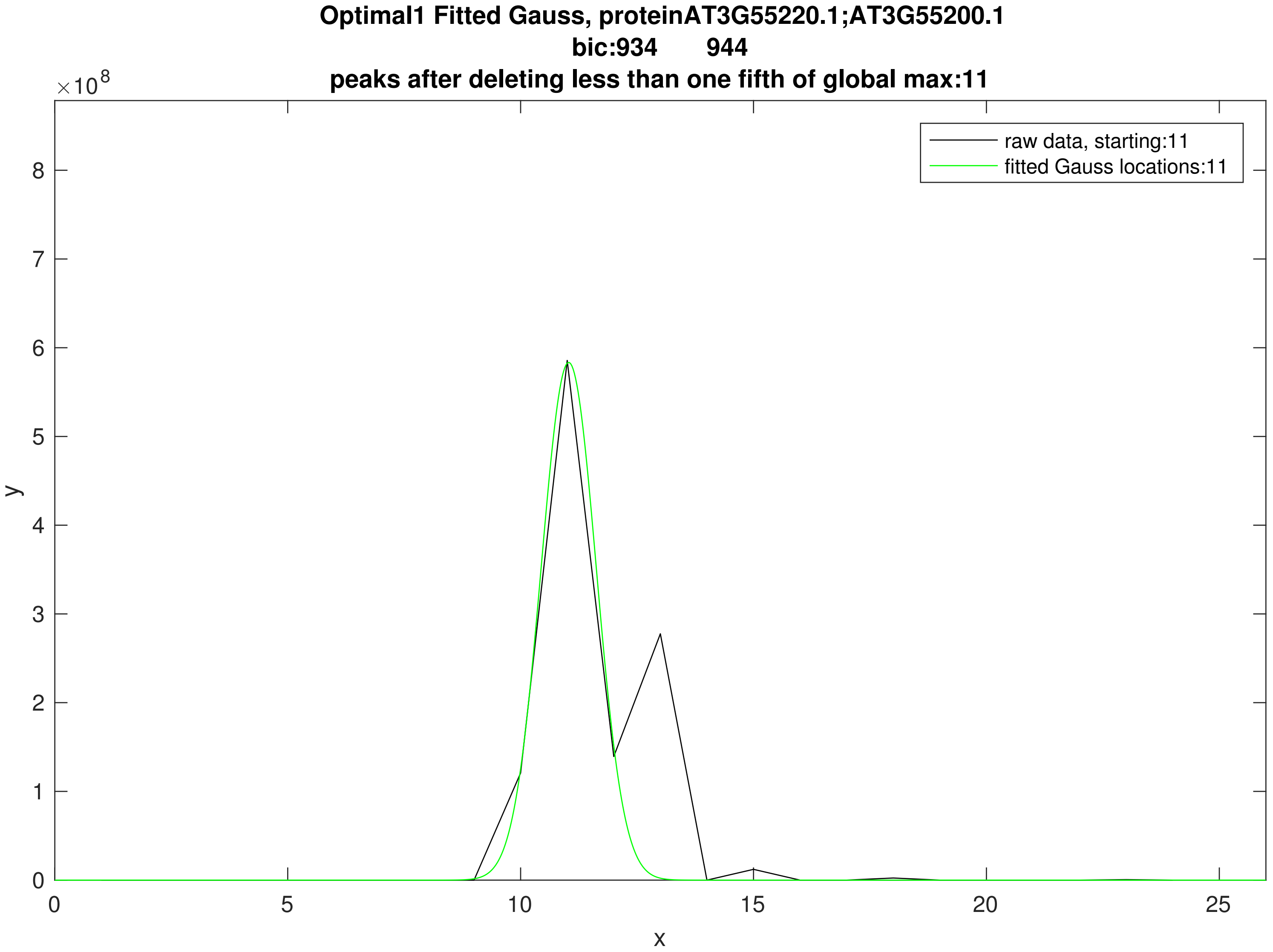
Optimal1 Fitted Gauss, proteinAT3G55220.1;AT3G55200.1
bic:934 944
peaks after deleting less than one fifth of global max:11
×10^8
raw data, starting:11
fitted Gauss locations:11
y
x
0
1
2
3
4
5
6
7
8
0
5
10
15
20
25

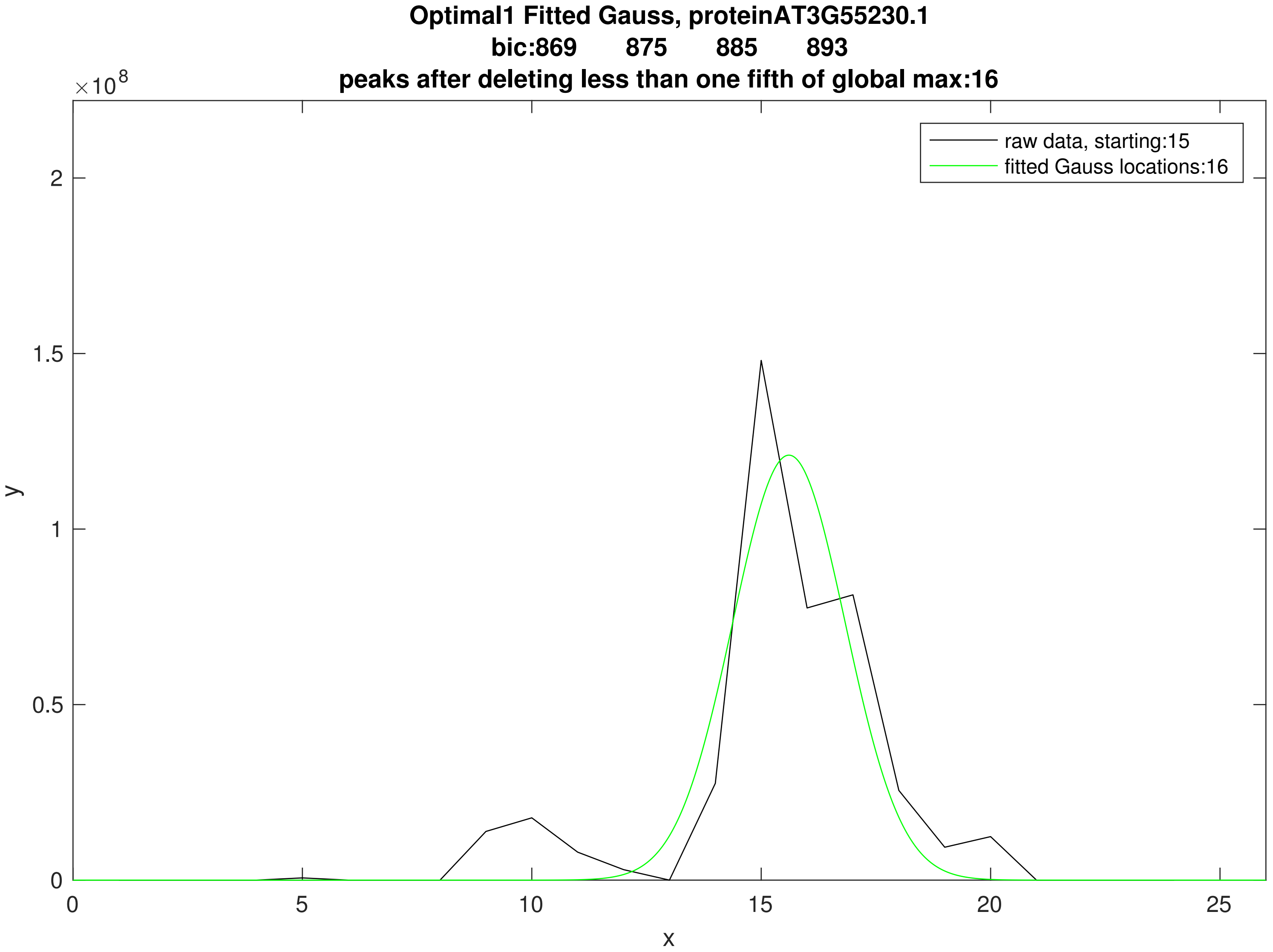
Optimal1 Fitted Gauss, proteinAT3G55230.1
bic:869 875 885 893
peaks after deleting less than one fifth of global max:16
×10^8
raw data, starting:15
fitted Gauss locations:16
y
x
0
0.5
1
1.5
2
5
10
15
20
25

#### Optimal1 Fitted Gauss, proteinAT3G55260.1
#### bic:1e+03  1.02e+03  1.03e+03
#### peaks after deleting less than one fifth of global max:12

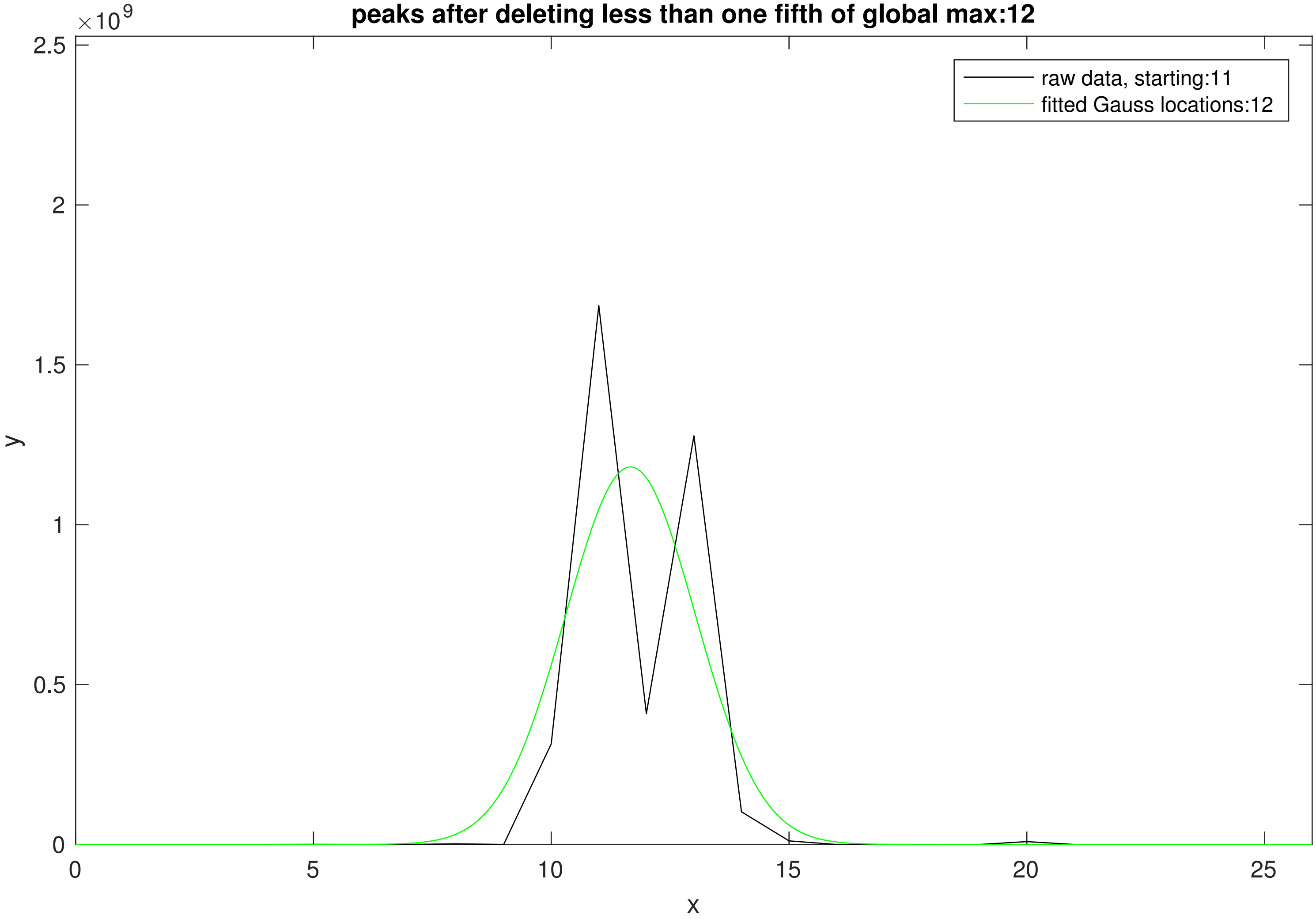

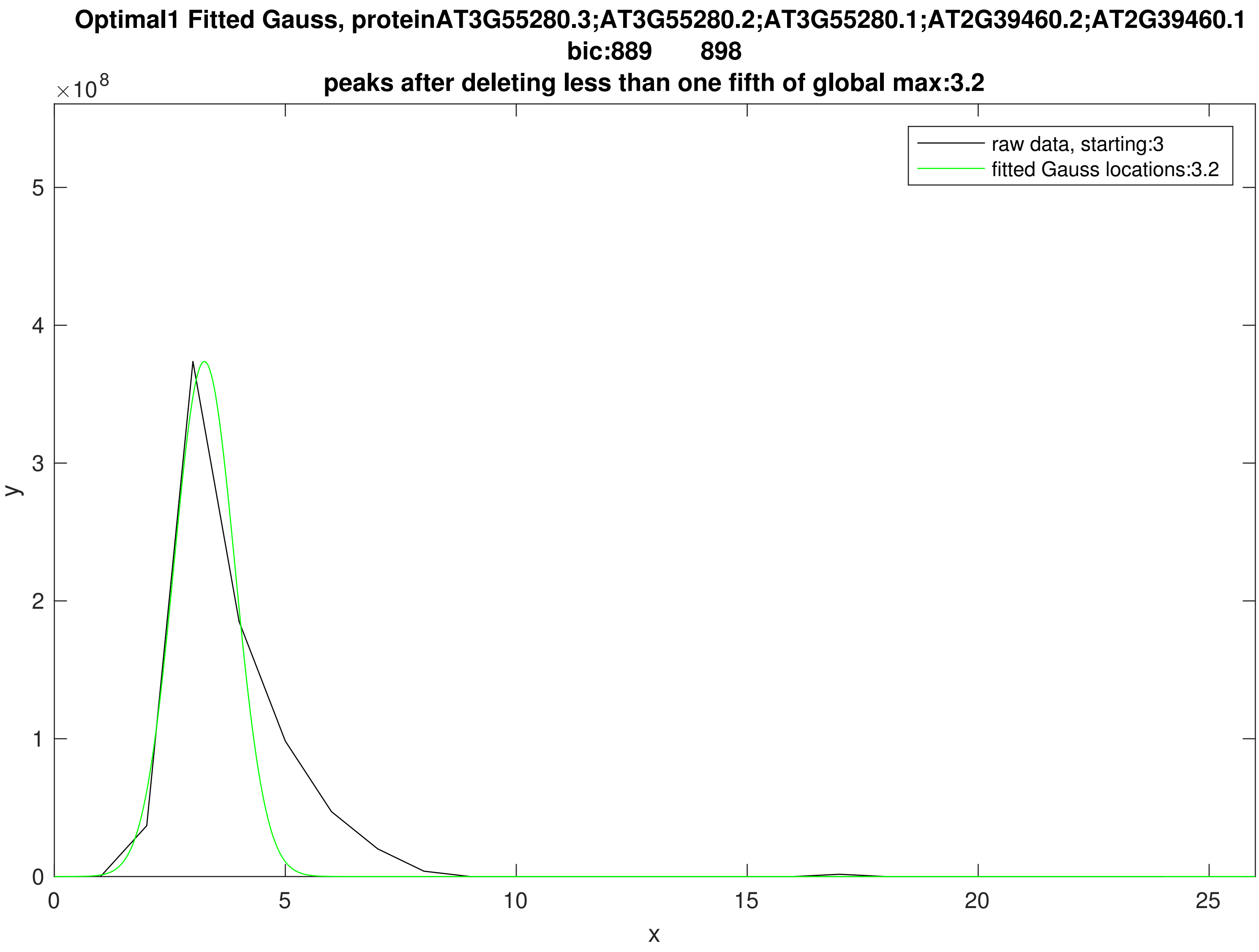

Optimal1 Fitted Gauss, proteinAT3G55280.3;AT3G55280.2;AT3G55280.1;AT2G39460.2;AT2G39460.1
bic:889 898
peaks after deleting less than one fifth of global max:3.2
×10^8
raw data, starting:3
fitted Gauss locations:3.2
y
x
0
1
2
3
4
5
0
5
10
15
20
25

#### Optimal2 Fitted Gauss, proteinAT3G55410.1
#### bic:980 959 1.03e+03 1.03e+03
#### peaks after deleting less than one fifth of global max:9.5

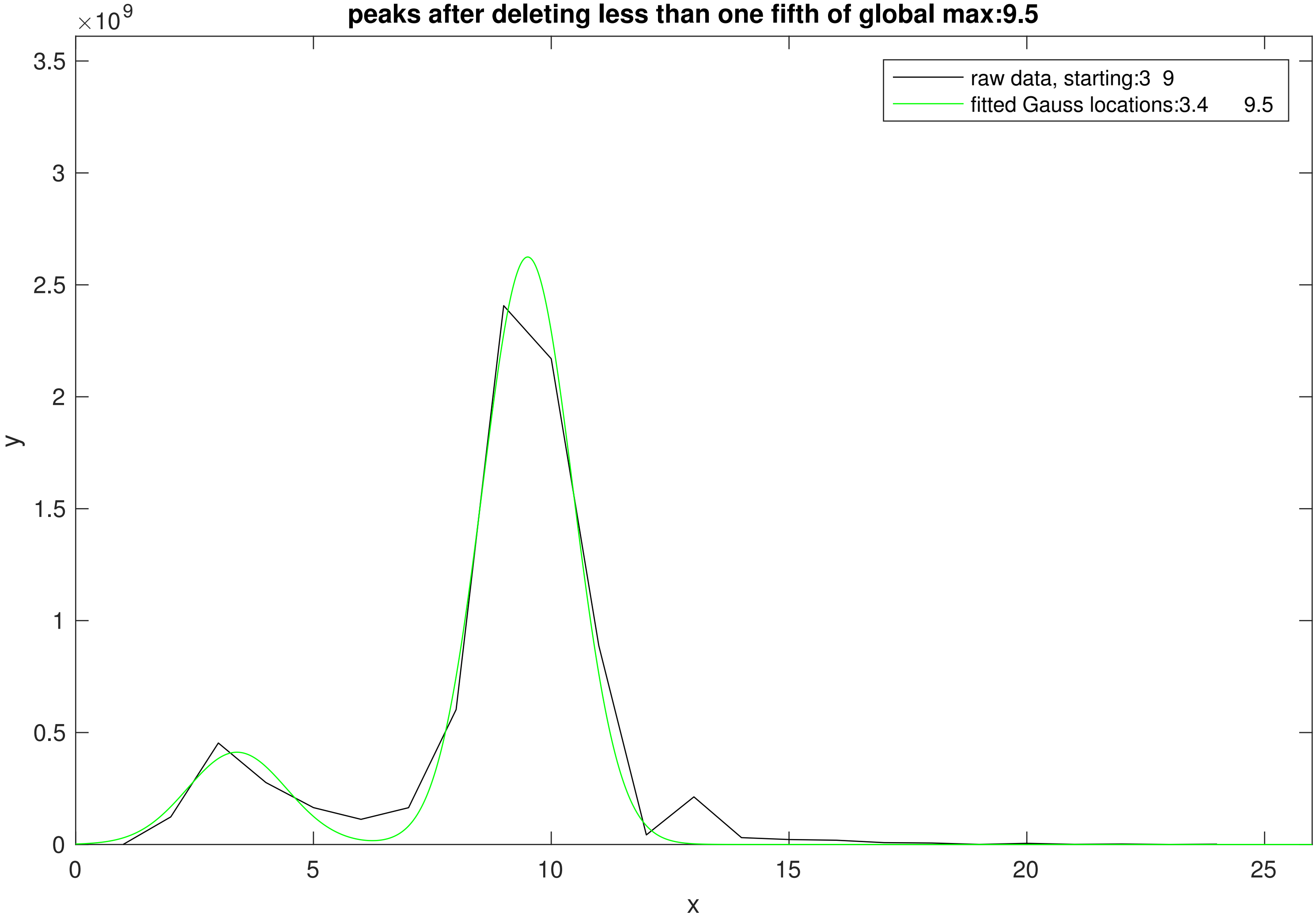

**Optimal1 Fitted Gauss, proteinAT3G55440.1**
**bic:1.11e+03 1.13e+03 1.14e+03**
**peaks after deleting less than one fifth of global max:16**

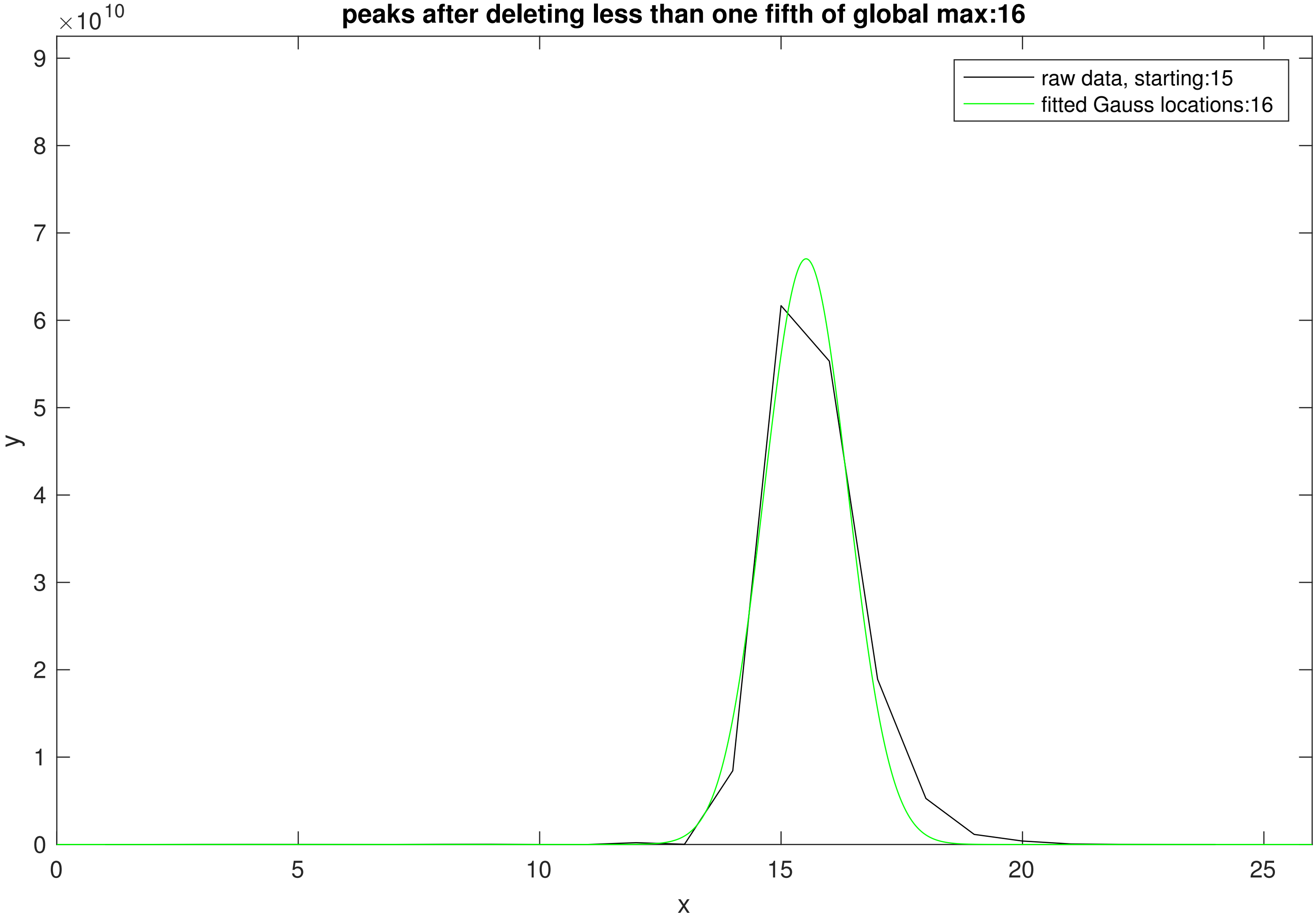

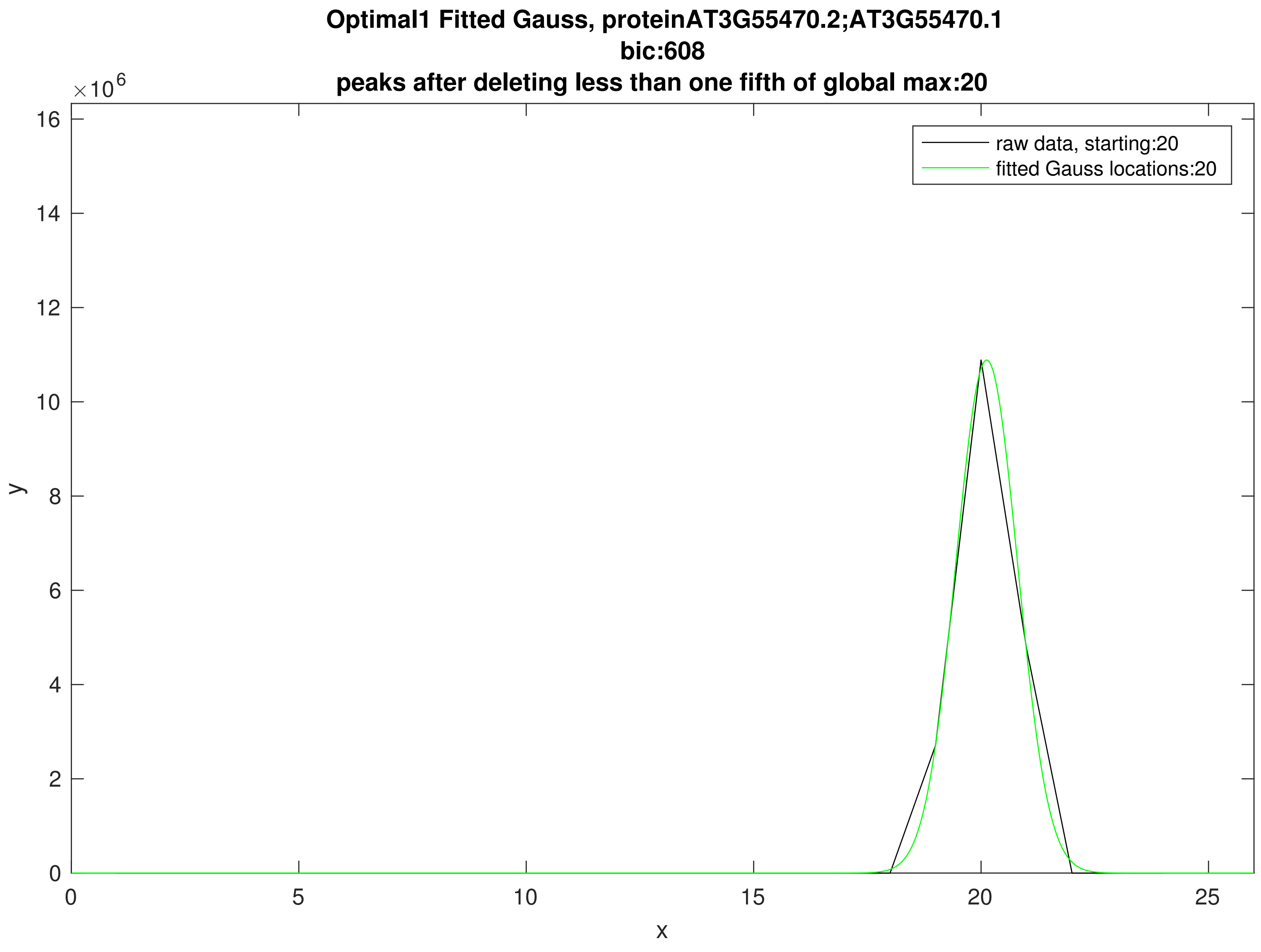

Optimal1 Fitted Gauss, proteinAT3G55470.2;AT3G55470.1
bic:608
peaks after deleting less than one fifth of global max:20
×10^6
raw data, starting:20
fitted Gauss locations:20
y
x
0
2
4
6
8
10
12
14
16
0
5
10
15
20
25

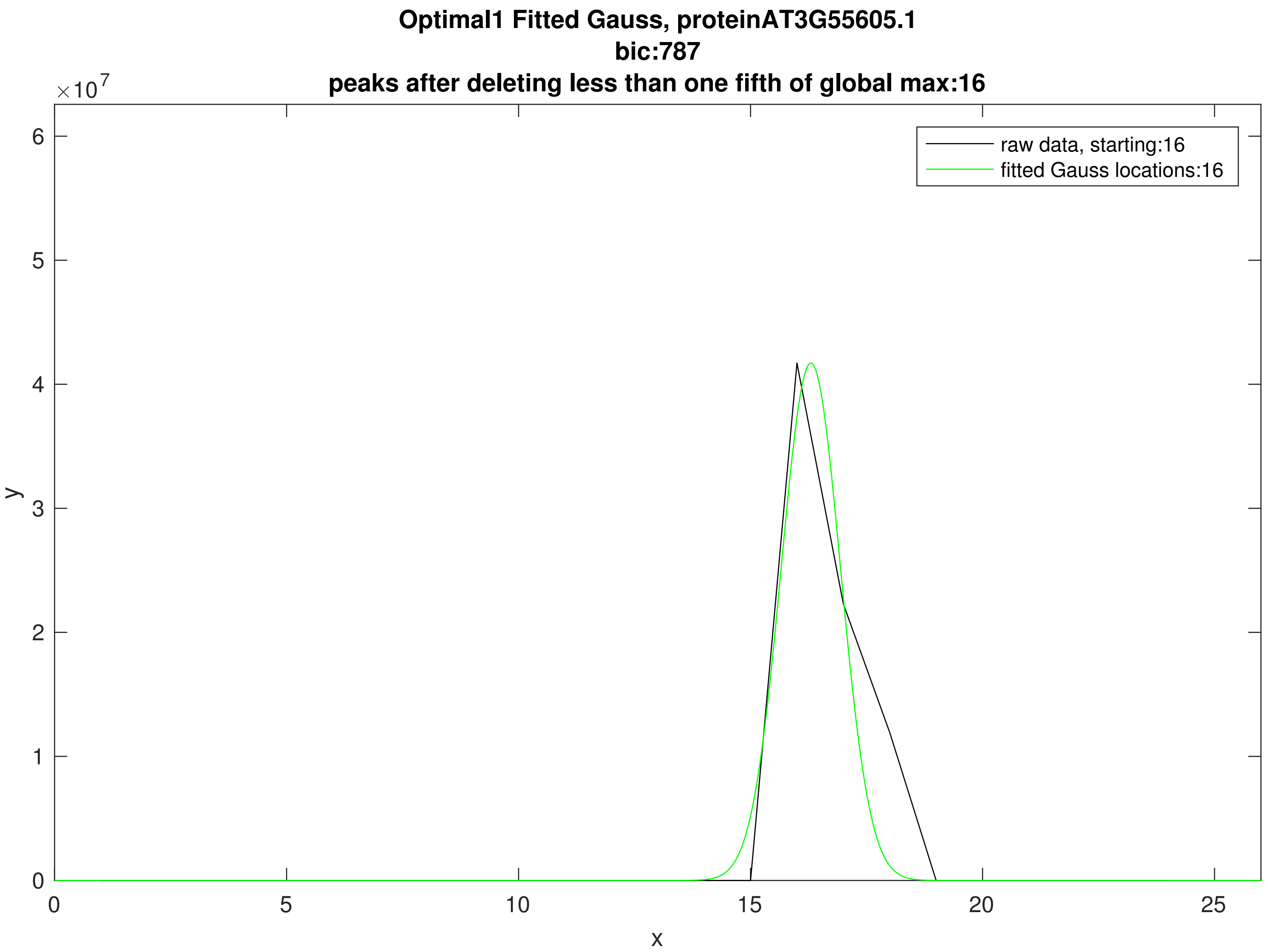
Optimal1 Fitted Gauss, proteinAT3G55605.1
bic:787
peaks after deleting less than one fifth of global max:16
×10^7
raw data, starting:16
fitted Gauss locations:16
y
x
0
1
2
3
4
5
6
0
5
10
15
20
25

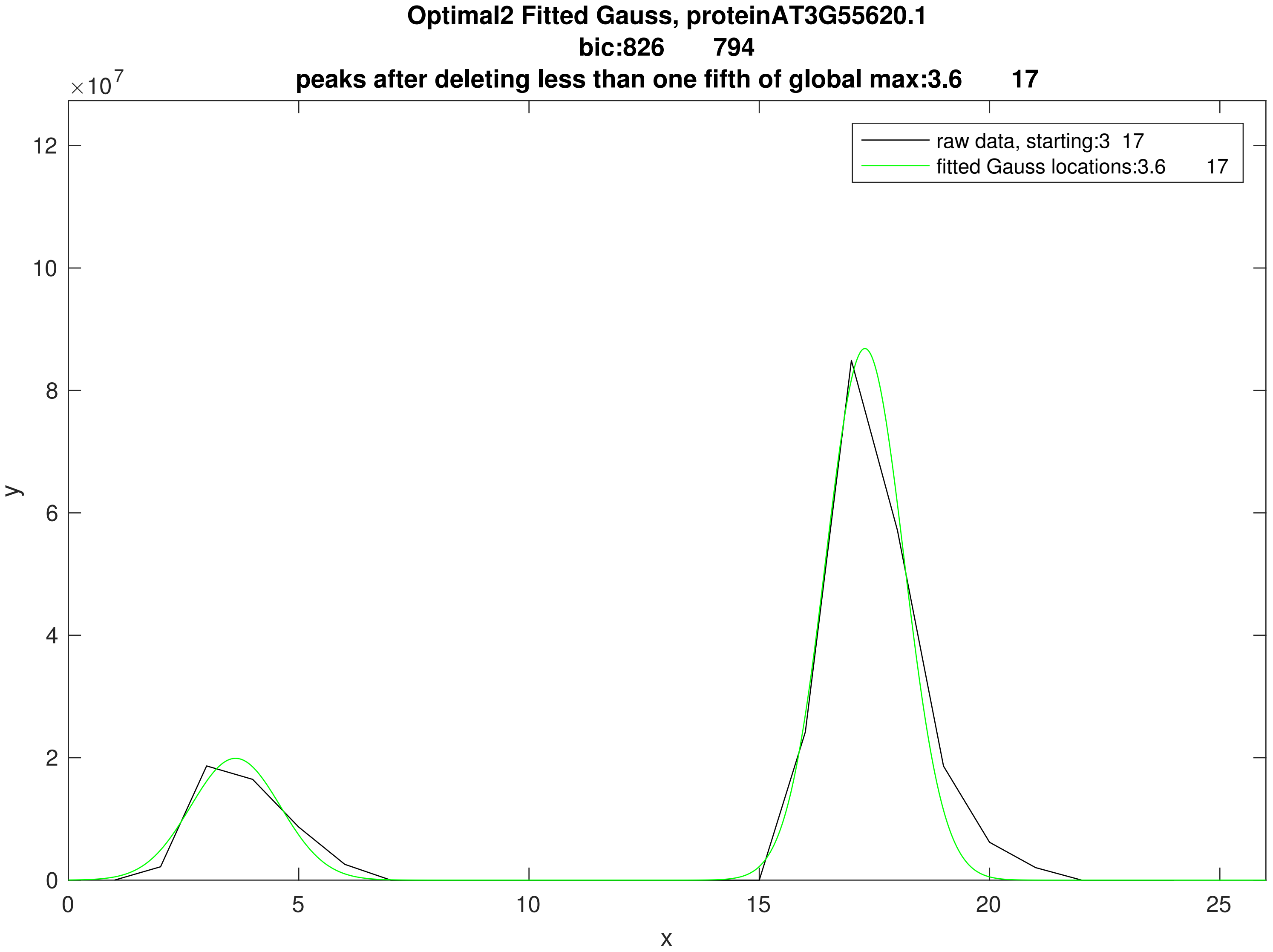
Optimal2 Fitted Gauss, proteinAT3G55620.1
bic:826 794
peaks after deleting less than one fifth of global max:3.6 17
×10^7
12
10
8
6
4
2
0
y
0
5
10
15
20
25
x
raw data, starting:3 17
fitted Gauss locations:3.6 17

#### Optimal1 Fitted Gauss, proteinAT3G55750.1;AT1G74270.1
#### bic:810 864
#### peaks after deleting less than one fifth of global max:3.1

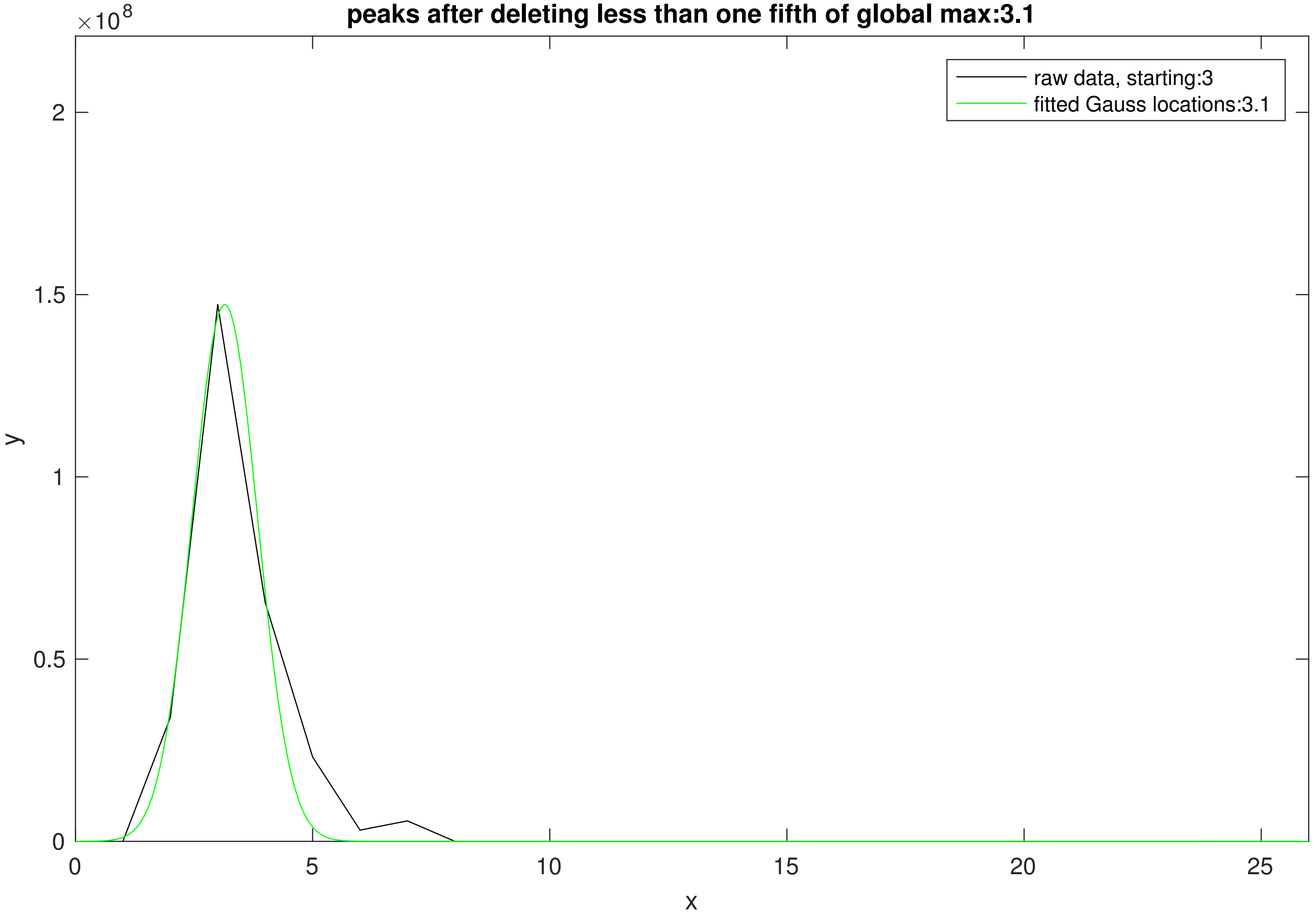

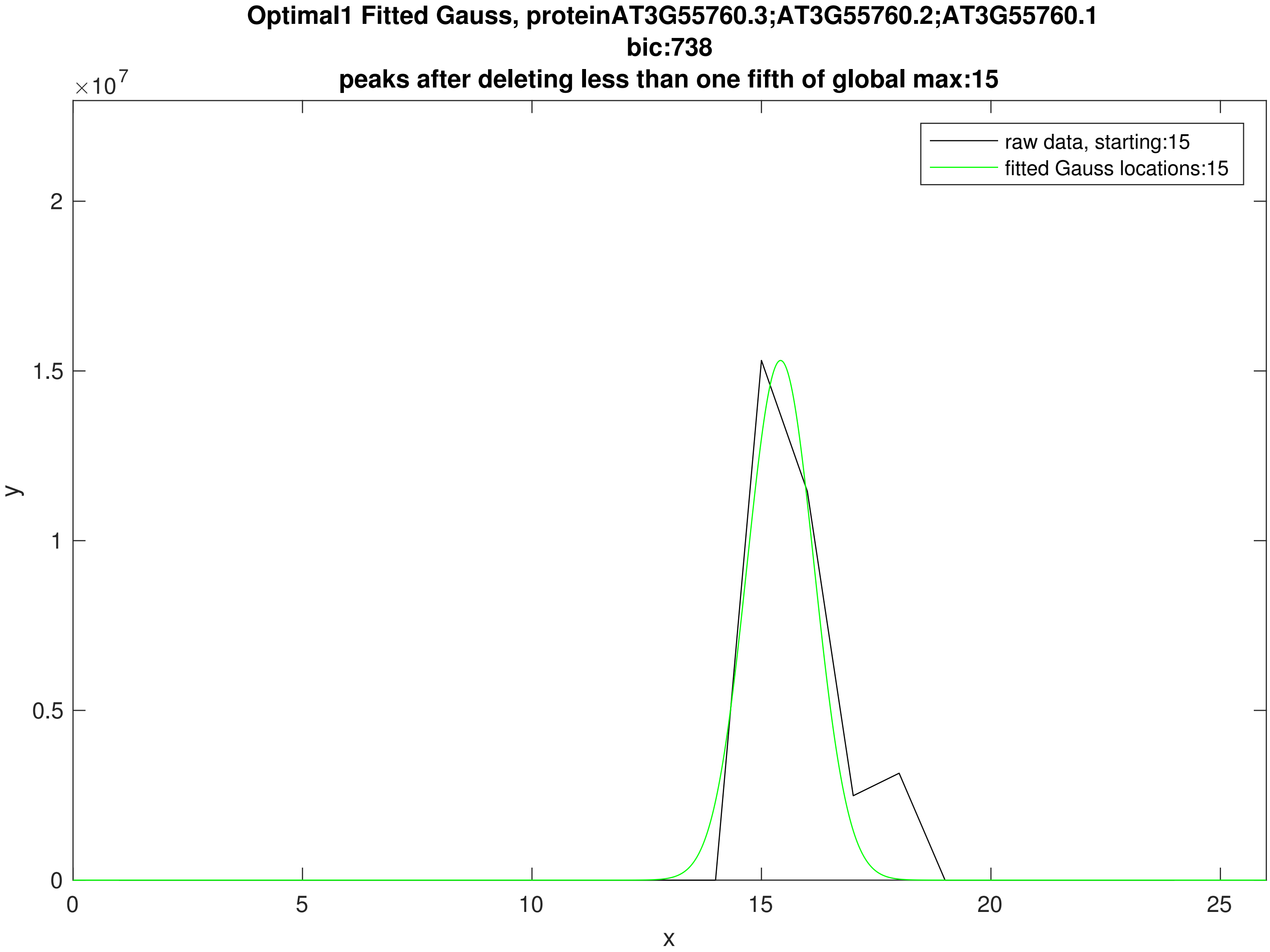

Optimal1 Fitted Gauss, proteinAT3G55760.3;AT3G55760.2;AT3G55760.1
bic:738
peaks after deleting less than one fifth of global max:15
×10^7
raw data, starting:15
fitted Gauss locations:15
2
1.5
1
0.5
0
0
5
10
15
20
25
x
y

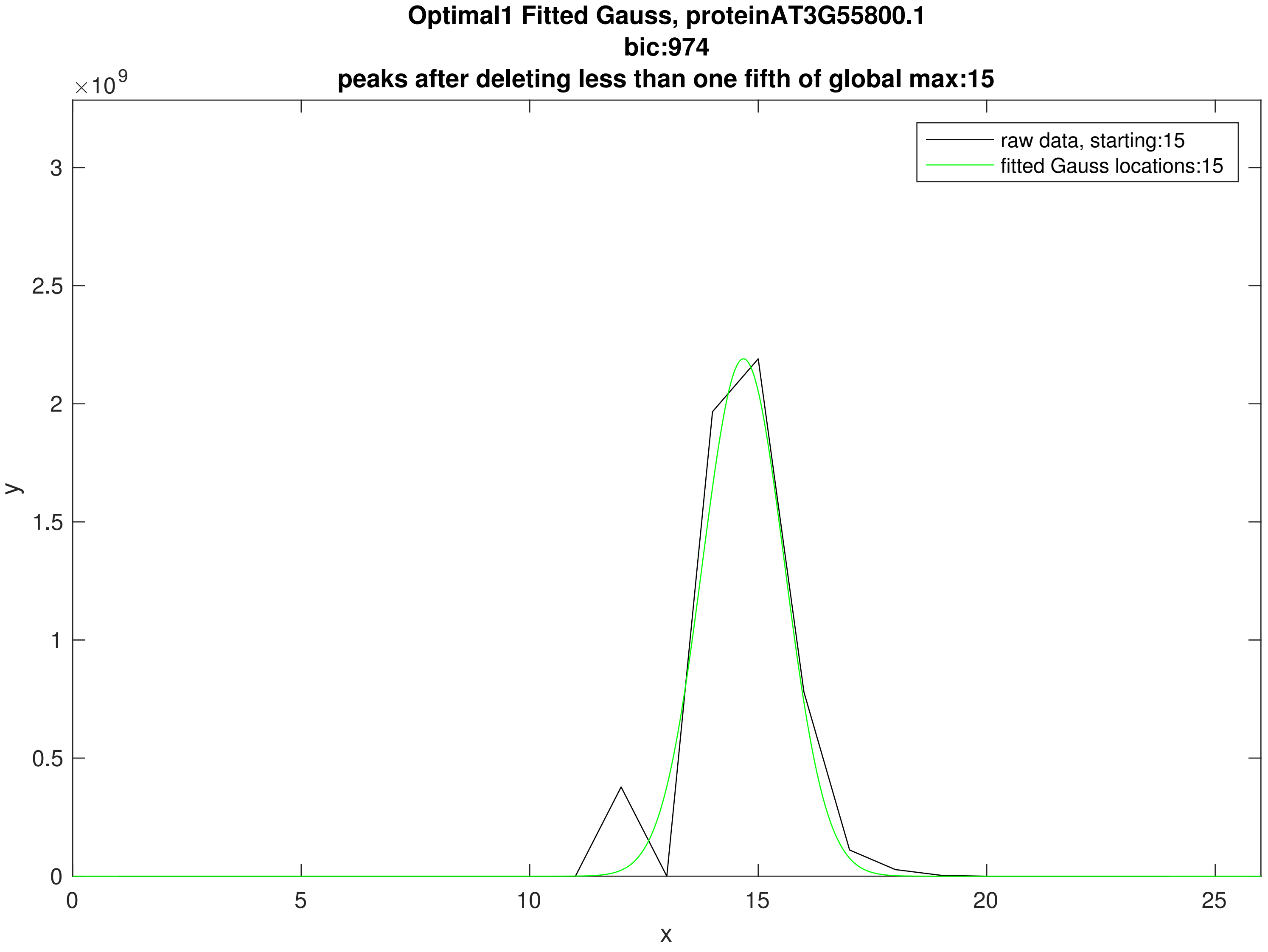
Optimal1 Fitted Gauss, proteinAT3G55800.1
bic:974
peaks after deleting less than one fifth of global max:15
×10^9
3
2.5
2
1.5
1
0.5
0
y
0
5
10
15
20
25
x
raw data, starting:15
fitted Gauss locations:15

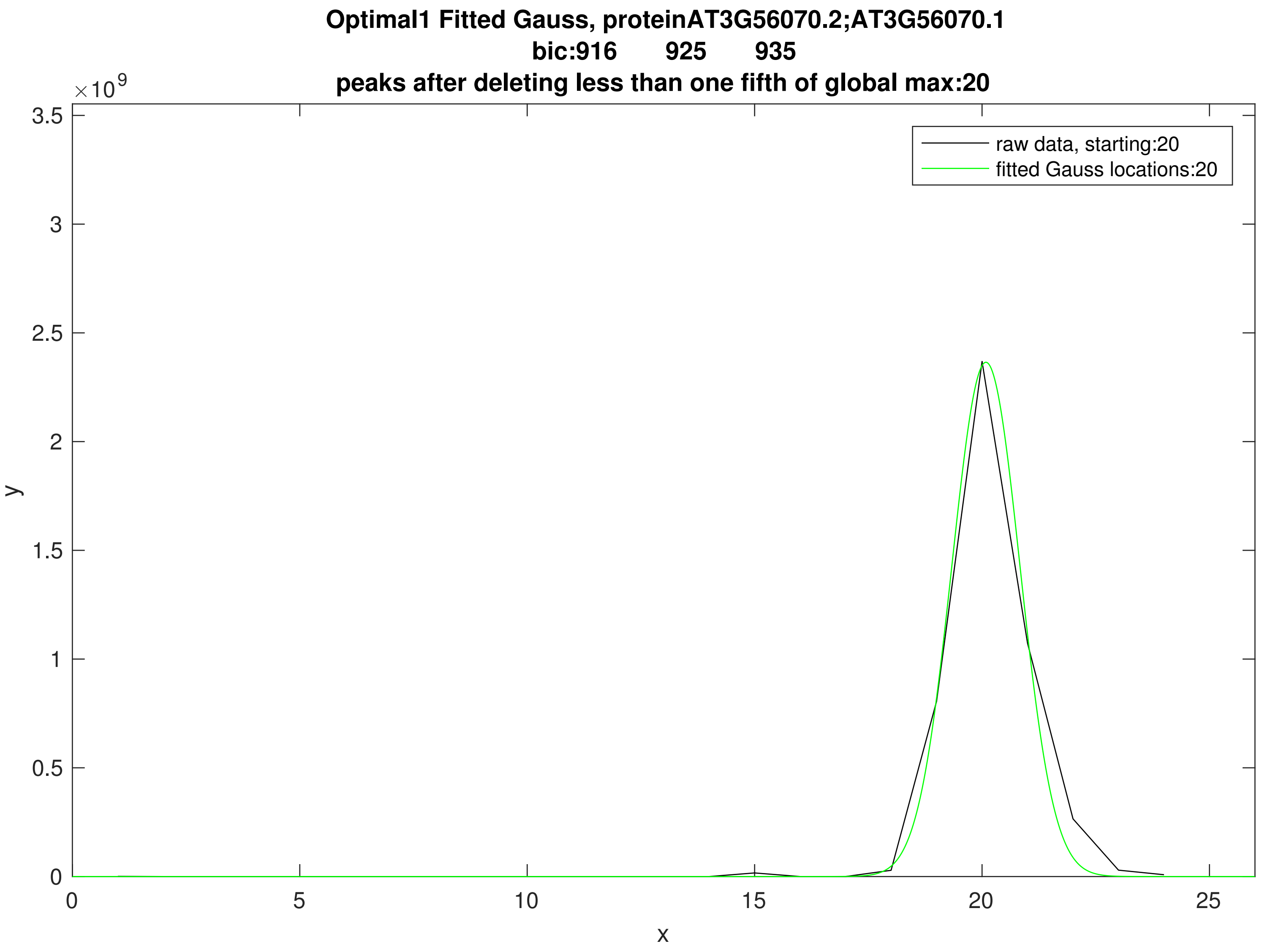

Optimal1 Fitted Gauss, proteinAT3G56070.2;AT3G56070.1
bic:916 925 935
peaks after deleting less than one fifth of global max:20
×10^9
3.5
3
2.5
2
1.5
1
0.5
0
y
0
5
10
15
20
25
x
raw data, starting:20
fitted Gauss locations:20

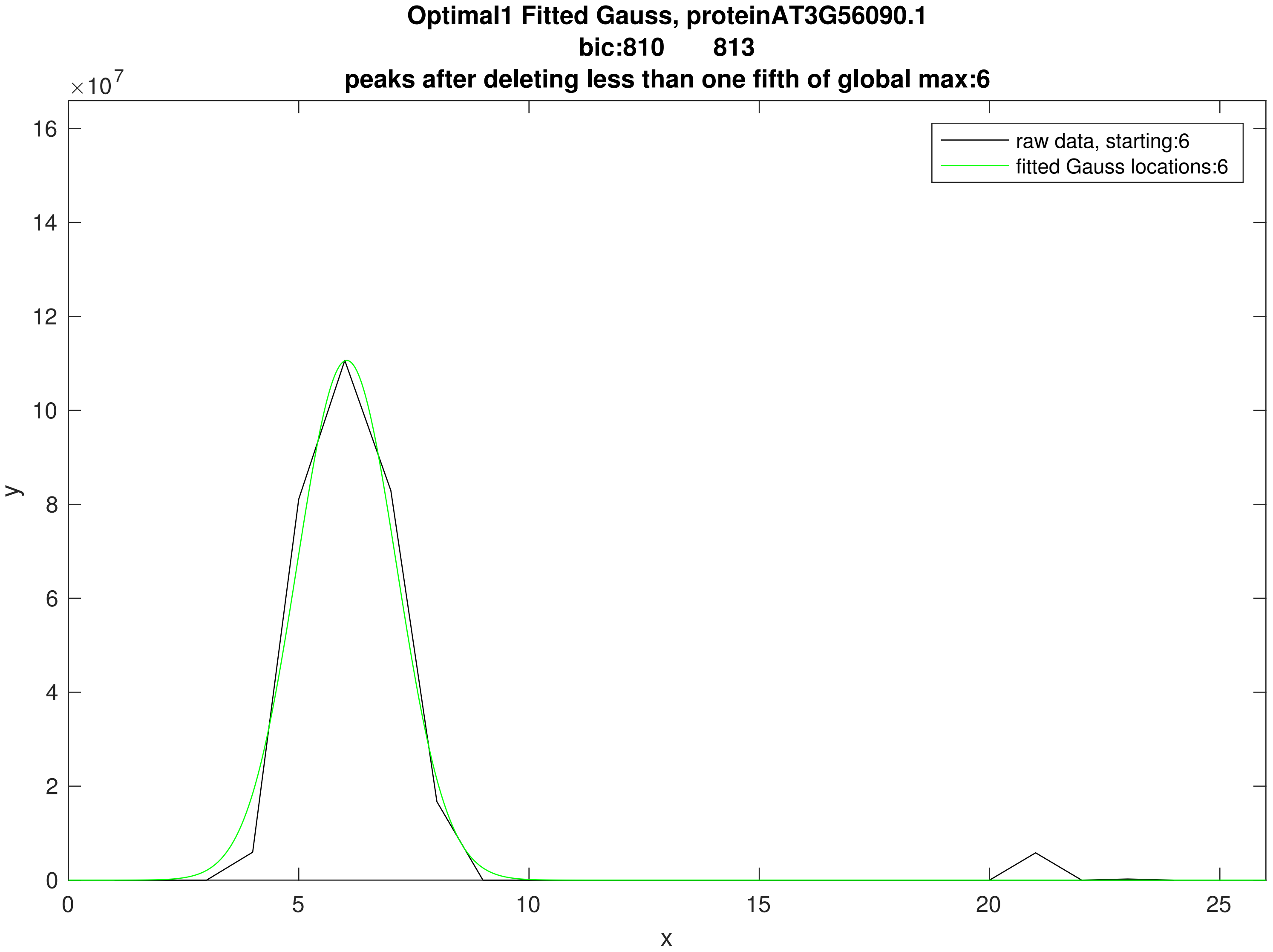
Optimal1 Fitted Gauss, proteinAT3G56090.1
bic:810 813
peaks after deleting less than one fifth of global max:6
×10^7
16
14
12
10
8
6
4
2
0
y
0
5
10
15
20
25
x
raw data, starting:6
fitted Gauss locations:6

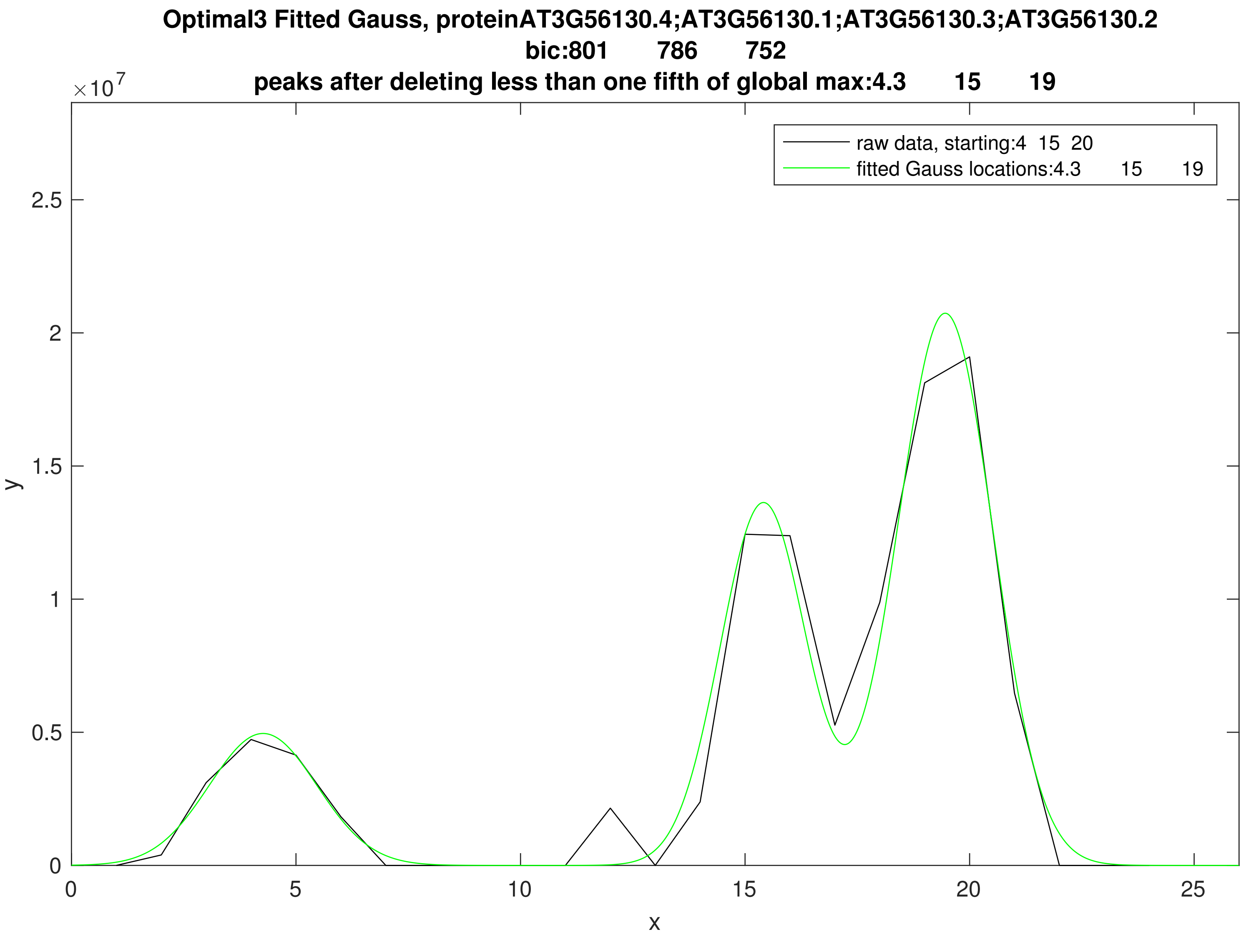
Optimal3 Fitted Gauss, proteinAT3G56130.4;AT3G56130.1;AT3G56130.3;AT3G56130.2
bic:801 786 752
peaks after deleting less than one fifth of global max:4.3 15 19
×10^7
raw data, starting:4 15 20
fitted Gauss locations:4.3 15 19
y
x
0
0.5
1
1.5
2
2.5
5
10
15
20
25

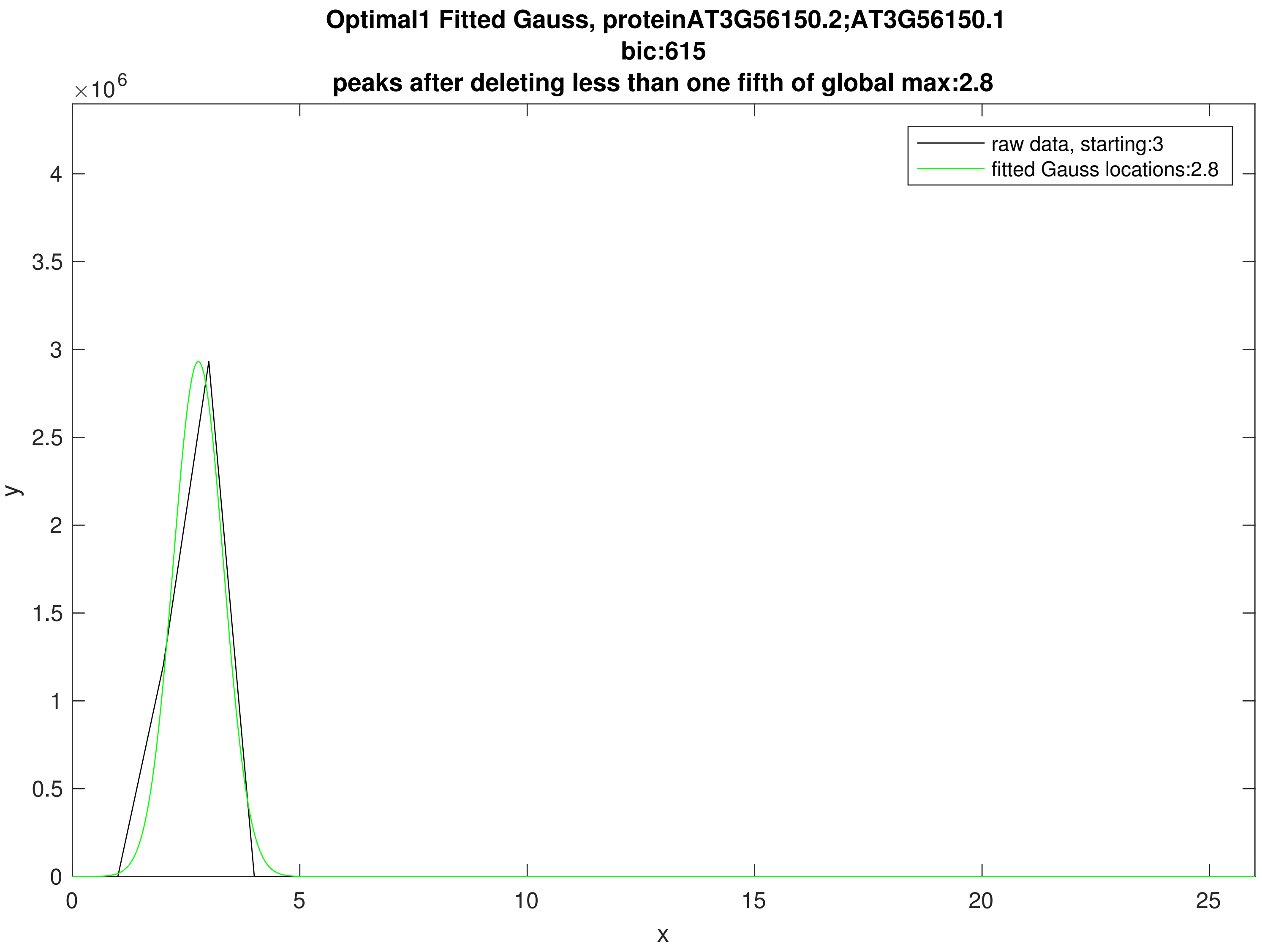
Optimal1 Fitted Gauss, proteinAT3G56150.2;AT3G56150.1
bic:615
peaks after deleting less than one fifth of global max:2.8
×10^6
raw data, starting:3
fitted Gauss locations:2.8
y
x
0
0.5
1
1.5
2
2.5
3
3.5
4
5
10
15
20
25

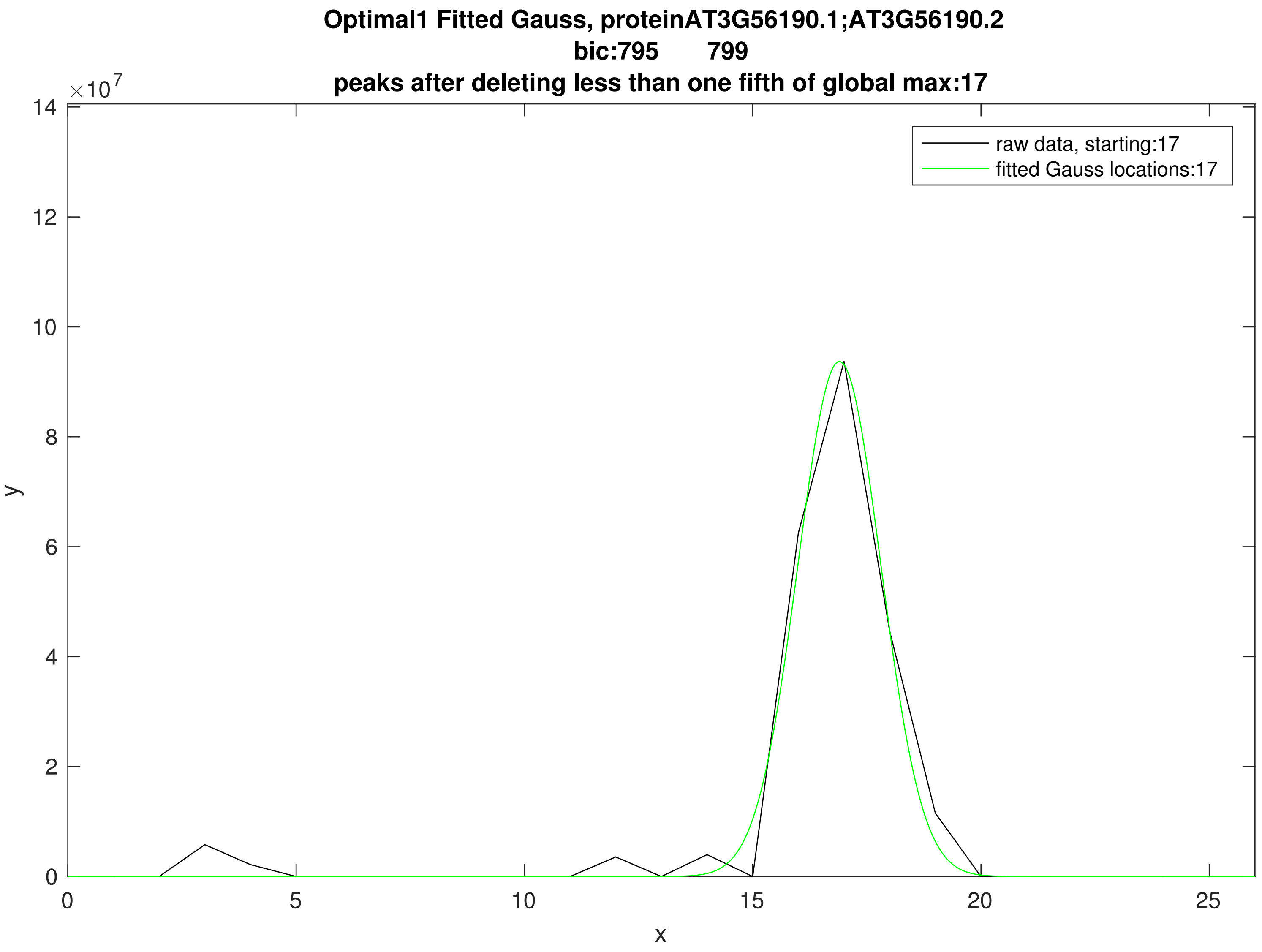
Optimal1 Fitted Gauss, proteinAT3G56190.1;AT3G56190.2
bic:795 799
peaks after deleting less than one fifth of global max:17
×10^7
raw data, starting:17
fitted Gauss locations:17
y
x
0
2
4
6
8
10
12
14
5
15
20
25

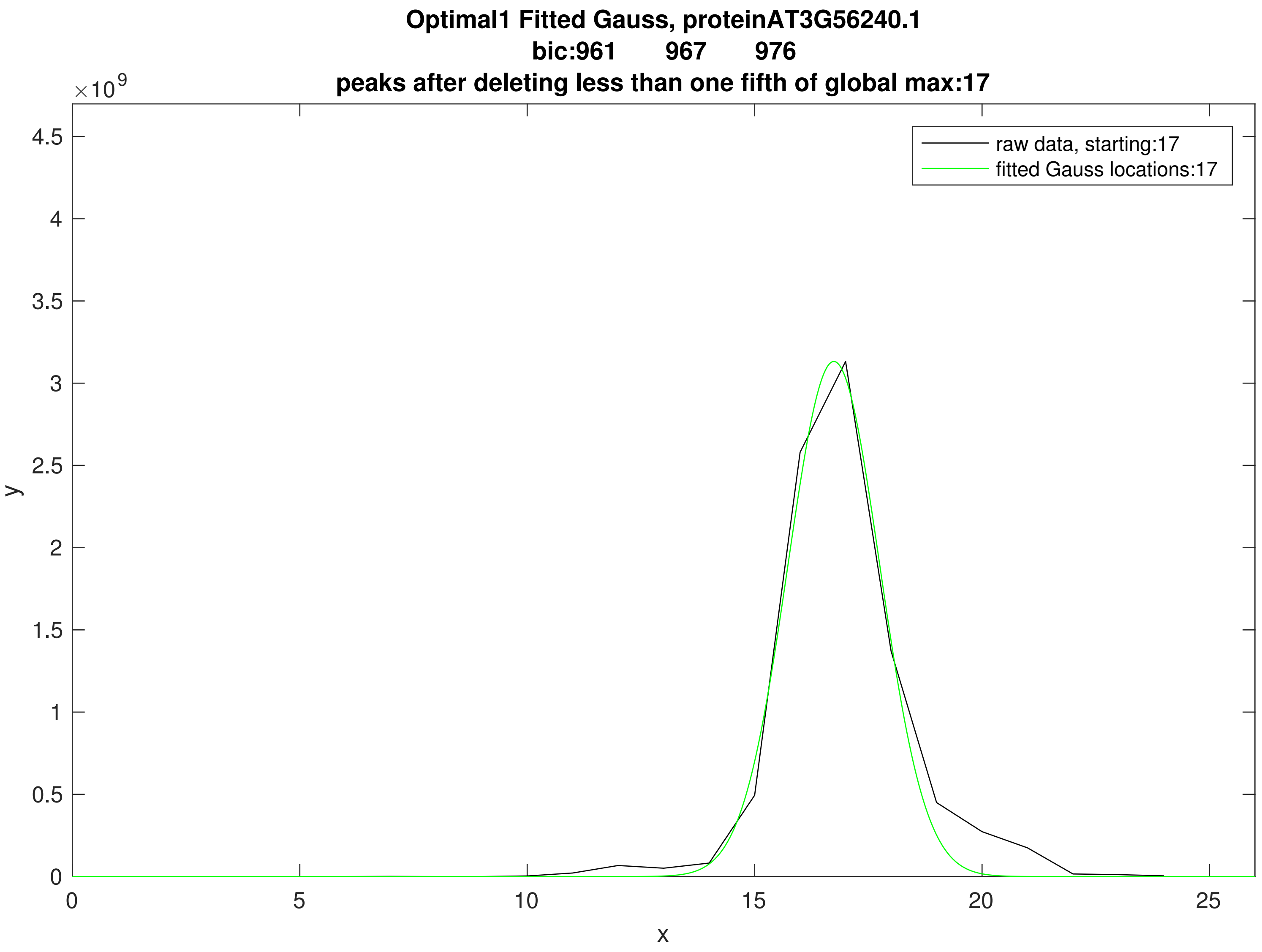

Optimal1 Fitted Gauss, proteinAT3G56240.1
bic:961 967 976
peaks after deleting less than one fifth of global max:17
×10^9
4.5
4
3.5
3
2.5
2
1.5
1
0.5
0
y
0
5
10
15
20
25
x
raw data, starting:17
fitted Gauss locations:17

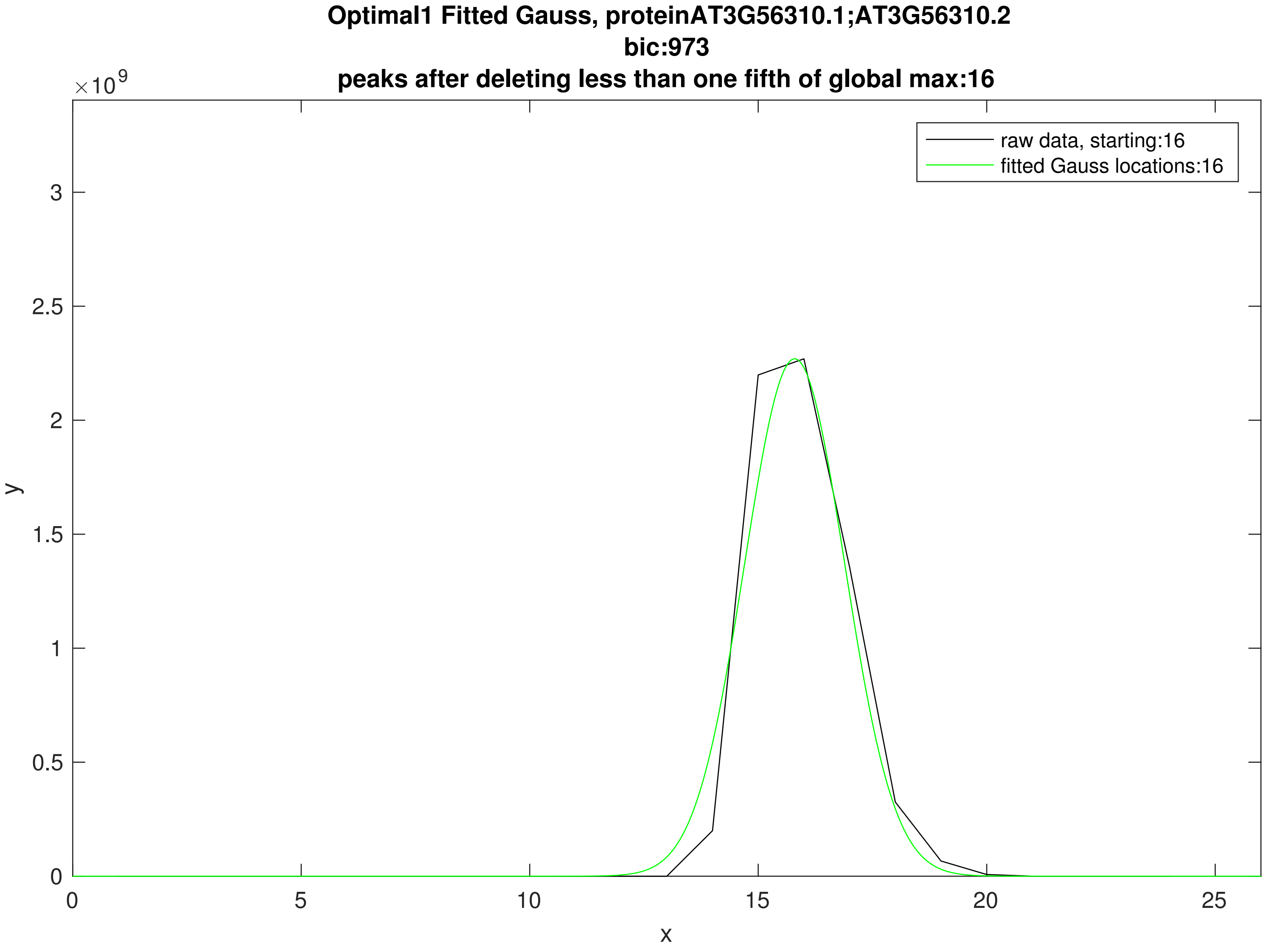
Optimal1 Fitted Gauss, proteinAT3G56310.1;AT3G56310.2
bic:973
peaks after deleting less than one fifth of global max:16
×10^9
raw data, starting:16
fitted Gauss locations:16
y
x
0
0.5
1
1.5
2
2.5
3
0
5
10
15
20
25

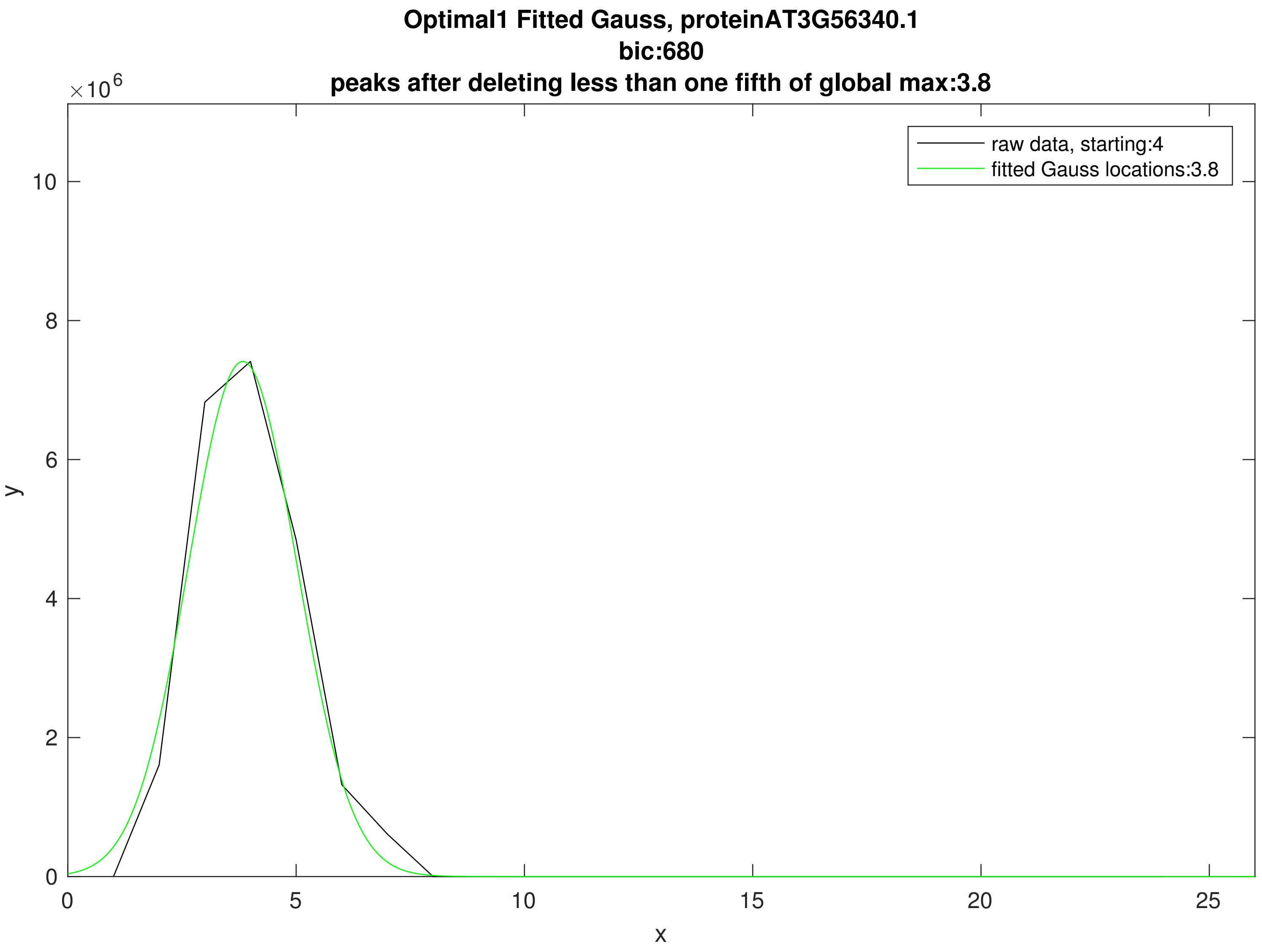
Optimal1 Fitted Gauss, proteinAT3G56340.1
bic:680
peaks after deleting less than one fifth of global max:3.8
×10^6
raw data, starting:4
fitted Gauss locations:3.8
10
8
6
4
2
0
y
0
5
10
15
20
25
x

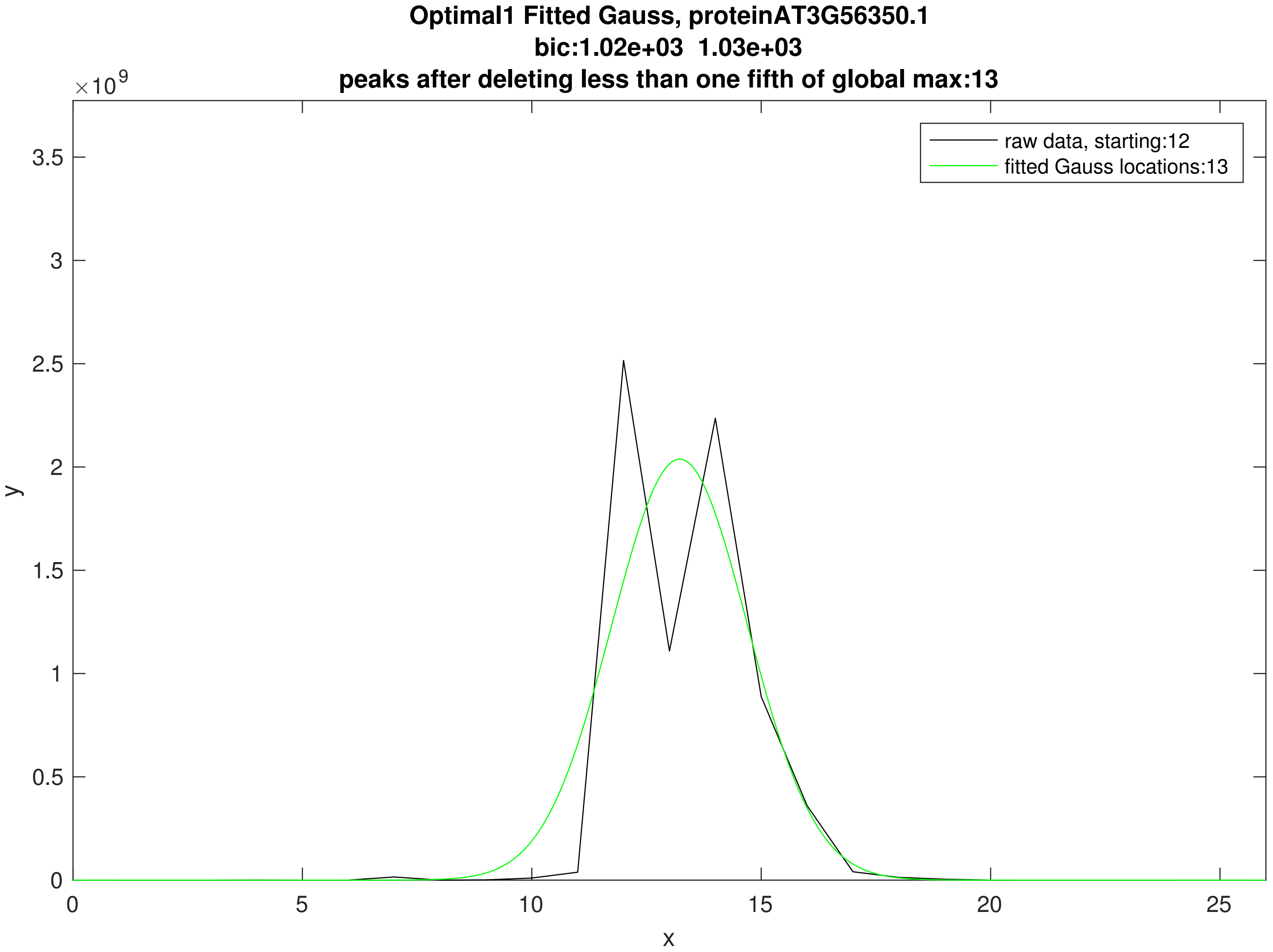
Optimal1 Fitted Gauss, proteinAT3G56350.1
bic:1.02e+03 1.03e+03
peaks after deleting less than one fifth of global max:13
×10^9
raw data, starting:12
fitted Gauss locations:13
y
x
0
0.5
1
1.5
2
2.5
3
3.5
0
5
10
15
20
25

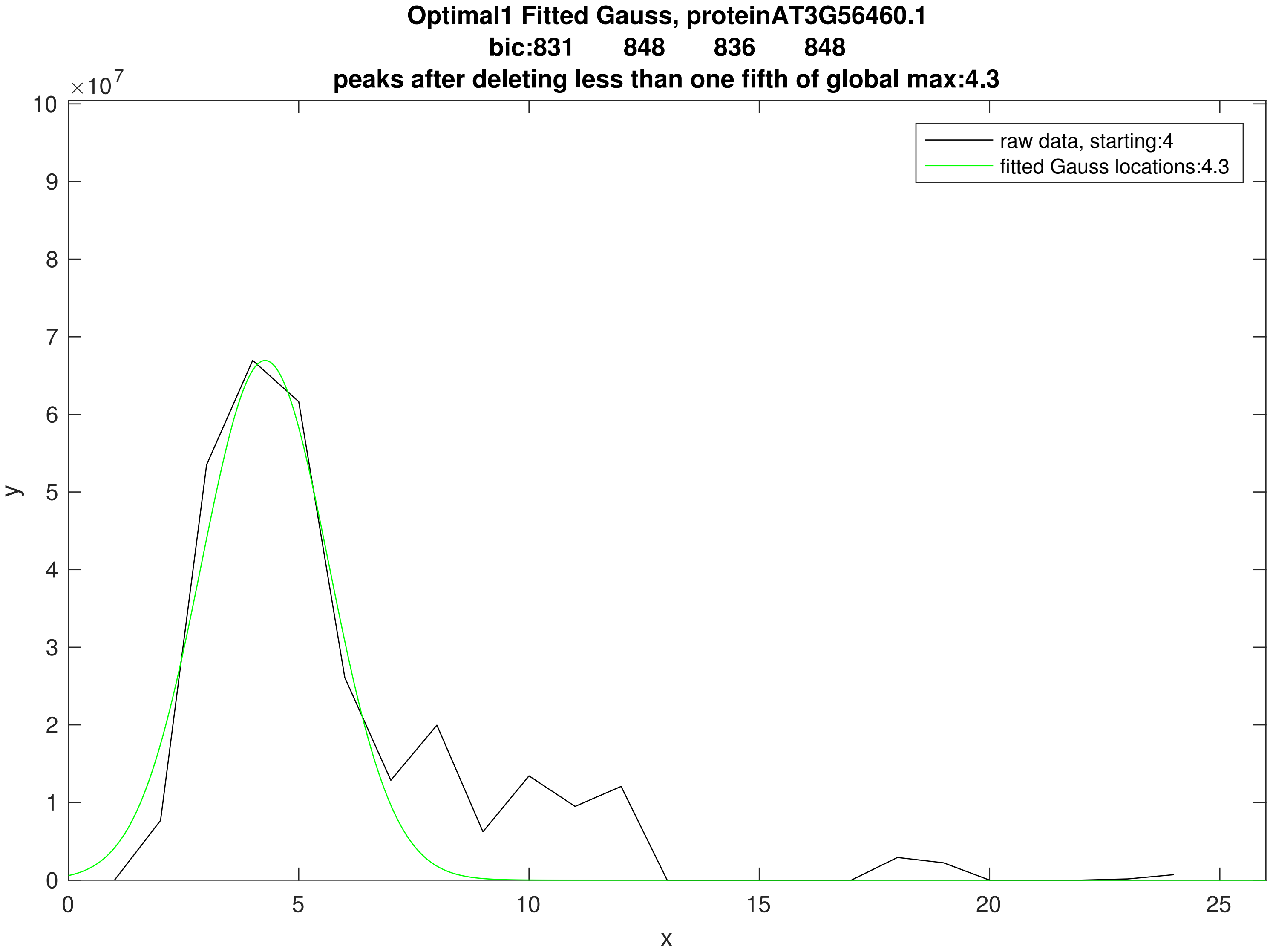

Optimal1 Fitted Gauss, proteinAT3G56460.1
bic:831 848 836 848
peaks after deleting less than one fifth of global max:4.3
×10^7
raw data, starting:4
fitted Gauss locations:4.3
y
x
0
1
2
3
4
5
6
7
8
9
10
0
5
10
15
20
25

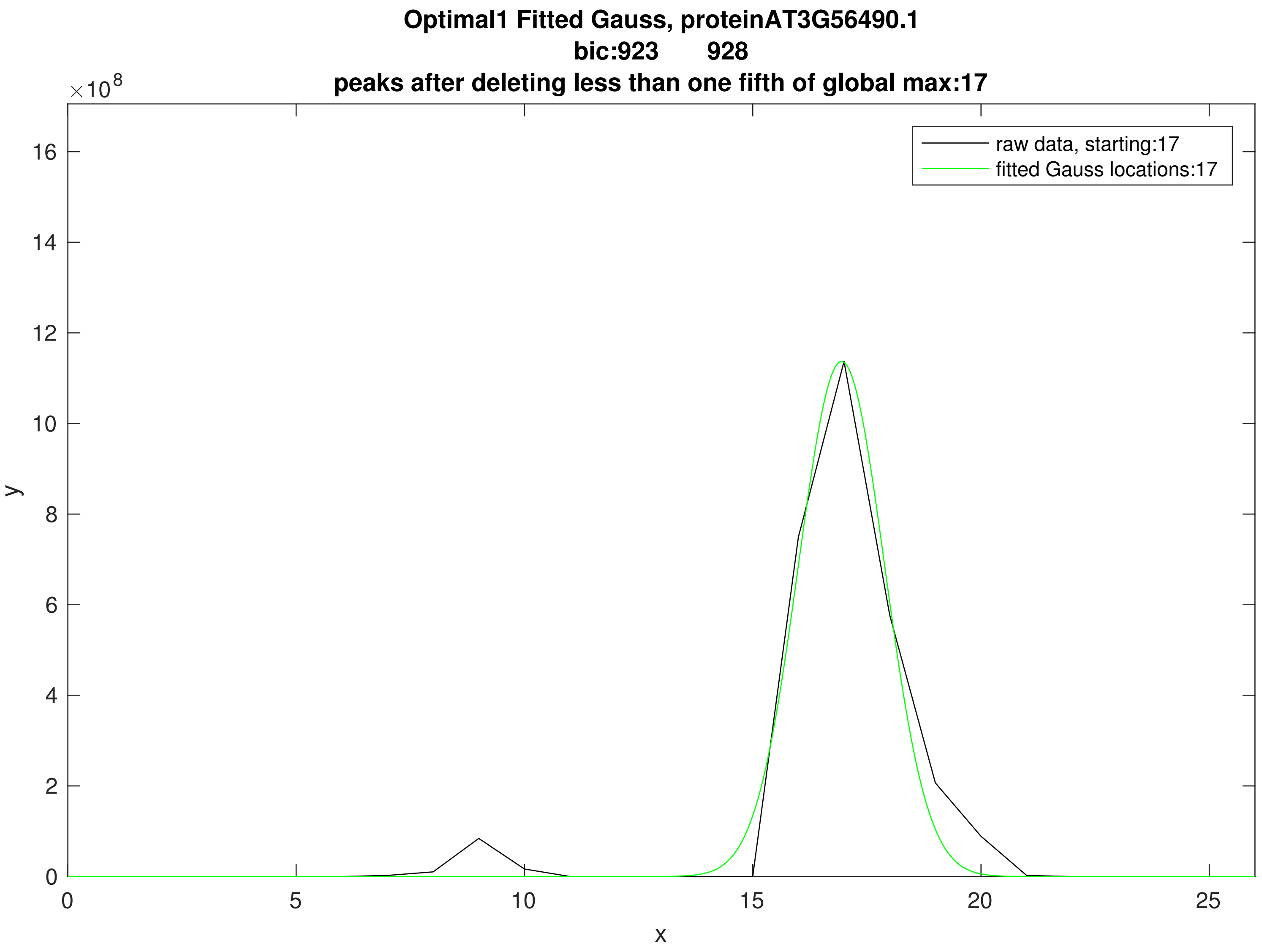
Optimal1 Fitted Gauss, proteinAT3G56490.1
bic:923 928
peaks after deleting less than one fifth of global max:17
×10^8
16
14
12
10
8
6
4
2
0
y
0
5
10
15
20
25
x
raw data, starting:17
fitted Gauss locations:17

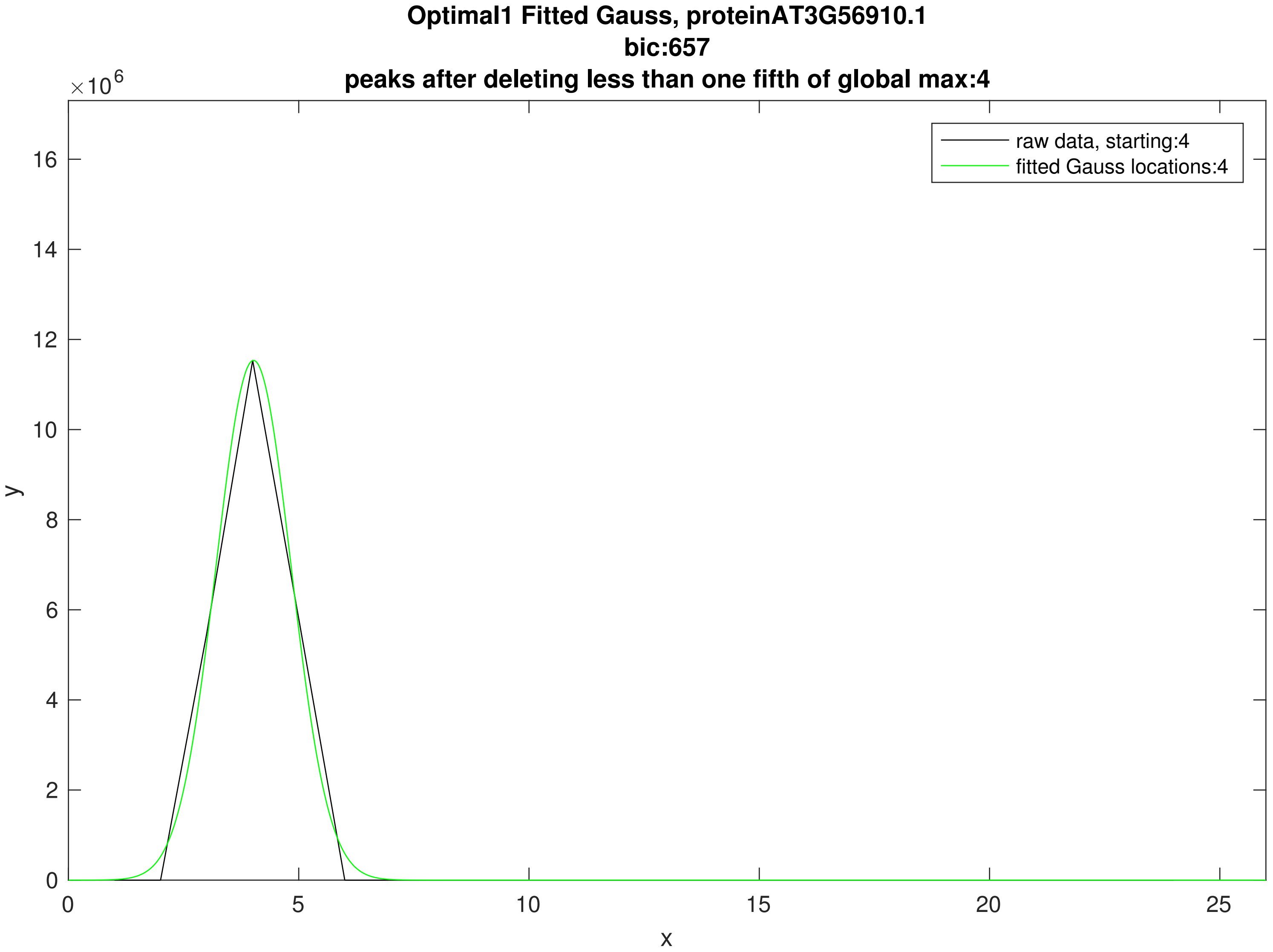

Optimal1 Fitted Gauss, proteinAT3G56910.1
bic:657
peaks after deleting less than one fifth of global max:4
×10^6
raw data, starting:4
fitted Gauss locations:4
y
x
0
2
4
6
8
10
12
14
16
0
5
10
15
20
25

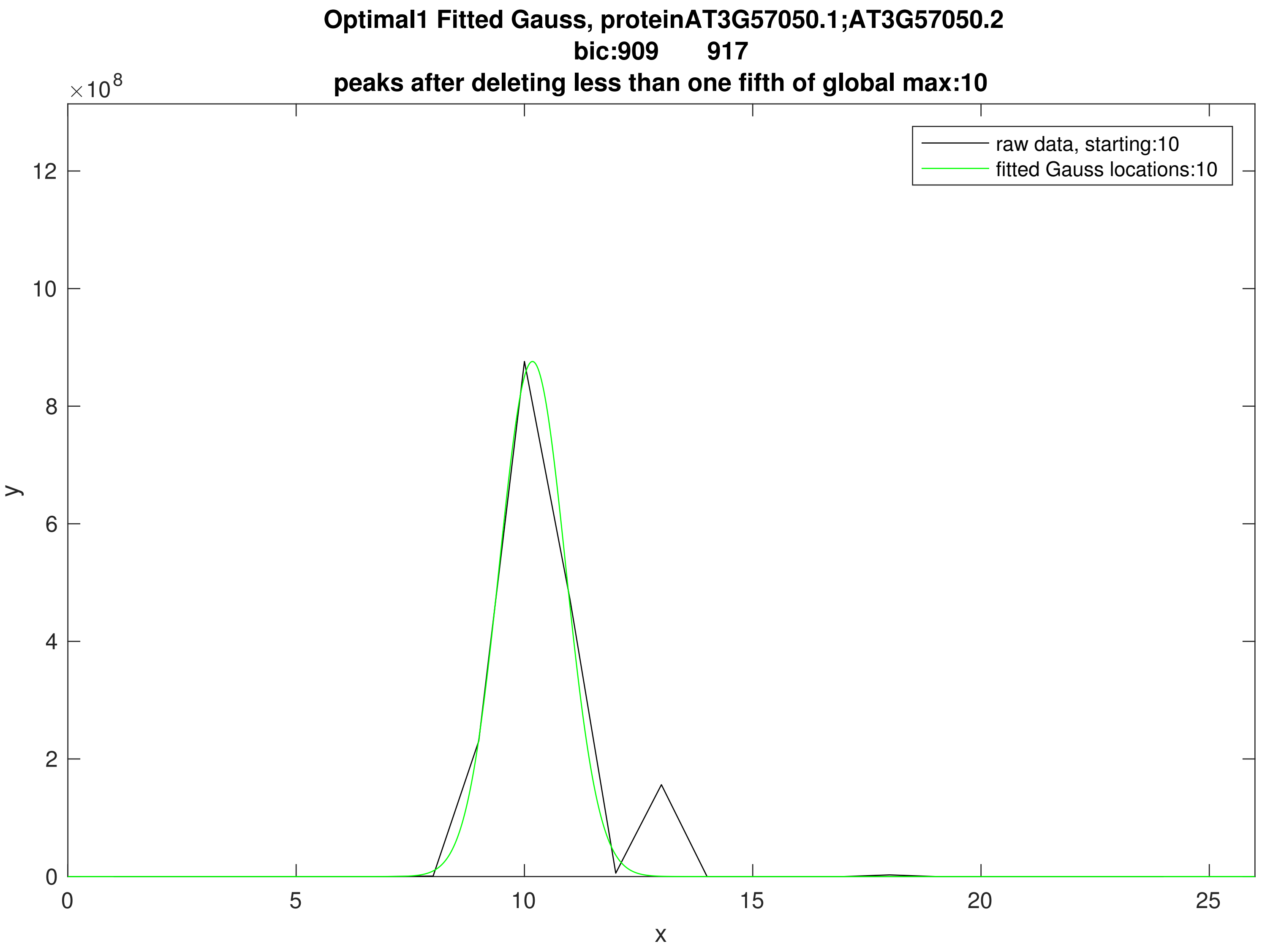
Optimal1 Fitted Gauss, proteinAT3G57050.1;AT3G57050.2
bic:909 917
peaks after deleting less than one fifth of global max:10
×10^8
raw data, starting:10
fitted Gauss locations:10
y
x
0
2
4
6
8
10
12
0
5
10
15
20
25

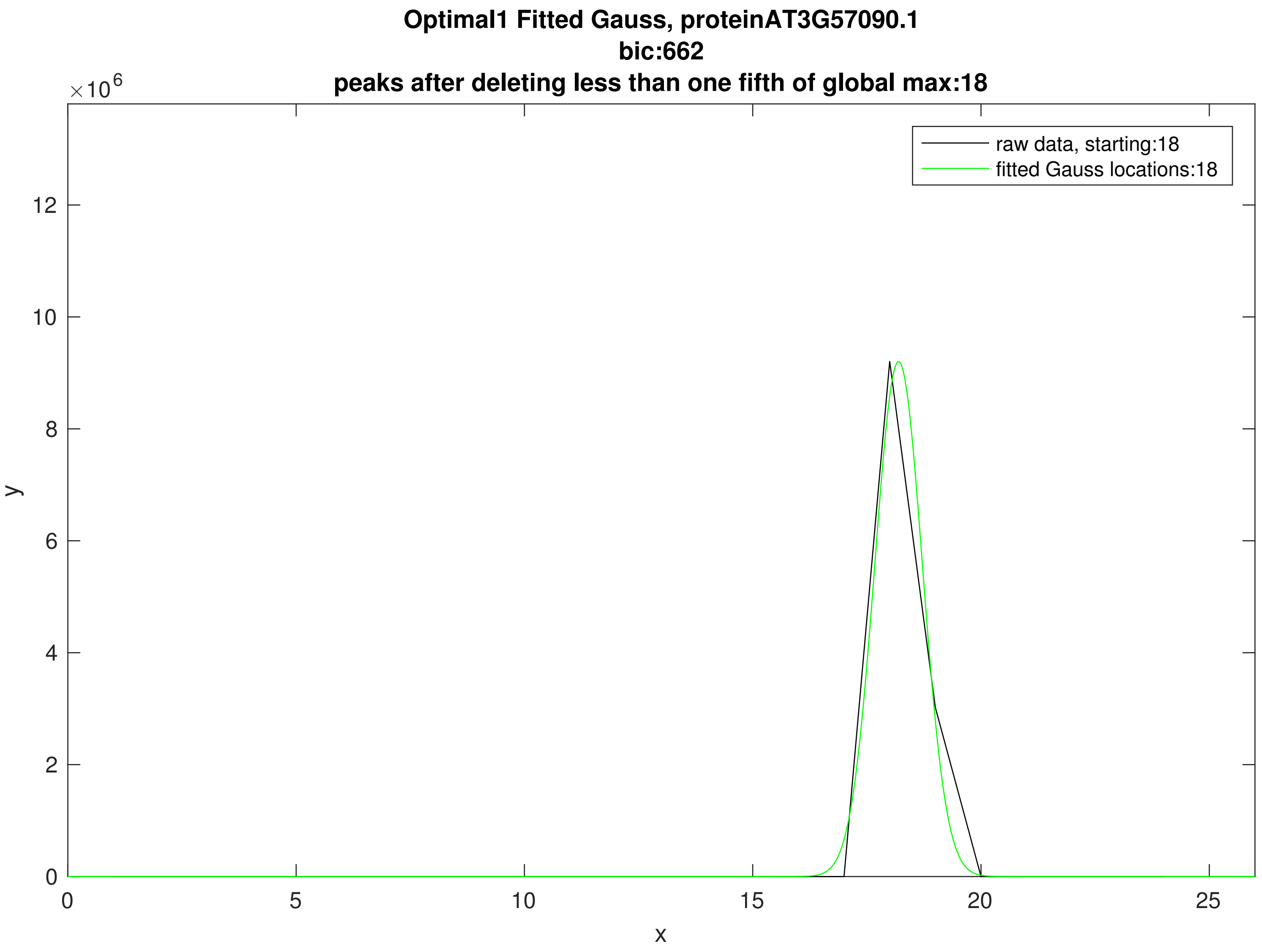
Optimal1 Fitted Gauss, proteinAT3G57090.1
bic:662
peaks after deleting less than one fifth of global max:18
×10^6
raw data, starting:18
fitted Gauss locations:18
y
x
0
2
4
6
8
10
12
0
5
10
15
20
25

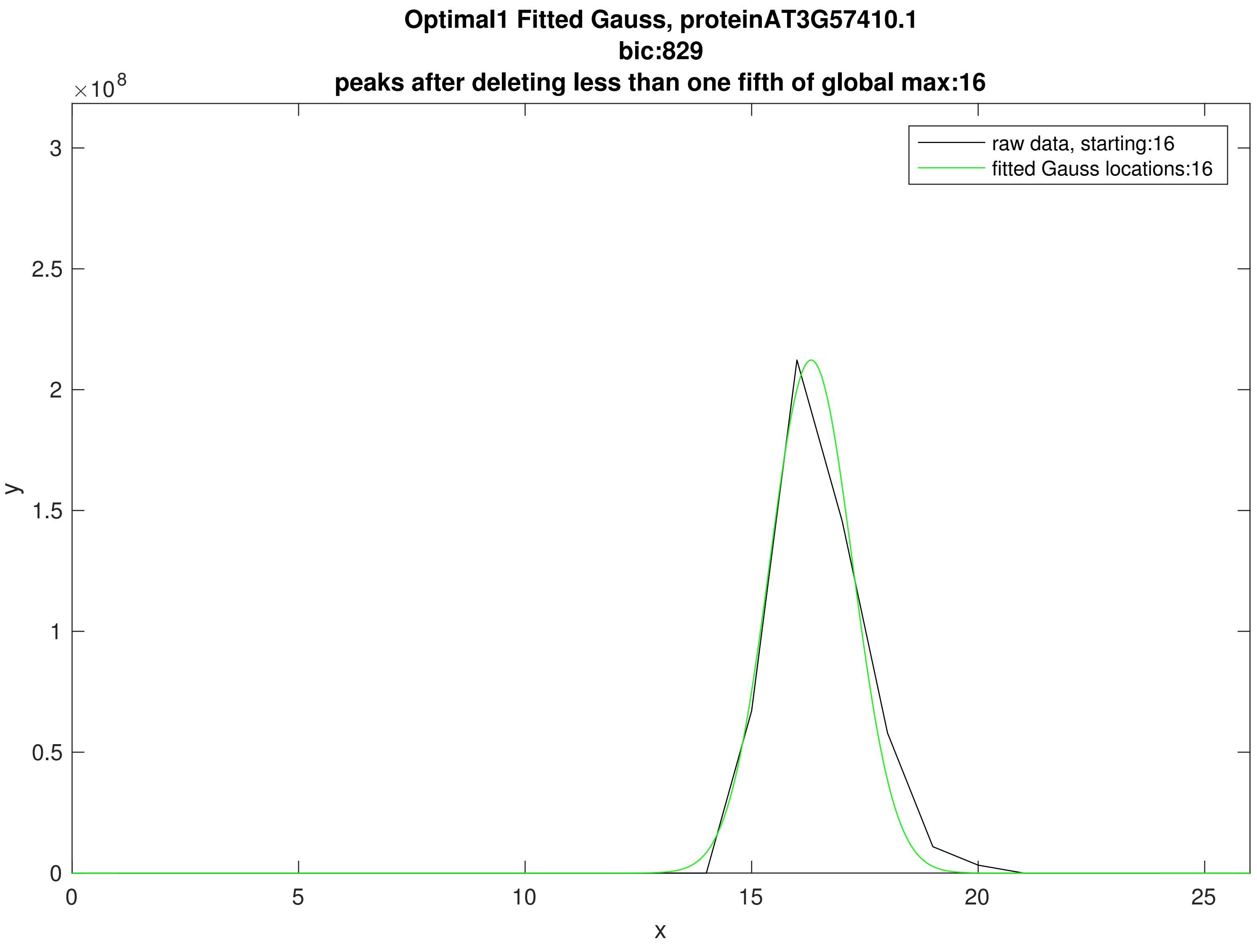
Optimal1 Fitted Gauss, proteinAT3G57410.1
bic:829
peaks after deleting less than one fifth of global max:16
$\times 10^8$
raw data, starting:16
fitted Gauss locations:16
y
x
0
0.5
1
1.5
2
2.5
3
0
5
10
15
20
25

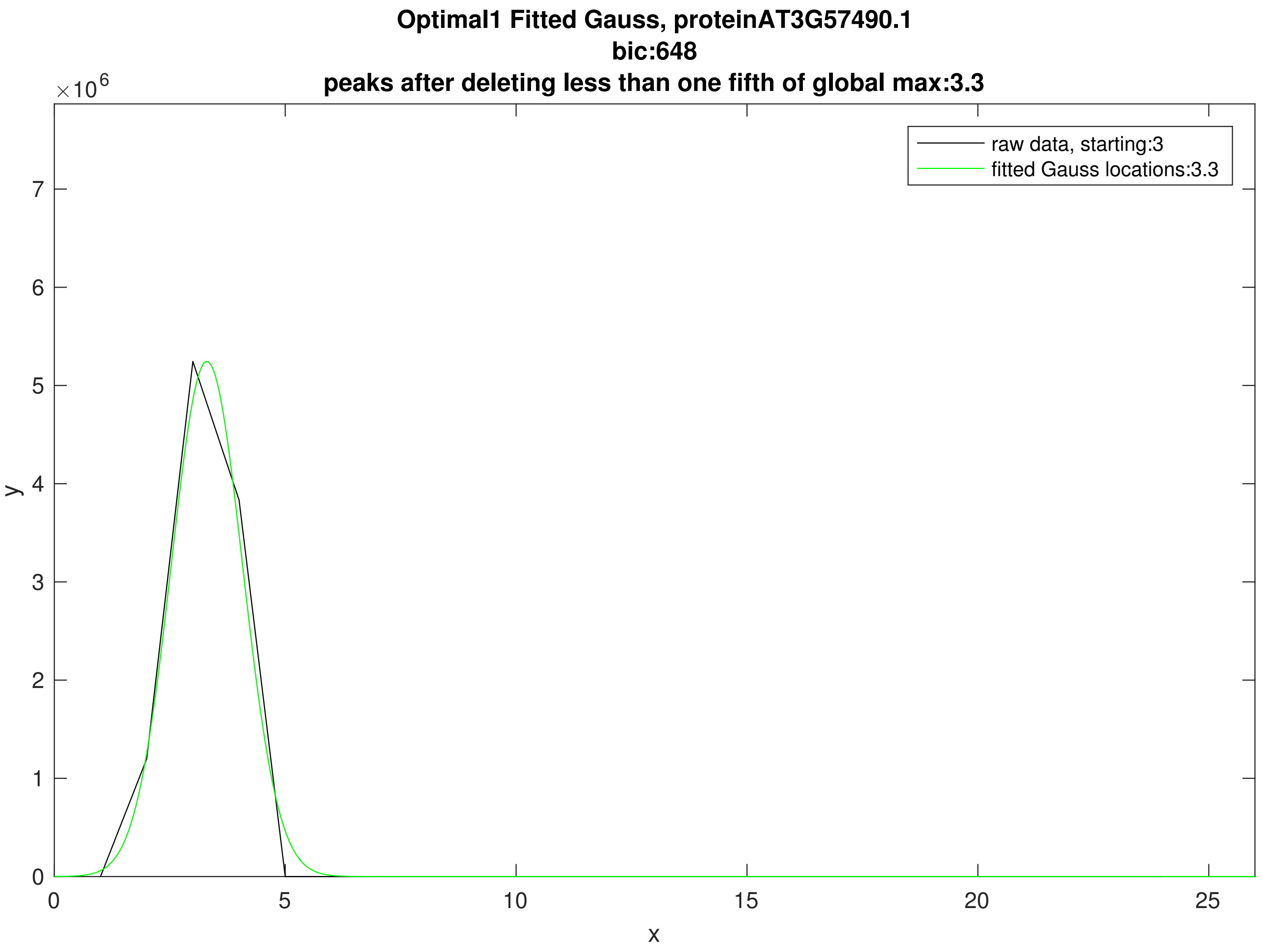

Optimal1 Fitted Gauss, proteinAT3G57490.1
bic:648
peaks after deleting less than one fifth of global max:3.3
×10^6
raw data, starting:3
fitted Gauss locations:3.3
y
x
0
1
2
3
4
5
6
7
0
5
10
15
20
25

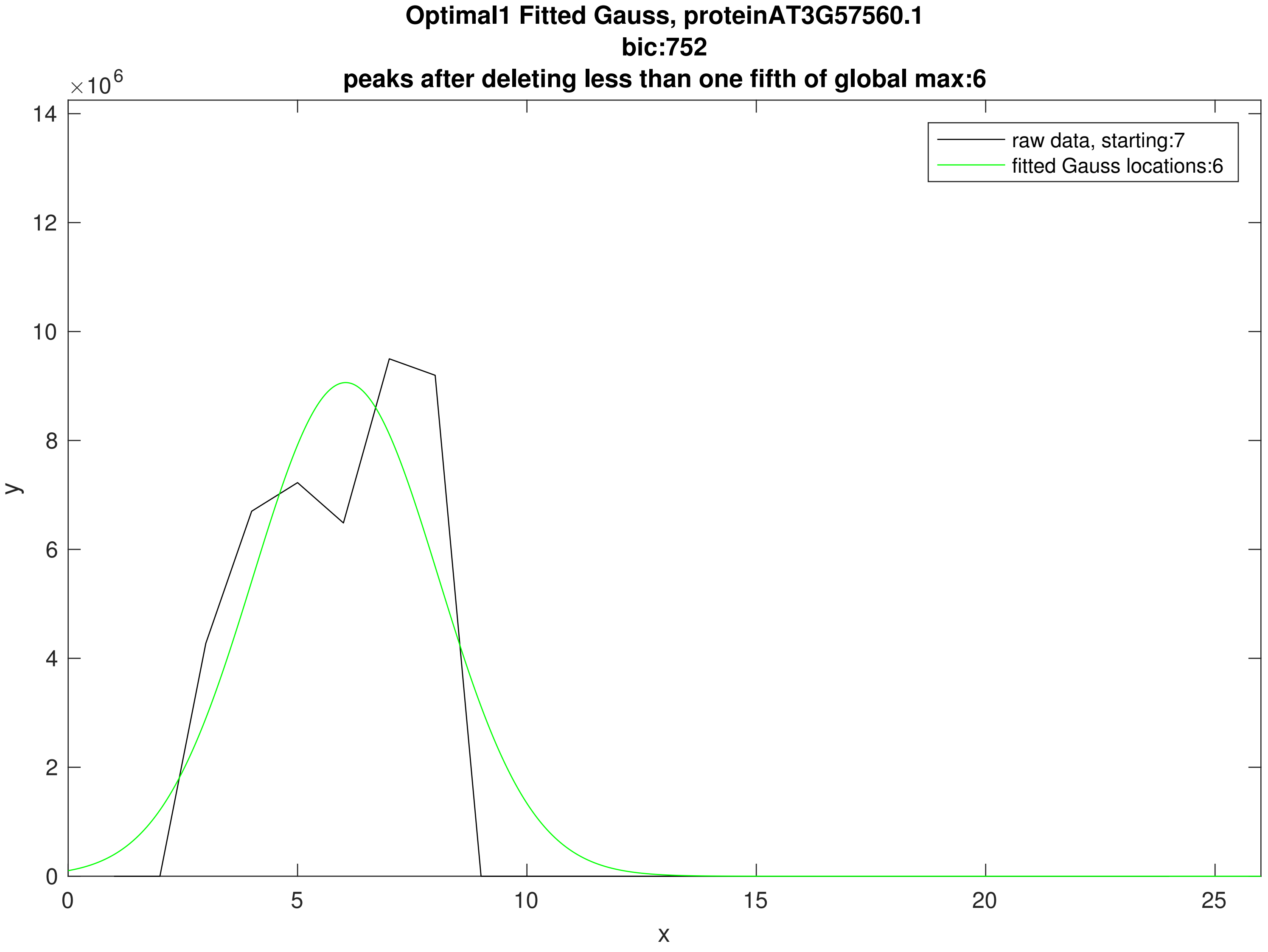

Optimal1 Fitted Gauss, proteinAT3G57560.1
bic:752
peaks after deleting less than one fifth of global max:6
×10^6
14
12
10
8
6
4
2
0
y
0
5
10
15
20
25
x
raw data, starting:7
fitted Gauss locations:6

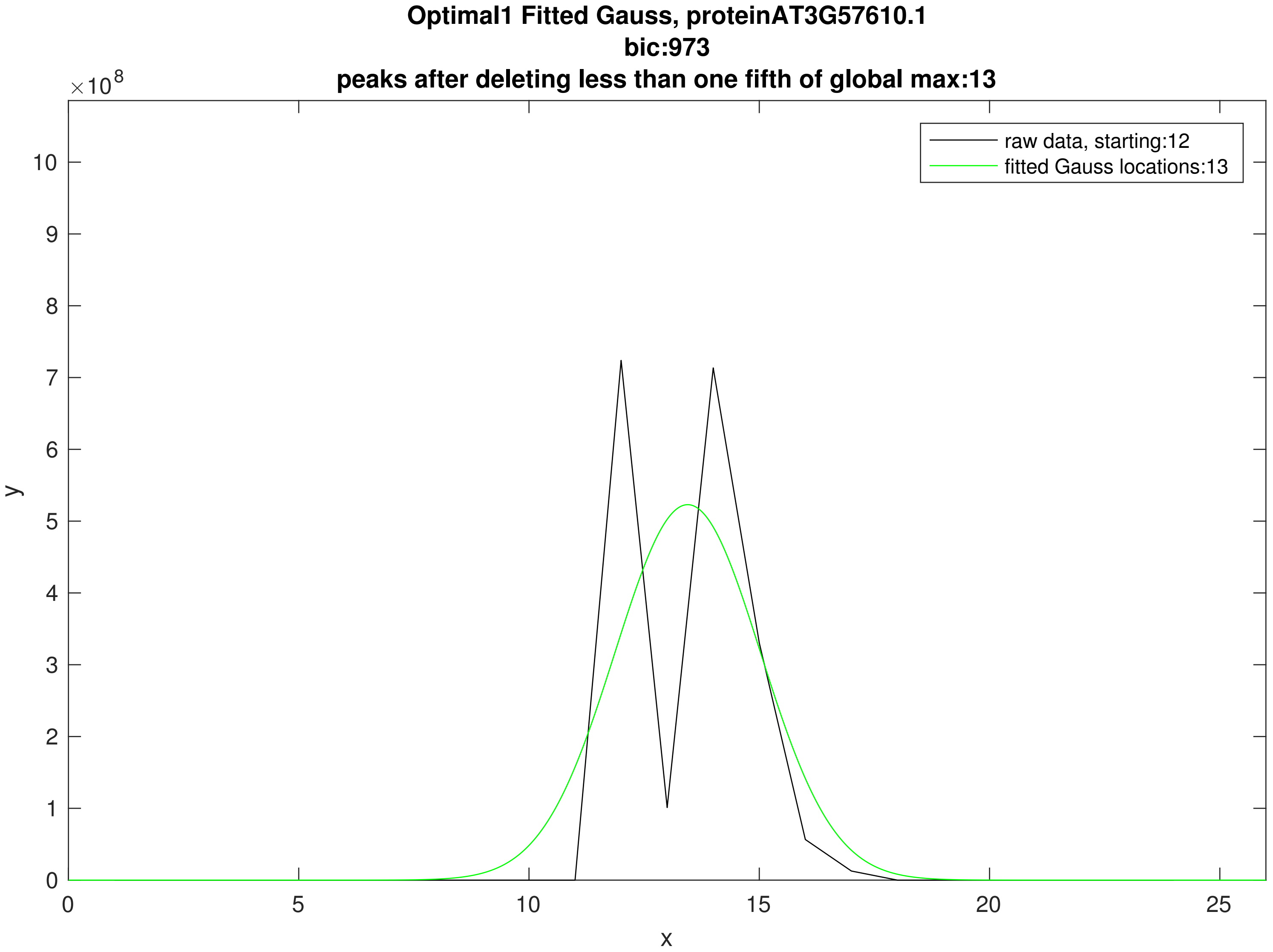
Optimal1 Fitted Gauss, proteinAT3G57610.1
bic:973
peaks after deleting less than one fifth of global max:13
×10^8
raw data, starting:12
fitted Gauss locations:13
10
9
8
7
6
5
4
3
2
1
0
y
0
5
10
15
20
25
x

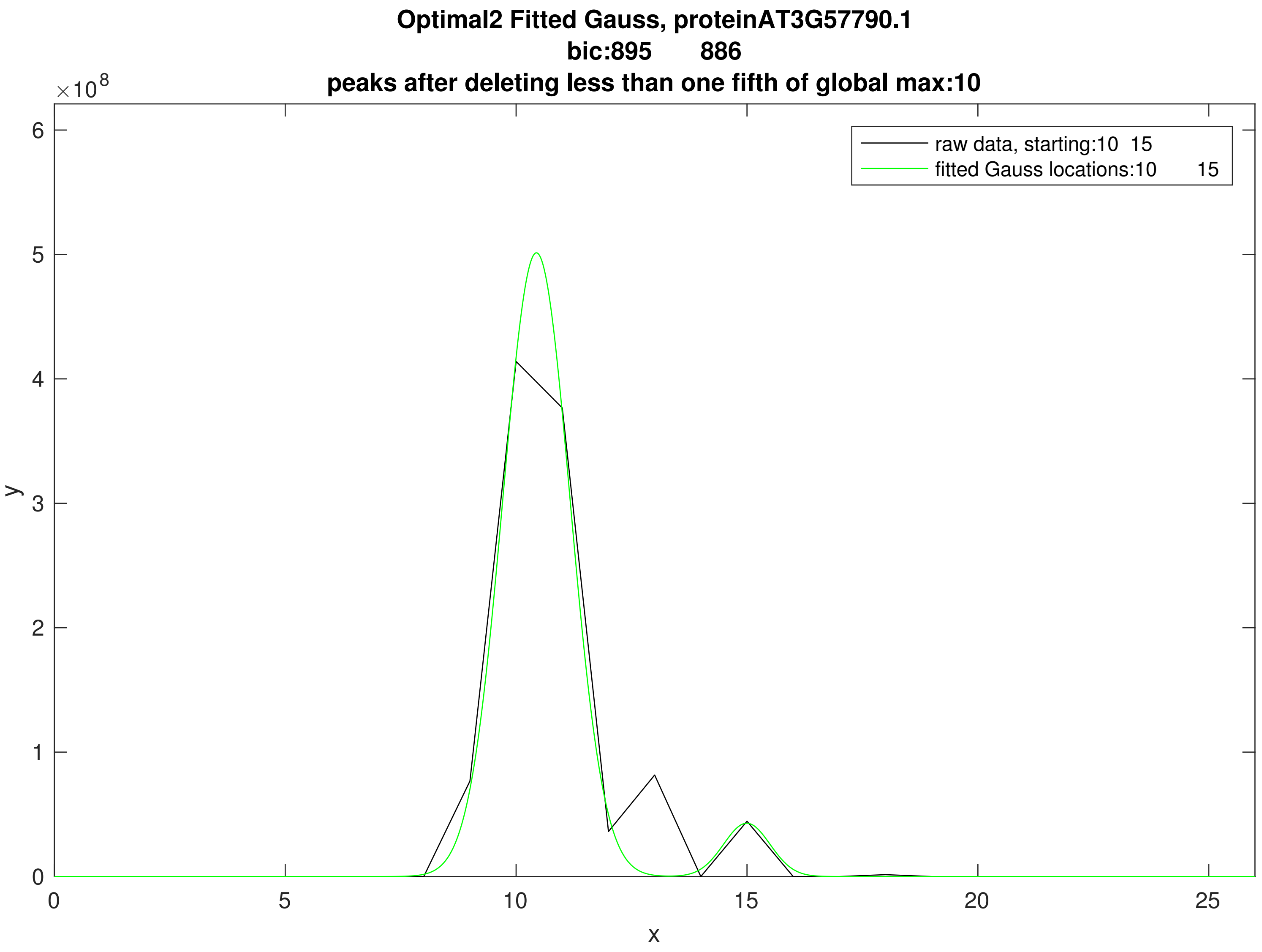
Optimal2 Fitted Gauss, proteinAT3G57790.1
bic:895 886
peaks after deleting less than one fifth of global max:10
×10^8
raw data, starting:10 15
fitted Gauss locations:10 15
y
x
0
1
2
3
4
5
6
10
15
20
25

Optimal1 Fitted Gauss, proteinAT3G57810.1;AT3G57810.3;AT3G57810.2
bic:661
peaks after deleting less than one fifth of global max:19

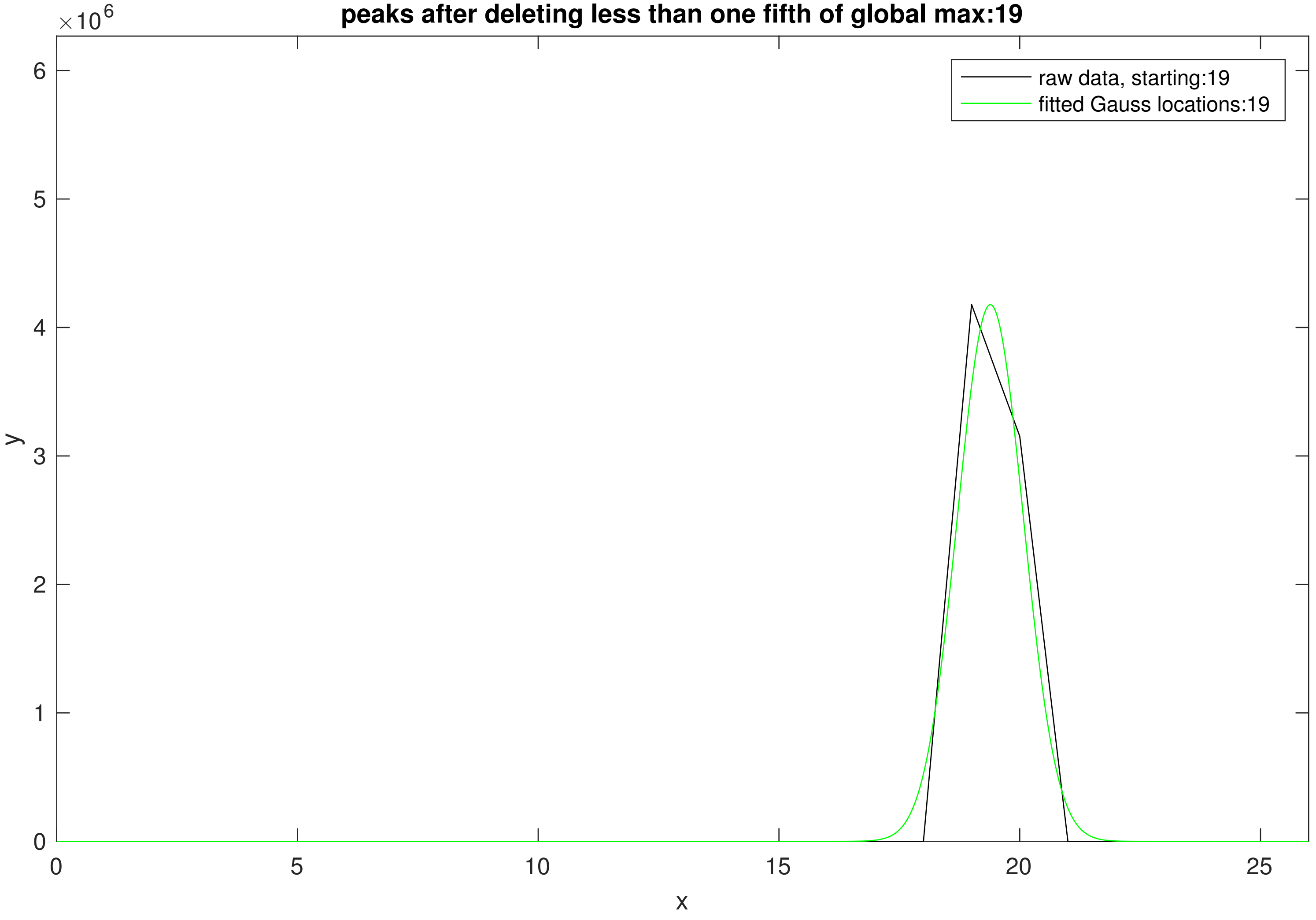

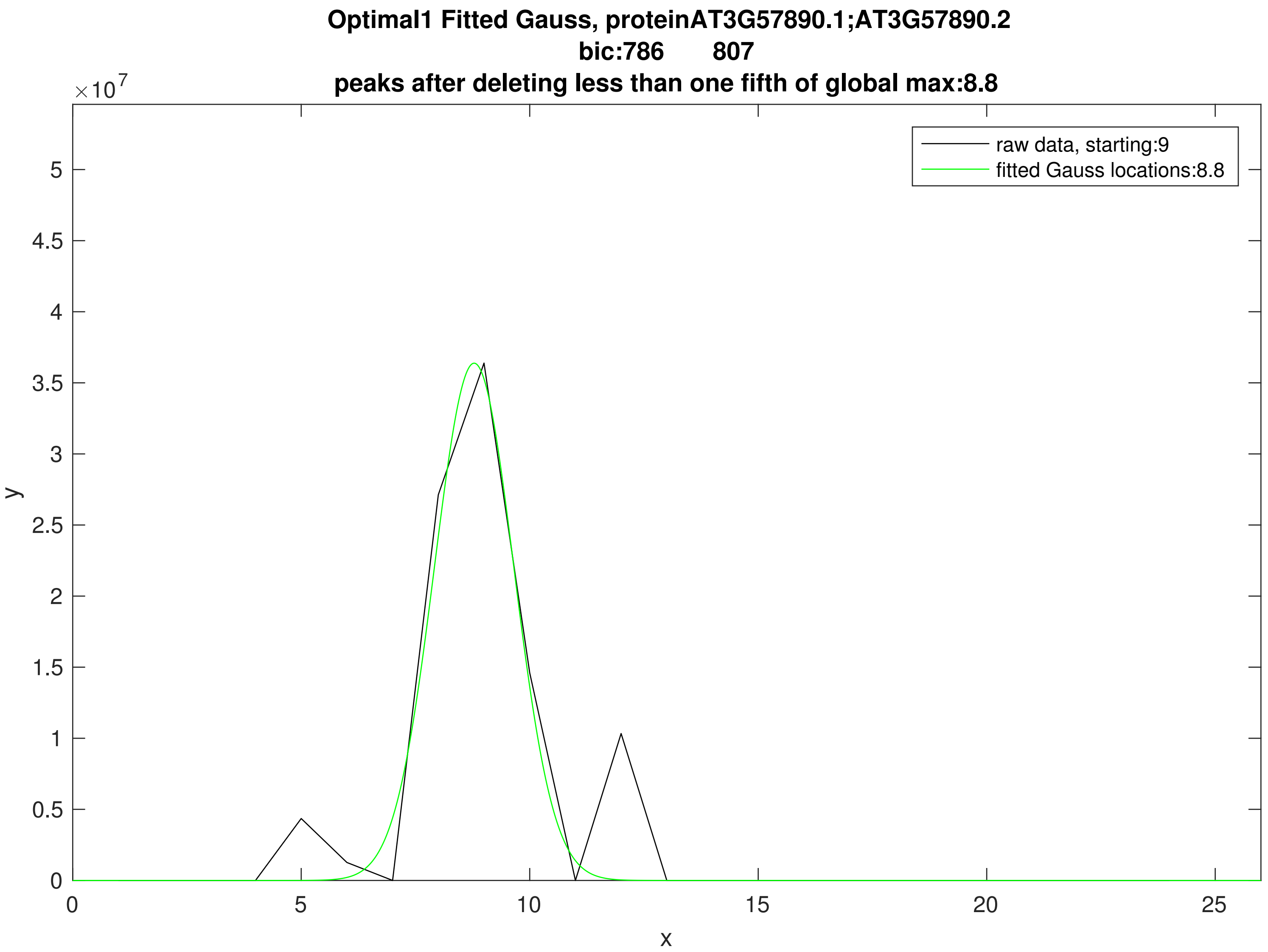
Optimal1 Fitted Gauss, proteinAT3G57890.1;AT3G57890.2
bic:786 807
peaks after deleting less than one fifth of global max:8.8
×10^7
raw data, starting:9
fitted Gauss locations:8.8
y
x
0
0.5
1
1.5
2
2.5
3
3.5
4
4.5
5
0
5
10
15
20
25

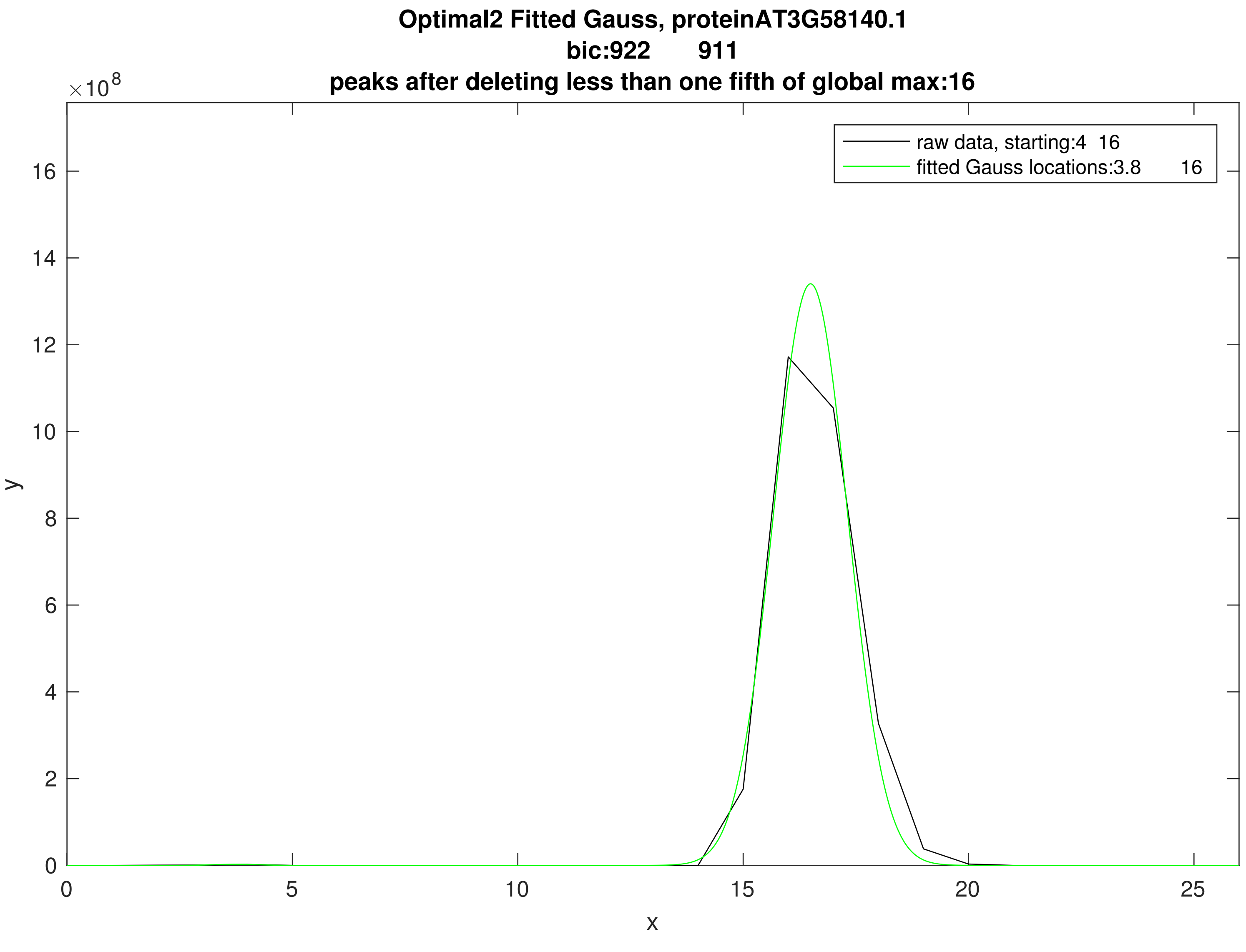

Optimal2 Fitted Gauss, proteinAT3G58140.1
bic:922 911
peaks after deleting less than one fifth of global max:16
$\times10^8$
raw data, starting:4 16
fitted Gauss locations:3.8 16
y
x
0
2
4
6
8
10
12
14
16
5
15
20
25

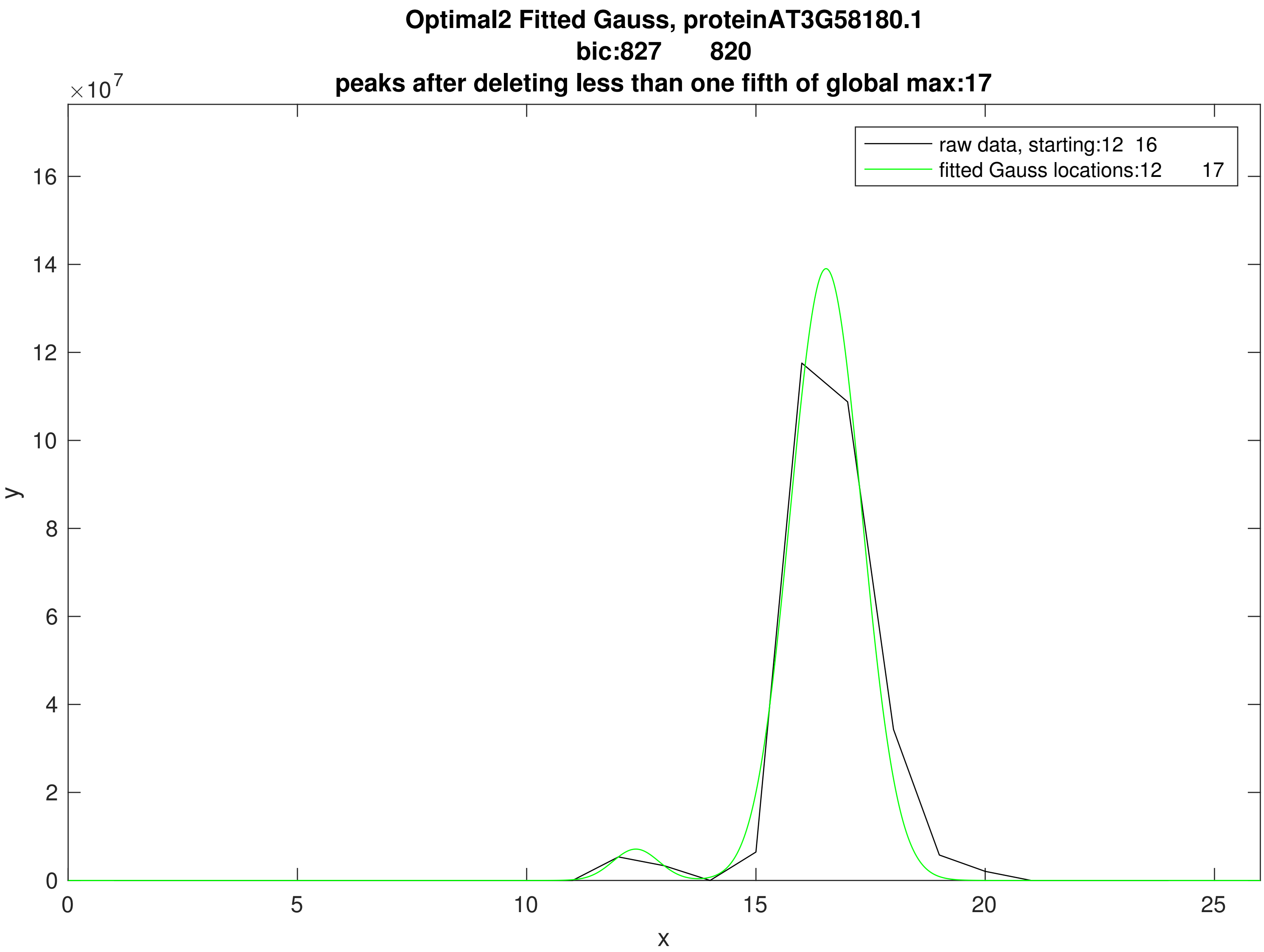

Optimal2 Fitted Gauss, proteinAT3G58180.1
bic:827 820
peaks after deleting less than one fifth of global max:17
×10^7
raw data, starting:12 16
fitted Gauss locations:12 17
y
x
0
2
4
6
8
10
12
14
16
0
5
10
15
20
25

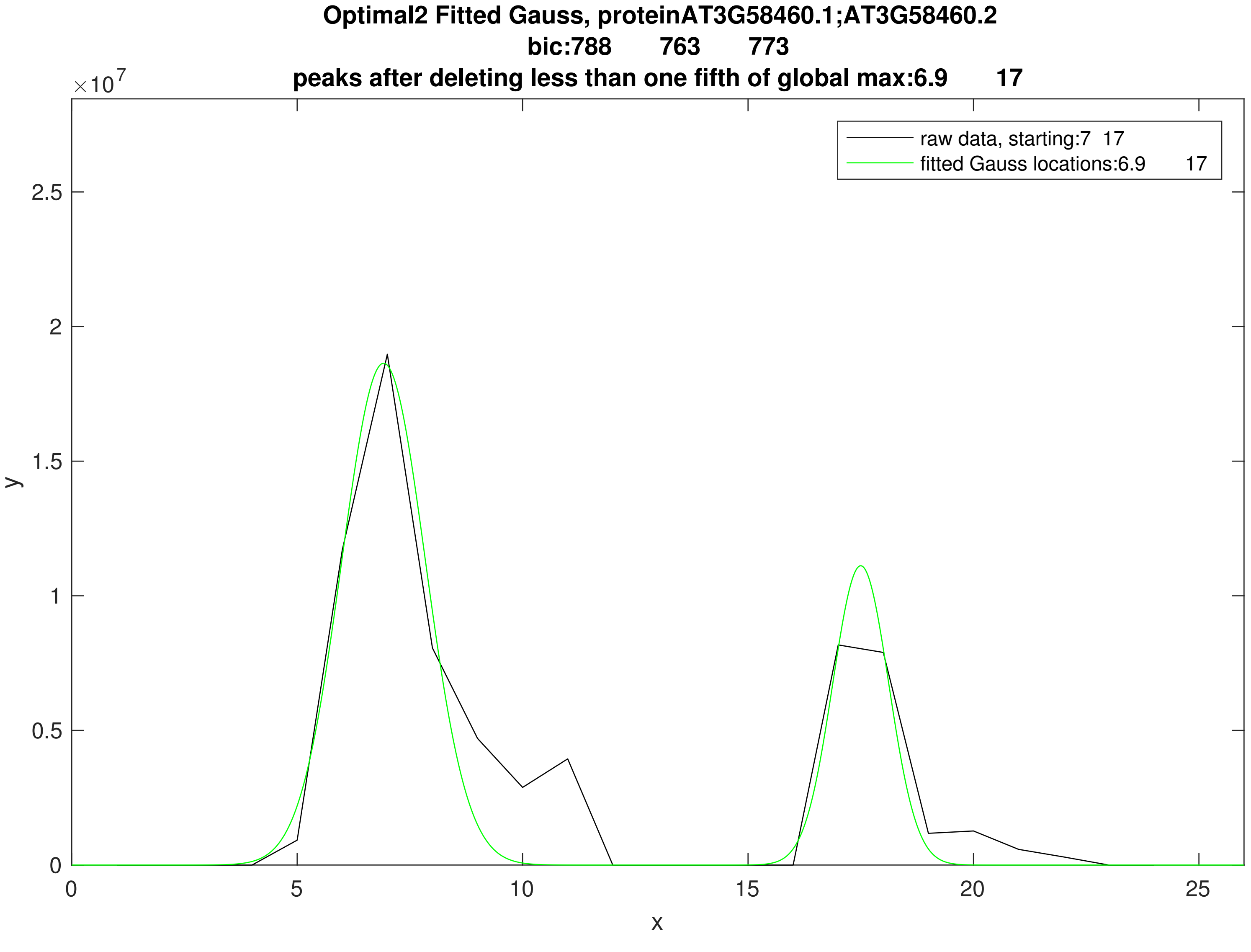

Optimal2 Fitted Gauss, proteinAT3G58460.1;AT3G58460.2
bic:788 763 773
peaks after deleting less than one fifth of global max:6.9 17
raw data, starting:7 17
fitted Gauss locations:6.9 17
×10^7
2.5
2
1.5
1
0.5
0
y
0
5
10
15
20
25
x

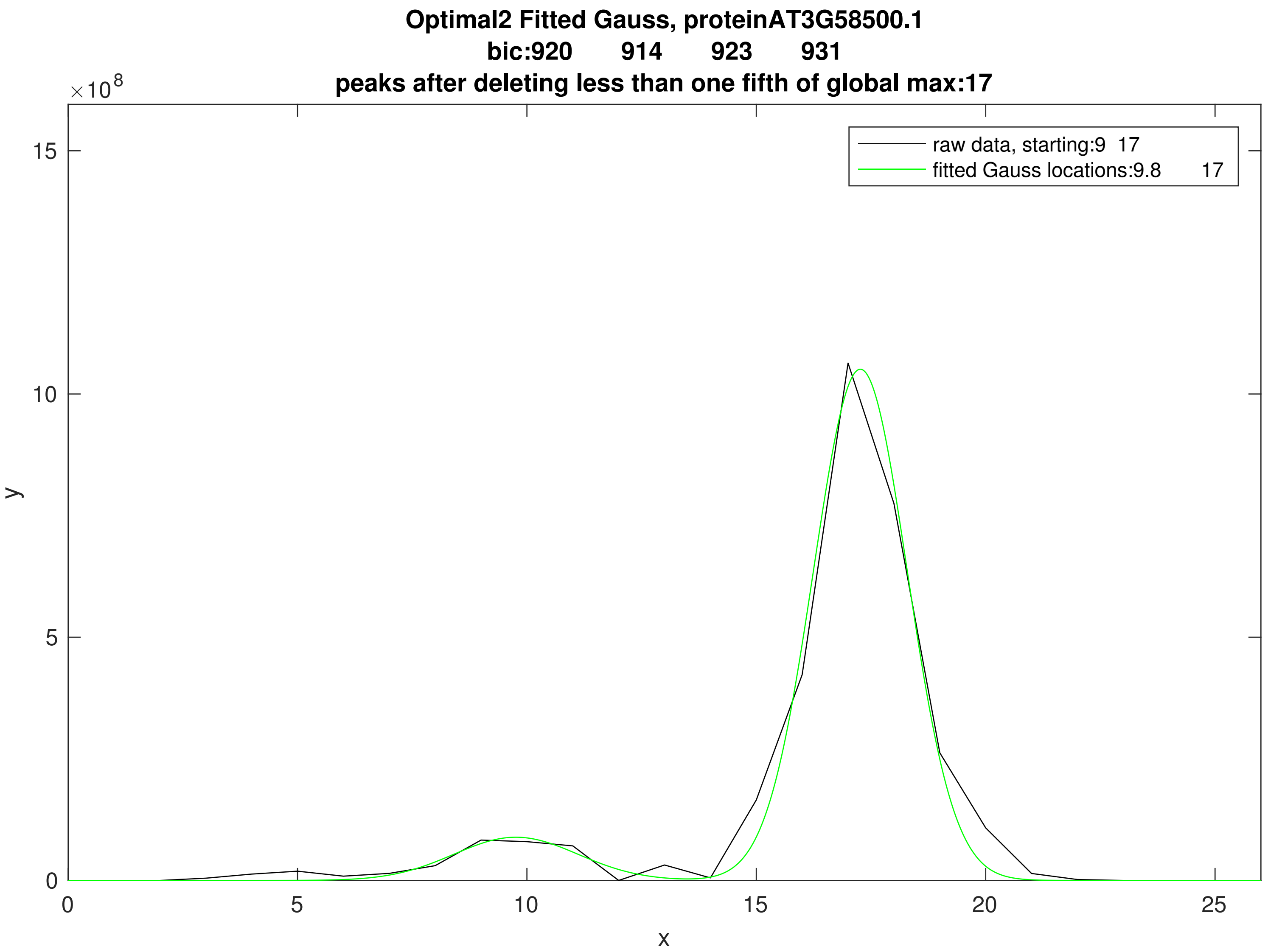
Optimal2 Fitted Gauss, proteinAT3G58500.1
bic:920 914 923 931
peaks after deleting less than one fifth of global max:17
×10^8
raw data, starting:9 17
fitted Gauss locations:9.8 17
y
x
0
5
10
15
20
25

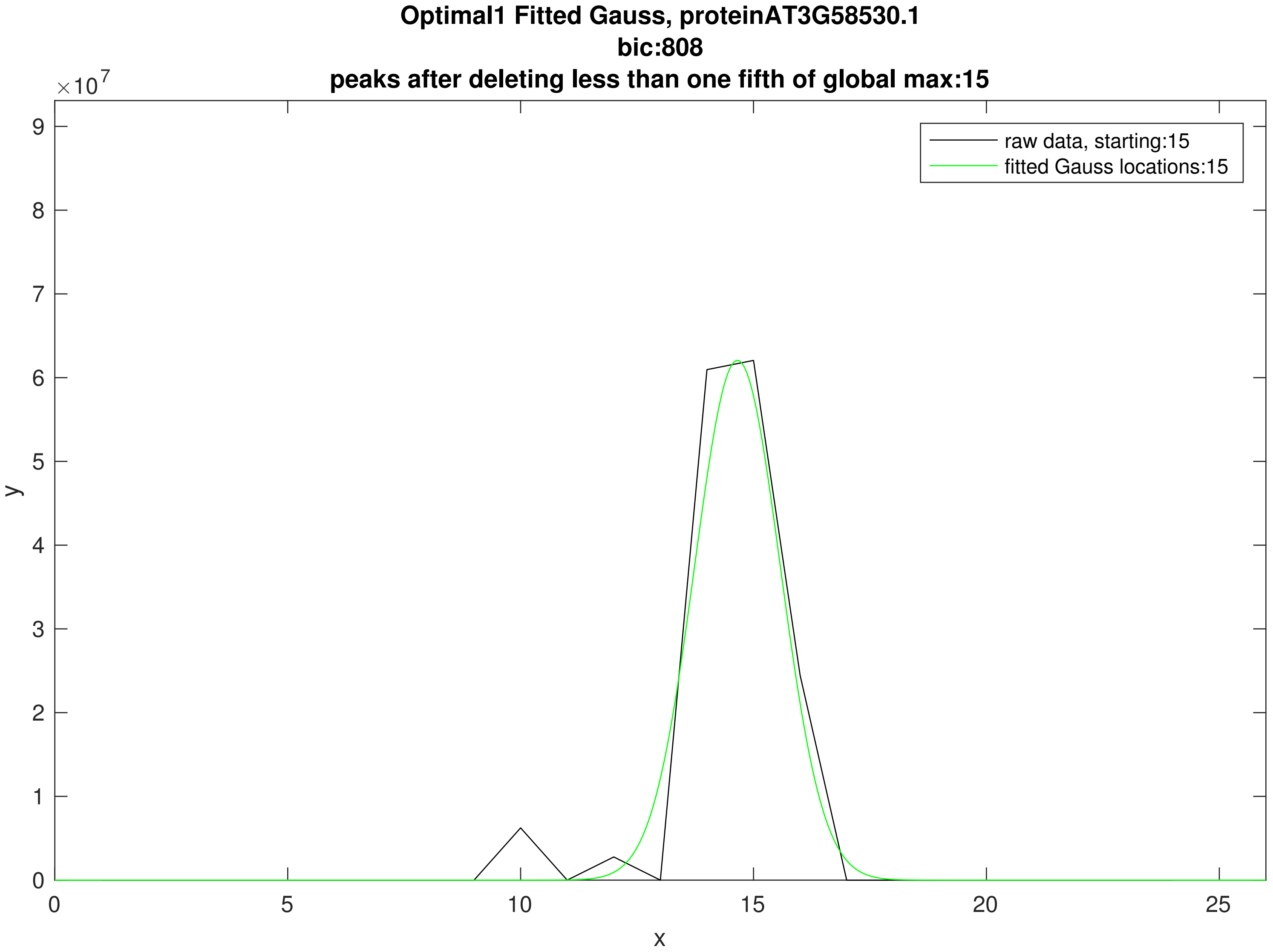
Optimal1 Fitted Gauss, proteinAT3G58530.1
bic:808
peaks after deleting less than one fifth of global max:15
×10^7
raw data, starting:15
fitted Gauss locations:15
y
x
0
1
2
3
4
5
6
7
8
9
0
5
10
15
20
25

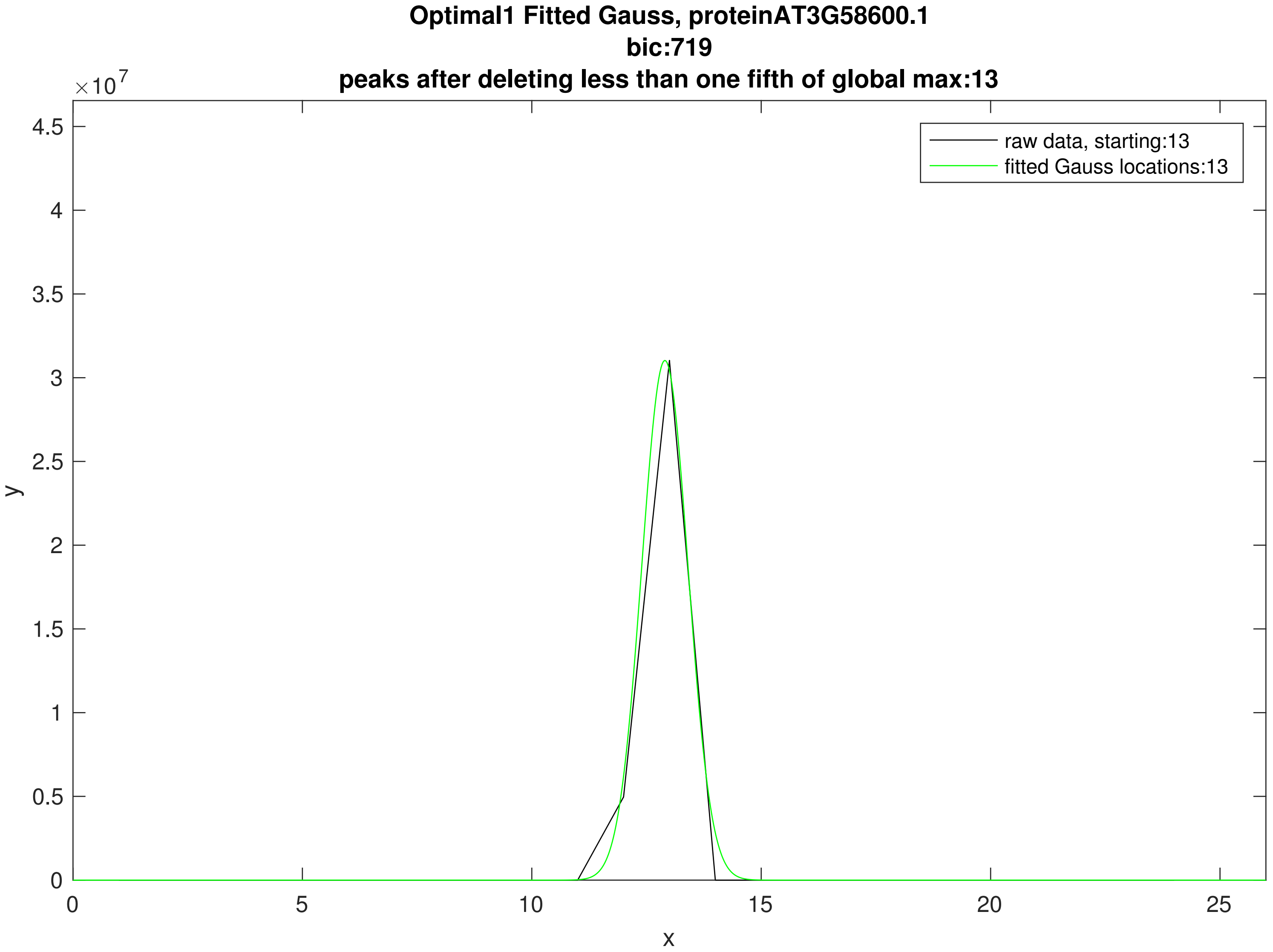
Optimal1 Fitted Gauss, proteinAT3G58600.1
bic:719
peaks after deleting less than one fifth of global max:13
×10^7
4.5
4
3.5
3
2.5
2
1.5
1
0.5
0
y
0
5
10
15
20
25
x
raw data, starting:13
fitted Gauss locations:13

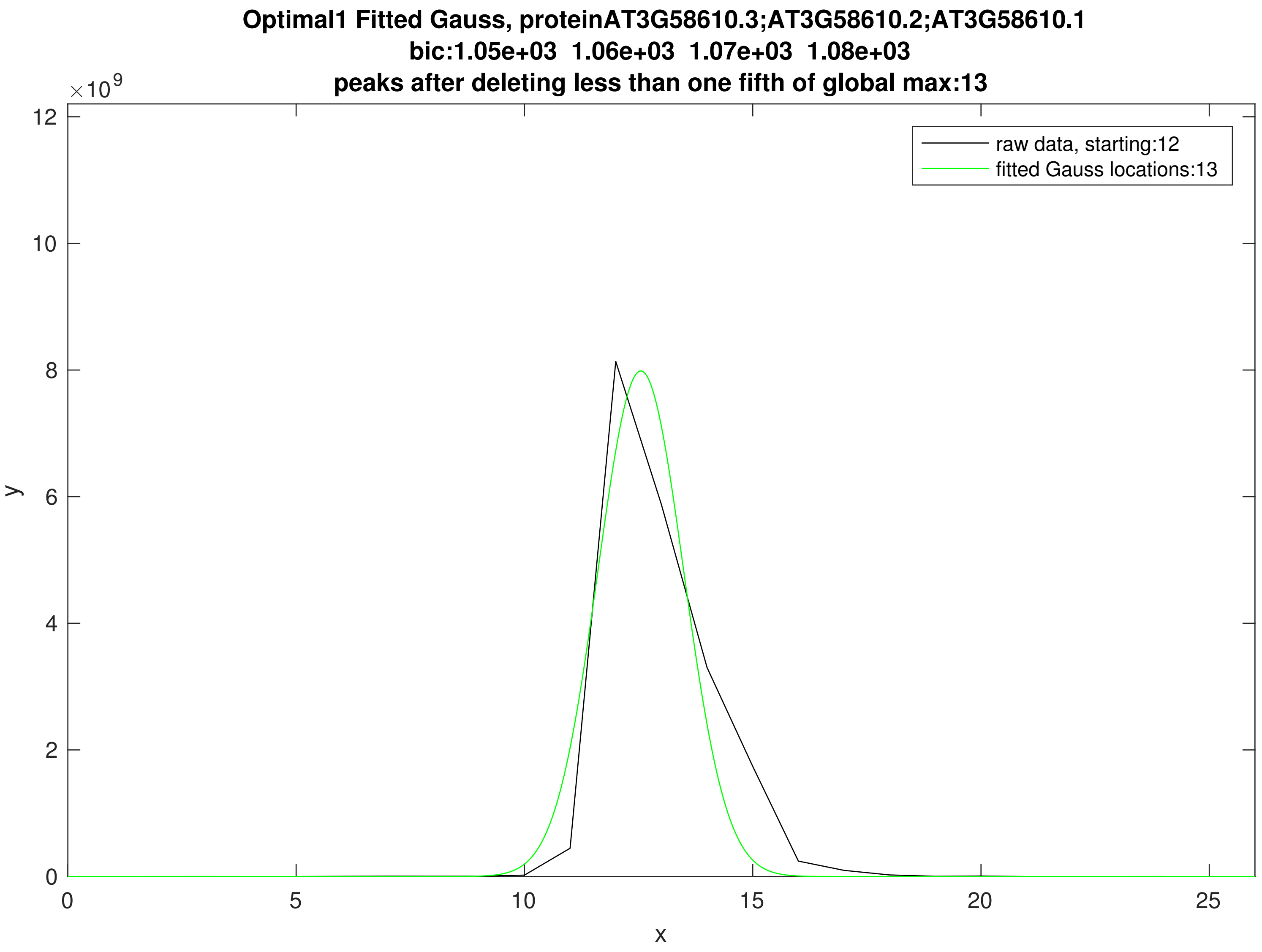

Optimal1 Fitted Gauss, proteinAT3G58610.3;AT3G58610.2;AT3G58610.1
bic:1.05e+03 1.06e+03 1.07e+03 1.08e+03
peaks after deleting less than one fifth of global max:13
×10^9
0
2
4
6
8
10
12
y
0
5
10
15
20
25
x
raw data, starting:12
fitted Gauss locations:13

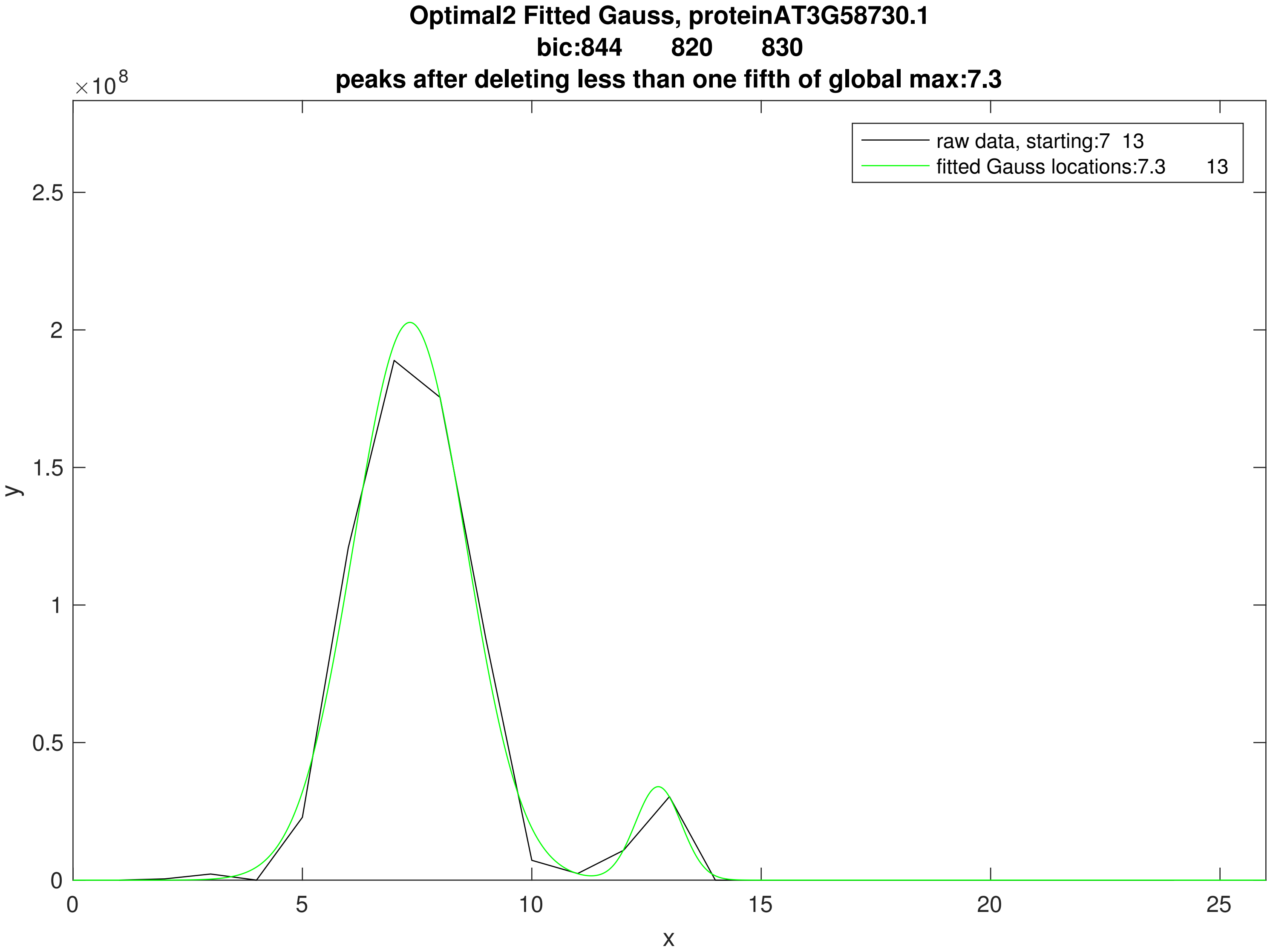
Optimal2 Fitted Gauss, proteinAT3G58730.1
bic:844 820 830
peaks after deleting less than one fifth of global max:7.3
×10^8
2.5
2
1.5
1
0.5
0
y
0
5
10
15
20
25
x
raw data, starting:7 13
fitted Gauss locations:7.3 13

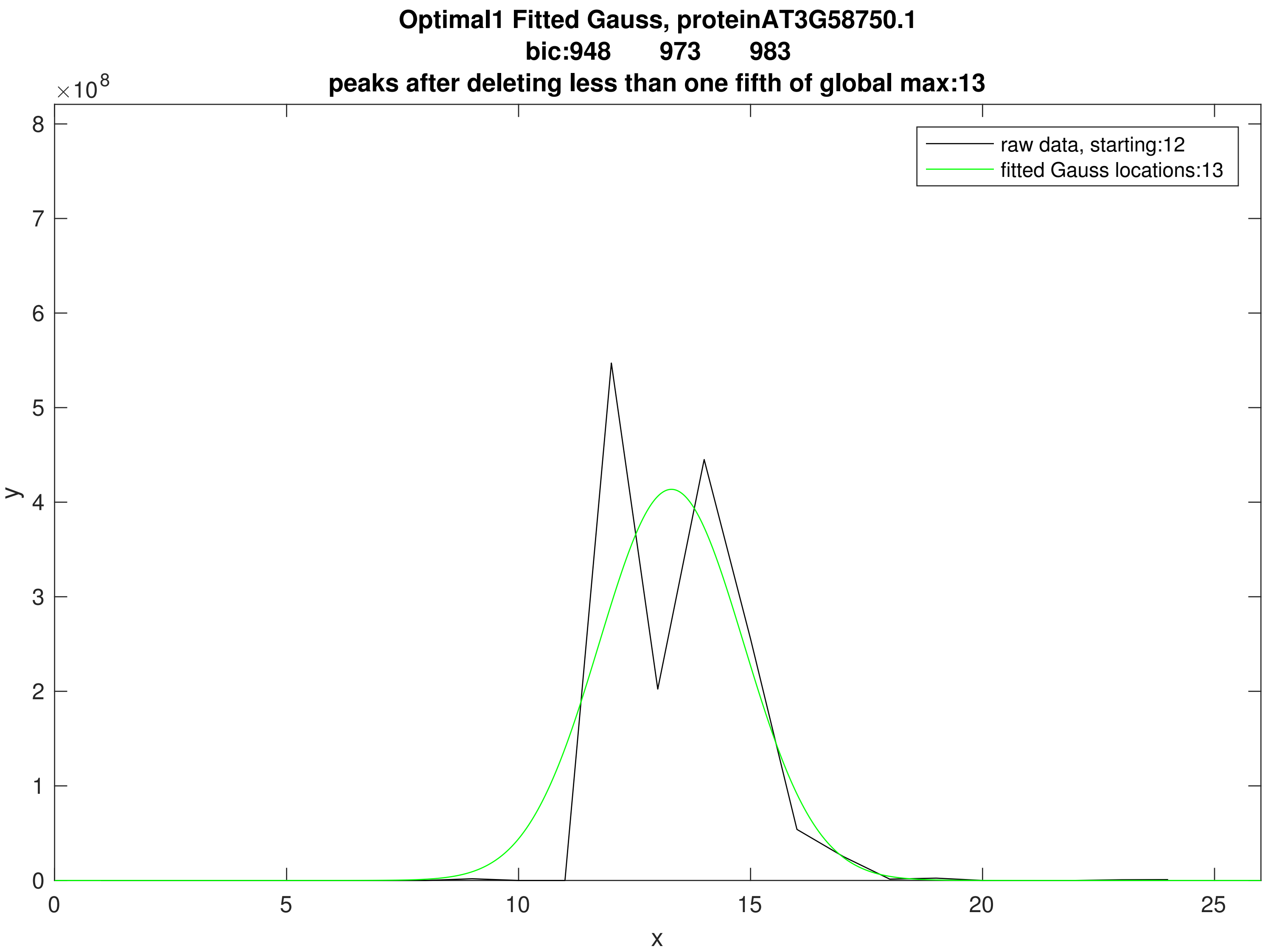

Optimal1 Fitted Gauss, proteinAT3G58750.1
bic:948 973 983
peaks after deleting less than one fifth of global max:13
×10^8
raw data, starting:12
fitted Gauss locations:13
y
x
0
1
2
3
4
5
6
7
8
0
5
10
15
20
25

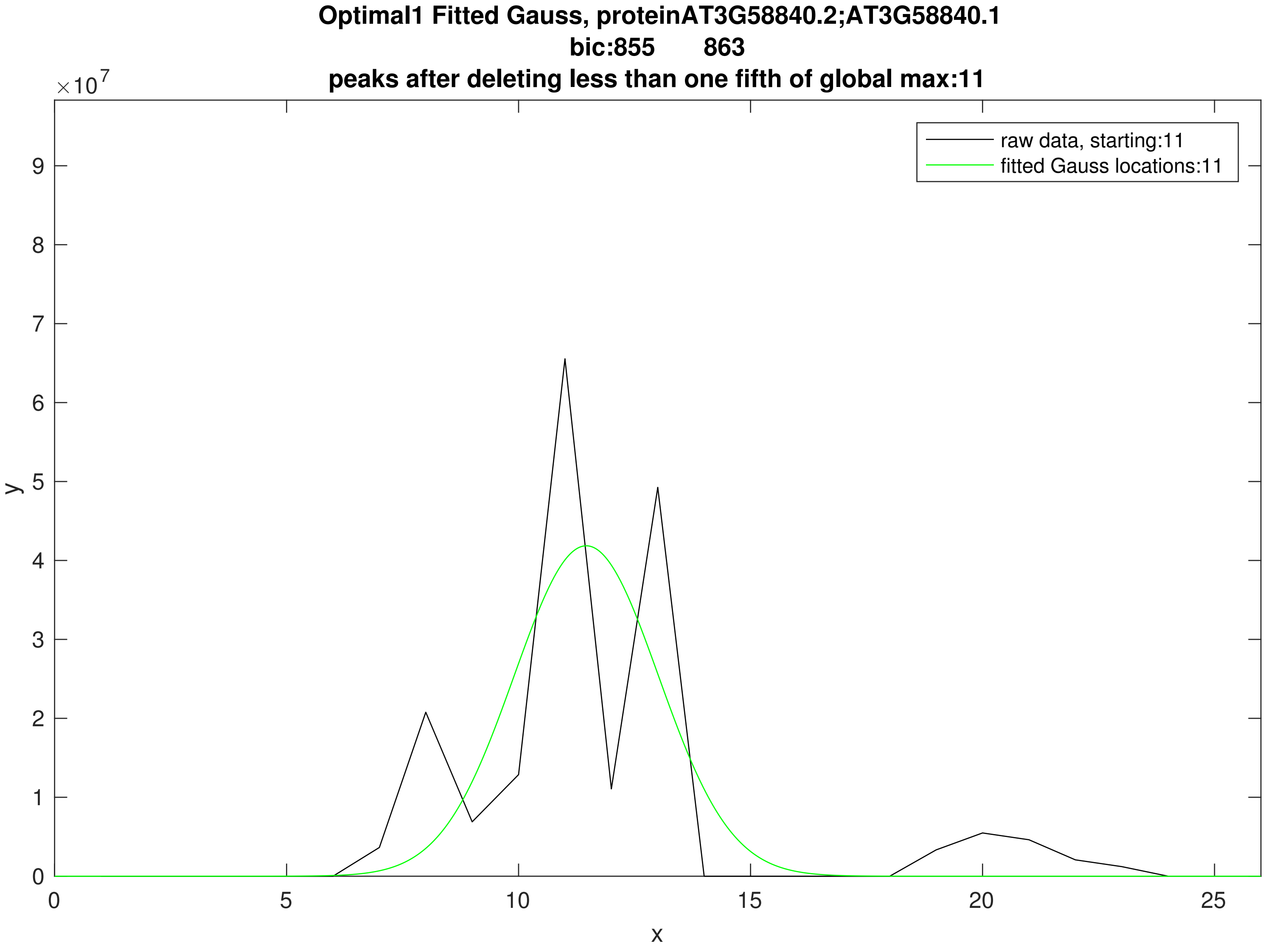

Optimal1 Fitted Gauss, proteinAT3G58840.2;AT3G58840.1
bic:855 863
peaks after deleting less than one fifth of global max:11
×10^7
raw data, starting:11
fitted Gauss locations:11
y
x
0
1
2
3
4
5
6
7
8
9
0
5
10
15
20
25

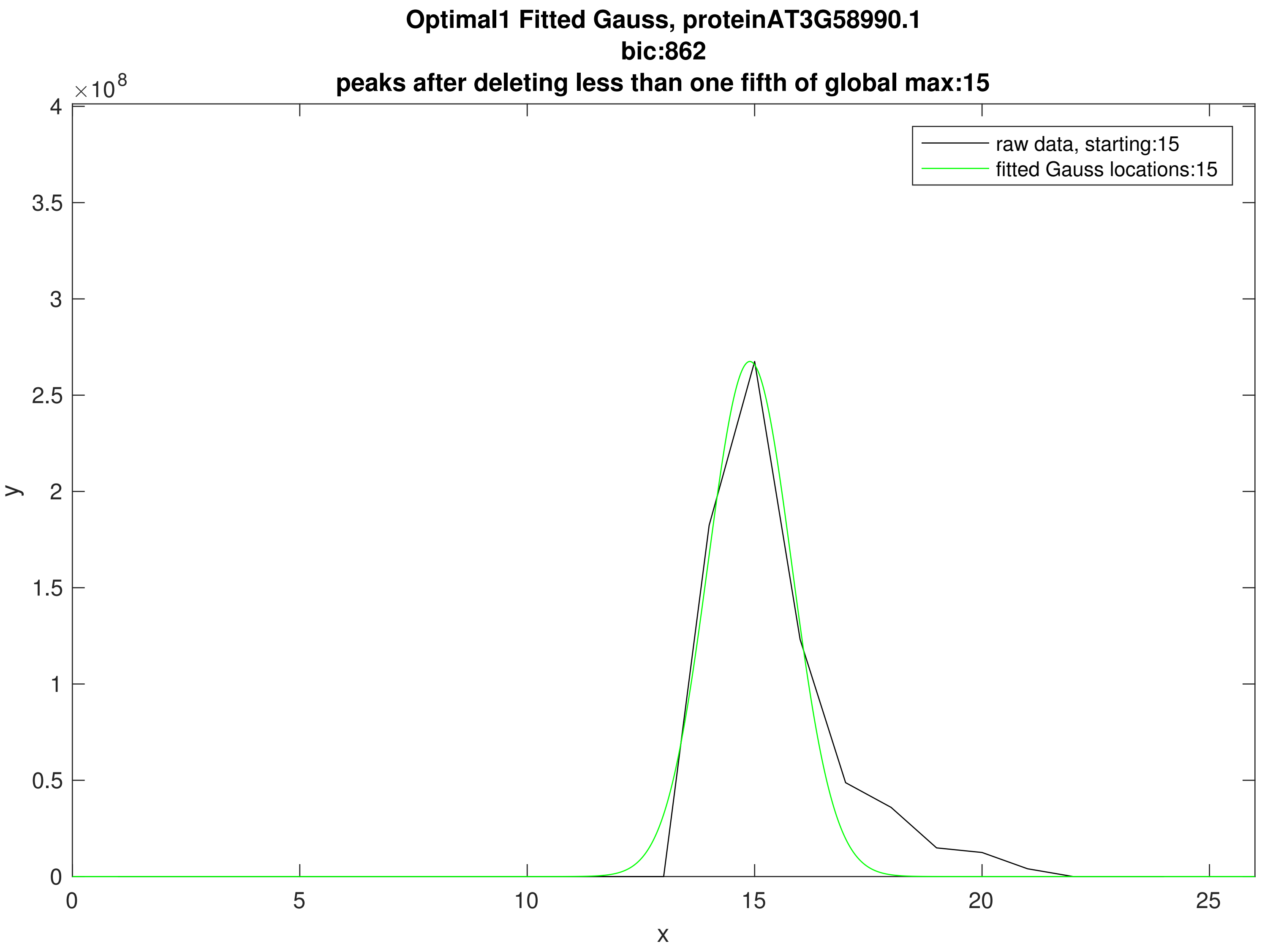
Optimal1 Fitted Gauss, proteinAT3G58990.1
bic:862
peaks after deleting less than one fifth of global max:15
×10^8
raw data, starting:15
fitted Gauss locations:15
y
x
0
0.5
1
1.5
2
2.5
3
3.5
4
0
5
10
15
20
25

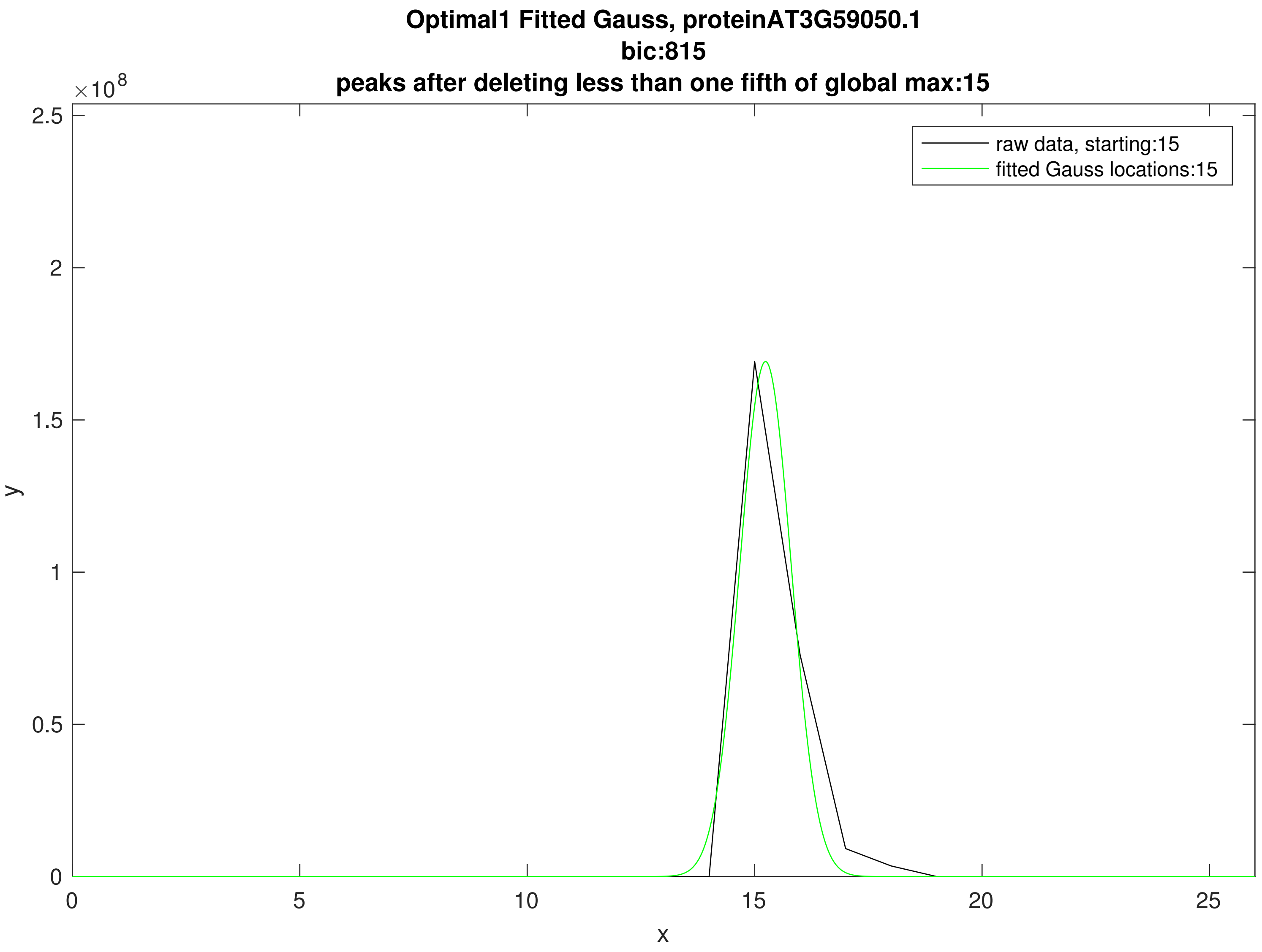

Optimal1 Fitted Gauss, proteinAT3G59050.1
bic:815
peaks after deleting less than one fifth of global max:15
×10^8
2.5
2
1.5
1
0.5
0
y
0
5
10
15
20
25
x
raw data, starting:15
fitted Gauss locations:15

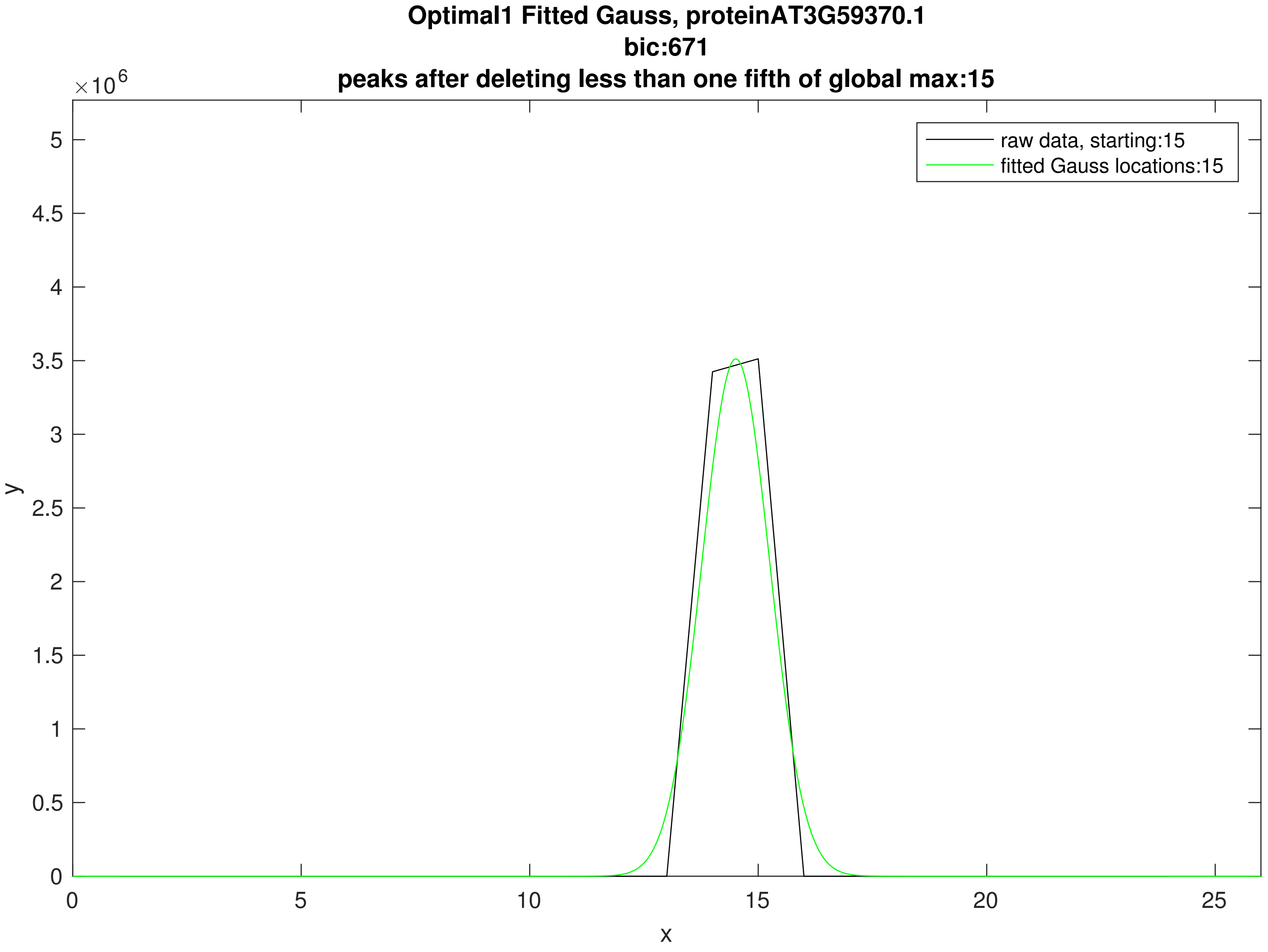
Optimal1 Fitted Gauss, proteinAT3G59370.1
bic:671
peaks after deleting less than one fifth of global max:15
×10^6
raw data, starting:15
fitted Gauss locations:15
y
x
0
0.5
1
1.5
2
2.5
3
3.5
4
4.5
5
0
5
10
15
20
25

Optimal1 Fitted Gauss, proteinAT3G59380.1
bic:911 919
peaks after deleting less than one fifth of global max:15

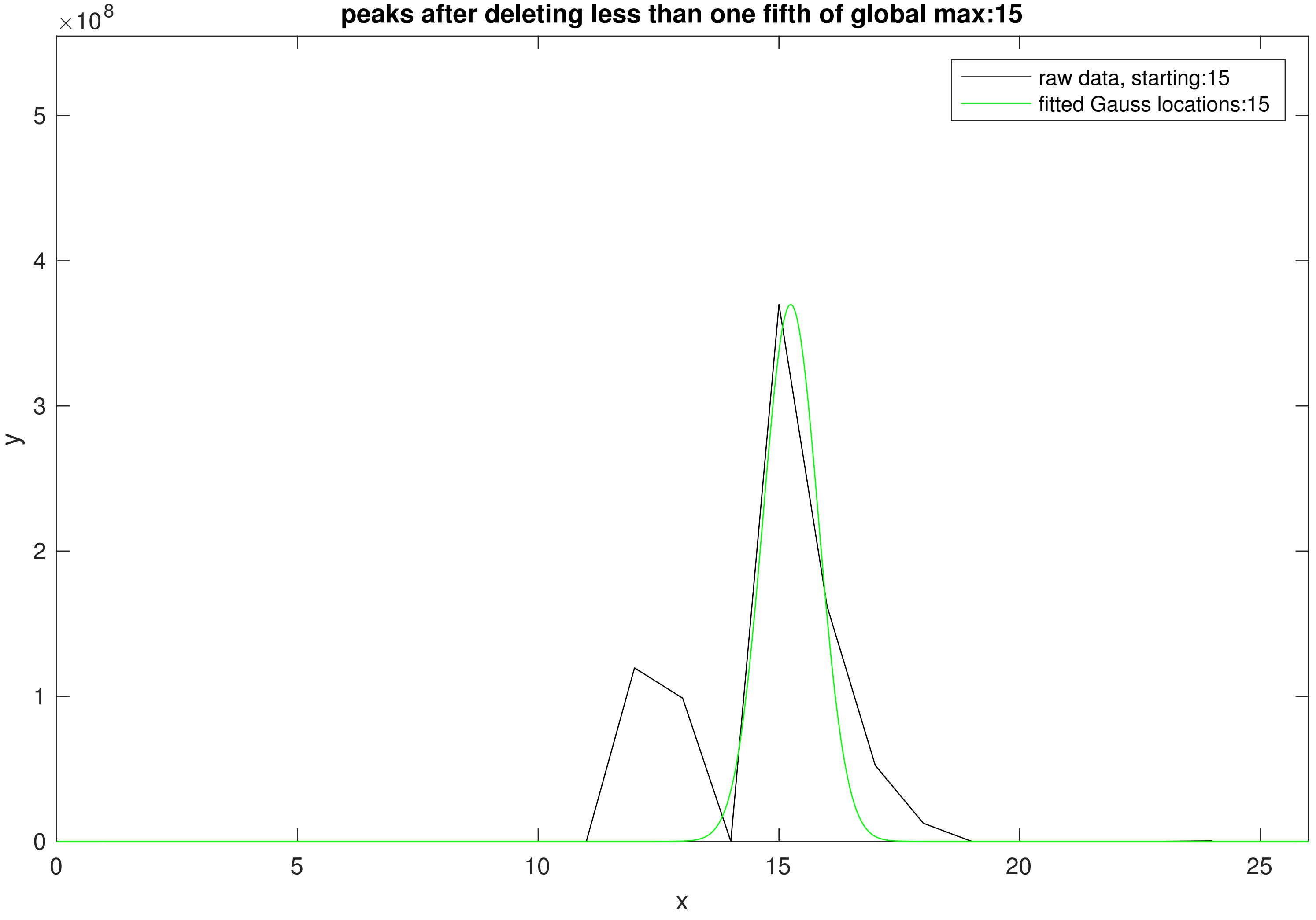

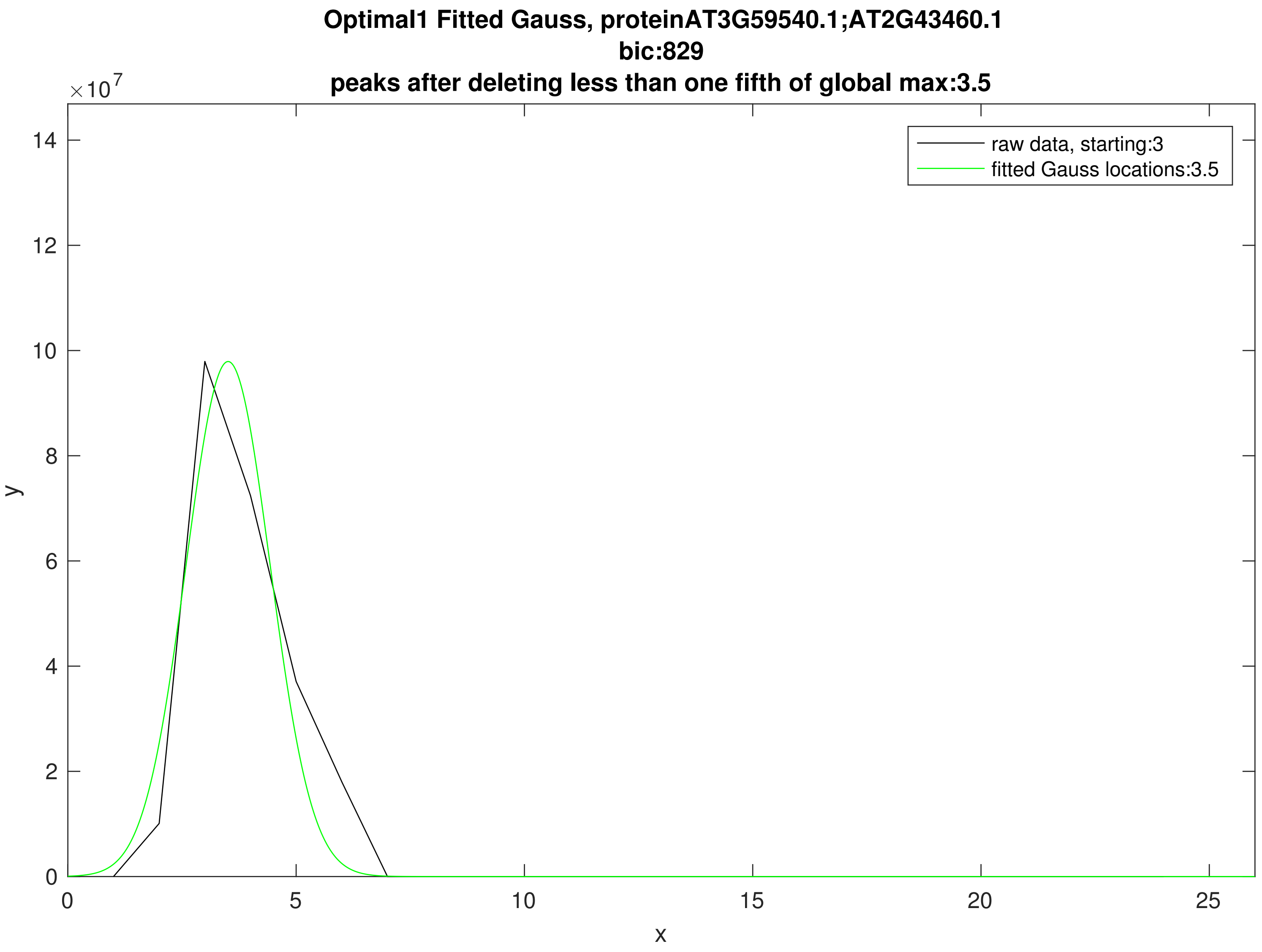

Optimal1 Fitted Gauss, proteinAT3G59540.1;AT2G43460.1
bic:829
peaks after deleting less than one fifth of global max:3.5
×10^7
raw data, starting:3
fitted Gauss locations:3.5
y
x
0
2
4
6
8
10
12
14
0
5
10
15
20
25

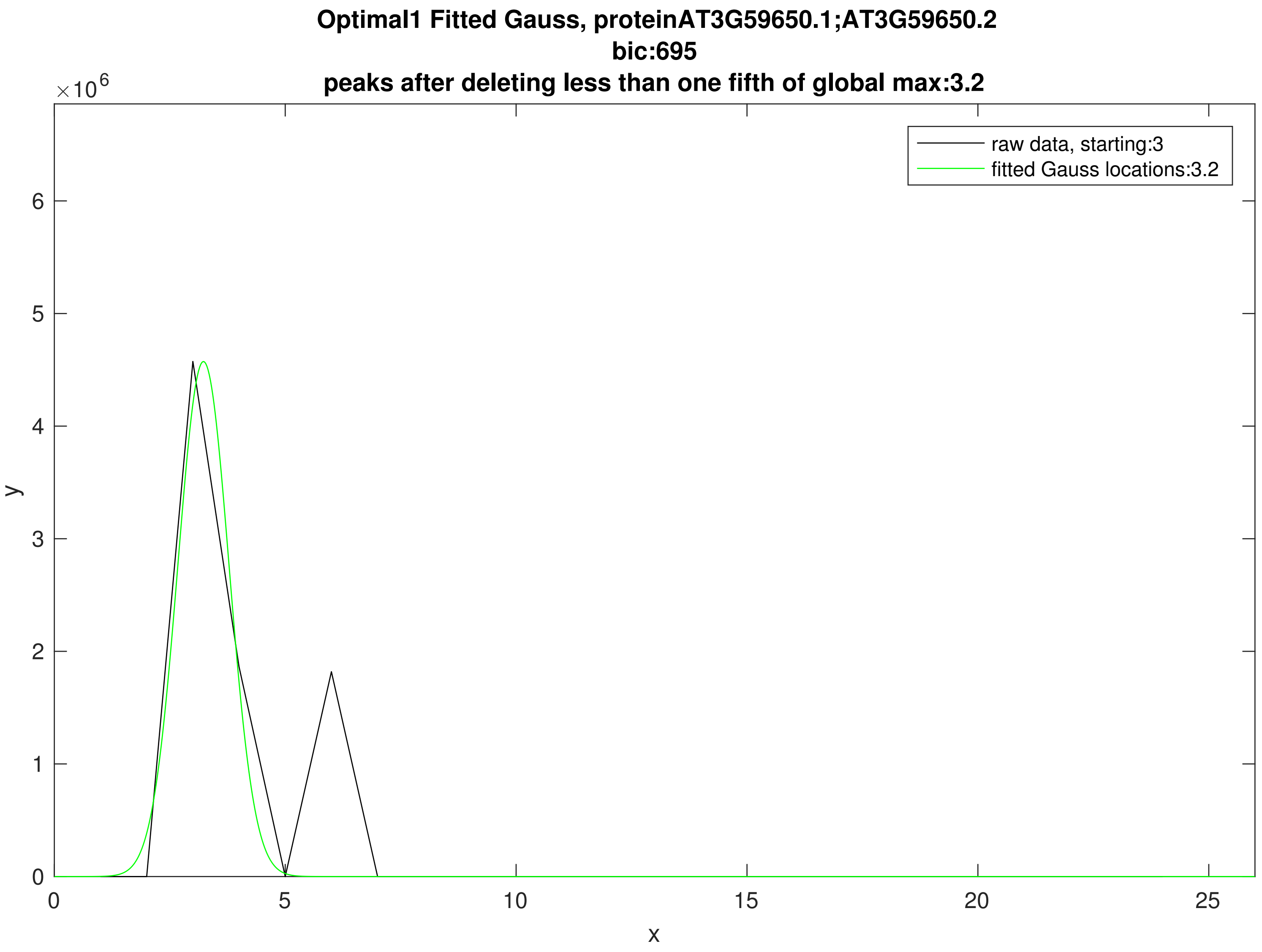
Optimal1 Fitted Gauss, proteinAT3G59650.1;AT3G59650.2
bic:695
peaks after deleting less than one fifth of global max:3.2
×10^6
raw data, starting:3
fitted Gauss locations:3.2
y
x
0
1
2
3
4
5
6
0
5
10
15
20
25

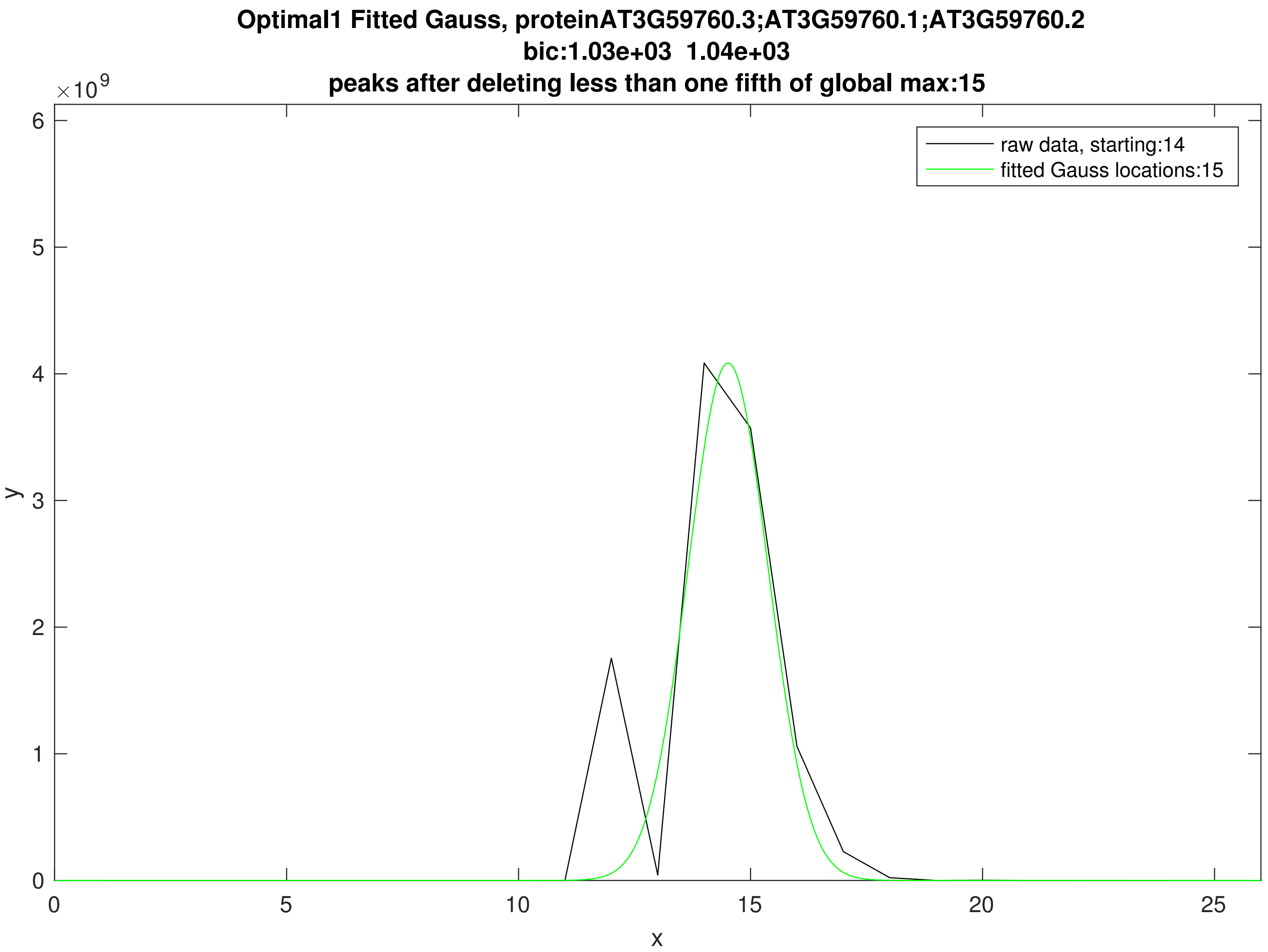

Optimal1 Fitted Gauss, proteinAT3G59760.3;AT3G59760.1;AT3G59760.2
bic:1.03e+03  1.04e+03
peaks after deleting less than one fifth of global max:15
×10^9
raw data, starting:14
fitted Gauss locations:15
y
x
0
1
2
3
4
5
6
0
5
10
15
20
25

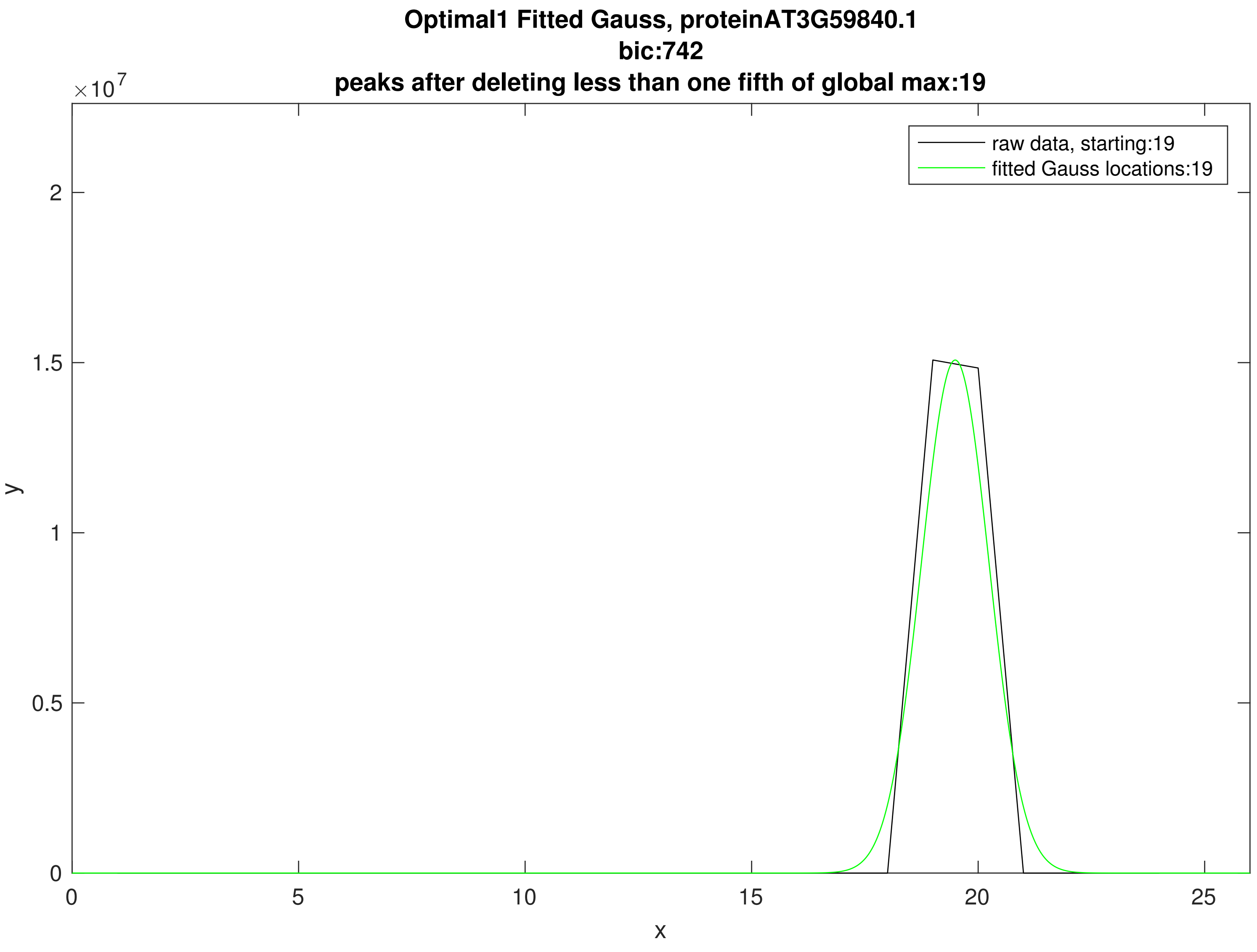

Optimal1 Fitted Gauss, proteinAT3G59840.1
bic:742
peaks after deleting less than one fifth of global max:19
×10^7
raw data, starting:19
fitted Gauss locations:19
y
x
0
0.5
1
1.5
2
0
5
10
15
20
25

#### Optimal1 Fitted Gauss, proteinAT3G59920.1
#### bic:915 924
#### peaks after deleting less than one fifth of global max:16

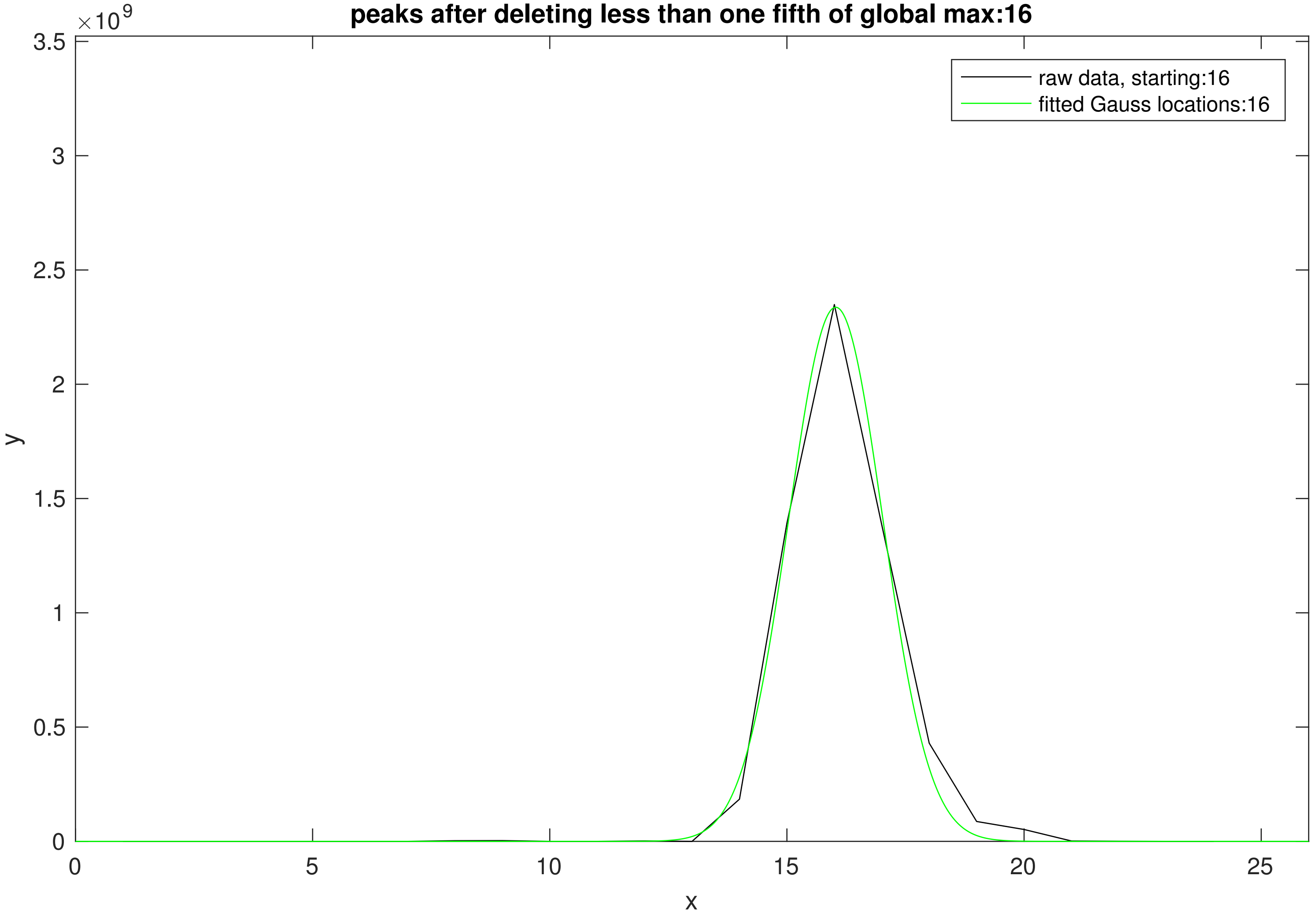

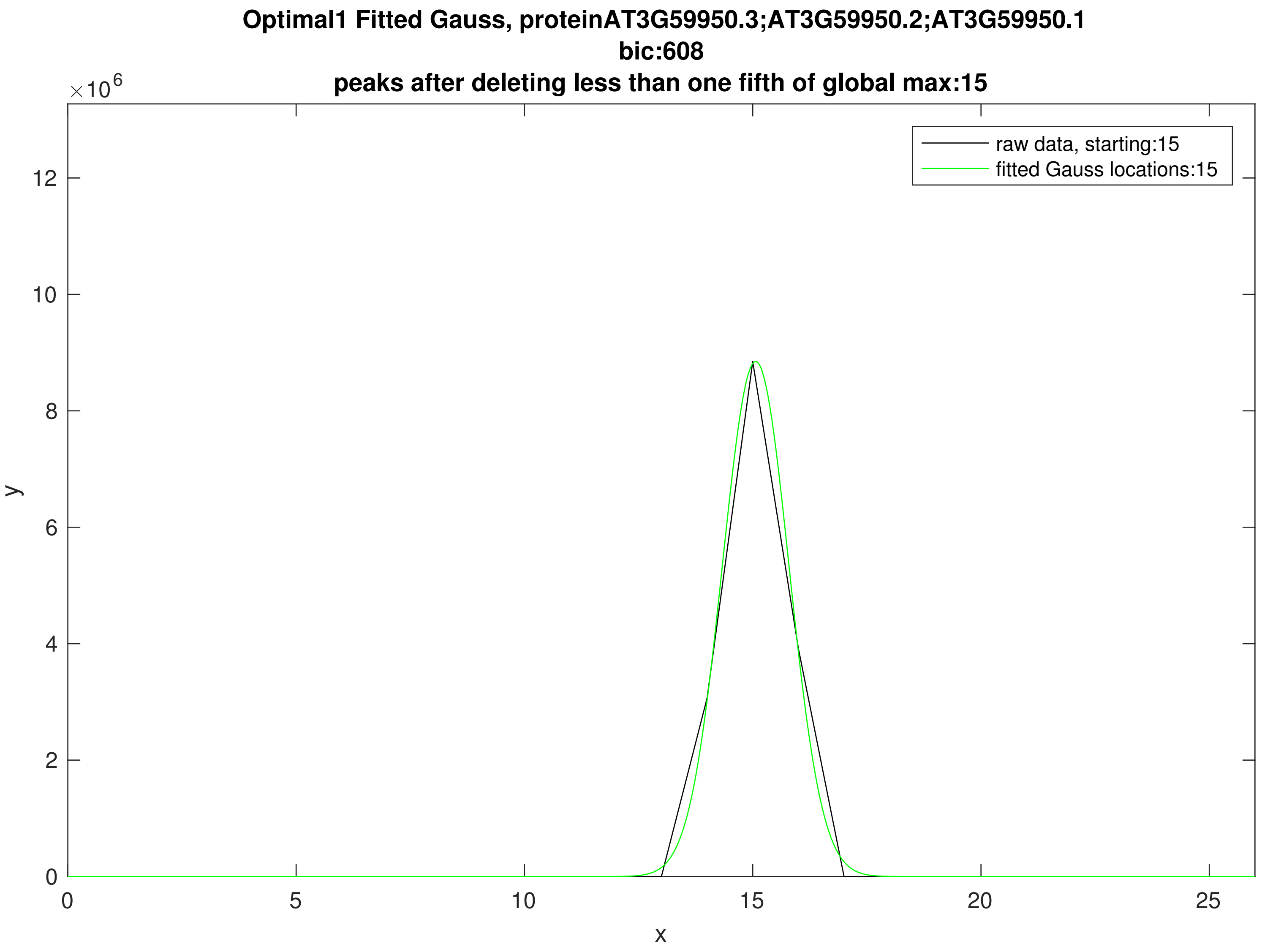
Optimal1 Fitted Gauss, proteinAT3G59950.3;AT3G59950.2;AT3G59950.1
bic:608
peaks after deleting less than one fifth of global max:15
×10^6
raw data, starting:15
fitted Gauss locations:15
y
x
0
2
4
6
8
10
12
0
5
10
15
20
25

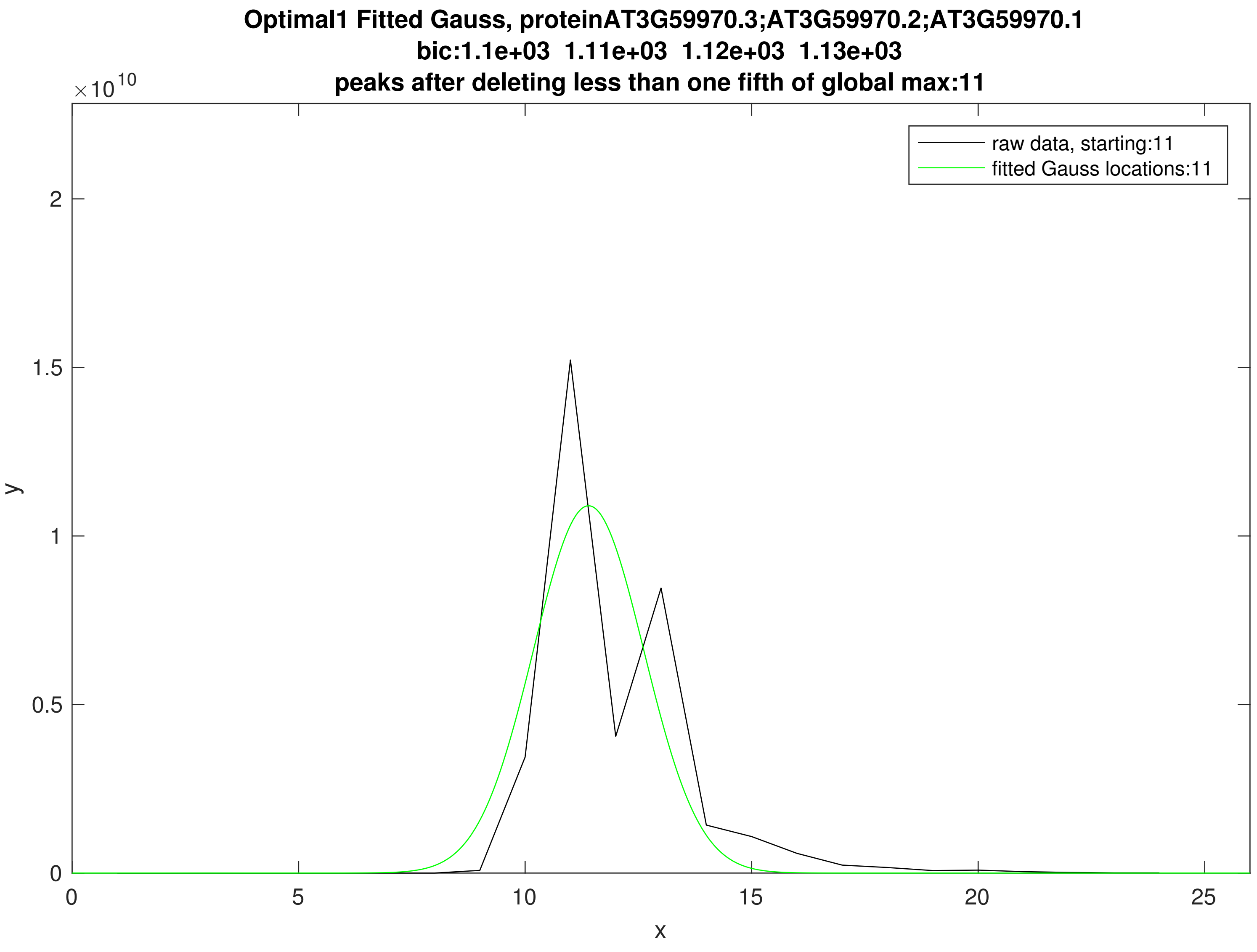
Optimal1 Fitted Gauss, proteinAT3G59970.3;AT3G59970.2;AT3G59970.1
bic:1.1e+03 1.11e+03 1.12e+03 1.13e+03
peaks after deleting less than one fifth of global max:11
×10^10
raw data, starting:11
fitted Gauss locations:11
y
x
0
0.5
1
1.5
2
5
10
15
20
25

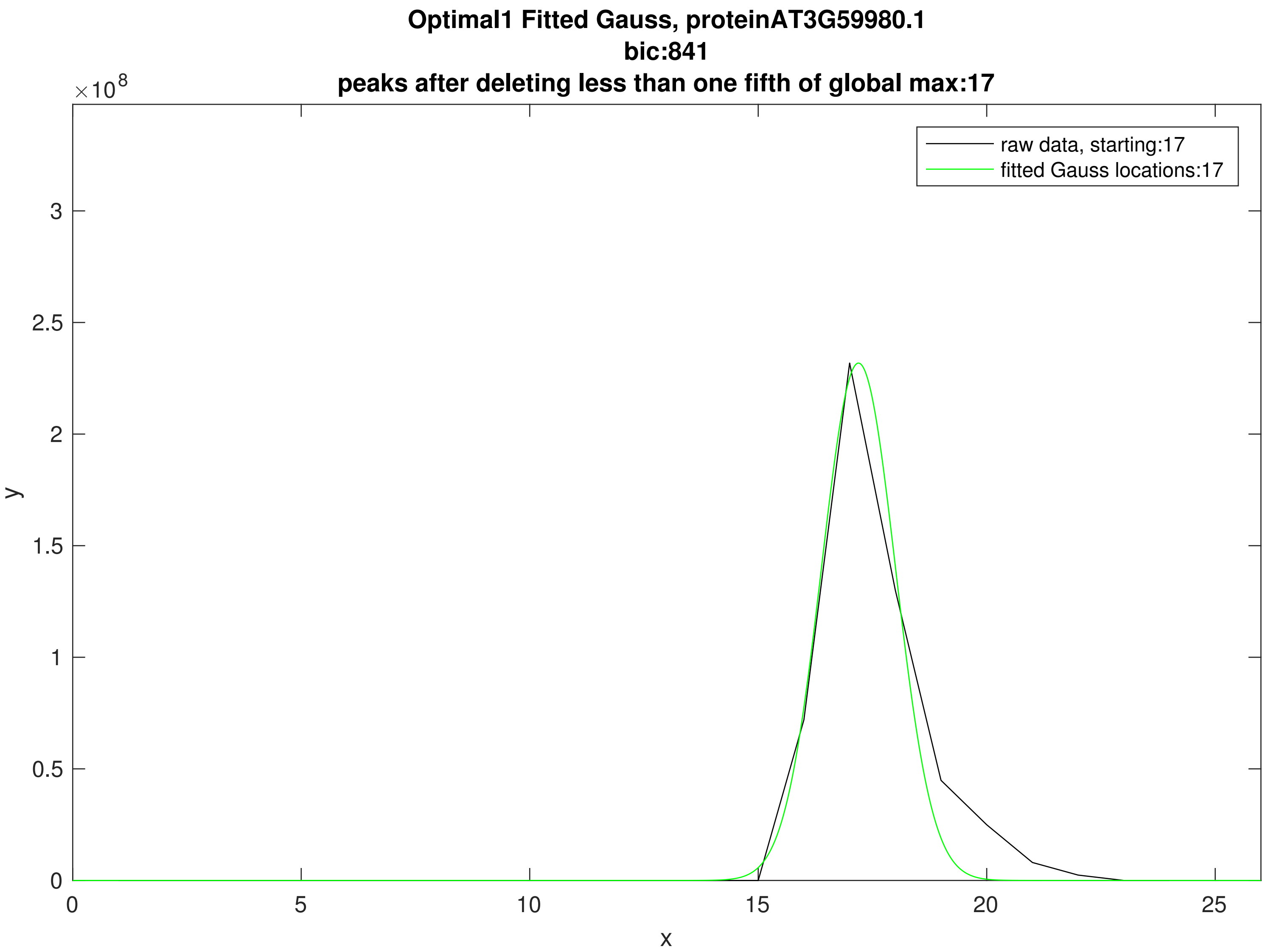

Optimal1 Fitted Gauss, proteinAT3G59980.1
bic:841
peaks after deleting less than one fifth of global max:17
×10^8
raw data, starting:17
fitted Gauss locations:17
y
x
0
0.5
1
1.5
2
2.5
3
0
5
10
15
20
25

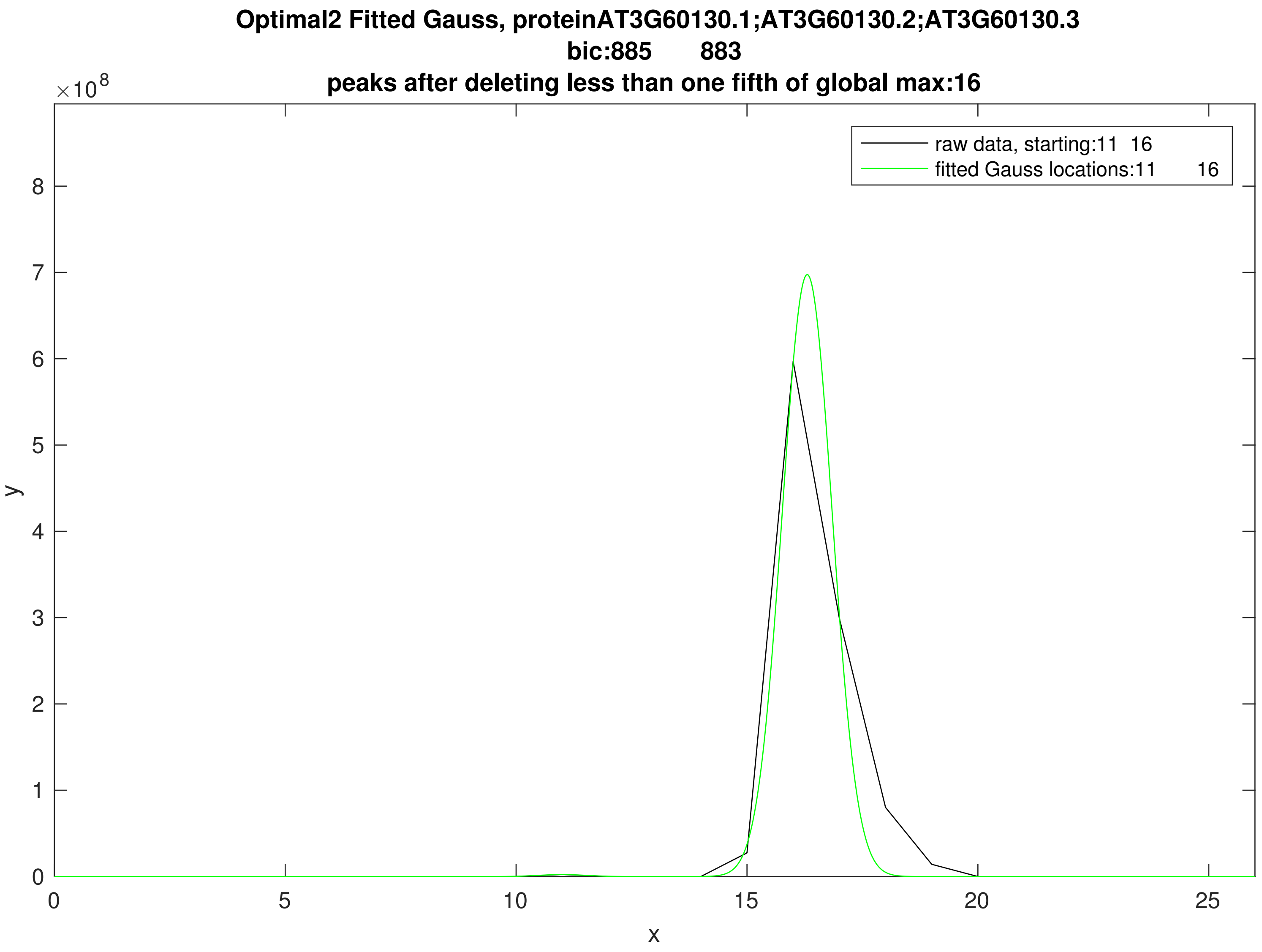
Optimal2 Fitted Gauss, proteinAT3G60130.1;AT3G60130.2;AT3G60130.3
bic:885 883
peaks after deleting less than one fifth of global max:16
×10^8
raw data, starting:11 16
fitted Gauss locations:11 16
y
x
0
1
2
3
4
5
6
7
8
0
5
10
15
20
25

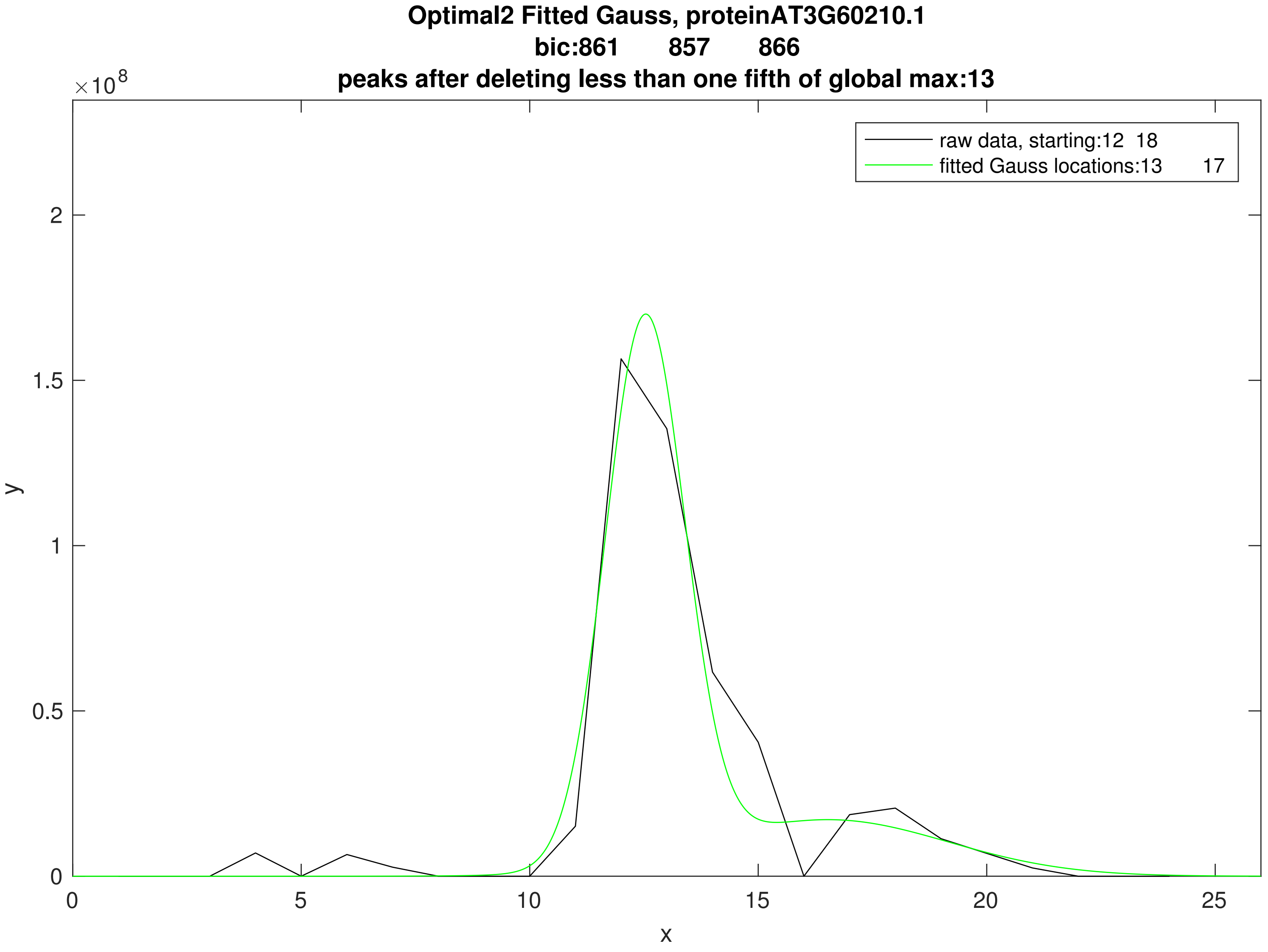

Optimal2 Fitted Gauss, proteinAT3G60210.1
bic:861 857 866
peaks after deleting less than one fifth of global max:13
×10^8
raw data, starting:12 18
fitted Gauss locations:13 17
y
x
0
0.5
1
1.5
2
5
10
15
20
25

**Optimal1 Fitted Gauss, proteinAT3G60245.1;AT3G10950.1**
**bic:852**
**peaks after deleting less than one fifth of global max:3.4**

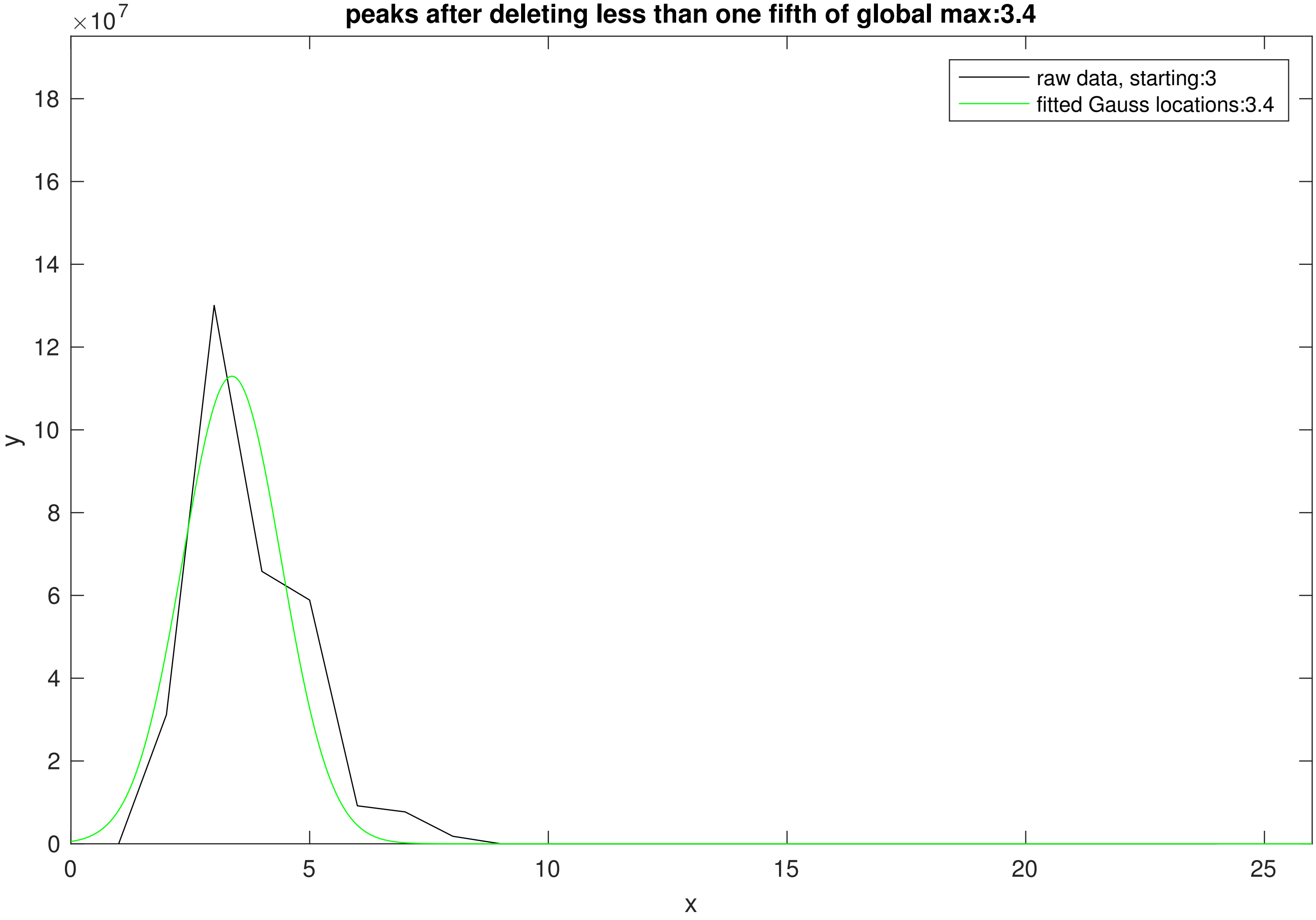

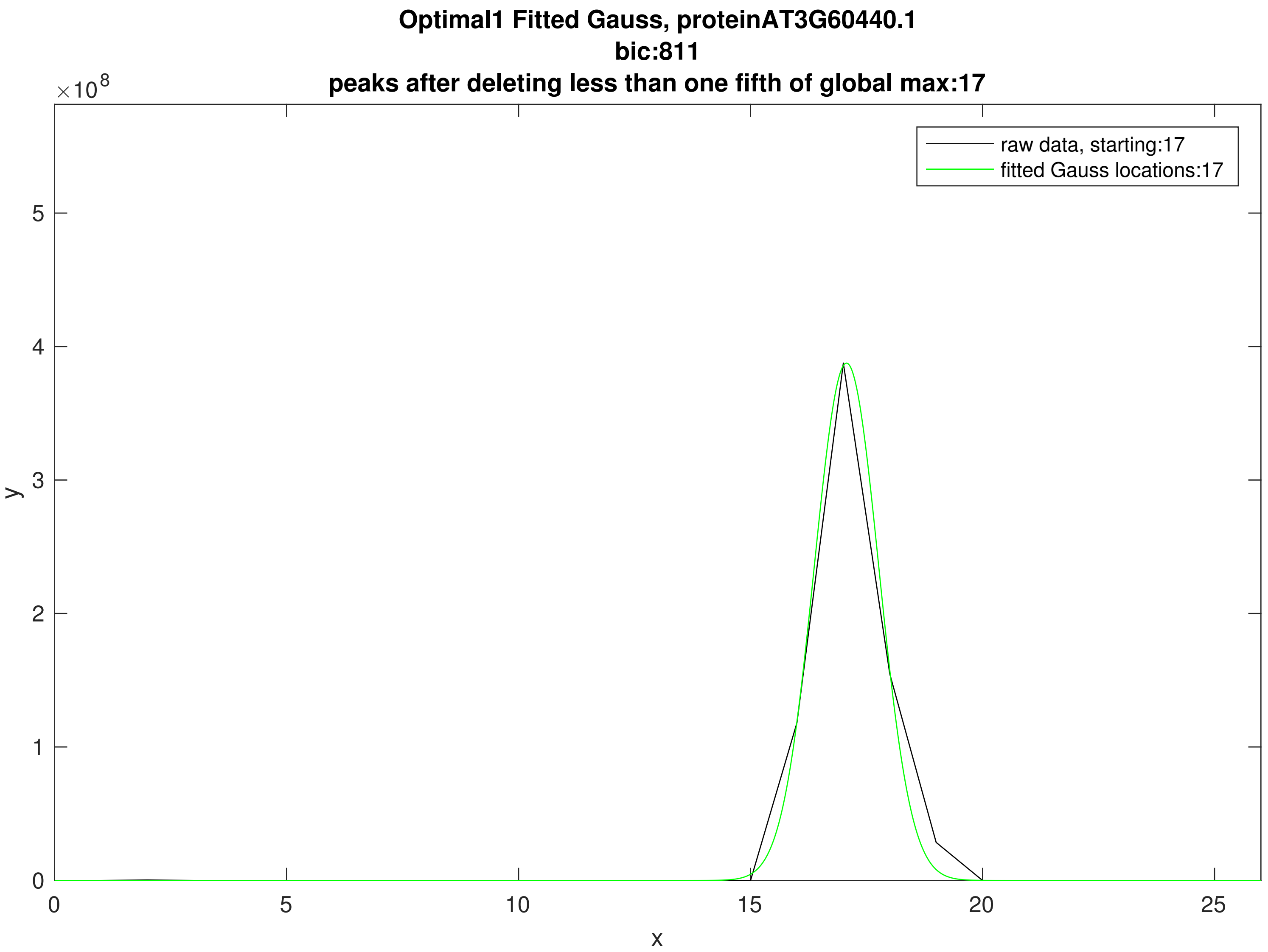

Optimal1 Fitted Gauss, proteinAT3G60440.1
bic:811
peaks after deleting less than one fifth of global max:17
×10^8
raw data, starting:17
fitted Gauss locations:17
y
x
0
1
2
3
4
5
0
5
10
15
20
25

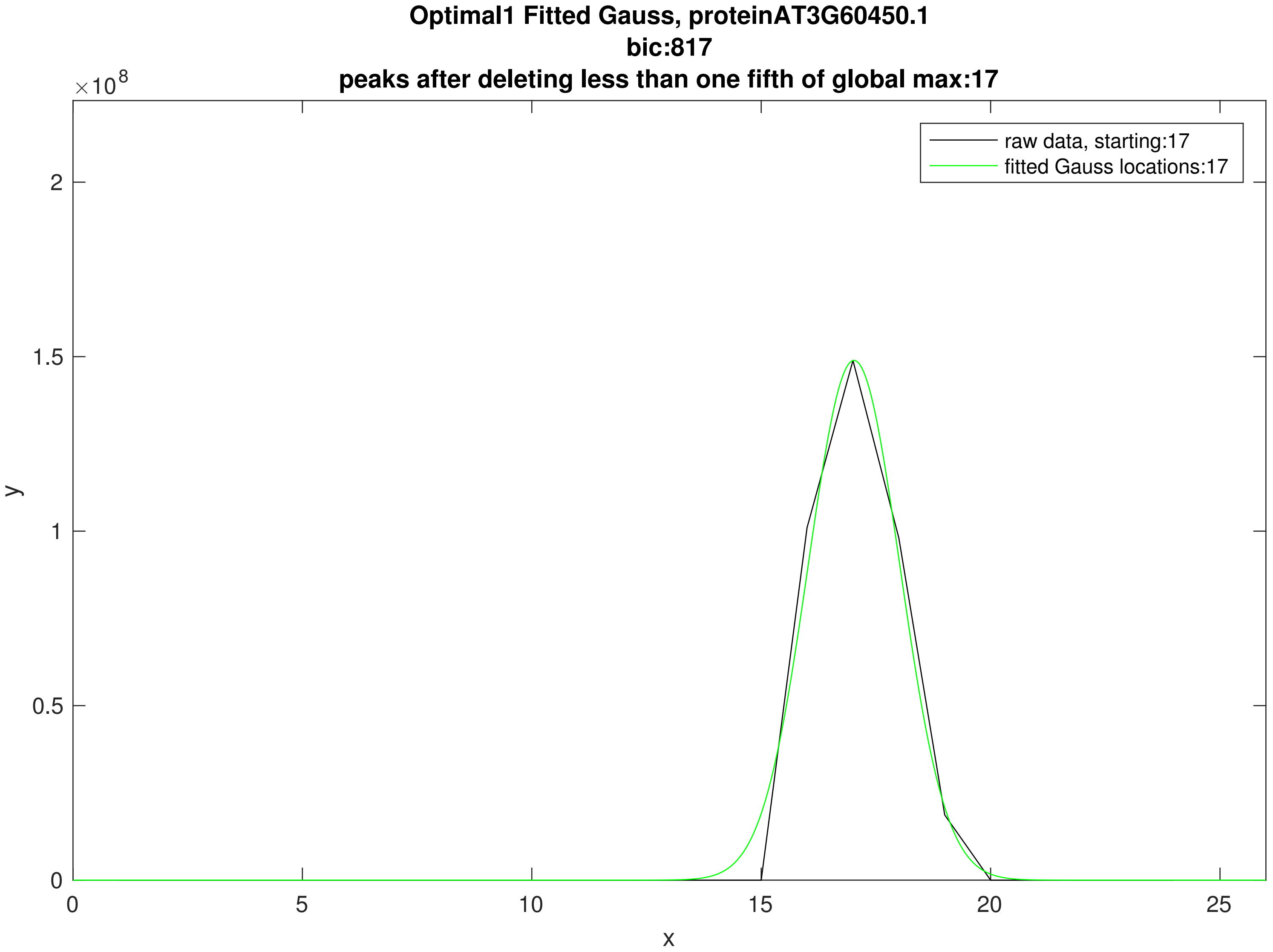
Optimal1 Fitted Gauss, proteinAT3G60450.1
bic:817
peaks after deleting less than one fifth of global max:17
×10^8
raw data, starting:17
fitted Gauss locations:17
2
1.5
1
0.5
0
y
0
5
10
15
20
25
x

**Optimal1 Fitted Gauss, proteinAT3G60750.1;AT3G60750.2**
**bic:1.14e+03 1.15e+03 1.16e+03**
**peaks after deleting less than one fifth of global max:11**

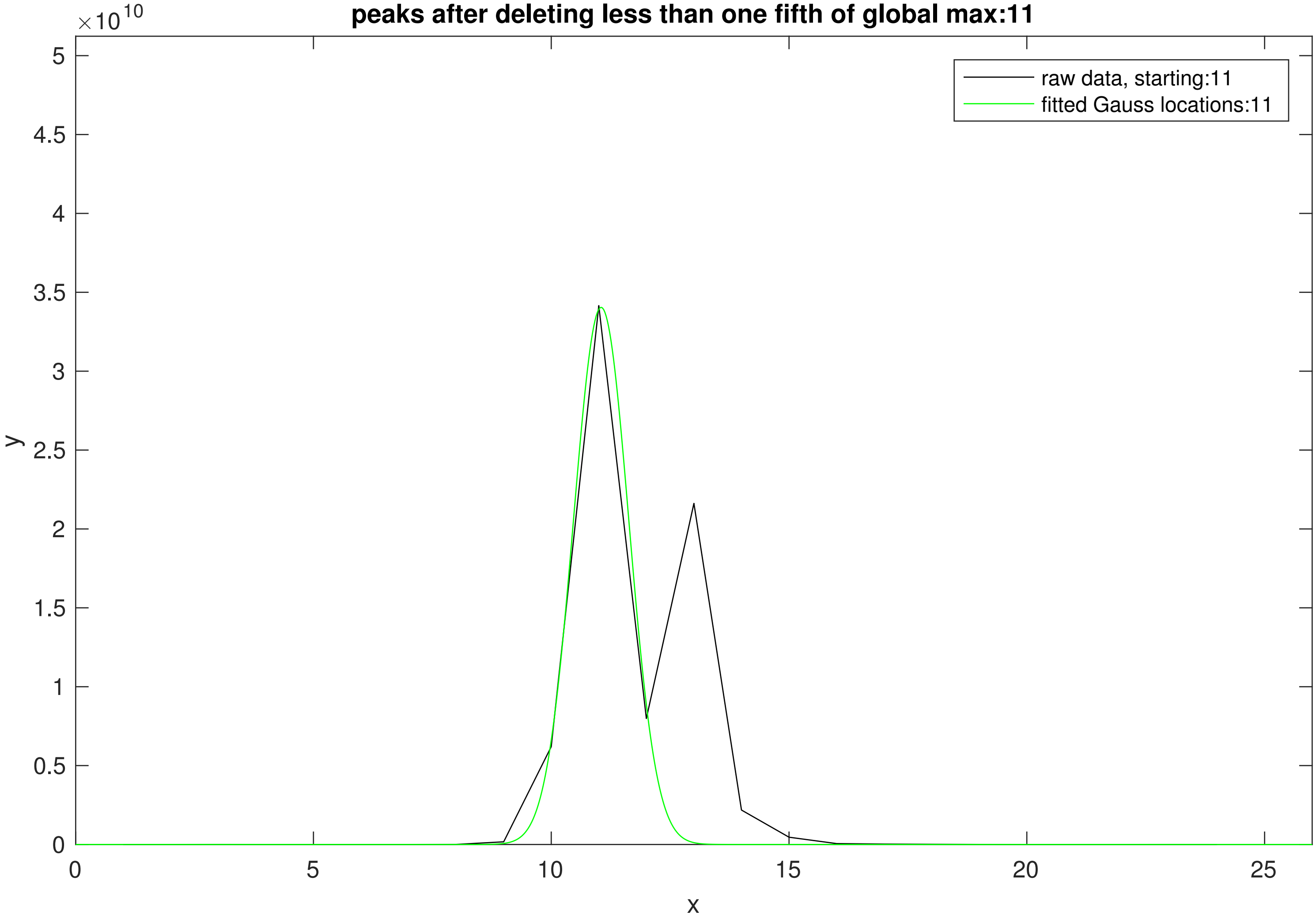

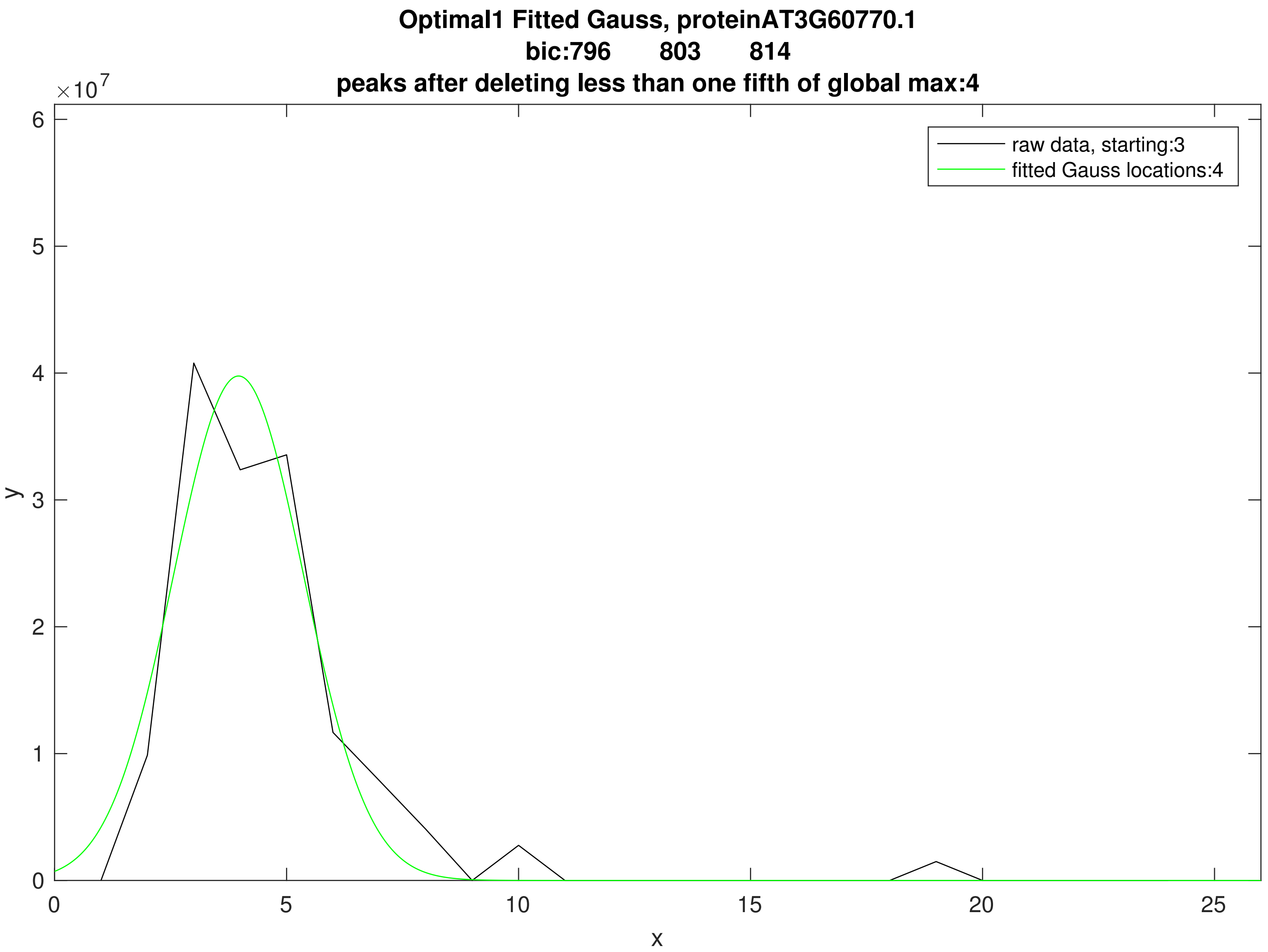
Optimal1 Fitted Gauss, proteinAT3G60770.1
bic:796 803 814
peaks after deleting less than one fifth of global max:4
×10^7
raw data, starting:3
fitted Gauss locations:4
y
x
0
1
2
3
4
5
6
0
5
10
15
20
25

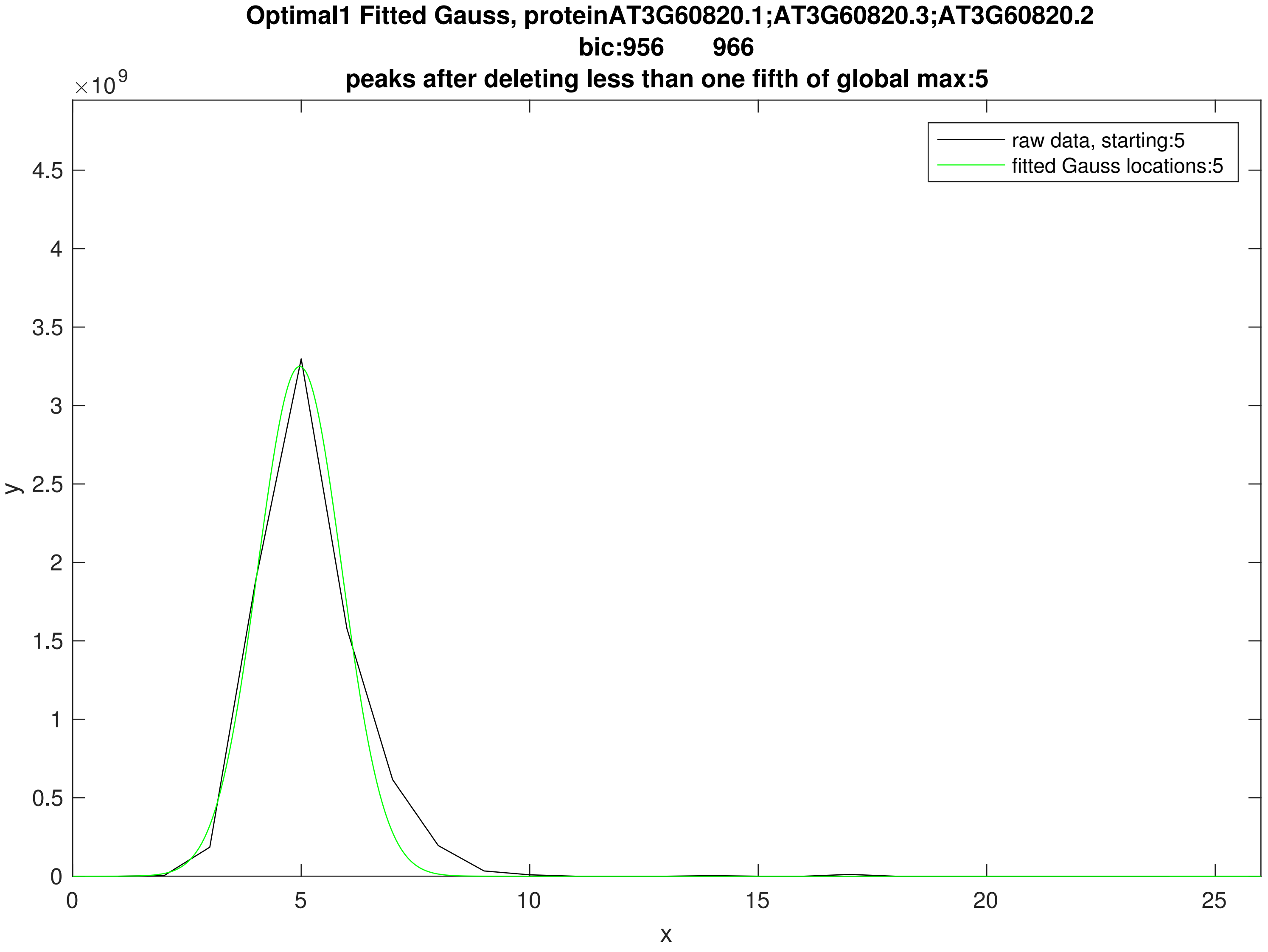

Optimal1 Fitted Gauss, proteinAT3G60820.1;AT3G60820.3;AT3G60820.2
bic:956 966
peaks after deleting less than one fifth of global max:5
×10^9
raw data, starting:5
fitted Gauss locations:5
y
x
0
0.5
1
1.5
2
2.5
3
3.5
4
4.5
5
10
15
20
25

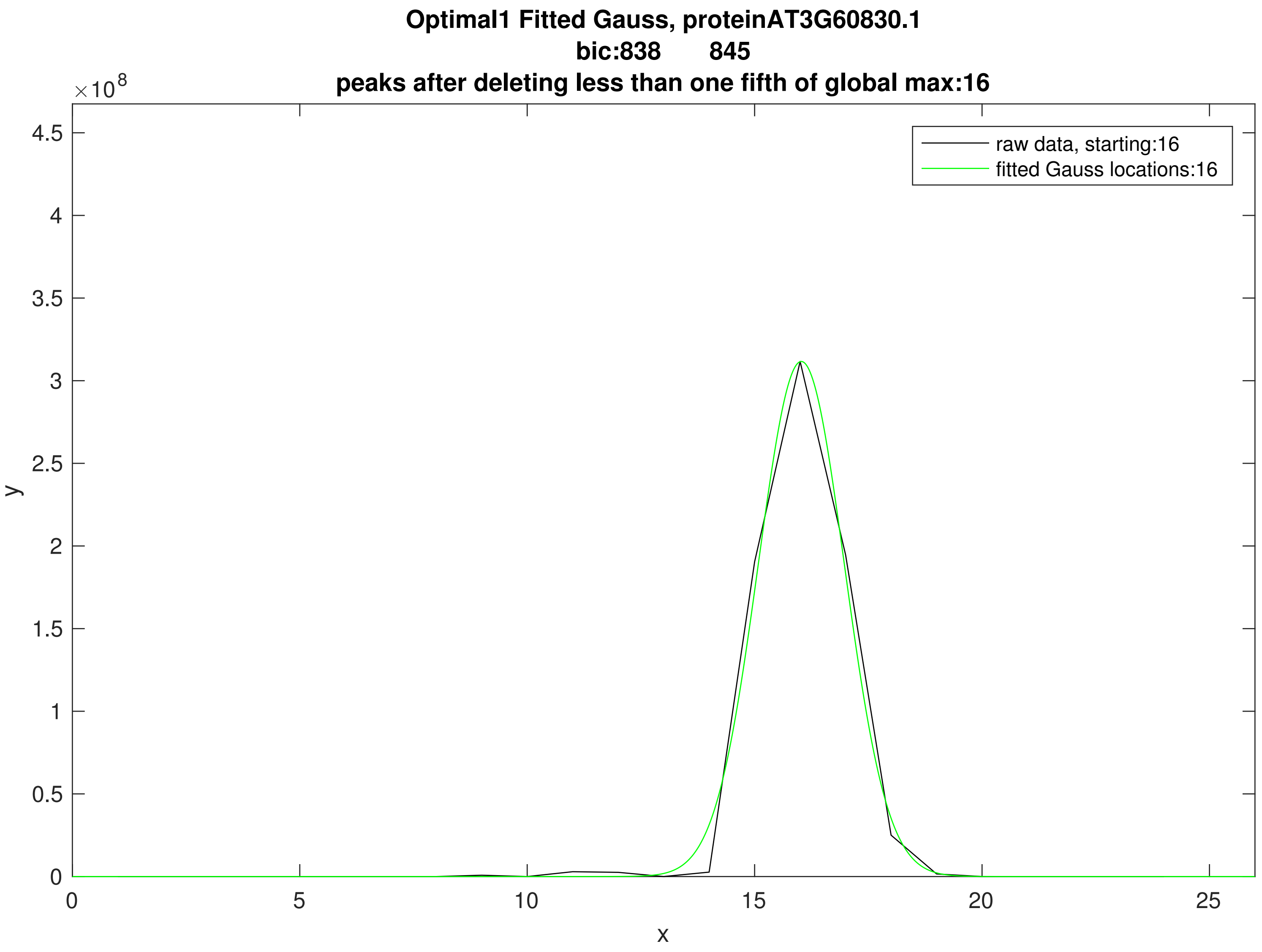
Optimal1 Fitted Gauss, proteinAT3G60830.1
bic:838 845
peaks after deleting less than one fifth of global max:16
×10^8
4.5
4
3.5
3
2.5
2
1.5
1
0.5
0
y
0
5
10
15
20
25
x
raw data, starting:16
fitted Gauss locations:16

#### Optimal1 Fitted Gauss, proteinAT3G60880.1;AT3G60880.2
#### bic:900
#### peaks after deleting less than one fifth of global max:13

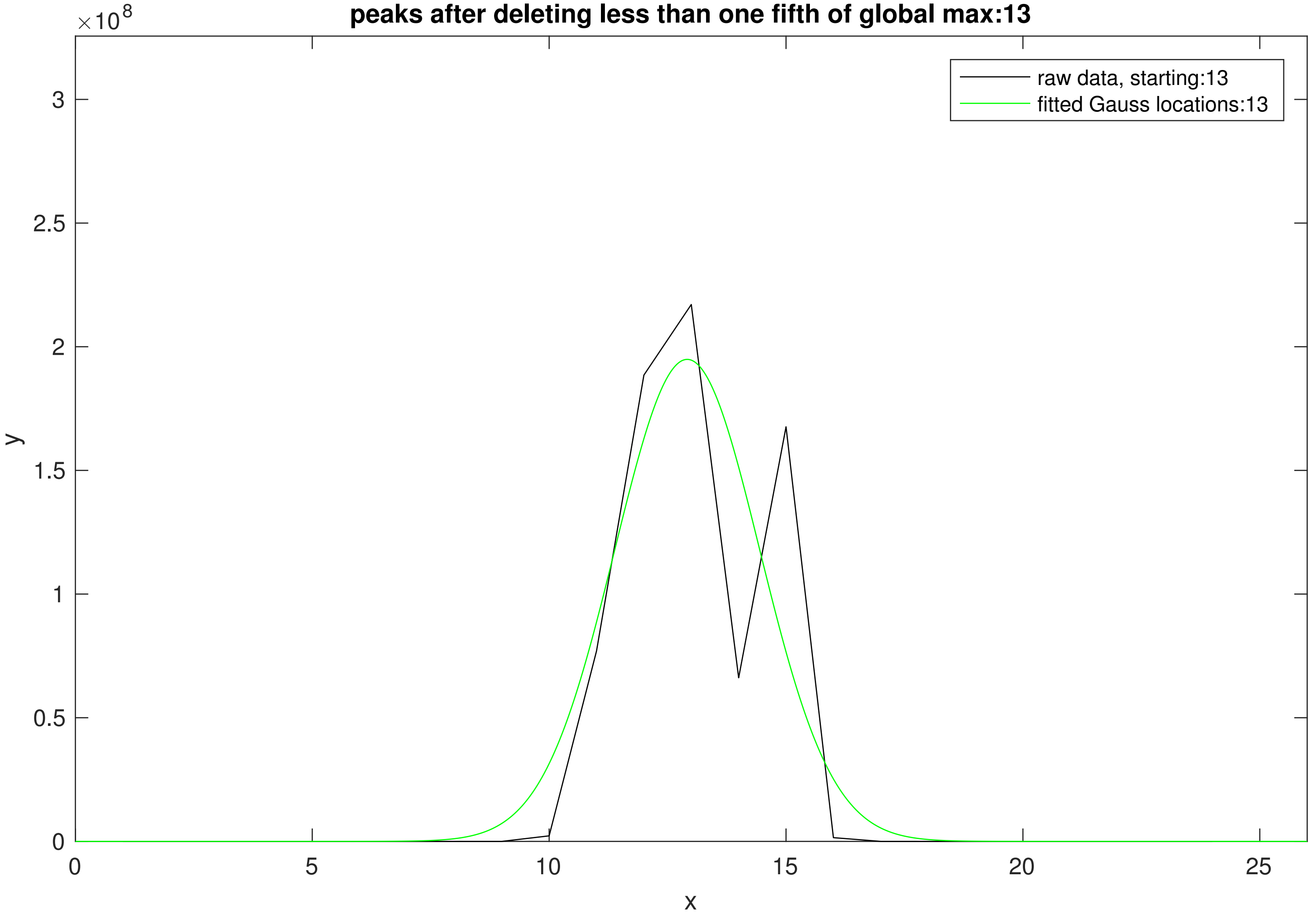

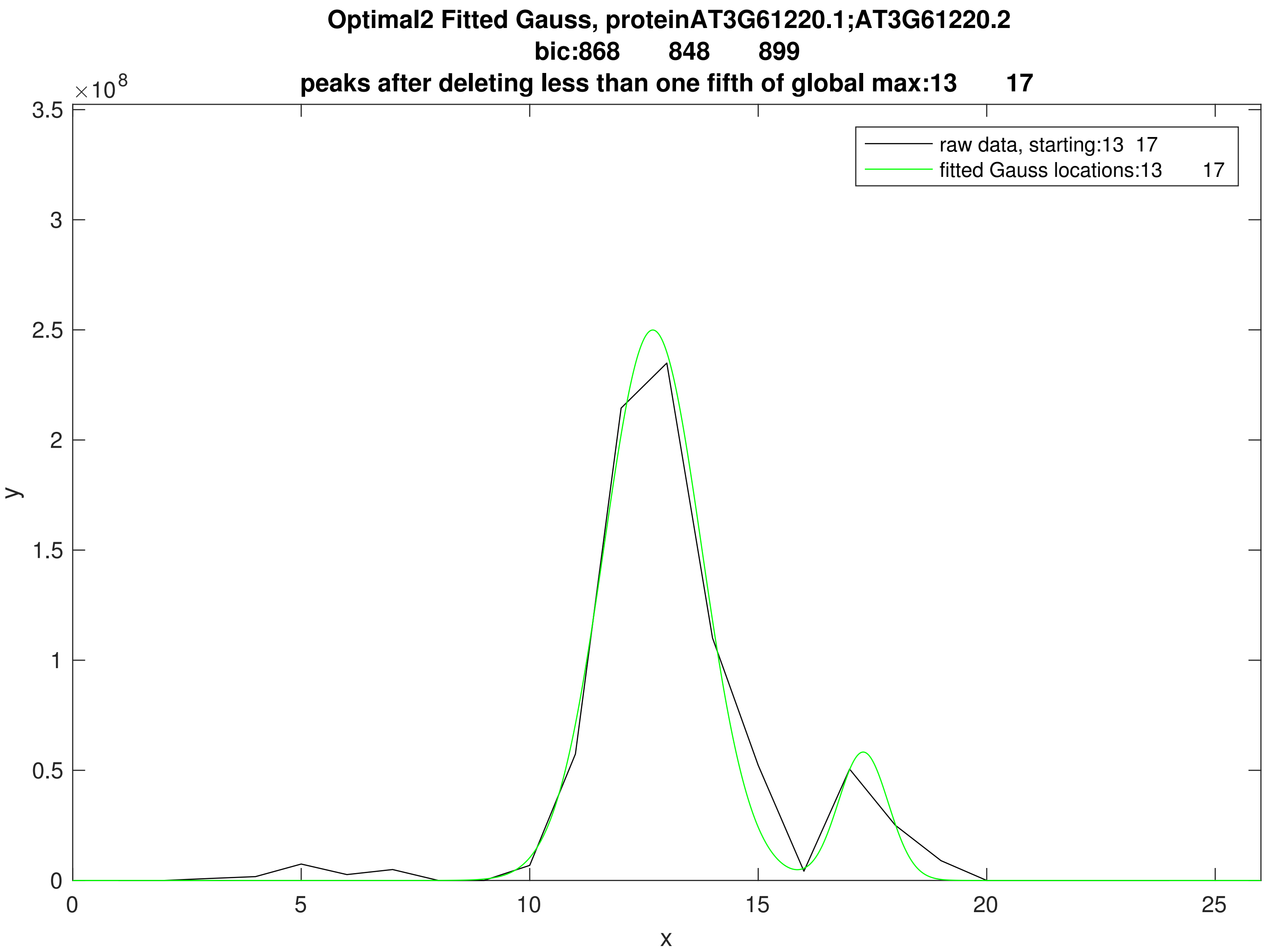
Optimal2 Fitted Gauss, proteinAT3G61220.1;AT3G61220.2
bic:868 848 899
peaks after deleting less than one fifth of global max:13 17
raw data, starting:13 17
fitted Gauss locations:13 17
×10^8
3.5
3
2.5
2
1.5
1
0.5
0
y
0
5
10
15
20
25
x

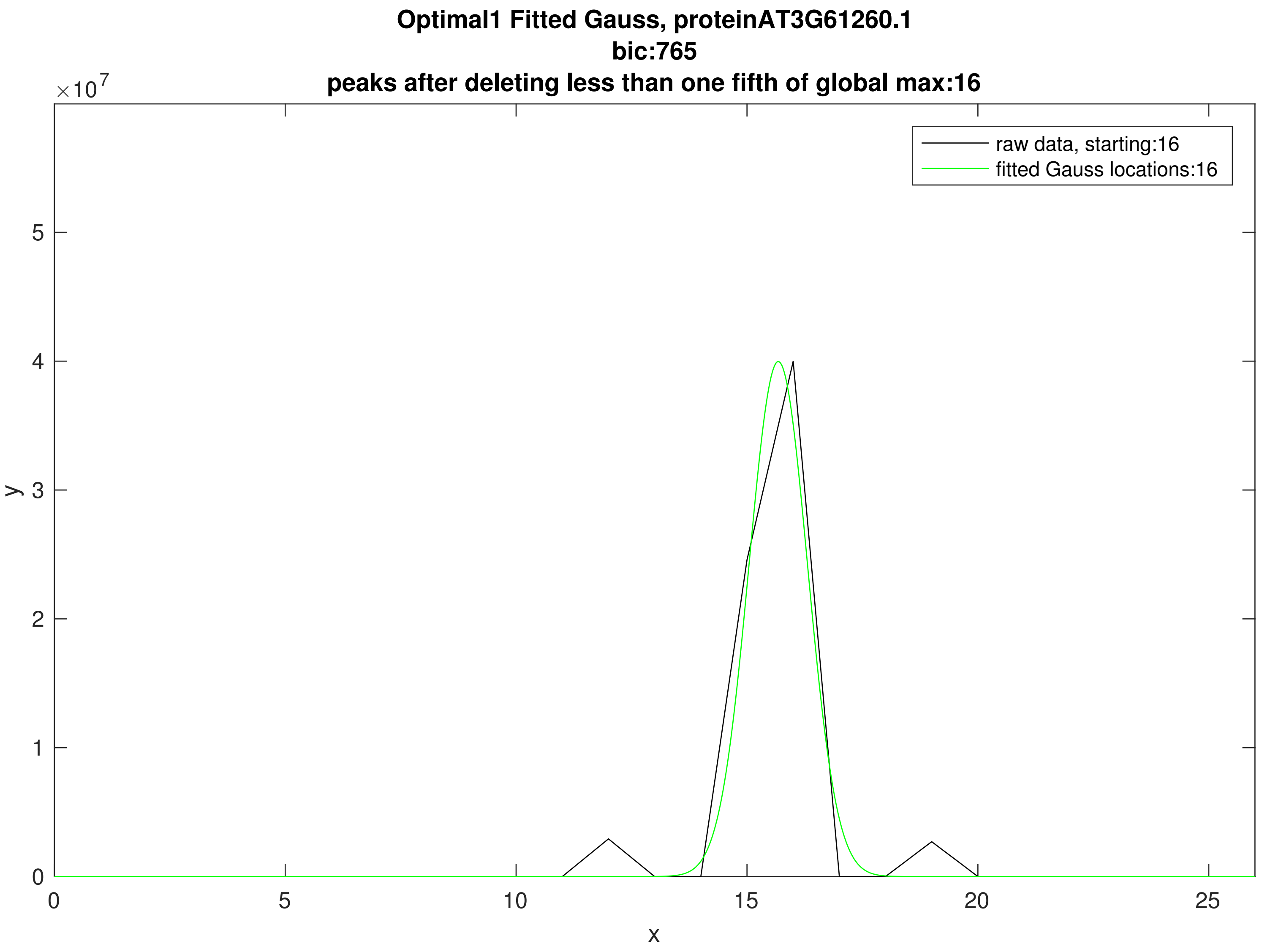
Optimal1 Fitted Gauss, proteinAT3G61260.1
bic:765
peaks after deleting less than one fifth of global max:16
×10^7
raw data, starting:16
fitted Gauss locations:16
5
4
3
2
1
0
y
0
5
10
15
20
25
x

**Optimal1 Fitted Gauss, proteinAT3G61440.1;AT3G61440.2**
**bic:1.01e+03  1.02e+03  1.03e+03**
**peaks after deleting less than one fifth of global max:15**

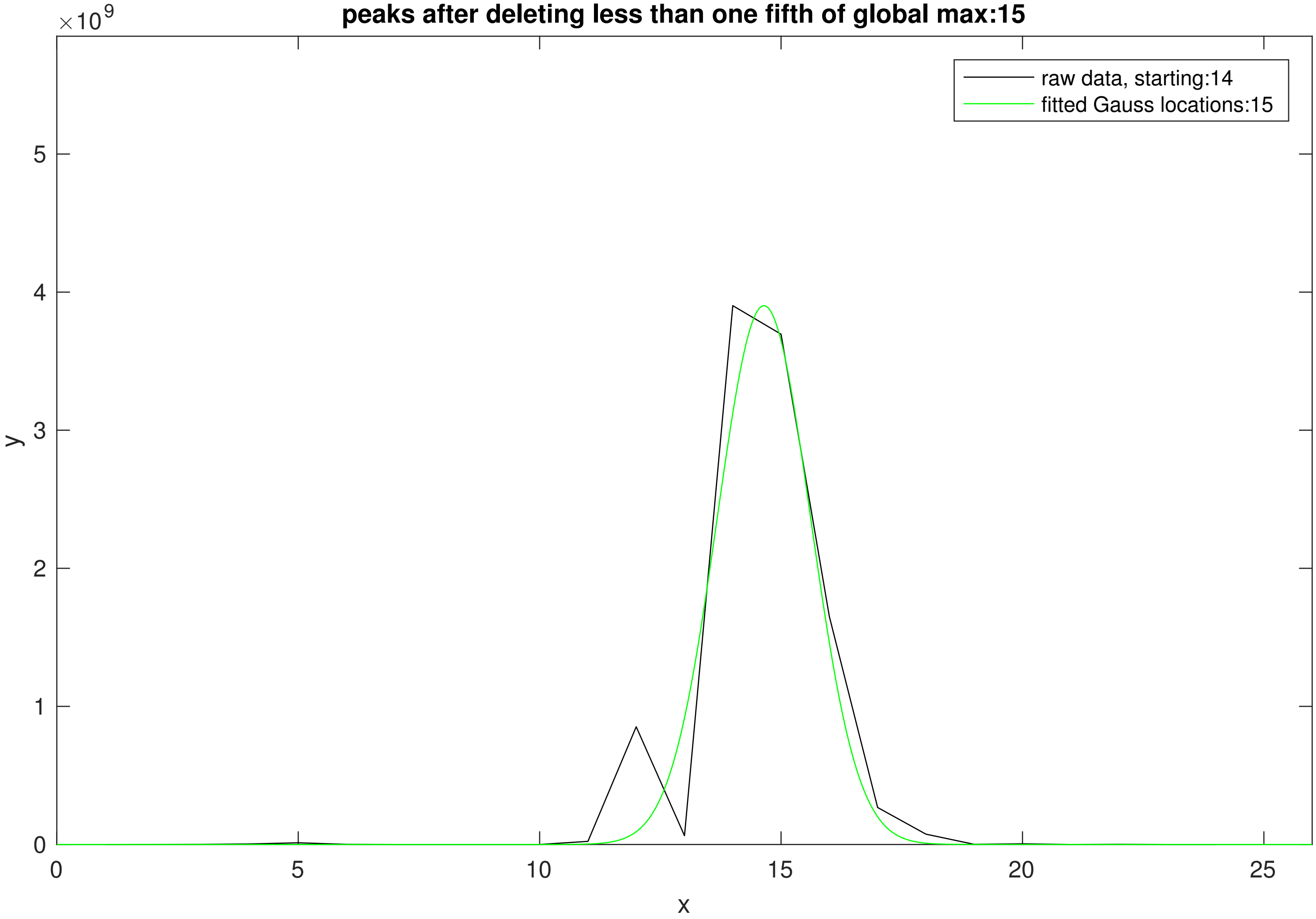

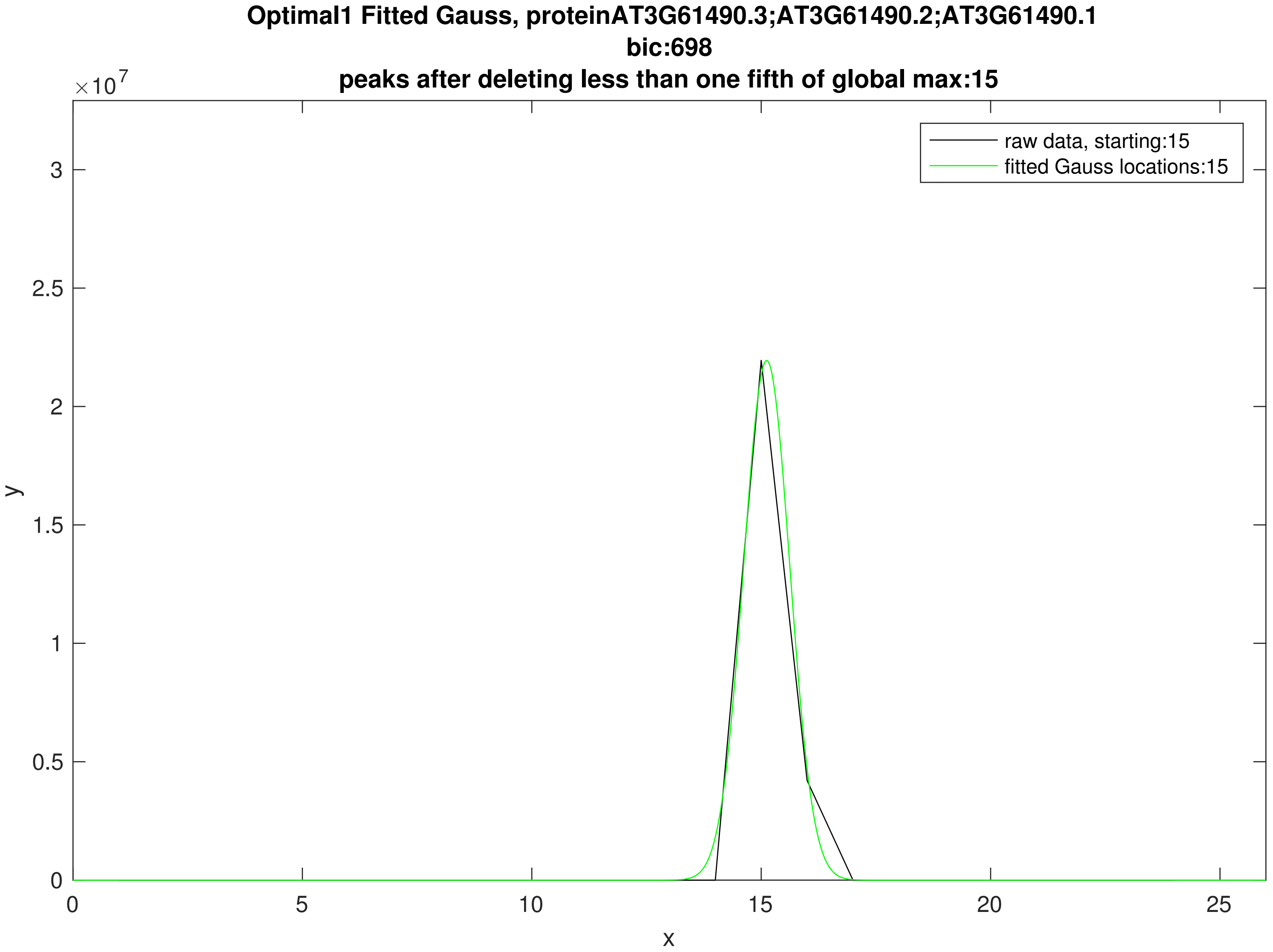
Optimal1 Fitted Gauss, proteinAT3G61490.3;AT3G61490.2;AT3G61490.1
bic:698
peaks after deleting less than one fifth of global max:15
×10^7
raw data, starting:15
fitted Gauss locations:15
y
x
0
0.5
1
1.5
2
2.5
3
0
5
10
15
20
25

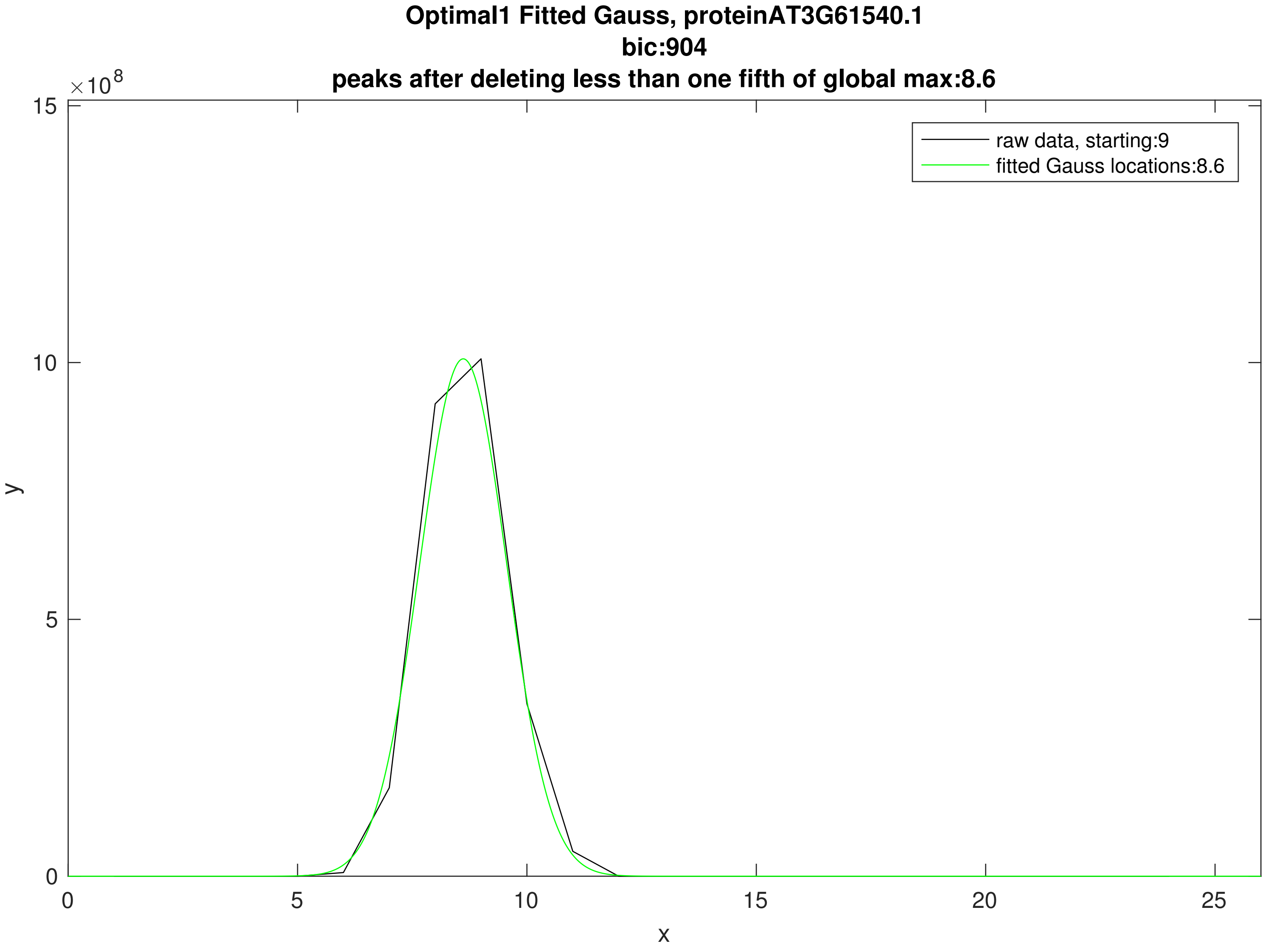

Optimal1 Fitted Gauss, proteinAT3G61540.1
bic:904
peaks after deleting less than one fifth of global max:8.6
×10^8
15
10
5
0
y
0
5
10
15
20
25
x
raw data, starting:9
fitted Gauss locations:8.6

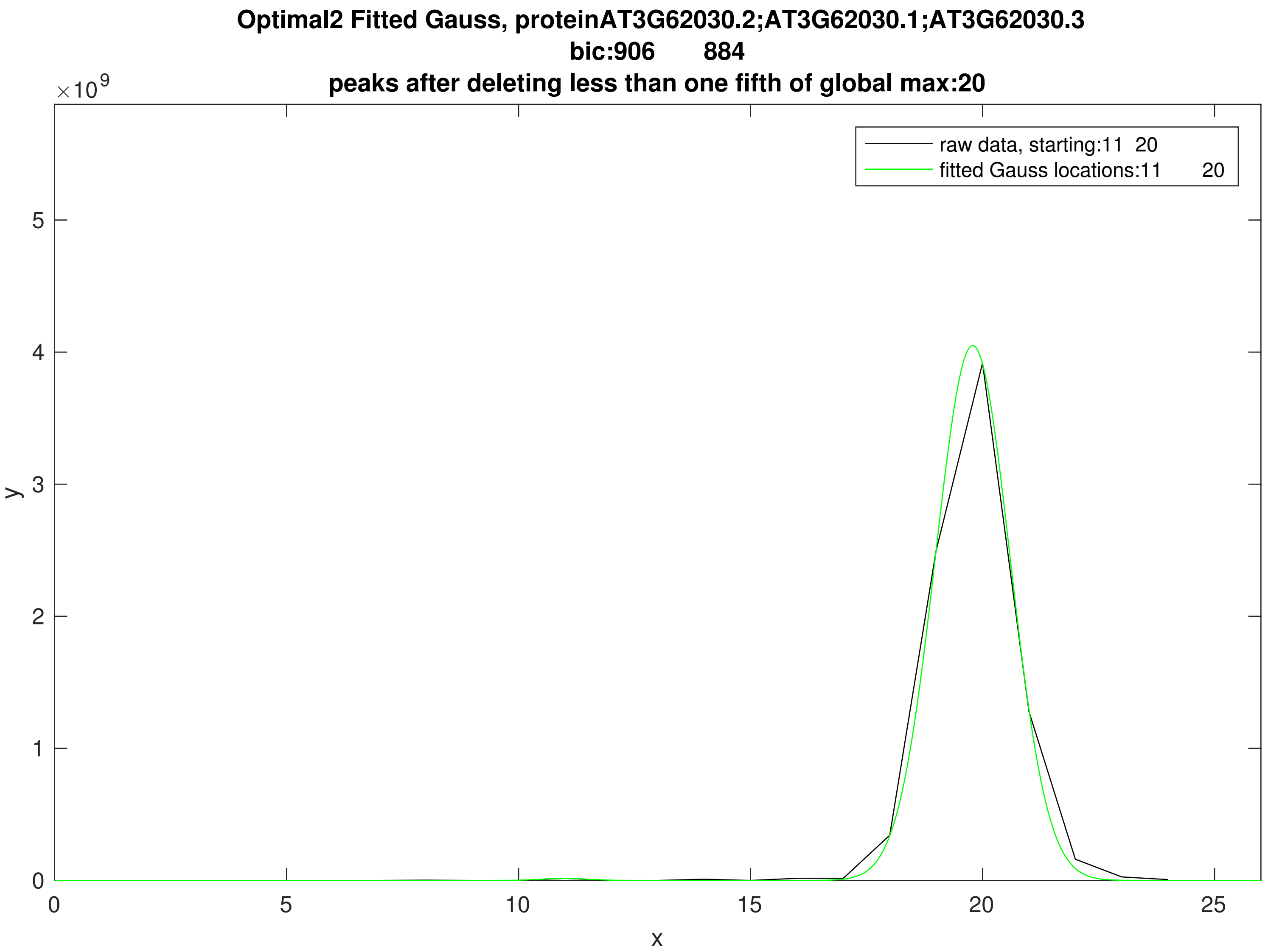

Optimal2 Fitted Gauss, proteinAT3G62030.2;AT3G62030.1;AT3G62030.3
bic:906 884
peaks after deleting less than one fifth of global max:20
×10^9
raw data, starting:11 20
fitted Gauss locations:11 20
y
x
0
1
2
3
4
5
0
5
10
15
20
25

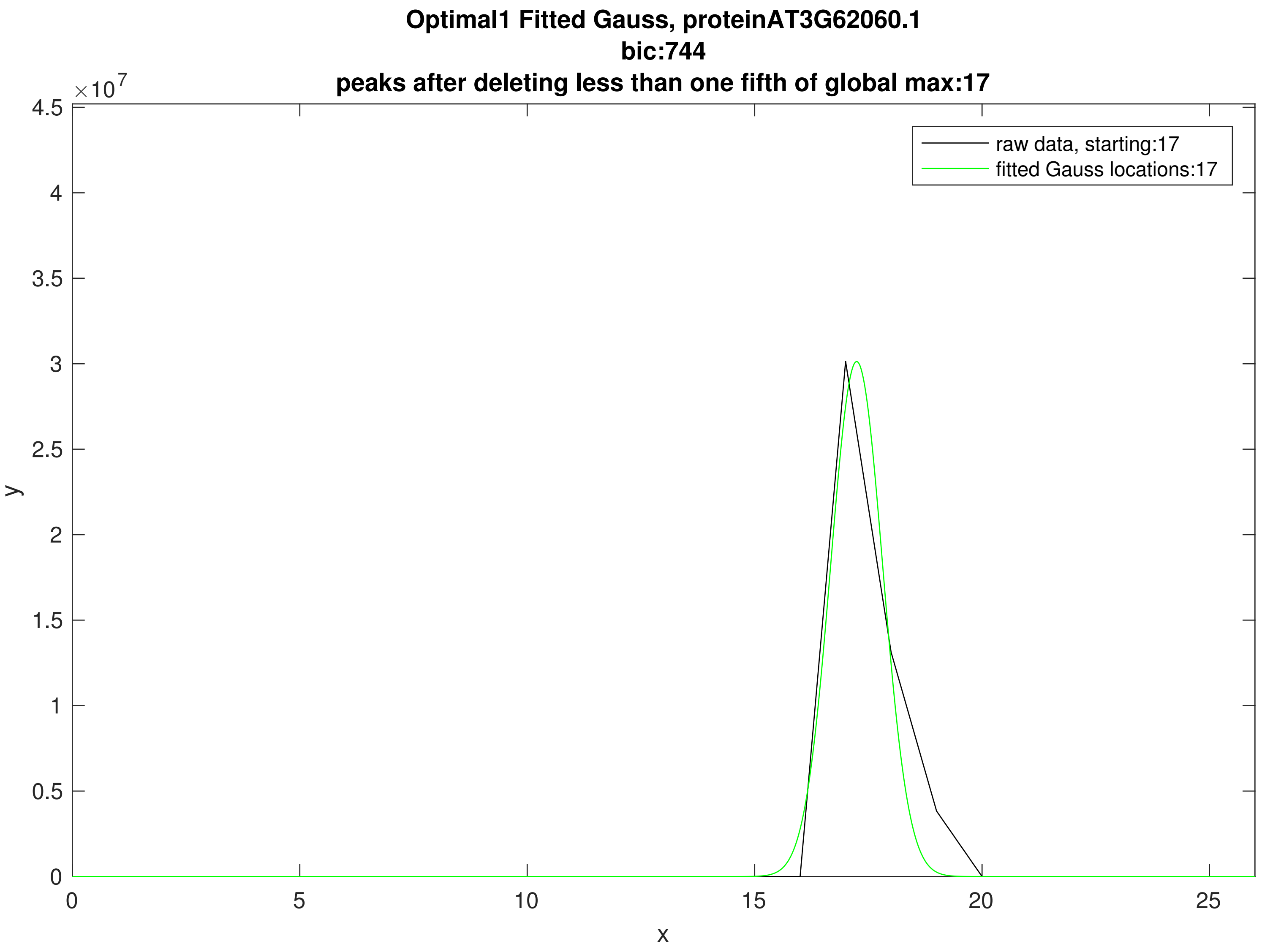
Optimal1 Fitted Gauss, proteinAT3G62060.1
bic:744
peaks after deleting less than one fifth of global max:17
×10^7
4.5
4
3.5
3
2.5
2
1.5
1
0.5
0
y
0
5
10
15
20
25
x
raw data, starting:17
fitted Gauss locations:17

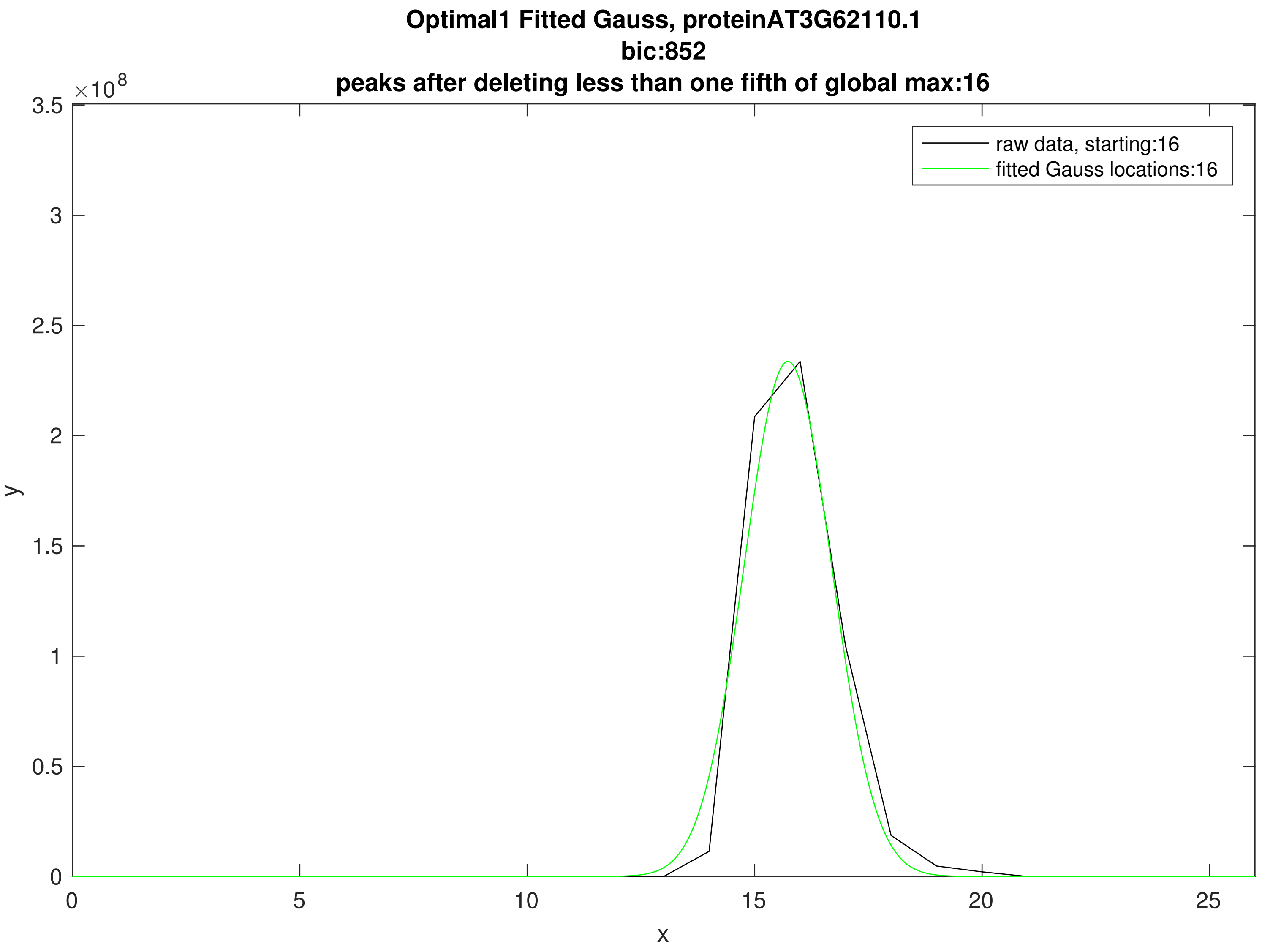
Optimal1 Fitted Gauss, proteinAT3G62110.1
bic:852
peaks after deleting less than one fifth of global max:16
×10^8
3.5
3
2.5
2
1.5
1
0.5
0
y
0
5
10
15
20
25
x
raw data, starting:16
fitted Gauss locations:16

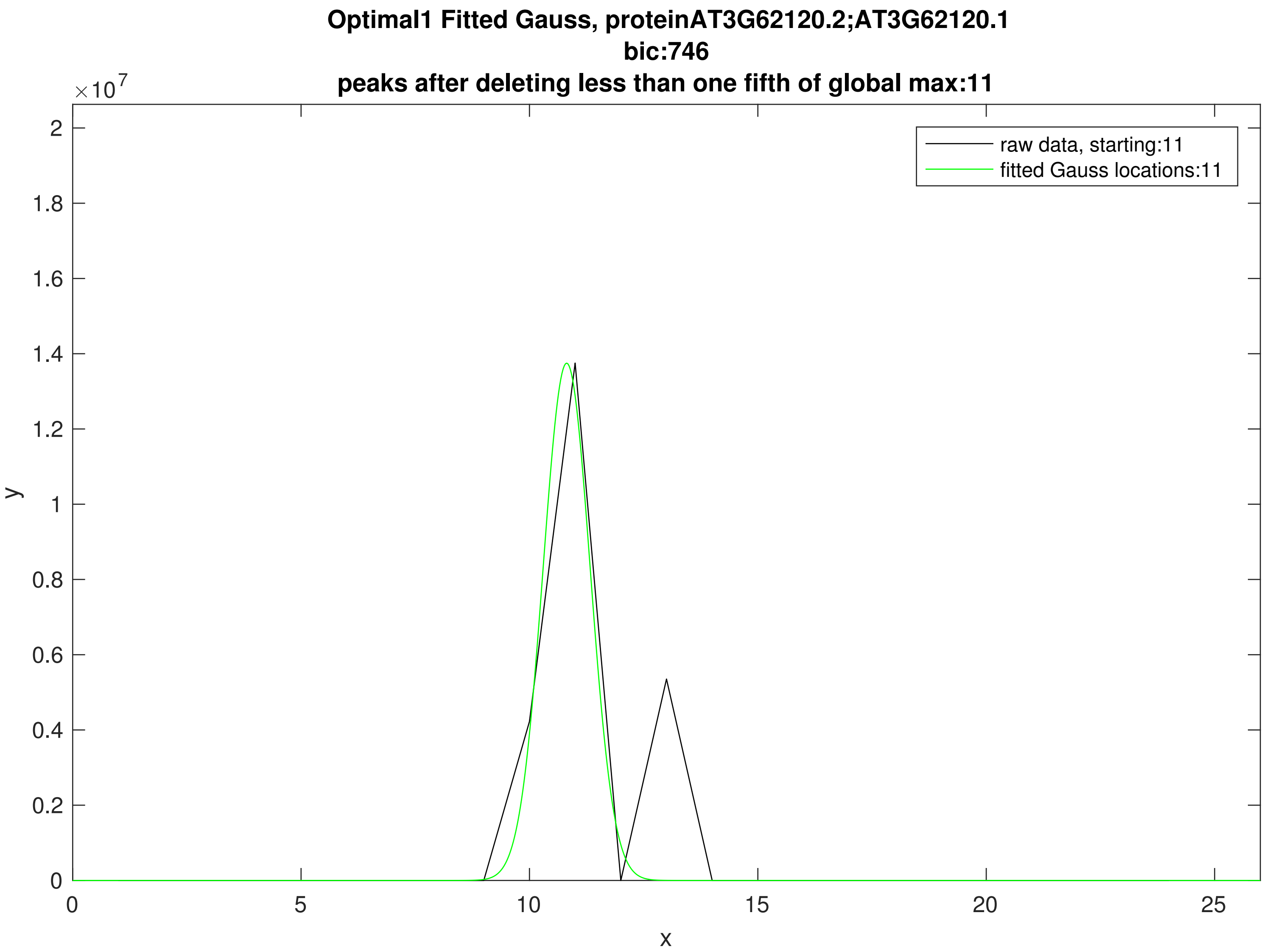
Optimal1 Fitted Gauss, proteinAT3G62120.2;AT3G62120.1
bic:746
peaks after deleting less than one fifth of global max:11
×10^7
2
1.8
1.6
1.4
1.2
1
0.8
0.6
0.4
0.2
0
y
0
5
10
15
20
25
x
raw data, starting:11
fitted Gauss locations:11

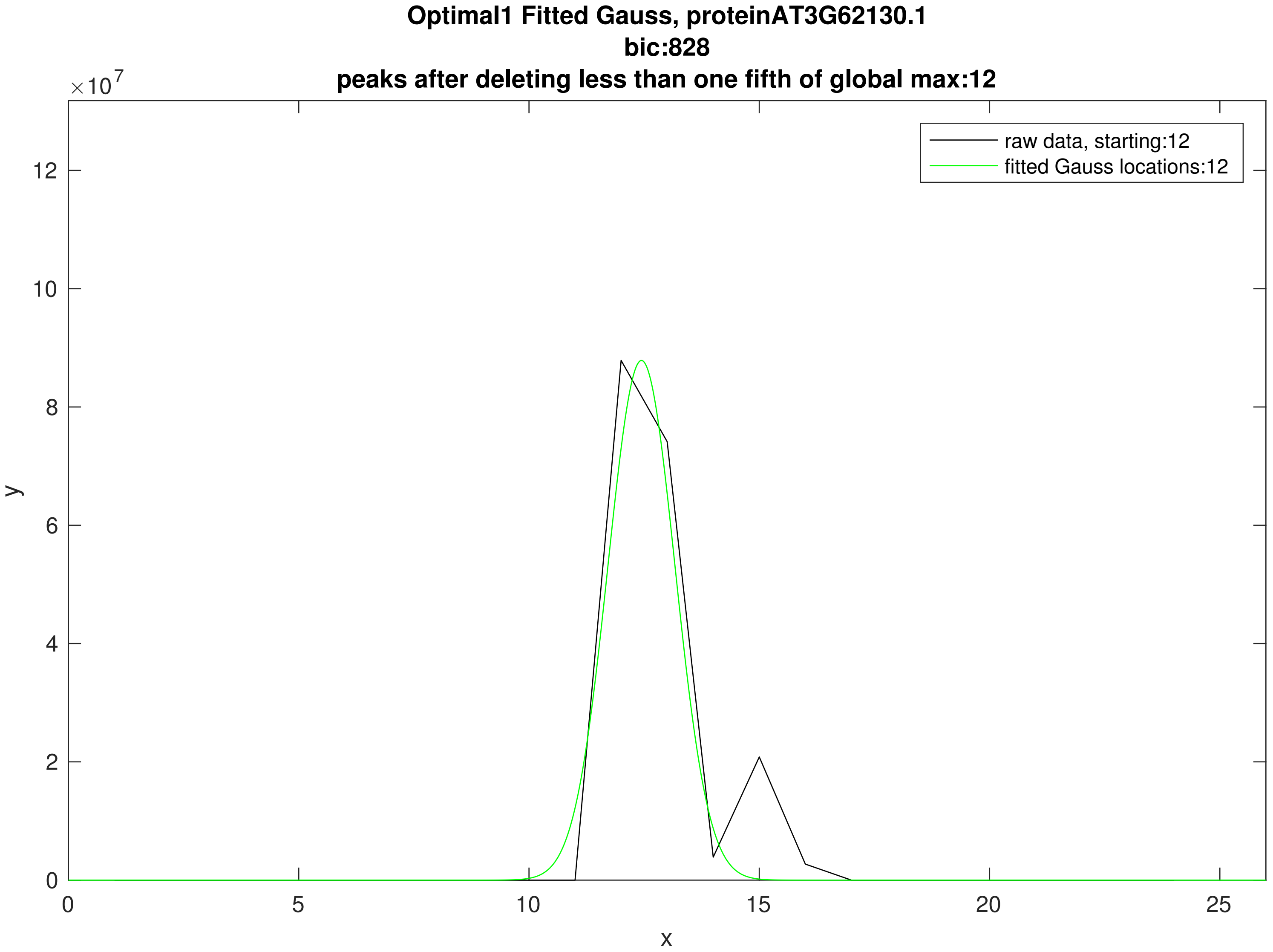
Optimal1 Fitted Gauss, proteinAT3G62130.1
bic:828
peaks after deleting less than one fifth of global max:12
×10^7
raw data, starting:12
fitted Gauss locations:12
y
x
0
2
4
6
8
10
12
0
5
10
15
20
25

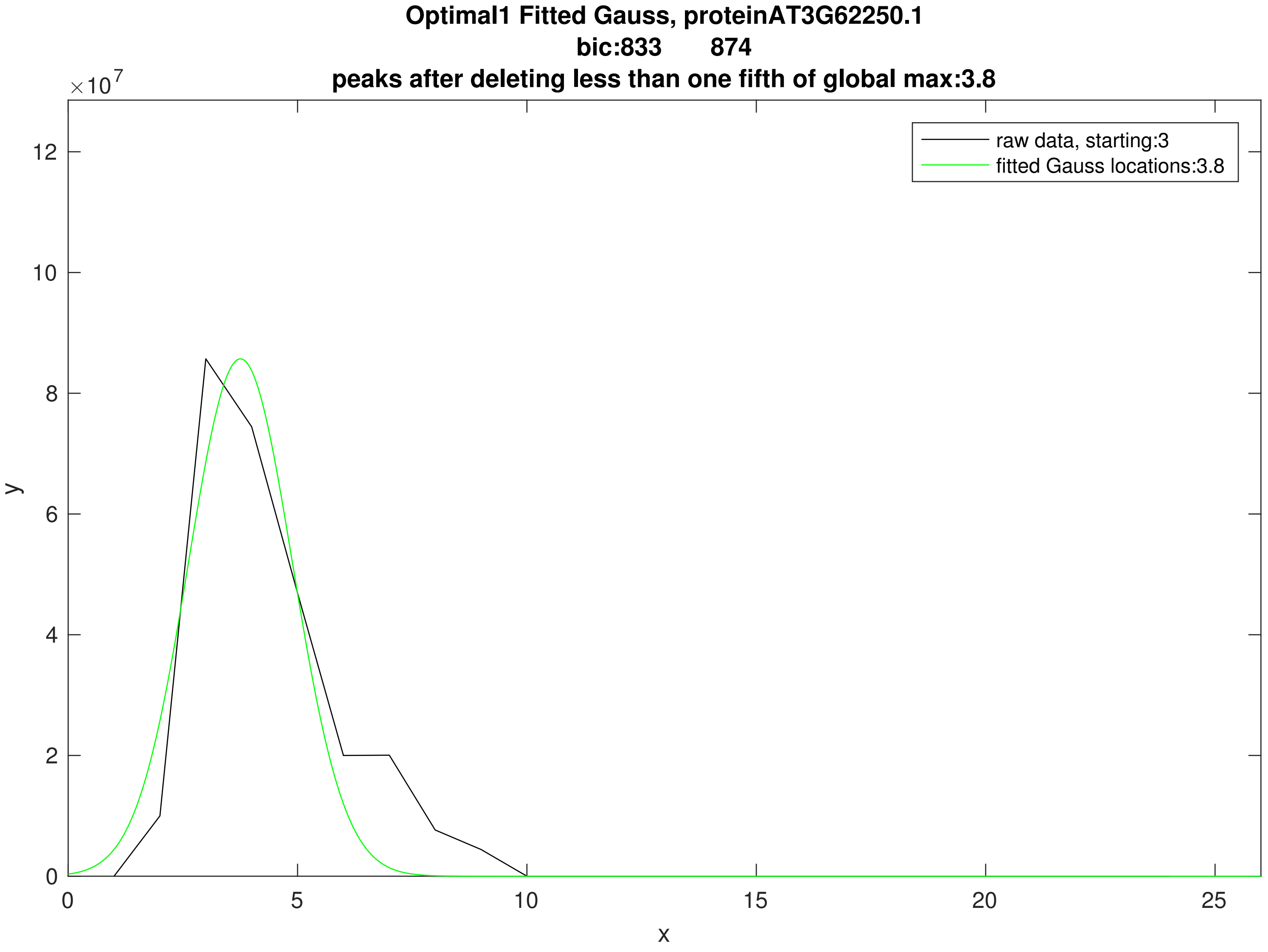
Optimal1 Fitted Gauss, proteinAT3G62250.1
bic:833 874
peaks after deleting less than one fifth of global max:3.8
×10^7
raw data, starting:3
fitted Gauss locations:3.8
y
x
0
2
4
6
8
10
12
0
5
10
15
20
25

, proteinAT3G62290.3;AT3G62290.2;AT3G62290.1;AT2G47170.1;AT1G70490.3;AT1G70490.2;AT1G70490.1;AT1G23490

bic:865 930 851

peaks after deleting less than one fifth of global max:20

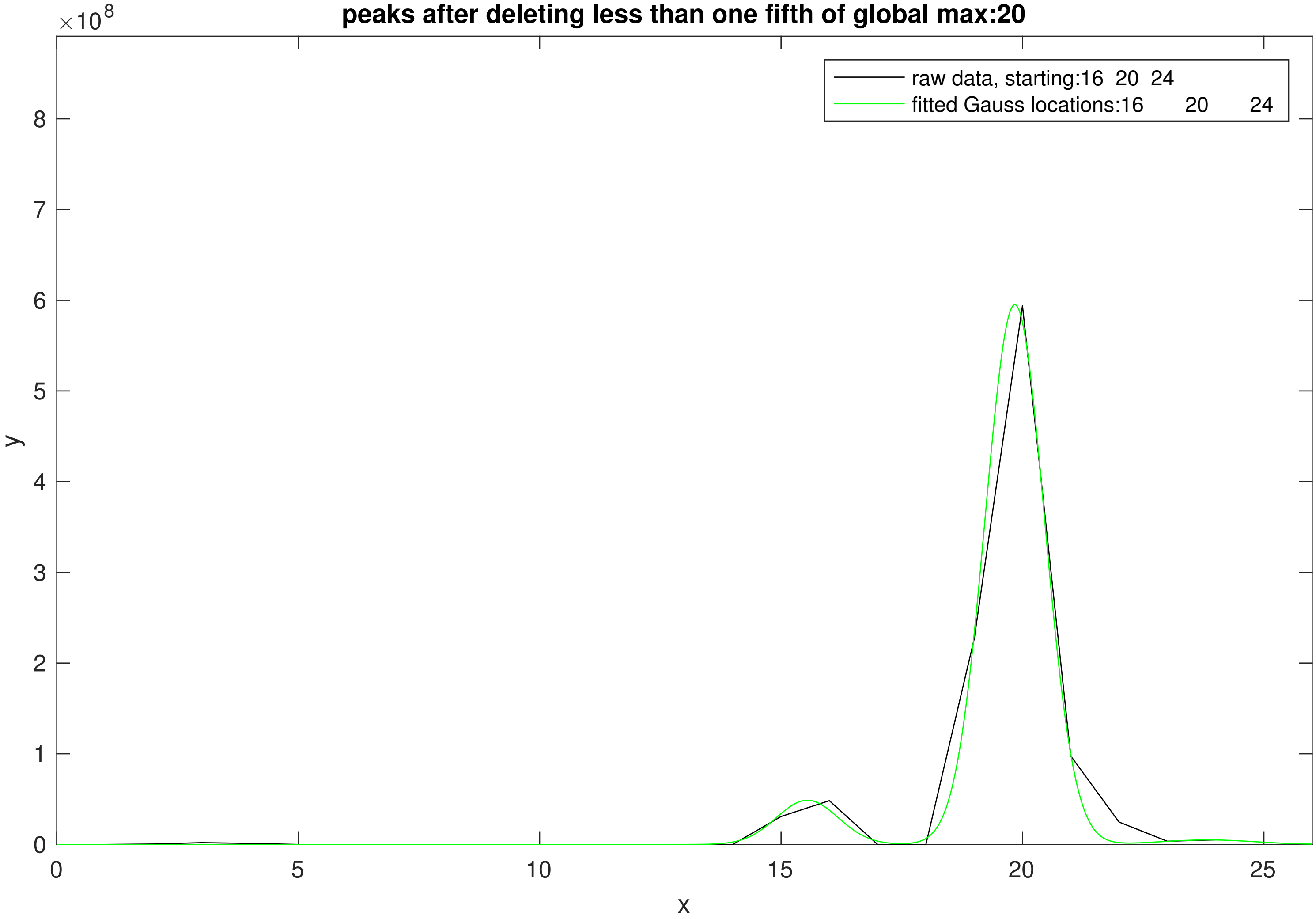

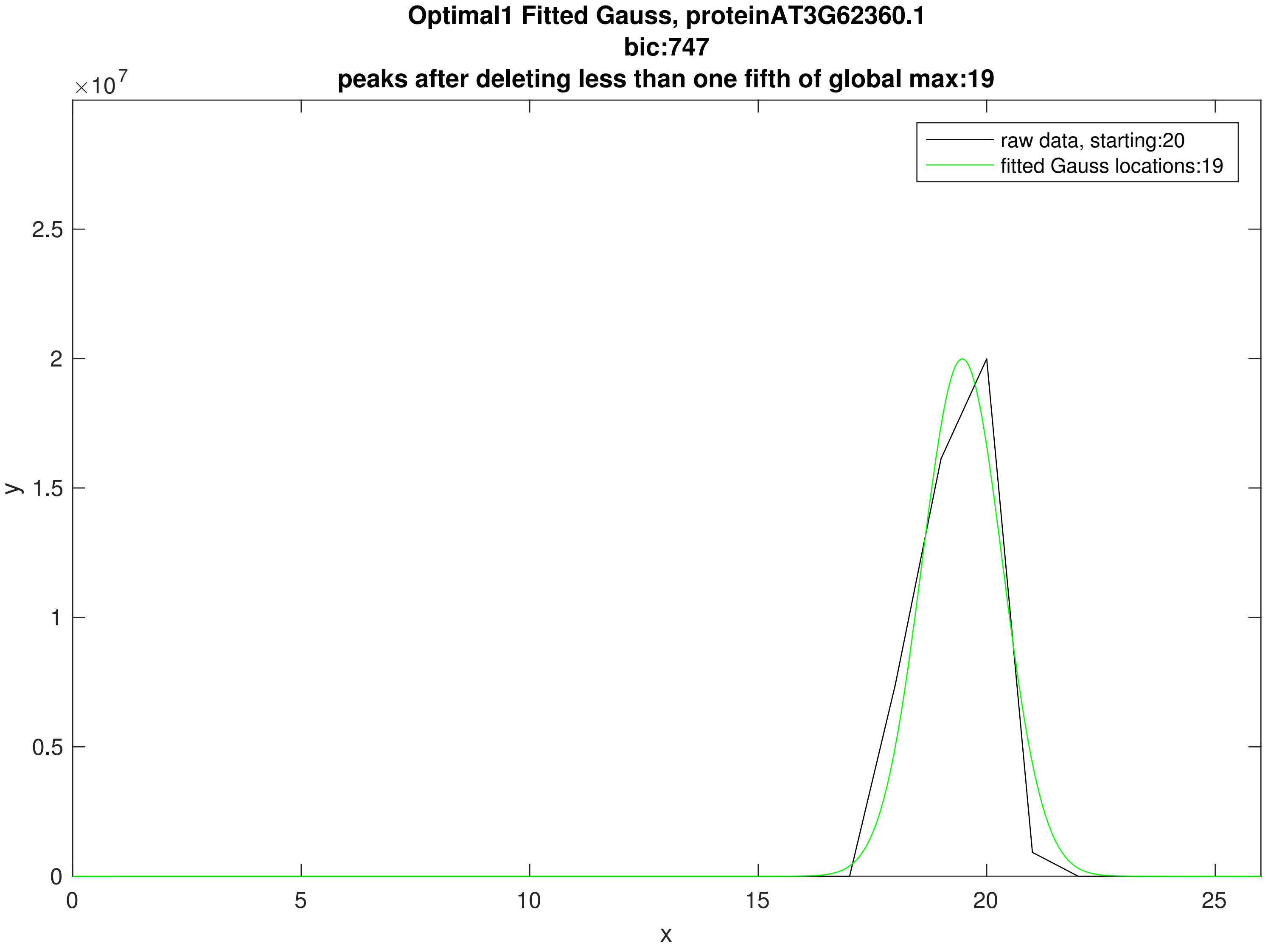
Optimal1 Fitted Gauss, proteinAT3G62360.1
bic:747
peaks after deleting less than one fifth of global max:19
×10^7
raw data, starting:20
fitted Gauss locations:19
2.5
2
1.5
1
0.5
0
y
0
5
10
15
20
25
x

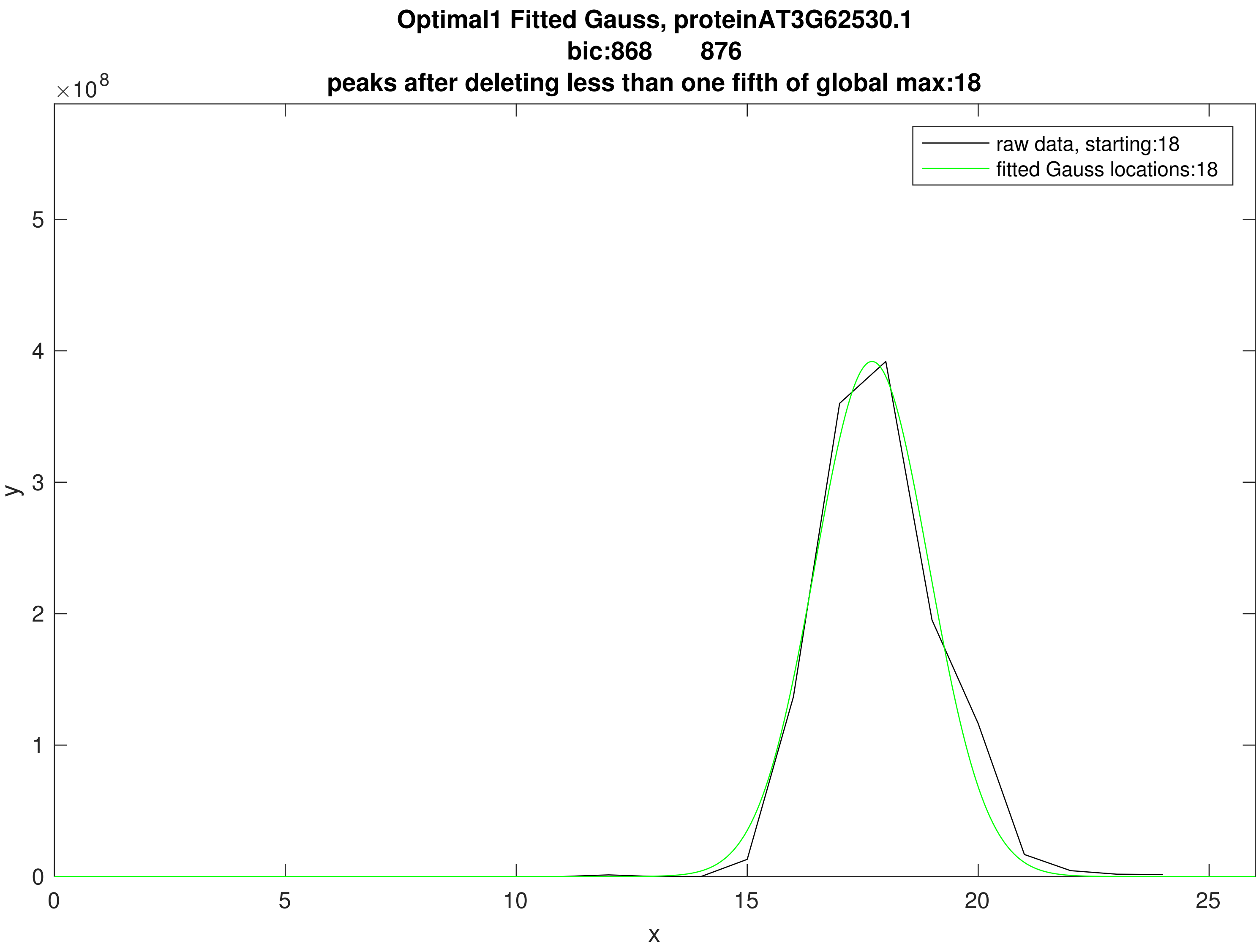
Optimal1 Fitted Gauss, proteinAT3G62530.1
bic:868 876
peaks after deleting less than one fifth of global max:18
×10^8
raw data, starting:18
fitted Gauss locations:18
5
4
3
2
1
0
y
0
5
10
15
20
25
x

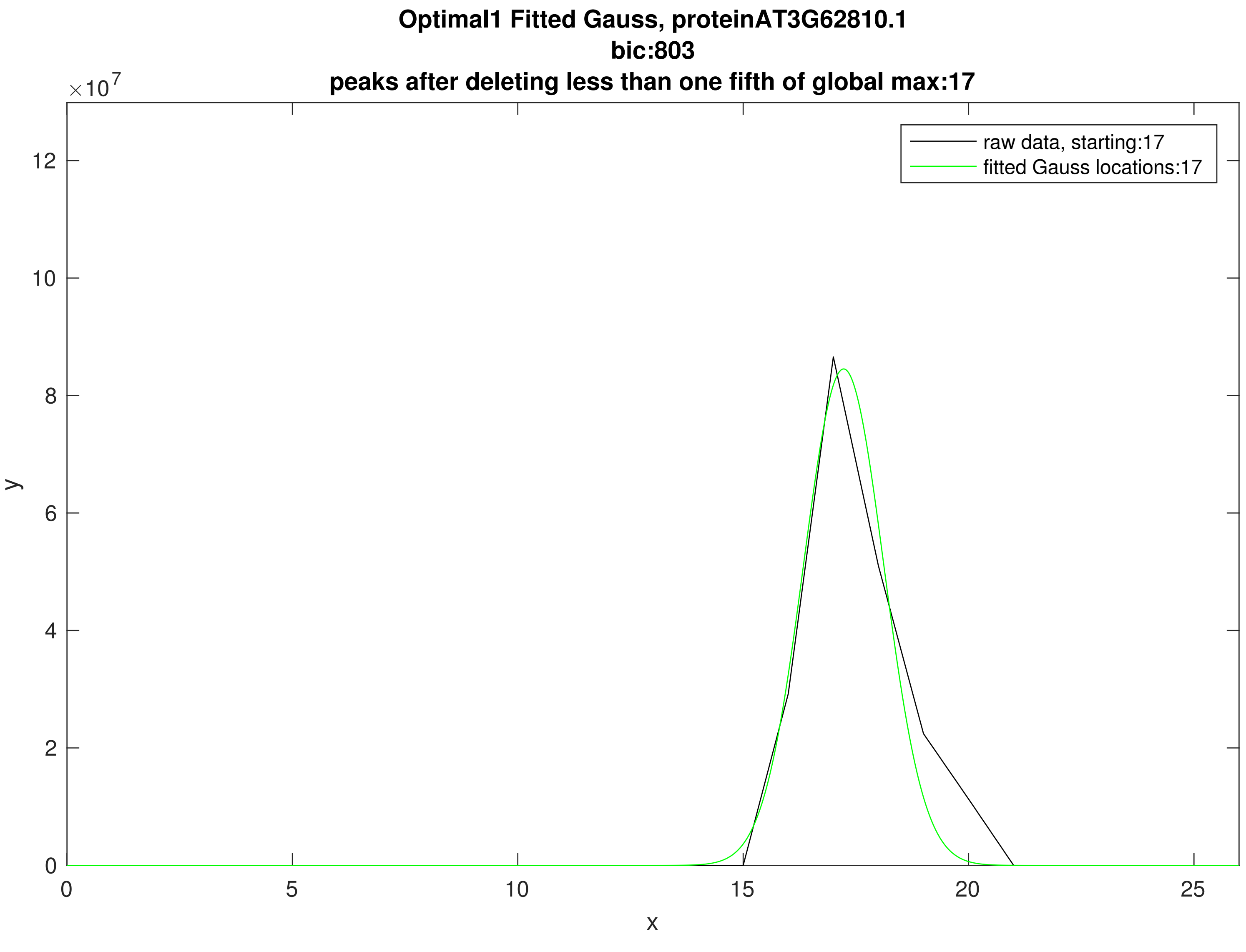

Optimal1 Fitted Gauss, proteinAT3G62810.1
bic:803
peaks after deleting less than one fifth of global max:17
×10^7
raw data, starting:17
fitted Gauss locations:17
12
10
8
6
4
2
0
y
0
5
10
15
20
25
x

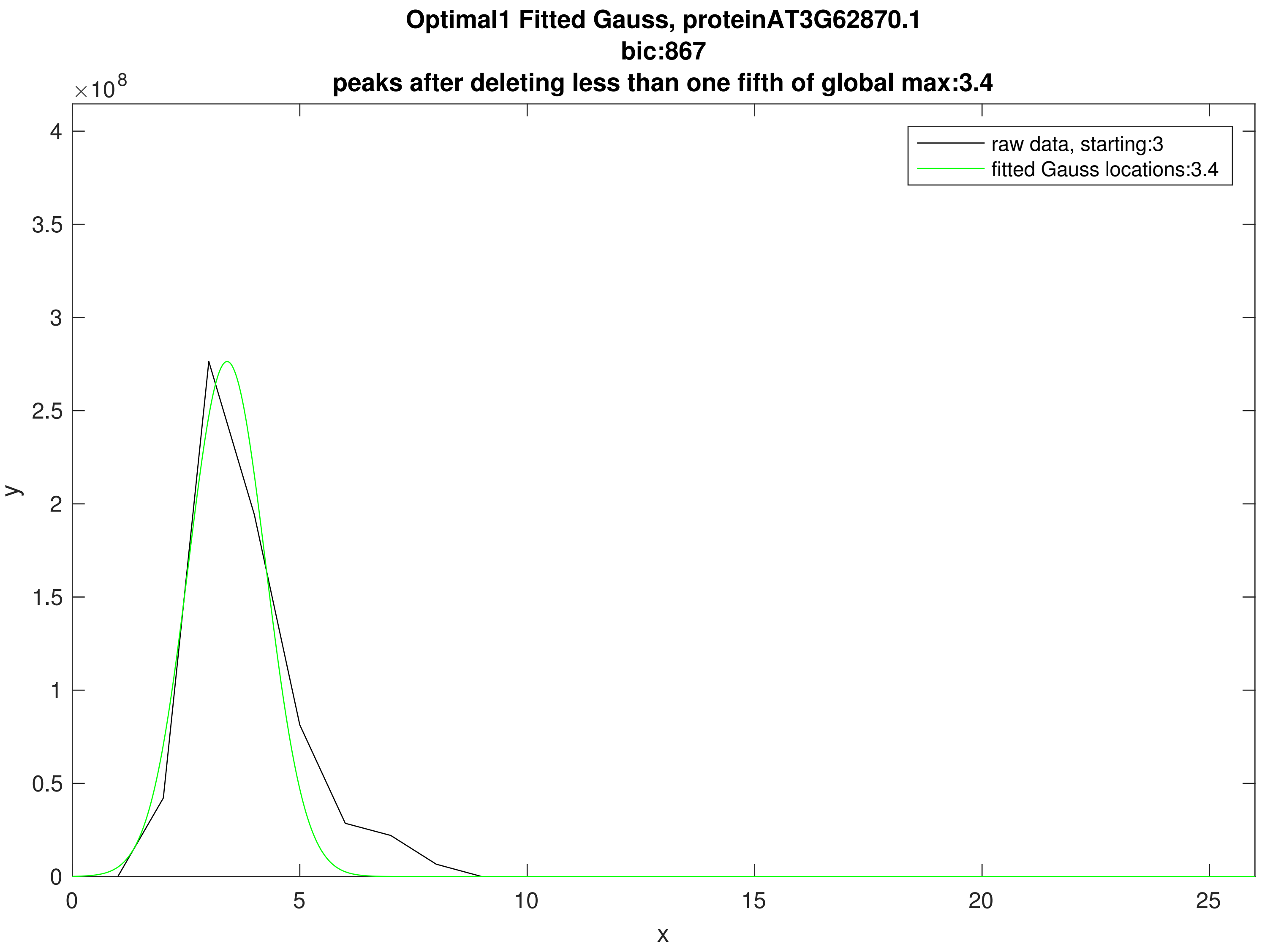
Optimal1 Fitted Gauss, proteinAT3G62870.1
bic:867
peaks after deleting less than one fifth of global max:3.4
×10^8
raw data, starting:3
fitted Gauss locations:3.4
y
x
0
0.5
1
1.5
2
2.5
3
3.5
4
0
5
10
15
20
25

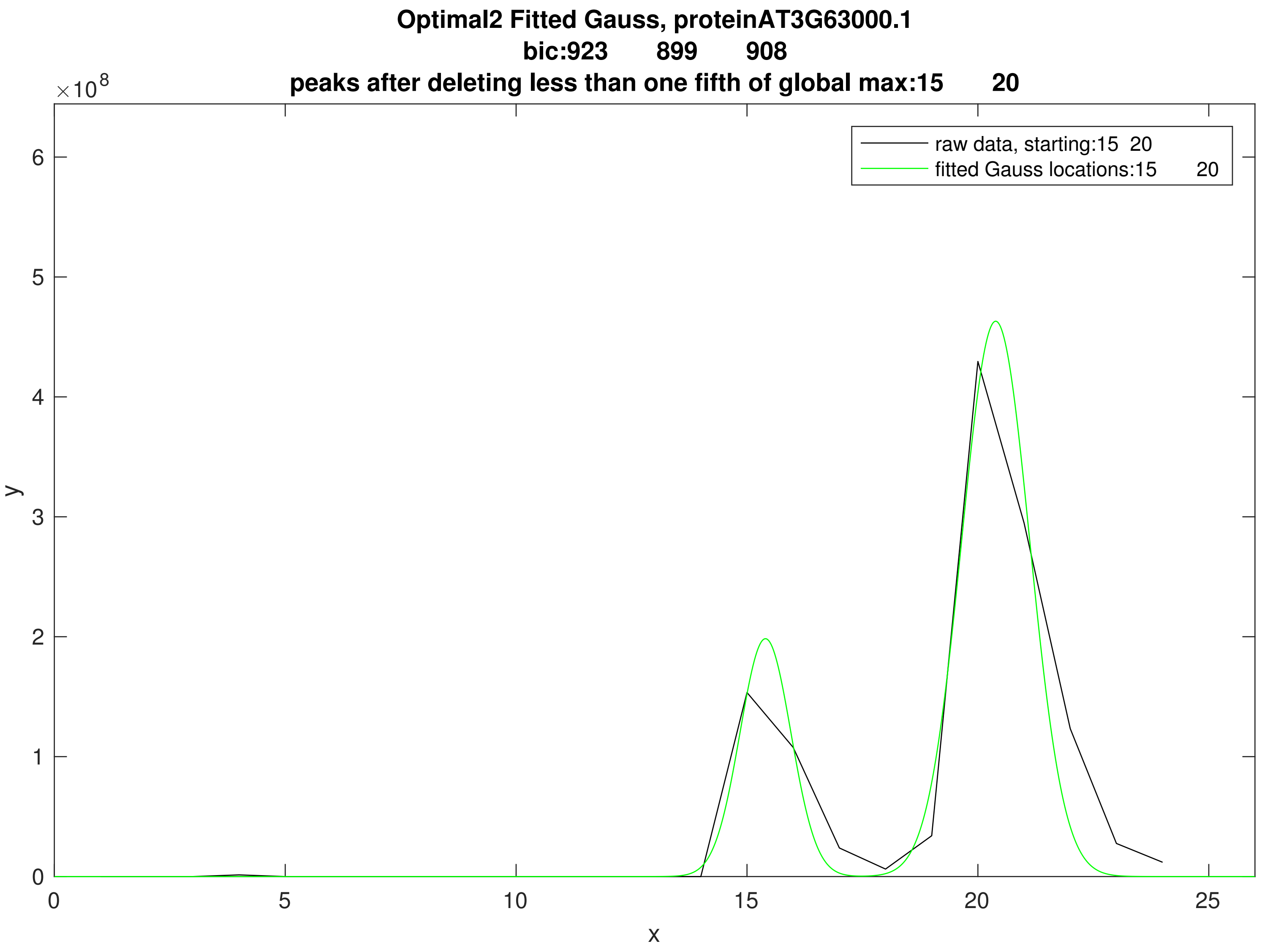
Optimal2 Fitted Gauss, proteinAT3G63000.1
bic:923 899 908
peaks after deleting less than one fifth of global max:15 20
×10^8
raw data, starting:15 20
fitted Gauss locations:15 20
y
x
0
1
2
3
4
5
6
0
5
10
15
20
25

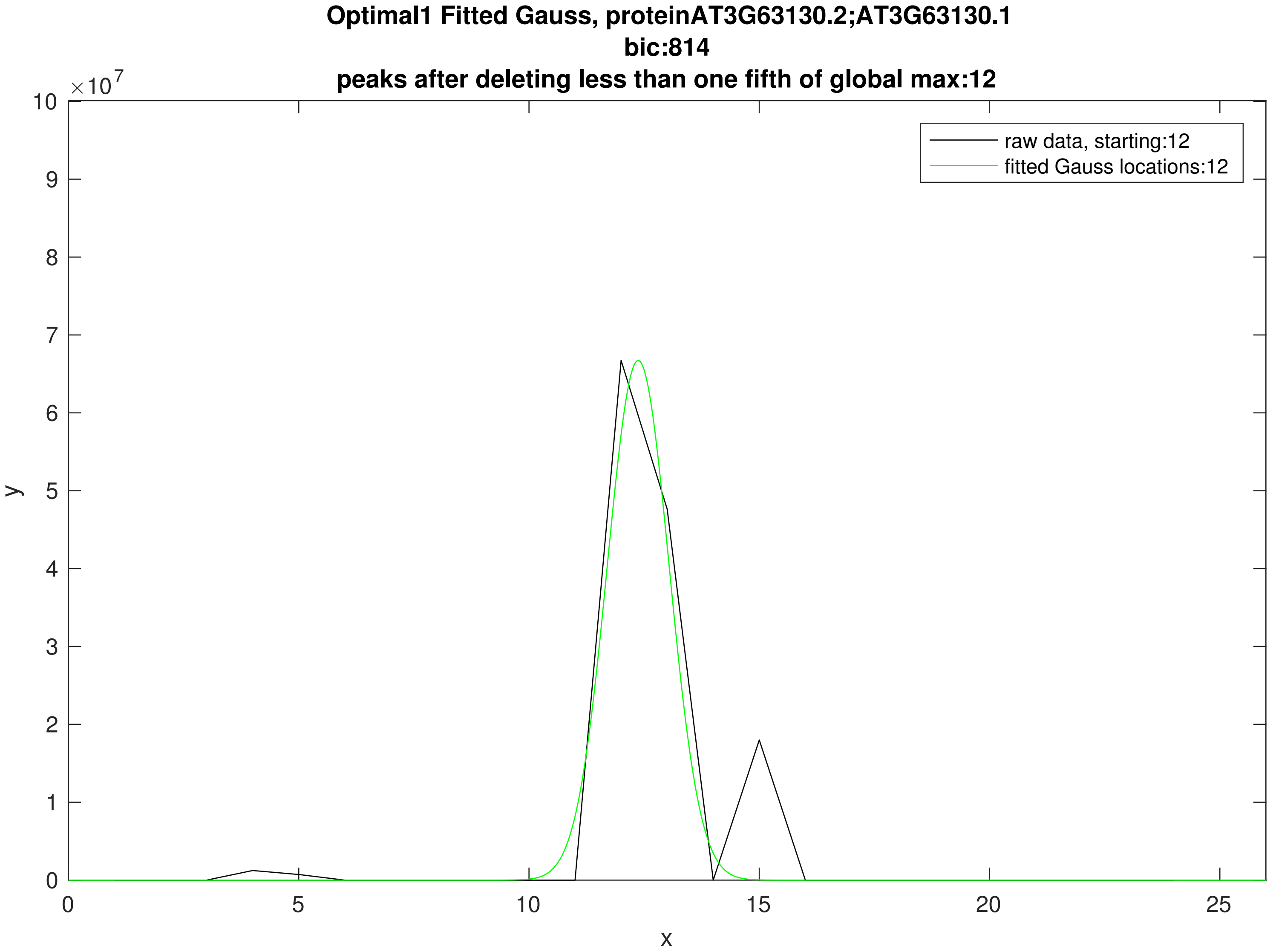
Optimal1 Fitted Gauss, proteinAT3G63130.2;AT3G63130.1
bic:814
peaks after deleting less than one fifth of global max:12
×10^7
raw data, starting:12
fitted Gauss locations:12
y
x
0
1
2
3
4
5
6
7
8
9
10
0
5
10
15
20
25

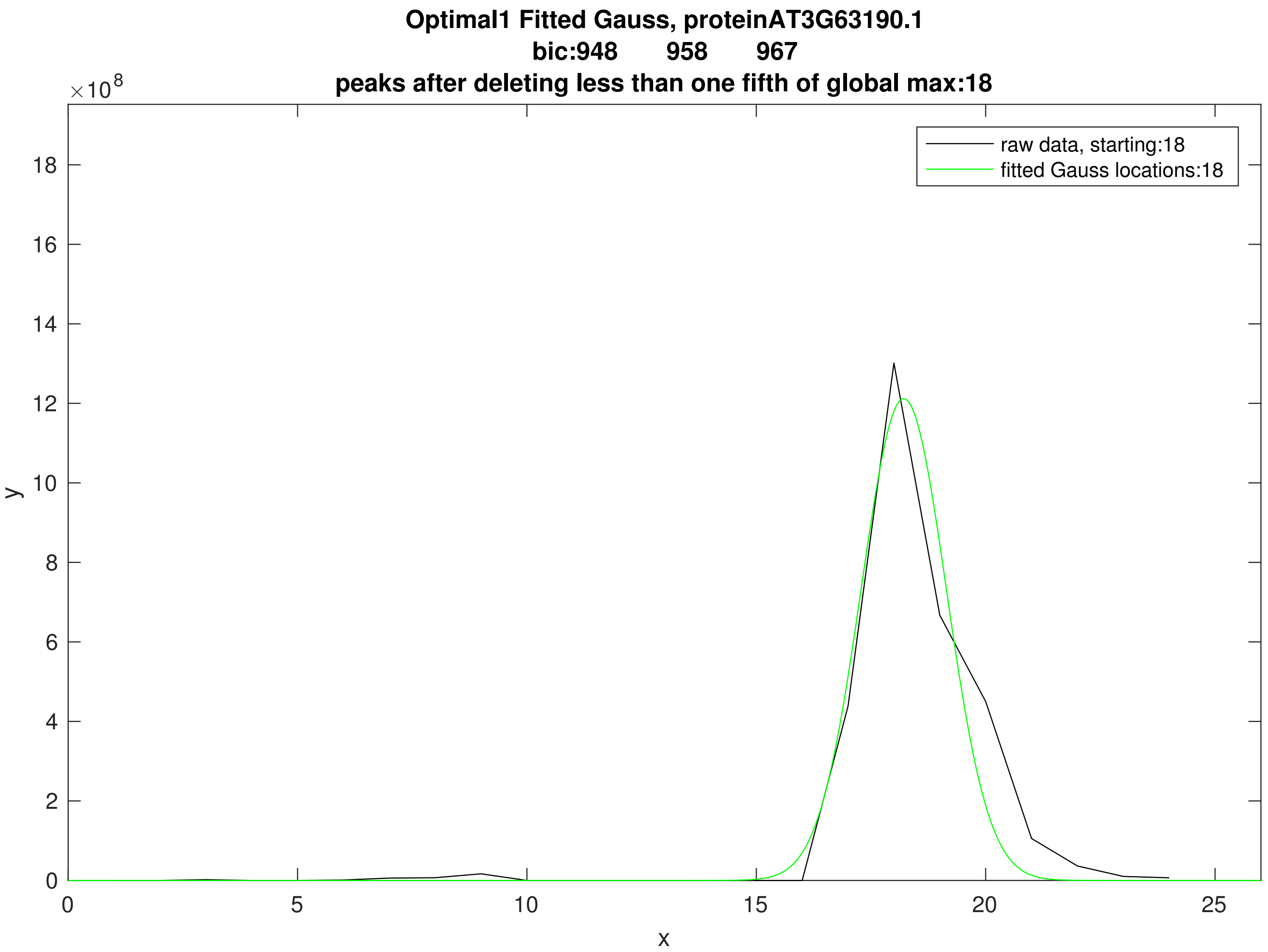
Optimal1 Fitted Gauss, proteinAT3G63190.1
bic:948 958 967
peaks after deleting less than one fifth of global max:18
×10^8
raw data, starting:18
fitted Gauss locations:18
y
x
0
2
4
6
8
10
12
14
16
18
0
5
10
15
20
25

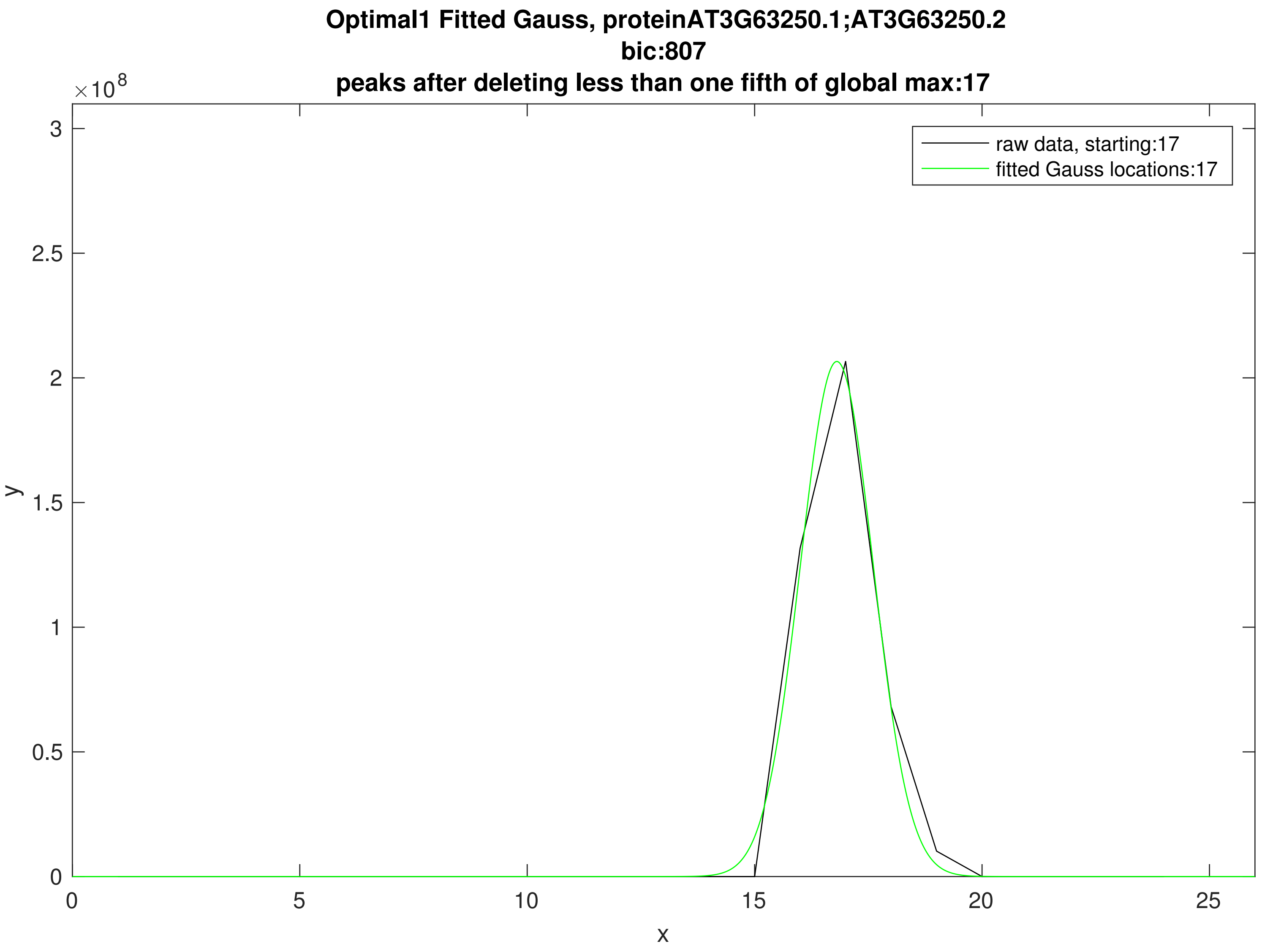

Optimal1 Fitted Gauss, proteinAT3G63250.1;AT3G63250.2
bic:807
peaks after deleting less than one fifth of global max:17
×10^8
raw data, starting:17
fitted Gauss locations:17
y
x
0
0.5
1
1.5
2
2.5
3
0
5
10
15
20
25

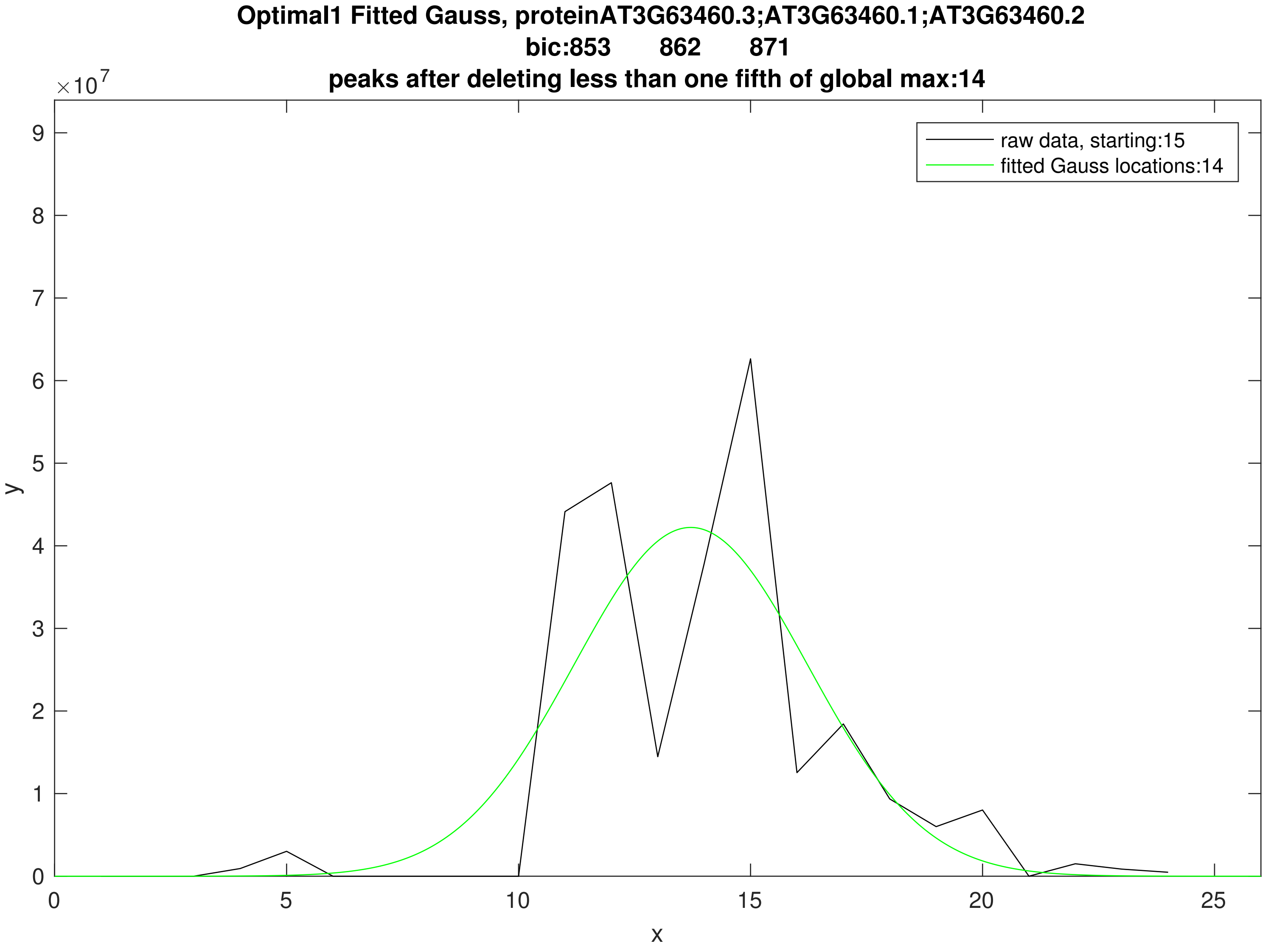

Optimal1 Fitted Gauss, proteinAT3G63460.3;AT3G63460.1;AT3G63460.2
bic:853 862 871
peaks after deleting less than one fifth of global max:14
×10^7
raw data, starting:15
fitted Gauss locations:14
y
x
0
1
2
3
4
5
6
7
8
9
0
5
10
15
20
25

#### Optimal1 Fitted Gauss, proteinAT3G63490.1;AT3G63490.2
#### bic:781 786
#### peaks after deleting less than one fifth of global max:4

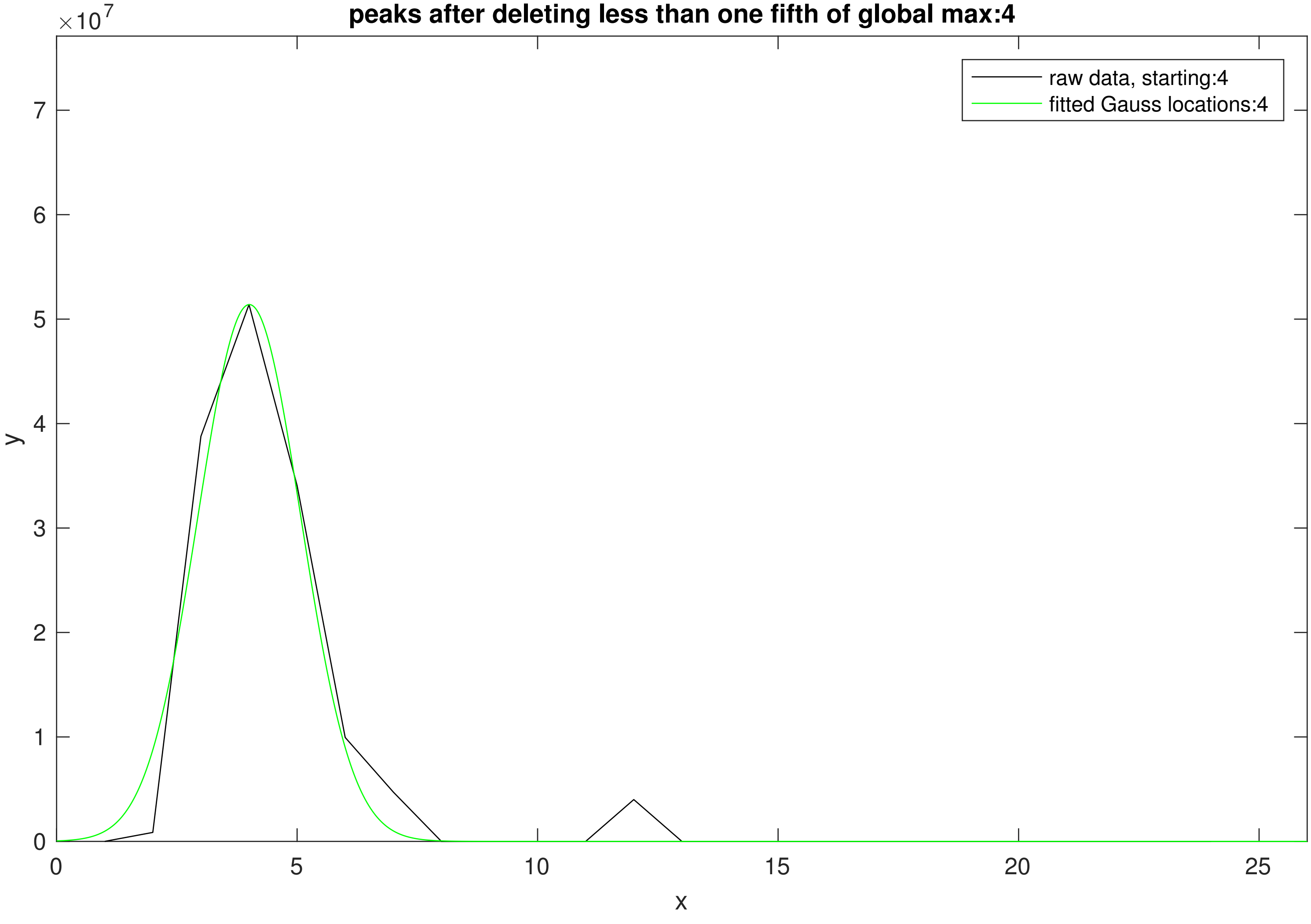

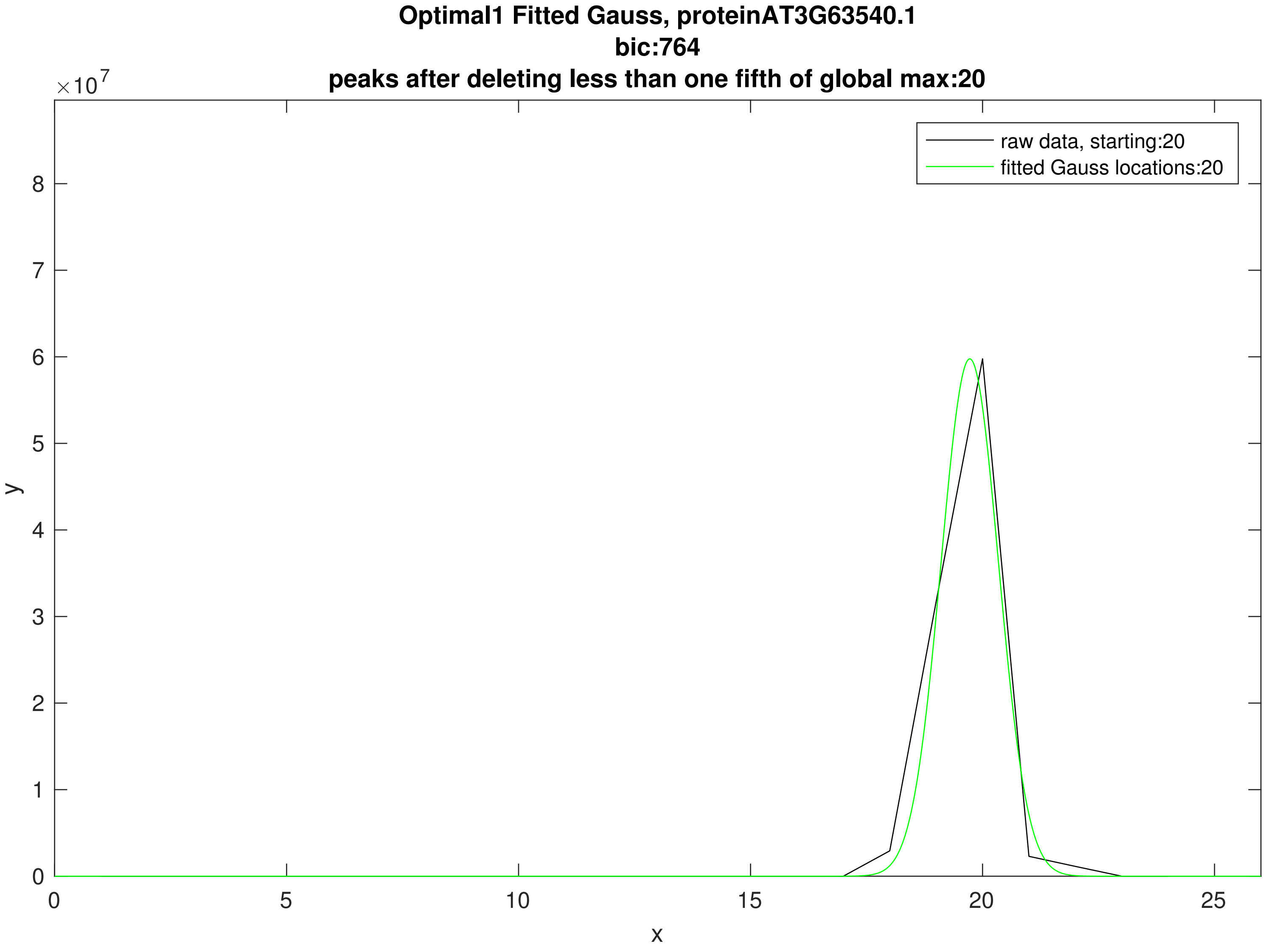

Optimal1 Fitted Gauss, proteinAT3G63540.1
bic:764
peaks after deleting less than one fifth of global max:20
×10^7
raw data, starting:20
fitted Gauss locations:20
8
7
6
5
4
3
2
1
0
y
0
5
10
15
20
25
x

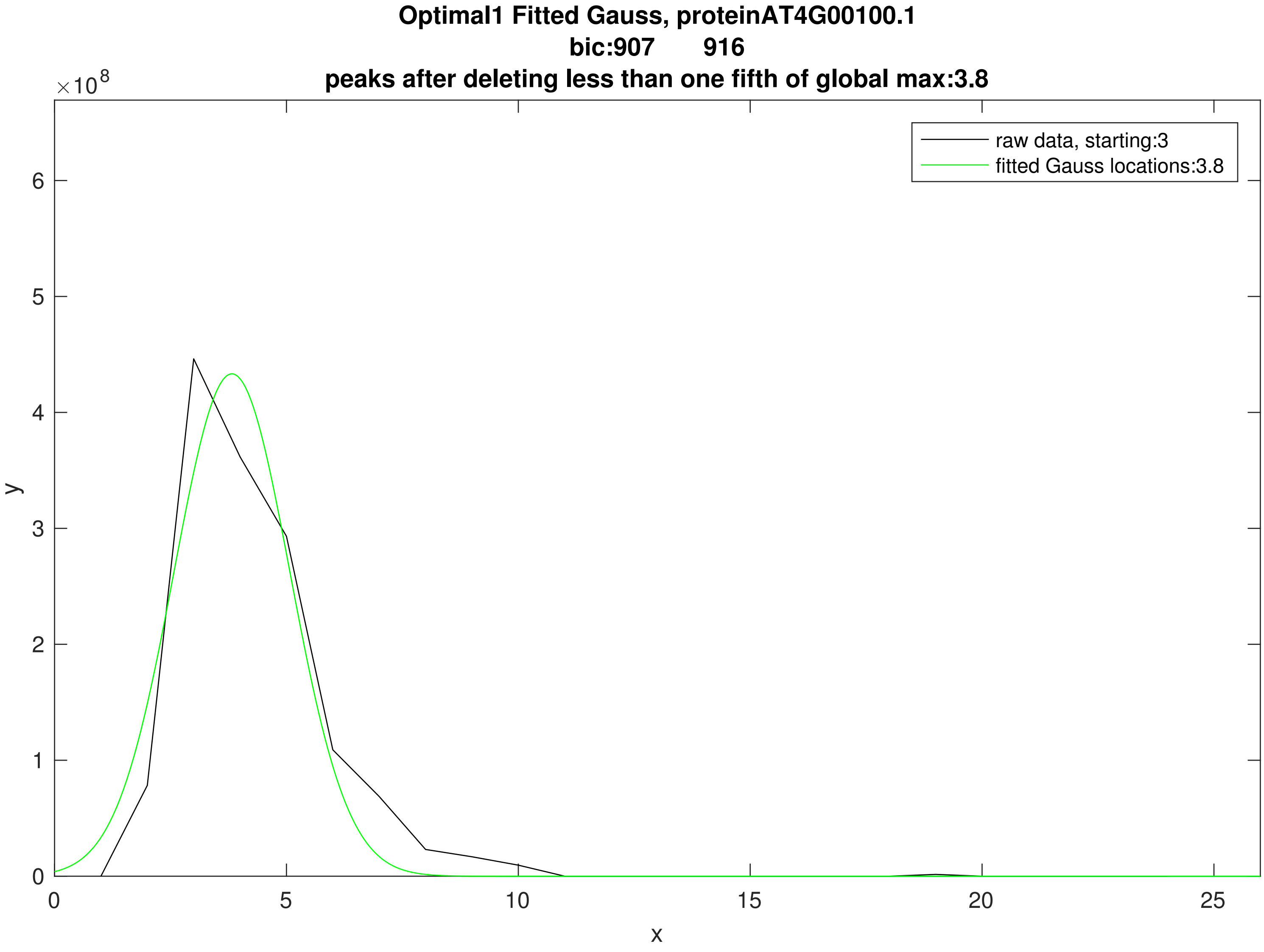

Optimal1 Fitted Gauss, proteinAT4G00100.1
bic:907 916
peaks after deleting less than one fifth of global max:3.8
×10^8
raw data, starting:3
fitted Gauss locations:3.8
y
x
0
1
2
3
4
5
6
0
5
10
15
20
25

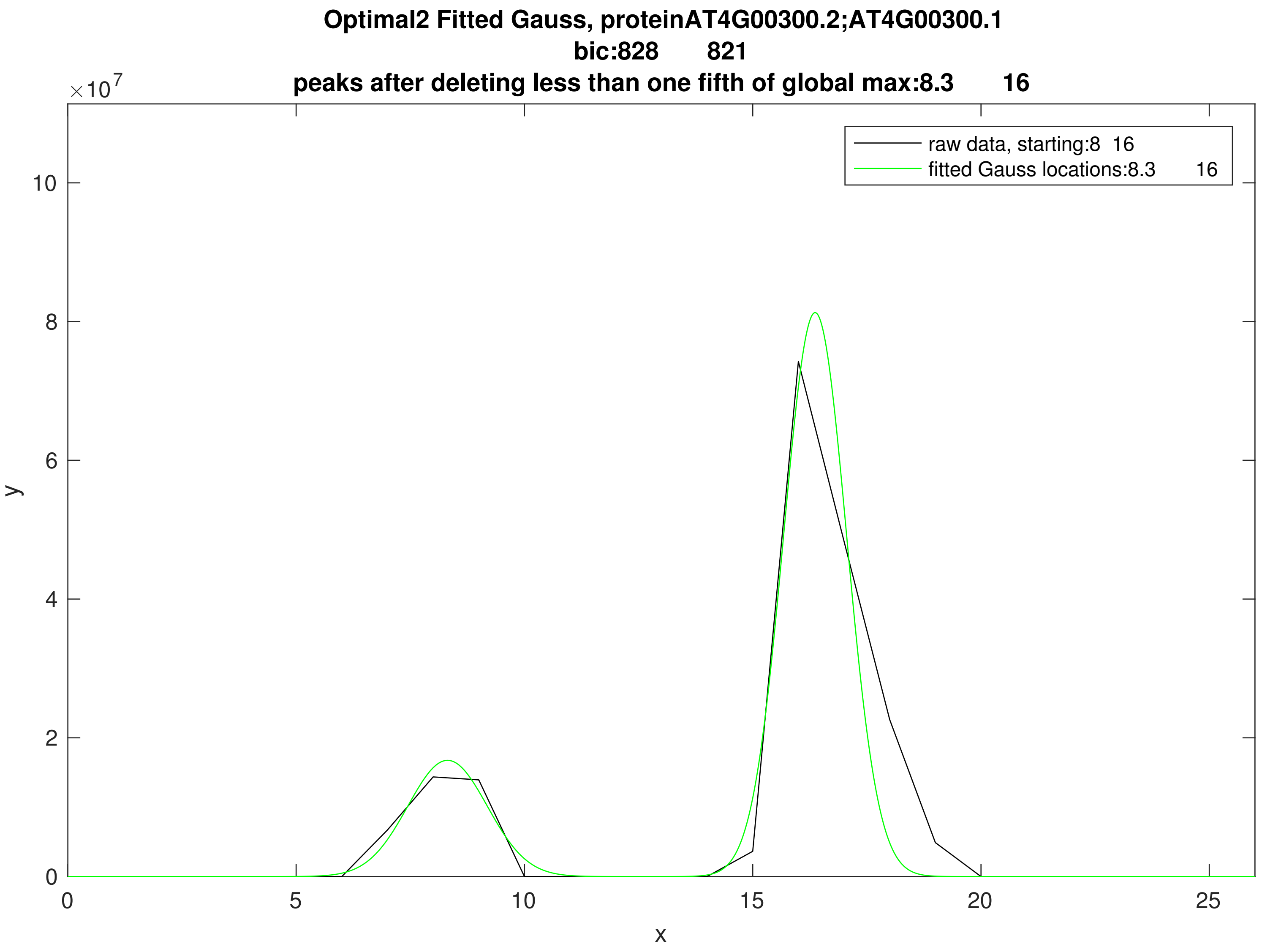
Optimal2 Fitted Gauss, proteinAT4G00300.2;AT4G00300.1
bic:828 821
peaks after deleting less than one fifth of global max:8.3 16
×10^7
raw data, starting:8 16
fitted Gauss locations:8.3 16
y
x
0
2
4
6
8
10
5
15
20
25

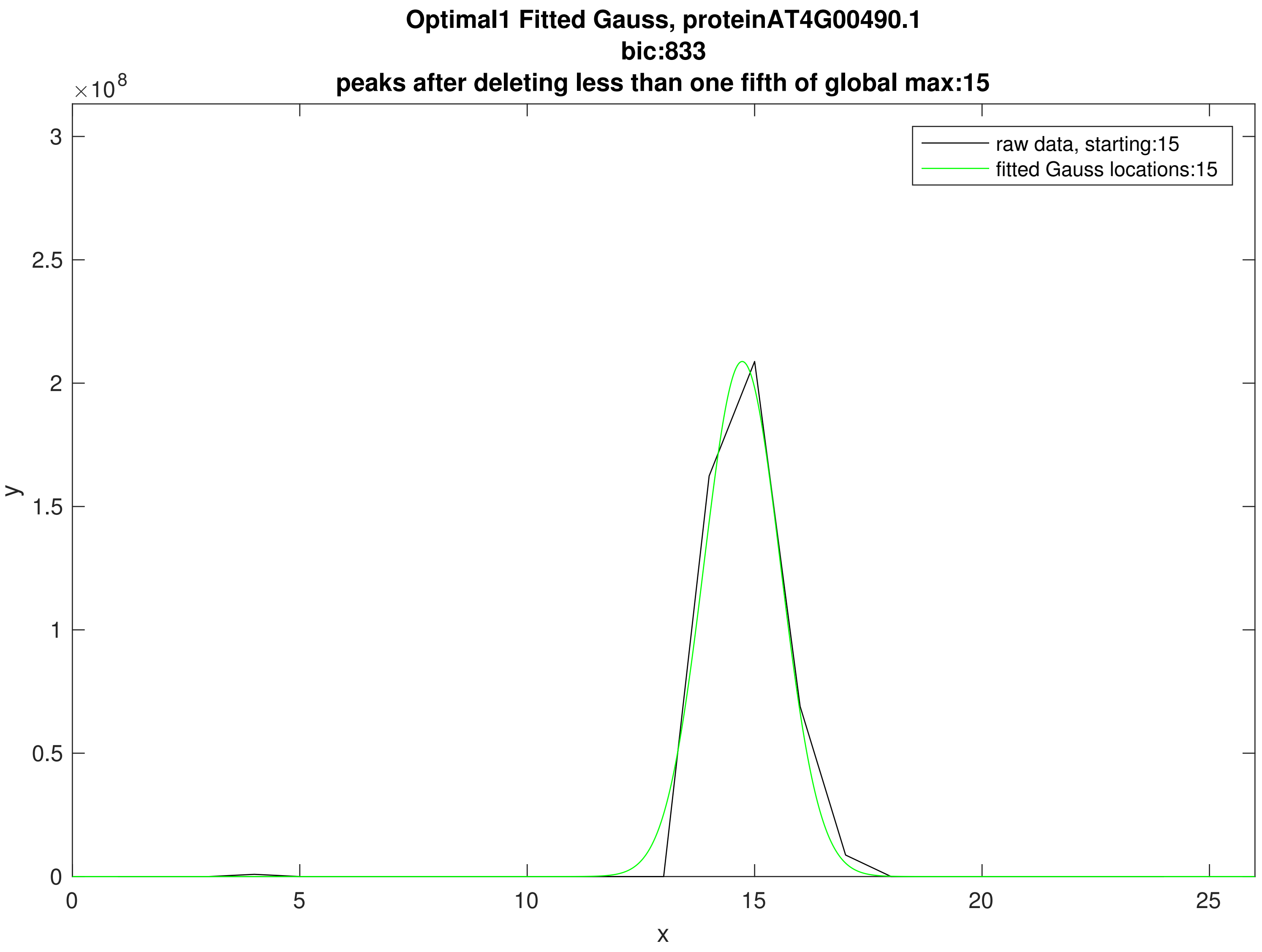
Optimal1 Fitted Gauss, proteinAT4G00490.1
bic:833
peaks after deleting less than one fifth of global max:15
×10^8
raw data, starting:15
fitted Gauss locations:15
3
2.5
2
1.5
1
0.5
0
y
0
5
10
15
20
25
x

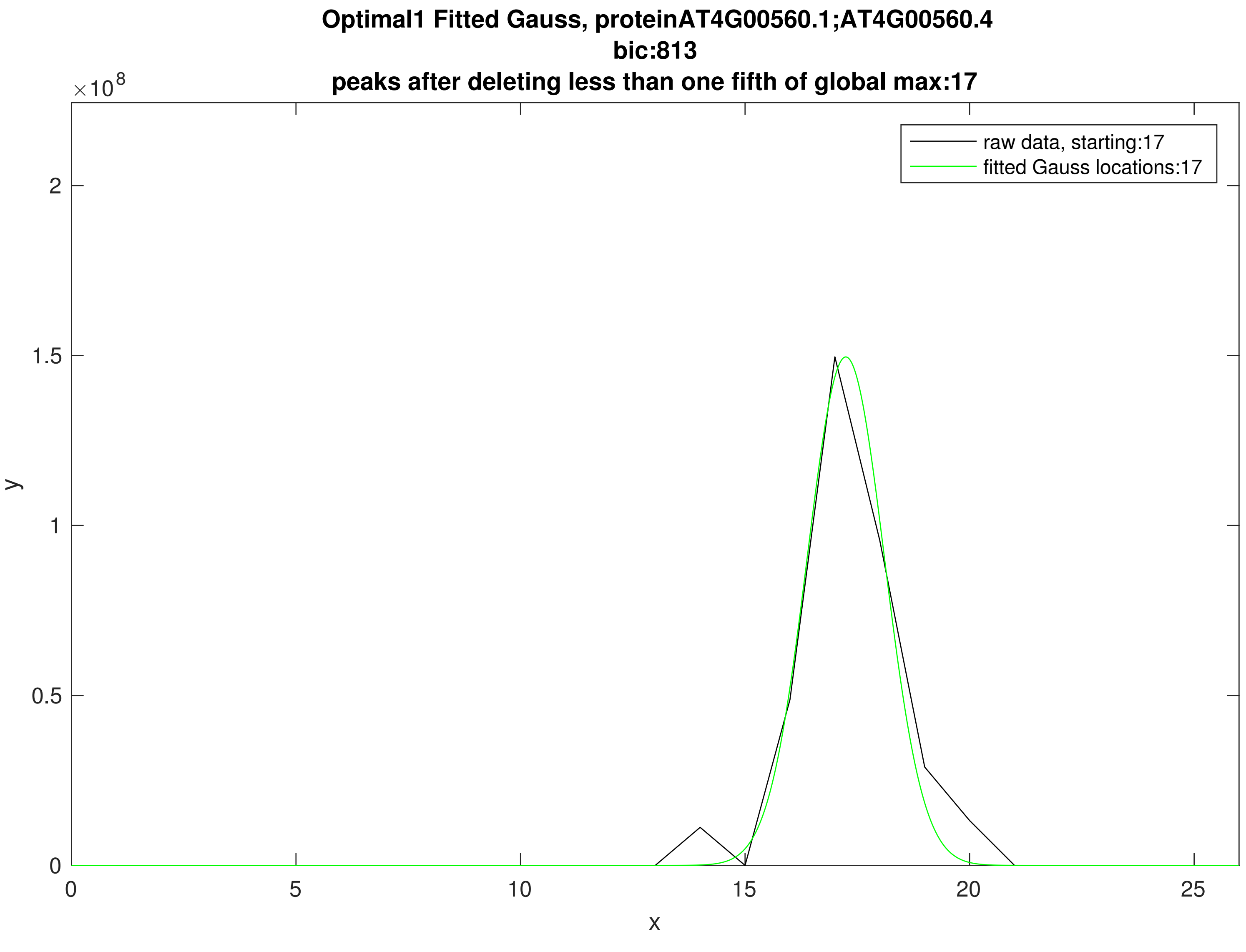
Optimal1 Fitted Gauss, proteinAT4G00560.1;AT4G00560.4
bic:813
peaks after deleting less than one fifth of global max:17
×10^8
raw data, starting:17
fitted Gauss locations:17
y
x
0
0.5
1
1.5
2
0
5
10
15
20
25

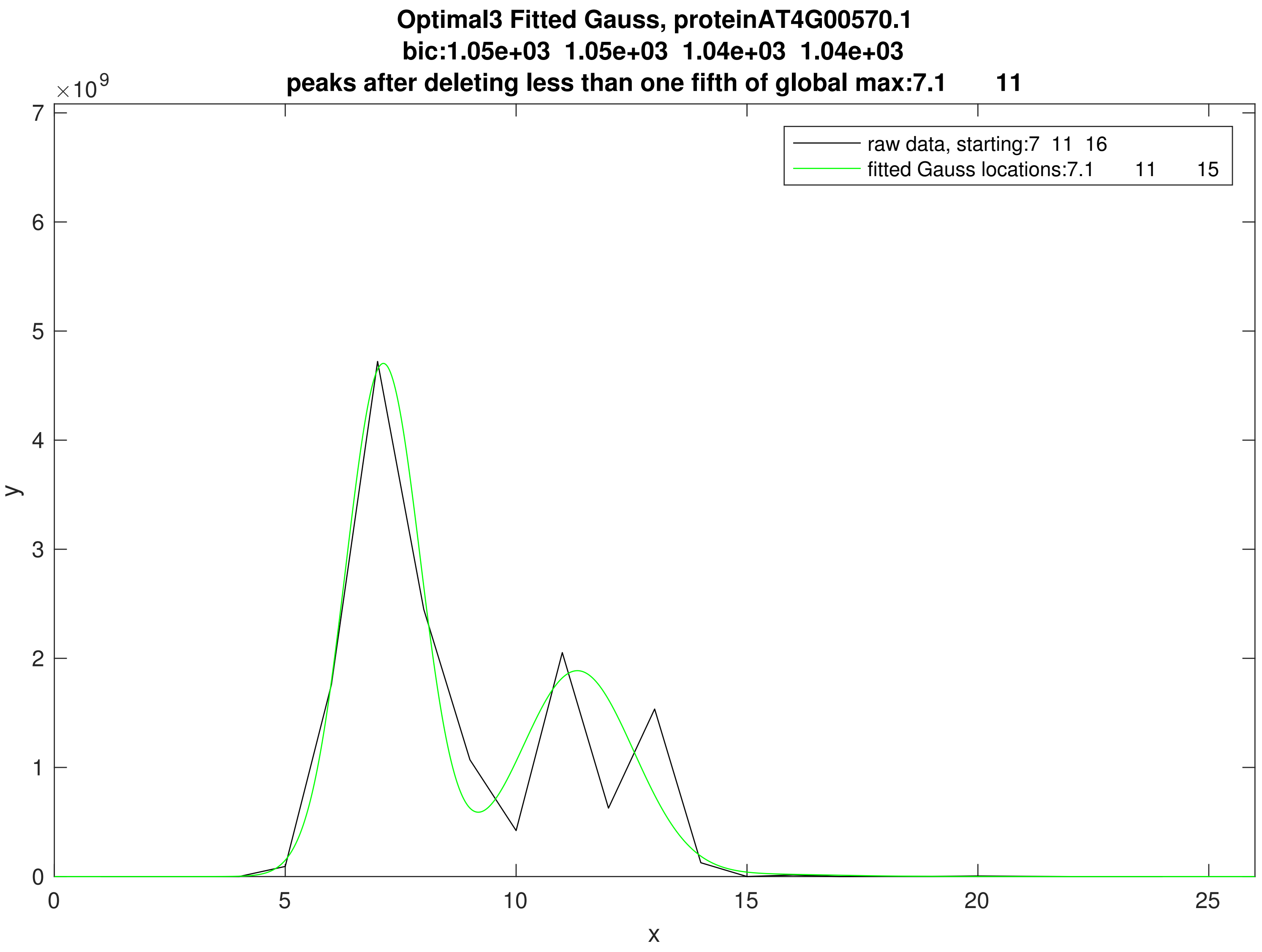
Optimal3 Fitted Gauss, proteinAT4G00570.1
bic:1.05e+03 1.05e+03 1.04e+03 1.04e+03
peaks after deleting less than one fifth of global max:7.1 11
×10^9
raw data, starting:7 11 16
fitted Gauss locations:7.1 11 15
y
x
0
1
2
3
4
5
6
7
0
5
10
15
20
25

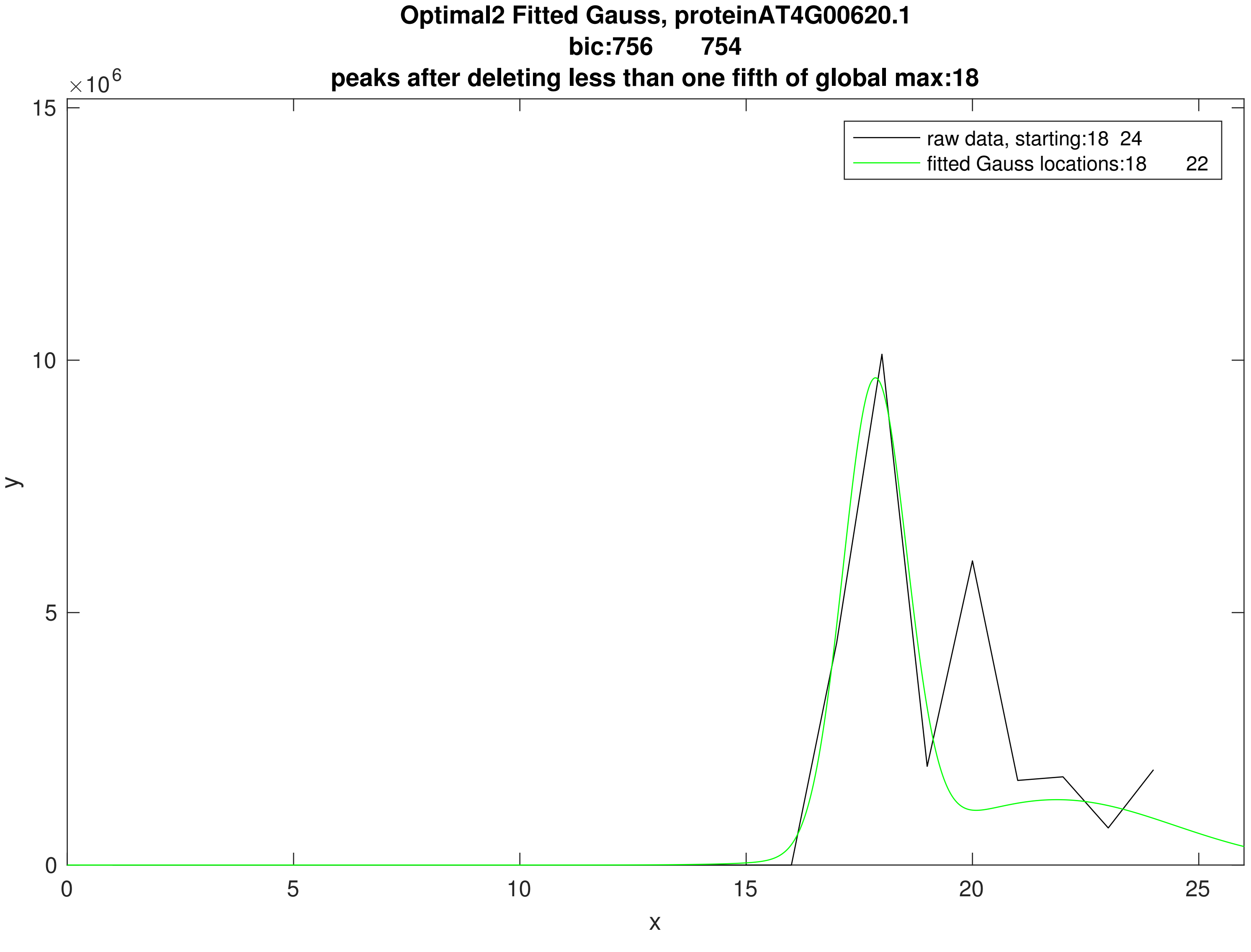

Optimal2 Fitted Gauss, proteinAT4G00620.1
bic:756 754
peaks after deleting less than one fifth of global max:18
raw data, starting:18 24
fitted Gauss locations:18 22
×10^6
y
x

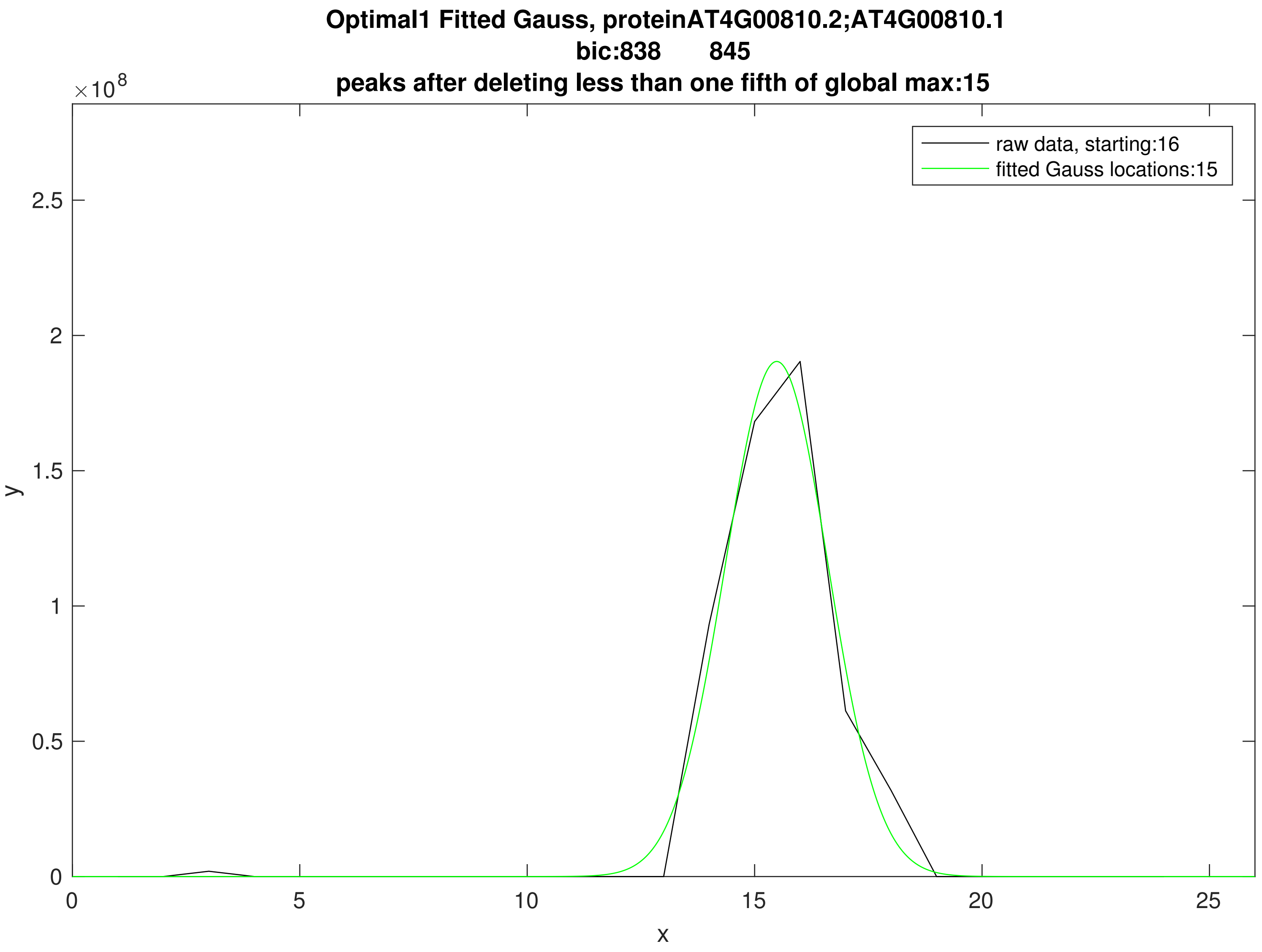
Optimal1 Fitted Gauss, proteinAT4G00810.2;AT4G00810.1
bic:838 845
peaks after deleting less than one fifth of global max:15
raw data, starting:16
fitted Gauss locations:15
×10^8
y
x
0
0.5
1
1.5
2
2.5
0
5
10
15
20
25

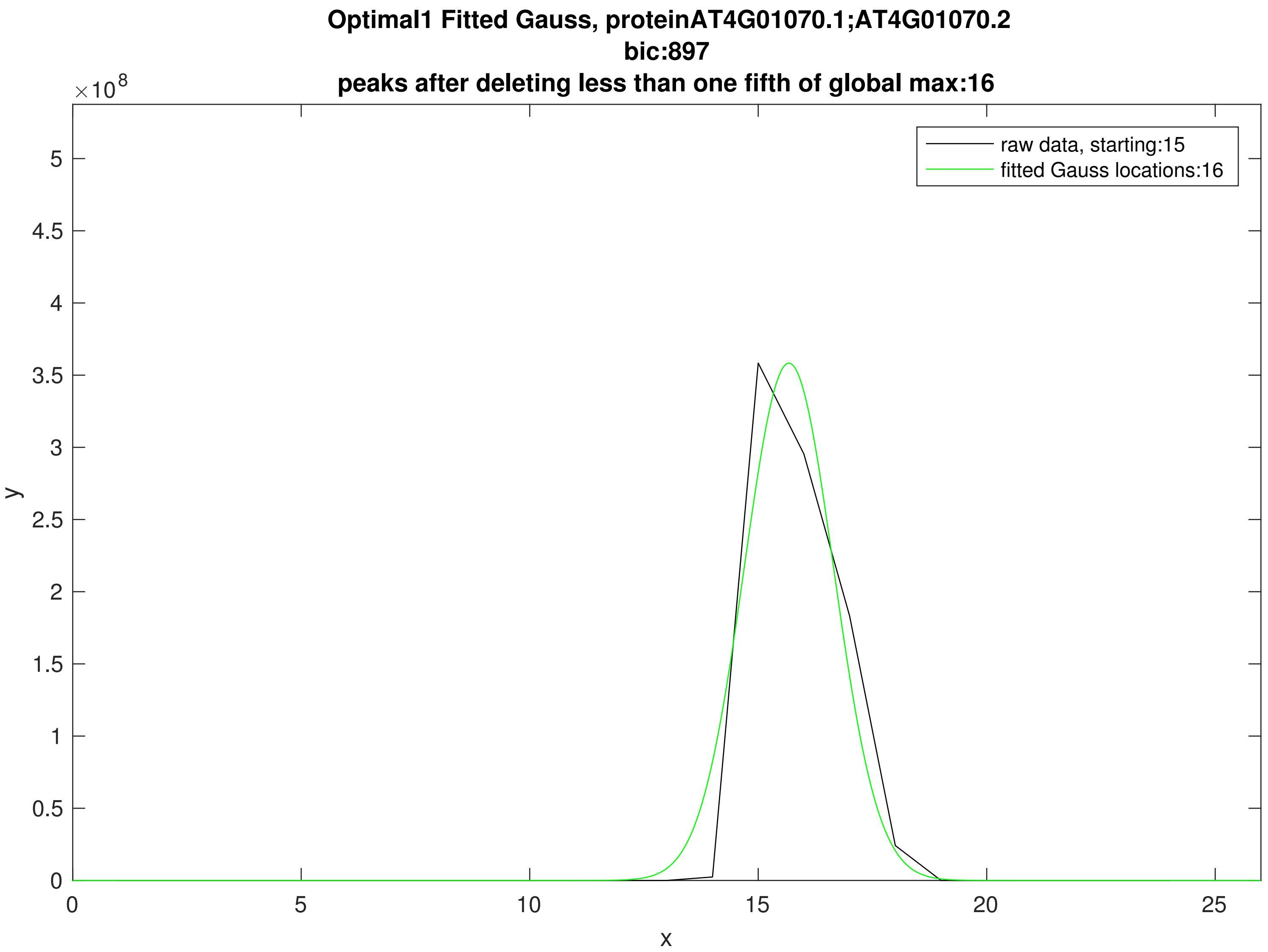
Optimal1 Fitted Gauss, proteinAT4G01070.1;AT4G01070.2
bic:897
peaks after deleting less than one fifth of global max:16
×10^8
raw data, starting:15
fitted Gauss locations:16
y
x
0
0.5
1
1.5
2
2.5
3
3.5
4
4.5
5
0
5
10
15
20
25

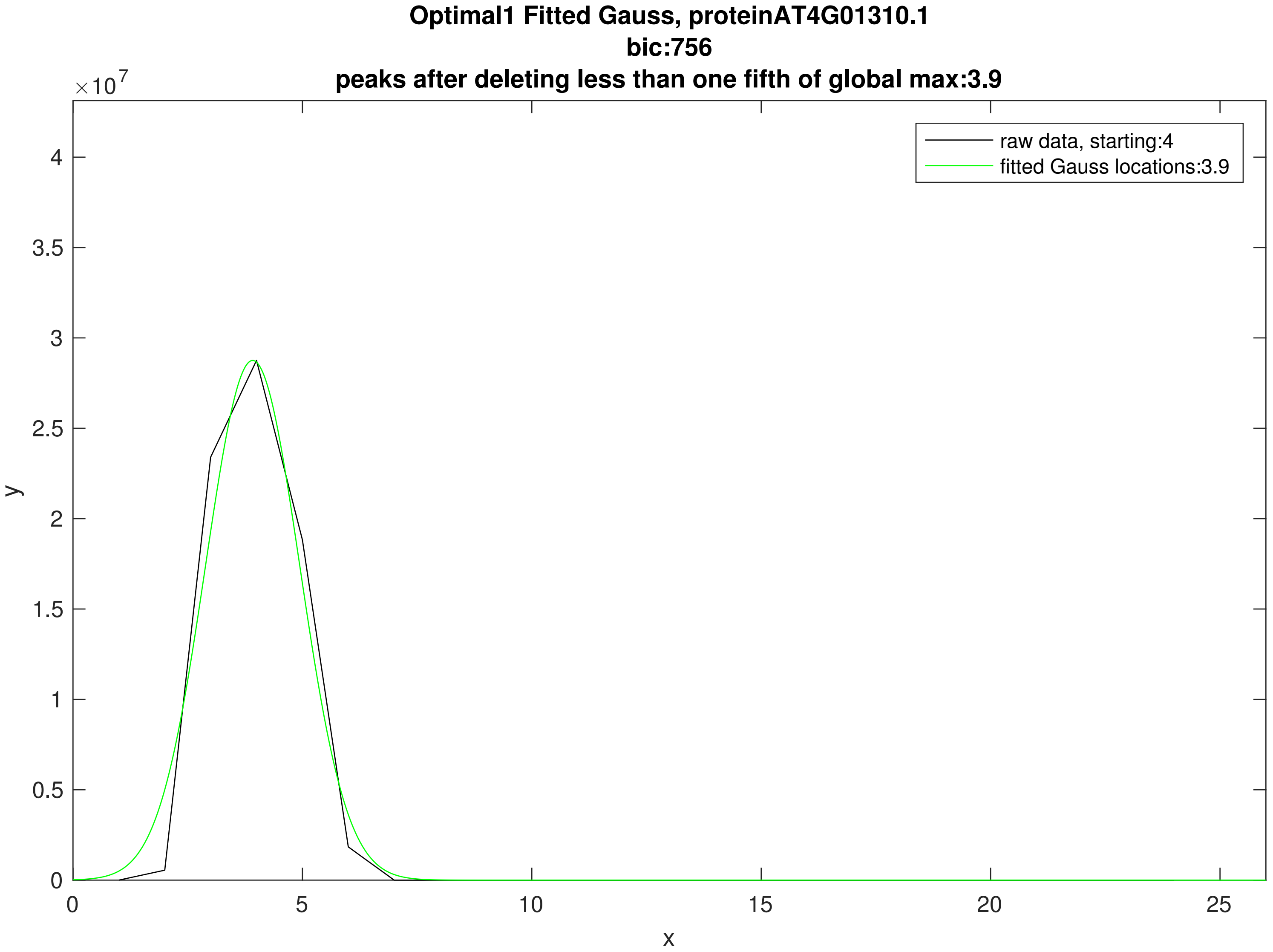

Optimal1 Fitted Gauss, proteinAT4G01310.1
bic:756
peaks after deleting less than one fifth of global max:3.9
×10^7
raw data, starting:4
fitted Gauss locations:3.9
y
x
0
0.5
1
1.5
2
2.5
3
3.5
4
0
5
10
15
20
25

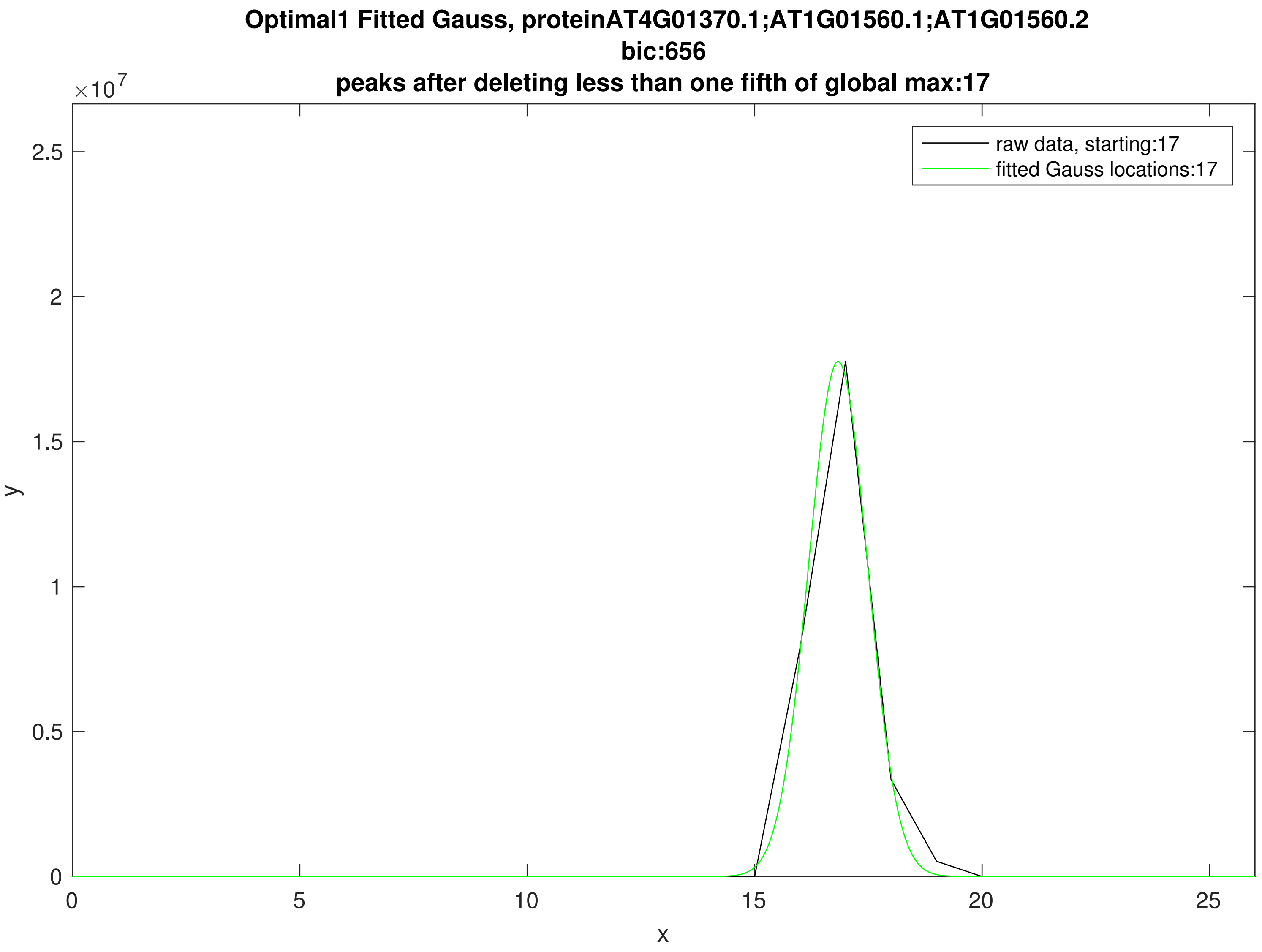
Optimal1 Fitted Gauss, proteinAT4G01370.1;AT1G01560.1;AT1G01560.2
bic:656
peaks after deleting less than one fifth of global max:17
×10^7
2.5
2
1.5
1
0.5
0
y
0
5
10
15
20
25
x
raw data, starting:17
fitted Gauss locations:17

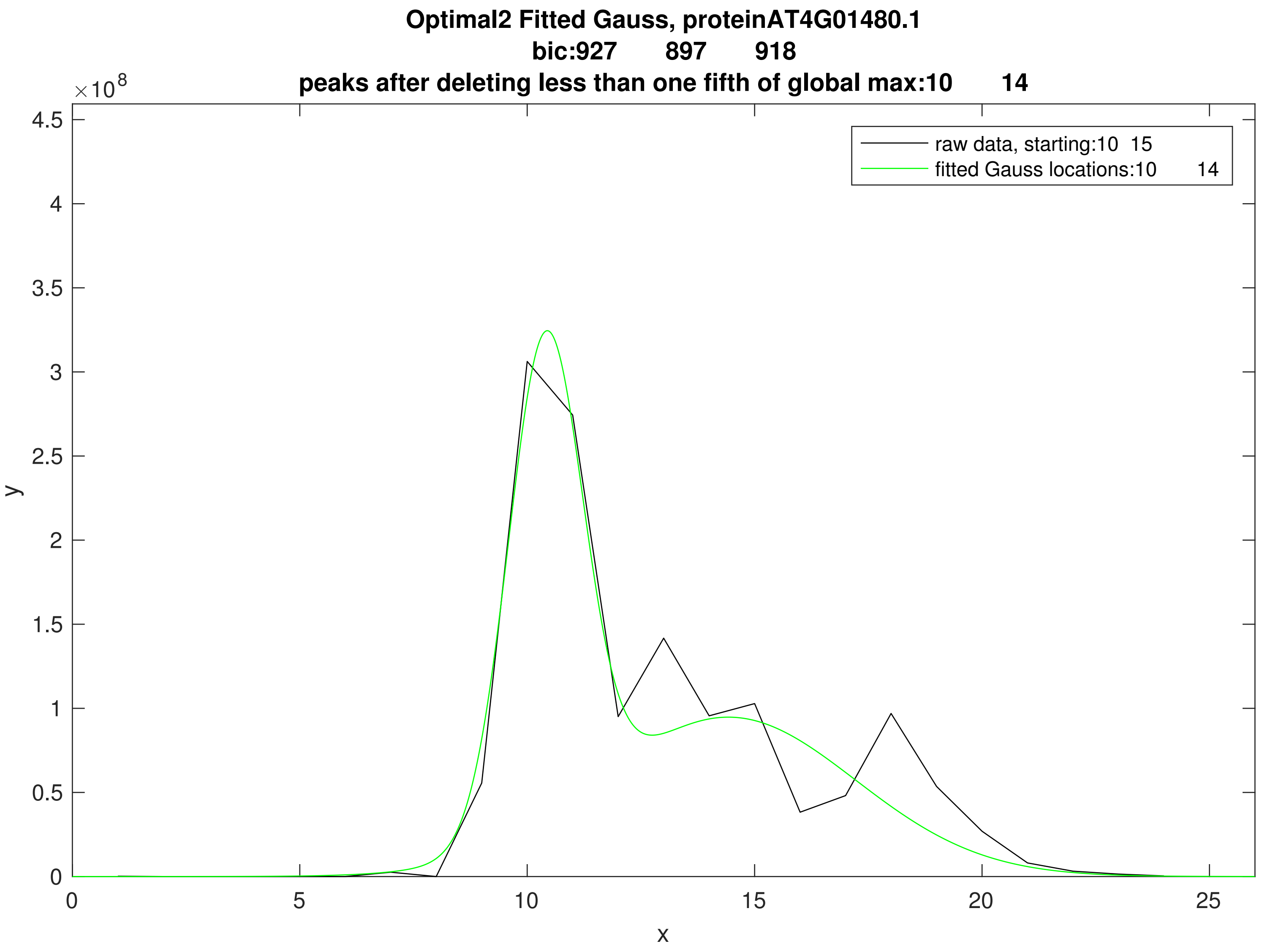
Optimal2 Fitted Gauss, proteinAT4G01480.1
bic:927 897 918
peaks after deleting less than one fifth of global max:10 14
×10^8
raw data, starting:10 15
fitted Gauss locations:10 14
y
x
0
0.5
1
1.5
2
2.5
3
3.5
4
4.5
0
5
10
15
20
25

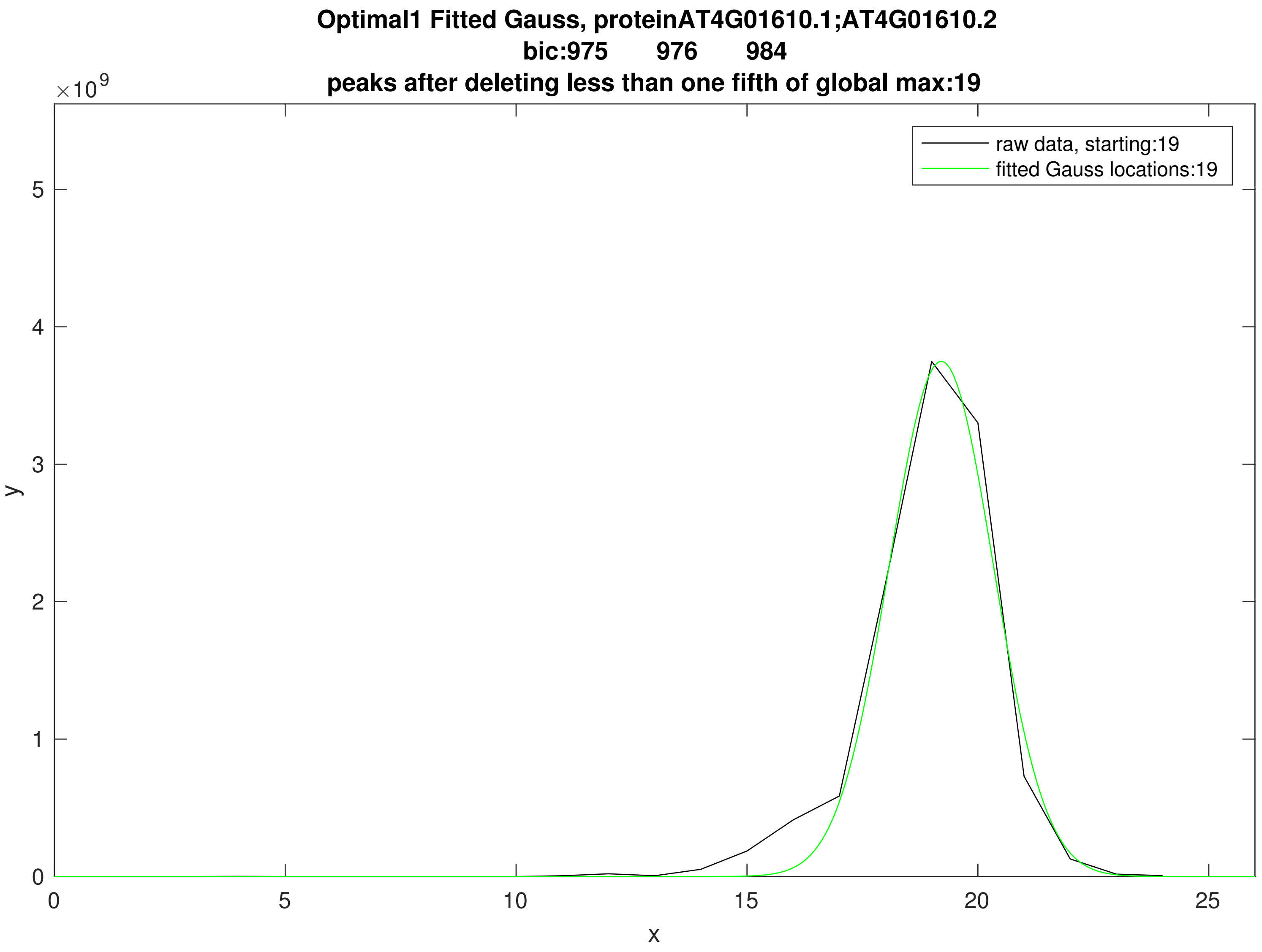
Optimal1 Fitted Gauss, proteinAT4G01610.1;AT4G01610.2
bic:975 976 984
peaks after deleting less than one fifth of global max:19
×10^9
raw data, starting:19
fitted Gauss locations:19
y
x
0
1
2
3
4
5
0
5
10
15
20
25

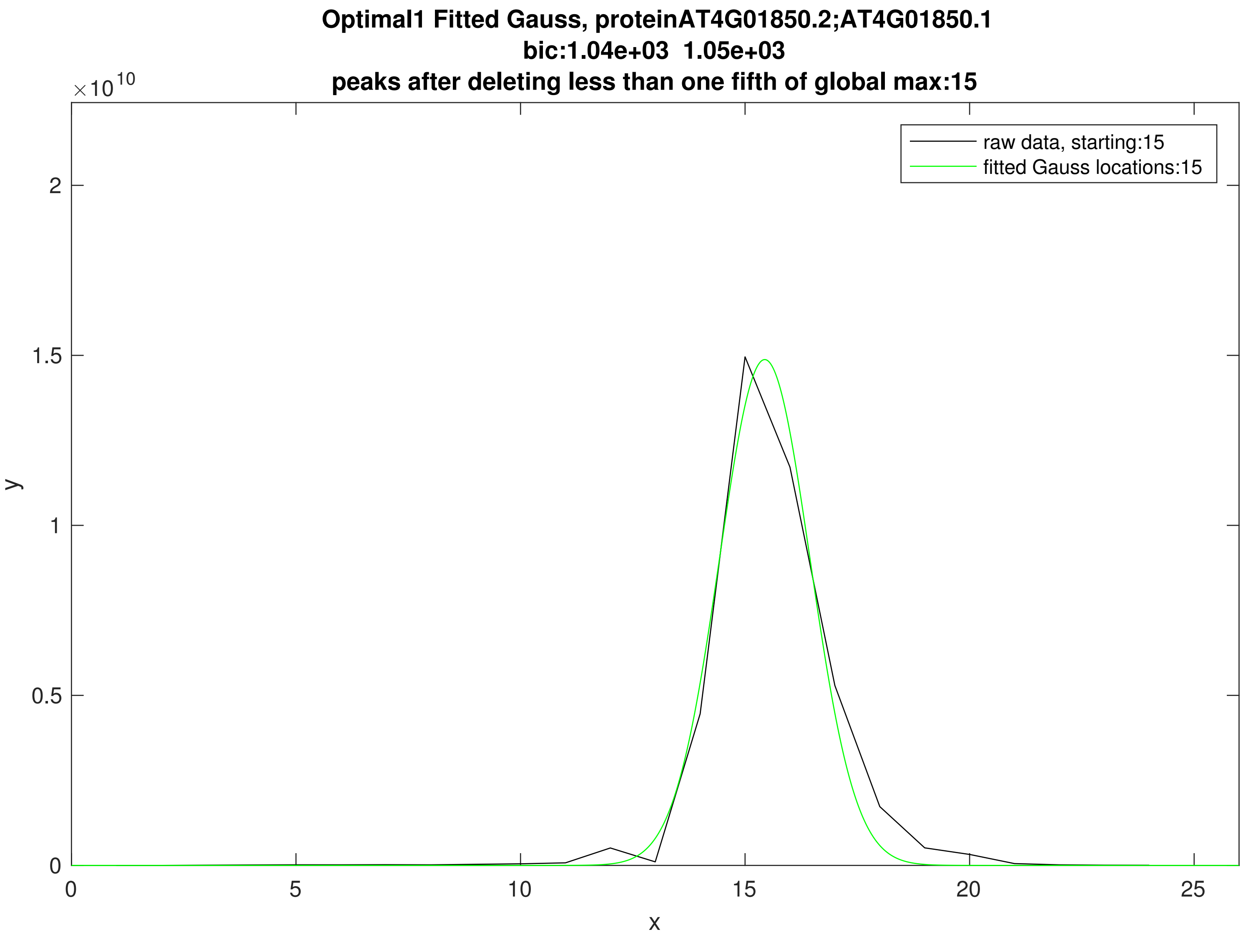
Optimal1 Fitted Gauss, proteinAT4G01850.2;AT4G01850.1
bic:1.04e+03  1.05e+03
peaks after deleting less than one fifth of global max:15
×10^10
raw data, starting:15
fitted Gauss locations:15
y
x
0
0.5
1
1.5
2
5
10
15
20
25

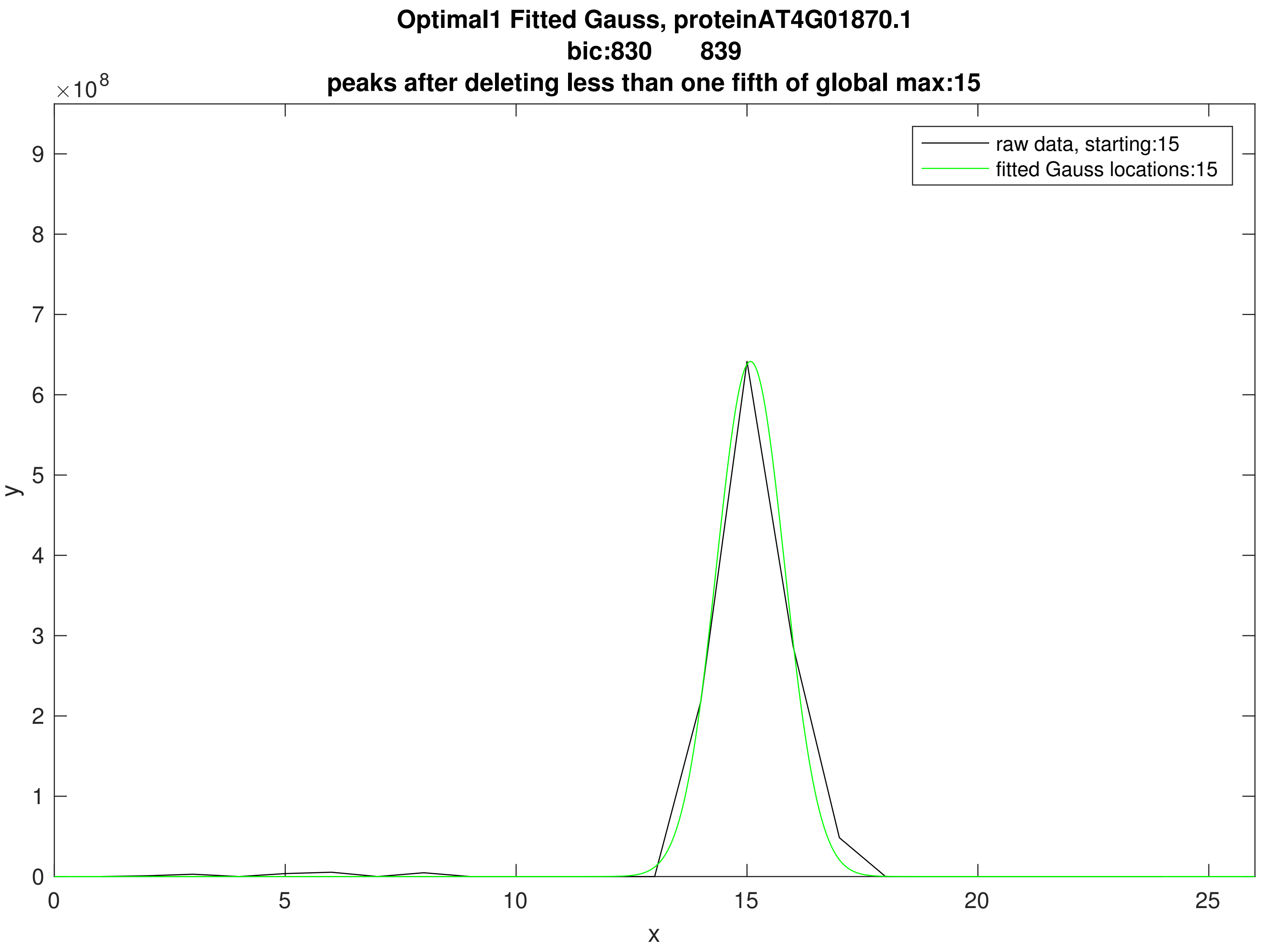
Optimal1 Fitted Gauss, proteinAT4G01870.1
bic:830 839
peaks after deleting less than one fifth of global max:15
×10^8
raw data, starting:15
fitted Gauss locations:15
y
x
0
1
2
3
4
5
6
7
8
9
0
5
10
15
20
25

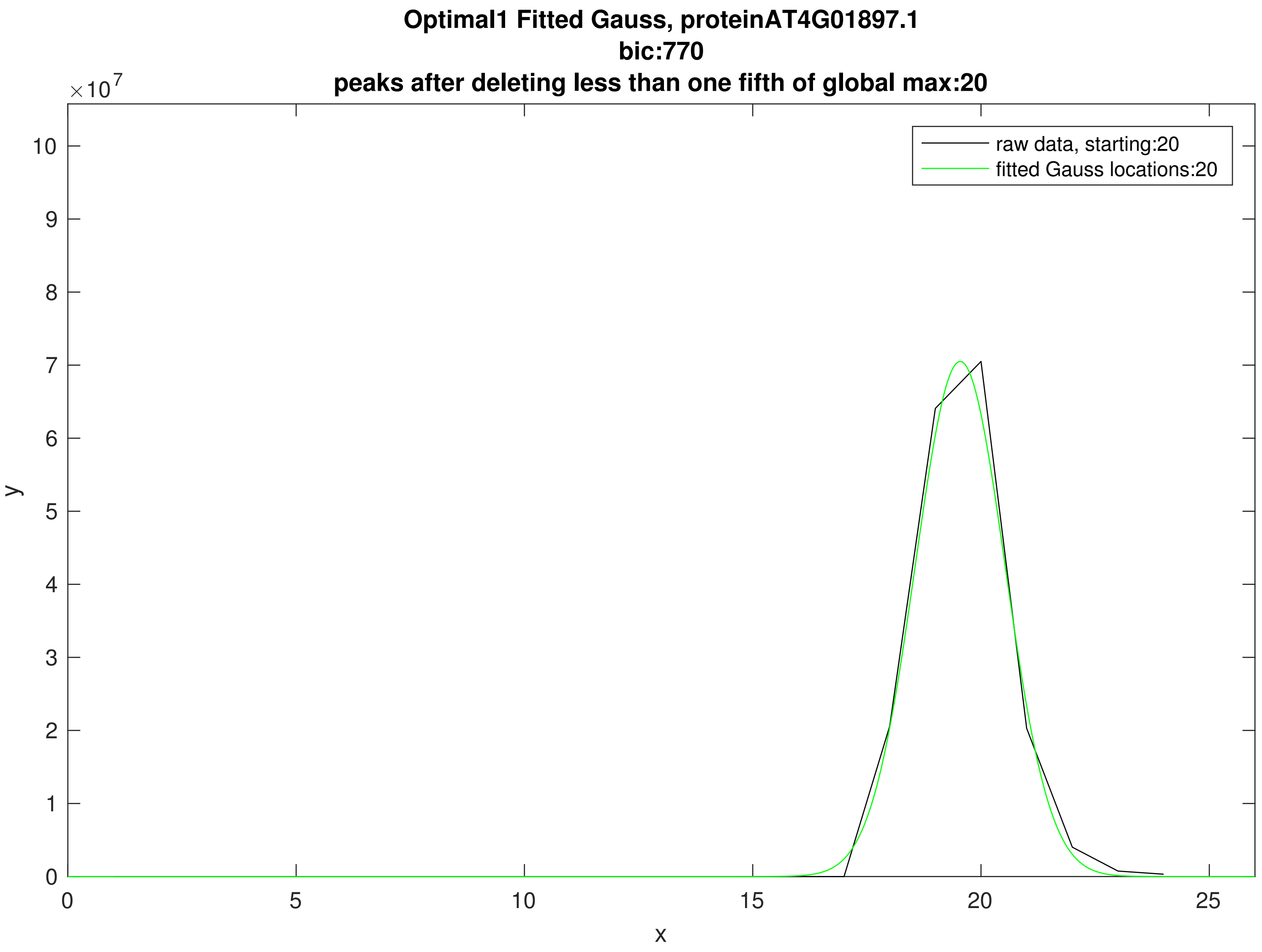
Optimal1 Fitted Gauss, proteinAT4G01897.1
bic:770
peaks after deleting less than one fifth of global max:20
×10^7
raw data, starting:20
fitted Gauss locations:20
y
x
0
1
2
3
4
5
6
7
8
9
10
0
5
10
15
20
25

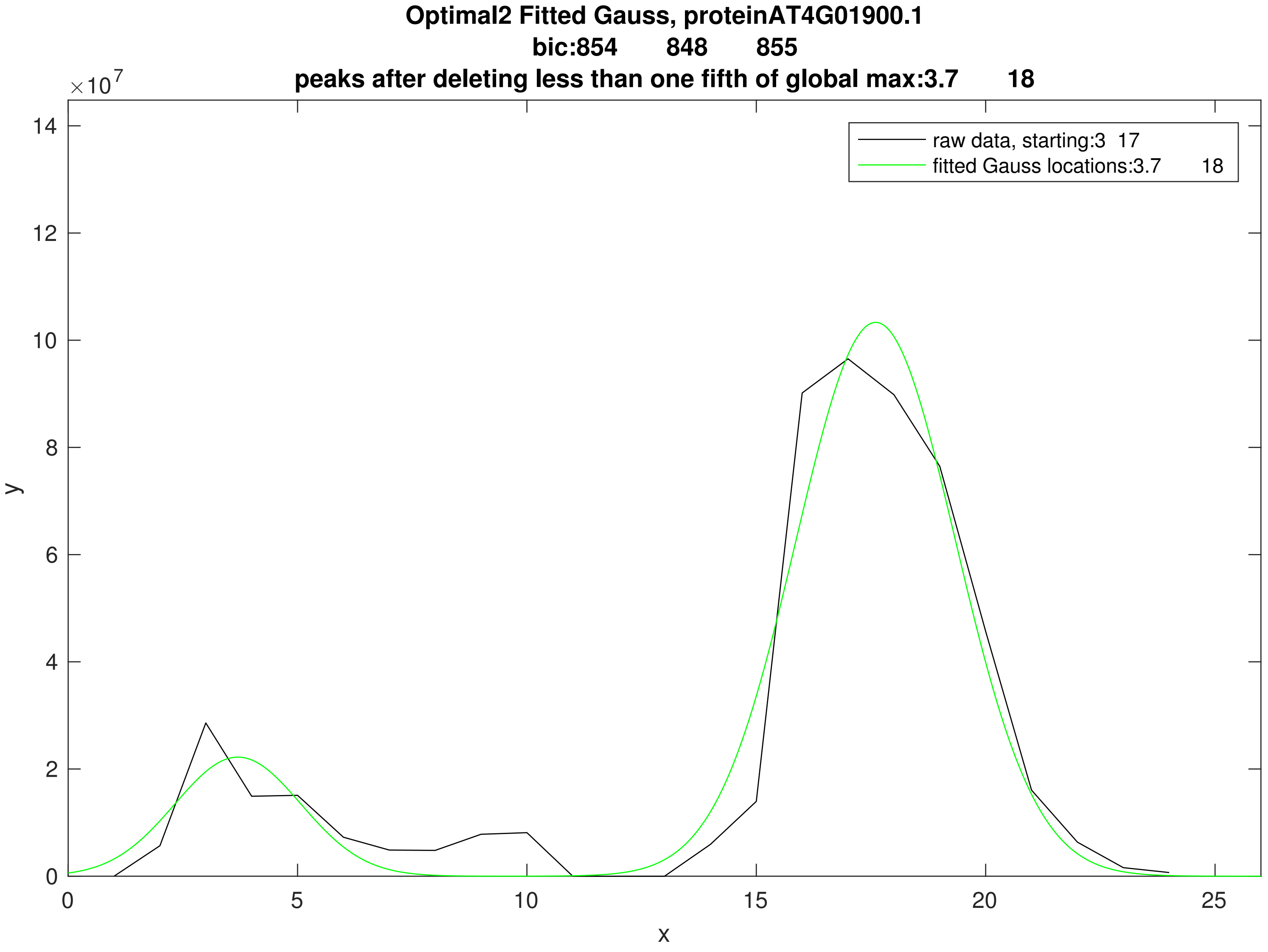
Optimal2 Fitted Gauss, proteinAT4G01900.1
bic:854 848 855
peaks after deleting less than one fifth of global max:3.7 18
×10^7
raw data, starting:3 17
fitted Gauss locations:3.7 18
y
x
0
2
4
6
8
10
12
14
5
15
20
25

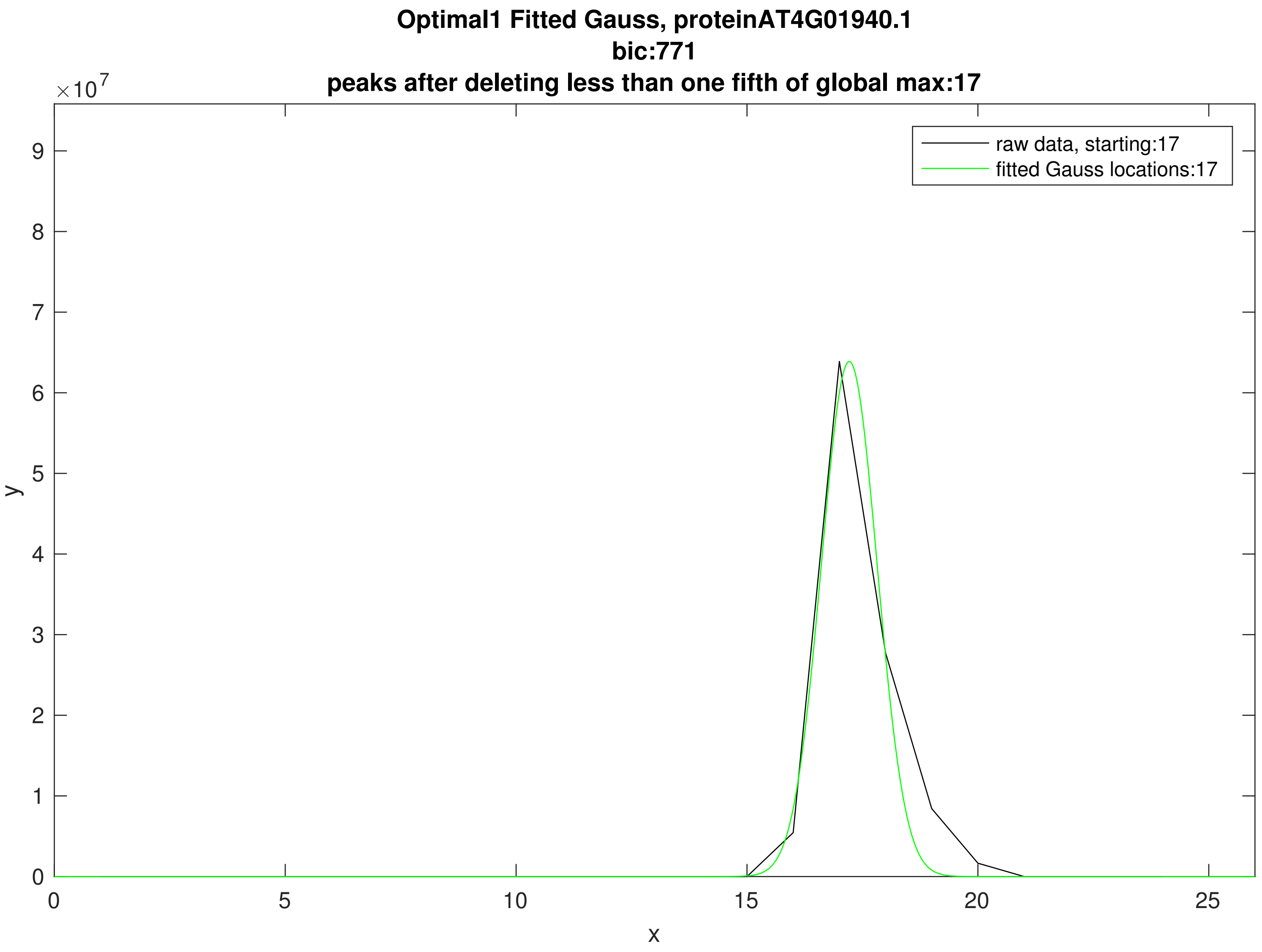
Optimal1 Fitted Gauss, proteinAT4G01940.1
bic:771
peaks after deleting less than one fifth of global max:17
×10^7
raw data, starting:17
fitted Gauss locations:17
y
x
0
1
2
3
4
5
6
7
8
9
0
5
10
15
20
25

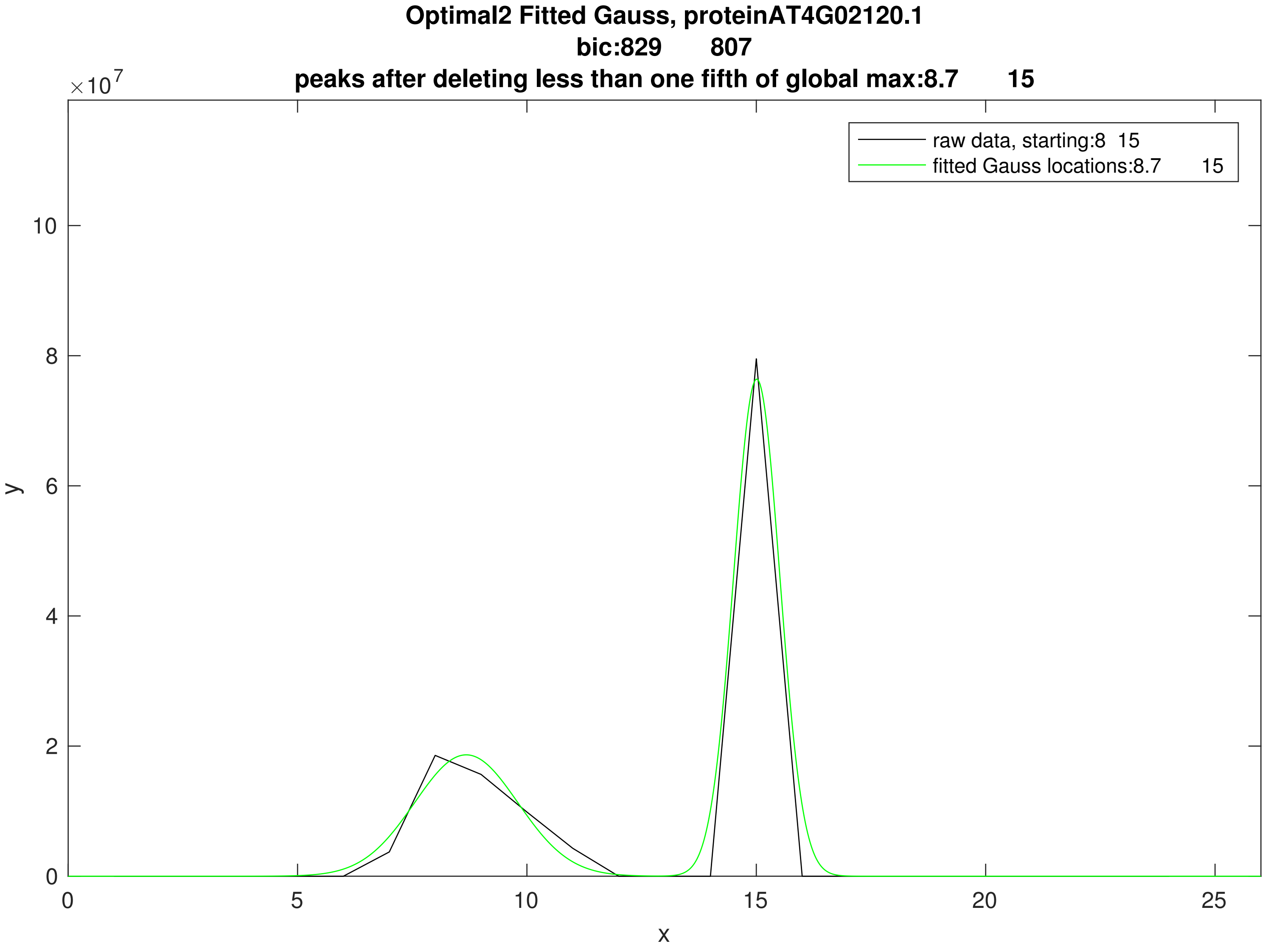
Optimal2 Fitted Gauss, proteinAT4G02120.1
bic:829 807
peaks after deleting less than one fifth of global max:8.7 15
×10^7
raw data, starting:8 15
fitted Gauss locations:8.7 15
y
x
0
2
4
6
8
10
0
5
10
15
20
25

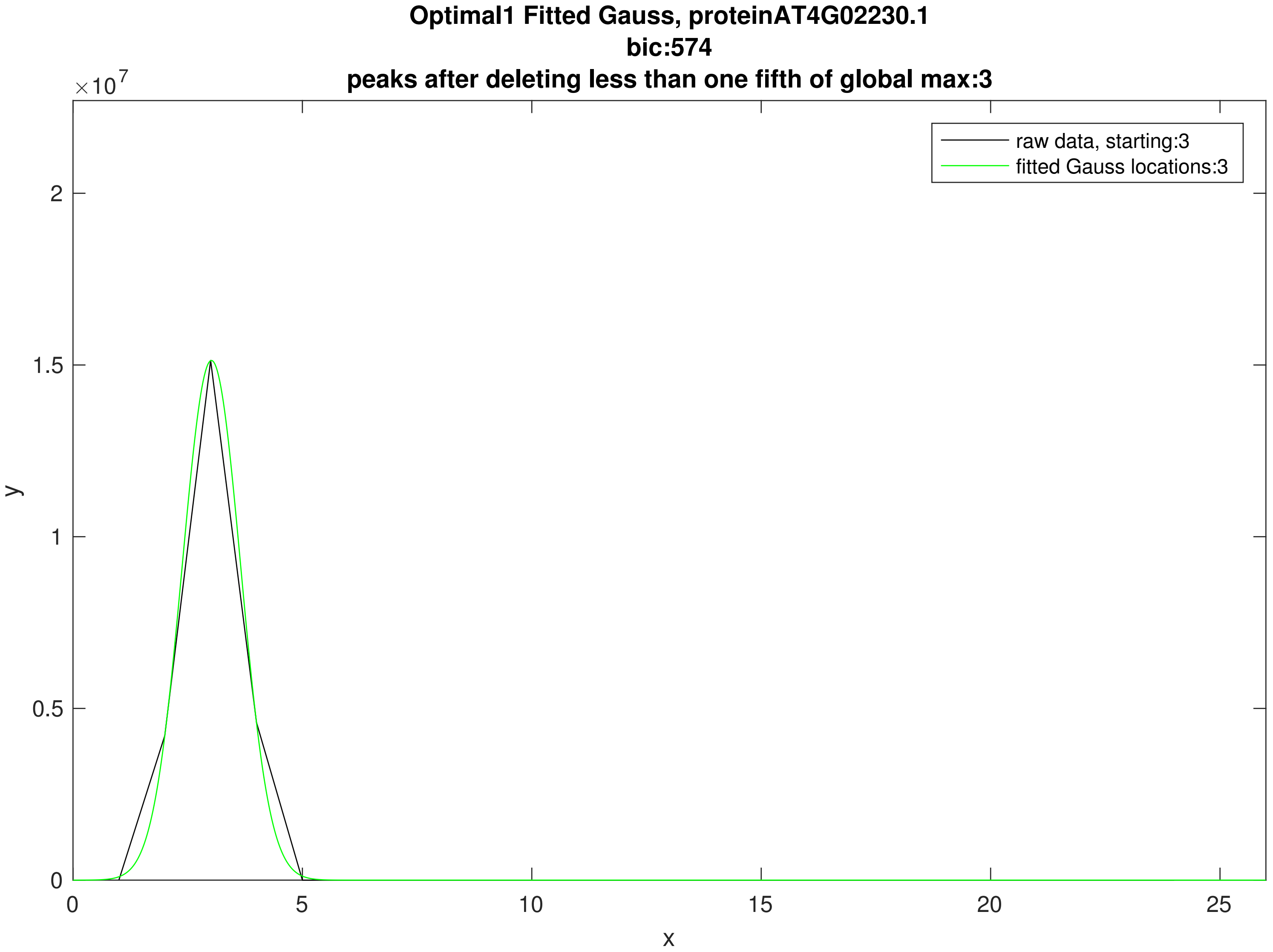
Optimal1 Fitted Gauss, proteinAT4G02230.1
bic:574
peaks after deleting less than one fifth of global max:3
×10^7
raw data, starting:3
fitted Gauss locations:3
y
x
0
0.5
1
1.5
2
0
5
10
15
20
25

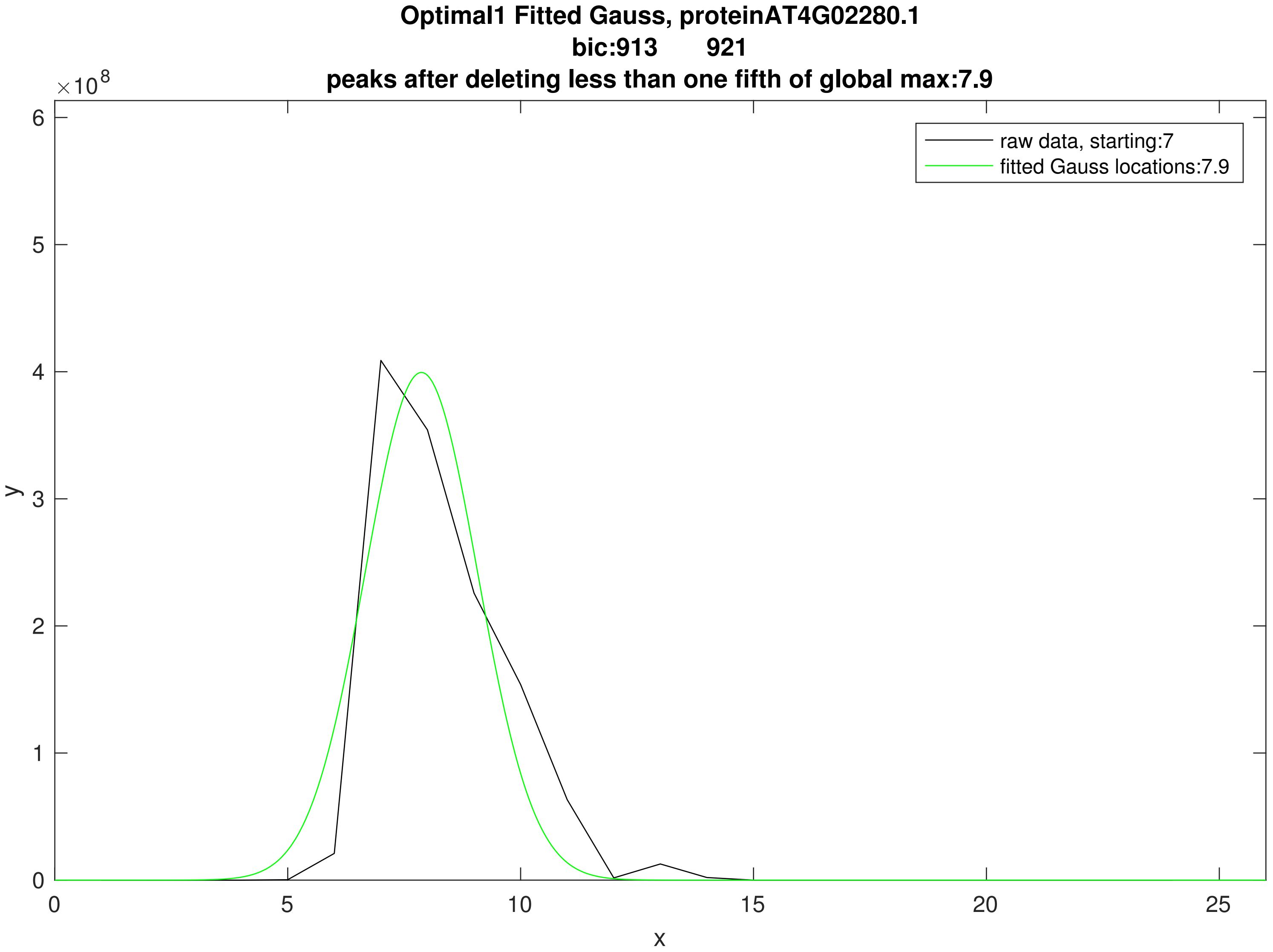
Optimal1 Fitted Gauss, proteinAT4G02280.1
bic:913 921
peaks after deleting less than one fifth of global max:7.9
×10^8
6
5
4
3
2
1
0
y
0
5
10
15
20
25
x
raw data, starting:7
fitted Gauss locations:7.9

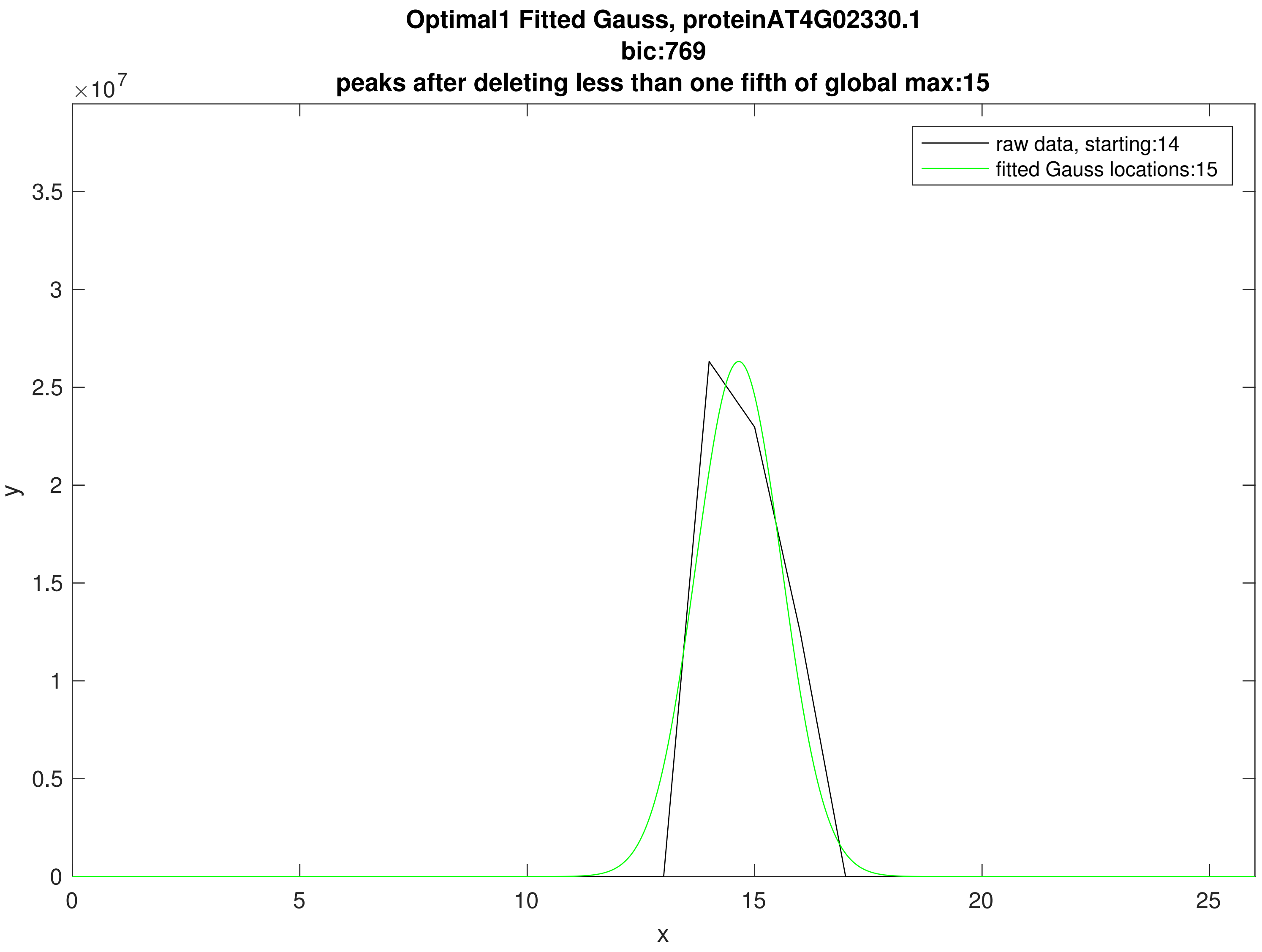

Optimal1 Fitted Gauss, proteinAT4G02330.1
bic:769
peaks after deleting less than one fifth of global max:15
×10^7
raw data, starting:14
fitted Gauss locations:15
3.5
3
2.5
2
1.5
1
0.5
0
y
0
5
10
15
20
25
x

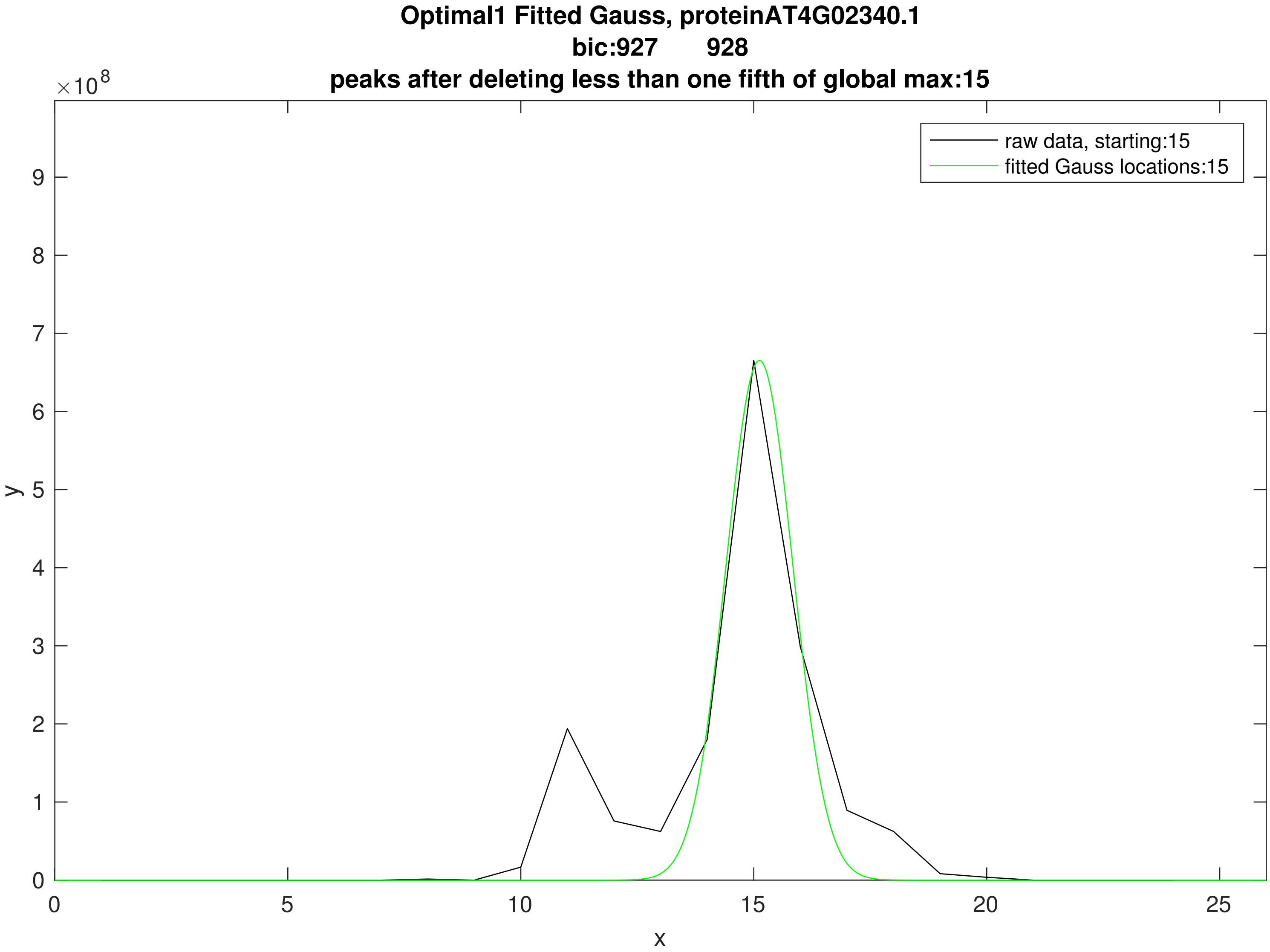

Optimal1 Fitted Gauss, proteinAT4G02340.1
bic:927 928
peaks after deleting less than one fifth of global max:15
×10^8
raw data, starting:15
fitted Gauss locations:15
y
x
0
1
2
3
4
5
6
7
8
9
0
5
10
15
20
25

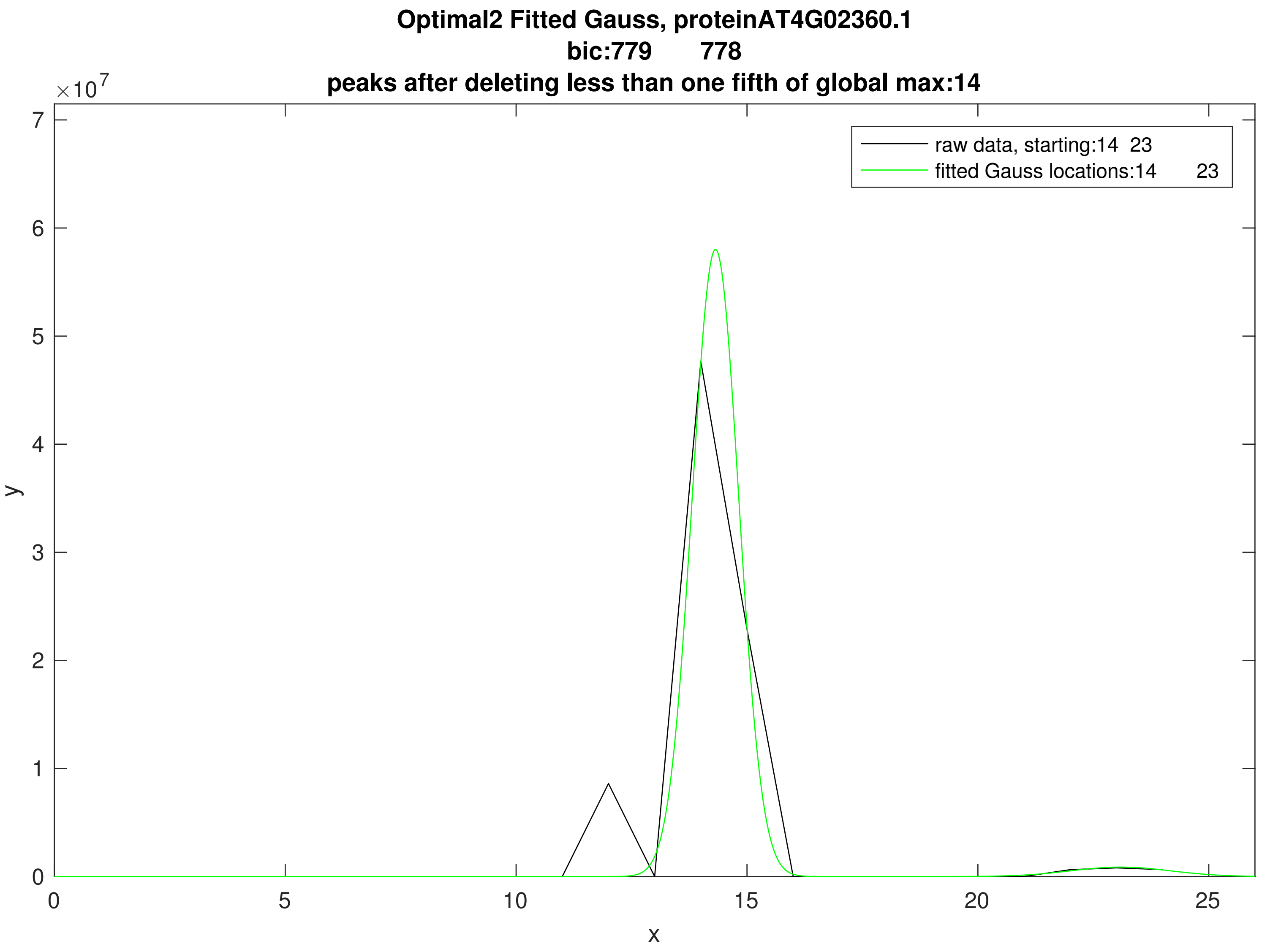

Optimal2 Fitted Gauss, proteinAT4G02360.1
bic:779 778
peaks after deleting less than one fifth of global max:14
×10^7
7
6
5
4
3
2
1
0
y
0
5
10
15
20
25
x
raw data, starting:14 23
fitted Gauss locations:14 23

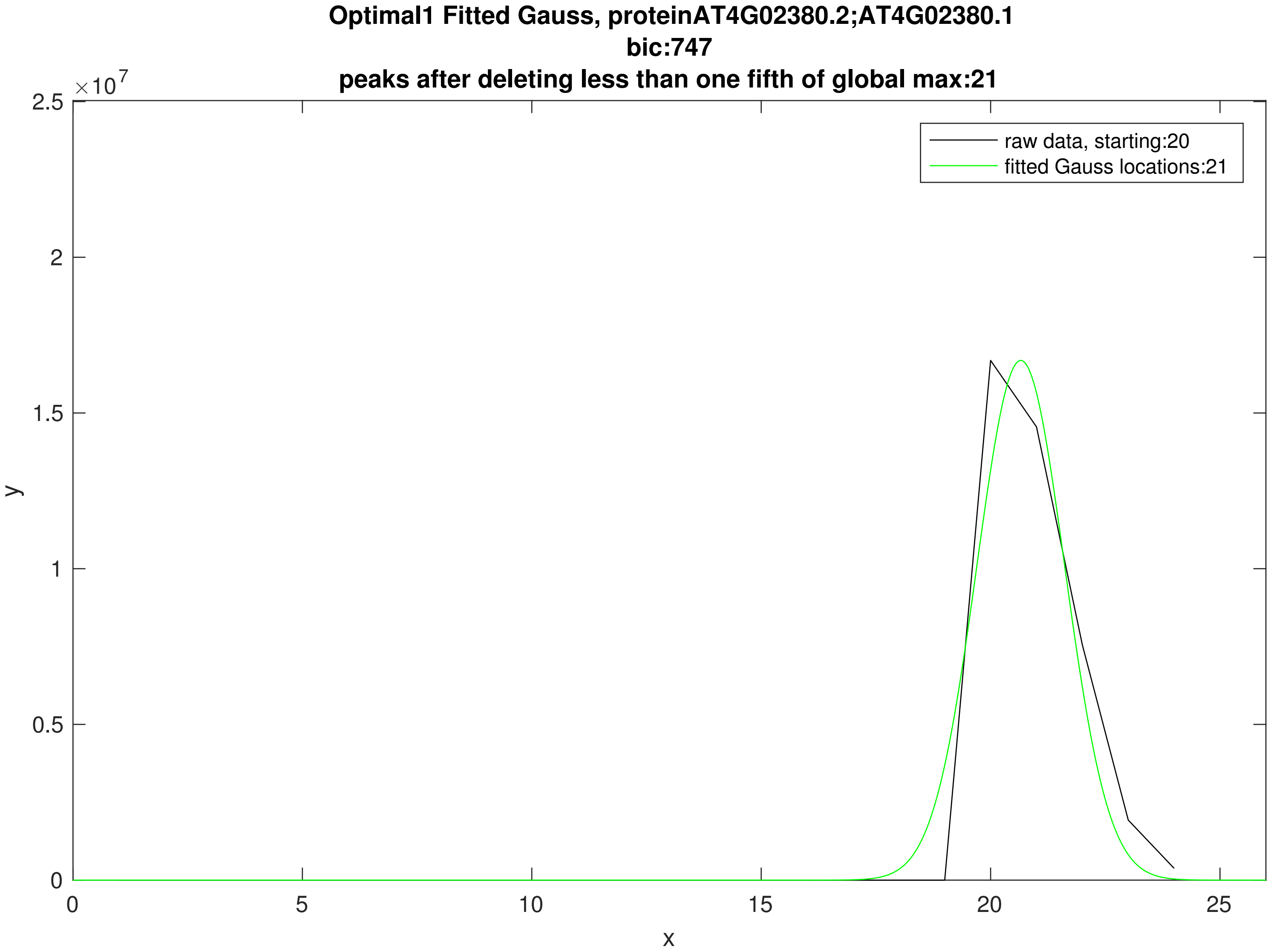

Optimal1 Fitted Gauss, proteinAT4G02380.2;AT4G02380.1
bic:747
peaks after deleting less than one fifth of global max:21
×10^7
2.5
2
1.5
1
0.5
0
y
0
5
10
15
20
25
x
raw data, starting:20
fitted Gauss locations:21

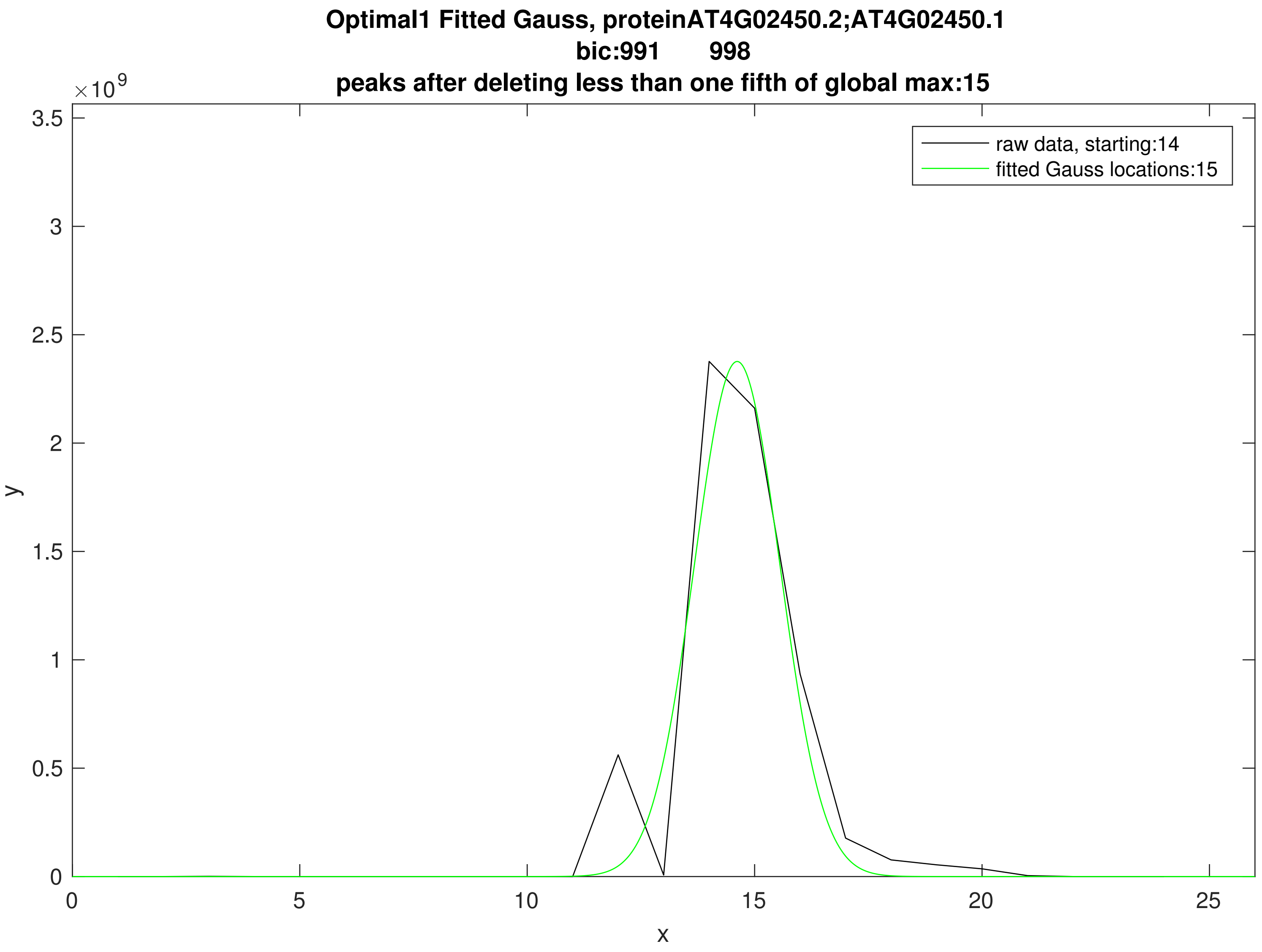

Optimal1 Fitted Gauss, proteinAT4G02450.2;AT4G02450.1
bic:991 998
peaks after deleting less than one fifth of global max:15
×10^9
3.5
3
2.5
2
1.5
1
0.5
0
y
0
5
10
15
20
25
x
raw data, starting:14
fitted Gauss locations:15

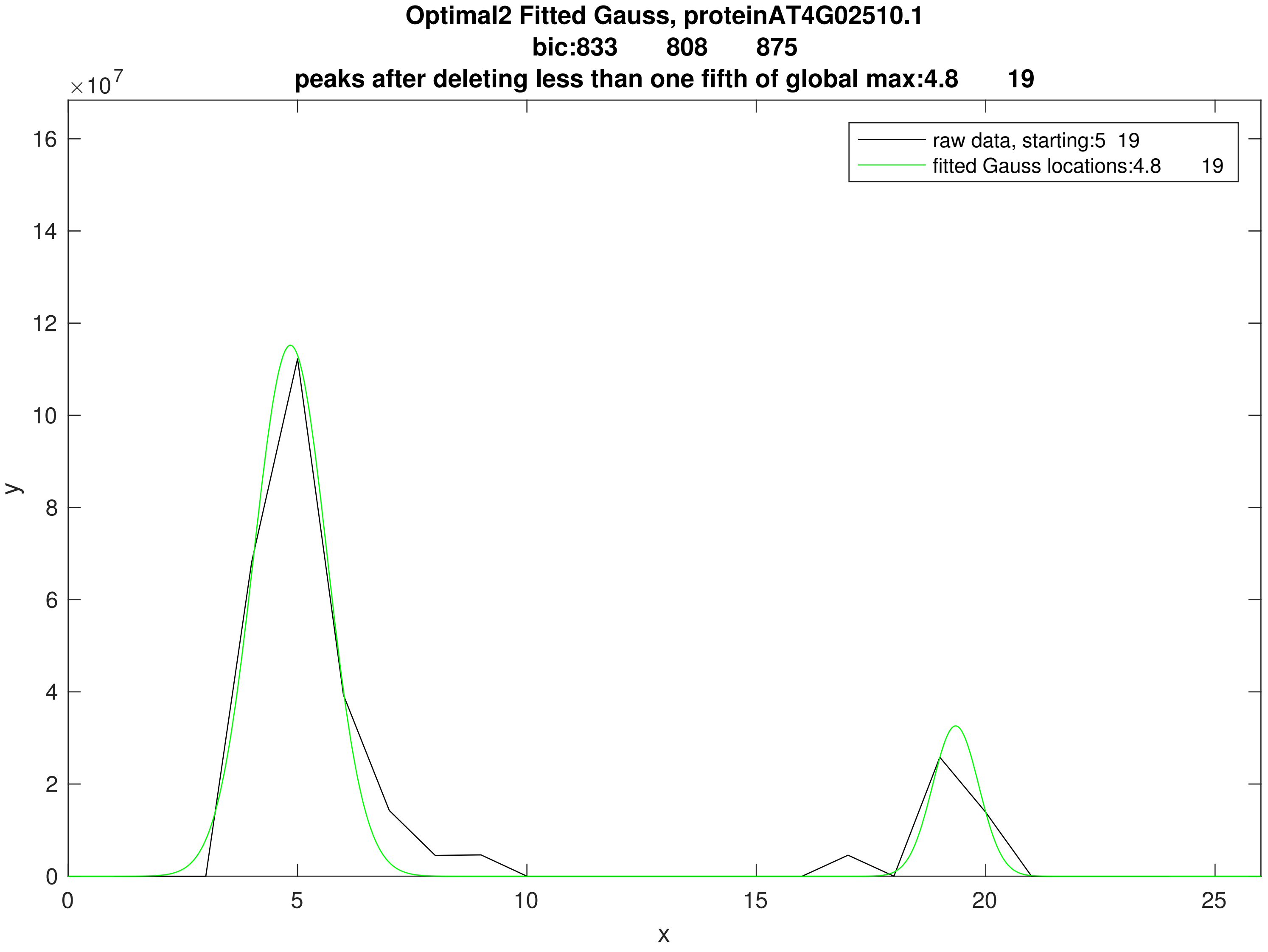
Optimal2 Fitted Gauss, proteinAT4G02510.1
bic:833 808 875
peaks after deleting less than one fifth of global max:4.8 19
×10^7
raw data, starting:5 19
fitted Gauss locations:4.8 19
y
x
0
2
4
6
8
10
12
14
16
0
5
10
15
20
25

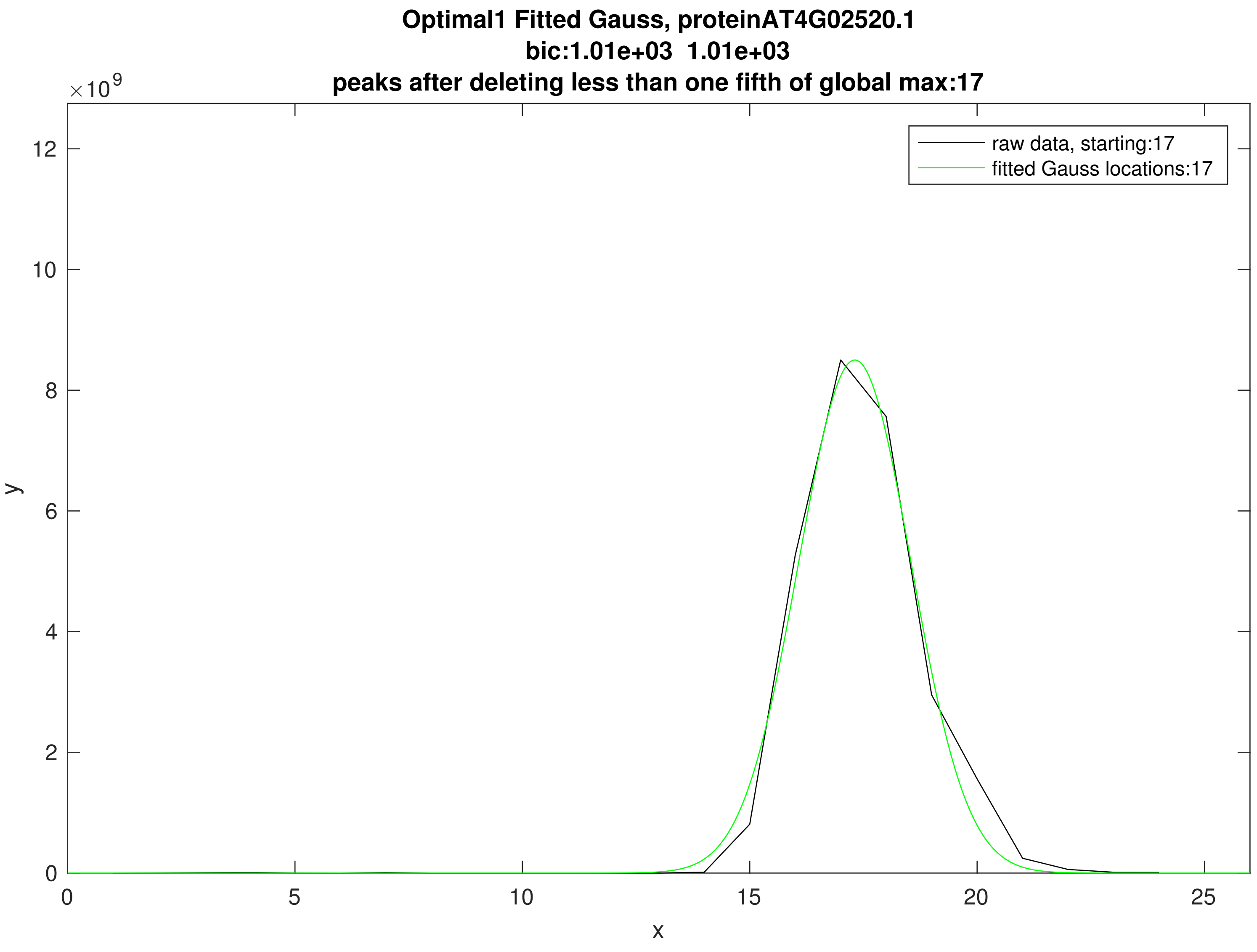
Optimal1 Fitted Gauss, proteinAT4G02520.1
bic:1.01e+03  1.01e+03
peaks after deleting less than one fifth of global max:17
×10^9
12
10
8
6
4
2
0
y
0
5
10
15
20
25
x
raw data, starting:17
fitted Gauss locations:17

#### Optimal1 Fitted Gauss, proteinAT4G02530.1
#### bic:848
#### peaks after deleting less than one fifth of global max:18

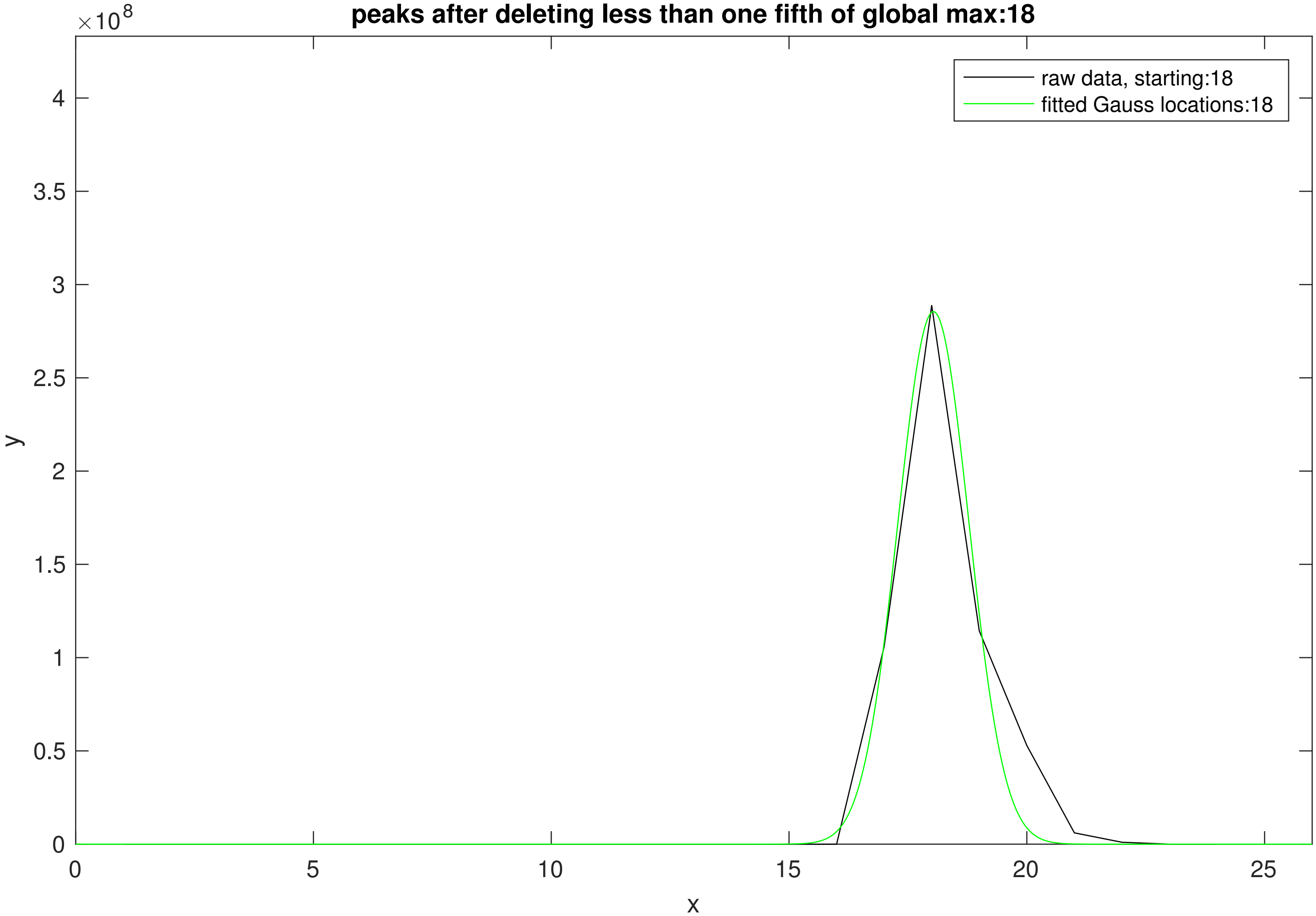

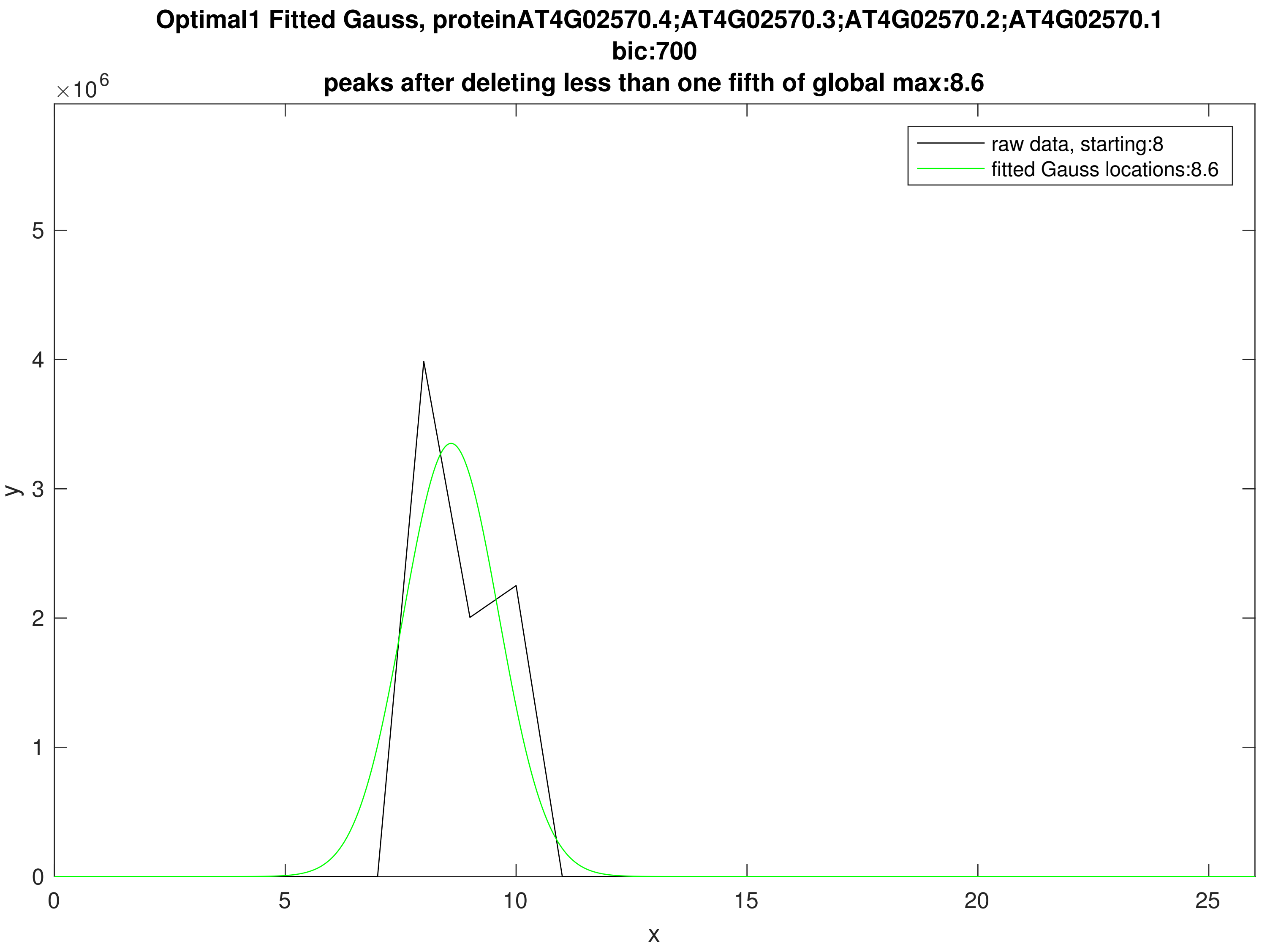
Optimal1 Fitted Gauss, proteinAT4G02570.4;AT4G02570.3;AT4G02570.2;AT4G02570.1
bic:700
peaks after deleting less than one fifth of global max:8.6
×10^6
raw data, starting:8
fitted Gauss locations:8.6
y
x
0
1
2
3
4
5
0
5
10
15
20
25

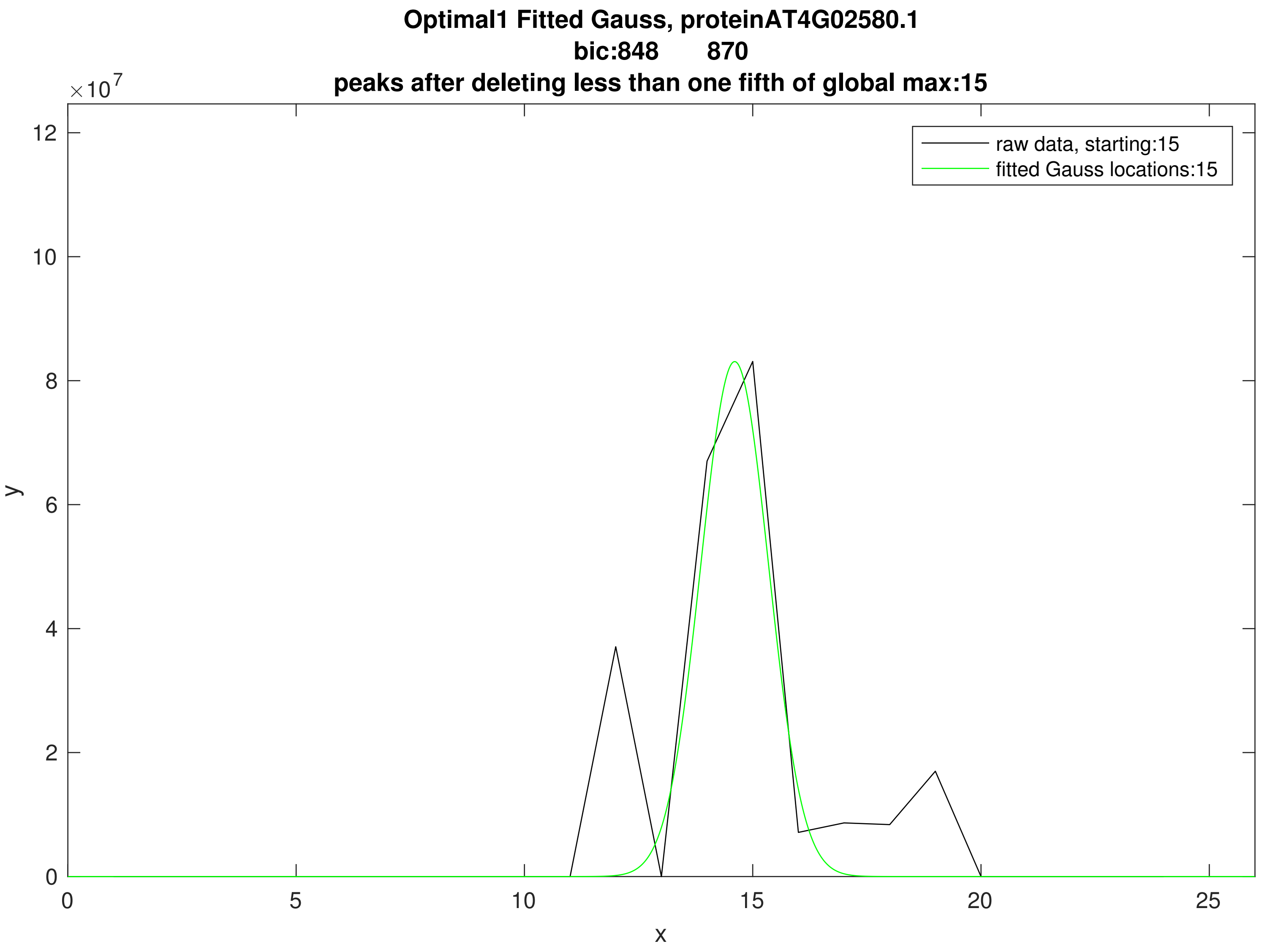

Optimal1 Fitted Gauss, proteinAT4G02580.1
bic:848 870
peaks after deleting less than one fifth of global max:15
×10^7
12
10
8
6
4
2
0
y
0
5
10
15
20
25
x
raw data, starting:15
fitted Gauss locations:15

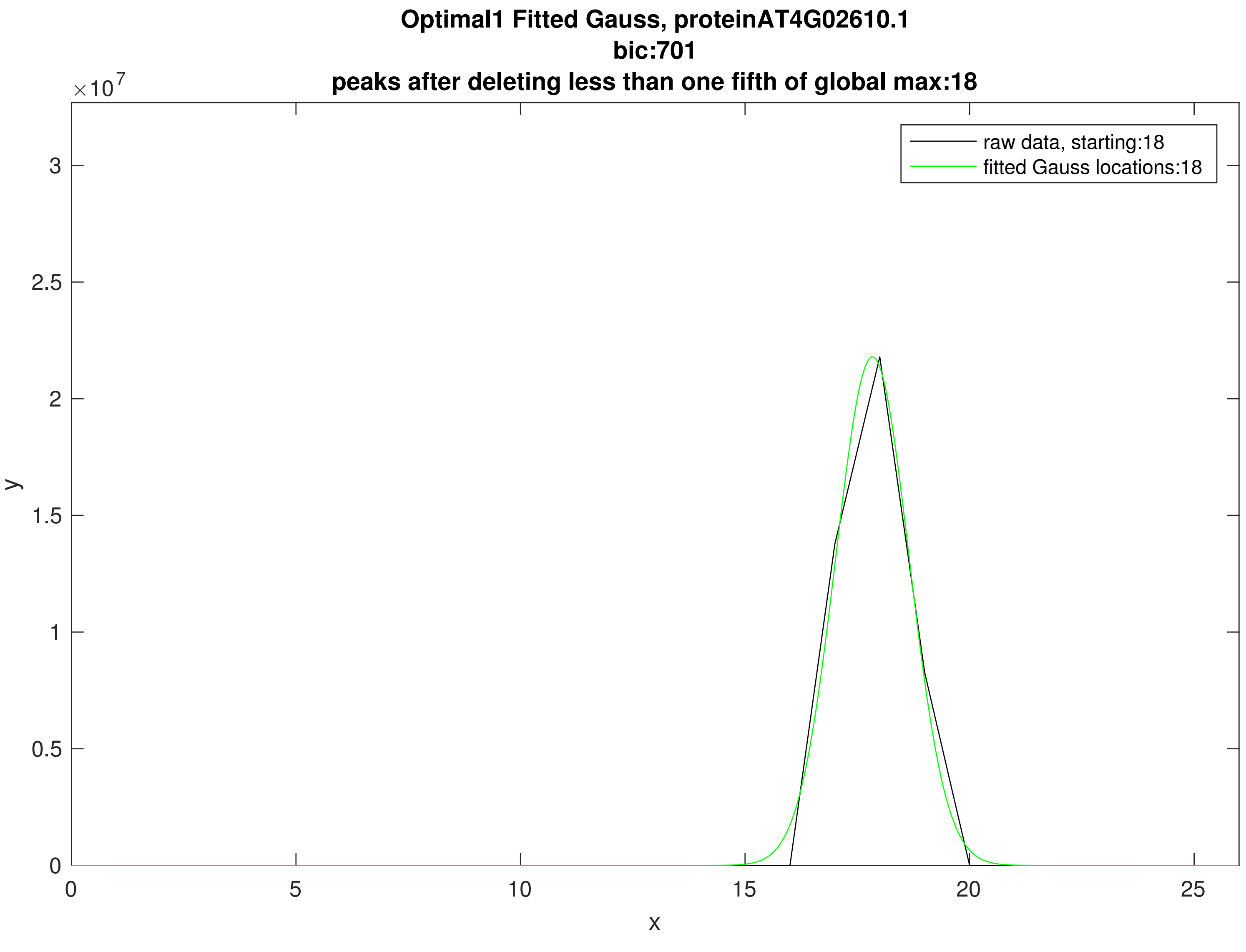
Optimal1 Fitted Gauss, proteinAT4G02610.1
bic:701
peaks after deleting less than one fifth of global max:18
×10^7
raw data, starting:18
fitted Gauss locations:18
y
x
0
0.5
1
1.5
2
2.5
3
0
5
10
15
20
25

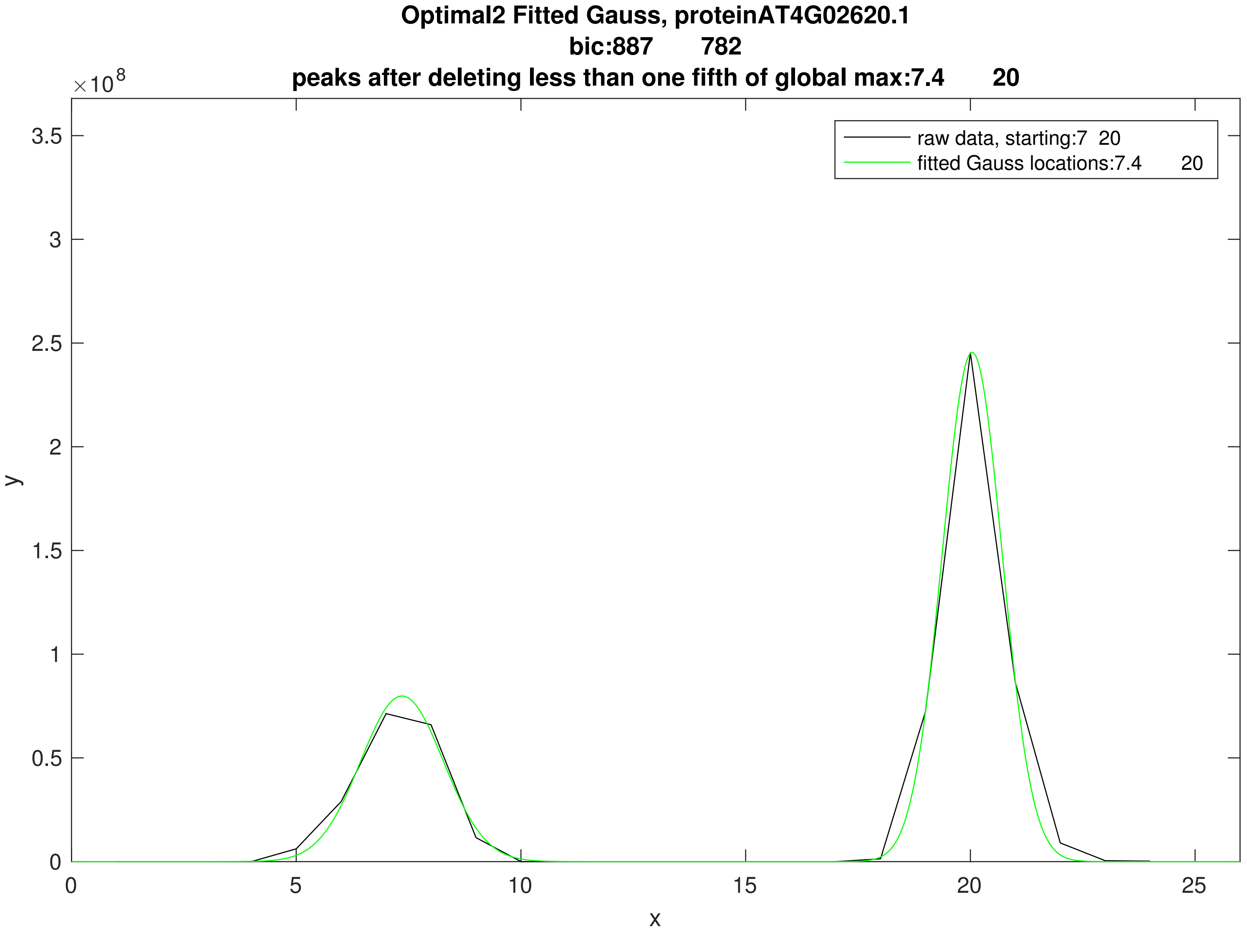

Optimal2 Fitted Gauss, proteinAT4G02620.1
bic:887 782
peaks after deleting less than one fifth of global max:7.4 20
×10^8
3.5
3
2.5
2
1.5
1
0.5
0
y
0
5
10
15
20
25
x
raw data, starting:7 20
fitted Gauss locations:7.4 20

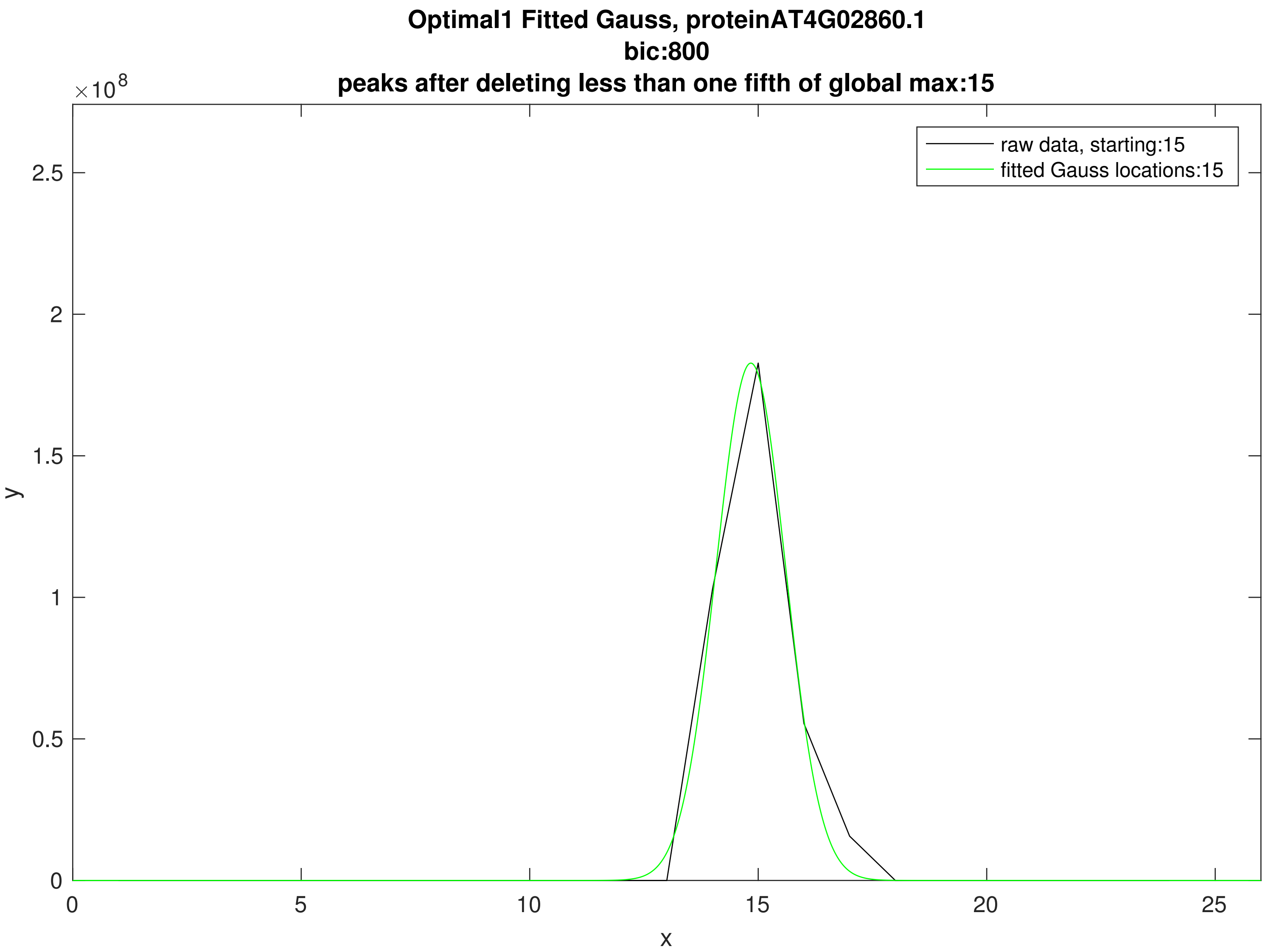

Optimal1 Fitted Gauss, proteinAT4G02860.1
bic:800
peaks after deleting less than one fifth of global max:15
×10^8
2.5
2
1.5
1
0.5
0
y
0
5
10
15
20
25
x
raw data, starting:15
fitted Gauss locations:15

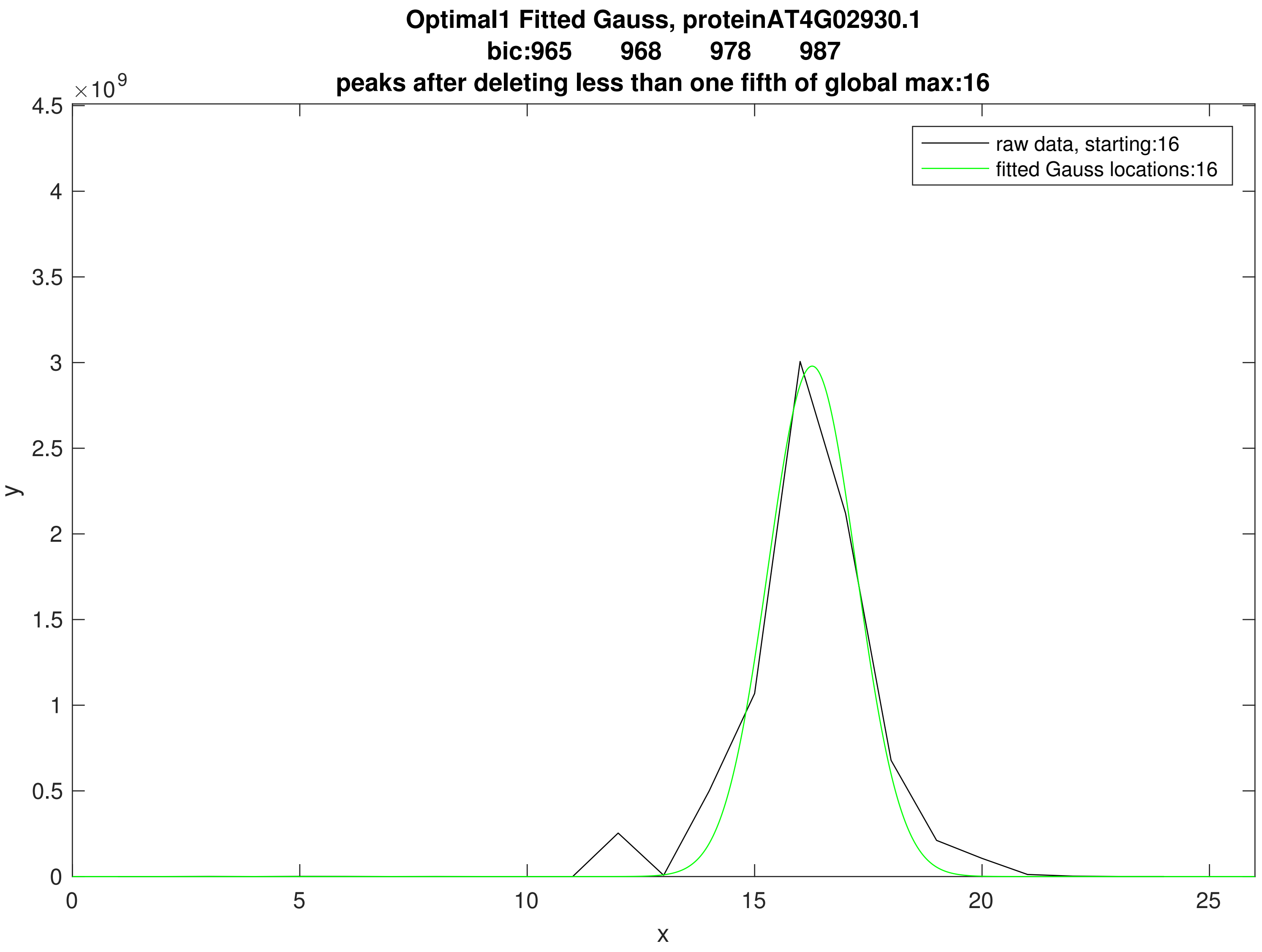

Optimal1 Fitted Gauss, proteinAT4G02930.1
bic:965 968 978 987
peaks after deleting less than one fifth of global max:16
×10^9
4.5
4
3.5
3
2.5
2
1.5
1
0.5
0
y
0
5
10
15
20
25
x
raw data, starting:16
fitted Gauss locations:16

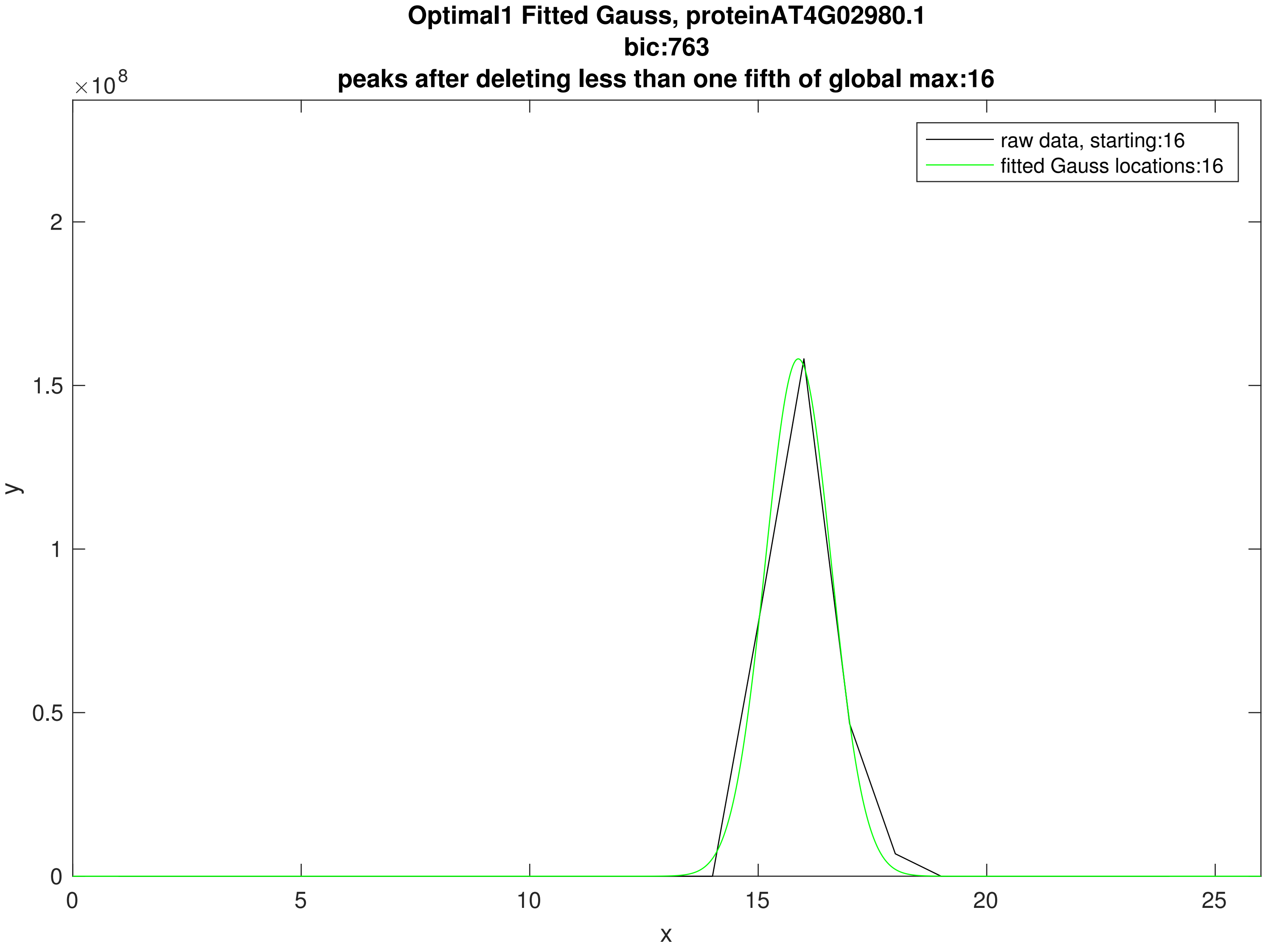
Optimal1 Fitted Gauss, proteinAT4G02980.1
bic:763
peaks after deleting less than one fifth of global max:16
×10^8
raw data, starting:16
fitted Gauss locations:16
y
x
0
0.5
1
1.5
2
0
5
10
15
20
25

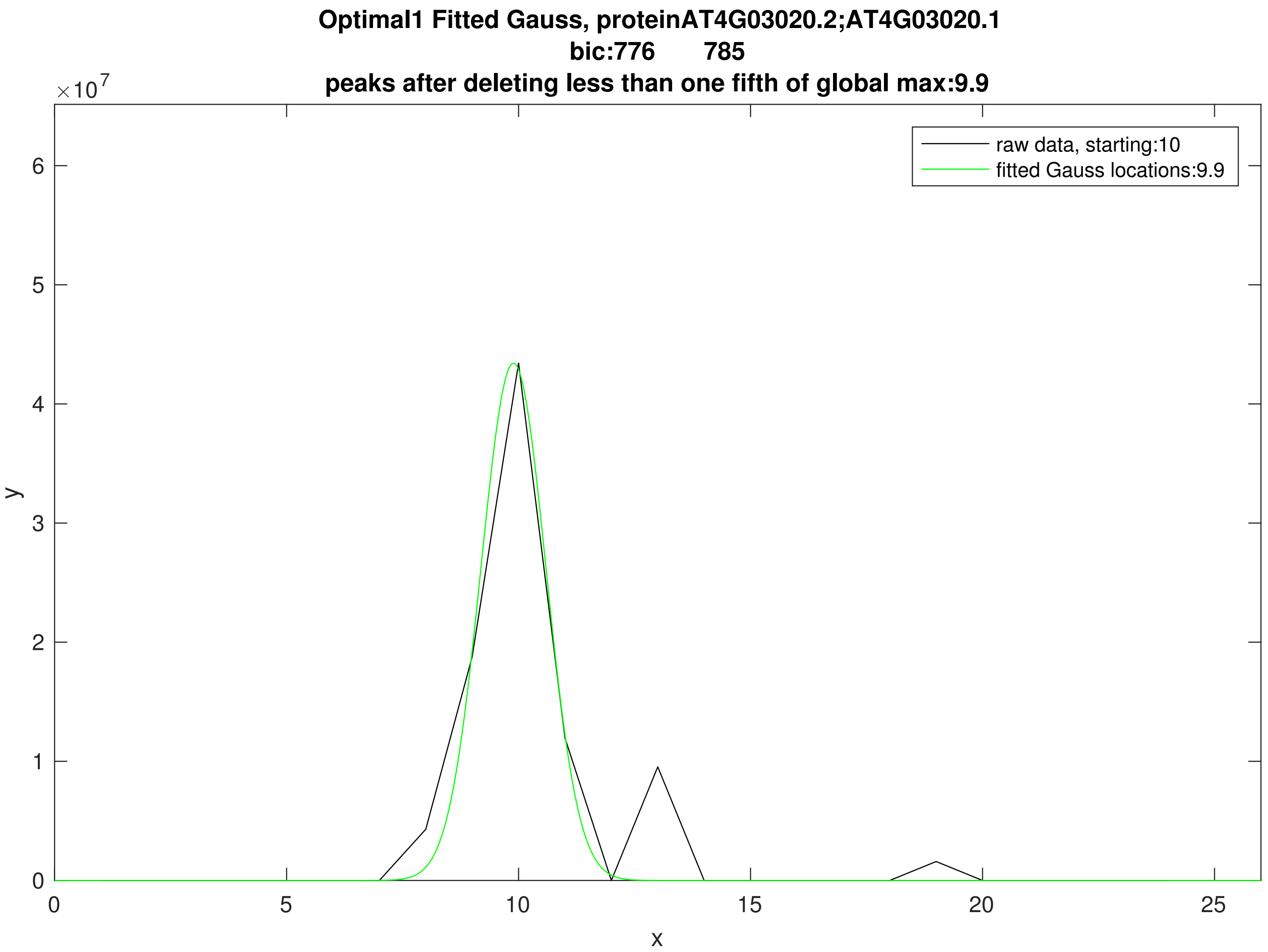

Optimal1 Fitted Gauss, proteinAT4G03020.2;AT4G03020.1
bic:776 785
peaks after deleting less than one fifth of global max:9.9
×10^7
6
5
4
3
2
1
0
y
0
5
10
15
20
25
x
raw data, starting:10
fitted Gauss locations:9.9

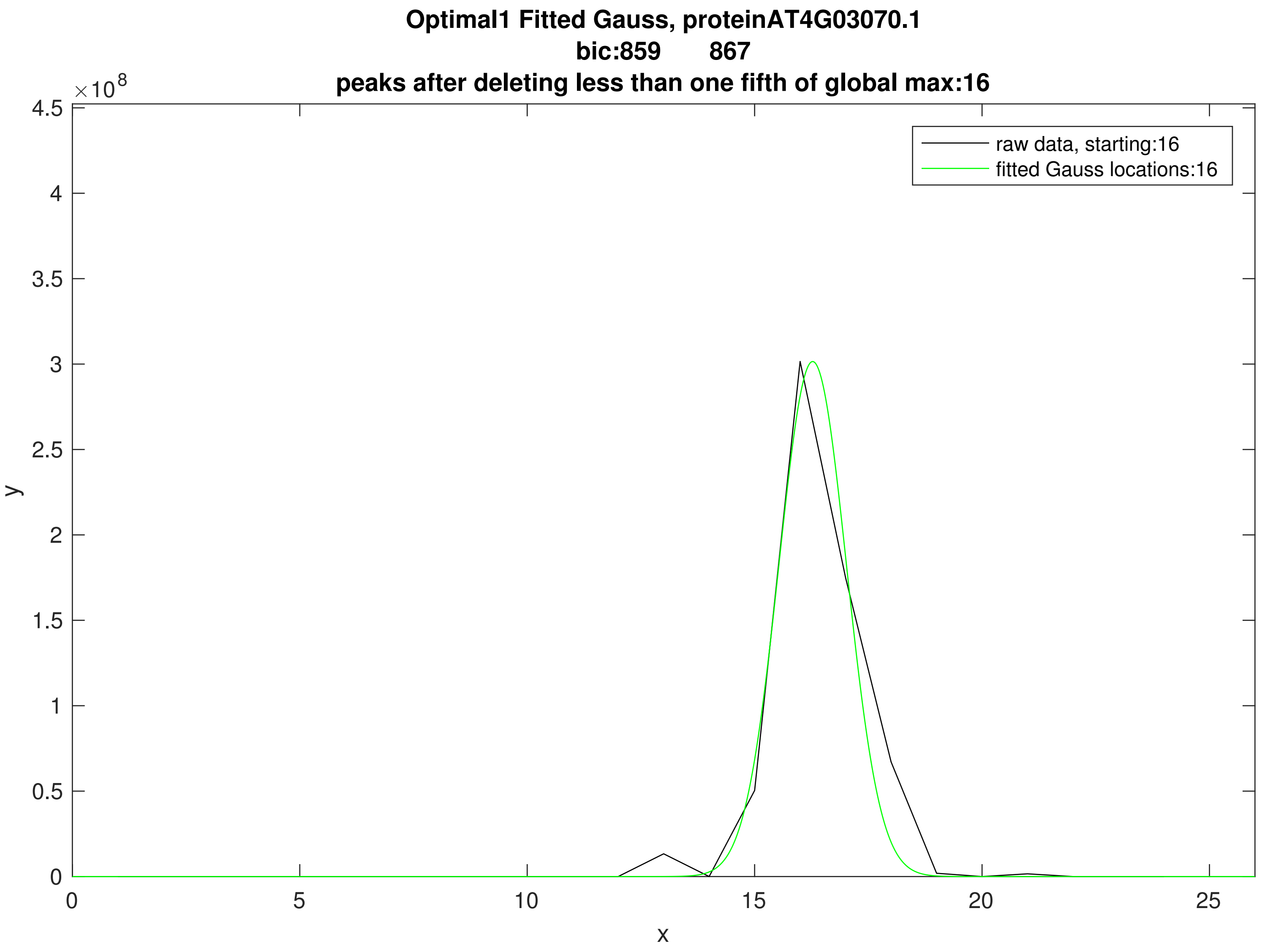

Optimal1 Fitted Gauss, proteinAT4G03070.1
bic:859 867
peaks after deleting less than one fifth of global max:16
×10^8
4.5
4
3.5
3
2.5
2
1.5
1
0.5
0
y
0
5
10
15
20
25
x
raw data, starting:16
fitted Gauss locations:16

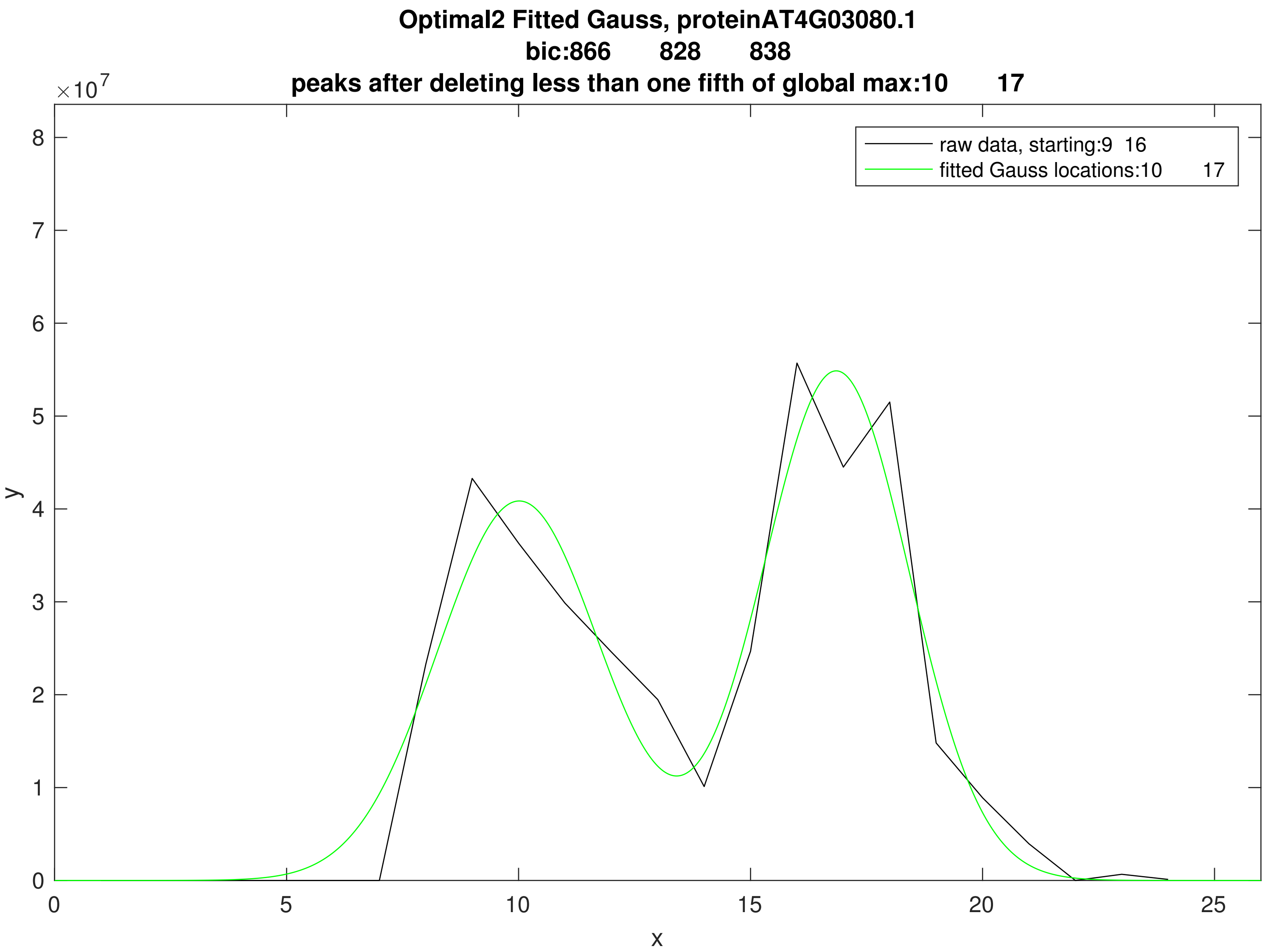

Optimal2 Fitted Gauss, proteinAT4G03080.1
bic:866 828 838
peaks after deleting less than one fifth of global max:10 17
×10^7
raw data, starting:9 16
fitted Gauss locations:10 17
y
x
0
1
2
3
4
5
6
7
8
0
5
10
15
20
25

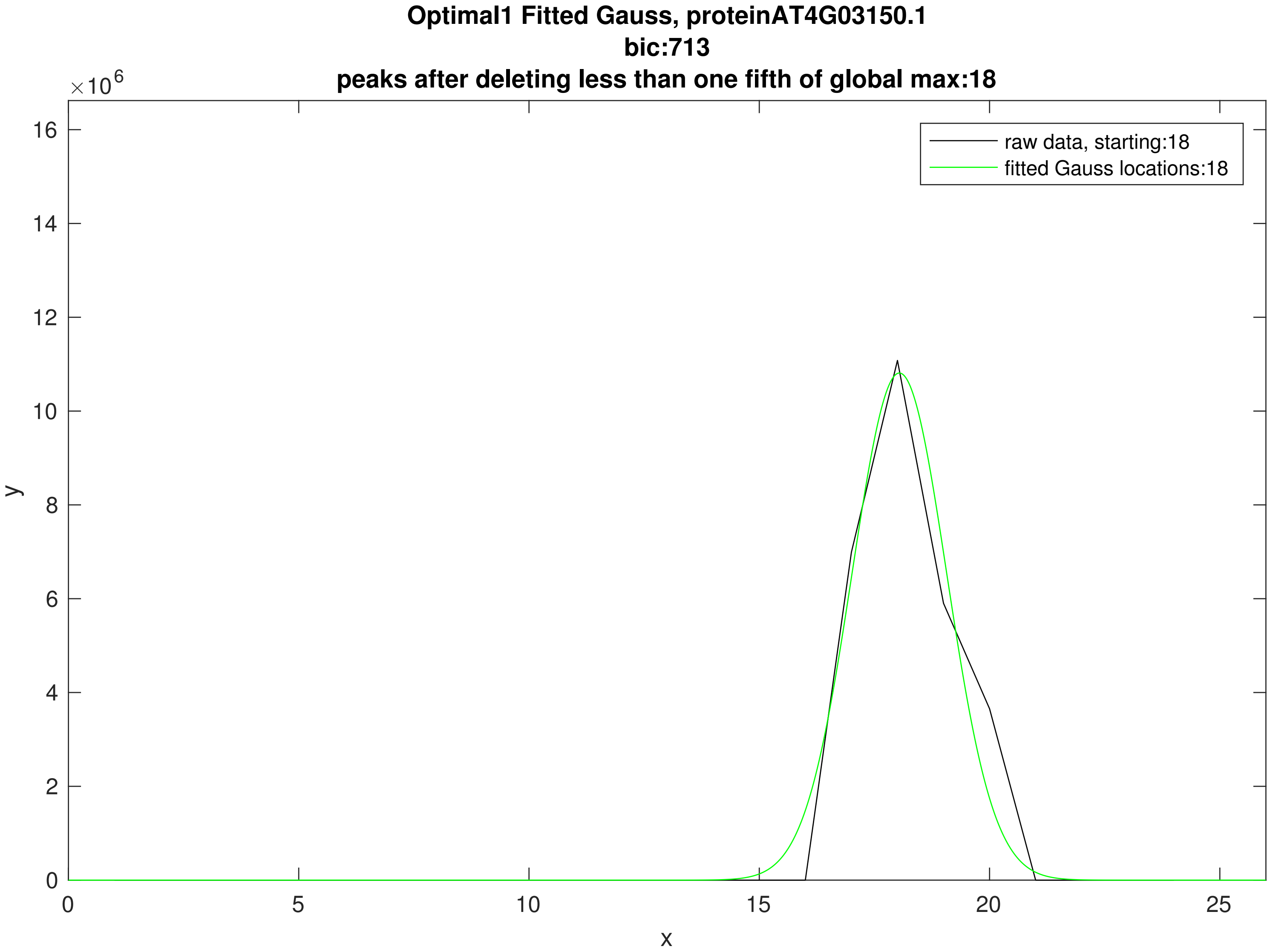
Optimal1 Fitted Gauss, proteinAT4G03150.1
bic:713
peaks after deleting less than one fifth of global max:18
×10^6
raw data, starting:18
fitted Gauss locations:18
y
x
0
2
4
6
8
10
12
14
16
0
5
10
15
20
25

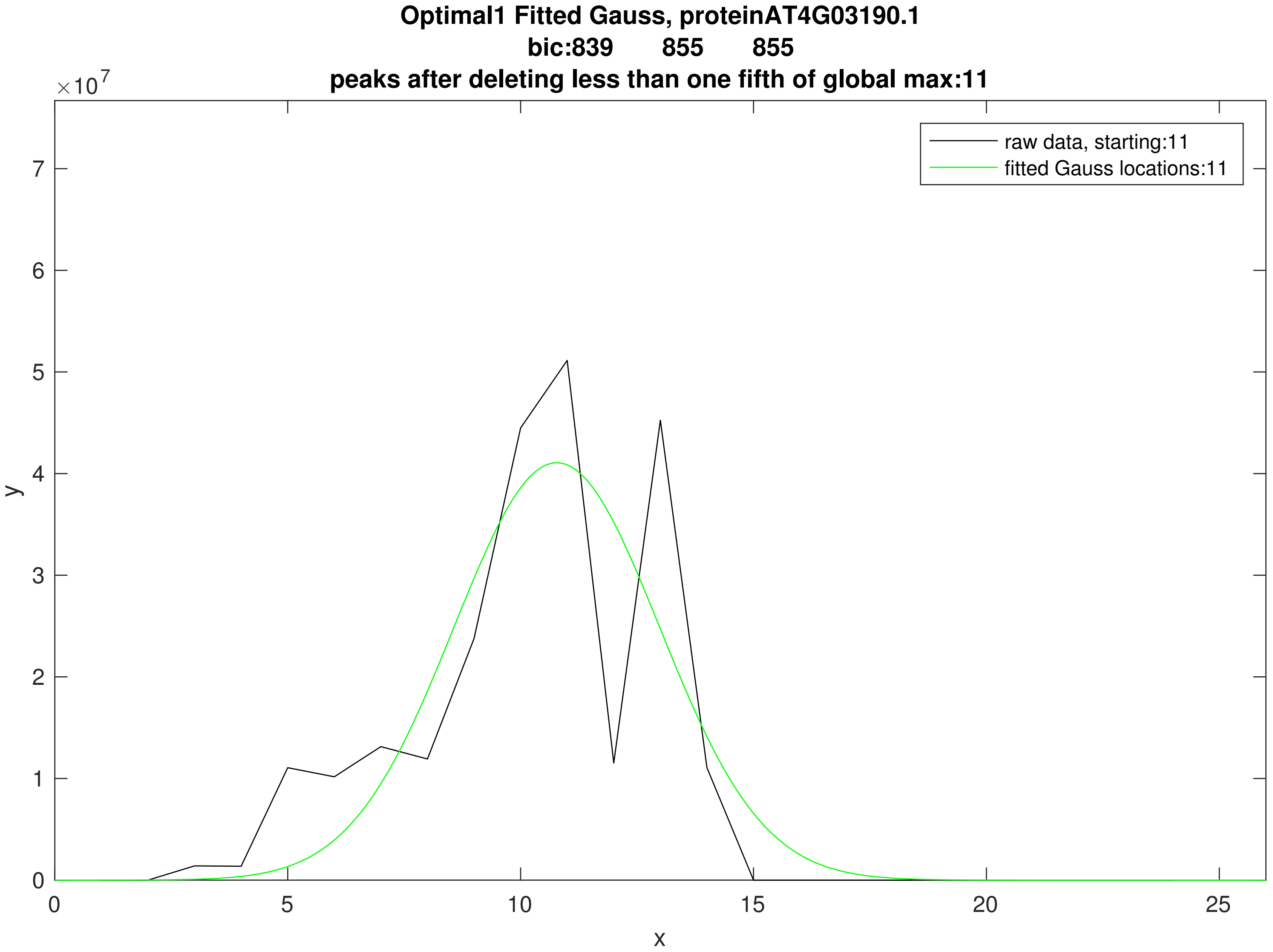

Optimal1 Fitted Gauss, proteinAT4G03190.1
bic:839 855 855
peaks after deleting less than one fifth of global max:11
×10^7
raw data, starting:11
fitted Gauss locations:11
y
x
0
1
2
3
4
5
6
7
0
5
10
15
20
25

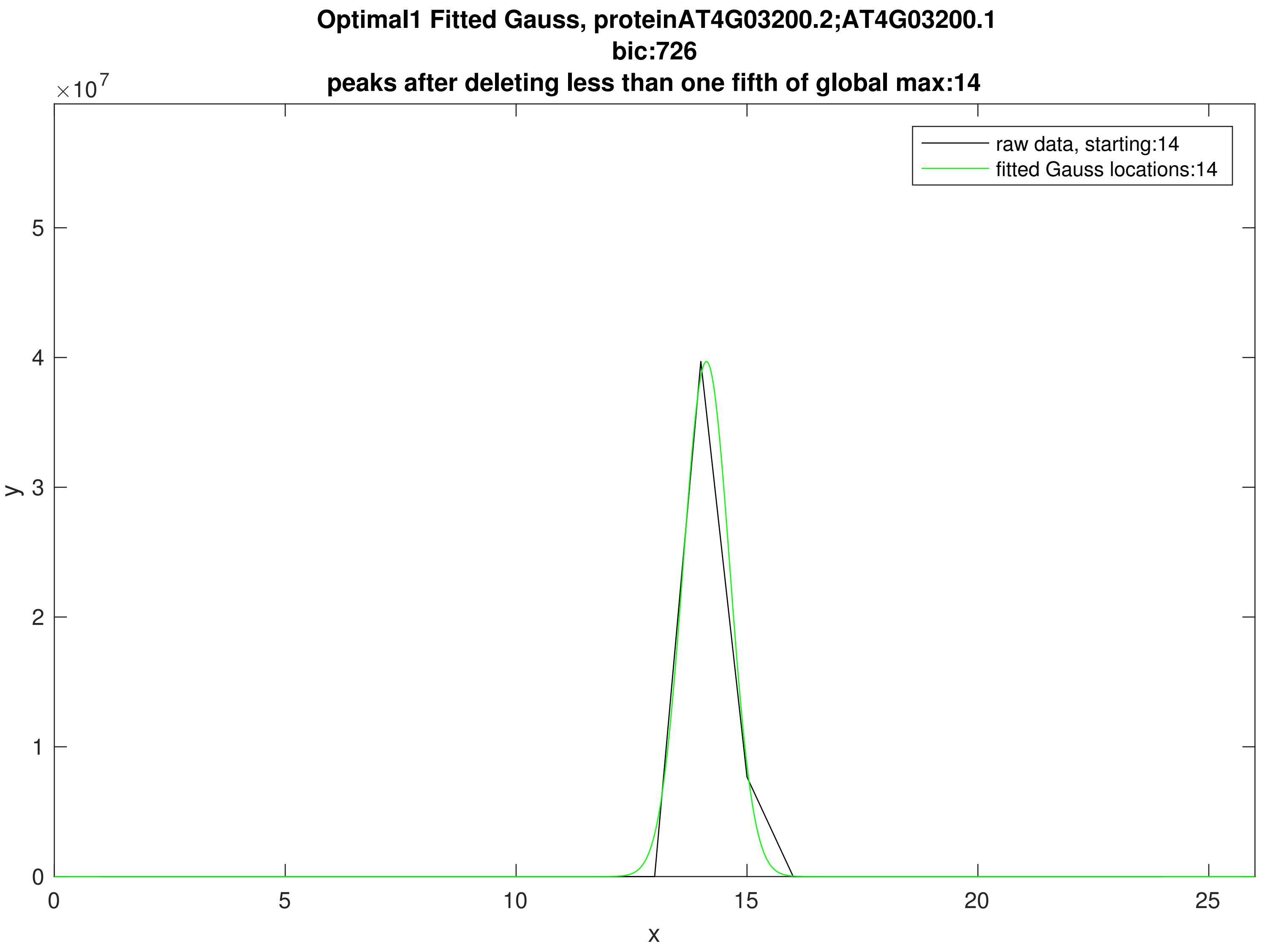
Optimal1 Fitted Gauss, proteinAT4G03200.2;AT4G03200.1
bic:726
peaks after deleting less than one fifth of global max:14
×10^7
raw data, starting:14
fitted Gauss locations:14
y
x
0
1
2
3
4
5
0
5
10
15
20
25

**Optimal1 Fitted Gauss, proteinAT4G03240.1**
**bic:805**
**peaks after deleting less than one fifth of global max:19**

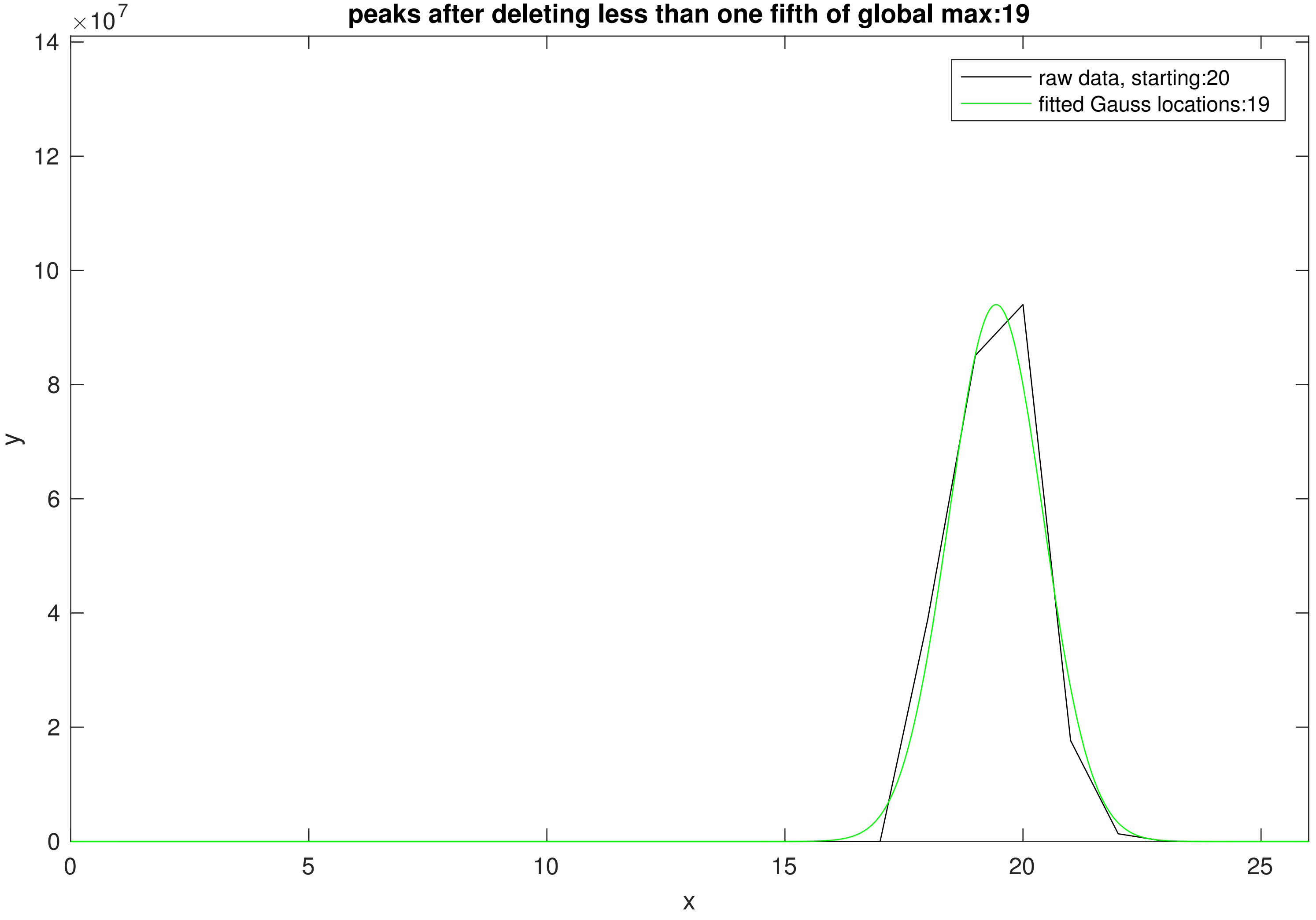

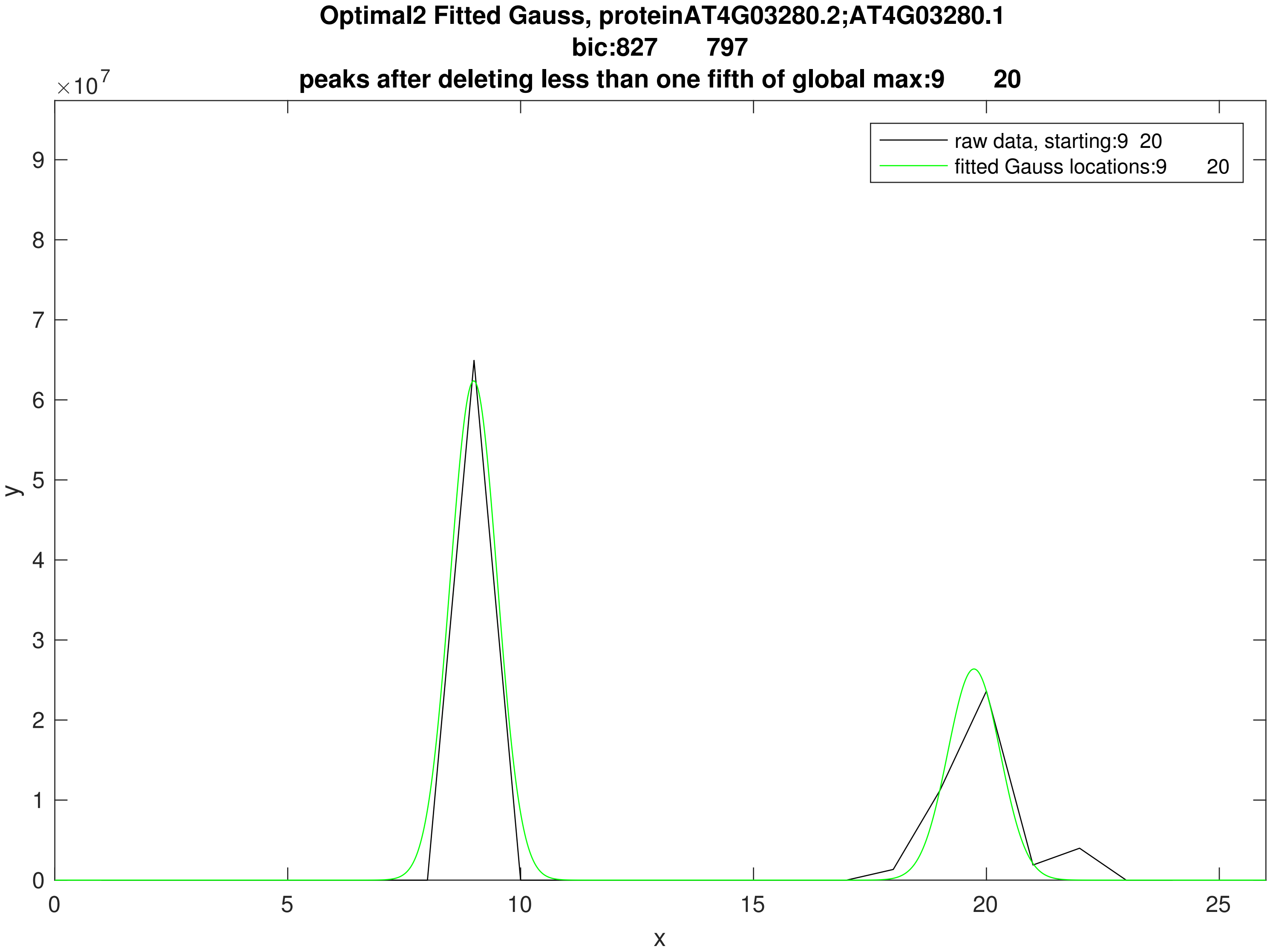

Optimal2 Fitted Gauss, proteinAT4G03280.2;AT4G03280.1
bic:827 797
peaks after deleting less than one fifth of global max:9 20
×10^7
raw data, starting:9 20
fitted Gauss locations:9 20
y
x
0
1
2
3
4
5
6
7
8
9
0
5
10
15
20
25

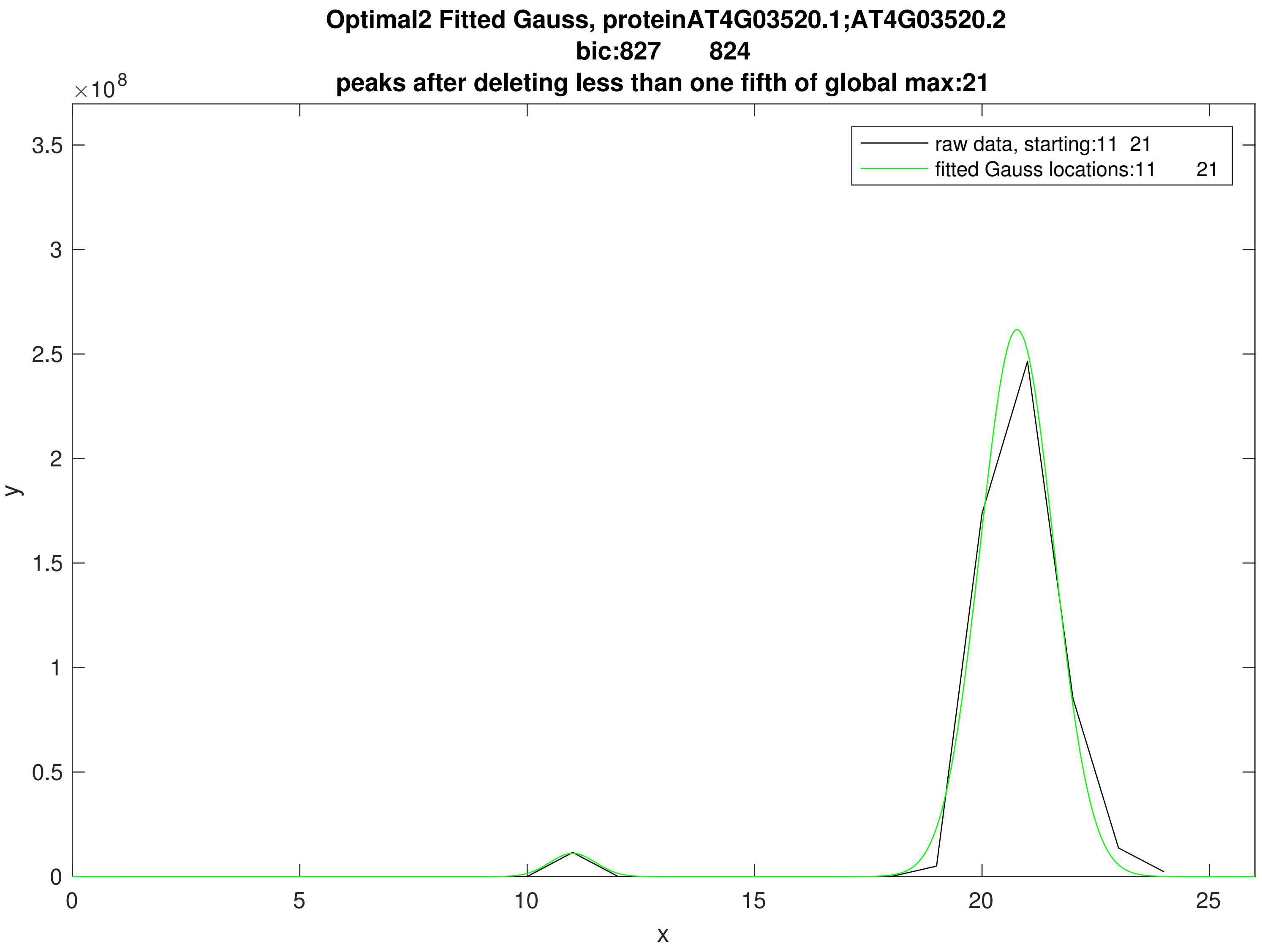

Optimal2 Fitted Gauss, proteinAT4G03520.1;AT4G03520.2
bic:827 824
peaks after deleting less than one fifth of global max:21
×10^8
3.5
3
2.5
2
1.5
1
0.5
0
y
0
5
10
15
20
25
x
raw data, starting:11 21
fitted Gauss locations:11 21

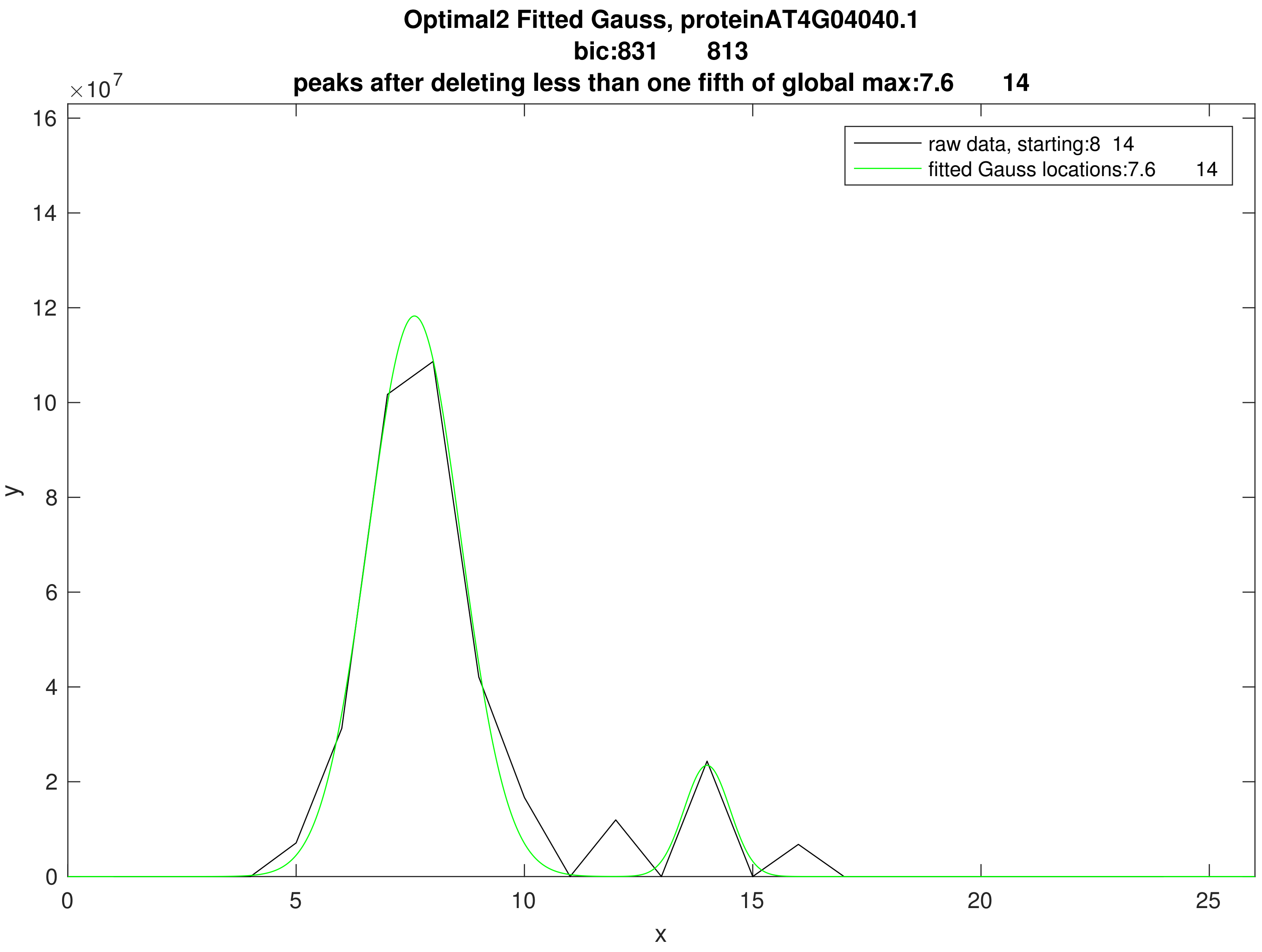

Optimal2 Fitted Gauss, proteinAT4G04040.1
bic:831 813
peaks after deleting less than one fifth of global max:7.6 14
×10^7
16
14
12
10
8
6
4
2
0
y
0
5
10
15
20
25
x
raw data, starting:8 14
fitted Gauss locations:7.6 14

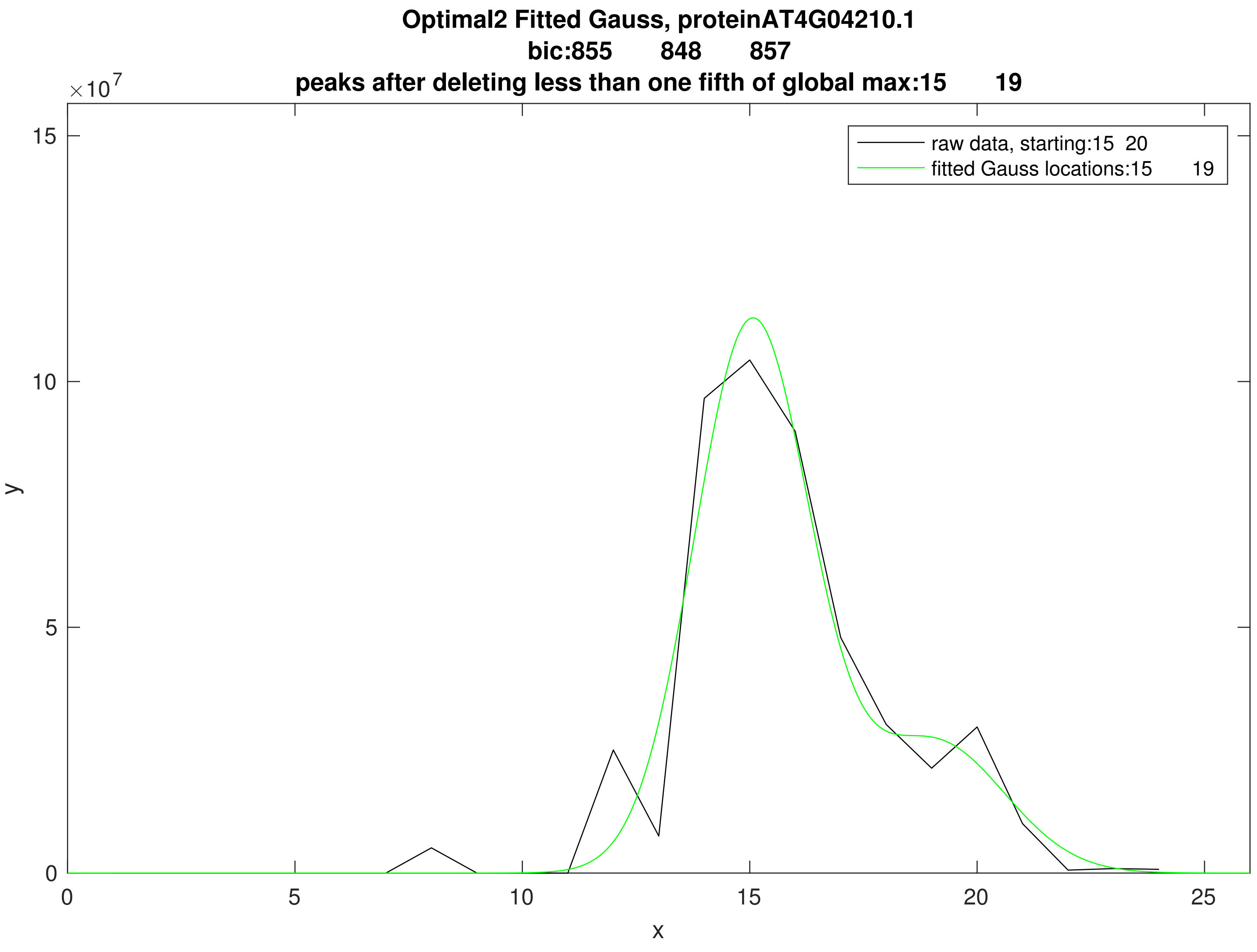
Optimal2 Fitted Gauss, proteinAT4G04210.1
bic:855 848 857
peaks after deleting less than one fifth of global max:15 19
×10^7
15
10
5
0
y
0
5
10
15
20
25
x
raw data, starting:15 20
fitted Gauss locations:15 19

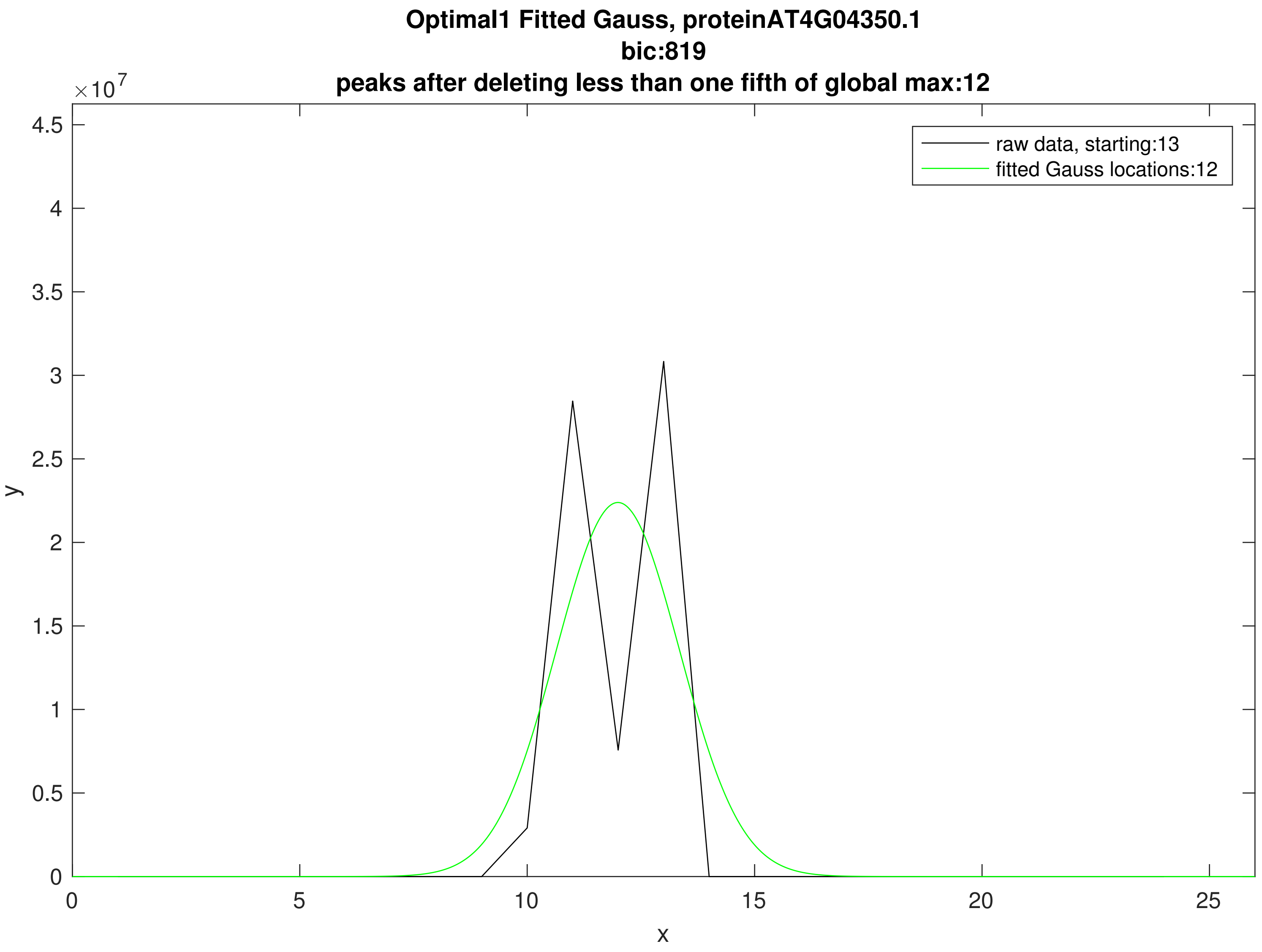
Optimal1 Fitted Gauss, proteinAT4G04350.1
bic:819
peaks after deleting less than one fifth of global max:12
×10^7
4.5
4
3.5
3
2.5
2
1.5
1
0.5
0
y
0
5
10
15
20
25
x
raw data, starting:13
fitted Gauss locations:12

**Optimal1 Fitted Gauss, proteinAT4G04640.1**
**bic:934 942**
**peaks after deleting less than one fifth of global max:7.5**

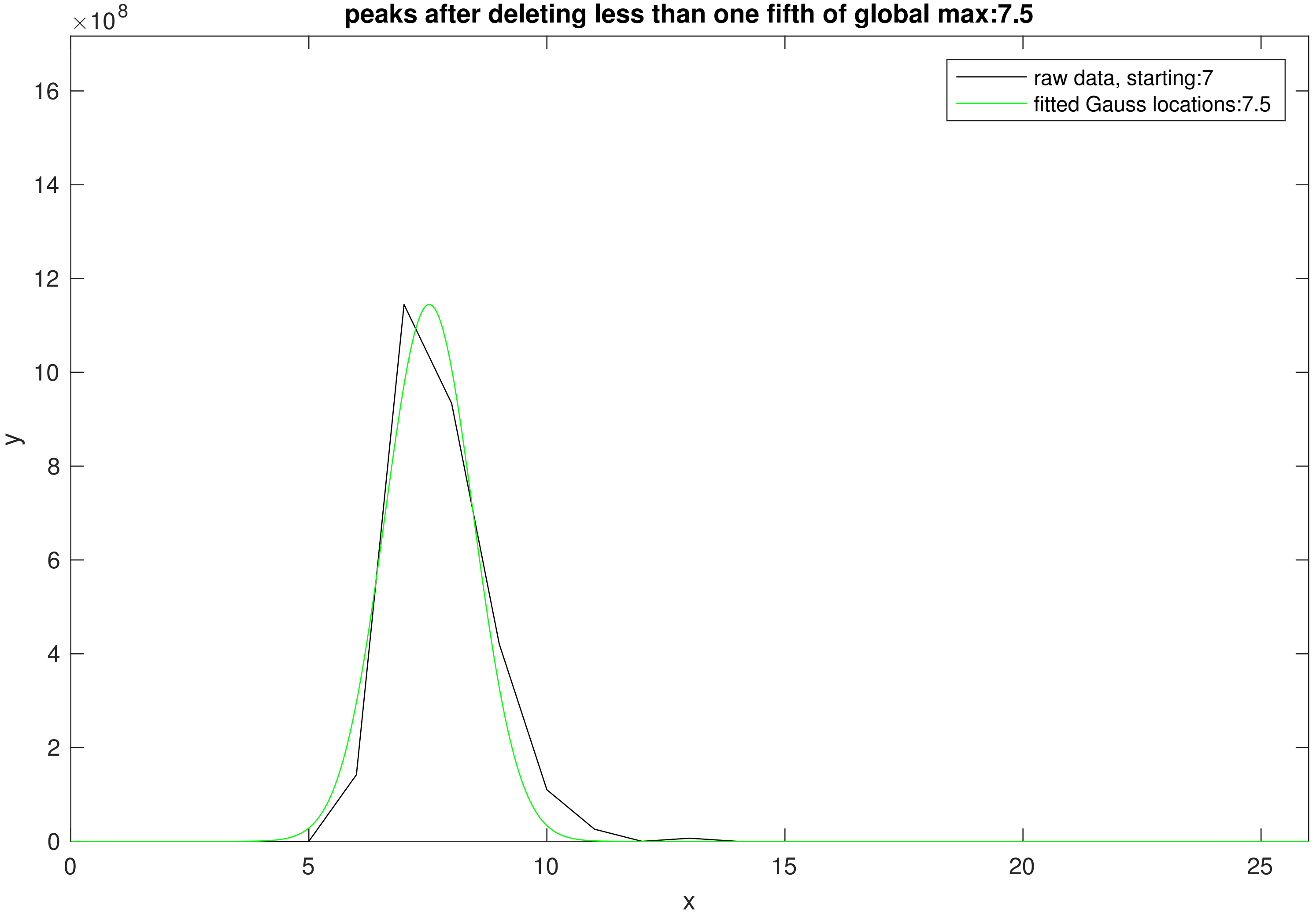

Optimal1 Fitted Gauss, proteinAT4G04800.1
bic:725
peaks after deleting less than one fifth of global max:20

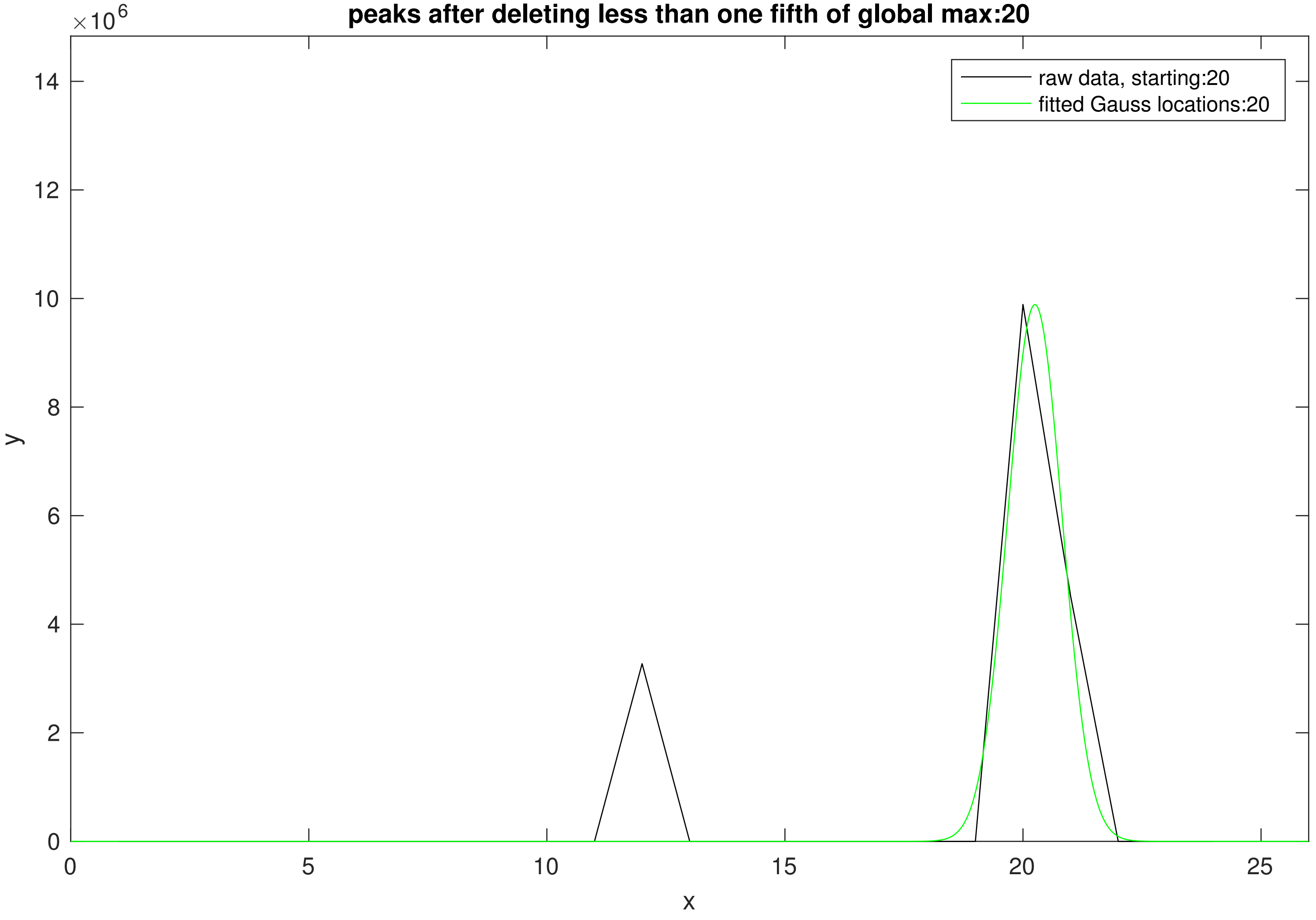

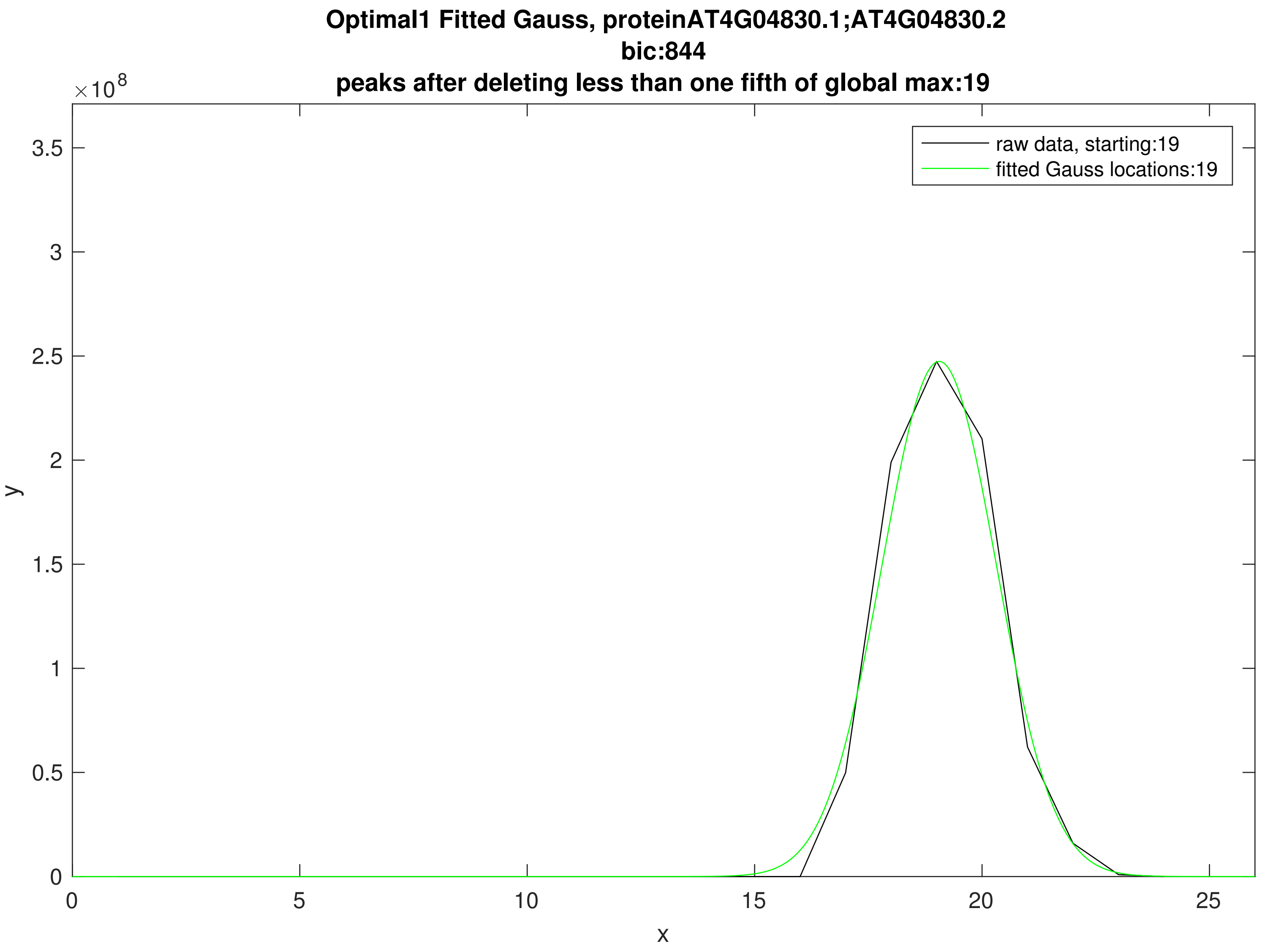

Optimal1 Fitted Gauss, proteinAT4G04830.1;AT4G04830.2
bic:844
peaks after deleting less than one fifth of global max:19
×10^8
raw data, starting:19
fitted Gauss locations:19
3.5
3
2.5
2
1.5
1
0.5
0
y
0
5
10
15
20
25
x

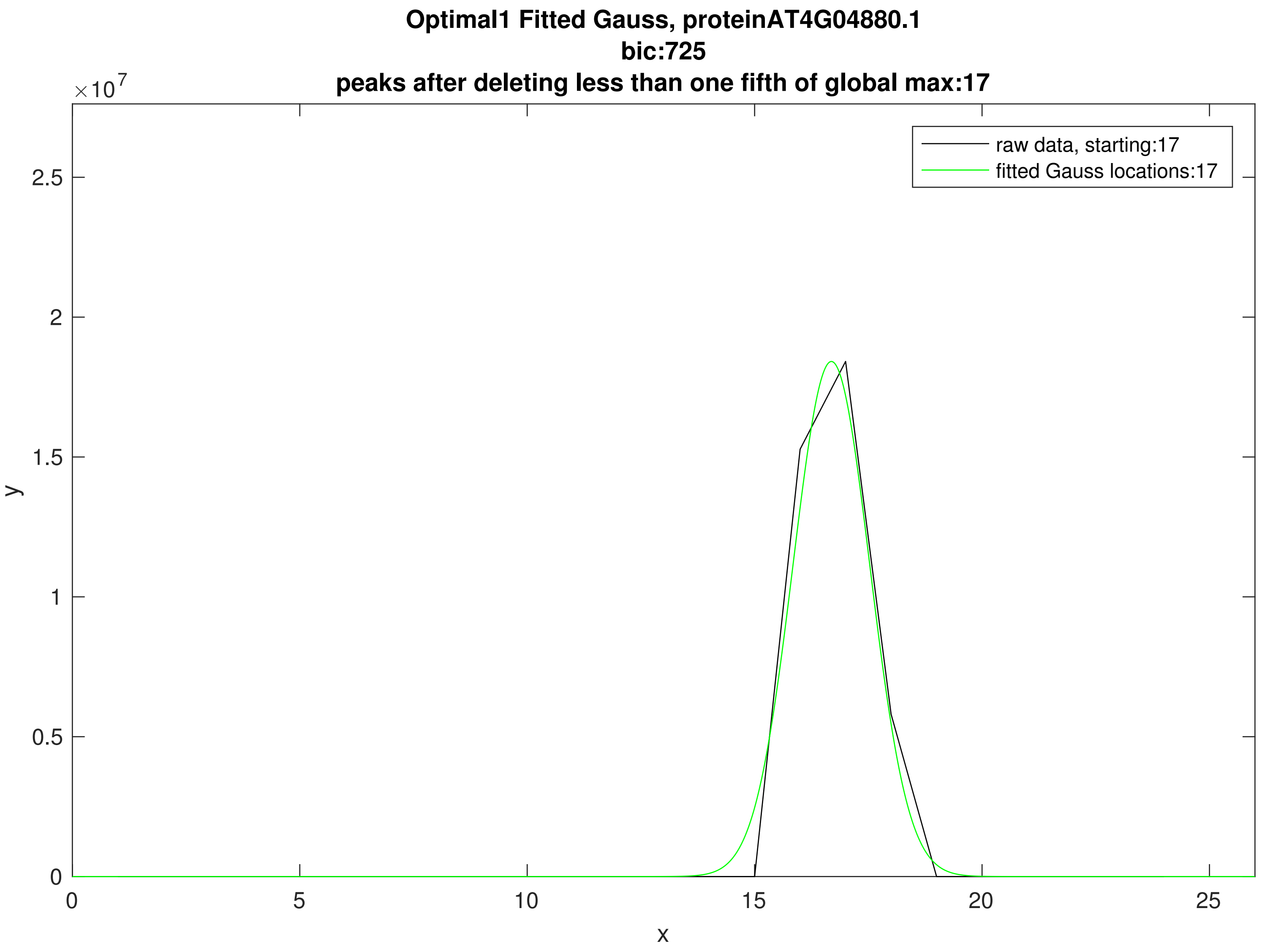

Optimal1 Fitted Gauss, proteinAT4G04880.1
bic:725
peaks after deleting less than one fifth of global max:17
×10^7
raw data, starting:17
fitted Gauss locations:17
2.5
2
1.5
1
0.5
0
y
0
5
10
15
20
25
x

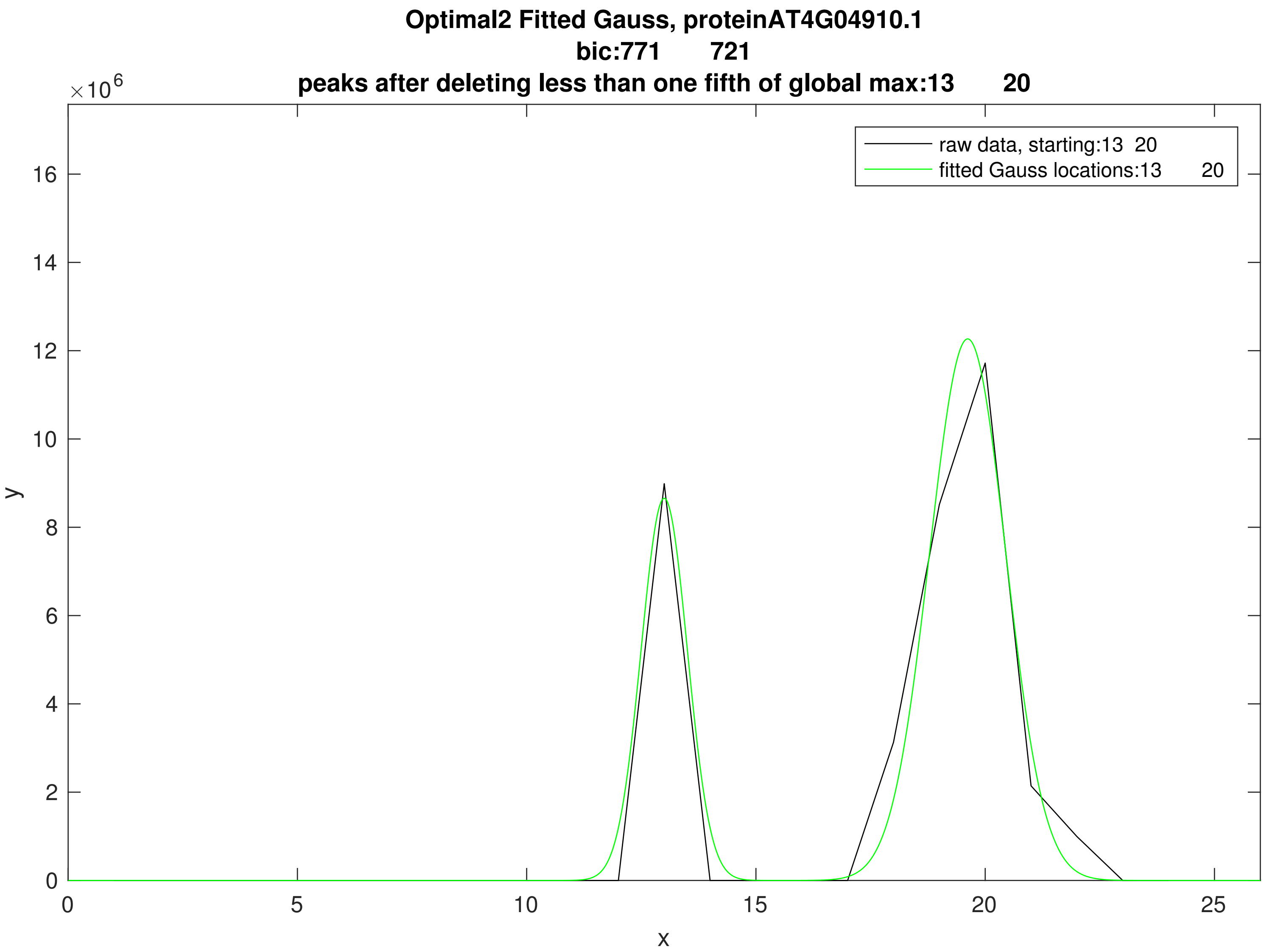
Optimal2 Fitted Gauss, proteinAT4G04910.1
bic:771 721
peaks after deleting less than one fifth of global max:13 20
raw data, starting:13 20
fitted Gauss locations:13 20
×10^6
y
x

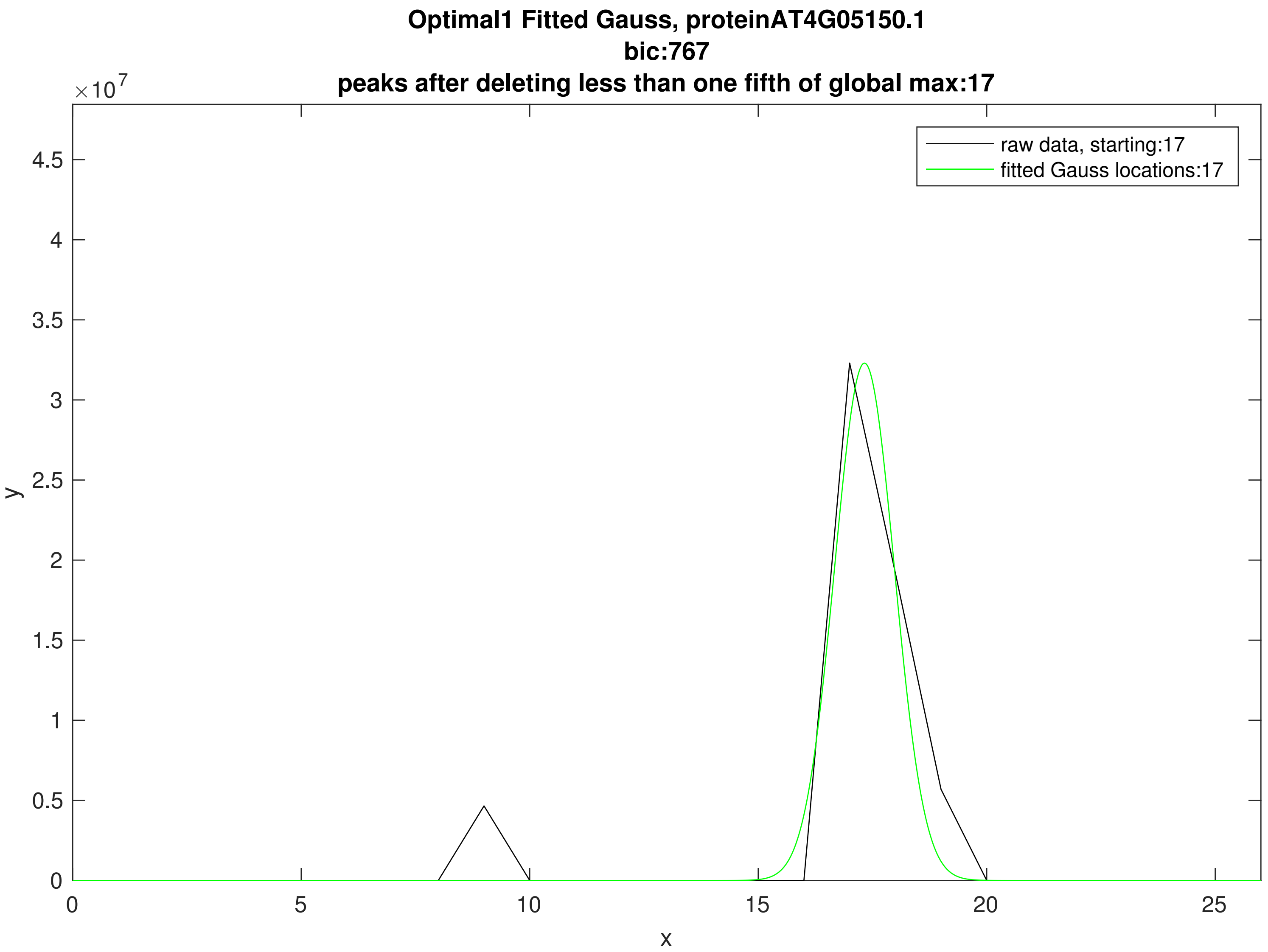
Optimal1 Fitted Gauss, proteinAT4G05150.1
bic:767
peaks after deleting less than one fifth of global max:17
×10^7
4.5
4
3.5
3
2.5
2
1.5
1
0.5
0
y
0
5
10
15
20
25
x
raw data, starting:17
fitted Gauss locations:17

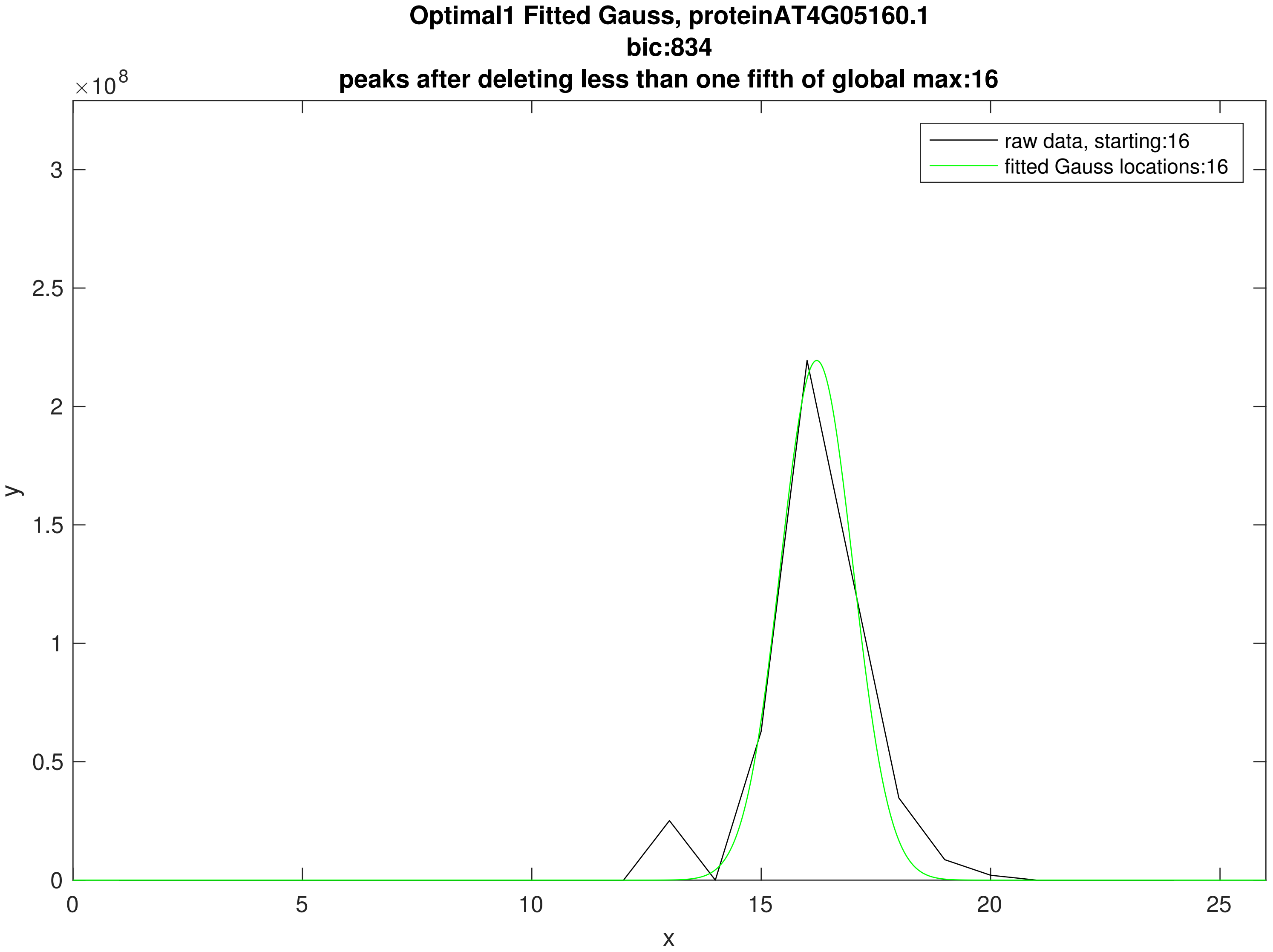
Optimal1 Fitted Gauss, proteinAT4G05160.1
bic:834
peaks after deleting less than one fifth of global max:16
×10^8
raw data, starting:16
fitted Gauss locations:16
y
x
0
0.5
1
1.5
2
2.5
3
0
5
10
15
20
25

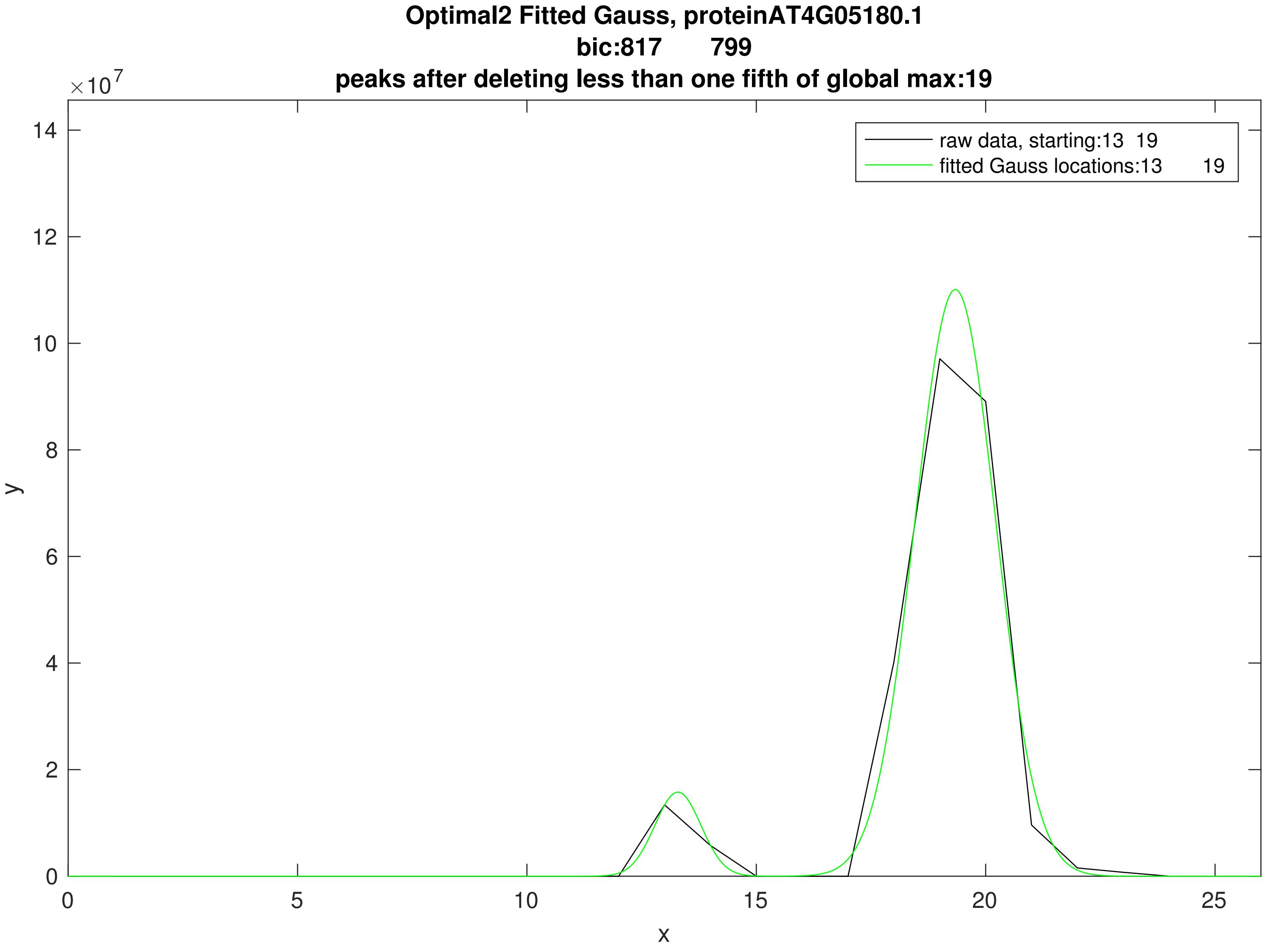

Optimal2 Fitted Gauss, proteinAT4G05180.1
bic:817 799
peaks after deleting less than one fifth of global max:19
×10^7
raw data, starting:13 19
fitted Gauss locations:13 19
y
x
0
2
4
6
8
10
12
14
0
5
10
15
20
25

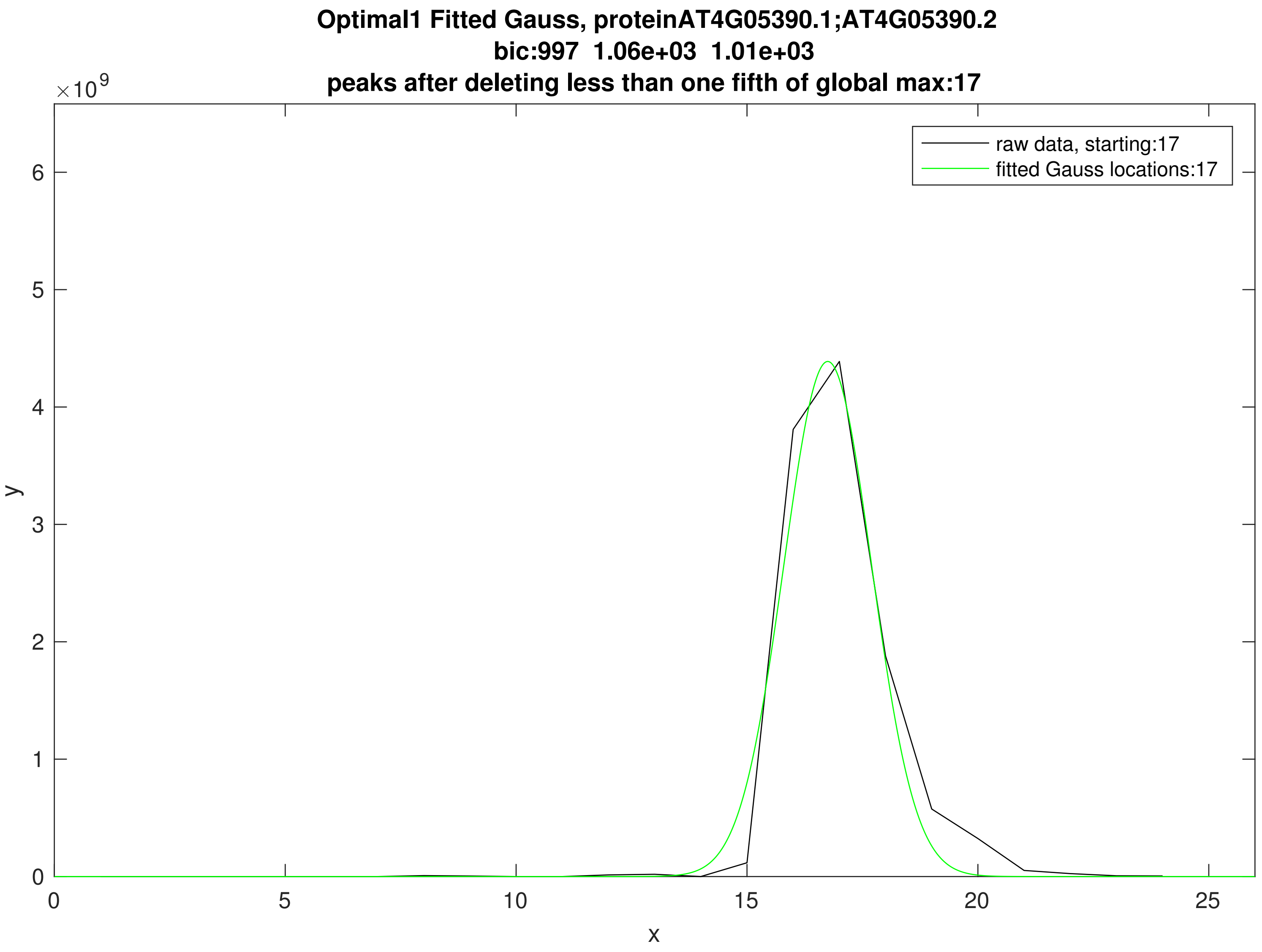
Optimal1 Fitted Gauss, proteinAT4G05390.1;AT4G05390.2
bic:997 1.06e+03 1.01e+03
peaks after deleting less than one fifth of global max:17
×10^9
raw data, starting:17
fitted Gauss locations:17
y
x
0
1
2
3
4
5
6
0
5
10
15
20
25

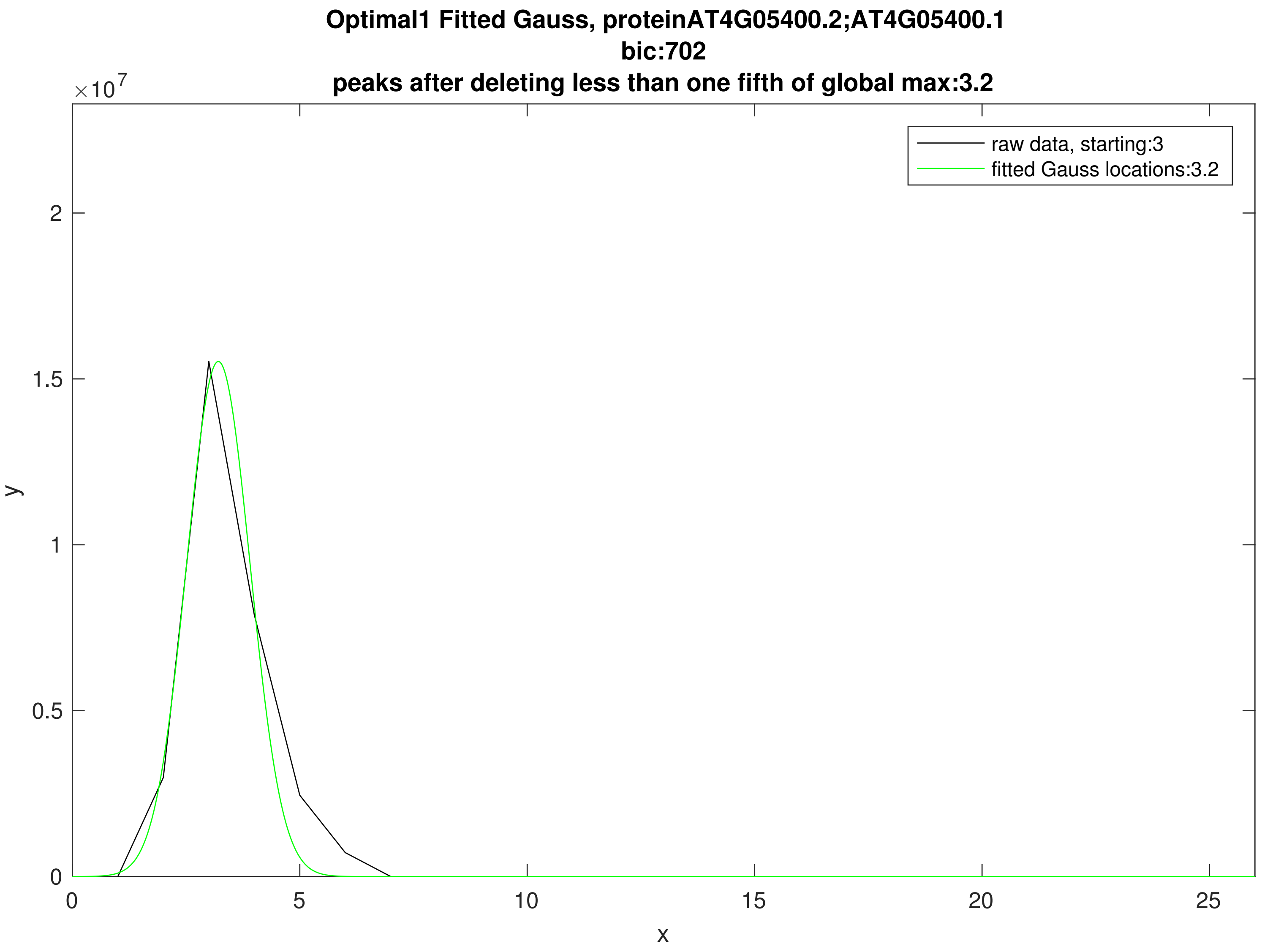

Optimal1 Fitted Gauss, proteinAT4G05400.2;AT4G05400.1
bic:702
peaks after deleting less than one fifth of global max:3.2
×10^7
raw data, starting:3
fitted Gauss locations:3.2
y
x
0
0.5
1
1.5
2
0
5
10
15
20
25

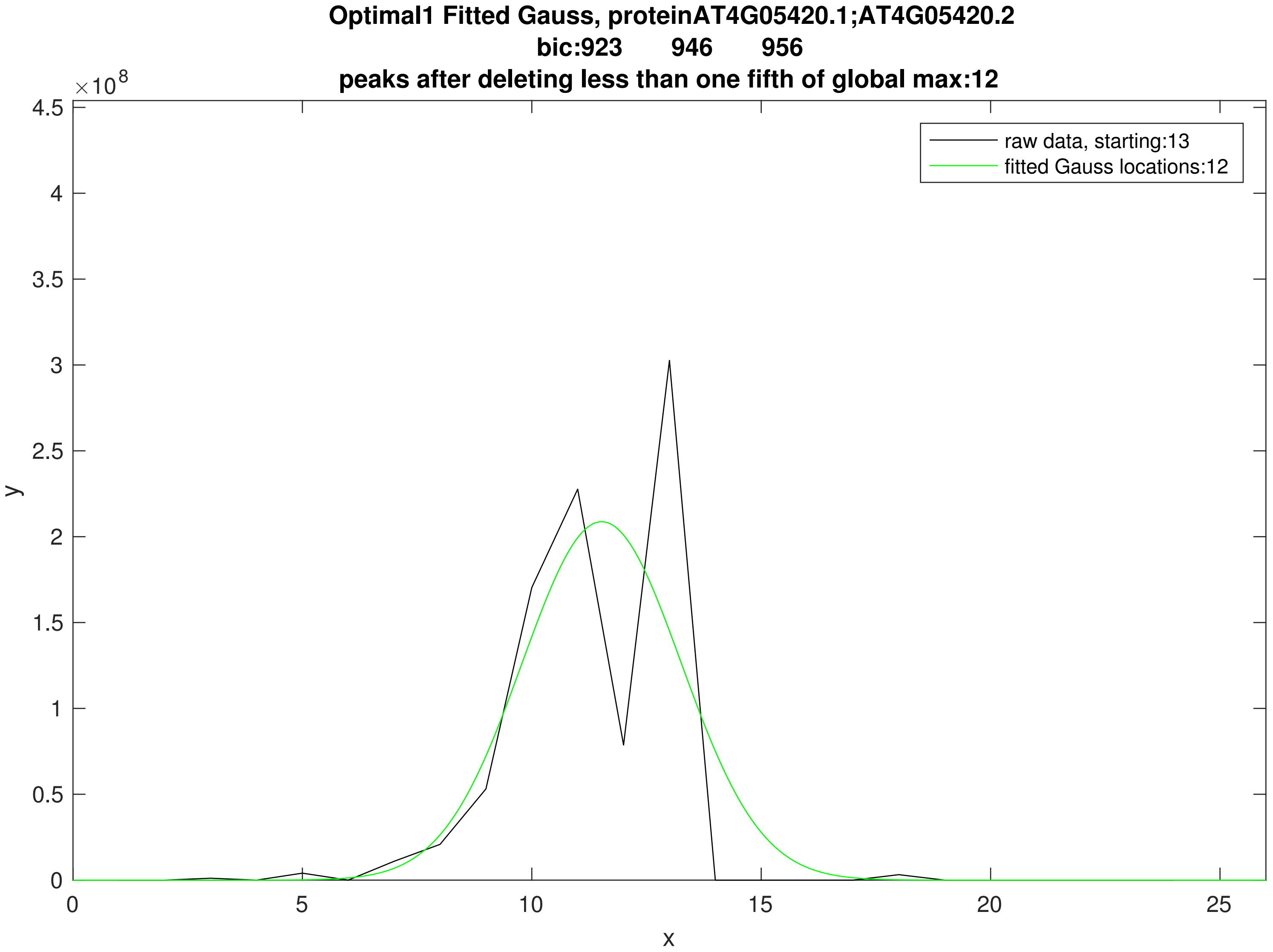
Optimal1 Fitted Gauss, proteinAT4G05420.1;AT4G05420.2
bic:923 946 956
peaks after deleting less than one fifth of global max:12
×10^8
raw data, starting:13
fitted Gauss locations:12
y
x
0
0.5
1
1.5
2
2.5
3
3.5
4
4.5
0
5
10
15
20
25

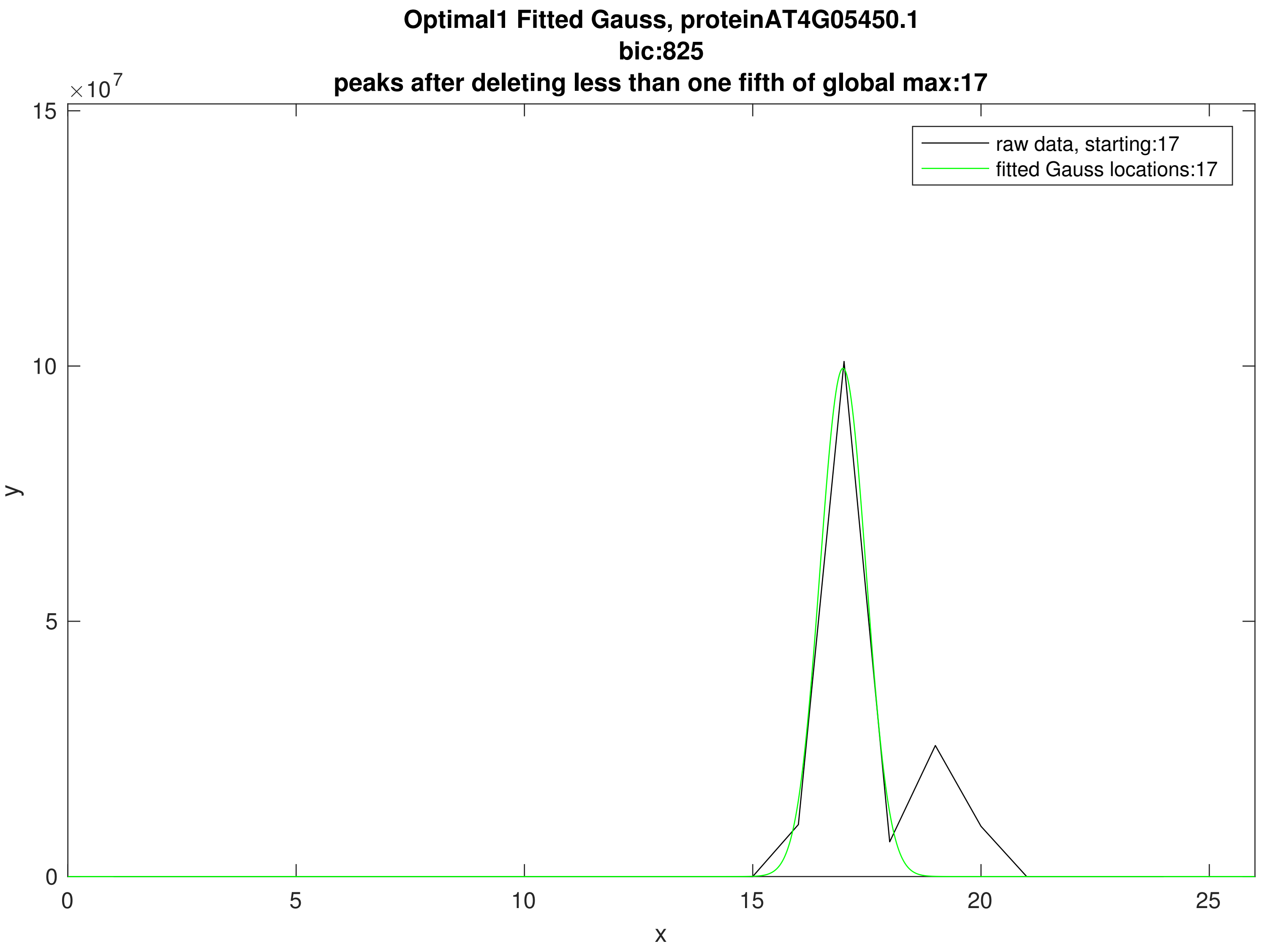
Optimal1 Fitted Gauss, proteinAT4G05450.1
bic:825
peaks after deleting less than one fifth of global max:17
×10^7
15
10
5
0
y
0
5
10
15
20
25
x
raw data, starting:17
fitted Gauss locations:17

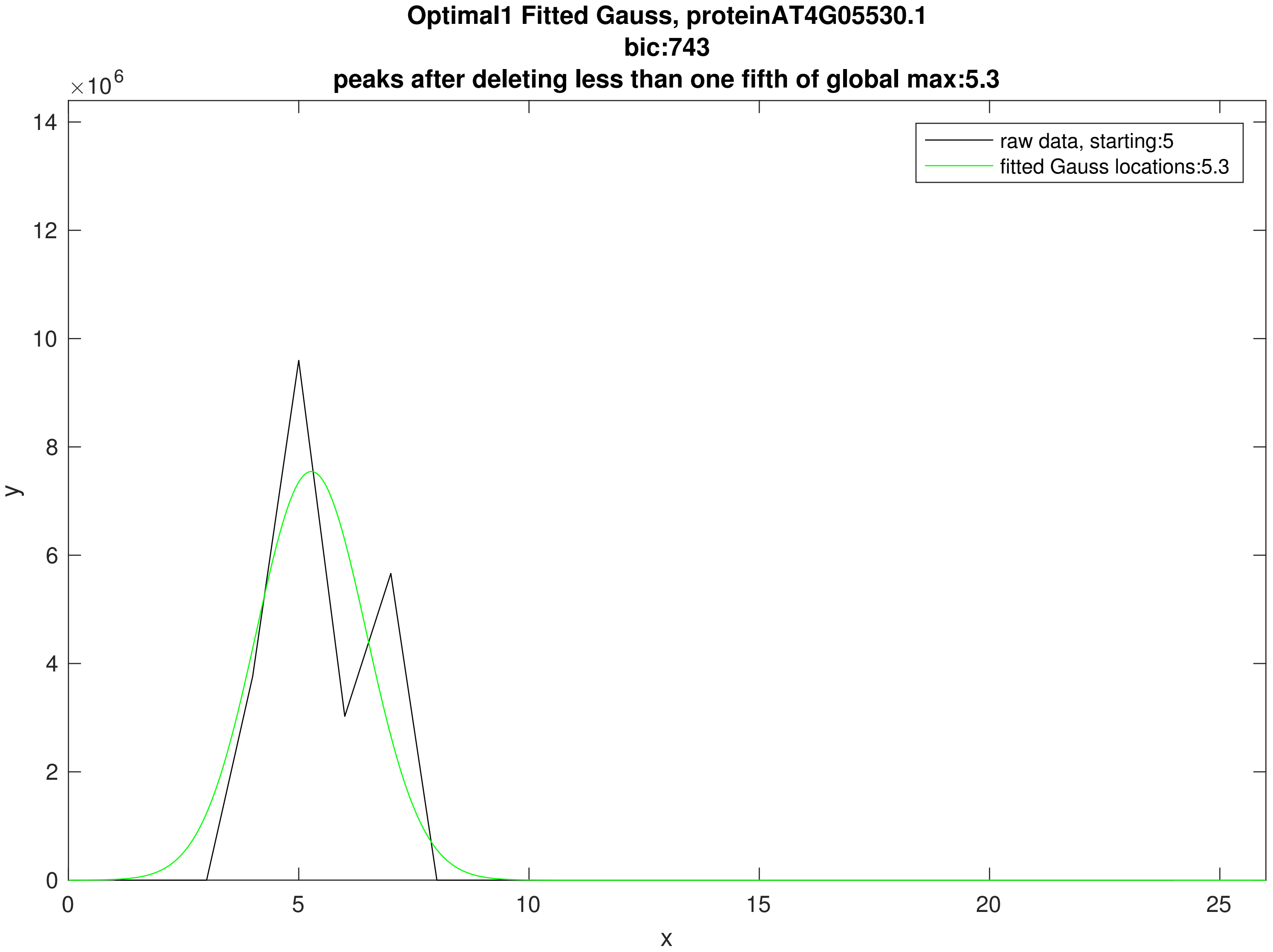

Optimal1 Fitted Gauss, proteinAT4G05530.1
bic:743
peaks after deleting less than one fifth of global max:5.3
×10^6
raw data, starting:5
fitted Gauss locations:5.3
y
x
0
2
4
6
8
10
12
14
0
5
10
15
20
25

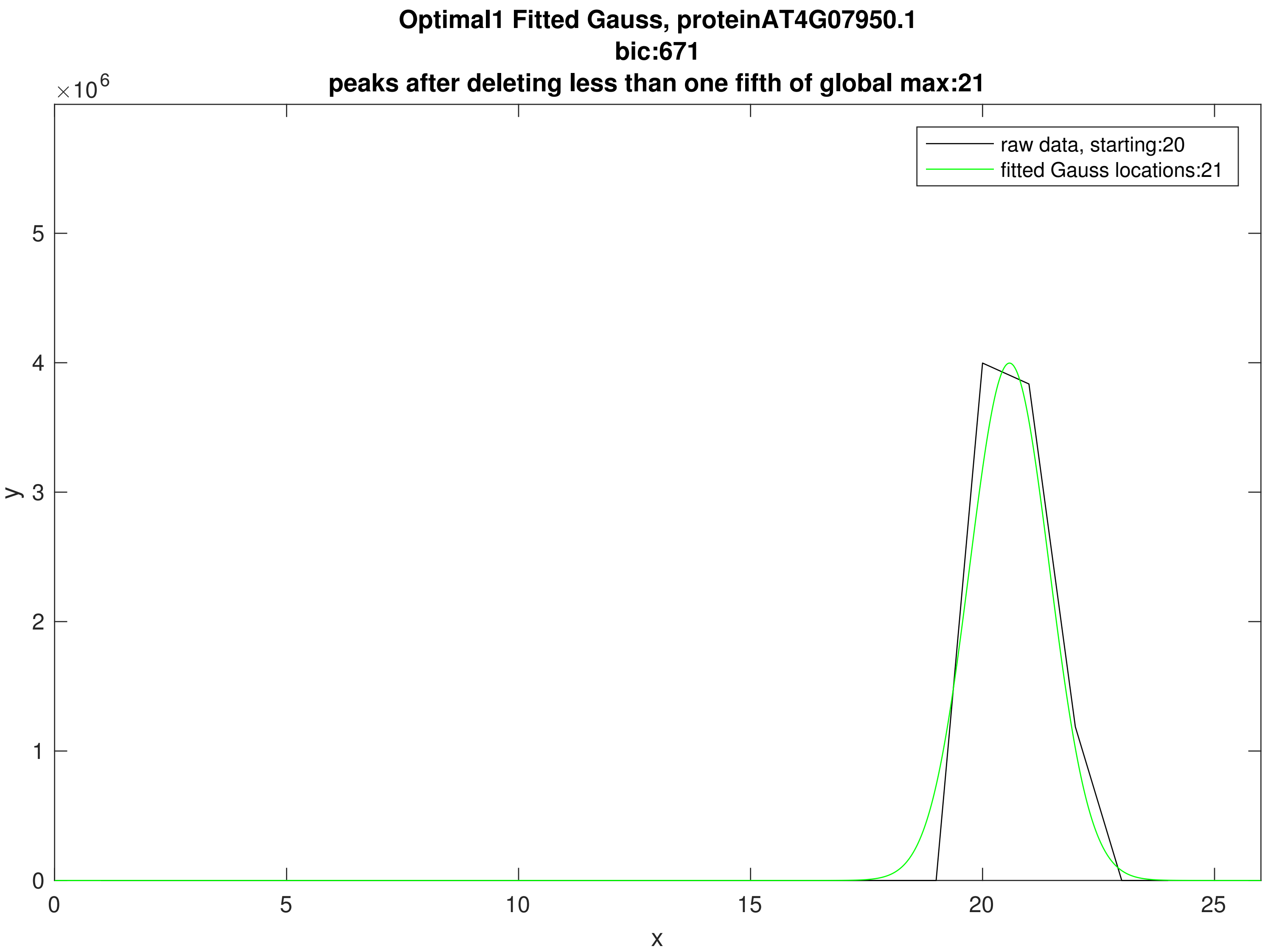
Optimal1 Fitted Gauss, proteinAT4G07950.1
bic:671
peaks after deleting less than one fifth of global max:21
×10^6
raw data, starting:20
fitted Gauss locations:21
y
x
0
1
2
3
4
5
0
5
10
15
20
25

Optimal1 Fitted Gauss, proteinAT4G08390.3;AT4G08390.2;AT4G08390.1;AT4G08390.4
bic:1e+03 1e+03 1.01e+03
peaks after deleting less than one fifth of global max:17

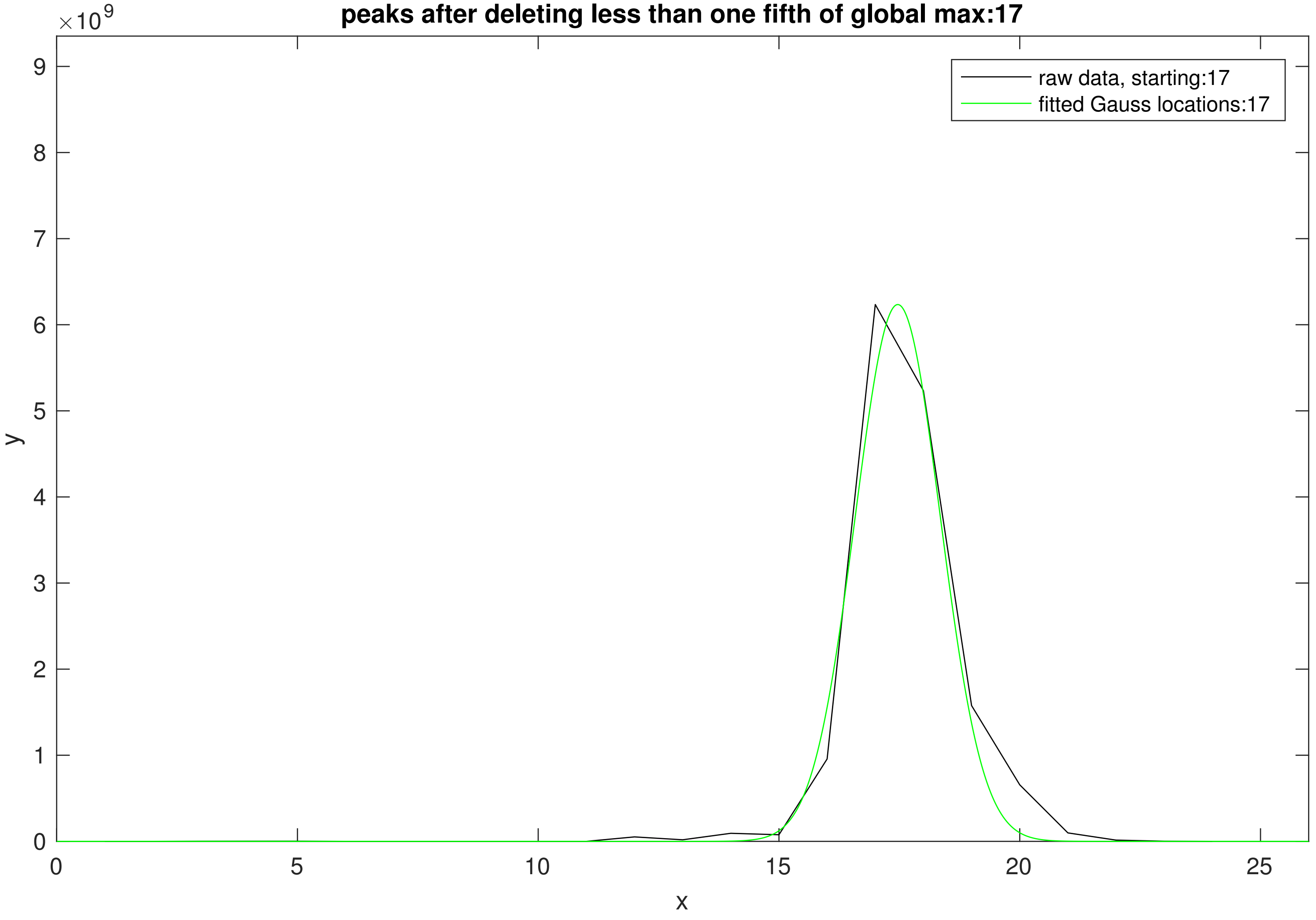

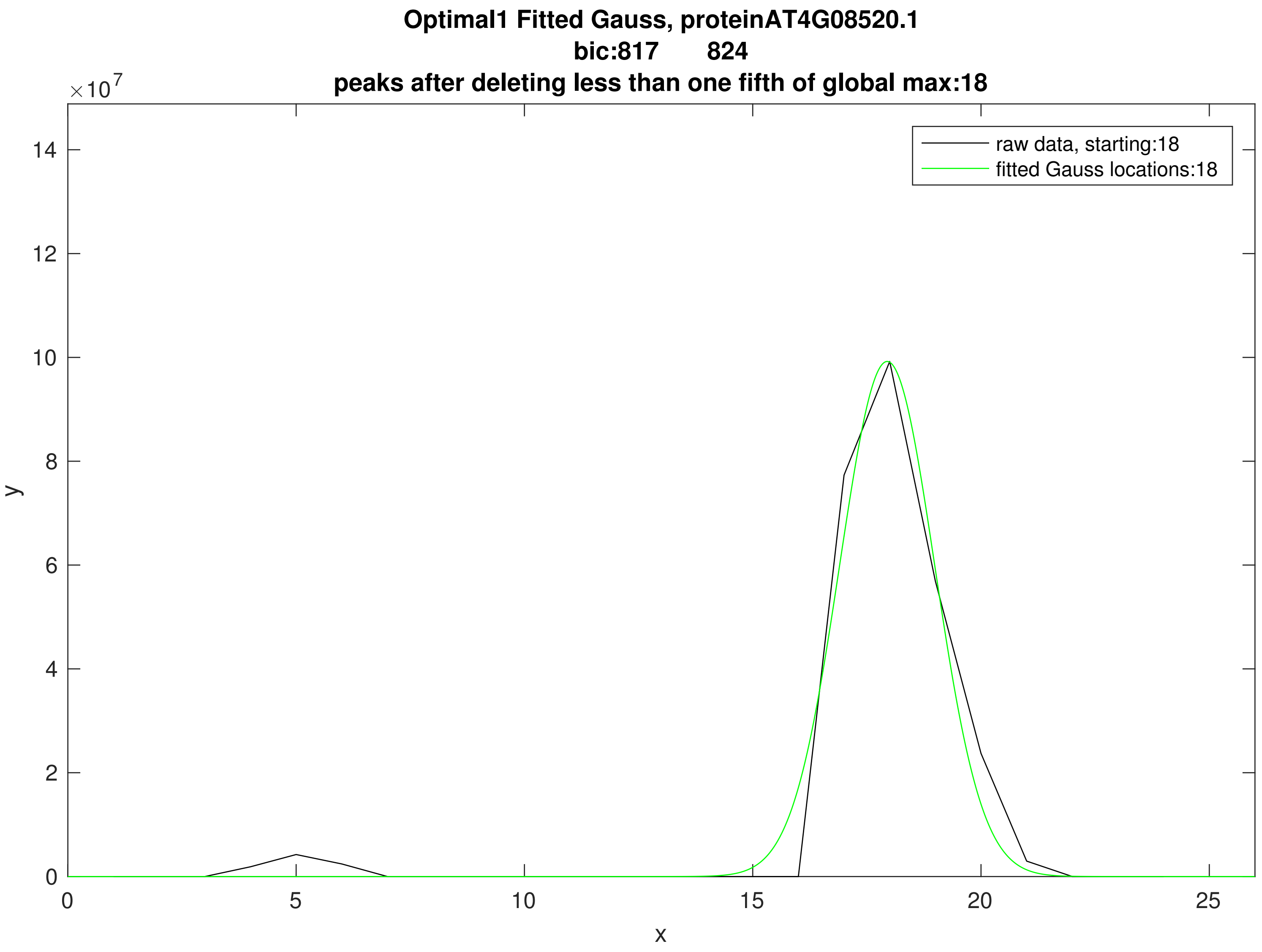
Optimal1 Fitted Gauss, proteinAT4G08520.1
bic:817 824
peaks after deleting less than one fifth of global max:18
×10^7
raw data, starting:18
fitted Gauss locations:18
y
x
0
2
4
6
8
10
12
14
5
15
20
25

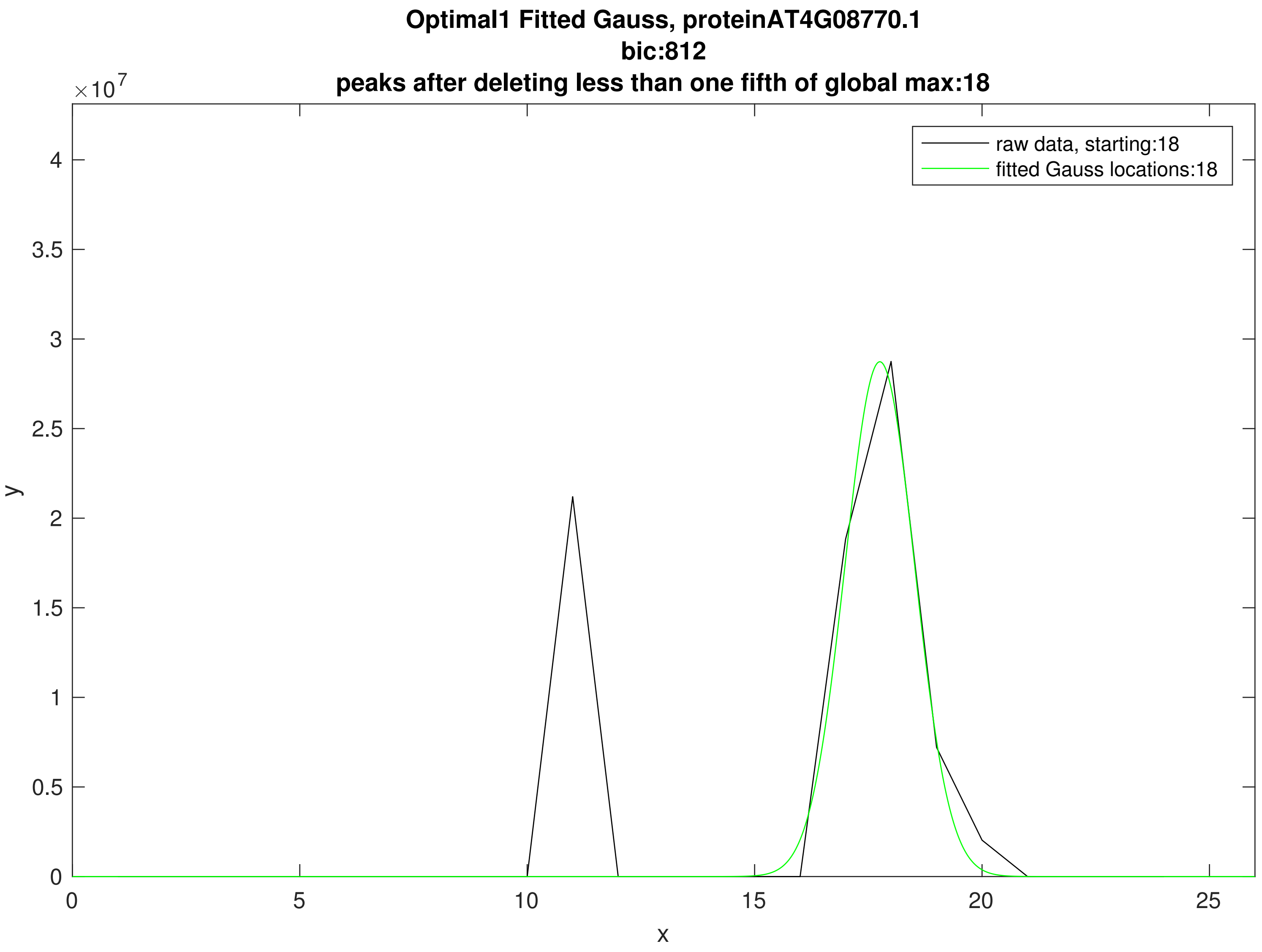

Optimal1 Fitted Gauss, proteinAT4G08770.1
bic:812
peaks after deleting less than one fifth of global max:18
×10^7
raw data, starting:18
fitted Gauss locations:18
y
x
0
0.5
1
1.5
2
2.5
3
3.5
4
0
5
10
15
20
25

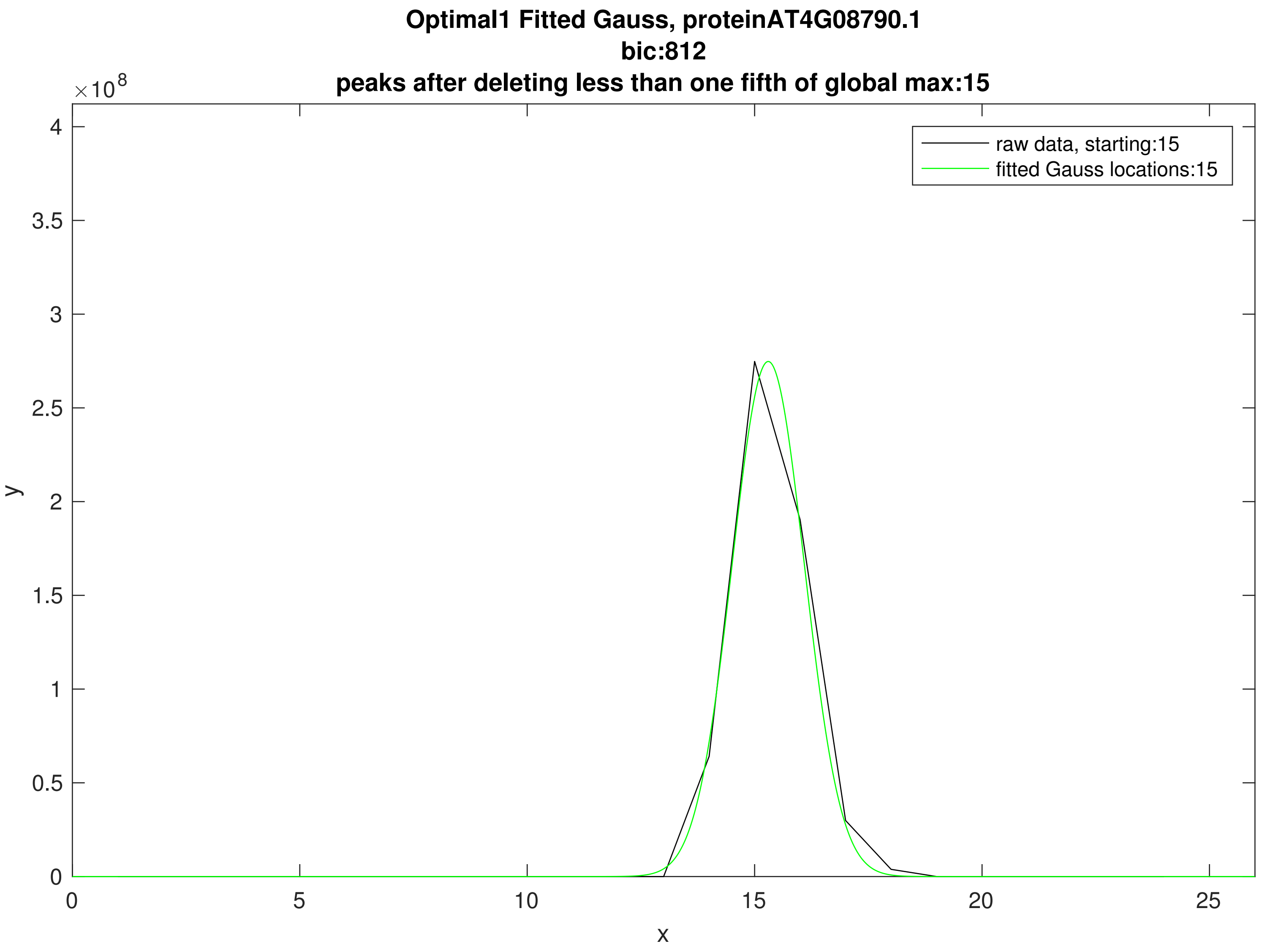

Optimal1 Fitted Gauss, proteinAT4G08790.1
bic:812
peaks after deleting less than one fifth of global max:15
×10^8
4
3.5
3
2.5
2
1.5
1
0.5
0
y
0
5
10
15
20
25
x
raw data, starting:15
fitted Gauss locations:15

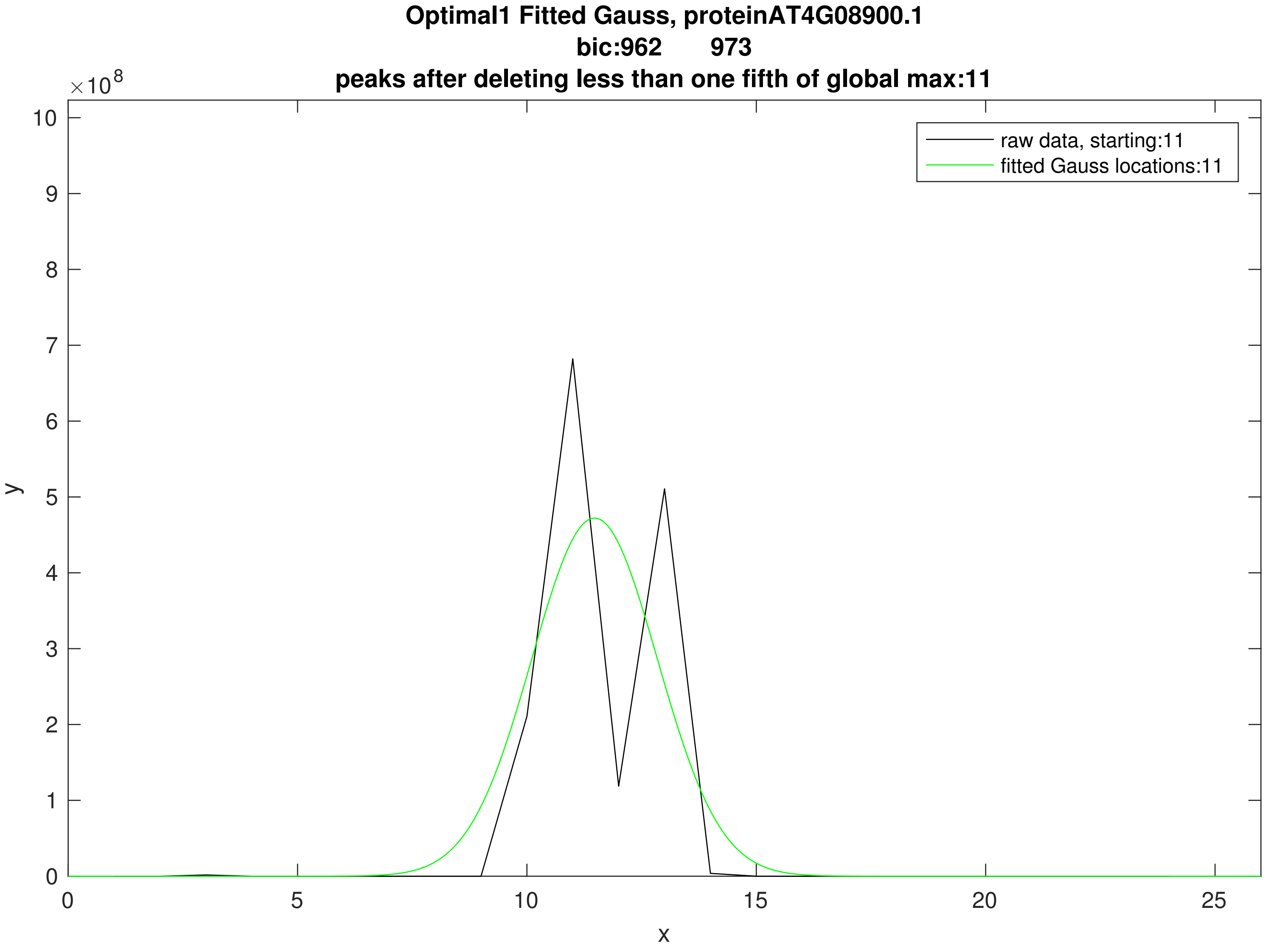

Optimal1 Fitted Gauss, proteinAT4G08900.1
bic:962 973
peaks after deleting less than one fifth of global max:11
×10^8
10
9
8
7
6
5
4
3
2
1
0
y
0
5
10
15
20
25
x
raw data, starting:11
fitted Gauss locations:11

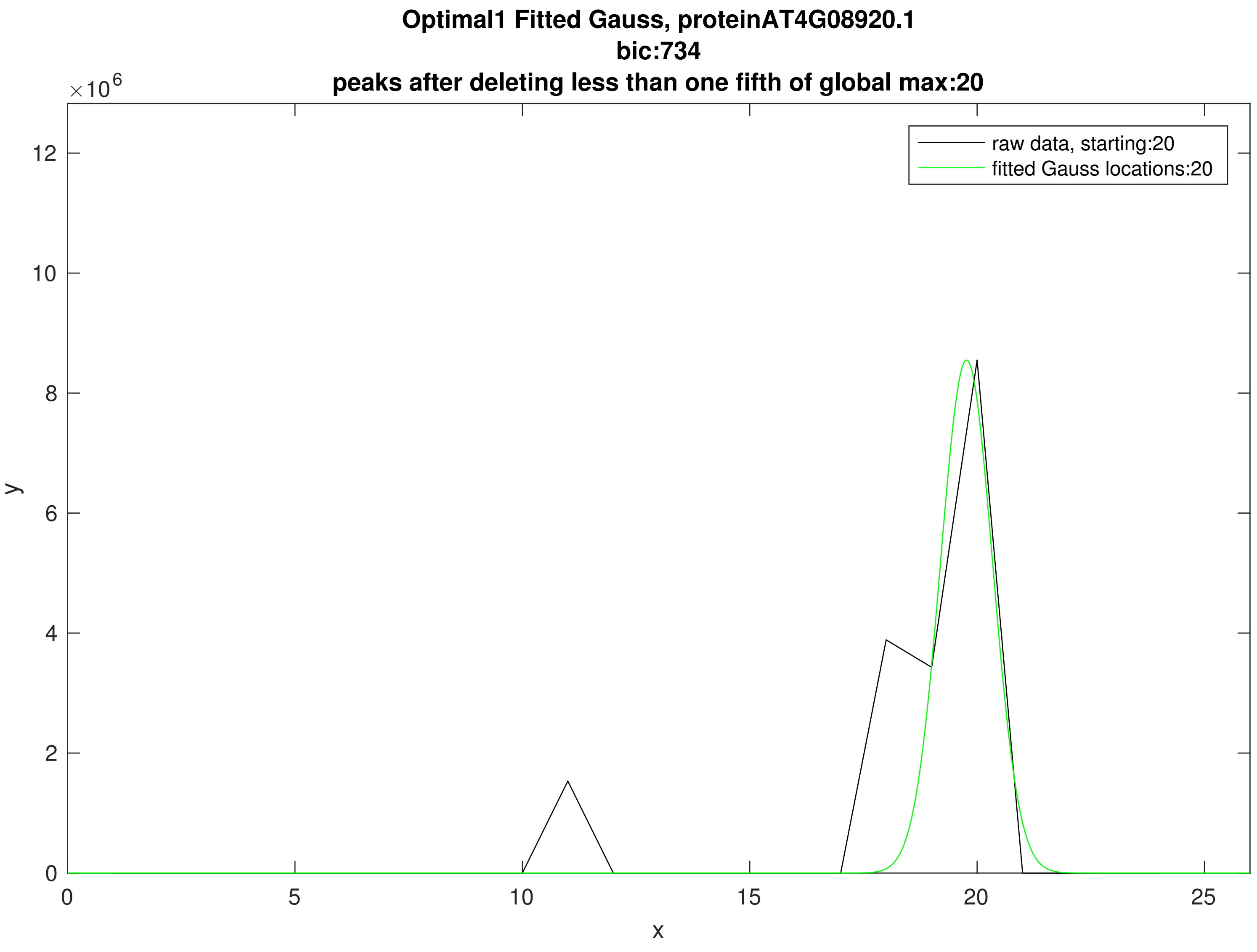

Optimal1 Fitted Gauss, proteinAT4G08920.1
bic:734
peaks after deleting less than one fifth of global max:20
×10^6
raw data, starting:20
fitted Gauss locations:20
y
x
0
2
4
6
8
10
12
0
5
10
15
20
25

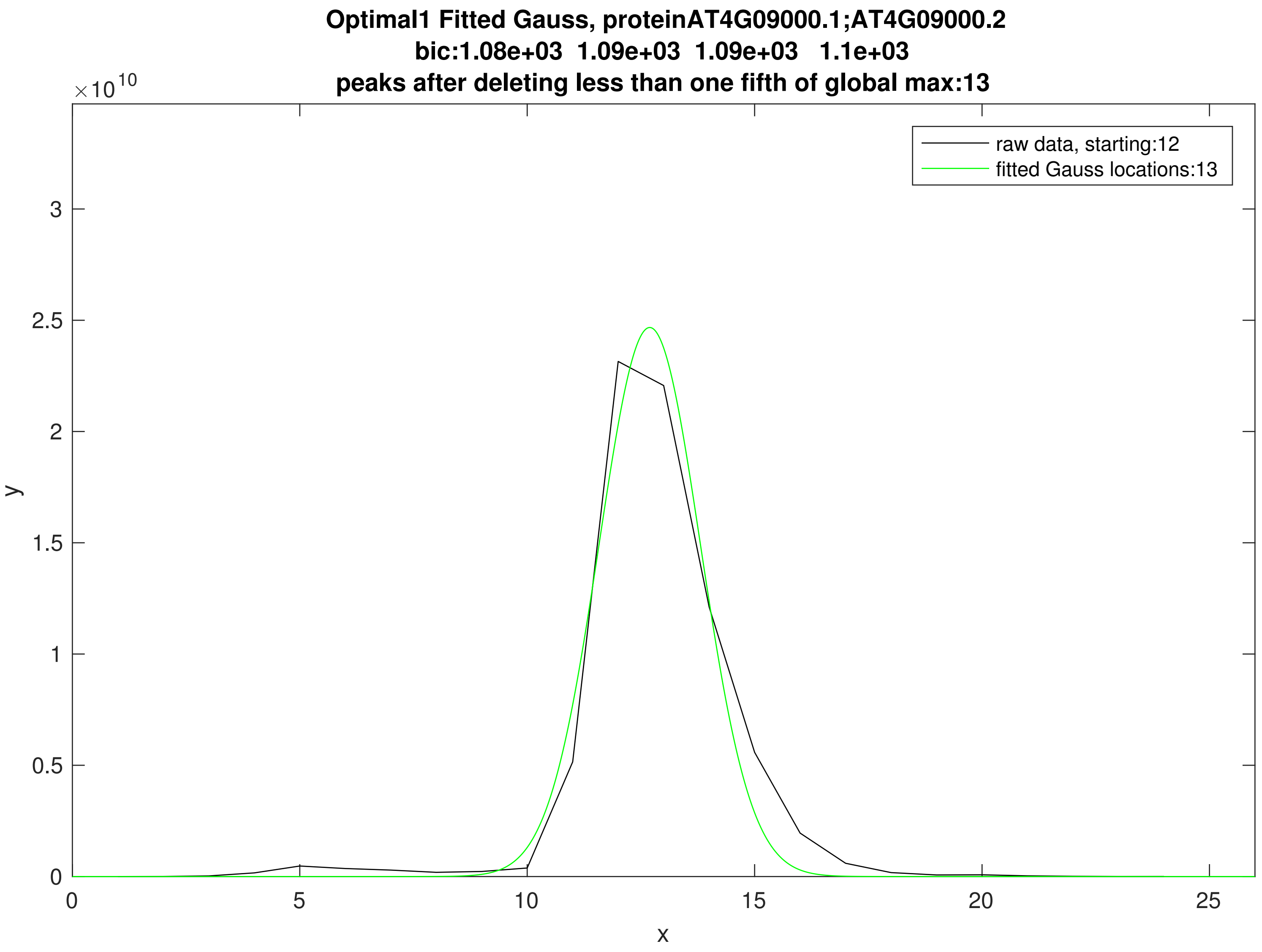
Optimal1 Fitted Gauss, proteinAT4G09000.1;AT4G09000.2
bic:1.08e+03 1.09e+03 1.09e+03 1.1e+03
peaks after deleting less than one fifth of global max:13
×10^10
raw data, starting:12
fitted Gauss locations:13
y
x
0
0.5
1
1.5
2
2.5
3
0
5
10
15
20
25

**Optimal1 Fitted Gauss, proteinAT4G09010.1**
**bic:753 760**
**peaks after deleting less than one fifth of global max:19**

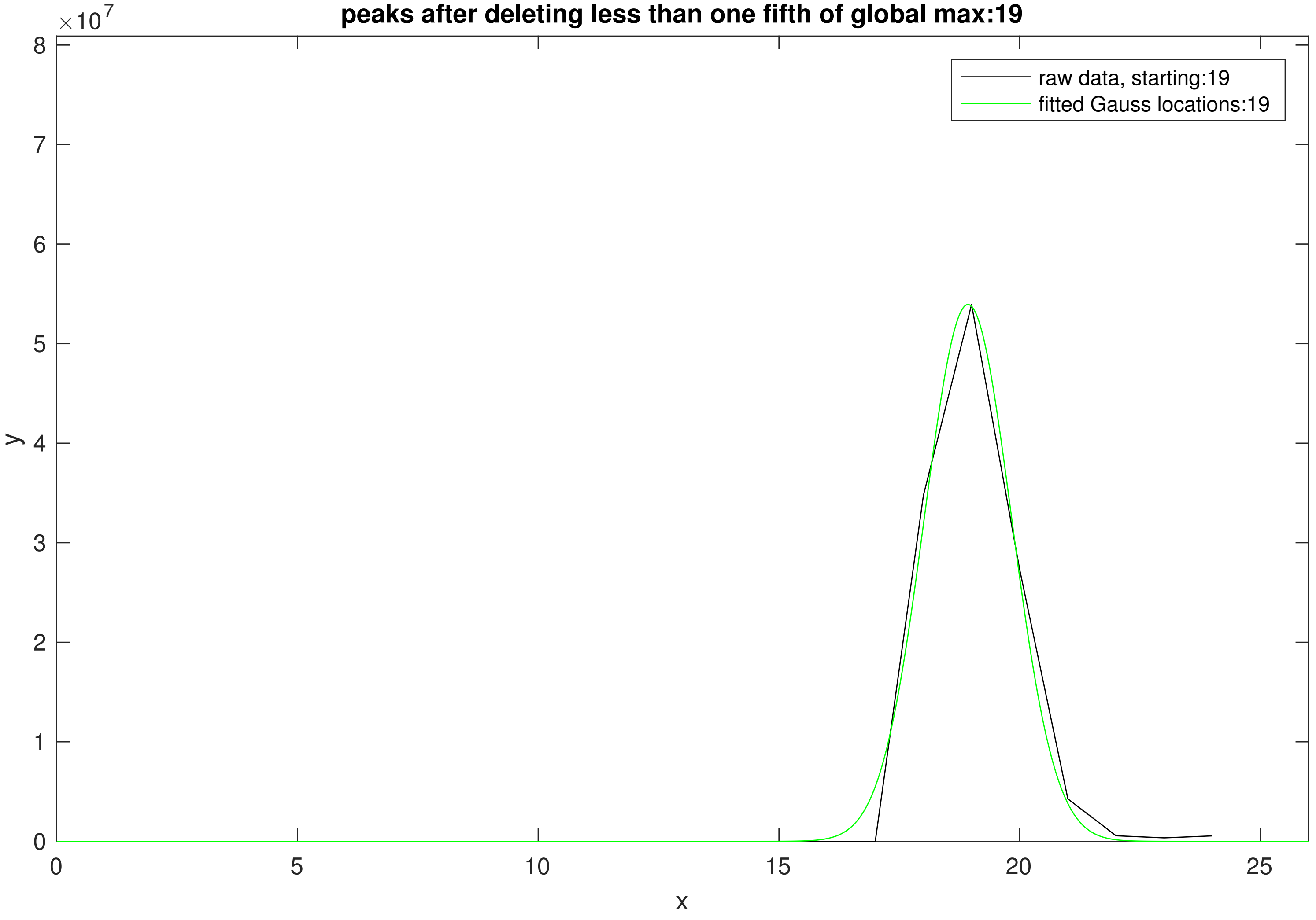

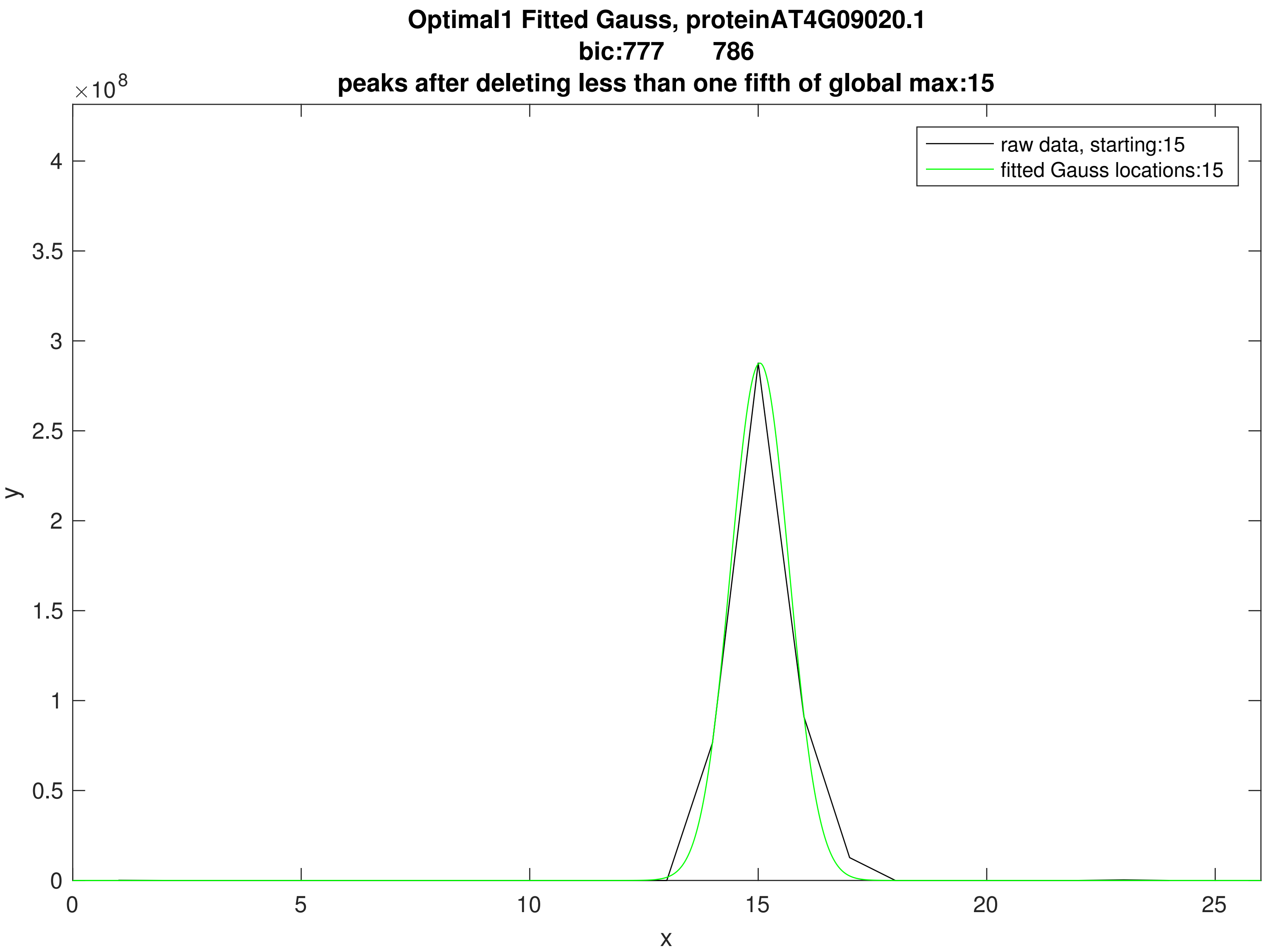
Optimal1 Fitted Gauss, proteinAT4G09020.1
bic:777 786
peaks after deleting less than one fifth of global max:15
raw data, starting:15
fitted Gauss locations:15
y
x
×10^8
0
0.5
1
1.5
2
2.5
3
3.5
4
0
5
10
15
20
25

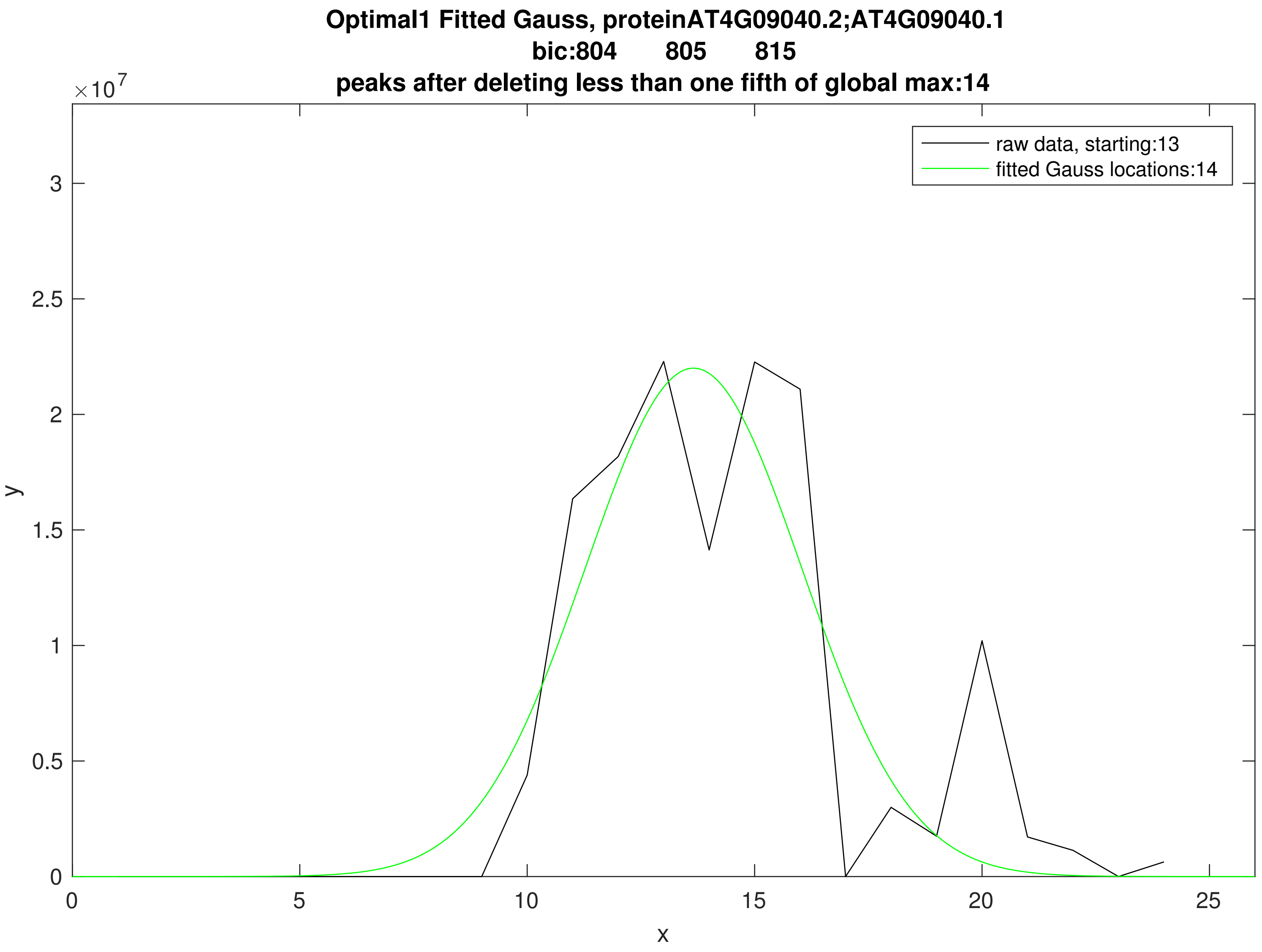
Optimal1 Fitted Gauss, proteinAT4G09040.2;AT4G09040.1
bic:804 805 815
peaks after deleting less than one fifth of global max:14
×10^7
raw data, starting:13
fitted Gauss locations:14
y
x
0
0.5
1
1.5
2
2.5
3
0
5
10
15
20
25

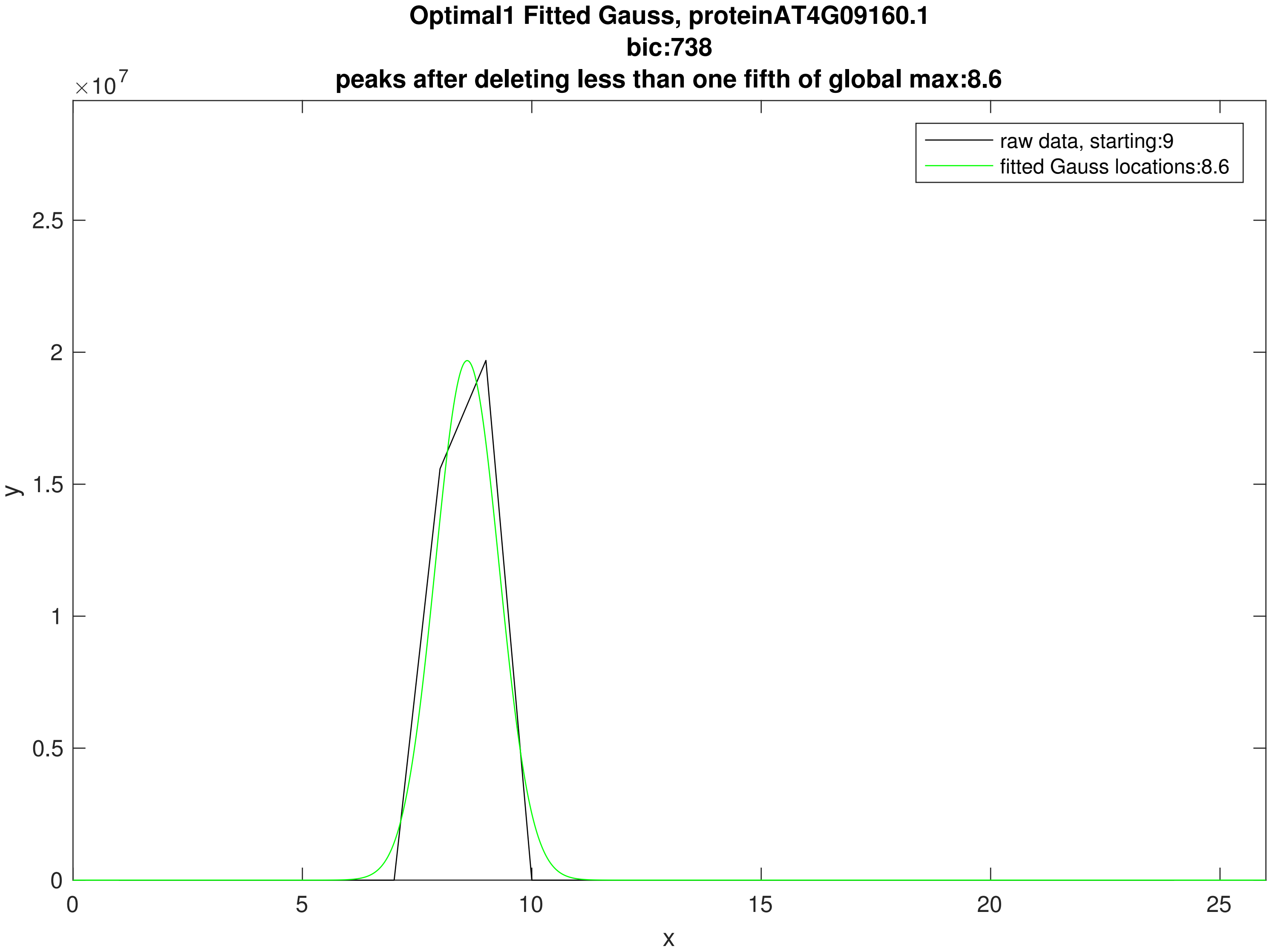
Optimal1 Fitted Gauss, proteinAT4G09160.1
bic:738
peaks after deleting less than one fifth of global max:8.6
×10^7
raw data, starting:9
fitted Gauss locations:8.6
2.5
2
1.5
1
0.5
0
y
0
5
10
15
20
25
x

**Optimal2 Fitted Gauss, proteinAT4G09320.1**
**bic:1.02e+03 1e+03 1.01e+03 1.02e+03**
**peaks after deleting less than one fifth of global max:16 20**

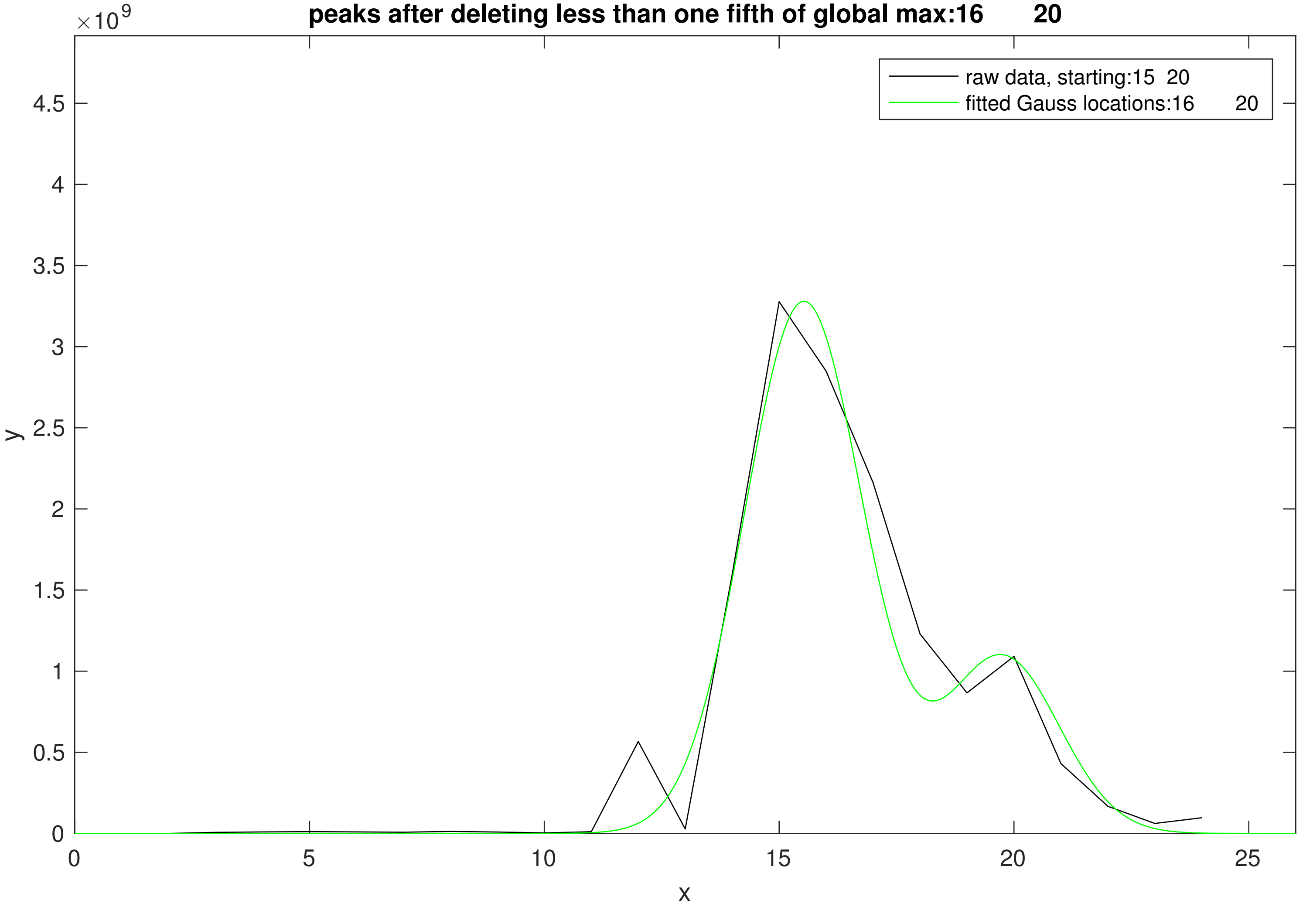

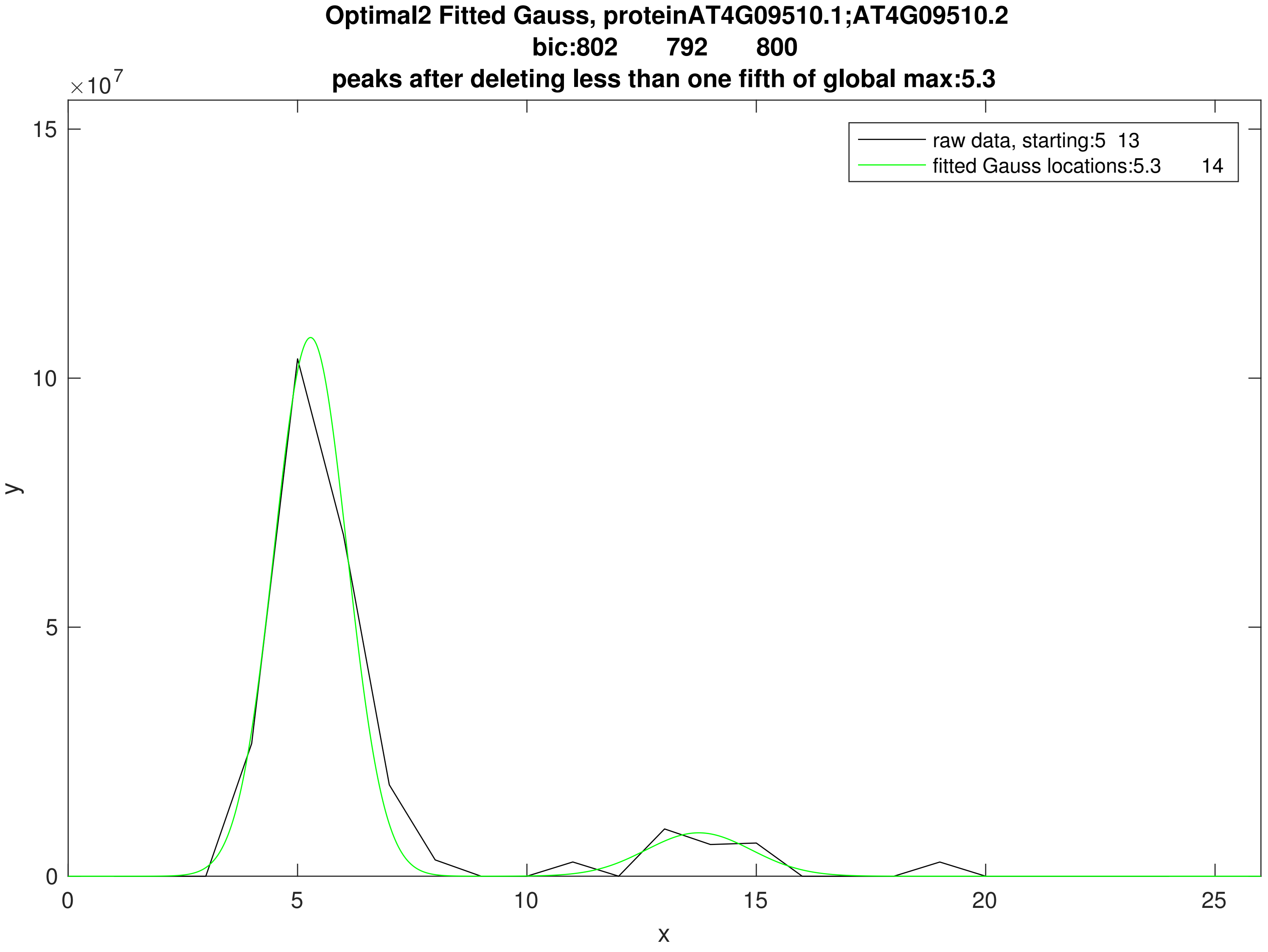
Optimal2 Fitted Gauss, proteinAT4G09510.1;AT4G09510.2
bic:802 792 800
peaks after deleting less than one fifth of global max:5.3
×10^7
15
10
5
0
y
0
5
10
15
20
25
x
raw data, starting:5 13
fitted Gauss locations:5.3 14

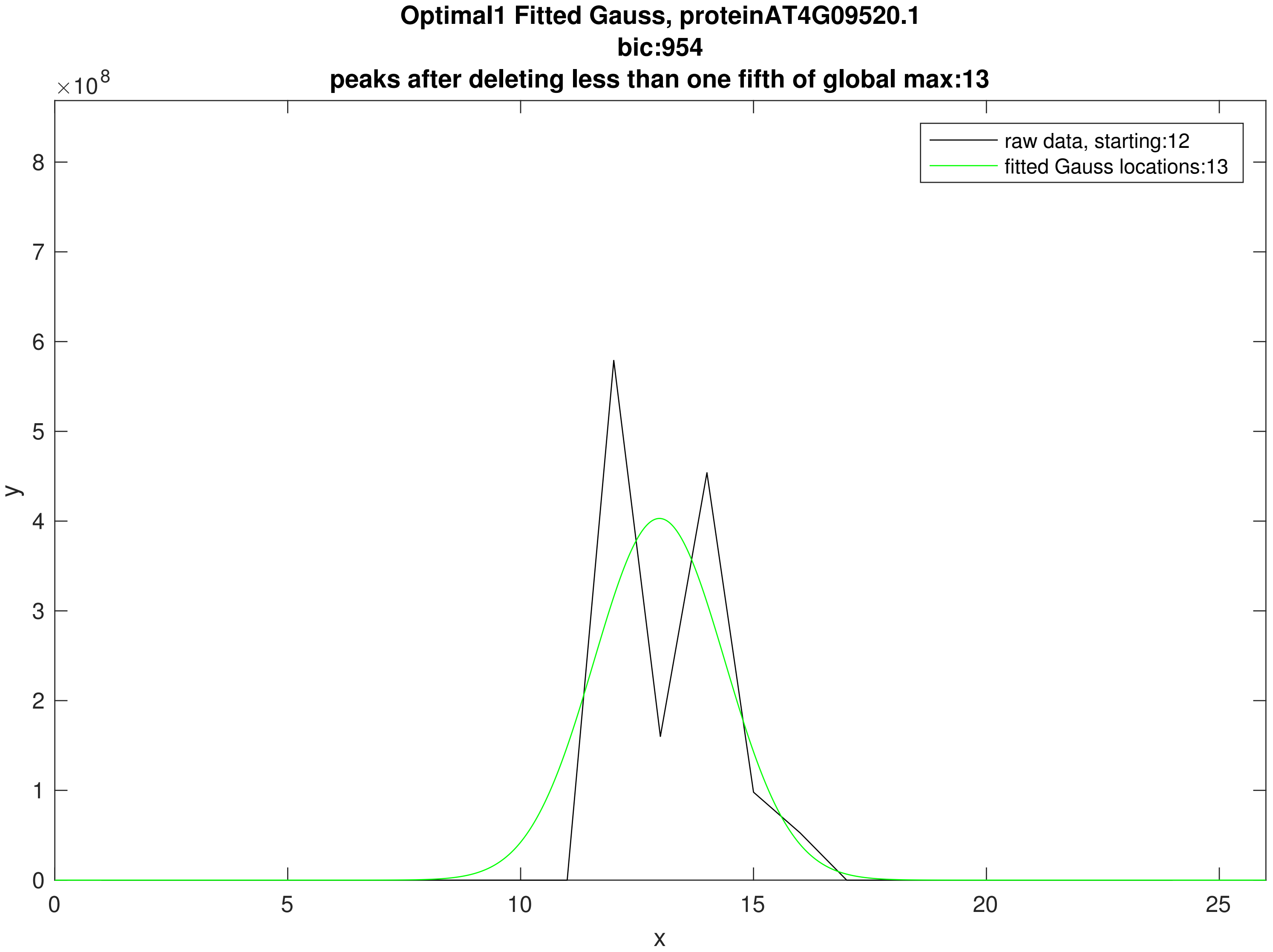

Optimal1 Fitted Gauss, proteinAT4G09520.1
bic:954
peaks after deleting less than one fifth of global max:13
×10^8
raw data, starting:12
fitted Gauss locations:13
y
x
0
1
2
3
4
5
6
7
8
0
5
10
15
20
25

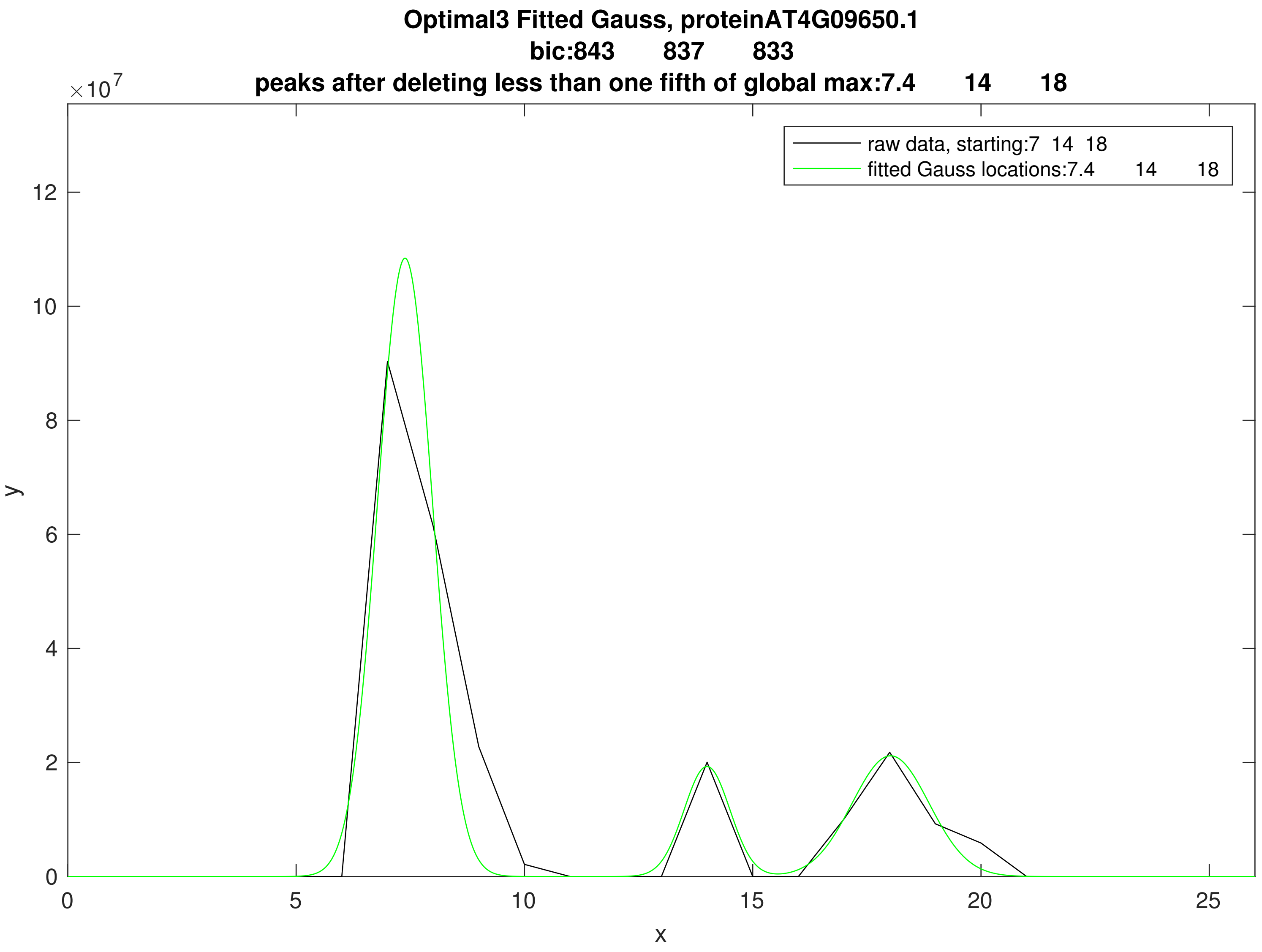
Optimal3 Fitted Gauss, proteinAT4G09650.1
bic:843 837 833
peaks after deleting less than one fifth of global max:7.4 14 18
×10^7
raw data, starting:7 14 18
fitted Gauss locations:7.4 14 18
y
x
0
2
4
6
8
10
12
0
5
10
15
20
25

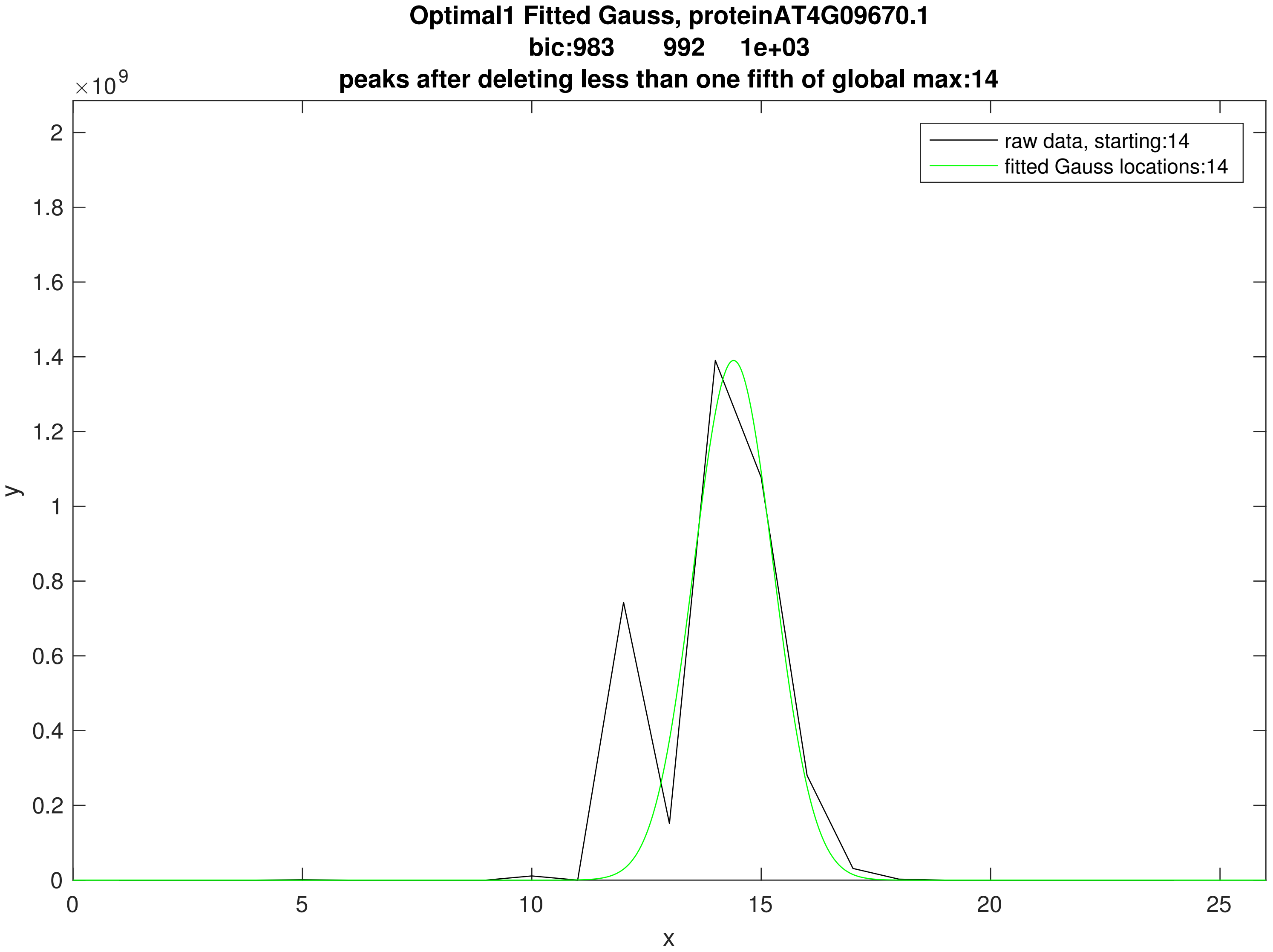

Optimal1 Fitted Gauss, proteinAT4G09670.1
bic:983 992 1e+03
peaks after deleting less than one fifth of global max:14
×10^9
2
1.8
1.6
1.4
1.2
1
0.8
0.6
0.4
0.2
0
y
0
5
10
15
20
25
x
raw data, starting:14
fitted Gauss locations:14

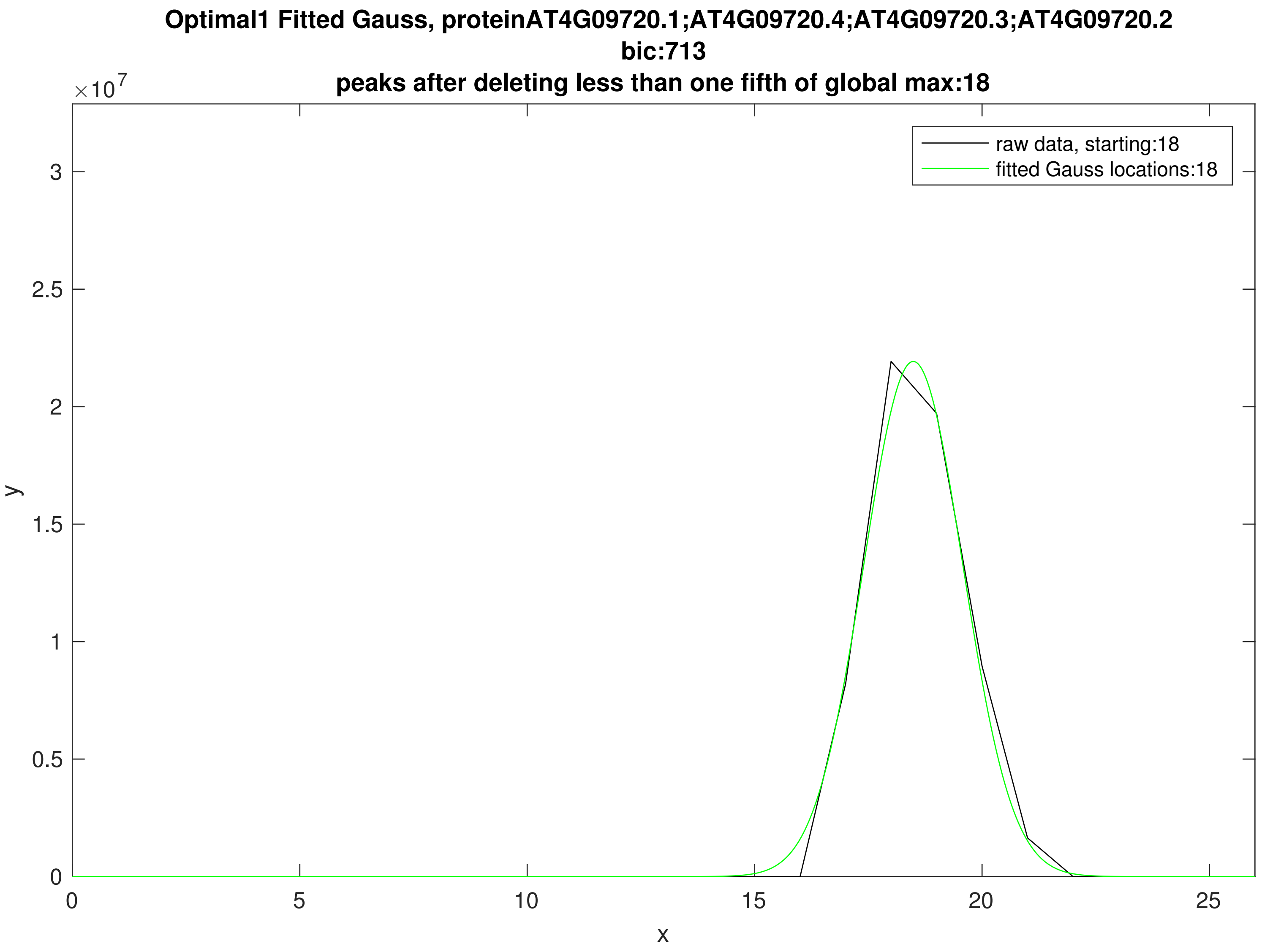

Optimal1 Fitted Gauss, proteinAT4G09720.1;AT4G09720.4;AT4G09720.3;AT4G09720.2
bic:713
peaks after deleting less than one fifth of global max:18
×10^7
raw data, starting:18
fitted Gauss locations:18
y
x
0
0.5
1
1.5
2
2.5
3
0
5
10
15
20
25

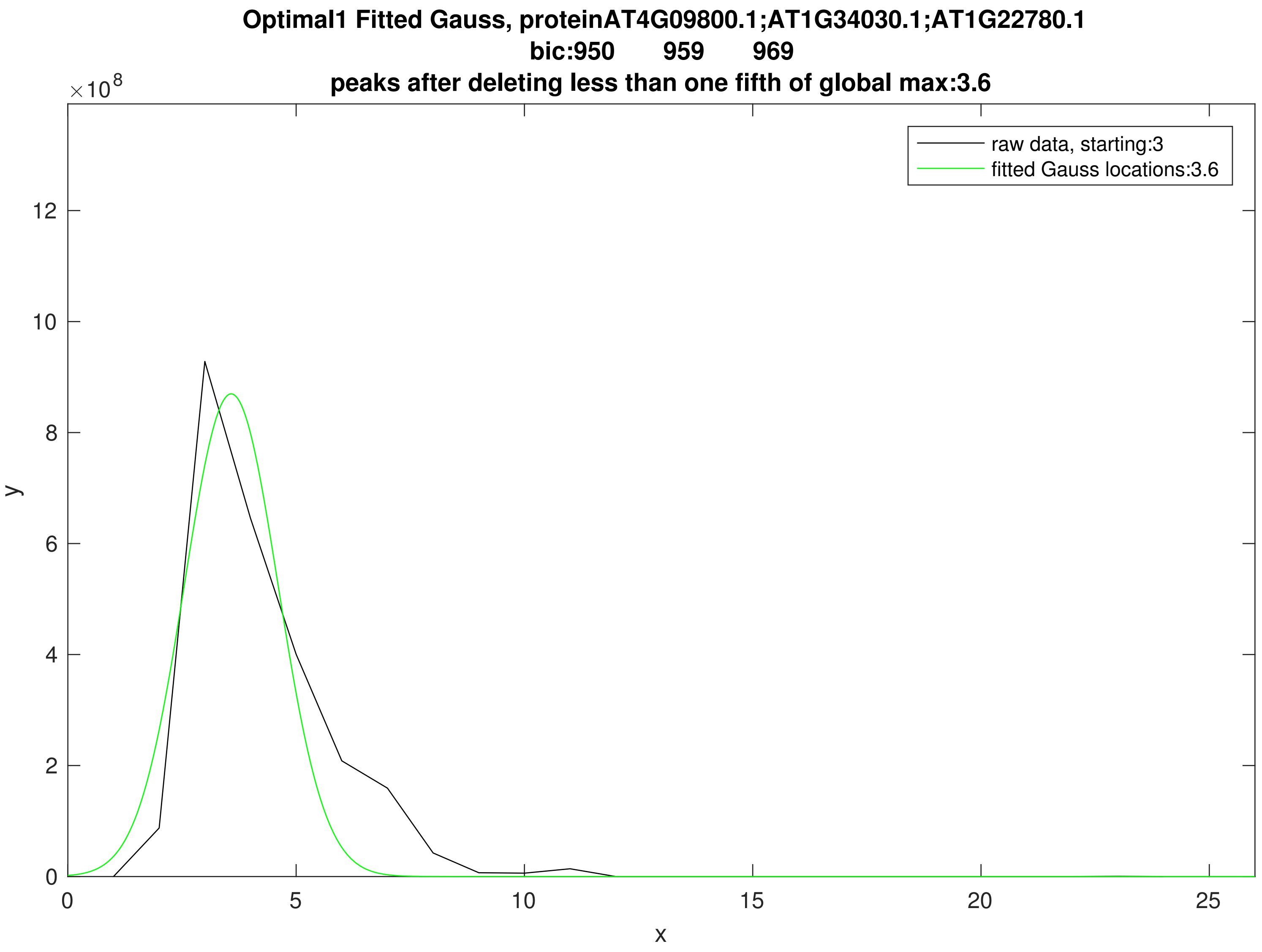

Optimal1 Fitted Gauss, proteinAT4G09800.1;AT1G34030.1;AT1G22780.1
bic:950 959 969
peaks after deleting less than one fifth of global max:3.6
×10^8
raw data, starting:3
fitted Gauss locations:3.6
y
x
0
2
4
6
8
10
12
0
5
10
15
20
25

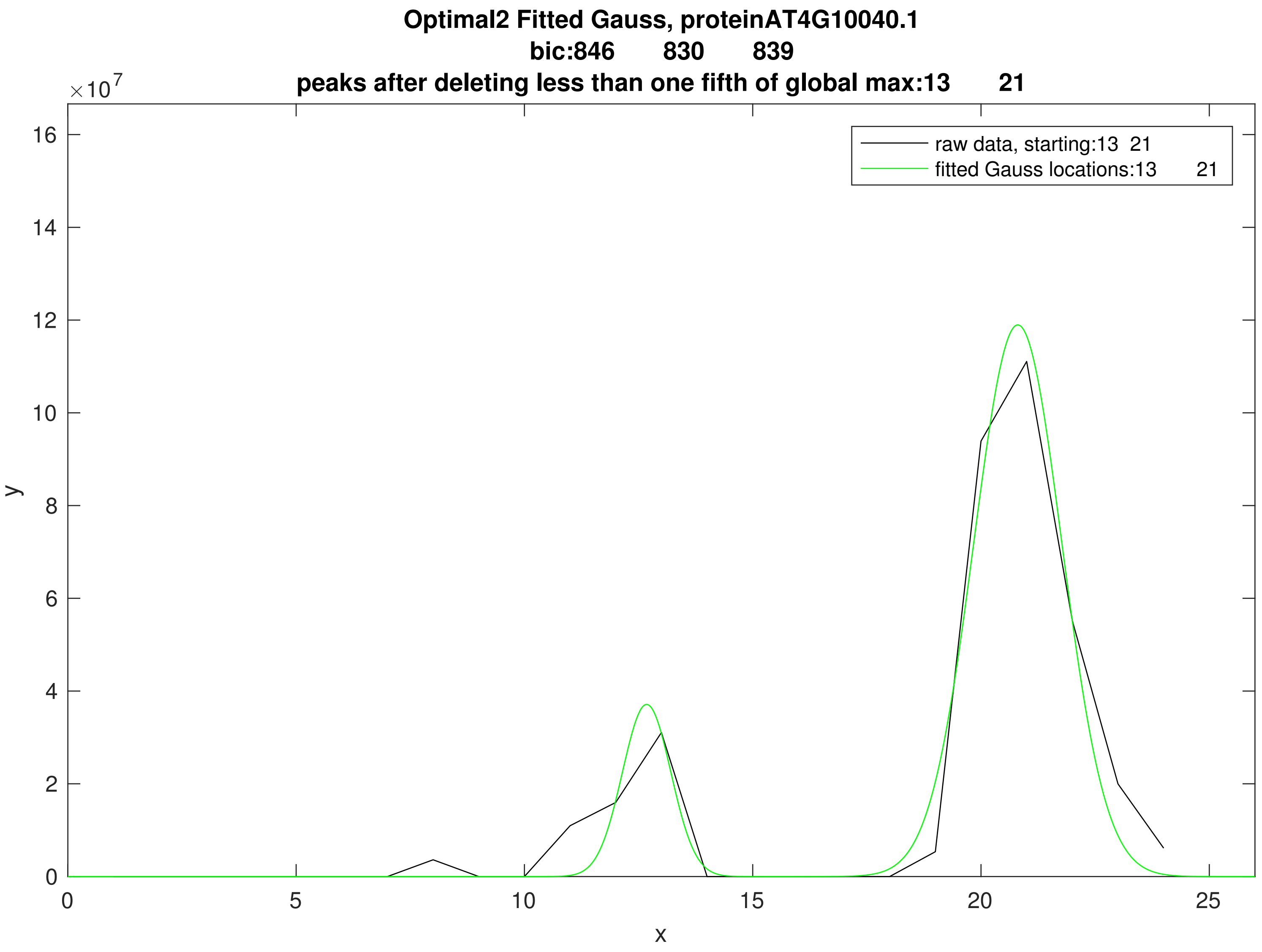

Optimal2 Fitted Gauss, proteinAT4G10040.1
bic:846 830 839
peaks after deleting less than one fifth of global max:13 21
×10^7
raw data, starting:13 21
fitted Gauss locations:13 21
y
x
0
2
4
6
8
10
12
14
16
0
5
10
15
20
25

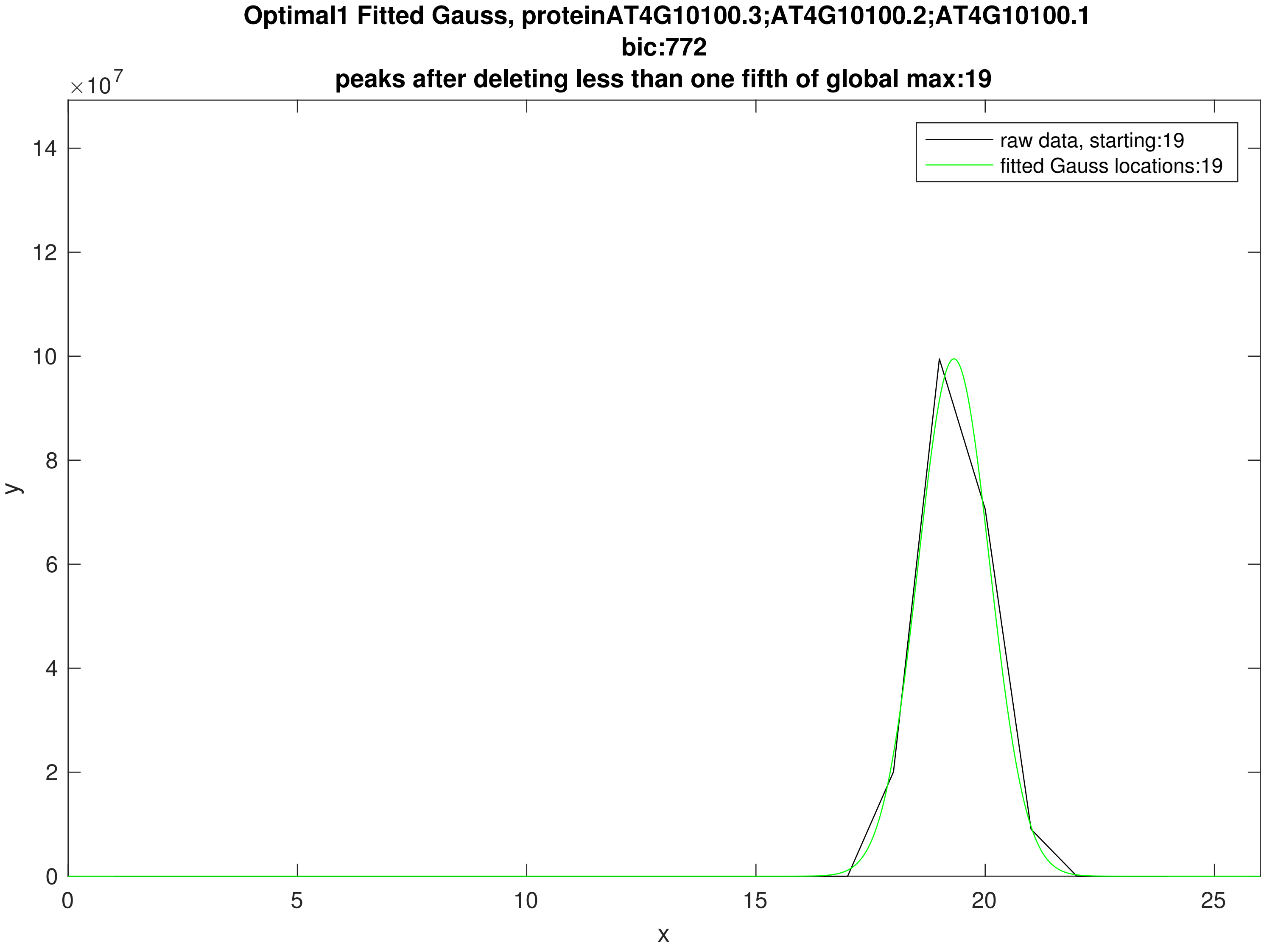
Optimal1 Fitted Gauss, proteinAT4G10100.3;AT4G10100.2;AT4G10100.1
bic:772
peaks after deleting less than one fifth of global max:19
$\times 10^7$
raw data, starting:19
fitted Gauss locations:19
y
x
0
2
4
6
8
10
12
14
5
15
20
25

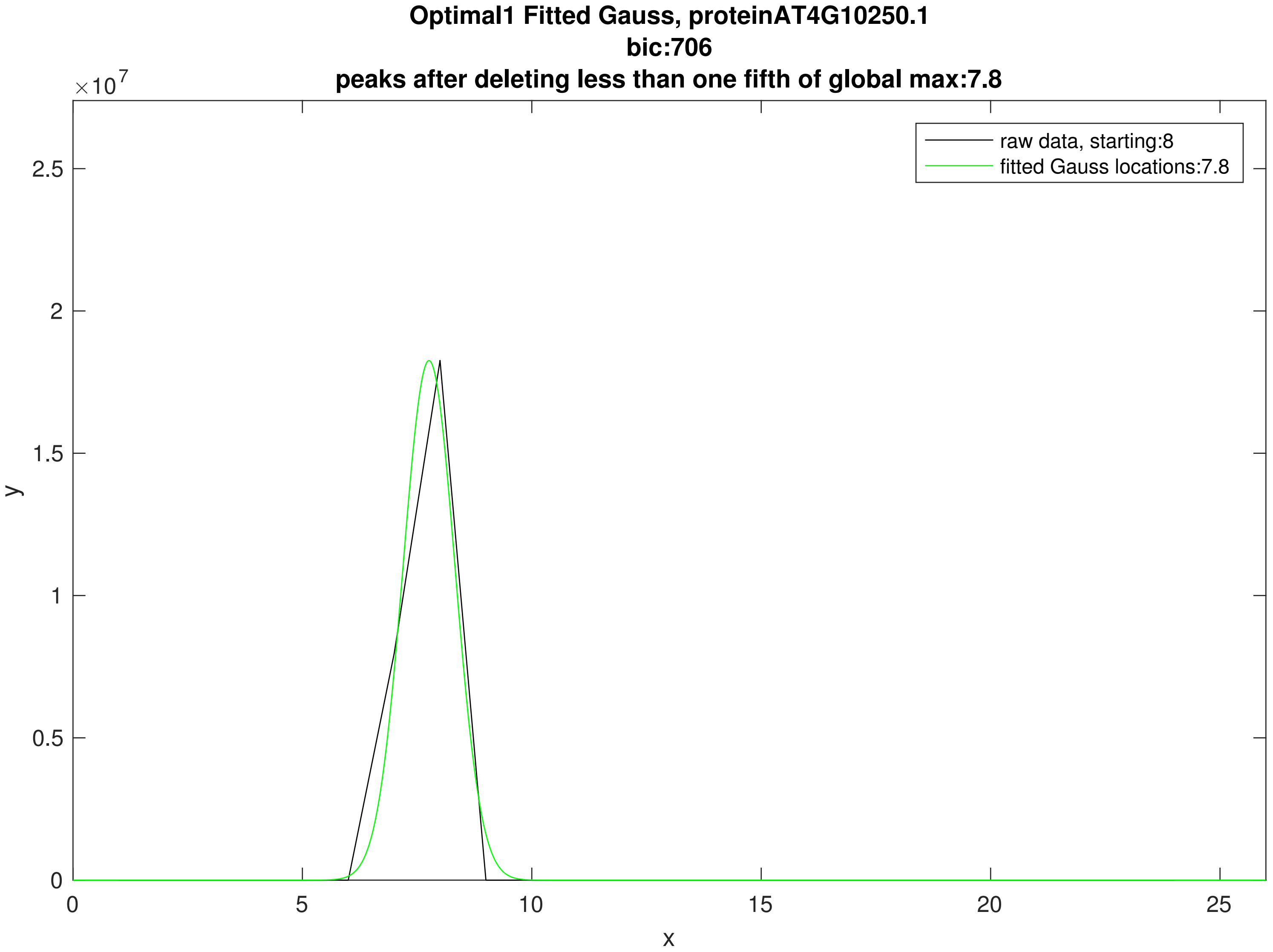
Optimal1 Fitted Gauss, proteinAT4G10250.1
bic:706
peaks after deleting less than one fifth of global max:7.8
×10^7
2.5
2
1.5
1
0.5
0
y
0
5
10
15
20
25
x
raw data, starting:8
fitted Gauss locations:7.8

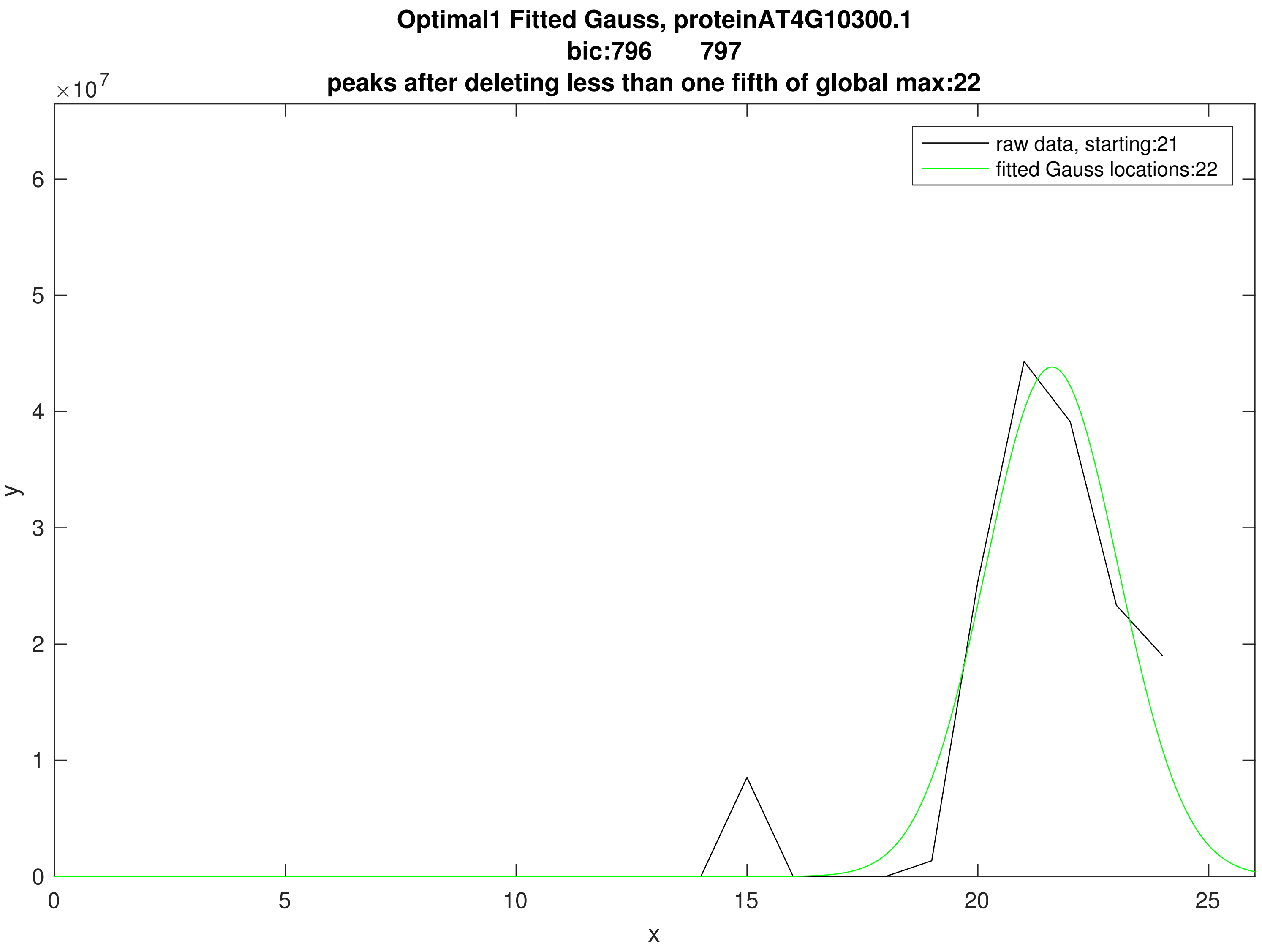
Optimal1 Fitted Gauss, proteinAT4G10300.1
bic:796 797
peaks after deleting less than one fifth of global max:22
×10^7
raw data, starting:21
fitted Gauss locations:22
y
x
0
1
2
3
4
5
6
0
5
10
15
20
25

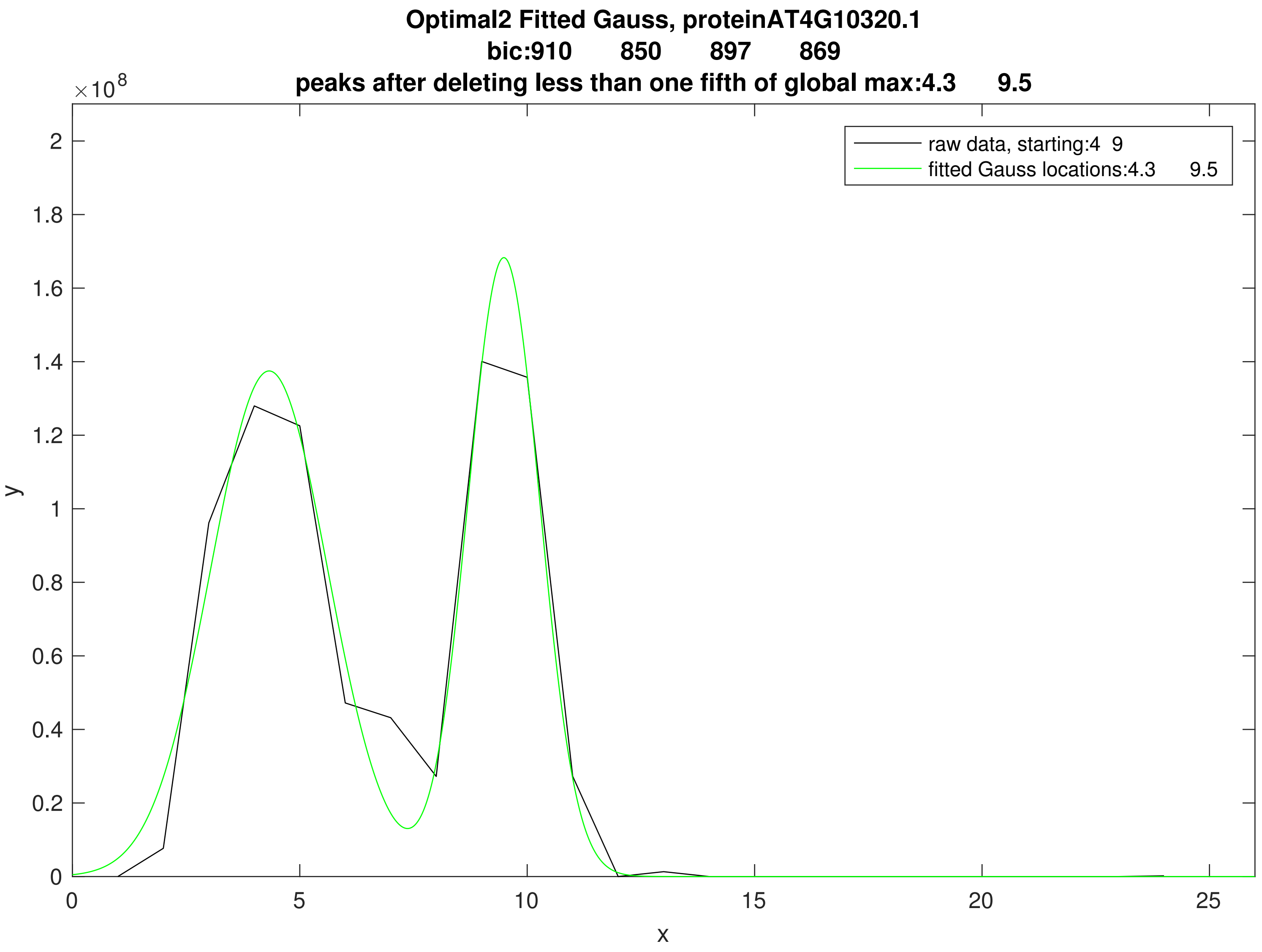

Optimal2 Fitted Gauss, proteinAT4G10320.1
bic:910 850 897 869
peaks after deleting less than one fifth of global max:4.3 9.5
×10^8
raw data, starting:4 9
fitted Gauss locations:4.3 9.5
y
x
0
0.2
0.4
0.6
0.8
1
1.2
1.4
1.6
1.8
2
5
10
15
20
25

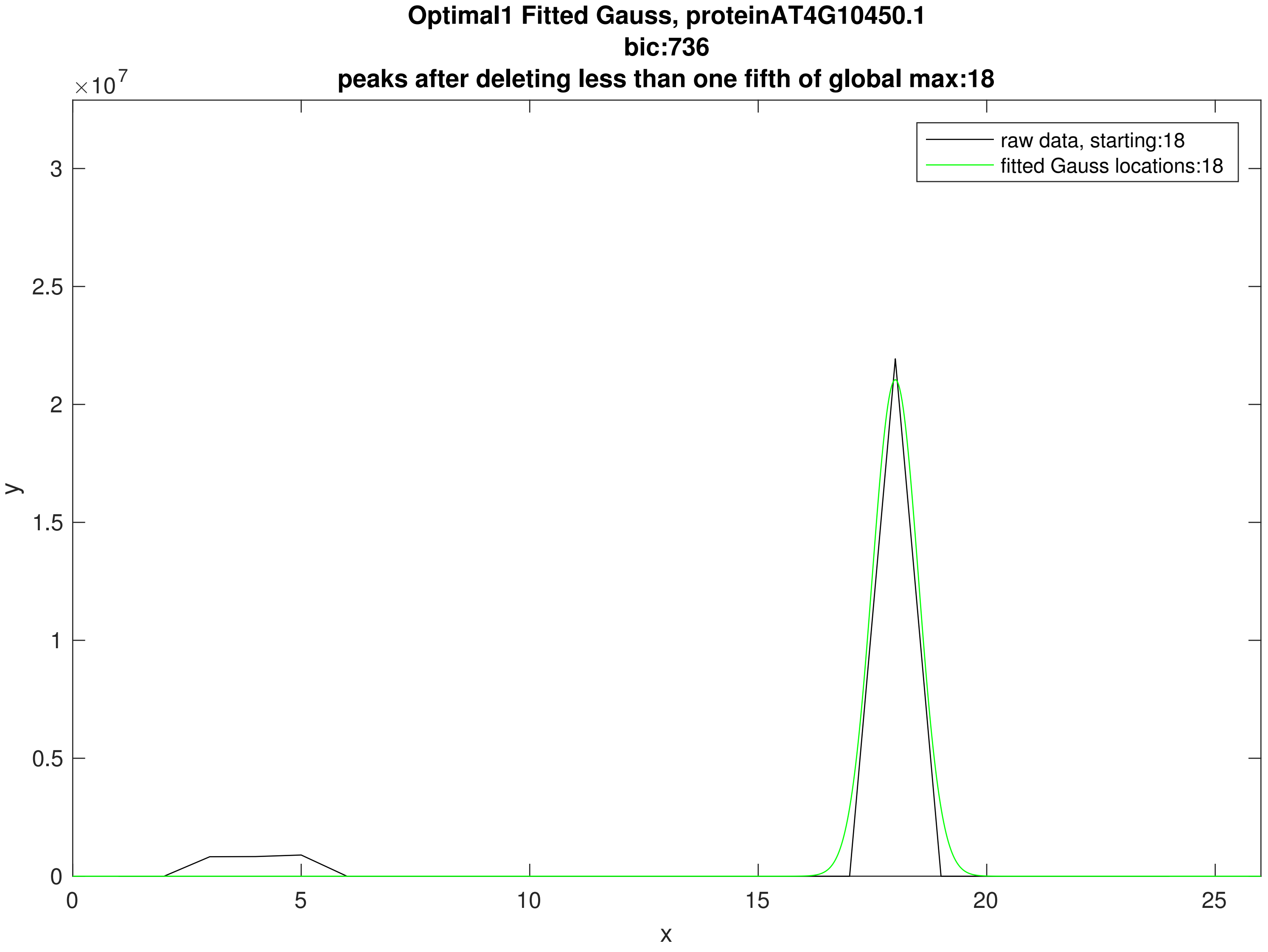

Optimal1 Fitted Gauss, proteinAT4G10450.1
bic:736
peaks after deleting less than one fifth of global max:18
×10^7
raw data, starting:18
fitted Gauss locations:18
y
x
0
0.5
1
1.5
2
2.5
3
0
5
10
15
20
25

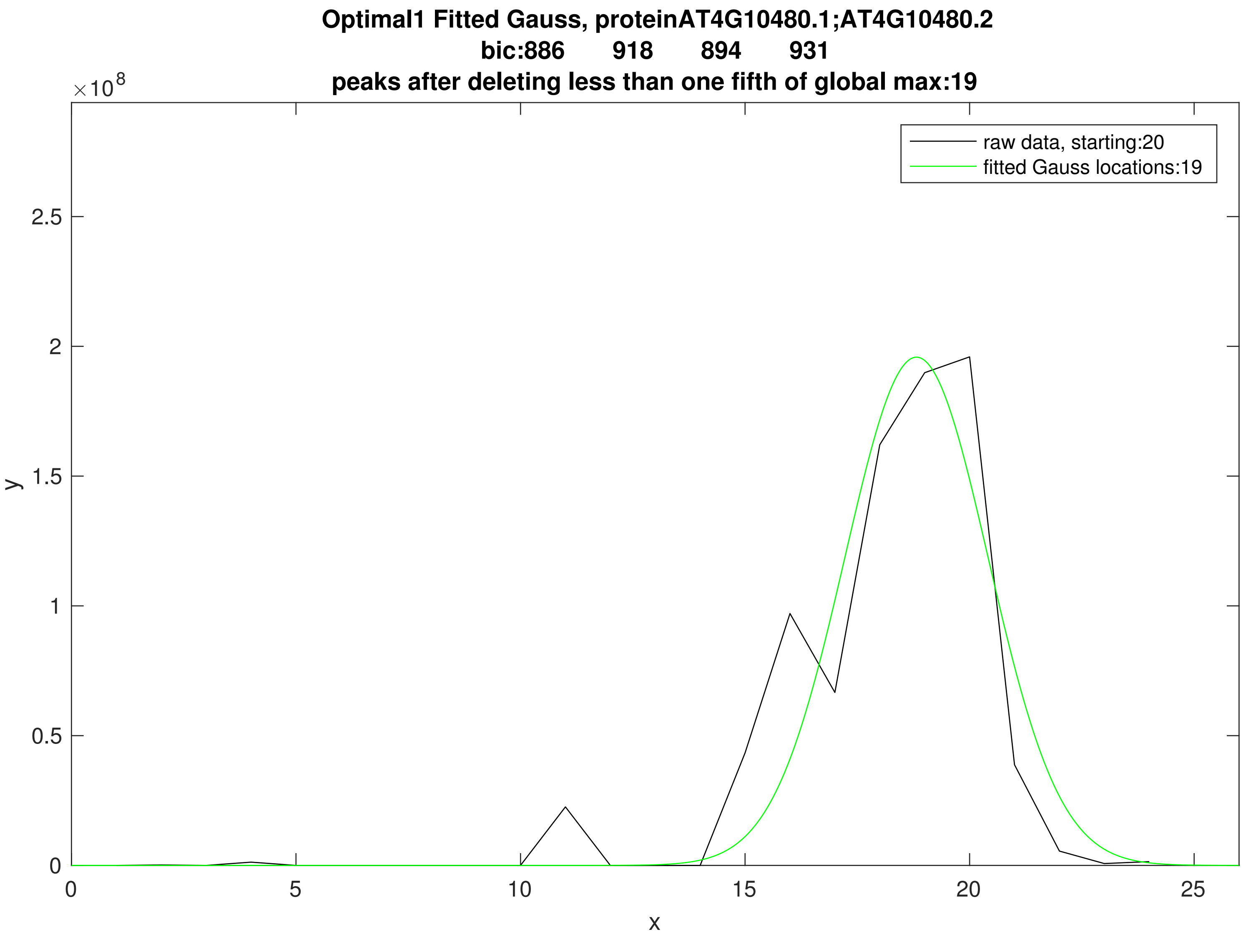

Optimal1 Fitted Gauss, proteinAT4G10480.1;AT4G10480.2
bic:886 918 894 931
peaks after deleting less than one fifth of global max:19
×10^8
raw data, starting:20
fitted Gauss locations:19
y
x
0
0.5
1
1.5
2
2.5
0
5
10
15
20
25

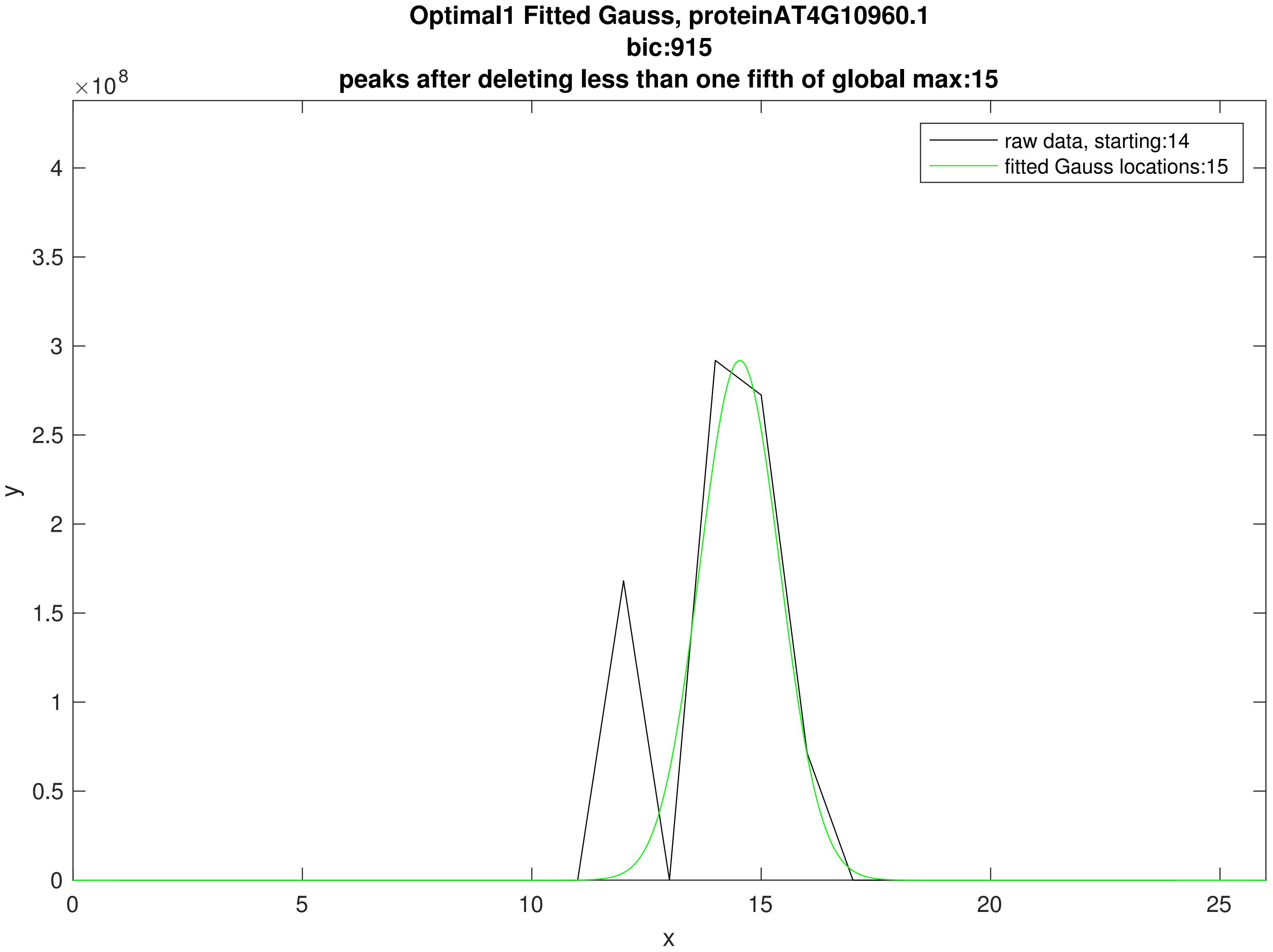

Optimal1 Fitted Gauss, proteinAT4G10960.1
bic:915
peaks after deleting less than one fifth of global max:15
×10^8
raw data, starting:14
fitted Gauss locations:15
4
3.5
3
2.5
2
1.5
1
0.5
0
y
0
5
10
15
20
25
x

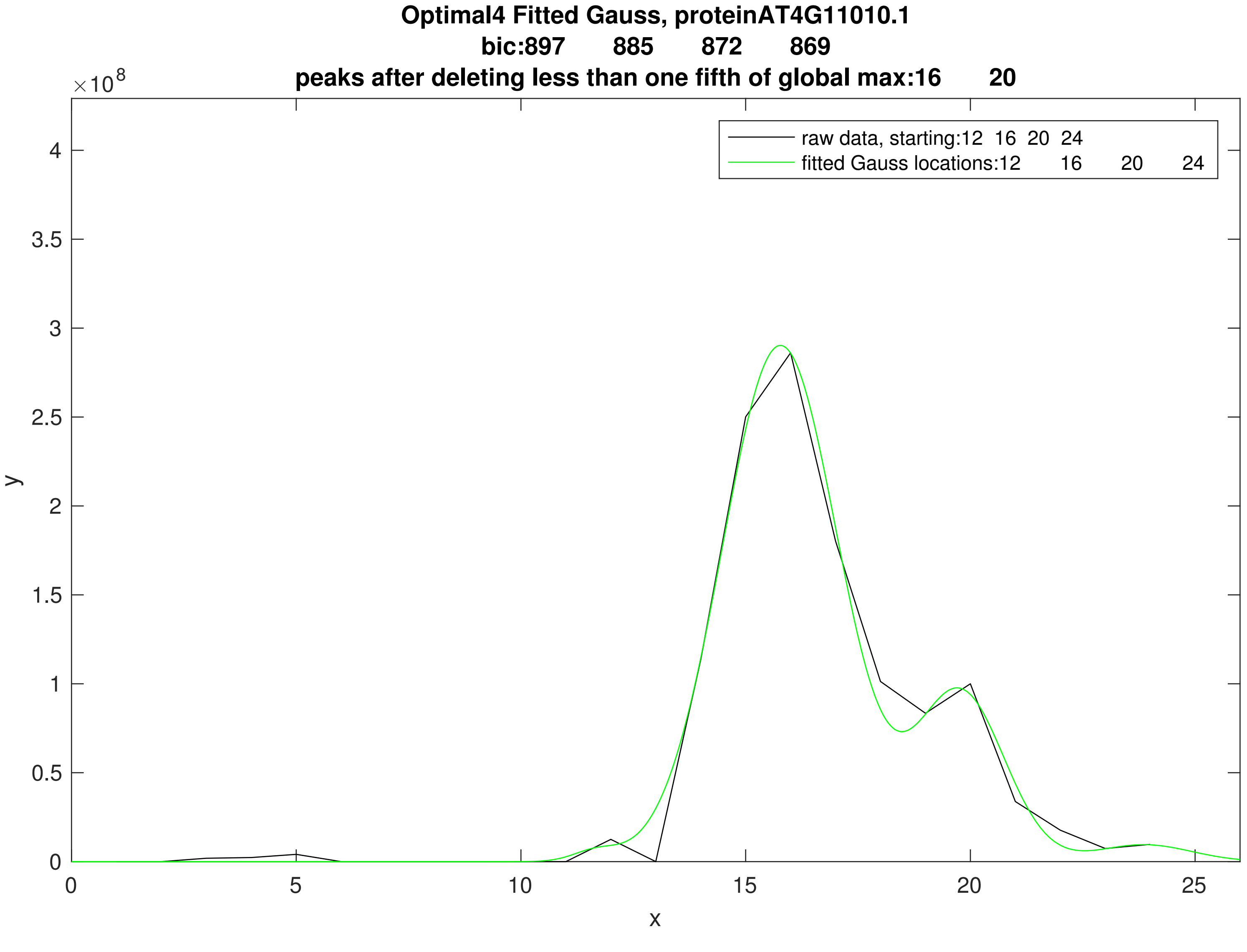

Optimal4 Fitted Gauss, proteinAT4G11010.1
bic:897 885 872 869
peaks after deleting less than one fifth of global max:16 20
×10^8
raw data, starting:12 16 20 24
fitted Gauss locations:12 16 20 24
y
x
0
0.5
1
1.5
2
2.5
3
3.5
4
5
10
15
20
25

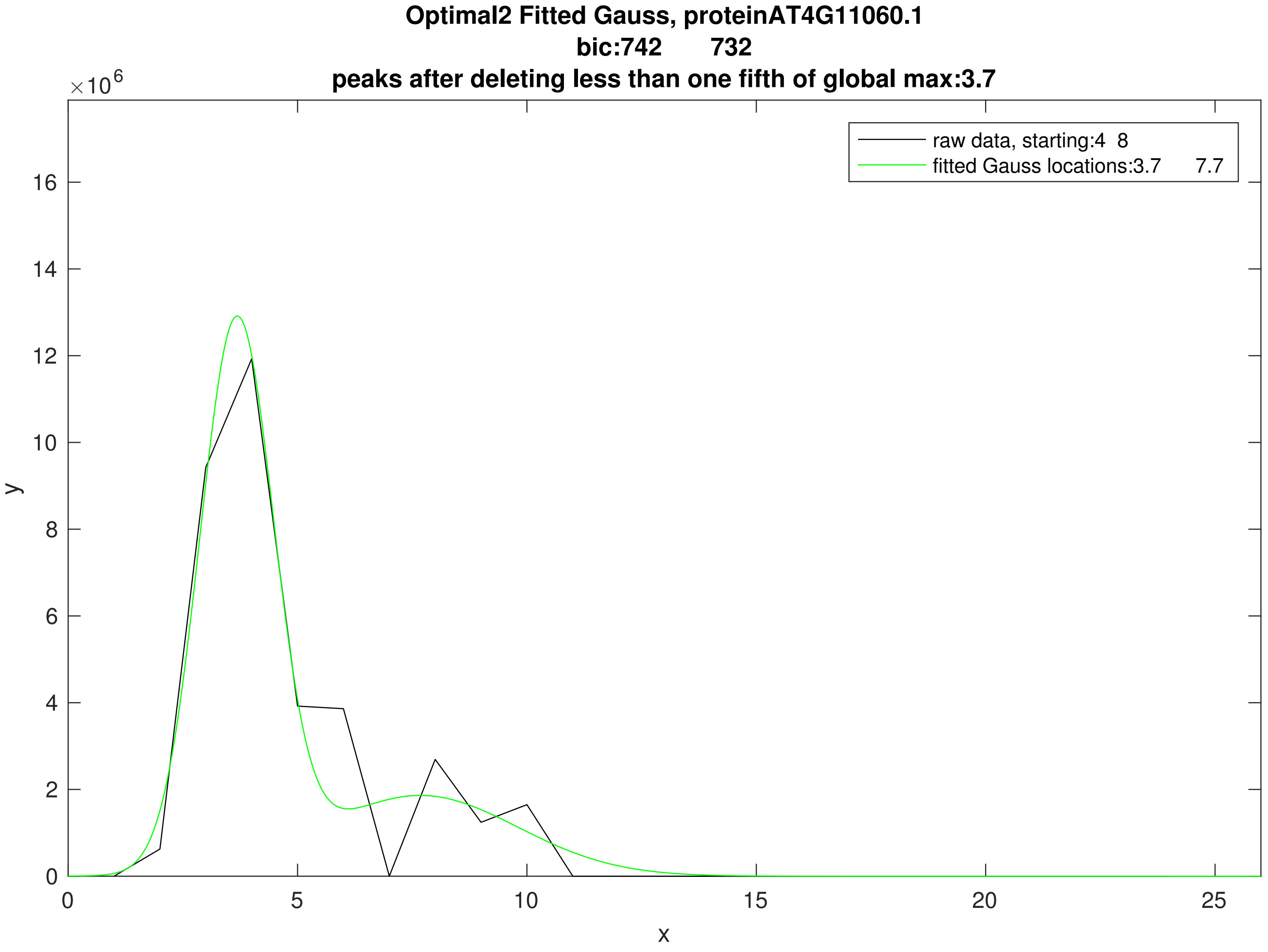

Optimal2 Fitted Gauss, proteinAT4G11060.1
bic:742 732
peaks after deleting less than one fifth of global max:3.7
×10^6
raw data, starting:4 8
fitted Gauss locations:3.7 7.7
y
x
0
2
4
6
8
10
12
14
16
5
15
20
25

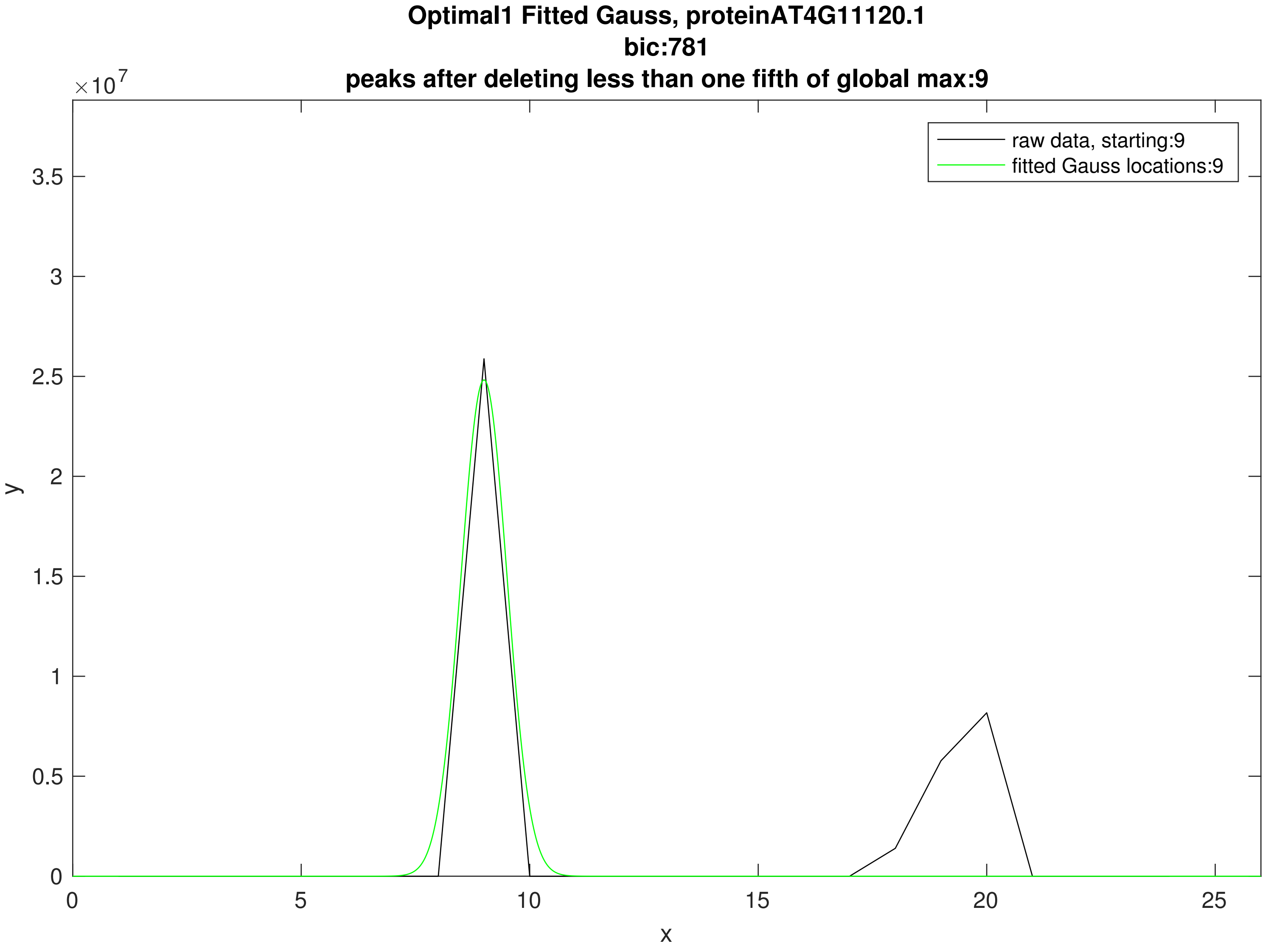
Optimal1 Fitted Gauss, proteinAT4G11120.1
bic:781
peaks after deleting less than one fifth of global max:9
×10^7
3.5
3
2.5
2
1.5
1
0.5
0
y
0
5
10
15
20
25
x
raw data, starting:9
fitted Gauss locations:9

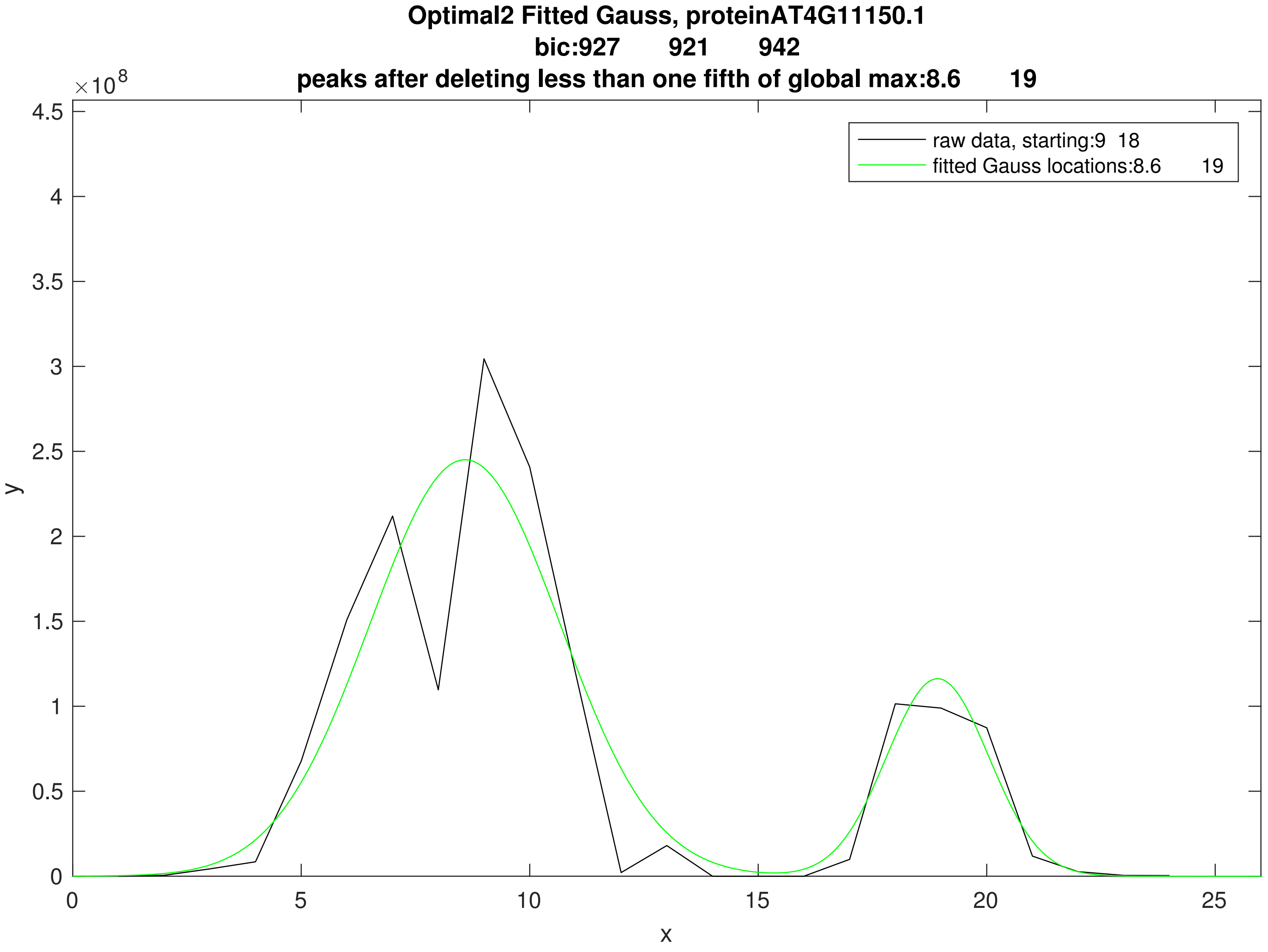
Optimal2 Fitted Gauss, proteinAT4G11150.1
bic:927 921 942
peaks after deleting less than one fifth of global max:8.6 19
×10^8
4.5
4
3.5
3
2.5
2
1.5
1
0.5
0
y
0
5
10
15
20
25
x
raw data, starting:9 18
fitted Gauss locations:8.6 19

**Optimal1 Fitted Gauss, proteinAT4G11190.1**
**bic:858 867 877**
**peaks after deleting less than one fifth of global max:18**

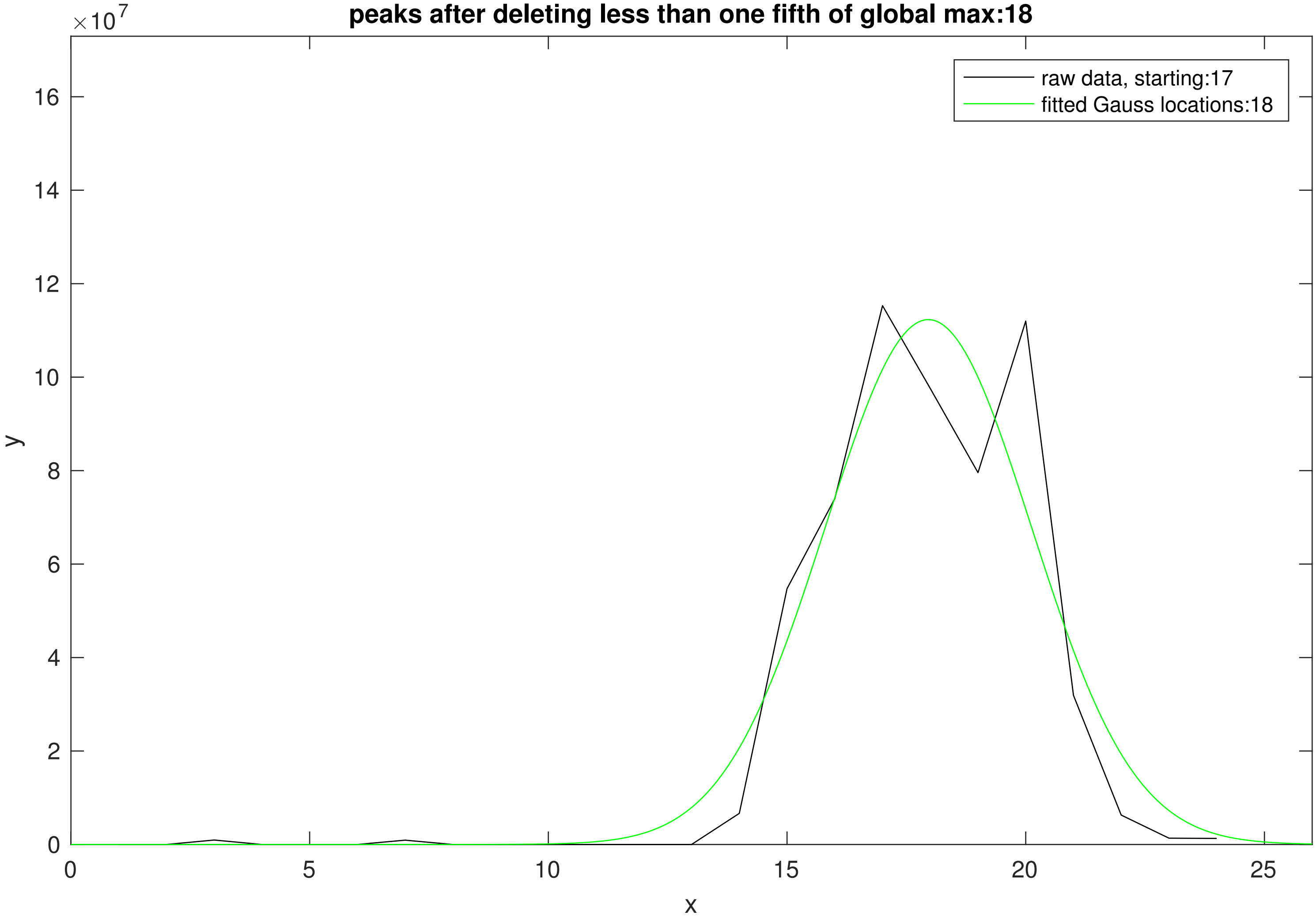

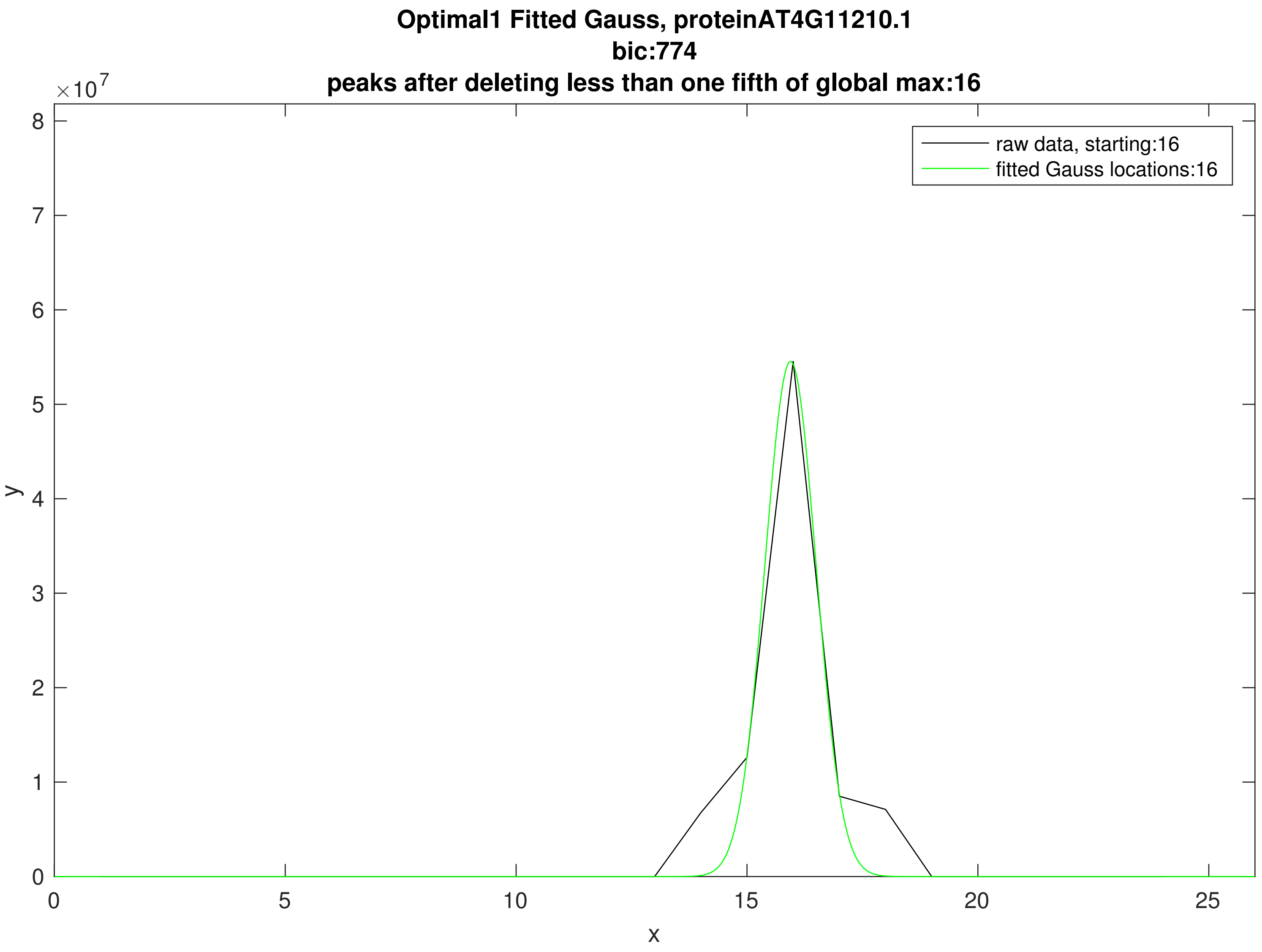
Optimal1 Fitted Gauss, proteinAT4G11210.1
bic:774
peaks after deleting less than one fifth of global max:16
×10^7
8
7
6
5
4
3
2
1
0
y
0
5
10
15
20
25
x
raw data, starting:16
fitted Gauss locations:16

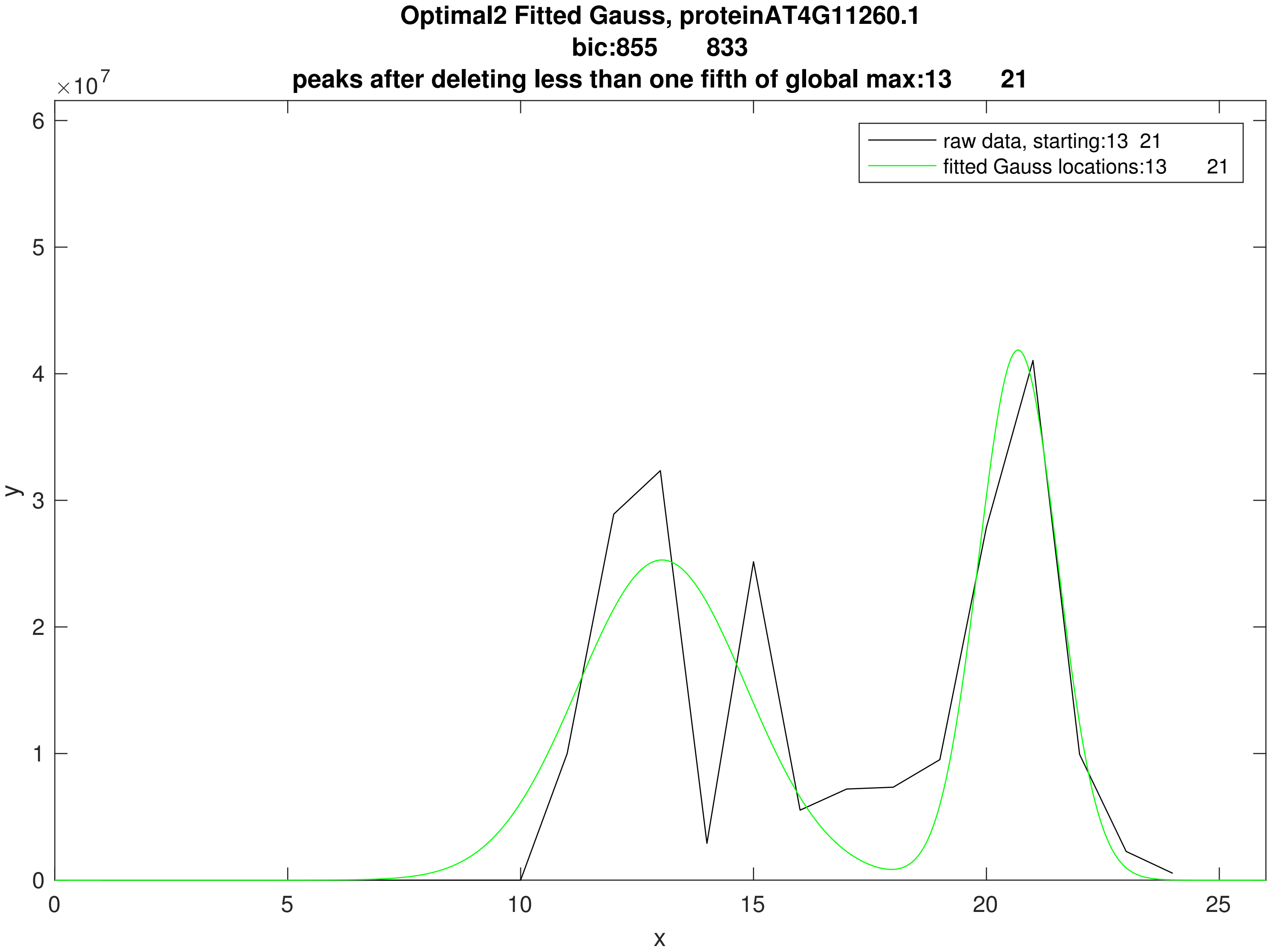

Optimal2 Fitted Gauss, proteinAT4G11260.1
bic:855 833
peaks after deleting less than one fifth of global max:13 21
×10^7
raw data, starting:13 21
fitted Gauss locations:13 21
0
1
2
3
4
5
6
y
0
5
10
15
20
25
x

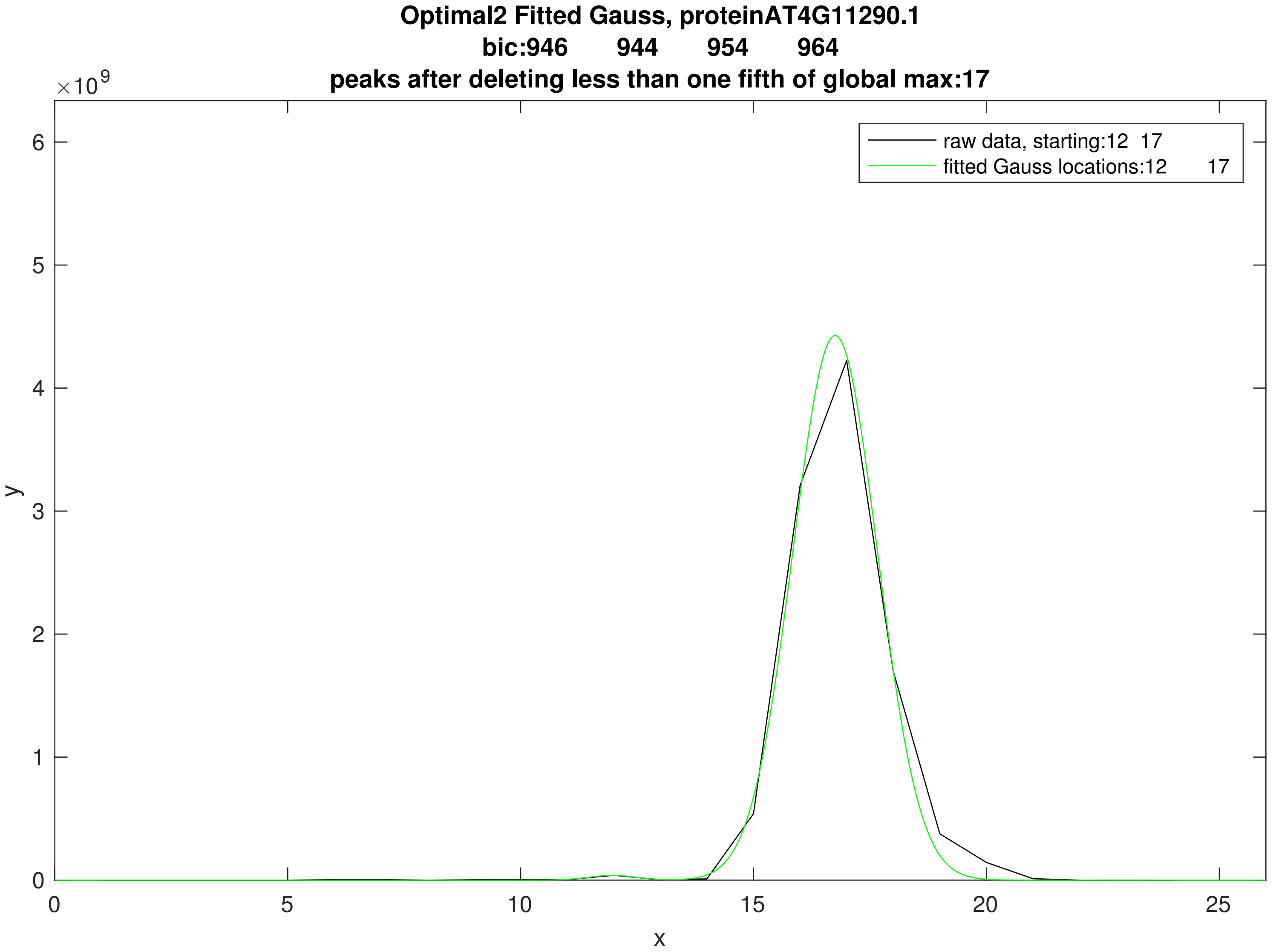
Optimal2 Fitted Gauss, proteinAT4G11290.1
bic:946 944 954 964
peaks after deleting less than one fifth of global max:17
raw data, starting:12 17
fitted Gauss locations:12 17
x
y

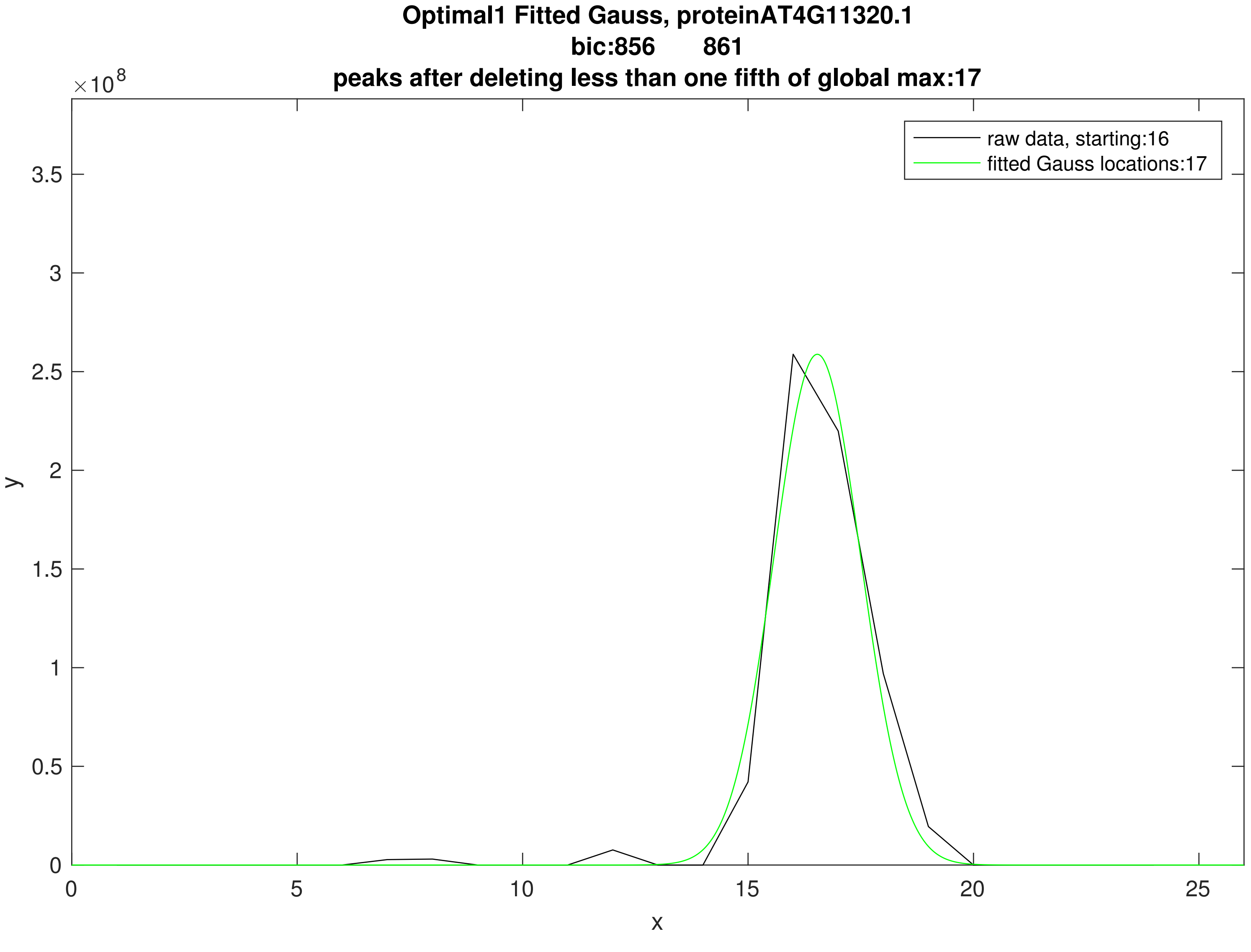
Optimal1 Fitted Gauss, proteinAT4G11320.1
bic:856 861
peaks after deleting less than one fifth of global max:17
raw data, starting:16
fitted Gauss locations:17
×10^8
3.5
3
2.5
2
1.5
1
0.5
0
y
0
5
10
15
20
25
x

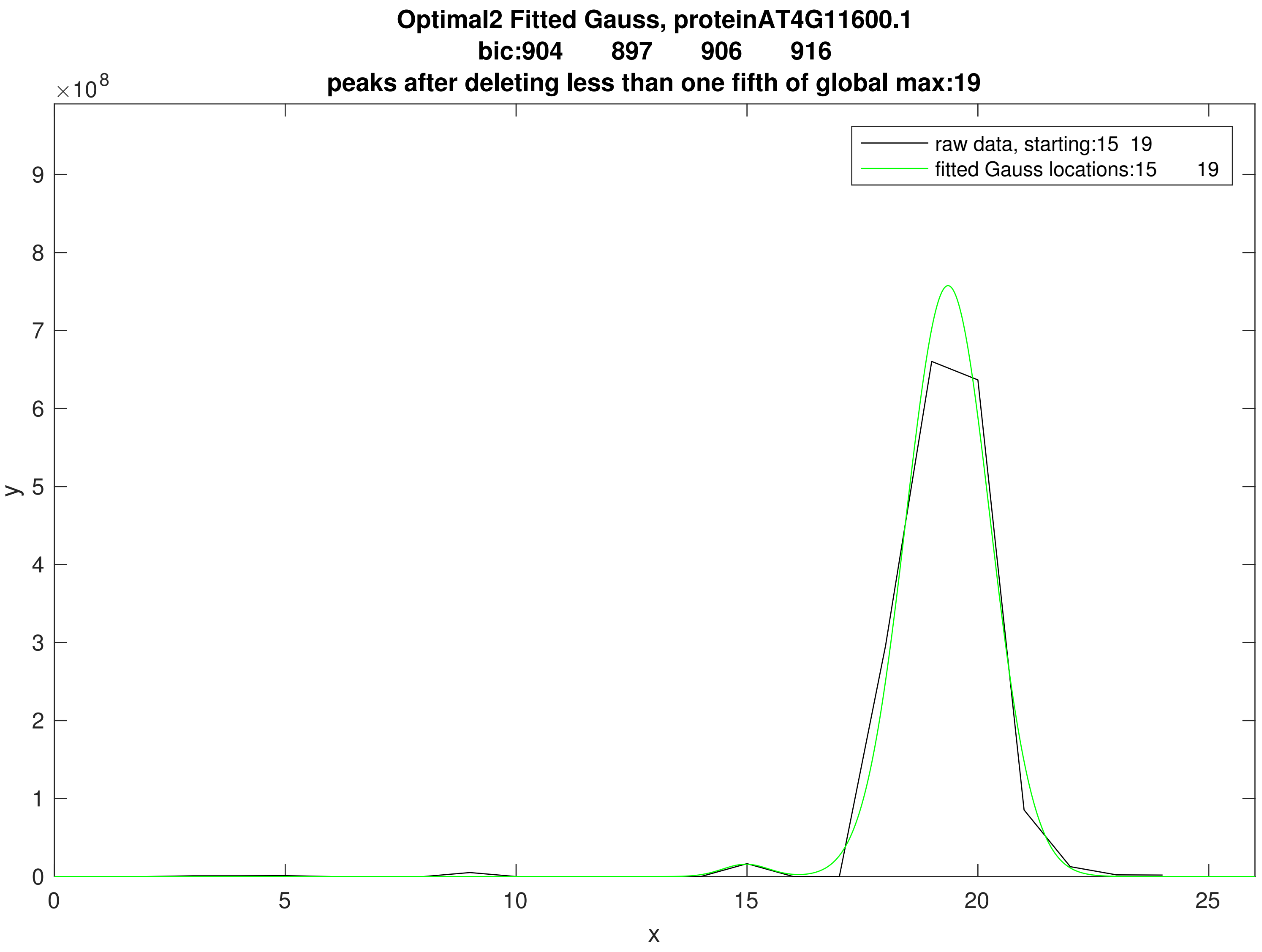
Optimal2 Fitted Gauss, proteinAT4G11600.1
bic:904 897 906 916
peaks after deleting less than one fifth of global max:19
×10^8
raw data, starting:15 19
fitted Gauss locations:15 19
y
x
0
1
2
3
4
5
6
7
8
9
0
5
10
15
20
25

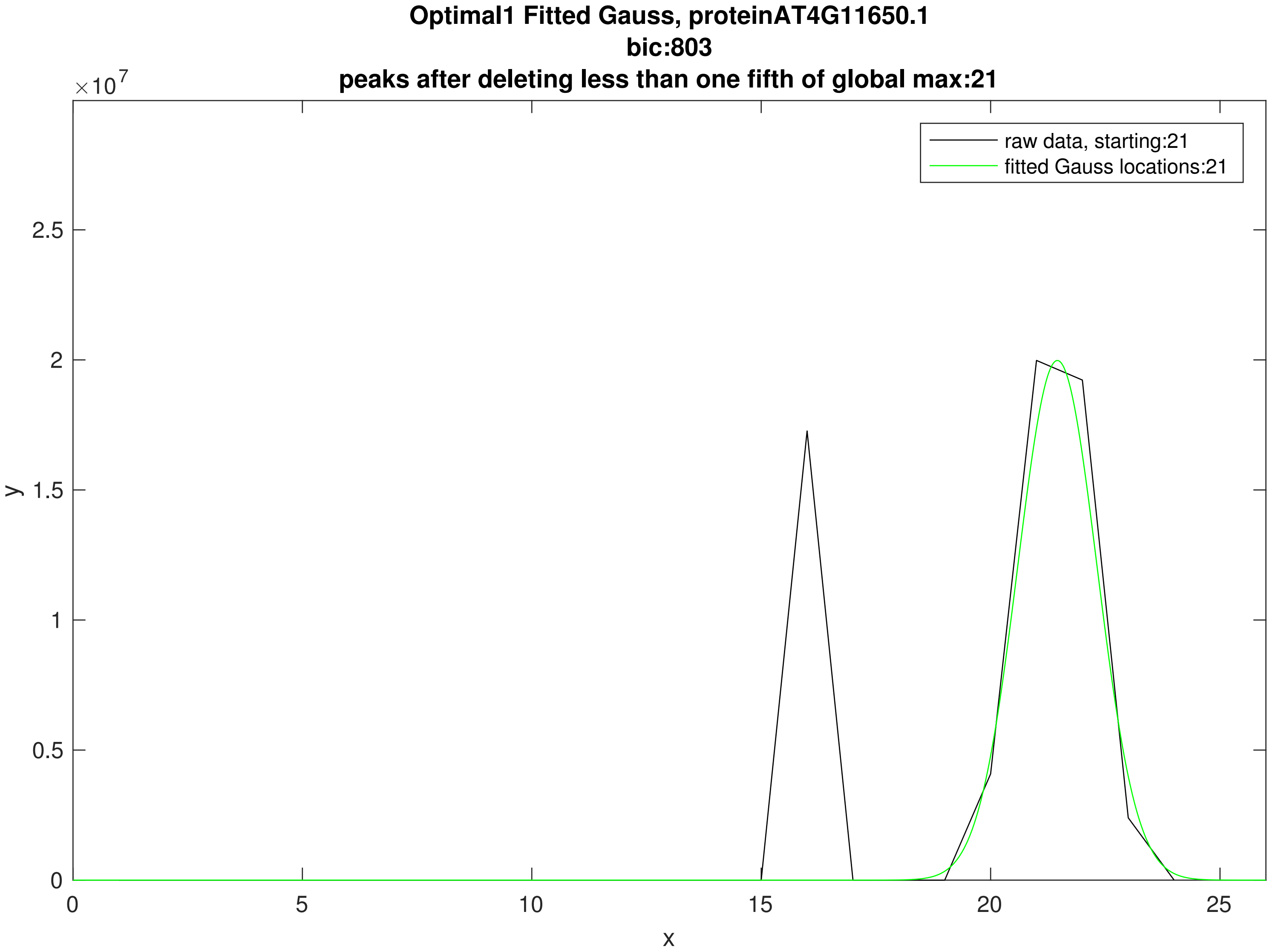
Optimal1 Fitted Gauss, proteinAT4G11650.1
bic:803
peaks after deleting less than one fifth of global max:21
×10^7
raw data, starting:21
fitted Gauss locations:21
y
x
0
0.5
1
1.5
2
2.5
0
5
10
15
20
25

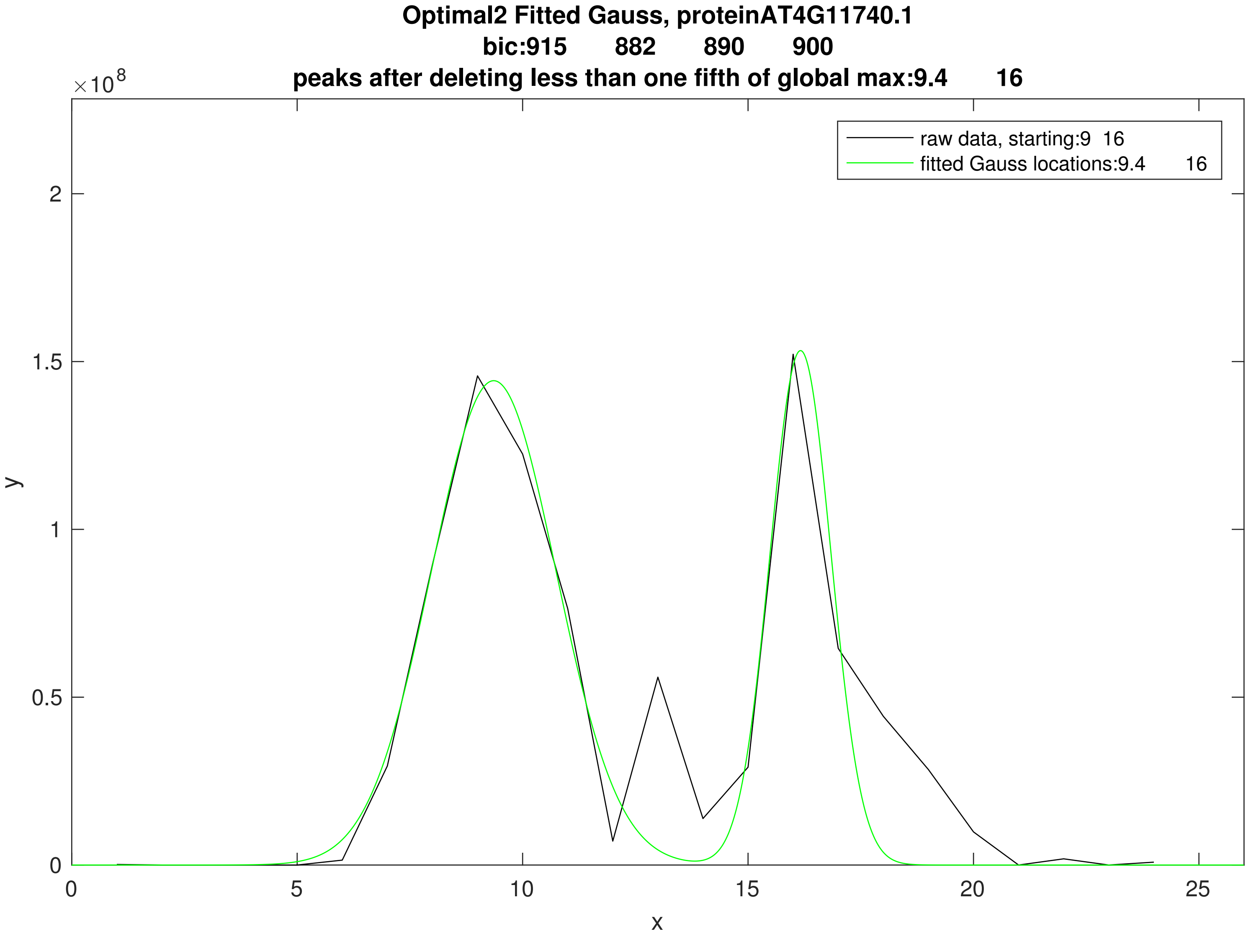
Optimal2 Fitted Gauss, proteinAT4G11740.1
bic:915 882 890 900
peaks after deleting less than one fifth of global max:9.4 16
×10^8
raw data, starting:9 16
fitted Gauss locations:9.4 16
y
x
0
0.5
1
1.5
2
5
10
15
20
25

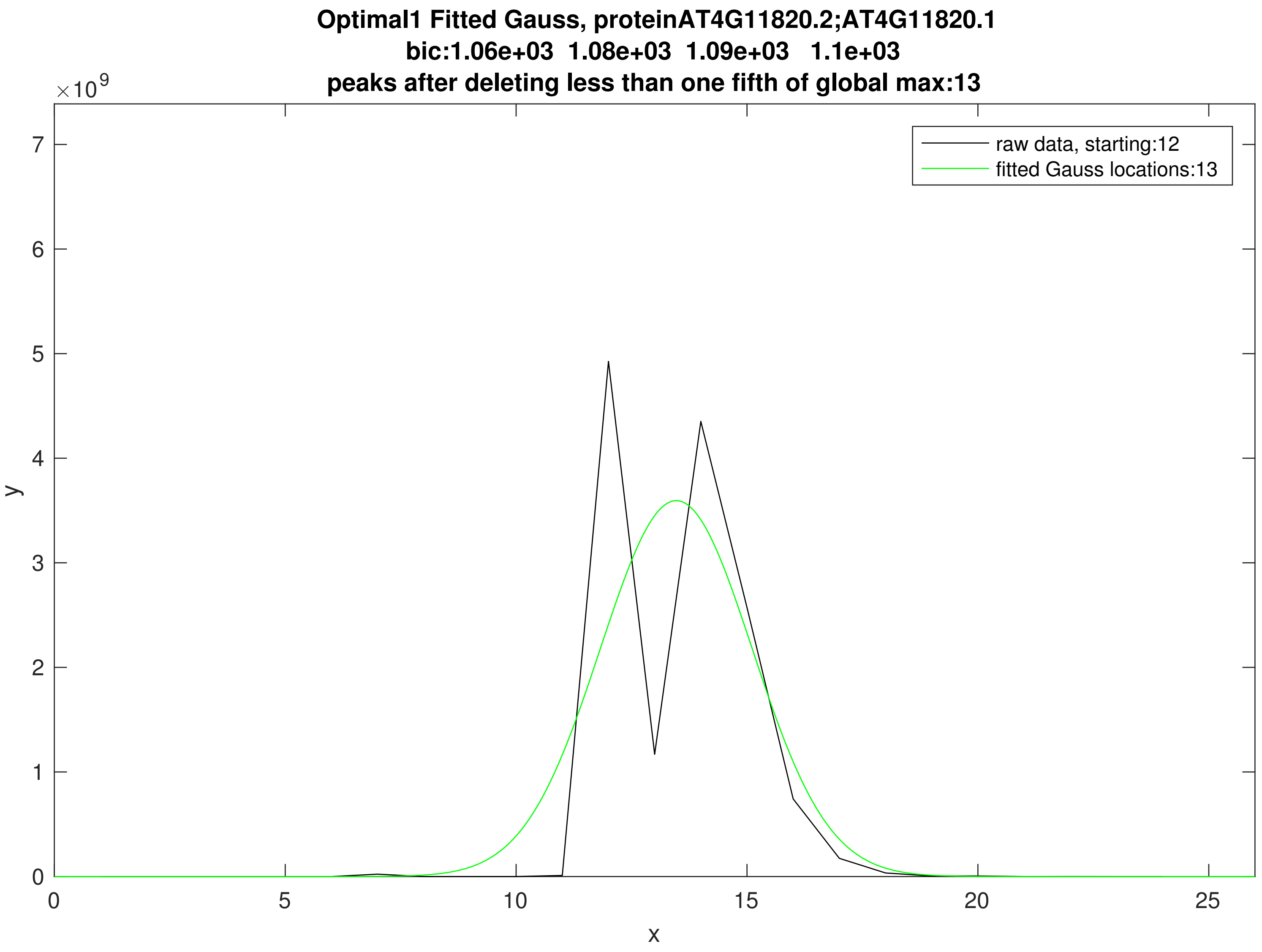
Optimal1 Fitted Gauss, proteinAT4G11820.2;AT4G11820.1
bic:1.06e+03 1.08e+03 1.09e+03 1.1e+03
peaks after deleting less than one fifth of global max:13
×10^9
raw data, starting:12
fitted Gauss locations:13
y
x
0
1
2
3
4
5
6
7
0
5
10
15
20
25

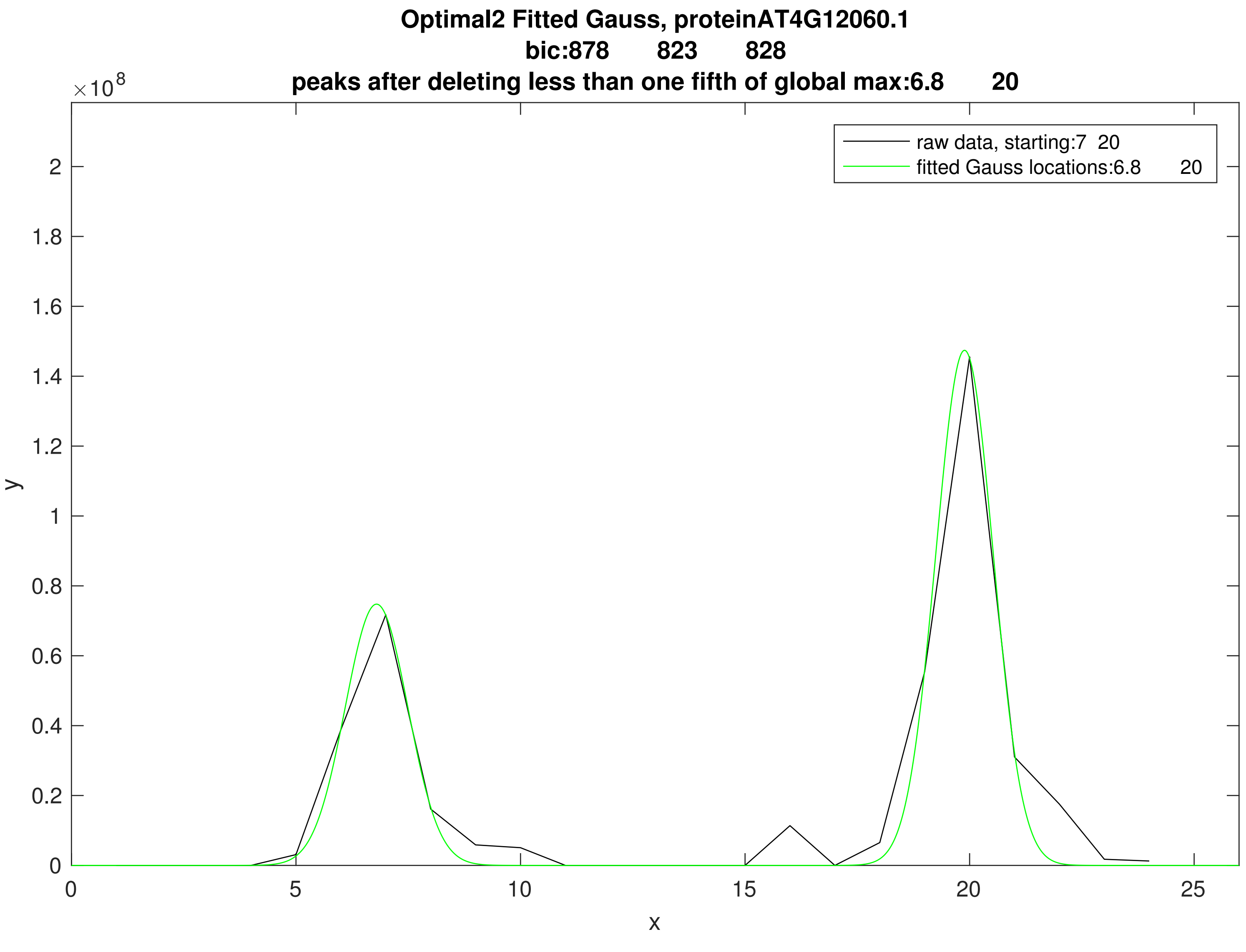
Optimal2 Fitted Gauss, proteinAT4G12060.1
bic:878 823 828
peaks after deleting less than one fifth of global max:6.8 20
×10^8
raw data, starting:7 20
fitted Gauss locations:6.8 20
y
x
0
0.2
0.4
0.6
0.8
1
1.2
1.4
1.6
1.8
2
5
10
15
20
25

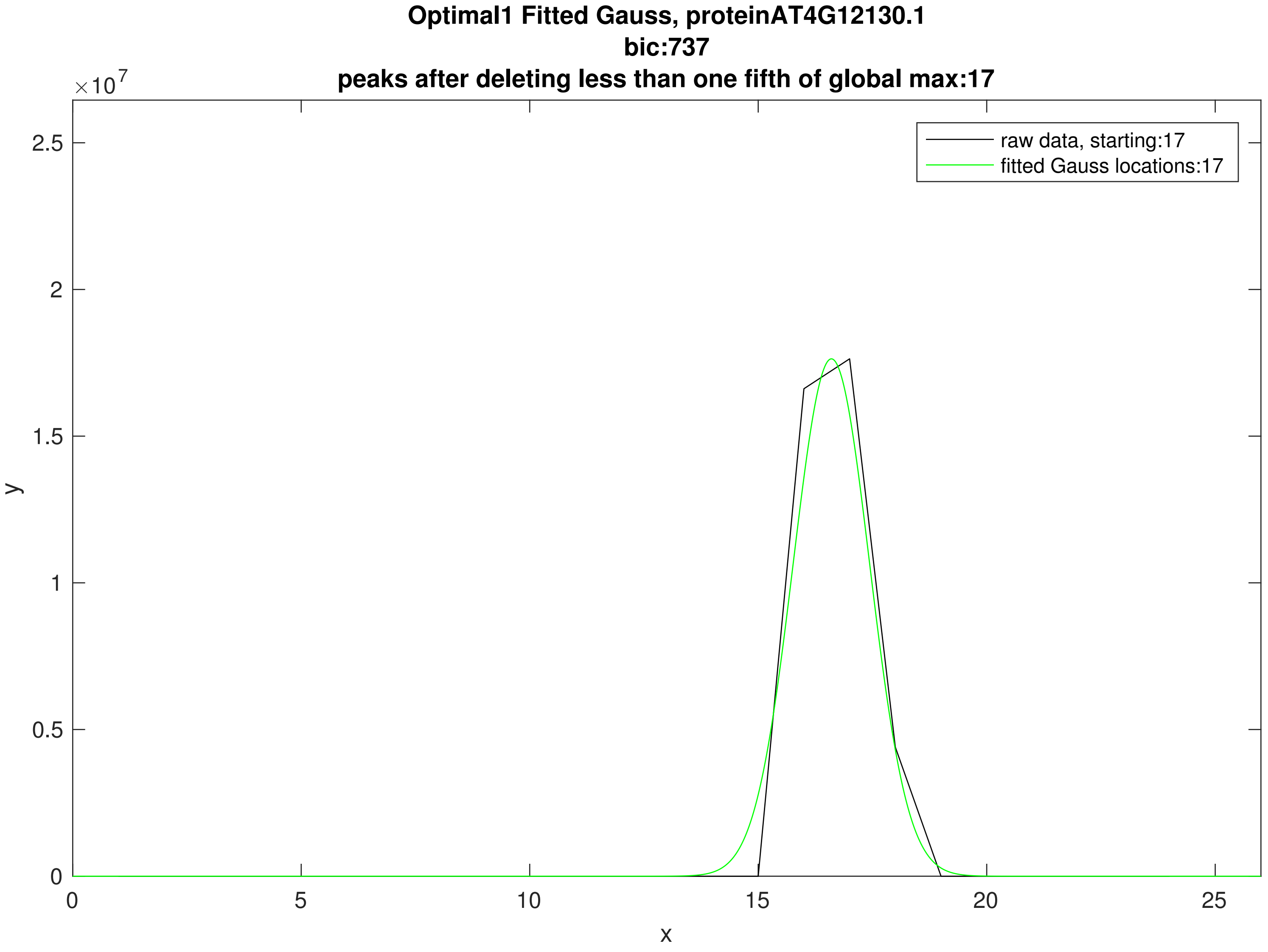
Optimal1 Fitted Gauss, proteinAT4G12130.1
bic:737
peaks after deleting less than one fifth of global max:17
×10^7
2.5
2
1.5
1
0.5
0
y
0
5
10
15
20
25
x
raw data, starting:17
fitted Gauss locations:17

**Optimal1 Fitted Gauss, proteinAT4G12290.1**
**bic:816**
**peaks after deleting less than one fifth of global max:8.8**

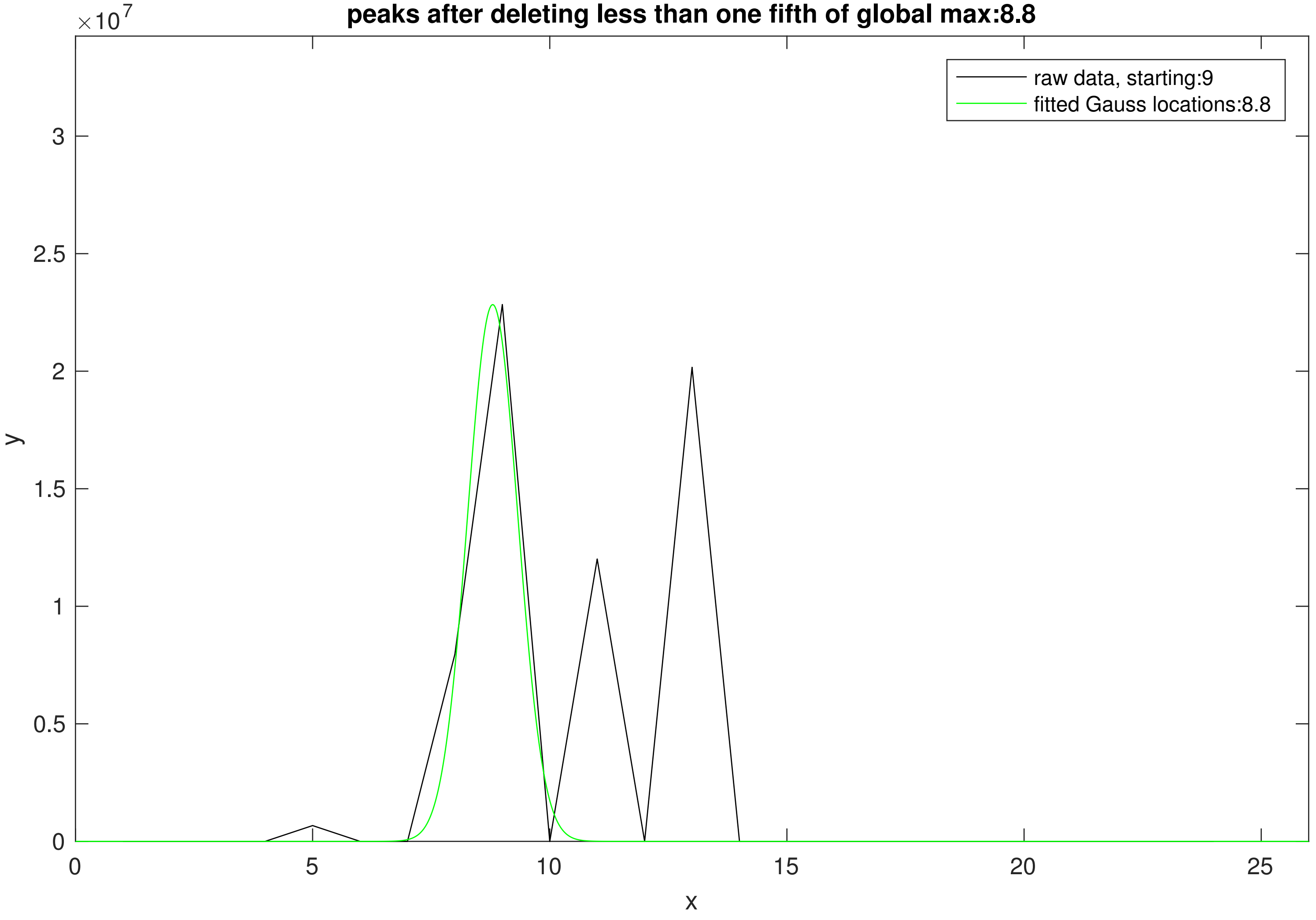

**Optimal1 Fitted Gauss, proteinAT4G12400.1;AT4G12400.2**
**bic:788 794**
**peaks after deleting less than one fifth of global max:8.7**

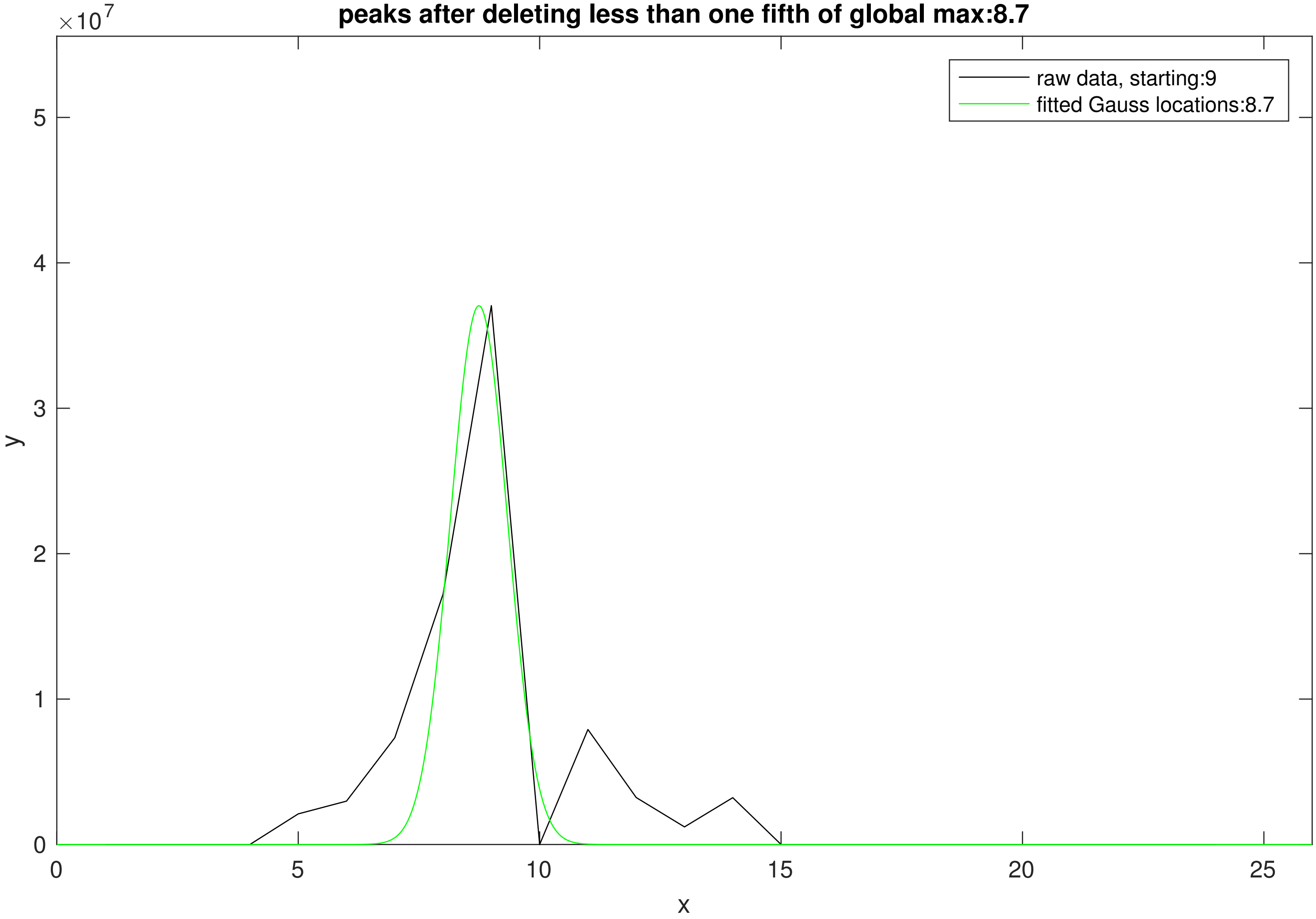

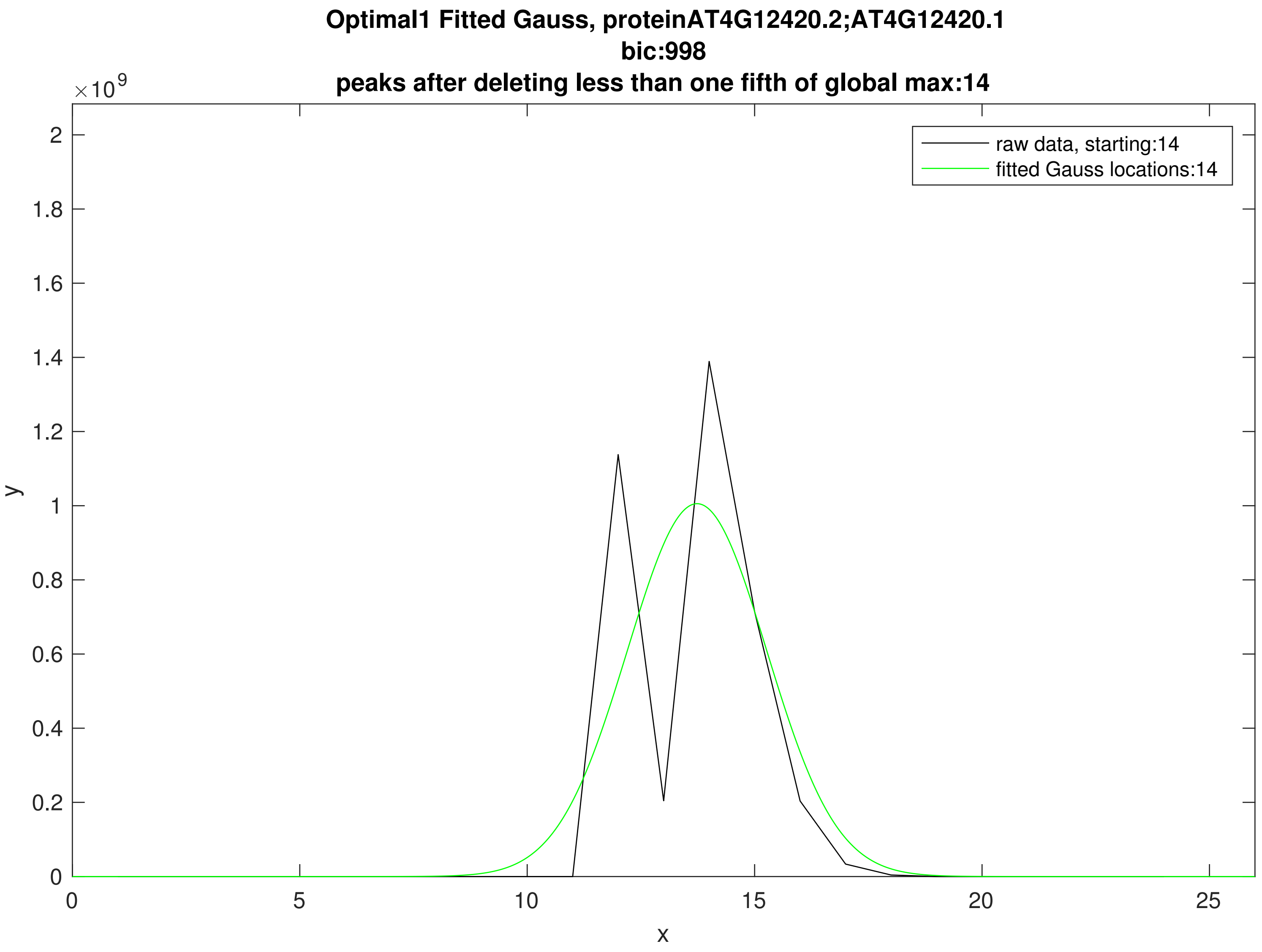
Optimal1 Fitted Gauss, proteinAT4G12420.2;AT4G12420.1
bic:998
peaks after deleting less than one fifth of global max:14
×10^9
raw data, starting:14
fitted Gauss locations:14
y
x
0
0.2
0.4
0.6
0.8
1
1.2
1.4
1.6
1.8
2
0
5
10
15
20
25

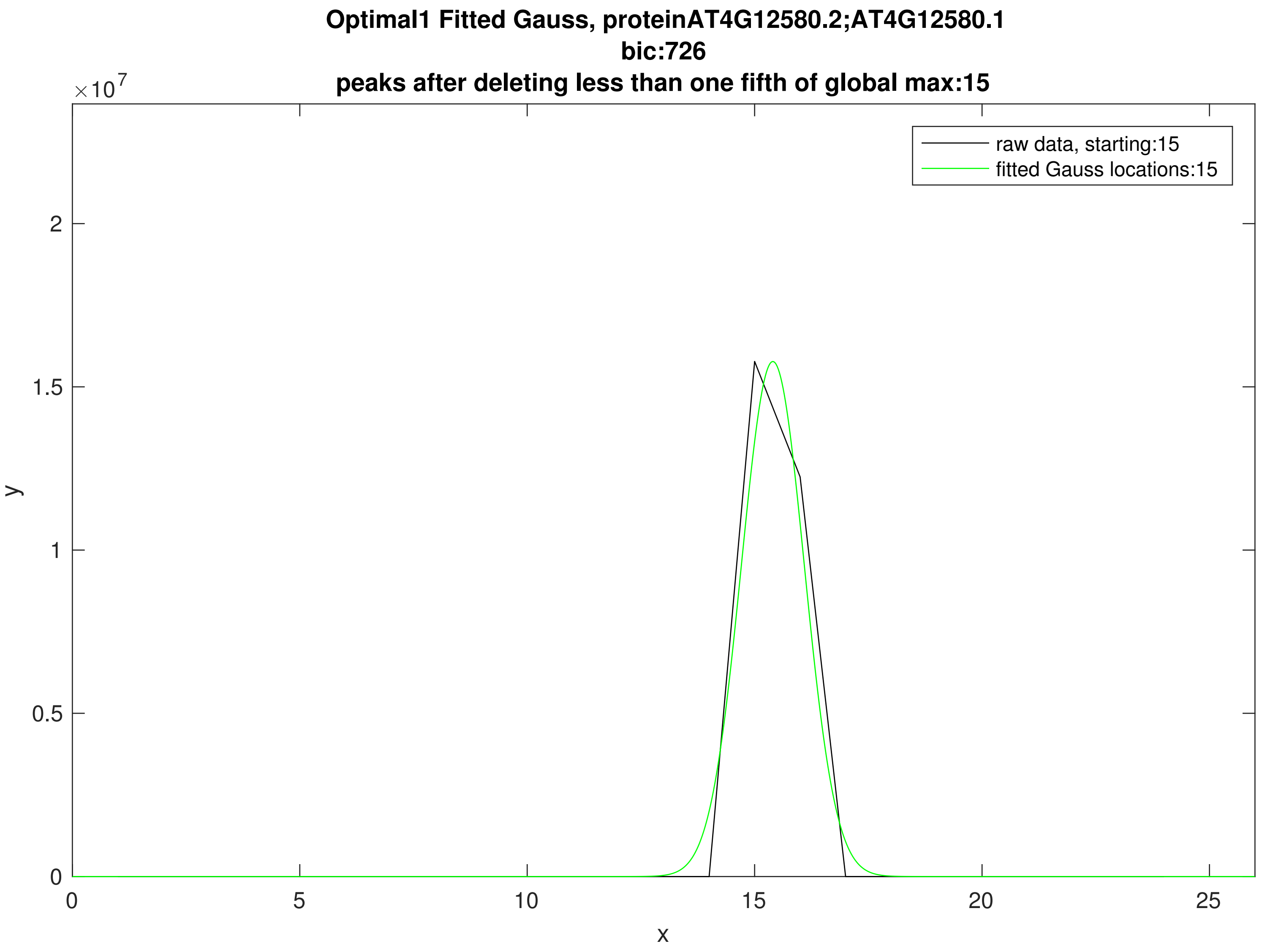
Optimal1 Fitted Gauss, proteinAT4G12580.2;AT4G12580.1
bic:726
peaks after deleting less than one fifth of global max:15
×10^7
raw data, starting:15
fitted Gauss locations:15
y
x
0
0.5
1
1.5
2
0
5
10
15
20
25

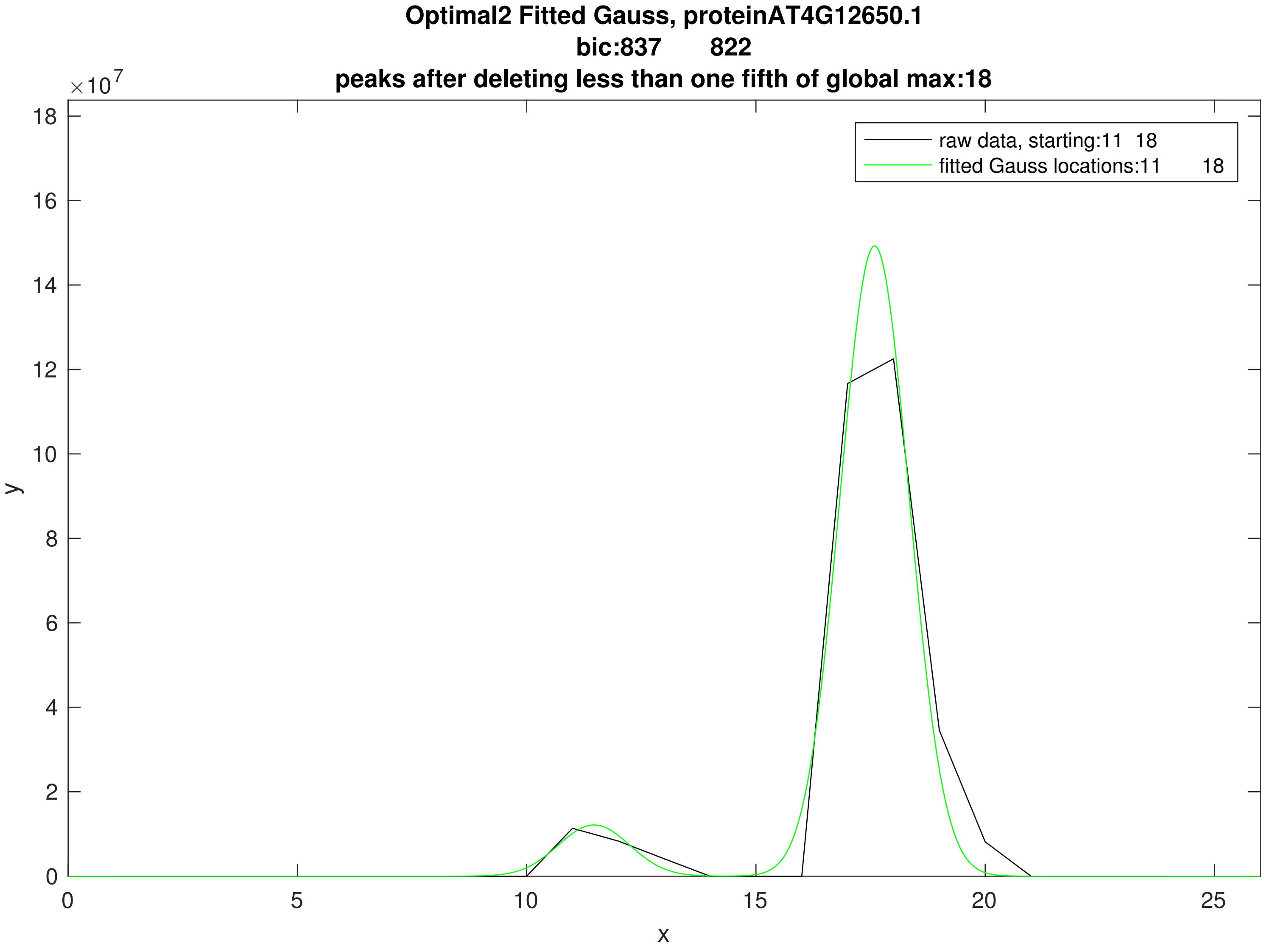
Optimal2 Fitted Gauss, proteinAT4G12650.1
bic:837 822
peaks after deleting less than one fifth of global max:18
×10^7
raw data, starting:11 18
fitted Gauss locations:11 18
y
x
0
2
4
6
8
10
12
14
16
18
0
5
10
15
20
25

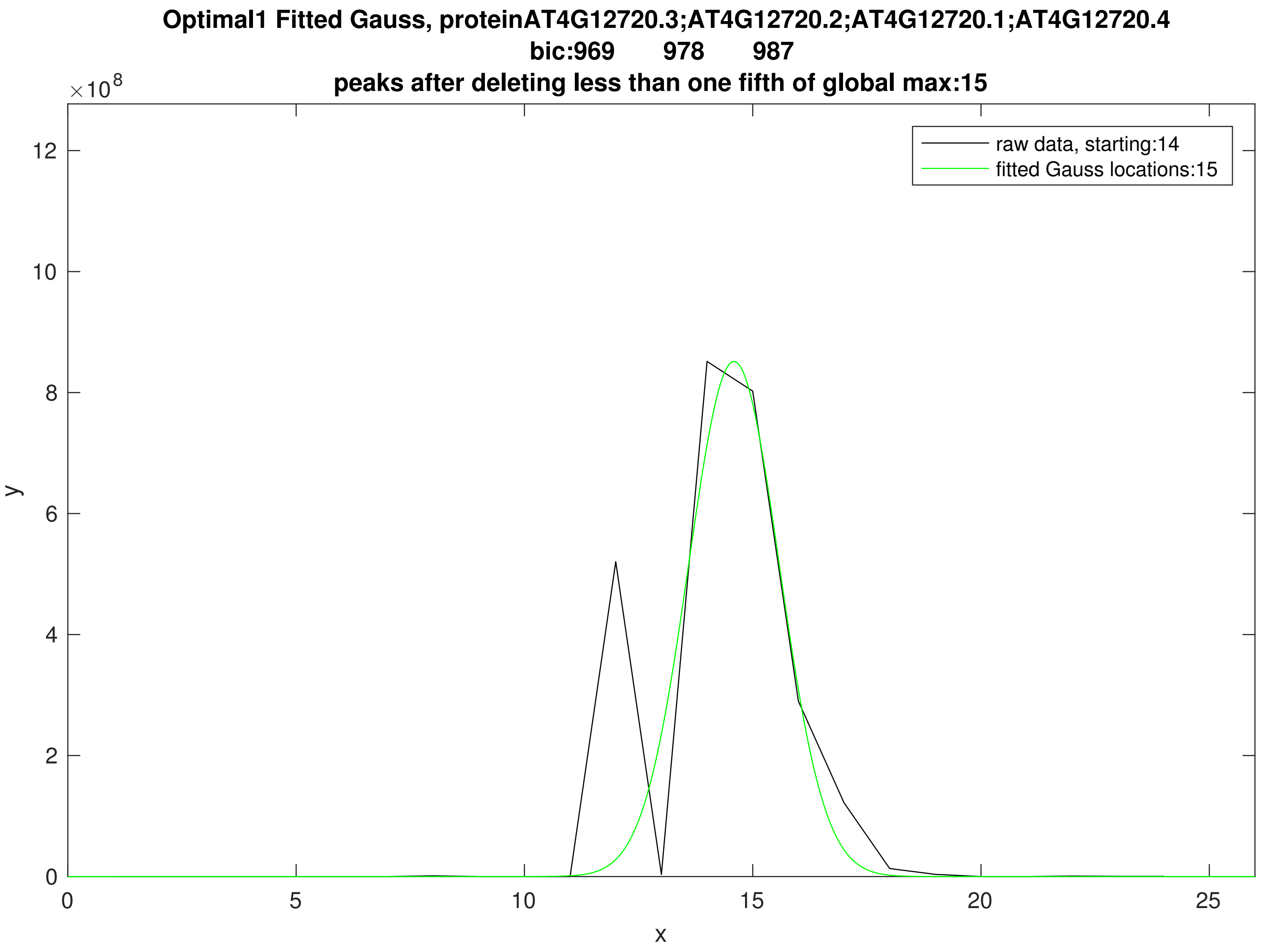
Optimal1 Fitted Gauss, proteinAT4G12720.3;AT4G12720.2;AT4G12720.1;AT4G12720.4
bic:969 978 987
peaks after deleting less than one fifth of global max:15
×10^8
12
10
8
6
4
2
0
y
0
5
10
15
20
25
x
raw data, starting:14
fitted Gauss locations:15

**Optimal2 Fitted Gauss, proteinAT4G12730.1**
**bic:1.06e+03  1.06e+03  1.07e+03**
**peaks after deleting less than one fifth of global max:12**

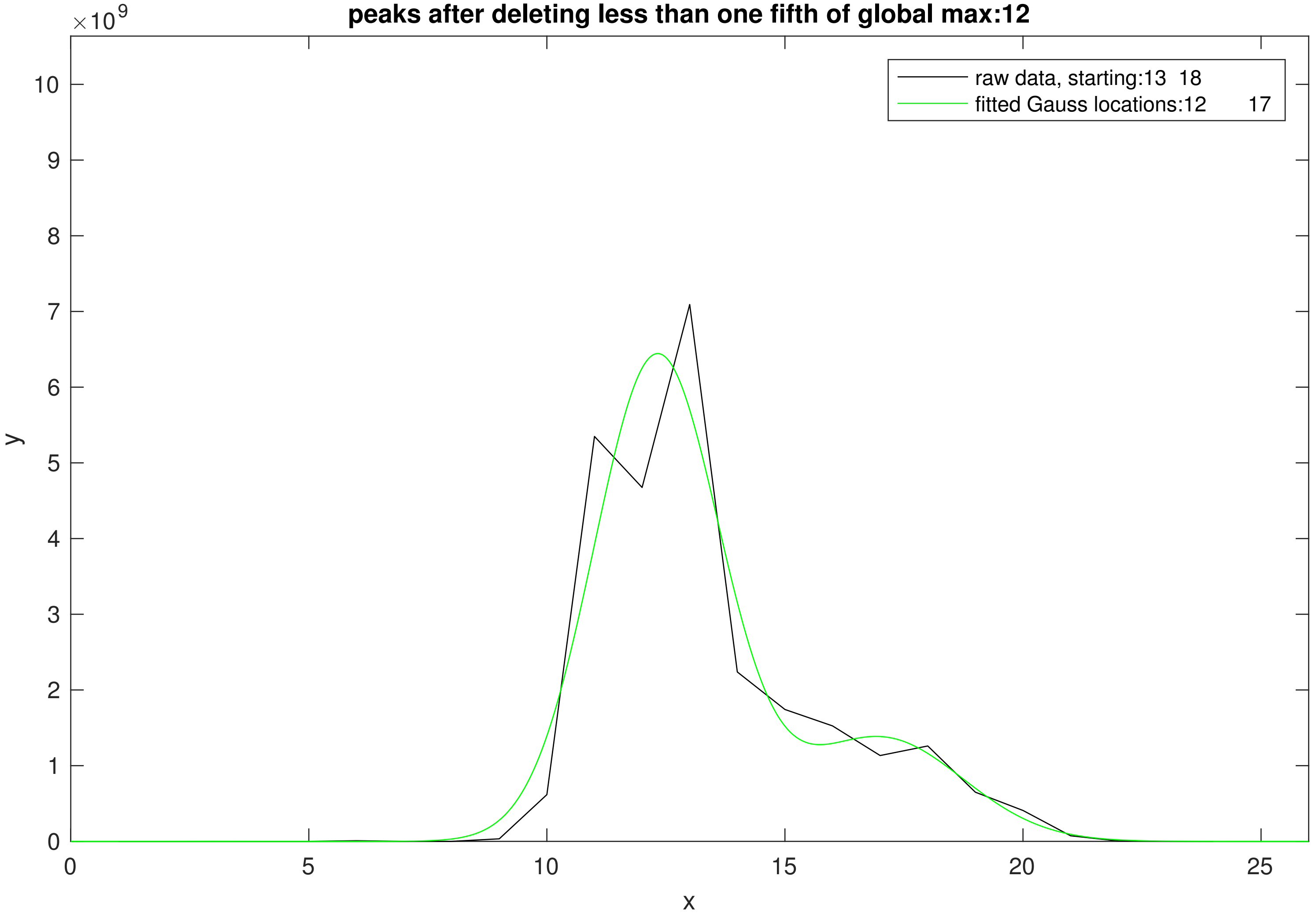

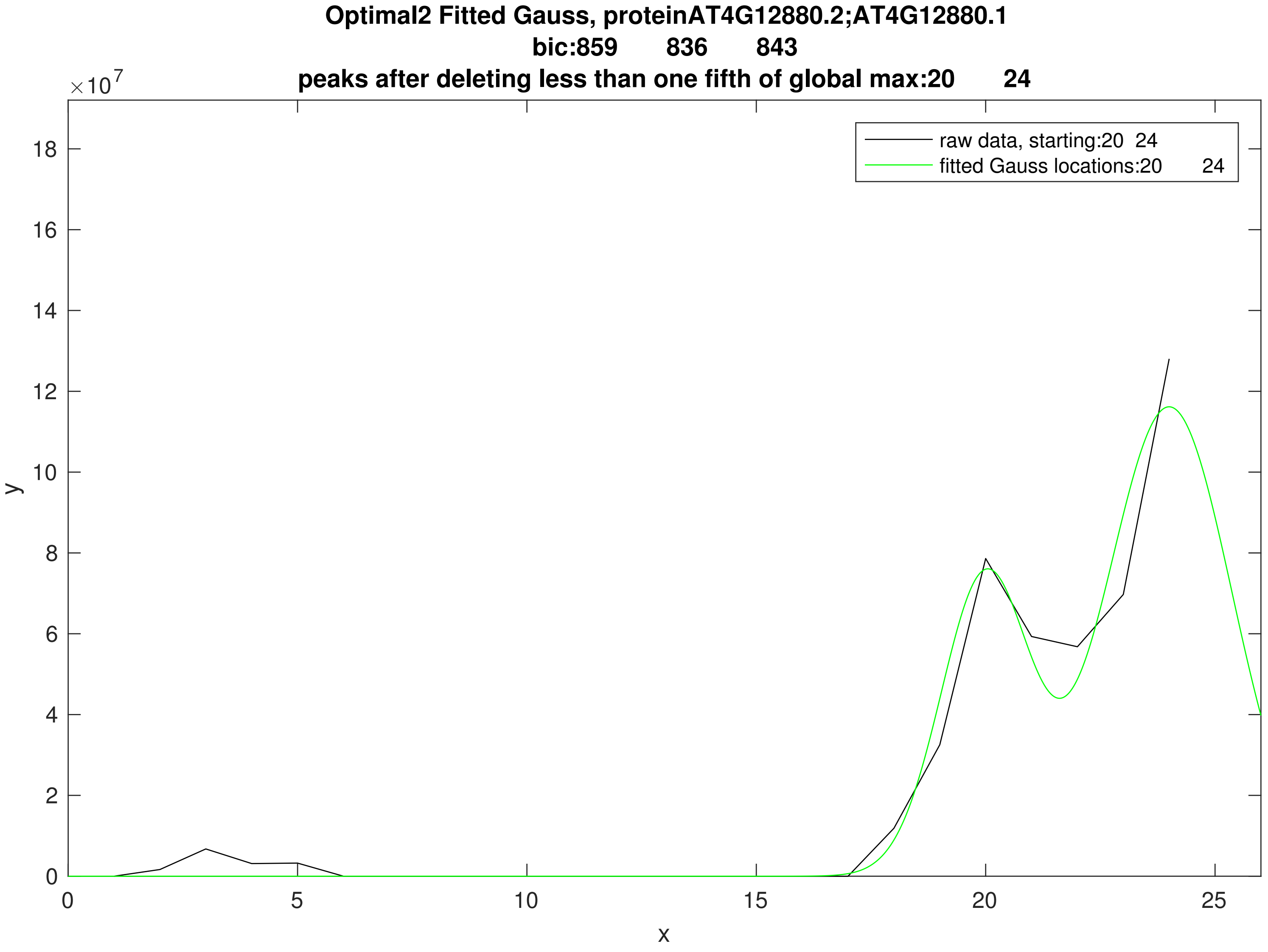

Optimal2 Fitted Gauss, proteinAT4G12880.2;AT4G12880.1
bic:859 836 843
peaks after deleting less than one fifth of global max:20 24
×10^7
raw data, starting:20 24
fitted Gauss locations:20 24
y
x
0
2
4
6
8
10
12
14
16
18
5
15
20
25

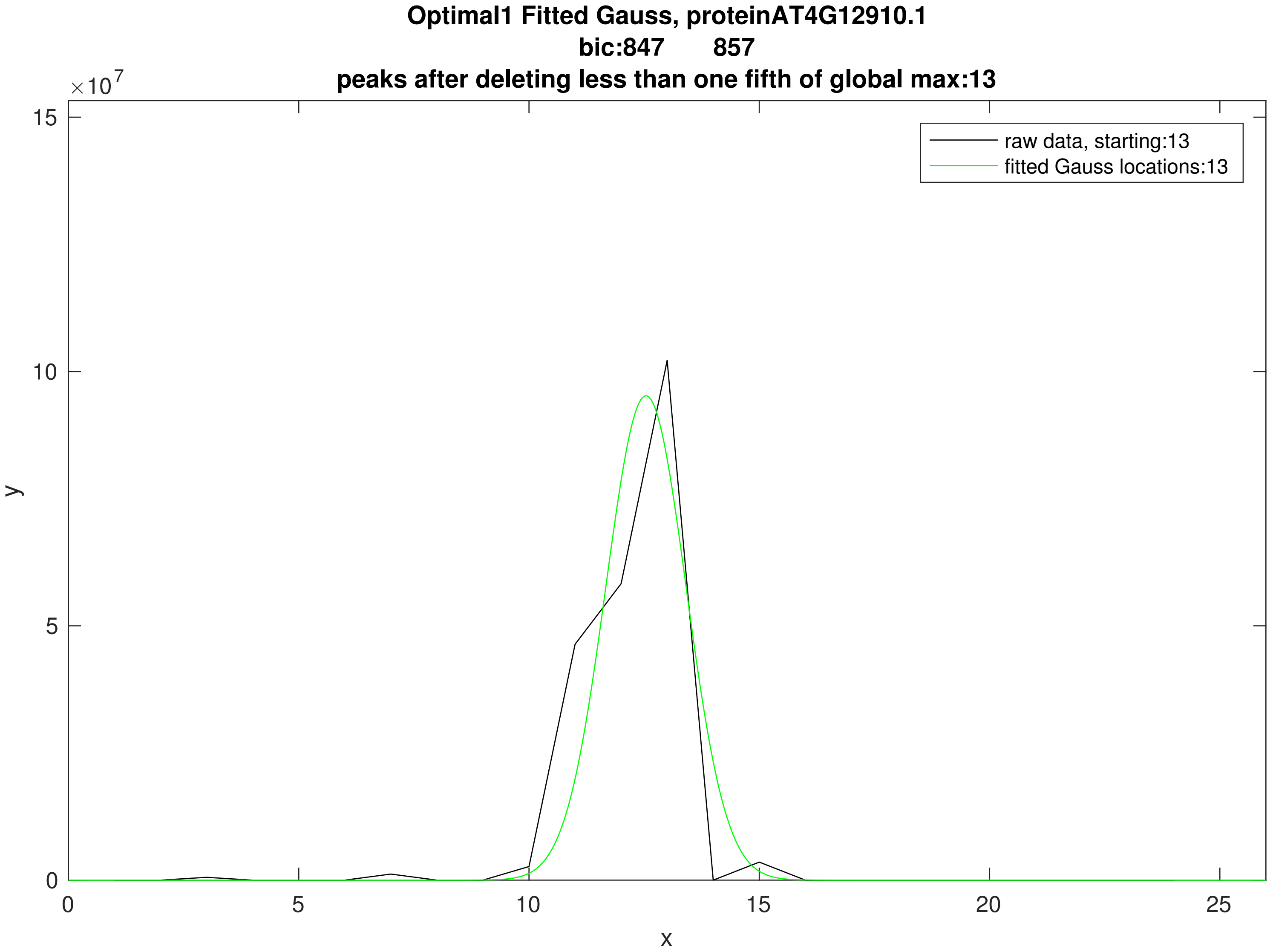
Optimal1 Fitted Gauss, proteinAT4G12910.1
bic:847 857
peaks after deleting less than one fifth of global max:13
×10^7
raw data, starting:13
fitted Gauss locations:13
y
x
0
5
10
15
20
25

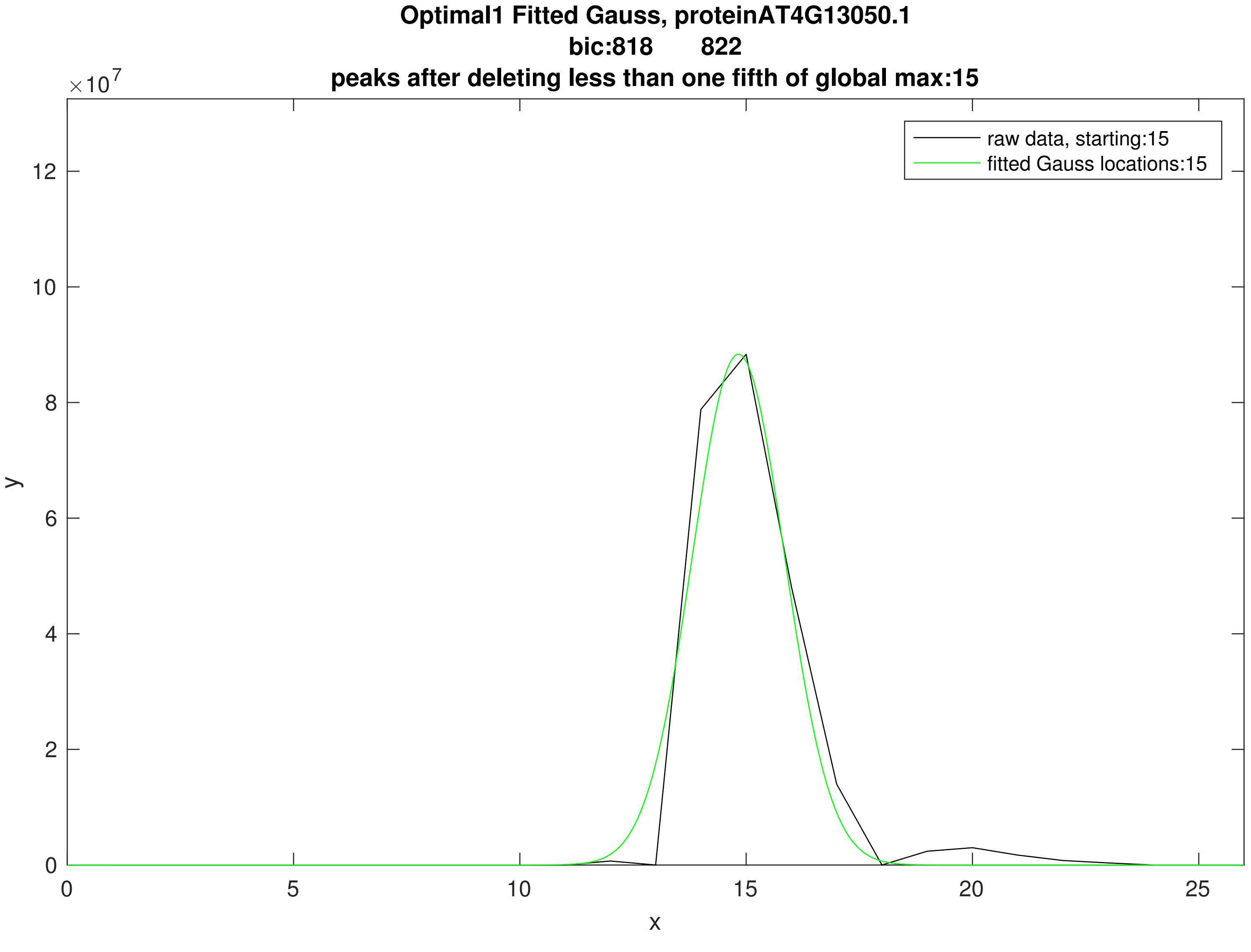
Optimal1 Fitted Gauss, proteinAT4G13050.1
bic:818 822
peaks after deleting less than one fifth of global max:15
×10^7
raw data, starting:15
fitted Gauss locations:15
12
10
8
6
4
2
0
y
0
5
10
15
20
25
x

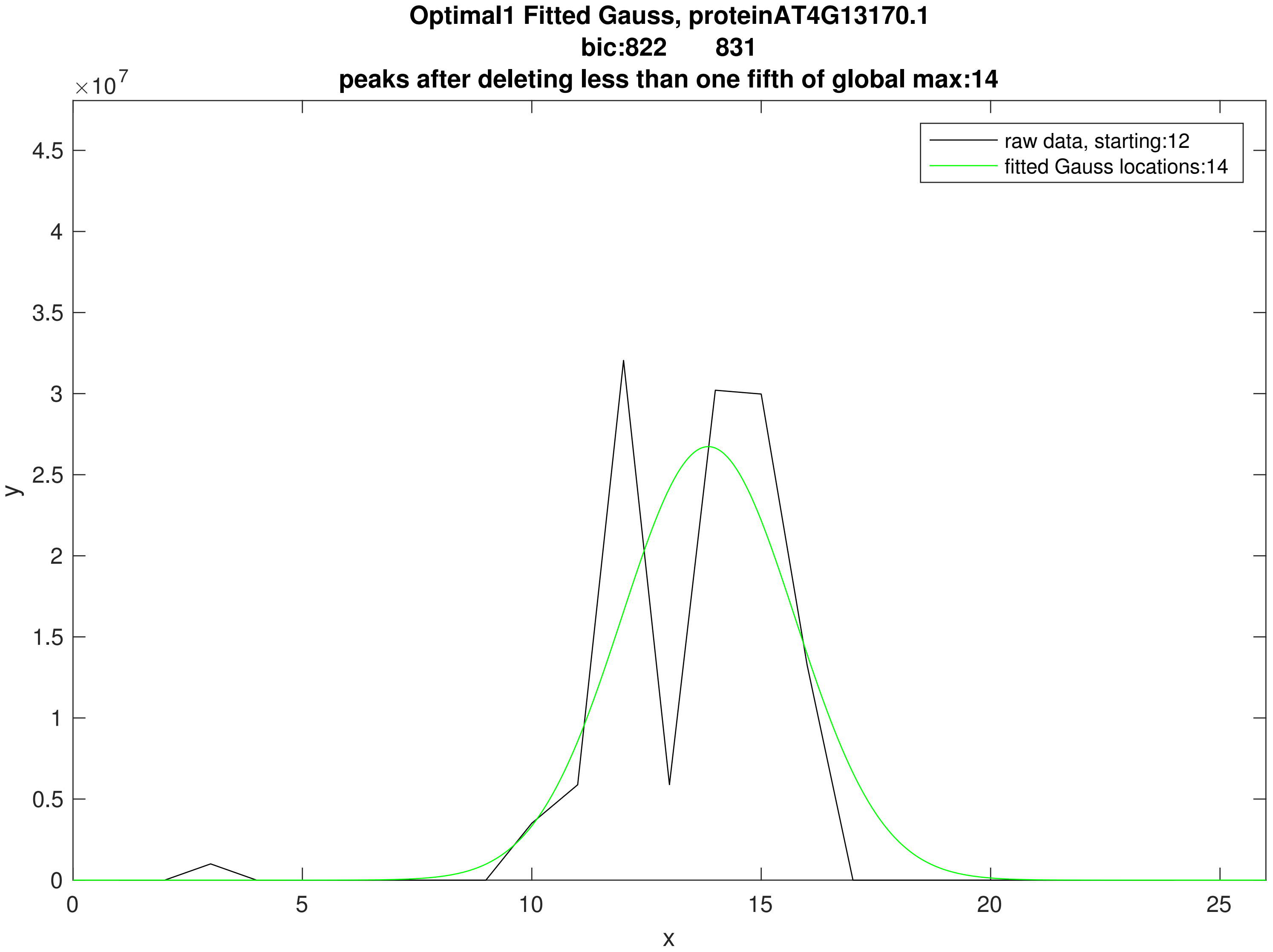
Optimal1 Fitted Gauss, proteinAT4G13170.1
bic:822 831
peaks after deleting less than one fifth of global max:14
×10^7
raw data, starting:12
fitted Gauss locations:14
y
x
0
0.5
1
1.5
2
2.5
3
3.5
4
4.5
5
10
15
20
25

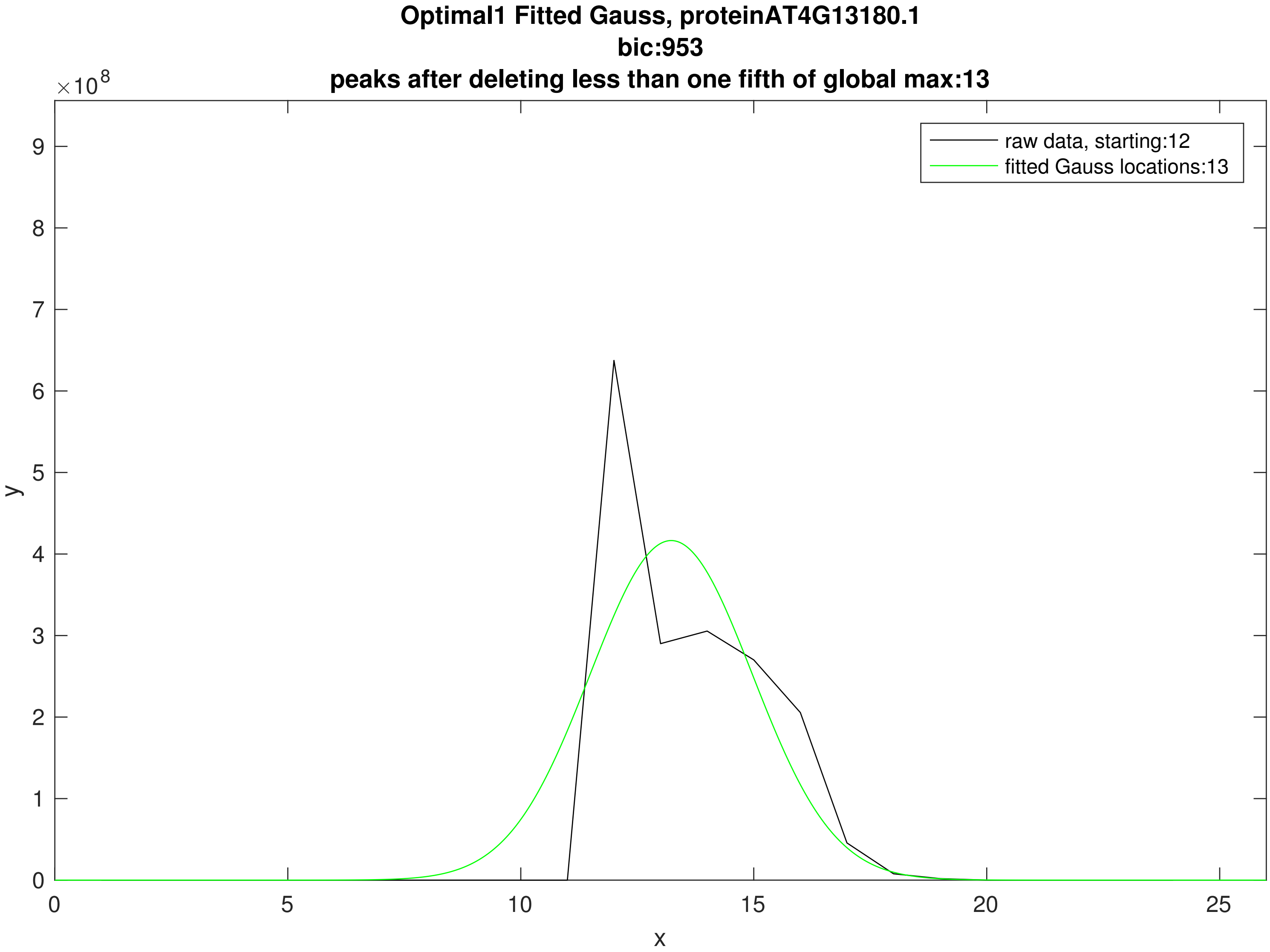
Optimal1 Fitted Gauss, proteinAT4G13180.1
bic:953
peaks after deleting less than one fifth of global max:13
×10^8
raw data, starting:12
fitted Gauss locations:13
y
x

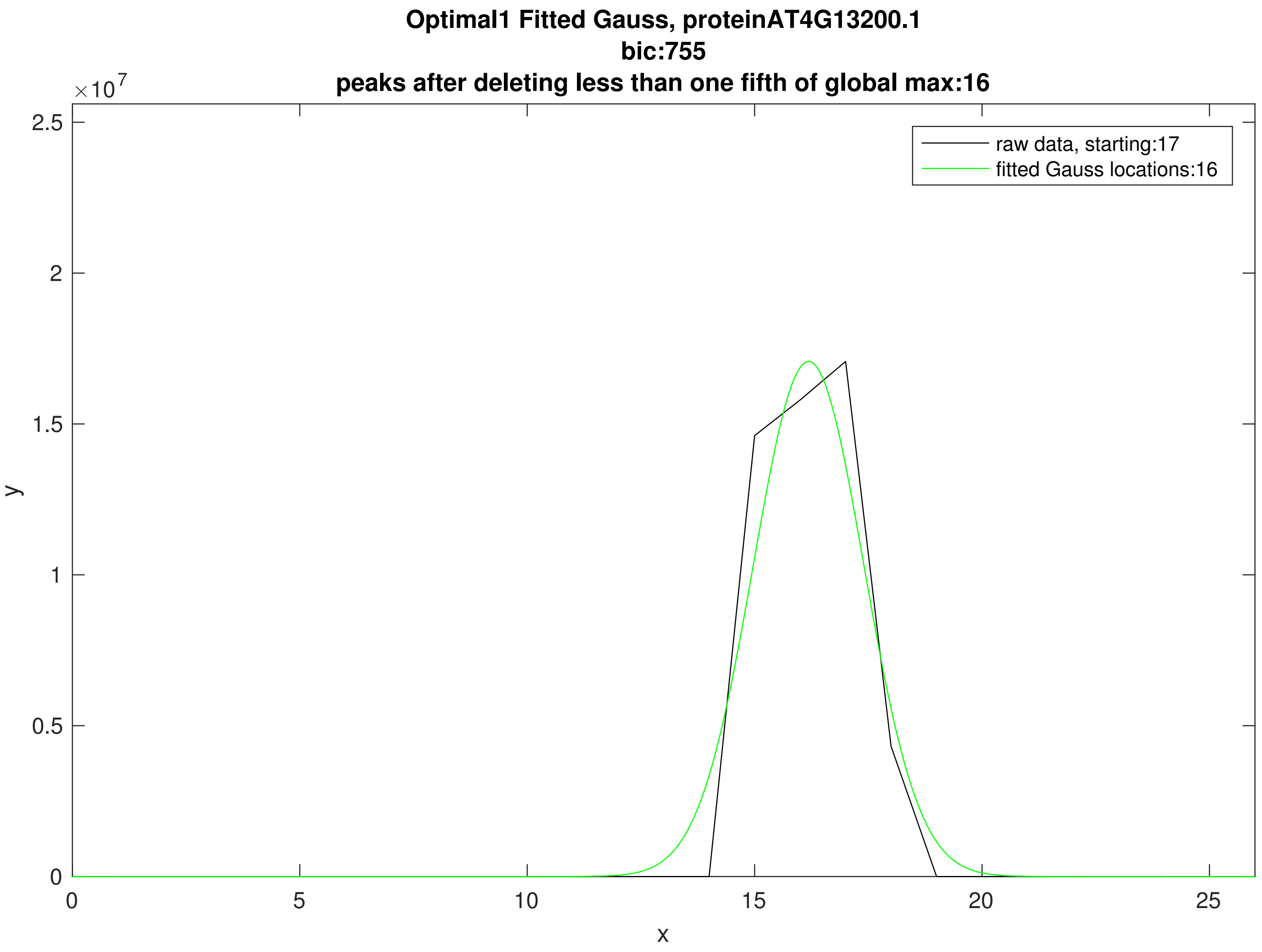

Optimal1 Fitted Gauss, proteinAT4G13200.1
bic:755
peaks after deleting less than one fifth of global max:16
×10^7
2.5
2
1.5
1
0.5
0
y
0
5
10
15
20
25
x
raw data, starting:17
fitted Gauss locations:16

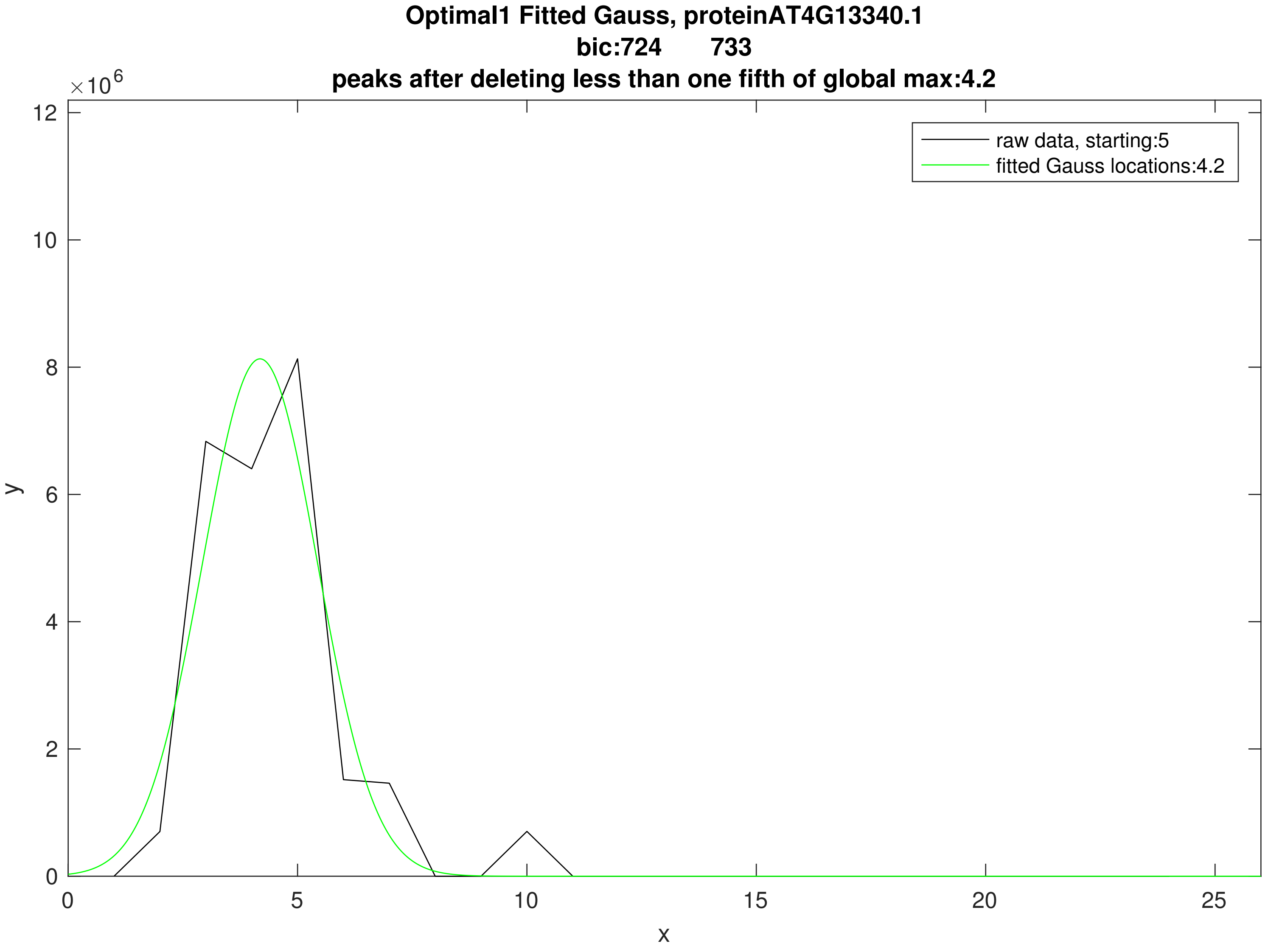

Optimal1 Fitted Gauss, proteinAT4G13340.1
bic:724 733
peaks after deleting less than one fifth of global max:4.2
raw data, starting:5
fitted Gauss locations:4.2
×10^6
y
x
0
2
4
6
8
10
12
5
15
20
25

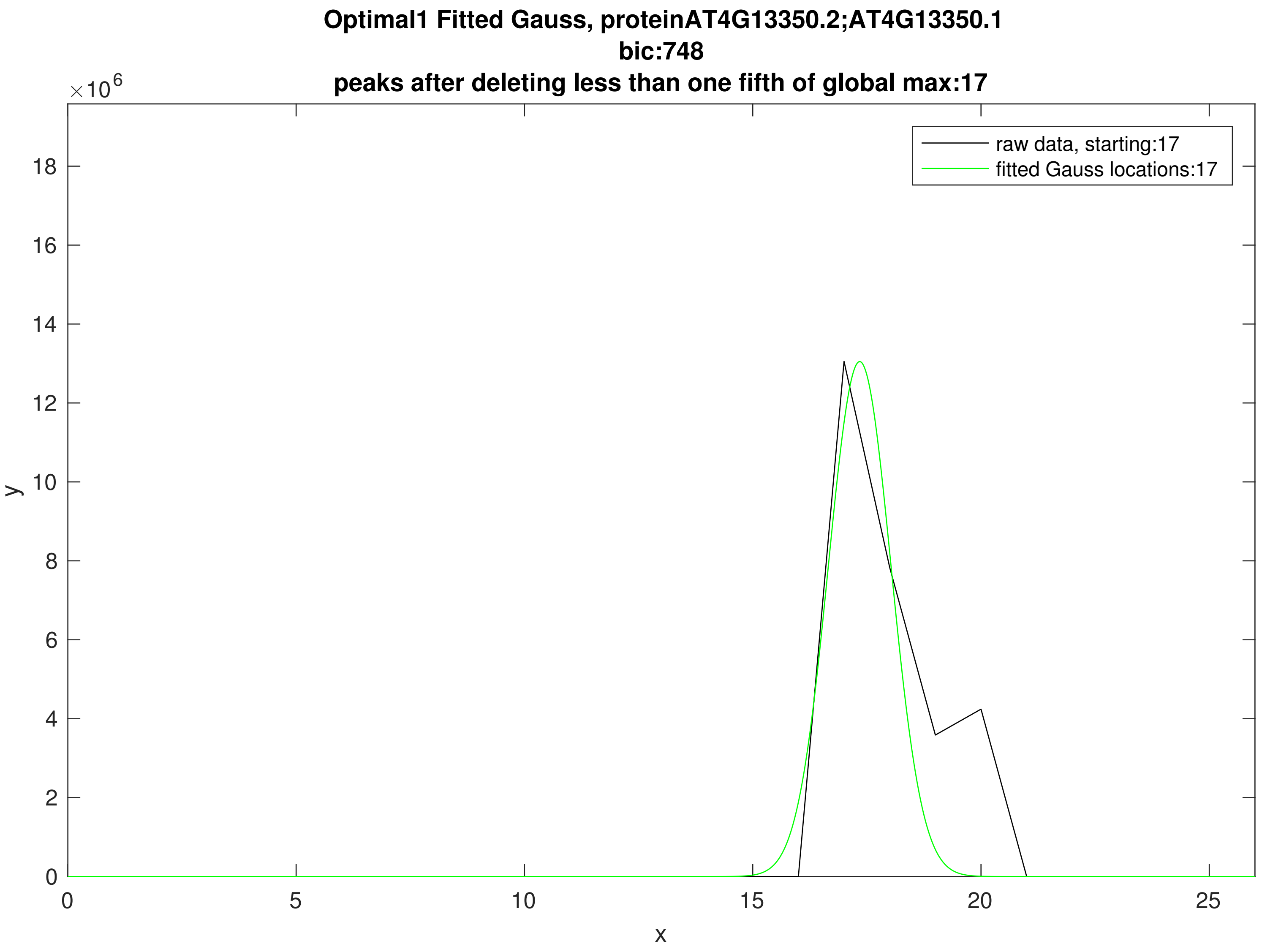

Optimal1 Fitted Gauss, proteinAT4G13350.2;AT4G13350.1
bic:748
peaks after deleting less than one fifth of global max:17
×10^6
raw data, starting:17
fitted Gauss locations:17
y
x
0
2
4
6
8
10
12
14
16
18
0
5
10
15
20
25

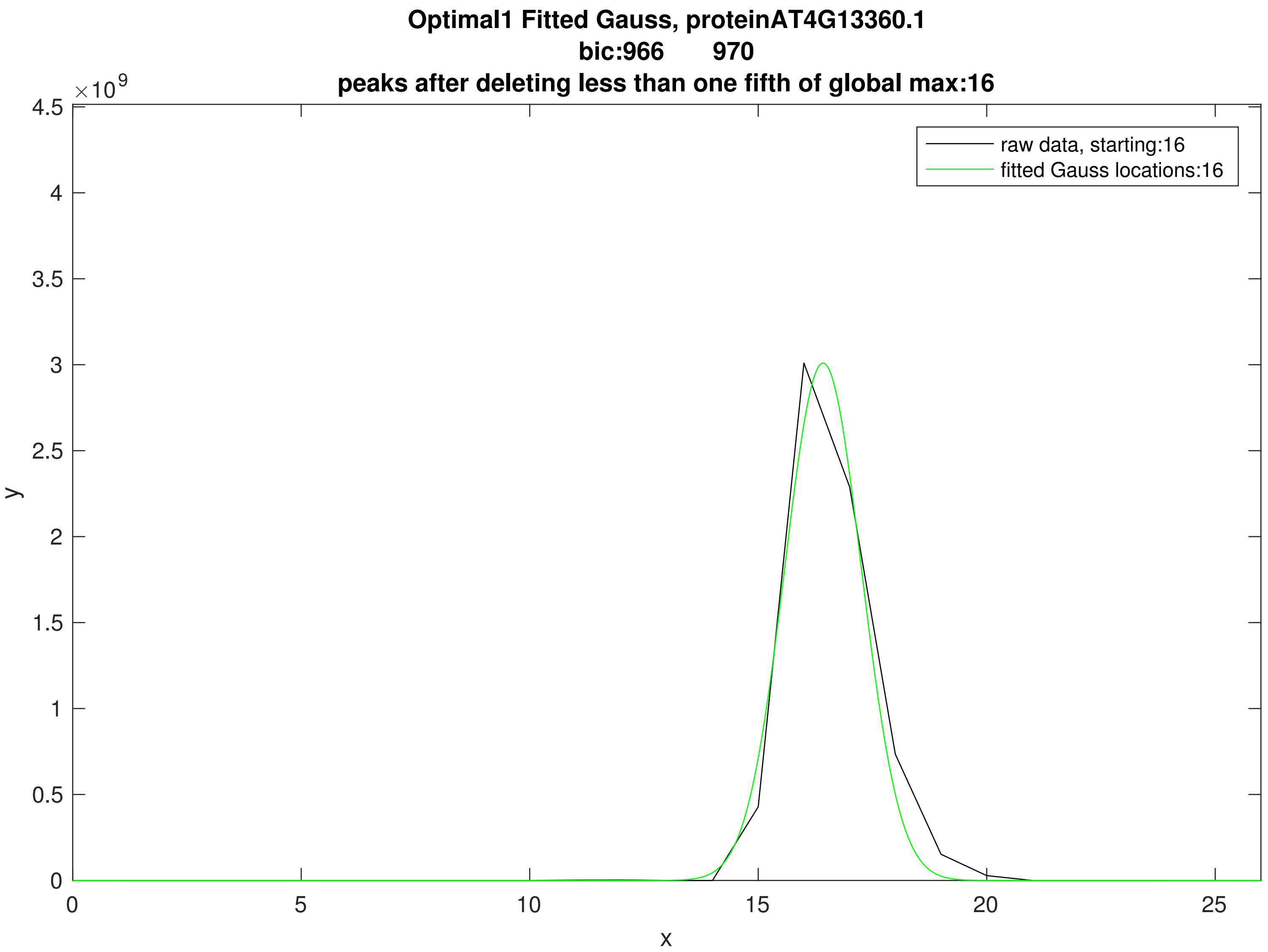

Optimal1 Fitted Gauss, proteinAT4G13360.1
bic:966 970
peaks after deleting less than one fifth of global max:16
×10^9
4.5
4
3.5
3
2.5
2
1.5
1
0.5
0
y
0
5
10
15
20
25
x
raw data, starting:16
fitted Gauss locations:16

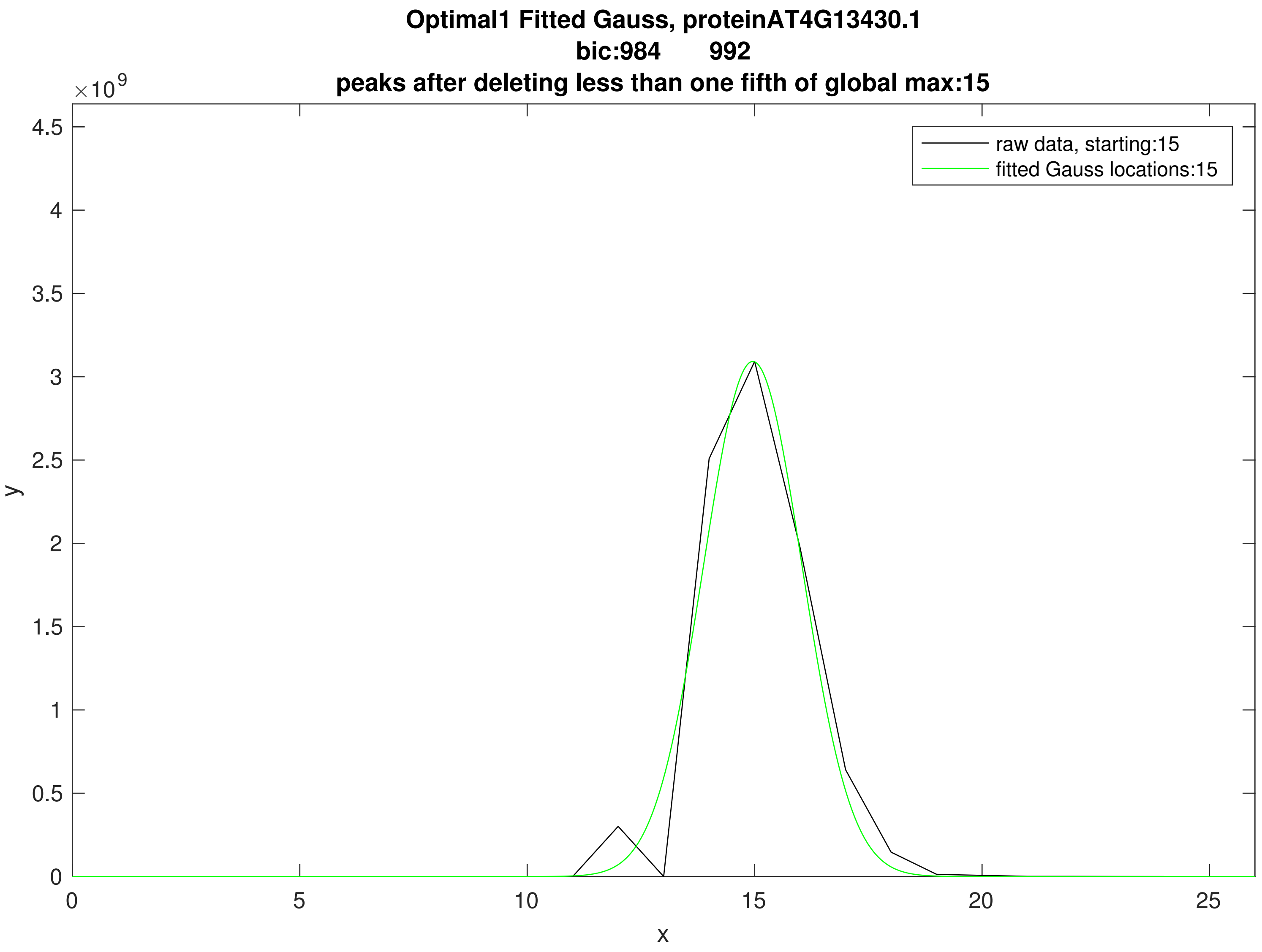

Optimal1 Fitted Gauss, proteinAT4G13430.1
bic:984 992
peaks after deleting less than one fifth of global max:15
×10^9
4.5
4
3.5
3
2.5
2
1.5
1
0.5
0
y
0
5
10
15
20
25
x
raw data, starting:15
fitted Gauss locations:15

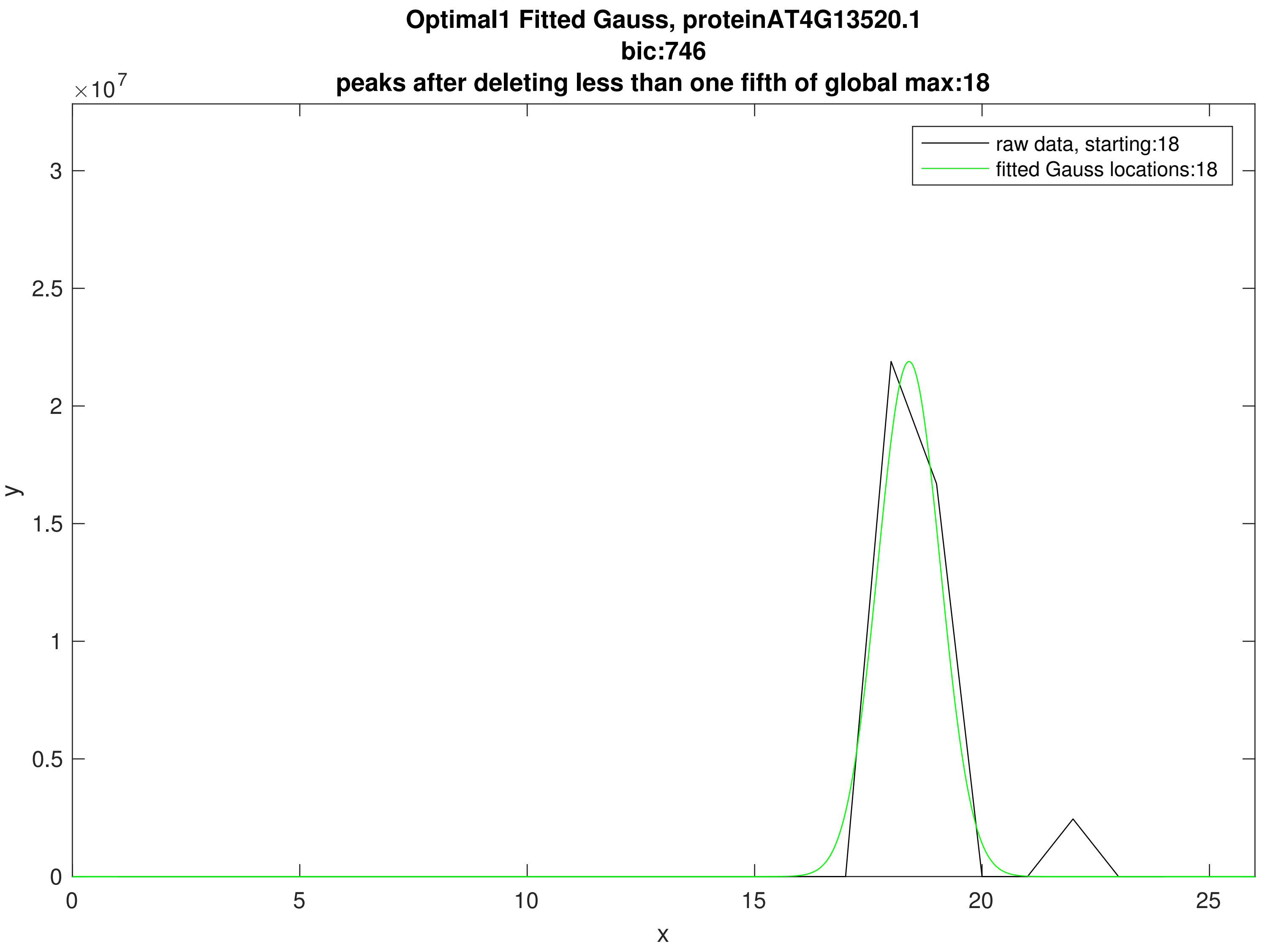
Optimal1 Fitted Gauss, proteinAT4G13520.1
bic:746
peaks after deleting less than one fifth of global max:18
×10^7
raw data, starting:18
fitted Gauss locations:18
y
x
0
0.5
1
1.5
2
2.5
3
0
5
10
15
20
25

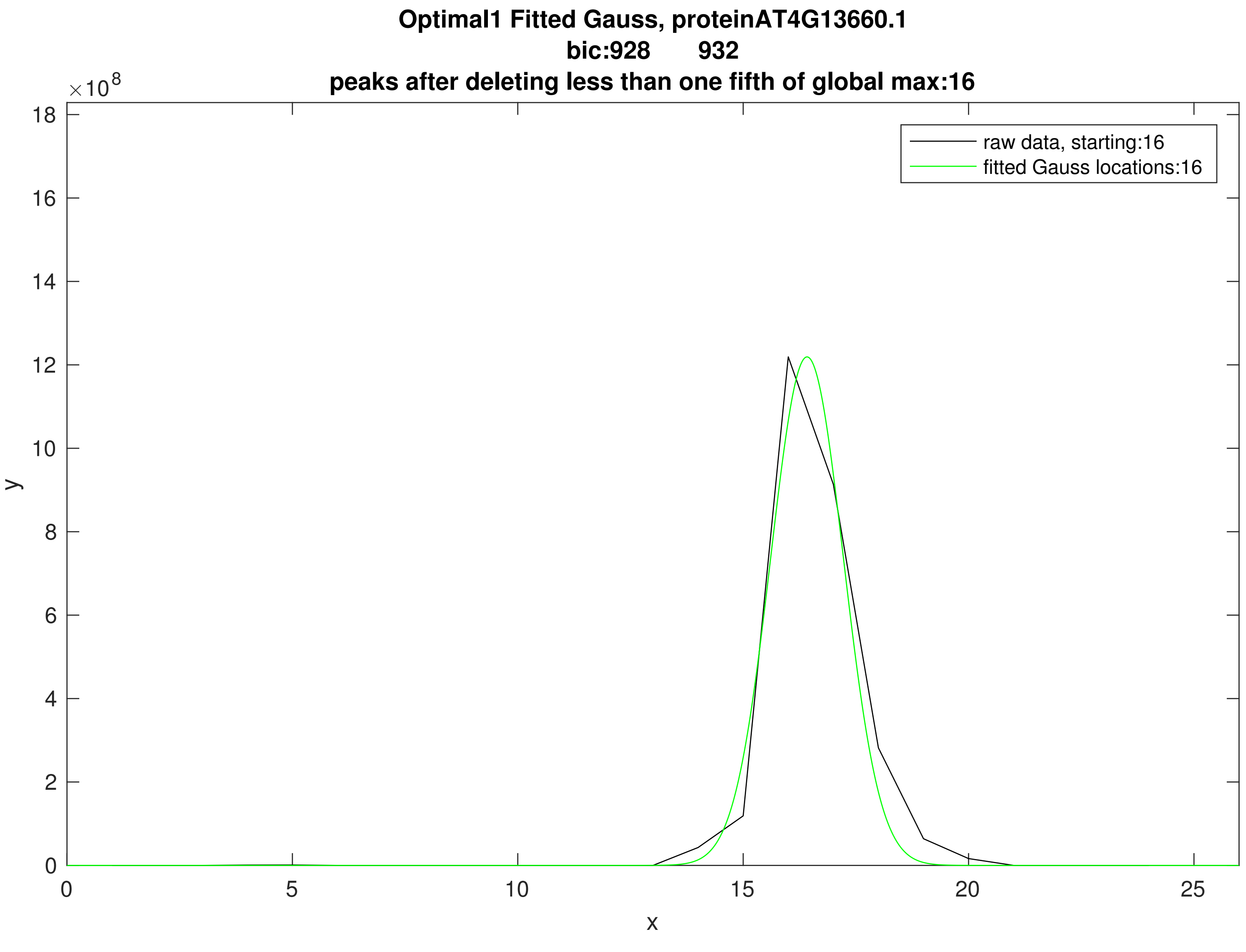

Optimal1 Fitted Gauss, proteinAT4G13660.1
bic:928 932
peaks after deleting less than one fifth of global max:16
×10^8
18
16
14
12
10
8
6
4
2
0
y
0
5
10
15
20
25
x
raw data, starting:16
fitted Gauss locations:16

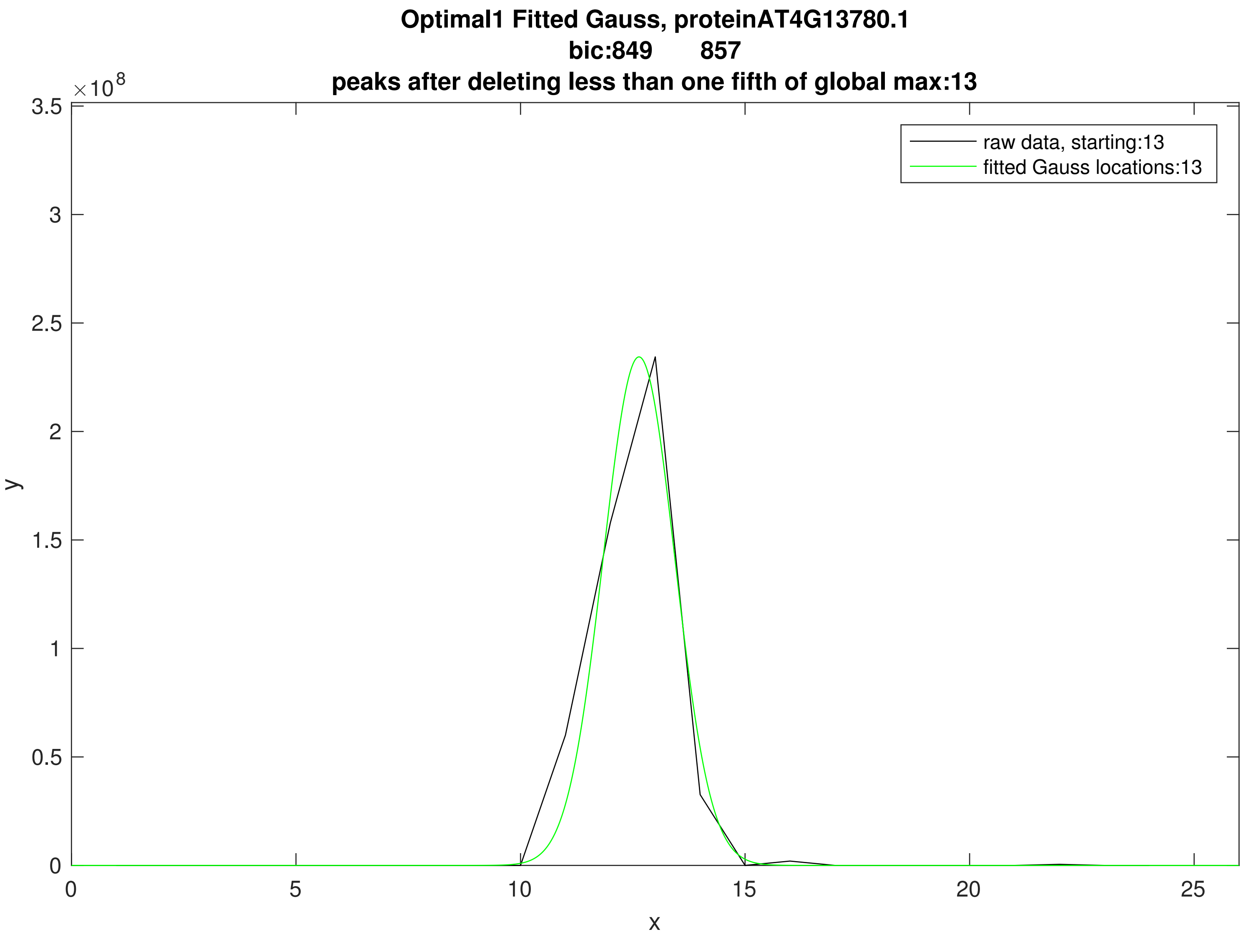

Optimal1 Fitted Gauss, proteinAT4G13780.1
bic:849 857
peaks after deleting less than one fifth of global max:13
×10^8
3.5
3
2.5
2
1.5
1
0.5
0
y
0
5
10
15
20
25
x
raw data, starting:13
fitted Gauss locations:13

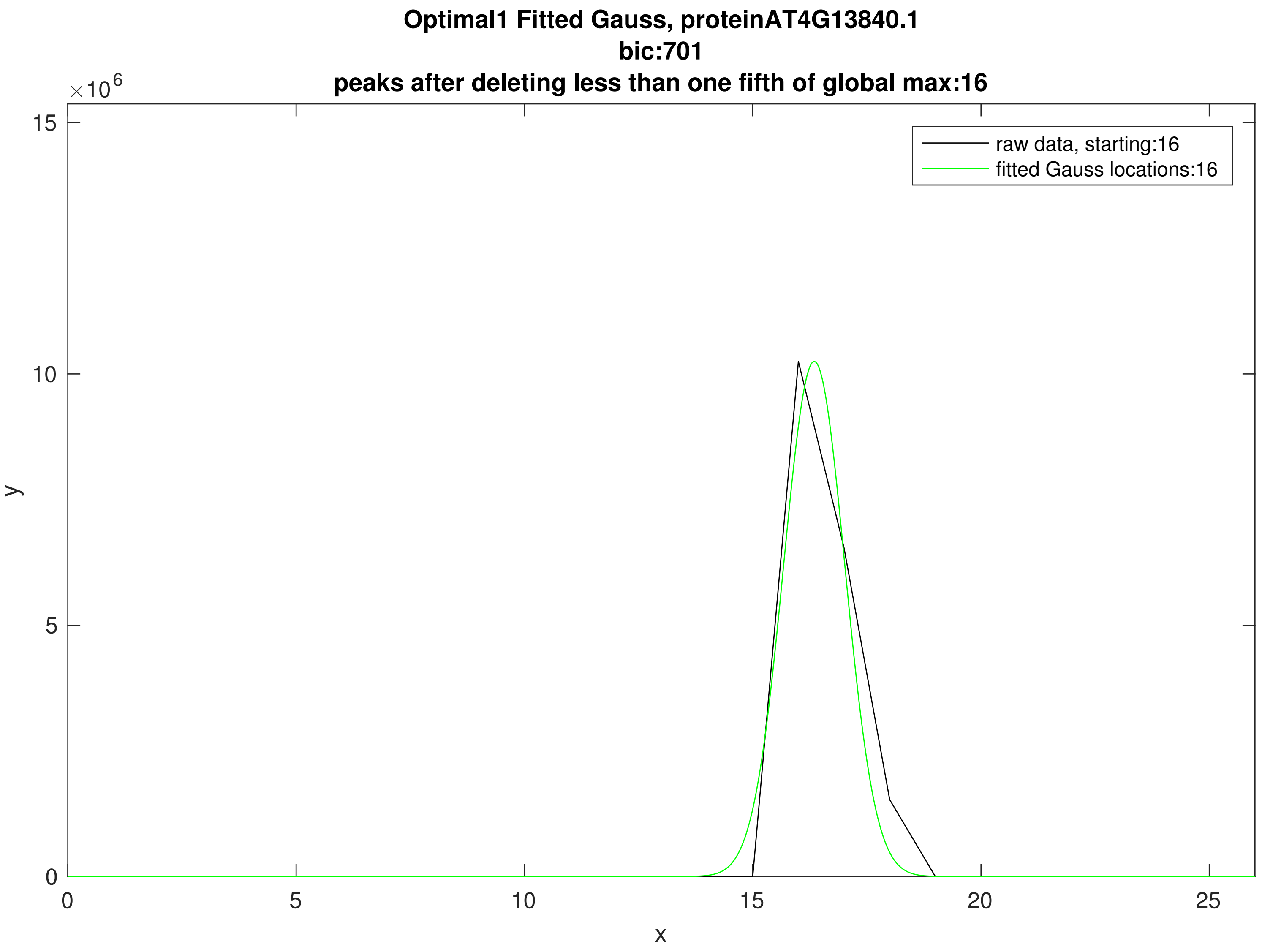
Optimal1 Fitted Gauss, proteinAT4G13840.1
bic:701
peaks after deleting less than one fifth of global max:16
×10^6
raw data, starting:16
fitted Gauss locations:16
y
x
0
5
10
15
20
25

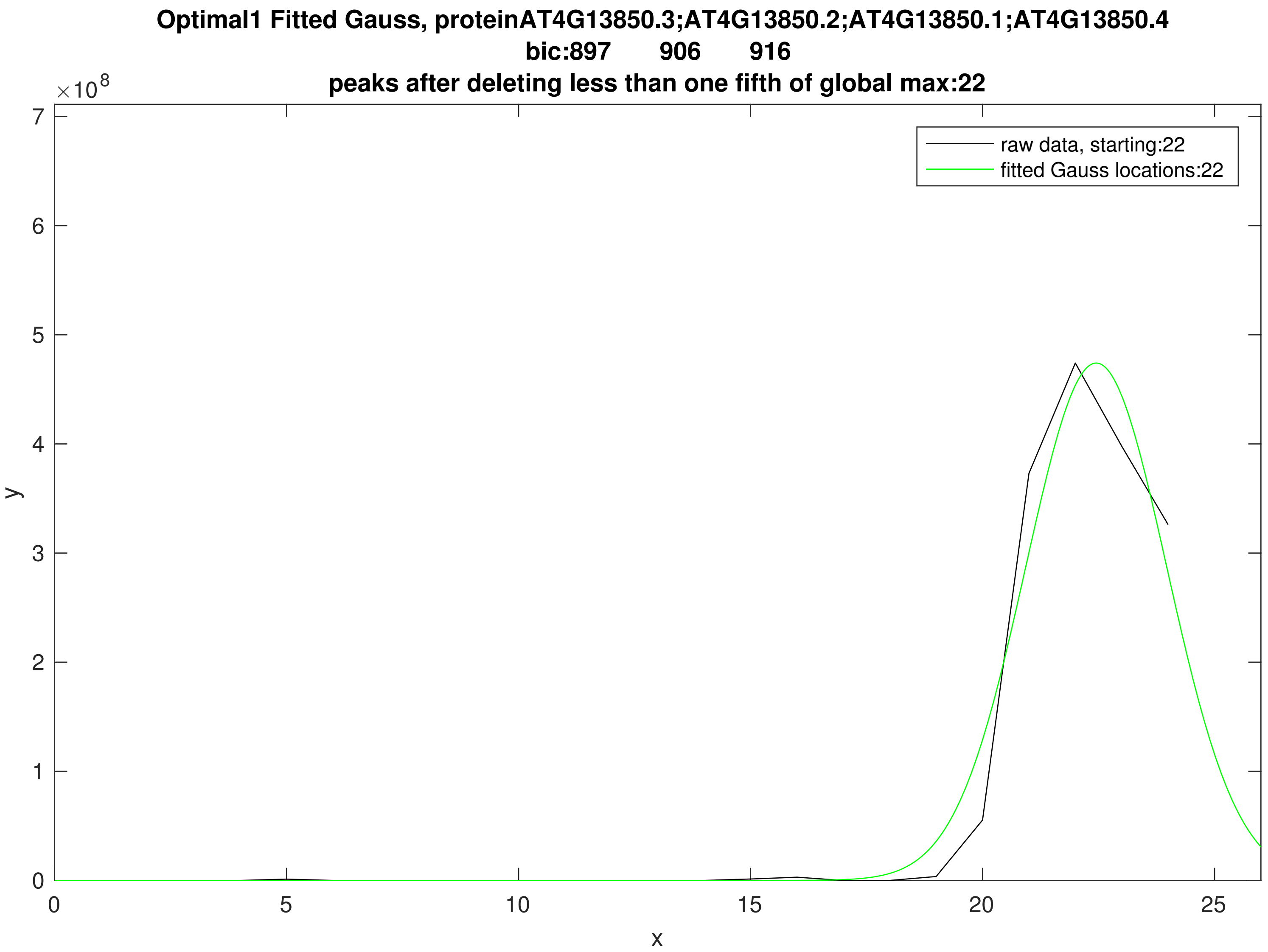
Optimal1 Fitted Gauss, proteinAT4G13850.3;AT4G13850.2;AT4G13850.1;AT4G13850.4
bic:897 906 916
peaks after deleting less than one fifth of global max:22
×10^8
7
6
5
4
3
2
1
0
y
0
5
10
15
20
25
x
raw data, starting:22
fitted Gauss locations:22

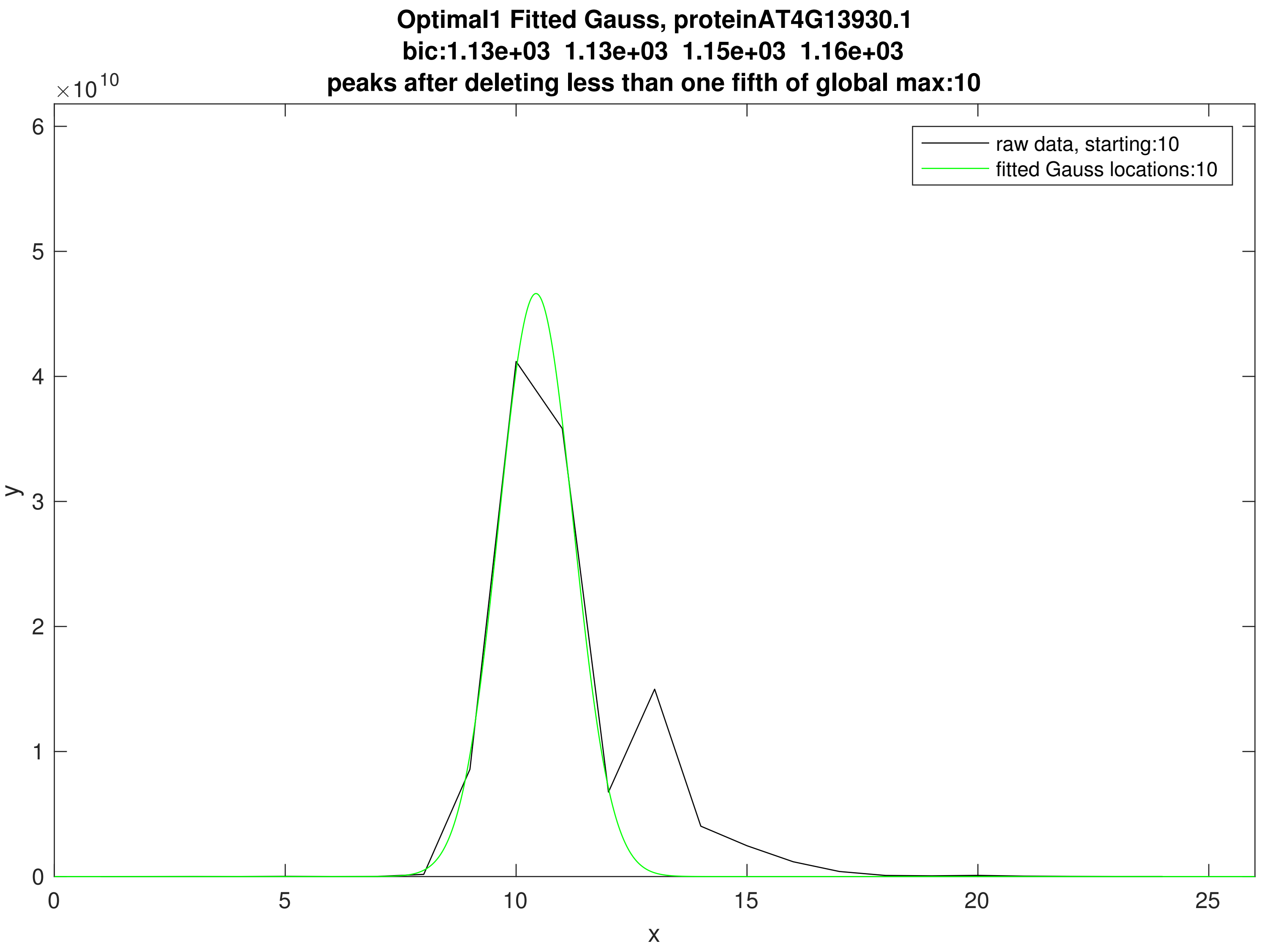
Optimal1 Fitted Gauss, proteinAT4G13930.1
bic:1.13e+03 1.13e+03 1.15e+03 1.16e+03
peaks after deleting less than one fifth of global max:10
×10^10
raw data, starting:10
fitted Gauss locations:10
y
x
0
1
2
3
4
5
6
0
5
10
15
20
25

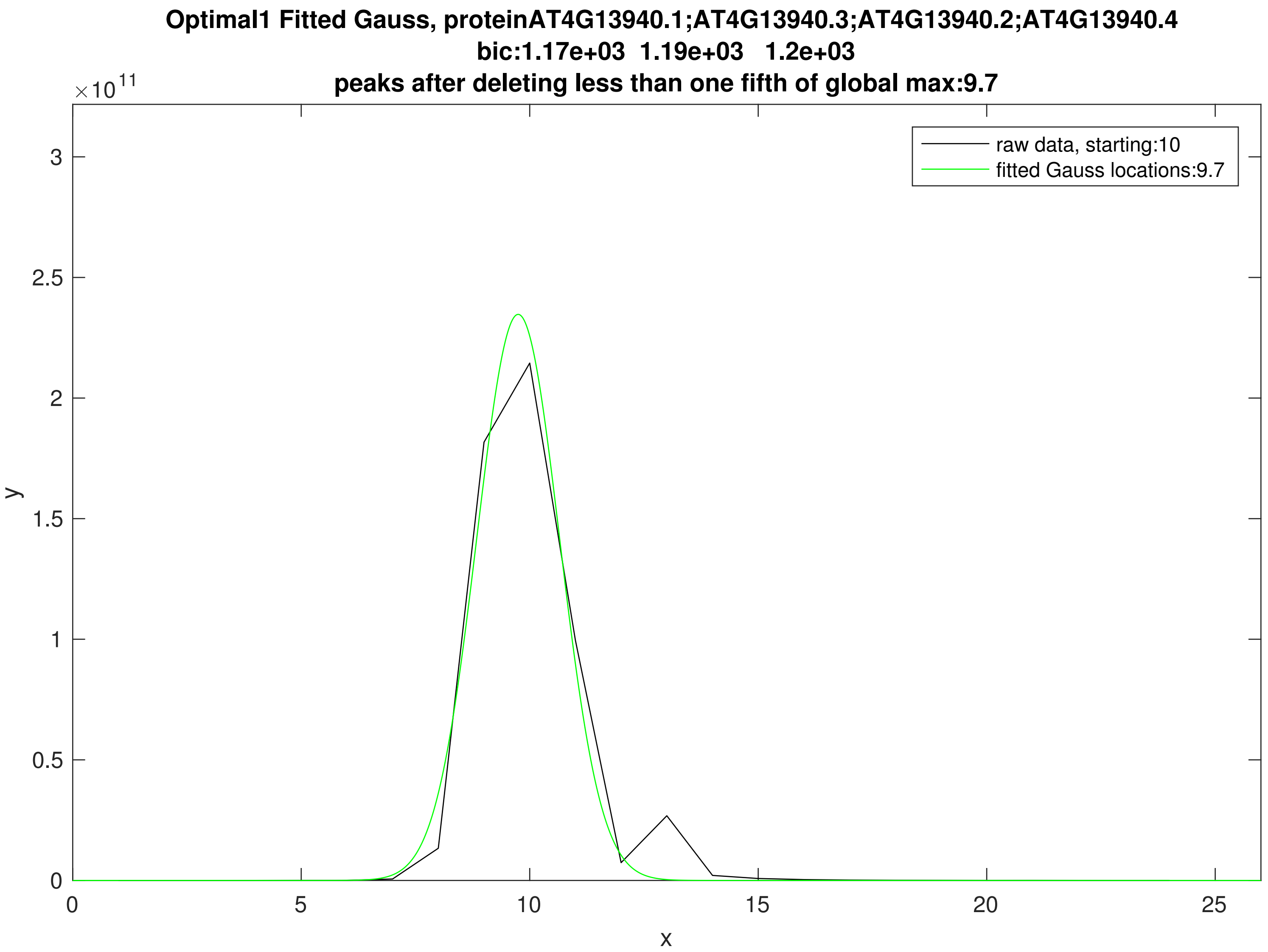
Optimal1 Fitted Gauss, proteinAT4G13940.1;AT4G13940.3;AT4G13940.2;AT4G13940.4
bic:1.17e+03 1.19e+03 1.2e+03
peaks after deleting less than one fifth of global max:9.7
×10^11
raw data, starting:10
fitted Gauss locations:9.7
3
2.5
2
1.5
1
0.5
0
y
0
5
10
15
20
25
x

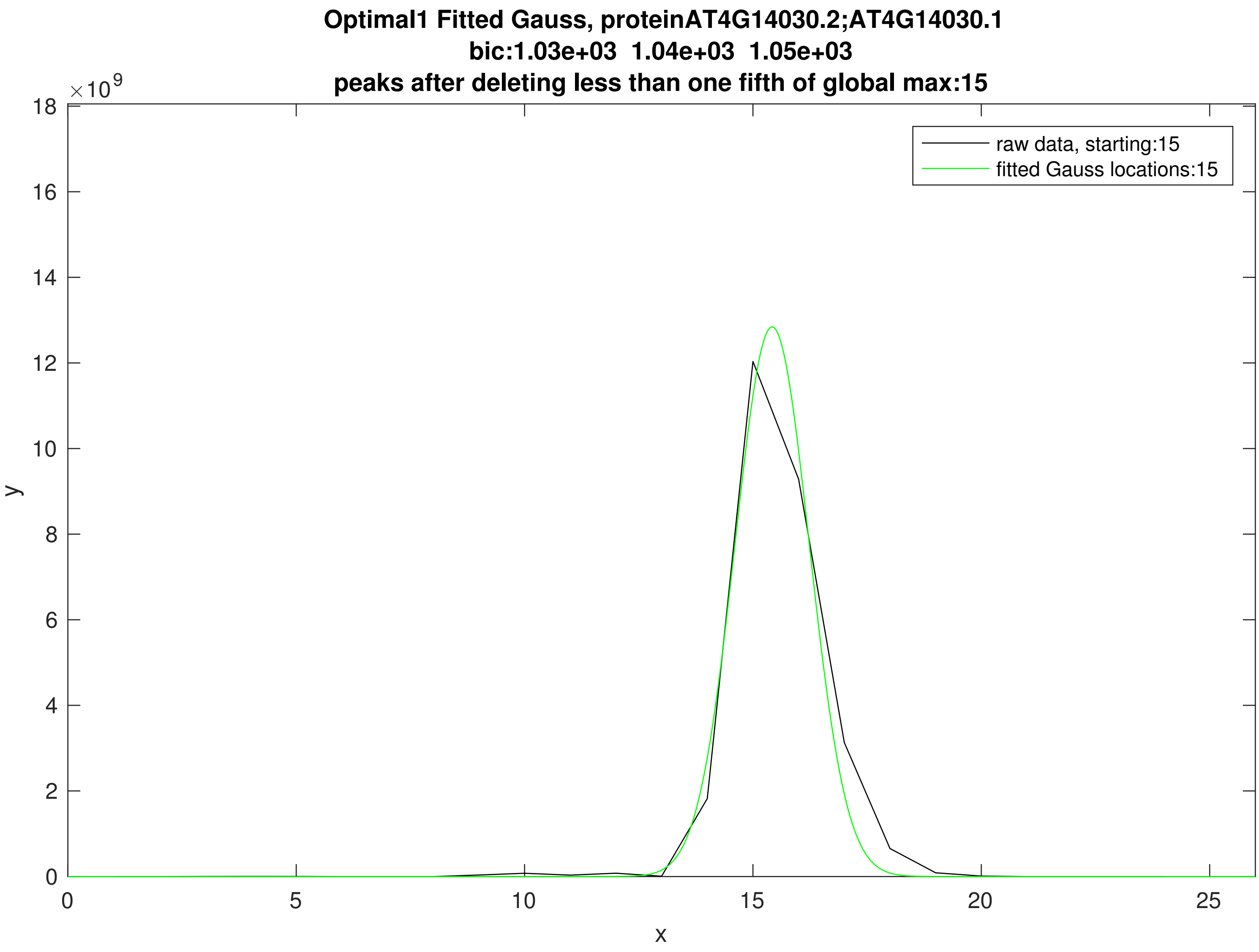

Optimal1 Fitted Gauss, proteinAT4G14030.2;AT4G14030.1
bic:1.03e+03 1.04e+03 1.05e+03
peaks after deleting less than one fifth of global max:15
×10^9
18
16
14
12
10
8
6
4
2
0
y
0
5
10
15
20
25
x
raw data, starting:15
fitted Gauss locations:15

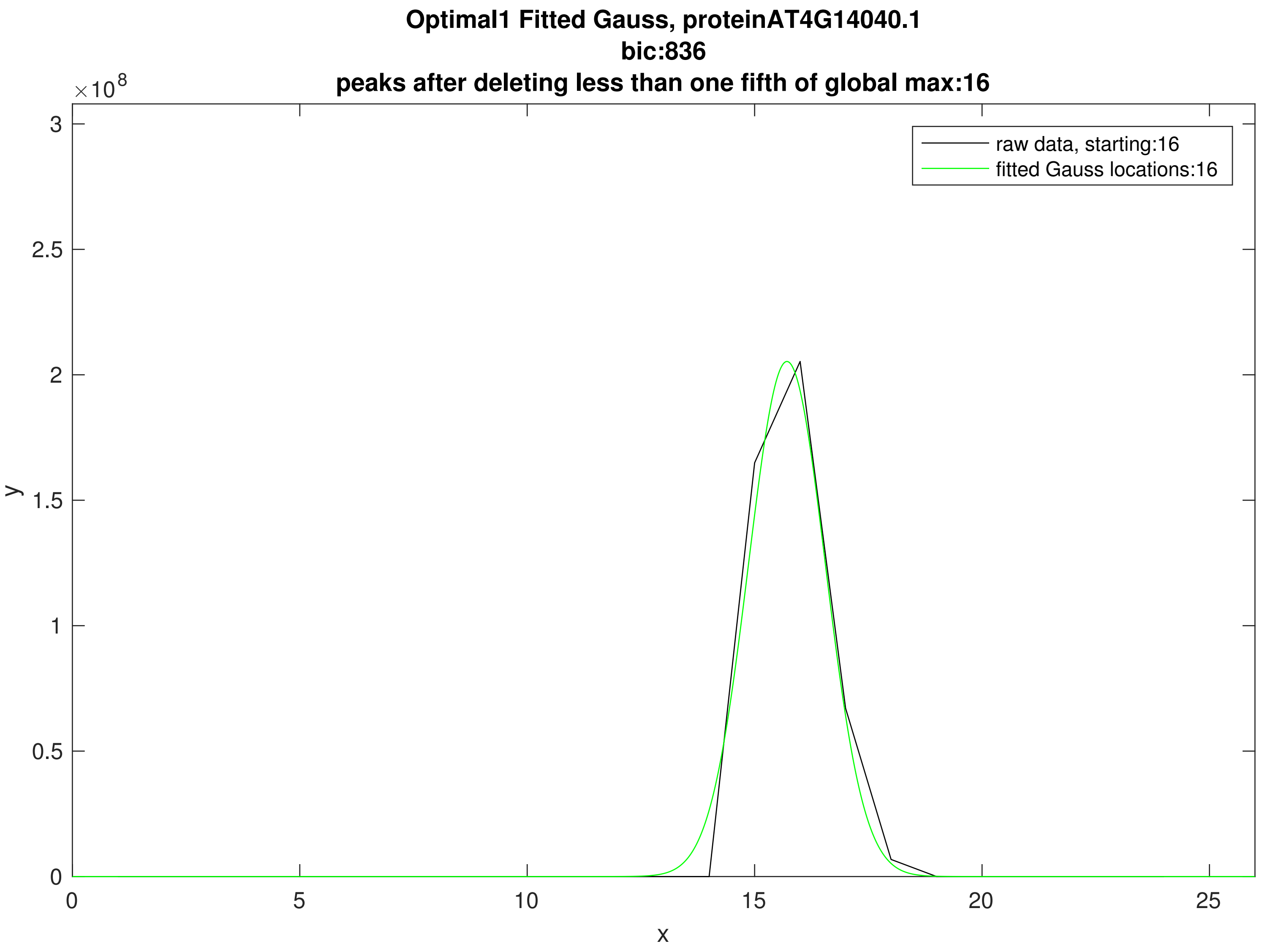

Optimal1 Fitted Gauss, proteinAT4G14040.1
bic:836
peaks after deleting less than one fifth of global max:16
×10^8
raw data, starting:16
fitted Gauss locations:16
y
x
0
0.5
1
1.5
2
2.5
3
5
10
15
20
25

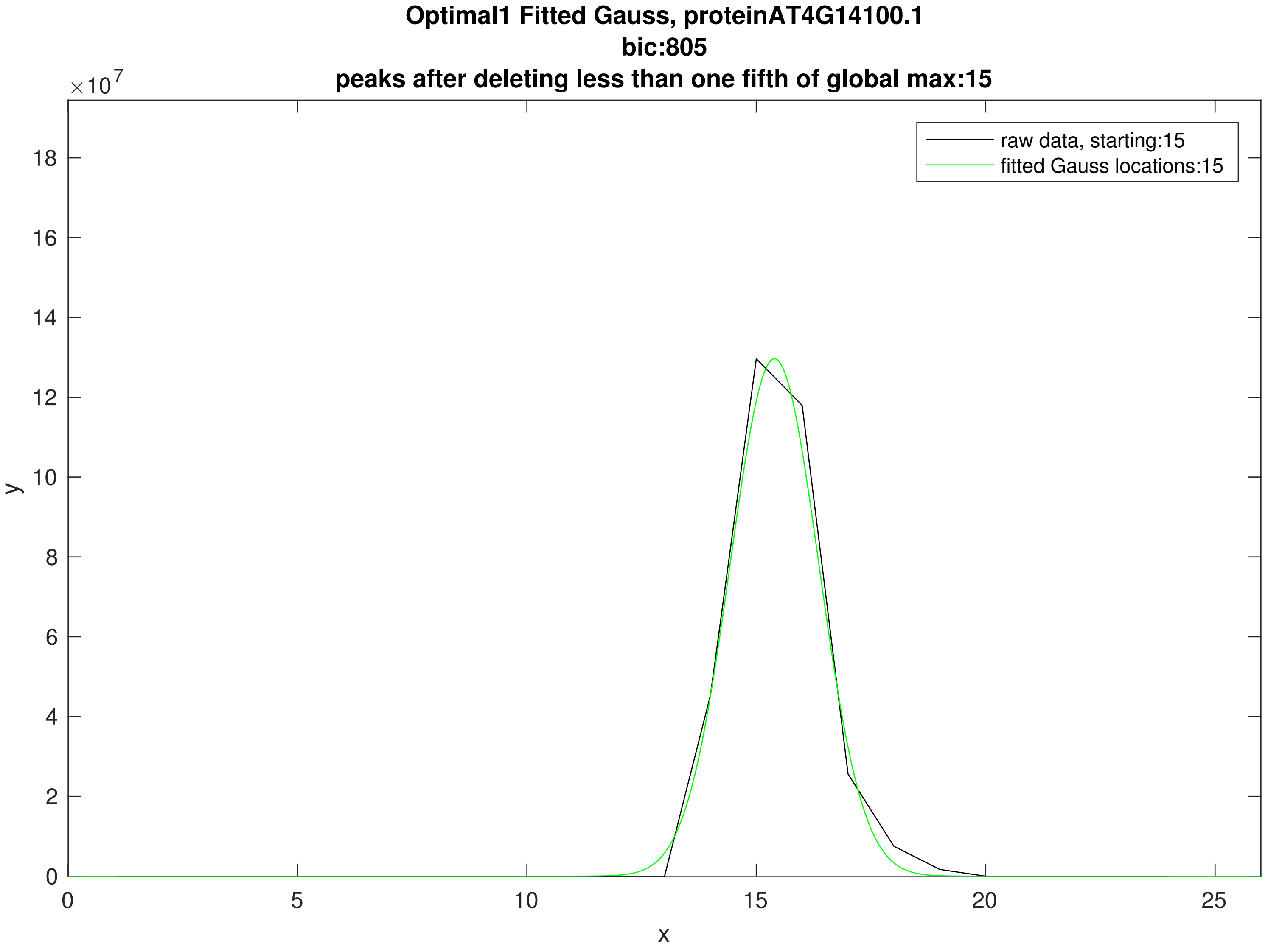
Optimal1 Fitted Gauss, proteinAT4G14100.1
bic:805
peaks after deleting less than one fifth of global max:15
×10^7
raw data, starting:15
fitted Gauss locations:15
y
x
0
2
4
6
8
10
12
14
16
18
0
5
10
15
20
25

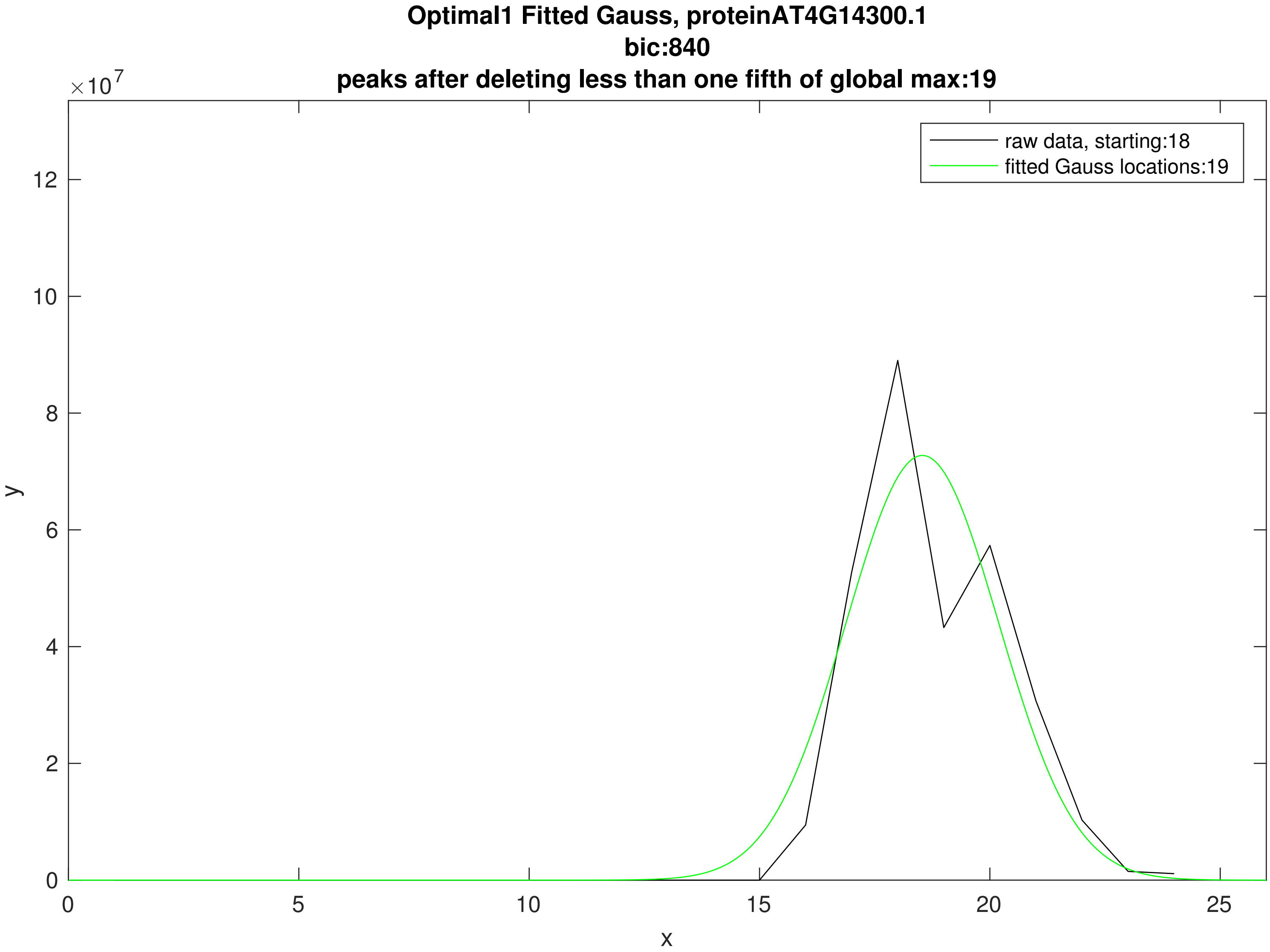
Optimal1 Fitted Gauss, proteinAT4G14300.1
bic:840
peaks after deleting less than one fifth of global max:19
×10^7
raw data, starting:18
fitted Gauss locations:19
12
10
8
6
4
2
0
y
0
5
10
15
20
25
x

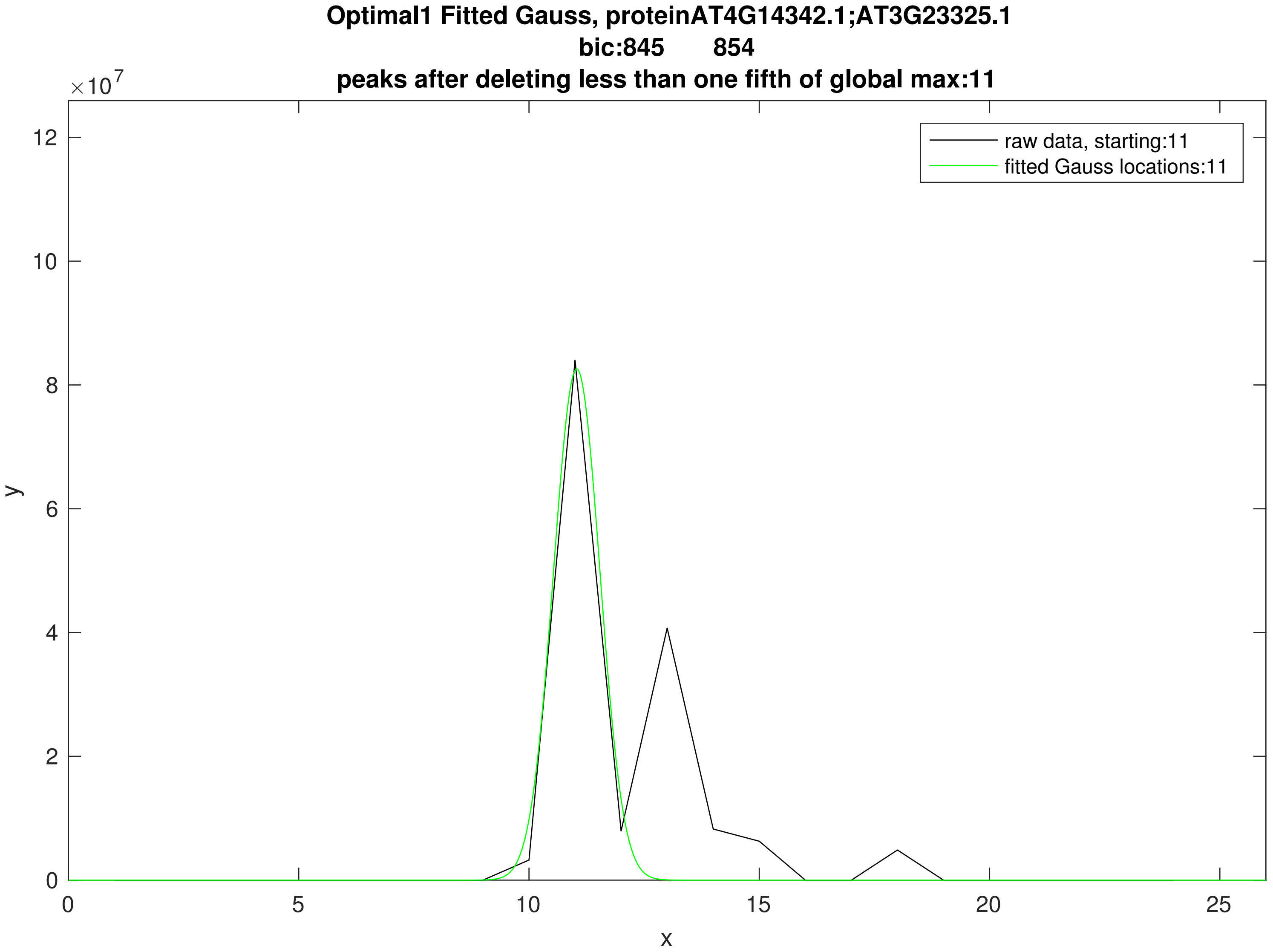
Optimal1 Fitted Gauss, proteinAT4G14342.1;AT3G23325.1
bic:845 854
peaks after deleting less than one fifth of global max:11
×10^7
12
10
8
6
4
2
0
y
0
5
10
15
20
25
x
raw data, starting:11
fitted Gauss locations:11

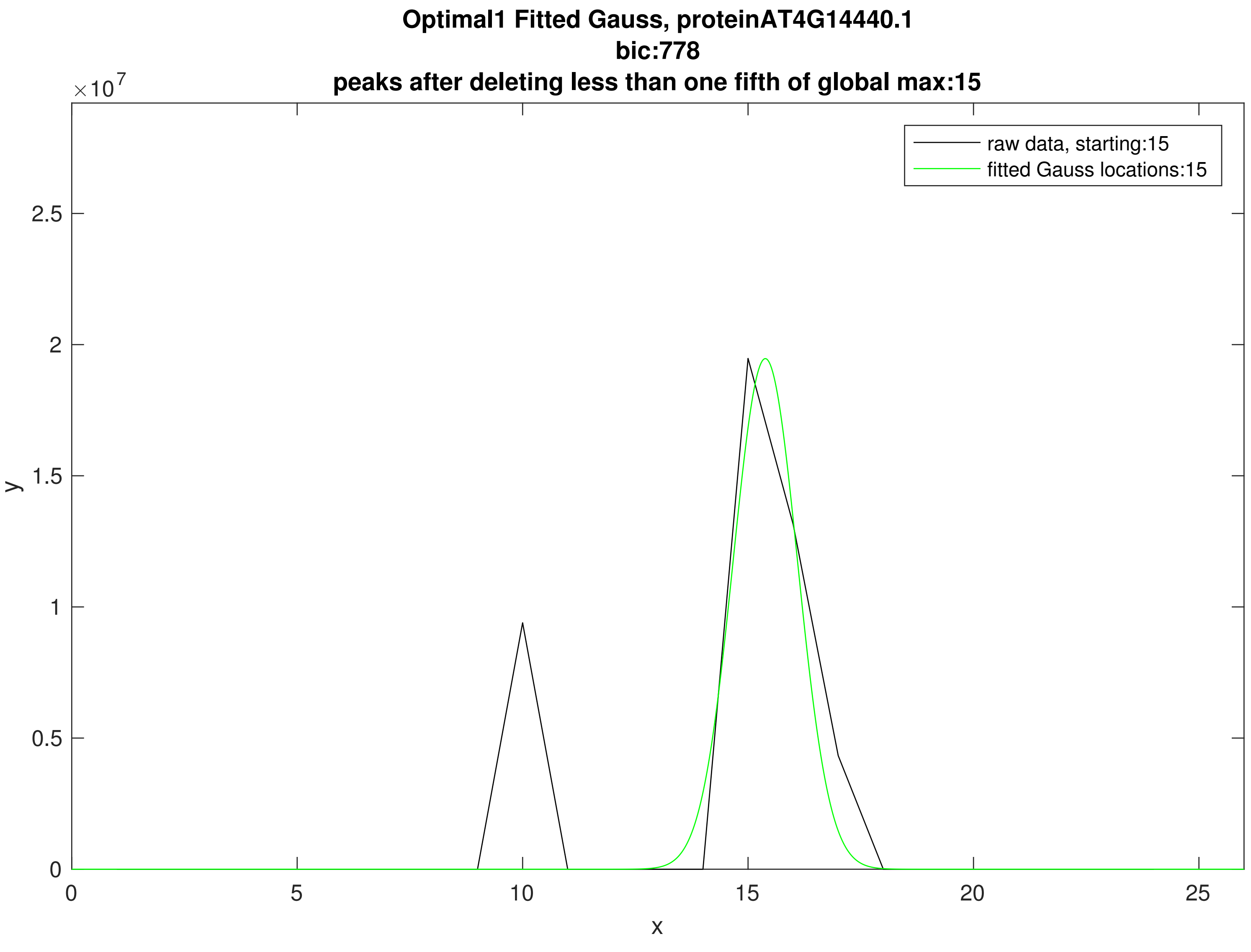
Optimal1 Fitted Gauss, proteinAT4G14440.1
bic:778
peaks after deleting less than one fifth of global max:15
×10^7
2.5
2
1.5
1
0.5
0
y
0
5
10
15
20
25
x
raw data, starting:15
fitted Gauss locations:15

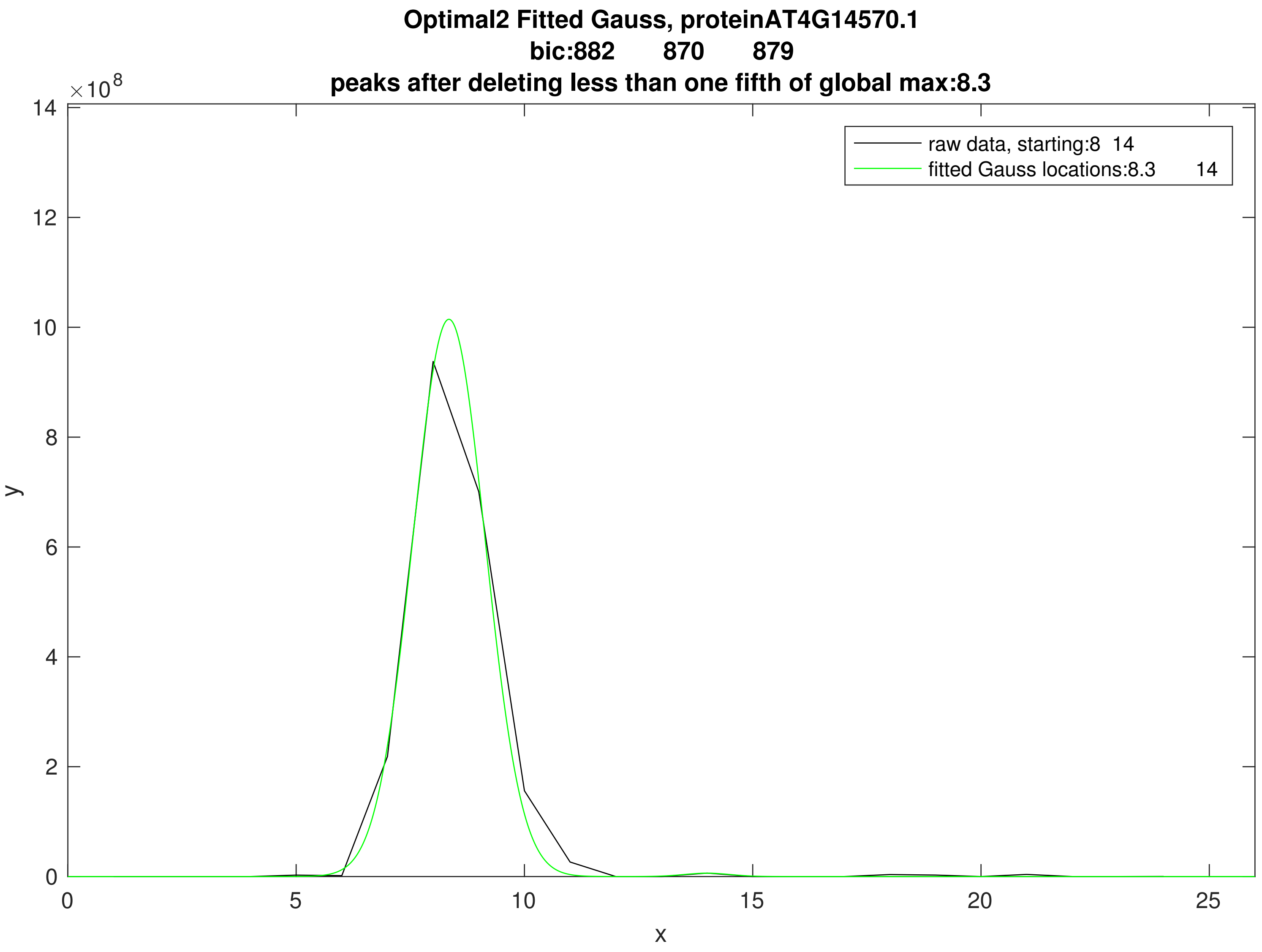

Optimal2 Fitted Gauss, proteinAT4G14570.1
bic:882 870 879
peaks after deleting less than one fifth of global max:8.3
×10^8
raw data, starting:8 14
fitted Gauss locations:8.3 14
y
x
0
2
4
6
8
10
12
14
0
5
10
15
20
25

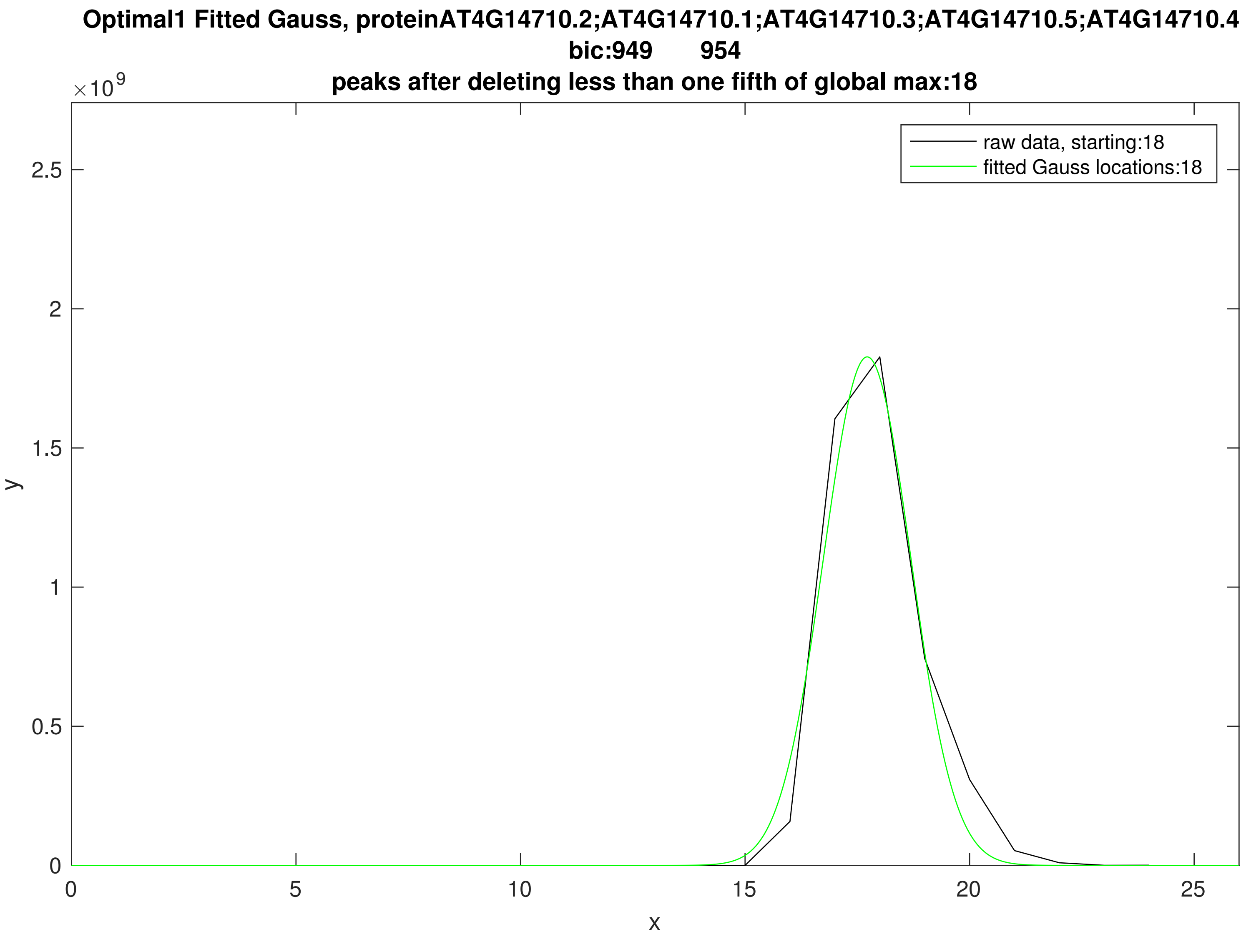

Optimal1 Fitted Gauss, proteinAT4G14710.2;AT4G14710.1;AT4G14710.3;AT4G14710.5;AT4G14710.4
bic:949 954
peaks after deleting less than one fifth of global max:18
×10^9
2.5
2
1.5
1
0.5
0
y
0
5
10
15
20
25
x
raw data, starting:18
fitted Gauss locations:18

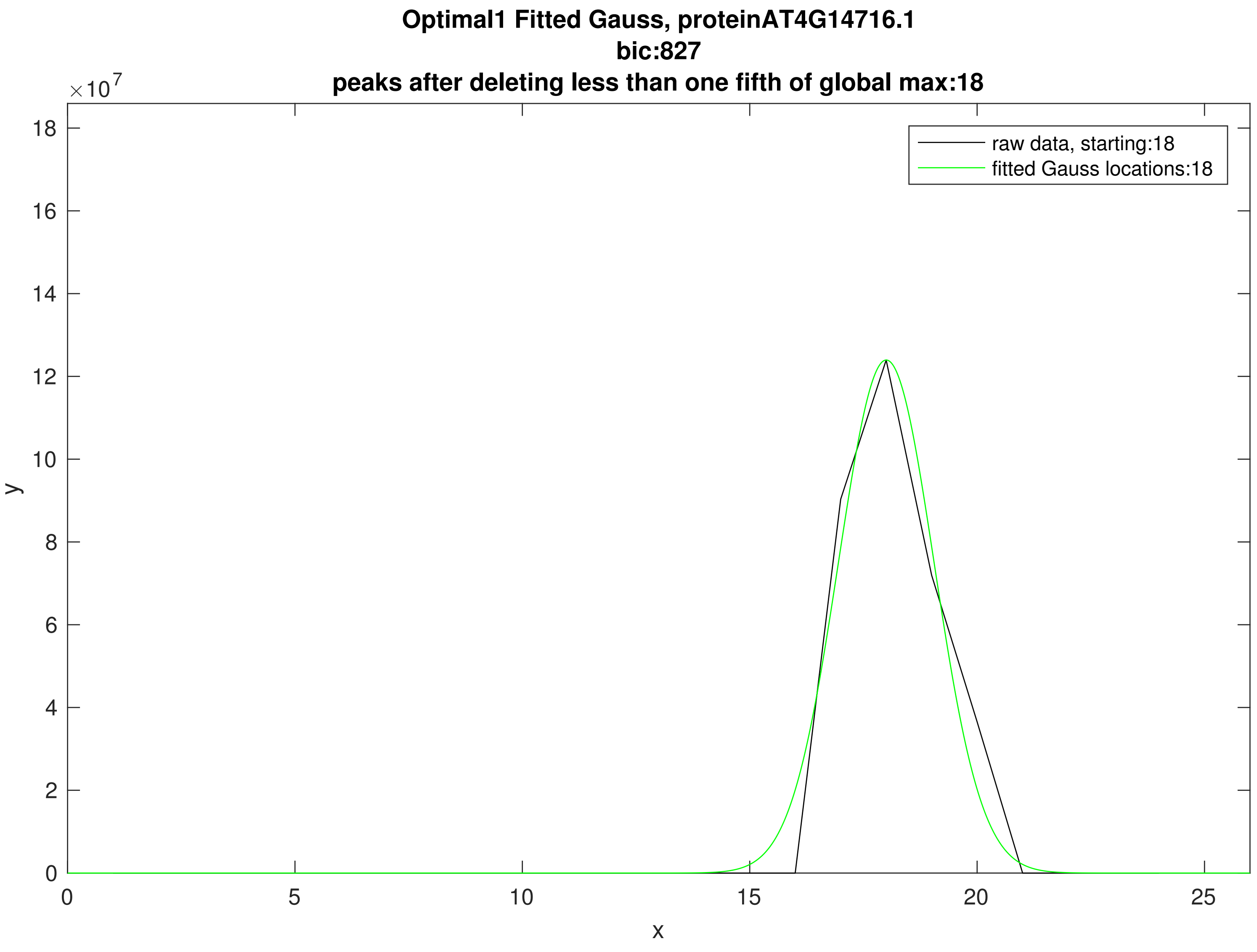
Optimal1 Fitted Gauss, proteinAT4G14716.1
bic:827
peaks after deleting less than one fifth of global max:18
×10^7
raw data, starting:18
fitted Gauss locations:18
y
x
0
2
4
6
8
10
12
14
16
18
0
5
10
15
20
25

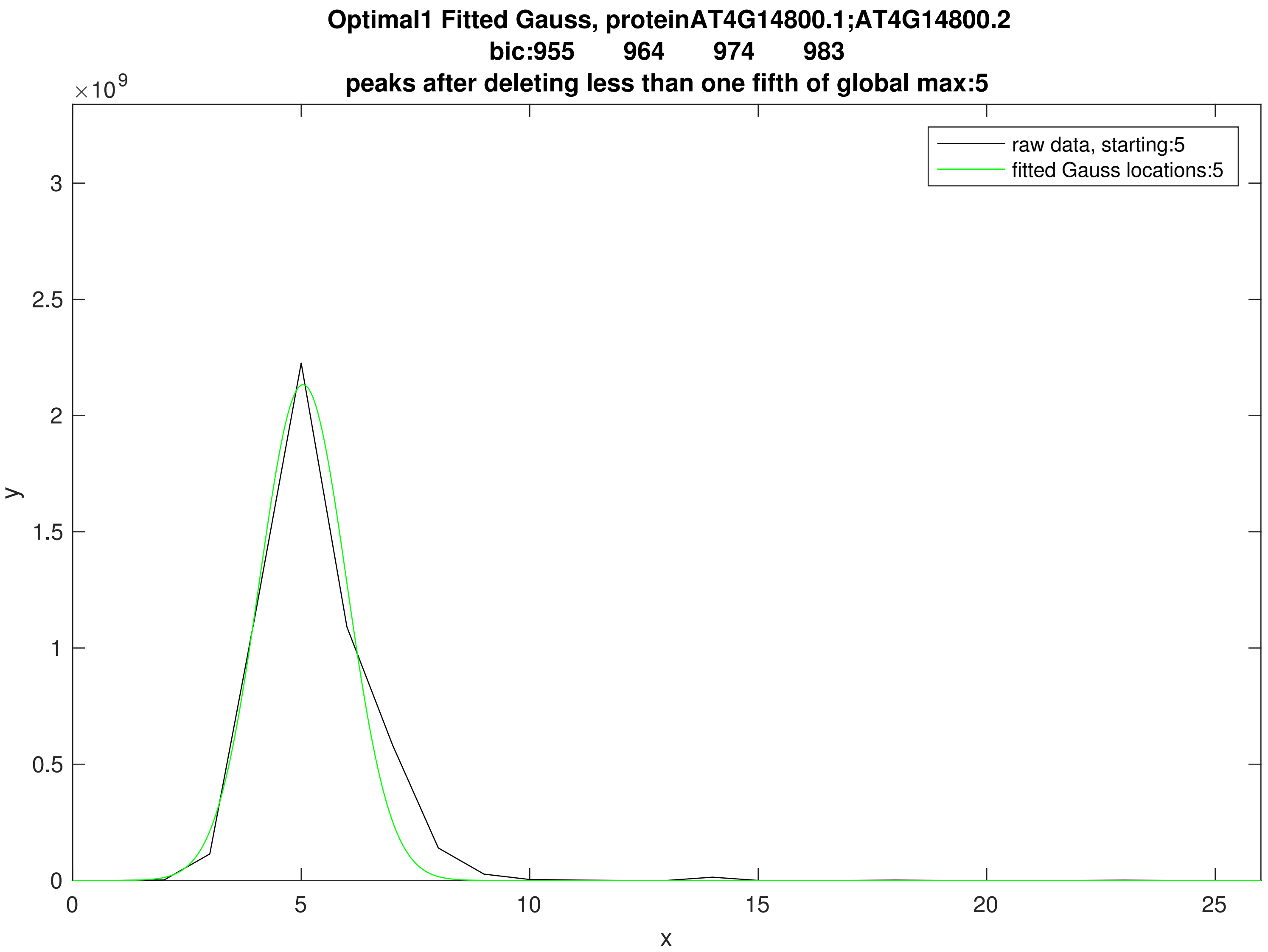

Optimal1 Fitted Gauss, proteinAT4G14800.1;AT4G14800.2
bic:955 964 974 983
peaks after deleting less than one fifth of global max:5
×10^9
raw data, starting:5
fitted Gauss locations:5
3
2.5
2
1.5
1
0.5
0
y
0
5
10
15
20
25
x

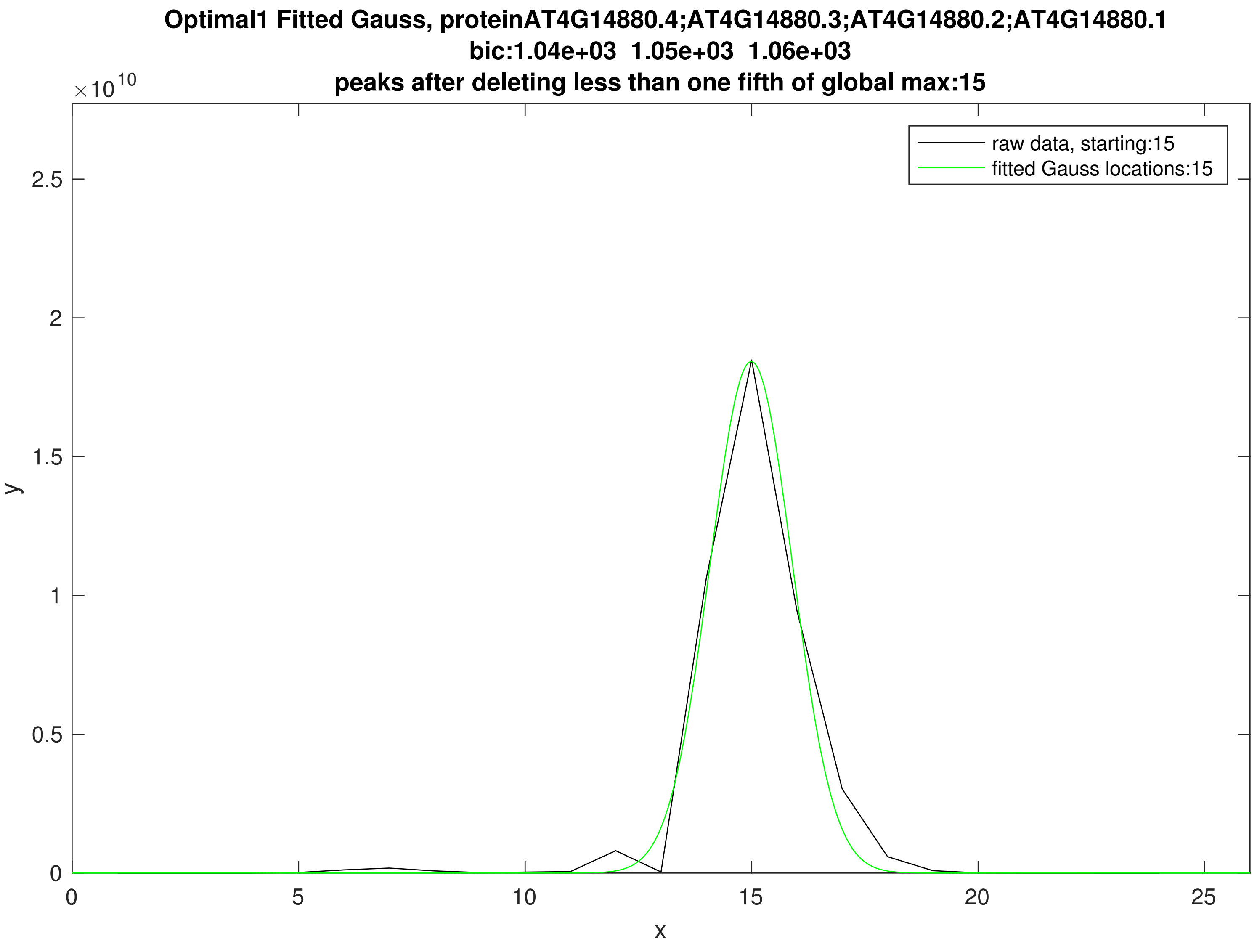

Optimal1 Fitted Gauss, proteinAT4G14880.4;AT4G14880.3;AT4G14880.2;AT4G14880.1
bic:1.04e+03 1.05e+03 1.06e+03
peaks after deleting less than one fifth of global max:15
×10^10
raw data, starting:15
fitted Gauss locations:15
y
x
0
0.5
1
1.5
2
2.5
5
10
15
20
25

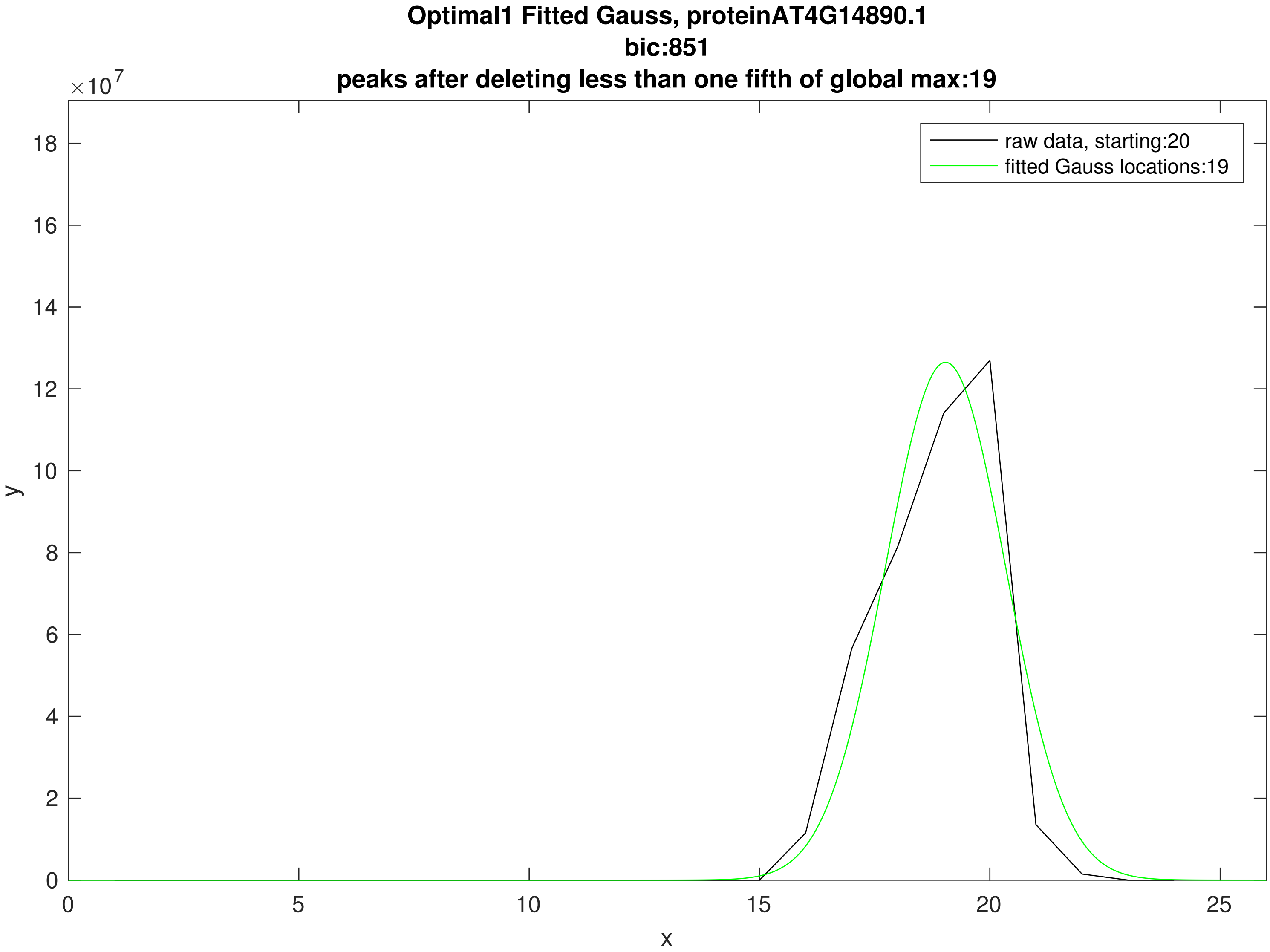

Optimal1 Fitted Gauss, proteinAT4G14890.1
bic:851
peaks after deleting less than one fifth of global max:19
×10^7
raw data, starting:20
fitted Gauss locations:19
18
16
14
12
10
8
6
4
2
0
y
0
5
10
15
20
25
x

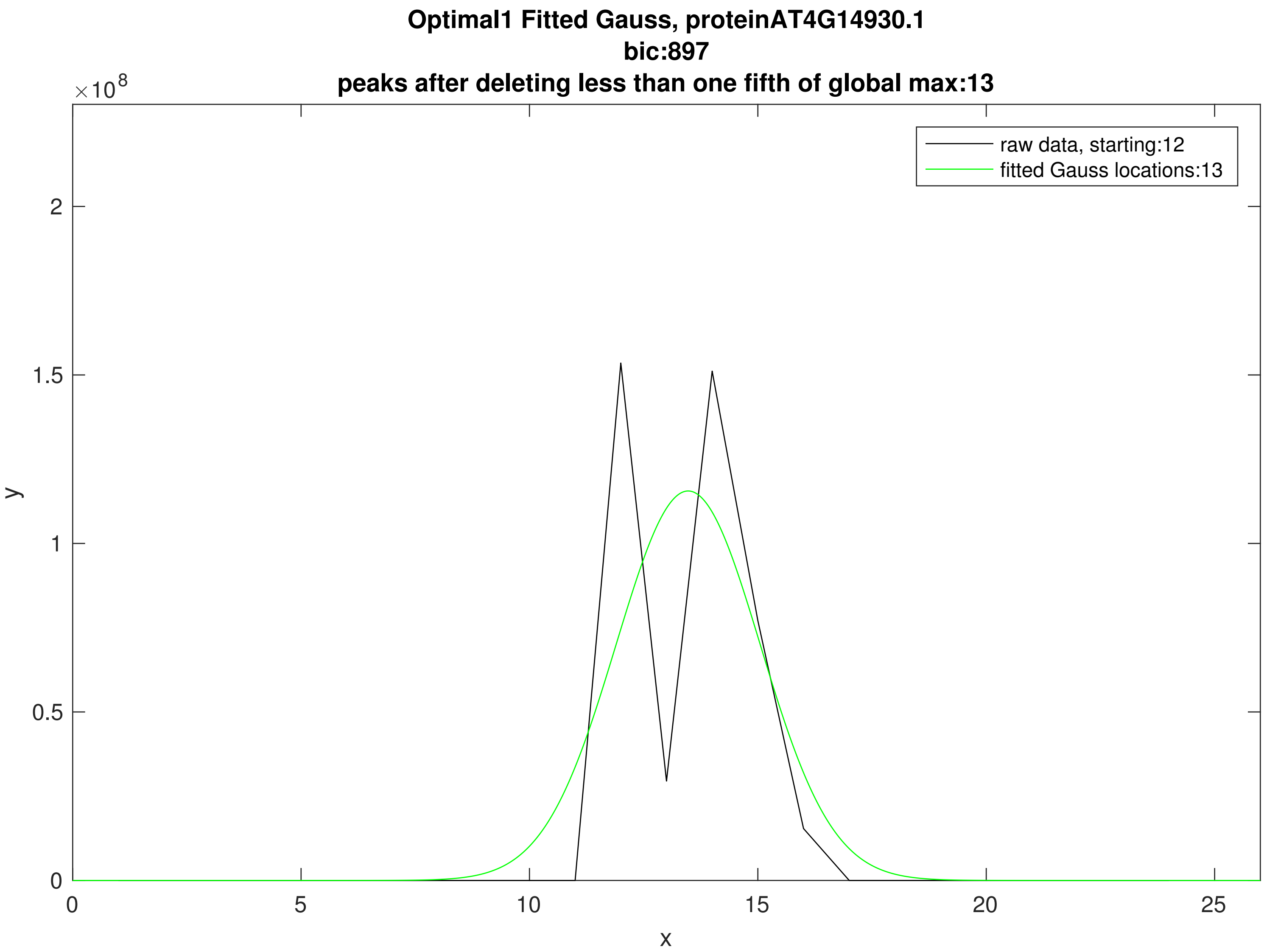

Optimal1 Fitted Gauss, proteinAT4G14930.1
bic:897
peaks after deleting less than one fifth of global max:13
×10^8
raw data, starting:12
fitted Gauss locations:13
2
1.5
1
0.5
0
y
0
5
10
15
20
25
x

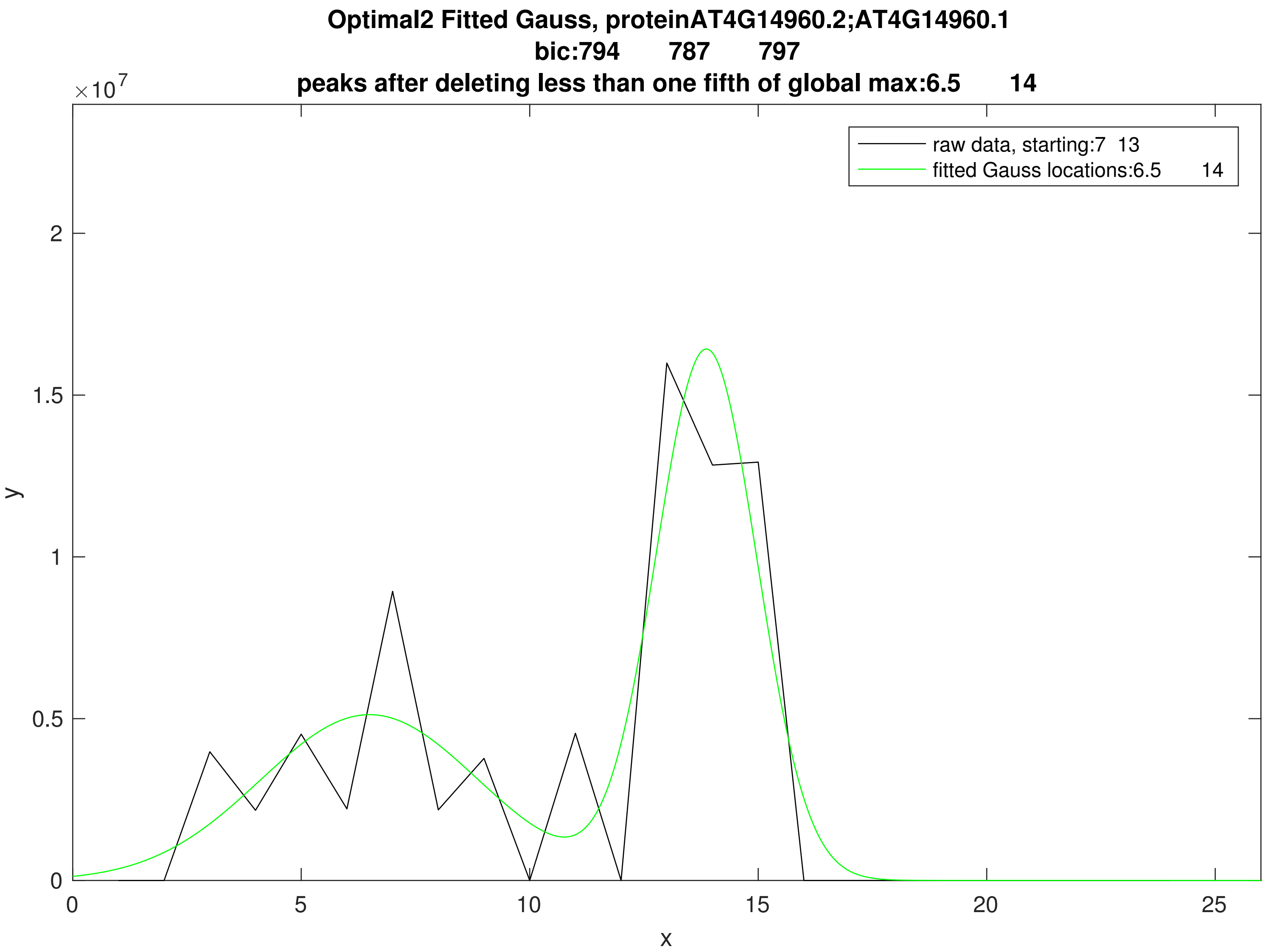
Optimal2 Fitted Gauss, proteinAT4G14960.2;AT4G14960.1
bic:794 787 797
peaks after deleting less than one fifth of global max:6.5 14
×10^7
raw data, starting:7 13
fitted Gauss locations:6.5 14
y
x
0
0.5
1
1.5
2
5
10
15
20
25

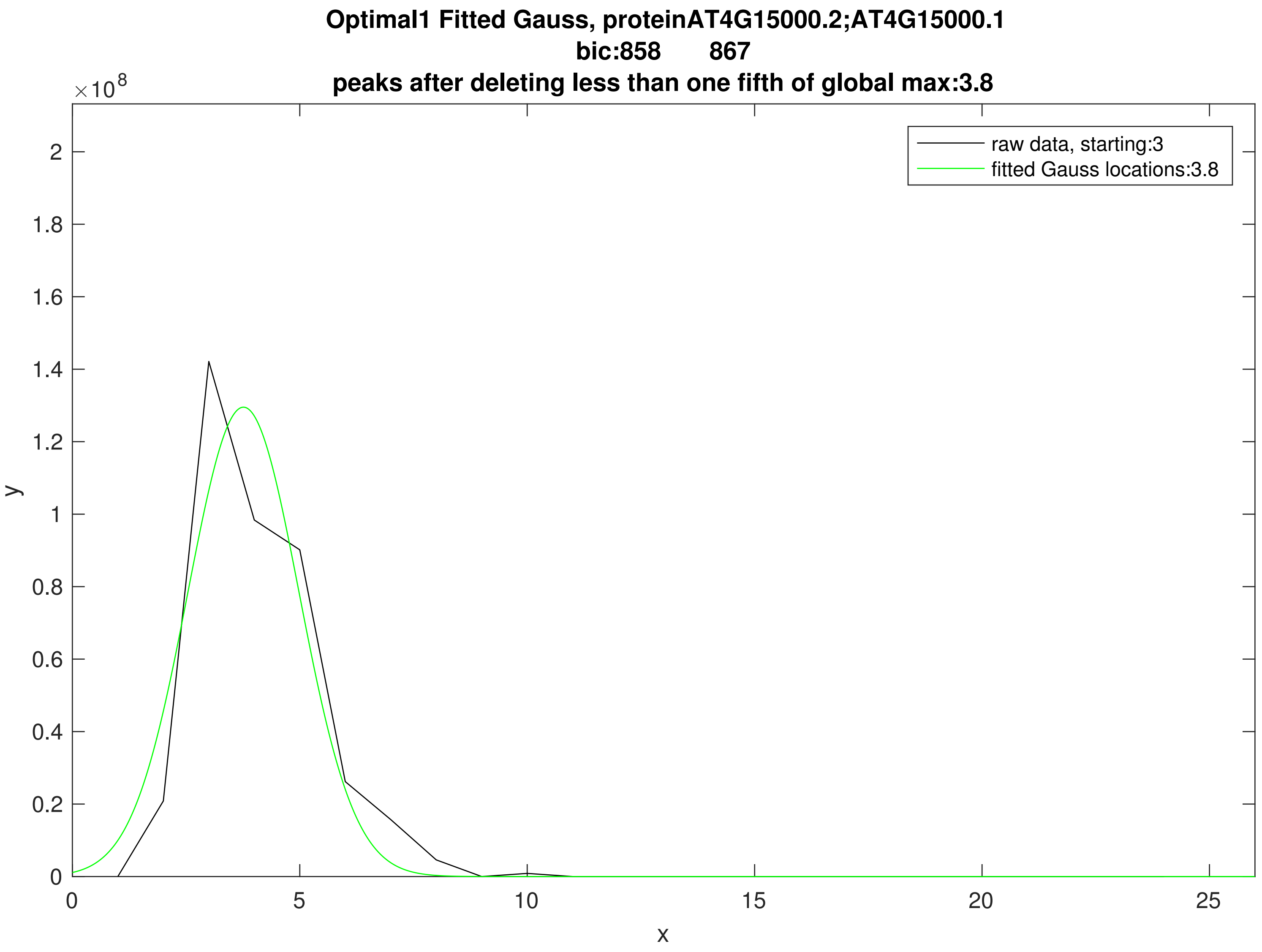

Optimal1 Fitted Gauss, proteinAT4G15000.2;AT4G15000.1
bic:858 867
peaks after deleting less than one fifth of global max:3.8
×10^8
raw data, starting:3
fitted Gauss locations:3.8
y
x
0
0.2
0.4
0.6
0.8
1
1.2
1.4
1.6
1.8
2
5
10
15
20
25

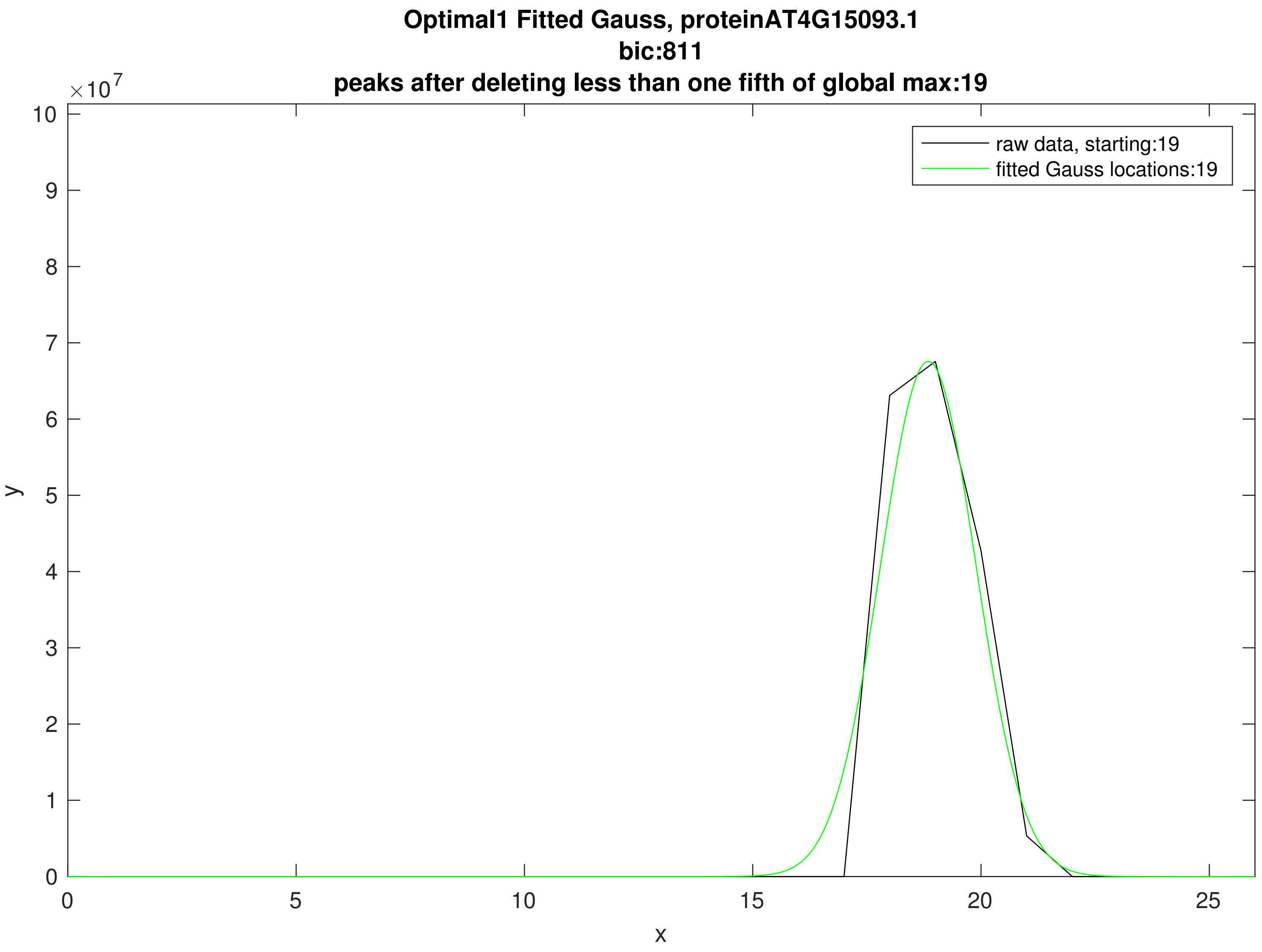
Optimal1 Fitted Gauss, proteinAT4G15093.1
bic:811
peaks after deleting less than one fifth of global max:19
×10^7
raw data, starting:19
fitted Gauss locations:19
y
x
0
1
2
3
4
5
6
7
8
9
10
0
5
10
15
20
25

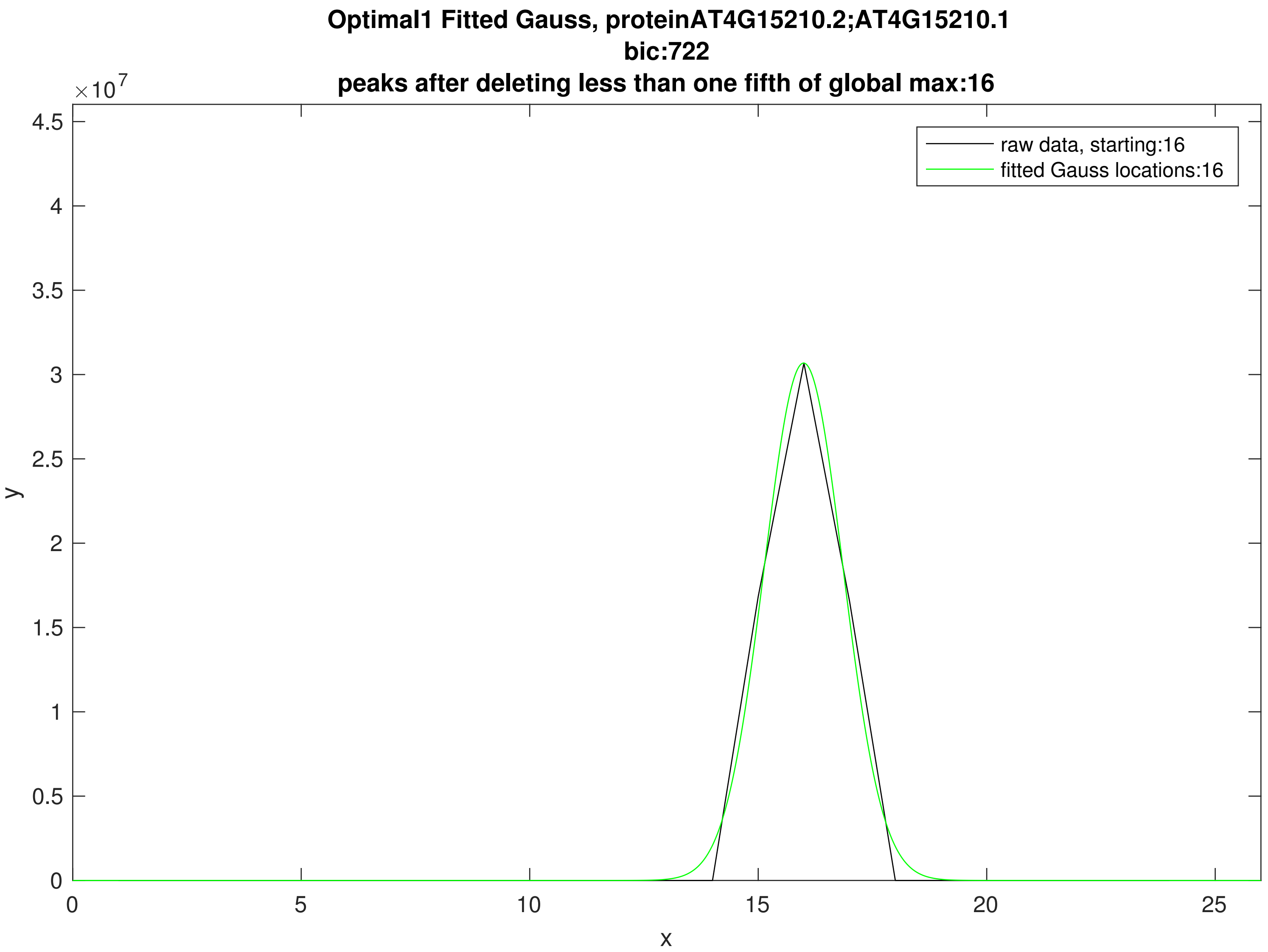
Optimal1 Fitted Gauss, proteinAT4G15210.2;AT4G15210.1
bic:722
peaks after deleting less than one fifth of global max:16
×10^7
4.5
4
3.5
3
2.5
2
1.5
1
0.5
0
y
0
5
10
15
20
25
x
raw data, starting:16
fitted Gauss locations:16

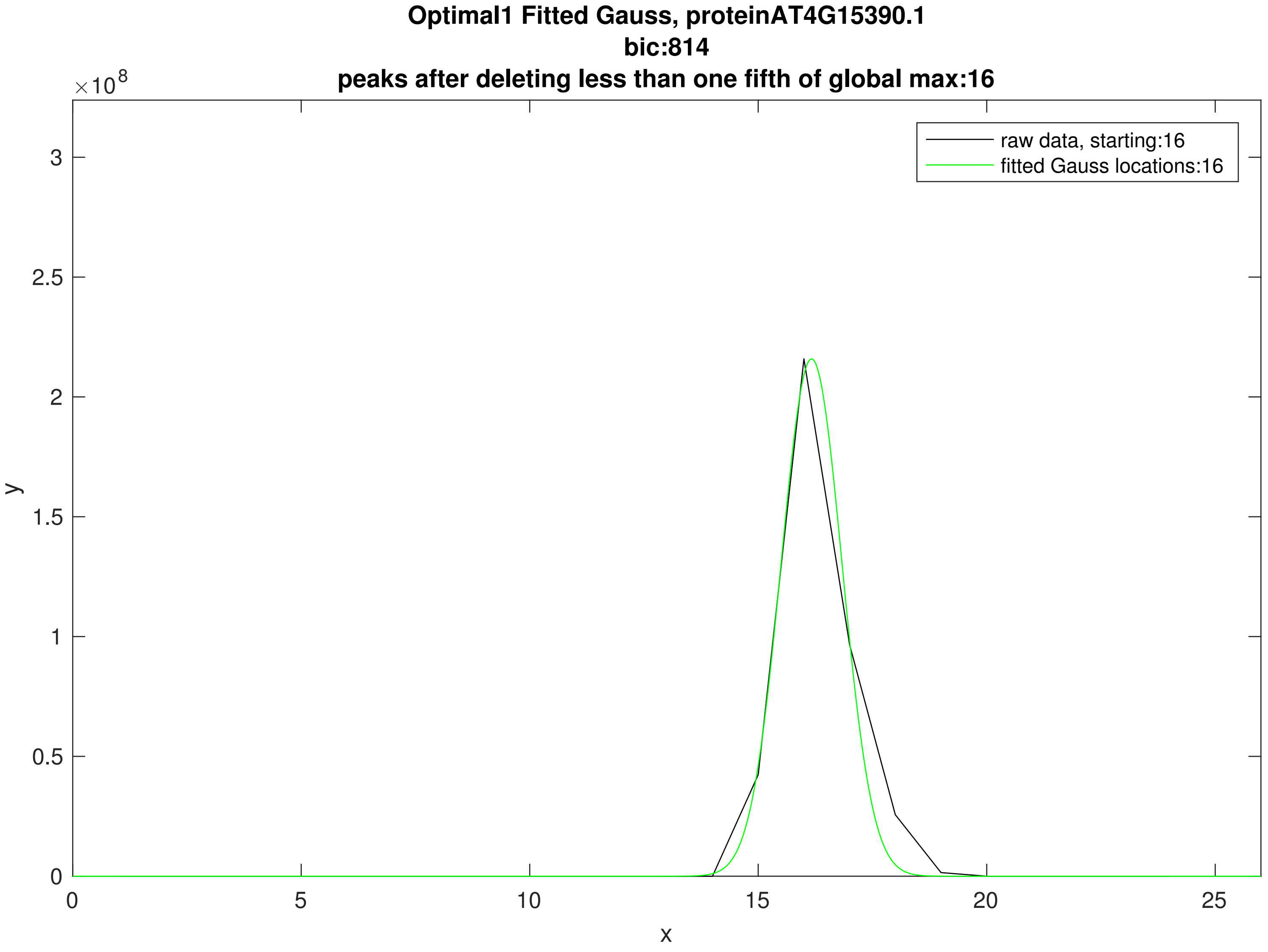
Optimal1 Fitted Gauss, proteinAT4G15390.1
bic:814
peaks after deleting less than one fifth of global max:16
×10^8
raw data, starting:16
fitted Gauss locations:16
y
x
0
0.5
1
1.5
2
2.5
3
0
5
10
15
20
25

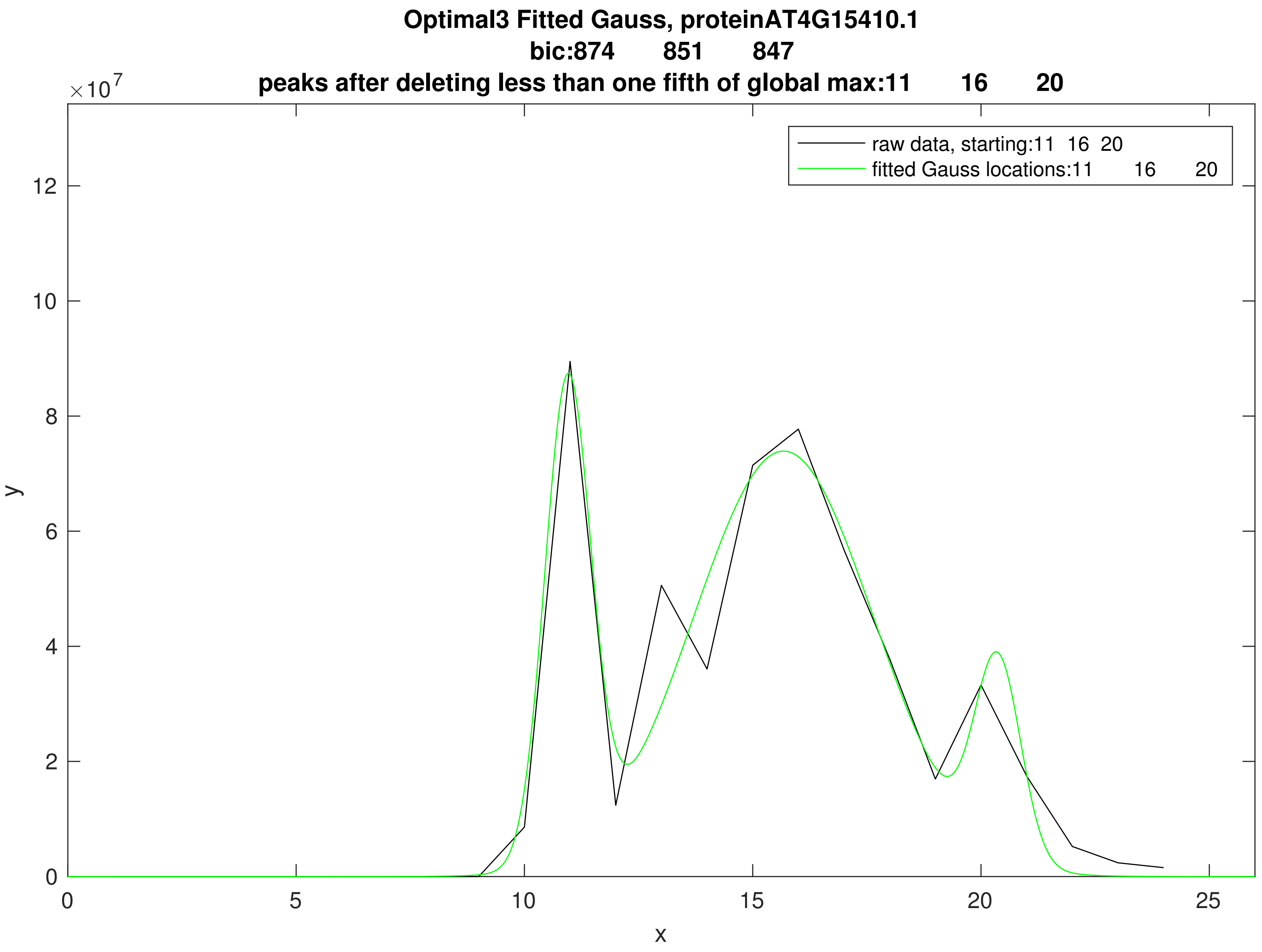
Optimal3 Fitted Gauss, proteinAT4G15410.1
bic:874 851 847
peaks after deleting less than one fifth of global max:11 16 20
×10^7
raw data, starting:11 16 20
fitted Gauss locations:11 16 20
y
x
0
2
4
6
8
10
12
5
15
20
25

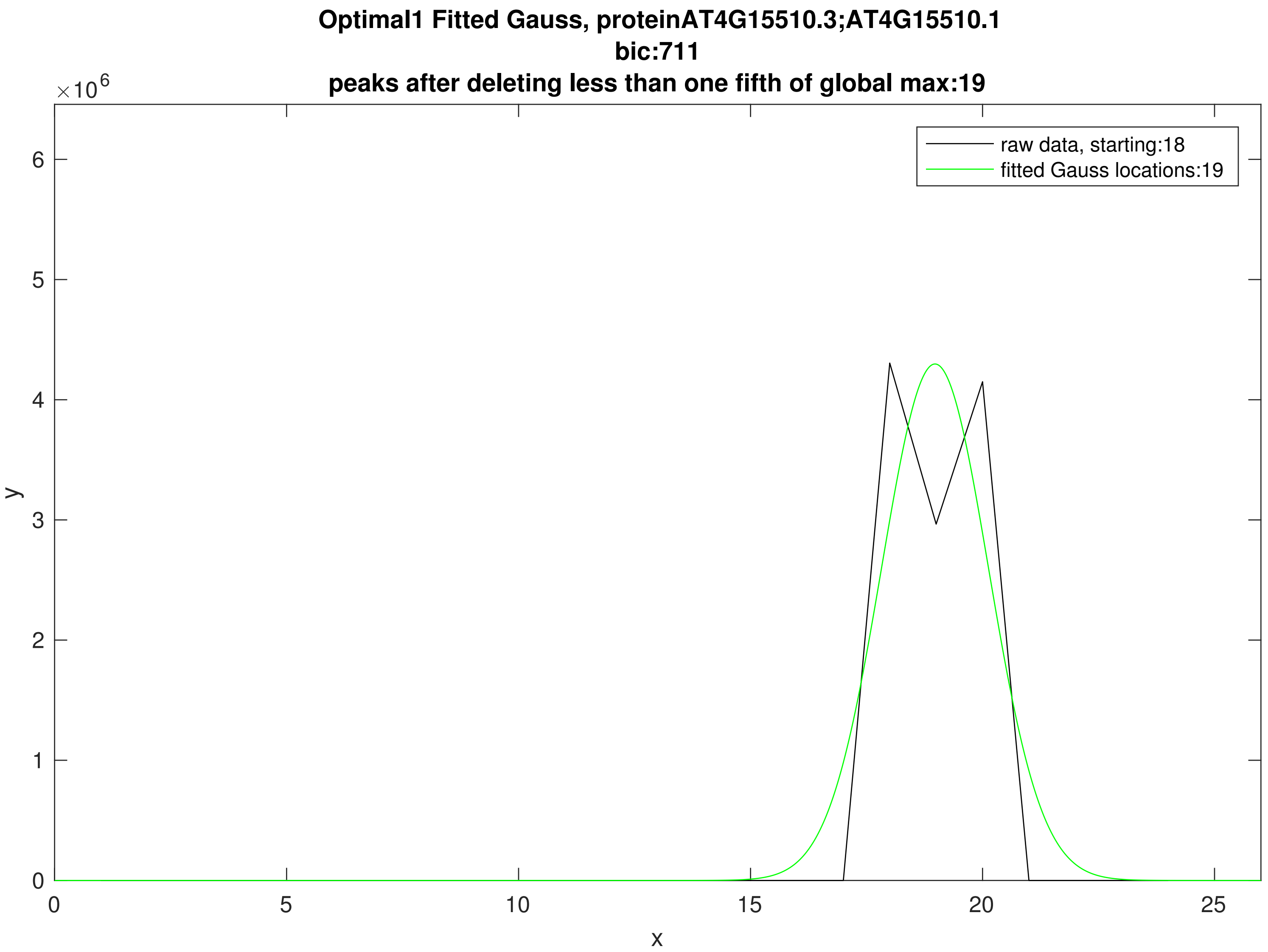
Optimal1 Fitted Gauss, proteinAT4G15510.3;AT4G15510.1
bic:711
peaks after deleting less than one fifth of global max:19
×10^6
raw data, starting:18
fitted Gauss locations:19
y
x
0
1
2
3
4
5
6
0
5
10
15
20
25

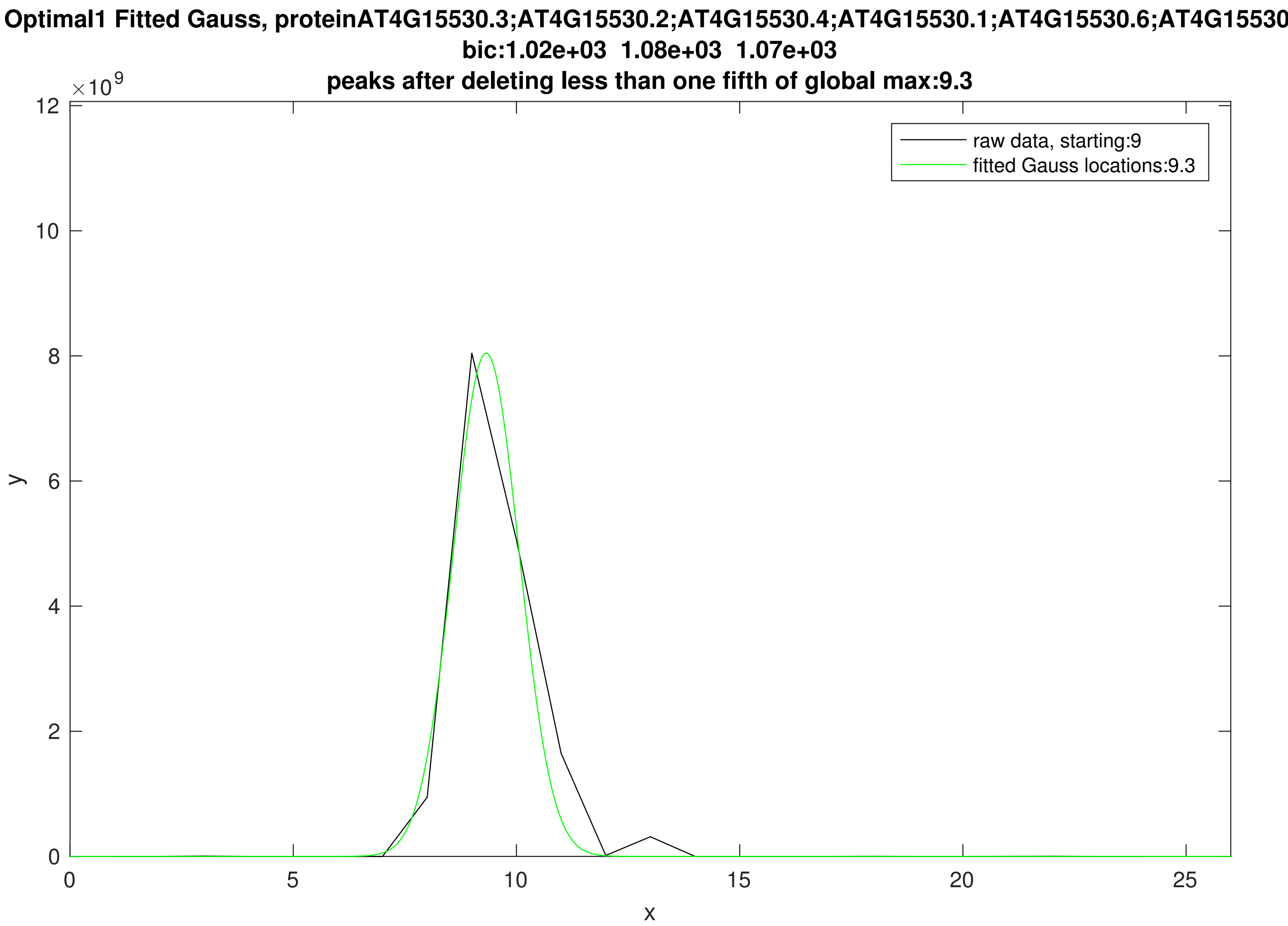

Optimal1 Fitted Gauss, proteinAT4G15530.3;AT4G15530.2;AT4G15530.4;AT4G15530.1;AT4G15530.6;AT4G15530
bic:1.02e+03 1.08e+03 1.07e+03
peaks after deleting less than one fifth of global max:9.3
raw data, starting:9
fitted Gauss locations:9.3
×10^9
12
10
8
6
4
2
0
y
0
5
10
15
20
25
x

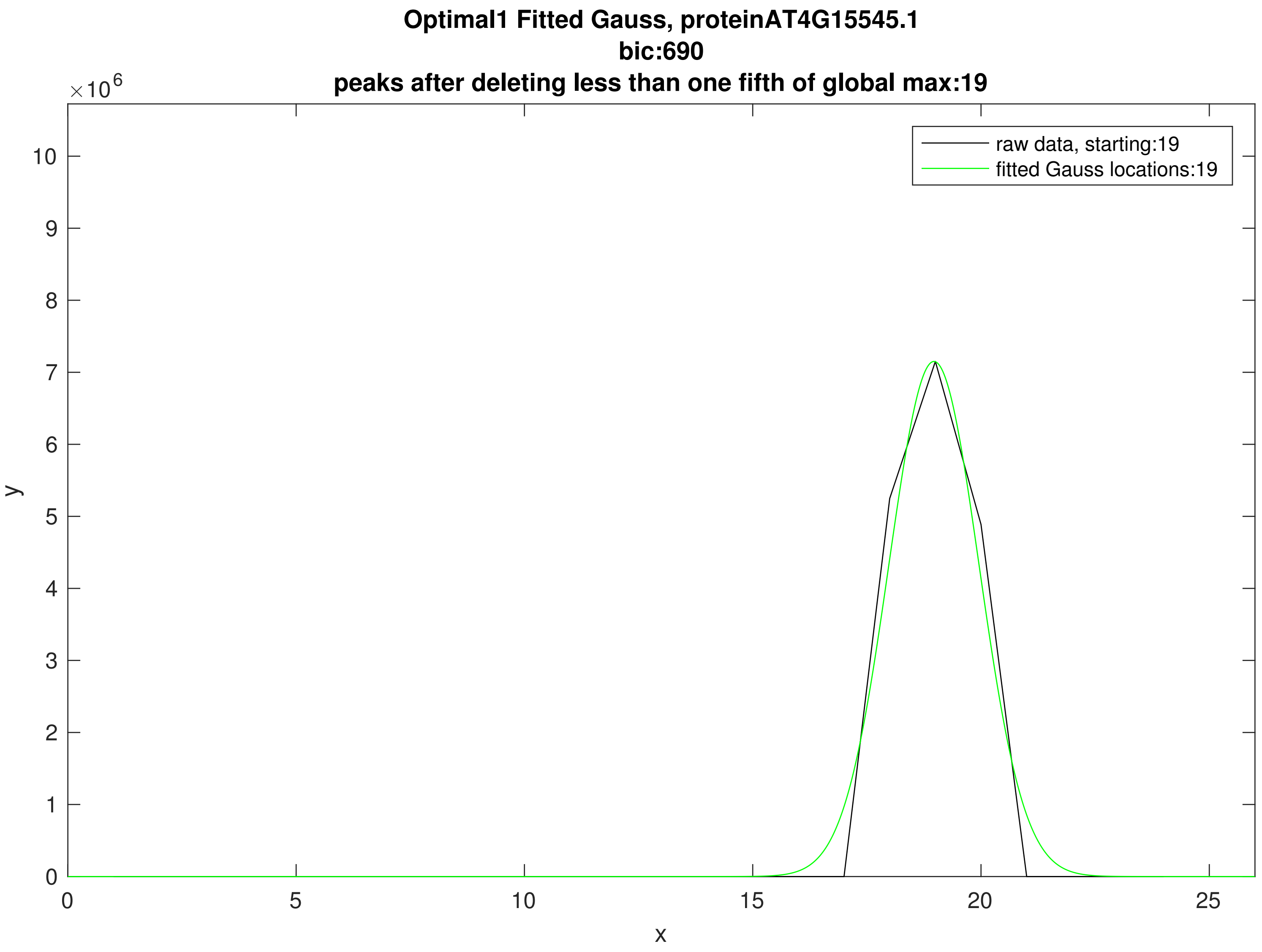
Optimal1 Fitted Gauss, proteinAT4G15545.1
bic:690
peaks after deleting less than one fifth of global max:19
$\times10^6$
raw data, starting:19
fitted Gauss locations:19
y
x
0
1
2
3
4
5
6
7
8
9
10
0
5
10
15
20
25

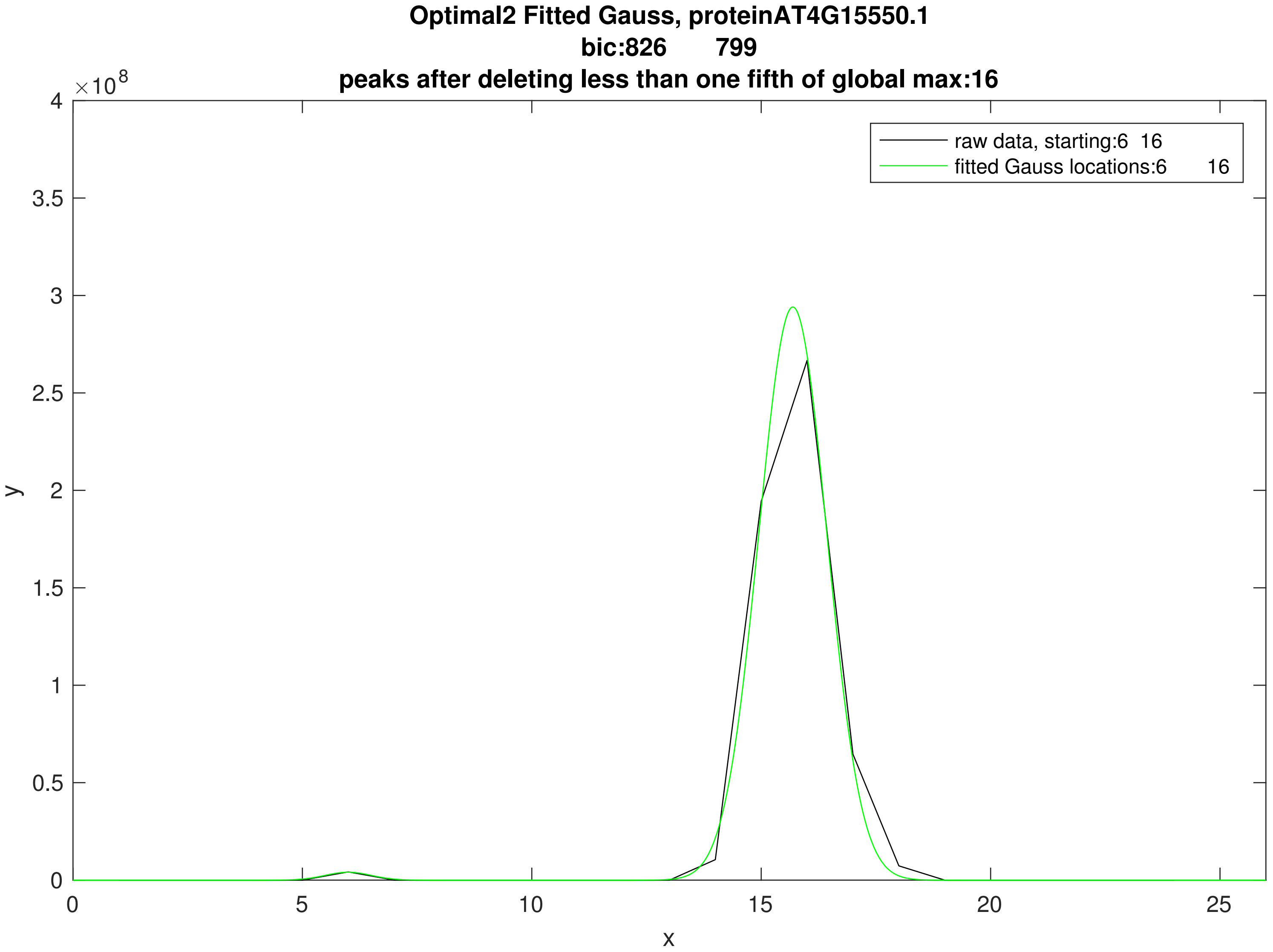

Optimal2 Fitted Gauss, proteinAT4G15550.1
bic:826 799
peaks after deleting less than one fifth of global max:16
×10^8
raw data, starting:6 16
fitted Gauss locations:6 16
y
x
0
0.5
1
1.5
2
2.5
3
3.5
4
5
10
15
20
25

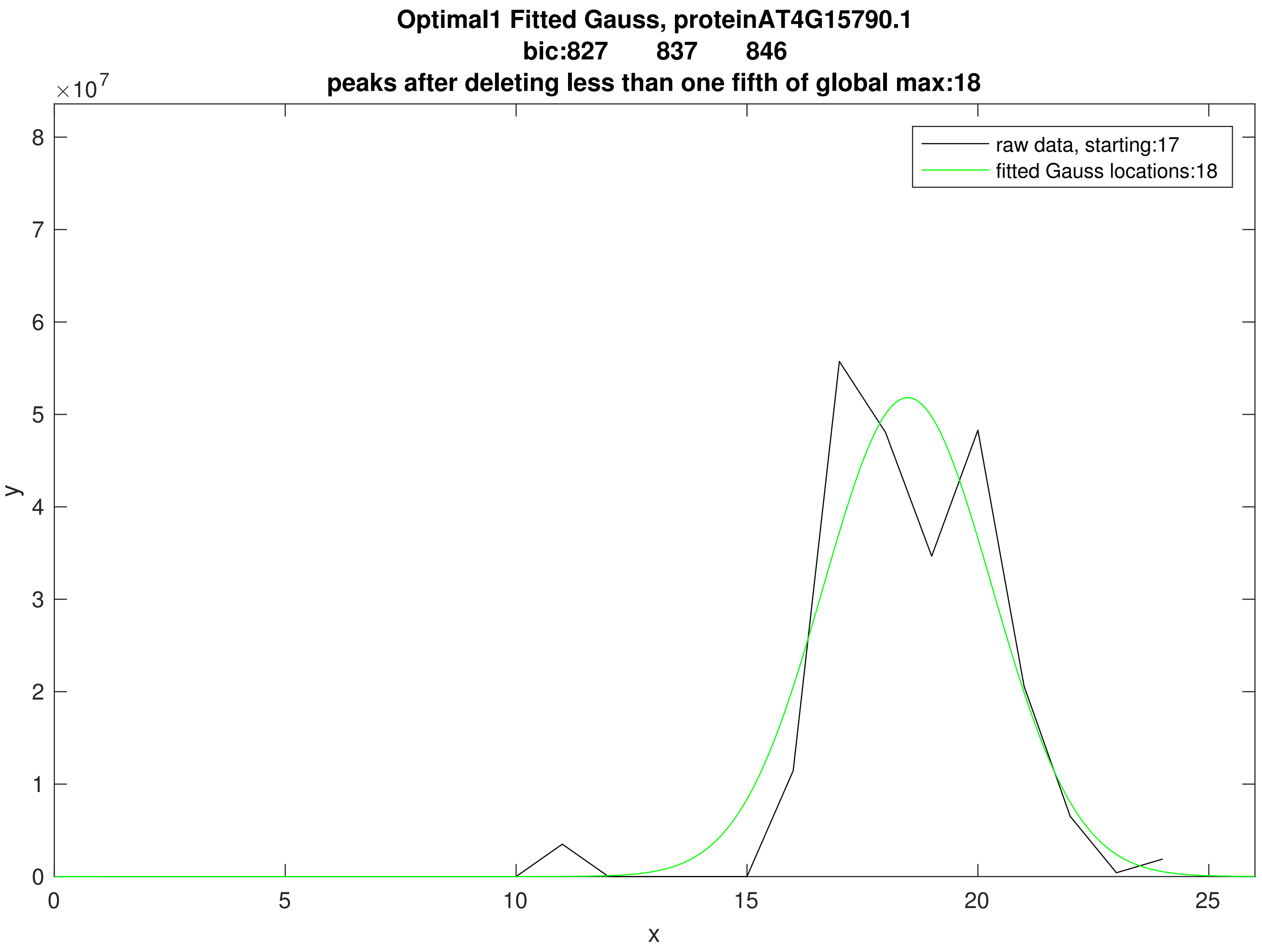
Optimal1 Fitted Gauss, proteinAT4G15790.1
bic:827 837 846
peaks after deleting less than one fifth of global max:18
×10^7
y
x
raw data, starting:17
fitted Gauss locations:18
0
1
2
3
4
5
6
7
8
0
5
10
15
20
25

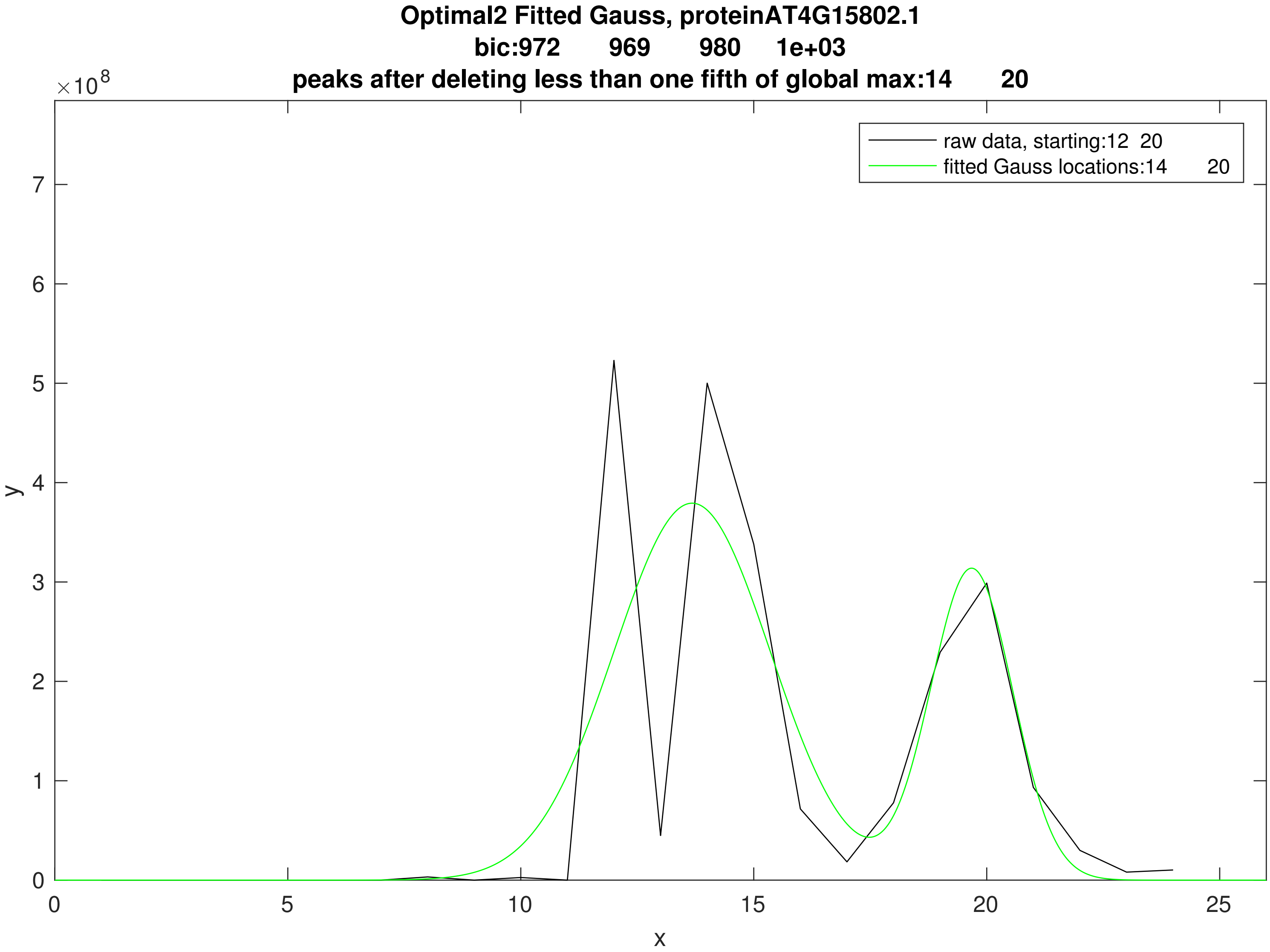

Optimal2 Fitted Gauss, proteinAT4G15802.1
bic:972 969 980 1e+03
peaks after deleting less than one fifth of global max:14 20
×10^8
raw data, starting:12 20
fitted Gauss locations:14 20
y
x

Optimal1 Fitted Gauss, proteinAT4G15930.1
bic:681
peaks after deleting less than one fifth of global max:20

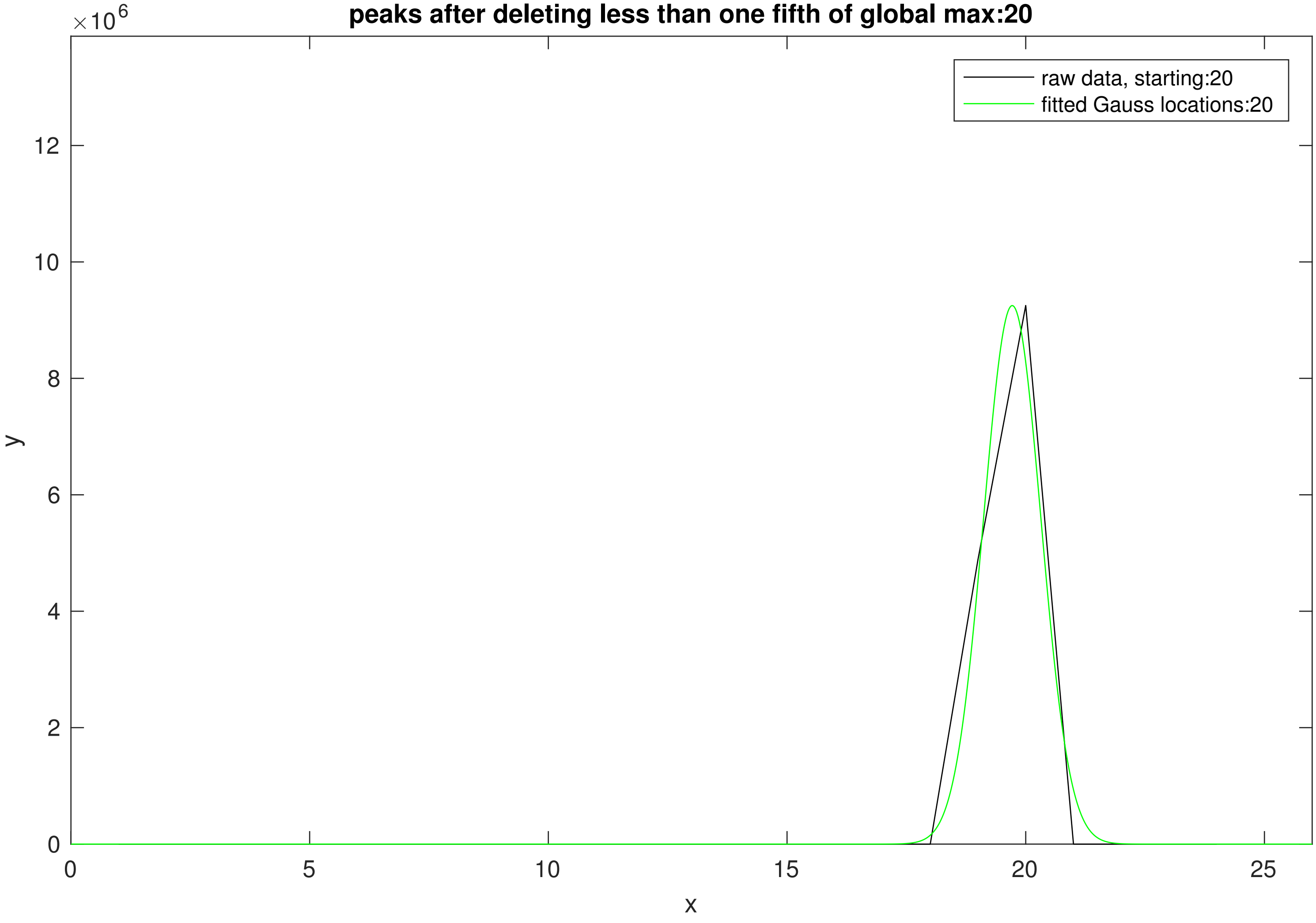

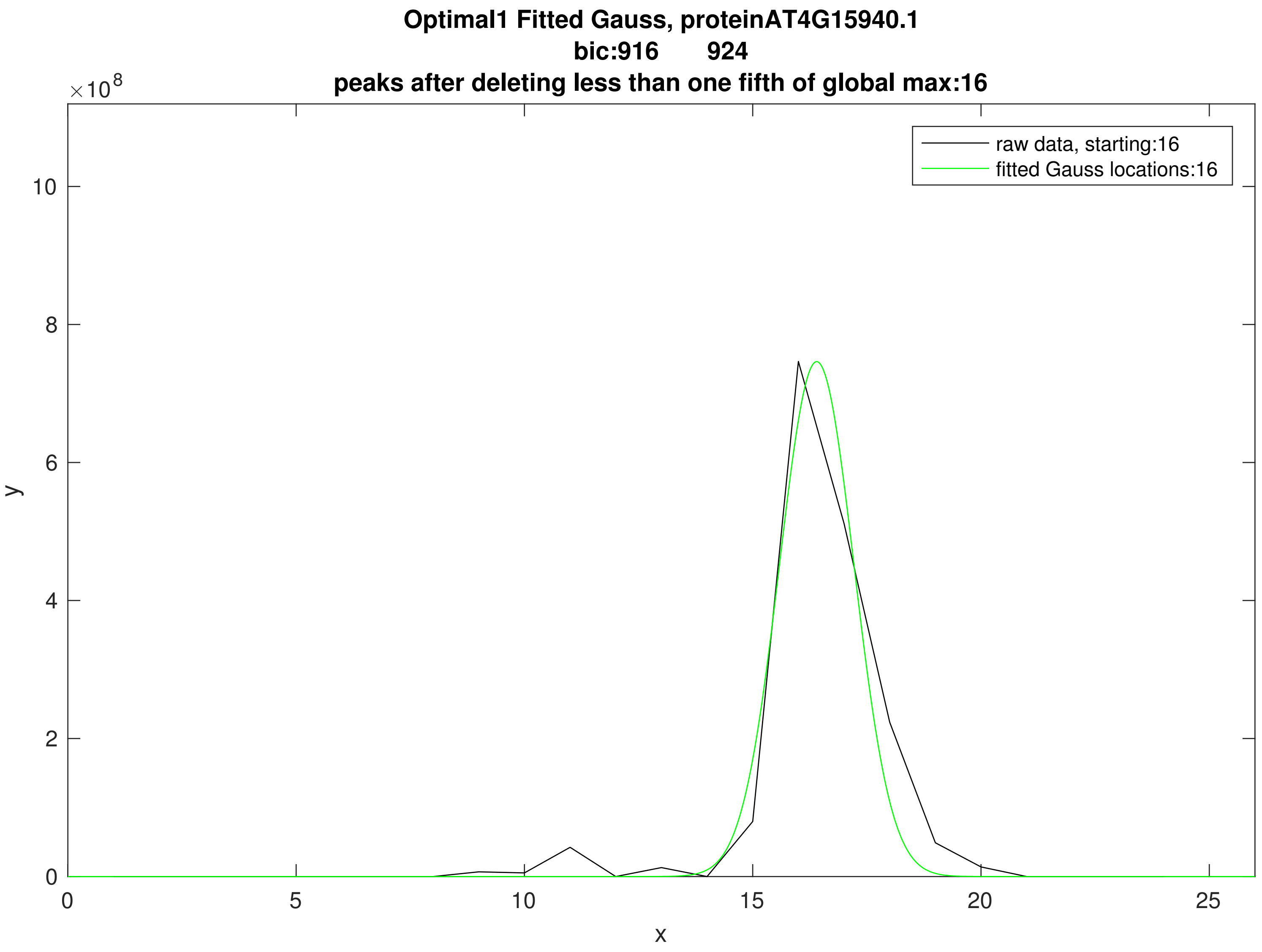
Optimal1 Fitted Gauss, proteinAT4G15940.1
bic:916 924
peaks after deleting less than one fifth of global max:16
×10^8
raw data, starting:16
fitted Gauss locations:16
y
x
0
2
4
6
8
10
0
5
10
15
20
25

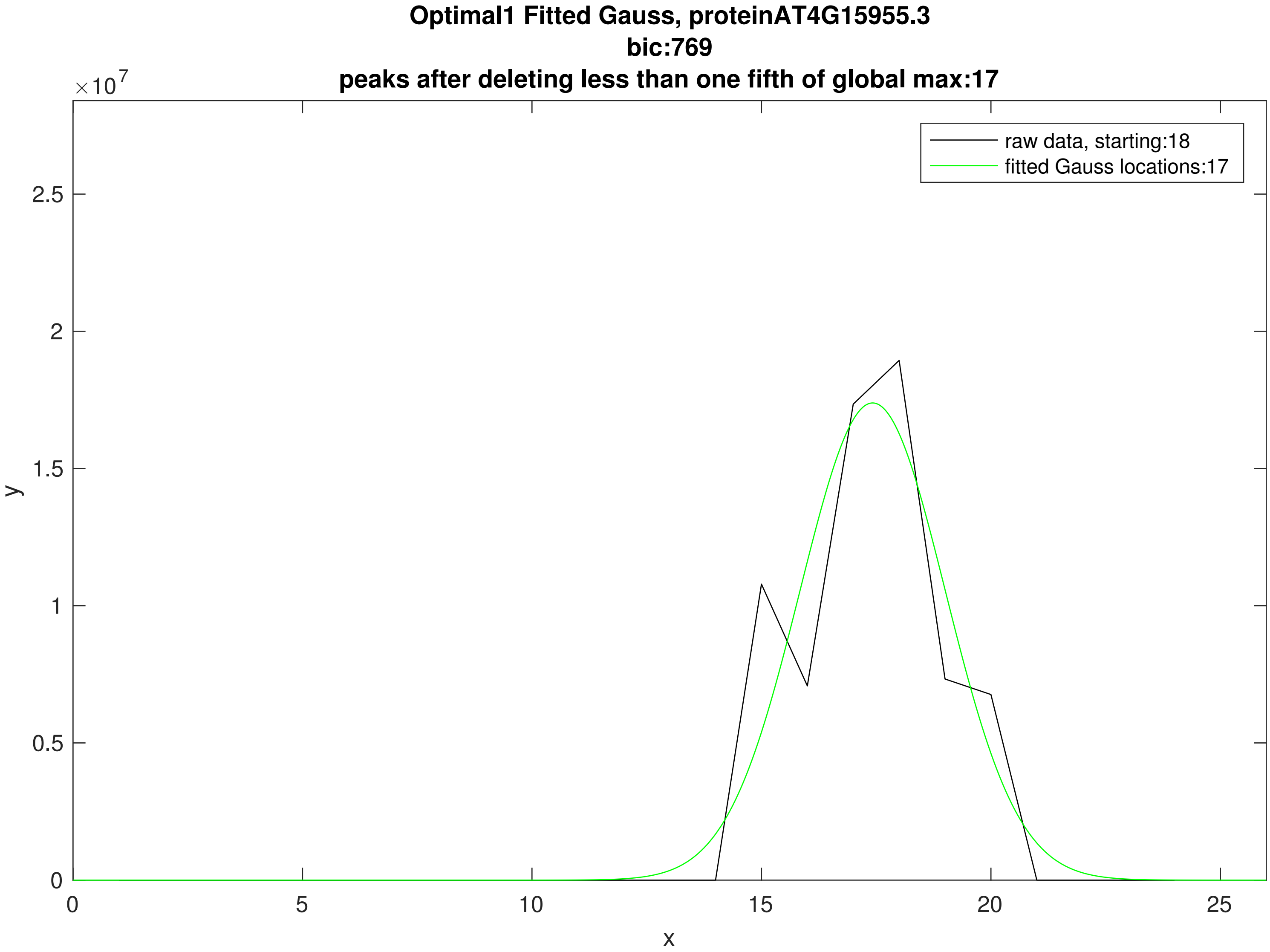

Optimal1 Fitted Gauss, proteinAT4G15955.3
bic:769
peaks after deleting less than one fifth of global max:17
×10^7
raw data, starting:18
fitted Gauss locations:17
2.5
2
1.5
1
0.5
0
y
0
5
10
15
20
25
x

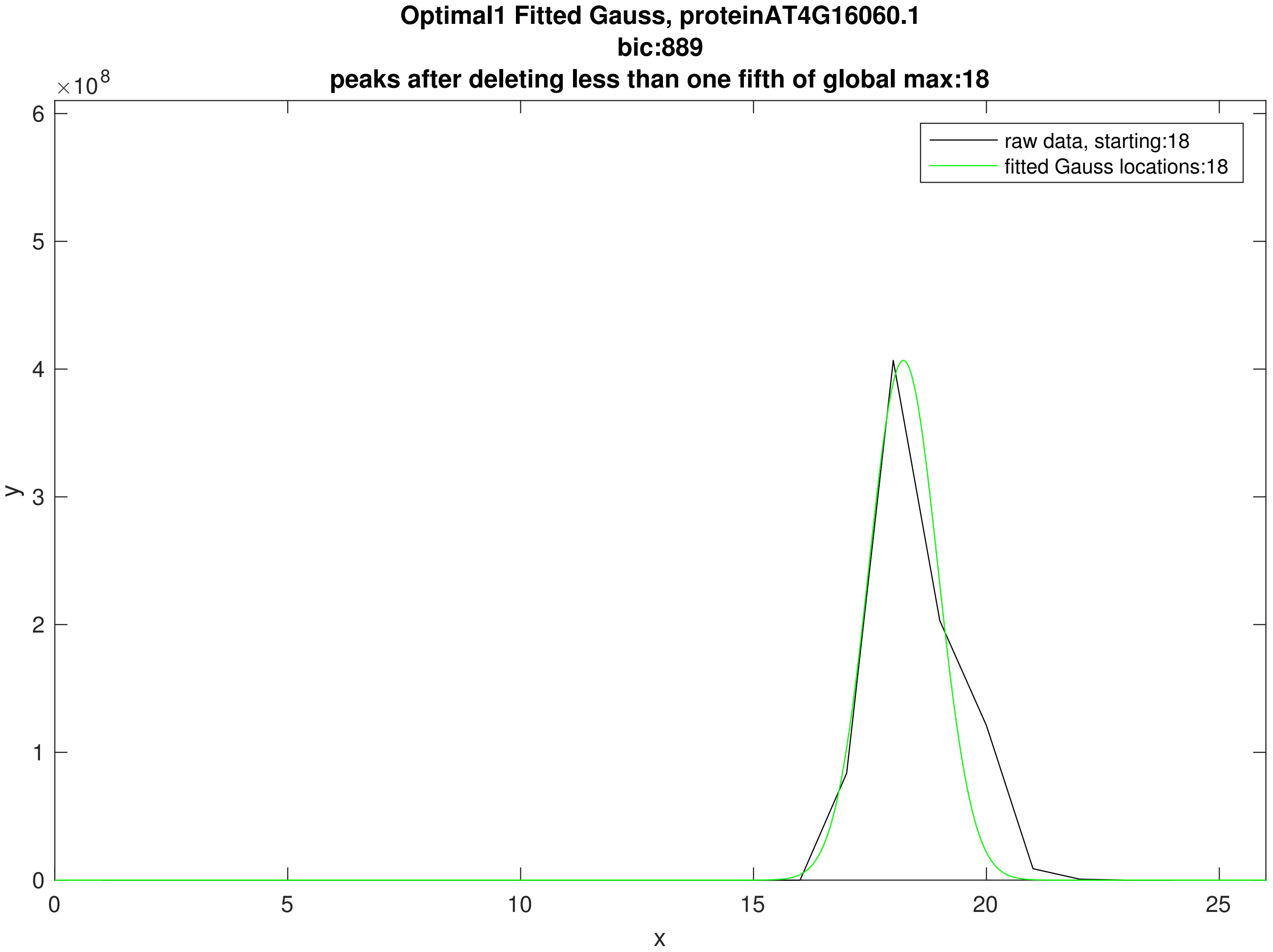

Optimal1 Fitted Gauss, proteinAT4G16060.1
bic:889
peaks after deleting less than one fifth of global max:18
×10^8
raw data, starting:18
fitted Gauss locations:18
y
x
0
1
2
3
4
5
6
0
5
10
15
20
25

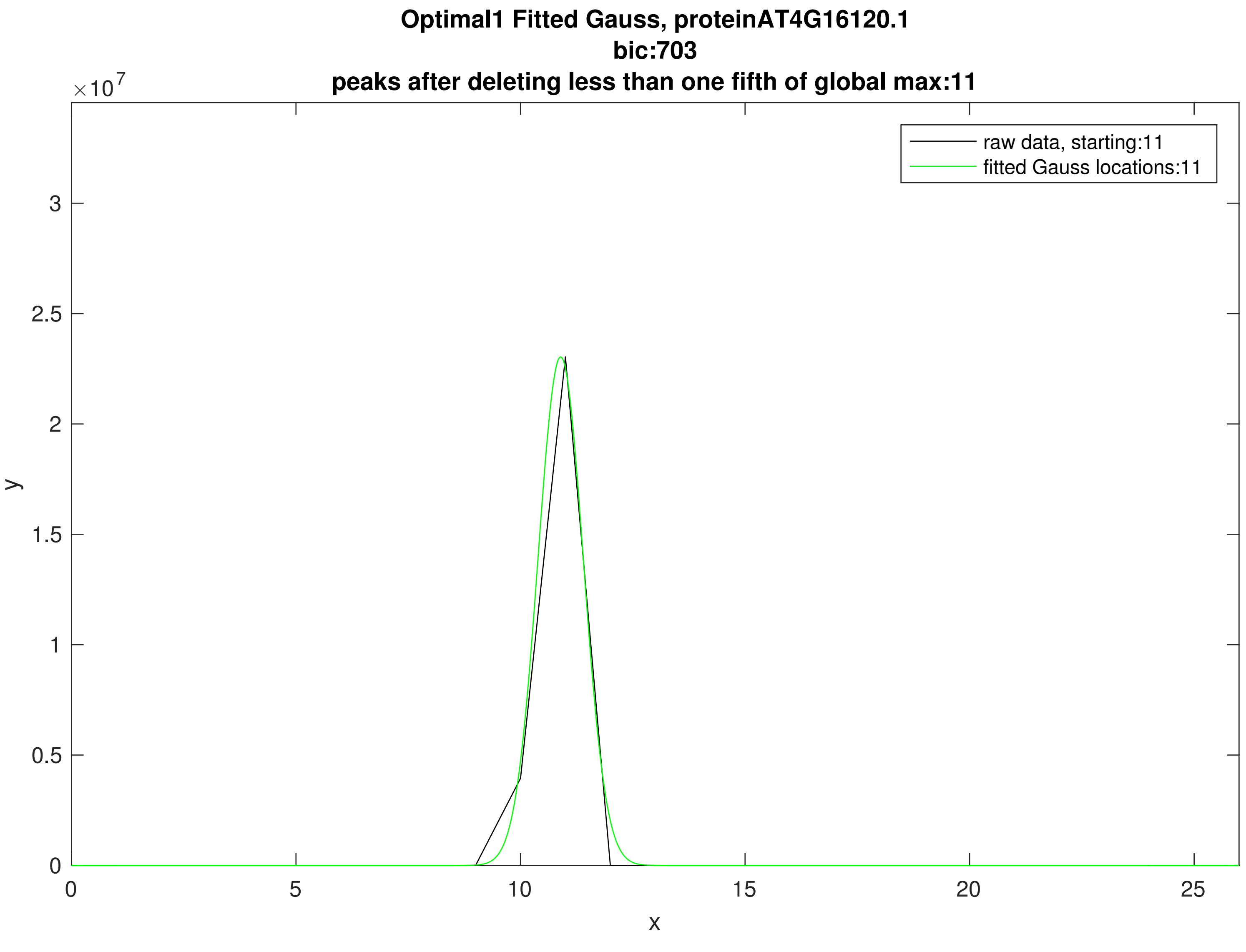
Optimal1 Fitted Gauss, proteinAT4G16120.1
bic:703
peaks after deleting less than one fifth of global max:11
×10^7
raw data, starting:11
fitted Gauss locations:11
y
x
0
0.5
1
1.5
2
2.5
3
0
5
10
15
20
25

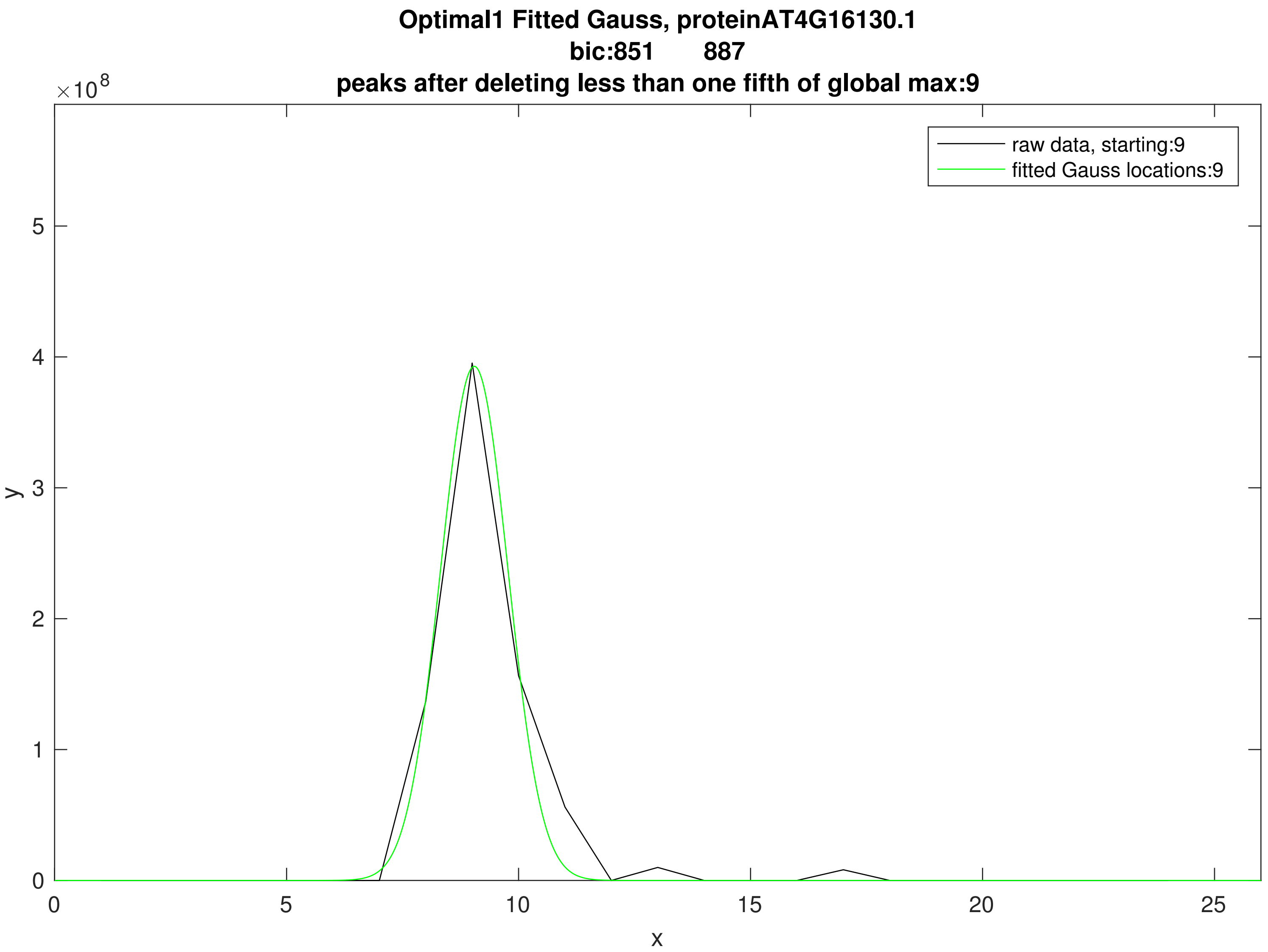
Optimal1 Fitted Gauss, proteinAT4G16130.1
bic:851 887
peaks after deleting less than one fifth of global max:9
×10^8
raw data, starting:9
fitted Gauss locations:9
y
x
0
1
2
3
4
5
0
5
10
15
20
25

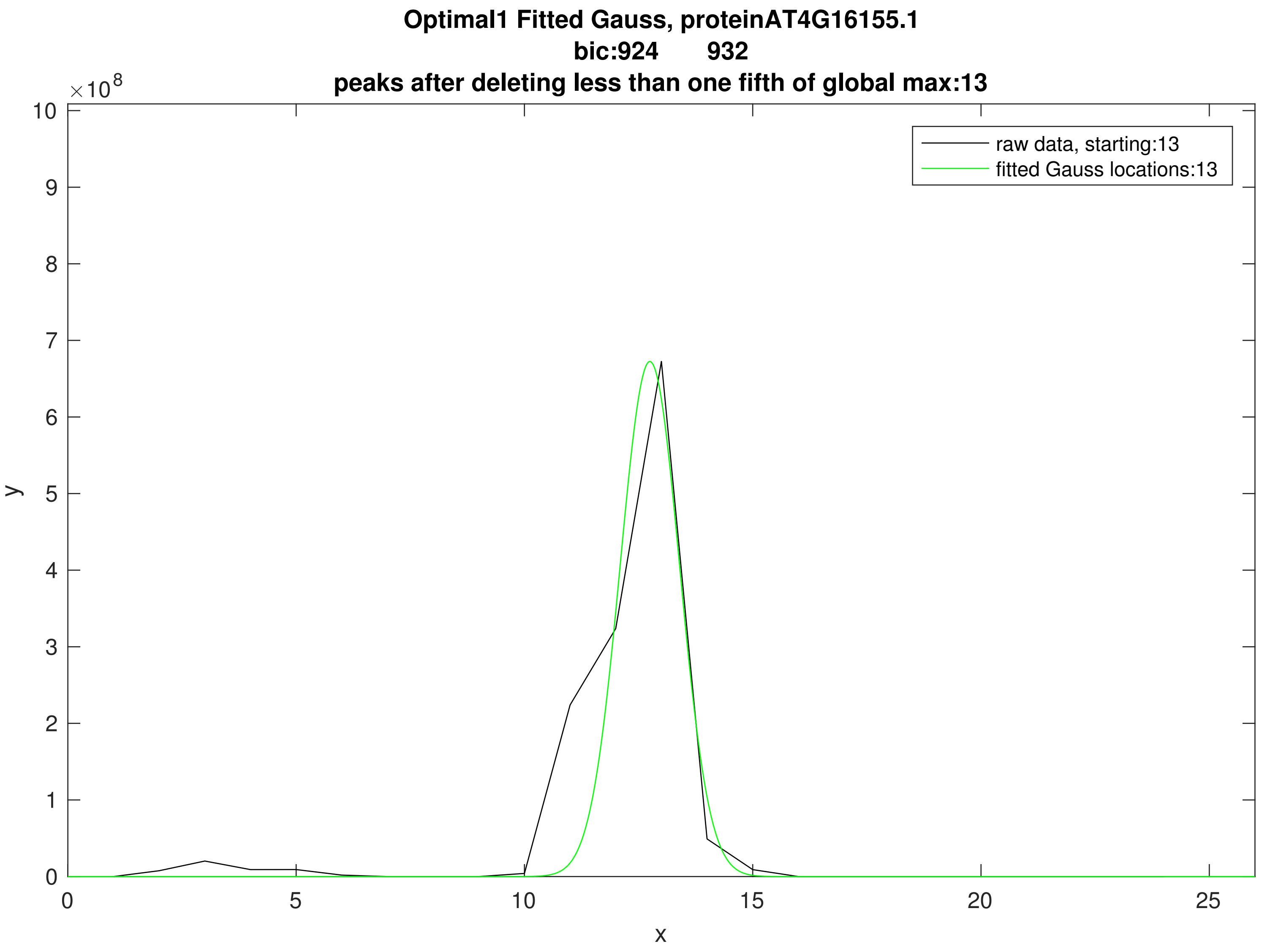
Optimal1 Fitted Gauss, proteinAT4G16155.1
bic:924 932
peaks after deleting less than one fifth of global max:13
×10^8
10
9
8
7
6
5
4
3
2
1
0
y
0
5
10
15
20
25
x
raw data, starting:13
fitted Gauss locations:13

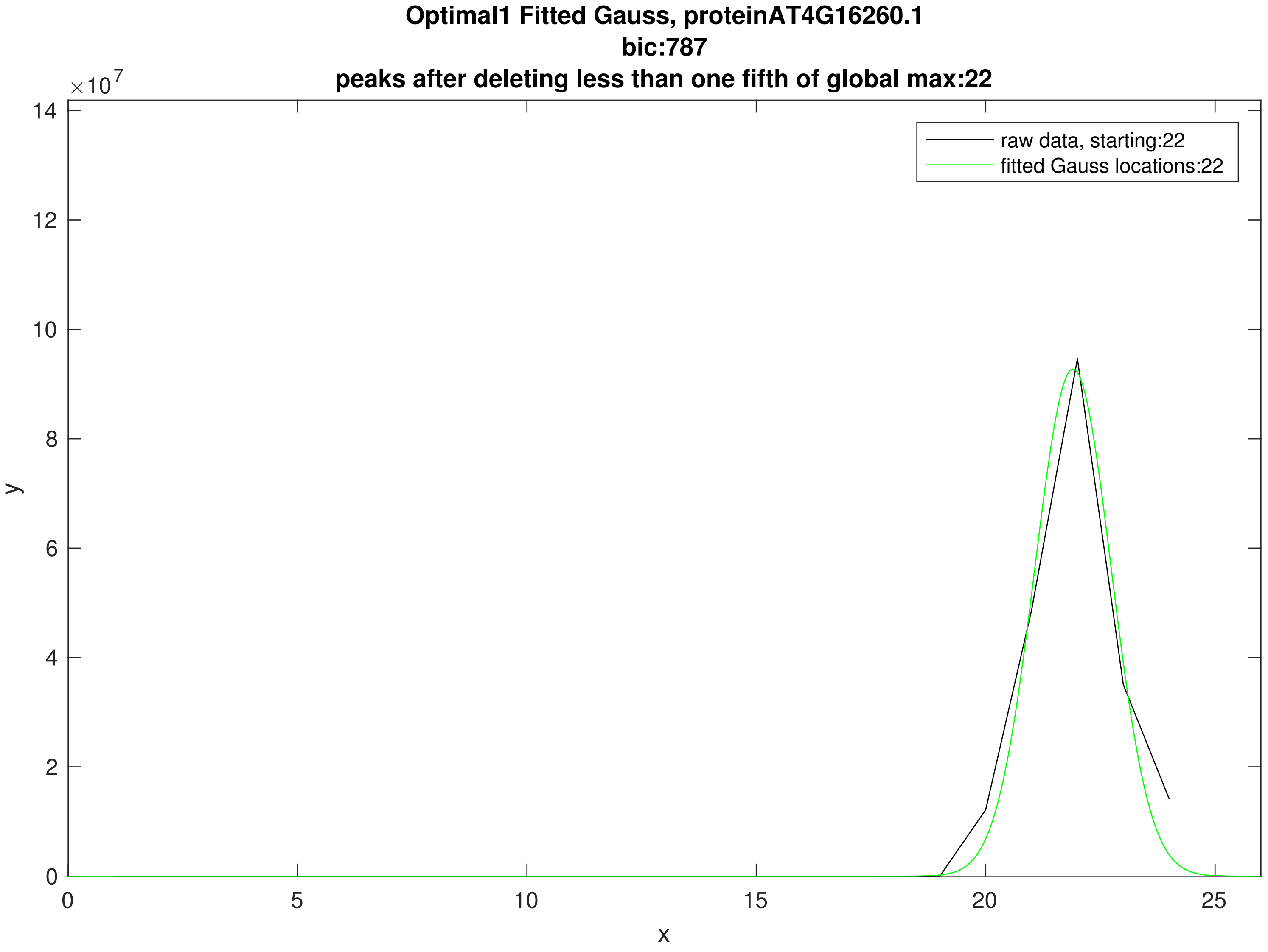
Optimal1 Fitted Gauss, proteinAT4G16260.1
bic:787
peaks after deleting less than one fifth of global max:22
×10^7
raw data, starting:22
fitted Gauss locations:22
y
x
0
2
4
6
8
10
12
14
0
5
10
15
20
25

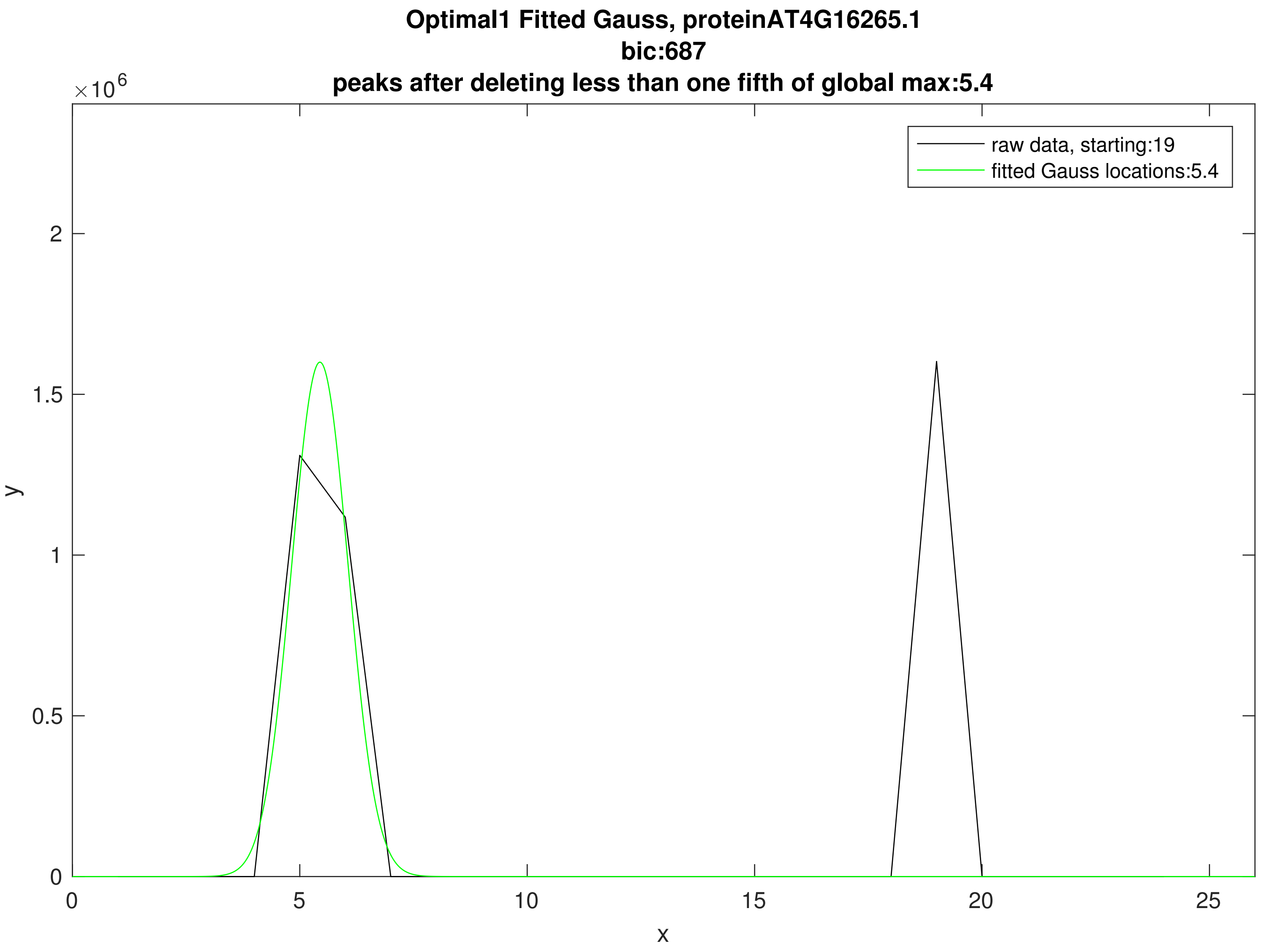
Optimal1 Fitted Gauss, proteinAT4G16265.1
bic:687
peaks after deleting less than one fifth of global max:5.4
×10^6
raw data, starting:19
fitted Gauss locations:5.4
y
x
0
0.5
1
1.5
2
0
5
10
15
20
25

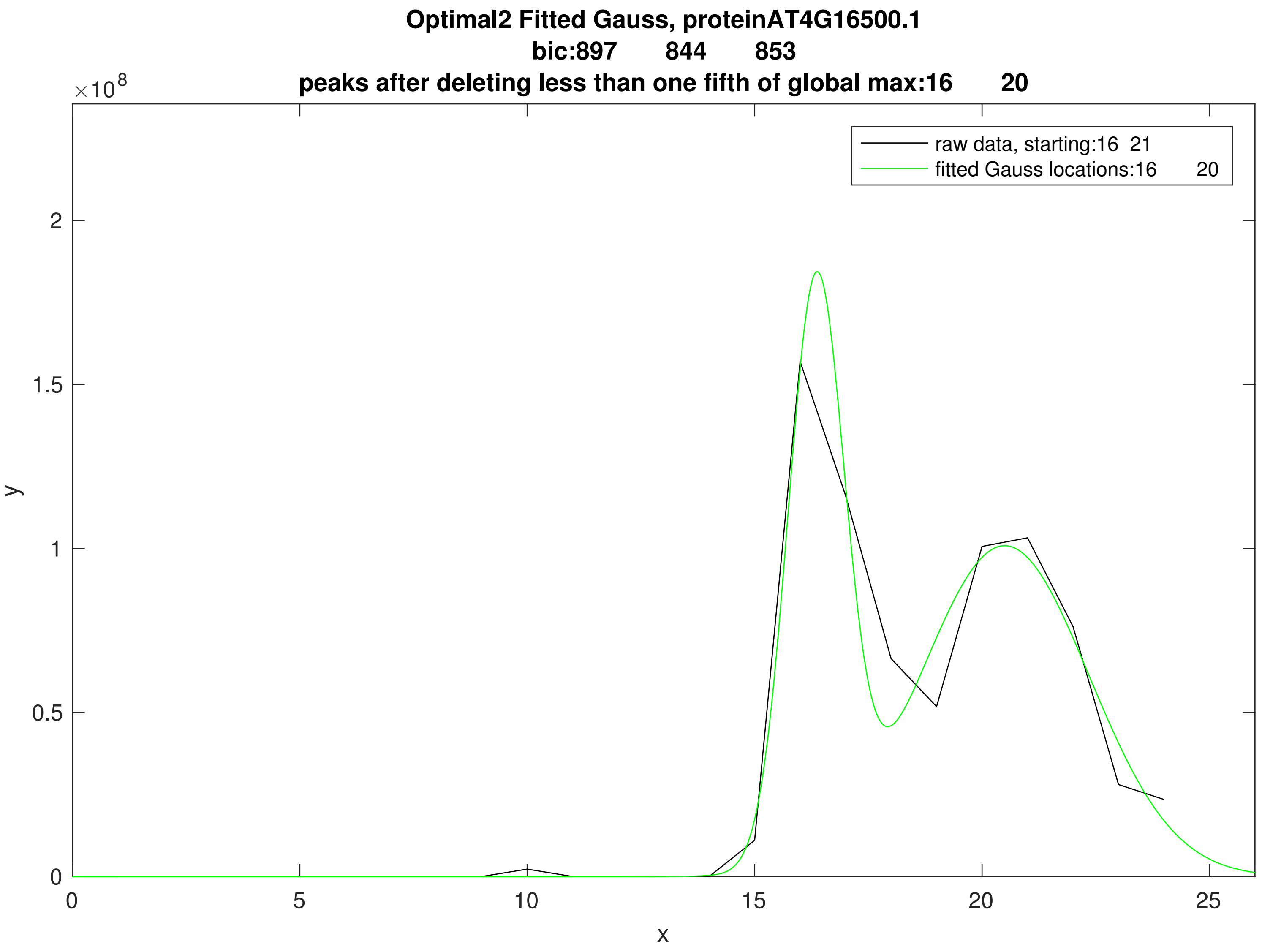
Optimal2 Fitted Gauss, proteinAT4G16500.1
bic:897 844 853
peaks after deleting less than one fifth of global max:16 20
×10^8
raw data, starting:16 21
fitted Gauss locations:16 20
2
1.5
1
0.5
0
y
0
5
10
15
20
25
x

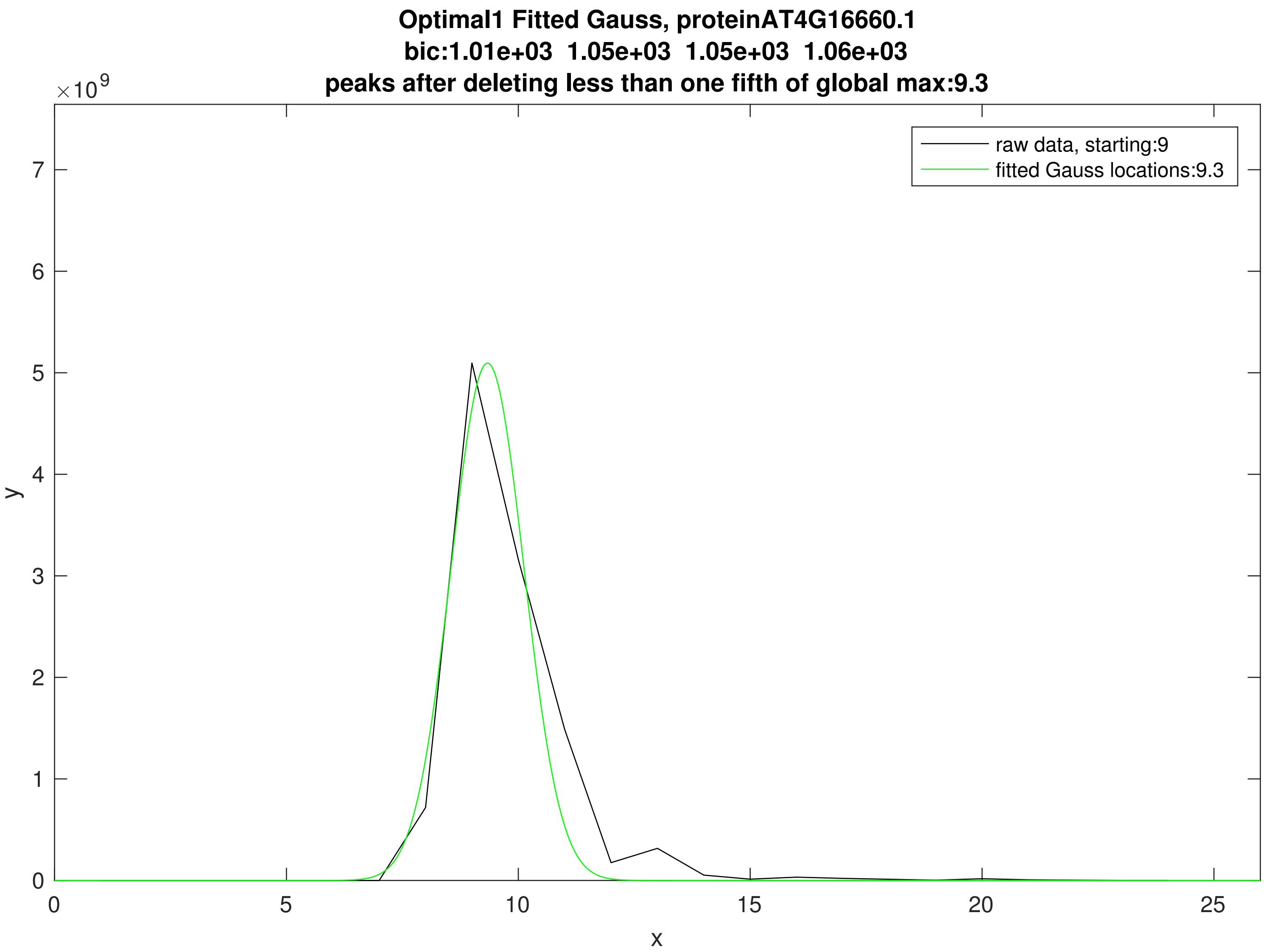

Optimal1 Fitted Gauss, proteinAT4G16660.1
bic:1.01e+03 1.05e+03 1.05e+03 1.06e+03
peaks after deleting less than one fifth of global max:9.3
×10^9
raw data, starting:9
fitted Gauss locations:9.3
y
x
0
1
2
3
4
5
6
7
0
5
10
15
20
25

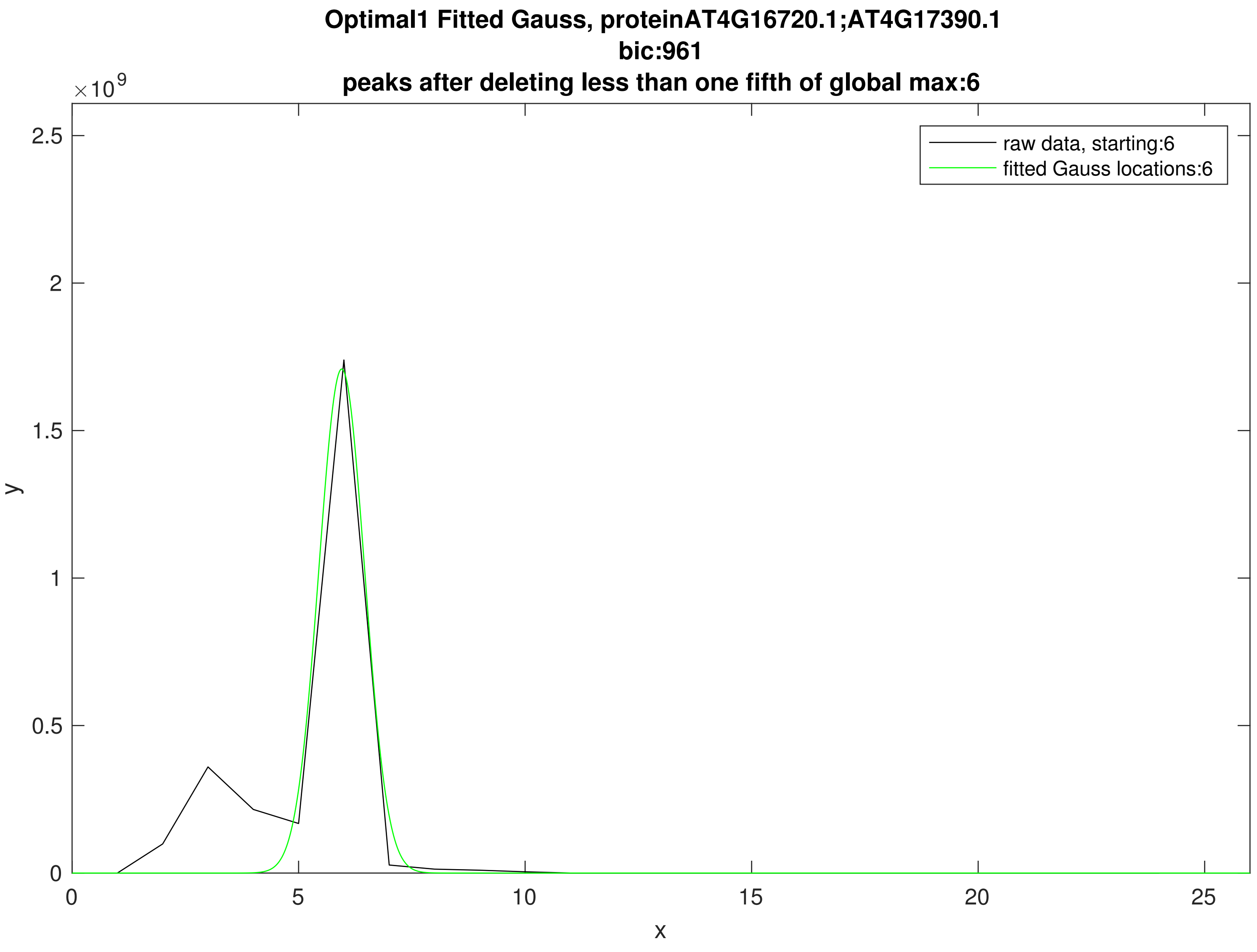
Optimal1 Fitted Gauss, proteinAT4G16720.1;AT4G17390.1
bic:961
peaks after deleting less than one fifth of global max:6
×10^9
raw data, starting:6
fitted Gauss locations:6
y
x
0
0.5
1
1.5
2
2.5
5
10
15
20
25

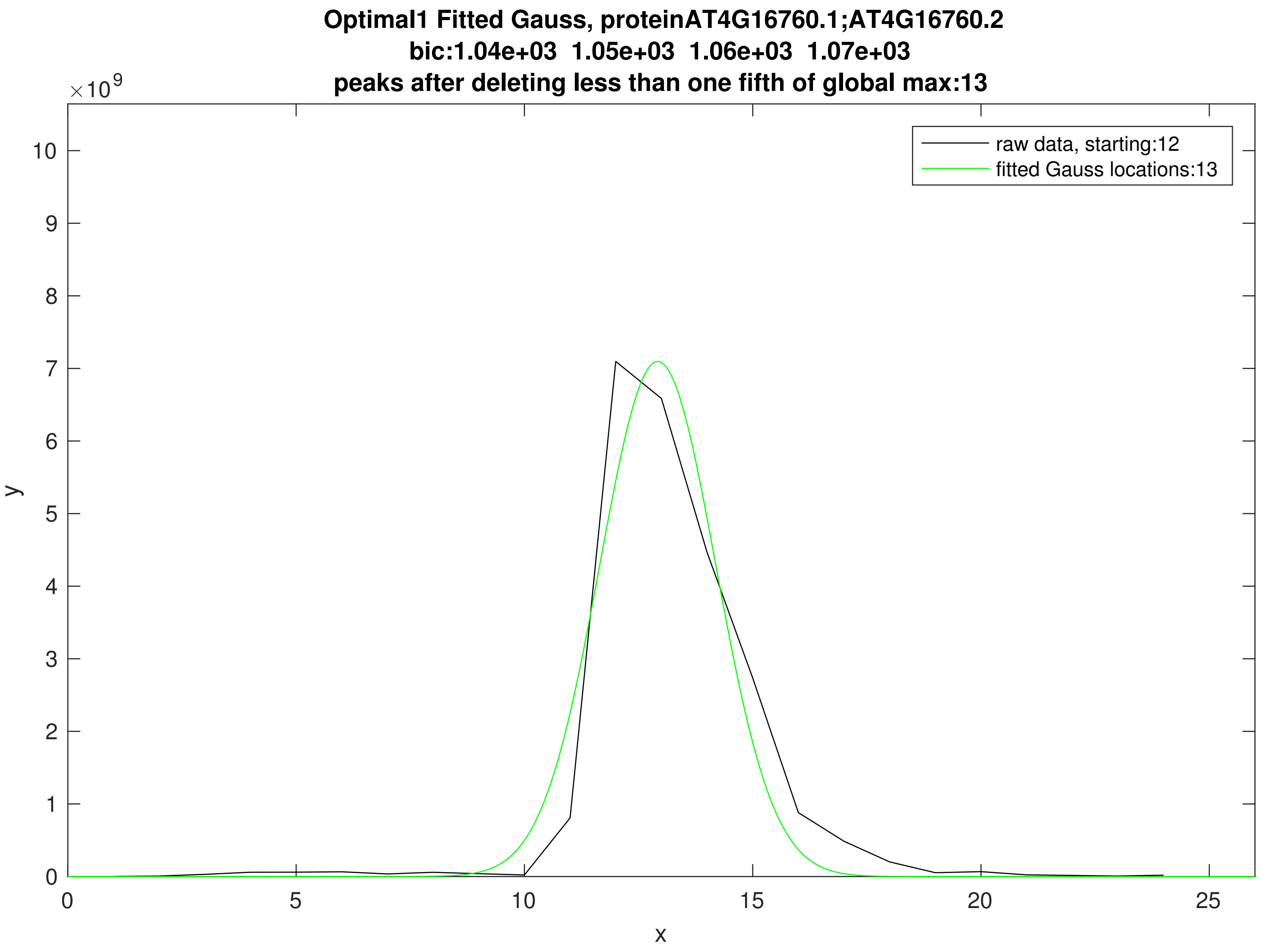

Optimal1 Fitted Gauss, proteinAT4G16760.1;AT4G16760.2
bic:1.04e+03 1.05e+03 1.06e+03 1.07e+03
peaks after deleting less than one fifth of global max:13
×10^9
raw data, starting:12
fitted Gauss locations:13
y
x
0
1
2
3
4
5
6
7
8
9
10
0
5
10
15
20
25

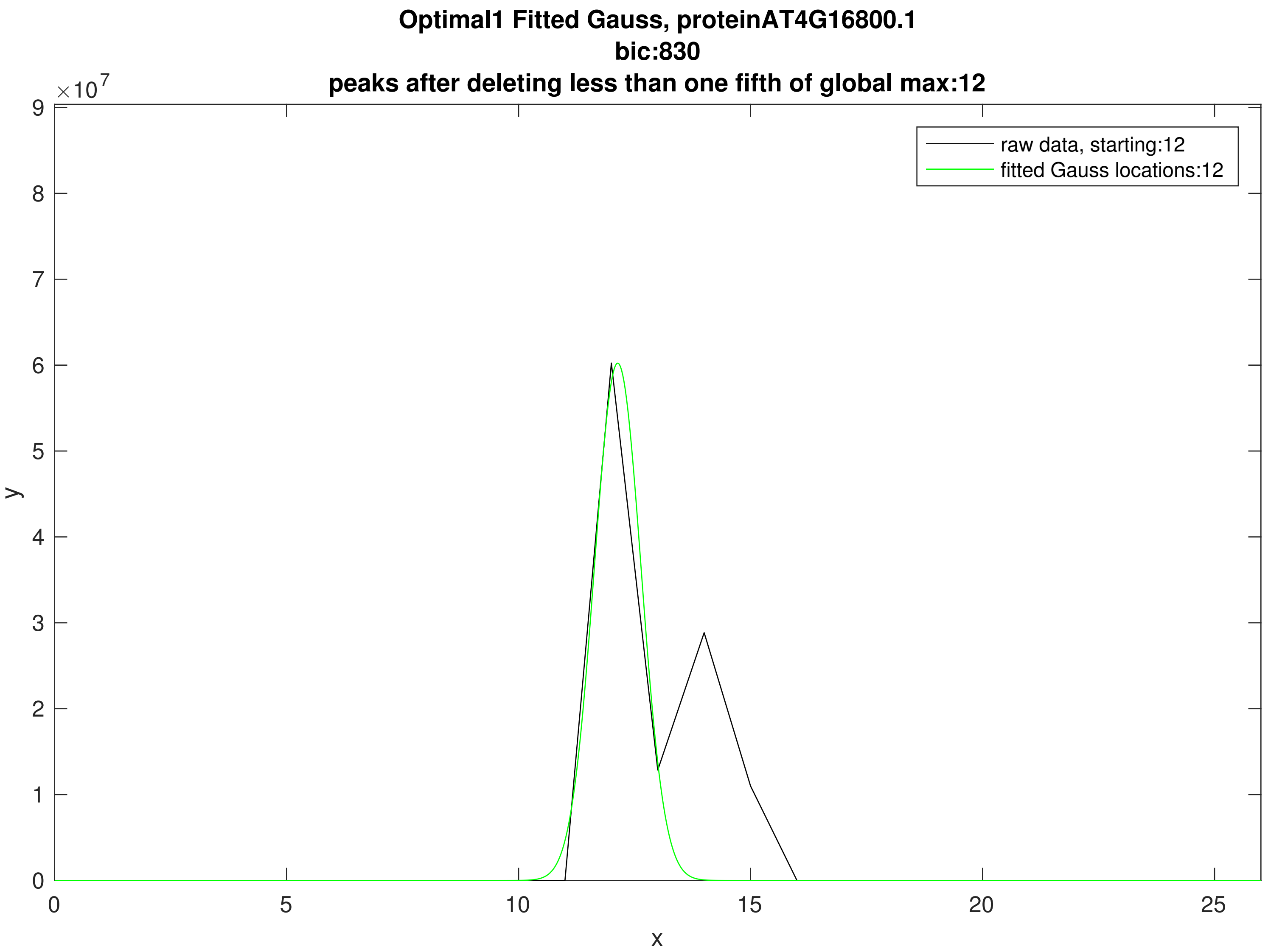

Optimal1 Fitted Gauss, proteinAT4G16800.1
bic:830
peaks after deleting less than one fifth of global max:12
×10^7
raw data, starting:12
fitted Gauss locations:12
y
x

**Optimal1 Fitted Gauss, proteinAT4G17030.1**
**bic:714**
**peaks after deleting less than one fifth of global max:20**

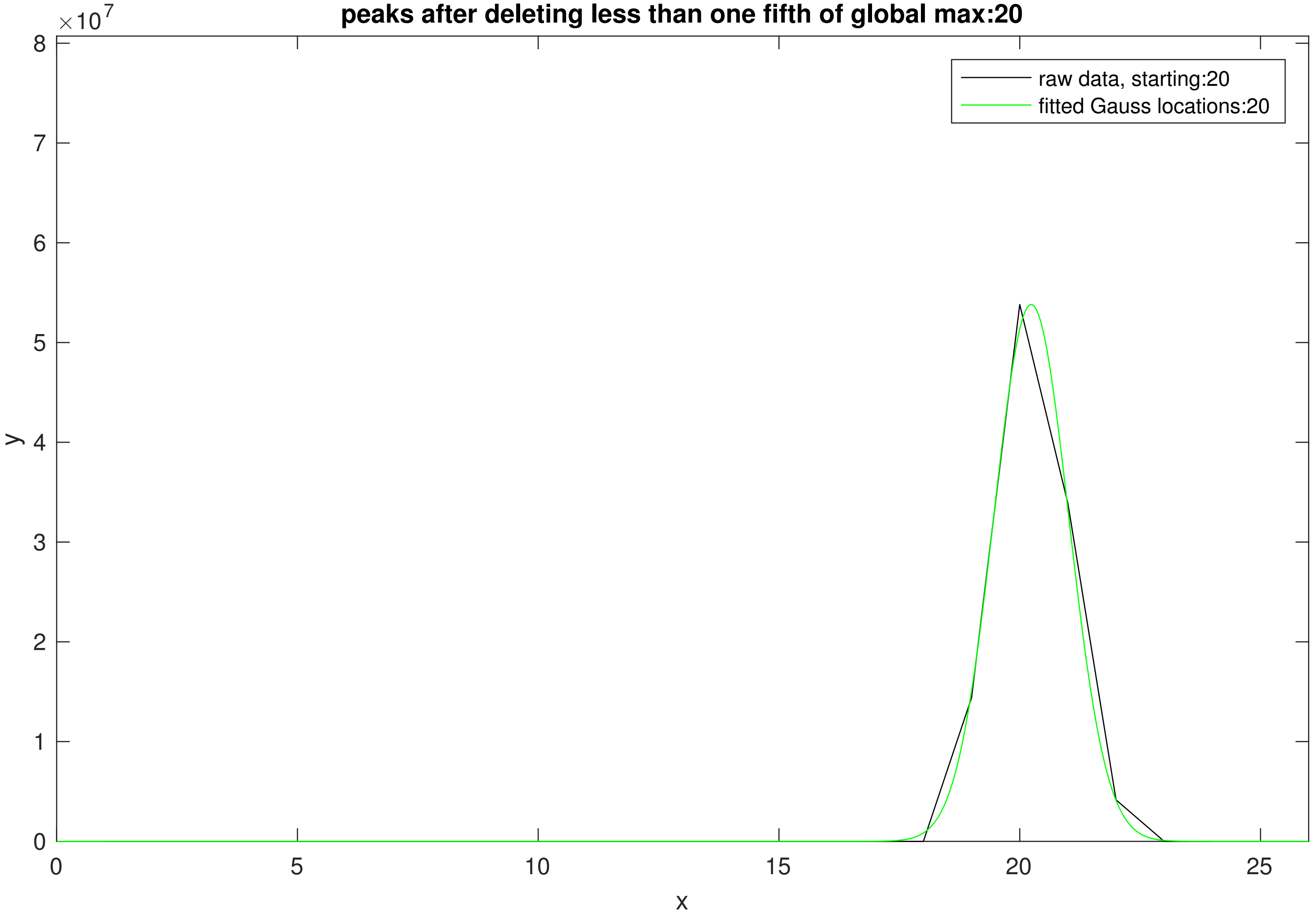

Optimal1 Fitted Gauss, proteinAT4G17040.1
bic:893 918 934
peaks after deleting less than one fifth of global max:8.6

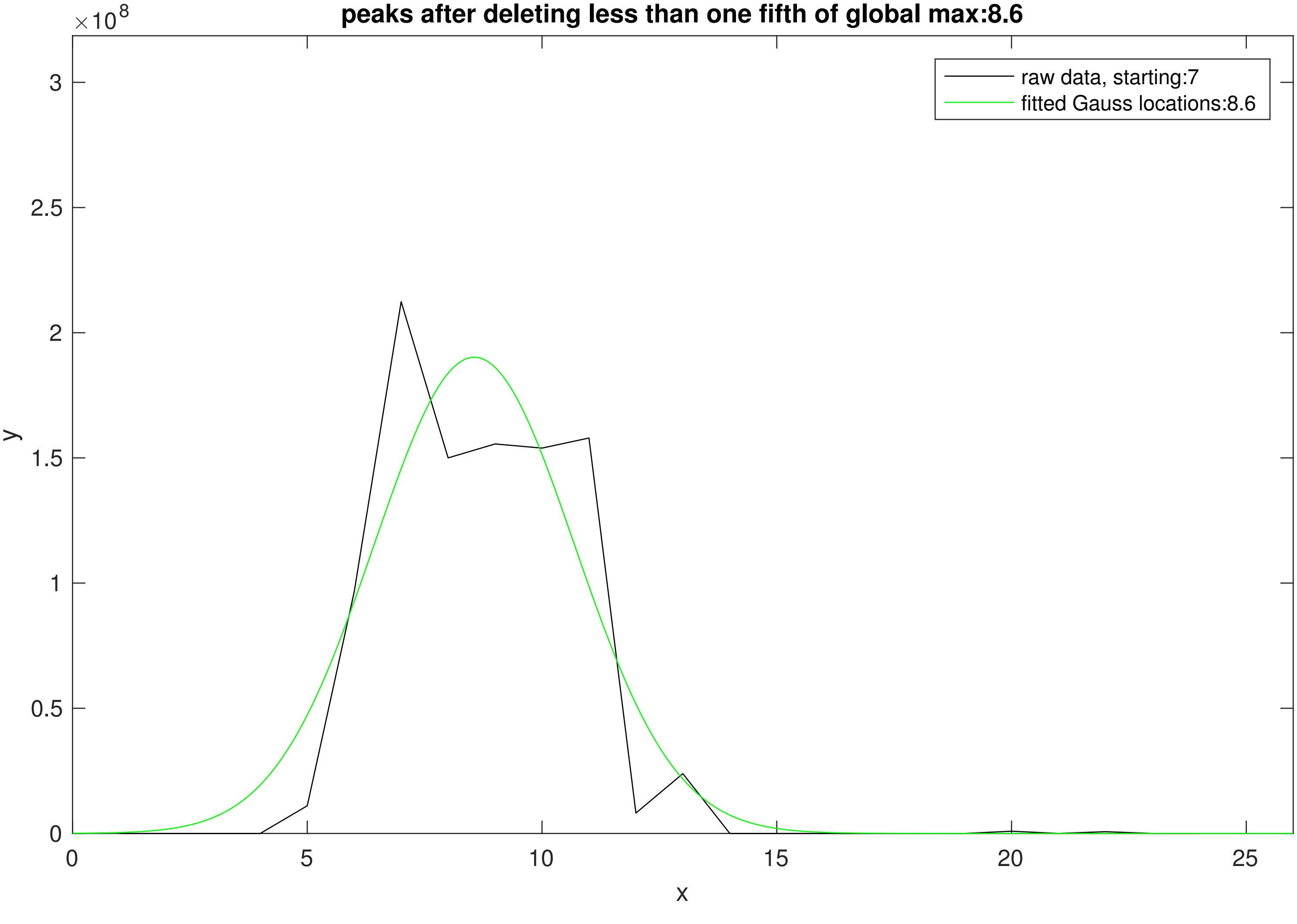

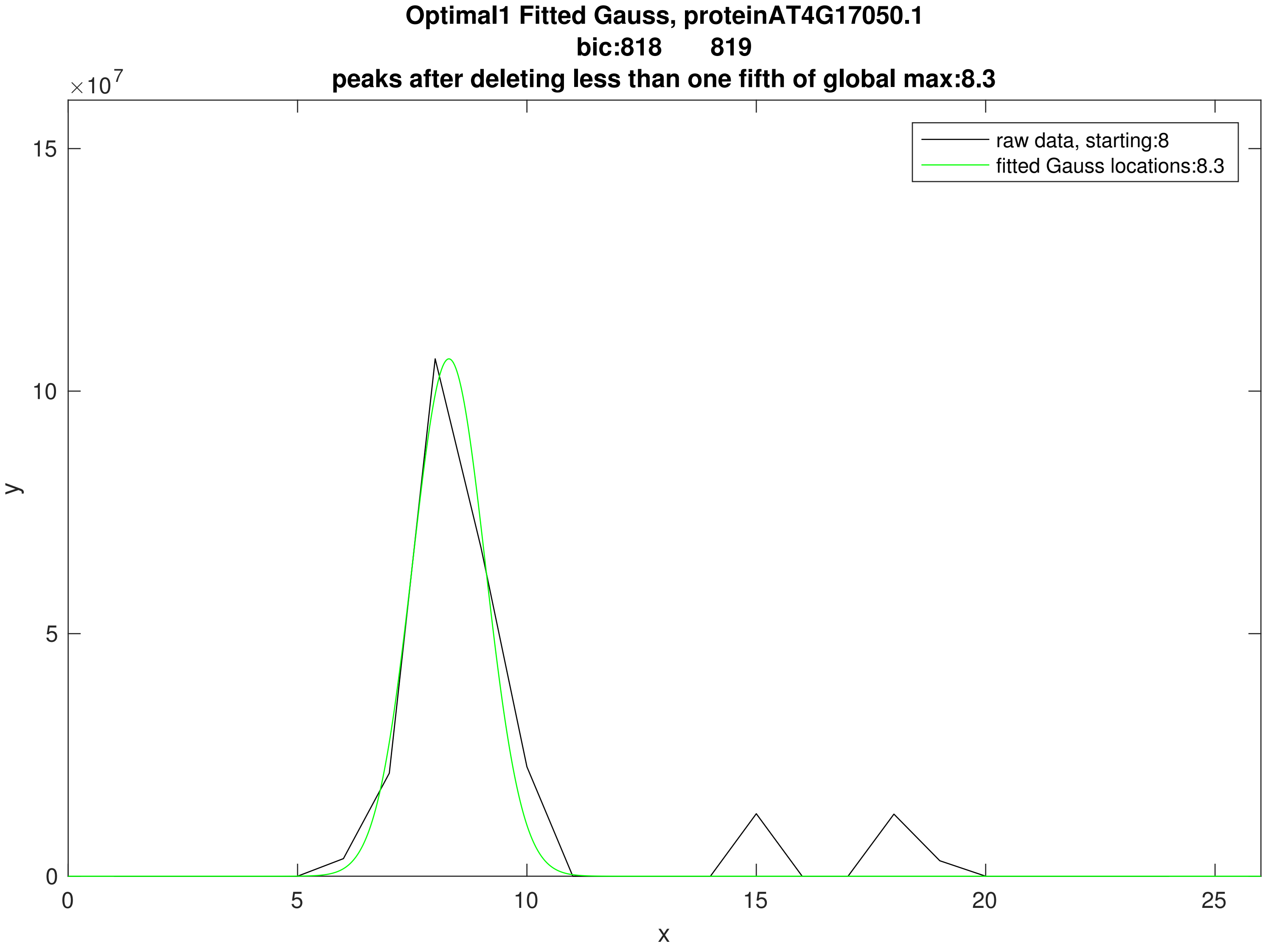
Optimal1 Fitted Gauss, proteinAT4G17050.1
bic:818 819
peaks after deleting less than one fifth of global max:8.3
×10^7
raw data, starting:8
fitted Gauss locations:8.3
y
x
0
5
10
15
20
25

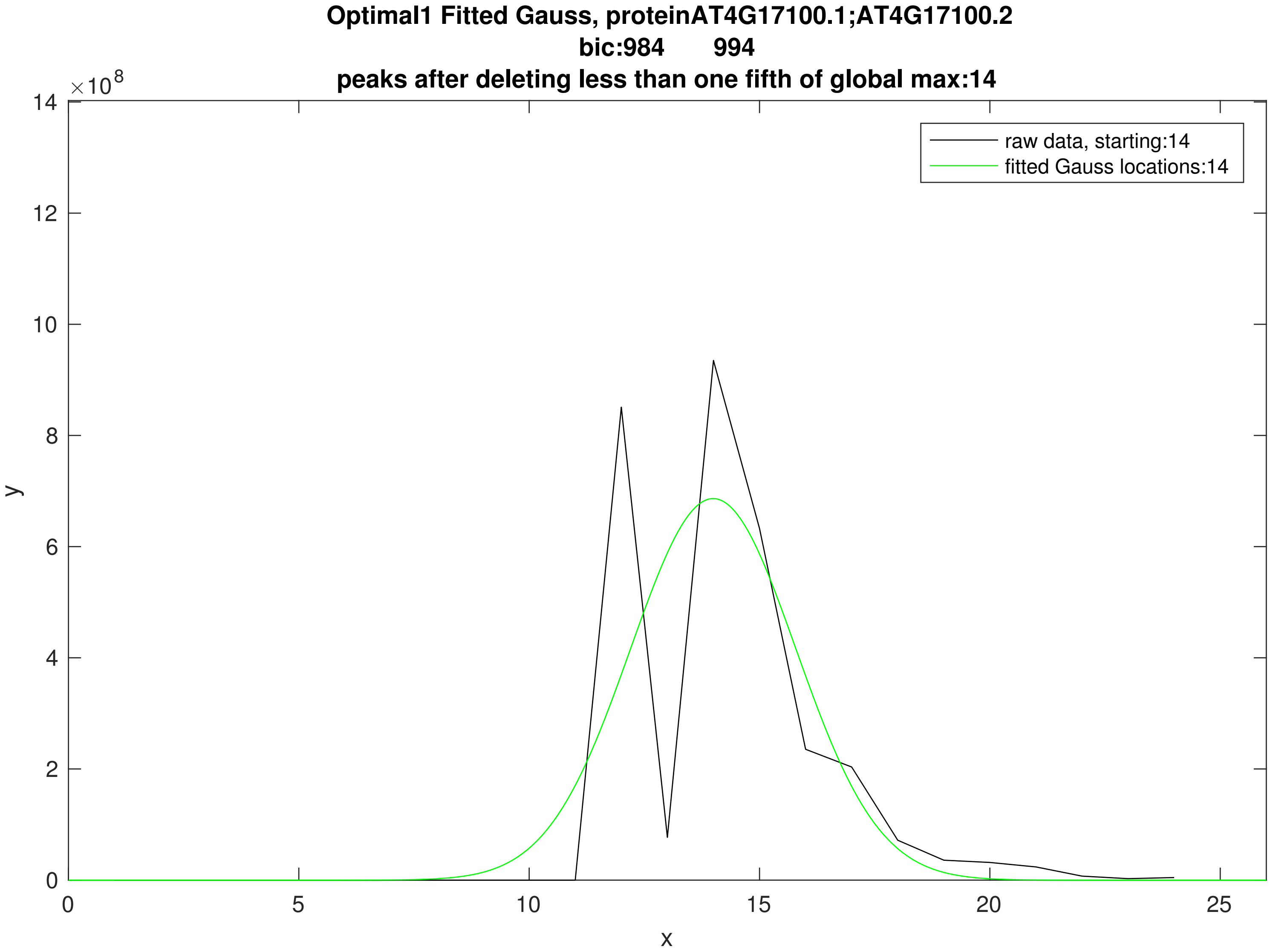
Optimal1 Fitted Gauss, proteinAT4G17100.1;AT4G17100.2
bic:984 994
peaks after deleting less than one fifth of global max:14
×10^8
14
12
10
8
6
4
2
0
y
0
5
10
15
20
25
x
raw data, starting:14
fitted Gauss locations:14

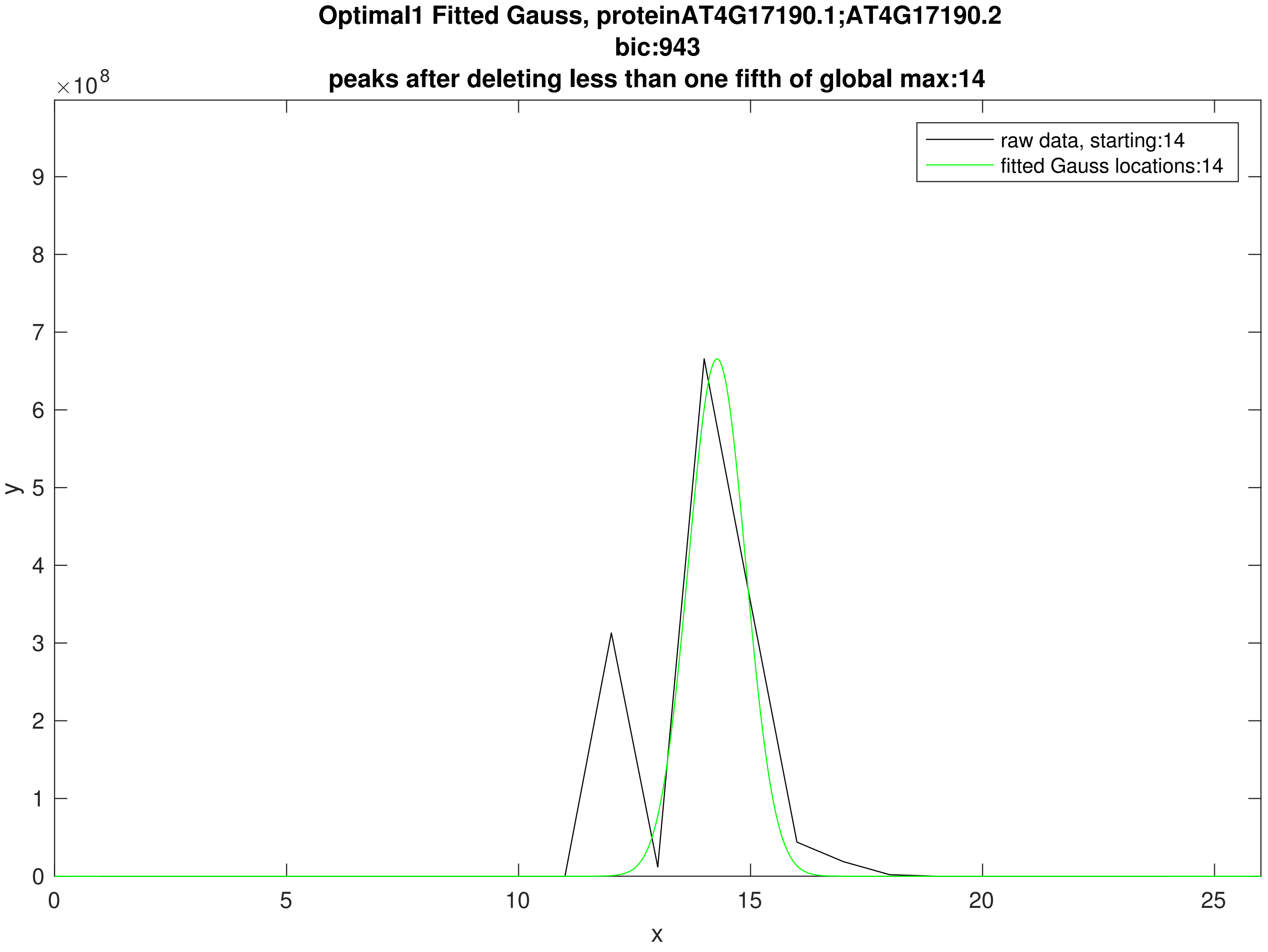
Optimal1 Fitted Gauss, proteinAT4G17190.1;AT4G17190.2
bic:943
peaks after deleting less than one fifth of global max:14
×10^8
raw data, starting:14
fitted Gauss locations:14
y
x
0
1
2
3
4
5
6
7
8
9
0
5
10
15
20
25

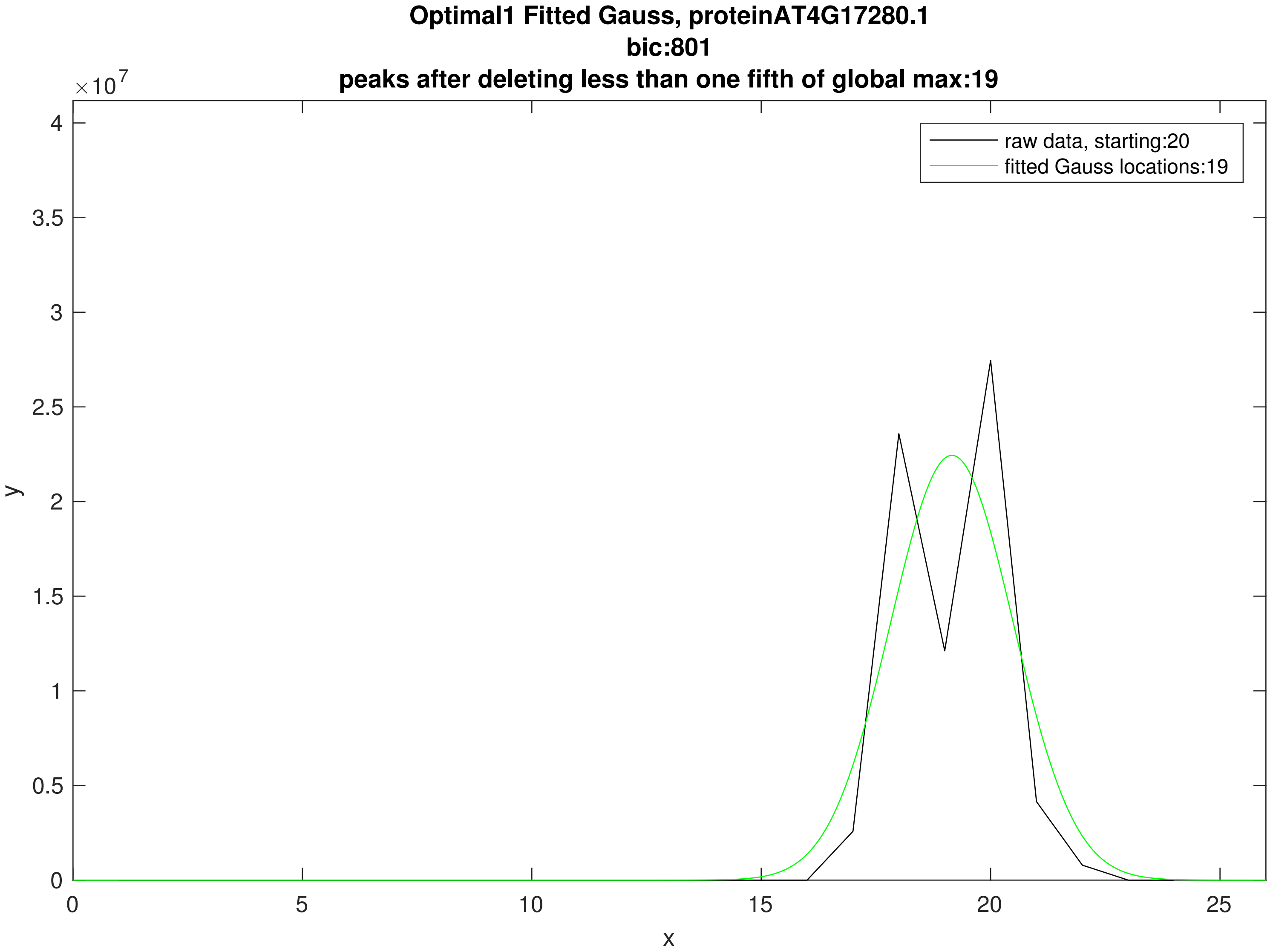

Optimal1 Fitted Gauss, proteinAT4G17280.1
bic:801
peaks after deleting less than one fifth of global max:19
×10^7
raw data, starting:20
fitted Gauss locations:19
y
x
0
0.5
1
1.5
2
2.5
3
3.5
4
5
10
15
20
25

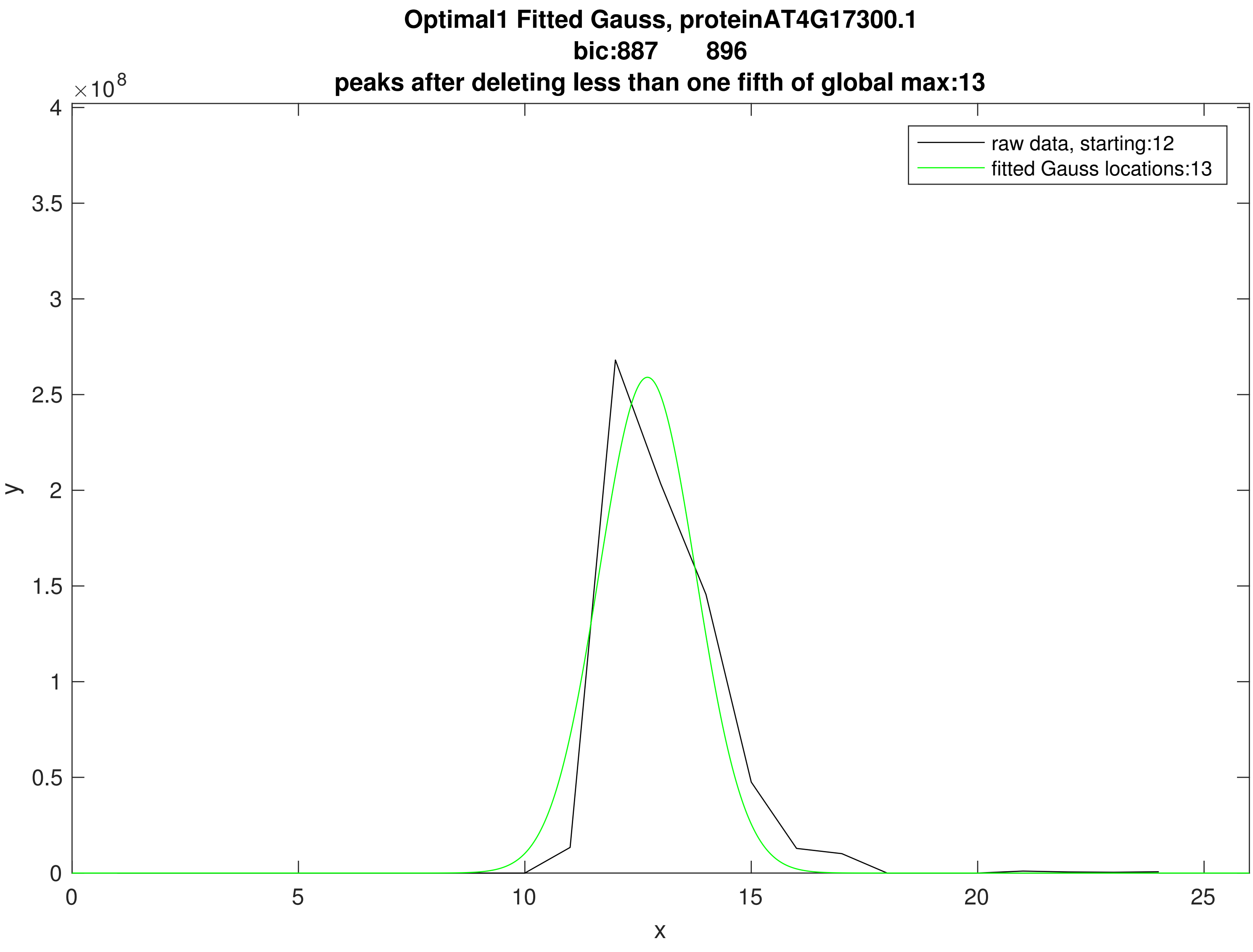

Optimal1 Fitted Gauss, proteinAT4G17300.1
bic:887 896
peaks after deleting less than one fifth of global max:13
×10^8
4
3.5
3
2.5
2
1.5
1
0.5
0
y
0
5
10
15
20
25
x
raw data, starting:12
fitted Gauss locations:13

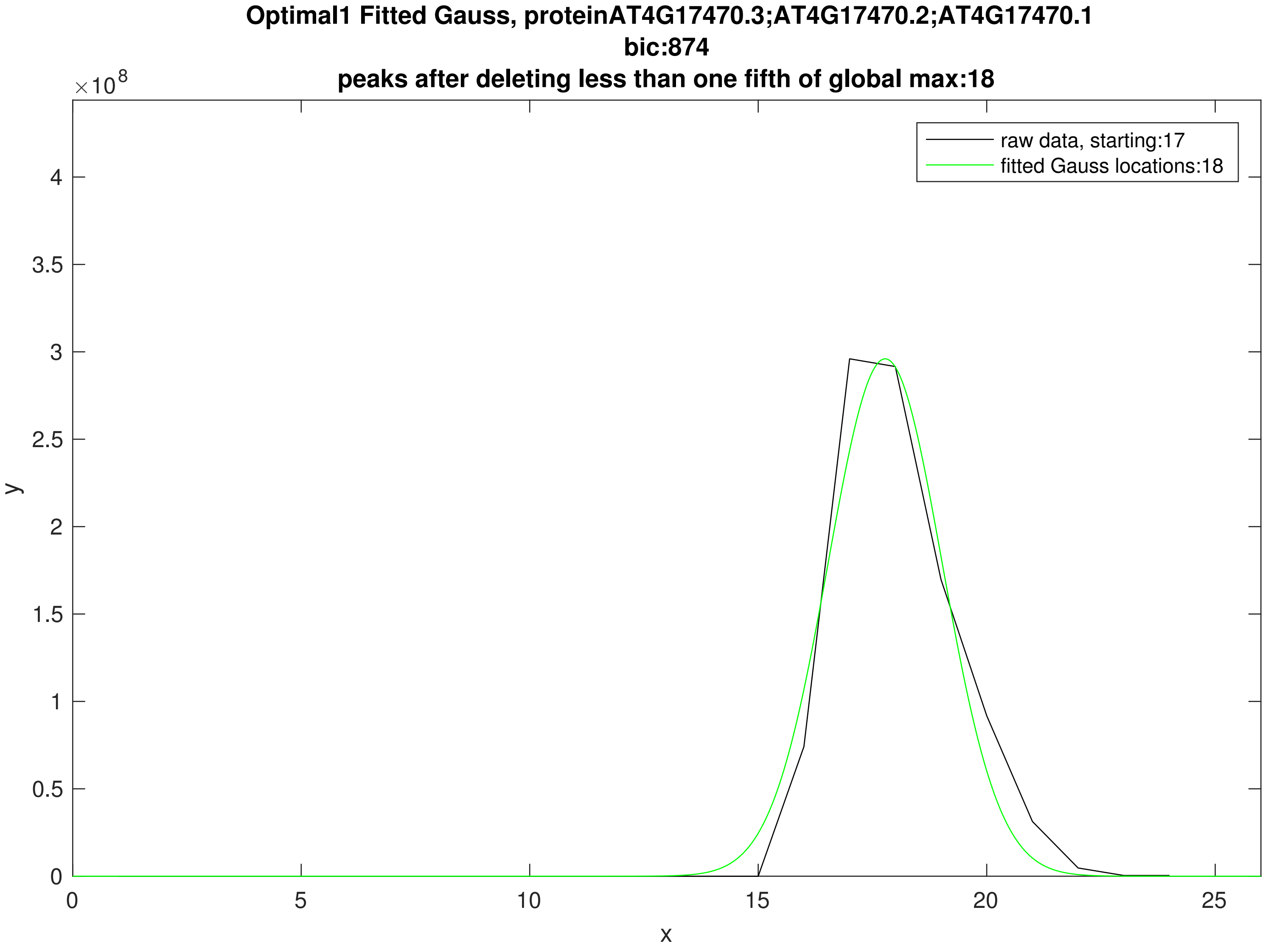

Optimal1 Fitted Gauss, proteinAT4G17470.3;AT4G17470.2;AT4G17470.1
bic:874
peaks after deleting less than one fifth of global max:18
×10^8
raw data, starting:17
fitted Gauss locations:18
y
x
0
0.5
1
1.5
2
2.5
3
3.5
4
0
5
10
15
20
25

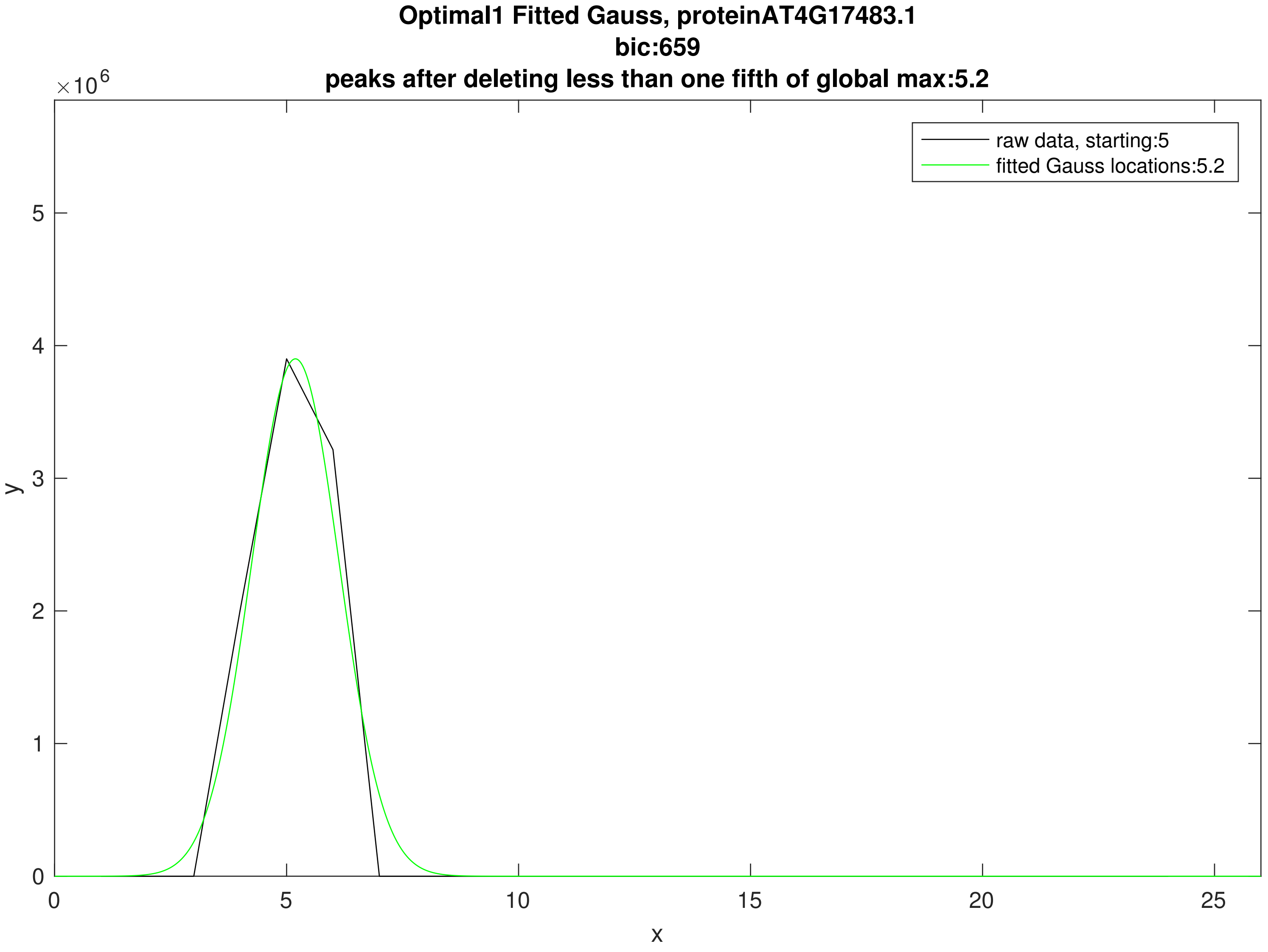
Optimal1 Fitted Gauss, proteinAT4G17483.1
bic:659
peaks after deleting less than one fifth of global max:5.2
×10^6
raw data, starting:5
fitted Gauss locations:5.2
y
x
0
1
2
3
4
5
0
5
10
15
20
25

**Optimal1 Fitted Gauss, proteinAT4G17510.1**
**bic:762**
**peaks after deleting less than one fifth of global max:17**

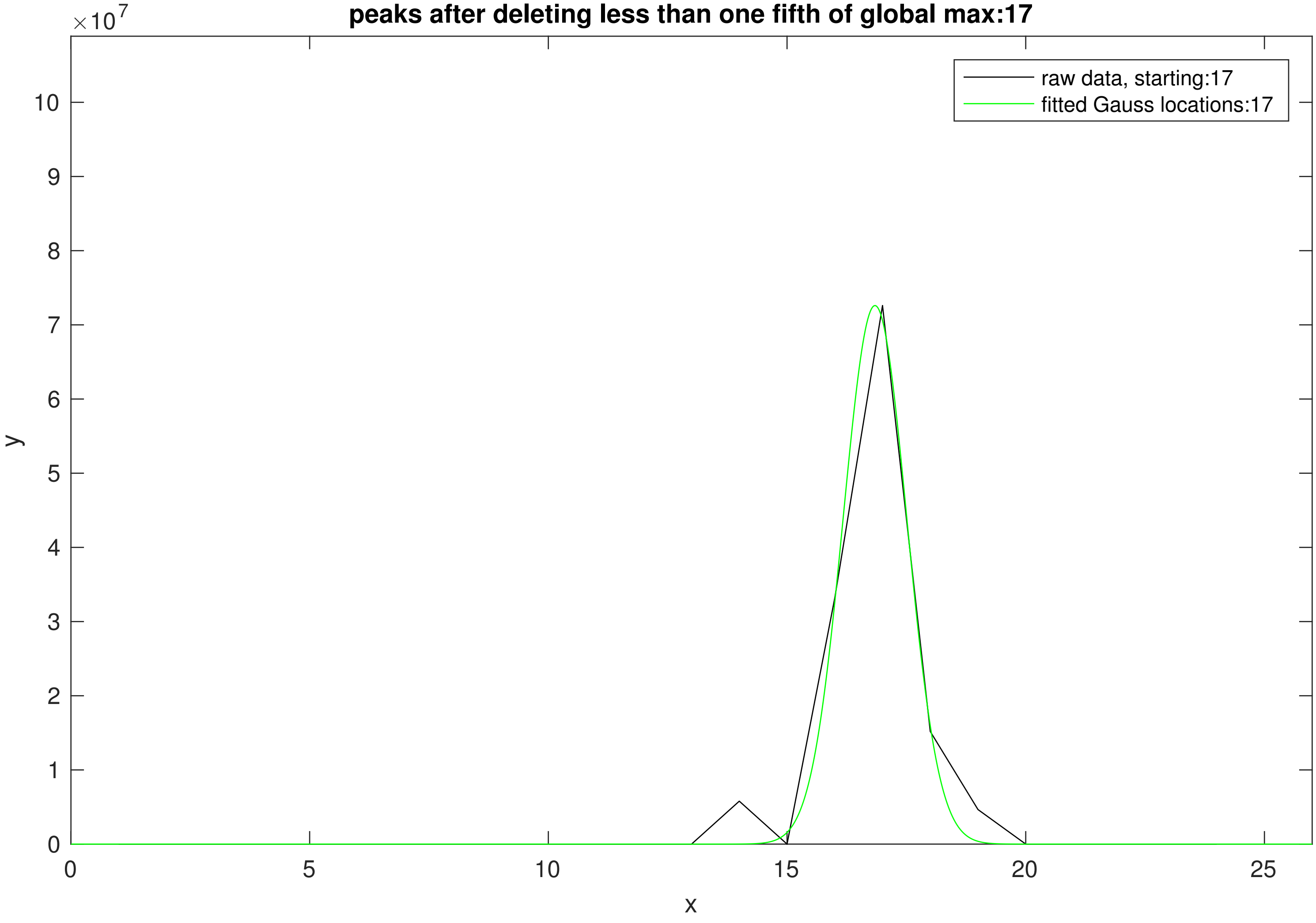

**Optimal1 Fitted Gauss, proteinAT4G17530.1**
**bic:790 806**
**peaks after deleting less than one fifth of global max:19**

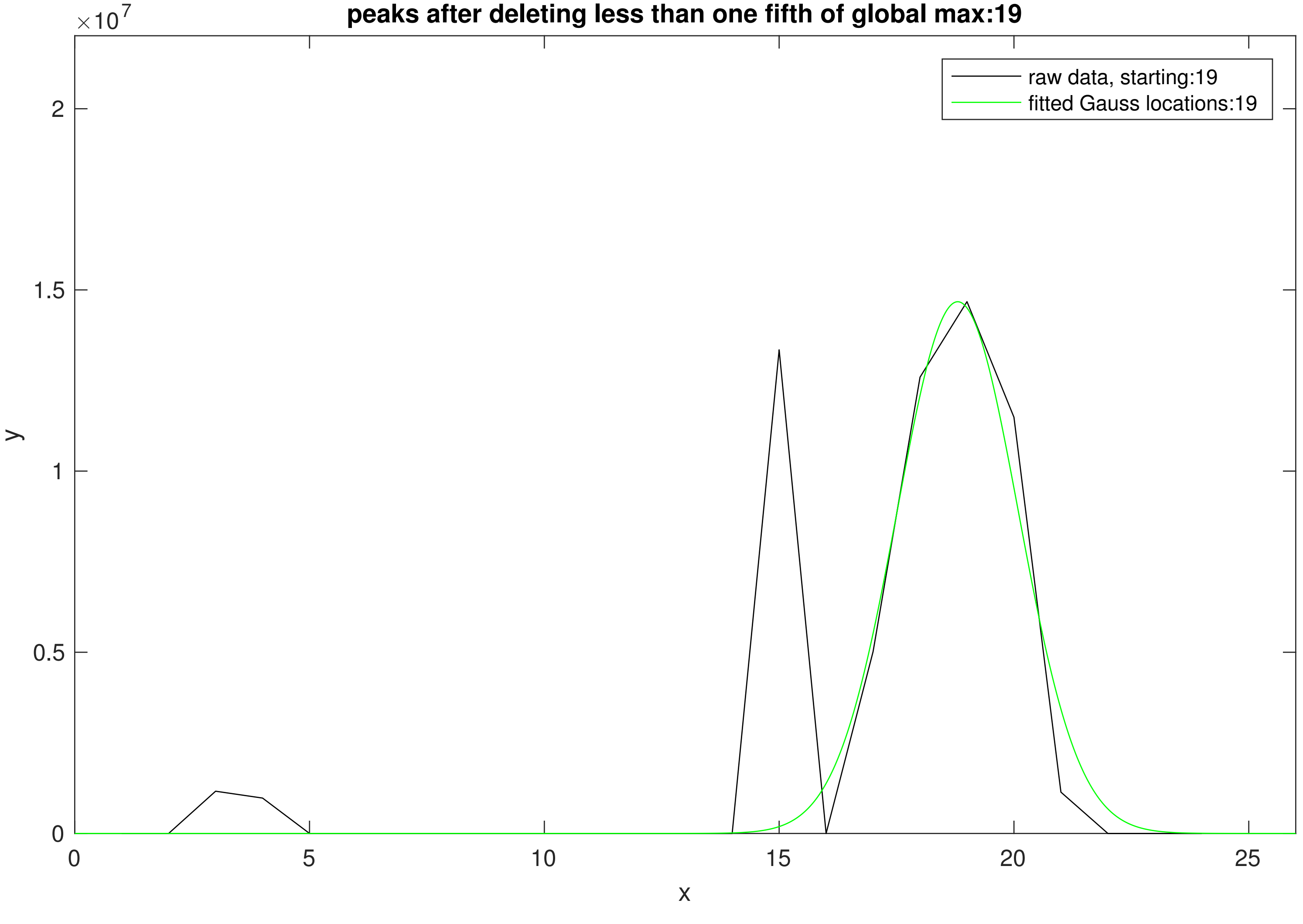

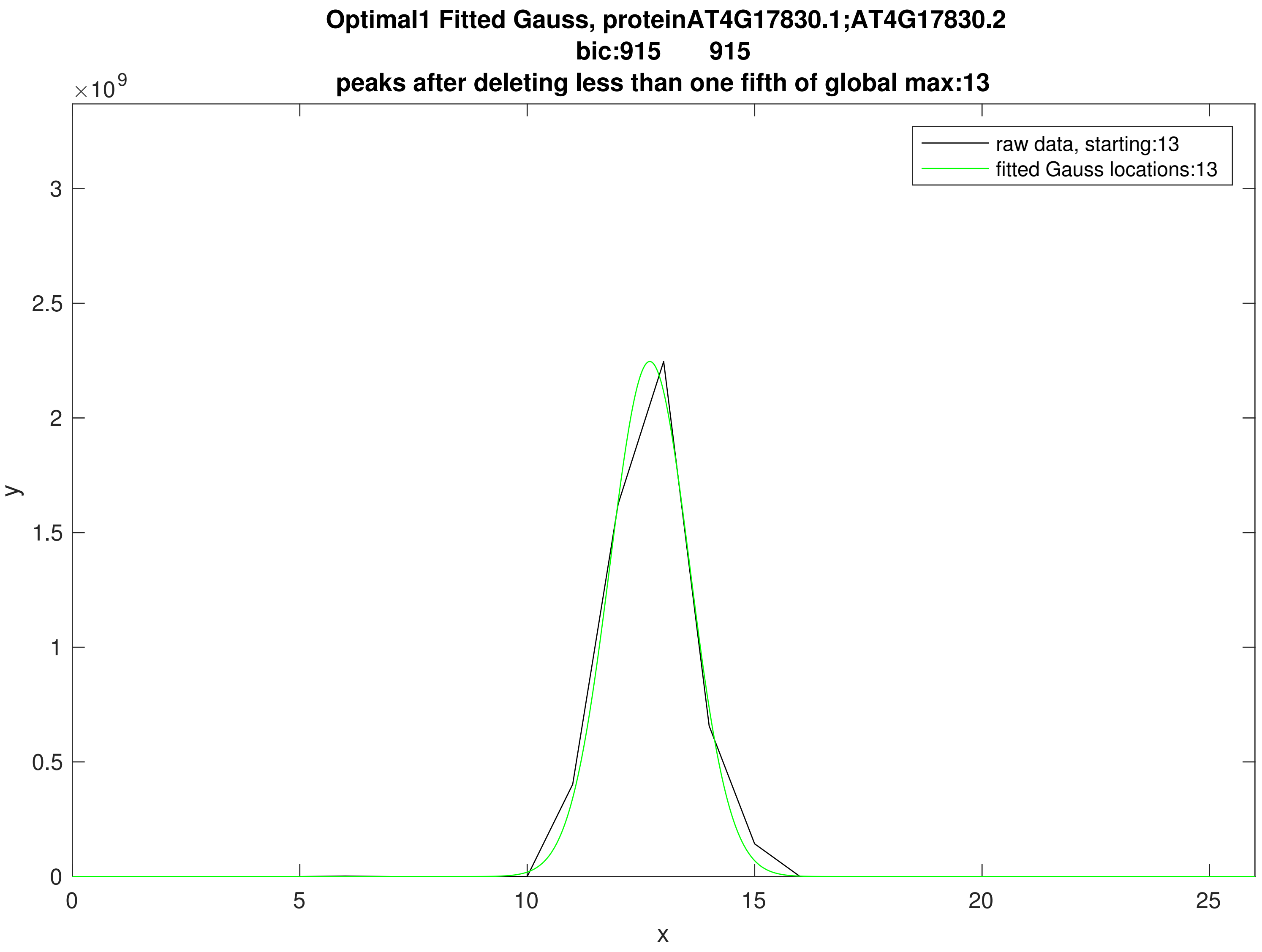
Optimal1 Fitted Gauss, proteinAT4G17830.1;AT4G17830.2
bic:915 915
peaks after deleting less than one fifth of global max:13
×10^9
raw data, starting:13
fitted Gauss locations:13
3
2.5
2
1.5
1
0.5
0
y
0
5
10
15
20
25
x

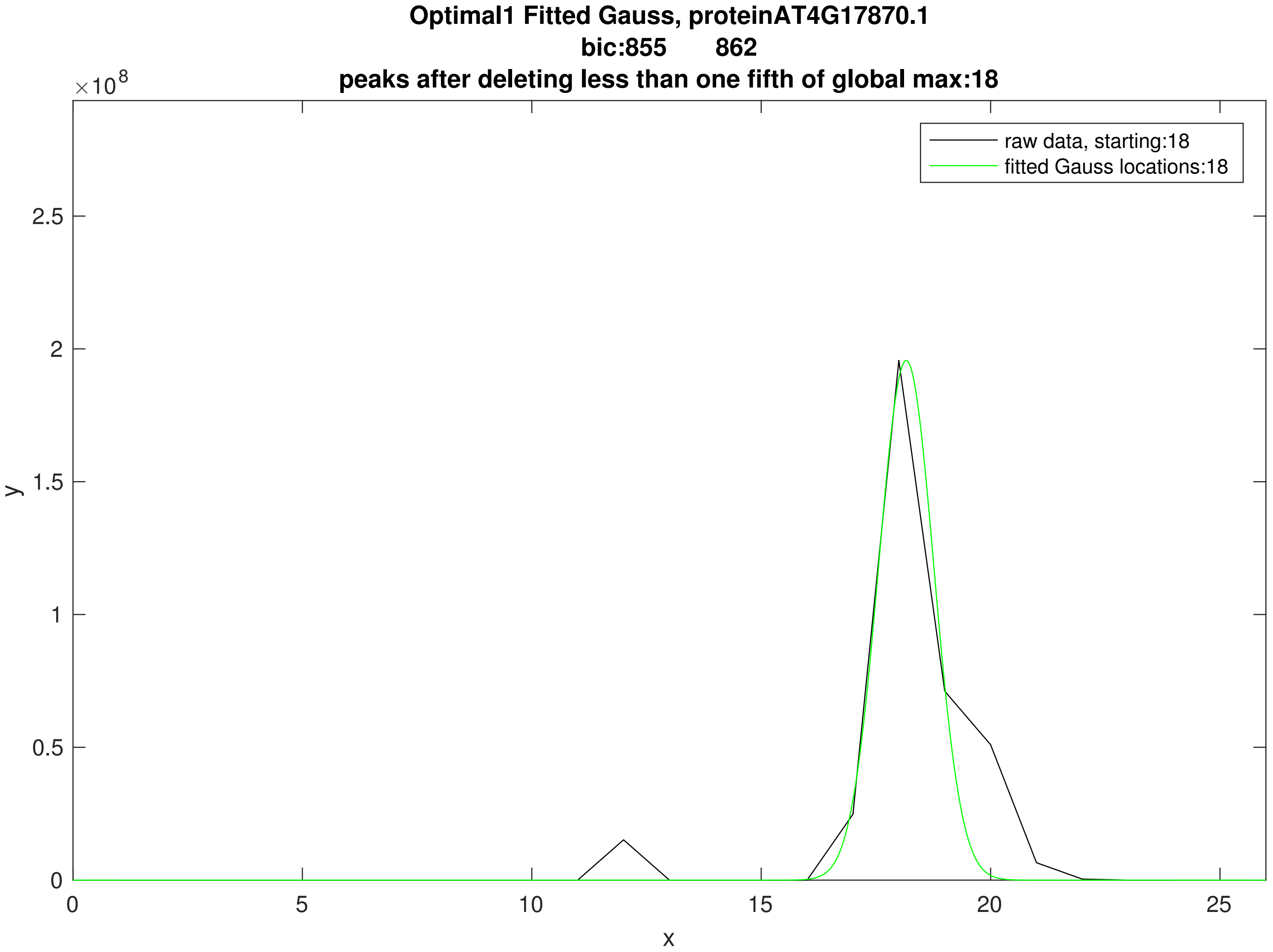
Optimal1 Fitted Gauss, proteinAT4G17870.1
bic:855 862
peaks after deleting less than one fifth of global max:18
×10^8
raw data, starting:18
fitted Gauss locations:18
y
x
0
0.5
1
1.5
2
2.5
0
5
10
15
20
25

Optimal1 Fitted Gauss, proteinAT4G17890.2;AT4G17890.1
bic:796
peaks after deleting less than one fifth of global max:16

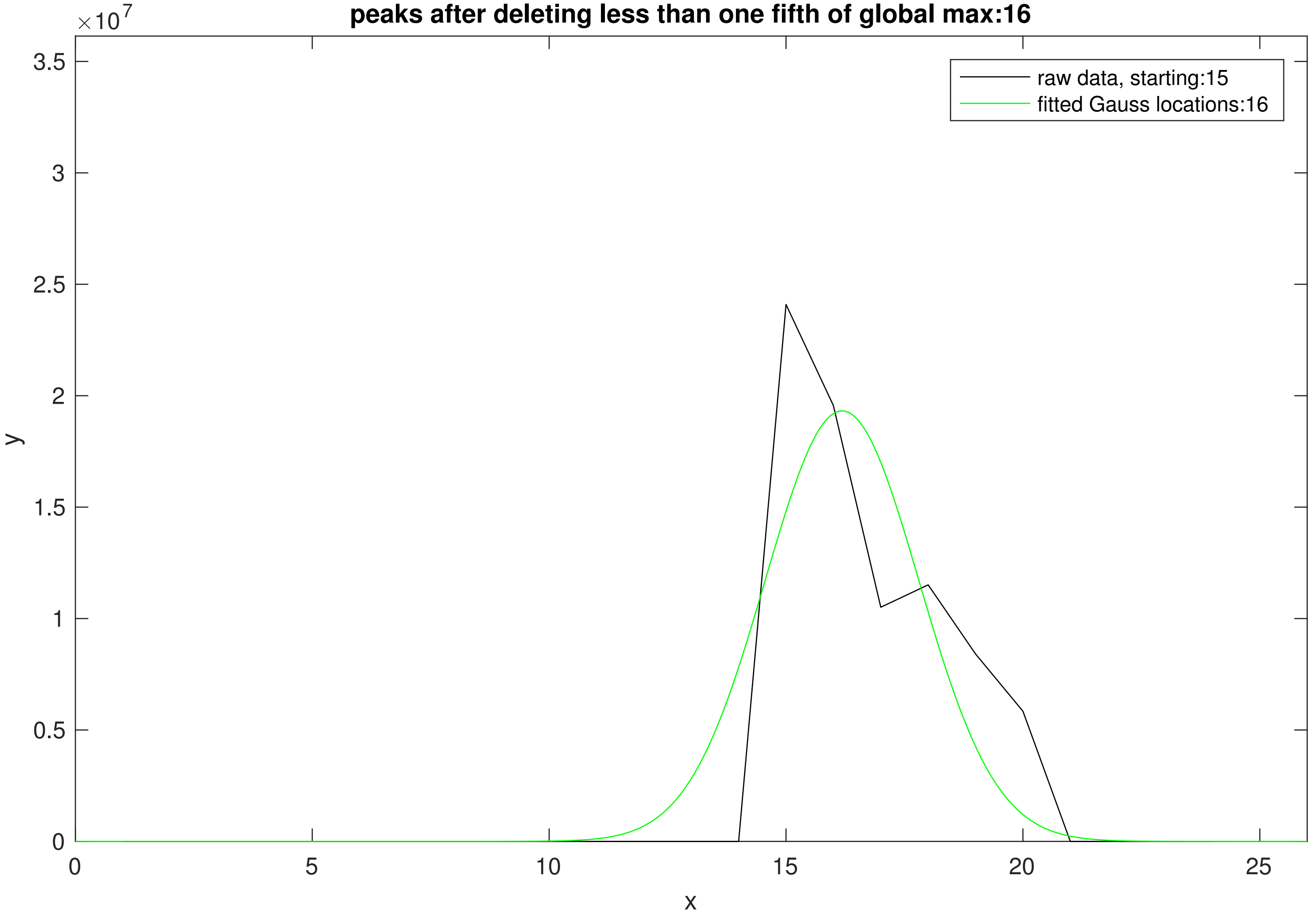

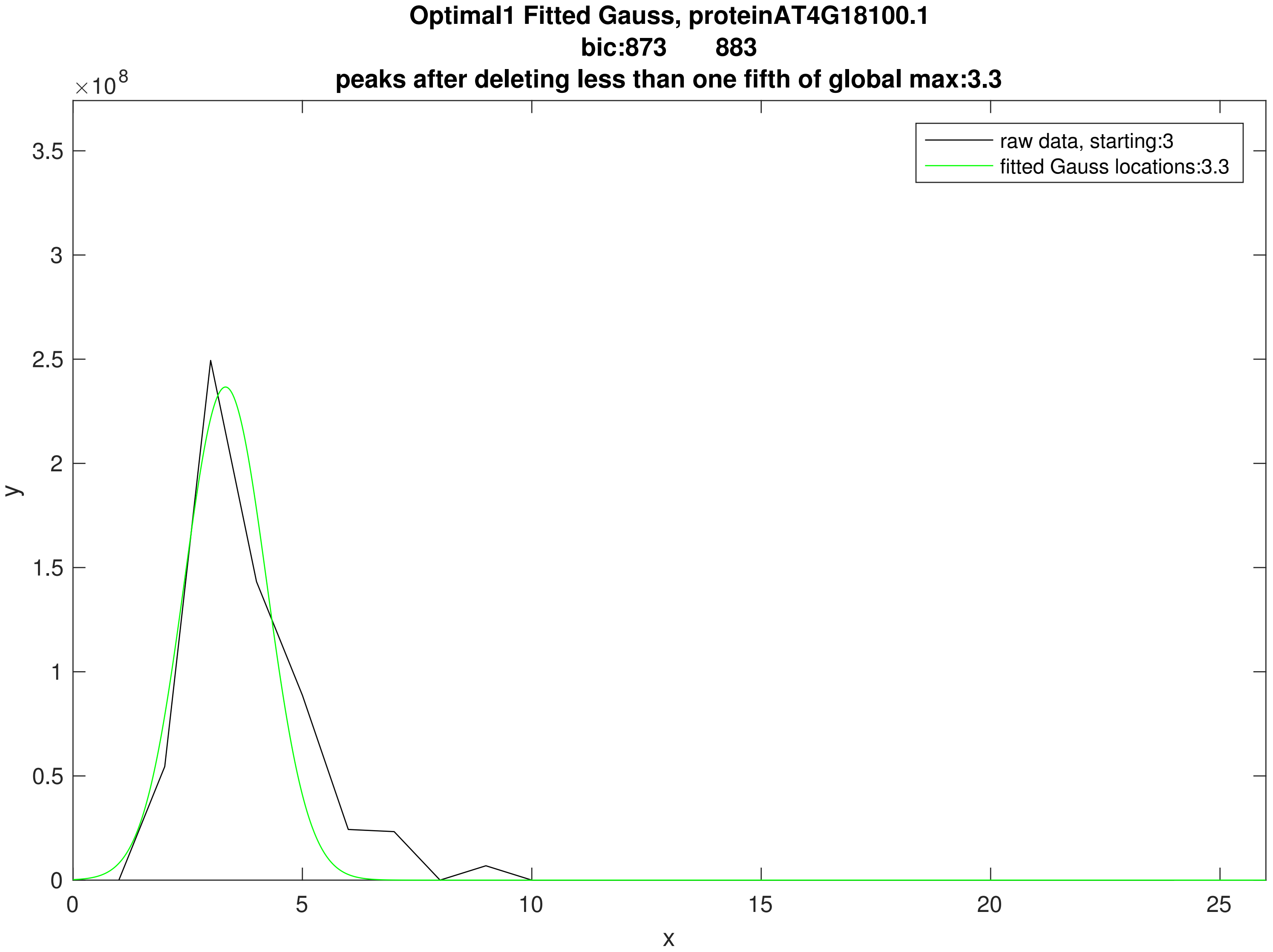
Optimal1 Fitted Gauss, proteinAT4G18100.1
bic:873 883
peaks after deleting less than one fifth of global max:3.3
×10^8
3.5
3
2.5
2
1.5
1
0.5
0
y
0
5
10
15
20
25
x
raw data, starting:3
fitted Gauss locations:3.3

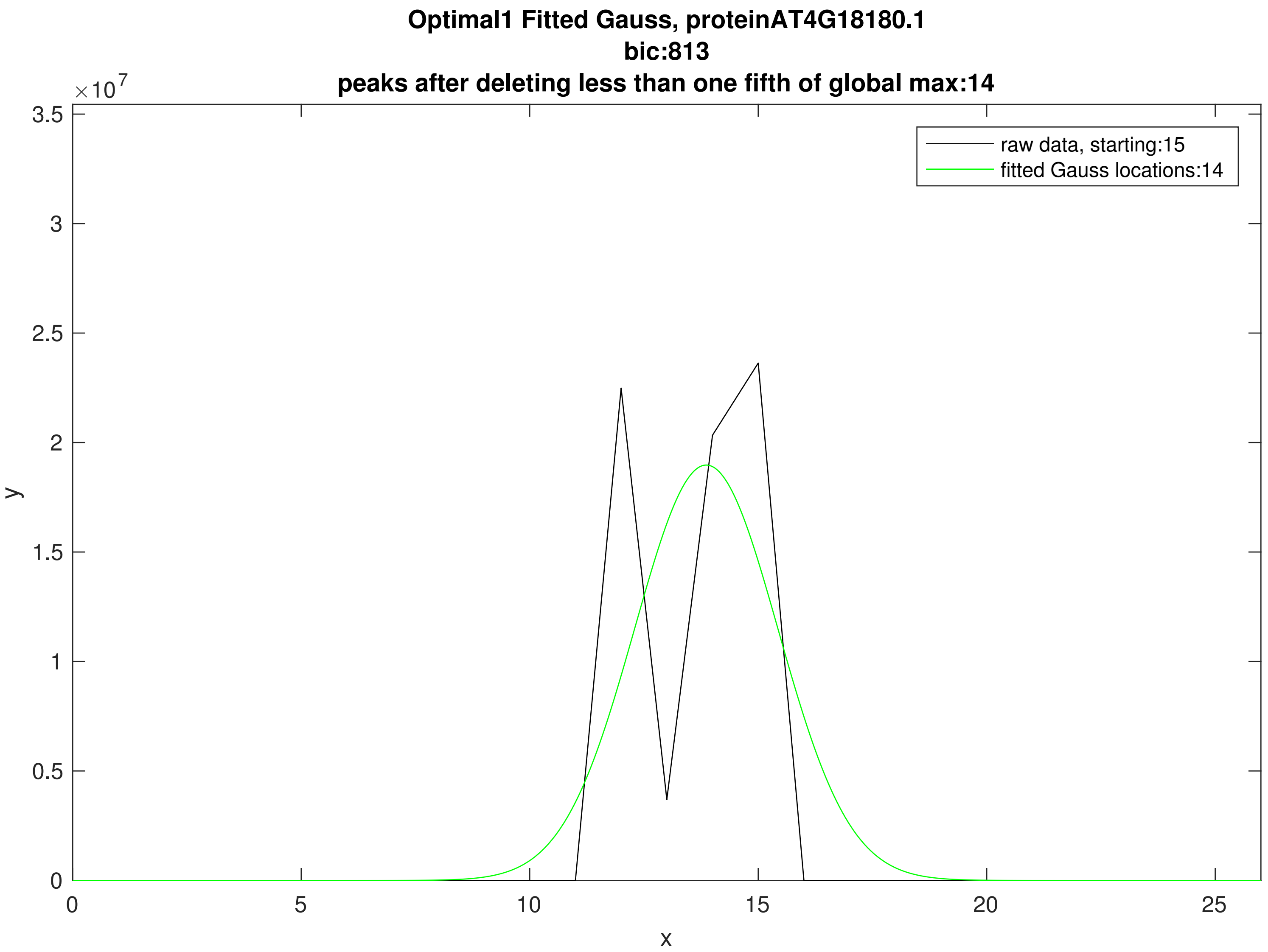
Optimal1 Fitted Gauss, proteinAT4G18180.1
bic:813
peaks after deleting less than one fifth of global max:14
×10^7
3.5
3
2.5
2
1.5
1
0.5
0
y
0
5
10
15
20
25
x
raw data, starting:15
fitted Gauss locations:14

**Optimal1 Fitted Gauss, proteinAT4G18360.2;AT4G18360.1**
**bic:817 823**
**peaks after deleting less than one fifth of global max:13**

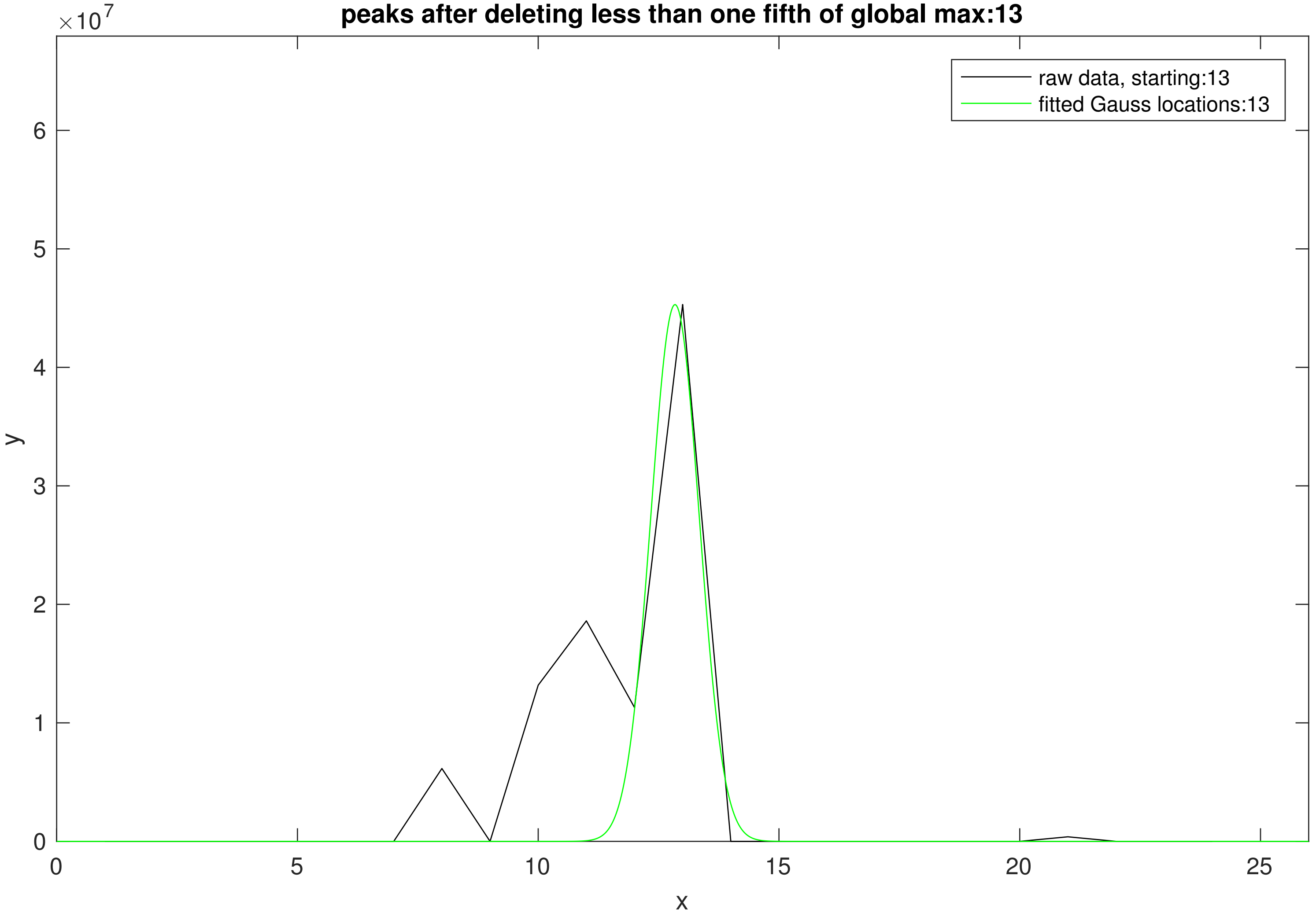

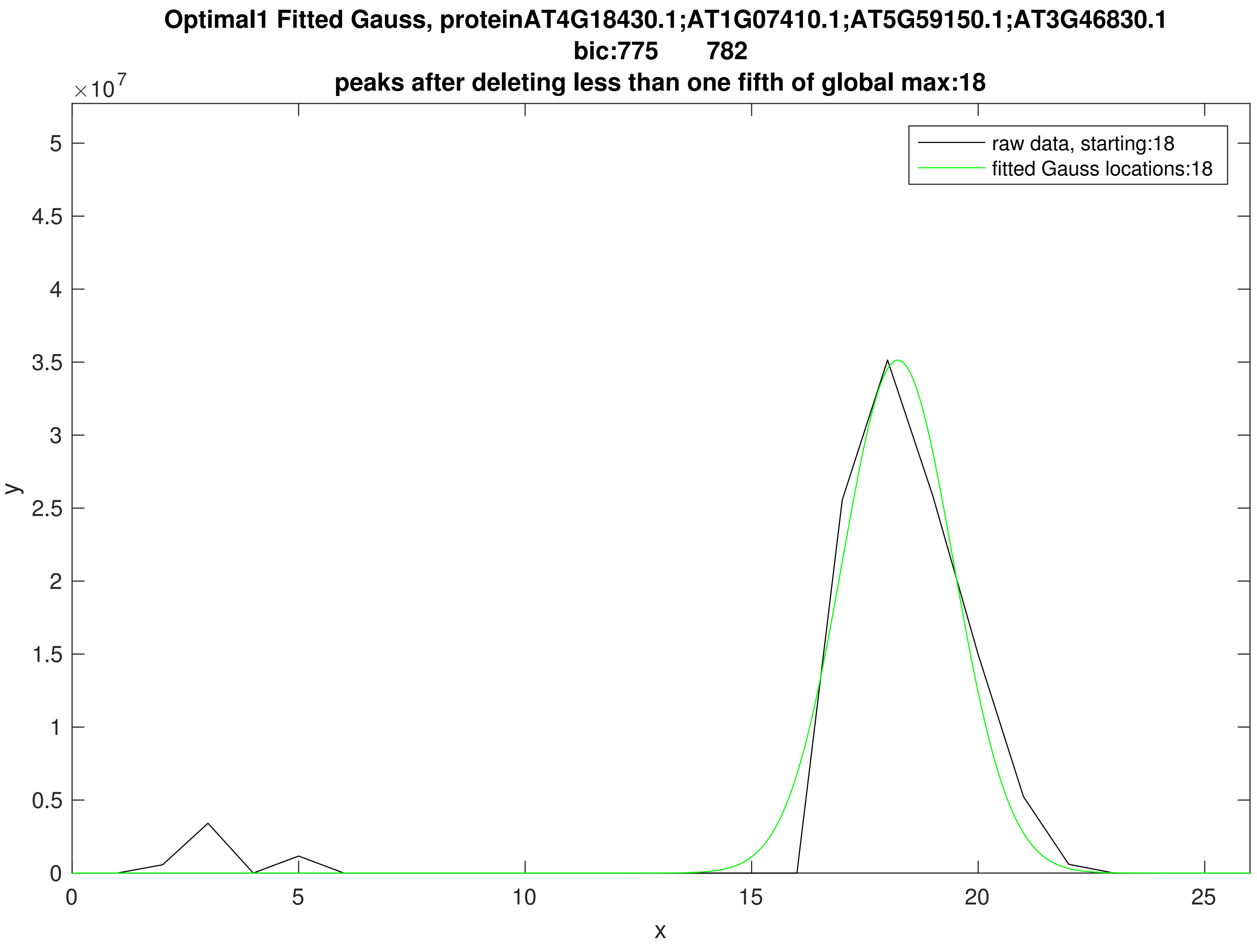
Optimal1 Fitted Gauss, proteinAT4G18430.1;AT1G07410.1;AT5G59150.1;AT3G46830.1
bic:775 782
peaks after deleting less than one fifth of global max:18
×10^7
5
4.5
4
3.5
3
2.5
2
1.5
1
0.5
0
y
0
5
10
15
20
25
x
raw data, starting:18
fitted Gauss locations:18

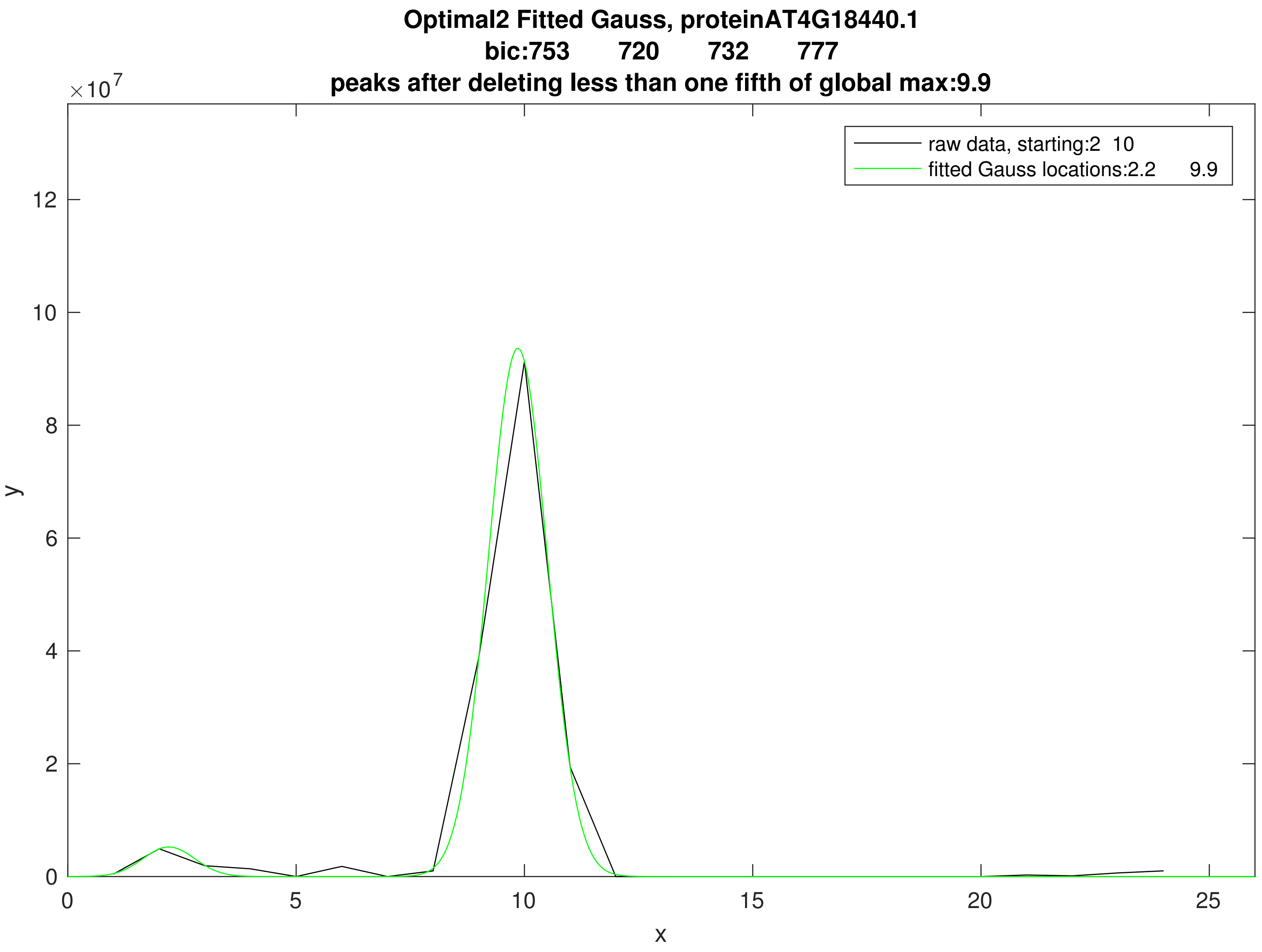

Optimal2 Fitted Gauss, proteinAT4G18440.1
bic:753 720 732 777
peaks after deleting less than one fifth of global max:9.9
×10^7
raw data, starting:2 10
fitted Gauss locations:2.2 9.9
y
x
0
2
4
6
8
10
12
0
5
10
15
20
25

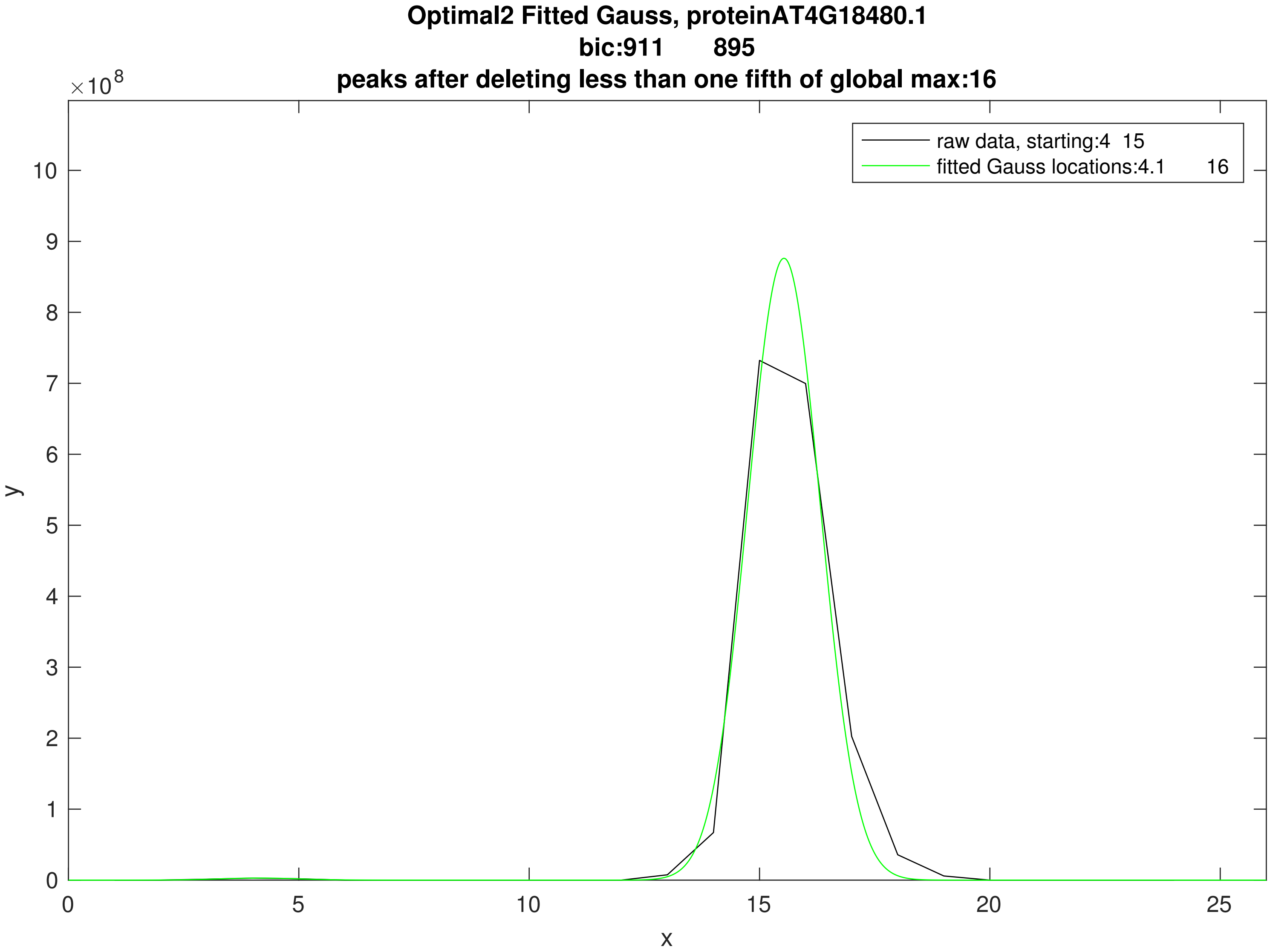
Optimal2 Fitted Gauss, proteinAT4G18480.1
bic:911 895
peaks after deleting less than one fifth of global max:16
×10^8
raw data, starting:4 15
fitted Gauss locations:4.1 16
y
x
0
1
2
3
4
5
6
7
8
9
10
0
5
10
15
20
25

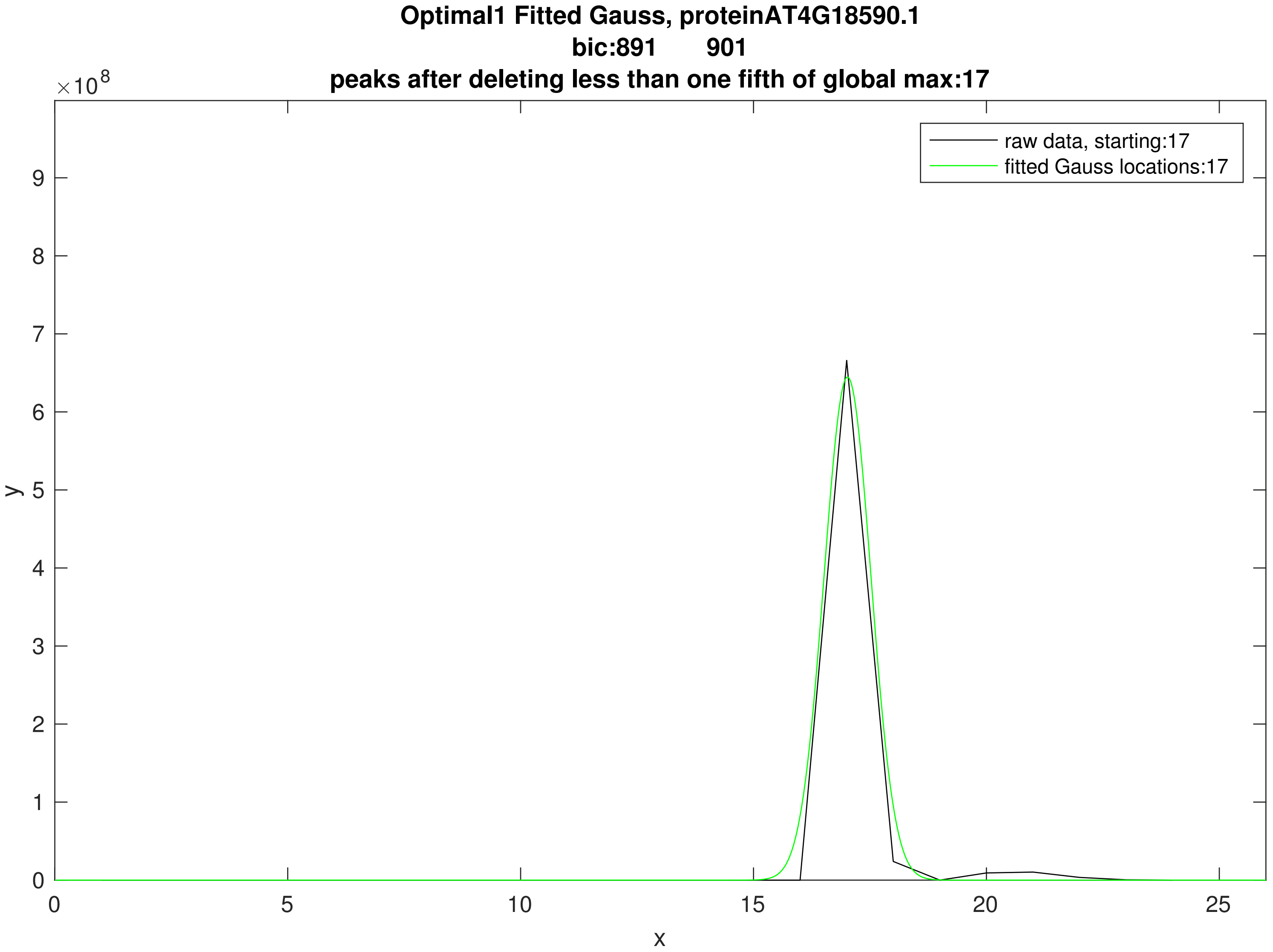

Optimal1 Fitted Gauss, proteinAT4G18590.1
bic:891 901
peaks after deleting less than one fifth of global max:17
×10^8
raw data, starting:17
fitted Gauss locations:17
y
x
0
1
2
3
4
5
6
7
8
9
0
5
10
15
20
25

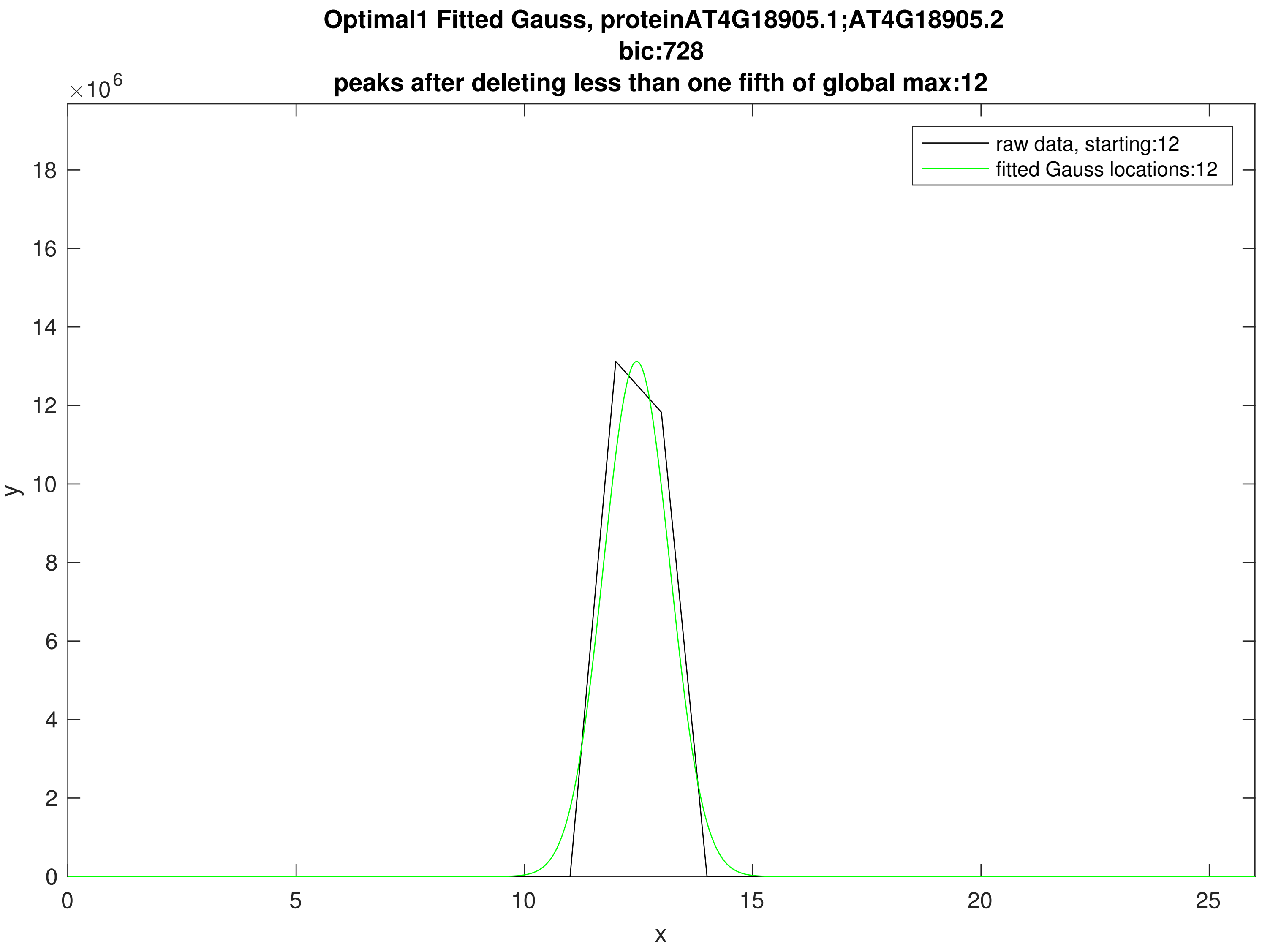

Optimal1 Fitted Gauss, proteinAT4G18905.1;AT4G18905.2
bic:728
peaks after deleting less than one fifth of global max:12
×10^6
raw data, starting:12
fitted Gauss locations:12
y
x
0
2
4
6
8
10
12
14
16
18
0
5
10
15
20
25

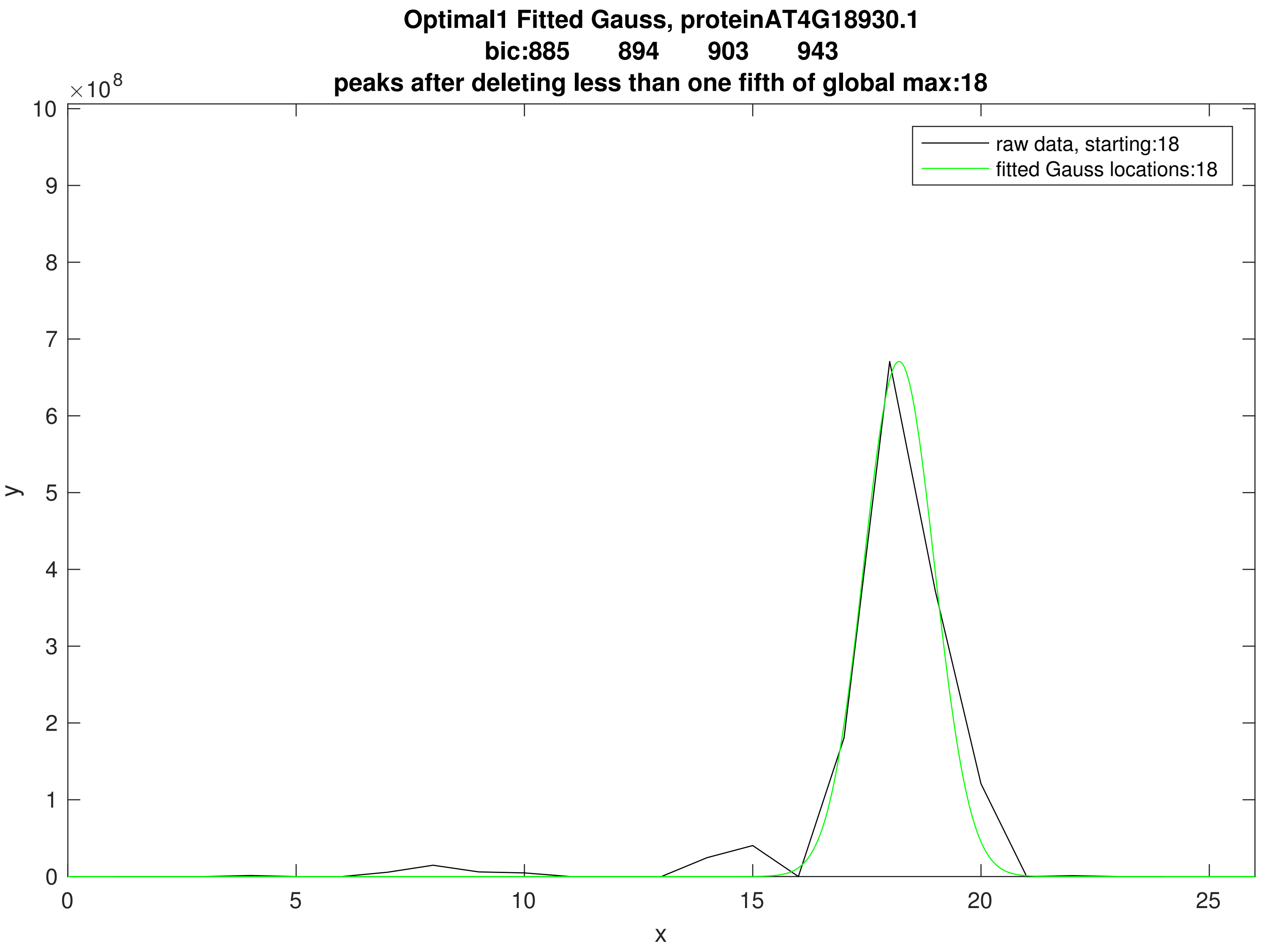
Optimal1 Fitted Gauss, proteinAT4G18930.1
bic:885 894 903 943
peaks after deleting less than one fifth of global max:18
×10^8
10
9
8
7
6
5
4
3
2
1
0
y
0
5
10
15
20
25
x
raw data, starting:18
fitted Gauss locations:18

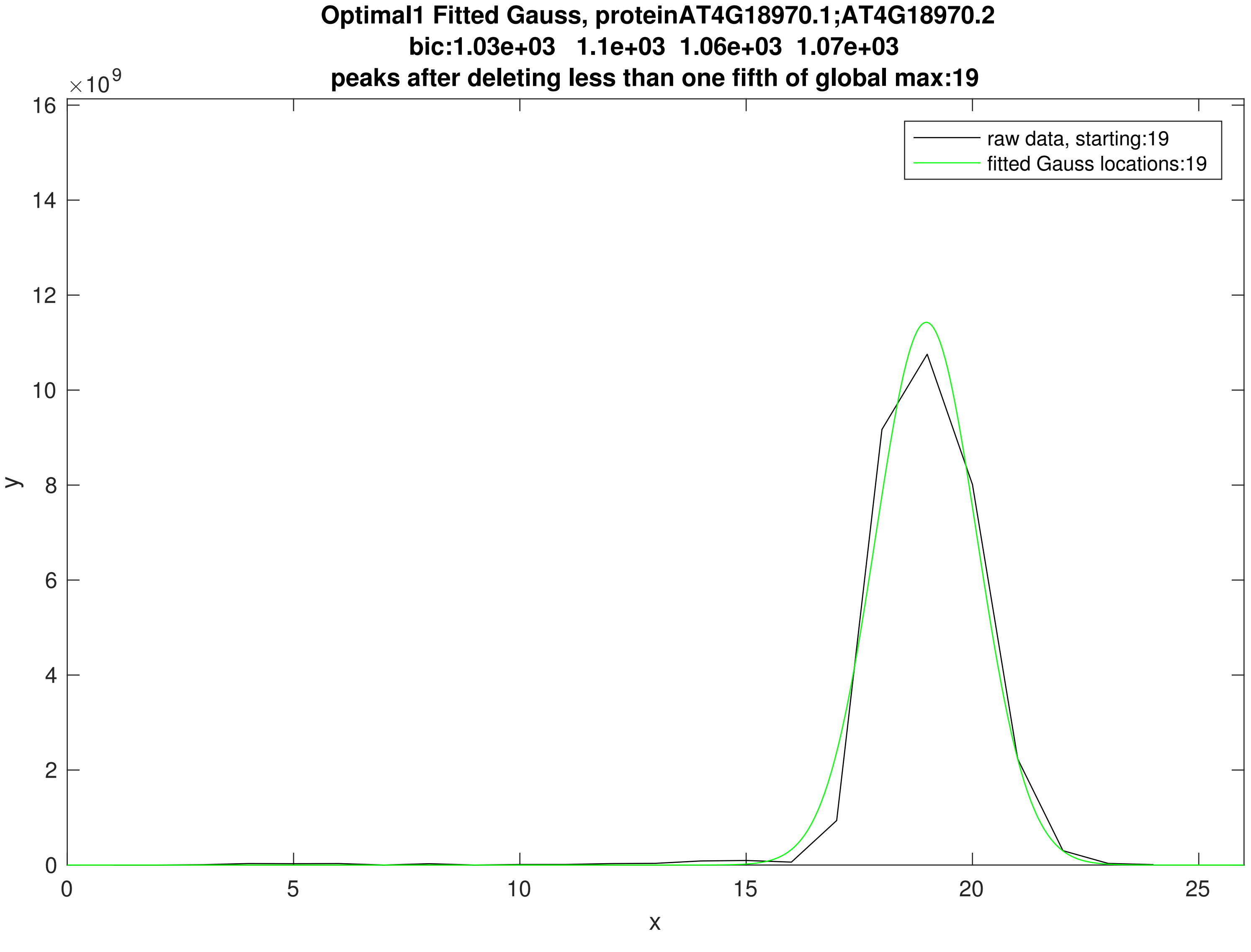

Optimal1 Fitted Gauss, proteinAT4G18970.1;AT4G18970.2
bic:1.03e+03   1.1e+03  1.06e+03  1.07e+03
peaks after deleting less than one fifth of global max:19
×10^9
16
14
12
10
8
6
4
2
0
y
0
5
10
15
20
25
x
raw data, starting:19
fitted Gauss locations:19

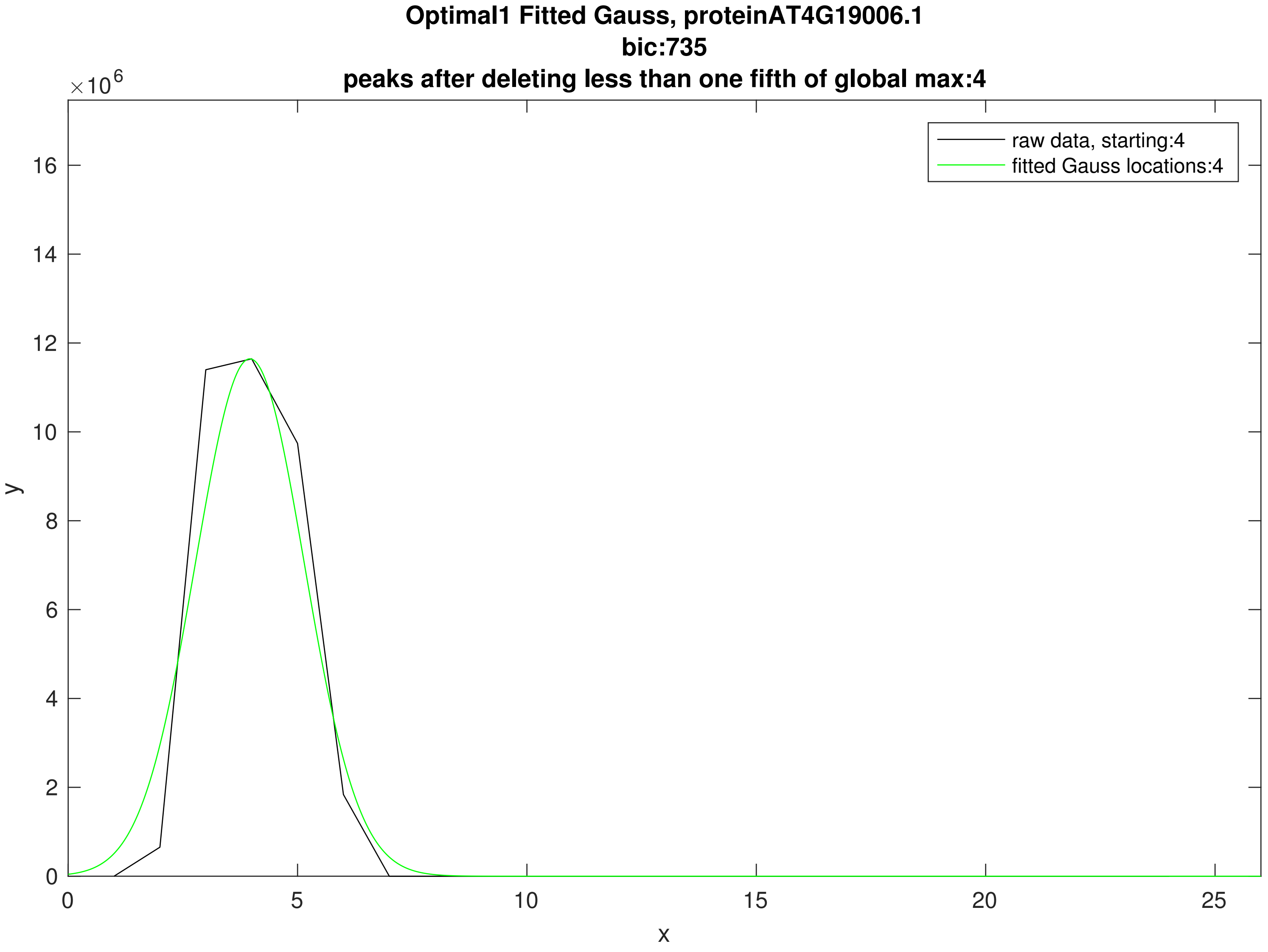
Optimal1 Fitted Gauss, proteinAT4G19006.1
bic:735
peaks after deleting less than one fifth of global max:4
×10^6
raw data, starting:4
fitted Gauss locations:4
16
14
12
10
8
6
4
2
0
y
0
5
10
15
20
25
x

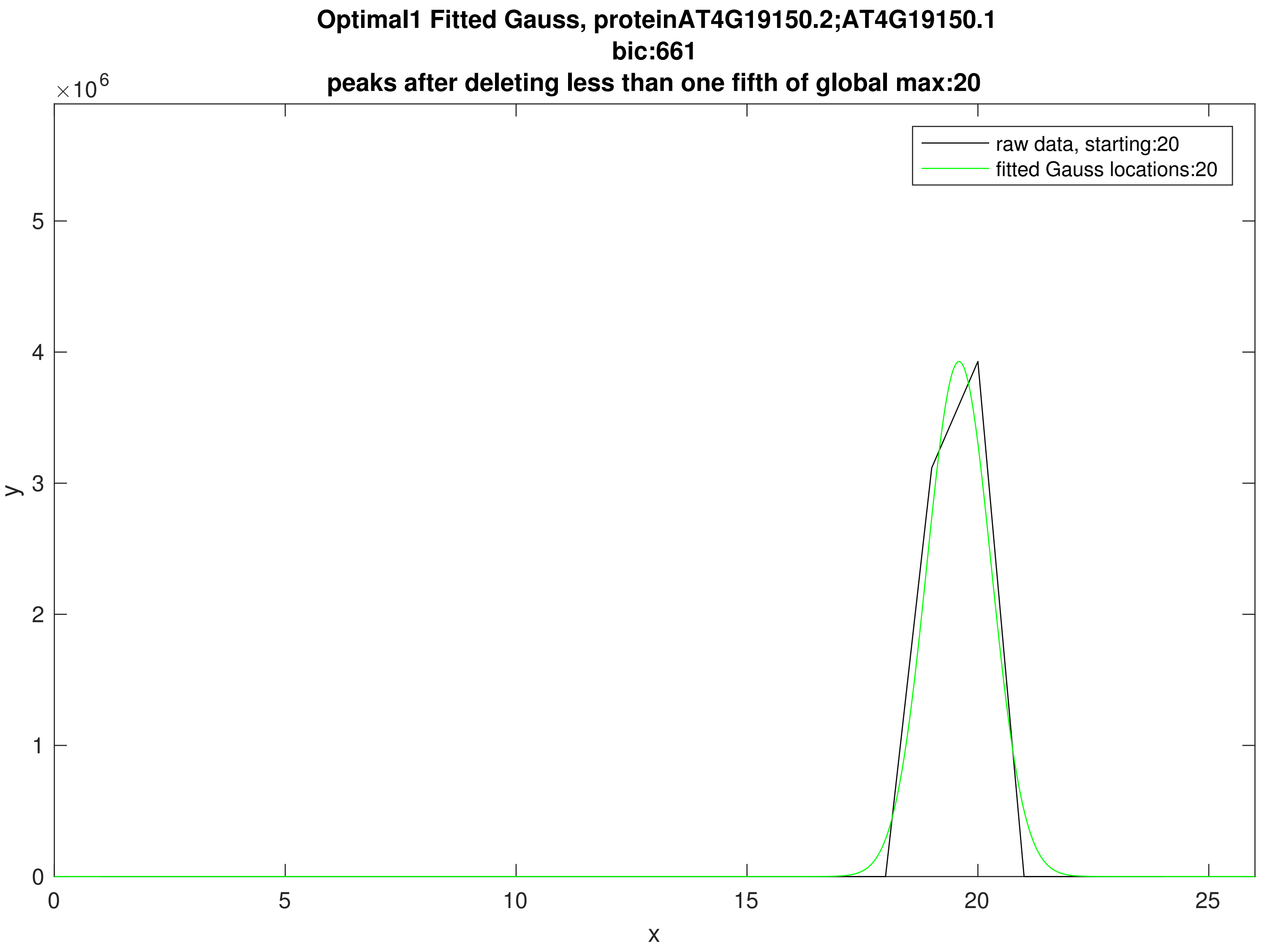
Optimal1 Fitted Gauss, proteinAT4G19150.2;AT4G19150.1
bic:661
peaks after deleting less than one fifth of global max:20
×10^6
raw data, starting:20
fitted Gauss locations:20
y
x
0
1
2
3
4
5
0
5
10
15
20
25

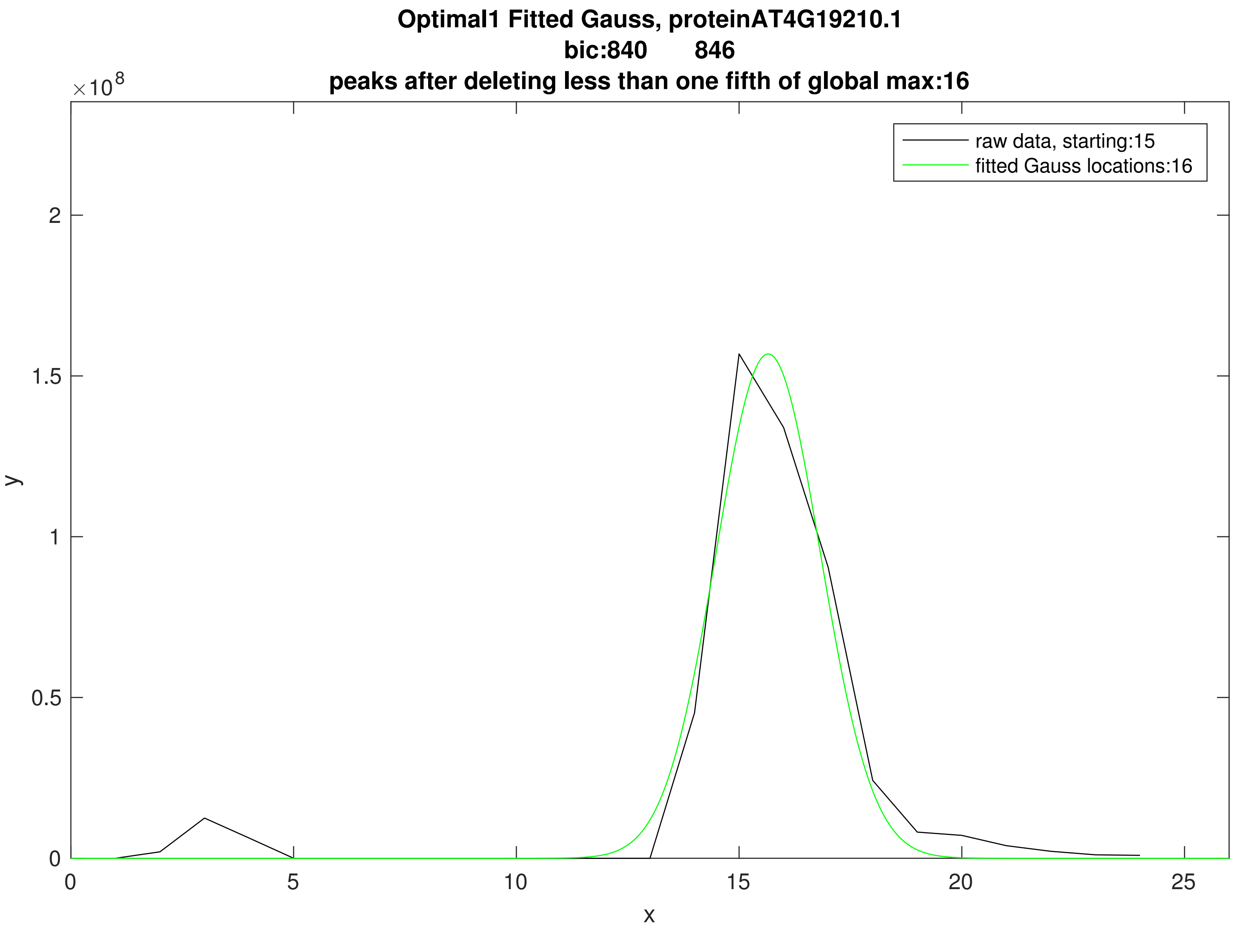

Optimal1 Fitted Gauss, proteinAT4G19210.1
bic:840 846
peaks after deleting less than one fifth of global max:16
×10^8
raw data, starting:15
fitted Gauss locations:16
y
x
0
0.5
1
1.5
2
0
5
10
15
20
25

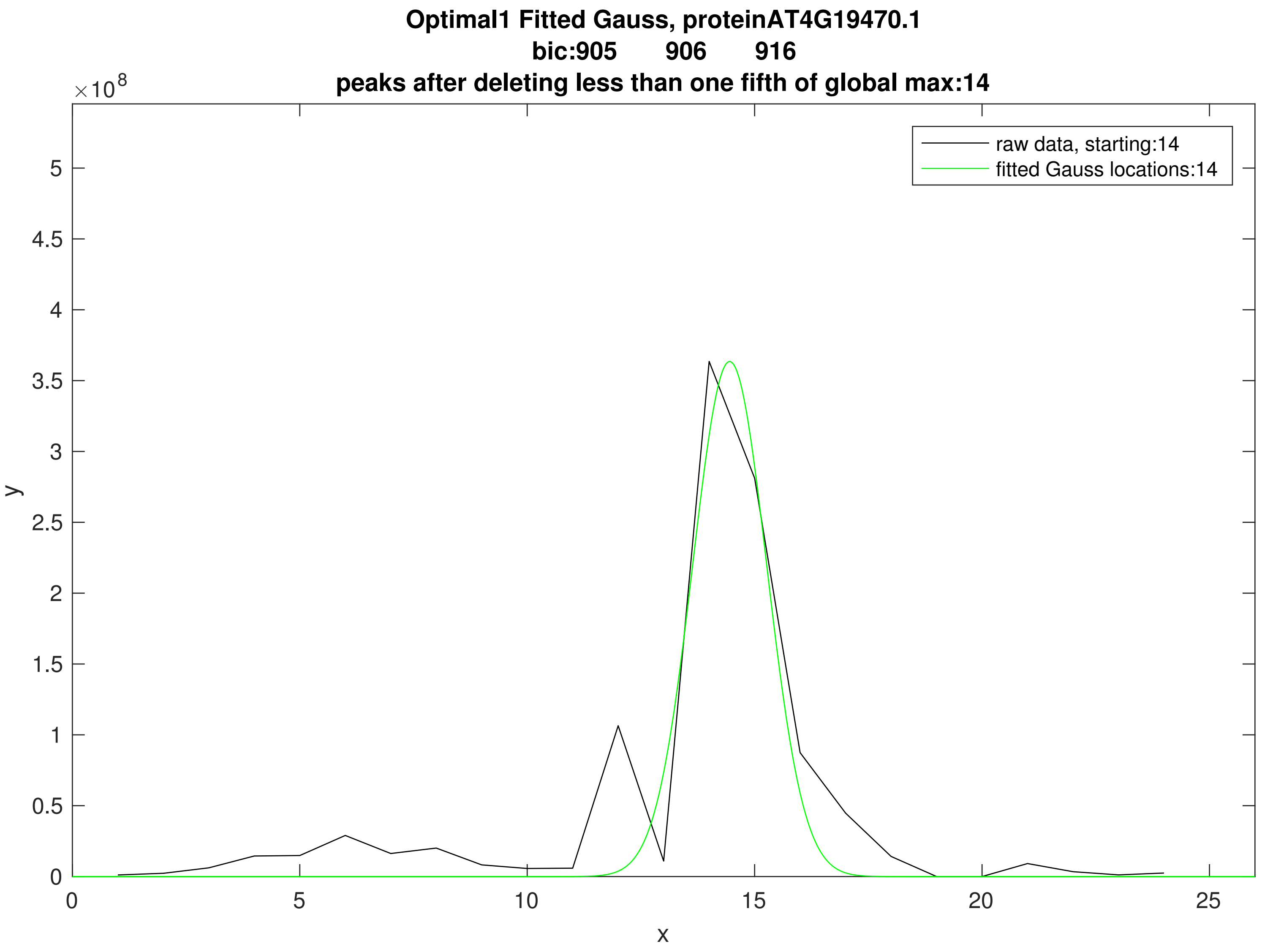

Optimal1 Fitted Gauss, proteinAT4G19470.1
bic:905 906 916
peaks after deleting less than one fifth of global max:14
×10^8
raw data, starting:14
fitted Gauss locations:14
y
x
0
0.5
1
1.5
2
2.5
3
3.5
4
4.5
5
0
5
10
15
20
25

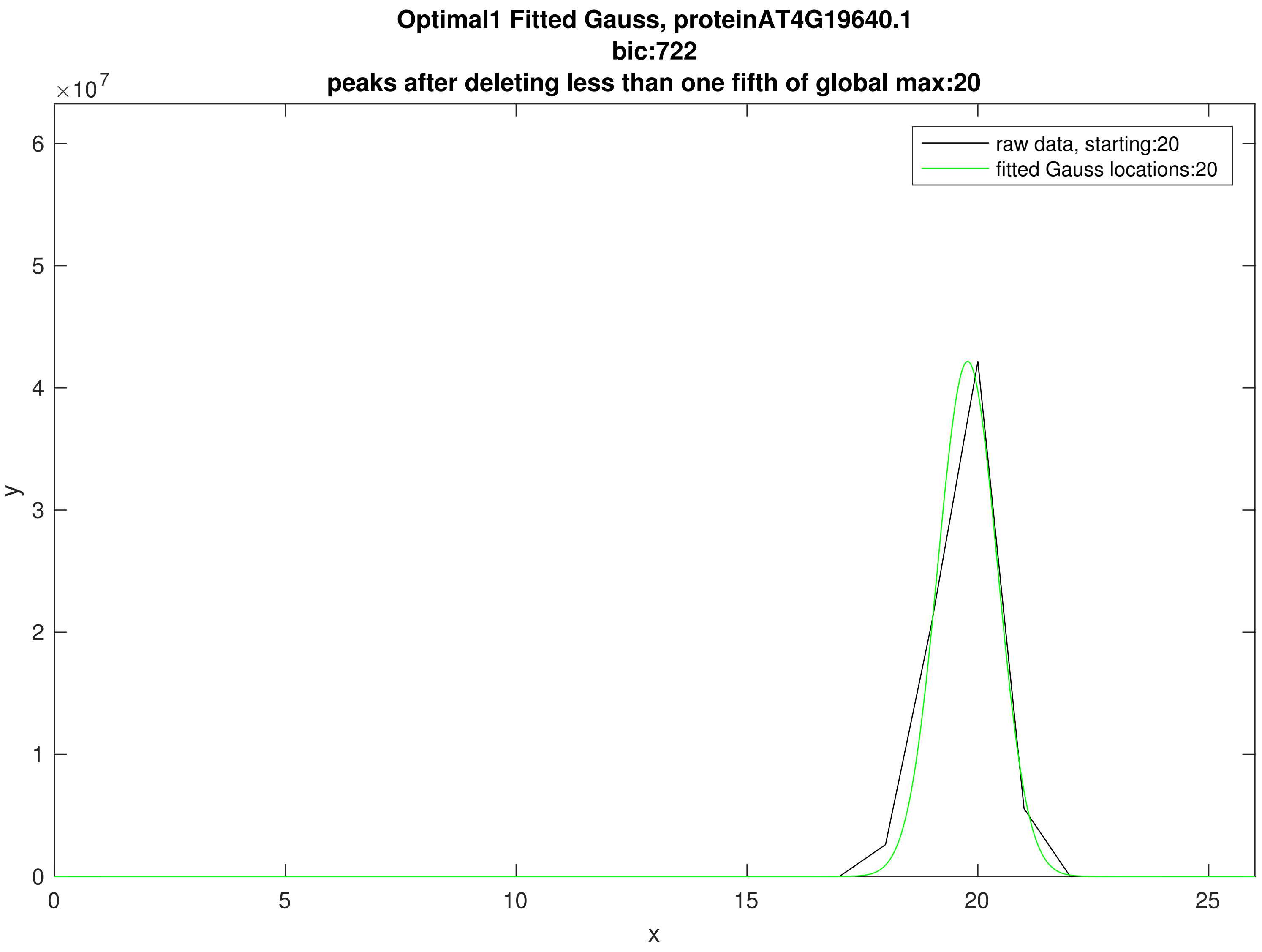
Optimal1 Fitted Gauss, proteinAT4G19640.1
bic:722
peaks after deleting less than one fifth of global max:20
×10^7
0
1
2
3
4
5
6
y
0
5
10
15
20
25
x
raw data, starting:20
fitted Gauss locations:20

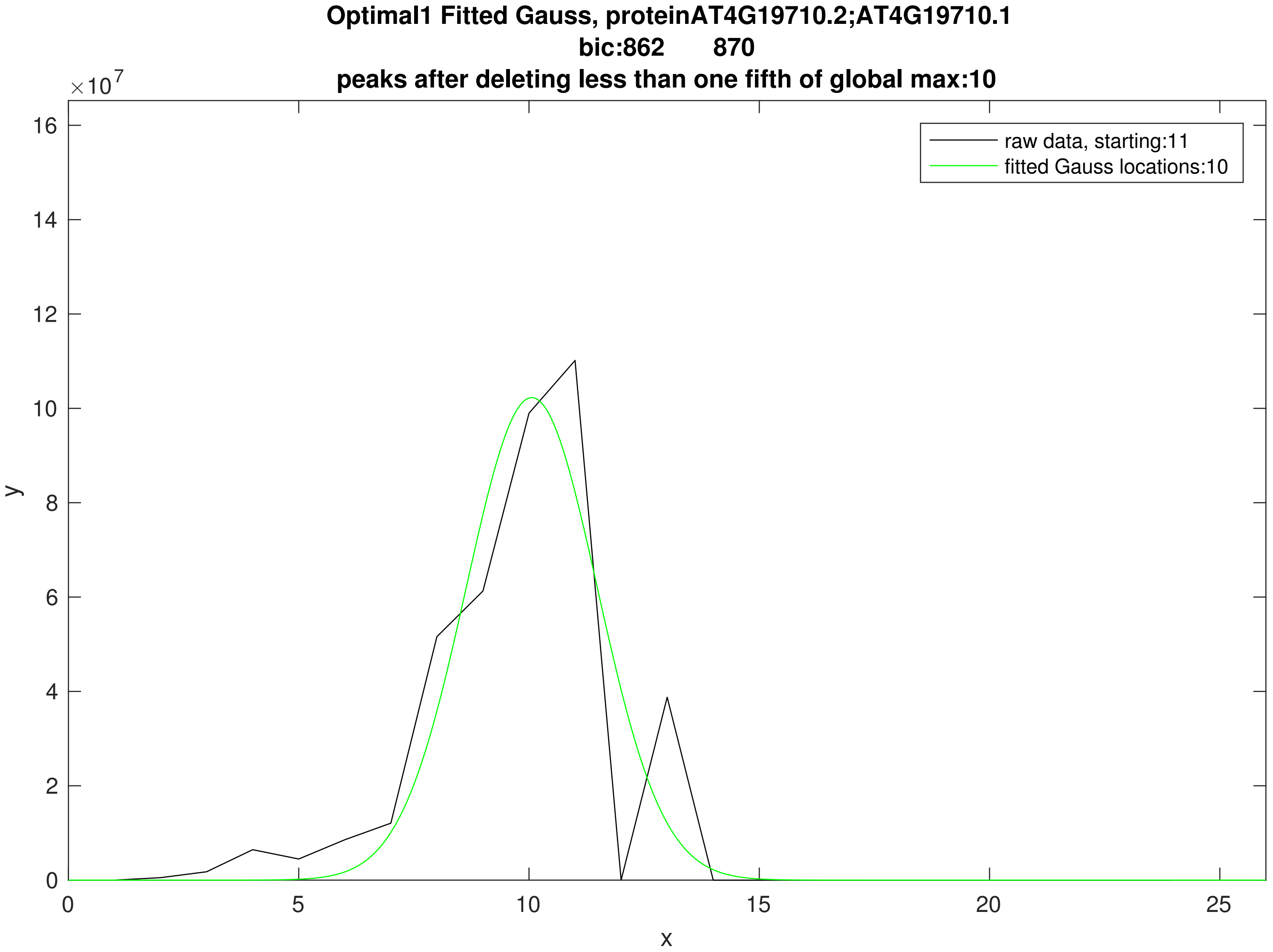

Optimal1 Fitted Gauss, proteinAT4G19710.2;AT4G19710.1
bic:862 870
peaks after deleting less than one fifth of global max:10
×10^7
raw data, starting:11
fitted Gauss locations:10
y
x
0
2
4
6
8
10
12
14
16
0
5
10
15
20
25

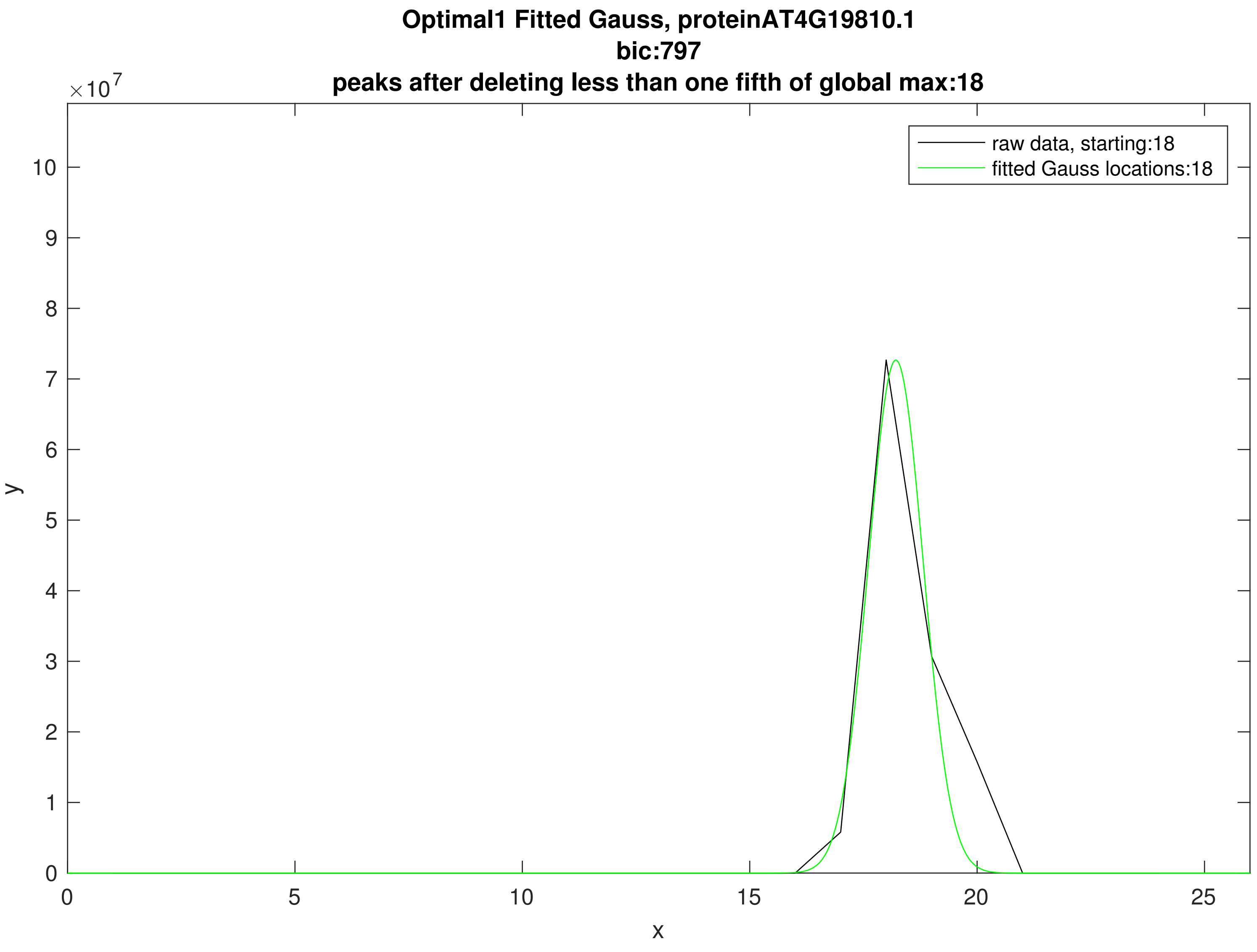

Optimal1 Fitted Gauss, proteinAT4G19810.1
bic:797
peaks after deleting less than one fifth of global max:18
×10^7
raw data, starting:18
fitted Gauss locations:18
y
x
0
1
2
3
4
5
6
7
8
9
10
0
5
10
15
20
25

Optimal1 Fitted Gauss, proteinAT4G19880.1;AT4G19880.2;AT4G19880.3
bic:1.02e+03  1.03e+03
peaks after deleting less than one fifth of global max:14

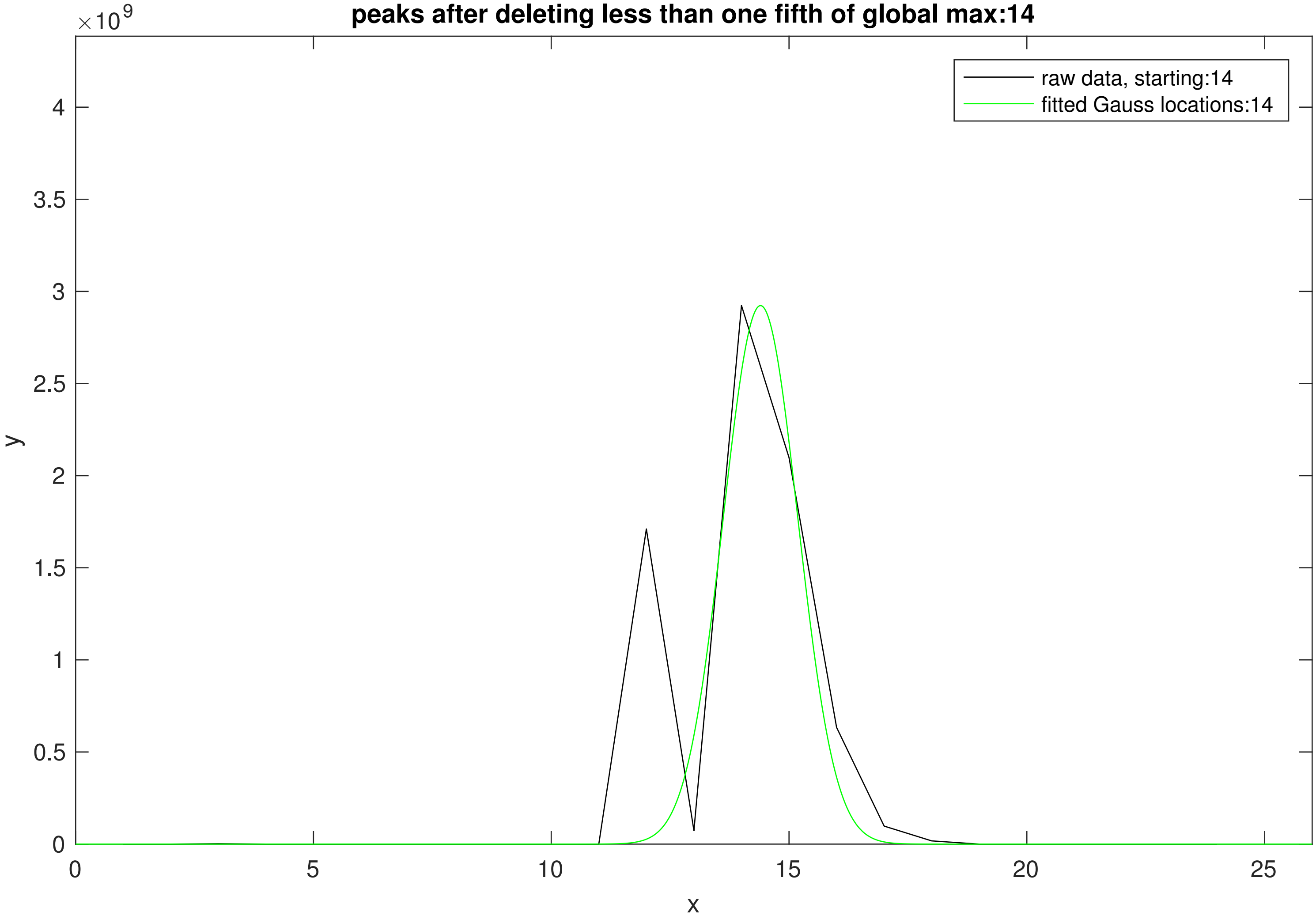

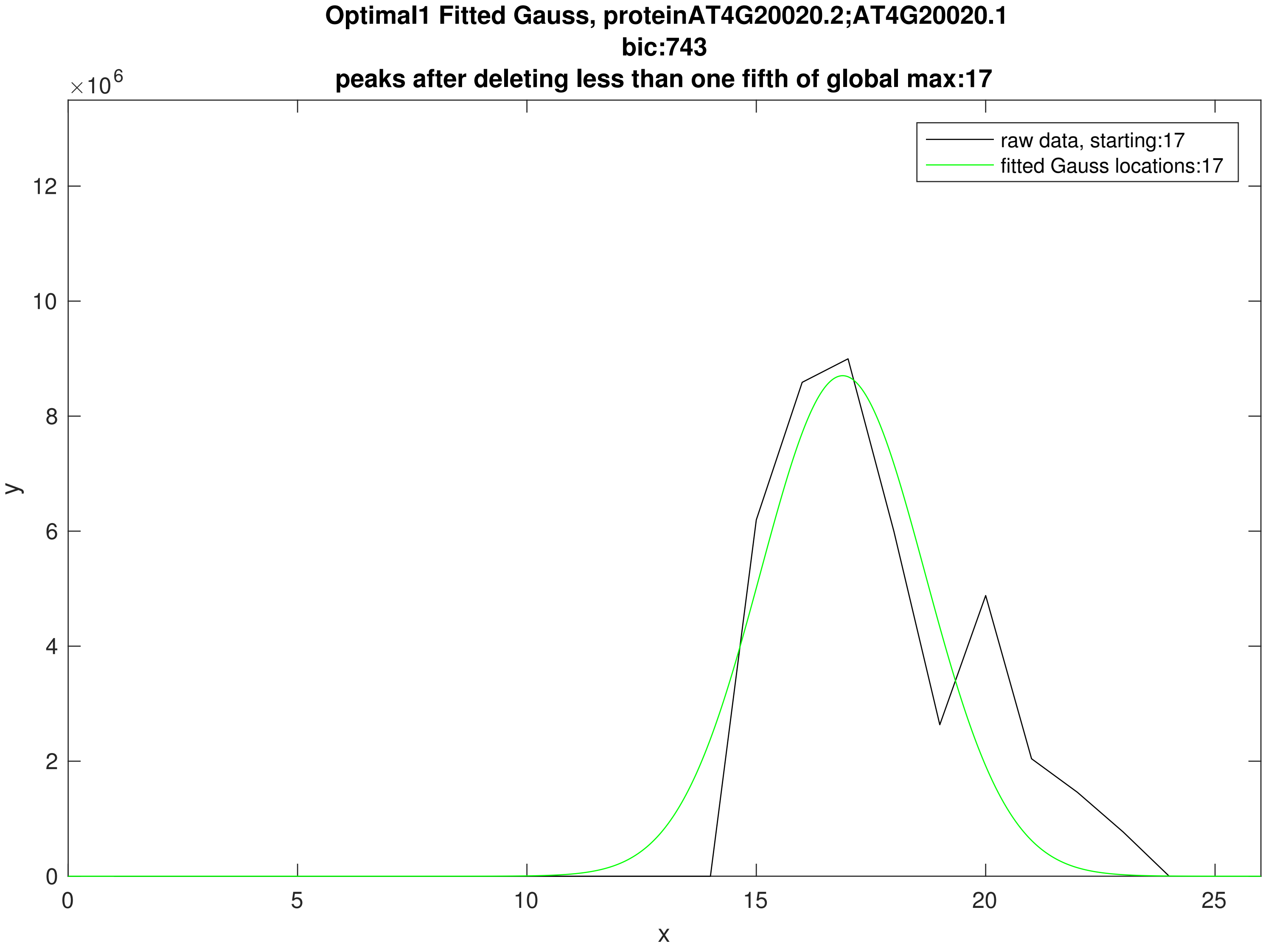

Optimal1 Fitted Gauss, proteinAT4G20020.2;AT4G20020.1
bic:743
peaks after deleting less than one fifth of global max:17
×10^6
raw data, starting:17
fitted Gauss locations:17
y
x
0
2
4
6
8
10
12
0
5
10
15
20
25

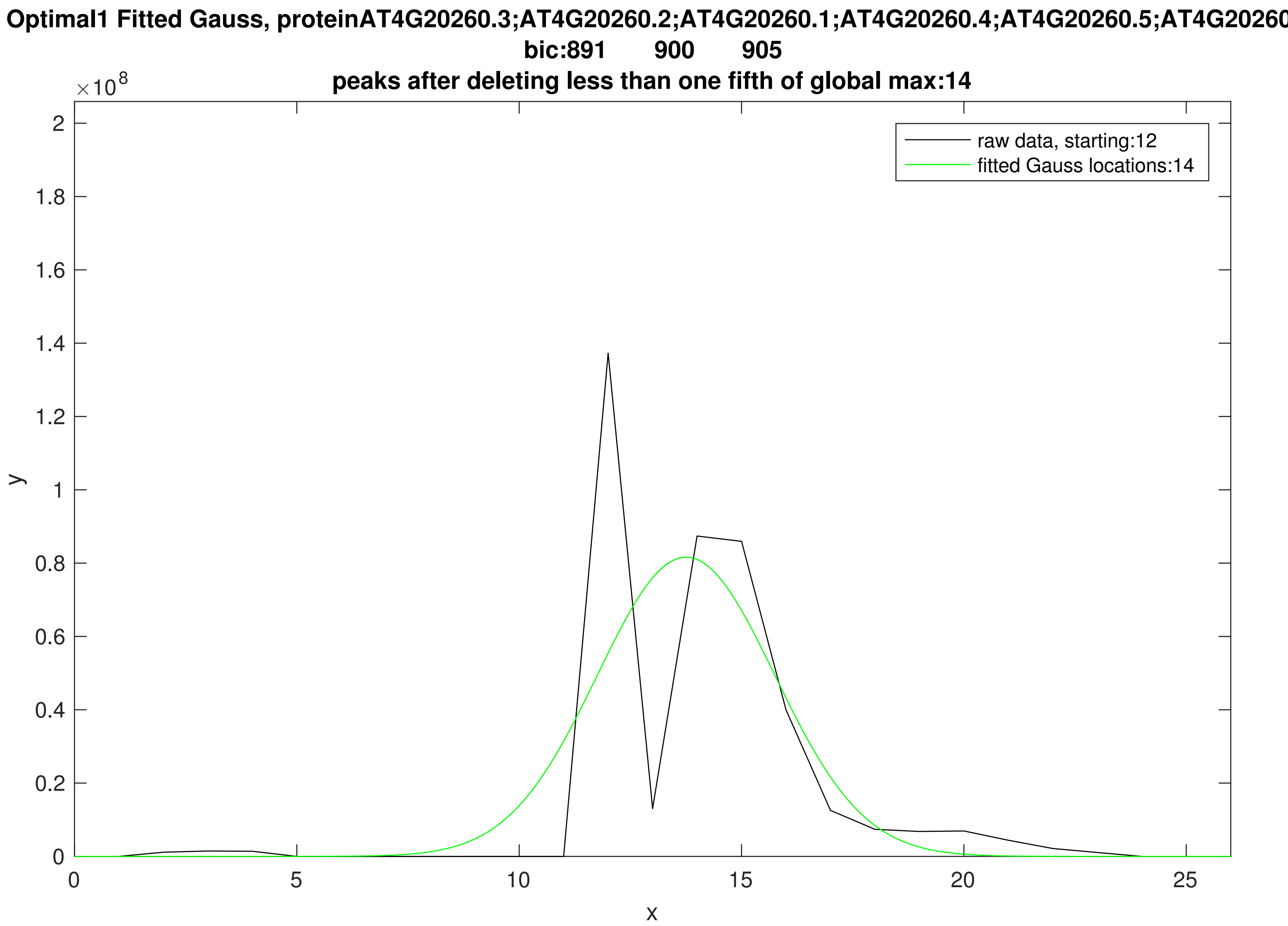

Optimal1 Fitted Gauss, proteinAT4G20260.3;AT4G20260.2;AT4G20260.1;AT4G20260.4;AT4G20260.5;AT4G20260
bic:891 900 905
peaks after deleting less than one fifth of global max:14
×10^8
raw data, starting:12
fitted Gauss locations:14
y
x
0
0.2
0.4
0.6
0.8
1
1.2
1.4
1.6
1.8
2
5
10
15
20
25

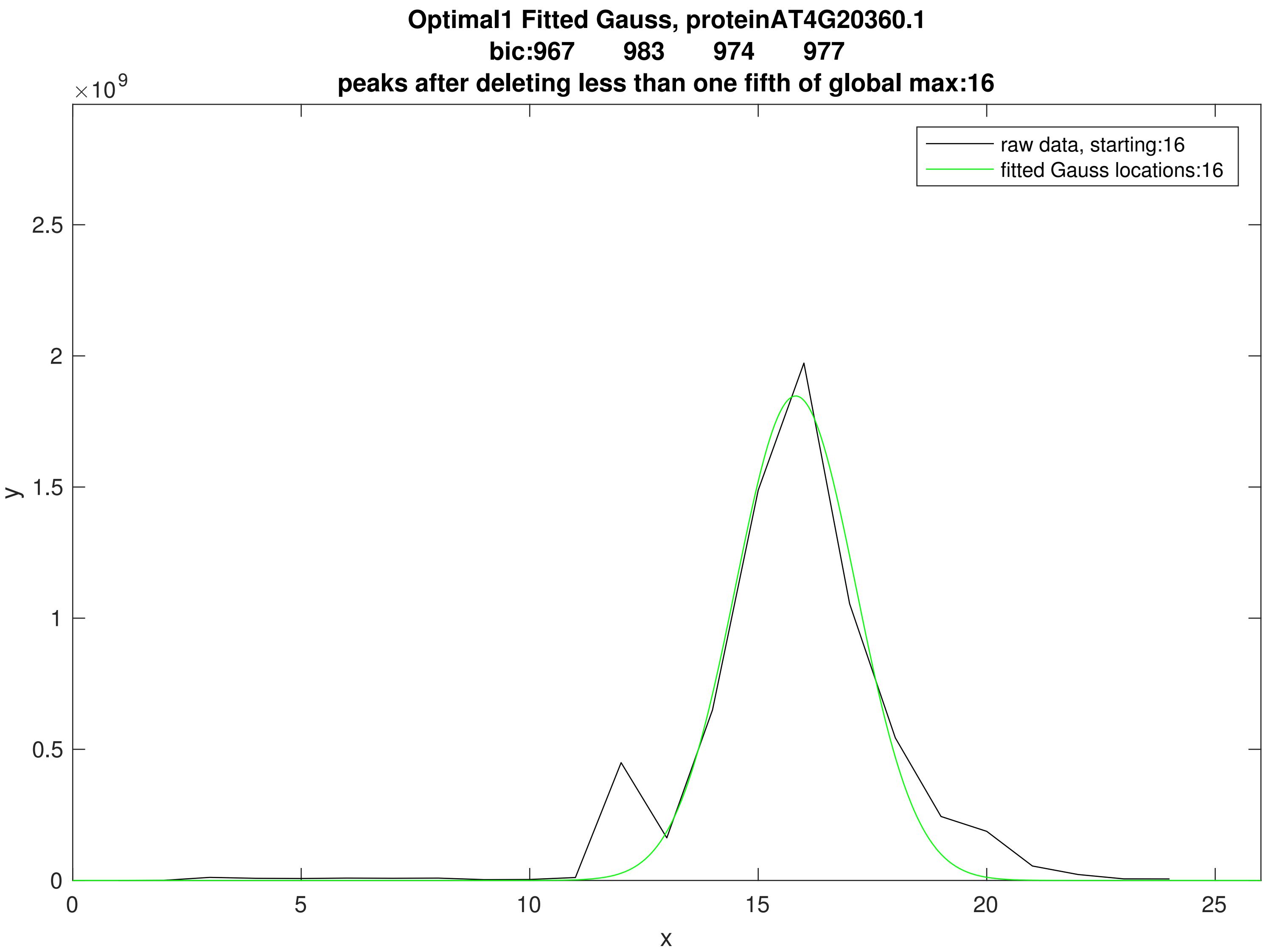
Optimal1 Fitted Gauss, proteinAT4G20360.1
bic:967 983 974 977
peaks after deleting less than one fifth of global max:16
×10^9
raw data, starting:16
fitted Gauss locations:16
y
x
0
0.5
1
1.5
2
2.5
0
5
10
15
20
25

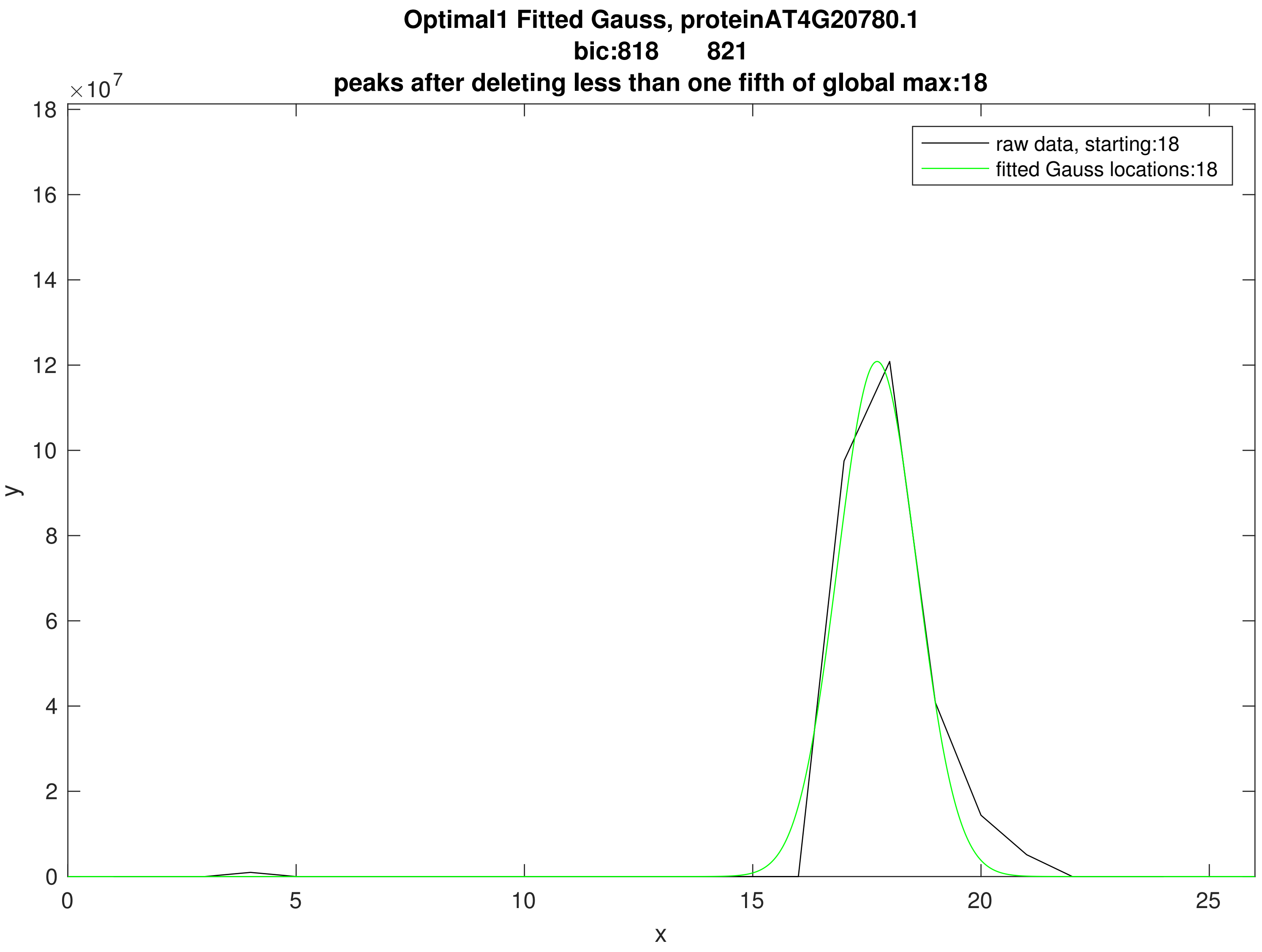

Optimal1 Fitted Gauss, proteinAT4G20780.1
bic:818 821
peaks after deleting less than one fifth of global max:18
×10^7
raw data, starting:18
fitted Gauss locations:18
y
x
0
2
4
6
8
10
12
14
16
18
0
5
10
15
20
25

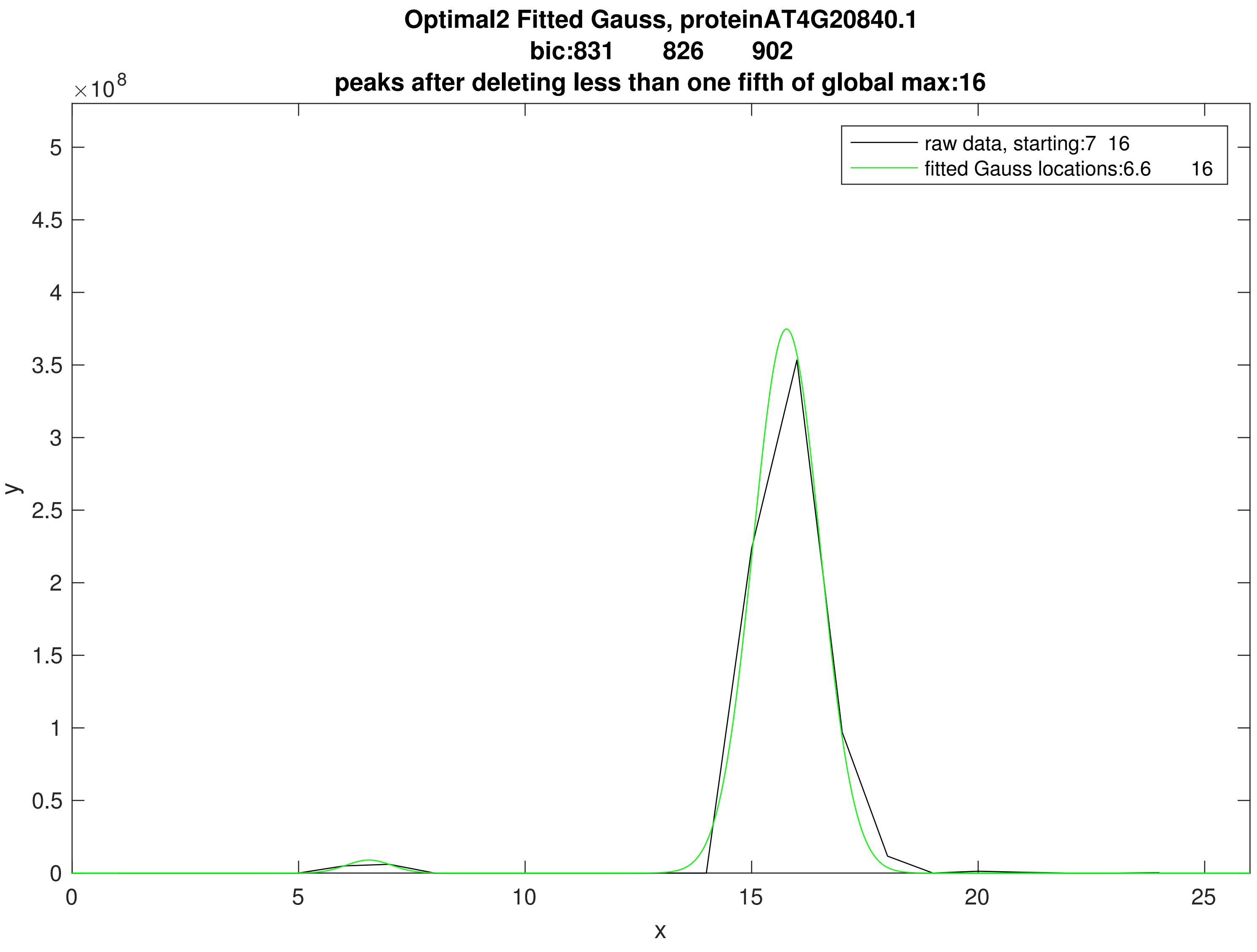
Optimal2 Fitted Gauss, proteinAT4G20840.1
bic:831 826 902
peaks after deleting less than one fifth of global max:16
×10^8
raw data, starting:7 16
fitted Gauss locations:6.6 16
y
x
0
0.5
1
1.5
2
2.5
3
3.5
4
4.5
5
0
5
10
15
20
25

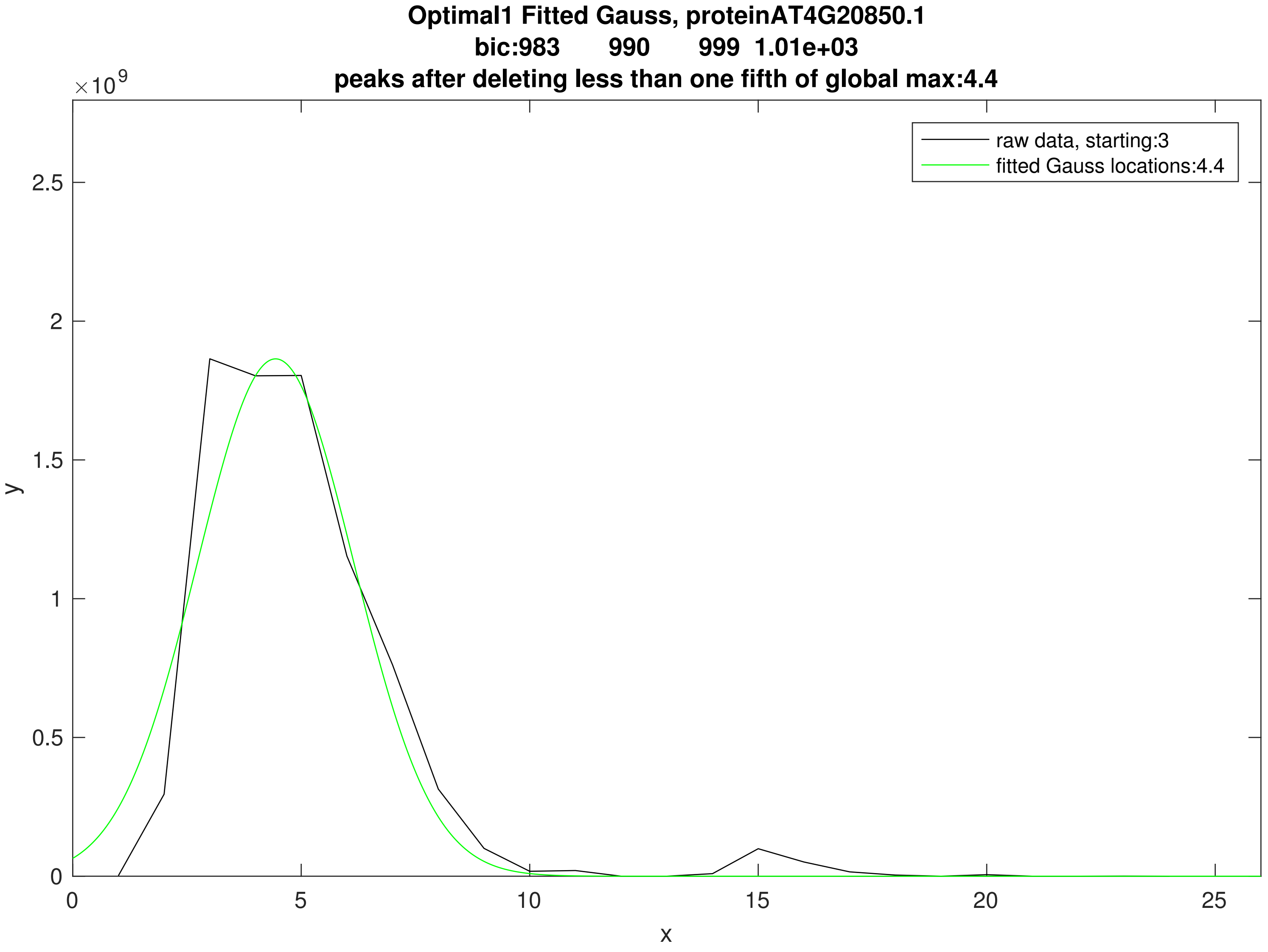
Optimal1 Fitted Gauss, proteinAT4G20850.1
bic:983 990 999 1.01e+03
peaks after deleting less than one fifth of global max:4.4
×10^9
raw data, starting:3
fitted Gauss locations:4.4
y
x
0
0.5
1
1.5
2
2.5
5
10
15
20
25

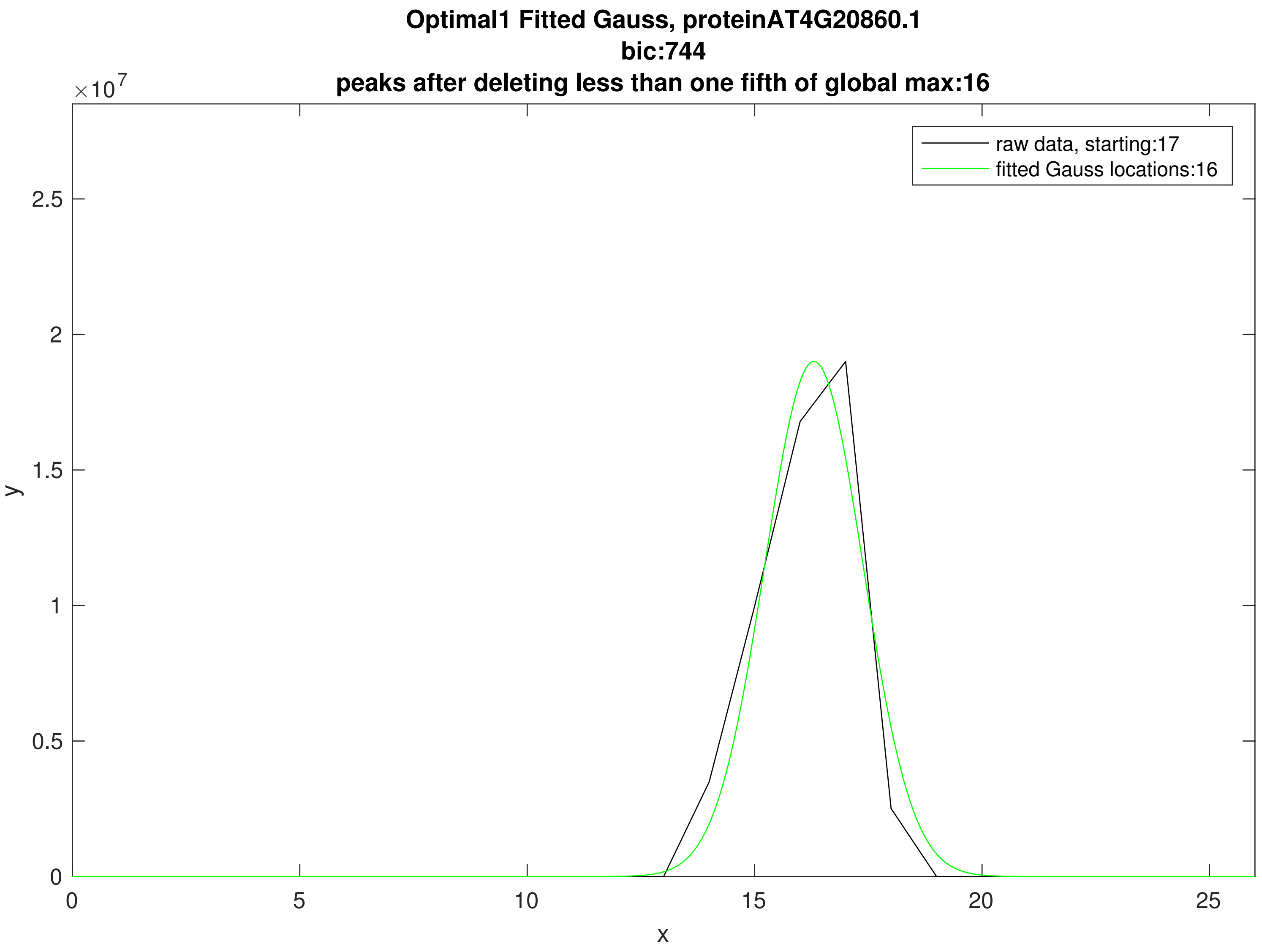
Optimal1 Fitted Gauss, proteinAT4G20860.1
bic:744
peaks after deleting less than one fifth of global max:16
×10^7
raw data, starting:17
fitted Gauss locations:16
y
x
0
0.5
1
1.5
2
2.5
0
5
10
15
20
25

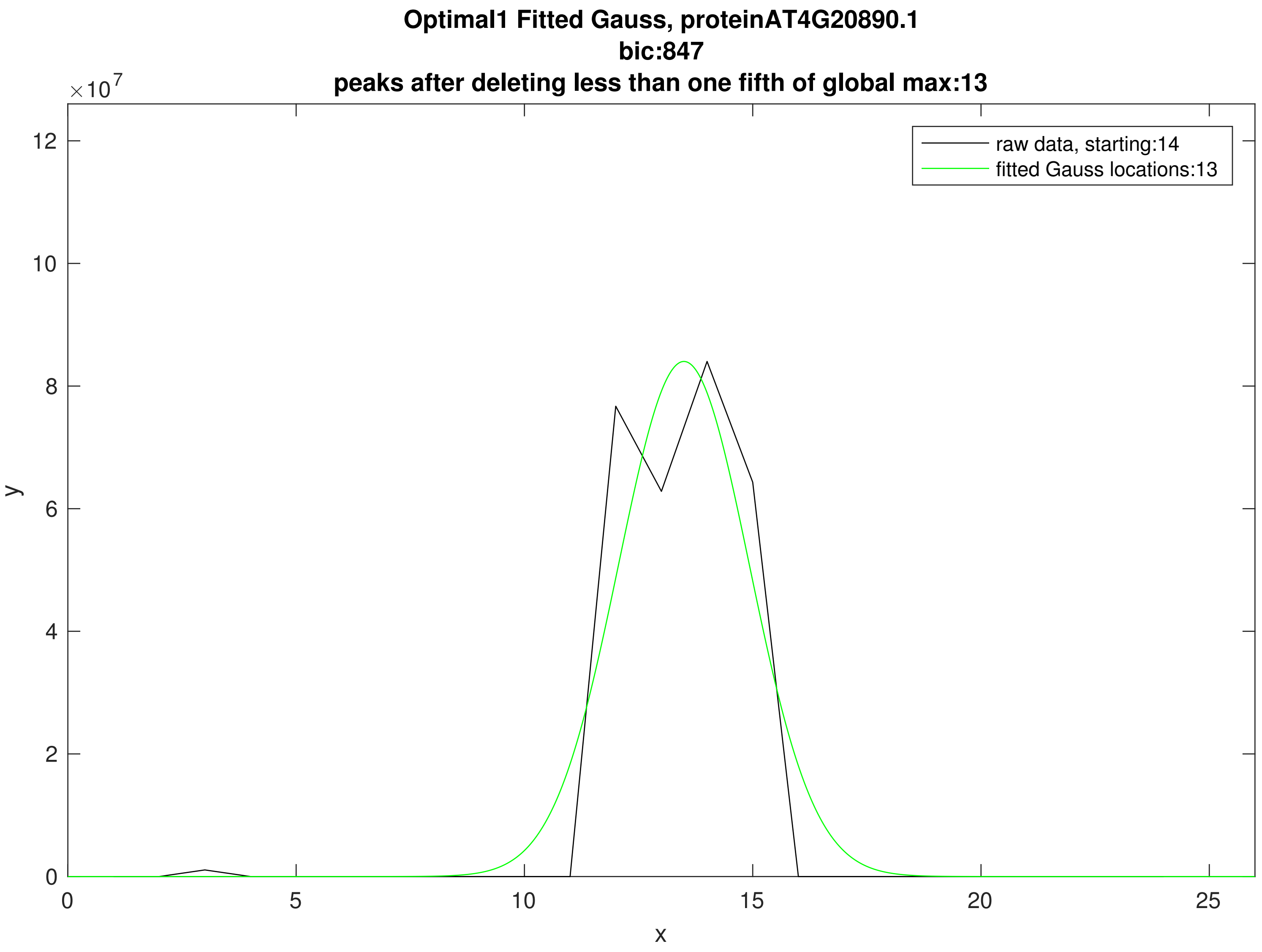
Optimal1 Fitted Gauss, proteinAT4G20890.1
bic:847
peaks after deleting less than one fifth of global max:13
×10^7
raw data, starting:14
fitted Gauss locations:13
y
x
0
2
4
6
8
10
12
0
5
10
15
20
25

**Optimal1 Fitted Gauss, proteinAT4G20960.1**
**bic:808**
**peaks after deleting less than one fifth of global max:13**

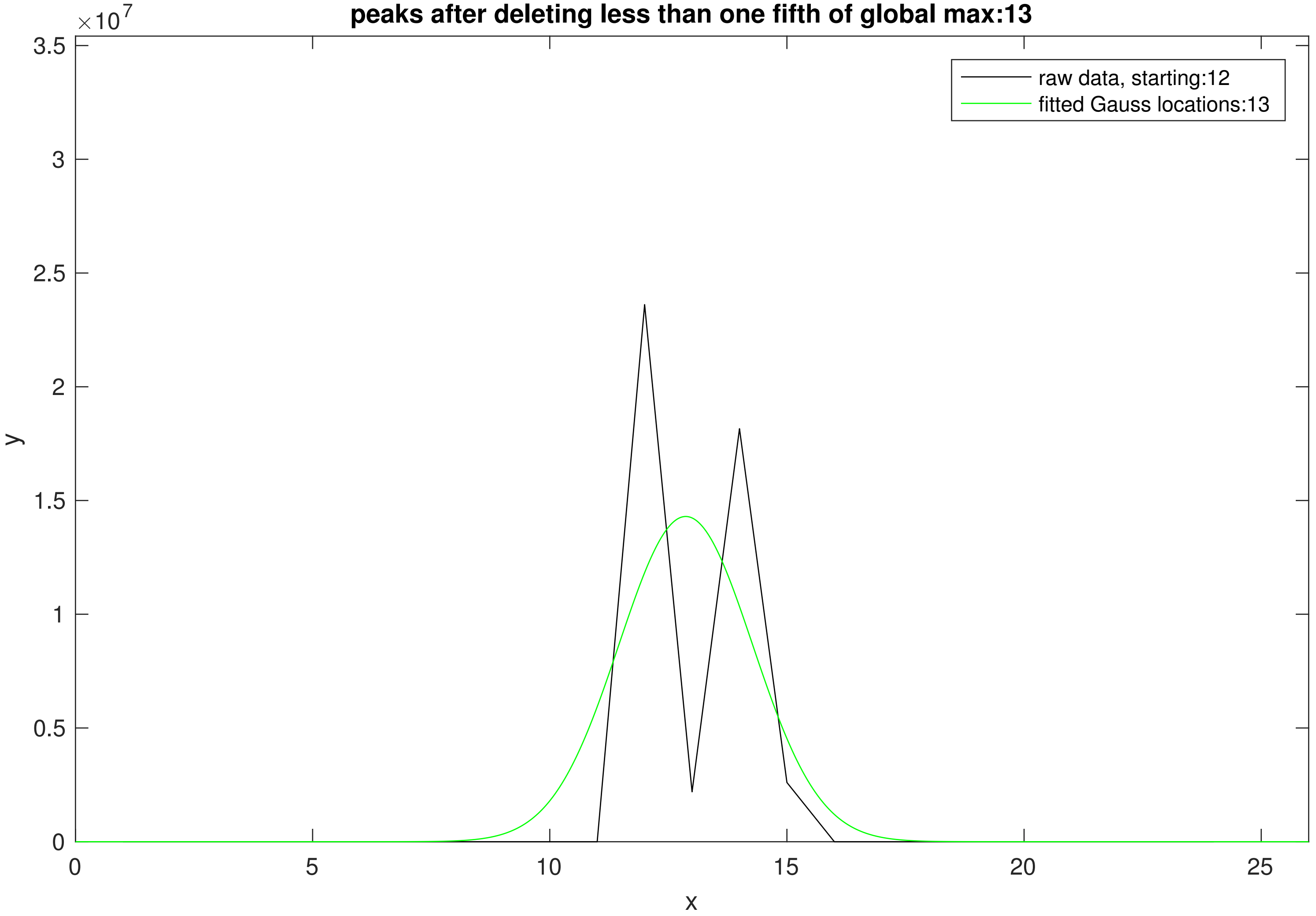

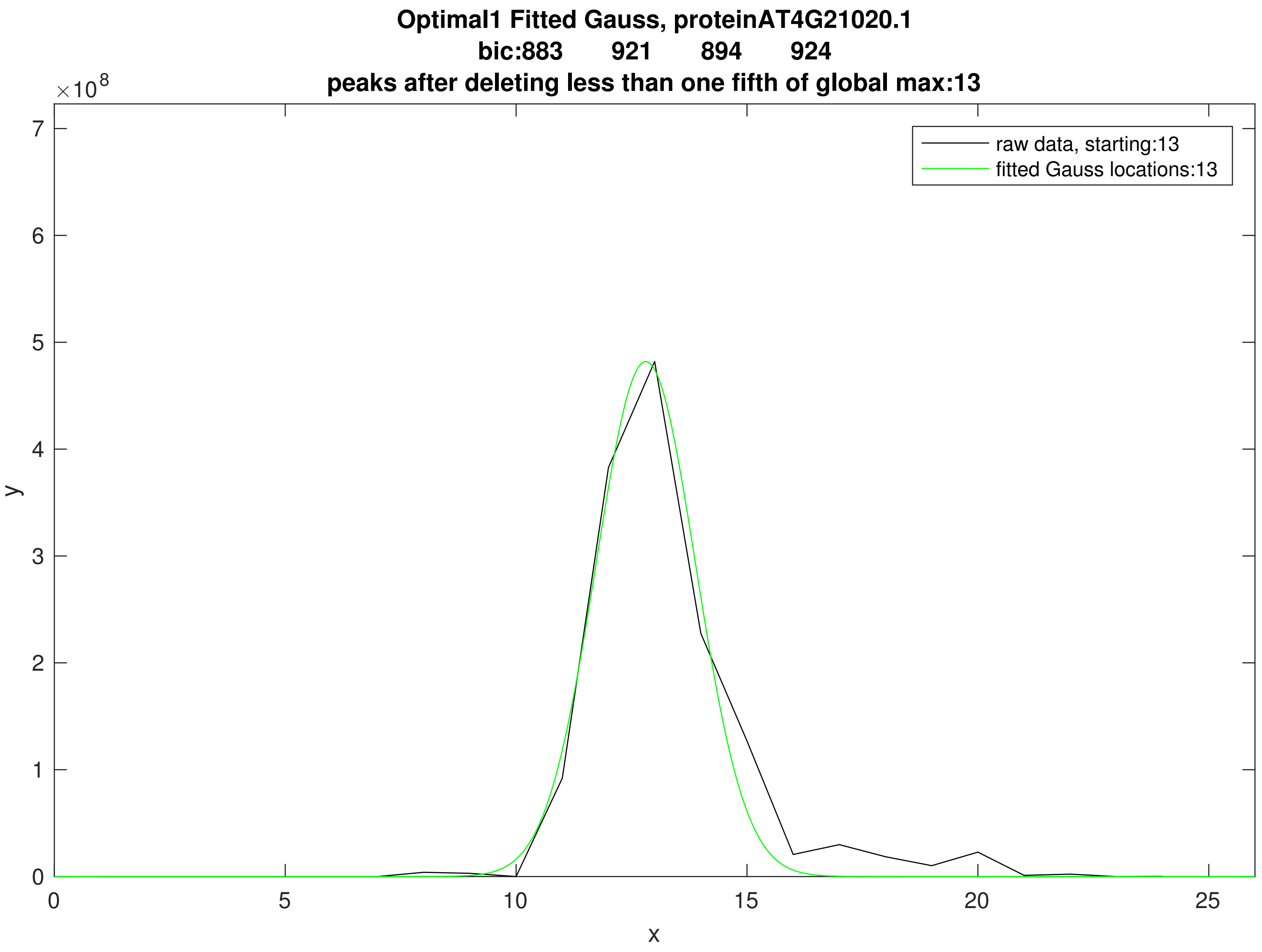

Optimal1 Fitted Gauss, proteinAT4G21020.1
bic:883 921 894 924
peaks after deleting less than one fifth of global max:13
×10^8
raw data, starting:13
fitted Gauss locations:13
y
x
0
1
2
3
4
5
6
7
0
5
10
15
20
25

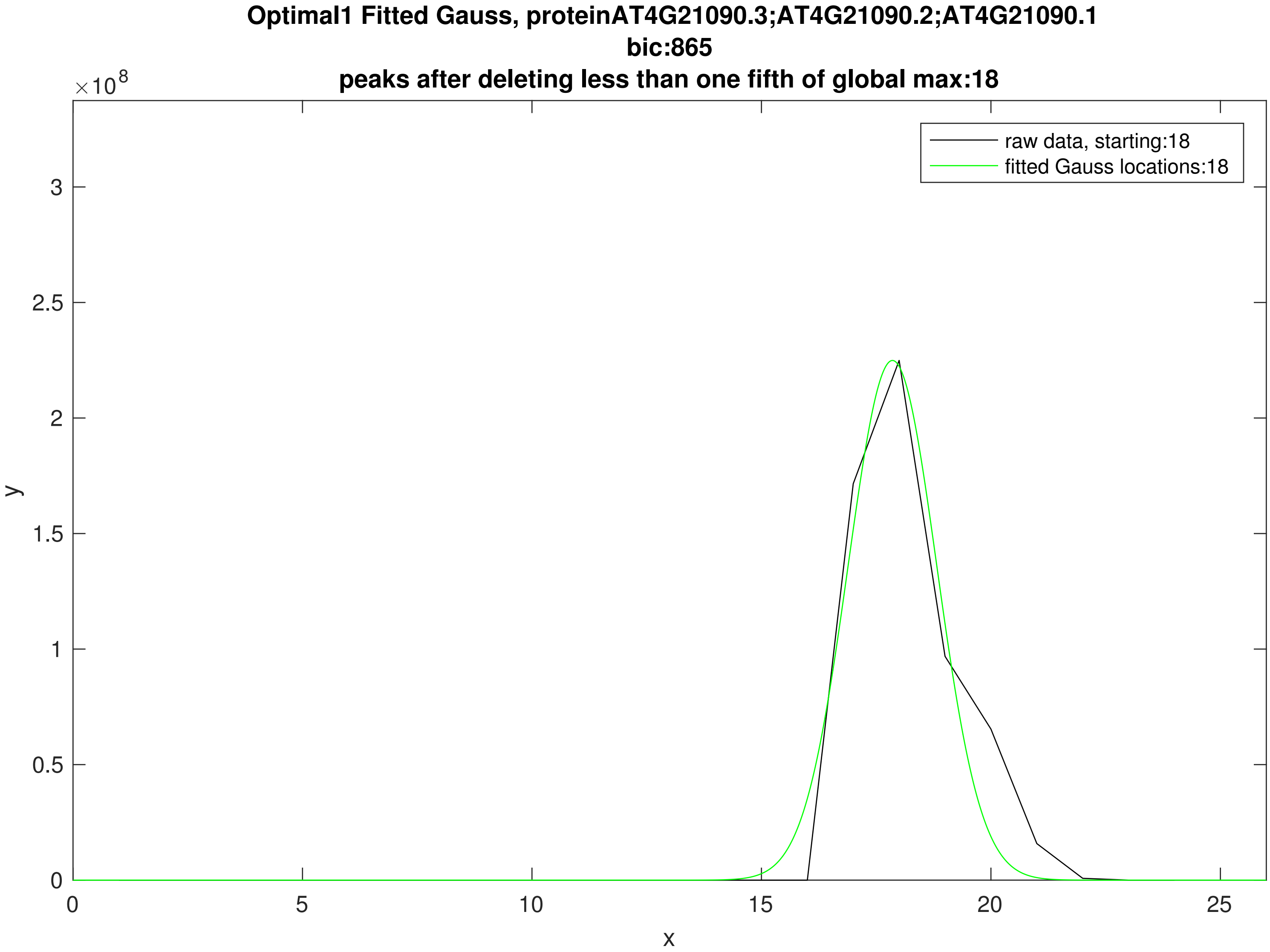

Optimal1 Fitted Gauss, proteinAT4G21090.3;AT4G21090.2;AT4G21090.1
bic:865
peaks after deleting less than one fifth of global max:18
×10^8
raw data, starting:18
fitted Gauss locations:18
3
2.5
2
1.5
1
0.5
0
y
0
5
10
15
20
25
x

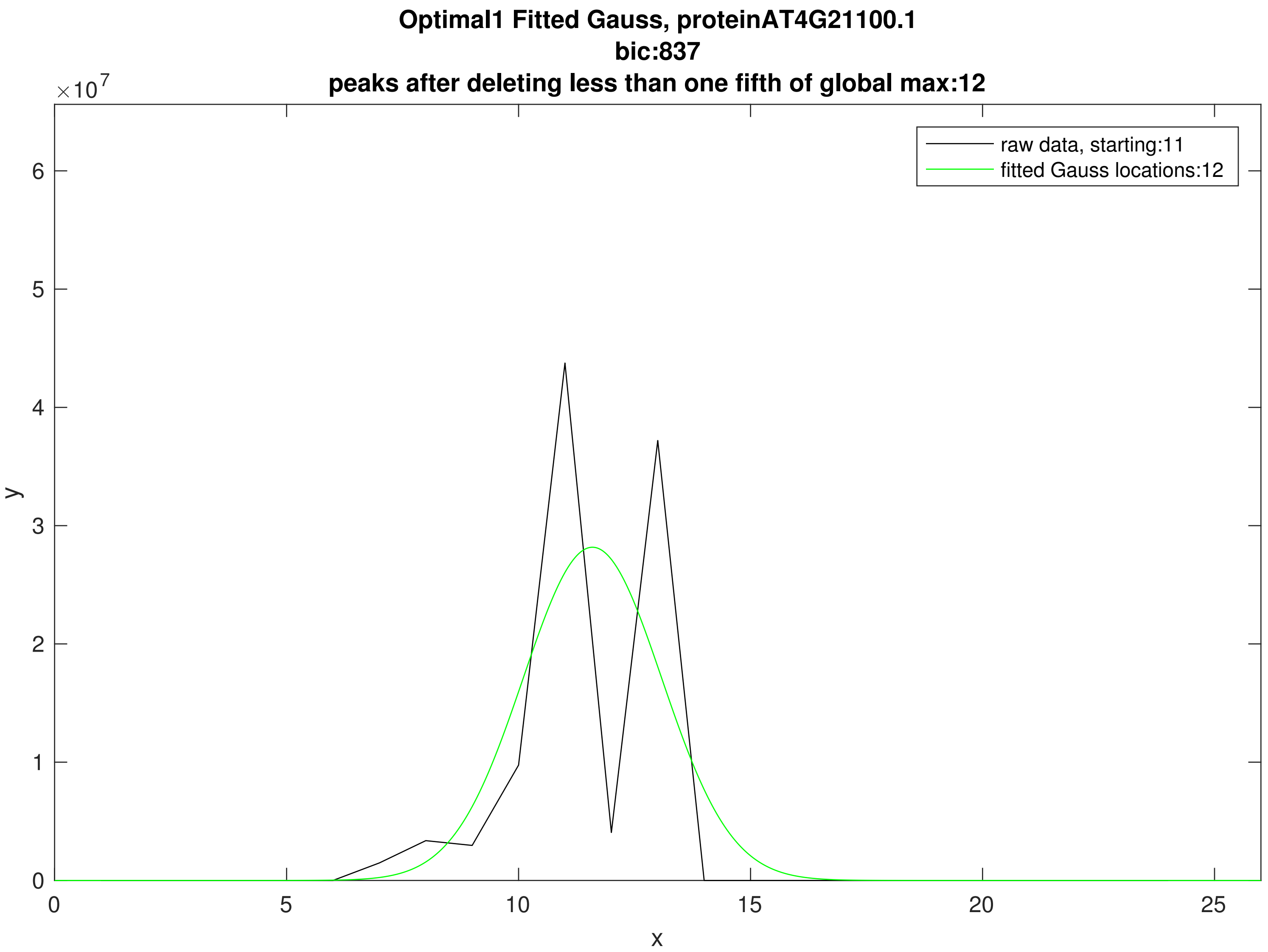

Optimal1 Fitted Gauss, proteinAT4G21100.1
bic:837
peaks after deleting less than one fifth of global max:12
×10^7
raw data, starting:11
fitted Gauss locations:12
6
5
4
3
2
1
0
y
0
5
10
15
20
25
x

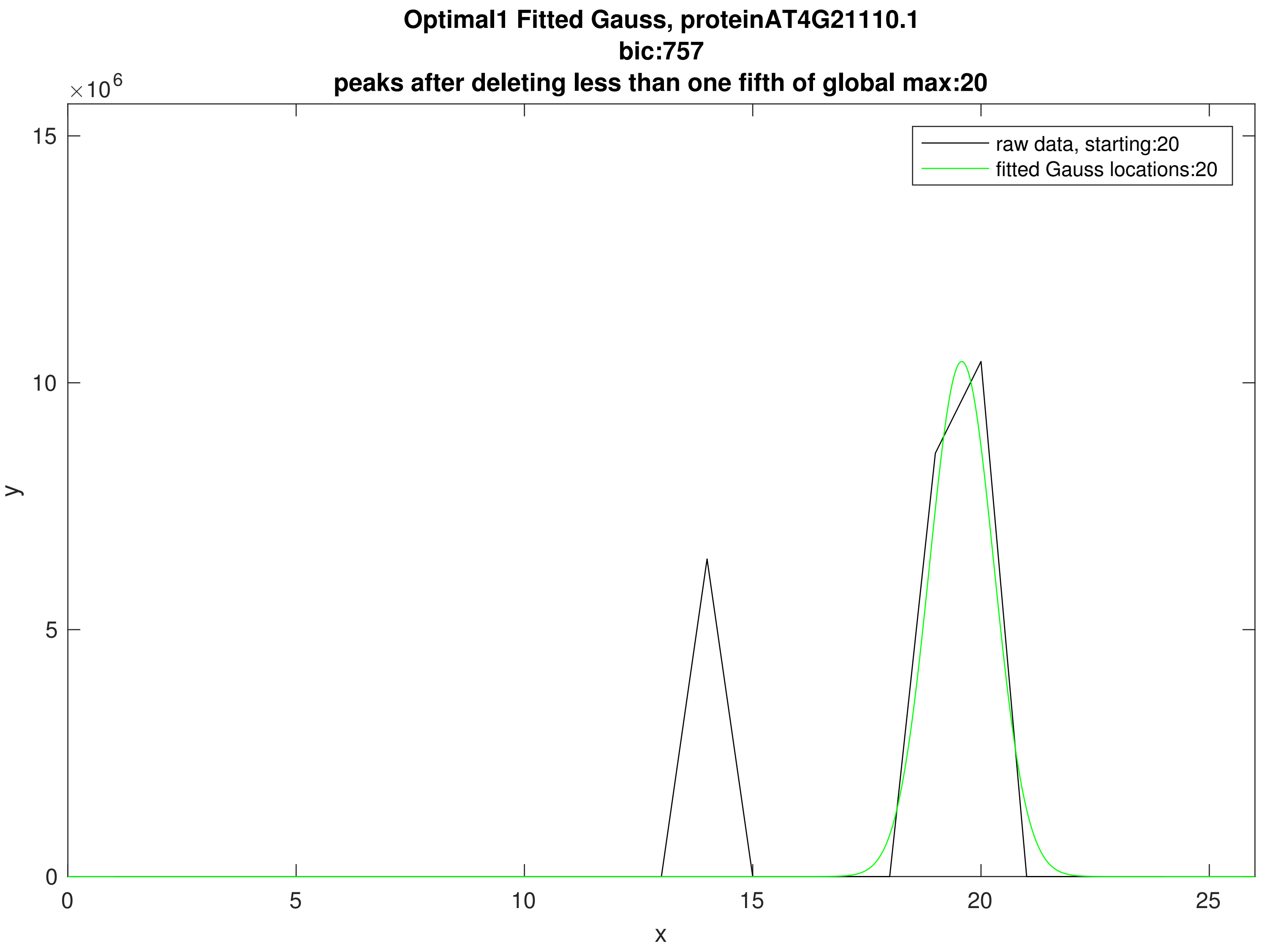
Optimal1 Fitted Gauss, proteinAT4G21110.1
bic:757
peaks after deleting less than one fifth of global max:20
×10^6
15
10
5
0
y
0
5
10
15
20
25
x
raw data, starting:20
fitted Gauss locations:20

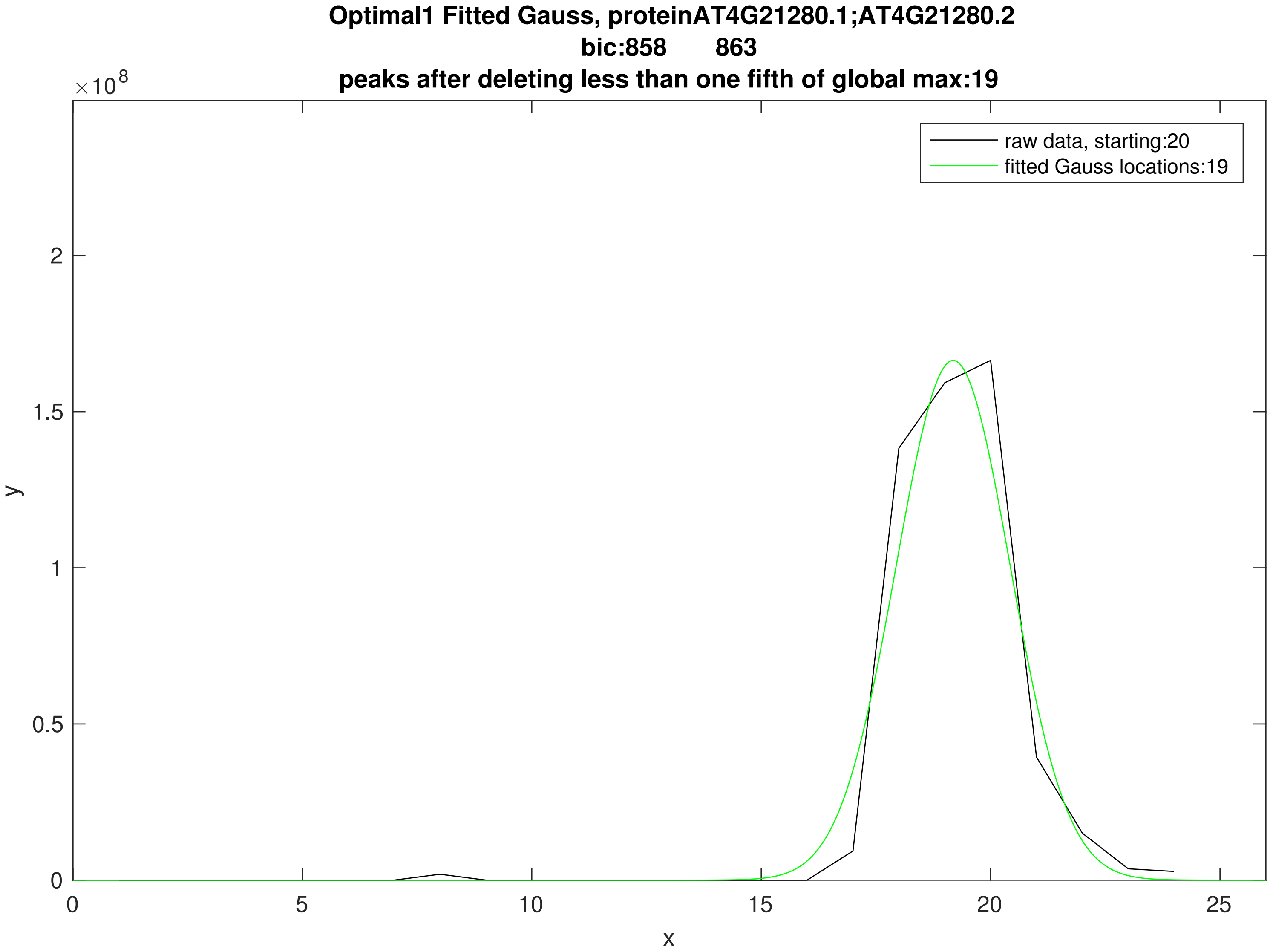
Optimal1 Fitted Gauss, proteinAT4G21280.1;AT4G21280.2
bic:858 863
peaks after deleting less than one fifth of global max:19
×10^8
raw data, starting:20
fitted Gauss locations:19
y
x
0
0.5
1
1.5
2
0
5
10
15
20
25

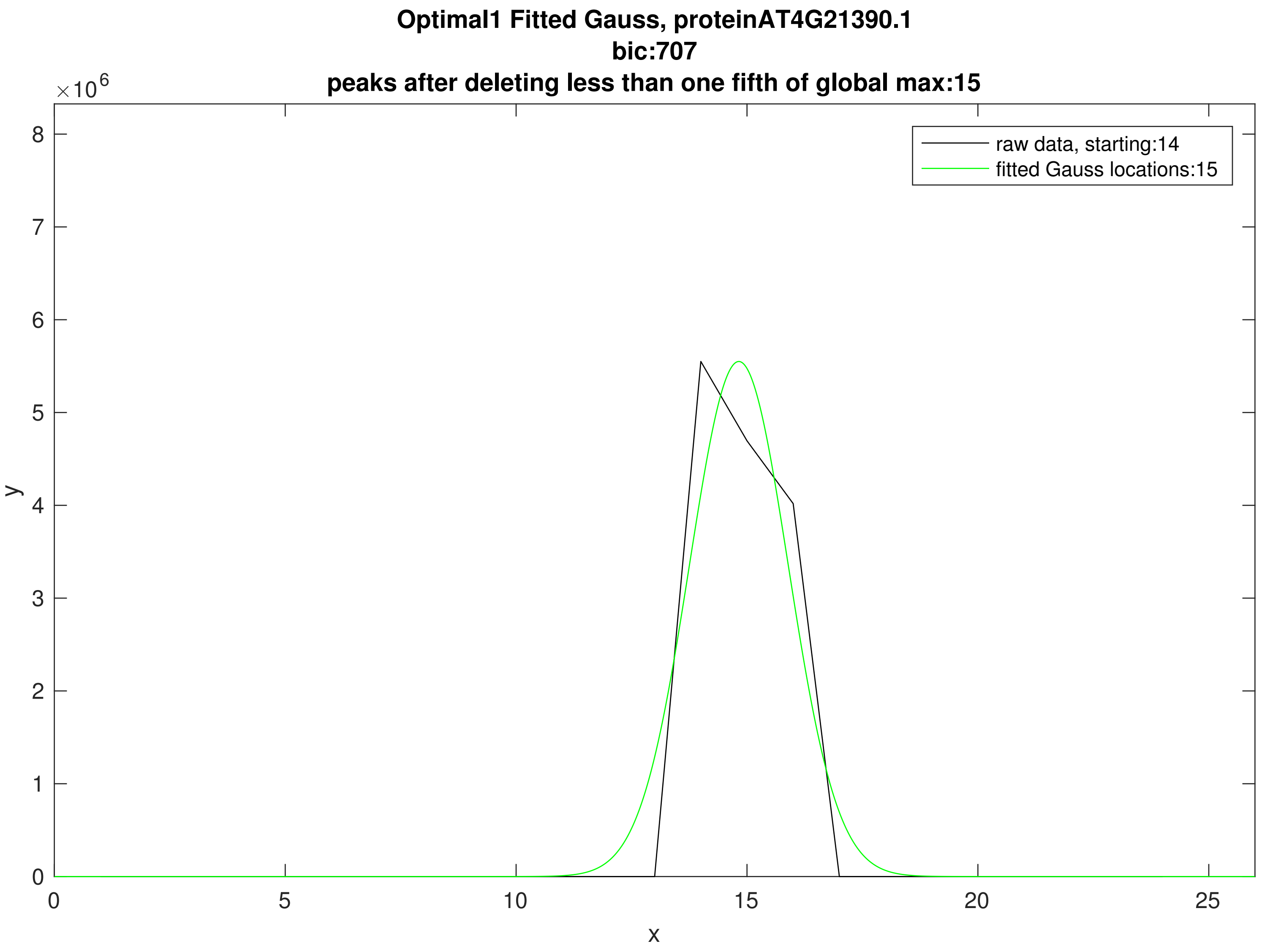
Optimal1 Fitted Gauss, proteinAT4G21390.1
bic:707
peaks after deleting less than one fifth of global max:15
×10^6
raw data, starting:14
fitted Gauss locations:15
y
x
0
1
2
3
4
5
6
7
8
0
5
10
15
20
25

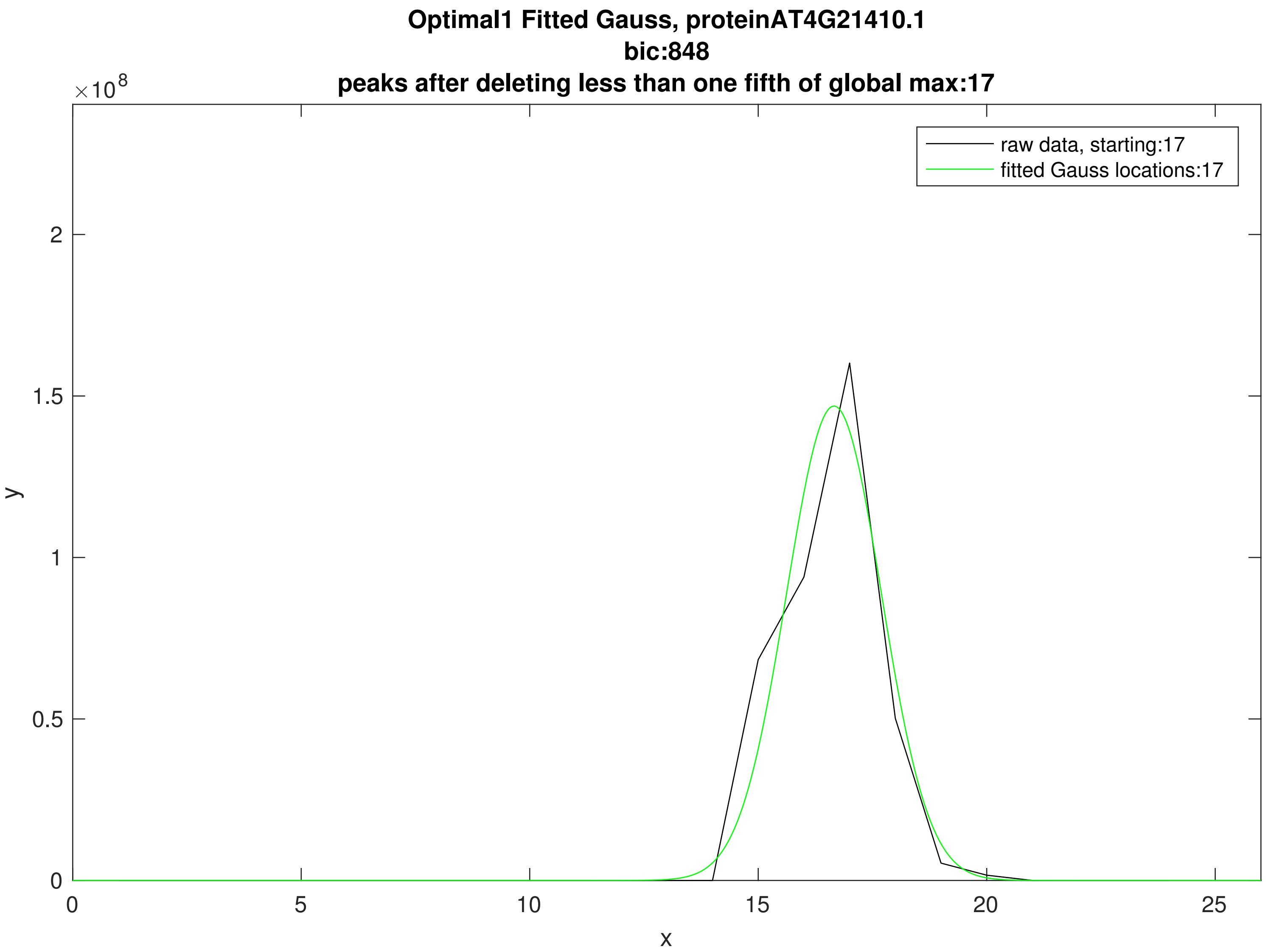

Optimal1 Fitted Gauss, proteinAT4G21410.1
bic:848
peaks after deleting less than one fifth of global max:17
×10^8
raw data, starting:17
fitted Gauss locations:17
2
1.5
1
0.5
0
y
0
5
10
15
20
25
x

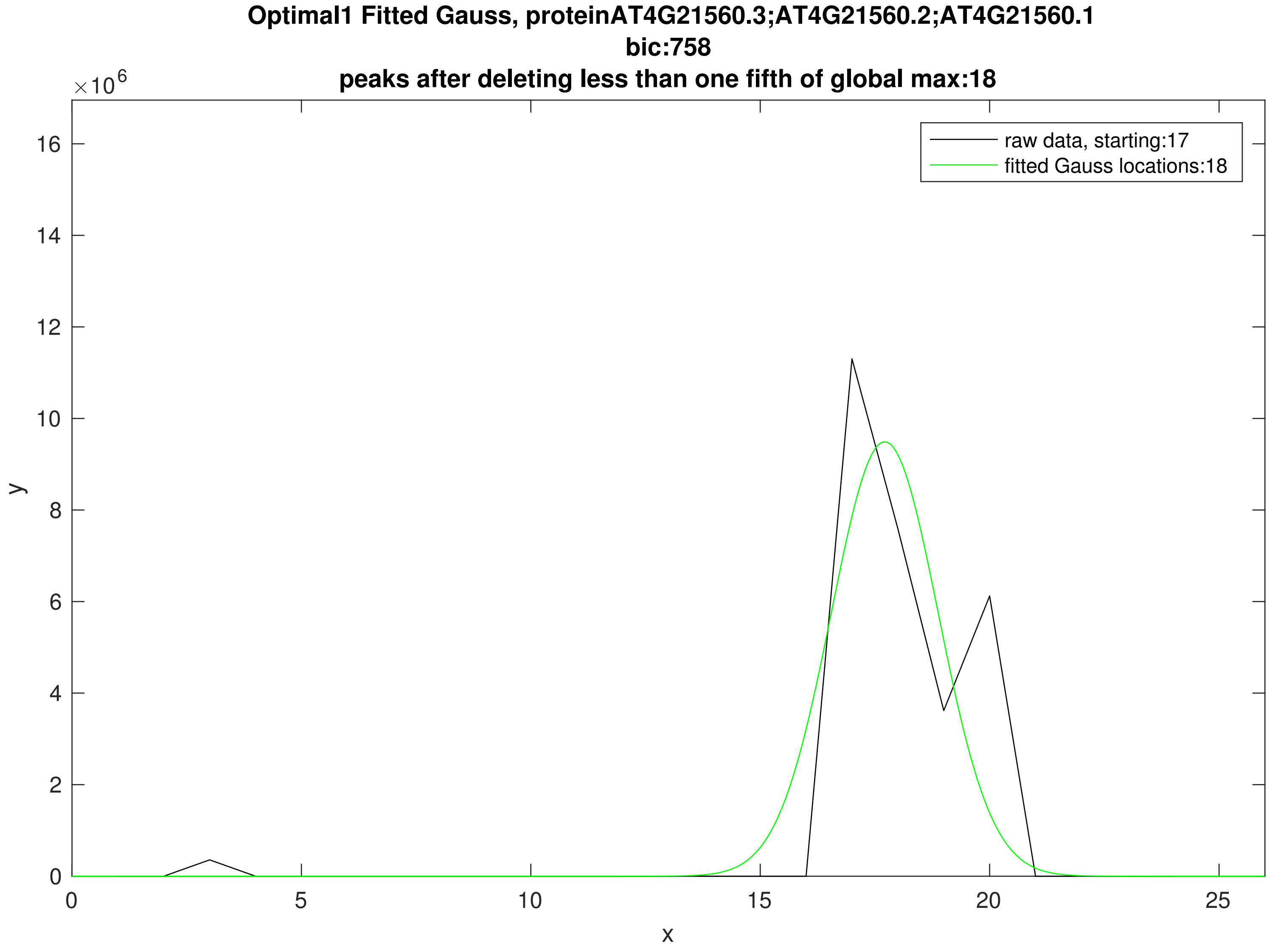
Optimal1 Fitted Gauss, proteinAT4G21560.3;AT4G21560.2;AT4G21560.1
bic:758
peaks after deleting less than one fifth of global max:18
×10^6
raw data, starting:17
fitted Gauss locations:18
y
x
0
2
4
6
8
10
12
14
16
0
5
10
15
20
25

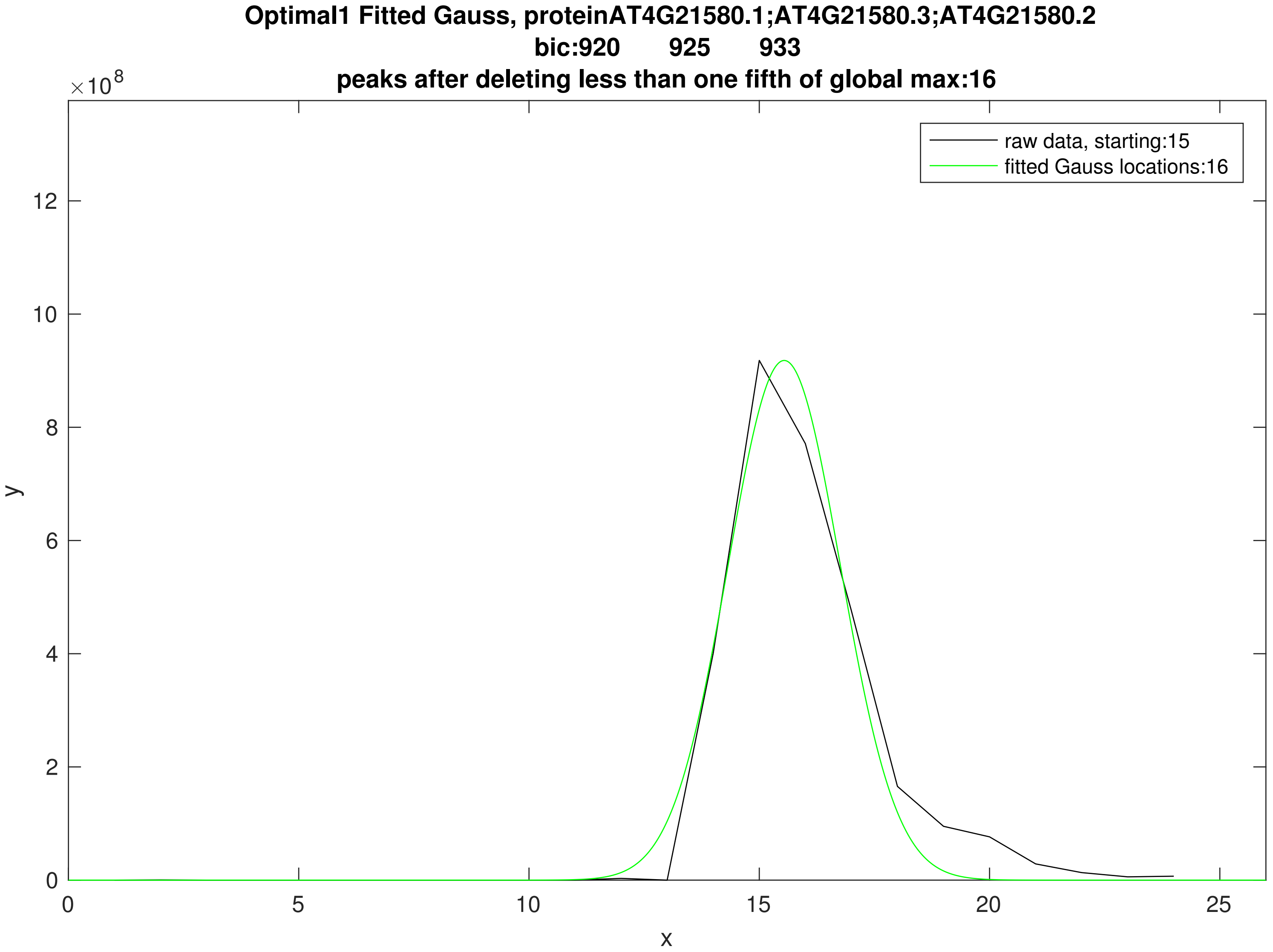

Optimal1 Fitted Gauss, proteinAT4G21580.1;AT4G21580.3;AT4G21580.2
bic:920 925 933
peaks after deleting less than one fifth of global max:16
×10^8
raw data, starting:15
fitted Gauss locations:16
y
x
0
2
4
6
8
10
12
0
5
10
15
20
25

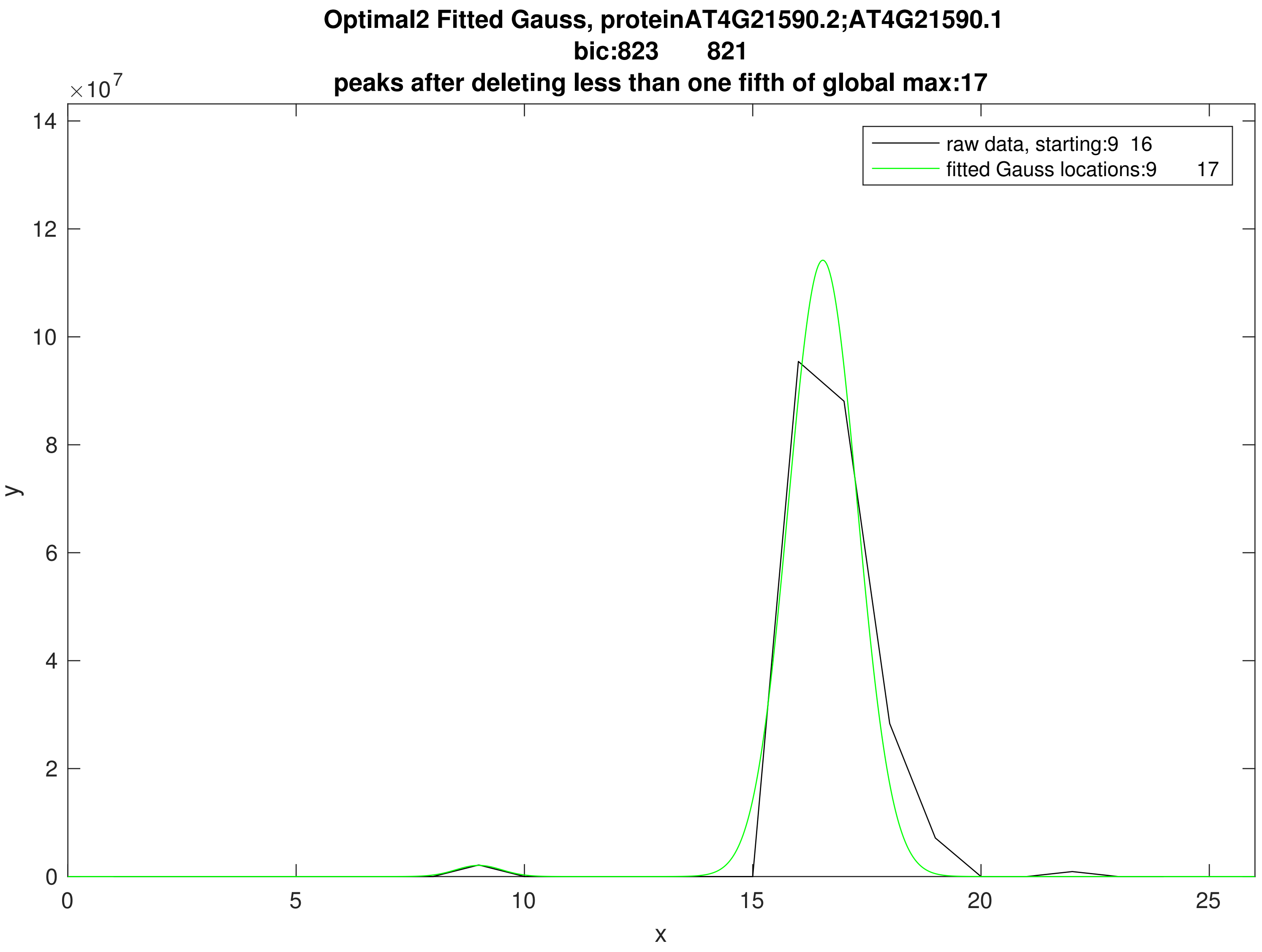
Optimal2 Fitted Gauss, proteinAT4G21590.2;AT4G21590.1
bic:823 821
peaks after deleting less than one fifth of global max:17
×10^7
14
12
10
8
6
4
2
0
y
0
5
10
15
20
25
x
raw data, starting:9 16
fitted Gauss locations:9 17

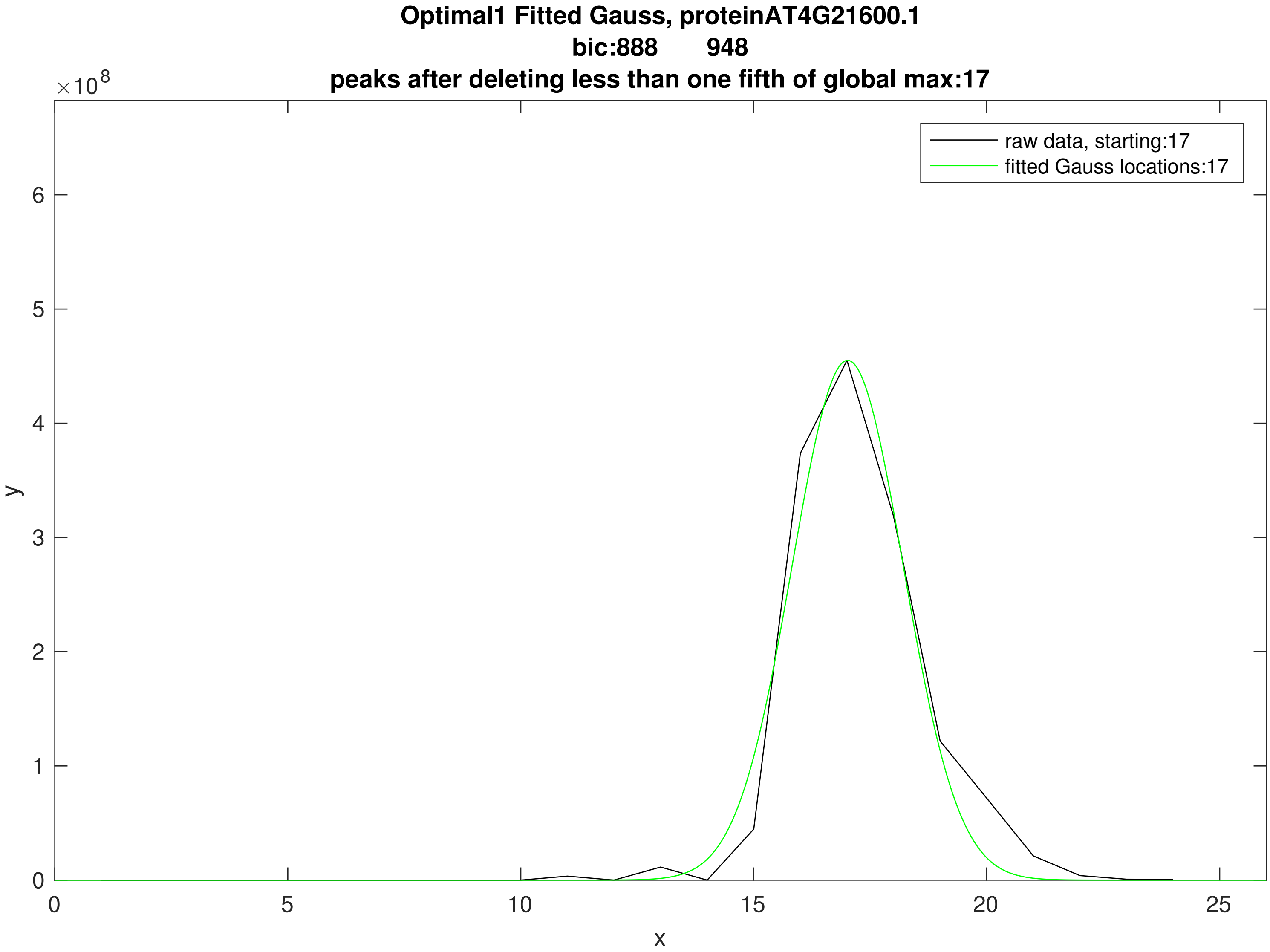
Optimal1 Fitted Gauss, proteinAT4G21600.1
bic:888 948
peaks after deleting less than one fifth of global max:17
×10^8
raw data, starting:17
fitted Gauss locations:17
y
x
0
1
2
3
4
5
6
0
5
10
15
20
25

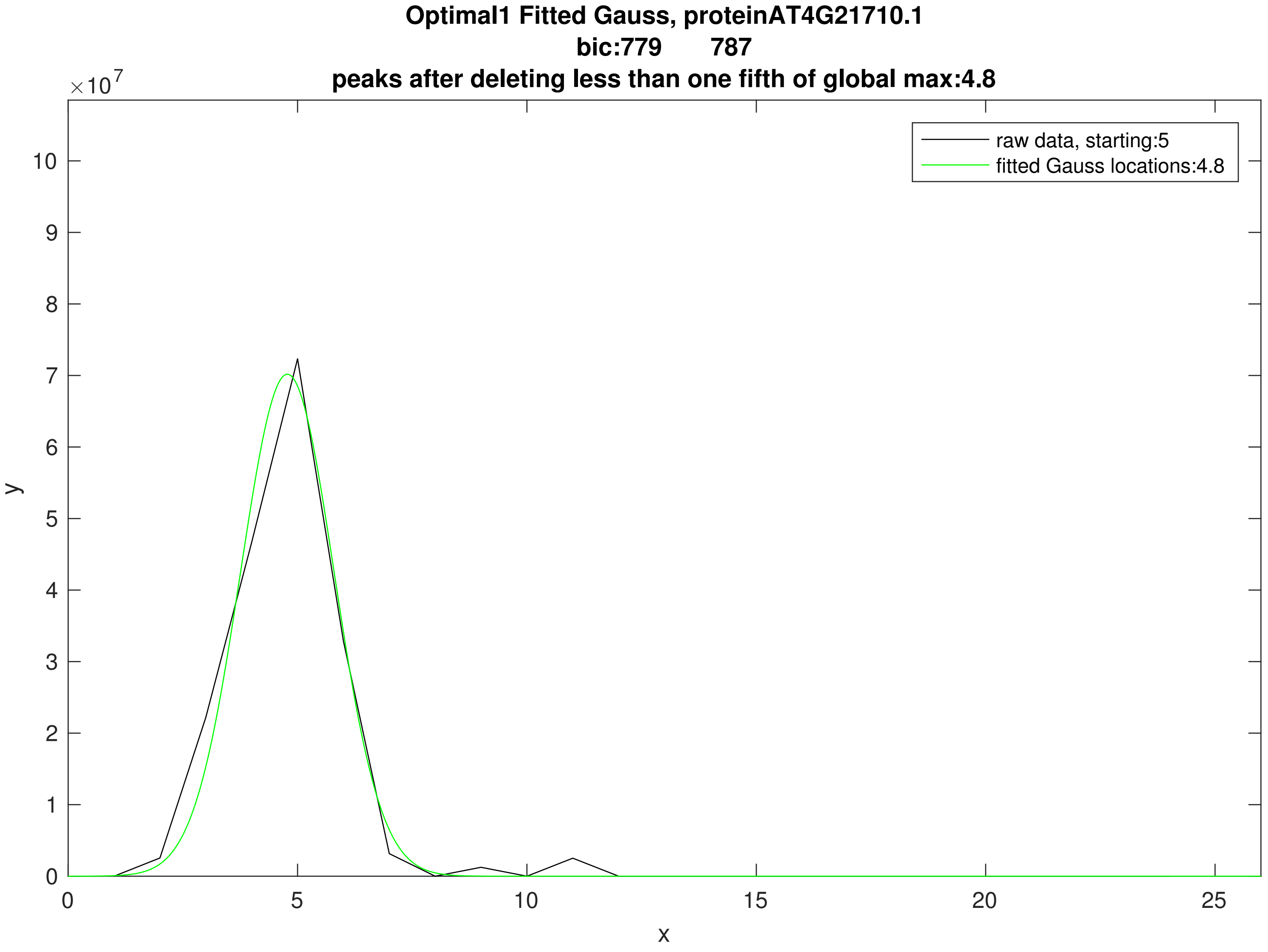
Optimal1 Fitted Gauss, proteinAT4G21710.1
bic:779 787
peaks after deleting less than one fifth of global max:4.8
×10^7
raw data, starting:5
fitted Gauss locations:4.8
y
x

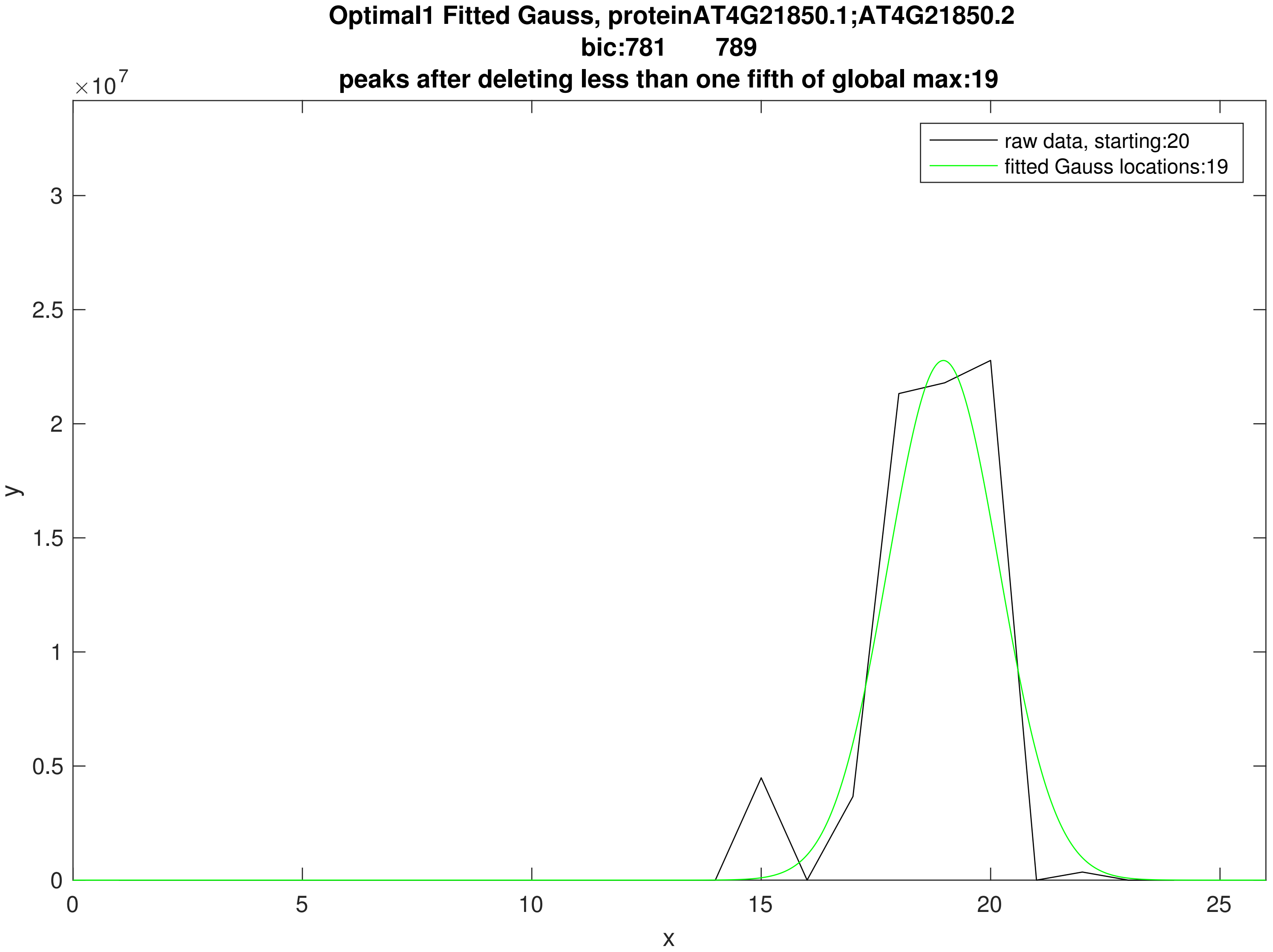
Optimal1 Fitted Gauss, proteinAT4G21850.1;AT4G21850.2
bic:781 789
peaks after deleting less than one fifth of global max:19
×10^7
raw data, starting:20
fitted Gauss locations:19
y
x
0
0.5
1
1.5
2
2.5
3
5
10
15
20
25

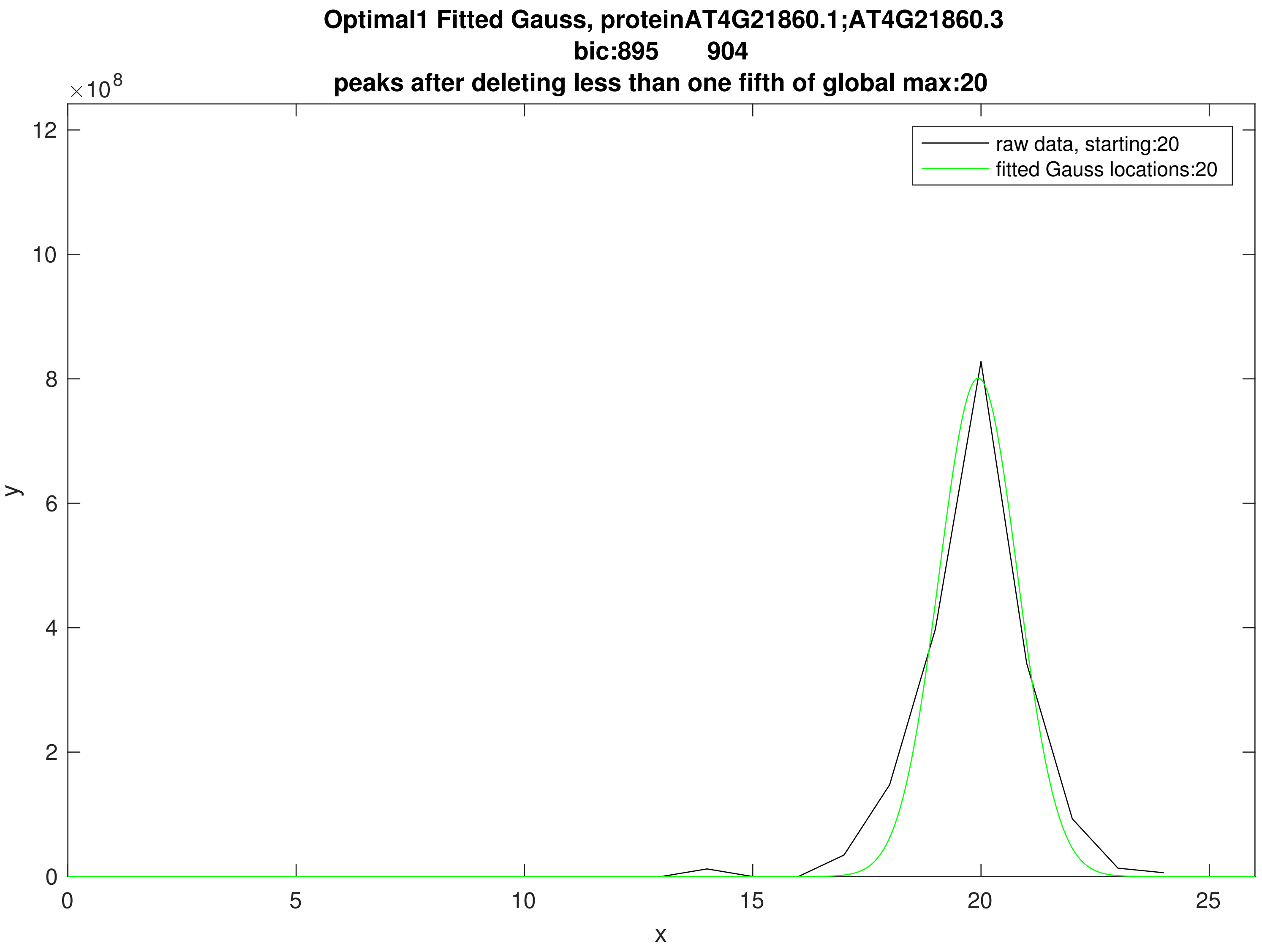
Optimal1 Fitted Gauss, proteinAT4G21860.1;AT4G21860.3
bic:895 904
peaks after deleting less than one fifth of global max:20
raw data, starting:20
fitted Gauss locations:20
×10^8
y
x
0
2
4
6
8
10
12
0
5
10
15
20
25

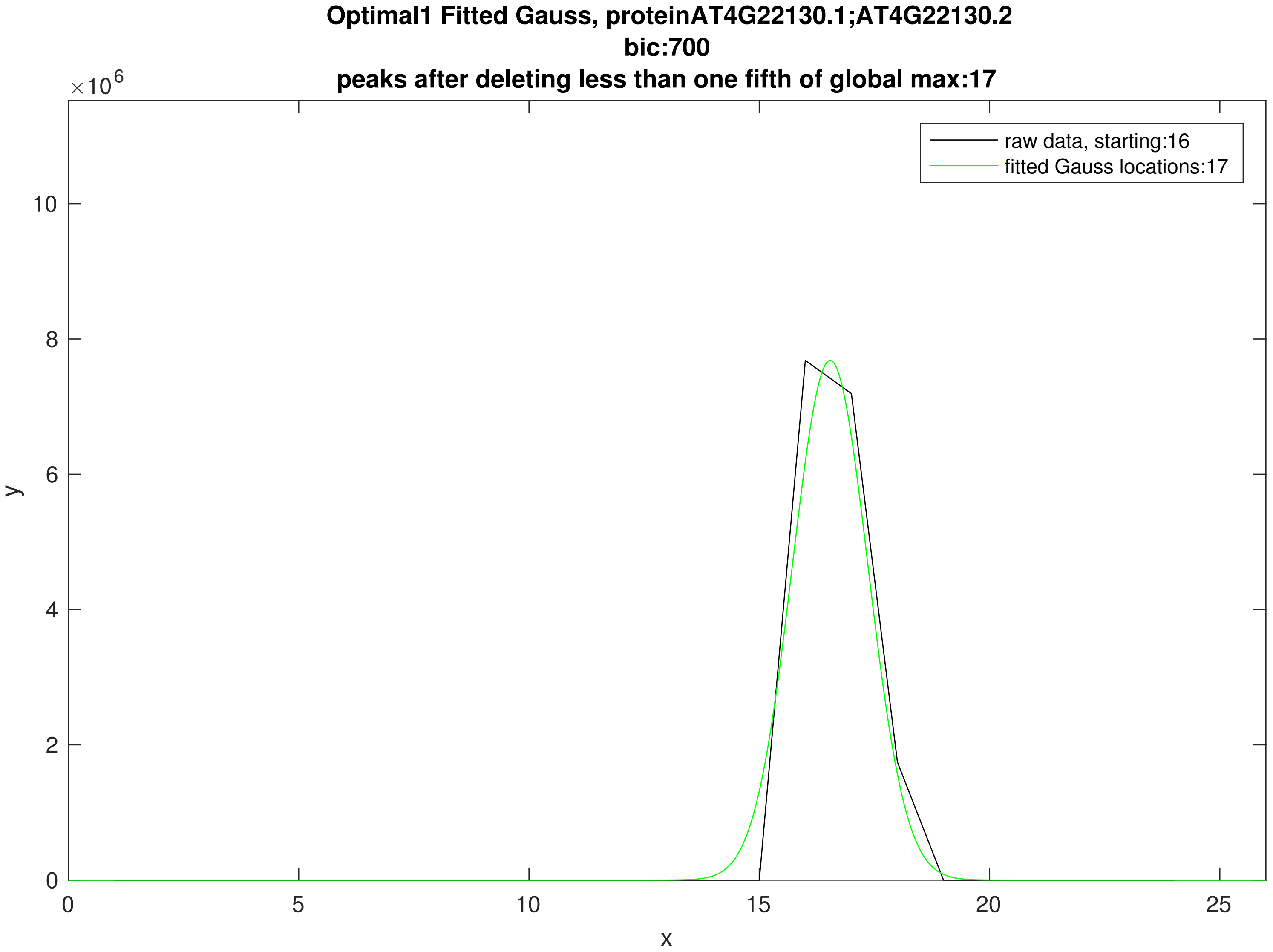
Optimal1 Fitted Gauss, proteinAT4G22130.1;AT4G22130.2
bic:700
peaks after deleting less than one fifth of global max:17
×10^6
raw data, starting:16
fitted Gauss locations:17
y
x
0
2
4
6
8
10
0
5
10
15
20
25

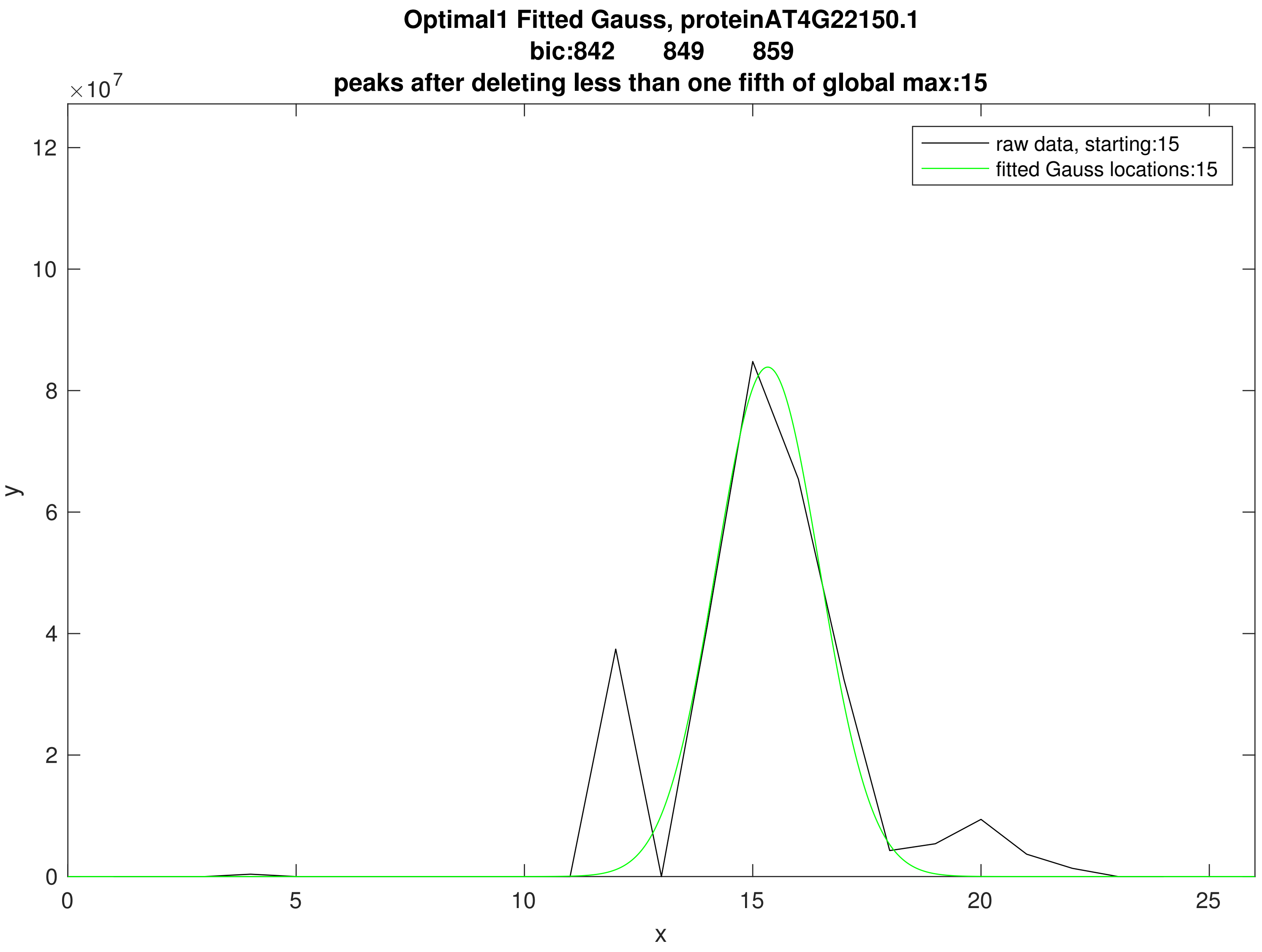
Optimal1 Fitted Gauss, proteinAT4G22150.1
bic:842 849 859
peaks after deleting less than one fifth of global max:15
×10^7
raw data, starting:15
fitted Gauss locations:15
y
x
0
2
4
6
8
10
12
0
5
10
15
20
25

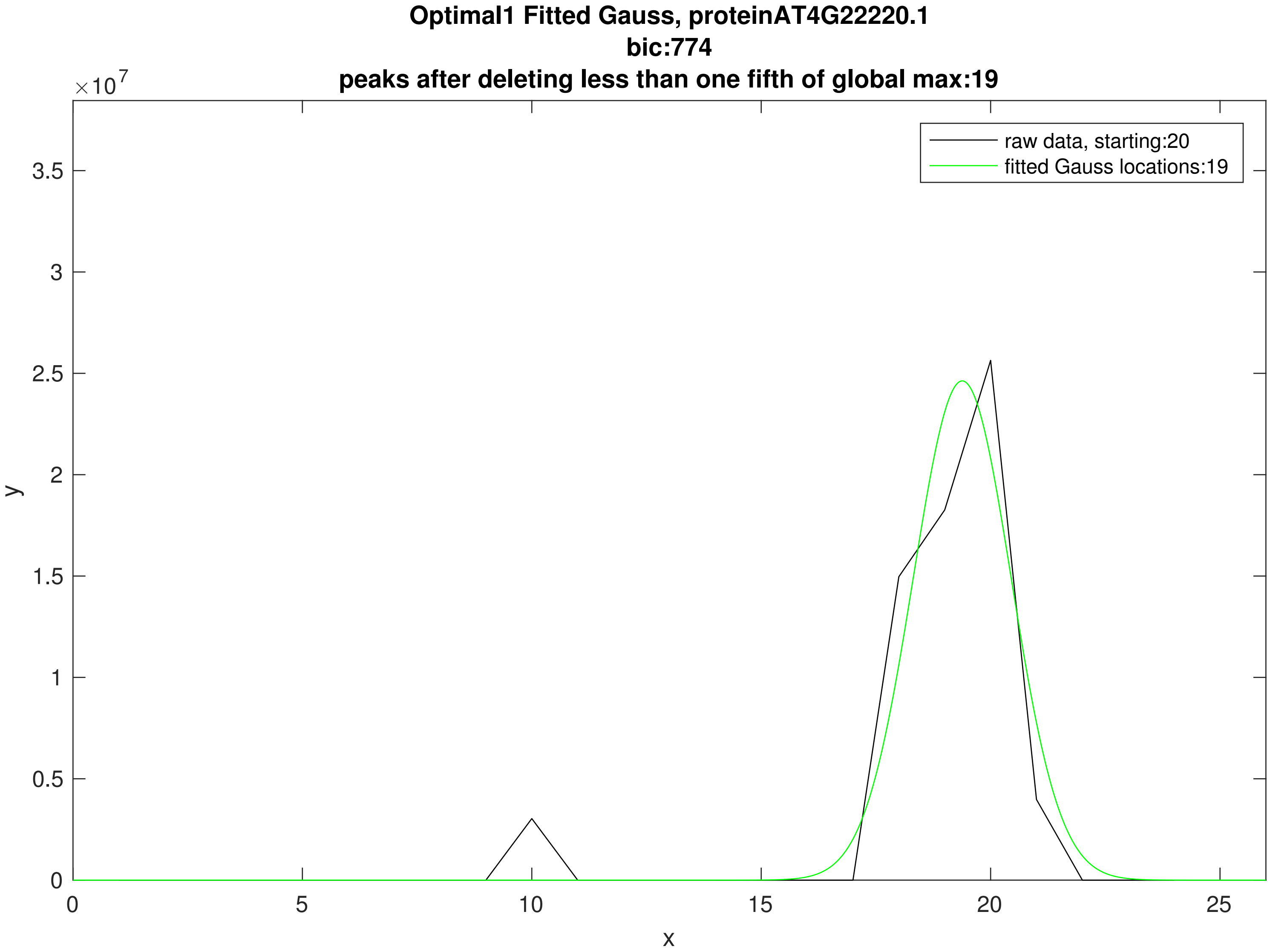

Optimal1 Fitted Gauss, proteinAT4G22220.1
bic:774
peaks after deleting less than one fifth of global max:19
×10^7
3.5
3
2.5
2
1.5
1
0.5
0
y
0
5
10
15
20
25
x
raw data, starting:20
fitted Gauss locations:19

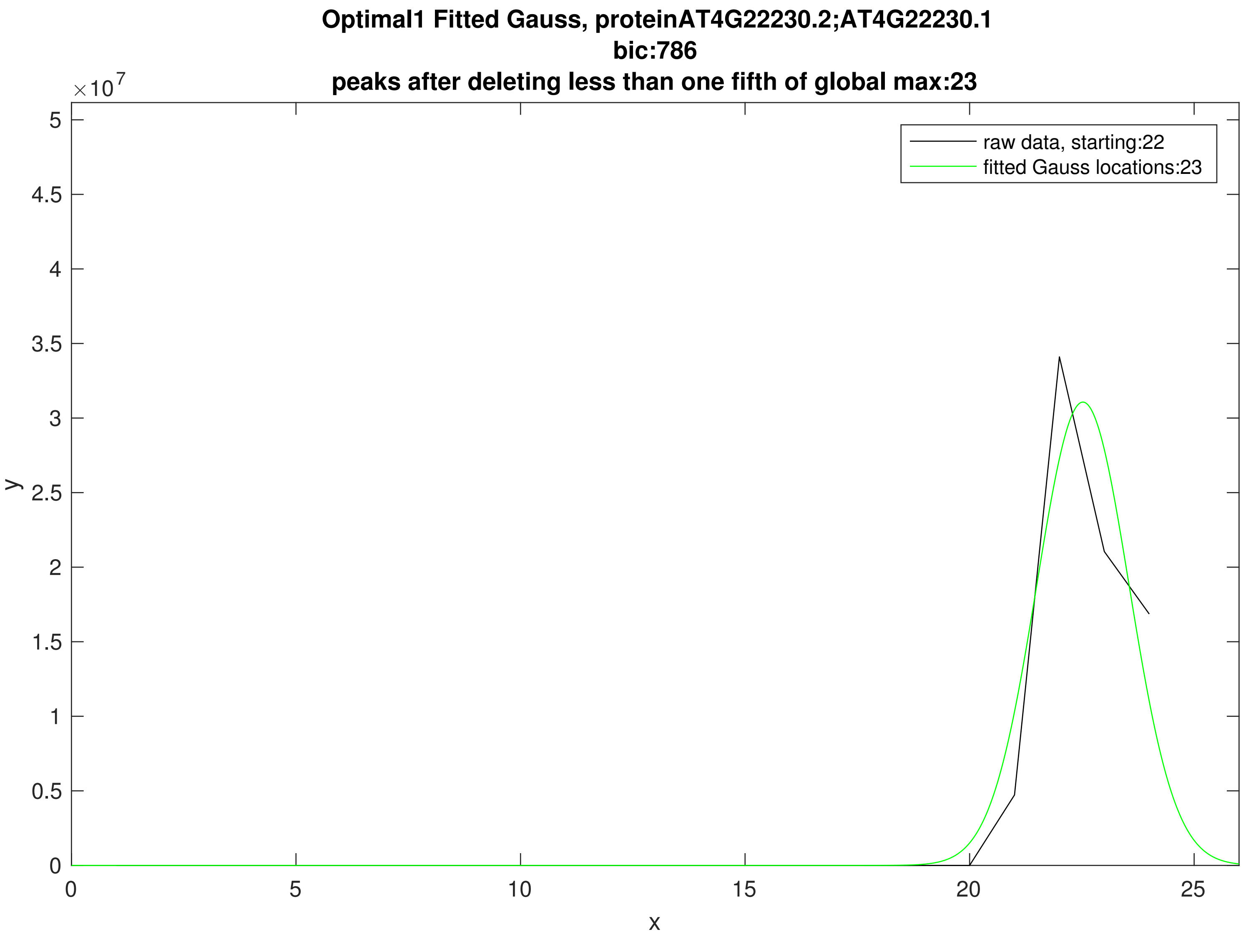
Optimal1 Fitted Gauss, proteinAT4G22230.2;AT4G22230.1
bic:786
peaks after deleting less than one fifth of global max:23
×10^7
raw data, starting:22
fitted Gauss locations:23
y
x
0
0.5
1
1.5
2
2.5
3
3.5
4
4.5
5
0
5
10
15
20
25

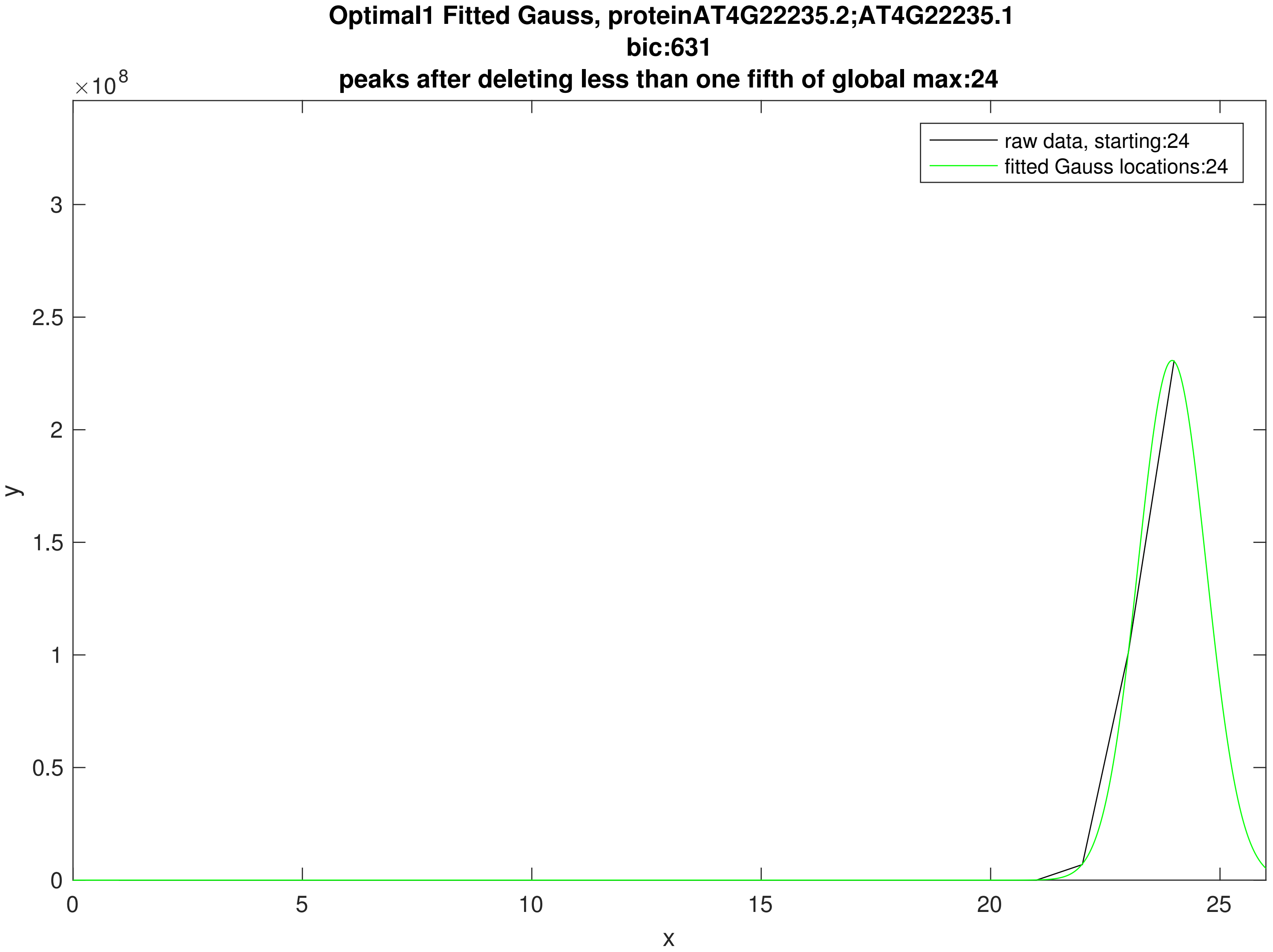

Optimal1 Fitted Gauss, proteinAT4G22235.2;AT4G22235.1
bic:631
peaks after deleting less than one fifth of global max:24
×10^8
raw data, starting:24
fitted Gauss locations:24
y
x
0
0.5
1
1.5
2
2.5
3
0
5
10
15
20
25

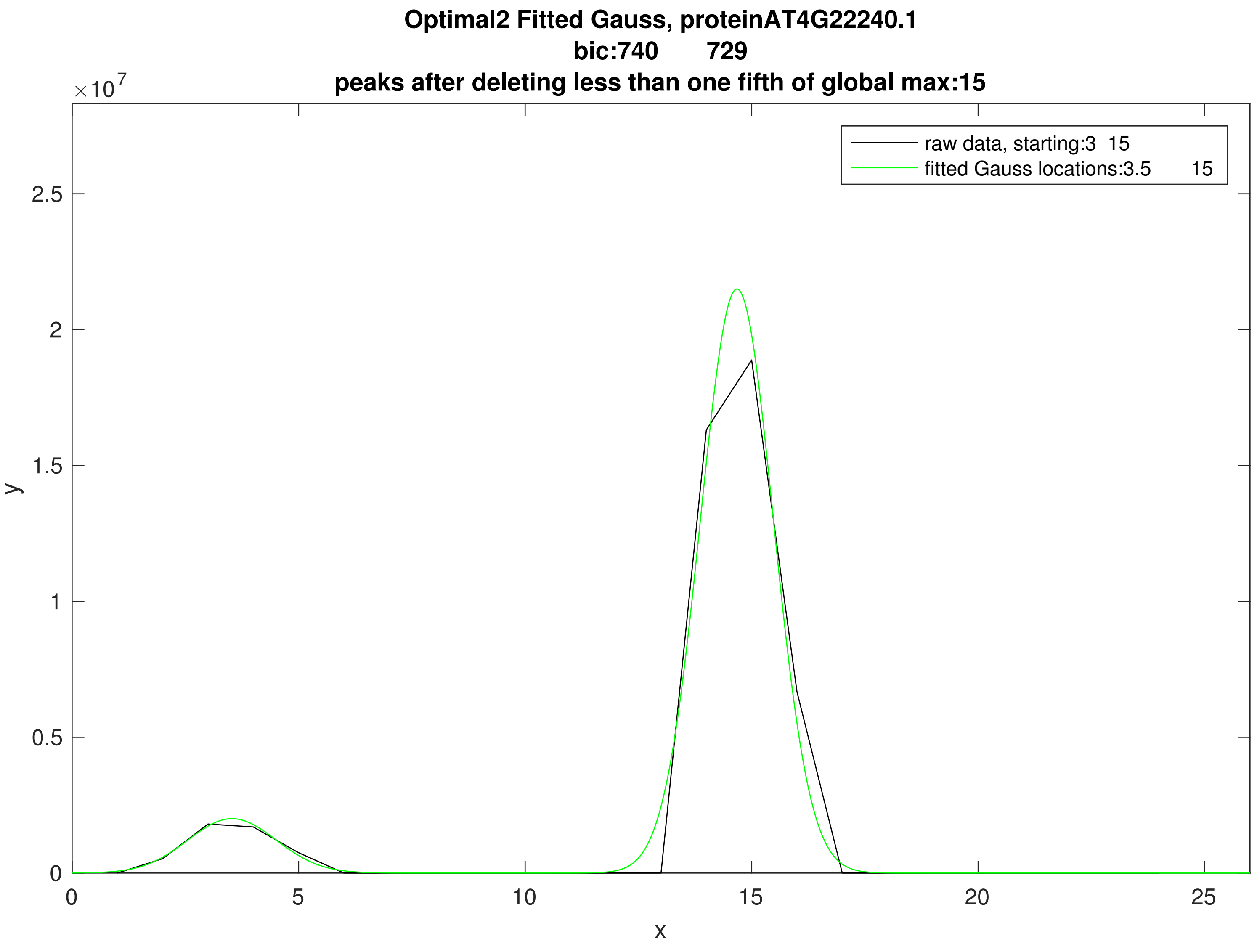

Optimal2 Fitted Gauss, proteinAT4G22240.1
bic:740 729
peaks after deleting less than one fifth of global max:15
×10^7
raw data, starting:3 15
fitted Gauss locations:3.5 15
y
x
0
0.5
1
1.5
2
2.5
0
5
10
15
20
25

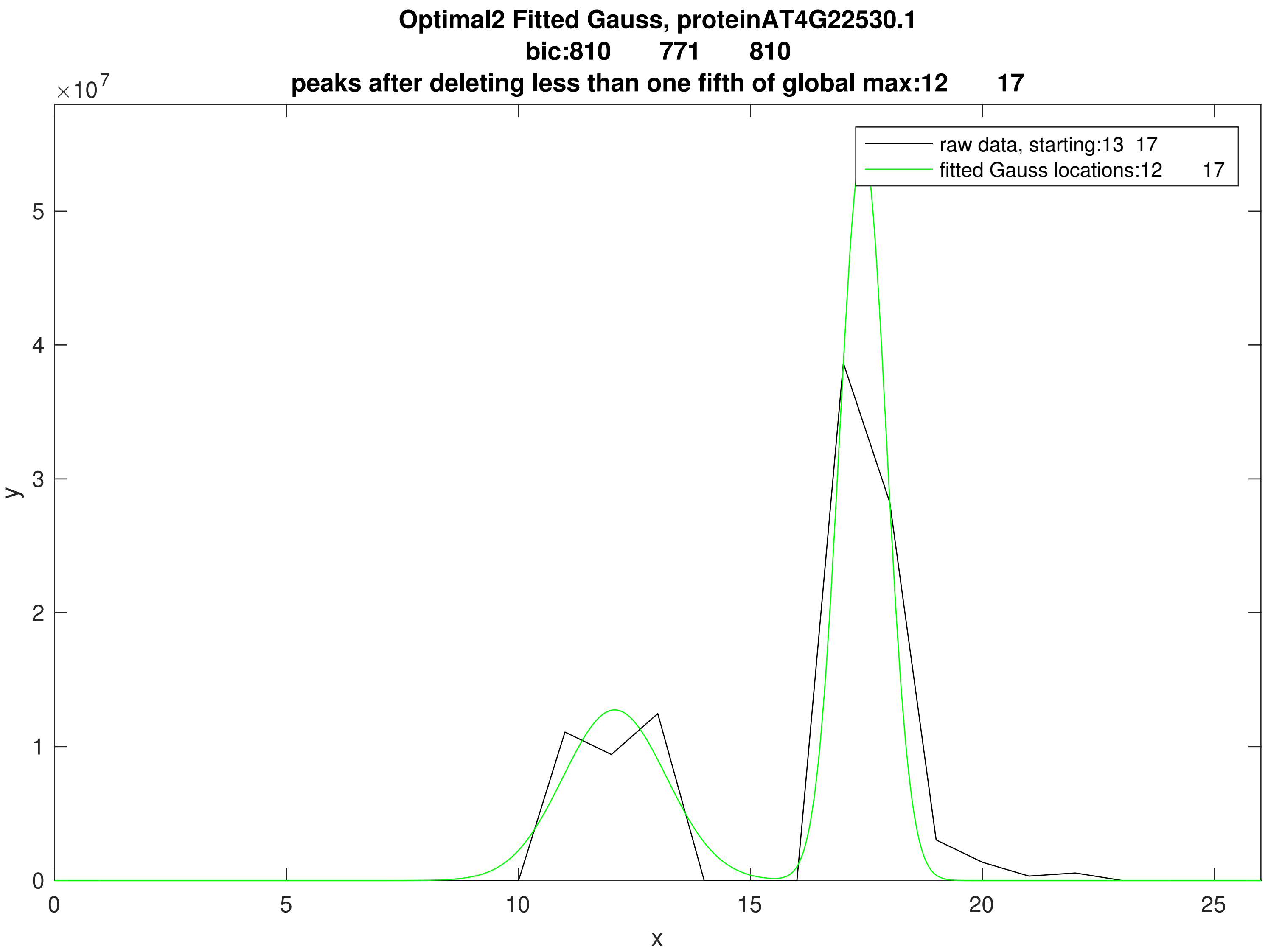
Optimal2 Fitted Gauss, proteinAT4G22530.1
bic:810 771 810
peaks after deleting less than one fifth of global max:12 17
×10^7
raw data, starting:13 17
fitted Gauss locations:12 17
y
x
0
1
2
3
4
5
0
5
10
15
20
25

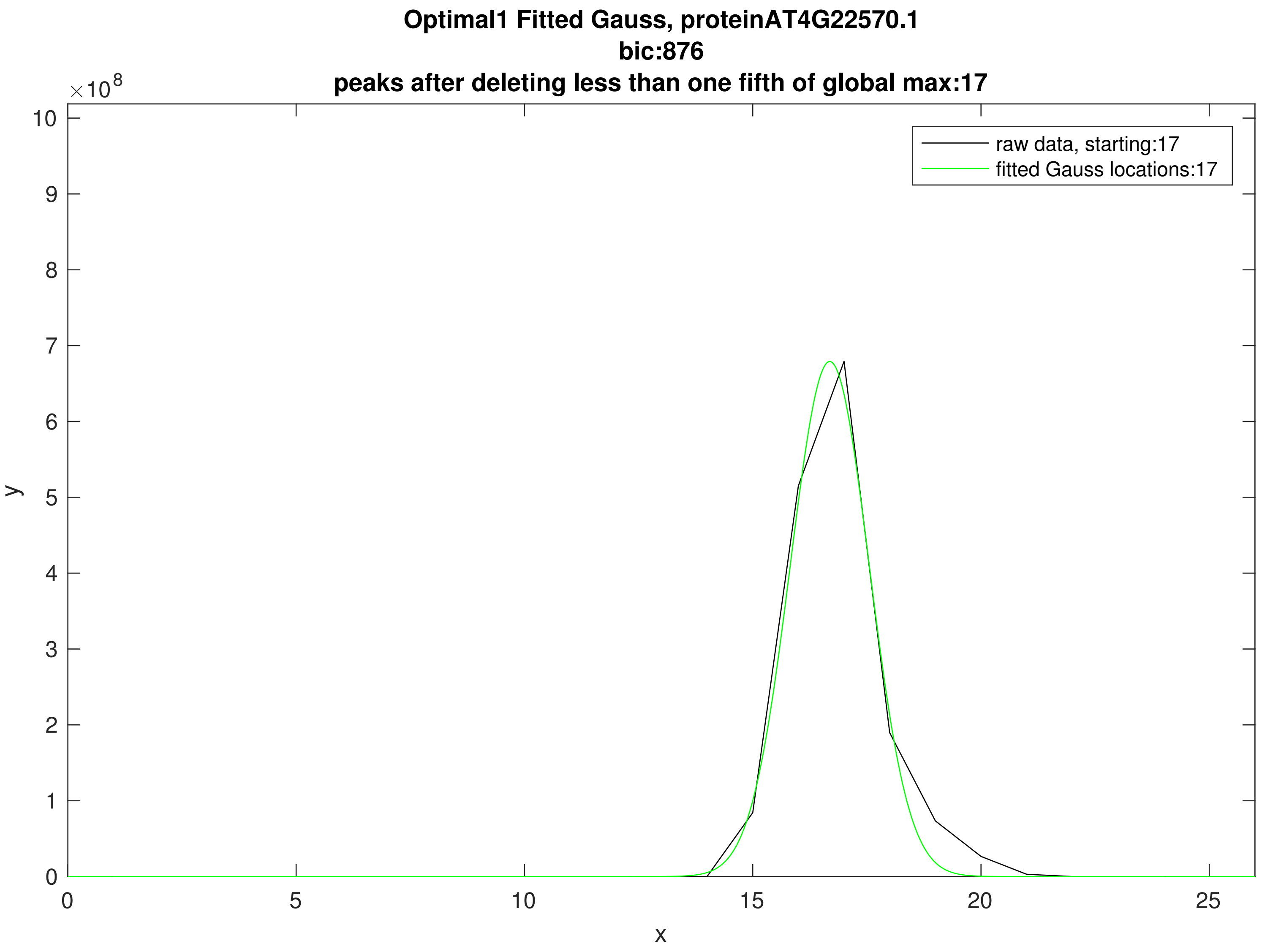

Optimal1 Fitted Gauss, proteinAT4G22570.1
bic:876
peaks after deleting less than one fifth of global max:17
×10^8
raw data, starting:17
fitted Gauss locations:17
y
x
0
1
2
3
4
5
6
7
8
9
10
0
5
10
15
20
25

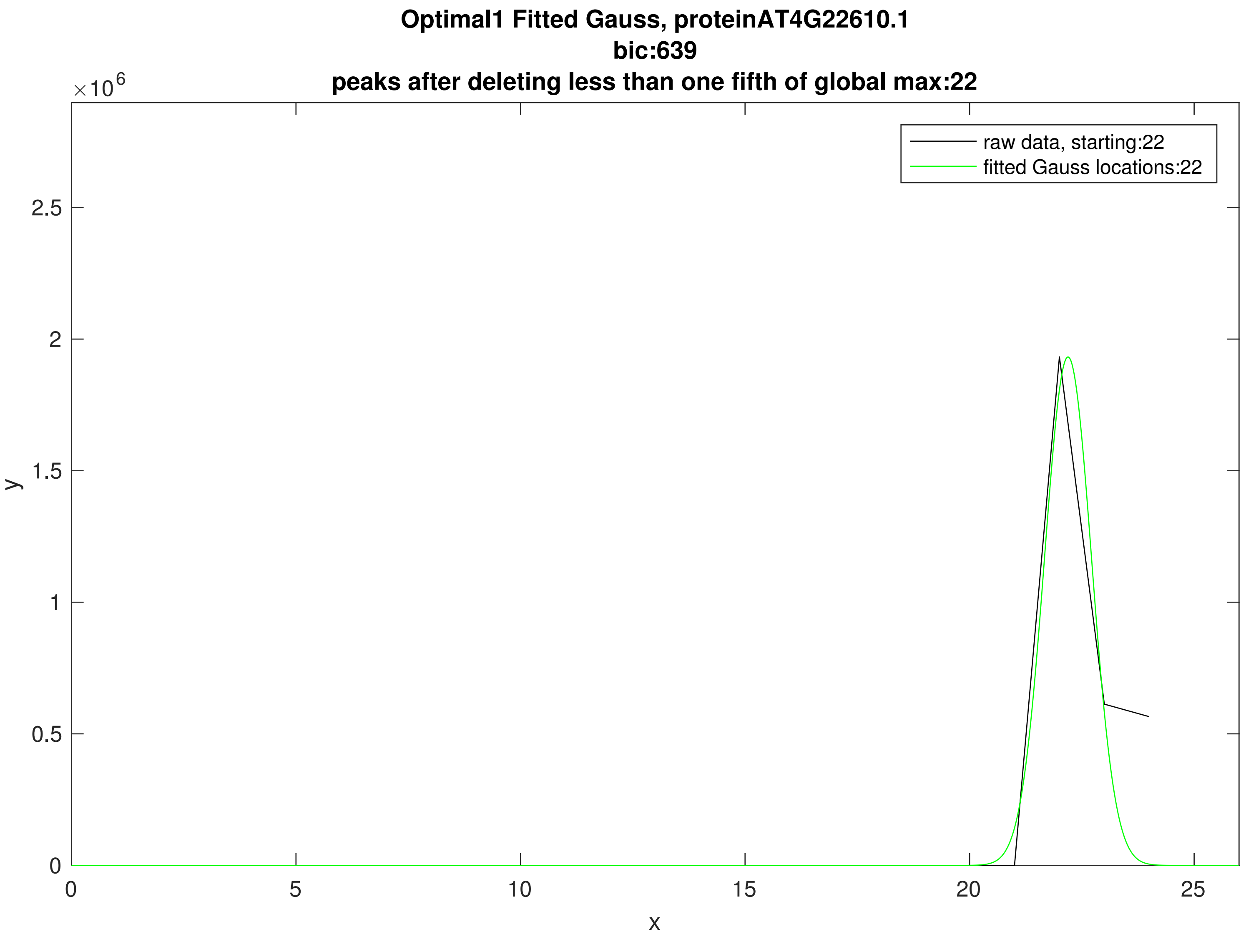

Optimal1 Fitted Gauss, proteinAT4G22610.1
bic:639
peaks after deleting less than one fifth of global max:22
×10^6
raw data, starting:22
fitted Gauss locations:22
y
x
0
0.5
1
1.5
2
2.5
0
5
10
15
20
25

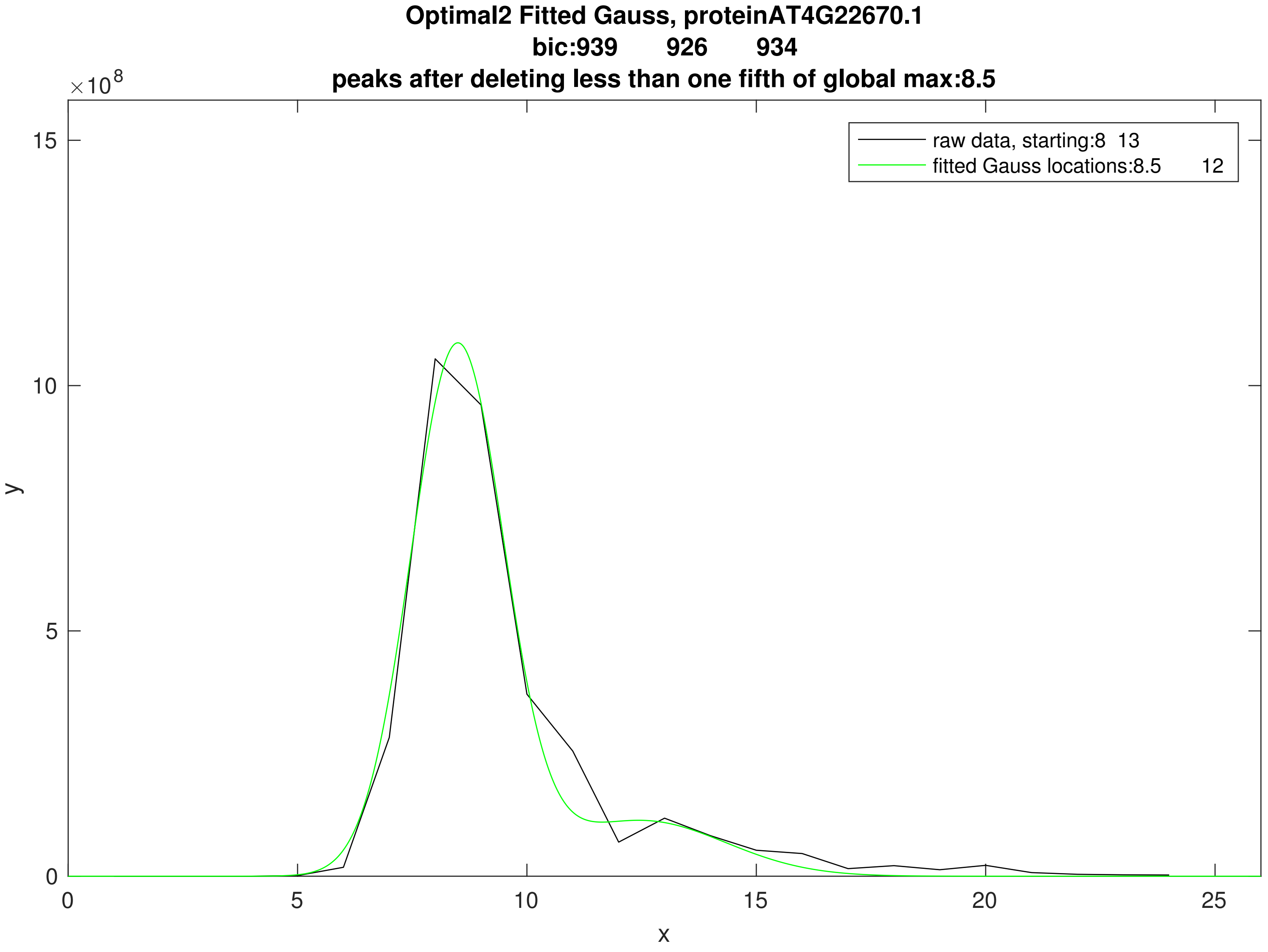

Optimal2 Fitted Gauss, proteinAT4G22670.1
bic:939 926 934
peaks after deleting less than one fifth of global max:8.5
×10^8
raw data, starting:8 13
fitted Gauss locations:8.5 12
y
x
0
5
10
15
20
25

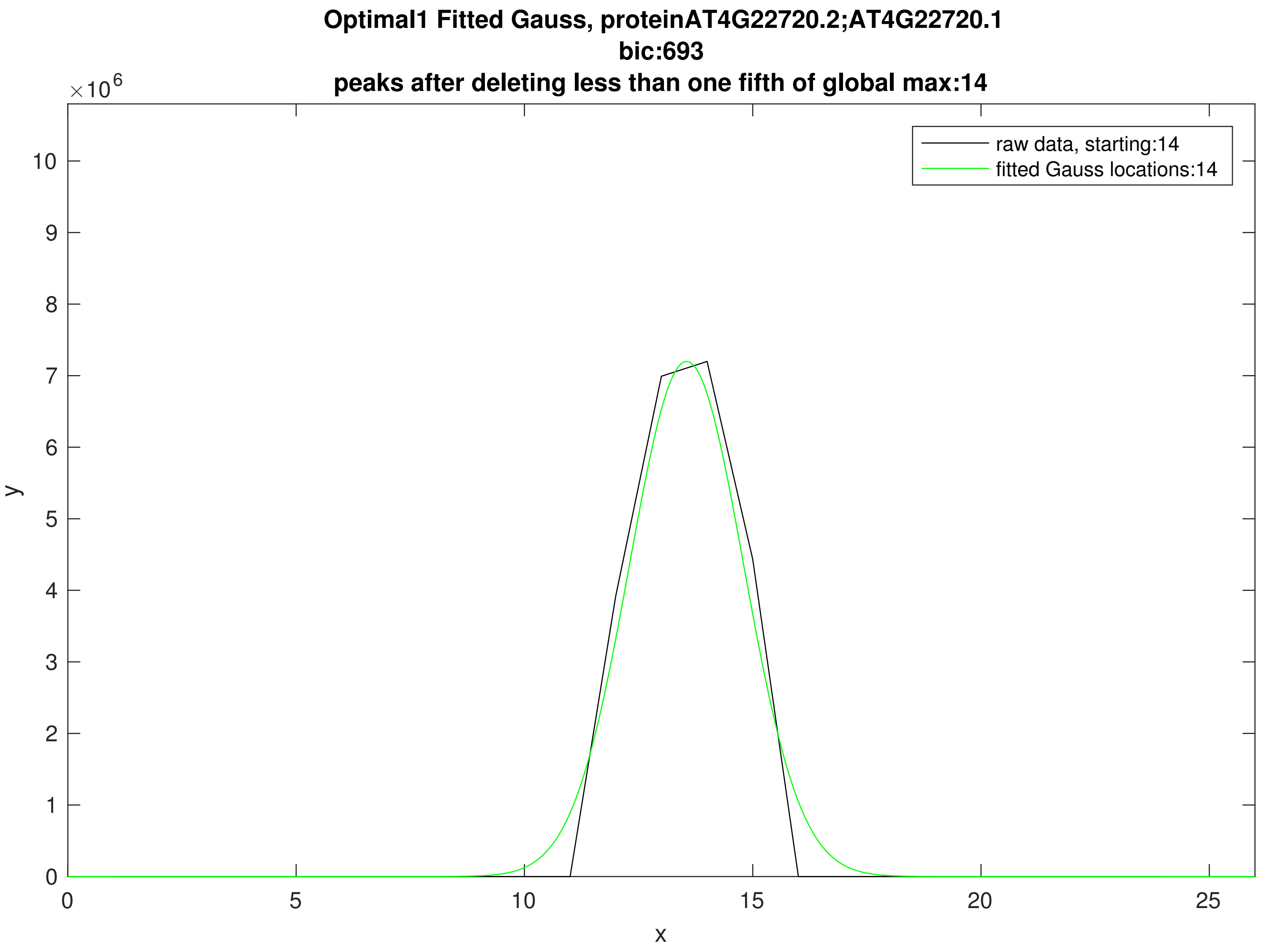

Optimal1 Fitted Gauss, proteinAT4G22720.2;AT4G22720.1
bic:693
peaks after deleting less than one fifth of global max:14
×10^6
raw data, starting:14
fitted Gauss locations:14
y
x
0
1
2
3
4
5
6
7
8
9
10
0
5
10
15
20
25

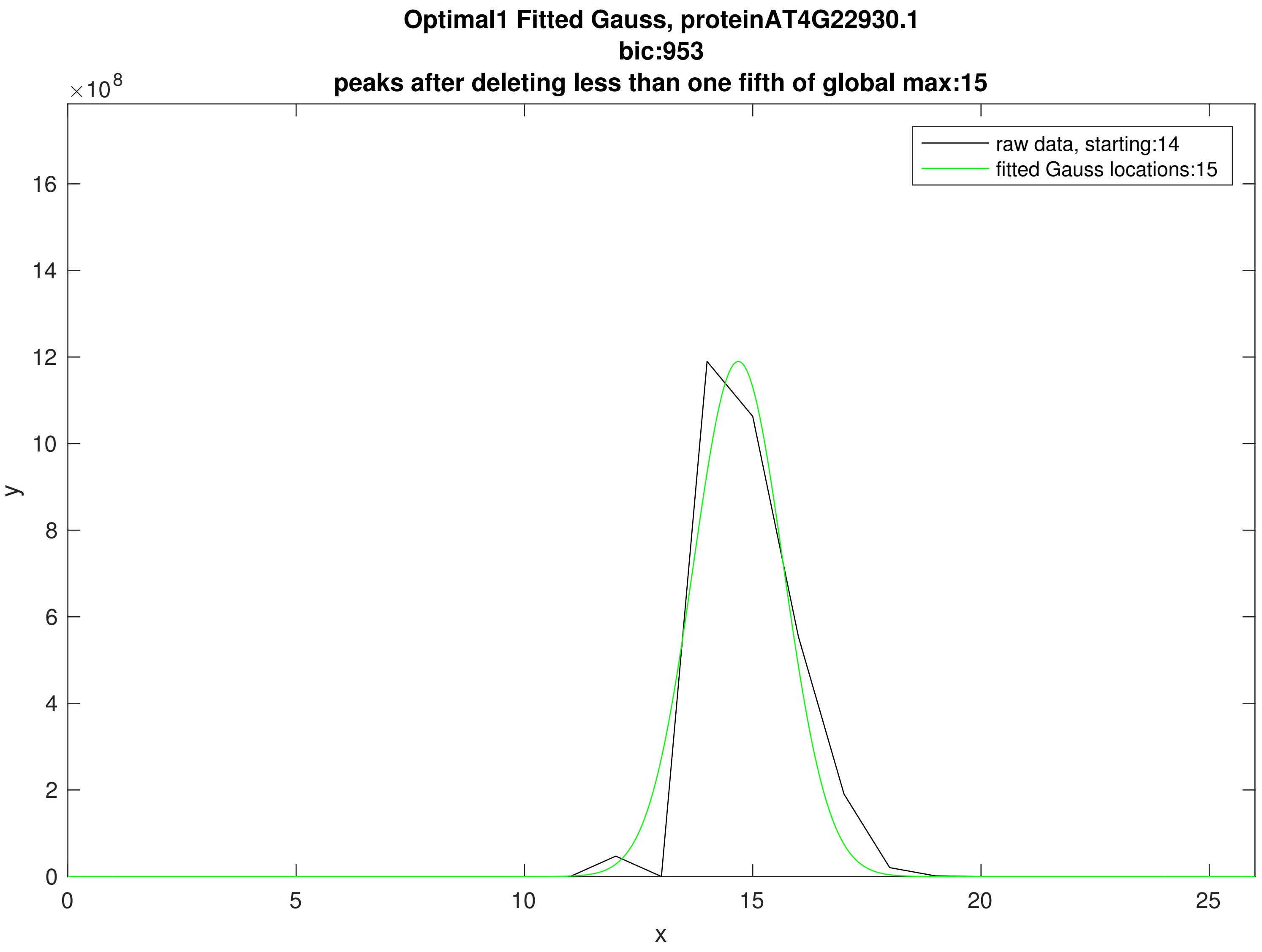

Optimal1 Fitted Gauss, proteinAT4G22930.1
bic:953
peaks after deleting less than one fifth of global max:15
×10^8
raw data, starting:14
fitted Gauss locations:15
16
14
12
10
8
6
4
2
0
y
0
5
10
15
20
25
x

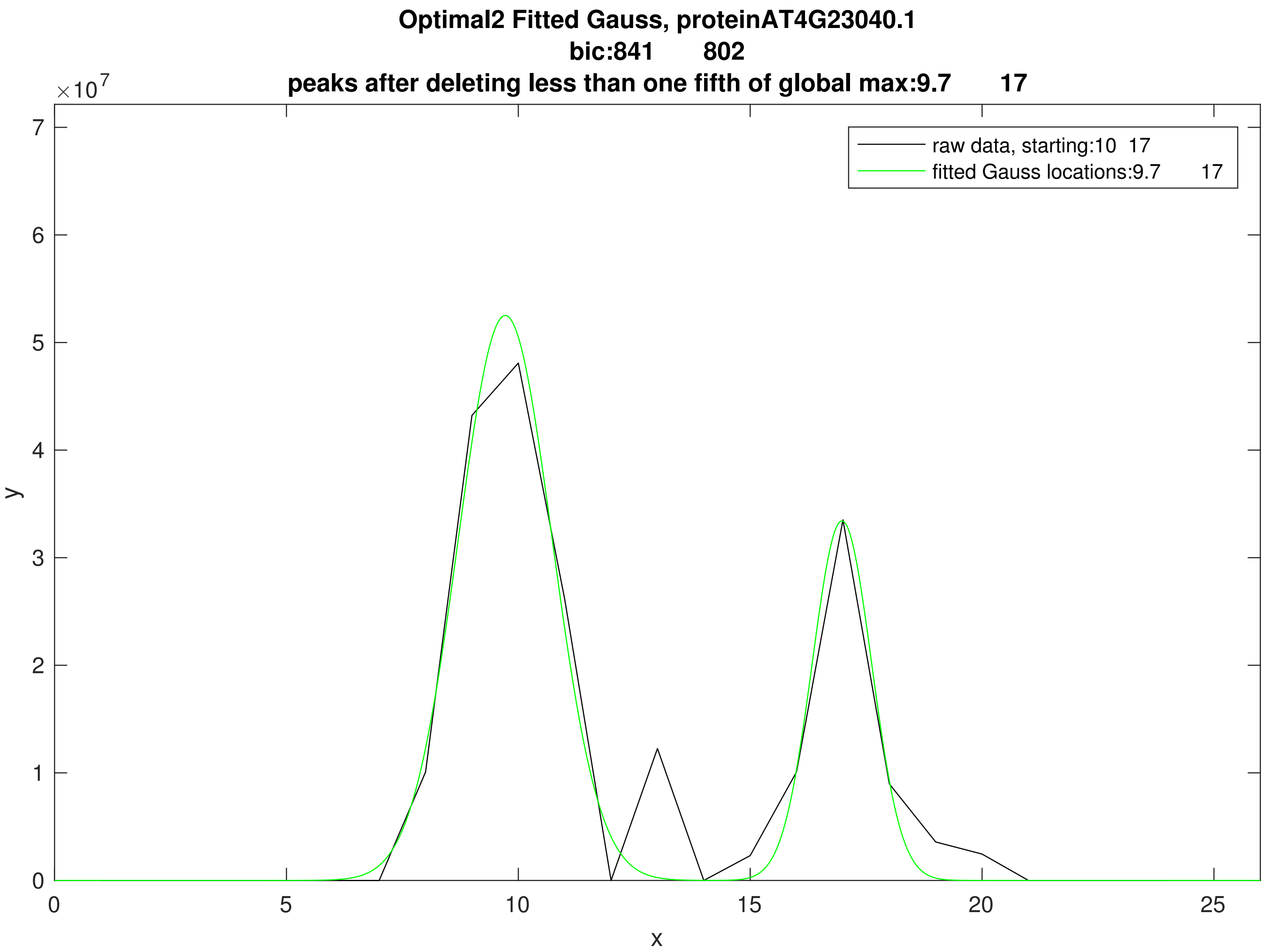
Optimal2 Fitted Gauss, proteinAT4G23040.1
bic:841 802
peaks after deleting less than one fifth of global max:9.7 17
×10^7
raw data, starting:10 17
fitted Gauss locations:9.7 17
y
x
0
1
2
3
4
5
6
7
0
5
10
15
20
25

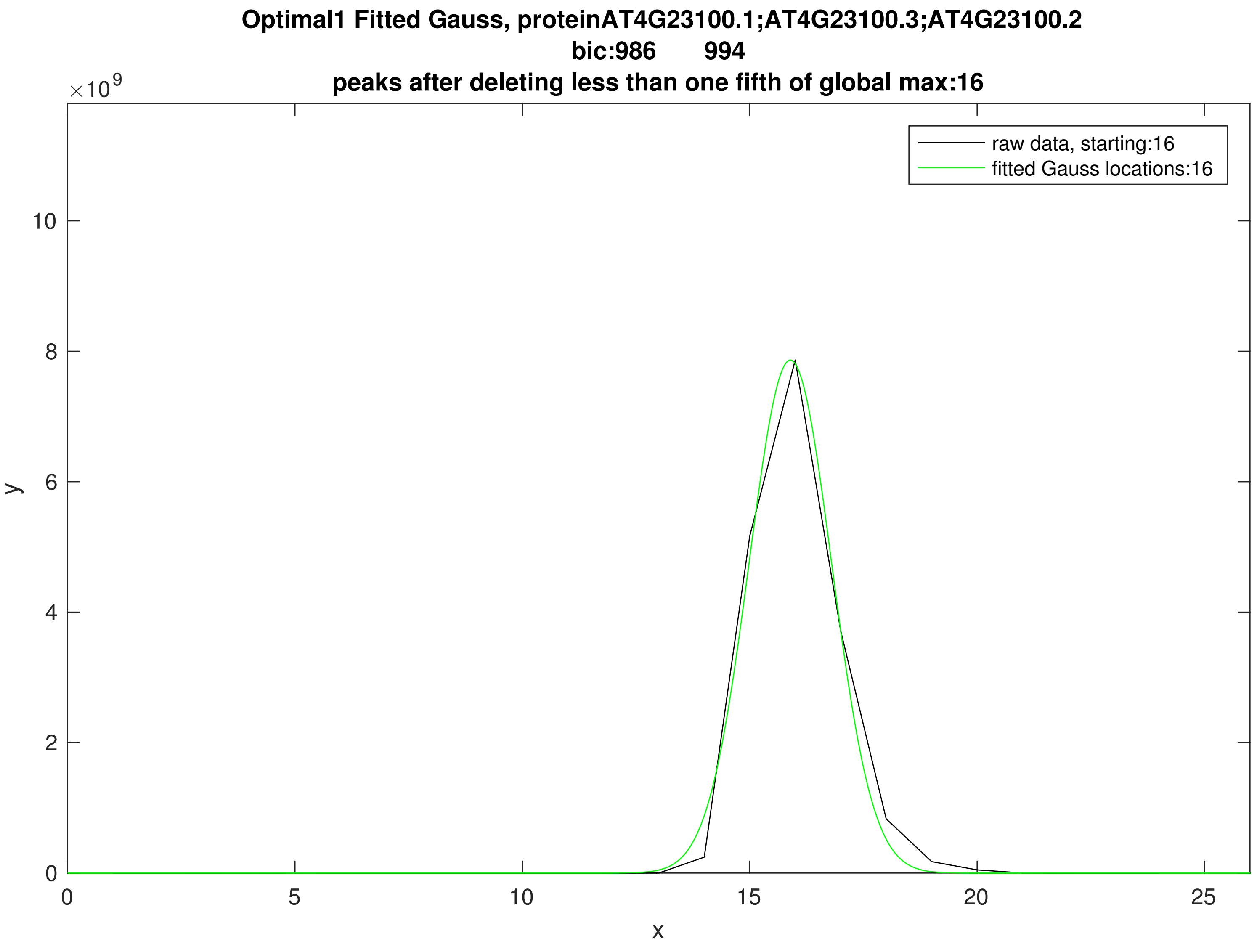

Optimal1 Fitted Gauss, proteinAT4G23100.1;AT4G23100.3;AT4G23100.2
bic:986 994
peaks after deleting less than one fifth of global max:16
×10^9
raw data, starting:16
fitted Gauss locations:16
10
8
6
4
2
0
y
0
5
10
15
20
25
x

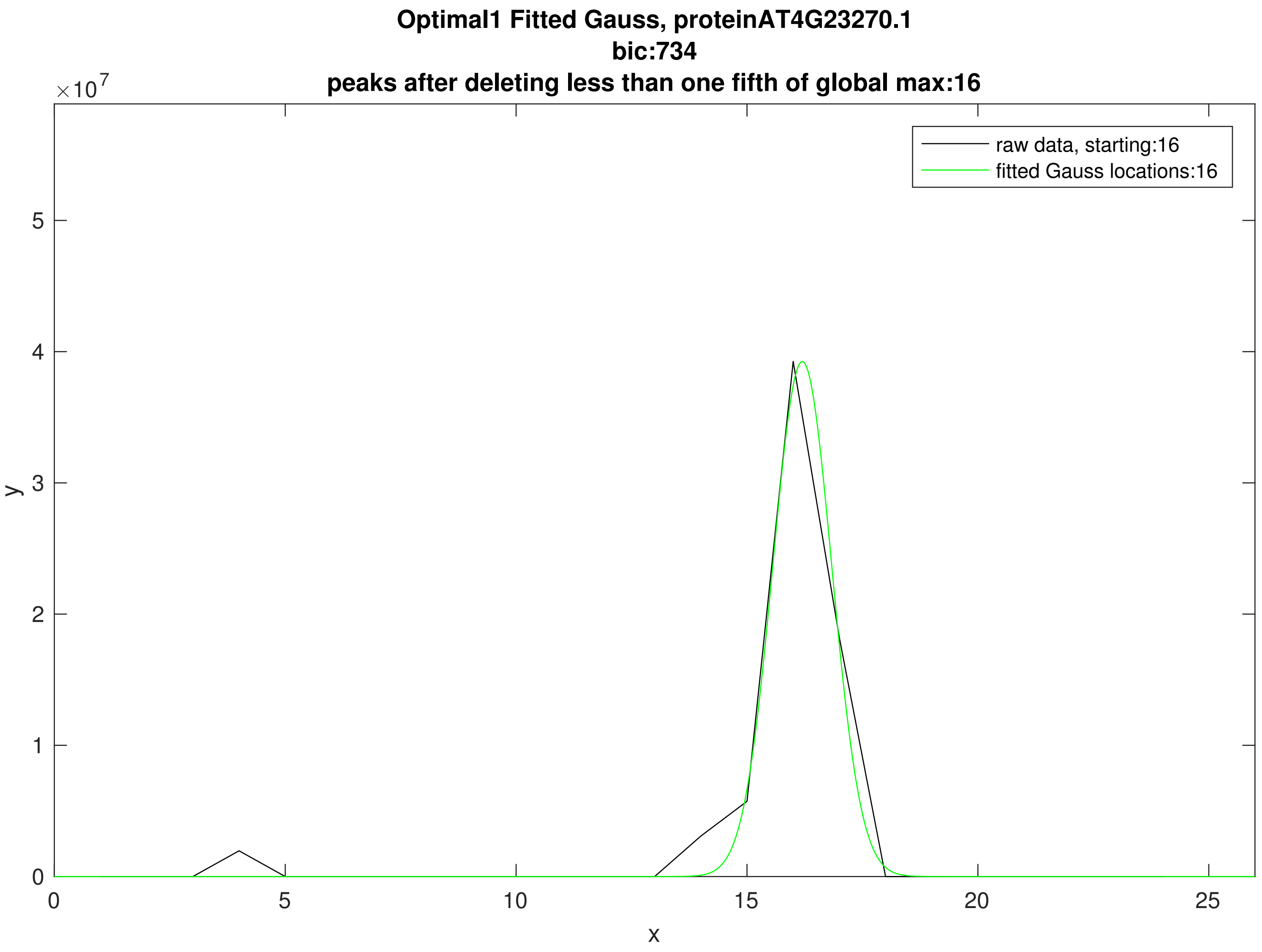

Optimal1 Fitted Gauss, proteinAT4G23270.1
bic:734
peaks after deleting less than one fifth of global max:16
×10^7
5
4
3
2
1
0
y
0
5
10
15
20
25
x
raw data, starting:16
fitted Gauss locations:16

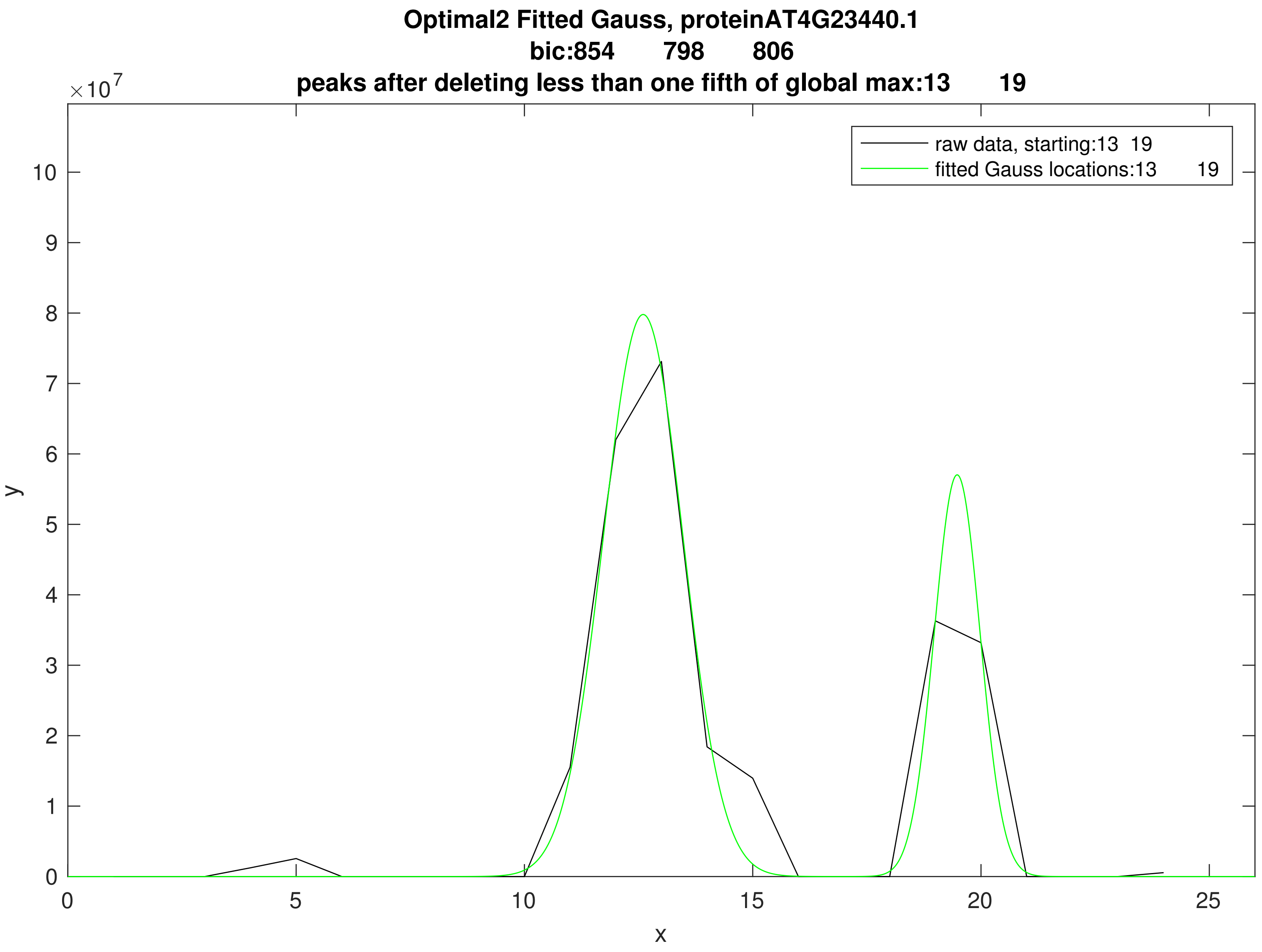
Optimal2 Fitted Gauss, proteinAT4G23440.1
bic:854 798 806
peaks after deleting less than one fifth of global max:13 19
×10^7
raw data, starting:13 19
fitted Gauss locations:13 19
y
x
0
1
2
3
4
5
6
7
8
9
10
0
5
10
15
20
25

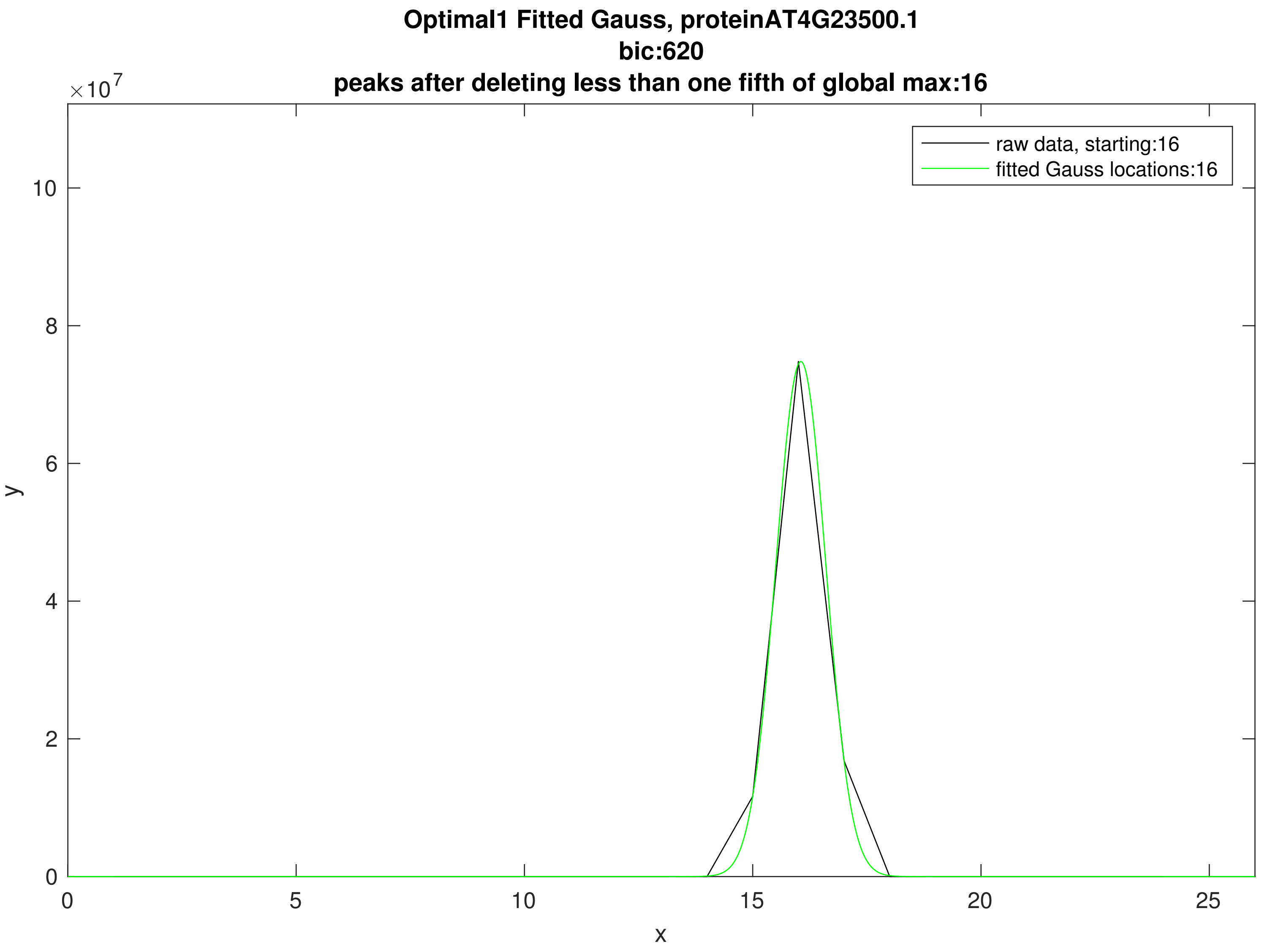
Optimal1 Fitted Gauss, proteinAT4G23500.1
bic:620
peaks after deleting less than one fifth of global max:16
×10^7
raw data, starting:16
fitted Gauss locations:16
y
x
0
2
4
6
8
10
0
5
10
15
20
25

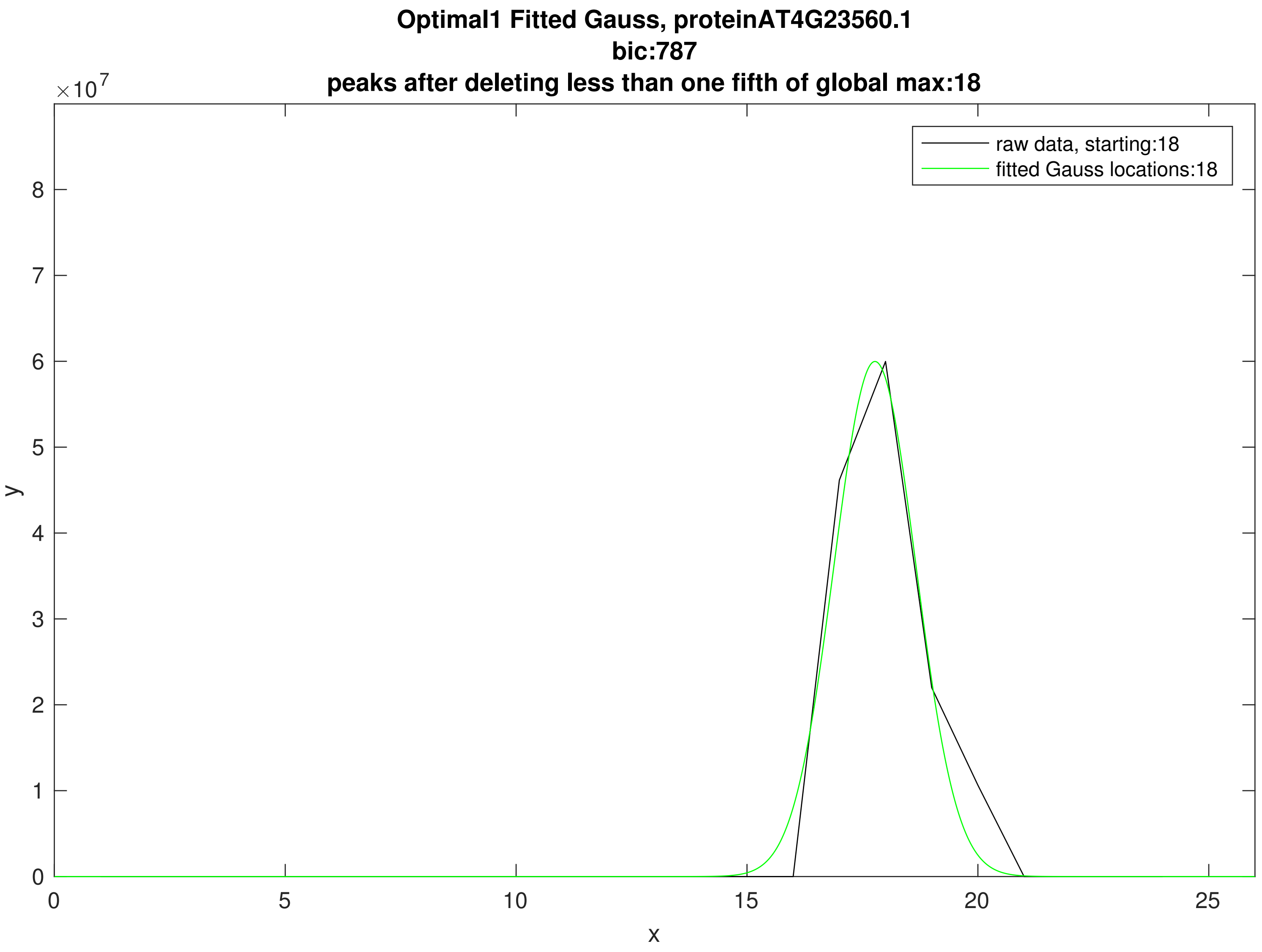

Optimal1 Fitted Gauss, proteinAT4G23560.1
bic:787
peaks after deleting less than one fifth of global max:18
×10^7
raw data, starting:18
fitted Gauss locations:18
y
x
0
1
2
3
4
5
6
7
8
0
5
10
15
20
25

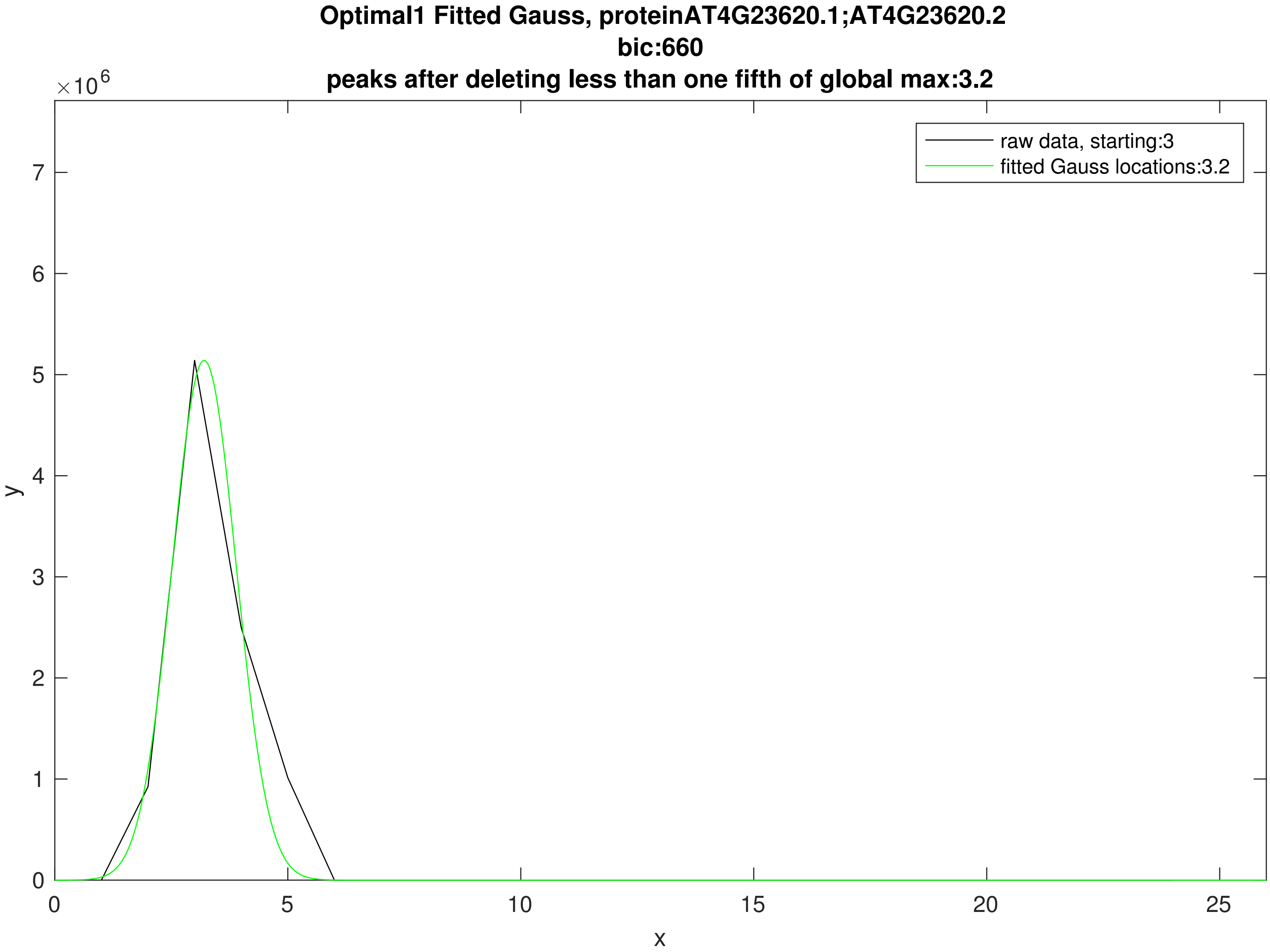

Optimal1 Fitted Gauss, proteinAT4G23620.1;AT4G23620.2
bic:660
peaks after deleting less than one fifth of global max:3.2
×10^6
raw data, starting:3
fitted Gauss locations:3.2
y
x
0
1
2
3
4
5
6
7
0
5
10
15
20
25

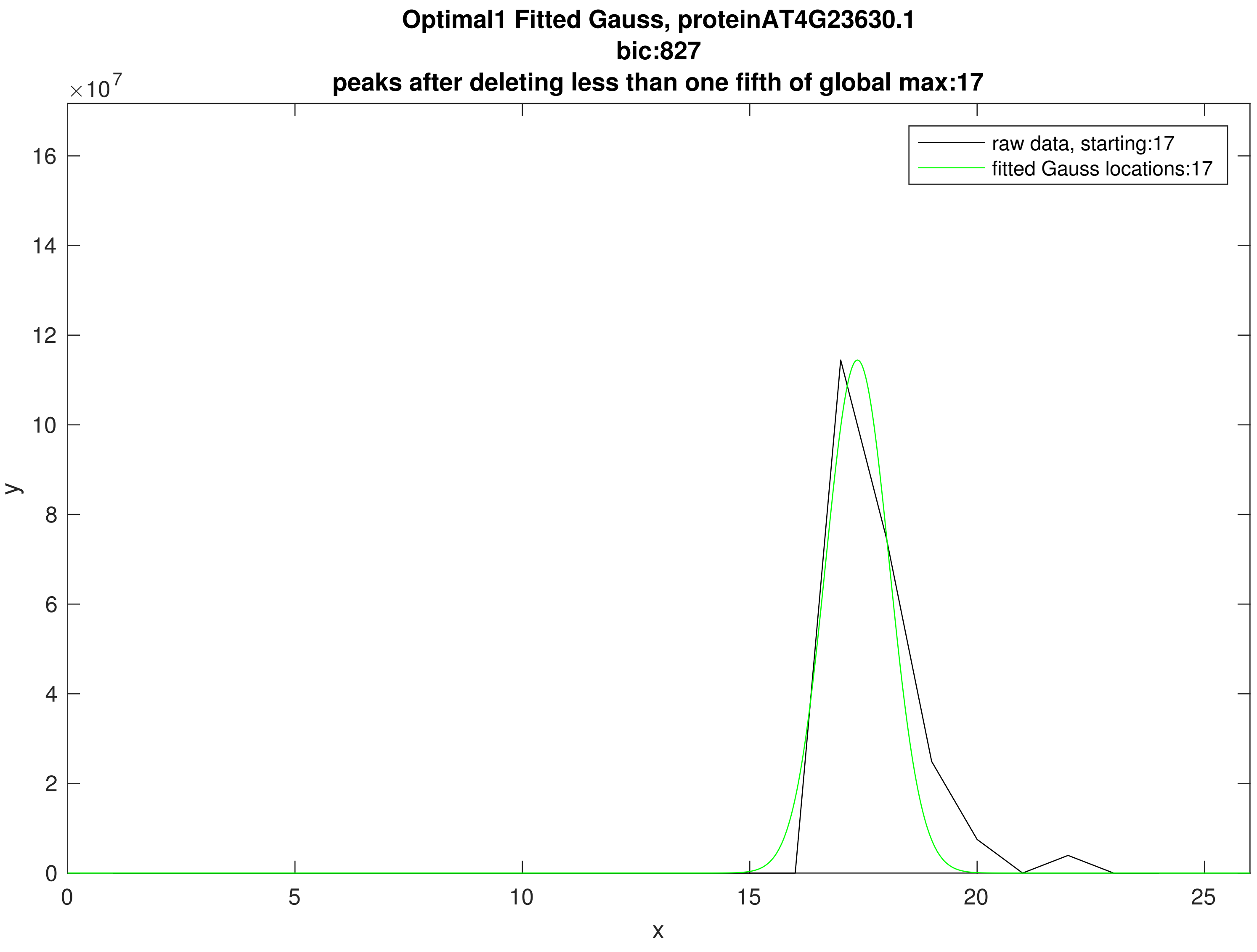
Optimal1 Fitted Gauss, proteinAT4G23630.1
bic:827
peaks after deleting less than one fifth of global max:17
×10^7
raw data, starting:17
fitted Gauss locations:17
y
x
0
2
4
6
8
10
12
14
16
0
5
10
15
20
25

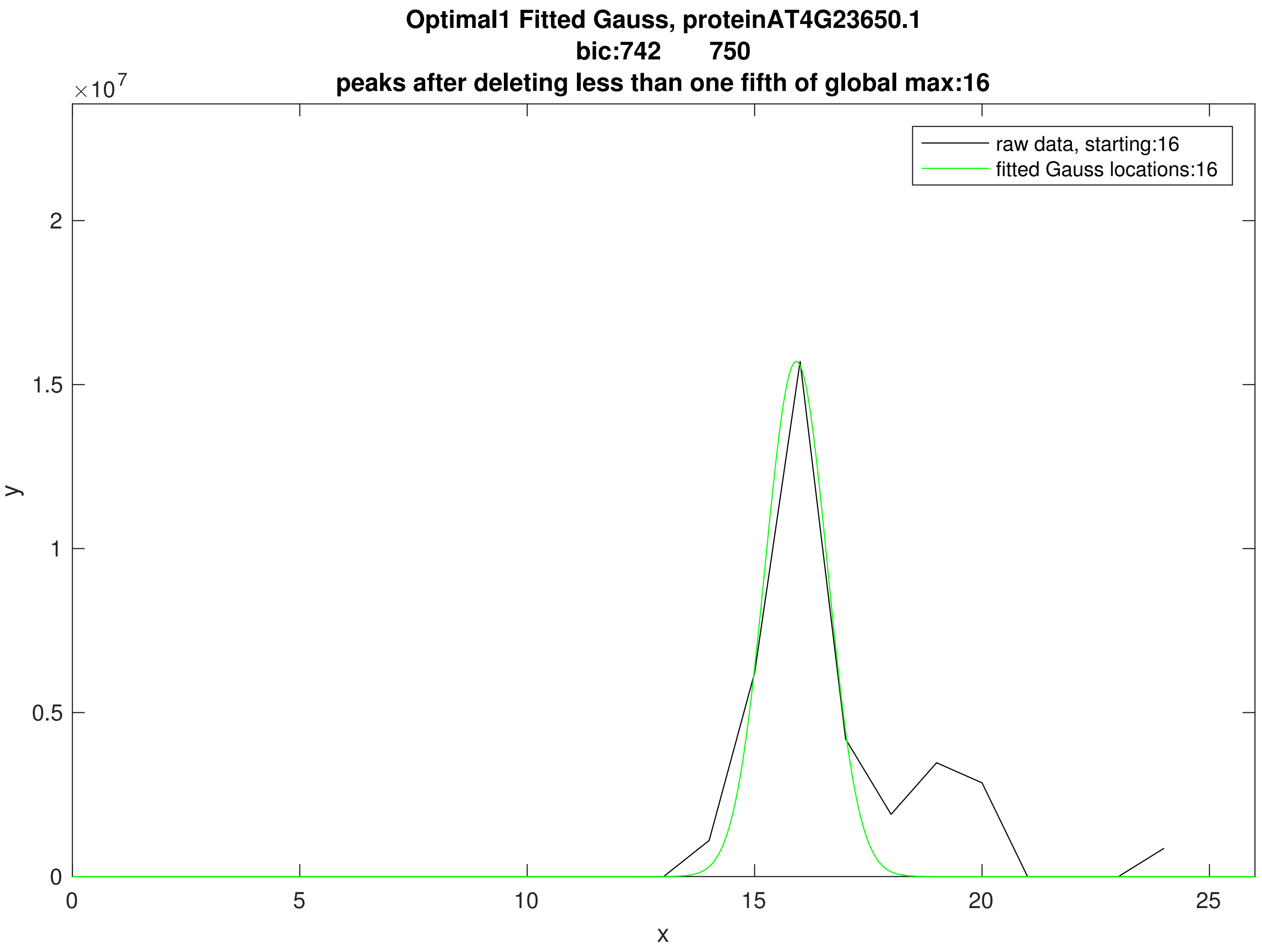

Optimal1 Fitted Gauss, proteinAT4G23650.1
bic:742 750
peaks after deleting less than one fifth of global max:16
×10^7
raw data, starting:16
fitted Gauss locations:16
y
x
0
0.5
1
1.5
2
0
5
10
15
20
25

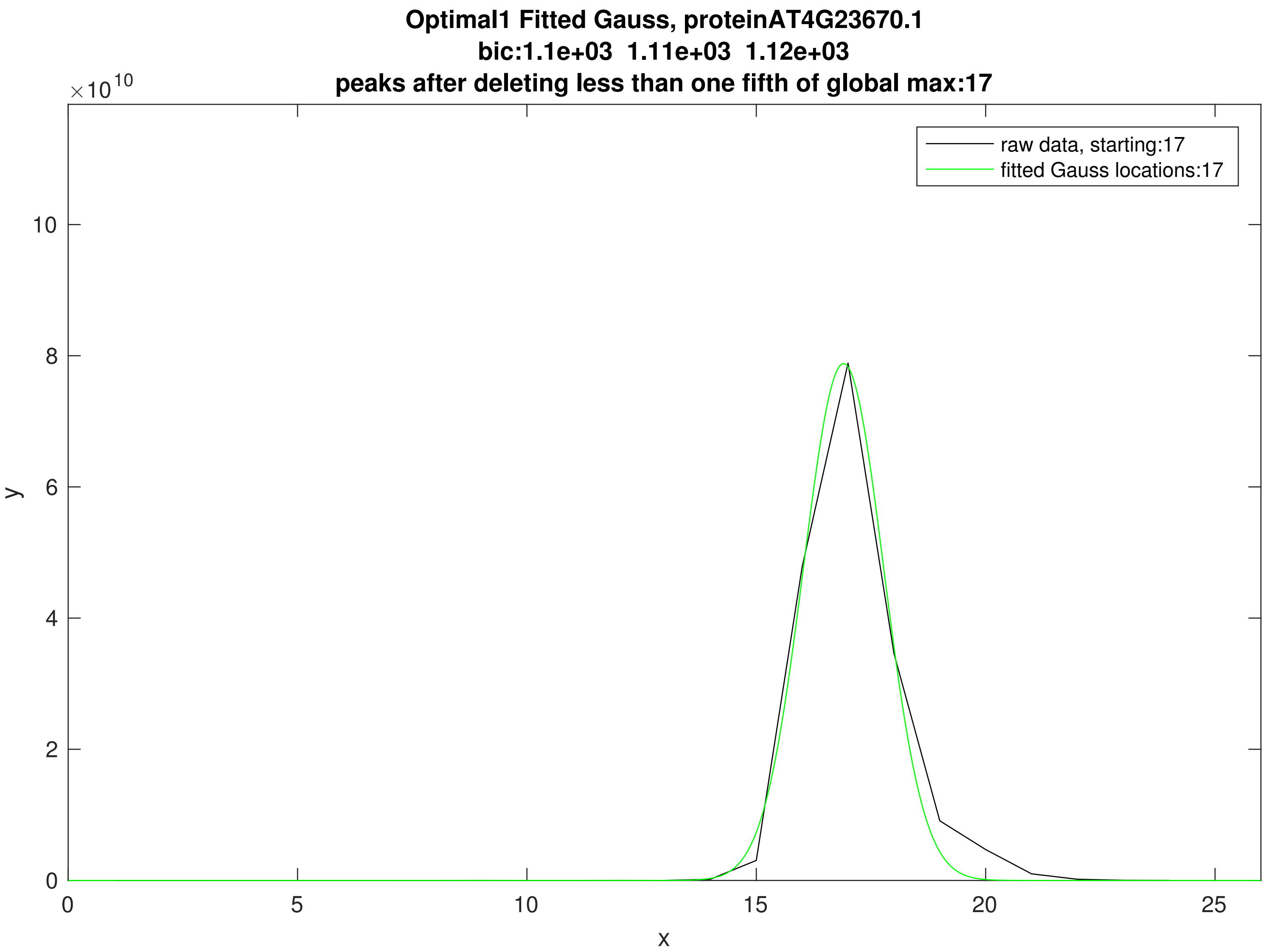
Optimal1 Fitted Gauss, proteinAT4G23670.1
bic:1.1e+03  1.11e+03  1.12e+03
peaks after deleting less than one fifth of global max:17
×10^10
raw data, starting:17
fitted Gauss locations:17
0
2
4
6
8
10
y
0
5
10
15
20
25
x

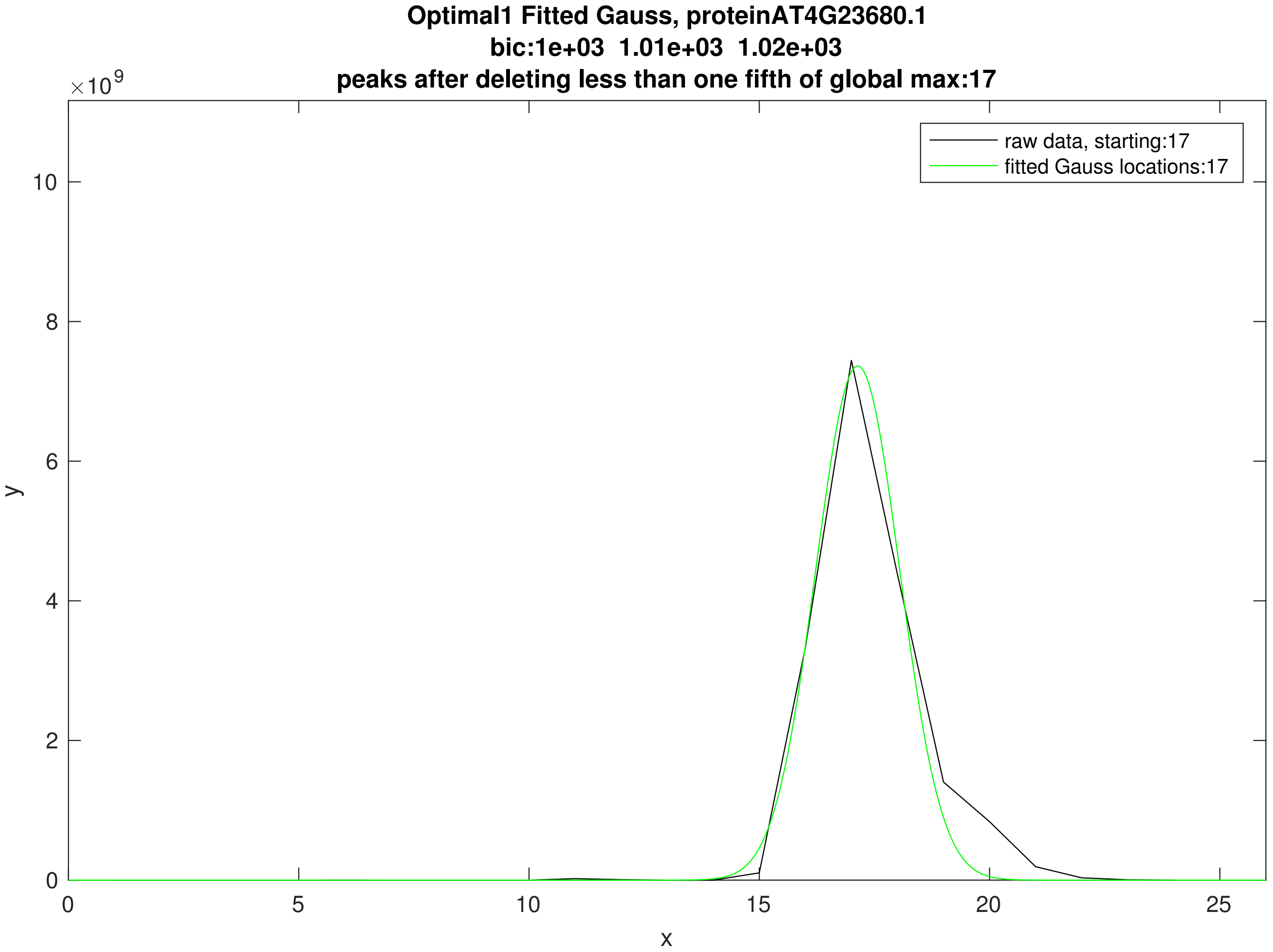

Optimal1 Fitted Gauss, proteinAT4G23680.1
bic:1e+03  1.01e+03  1.02e+03
peaks after deleting less than one fifth of global max:17
×10^9
raw data, starting:17
fitted Gauss locations:17
10
8
6
4
2
0
y
0
5
10
15
20
25
x

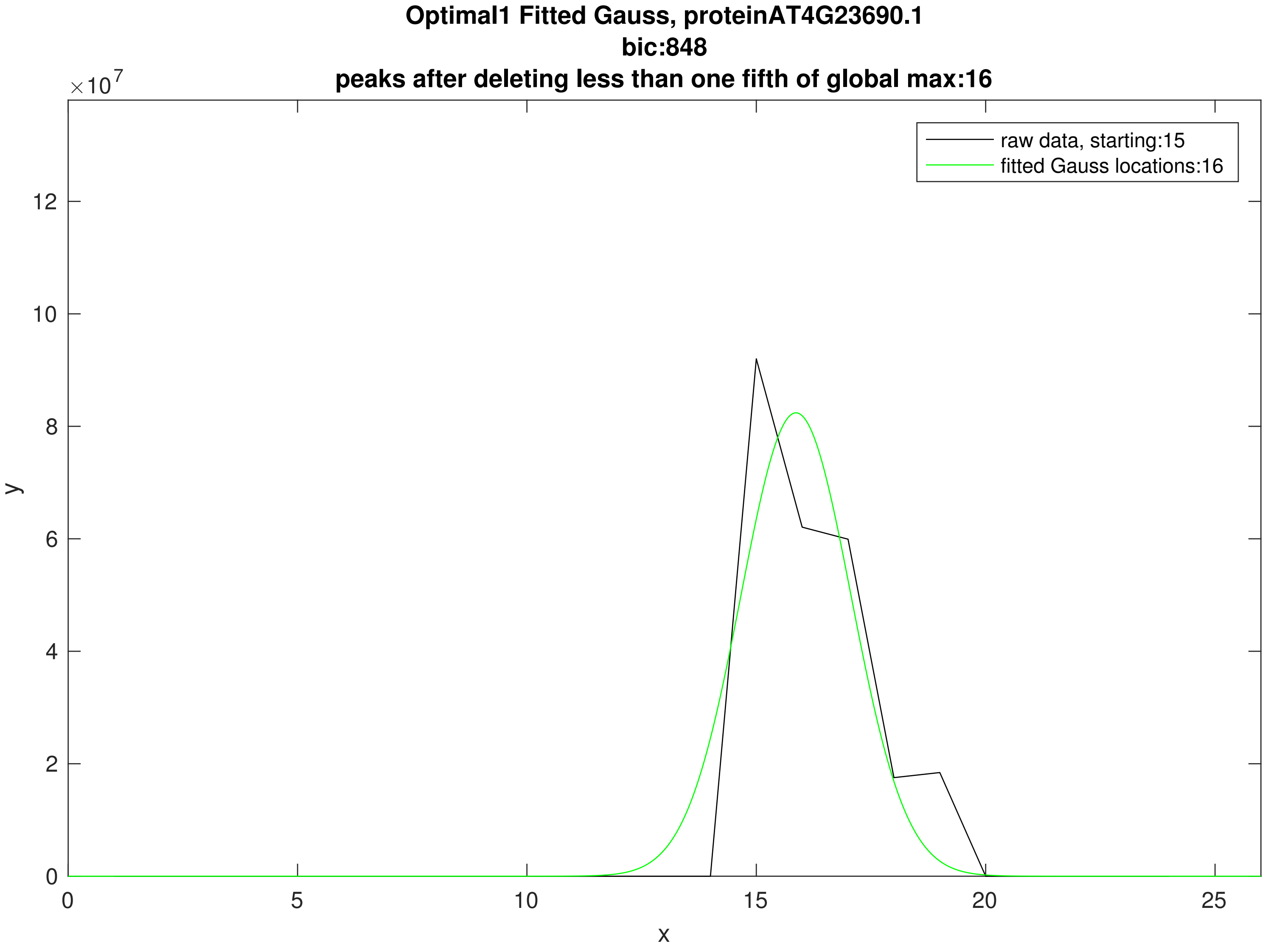
Optimal1 Fitted Gauss, proteinAT4G23690.1
bic:848
peaks after deleting less than one fifth of global max:16
×10^7
raw data, starting:15
fitted Gauss locations:16
12
10
8
6
4
2
0
y
0
5
10
15
20
25
x

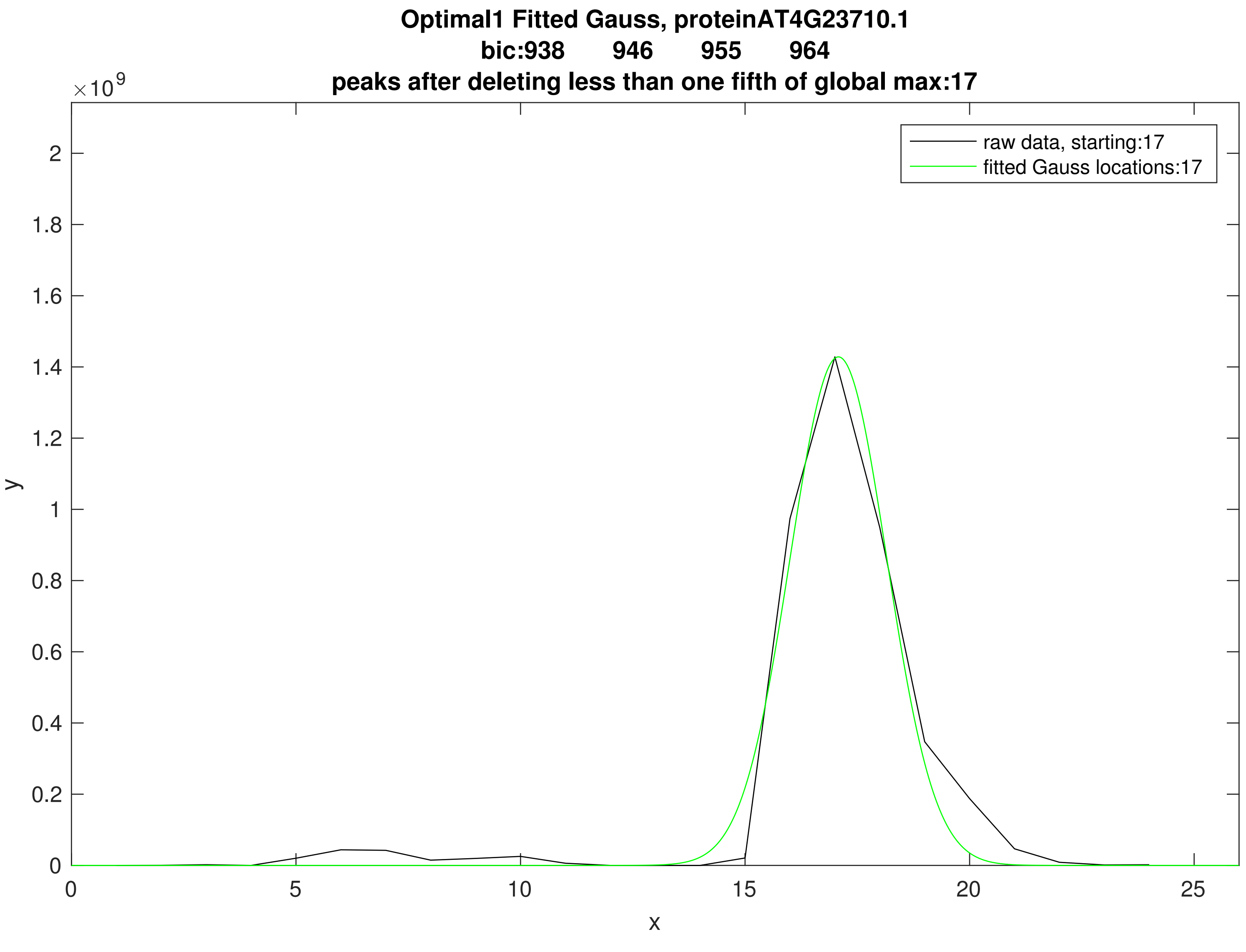
Optimal1 Fitted Gauss, proteinAT4G23710.1
bic:938 946 955 964
peaks after deleting less than one fifth of global max:17
×10^9
raw data, starting:17
fitted Gauss locations:17
y
x
0
0.2
0.4
0.6
0.8
1
1.2
1.4
1.6
1.8
2
5
10
15
20
25

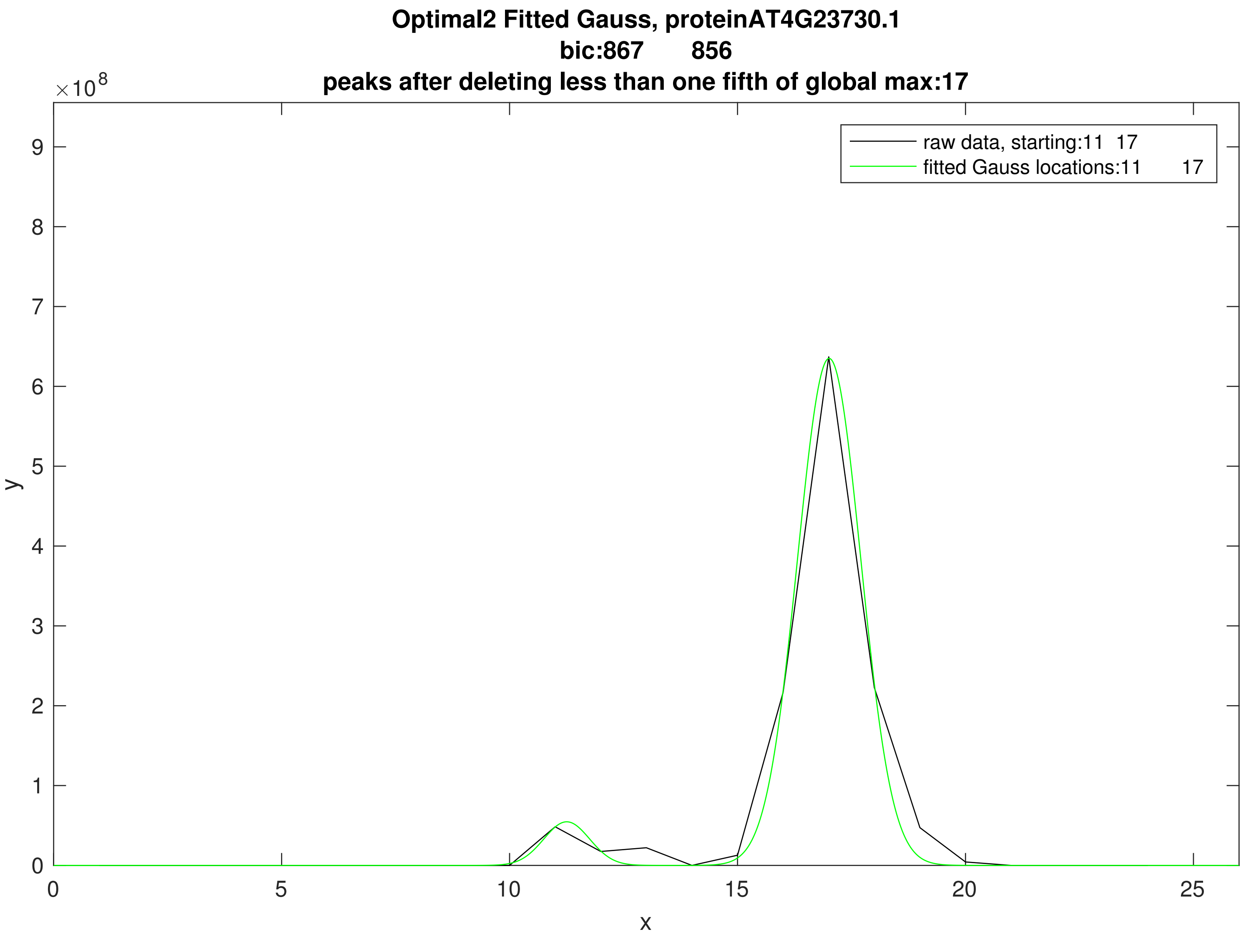

Optimal2 Fitted Gauss, proteinAT4G23730.1
bic:867 856
peaks after deleting less than one fifth of global max:17
×10^8
raw data, starting:11 17
fitted Gauss locations:11 17
y
x
0
1
2
3
4
5
6
7
8
9
0
5
10
15
20
25

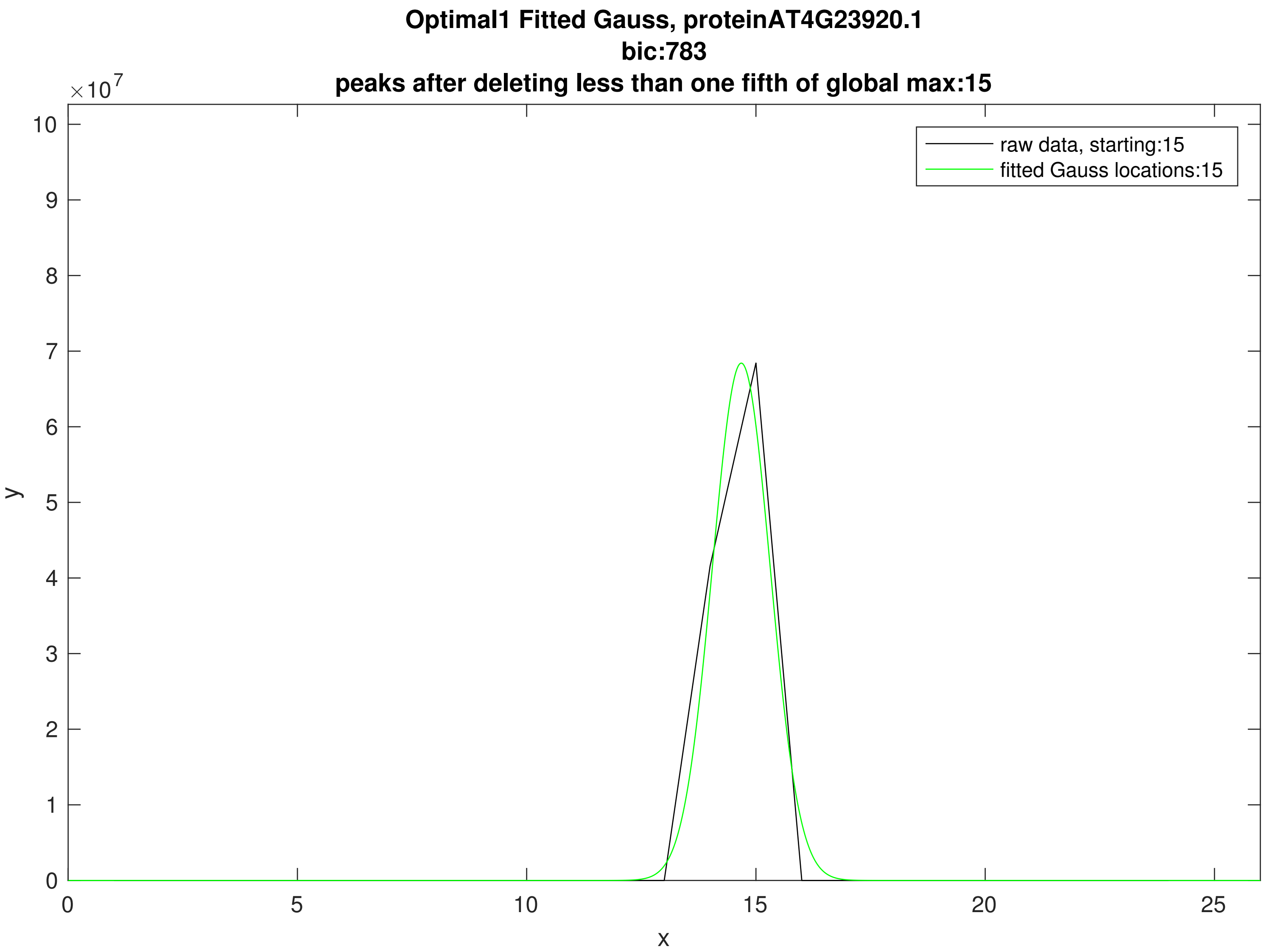

Optimal1 Fitted Gauss, proteinAT4G23920.1
bic:783
peaks after deleting less than one fifth of global max:15
×10^7
10
9
8
7
6
5
4
3
2
1
0
y
0
5
10
15
20
25
x
raw data, starting:15
fitted Gauss locations:15

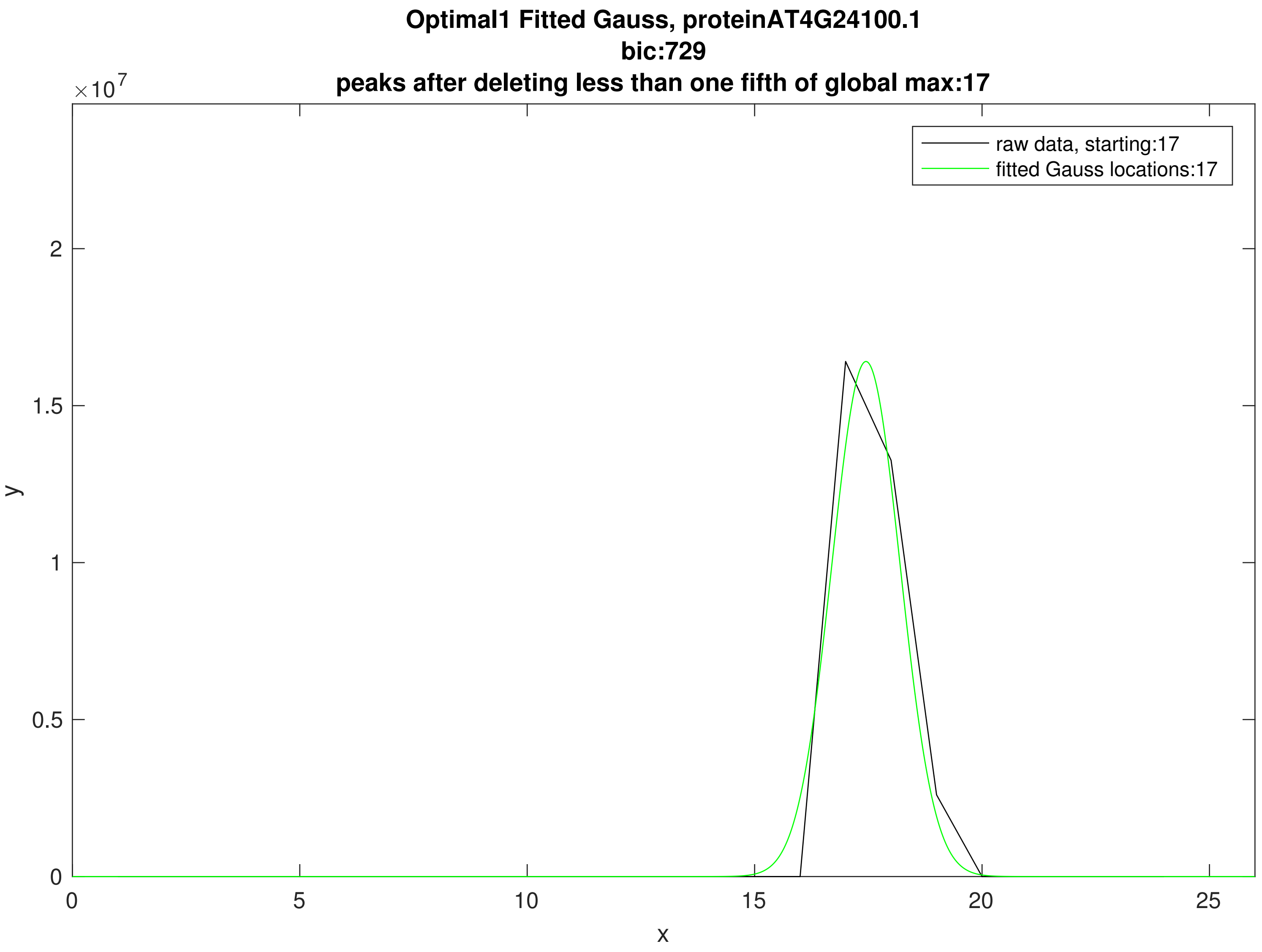

Optimal1 Fitted Gauss, proteinAT4G24100.1
bic:729
peaks after deleting less than one fifth of global max:17
×10^7
raw data, starting:17
fitted Gauss locations:17
y
x
0
0.5
1
1.5
2
0
5
10
15
20
25

**Optimal4 Fitted Gauss, proteinAT4G24190.2;AT4G24190.1**
**bic:874 902 866 865**
**peaks after deleting less than one fifth of global max:6.8 11**

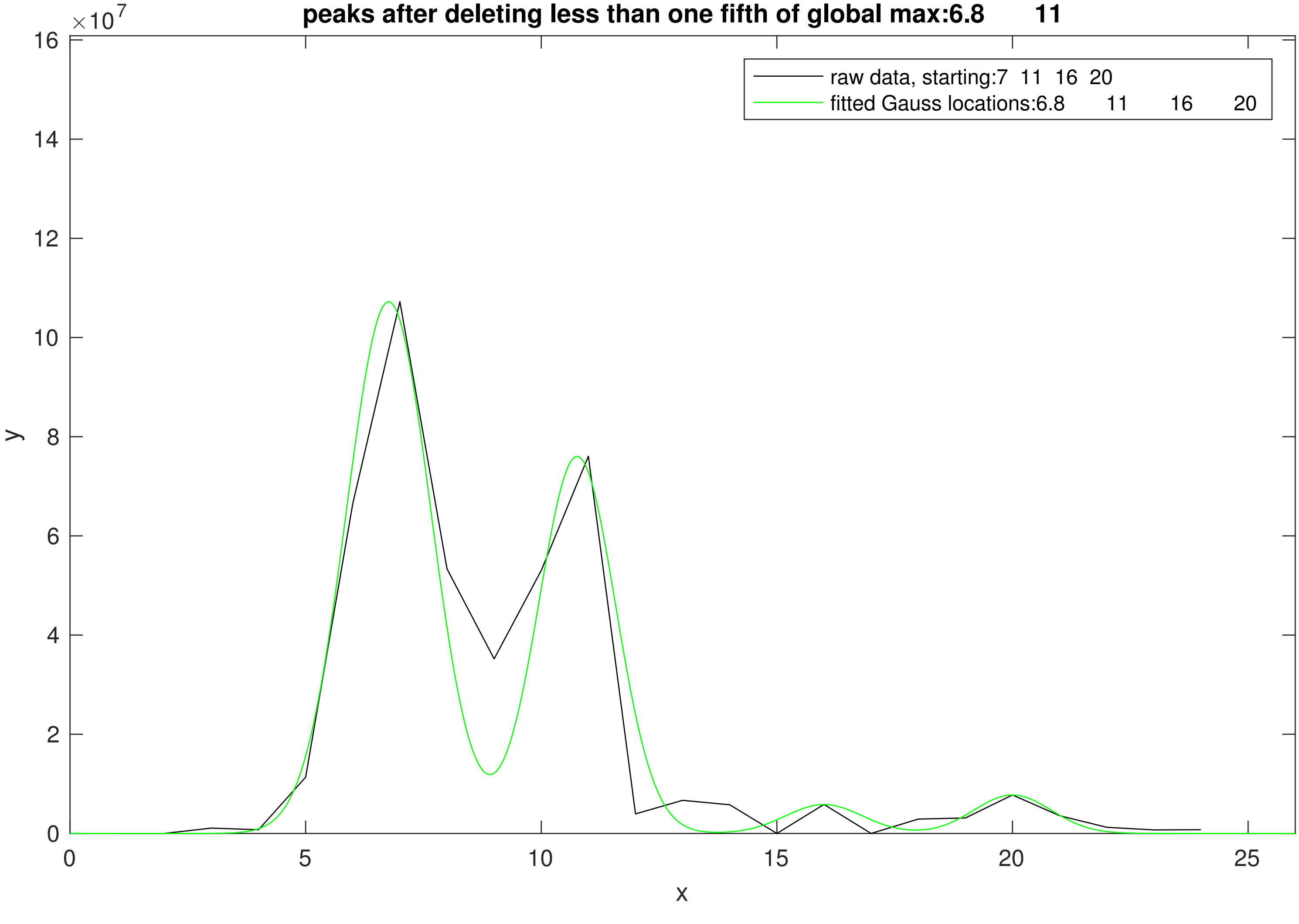

Optimal1 Fitted Gauss, proteinAT4G24220.2;AT4G24220.1
bic:1.06e+03  1.07e+03  1.08e+03
peaks after deleting less than one fifth of global max:13

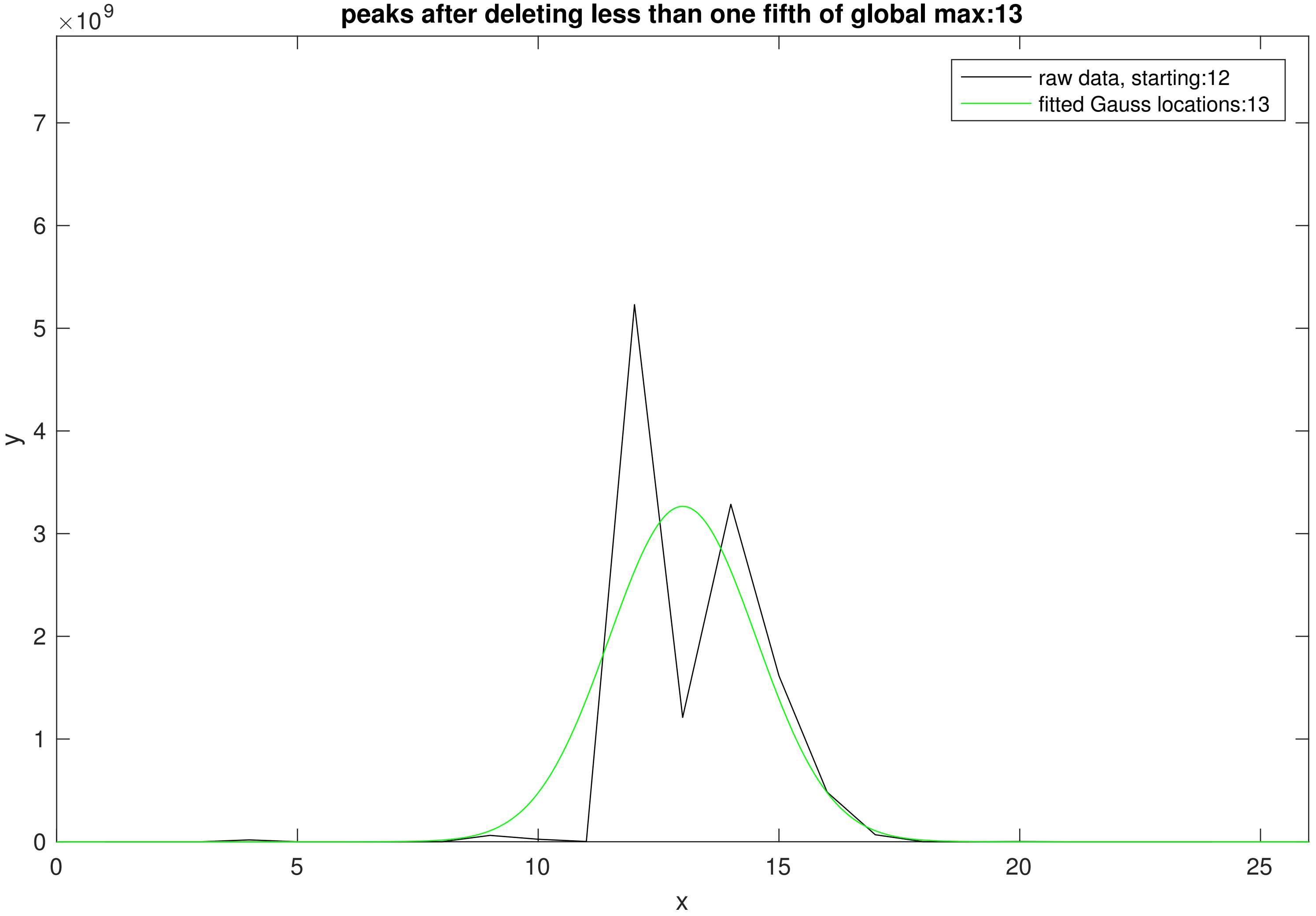

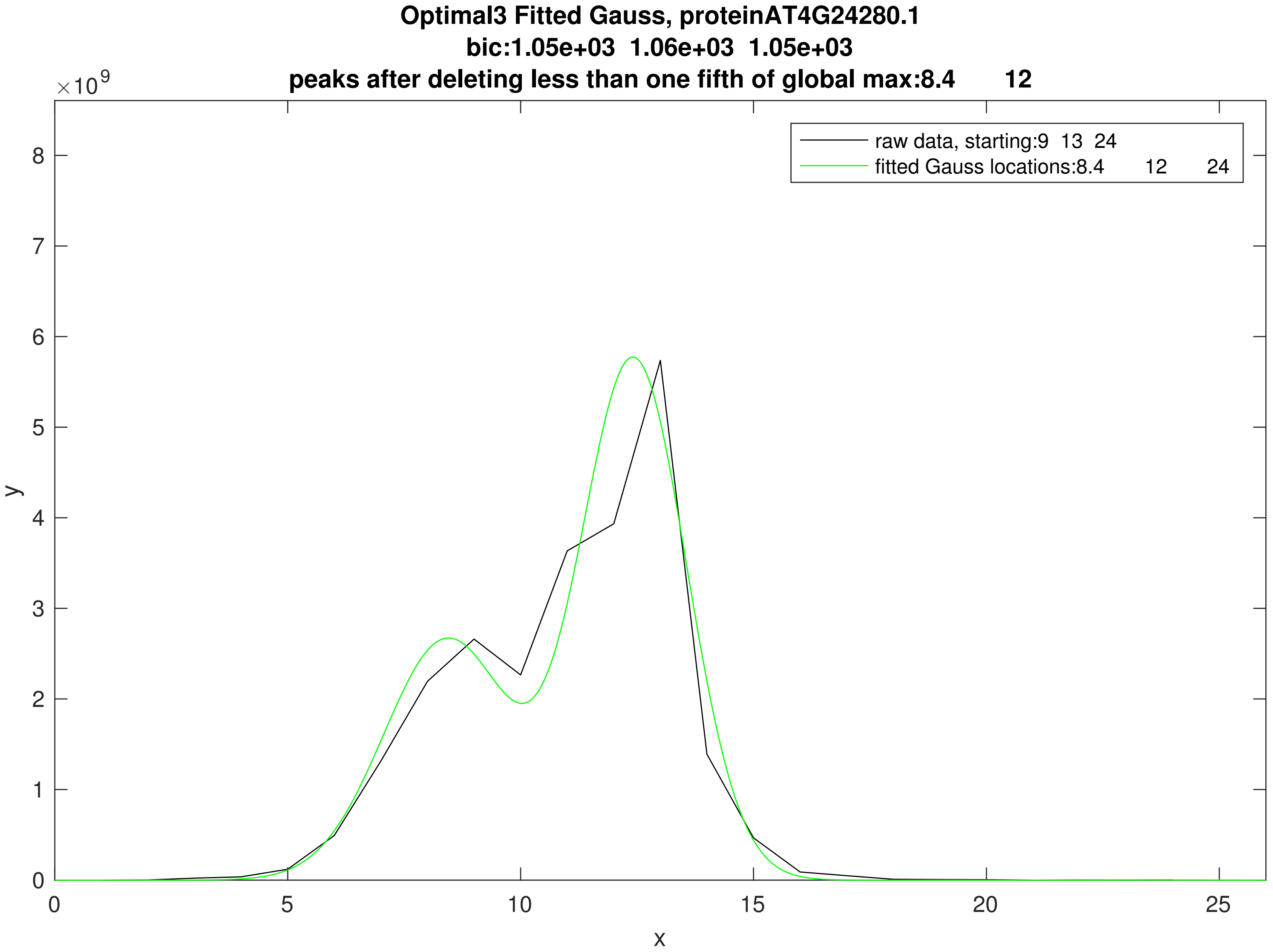
Optimal3 Fitted Gauss, proteinAT4G24280.1
bic:1.05e+03 1.06e+03 1.05e+03
peaks after deleting less than one fifth of global max:8.4 12
×10^9
raw data, starting:9 13 24
fitted Gauss locations:8.4 12 24
y
x
0
1
2
3
4
5
6
7
8
0
5
10
15
20
25

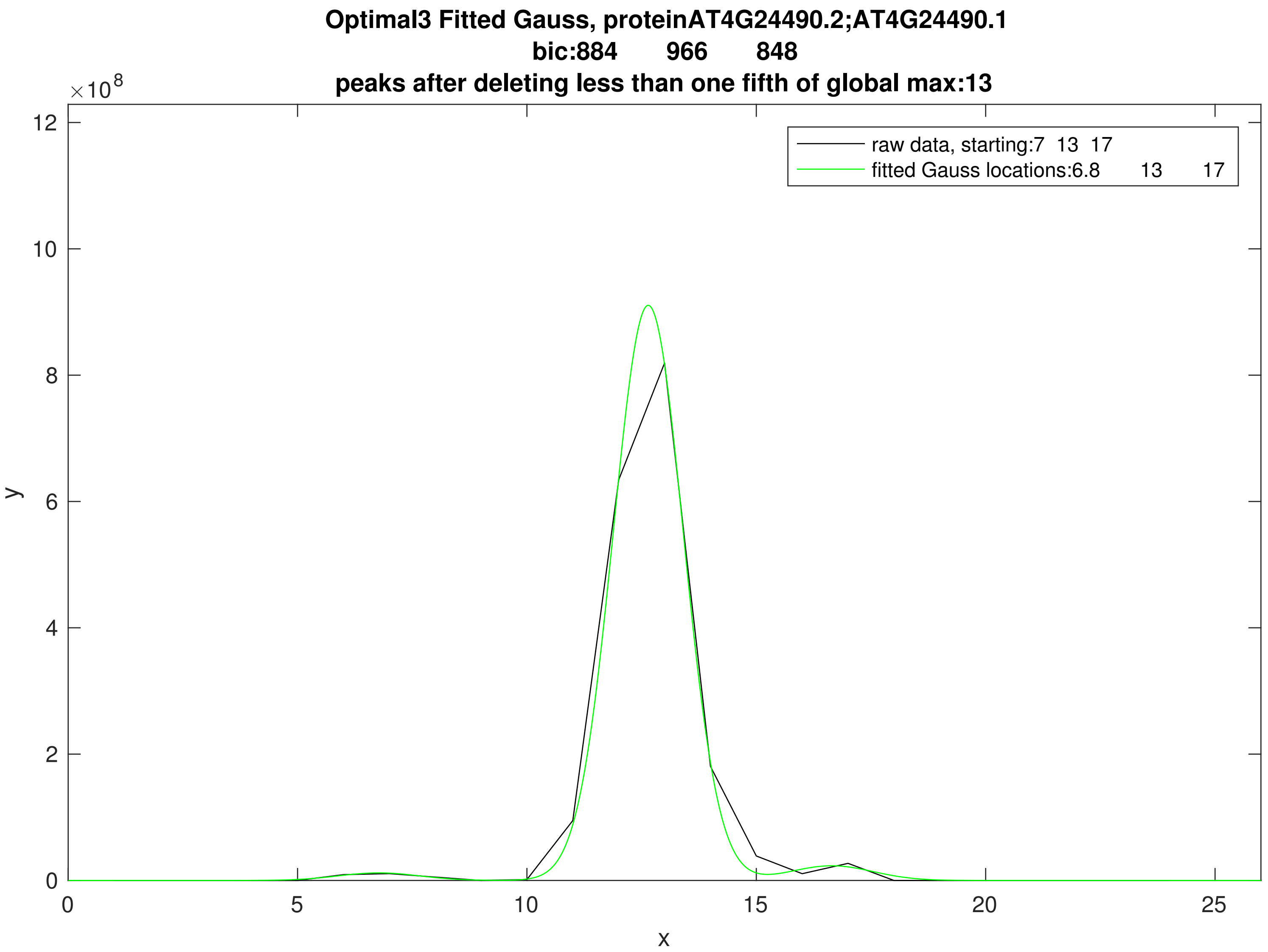
Optimal3 Fitted Gauss, proteinAT4G24490.2;AT4G24490.1
bic:884 966 848
peaks after deleting less than one fifth of global max:13
×10^8
raw data, starting:7 13 17
fitted Gauss locations:6.8 13 17
y
x
0
2
4
6
8
10
12
0
5
10
15
20
25

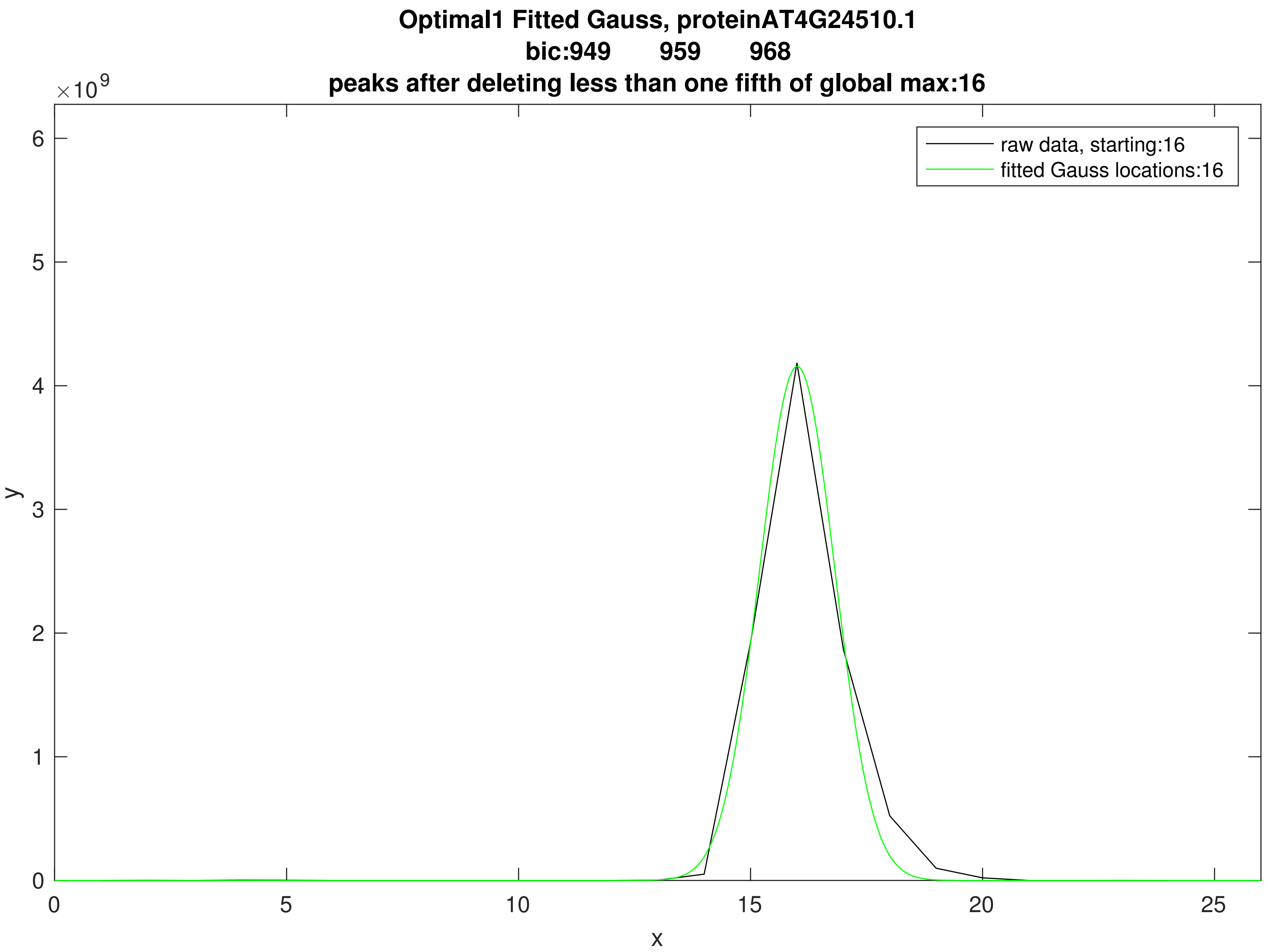

Optimal1 Fitted Gauss, proteinAT4G24510.1
bic:949 959 968
peaks after deleting less than one fifth of global max:16
×10^9
6
5
4
3
2
1
0
y
0
5
10
15
20
25
x
raw data, starting:16
fitted Gauss locations:16

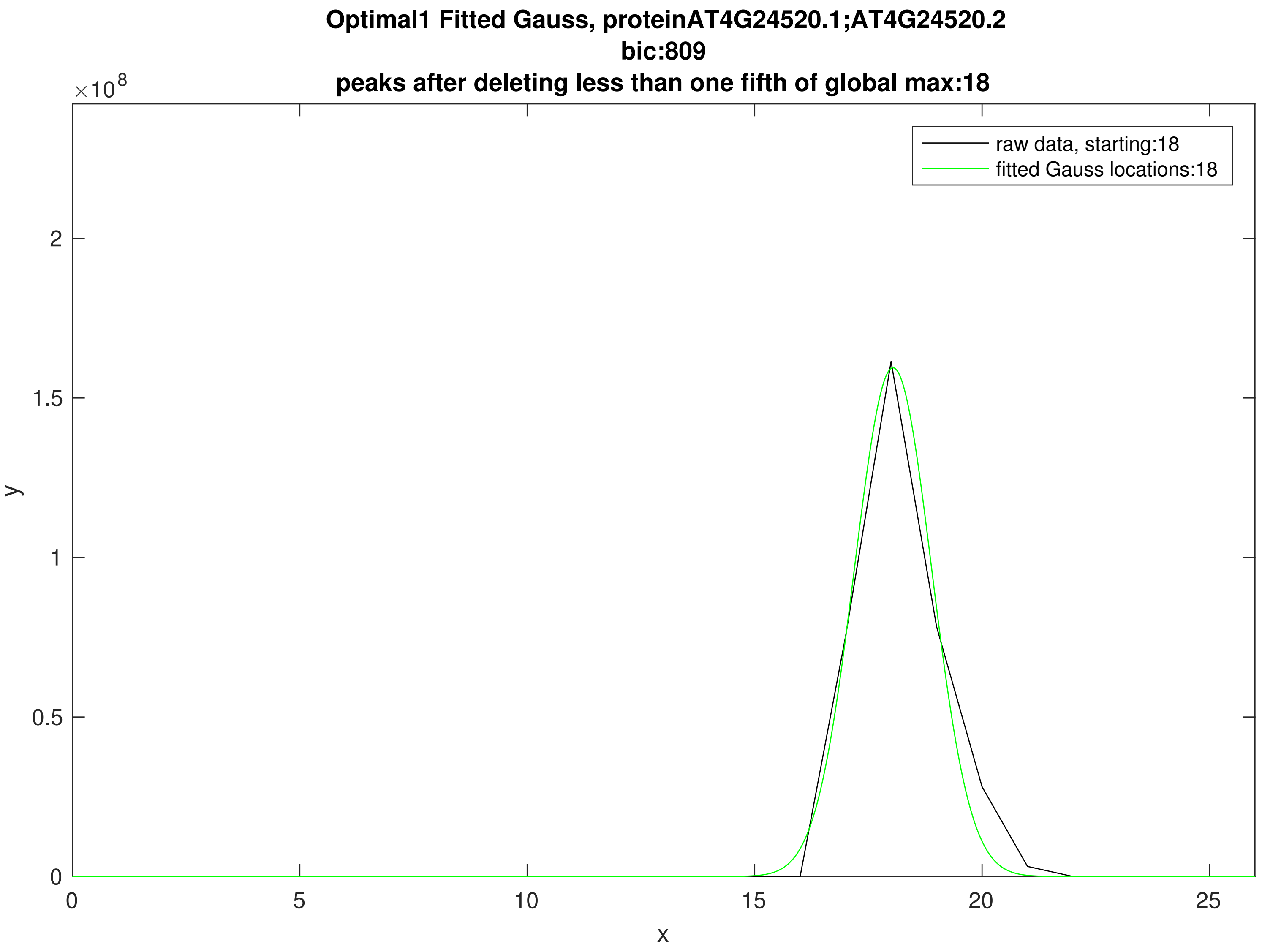

Optimal1 Fitted Gauss, proteinAT4G24520.1;AT4G24520.2
bic:809
peaks after deleting less than one fifth of global max:18
×10^8
raw data, starting:18
fitted Gauss locations:18
y
x
0
0.5
1
1.5
2
0
5
10
15
20
25

#### Optimal1 Fitted Gauss, proteinAT4G24620.1;AT4G24620.2
#### bic:997  1.01e+03
#### peaks after deleting less than one fifth of global max:13

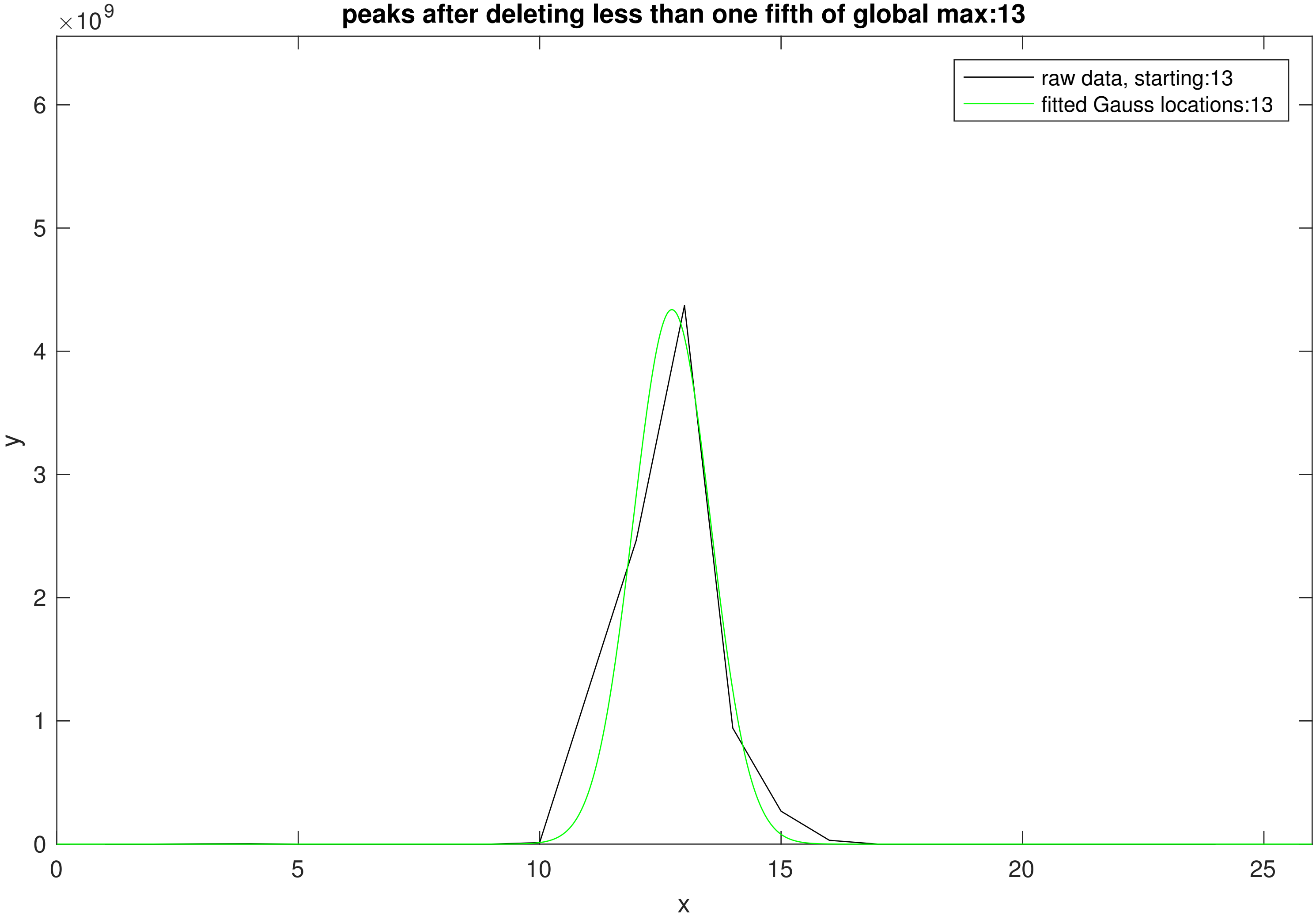

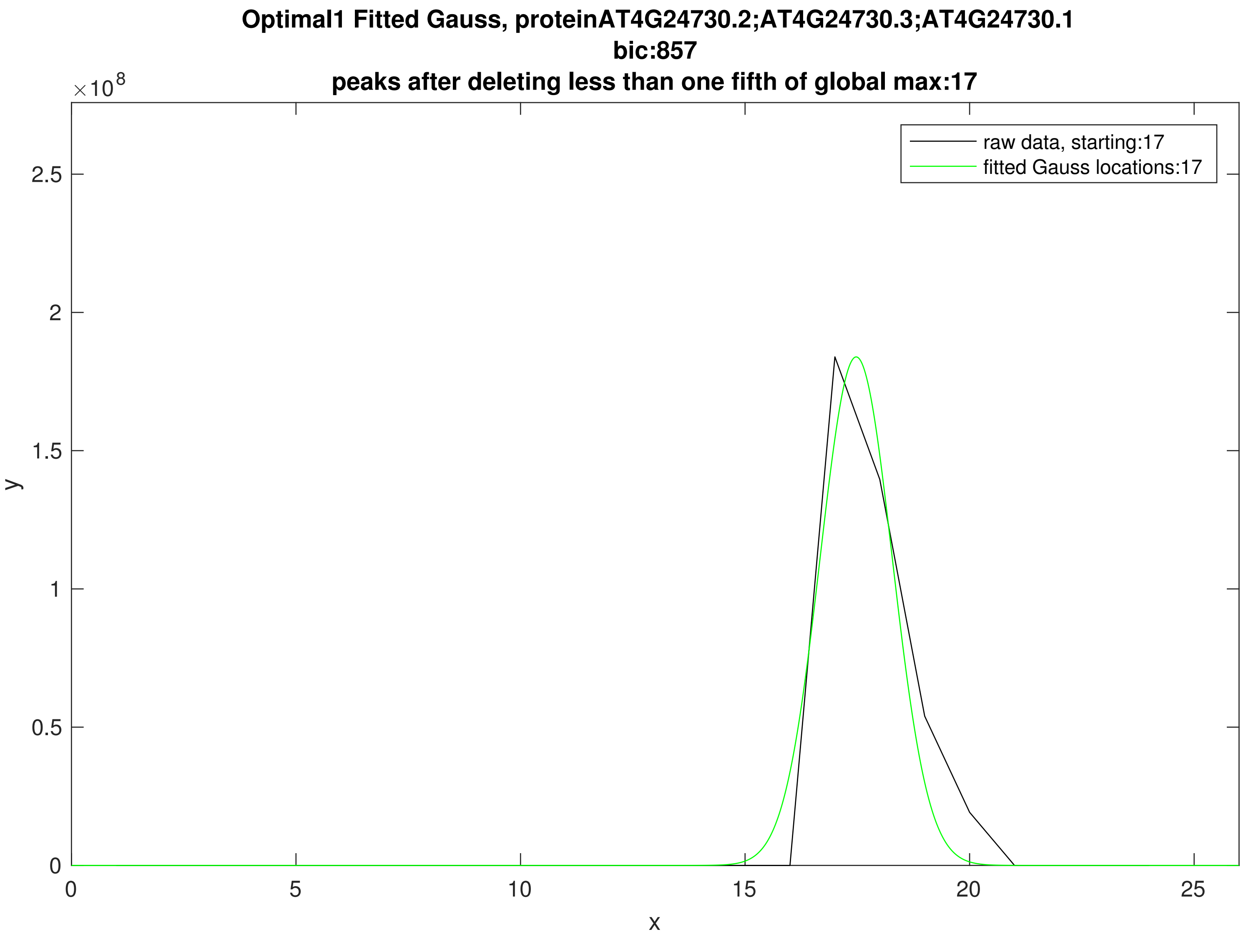
Optimal1 Fitted Gauss, proteinAT4G24730.2;AT4G24730.3;AT4G24730.1
bic:857
peaks after deleting less than one fifth of global max:17
×10^8
2.5
2
1.5
1
0.5
0
y
0
5
10
15
20
25
x
raw data, starting:17
fitted Gauss locations:17

Optimal1 Fitted Gauss, proteinAT4G24770.1
bic:984 988 996
peaks after deleting less than one fifth of global max:15

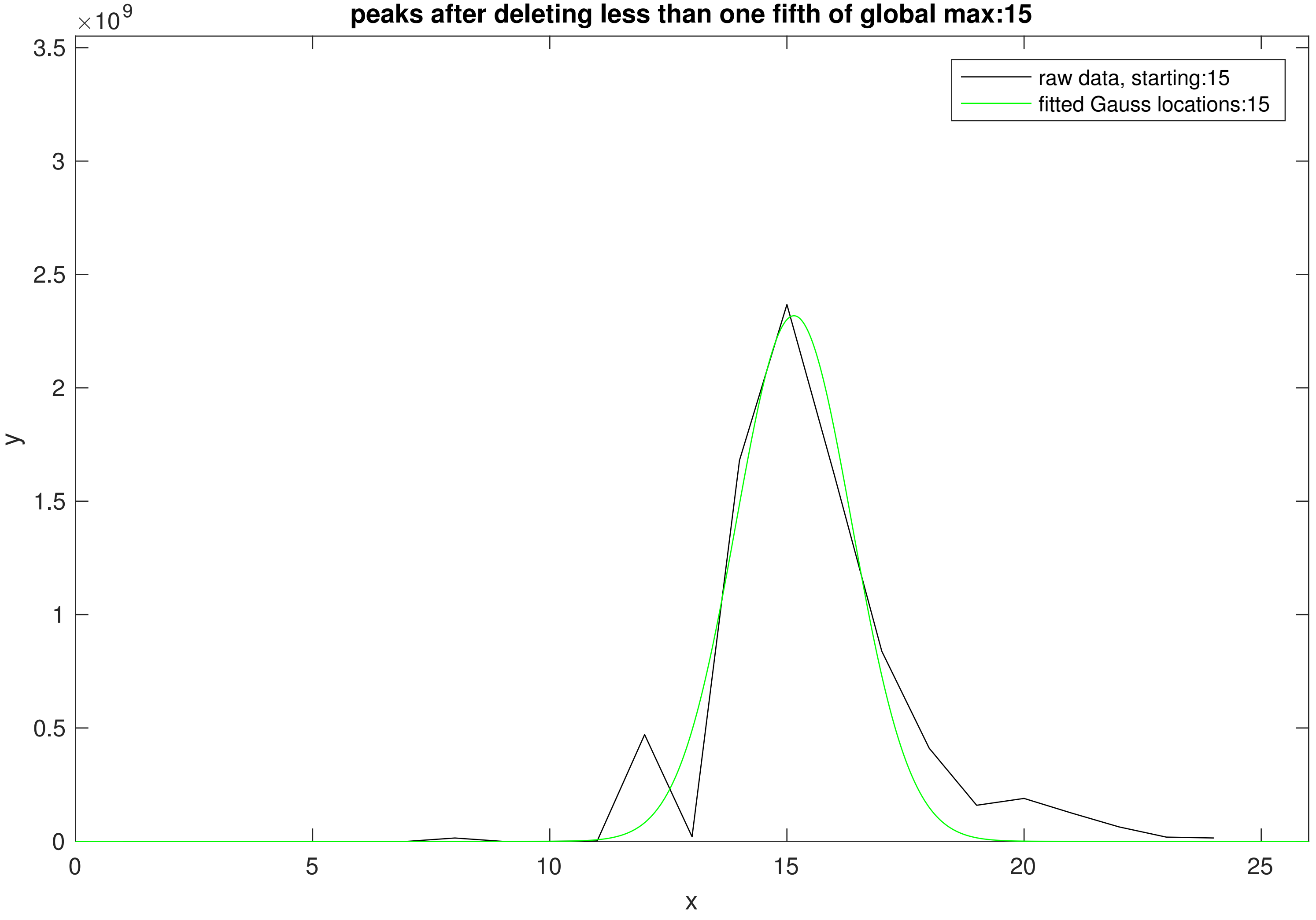

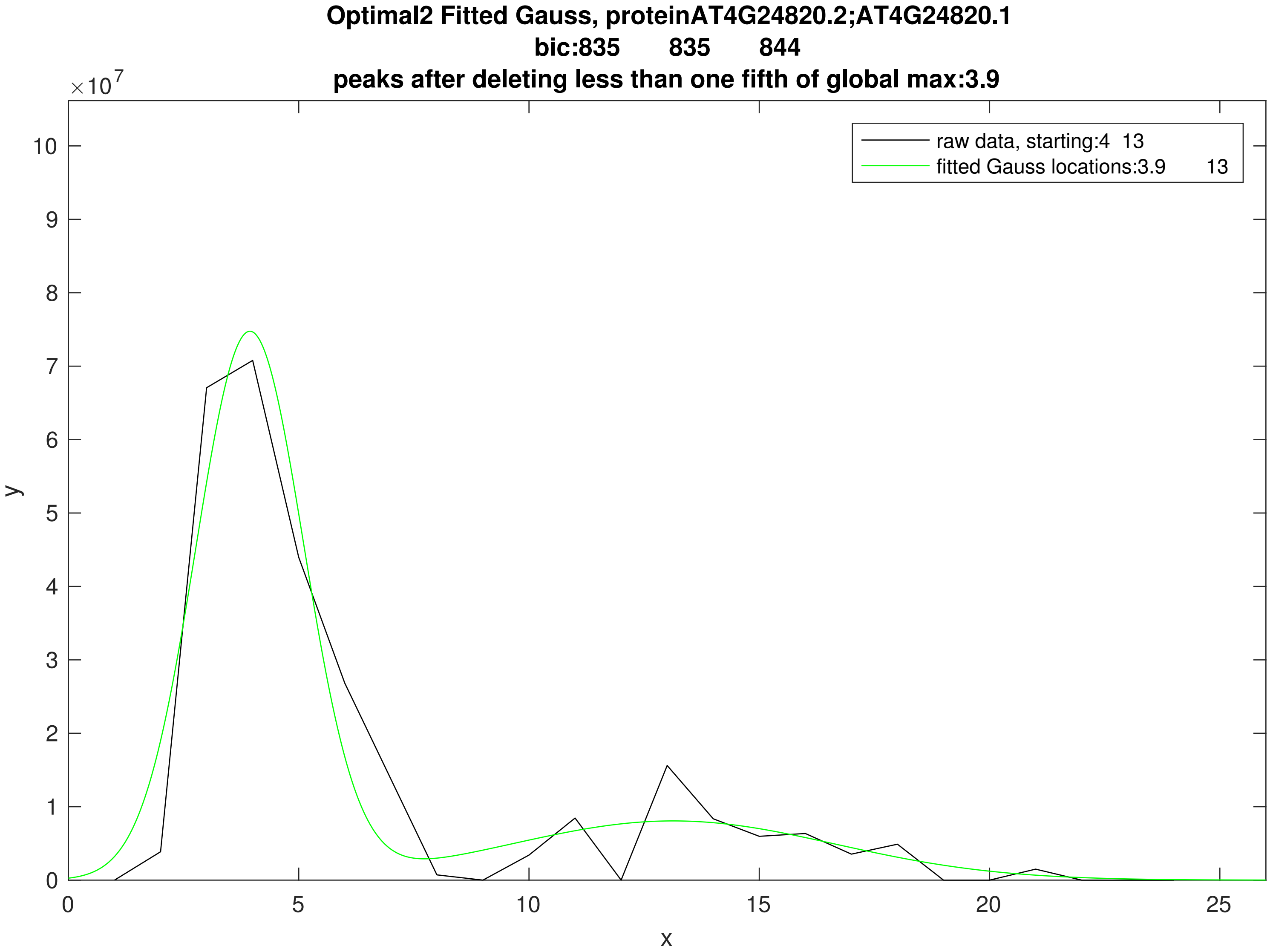

Optimal2 Fitted Gauss, proteinAT4G24820.2;AT4G24820.1
bic:835 835 844
peaks after deleting less than one fifth of global max:3.9
×10^7
raw data, starting:4 13
fitted Gauss locations:3.9 13
y
x
0
1
2
3
4
5
6
7
8
9
10
0
5
10
15
20
25

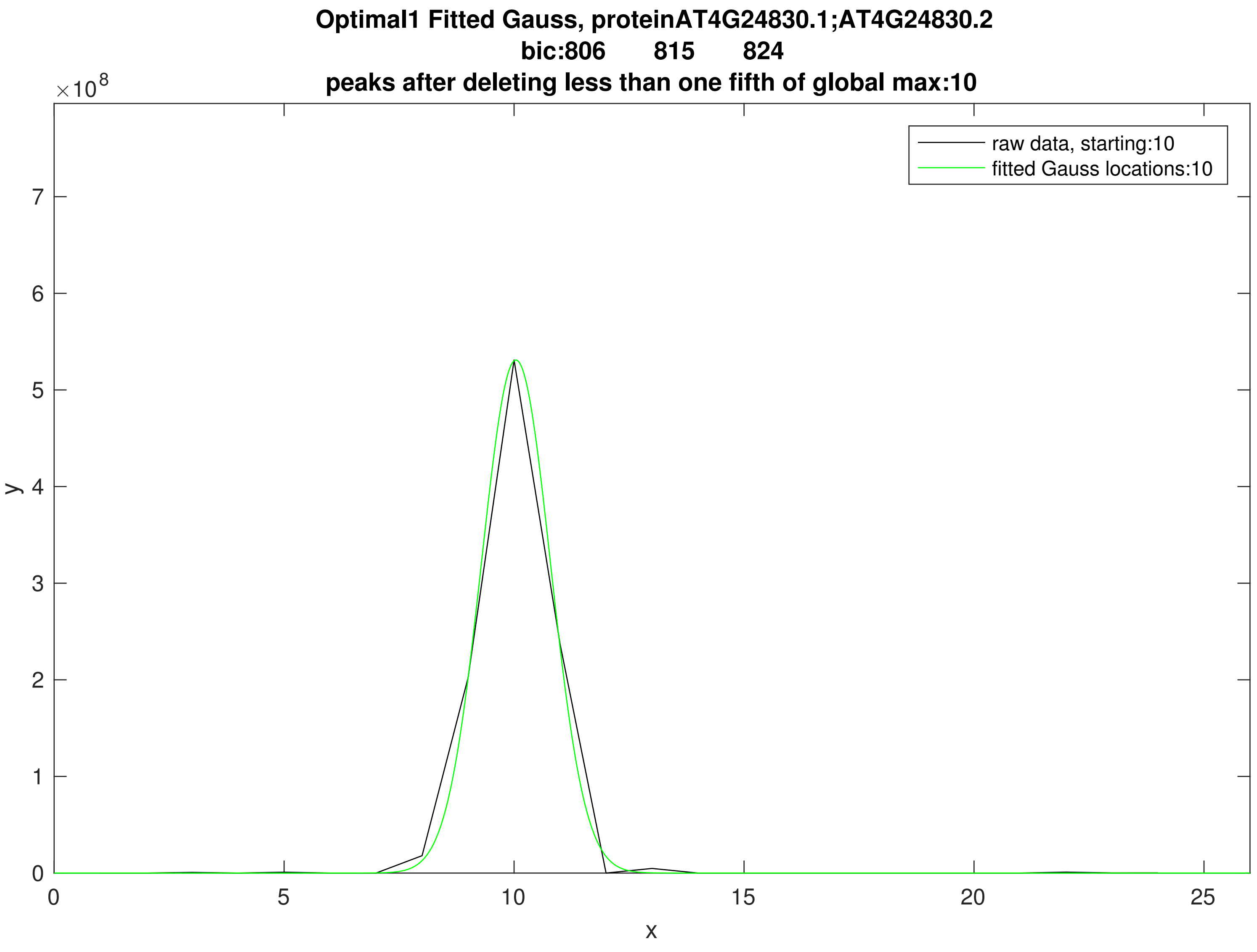
Optimal1 Fitted Gauss, proteinAT4G24830.1;AT4G24830.2
bic:806 815 824
peaks after deleting less than one fifth of global max:10
×10^8
raw data, starting:10
fitted Gauss locations:10
y
x
0
1
2
3
4
5
6
7
0
5
10
15
20
25

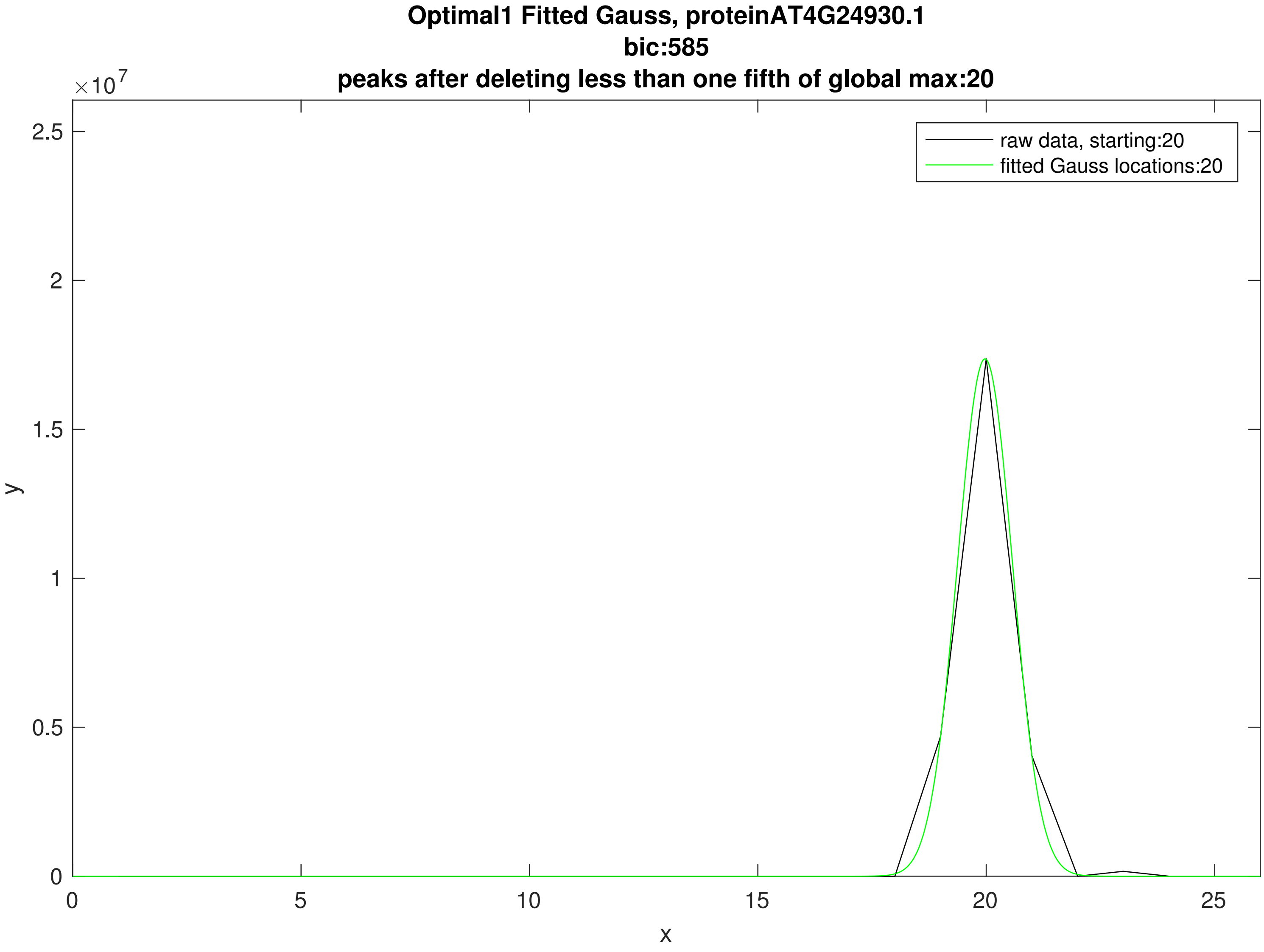

Optimal1 Fitted Gauss, proteinAT4G24930.1
bic:585
peaks after deleting less than one fifth of global max:20
×10^7
2.5
2
1.5
1
0.5
0
y
0
5
10
15
20
25
x
raw data, starting:20
fitted Gauss locations:20

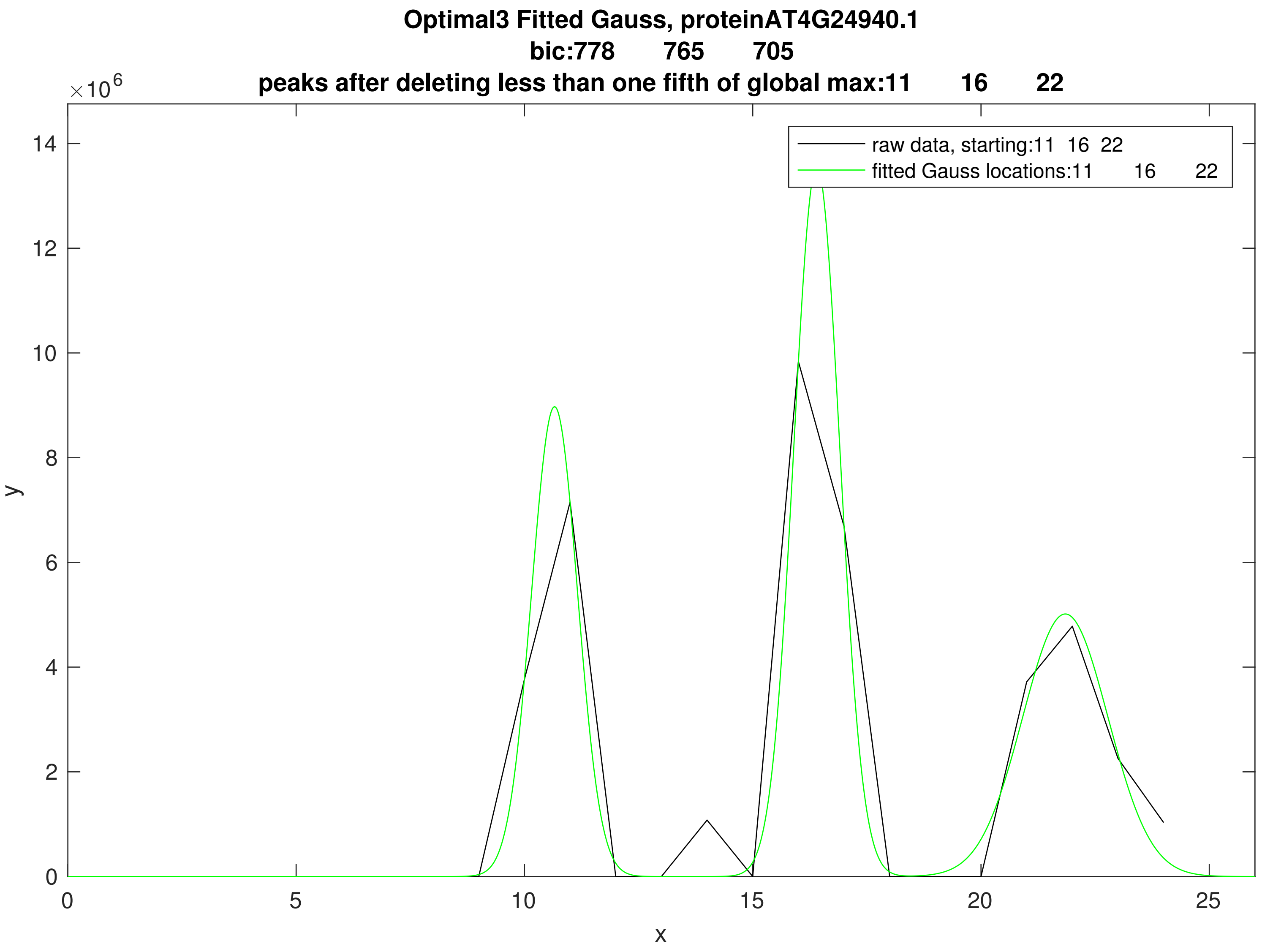

Optimal3 Fitted Gauss, proteinAT4G24940.1
bic:778 765 705
peaks after deleting less than one fifth of global max:11 16 22
×10^6
raw data, starting:11 16 22
fitted Gauss locations:11 16 22
y
x
0
2
4
6
8
10
12
14
0
5
10
15
20
25

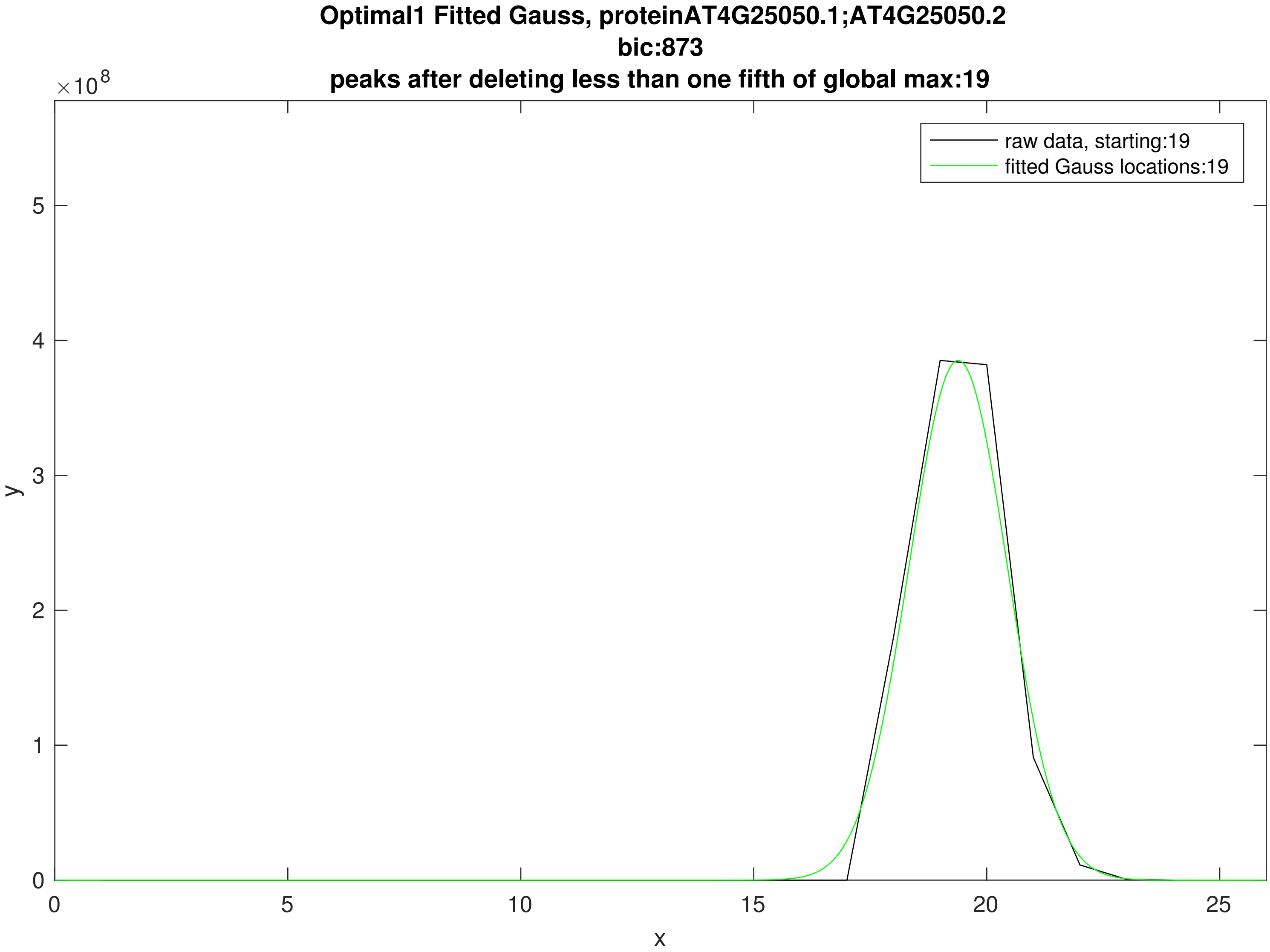

Optimal1 Fitted Gauss, proteinAT4G25050.1;AT4G25050.2
bic:873
peaks after deleting less than one fifth of global max:19
×10^8
raw data, starting:19
fitted Gauss locations:19
y
x
0
1
2
3
4
5
0
5
10
15
20
25

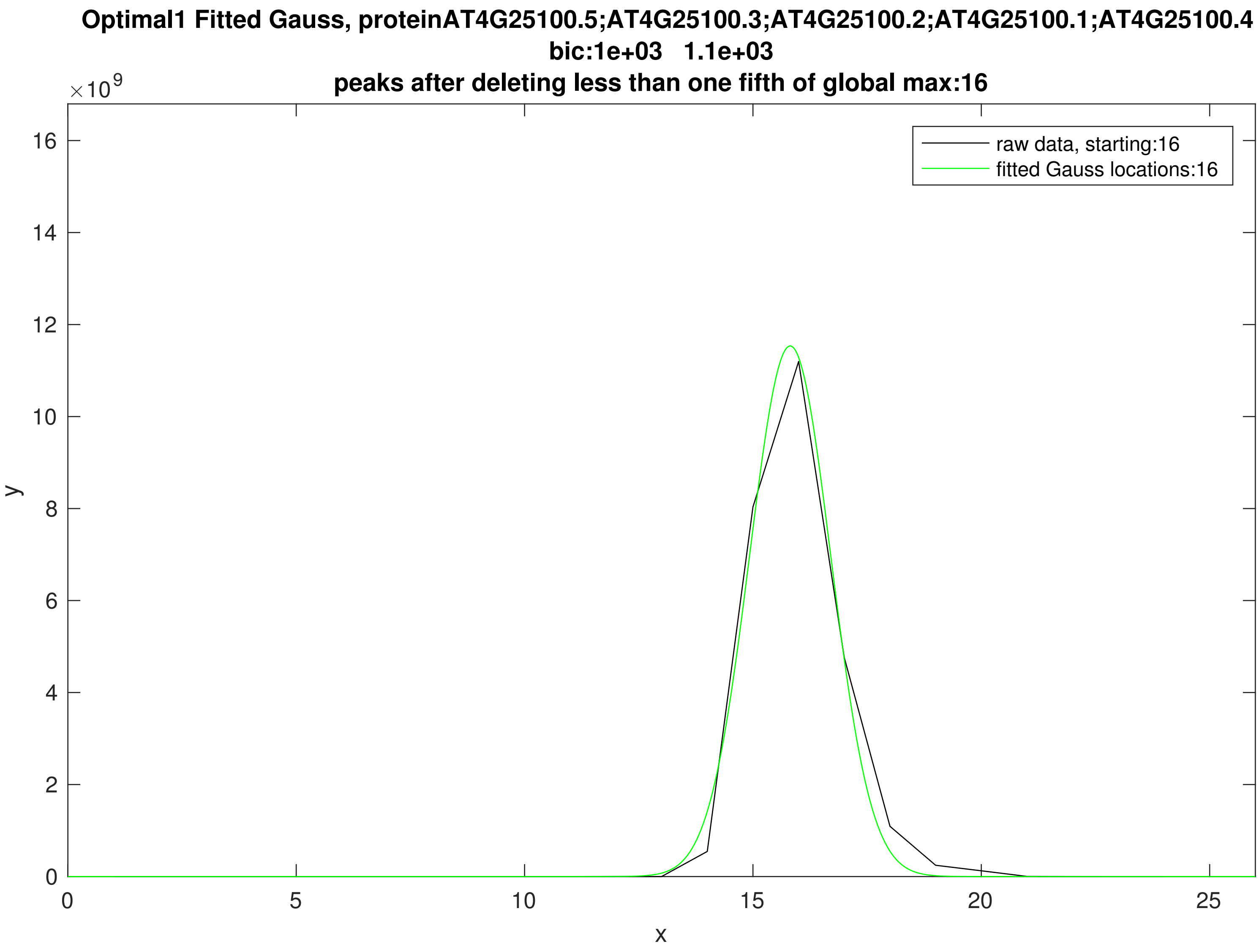
Optimal1 Fitted Gauss, proteinAT4G25100.5;AT4G25100.3;AT4G25100.2;AT4G25100.1;AT4G25100.4
bic:1e+03 1.1e+03
peaks after deleting less than one fifth of global max:16
×10^9
raw data, starting:16
fitted Gauss locations:16
y
x
0
2
4
6
8
10
12
14
16
0
5
10
15
20
25

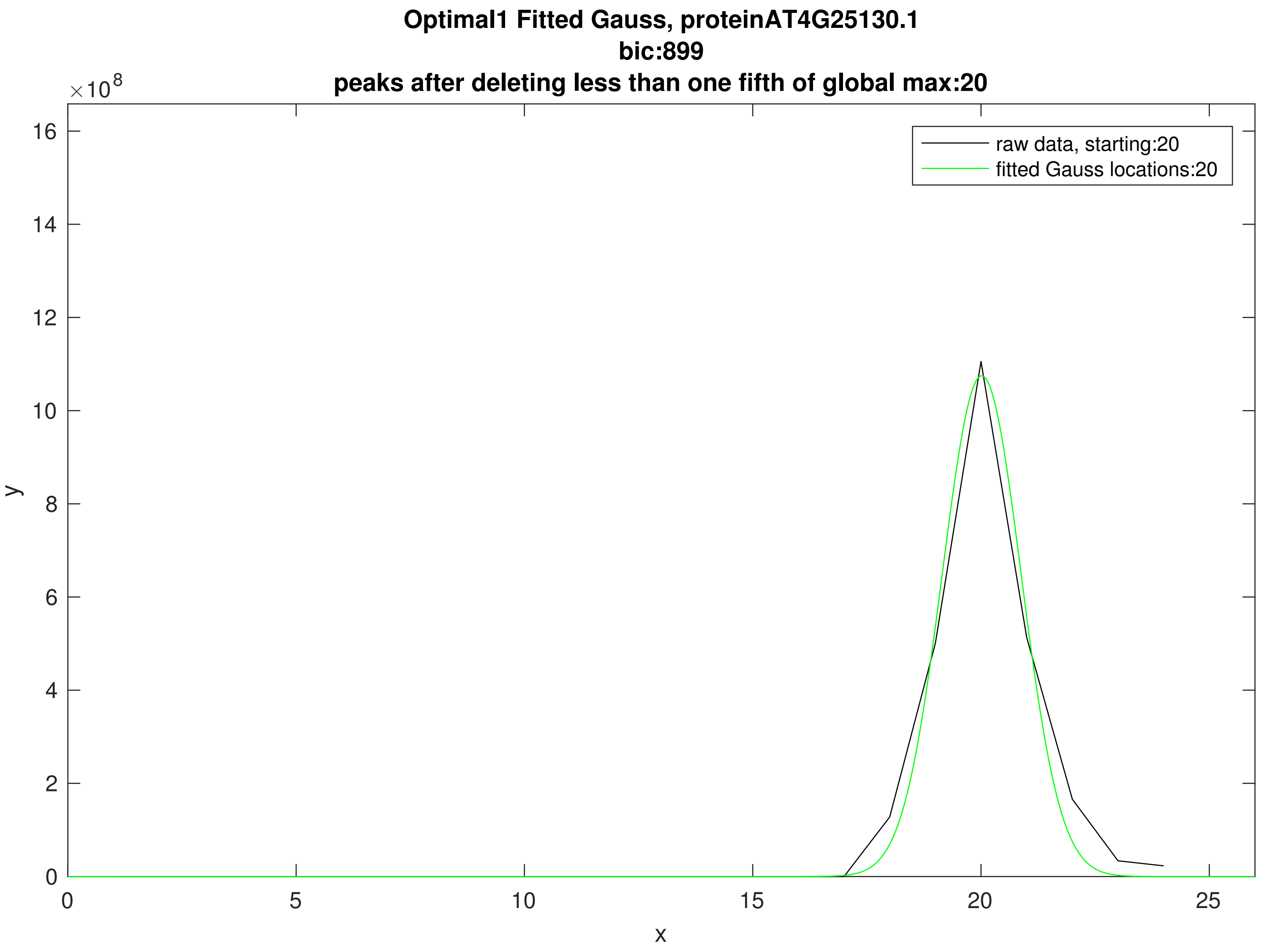
Optimal1 Fitted Gauss, proteinAT4G25130.1
bic:899
peaks after deleting less than one fifth of global max:20
$\times 10^8$
16
14
12
10
8
6
4
2
0
y
0
5
10
15
20
25
x
raw data, starting:20
fitted Gauss locations:20

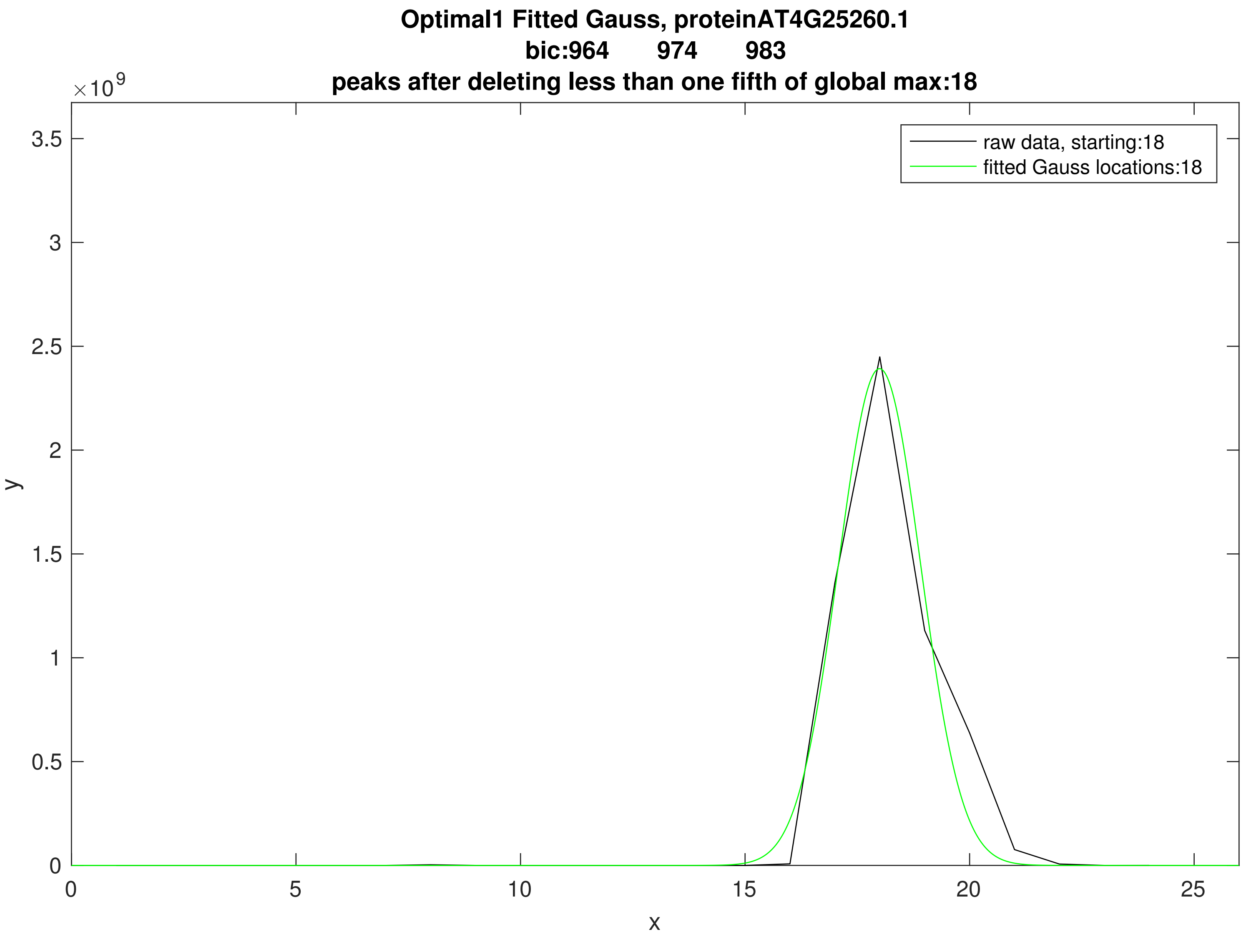

Optimal1 Fitted Gauss, proteinAT4G25260.1
bic:964 974 983
peaks after deleting less than one fifth of global max:18
×10^9
3.5
3
2.5
2
1.5
1
0.5
0
y
0
5
10
15
20
25
x
raw data, starting:18
fitted Gauss locations:18

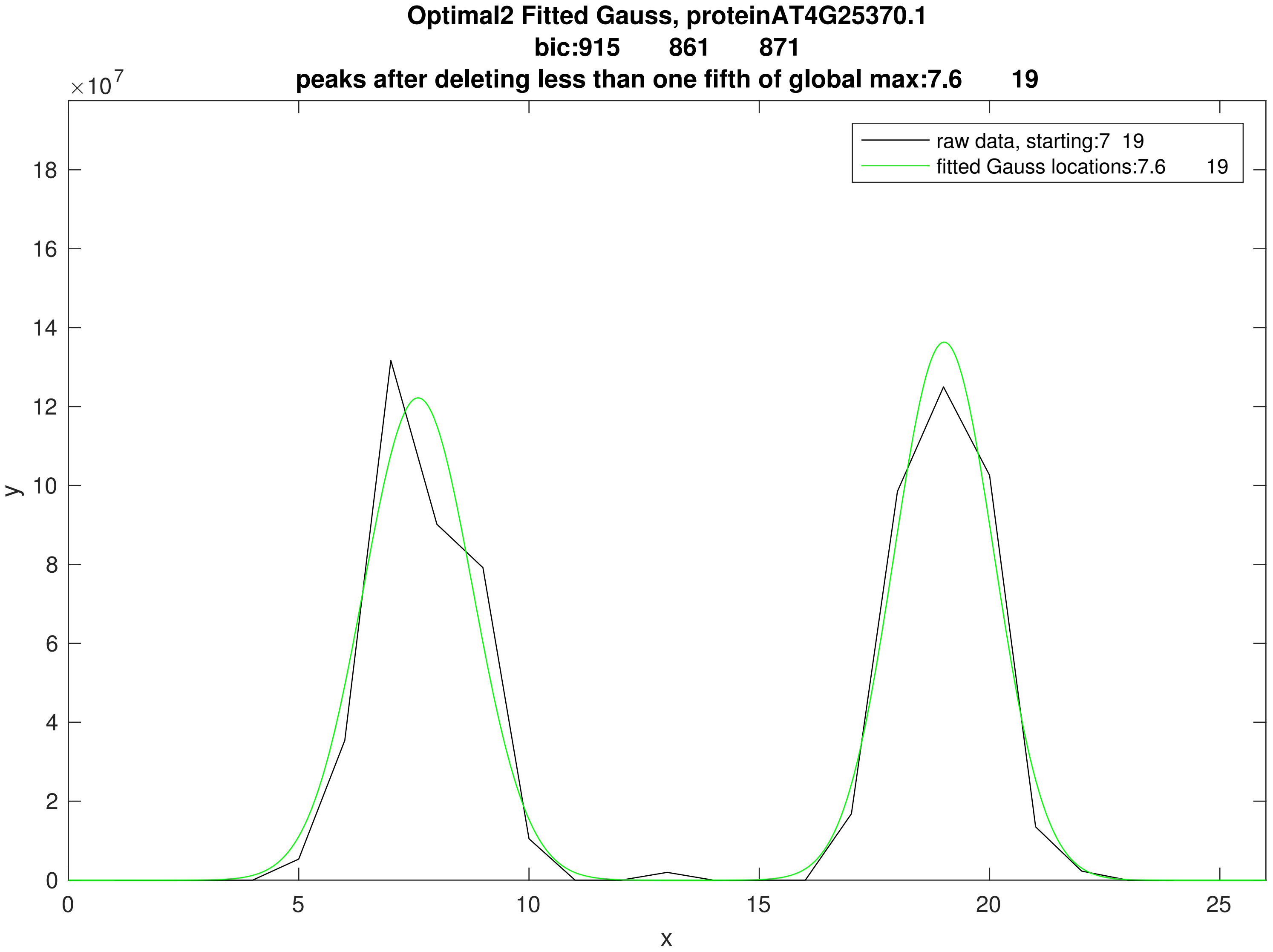
Optimal2 Fitted Gauss, proteinAT4G25370.1
bic:915 861 871
peaks after deleting less than one fifth of global max:7.6 19
×10^7
raw data, starting:7 19
fitted Gauss locations:7.6 19
18
16
14
12
10
8
6
4
2
0
y
0
5
10
15
20
25
x

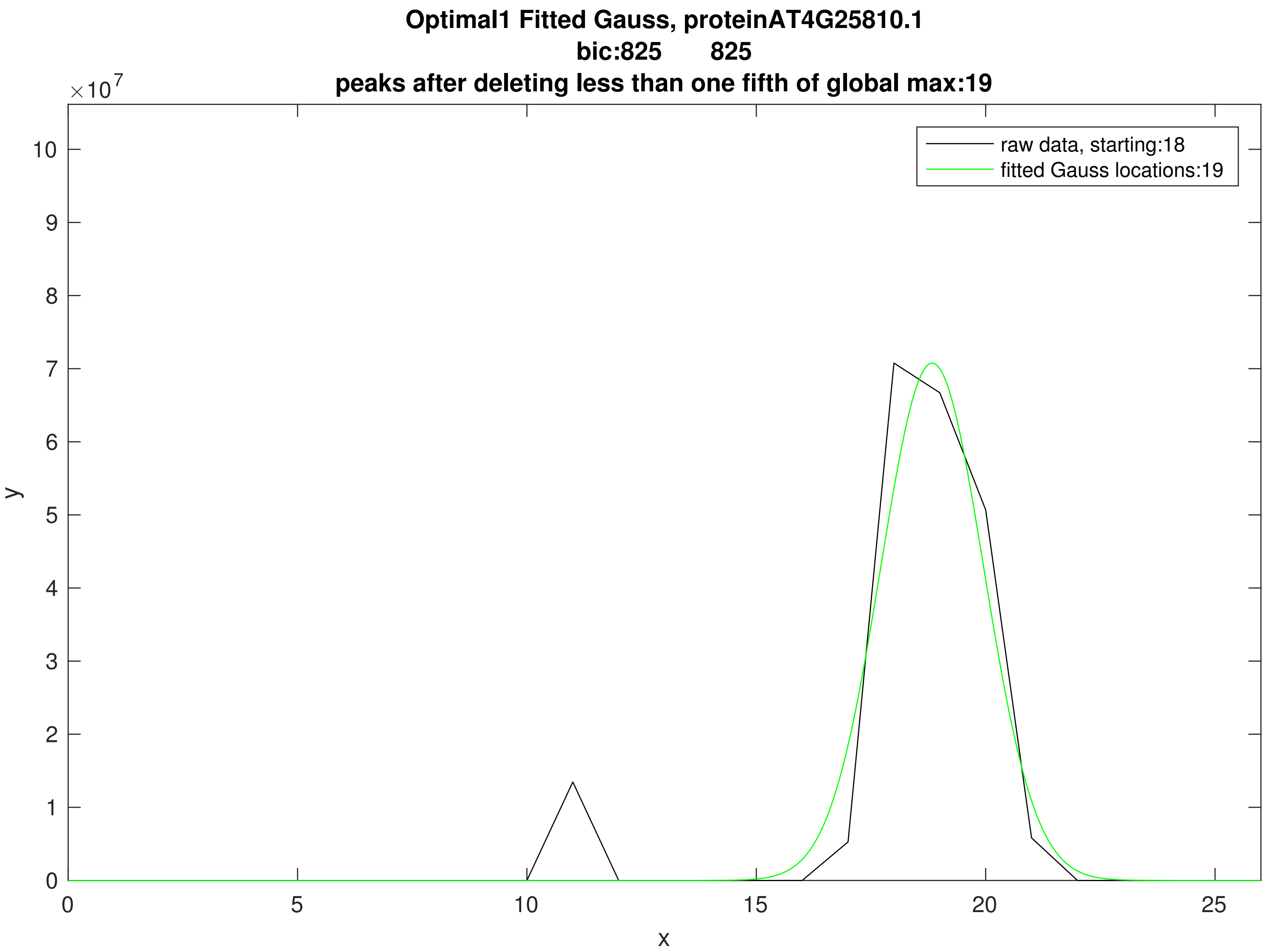
Optimal1 Fitted Gauss, proteinAT4G25810.1
bic:825 825
peaks after deleting less than one fifth of global max:19
×10^7
10
9
8
7
6
5
4
3
2
1
0
y
0
5
10
15
20
25
x
raw data, starting:18
fitted Gauss locations:19

#### Optimal1 Fitted Gauss, proteinAT4G26110.2;AT4G26110.1
#### bic:761
#### peaks after deleting less than one fifth of global max:10

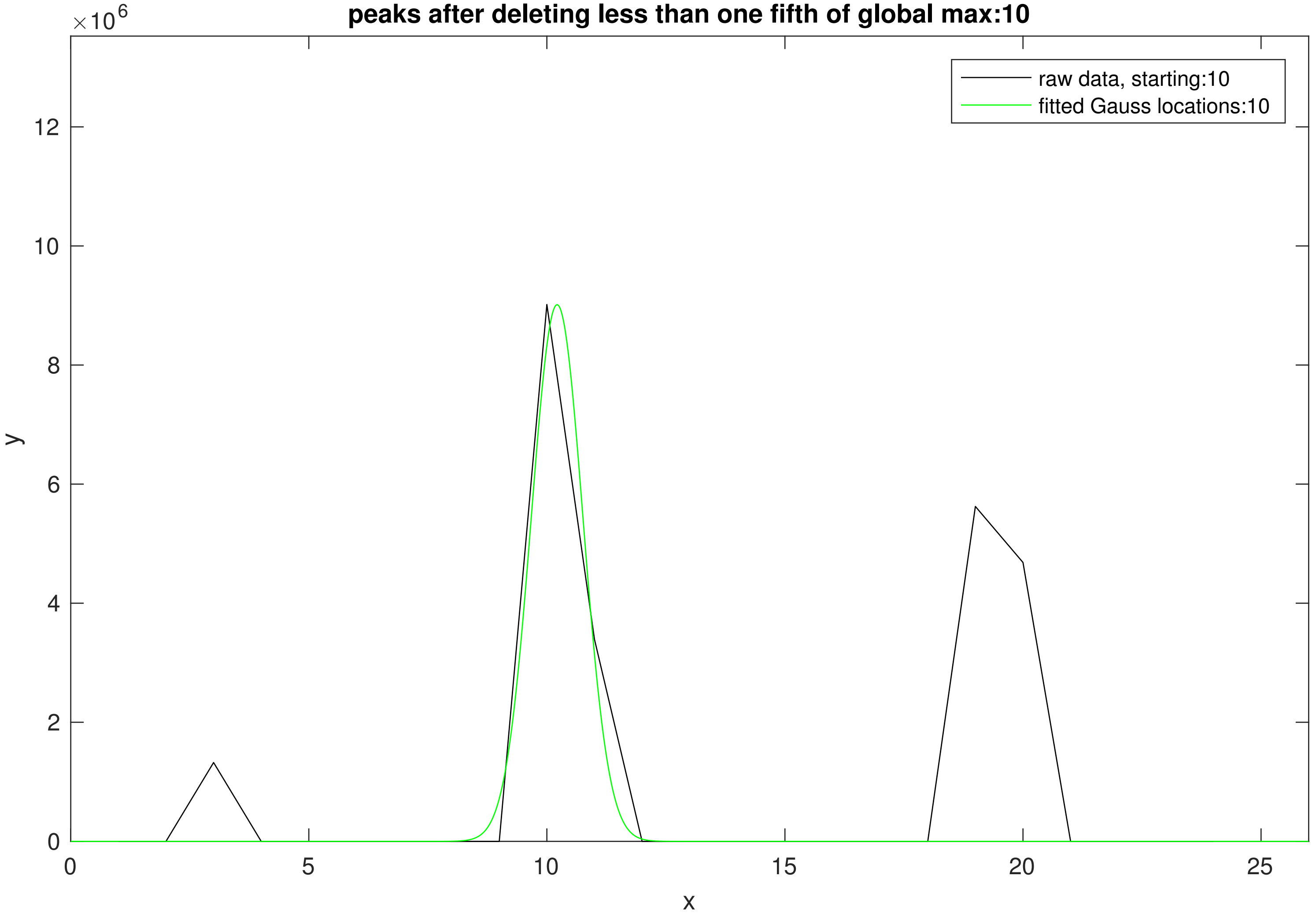

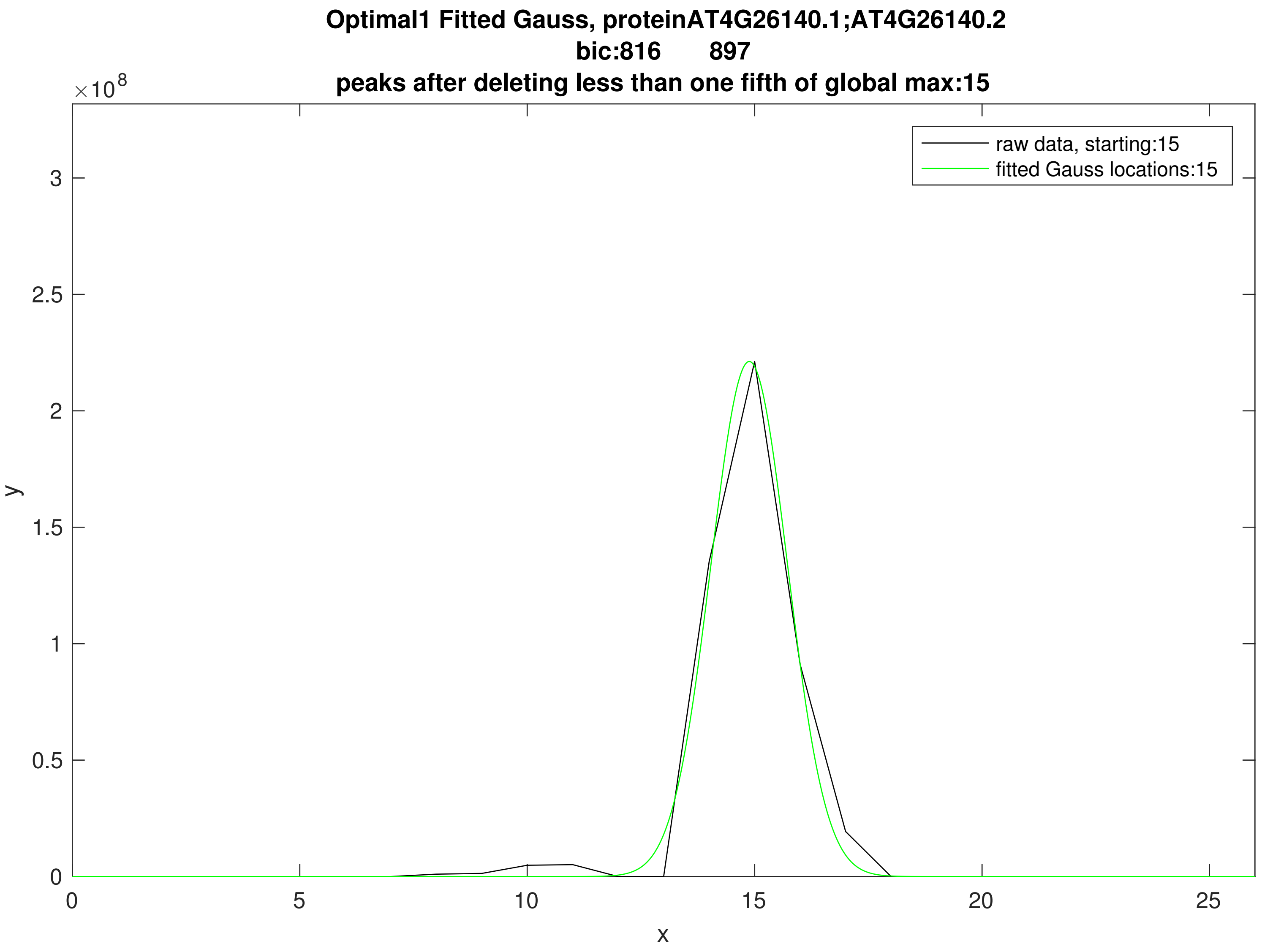
Optimal1 Fitted Gauss, proteinAT4G26140.1;AT4G26140.2
bic:816 897
peaks after deleting less than one fifth of global max:15
×10^8
raw data, starting:15
fitted Gauss locations:15
y
x
0
0.5
1
1.5
2
2.5
3
0
5
10
15
20
25

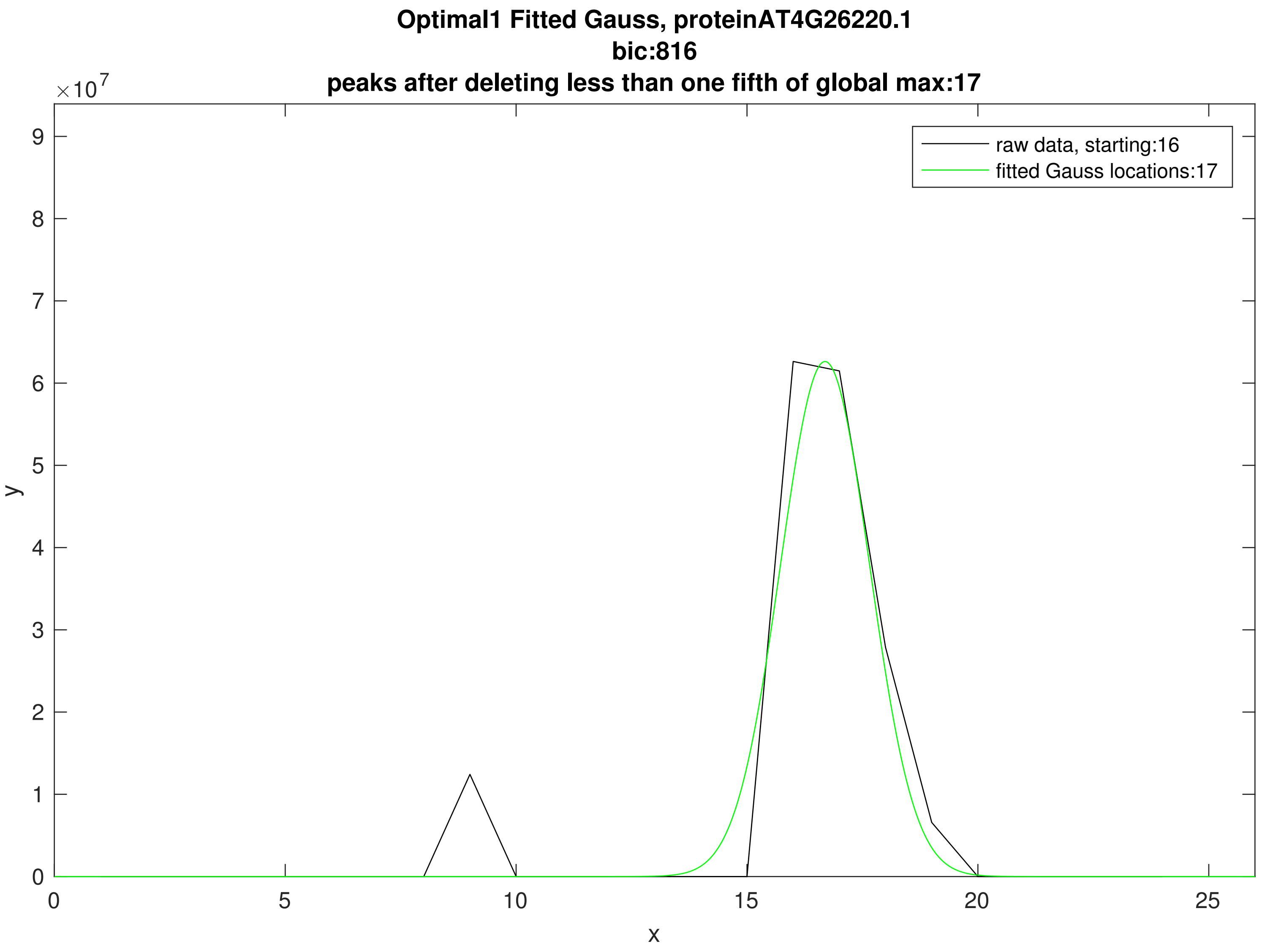
Optimal1 Fitted Gauss, proteinAT4G26220.1
bic:816
peaks after deleting less than one fifth of global max:17
×10^7
raw data, starting:16
fitted Gauss locations:17
y
x
0
1
2
3
4
5
6
7
8
9
0
5
10
15
20
25

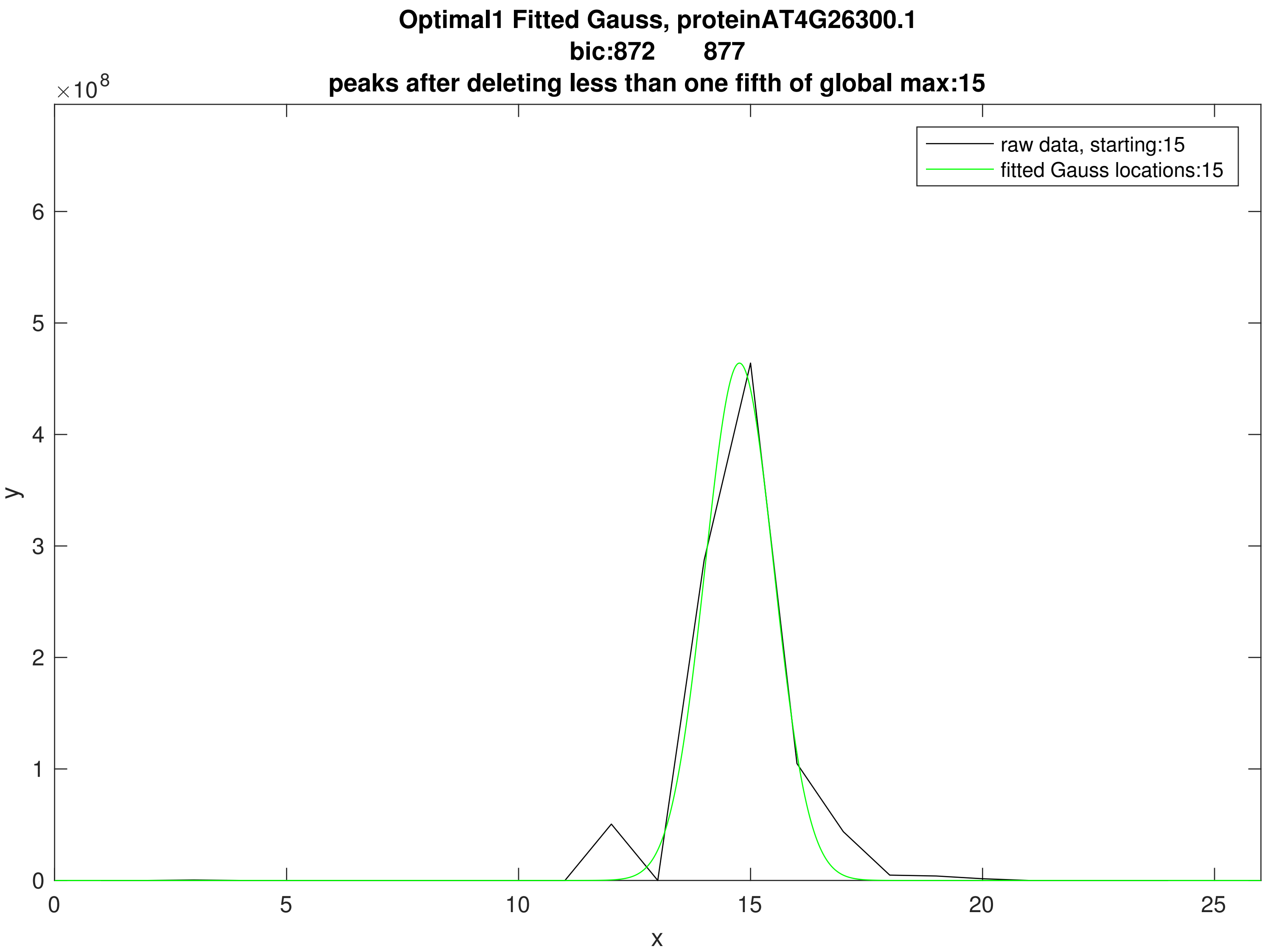
Optimal1 Fitted Gauss, proteinAT4G26300.1
bic:872 877
peaks after deleting less than one fifth of global max:15
×10^8
raw data, starting:15
fitted Gauss locations:15
y
x
0
1
2
3
4
5
6
0
5
10
15
20
25

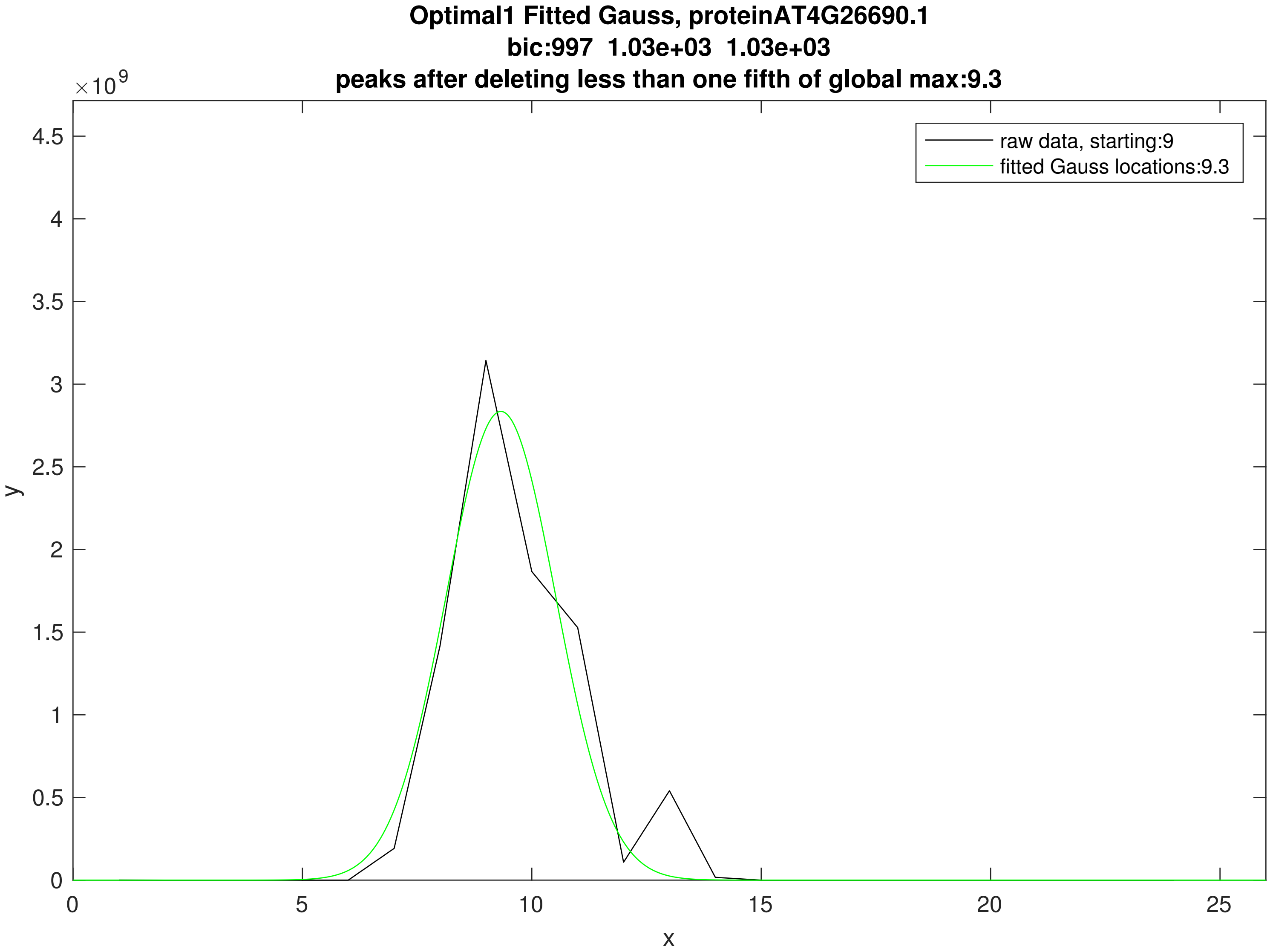
Optimal1 Fitted Gauss, proteinAT4G26690.1
bic:997  1.03e+03  1.03e+03
peaks after deleting less than one fifth of global max:9.3
×10^9
raw data, starting:9
fitted Gauss locations:9.3
y
x
0
0.5
1
1.5
2
2.5
3
3.5
4
4.5
0
5
10
15
20
25

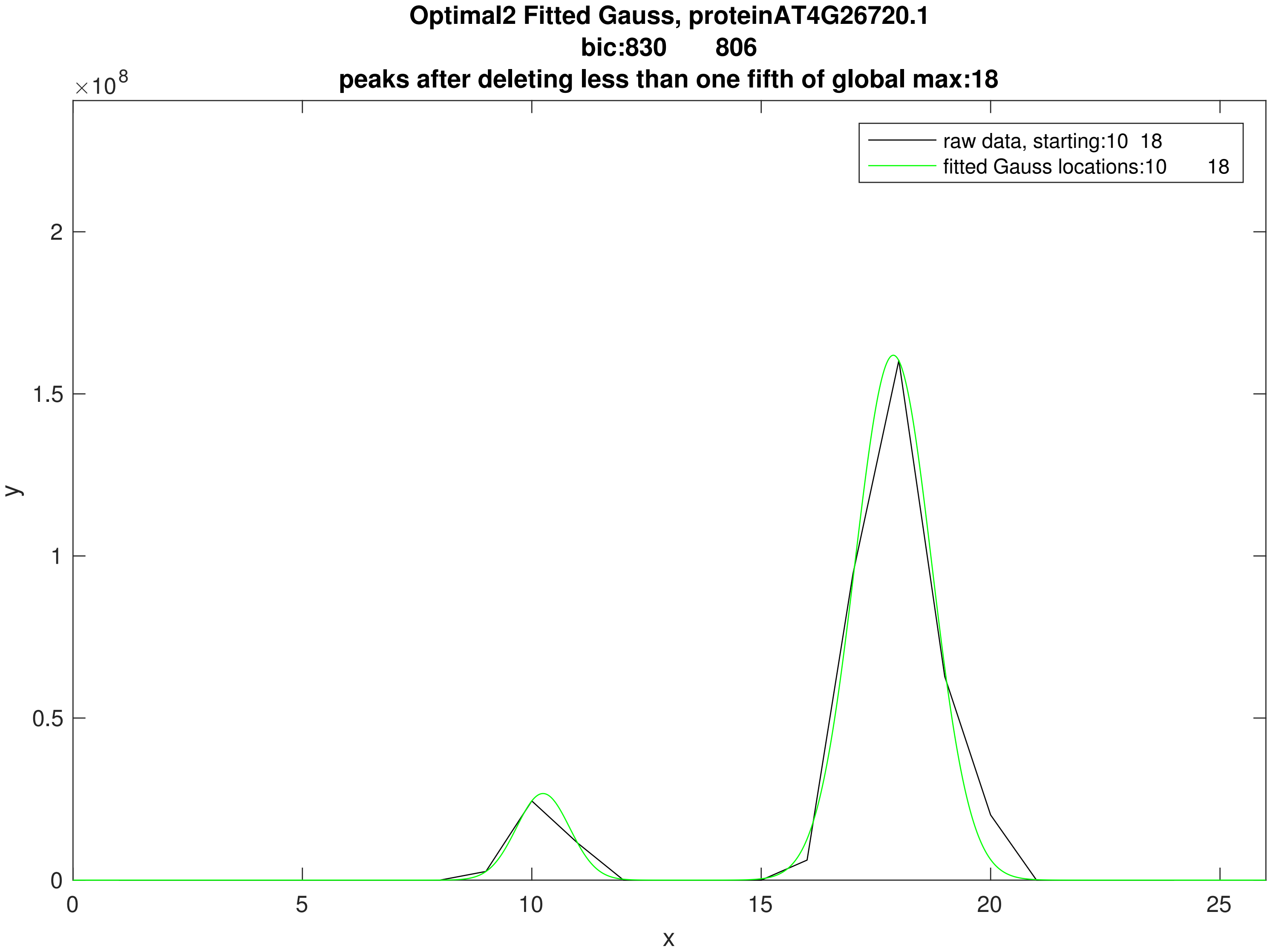

Optimal2 Fitted Gauss, proteinAT4G26720.1
bic:830 806
peaks after deleting less than one fifth of global max:18
×10^8
raw data, starting:10 18
fitted Gauss locations:10 18
2
1.5
1
0.5
0
y
0
5
10
15
20
25
x

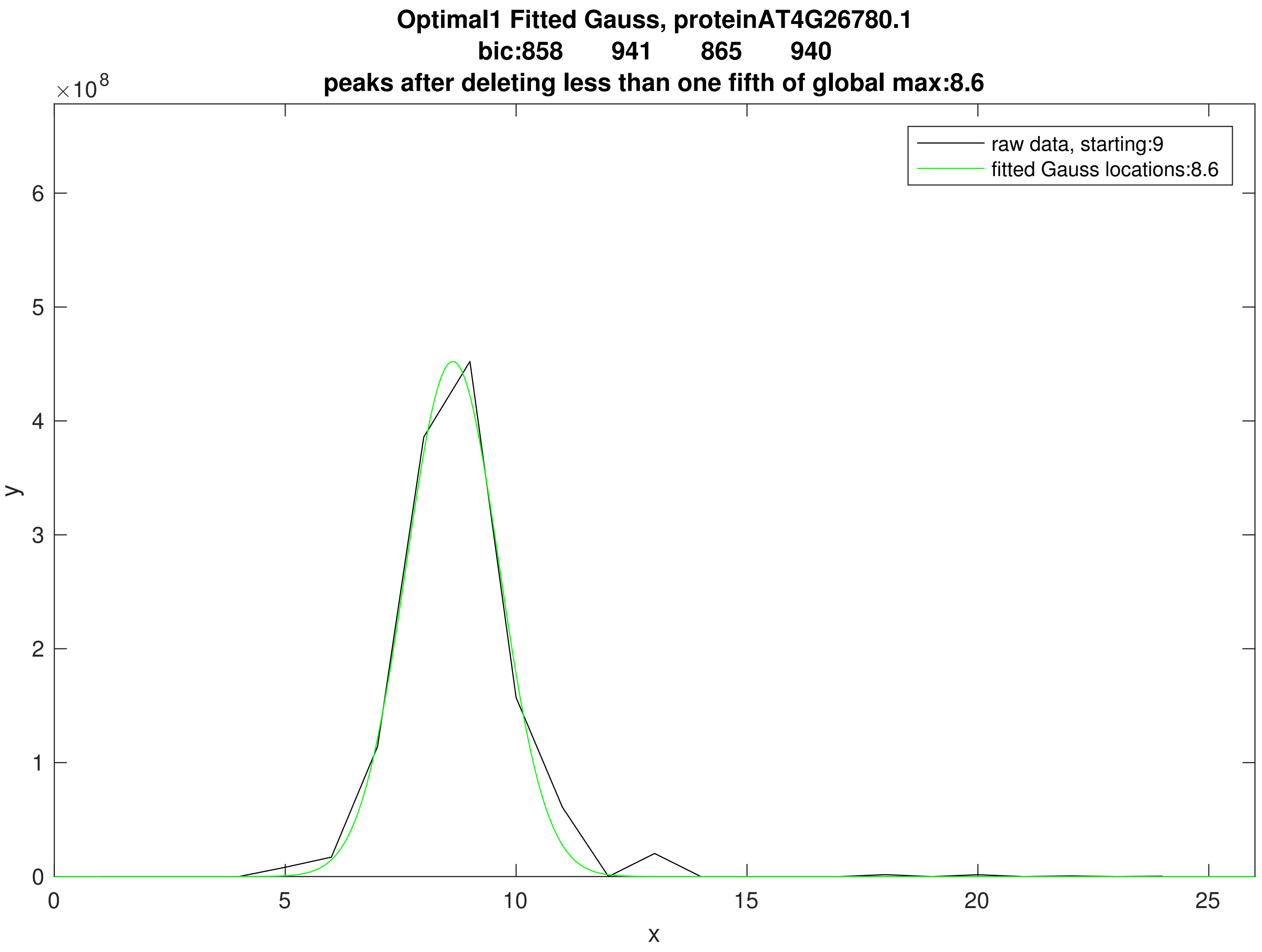
Optimal1 Fitted Gauss, proteinAT4G26780.1
bic:858 941 865 940
peaks after deleting less than one fifth of global max:8.6
×10^8
raw data, starting:9
fitted Gauss locations:8.6
y
x
0
1
2
3
4
5
6
0
5
10
15
20
25

Optimal1 Fitted Gauss, proteinAT4G26840.1
bic:888
peaks after deleting less than one fifth of global max:20

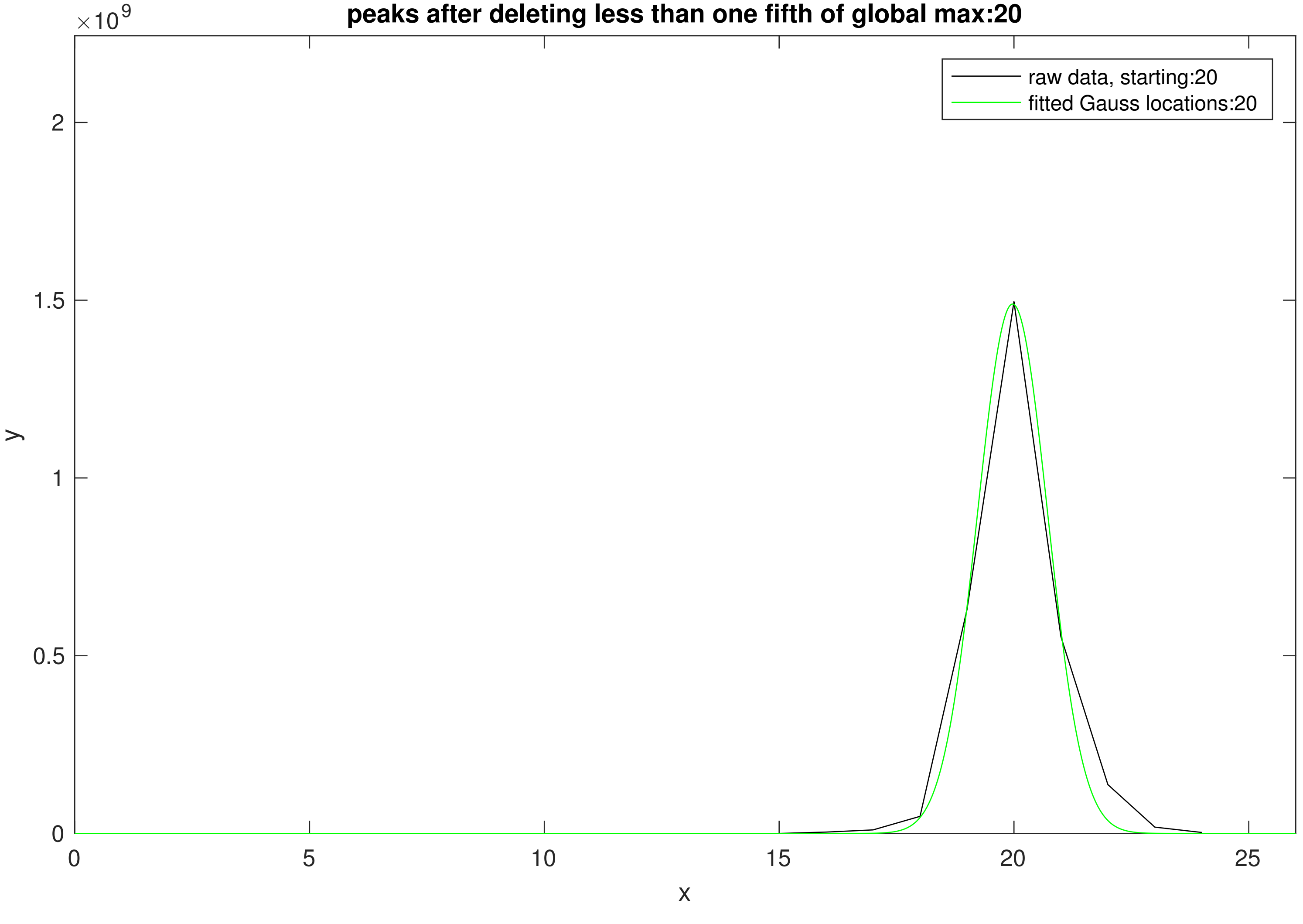

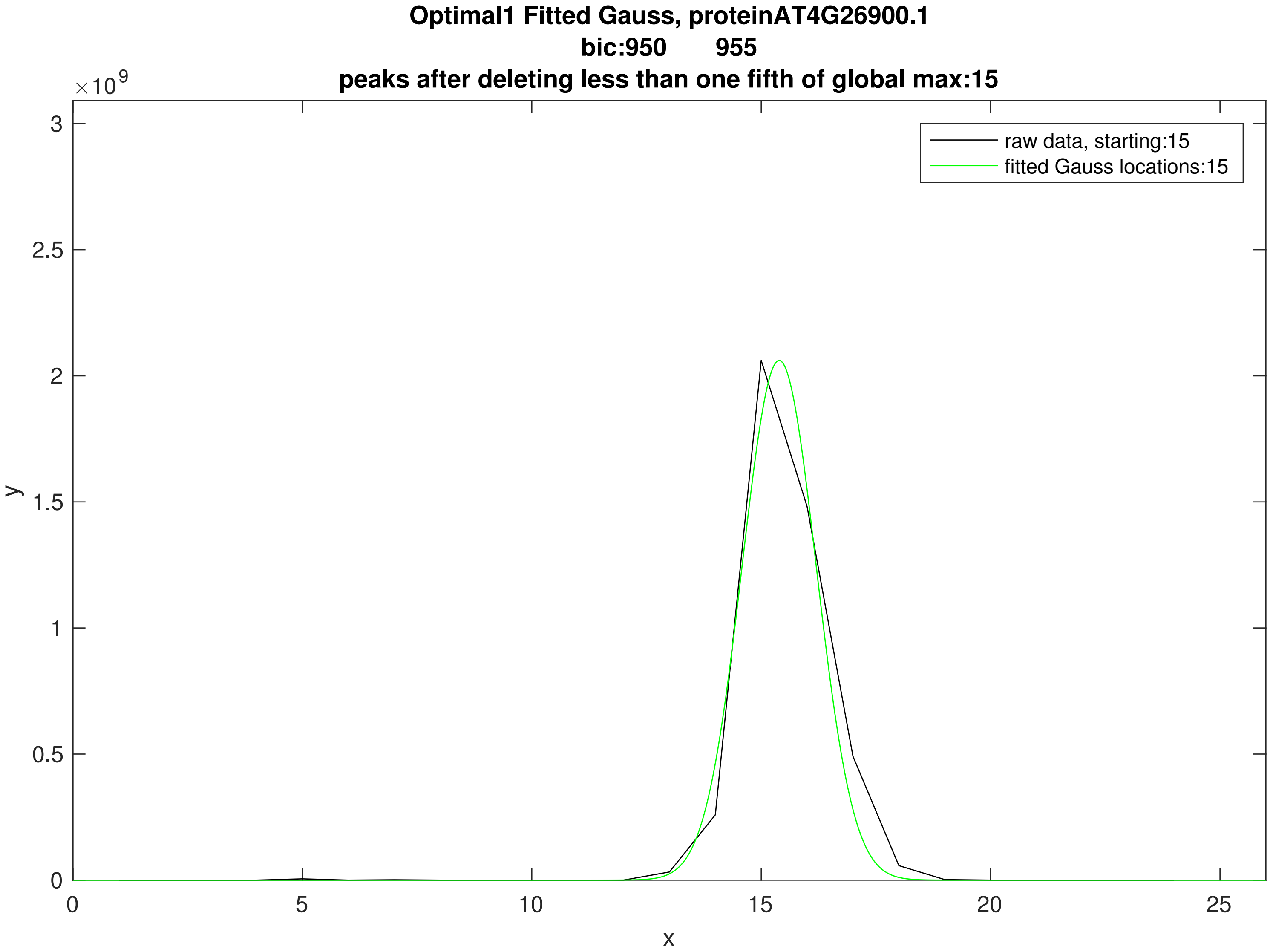
Optimal1 Fitted Gauss, proteinAT4G26900.1
bic:950 955
peaks after deleting less than one fifth of global max:15
×10^9
raw data, starting:15
fitted Gauss locations:15
y
x
0
0.5
1
1.5
2
2.5
3
0
5
10
15
20
25

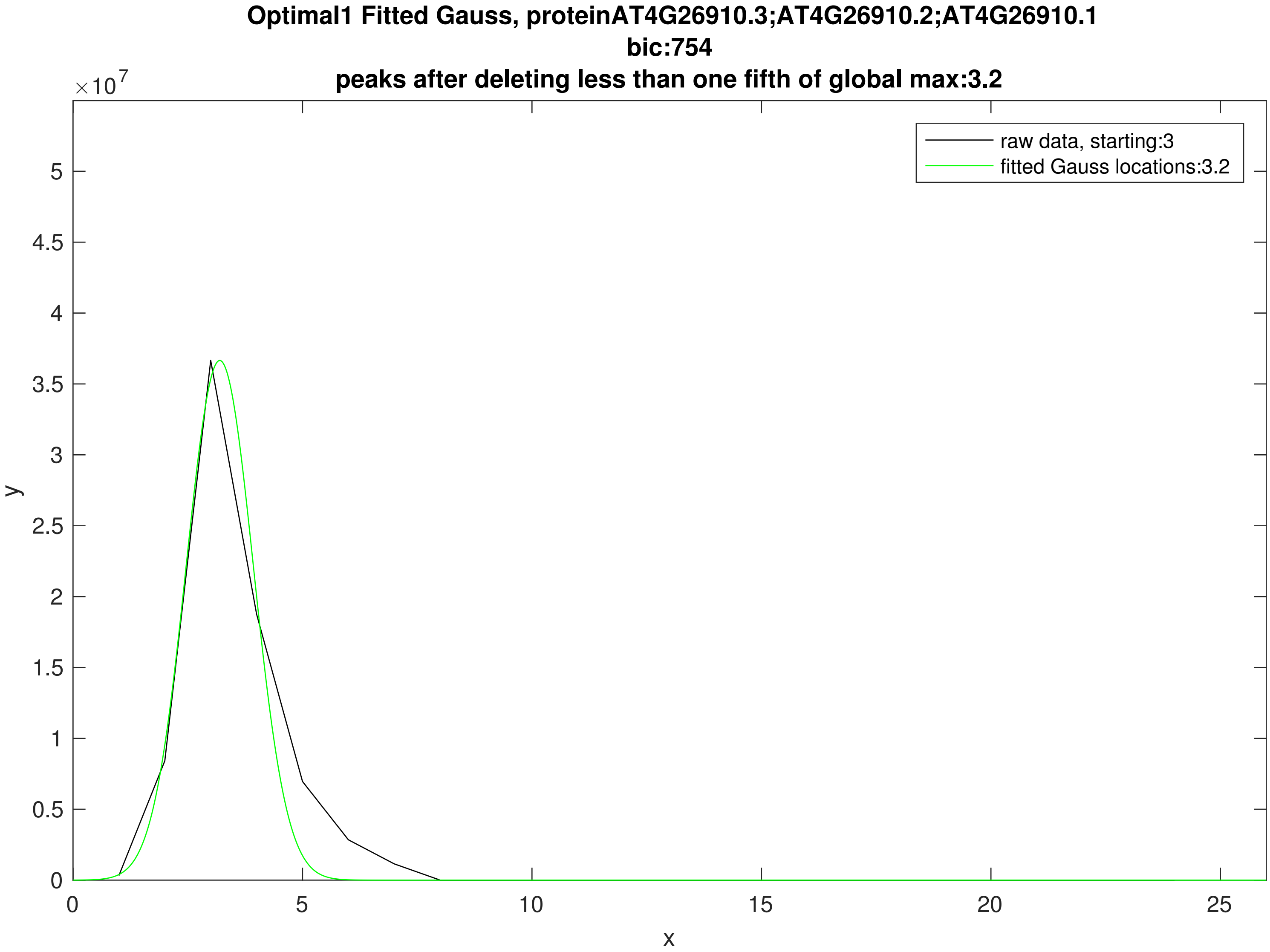
Optimal1 Fitted Gauss, proteinAT4G26910.3;AT4G26910.2;AT4G26910.1
bic:754
peaks after deleting less than one fifth of global max:3.2
×10^7
raw data, starting:3
fitted Gauss locations:3.2
y
x
0
0.5
1
1.5
2
2.5
3
3.5
4
4.5
5
0
5
10
15
20
25

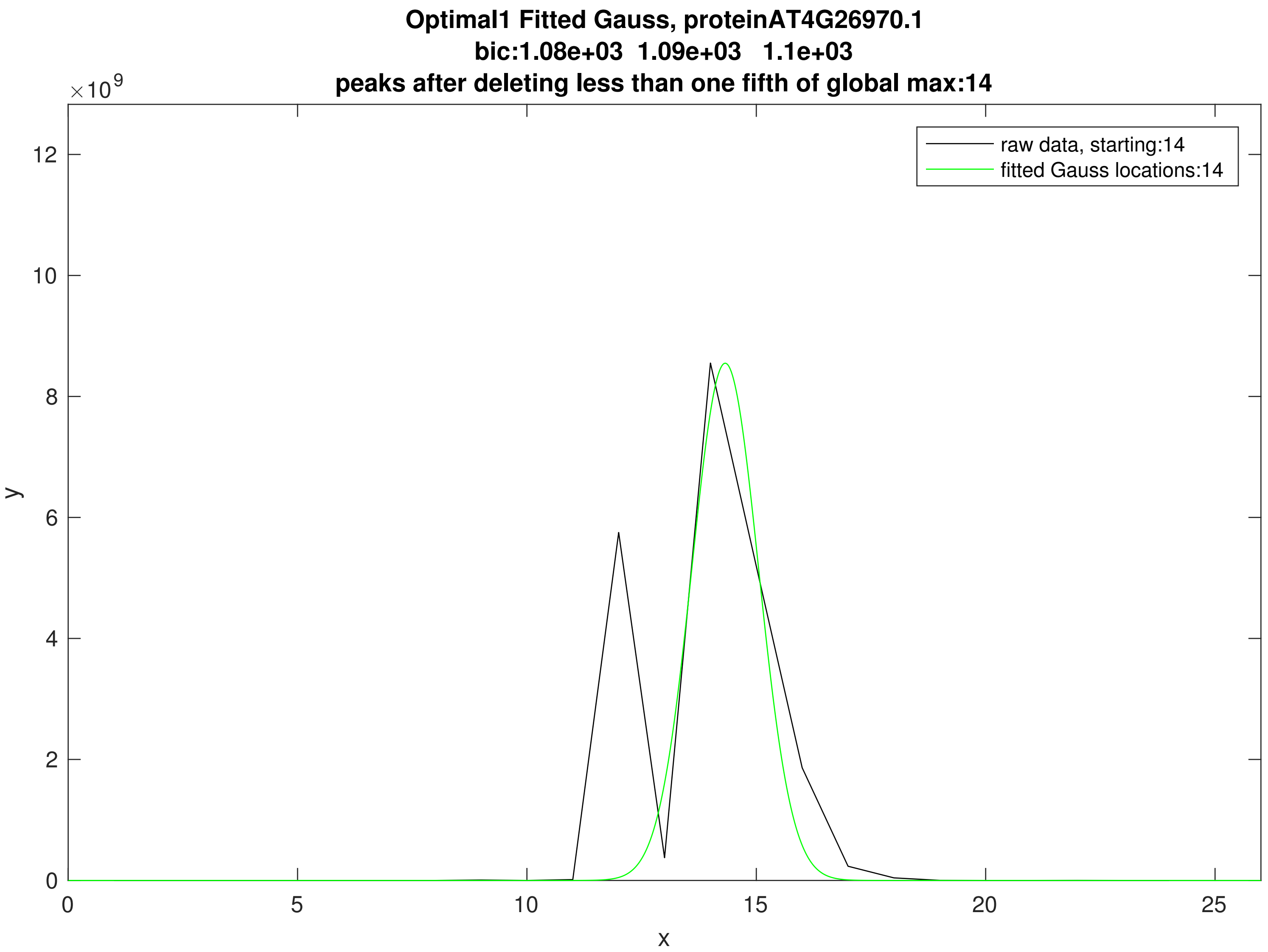
Optimal1 Fitted Gauss, proteinAT4G26970.1
bic:1.08e+03 1.09e+03 1.1e+03
peaks after deleting less than one fifth of global max:14
×10^9
12
10
8
6
4
2
0
y
0
5
10
15
20
25
x
raw data, starting:14
fitted Gauss locations:14

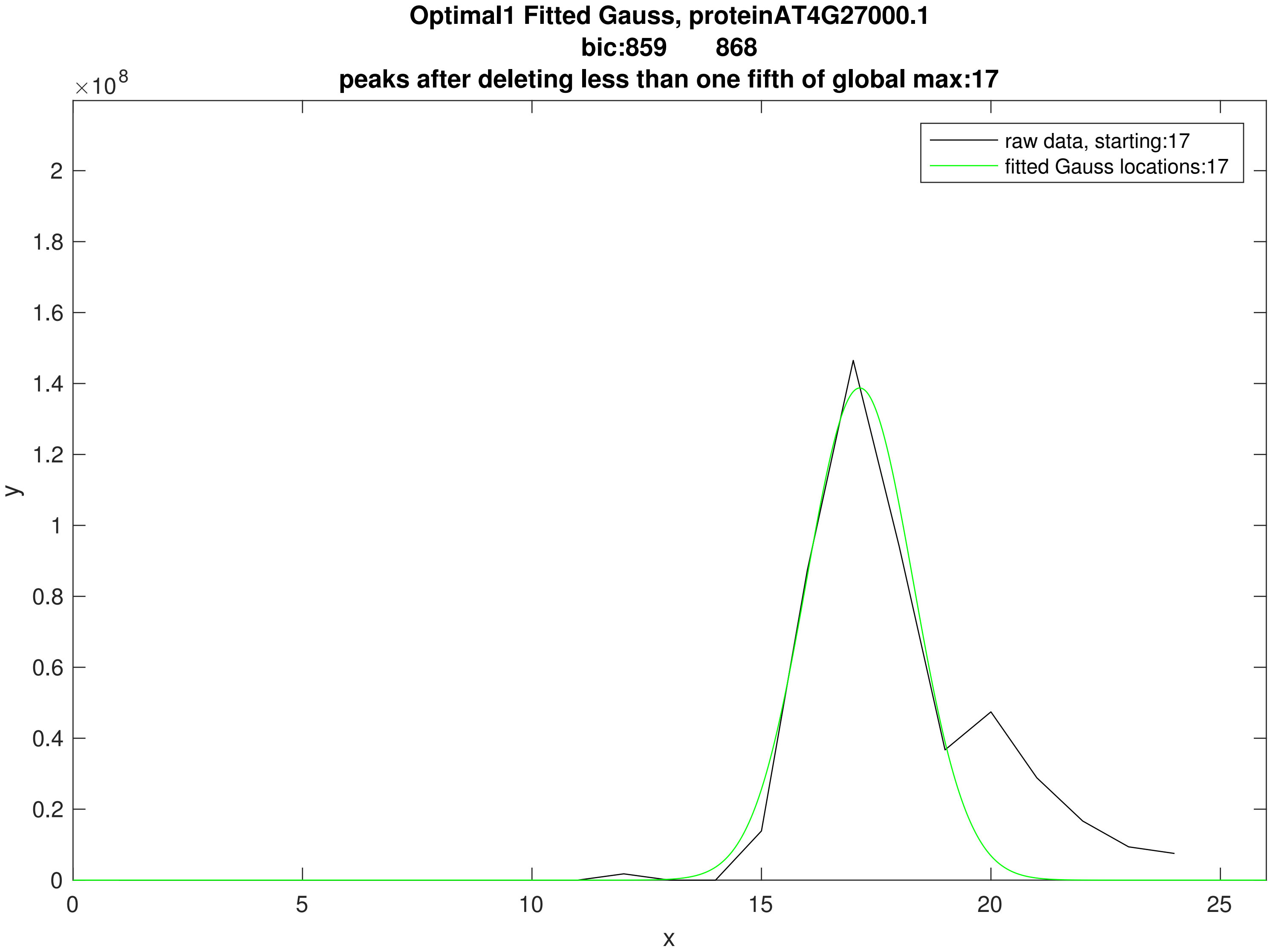
Optimal1 Fitted Gauss, proteinAT4G27000.1
bic:859 868
peaks after deleting less than one fifth of global max:17
×10^8
raw data, starting:17
fitted Gauss locations:17
y
x
0
0.2
0.4
0.6
0.8
1
1.2
1.4
1.6
1.8
2
5
10
15
20
25

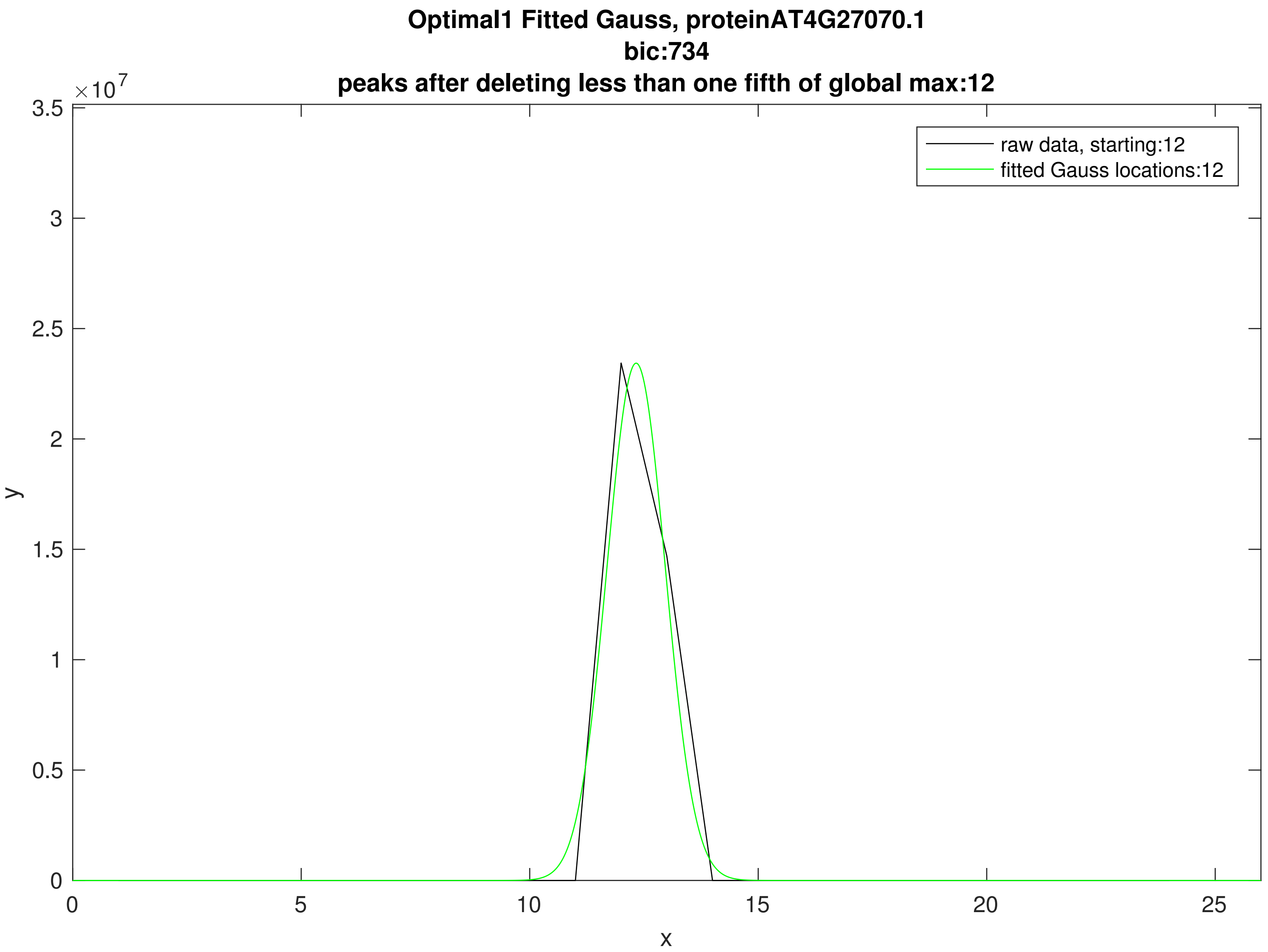
Optimal1 Fitted Gauss, proteinAT4G27070.1
bic:734
peaks after deleting less than one fifth of global max:12
×10^7
3.5
3
2.5
2
1.5
1
0.5
0
y
0
5
10
15
20
25
x
raw data, starting:12
fitted Gauss locations:12

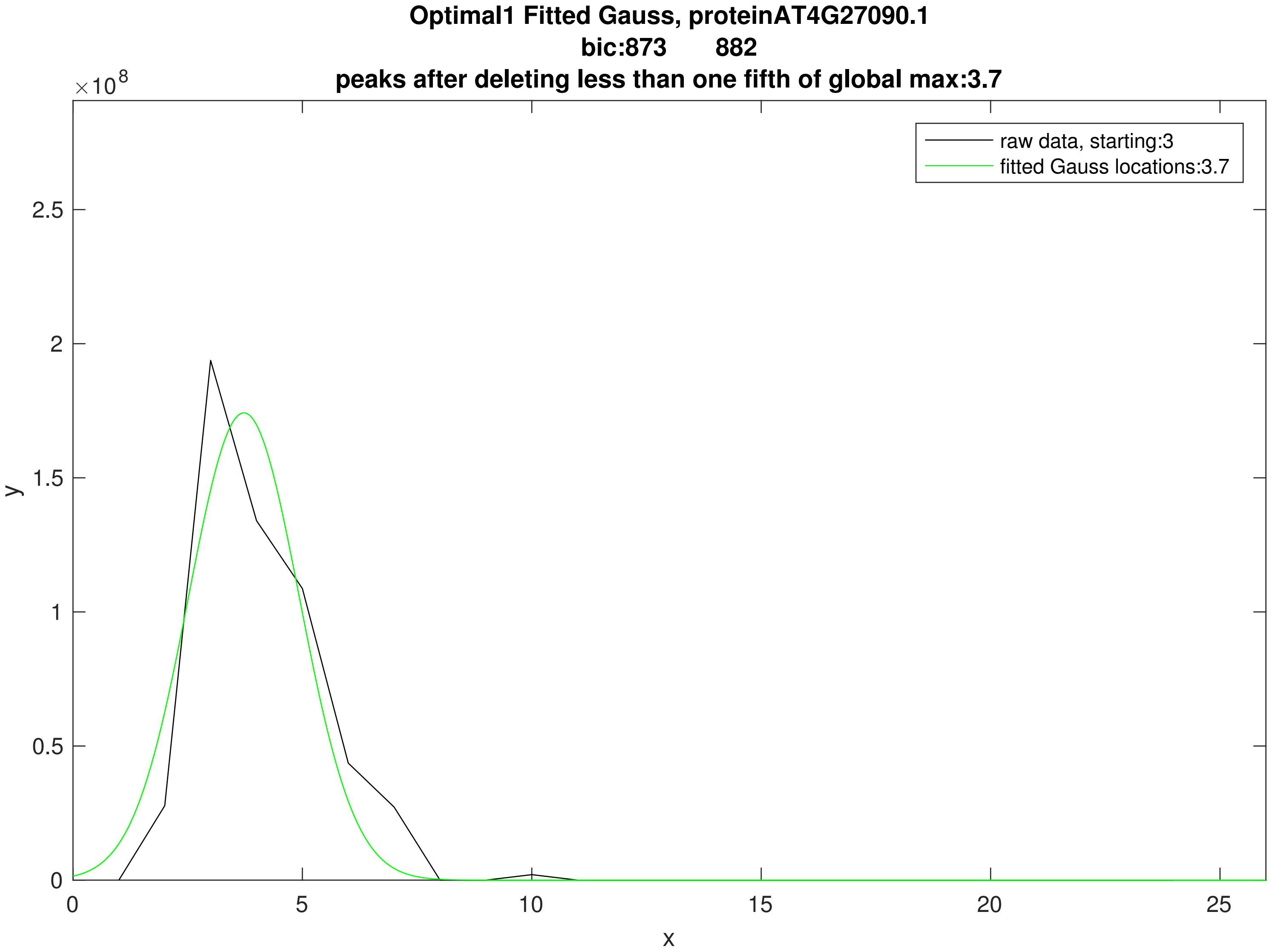
Optimal1 Fitted Gauss, proteinAT4G27090.1
bic:873 882
peaks after deleting less than one fifth of global max:3.7
×10^8
raw data, starting:3
fitted Gauss locations:3.7
y
0
0.5
1
1.5
2
2.5
x
0
5
10
15
20
25

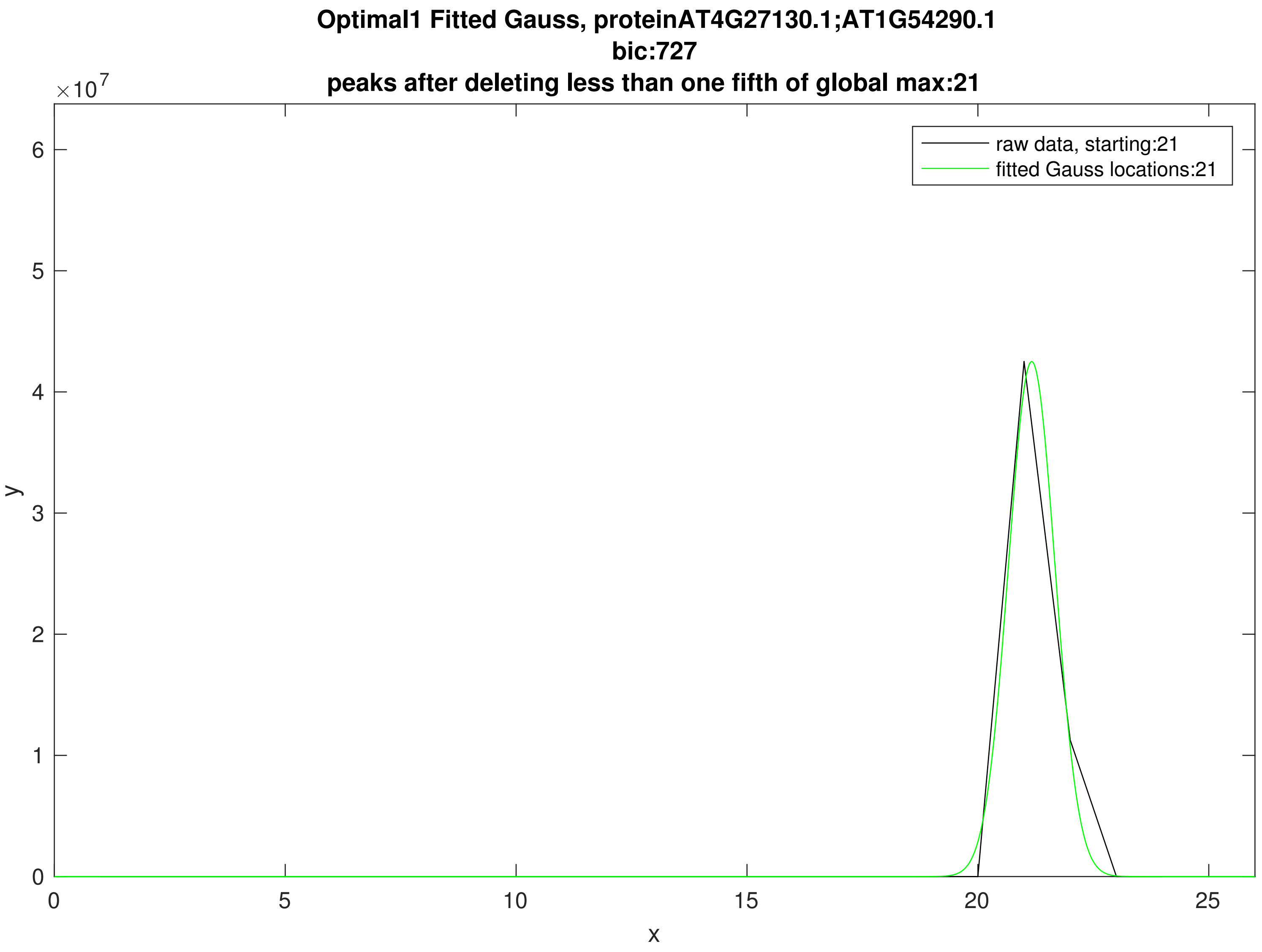
Optimal1 Fitted Gauss, proteinAT4G27130.1;AT1G54290.1
bic:727
peaks after deleting less than one fifth of global max:21
×10^7
6
5
4
3
2
1
0
y
0
5
10
15
20
25
x
raw data, starting:21
fitted Gauss locations:21

**Optimal1 Fitted Gauss, proteinAT4G27140.1**
**bic:794 802 811 819**
**peaks after deleting less than one fifth of global max:20**

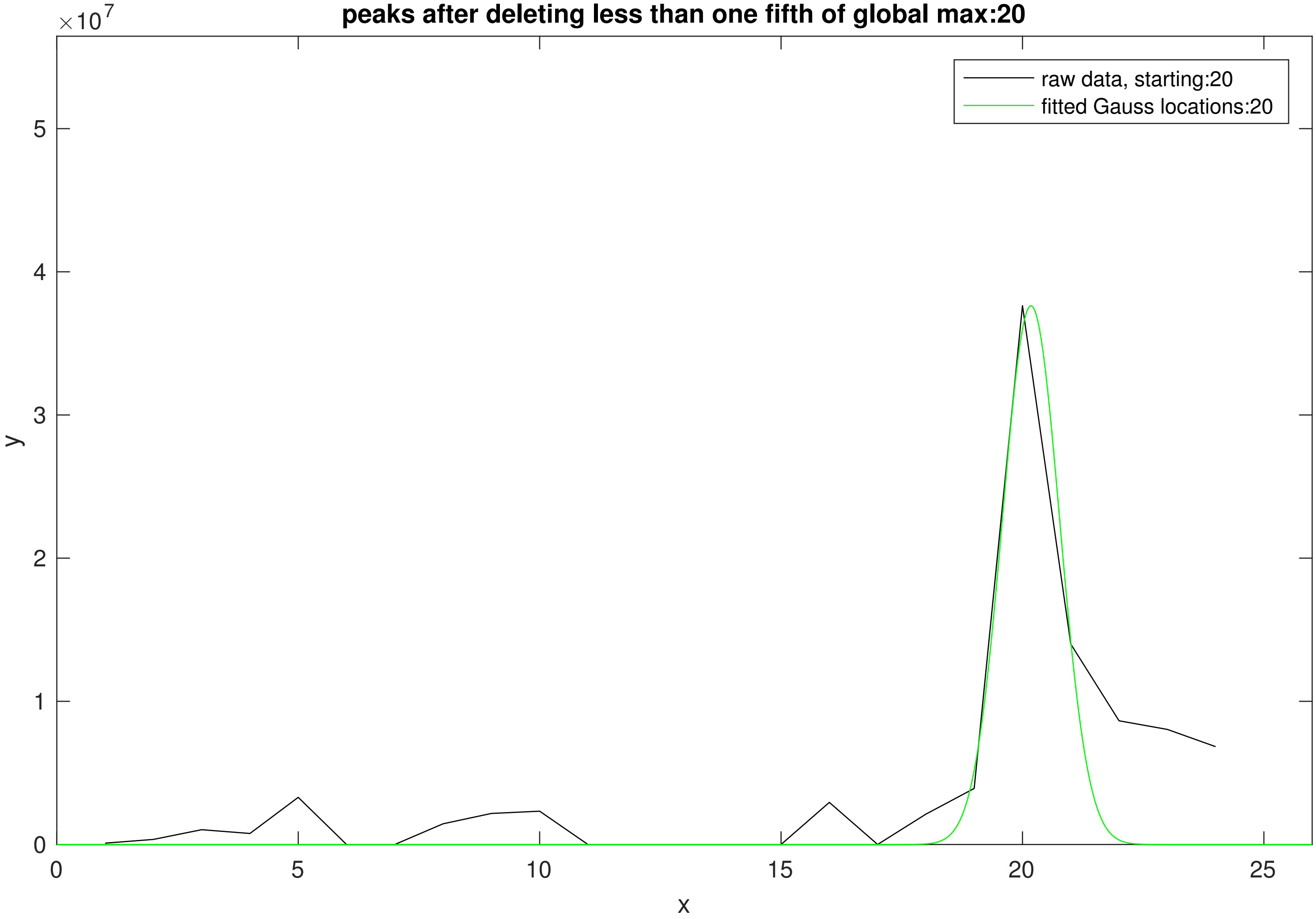

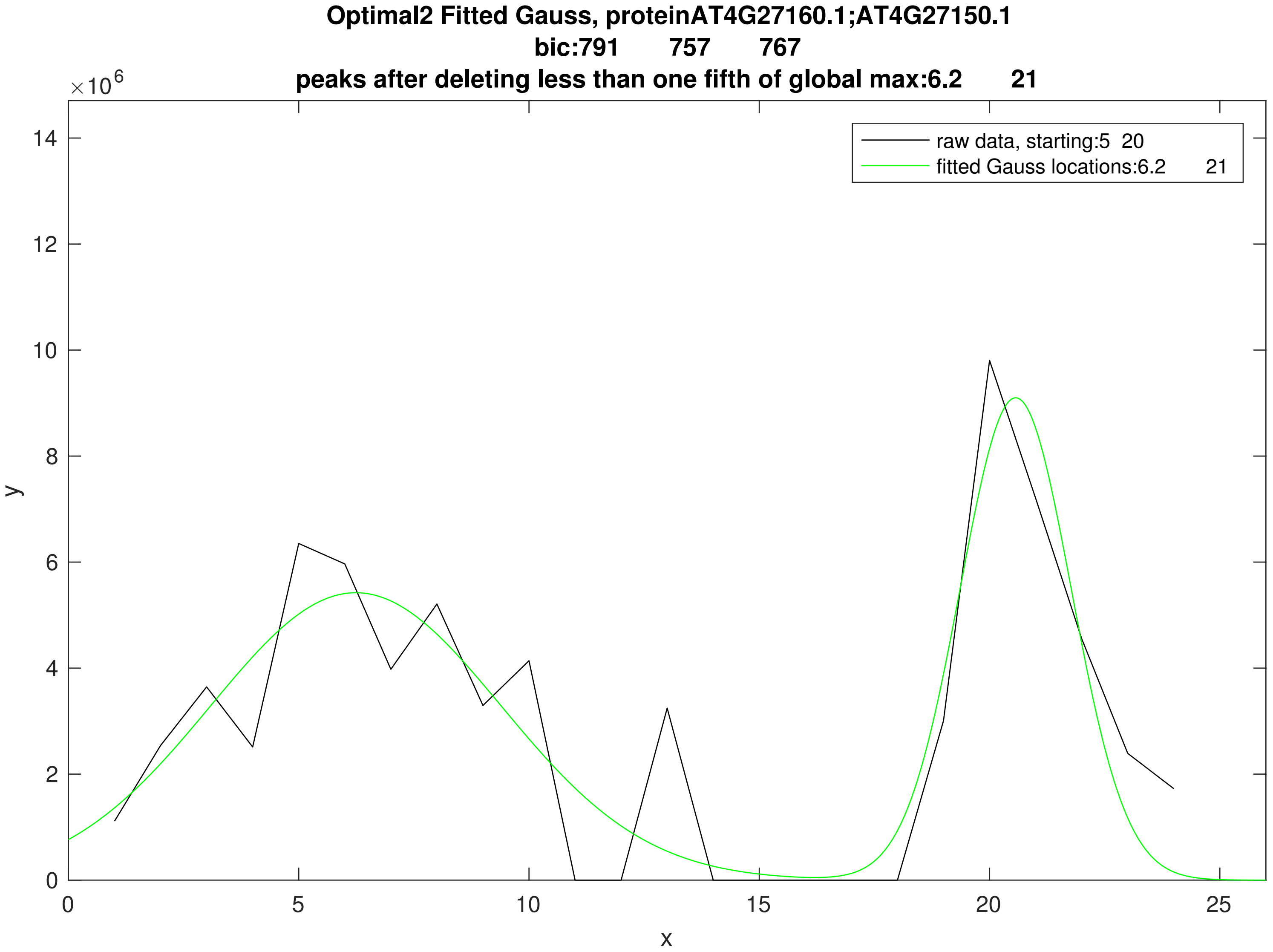
Optimal2 Fitted Gauss, proteinAT4G27160.1;AT4G27150.1
bic:791 757 767
peaks after deleting less than one fifth of global max:6.2 21
×10^6
raw data, starting:5 20
fitted Gauss locations:6.2 21
y
x
0
2
4
6
8
10
12
14
0
5
10
15
20
25

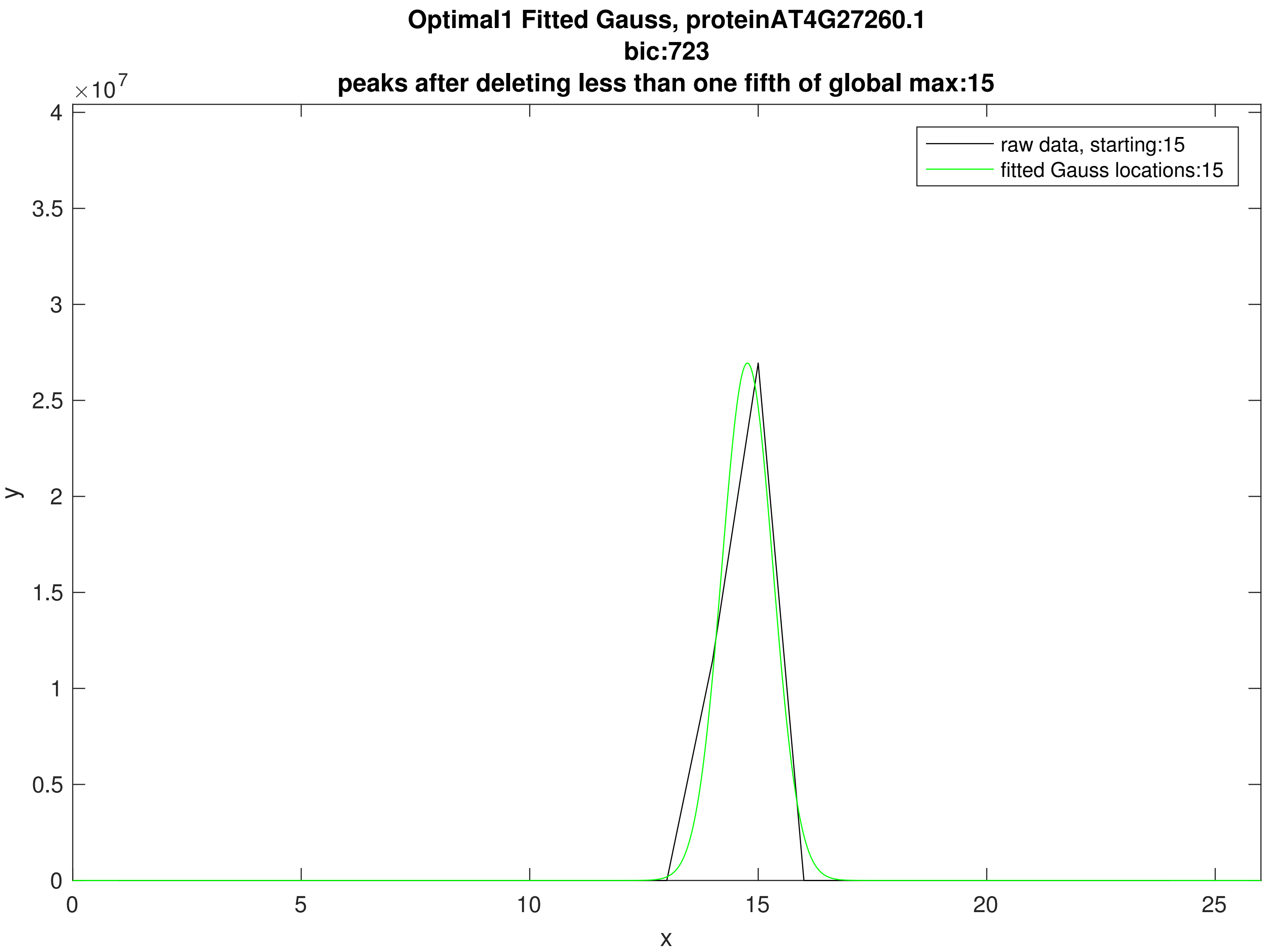

Optimal1 Fitted Gauss, proteinAT4G27260.1
bic:723
peaks after deleting less than one fifth of global max:15
×10^7
4
3.5
3
2.5
2
1.5
1
0.5
0
y
0
5
10
15
20
25
x
raw data, starting:15
fitted Gauss locations:15

**Optimal1 Fitted Gauss, proteinAT4G27320.1**
**bic:857 862**
**peaks after deleting less than one fifth of global max:15**

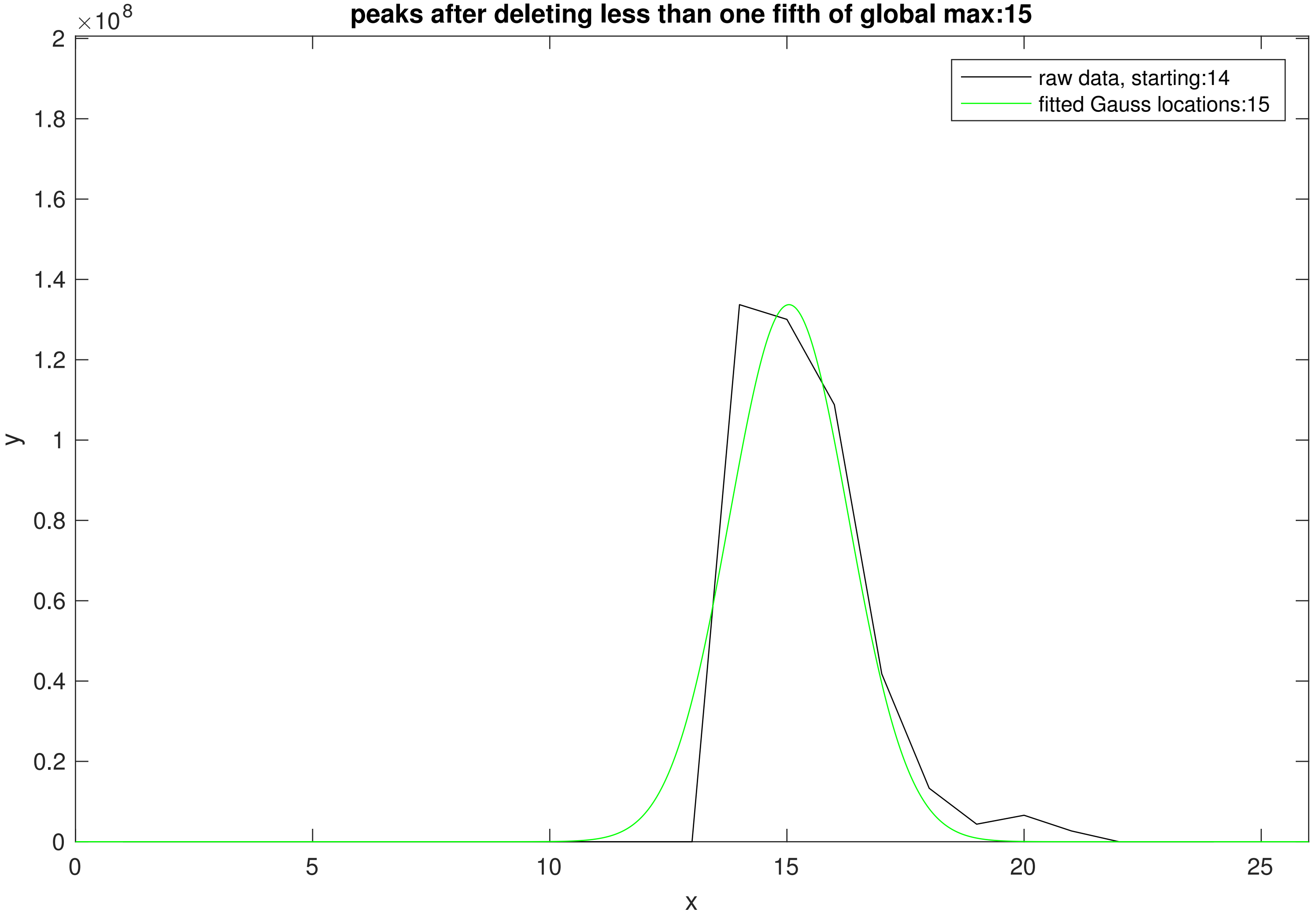

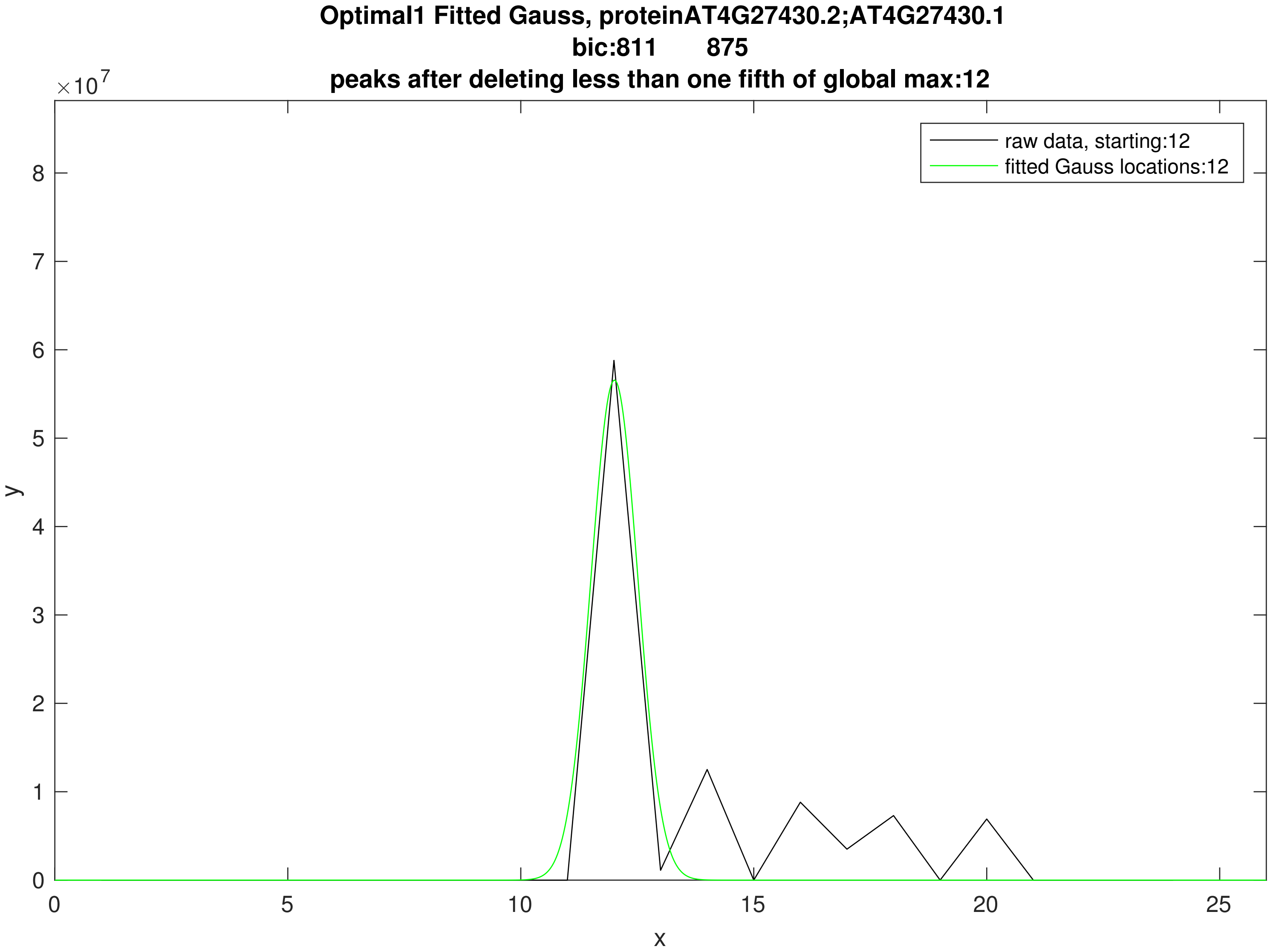

Optimal1 Fitted Gauss, proteinAT4G27430.2;AT4G27430.1
bic:811 875
peaks after deleting less than one fifth of global max:12
×10^7
raw data, starting:12
fitted Gauss locations:12
y
x
0
1
2
3
4
5
6
7
8
0
5
10
15
20
25

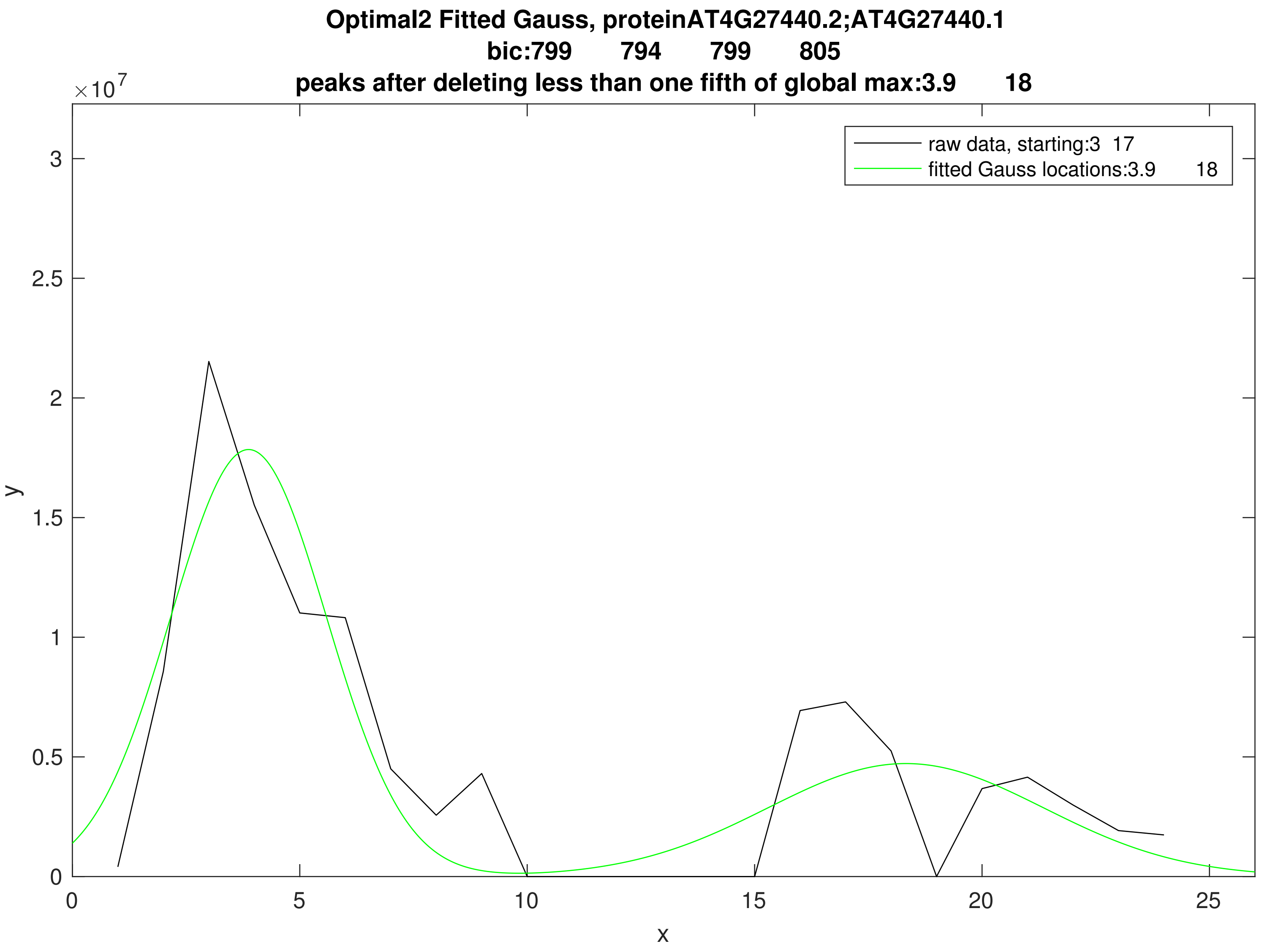

Optimal2 Fitted Gauss, proteinAT4G27440.2;AT4G27440.1
bic:799 794 799 805
peaks after deleting less than one fifth of global max:3.9 18
×10^7
raw data, starting:3 17
fitted Gauss locations:3.9 18
y
x
0
0.5
1
1.5
2
2.5
3
5
10
15
20
25

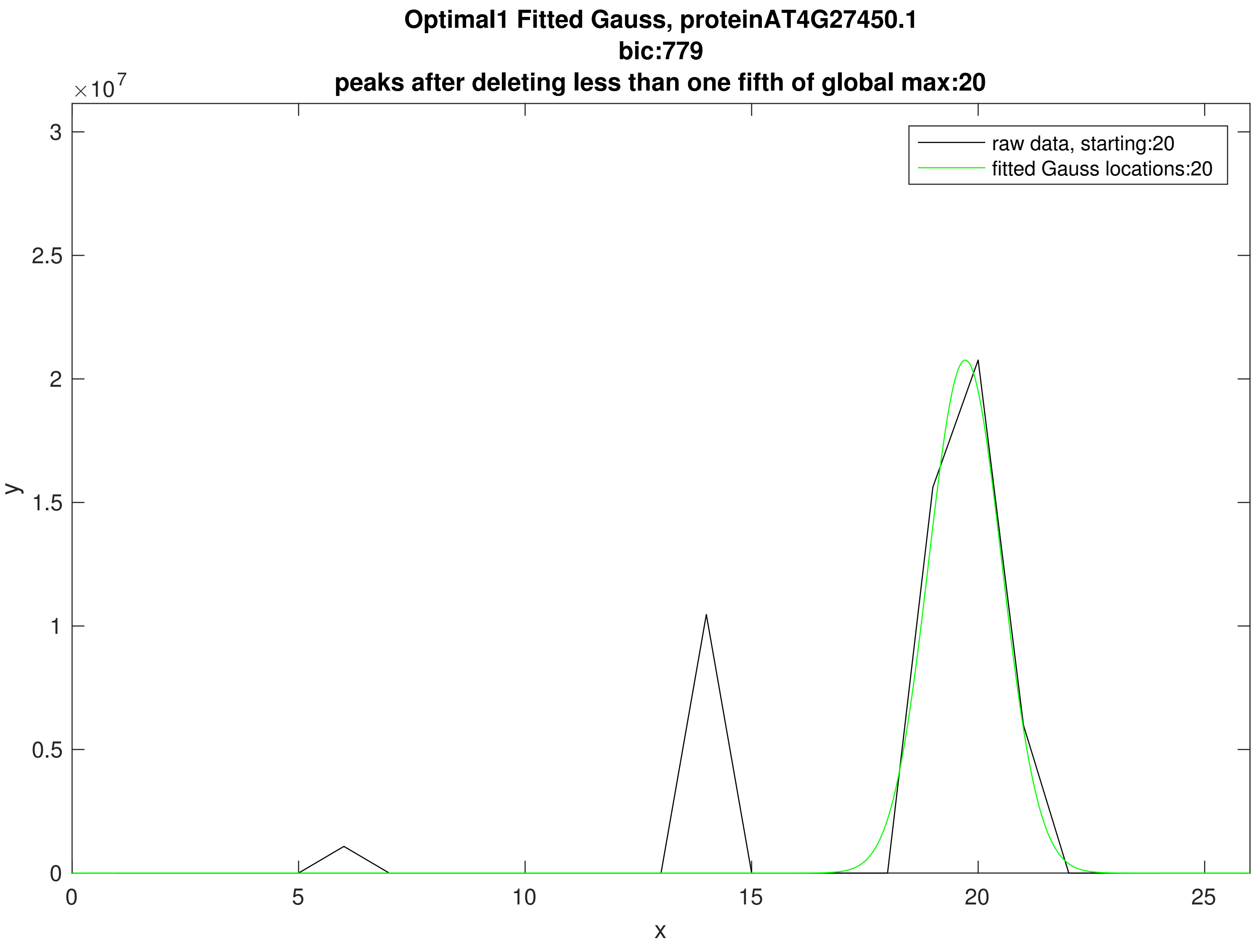

Optimal1 Fitted Gauss, proteinAT4G27450.1
bic:779
peaks after deleting less than one fifth of global max:20
×10^7
3
2.5
2
1.5
1
0.5
0
y
0
5
10
15
20
25
x
raw data, starting:20
fitted Gauss locations:20

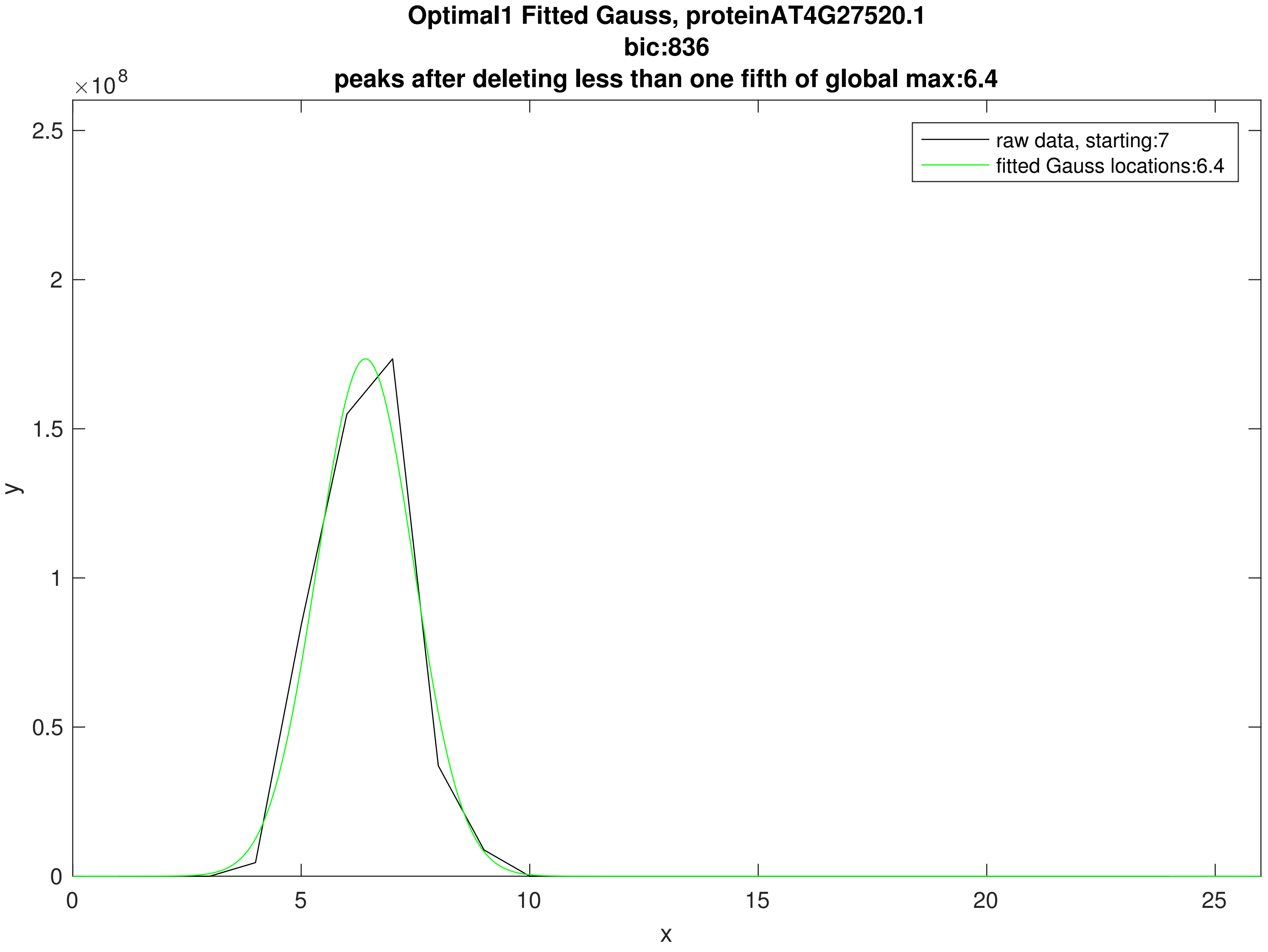
Optimal1 Fitted Gauss, proteinAT4G27520.1
bic:836
peaks after deleting less than one fifth of global max:6.4
×10^8
2.5
2
1.5
1
0.5
0
y
0
5
10
15
20
25
x
raw data, starting:7
fitted Gauss locations:6.4

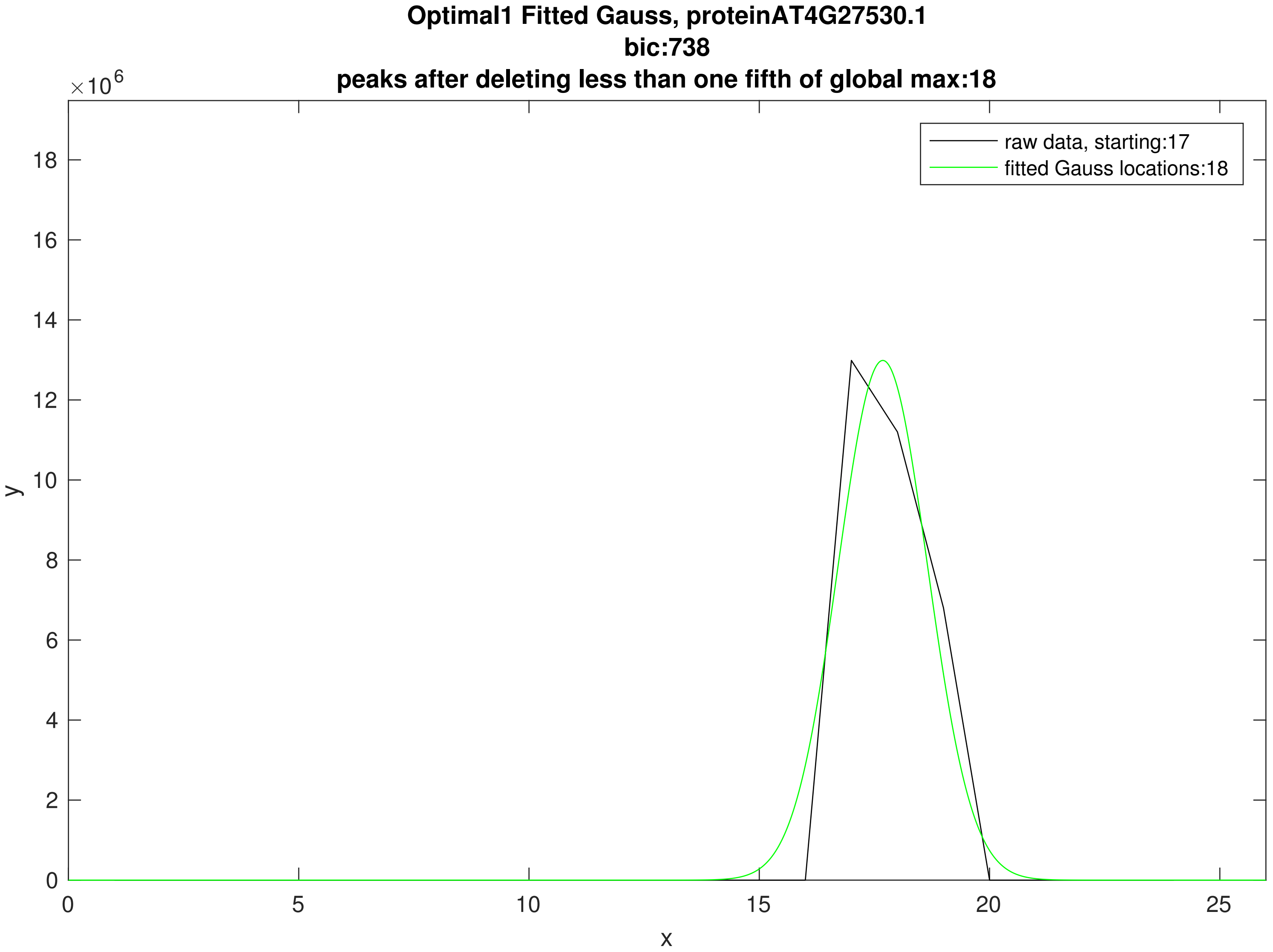
Optimal1 Fitted Gauss, proteinAT4G27530.1
bic:738
peaks after deleting less than one fifth of global max:18
×10^6
raw data, starting:17
fitted Gauss locations:18
y
x
0
2
4
6
8
10
12
14
16
18
0
5
10
15
20
25

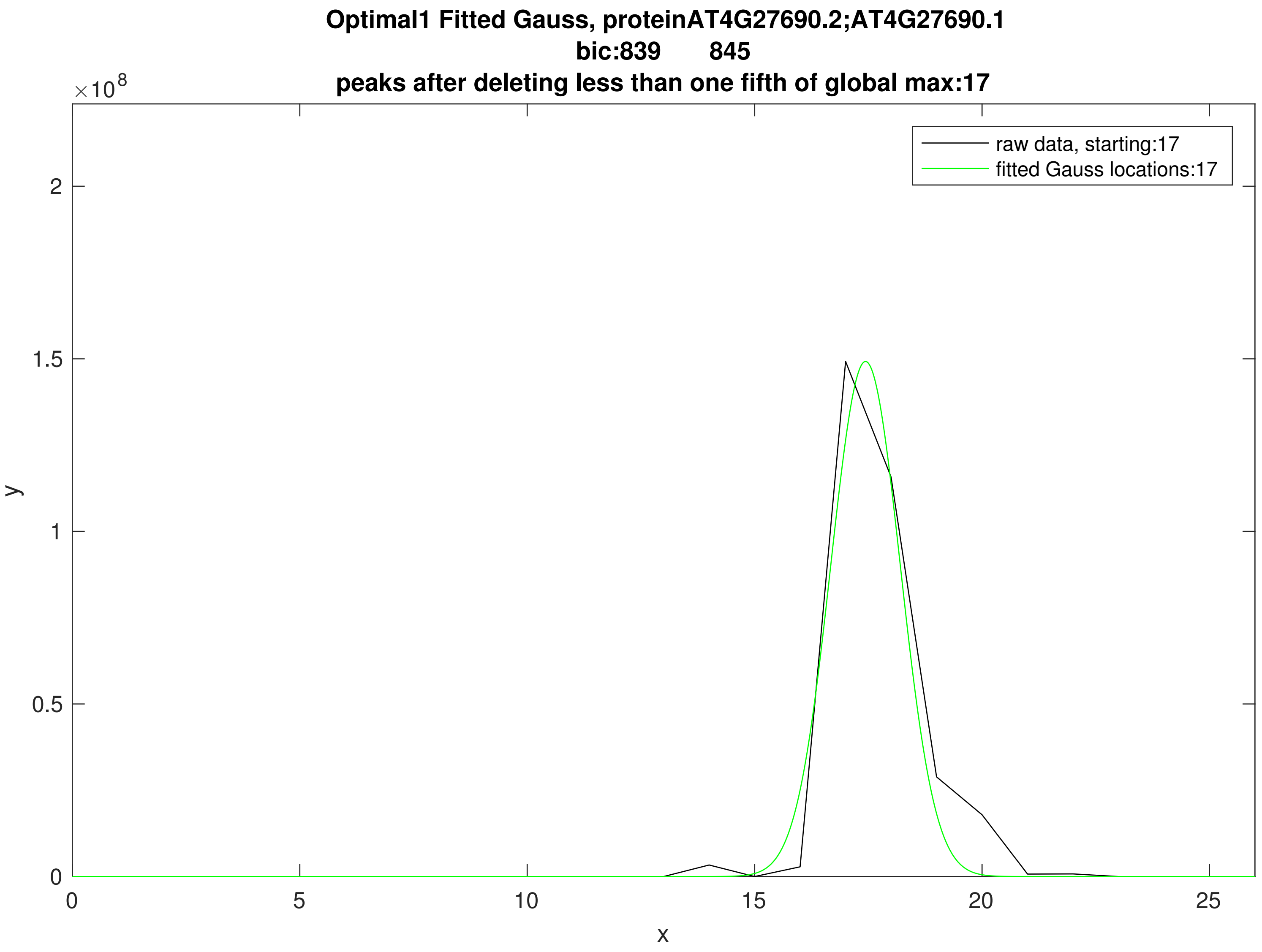

Optimal1 Fitted Gauss, proteinAT4G27690.2;AT4G27690.1
bic:839 845
peaks after deleting less than one fifth of global max:17
×10^8
raw data, starting:17
fitted Gauss locations:17
y
x
0
0.5
1
1.5
2
0
5
10
15
20
25

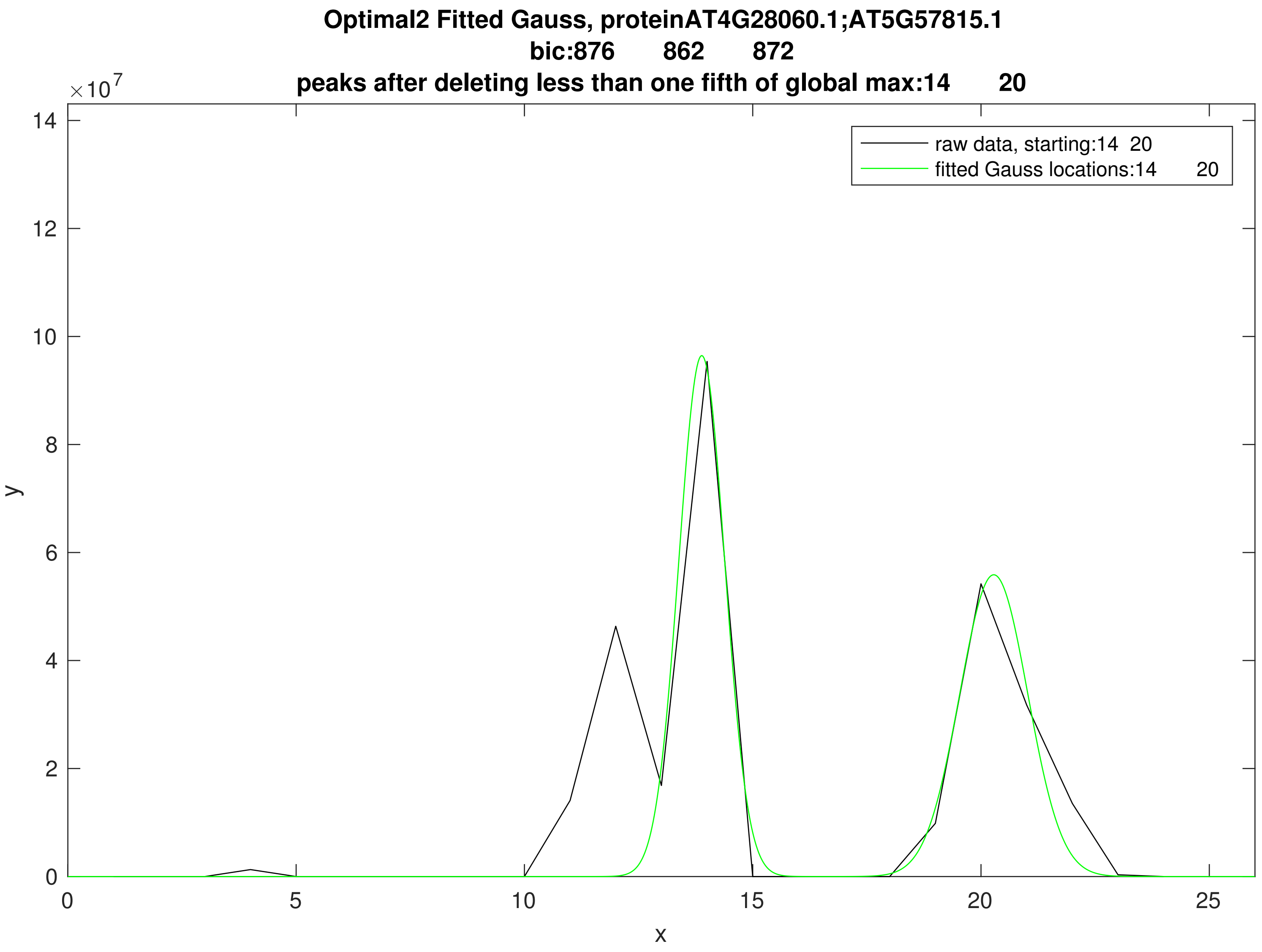
Optimal2 Fitted Gauss, proteinAT4G28060.1;AT5G57815.1
bic:876 862 872
peaks after deleting less than one fifth of global max:14 20
×10^7
raw data, starting:14 20
fitted Gauss locations:14 20
y
x
0
2
4
6
8
10
12
14
0
5
10
15
20
25

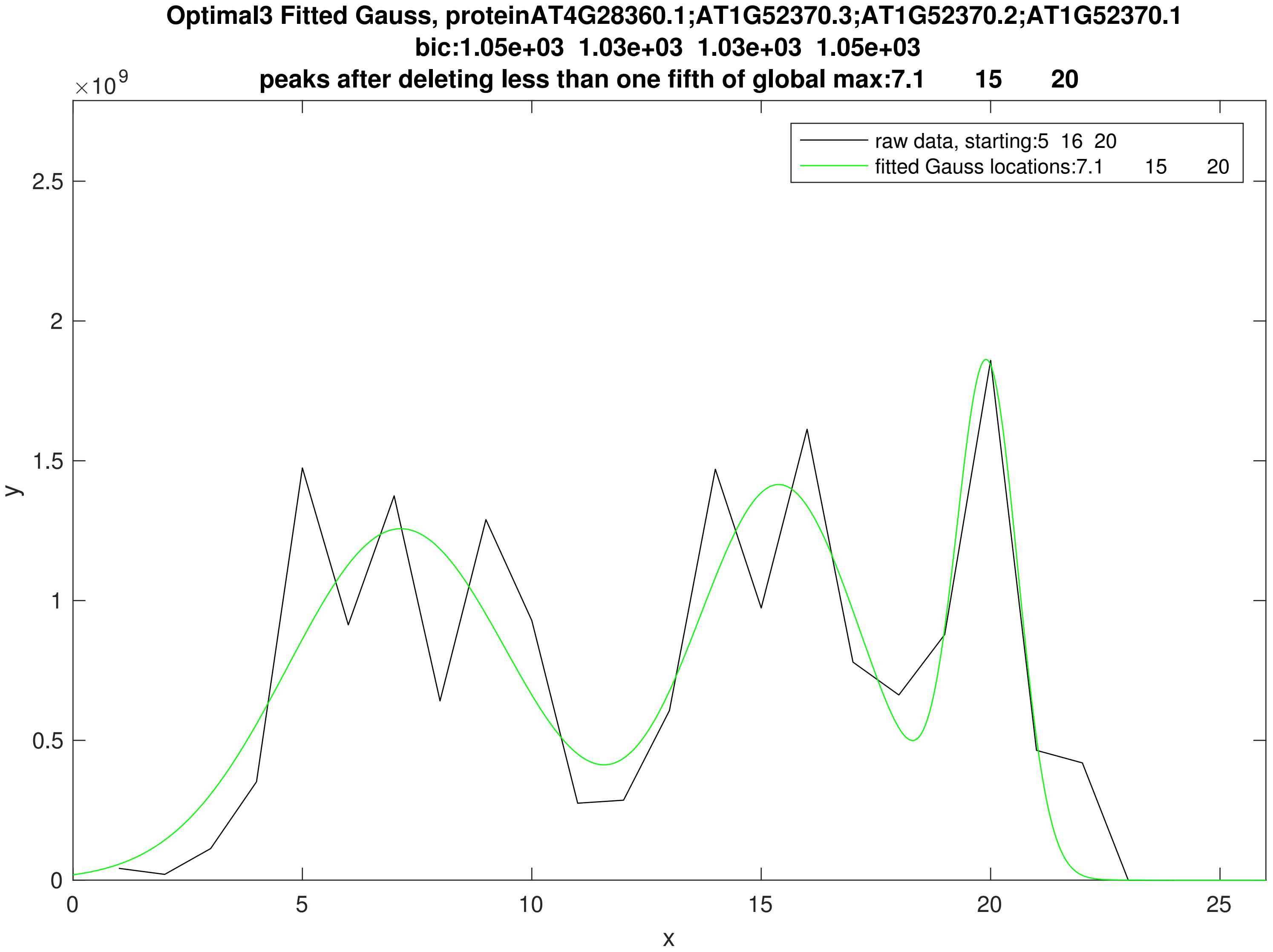

Optimal3 Fitted Gauss, proteinAT4G28360.1;AT1G52370.3;AT1G52370.2;AT1G52370.1
bic:1.05e+03 1.03e+03 1.03e+03 1.05e+03
peaks after deleting less than one fifth of global max:7.1 15 20
×10^9
raw data, starting:5 16 20
fitted Gauss locations:7.1 15 20
y
x
0
0.5
1
1.5
2
2.5
5
10
15
20
25

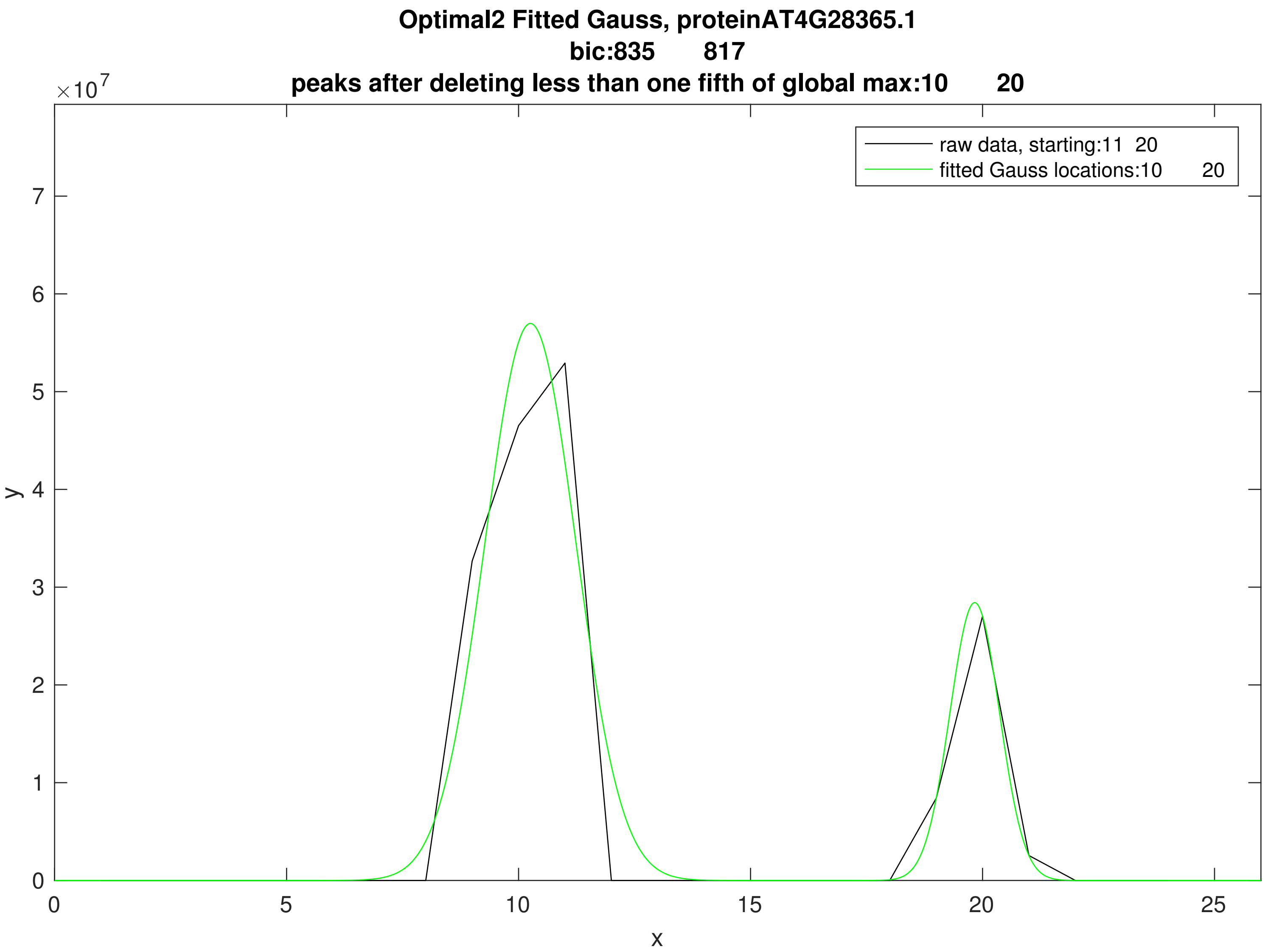

Optimal2 Fitted Gauss, proteinAT4G28365.1
bic:835 817
peaks after deleting less than one fifth of global max:10 20
×10^7
raw data, starting:11 20
fitted Gauss locations:10 20
y
x
0
1
2
3
4
5
6
7
0
5
10
15
20
25

#### Optimal1 Fitted Gauss, proteinAT4G28410.1
#### bic:844
#### peaks after deleting less than one fifth of global max:13

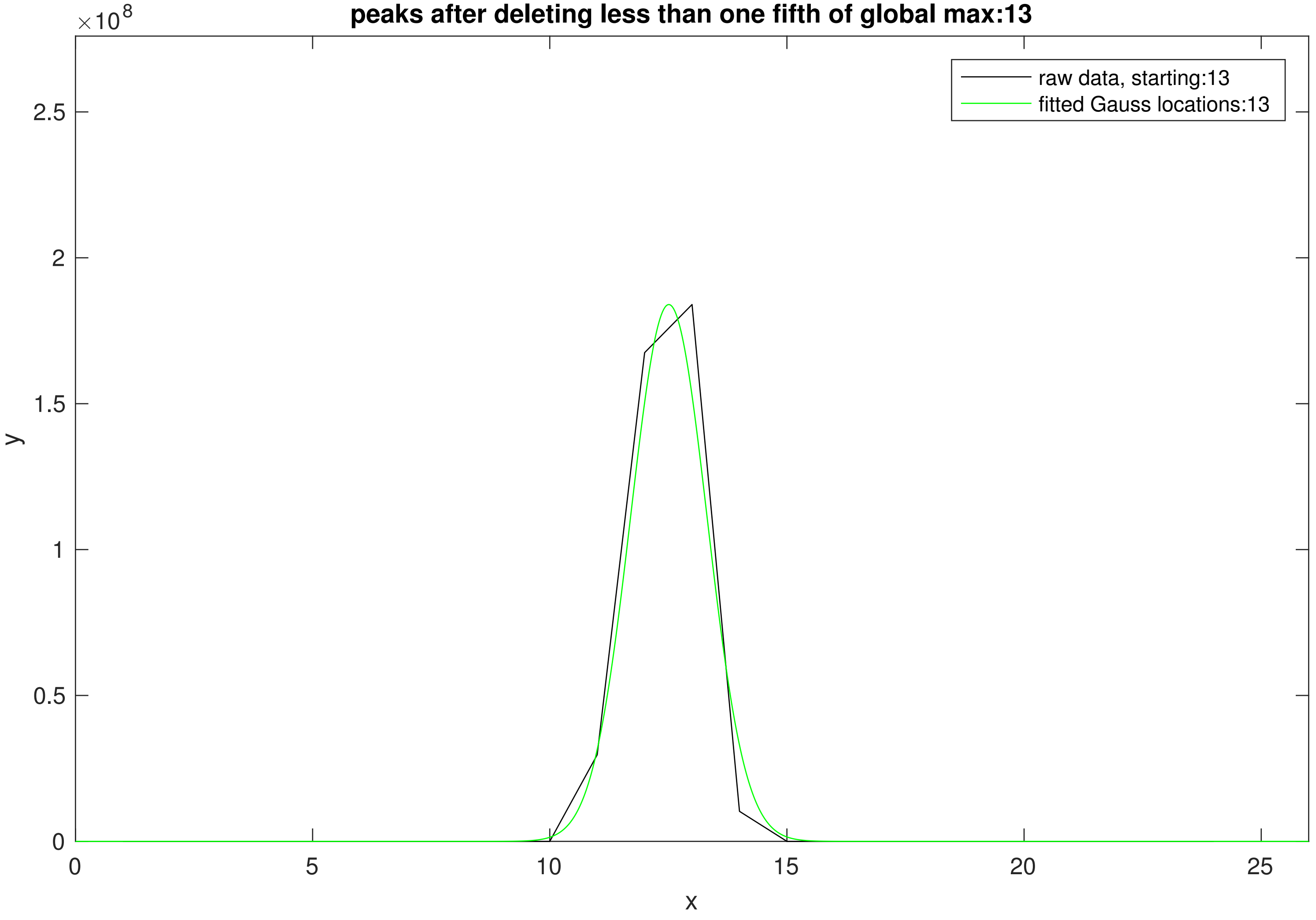

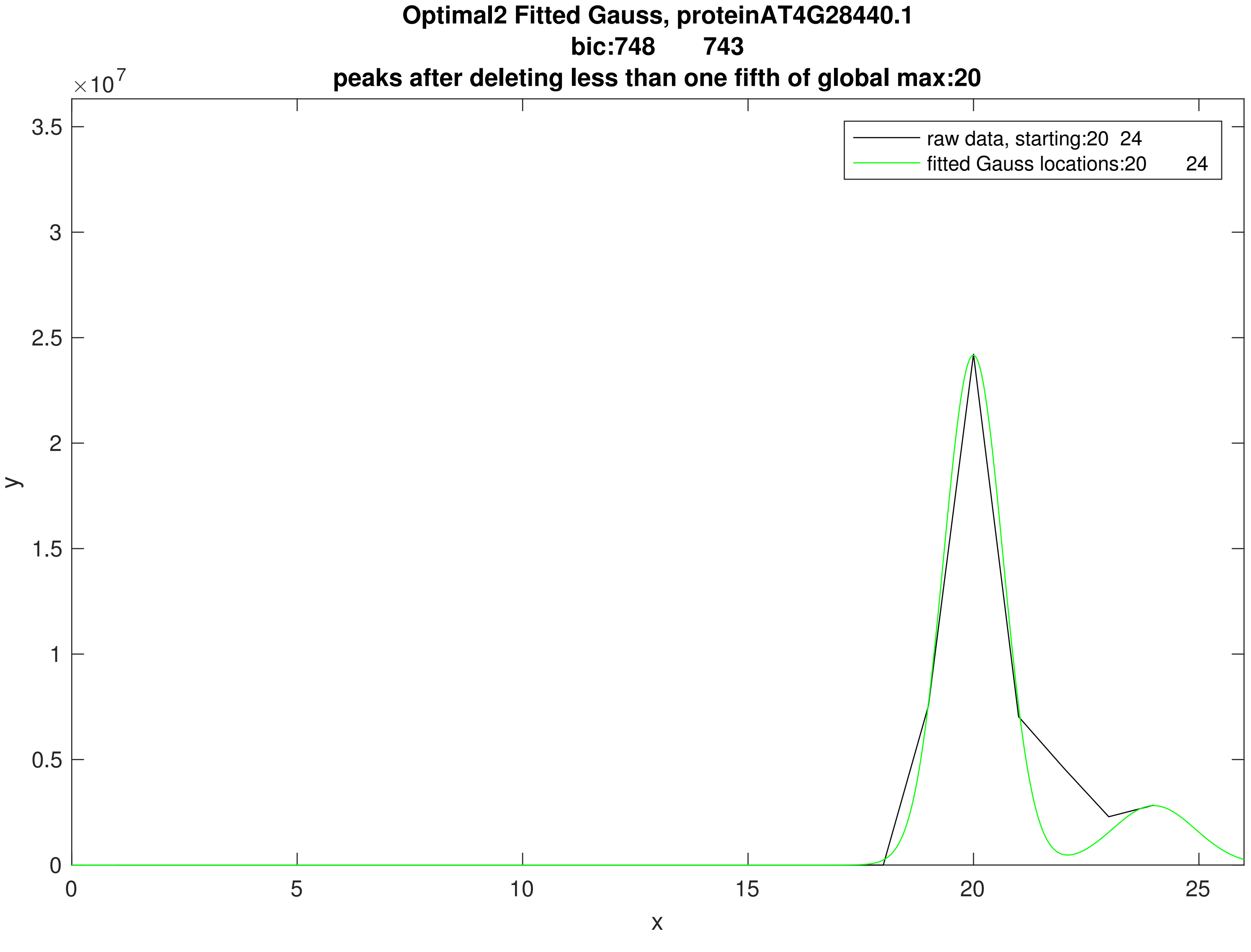

Optimal2 Fitted Gauss, proteinAT4G28440.1
bic:748 743
peaks after deleting less than one fifth of global max:20
×10^7
3.5
3
2.5
2
1.5
1
0.5
0
y
0
5
10
15
20
25
x
raw data, starting:20 24
fitted Gauss locations:20 24

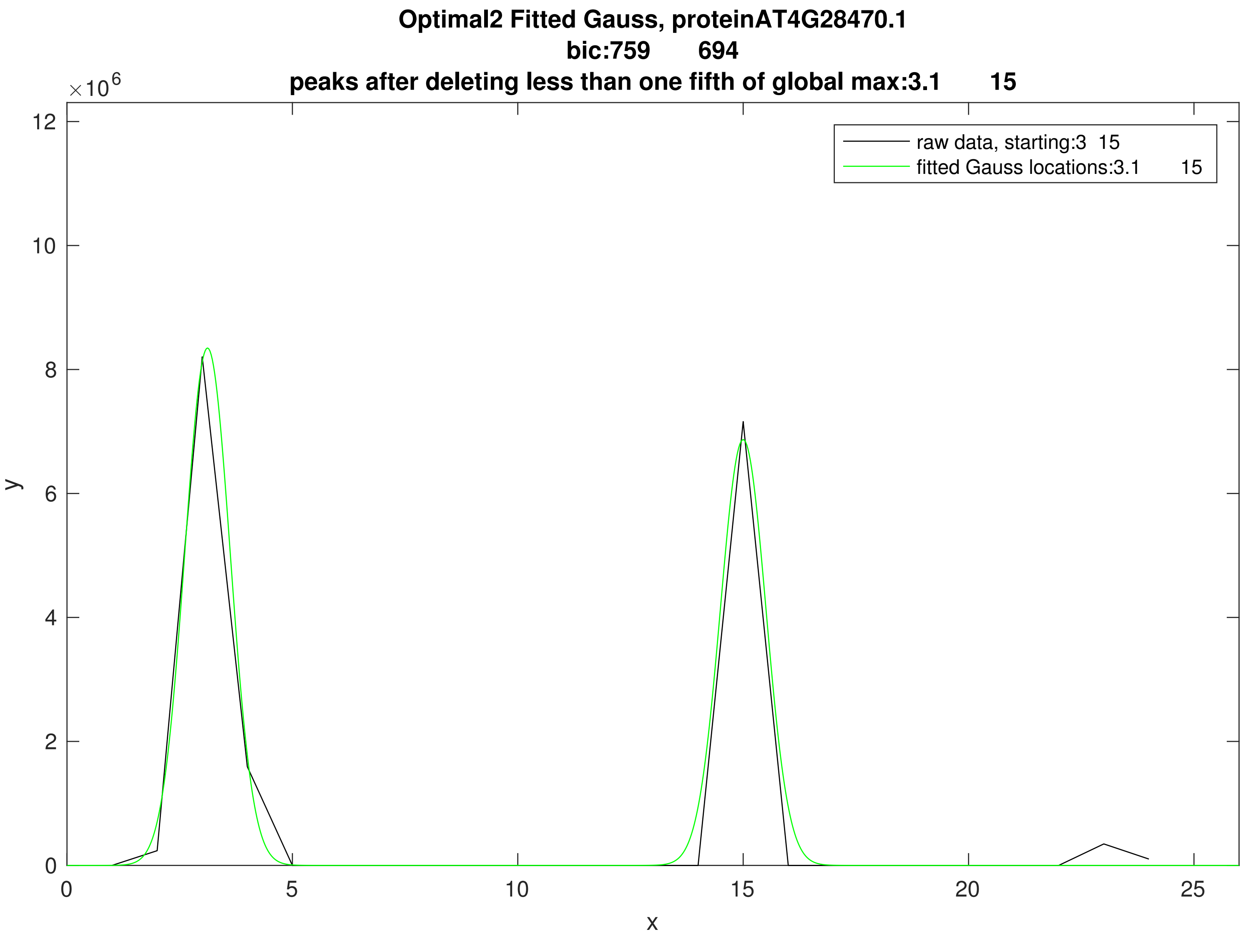
Optimal2 Fitted Gauss, proteinAT4G28470.1
bic:759 694
peaks after deleting less than one fifth of global max:3.1 15
×10^6
raw data, starting:3 15
fitted Gauss locations:3.1 15
y
x
0
2
4
6
8
10
12
5
15
20
25

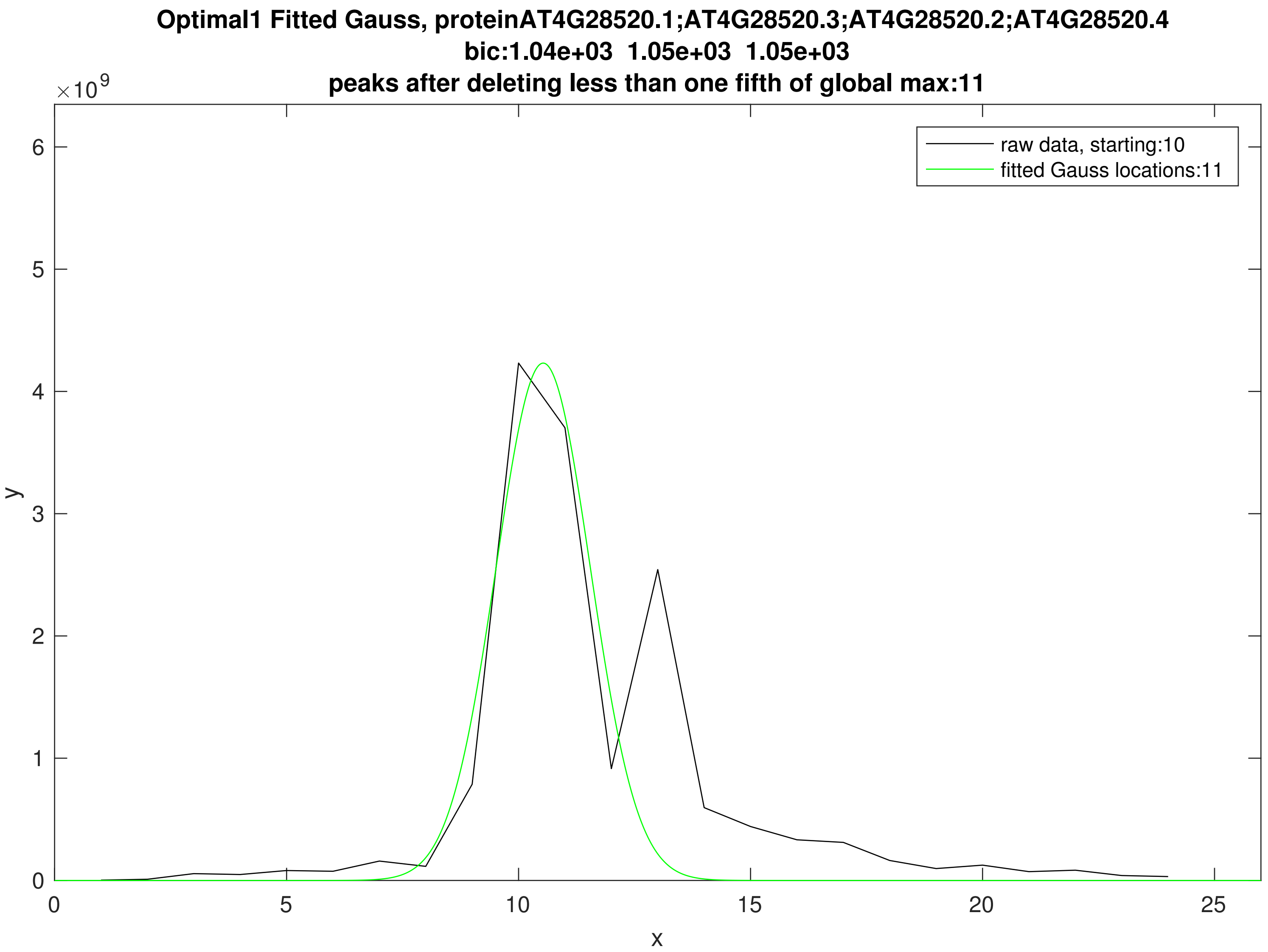

Optimal1 Fitted Gauss, proteinAT4G28520.1;AT4G28520.3;AT4G28520.2;AT4G28520.4
bic:1.04e+03 1.05e+03 1.05e+03
peaks after deleting less than one fifth of global max:11
×10^9
raw data, starting:10
fitted Gauss locations:11
y
x
0
1
2
3
4
5
6
0
5
10
15
20
25

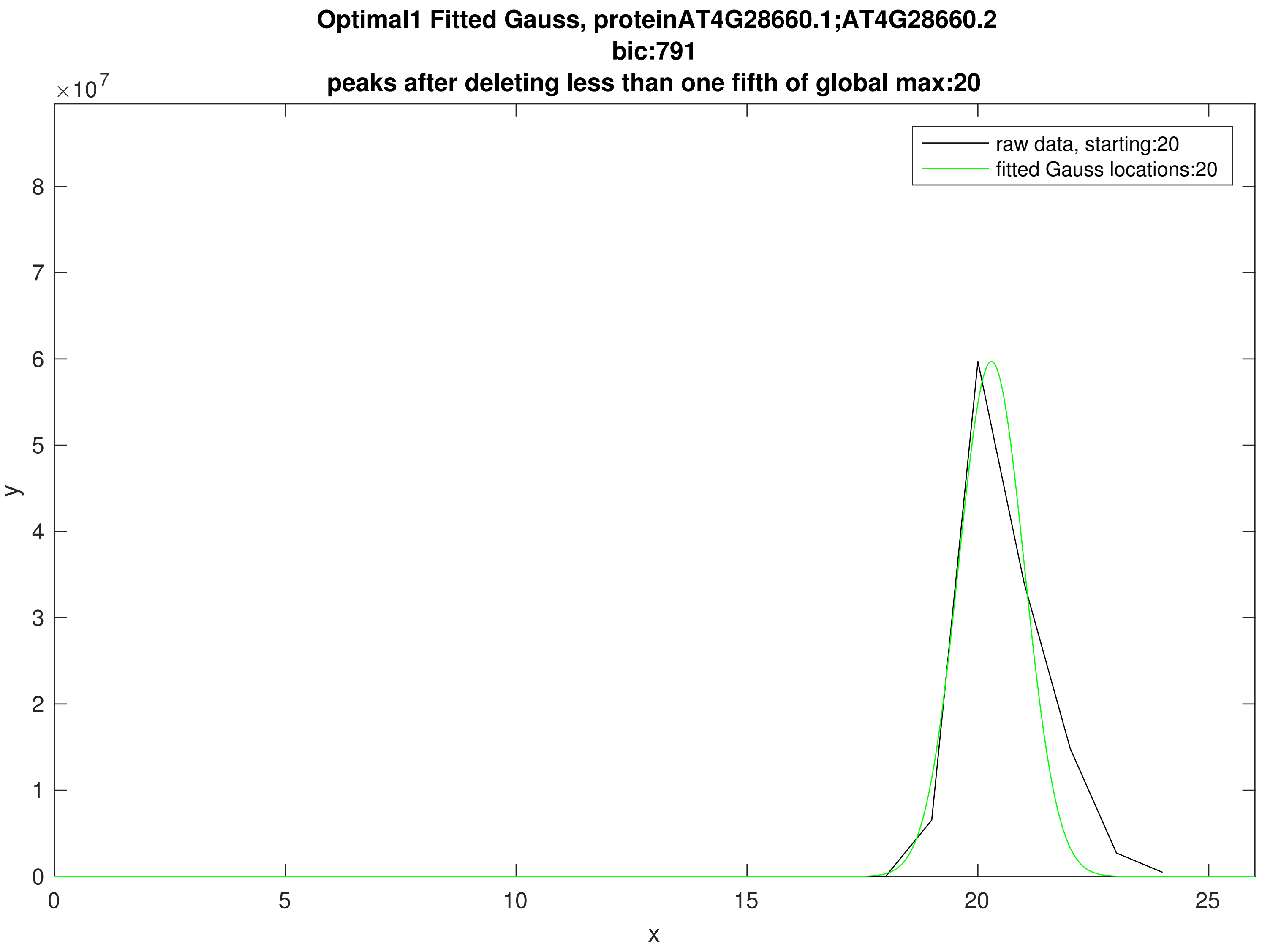
Optimal1 Fitted Gauss, proteinAT4G28660.1;AT4G28660.2
bic:791
peaks after deleting less than one fifth of global max:20
×10^7
raw data, starting:20
fitted Gauss locations:20
y
x
0
1
2
3
4
5
6
7
8
0
5
10
15
20
25

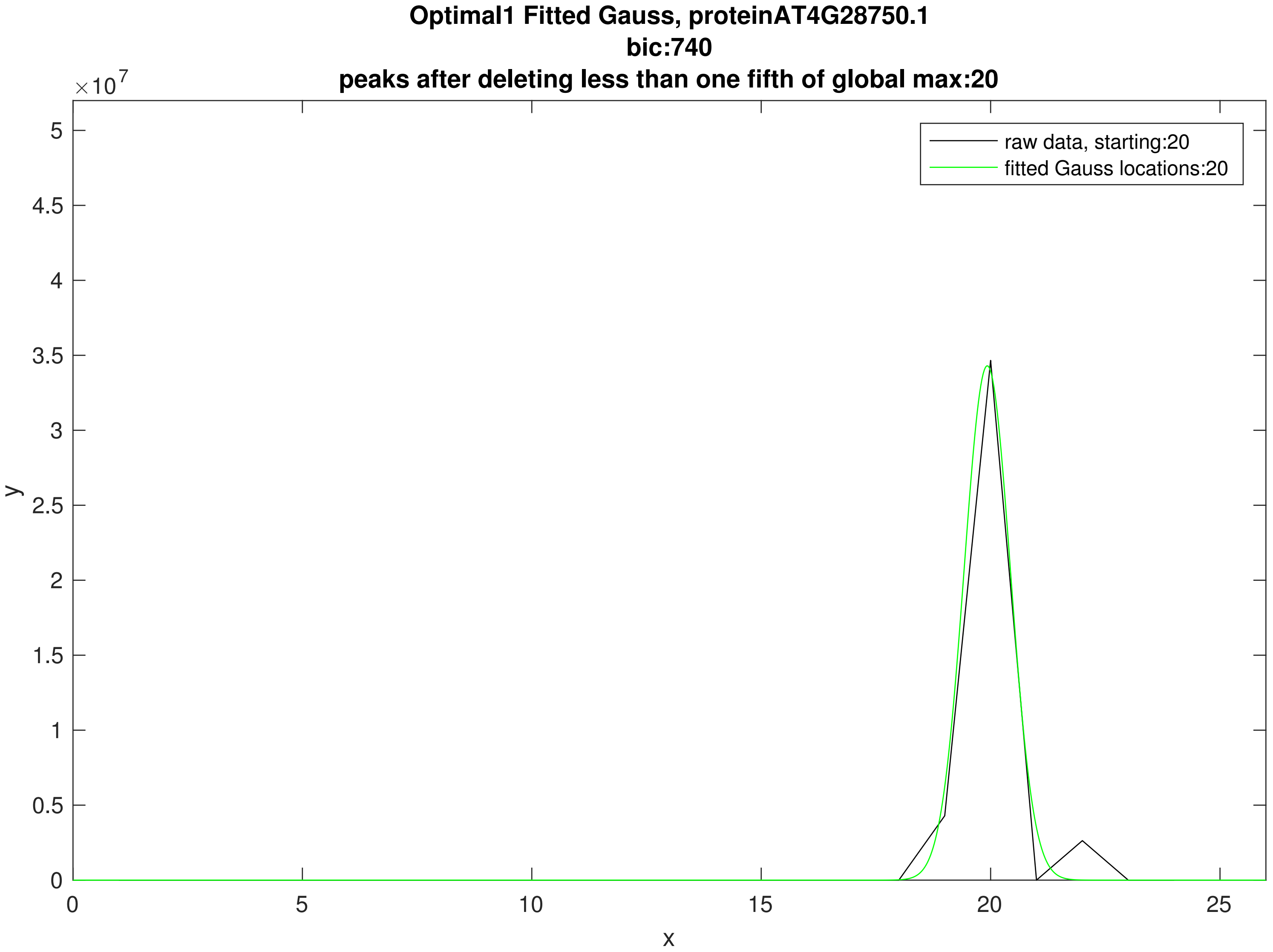

Optimal1 Fitted Gauss, proteinAT4G28750.1
bic:740
peaks after deleting less than one fifth of global max:20
×10^7
raw data, starting:20
fitted Gauss locations:20
y
x
0
0.5
1
1.5
2
2.5
3
3.5
4
4.5
5
0
5
10
15
20
25

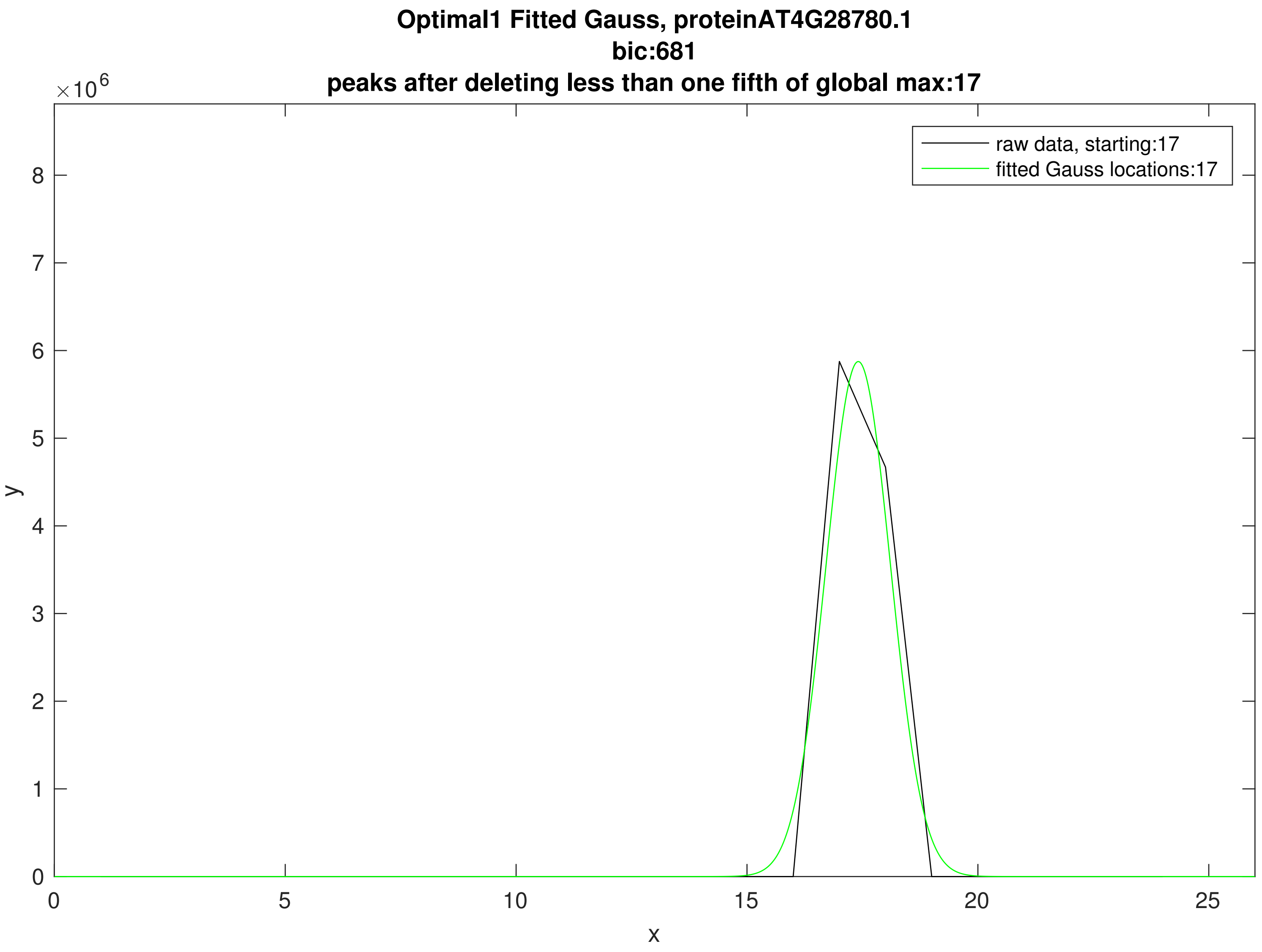
Optimal1 Fitted Gauss, proteinAT4G28780.1
bic:681
peaks after deleting less than one fifth of global max:17
×10^6
raw data, starting:17
fitted Gauss locations:17
y
x
0
1
2
3
4
5
6
7
8
0
5
10
15
20
25

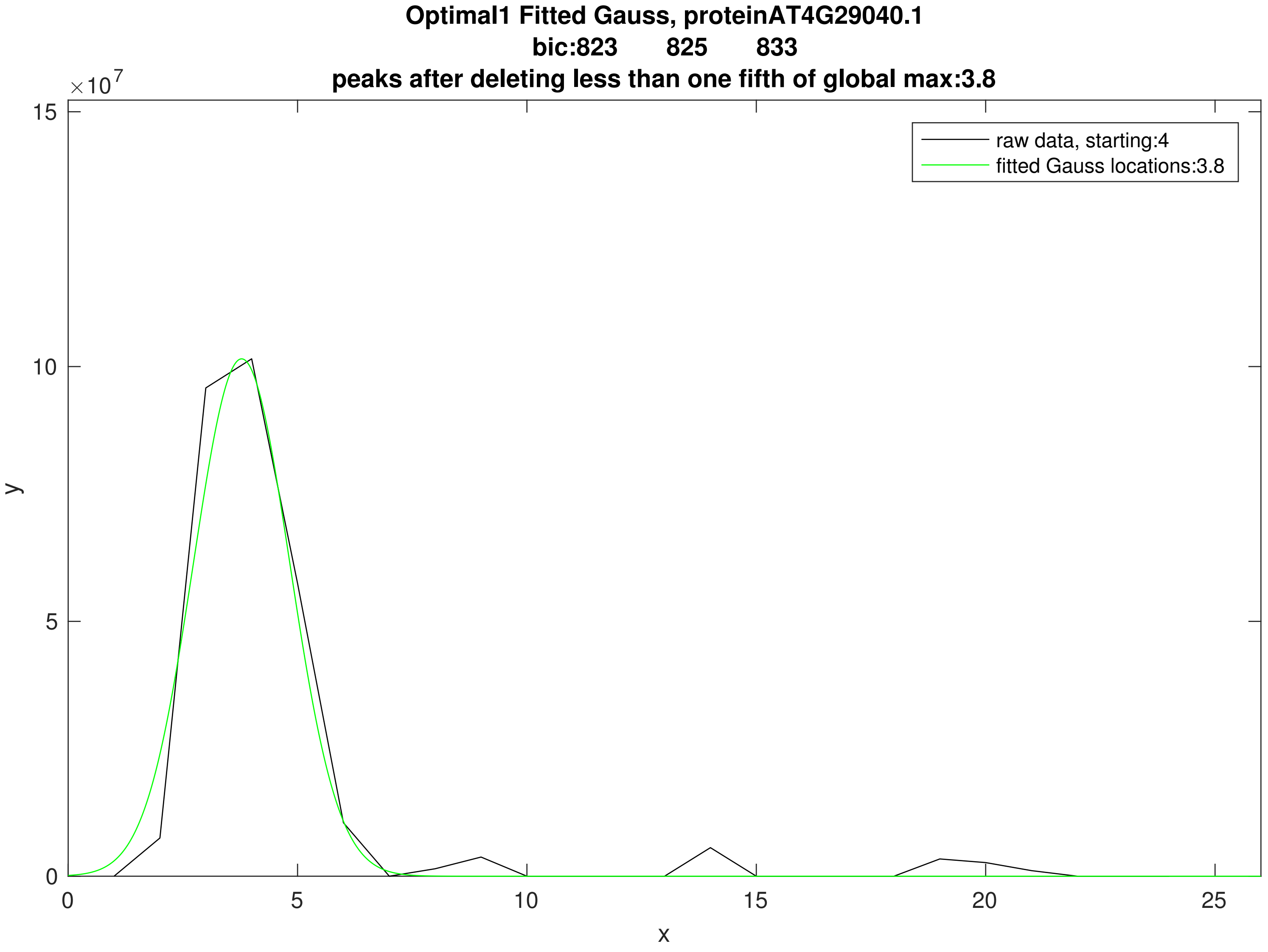
Optimal1 Fitted Gauss, proteinAT4G29040.1
bic:823 825 833
peaks after deleting less than one fifth of global max:3.8
×10^7
15
10
5
0
y
0
5
10
15
20
25
x
raw data, starting:4
fitted Gauss locations:3.8

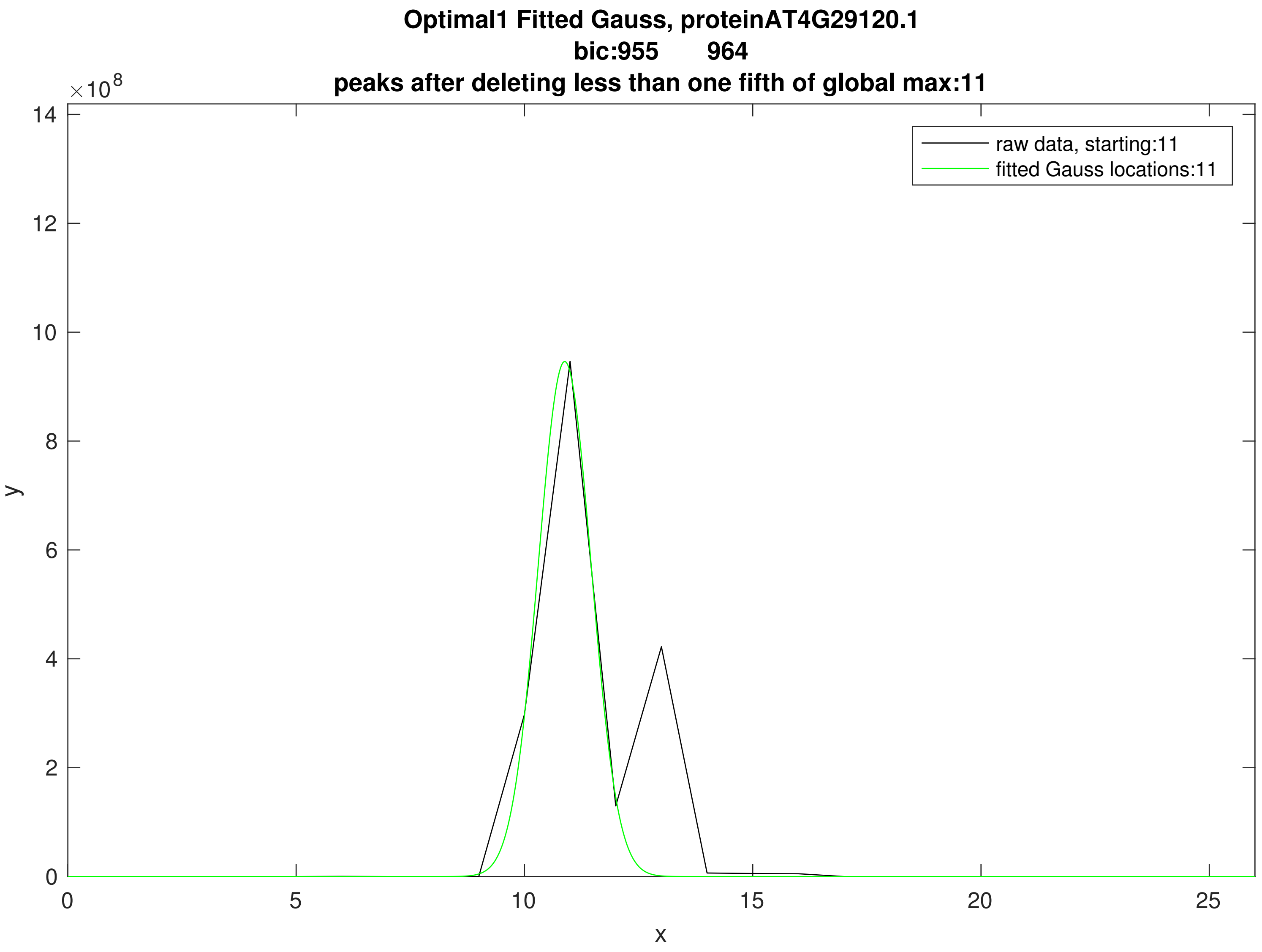
Optimal1 Fitted Gauss, proteinAT4G29120.1
bic:955 964
peaks after deleting less than one fifth of global max:11
×10^8
raw data, starting:11
fitted Gauss locations:11
y
x
0
2
4
6
8
10
12
14
0
5
10
15
20
25

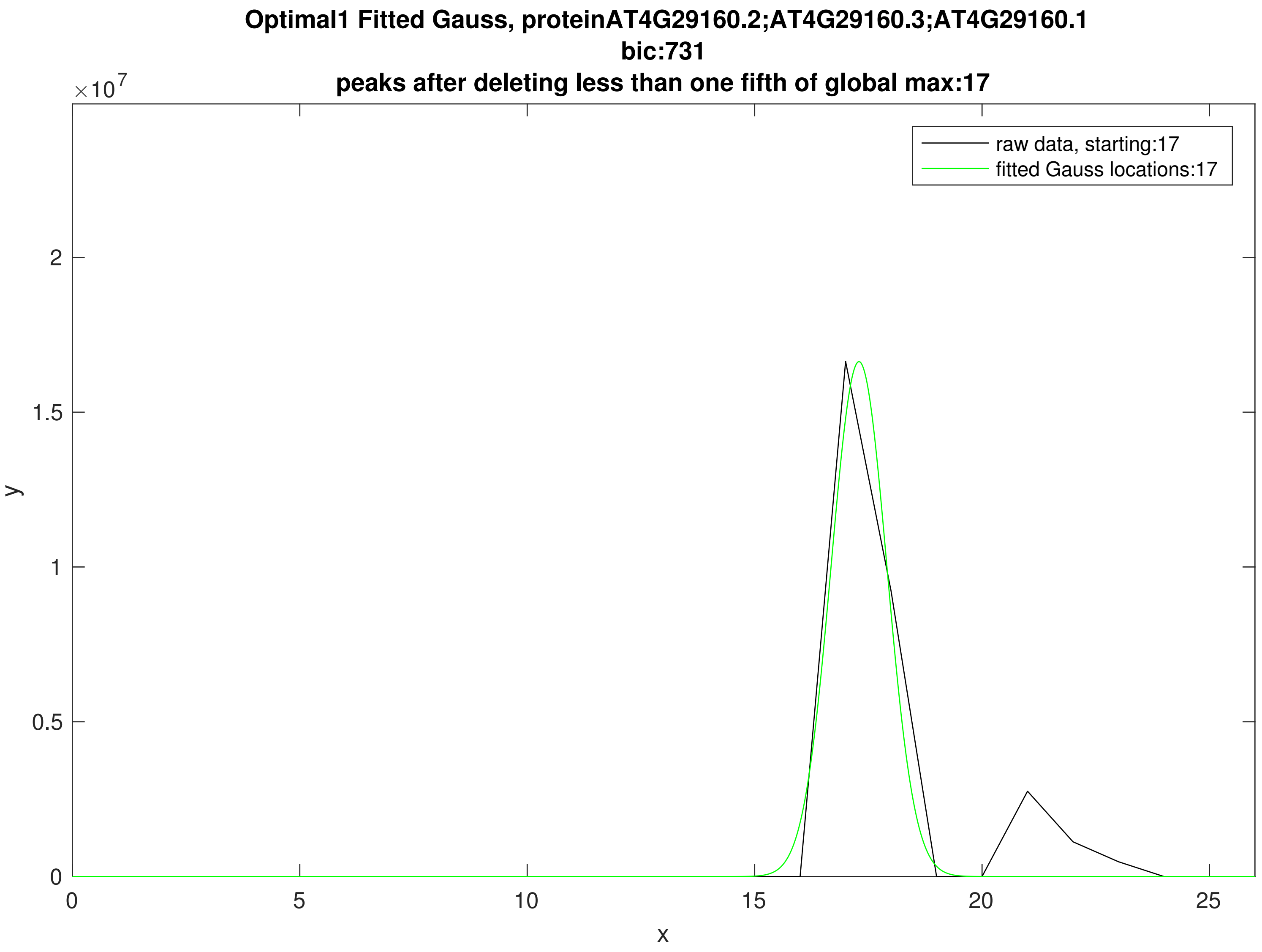
Optimal1 Fitted Gauss, proteinAT4G29160.2;AT4G29160.3;AT4G29160.1
bic:731
peaks after deleting less than one fifth of global max:17
×10^7
raw data, starting:17
fitted Gauss locations:17
2
1.5
1
0.5
0
y
0
5
10
15
20
25
x

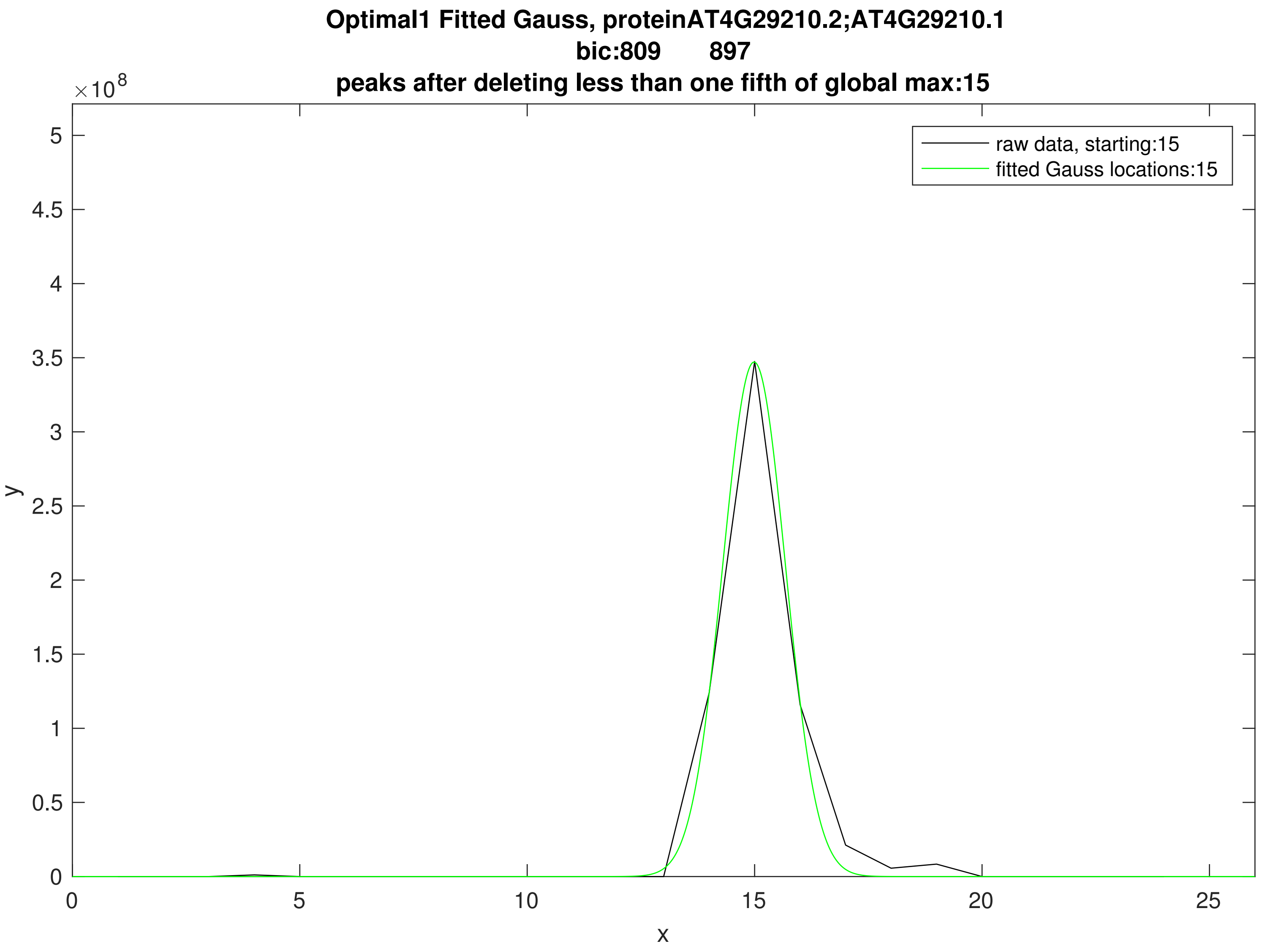
Optimal1 Fitted Gauss, proteinAT4G29210.2;AT4G29210.1
bic:809 897
peaks after deleting less than one fifth of global max:15
×10^8
raw data, starting:15
fitted Gauss locations:15
y
x
0
0.5
1
1.5
2
2.5
3
3.5
4
4.5
5
0
5
10
15
20
25

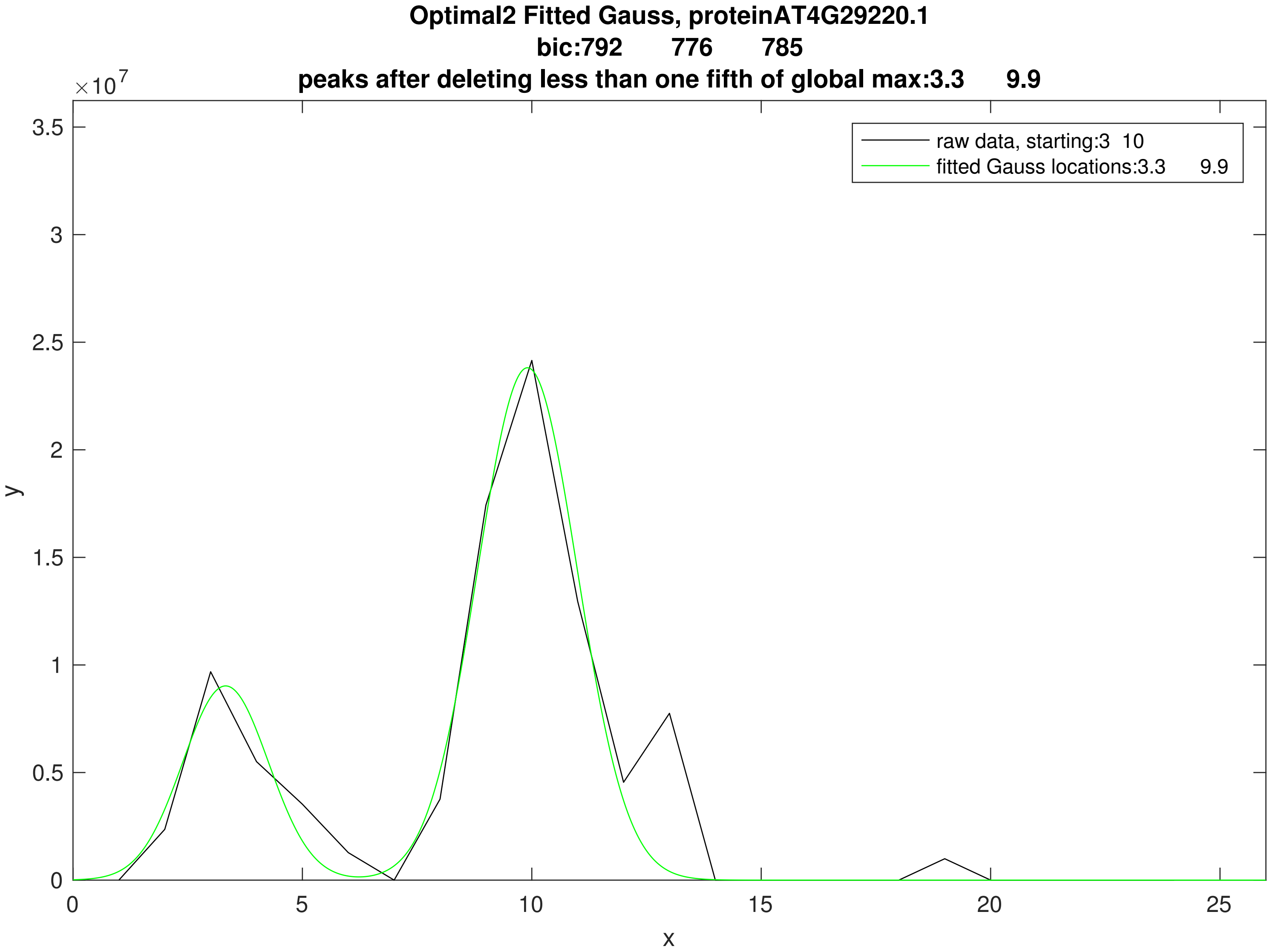
Optimal2 Fitted Gauss, proteinAT4G29220.1
bic:792 776 785
peaks after deleting less than one fifth of global max:3.3 9.9
×10^7
raw data, starting:3 10
fitted Gauss locations:3.3 9.9
y
x
0
0.5
1
1.5
2
2.5
3
3.5
5
10
15
20
25

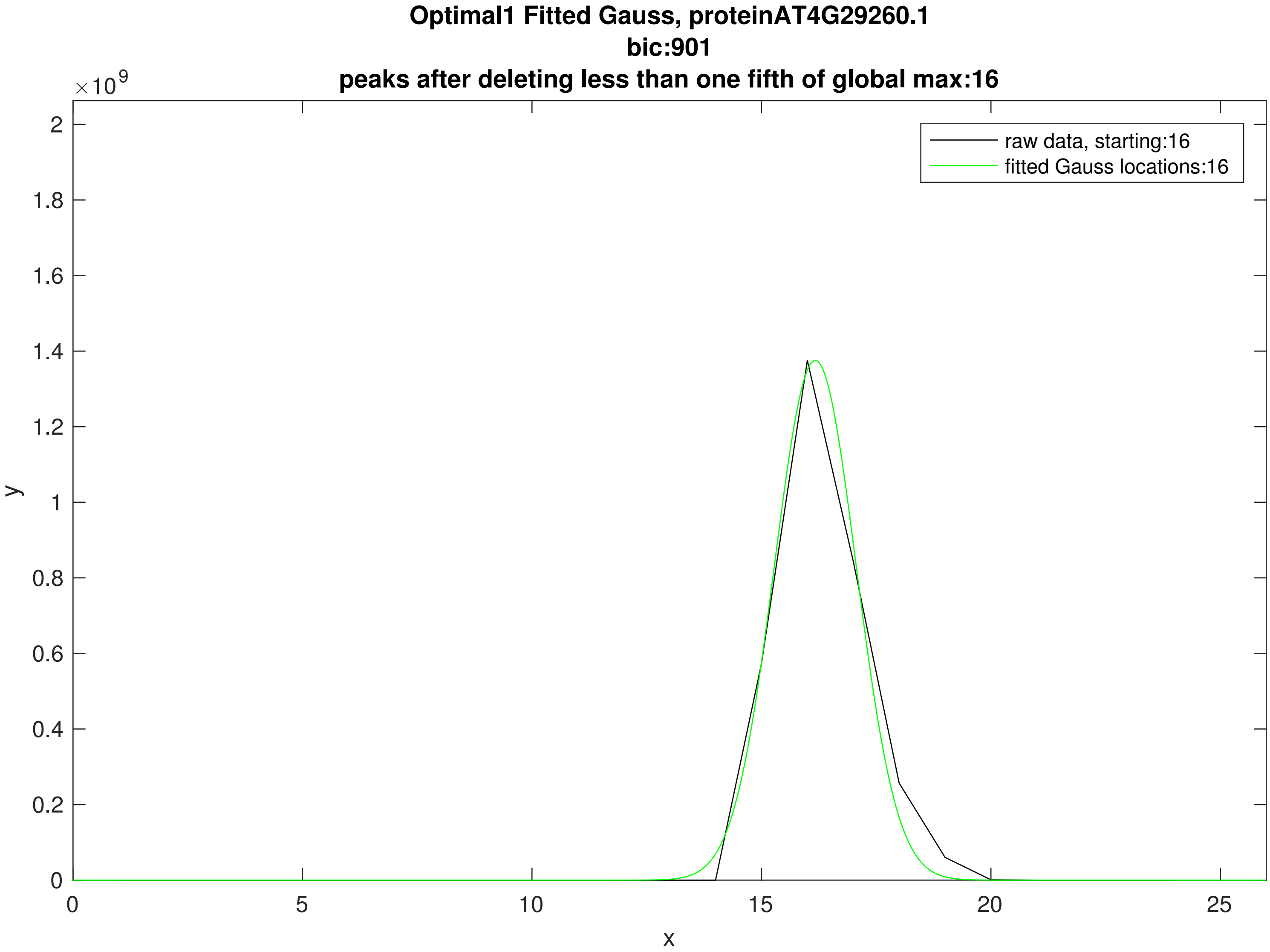
Optimal1 Fitted Gauss, proteinAT4G29260.1
bic:901
peaks after deleting less than one fifth of global max:16
×10^9
raw data, starting:16
fitted Gauss locations:16
y
x
0
0.2
0.4
0.6
0.8
1
1.2
1.4
1.6
1.8
2
0
5
10
15
20
25

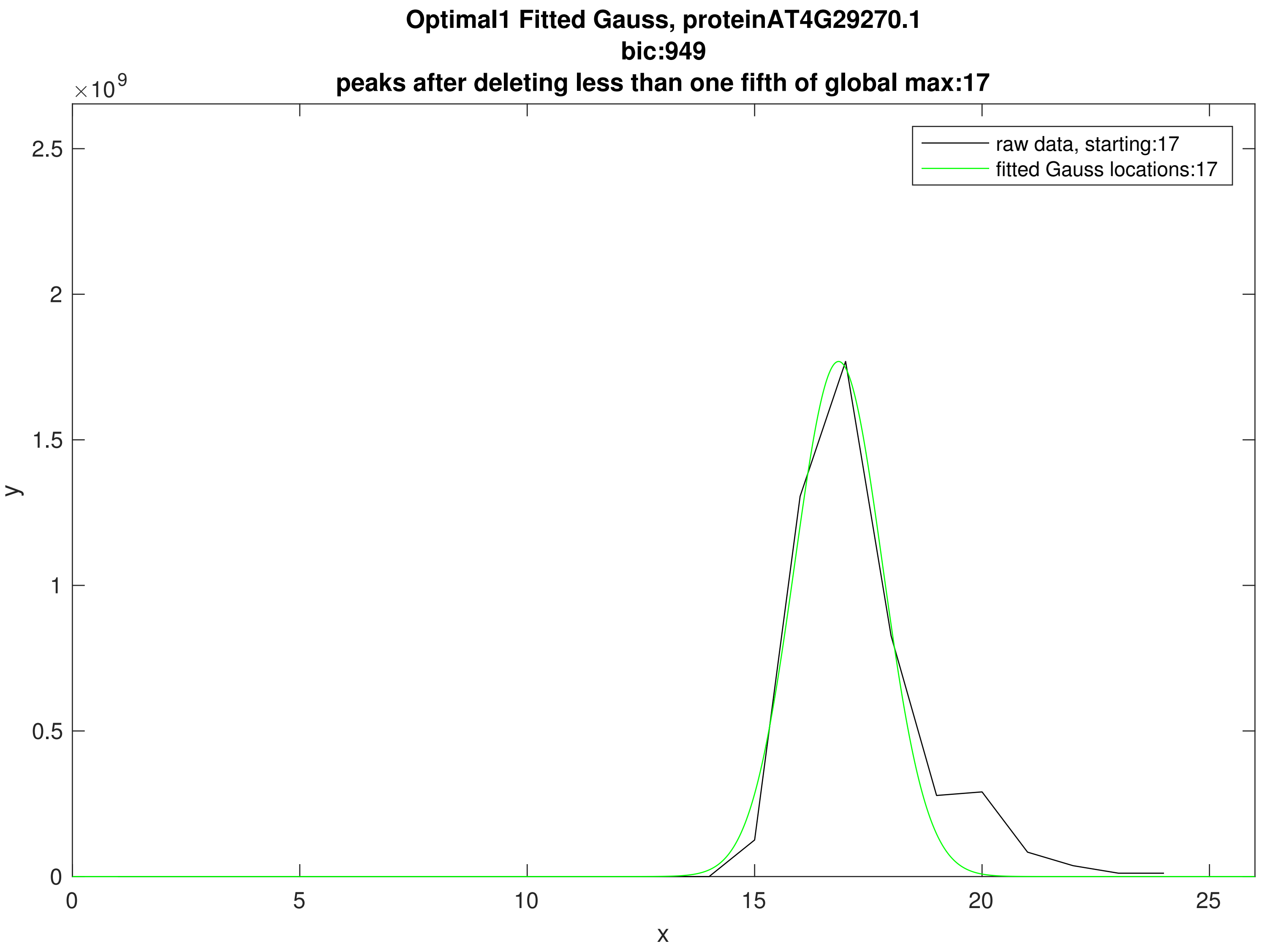
Optimal1 Fitted Gauss, proteinAT4G29270.1
bic:949
peaks after deleting less than one fifth of global max:17
×10^9
2.5
2
1.5
1
0.5
0
y
0
5
10
15
20
25
x
raw data, starting:17
fitted Gauss locations:17

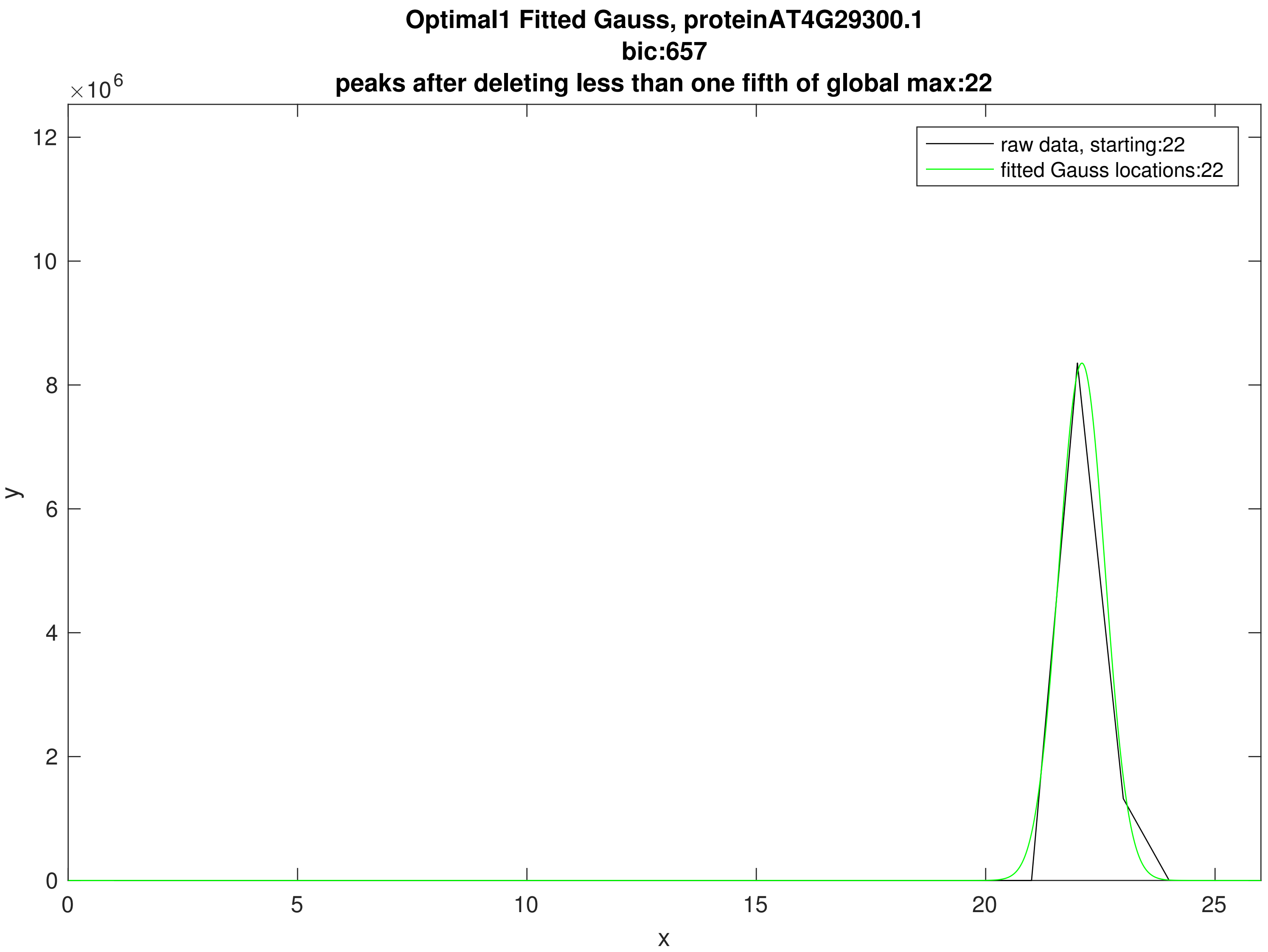
Optimal1 Fitted Gauss, proteinAT4G29300.1
bic:657
peaks after deleting less than one fifth of global max:22
×10^6
12
10
8
6
4
2
0
y
0
5
10
15
20
25
x
raw data, starting:22
fitted Gauss locations:22

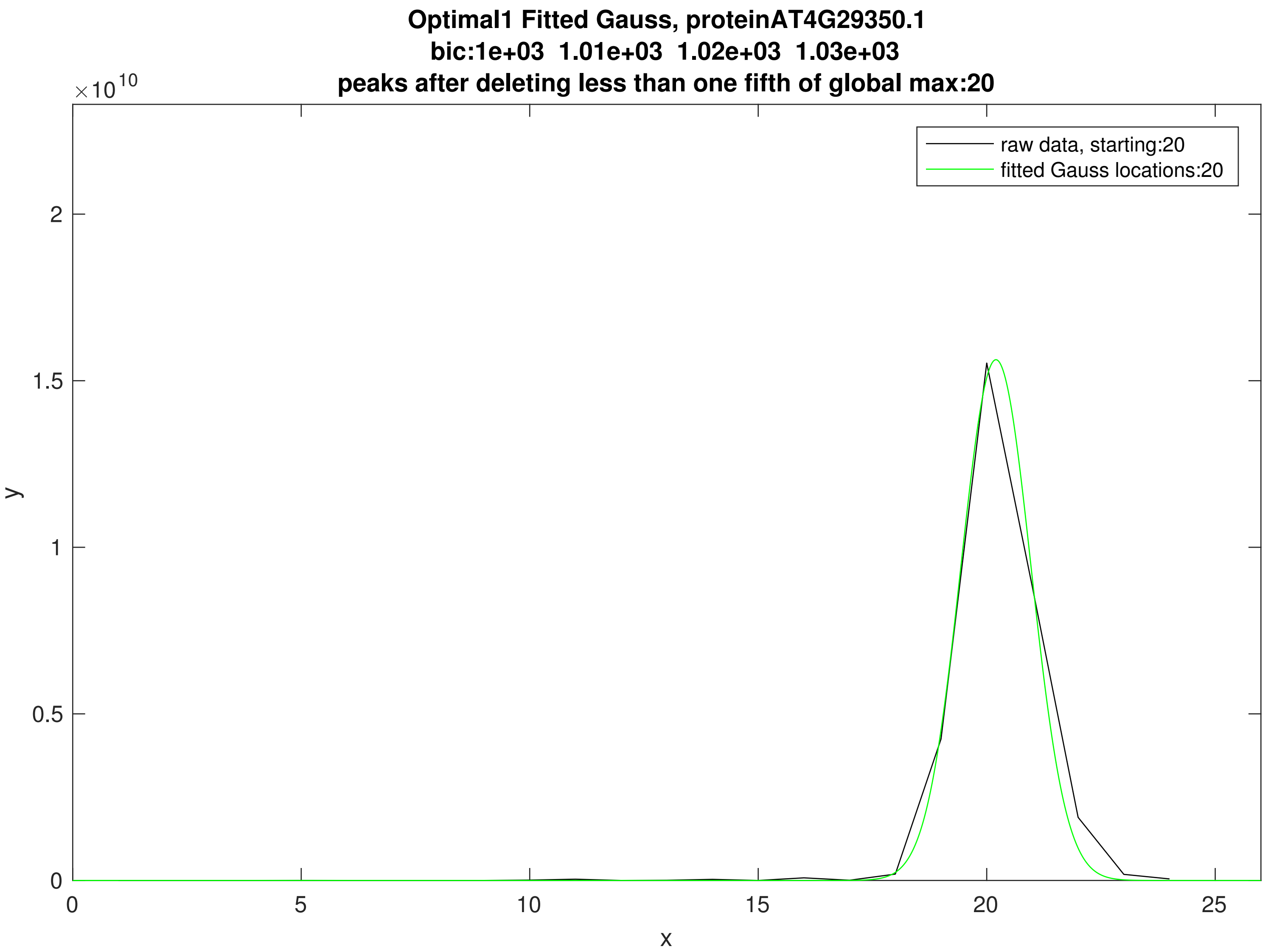

Optimal1 Fitted Gauss, proteinAT4G29350.1
bic:1e+03 1.01e+03 1.02e+03 1.03e+03
peaks after deleting less than one fifth of global max:20
×10^10
raw data, starting:20
fitted Gauss locations:20
2
1.5
1
0.5
0
y
0
5
10
15
20
25
x

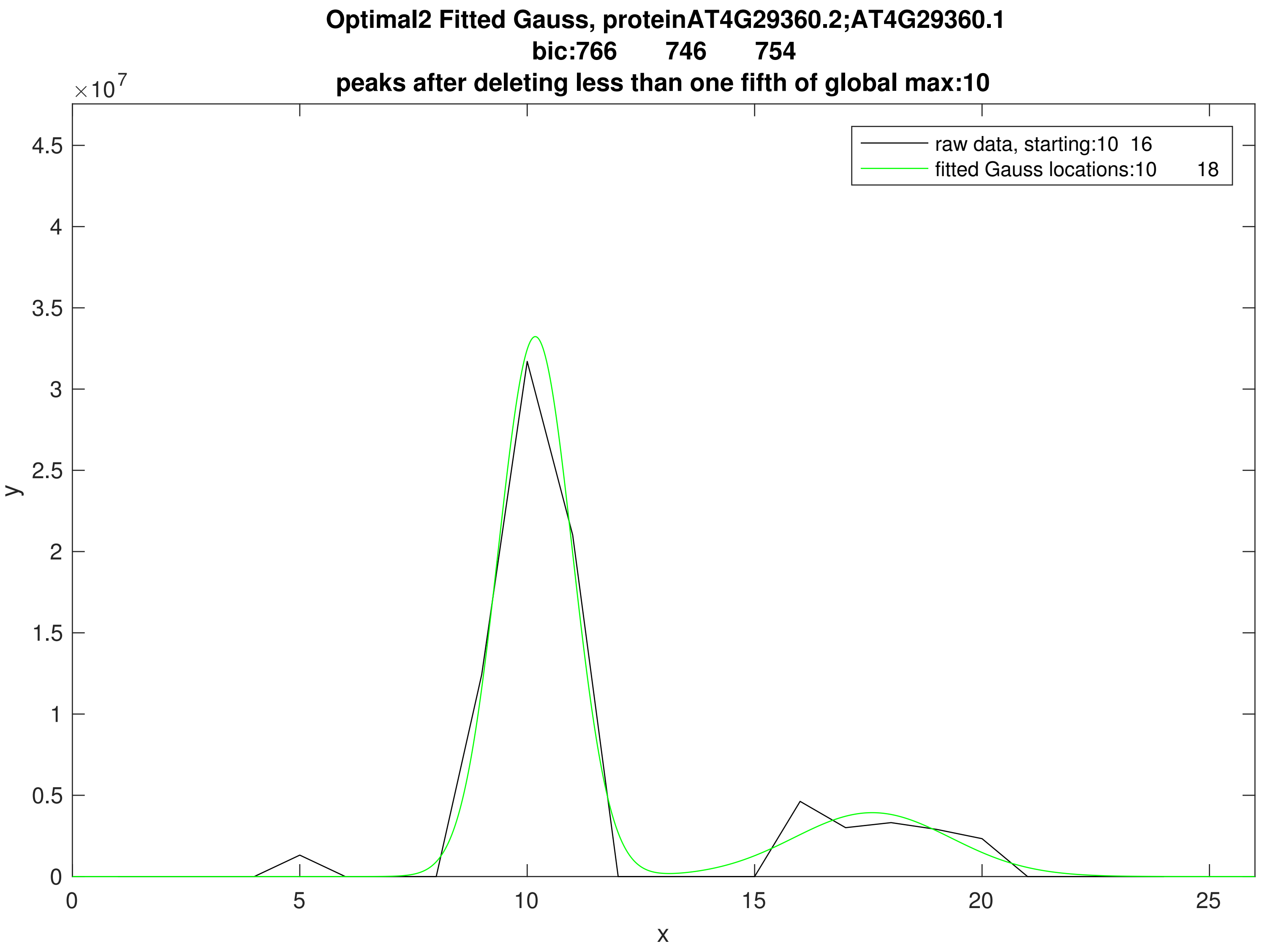
Optimal2 Fitted Gauss, proteinAT4G29360.2;AT4G29360.1
bic:766 746 754
peaks after deleting less than one fifth of global max:10
×10^7
raw data, starting:10 16
fitted Gauss locations:10 18
y
x
0
0.5
1
1.5
2
2.5
3
3.5
4
4.5
5
10
15
20
25

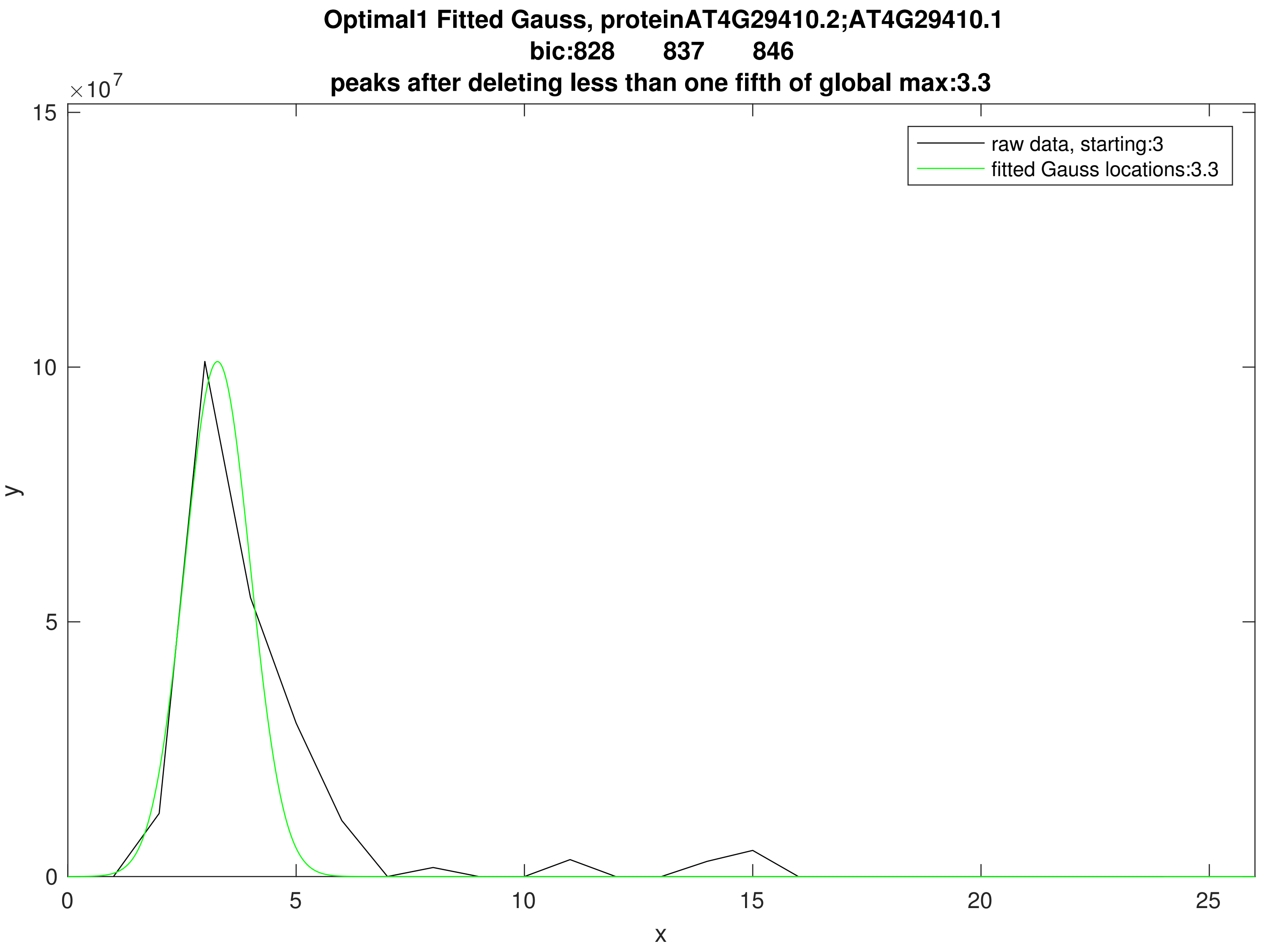
Optimal1 Fitted Gauss, proteinAT4G29410.2;AT4G29410.1
bic:828 837 846
peaks after deleting less than one fifth of global max:3.3
×10^7
raw data, starting:3
fitted Gauss locations:3.3
y
x
0
5
10
15
20
25

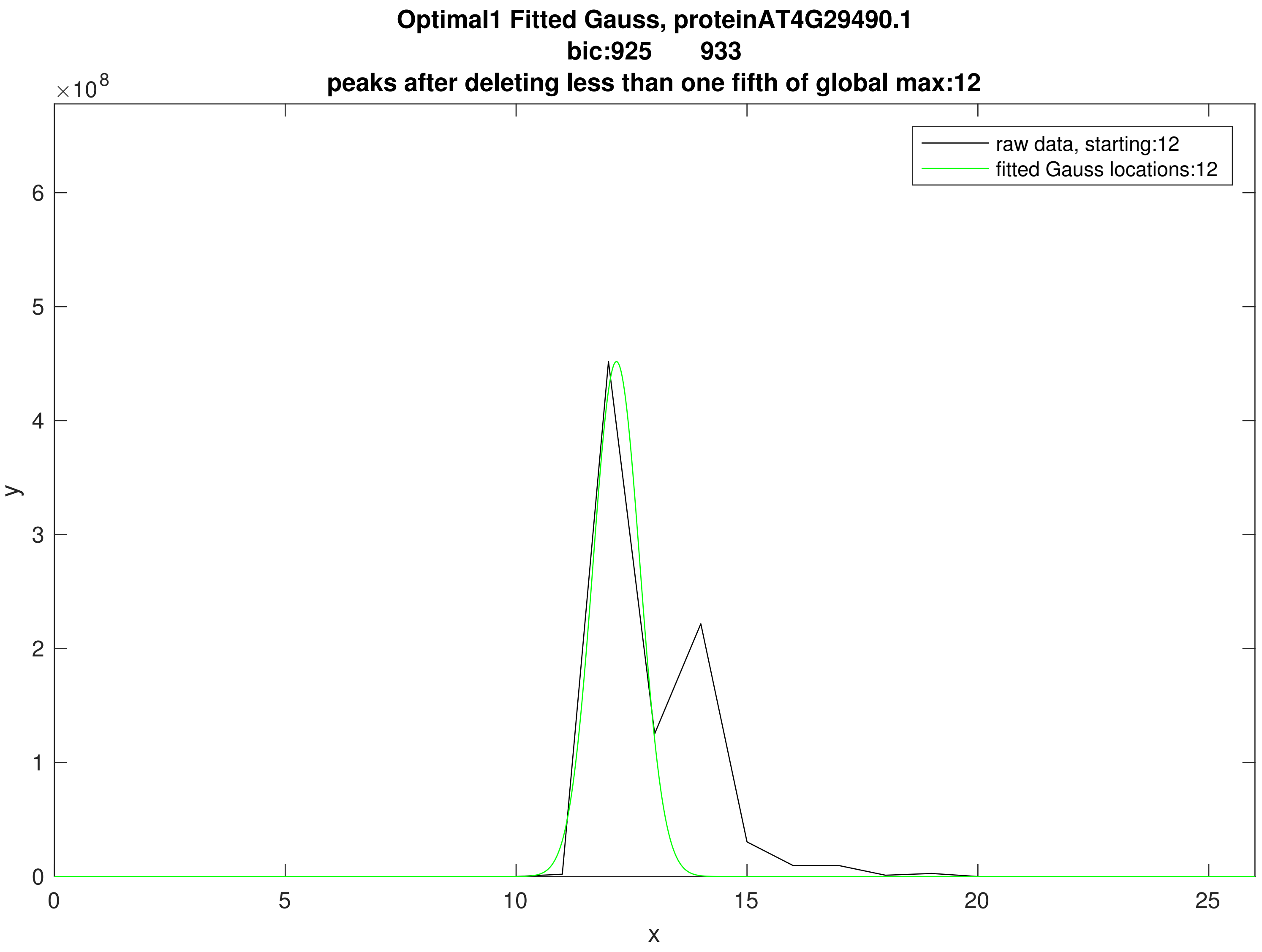
Optimal1 Fitted Gauss, proteinAT4G29490.1
bic:925 933
peaks after deleting less than one fifth of global max:12
×10^8
raw data, starting:12
fitted Gauss locations:12
y
x
0
1
2
3
4
5
6
0
5
10
15
20
25

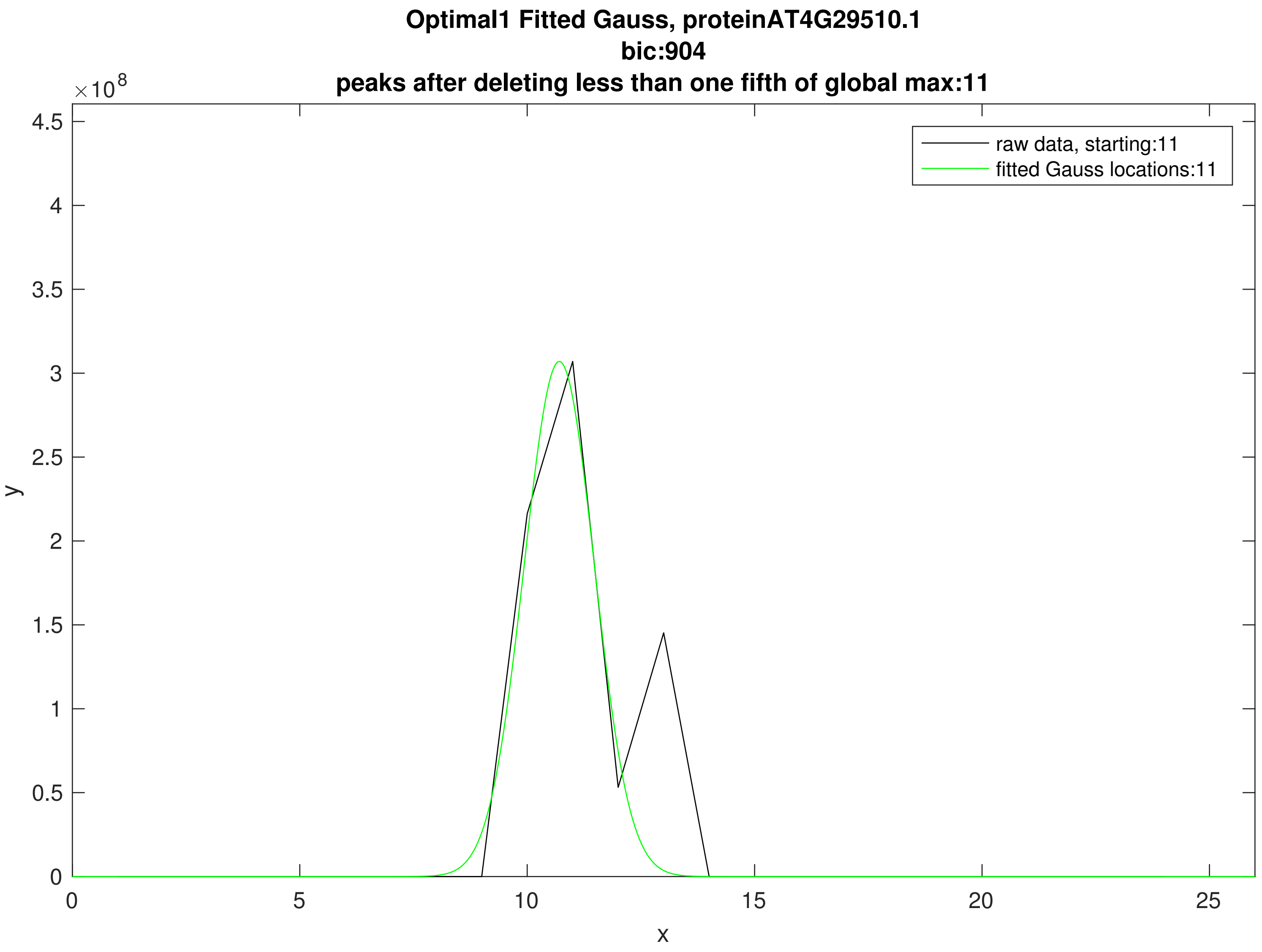

Optimal1 Fitted Gauss, proteinAT4G29510.1
bic:904
peaks after deleting less than one fifth of global max:11
×10^8
4.5
4
3.5
3
2.5
2
1.5
1
0.5
0
y
0
5
10
15
20
25
x
raw data, starting:11
fitted Gauss locations:11

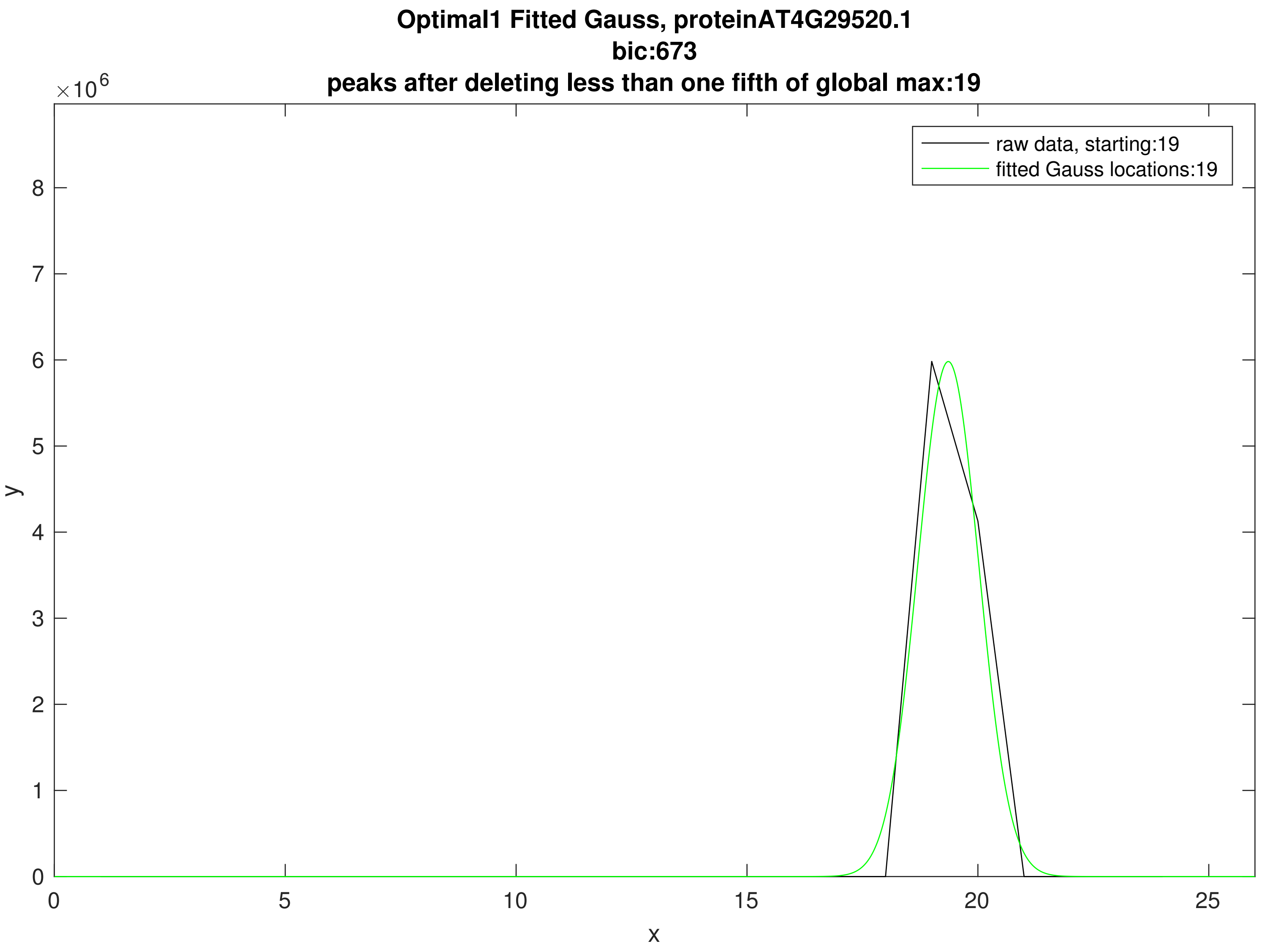

Optimal1 Fitted Gauss, proteinAT4G29520.1
bic:673
peaks after deleting less than one fifth of global max:19
×10^6
raw data, starting:19
fitted Gauss locations:19
y
x
0
1
2
3
4
5
6
7
8
0
5
10
15
20
25

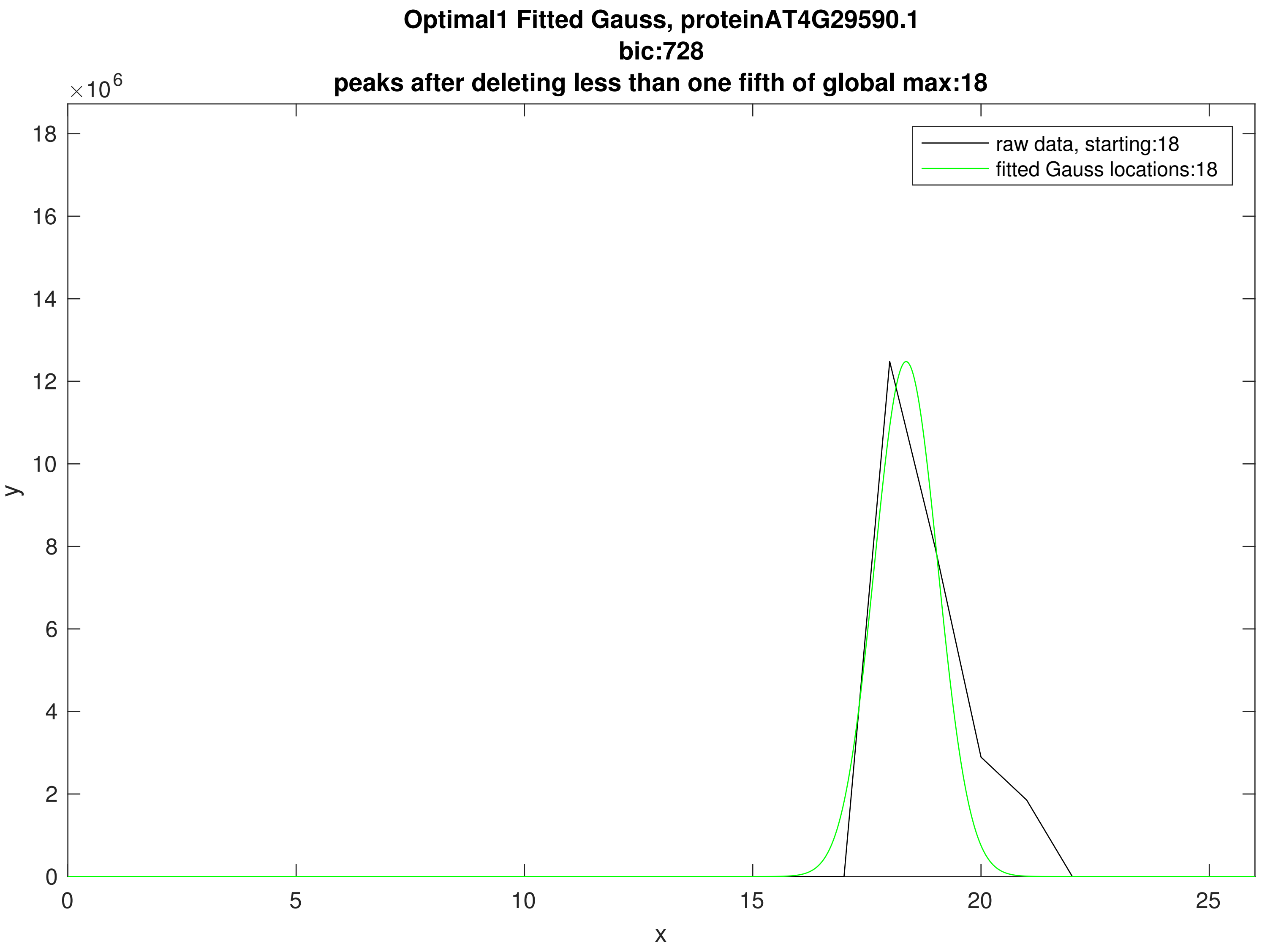
Optimal1 Fitted Gauss, proteinAT4G29590.1
bic:728
peaks after deleting less than one fifth of global max:18
×10^6
18
16
14
12
10
8
6
4
2
0
y
0
5
10
15
20
25
x
raw data, starting:18
fitted Gauss locations:18

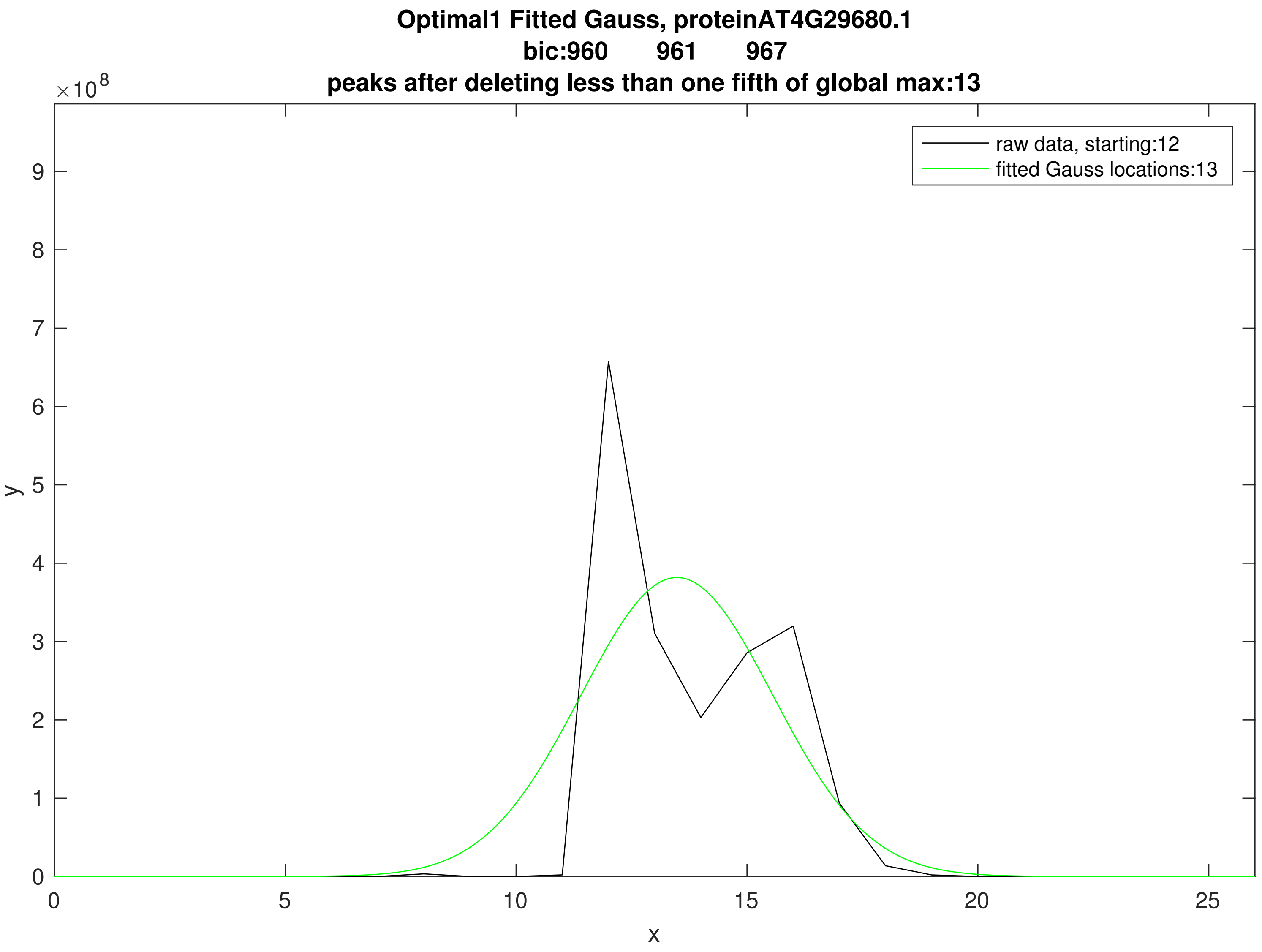
Optimal1 Fitted Gauss, proteinAT4G29680.1
bic:960 961 967
peaks after deleting less than one fifth of global max:13
×10^8
raw data, starting:12
fitted Gauss locations:13
y
x
0
1
2
3
4
5
6
7
8
9
0
5
10
15
20
25

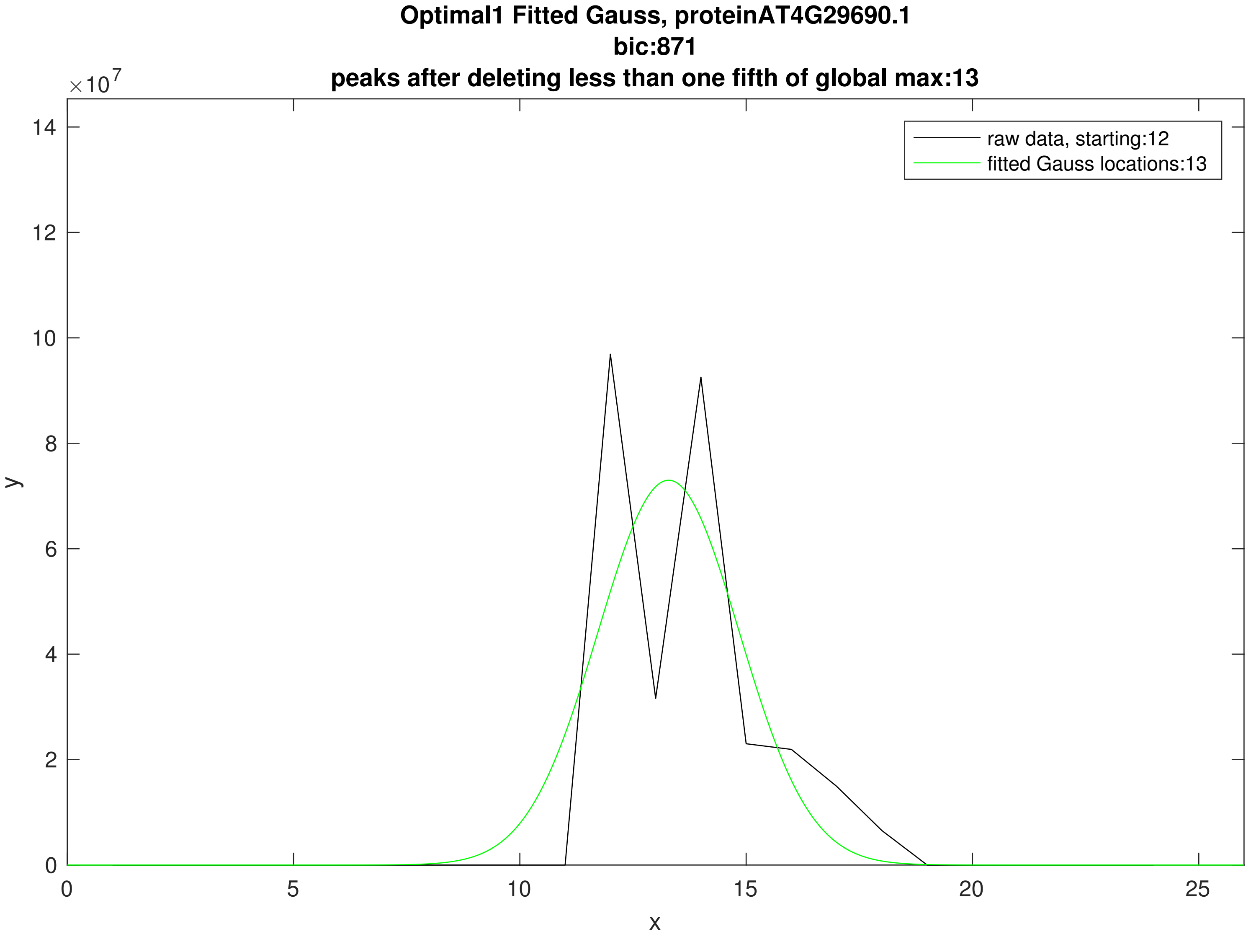
Optimal1 Fitted Gauss, proteinAT4G29690.1
bic:871
peaks after deleting less than one fifth of global max:13
×10^7
raw data, starting:12
fitted Gauss locations:13
y
x
0
2
4
6
8
10
12
14
0
5
10
15
20
25

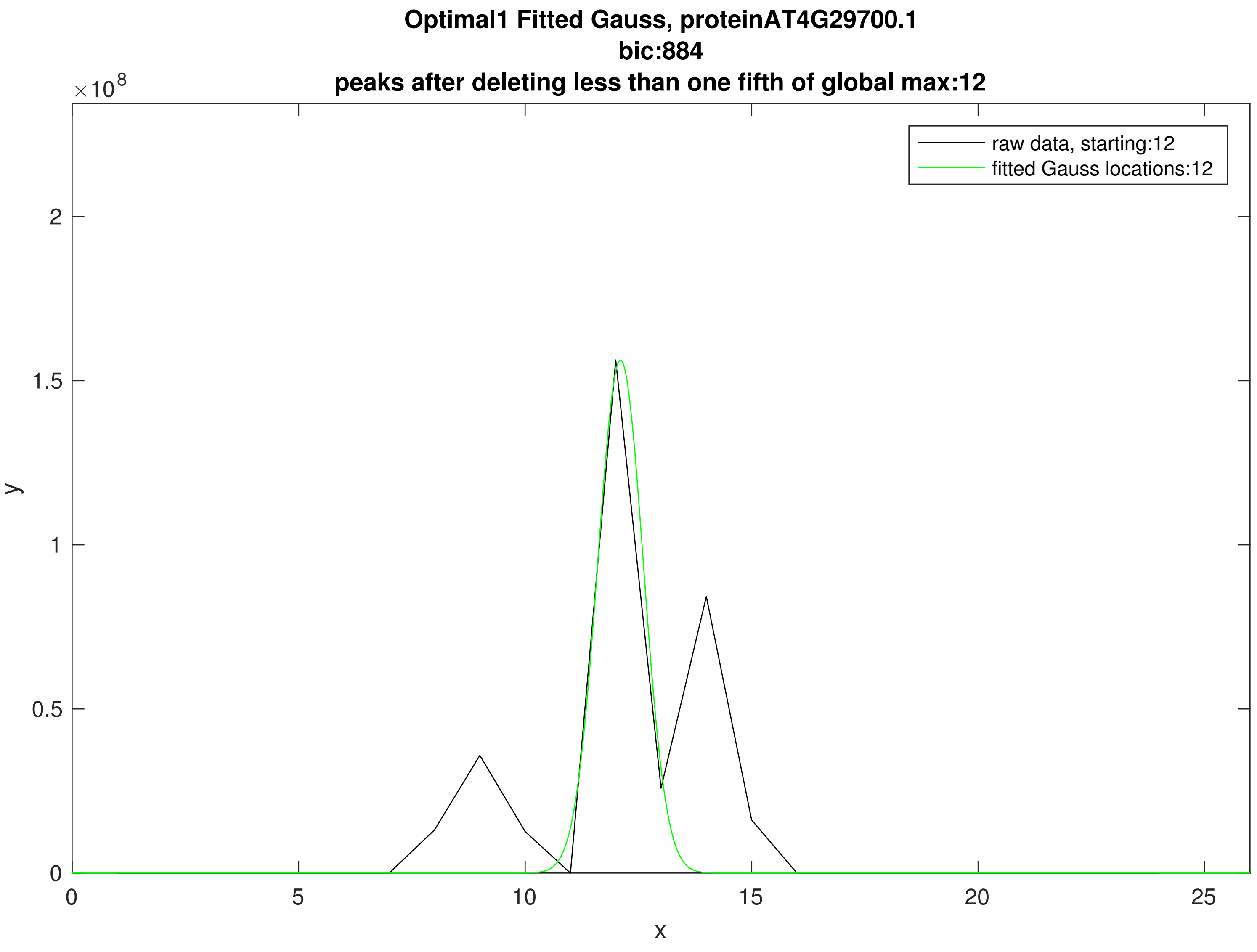

Optimal1 Fitted Gauss, proteinAT4G29700.1
bic:884
peaks after deleting less than one fifth of global max:12
×10^8
raw data, starting:12
fitted Gauss locations:12
2
1.5
1
0.5
0
y
0
5
10
15
20
25
x

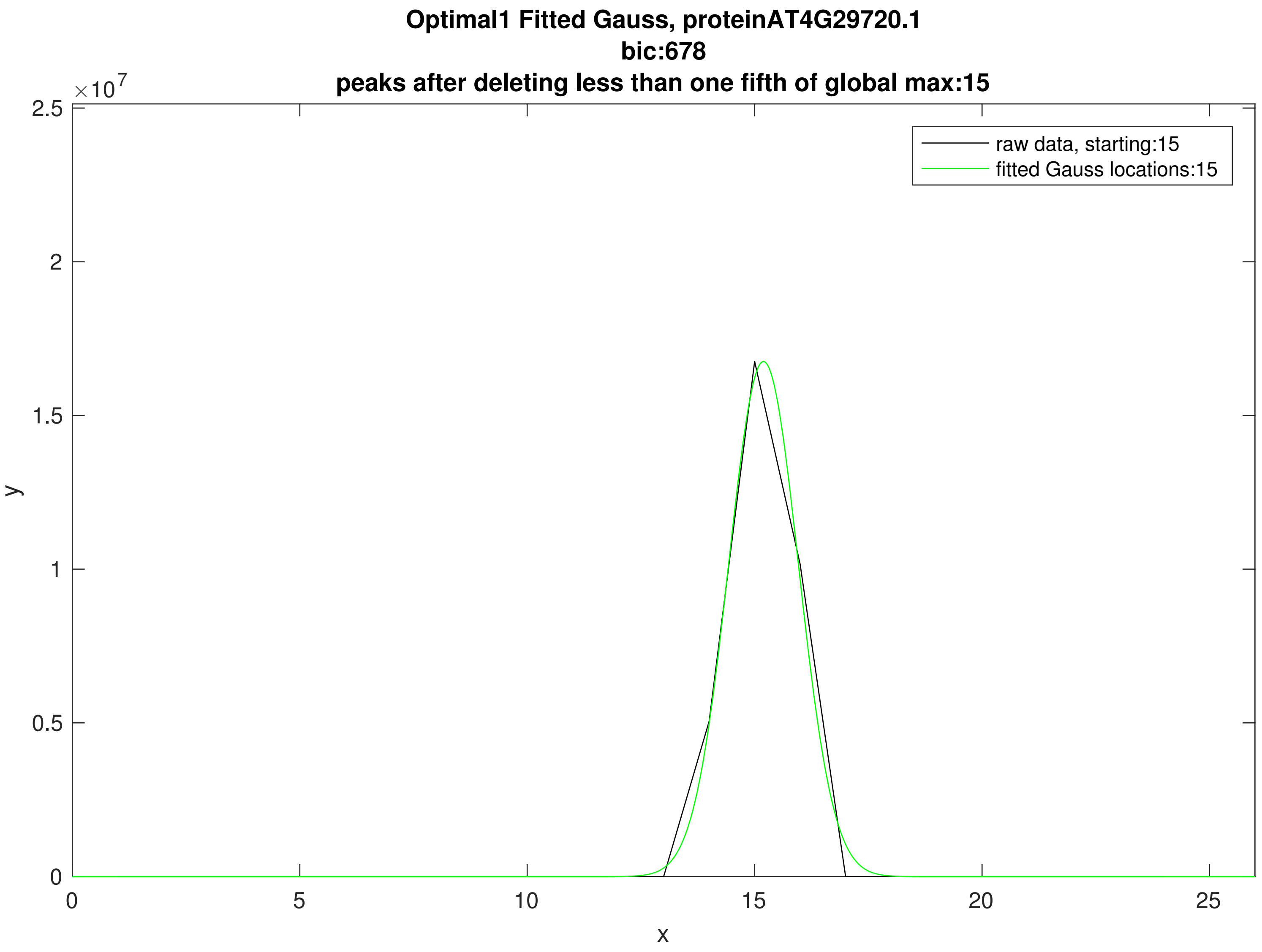

Optimal1 Fitted Gauss, proteinAT4G29720.1
bic:678
peaks after deleting less than one fifth of global max:15
×10^7
2.5
2
1.5
1
0.5
0
y
0
5
10
15
20
25
x
raw data, starting:15
fitted Gauss locations:15

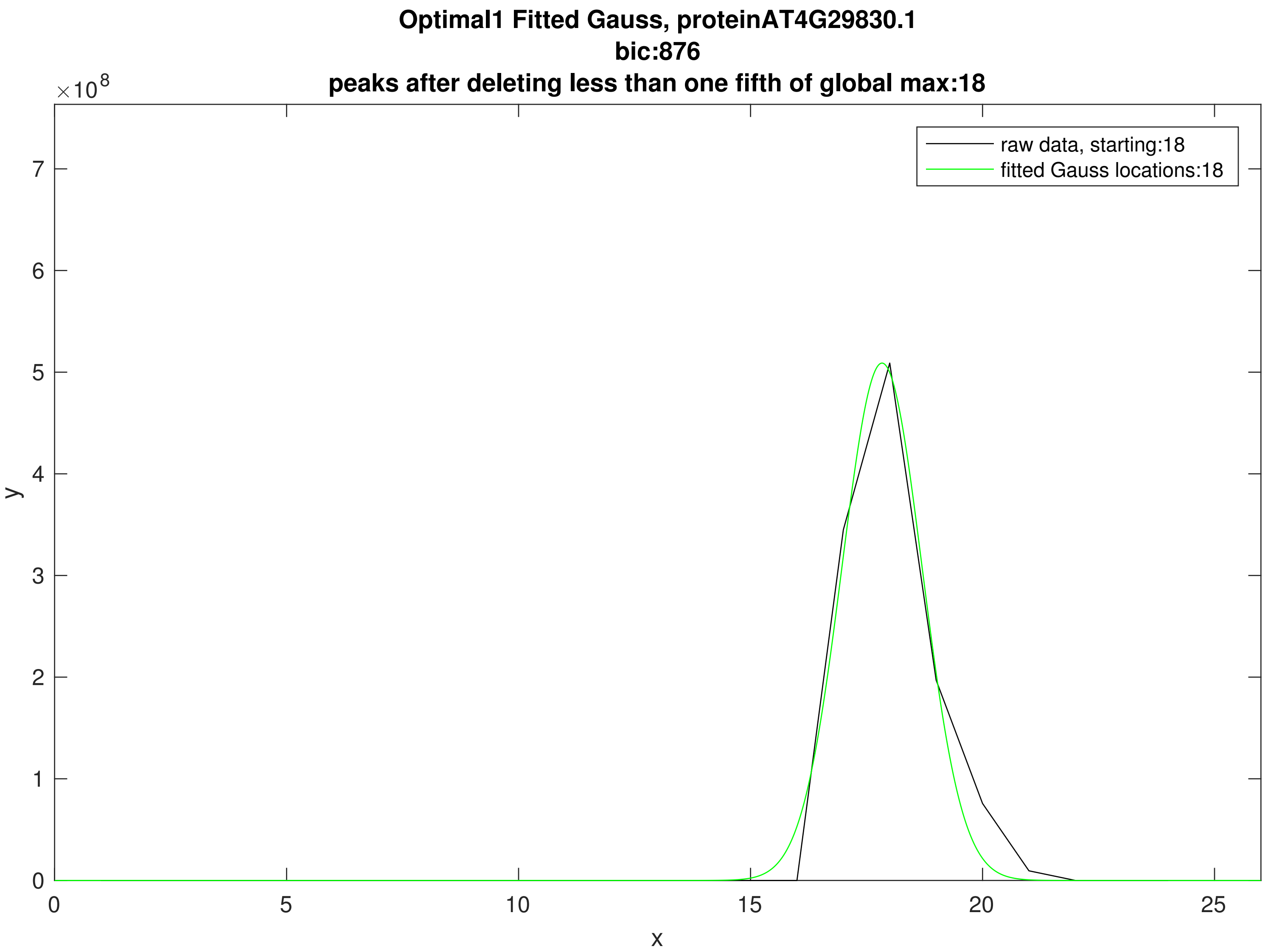
Optimal1 Fitted Gauss, proteinAT4G29830.1
bic:876
peaks after deleting less than one fifth of global max:18
×10^8
raw data, starting:18
fitted Gauss locations:18
y
x
0
1
2
3
4
5
6
7
0
5
10
15
20
25

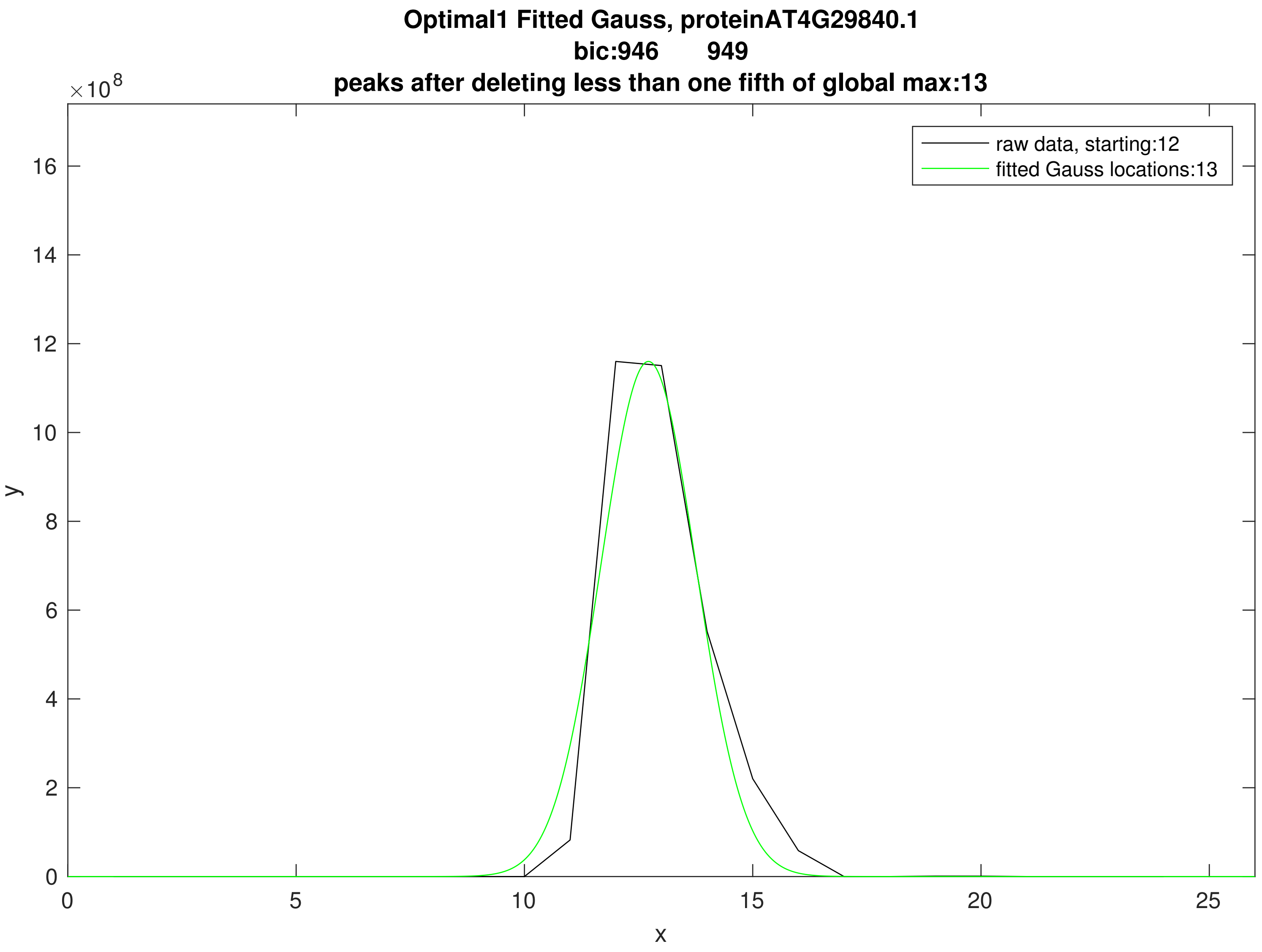

Optimal1 Fitted Gauss, proteinAT4G29840.1
bic:946 949
peaks after deleting less than one fifth of global max:13
×10^8
16
14
12
10
8
6
4
2
0
y
0
5
10
15
20
25
x
raw data, starting:12
fitted Gauss locations:13

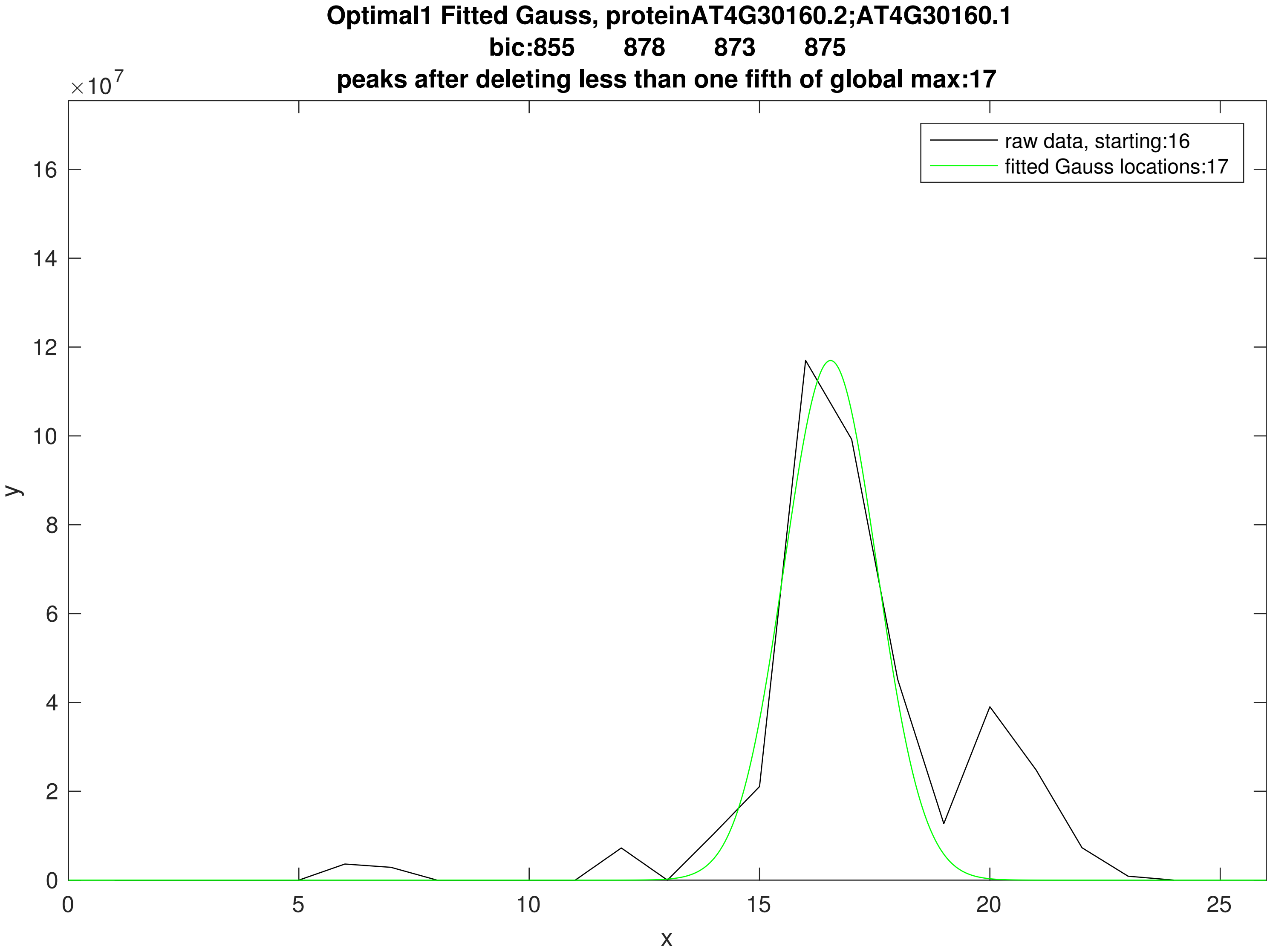
Optimal1 Fitted Gauss, proteinAT4G30160.2;AT4G30160.1
bic:855 878 873 875
peaks after deleting less than one fifth of global max:17
×10^7
raw data, starting:16
fitted Gauss locations:17
y
x
0
2
4
6
8
10
12
14
16
0
5
10
15
20
25

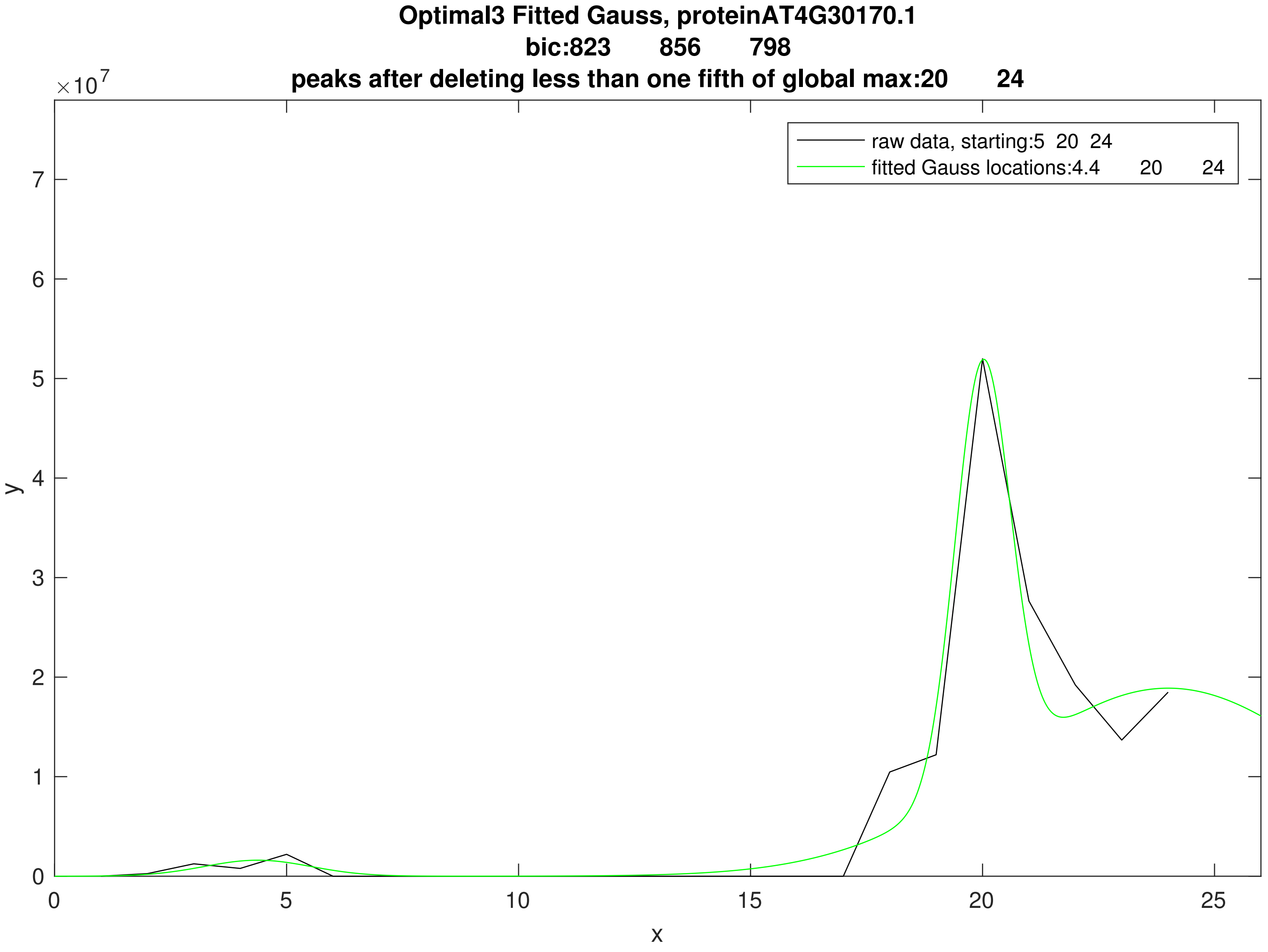

Optimal3 Fitted Gauss, proteinAT4G30170.1
bic:823 856 798
peaks after deleting less than one fifth of global max:20 24
×10^7
raw data, starting:5 20 24
fitted Gauss locations:4.4 20 24
y
x
0
1
2
3
4
5
6
7
0
5
10
15
20
25

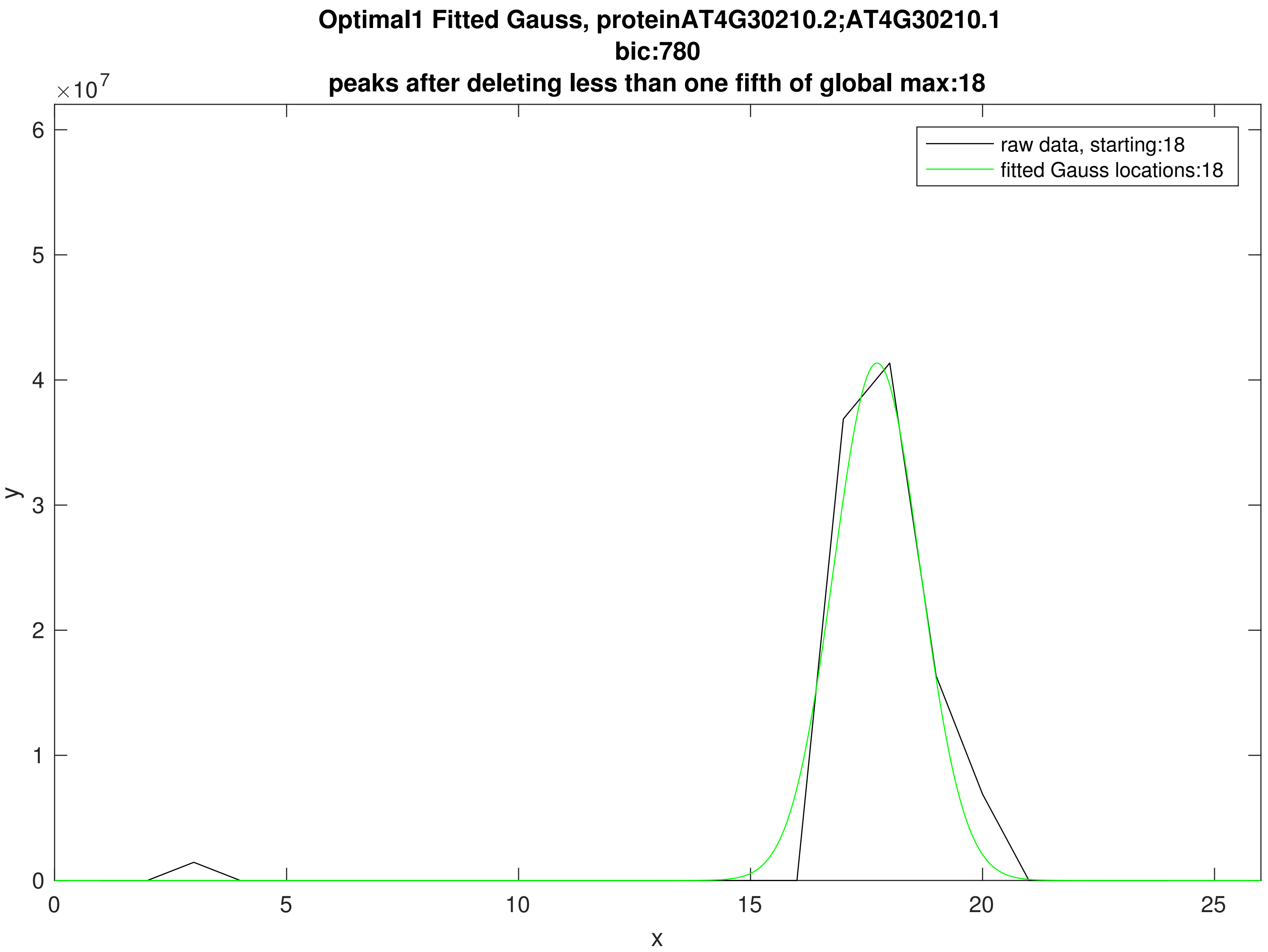
Optimal1 Fitted Gauss, proteinAT4G30210.2;AT4G30210.1
bic:780
peaks after deleting less than one fifth of global max:18
×10^7
6
5
4
3
2
1
0
y
0
5
10
15
20
25
x
raw data, starting:18
fitted Gauss locations:18

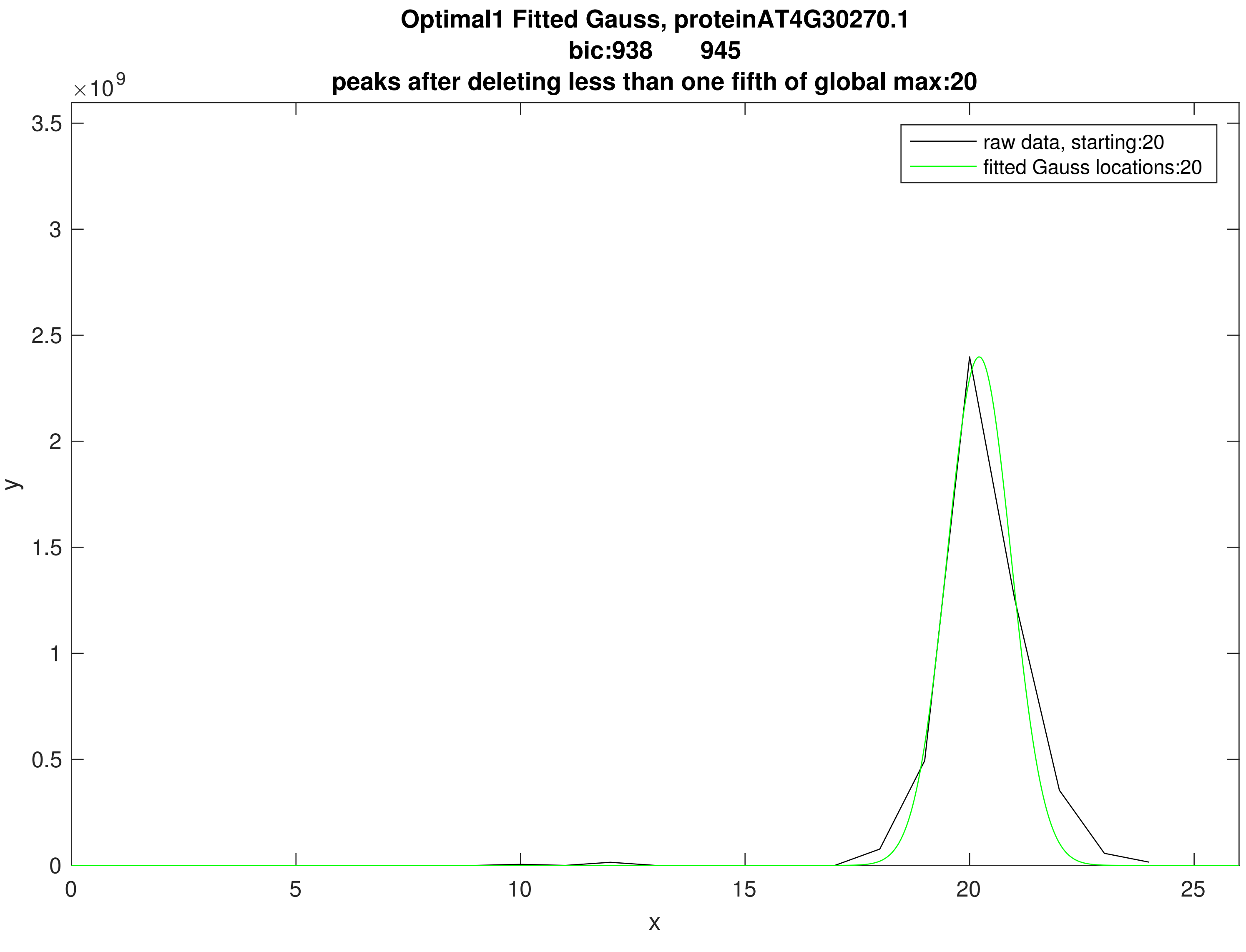

Optimal1 Fitted Gauss, proteinAT4G30270.1
bic:938 945
peaks after deleting less than one fifth of global max:20
×10^9
3.5
3
2.5
2
1.5
1
0.5
0
y
0
5
10
15
20
25
x
raw data, starting:20
fitted Gauss locations:20

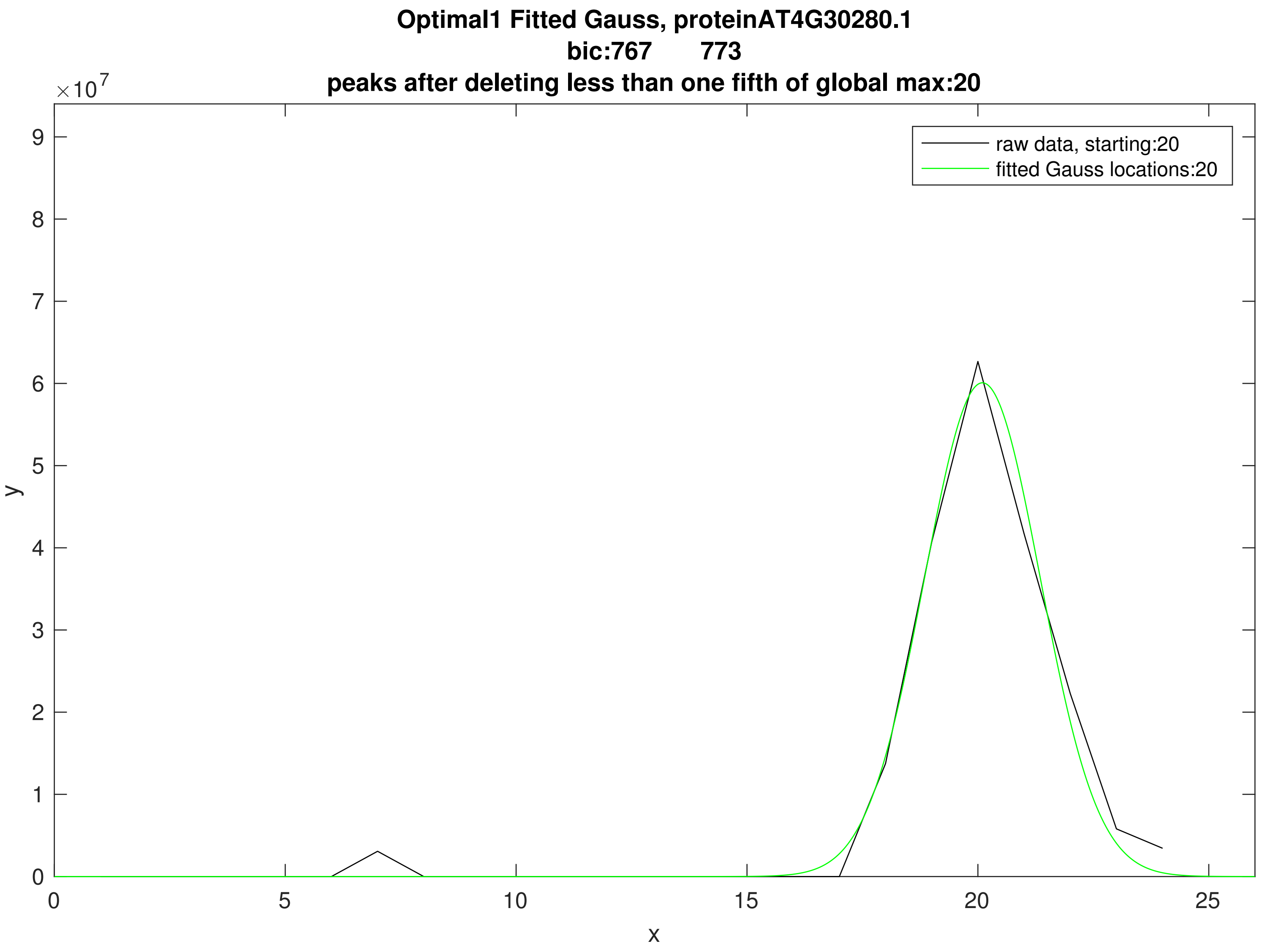
Optimal1 Fitted Gauss, proteinAT4G30280.1
bic:767 773
peaks after deleting less than one fifth of global max:20
×10^7
raw data, starting:20
fitted Gauss locations:20
y
x
0
1
2
3
4
5
6
7
8
9
0
5
10
15
20
25

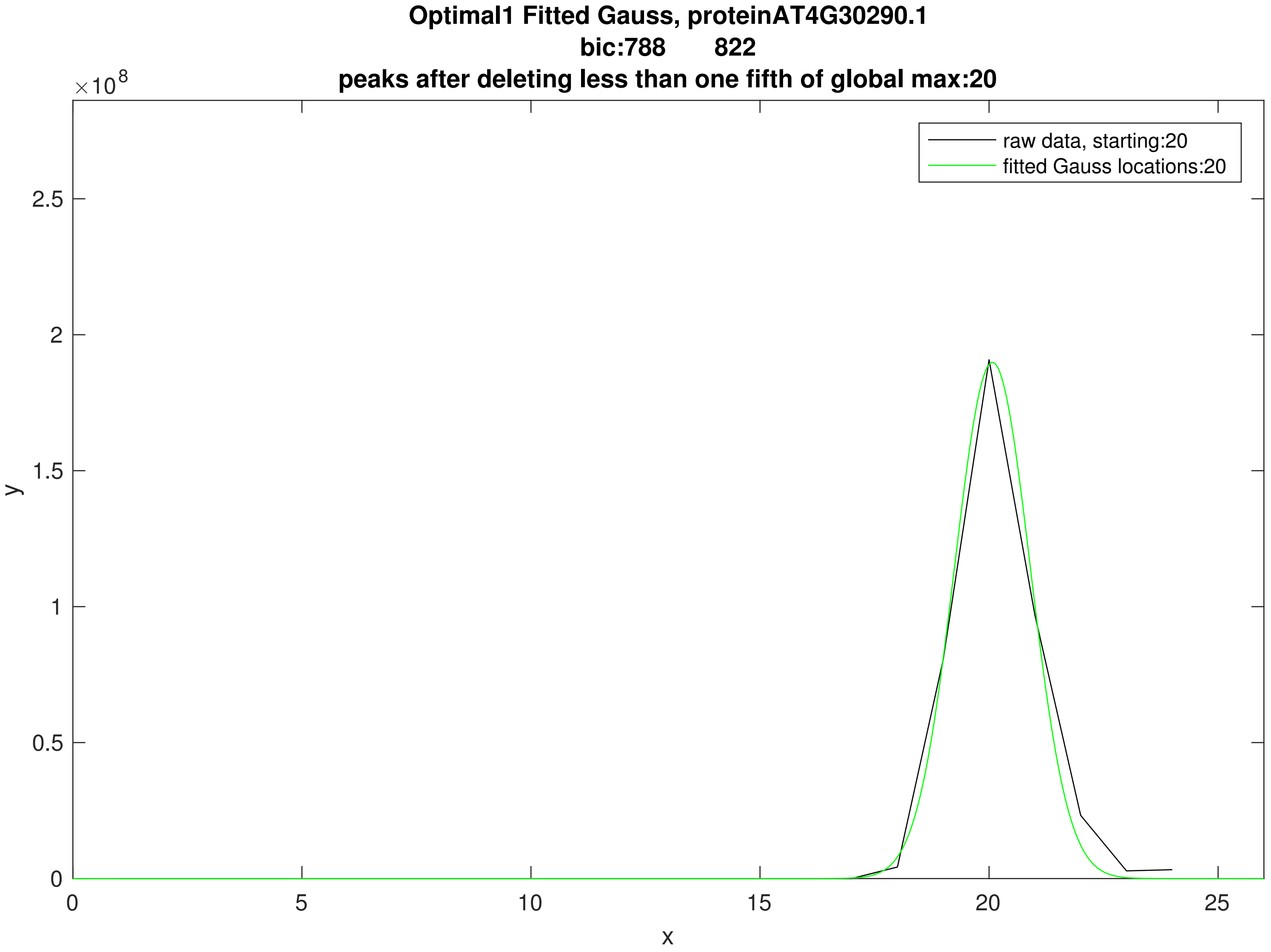
Optimal1 Fitted Gauss, proteinAT4G30290.1
bic:788 822
peaks after deleting less than one fifth of global max:20
×10^8
raw data, starting:20
fitted Gauss locations:20
2.5
2
1.5
1
0.5
0
y
0
5
10
15
20
25
x

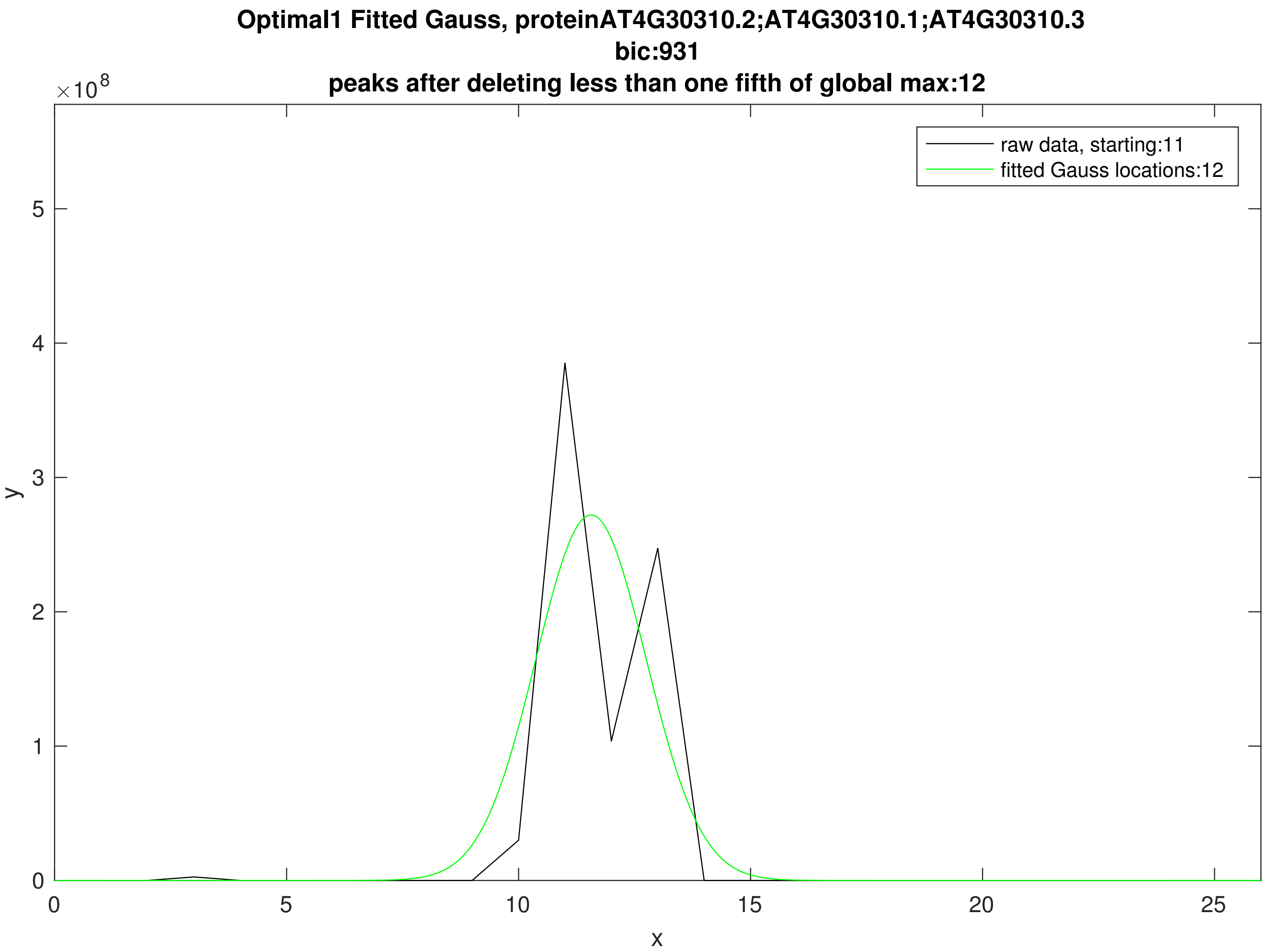

Optimal1 Fitted Gauss, proteinAT4G30310.2;AT4G30310.1;AT4G30310.3
bic:931
peaks after deleting less than one fifth of global max:12
×10^8
raw data, starting:11
fitted Gauss locations:12
0
1
2
3
4
5
y
0
5
10
15
20
25
x

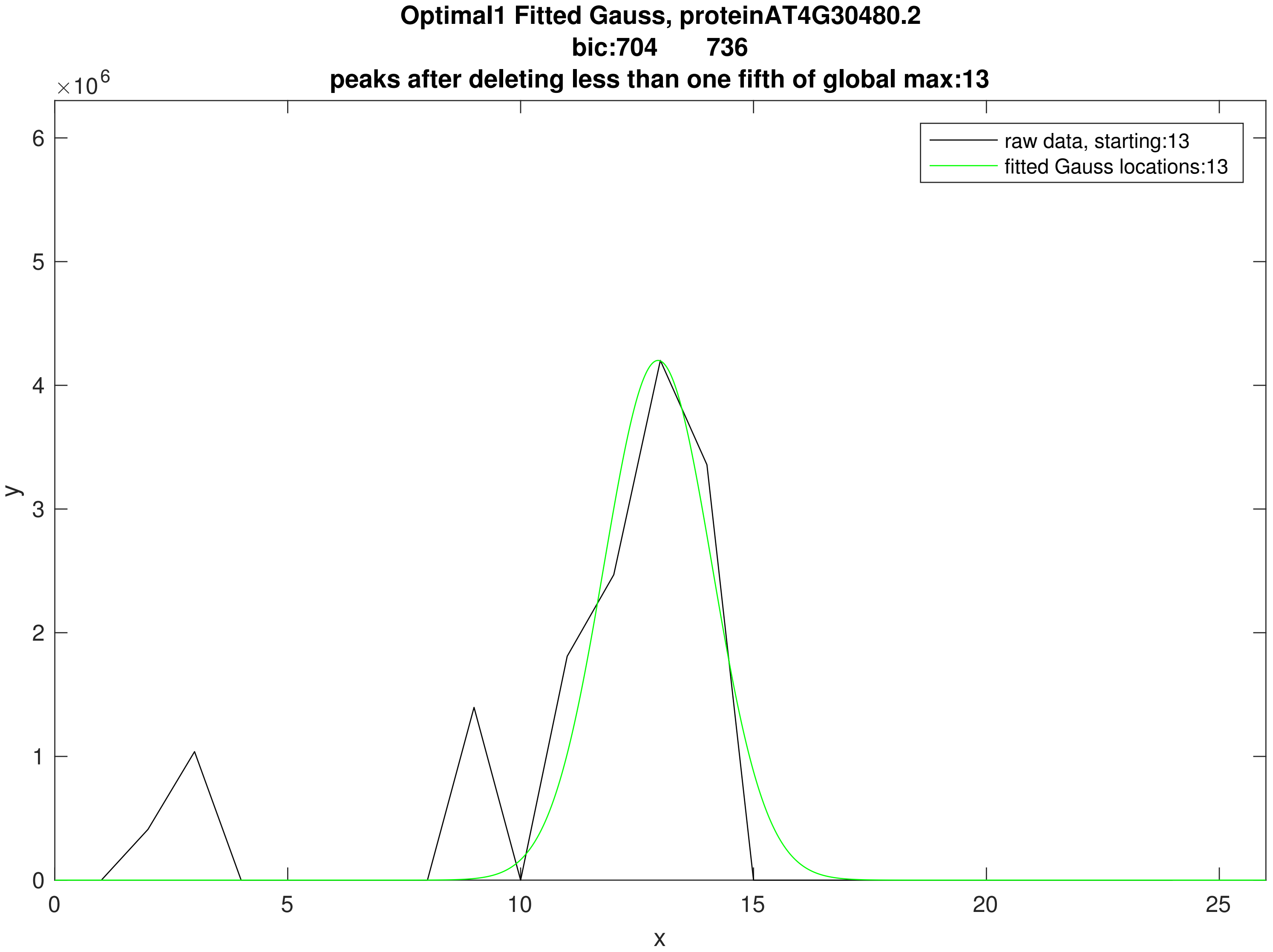
Optimal1 Fitted Gauss, proteinAT4G30480.2
bic:704 736
peaks after deleting less than one fifth of global max:13
×10^6
raw data, starting:13
fitted Gauss locations:13
y
x
0
1
2
3
4
5
6
0
5
10
15
20
25

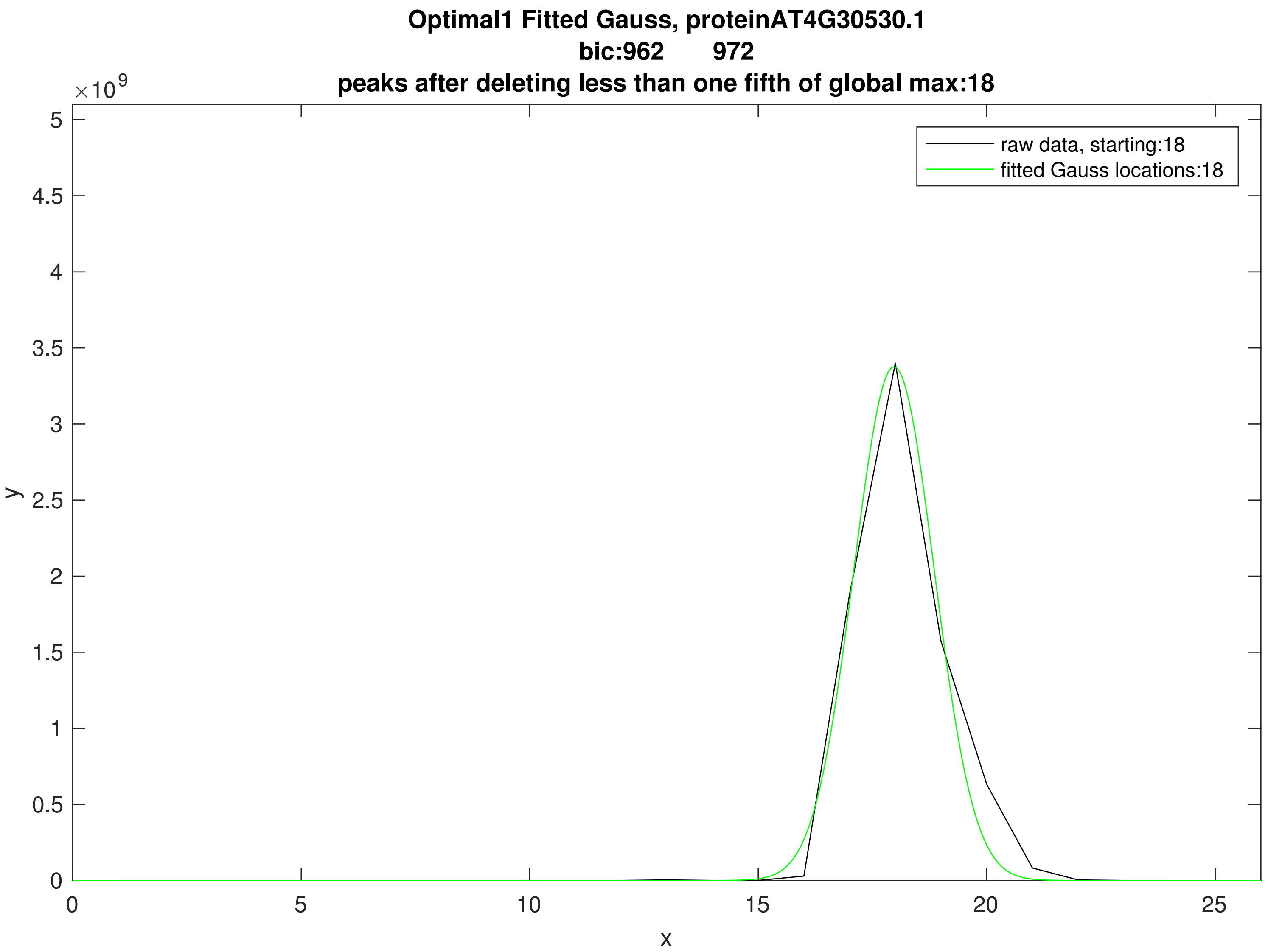

Optimal1 Fitted Gauss, proteinAT4G30530.1
bic:962 972
peaks after deleting less than one fifth of global max:18
×10^9
raw data, starting:18
fitted Gauss locations:18
y
x
0
0.5
1
1.5
2
2.5
3
3.5
4
4.5
5
0
5
10
15
20
25

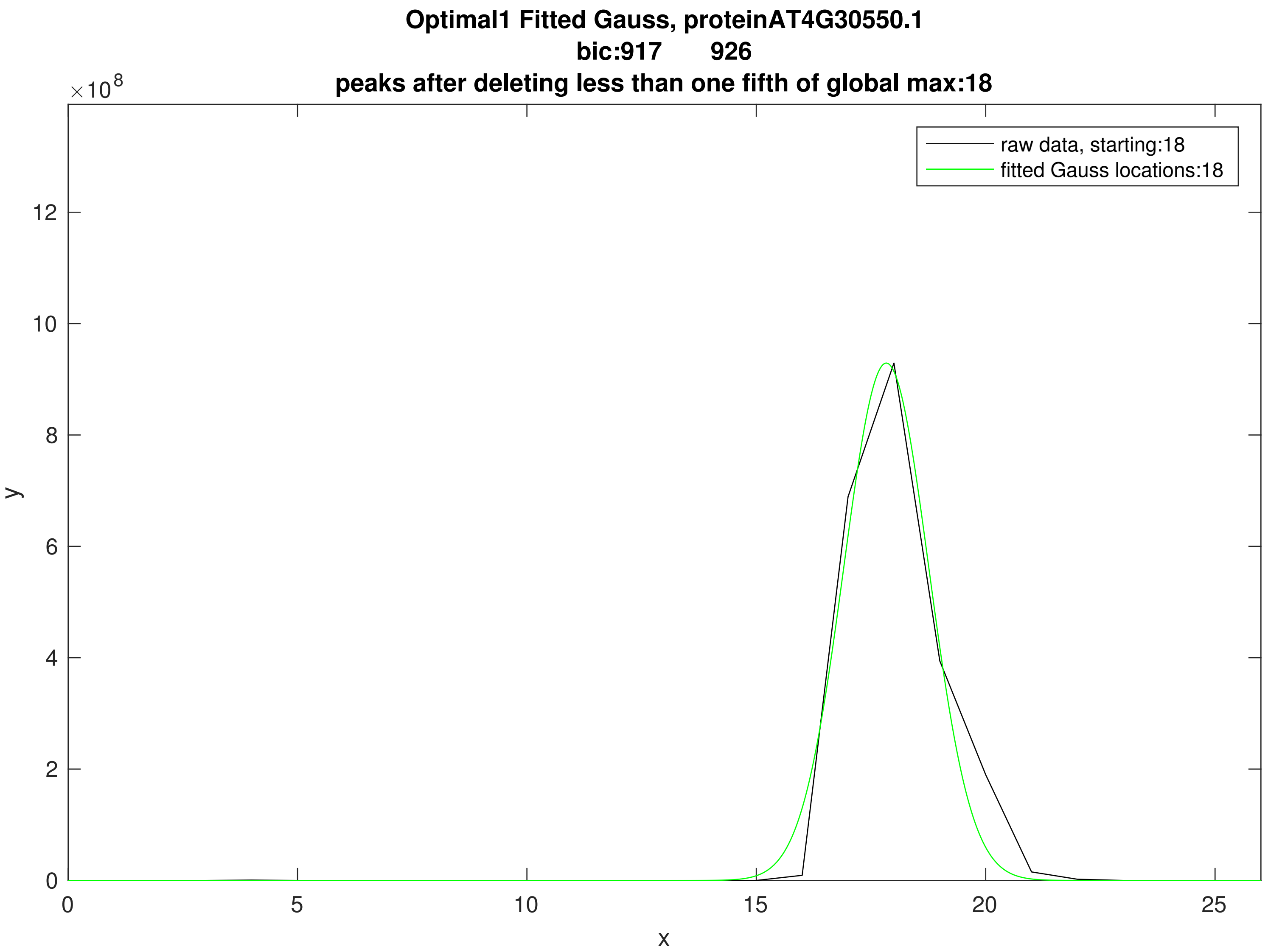
Optimal1 Fitted Gauss, proteinAT4G30550.1
bic:917 926
peaks after deleting less than one fifth of global max:18
×10^8
raw data, starting:18
fitted Gauss locations:18
y
x
0
2
4
6
8
10
12
0
5
10
15
20
25

Optimal1 Fitted Gauss, proteinAT4G30810.1
bic:884 897
peaks after deleting less than one fifth of global max:13

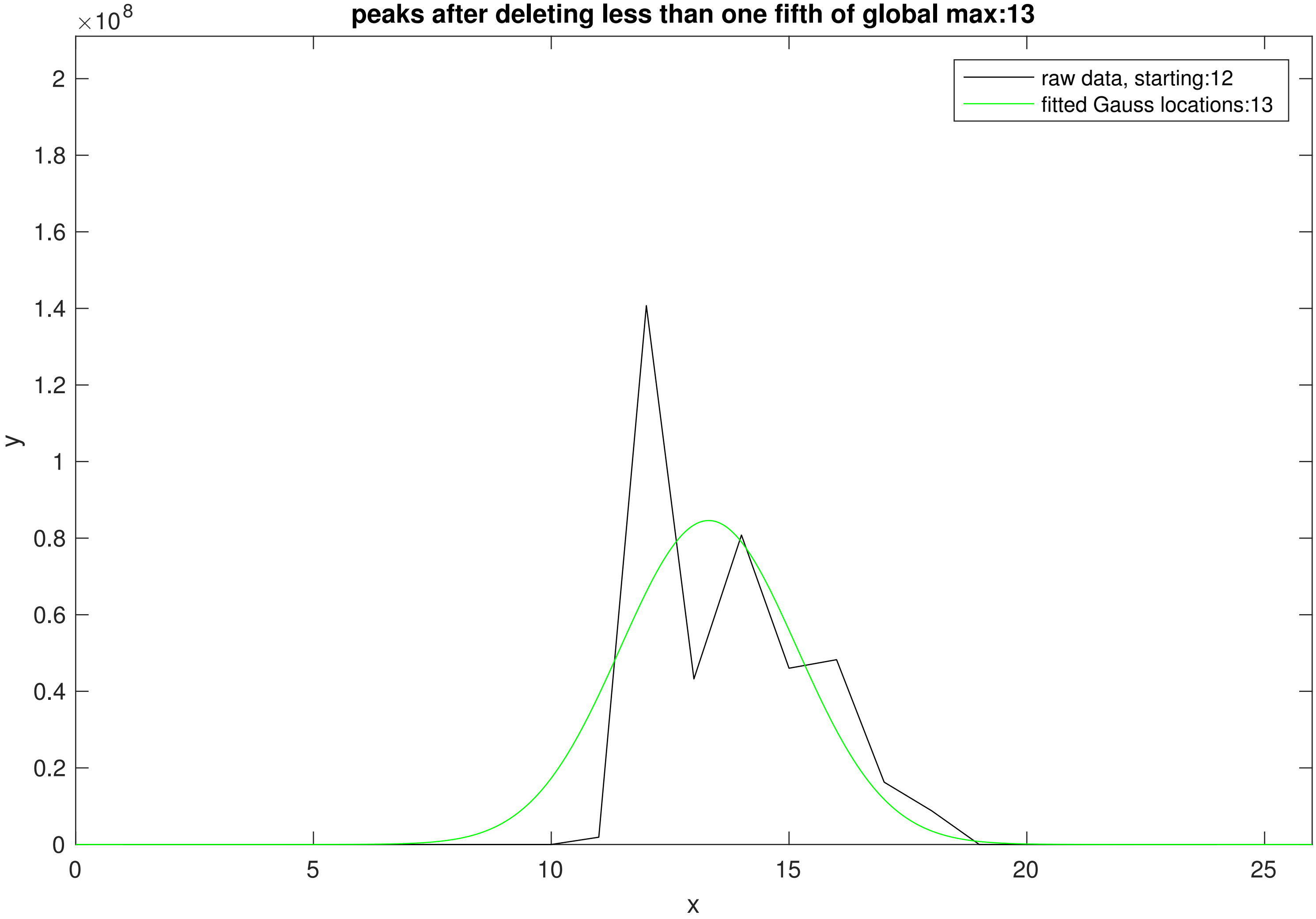

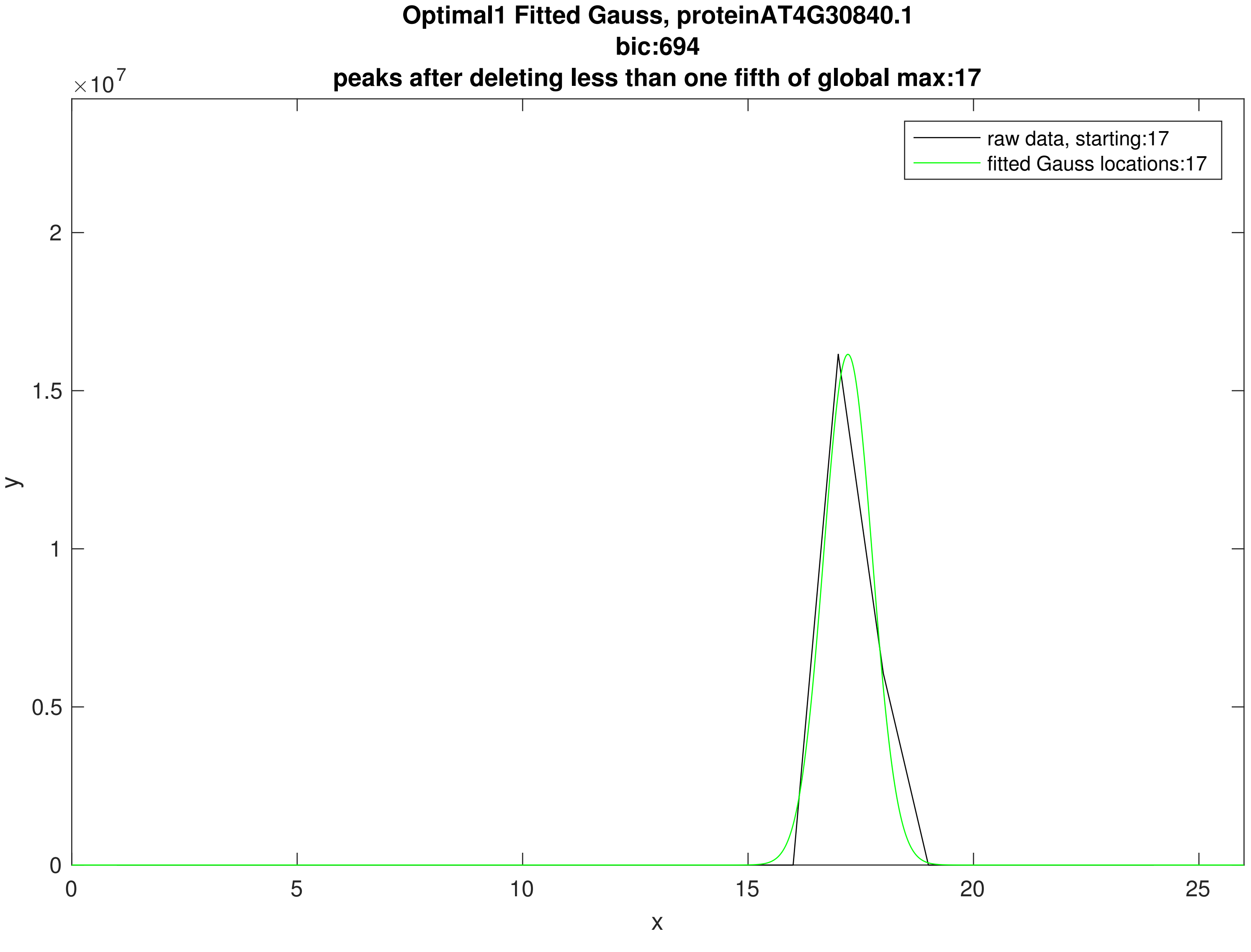
Optimal1 Fitted Gauss, proteinAT4G30840.1
bic:694
peaks after deleting less than one fifth of global max:17
×10^7
raw data, starting:17
fitted Gauss locations:17
2
1.5
1
0.5
0
y
0
5
10
15
20
25
x

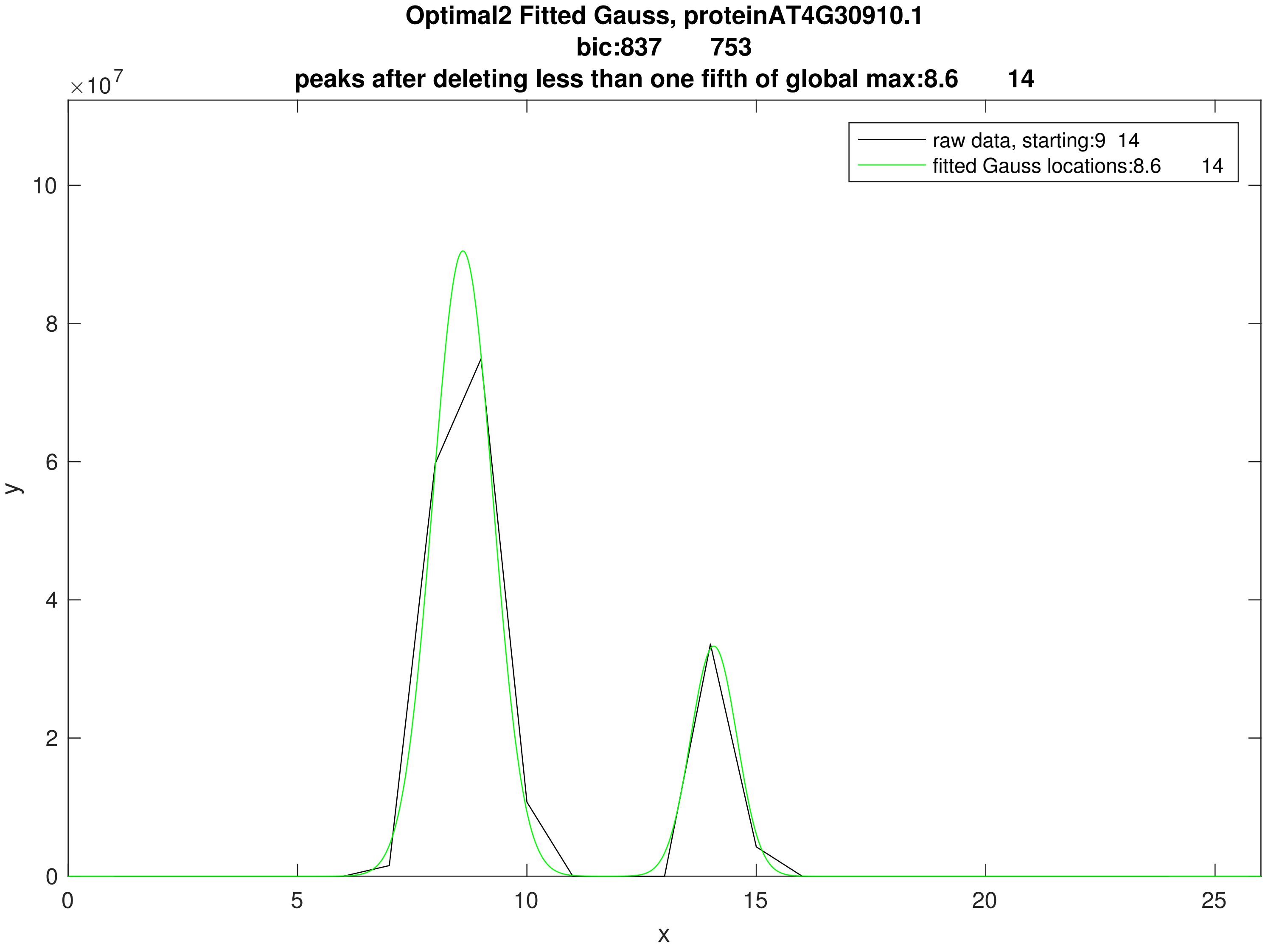
Optimal2 Fitted Gauss, proteinAT4G30910.1
bic:837 753
peaks after deleting less than one fifth of global max:8.6 14
×10^7
raw data, starting:9 14
fitted Gauss locations:8.6 14
10
8
6
4
2
0
y
0
5
10
15
20
25
x

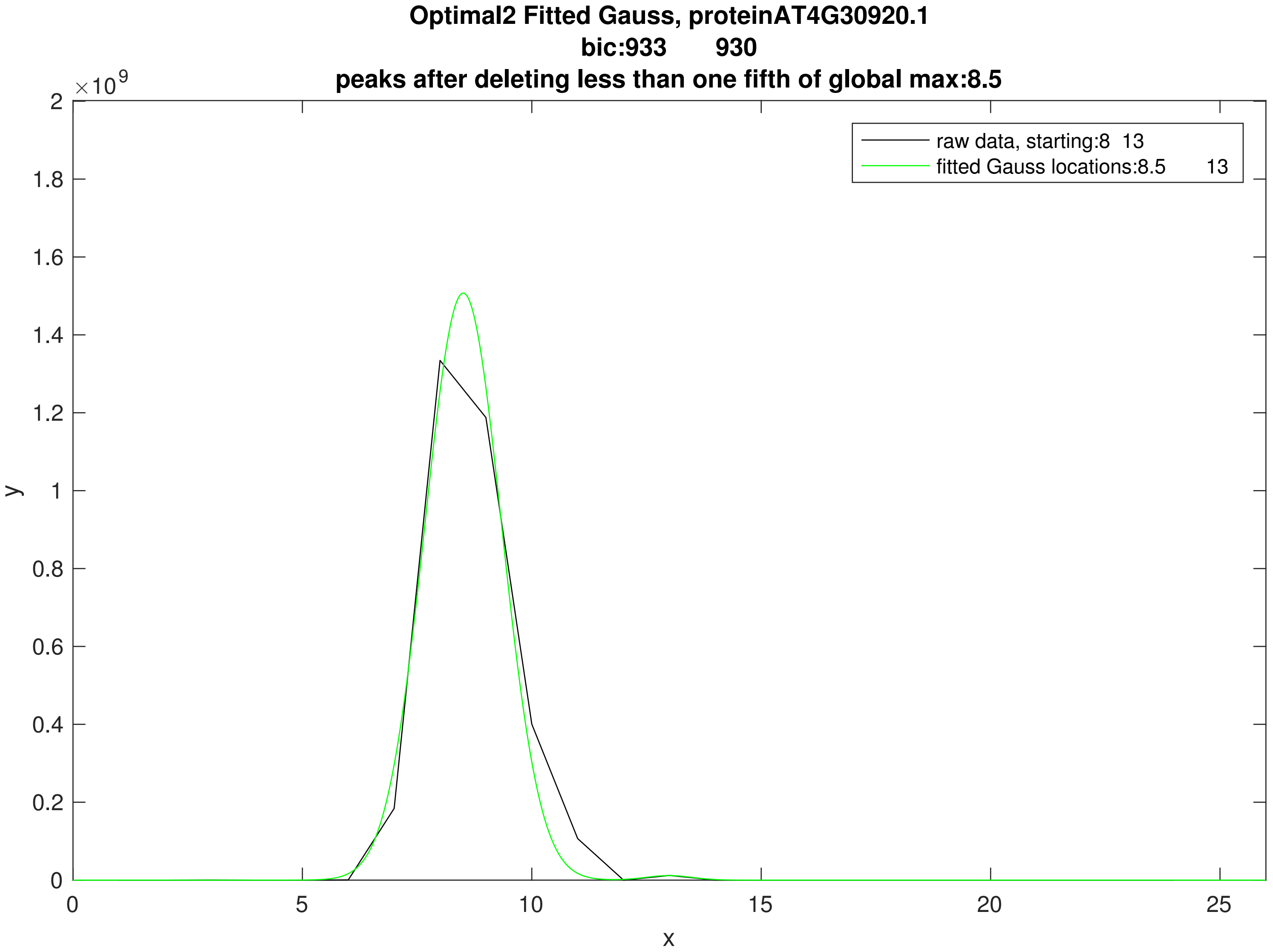
Optimal2 Fitted Gauss, proteinAT4G30920.1
bic:933 930
peaks after deleting less than one fifth of global max:8.5
$\times 10^9$
raw data, starting:8 13
fitted Gauss locations:8.5 13
y
x
0
0.2
0.4
0.6
0.8
1
1.2
1.4
1.6
1.8
2
0
5
10
15
20
25

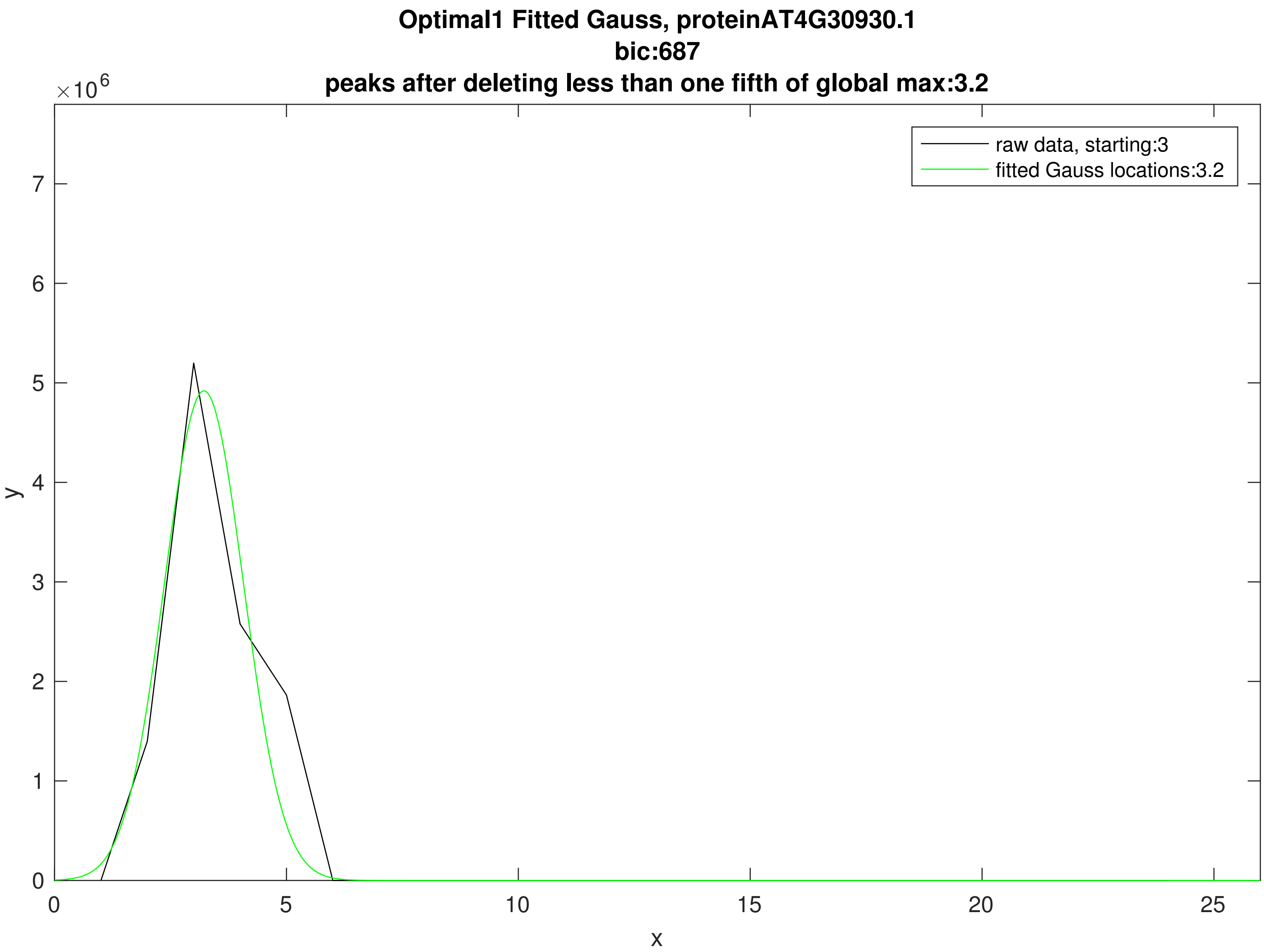

Optimal1 Fitted Gauss, proteinAT4G30930.1
bic:687
peaks after deleting less than one fifth of global max:3.2
×10^6
raw data, starting:3
fitted Gauss locations:3.2
y
x
0
1
2
3
4
5
6
7
0
5
10
15
20
25

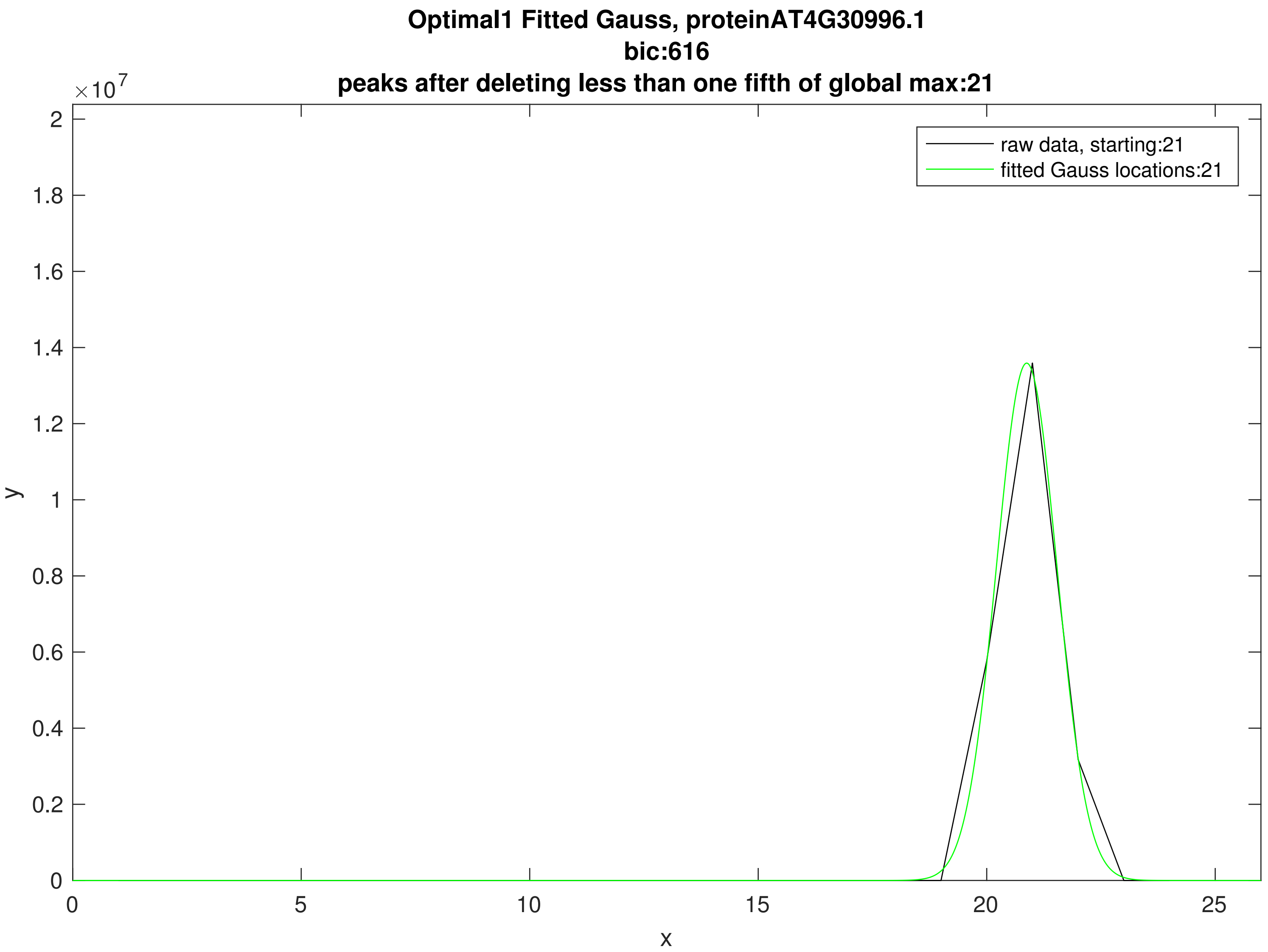
Optimal1 Fitted Gauss, proteinAT4G30996.1
bic:616
peaks after deleting less than one fifth of global max:21
×10^7
raw data, starting:21
fitted Gauss locations:21
y
x
0
0.2
0.4
0.6
0.8
1
1.2
1.4
1.6
1.8
2
0
5
10
15
20
25

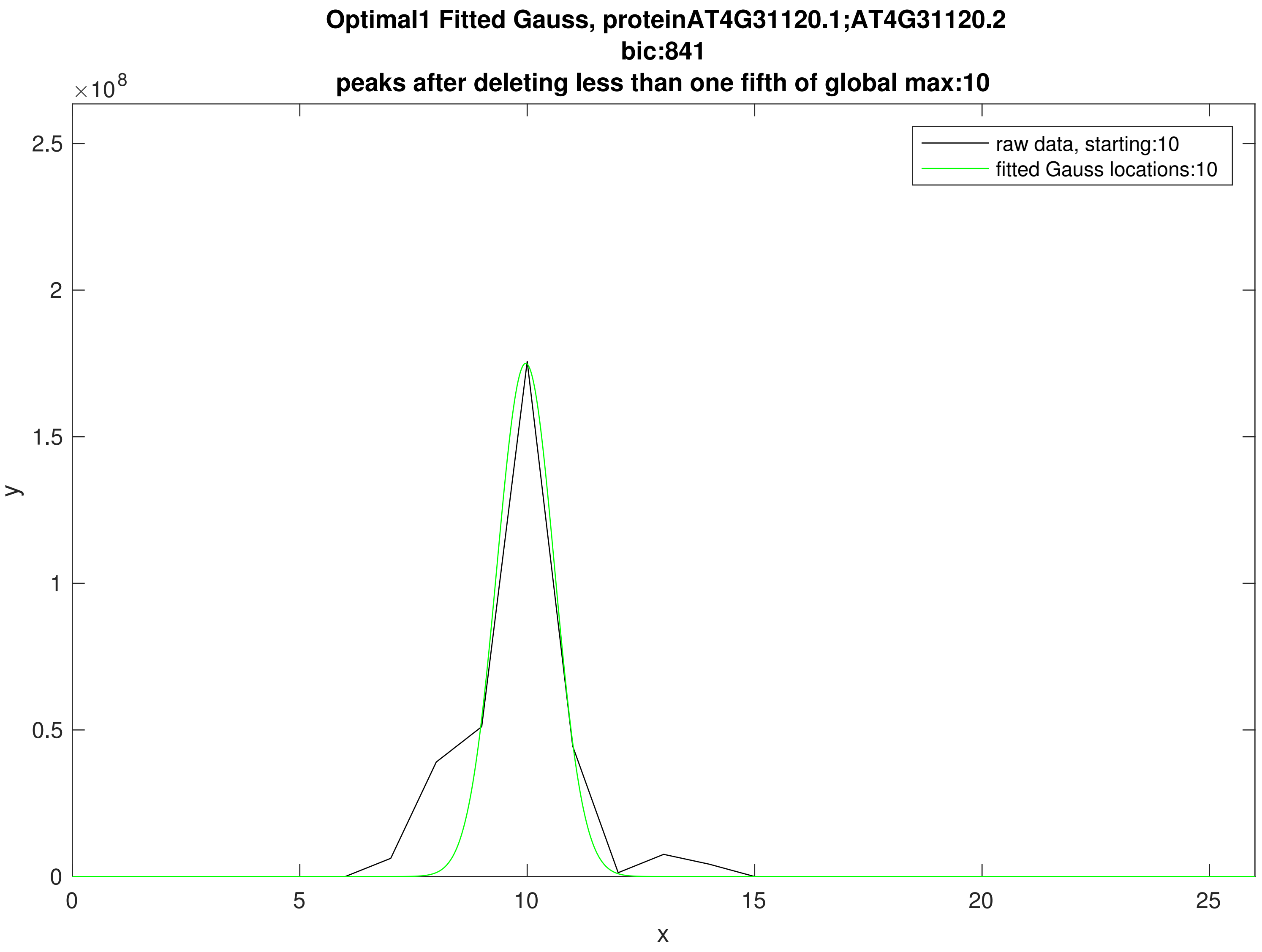

Optimal1 Fitted Gauss, proteinAT4G31120.1;AT4G31120.2
bic:841
peaks after deleting less than one fifth of global max:10
×10^8
2.5
2
1.5
1
0.5
0
y
0
5
10
15
20
25
x
raw data, starting:10
fitted Gauss locations:10

#### Optimal1 Fitted Gauss, proteinAT4G31140.1
#### bic:837 842
#### peaks after deleting less than one fifth of global max:16

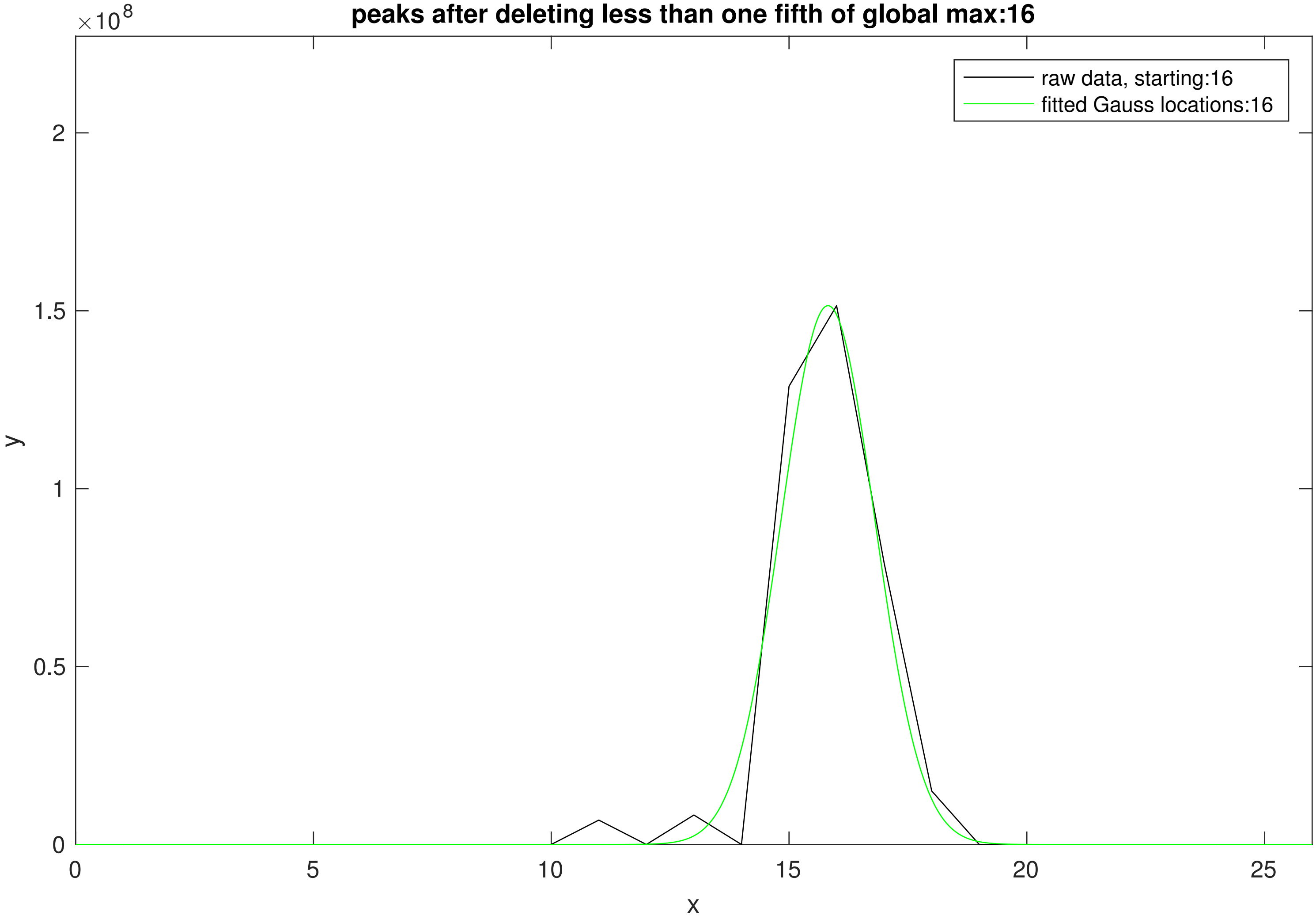

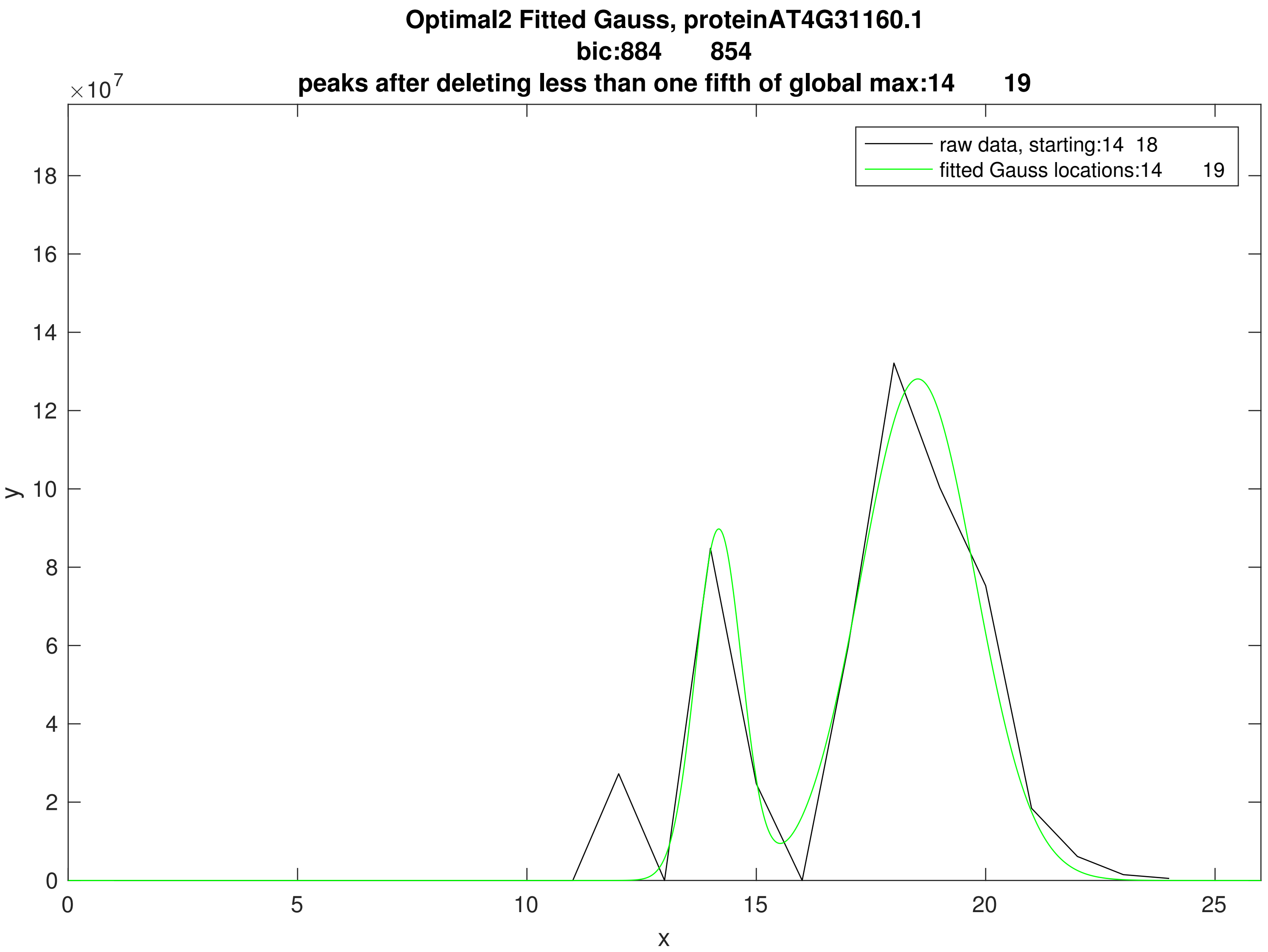
Optimal2 Fitted Gauss, proteinAT4G31160.1
bic:884 854
peaks after deleting less than one fifth of global max:14 19
×10^7
raw data, starting:14 18
fitted Gauss locations:14 19
y
x
0
2
4
6
8
10
12
14
16
18
0
5
10
15
20
25

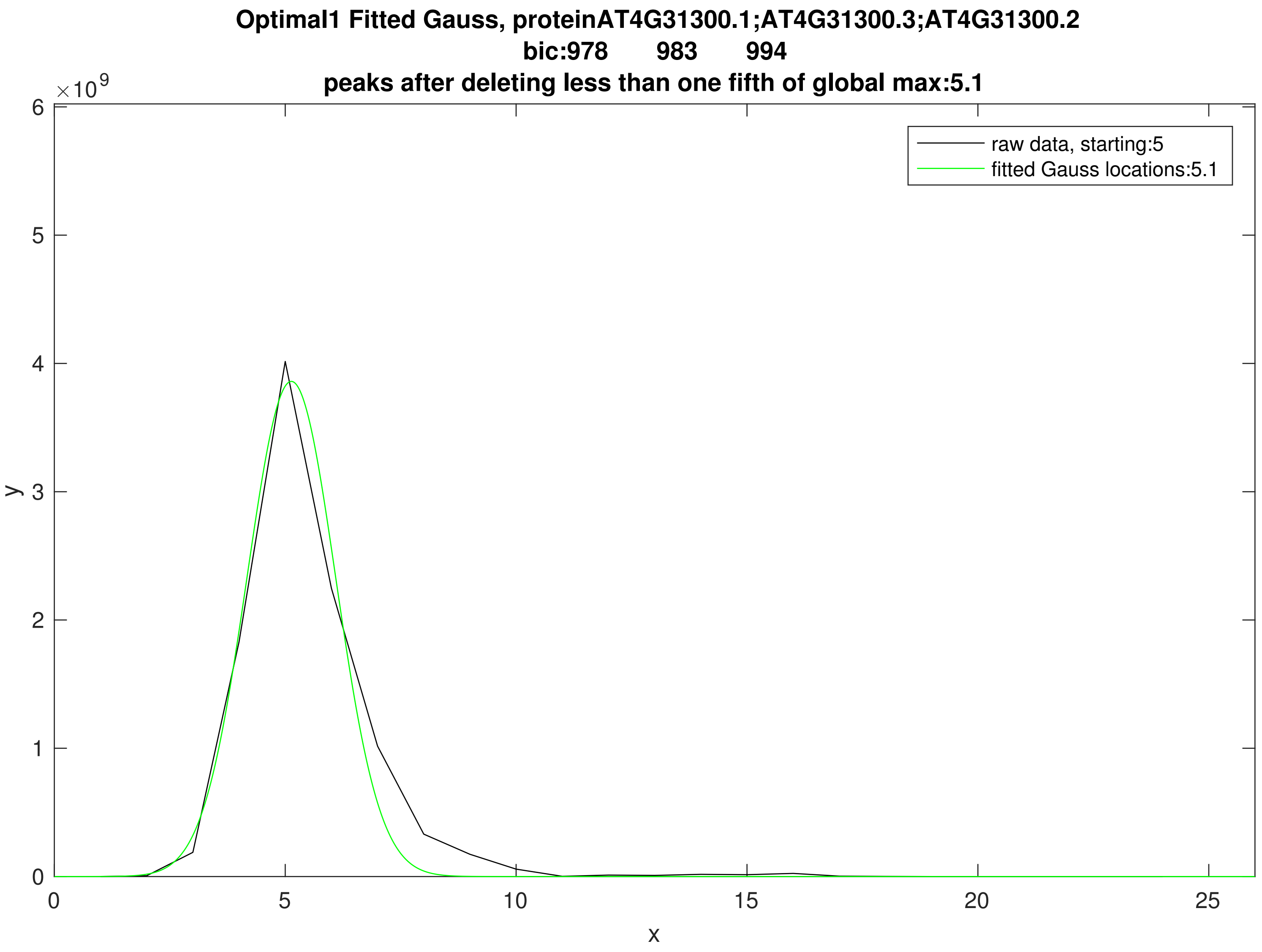
Optimal1 Fitted Gauss, proteinAT4G31300.1;AT4G31300.3;AT4G31300.2
bic:978 983 994
peaks after deleting less than one fifth of global max:5.1
×10^9
raw data, starting:5
fitted Gauss locations:5.1
y
x
0
1
2
3
4
5
6
10
15
20
25

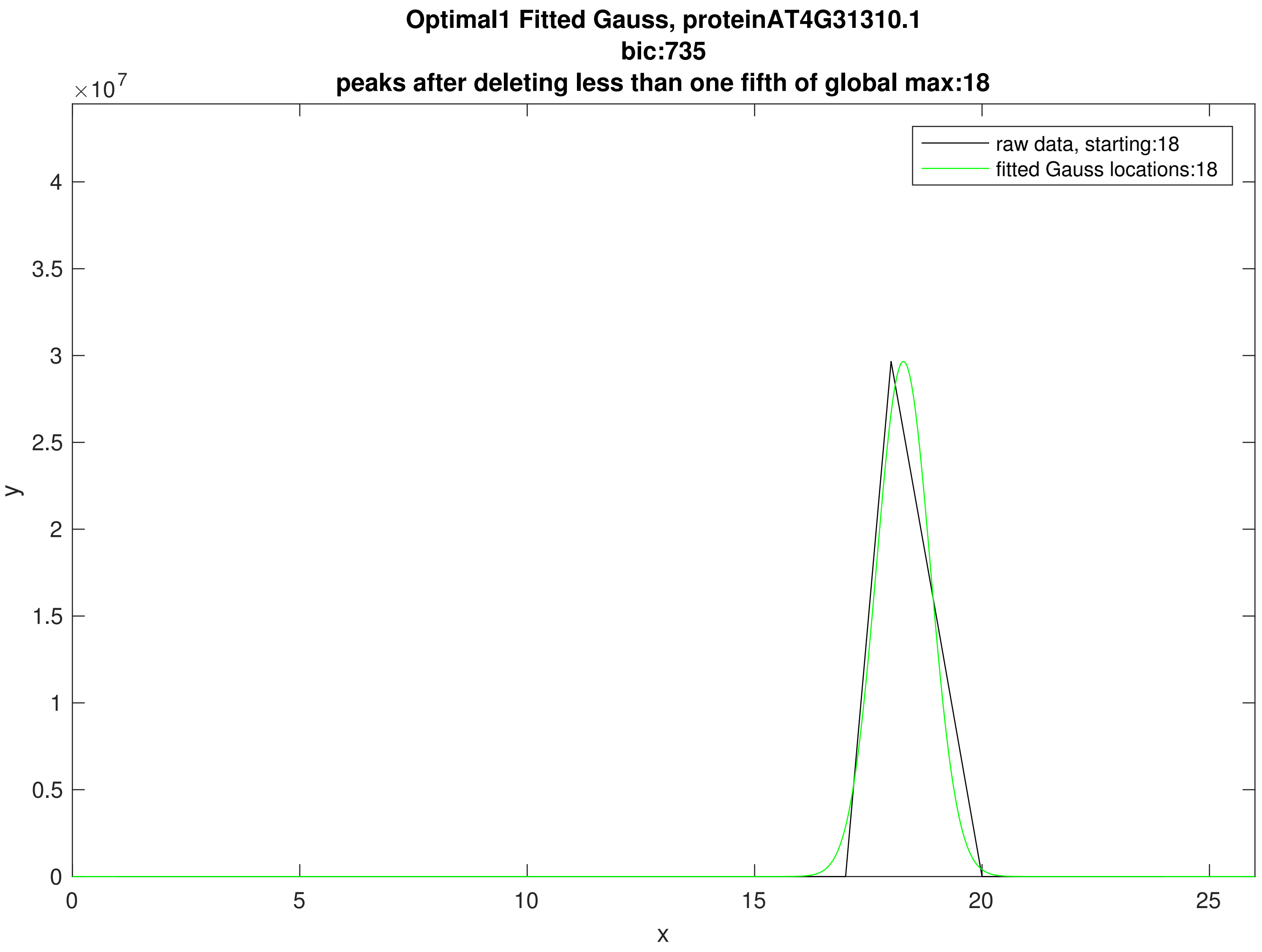

Optimal1 Fitted Gauss, proteinAT4G31310.1
bic:735
peaks after deleting less than one fifth of global max:18
×10^7
4
3.5
3
2.5
2
1.5
1
0.5
0
y
0
5
10
15
20
25
x
raw data, starting:18
fitted Gauss locations:18

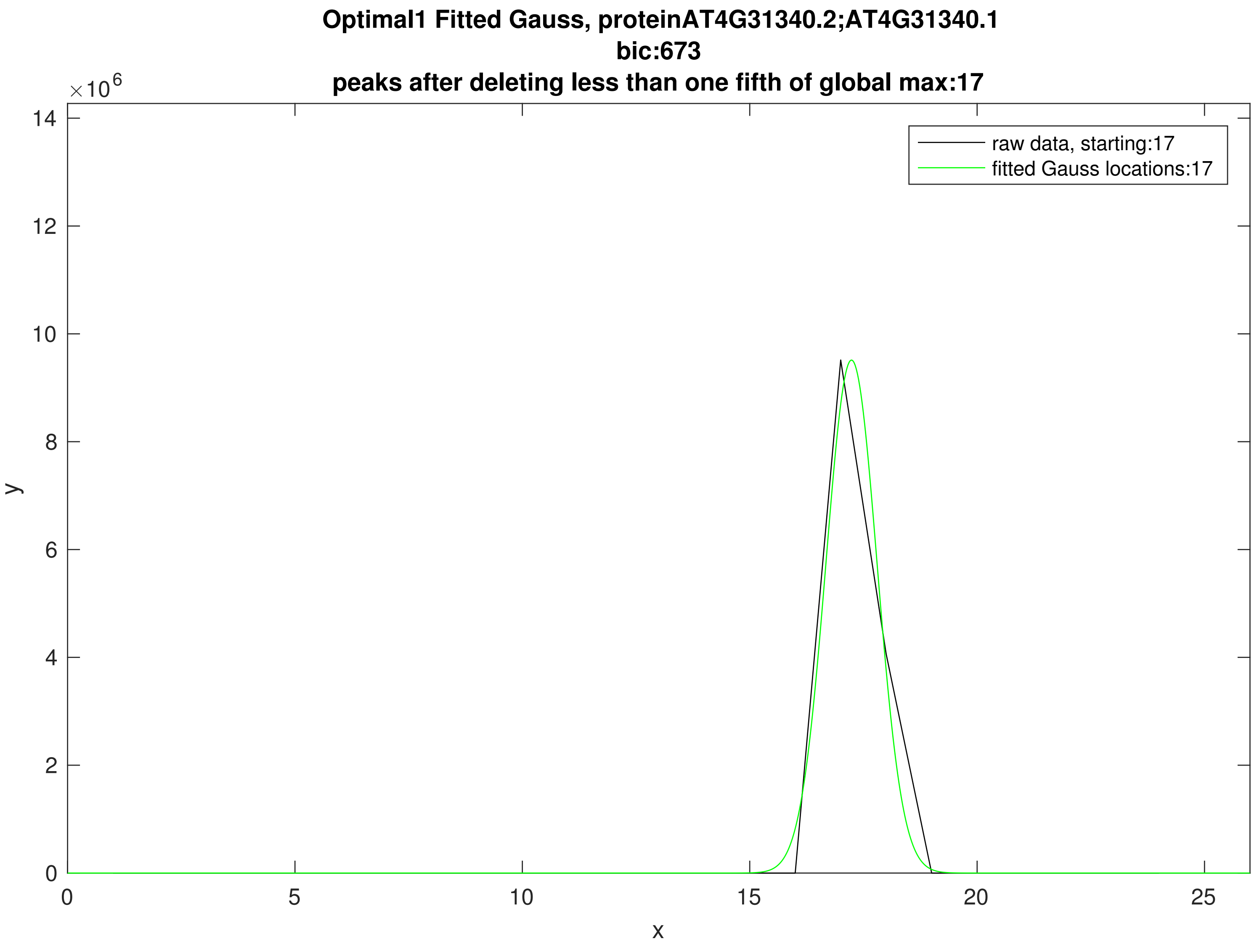

Optimal1 Fitted Gauss, proteinAT4G31340.2;AT4G31340.1
bic:673
peaks after deleting less than one fifth of global max:17
×10^6
14
12
10
8
6
4
2
0
y
0
5
10
15
20
25
x
raw data, starting:17
fitted Gauss locations:17

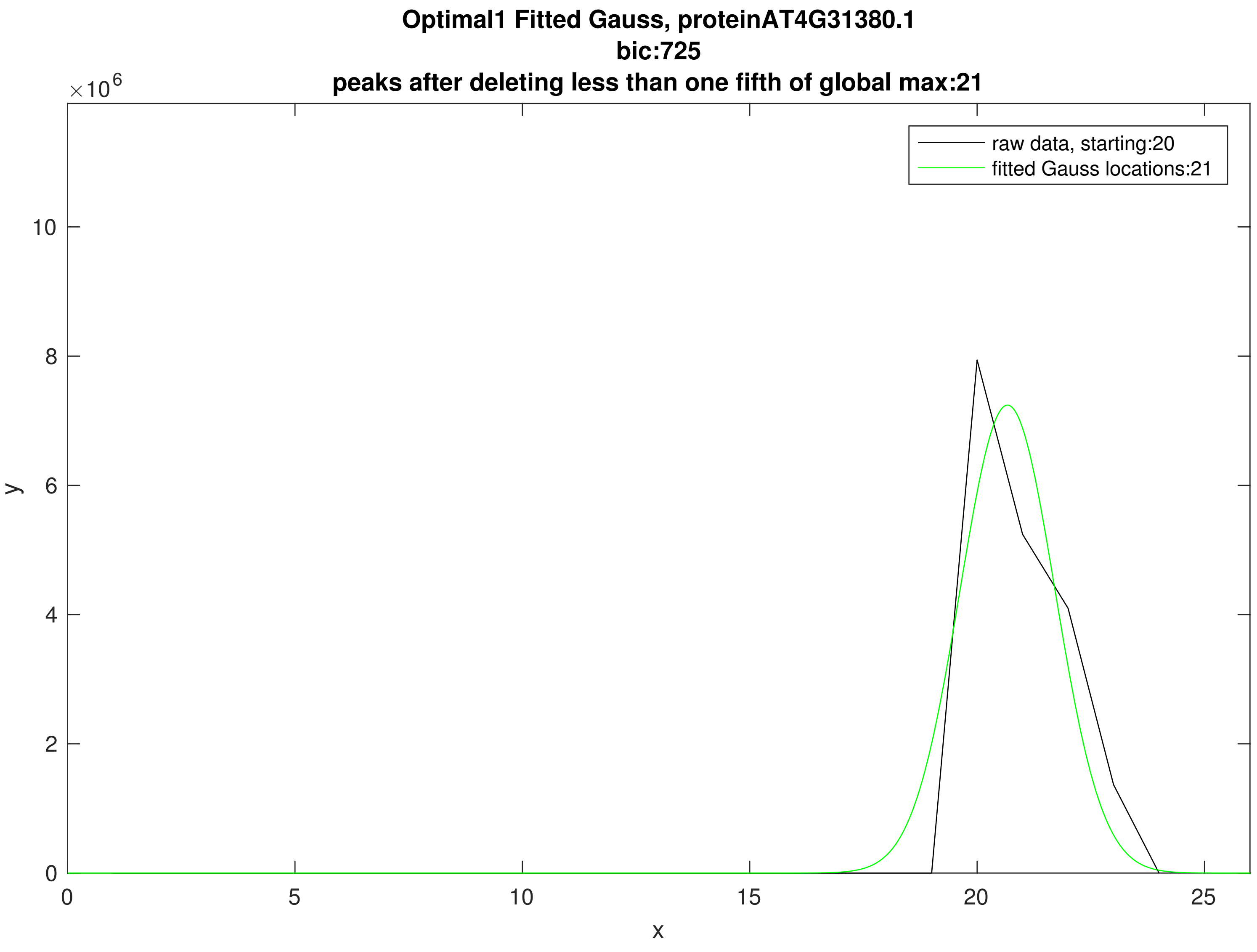

Optimal1 Fitted Gauss, proteinAT4G31380.1
bic:725
peaks after deleting less than one fifth of global max:21
×10^6
raw data, starting:20
fitted Gauss locations:21
0
2
4
6
8
10
y
0
5
10
15
20
25
x

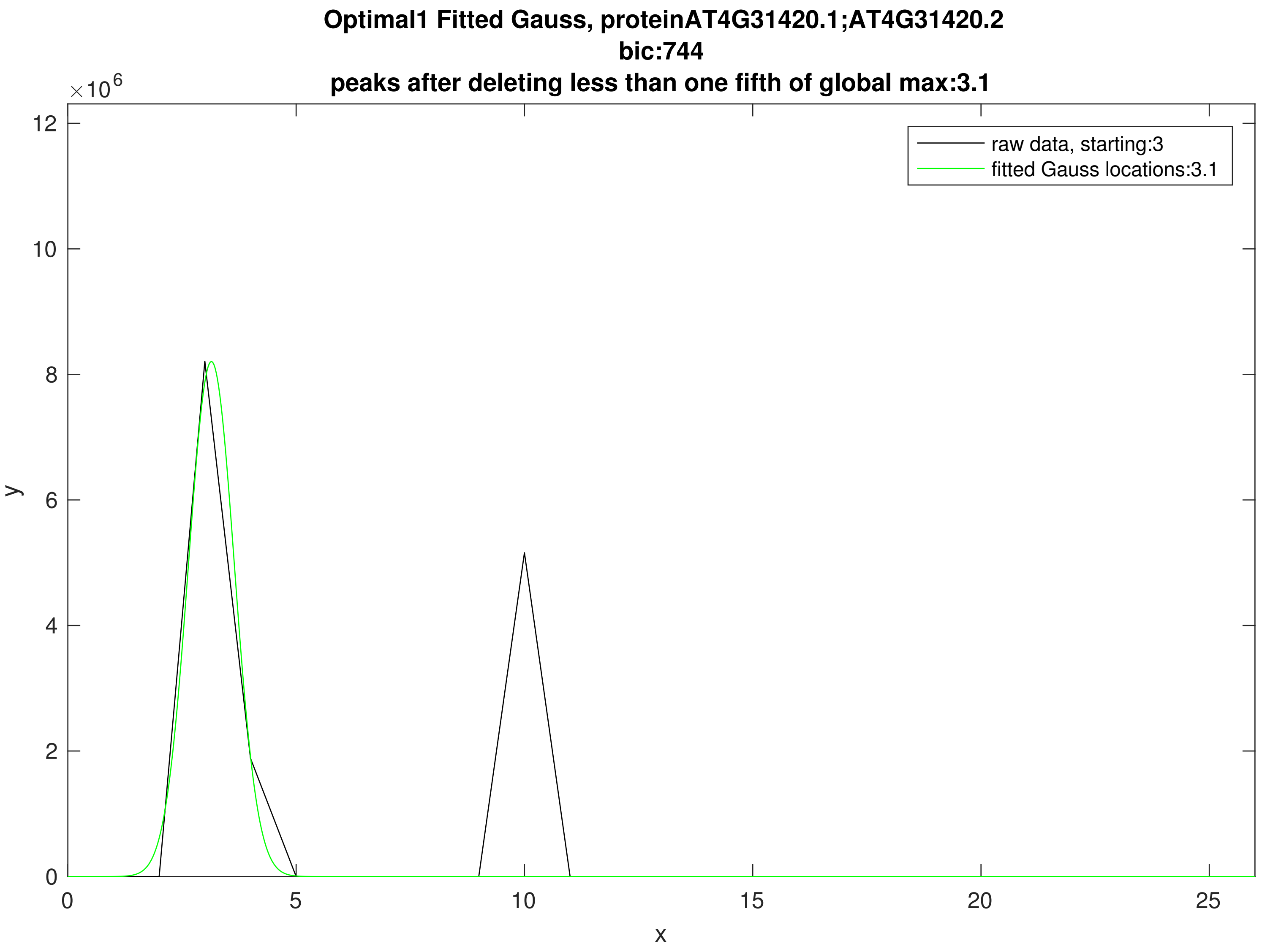

Optimal1 Fitted Gauss, proteinAT4G31420.1;AT4G31420.2
bic:744
peaks after deleting less than one fifth of global max:3.1
×10^6
raw data, starting:3
fitted Gauss locations:3.1
y
x
0
2
4
6
8
10
12
0
5
10
15
20
25

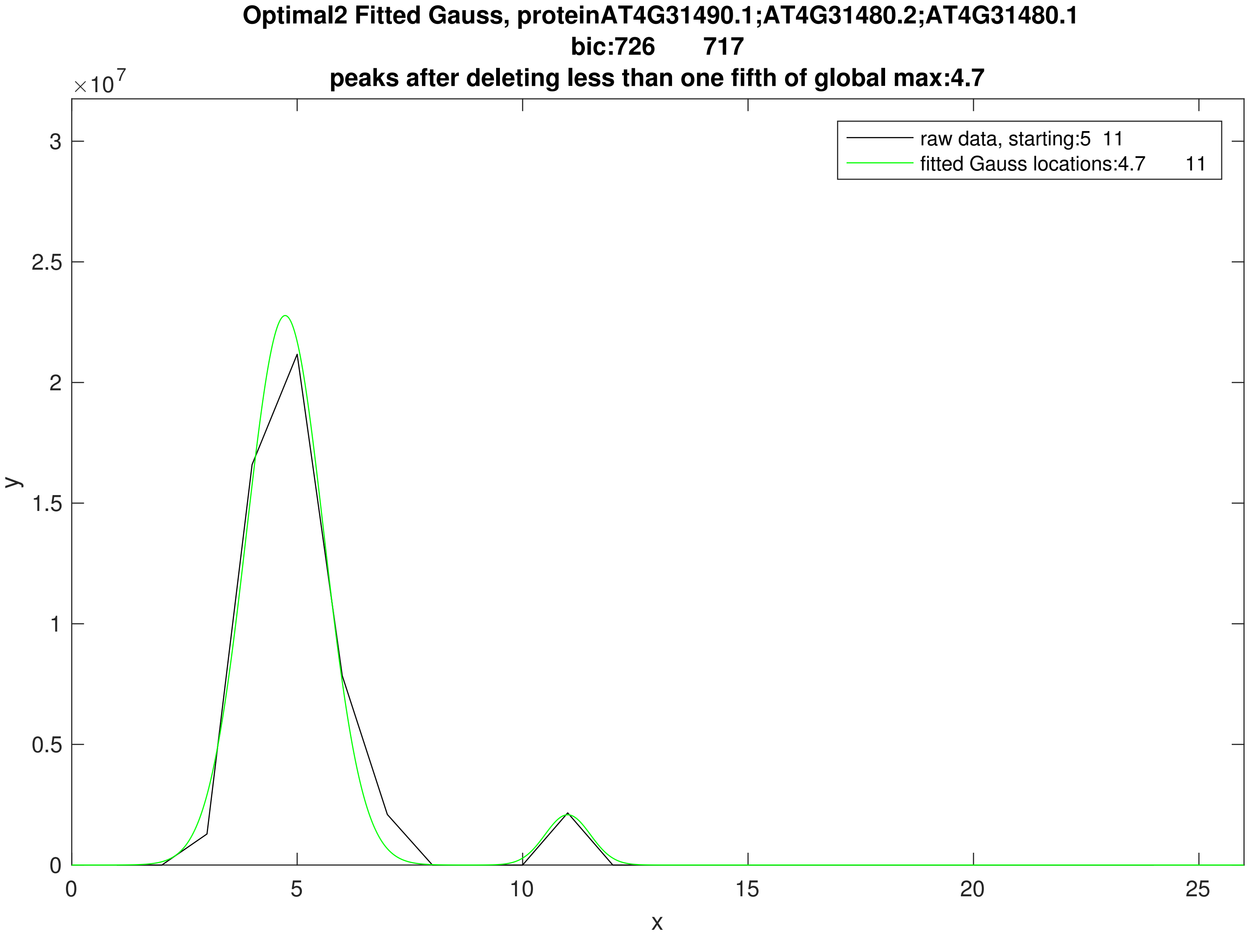

Optimal2 Fitted Gauss, proteinAT4G31490.1;AT4G31480.2;AT4G31480.1
bic:726 717
peaks after deleting less than one fifth of global max:4.7
×10^7
raw data, starting:5 11
fitted Gauss locations:4.7 11
y
x
0
0.5
1
1.5
2
2.5
3
0
5
10
15
20
25

#### Optimal1 Fitted Gauss, proteinAT4G31700.1;AT4G31700.2
#### bic:864 873
#### peaks after deleting less than one fifth of global max:3.7

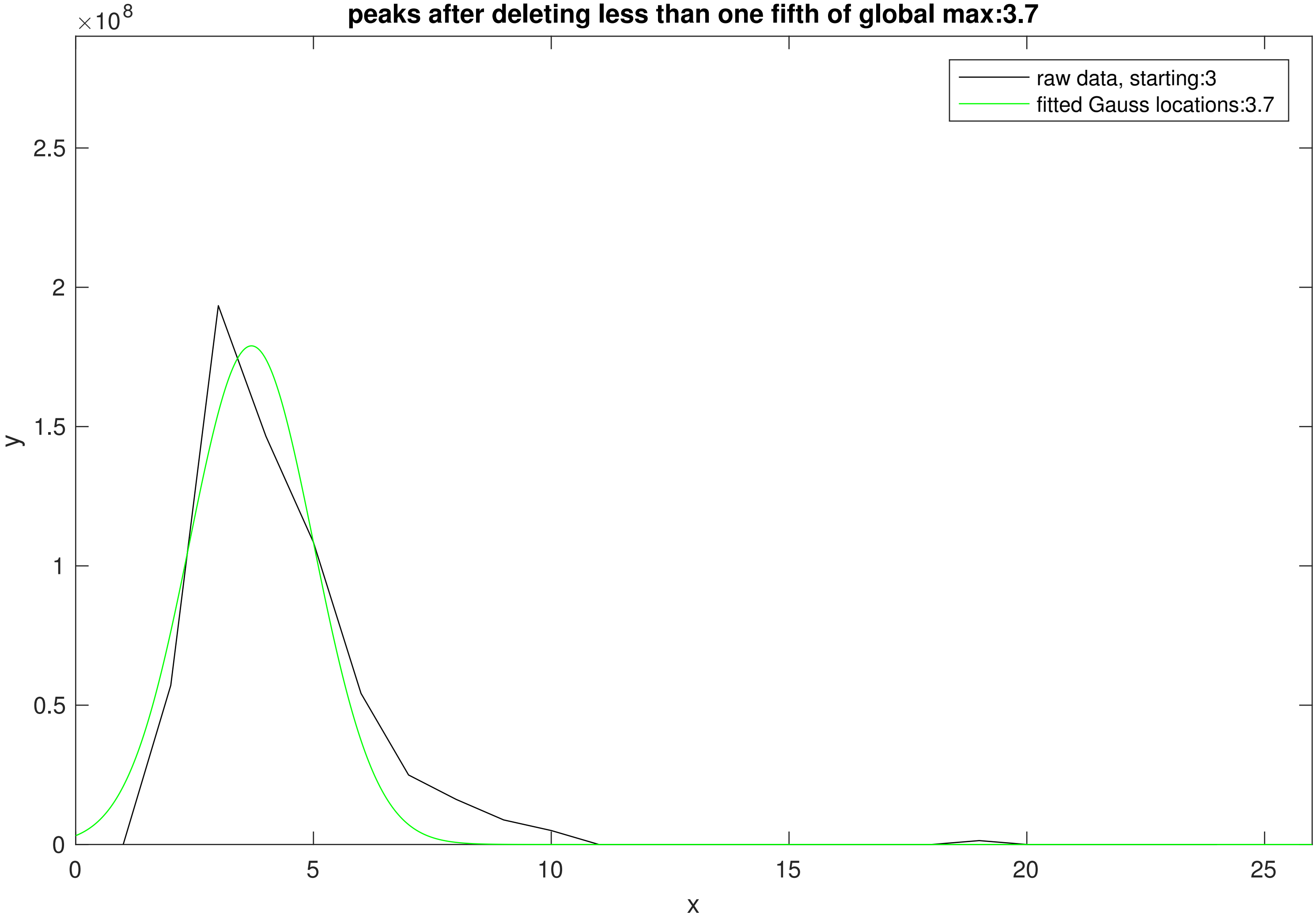

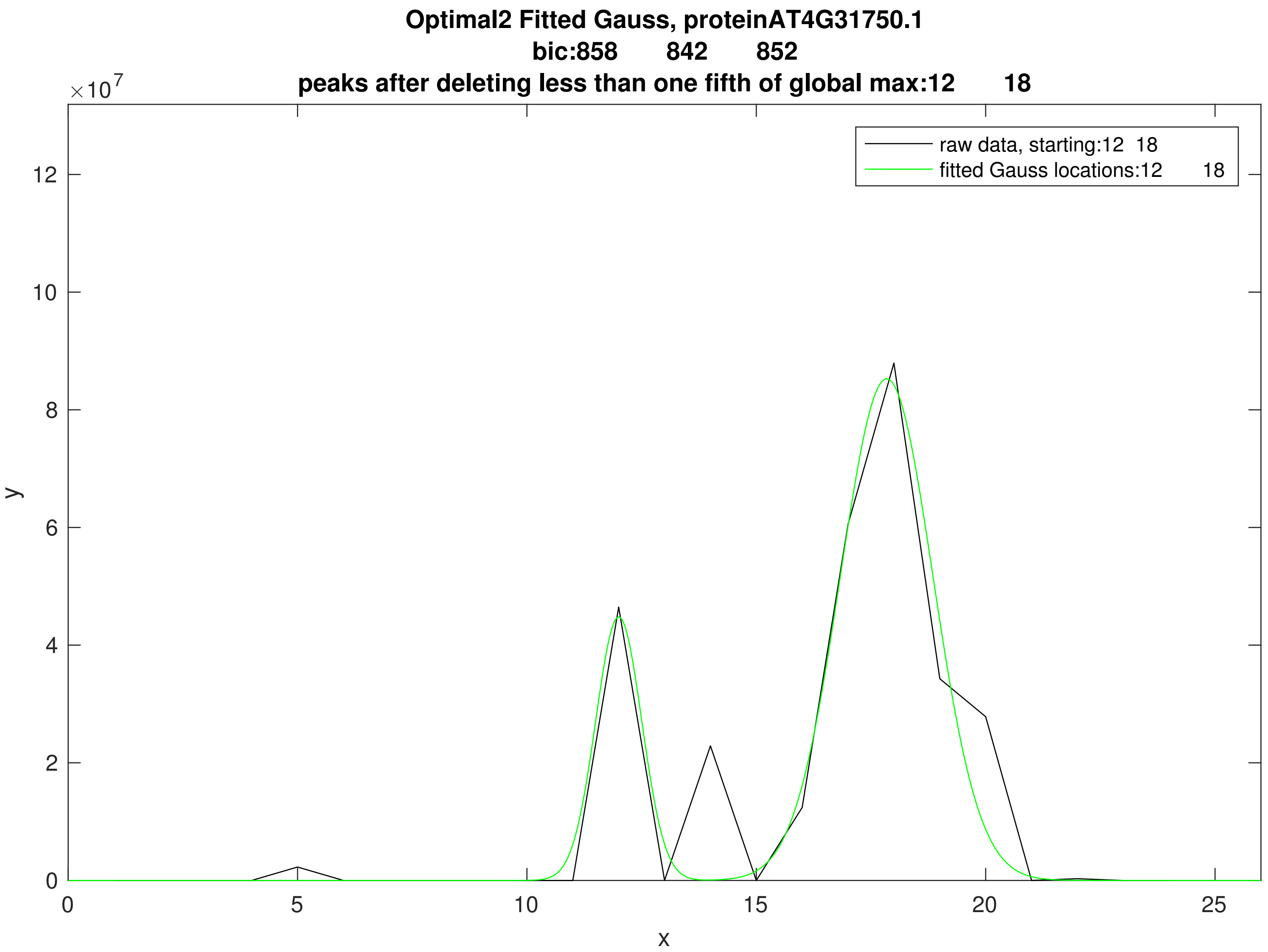
Optimal2 Fitted Gauss, proteinAT4G31750.1
bic:858 842 852
peaks after deleting less than one fifth of global max:12 18
×10^7
raw data, starting:12 18
fitted Gauss locations:12 18
y
x
0
2
4
6
8
10
12
0
5
10
15
20
25

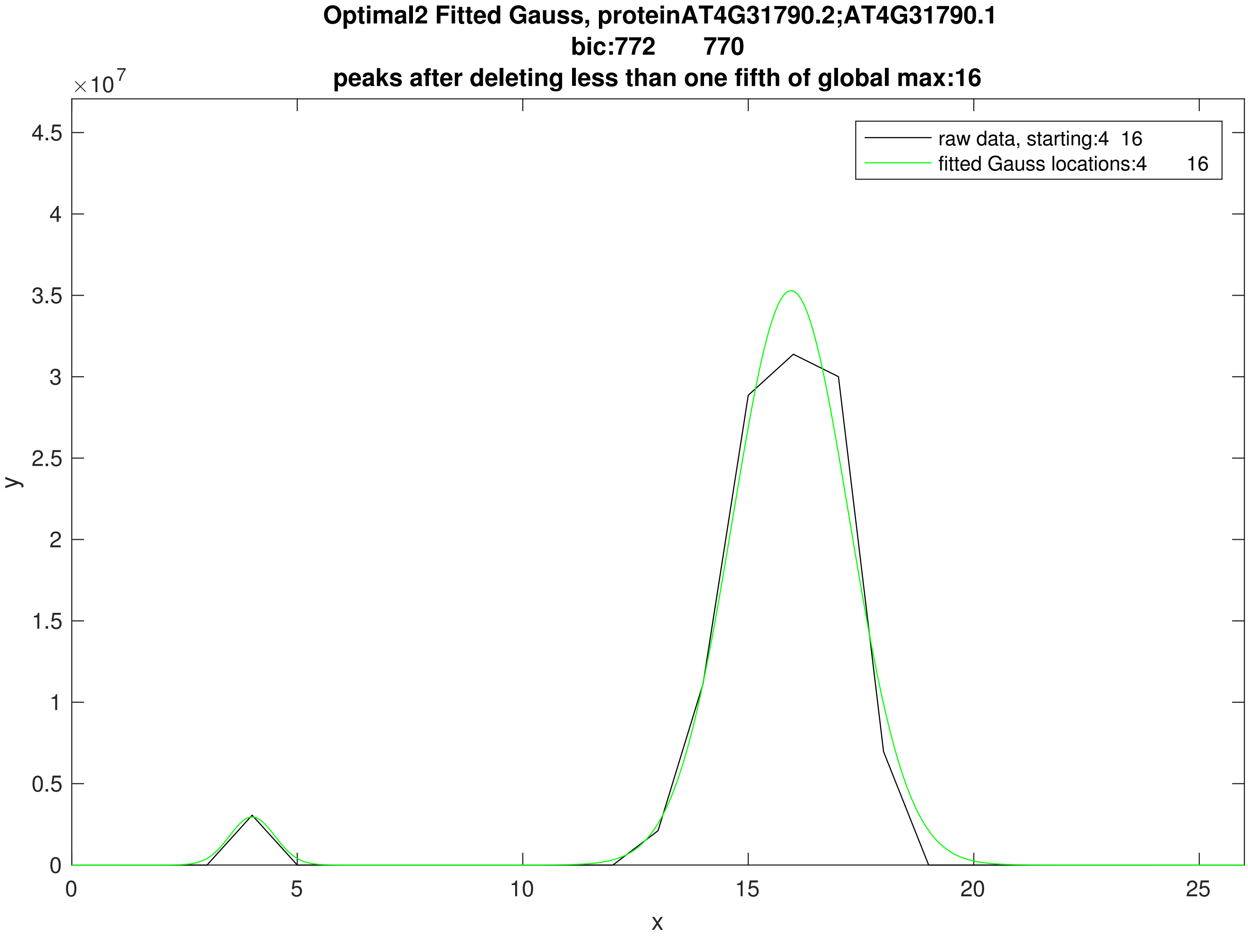

Optimal2 Fitted Gauss, proteinAT4G31790.2;AT4G31790.1
bic:772 770
peaks after deleting less than one fifth of global max:16
×10^7
raw data, starting:4 16
fitted Gauss locations:4 16
y
x
0
0.5
1
1.5
2
2.5
3
3.5
4
4.5
5
10
15
20
25

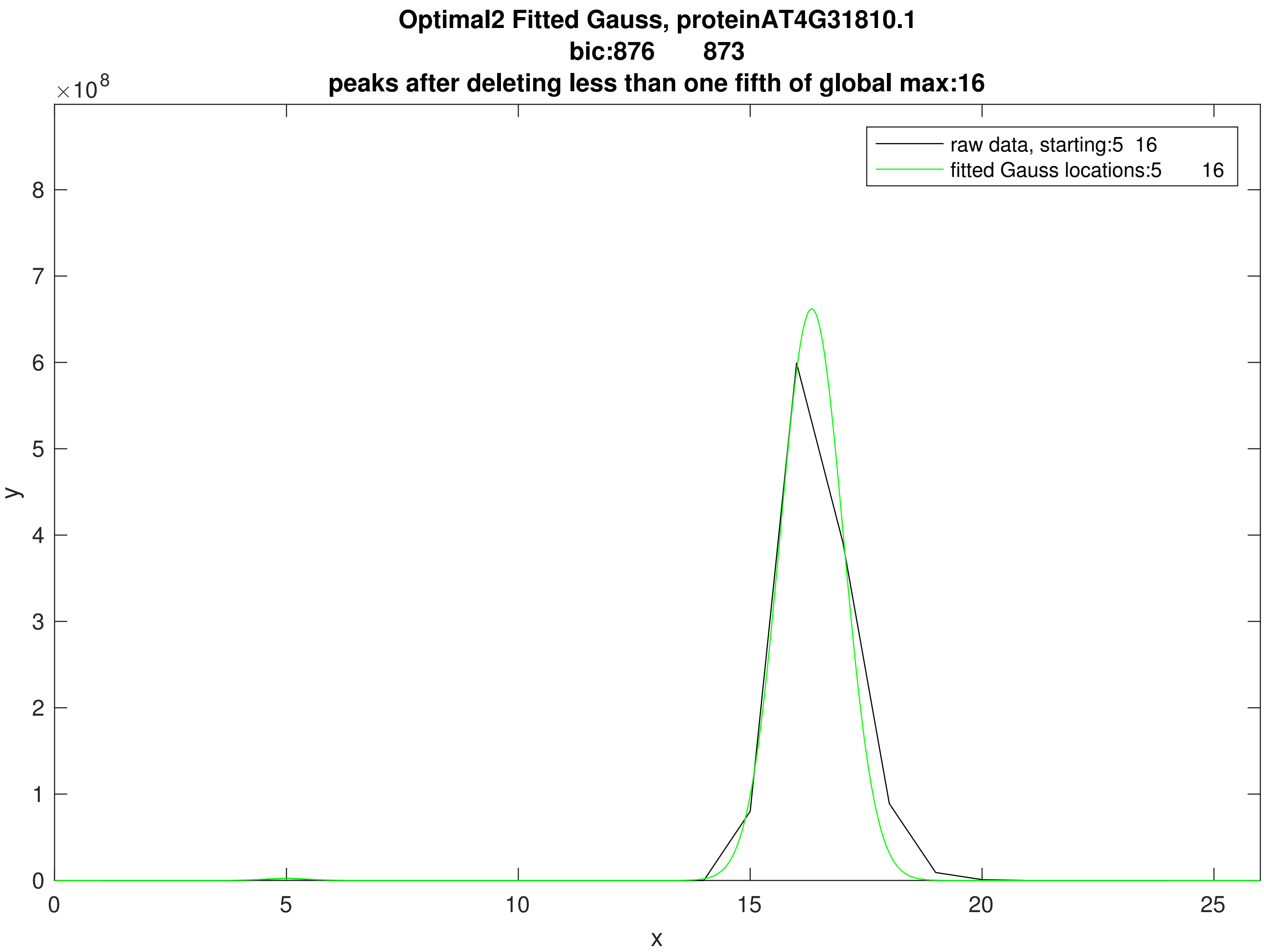

Optimal2 Fitted Gauss, proteinAT4G31810.1
bic:876 873
peaks after deleting less than one fifth of global max:16
×10^8
raw data, starting:5 16
fitted Gauss locations:5 16
y
x
0
1
2
3
4
5
6
7
8
0
5
10
15
20
25

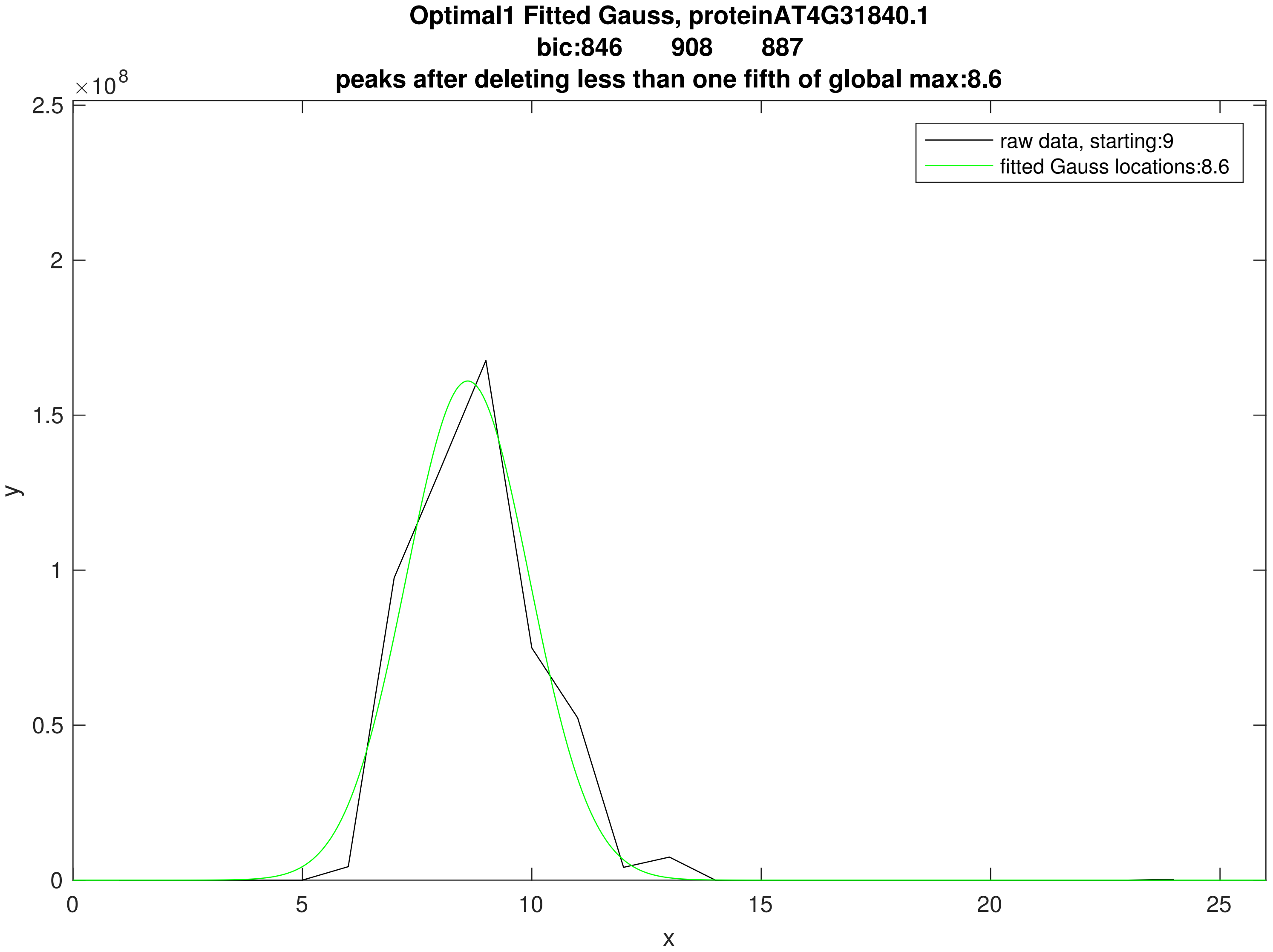
Optimal1 Fitted Gauss, proteinAT4G31840.1
bic:846 908 887
peaks after deleting less than one fifth of global max:8.6
raw data, starting:9
fitted Gauss locations:8.6
×10^8
2.5
2
1.5
1
0.5
0
y
0
5
10
15
20
25
x

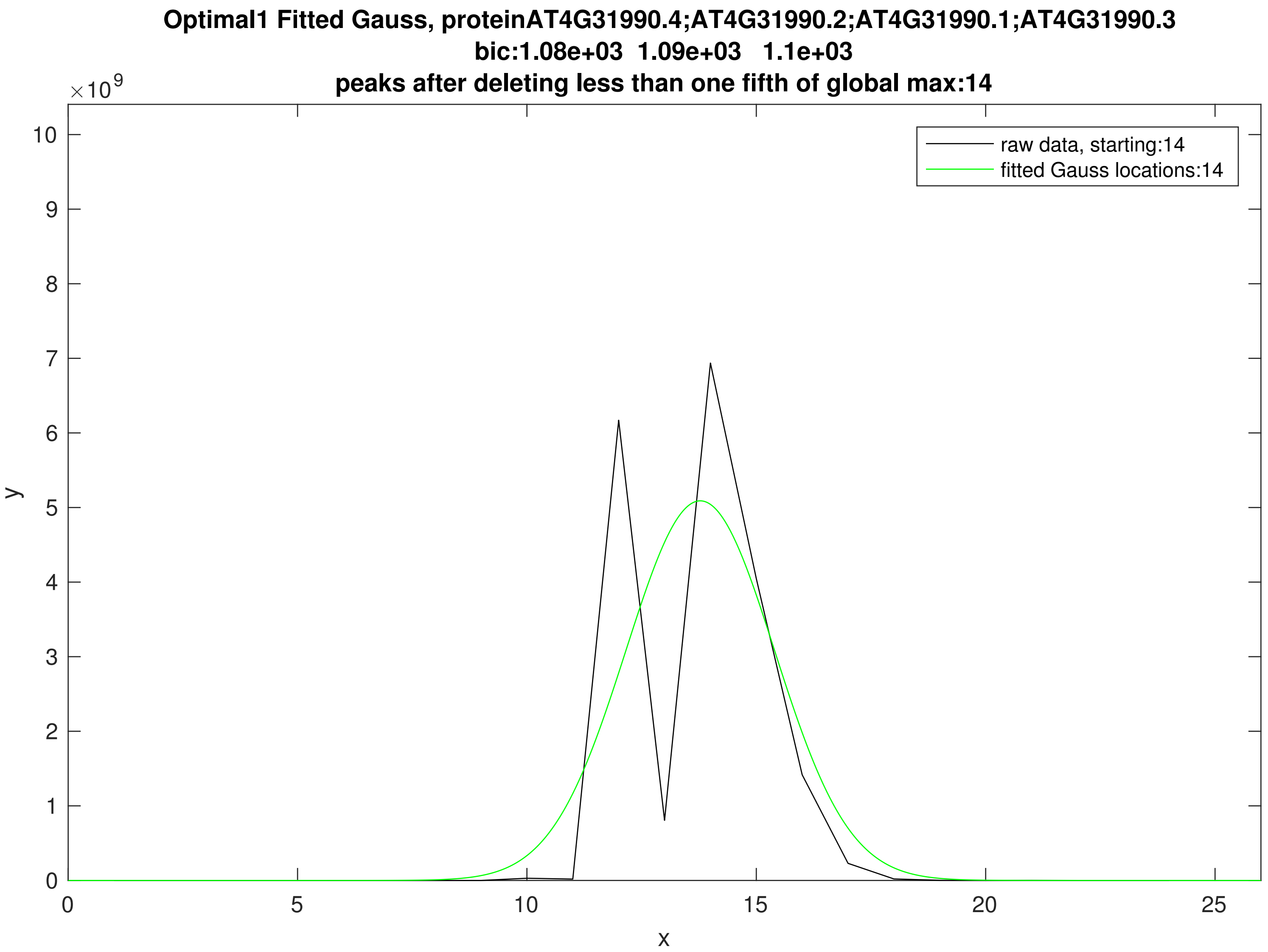
Optimal1 Fitted Gauss, proteinAT4G31990.4;AT4G31990.2;AT4G31990.1;AT4G31990.3
bic:1.08e+03 1.09e+03 1.1e+03
peaks after deleting less than one fifth of global max:14
×10^9
raw data, starting:14
fitted Gauss locations:14
y
x
0
1
2
3
4
5
6
7
8
9
10
0
5
10
15
20
25

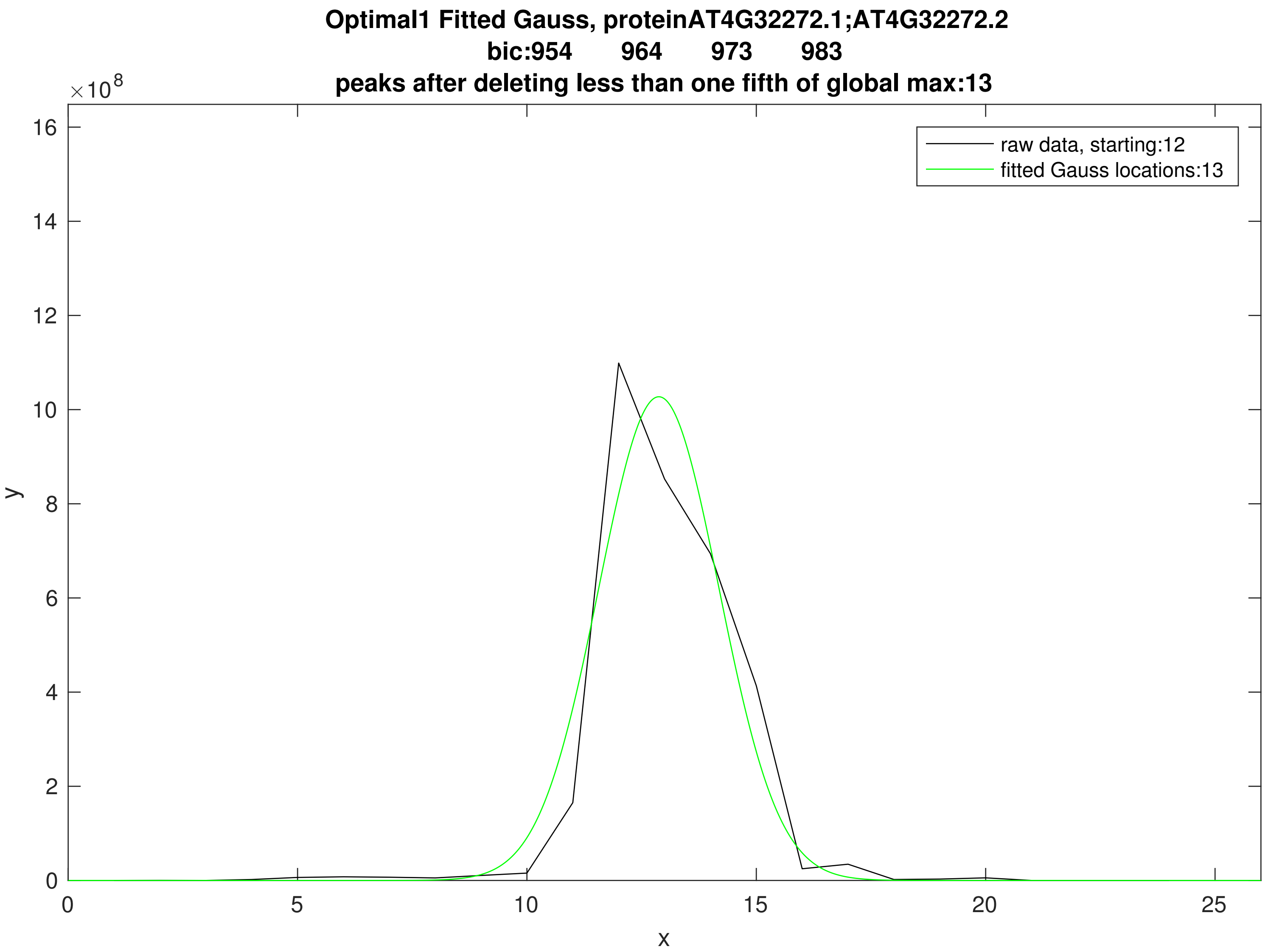
Optimal1 Fitted Gauss, proteinAT4G32272.1;AT4G32272.2
bic:954 964 973 983
peaks after deleting less than one fifth of global max:13
×10^8
raw data, starting:12
fitted Gauss locations:13
y
x
0
2
4
6
8
10
12
14
16
0
5
10
15
20
25

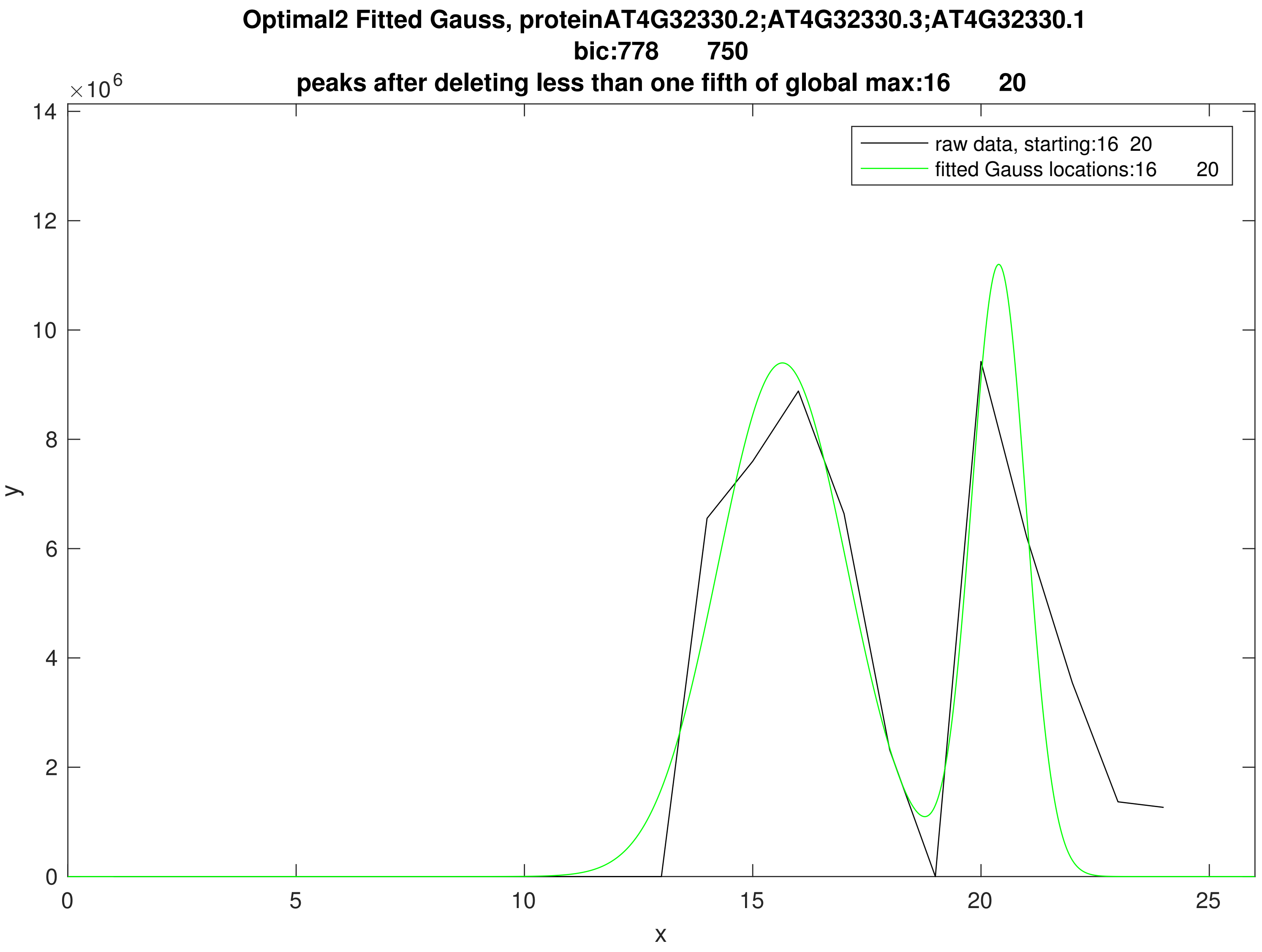
Optimal2 Fitted Gauss, proteinAT4G32330.2;AT4G32330.3;AT4G32330.1
bic:778 750
peaks after deleting less than one fifth of global max:16 20
×10^6
raw data, starting:16 20
fitted Gauss locations:16 20
y
x
0
2
4
6
8
10
12
14
0
5
10
15
20
25

Optimal1 Fitted Gauss, proteinAT4G32520.2;AT4G32520.1
bic:914
peaks after deleting less than one fifth of global max:9.9

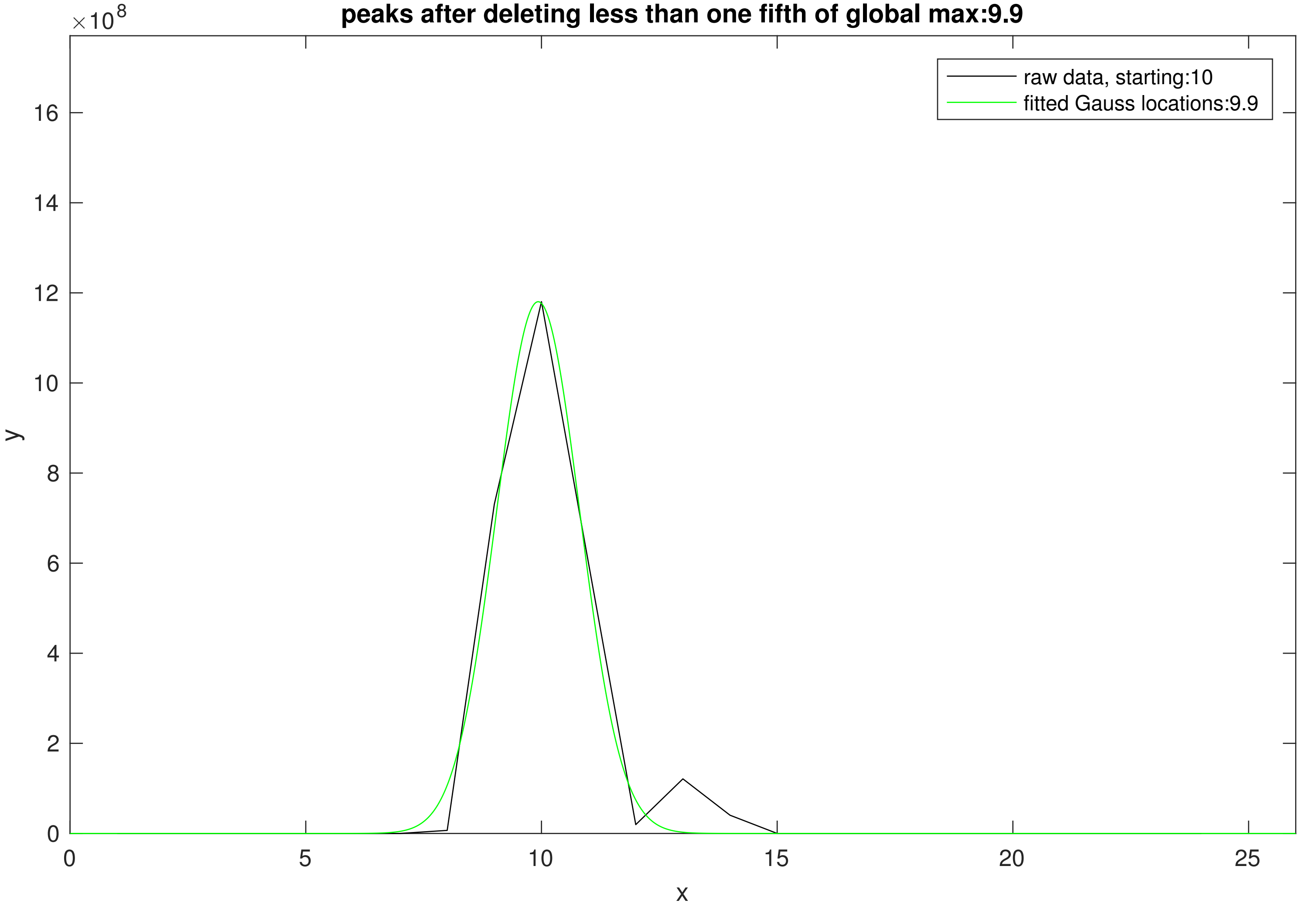

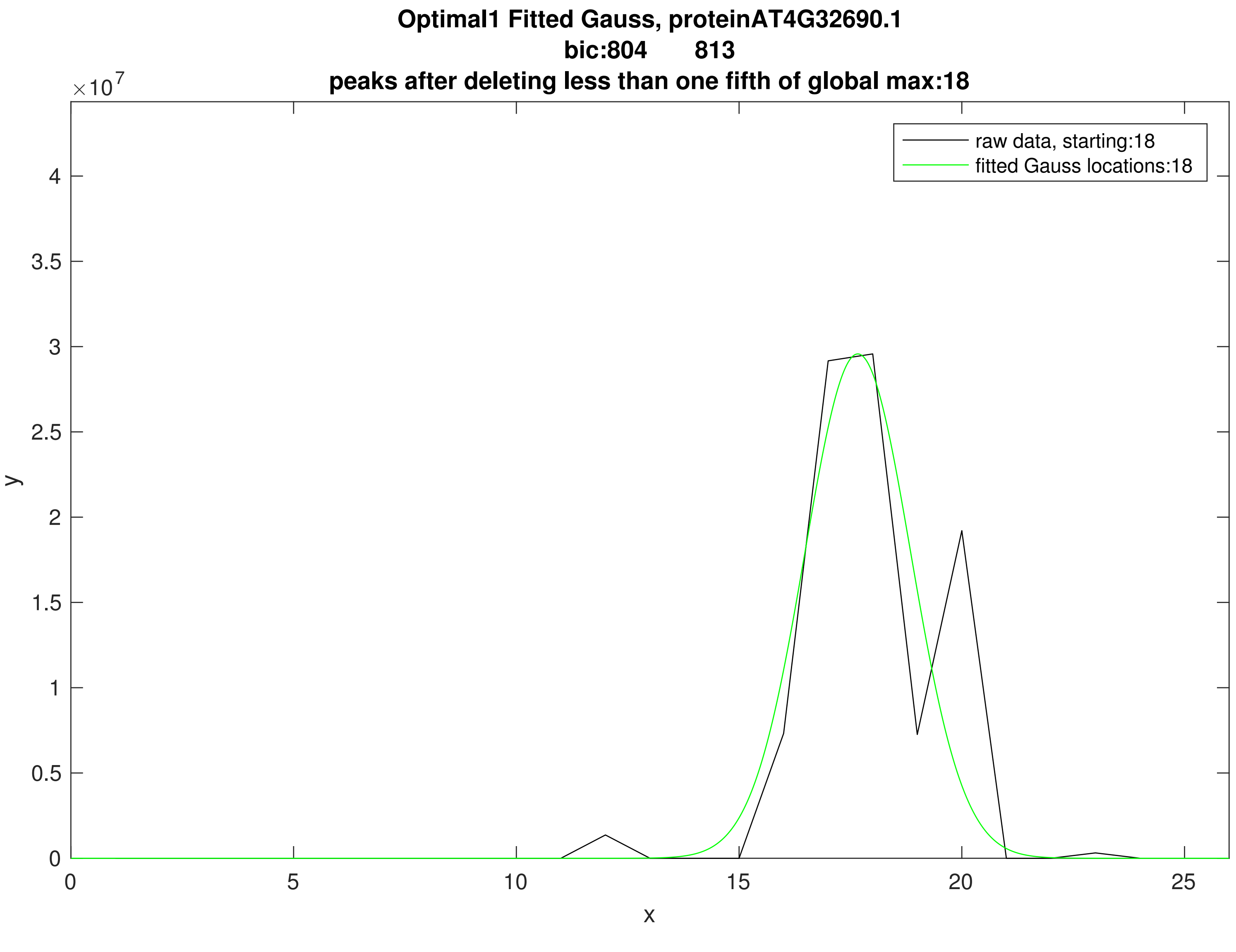

Optimal1 Fitted Gauss, proteinAT4G32690.1
bic:804 813
peaks after deleting less than one fifth of global max:18
×10^7
raw data, starting:18
fitted Gauss locations:18
y
x
0
0.5
1
1.5
2
2.5
3
3.5
4
5
10
15
20
25

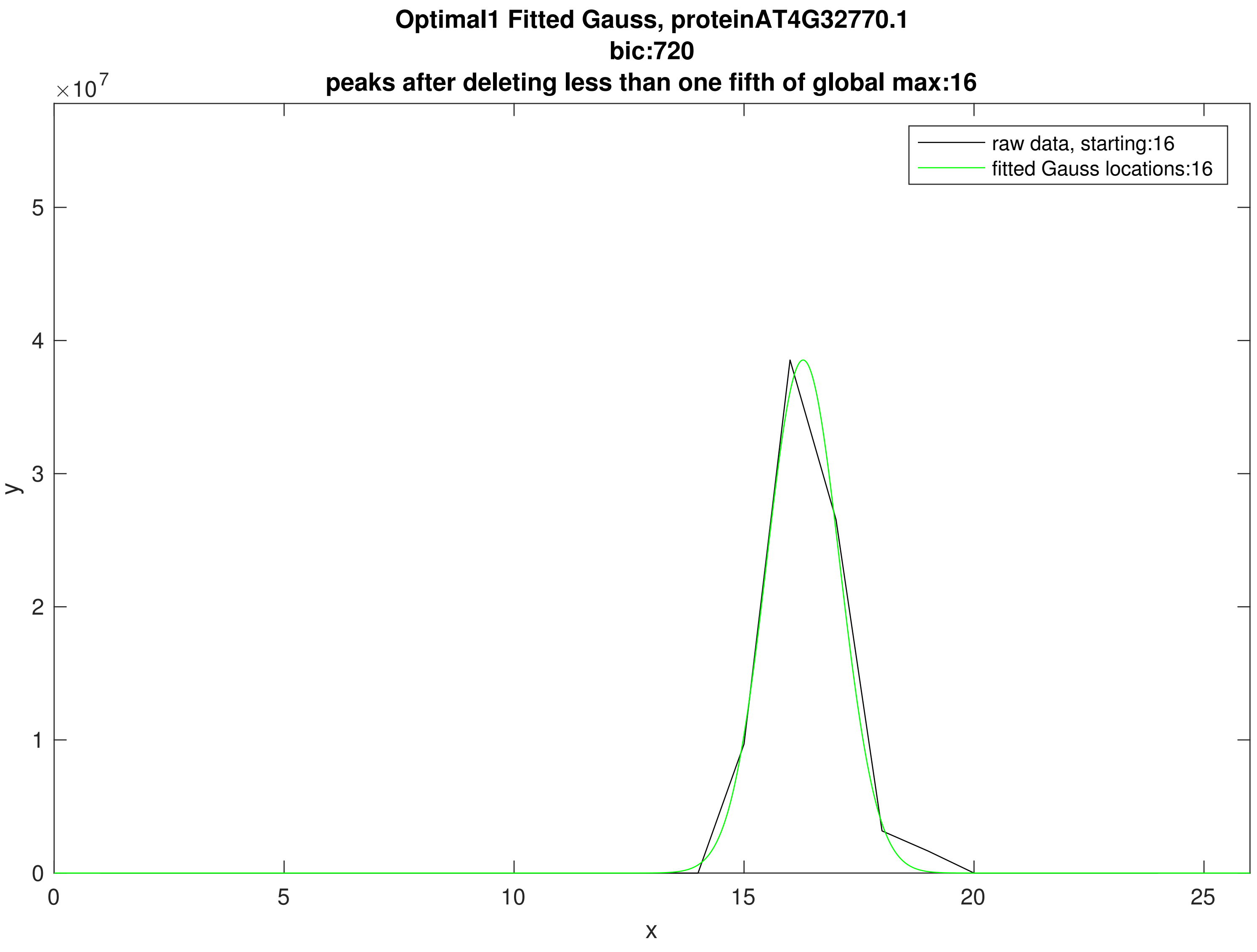

Optimal1 Fitted Gauss, proteinAT4G32770.1
bic:720
peaks after deleting less than one fifth of global max:16
×10^7
raw data, starting:16
fitted Gauss locations:16
y
x
0
1
2
3
4
5
0
5
10
15
20
25

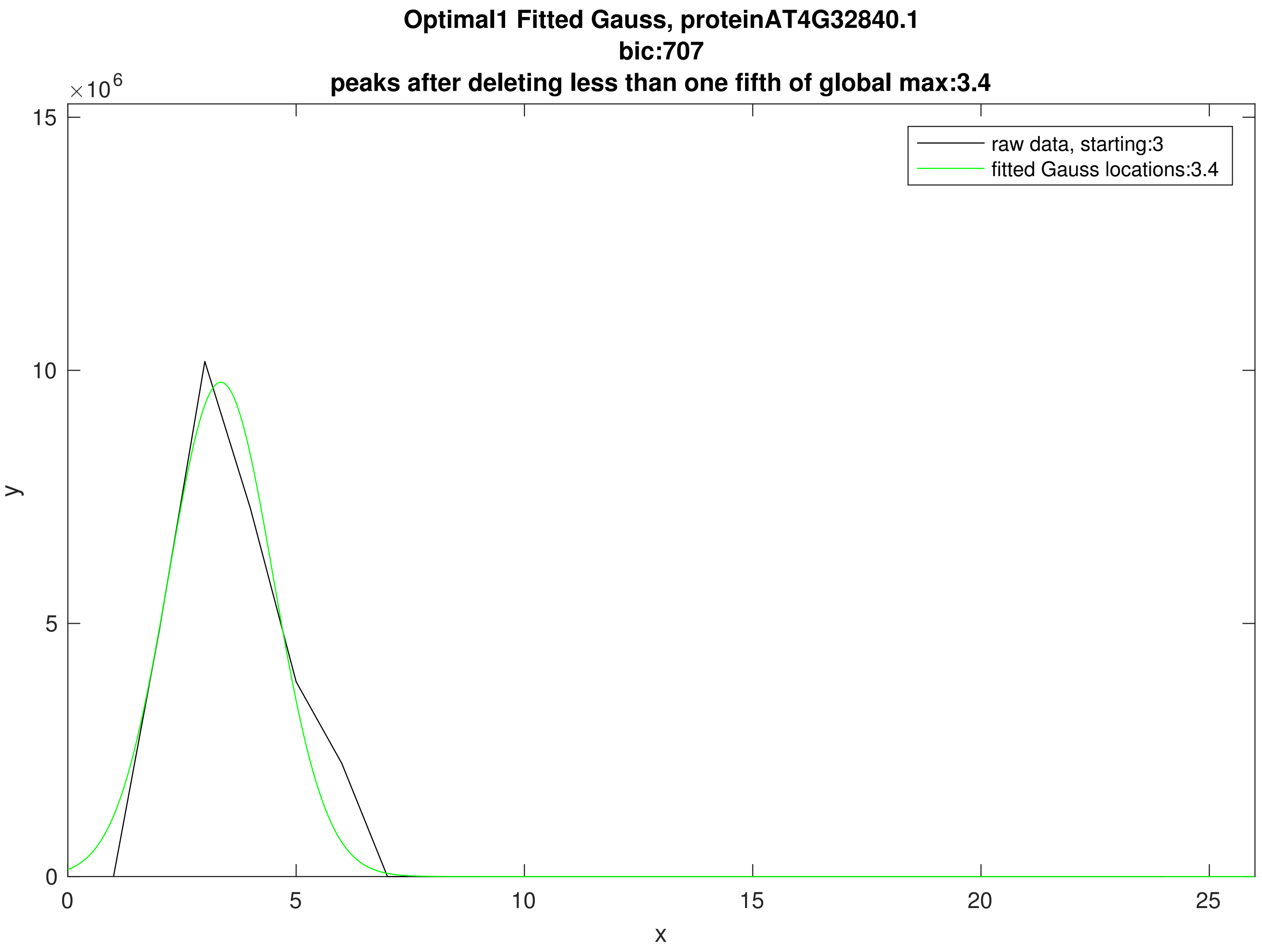
Optimal1 Fitted Gauss, proteinAT4G32840.1
bic:707
peaks after deleting less than one fifth of global max:3.4
×10^6
15
10
5
0
y
0
5
10
15
20
25
x
raw data, starting:3
fitted Gauss locations:3.4

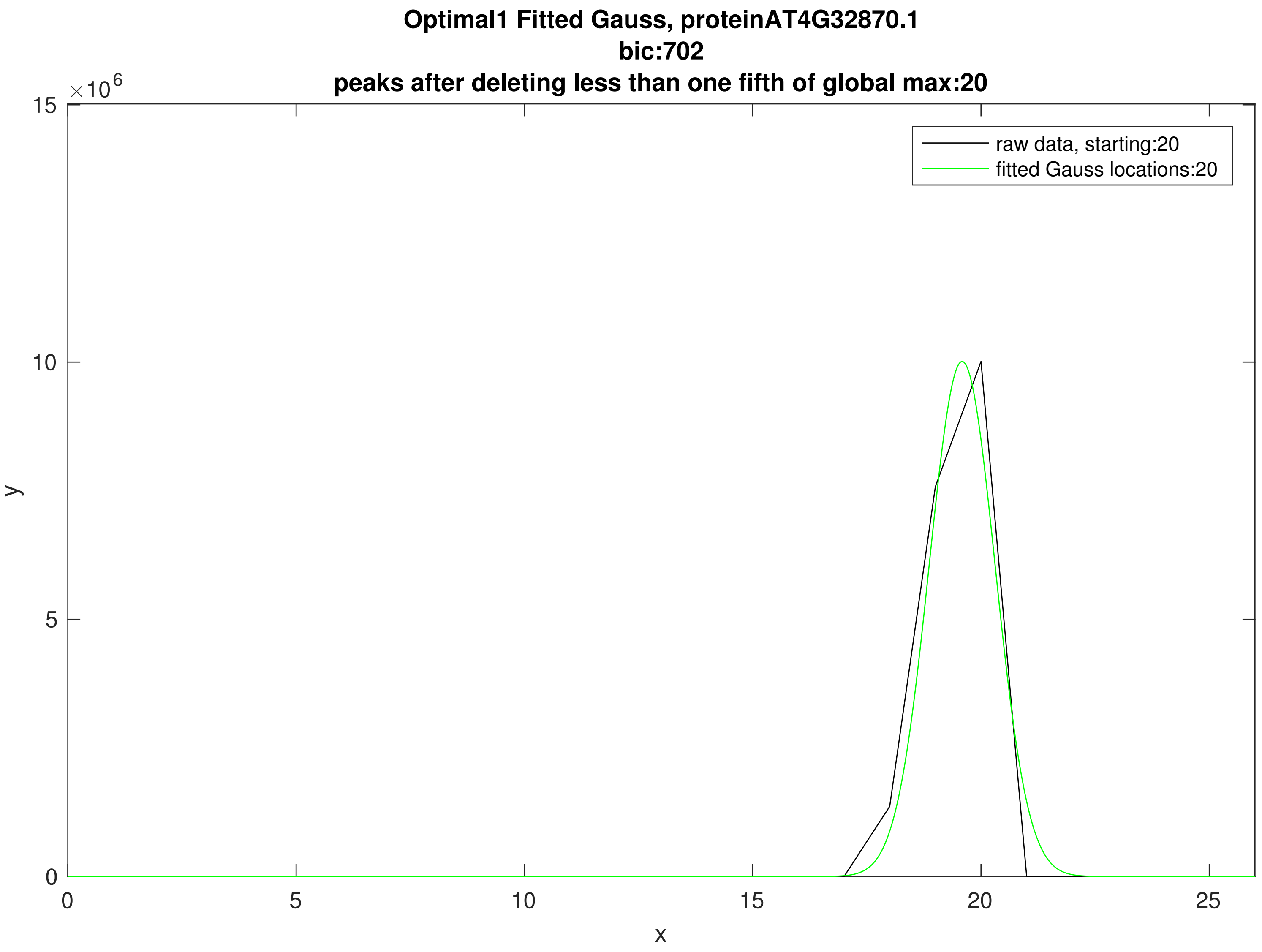
Optimal1 Fitted Gauss, proteinAT4G32870.1
bic:702
peaks after deleting less than one fifth of global max:20
×10^6
15
10
5
0
0
5
10
15
20
25
x
y
raw data, starting:20
fitted Gauss locations:20

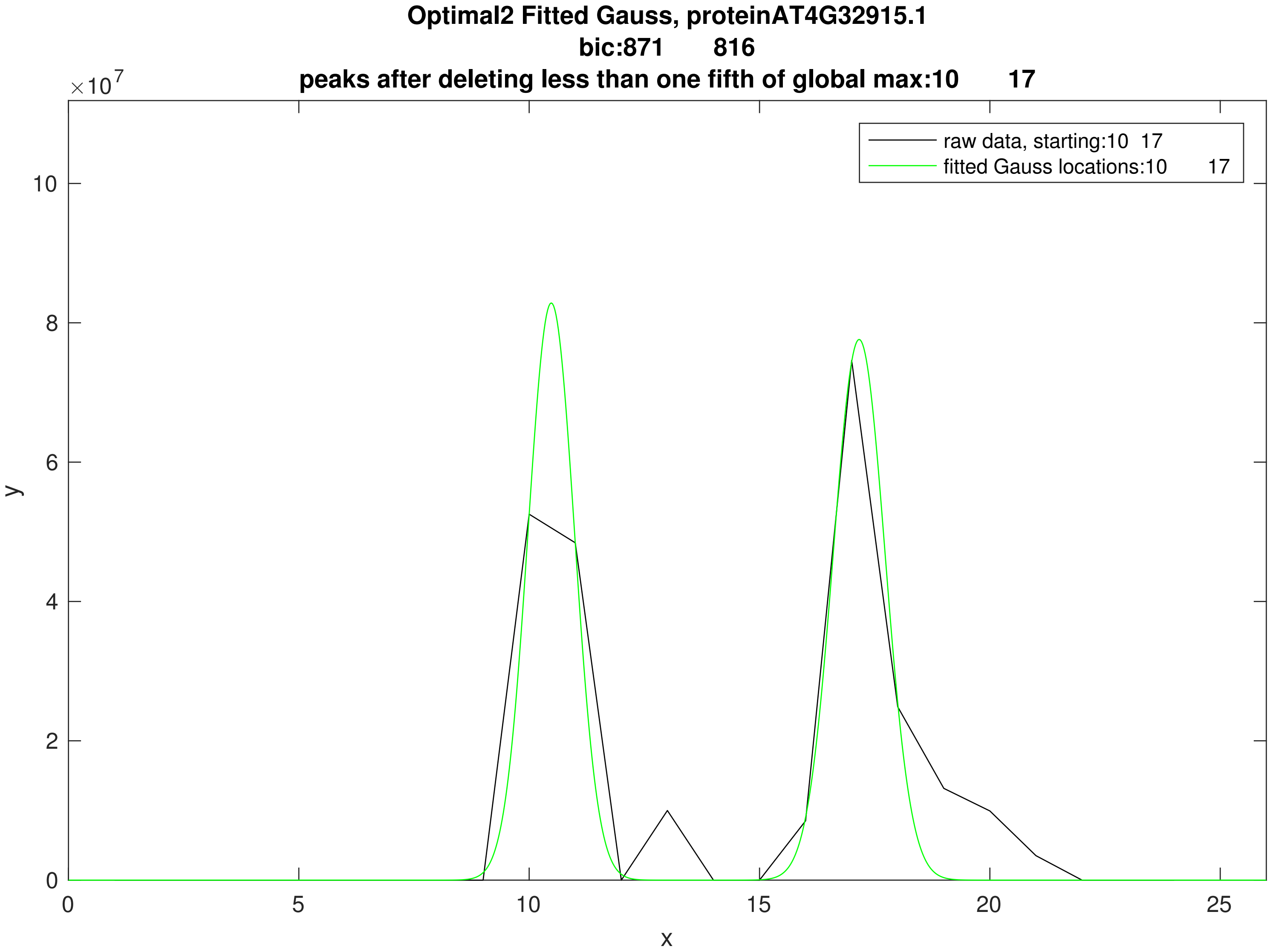
Optimal2 Fitted Gauss, proteinAT4G32915.1
bic:871 816
peaks after deleting less than one fifth of global max:10 17
×10^7
raw data, starting:10 17
fitted Gauss locations:10 17
10
8
6
4
2
0
y
0
5
10
15
20
25
x

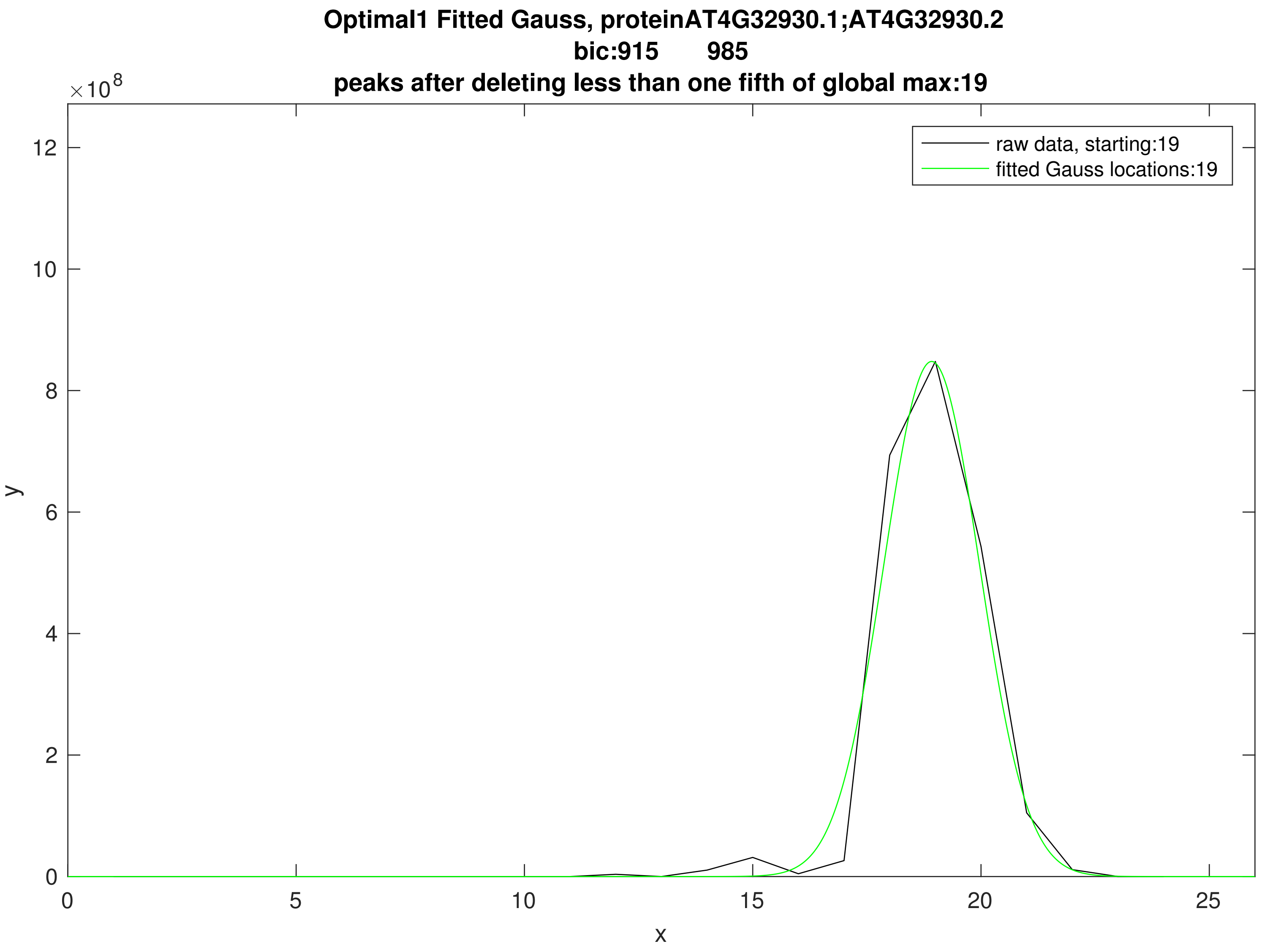

Optimal1 Fitted Gauss, proteinAT4G32930.1;AT4G32930.2
bic:915 985
peaks after deleting less than one fifth of global max:19
×10^8
raw data, starting:19
fitted Gauss locations:19
y
x
0
2
4
6
8
10
12
0
5
10
15
20
25

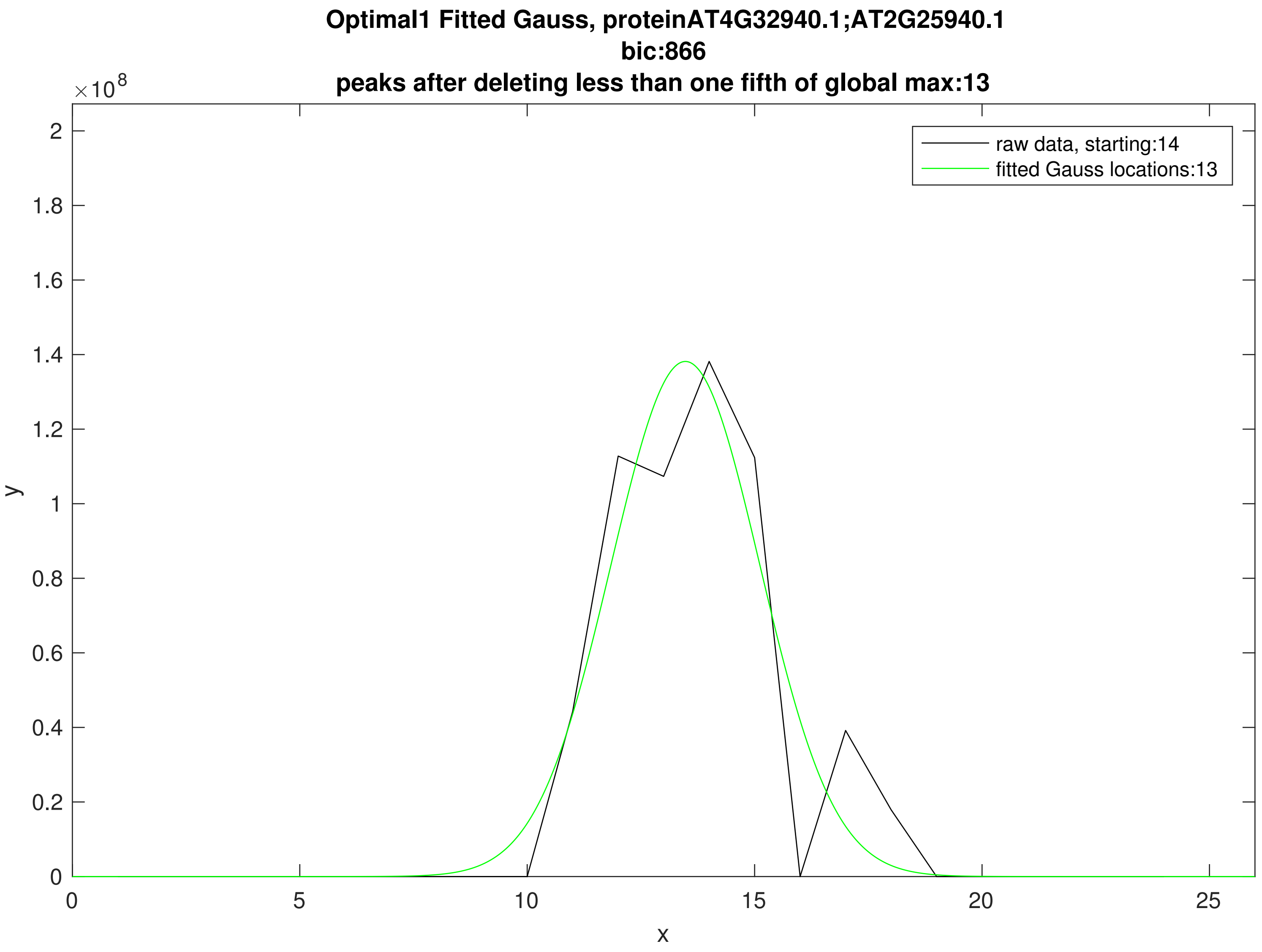
Optimal1 Fitted Gauss, proteinAT4G32940.1;AT2G25940.1
bic:866
peaks after deleting less than one fifth of global max:13
×10^8
raw data, starting:14
fitted Gauss locations:13
y
x
0
0.2
0.4
0.6
0.8
1
1.2
1.4
1.6
1.8
2
0
5
10
15
20
25

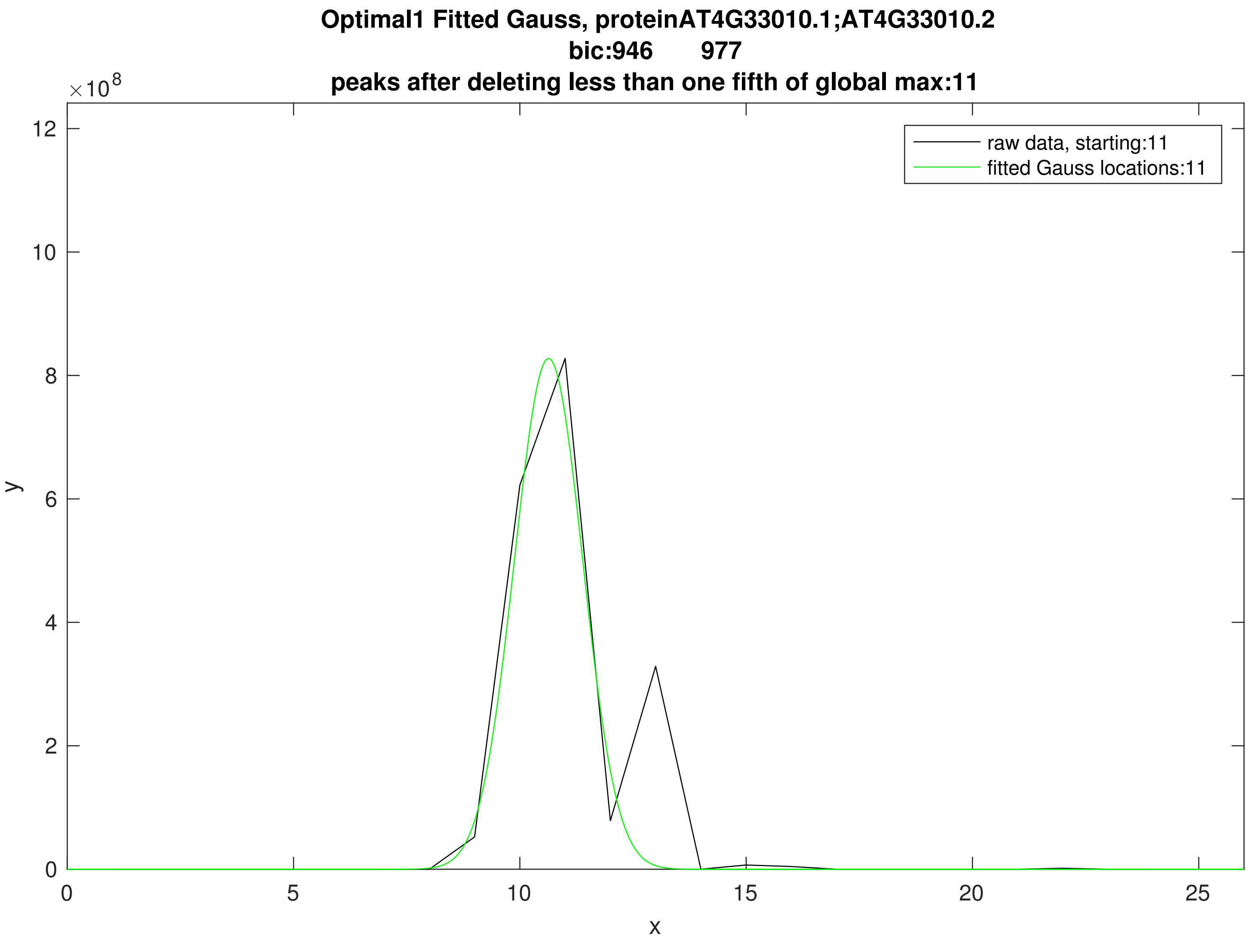
Optimal1 Fitted Gauss, proteinAT4G33010.1;AT4G33010.2
bic:946 977
peaks after deleting less than one fifth of global max:11
×10^8
raw data, starting:11
fitted Gauss locations:11
y
x
0
2
4
6
8
10
12
0
5
10
15
20
25

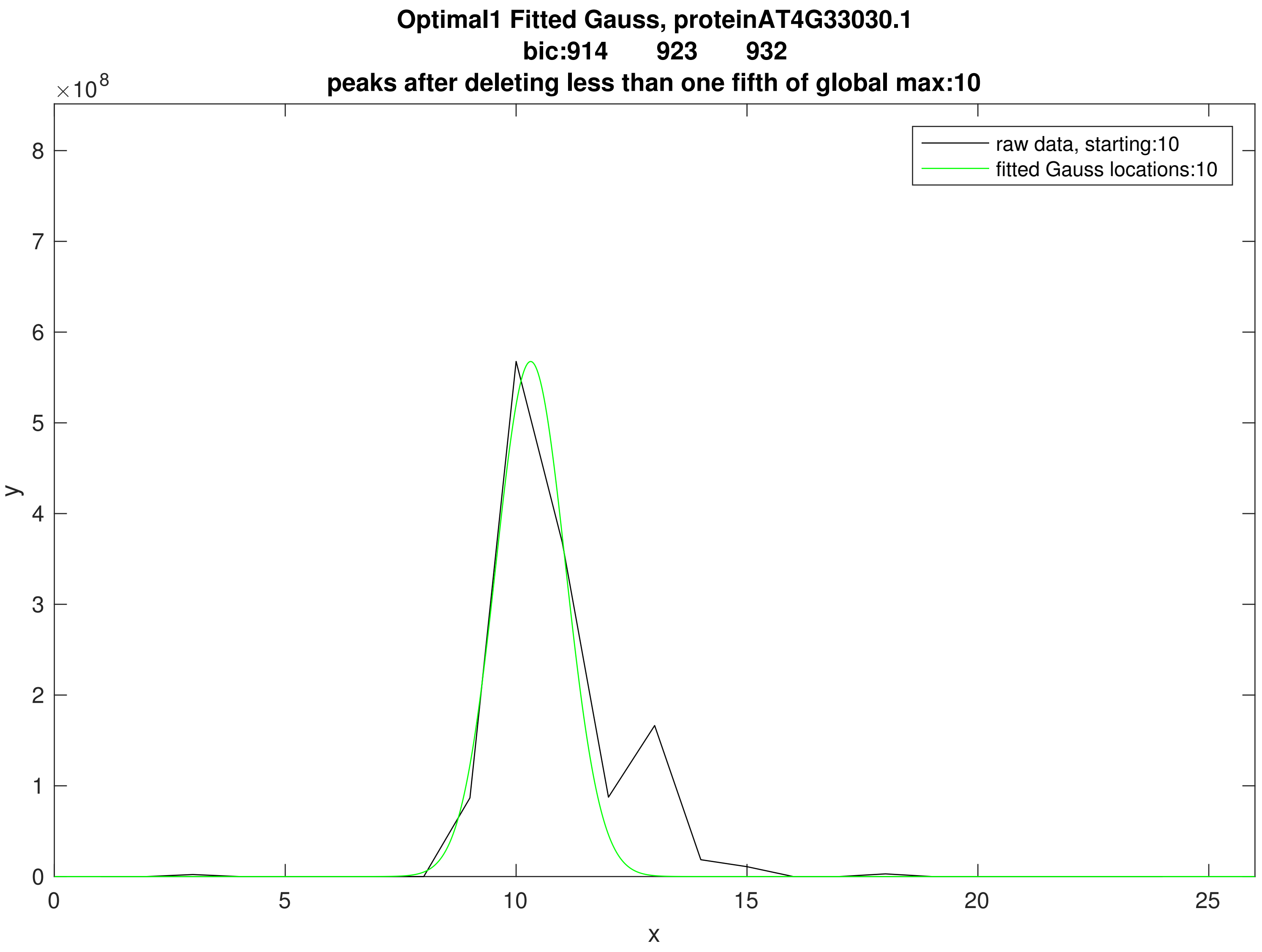
Optimal1 Fitted Gauss, proteinAT4G33030.1
bic:914 923 932
peaks after deleting less than one fifth of global max:10
×10^8
raw data, starting:10
fitted Gauss locations:10
y
x
0
1
2
3
4
5
6
7
8
0
5
10
15
20
25

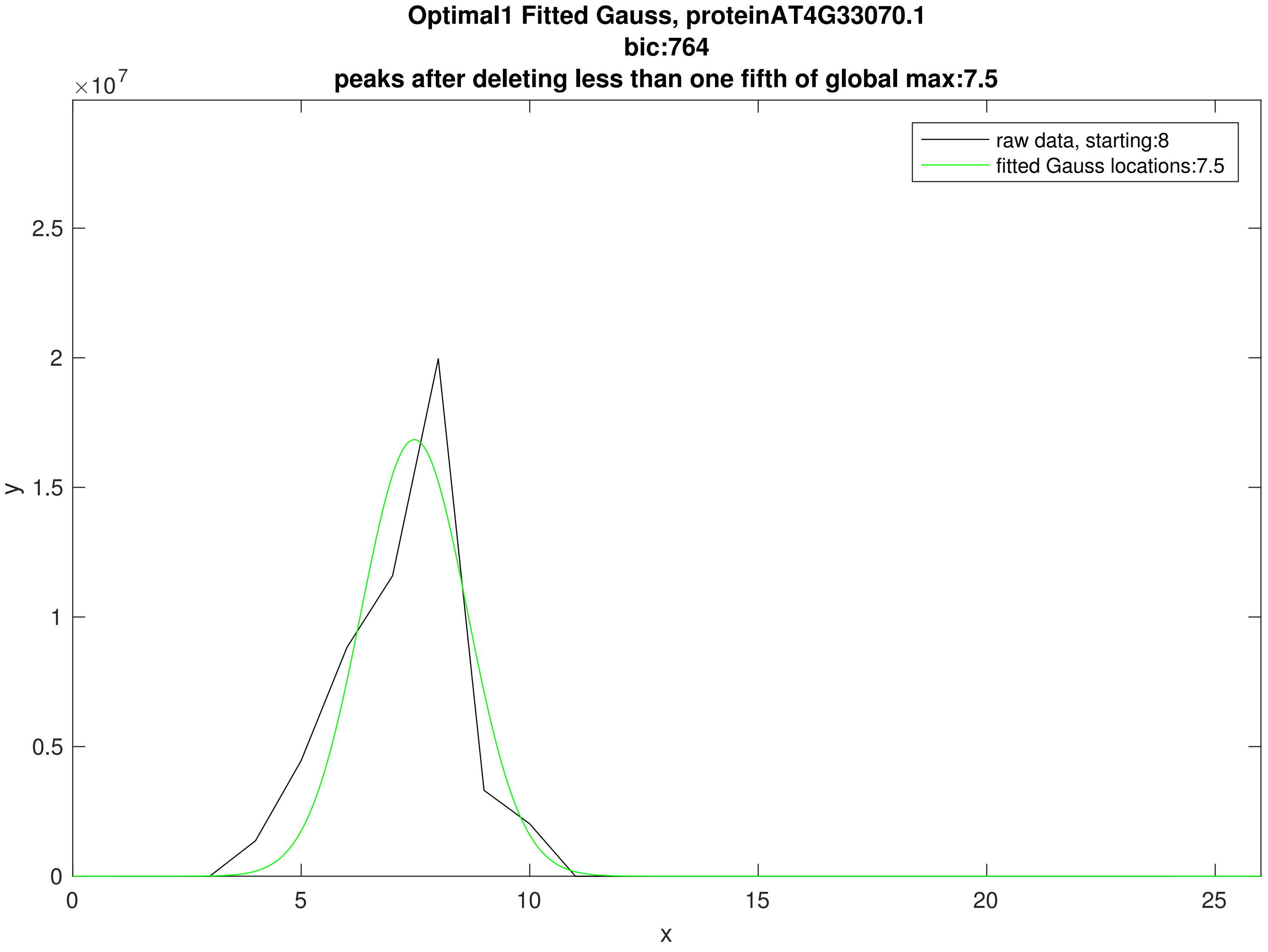
Optimal1 Fitted Gauss, proteinAT4G33070.1
bic:764
peaks after deleting less than one fifth of global max:7.5
×10^7
raw data, starting:8
fitted Gauss locations:7.5
y
x
0
0.5
1
1.5
2
2.5
0
5
10
15
20
25

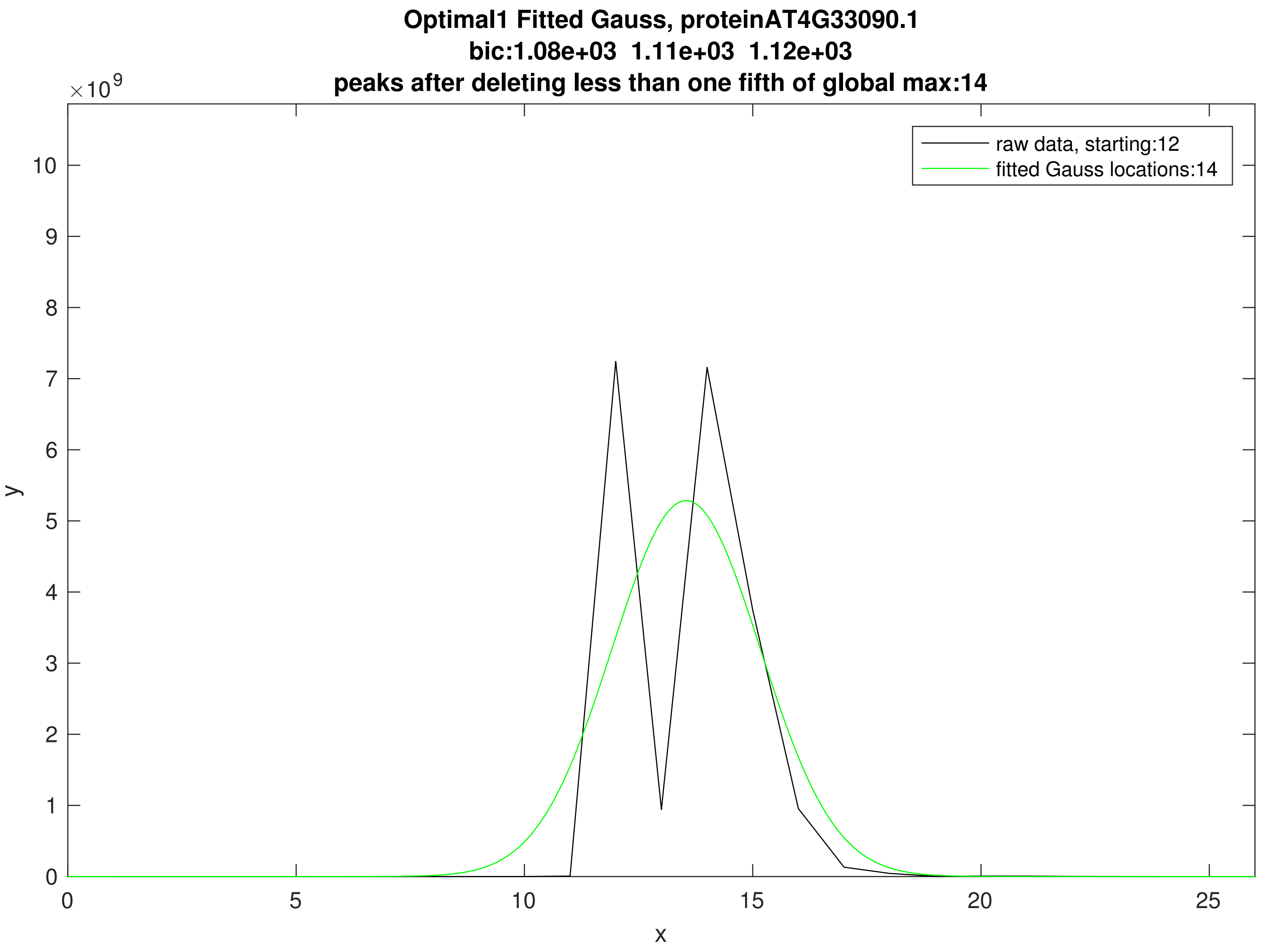

Optimal1 Fitted Gauss, proteinAT4G33090.1
bic:1.08e+03 1.11e+03 1.12e+03
peaks after deleting less than one fifth of global max:14
×10^9
raw data, starting:12
fitted Gauss locations:14
y
x
0
1
2
3
4
5
6
7
8
9
10
0
5
10
15
20
25

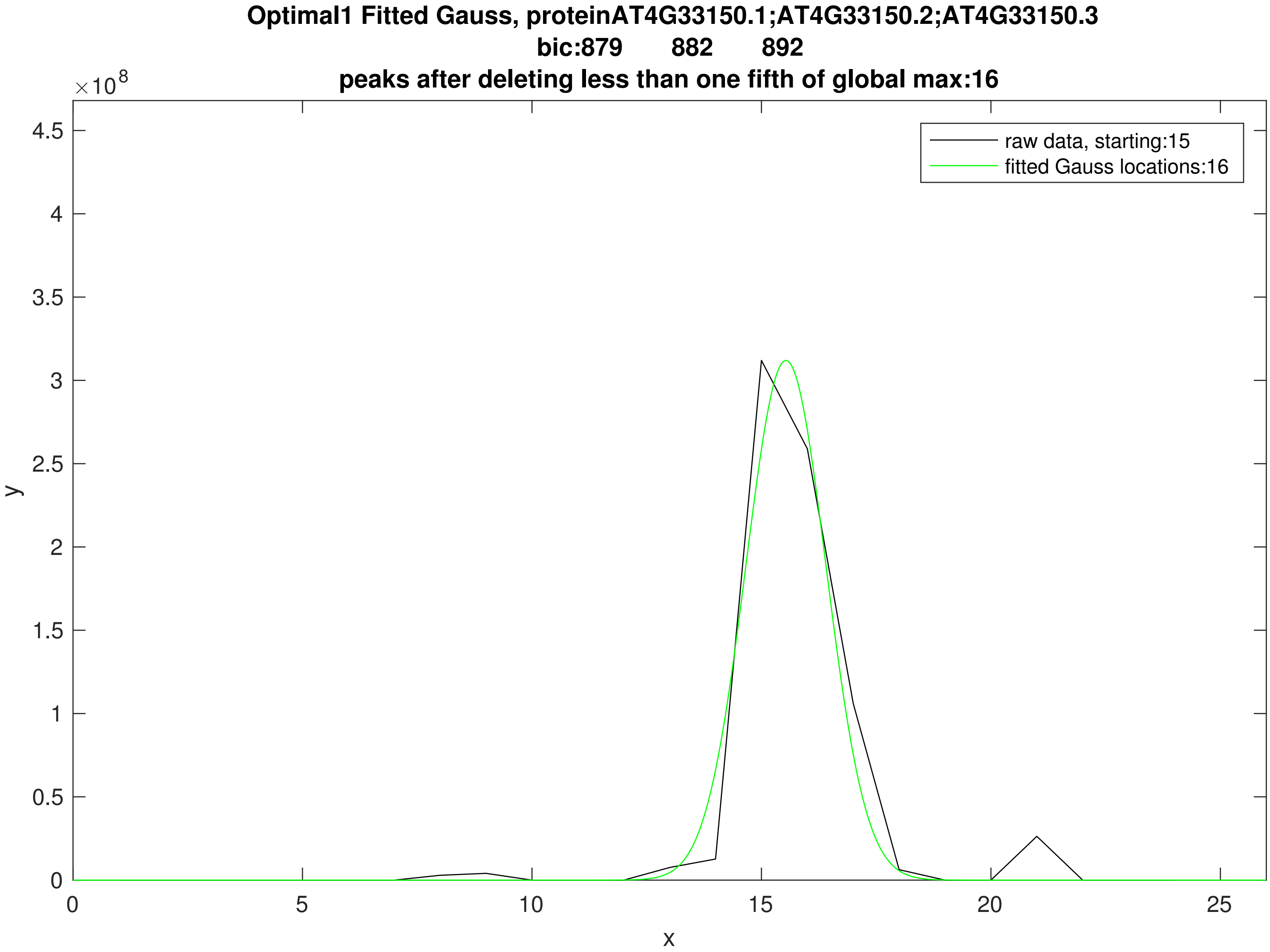
Optimal1 Fitted Gauss, proteinAT4G33150.1;AT4G33150.2;AT4G33150.3
bic:879 882 892
peaks after deleting less than one fifth of global max:16
raw data, starting:15
fitted Gauss locations:16
y
x
×10^8

Optimal1 Fitted Gauss, proteinAT4G33220.1
bic:904 913
peaks after deleting less than one fifth of global max:15

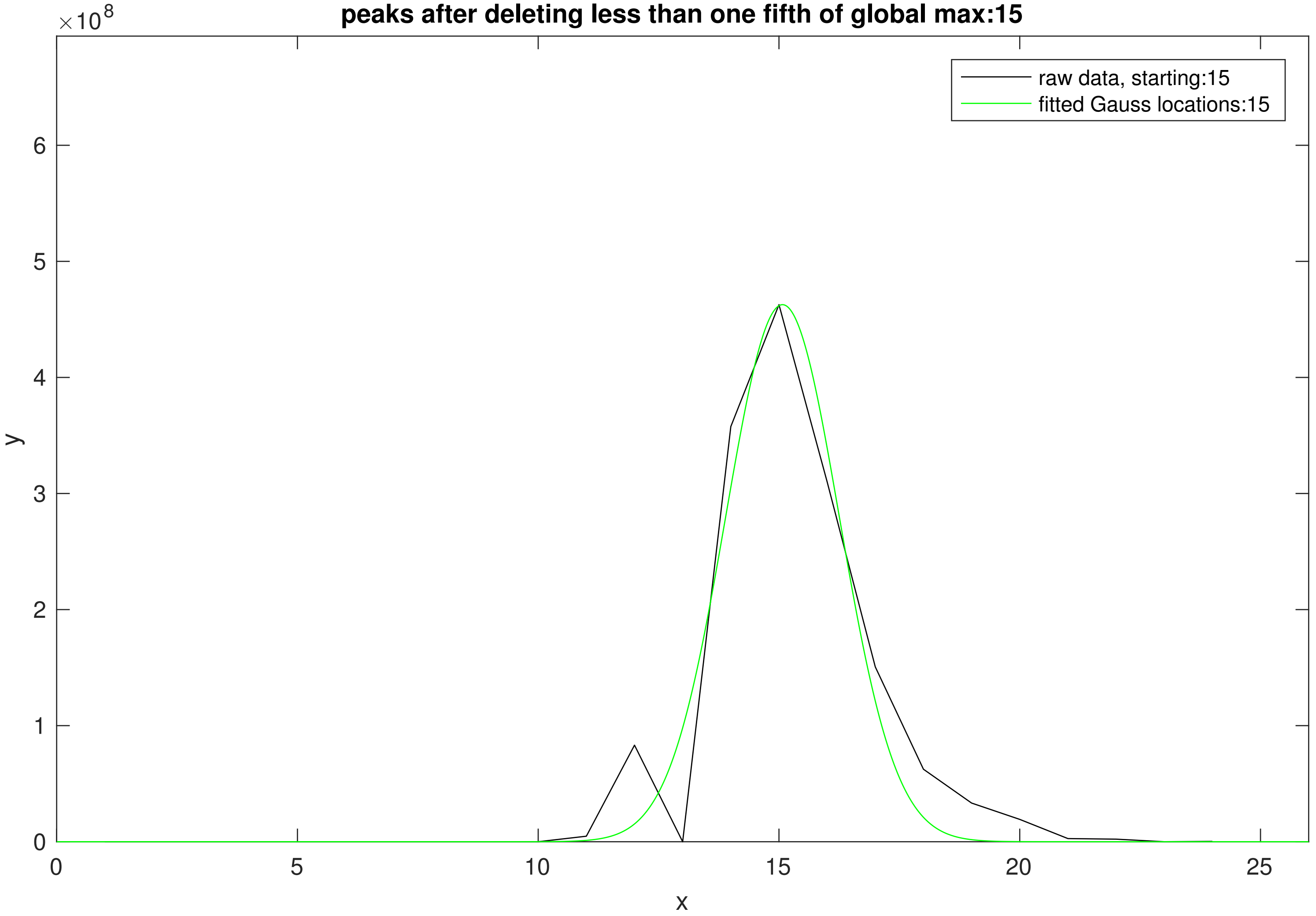

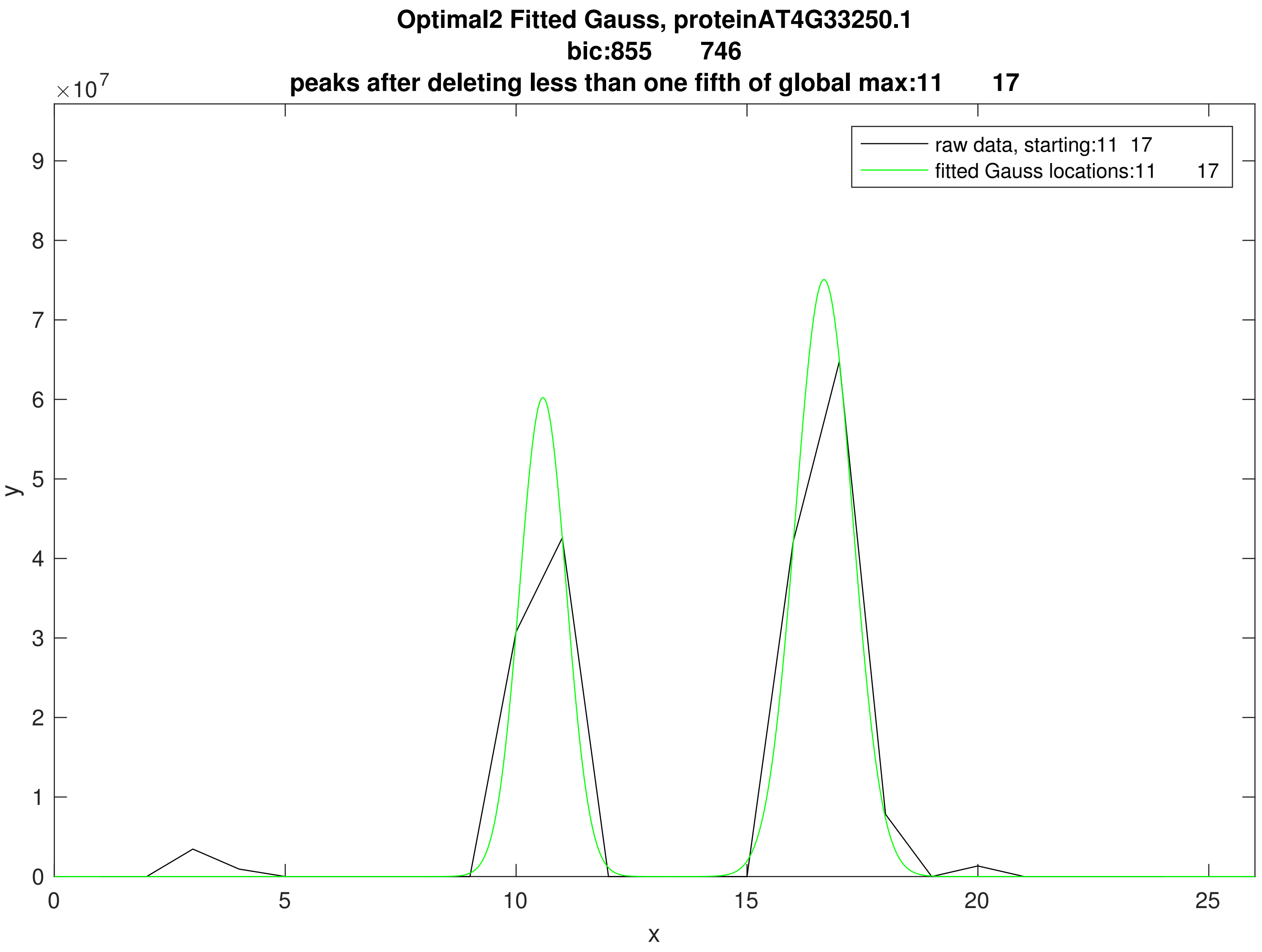
Optimal2 Fitted Gauss, proteinAT4G33250.1
bic:855 746
peaks after deleting less than one fifth of global max:11 17
×10^7
raw data, starting:11 17
fitted Gauss locations:11 17
y
x
0
1
2
3
4
5
6
7
8
9
0
5
10
15
20
25

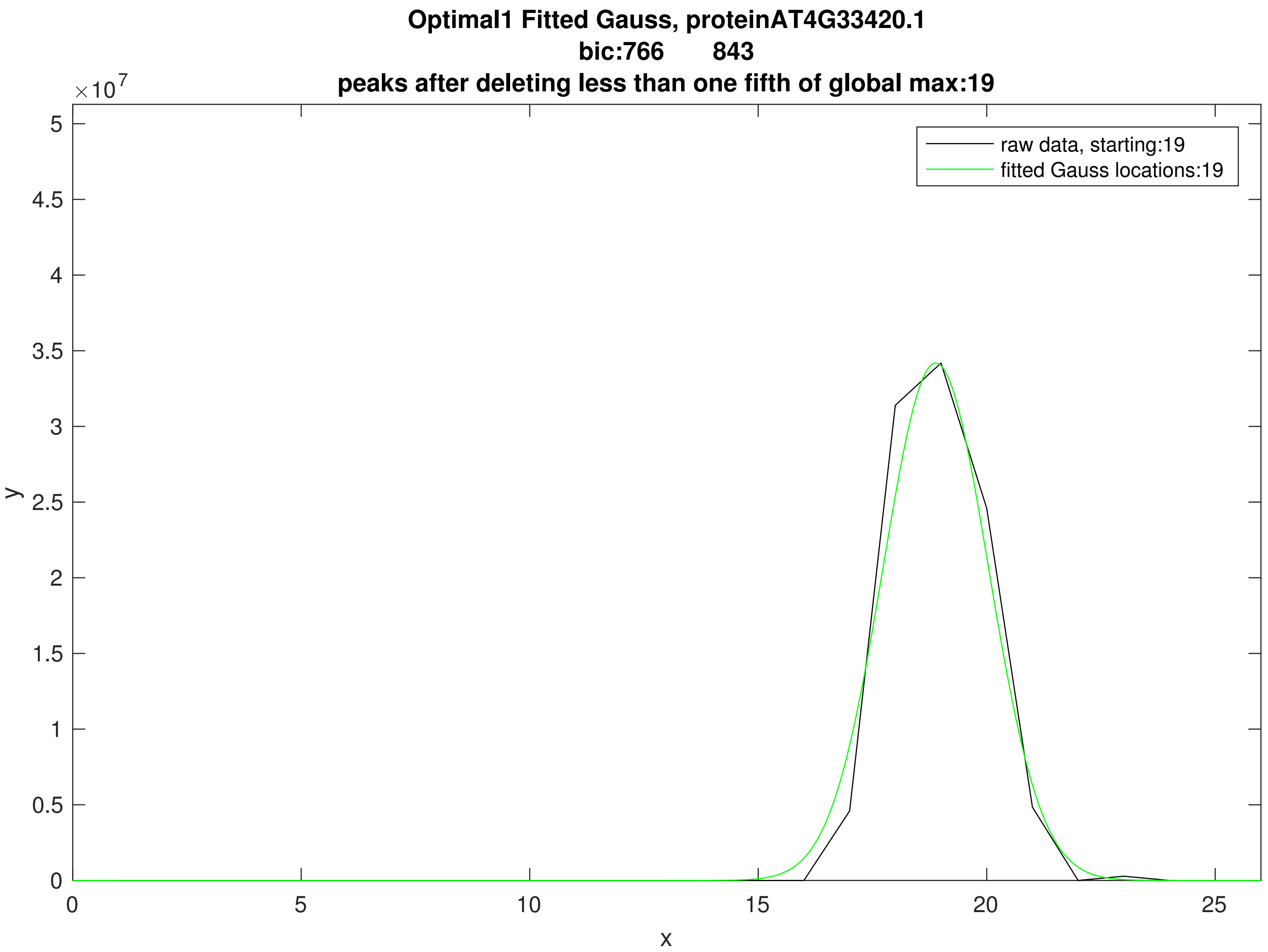

Optimal1 Fitted Gauss, proteinAT4G33420.1
bic:766 843
peaks after deleting less than one fifth of global max:19
×10^7
5
4.5
4
3.5
3
2.5
2
1.5
1
0.5
0
y
0
5
10
15
20
25
x
raw data, starting:19
fitted Gauss locations:19

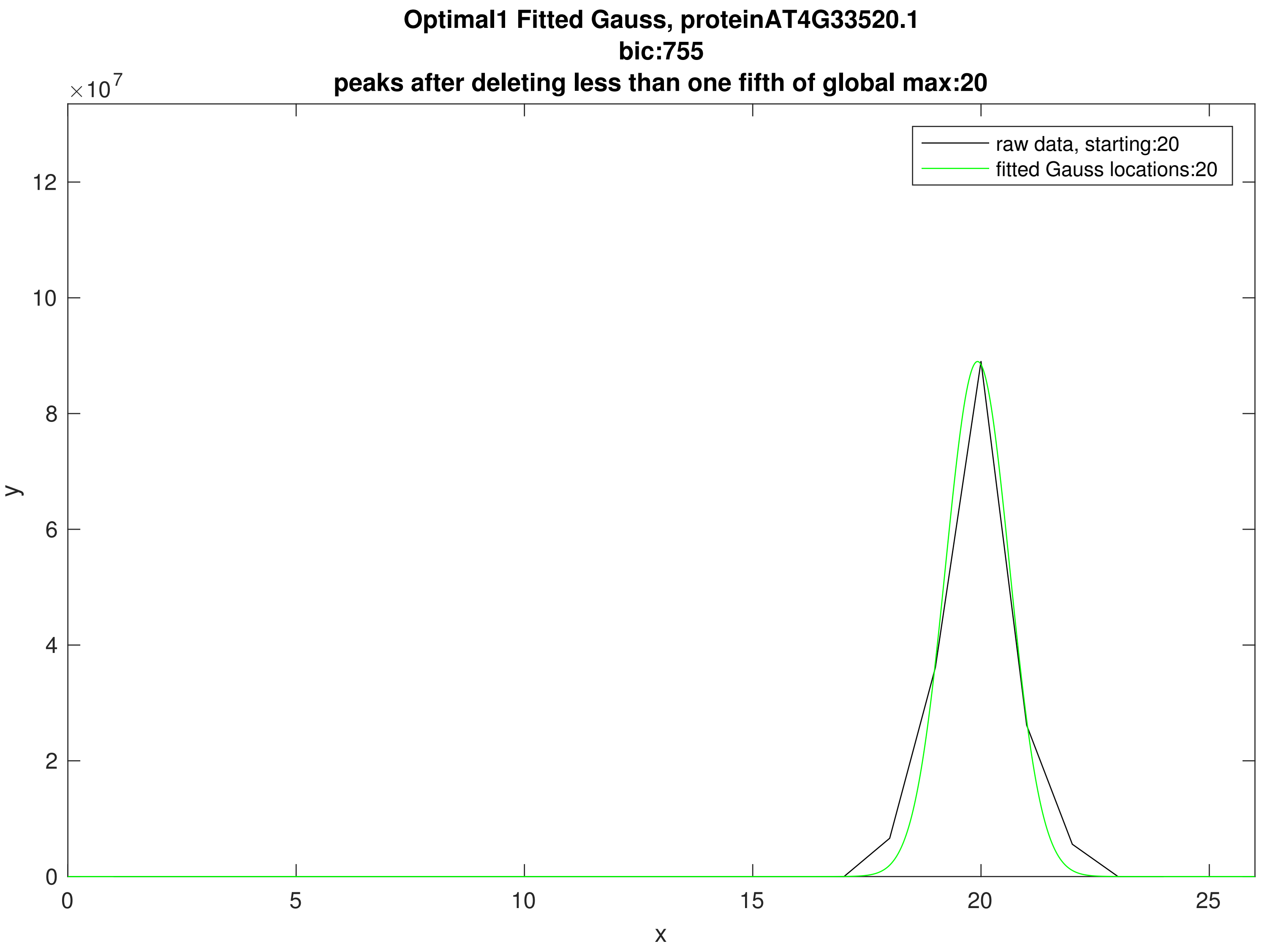

Optimal1 Fitted Gauss, proteinAT4G33520.1
bic:755
peaks after deleting less than one fifth of global max:20
×10^7
raw data, starting:20
fitted Gauss locations:20
y
x
0
2
4
6
8
10
12
0
5
10
15
20
25

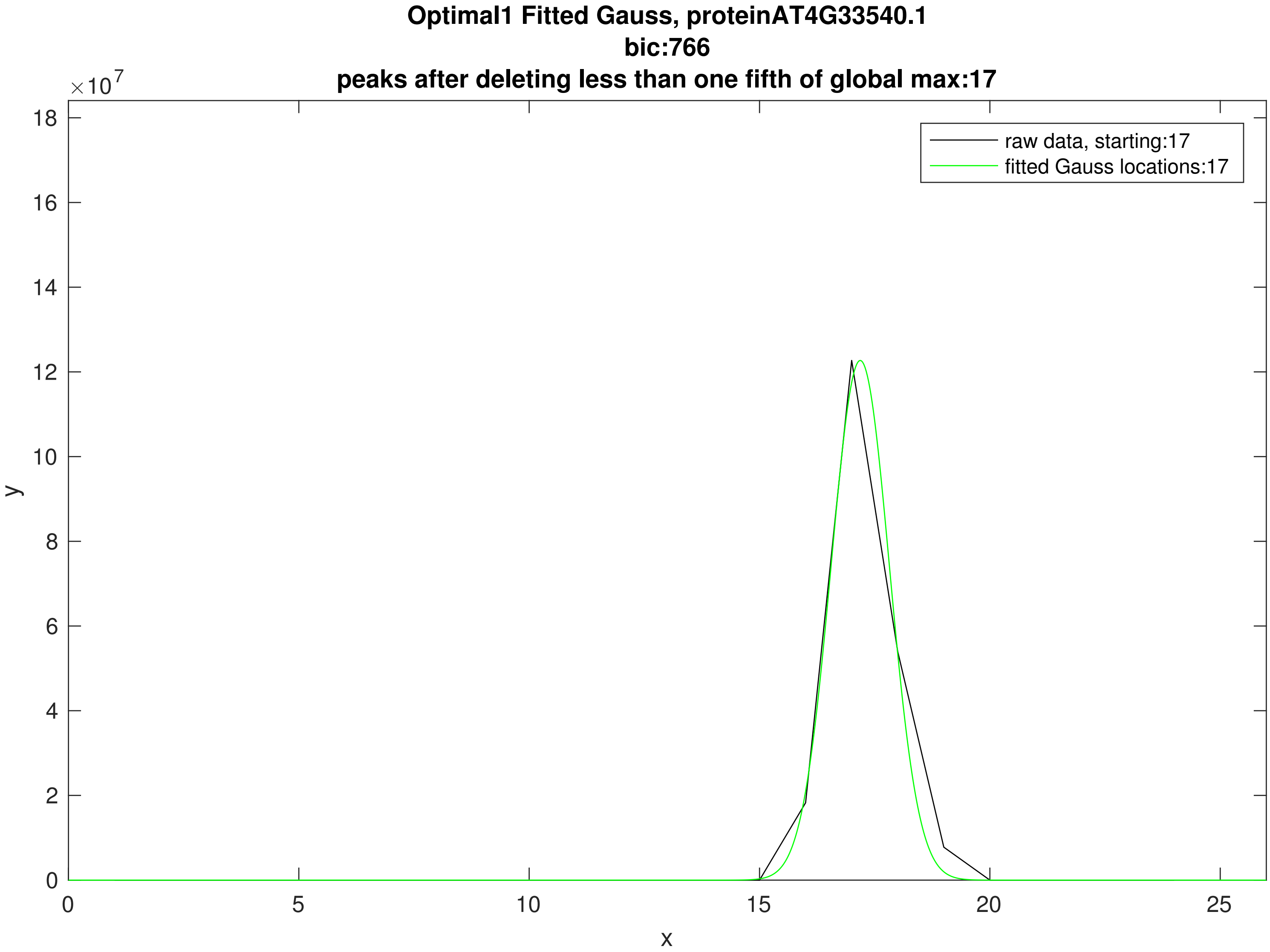

Optimal1 Fitted Gauss, proteinAT4G33540.1
bic:766
peaks after deleting less than one fifth of global max:17
×10^7
18
16
14
12
10
8
6
4
2
0
y
0
5
10
15
20
25
x
raw data, starting:17
fitted Gauss locations:17

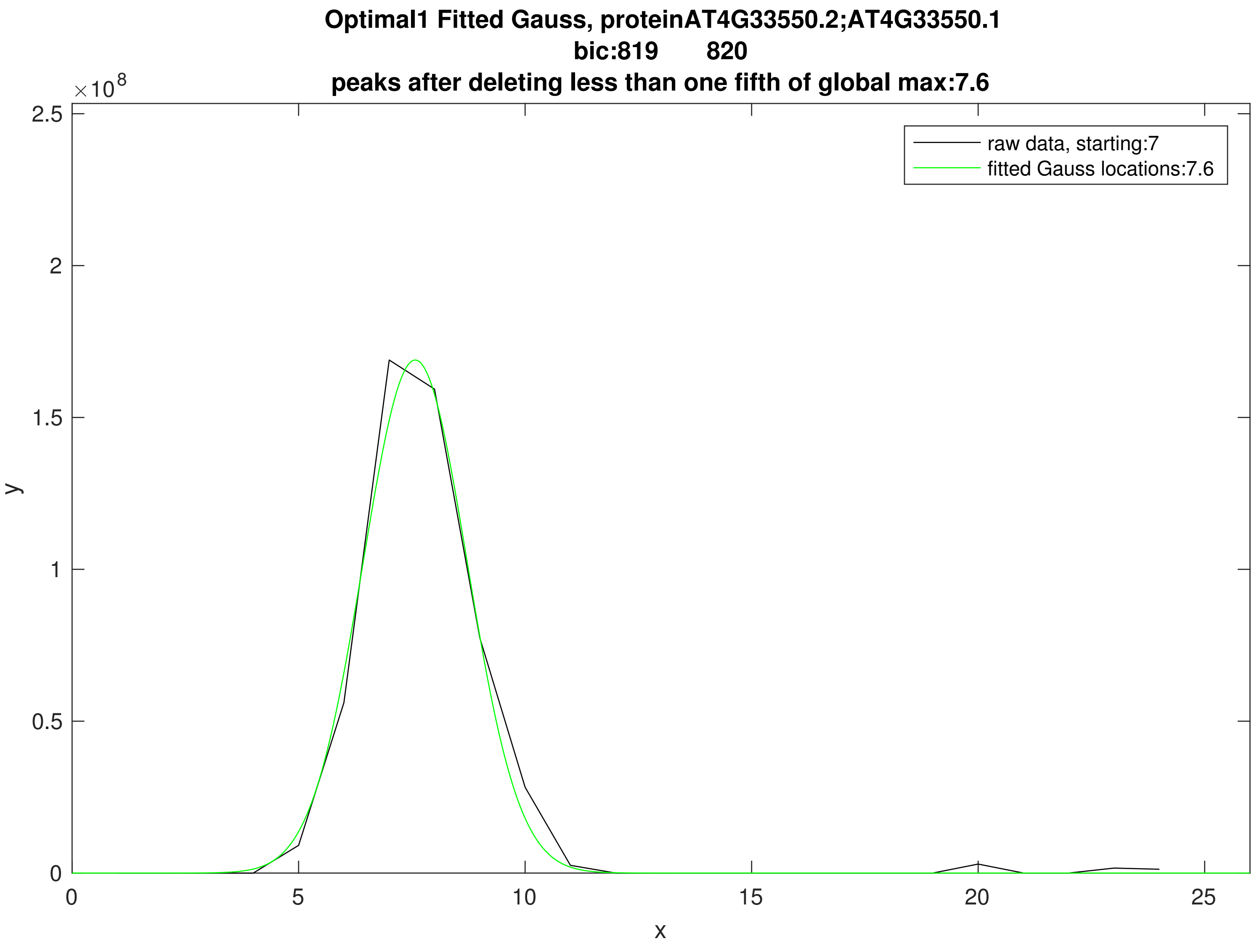
Optimal1 Fitted Gauss, proteinAT4G33550.2;AT4G33550.1
bic:819 820
peaks after deleting less than one fifth of global max:7.6
×10^8
2.5
2
1.5
1
0.5
0
y
0
5
10
15
20
25
x
raw data, starting:7
fitted Gauss locations:7.6

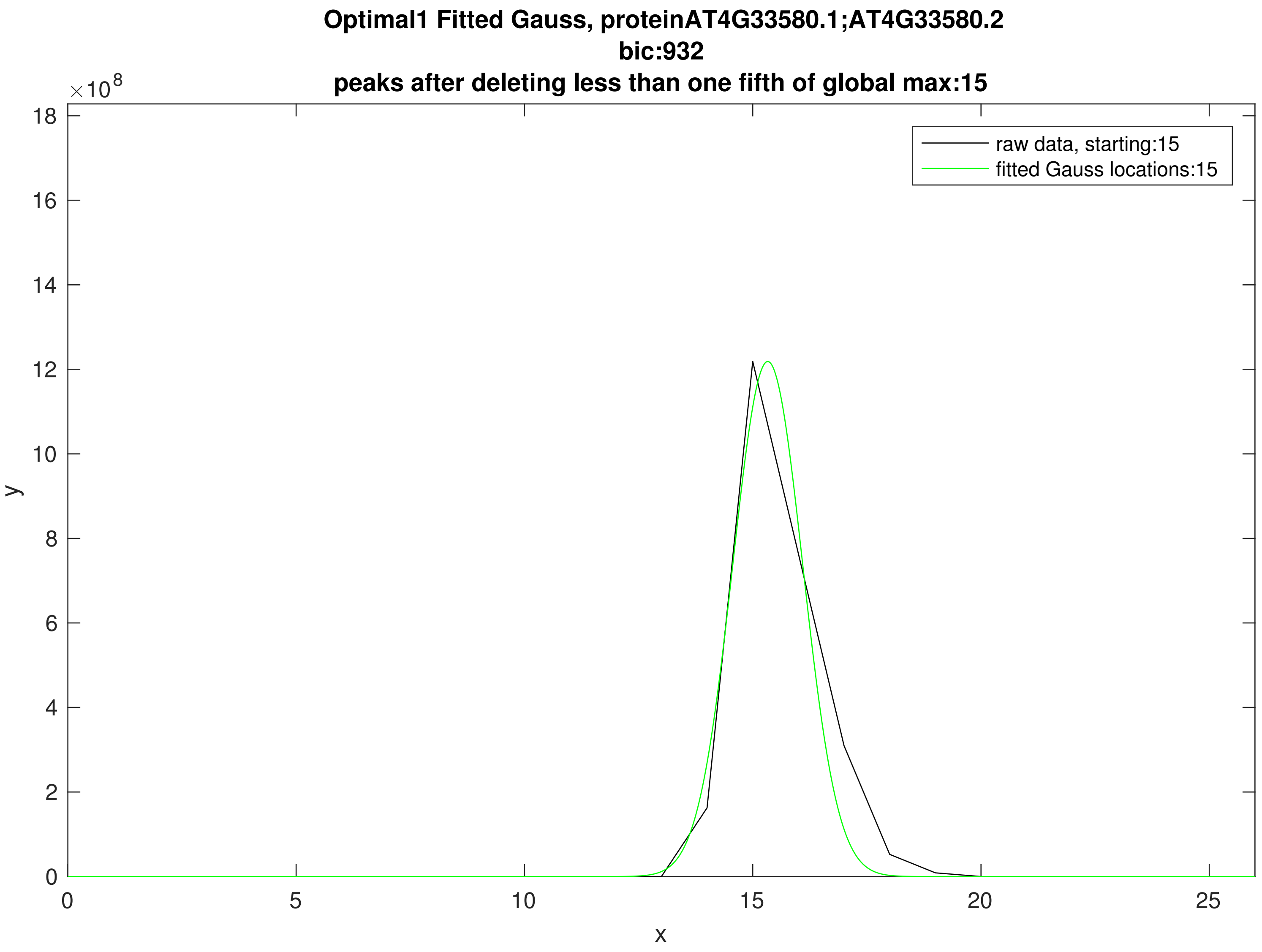
Optimal1 Fitted Gauss, proteinAT4G33580.1;AT4G33580.2
bic:932
peaks after deleting less than one fifth of global max:15
×10^8
raw data, starting:15
fitted Gauss locations:15
y
x
0
2
4
6
8
10
12
14
16
18
0
5
10
15
20
25

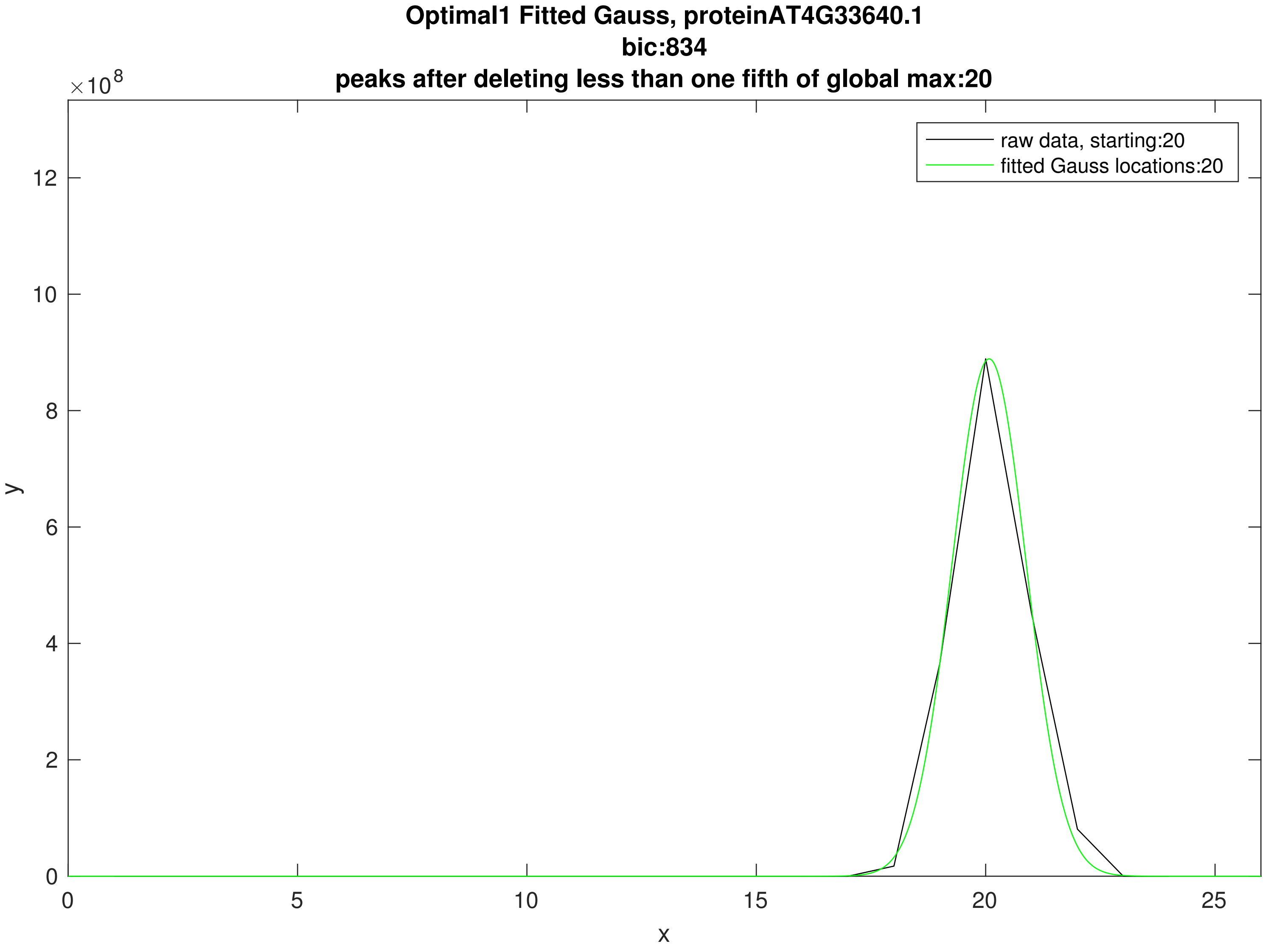
Optimal1 Fitted Gauss, proteinAT4G33640.1
bic:834
peaks after deleting less than one fifth of global max:20
×10^8
raw data, starting:20
fitted Gauss locations:20
y
x
0
2
4
6
8
10
12
0
5
10
15
20
25

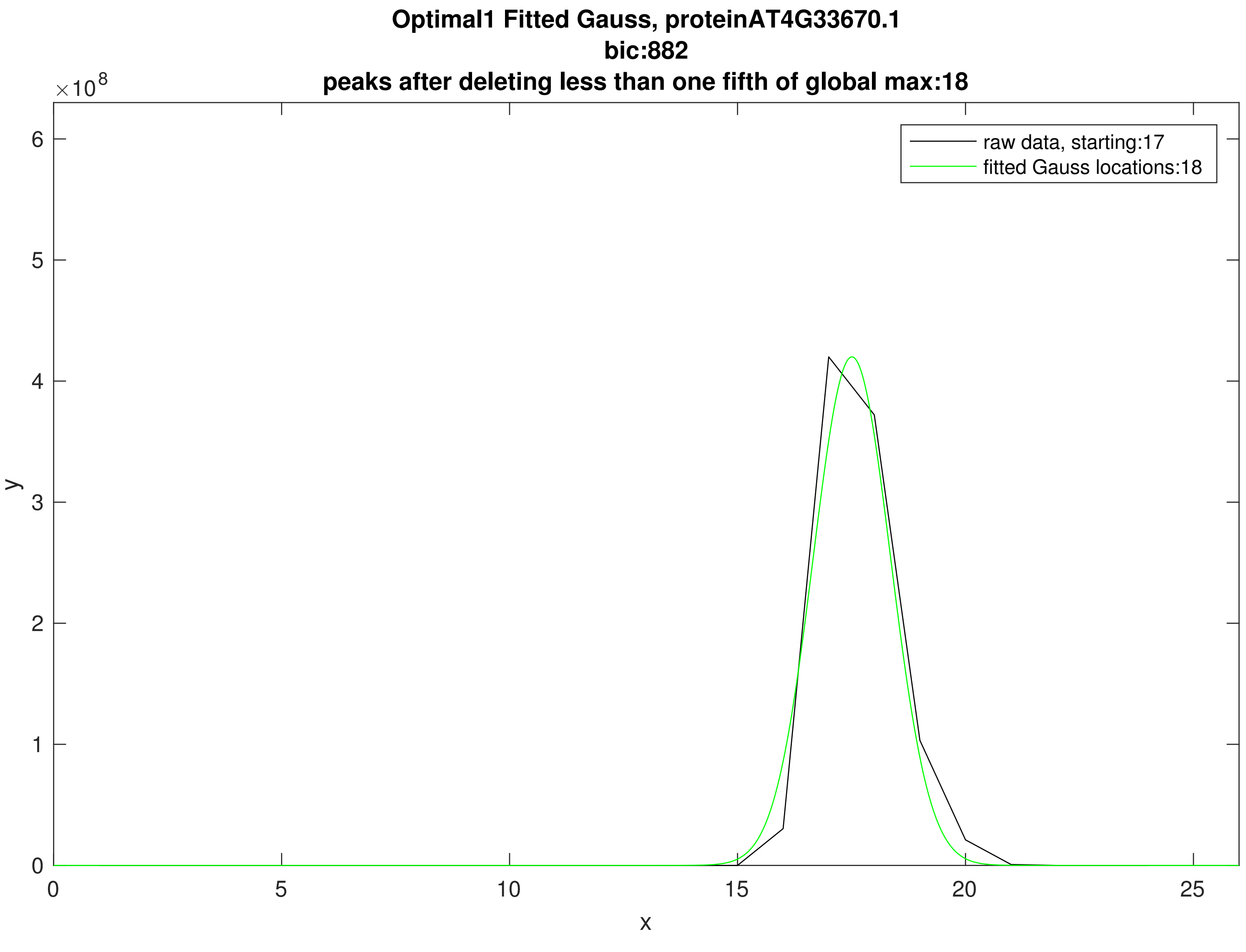
Optimal1 Fitted Gauss, proteinAT4G33670.1
bic:882
peaks after deleting less than one fifth of global max:18
×10^8
raw data, starting:17
fitted Gauss locations:18
y
x
0
1
2
3
4
5
6
0
5
10
15
20
25

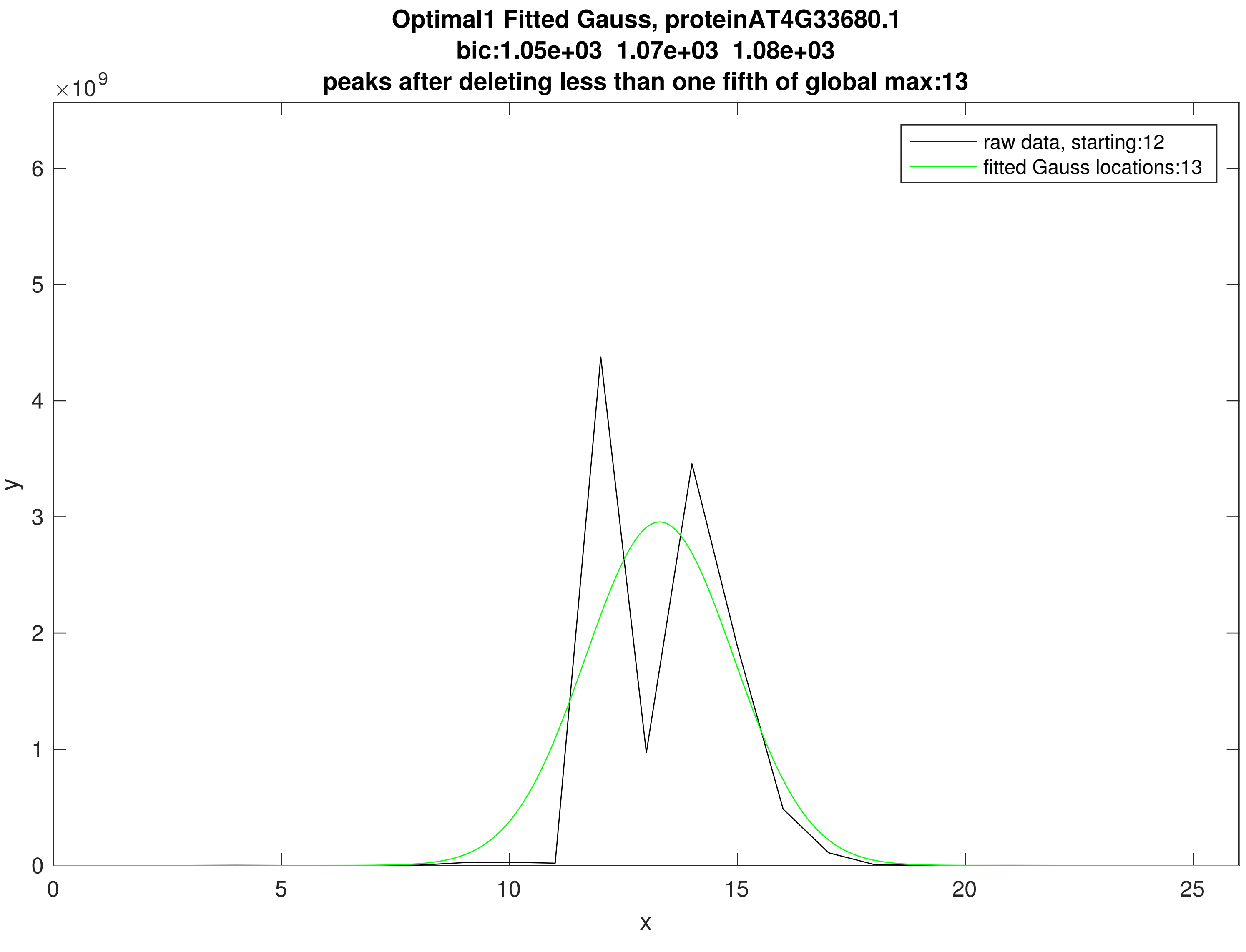
Optimal1 Fitted Gauss, proteinAT4G33680.1
bic:1.05e+03  1.07e+03  1.08e+03
peaks after deleting less than one fifth of global max:13
×10^9
raw data, starting:12
fitted Gauss locations:13
y
x
0
1
2
3
4
5
6
0
5
10
15
20
25

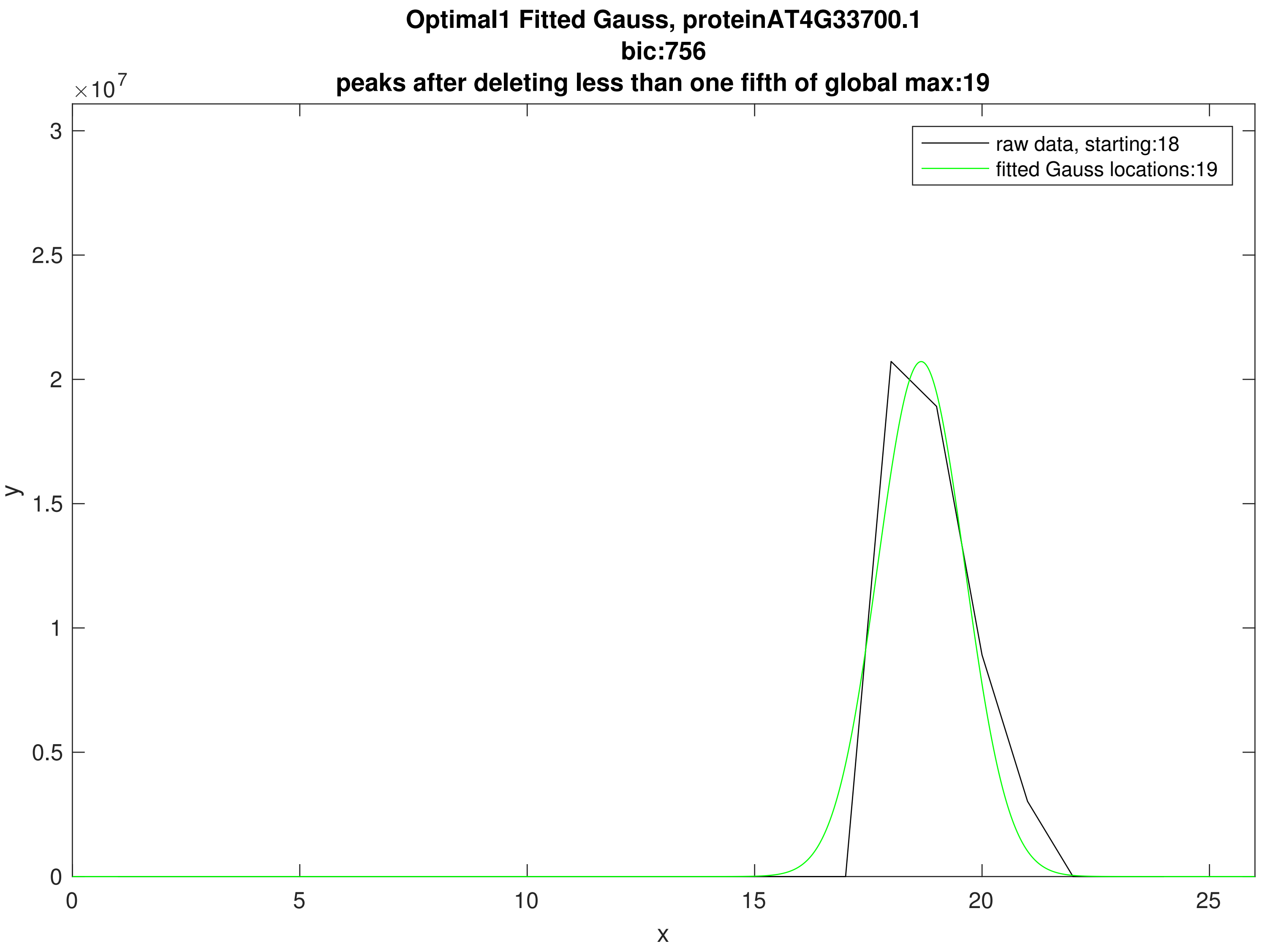

Optimal1 Fitted Gauss, proteinAT4G33700.1
bic:756
peaks after deleting less than one fifth of global max:19
×10^7
raw data, starting:18
fitted Gauss locations:19
y
x
0
0.5
1
1.5
2
2.5
3
0
5
10
15
20
25

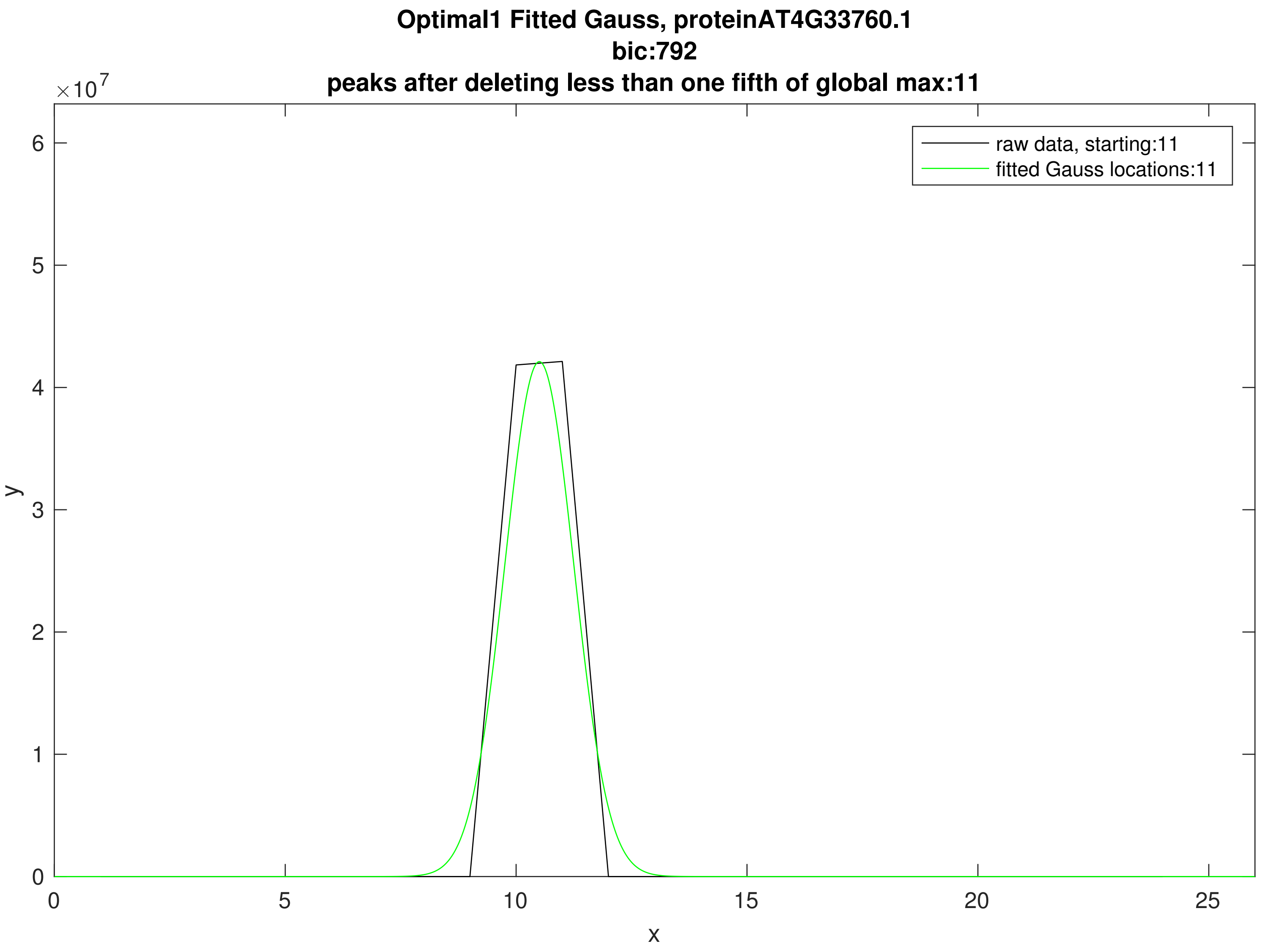

Optimal1 Fitted Gauss, proteinAT4G33760.1
bic:792
peaks after deleting less than one fifth of global max:11
×10^7
raw data, starting:11
fitted Gauss locations:11
y
x
0
1
2
3
4
5
6
0
5
10
15
20
25

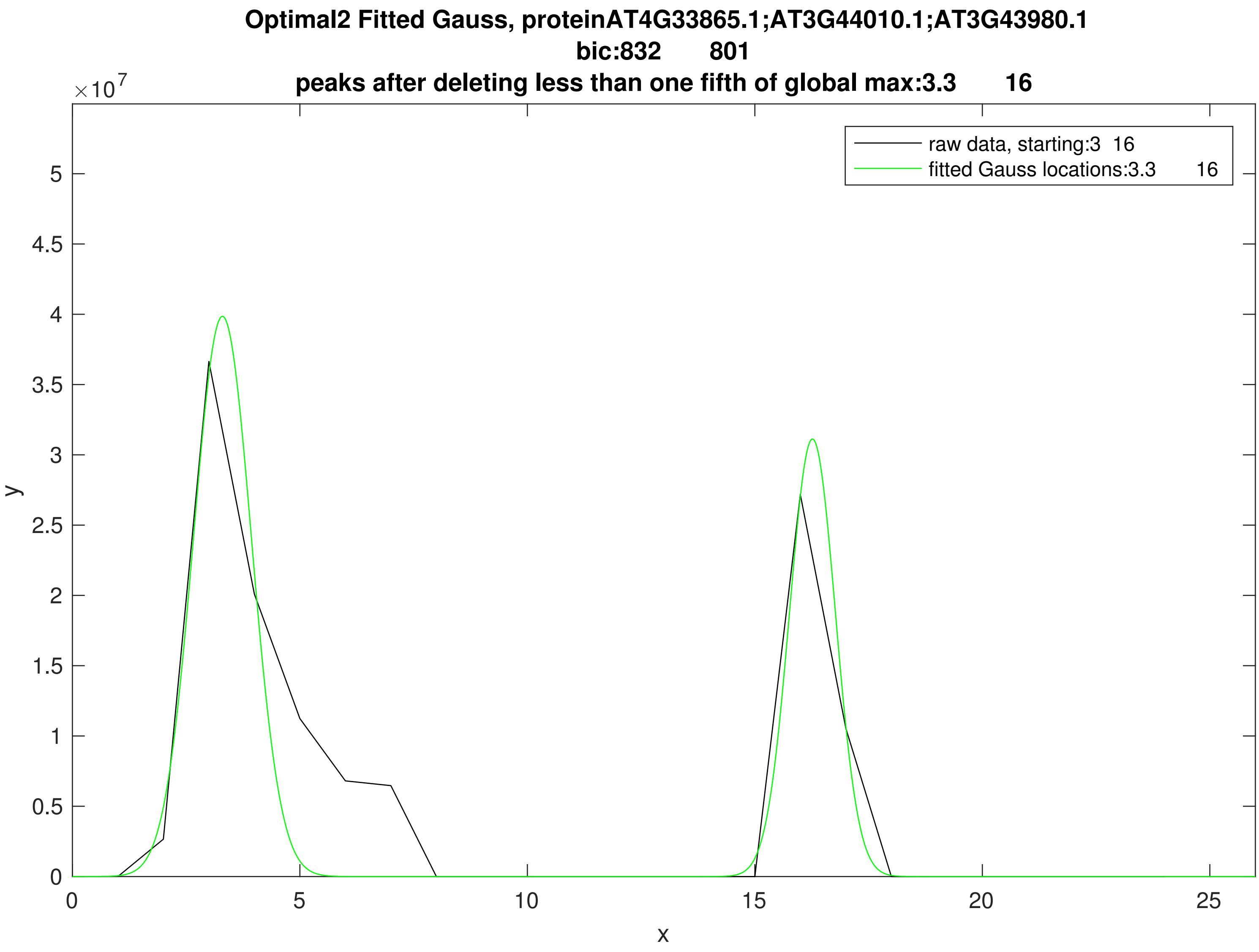
Optimal2 Fitted Gauss, proteinAT4G33865.1;AT3G44010.1;AT3G43980.1
bic:832 801
peaks after deleting less than one fifth of global max:3.3 16
×10^7
raw data, starting:3 16
fitted Gauss locations:3.3 16
5
4.5
4
3.5
3
2.5
2
1.5
1
0.5
0
y
0
5
10
15
20
25
x

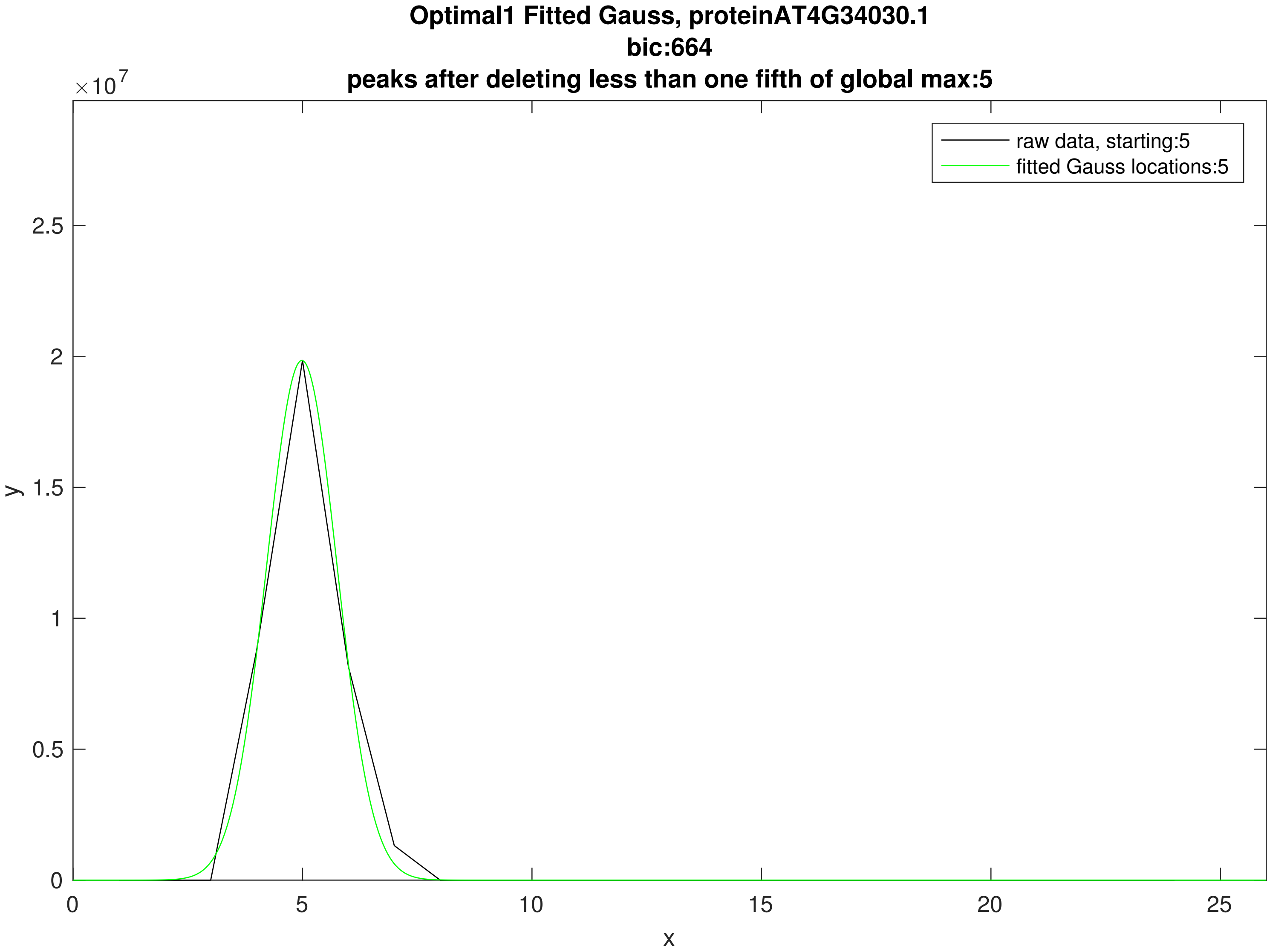

Optimal1 Fitted Gauss, proteinAT4G34030.1
bic:664
peaks after deleting less than one fifth of global max:5
×10^7
raw data, starting:5
fitted Gauss locations:5
y
x
0
0.5
1
1.5
2
2.5
0
5
10
15
20
25

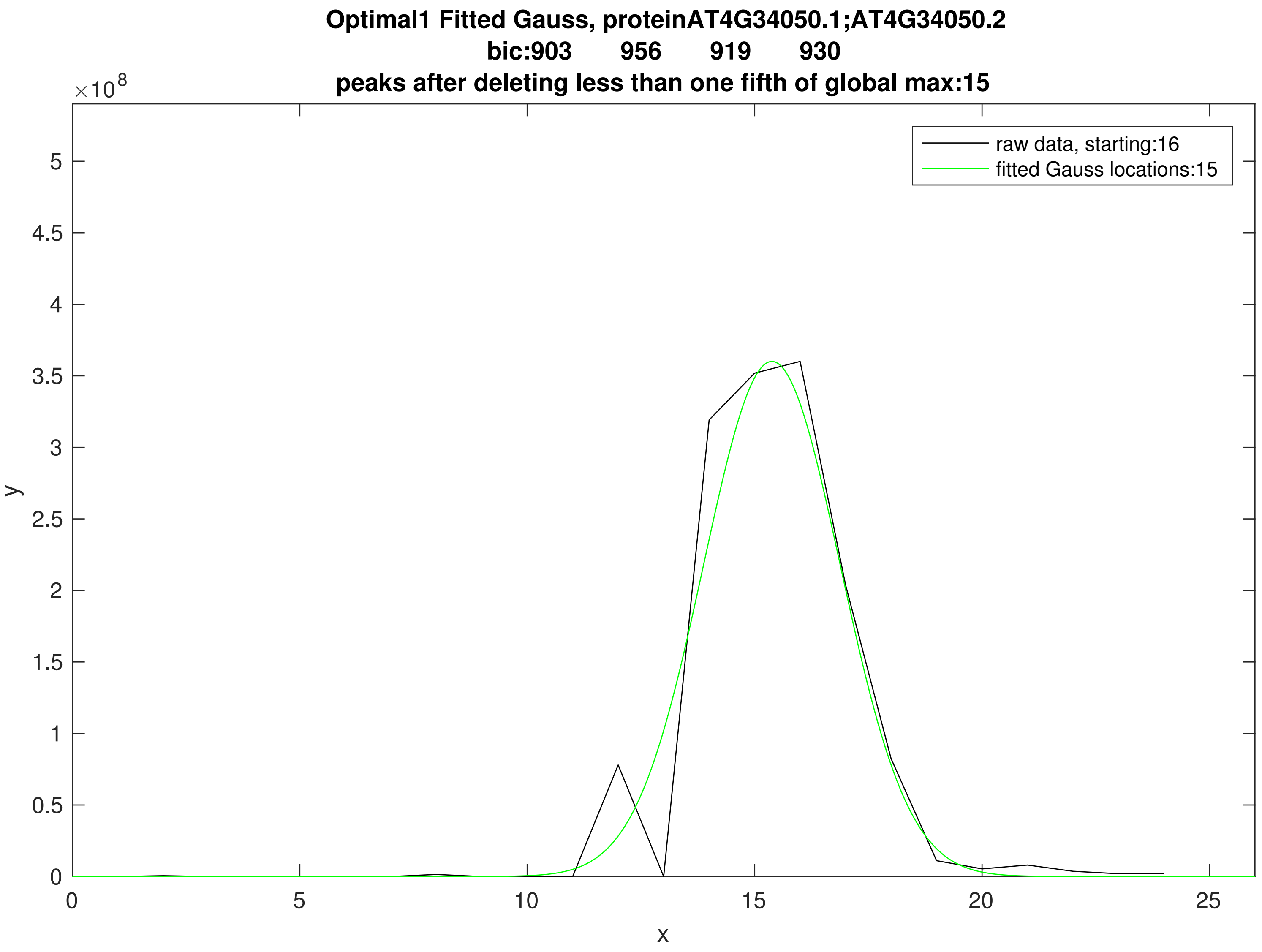
Optimal1 Fitted Gauss, proteinAT4G34050.1;AT4G34050.2
bic:903 956 919 930
peaks after deleting less than one fifth of global max:15
×10^8
raw data, starting:16
fitted Gauss locations:15
y
x
0
0.5
1
1.5
2
2.5
3
3.5
4
4.5
5
0
5
10
15
20
25

**Optimal1 Fitted Gauss, proteinAT4G34110.1**
**bic:868 872 874**
**peaks after deleting less than one fifth of global max:13**

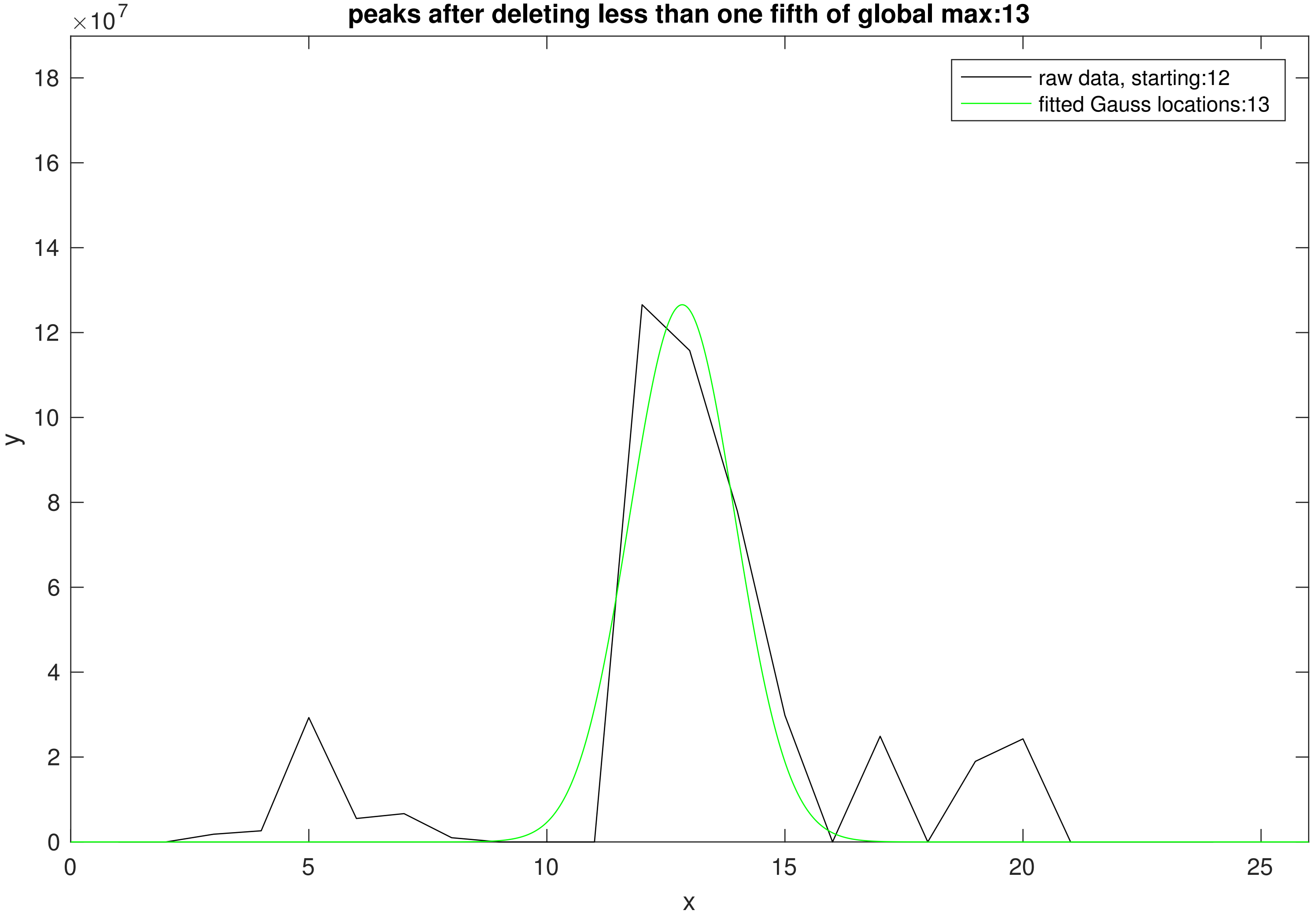

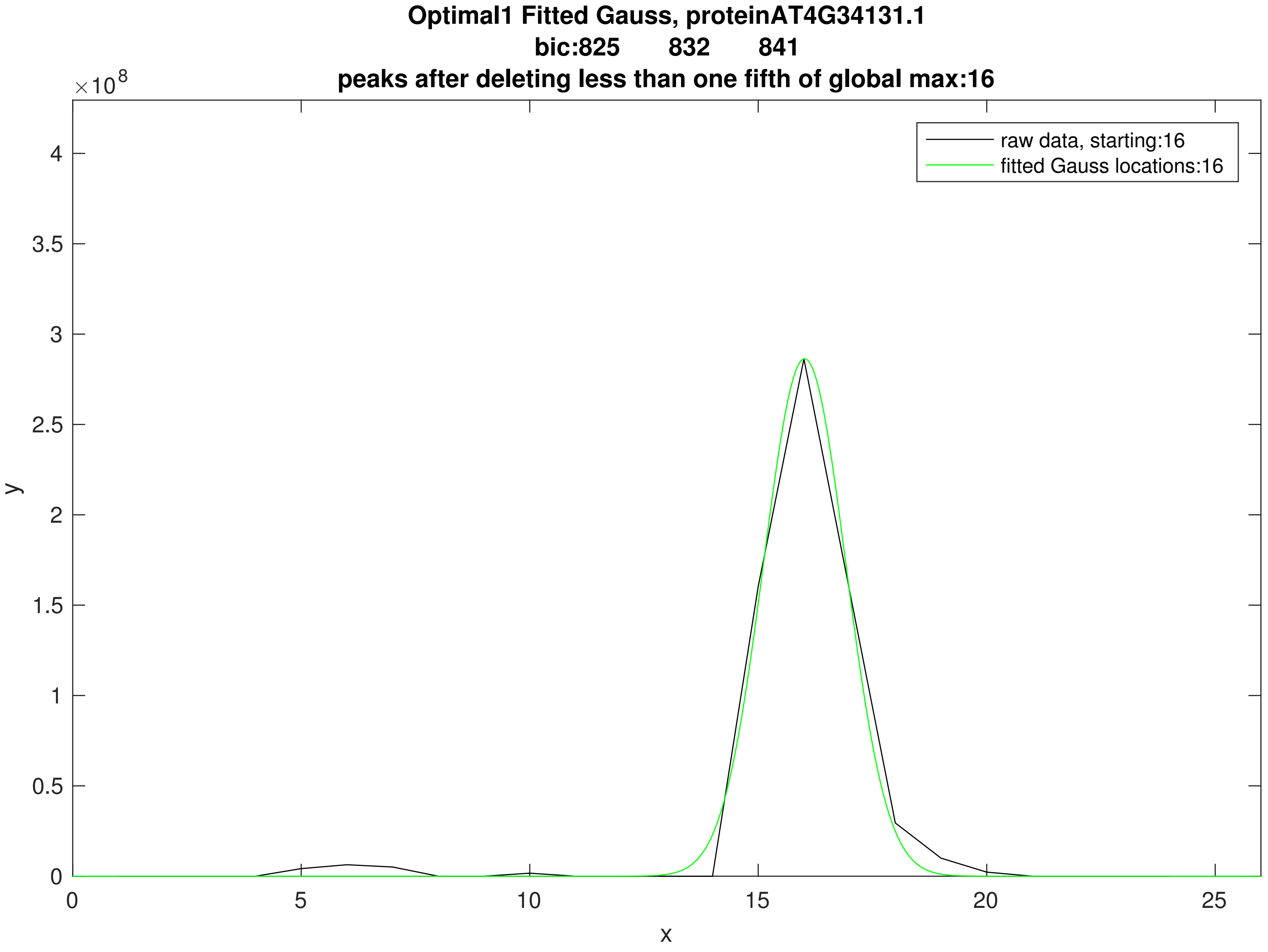

Optimal1 Fitted Gauss, proteinAT4G34131.1
bic:825 832 841
peaks after deleting less than one fifth of global max:16
×10^8
raw data, starting:16
fitted Gauss locations:16
y
x
0
0.5
1
1.5
2
2.5
3
3.5
4
5
10
15
20
25

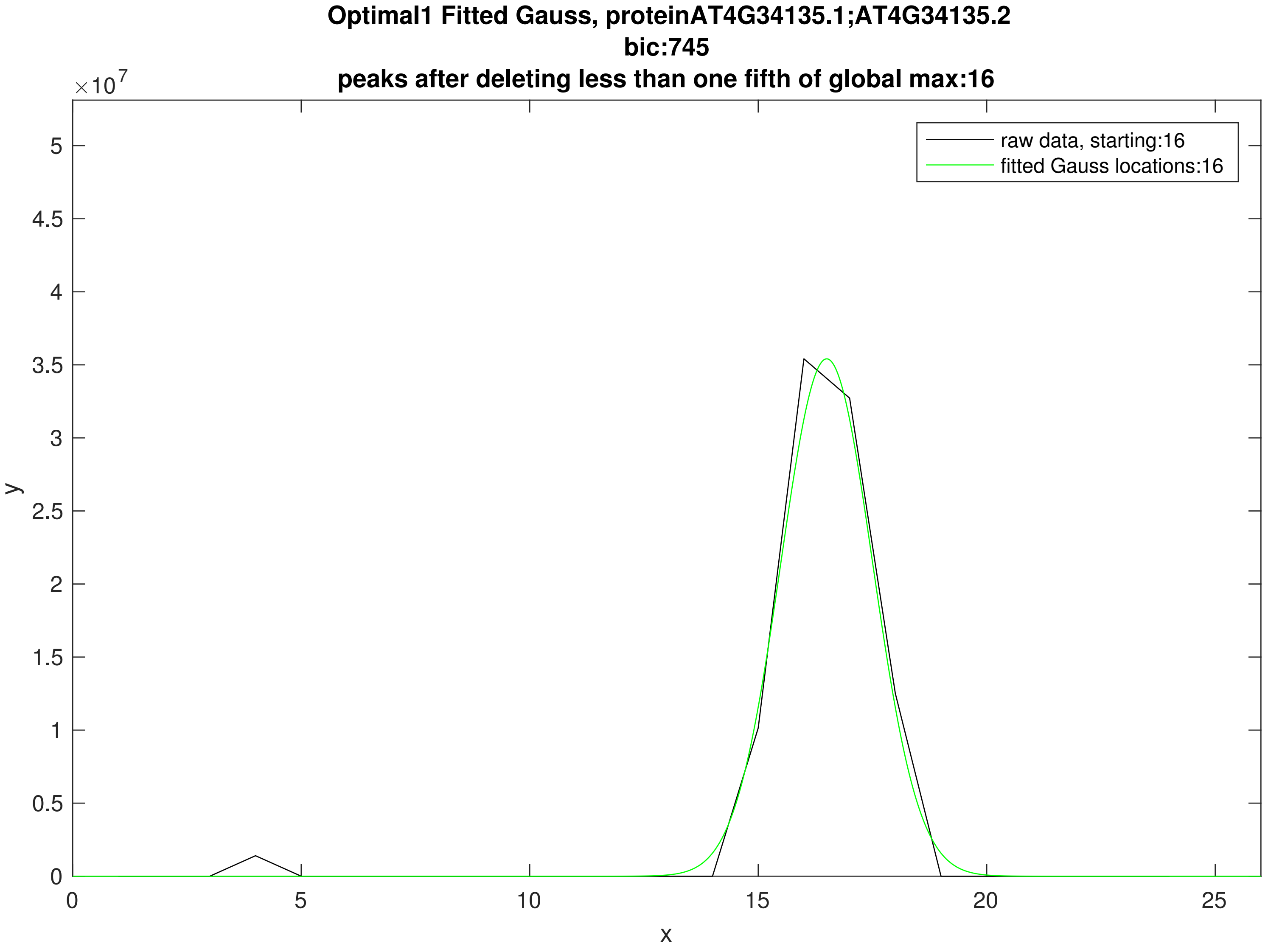

Optimal1 Fitted Gauss, proteinAT4G34135.1;AT4G34135.2
bic:745
peaks after deleting less than one fifth of global max:16
×10^7
raw data, starting:16
fitted Gauss locations:16
y
x
0
0.5
1
1.5
2
2.5
3
3.5
4
4.5
5
0
5
10
15
20
25

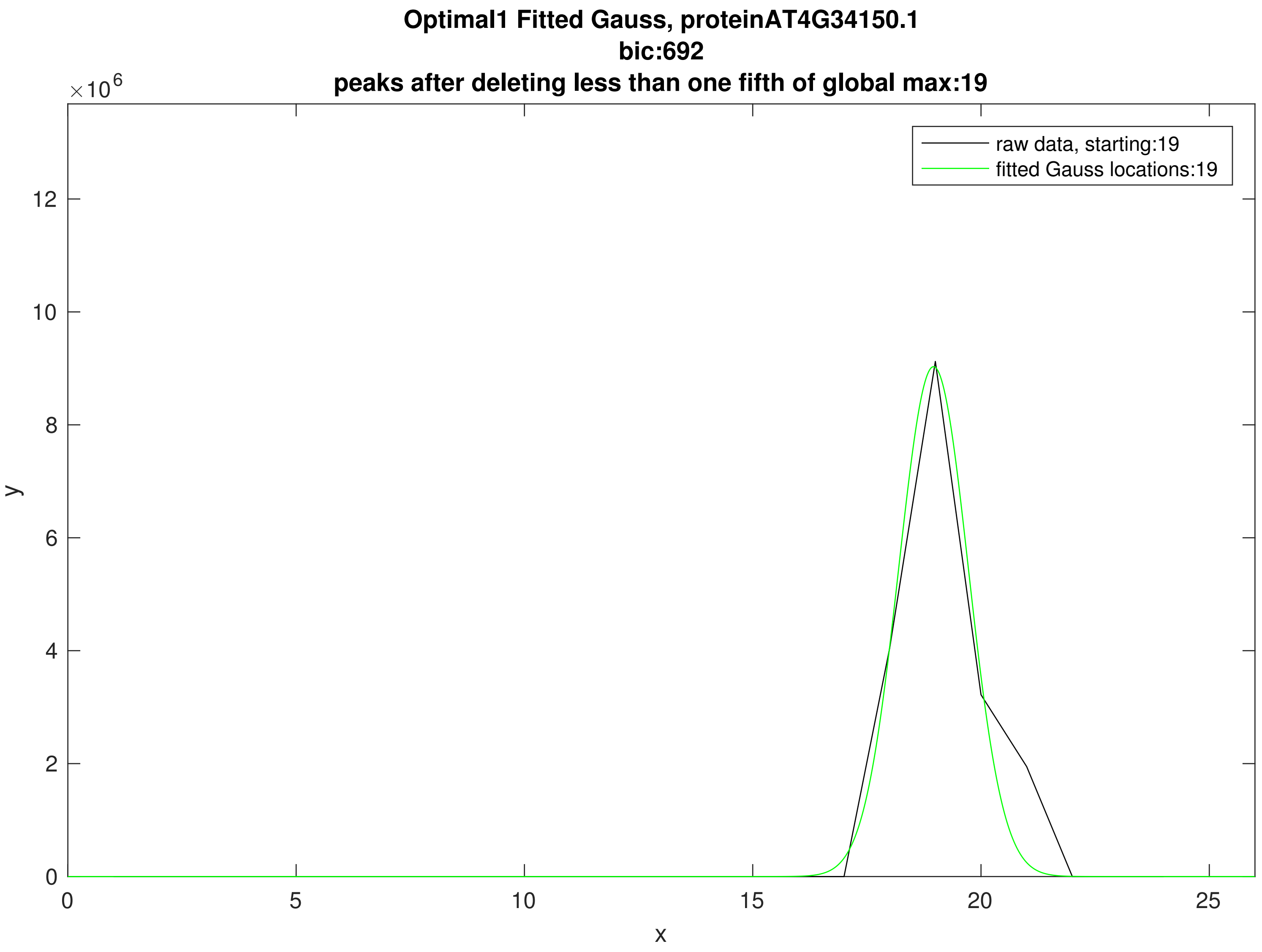

Optimal1 Fitted Gauss, proteinAT4G34150.1
bic:692
peaks after deleting less than one fifth of global max:19
×10^6
raw data, starting:19
fitted Gauss locations:19
12
10
8
6
4
2
0
y
0
5
10
15
20
25
x

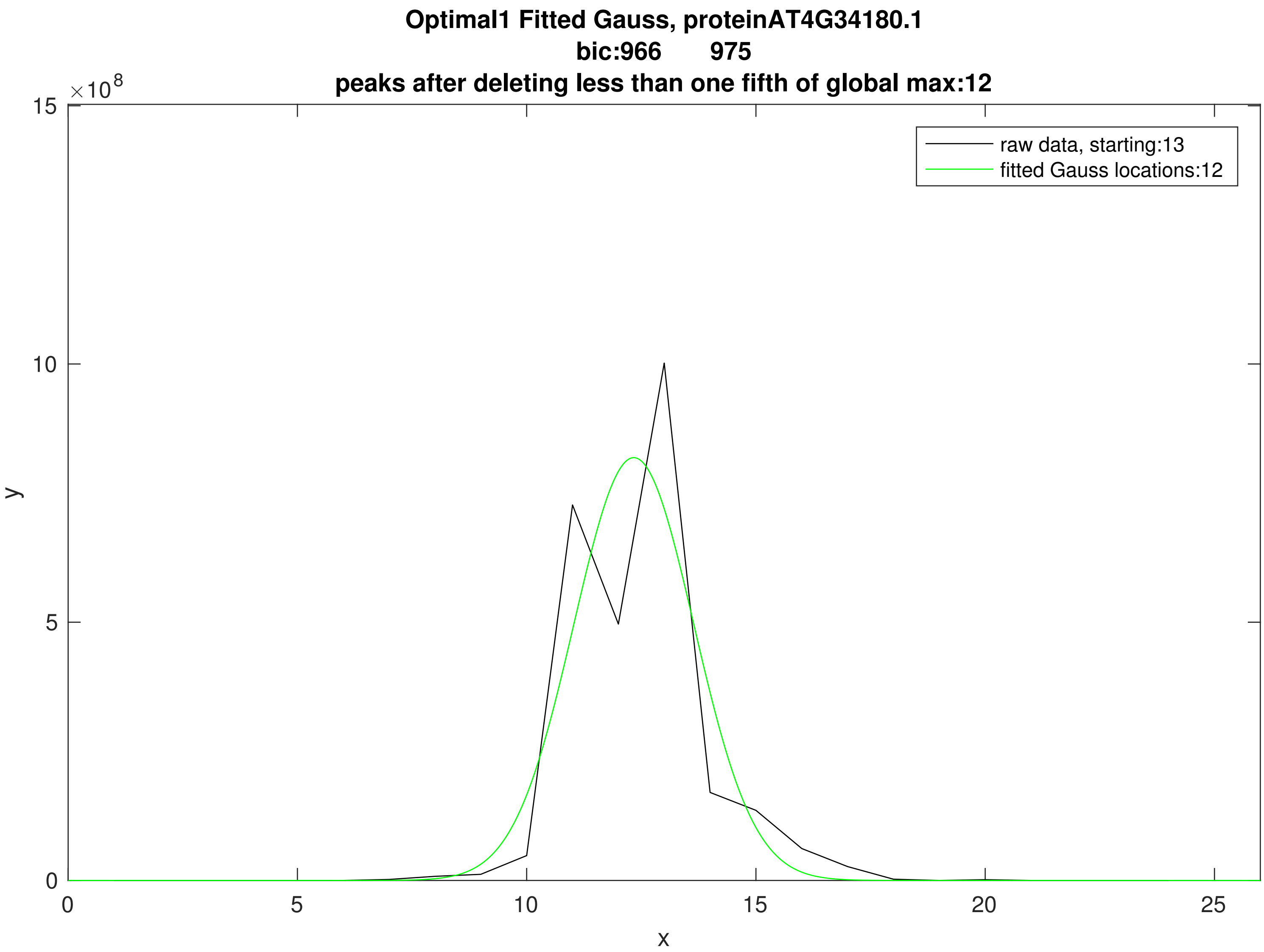
Optimal1 Fitted Gauss, proteinAT4G34180.1
bic:966 975
peaks after deleting less than one fifth of global max:12
×10^8
15
10
5
0
y
0
5
10
15
20
25
x
raw data, starting:13
fitted Gauss locations:12

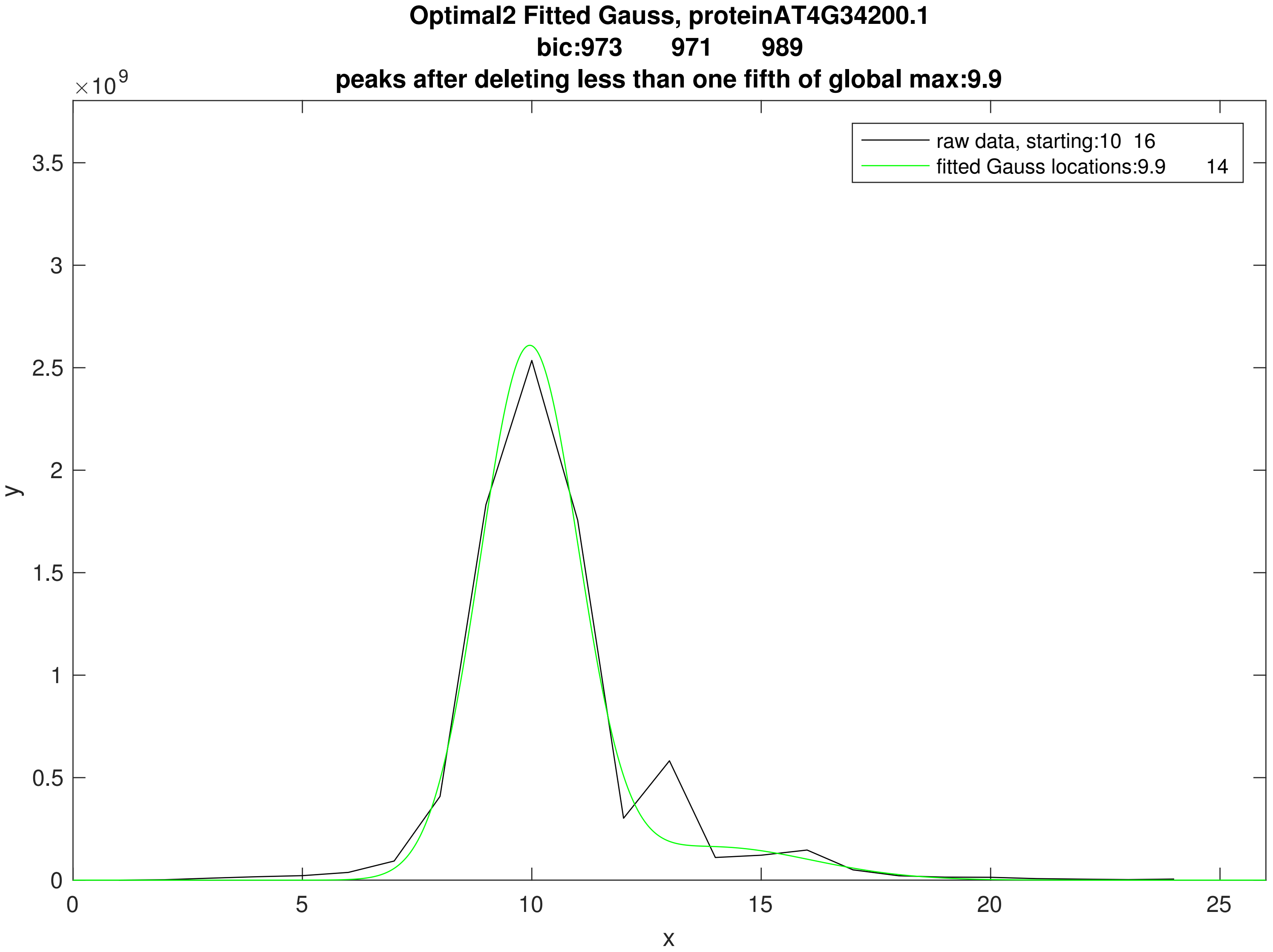
Optimal2 Fitted Gauss, proteinAT4G34200.1
bic:973 971 989
peaks after deleting less than one fifth of global max:9.9
×10^9
3.5
3
2.5
2
1.5
1
0.5
0
y
0
5
10
15
20
25
x
raw data, starting:10 16
fitted Gauss locations:9.9 14

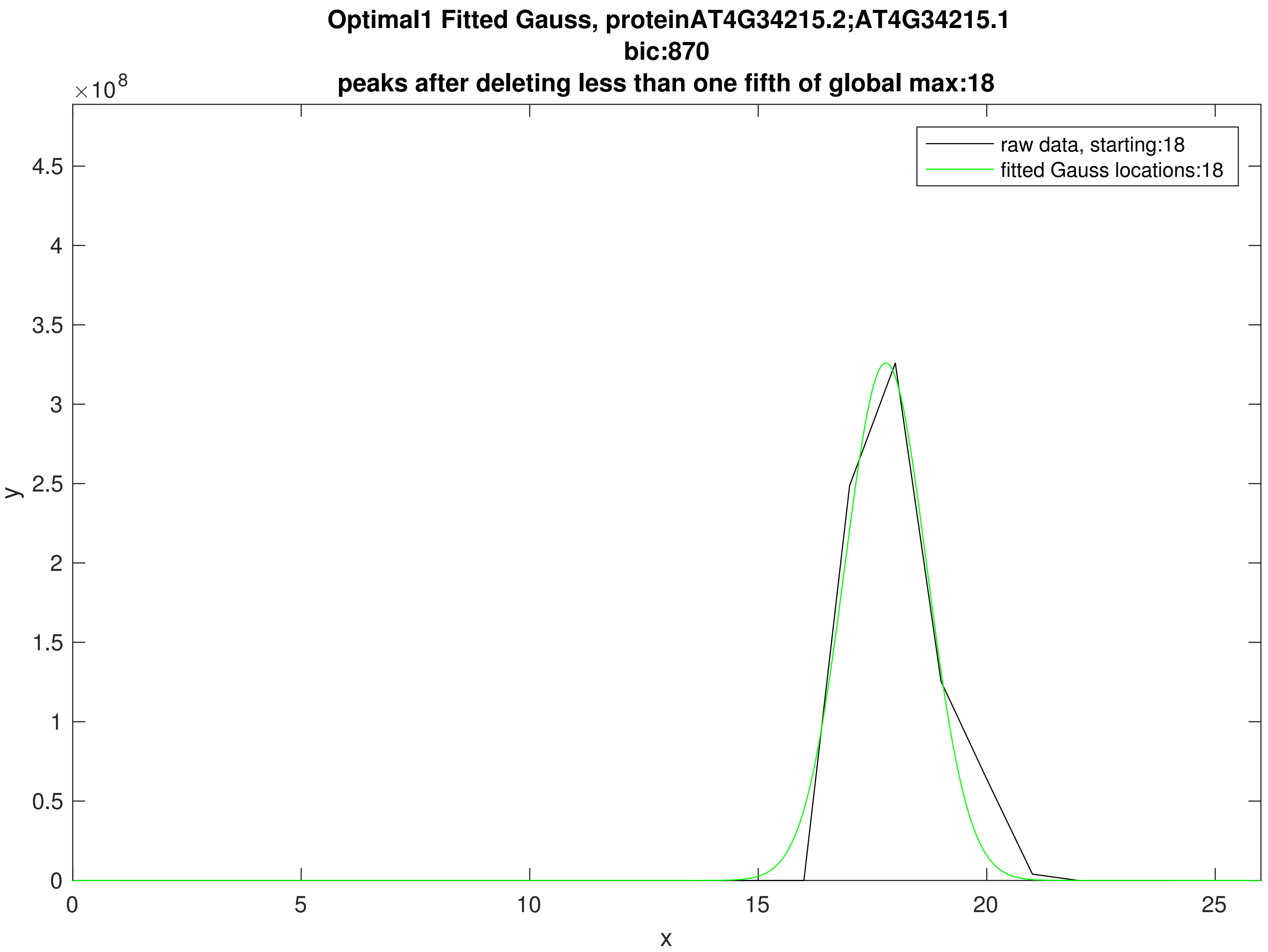

Optimal1 Fitted Gauss, proteinAT4G34215.2;AT4G34215.1
bic:870
peaks after deleting less than one fifth of global max:18
×10^8
raw data, starting:18
fitted Gauss locations:18
y
x
0
0.5
1
1.5
2
2.5
3
3.5
4
4.5
0
5
10
15
20
25

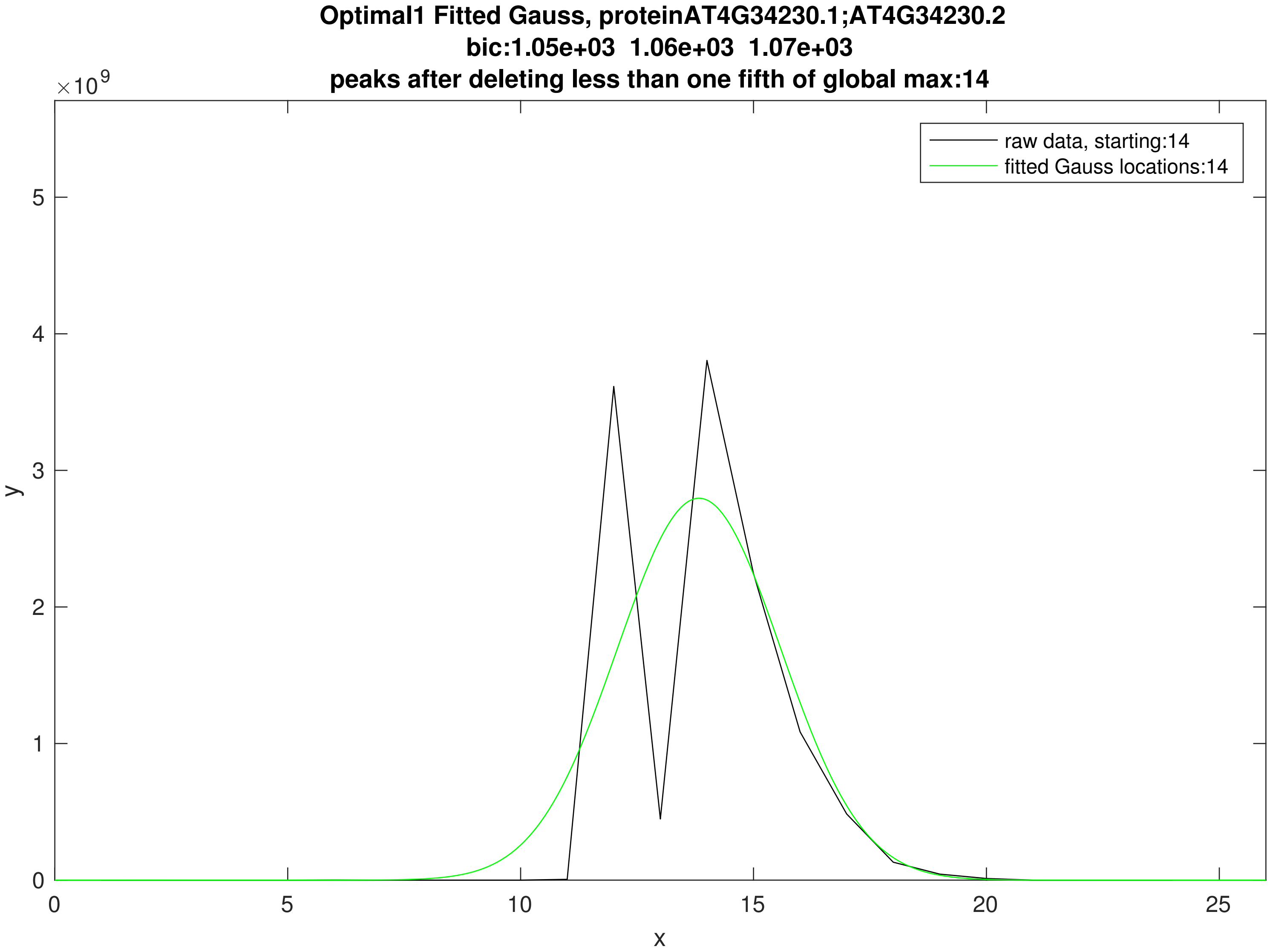
Optimal1 Fitted Gauss, proteinAT4G34230.1;AT4G34230.2
bic:1.05e+03 1.06e+03 1.07e+03
peaks after deleting less than one fifth of global max:14
×10^9
raw data, starting:14
fitted Gauss locations:14
y
x
0
1
2
3
4
5
0
5
10
15
20
25

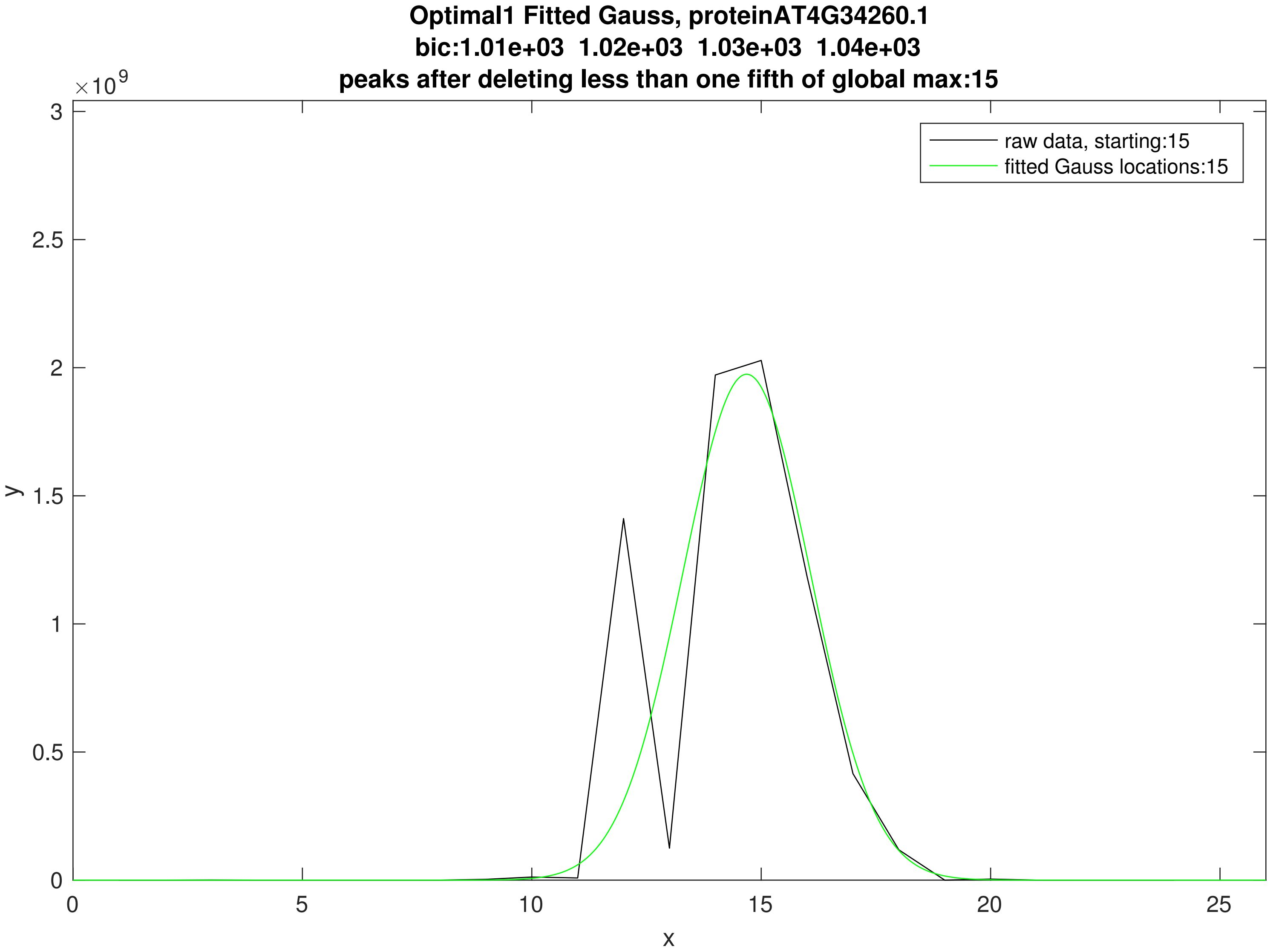
Optimal1 Fitted Gauss, proteinAT4G34260.1
bic:1.01e+03  1.02e+03  1.03e+03  1.04e+03
peaks after deleting less than one fifth of global max:15
×10^9
3
2.5
2
1.5
1
0.5
0
y
0
5
10
15
20
25
x
raw data, starting:15
fitted Gauss locations:15

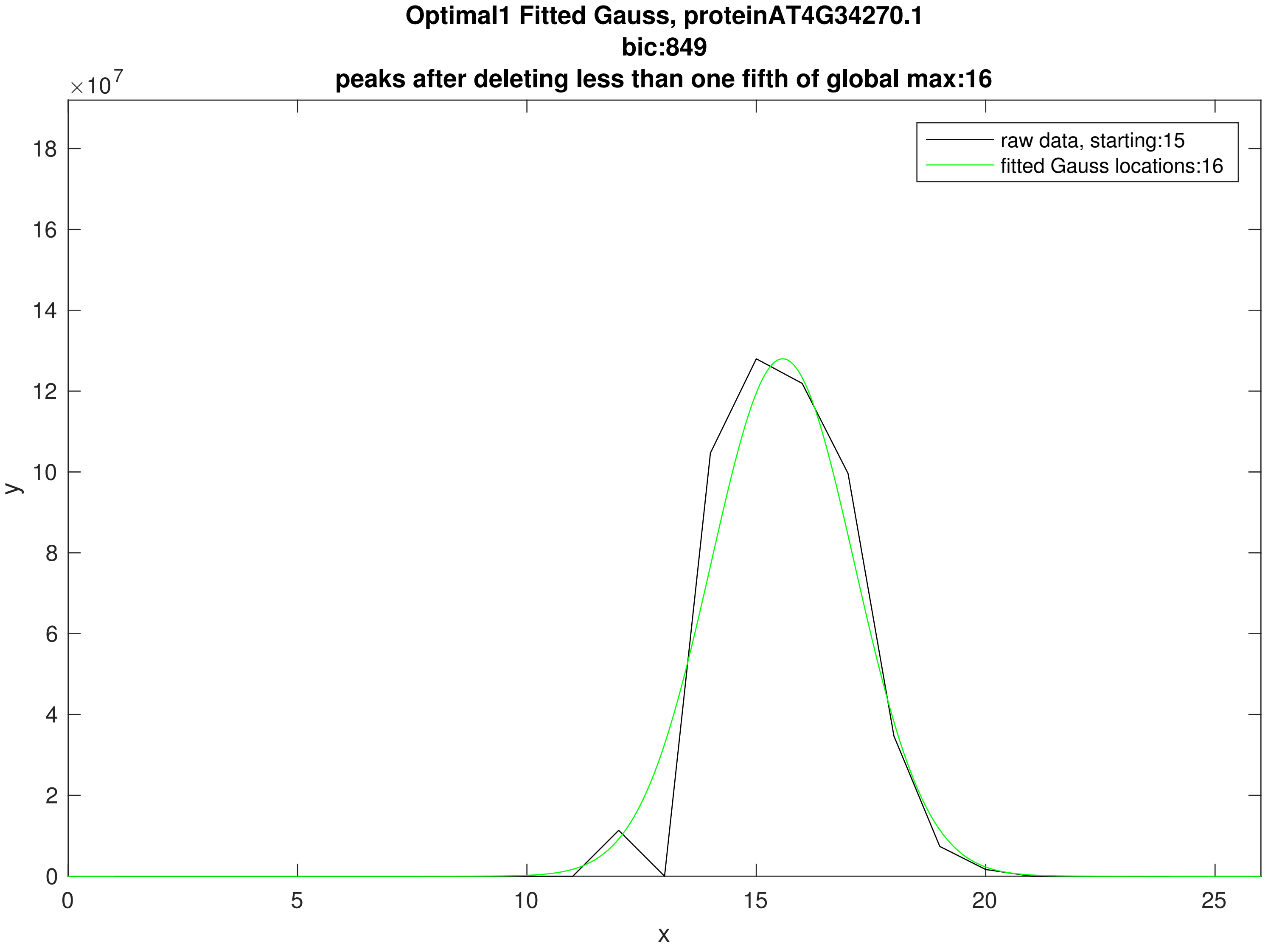
Optimal1 Fitted Gauss, proteinAT4G34270.1
bic:849
peaks after deleting less than one fifth of global max:16
×10^7
18
16
14
12
10
8
6
4
2
0
y
0
5
10
15
20
25
x
raw data, starting:15
fitted Gauss locations:16

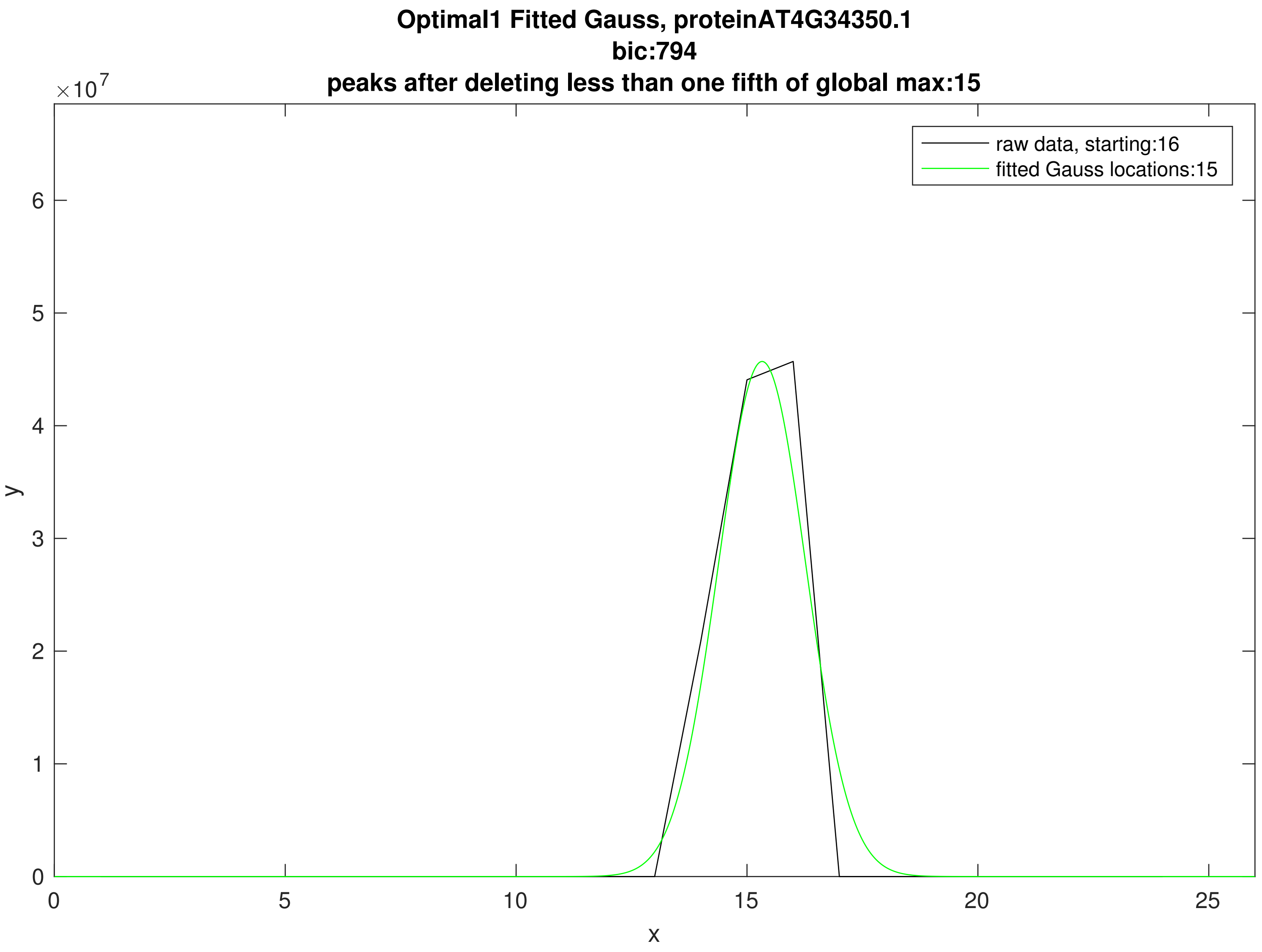
Optimal1 Fitted Gauss, proteinAT4G34350.1
bic:794
peaks after deleting less than one fifth of global max:15
×10^7
raw data, starting:16
fitted Gauss locations:15
y
x
0
1
2
3
4
5
6
0
5
10
15
20
25

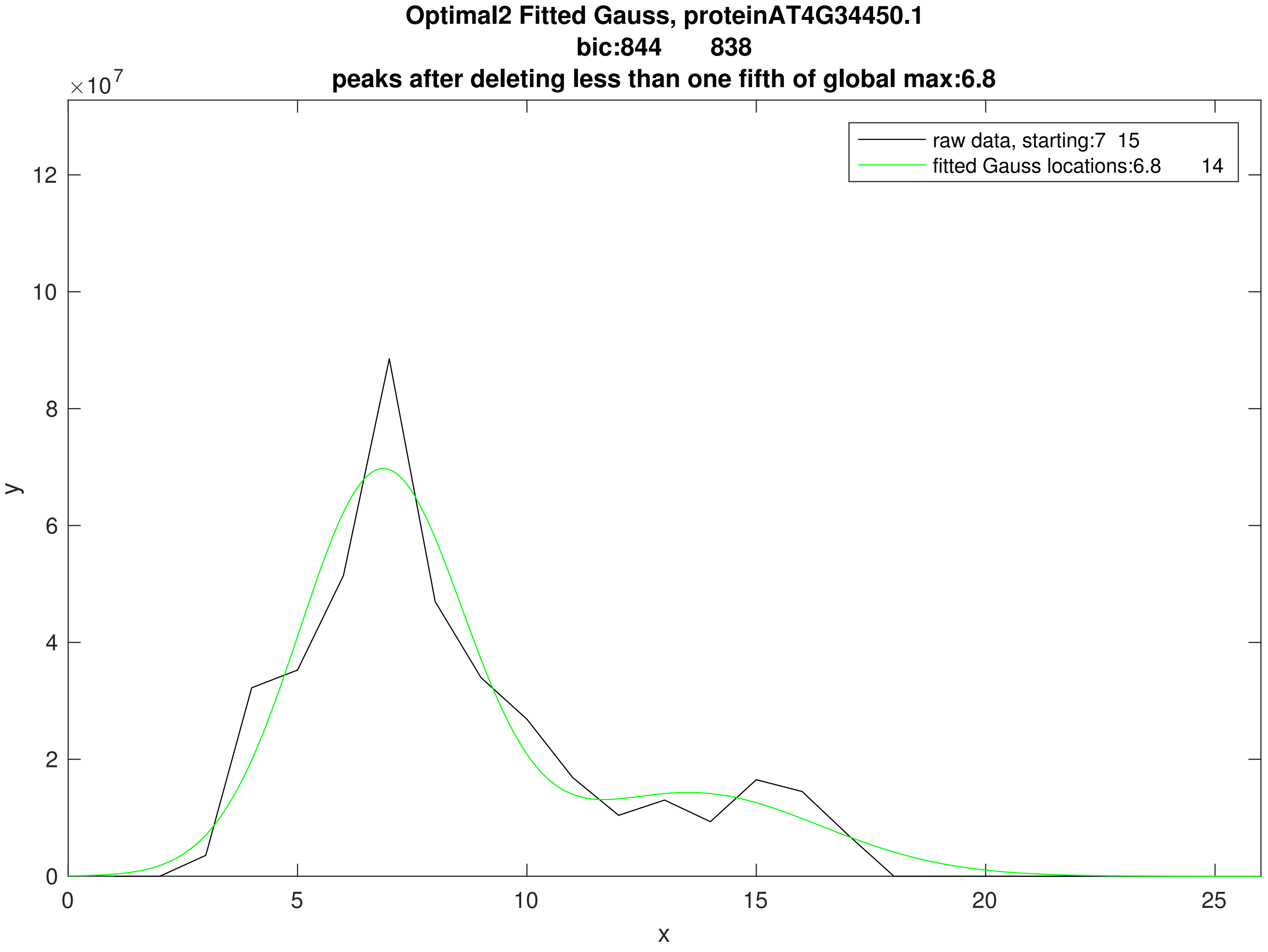
Optimal2 Fitted Gauss, proteinAT4G34450.1
bic:844 838
peaks after deleting less than one fifth of global max:6.8
×10^7
raw data, starting:7 15
fitted Gauss locations:6.8 14
y
x
0
2
4
6
8
10
12
0
5
10
15
20
25

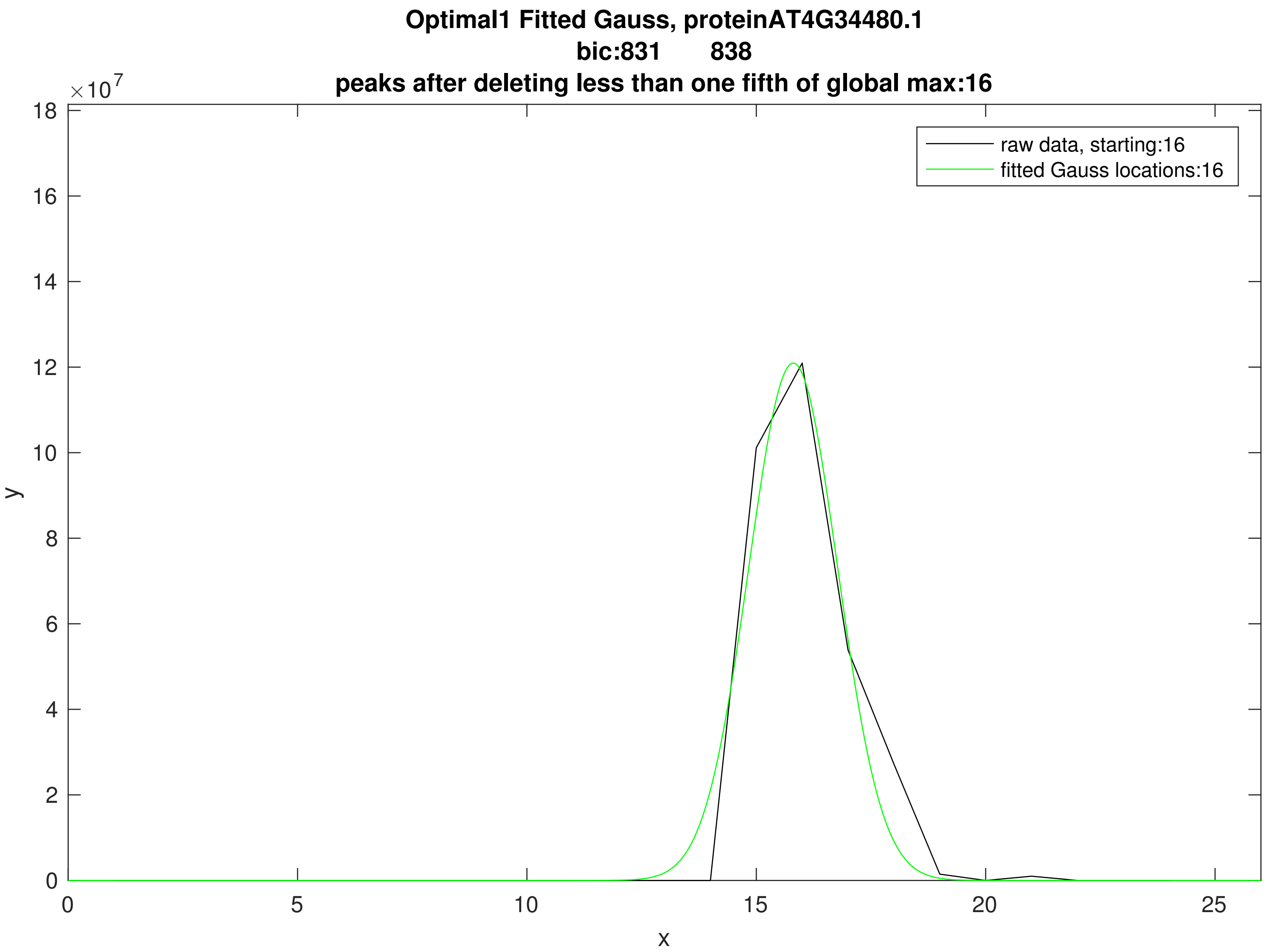
Optimal1 Fitted Gauss, proteinAT4G34480.1
bic:831 838
peaks after deleting less than one fifth of global max:16
×10^7
18
16
14
12
10
8
6
4
2
0
y
0
5
10
15
20
25
x
raw data, starting:16
fitted Gauss locations:16

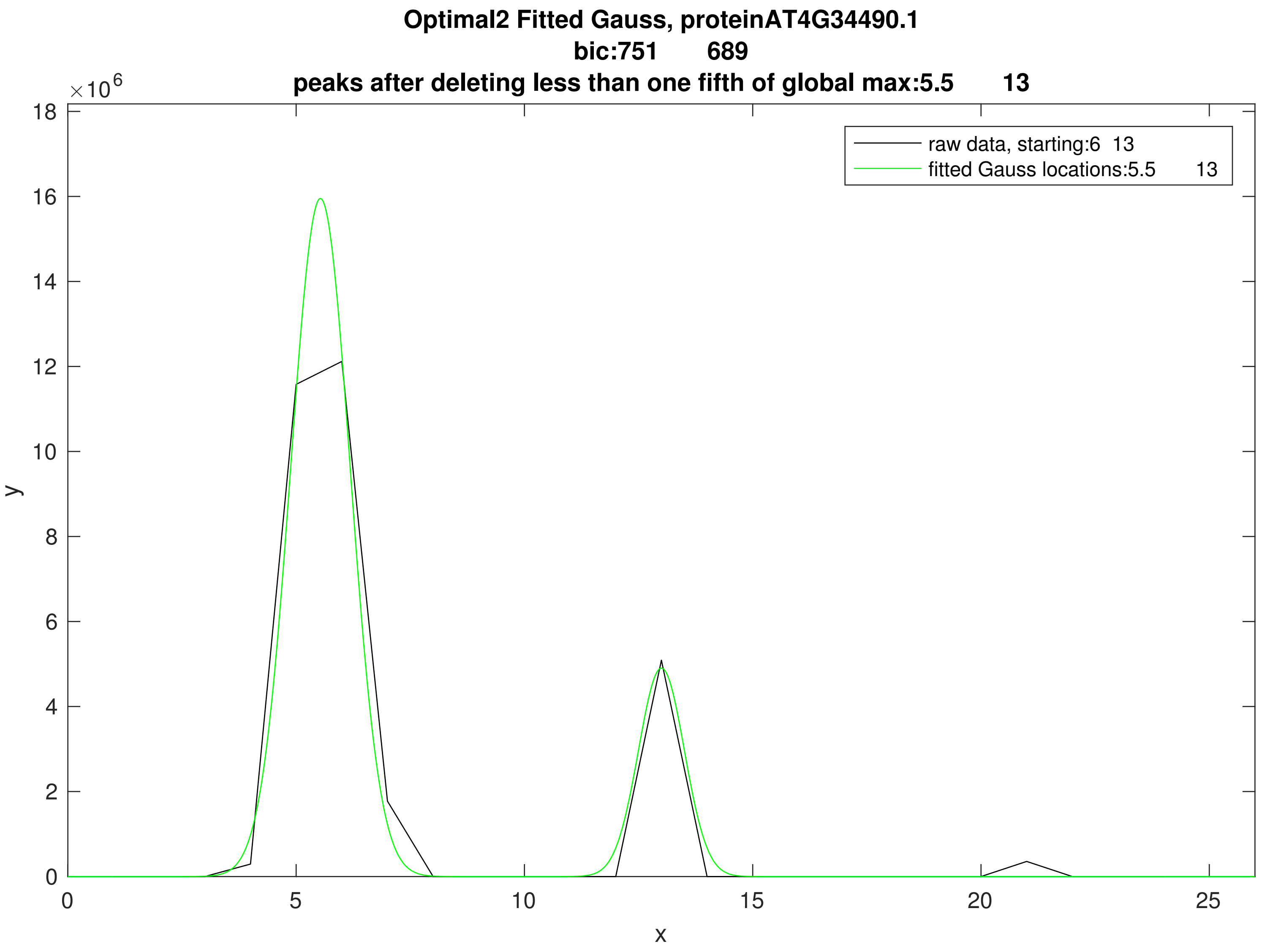

Optimal2 Fitted Gauss, proteinAT4G34490.1
bic:751 689
peaks after deleting less than one fifth of global max:5.5 13
×10^6
raw data, starting:6 13
fitted Gauss locations:5.5 13
y
x
0
2
4
6
8
10
12
14
16
18
0
5
10
15
20
25

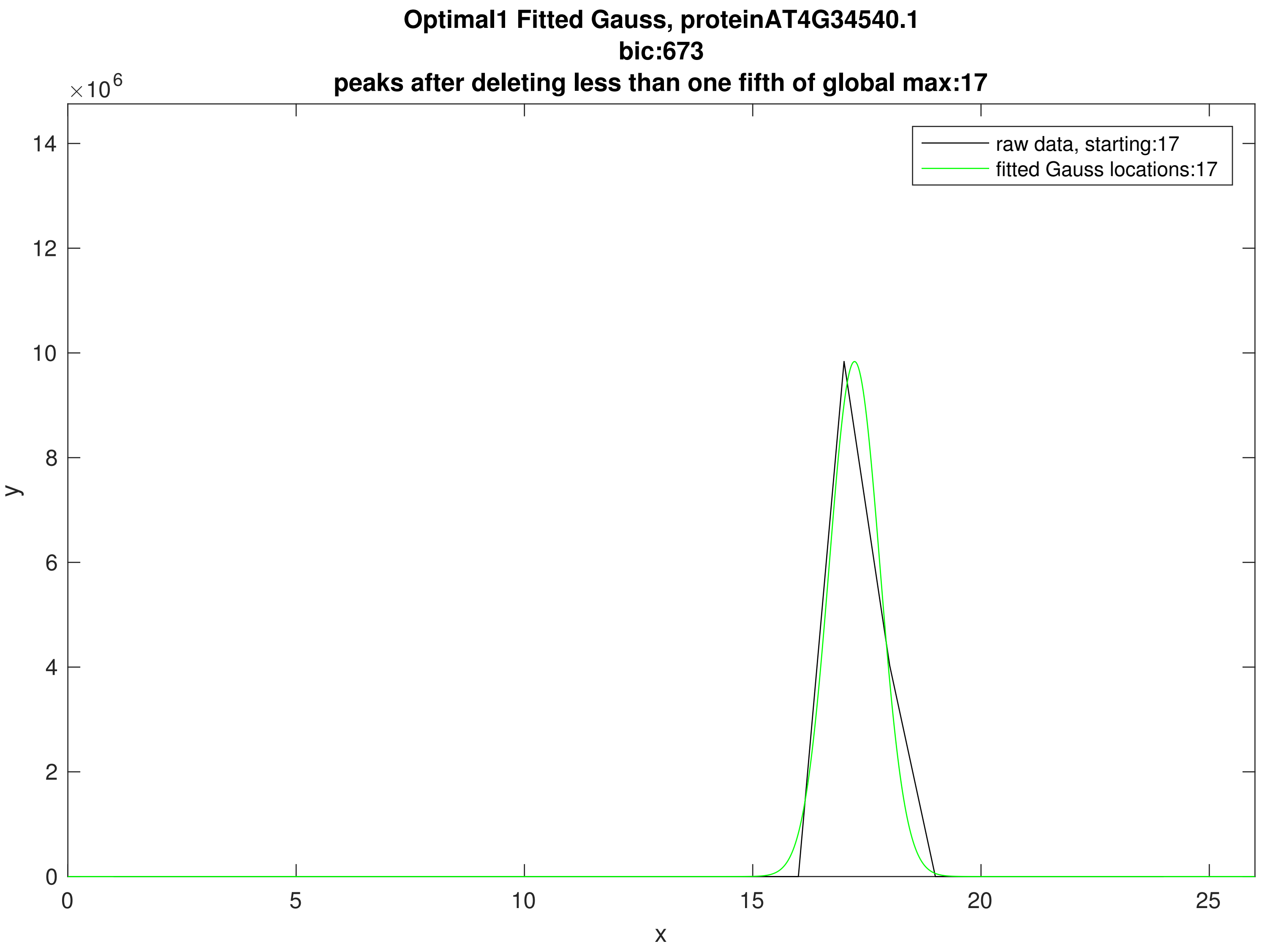
Optimal1 Fitted Gauss, proteinAT4G34540.1
bic:673
peaks after deleting less than one fifth of global max:17
×10^6
14
12
10
8
6
4
2
0
y
0
5
10
15
20
25
x
raw data, starting:17
fitted Gauss locations:17

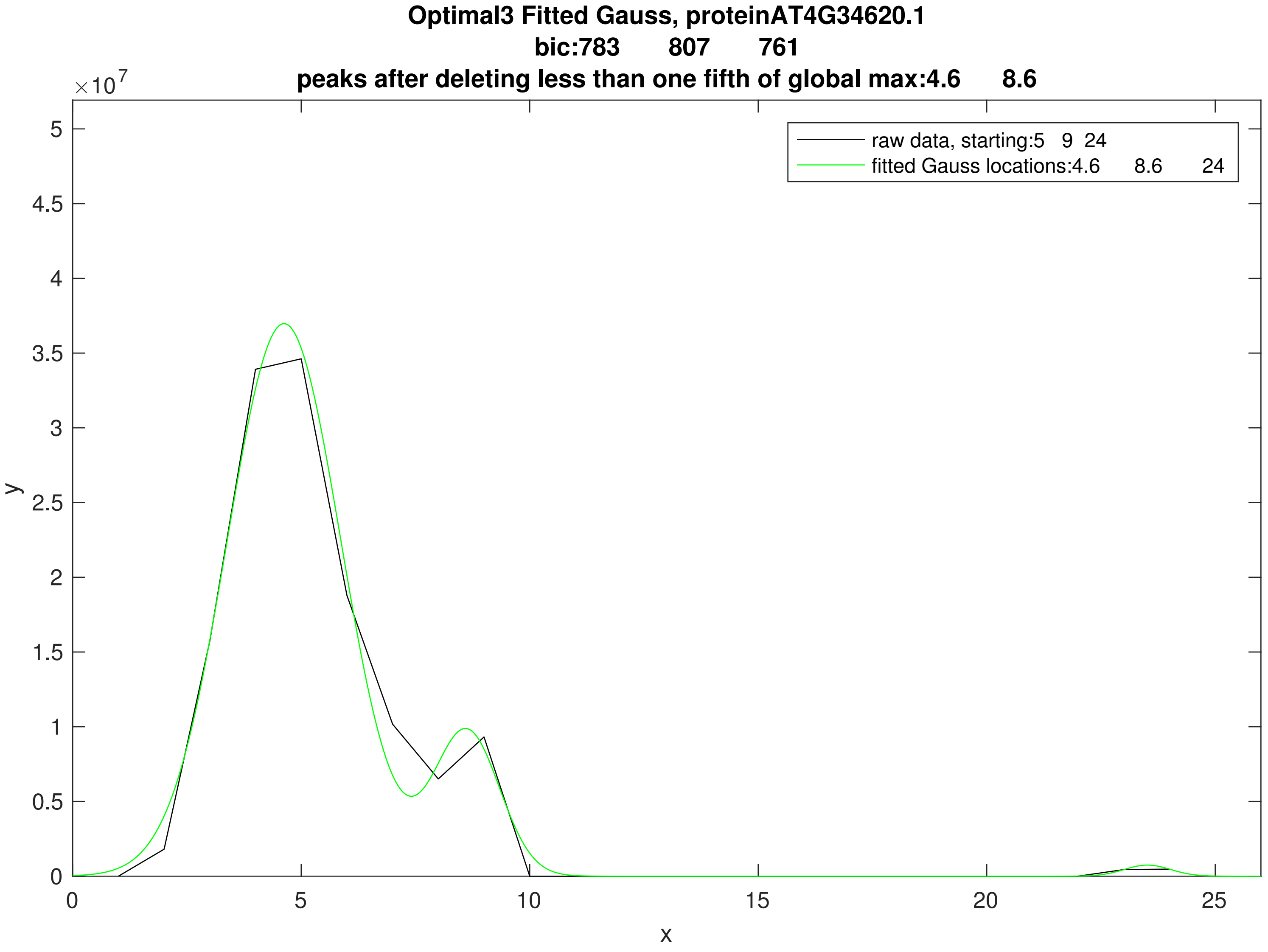
Optimal3 Fitted Gauss, proteinAT4G34620.1
bic:783 807 761
peaks after deleting less than one fifth of global max:4.6 8.6
raw data, starting:5 9 24
fitted Gauss locations:4.6 8.6 24
×10^7
y
x

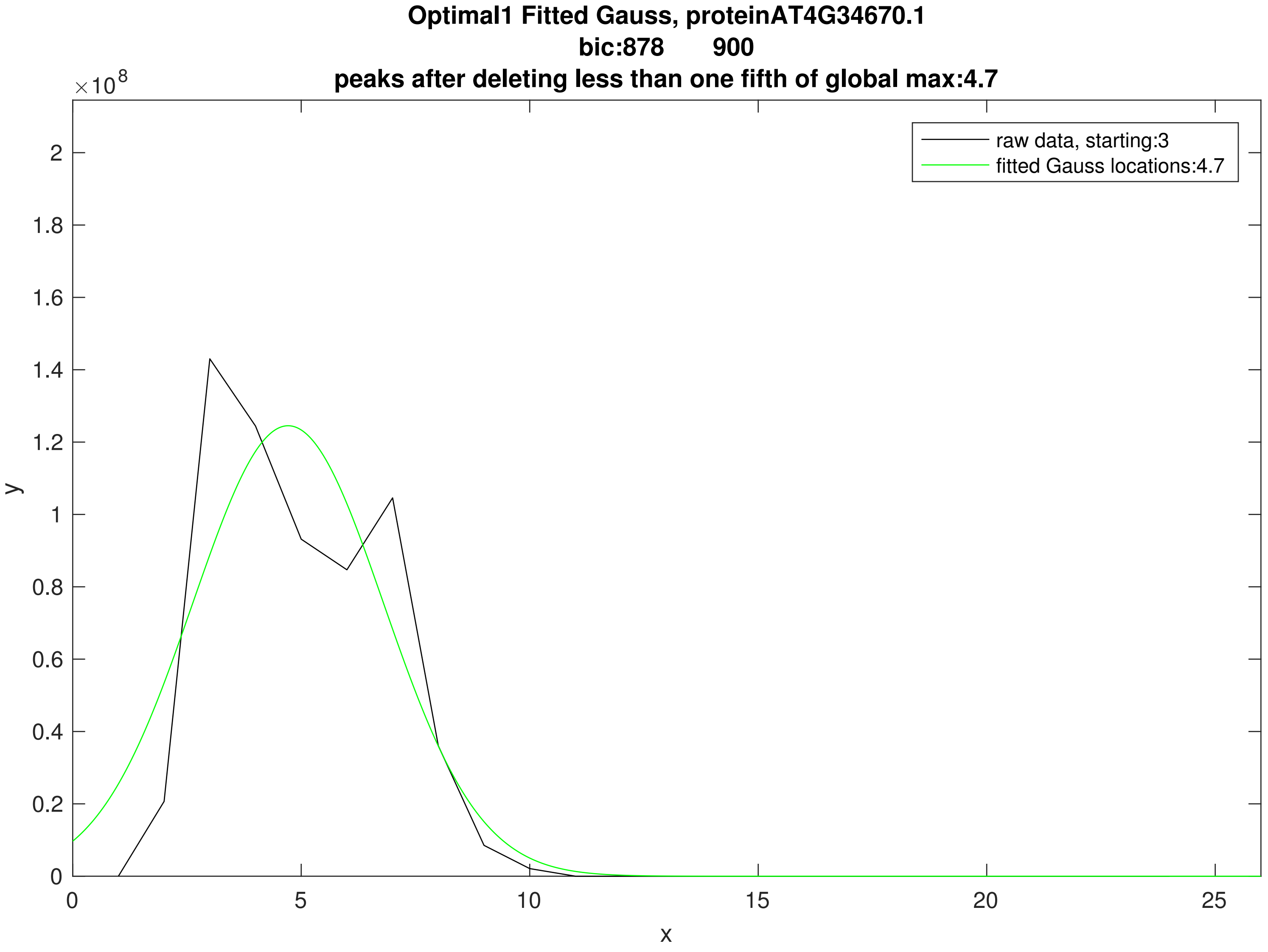
Optimal1 Fitted Gauss, proteinAT4G34670.1
bic:878 900
peaks after deleting less than one fifth of global max:4.7
×10^8
raw data, starting:3
fitted Gauss locations:4.7
y
x
0
0.2
0.4
0.6
0.8
1
1.2
1.4
1.6
1.8
2
5
10
15
20
25

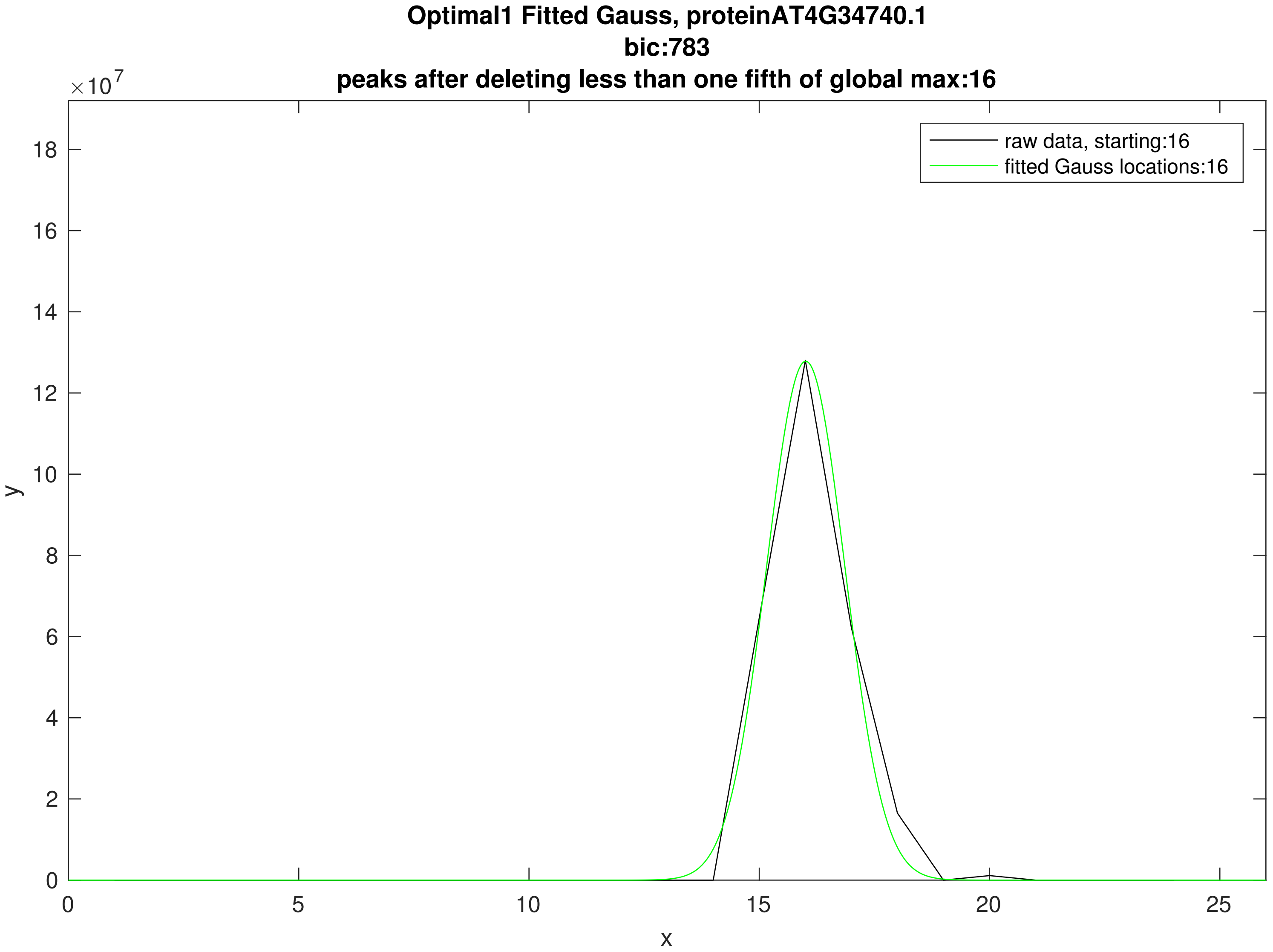

Optimal1 Fitted Gauss, proteinAT4G34740.1
bic:783
peaks after deleting less than one fifth of global max:16
×10^7
raw data, starting:16
fitted Gauss locations:16
y
x
0
2
4
6
8
10
12
14
16
18
0
5
10
15
20
25

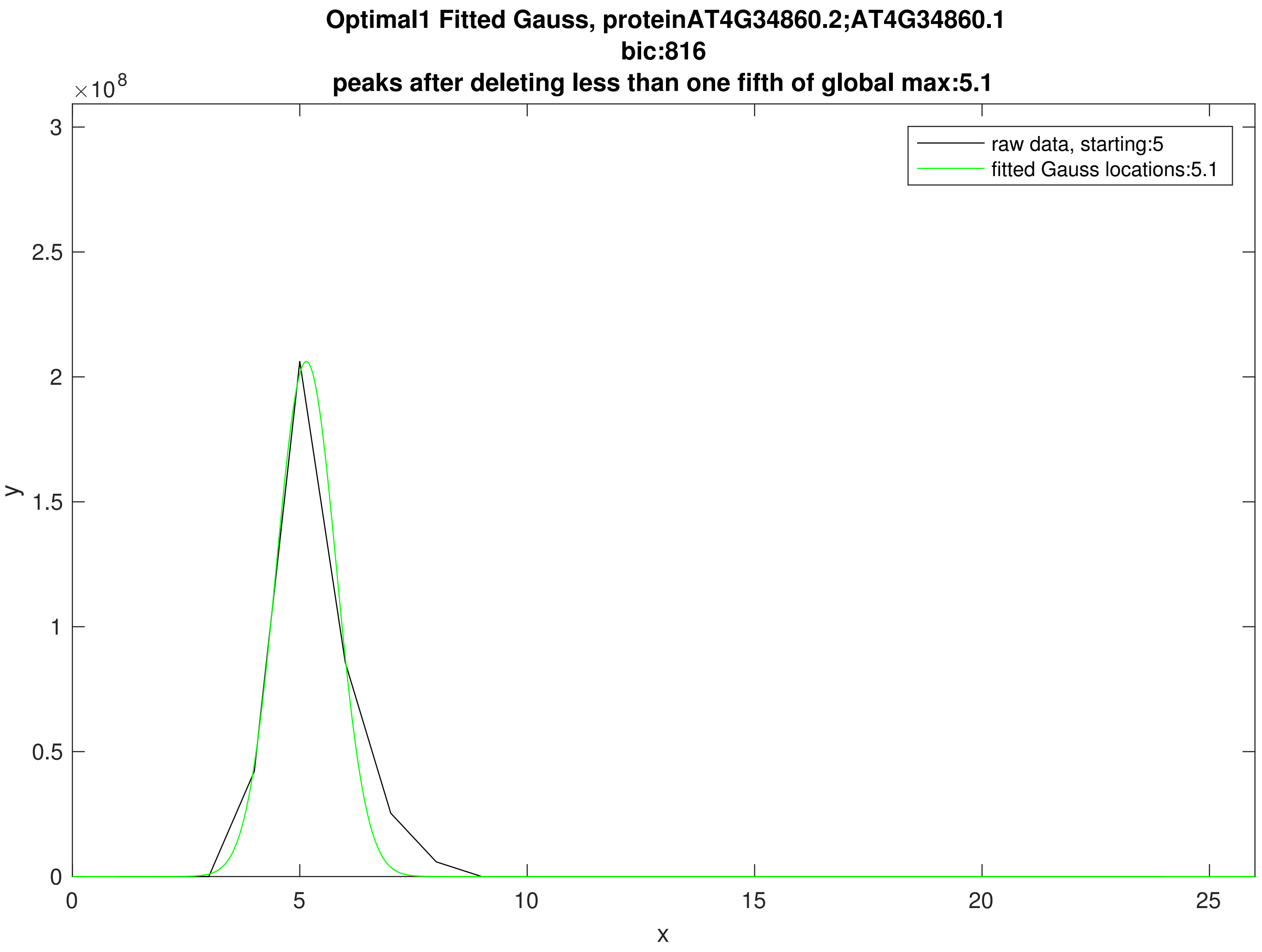
Optimal1 Fitted Gauss, proteinAT4G34860.2;AT4G34860.1
bic:816
peaks after deleting less than one fifth of global max:5.1
×10^8
raw data, starting:5
fitted Gauss locations:5.1
y
x
0
0.5
1
1.5
2
2.5
3
5
10
15
20
25

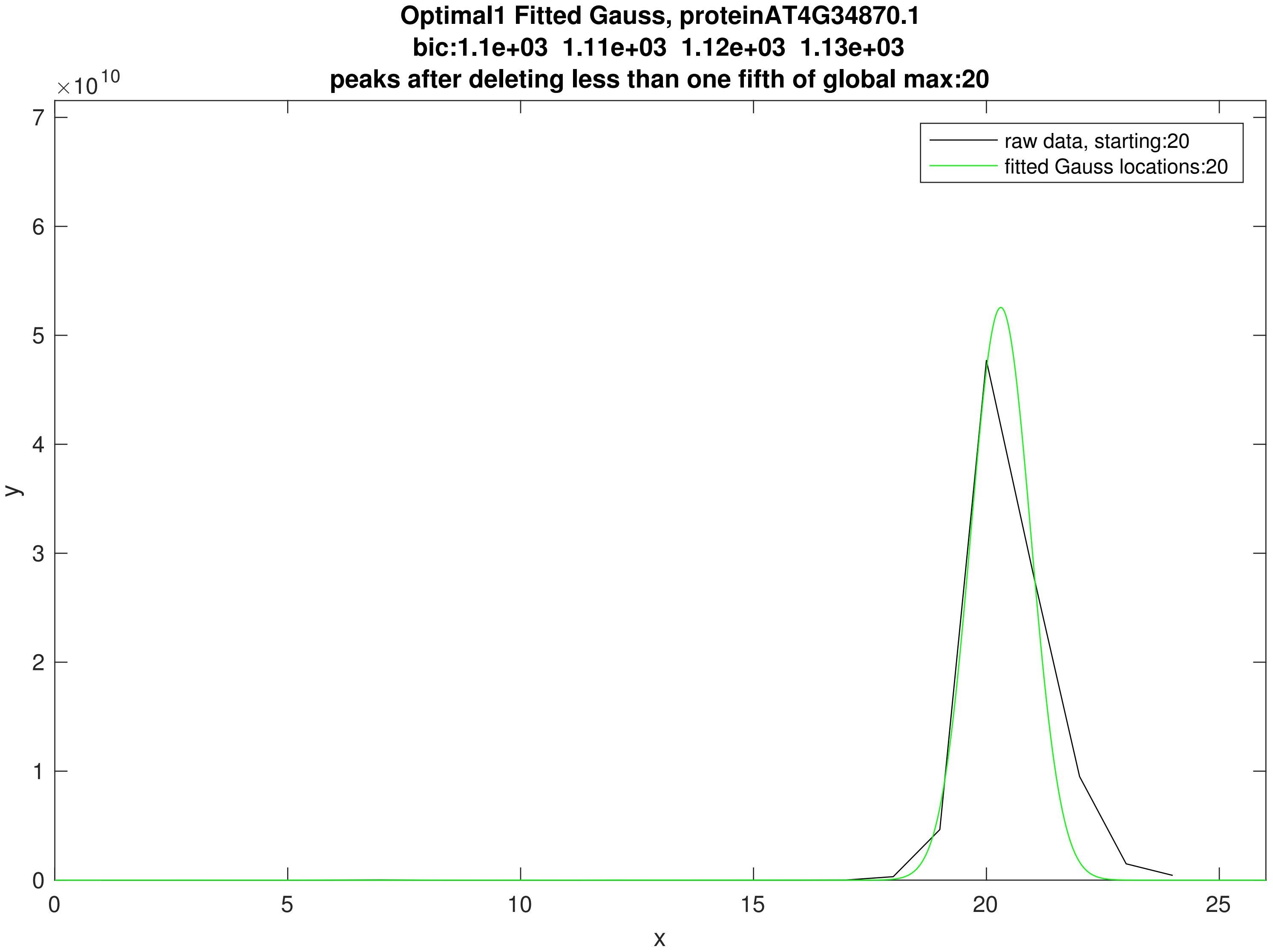

Optimal1 Fitted Gauss, proteinAT4G34870.1
bic:1.1e+03 1.11e+03 1.12e+03 1.13e+03
peaks after deleting less than one fifth of global max:20
×10^10
raw data, starting:20
fitted Gauss locations:20
y
x
0
1
2
3
4
5
6
7
0
5
10
15
20
25

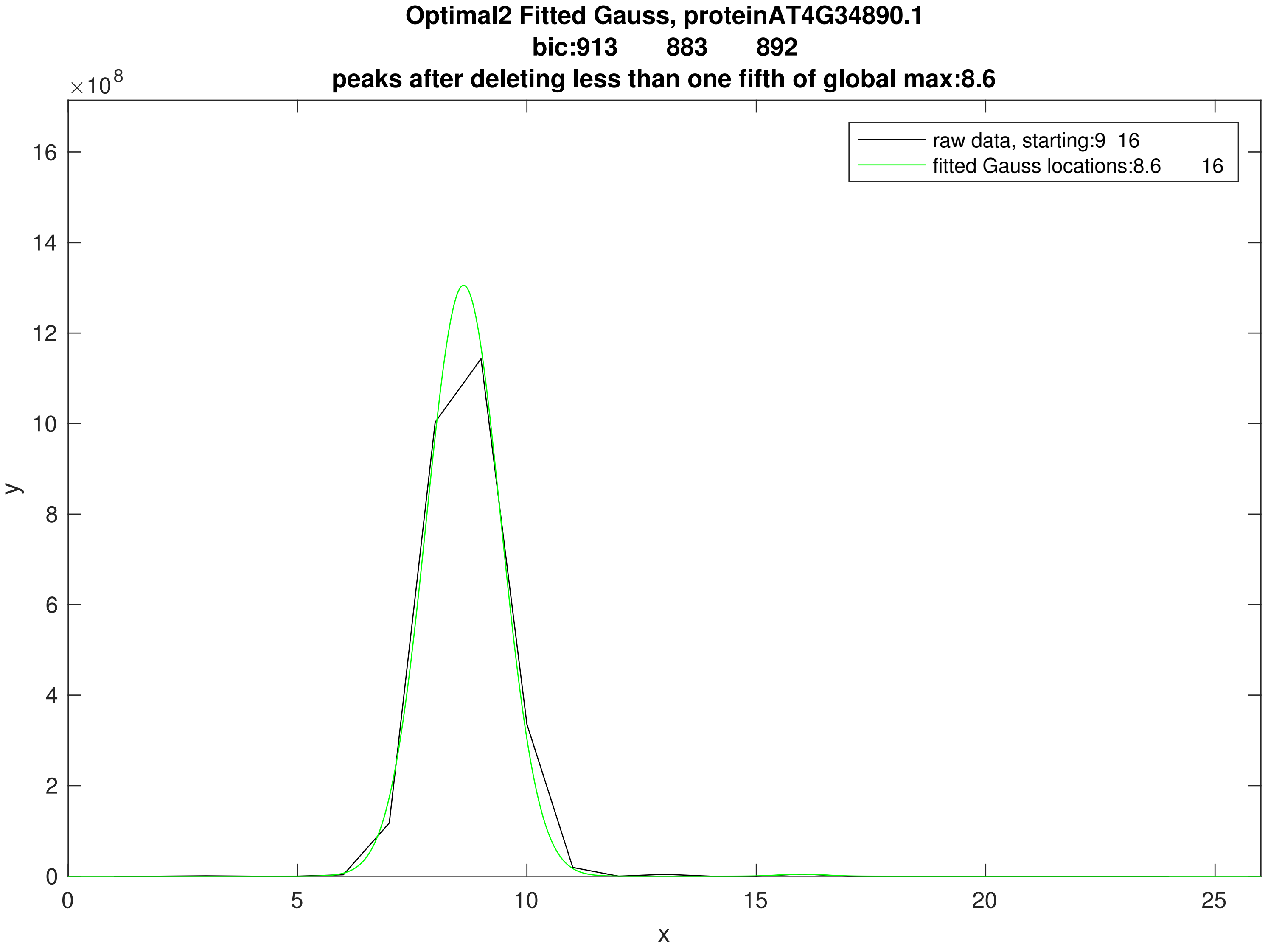
Optimal2 Fitted Gauss, proteinAT4G34890.1
bic:913 883 892
peaks after deleting less than one fifth of global max:8.6
×10^8
raw data, starting:9 16
fitted Gauss locations:8.6 16
y
x
0
2
4
6
8
10
12
14
16
0
5
10
15
20
25

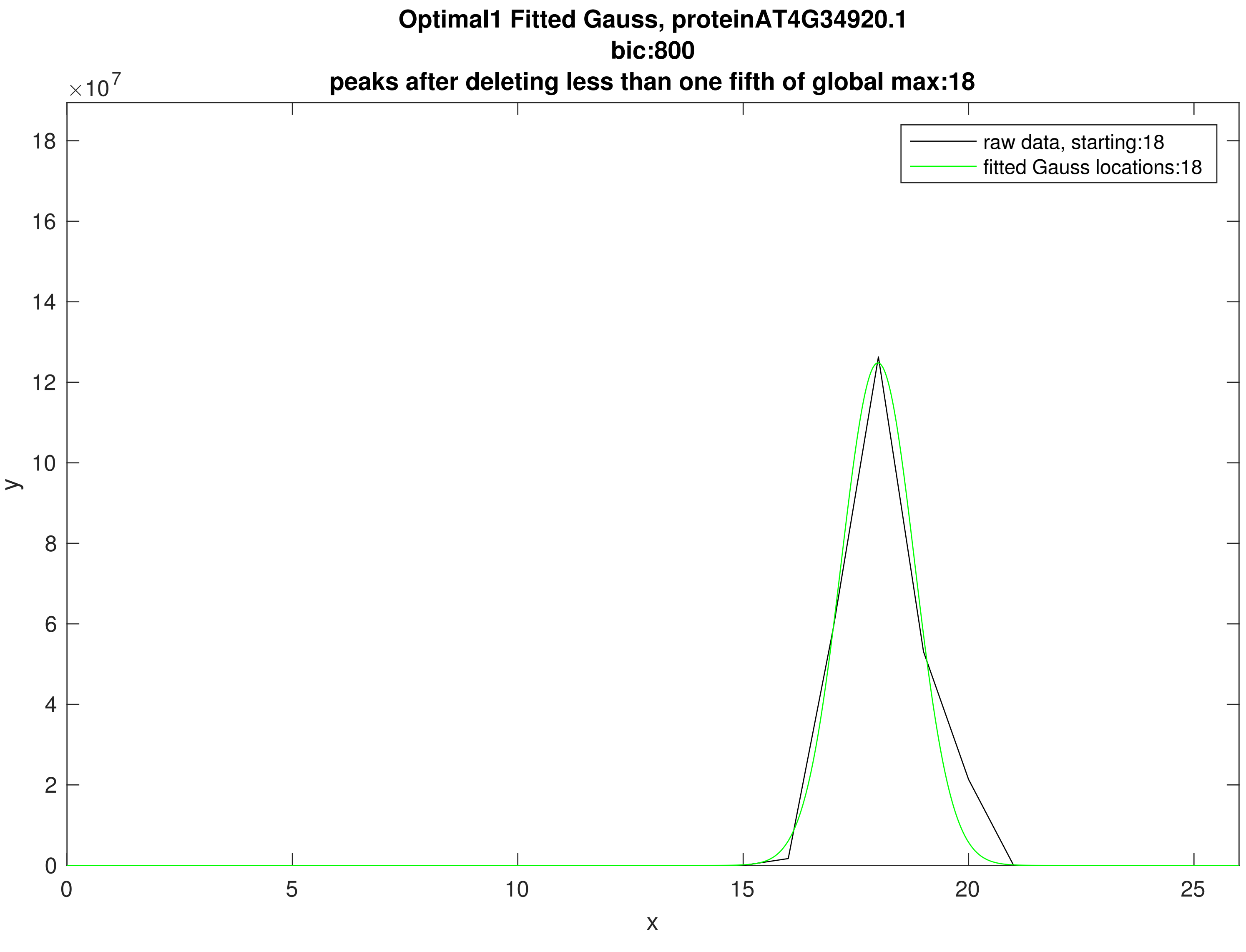
Optimal1 Fitted Gauss, proteinAT4G34920.1
bic:800
peaks after deleting less than one fifth of global max:18
×10^7
raw data, starting:18
fitted Gauss locations:18
y
x
0
2
4
6
8
10
12
14
16
18
0
5
10
15
20
25

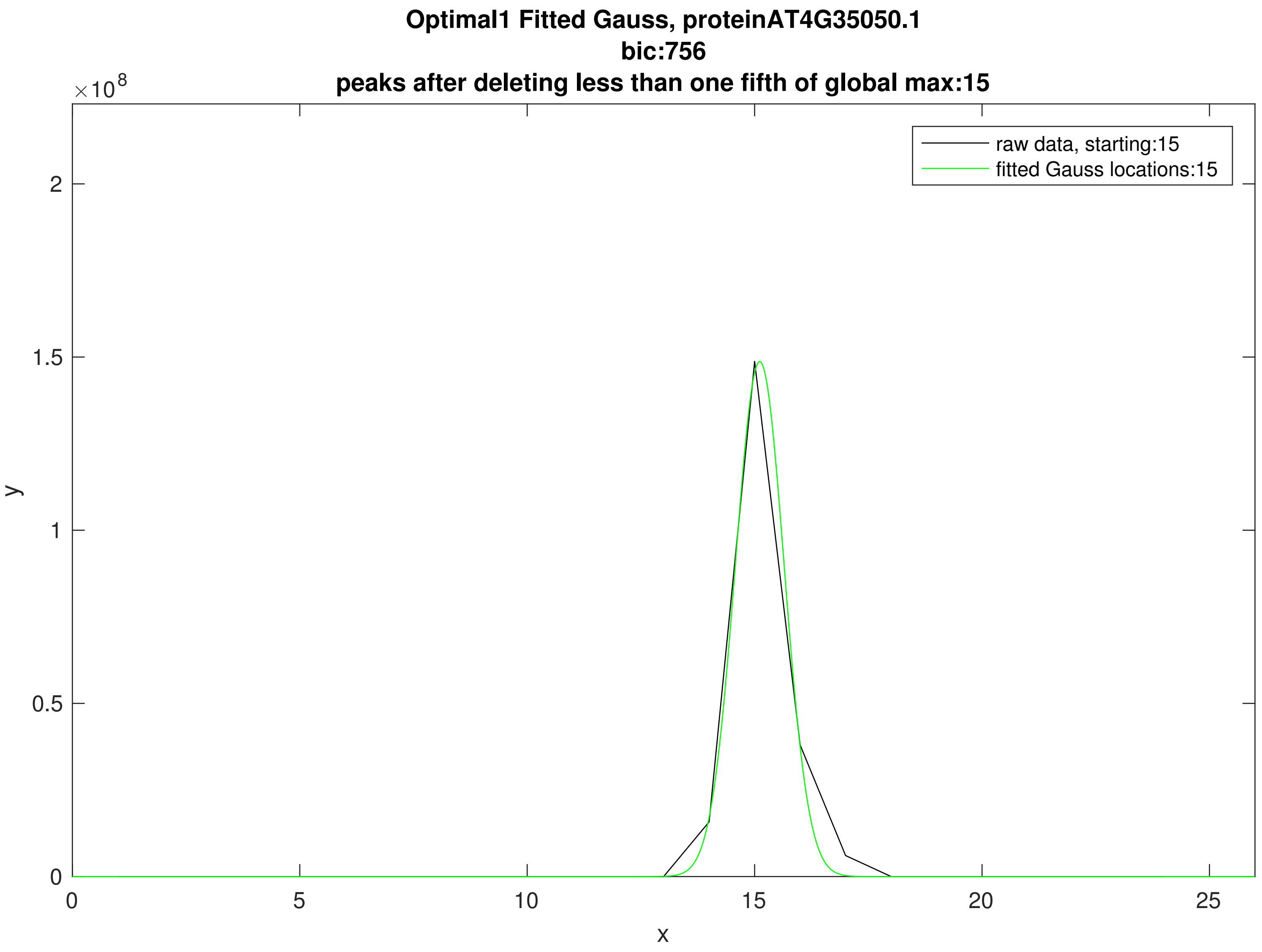
Optimal1 Fitted Gauss, proteinAT4G35050.1
bic:756
peaks after deleting less than one fifth of global max:15
×10^8
raw data, starting:15
fitted Gauss locations:15
y
x
0
0.5
1
1.5
2
0
5
10
15
20
25

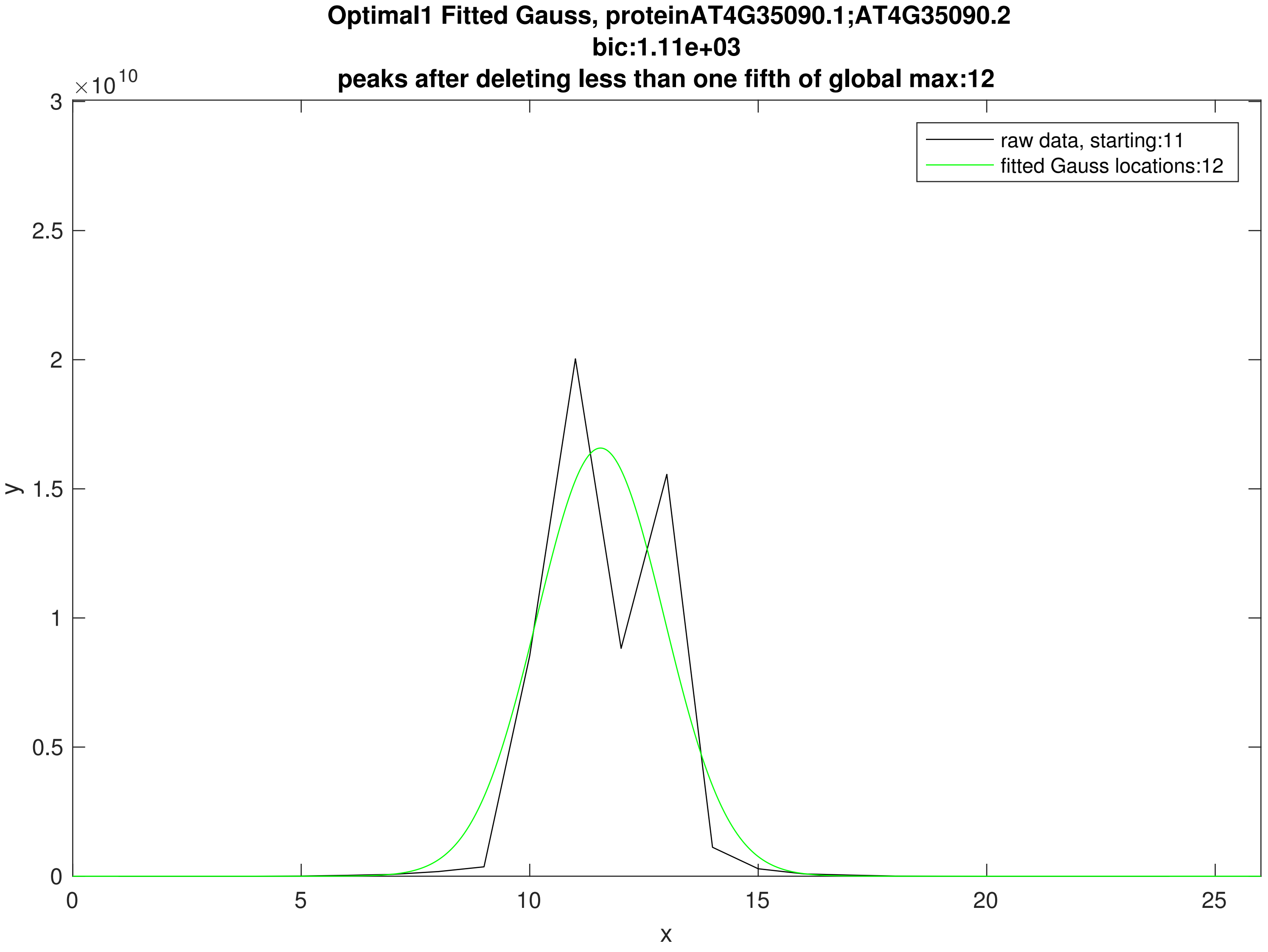

Optimal1 Fitted Gauss, proteinAT4G35090.1;AT4G35090.2
bic:1.11e+03
peaks after deleting less than one fifth of global max:12
×10^10
3
2.5
2
1.5
1
0.5
0
y
0
5
10
15
20
25
x
raw data, starting:11
fitted Gauss locations:12

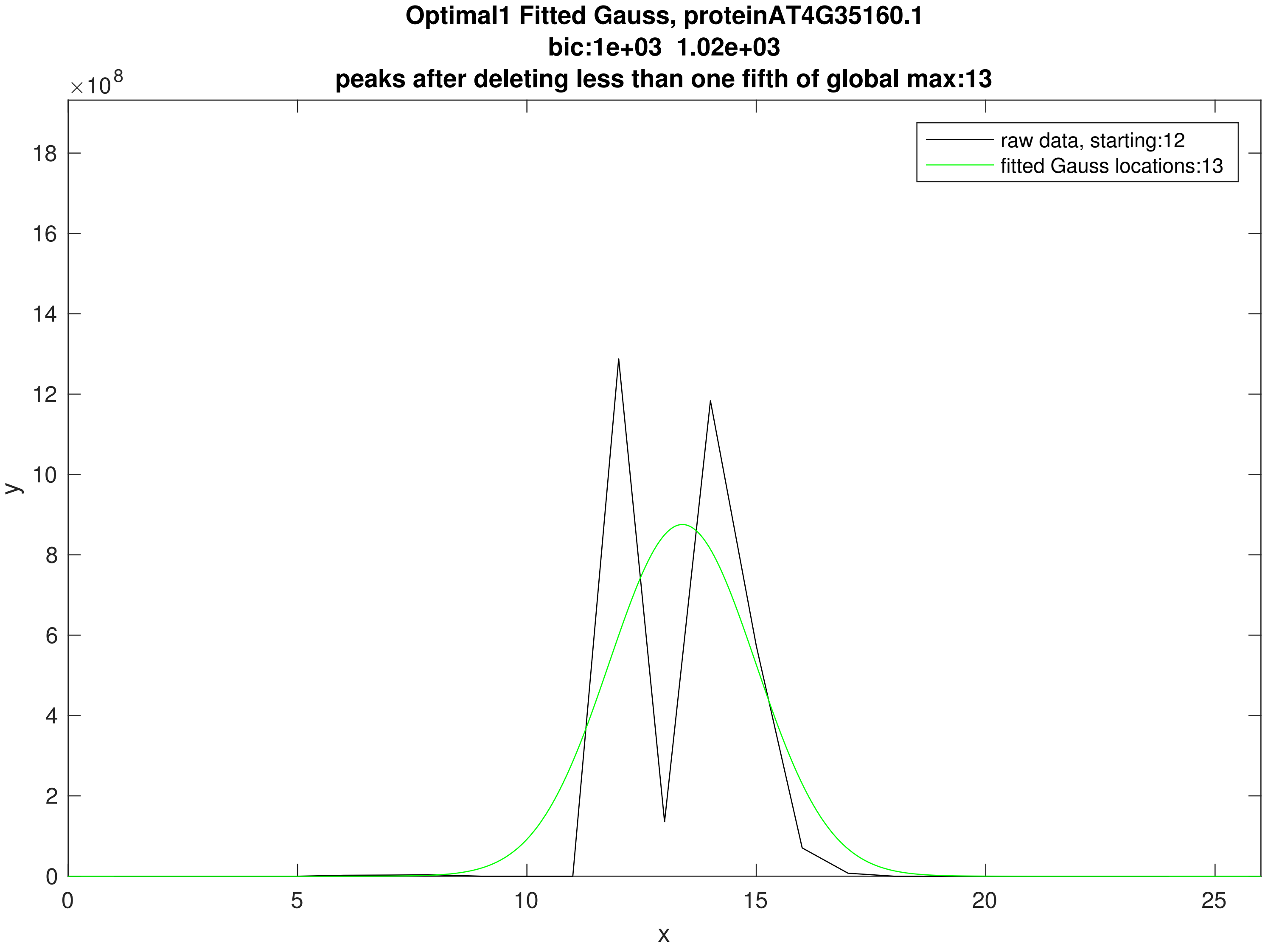

Optimal1 Fitted Gauss, proteinAT4G35160.1
bic:1e+03 1.02e+03
peaks after deleting less than one fifth of global max:13
×10^8
raw data, starting:12
fitted Gauss locations:13
y
x
0
2
4
6
8
10
12
14
16
18
0
5
10
15
20
25

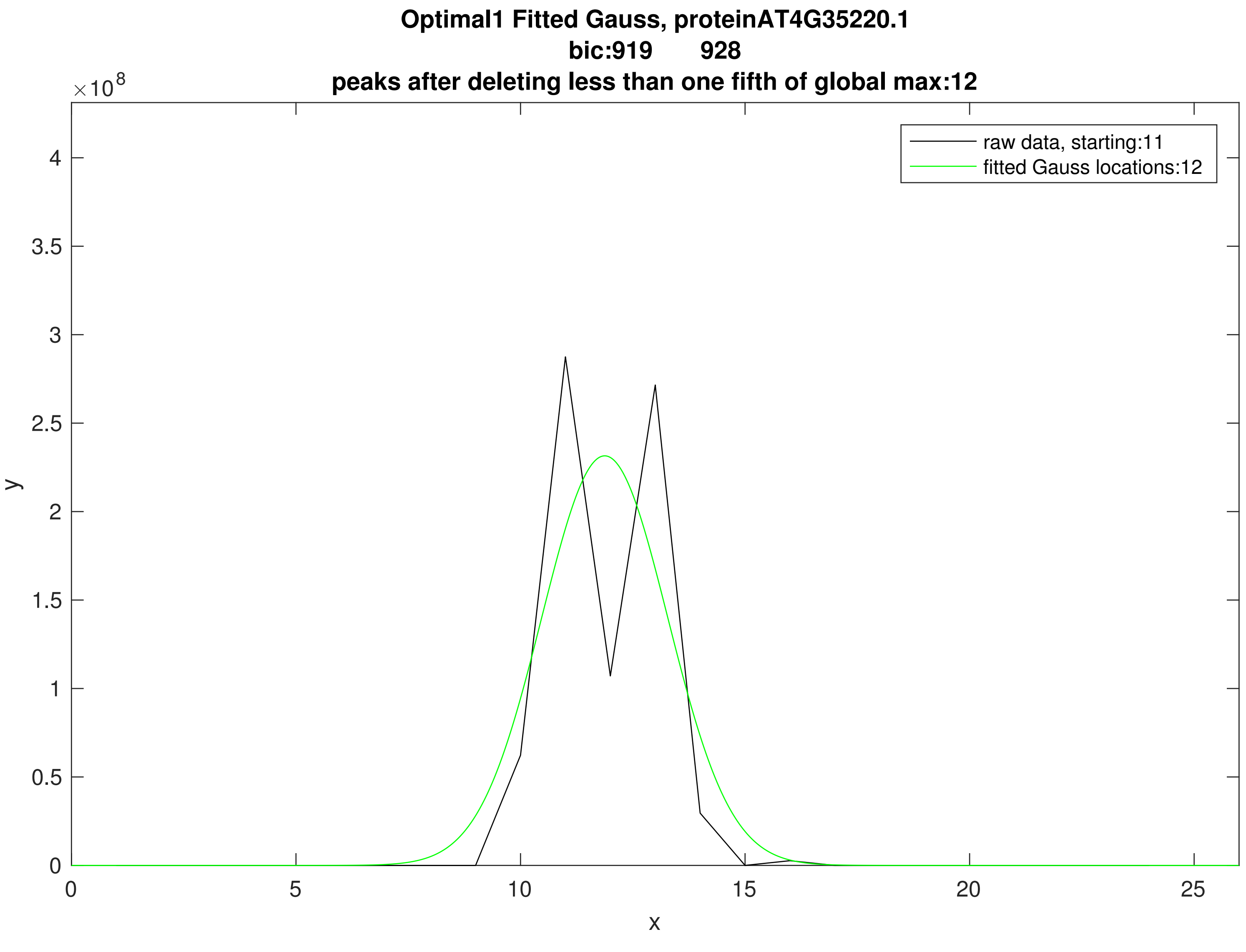
Optimal1 Fitted Gauss, proteinAT4G35220.1
bic:919 928
peaks after deleting less than one fifth of global max:12
×10^8
raw data, starting:11
fitted Gauss locations:12
y
x
0
0.5
1
1.5
2
2.5
3
3.5
4
5
10
15
20
25

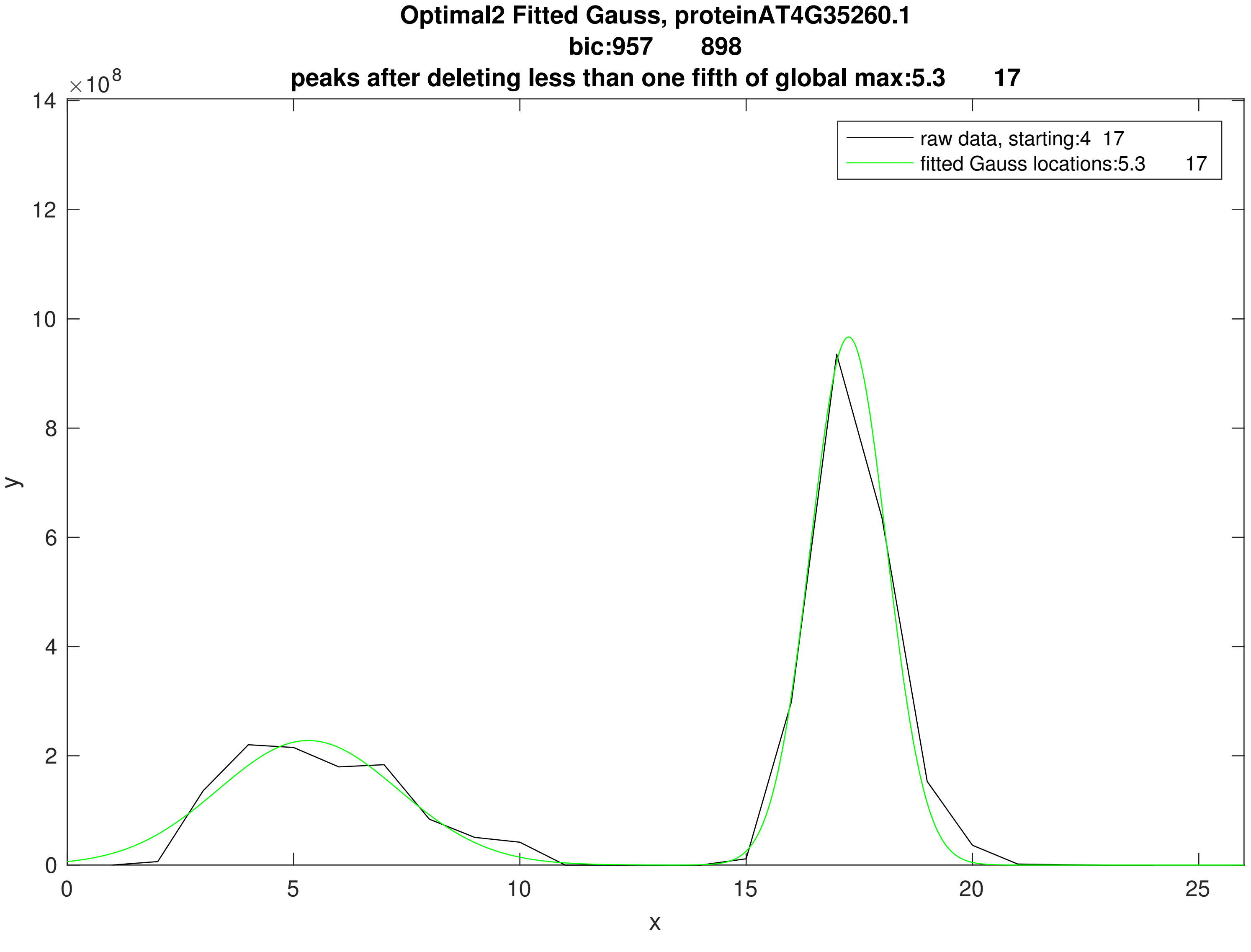

Optimal2 Fitted Gauss, proteinAT4G35260.1
bic:957 898
peaks after deleting less than one fifth of global max:5.3 17
×10^8
14
12
10
8
6
4
2
0
y
0
5
10
15
20
25
x
raw data, starting:4 17
fitted Gauss locations:5.3 17

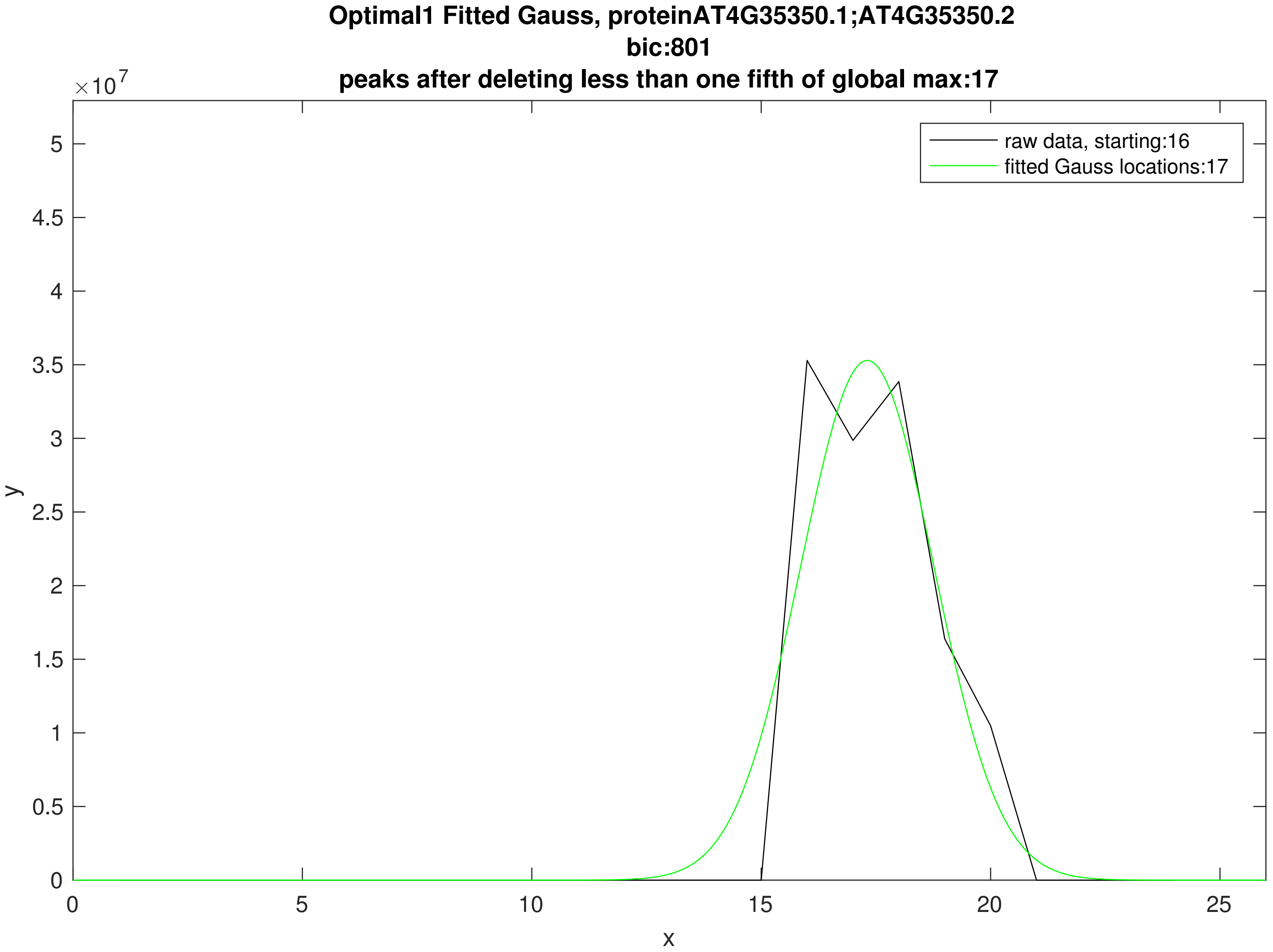
Optimal1 Fitted Gauss, proteinAT4G35350.1;AT4G35350.2
bic:801
peaks after deleting less than one fifth of global max:17
×10^7
raw data, starting:16
fitted Gauss locations:17
y
x
0
0.5
1
1.5
2
2.5
3
3.5
4
4.5
5
0
5
10
15
20
25

**Optimal2 Fitted Gauss, proteinAT4G35360.1**
**bic:880 865**
**peaks after deleting less than one fifth of global max:16**

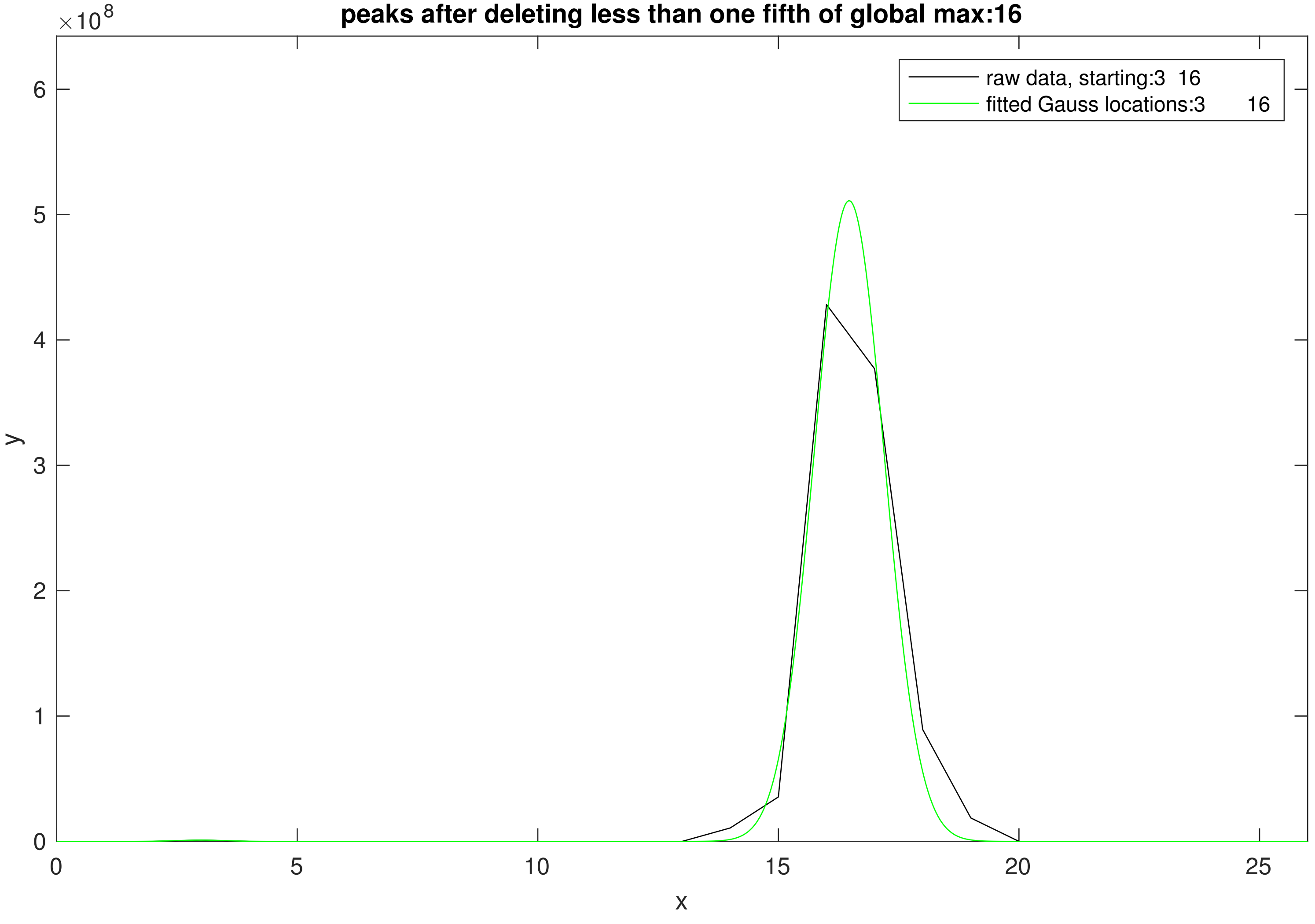

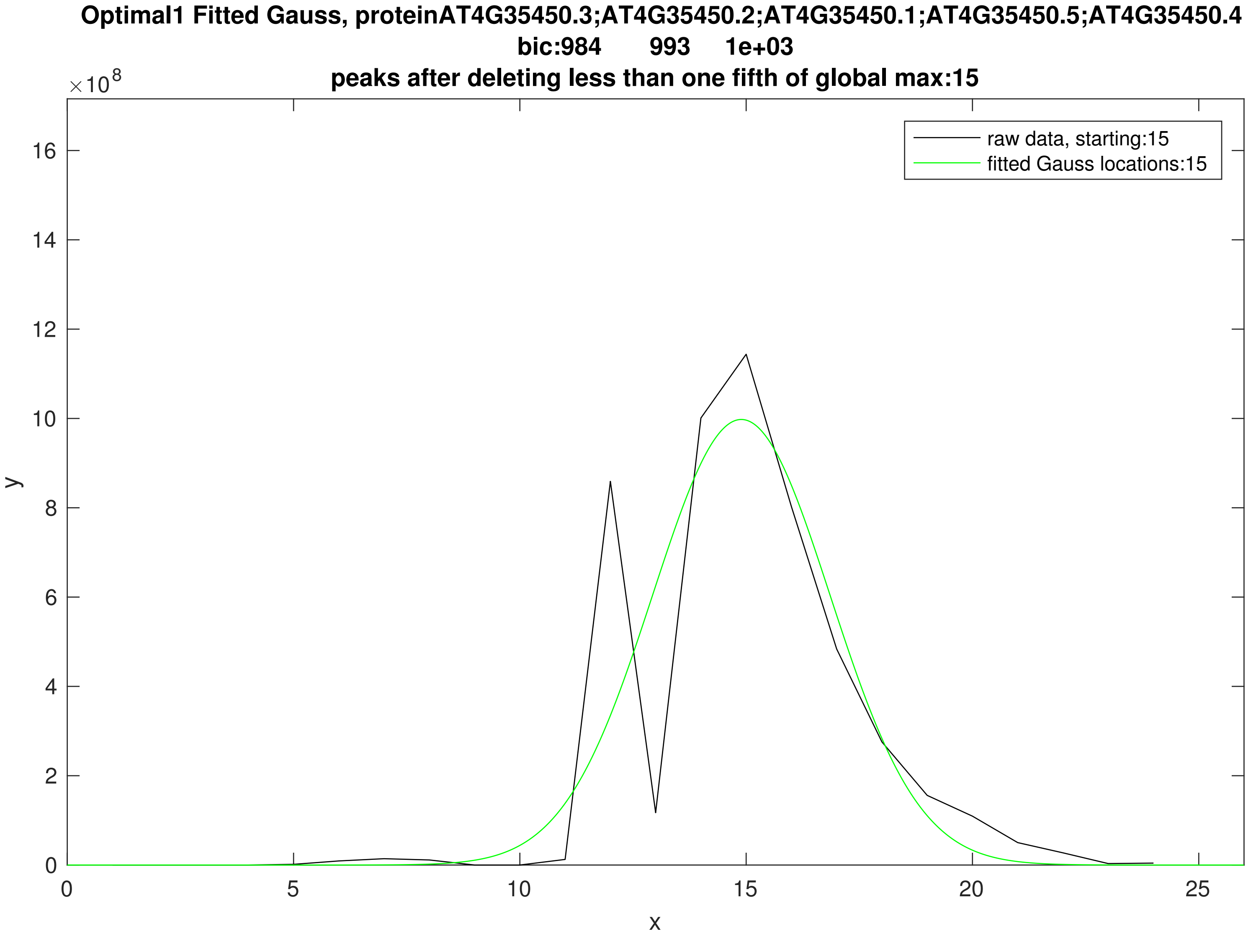

Optimal1 Fitted Gauss, proteinAT4G35450.3;AT4G35450.2;AT4G35450.1;AT4G35450.5;AT4G35450.4
bic:984 993 1e+03
peaks after deleting less than one fifth of global max:15
×10^8
raw data, starting:15
fitted Gauss locations:15
y
x

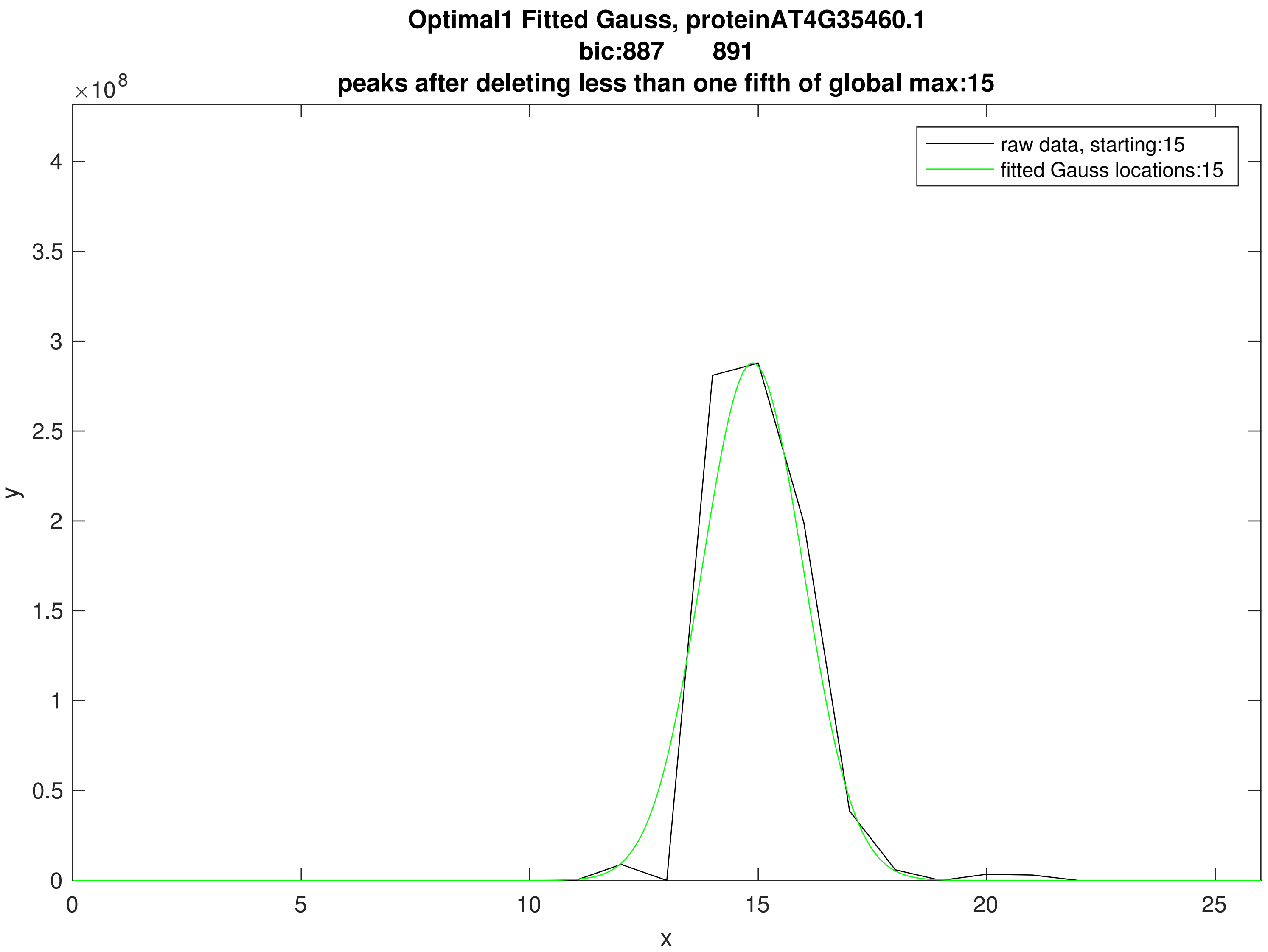
Optimal1 Fitted Gauss, proteinAT4G35460.1
bic:887 891
peaks after deleting less than one fifth of global max:15
×10^8
raw data, starting:15
fitted Gauss locations:15
y
x
0
0.5
1
1.5
2
2.5
3
3.5
4
5
10
15
20
25

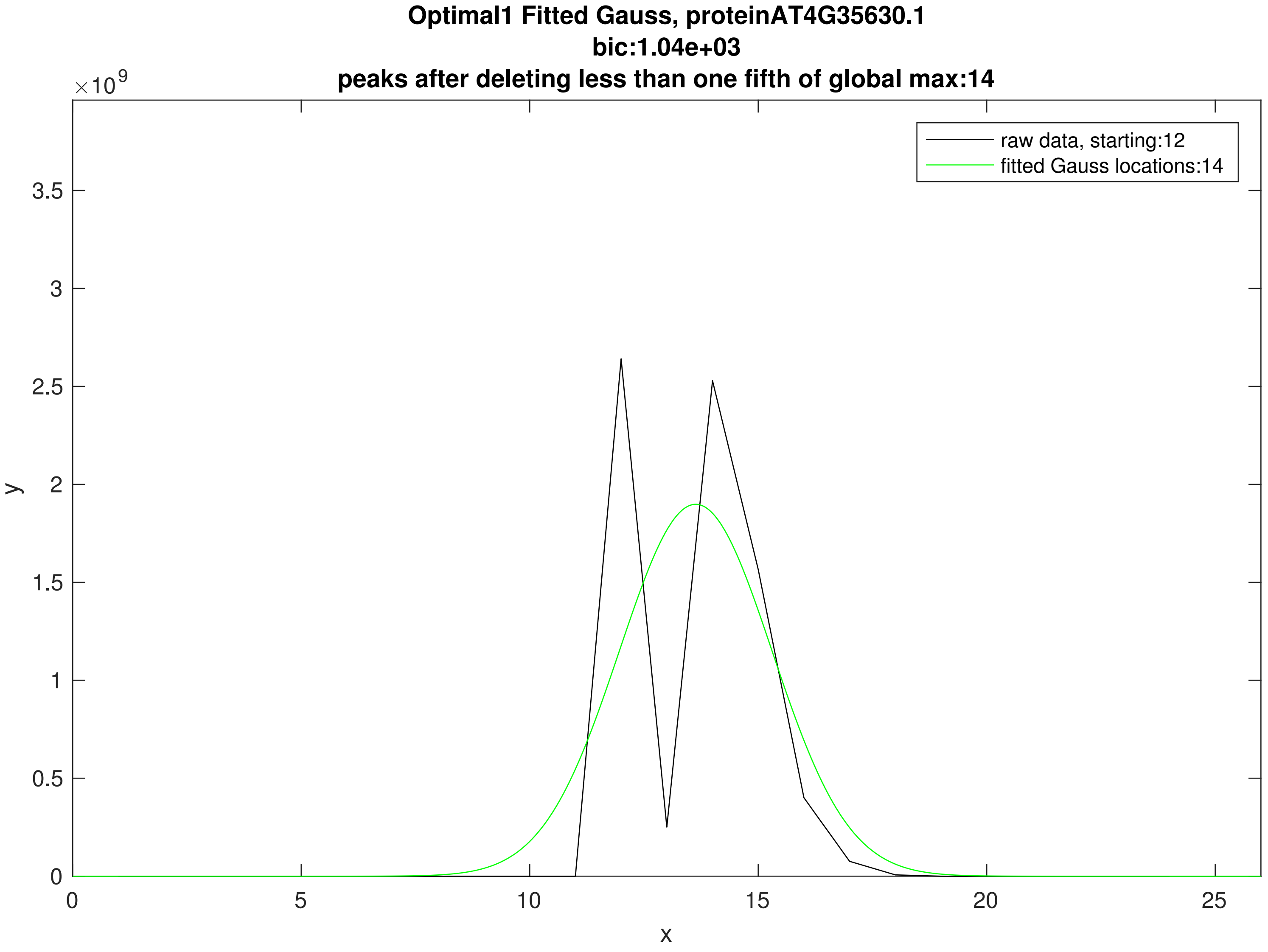

Optimal1 Fitted Gauss, proteinAT4G35630.1
bic:1.04e+03
peaks after deleting less than one fifth of global max:14
×10^9
3.5
3
2.5
2
1.5
1
0.5
0
y
0
5
10
15
20
25
x
raw data, starting:12
fitted Gauss locations:14

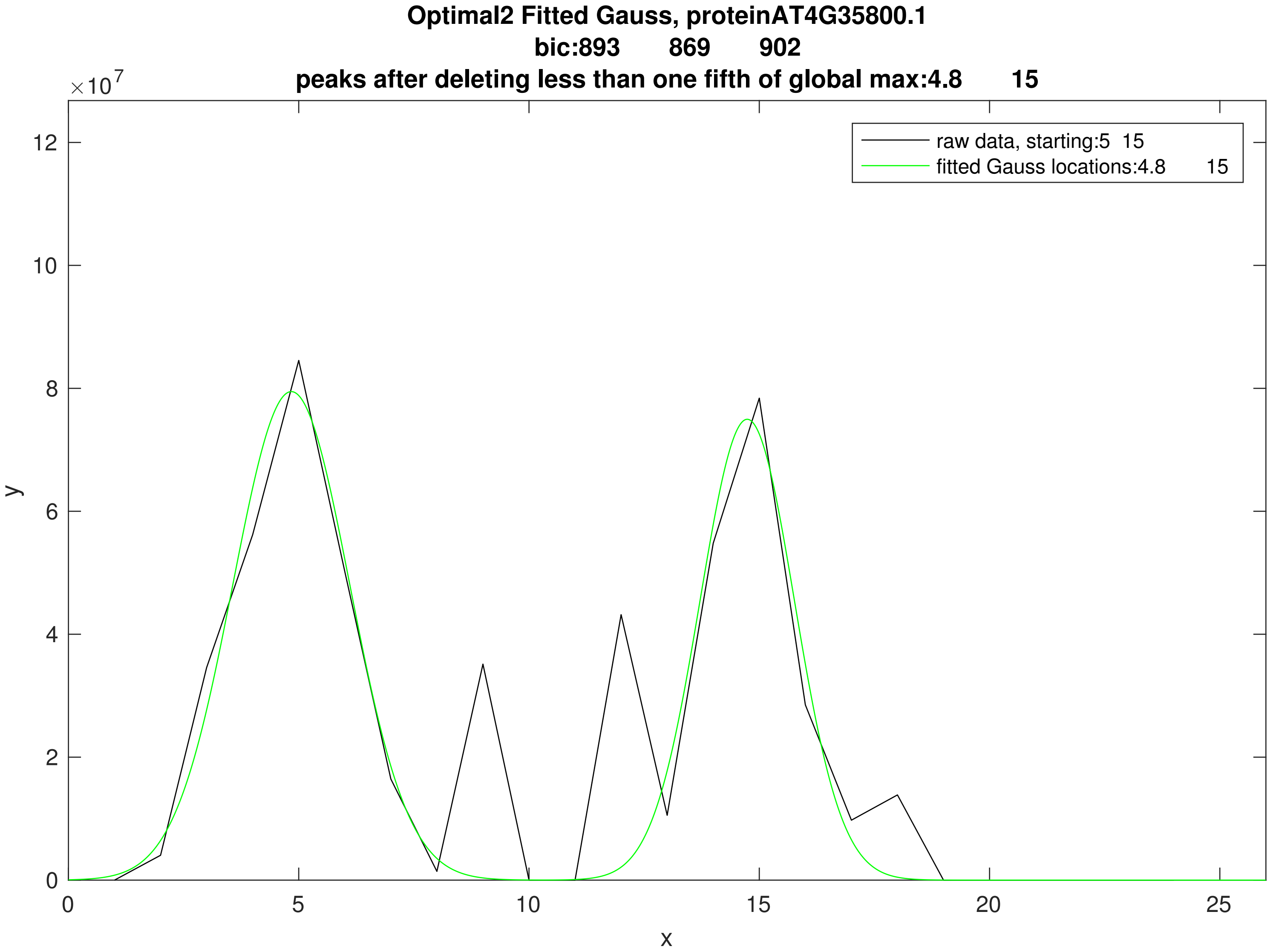
Optimal2 Fitted Gauss, proteinAT4G35800.1
bic:893 869 902
peaks after deleting less than one fifth of global max:4.8 15
×10^7
raw data, starting:5 15
fitted Gauss locations:4.8 15
y
x
0
2
4
6
8
10
12
0
5
10
15
20
25

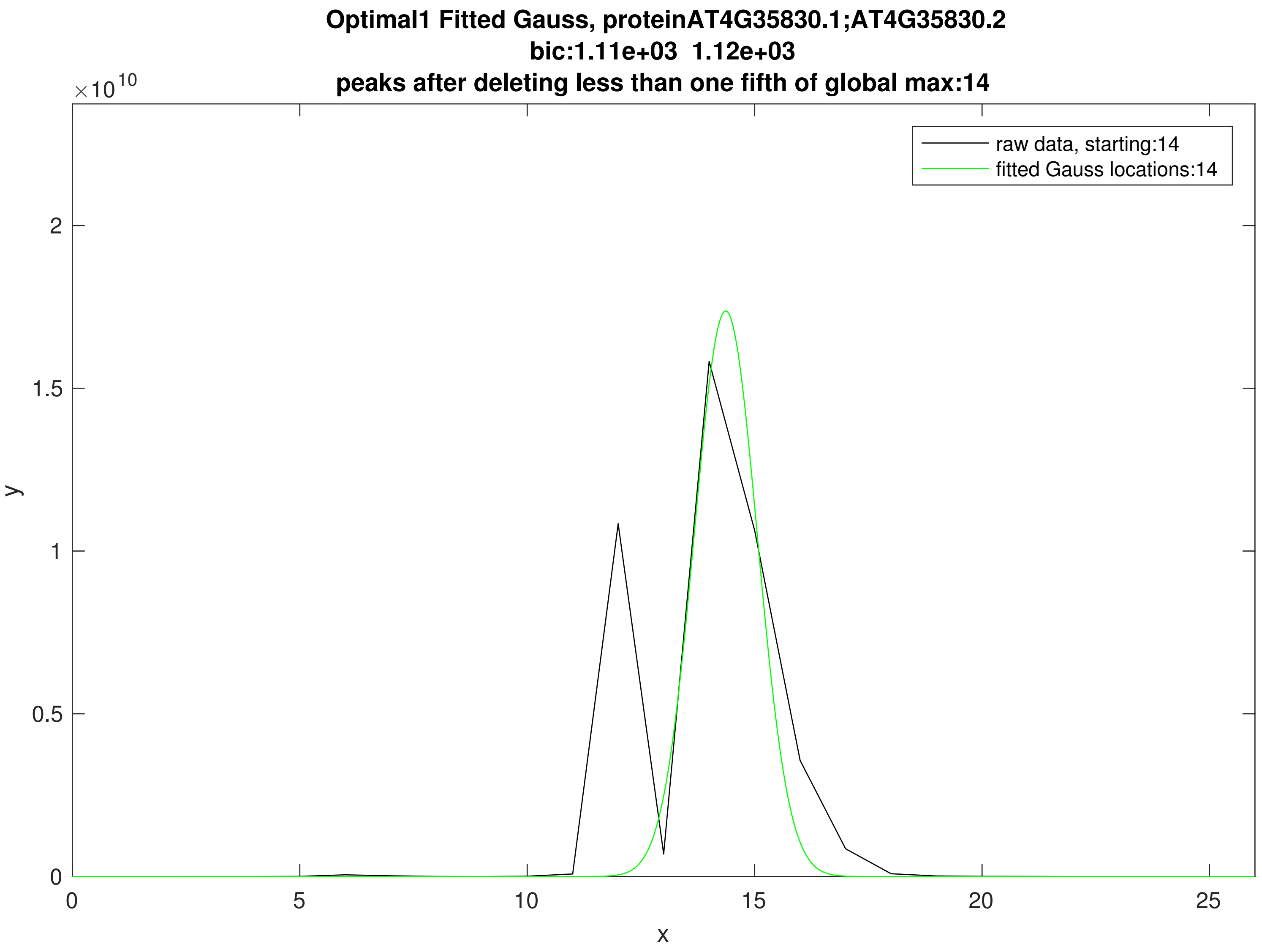
Optimal1 Fitted Gauss, proteinAT4G35830.1;AT4G35830.2
bic:1.11e+03  1.12e+03
peaks after deleting less than one fifth of global max:14
×10^10
raw data, starting:14
fitted Gauss locations:14
y
x
0
0.5
1
1.5
2
5
10
15
20
25

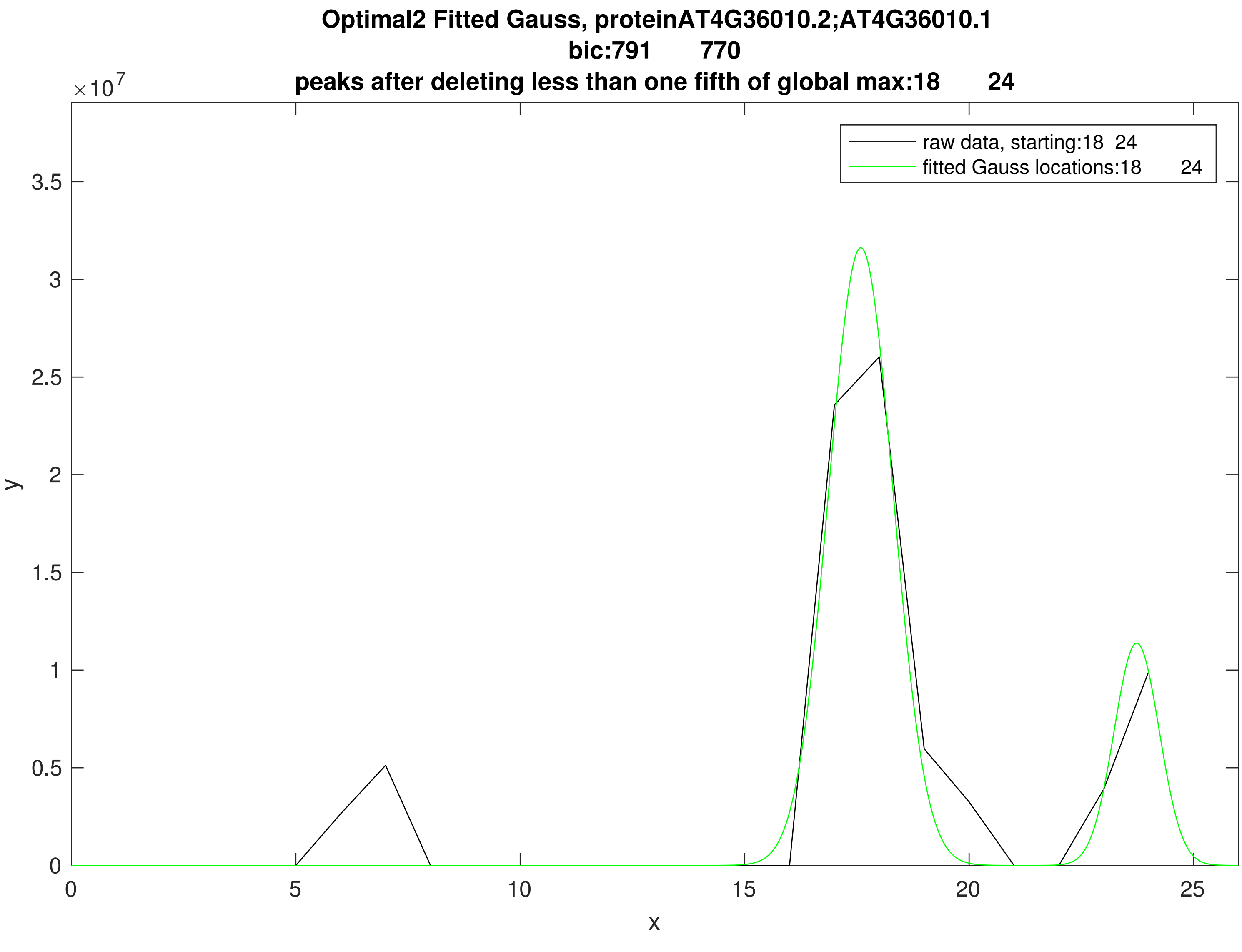

Optimal2 Fitted Gauss, proteinAT4G36010.2;AT4G36010.1
bic:791 770
peaks after deleting less than one fifth of global max:18 24
×10^7
raw data, starting:18 24
fitted Gauss locations:18 24
3.5
3
2.5
2
1.5
1
0.5
0
y
0
5
10
15
20
25
x

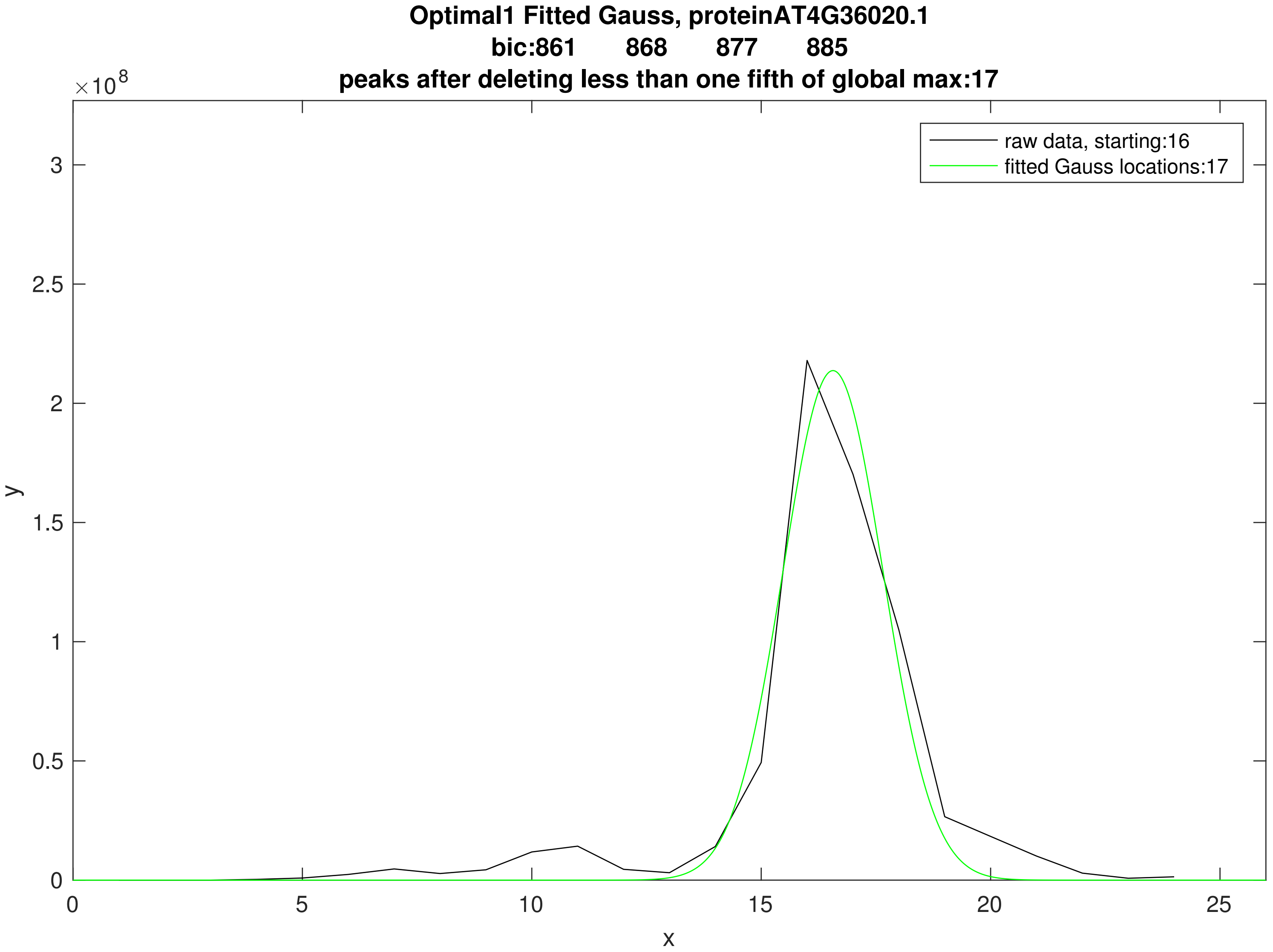
Optimal1 Fitted Gauss, proteinAT4G36020.1
bic:861 868 877 885
peaks after deleting less than one fifth of global max:17
×10^8
3
2.5
2
1.5
1
0.5
0
y
0
5
10
15
20
25
x
raw data, starting:16
fitted Gauss locations:17

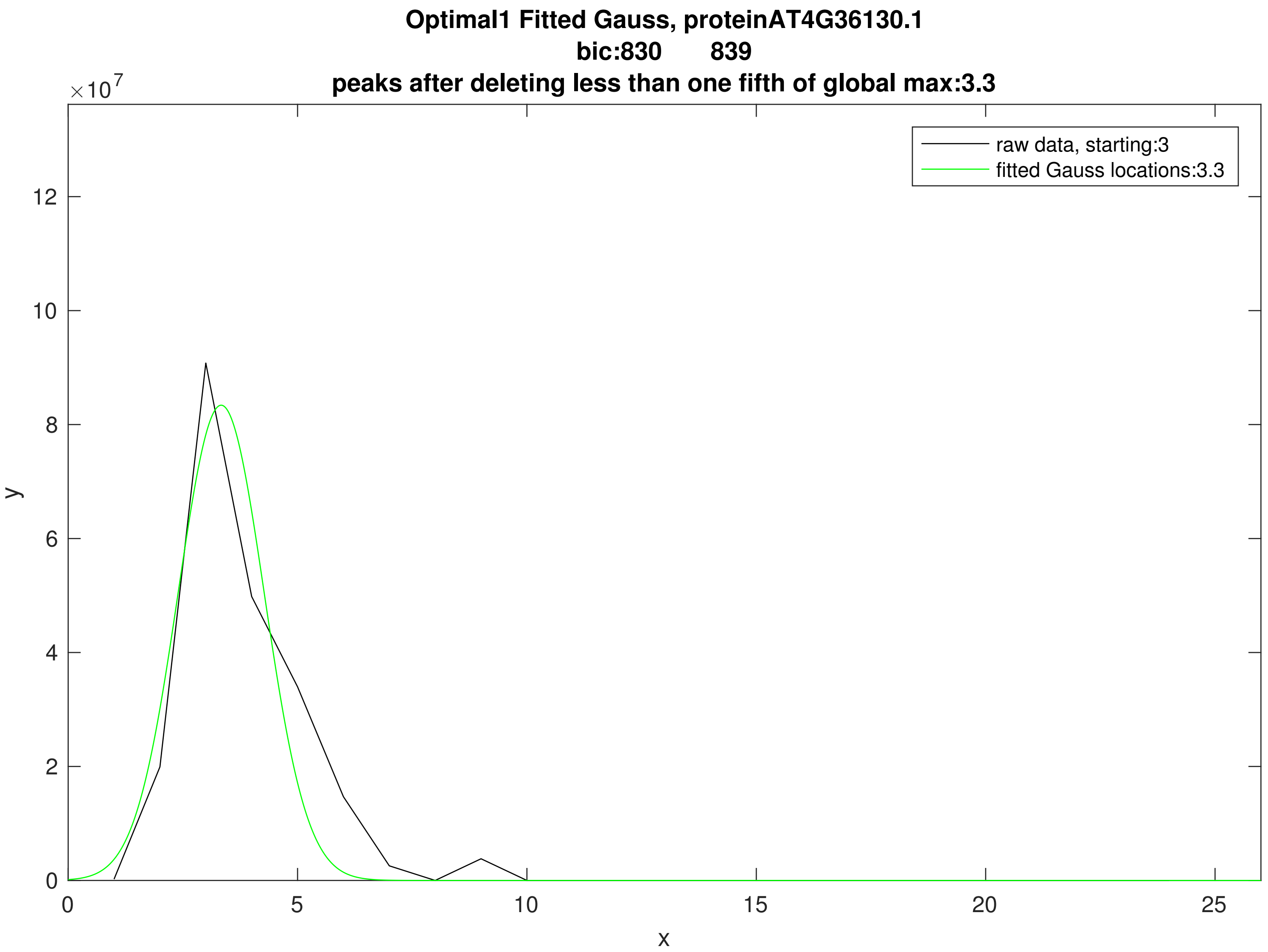
Optimal1 Fitted Gauss, proteinAT4G36130.1
bic:830 839
peaks after deleting less than one fifth of global max:3.3
×10^7
raw data, starting:3
fitted Gauss locations:3.3
y
x
0
2
4
6
8
10
12
5
15
20
25

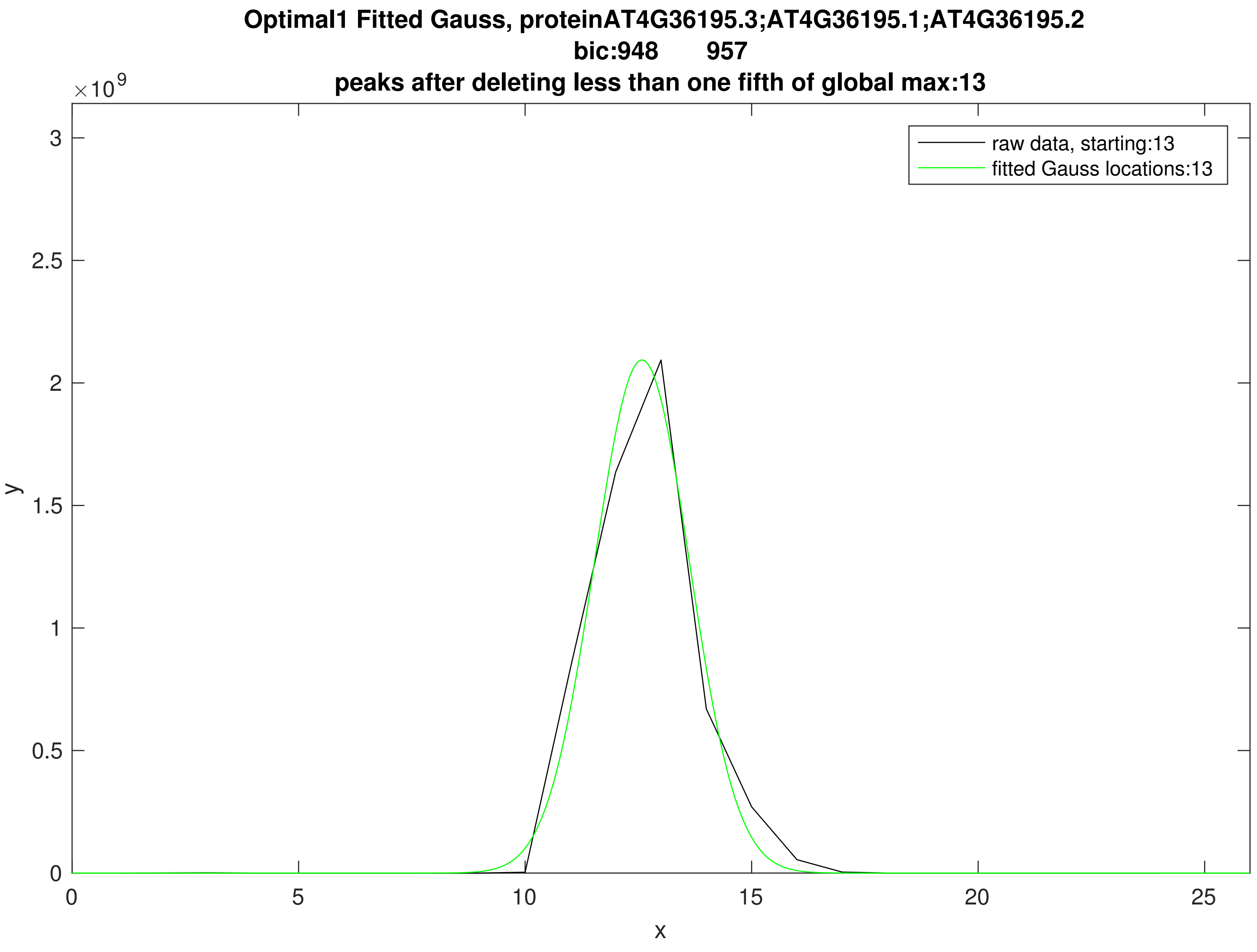
Optimal1 Fitted Gauss, proteinAT4G36195.3;AT4G36195.1;AT4G36195.2
bic:948 957
peaks after deleting less than one fifth of global max:13
×10^9
3
2.5
2
1.5
1
0.5
0
y
0
5
10
15
20
25
x
raw data, starting:13
fitted Gauss locations:13

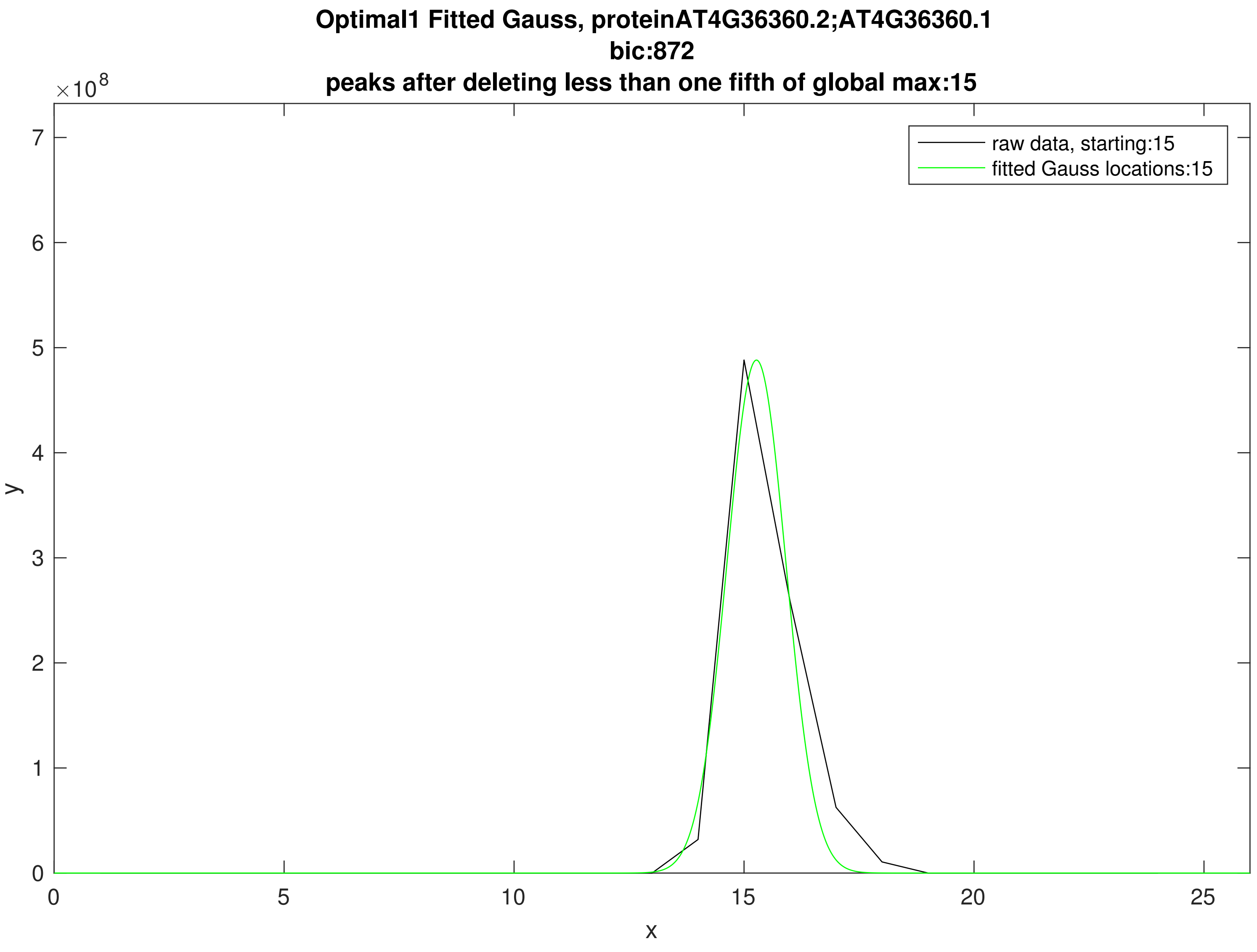
Optimal1 Fitted Gauss, proteinAT4G36360.2;AT4G36360.1
bic:872
peaks after deleting less than one fifth of global max:15
×10^8
raw data, starting:15
fitted Gauss locations:15
y
x
0
1
2
3
4
5
6
7
0
5
10
15
20
25

#### Optimal1 Fitted Gauss, proteinAT4G36400.2;AT4G36400.1
#### bic:816
#### peaks after deleting less than one fifth of global max:12

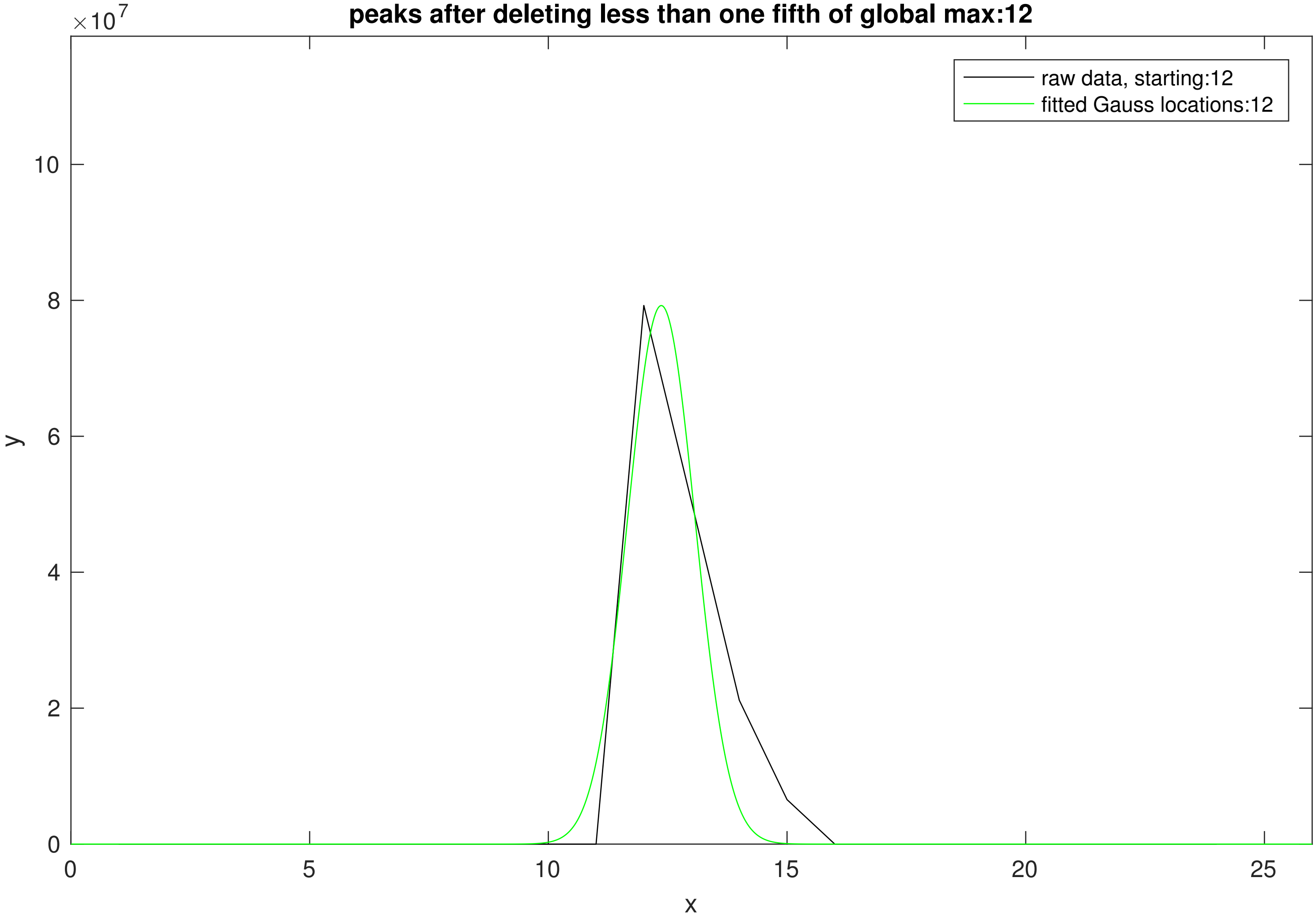

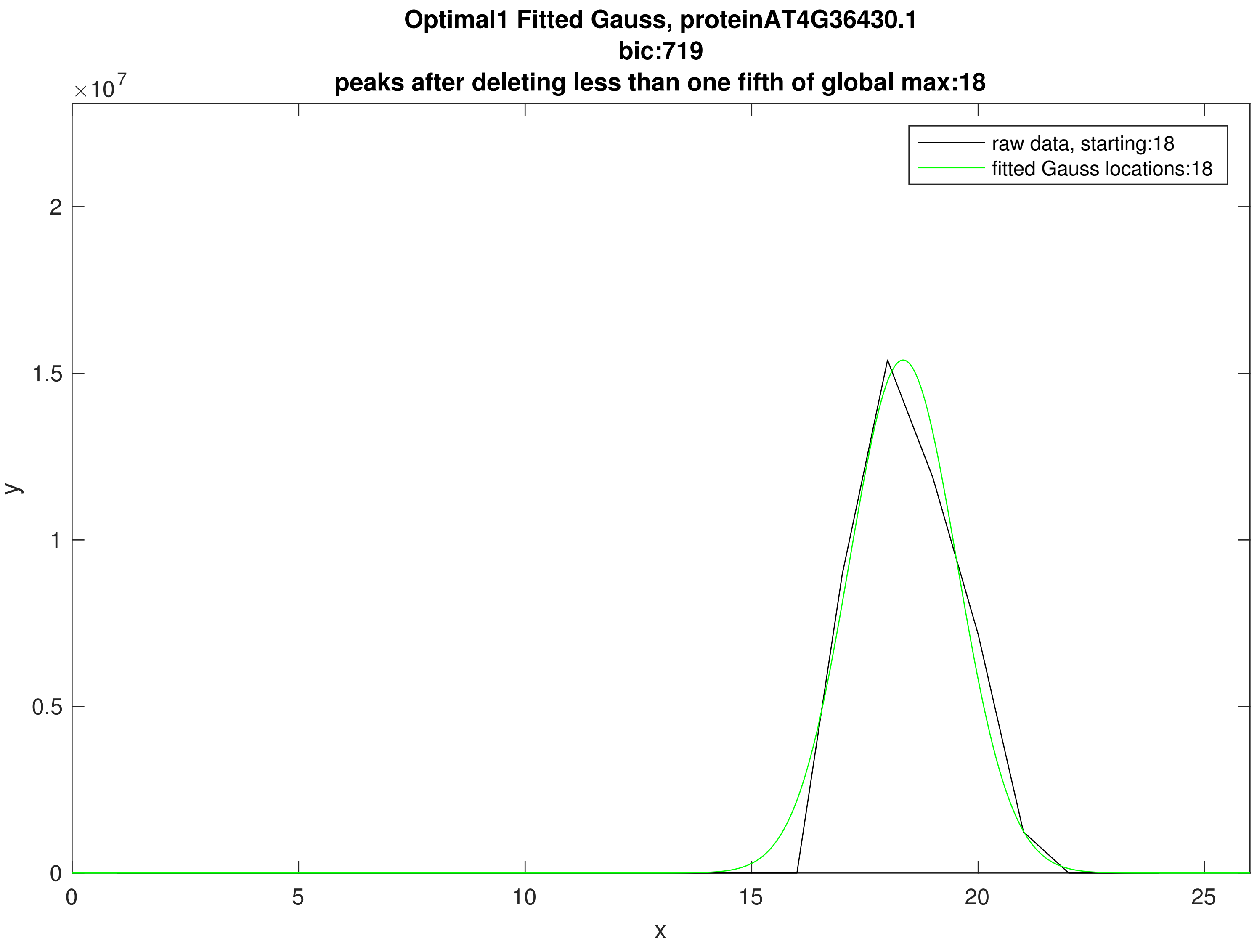
Optimal1 Fitted Gauss, proteinAT4G36430.1
bic:719
peaks after deleting less than one fifth of global max:18
×10^7
raw data, starting:18
fitted Gauss locations:18
y
x
0
0.5
1
1.5
2
0
5
10
15
20
25

**Optimal1 Fitted Gauss, proteinAT4G36680.1**
**bic:635**
**peaks after deleting less than one fifth of global max:3.2**

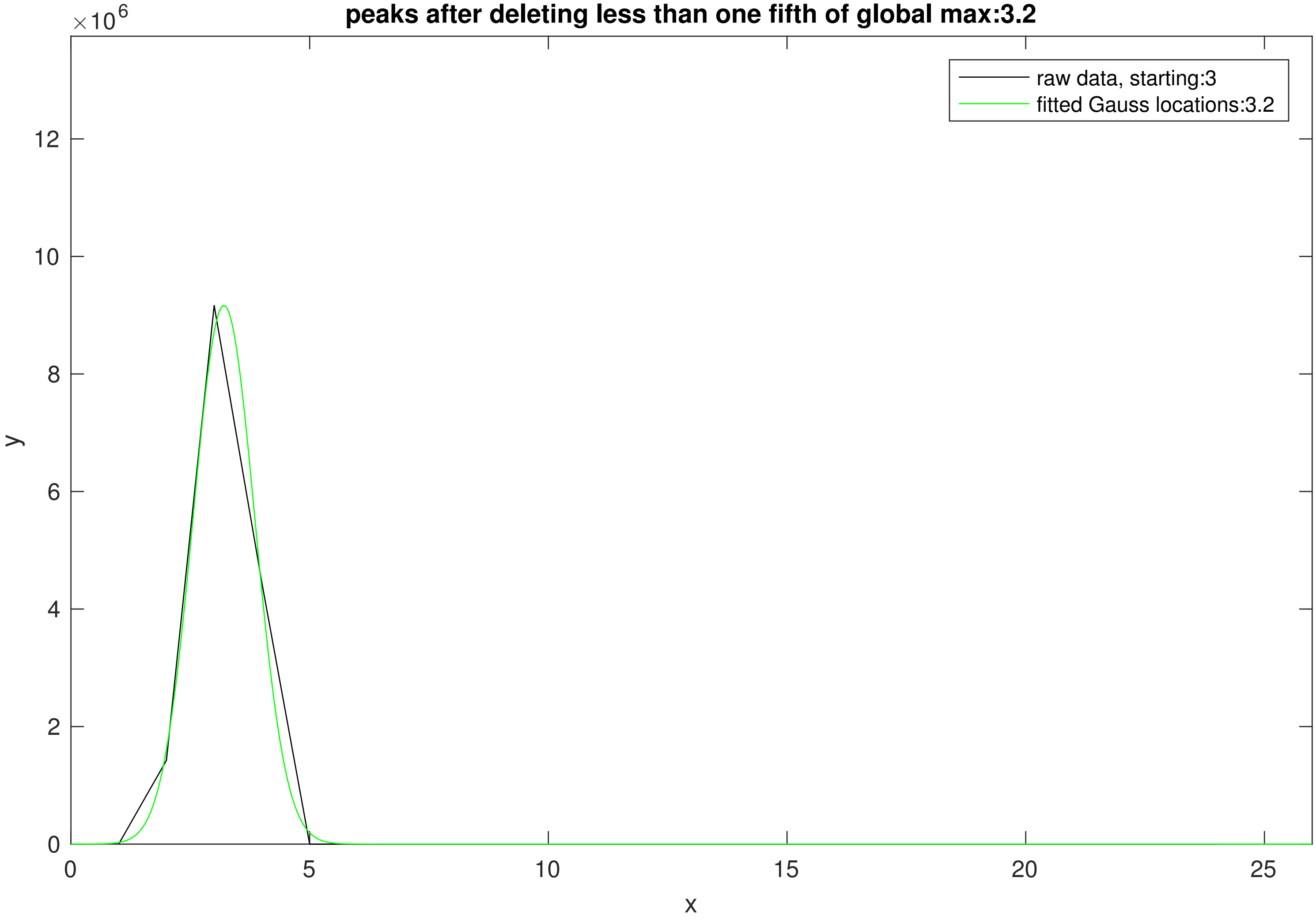

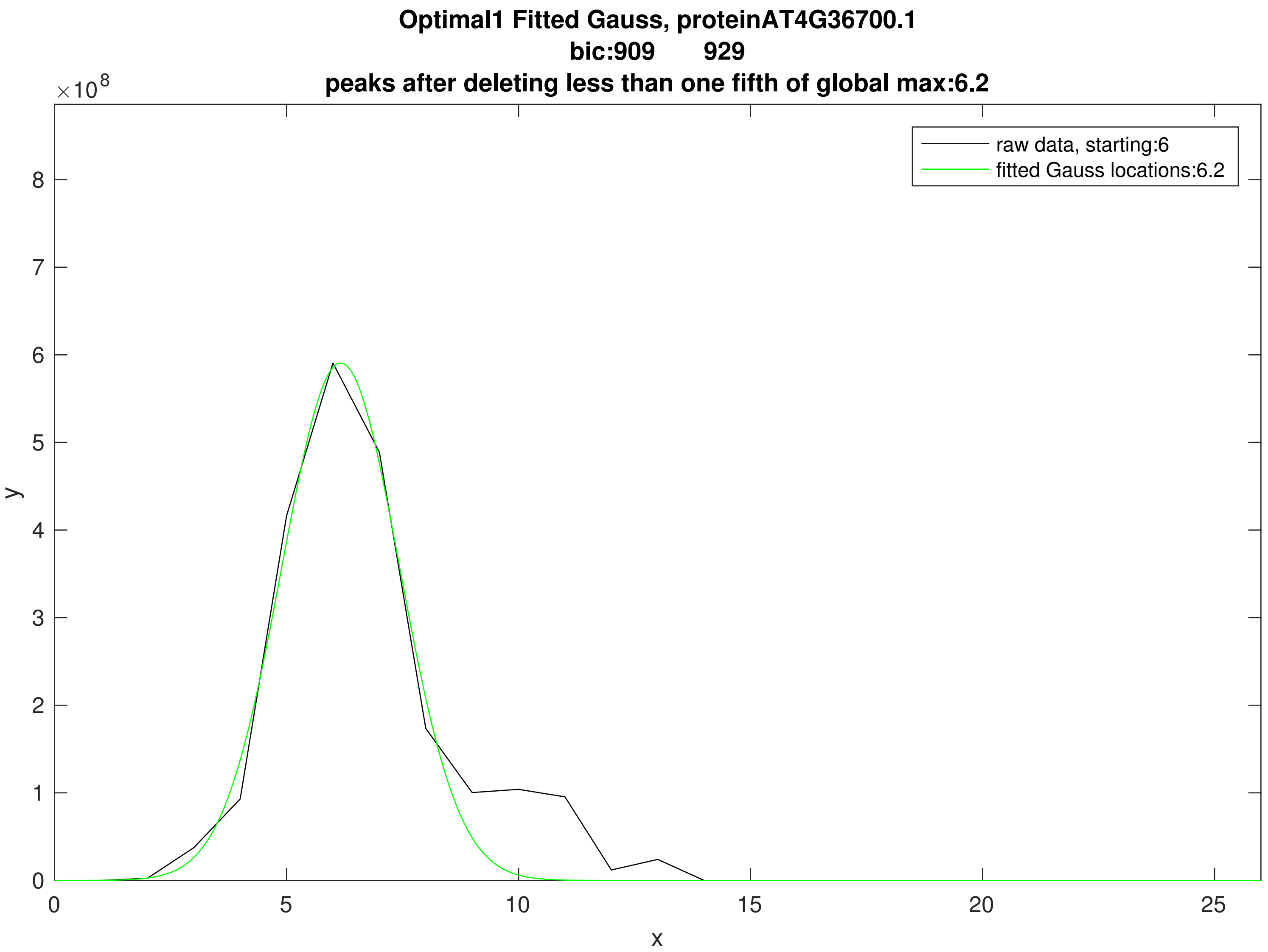
Optimal1 Fitted Gauss, proteinAT4G36700.1
bic:909 929
peaks after deleting less than one fifth of global max:6.2
×10^8
raw data, starting:6
fitted Gauss locations:6.2
y
x
0
1
2
3
4
5
6
7
8
0
5
10
15
20
25

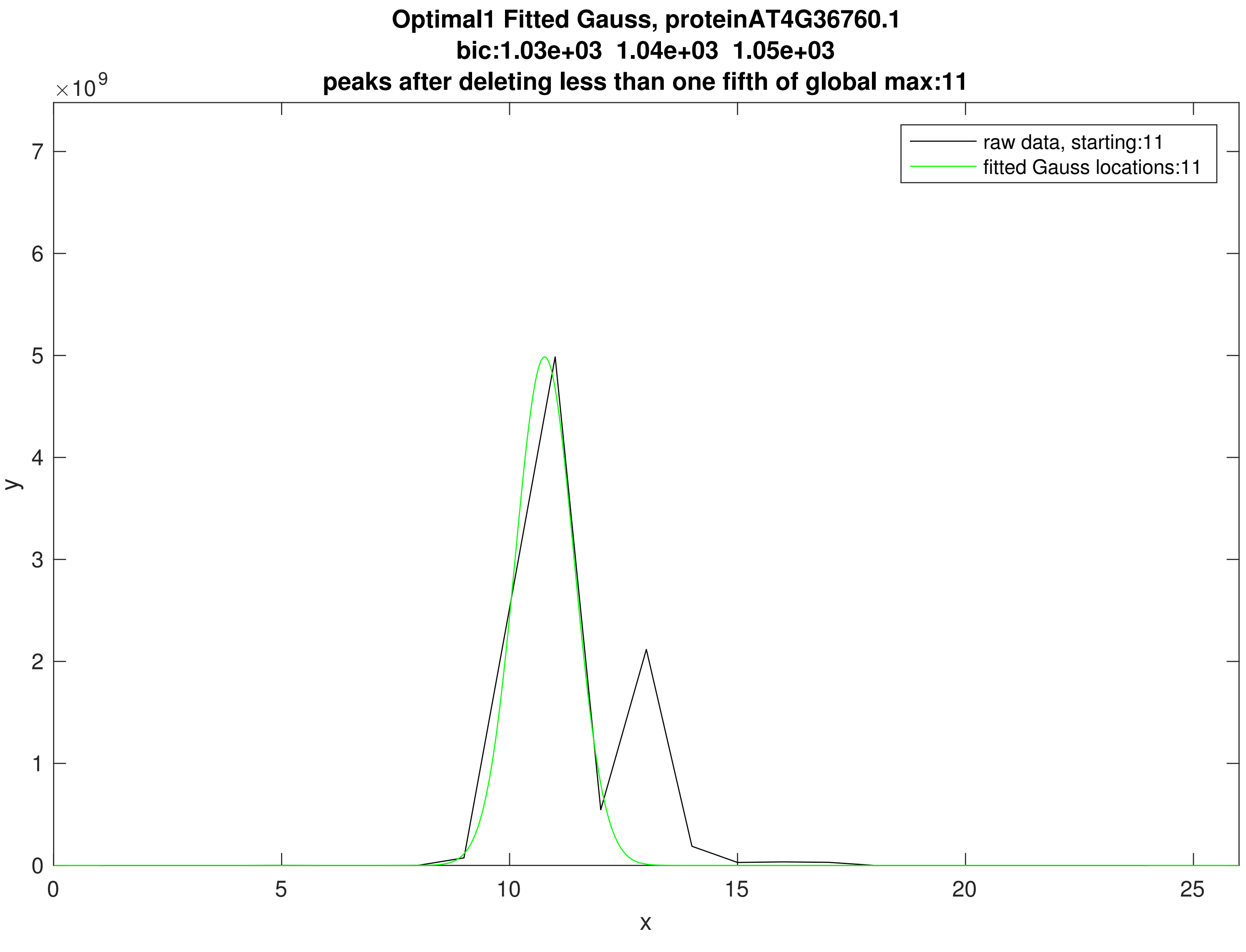
Optimal1 Fitted Gauss, proteinAT4G36760.1
bic:1.03e+03  1.04e+03  1.05e+03
peaks after deleting less than one fifth of global max:11
×10^9
raw data, starting:11
fitted Gauss locations:11
y
x
0
1
2
3
4
5
6
7
0
5
10
15
20
25

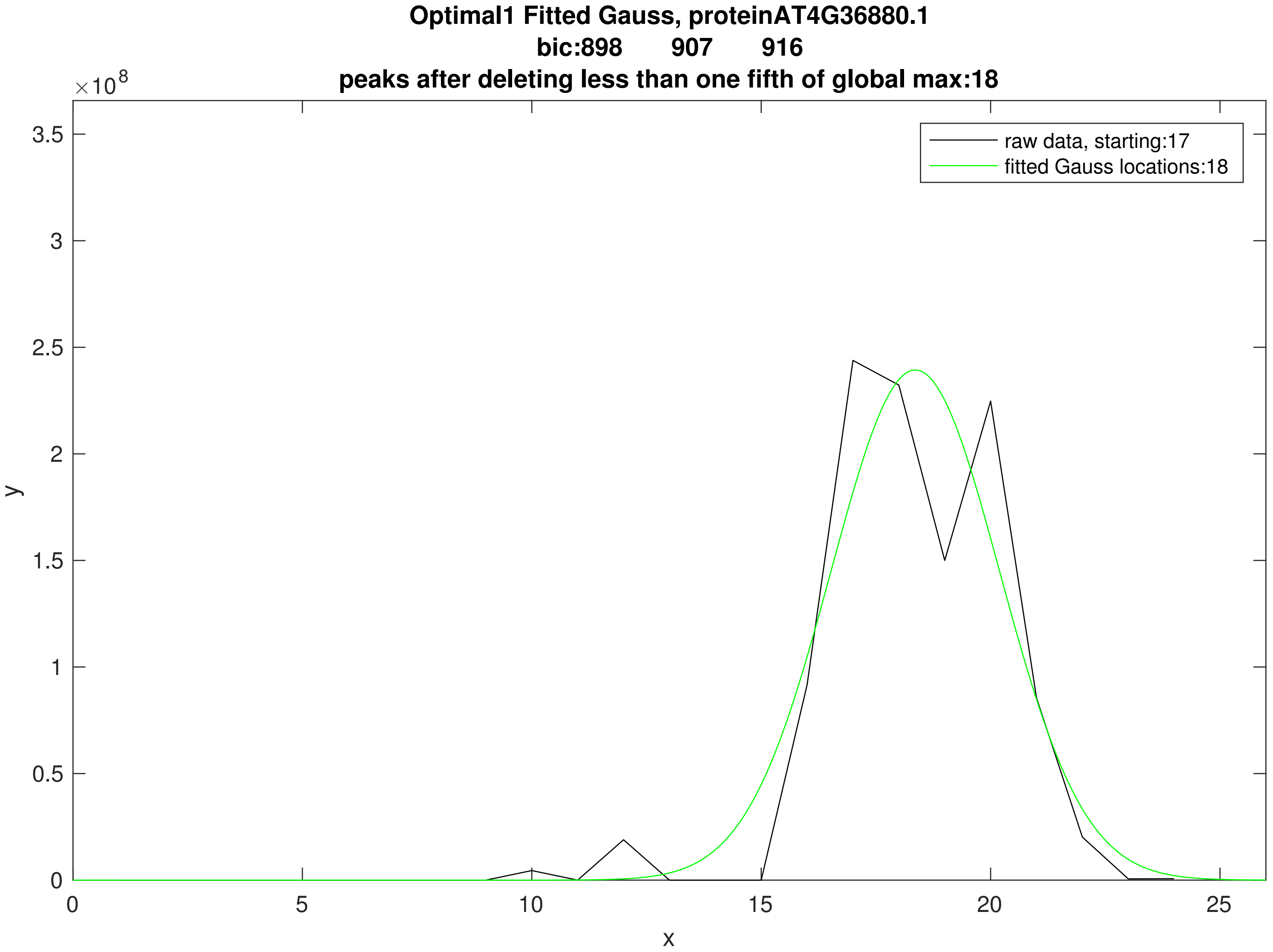
Optimal1 Fitted Gauss, proteinAT4G36880.1
bic:898 907 916
peaks after deleting less than one fifth of global max:18
×10^8
raw data, starting:17
fitted Gauss locations:18
y
x
0
0.5
1
1.5
2
2.5
3
3.5
0
5
10
15
20
25

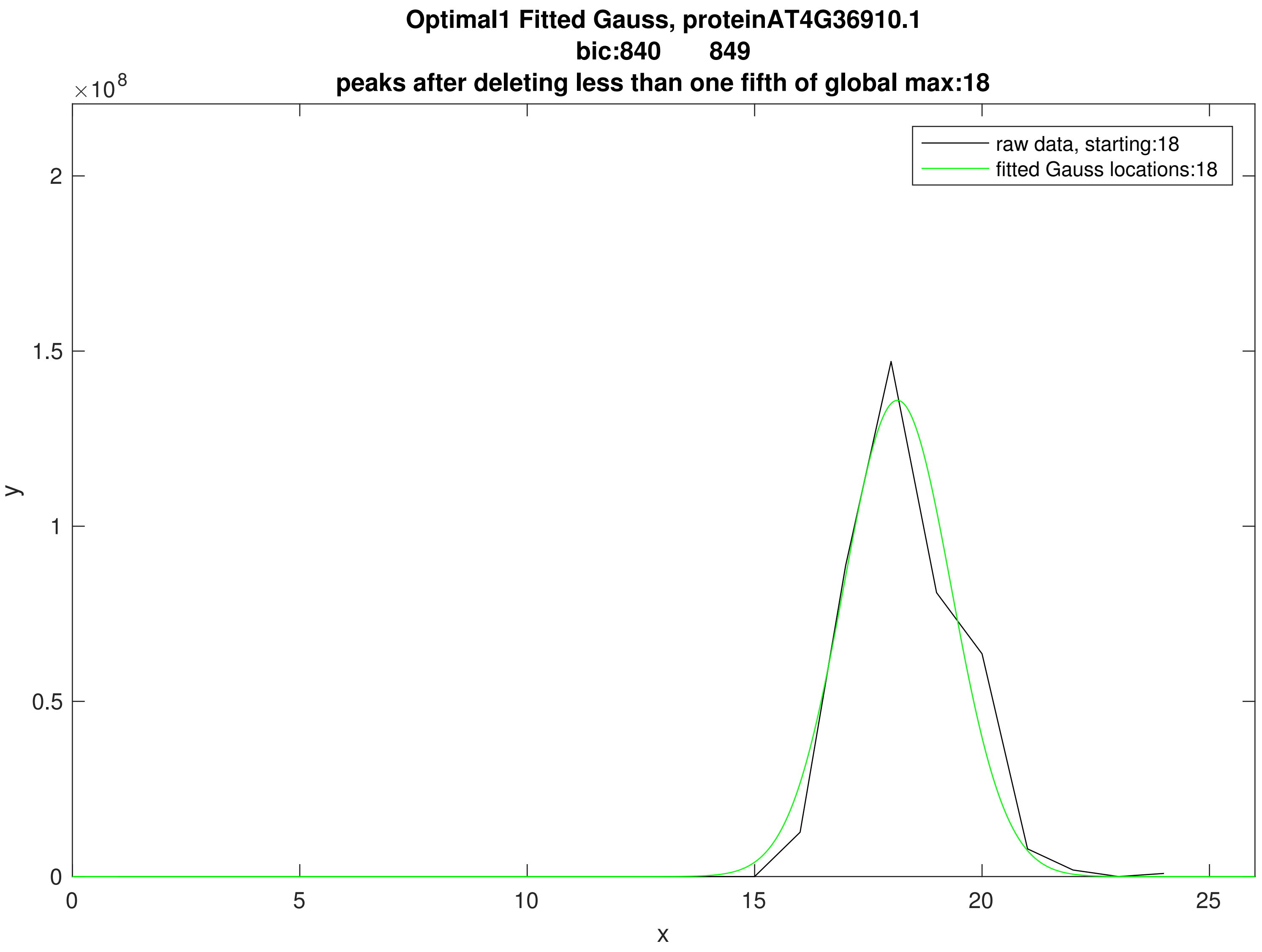

Optimal1 Fitted Gauss, proteinAT4G36910.1
bic:840 849
peaks after deleting less than one fifth of global max:18
×10^8
raw data, starting:18
fitted Gauss locations:18
y
x
0
0.5
1
1.5
2
0
5
10
15
20
25

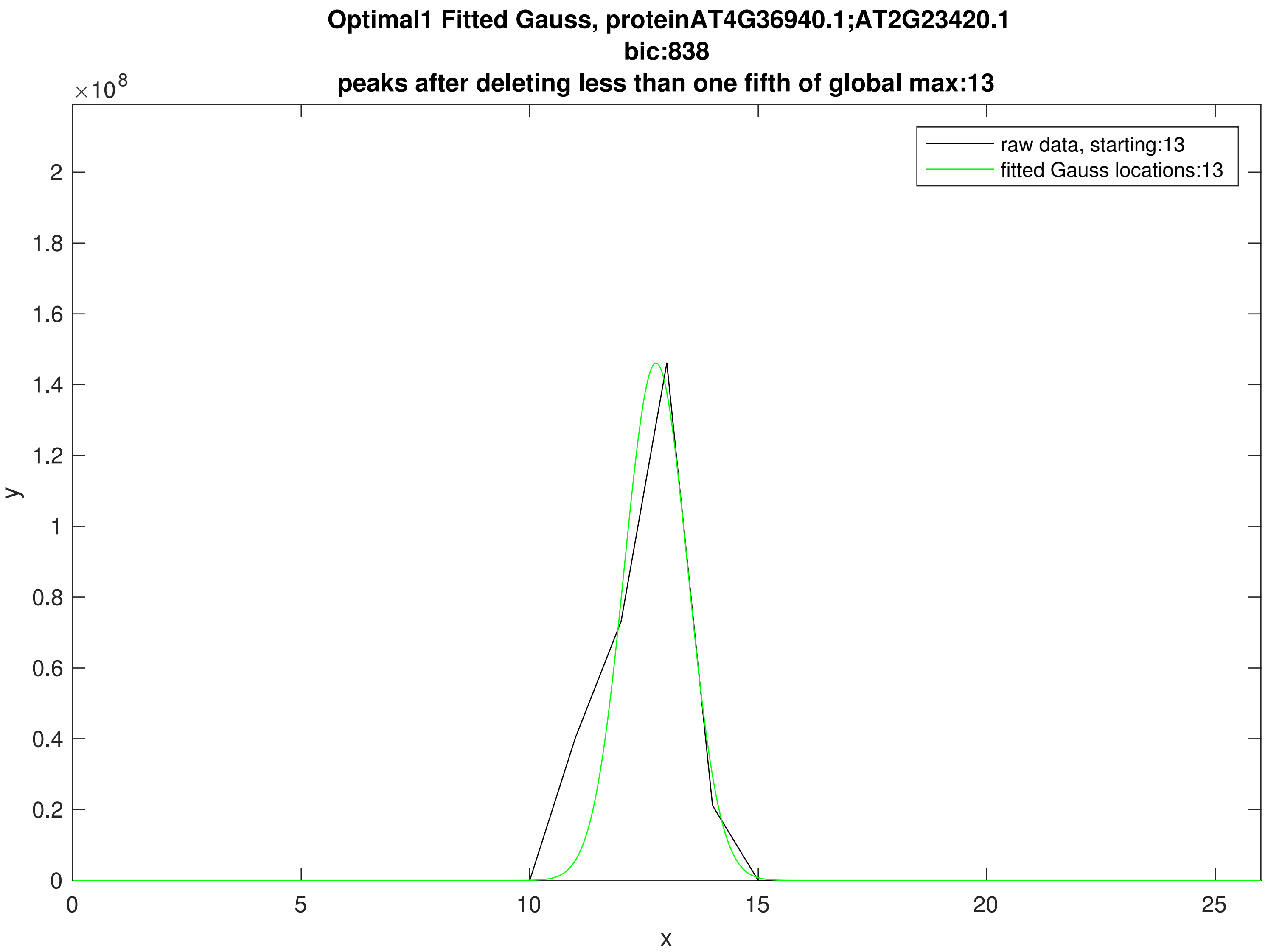

Optimal1 Fitted Gauss, proteinAT4G36940.1;AT2G23420.1
bic:838
peaks after deleting less than one fifth of global max:13
×10^8
raw data, starting:13
fitted Gauss locations:13
y
x
0
0.2
0.4
0.6
0.8
1
1.2
1.4
1.6
1.8
2
0
5
10
15
20
25

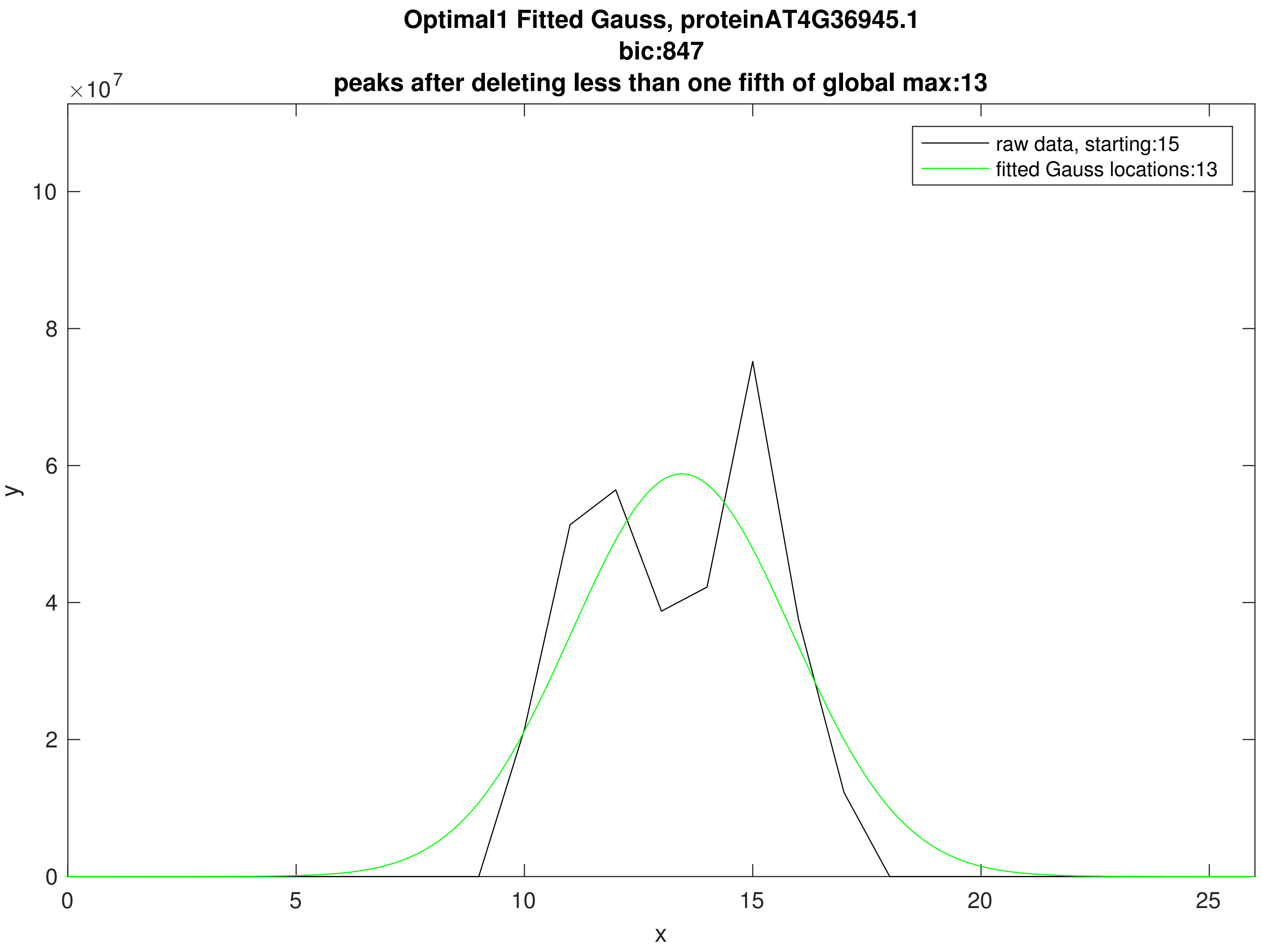
Optimal1 Fitted Gauss, proteinAT4G36945.1
bic:847
peaks after deleting less than one fifth of global max:13
×10^7
raw data, starting:15
fitted Gauss locations:13
10
8
6
4
2
0
y
0
5
10
15
20
25
x

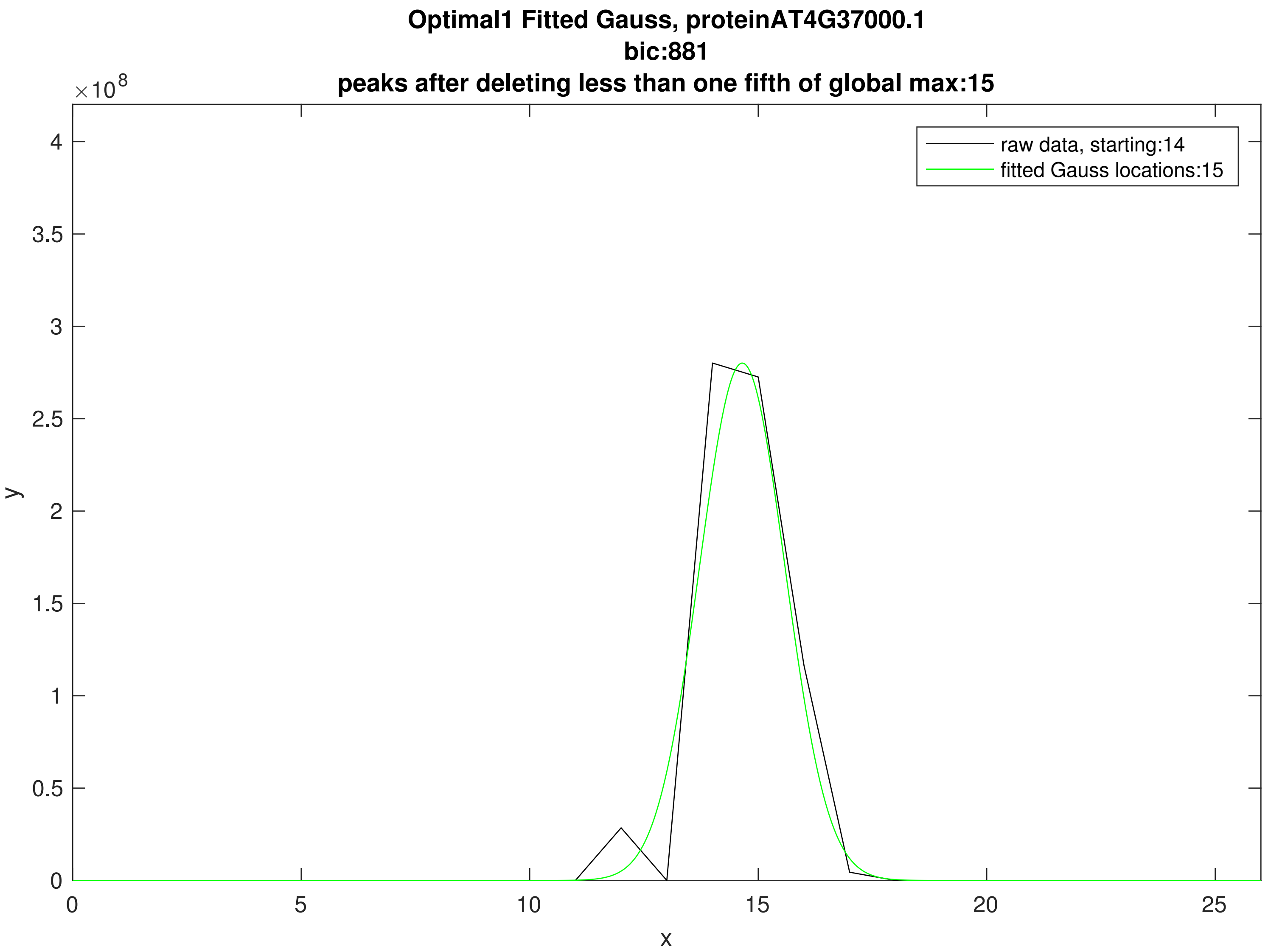

Optimal1 Fitted Gauss, proteinAT4G37000.1
bic:881
peaks after deleting less than one fifth of global max:15
×10^8
raw data, starting:14
fitted Gauss locations:15
4
3.5
3
2.5
2
1.5
1
0.5
0
y
0
5
10
15
20
25
x

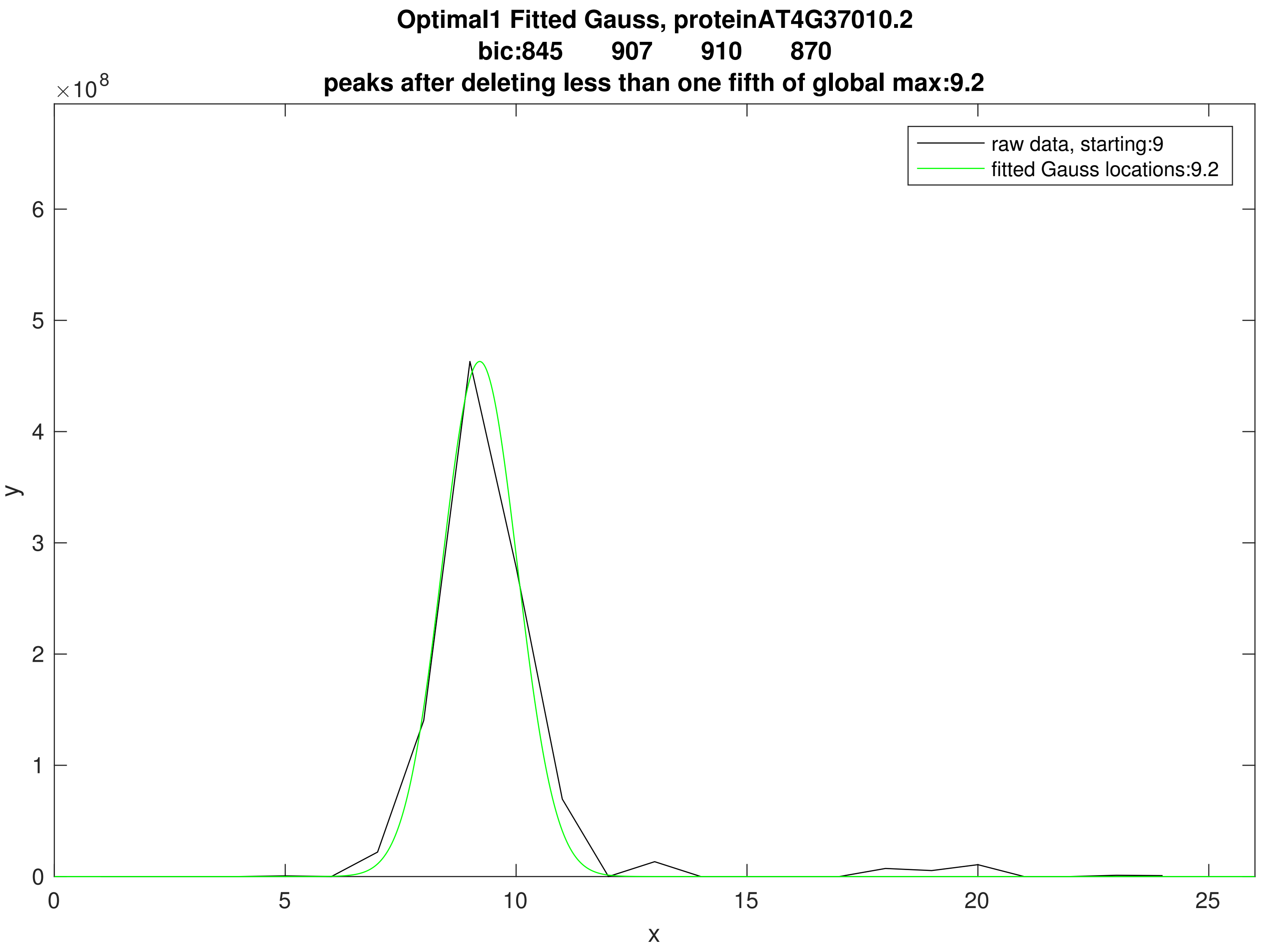

Optimal1 Fitted Gauss, proteinAT4G37010.2
bic:845 907 910 870
peaks after deleting less than one fifth of global max:9.2
×10^8
raw data, starting:9
fitted Gauss locations:9.2
y
x
0
1
2
3
4
5
6
0
5
10
15
20
25

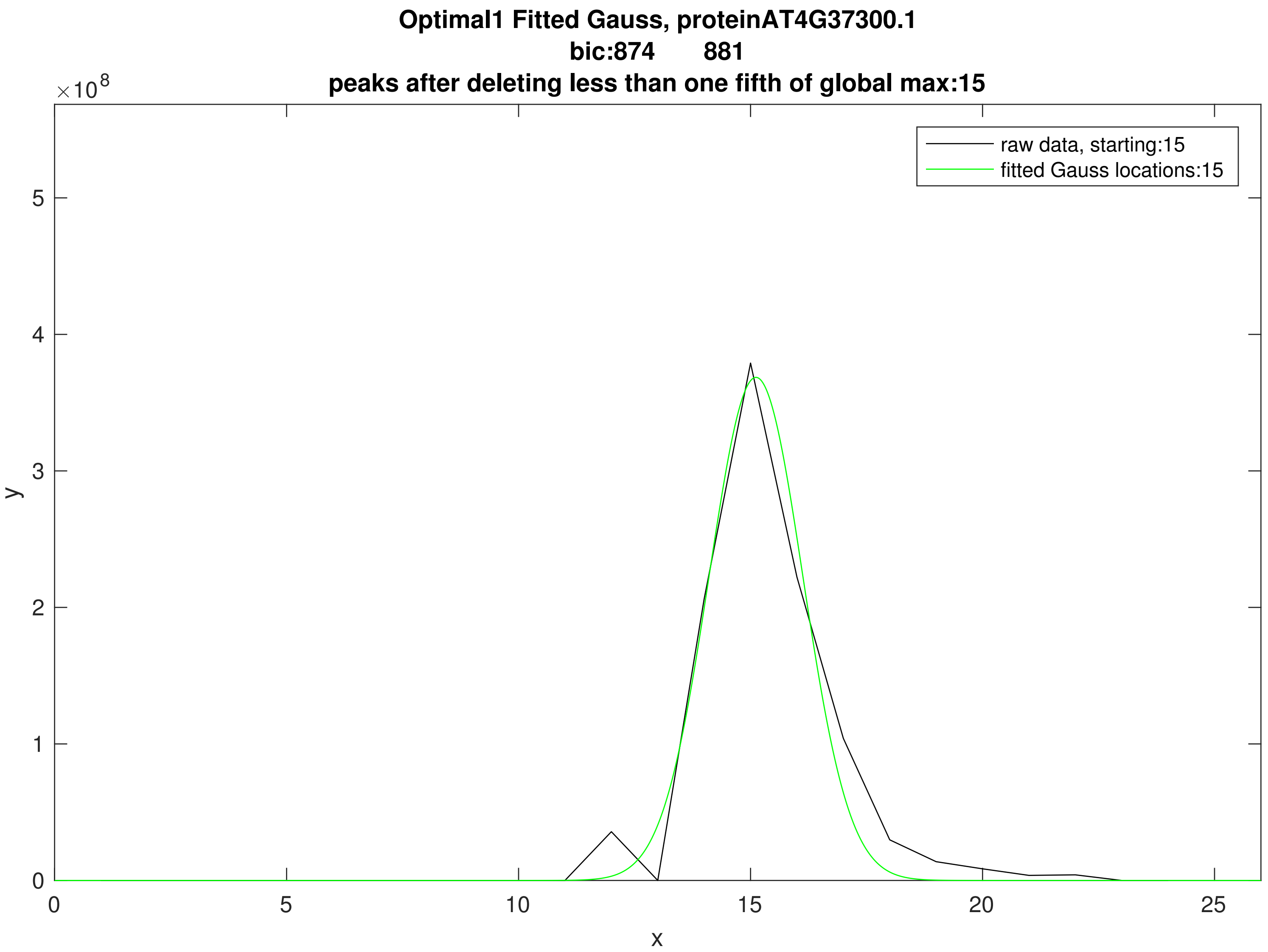

Optimal1 Fitted Gauss, proteinAT4G37300.1
bic:874 881
peaks after deleting less than one fifth of global max:15
×10^8
raw data, starting:15
fitted Gauss locations:15
y
x
0
1
2
3
4
5
0
5
10
15
20
25

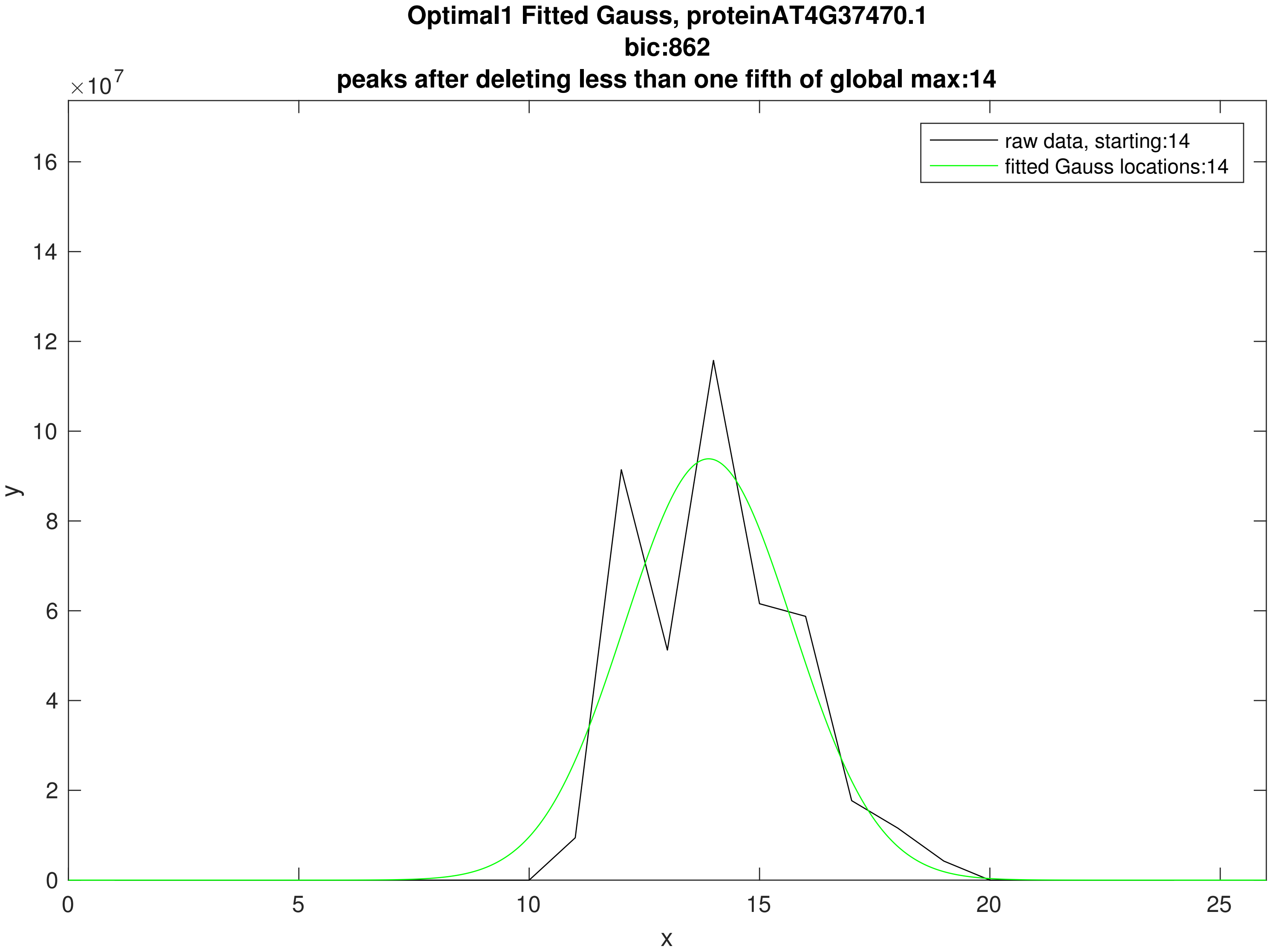

Optimal1 Fitted Gauss, proteinAT4G37470.1
bic:862
peaks after deleting less than one fifth of global max:14
×10^7
16
14
12
10
8
6
4
2
0
y
0
5
10
15
20
25
x
raw data, starting:14
fitted Gauss locations:14

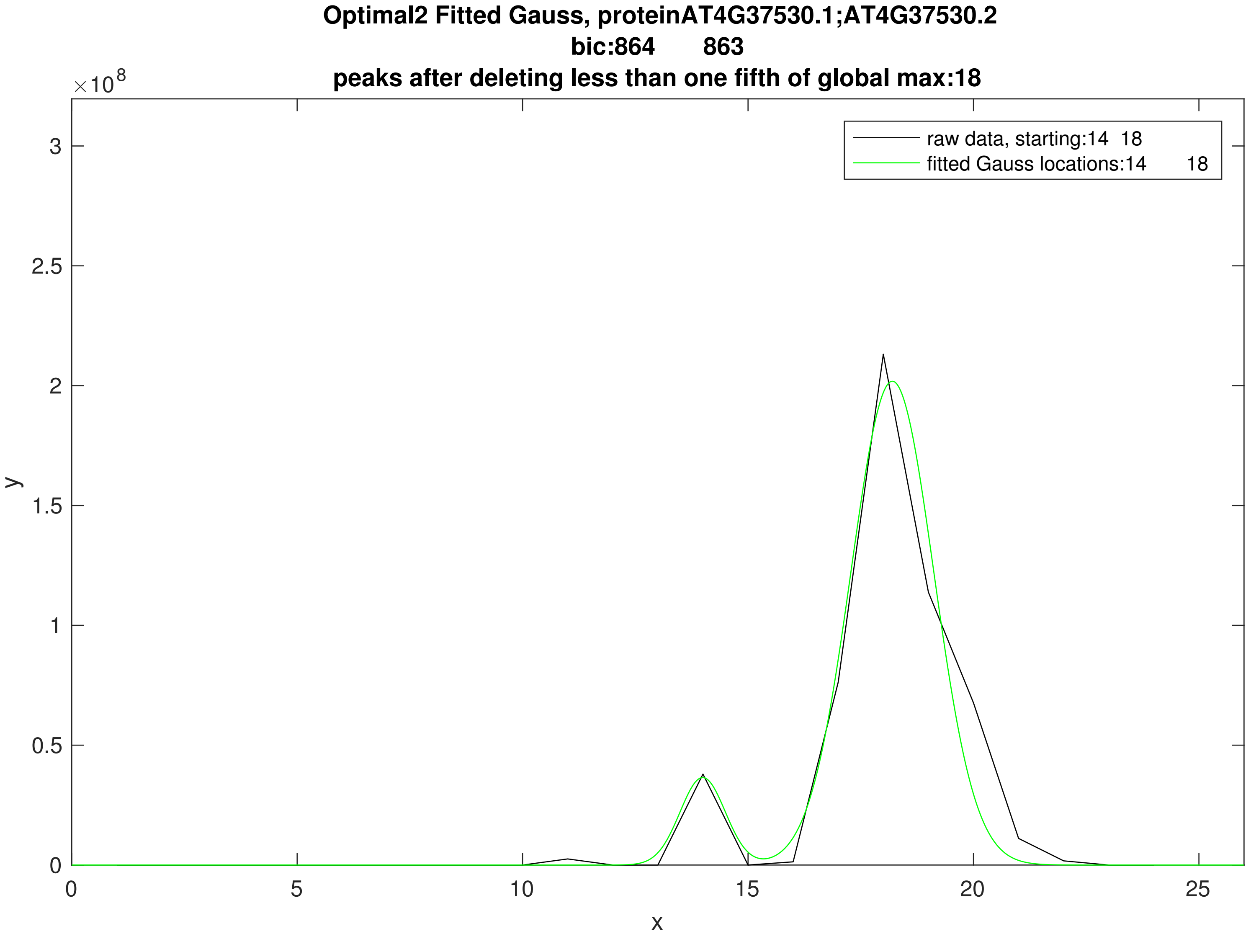

Optimal2 Fitted Gauss, proteinAT4G37530.1;AT4G37530.2
bic:864 863
peaks after deleting less than one fifth of global max:18
×10^8
raw data, starting:14 18
fitted Gauss locations:14 18
y
x
0
0.5
1
1.5
2
2.5
3
0
5
10
15
20
25

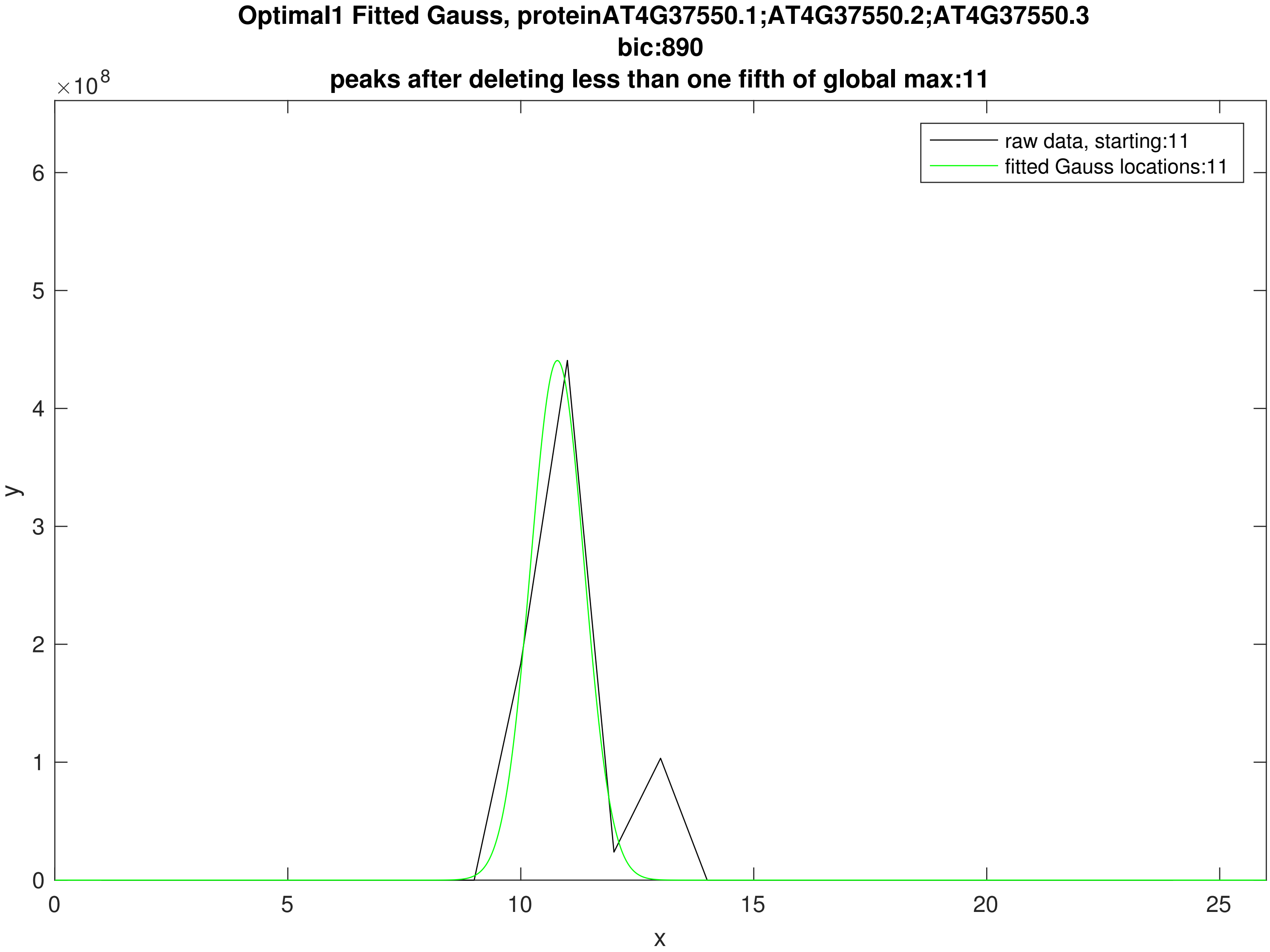

Optimal1 Fitted Gauss, proteinAT4G37550.1;AT4G37550.2;AT4G37550.3
bic:890
peaks after deleting less than one fifth of global max:11
×10^8
raw data, starting:11
fitted Gauss locations:11
y
x
0
1
2
3
4
5
6
0
5
10
15
20
25

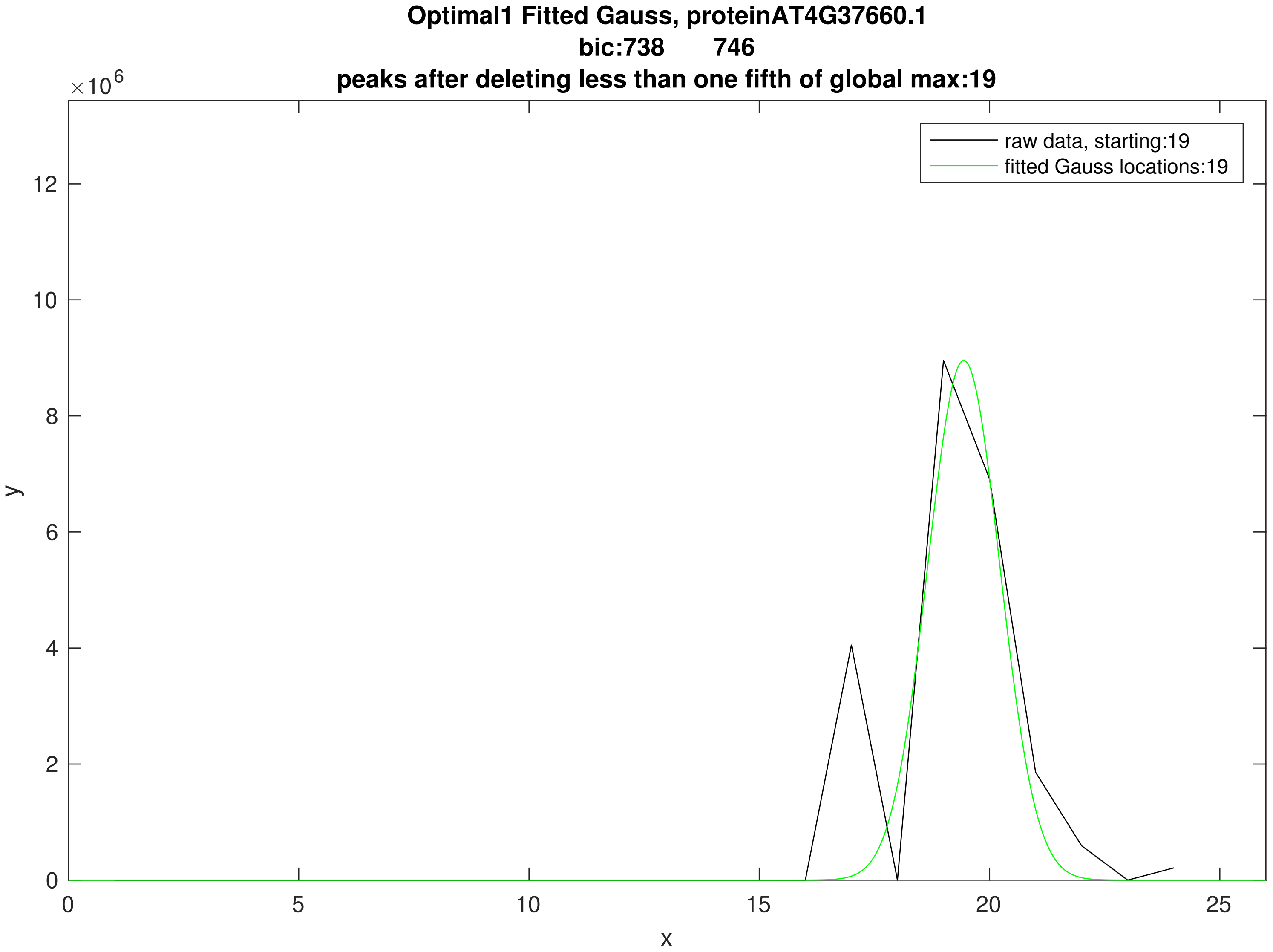
Optimal1 Fitted Gauss, proteinAT4G37660.1
bic:738 746
peaks after deleting less than one fifth of global max:19
×10^6
raw data, starting:19
fitted Gauss locations:19
y
x
0
2
4
6
8
10
12
0
5
10
15
20
25

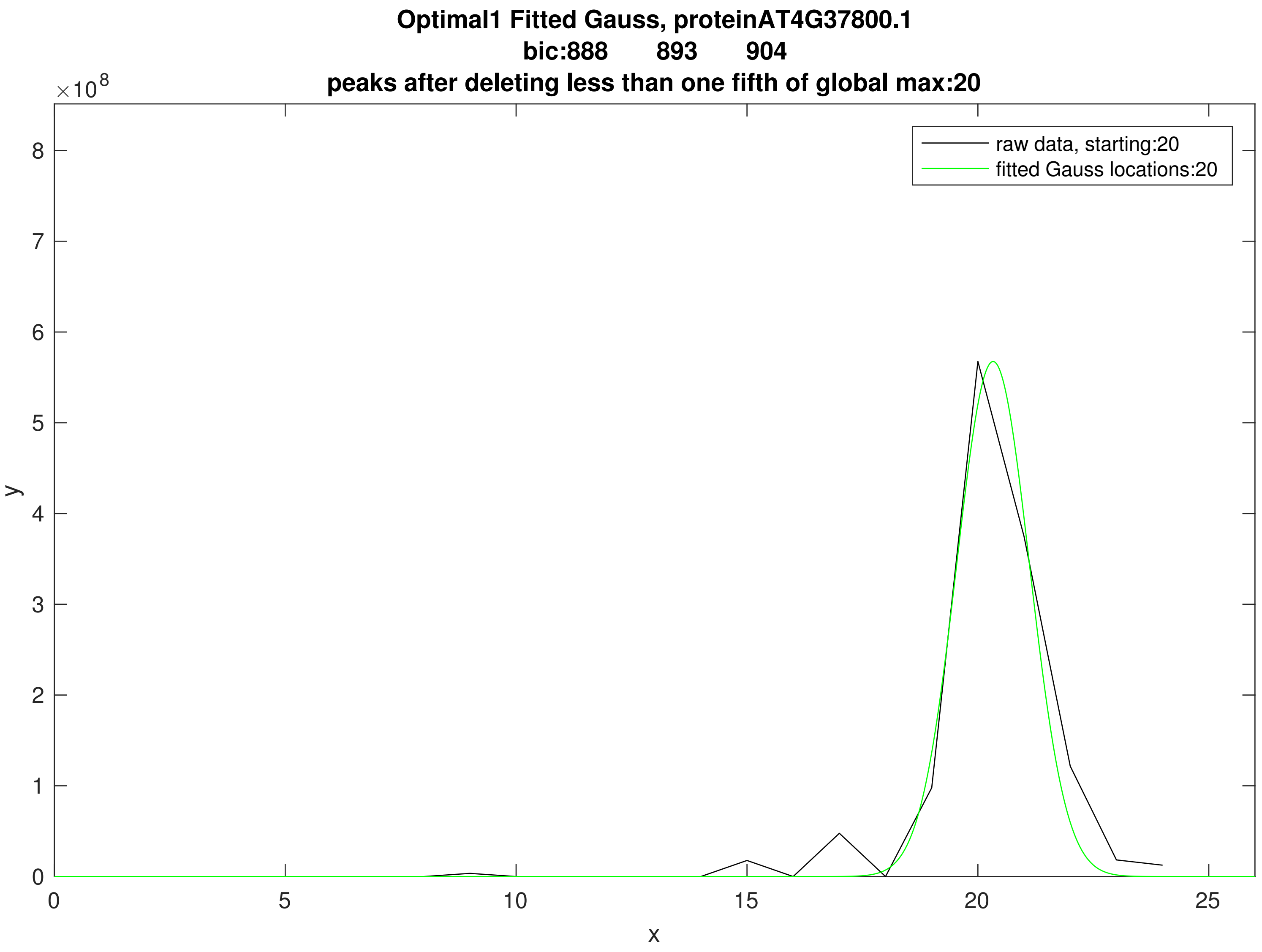

Optimal1 Fitted Gauss, proteinAT4G37800.1
bic:888 893 904
peaks after deleting less than one fifth of global max:20
×10^8
8
7
6
5
4
3
2
1
0
y
0
5
10
15
20
25
x
raw data, starting:20
fitted Gauss locations:20

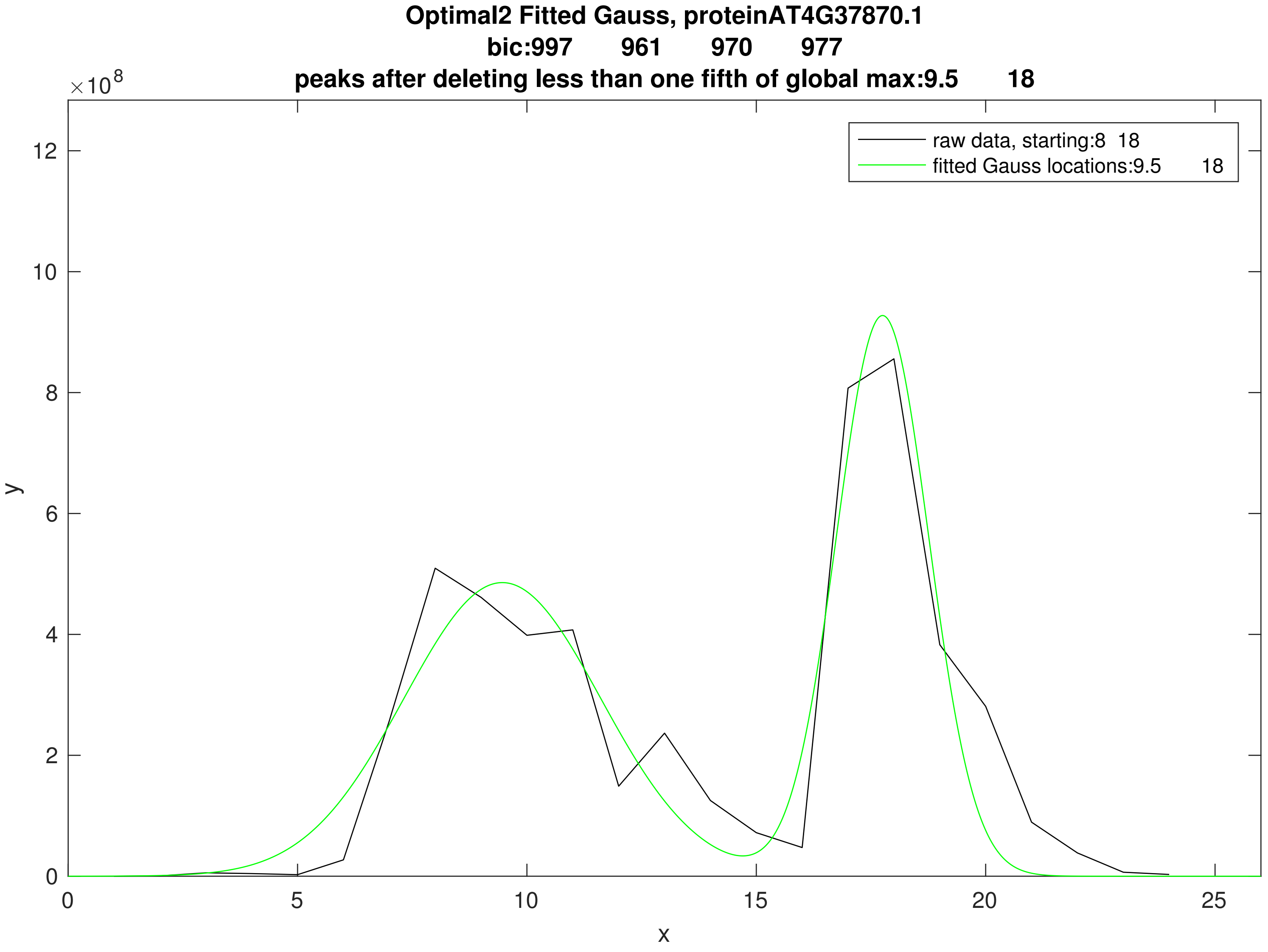

Optimal2 Fitted Gauss, proteinAT4G37870.1
bic:997 961 970 977
peaks after deleting less than one fifth of global max:9.5 18
×10^8
raw data, starting:8 18
fitted Gauss locations:9.5 18
y
x
0
2
4
6
8
10
12
0
5
10
15
20
25

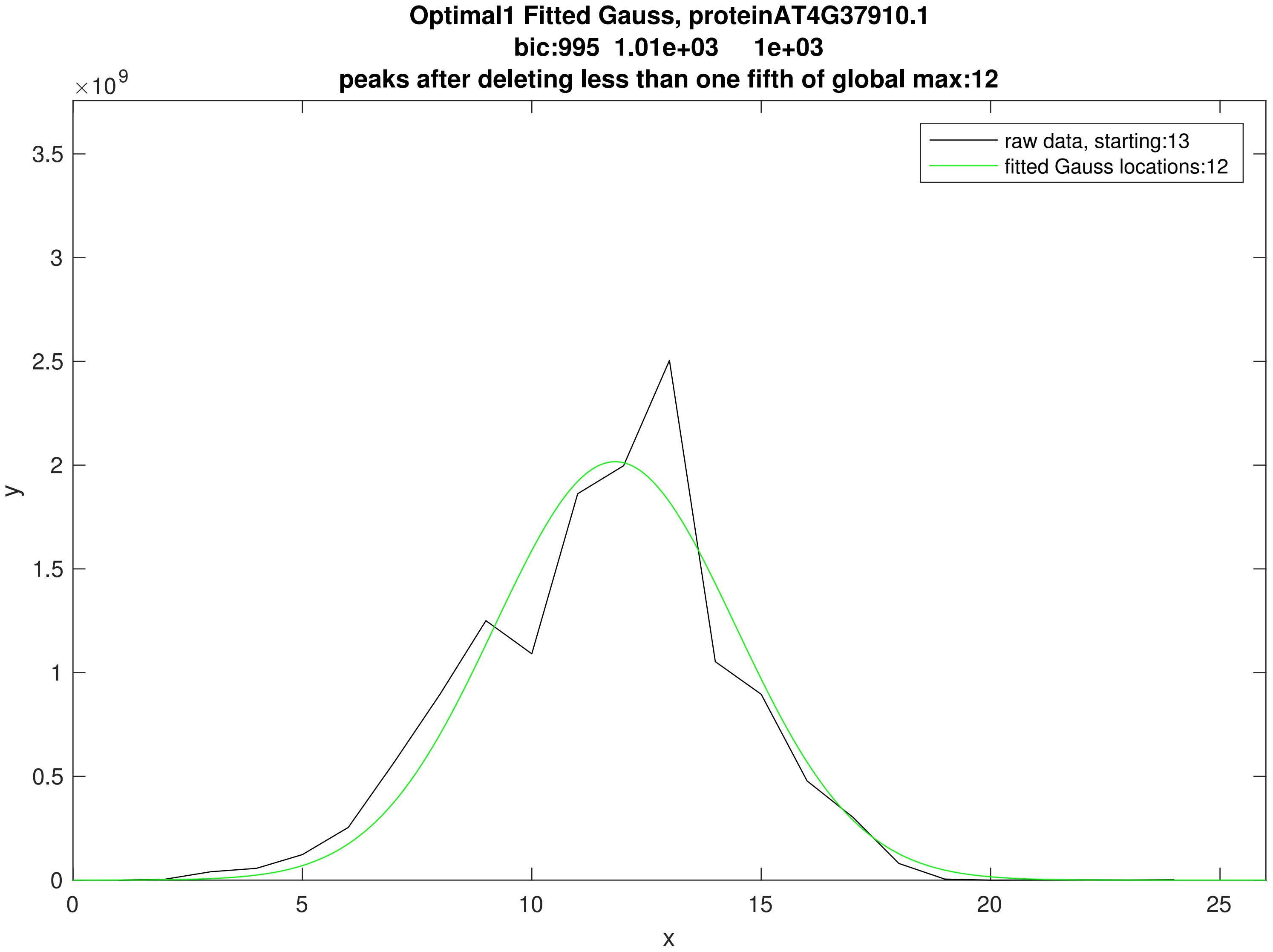
Optimal1 Fitted Gauss, proteinAT4G37910.1
bic:995  1.01e+03     1e+03
peaks after deleting less than one fifth of global max:12
×10^9
3.5
3
2.5
2
1.5
1
0.5
0
y
0
5
10
15
20
25
x
raw data, starting:13
fitted Gauss locations:12

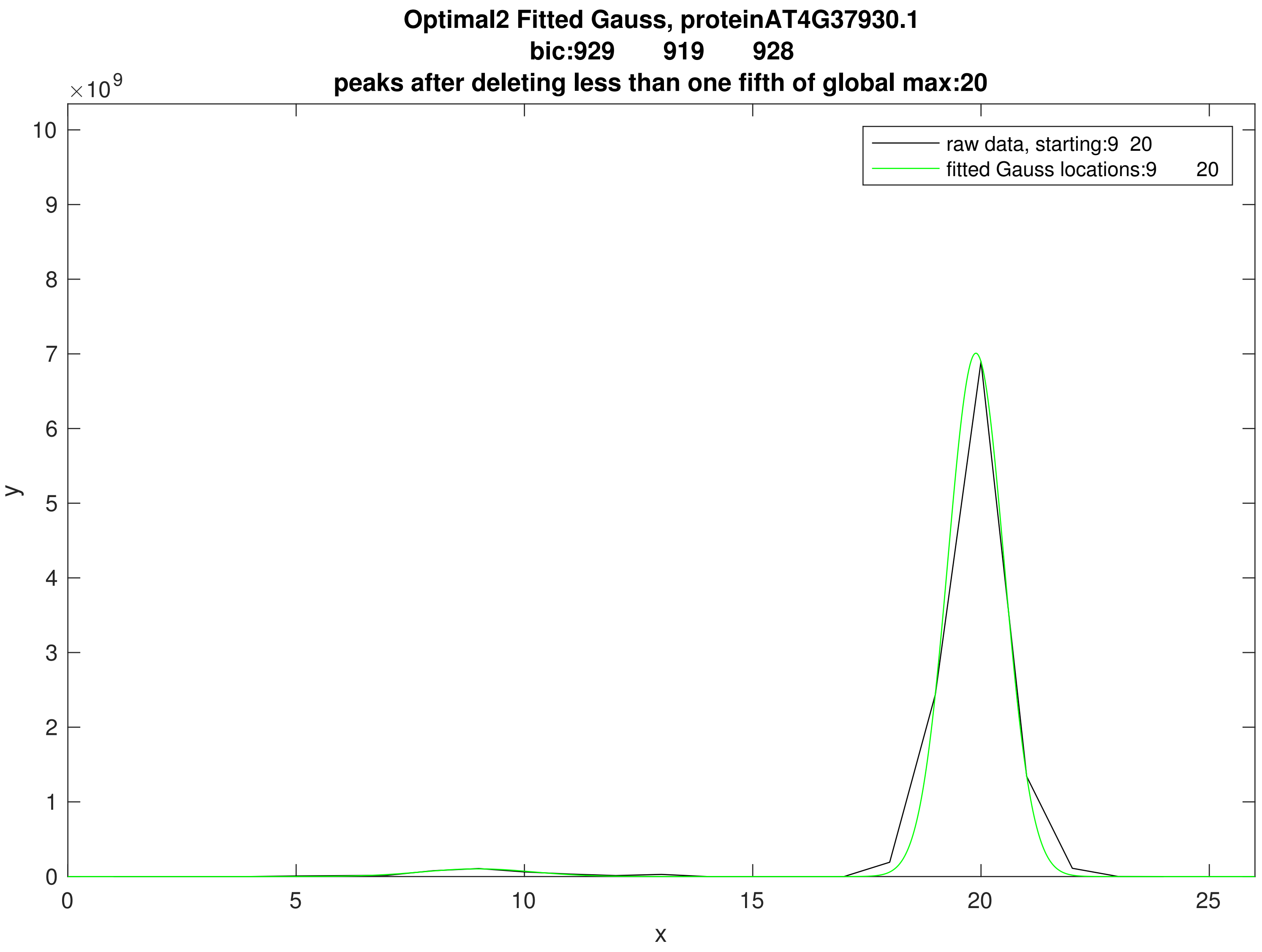
Optimal2 Fitted Gauss, proteinAT4G37930.1
bic:929 919 928
peaks after deleting less than one fifth of global max:20
×10^9
raw data, starting:9 20
fitted Gauss locations:9 20
y
x

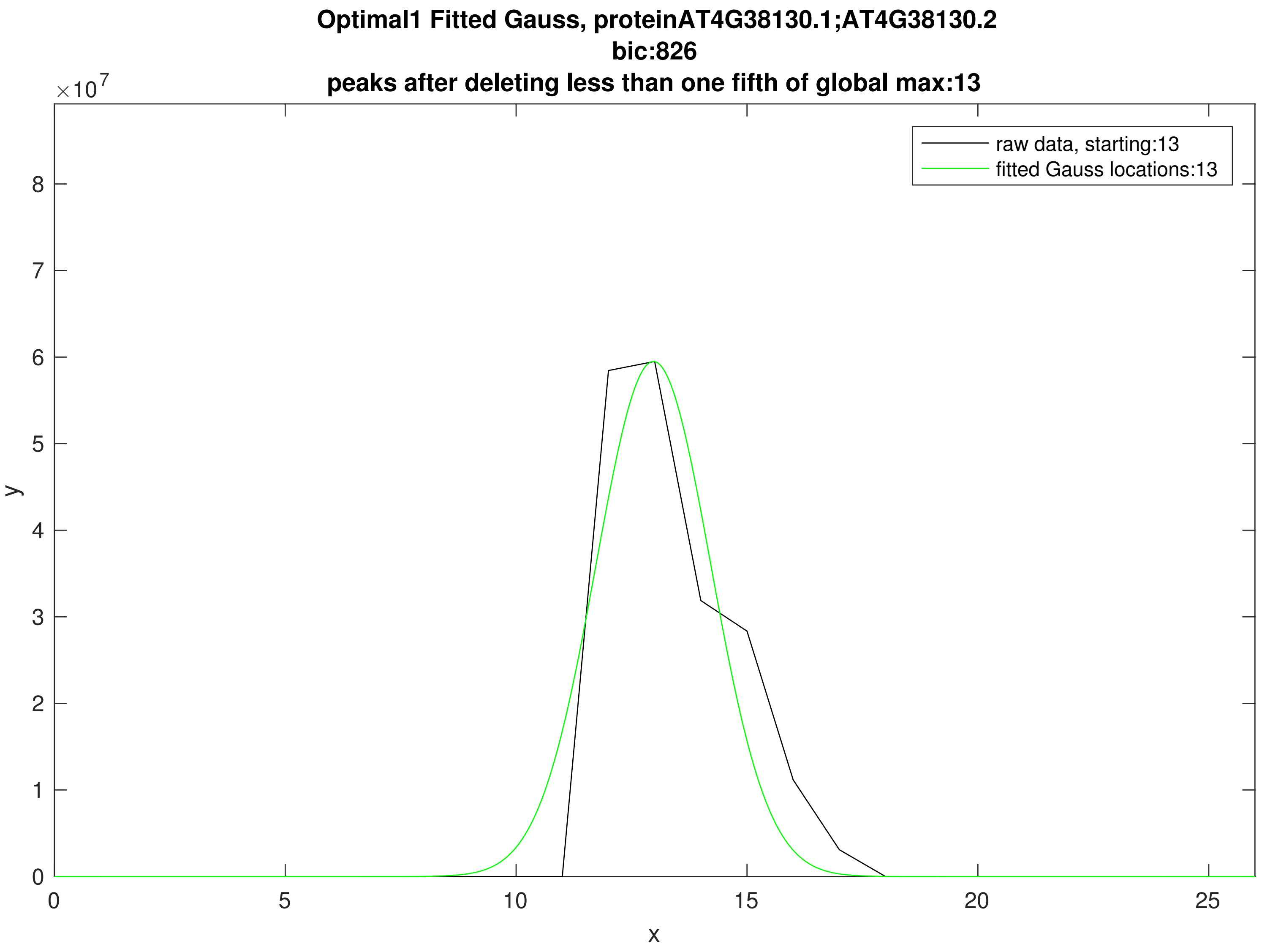

Optimal1 Fitted Gauss, proteinAT4G38130.1;AT4G38130.2
bic:826
peaks after deleting less than one fifth of global max:13
×10^7
raw data, starting:13
fitted Gauss locations:13
y
x
0
1
2
3
4
5
6
7
8
0
5
10
15
20
25

Optimal1 Fitted Gauss, proteinAT4G38220.1;AT4G38220.2
bic:984 1.03e+03 1e+03
peaks after deleting less than one fifth of global max:9.6

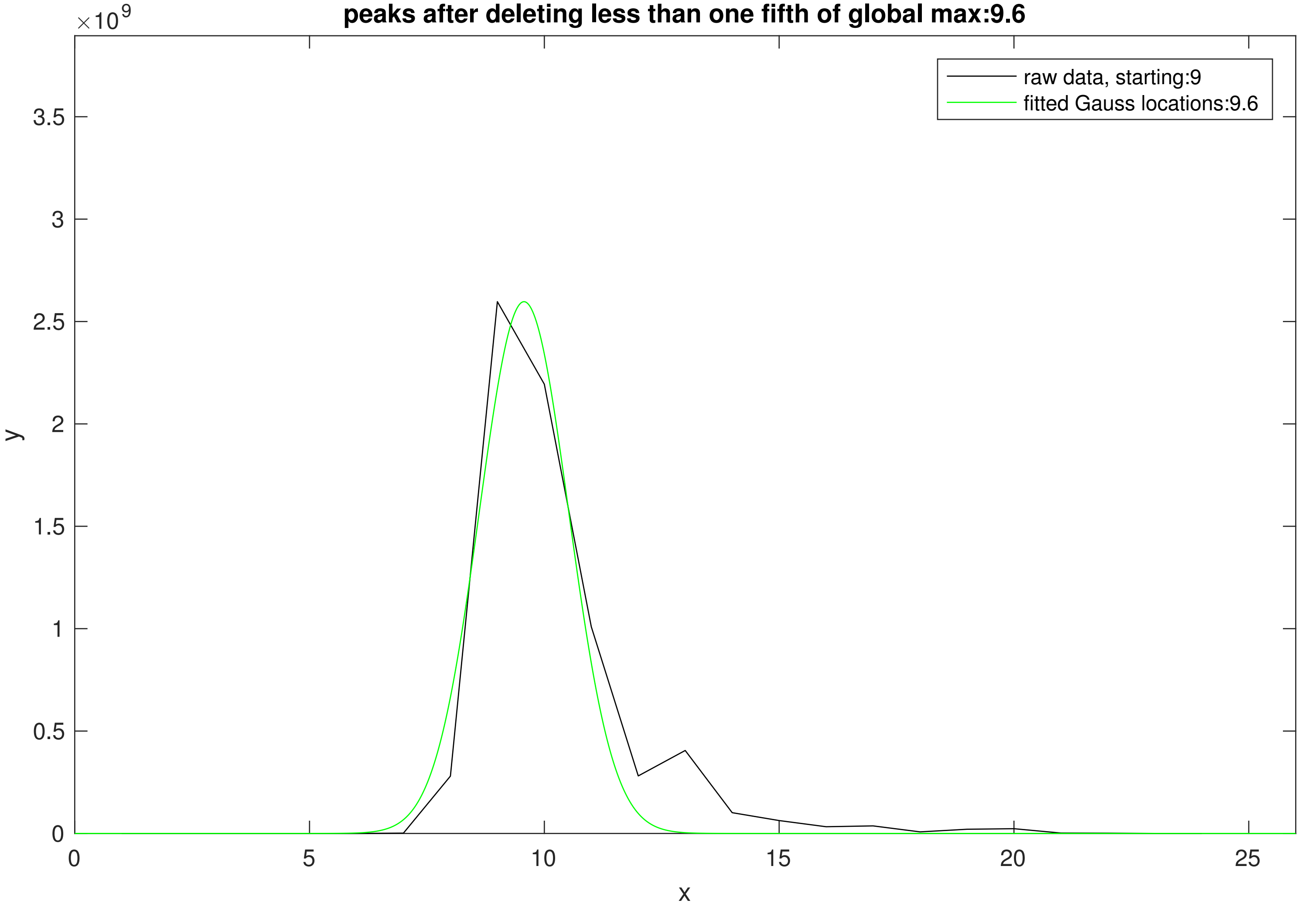

Optimal1 Fitted Gauss, proteinAT4G38225.2;AT4G38225.1;AT4G38225.3
bic:768
peaks after deleting less than one fifth of global max:17

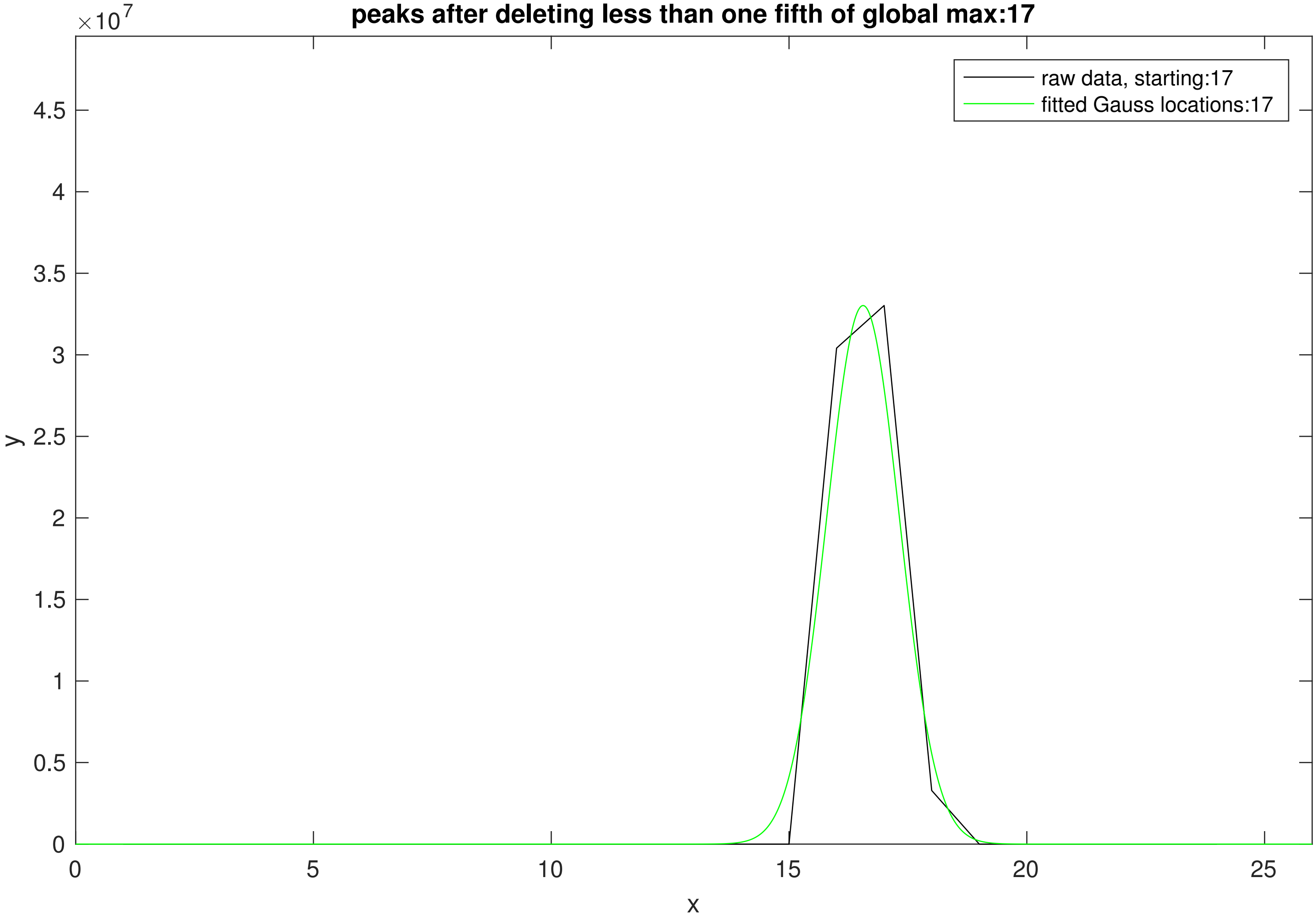

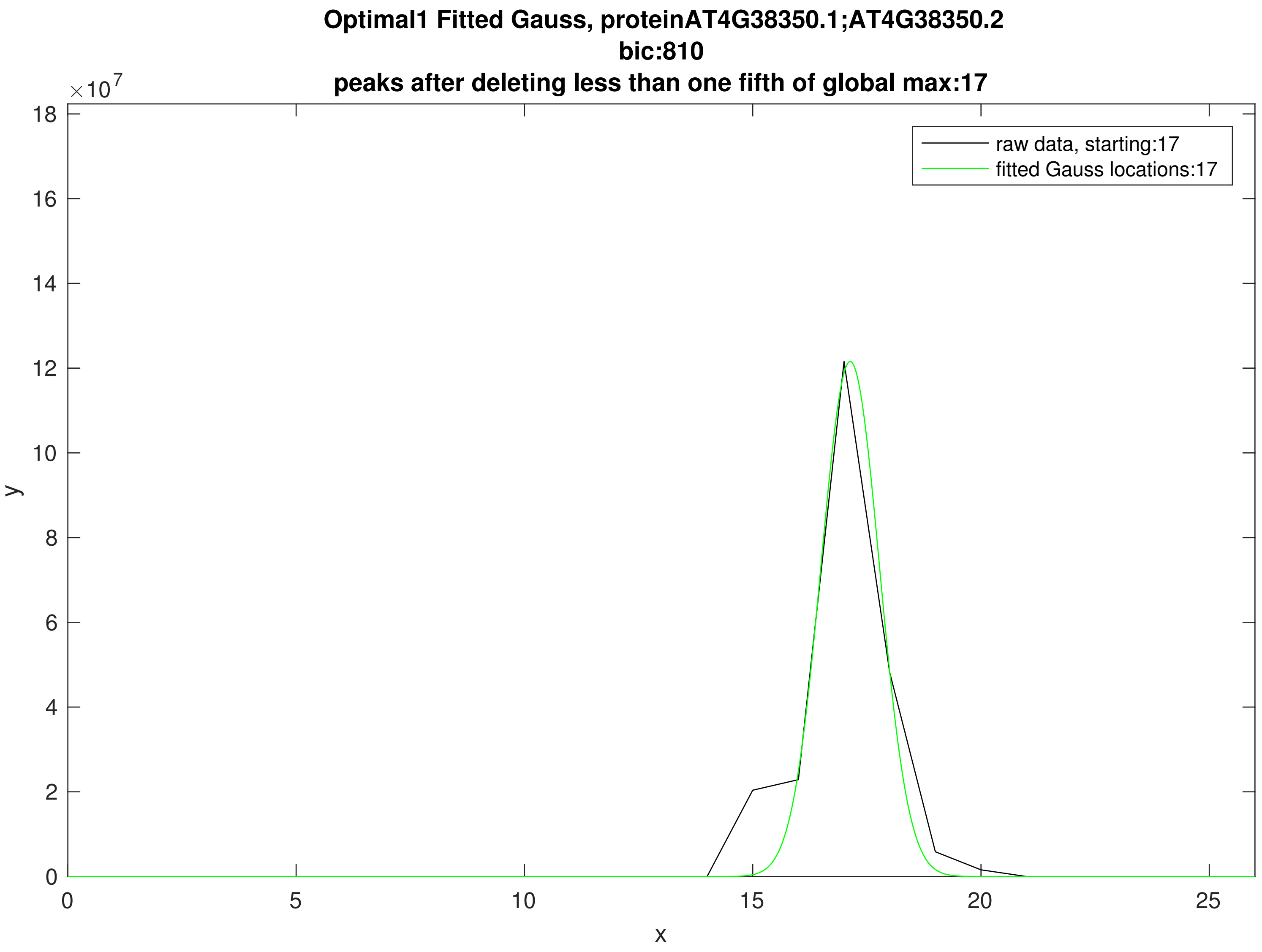
Optimal1 Fitted Gauss, proteinAT4G38350.1;AT4G38350.2
bic:810
peaks after deleting less than one fifth of global max:17
×10^7
raw data, starting:17
fitted Gauss locations:17
y
x
0
2
4
6
8
10
12
14
16
18
0
5
10
15
20
25

**Optimal1 Fitted Gauss, proteinAT4G38370.1**
**bic:828**
**peaks after deleting less than one fifth of global max:18**

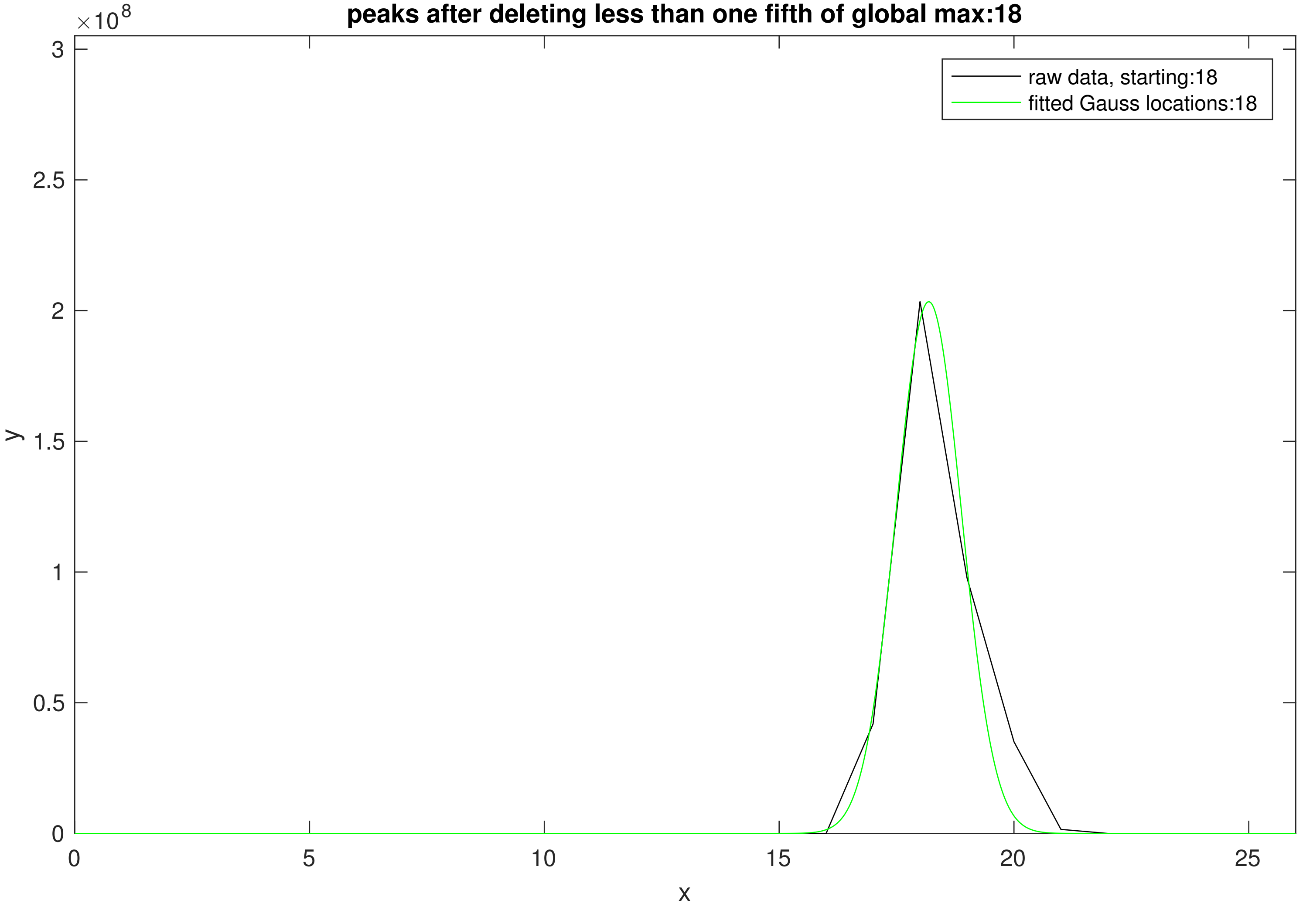

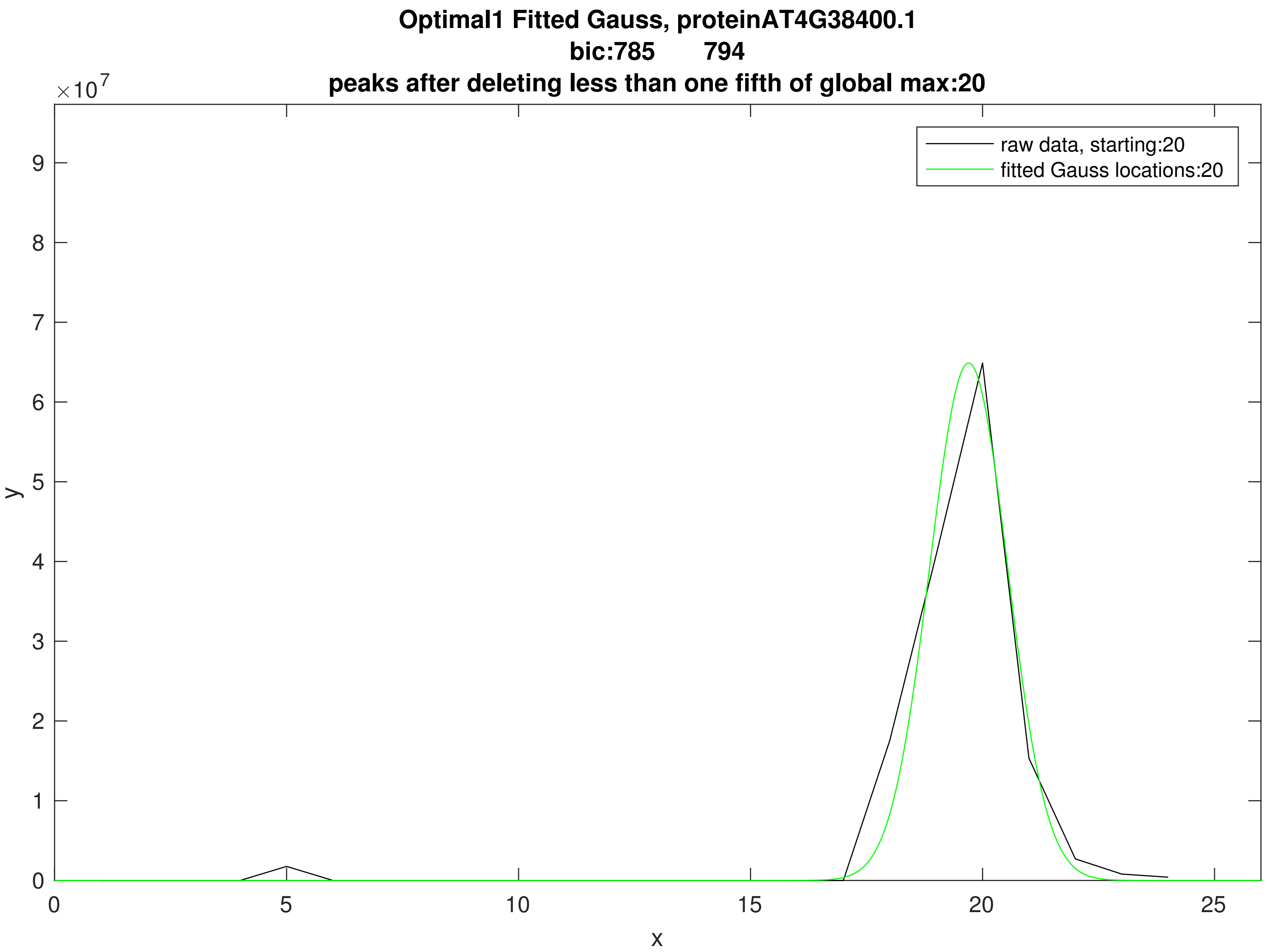
Optimal1 Fitted Gauss, proteinAT4G38400.1
bic:785 794
peaks after deleting less than one fifth of global max:20
×10^7
raw data, starting:20
fitted Gauss locations:20
y
x
0
1
2
3
4
5
6
7
8
9
0
5
10
15
20
25

**Optimal1 Fitted Gauss, proteinAT4G38510.4;AT4G38510.3;AT4G38510.2;AT4G38510.1;AT4G38510.5**
**bic:929 937 947**
**peaks after deleting less than one fifth of global max:12**

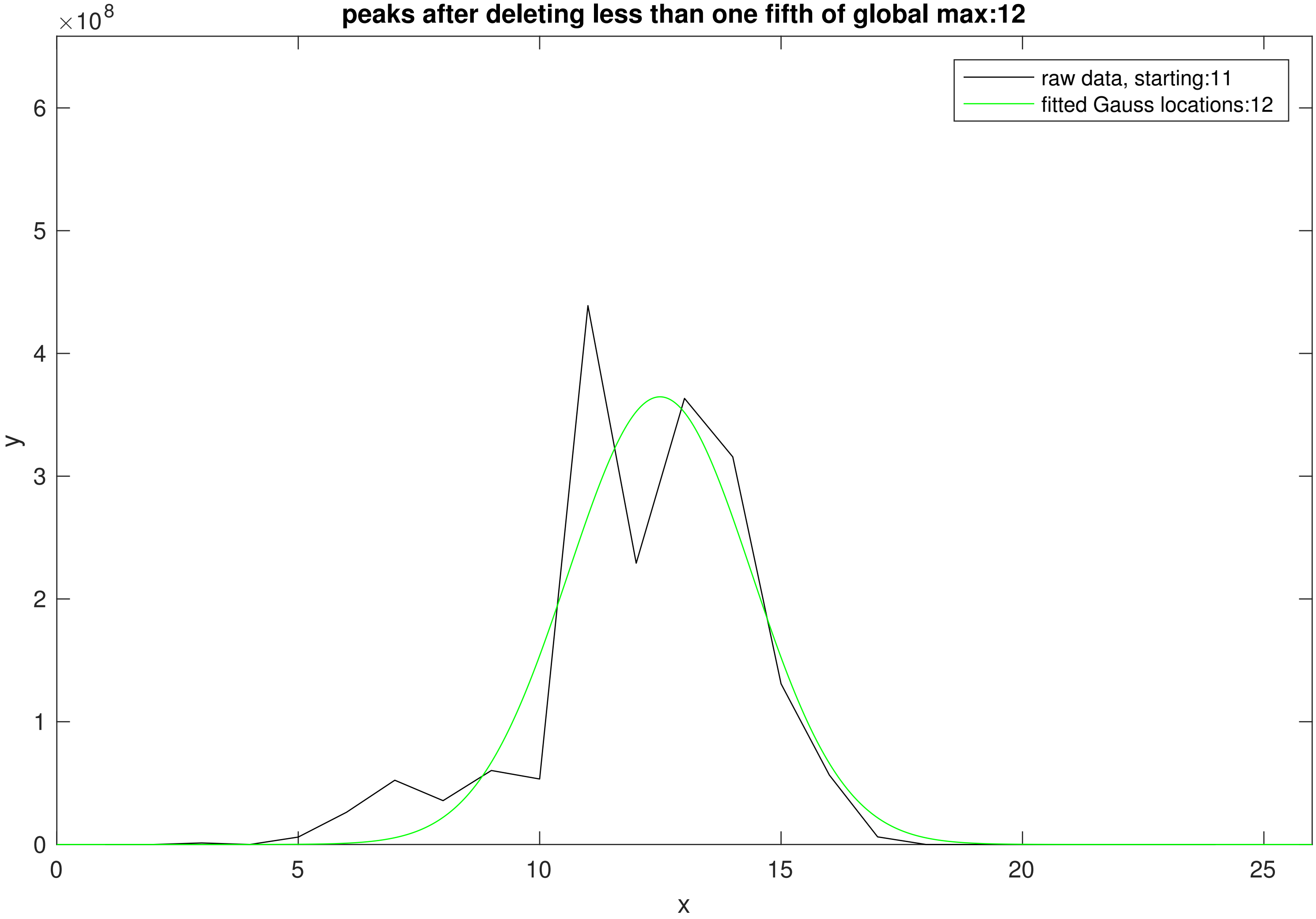

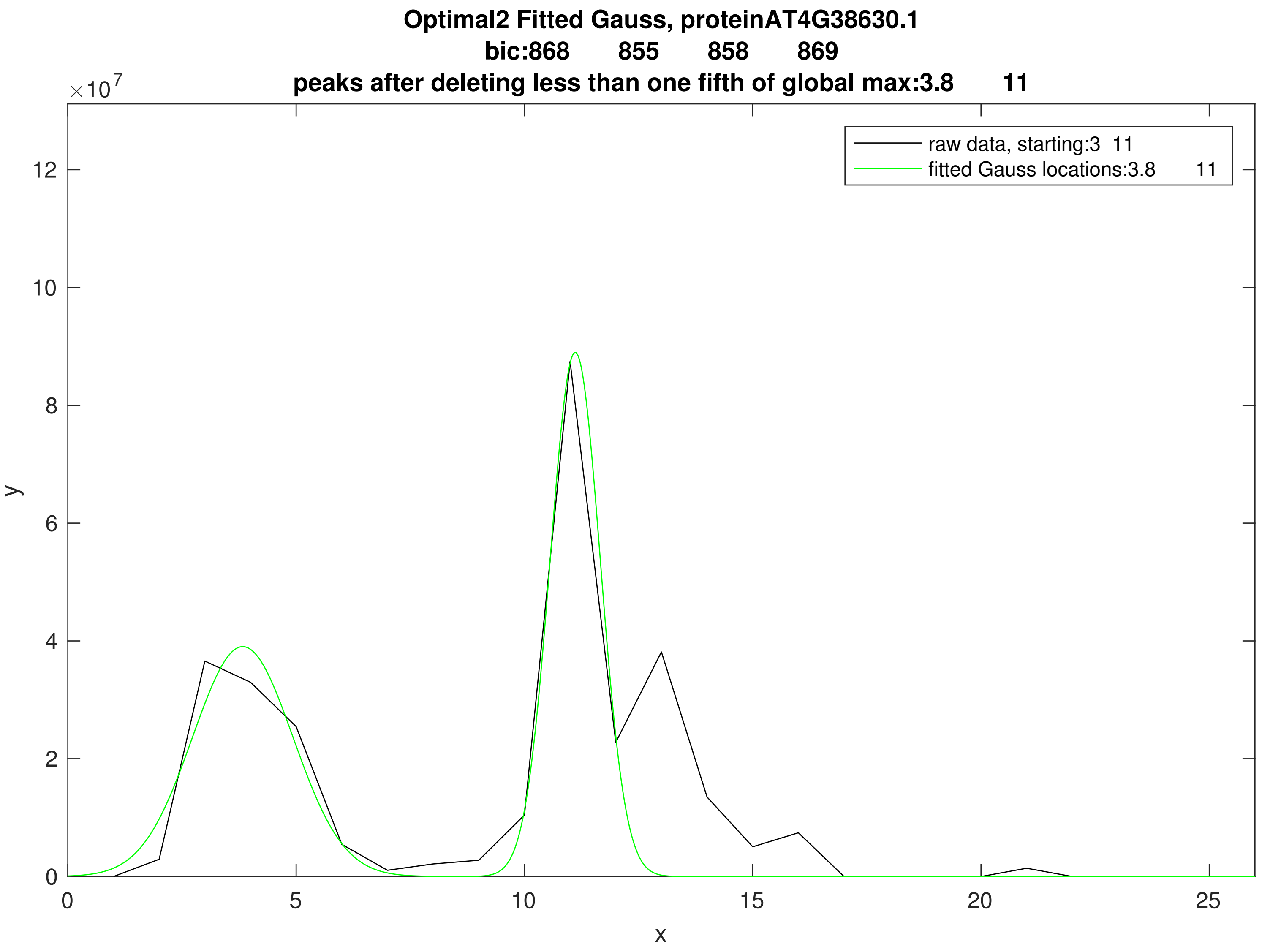

Optimal2 Fitted Gauss, proteinAT4G38630.1
bic:868 855 858 869
peaks after deleting less than one fifth of global max:3.8 11
$\times 10^7$
raw data, starting:3 11
fitted Gauss locations:3.8 11
y
x
0
2
4
6
8
10
12
0
5
10
15
20
25

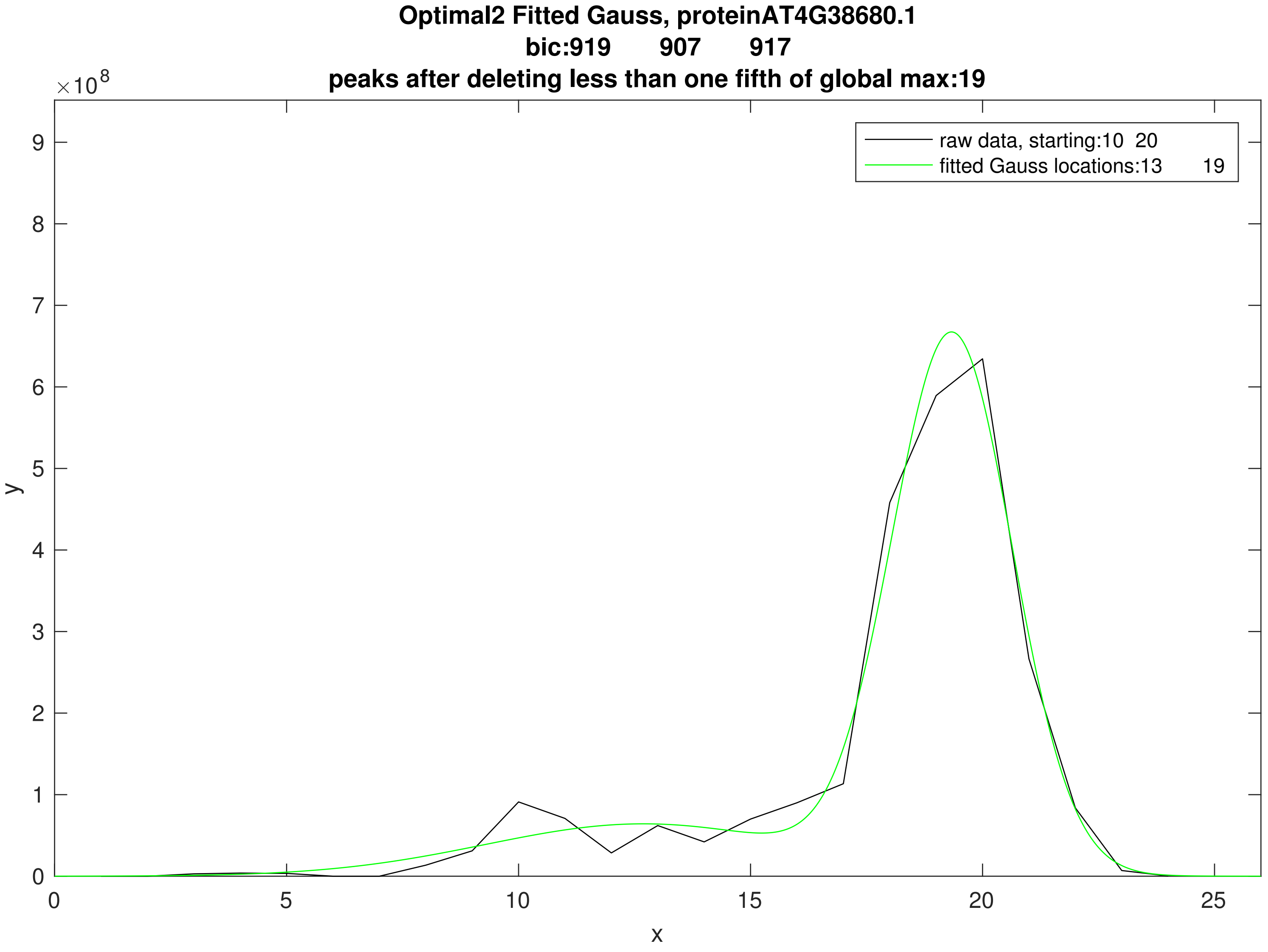

Optimal2 Fitted Gauss, proteinAT4G38680.1
bic:919 907 917
peaks after deleting less than one fifth of global max:19
×10^8
raw data, starting:10 20
fitted Gauss locations:13 19
y
x
0
1
2
3
4
5
6
7
8
9
0
5
10
15
20
25

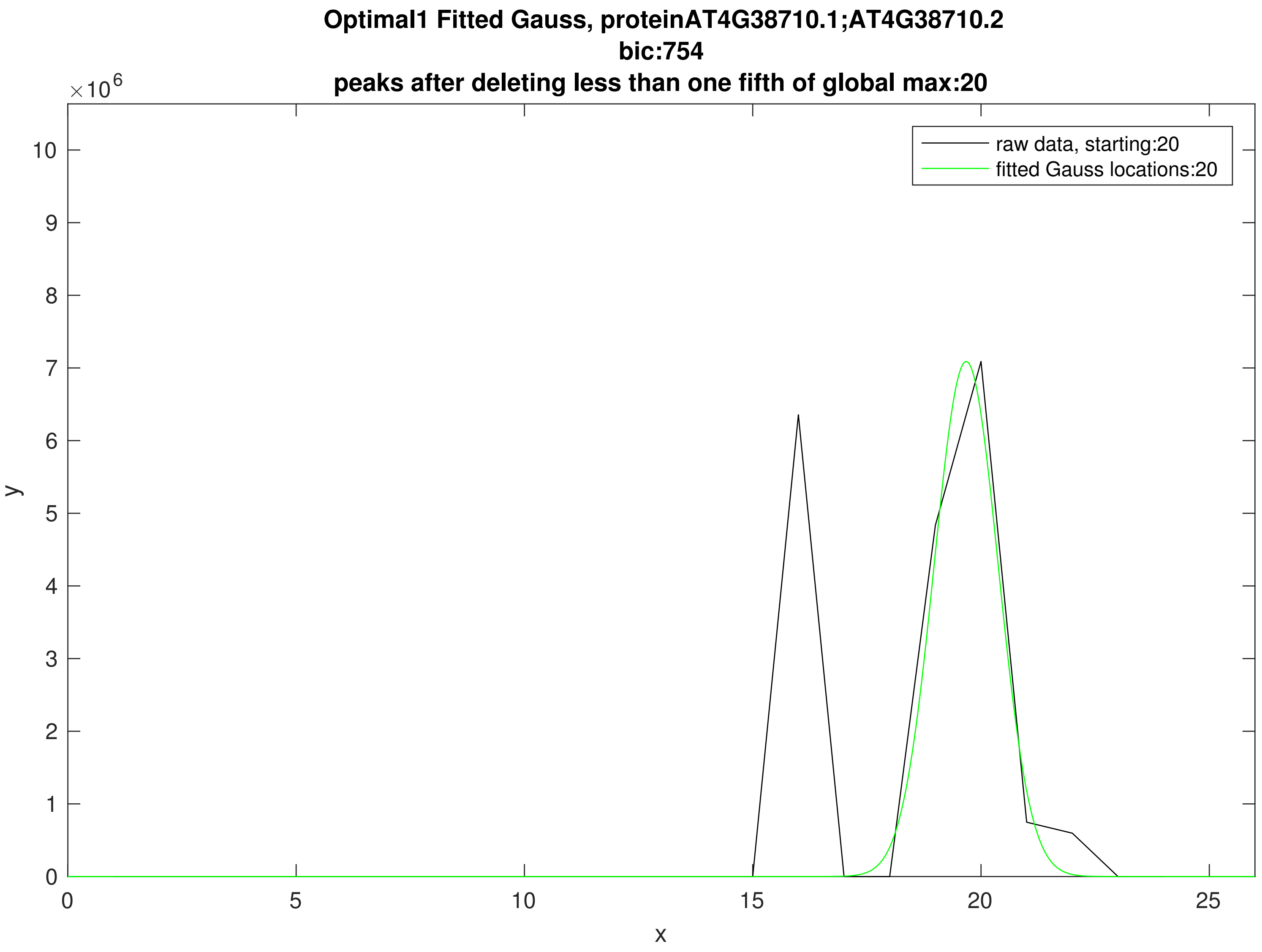
Optimal1 Fitted Gauss, proteinAT4G38710.1;AT4G38710.2
bic:754
peaks after deleting less than one fifth of global max:20
×10^6
raw data, starting:20
fitted Gauss locations:20
y
x
0
1
2
3
4
5
6
7
8
9
10
0
5
10
15
20
25

**Optimal1 Fitted Gauss, proteinAT4G38740.1**
**bic:1.03e+03  1.04e+03**
**peaks after deleting less than one fifth of global max:20**

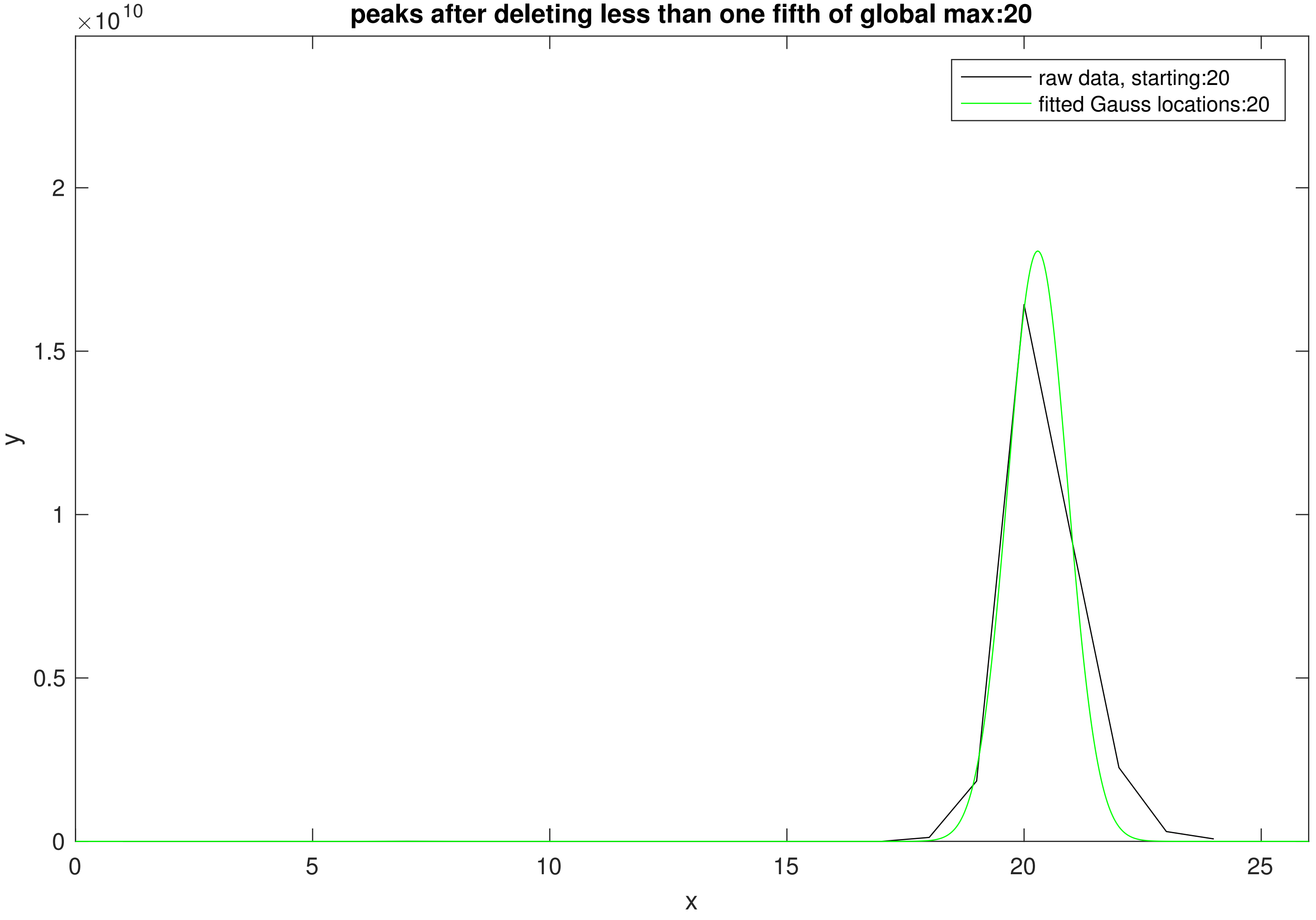

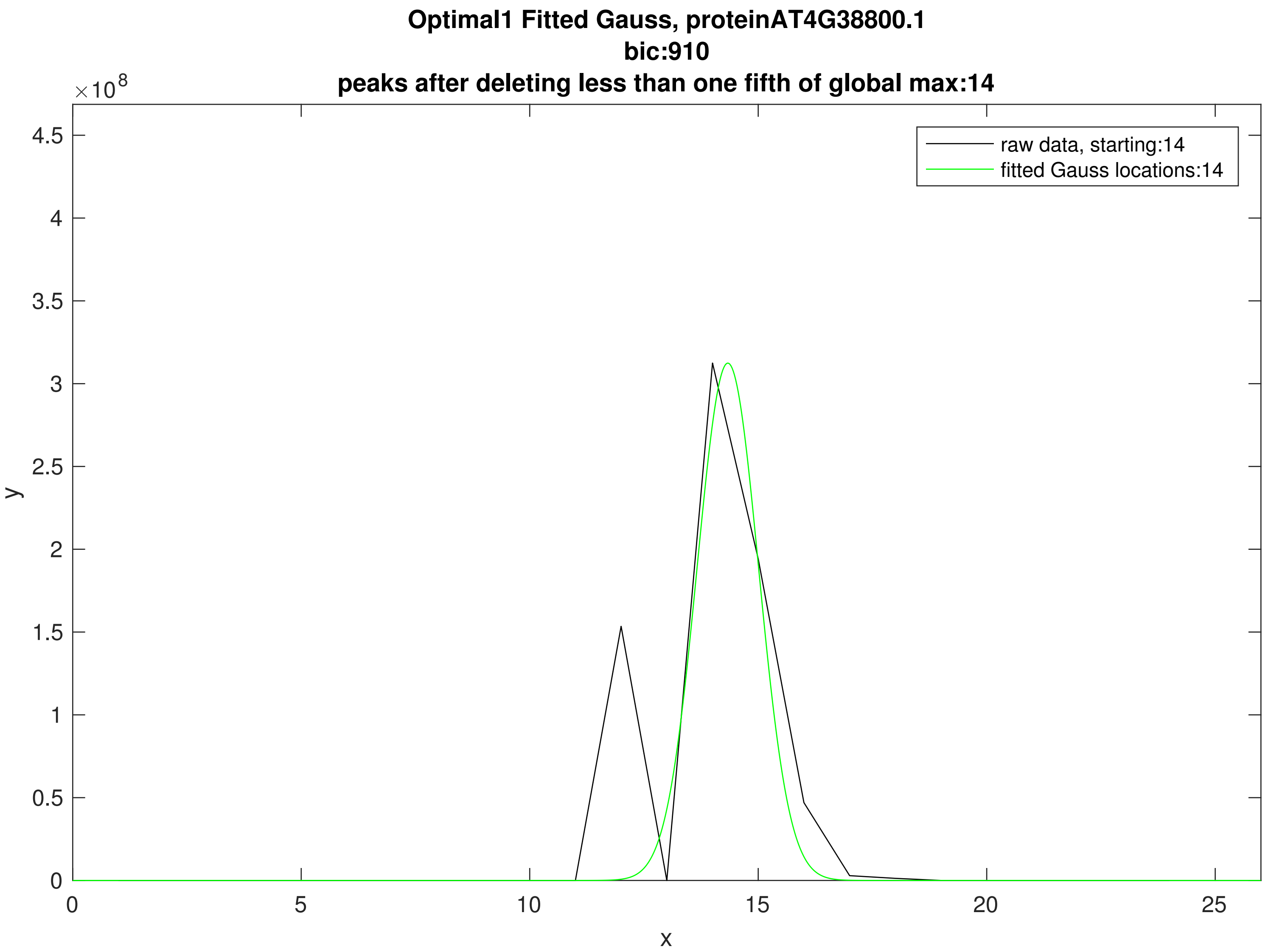
Optimal1 Fitted Gauss, proteinAT4G38800.1
bic:910
peaks after deleting less than one fifth of global max:14
×10^8
4.5
4
3.5
3
2.5
2
1.5
1
0.5
0
y
0
5
10
15
20
25
x
raw data, starting:14
fitted Gauss locations:14

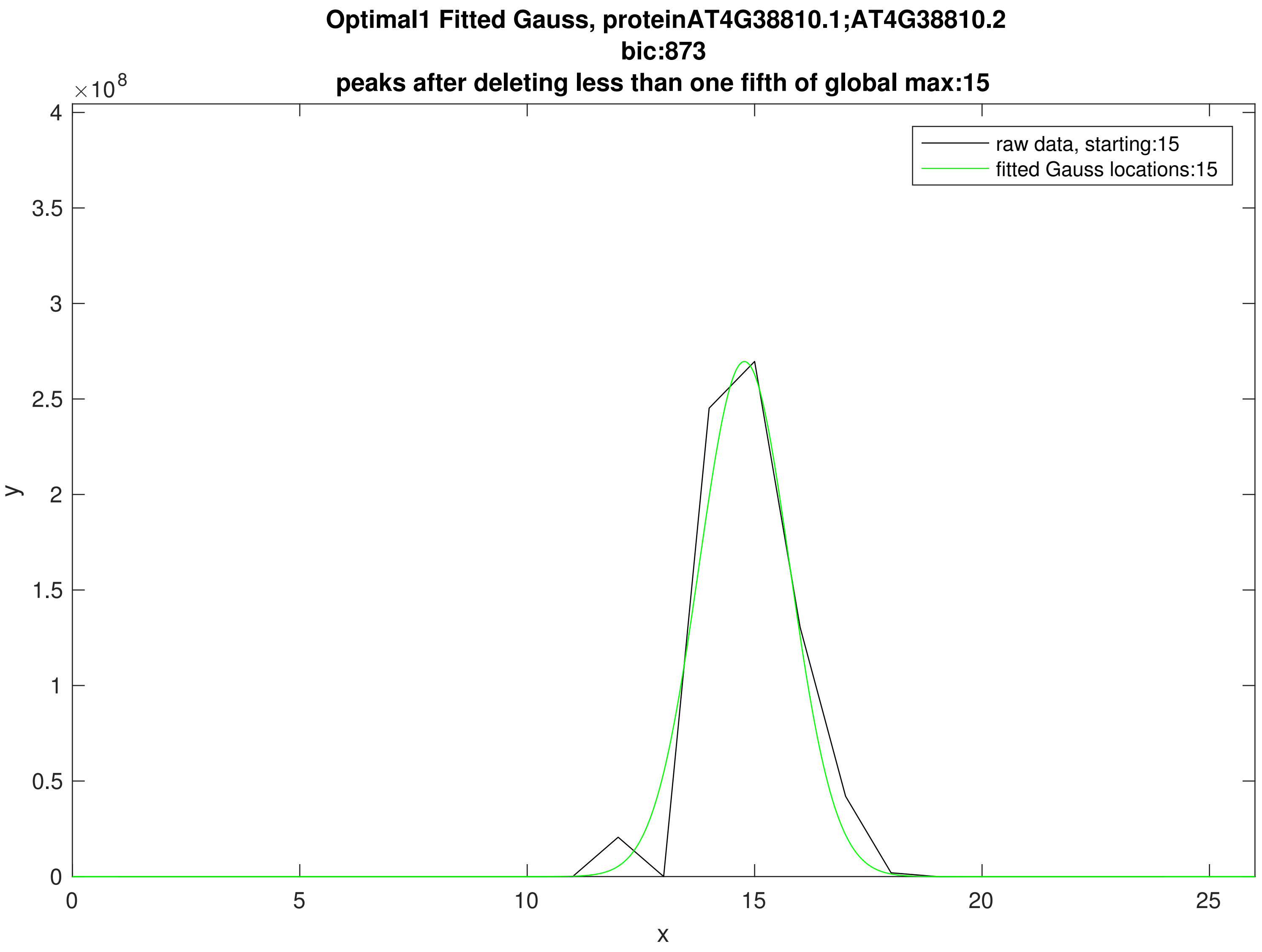

Optimal1 Fitted Gauss, proteinAT4G38810.1;AT4G38810.2
bic:873
peaks after deleting less than one fifth of global max:15
×10^8
raw data, starting:15
fitted Gauss locations:15
y
x
0
0.5
1
1.5
2
2.5
3
3.5
4
5
10
15
20
25

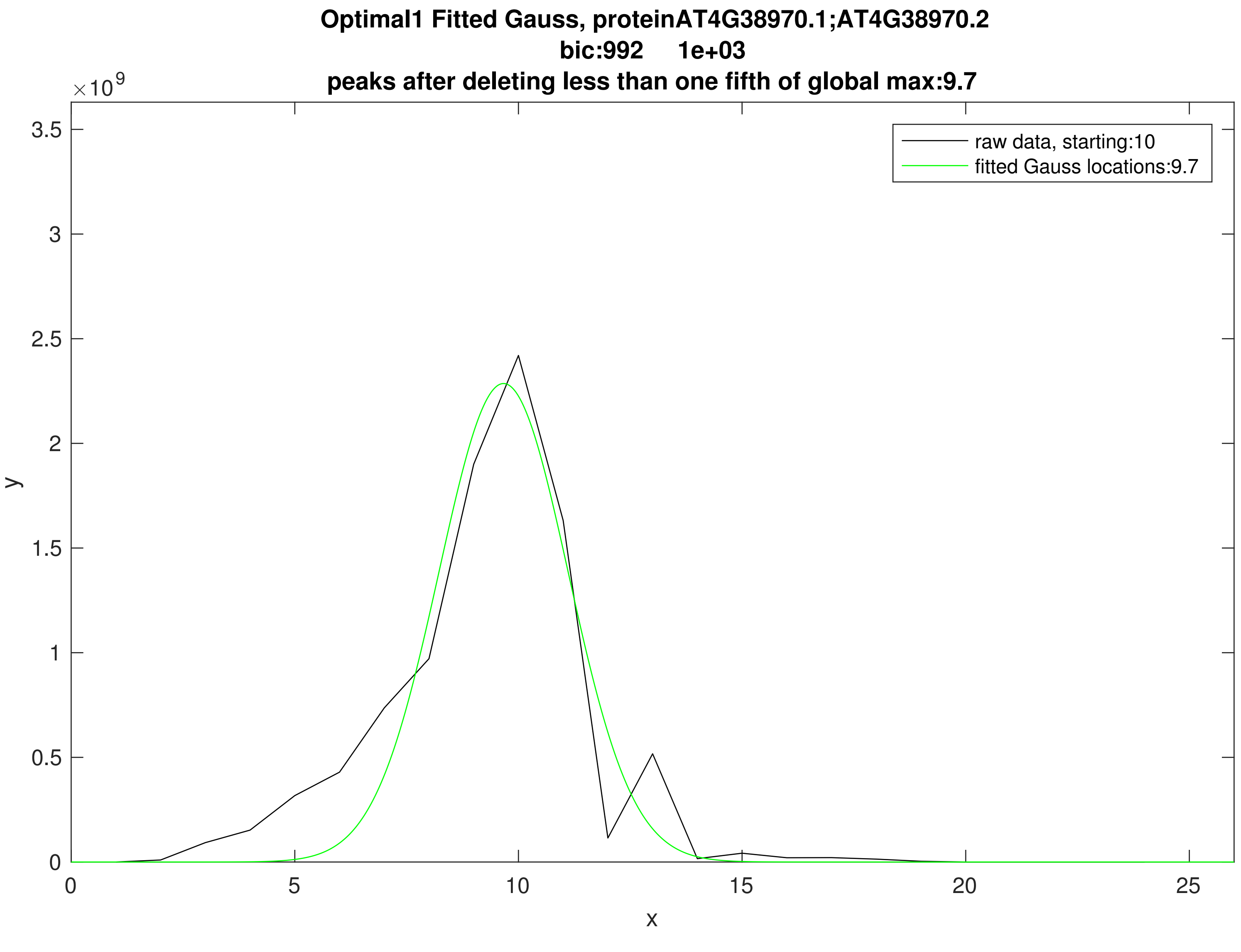
Optimal1 Fitted Gauss, proteinAT4G38970.1;AT4G38970.2
bic:992 1e+03
peaks after deleting less than one fifth of global max:9.7
×10^9
3.5
3
2.5
2
1.5
1
0.5
0
y
0
5
10
15
20
25
x
raw data, starting:10
fitted Gauss locations:9.7

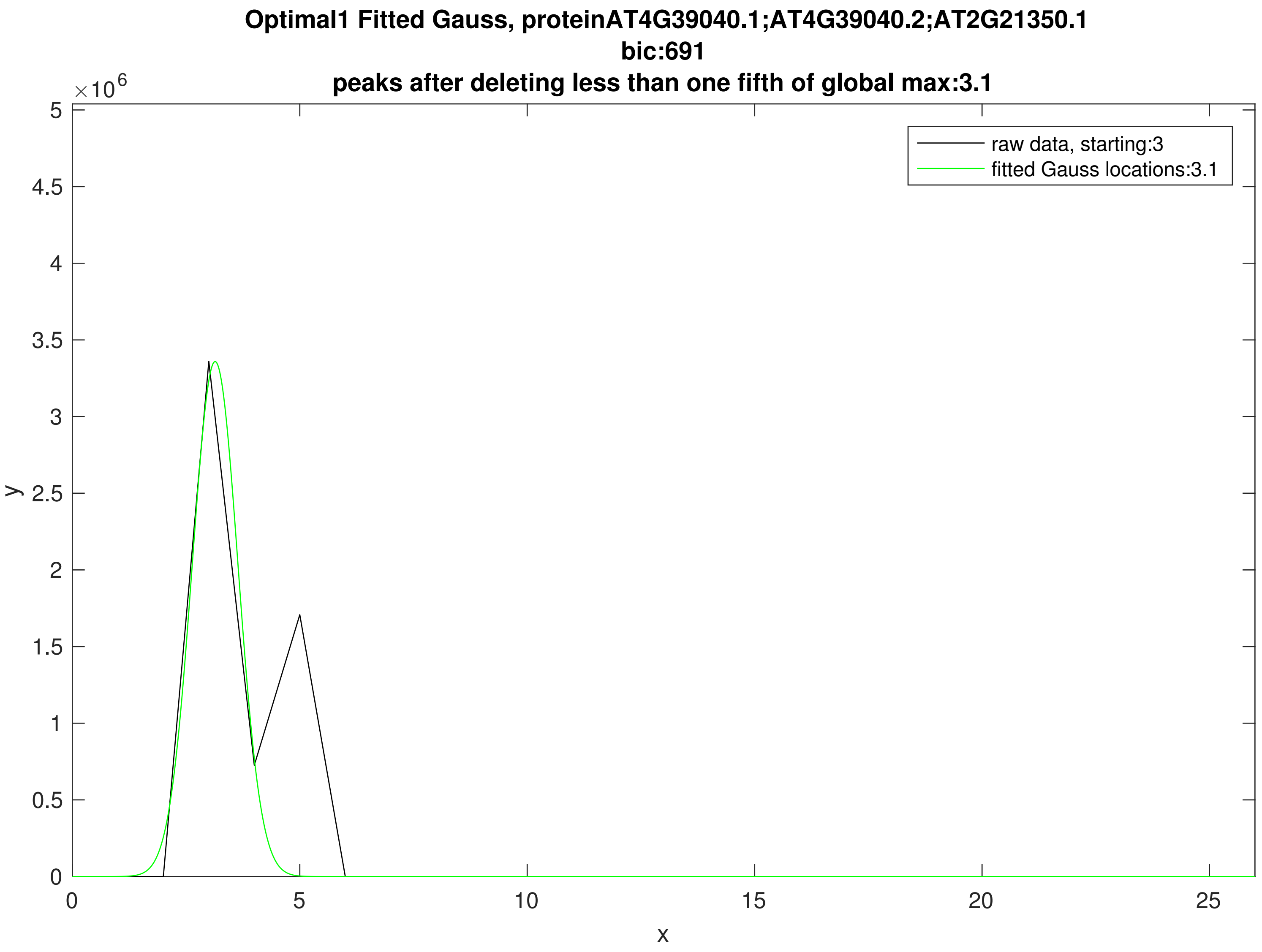

Optimal1 Fitted Gauss, proteinAT4G39040.1;AT4G39040.2;AT2G21350.1
bic:691
peaks after deleting less than one fifth of global max:3.1
×10^6
raw data, starting:3
fitted Gauss locations:3.1
y
x
0
0.5
1
1.5
2
2.5
3
3.5
4
4.5
5
0
5
10
15
20
25

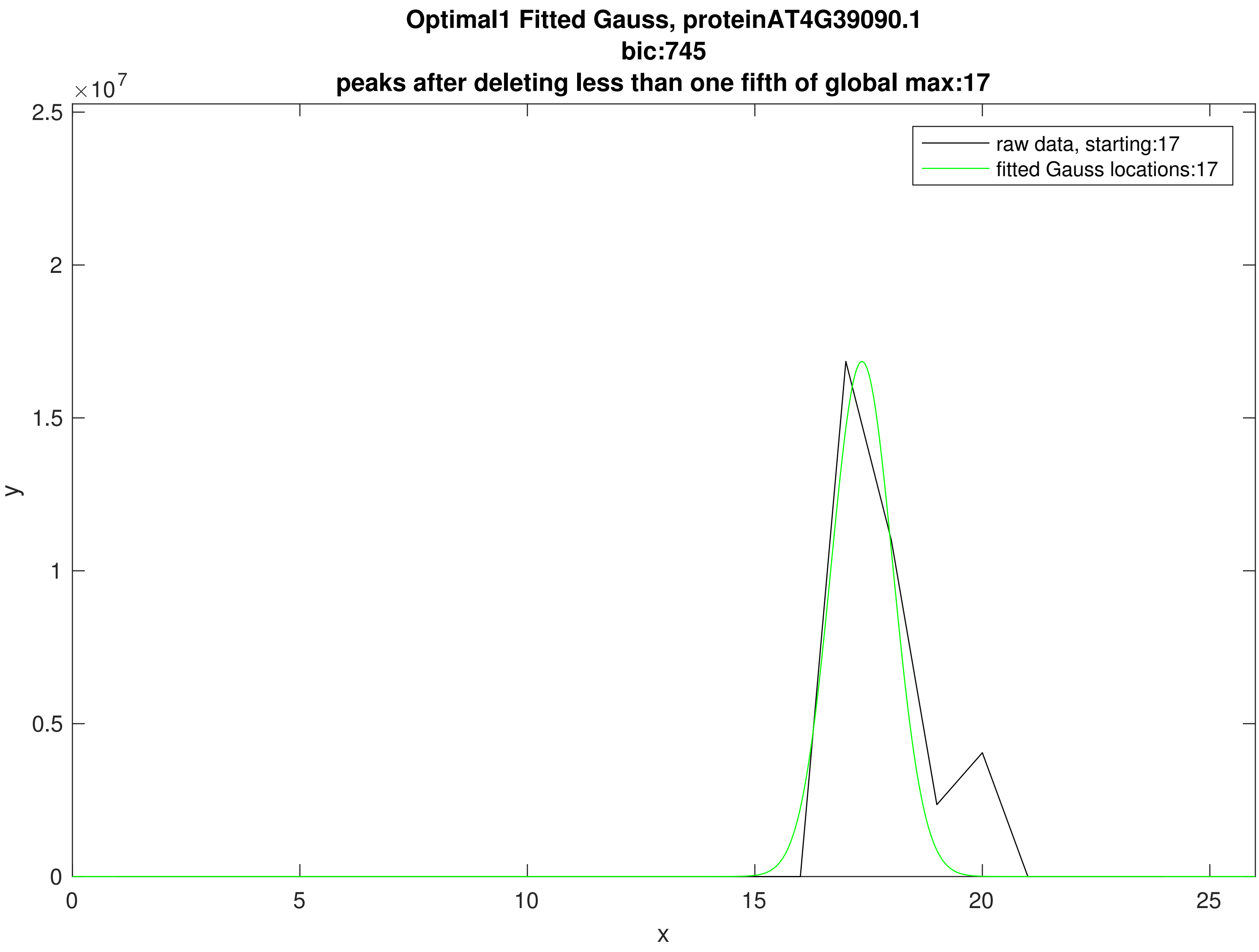
Optimal1 Fitted Gauss, proteinAT4G39090.1
bic:745
peaks after deleting less than one fifth of global max:17
×10^7
2.5
2
1.5
1
0.5
0
y
0
5
10
15
20
25
x
raw data, starting:17
fitted Gauss locations:17

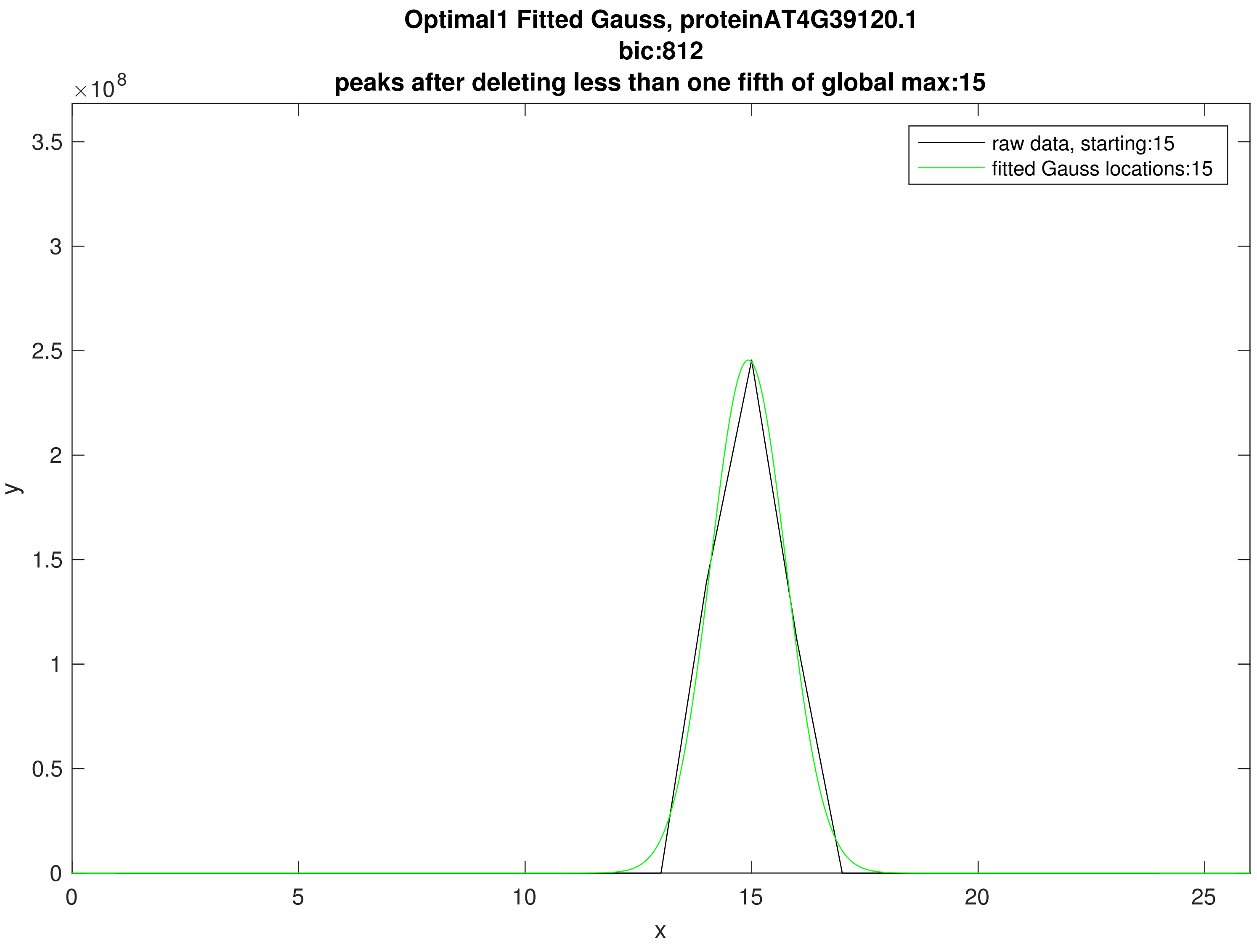

Optimal1 Fitted Gauss, proteinAT4G39120.1
bic:812
peaks after deleting less than one fifth of global max:15
×10^8
3.5
3
2.5
2
1.5
1
0.5
0
y
0
5
10
15
20
25
x
raw data, starting:15
fitted Gauss locations:15

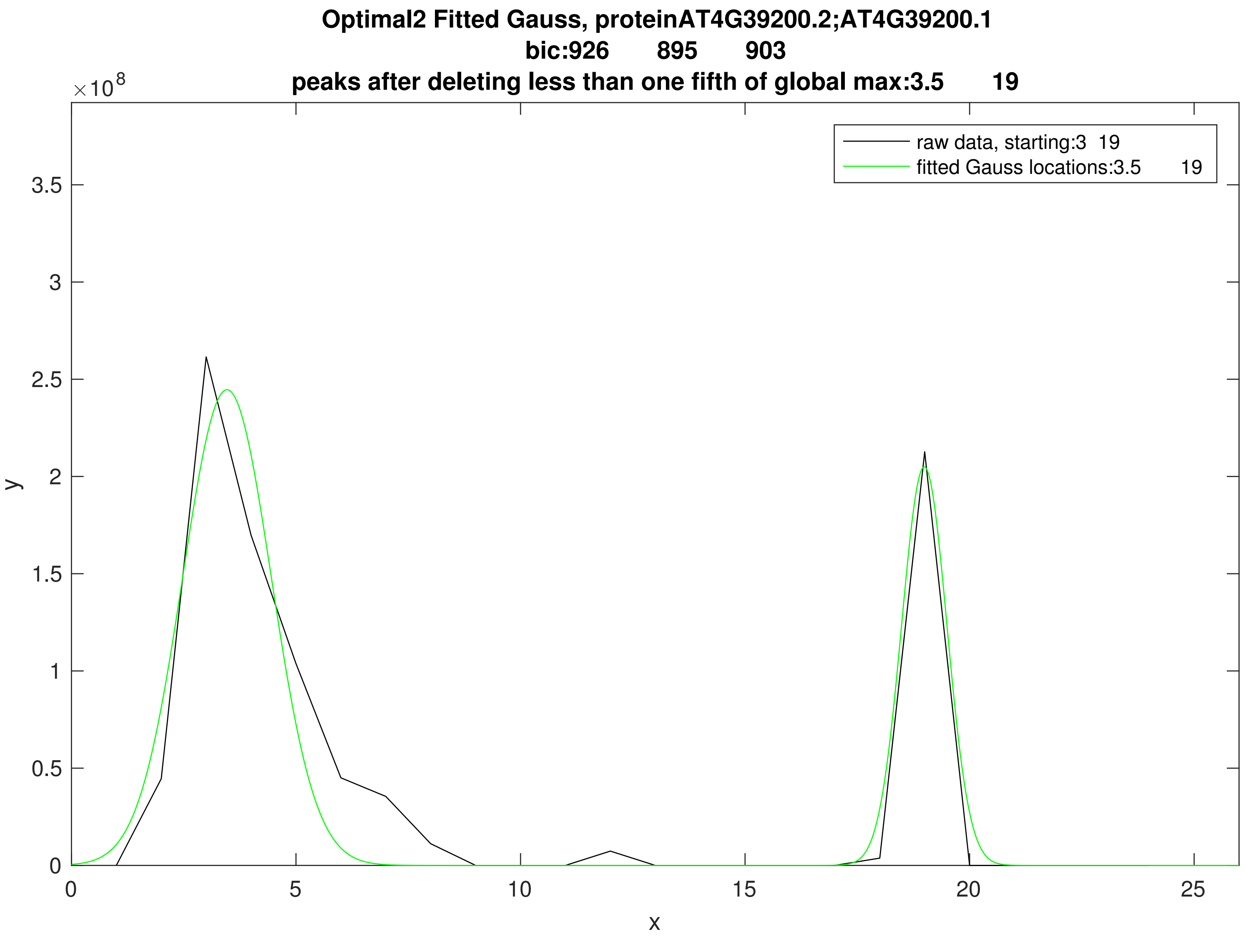
Optimal2 Fitted Gauss, proteinAT4G39200.2;AT4G39200.1
bic:926 895 903
peaks after deleting less than one fifth of global max:3.5 19
×10^8
raw data, starting:3 19
fitted Gauss locations:3.5 19
y
x
0
0.5
1
1.5
2
2.5
3
3.5
5
10
15
20
25

**Optimal1 Fitted Gauss, proteinAT4G39210.1**
**bic:806**
**peaks after deleting less than one fifth of global max:10**

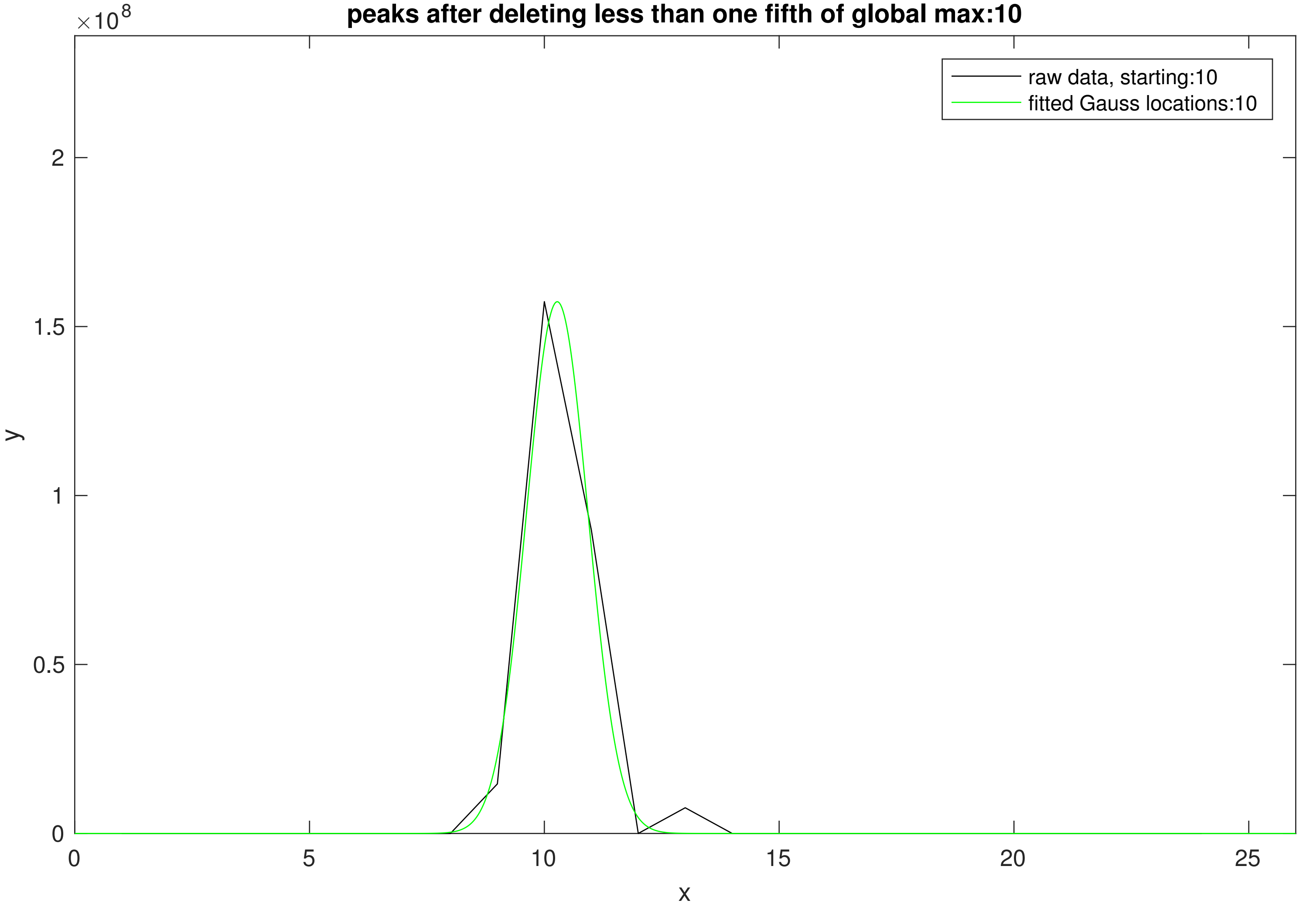

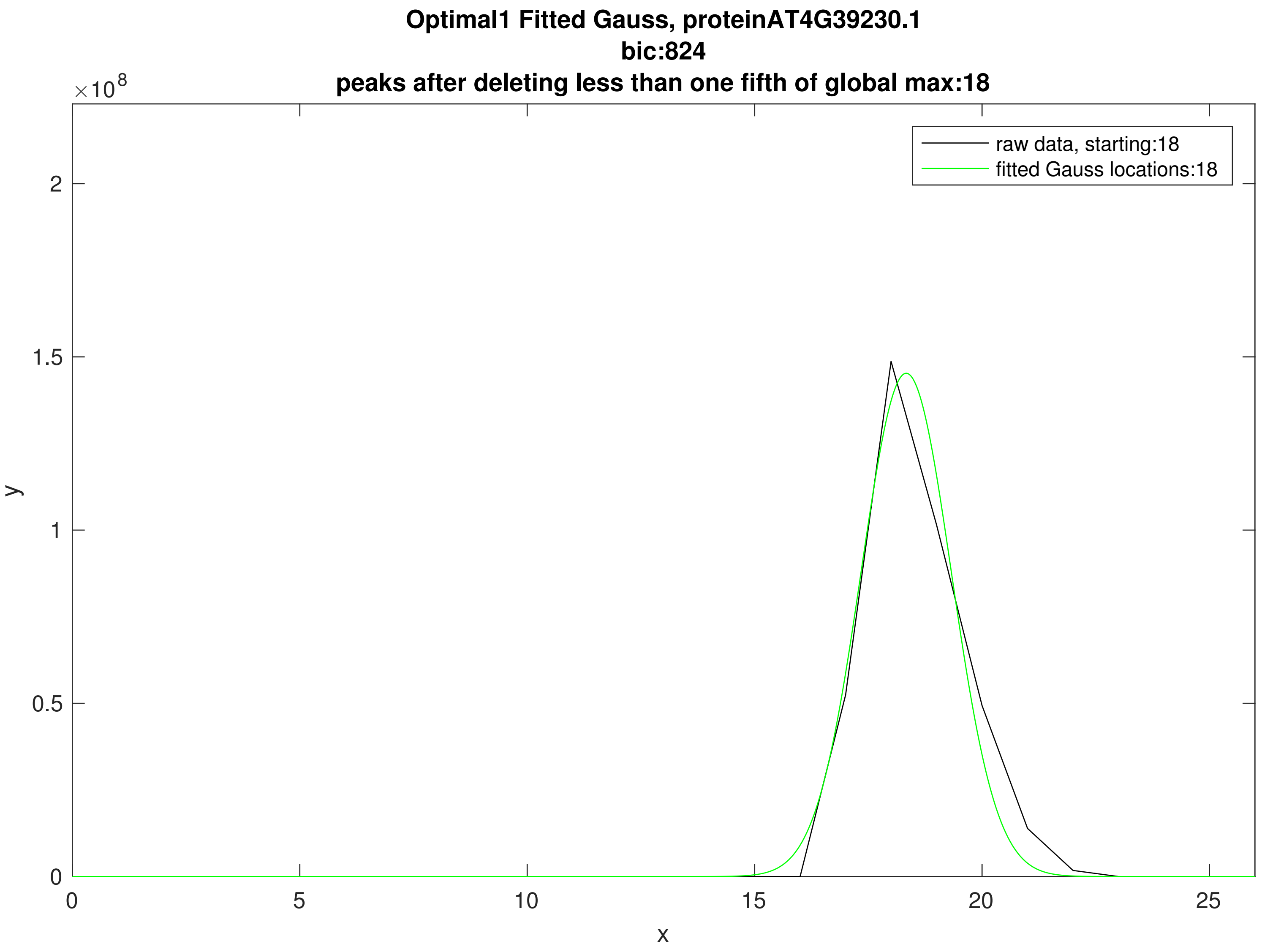
Optimal1 Fitted Gauss, proteinAT4G39230.1
bic:824
peaks after deleting less than one fifth of global max:18
×10^8
raw data, starting:18
fitted Gauss locations:18
y
x
0
0.5
1
1.5
2
0
5
10
15
20
25

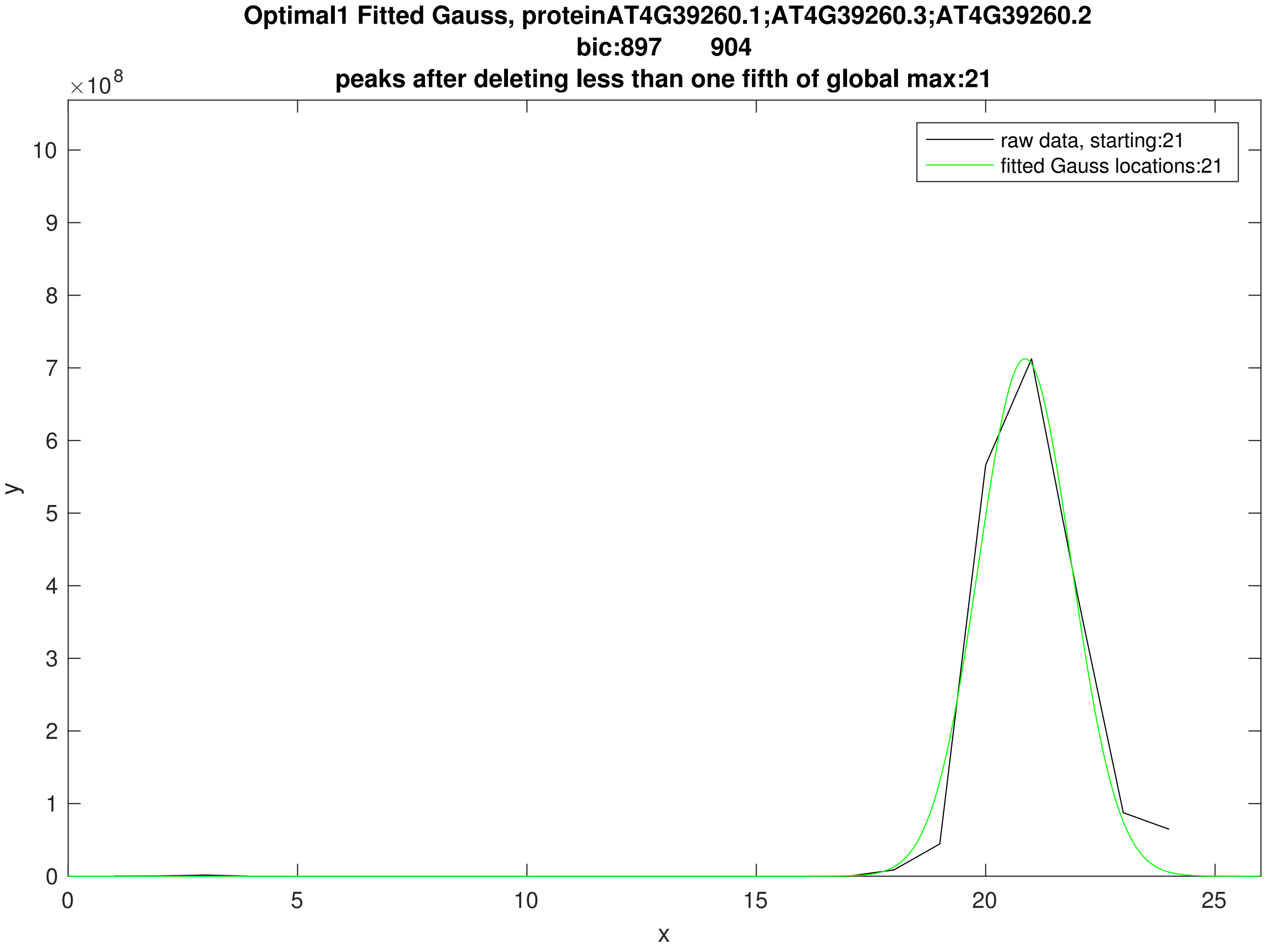
Optimal1 Fitted Gauss, proteinAT4G39260.1;AT4G39260.3;AT4G39260.2
bic:897 904
peaks after deleting less than one fifth of global max:21
×10^8
raw data, starting:21
fitted Gauss locations:21
y
x
0
1
2
3
4
5
6
7
8
9
10
0
5
10
15
20
25

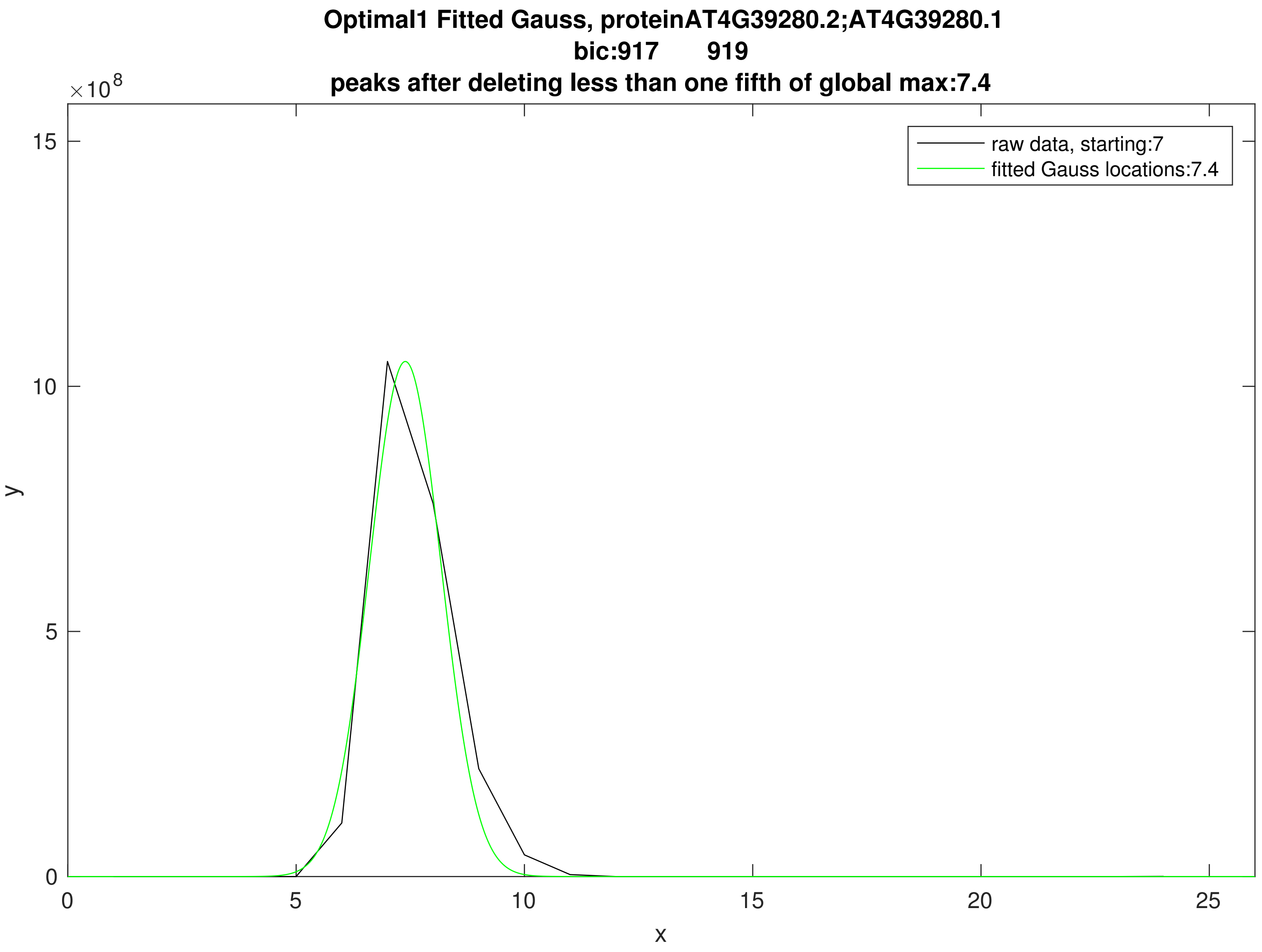
Optimal1 Fitted Gauss, proteinAT4G39280.2;AT4G39280.1
bic:917 919
peaks after deleting less than one fifth of global max:7.4
×10^8
15
10
5
0
y
0
5
10
15
20
25
x
raw data, starting:7
fitted Gauss locations:7.4

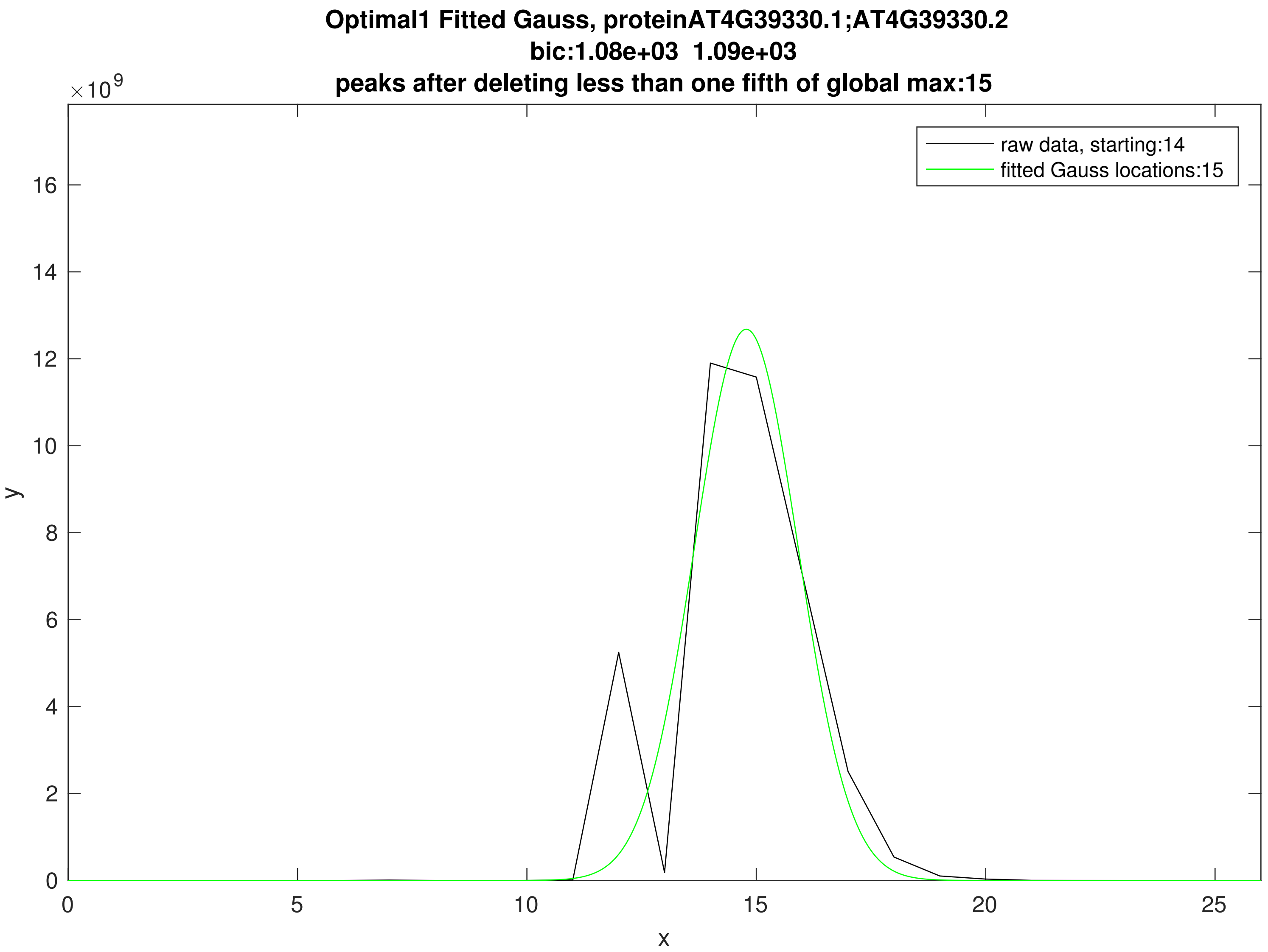
Optimal1 Fitted Gauss, proteinAT4G39330.1;AT4G39330.2
bic:1.08e+03 1.09e+03
peaks after deleting less than one fifth of global max:15
×10^9
raw data, starting:14
fitted Gauss locations:15
y
x

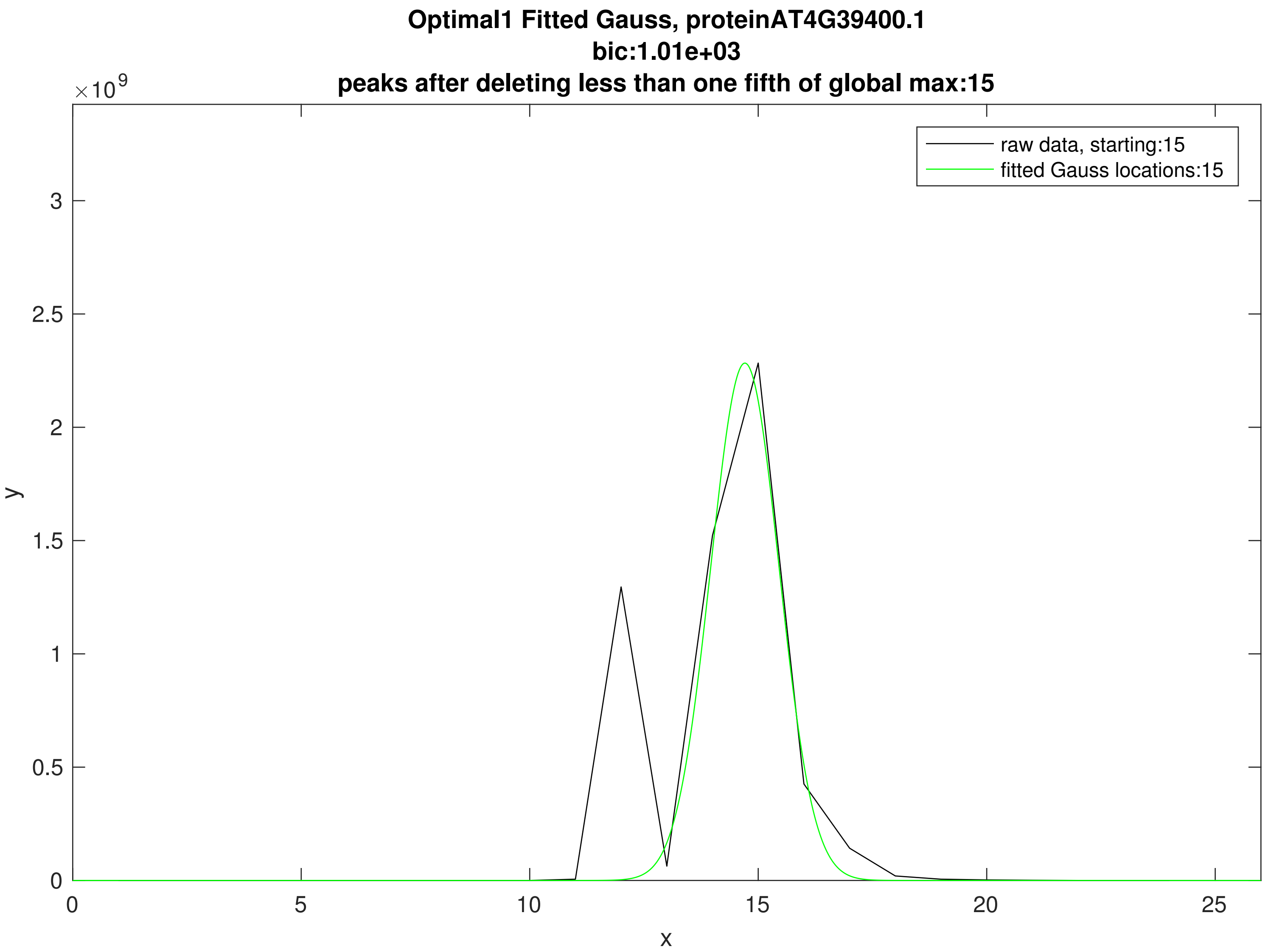

Optimal1 Fitted Gauss, proteinAT4G39400.1
bic:1.01e+03
peaks after deleting less than one fifth of global max:15
×10^9
raw data, starting:15
fitted Gauss locations:15
y
x
0
0.5
1
1.5
2
2.5
3
0
5
10
15
20
25

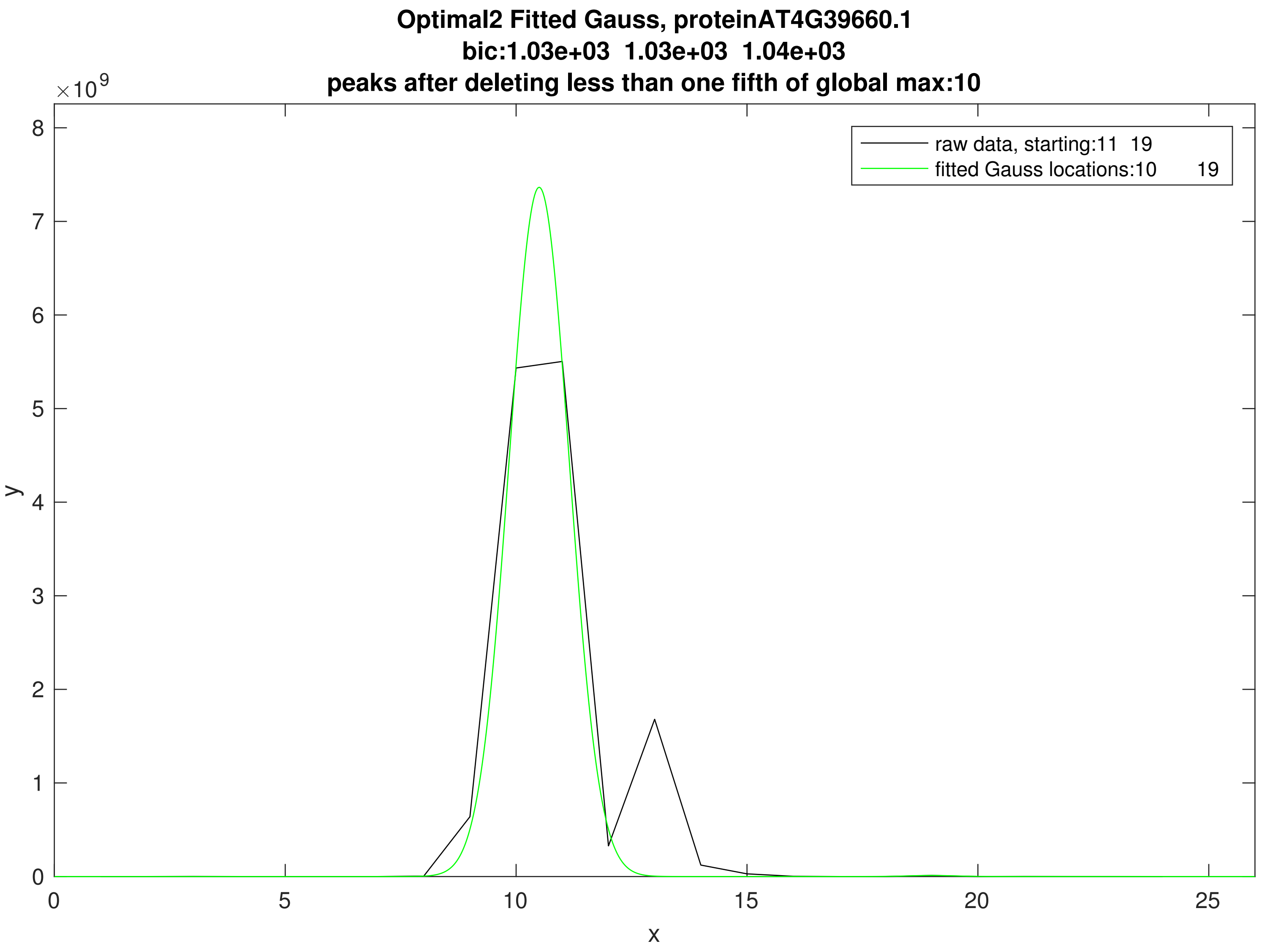

Optimal2 Fitted Gauss, proteinAT4G39660.1
bic:1.03e+03  1.03e+03  1.04e+03
peaks after deleting less than one fifth of global max:10
×10^9
raw data, starting:11  19
fitted Gauss locations:10  19
y
x
0
1
2
3
4
5
6
7
8
0
5
10
15
20
25

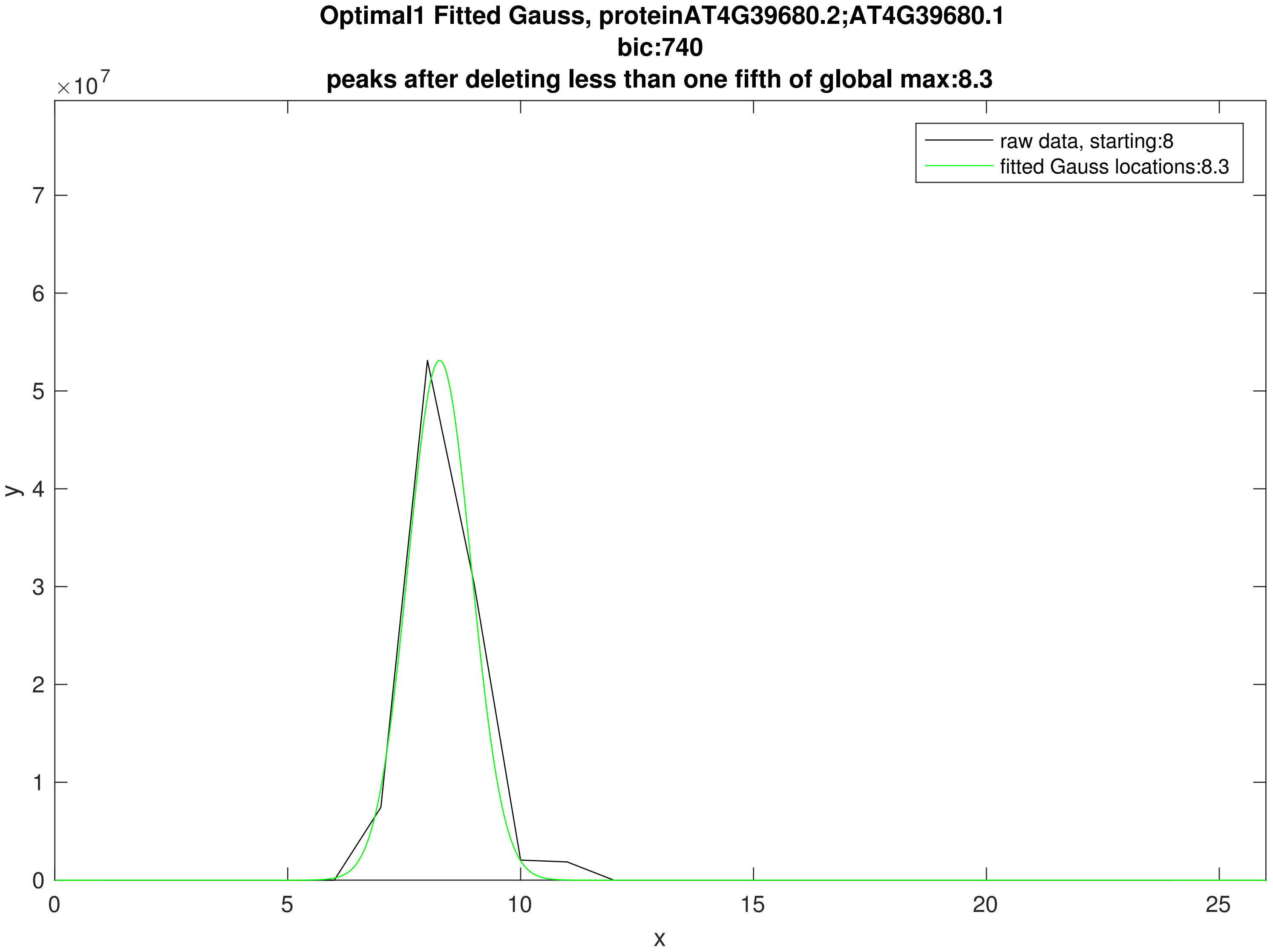
Optimal1 Fitted Gauss, proteinAT4G39680.2;AT4G39680.1
bic:740
peaks after deleting less than one fifth of global max:8.3
×10^7
raw data, starting:8
fitted Gauss locations:8.3
y
x
0
1
2
3
4
5
6
7
0
5
10
15
20
25

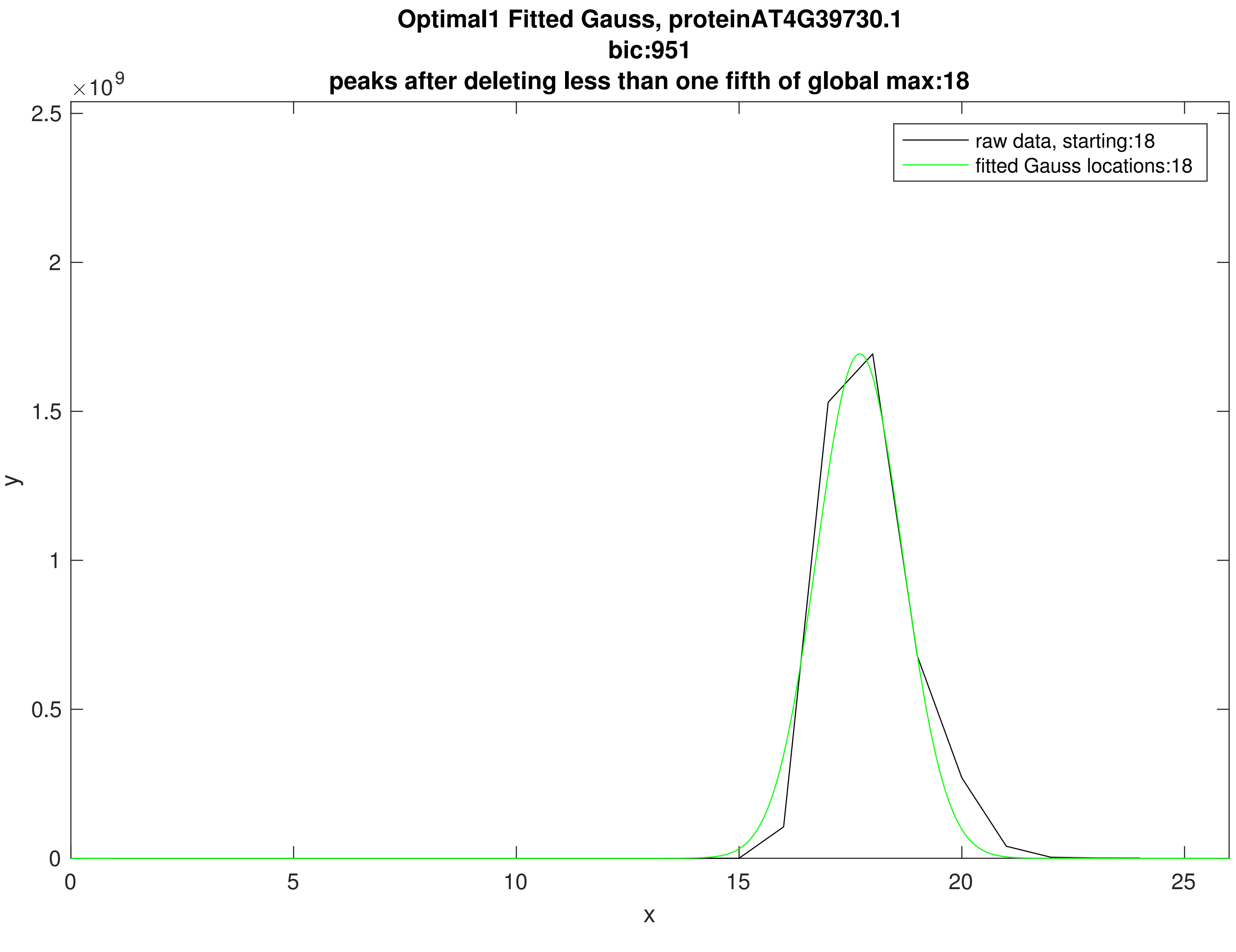
Optimal1 Fitted Gauss, proteinAT4G39730.1
bic:951
peaks after deleting less than one fifth of global max:18
×10^9
2.5
2
1.5
1
0.5
0
y
0
5
10
15
20
25
x
raw data, starting:18
fitted Gauss locations:18

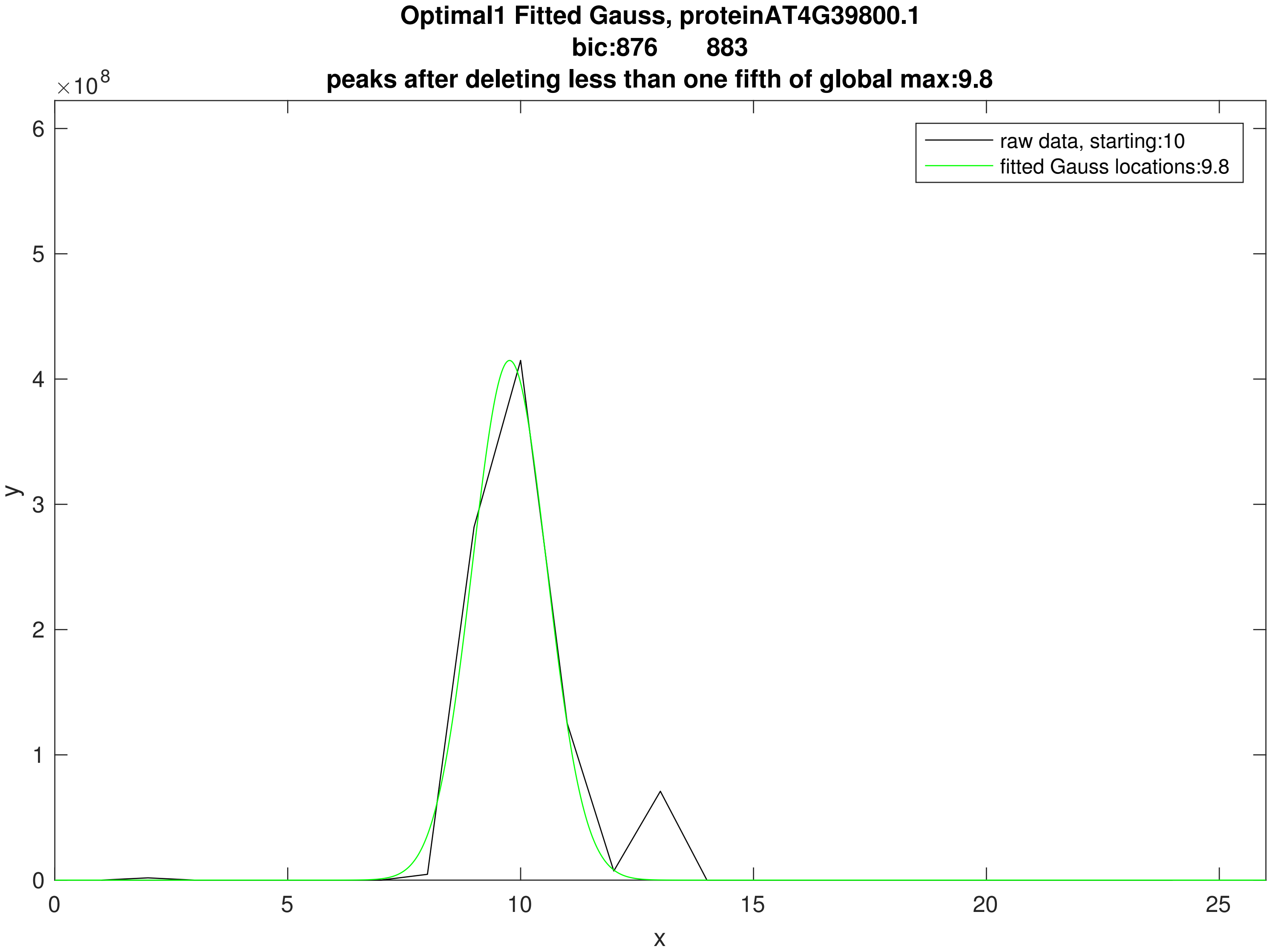
Optimal1 Fitted Gauss, proteinAT4G39800.1
bic:876 883
peaks after deleting less than one fifth of global max:9.8
×10^8
6
5
4
3
2
1
0
y
0
5
10
15
20
25
x
raw data, starting:10
fitted Gauss locations:9.8

#### Optimal1 Fitted Gauss, proteinAT4G39860.2;AT4G39860.1
#### bic:836 846 855
#### peaks after deleting less than one fifth of global max:18

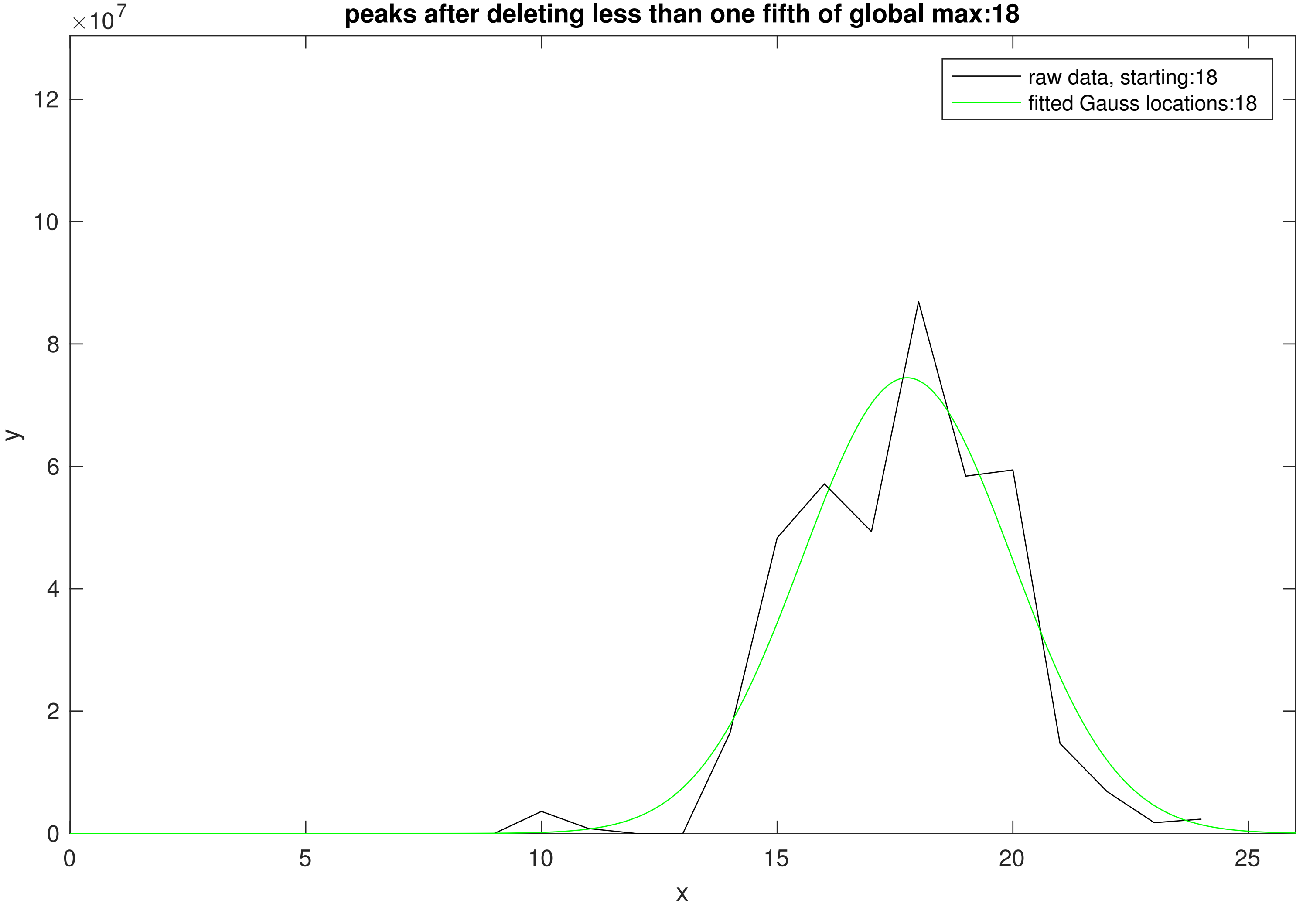

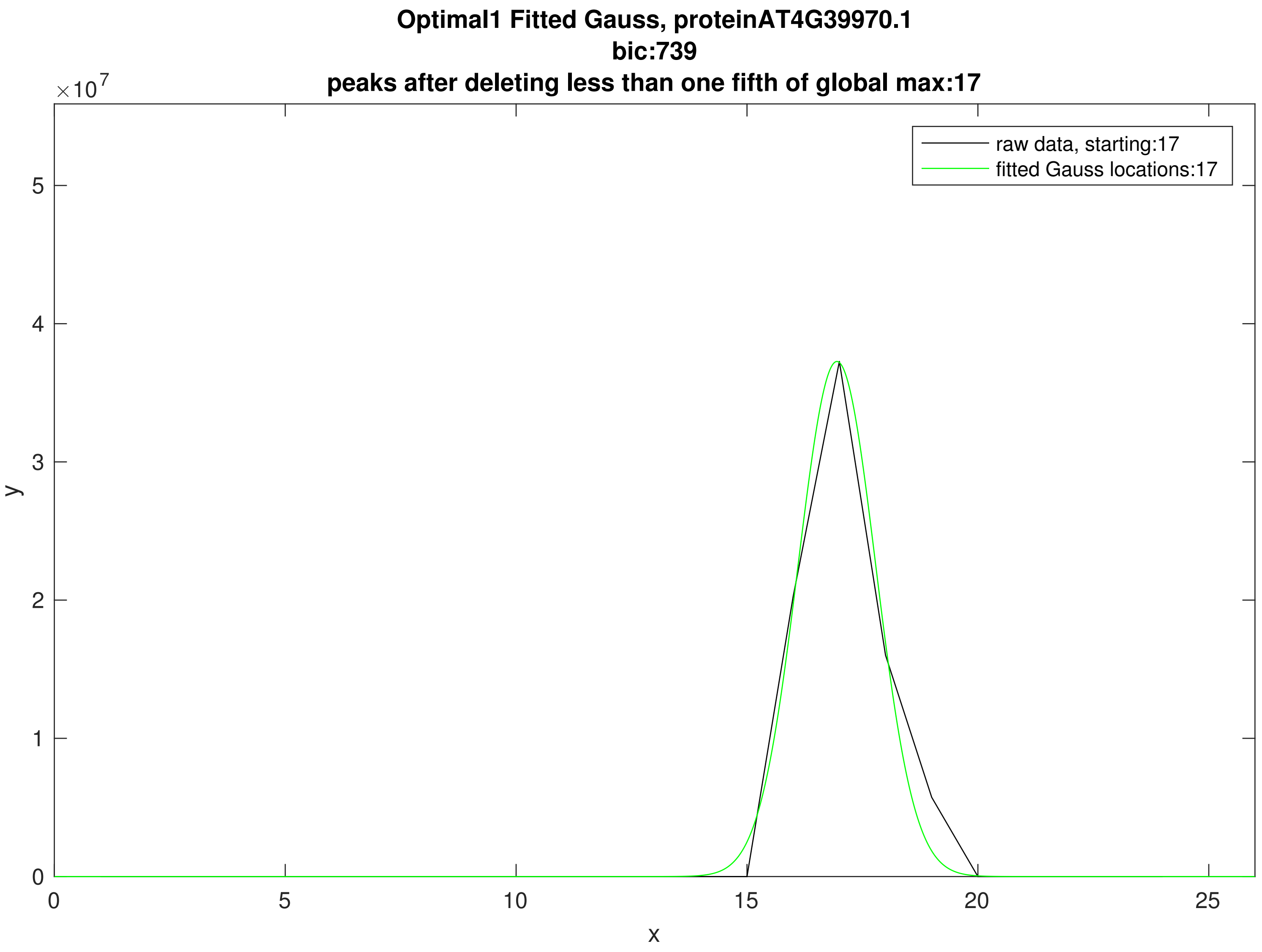

Optimal1 Fitted Gauss, proteinAT4G39970.1
bic:739
peaks after deleting less than one fifth of global max:17
×10^7
raw data, starting:17
fitted Gauss locations:17
y
x
0
1
2
3
4
5
0
5
10
15
20
25

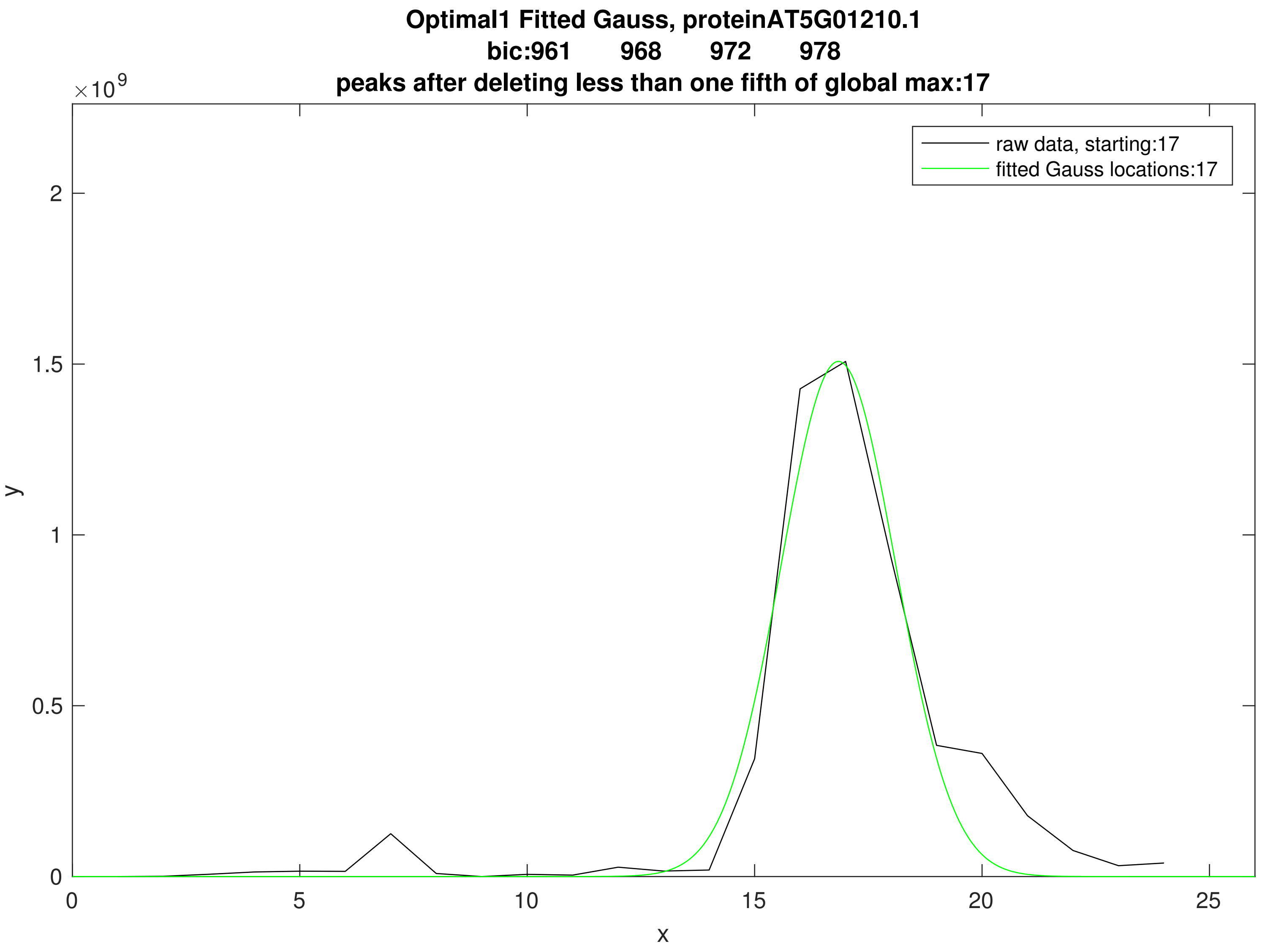
Optimal1 Fitted Gauss, proteinAT5G01210.1
bic:961 968 972 978
peaks after deleting less than one fifth of global max:17
×10^9
raw data, starting:17
fitted Gauss locations:17
y
x
0
0.5
1
1.5
2
0
5
10
15
20
25

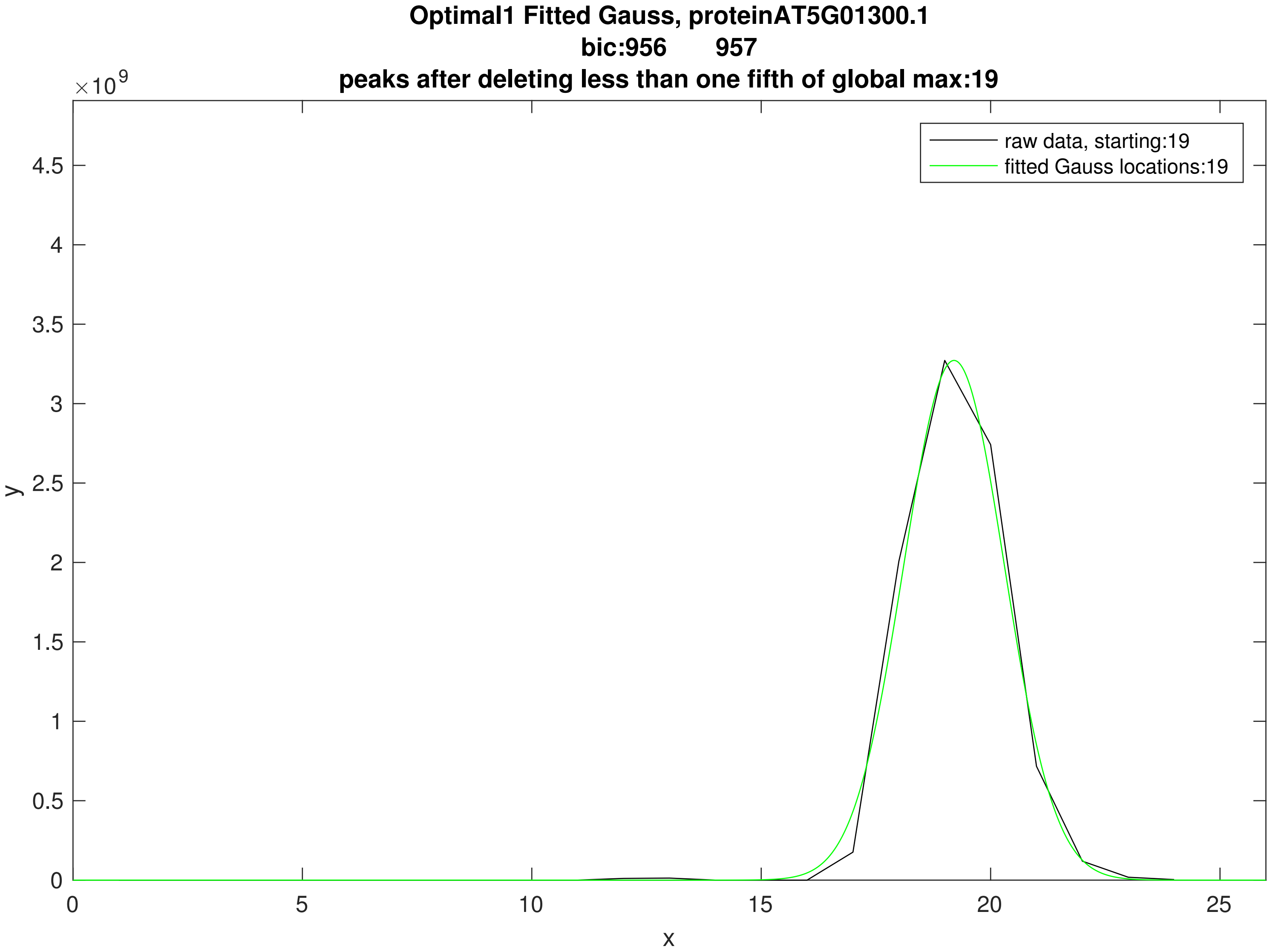

Optimal1 Fitted Gauss, proteinAT5G01300.1
bic:956 957
peaks after deleting less than one fifth of global max:19
×10^9
4.5
4
3.5
3
2.5
2
1.5
1
0.5
0
y
0
5
10
15
20
25
x
raw data, starting:19
fitted Gauss locations:19

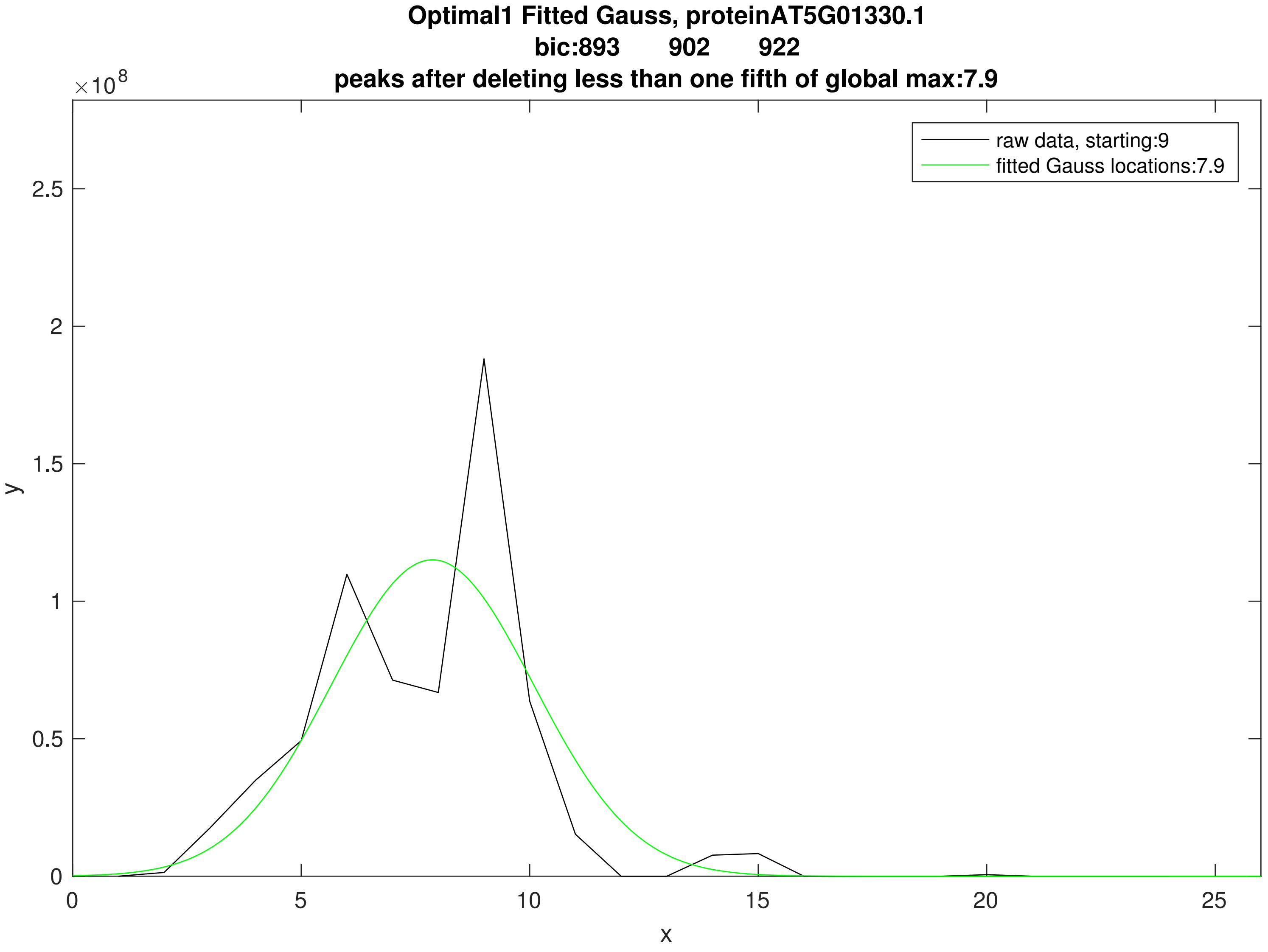
Optimal1 Fitted Gauss, proteinAT5G01330.1
bic:893 902 922
peaks after deleting less than one fifth of global max:7.9
×10^8
y
x
raw data, starting:9
fitted Gauss locations:7.9
0
0.5
1
1.5
2
2.5
5
10
15
20
25

Optimal1 Fitted Gauss, proteinAT5G01390.3;AT5G01390.2;AT5G01390.4;AT5G01390.1
bic:705
peaks after deleting less than one fifth of global max:20

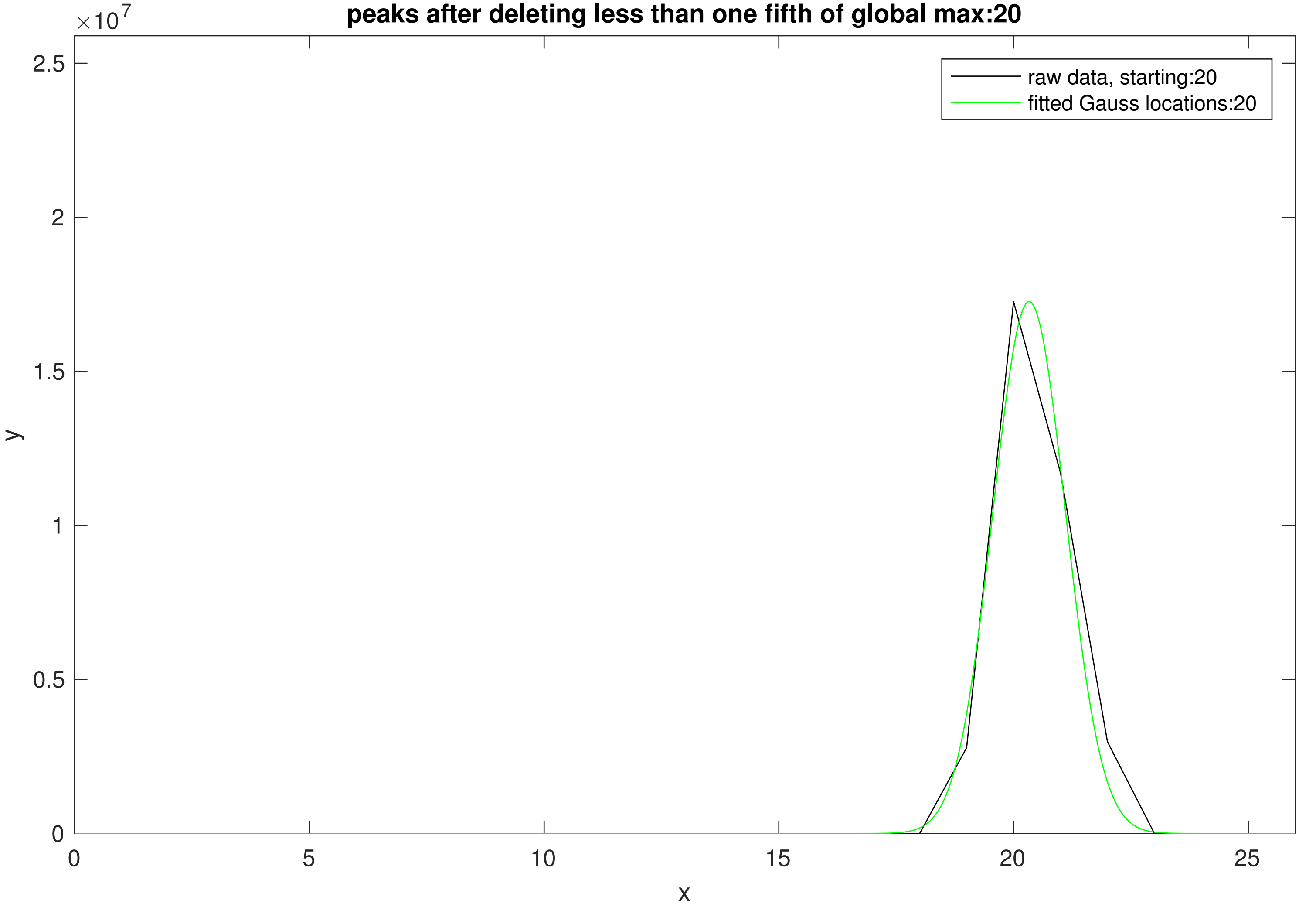

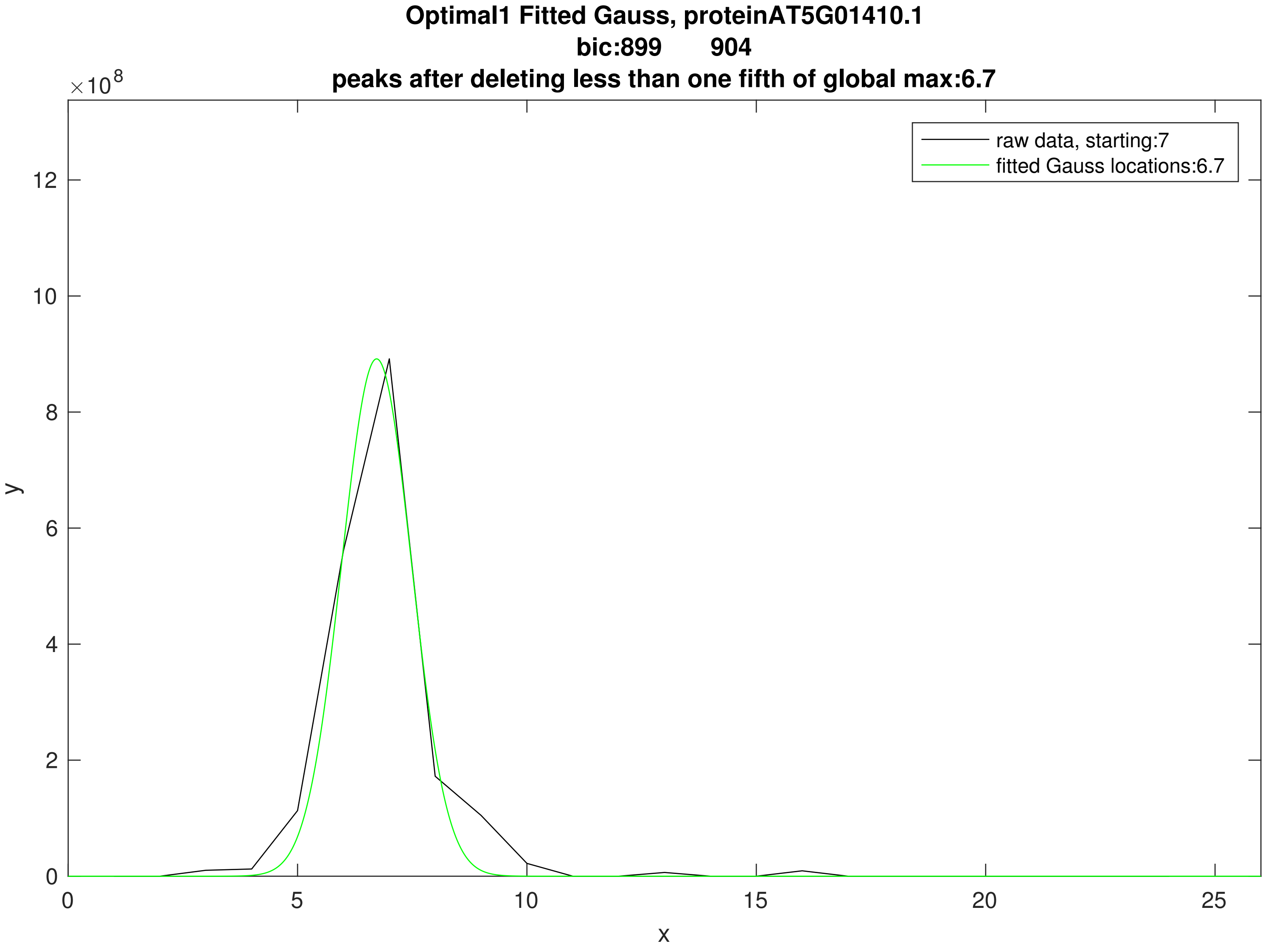

Optimal1 Fitted Gauss, proteinAT5G01410.1
bic:899 904
peaks after deleting less than one fifth of global max:6.7
×10^8
raw data, starting:7
fitted Gauss locations:6.7
12
10
8
6
4
2
0
y
0
5
10
15
20
25
x

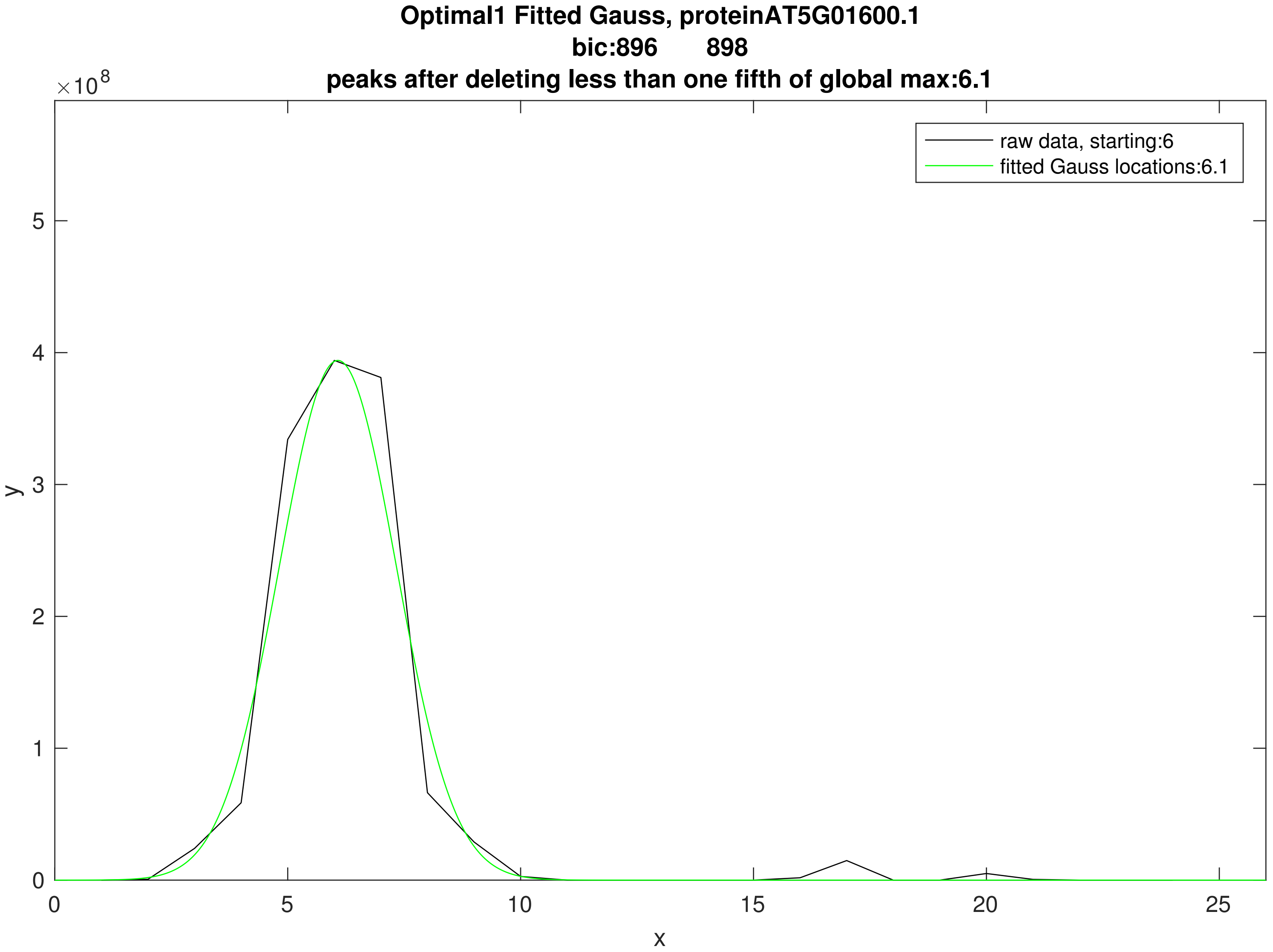
Optimal1 Fitted Gauss, proteinAT5G01600.1
bic:896 898
peaks after deleting less than one fifth of global max:6.1
×10^8
raw data, starting:6
fitted Gauss locations:6.1
y
x
0
1
2
3
4
5
10
15
20
25

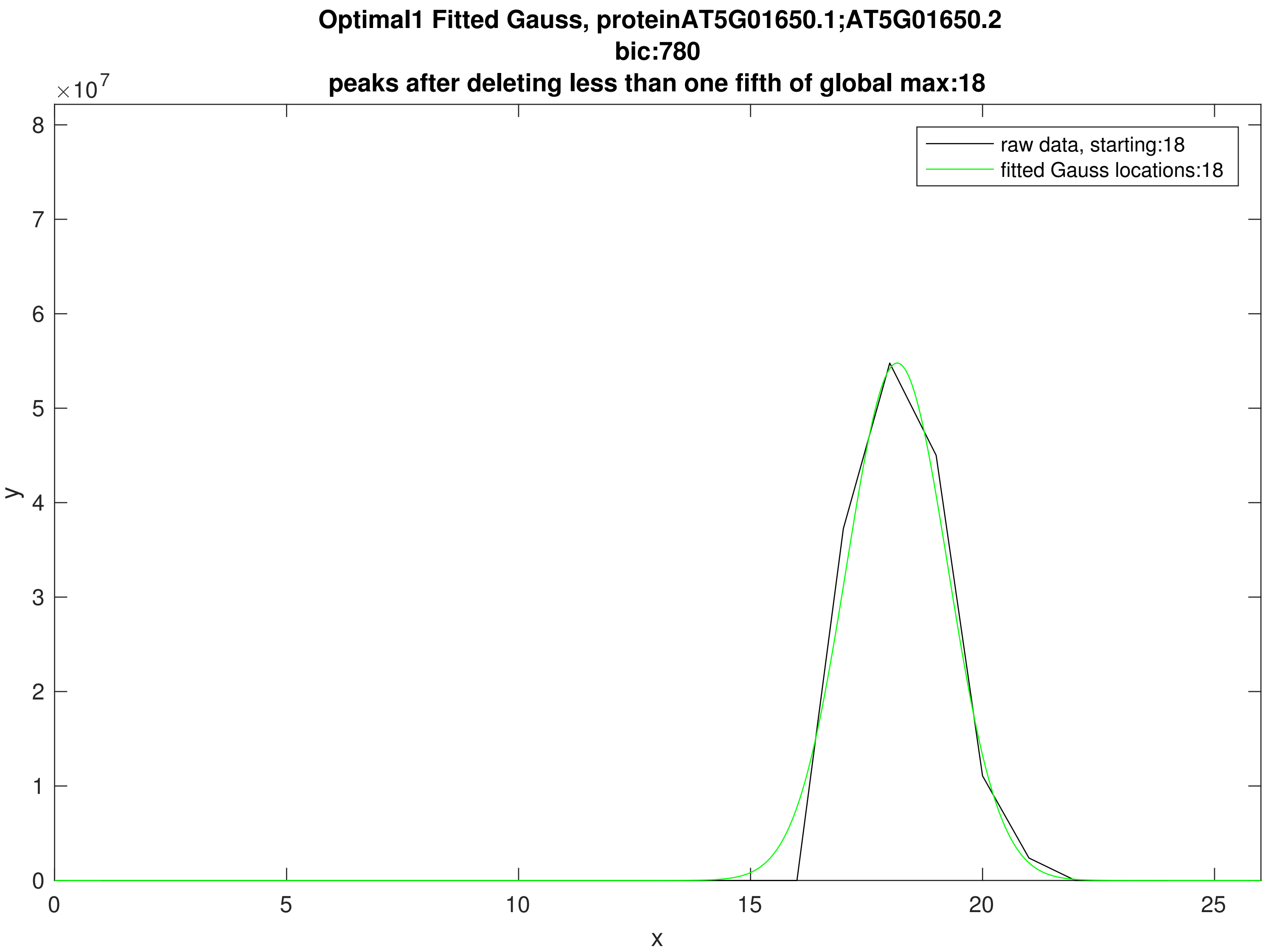
Optimal1 Fitted Gauss, proteinAT5G01650.1;AT5G01650.2
bic:780
peaks after deleting less than one fifth of global max:18
×10^7
raw data, starting:18
fitted Gauss locations:18
y
x
0
1
2
3
4
5
6
7
8
0
5
10
15
20
25

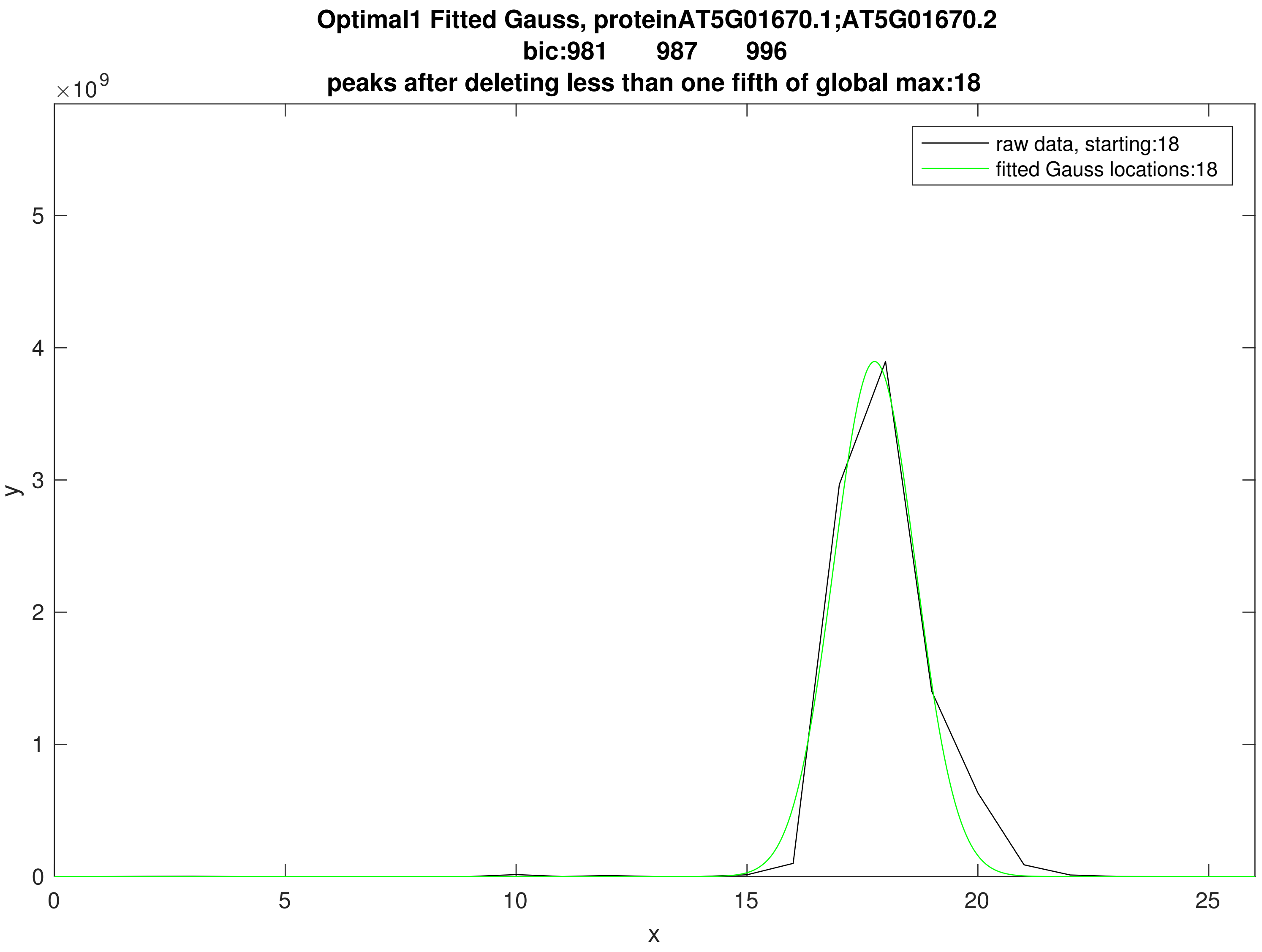

Optimal1 Fitted Gauss, proteinAT5G01670.1;AT5G01670.2
bic:981 987 996
peaks after deleting less than one fifth of global max:18
×10^9
raw data, starting:18
fitted Gauss locations:18
y
x
0
1
2
3
4
5
0
5
10
15
20
25

**Optimal1 Fitted Gauss, proteinAT5G01750.2;AT5G01750.1**
**bic:764 768**
**peaks after deleting less than one fifth of global max:20**

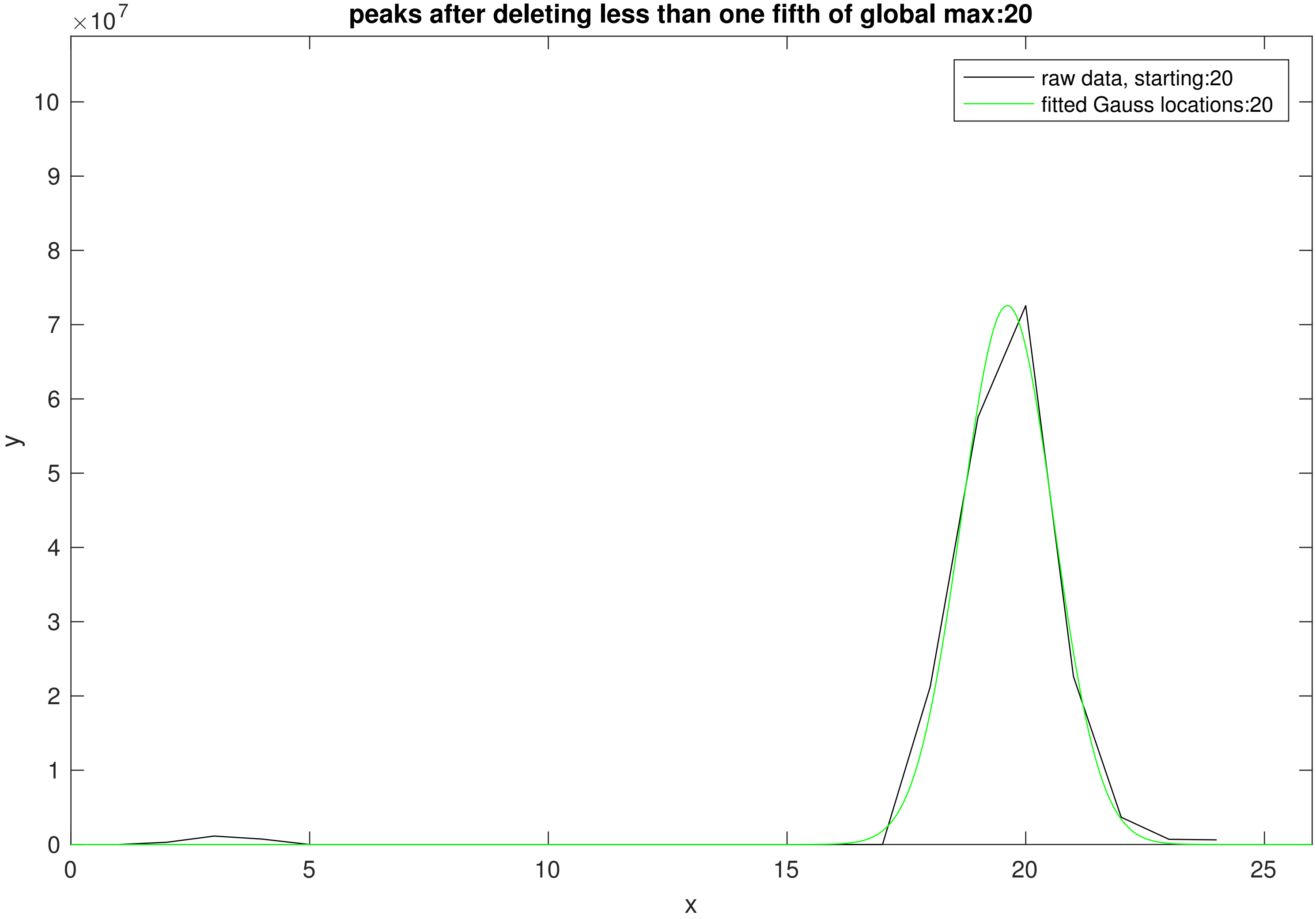

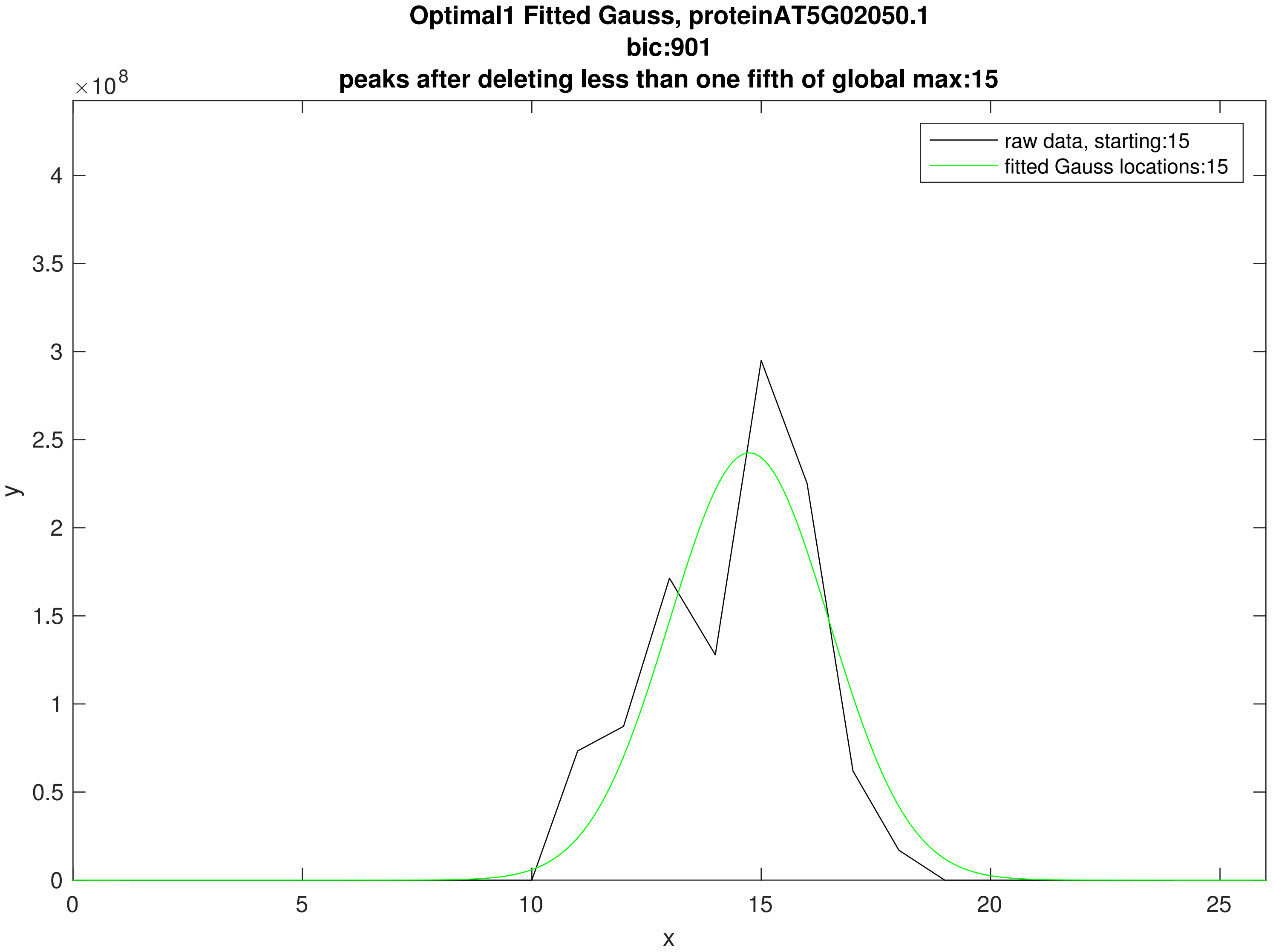
Optimal1 Fitted Gauss, proteinAT5G02050.1
bic:901
peaks after deleting less than one fifth of global max:15
×10^8
raw data, starting:15
fitted Gauss locations:15
y
x
0
0.5
1
1.5
2
2.5
3
3.5
4
0
5
10
15
20
25

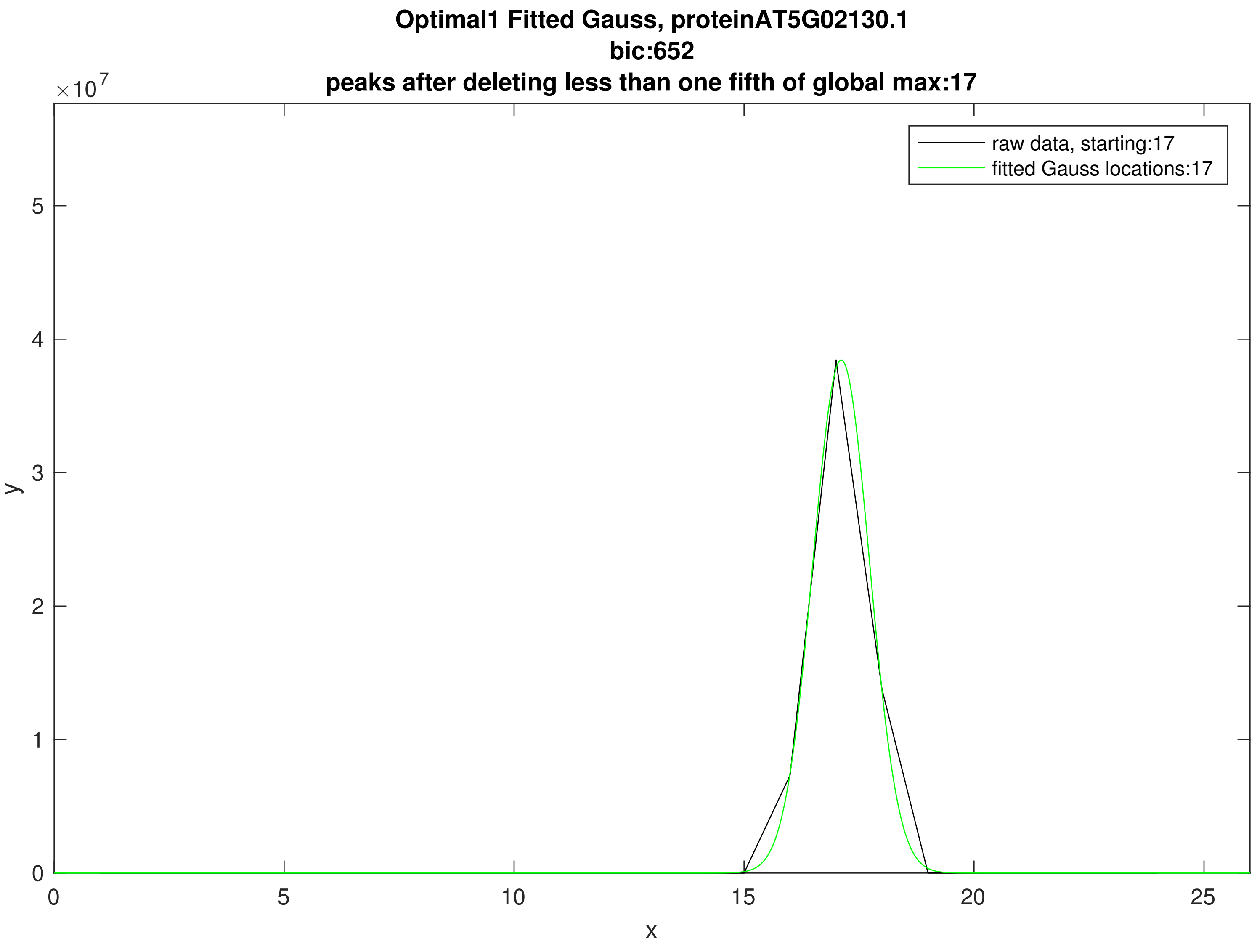
Optimal1 Fitted Gauss, proteinAT5G02130.1
bic:652
peaks after deleting less than one fifth of global max:17
×10^7
raw data, starting:17
fitted Gauss locations:17
y
x
0
1
2
3
4
5
0
5
10
15
20
25

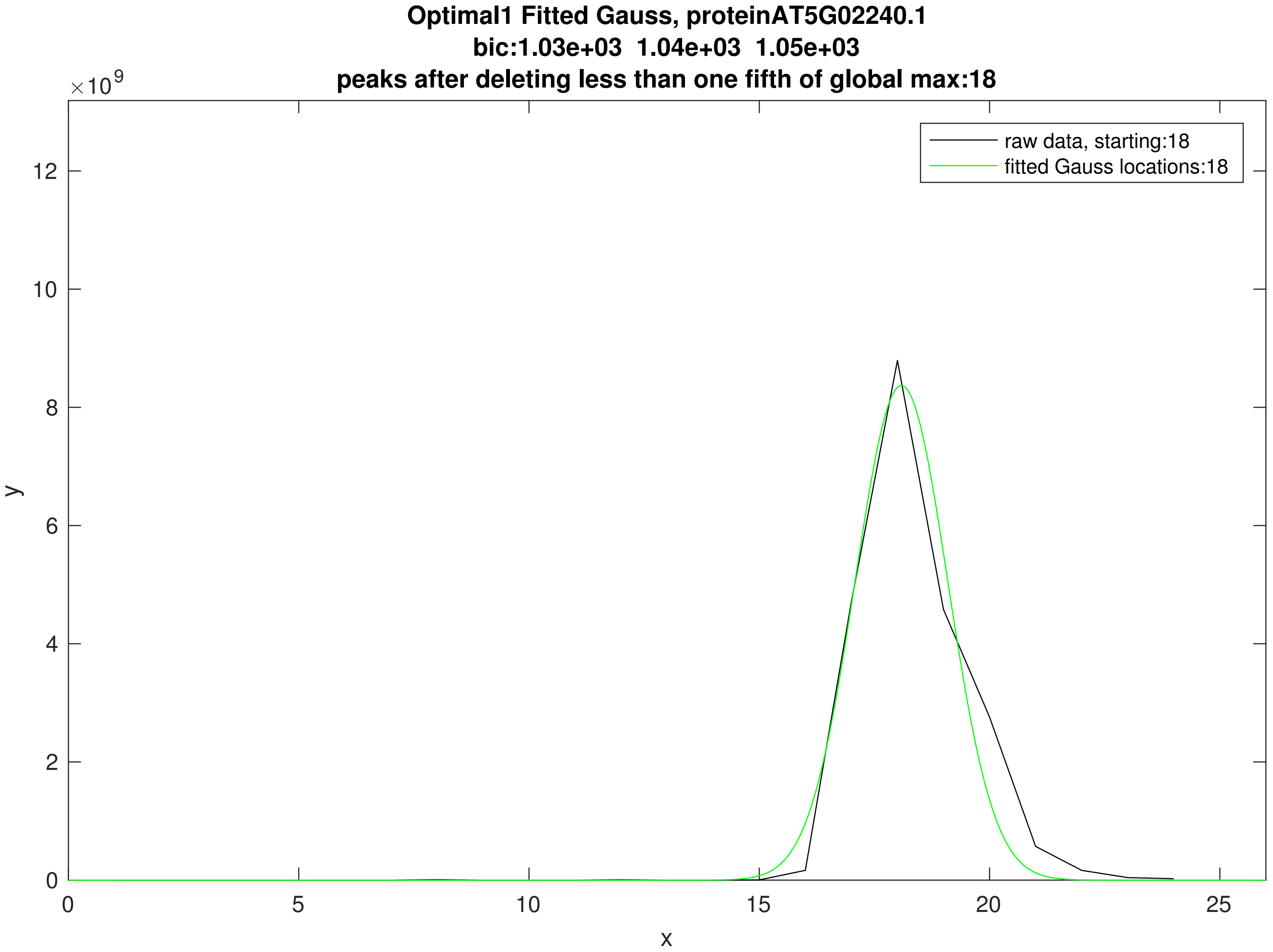
Optimal1 Fitted Gauss, proteinAT5G02240.1
bic:1.03e+03 1.04e+03 1.05e+03
peaks after deleting less than one fifth of global max:18
×10^9
raw data, starting:18
fitted Gauss locations:18
y
x
0
2
4
6
8
10
12
0
5
10
15
20
25

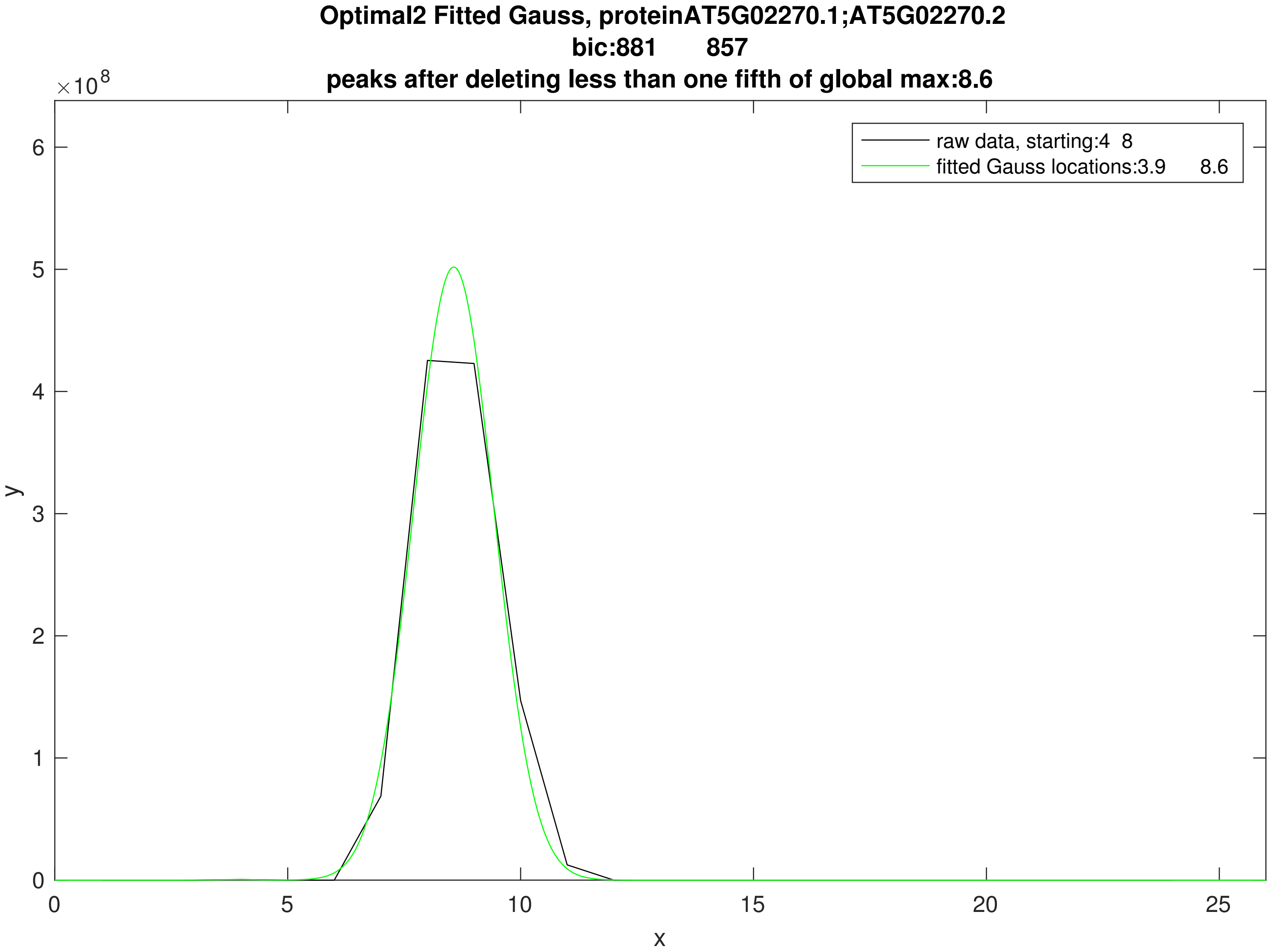
Optimal2 Fitted Gauss, proteinAT5G02270.1;AT5G02270.2
bic:881 857
peaks after deleting less than one fifth of global max:8.6
×10^8
raw data, starting:4 8
fitted Gauss locations:3.9 8.6
y
x
0
1
2
3
4
5
6
0
5
10
15
20
25

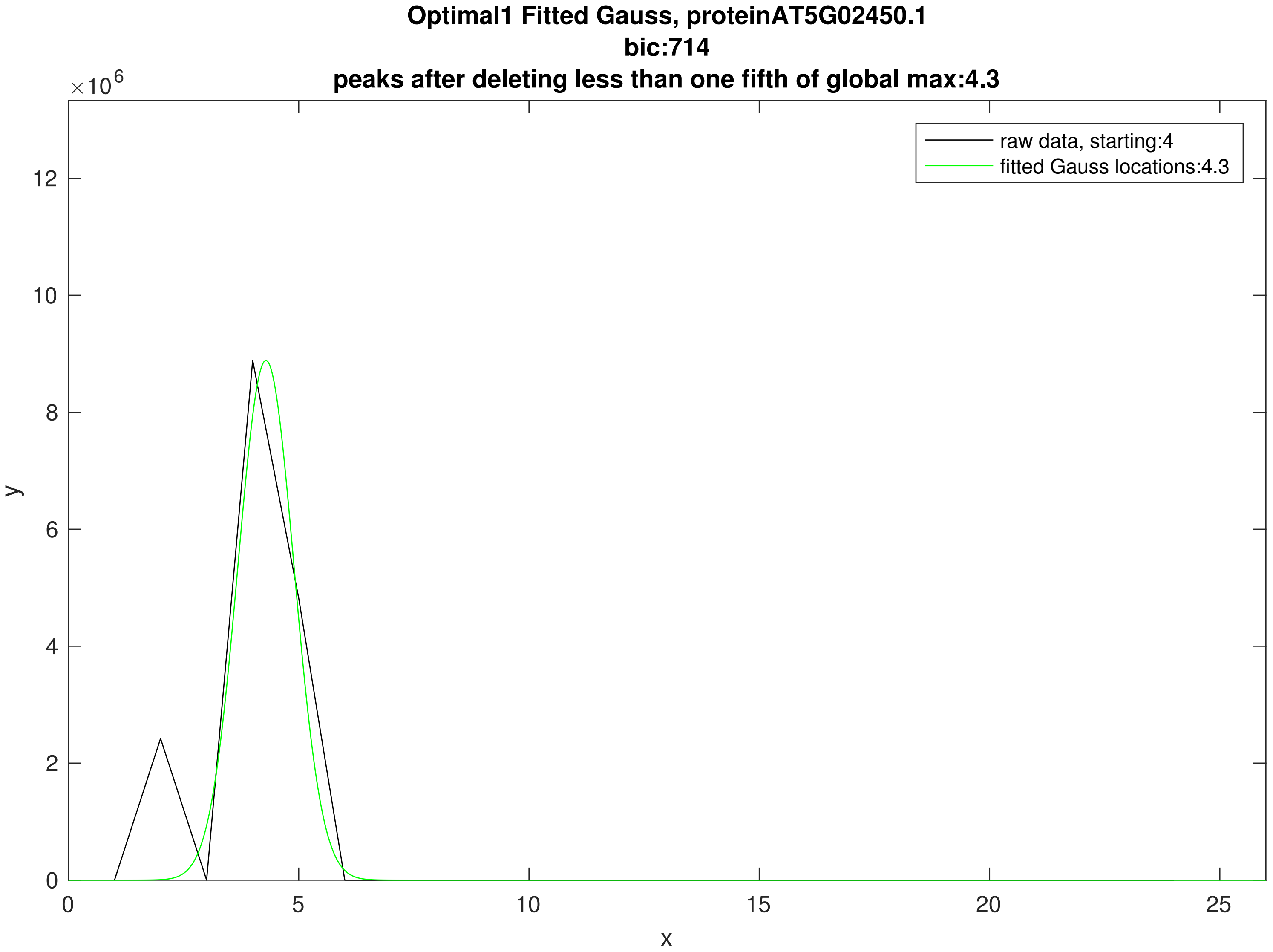
Optimal1 Fitted Gauss, proteinAT5G02450.1
bic:714
peaks after deleting less than one fifth of global max:4.3
×10^6
raw data, starting:4
fitted Gauss locations:4.3
y
x
0
2
4
6
8
10
12
0
5
10
15
20
25

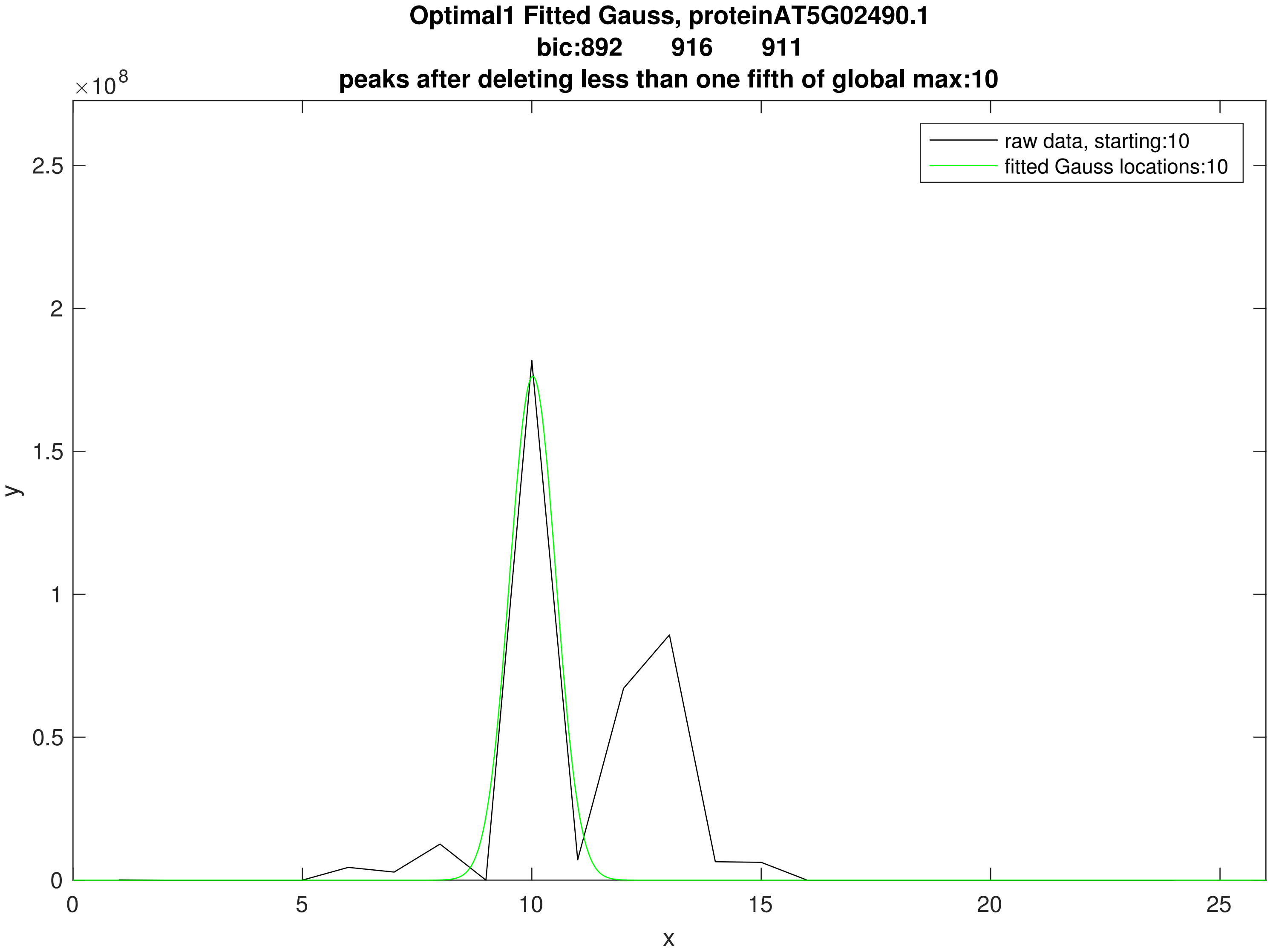
Optimal1 Fitted Gauss, proteinAT5G02490.1
bic:892 916 911
peaks after deleting less than one fifth of global max:10
×10^8
raw data, starting:10
fitted Gauss locations:10
y
x
0
0.5
1
1.5
2
2.5
0
5
10
15
20
25

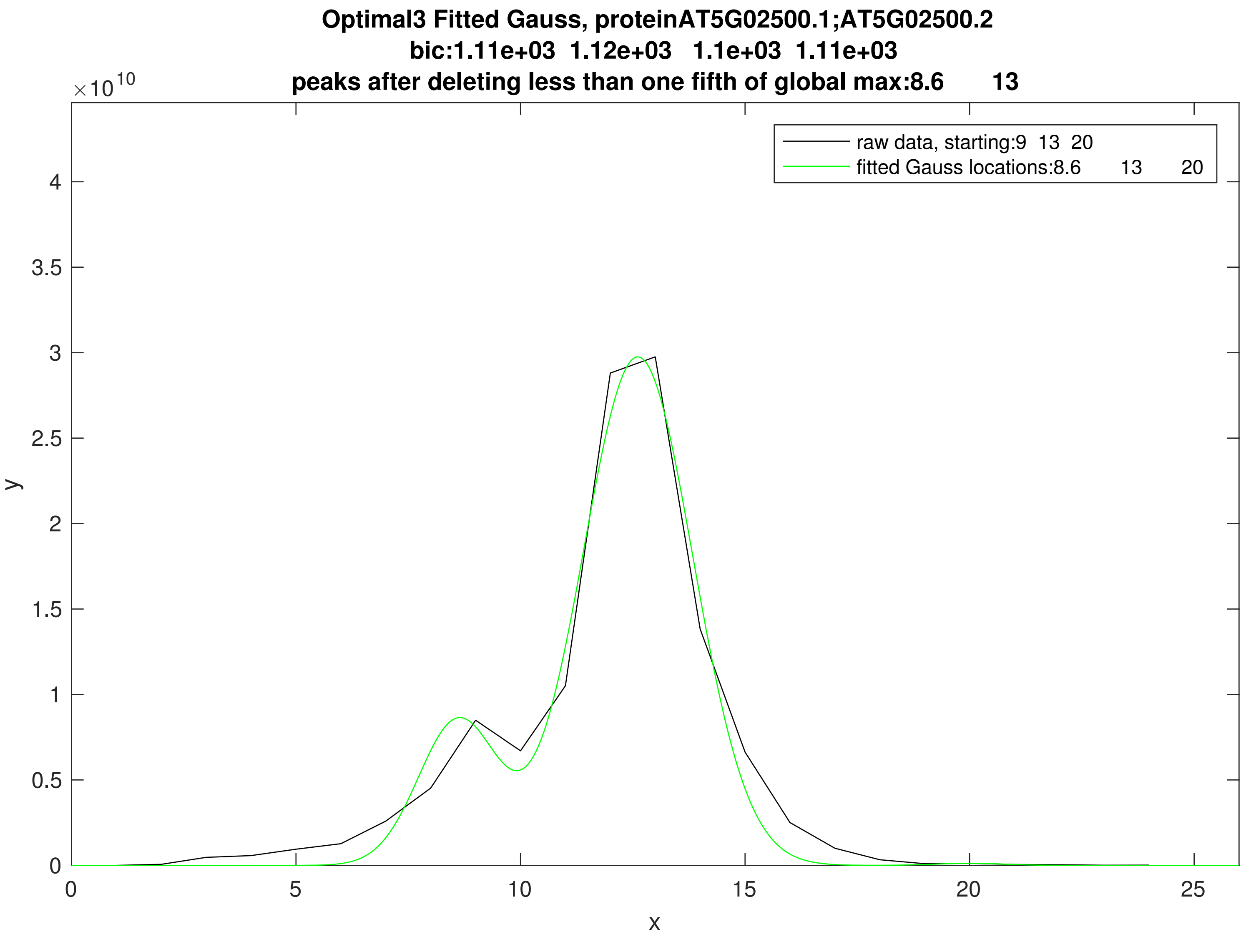
Optimal3 Fitted Gauss, proteinAT5G02500.1;AT5G02500.2
bic:1.11e+03 1.12e+03 1.1e+03 1.11e+03
peaks after deleting less than one fifth of global max:8.6 13
×10^10
raw data, starting:9 13 20
fitted Gauss locations:8.6 13 20
4
3.5
3
2.5
2
1.5
1
0.5
0
y
0
5
10
15
20
25
x

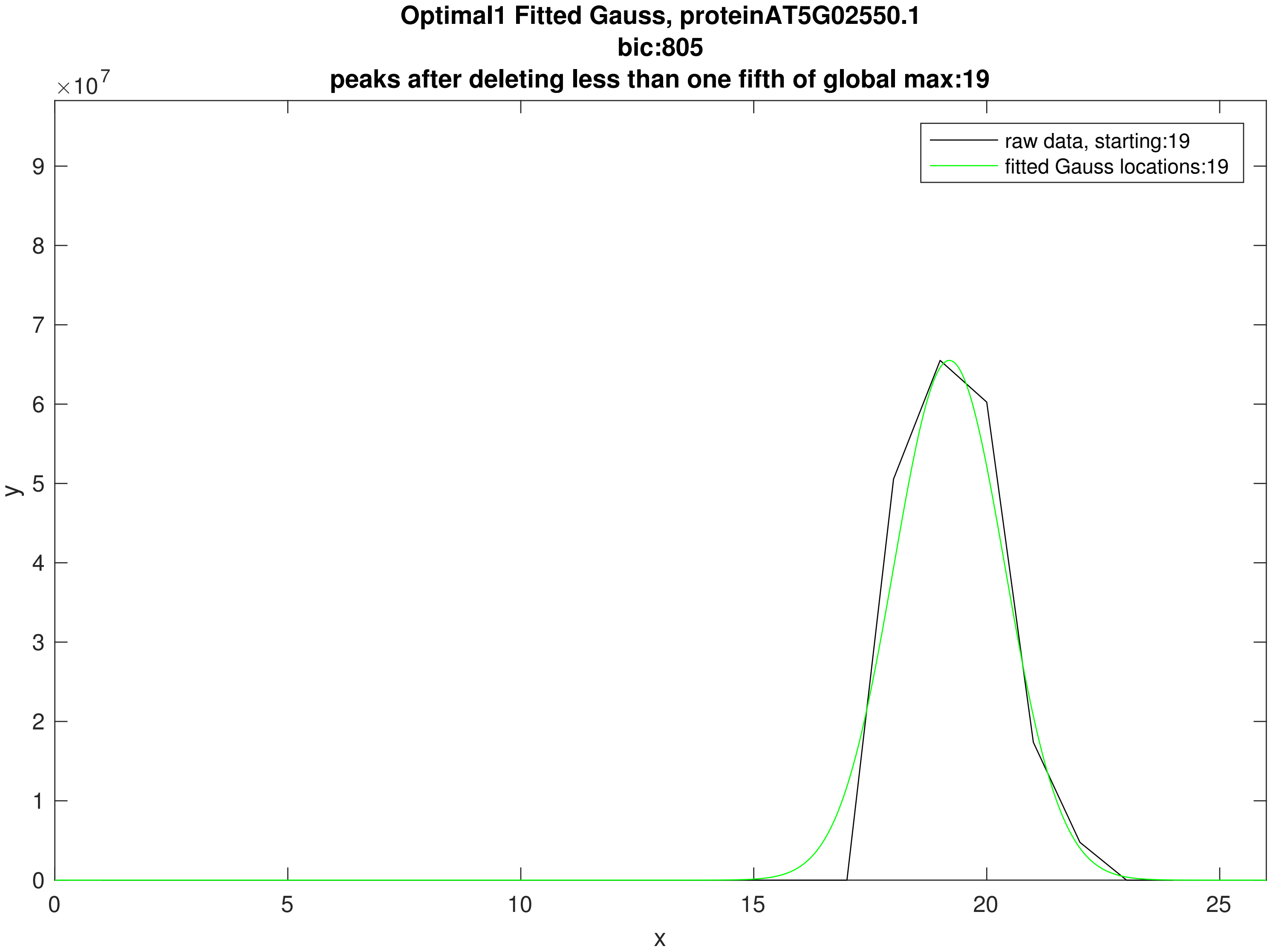

Optimal1 Fitted Gauss, proteinAT5G02550.1
bic:805
peaks after deleting less than one fifth of global max:19
×10^7
raw data, starting:19
fitted Gauss locations:19
y
x
0
1
2
3
4
5
6
7
8
9
0
5
10
15
20
25

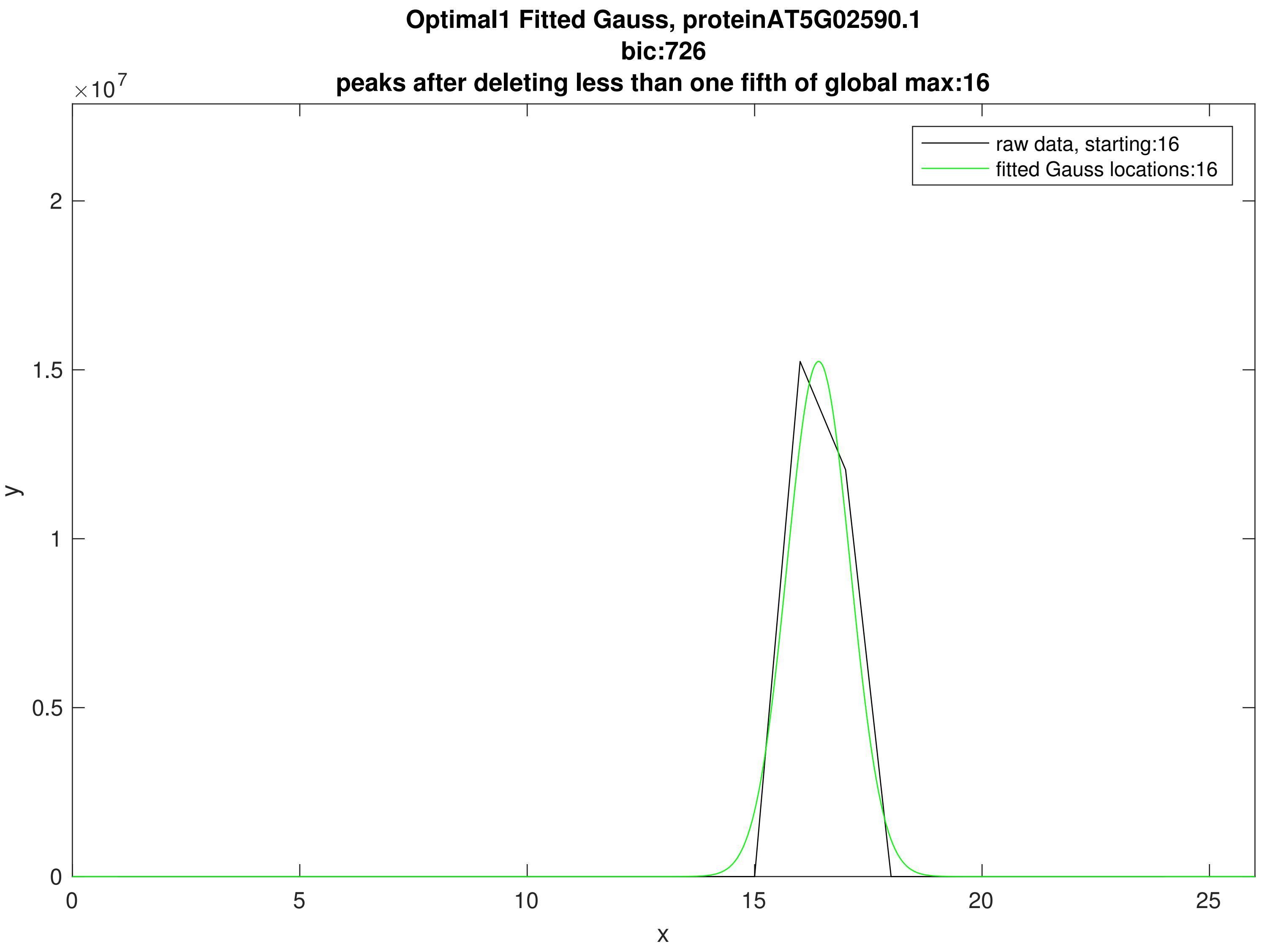
Optimal1 Fitted Gauss, proteinAT5G02590.1
bic:726
peaks after deleting less than one fifth of global max:16
×10^7
raw data, starting:16
fitted Gauss locations:16
2
1.5
1
0.5
0
y
0
5
10
15
20
25
x

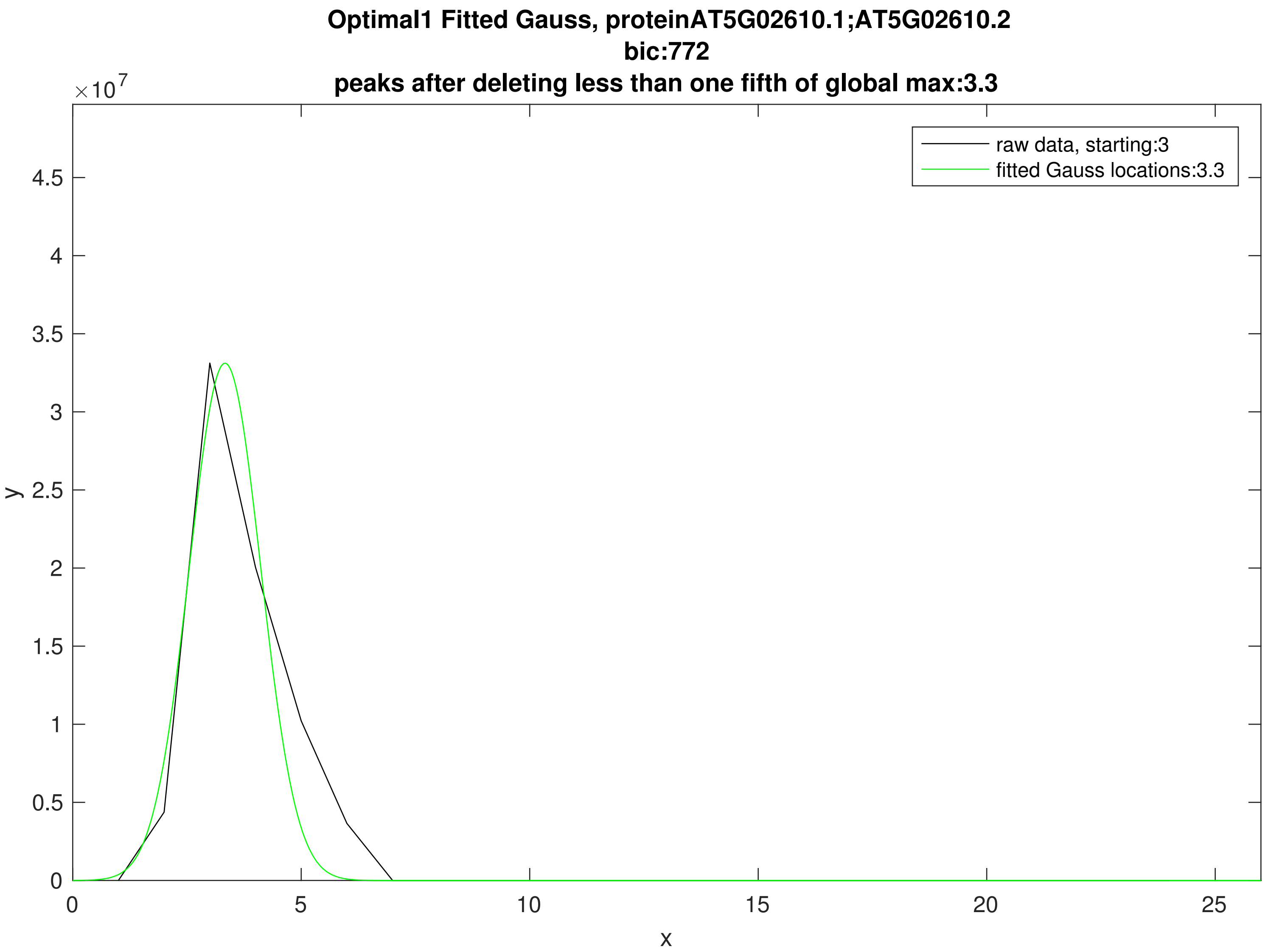

Optimal1 Fitted Gauss, proteinAT5G02610.1;AT5G02610.2
bic:772
peaks after deleting less than one fifth of global max:3.3
×10^7
raw data, starting:3
fitted Gauss locations:3.3
4.5
4
3.5
3
2.5
2
1.5
1
0.5
0
y
0
5
10
15
20
25
x

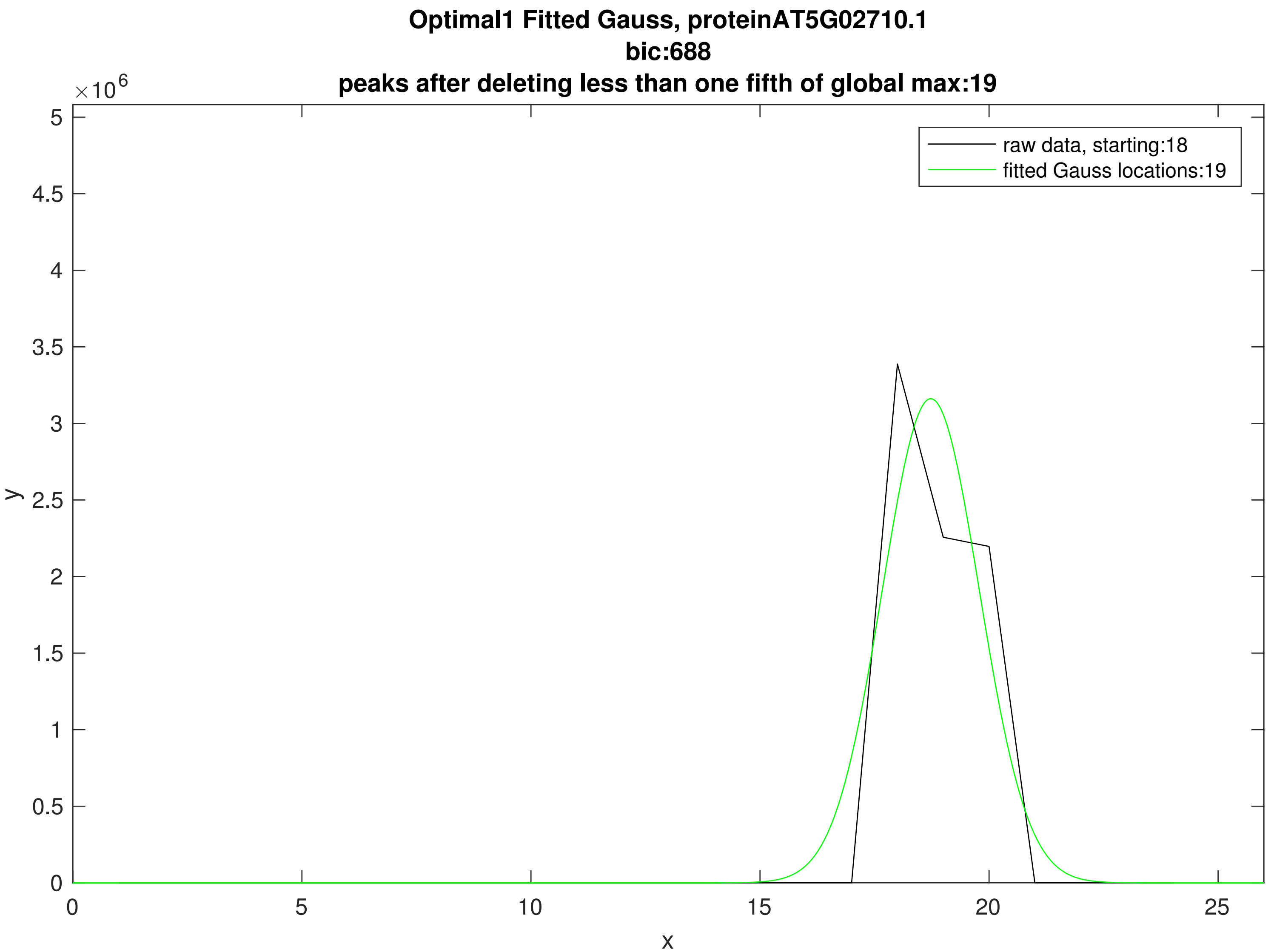

Optimal1 Fitted Gauss, proteinAT5G02710.1
bic:688
peaks after deleting less than one fifth of global max:19
×10^6
raw data, starting:18
fitted Gauss locations:19
y
x
0
0.5
1
1.5
2
2.5
3
3.5
4
4.5
5
0
5
10
15
20
25

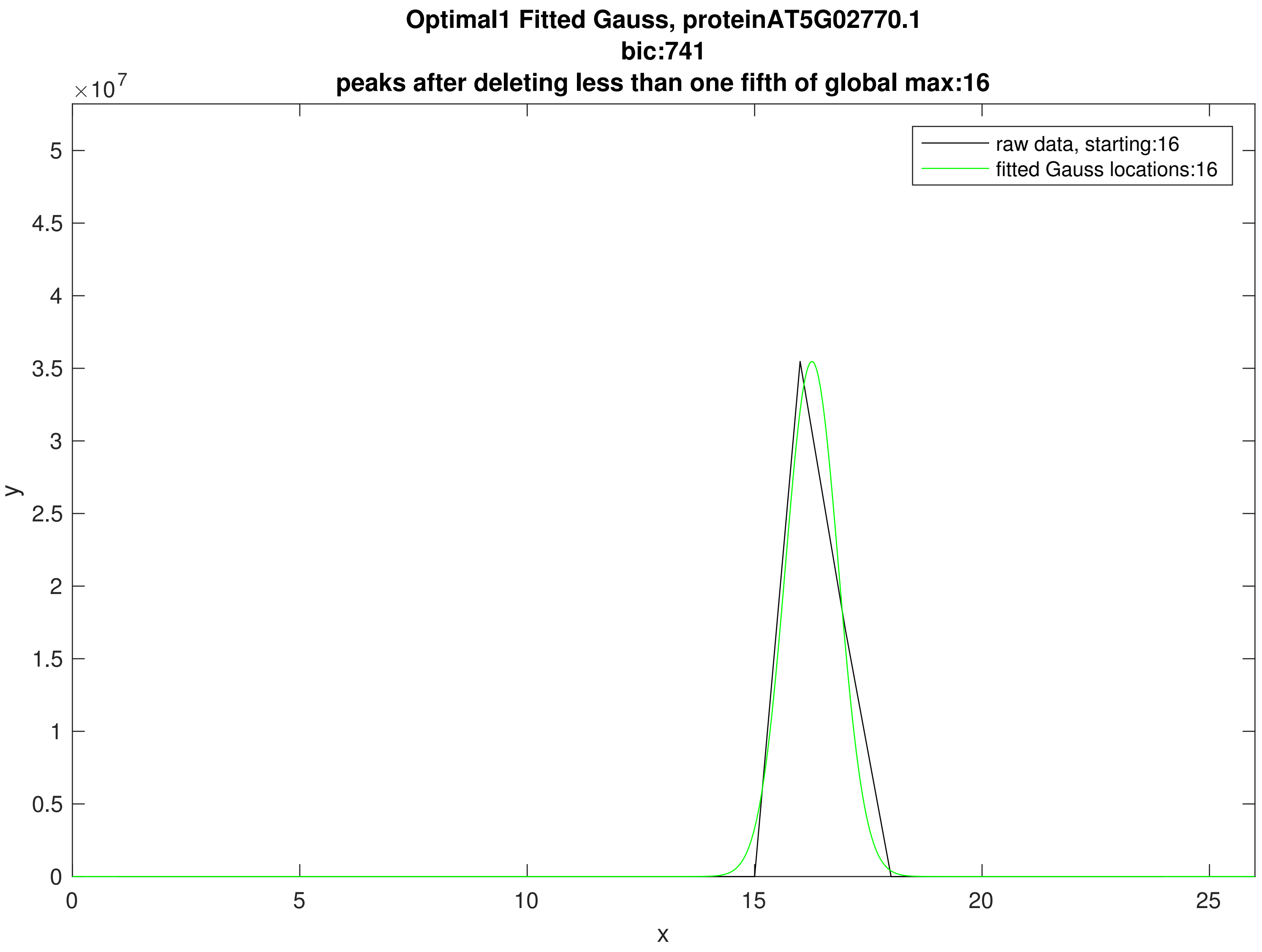
Optimal1 Fitted Gauss, proteinAT5G02770.1
bic:741
peaks after deleting less than one fifth of global max:16
×10^7
raw data, starting:16
fitted Gauss locations:16
y
x
0
0.5
1
1.5
2
2.5
3
3.5
4
4.5
5
0
5
10
15
20
25

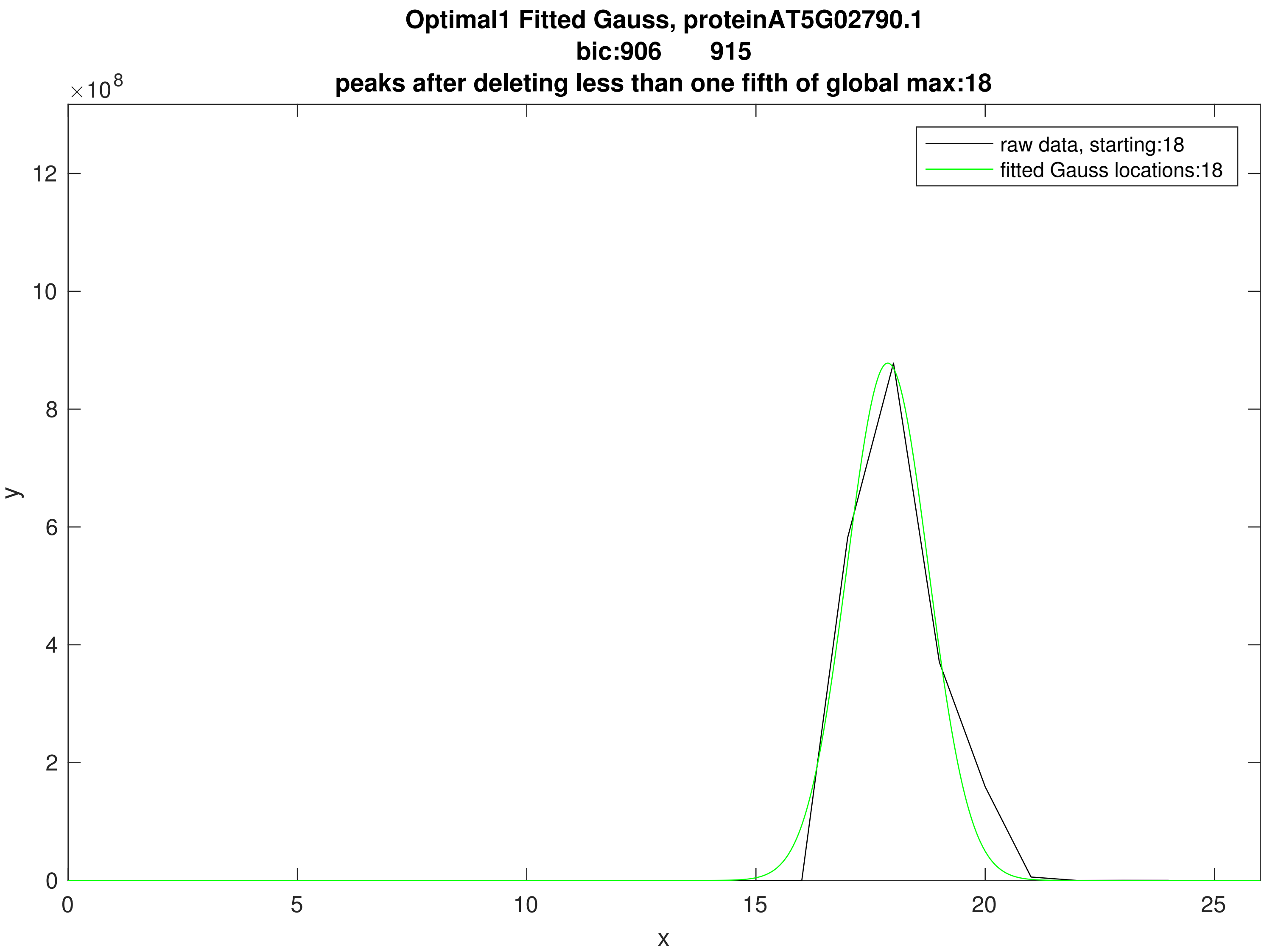

Optimal1 Fitted Gauss, proteinAT5G02790.1
bic:906 915
peaks after deleting less than one fifth of global max:18
×10^8
12
10
8
6
4
2
0
y
0
5
10
15
20
25
x
raw data, starting:18
fitted Gauss locations:18

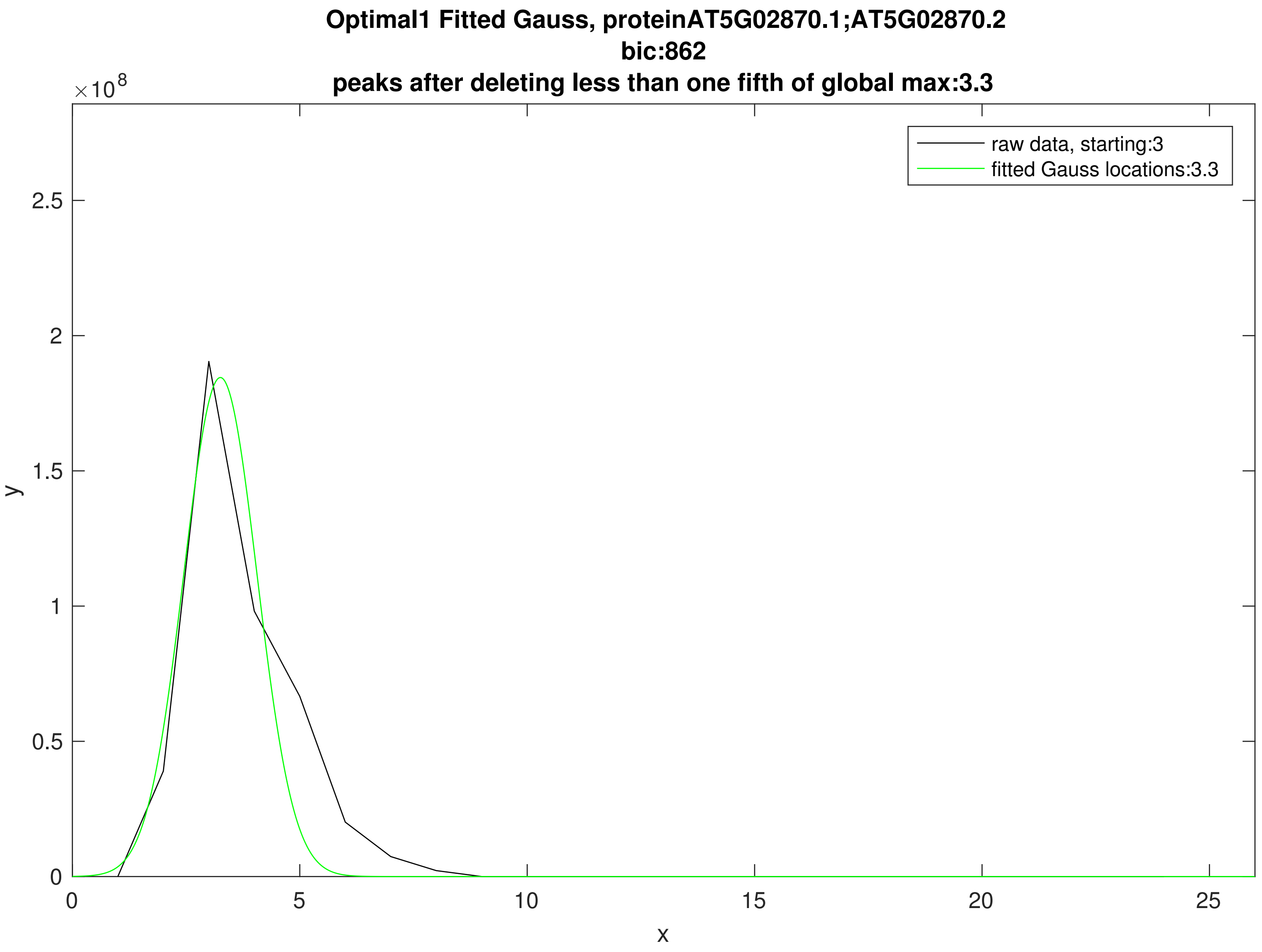
Optimal1 Fitted Gauss, proteinAT5G02870.1;AT5G02870.2
bic:862
peaks after deleting less than one fifth of global max:3.3
×10^8
raw data, starting:3
fitted Gauss locations:3.3
y
x
0
0.5
1
1.5
2
2.5
5
10
15
20
25

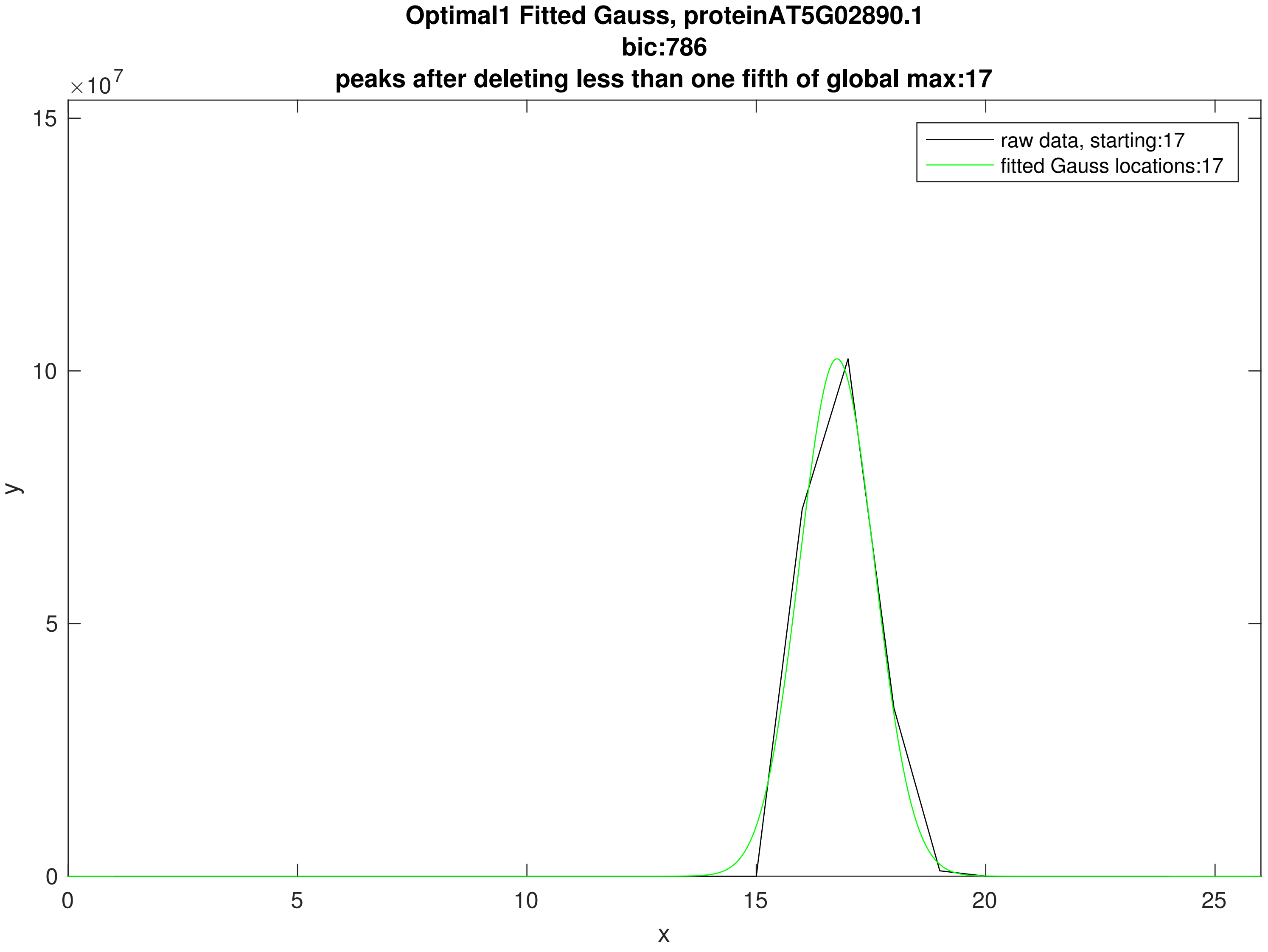

Optimal1 Fitted Gauss, proteinAT5G02890.1
bic:786
peaks after deleting less than one fifth of global max:17
×10^7
15
10
5
0
y
0
5
10
15
20
25
x
raw data, starting:17
fitted Gauss locations:17

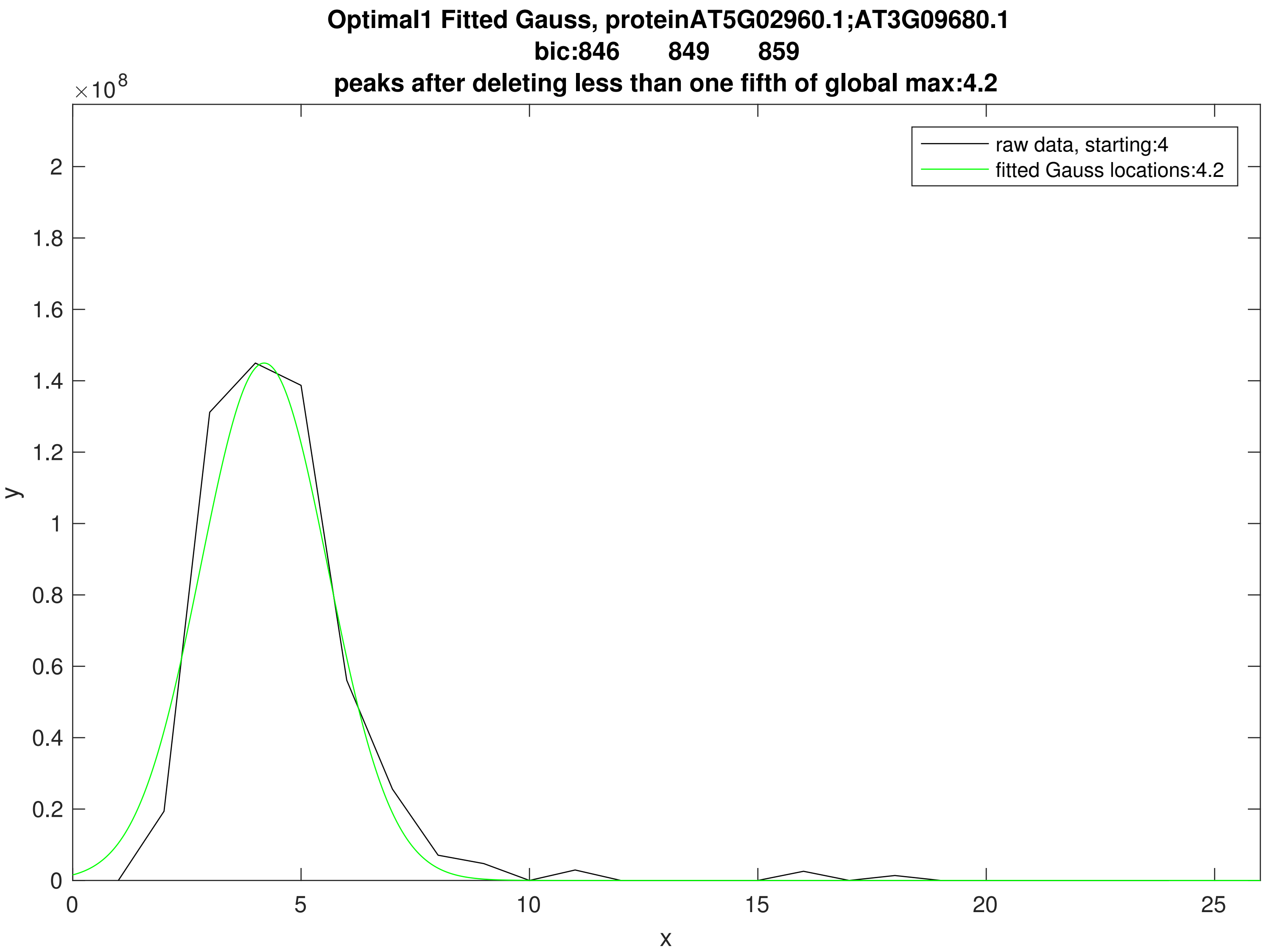
Optimal1 Fitted Gauss, proteinAT5G02960.1;AT3G09680.1
bic:846 849 859
peaks after deleting less than one fifth of global max:4.2
×10^8
raw data, starting:4
fitted Gauss locations:4.2
y
x
0
0.2
0.4
0.6
0.8
1
1.2
1.4
1.6
1.8
2
0
5
10
15
20
25

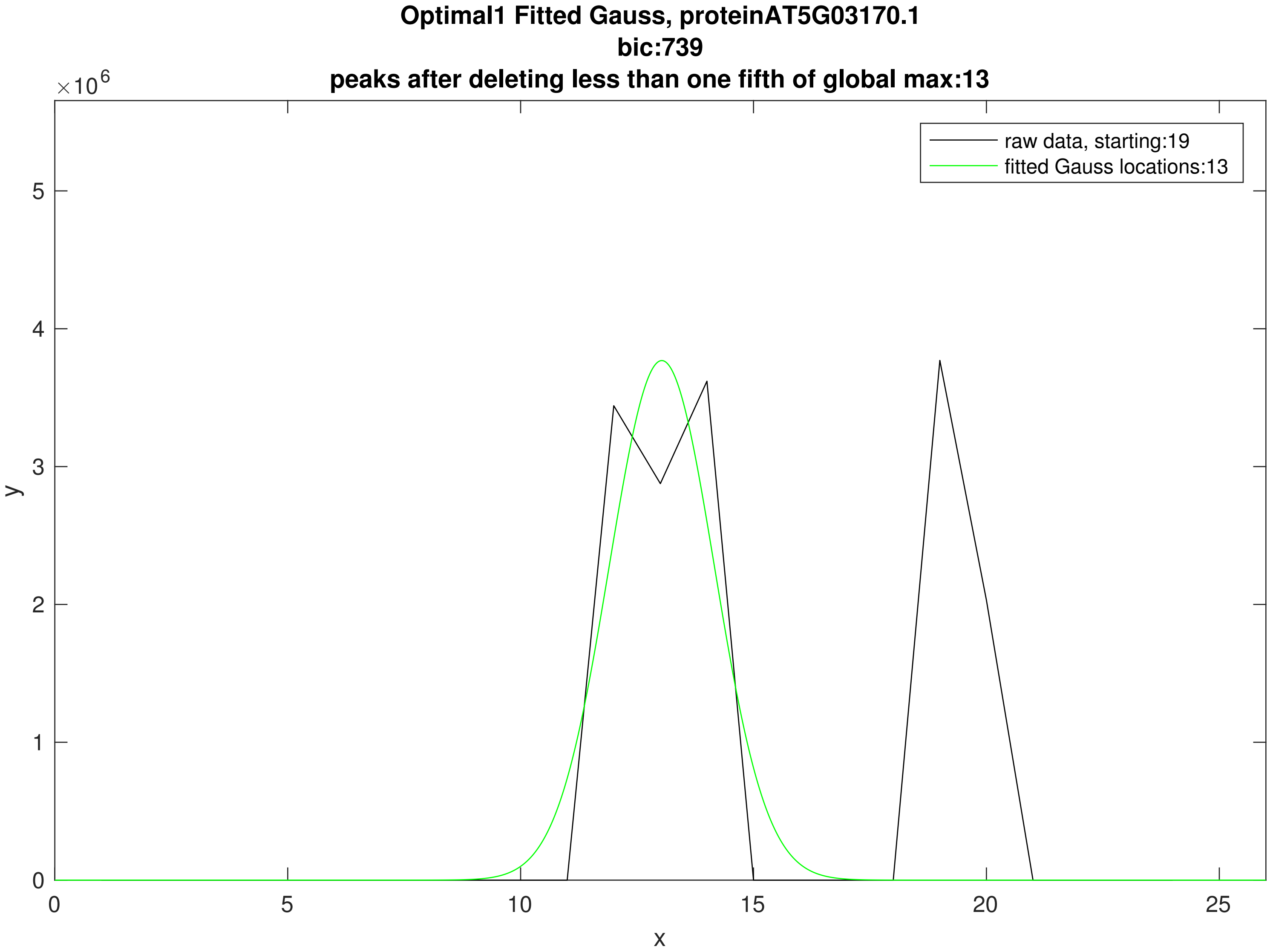
Optimal1 Fitted Gauss, proteinAT5G03170.1
bic:739
peaks after deleting less than one fifth of global max:13
×10^6
raw data, starting:19
fitted Gauss locations:13
5
4
3
2
1
0
y
0
5
10
15
20
25
x

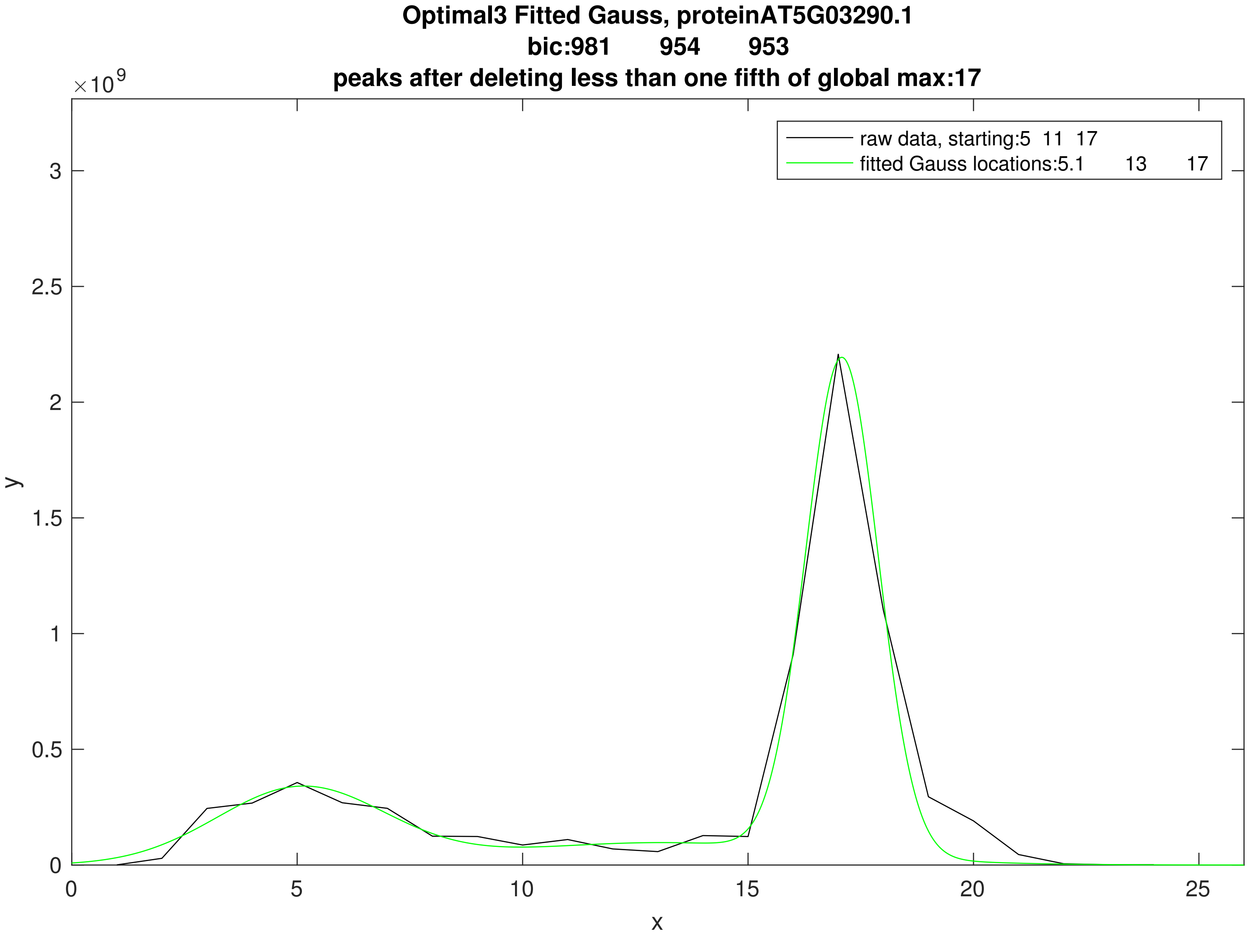

Optimal3 Fitted Gauss, proteinAT5G03290.1
bic:981 954 953
peaks after deleting less than one fifth of global max:17
×10^9
raw data, starting:5 11 17
fitted Gauss locations:5.1 13 17
y
x
0
0.5
1
1.5
2
2.5
3
0
5
10
15
20
25

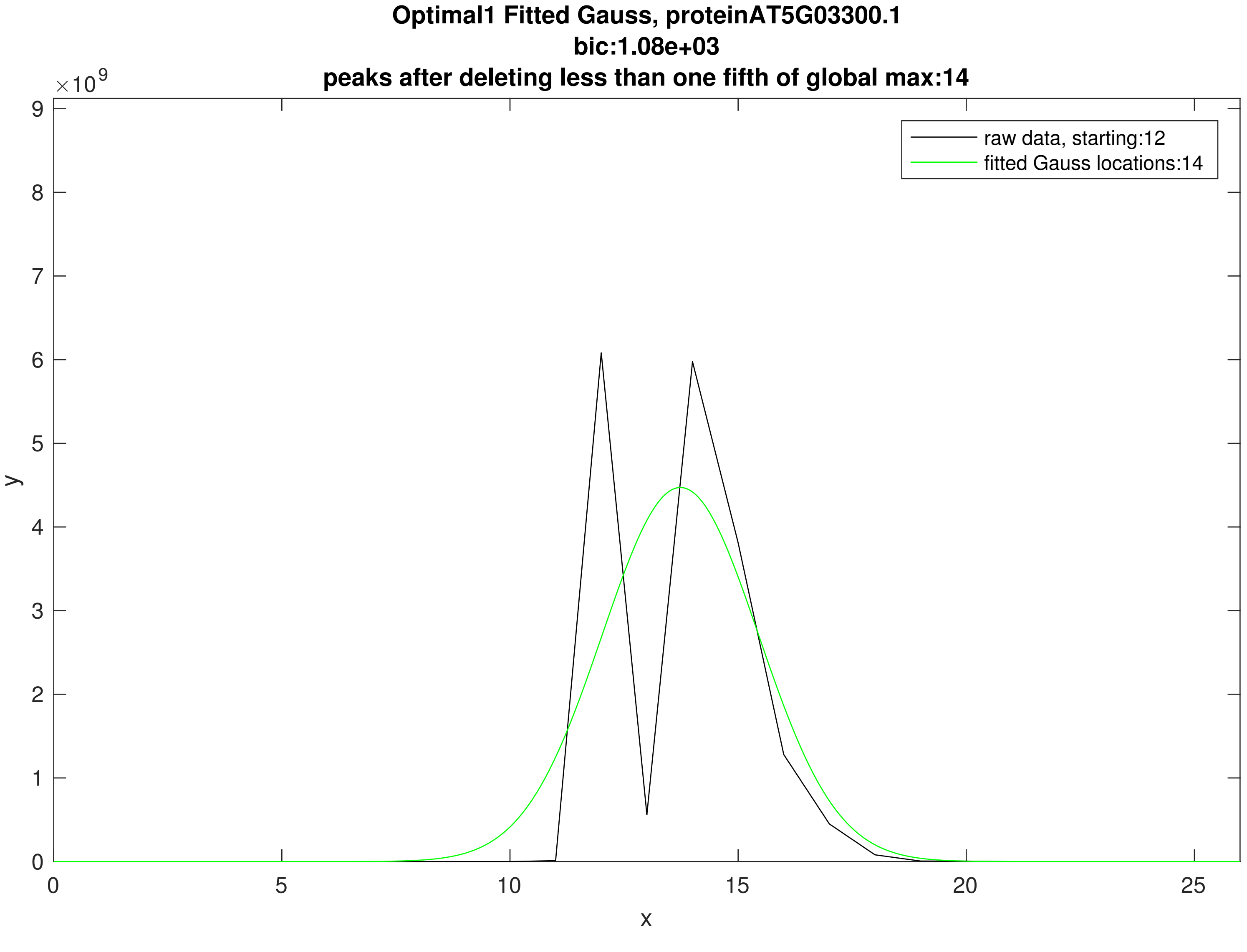
Optimal1 Fitted Gauss, proteinAT5G03300.1
bic:1.08e+03
peaks after deleting less than one fifth of global max:14
×10^9
raw data, starting:12
fitted Gauss locations:14
y
x
0
1
2
3
4
5
6
7
8
9
0
5
10
15
20
25

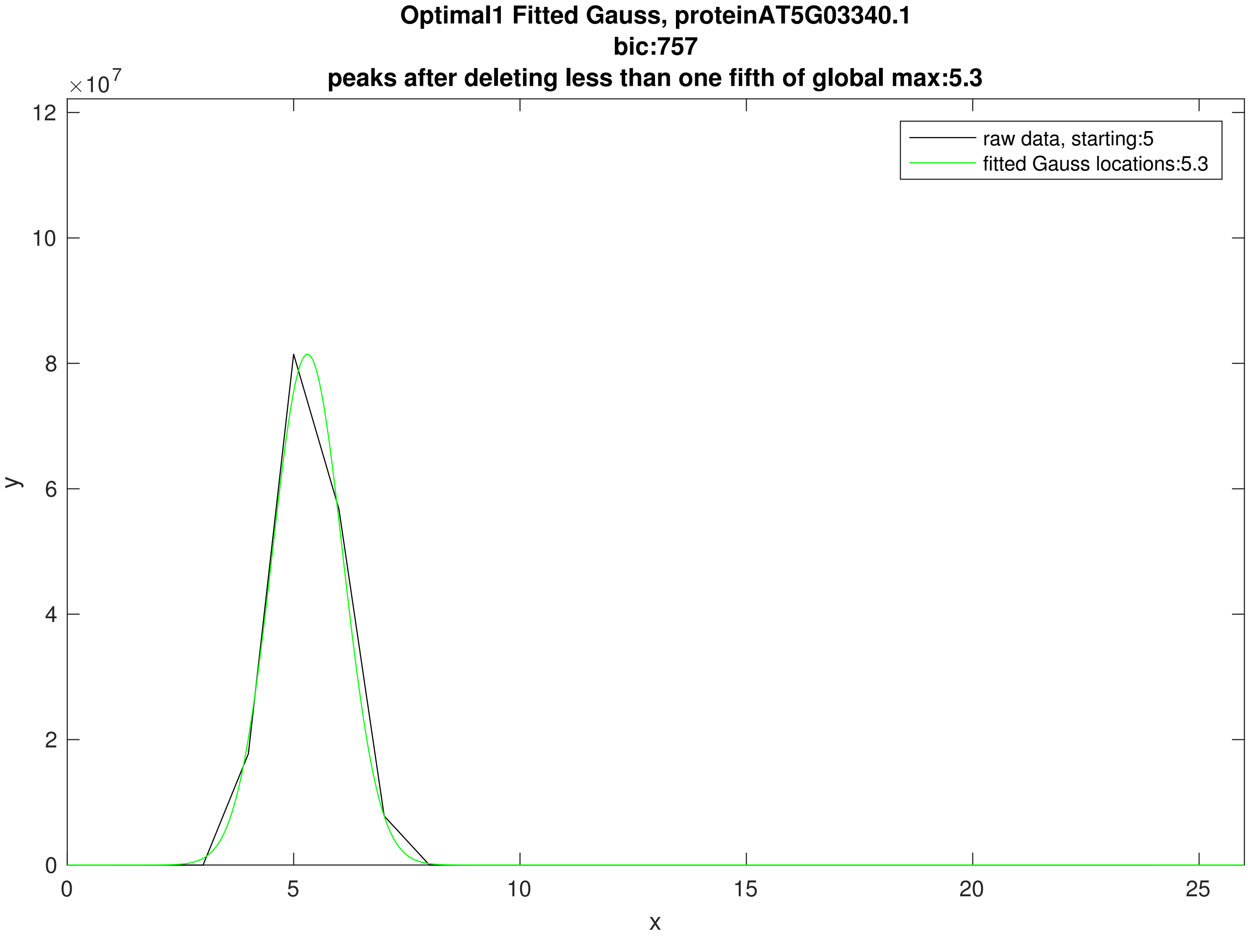

Optimal1 Fitted Gauss, proteinAT5G03340.1
bic:757
peaks after deleting less than one fifth of global max:5.3
×10^7
12
10
8
6
4
2
0
y
0
5
10
15
20
25
x
raw data, starting:5
fitted Gauss locations:5.3

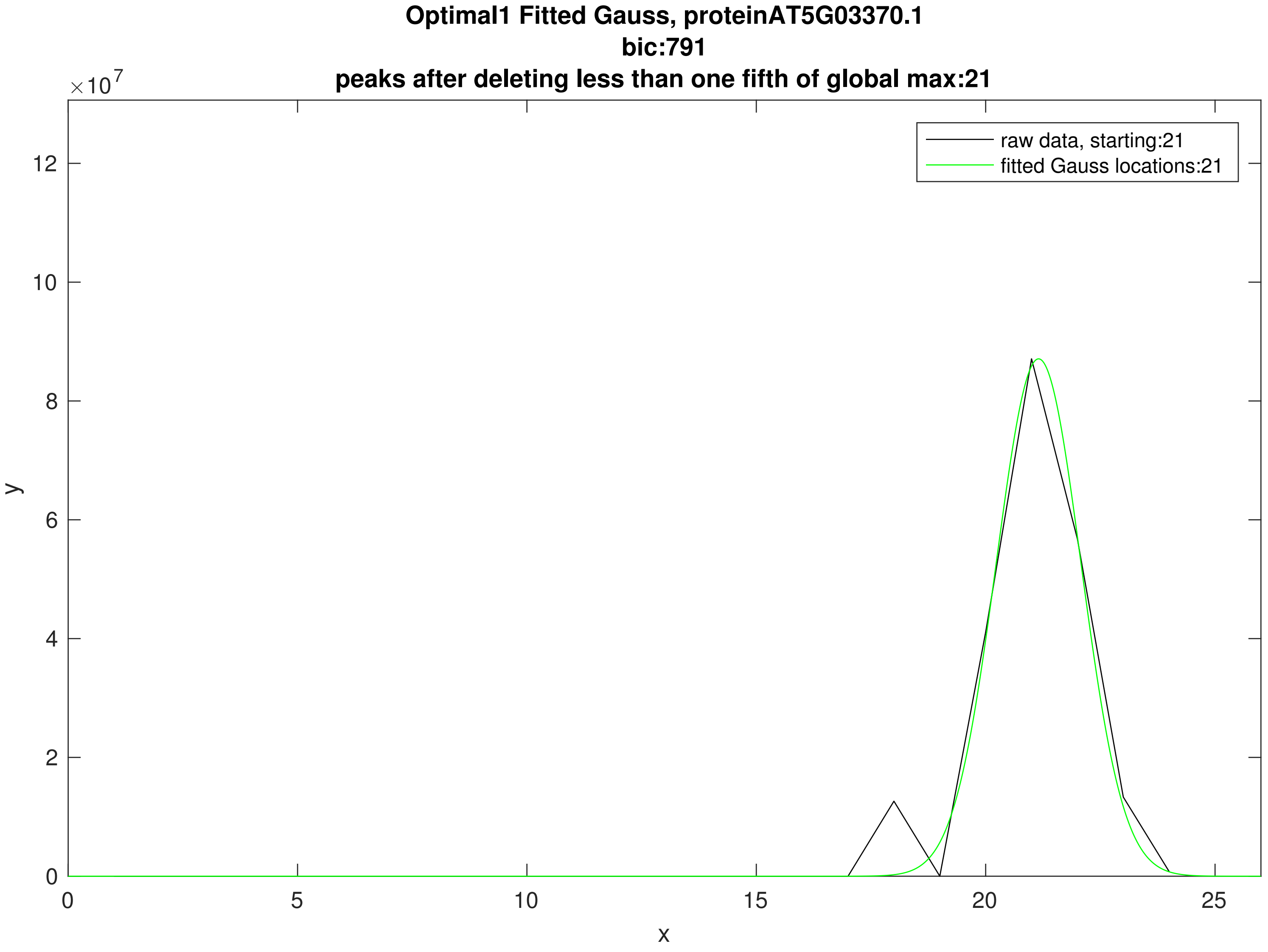

Optimal1 Fitted Gauss, proteinAT5G03370.1
bic:791
peaks after deleting less than one fifth of global max:21
×10^7
raw data, starting:21
fitted Gauss locations:21
12
10
8
6
4
2
0
y
0
5
10
15
20
25
x

Optimal1 Fitted Gauss, proteinAT5G03455.1
bic:811 820
peaks after deleting less than one fifth of global max:20

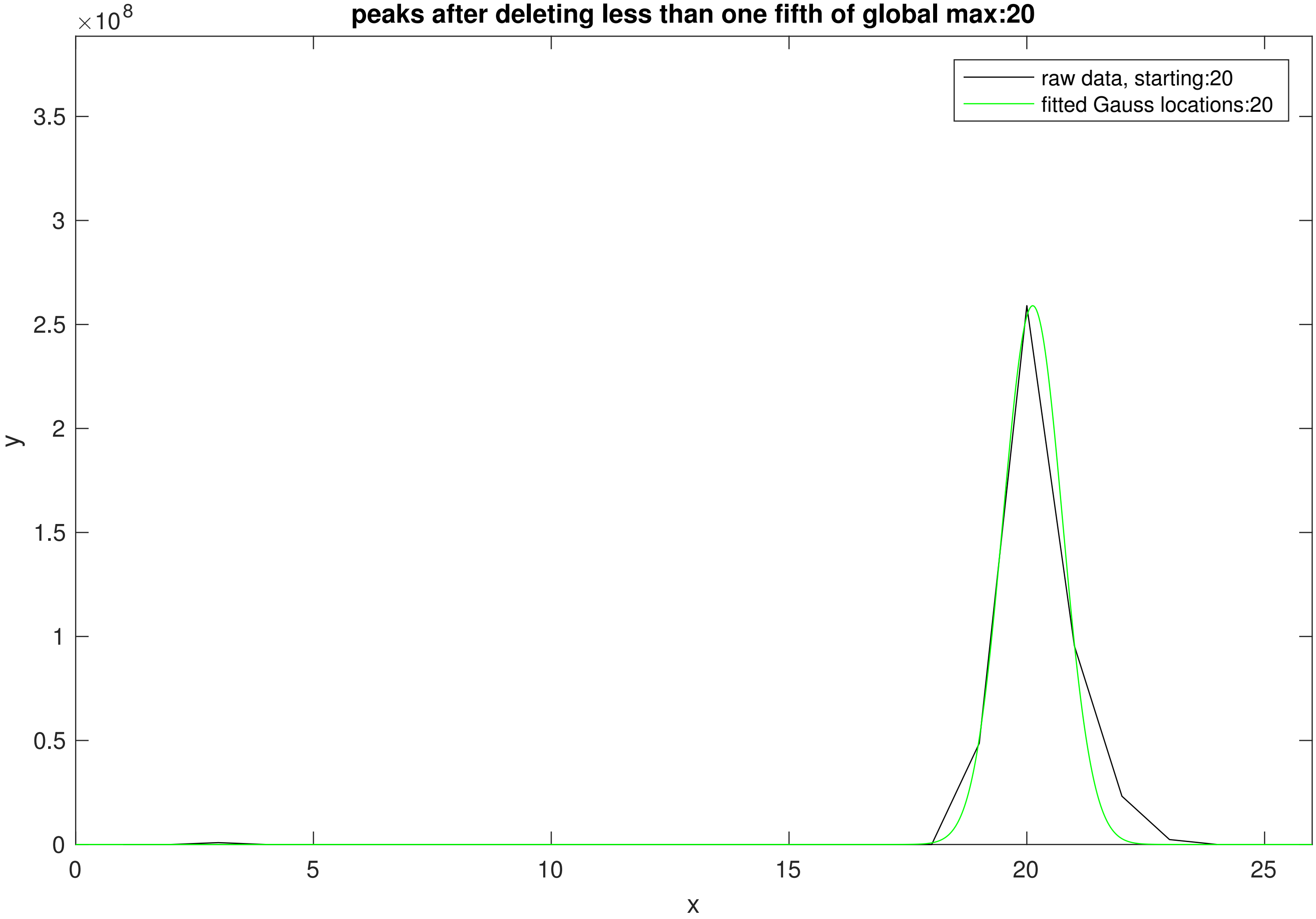

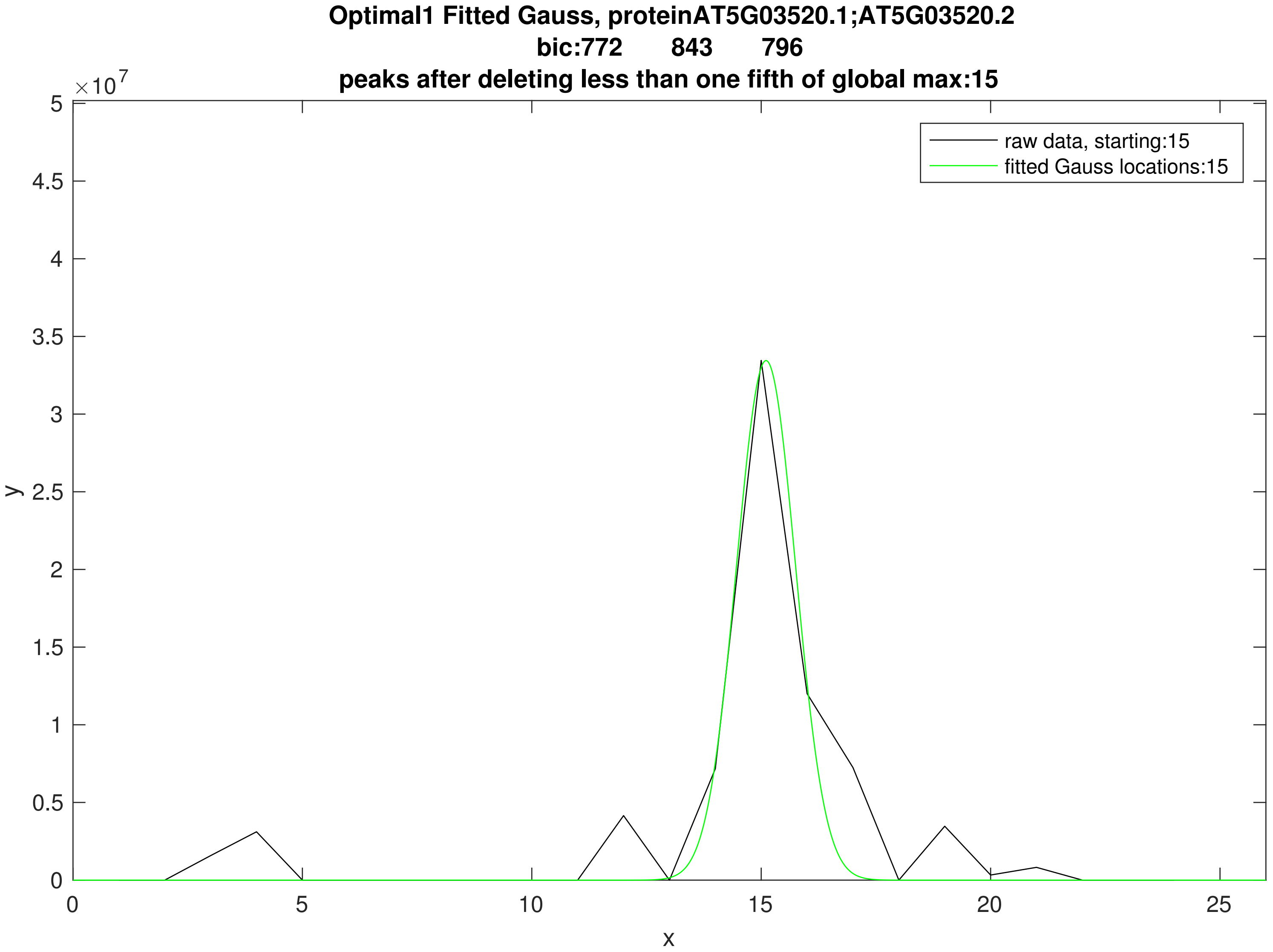
Optimal1 Fitted Gauss, proteinAT5G03520.1;AT5G03520.2
bic:772 843 796
peaks after deleting less than one fifth of global max:15
×10^7
raw data, starting:15
fitted Gauss locations:15
y
x
0
0.5
1
1.5
2
2.5
3
3.5
4
4.5
5
0
5
10
15
20
25

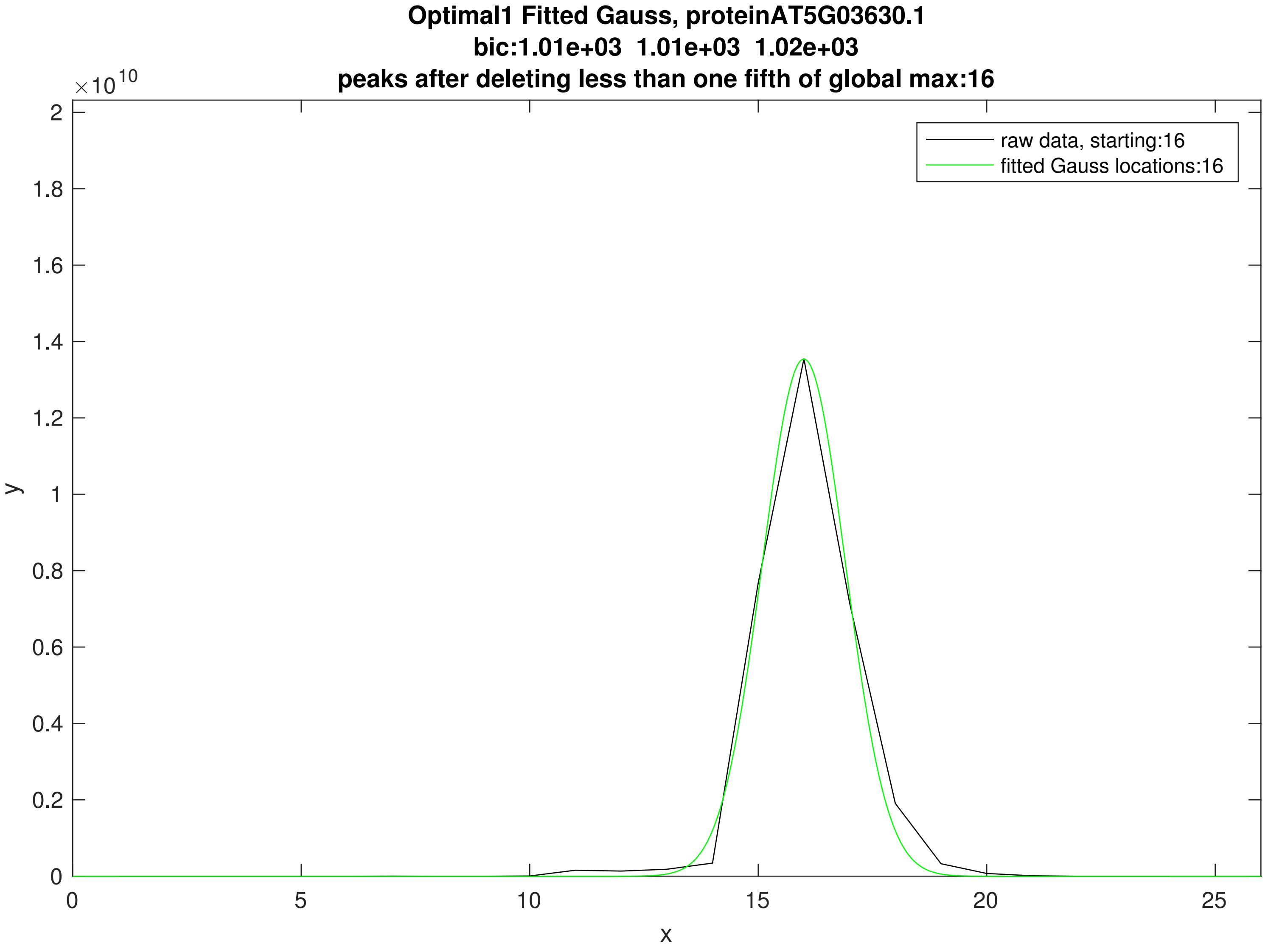
Optimal1 Fitted Gauss, proteinAT5G03630.1
bic:1.01e+03 1.01e+03 1.02e+03
peaks after deleting less than one fifth of global max:16
×10^10
raw data, starting:16
fitted Gauss locations:16
y
x
0
0.2
0.4
0.6
0.8
1
1.2
1.4
1.6
1.8
2
0
5
10
15
20
25

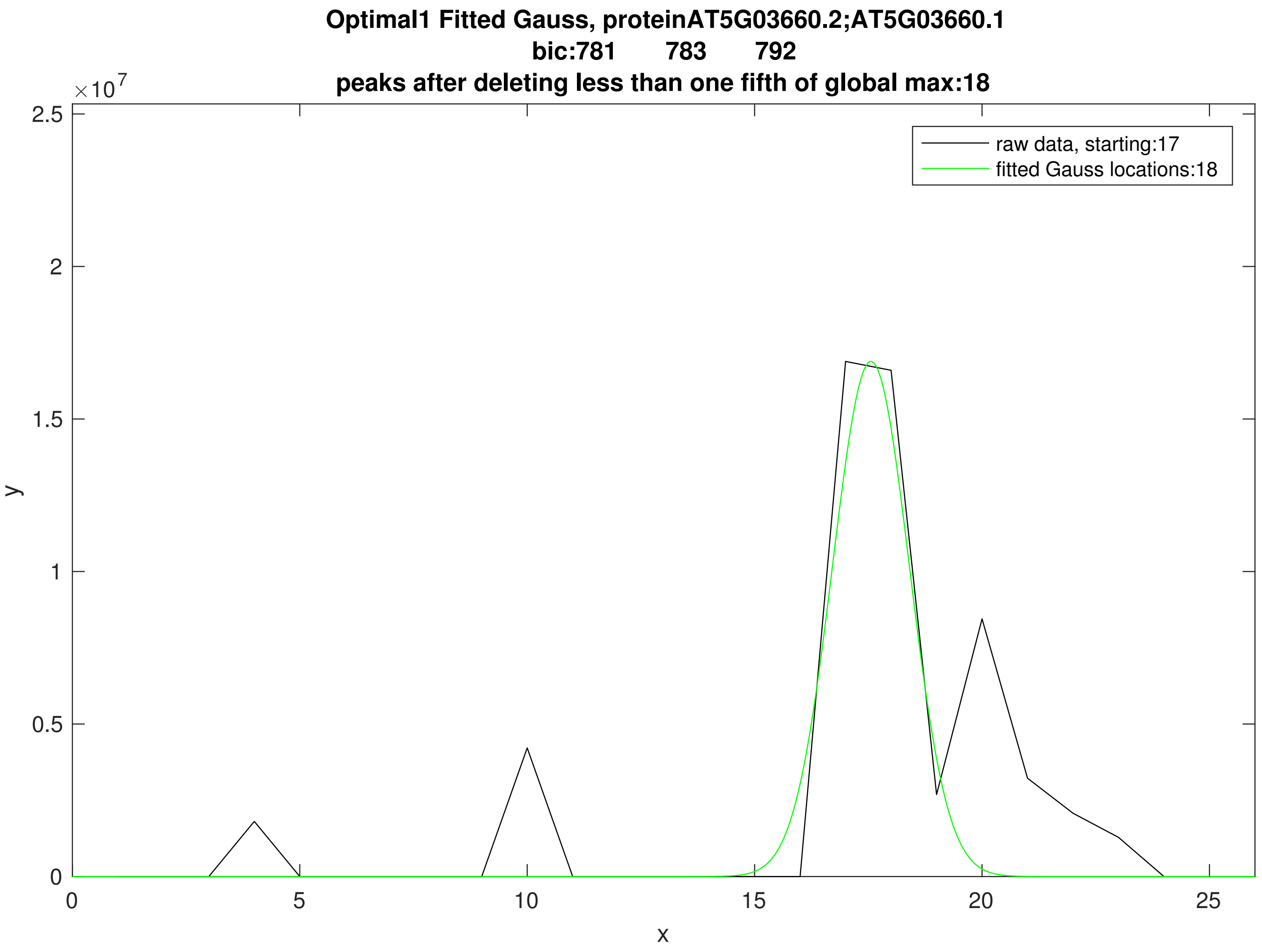

Optimal1 Fitted Gauss, proteinAT5G03660.2;AT5G03660.1
bic:781 783 792
peaks after deleting less than one fifth of global max:18
×10^7
2.5
2
1.5
1
0.5
0
y
0
5
10
15
20
25
x
raw data, starting:17
fitted Gauss locations:18

Optimal3 Fitted Gauss, proteinAT5G03690.2;AT5G03690.1
bic:1e+03 978 978
peaks after deleting less than one fifth of global max:5.2 9.9

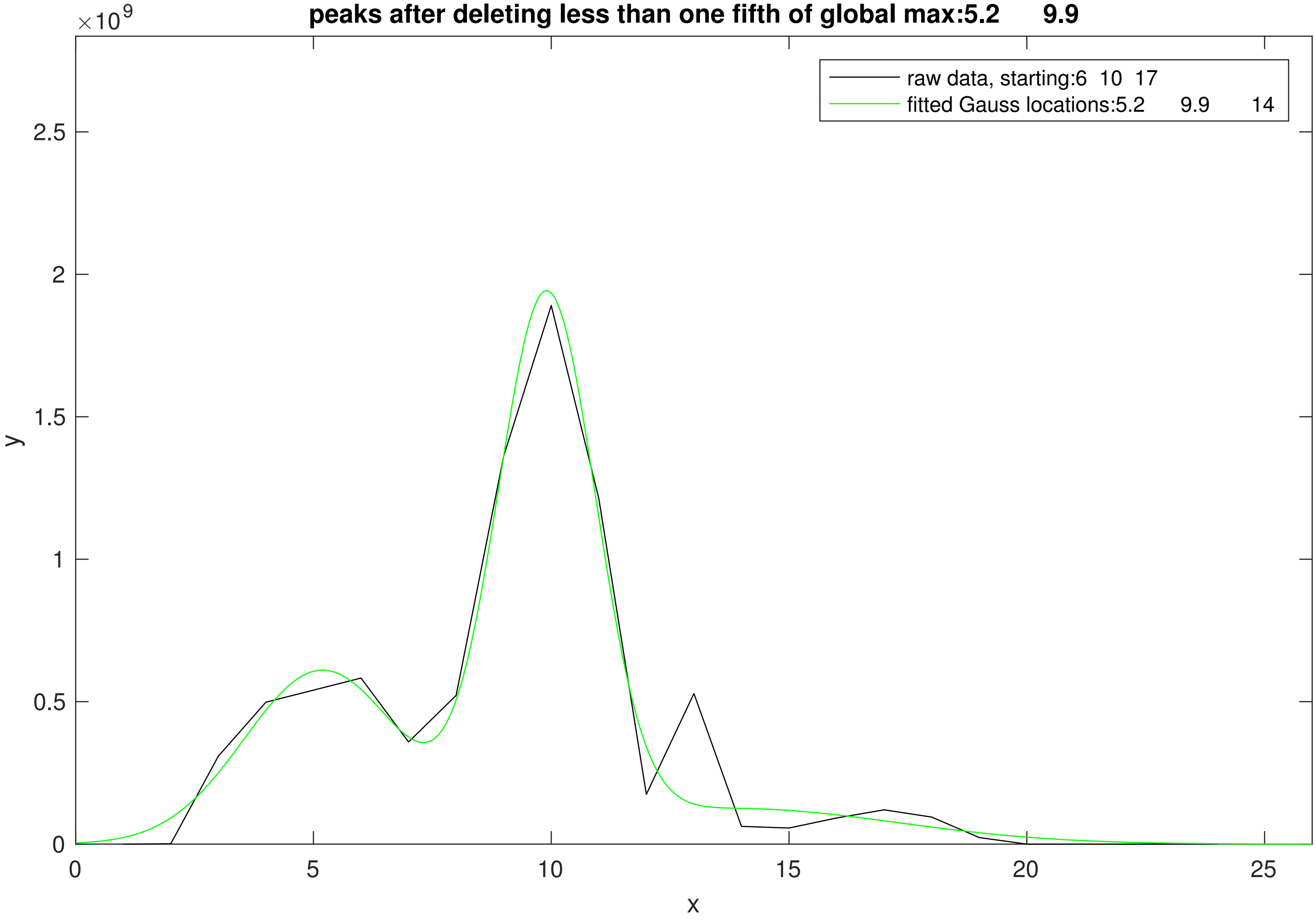

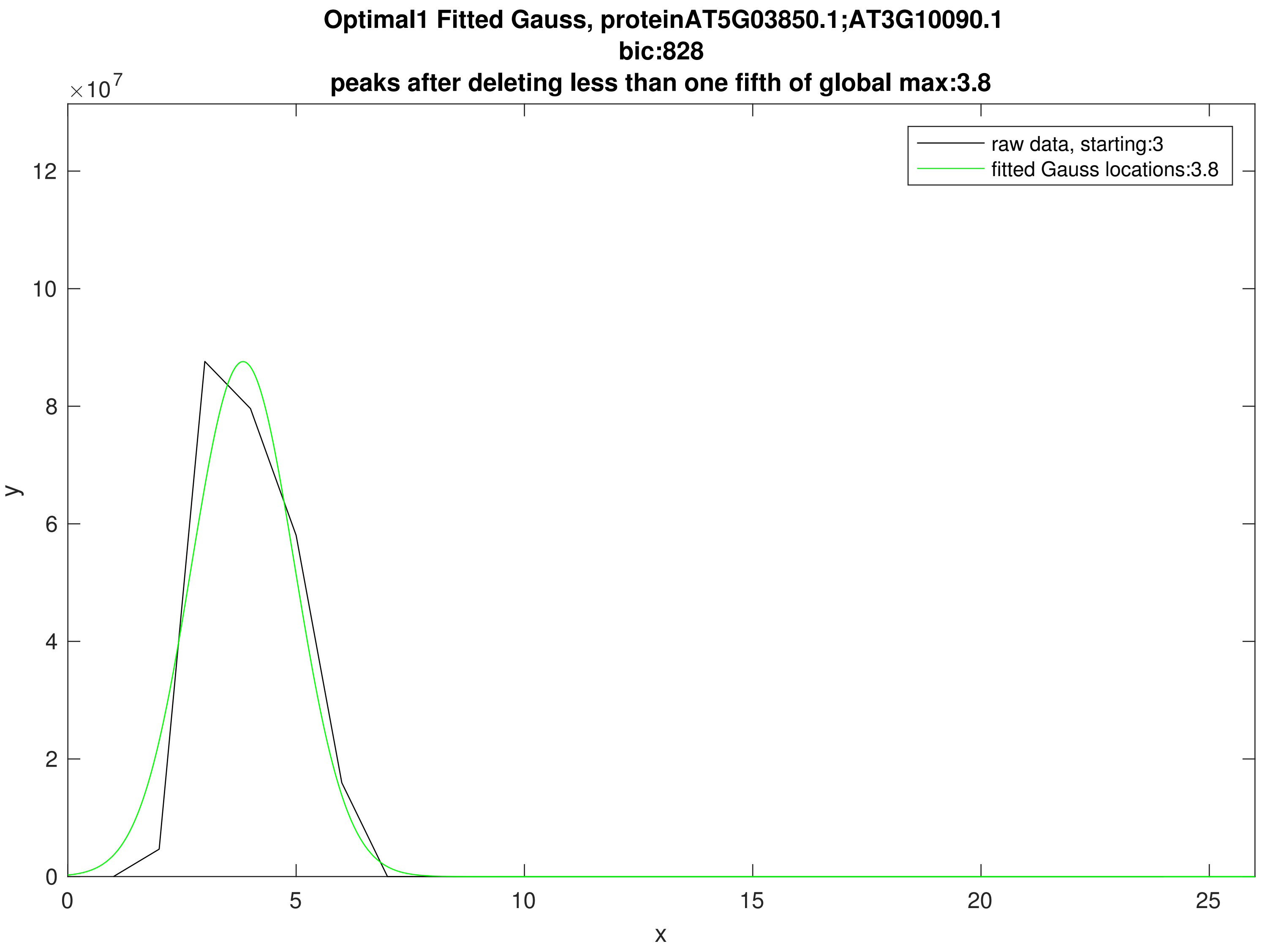

Optimal1 Fitted Gauss, proteinAT5G03850.1;AT3G10090.1
bic:828
peaks after deleting less than one fifth of global max:3.8
×10^7
raw data, starting:3
fitted Gauss locations:3.8
y
x
0
2
4
6
8
10
12
0
5
10
15
20
25

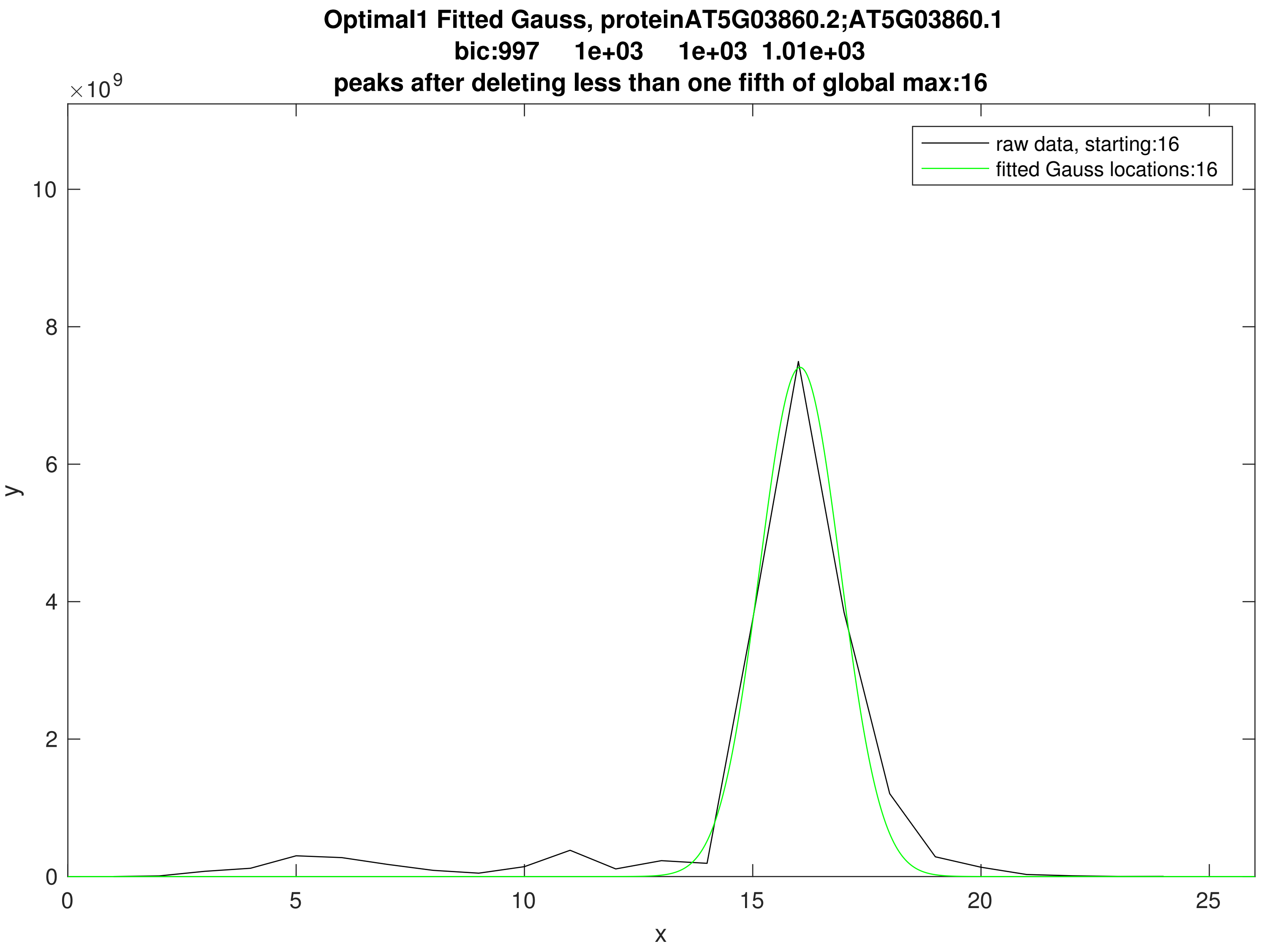
Optimal1 Fitted Gauss, proteinAT5G03860.2;AT5G03860.1
bic:997 1e+03 1e+03 1.01e+03
peaks after deleting less than one fifth of global max:16
$\times 10^9$
raw data, starting:16
fitted Gauss locations:16
y
x
0
2
4
6
8
10
0
5
10
15
20
25

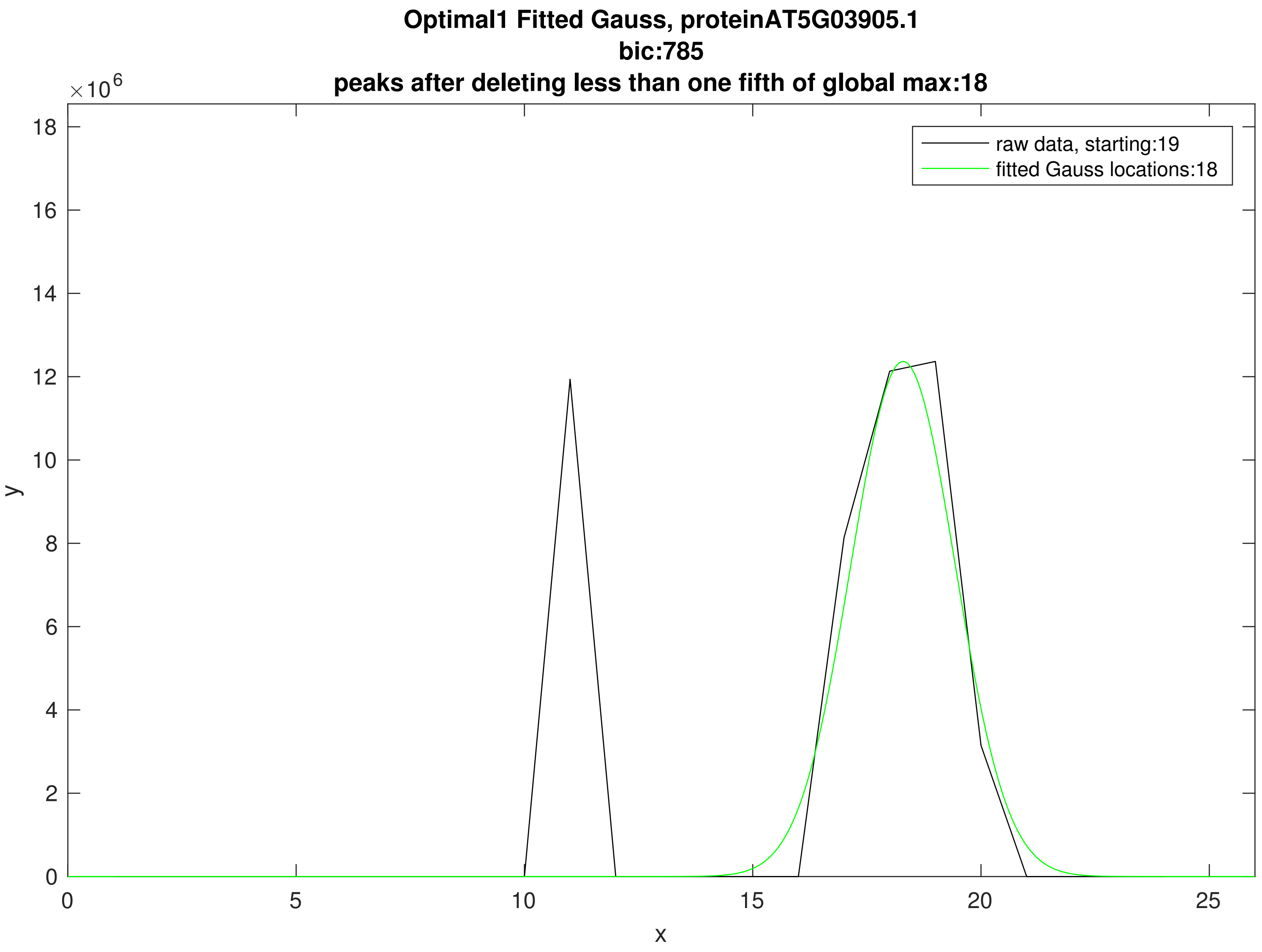

Optimal1 Fitted Gauss, proteinAT5G03905.1
bic:785
peaks after deleting less than one fifth of global max:18
×10^6
raw data, starting:19
fitted Gauss locations:18
y
x
0
2
4
6
8
10
12
14
16
18
0
5
10
15
20
25

Optimal1 Fitted Gauss, proteinAT5G04000.1;AT5G04000.2
bic:707
peaks after deleting less than one fifth of global max:16

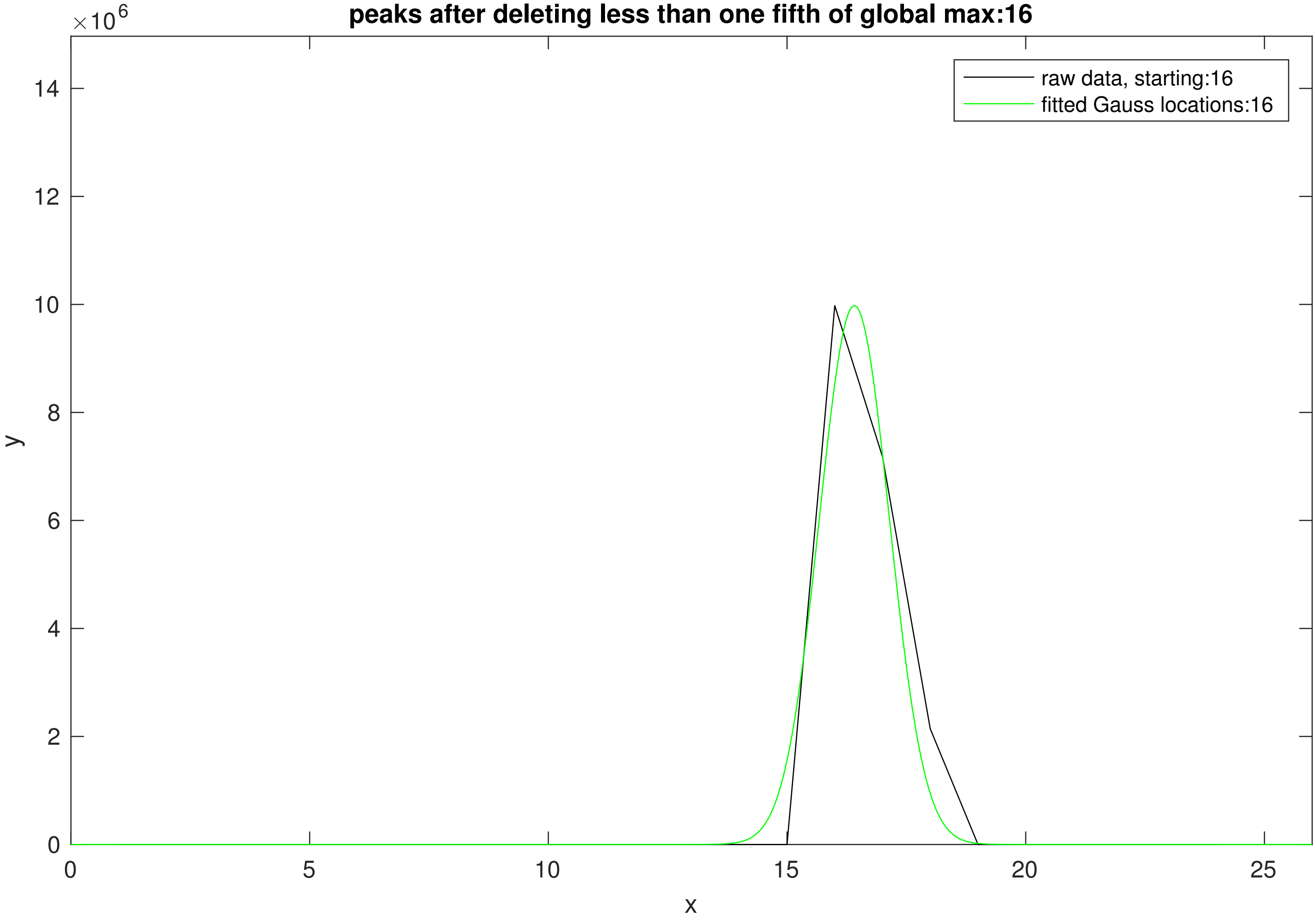

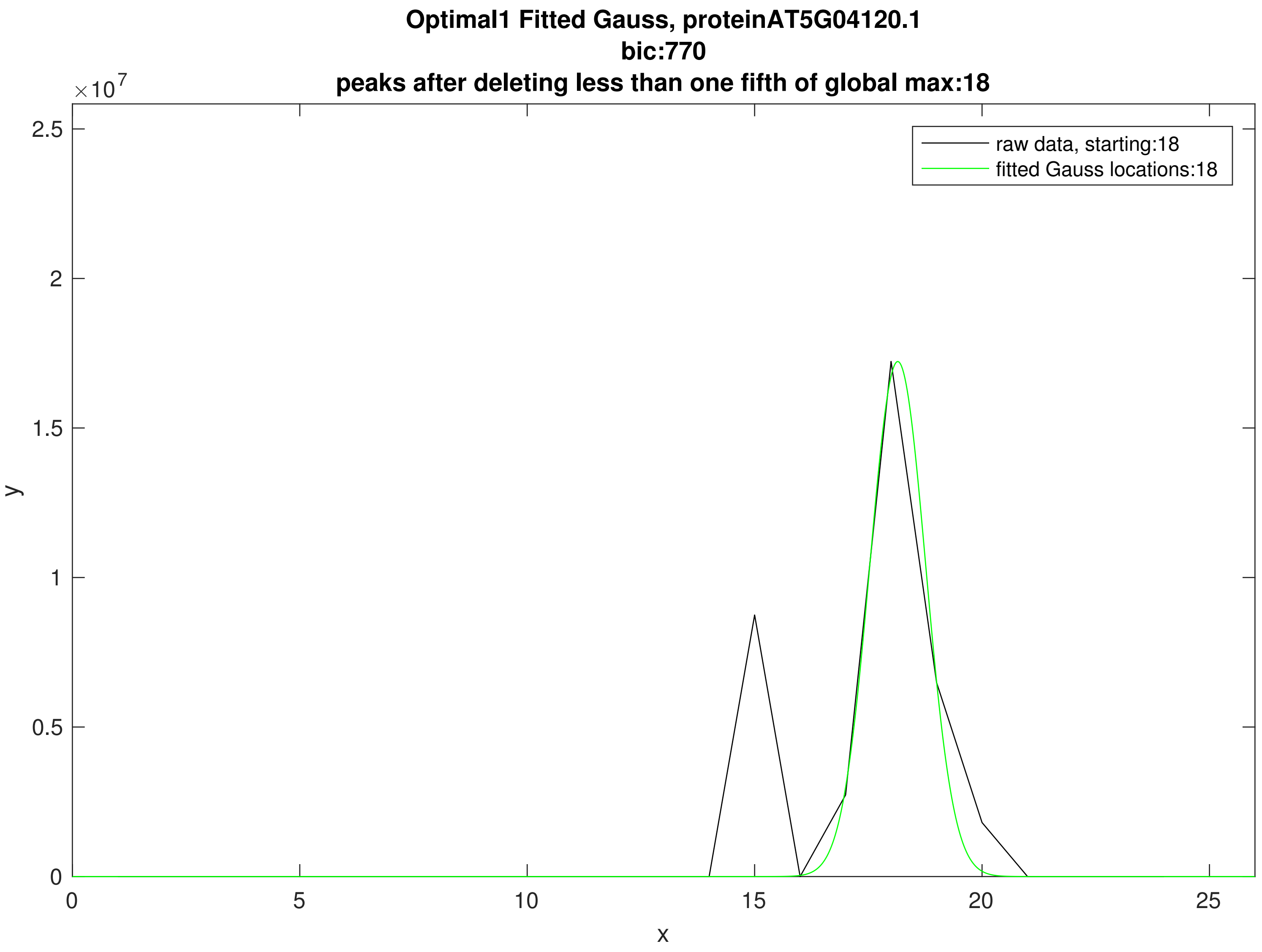

Optimal1 Fitted Gauss, proteinAT5G04120.1
bic:770
peaks after deleting less than one fifth of global max:18
×10^7
2.5
2
1.5
1
0.5
0
y
0
5
10
15
20
25
x
raw data, starting:18
fitted Gauss locations:18

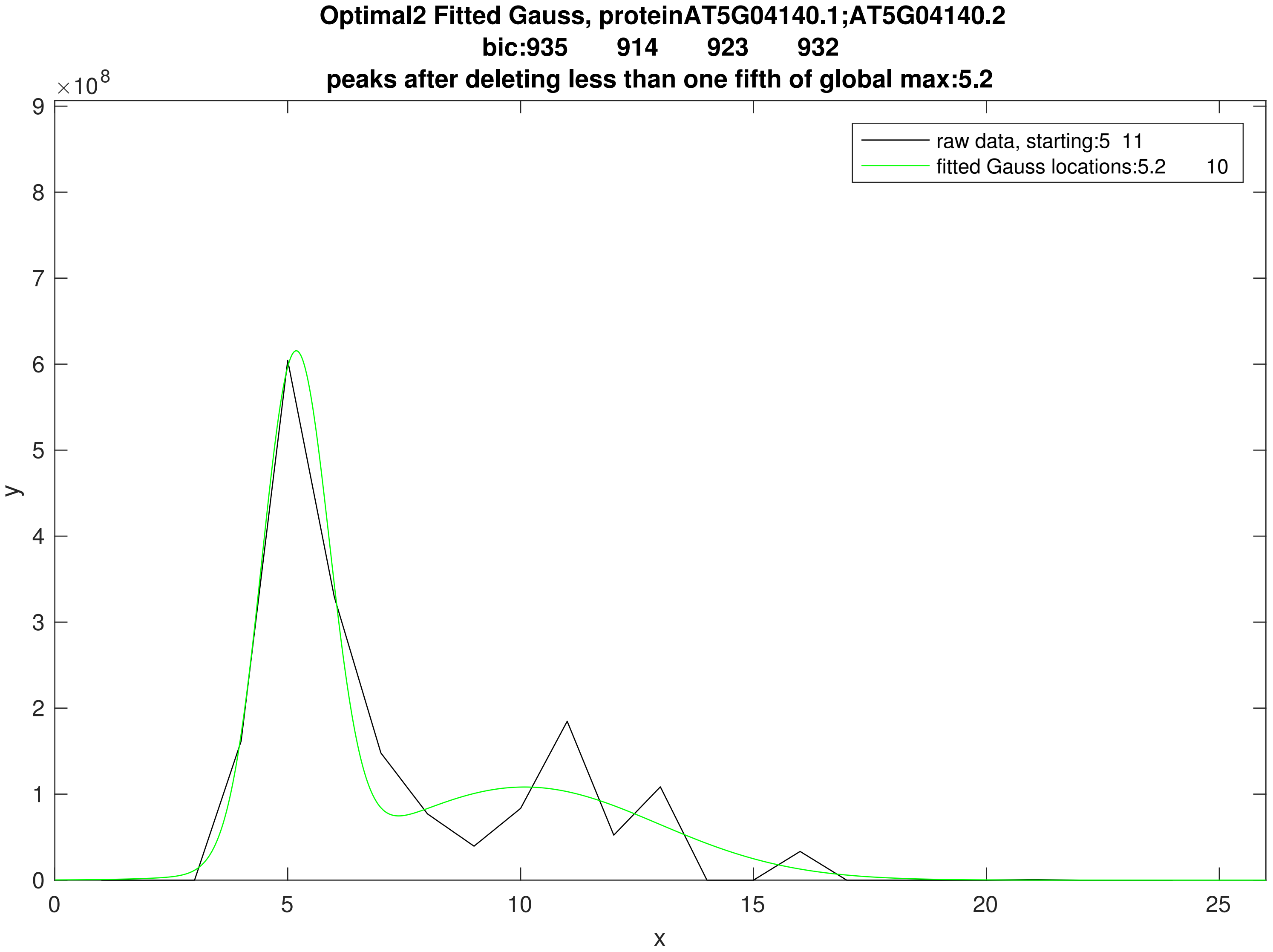
Optimal2 Fitted Gauss, proteinAT5G04140.1;AT5G04140.2
bic:935 914 923 932
peaks after deleting less than one fifth of global max:5.2
×10^8
raw data, starting:5 11
fitted Gauss locations:5.2 10
y
x
0
1
2
3
4
5
6
7
8
9
0
5
10
15
20
25

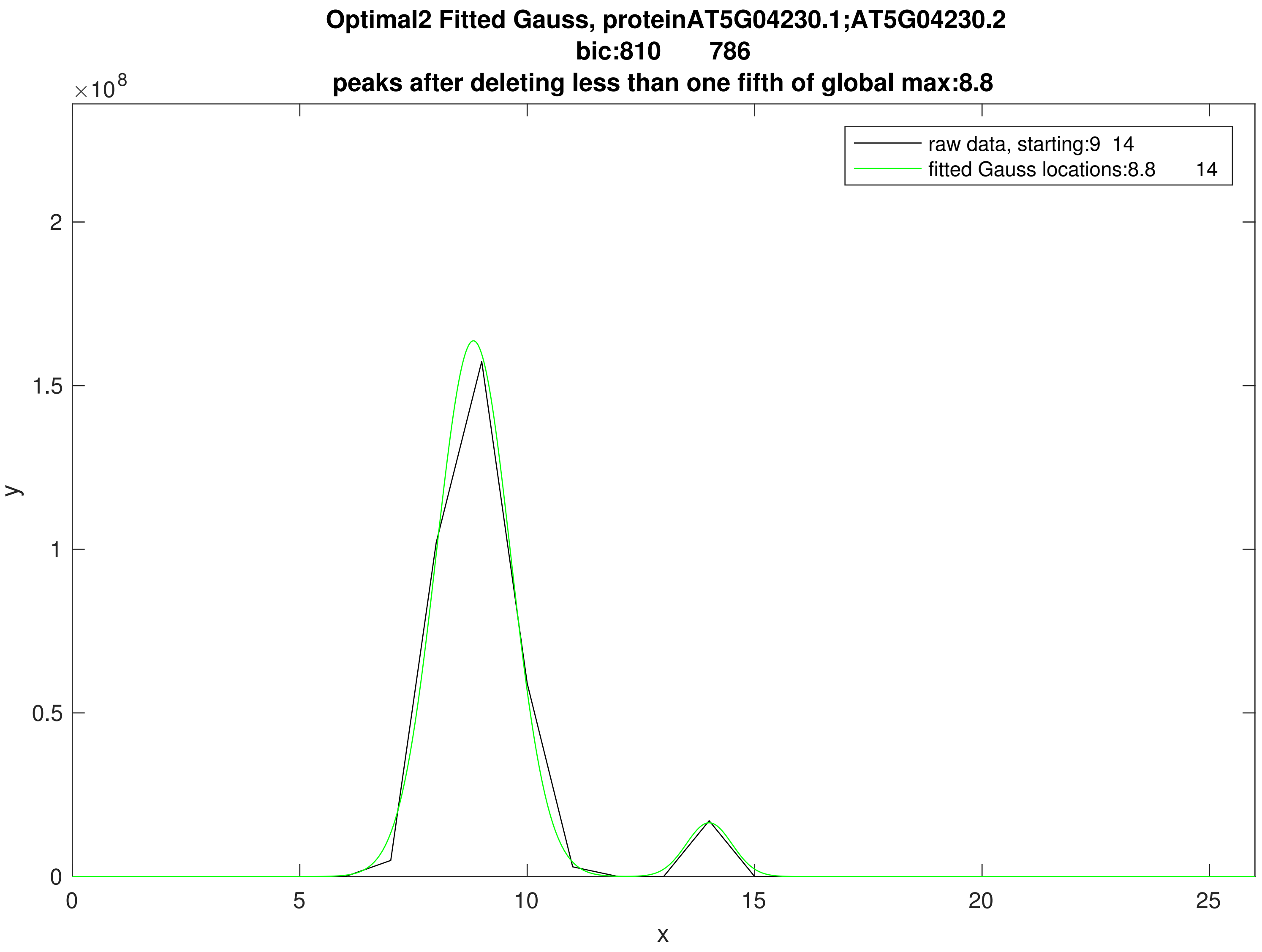
Optimal2 Fitted Gauss, proteinAT5G04230.1;AT5G04230.2
bic:810 786
peaks after deleting less than one fifth of global max:8.8
×10^8
raw data, starting:9 14
fitted Gauss locations:8.8 14
y
x
0
0.5
1
1.5
2
0
5
10
15
20
25

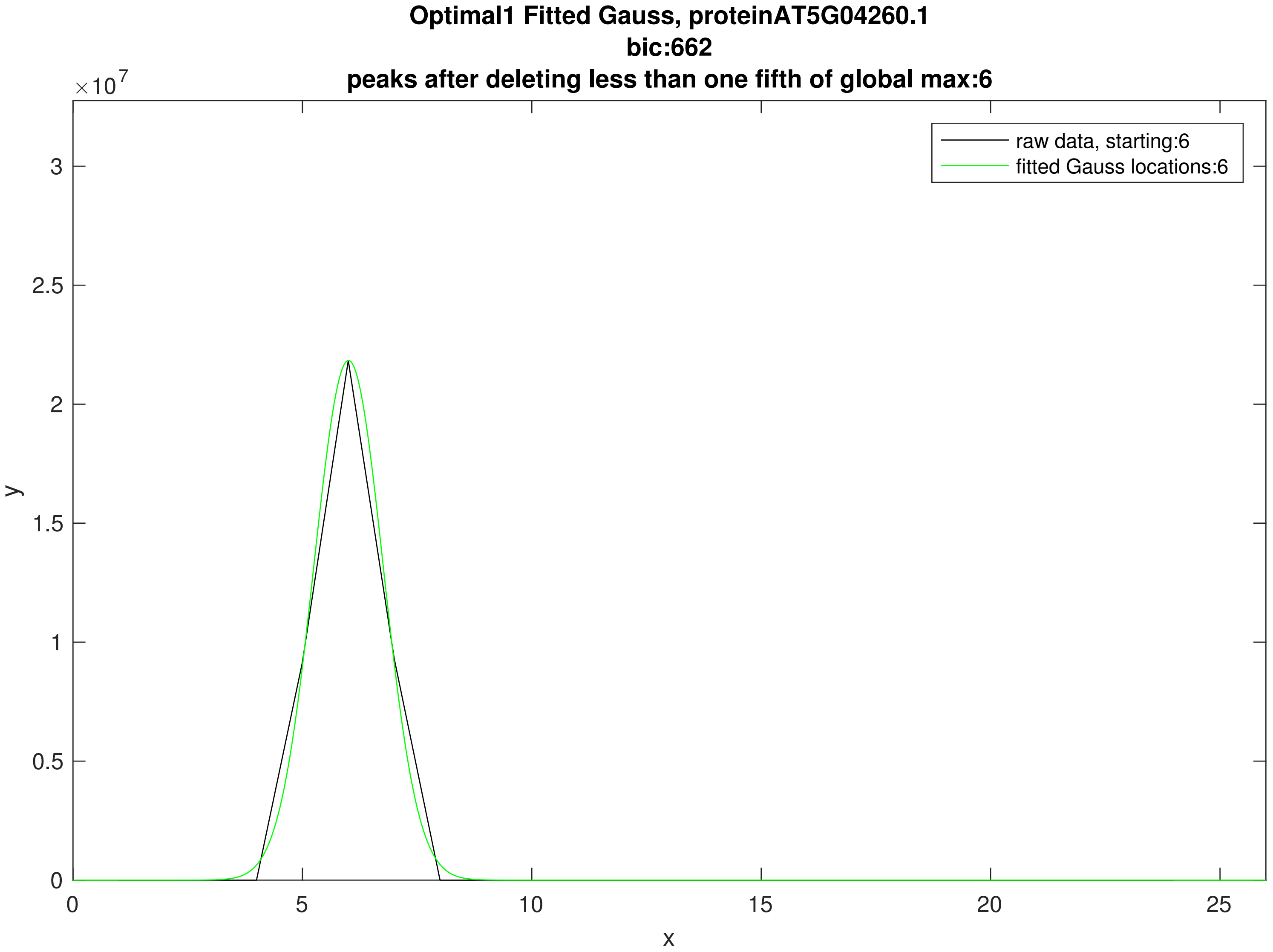

Optimal1 Fitted Gauss, proteinAT5G04260.1
bic:662
peaks after deleting less than one fifth of global max:6
×10^7
raw data, starting:6
fitted Gauss locations:6
3
2.5
2
1.5
1
0.5
0
y
0
5
10
15
20
25
x

**Optimal1 Fitted Gauss, proteinAT5G04420.3;AT5G04420.2;AT5G04420.1**
**bic:944 957 965 959**
**peaks after deleting less than one fifth of global max:13**

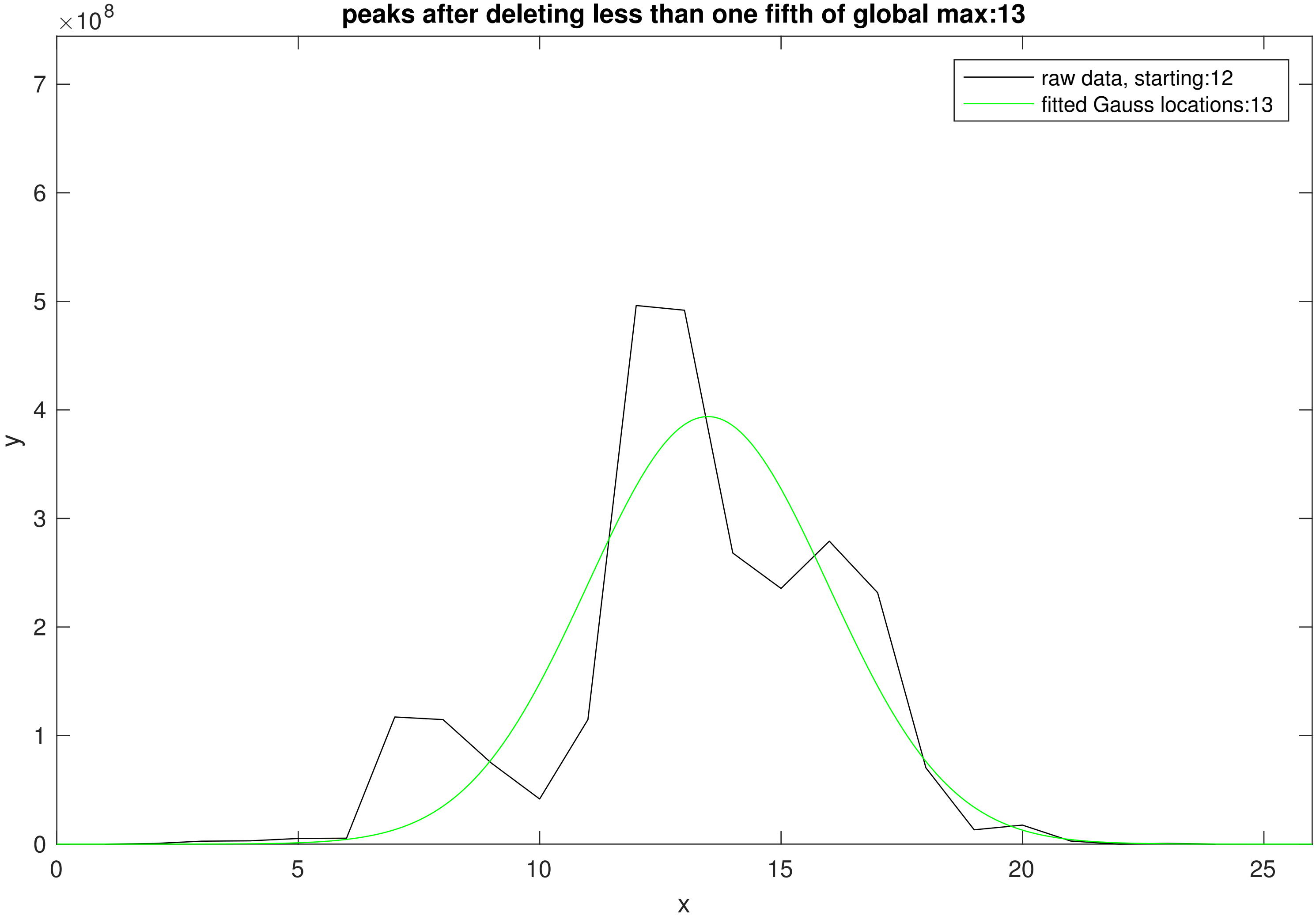

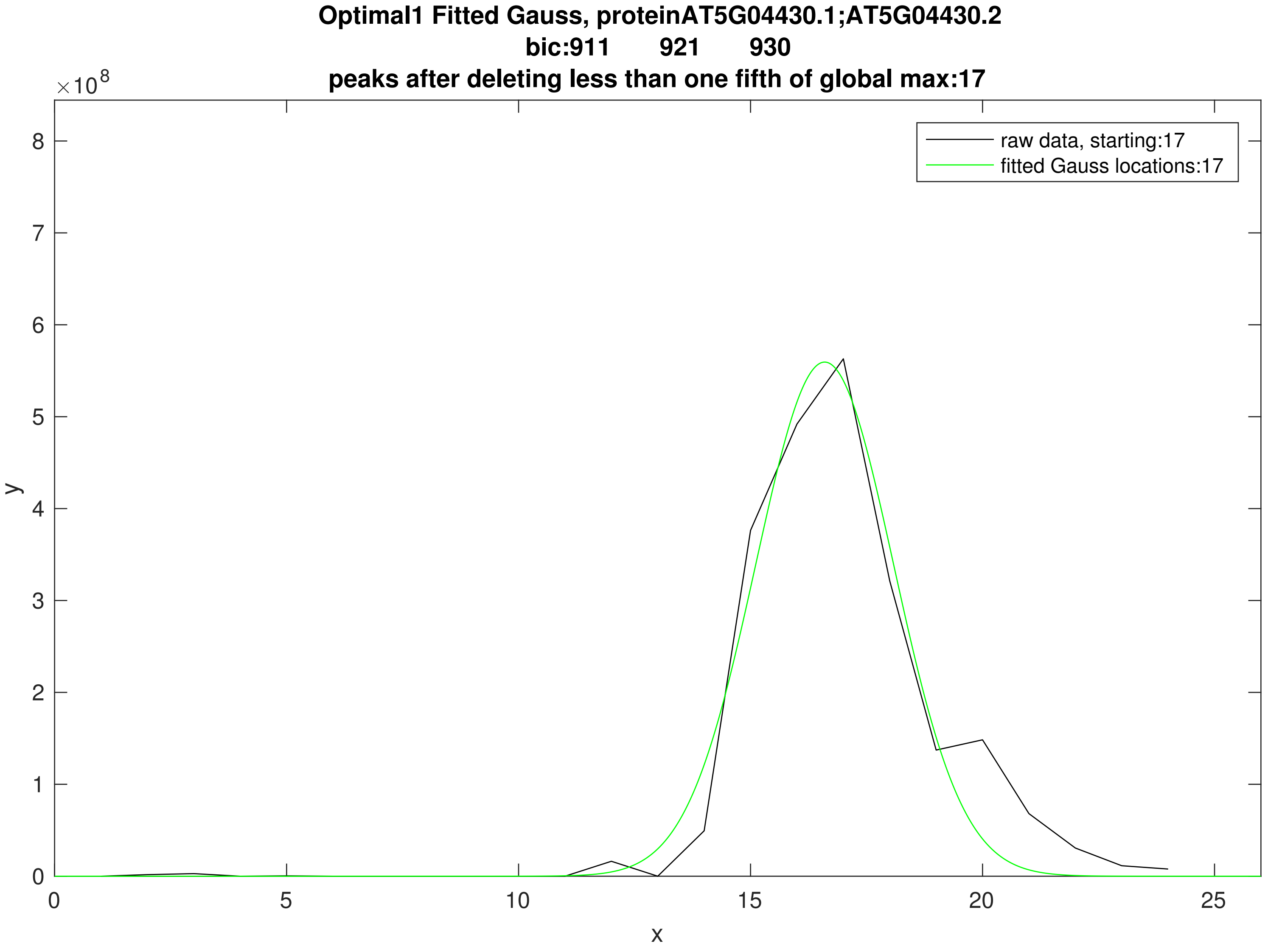
Optimal1 Fitted Gauss, proteinAT5G04430.1;AT5G04430.2
bic:911 921 930
peaks after deleting less than one fifth of global max:17
×10^8
raw data, starting:17
fitted Gauss locations:17
y
x
0
1
2
3
4
5
6
7
8
0
5
10
15
20
25

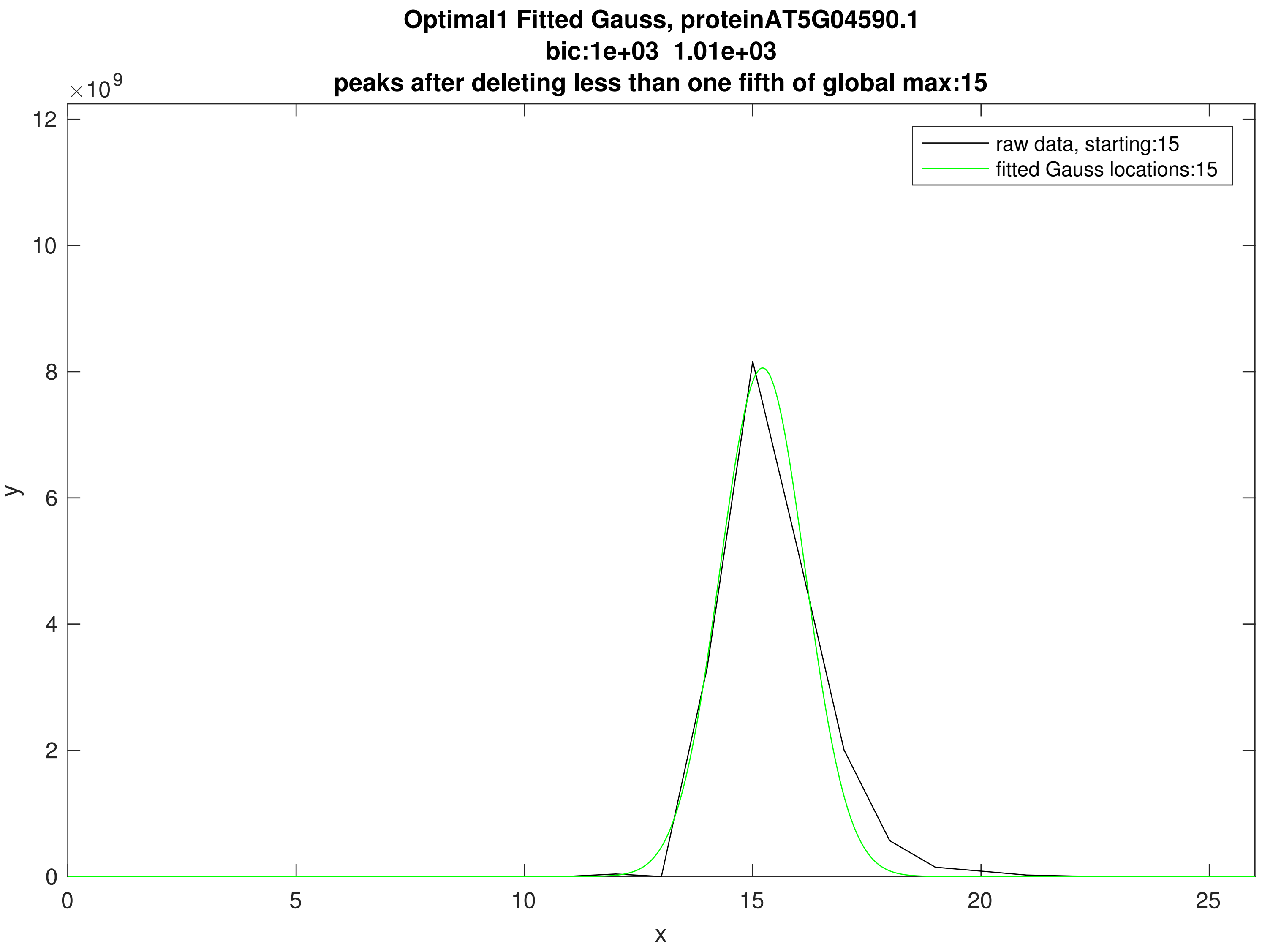
Optimal1 Fitted Gauss, proteinAT5G04590.1
bic:1e+03 1.01e+03
peaks after deleting less than one fifth of global max:15
×10^9
12
10
8
6
4
2
0
y
0
5
10
15
20
25
x
raw data, starting:15
fitted Gauss locations:15

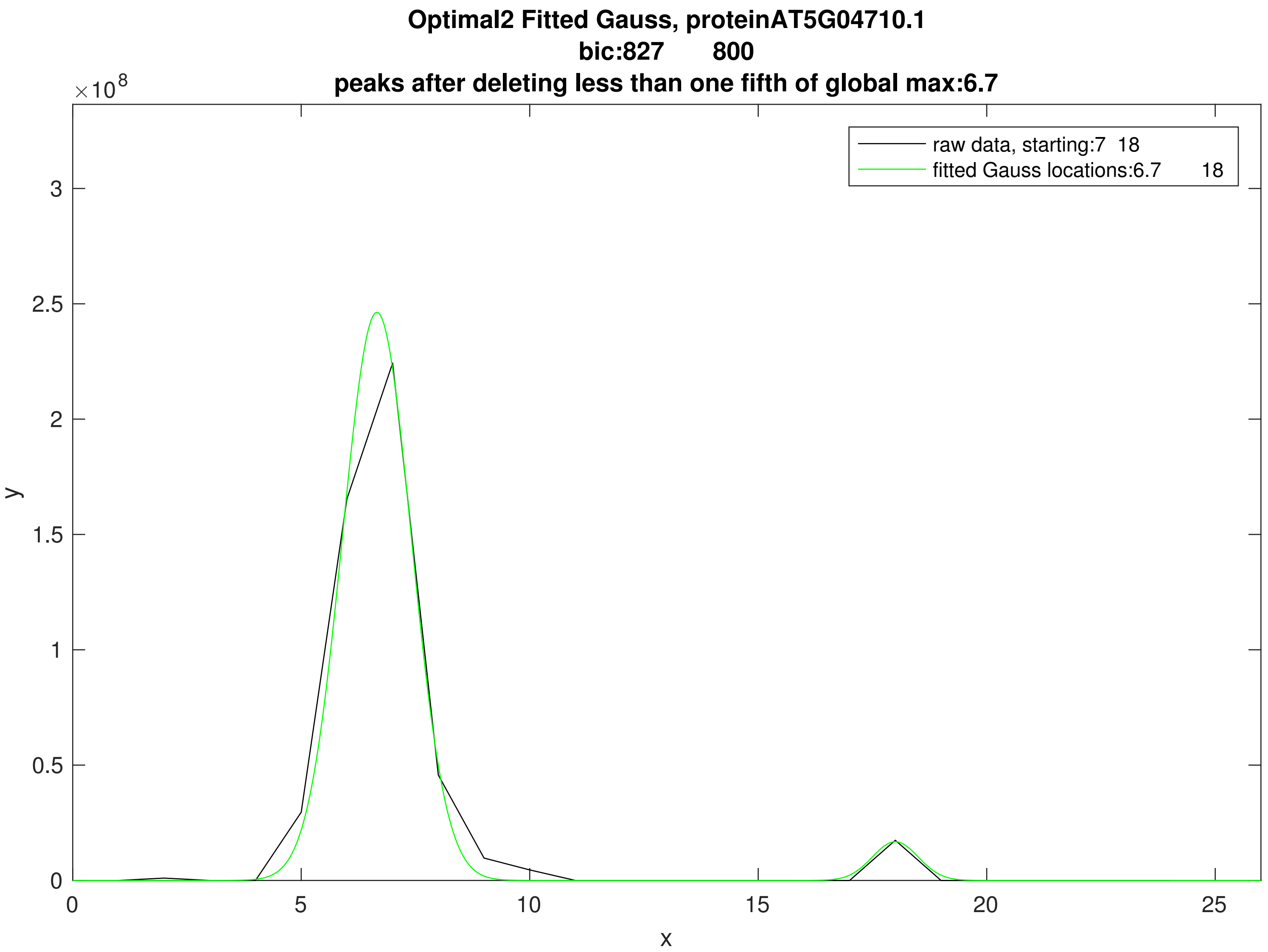
Optimal2 Fitted Gauss, proteinAT5G04710.1
bic:827 800
peaks after deleting less than one fifth of global max:6.7
×10^8
raw data, starting:7 18
fitted Gauss locations:6.7 18
y
x
0
0.5
1
1.5
2
2.5
3
0
5
10
15
20
25

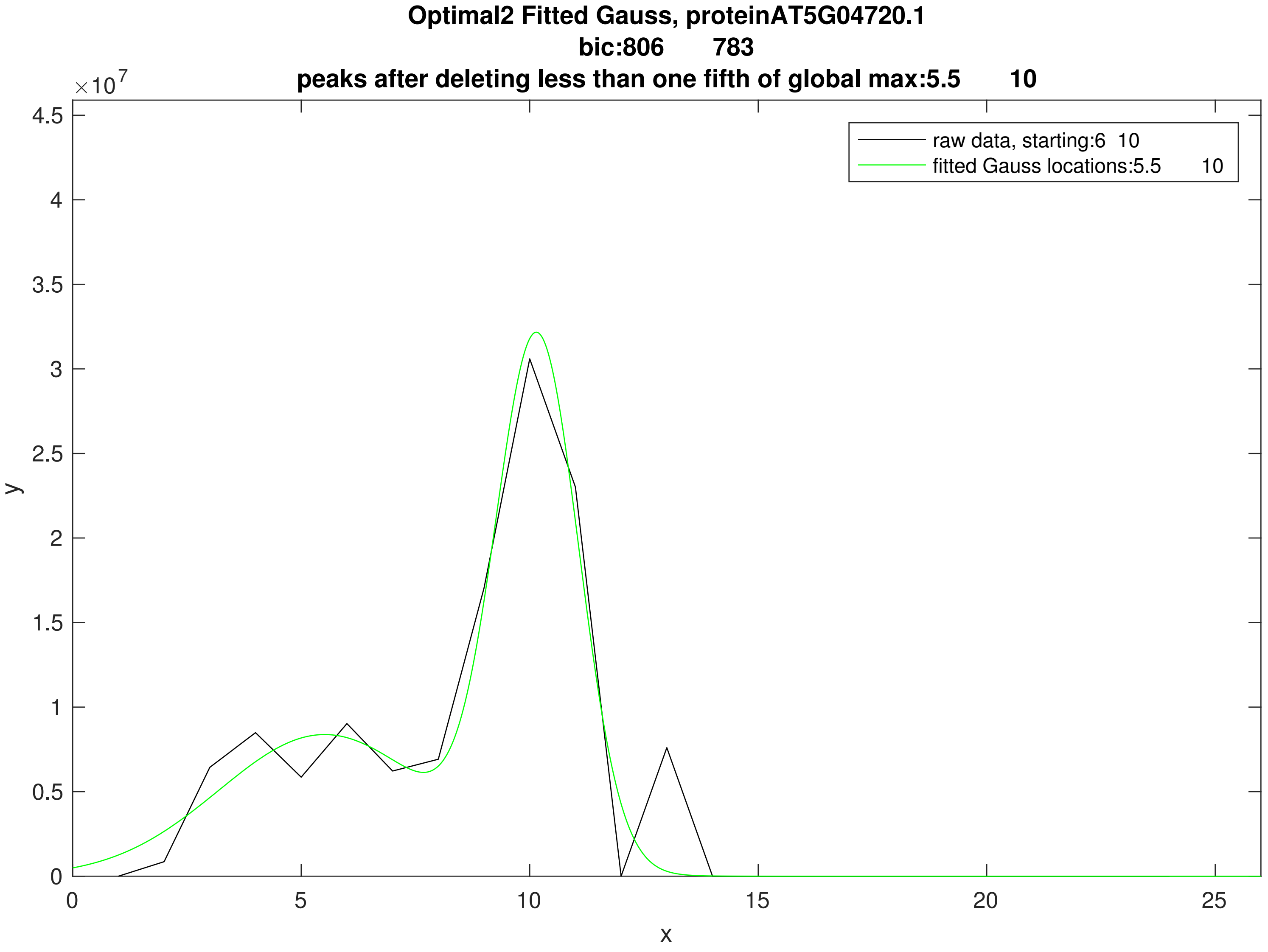
Optimal2 Fitted Gauss, proteinAT5G04720.1
bic:806 783
peaks after deleting less than one fifth of global max:5.5 10
×10^7
4.5
4
3.5
3
2.5
2
1.5
1
0.5
0
y
0
5
10
15
20
25
x
raw data, starting:6 10
fitted Gauss locations:5.5 10

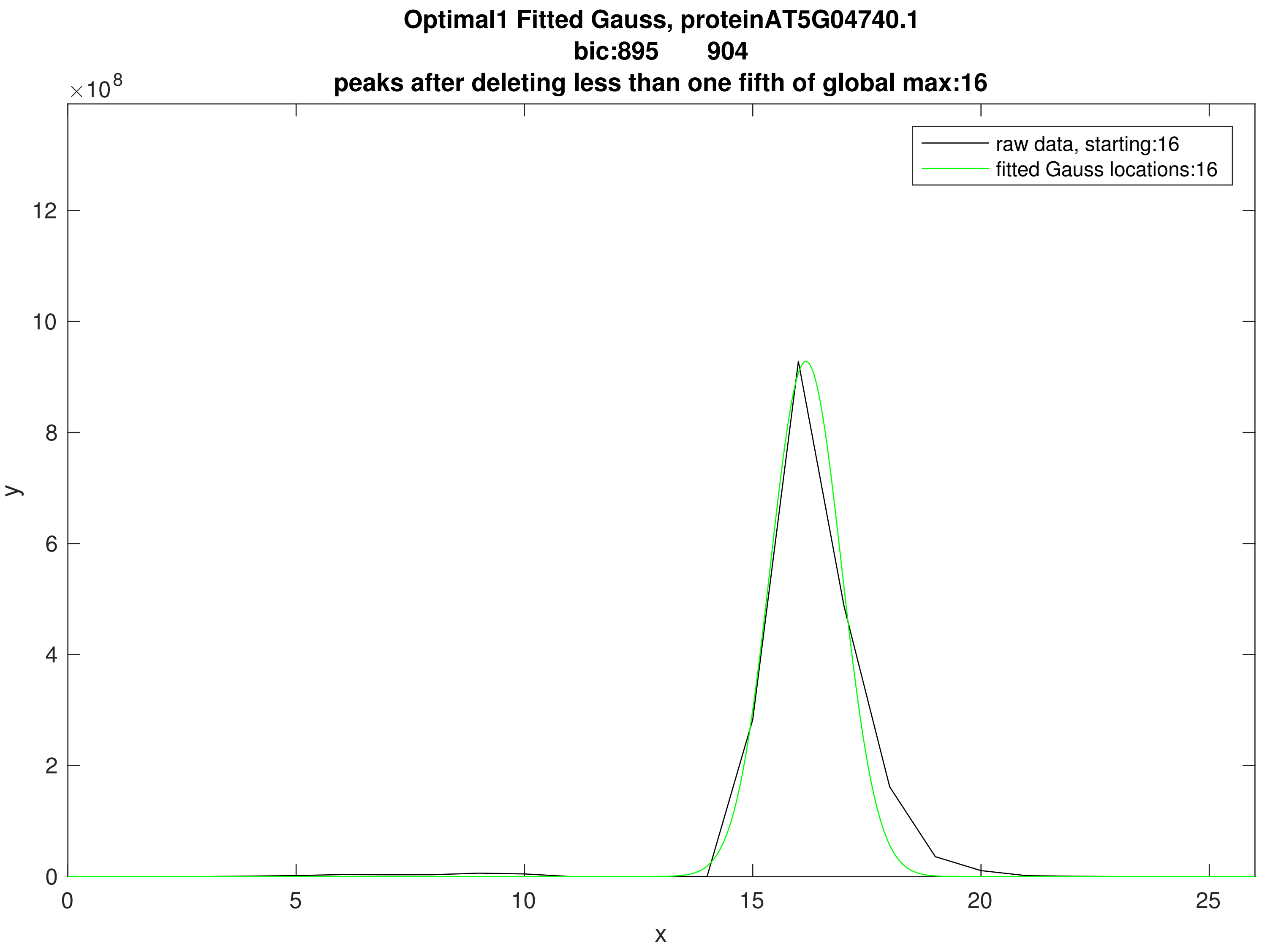

Optimal1 Fitted Gauss, proteinAT5G04740.1
bic:895 904
peaks after deleting less than one fifth of global max:16
×10^8
raw data, starting:16
fitted Gauss locations:16
12
10
8
6
4
2
0
y
0
5
10
15
20
25
x

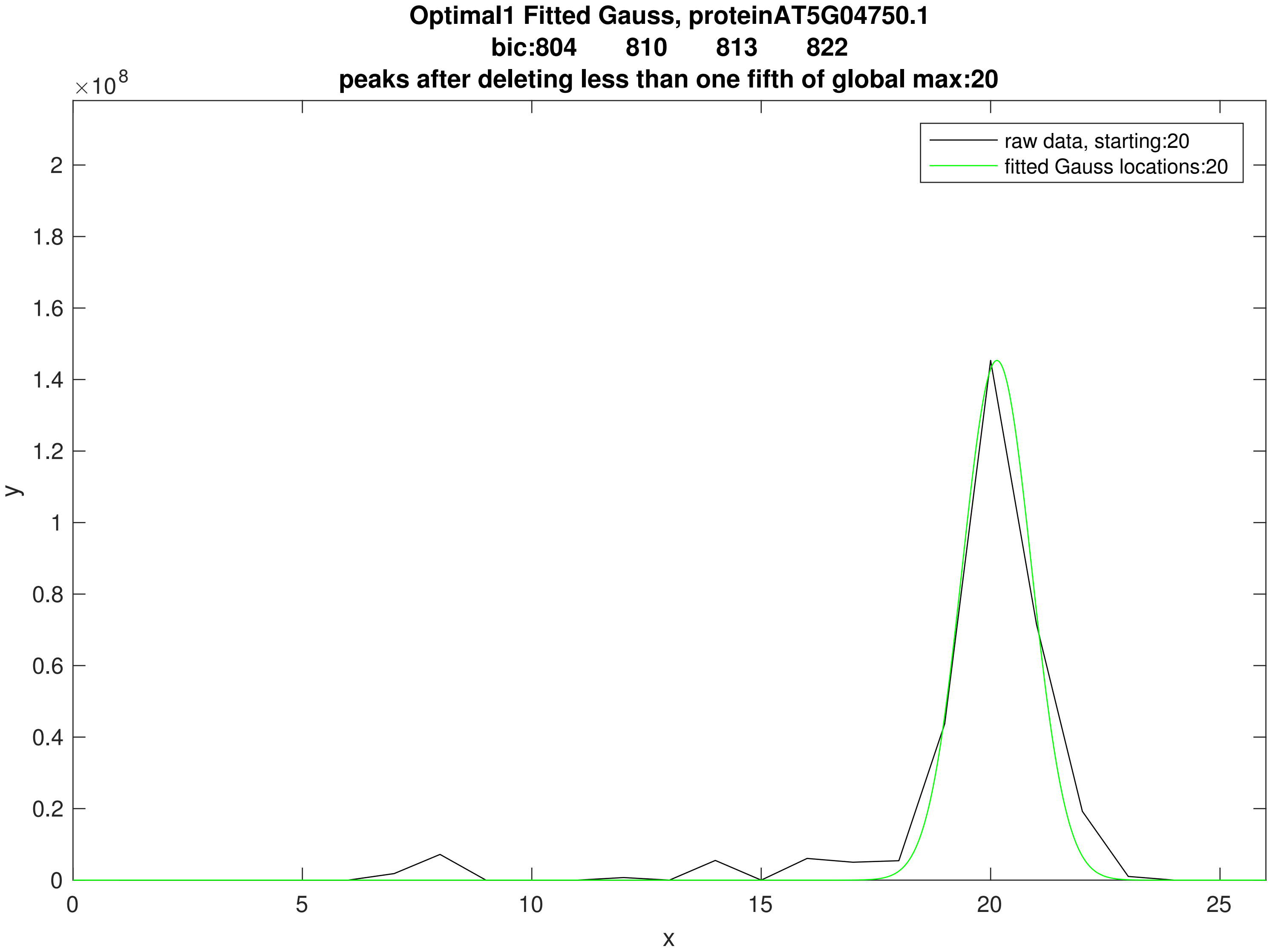

Optimal1 Fitted Gauss, proteinAT5G04750.1
bic:804 810 813 822
peaks after deleting less than one fifth of global max:20
×10^8
raw data, starting:20
fitted Gauss locations:20
y
x
0
0.2
0.4
0.6
0.8
1
1.2
1.4
1.6
1.8
2
0
5
10
15
20
25

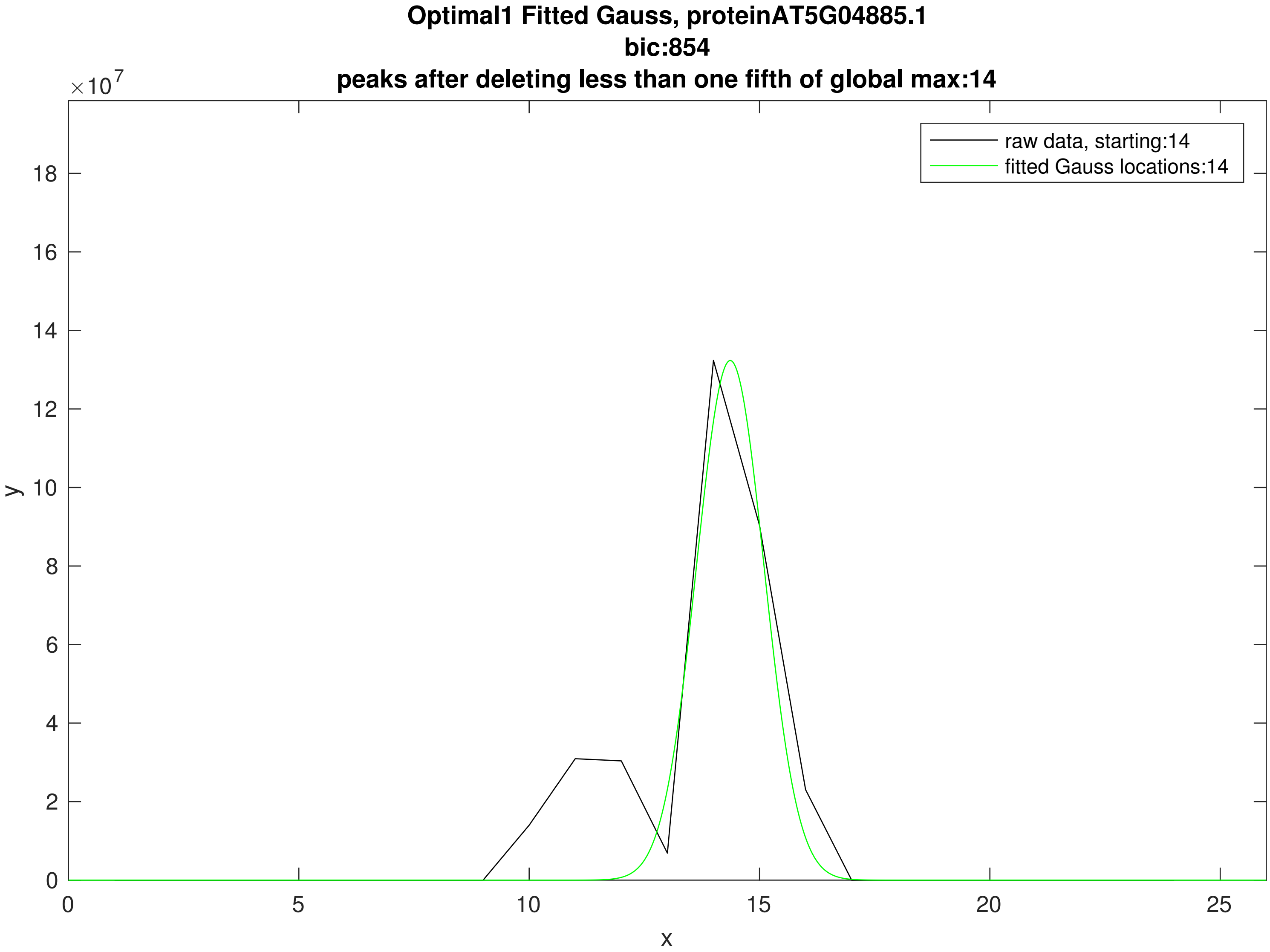

Optimal1 Fitted Gauss, proteinAT5G04885.1
bic:854
peaks after deleting less than one fifth of global max:14
×10^7
raw data, starting:14
fitted Gauss locations:14
18
16
14
12
10
8
6
4
2
0
y
0
5
10
15
20
25
x

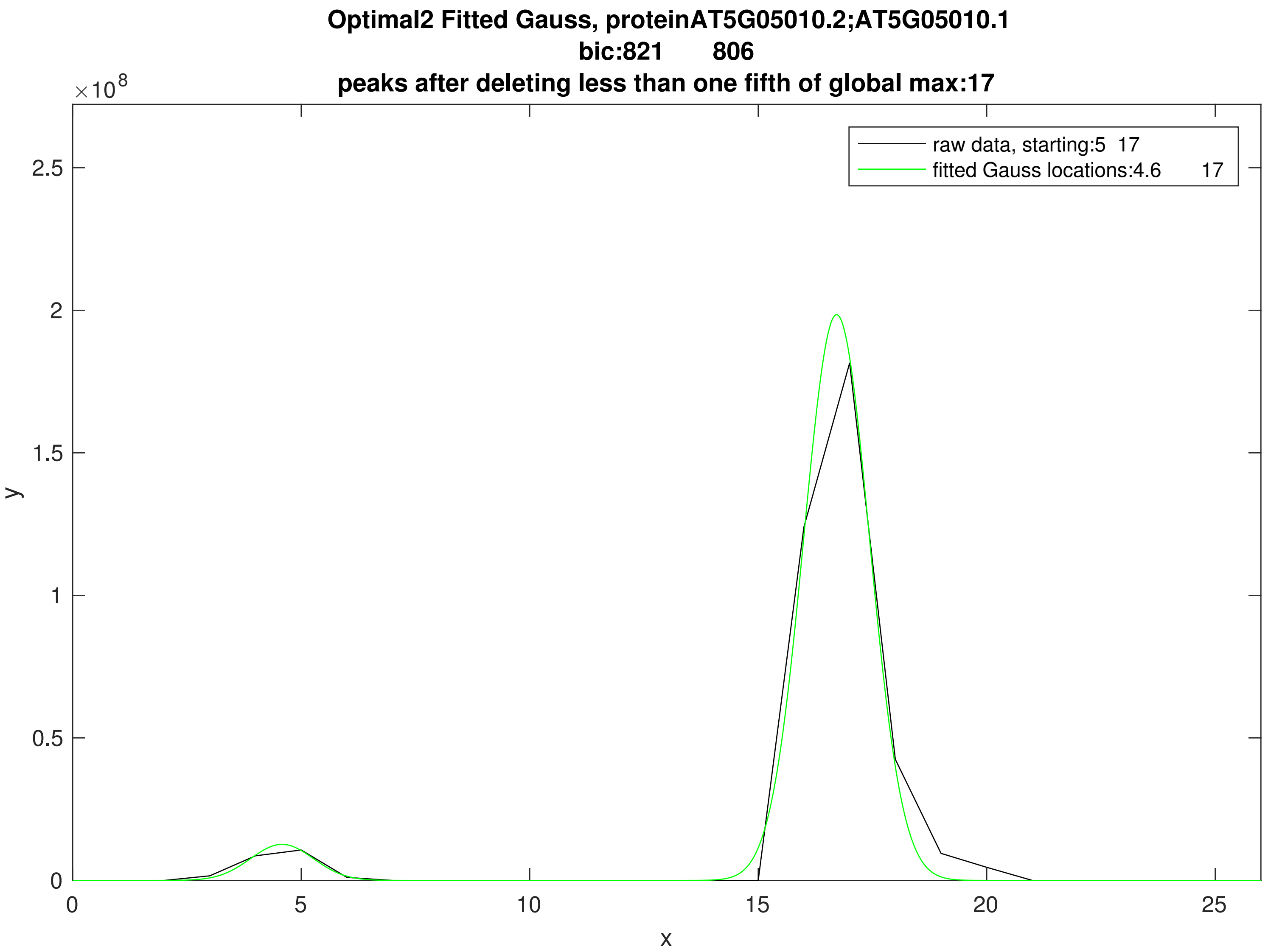
Optimal2 Fitted Gauss, proteinAT5G05010.2;AT5G05010.1
bic:821 806
peaks after deleting less than one fifth of global max:17
×10^8
2.5
2
1.5
1
0.5
0
y
0
5
10
15
20
25
x
raw data, starting:5 17
fitted Gauss locations:4.6 17

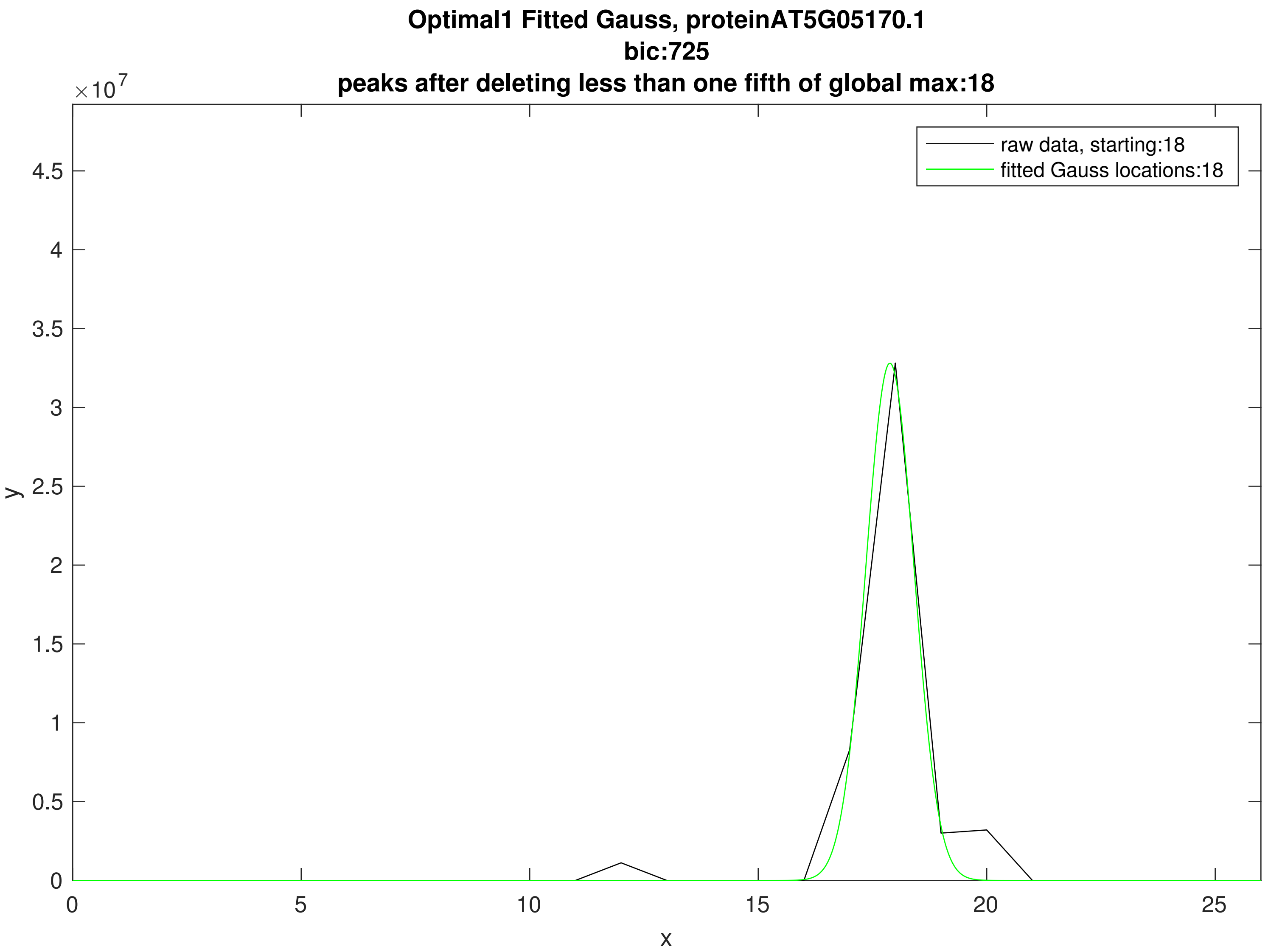
Optimal1 Fitted Gauss, proteinAT5G05170.1
bic:725
peaks after deleting less than one fifth of global max:18
×10^7
4.5
4
3.5
3
2.5
2
1.5
1
0.5
0
y
0
5
10
15
20
25
x
raw data, starting:18
fitted Gauss locations:18

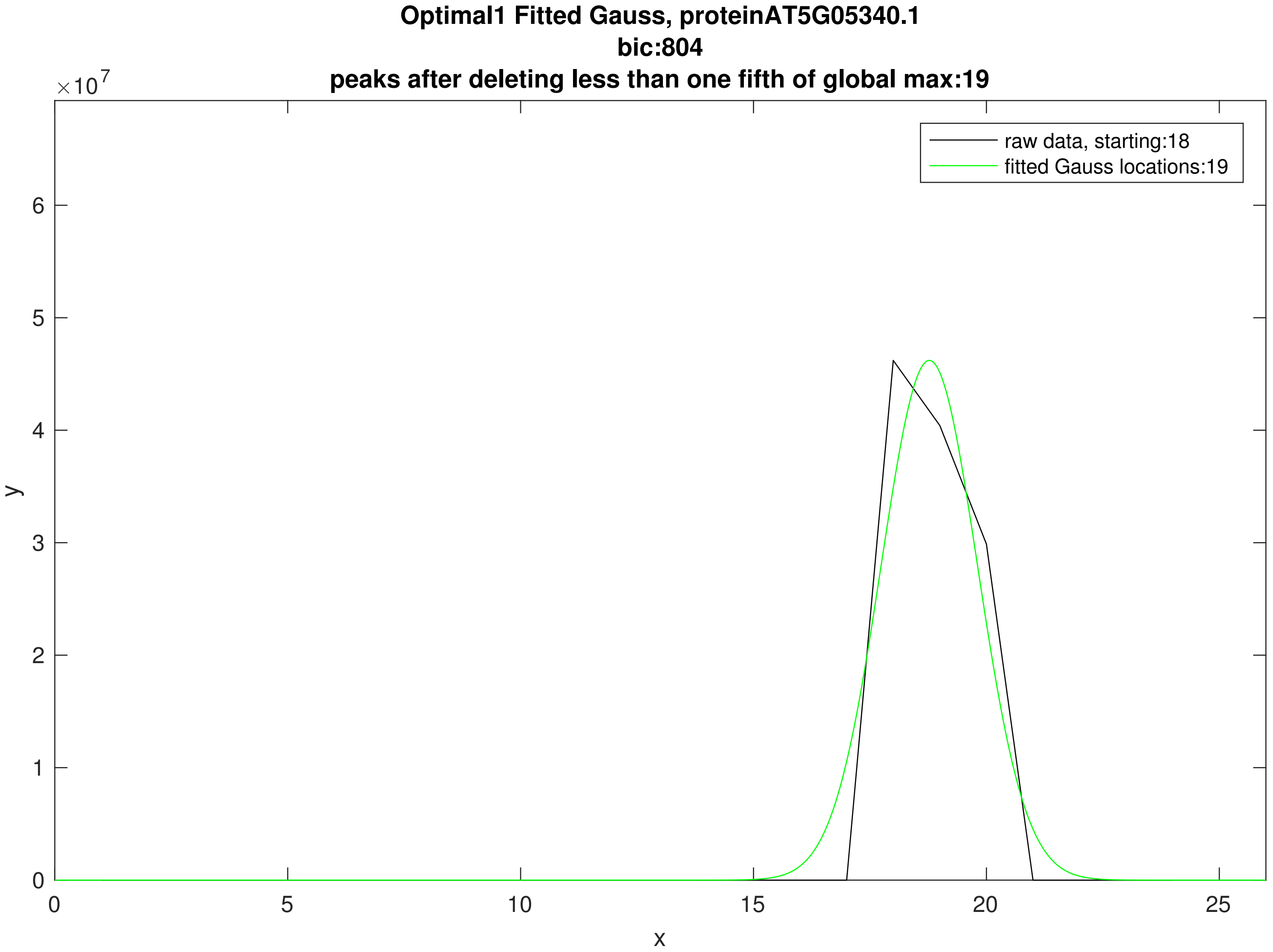

Optimal1 Fitted Gauss, proteinAT5G05340.1
bic:804
peaks after deleting less than one fifth of global max:19
×10^7
raw data, starting:18
fitted Gauss locations:19
y
x
0
1
2
3
4
5
6
0
5
10
15
20
25

Optimal1 Fitted Gauss, proteinAT5G05460.1
bic:836
peaks after deleting less than one fifth of global max:12

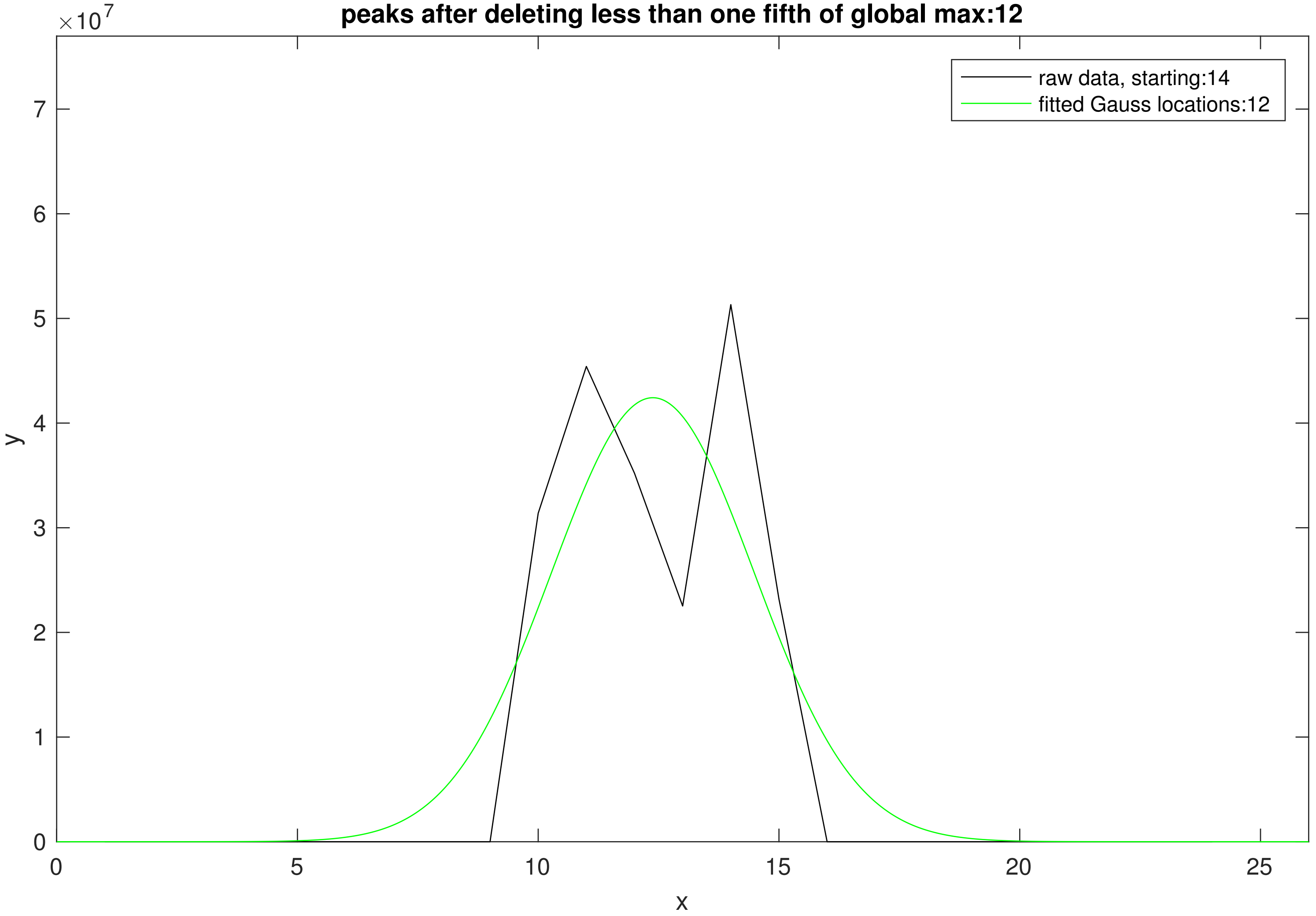

**Optimal1 Fitted Gauss, proteinAT5G05480.1**
**bic:899 906**
**peaks after deleting less than one fifth of global max:11**

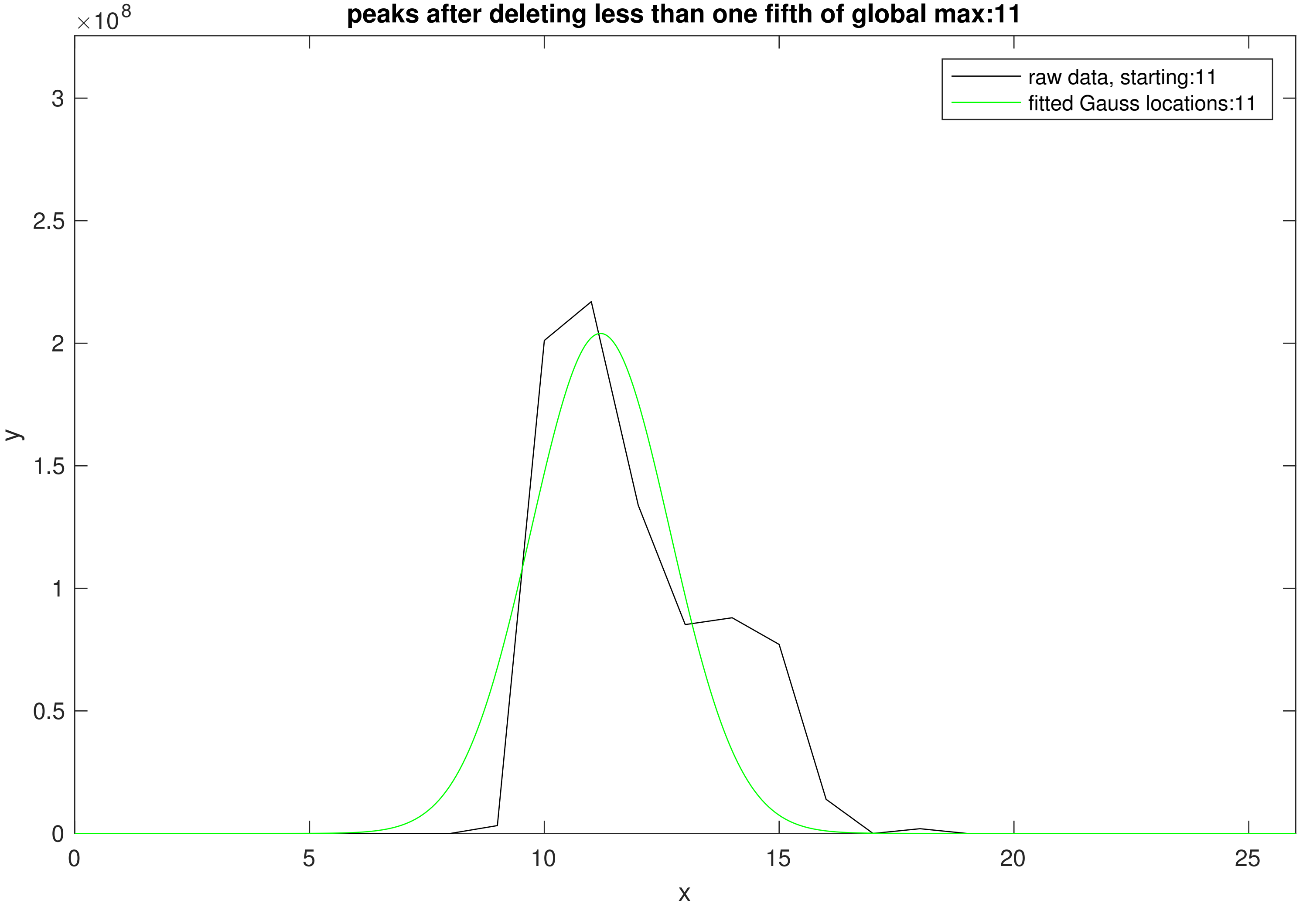

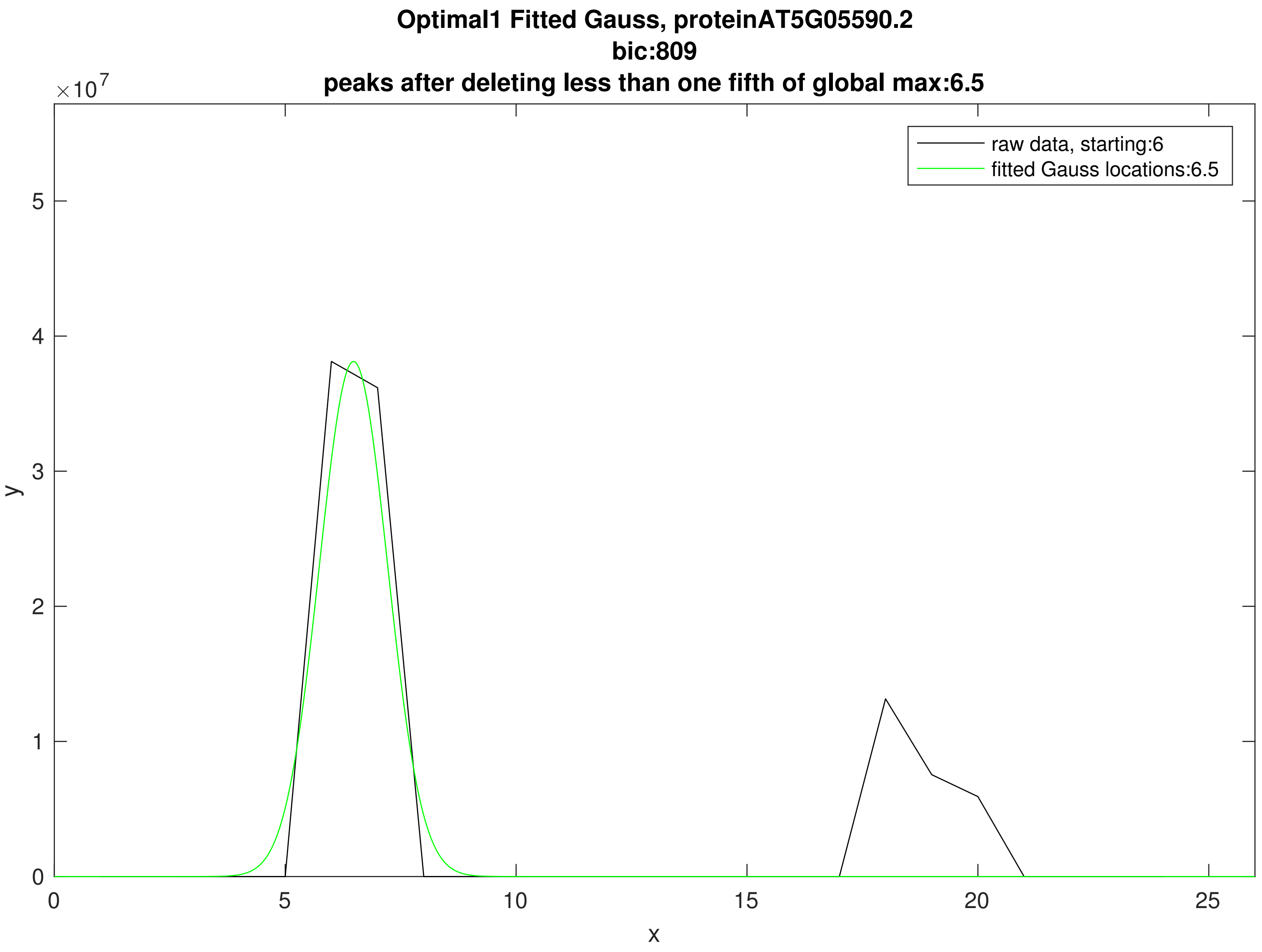
Optimal1 Fitted Gauss, proteinAT5G05590.2
bic:809
peaks after deleting less than one fifth of global max:6.5
×10^7
raw data, starting:6
fitted Gauss locations:6.5
y
x
0
1
2
3
4
5
0
5
10
15
20
25

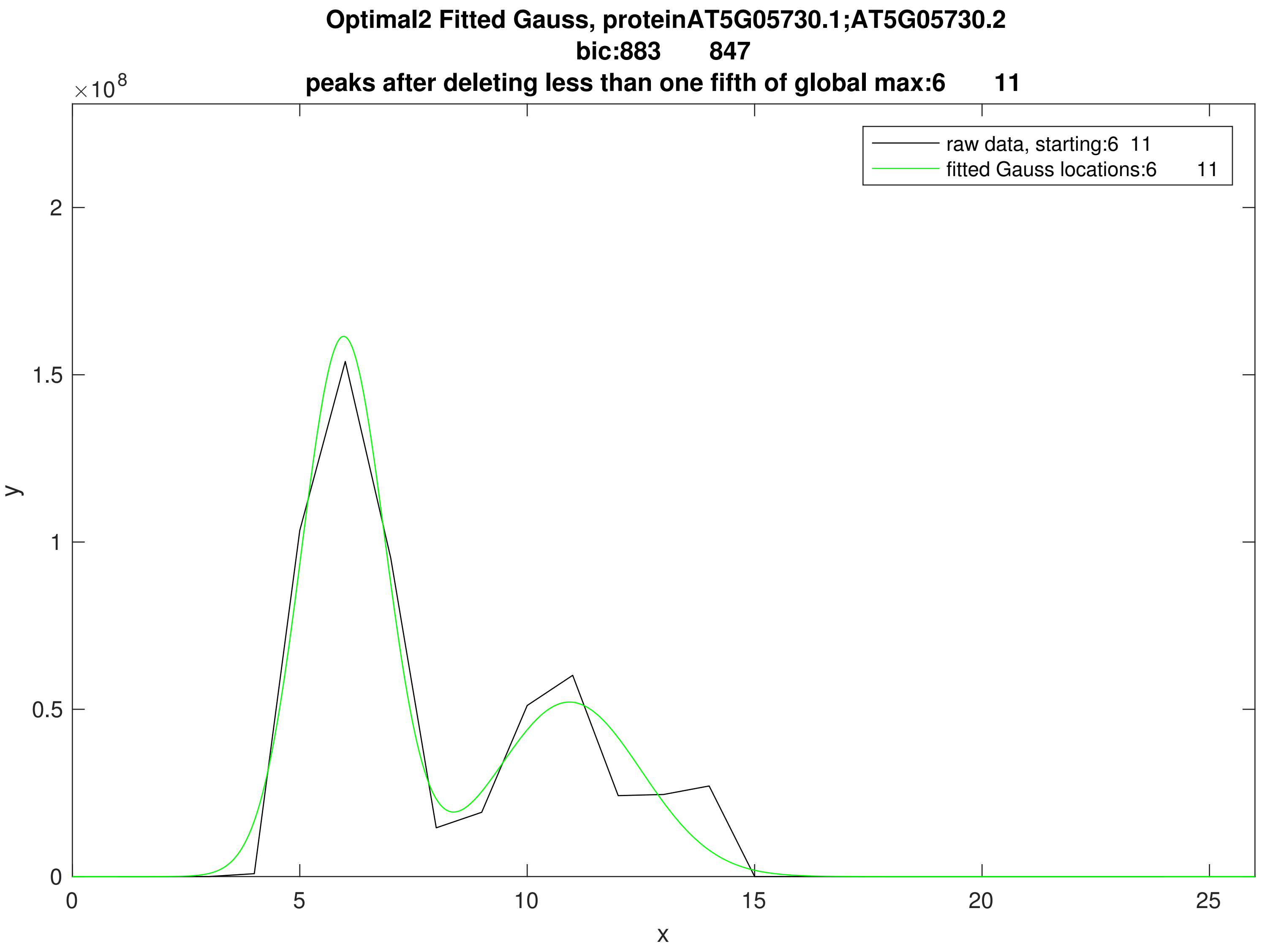
Optimal2 Fitted Gauss, proteinAT5G05730.1;AT5G05730.2
bic:883 847
peaks after deleting less than one fifth of global max:6 11
×10^8
raw data, starting:6 11
fitted Gauss locations:6 11
y
x
0
0.5
1
1.5
2
5
10
15
20
25

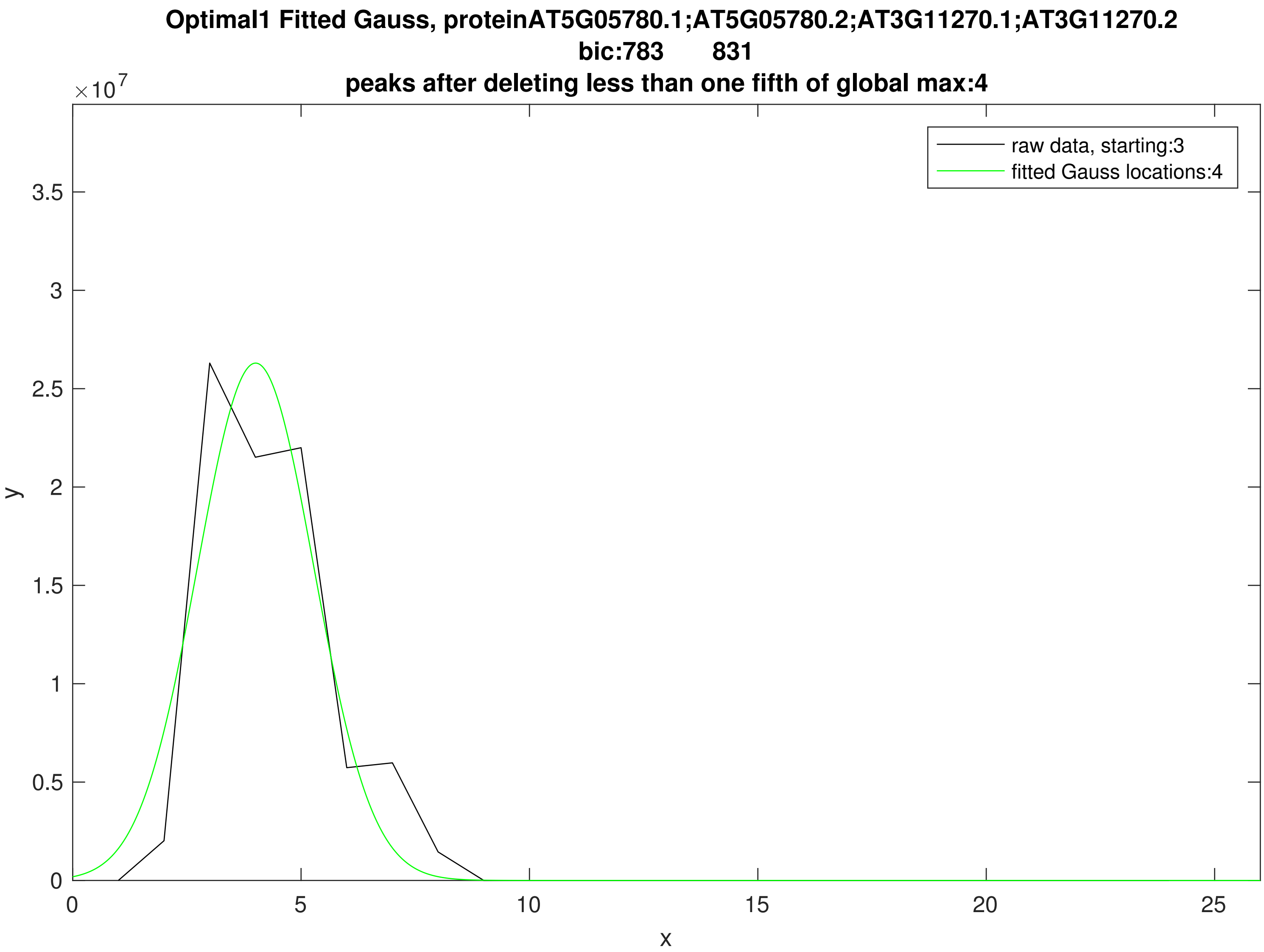

Optimal1 Fitted Gauss, proteinAT5G05780.1;AT5G05780.2;AT3G11270.1;AT3G11270.2
bic:783 831
peaks after deleting less than one fifth of global max:4
$\times 10^7$
raw data, starting:3
fitted Gauss locations:4
y
x
0
0.5
1
1.5
2
2.5
3
3.5
0
5
10
15
20
25

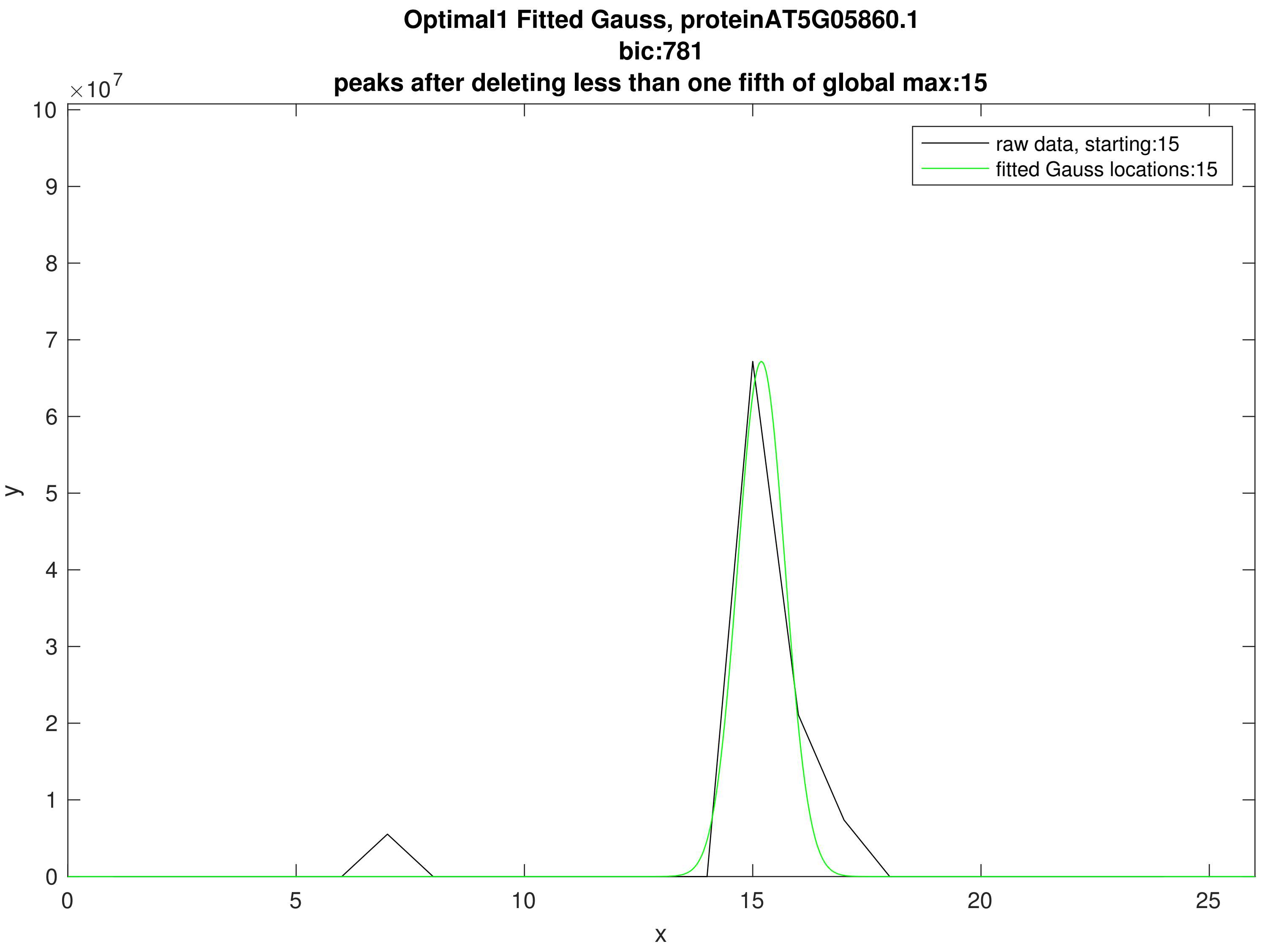

Optimal1 Fitted Gauss, proteinAT5G05860.1
bic:781
peaks after deleting less than one fifth of global max:15
×10^7
10
9
8
7
6
5
4
3
2
1
0
y
0
5
10
15
20
25
x
raw data, starting:15
fitted Gauss locations:15

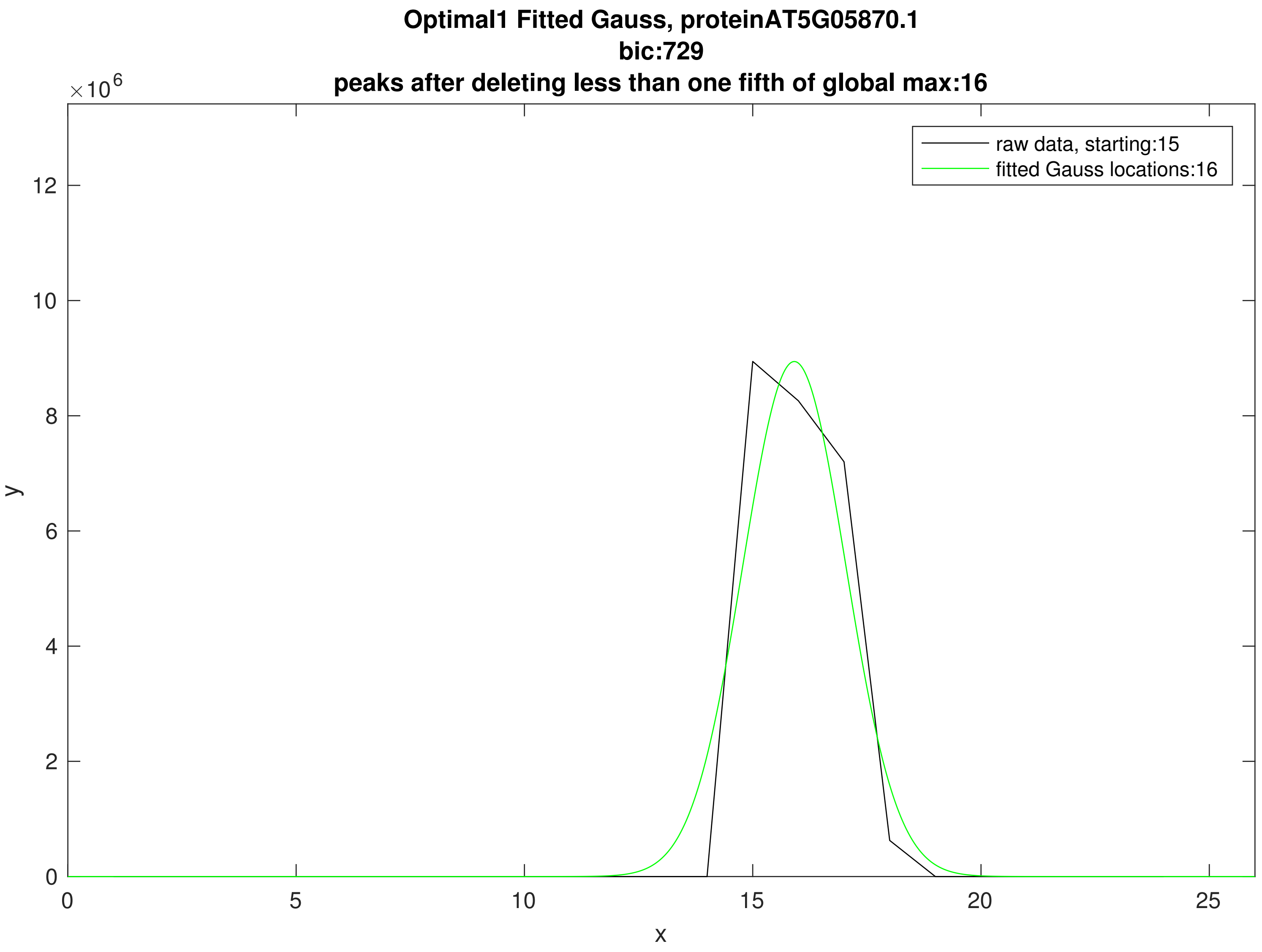

Optimal1 Fitted Gauss, proteinAT5G05870.1
bic:729
peaks after deleting less than one fifth of global max:16
×10^6
raw data, starting:15
fitted Gauss locations:16
y
x
0
2
4
6
8
10
12
0
5
10
15
20
25

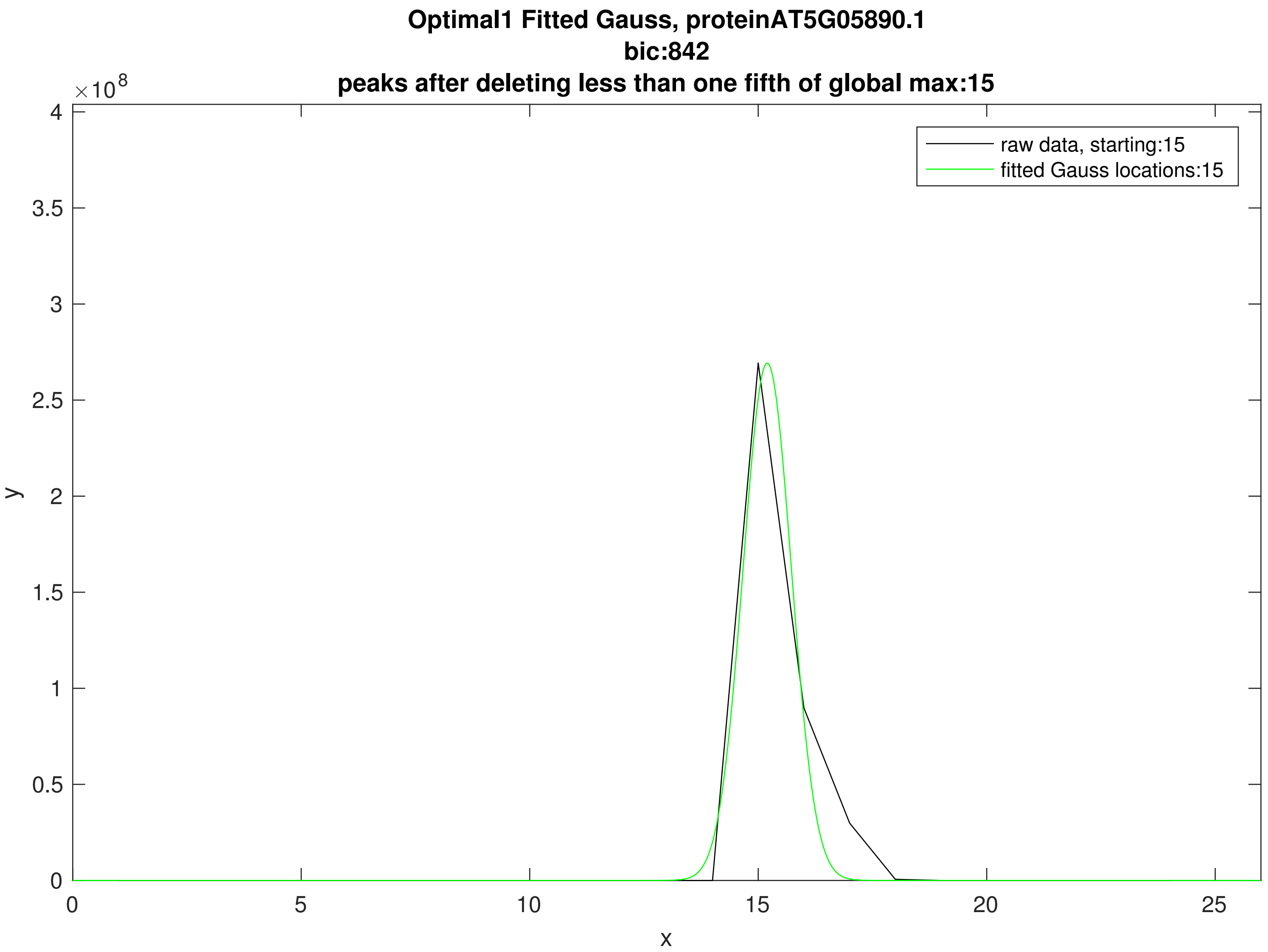
Optimal1 Fitted Gauss, proteinAT5G05890.1
bic:842
peaks after deleting less than one fifth of global max:15
×10^8
raw data, starting:15
fitted Gauss locations:15
y
x
0
0.5
1
1.5
2
2.5
3
3.5
4
5
10
15
20
25

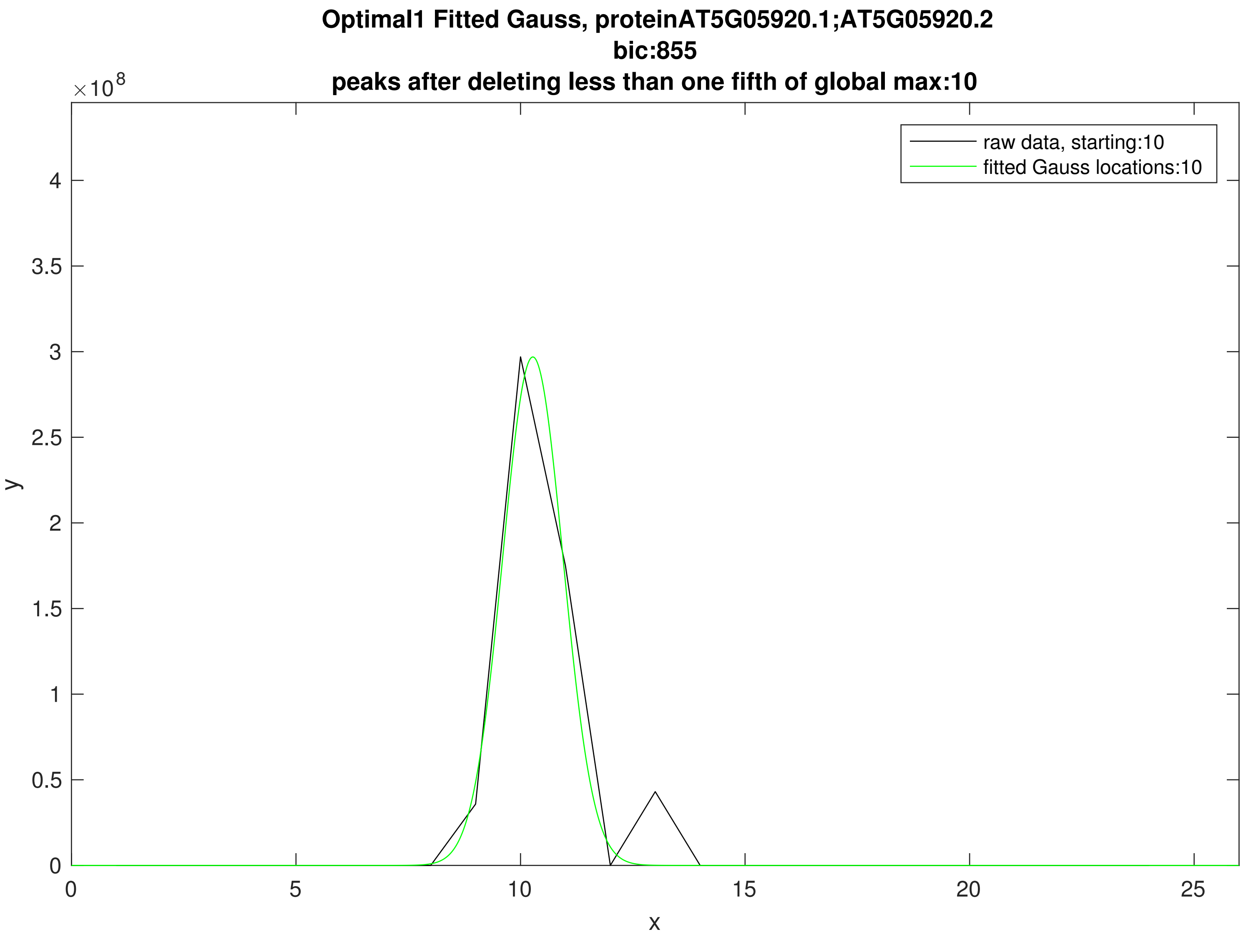

Optimal1 Fitted Gauss, proteinAT5G05920.1;AT5G05920.2
bic:855
peaks after deleting less than one fifth of global max:10
×10^8
raw data, starting:10
fitted Gauss locations:10
4
3.5
3
2.5
2
1.5
1
0.5
0
y
0
5
10
15
20
25
x

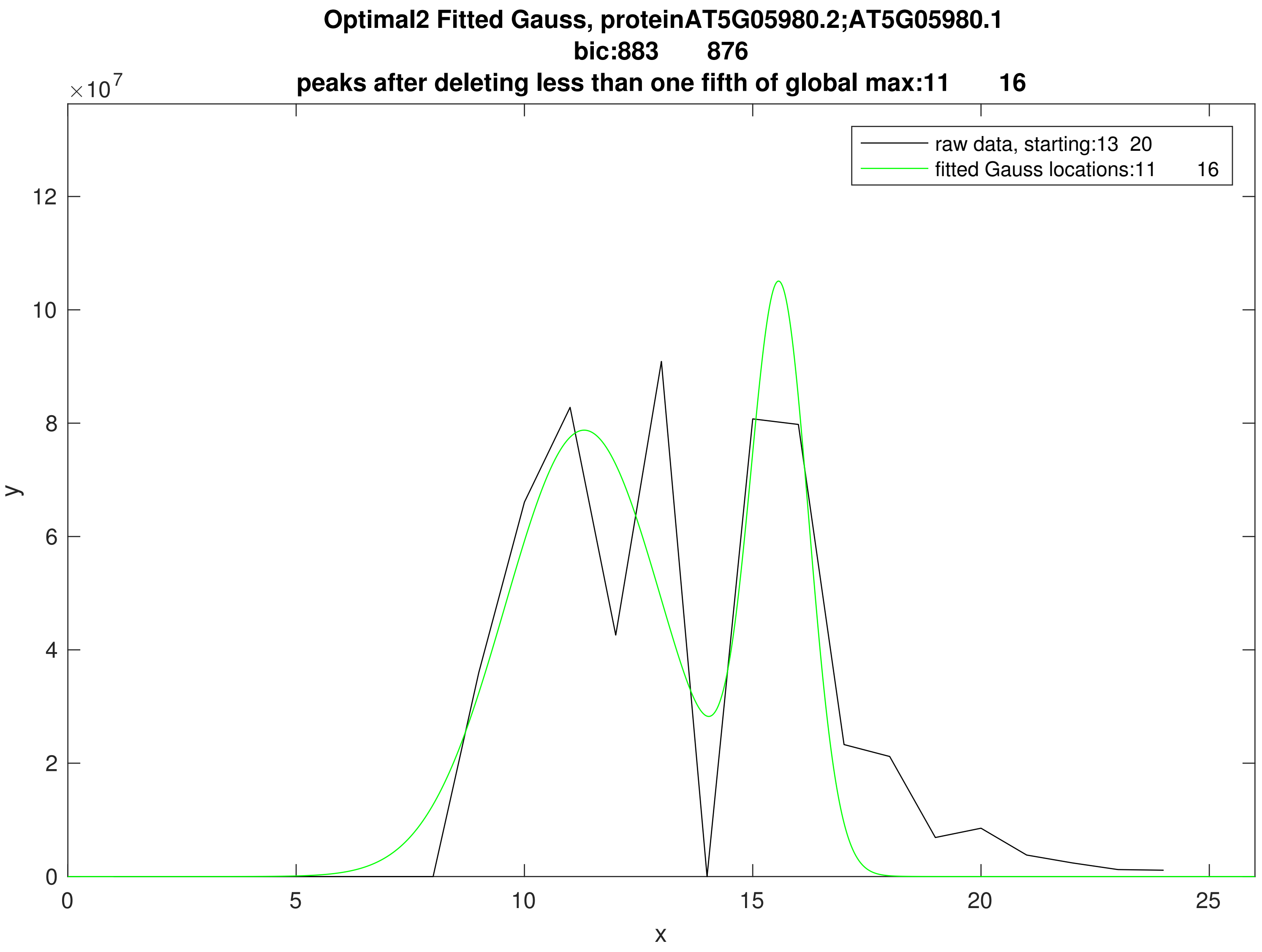

Optimal2 Fitted Gauss, proteinAT5G05980.2;AT5G05980.1
bic:883 876
peaks after deleting less than one fifth of global max:11 16
×10^7
raw data, starting:13 20
fitted Gauss locations:11 16
y
x
0
2
4
6
8
10
12
0
5
10
15
20
25

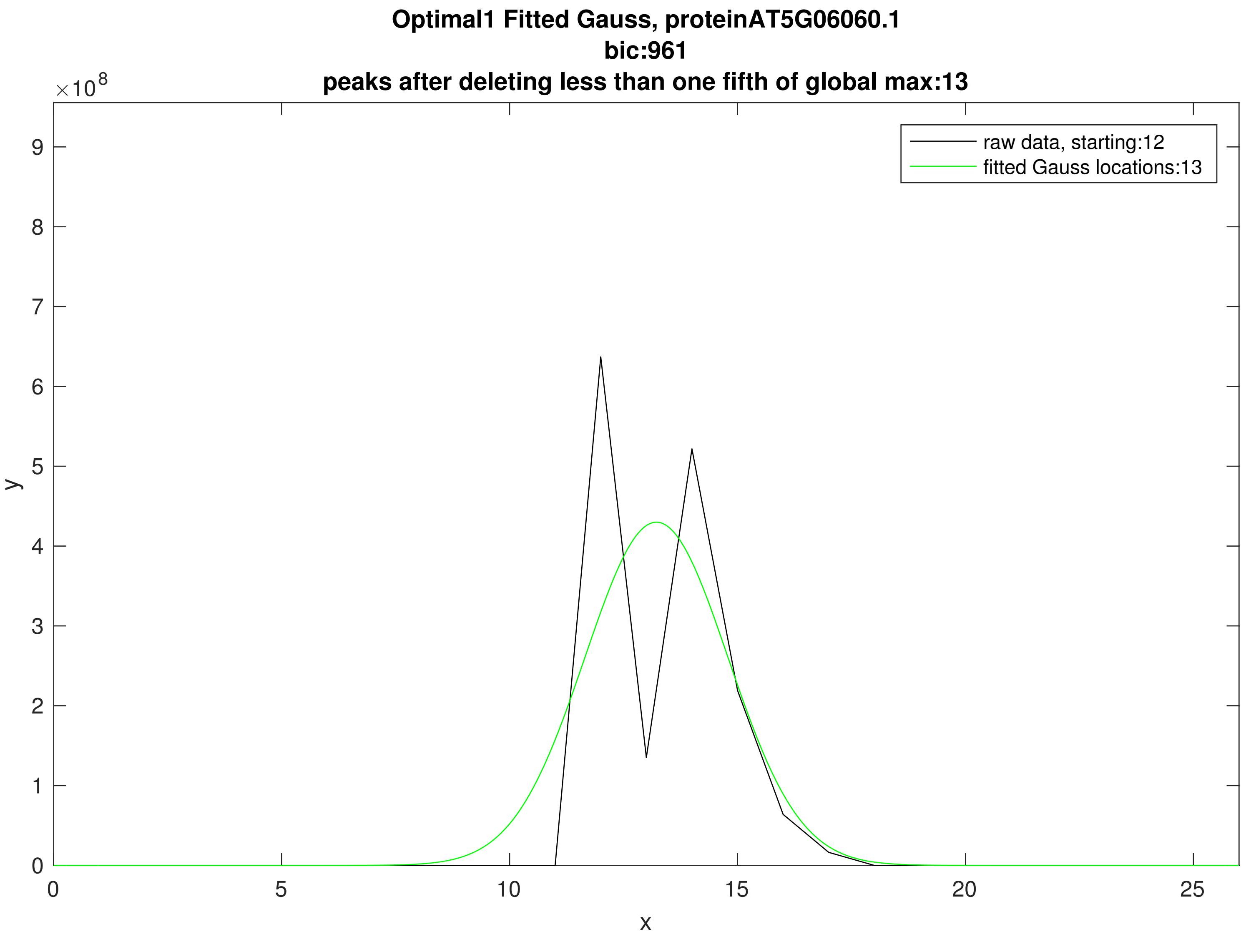
Optimal1 Fitted Gauss, proteinAT5G06060.1
bic:961
peaks after deleting less than one fifth of global max:13
×10^8
raw data, starting:12
fitted Gauss locations:13
y
x
0
1
2
3
4
5
6
7
8
9
0
5
10
15
20
25

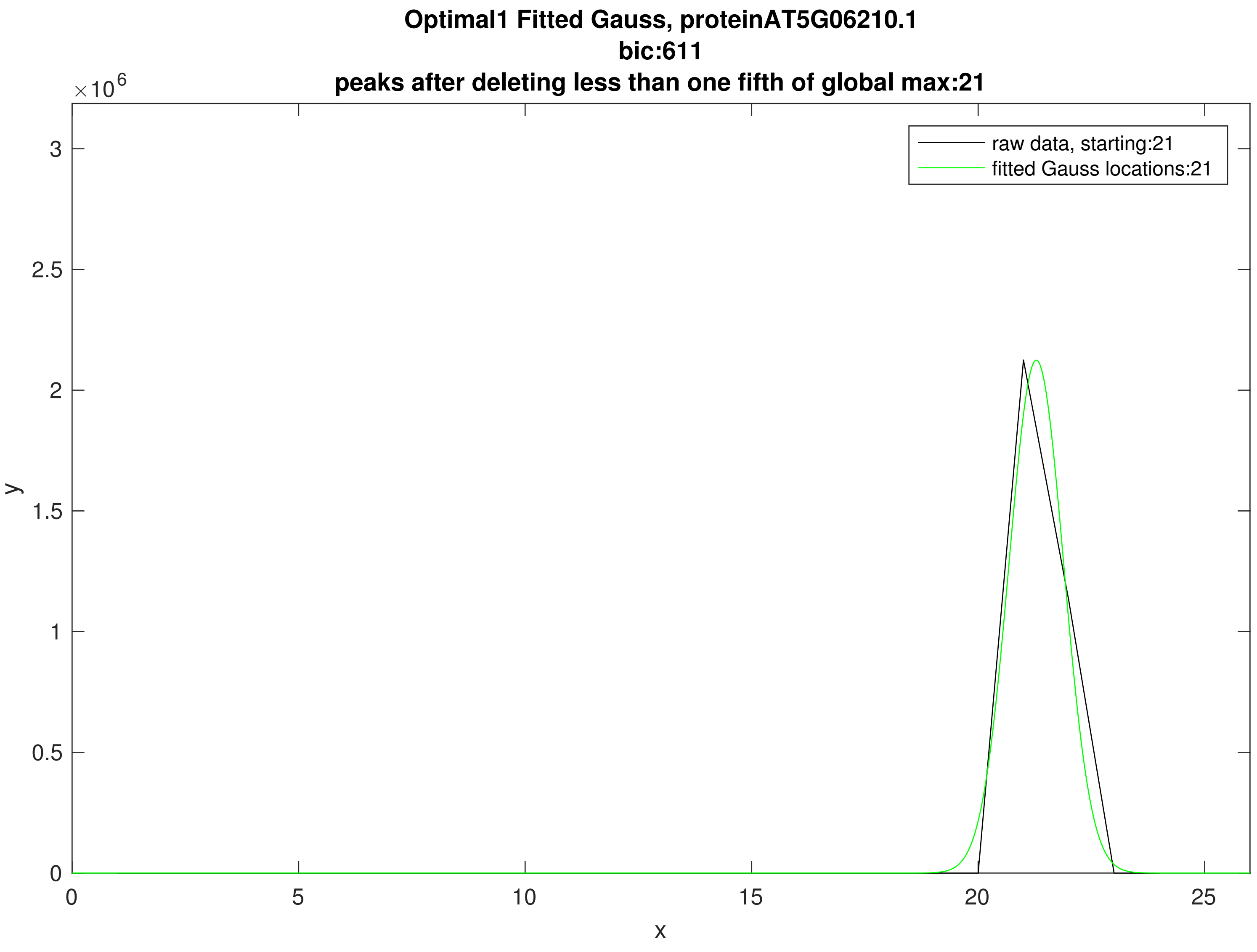
Optimal1 Fitted Gauss, proteinAT5G06210.1
bic:611
peaks after deleting less than one fifth of global max:21
×10^6
3
2.5
2
1.5
1
0.5
0
y
0
5
10
15
20
25
x
raw data, starting:21
fitted Gauss locations:21

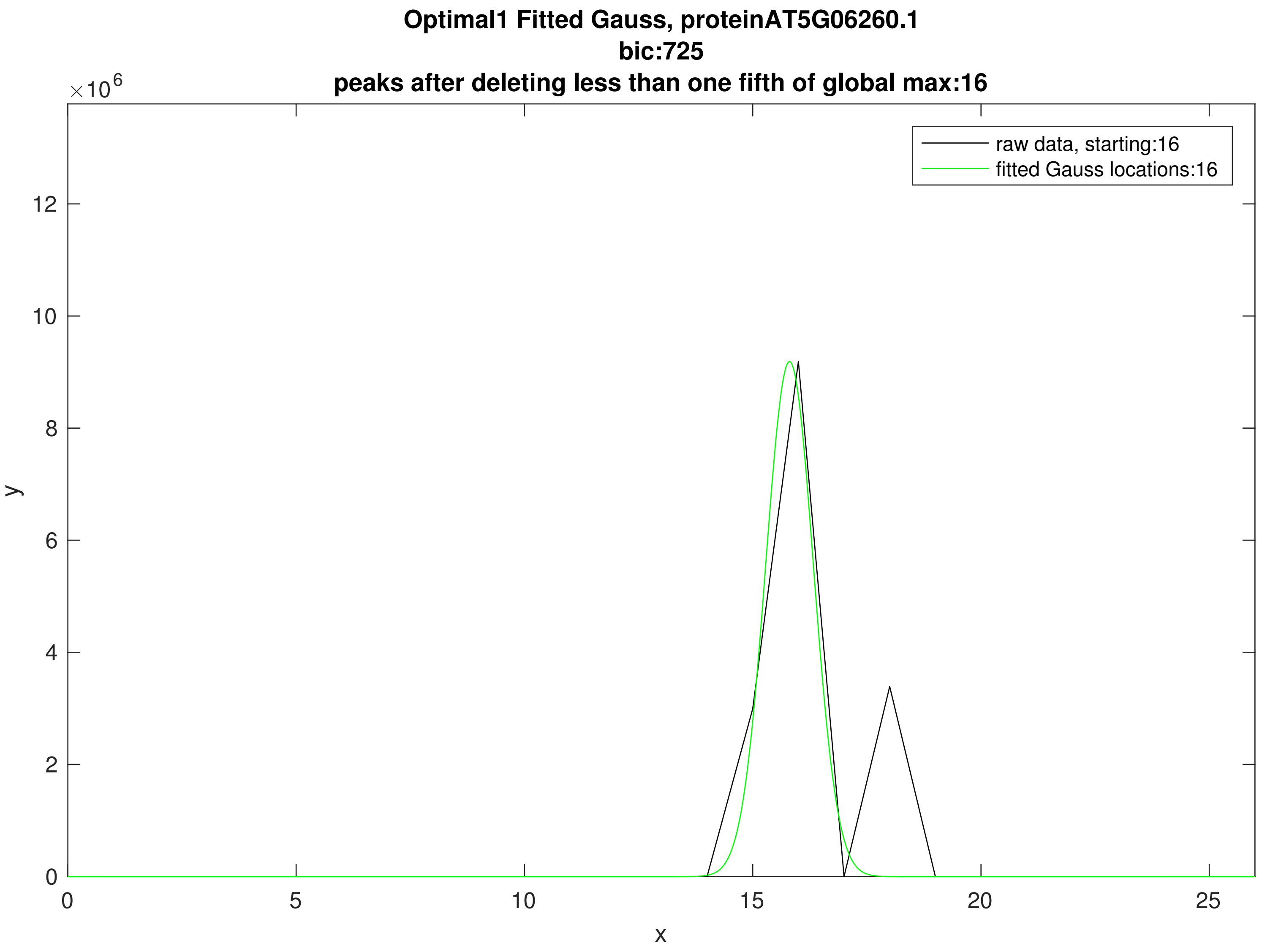
Optimal1 Fitted Gauss, proteinAT5G06260.1
bic:725
peaks after deleting less than one fifth of global max:16
×10^6
raw data, starting:16
fitted Gauss locations:16
12
10
8
6
4
2
0
y
0
5
10
15
20
25
x

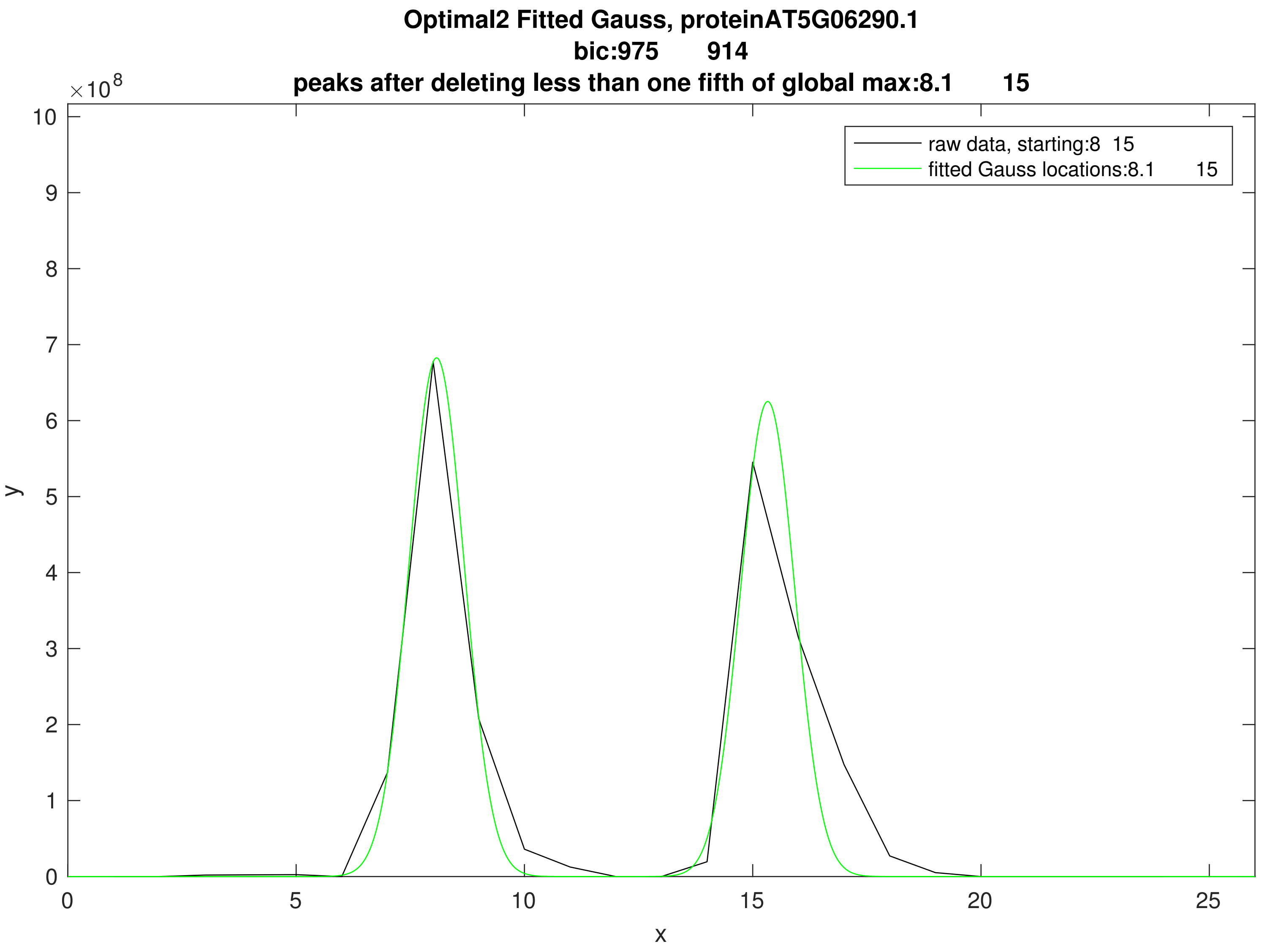

Optimal2 Fitted Gauss, proteinAT5G06290.1
bic:975 914
peaks after deleting less than one fifth of global max:8.1 15
×10^8
raw data, starting:8 15
fitted Gauss locations:8.1 15
y
x
0
1
2
3
4
5
6
7
8
9
10
0
5
10
15
20
25

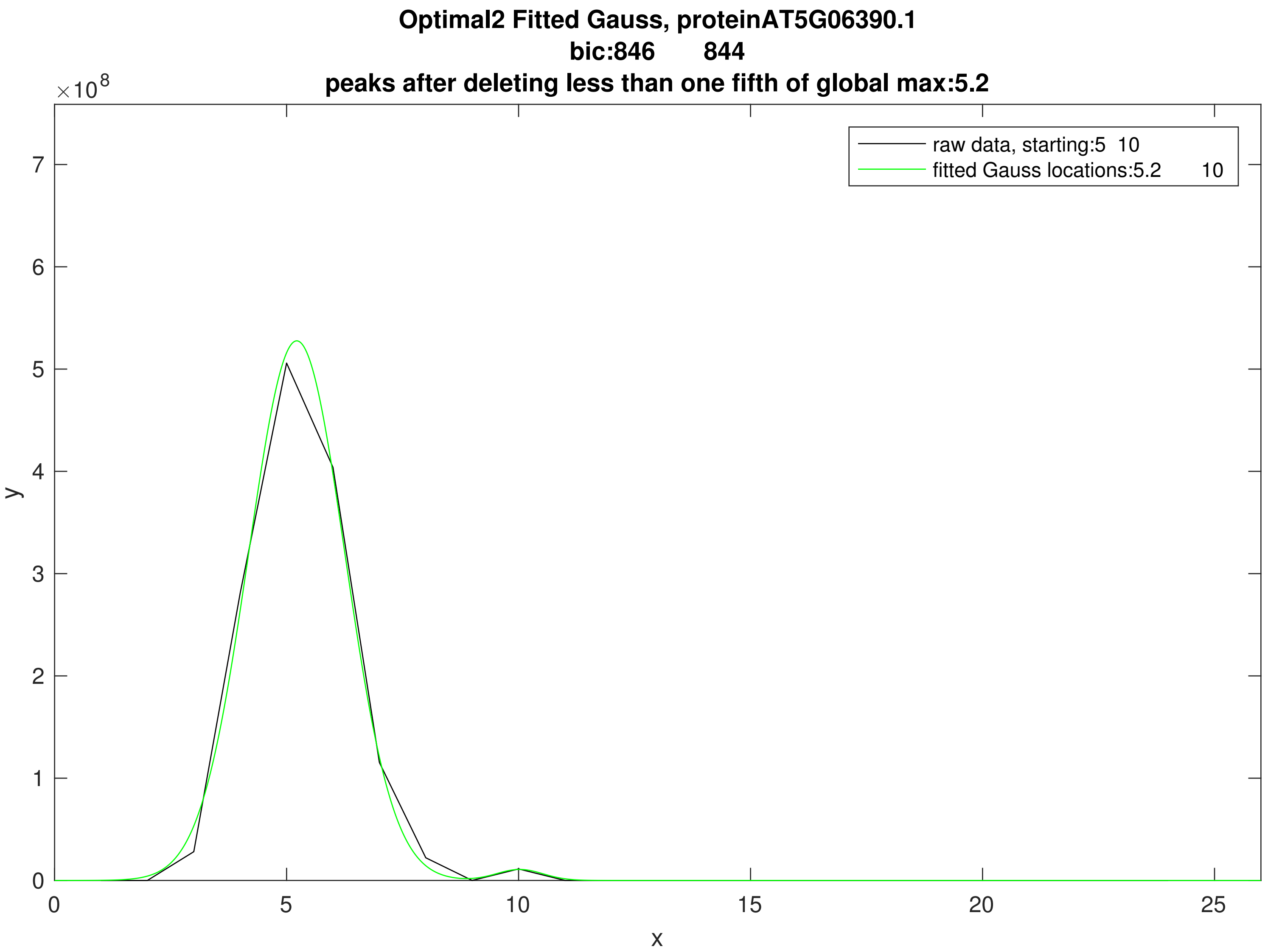
Optimal2 Fitted Gauss, proteinAT5G06390.1
bic:846 844
peaks after deleting less than one fifth of global max:5.2
×10^8
raw data, starting:5 10
fitted Gauss locations:5.2 10
y
x
0
1
2
3
4
5
6
7
0
5
10
15
20
25

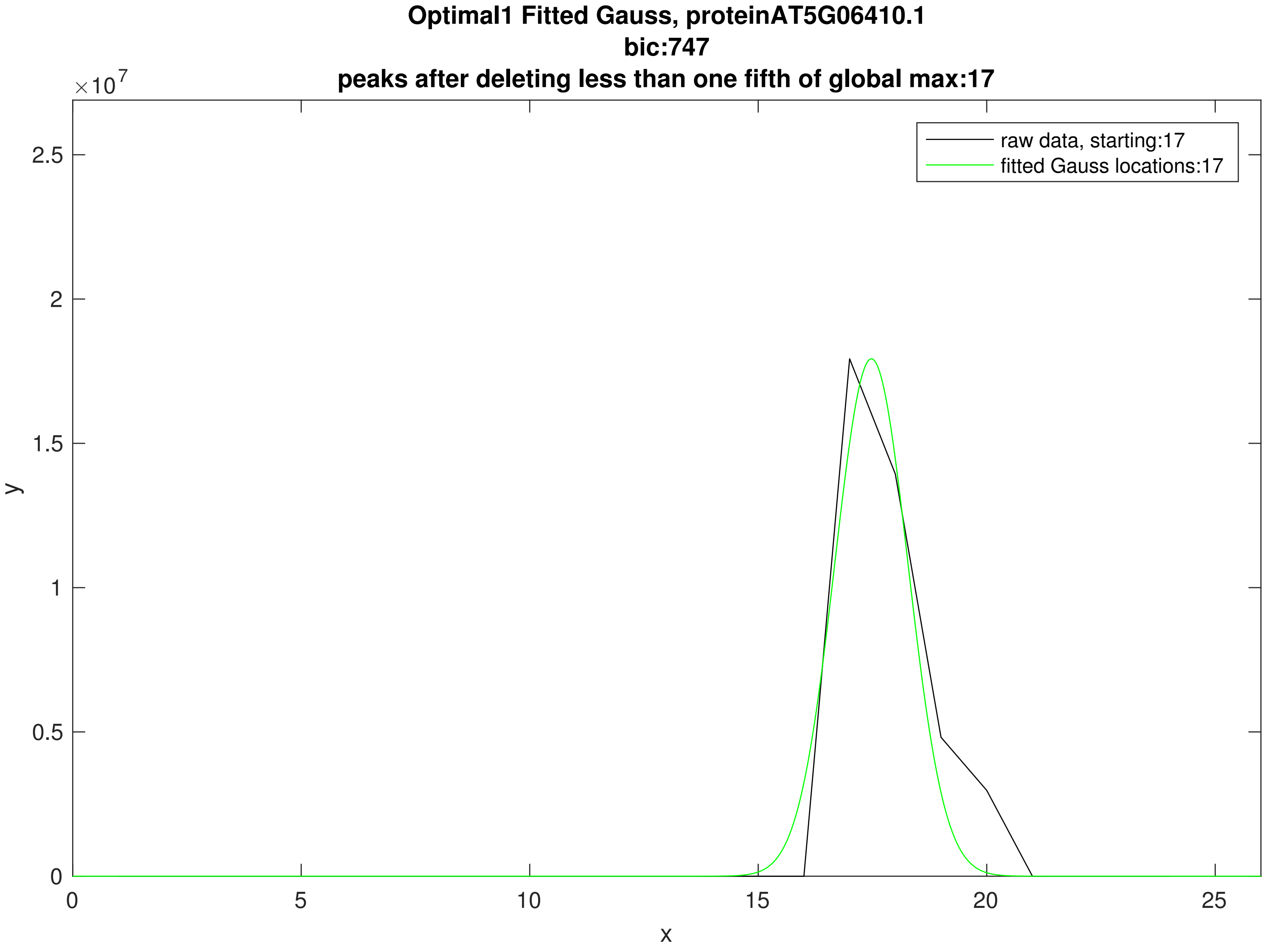

Optimal1 Fitted Gauss, proteinAT5G06410.1
bic:747
peaks after deleting less than one fifth of global max:17
×10^7
2.5
2
1.5
1
0.5
0
y
0
5
10
15
20
25
x
raw data, starting:17
fitted Gauss locations:17

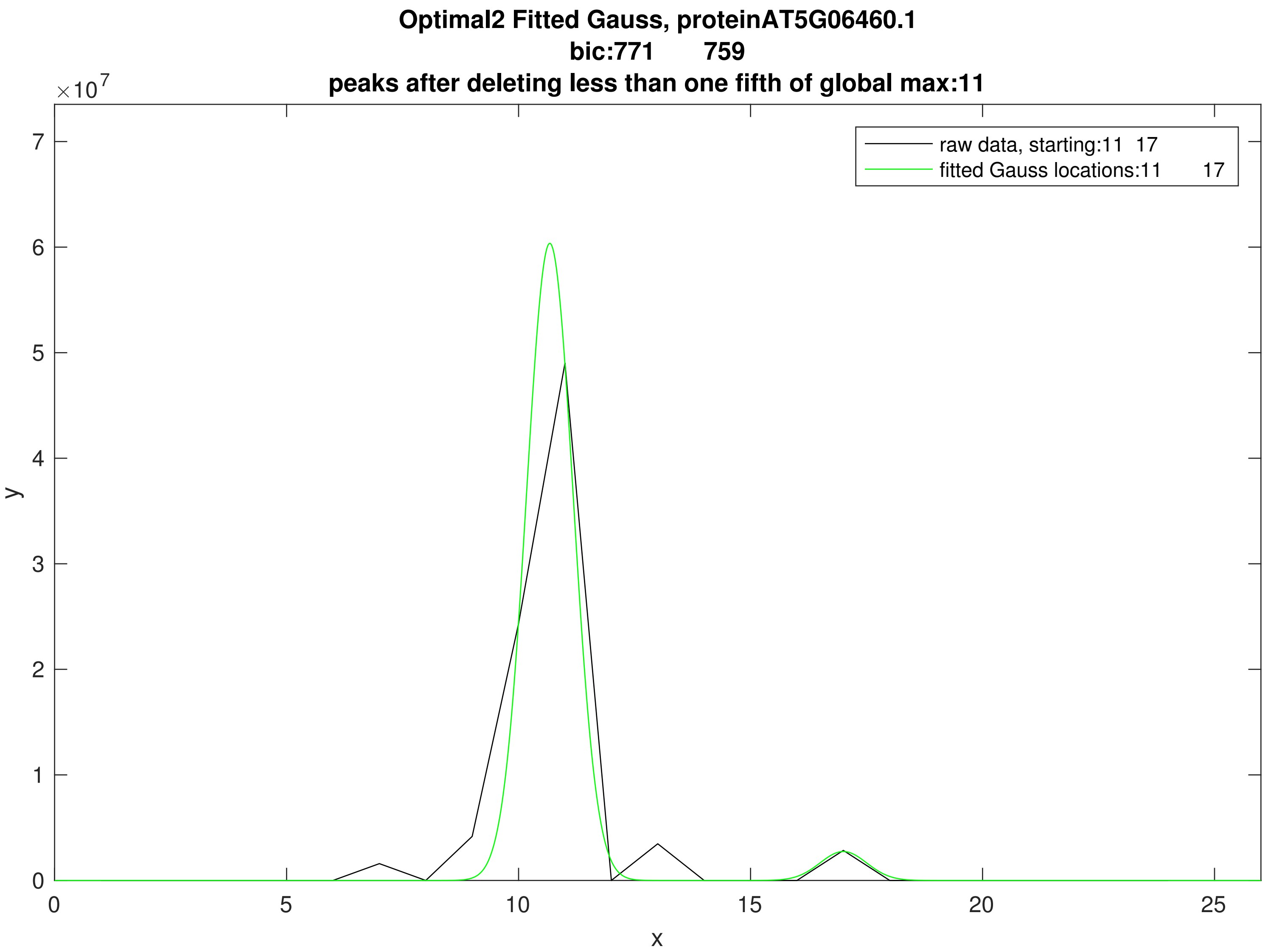
Optimal2 Fitted Gauss, proteinAT5G06460.1
bic:771 759
peaks after deleting less than one fifth of global max:11
×10^7
raw data, starting:11 17
fitted Gauss locations:11 17
y
x
0
1
2
3
4
5
6
7
0
5
10
15
20
25

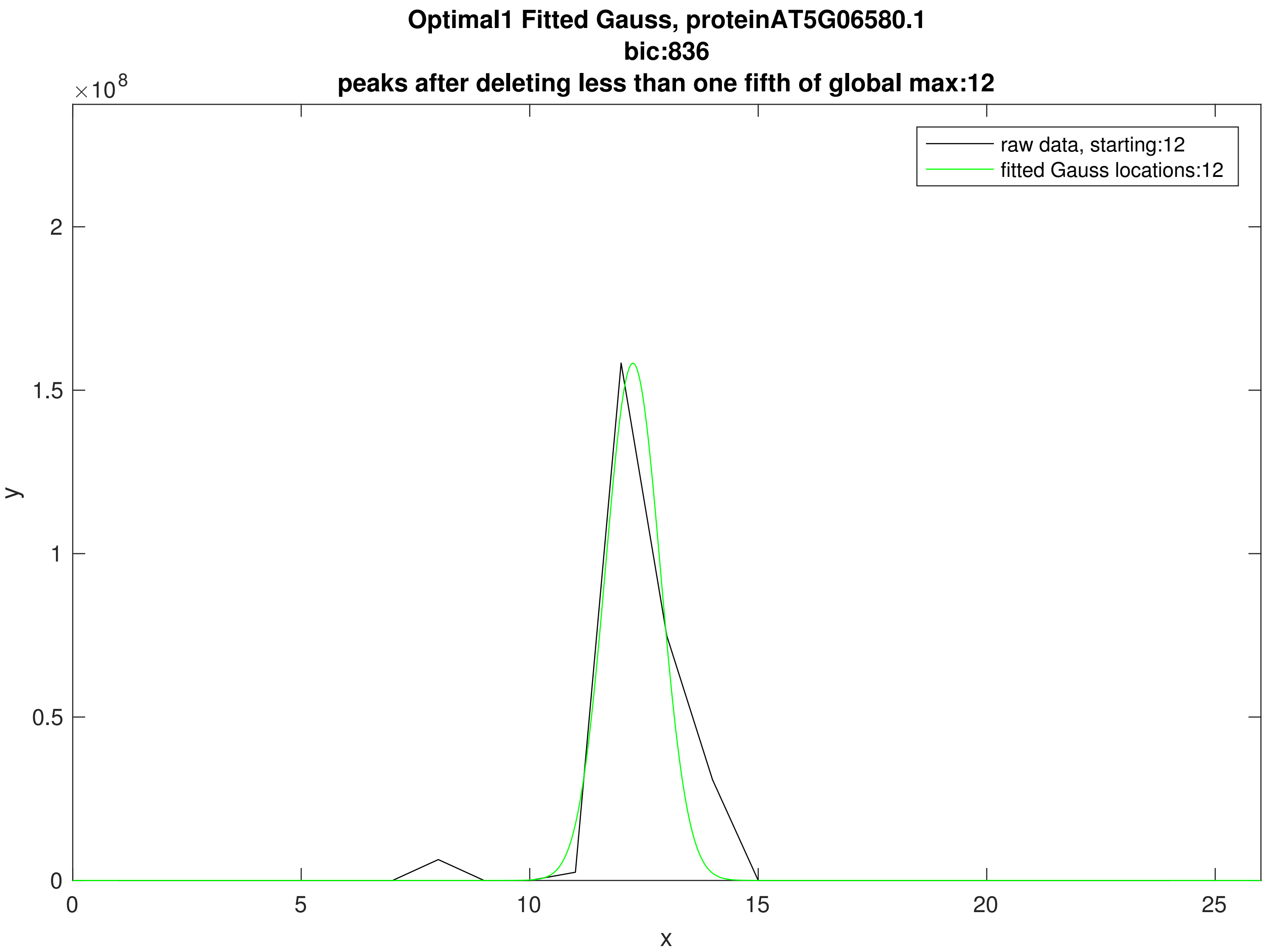

Optimal1 Fitted Gauss, proteinAT5G06580.1
bic:836
peaks after deleting less than one fifth of global max:12
×10^8
raw data, starting:12
fitted Gauss locations:12
y
x
0
0.5
1
1.5
2
0
5
10
15
20
25

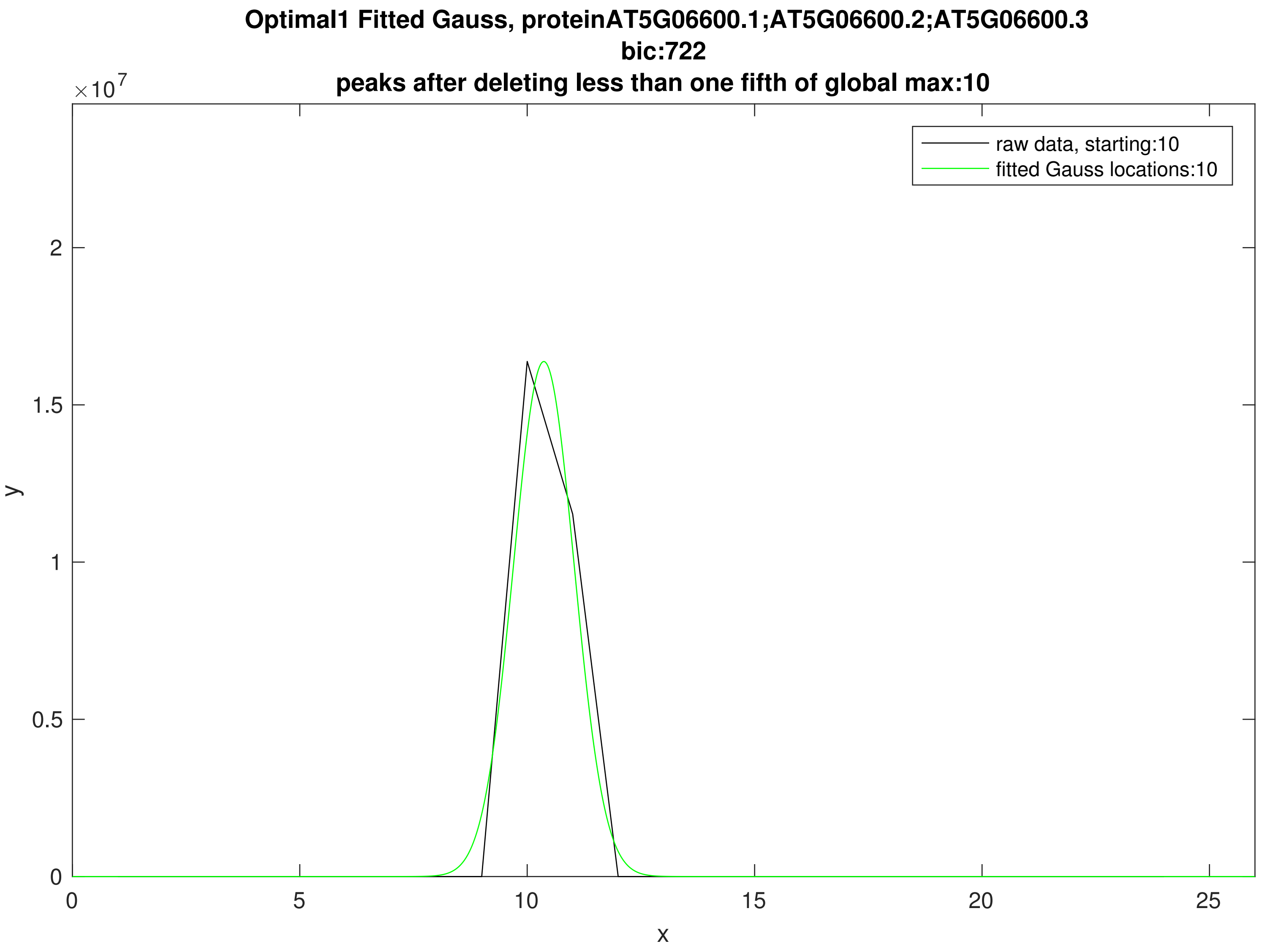

Optimal1 Fitted Gauss, proteinAT5G06600.1;AT5G06600.2;AT5G06600.3
bic:722
peaks after deleting less than one fifth of global max:10
×10^7
raw data, starting:10
fitted Gauss locations:10
2
1.5
1
0.5
0
y
0
5
10
15
20
25
x

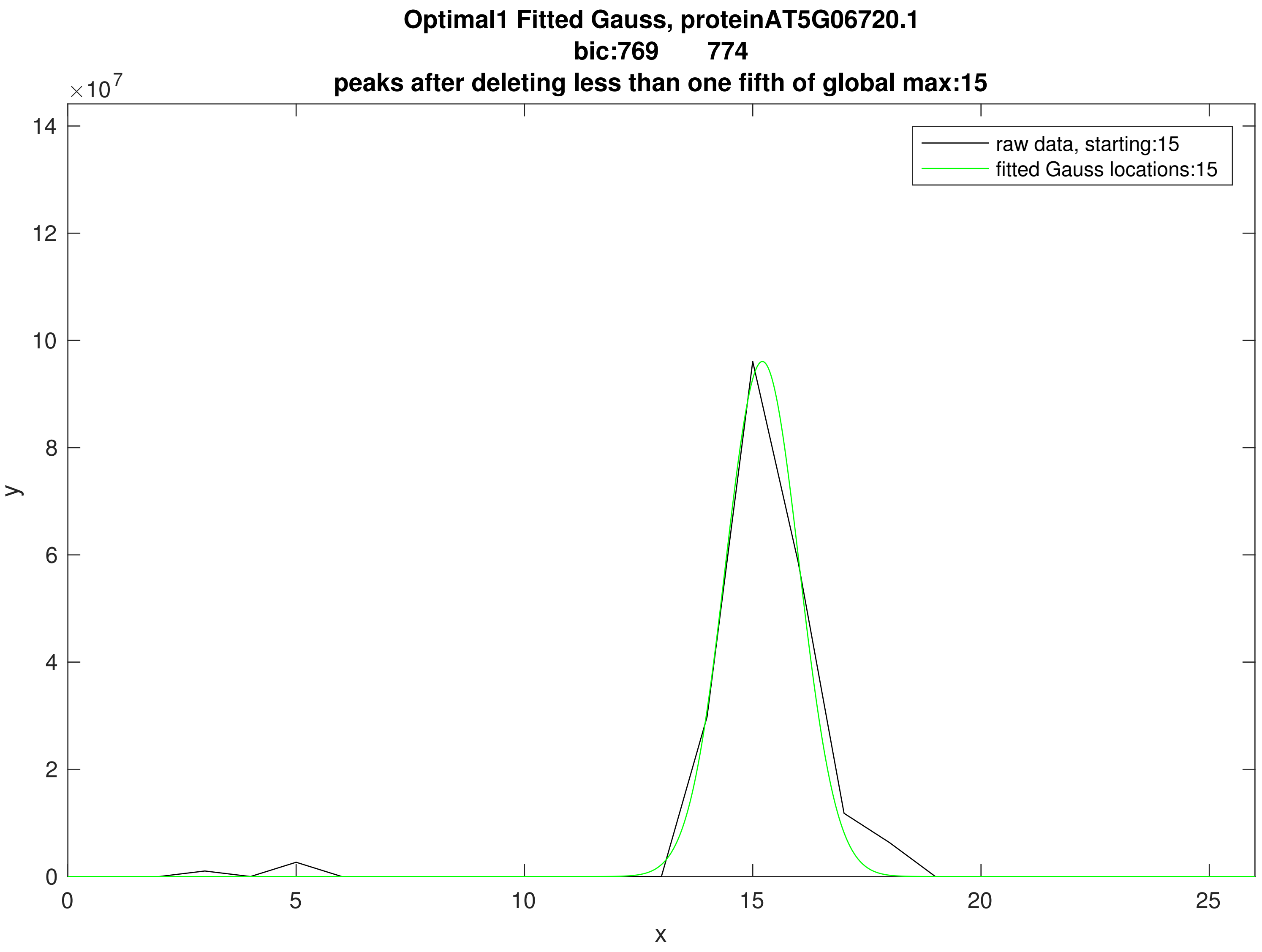
Optimal1 Fitted Gauss, proteinAT5G06720.1
bic:769 774
peaks after deleting less than one fifth of global max:15
×10^7
raw data, starting:15
fitted Gauss locations:15
y
x
0
2
4
6
8
10
12
14
0
5
10
15
20
25

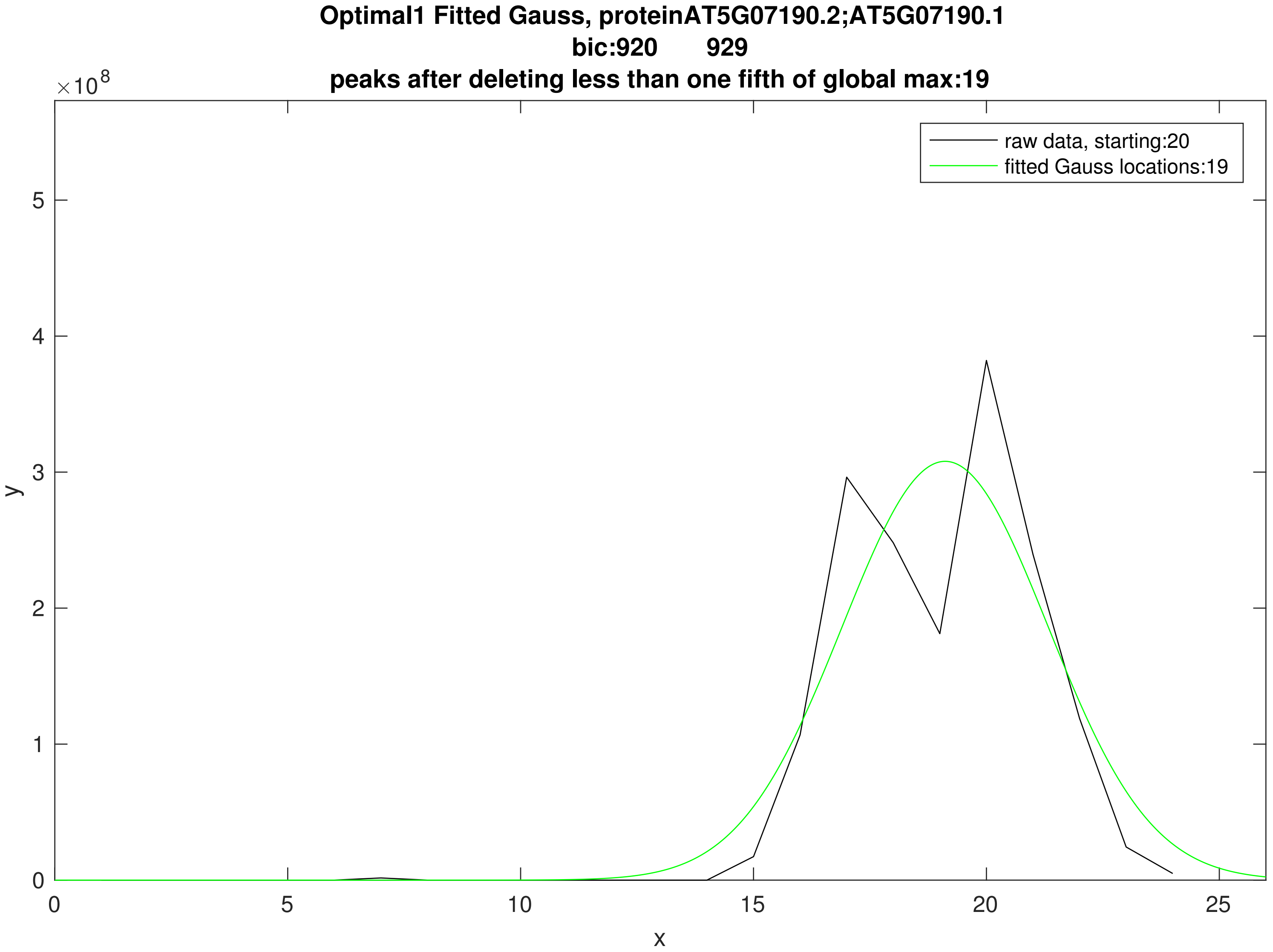
Optimal1 Fitted Gauss, proteinAT5G07190.2;AT5G07190.1
bic:920 929
peaks after deleting less than one fifth of global max:19
×10^8
raw data, starting:20
fitted Gauss locations:19
y
x
0
1
2
3
4
5
0
5
10
15
20
25

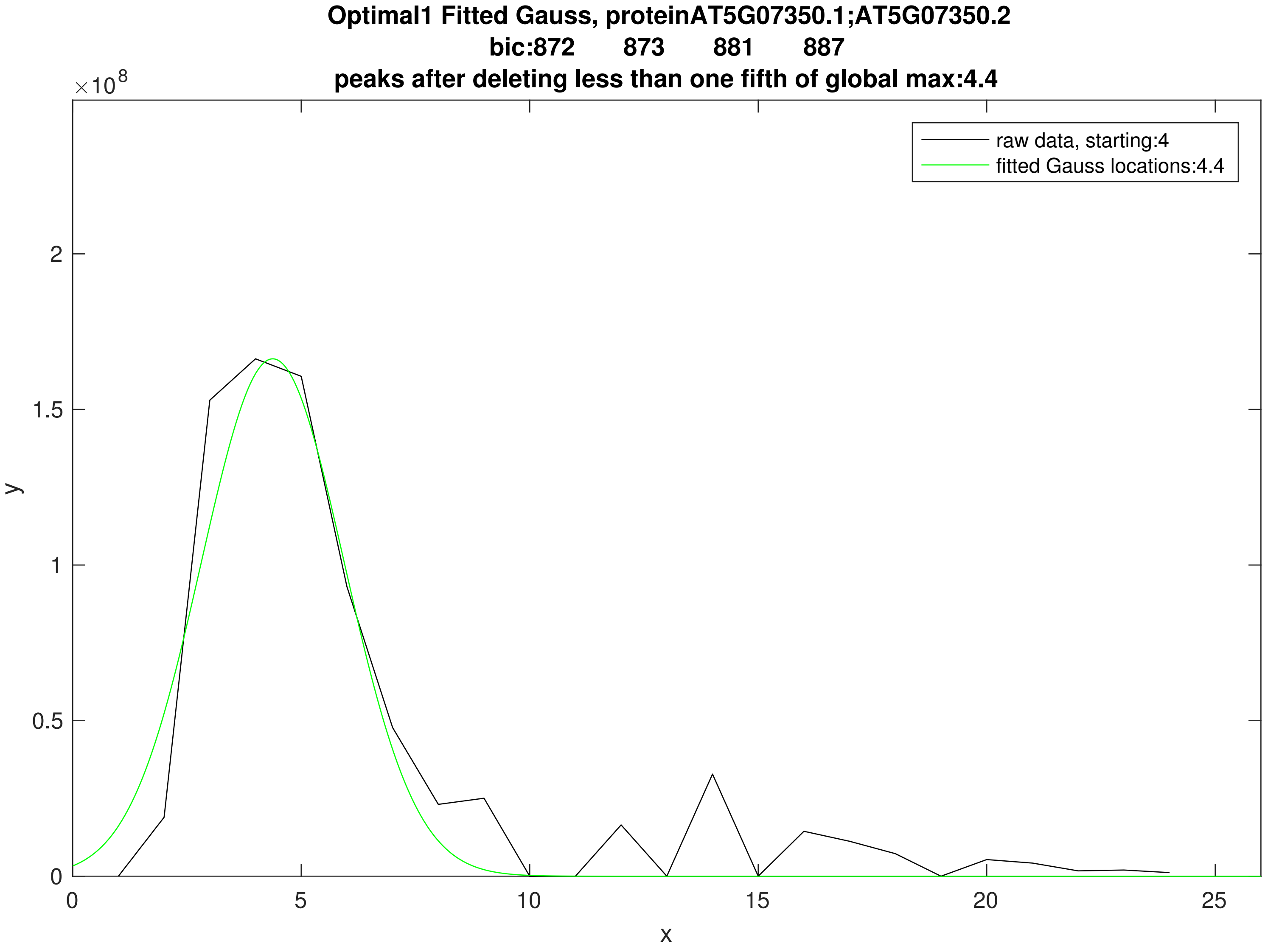
Optimal1 Fitted Gauss, proteinAT5G07350.1;AT5G07350.2
bic:872 873 881 887
peaks after deleting less than one fifth of global max:4.4
×10^8
raw data, starting:4
fitted Gauss locations:4.4
y
x
0
0.5
1
1.5
2
5
10
15
20
25

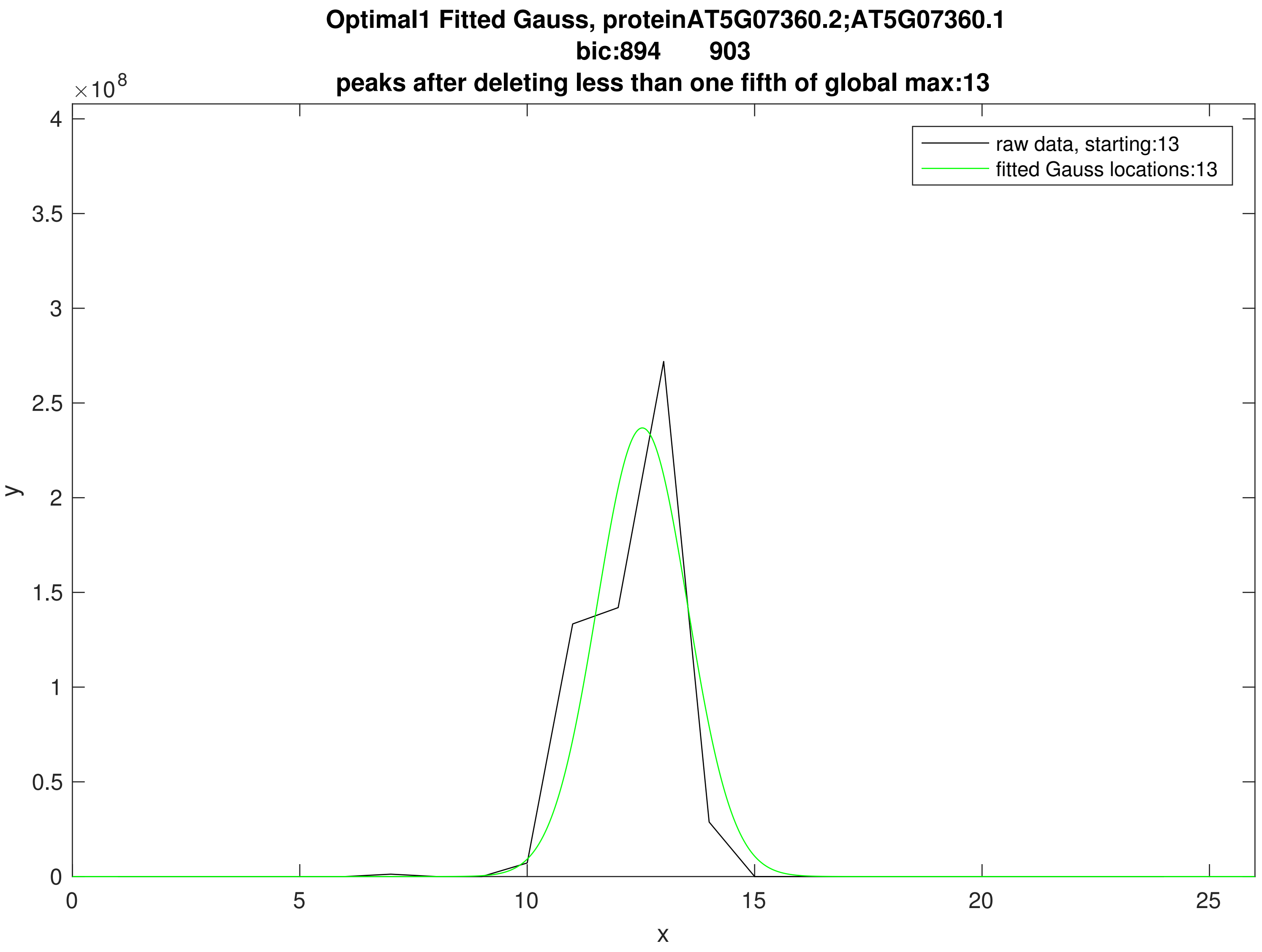

Optimal1 Fitted Gauss, proteinAT5G07360.2;AT5G07360.1
bic:894 903
peaks after deleting less than one fifth of global max:13
×10^8
raw data, starting:13
fitted Gauss locations:13
y
x
0
0.5
1
1.5
2
2.5
3
3.5
4
0
5
10
15
20
25

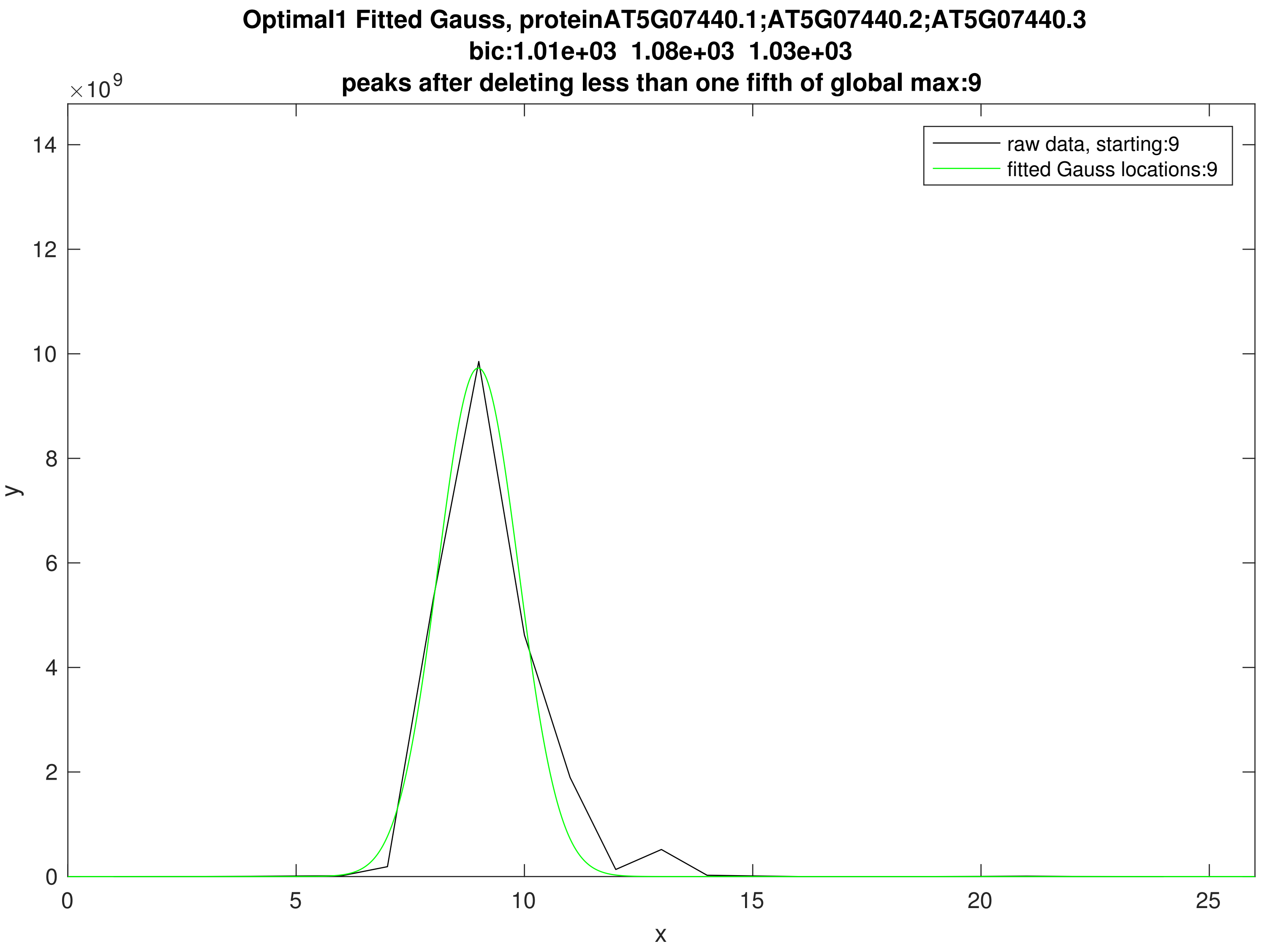
Optimal1 Fitted Gauss, proteinAT5G07440.1;AT5G07440.2;AT5G07440.3
bic:1.01e+03 1.08e+03 1.03e+03
peaks after deleting less than one fifth of global max:9
×10^9
raw data, starting:9
fitted Gauss locations:9
y
x
0
2
4
6
8
10
12
14
0
5
10
15
20
25

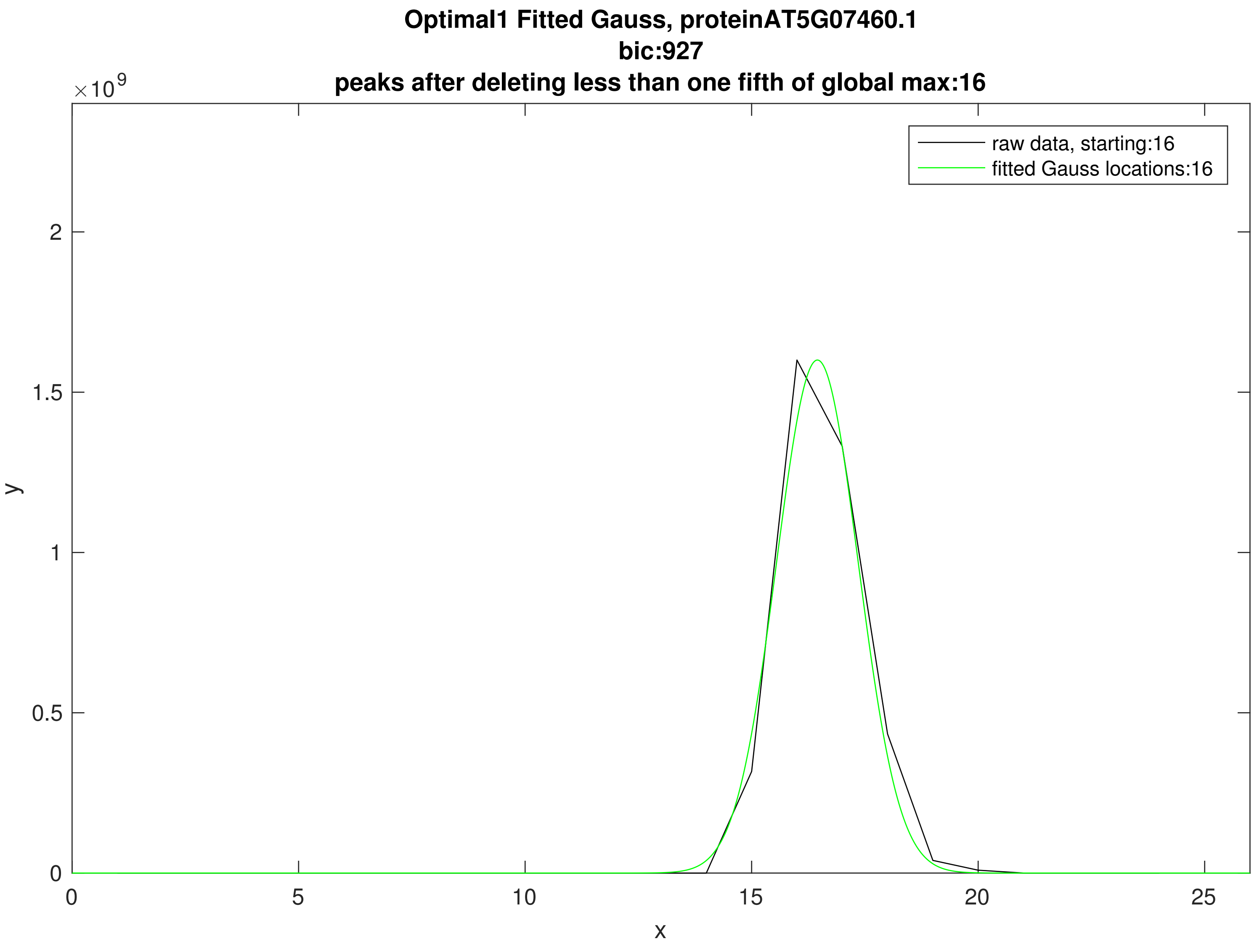
Optimal1 Fitted Gauss, proteinAT5G07460.1
bic:927
peaks after deleting less than one fifth of global max:16
×10^9
raw data, starting:16
fitted Gauss locations:16
y
x
0
0.5
1
1.5
2
5
10
15
20
25

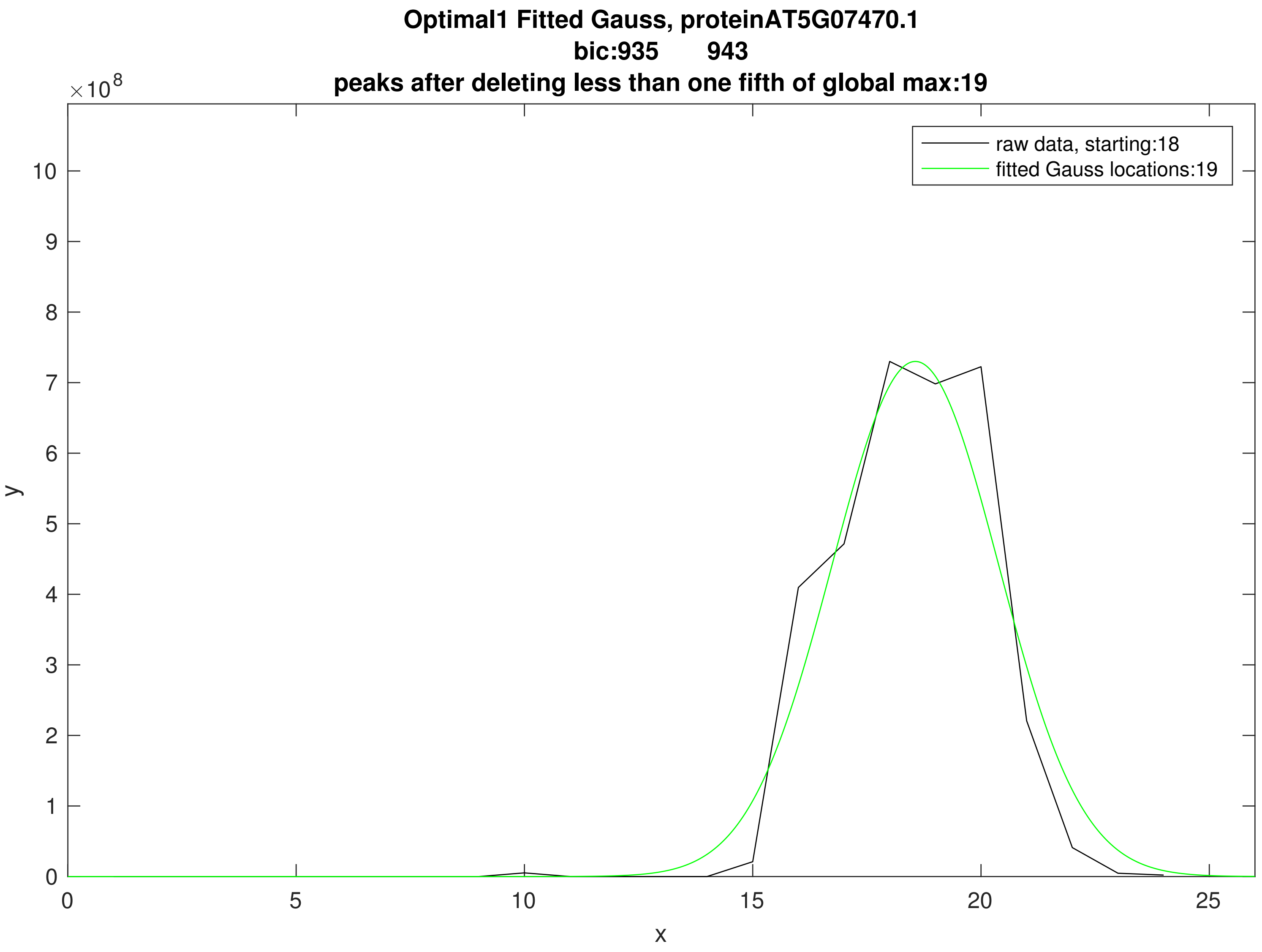
Optimal1 Fitted Gauss, proteinAT5G07470.1
bic:935 943
peaks after deleting less than one fifth of global max:19
×10^8
raw data, starting:18
fitted Gauss locations:19
y
x

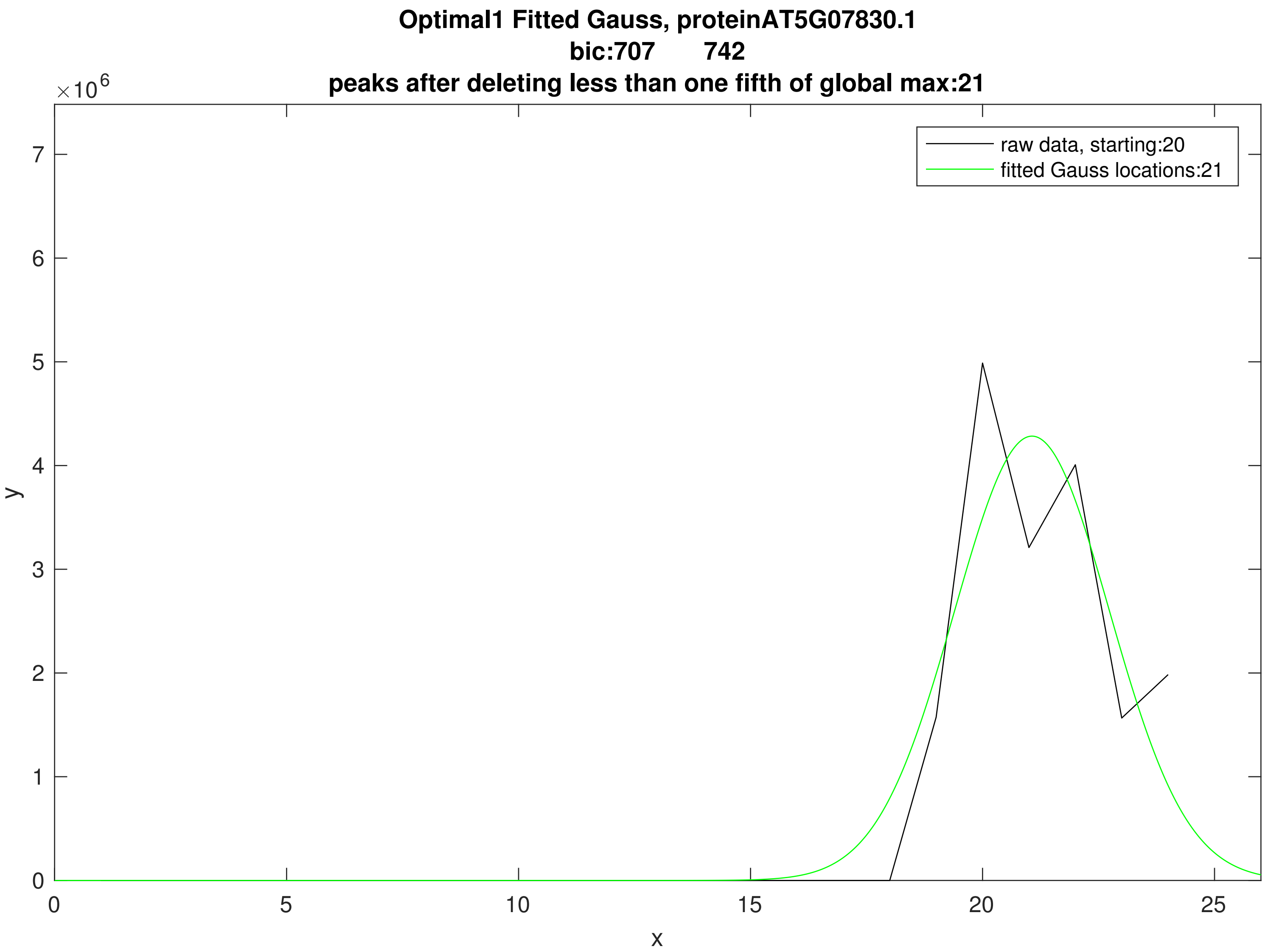
Optimal1 Fitted Gauss, proteinAT5G07830.1
bic:707 742
peaks after deleting less than one fifth of global max:21
$\times 10^6$
raw data, starting:20
fitted Gauss locations:21
y
x
0
1
2
3
4
5
6
7
0
5
10
15
20
25

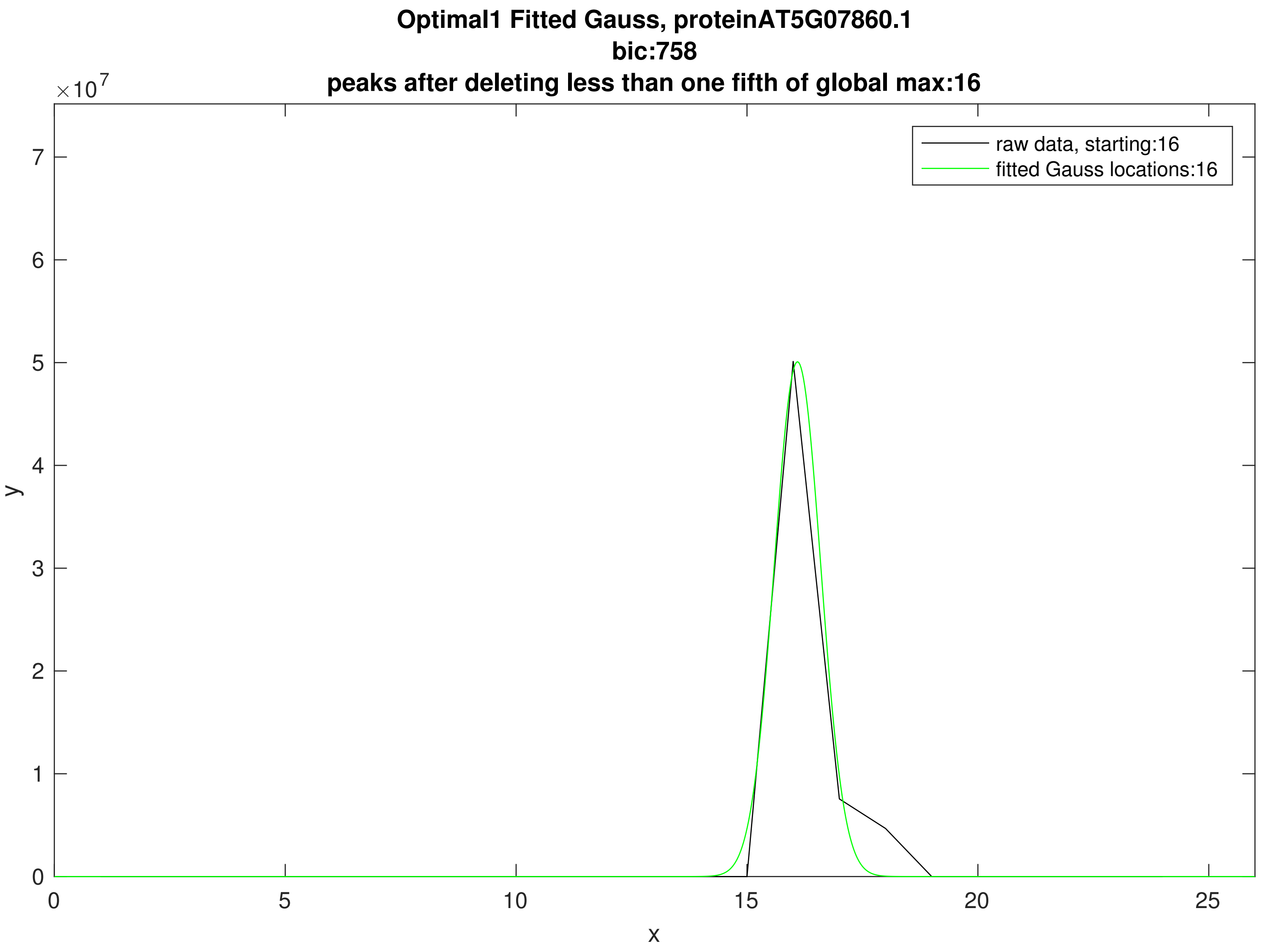
Optimal1 Fitted Gauss, proteinAT5G07860.1
bic:758
peaks after deleting less than one fifth of global max:16
×10^7
raw data, starting:16
fitted Gauss locations:16
y
x
0
1
2
3
4
5
6
7
0
5
10
15
20
25

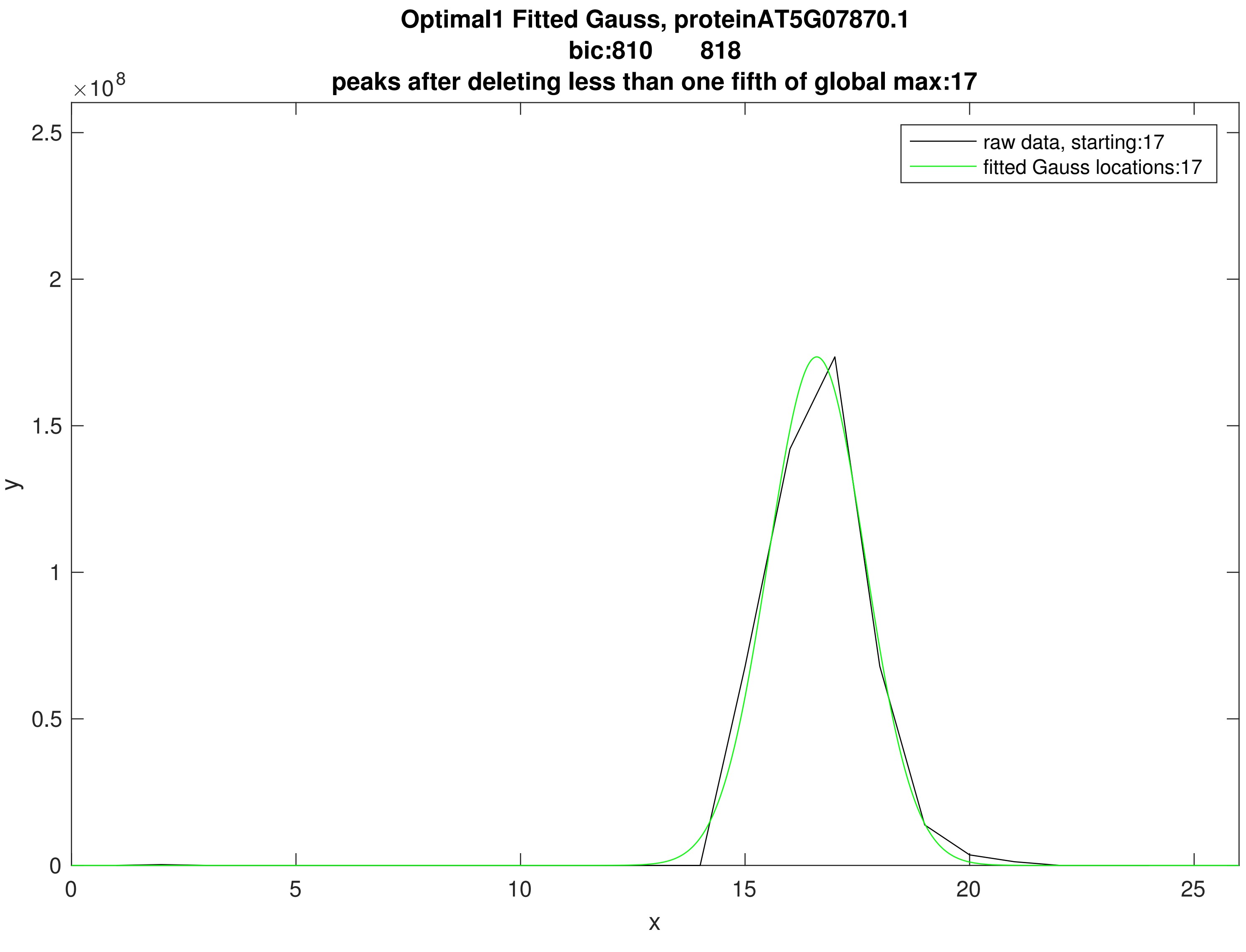
Optimal1 Fitted Gauss, proteinAT5G07870.1
bic:810 818
peaks after deleting less than one fifth of global max:17
×10^8
2.5
2
1.5
1
0.5
0
y
0
5
10
15
20
25
x
raw data, starting:17
fitted Gauss locations:17

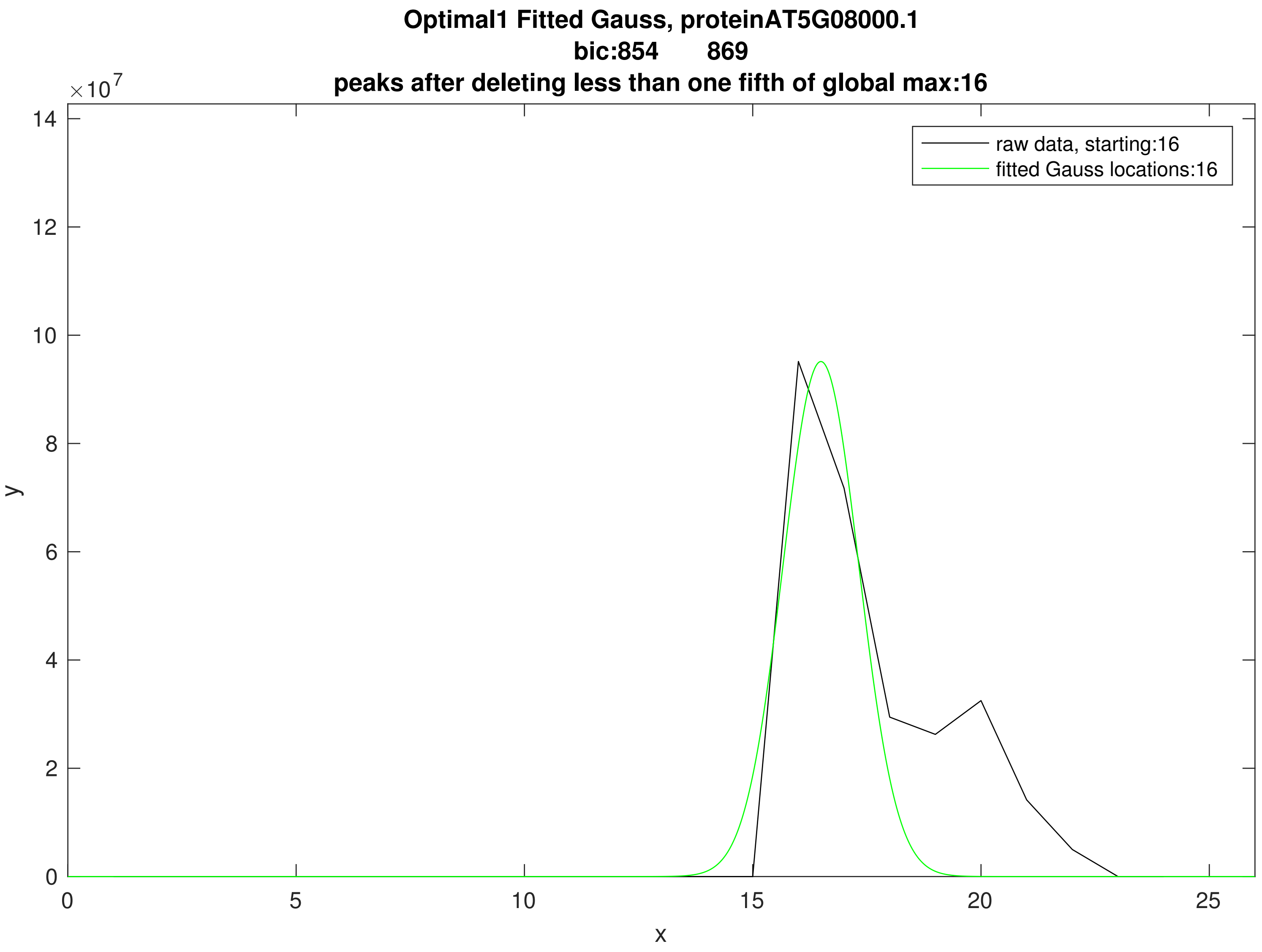
Optimal1 Fitted Gauss, proteinAT5G08000.1
bic:854 869
peaks after deleting less than one fifth of global max:16
×10^7
raw data, starting:16
fitted Gauss locations:16
y
x
0
2
4
6
8
10
12
14
5
15
20
25

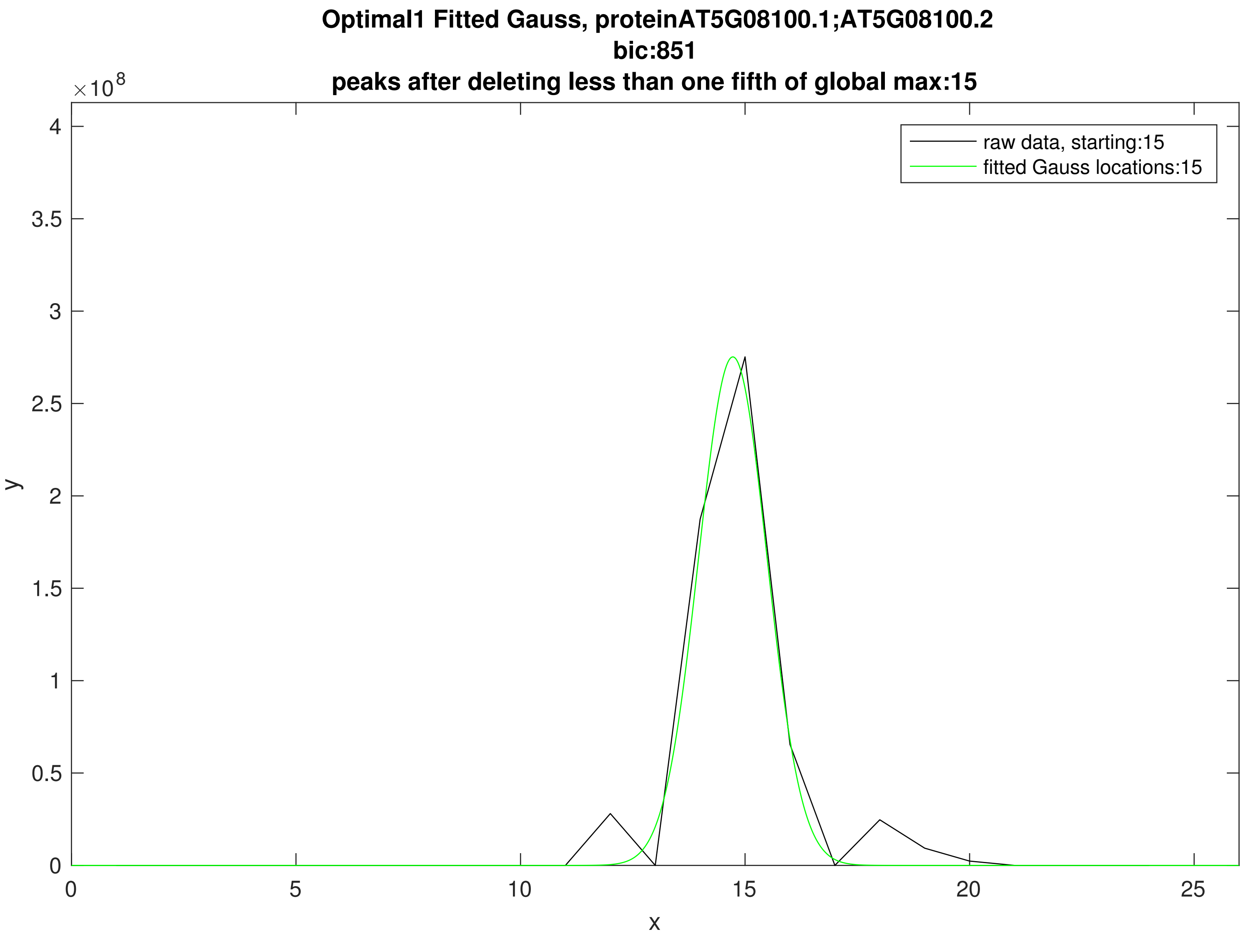
Optimal1 Fitted Gauss, proteinAT5G08100.1;AT5G08100.2
bic:851
peaks after deleting less than one fifth of global max:15
×10^8
4
3.5
3
2.5
2
1.5
1
0.5
0
y
0
5
10
15
20
25
x
raw data, starting:15
fitted Gauss locations:15

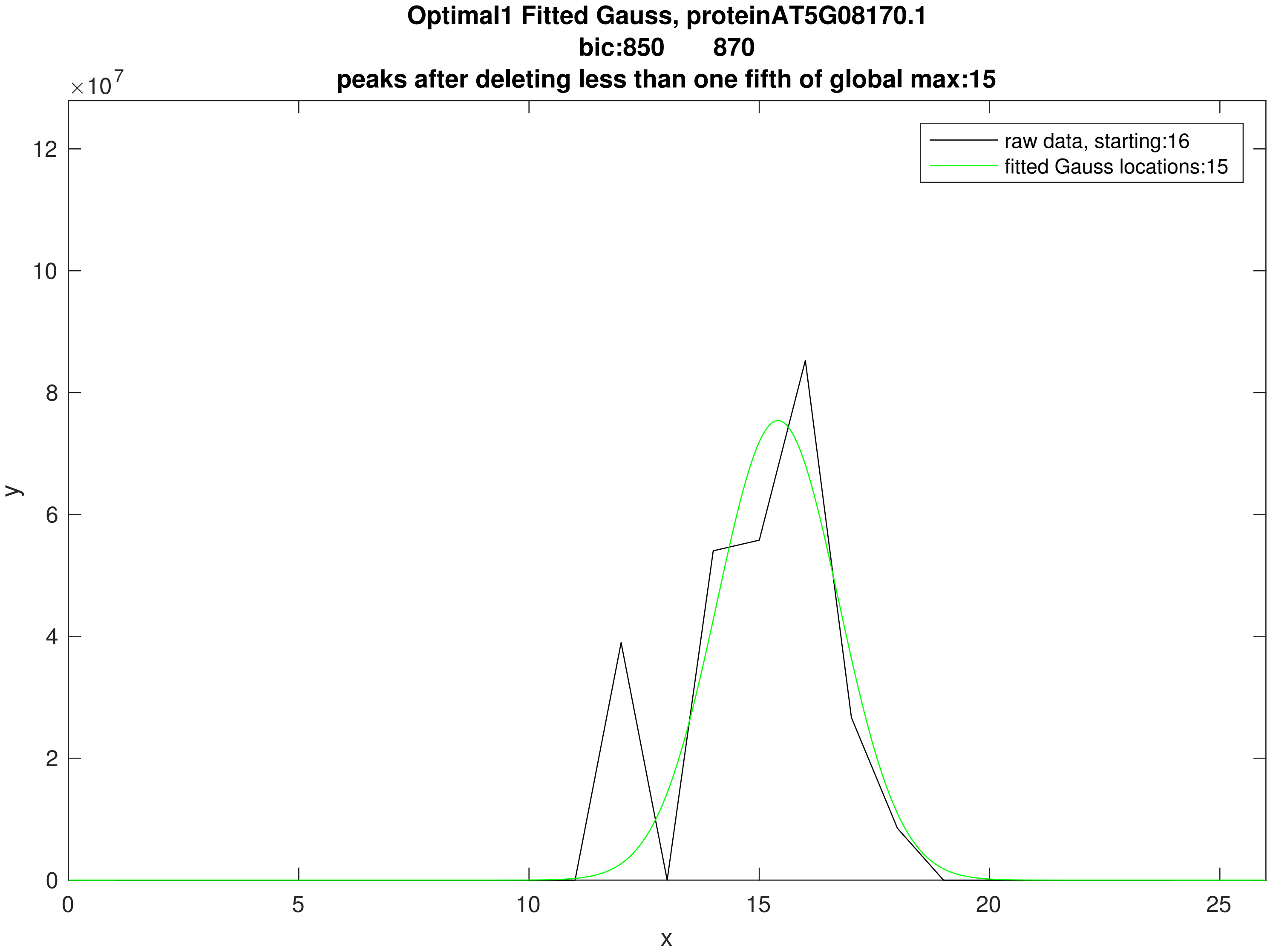

Optimal1 Fitted Gauss, proteinAT5G08170.1
bic:850 870
peaks after deleting less than one fifth of global max:15
×10^7
raw data, starting:16
fitted Gauss locations:15
y
x
0
2
4
6
8
10
12
0
5
10
15
20
25

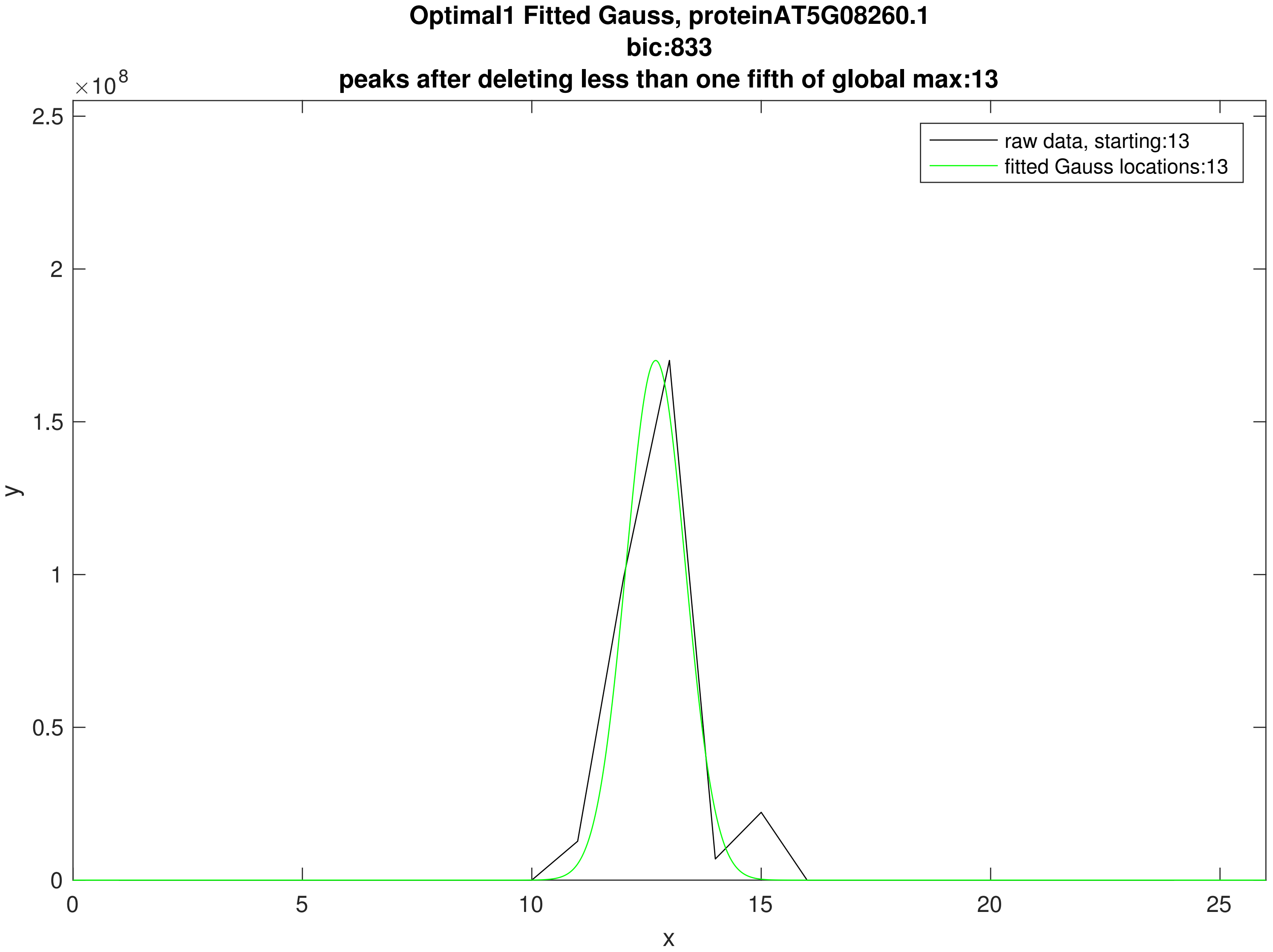

Optimal1 Fitted Gauss, proteinAT5G08260.1
bic:833
peaks after deleting less than one fifth of global max:13
×10^8
2.5
2
1.5
1
0.5
0
y
0
5
10
15
20
25
x
raw data, starting:13
fitted Gauss locations:13

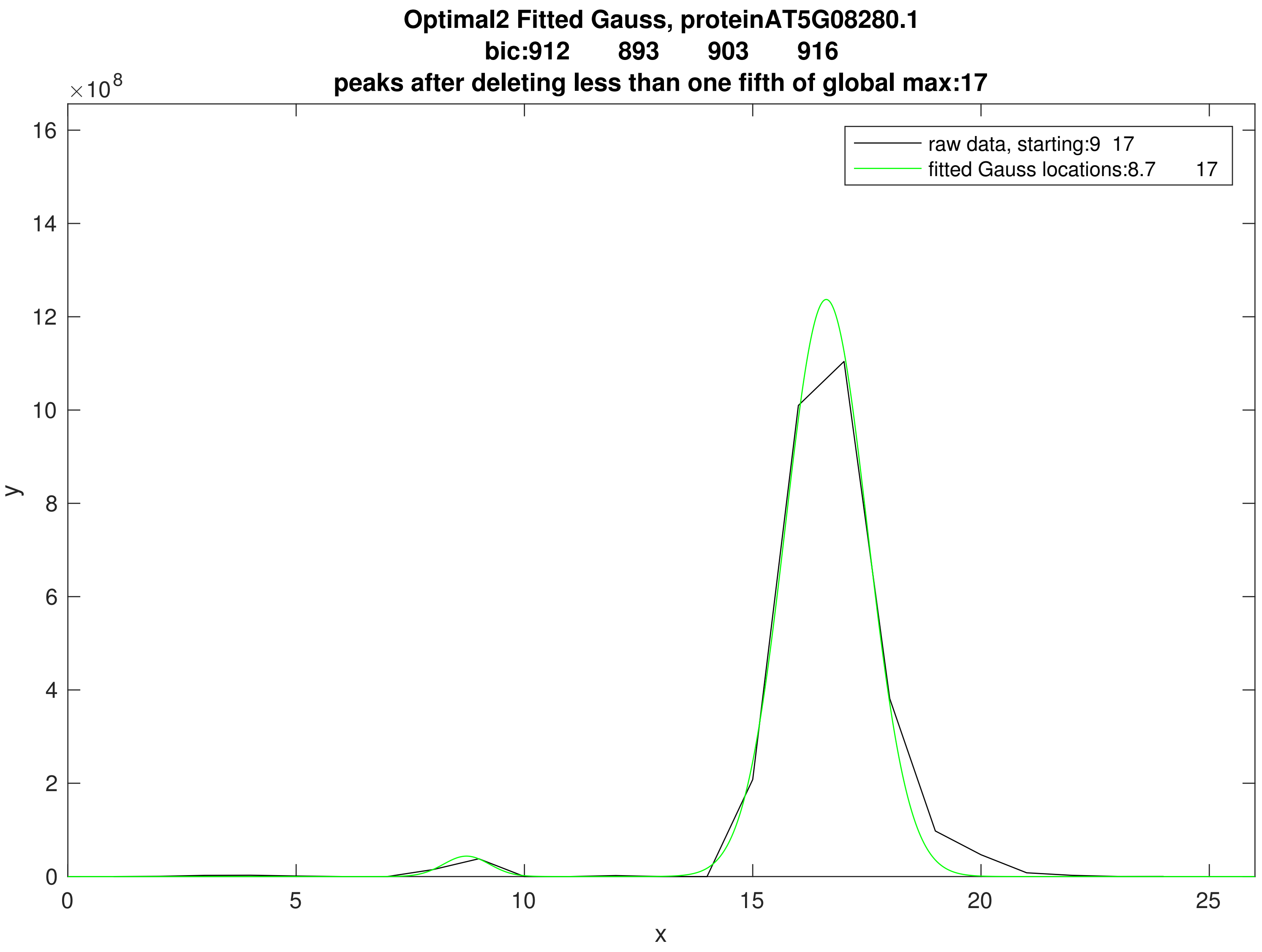

Optimal2 Fitted Gauss, proteinAT5G08280.1
bic:912 893 903 916
peaks after deleting less than one fifth of global max:17
×10^8
raw data, starting:9 17
fitted Gauss locations:8.7 17
y
x
0
2
4
6
8
10
12
14
16
0
5
10
15
20
25

**Optimal1 Fitted Gauss, proteinAT5G08300.1**
**bic:1.06e+03  1.07e+03**
**peaks after deleting less than one fifth of global max:15**

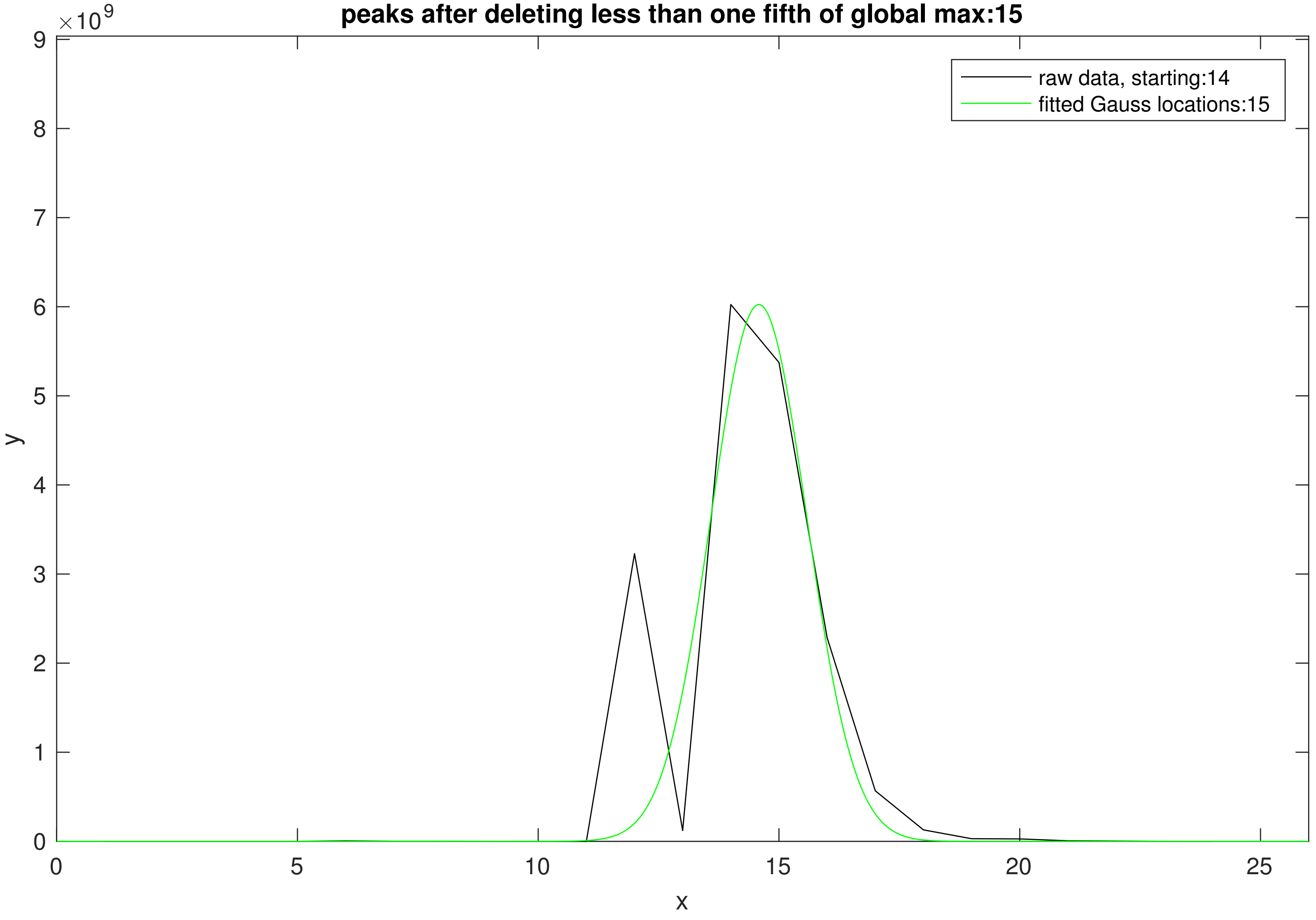

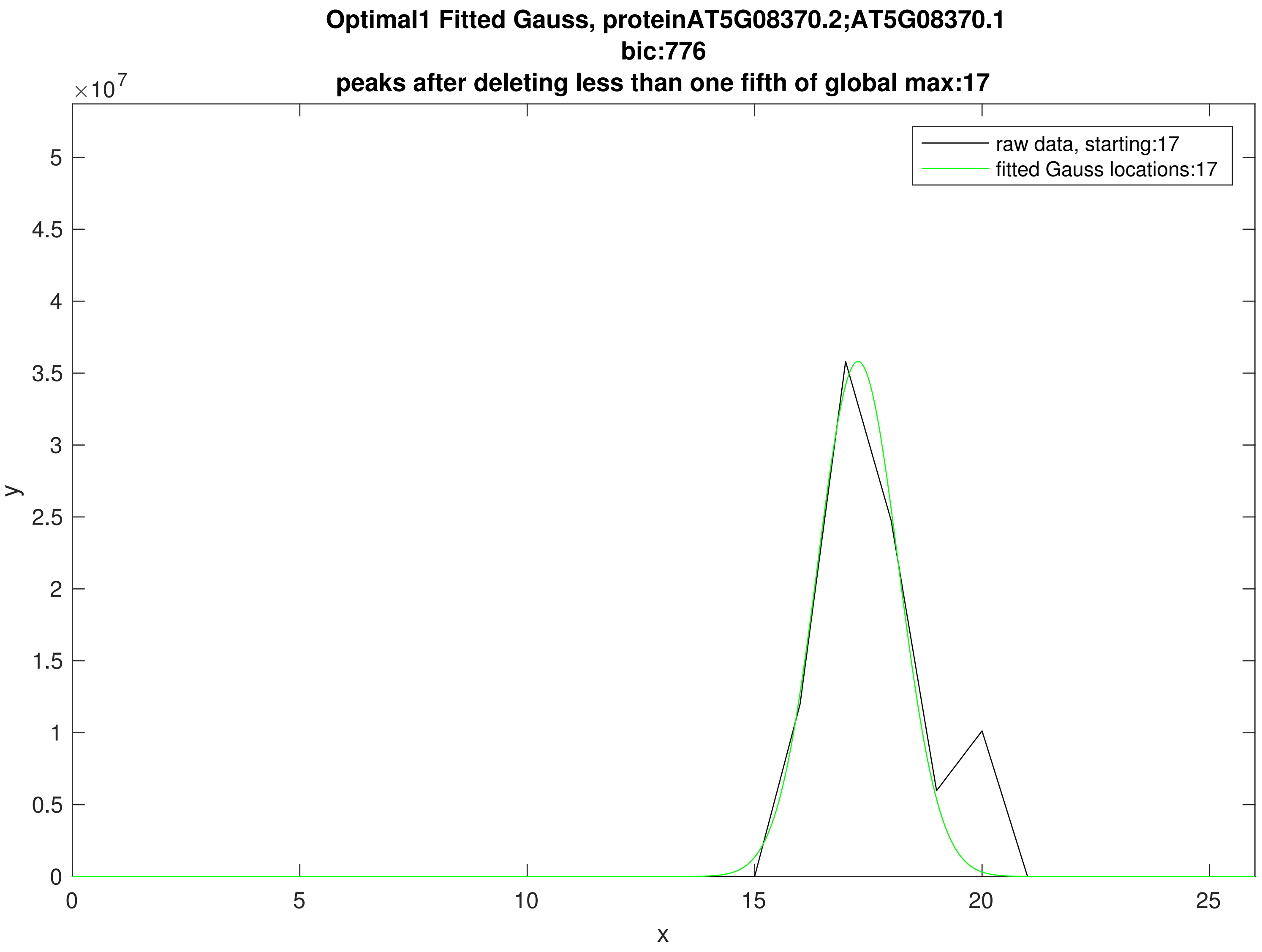
Optimal1 Fitted Gauss, proteinAT5G08370.2;AT5G08370.1
bic:776
peaks after deleting less than one fifth of global max:17
$\times 10^7$
raw data, starting:17
fitted Gauss locations:17
y
x
0
0.5
1
1.5
2
2.5
3
3.5
4
4.5
5
0
5
10
15
20
25

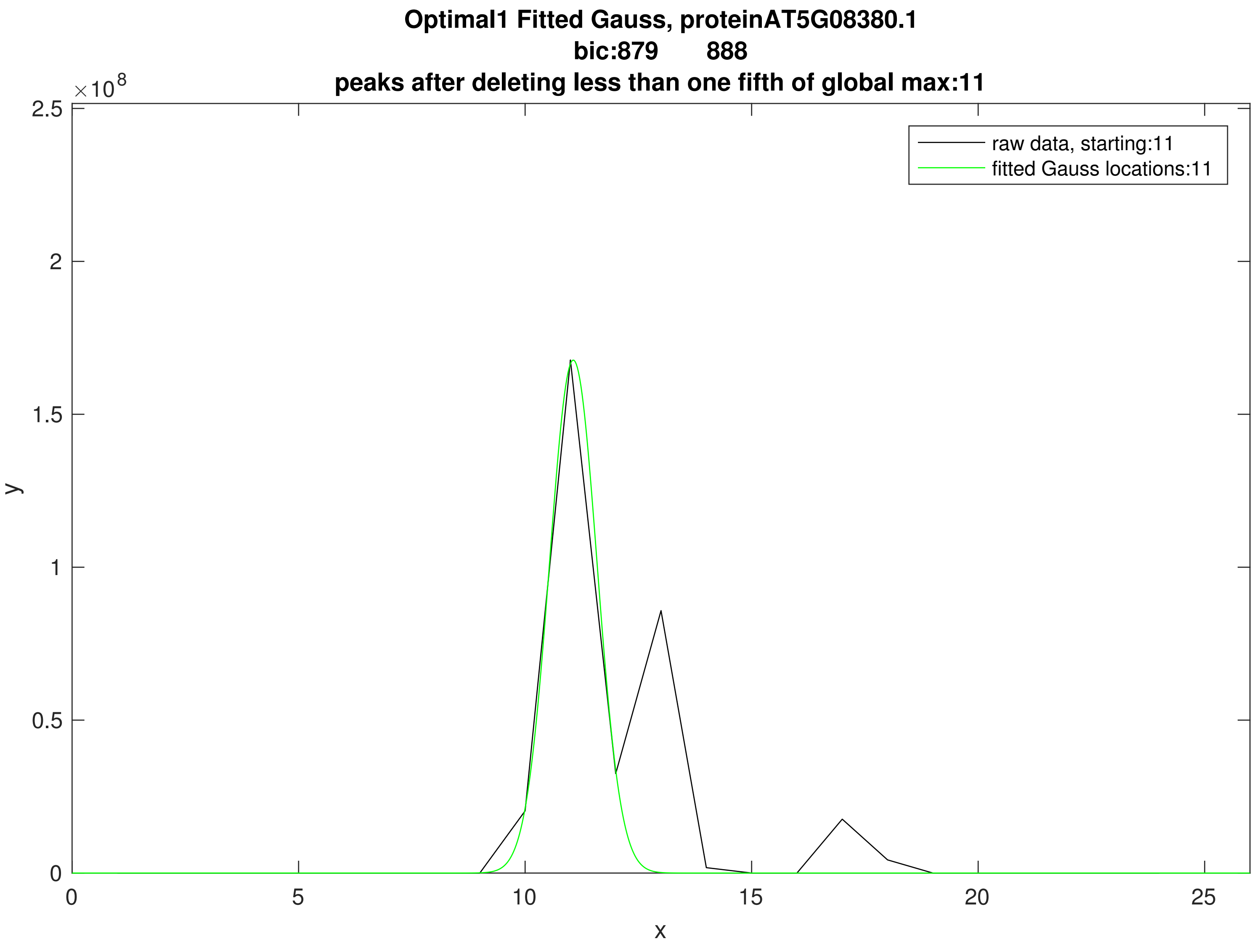
Optimal1 Fitted Gauss, proteinAT5G08380.1
bic:879 888
peaks after deleting less than one fifth of global max:11
×10^8
2.5
2
1.5
1
0.5
0
y
0
5
10
15
20
25
x
raw data, starting:11
fitted Gauss locations:11

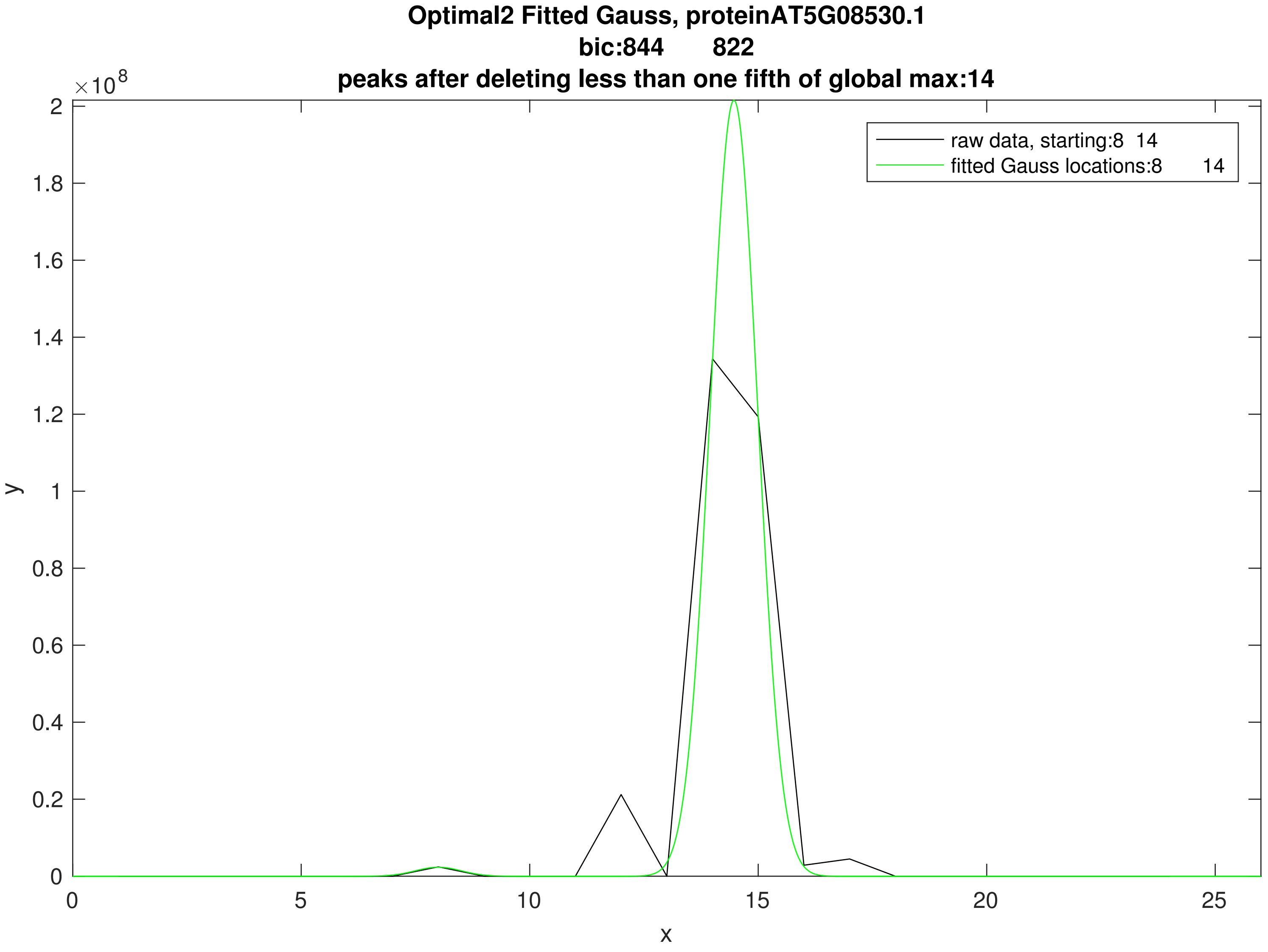

Optimal2 Fitted Gauss, proteinAT5G08530.1
bic:844 822
peaks after deleting less than one fifth of global max:14
×10^8
2
1.8
1.6
1.4
1.2
1
0.8
0.6
0.4
0.2
0
y
raw data, starting:8 14
fitted Gauss locations:8 14
0
5
10
15
20
25
x

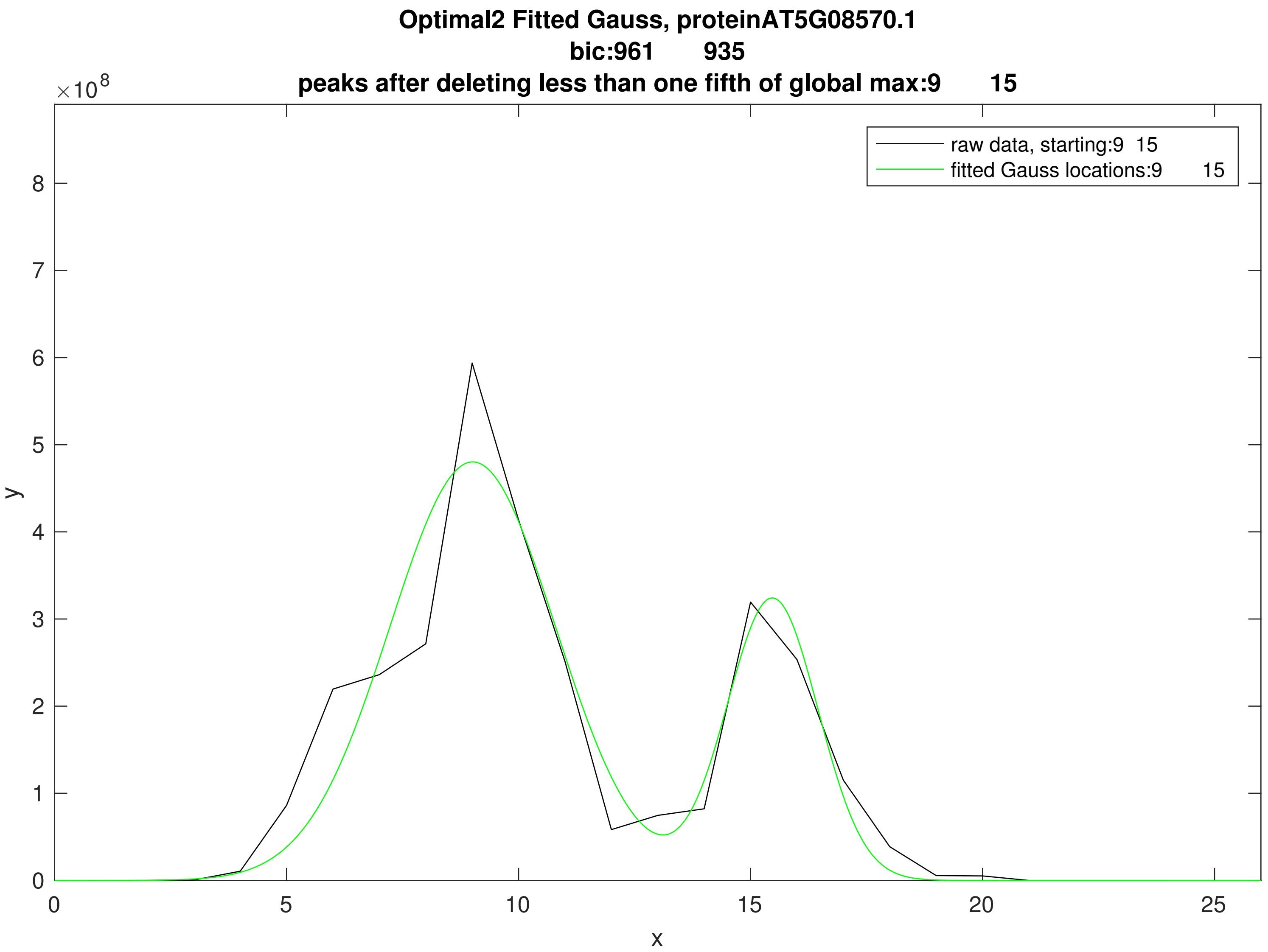
Optimal2 Fitted Gauss, proteinAT5G08570.1
bic:961 935
peaks after deleting less than one fifth of global max:9 15
×10^8
raw data, starting:9 15
fitted Gauss locations:9 15
y
x
0
1
2
3
4
5
6
7
8
0
5
10
15
20
25

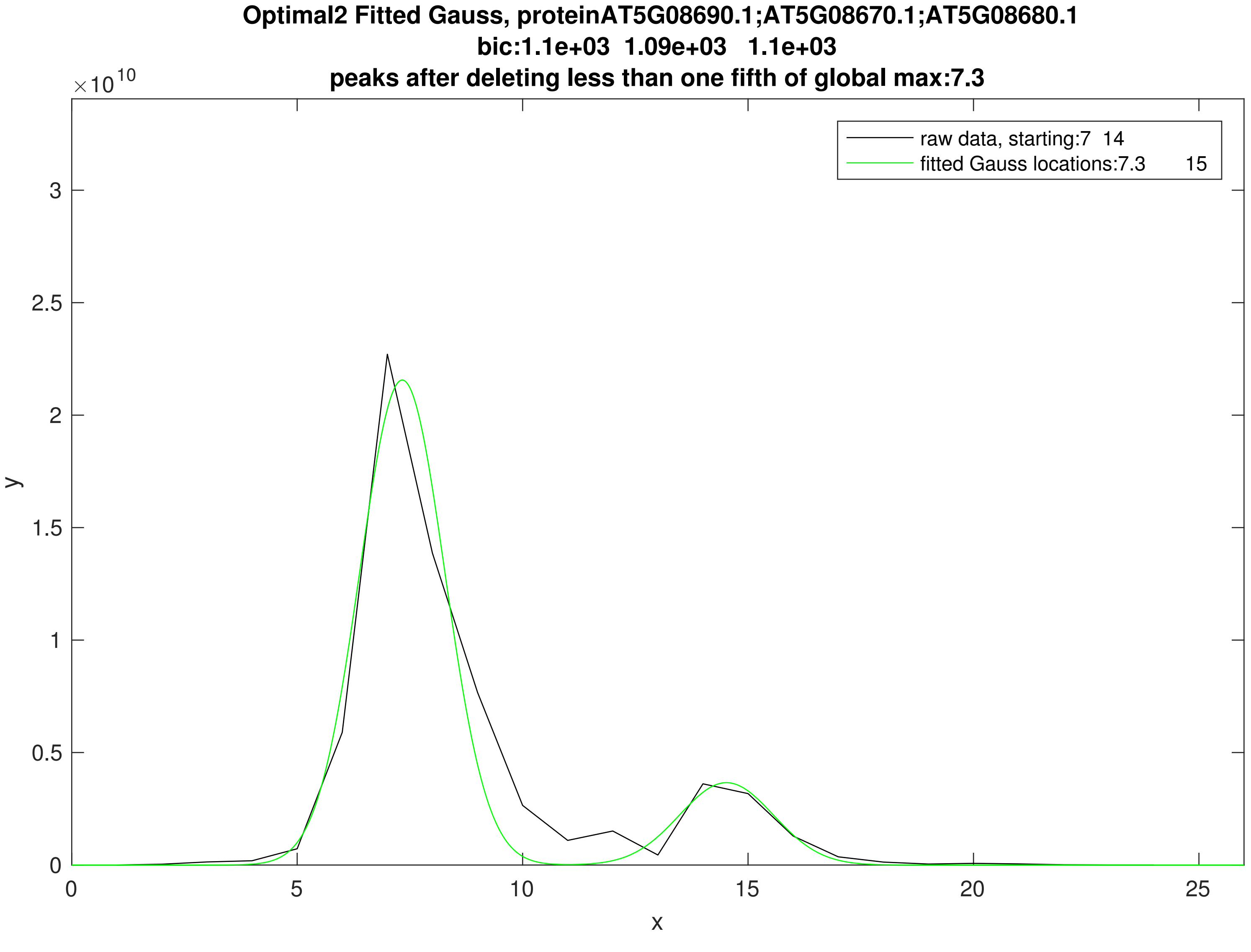

Optimal2 Fitted Gauss, proteinAT5G08690.1;AT5G08670.1;AT5G08680.1
bic:1.1e+03 1.09e+03 1.1e+03
peaks after deleting less than one fifth of global max:7.3
×10^10
raw data, starting:7 14
fitted Gauss locations:7.3 15
y
x
0
0.5
1
1.5
2
2.5
3
5
10
15
20
25

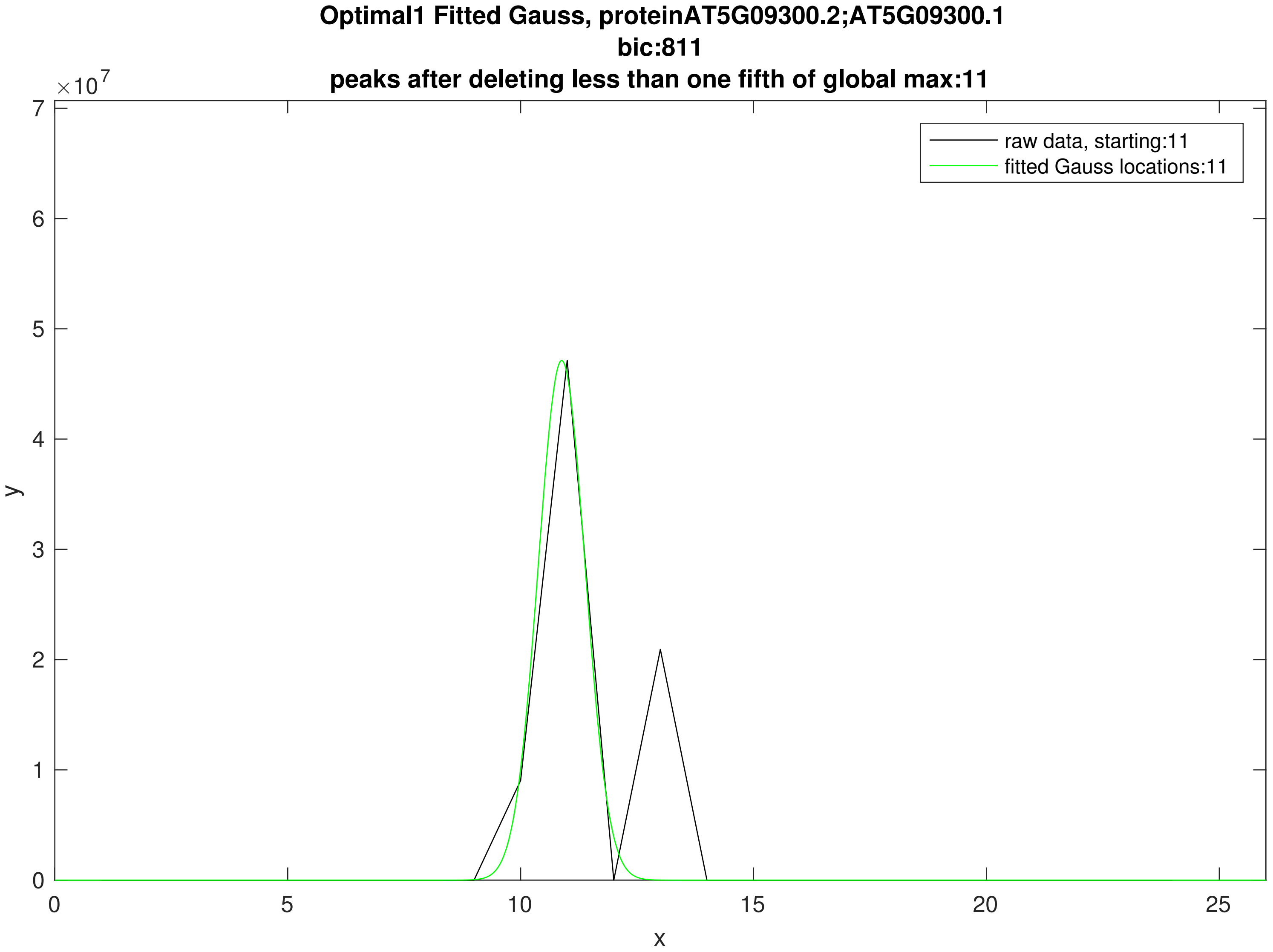

Optimal1 Fitted Gauss, proteinAT5G09300.2;AT5G09300.1
bic:811
peaks after deleting less than one fifth of global max:11
×10^7
raw data, starting:11
fitted Gauss locations:11
y
x
0
1
2
3
4
5
6
7
0
5
10
15
20
25

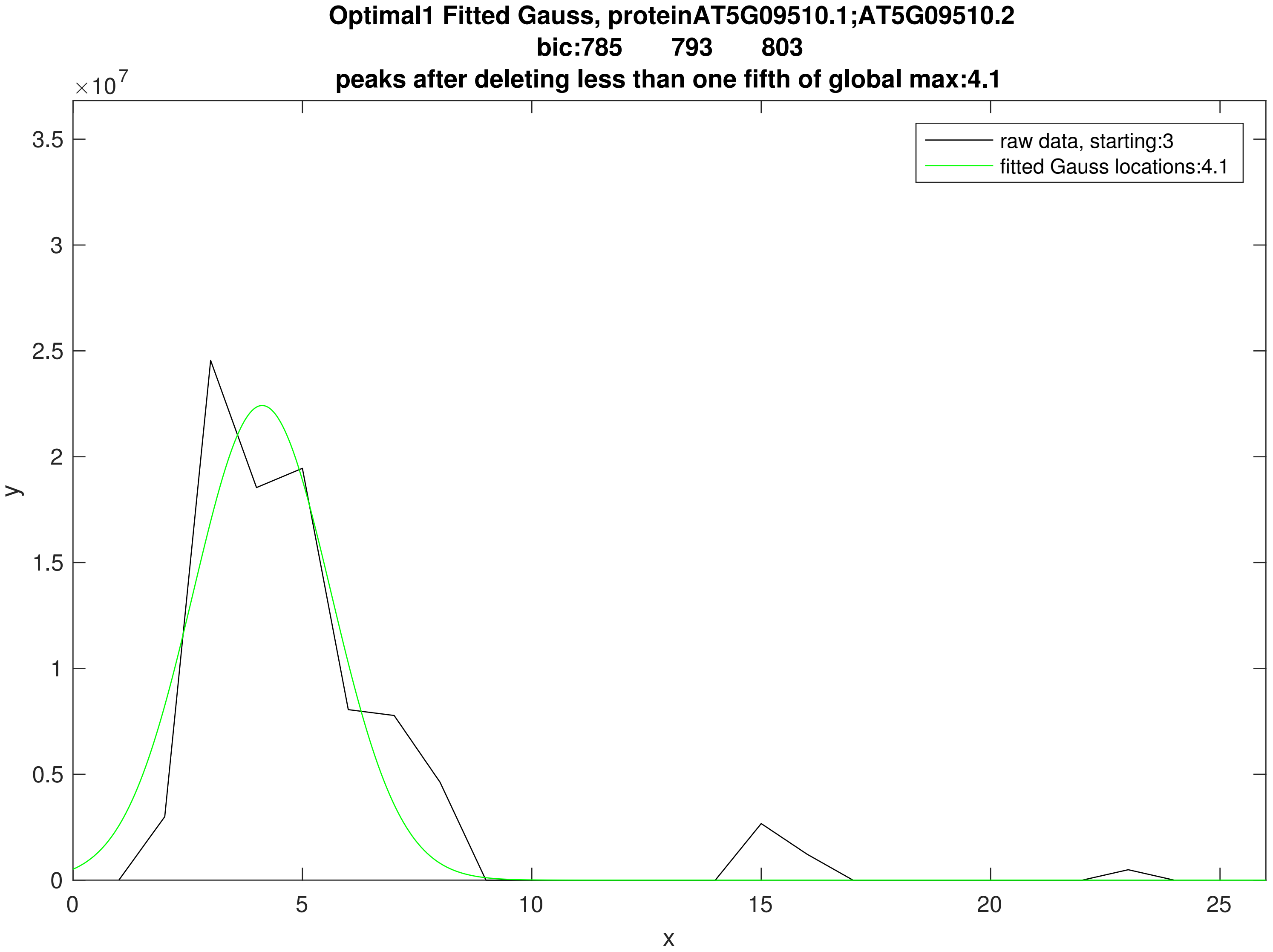

Optimal1 Fitted Gauss, proteinAT5G09510.1;AT5G09510.2
bic:785 793 803
peaks after deleting less than one fifth of global max:4.1
×10^7
3.5
3
2.5
2
1.5
1
0.5
0
y
0
5
10
15
20
25
x
raw data, starting:3
fitted Gauss locations:4.1

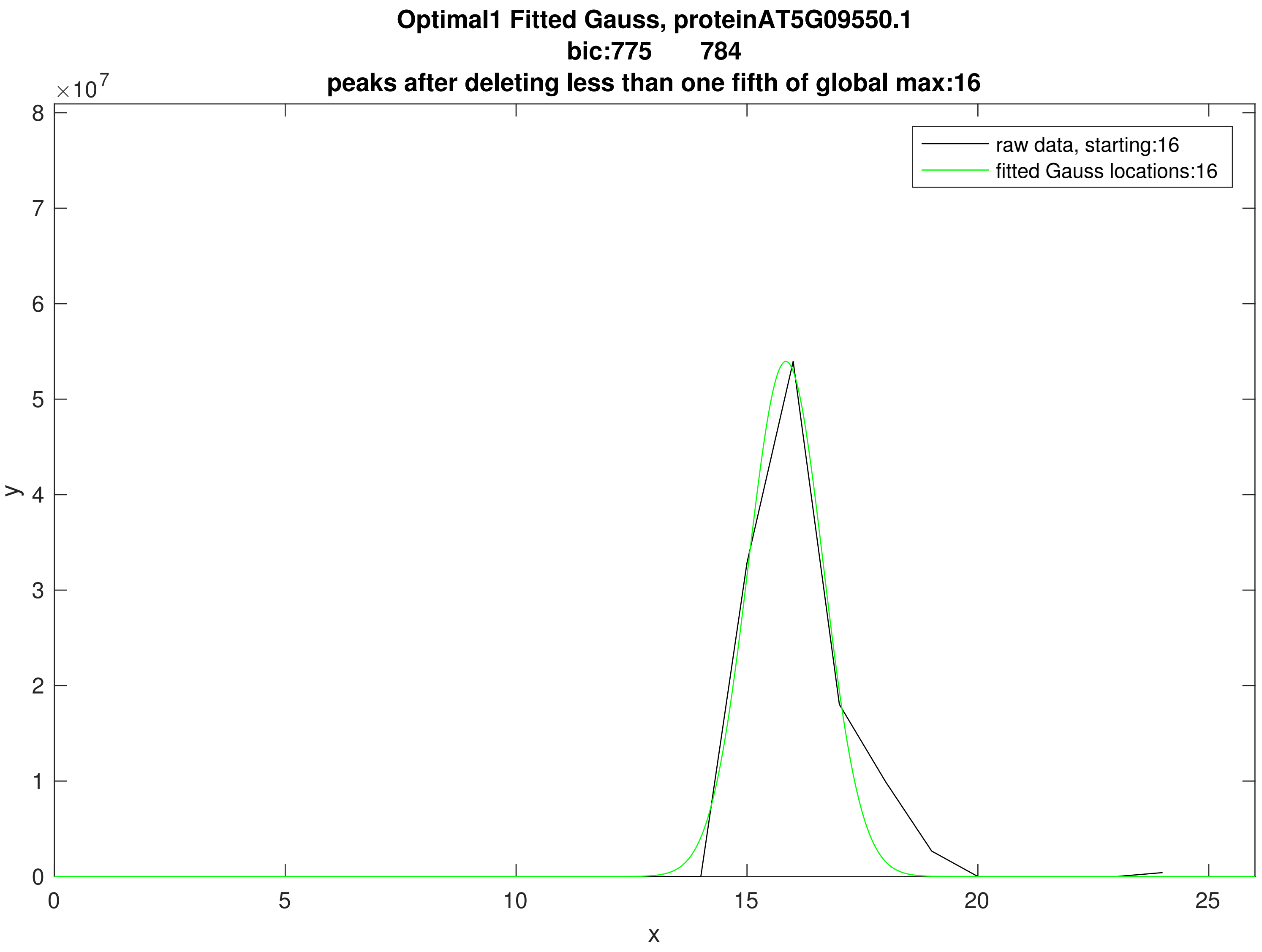
Optimal1 Fitted Gauss, proteinAT5G09550.1
bic:775 784
peaks after deleting less than one fifth of global max:16
×10^7
8
7
6
5
4
3
2
1
0
y
0
5
10
15
20
25
x
raw data, starting:16
fitted Gauss locations:16

**Optimal3 Fitted Gauss, proteinAT5G09590.1**
**bic:972 963 931 940**
**peaks after deleting less than one fifth of global max:8.6 13**

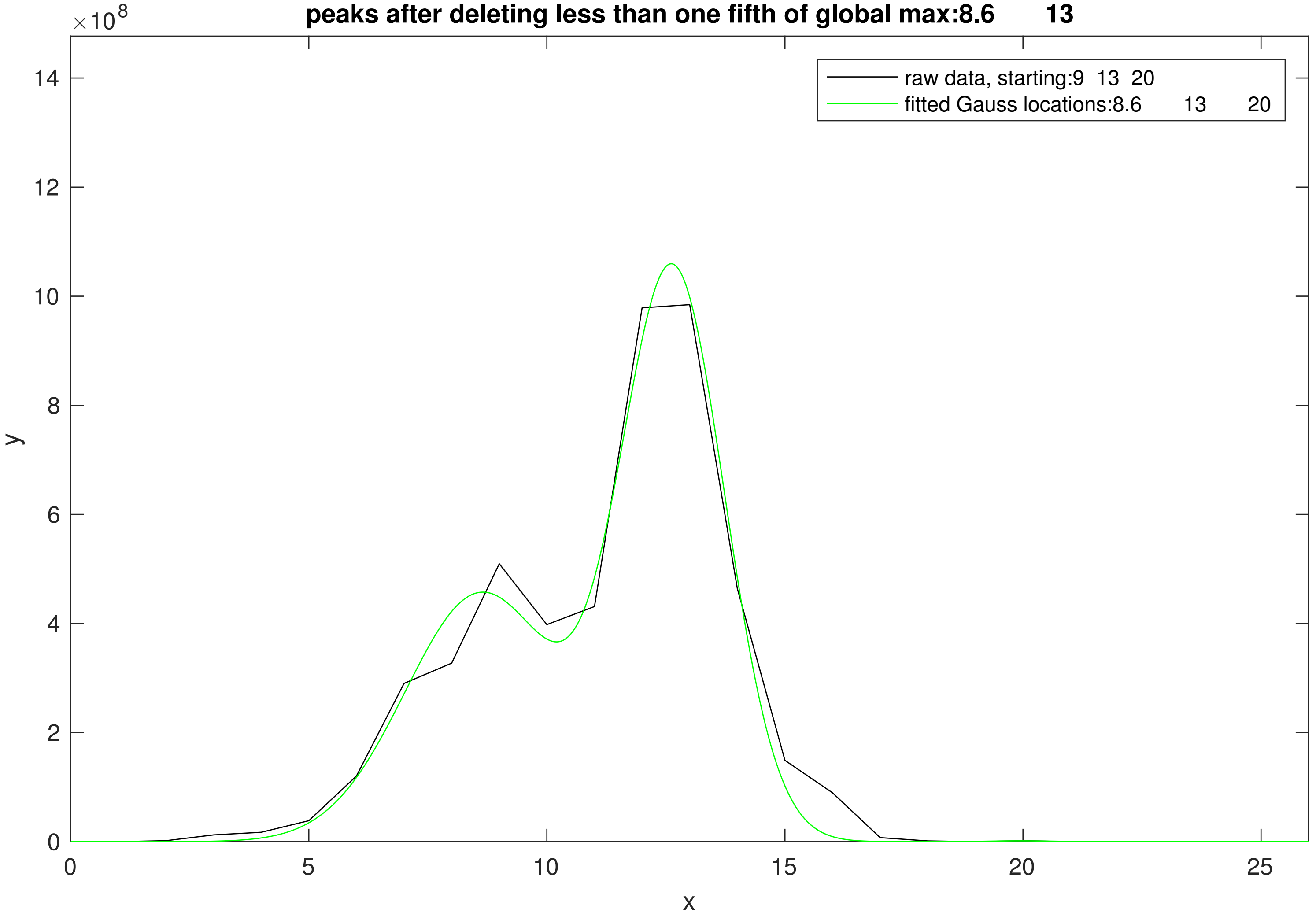

**Optimal1 Fitted Gauss, proteinAT5G09620.1**
**bic:821 828**
**peaks after deleting less than one fifth of global max:11**

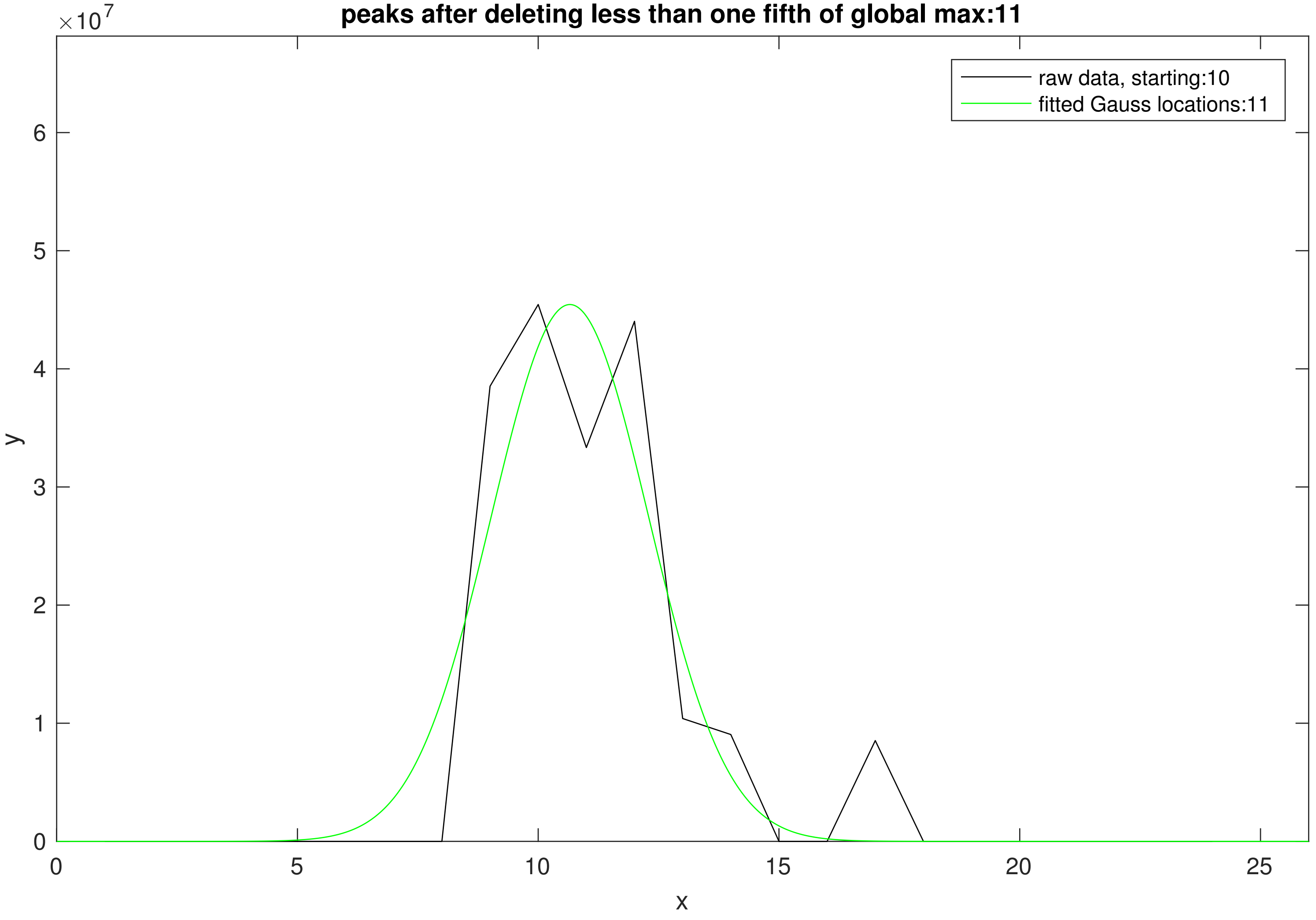

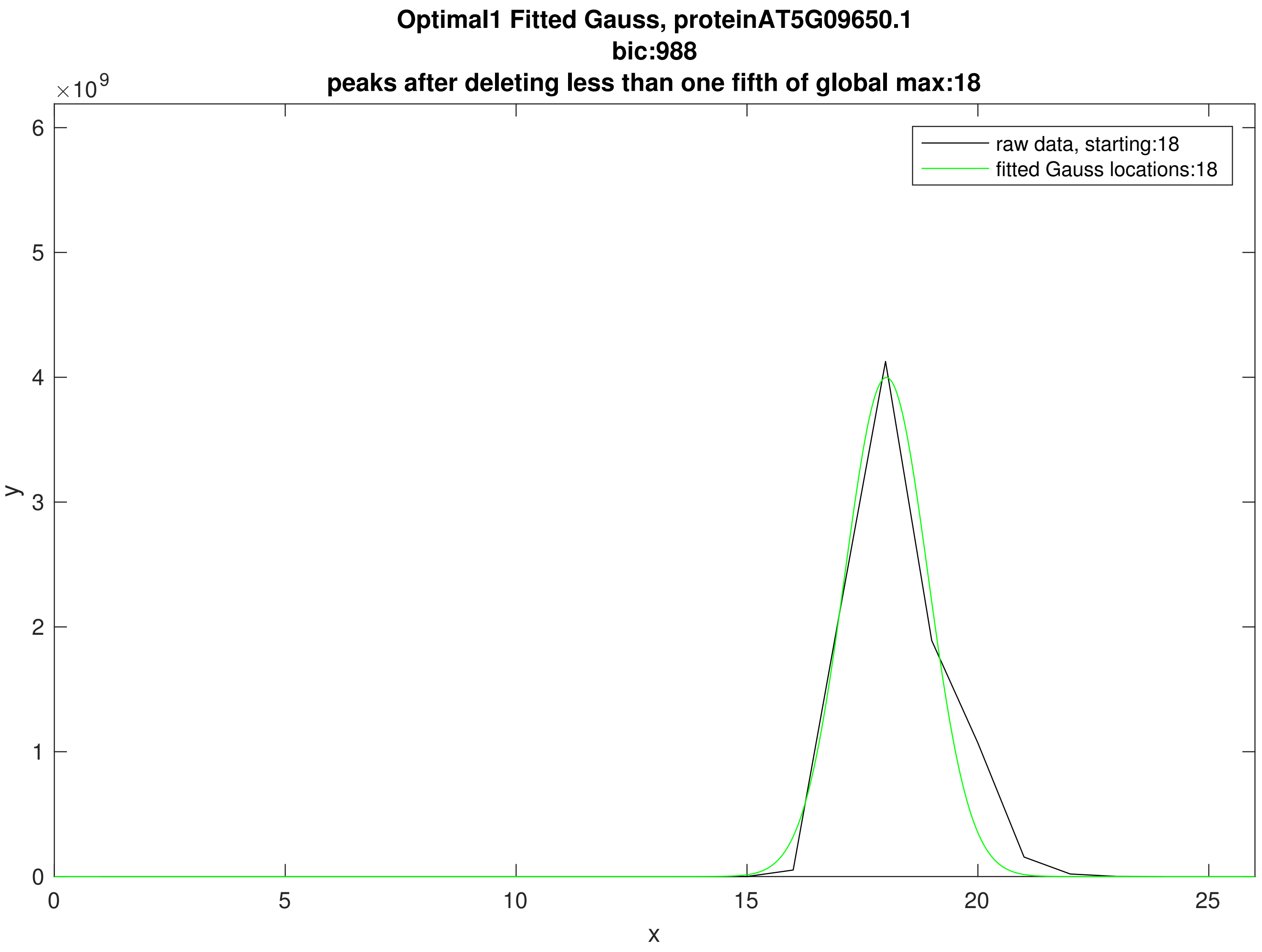
Optimal1 Fitted Gauss, proteinAT5G09650.1
bic:988
peaks after deleting less than one fifth of global max:18
×10^9
raw data, starting:18
fitted Gauss locations:18
y
x
0
1
2
3
4
5
6
0
5
10
15
20
25

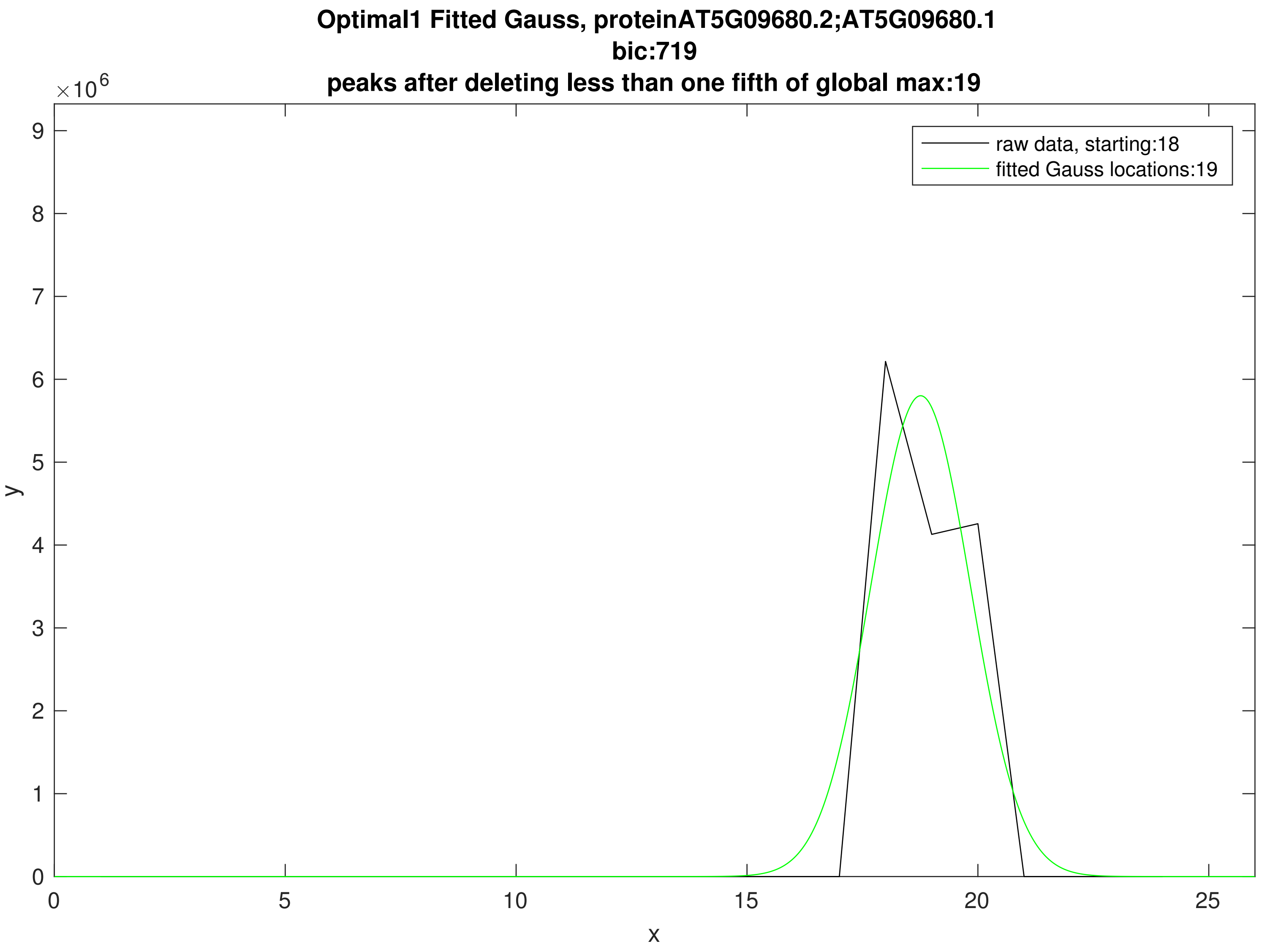
Optimal1 Fitted Gauss, proteinAT5G09680.2;AT5G09680.1
bic:719
peaks after deleting less than one fifth of global max:19
×10^6
raw data, starting:18
fitted Gauss locations:19
y
x
0
1
2
3
4
5
6
7
8
9
0
5
10
15
20
25

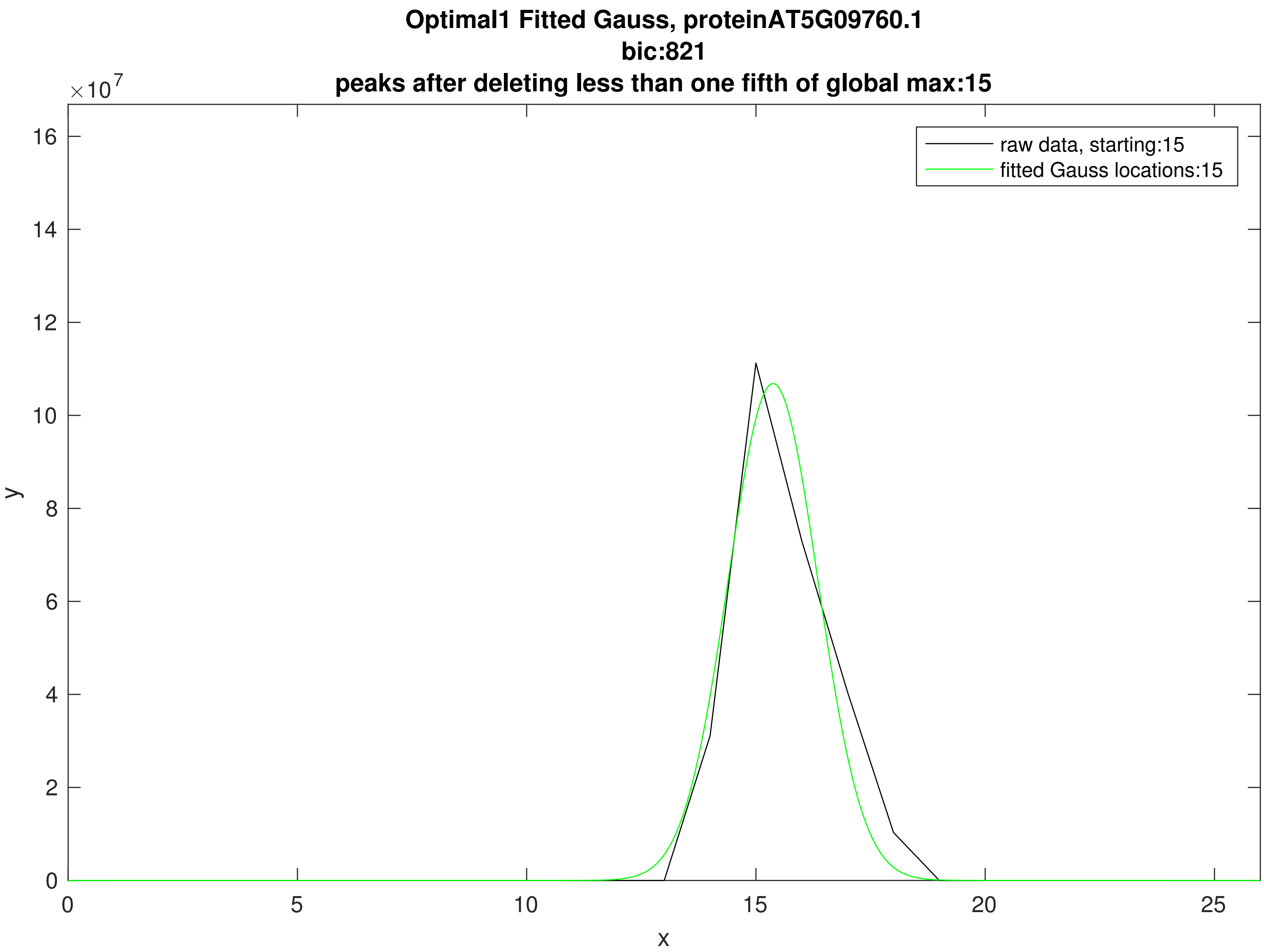

Optimal1 Fitted Gauss, proteinAT5G09760.1
bic:821
peaks after deleting less than one fifth of global max:15
×10^7
16
14
12
10
8
6
4
2
0
y
0
5
10
15
20
25
x
raw data, starting:15
fitted Gauss locations:15

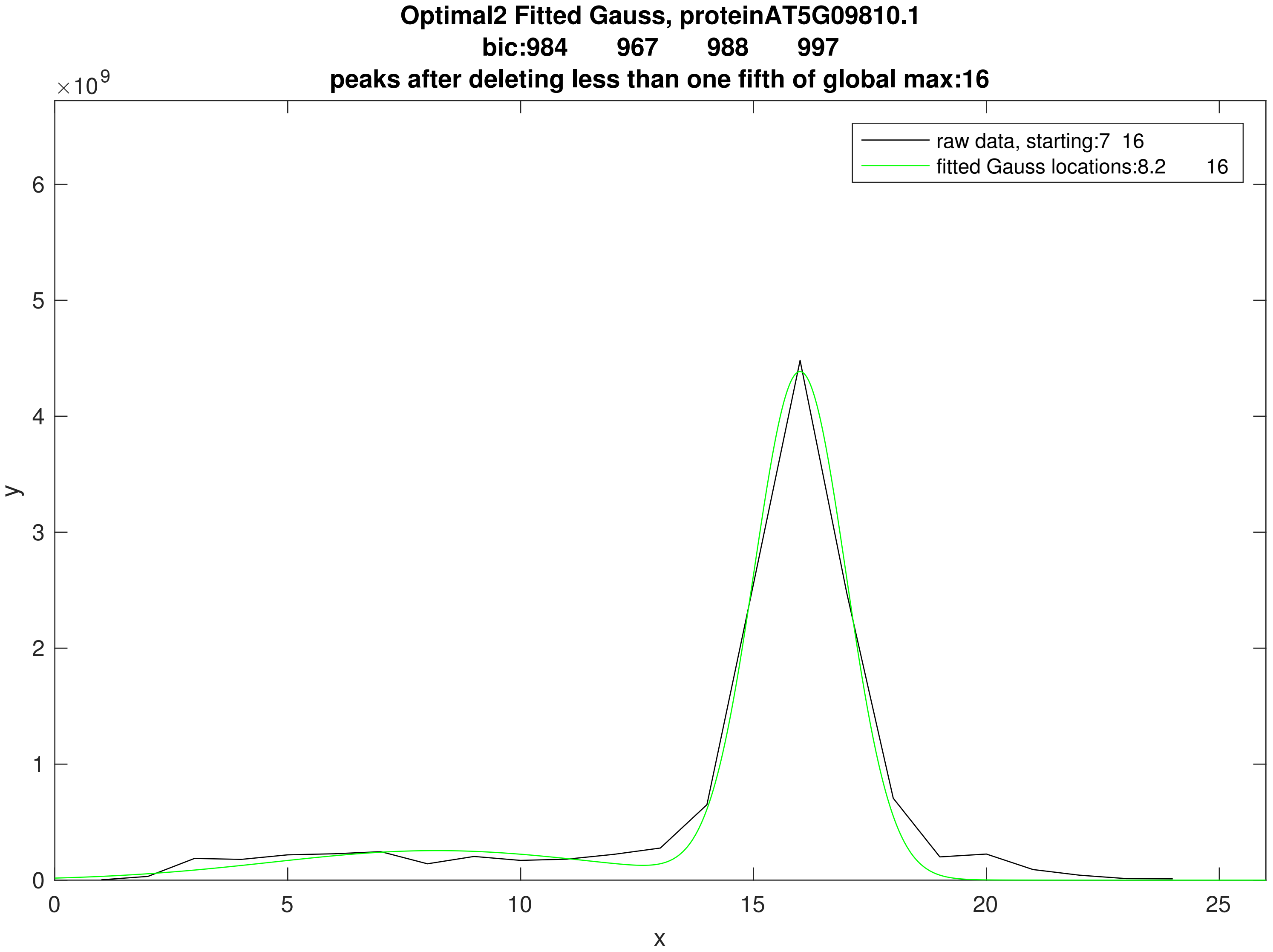
Optimal2 Fitted Gauss, proteinAT5G09810.1
bic:984 967 988 997
peaks after deleting less than one fifth of global max:16
raw data, starting:7 16
fitted Gauss locations:8.2 16
×10^9
y
x
0
1
2
3
4
5
6
0
5
10
15
20
25

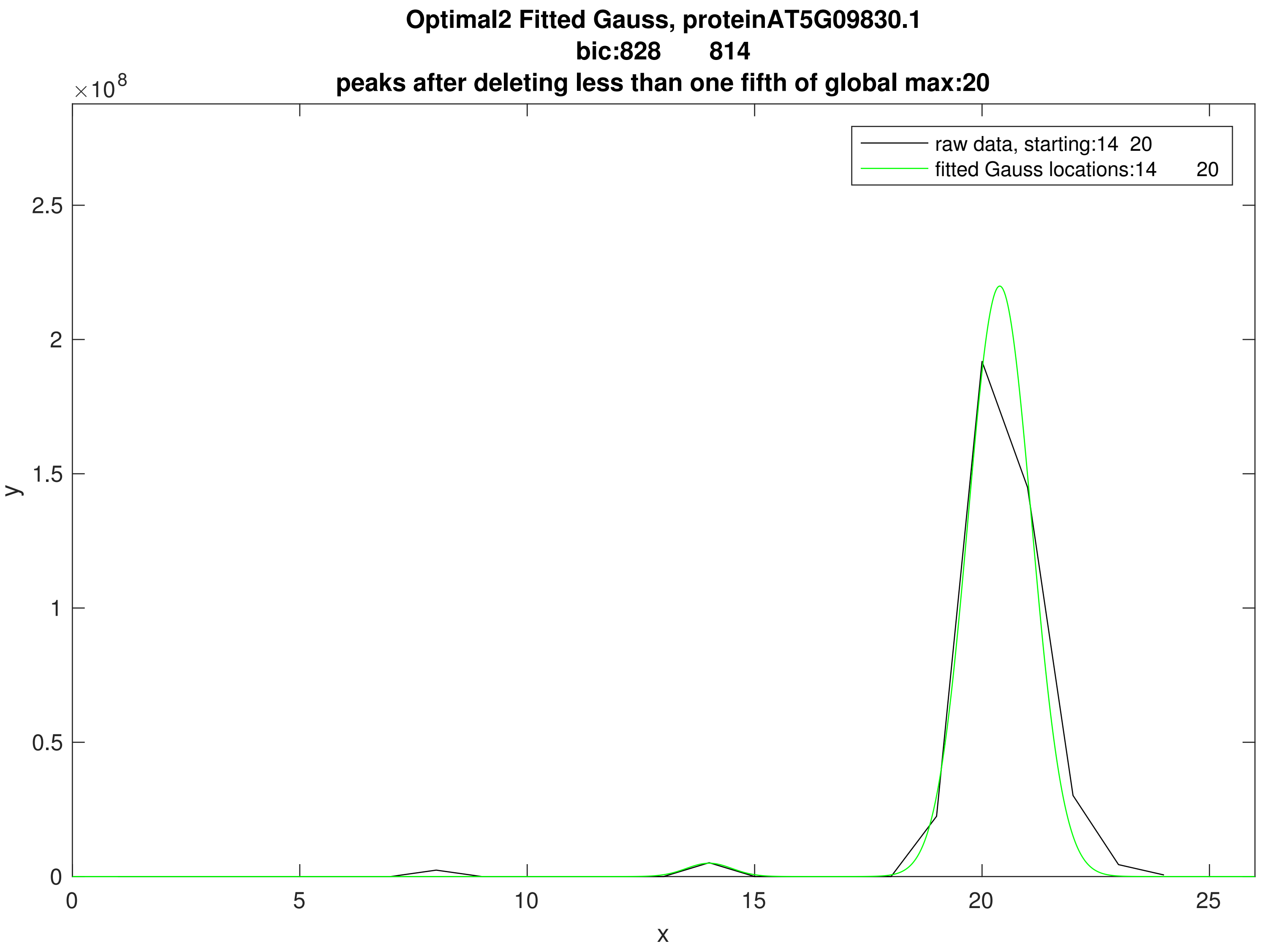
Optimal2 Fitted Gauss, proteinAT5G09830.1
bic:828 814
peaks after deleting less than one fifth of global max:20
raw data, starting:14 20
fitted Gauss locations:14 20
×10^8
2.5
2
1.5
1
0.5
0
y
0
5
10
15
20
25
x

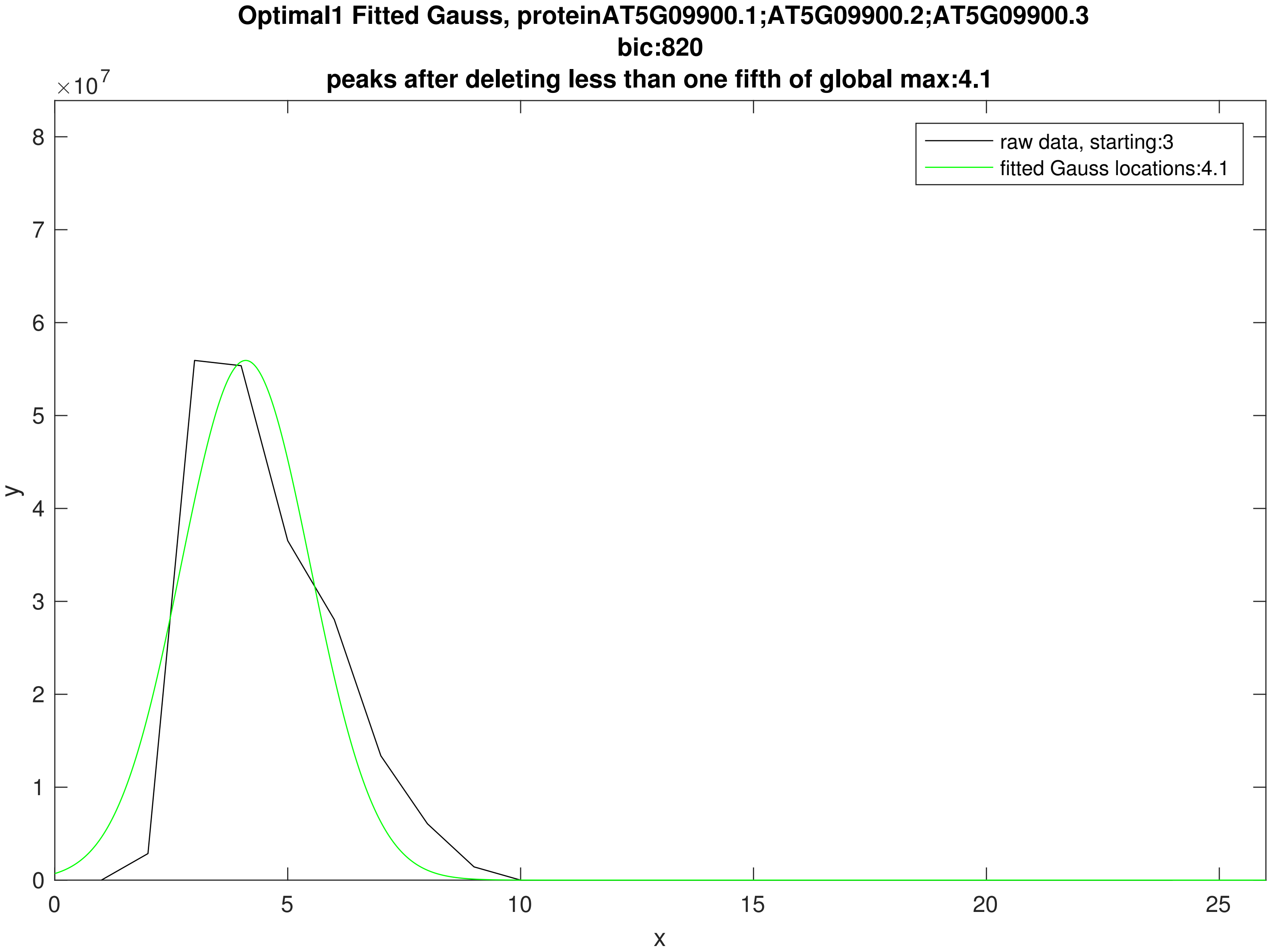
Optimal1 Fitted Gauss, proteinAT5G09900.1;AT5G09900.2;AT5G09900.3
bic:820
peaks after deleting less than one fifth of global max:4.1
×10^7
raw data, starting:3
fitted Gauss locations:4.1
y
x
0
1
2
3
4
5
6
7
8
0
5
10
15
20
25

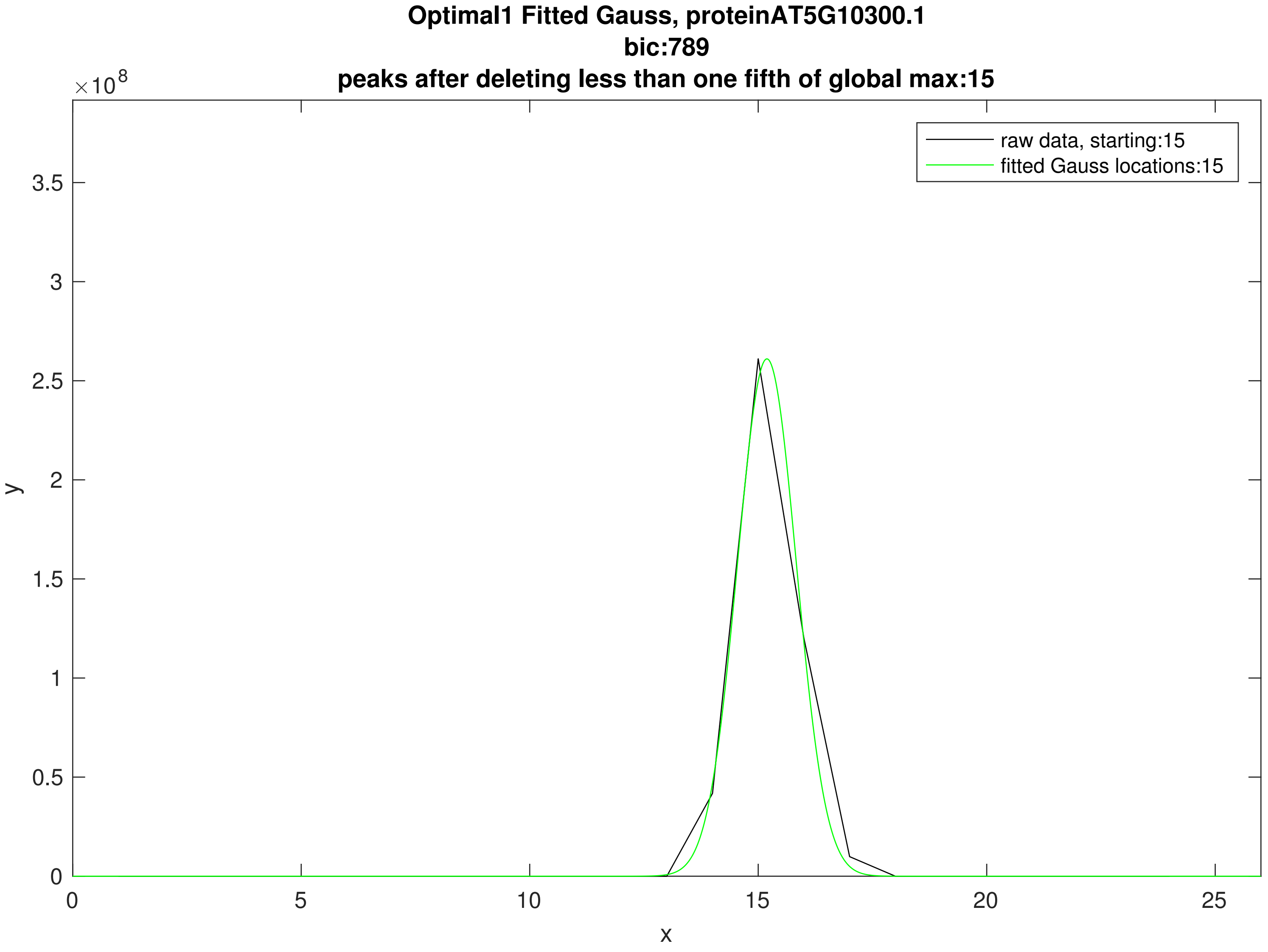
Optimal1 Fitted Gauss, proteinAT5G10300.1
bic:789
peaks after deleting less than one fifth of global max:15
×10^8
raw data, starting:15
fitted Gauss locations:15
3.5
3
2.5
2
1.5
1
0.5
0
y
0
5
10
15
20
25
x

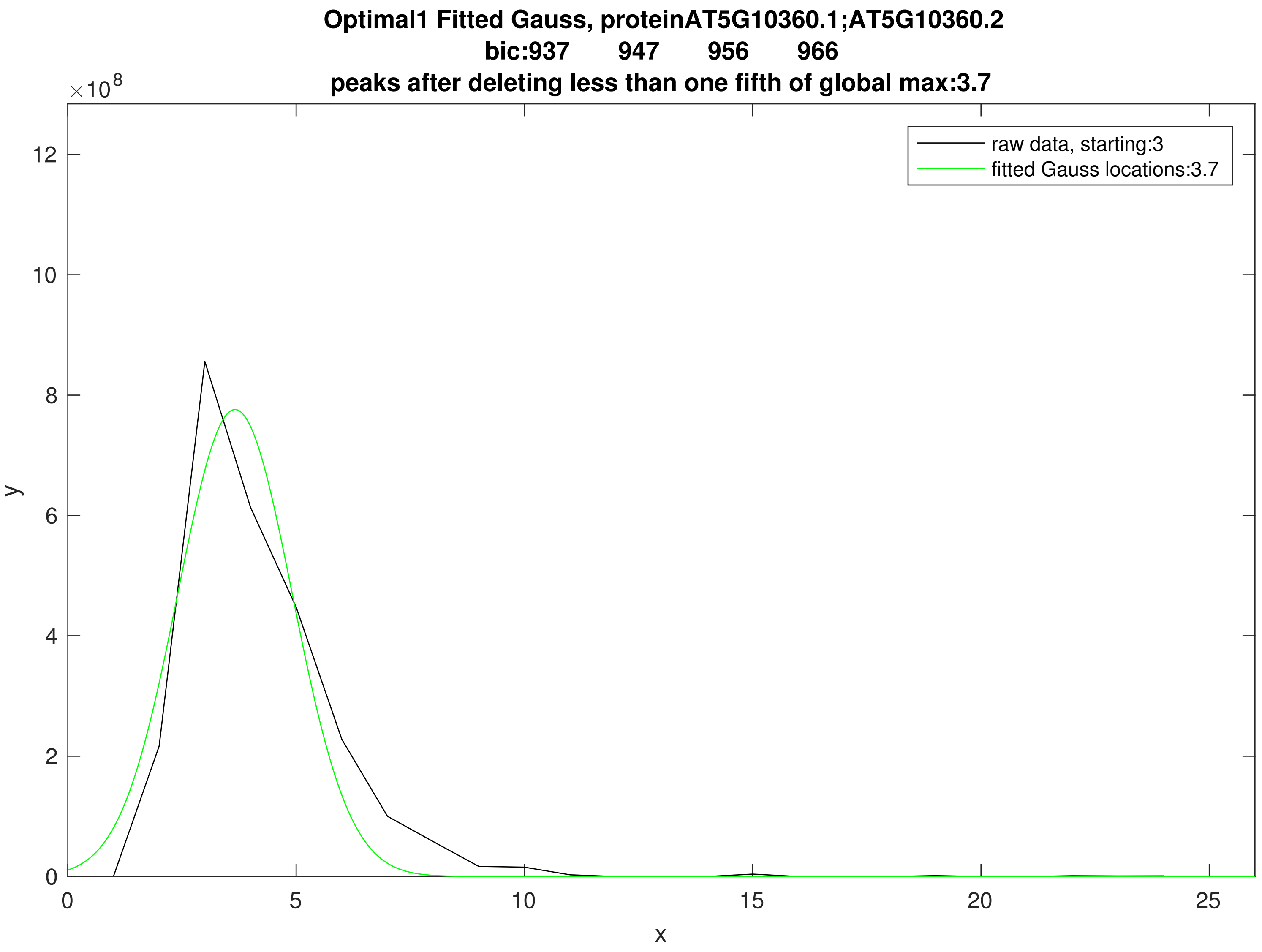
Optimal1 Fitted Gauss, proteinAT5G10360.1;AT5G10360.2
bic:937 947 956 966
peaks after deleting less than one fifth of global max:3.7
×10^8
raw data, starting:3
fitted Gauss locations:3.7
y
x
0
2
4
6
8
10
12
5
15
20
25

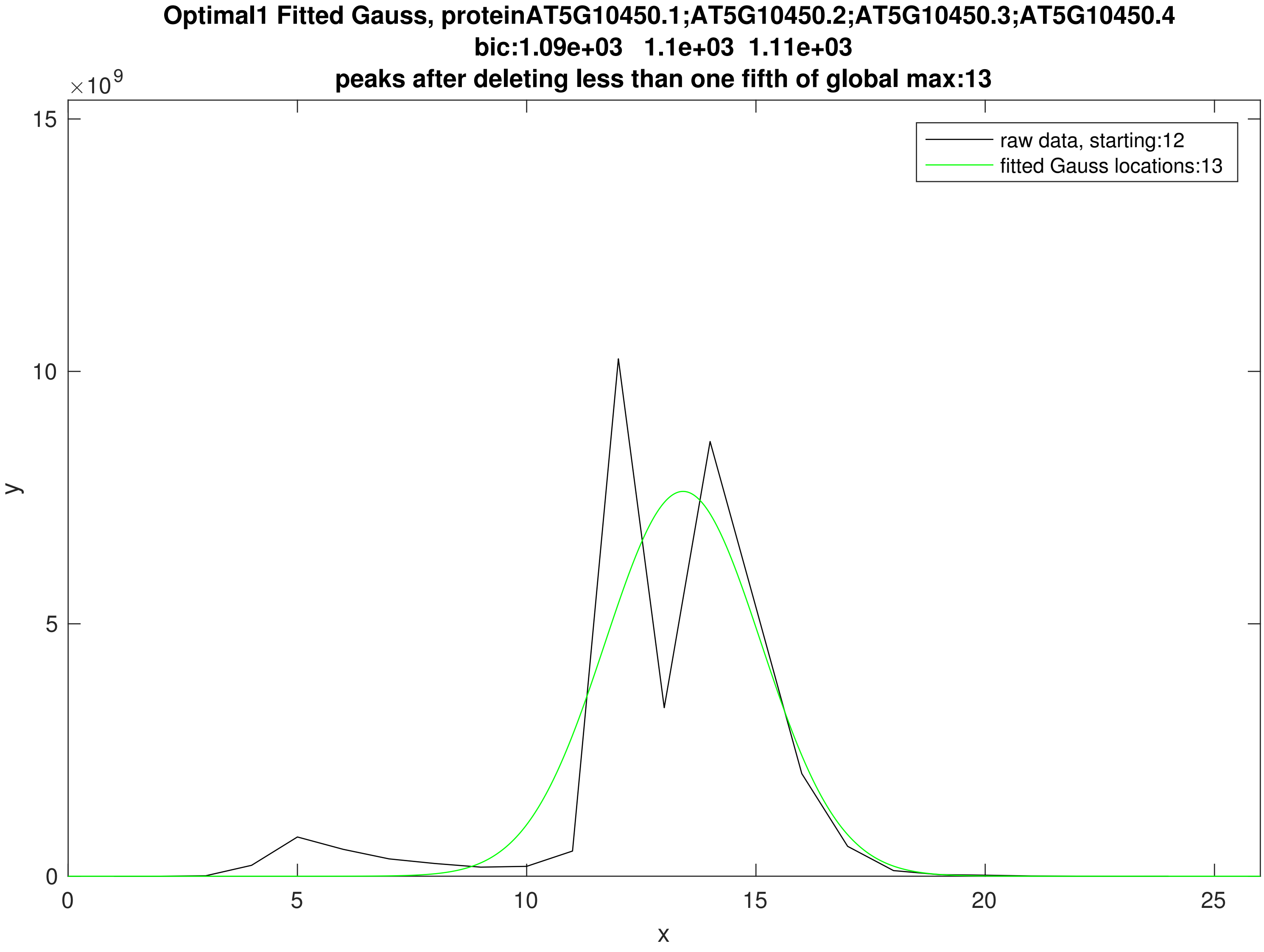

Optimal1 Fitted Gauss, proteinAT5G10450.1;AT5G10450.2;AT5G10450.3;AT5G10450.4
bic:1.09e+03 1.1e+03 1.11e+03
peaks after deleting less than one fifth of global max:13
×10^9
15
10
5
0
y
0
5
10
15
20
25
x
raw data, starting:12
fitted Gauss locations:13

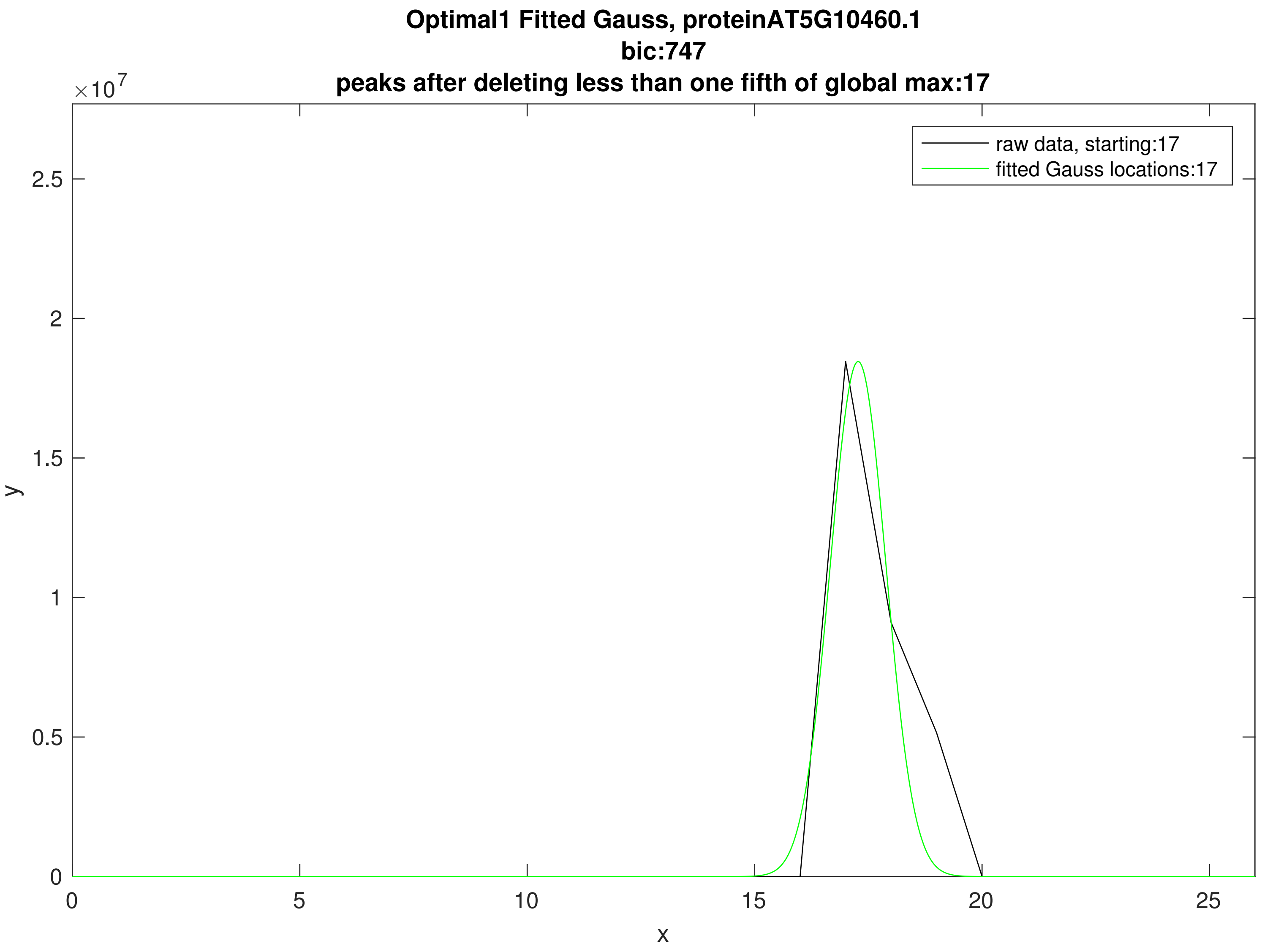

Optimal1 Fitted Gauss, proteinAT5G10460.1
bic:747
peaks after deleting less than one fifth of global max:17
×10^7
2.5
2
1.5
1
0.5
0
y
0
5
10
15
20
25
x
raw data, starting:17
fitted Gauss locations:17

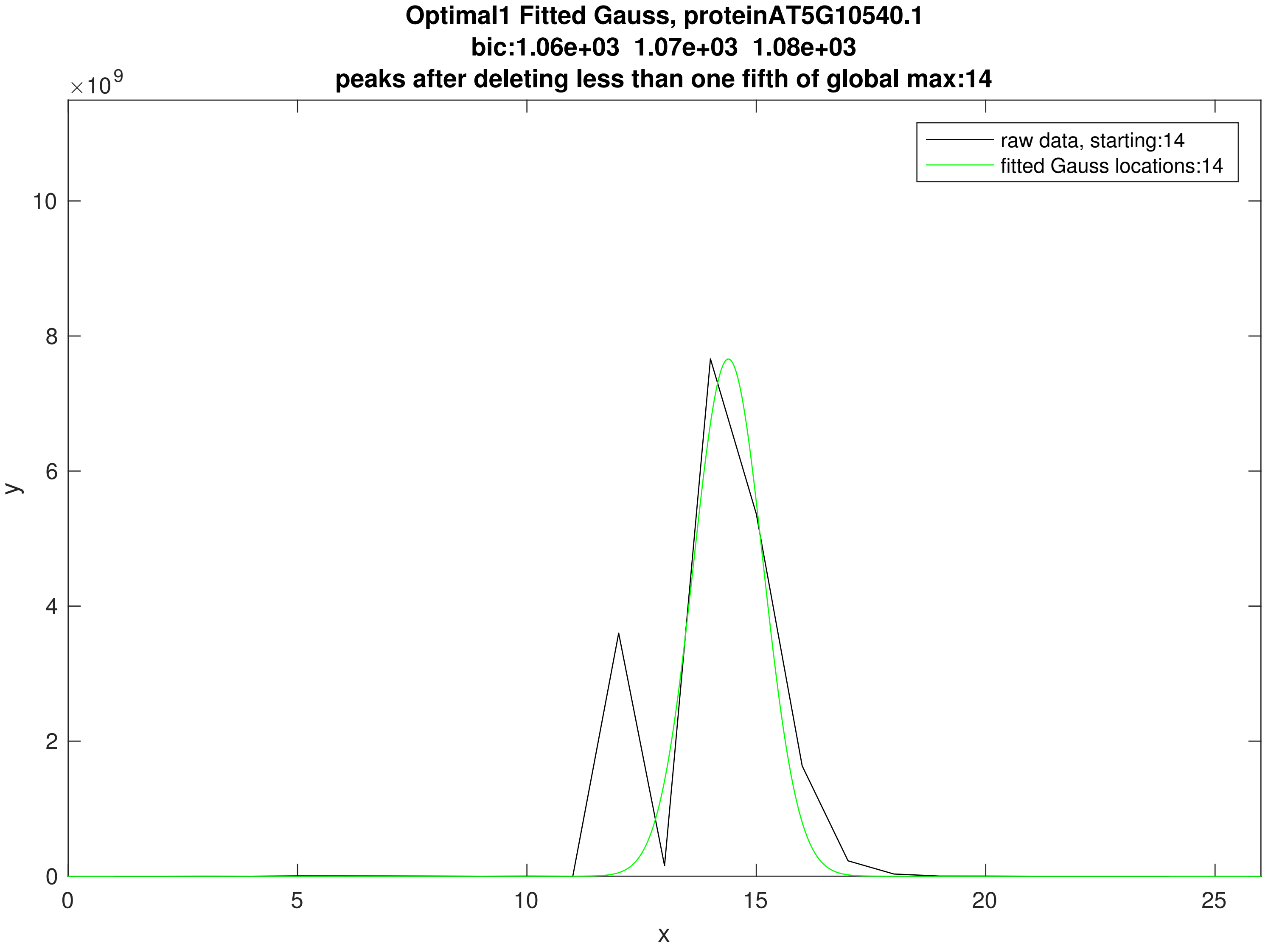
Optimal1 Fitted Gauss, proteinAT5G10540.1
bic:1.06e+03  1.07e+03  1.08e+03
peaks after deleting less than one fifth of global max:14
×10^9
raw data, starting:14
fitted Gauss locations:14
10
8
6
4
2
0
y
0
5
10
15
20
25
x

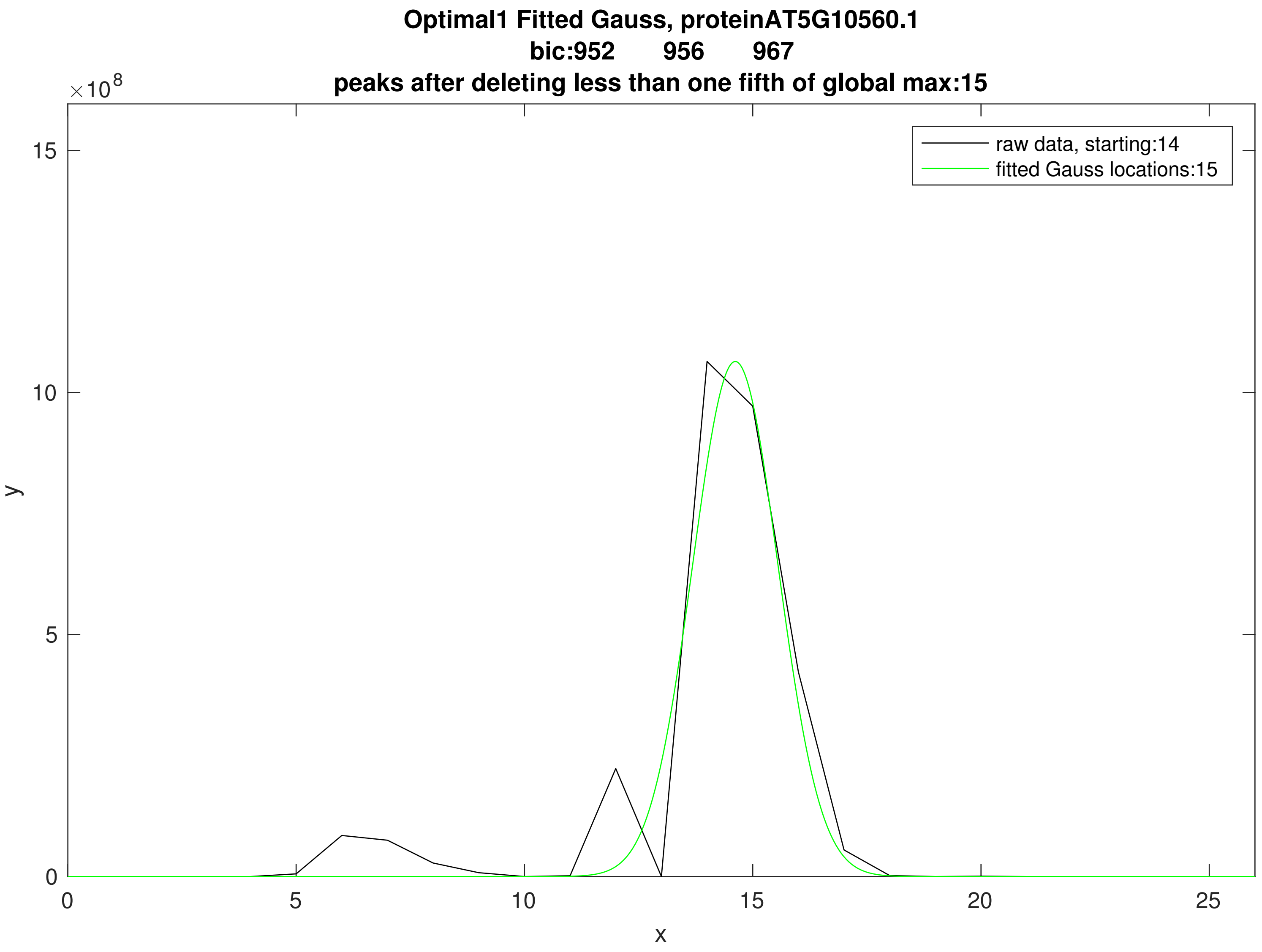
Optimal1 Fitted Gauss, proteinAT5G10560.1
bic:952 956 967
peaks after deleting less than one fifth of global max:15
×10^8
15
10
5
0
y
0
5
10
15
20
25
x
raw data, starting:14
fitted Gauss locations:15

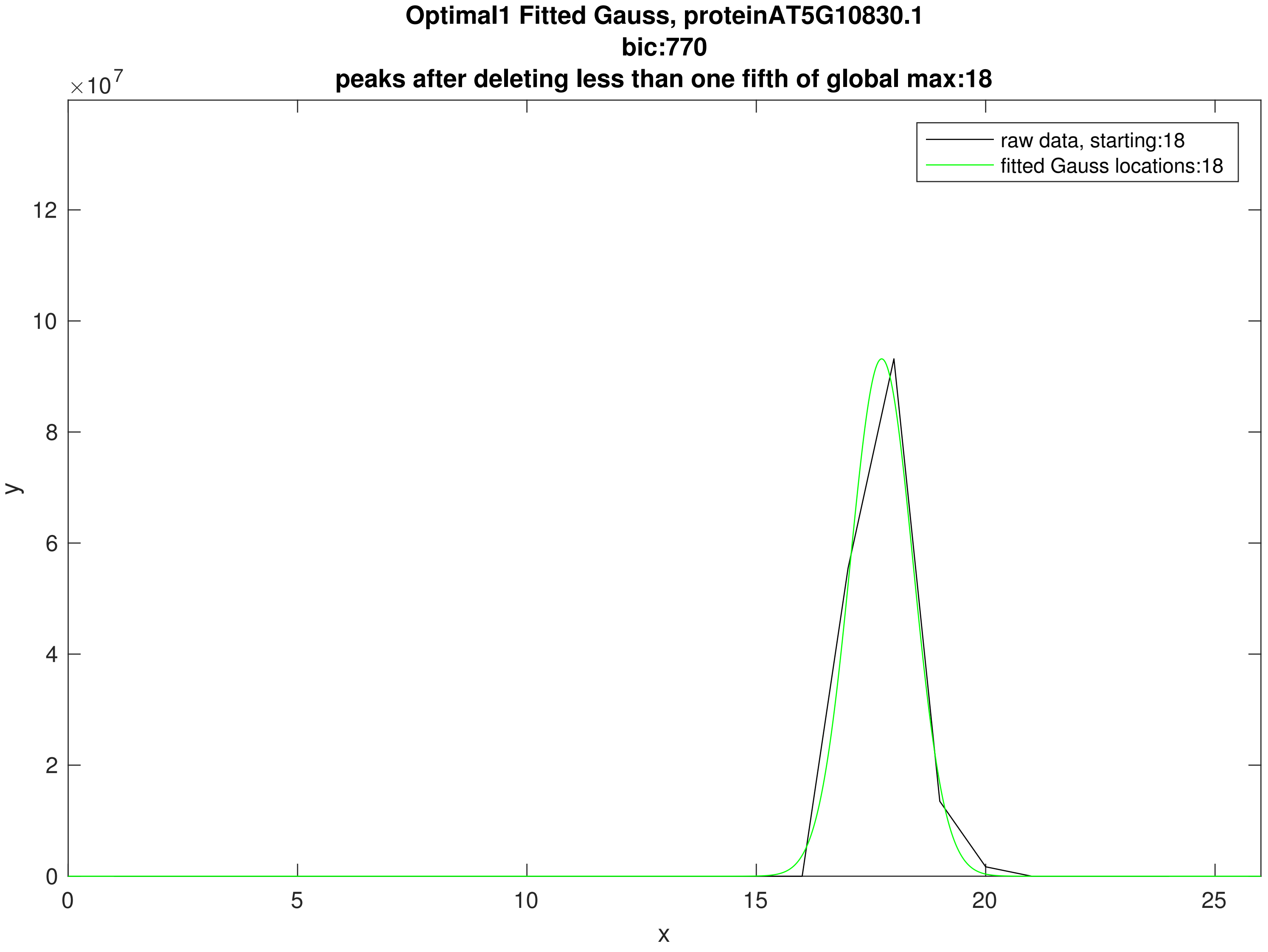
Optimal1 Fitted Gauss, proteinAT5G10830.1
bic:770
peaks after deleting less than one fifth of global max:18
×10^7
raw data, starting:18
fitted Gauss locations:18
y
x
0
2
4
6
8
10
12
0
5
10
15
20
25

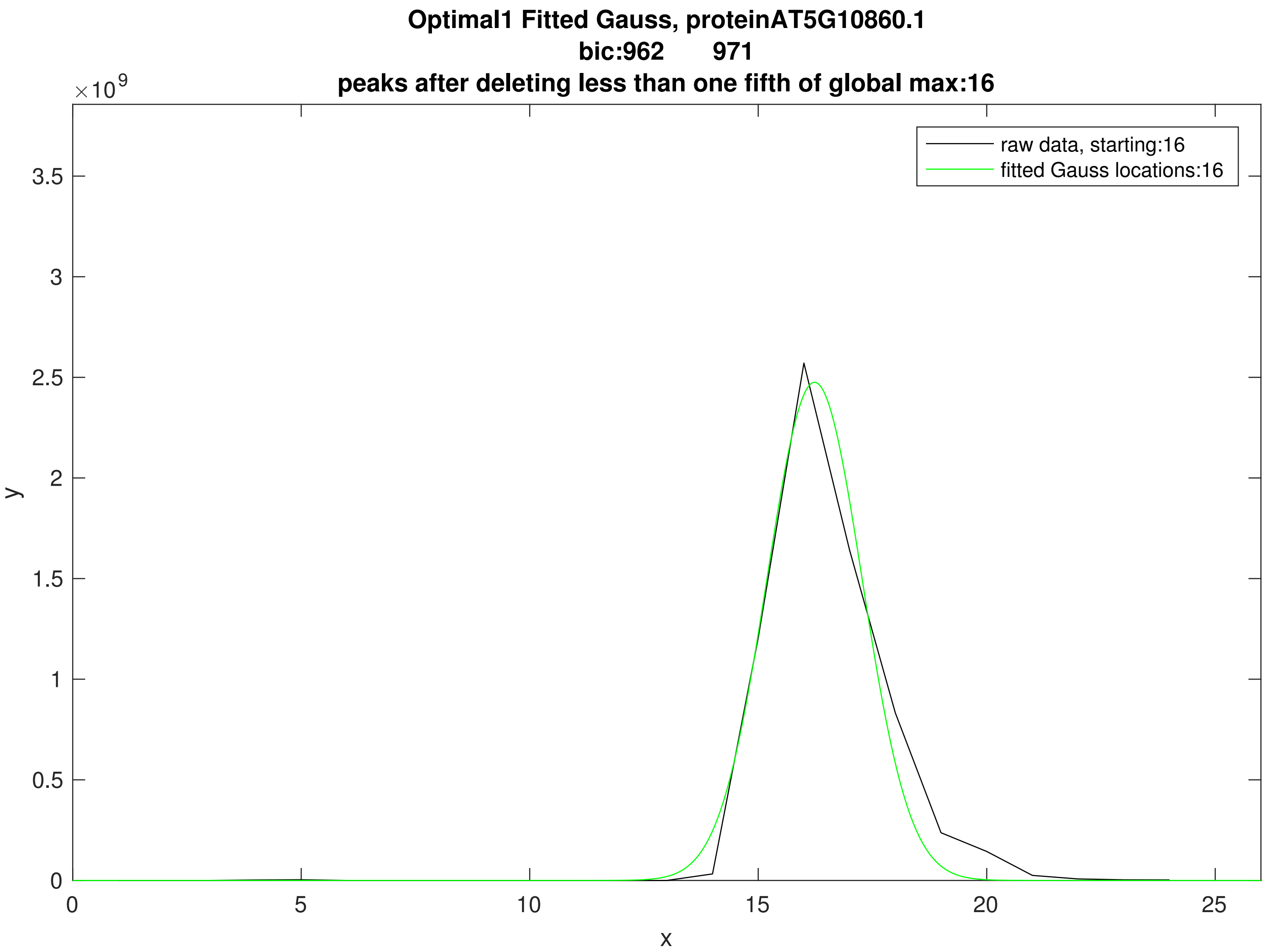

Optimal1 Fitted Gauss, proteinAT5G10860.1
bic:962 971
peaks after deleting less than one fifth of global max:16
×10^9
raw data, starting:16
fitted Gauss locations:16
3.5
3
2.5
2
1.5
1
0.5
0
y
0
5
10
15
20
25
x

#### Optimal1 Fitted Gauss, proteinAT5G10920.1
#### bic:963
#### peaks after deleting less than one fifth of global max:10

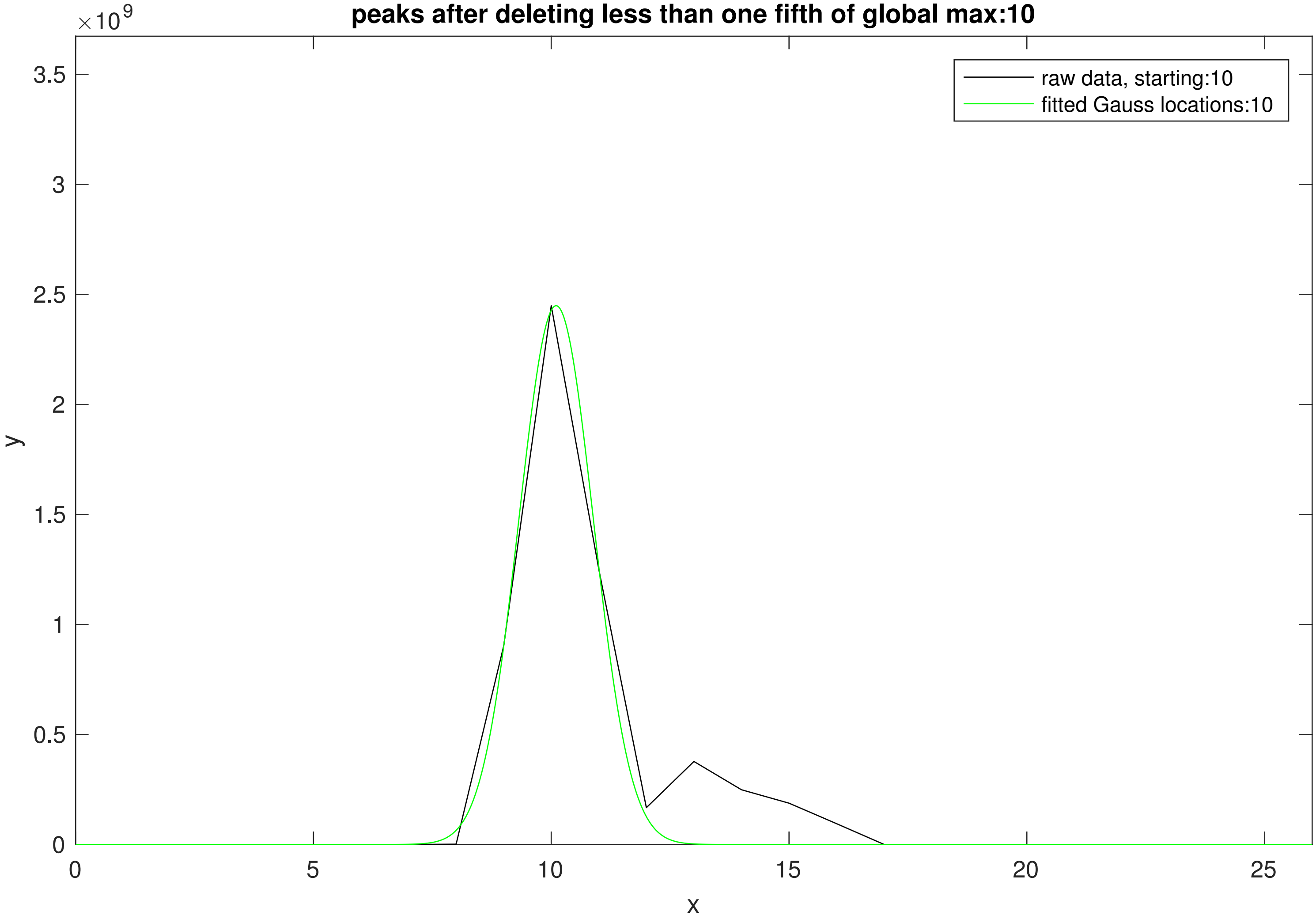

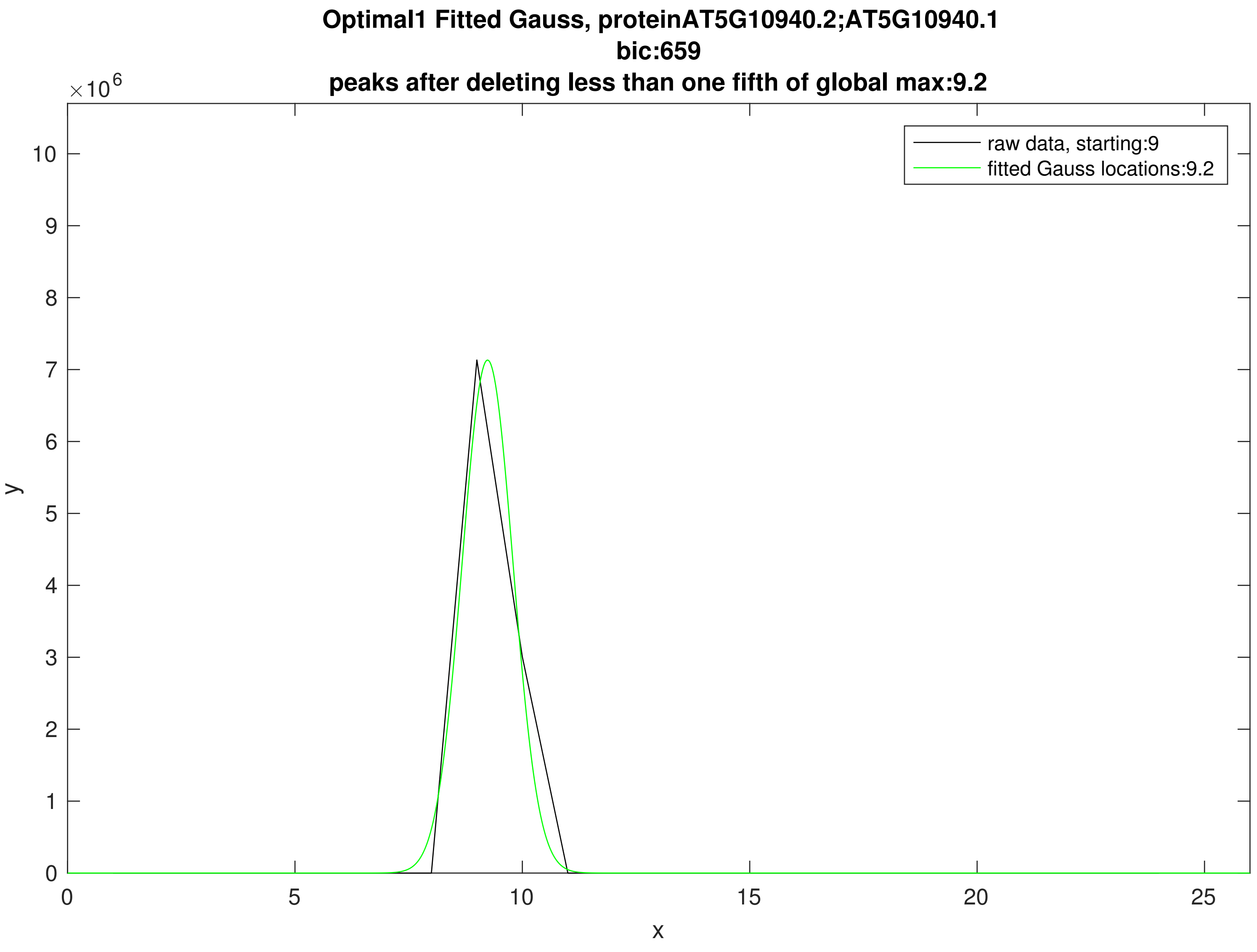

Optimal1 Fitted Gauss, proteinAT5G10940.2;AT5G10940.1
bic:659
peaks after deleting less than one fifth of global max:9.2
×10^6
raw data, starting:9
fitted Gauss locations:9.2
y
x
0
1
2
3
4
5
6
7
8
9
10
0
5
10
15
20
25

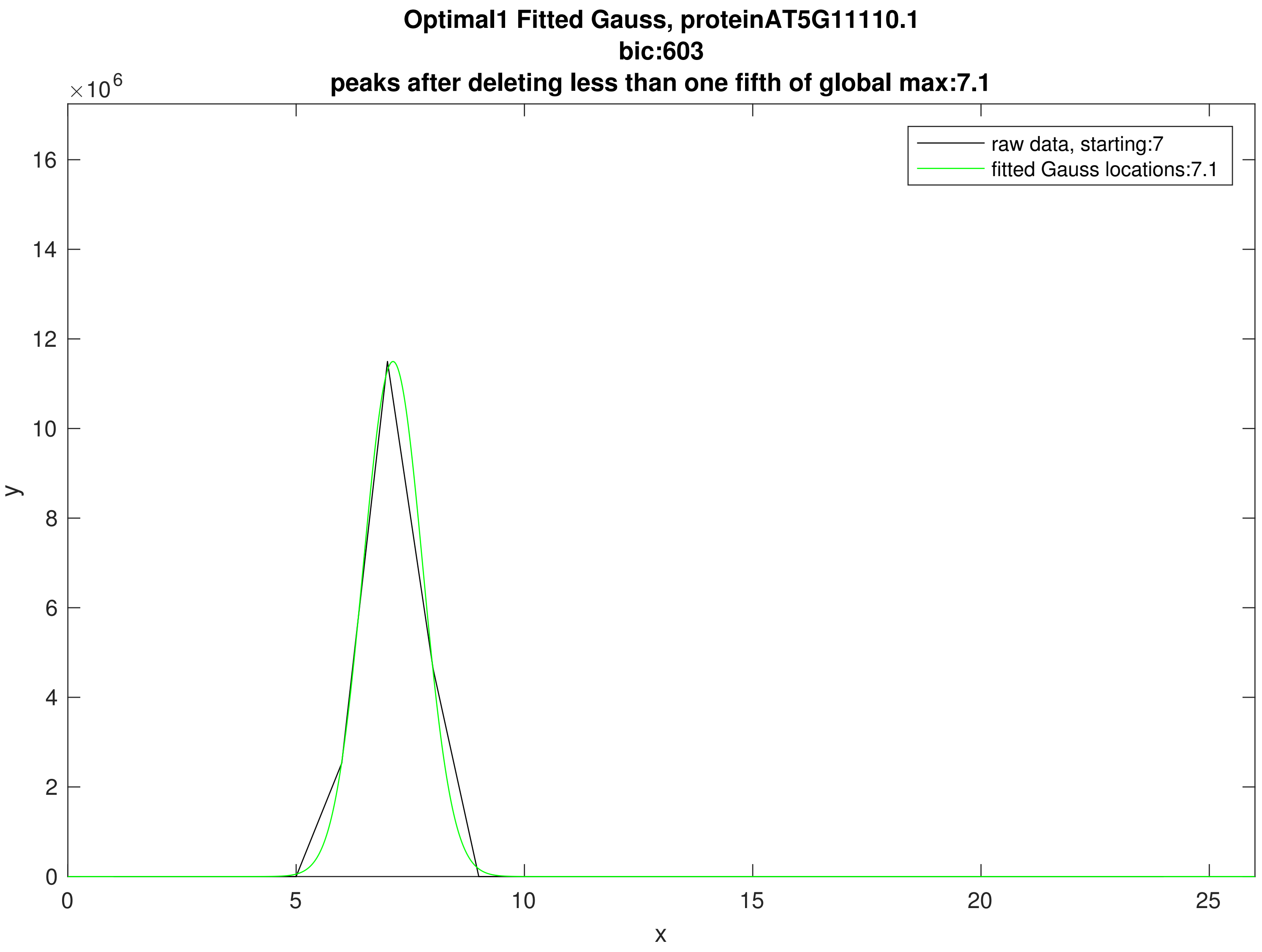

Optimal1 Fitted Gauss, proteinAT5G11110.1
bic:603
peaks after deleting less than one fifth of global max:7.1
×10^6
raw data, starting:7
fitted Gauss locations:7.1
y
x
0
2
4
6
8
10
12
14
16
0
5
10
15
20
25

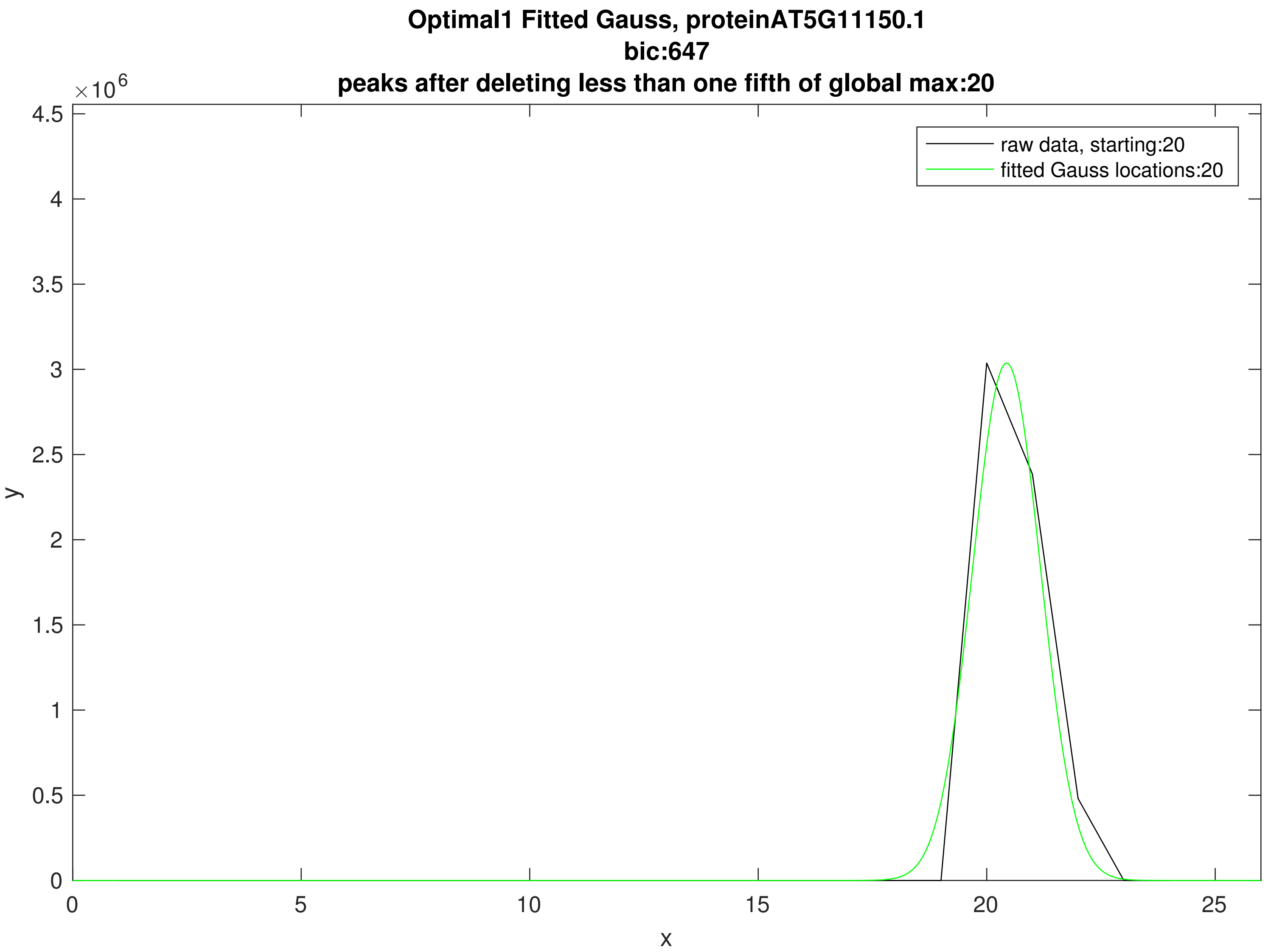
Optimal1 Fitted Gauss, proteinAT5G11150.1
bic:647
peaks after deleting less than one fifth of global max:20
×10^6
4.5
4
3.5
3
2.5
2
1.5
1
0.5
0
y
0
5
10
15
20
25
x
raw data, starting:20
fitted Gauss locations:20

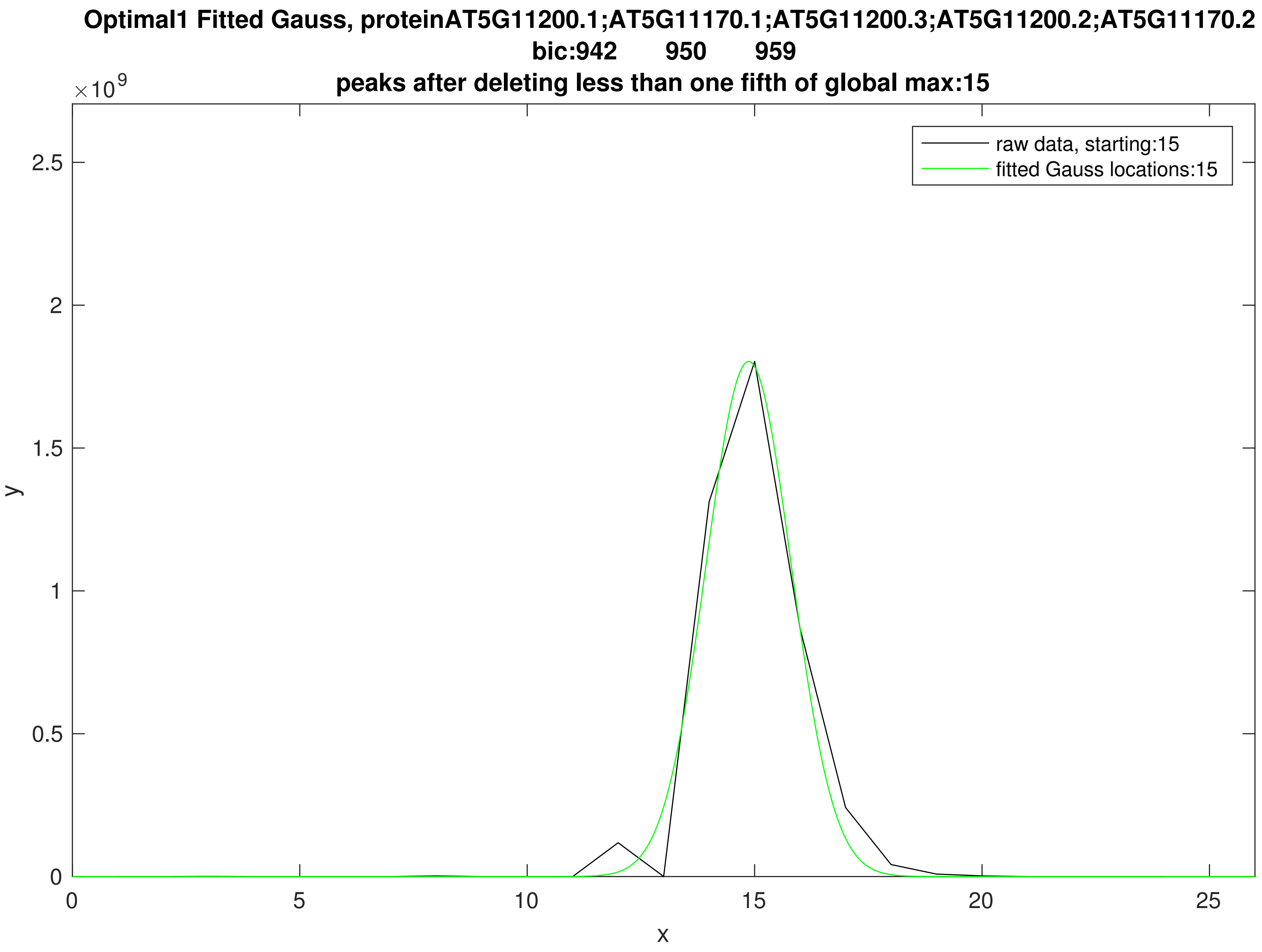

Optimal1 Fitted Gauss, proteinAT5G11200.1;AT5G11170.1;AT5G11200.3;AT5G11200.2;AT5G11170.2
bic:942 950 959
peaks after deleting less than one fifth of global max:15
×10^9
2.5
2
1.5
1
0.5
0
y
0
5
10
15
20
25
x
raw data, starting:15
fitted Gauss locations:15

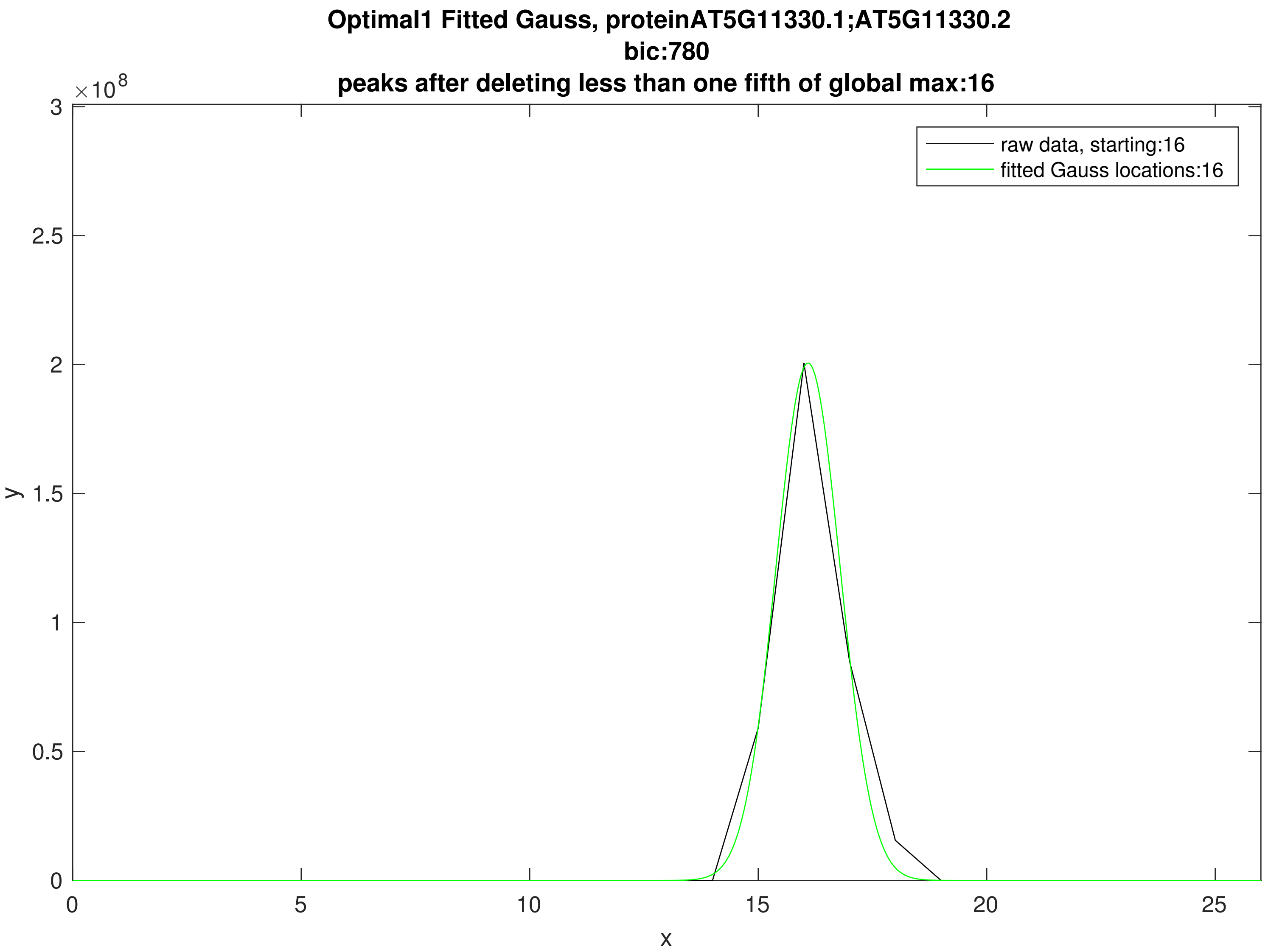
Optimal1 Fitted Gauss, proteinAT5G11330.1;AT5G11330.2
bic:780
peaks after deleting less than one fifth of global max:16
×10^8
3
2.5
2
1.5
1
0.5
0
y
0
5
10
15
20
25
x
raw data, starting:16
fitted Gauss locations:16

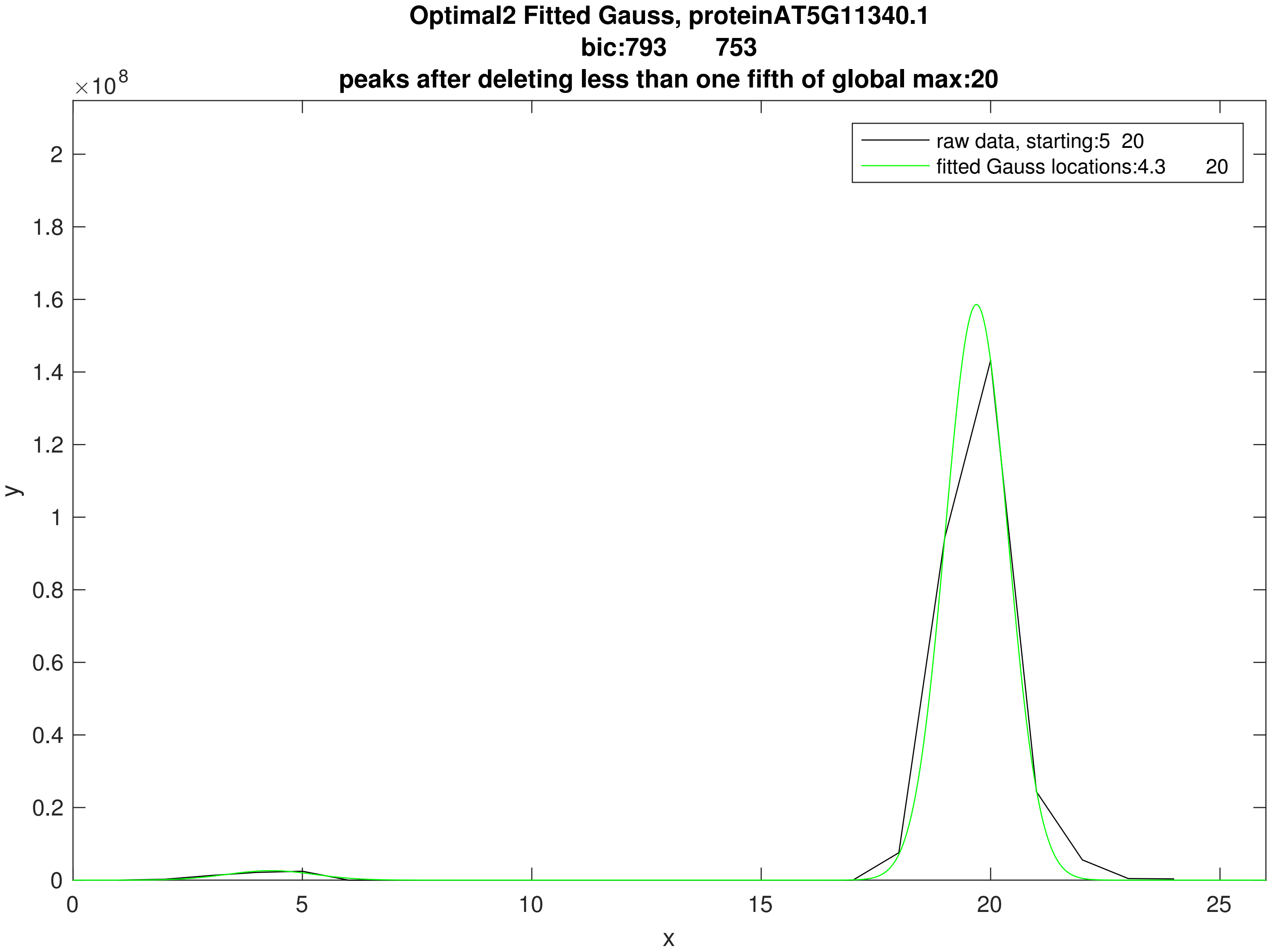
Optimal2 Fitted Gauss, proteinAT5G11340.1
bic:793 753
peaks after deleting less than one fifth of global max:20
×10^8
raw data, starting:5 20
fitted Gauss locations:4.3 20
2
1.8
1.6
1.4
1.2
1
0.8
0.6
0.4
0.2
0
y
0
5
10
15
20
25
x

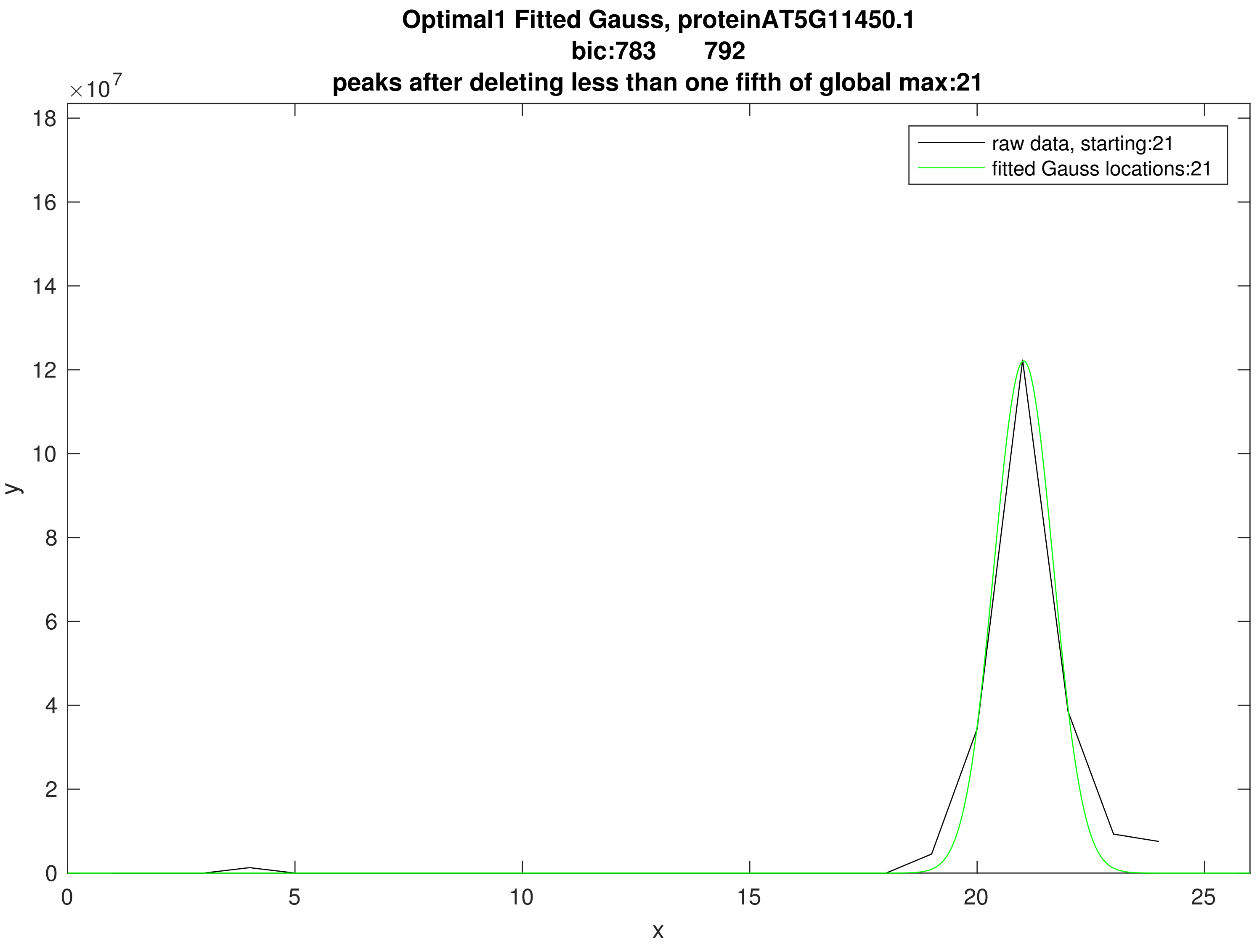

Optimal1 Fitted Gauss, proteinAT5G11450.1
bic:783 792
peaks after deleting less than one fifth of global max:21
×10^7
18
16
14
12
10
8
6
4
2
0
y
0
5
10
15
20
25
x
raw data, starting:21
fitted Gauss locations:21

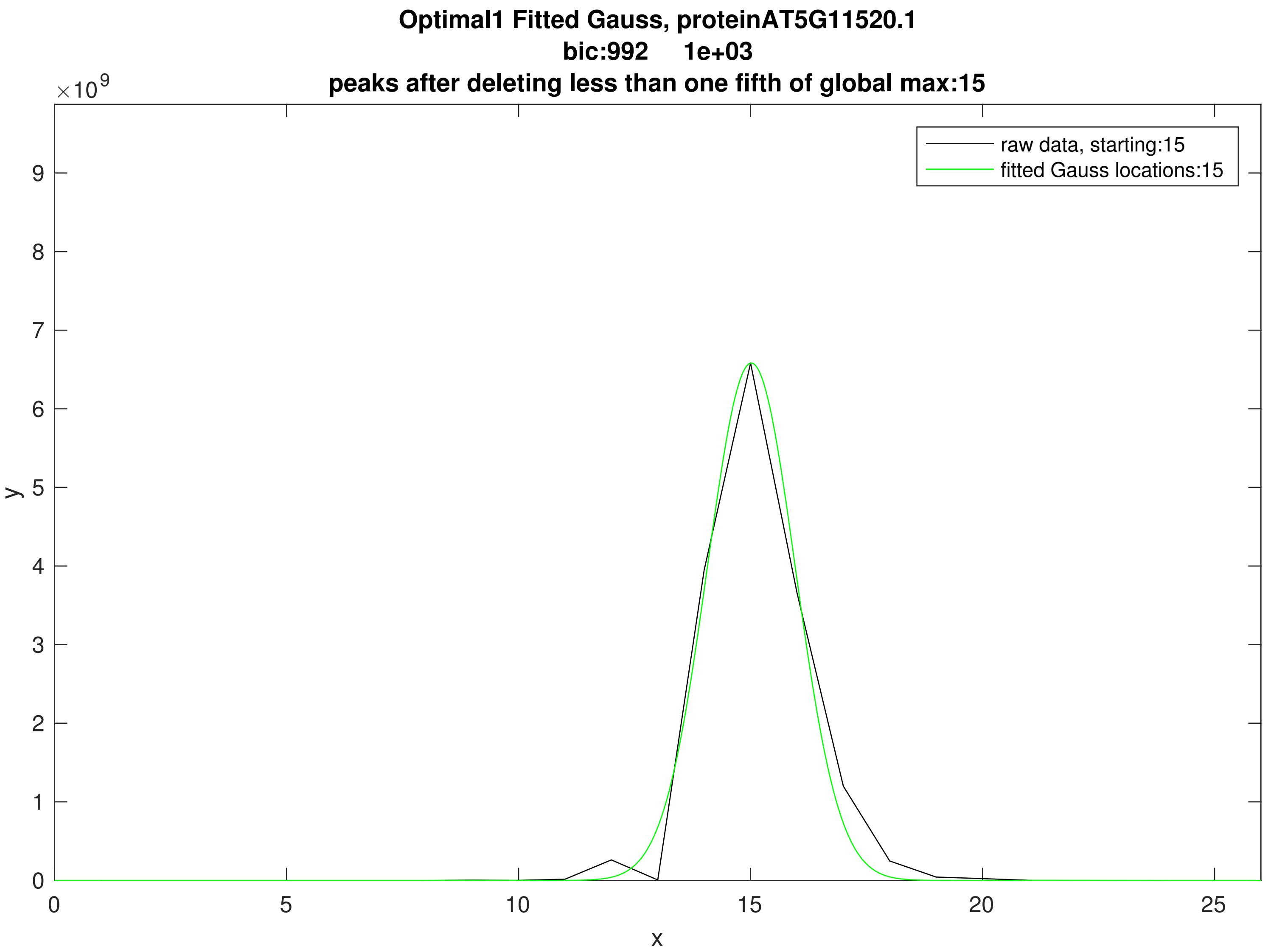
Optimal1 Fitted Gauss, proteinAT5G11520.1
bic:992 1e+03
peaks after deleting less than one fifth of global max:15
×10^9
raw data, starting:15
fitted Gauss locations:15
y
x
0
1
2
3
4
5
6
7
8
9
0
5
10
15
20
25

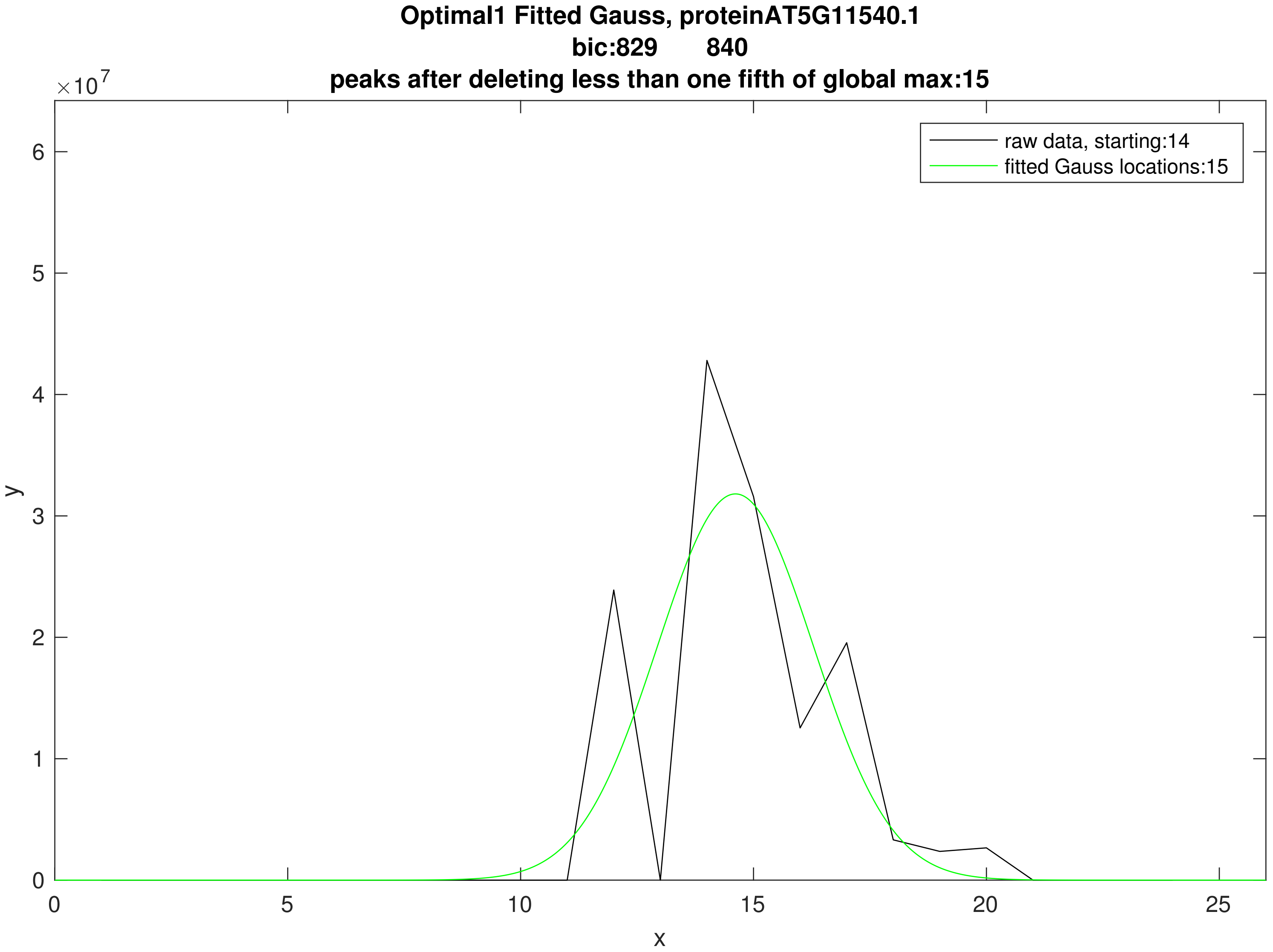
Optimal1 Fitted Gauss, proteinAT5G11540.1
bic:829 840
peaks after deleting less than one fifth of global max:15
×10^7
raw data, starting:14
fitted Gauss locations:15
y
x
0
1
2
3
4
5
6
0
5
10
15
20
25

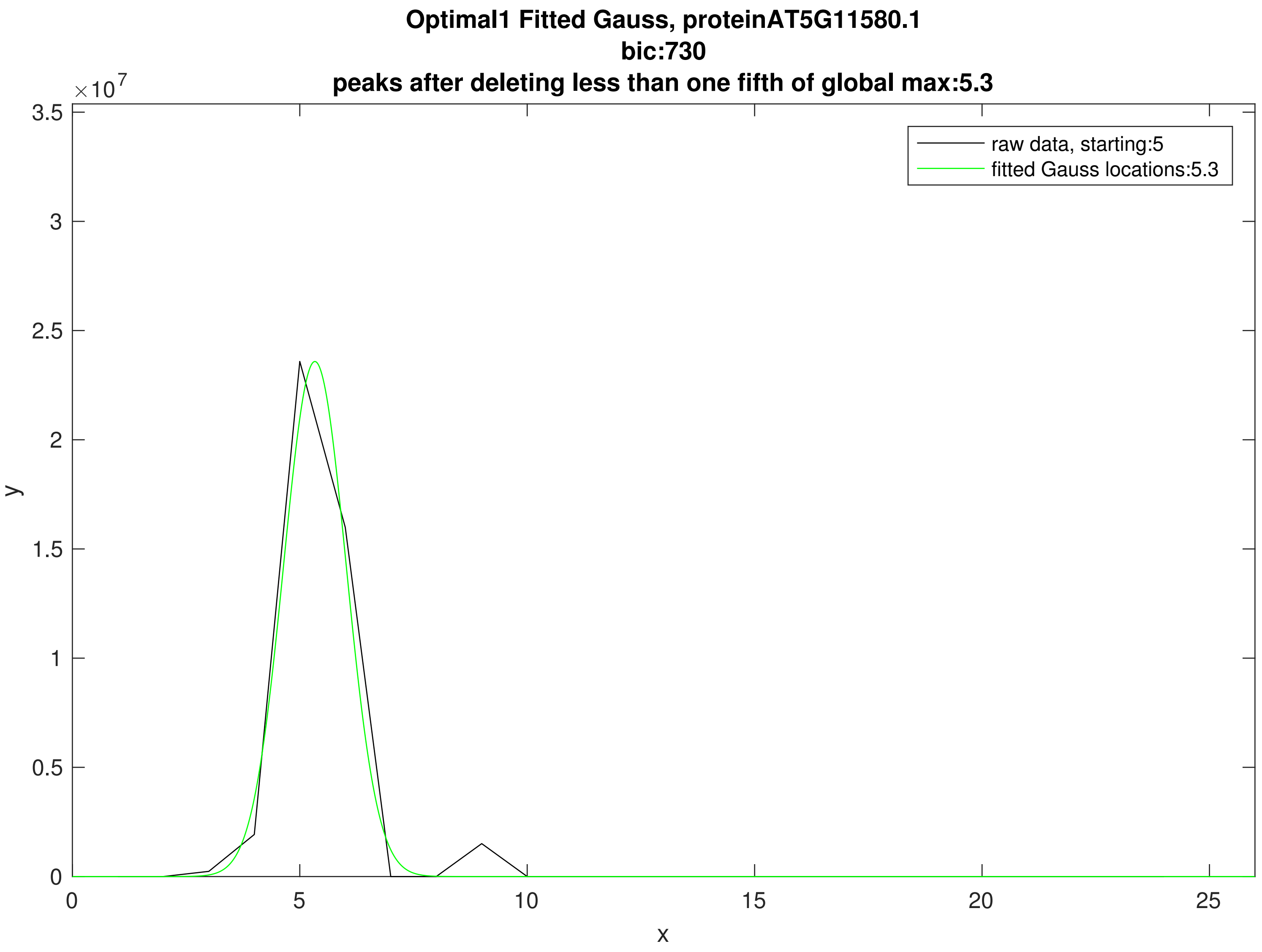
Optimal1 Fitted Gauss, proteinAT5G11580.1
bic:730
peaks after deleting less than one fifth of global max:5.3
×10^7
3.5
3
2.5
2
1.5
1
0.5
0
y
0
5
10
15
20
25
x
raw data, starting:5
fitted Gauss locations:5.3

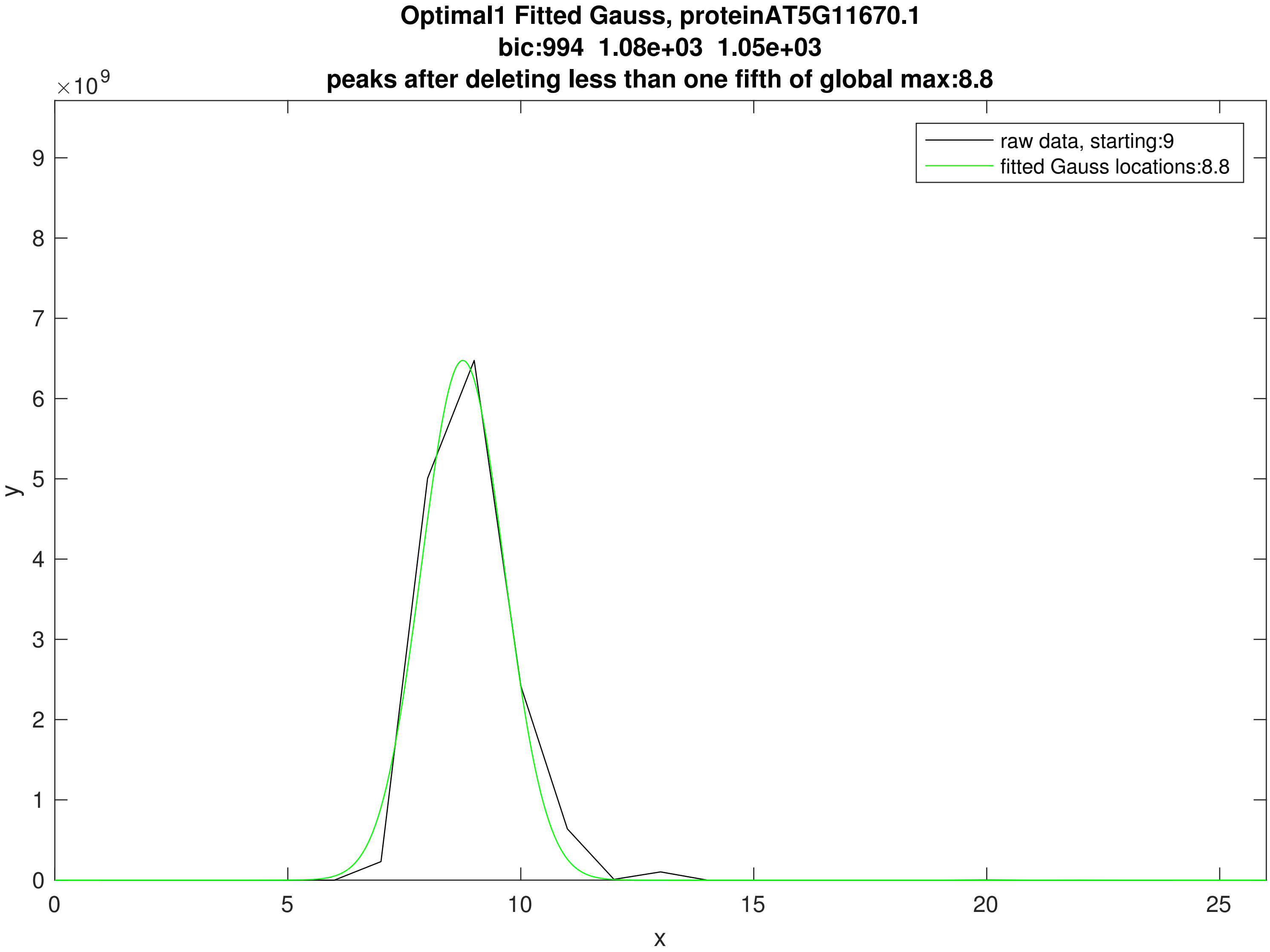

Optimal1 Fitted Gauss, proteinAT5G11670.1
bic:994 1.08e+03 1.05e+03
peaks after deleting less than one fifth of global max:8.8
×10^9
raw data, starting:9
fitted Gauss locations:8.8
y
x

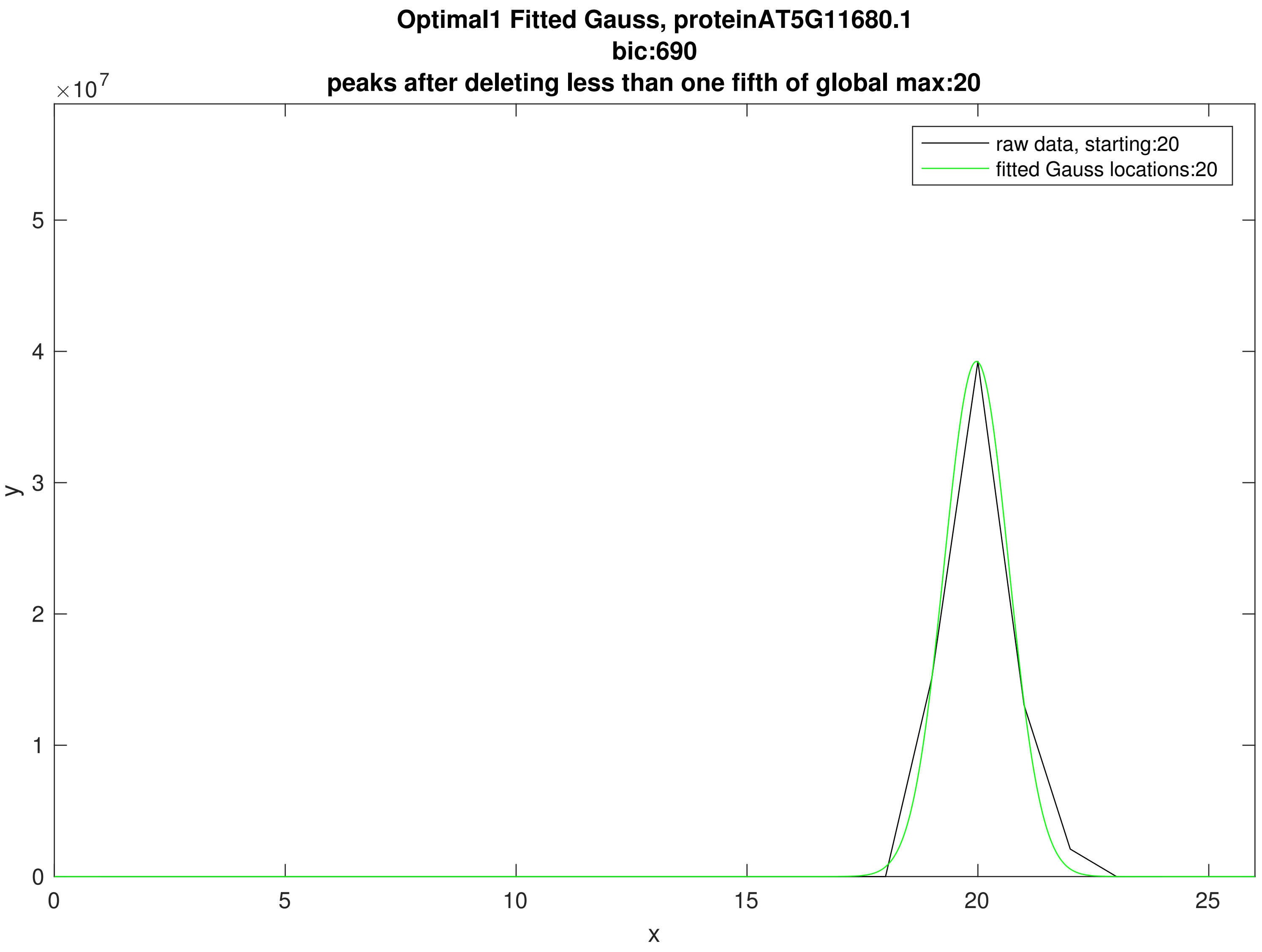

Optimal1 Fitted Gauss, proteinAT5G11680.1
bic:690
peaks after deleting less than one fifth of global max:20
×10^7
raw data, starting:20
fitted Gauss locations:20
y
x
0
1
2
3
4
5
0
5
10
15
20
25

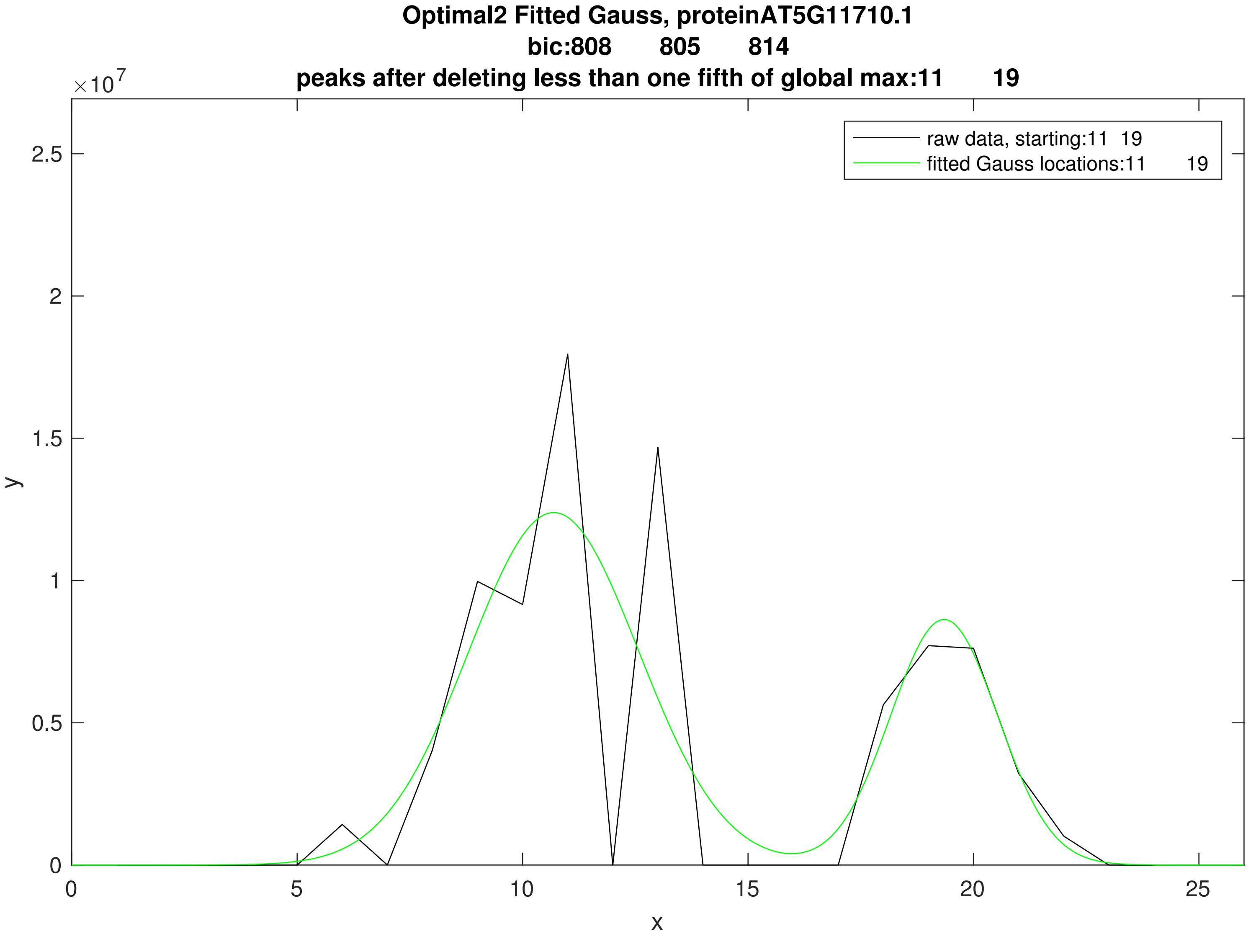
Optimal2 Fitted Gauss, proteinAT5G11710.1
bic:808 805 814
peaks after deleting less than one fifth of global max:11 19
×10^7
raw data, starting:11 19
fitted Gauss locations:11 19
y
x
0
0.5
1
1.5
2
2.5
5
10
15
20
25

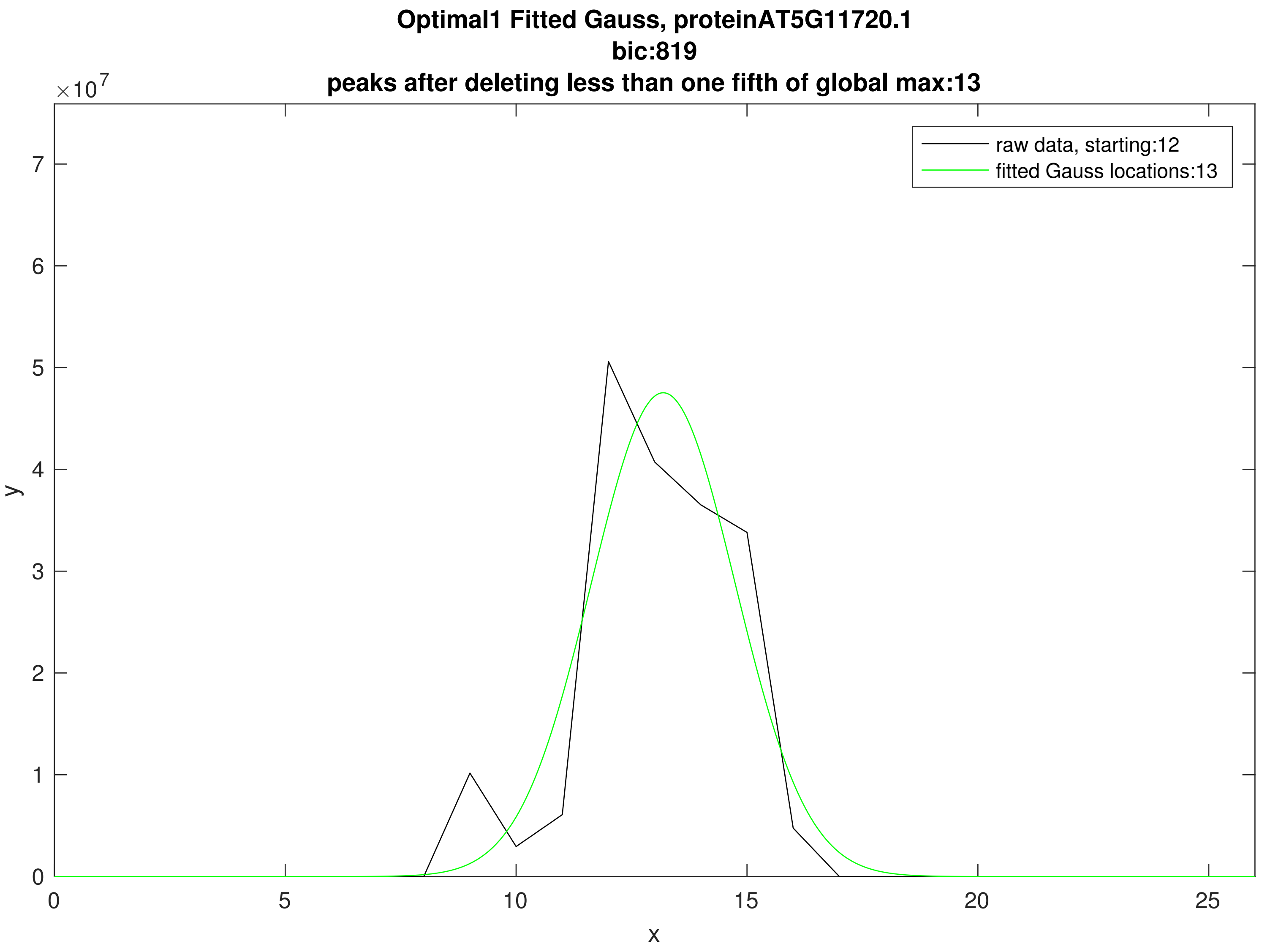
Optimal1 Fitted Gauss, proteinAT5G11720.1
bic:819
peaks after deleting less than one fifth of global max:13
×10^7
raw data, starting:12
fitted Gauss locations:13
y
x
0
1
2
3
4
5
6
7
0
5
10
15
20
25

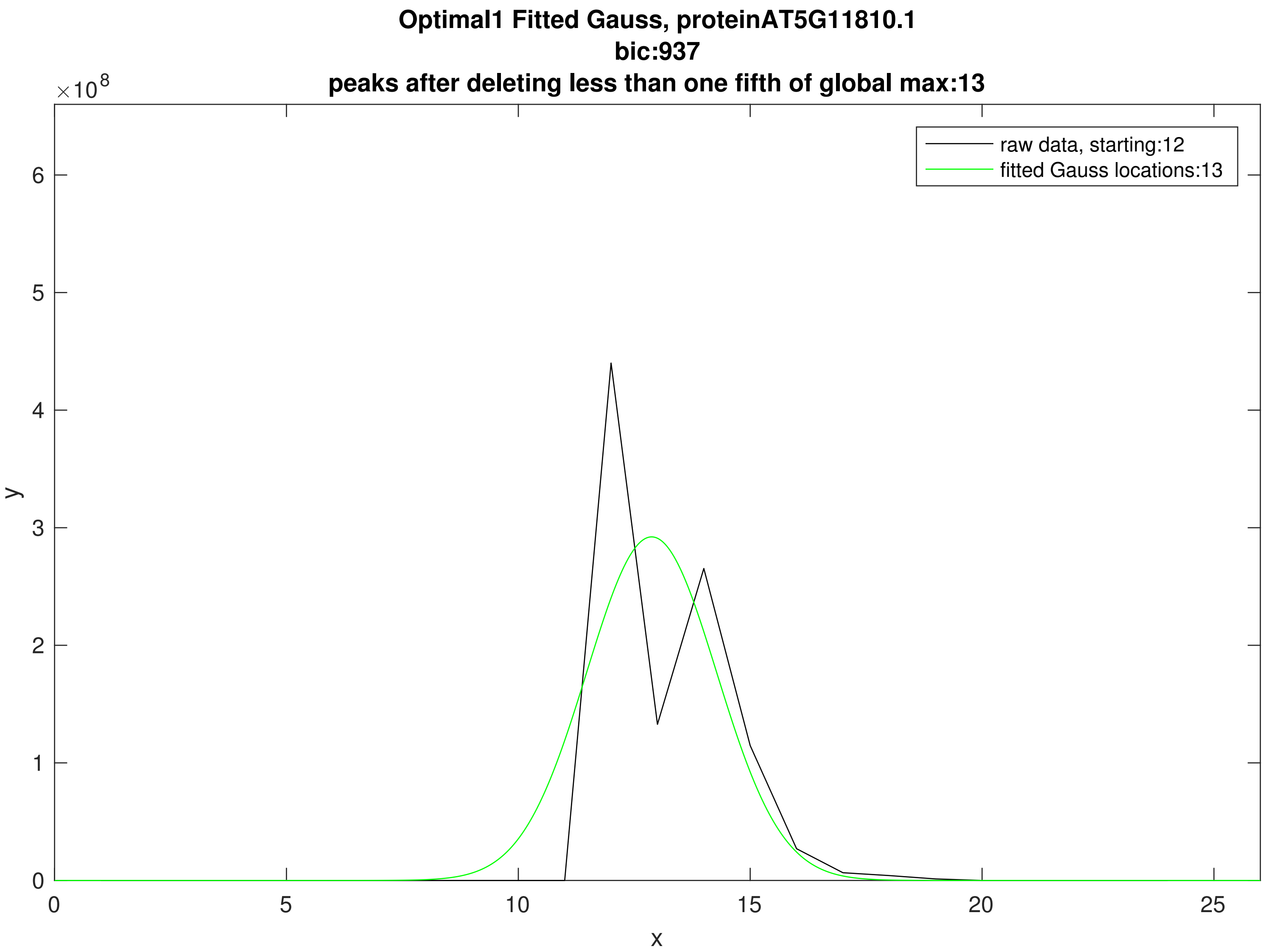
Optimal1 Fitted Gauss, proteinAT5G11810.1
bic:937
peaks after deleting less than one fifth of global max:13
×10^8
raw data, starting:12
fitted Gauss locations:13
y
x
0
1
2
3
4
5
6
0
5
10
15
20
25

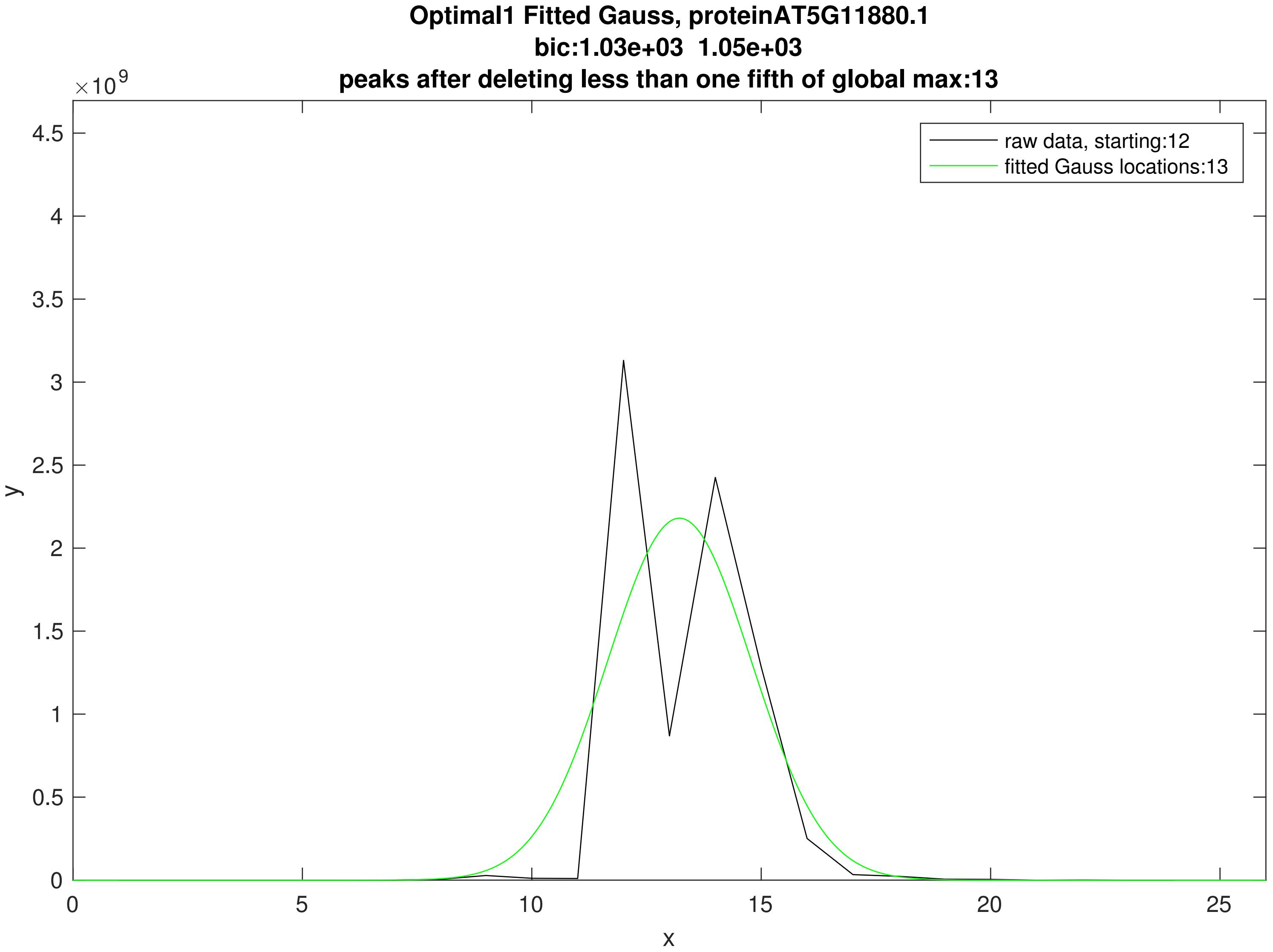
Optimal1 Fitted Gauss, proteinAT5G11880.1
bic:1.03e+03  1.05e+03
peaks after deleting less than one fifth of global max:13
×10^9
raw data, starting:12
fitted Gauss locations:13
y
x

Optimal1 Fitted Gauss, proteinAT5G11920.2;AT5G11920.1
bic:762
peaks after deleting less than one fifth of global max:13

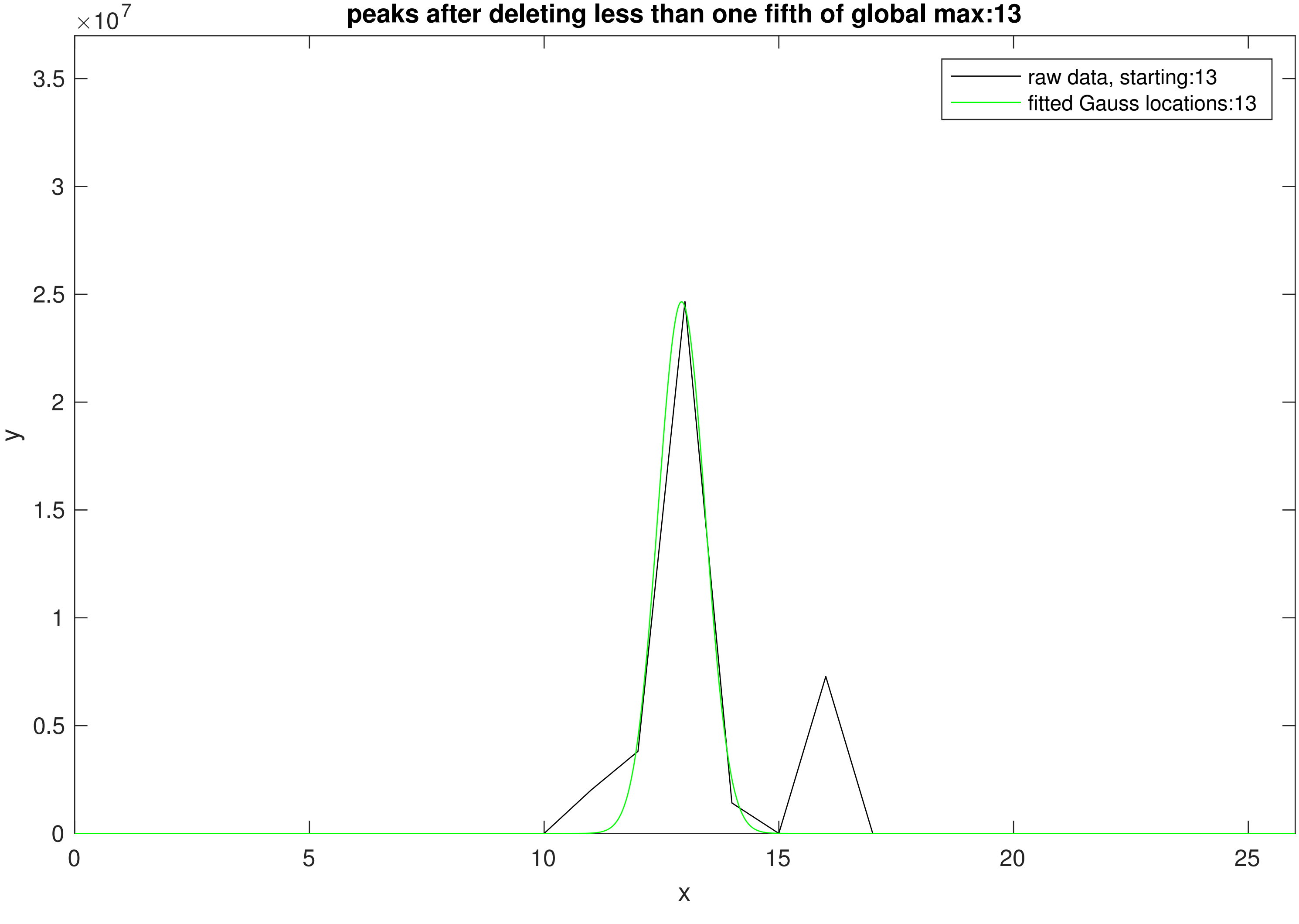

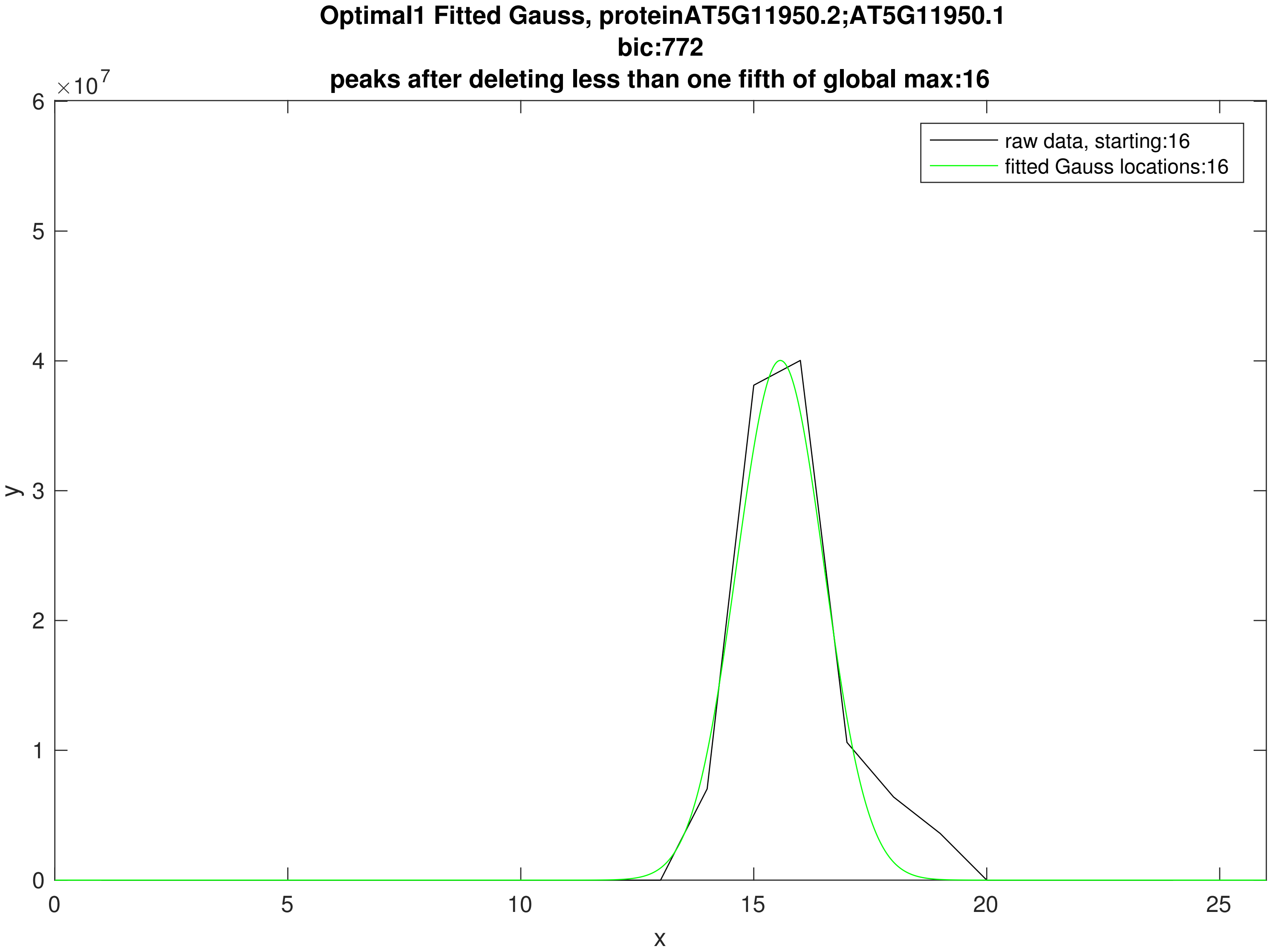

Optimal1 Fitted Gauss, proteinAT5G11950.2;AT5G11950.1
bic:772
peaks after deleting less than one fifth of global max:16
×10^7
raw data, starting:16
fitted Gauss locations:16
y
x
0
1
2
3
4
5
6
0
5
10
15
20
25

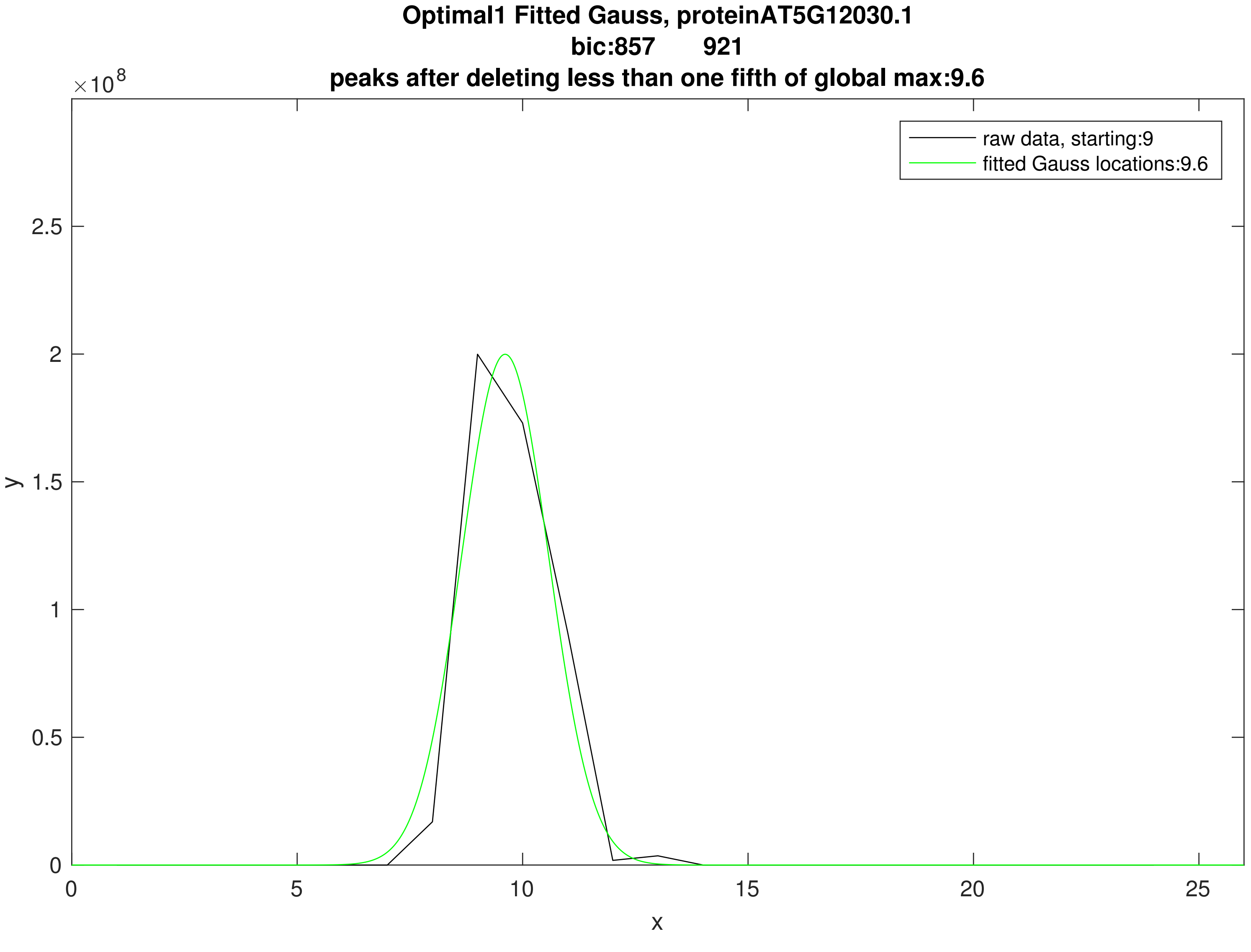
Optimal1 Fitted Gauss, proteinAT5G12030.1
bic:857 921
peaks after deleting less than one fifth of global max:9.6
×10^8
2.5
2
1.5
1
0.5
0
y
0
5
10
15
20
25
x
raw data, starting:9
fitted Gauss locations:9.6

**Optimal1 Fitted Gauss, proteinAT5G12040.1;AT5G12040.2**
**bic:973 982 992**
**peaks after deleting less than one fifth of global max:15**

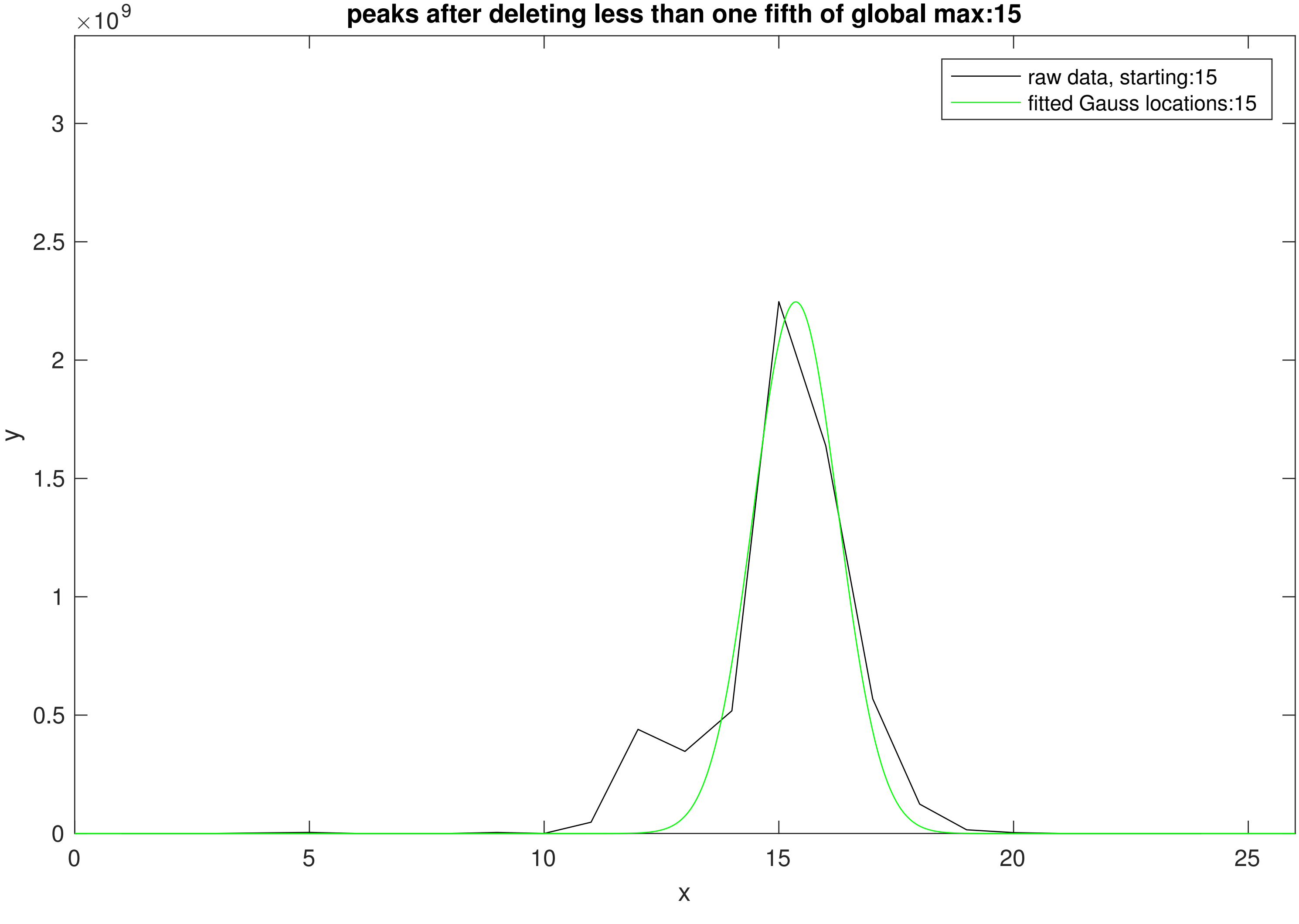

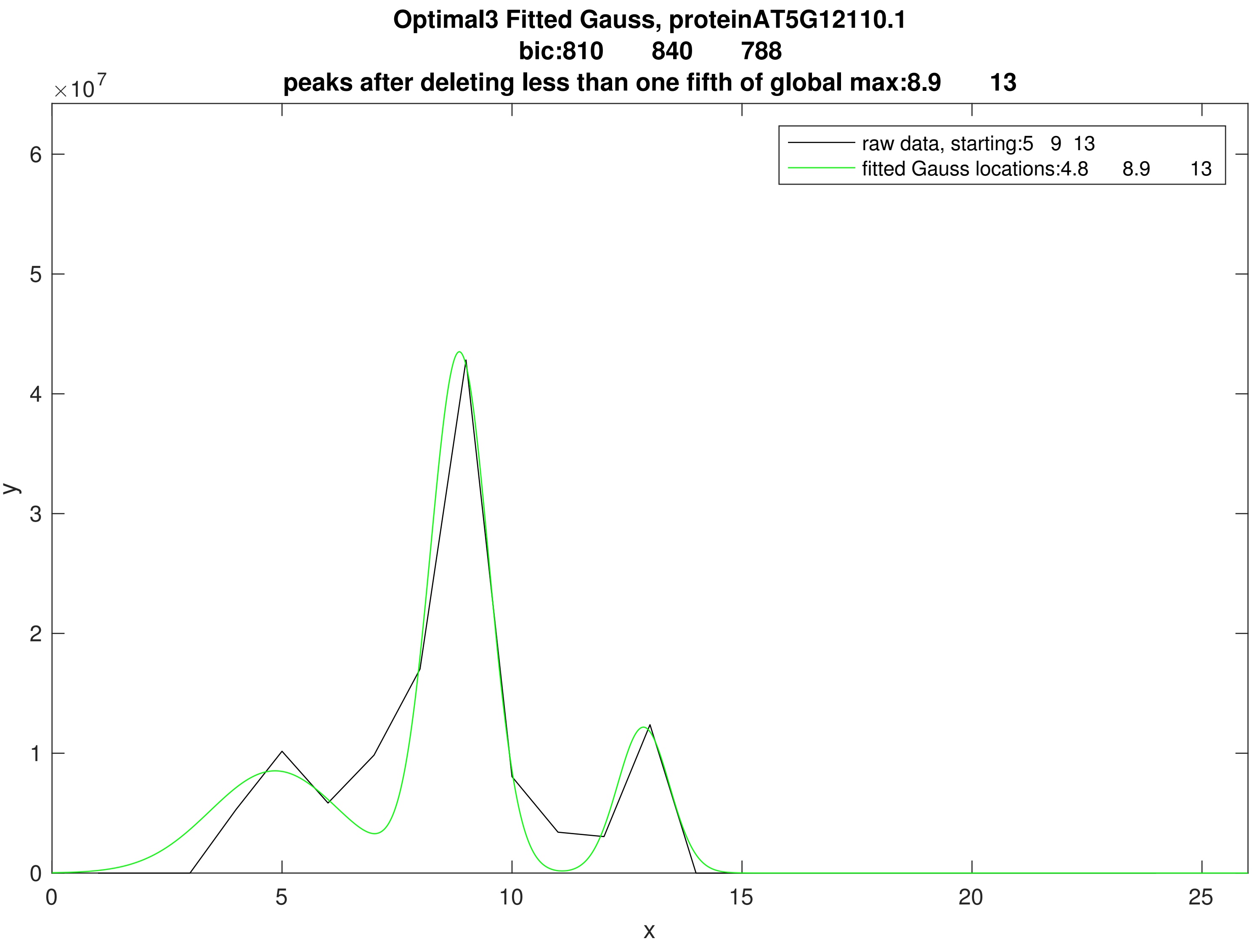

Optimal3 Fitted Gauss, proteinAT5G12110.1
bic:810 840 788
peaks after deleting less than one fifth of global max:8.9 13
×10^7
raw data, starting:5 9 13
fitted Gauss locations:4.8 8.9 13
y
x
0
1
2
3
4
5
6
0
5
10
15
20
25

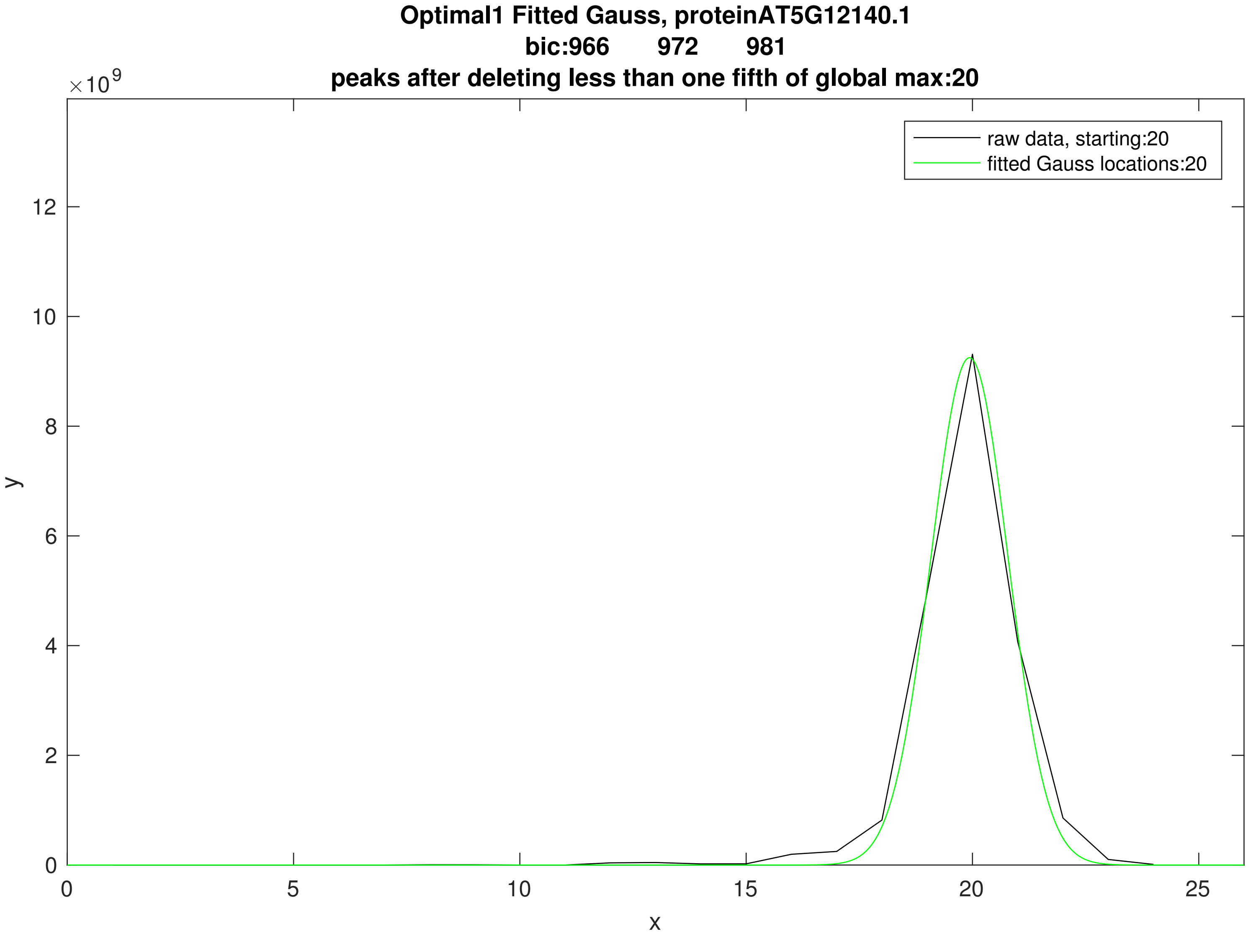

Optimal1 Fitted Gauss, proteinAT5G12140.1
bic:966 972 981
peaks after deleting less than one fifth of global max:20
×10^9
raw data, starting:20
fitted Gauss locations:20
y
x
0
2
4
6
8
10
12
0
5
10
15
20
25

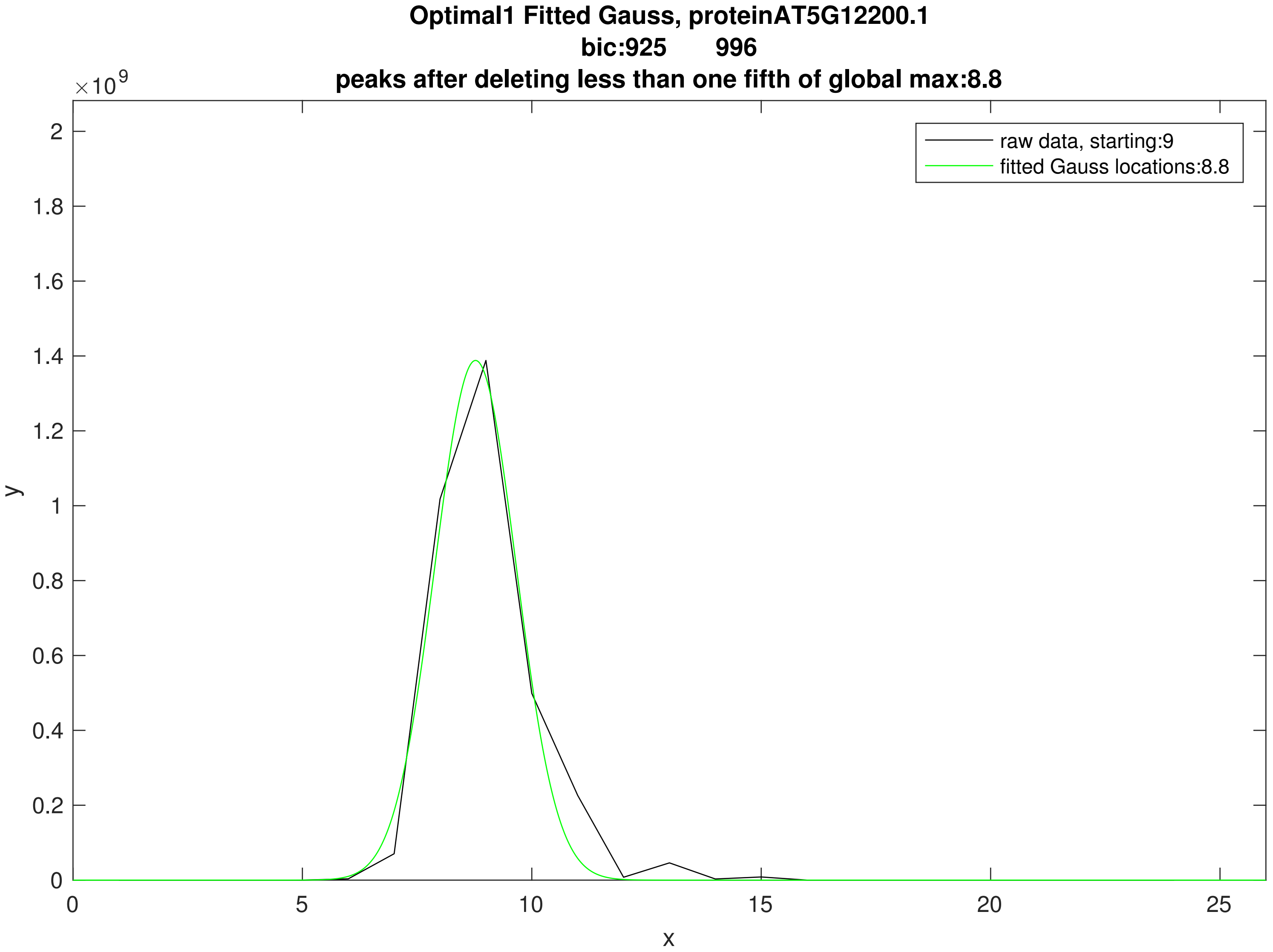
Optimal1 Fitted Gauss, proteinAT5G12200.1
bic:925 996
peaks after deleting less than one fifth of global max:8.8
×10^9
raw data, starting:9
fitted Gauss locations:8.8
y
x
0
0.2
0.4
0.6
0.8
1
1.2
1.4
1.6
1.8
2
5
10
15
20
25

**Optimal1 Fitted Gauss, proteinAT5G12210.2;AT5G12210.1**
**bic:835 887 837**
**peaks after deleting less than one fifth of global max:13**

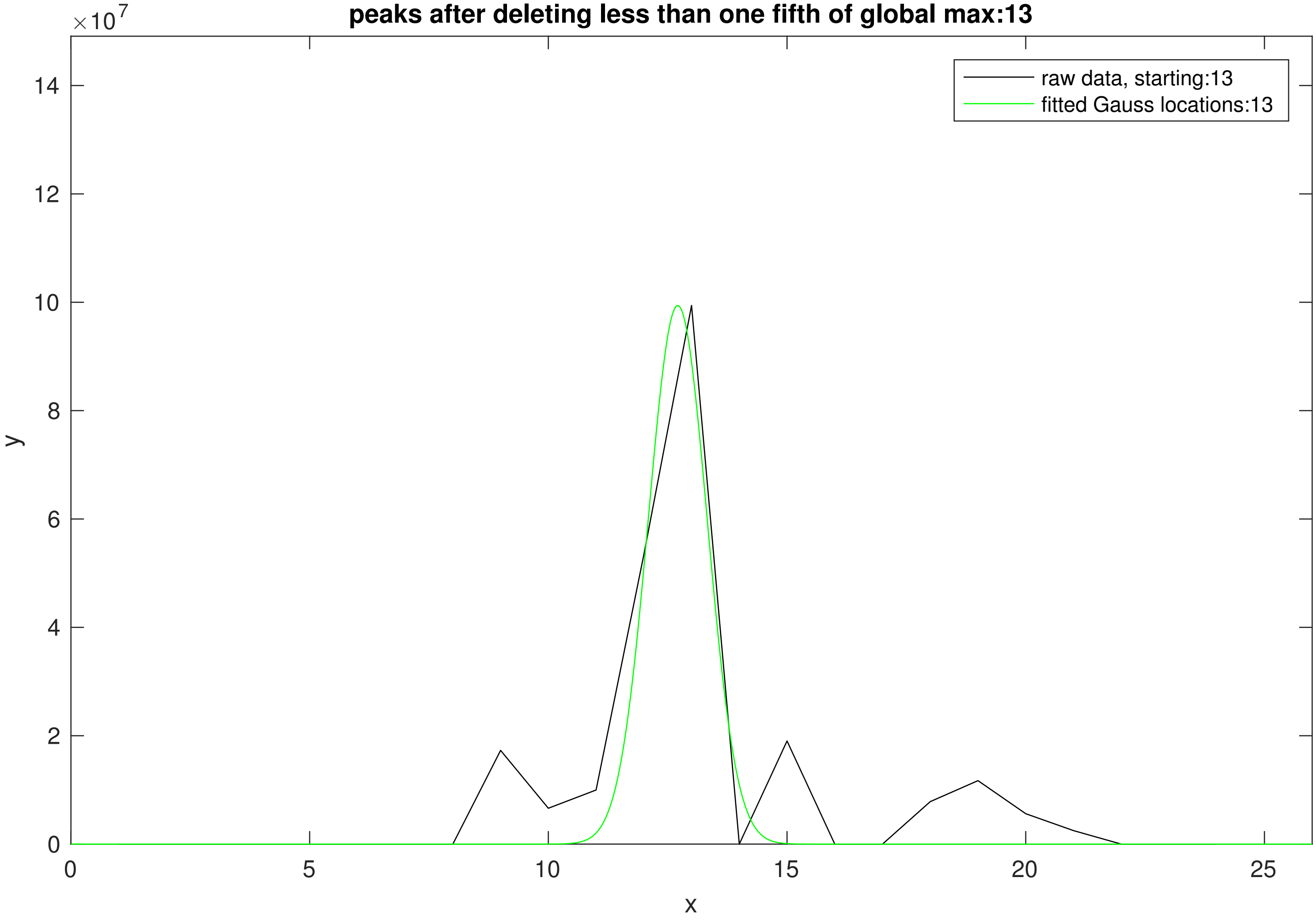

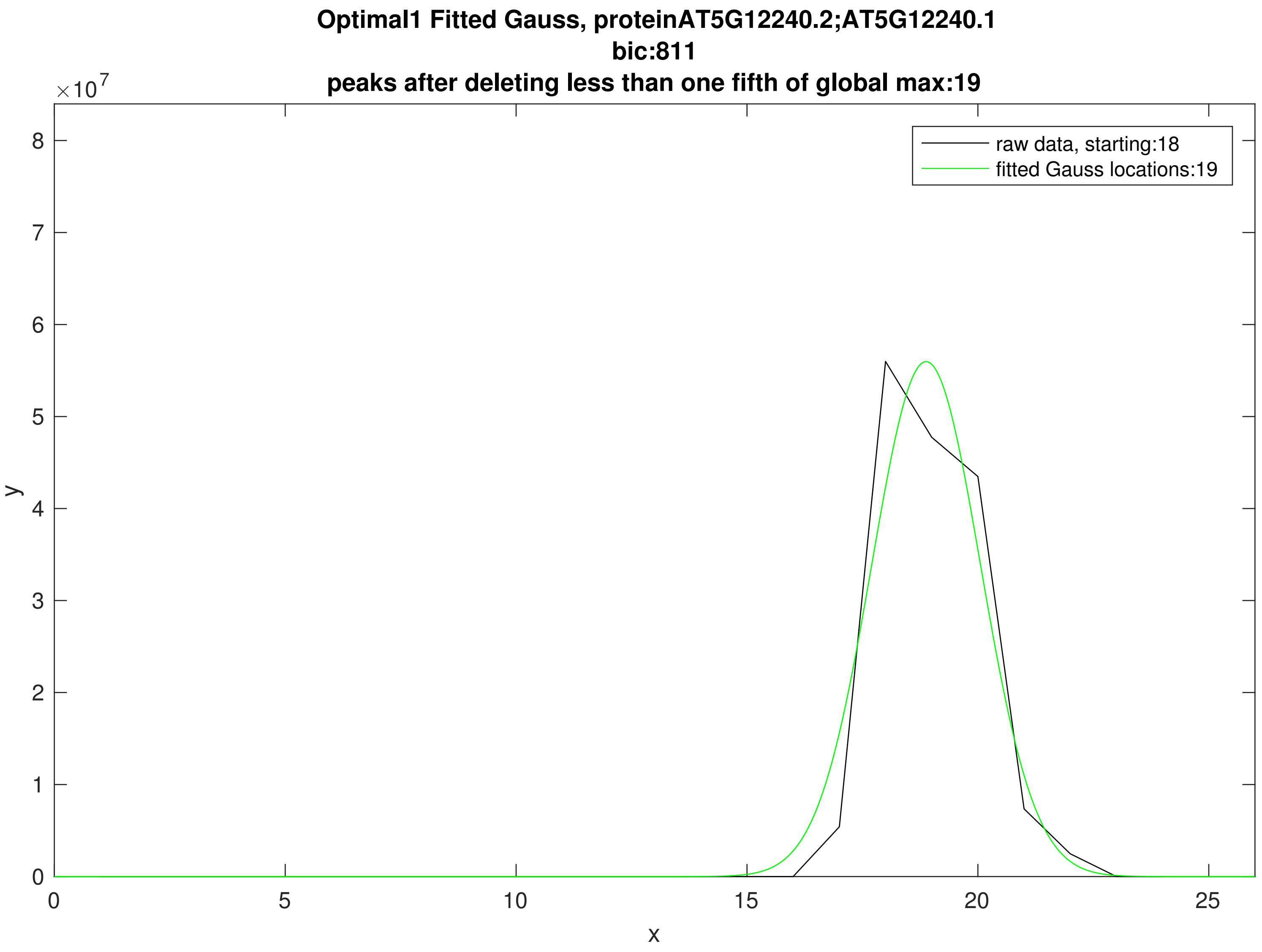
Optimal1 Fitted Gauss, proteinAT5G12240.2;AT5G12240.1
bic:811
peaks after deleting less than one fifth of global max:19
×10^7
raw data, starting:18
fitted Gauss locations:19
y
x
0
1
2
3
4
5
6
7
8
0
5
10
15
20
25

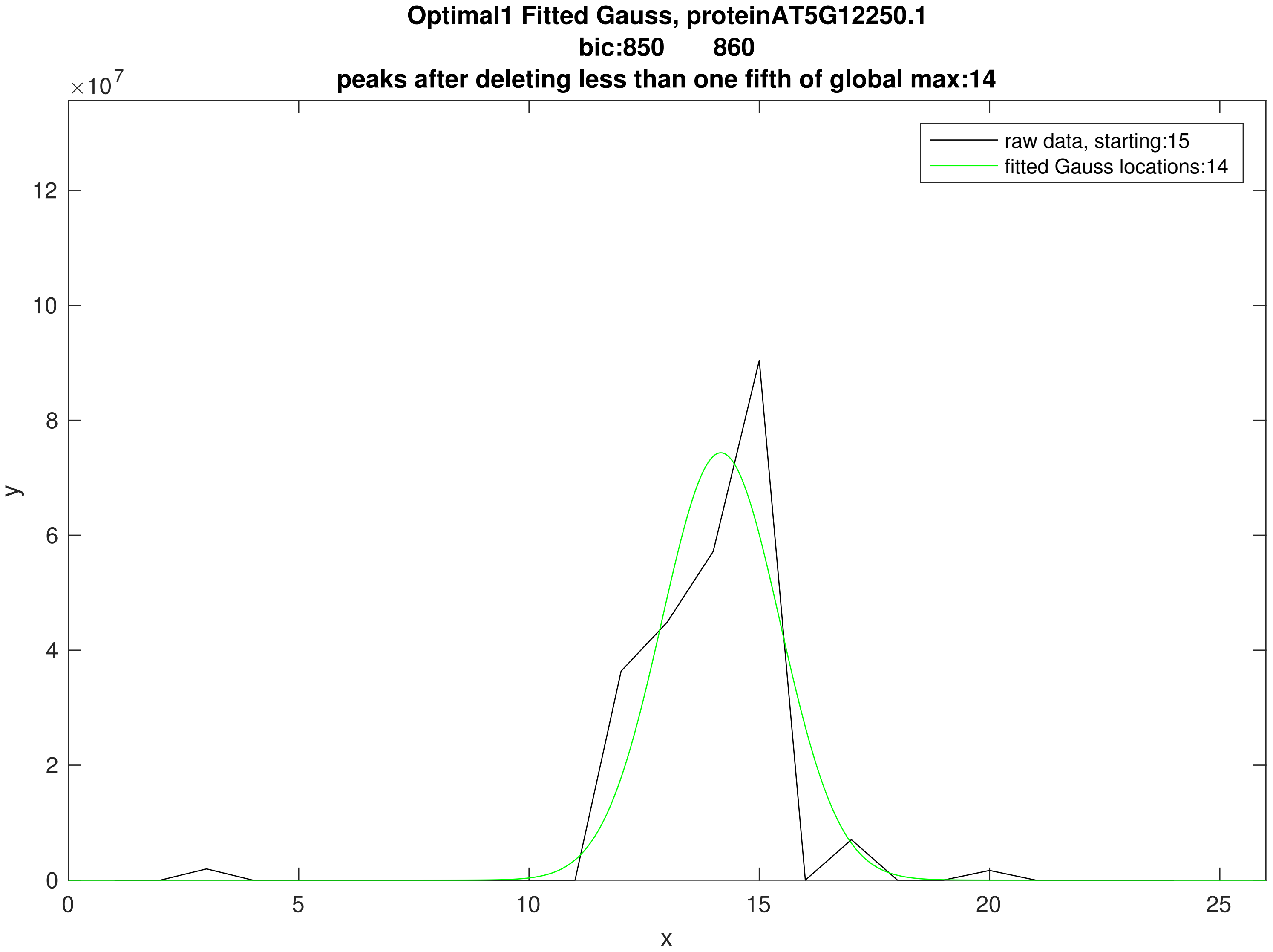

Optimal1 Fitted Gauss, proteinAT5G12250.1
bic:850 860
peaks after deleting less than one fifth of global max:14
×10^7
raw data, starting:15
fitted Gauss locations:14
y
x
0
2
4
6
8
10
12
0
5
10
15
20
25

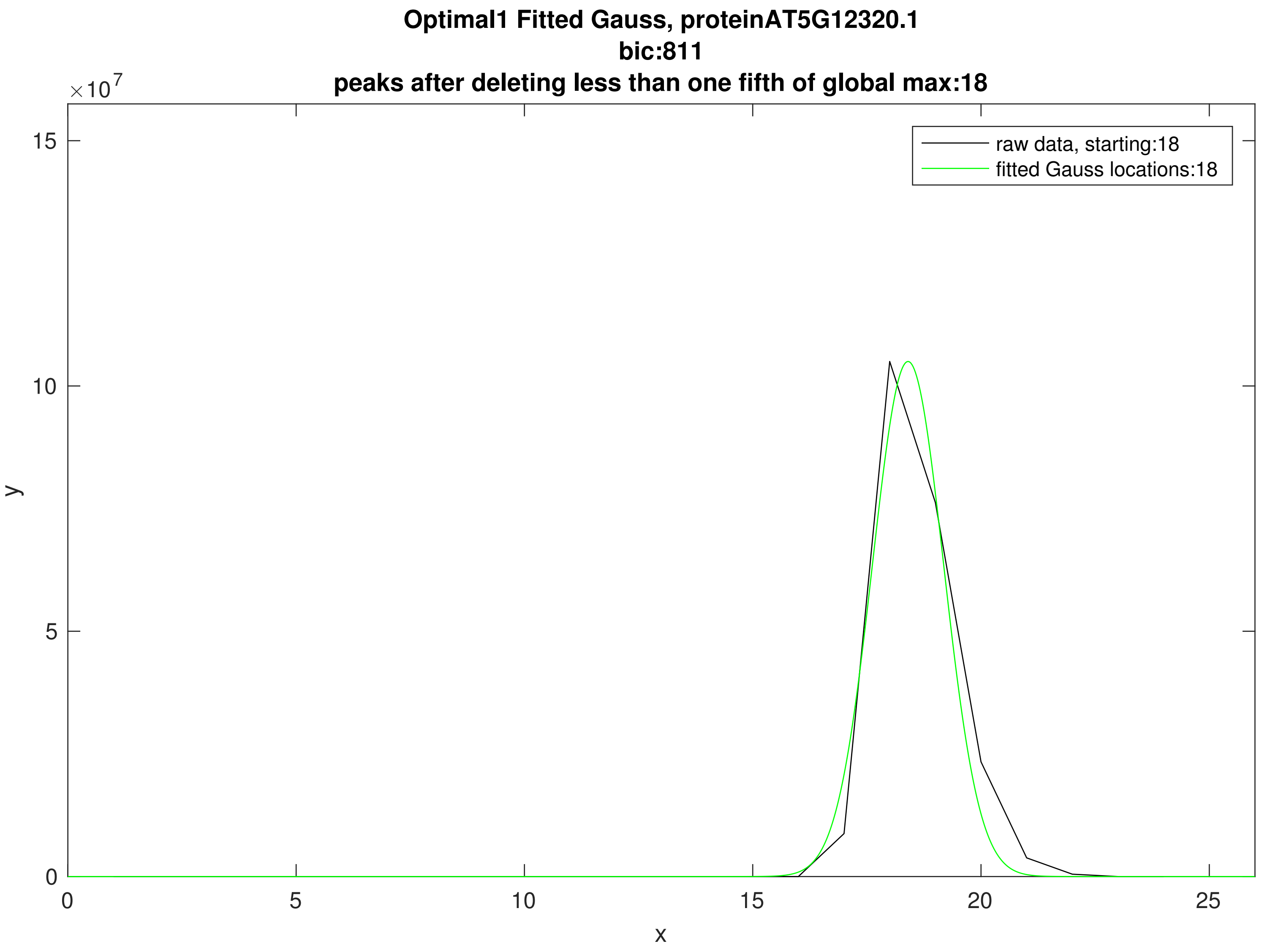

Optimal1 Fitted Gauss, proteinAT5G12320.1
bic:811
peaks after deleting less than one fifth of global max:18
×10^7
15
10
5
0
y
0
5
10
15
20
25
x
raw data, starting:18
fitted Gauss locations:18

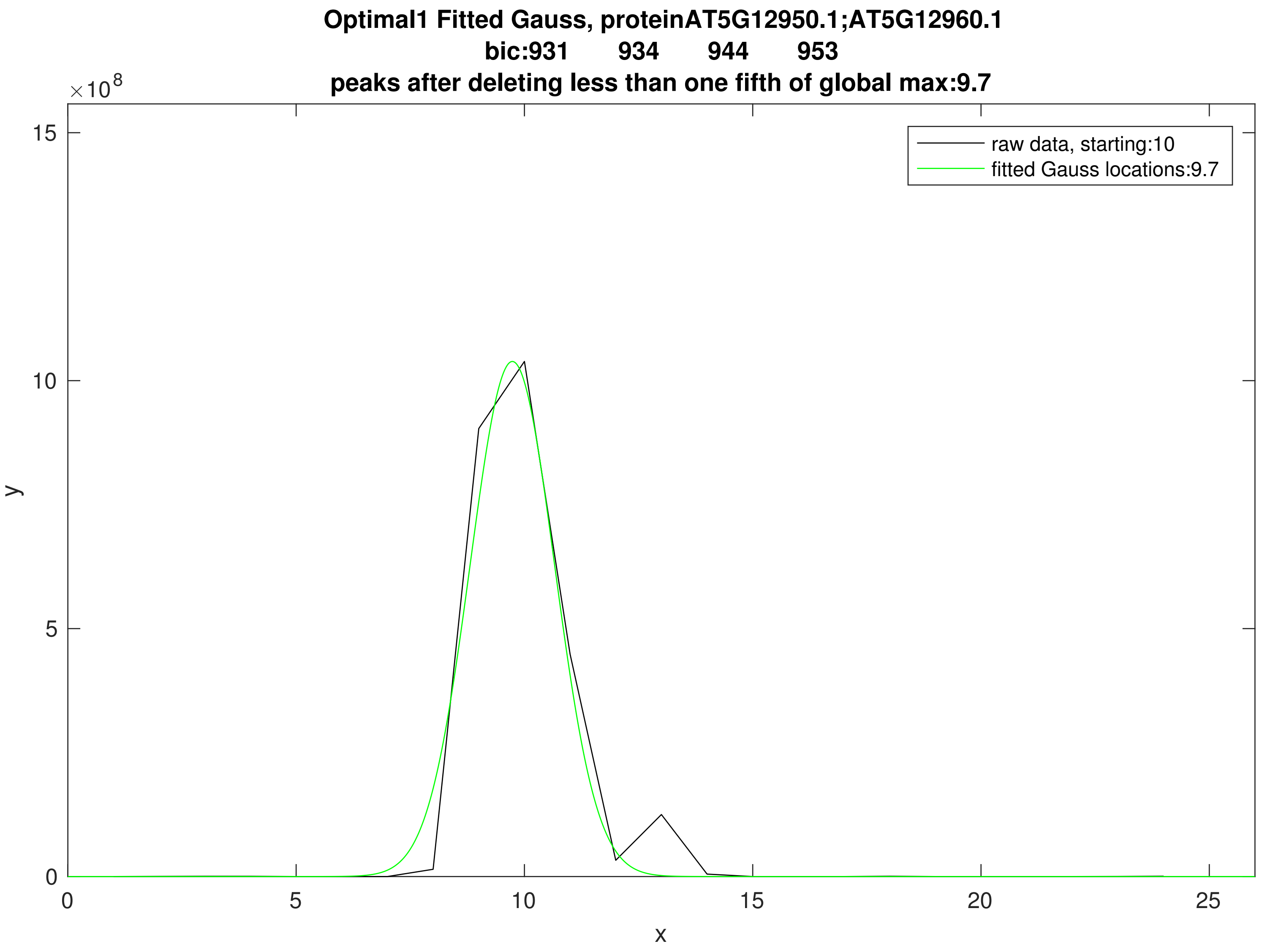

Optimal1 Fitted Gauss, proteinAT5G12950.1;AT5G12960.1
bic:931 934 944 953
peaks after deleting less than one fifth of global max:9.7
×10^8
15
10
5
0
y
0
5
10
15
20
25
x
raw data, starting:10
fitted Gauss locations:9.7

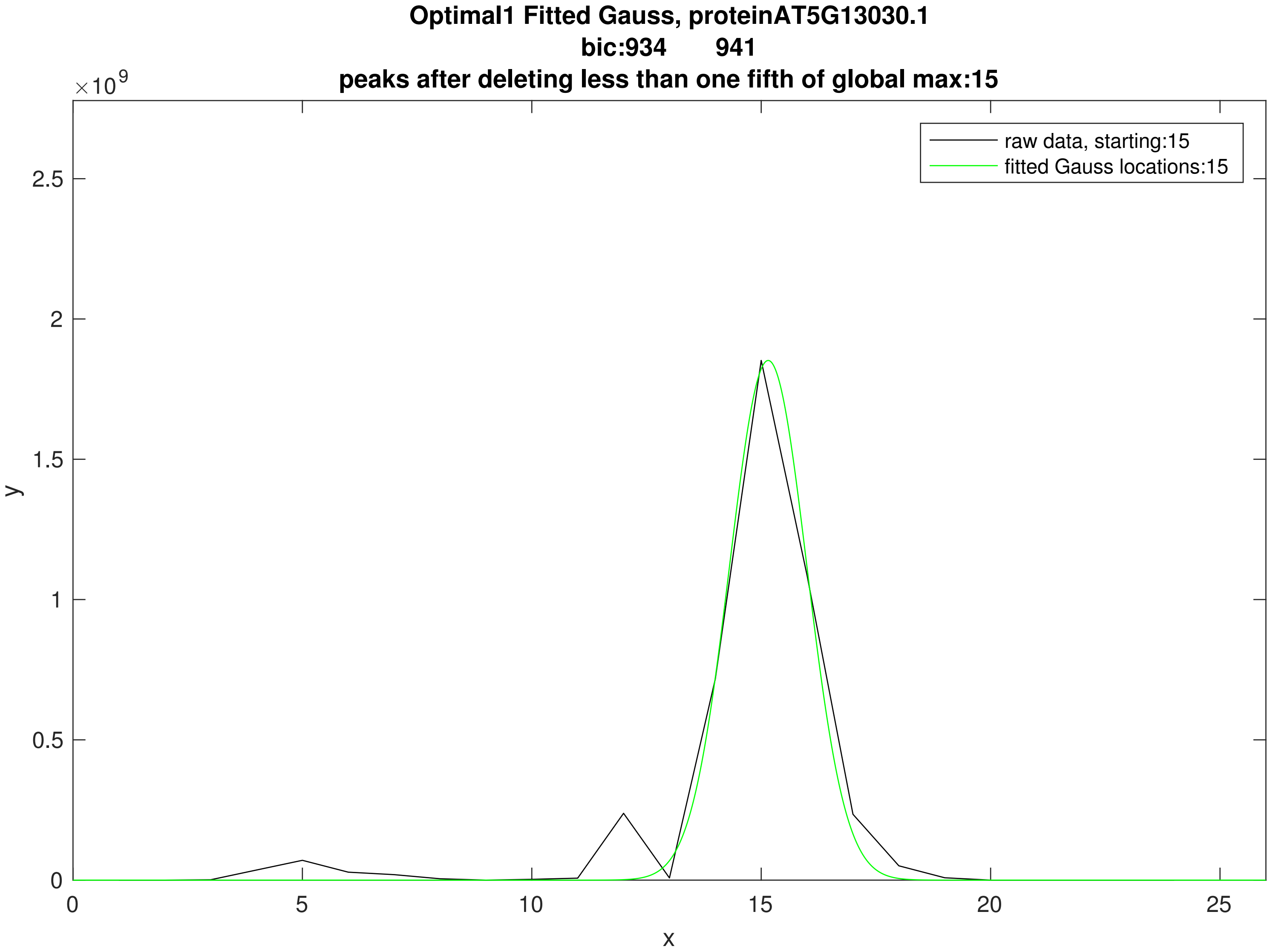
Optimal1 Fitted Gauss, proteinAT5G13030.1
bic:934 941
peaks after deleting less than one fifth of global max:15
×10^9
raw data, starting:15
fitted Gauss locations:15
y
x
0
0.5
1
1.5
2
2.5
0
5
10
15
20
25

#### Optimal2 Fitted Gauss, proteinAT5G13110.1;AT1G24280.1
bic:809 808 818
peaks after deleting less than one fifth of global max:9.6 18

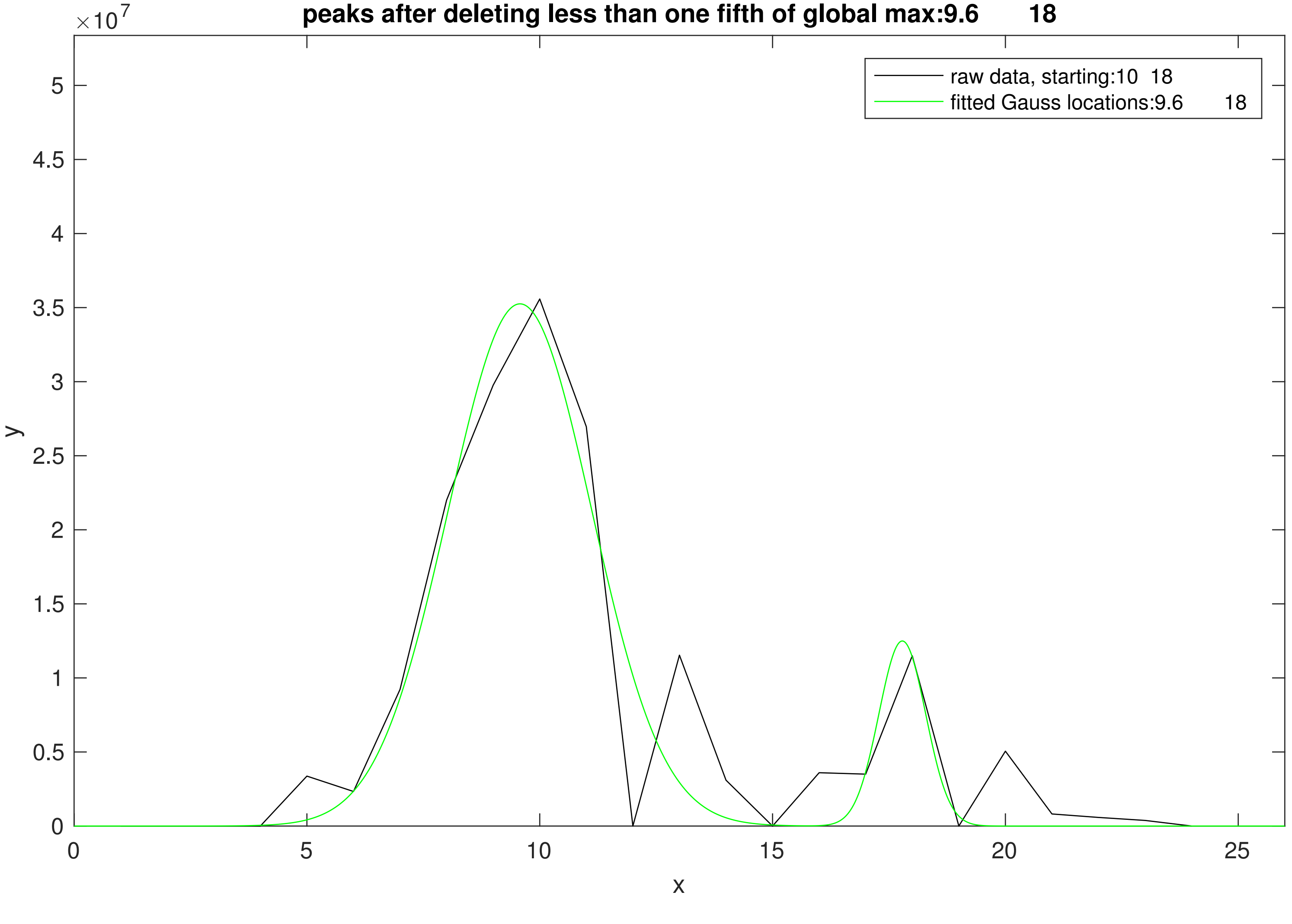

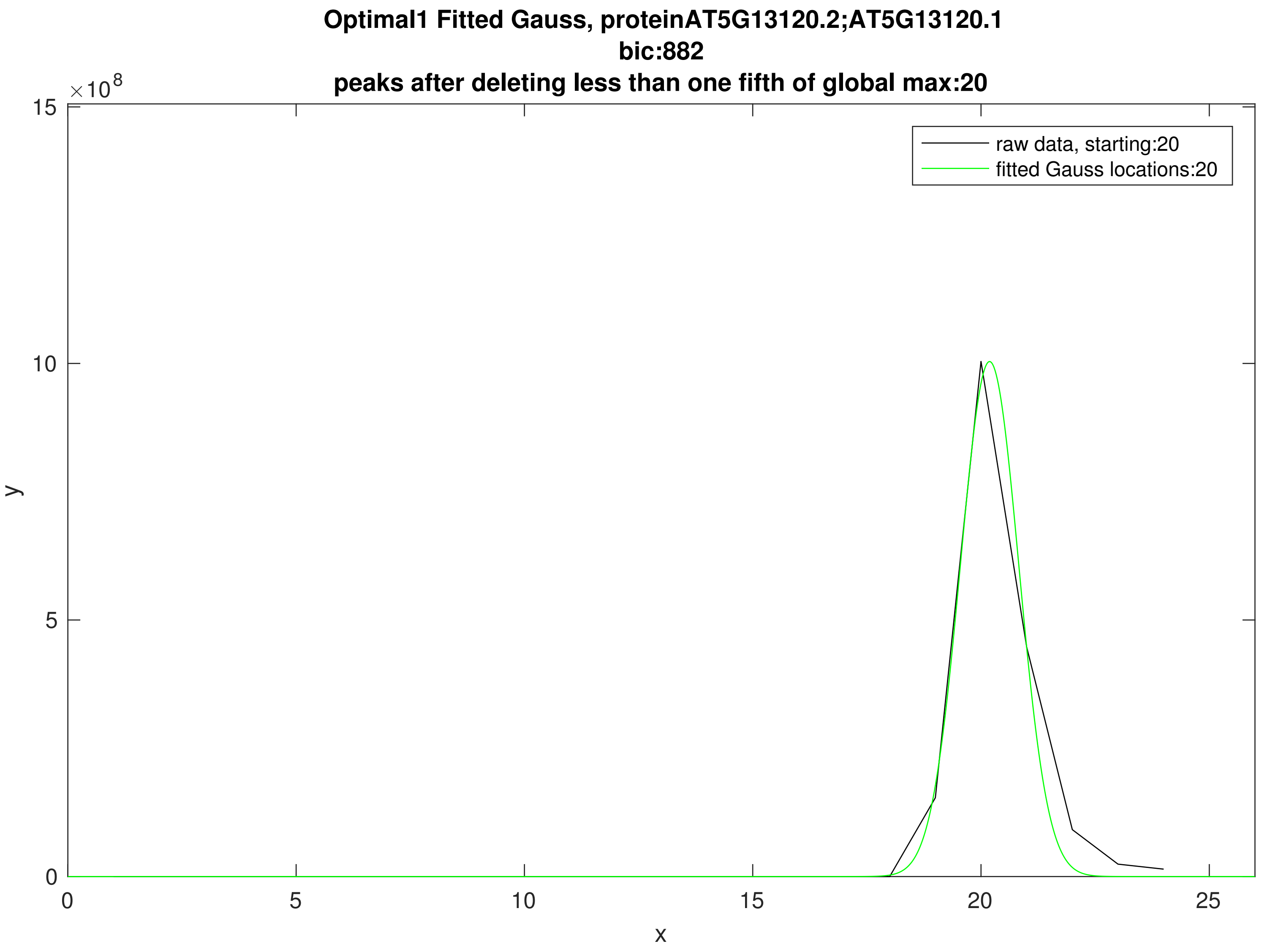

Optimal1 Fitted Gauss, proteinAT5G13120.2;AT5G13120.1
bic:882
peaks after deleting less than one fifth of global max:20
×10^8
15
10
5
0
y
raw data, starting:20
fitted Gauss locations:20
0
5
10
15
20
25
x

**Optimal1 Fitted Gauss, proteinAT5G13360.1;AT5G13360.2;AT5G13360.3**
**bic:906**
**peaks after deleting less than one fifth of global max:15**

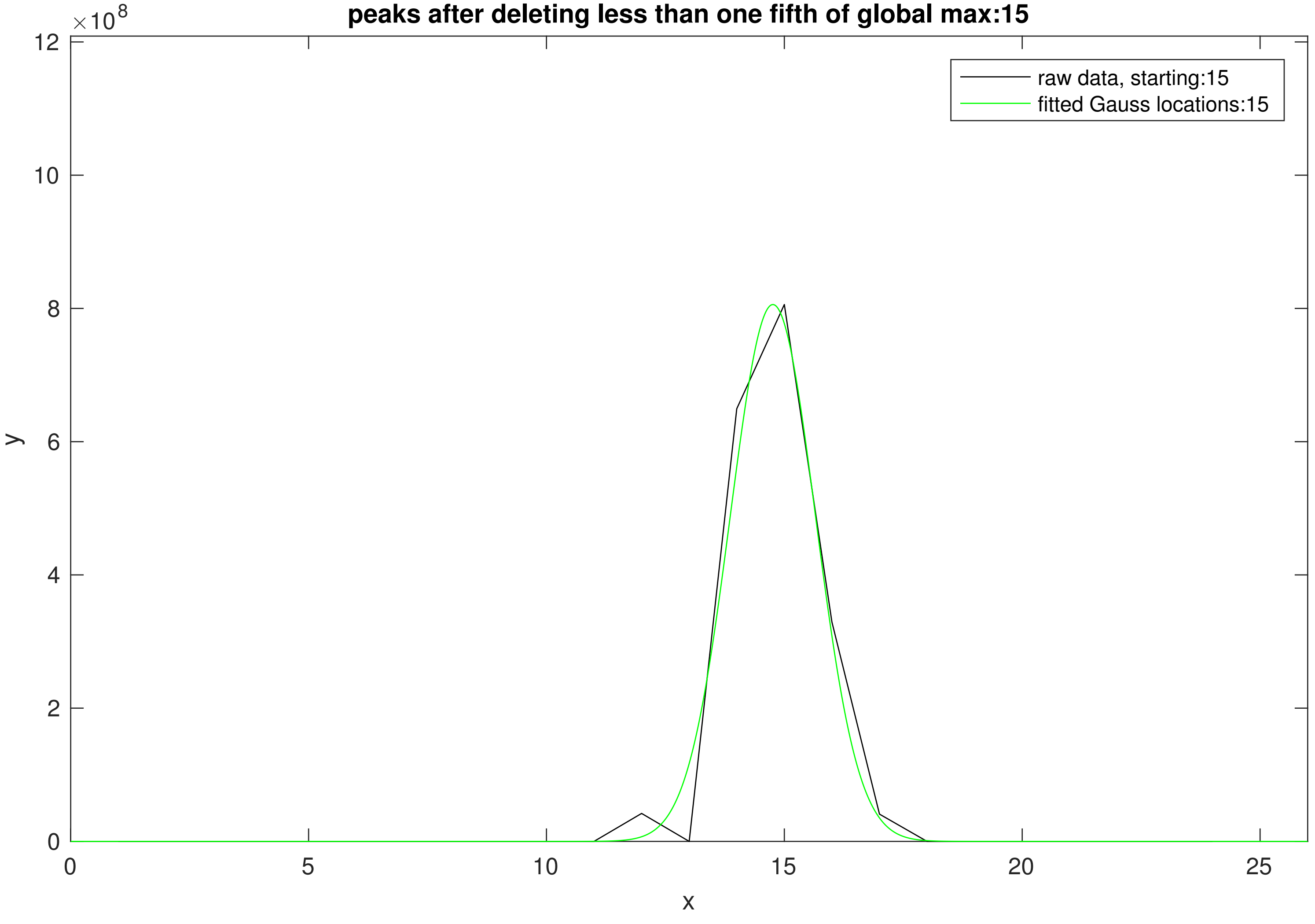

**Optimal1 Fitted Gauss, proteinAT5G13370.1**
**bic:902**
**peaks after deleting less than one fifth of global max:15**

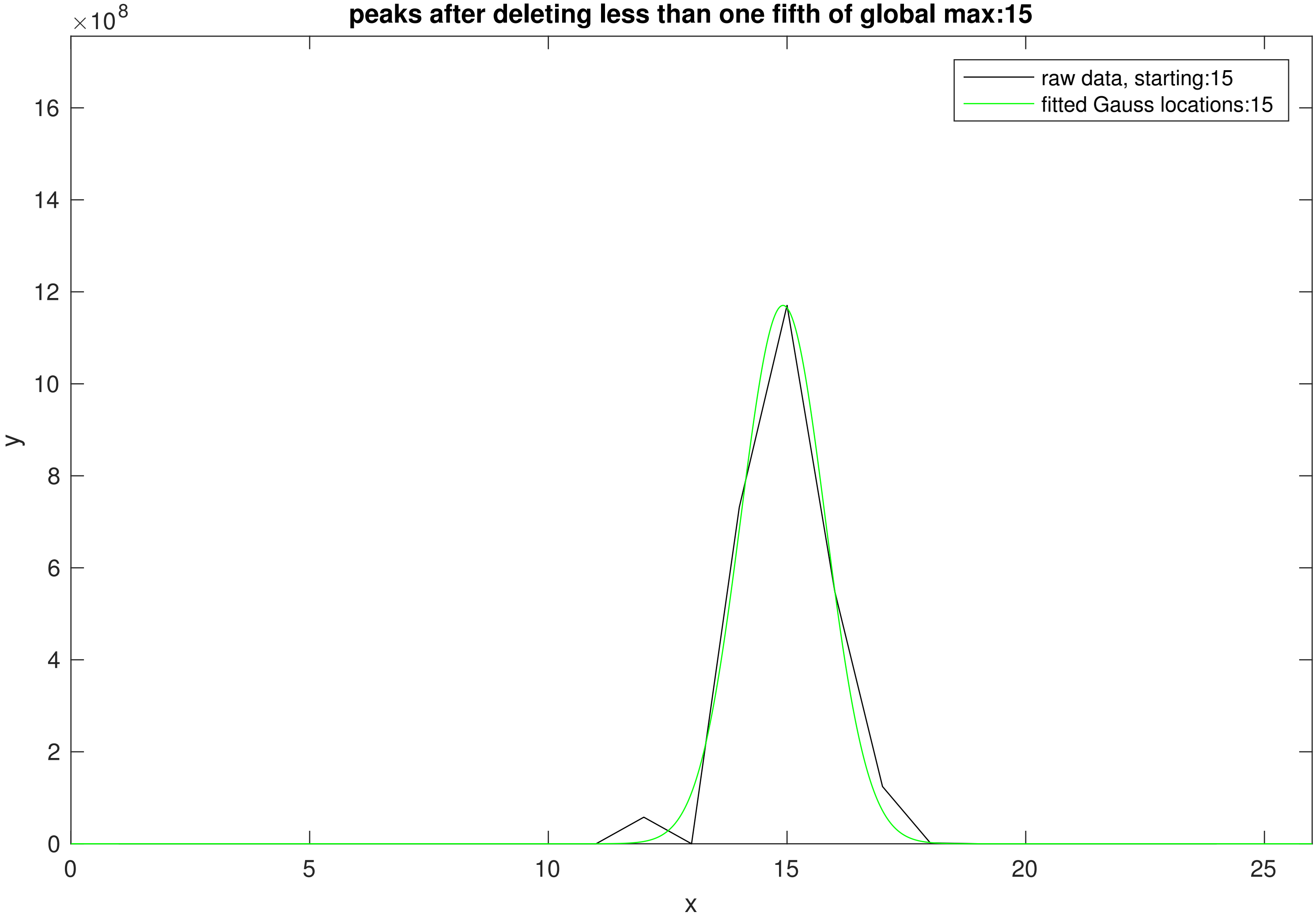

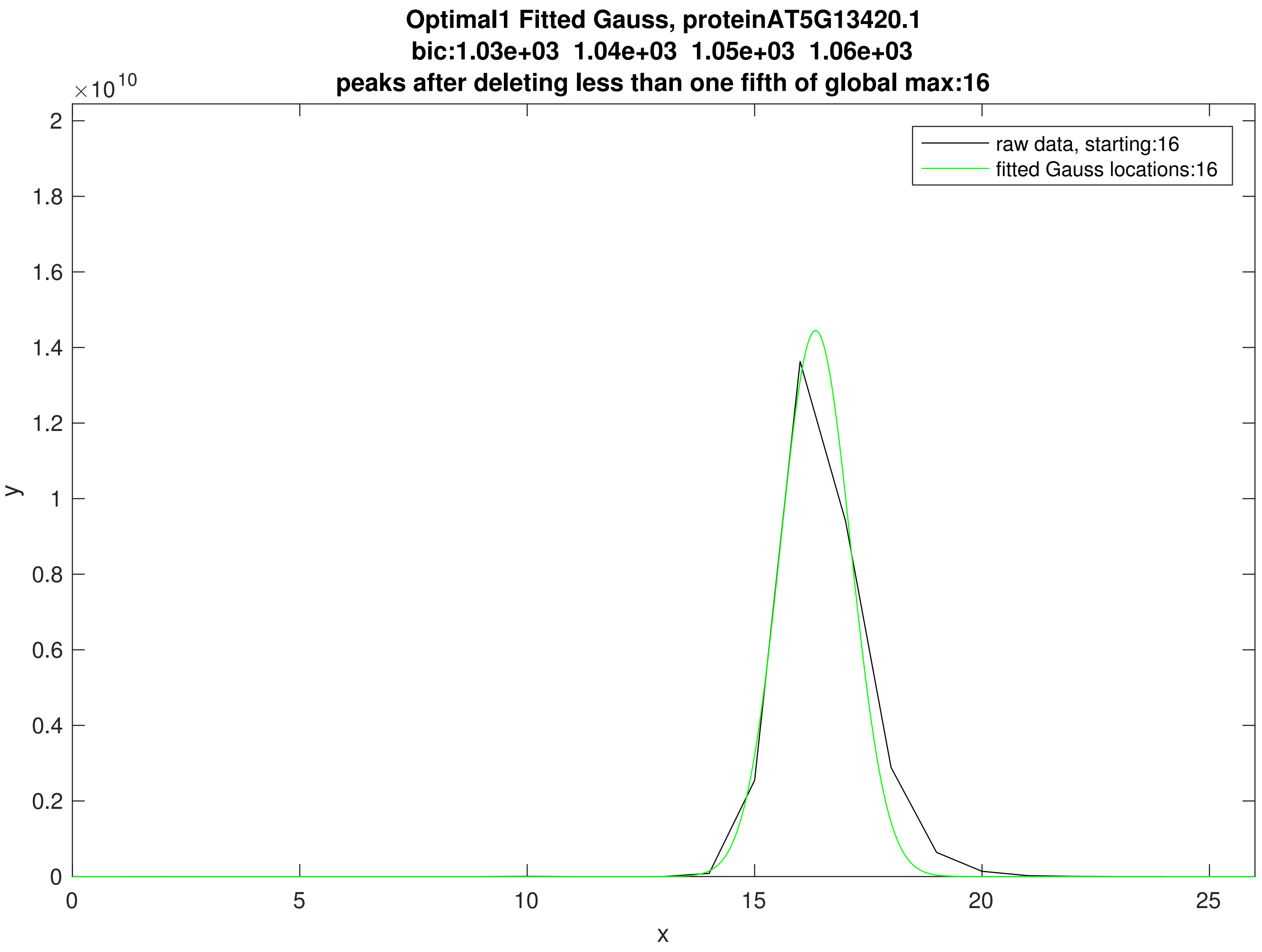

Optimal1 Fitted Gauss, proteinAT5G13420.1
bic:1.03e+03 1.04e+03 1.05e+03 1.06e+03
peaks after deleting less than one fifth of global max:16
×10^10
raw data, starting:16
fitted Gauss locations:16
y
x
0
0.2
0.4
0.6
0.8
1
1.2
1.4
1.6
1.8
2
0
5
10
15
20
25

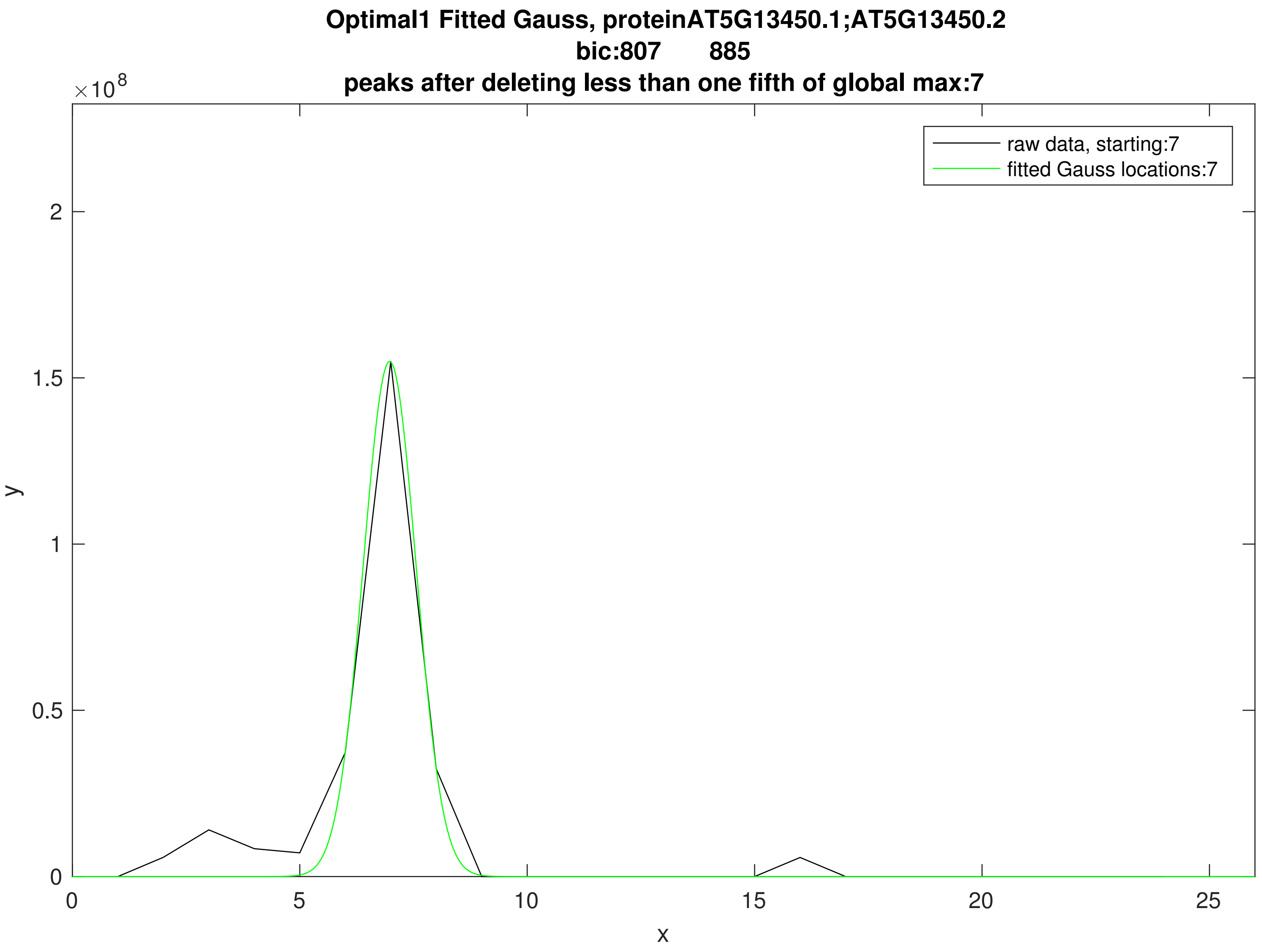

Optimal1 Fitted Gauss, proteinAT5G13450.1;AT5G13450.2
bic:807 885
peaks after deleting less than one fifth of global max:7
×10^8
raw data, starting:7
fitted Gauss locations:7
y
x
0
0.5
1
1.5
2
5
10
15
20
25

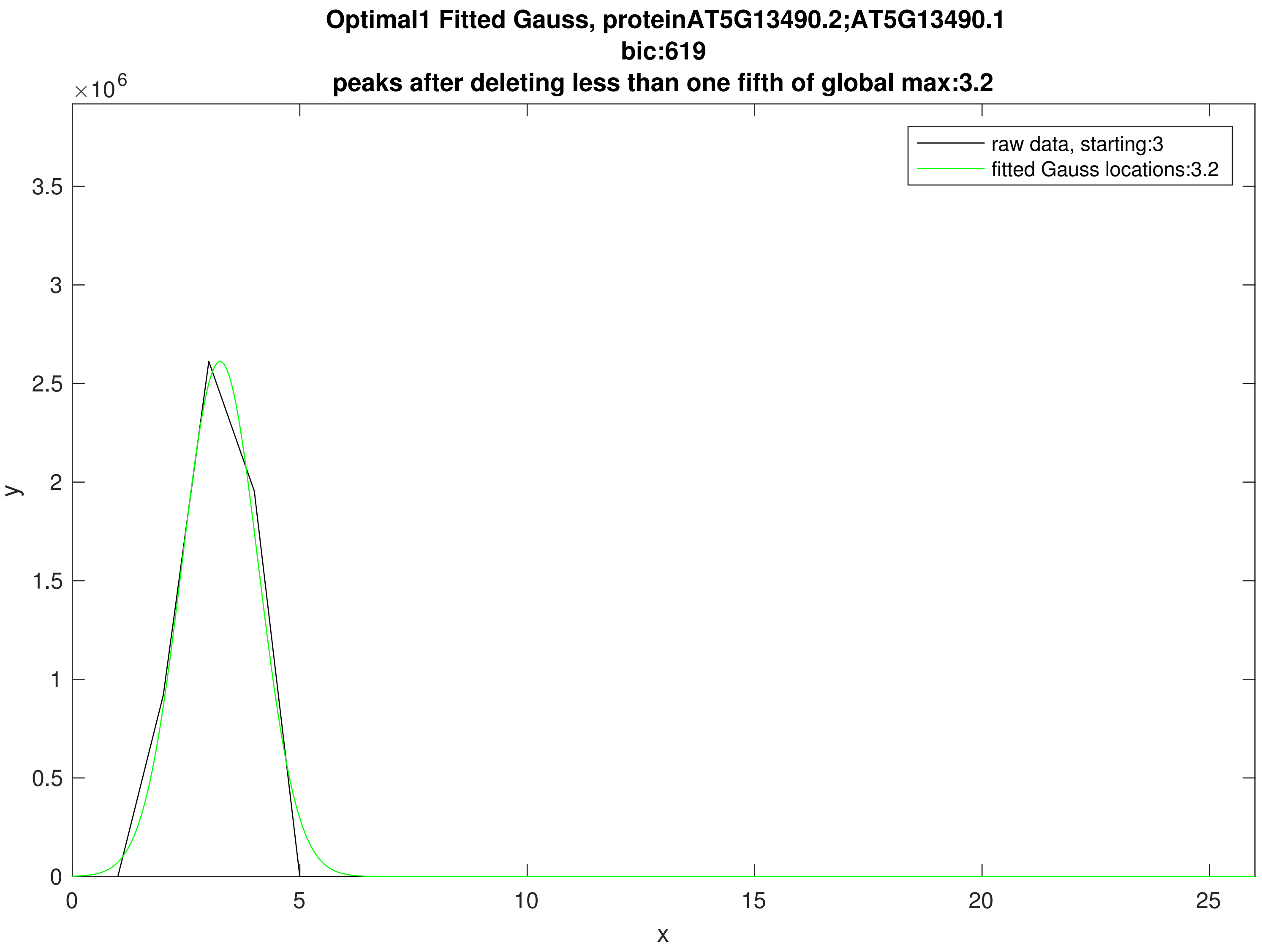
Optimal1 Fitted Gauss, proteinAT5G13490.2;AT5G13490.1
bic:619
peaks after deleting less than one fifth of global max:3.2
×10^6
raw data, starting:3
fitted Gauss locations:3.2
y
x
0
0.5
1
1.5
2
2.5
3
3.5
0
5
10
15
20
25

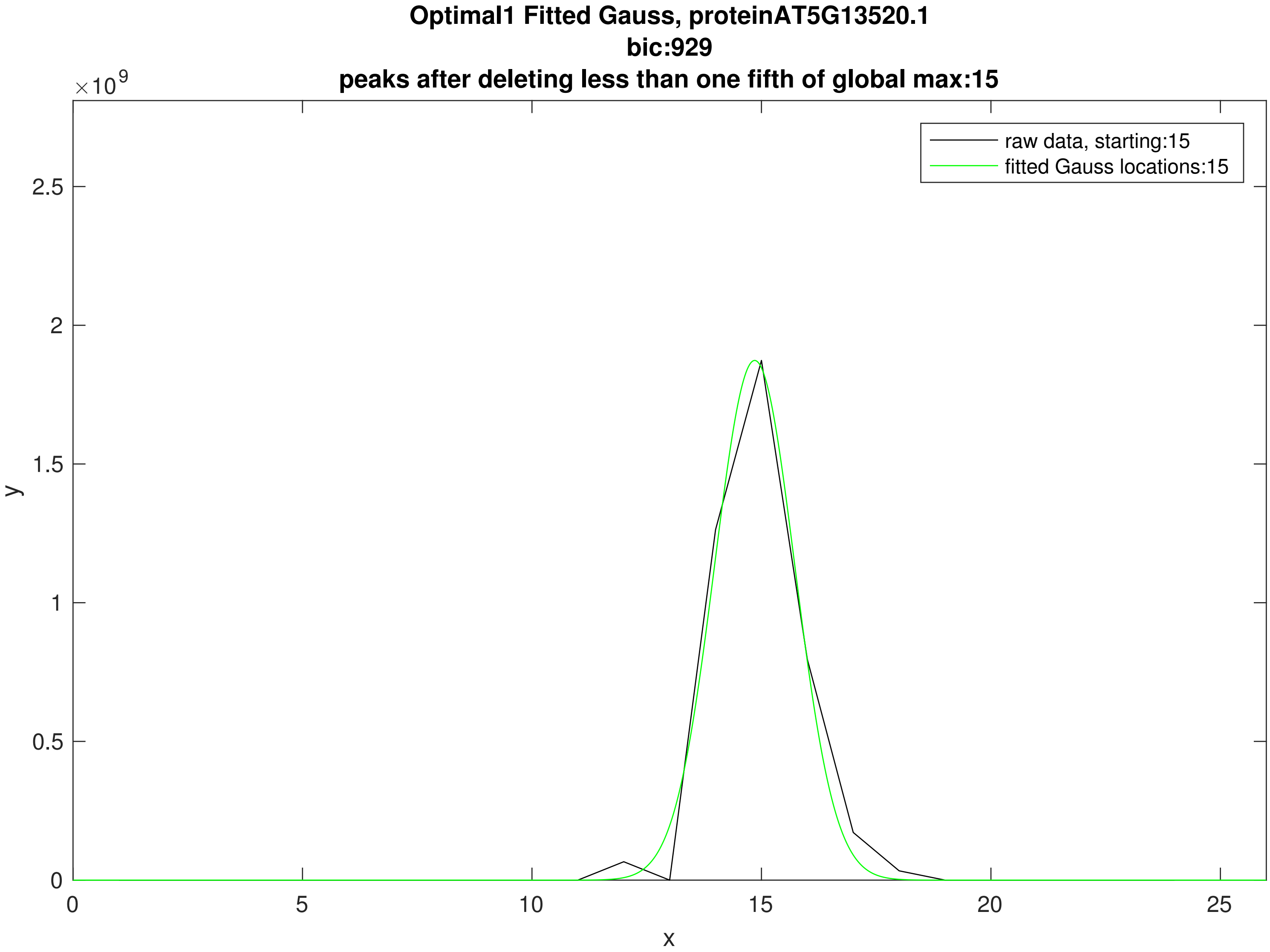

Optimal1 Fitted Gauss, proteinAT5G13520.1
bic:929
peaks after deleting less than one fifth of global max:15
×10^9
raw data, starting:15
fitted Gauss locations:15
2.5
2
1.5
1
0.5
0
y
0
5
10
15
20
25
x

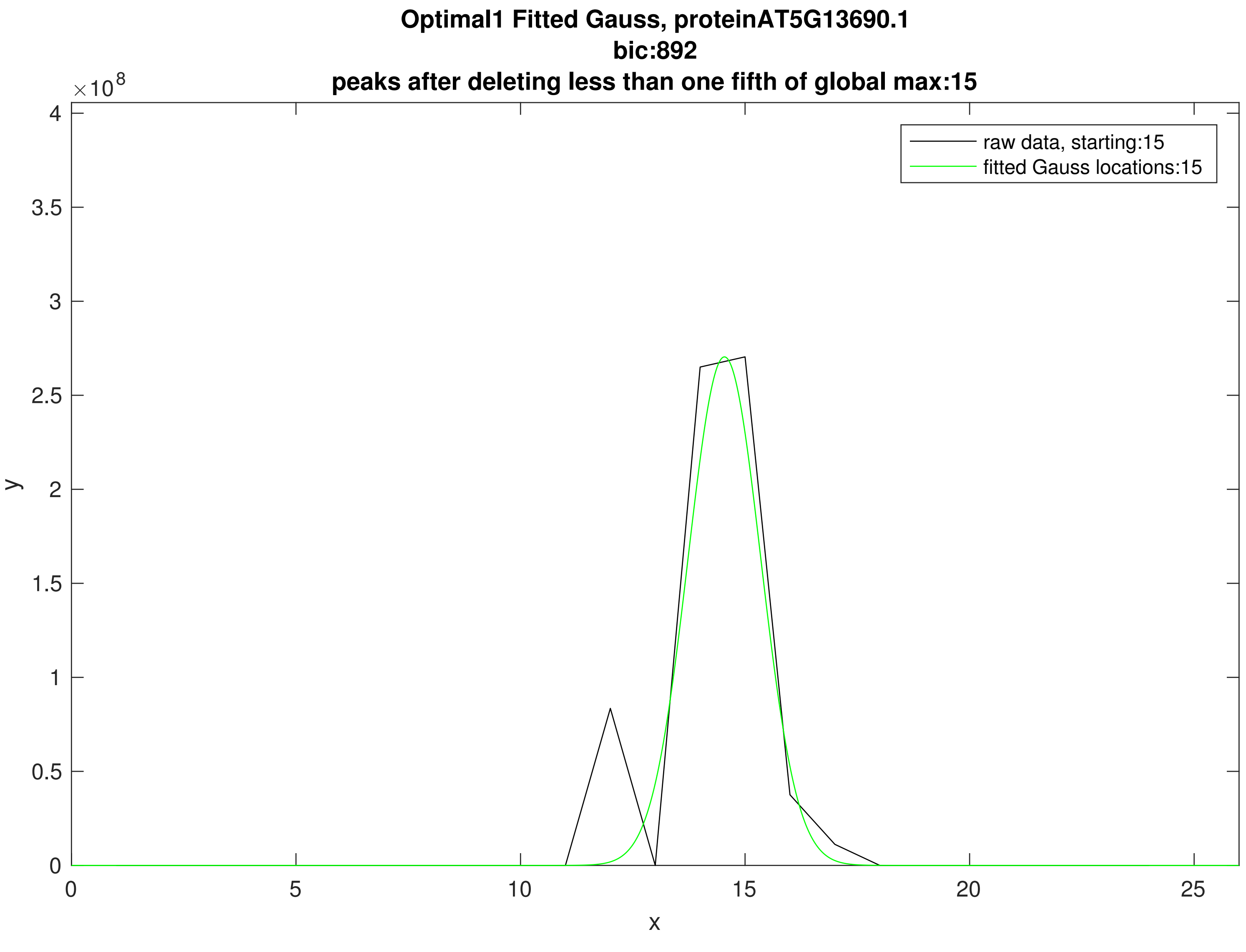
Optimal1 Fitted Gauss, proteinAT5G13690.1
bic:892
peaks after deleting less than one fifth of global max:15
×10^8
4
3.5
3
2.5
2
1.5
1
0.5
0
y
0
5
10
15
20
25
x
raw data, starting:15
fitted Gauss locations:15

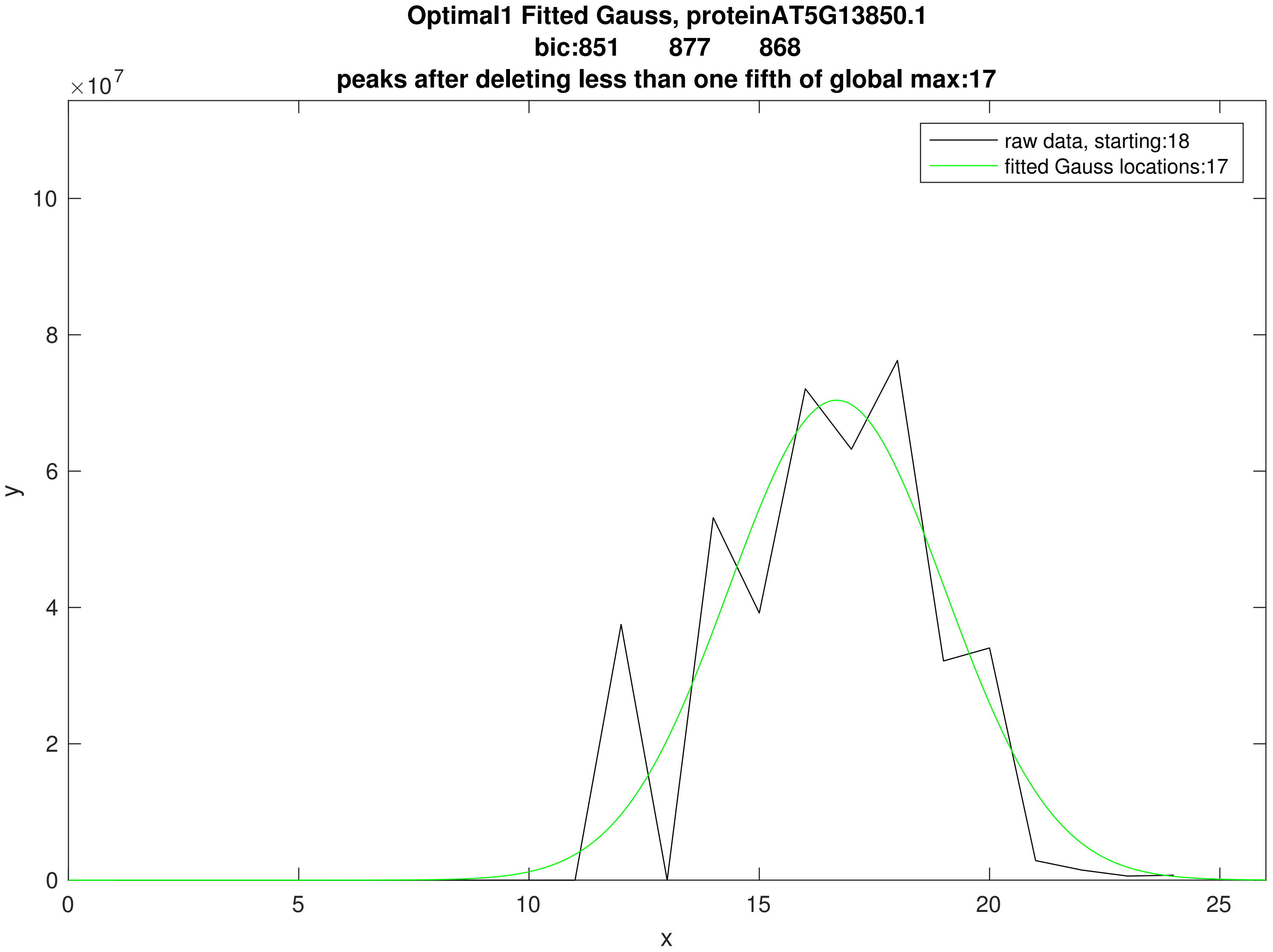
Optimal1 Fitted Gauss, proteinAT5G13850.1
bic:851 877 868
peaks after deleting less than one fifth of global max:17
×10^7
raw data, starting:18
fitted Gauss locations:17
y
x
0
2
4
6
8
10
0
5
10
15
20
25

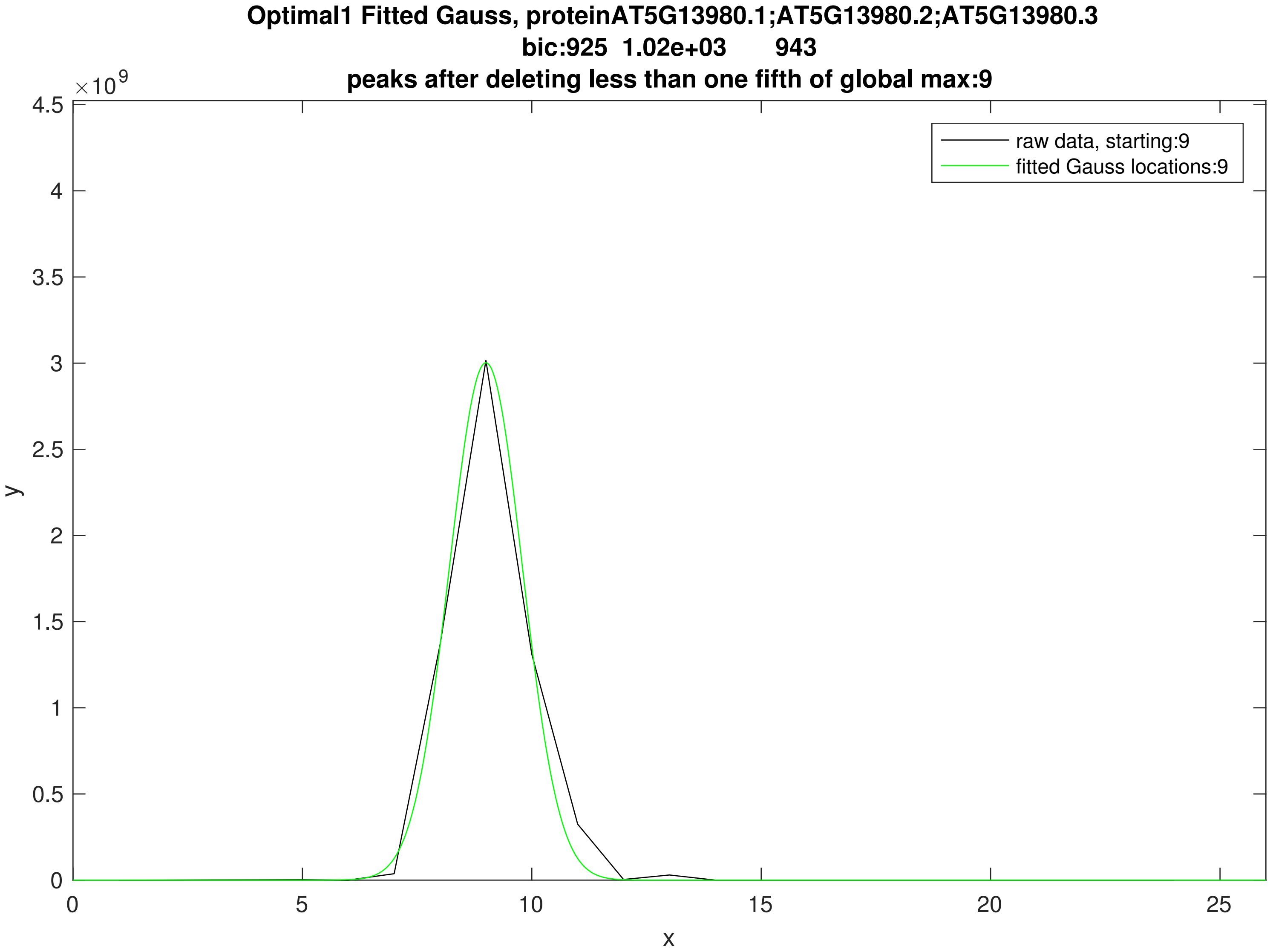
Optimal1 Fitted Gauss, proteinAT5G13980.1;AT5G13980.2;AT5G13980.3
bic:925 1.02e+03 943
peaks after deleting less than one fifth of global max:9
×10^9
4.5
4
3.5
3
2.5
2
1.5
1
0.5
0
y
0
5
10
15
20
25
x
raw data, starting:9
fitted Gauss locations:9

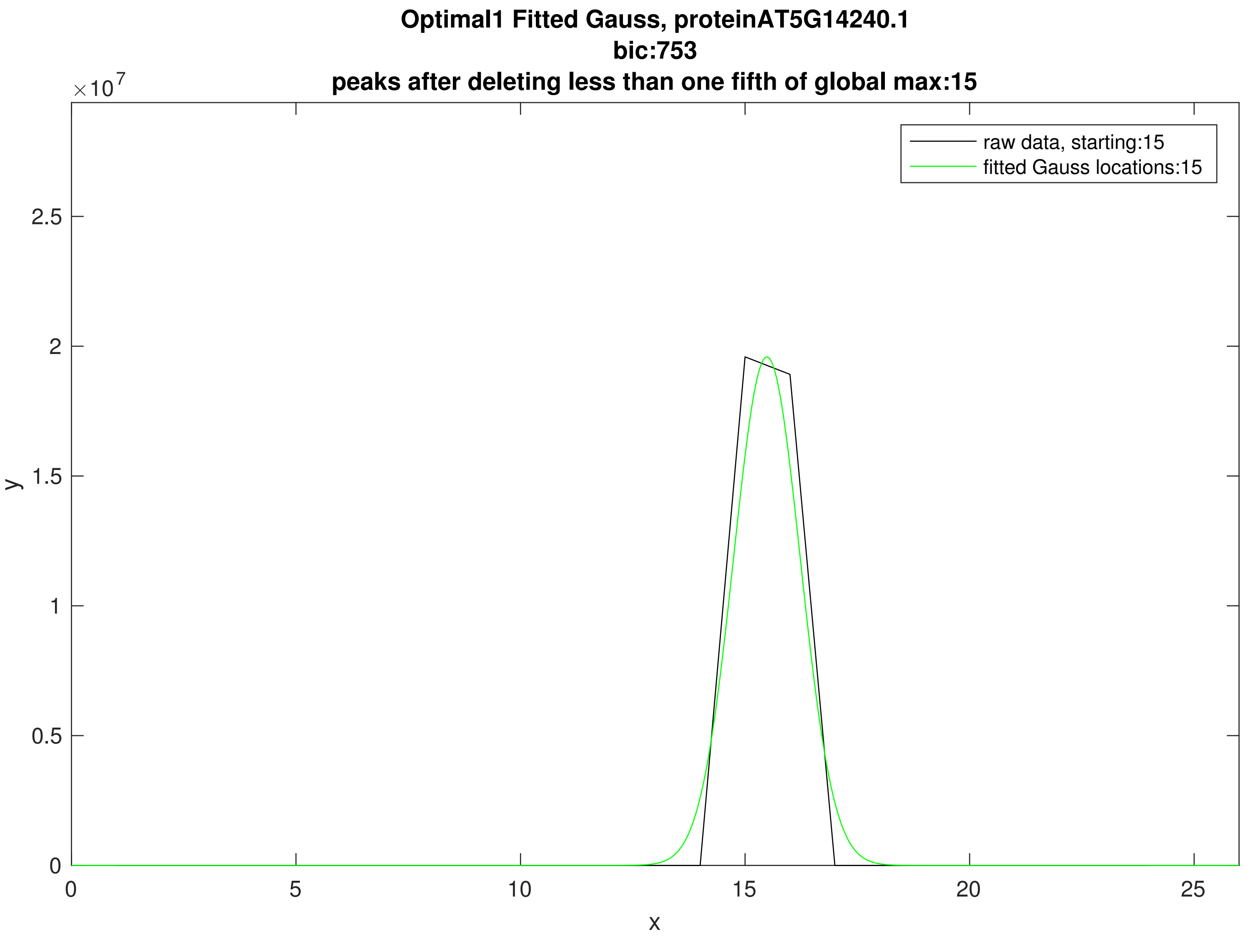
Optimal1 Fitted Gauss, proteinAT5G14240.1
bic:753
peaks after deleting less than one fifth of global max:15
×10^7
raw data, starting:15
fitted Gauss locations:15
y
x
0
0.5
1
1.5
2
2.5
0
5
10
15
20
25

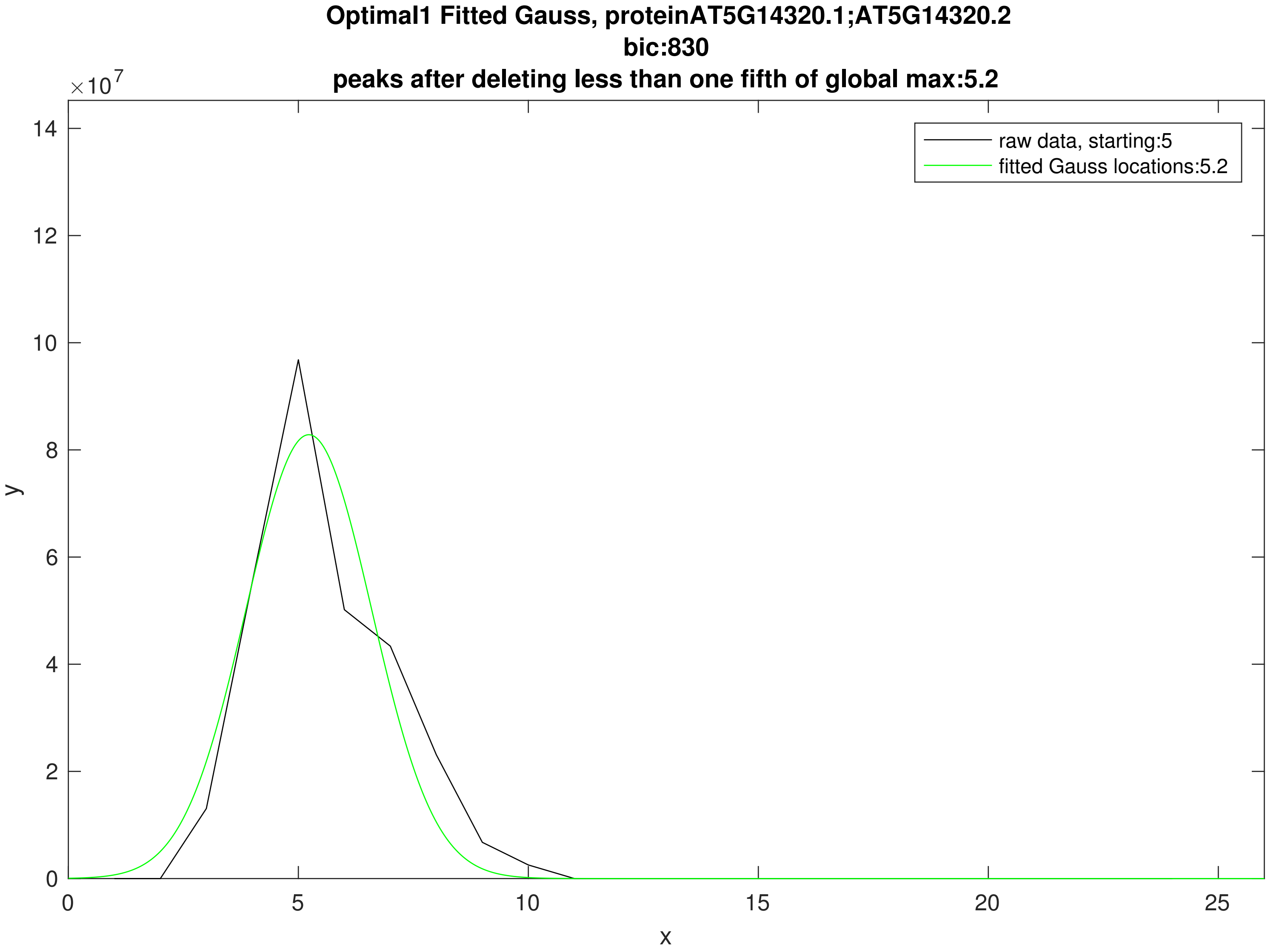
Optimal1 Fitted Gauss, proteinAT5G14320.1;AT5G14320.2
bic:830
peaks after deleting less than one fifth of global max:5.2
×10^7
raw data, starting:5
fitted Gauss locations:5.2
y
x
0
2
4
6
8
10
12
14
0
5
10
15
20
25

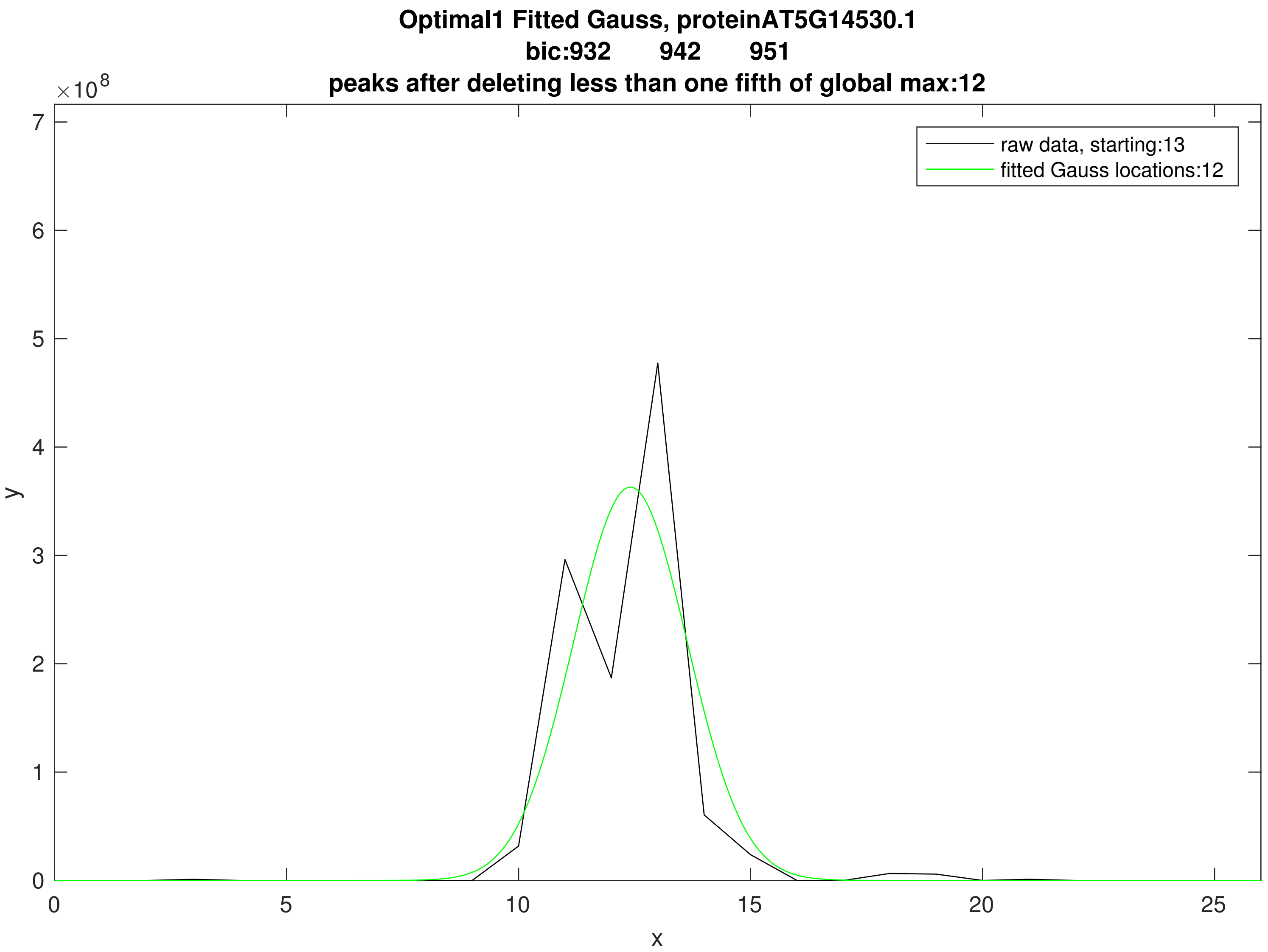
Optimal1 Fitted Gauss, proteinAT5G14530.1
bic:932 942 951
peaks after deleting less than one fifth of global max:12
×10^8
raw data, starting:13
fitted Gauss locations:12
y
x
0
1
2
3
4
5
6
7
0
5
10
15
20
25

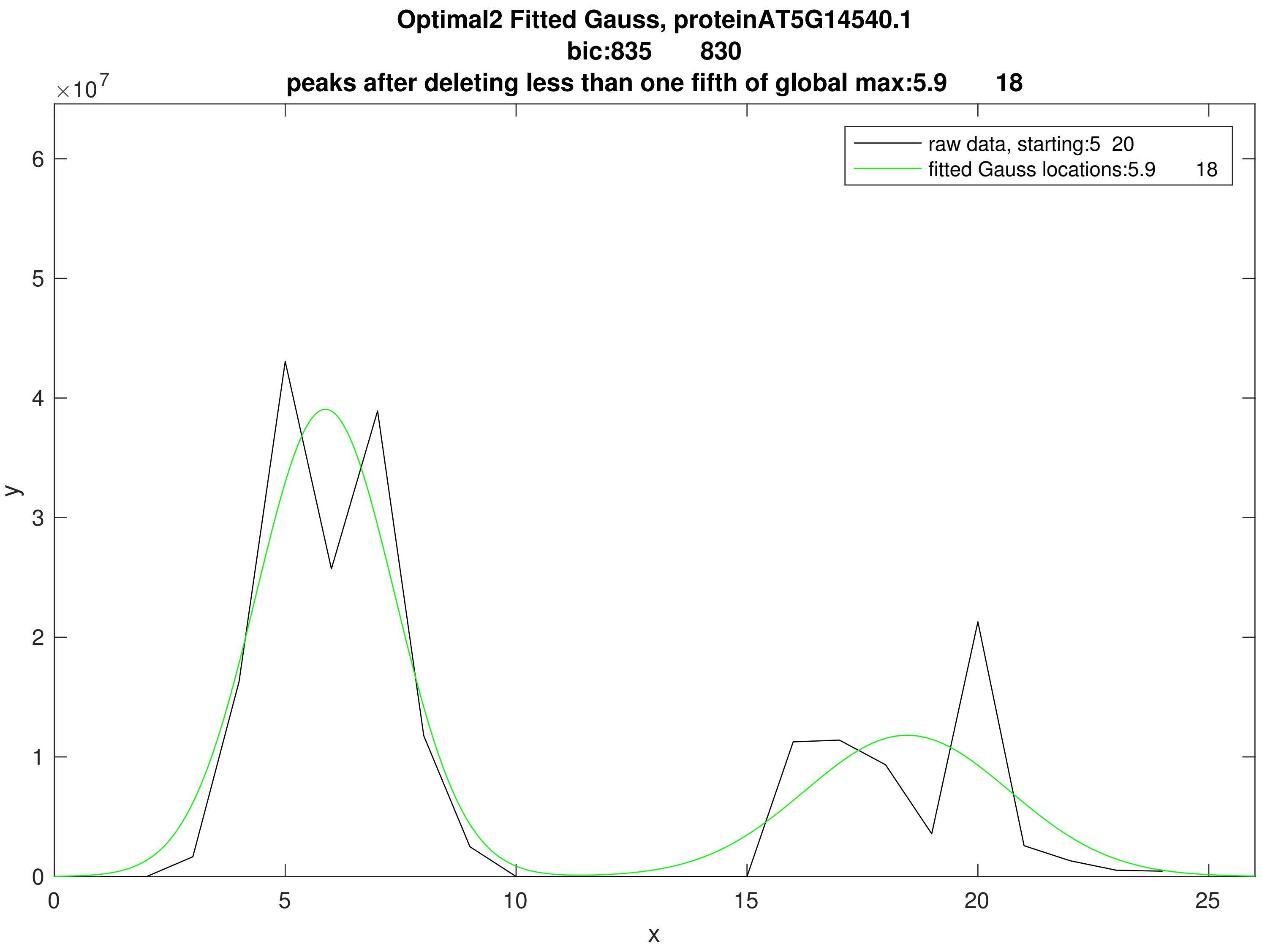

Optimal2 Fitted Gauss, proteinAT5G14540.1
bic:835 830
peaks after deleting less than one fifth of global max:5.9 18
×10^7
raw data, starting:5 20
fitted Gauss locations:5.9 18
y
x
0
1
2
3
4
5
6
0
5
10
15
20
25

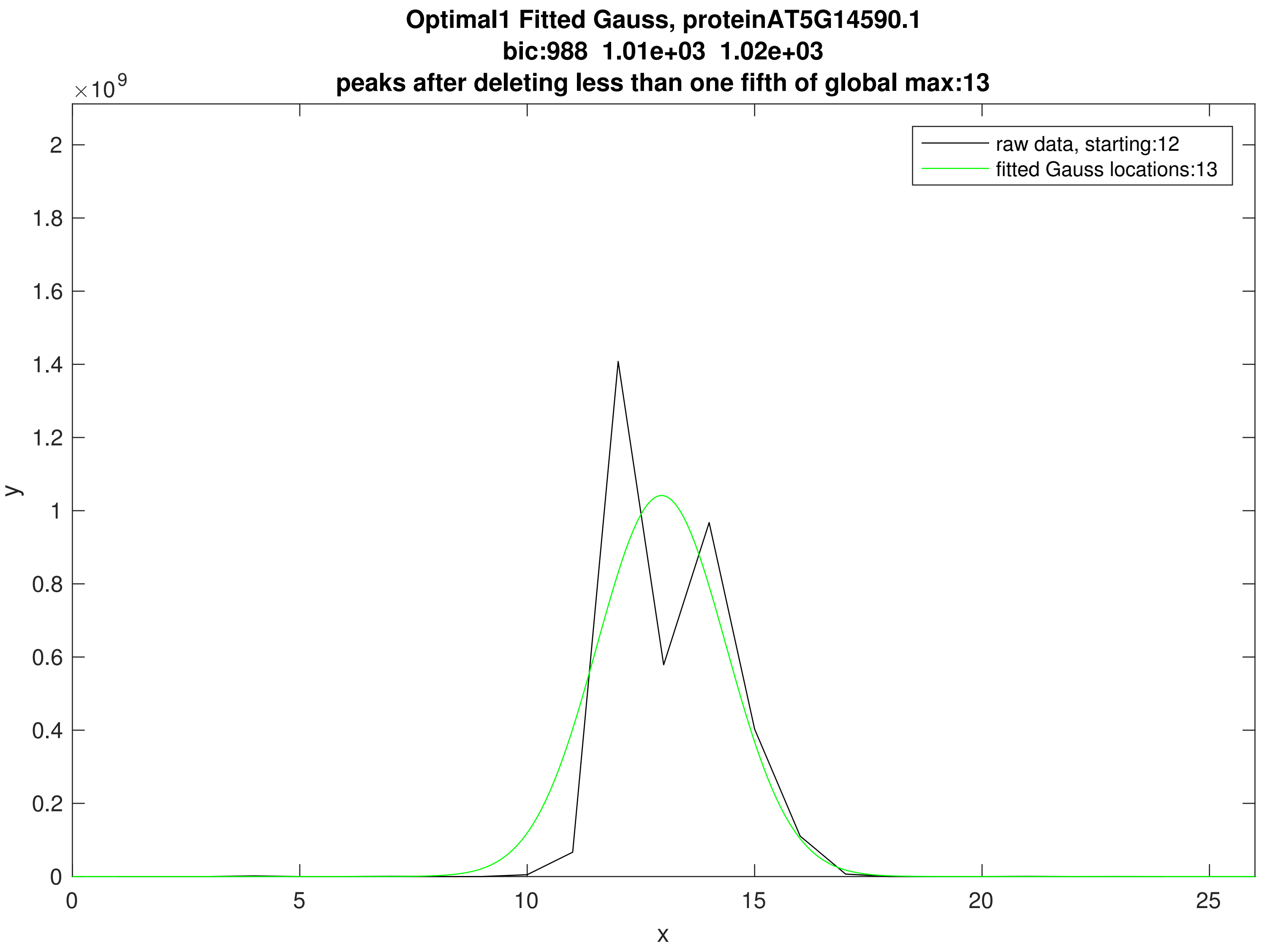
Optimal1 Fitted Gauss, proteinAT5G14590.1
bic:988 1.01e+03 1.02e+03
peaks after deleting less than one fifth of global max:13
×10^9
raw data, starting:12
fitted Gauss locations:13
y
x
0
0.2
0.4
0.6
0.8
1
1.2
1.4
1.6
1.8
2
5
10
15
20
25

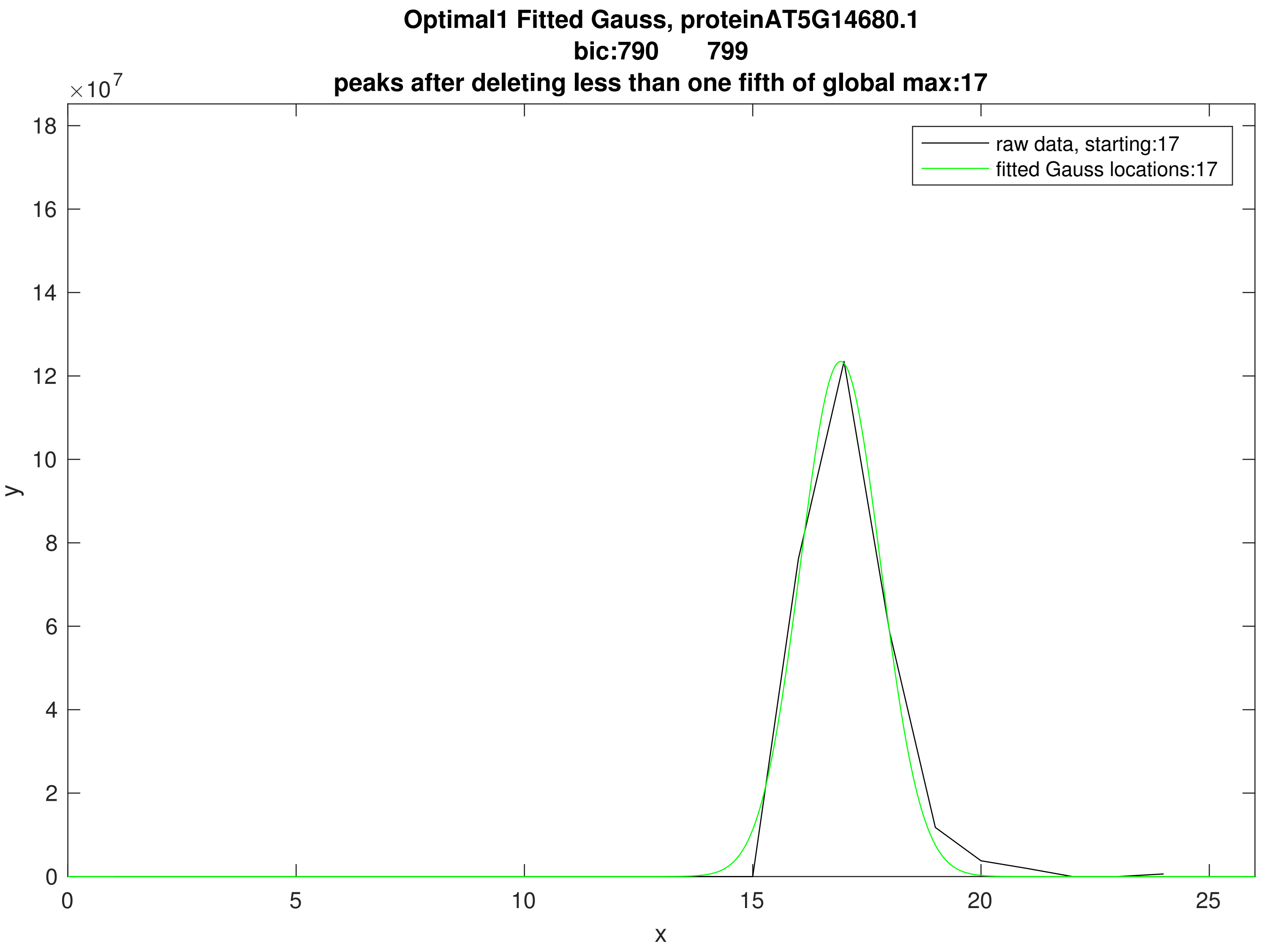

Optimal1 Fitted Gauss, proteinAT5G14680.1
bic:790 799
peaks after deleting less than one fifth of global max:17
×10^7
raw data, starting:17
fitted Gauss locations:17
y
x

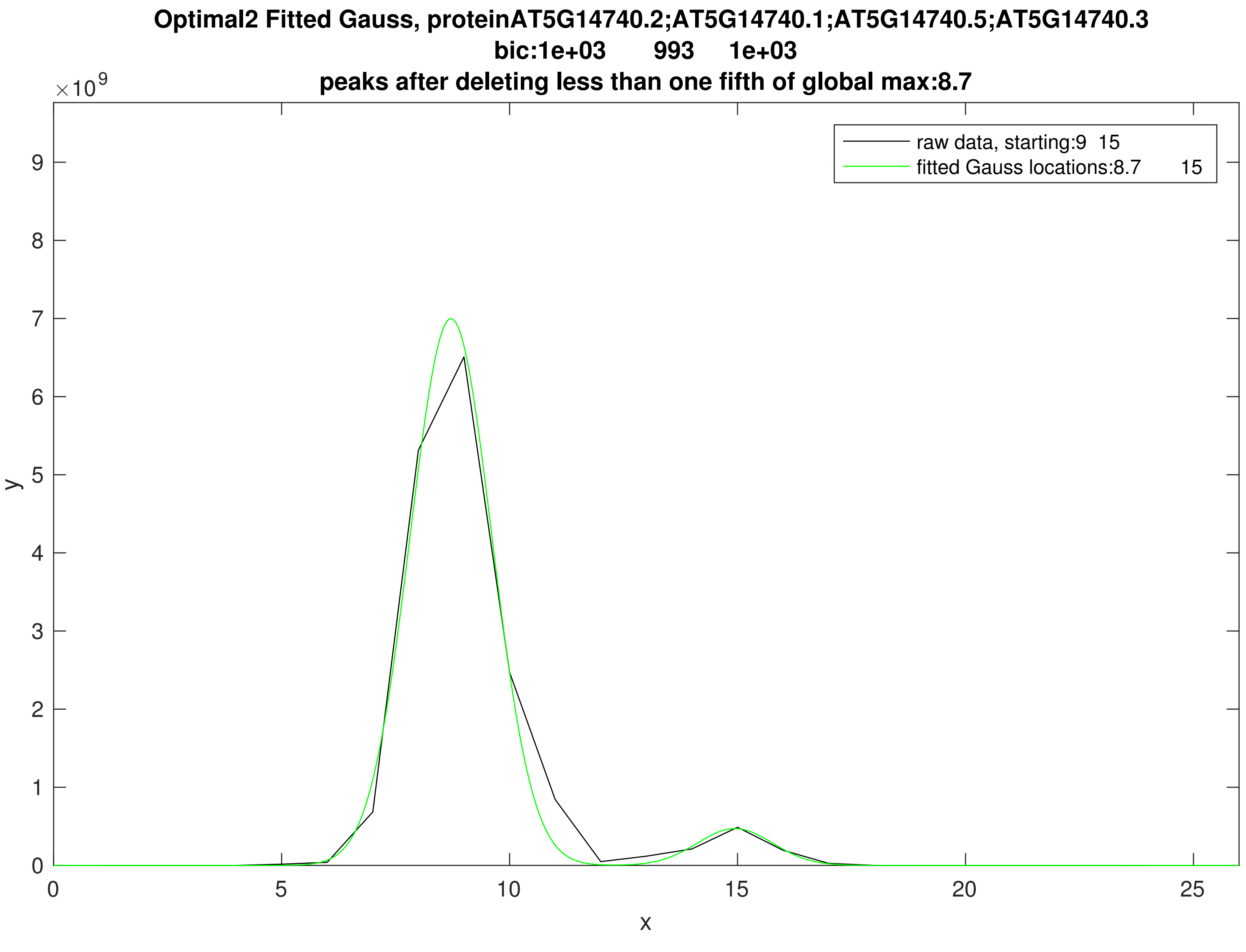
Optimal2 Fitted Gauss, proteinAT5G14740.2;AT5G14740.1;AT5G14740.5;AT5G14740.3
bic:1e+03 993 1e+03
peaks after deleting less than one fifth of global max:8.7
×10^9
raw data, starting:9 15
fitted Gauss locations:8.7 15
y
x
0
1
2
3
4
5
6
7
8
9
0
5
10
15
20
25

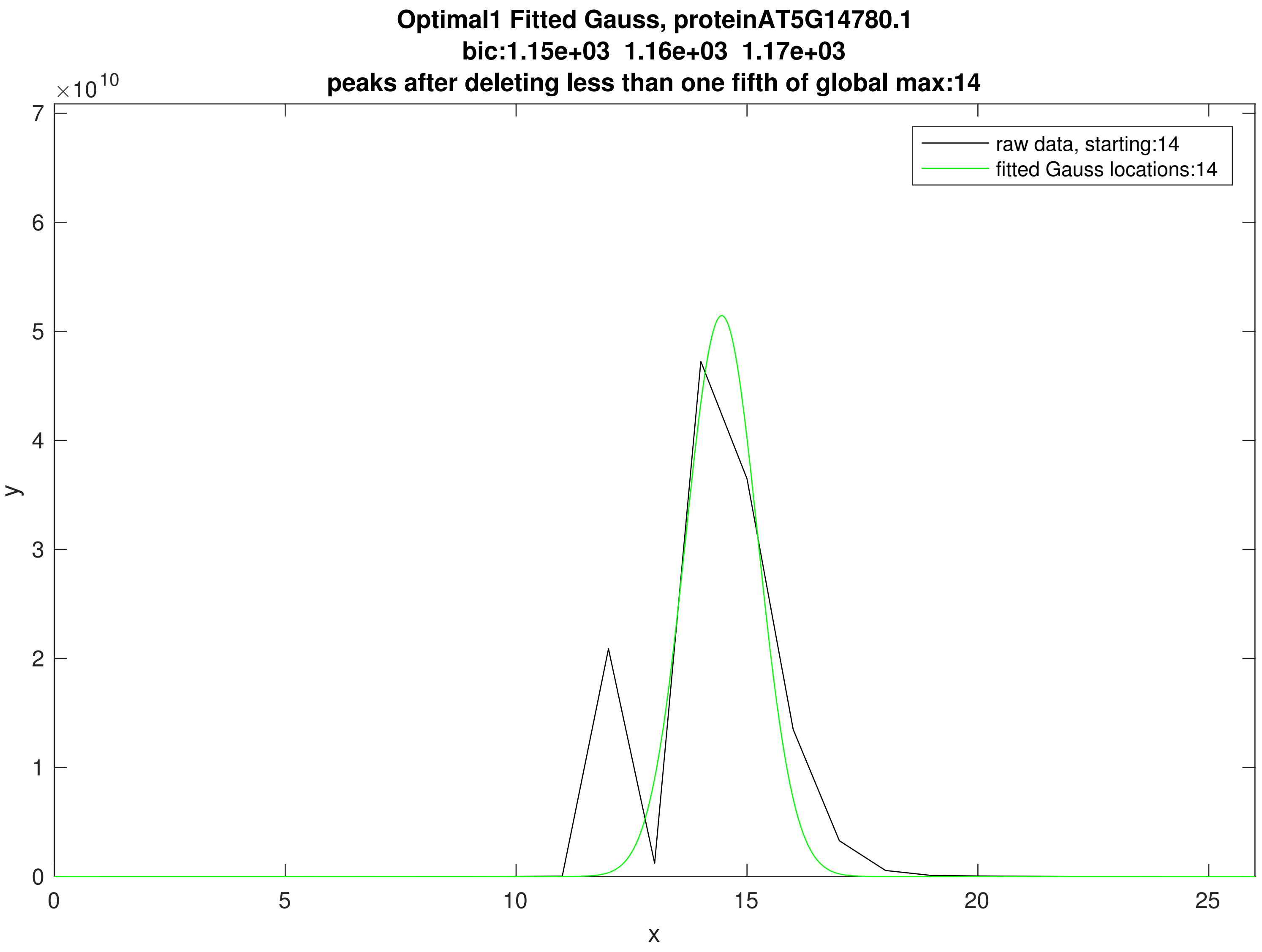

Optimal1 Fitted Gauss, proteinAT5G14780.1
bic:1.15e+03 1.16e+03 1.17e+03
peaks after deleting less than one fifth of global max:14
×10^10
raw data, starting:14
fitted Gauss locations:14
y
x
0
1
2
3
4
5
6
7
0
5
10
15
20
25

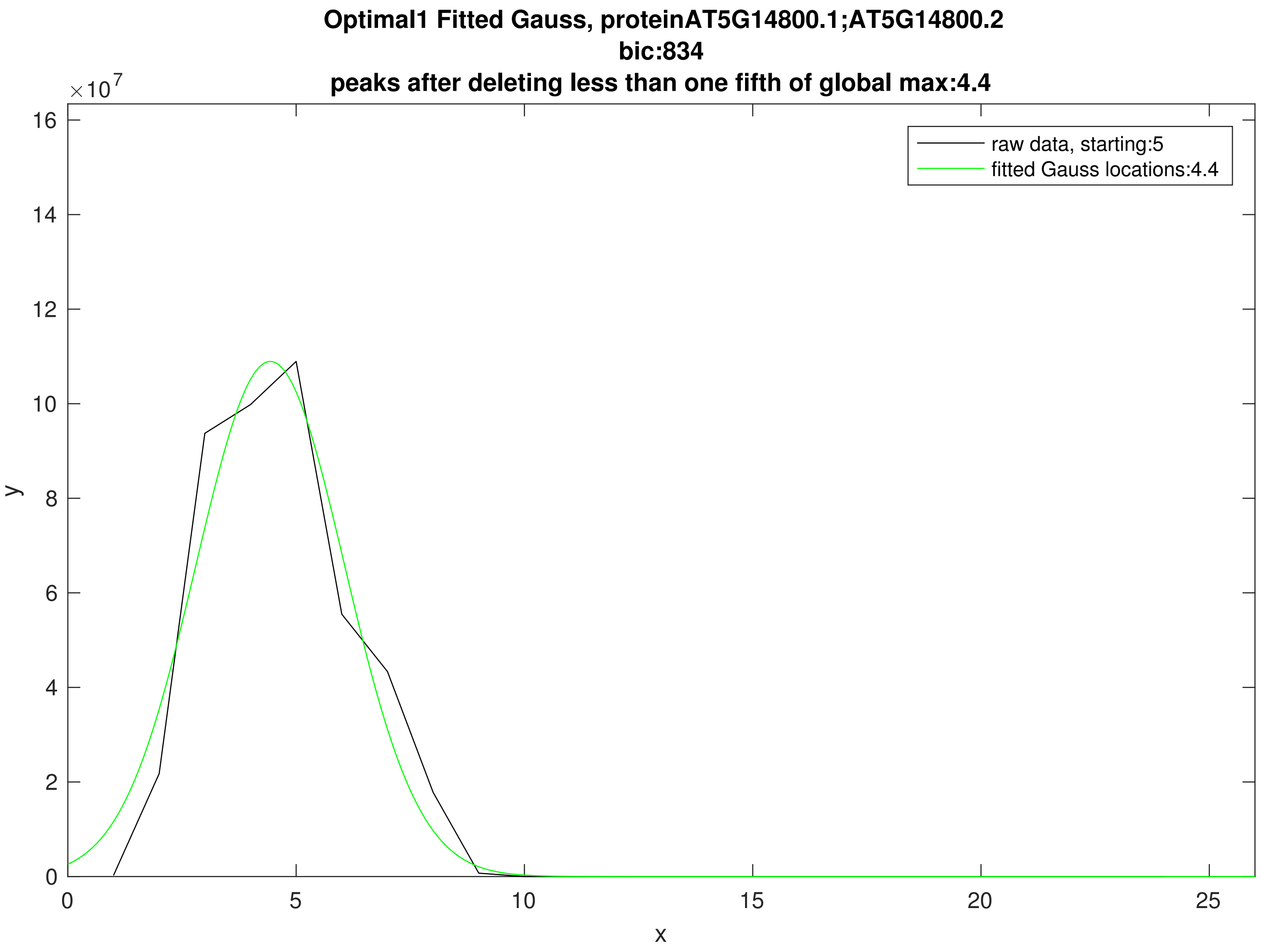

Optimal1 Fitted Gauss, proteinAT5G14800.1;AT5G14800.2
bic:834
peaks after deleting less than one fifth of global max:4.4
×10^7
raw data, starting:5
fitted Gauss locations:4.4
y
x
0
2
4
6
8
10
12
14
16
0
5
10
15
20
25

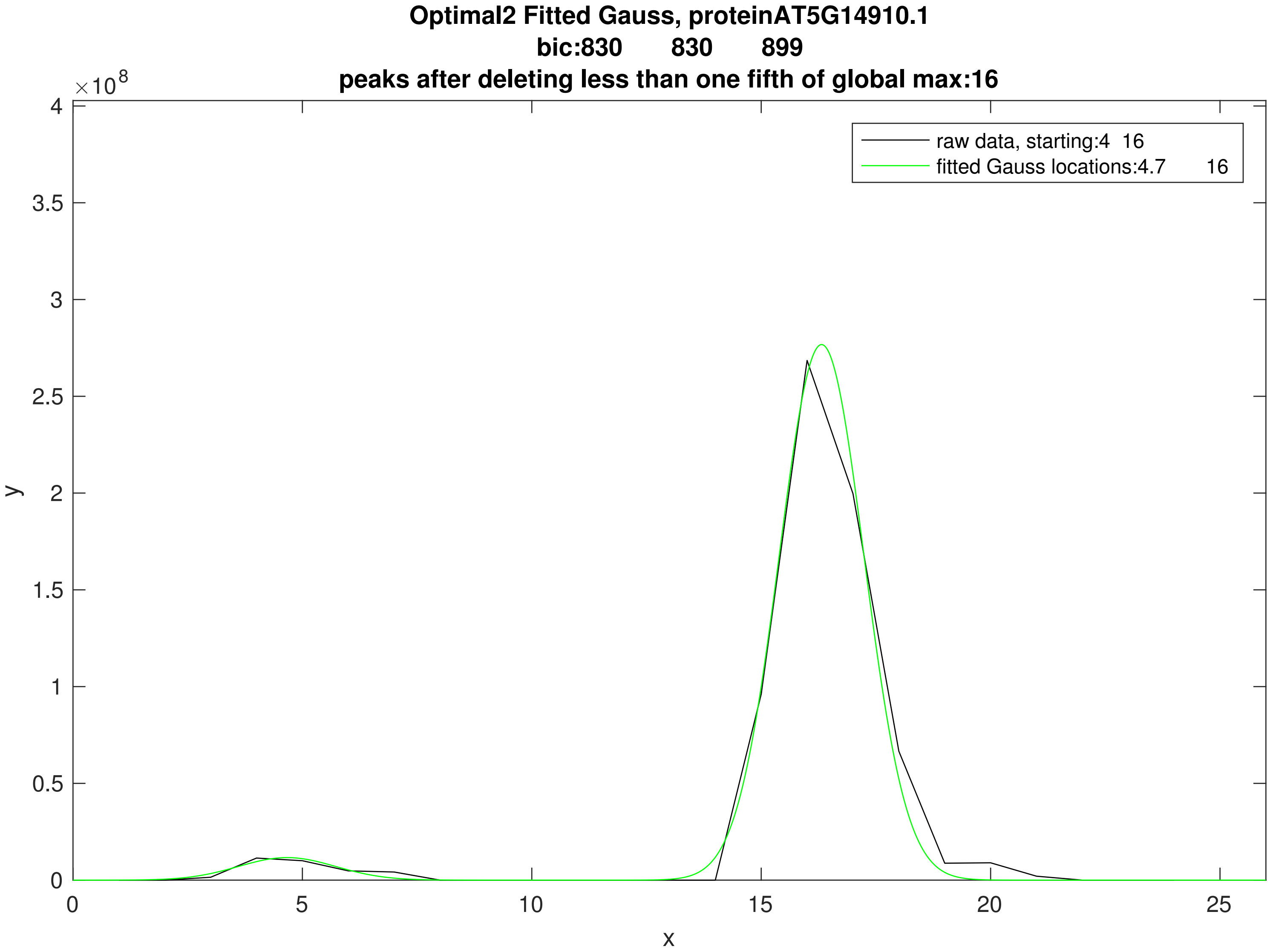
Optimal2 Fitted Gauss, proteinAT5G14910.1
bic:830 830 899
peaks after deleting less than one fifth of global max:16
raw data, starting:4 16
fitted Gauss locations:4.7 16
×10^8
y
x
0
0.5
1
1.5
2
2.5
3
3.5
4
5
10
15
20
25

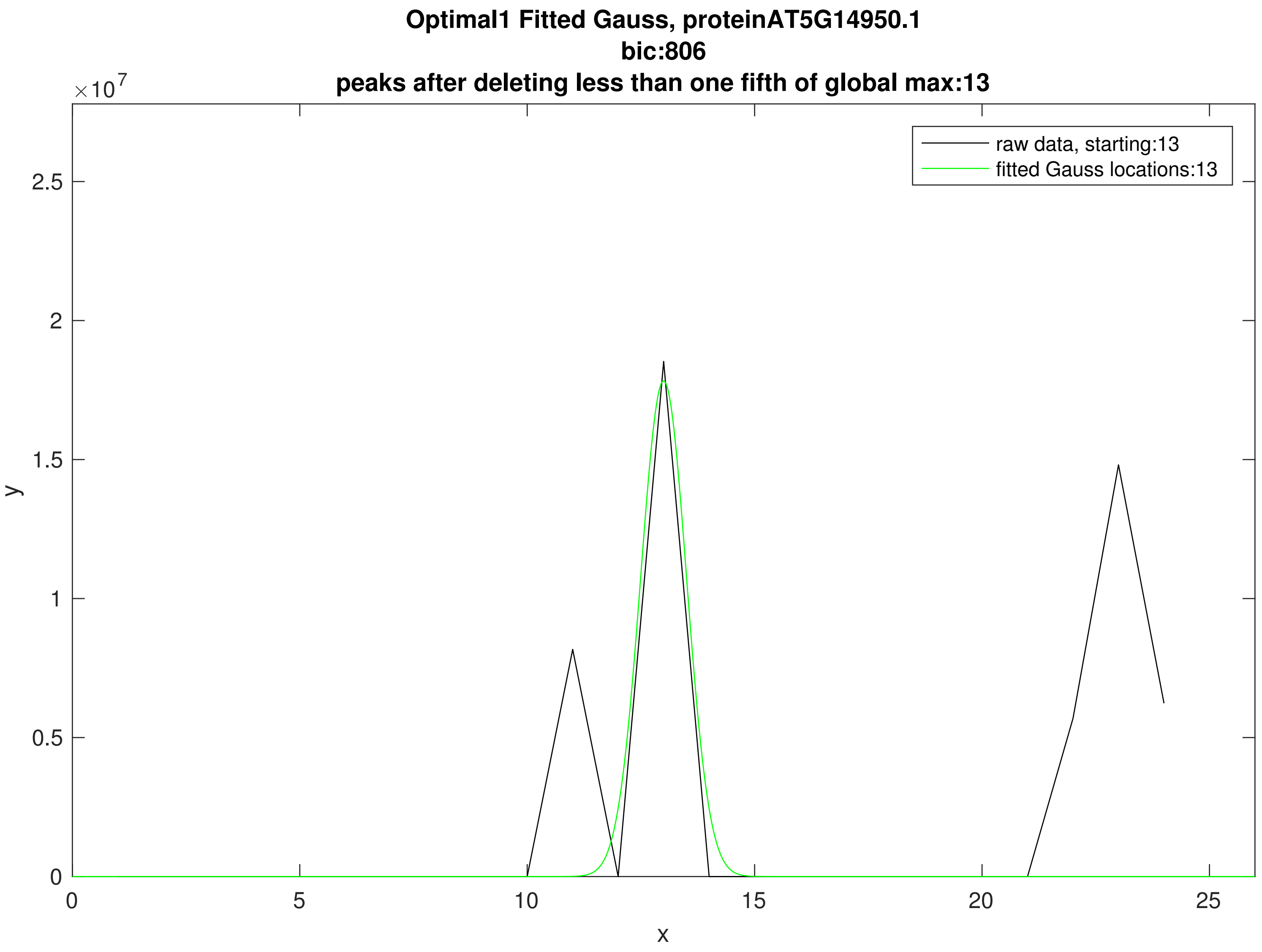
Optimal1 Fitted Gauss, proteinAT5G14950.1
bic:806
peaks after deleting less than one fifth of global max:13
×10^7
y
x
raw data, starting:13
fitted Gauss locations:13
0
0.5
1
1.5
2
2.5
0
5
10
15
20
25

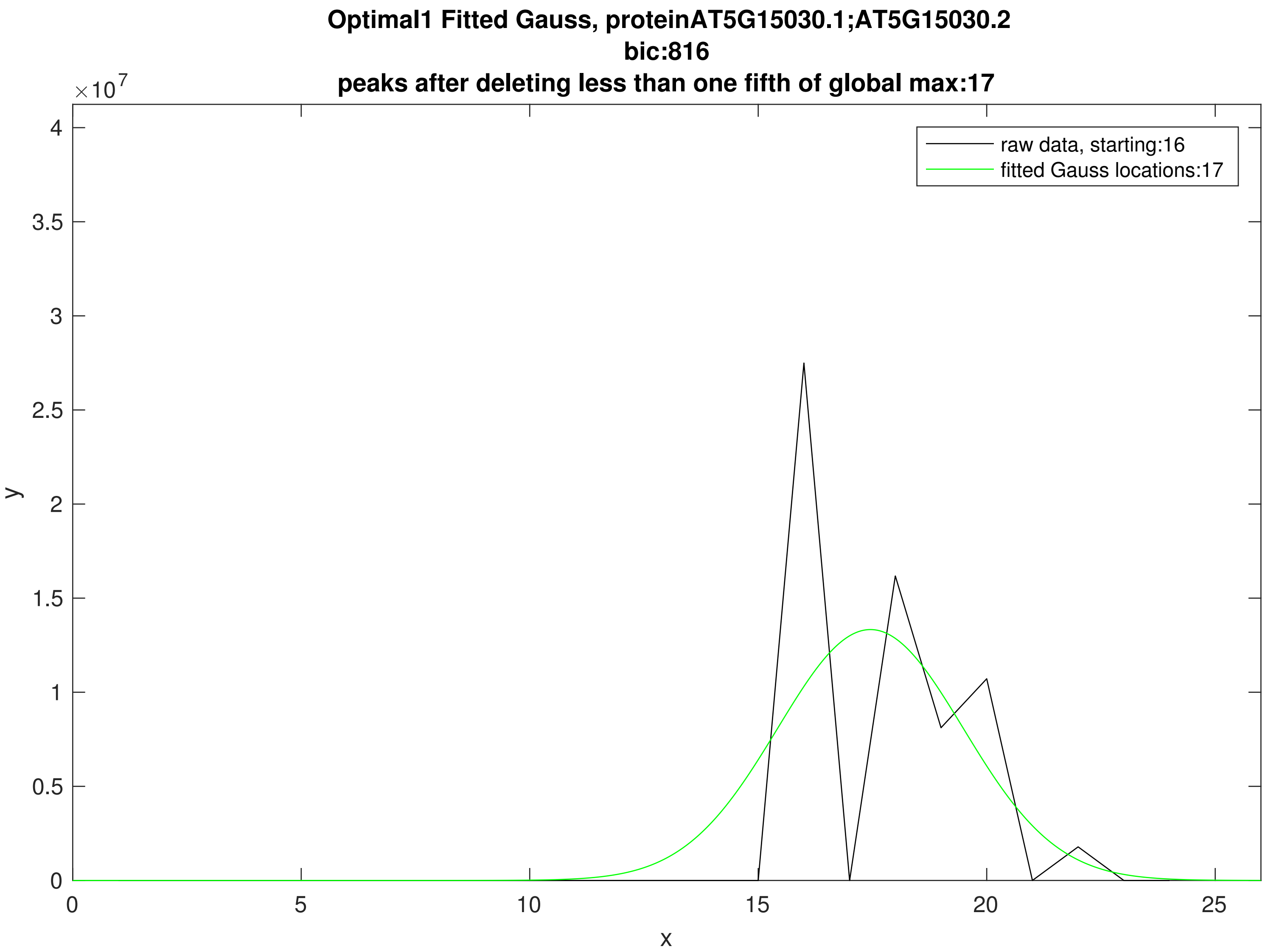

Optimal1 Fitted Gauss, proteinAT5G15030.1;AT5G15030.2
bic:816
peaks after deleting less than one fifth of global max:17
×10^7
raw data, starting:16
fitted Gauss locations:17
y
x
0
0.5
1
1.5
2
2.5
3
3.5
4
5
10
15
20
25

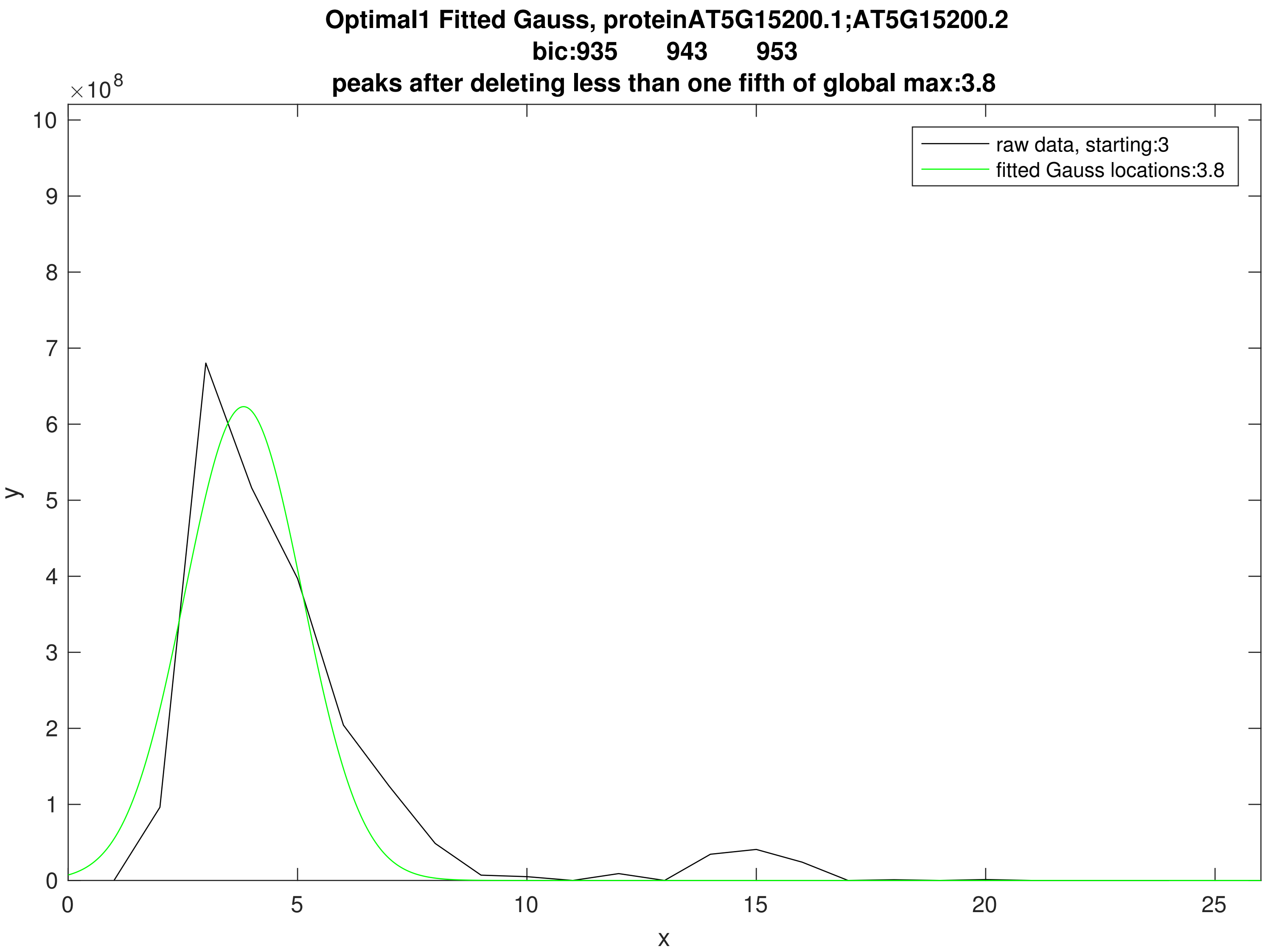
Optimal1 Fitted Gauss, proteinAT5G15200.1;AT5G15200.2
bic:935 943 953
peaks after deleting less than one fifth of global max:3.8
×10^8
raw data, starting:3
fitted Gauss locations:3.8
y
x
0
1
2
3
4
5
6
7
8
9
10
0
5
10
15
20
25

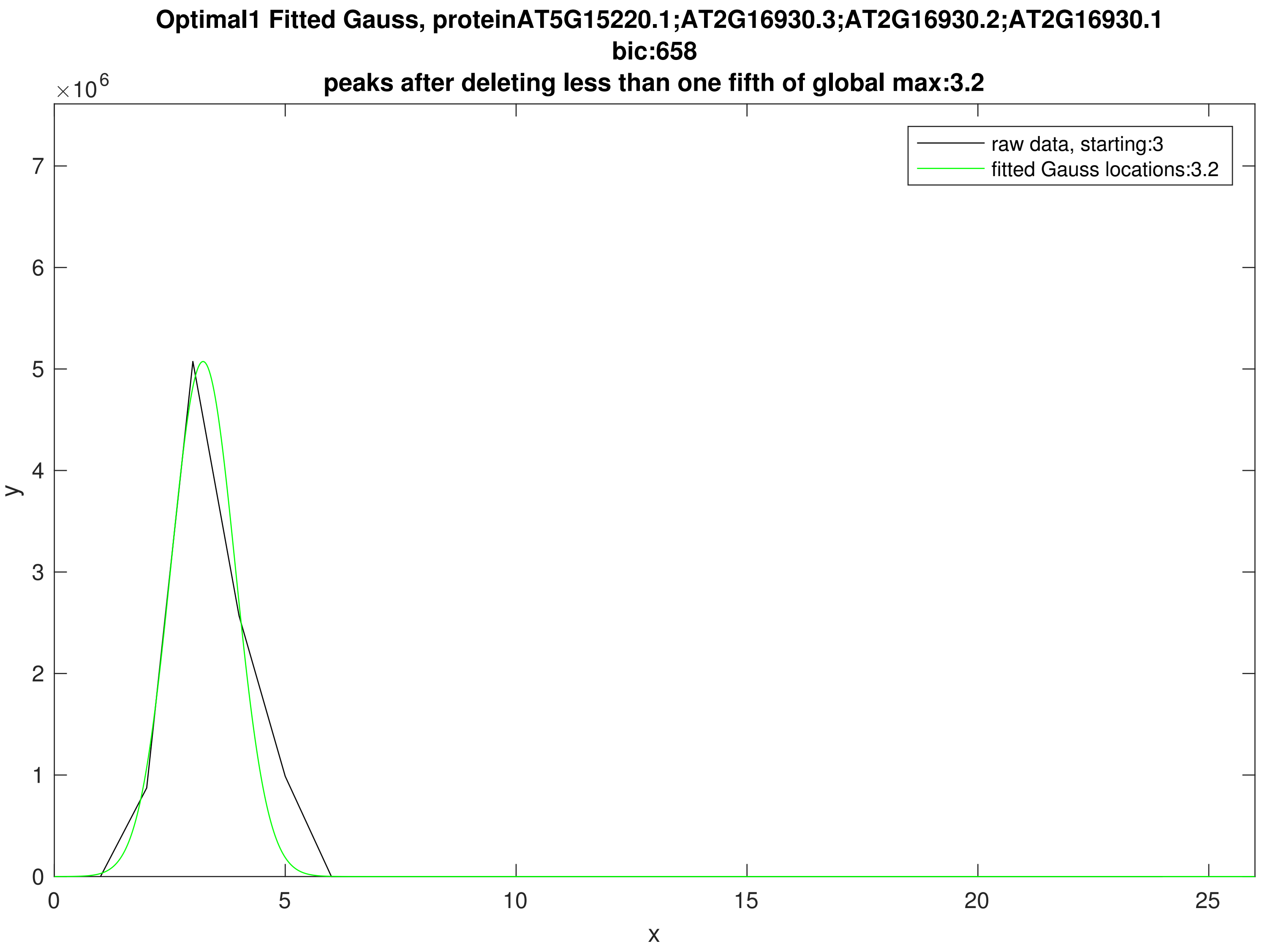
Optimal1 Fitted Gauss, proteinAT5G15220.1;AT2G16930.3;AT2G16930.2;AT2G16930.1
bic:658
peaks after deleting less than one fifth of global max:3.2
×10^6
raw data, starting:3
fitted Gauss locations:3.2
y
x
0
1
2
3
4
5
6
7
0
5
10
15
20
25

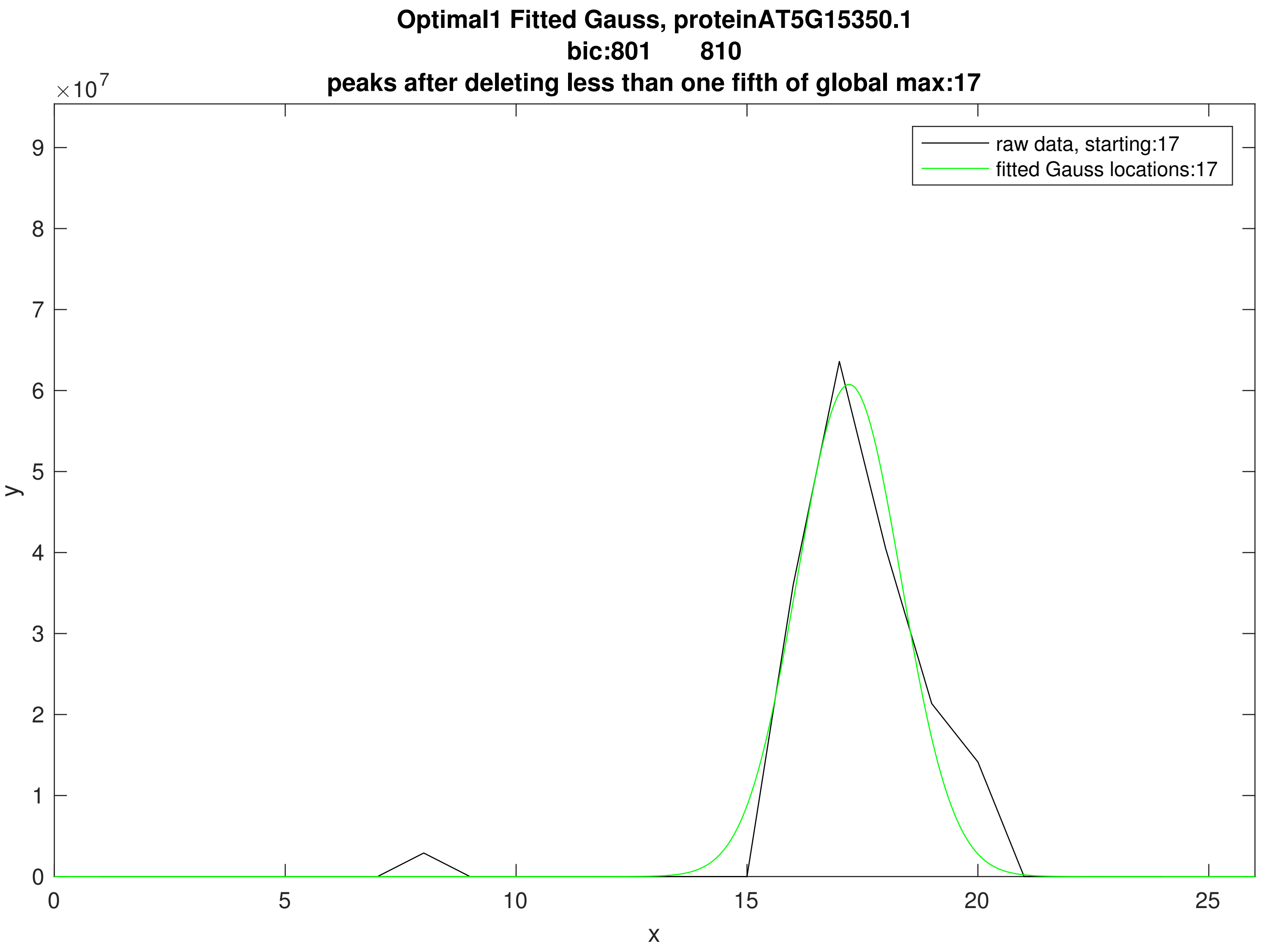
Optimal1 Fitted Gauss, proteinAT5G15350.1
bic:801 810
peaks after deleting less than one fifth of global max:17
×10^7
raw data, starting:17
fitted Gauss locations:17
y
x
0
1
2
3
4
5
6
7
8
9
0
5
10
15
20
25

**Optimal1 Fitted Gauss, proteinAT5G15400.1**
**bic:833 842 851**
**peaks after deleting less than one fifth of global max:17**

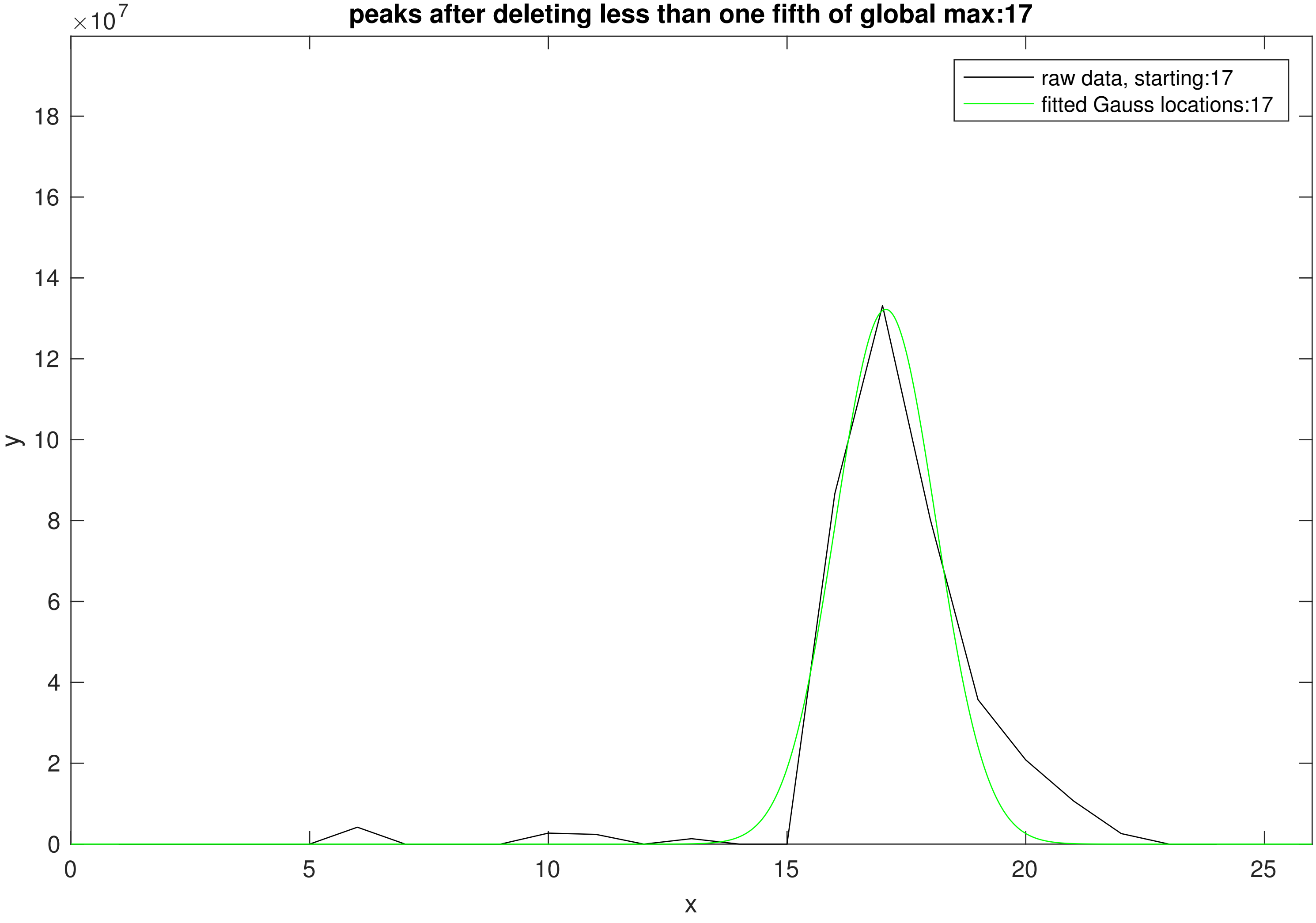

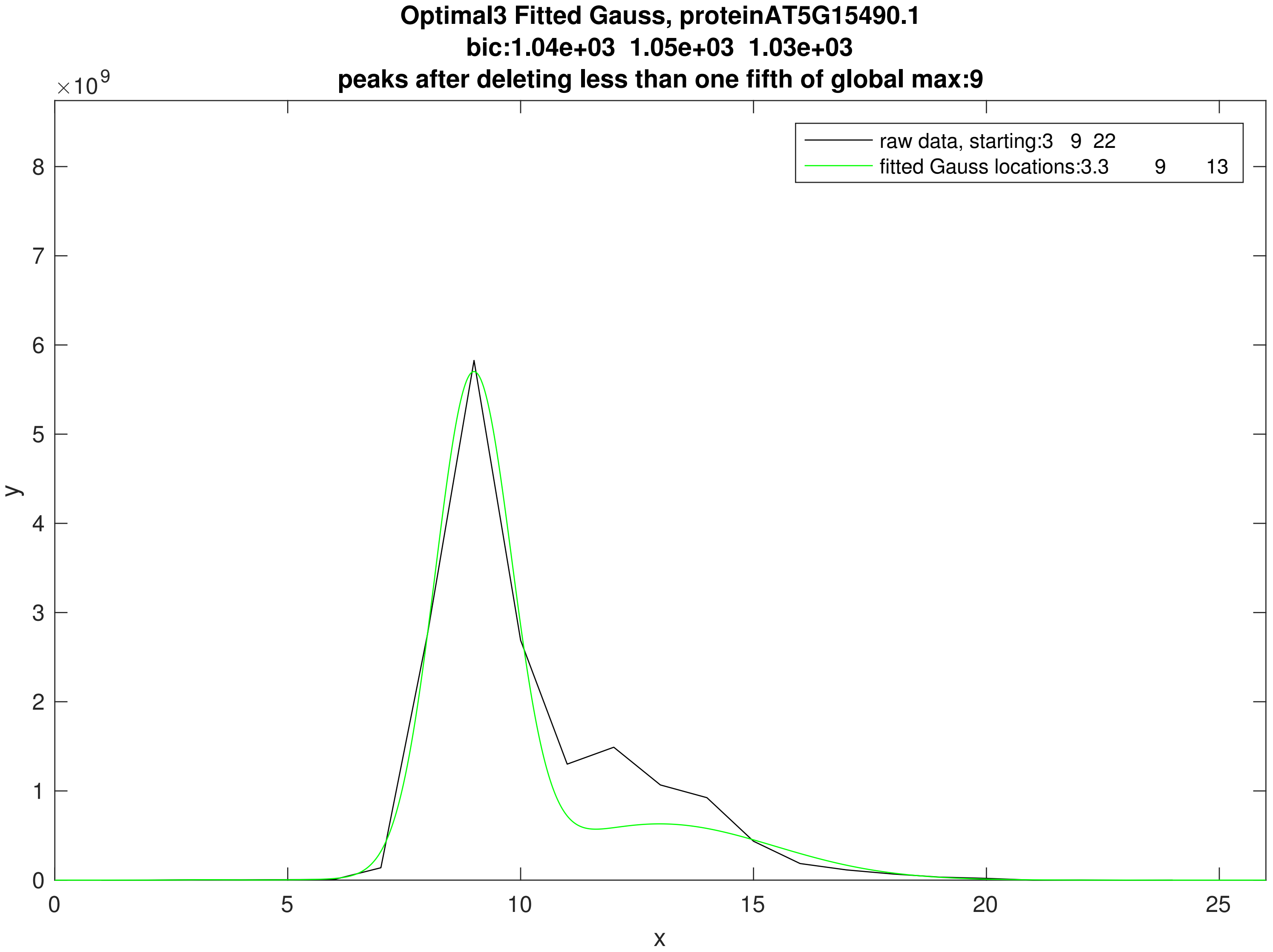

Optimal3 Fitted Gauss, proteinAT5G15490.1
bic:1.04e+03 1.05e+03 1.03e+03
peaks after deleting less than one fifth of global max:9
×10^9
raw data, starting:3 9 22
fitted Gauss locations:3.3 9 13
y
x
0
1
2
3
4
5
6
7
8
0
5
10
15
20
25

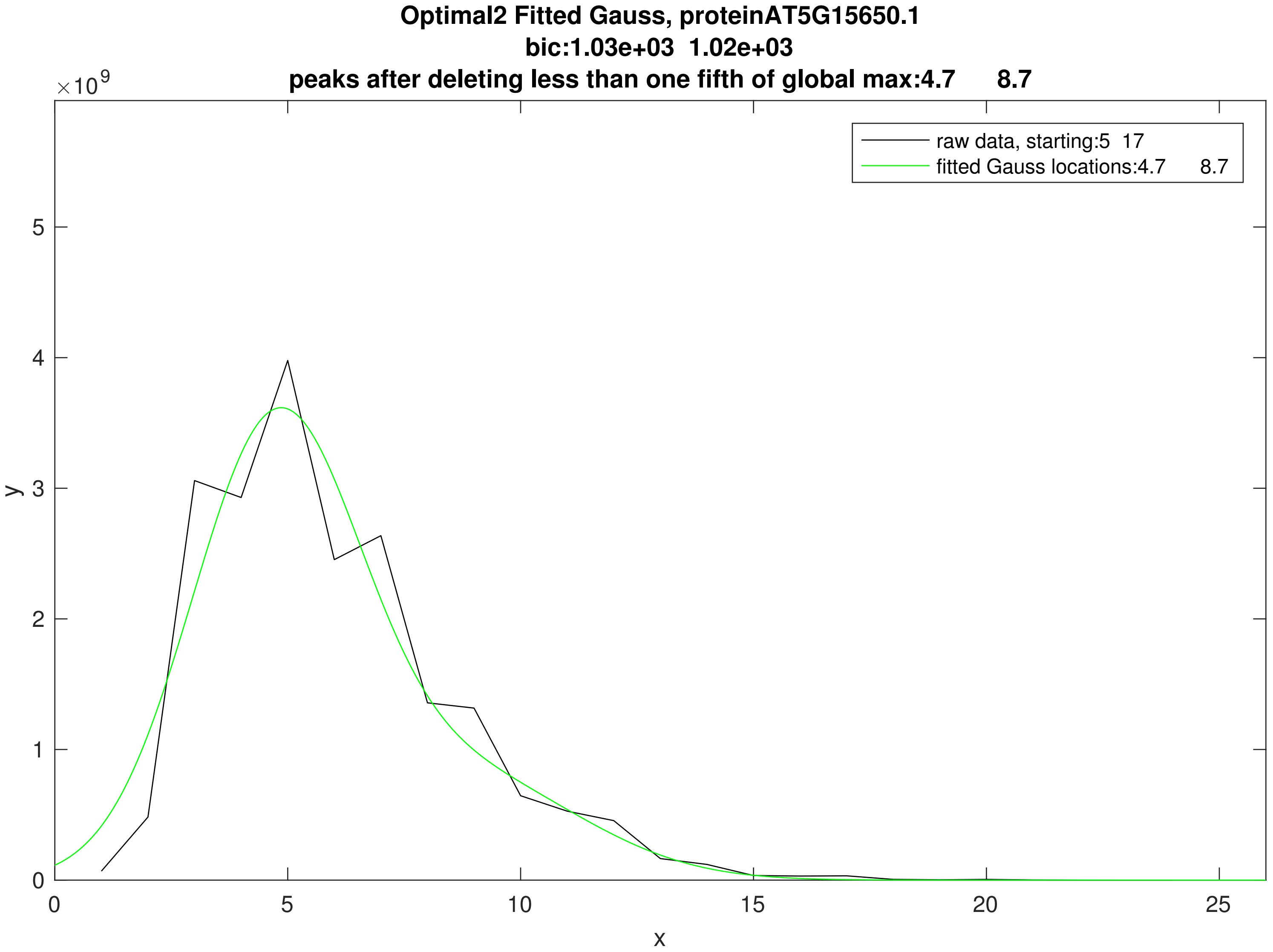
Optimal2 Fitted Gauss, proteinAT5G15650.1
bic:1.03e+03  1.02e+03
peaks after deleting less than one fifth of global max:4.7      8.7
×10^9
raw data, starting:5  17
fitted Gauss locations:4.7      8.7
y
x
0
1
2
3
4
5
0
5
10
15
20
25

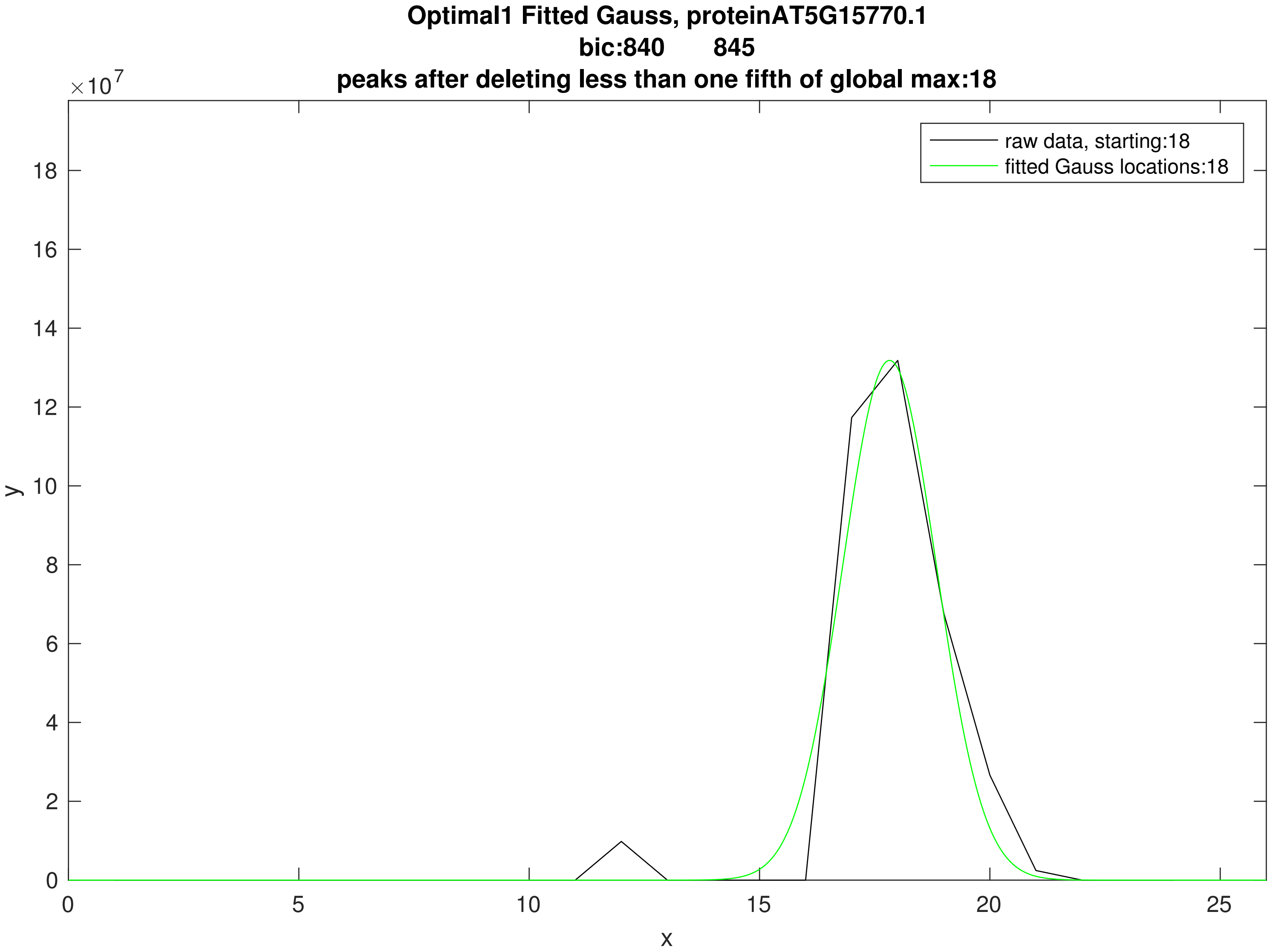

Optimal1 Fitted Gauss, proteinAT5G15770.1
bic:840 845
peaks after deleting less than one fifth of global max:18
×10^7
raw data, starting:18
fitted Gauss locations:18
y
x
0
2
4
6
8
10
12
14
16
18
0
5
10
15
20
25

**Optimal1 Fitted Gauss, proteinAT5G15960.1**
**bic:716**
**peaks after deleting less than one fifth of global max:20**

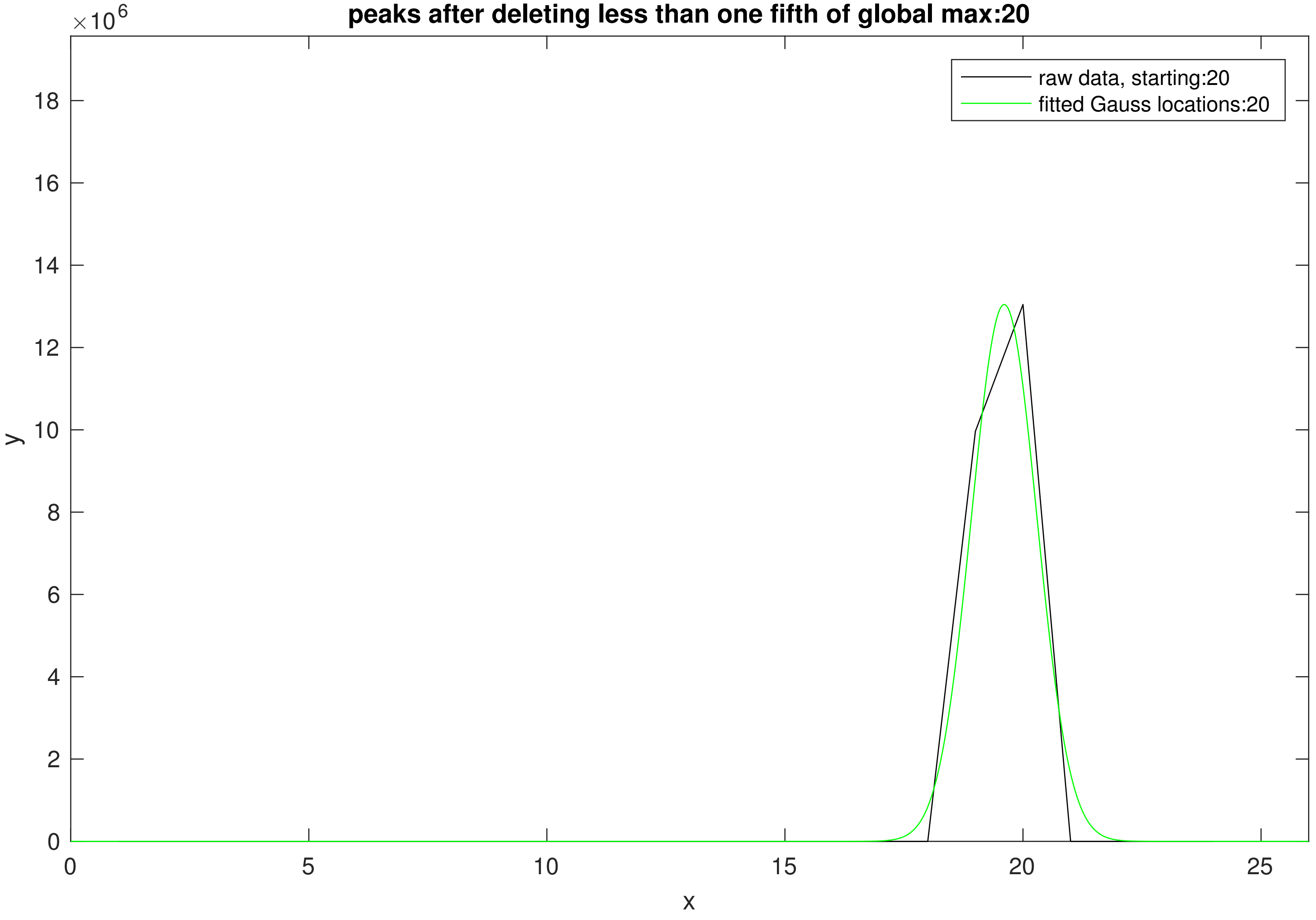

**Optimal1 Fitted Gauss, proteinAT5G15970.1**
**bic:910 976 928**
**peaks after deleting less than one fifth of global max:20**

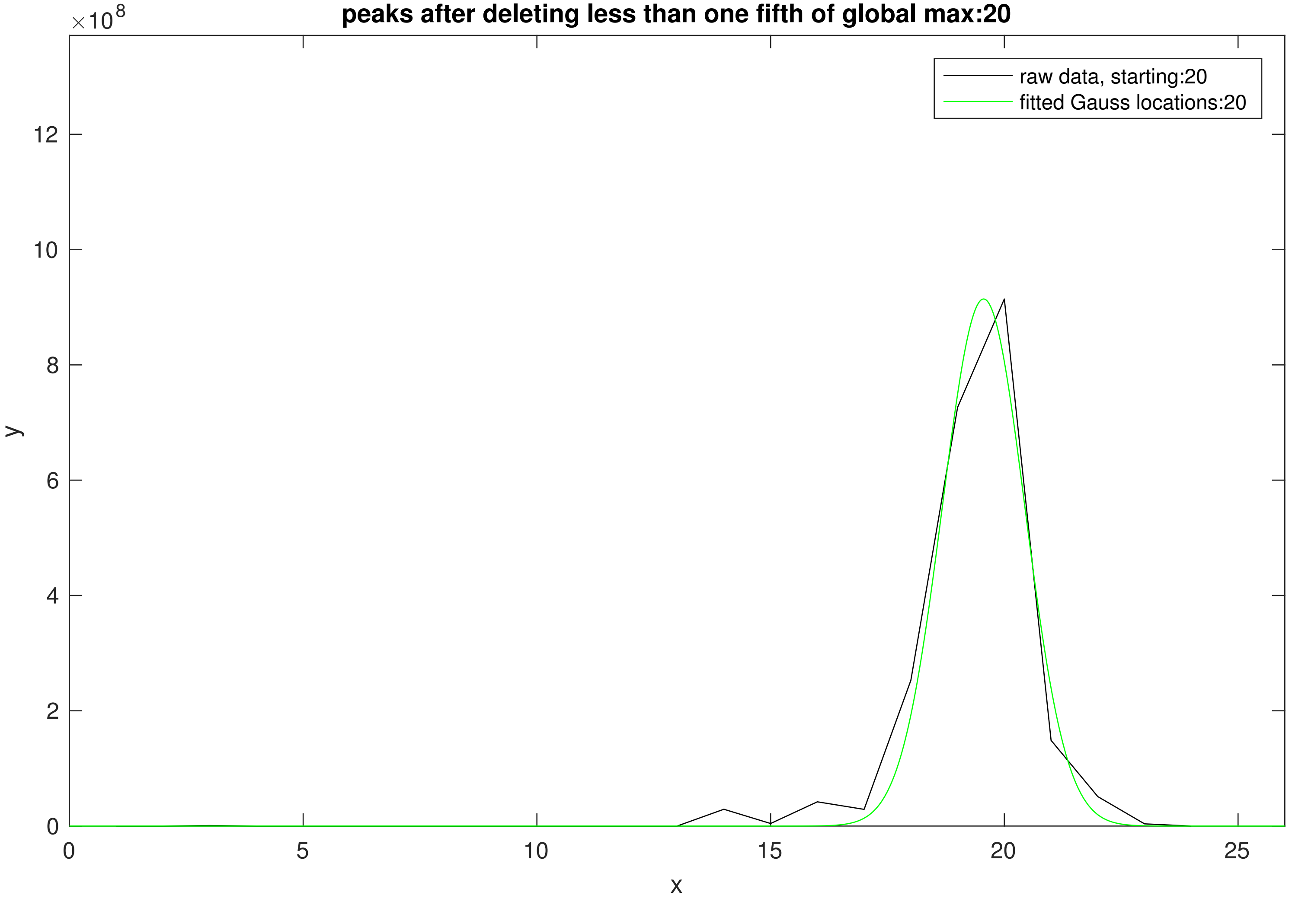

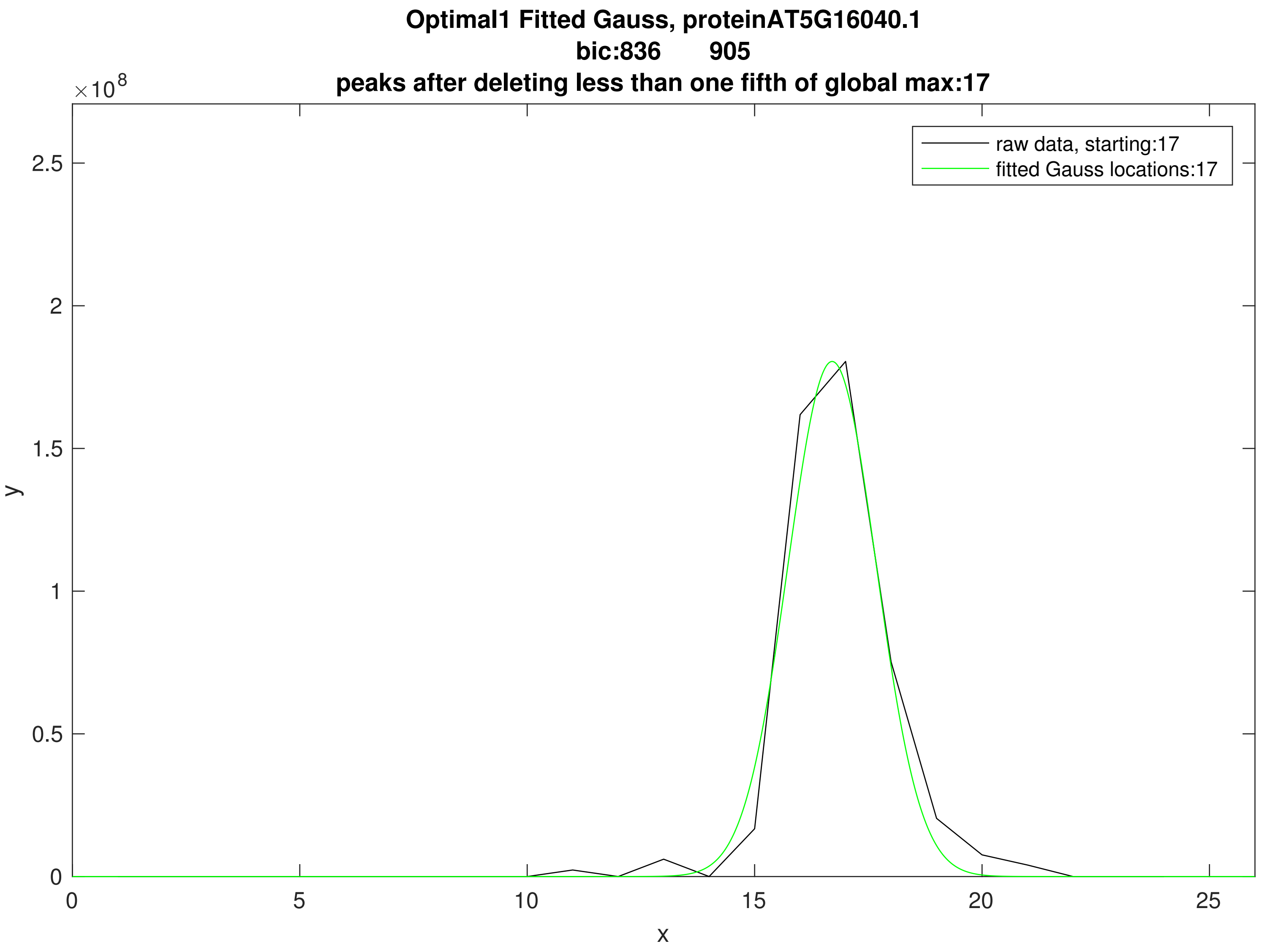
Optimal1 Fitted Gauss, proteinAT5G16040.1
bic:836 905
peaks after deleting less than one fifth of global max:17
×10^8
raw data, starting:17
fitted Gauss locations:17
y
x
0
0.5
1
1.5
2
2.5
5
10
15
20
25

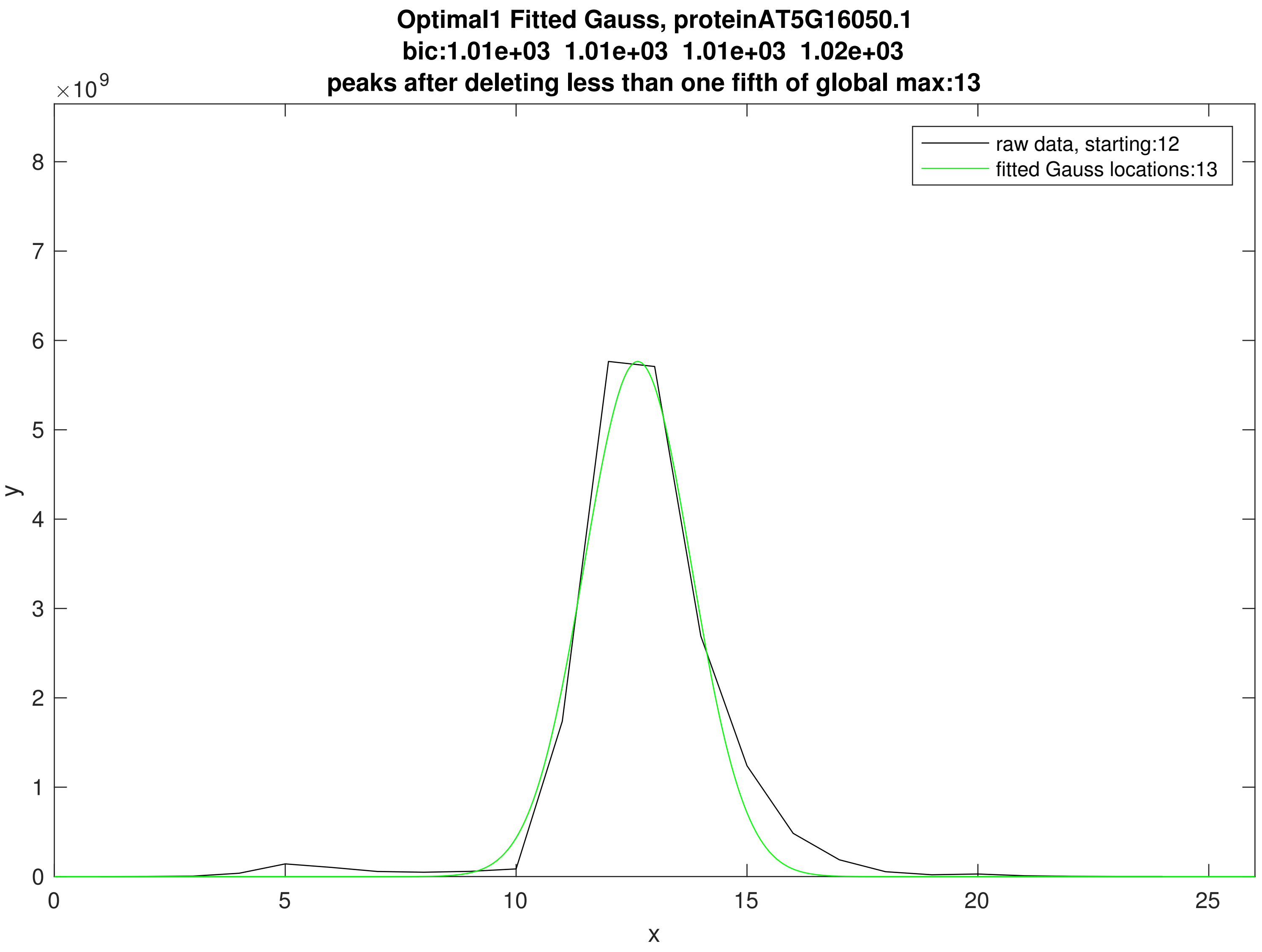

Optimal1 Fitted Gauss, proteinAT5G16050.1
bic:1.01e+03 1.01e+03 1.01e+03 1.02e+03
peaks after deleting less than one fifth of global max:13
×10^9
raw data, starting:12
fitted Gauss locations:13
y
x
0
1
2
3
4
5
6
7
8
0
5
10
15
20
25

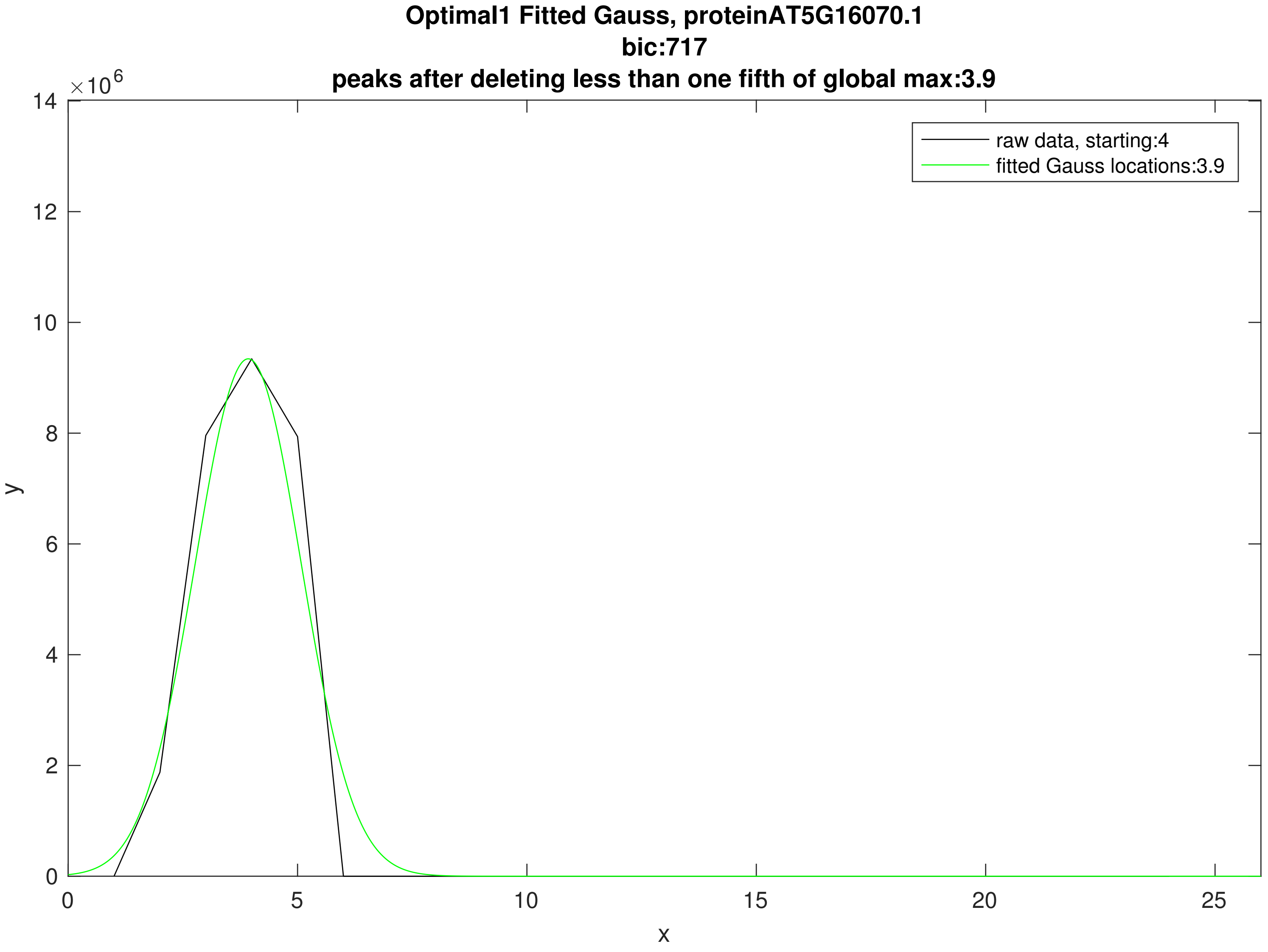
Optimal1 Fitted Gauss, proteinAT5G16070.1
bic:717
peaks after deleting less than one fifth of global max:3.9
×10^6
raw data, starting:4
fitted Gauss locations:3.9
y
x
0
2
4
6
8
10
12
14
5
15
20
25

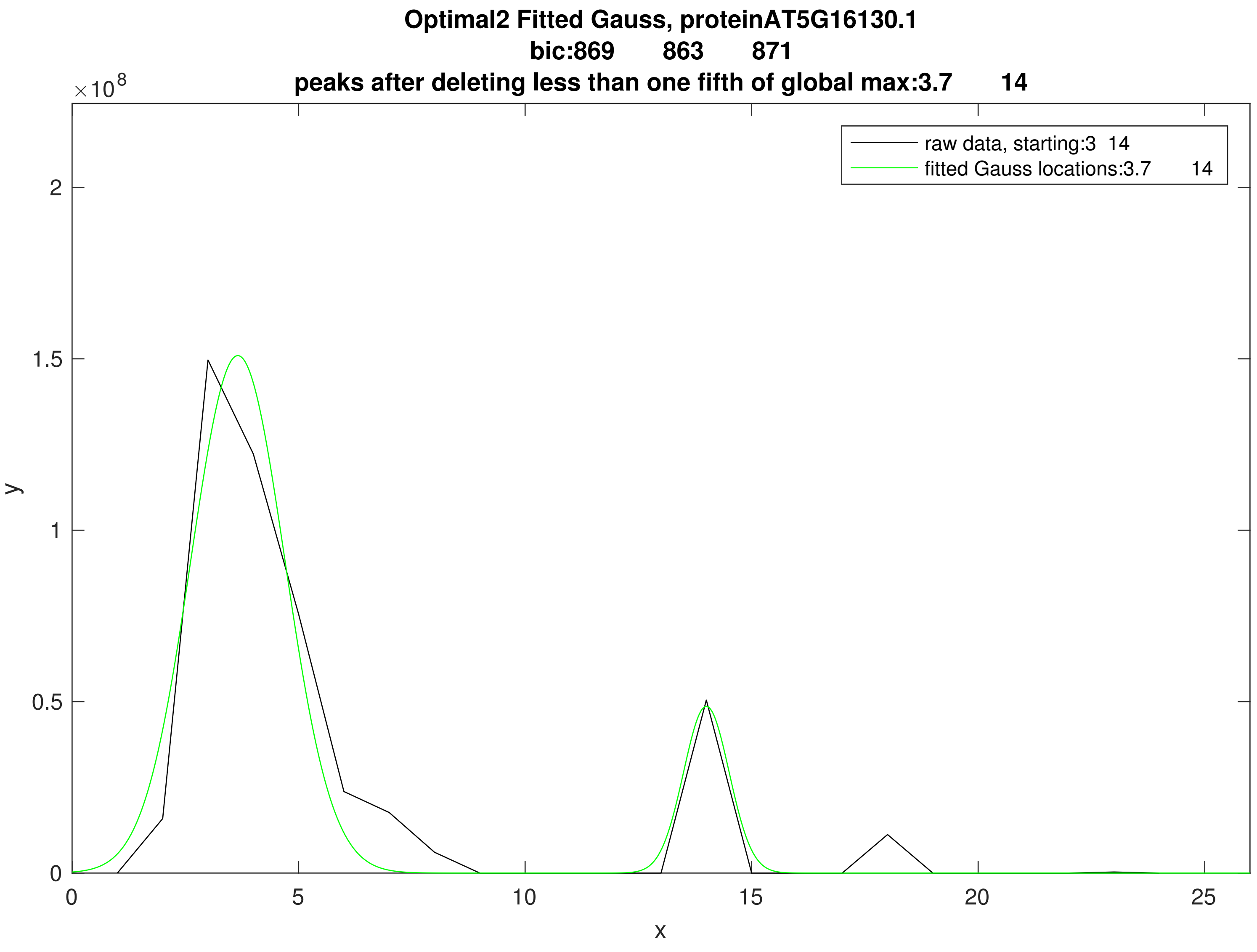

Optimal2 Fitted Gauss, proteinAT5G16130.1
bic:869 863 871
peaks after deleting less than one fifth of global max:3.7 14
×10^8
raw data, starting:3 14
fitted Gauss locations:3.7 14
2
1.5
1
0.5
0
y
0
5
10
15
20
25
x

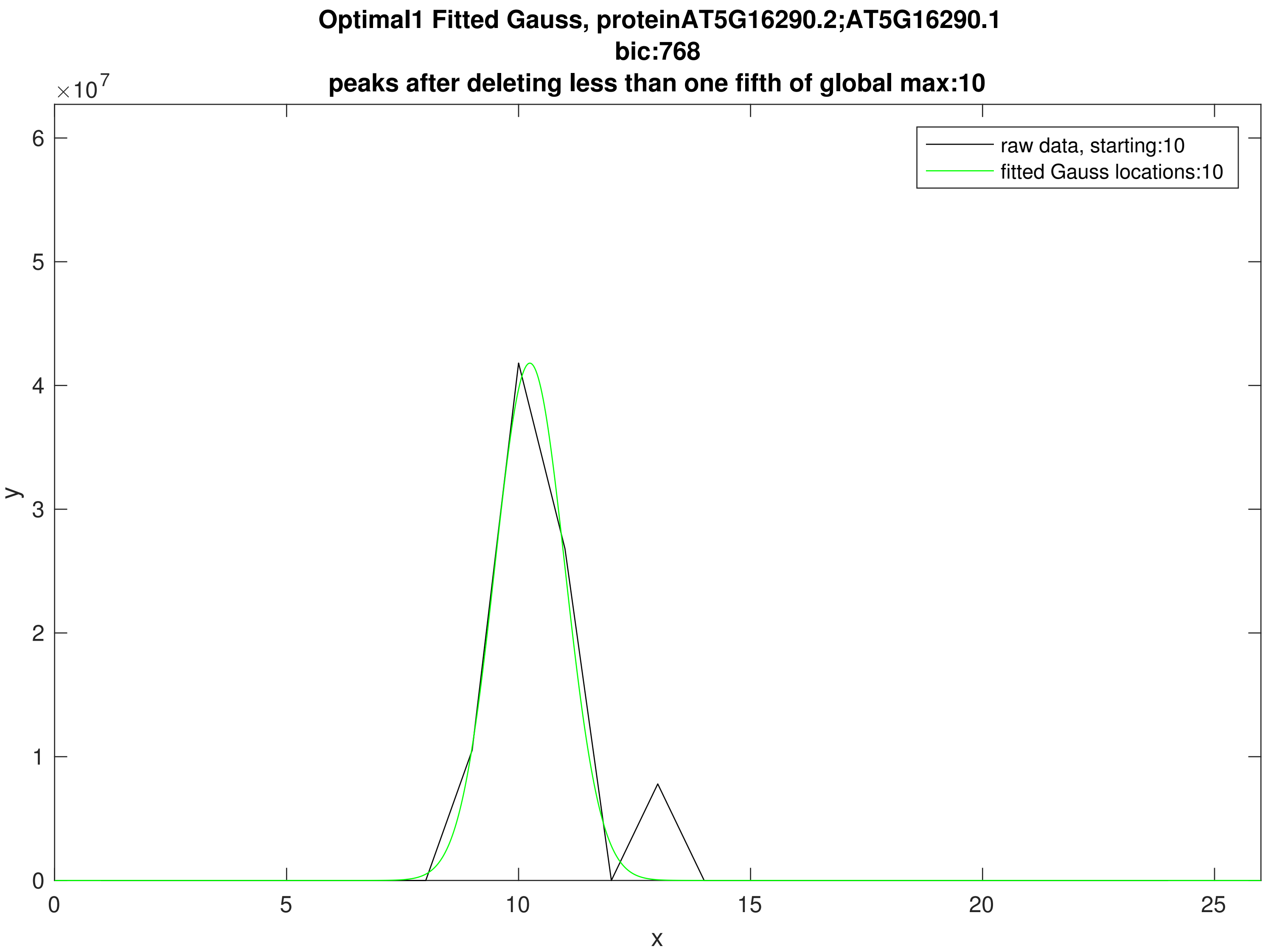
Optimal1 Fitted Gauss, proteinAT5G16290.2;AT5G16290.1
bic:768
peaks after deleting less than one fifth of global max:10
×10^7
raw data, starting:10
fitted Gauss locations:10
y
x
0
1
2
3
4
5
6
0
5
10
15
20
25

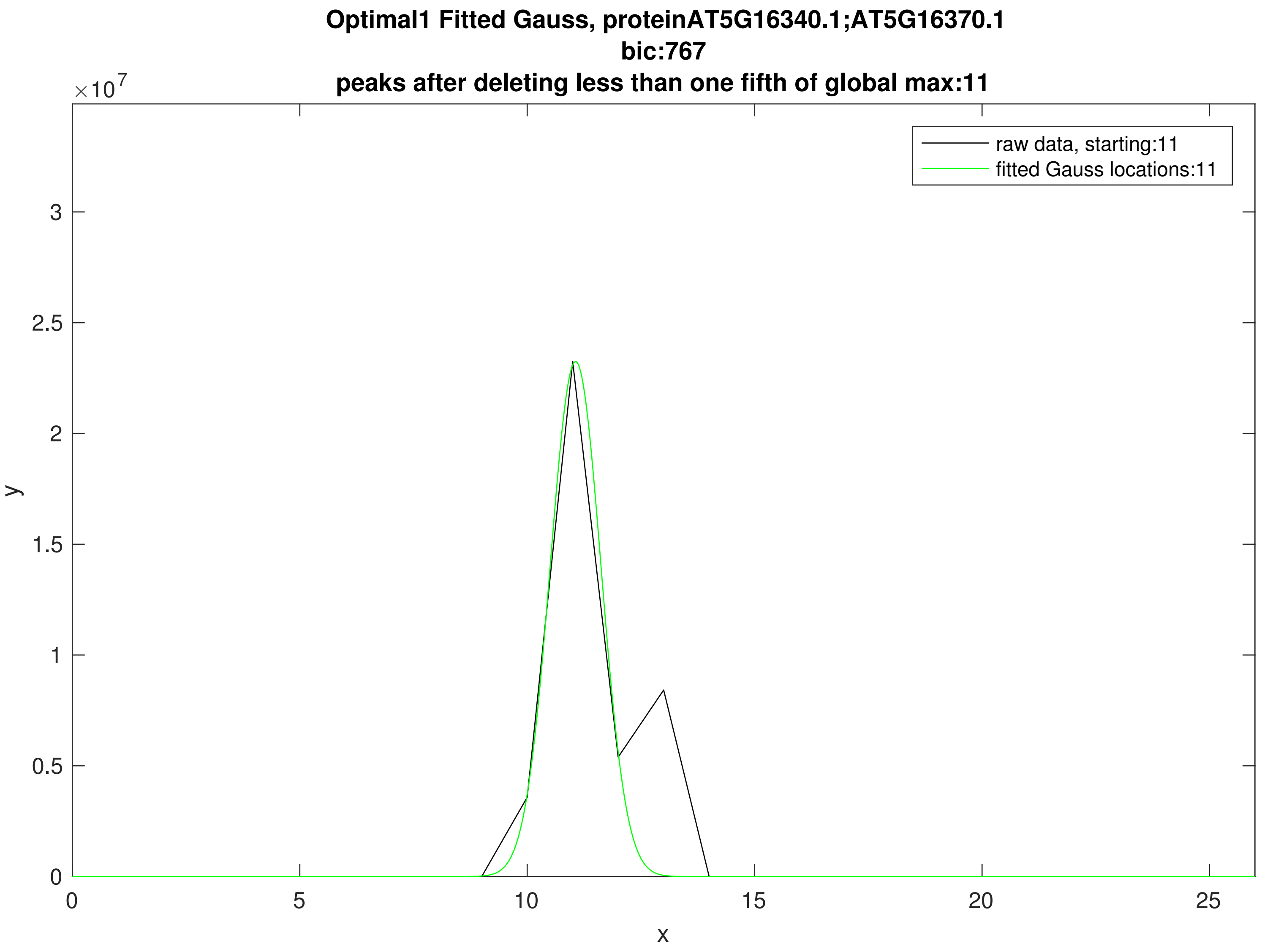
Optimal1 Fitted Gauss, proteinAT5G16340.1;AT5G16370.1
bic:767
peaks after deleting less than one fifth of global max:11
×10^7
raw data, starting:11
fitted Gauss locations:11
3
2.5
2
1.5
1
0.5
0
y
0
5
10
15
20
25
x

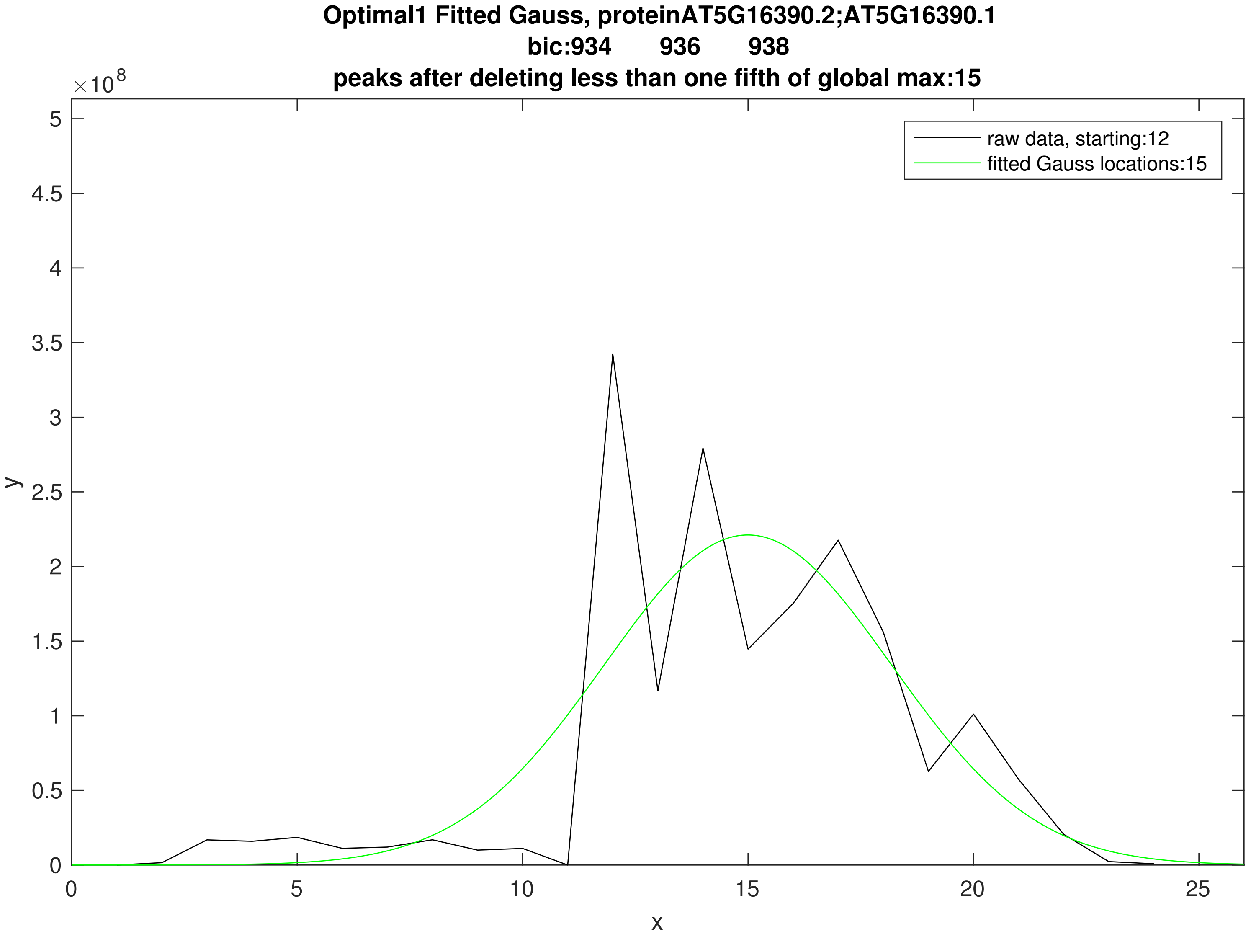

Optimal1 Fitted Gauss, proteinAT5G16390.2;AT5G16390.1
bic:934 936 938
peaks after deleting less than one fifth of global max:15
×10^8
raw data, starting:12
fitted Gauss locations:15
y
x
0
0.5
1
1.5
2
2.5
3
3.5
4
4.5
5
0
5
10
15
20
25

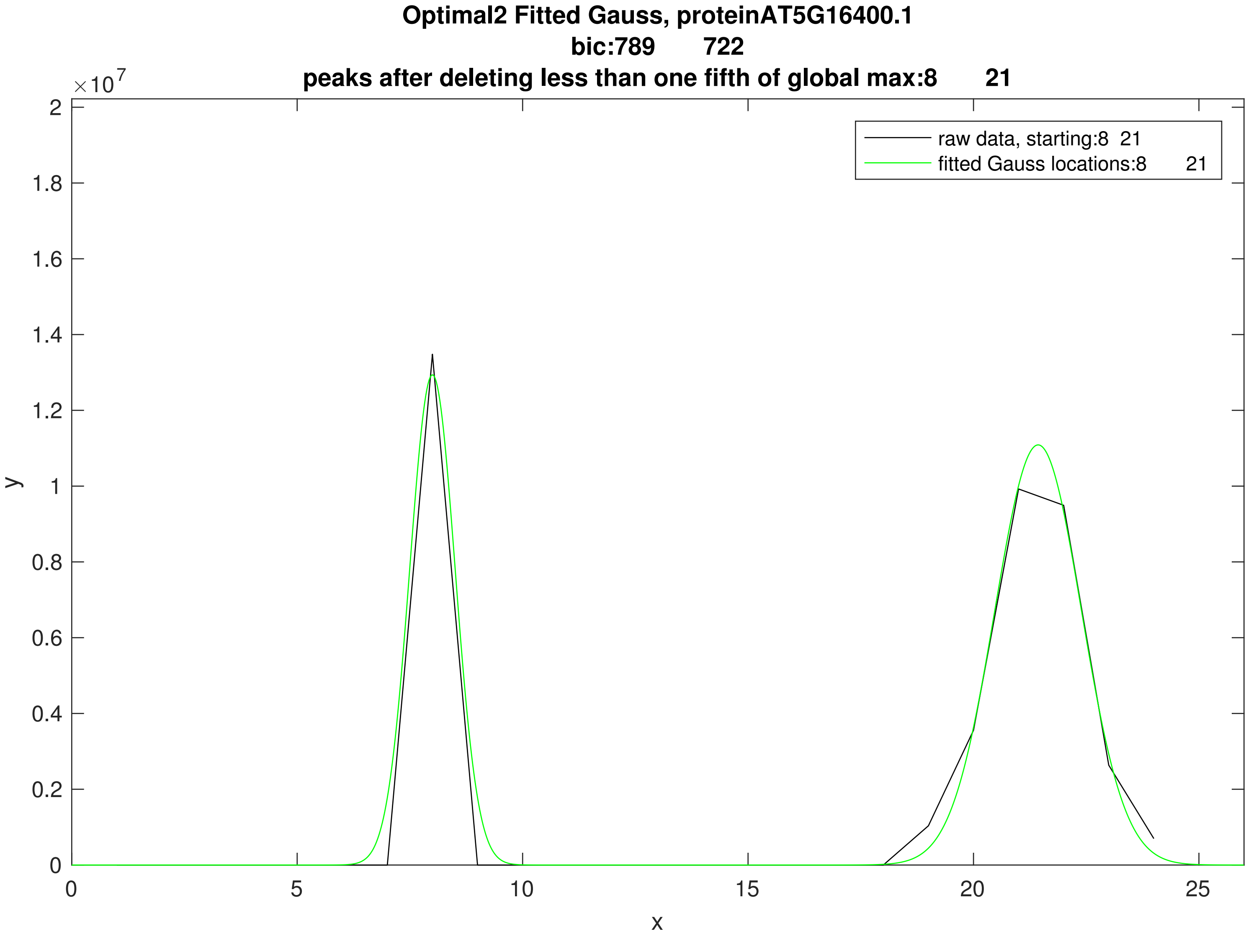

Optimal2 Fitted Gauss, proteinAT5G16400.1
bic:789 722
peaks after deleting less than one fifth of global max:8 21
×10^7
raw data, starting:8 21
fitted Gauss locations:8 21
y
x
0
0.2
0.4
0.6
0.8
1
1.2
1.4
1.6
1.8
2
0
5
10
15
20
25

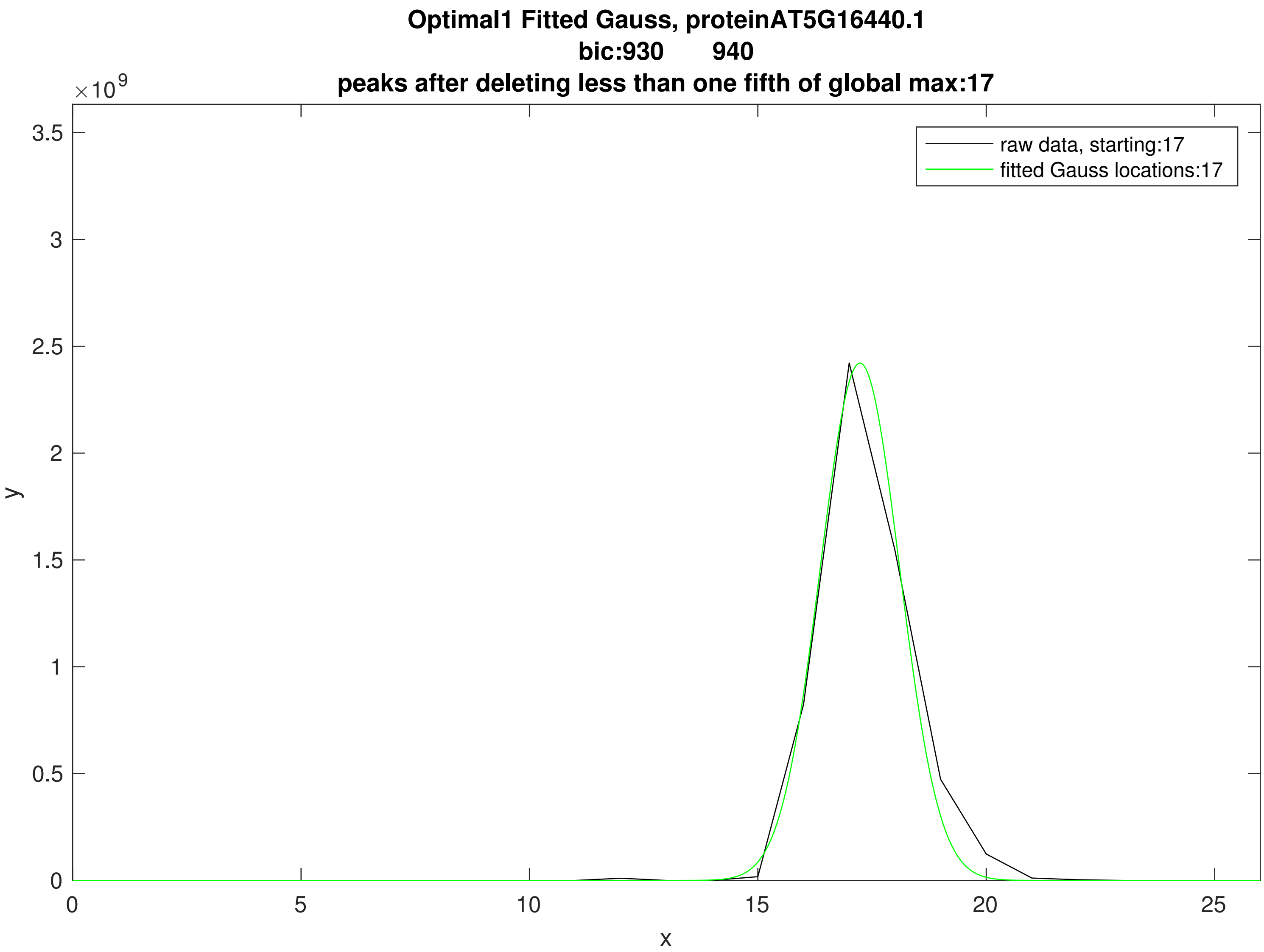
Optimal1 Fitted Gauss, proteinAT5G16440.1
bic:930 940
peaks after deleting less than one fifth of global max:17
×10^9
3.5
3
2.5
2
1.5
1
0.5
0
y
0
5
10
15
20
25
x
raw data, starting:17
fitted Gauss locations:17

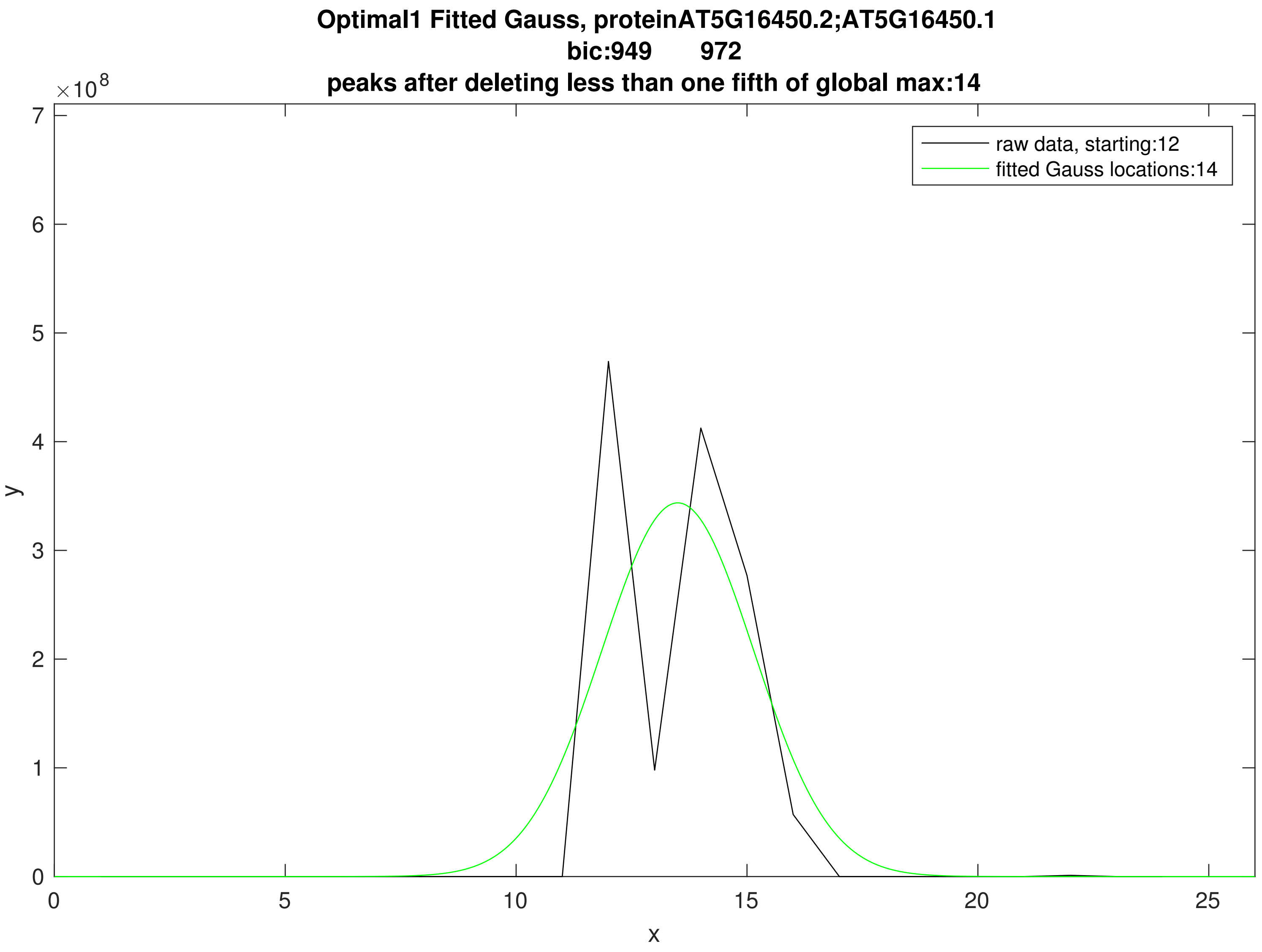

Optimal1 Fitted Gauss, proteinAT5G16450.2;AT5G16450.1
bic:949 972
peaks after deleting less than one fifth of global max:14
×10^8
raw data, starting:12
fitted Gauss locations:14
y
x

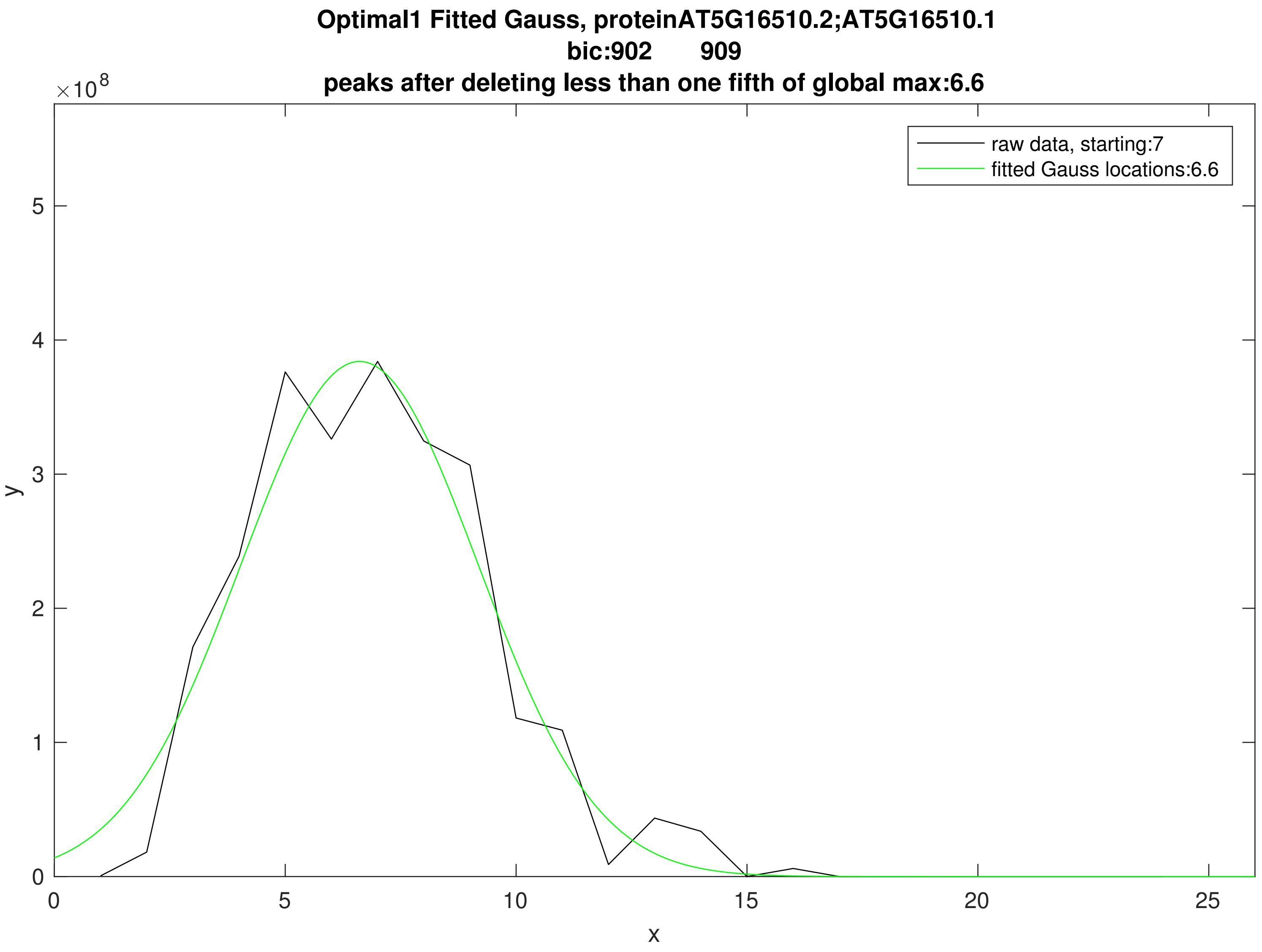
Optimal1 Fitted Gauss, proteinAT5G16510.2;AT5G16510.1
bic:902 909
peaks after deleting less than one fifth of global max:6.6
×10^8
raw data, starting:7
fitted Gauss locations:6.6
y
x
0
1
2
3
4
5
0
5
10
15
20
25

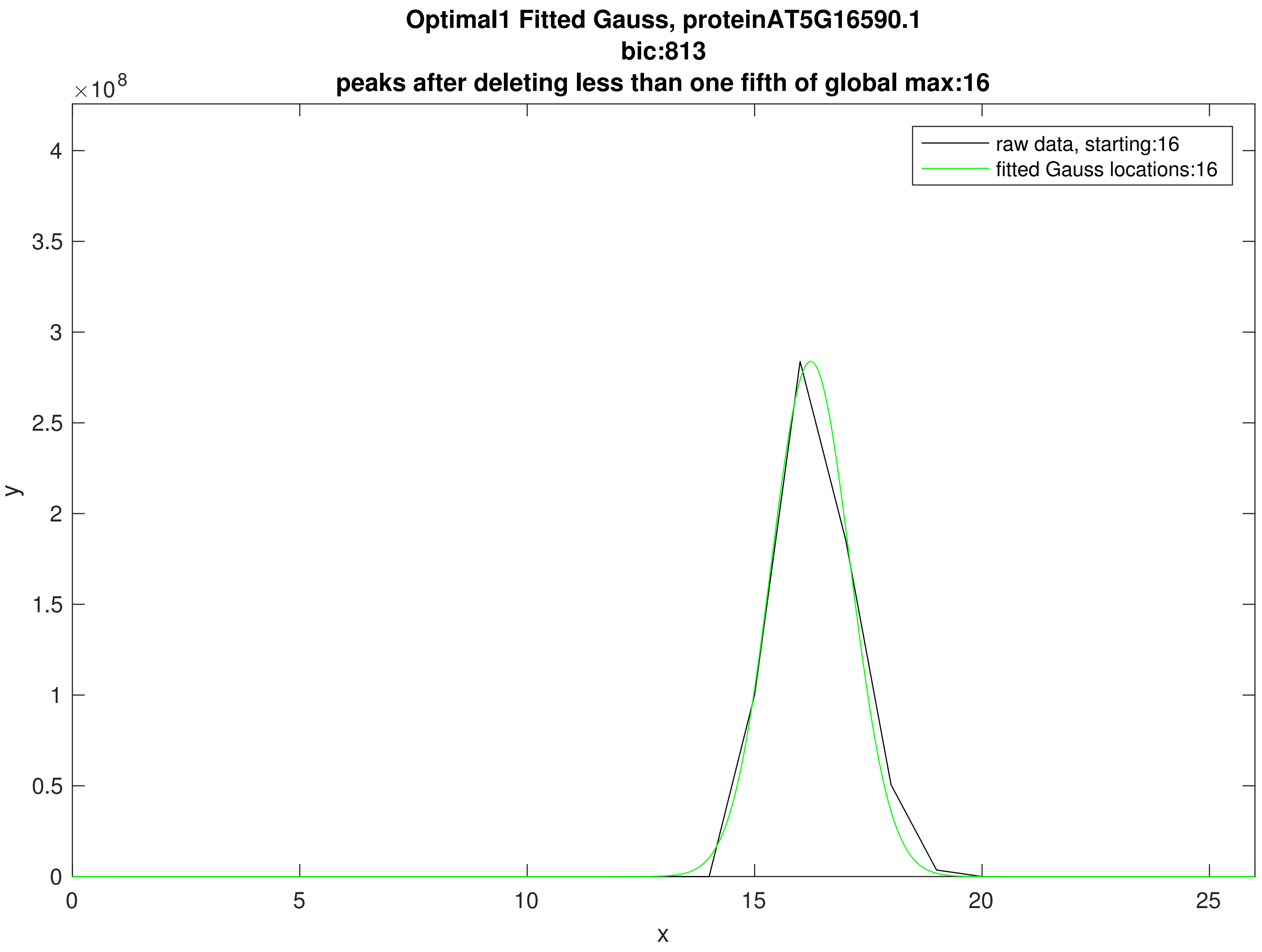

Optimal1 Fitted Gauss, proteinAT5G16590.1
bic:813
peaks after deleting less than one fifth of global max:16
×10^8
raw data, starting:16
fitted Gauss locations:16
4
3.5
3
2.5
2
1.5
1
0.5
0
y
0
5
10
15
20
25
x

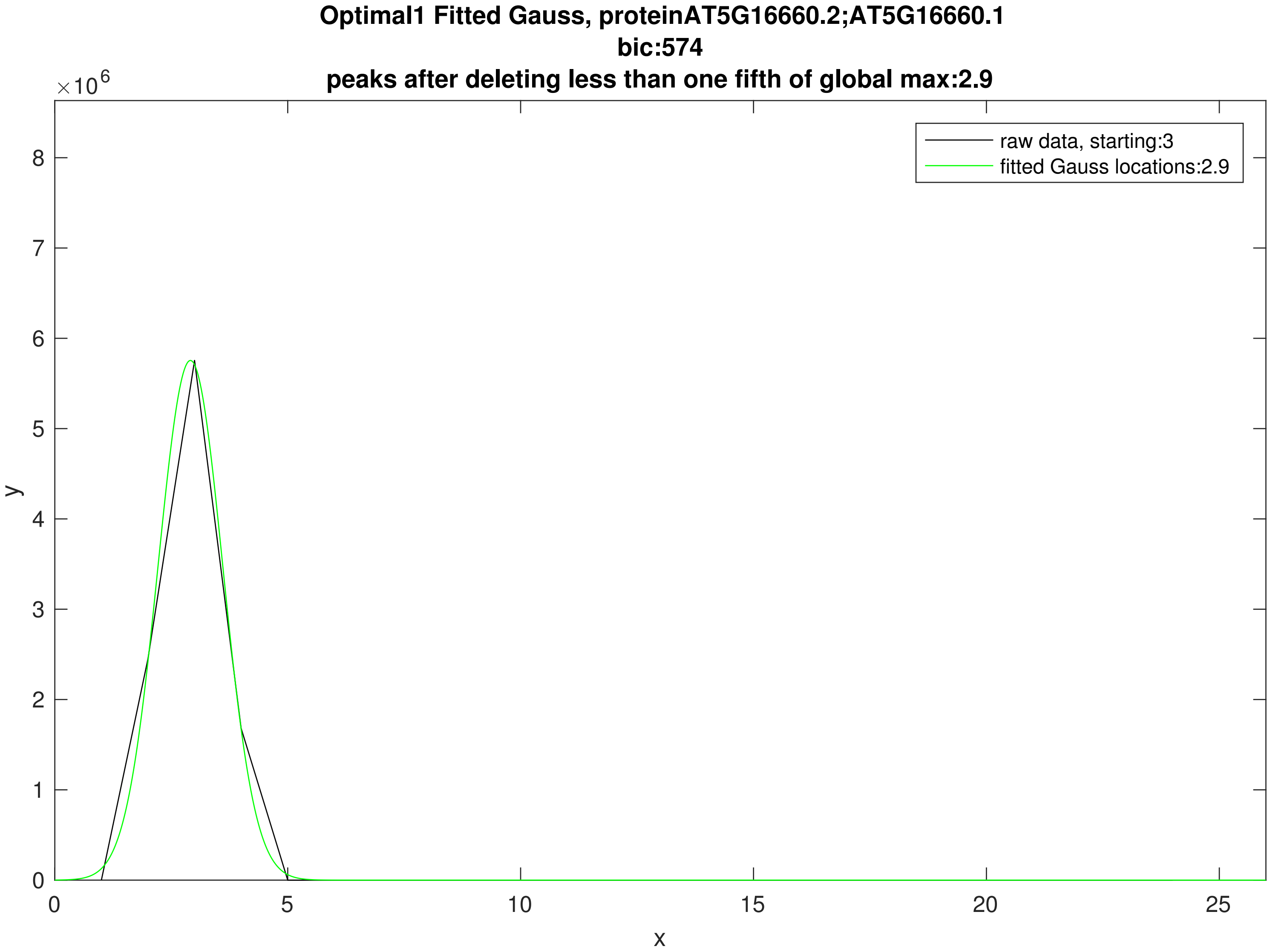

Optimal1 Fitted Gauss, proteinAT5G16660.2;AT5G16660.1
bic:574
peaks after deleting less than one fifth of global max:2.9
×10^6
raw data, starting:3
fitted Gauss locations:2.9
y
x
0
1
2
3
4
5
6
7
8
0
5
10
15
20
25

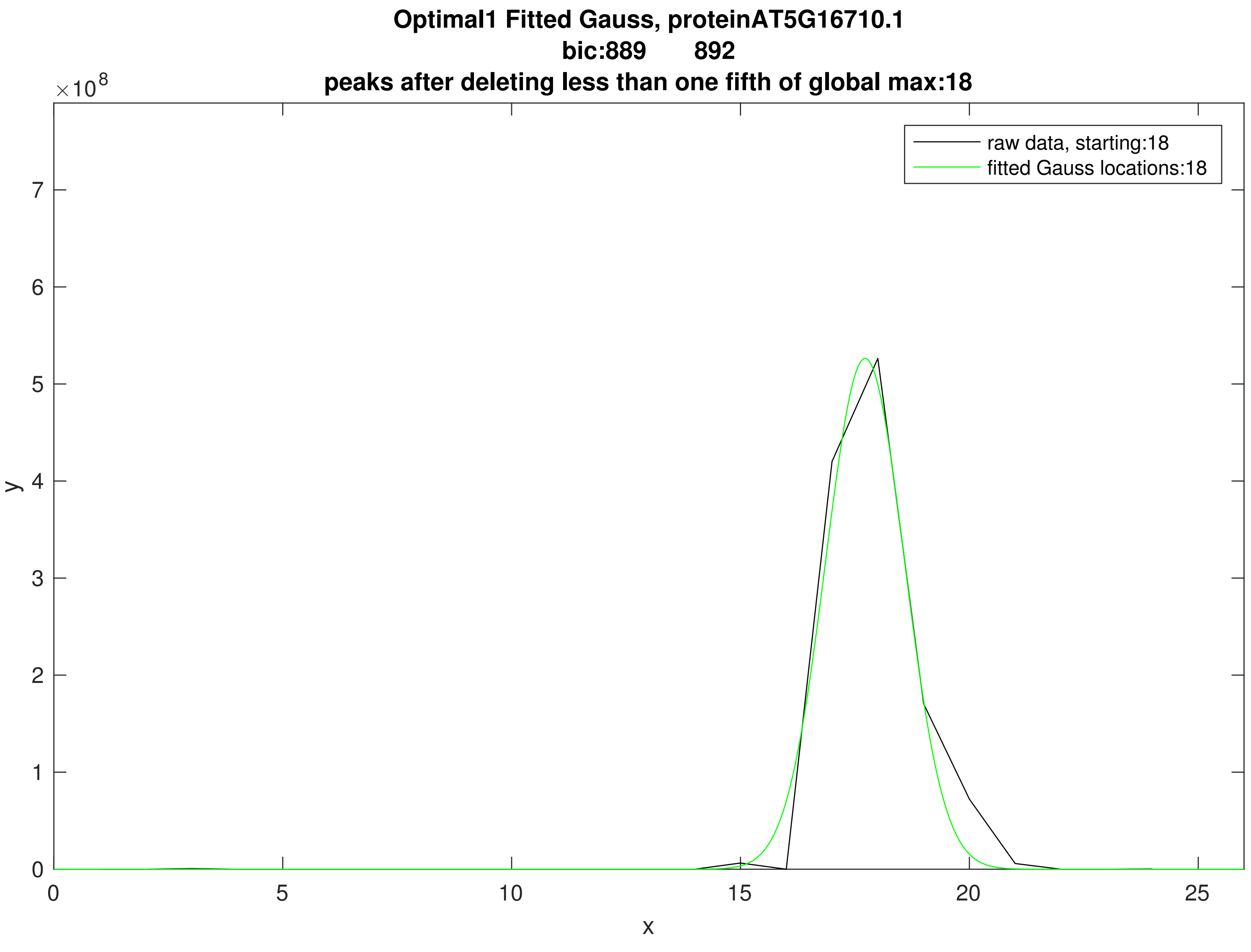

Optimal1 Fitted Gauss, proteinAT5G16710.1
bic:889 892
peaks after deleting less than one fifth of global max:18
×10^8
raw data, starting:18
fitted Gauss locations:18
y
x
0
1
2
3
4
5
6
7
0
5
10
15
20
25

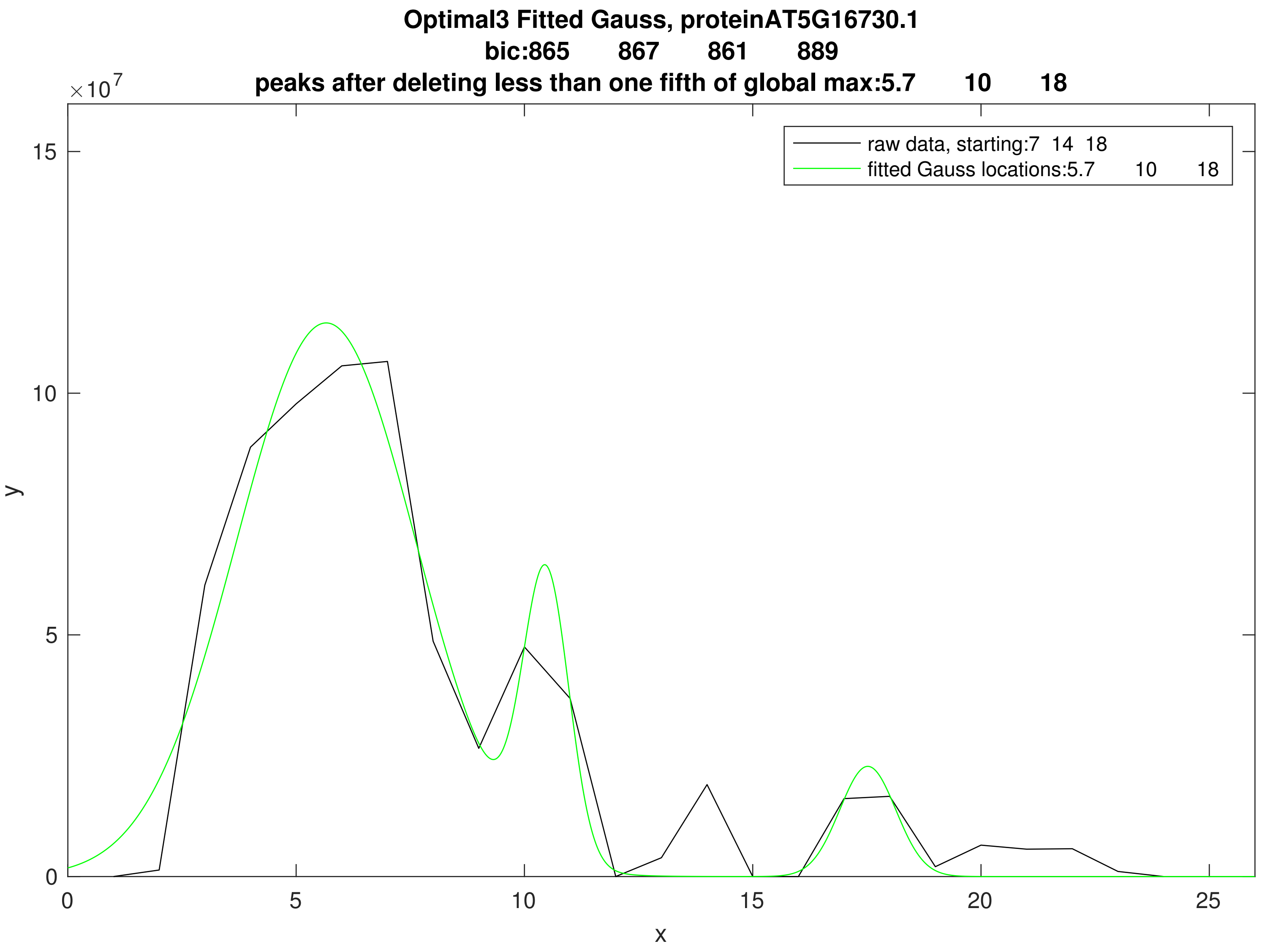
Optimal3 Fitted Gauss, proteinAT5G16730.1
bic:865 867 861 889
peaks after deleting less than one fifth of global max:5.7 10 18
×10^7
raw data, starting:7 14 18
fitted Gauss locations:5.7 10 18
y
x
0
5
10
15
20
25

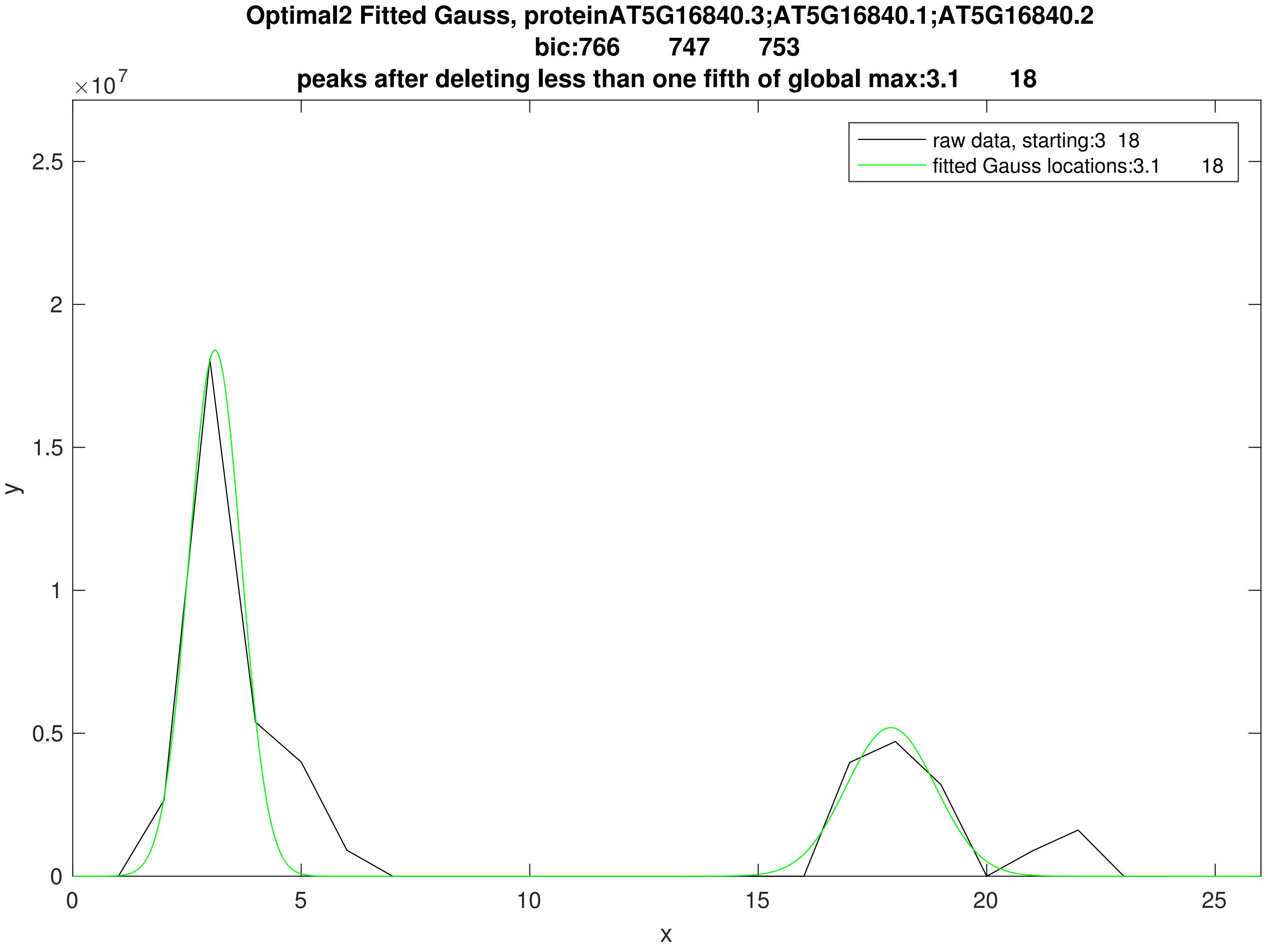

Optimal2 Fitted Gauss, proteinAT5G16840.3;AT5G16840.1;AT5G16840.2
bic:766 747 753
peaks after deleting less than one fifth of global max:3.1 18
×10^7
raw data, starting:3 18
fitted Gauss locations:3.1 18
y
x
0
0.5
1
1.5
2
2.5
0
5
10
15
20
25

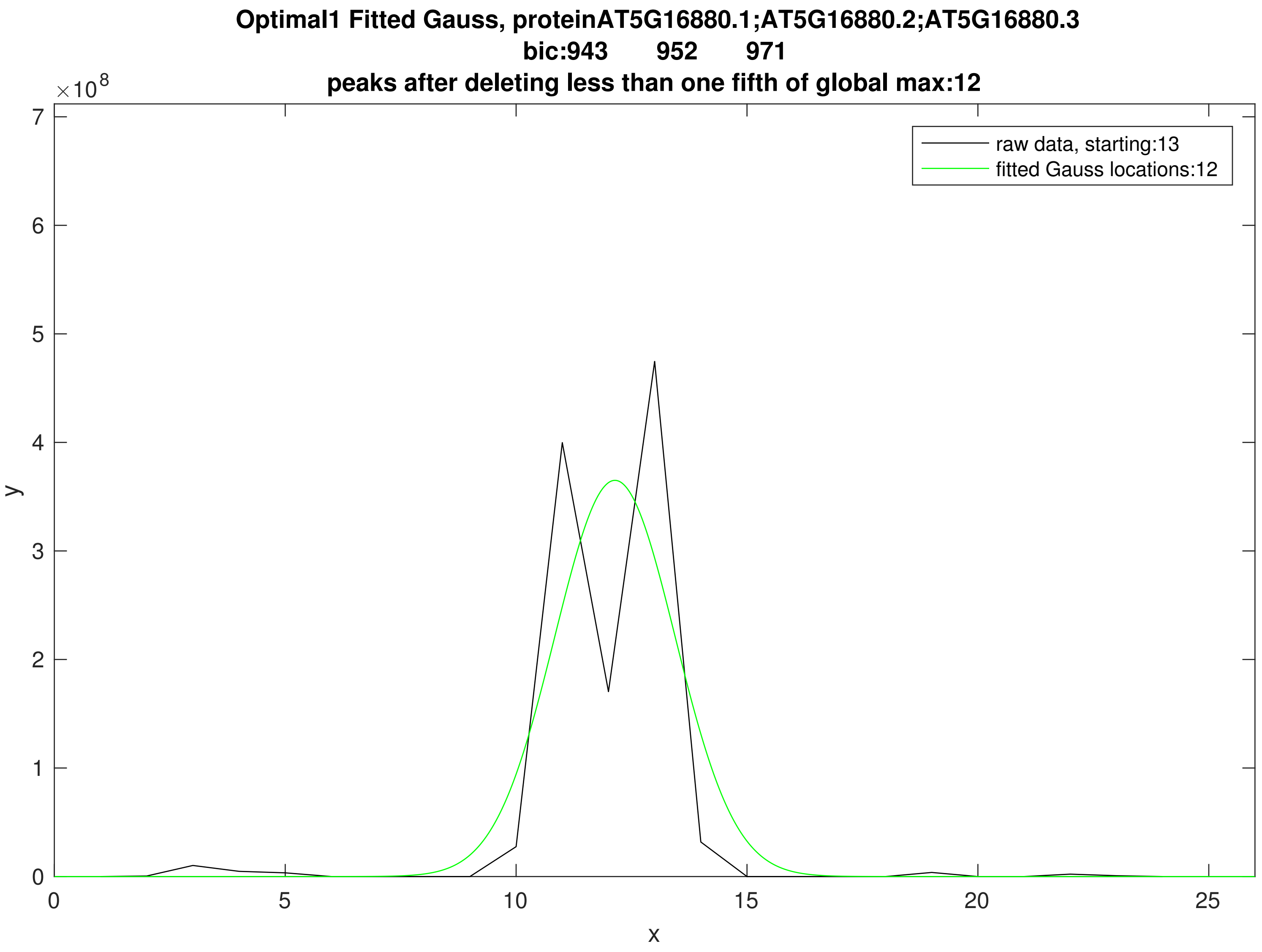
Optimal1 Fitted Gauss, proteinAT5G16880.1;AT5G16880.2;AT5G16880.3
bic:943 952 971
peaks after deleting less than one fifth of global max:12
×10^8
raw data, starting:13
fitted Gauss locations:12
y
x
0
1
2
3
4
5
6
7
10
15
20
25

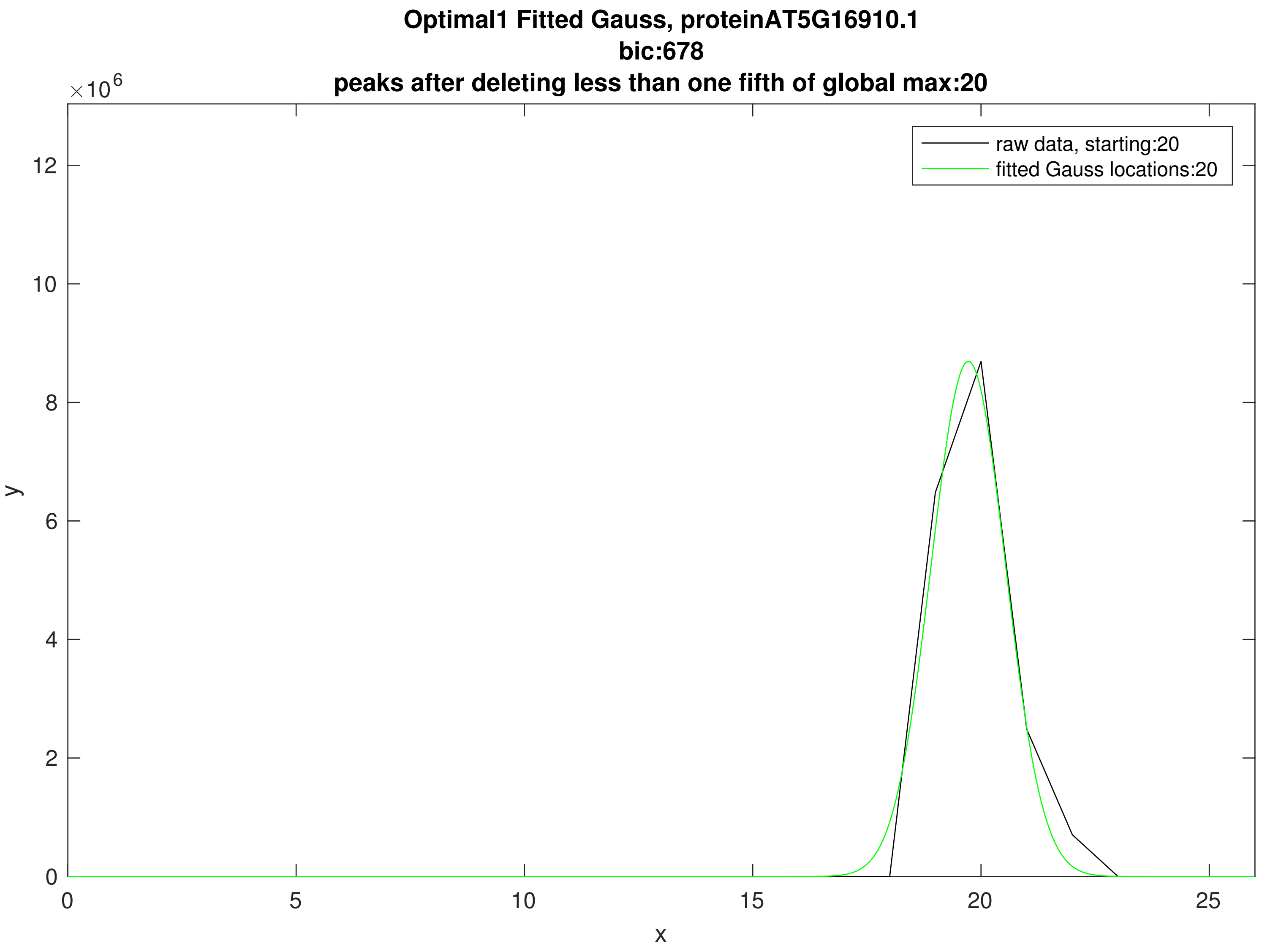

Optimal1 Fitted Gauss, proteinAT5G16910.1
bic:678
peaks after deleting less than one fifth of global max:20
×10^6
raw data, starting:20
fitted Gauss locations:20
12
10
8
6
4
2
0
y
0
5
10
15
20
25
x

**Optimal2 Fitted Gauss, proteinAT5G16940.2;AT5G16940.1**
**bic:822 816**
**peaks after deleting less than one fifth of global max:12**

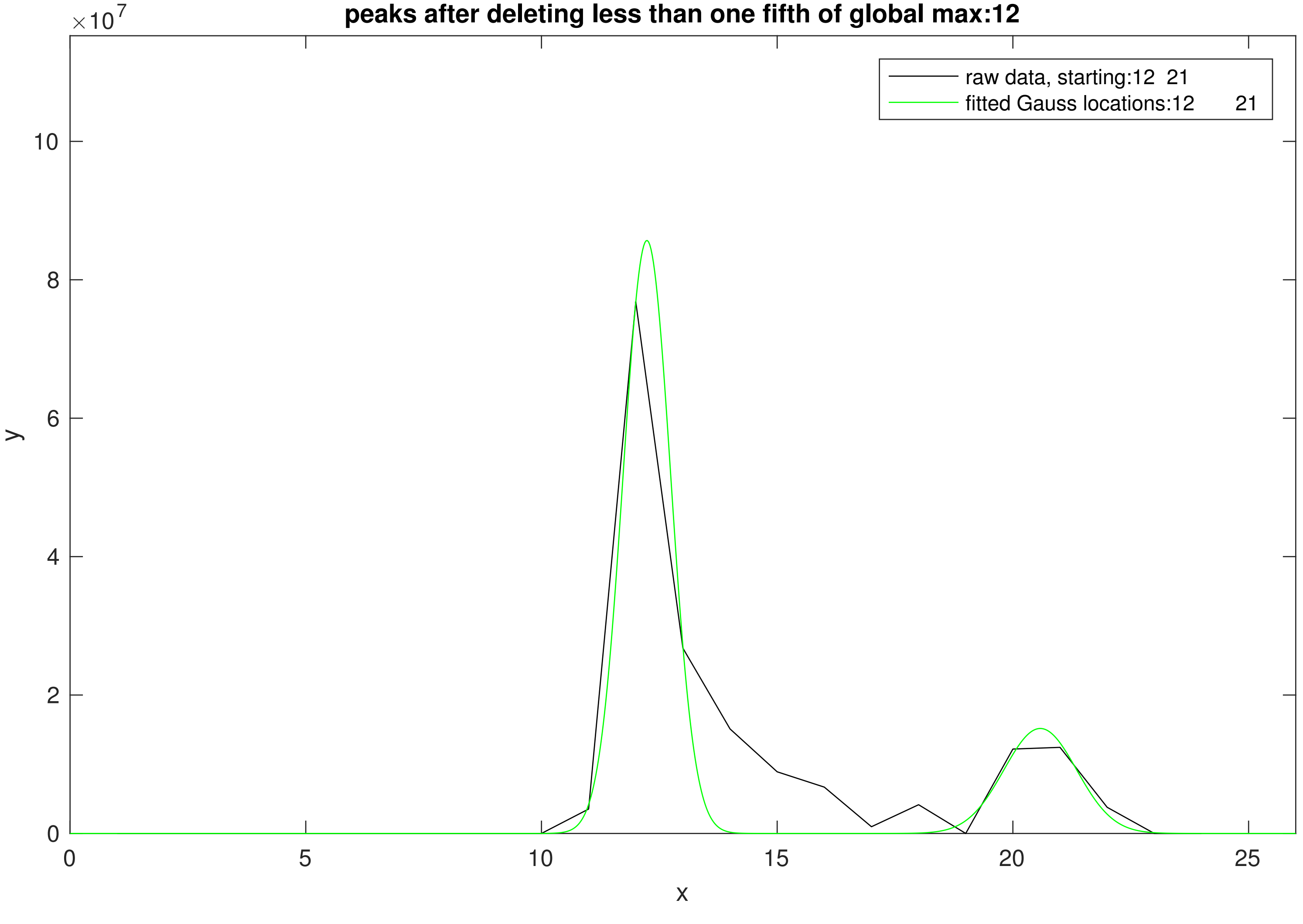

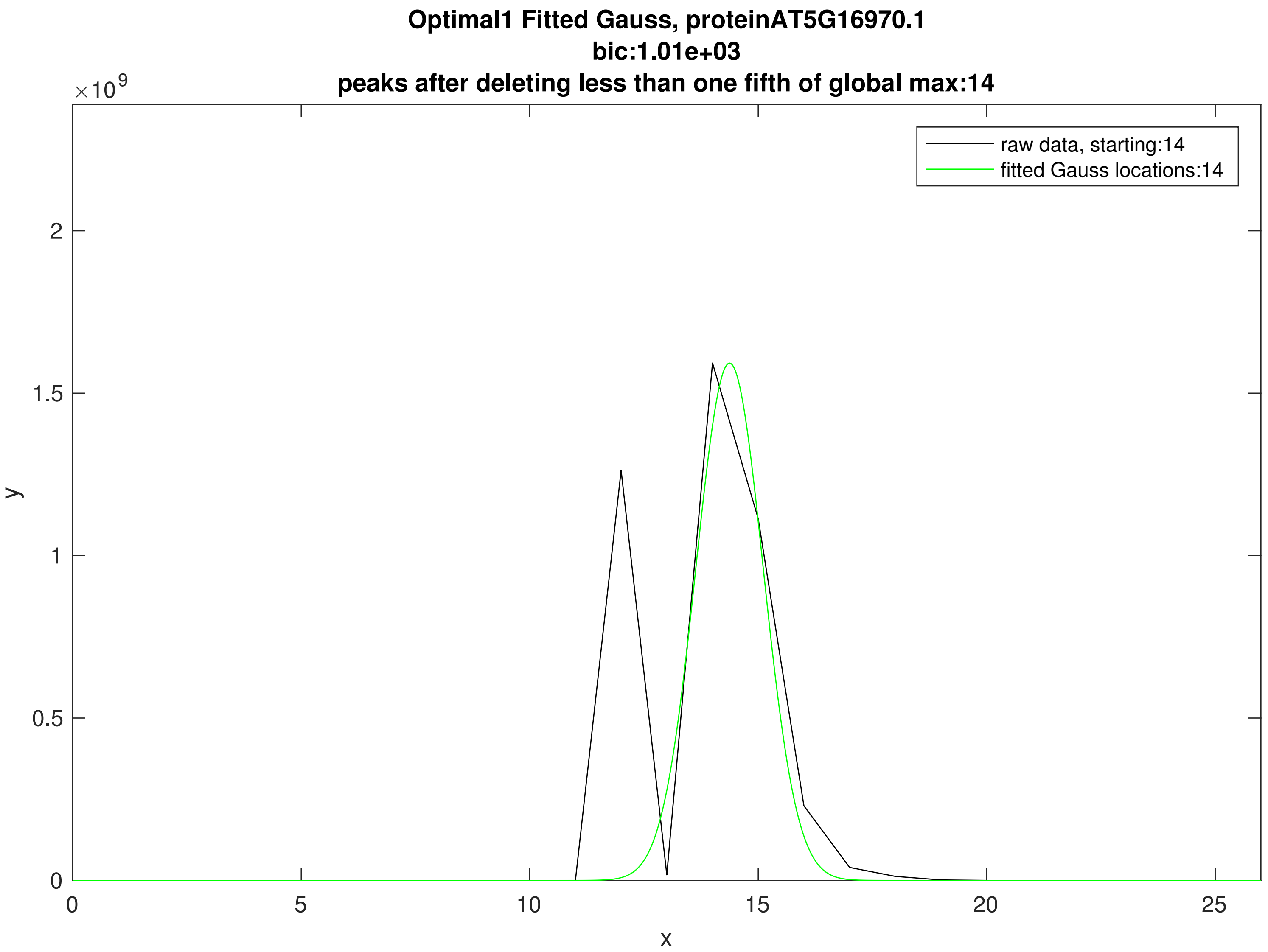
Optimal1 Fitted Gauss, proteinAT5G16970.1
bic:1.01e+03
peaks after deleting less than one fifth of global max:14
×10^9
raw data, starting:14
fitted Gauss locations:14
y
x
0
0.5
1
1.5
2
0
5
10
15
20
25

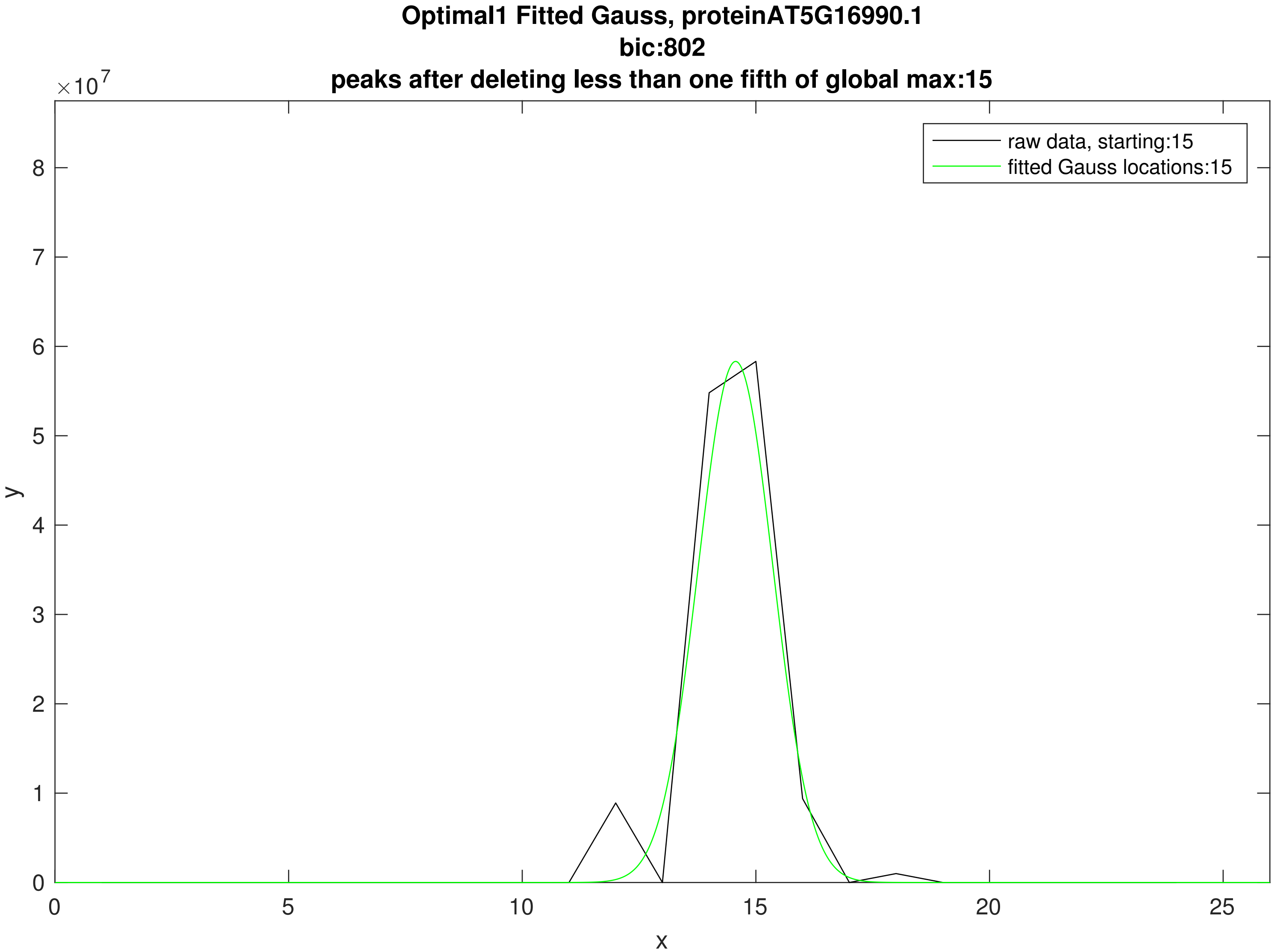
Optimal1 Fitted Gauss, proteinAT5G16990.1
bic:802
peaks after deleting less than one fifth of global max:15
×10^7
raw data, starting:15
fitted Gauss locations:15
y
x
0
1
2
3
4
5
6
7
8
0
5
10
15
20
25

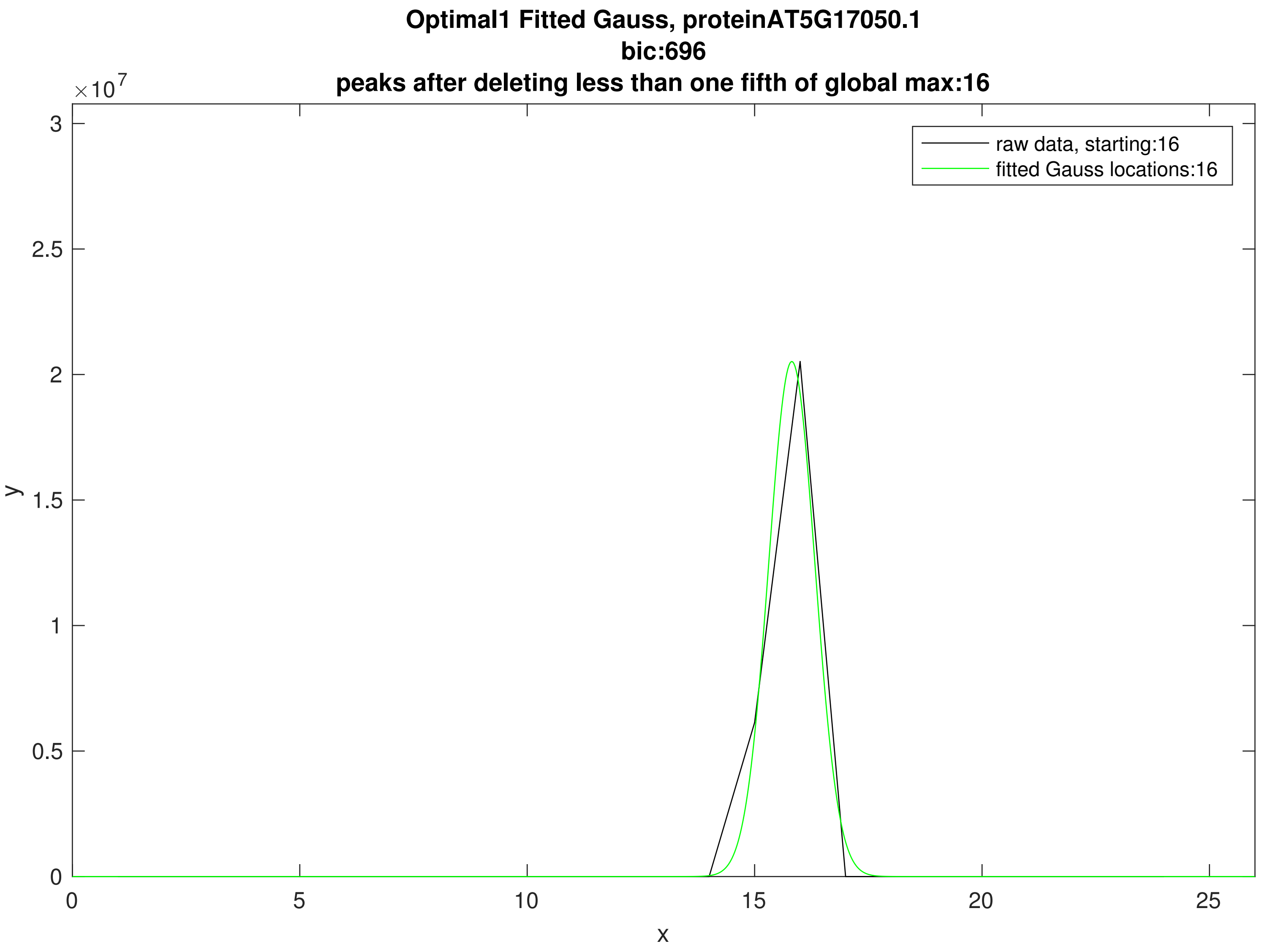
Optimal1 Fitted Gauss, proteinAT5G17050.1
bic:696
peaks after deleting less than one fifth of global max:16
×10^7
3
2.5
2
1.5
1
0.5
0
y
0
5
10
15
20
25
x
raw data, starting:16
fitted Gauss locations:16

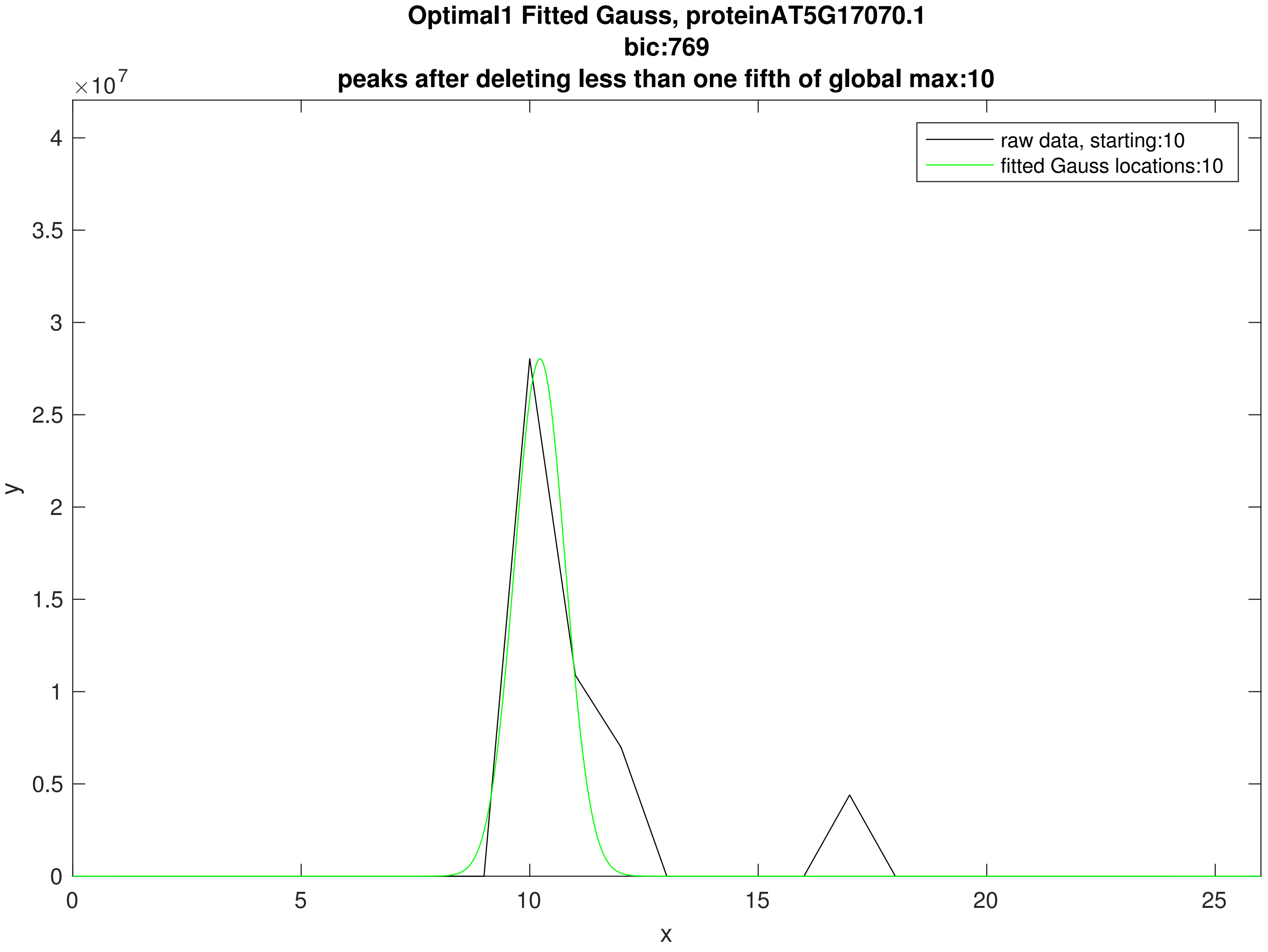
Optimal1 Fitted Gauss, proteinAT5G17070.1
bic:769
peaks after deleting less than one fifth of global max:10
×10^7
4
3.5
3
2.5
2
1.5
1
0.5
0
y
0
5
10
15
20
25
x
raw data, starting:10
fitted Gauss locations:10

**Optimal1 Fitted Gauss, proteinAT5G17310.2;AT5G17310.1**
**bic:993 1.05e+03 996 1.01e+03**
**peaks after deleting less than one fifth of global max:16**

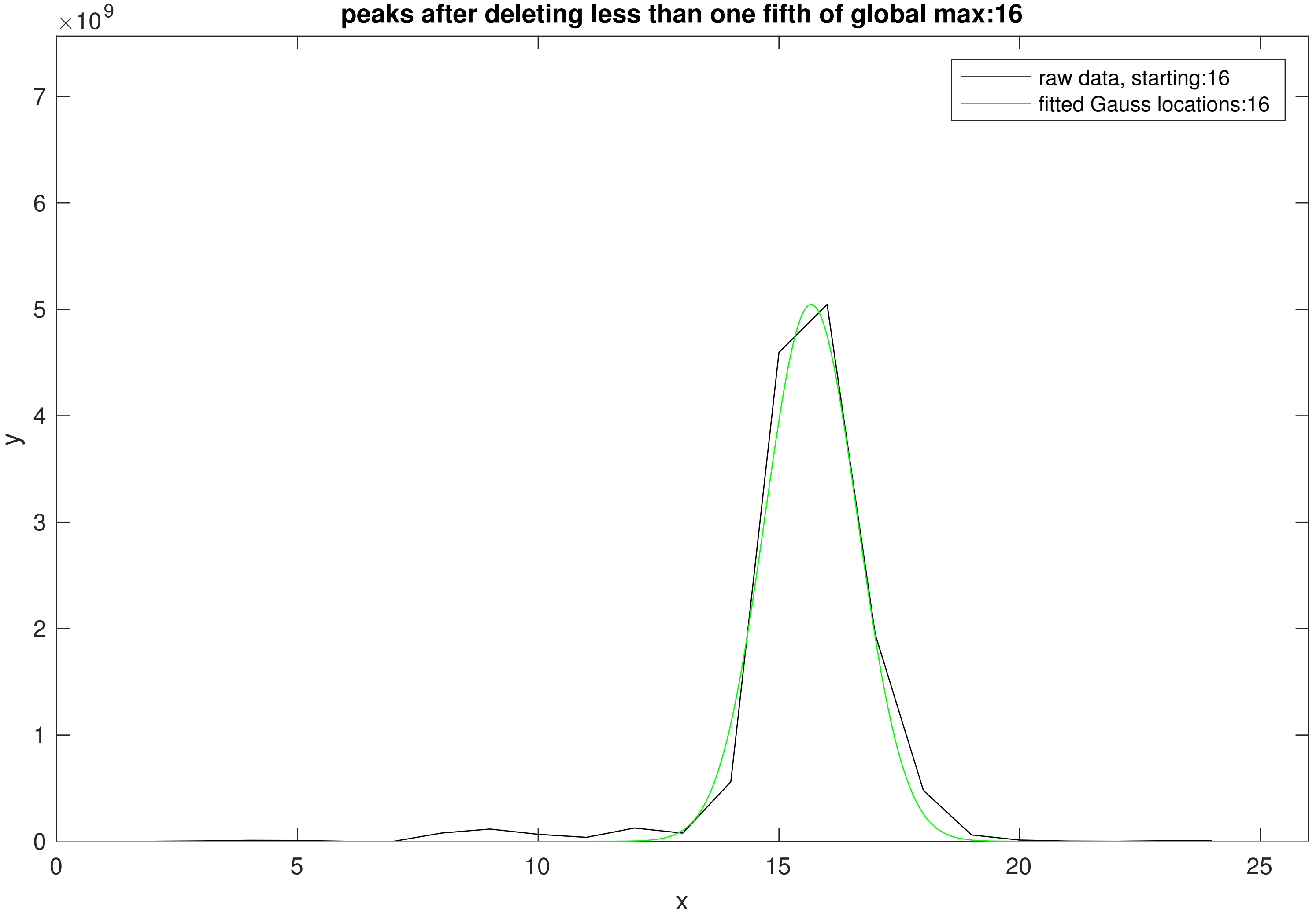

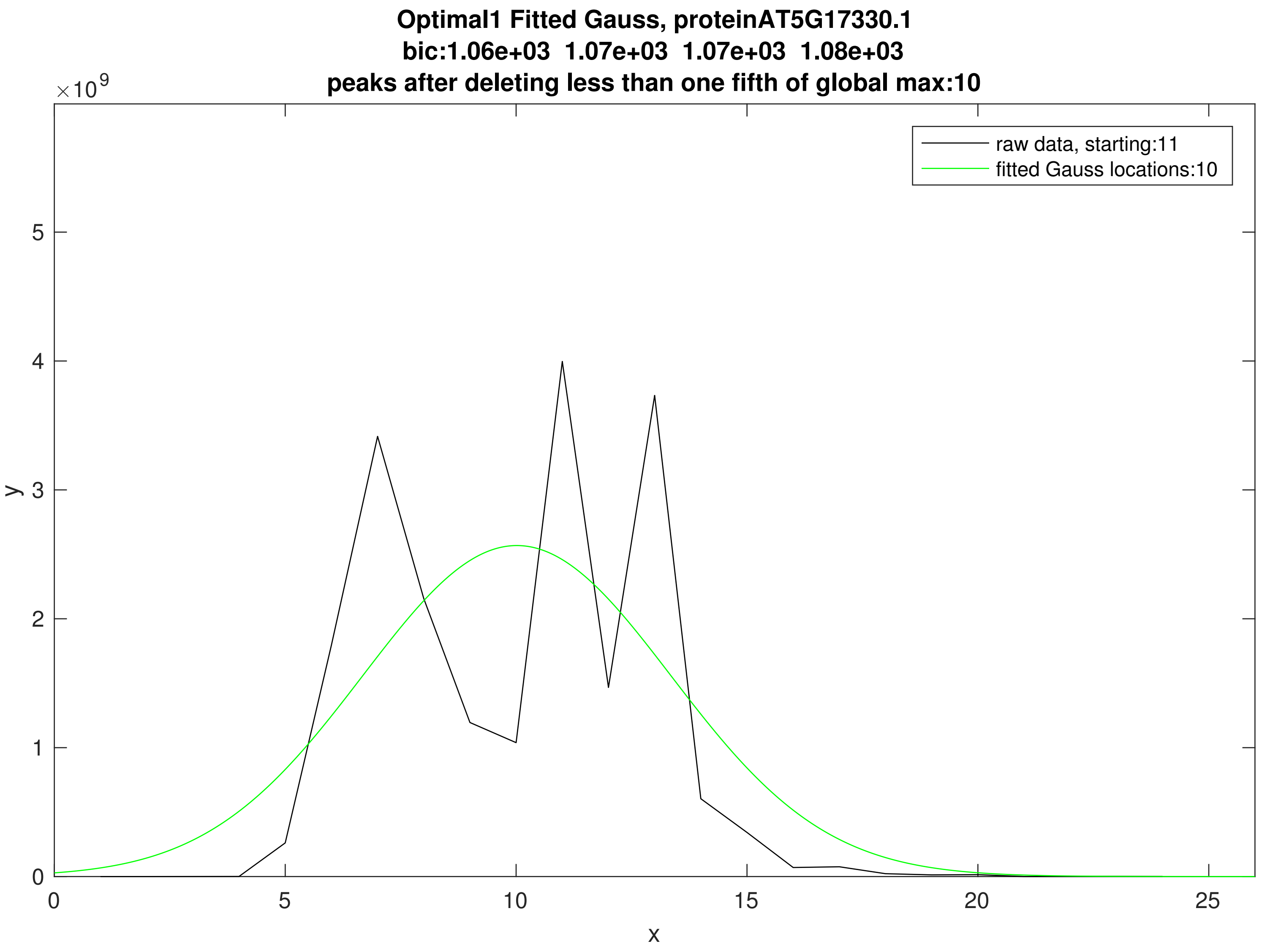
Optimal1 Fitted Gauss, proteinAT5G17330.1
bic:1.06e+03 1.07e+03 1.07e+03 1.08e+03
peaks after deleting less than one fifth of global max:10
×10^9
raw data, starting:11
fitted Gauss locations:10
y
x
0
1
2
3
4
5
0
5
10
15
20
25

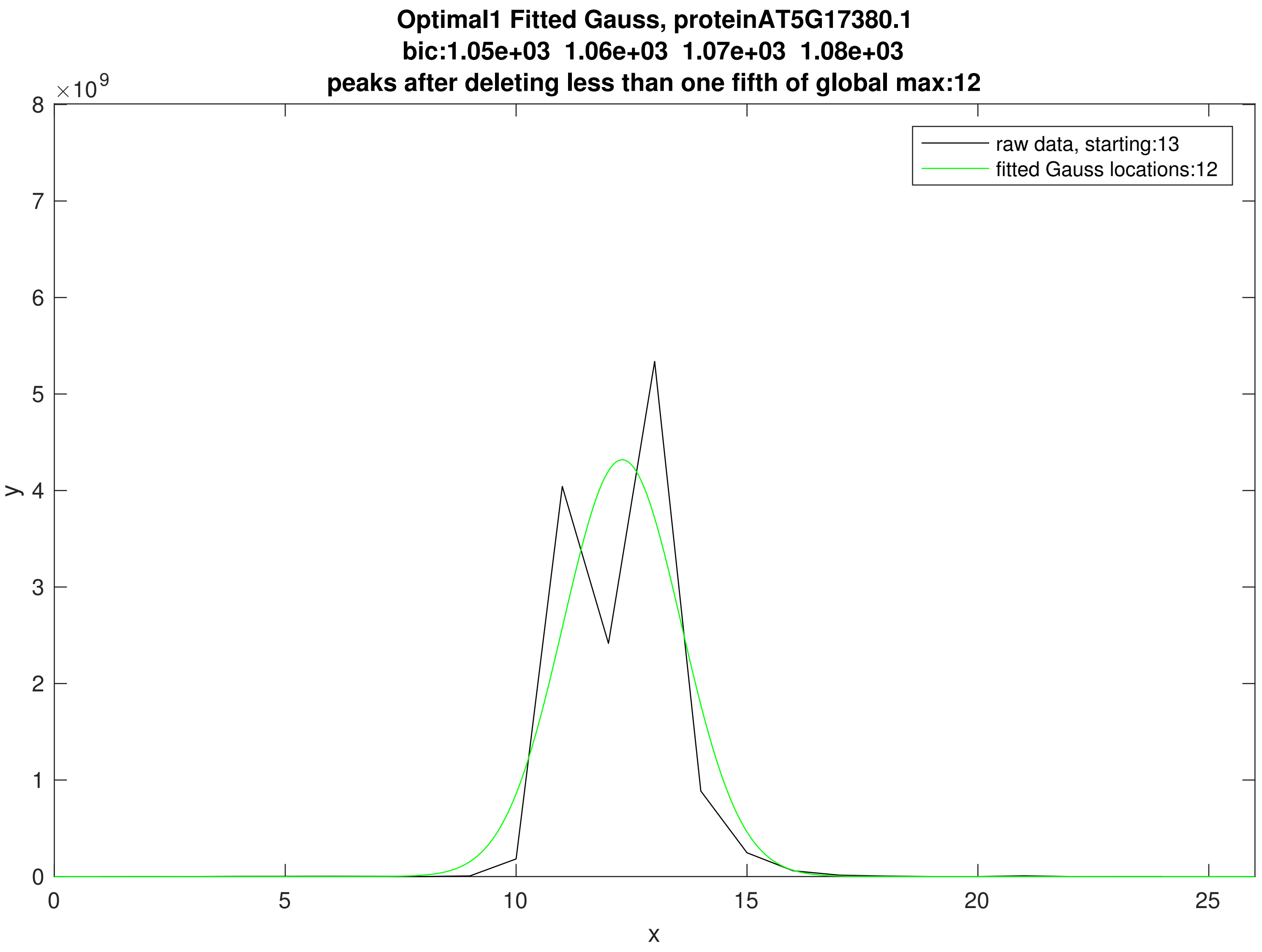

Optimal1 Fitted Gauss, proteinAT5G17380.1
bic:1.05e+03 1.06e+03 1.07e+03 1.08e+03
peaks after deleting less than one fifth of global max:12
×10^9
raw data, starting:13
fitted Gauss locations:12
y
x
0
1
2
3
4
5
6
7
8
0
5
10
15
20
25

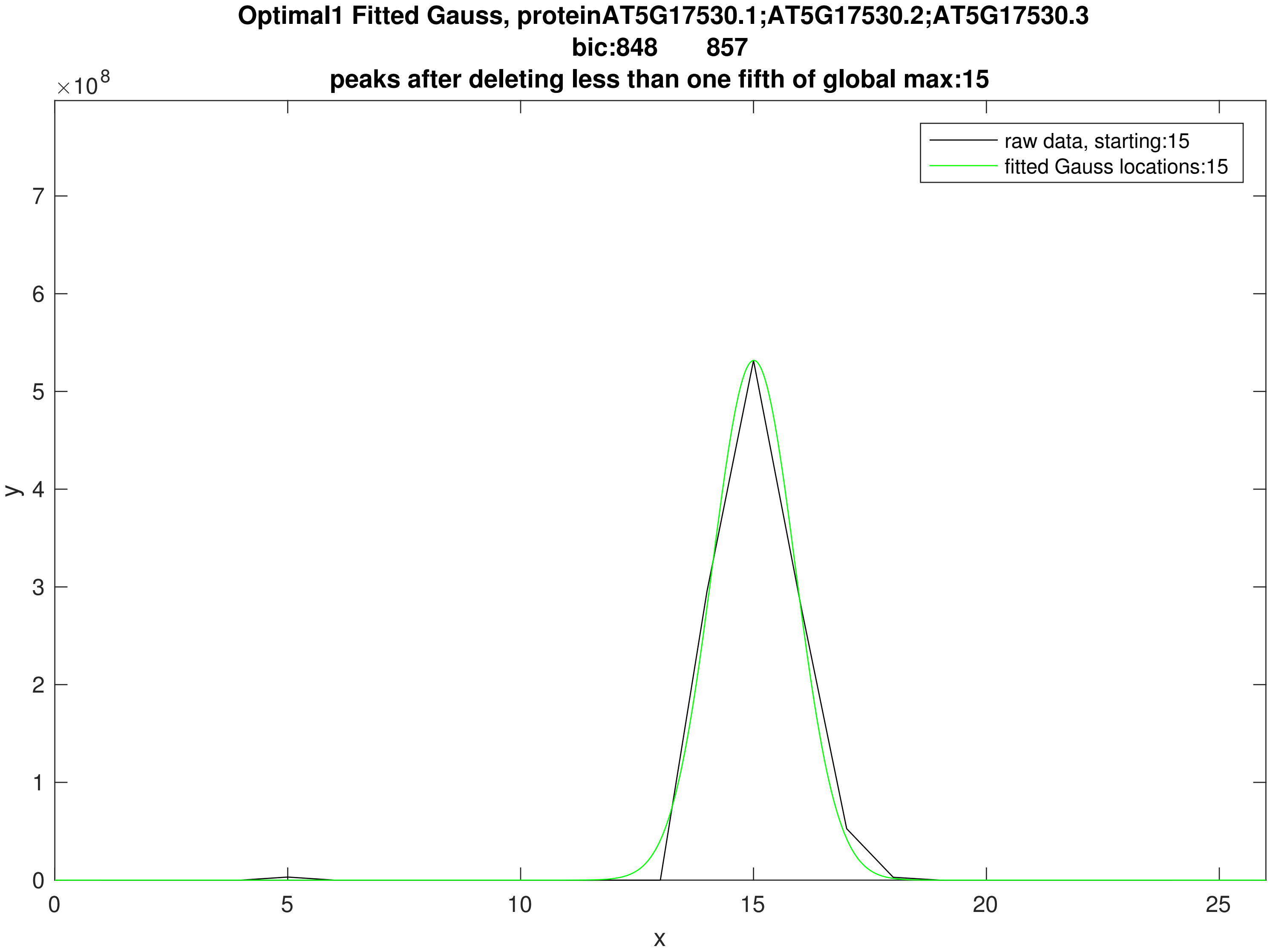
Optimal1 Fitted Gauss, proteinAT5G17530.1;AT5G17530.2;AT5G17530.3
bic:848 857
peaks after deleting less than one fifth of global max:15
×10^8
raw data, starting:15
fitted Gauss locations:15
y
x
0
1
2
3
4
5
6
7
0
5
10
15
20
25

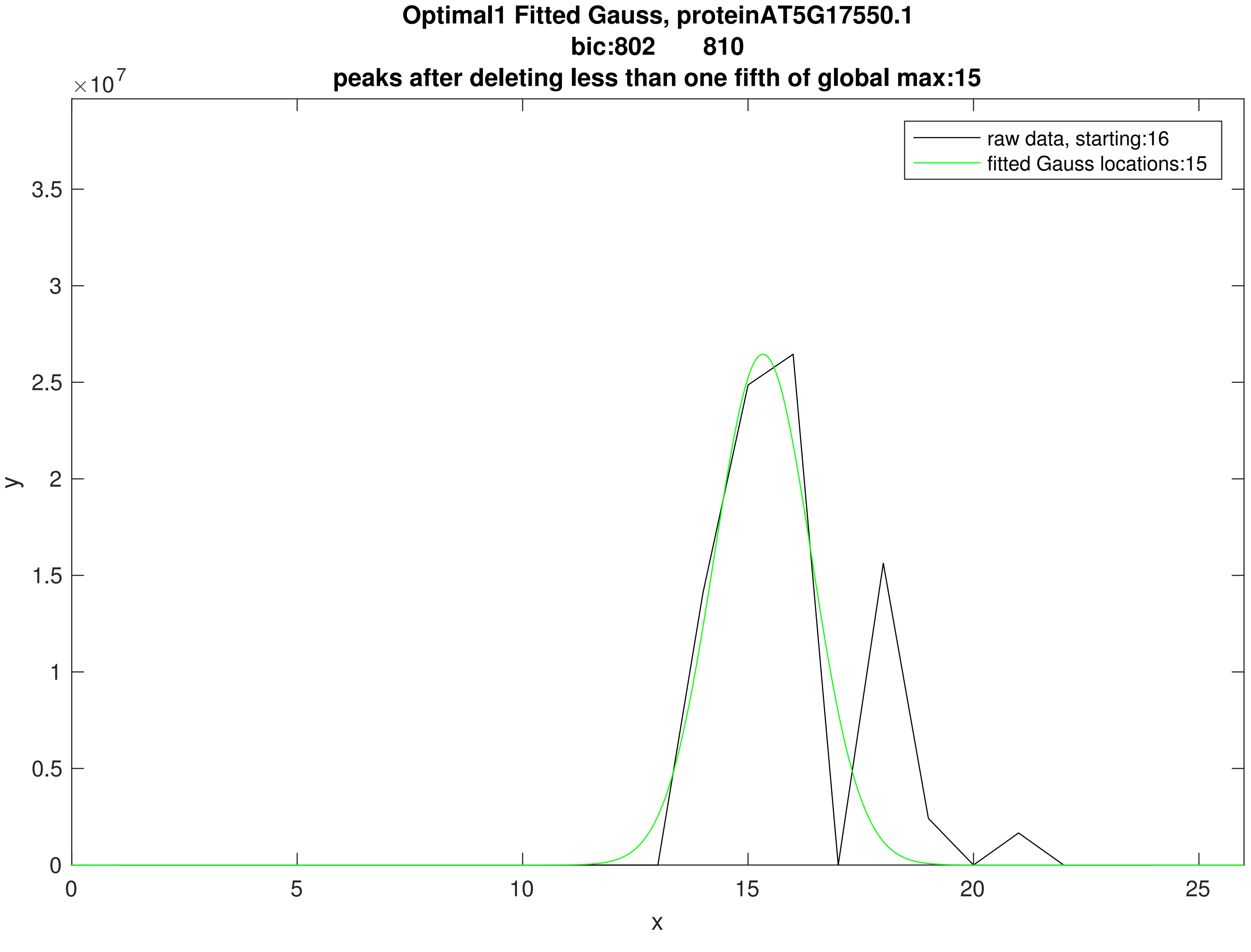

Optimal1 Fitted Gauss, proteinAT5G17550.1
bic:802 810
peaks after deleting less than one fifth of global max:15
×10^7
raw data, starting:16
fitted Gauss locations:15
3.5
3
2.5
2
1.5
1
0.5
0
y
0
5
10
15
20
25
x

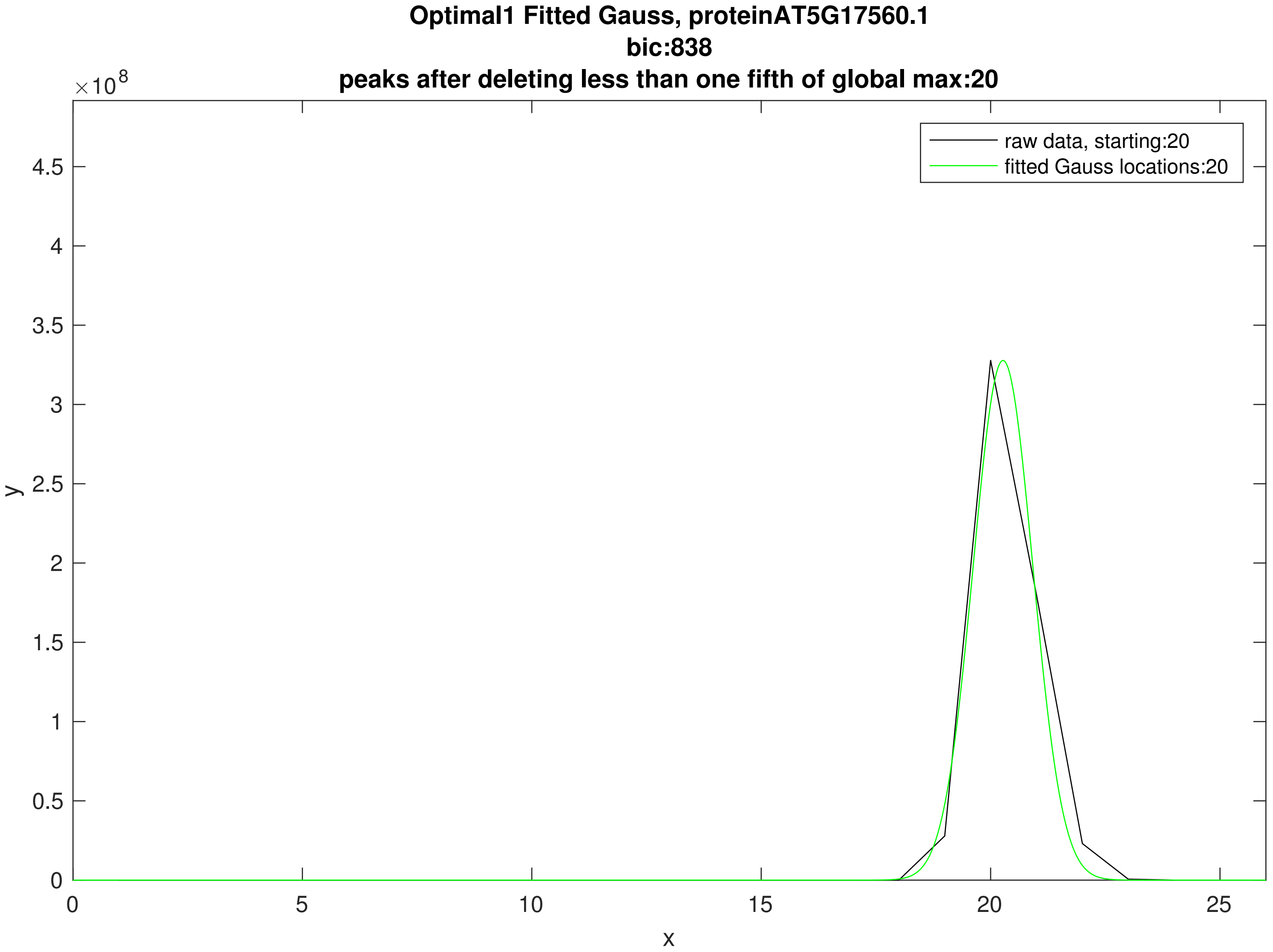
Optimal1 Fitted Gauss, proteinAT5G17560.1
bic:838
peaks after deleting less than one fifth of global max:20
×10^8
raw data, starting:20
fitted Gauss locations:20
y
x
0
0.5
1
1.5
2
2.5
3
3.5
4
4.5
5
10
15
20
25

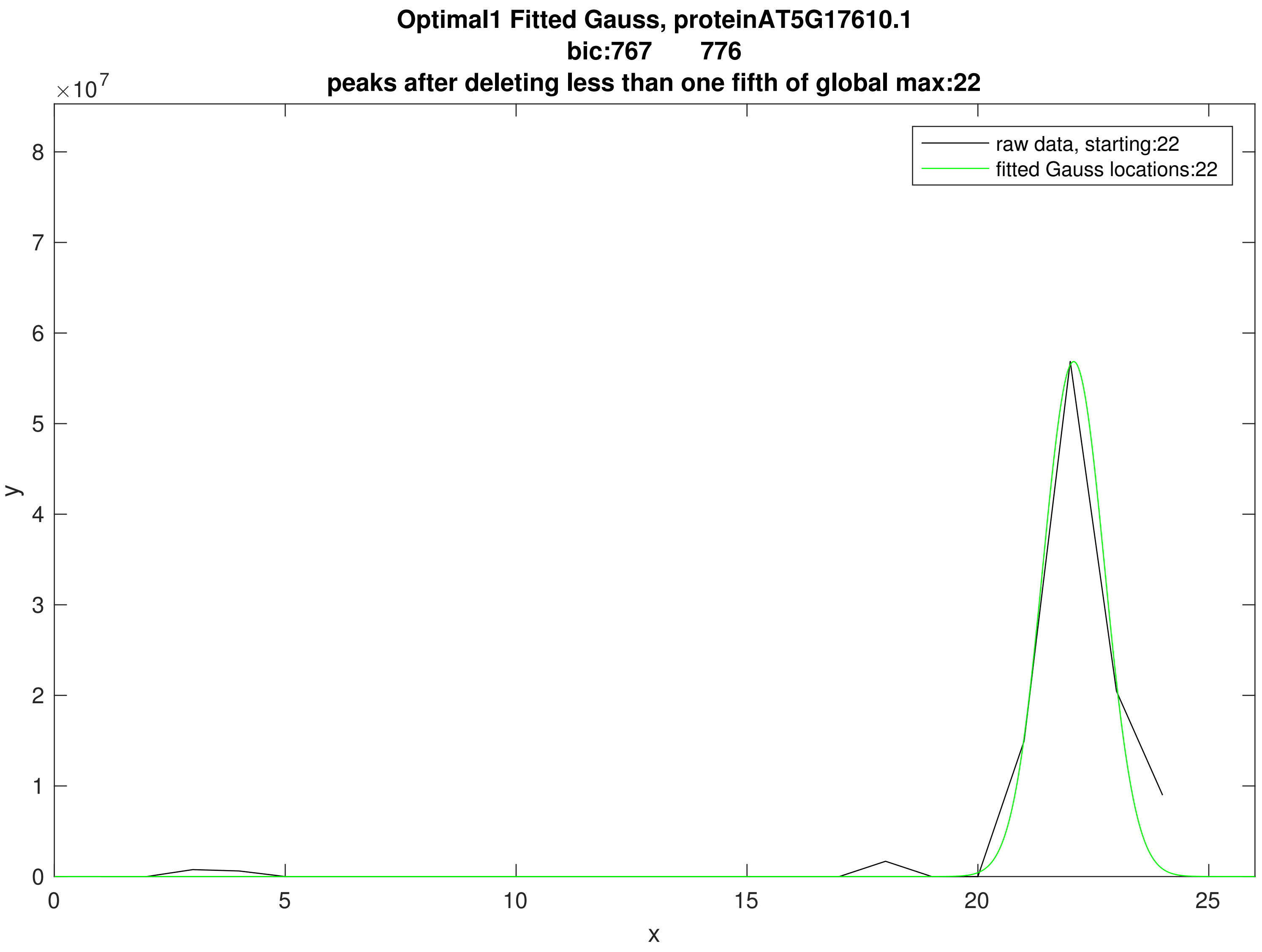
Optimal1 Fitted Gauss, proteinAT5G17610.1
bic:767 776
peaks after deleting less than one fifth of global max:22
×10^7
raw data, starting:22
fitted Gauss locations:22
y
x
0
1
2
3
4
5
6
7
8
0
5
10
15
20
25

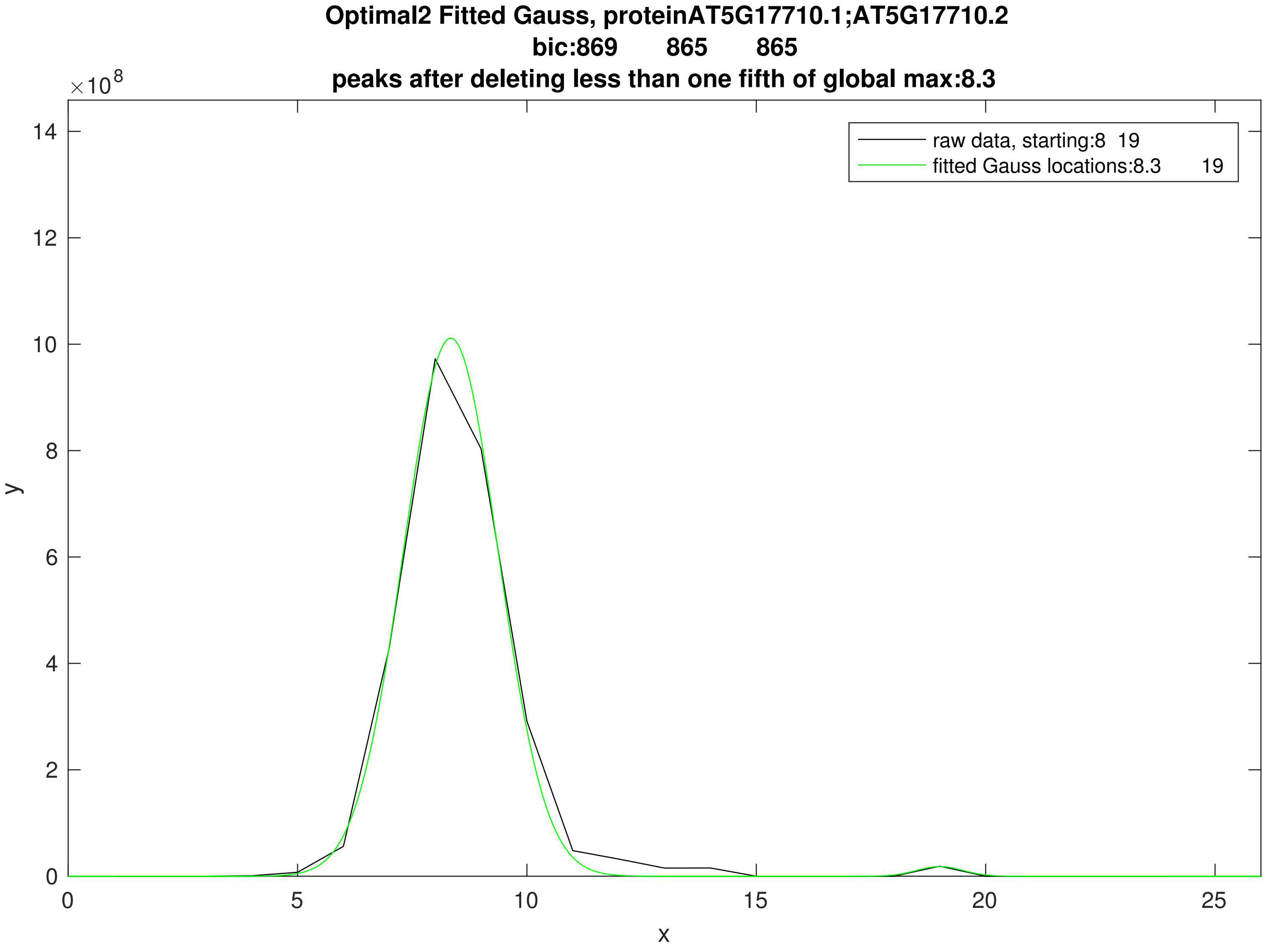

Optimal2 Fitted Gauss, proteinAT5G17710.1;AT5G17710.2
bic:869 865 865
peaks after deleting less than one fifth of global max:8.3
×10^8
raw data, starting:8 19
fitted Gauss locations:8.3 19
y
x
0
2
4
6
8
10
12
14
0
5
10
15
20
25

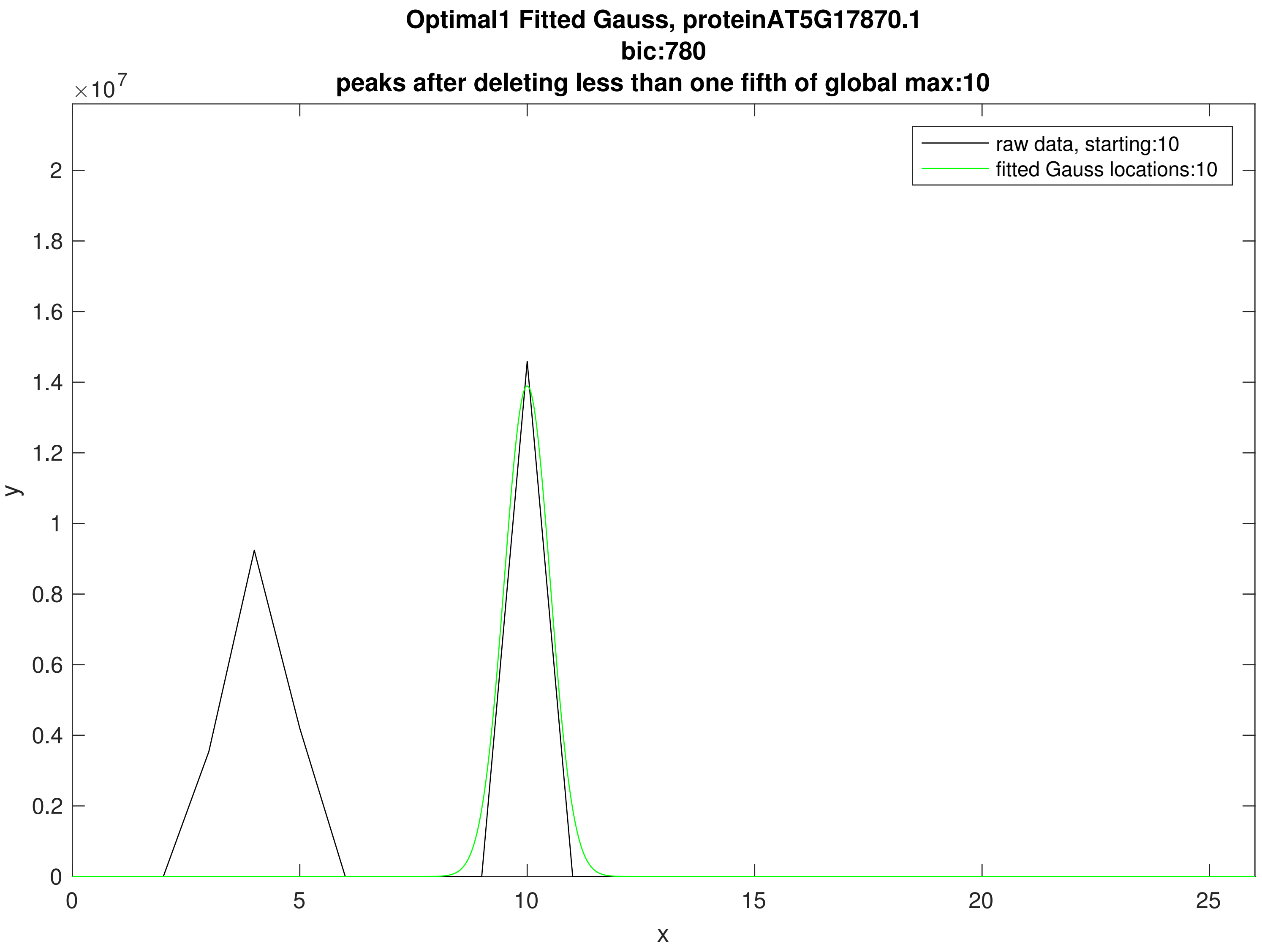

Optimal1 Fitted Gauss, proteinAT5G17870.1
bic:780
peaks after deleting less than one fifth of global max:10
×10^7
raw data, starting:10
fitted Gauss locations:10
2
1.8
1.6
1.4
1.2
1
0.8
0.6
0.4
0.2
0
y
0
5
10
15
20
25
x

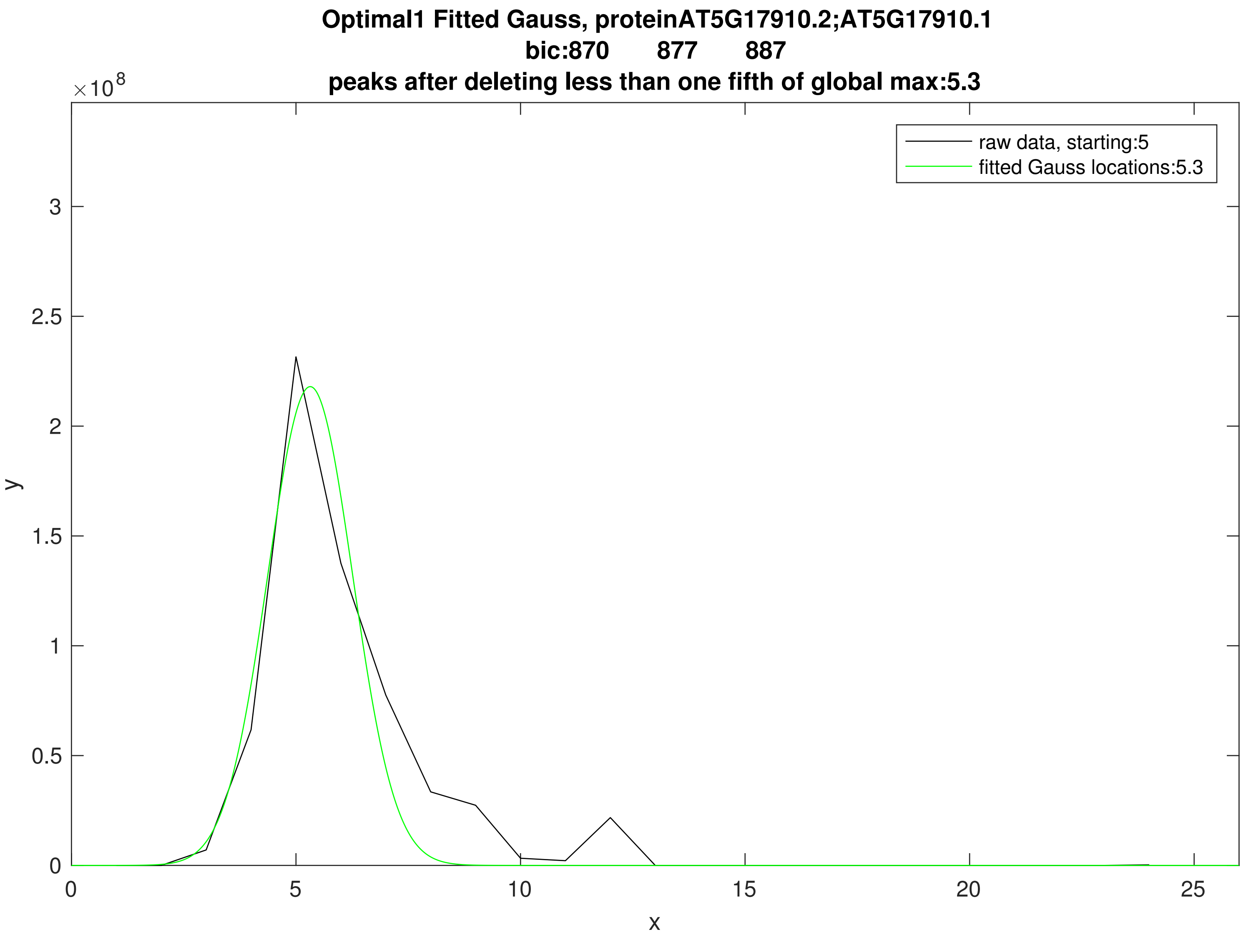

Optimal1 Fitted Gauss, proteinAT5G17910.2;AT5G17910.1
bic:870 877 887
peaks after deleting less than one fifth of global max:5.3
×10^8
raw data, starting:5
fitted Gauss locations:5.3
y
x
0
0.5
1
1.5
2
2.5
3
5
10
15
20
25

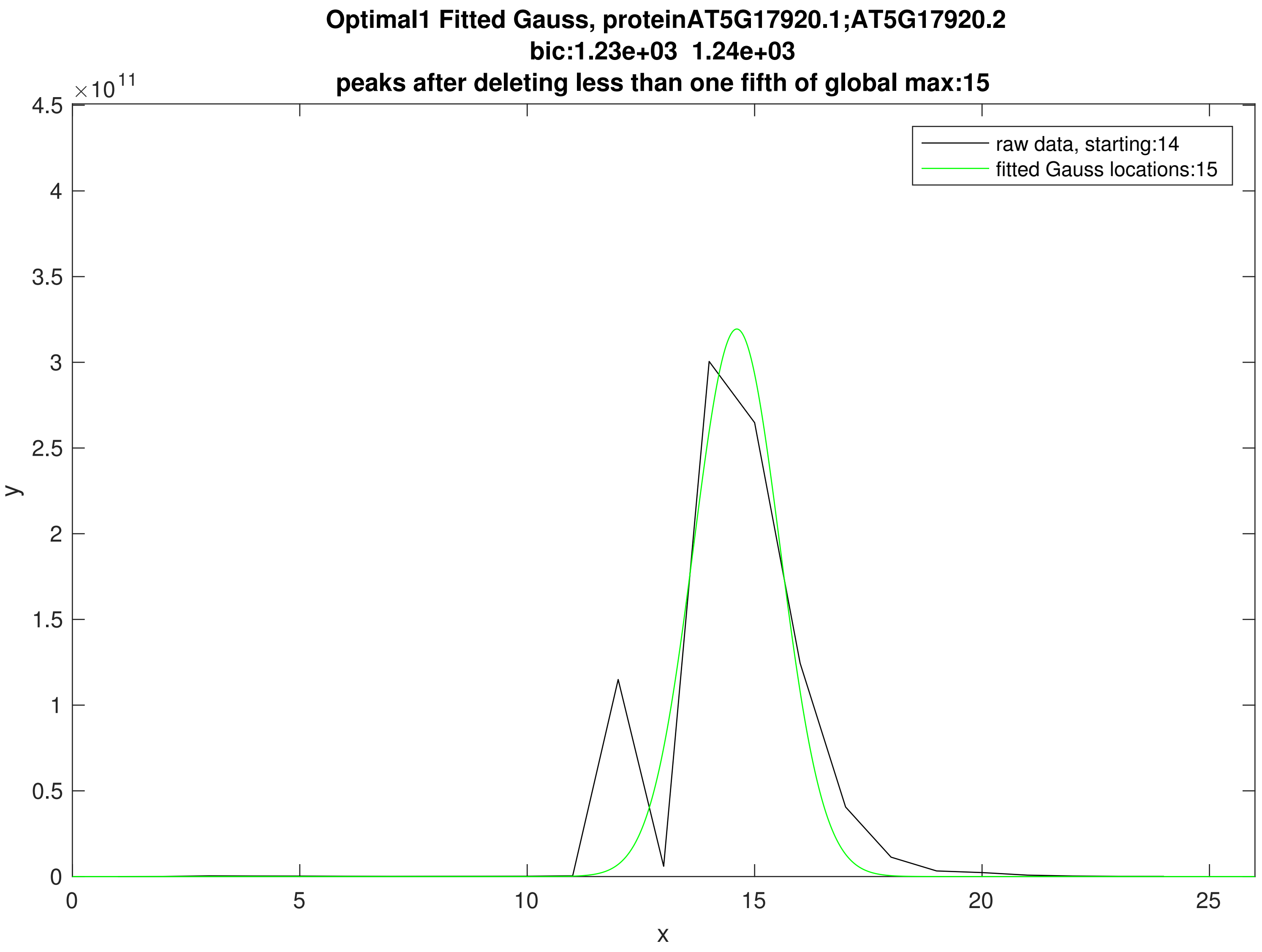

Optimal1 Fitted Gauss, proteinAT5G17920.1;AT5G17920.2
bic:1.23e+03  1.24e+03
peaks after deleting less than one fifth of global max:15
×10^11
4.5
4
3.5
3
2.5
2
1.5
1
0.5
0
y
0
5
10
15
20
25
x
raw data, starting:14
fitted Gauss locations:15

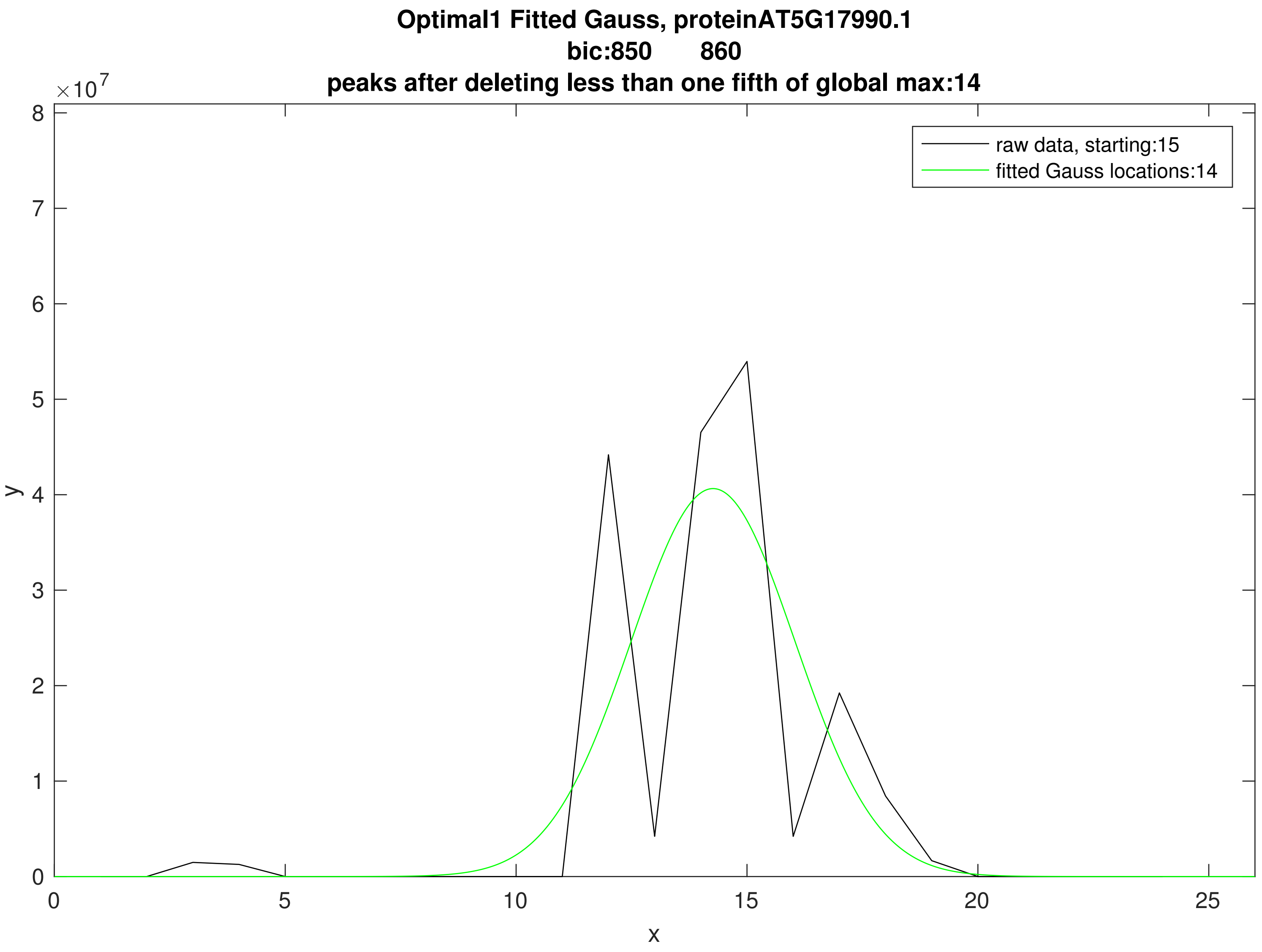
Optimal1 Fitted Gauss, proteinAT5G17990.1
bic:850 860
peaks after deleting less than one fifth of global max:14
×10^7
y
x
raw data, starting:15
fitted Gauss locations:14
0
1
2
3
4
5
6
7
8
0
5
10
15
20
25

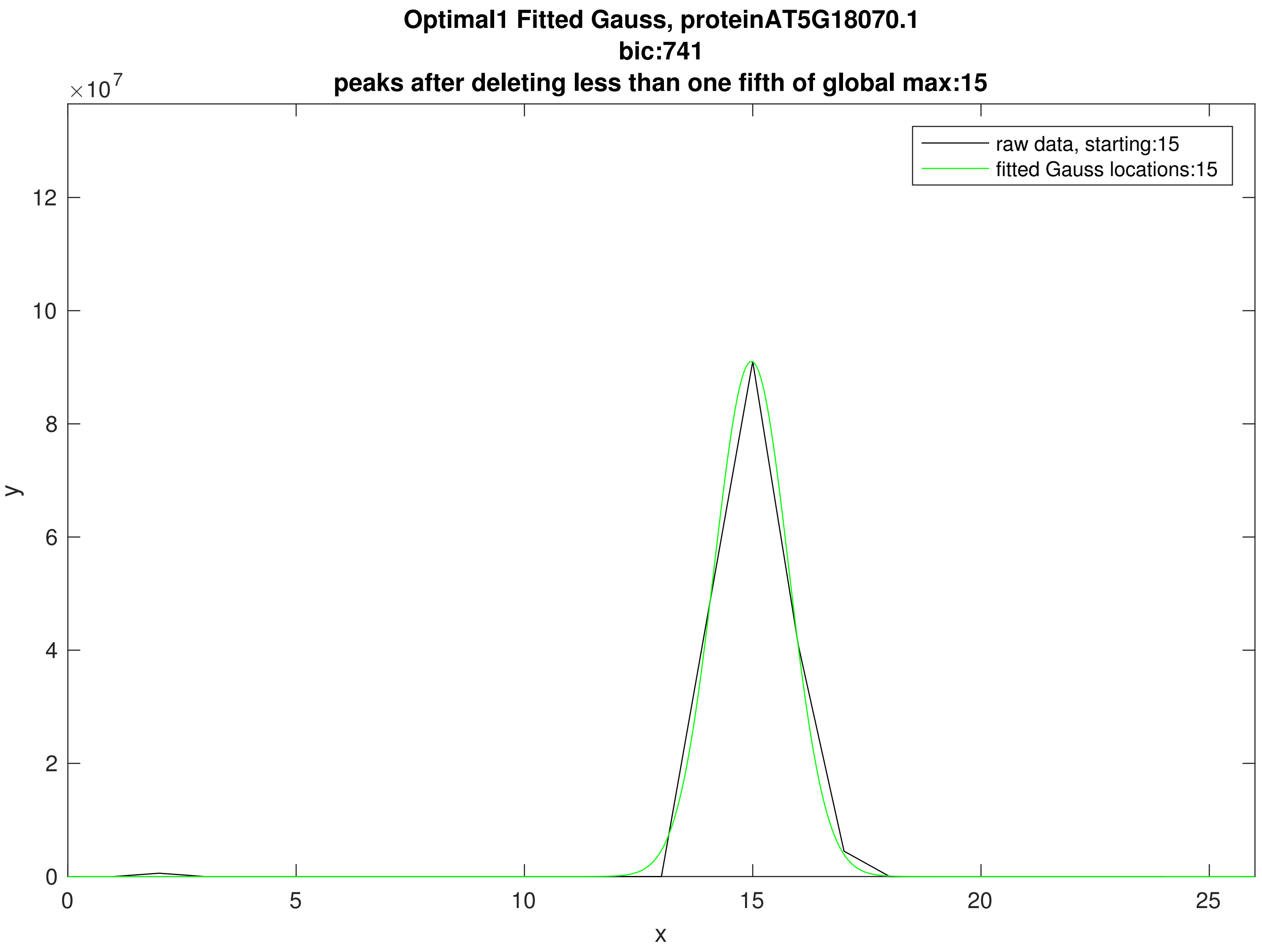
Optimal1 Fitted Gauss, proteinAT5G18070.1
bic:741
peaks after deleting less than one fifth of global max:15
×10^7
raw data, starting:15
fitted Gauss locations:15
y
x
0
2
4
6
8
10
12
0
5
10
15
20
25

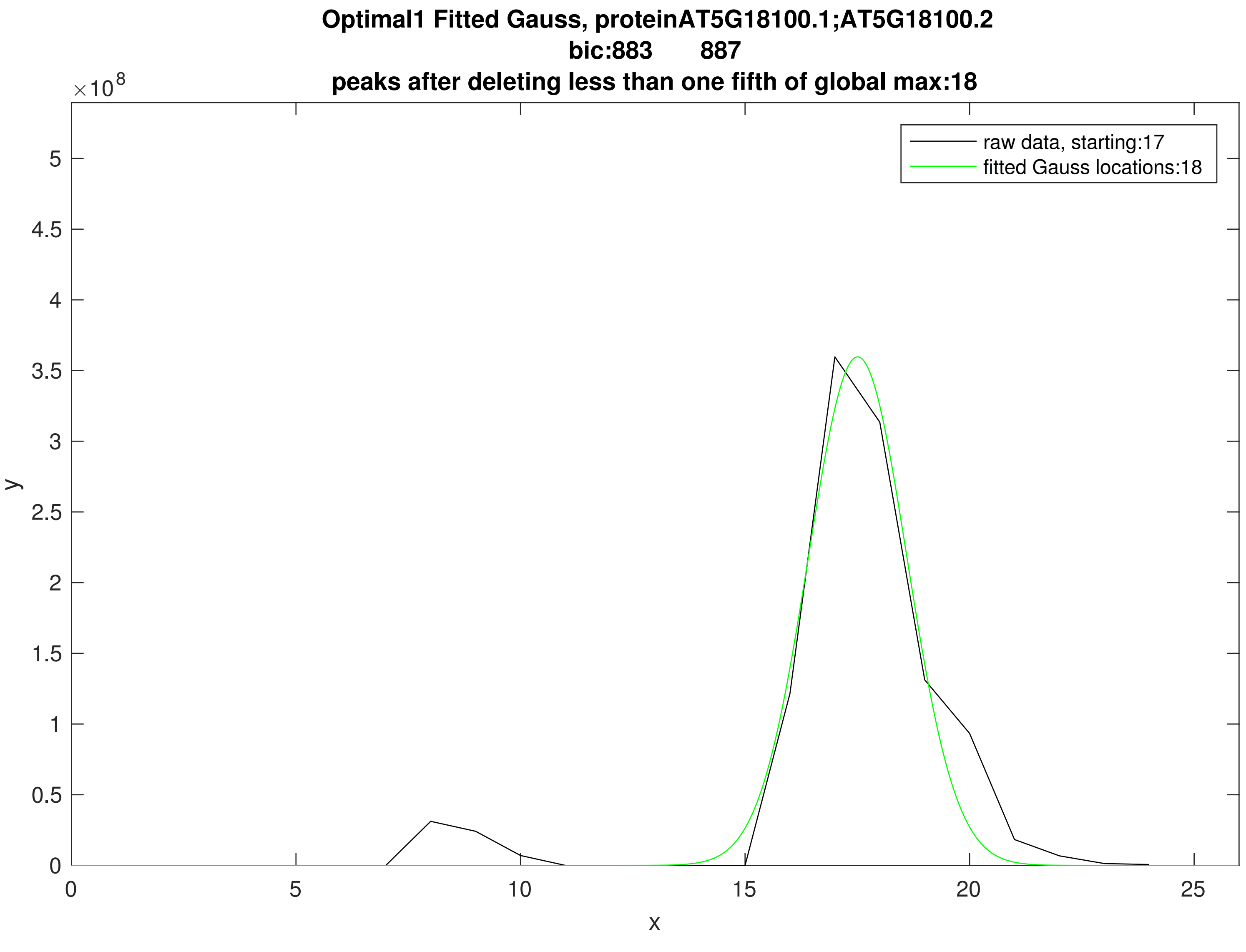

Optimal1 Fitted Gauss, proteinAT5G18100.1;AT5G18100.2
bic:883 887
peaks after deleting less than one fifth of global max:18
×10^8
raw data, starting:17
fitted Gauss locations:18
y
x
0
0.5
1
1.5
2
2.5
3
3.5
4
4.5
5
0
5
10
15
20
25

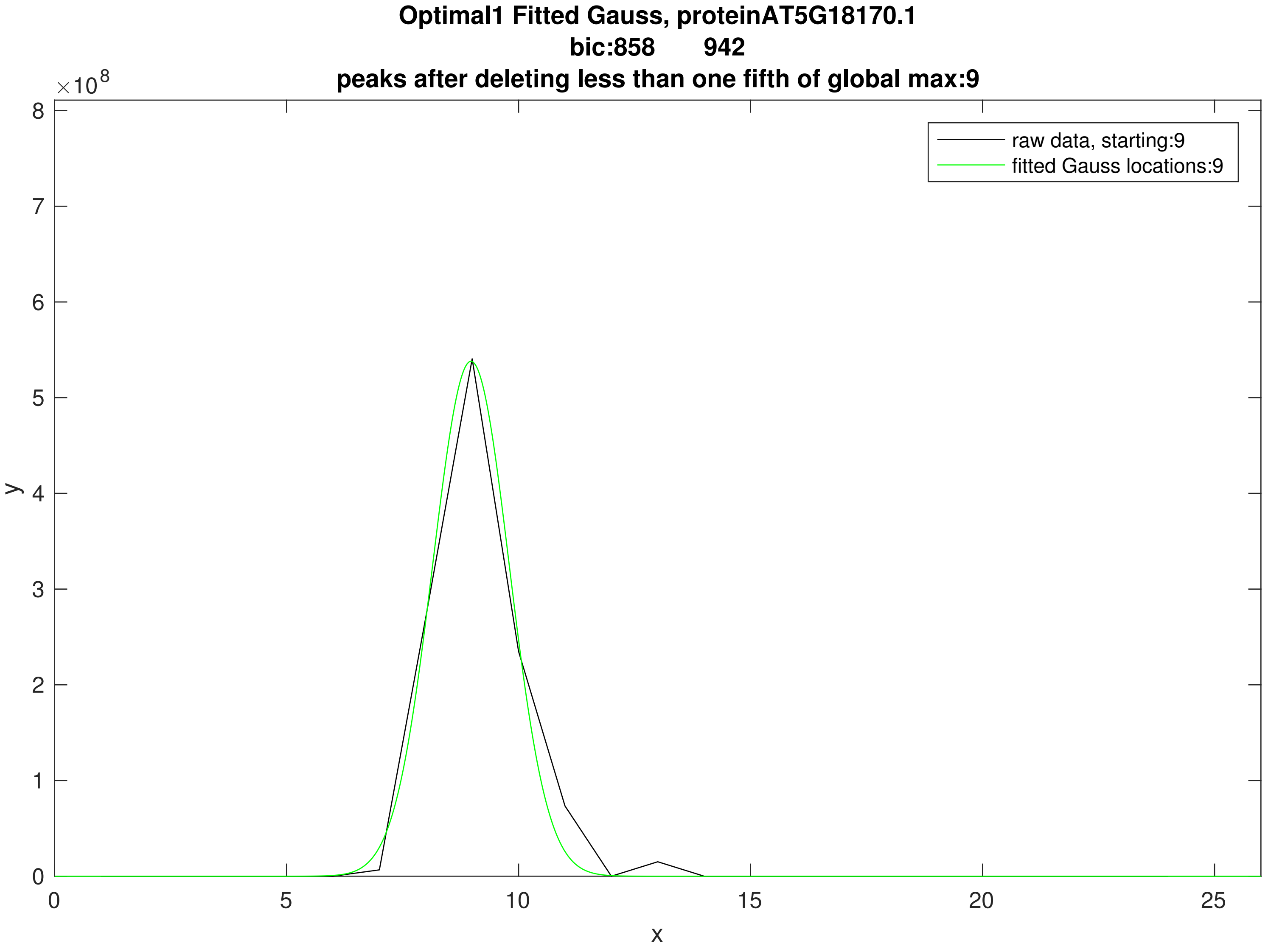

Optimal1 Fitted Gauss, proteinAT5G18170.1
bic:858 942
peaks after deleting less than one fifth of global max:9
×10^8
8
7
6
5
4
3
2
1
0
y
0
5
10
15
20
25
x
raw data, starting:9
fitted Gauss locations:9

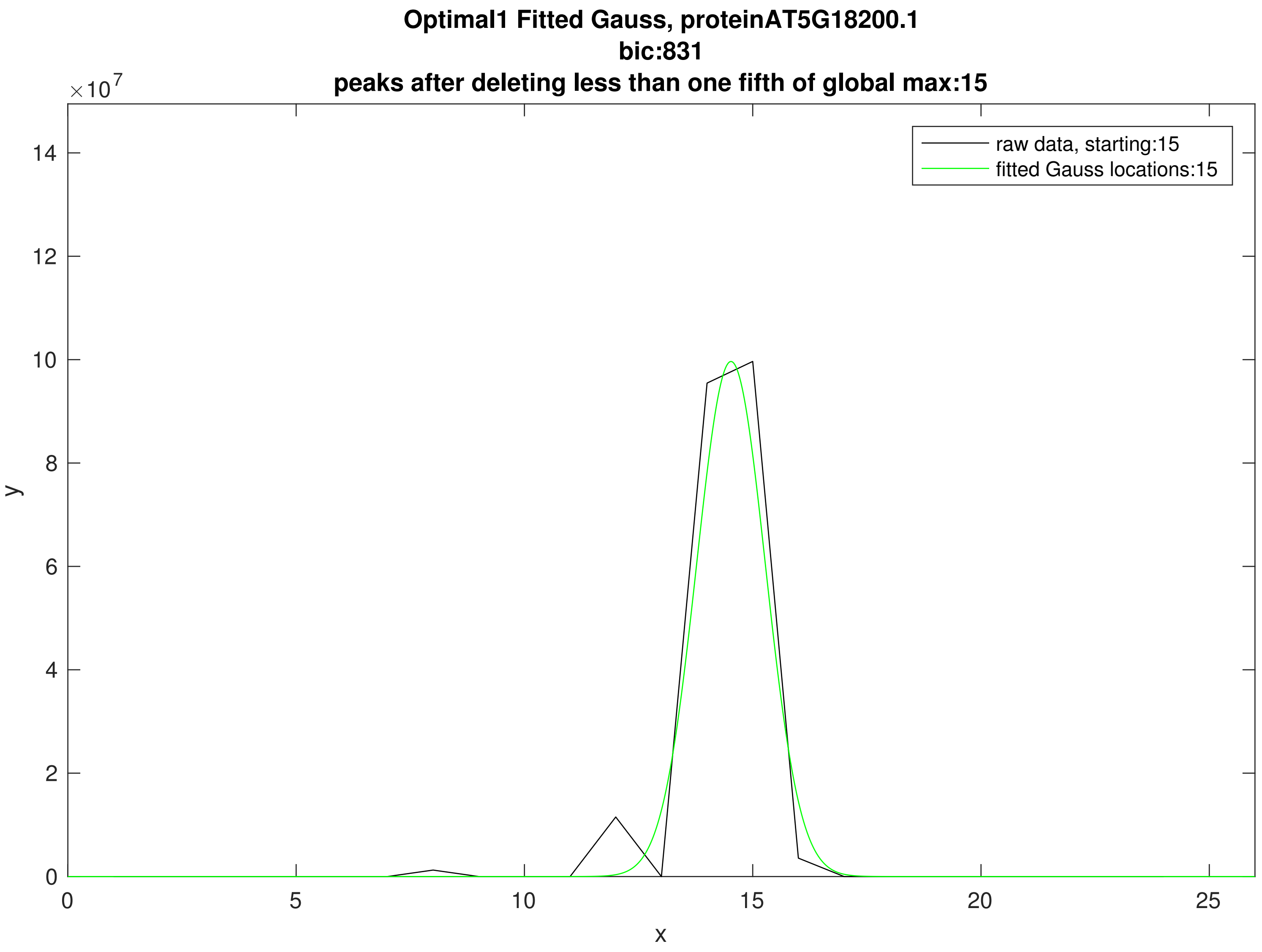
Optimal1 Fitted Gauss, proteinAT5G18200.1
bic:831
peaks after deleting less than one fifth of global max:15
×10^7
raw data, starting:15
fitted Gauss locations:15
y
x
0
2
4
6
8
10
12
14
0
5
10
15
20
25

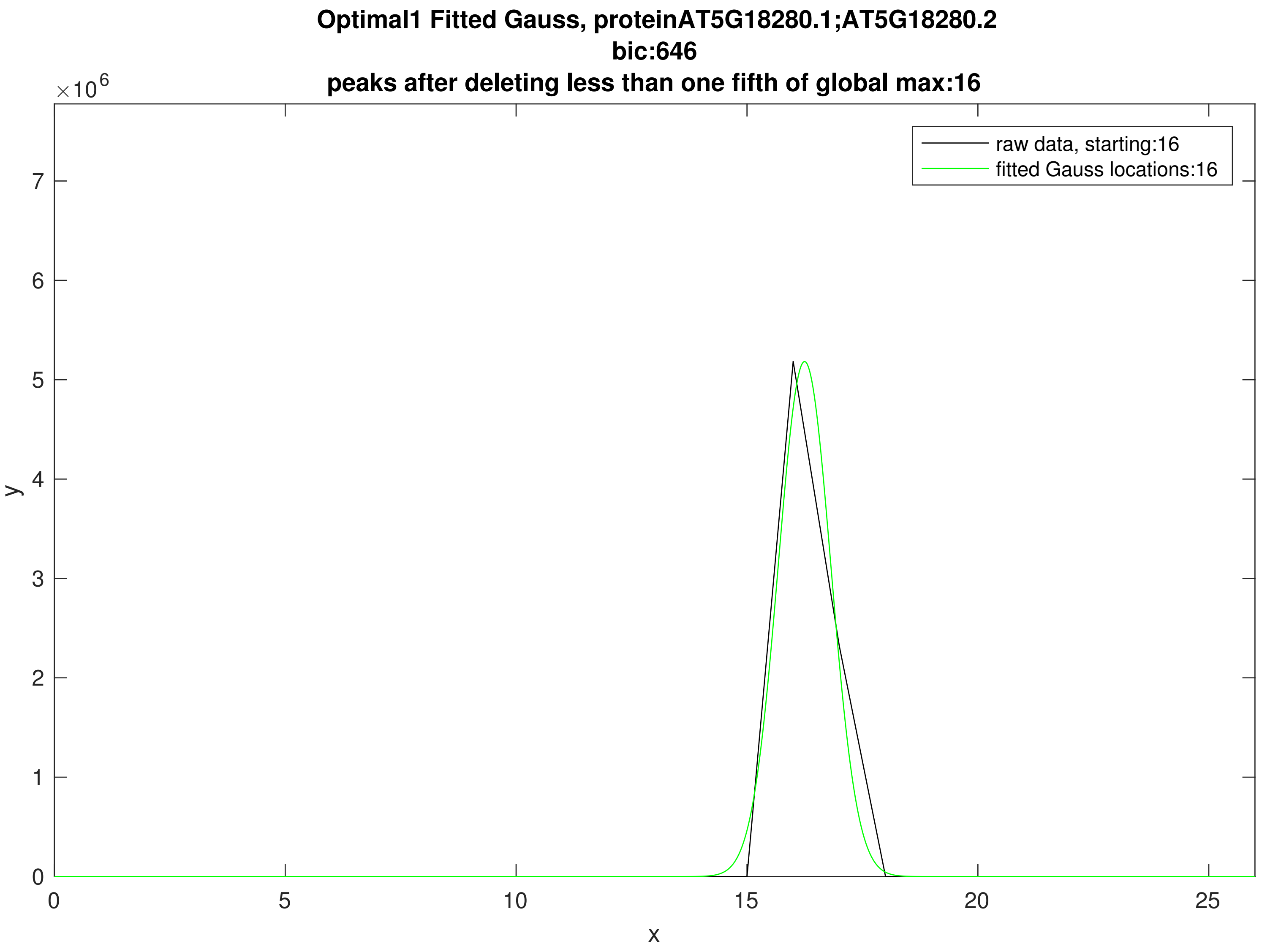
Optimal1 Fitted Gauss, proteinAT5G18280.1;AT5G18280.2
bic:646
peaks after deleting less than one fifth of global max:16
×10^6
raw data, starting:16
fitted Gauss locations:16
y
x
0
1
2
3
4
5
6
7
0
5
10
15
20
25

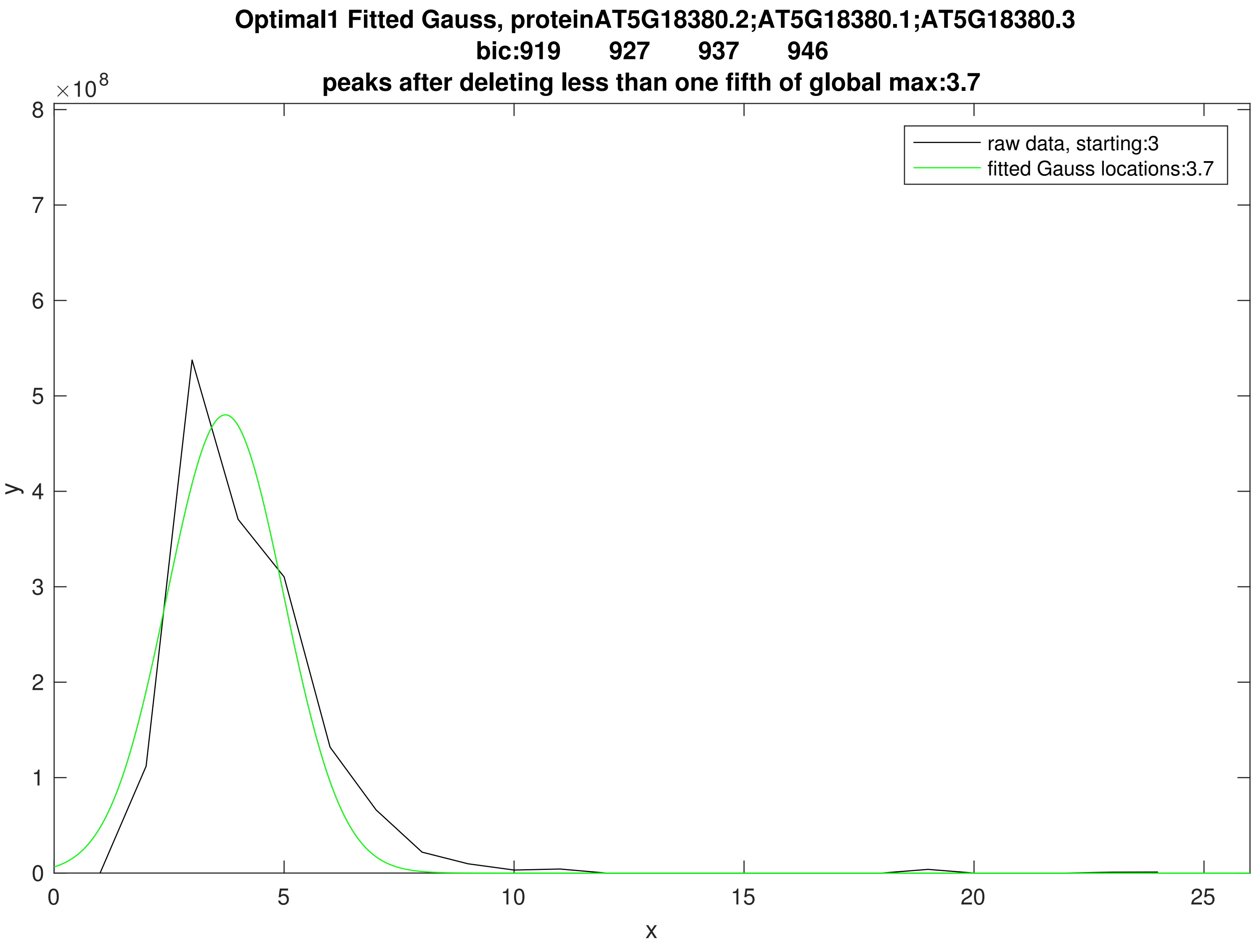

Optimal1 Fitted Gauss, proteinAT5G18380.2;AT5G18380.1;AT5G18380.3
bic:919 927 937 946
peaks after deleting less than one fifth of global max:3.7
×10^8
raw data, starting:3
fitted Gauss locations:3.7
y
x

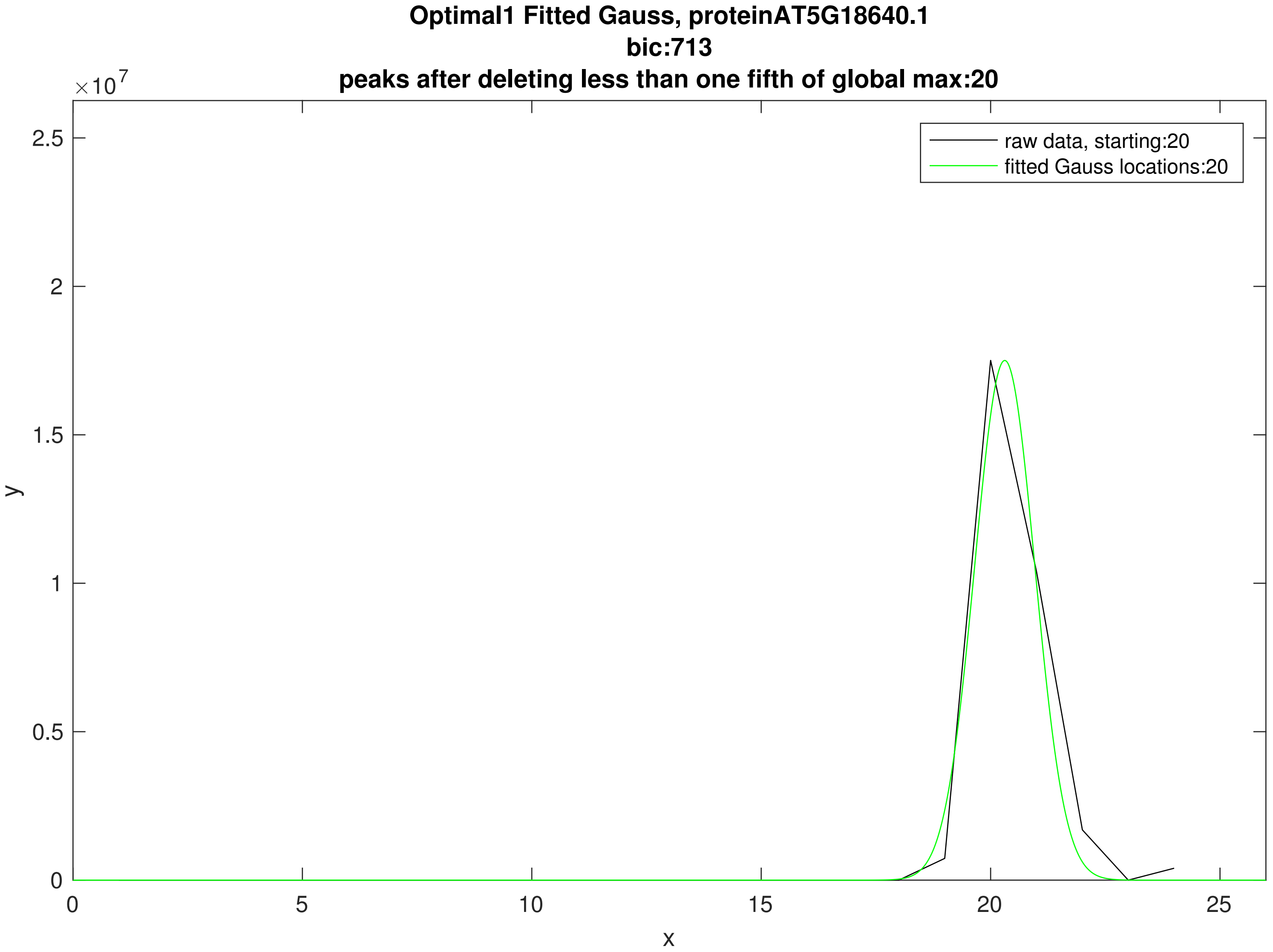

Optimal1 Fitted Gauss, proteinAT5G18640.1
bic:713
peaks after deleting less than one fifth of global max:20
×10^7
2.5
2
1.5
1
0.5
0
y
0
5
10
15
20
25
x
raw data, starting:20
fitted Gauss locations:20

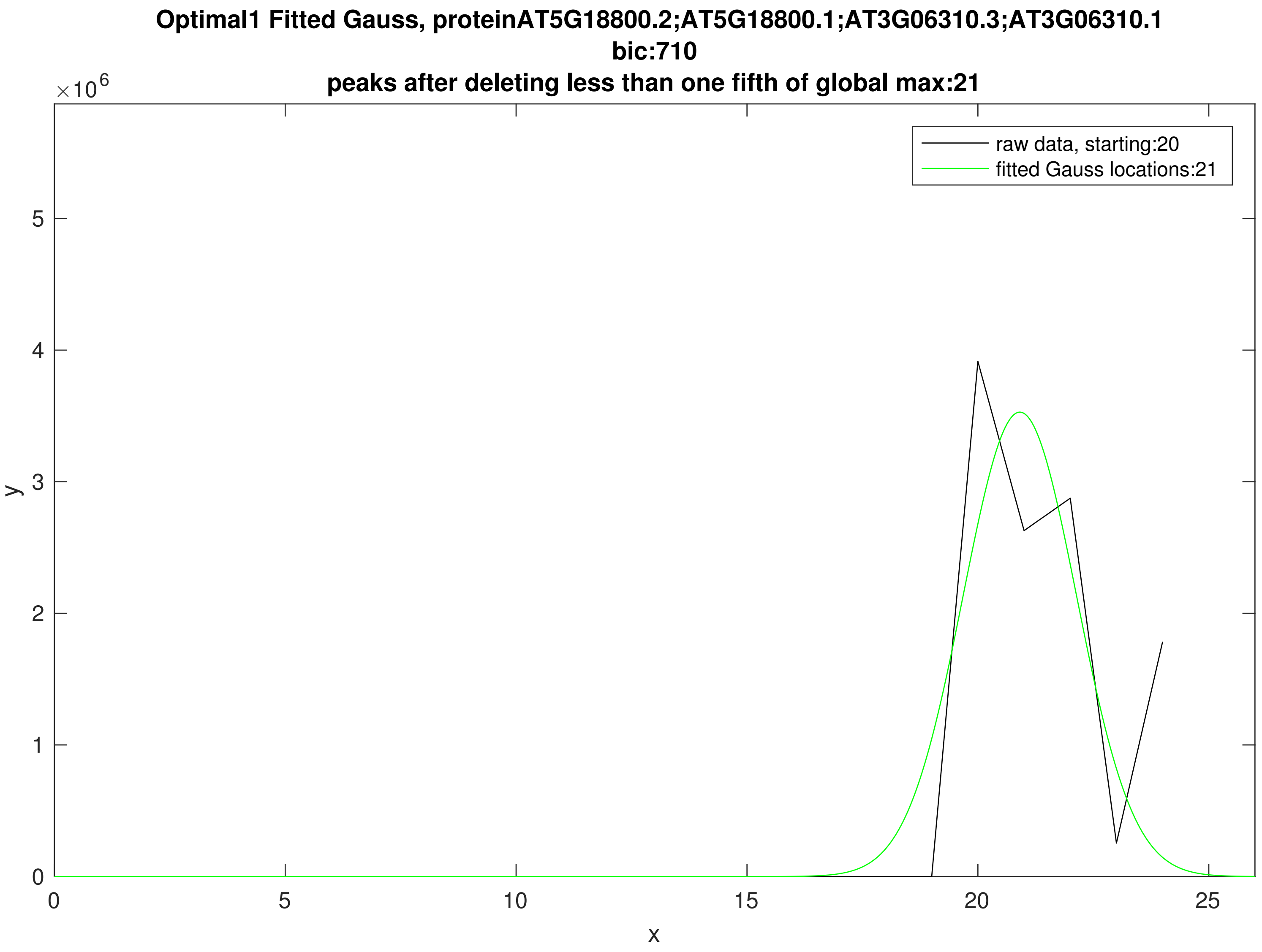
Optimal1 Fitted Gauss, proteinAT5G18800.2;AT5G18800.1;AT3G06310.3;AT3G06310.1
bic:710
peaks after deleting less than one fifth of global max:21
×10^6
raw data, starting:20
fitted Gauss locations:21
y
x
0
1
2
3
4
5
0
5
10
15
20
25

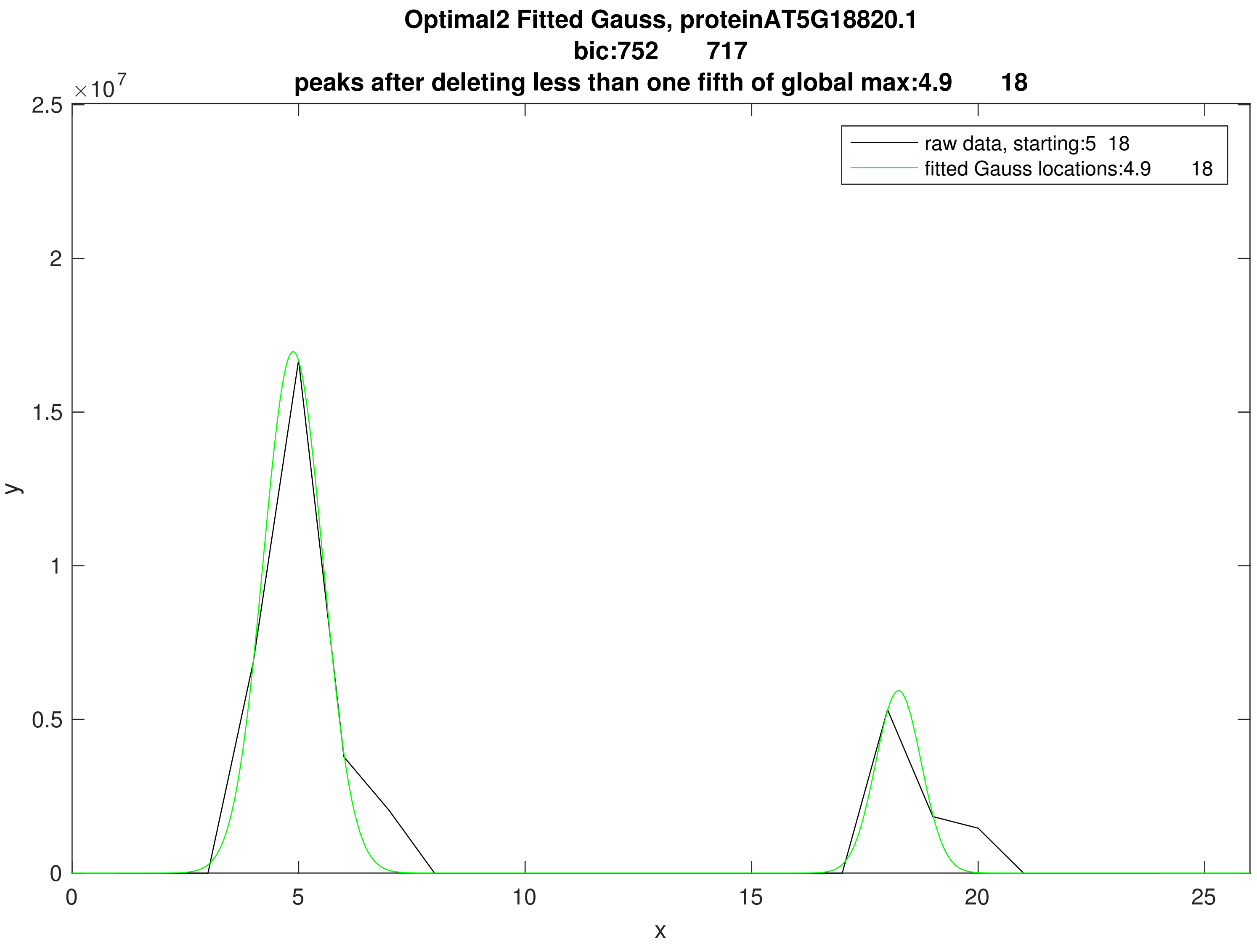

Optimal2 Fitted Gauss, proteinAT5G18820.1
bic:752 717
peaks after deleting less than one fifth of global max:4.9 18
×10^7
2.5
2
1.5
1
0.5
0
y
0
5
10
15
20
25
x
raw data, starting:5 18
fitted Gauss locations:4.9 18

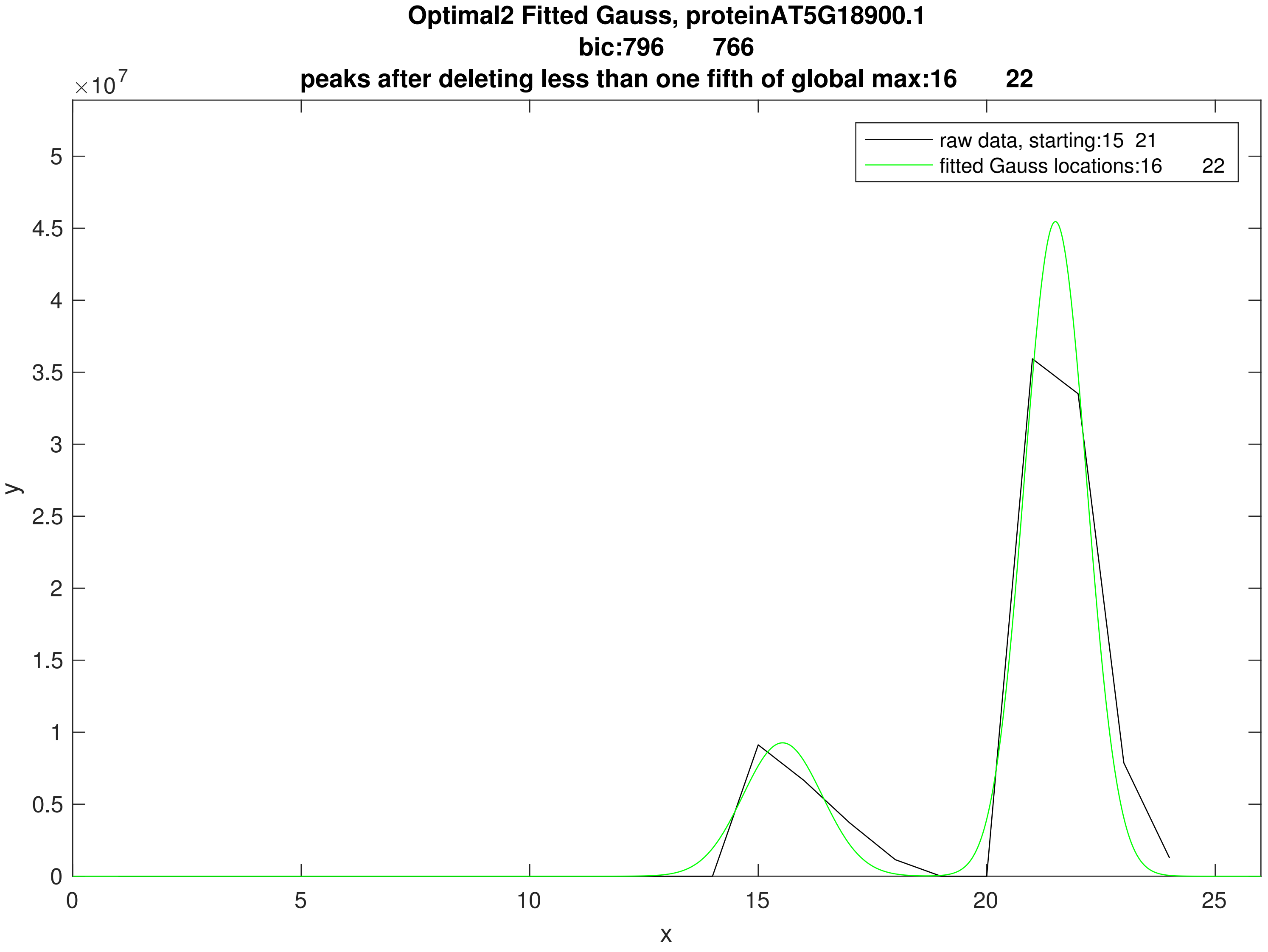
Optimal2 Fitted Gauss, proteinAT5G18900.1
bic:796 766
peaks after deleting less than one fifth of global max:16 22
×10^7
raw data, starting:15 21
fitted Gauss locations:16 22
5
4.5
4
3.5
3
2.5
2
1.5
1
0.5
0
y
0
5
10
15
20
25
x

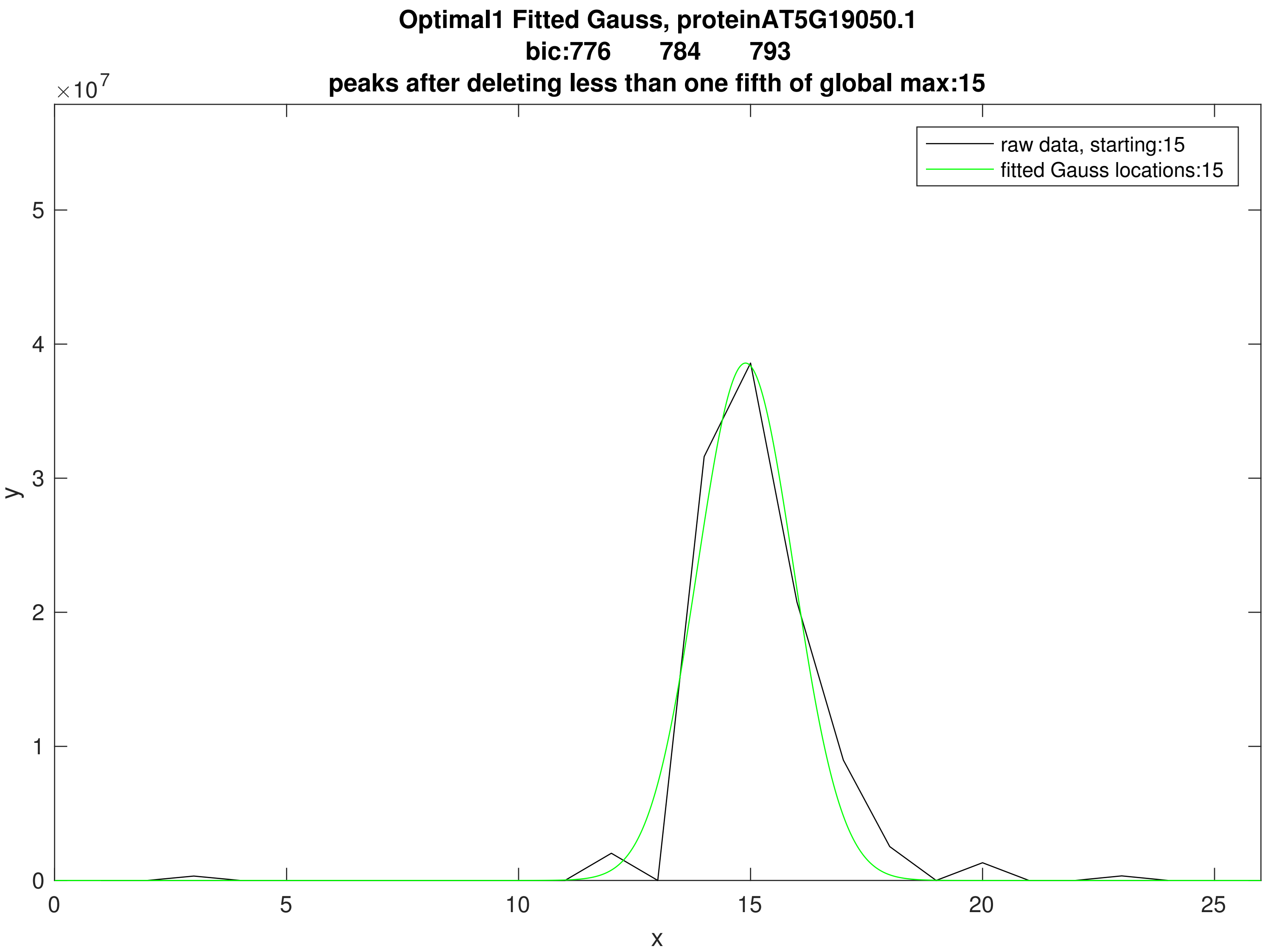
Optimal1 Fitted Gauss, proteinAT5G19050.1
bic:776 784 793
peaks after deleting less than one fifth of global max:15
×10^7
raw data, starting:15
fitted Gauss locations:15
y
x
0
1
2
3
4
5
0
5
10
15
20
25

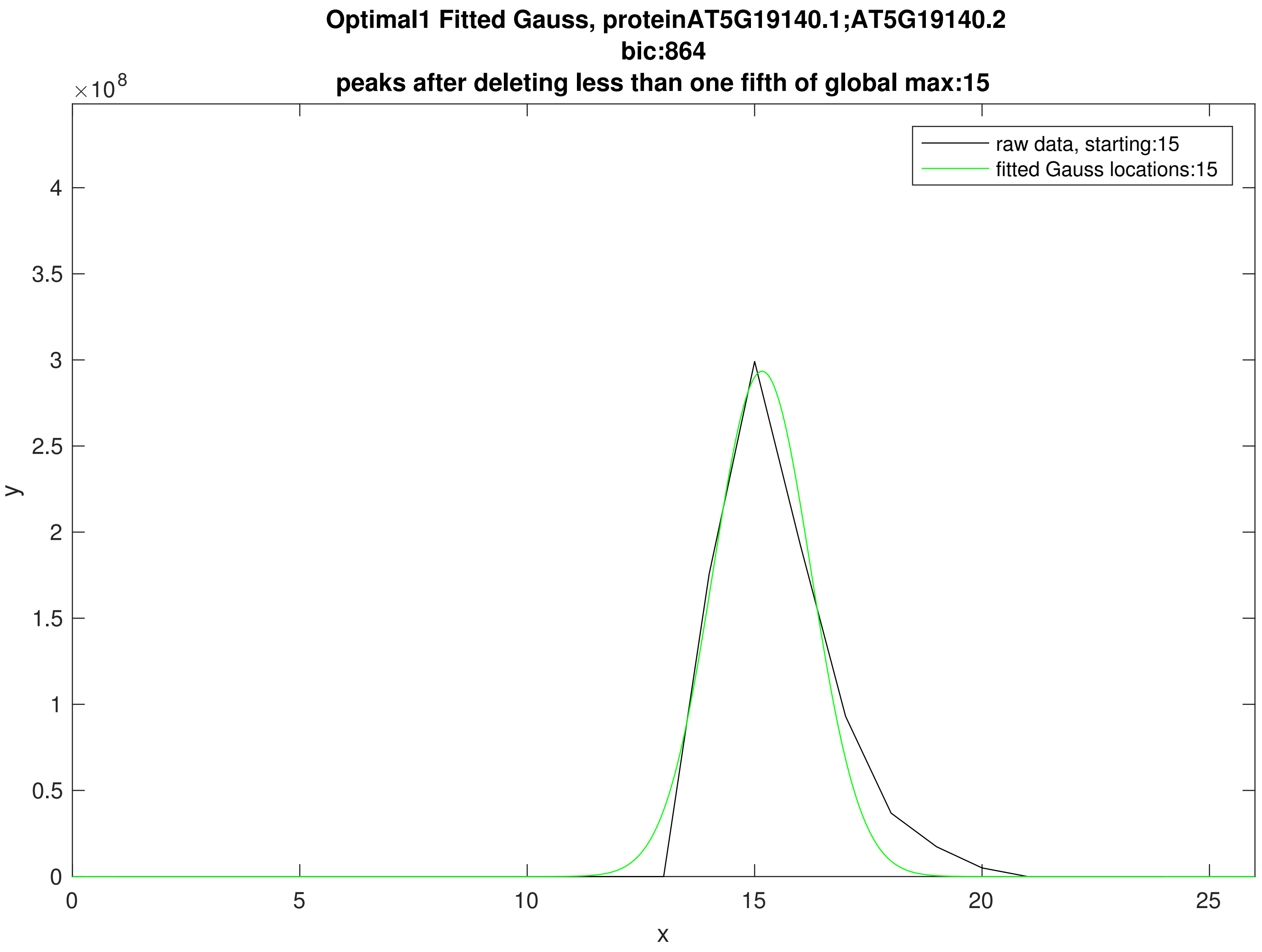
Optimal1 Fitted Gauss, proteinAT5G19140.1;AT5G19140.2
bic:864
peaks after deleting less than one fifth of global max:15
×10^8
raw data, starting:15
fitted Gauss locations:15
y
x
0
0.5
1
1.5
2
2.5
3
3.5
4
0
5
10
15
20
25

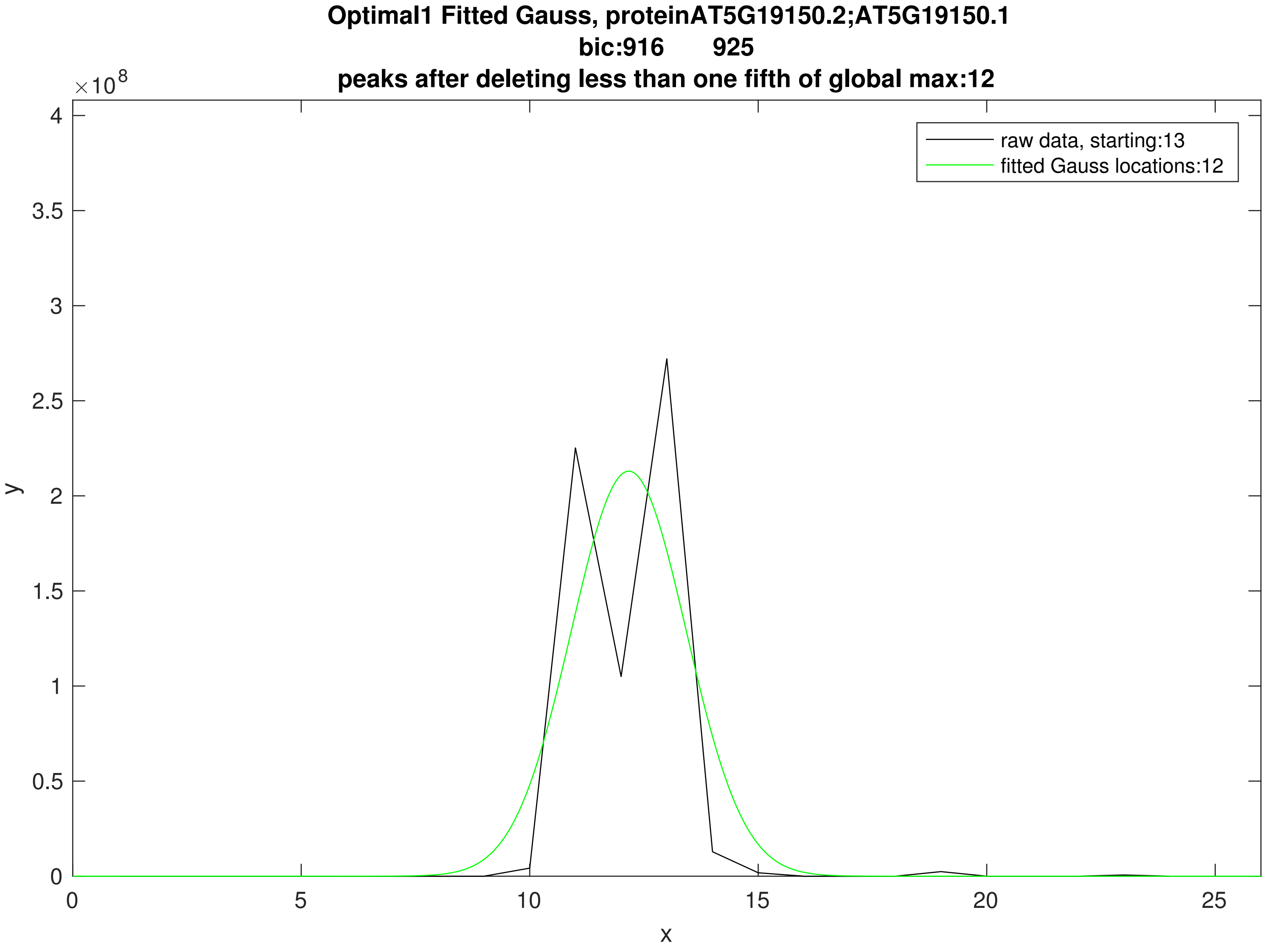
Optimal1 Fitted Gauss, proteinAT5G19150.2;AT5G19150.1
bic:916 925
peaks after deleting less than one fifth of global max:12
×10^8
raw data, starting:13
fitted Gauss locations:12
y
x
0
0.5
1
1.5
2
2.5
3
3.5
4
5
10
15
20
25

#### Optimal1 Fitted Gauss, proteinAT5G19180.1
#### bic:972 988 997
#### peaks after deleting less than one fifth of global max:12

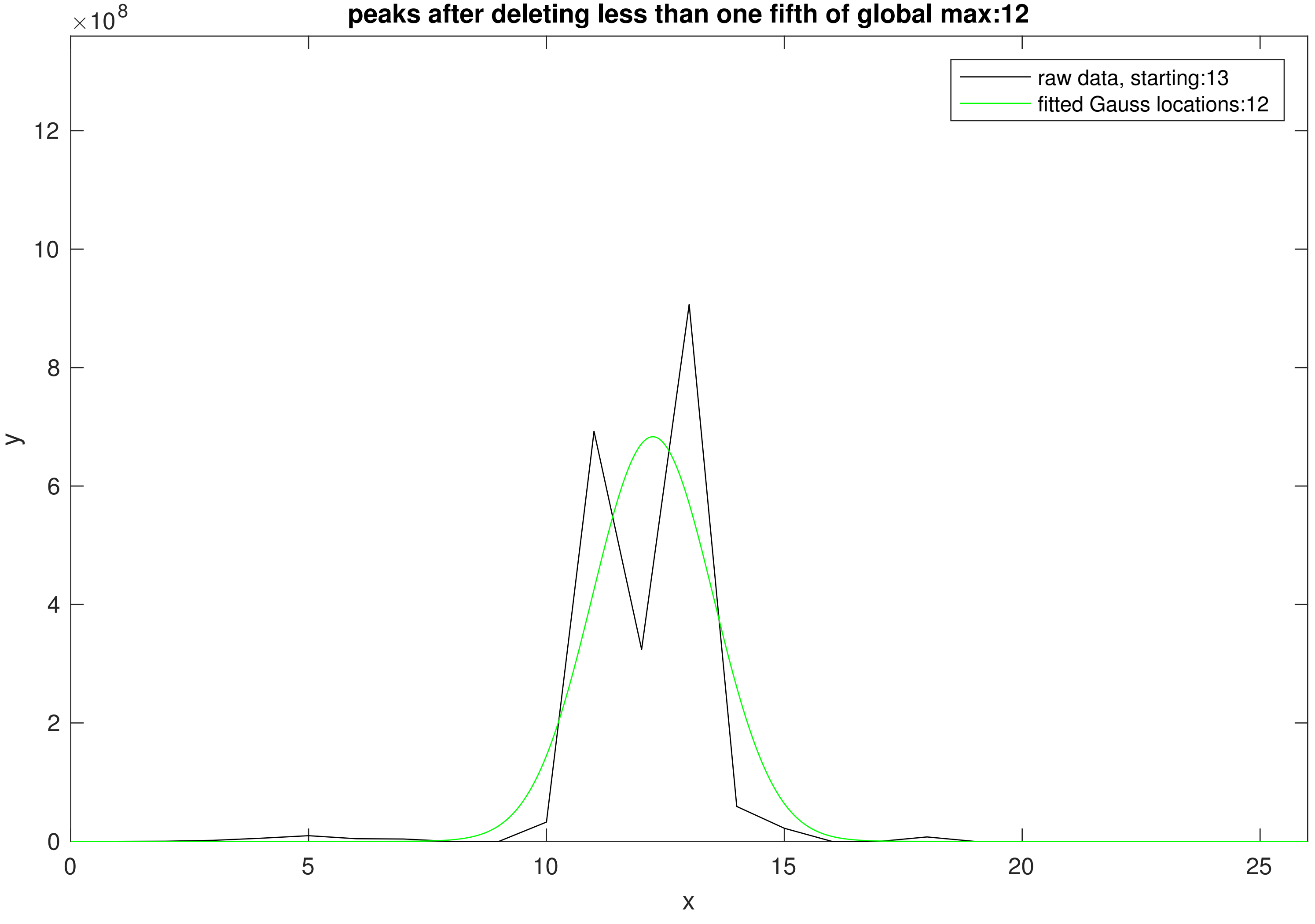

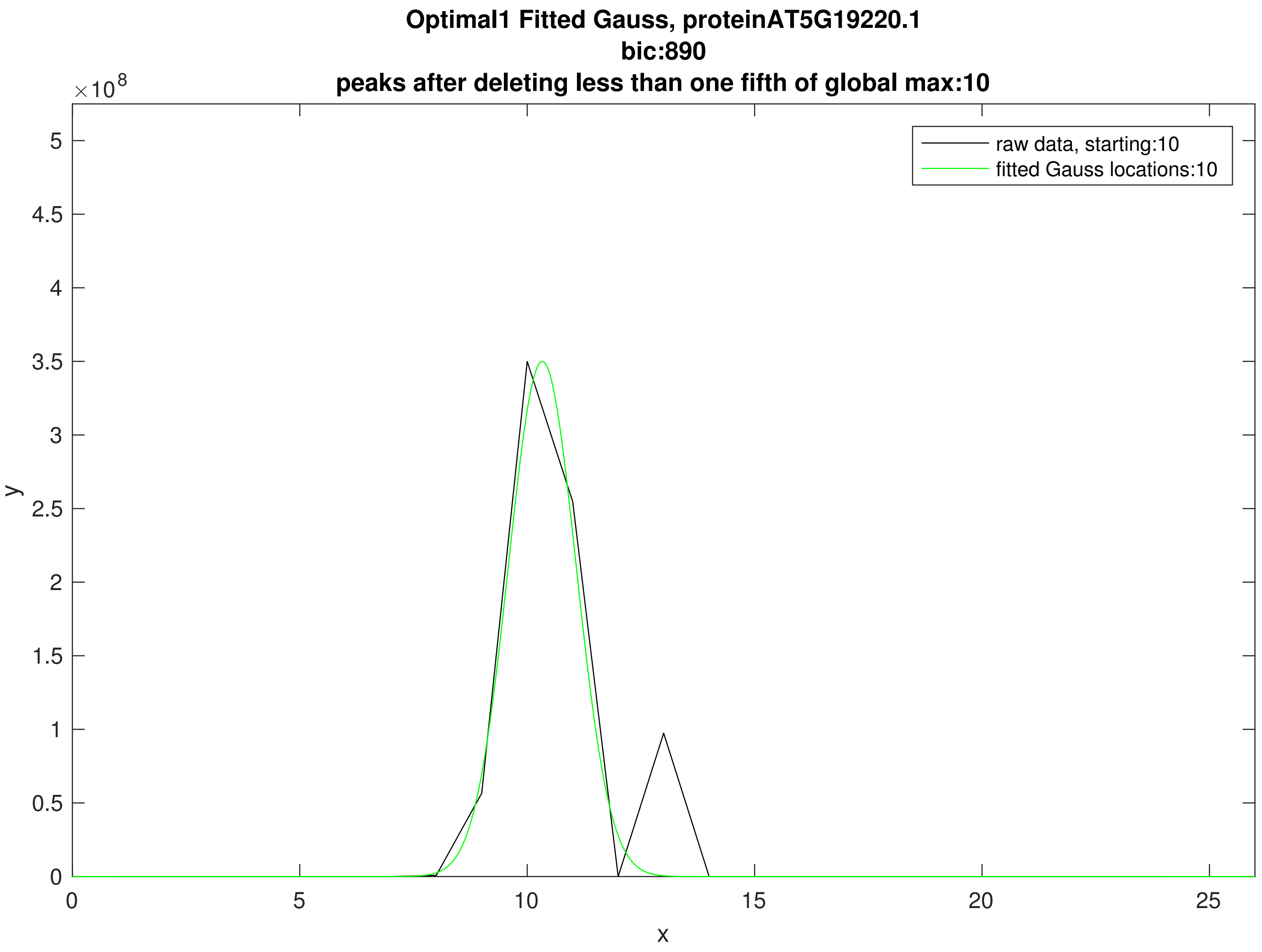
Optimal1 Fitted Gauss, proteinAT5G19220.1
bic:890
peaks after deleting less than one fifth of global max:10
×10^8
raw data, starting:10
fitted Gauss locations:10
5
4.5
4
3.5
3
2.5
2
1.5
1
0.5
0
y
0
5
10
15
20
25
x

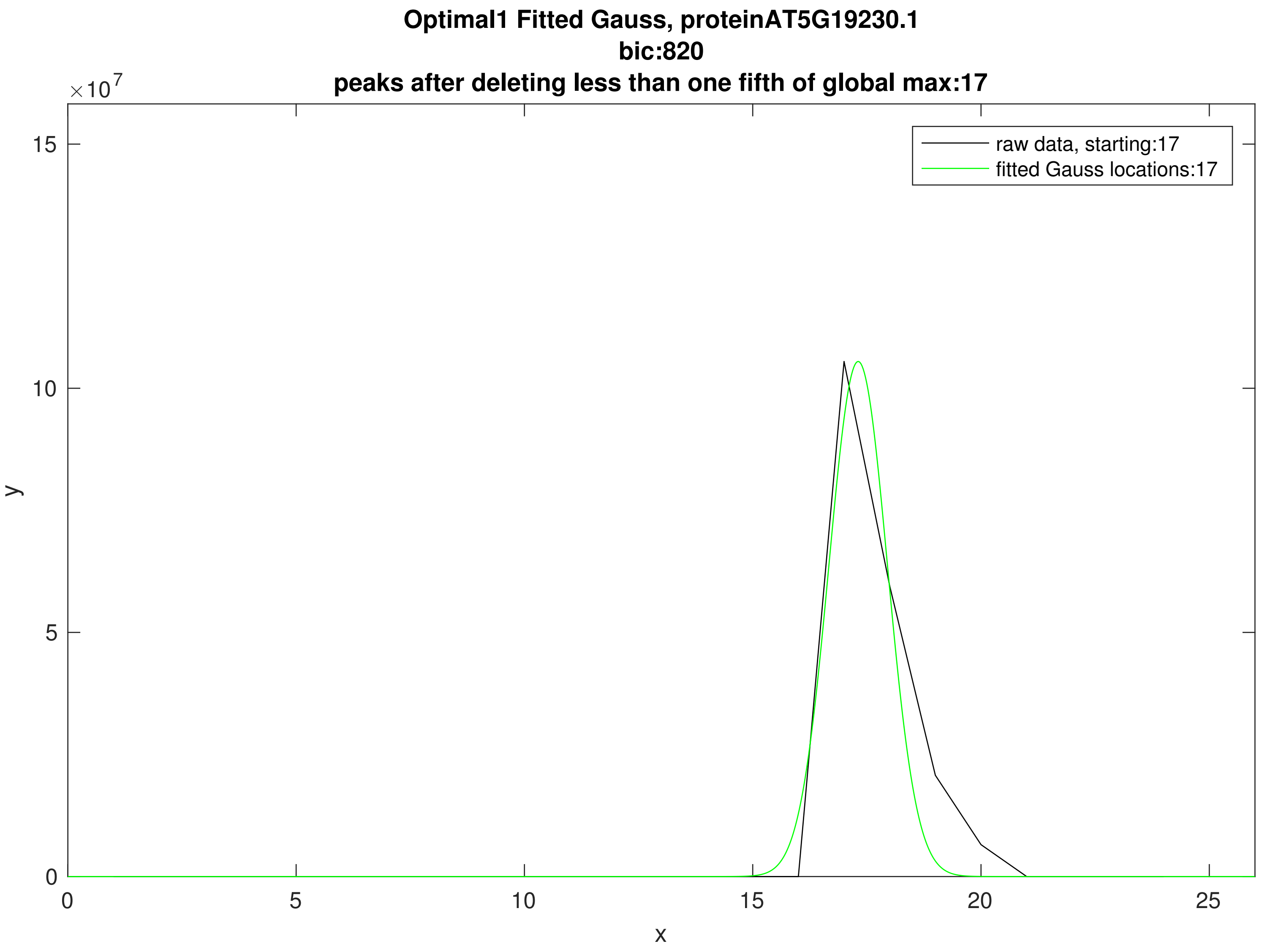
Optimal1 Fitted Gauss, proteinAT5G19230.1
bic:820
peaks after deleting less than one fifth of global max:17
×10^7
15
10
5
0
y
0
5
10
15
20
25
x
raw data, starting:17
fitted Gauss locations:17

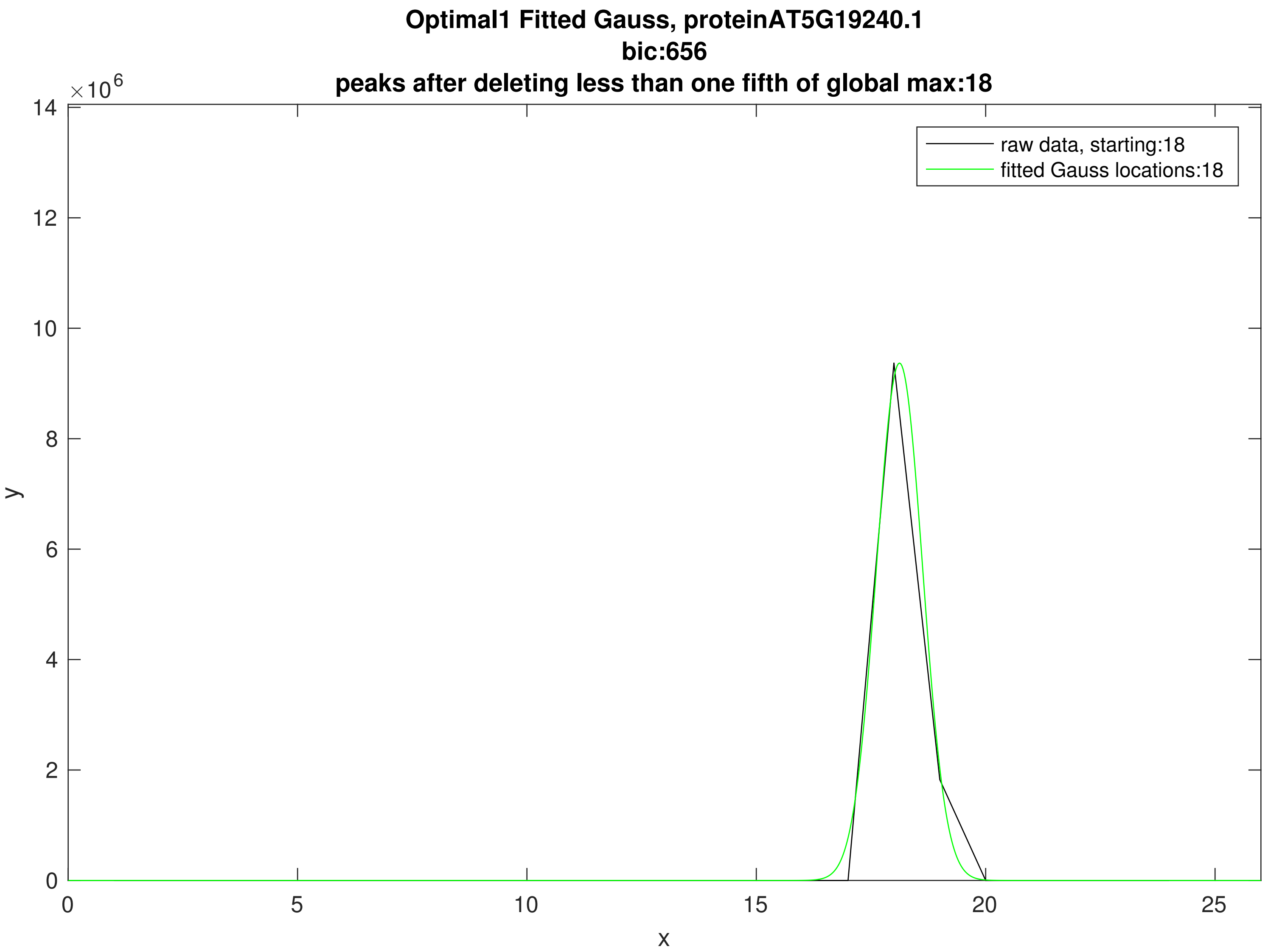
Optimal1 Fitted Gauss, proteinAT5G19240.1
bic:656
peaks after deleting less than one fifth of global max:18
×10^6
raw data, starting:18
fitted Gauss locations:18
y
x
0
2
4
6
8
10
12
14
0
5
10
15
20
25

#### Optimal1 Fitted Gauss, proteinAT5G19320.1
#### bic:733
#### peaks after deleting less than one fifth of global max:13

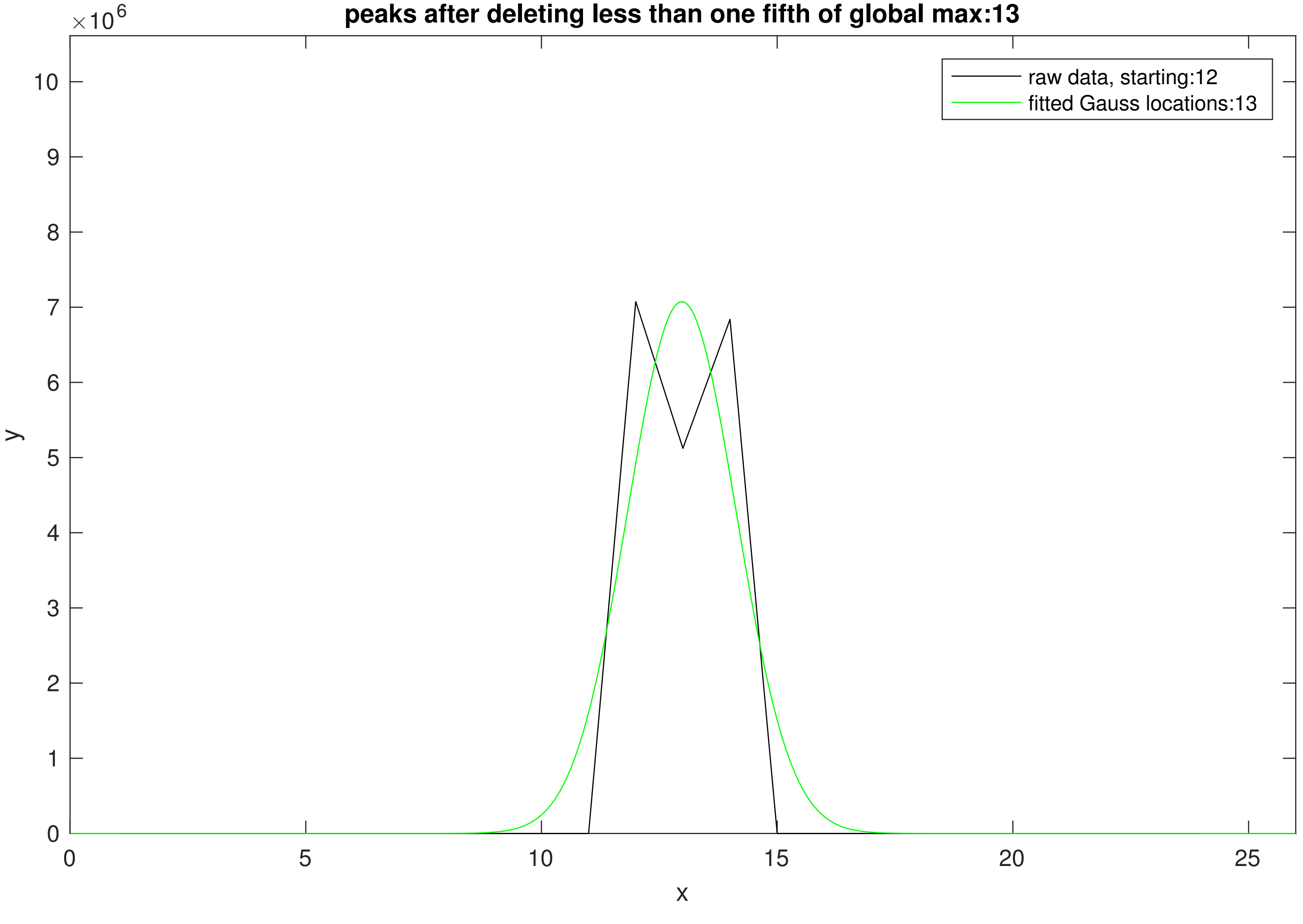

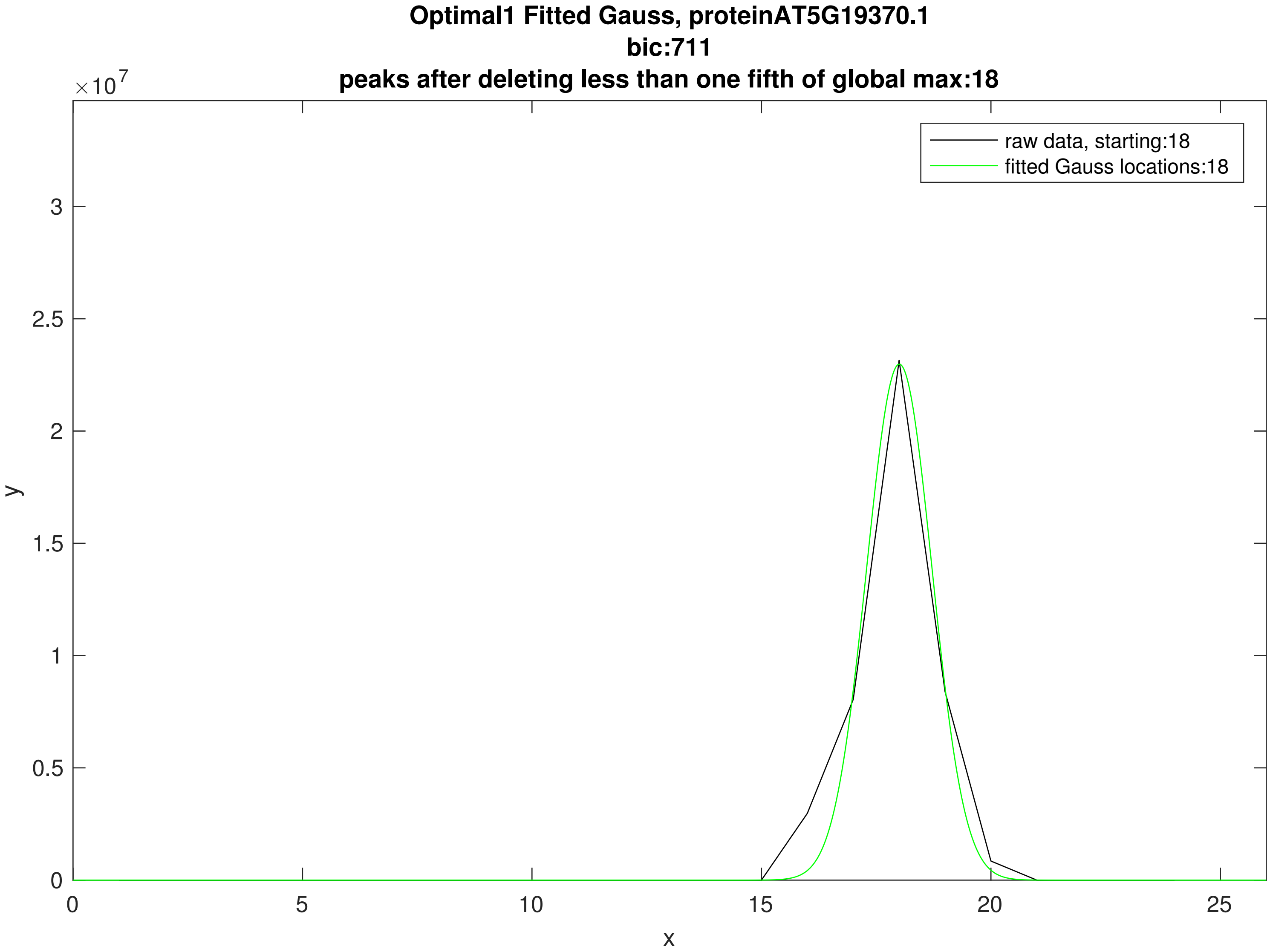
Optimal1 Fitted Gauss, proteinAT5G19370.1
bic:711
peaks after deleting less than one fifth of global max:18
×10^7
raw data, starting:18
fitted Gauss locations:18
y
x
0
0.5
1
1.5
2
2.5
3
0
5
10
15
20
25

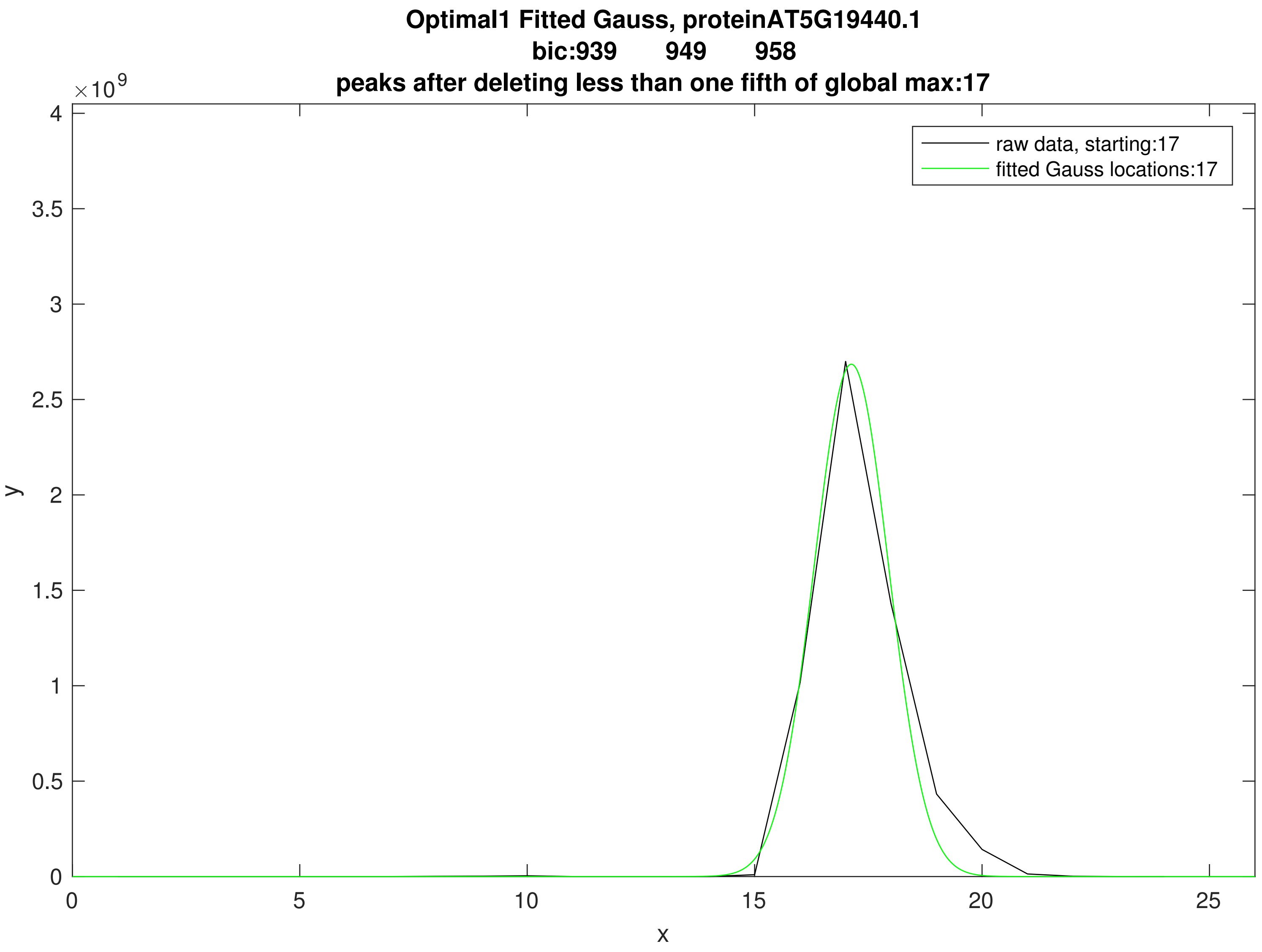

Optimal1 Fitted Gauss, proteinAT5G19440.1
bic:939 949 958
peaks after deleting less than one fifth of global max:17
×10^9
raw data, starting:17
fitted Gauss locations:17
y
x
0
0.5
1
1.5
2
2.5
3
3.5
4
5
10
15
20
25

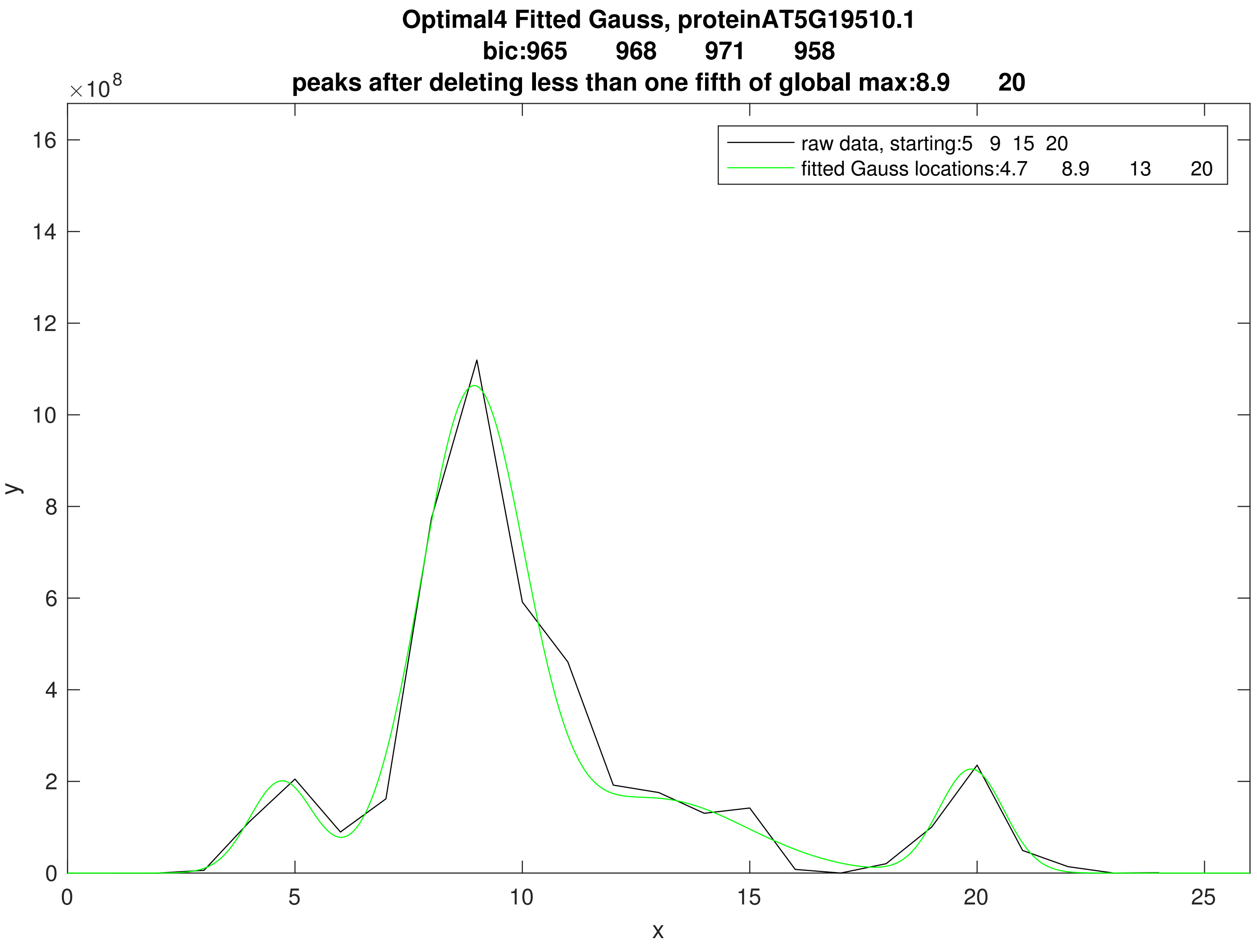
Optimal4 Fitted Gauss, proteinAT5G19510.1
bic:965 968 971 958
peaks after deleting less than one fifth of global max:8.9 20
×10^8
raw data, starting:5 9 15 20
fitted Gauss locations:4.7 8.9 13 20
y
x
0
2
4
6
8
10
12
14
16
0
5
10
15
20
25

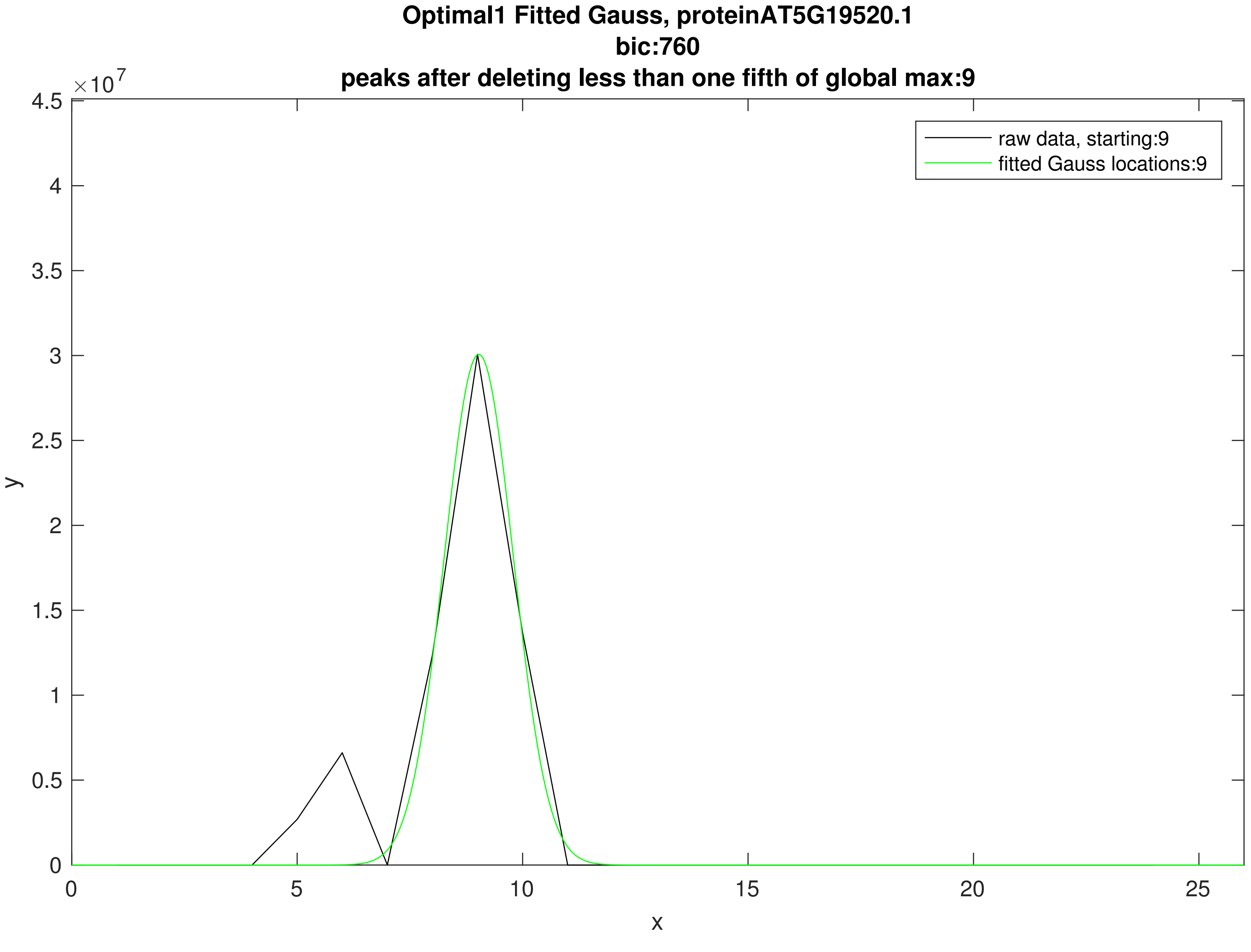

Optimal1 Fitted Gauss, proteinAT5G19520.1
bic:760
peaks after deleting less than one fifth of global max:9
×10^7
4.5
4
3.5
3
2.5
2
1.5
1
0.5
0
y
0
5
10
15
20
25
x
raw data, starting:9
fitted Gauss locations:9

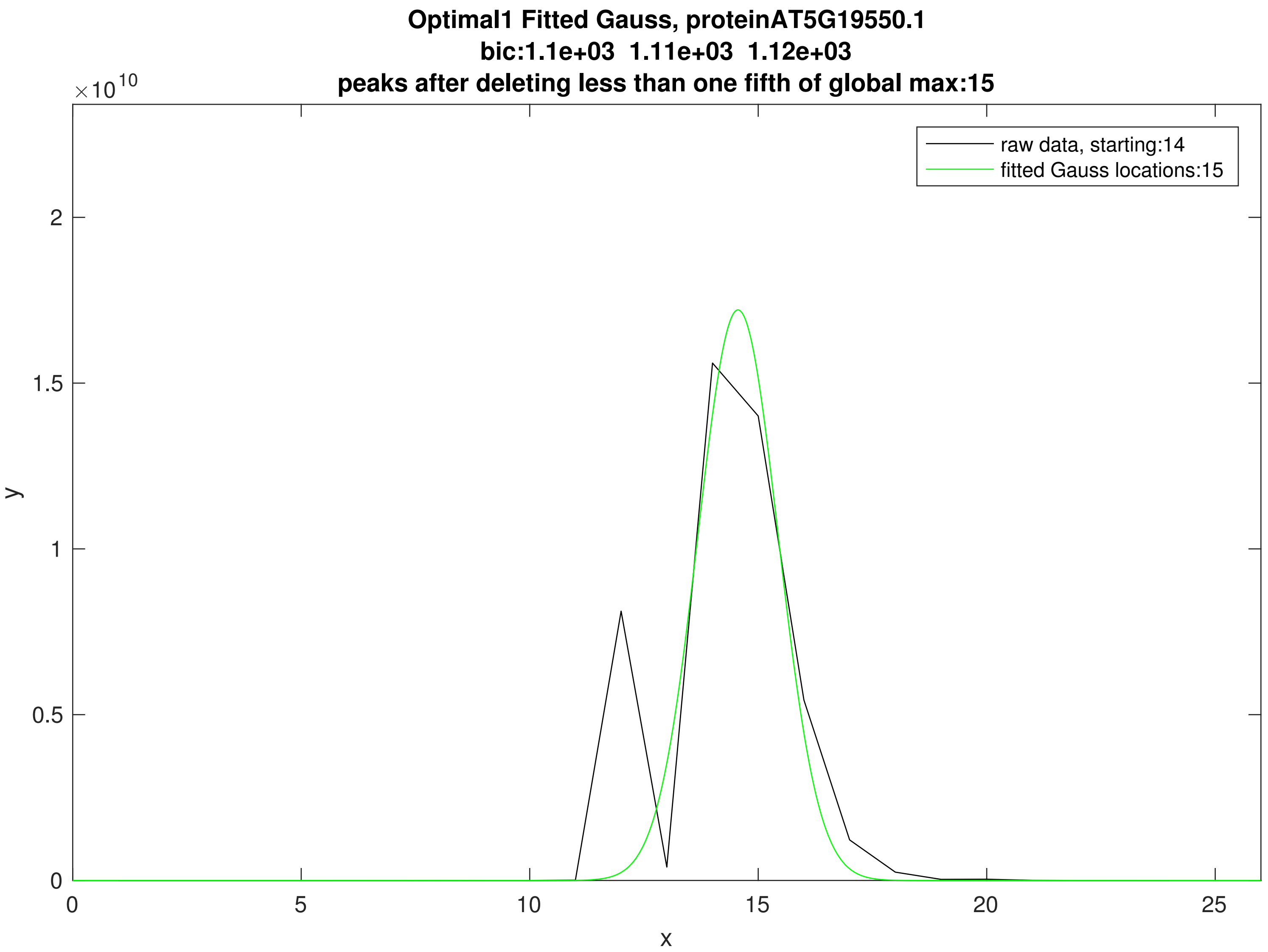
Optimal1 Fitted Gauss, proteinAT5G19550.1
bic:1.1e+03  1.11e+03  1.12e+03
peaks after deleting less than one fifth of global max:15
×10^10
raw data, starting:14
fitted Gauss locations:15
2
1.5
1
0.5
0
y
0
5
10
15
20
25
x

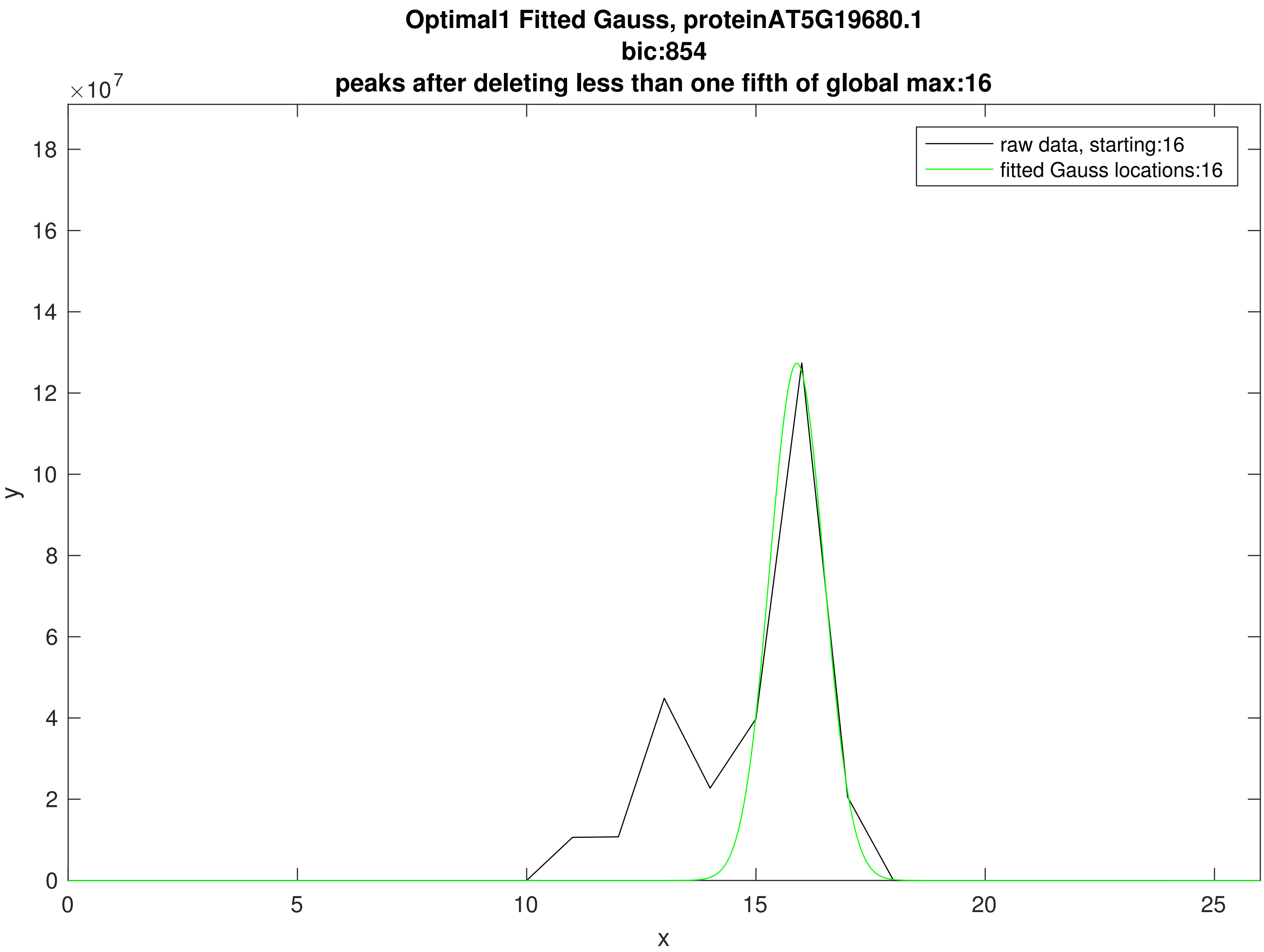
Optimal1 Fitted Gauss, proteinAT5G19680.1
bic:854
peaks after deleting less than one fifth of global max:16
×10^7
raw data, starting:16
fitted Gauss locations:16
y
x
0
2
4
6
8
10
12
14
16
18
0
5
10
15
20
25

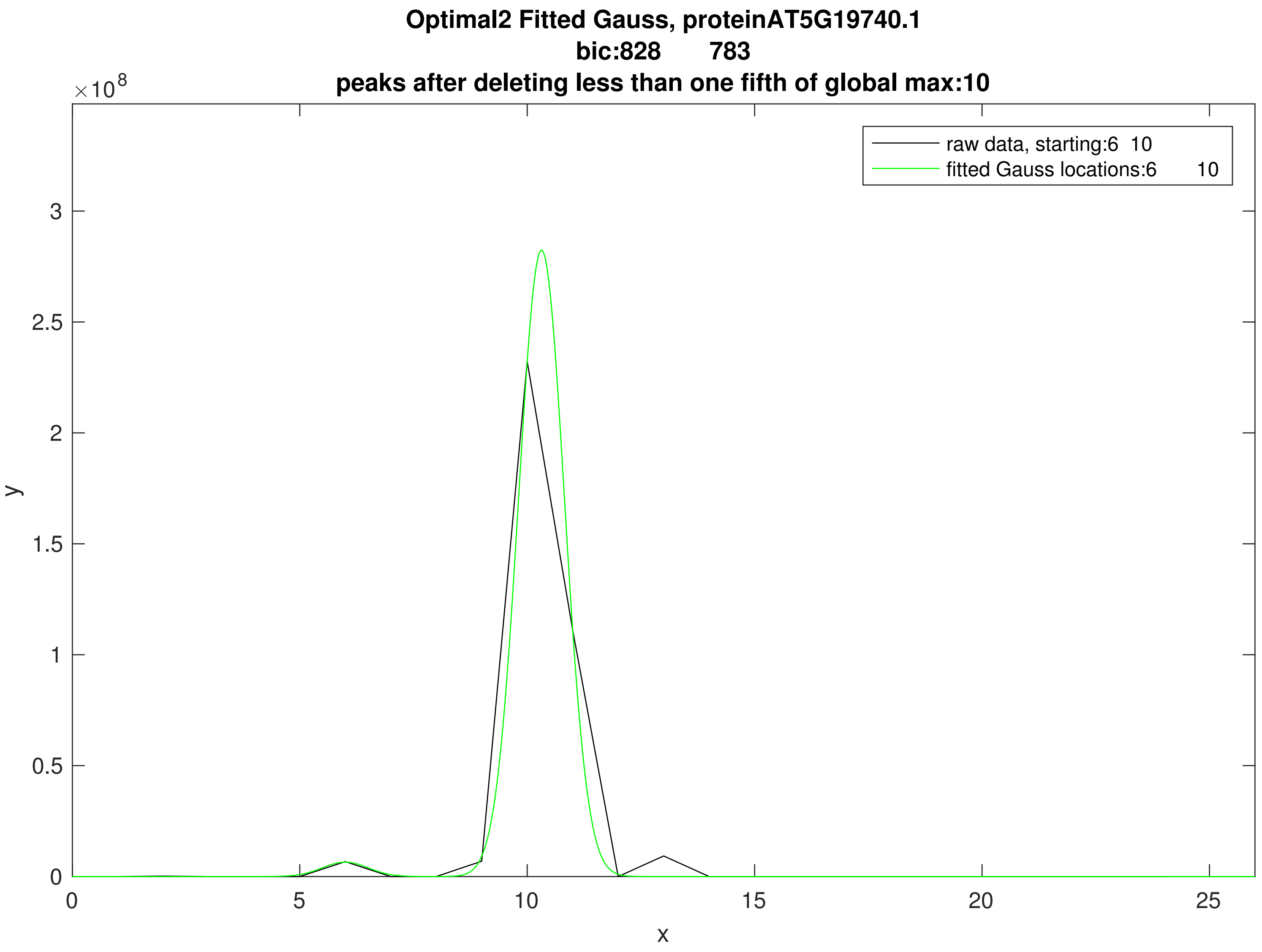

Optimal2 Fitted Gauss, proteinAT5G19740.1
bic:828 783
peaks after deleting less than one fifth of global max:10
×10^8
raw data, starting:6 10
fitted Gauss locations:6 10
3
2.5
2
1.5
1
0.5
0
y
0
5
10
15
20
25
x

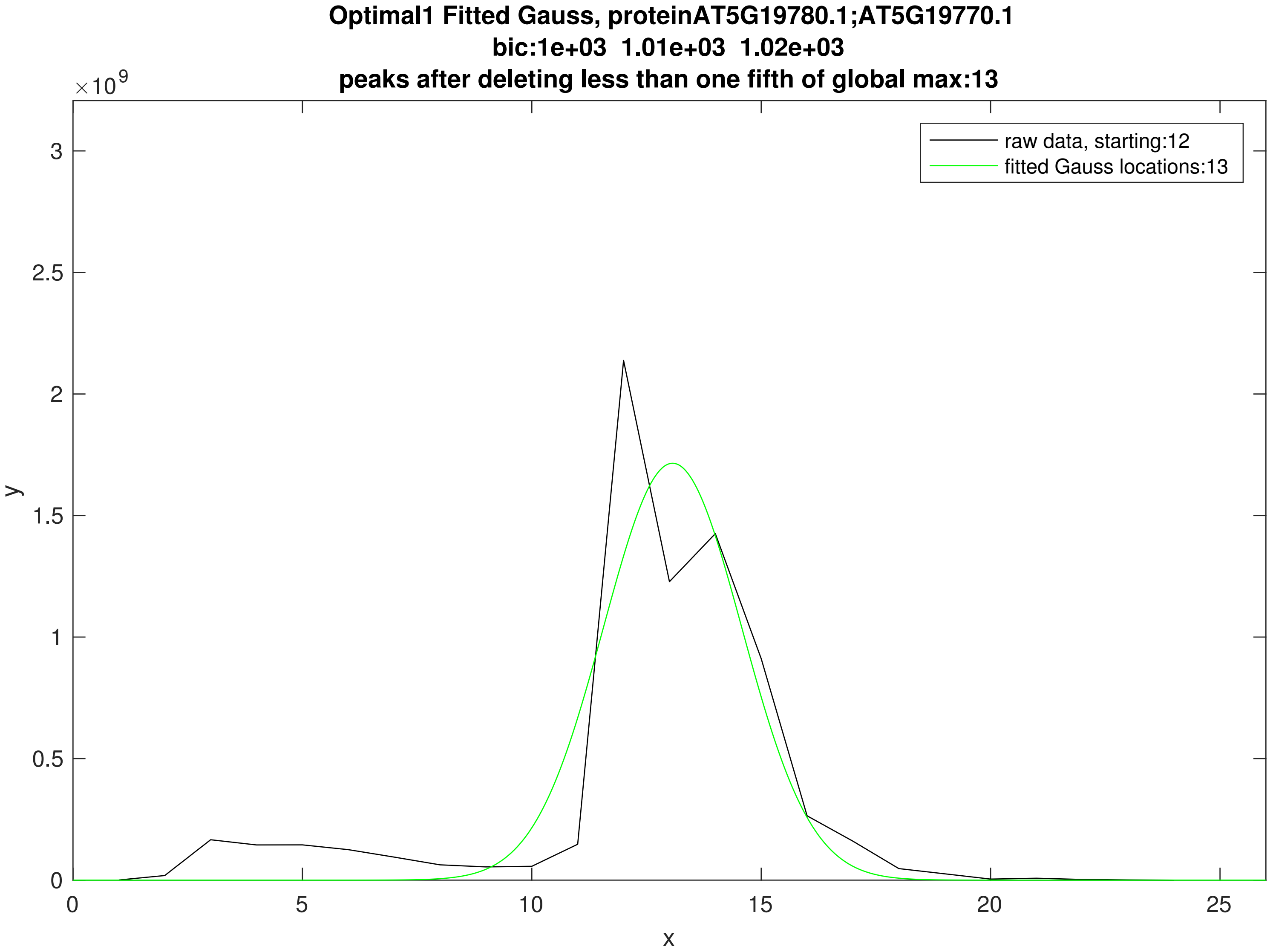

Optimal1 Fitted Gauss, proteinAT5G19780.1;AT5G19770.1
bic:1e+03 1.01e+03 1.02e+03
peaks after deleting less than one fifth of global max:13
×10^9
raw data, starting:12
fitted Gauss locations:13
y
x
0
0.5
1
1.5
2
2.5
3
5
10
15
20
25

**Optimal1 Fitted Gauss, proteinAT5G19860.1**
**bic:876**
**peaks after deleting less than one fifth of global max:20**

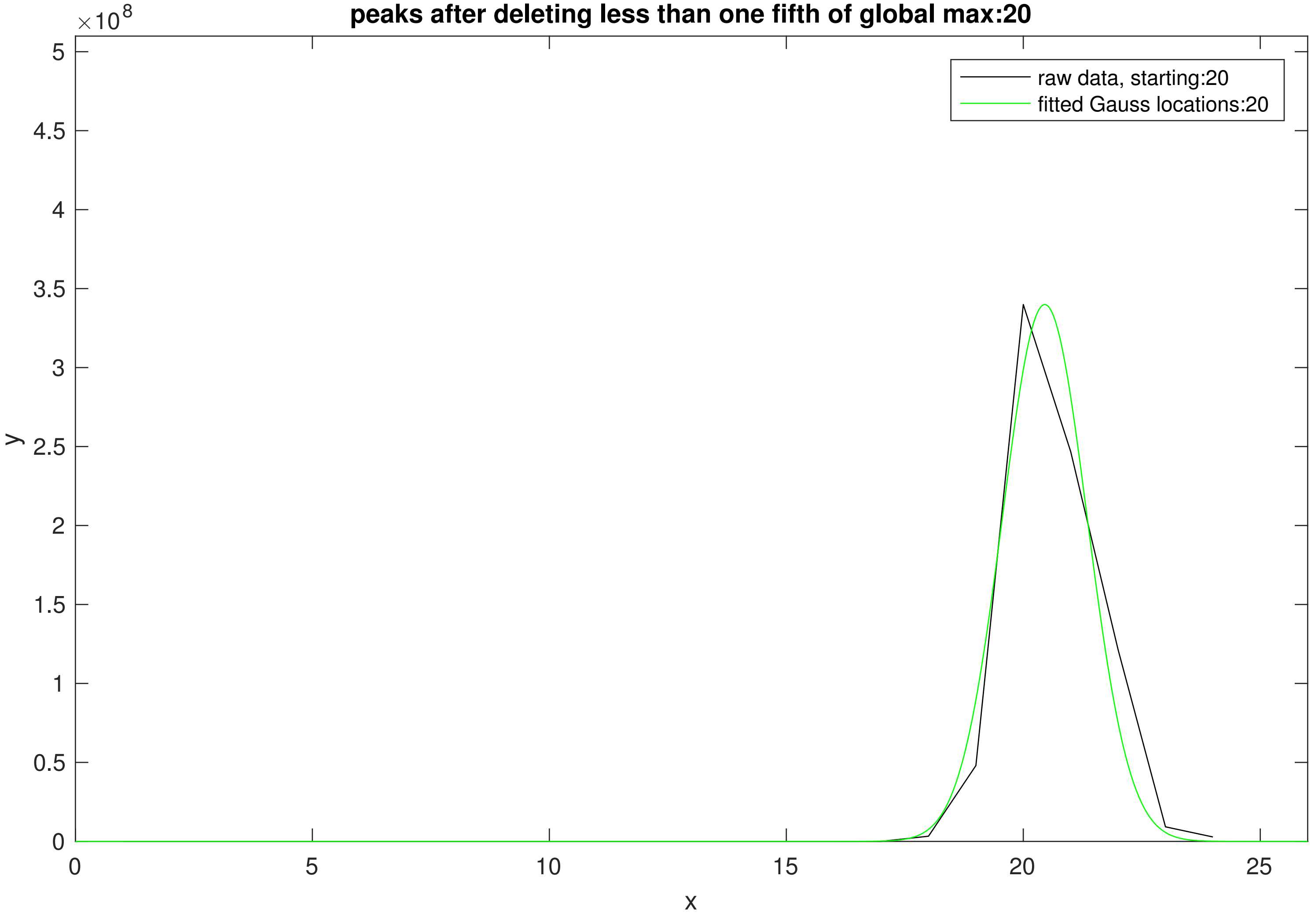

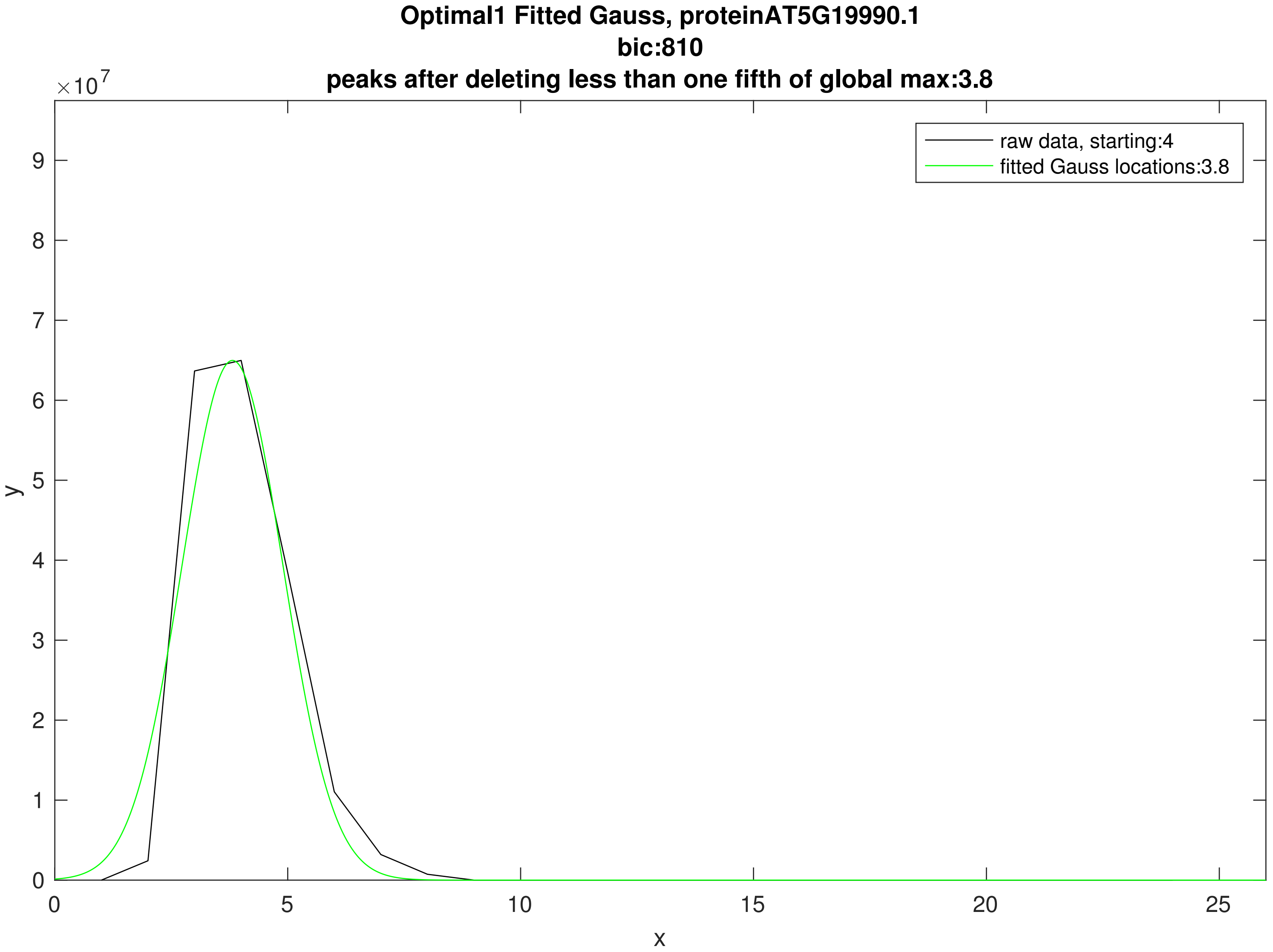
Optimal1 Fitted Gauss, proteinAT5G19990.1
bic:810
peaks after deleting less than one fifth of global max:3.8
×10^7
raw data, starting:4
fitted Gauss locations:3.8
y
x
0
1
2
3
4
5
6
7
8
9
0
5
10
15
20
25

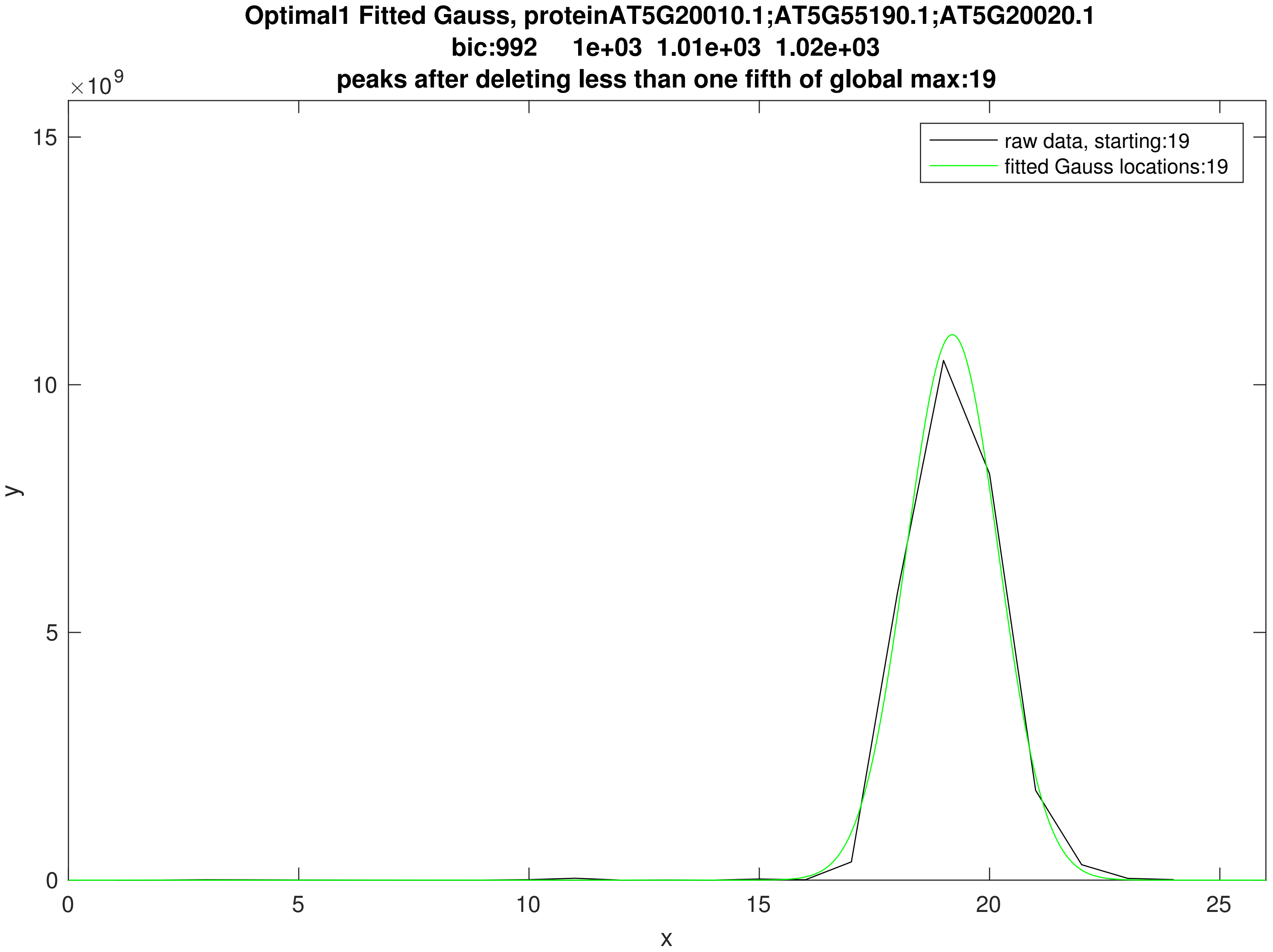
Optimal1 Fitted Gauss, proteinAT5G20010.1;AT5G55190.1;AT5G20020.1
bic:992 1e+03 1.01e+03 1.02e+03
peaks after deleting less than one fifth of global max:19
$\times 10^9$
15
10
5
0
y
0
5
10
15
20
25
x
raw data, starting:19
fitted Gauss locations:19

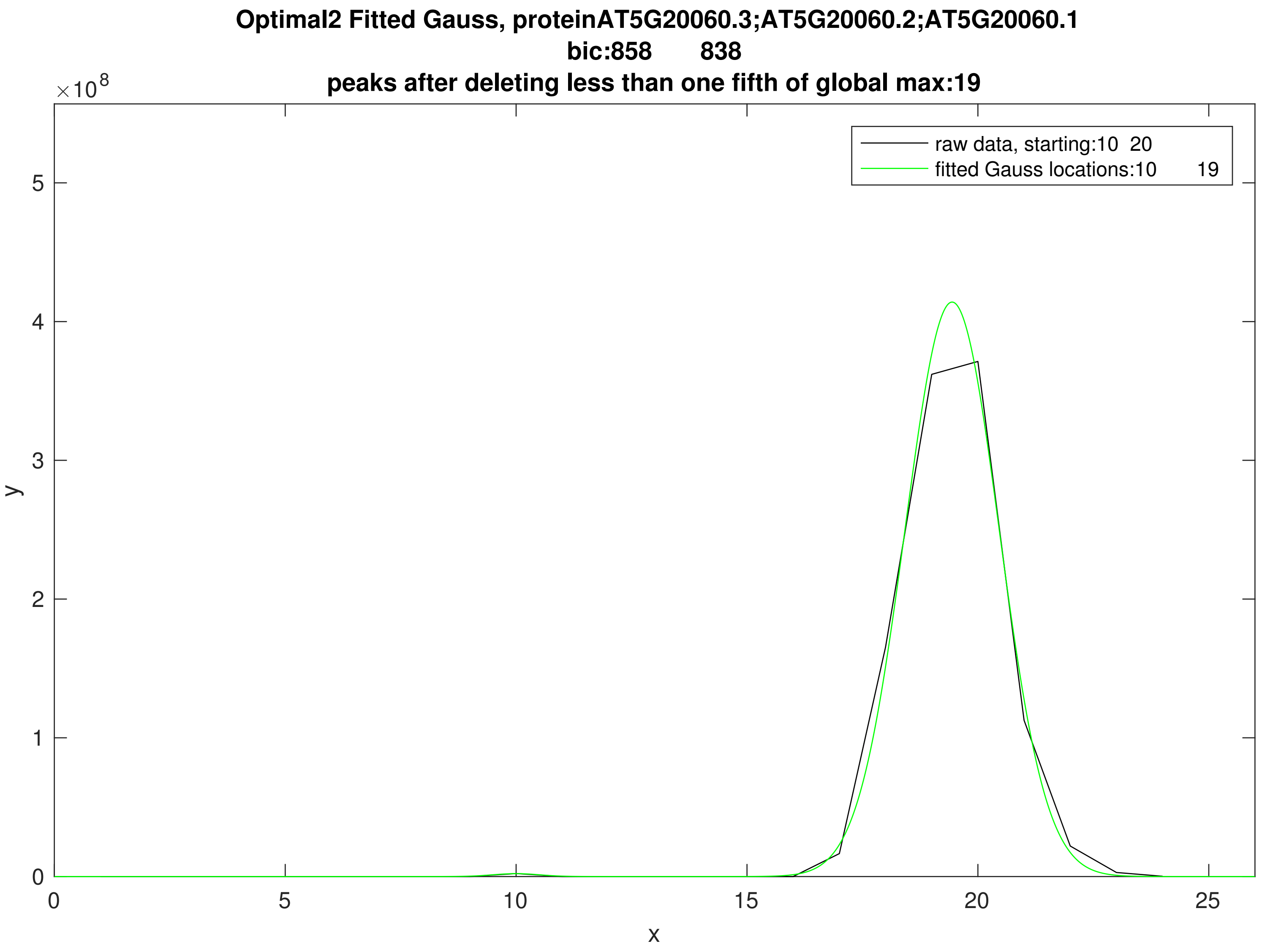
Optimal2 Fitted Gauss, proteinAT5G20060.3;AT5G20060.2;AT5G20060.1
bic:858 838
peaks after deleting less than one fifth of global max:19
×10^8
raw data, starting:10 20
fitted Gauss locations:10 19
y
x
0
1
2
3
4
5
0
5
10
15
20
25

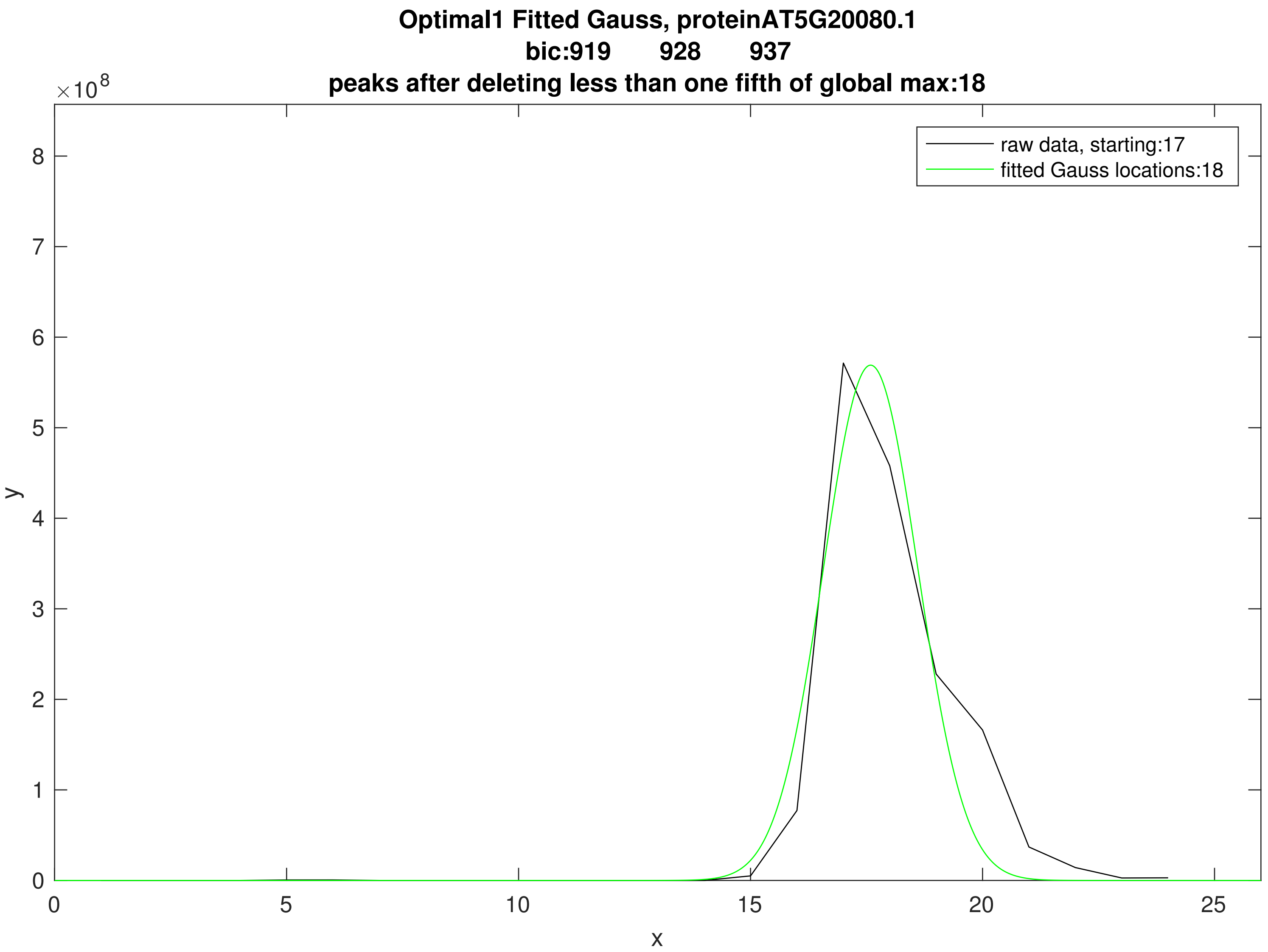

Optimal1 Fitted Gauss, proteinAT5G20080.1
bic:919 928 937
peaks after deleting less than one fifth of global max:18
×10^8
raw data, starting:17
fitted Gauss locations:18
y
x
0
1
2
3
4
5
6
7
8
0
5
10
15
20
25

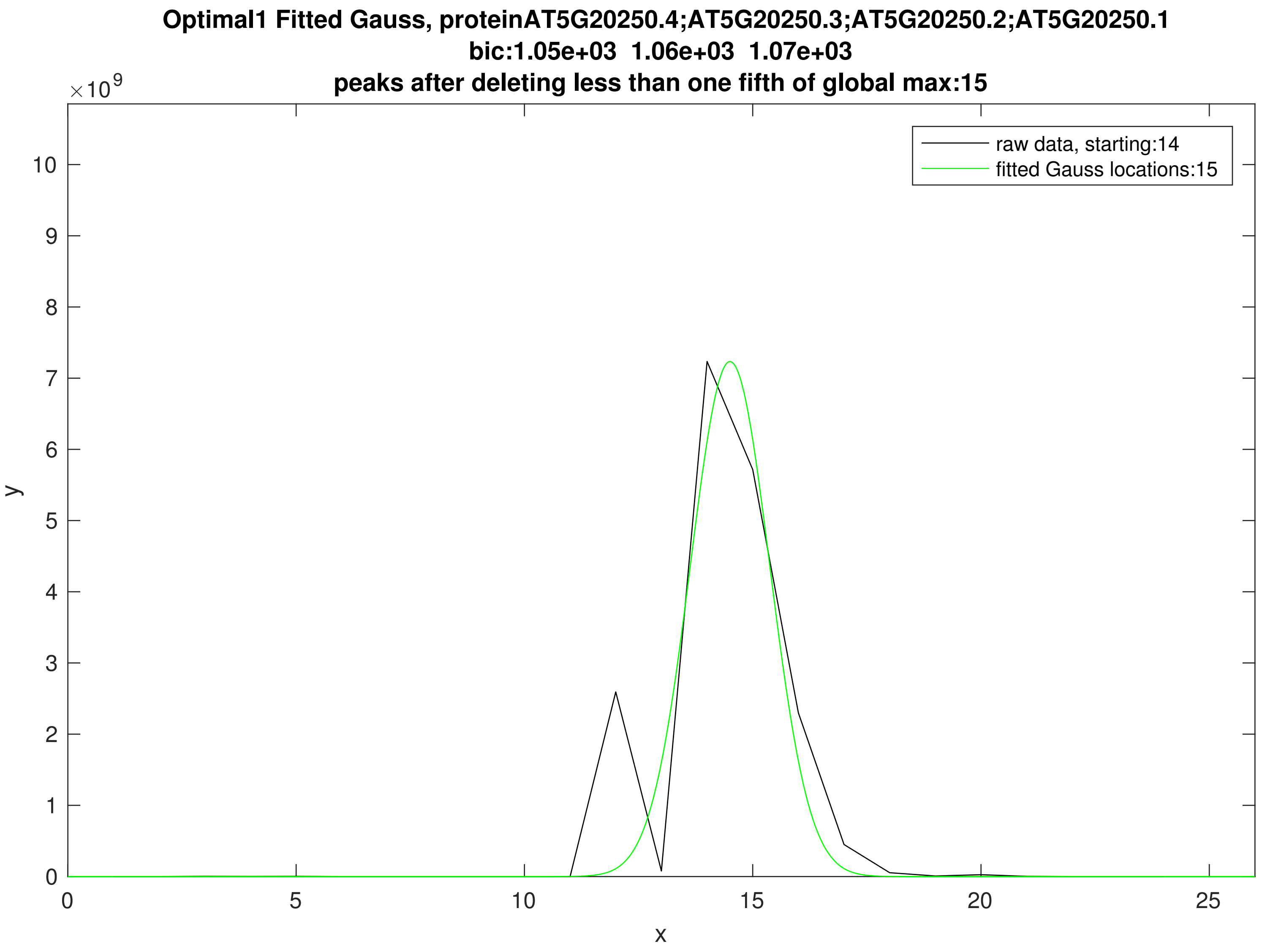
Optimal1 Fitted Gauss, proteinAT5G20250.4;AT5G20250.3;AT5G20250.2;AT5G20250.1
bic:1.05e+03  1.06e+03  1.07e+03
peaks after deleting less than one fifth of global max:15
×10^9
raw data, starting:14
fitted Gauss locations:15
y
x
0
1
2
3
4
5
6
7
8
9
10
0
5
10
15
20
25

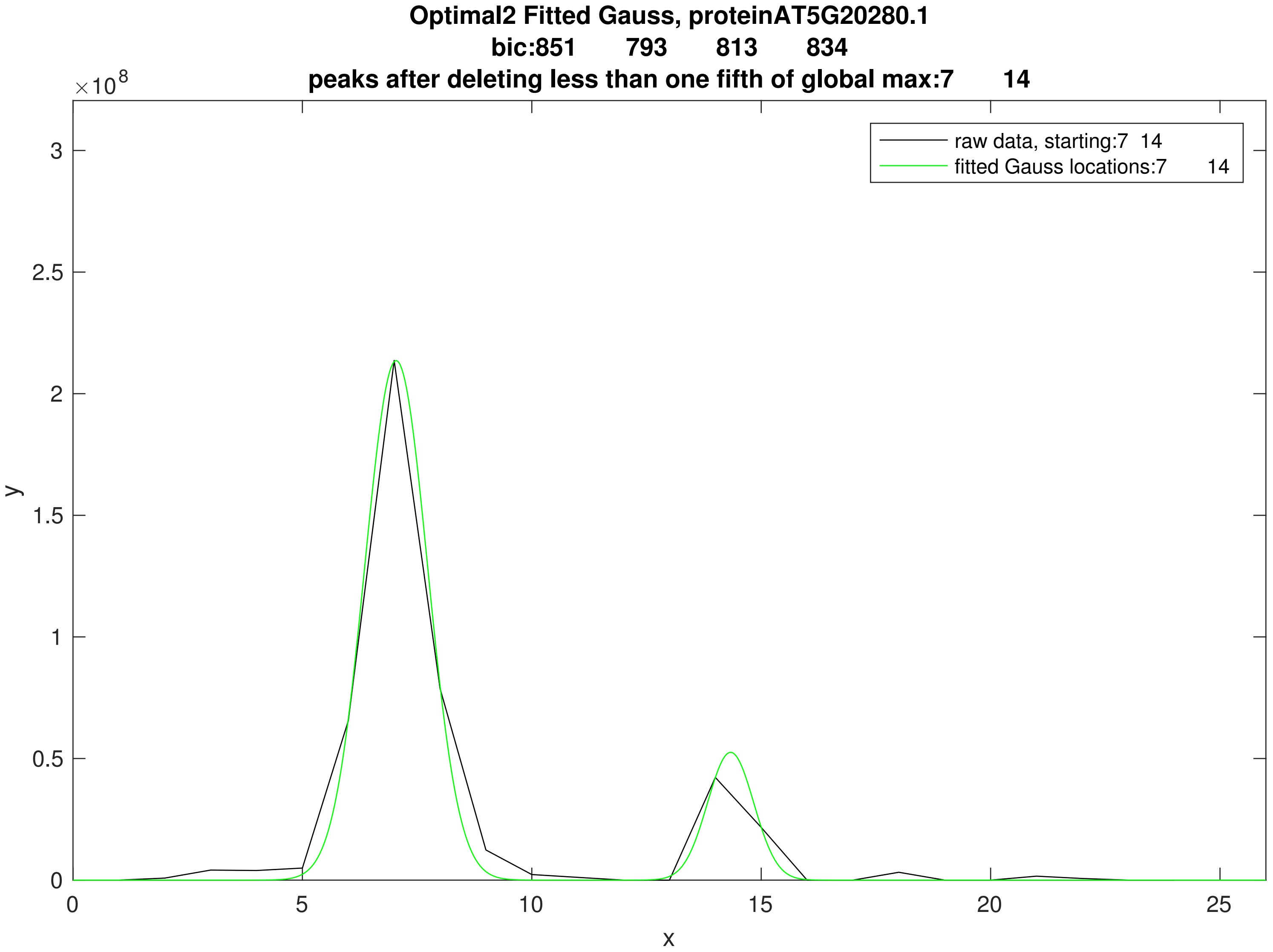

Optimal2 Fitted Gauss, proteinAT5G20280.1
bic:851 793 813 834
peaks after deleting less than one fifth of global max:7 14
raw data, starting:7 14
fitted Gauss locations:7 14
×10^8
y
x

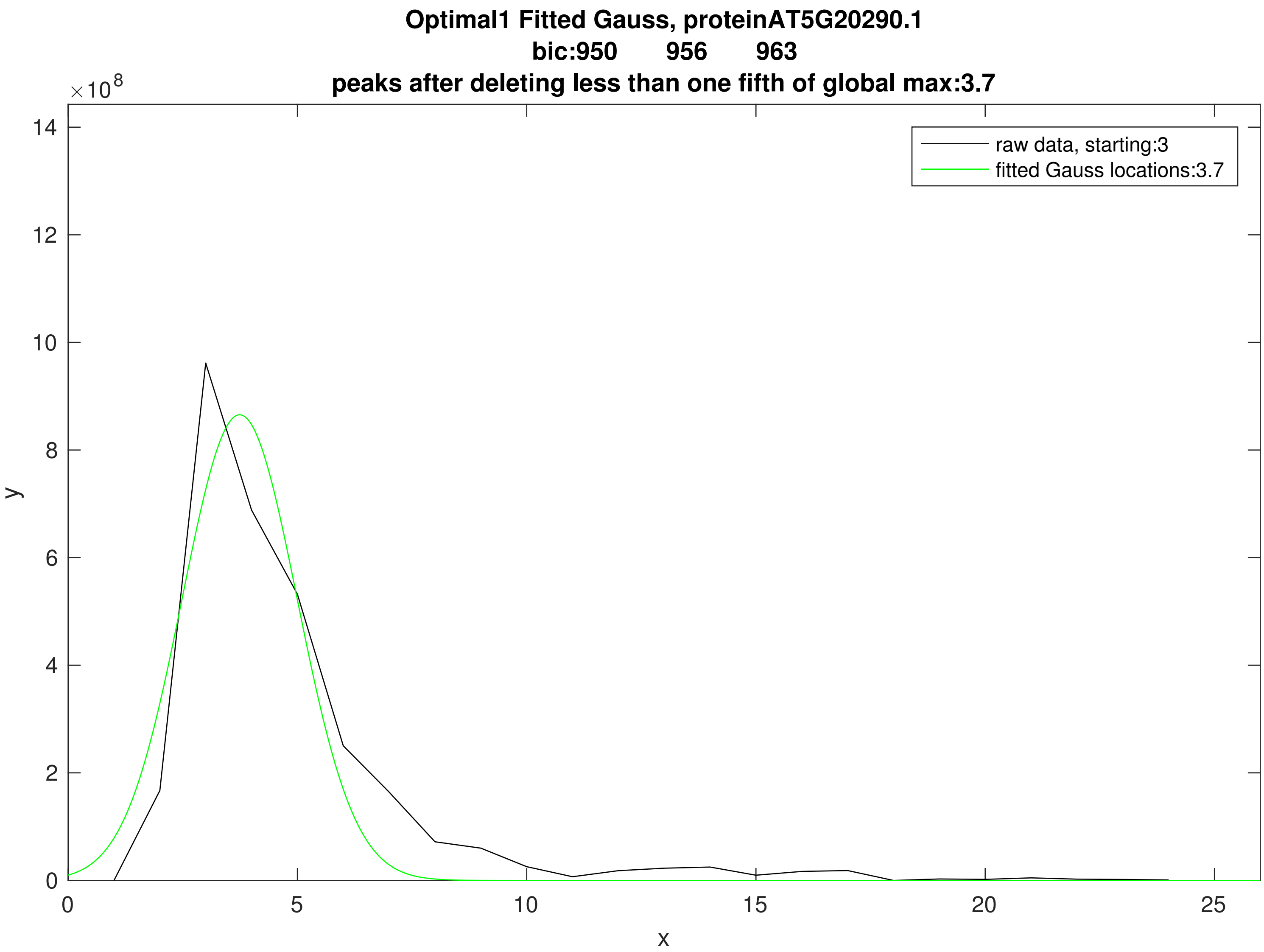

Optimal1 Fitted Gauss, proteinAT5G20290.1
bic:950 956 963
peaks after deleting less than one fifth of global max:3.7
×10^8
raw data, starting:3
fitted Gauss locations:3.7
y
x
0
2
4
6
8
10
12
14
0
5
10
15
20
25

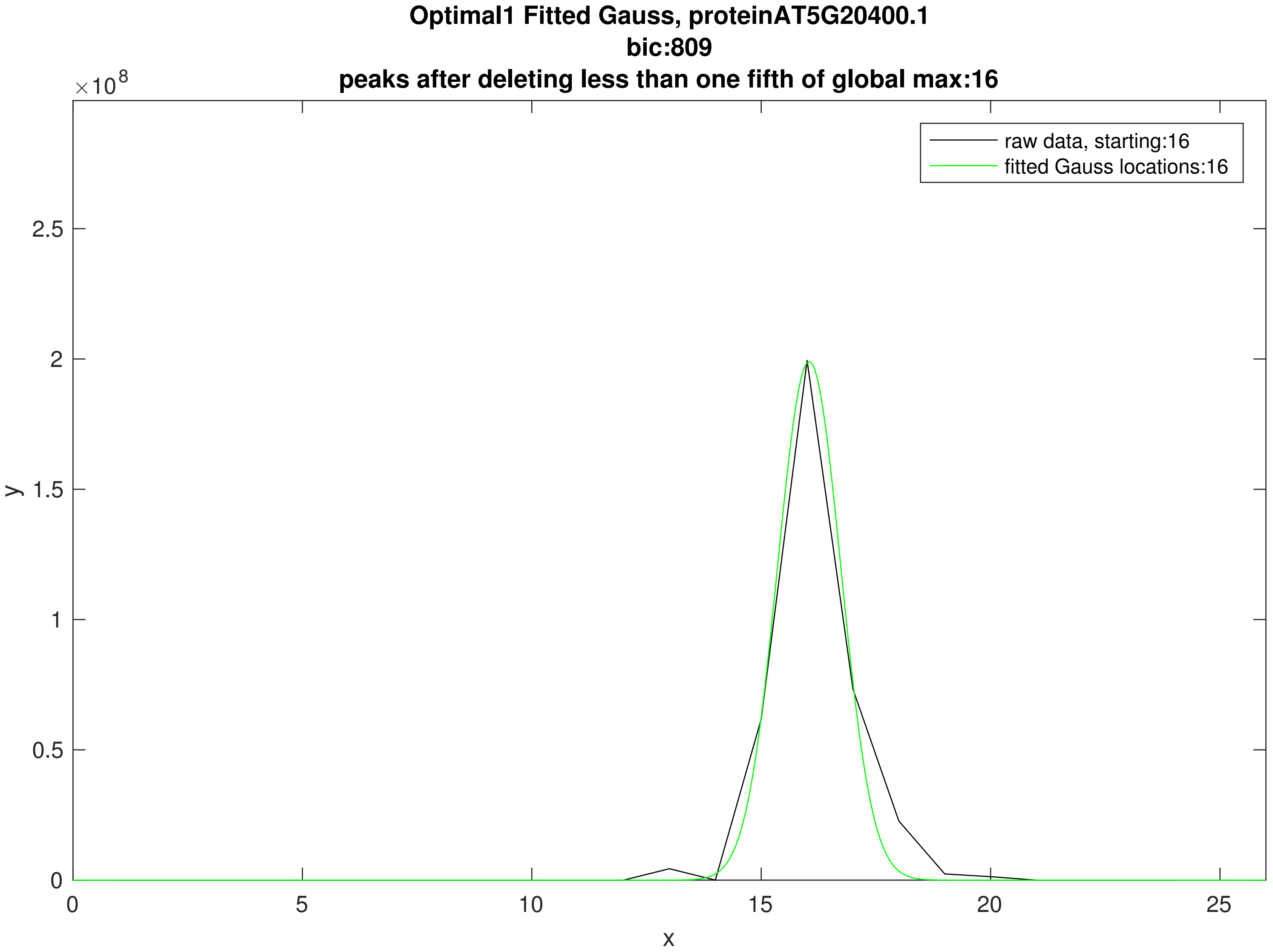
Optimal1 Fitted Gauss, proteinAT5G20400.1
bic:809
peaks after deleting less than one fifth of global max:16
×10^8
raw data, starting:16
fitted Gauss locations:16
y
x
0
0.5
1
1.5
2
2.5
0
5
10
15
20
25

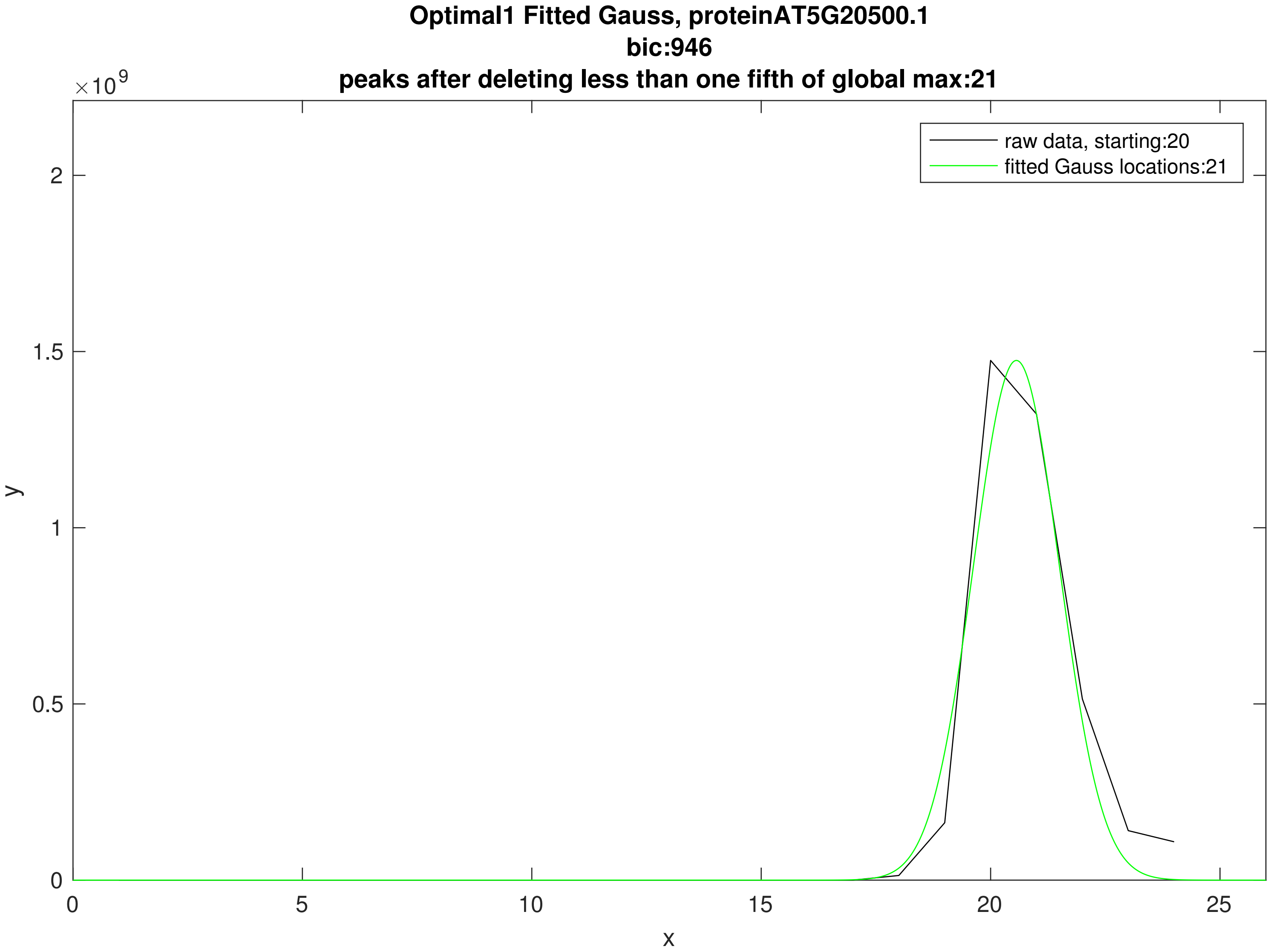

Optimal1 Fitted Gauss, proteinAT5G20500.1
bic:946
peaks after deleting less than one fifth of global max:21
×10^9
raw data, starting:20
fitted Gauss locations:21
2
1.5
1
0.5
0
y
0
5
10
15
20
25
x

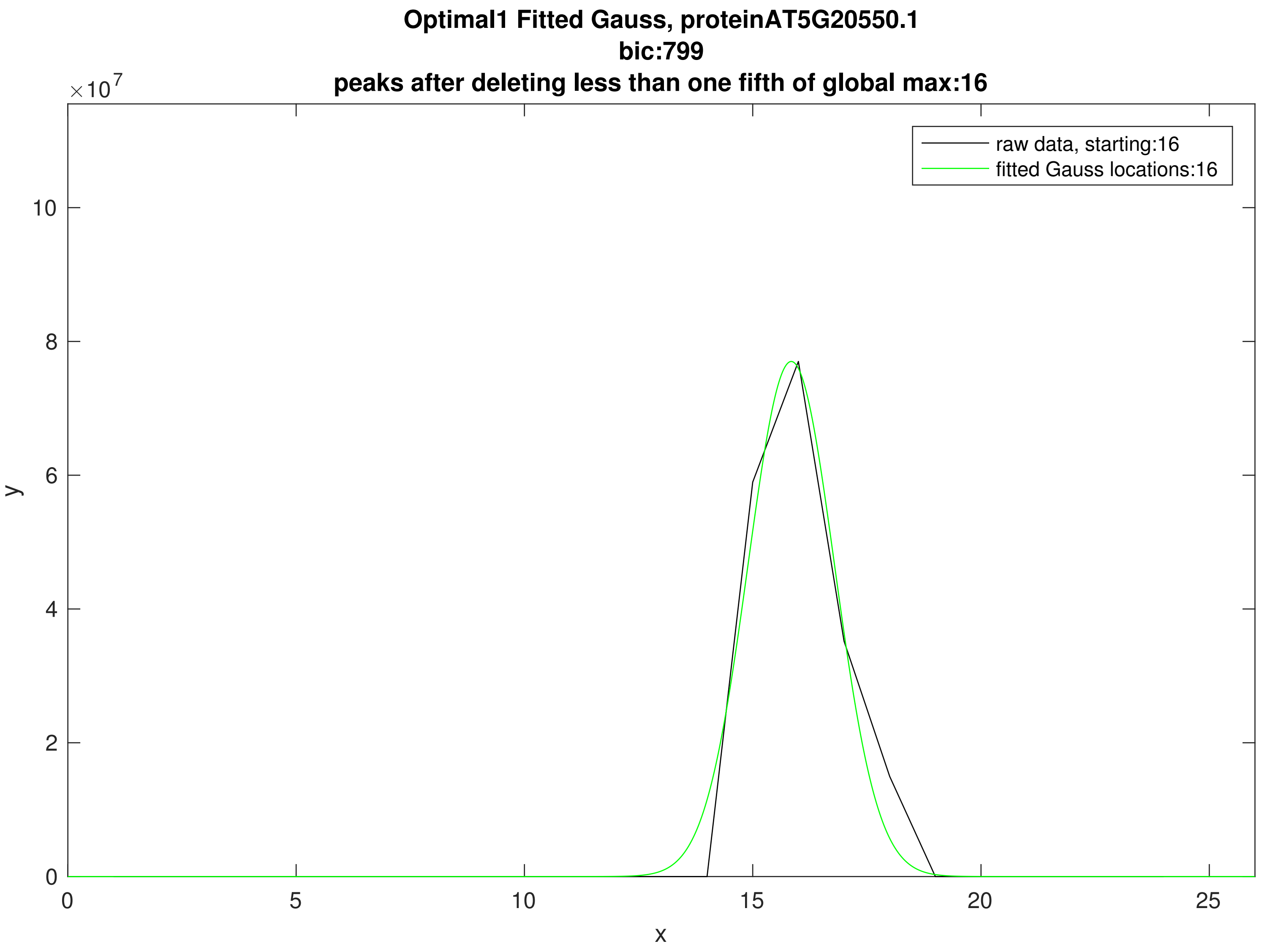
Optimal1 Fitted Gauss, proteinAT5G20550.1
bic:799
peaks after deleting less than one fifth of global max:16
×10^7
raw data, starting:16
fitted Gauss locations:16
y
x
0
2
4
6
8
10
0
5
10
15
20
25

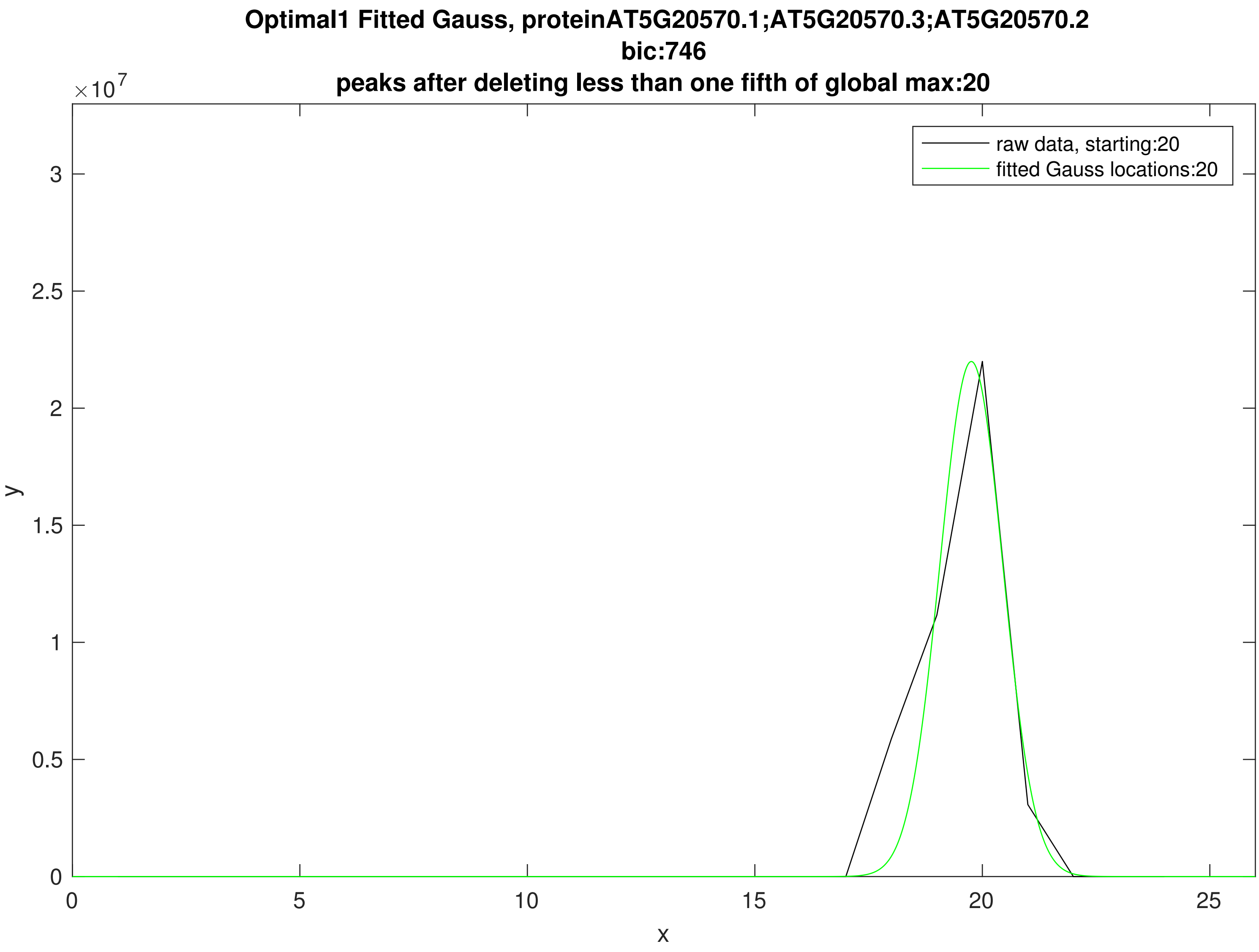
Optimal1 Fitted Gauss, proteinAT5G20570.1;AT5G20570.3;AT5G20570.2
bic:746
peaks after deleting less than one fifth of global max:20
×10^7
raw data, starting:20
fitted Gauss locations:20
y
x
0
0.5
1
1.5
2
2.5
3
5
10
15
20
25

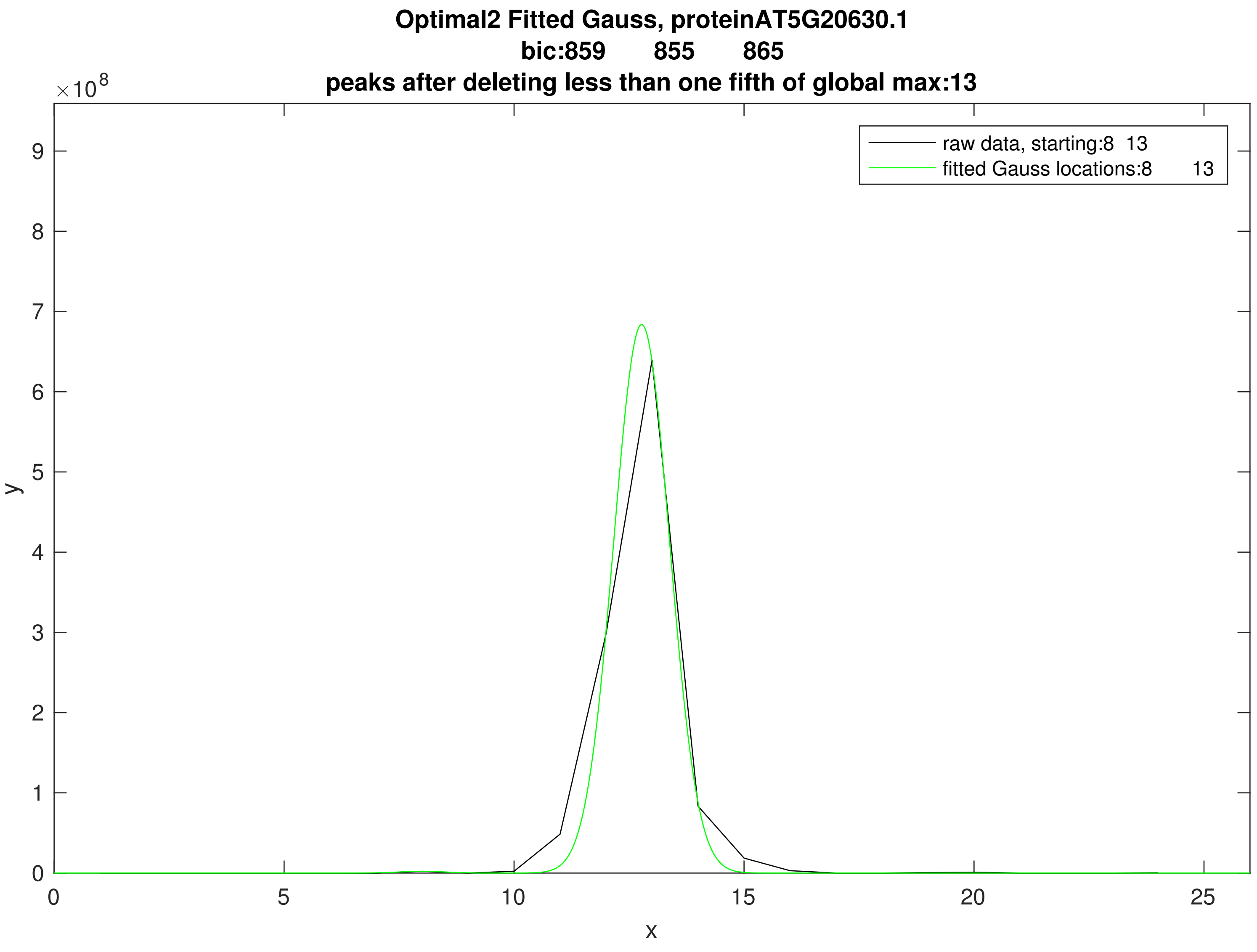

Optimal2 Fitted Gauss, proteinAT5G20630.1
bic:859 855 865
peaks after deleting less than one fifth of global max:13
×10^8
raw data, starting:8 13
fitted Gauss locations:8 13
y
x
0
1
2
3
4
5
6
7
8
9
0
5
10
15
20
25

**Optimal1 Fitted Gauss, proteinAT5G20720.3;AT5G20720.2;AT5G20720.1**
**bic:1.03e+03  1.04e+03**
**peaks after deleting less than one fifth of global max:13**

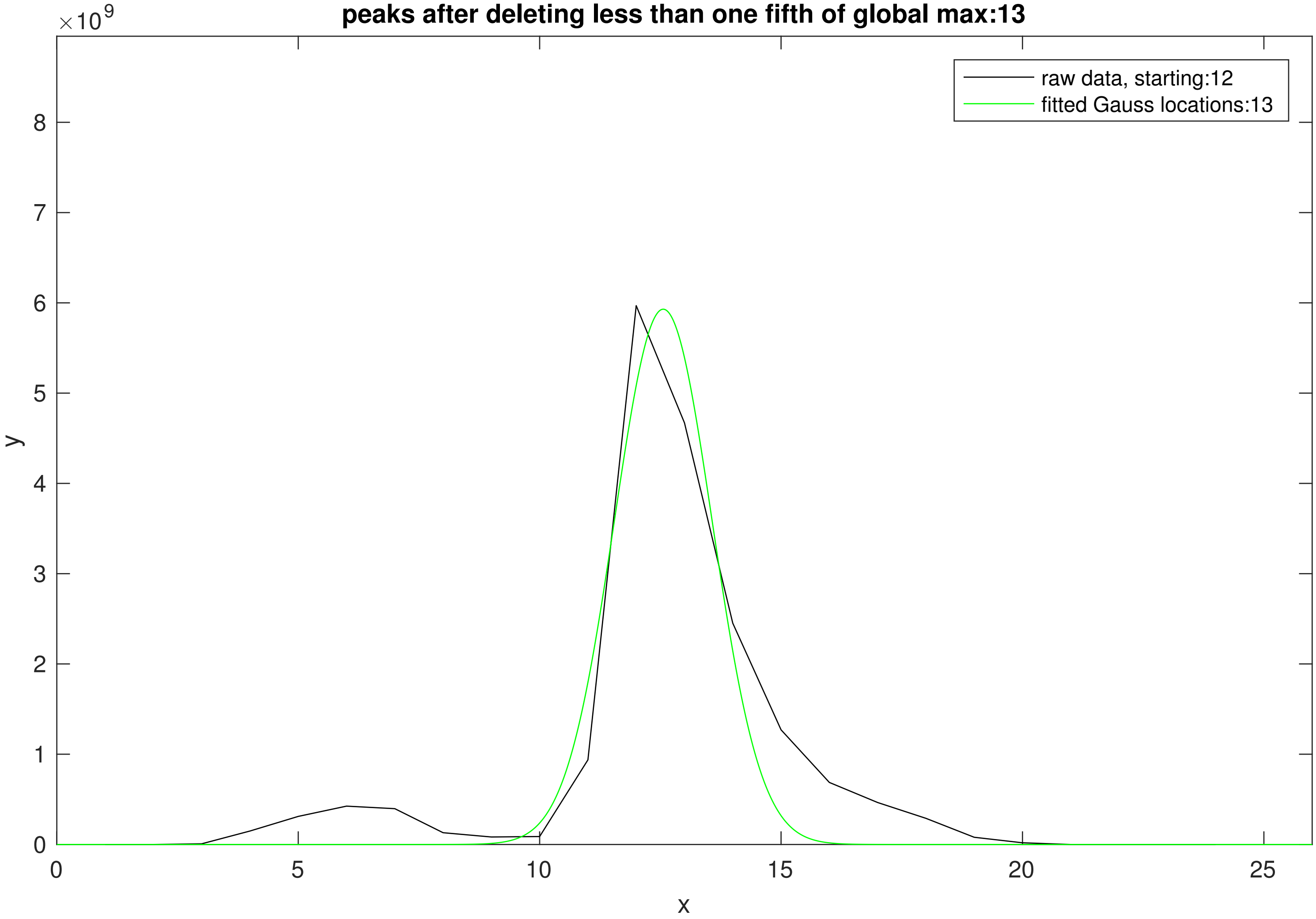

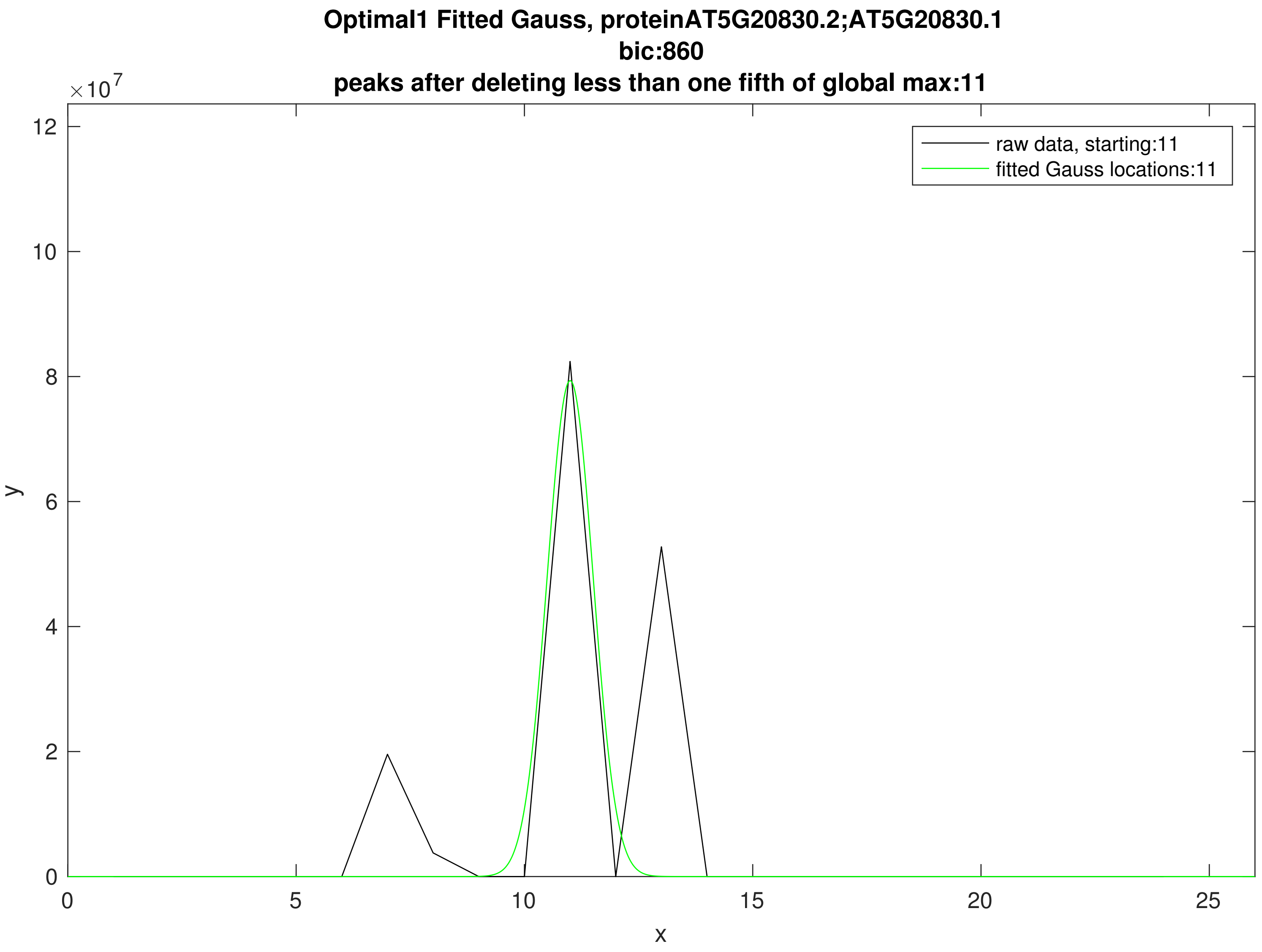

Optimal1 Fitted Gauss, proteinAT5G20830.2;AT5G20830.1
bic:860
peaks after deleting less than one fifth of global max:11
×10^7
12
10
8
6
4
2
0
y
0
5
10
15
20
25
x
raw data, starting:11
fitted Gauss locations:11

**Optimal2 Fitted Gauss, proteinAT5G20890.1**
**bic:862 853 863**
**peaks after deleting less than one fifth of global max:4.4**

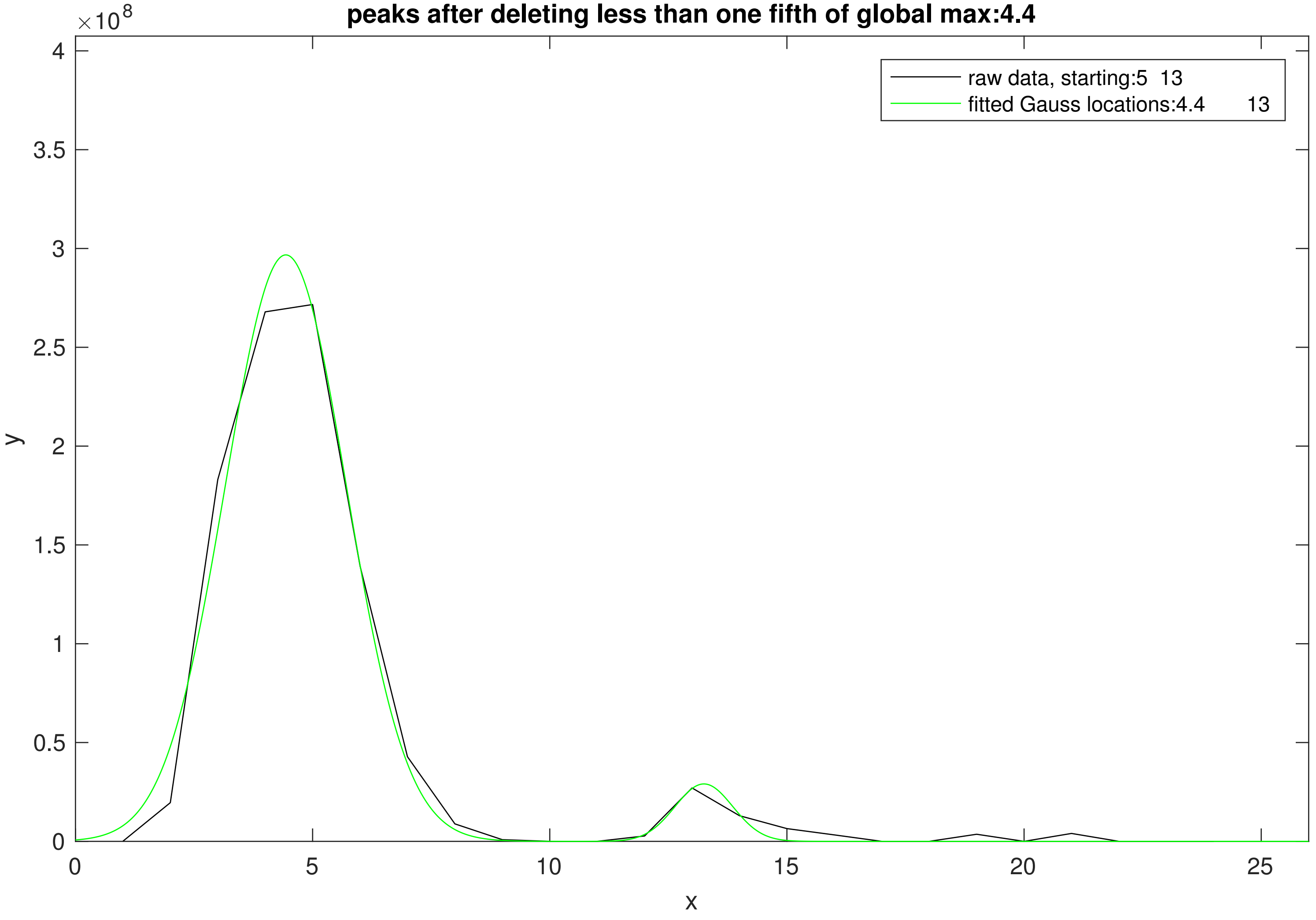

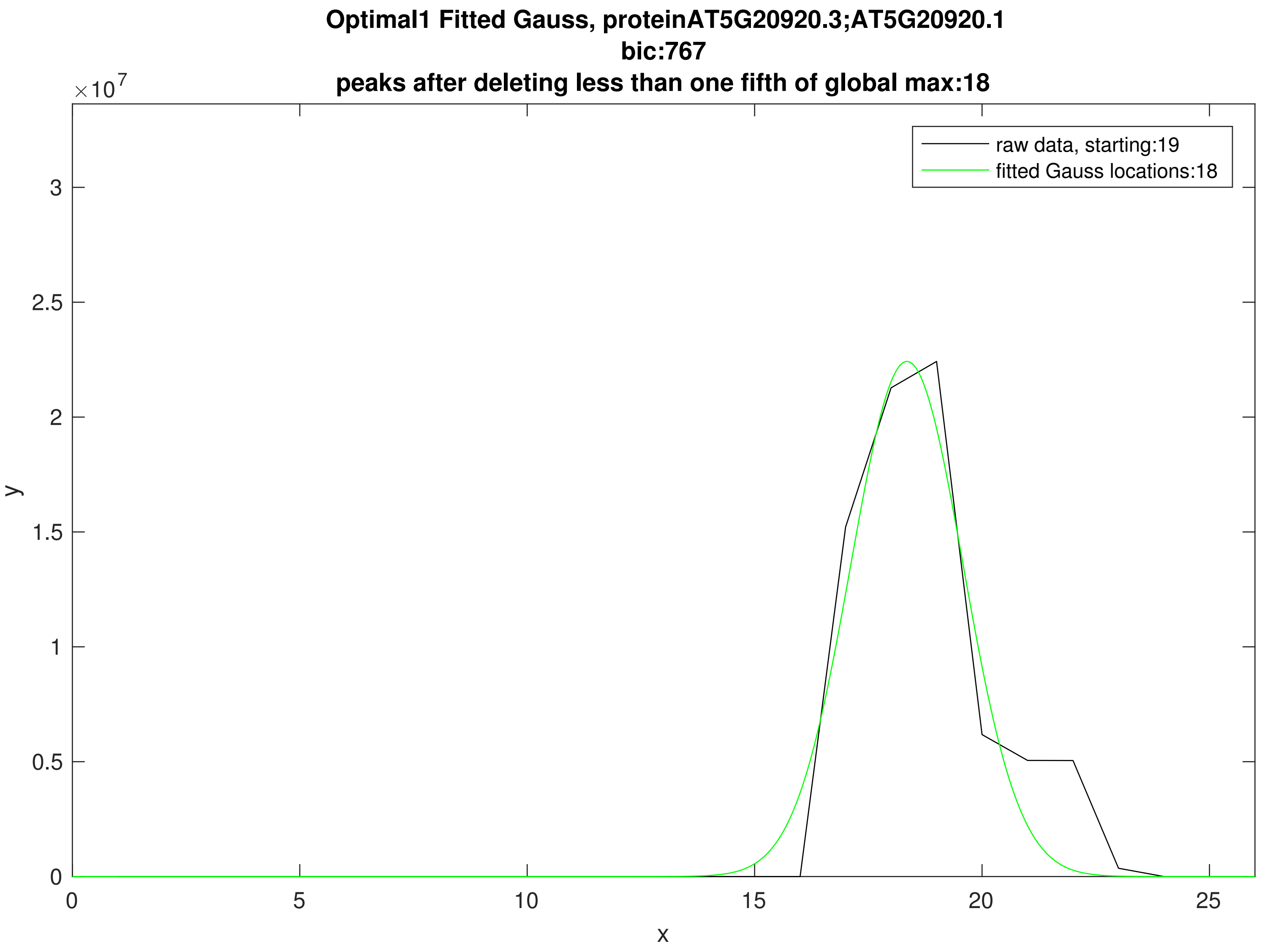
Optimal1 Fitted Gauss, proteinAT5G20920.3;AT5G20920.1
bic:767
peaks after deleting less than one fifth of global max:18
×10^7
raw data, starting:19
fitted Gauss locations:18
y
x
0
0.5
1
1.5
2
2.5
3
0
5
10
15
20
25

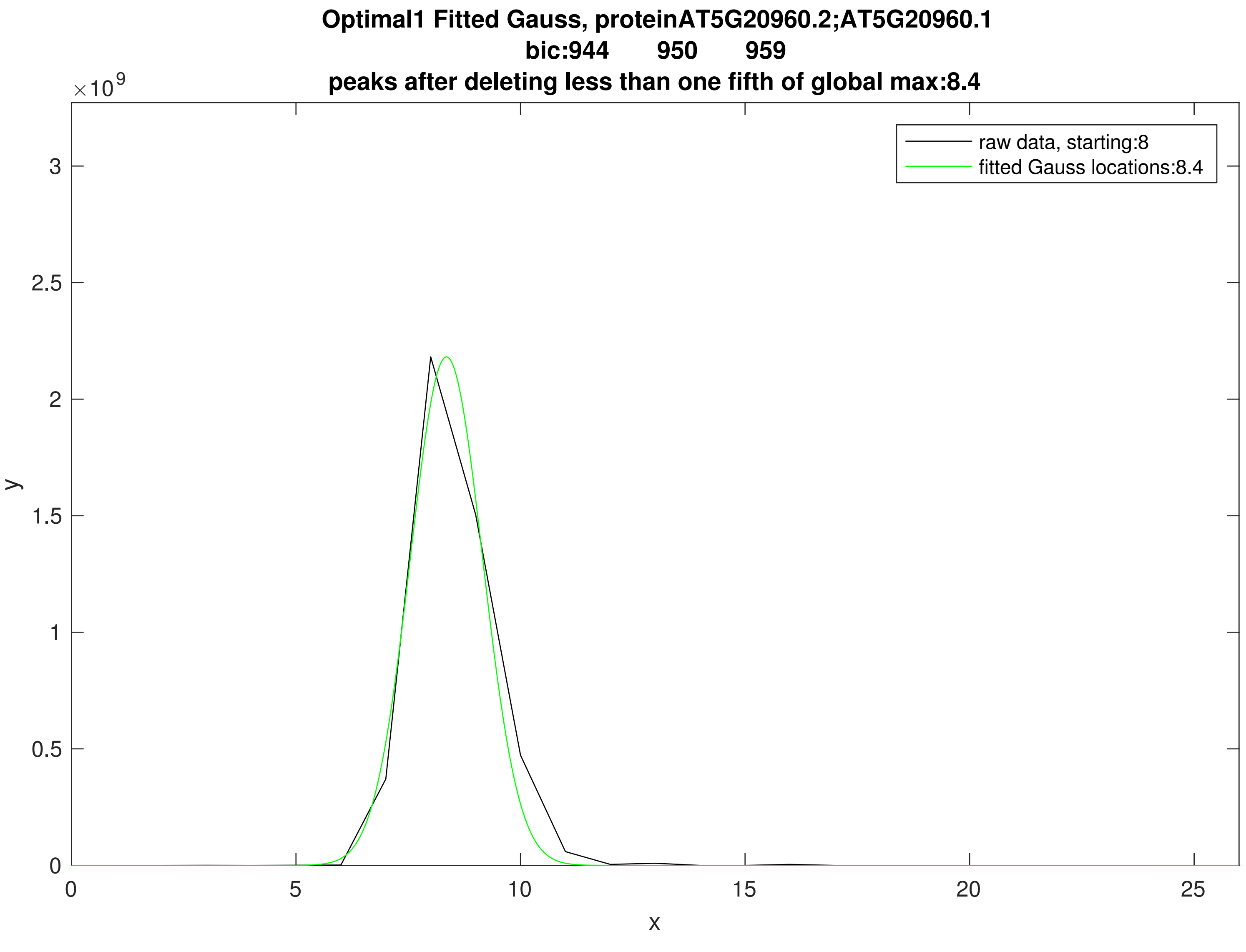
Optimal1 Fitted Gauss, proteinAT5G20960.2;AT5G20960.1
bic:944 950 959
peaks after deleting less than one fifth of global max:8.4
×10^9
raw data, starting:8
fitted Gauss locations:8.4
y
x
0
0.5
1
1.5
2
2.5
3
5
10
15
20
25

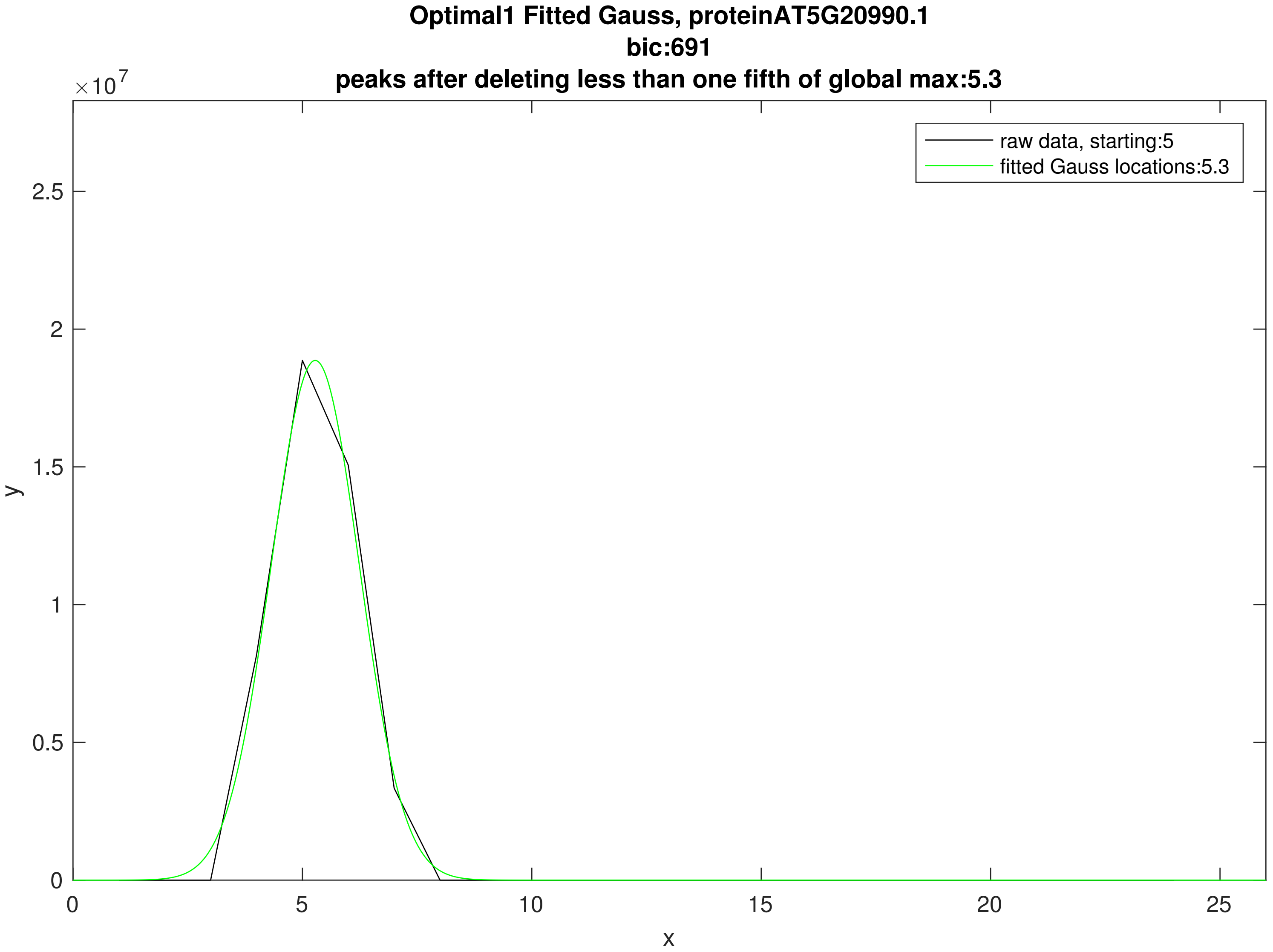

Optimal1 Fitted Gauss, proteinAT5G20990.1
bic:691
peaks after deleting less than one fifth of global max:5.3
×10^7
2.5
2
1.5
1
0.5
0
y
0
5
10
15
20
25
x
raw data, starting:5
fitted Gauss locations:5.3

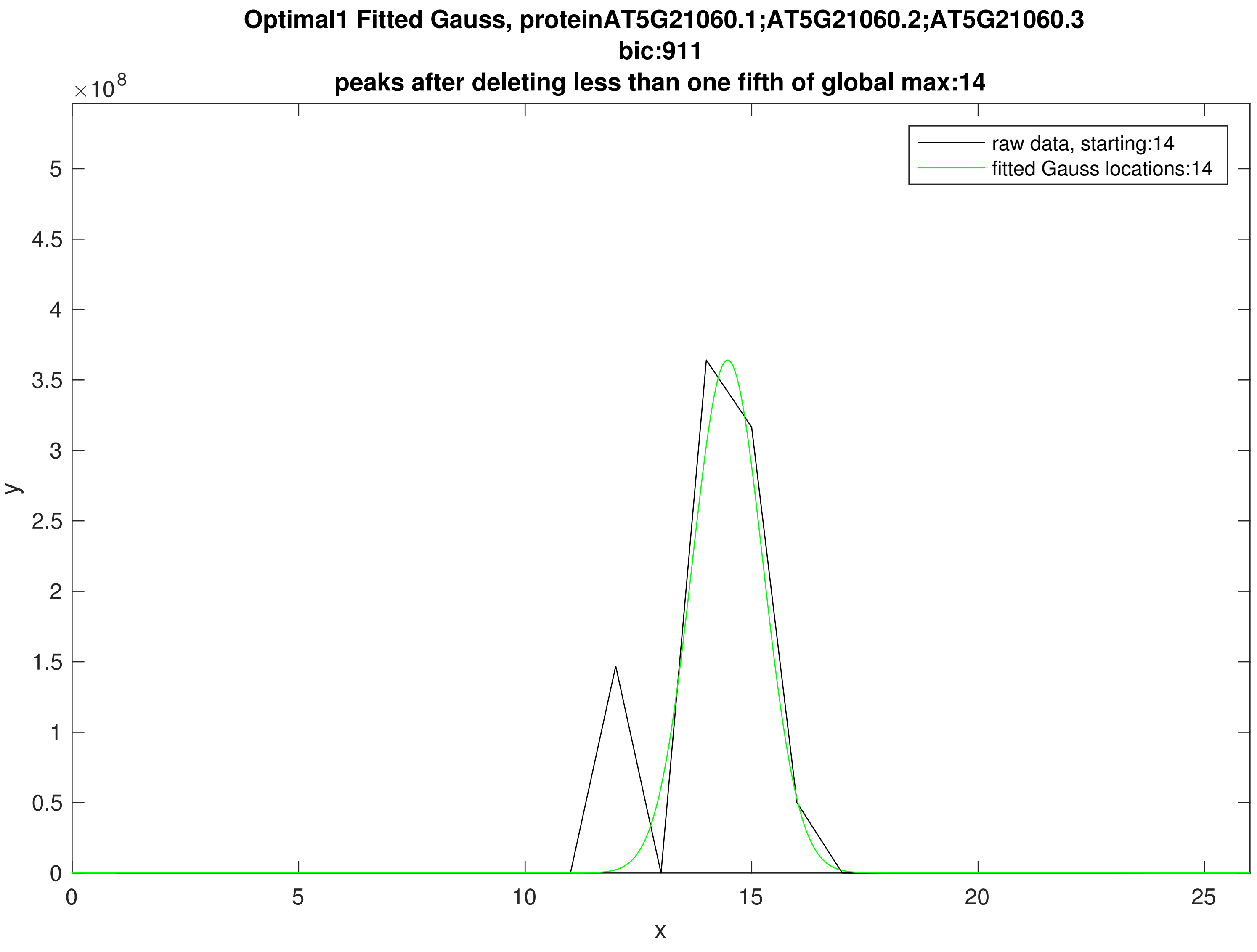

Optimal1 Fitted Gauss, proteinAT5G21060.1;AT5G21060.2;AT5G21060.3
bic:911
peaks after deleting less than one fifth of global max:14
×10^8
raw data, starting:14
fitted Gauss locations:14
y
x
0
0.5
1
1.5
2
2.5
3
3.5
4
4.5
5
0
5
10
15
20
25

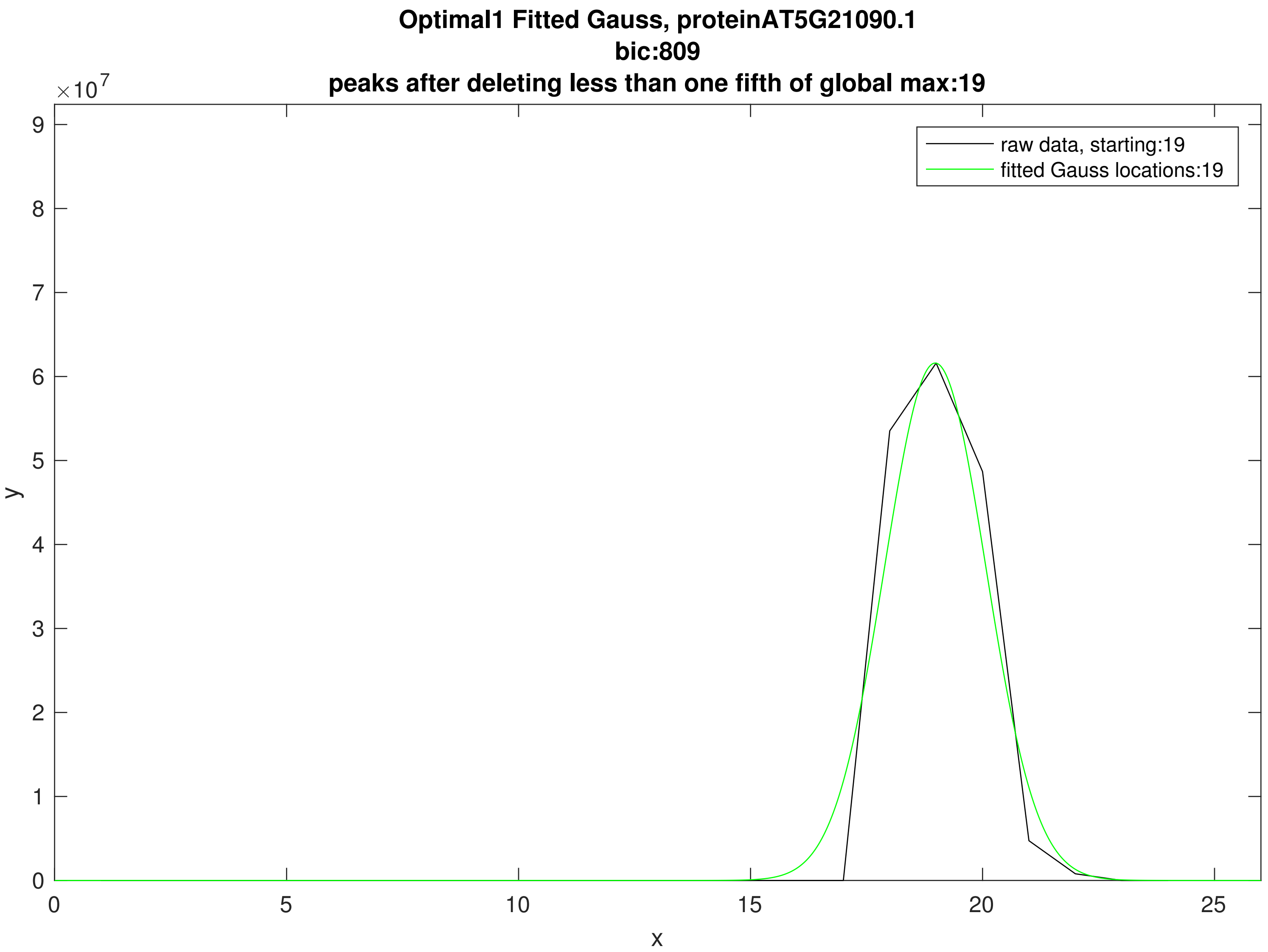

Optimal1 Fitted Gauss, proteinAT5G21090.1
bic:809
peaks after deleting less than one fifth of global max:19
×10^7
raw data, starting:19
fitted Gauss locations:19
y
x
0
1
2
3
4
5
6
7
8
9
0
5
10
15
20
25

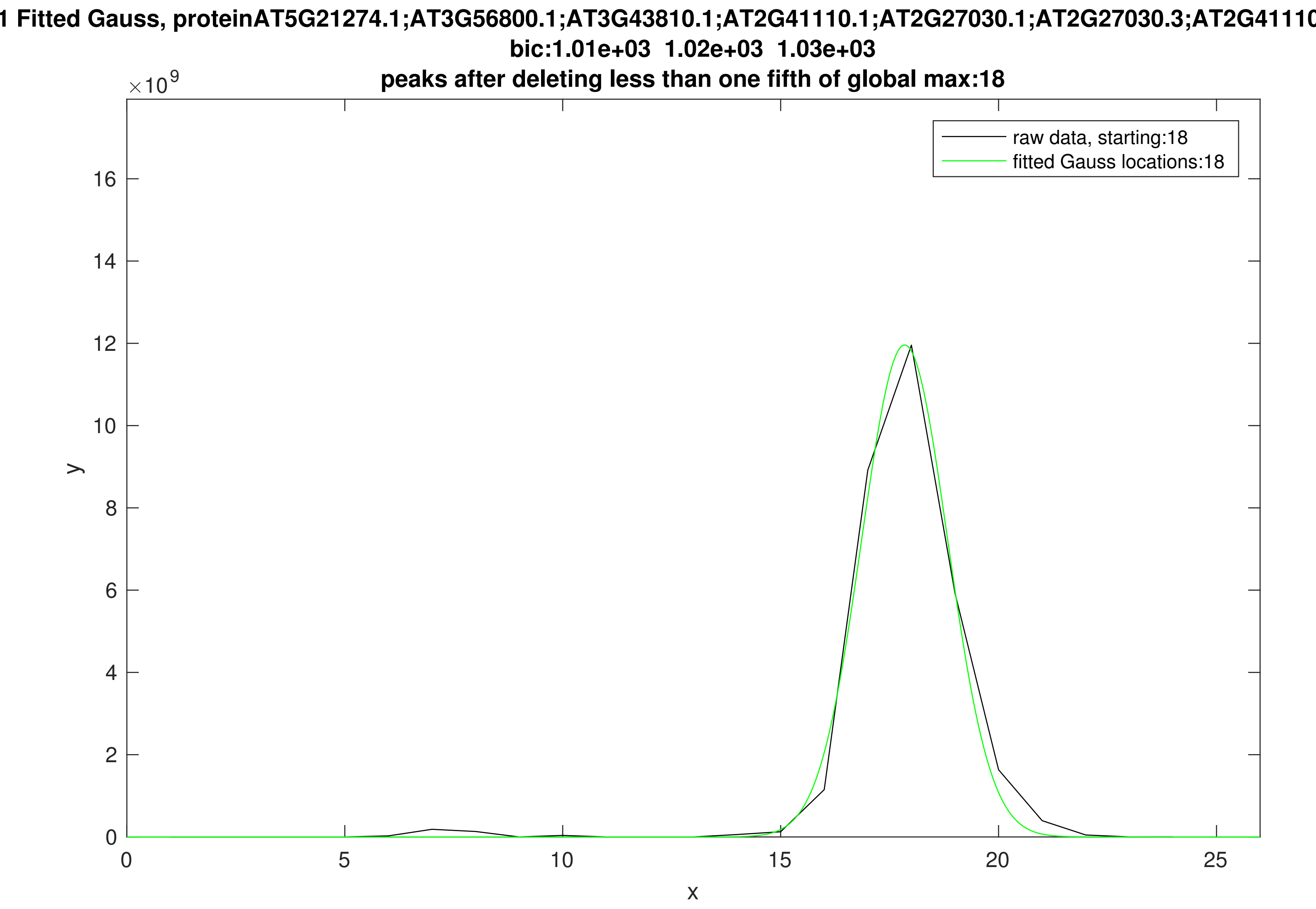
1 Fitted Gauss, proteinAT5G21274.1;AT3G56800.1;AT3G43810.1;AT2G41110.1;AT2G27030.1;AT2G27030.3;AT2G4111
bic:1.01e+03 1.02e+03 1.03e+03
peaks after deleting less than one fifth of global max:18
×10^9
raw data, starting:18
fitted Gauss locations:18
y
x
0
2
4
6
8
10
12
14
16
0
5
10
15
20
25

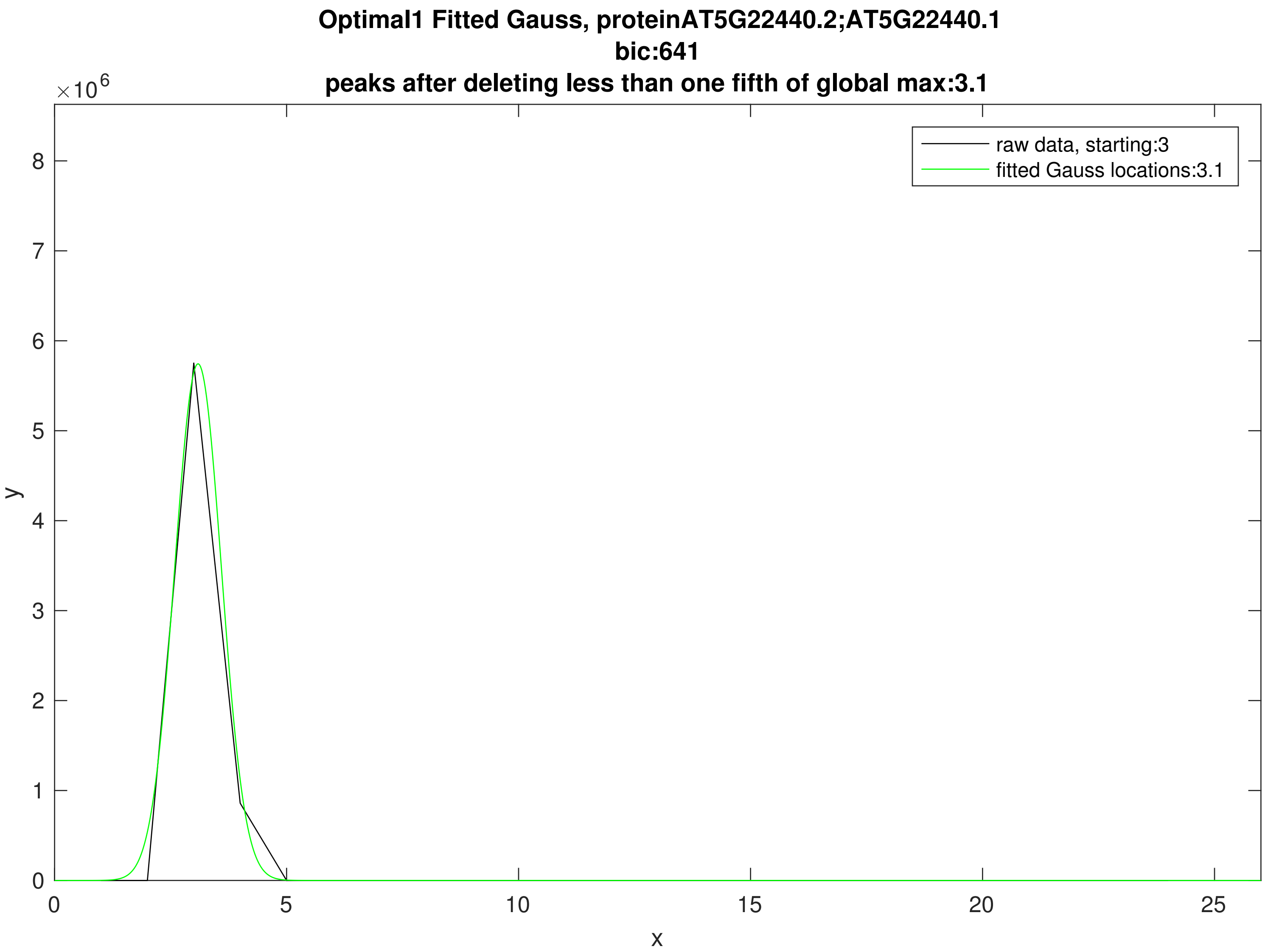

Optimal1 Fitted Gauss, proteinAT5G22440.2;AT5G22440.1
bic:641
peaks after deleting less than one fifth of global max:3.1
×10^6
raw data, starting:3
fitted Gauss locations:3.1
y
x
0
1
2
3
4
5
6
7
8
0
5
10
15
20
25

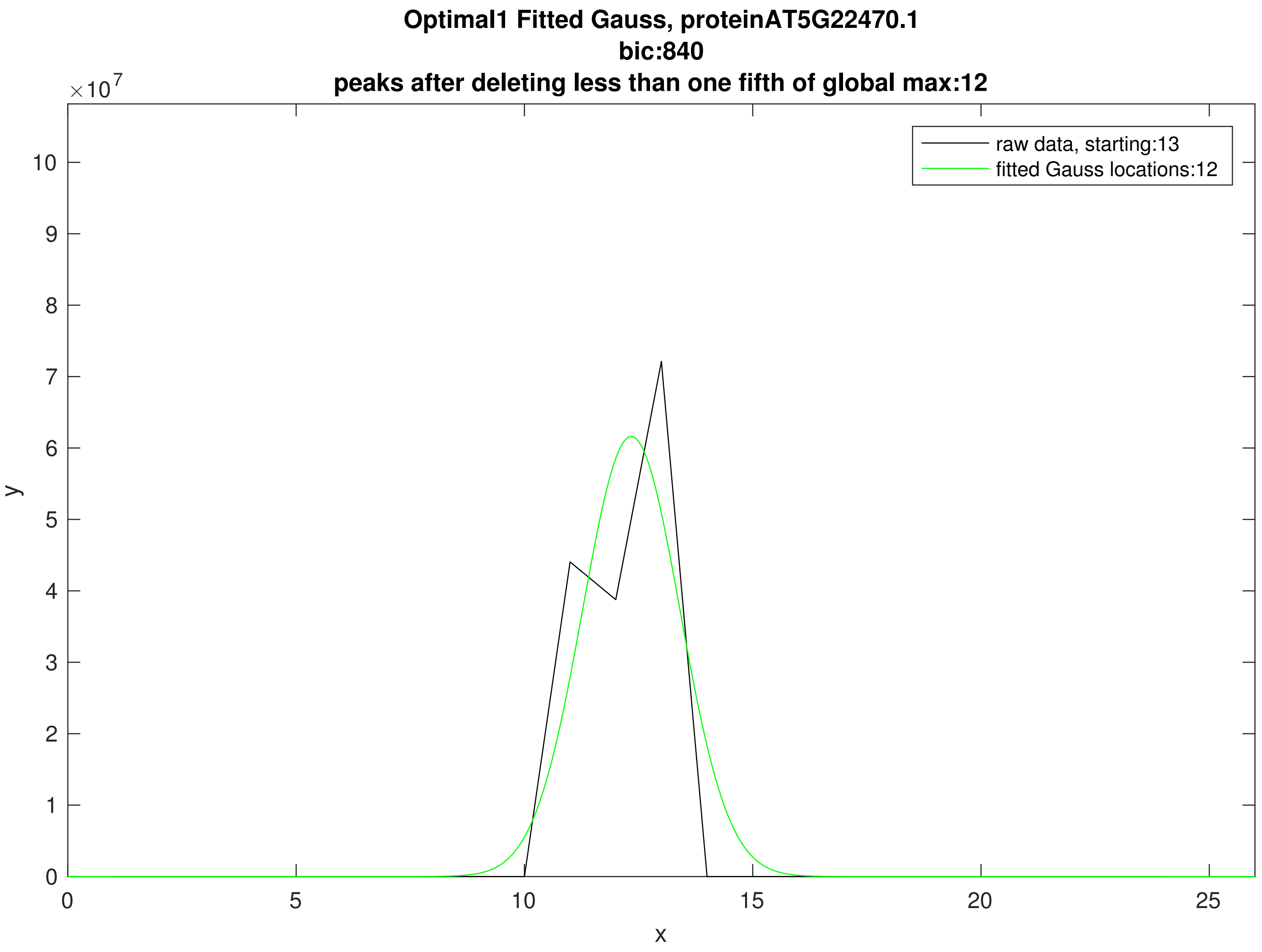

Optimal1 Fitted Gauss, proteinAT5G22470.1
bic:840
peaks after deleting less than one fifth of global max:12
×10^7
raw data, starting:13
fitted Gauss locations:12
0
1
2
3
4
5
6
7
8
9
10
y
0
5
10
15
20
25
x

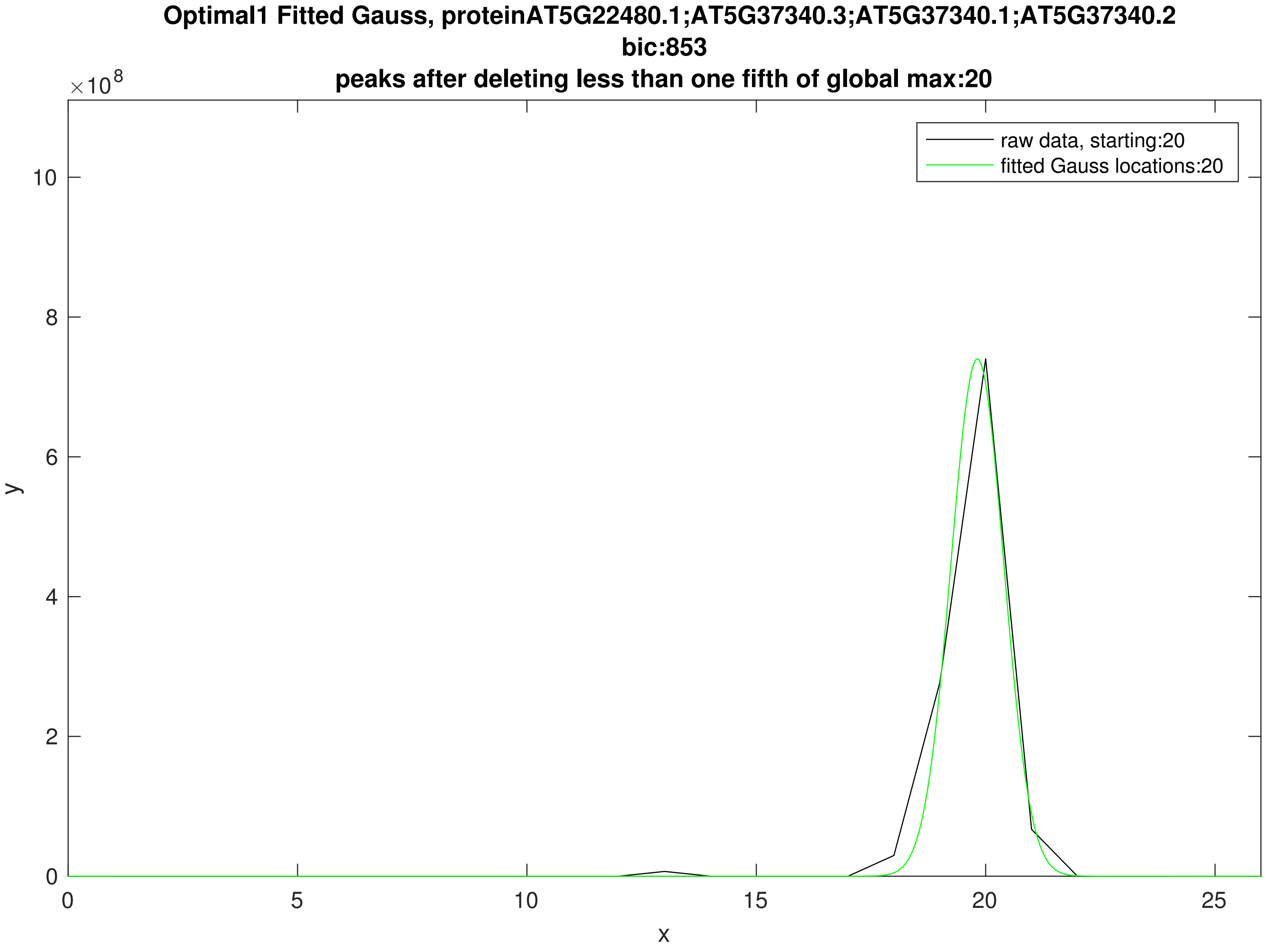
Optimal1 Fitted Gauss, proteinAT5G22480.1;AT5G37340.3;AT5G37340.1;AT5G37340.2
bic:853
peaks after deleting less than one fifth of global max:20
×10^8
raw data, starting:20
fitted Gauss locations:20
y
x
0
2
4
6
8
10
0
5
10
15
20
25

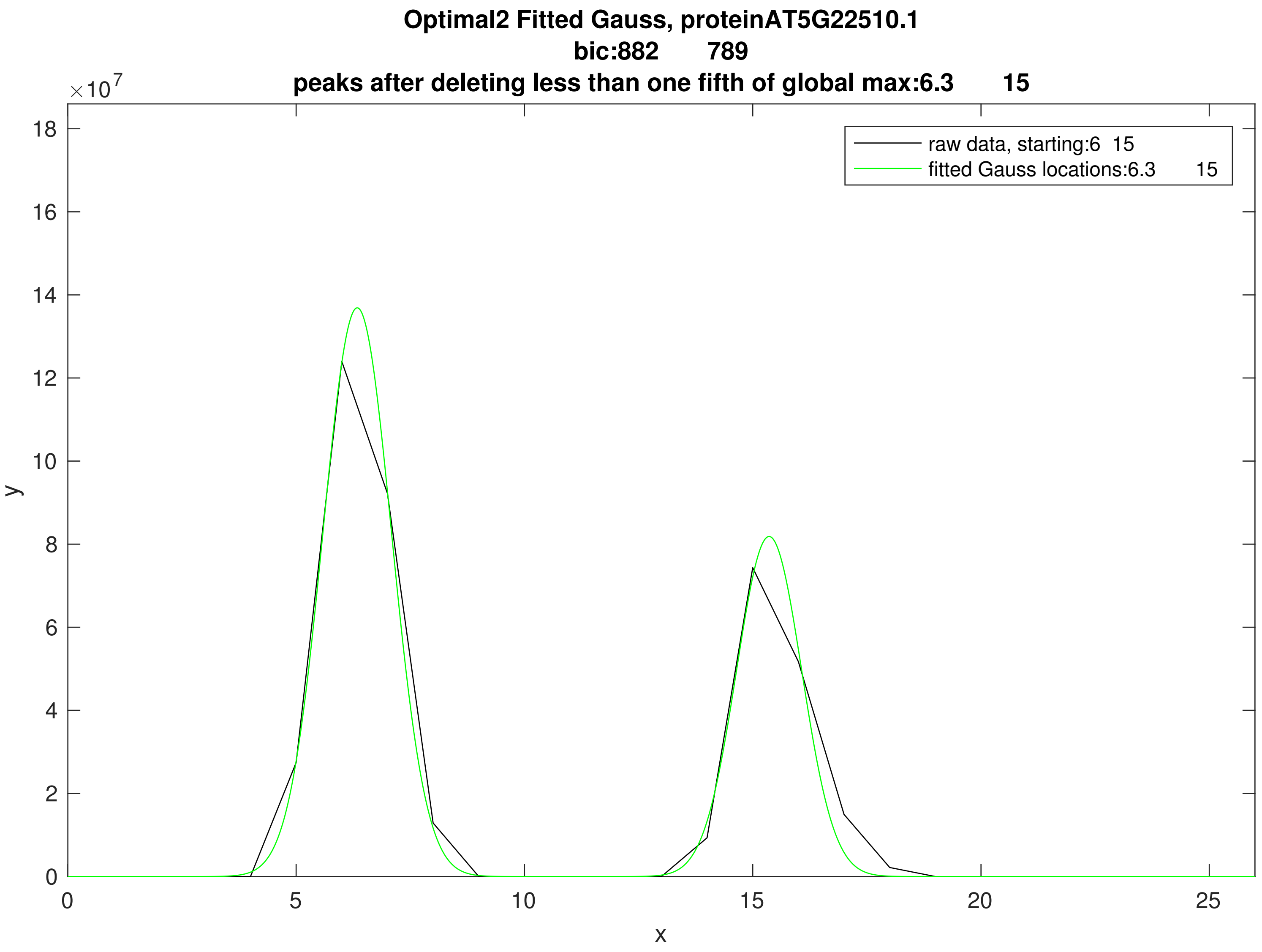

Optimal2 Fitted Gauss, proteinAT5G22510.1
bic:882 789
peaks after deleting less than one fifth of global max:6.3 15
×10^7
raw data, starting:6 15
fitted Gauss locations:6.3 15
y
x
0
2
4
6
8
10
12
14
16
18
0
5
10
15
20
25

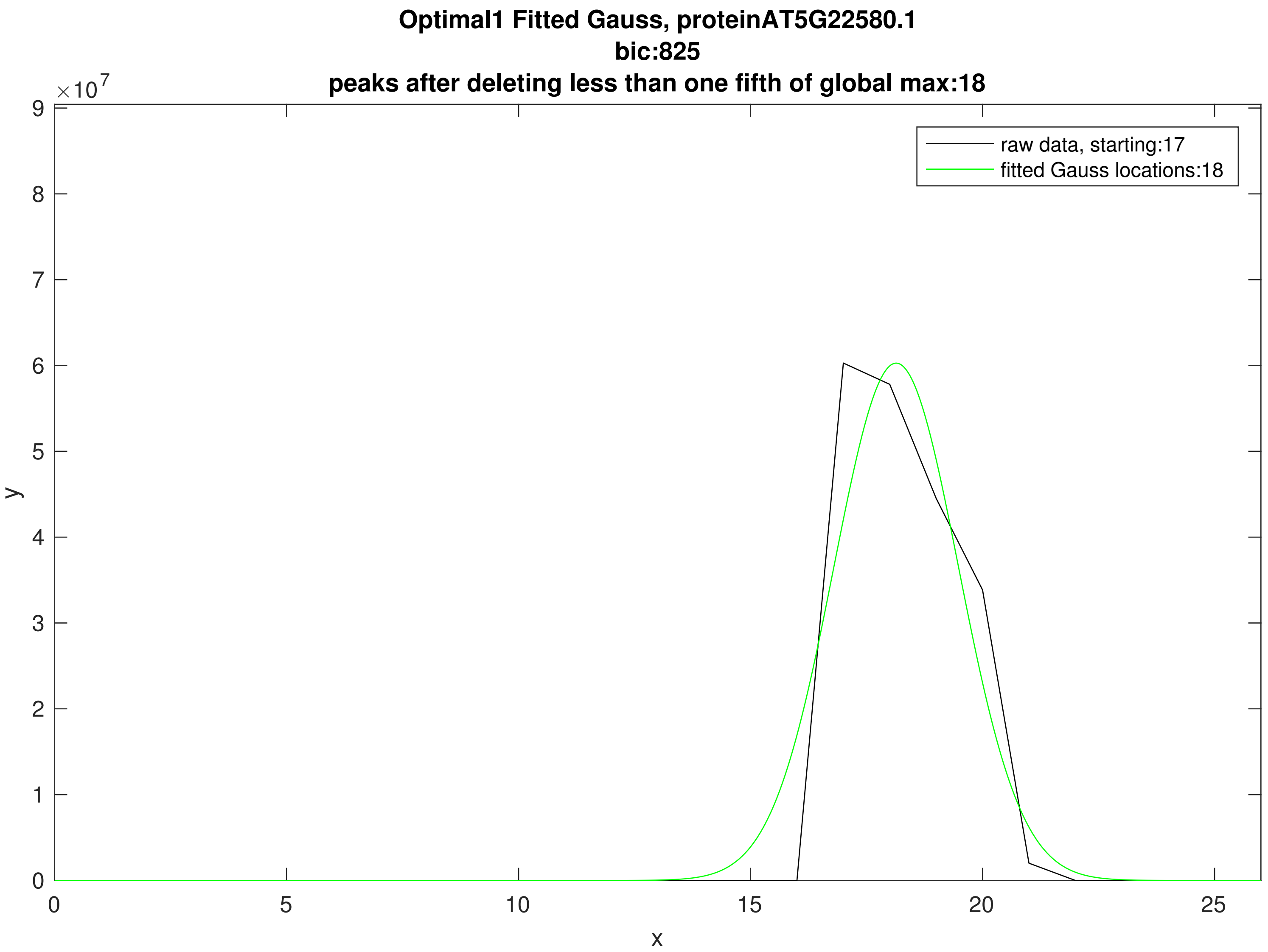

Optimal1 Fitted Gauss, proteinAT5G22580.1
bic:825
peaks after deleting less than one fifth of global max:18
×10^7
9
8
7
6
5
4
3
2
1
0
y
0
5
10
15
20
25
x
raw data, starting:17
fitted Gauss locations:18

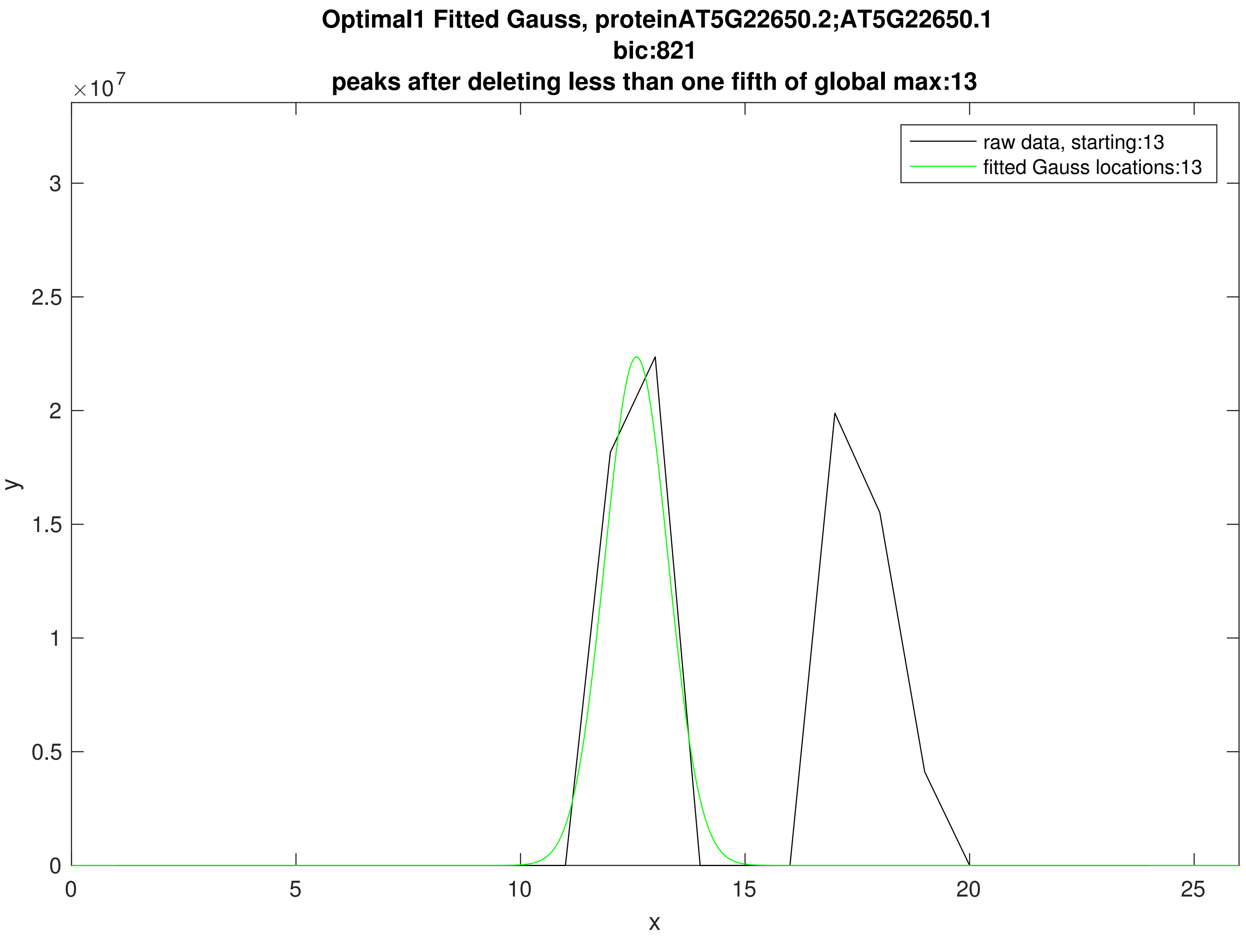
Optimal1 Fitted Gauss, proteinAT5G22650.2;AT5G22650.1
bic:821
peaks after deleting less than one fifth of global max:13
×10^7
raw data, starting:13
fitted Gauss locations:13
3
2.5
2
1.5
1
0.5
0
y
0
5
10
15
20
25
x

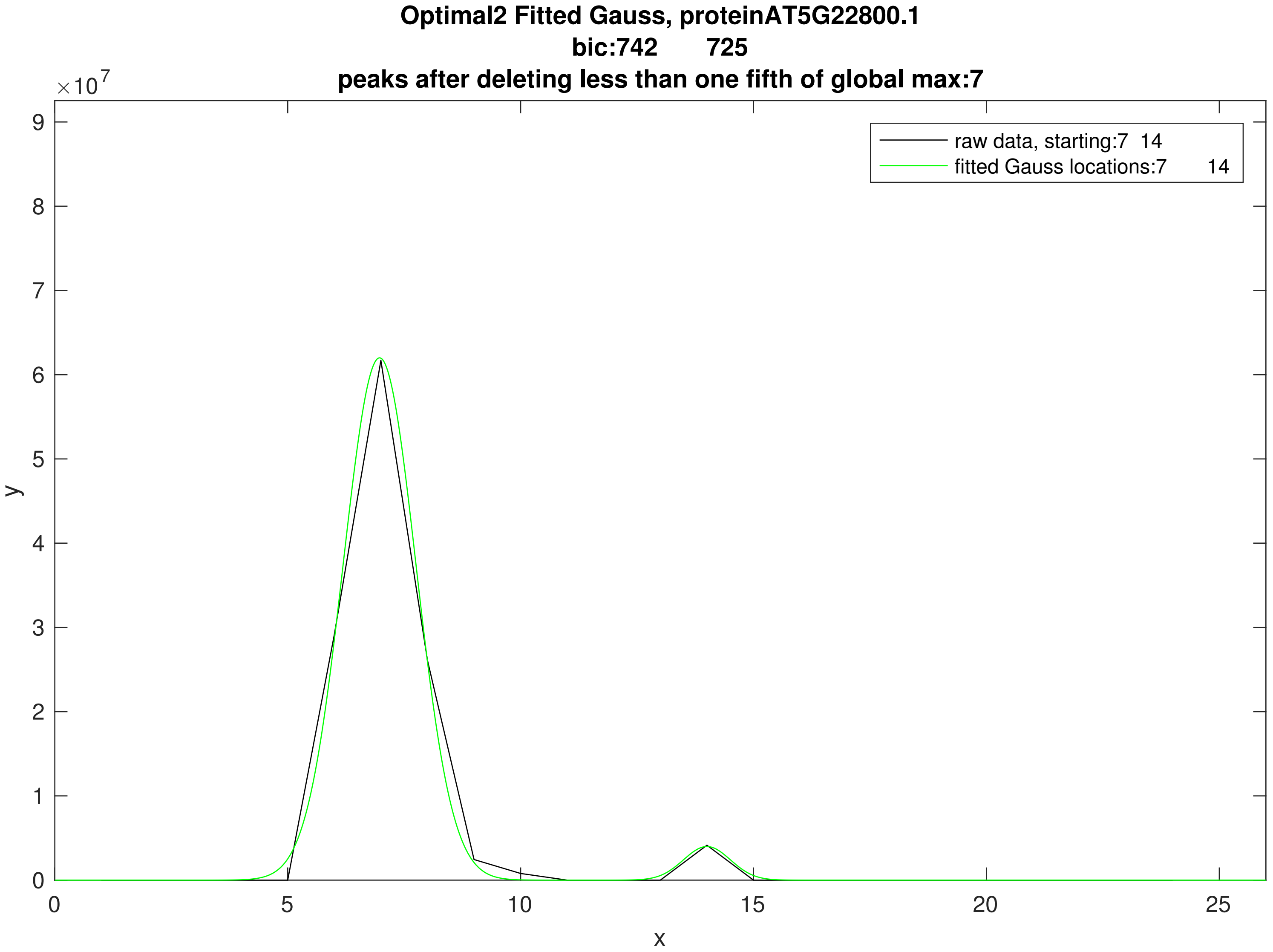
Optimal2 Fitted Gauss, proteinAT5G22800.1
bic:742 725
peaks after deleting less than one fifth of global max:7
×10^7
raw data, starting:7 14
fitted Gauss locations:7 14
y
x
0
1
2
3
4
5
6
7
8
9
0
5
10
15
20
25

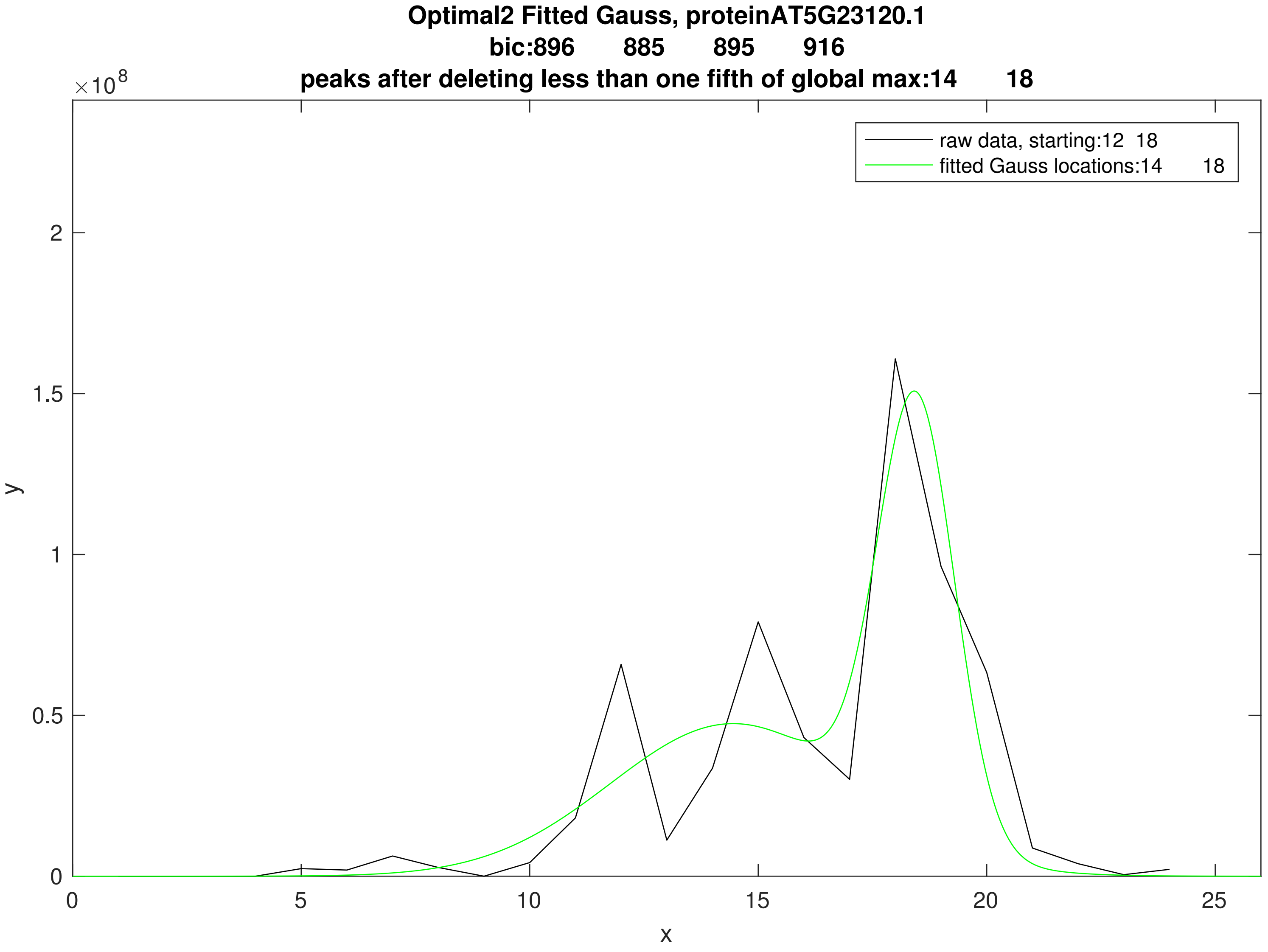
Optimal2 Fitted Gauss, proteinAT5G23120.1
bic:896 885 895 916
peaks after deleting less than one fifth of global max:14 18
raw data, starting:12 18
fitted Gauss locations:14 18
$\times 10^8$
y
x
0
0.5
1
1.5
2
5
10
15
20
25

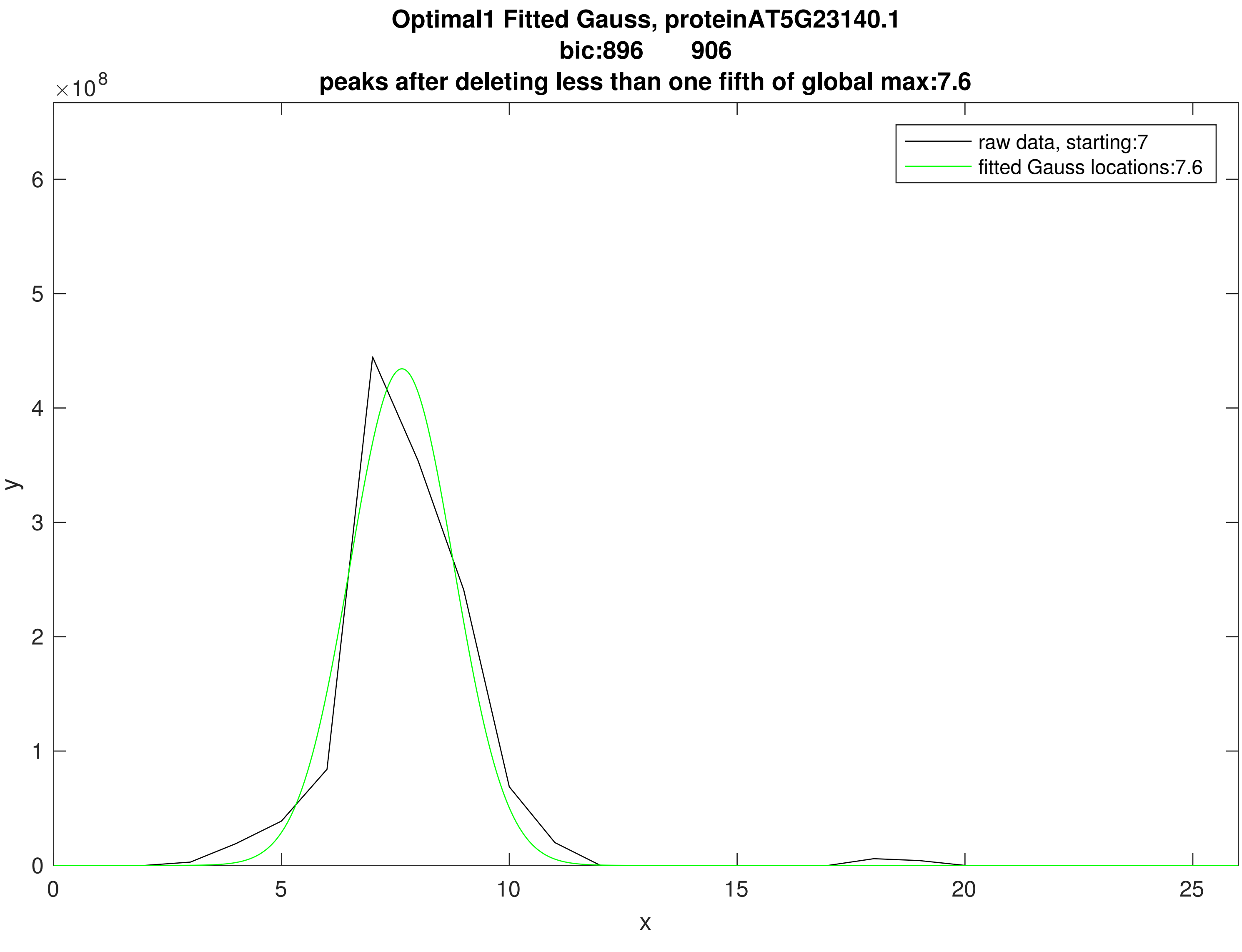

Optimal1 Fitted Gauss, proteinAT5G23140.1
bic:896 906
peaks after deleting less than one fifth of global max:7.6
×10^8
raw data, starting:7
fitted Gauss locations:7.6
y
x
0
1
2
3
4
5
6
0
5
10
15
20
25

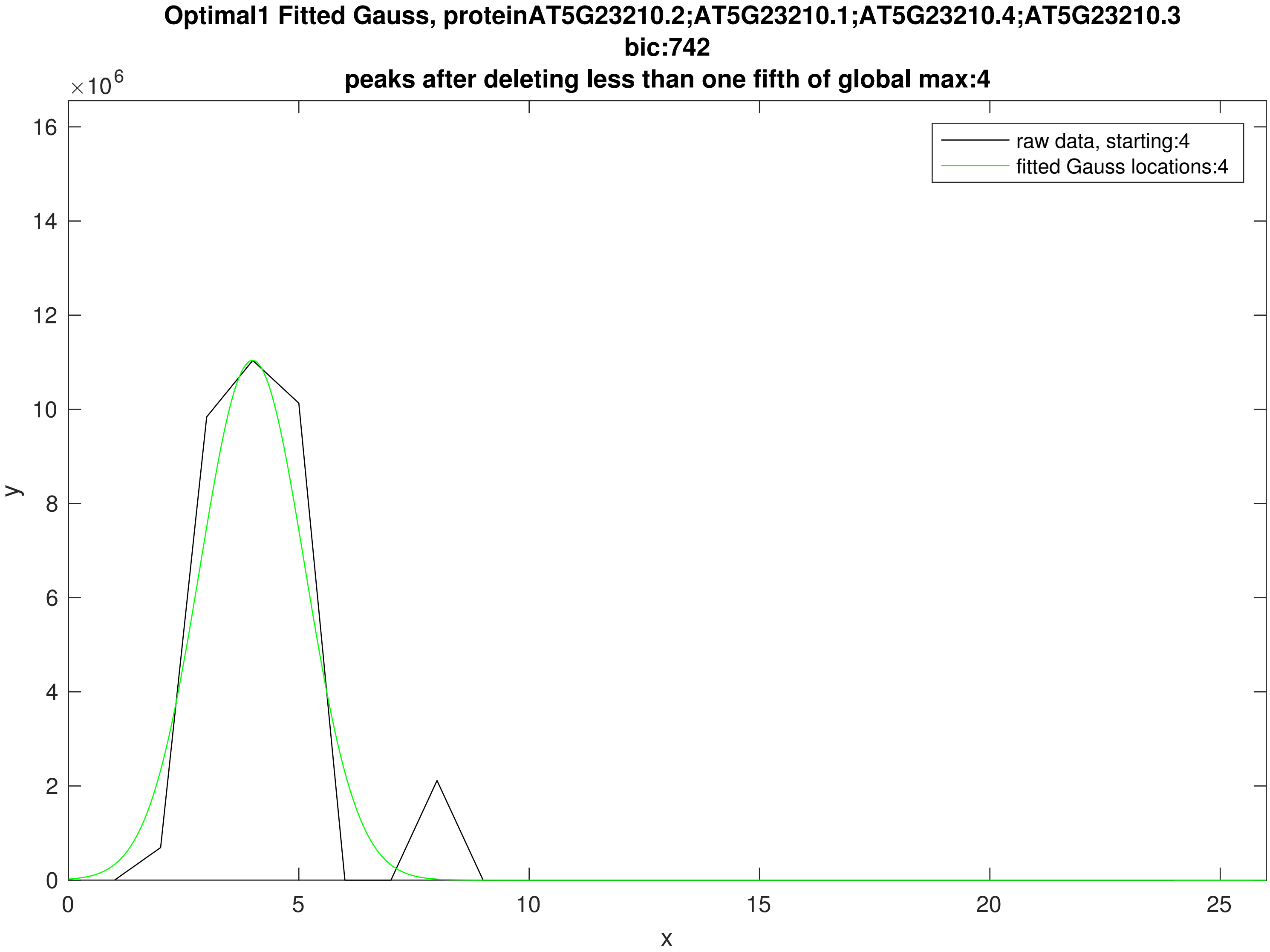
Optimal1 Fitted Gauss, proteinAT5G23210.2;AT5G23210.1;AT5G23210.4;AT5G23210.3
bic:742
peaks after deleting less than one fifth of global max:4
×10^6
raw data, starting:4
fitted Gauss locations:4
y
x
0
2
4
6
8
10
12
14
16
0
5
10
15
20
25

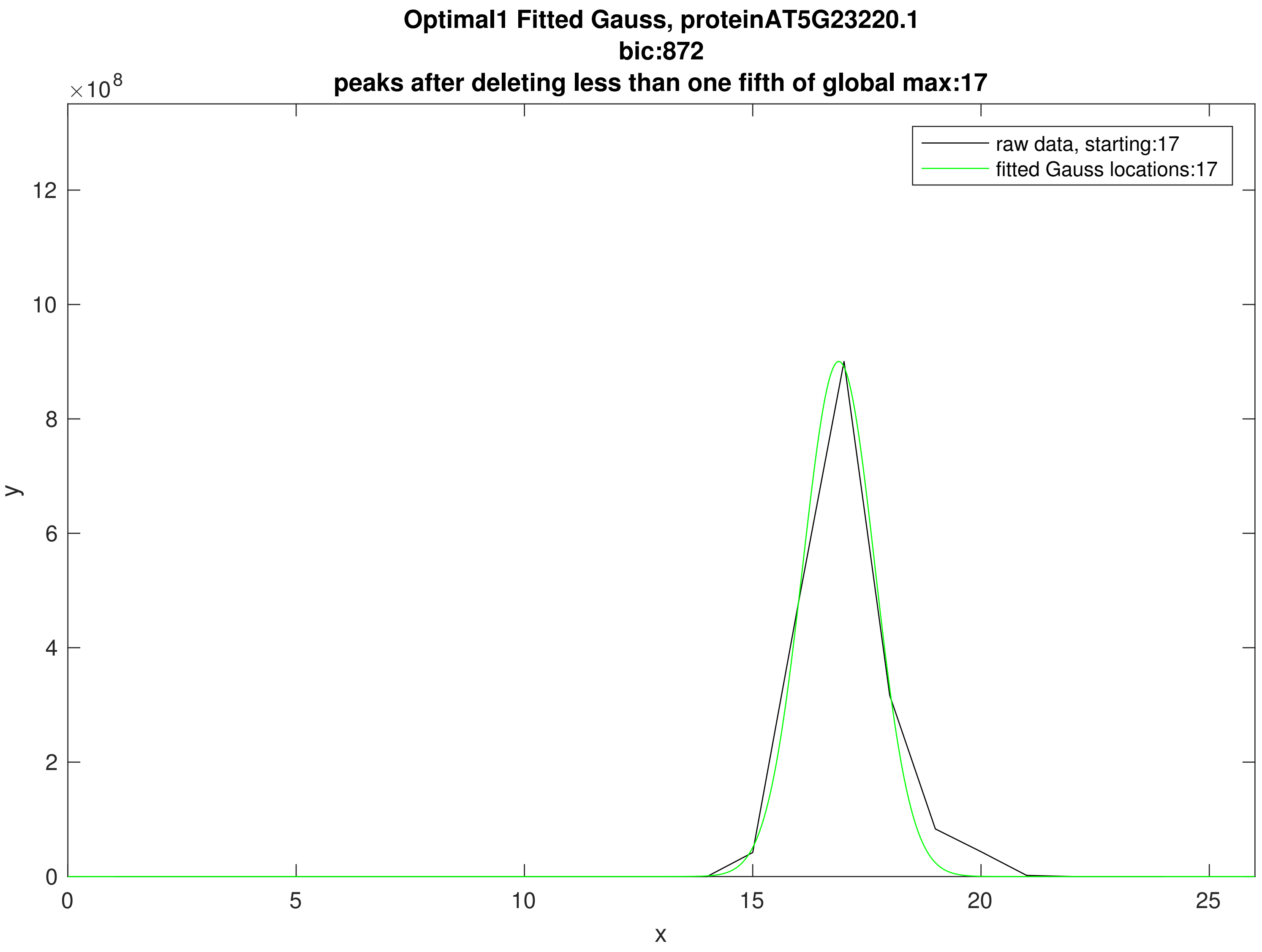

Optimal1 Fitted Gauss, proteinAT5G23220.1
bic:872
peaks after deleting less than one fifth of global max:17
$\times 10^8$
raw data, starting:17
fitted Gauss locations:17
y
x
0
2
4
6
8
10
12
0
5
10
15
20
25

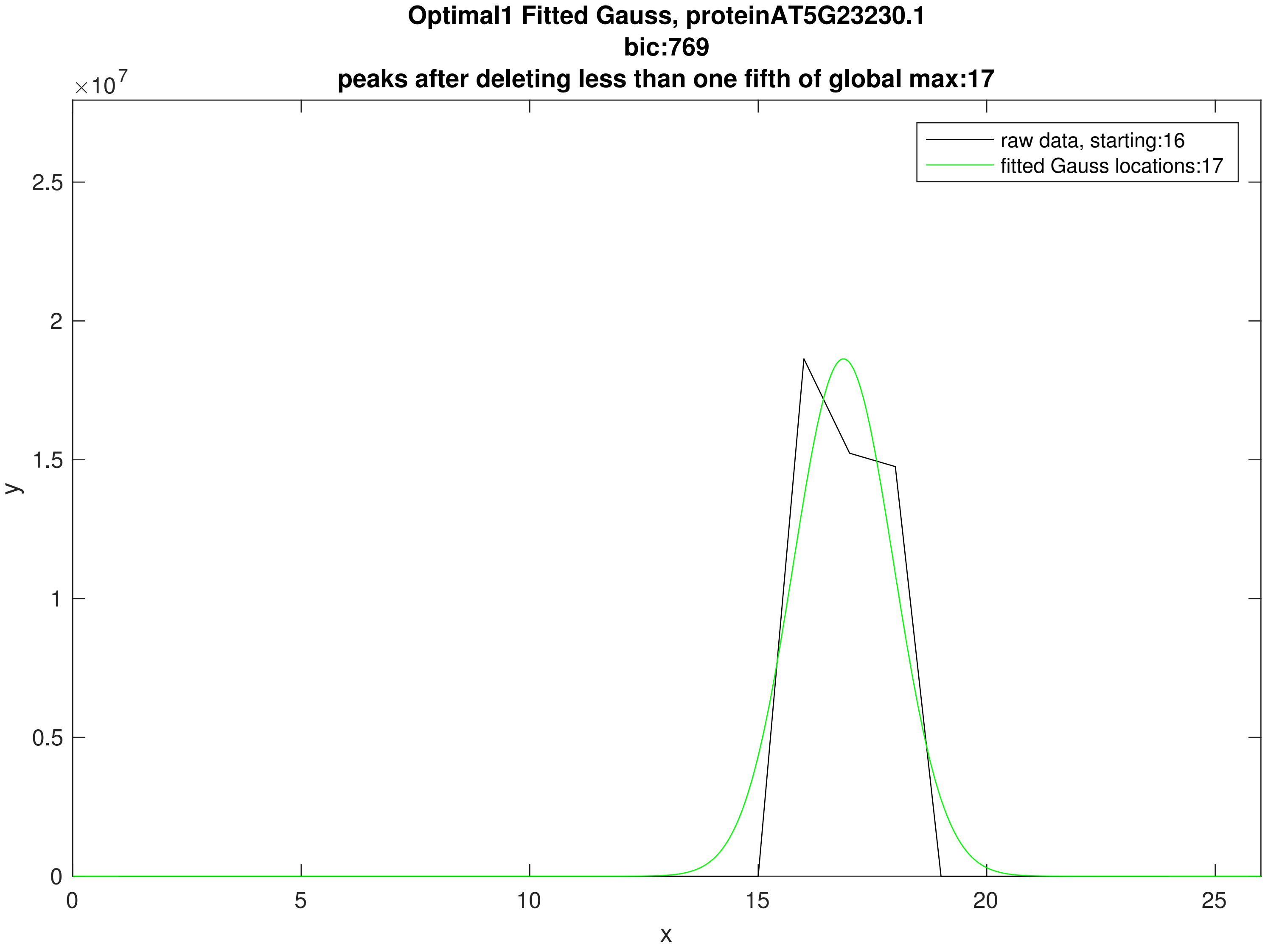

Optimal1 Fitted Gauss, proteinAT5G23230.1
bic:769
peaks after deleting less than one fifth of global max:17
×10^7
raw data, starting:16
fitted Gauss locations:17
2.5
2
1.5
1
0.5
0
y
0
5
10
15
20
25
x

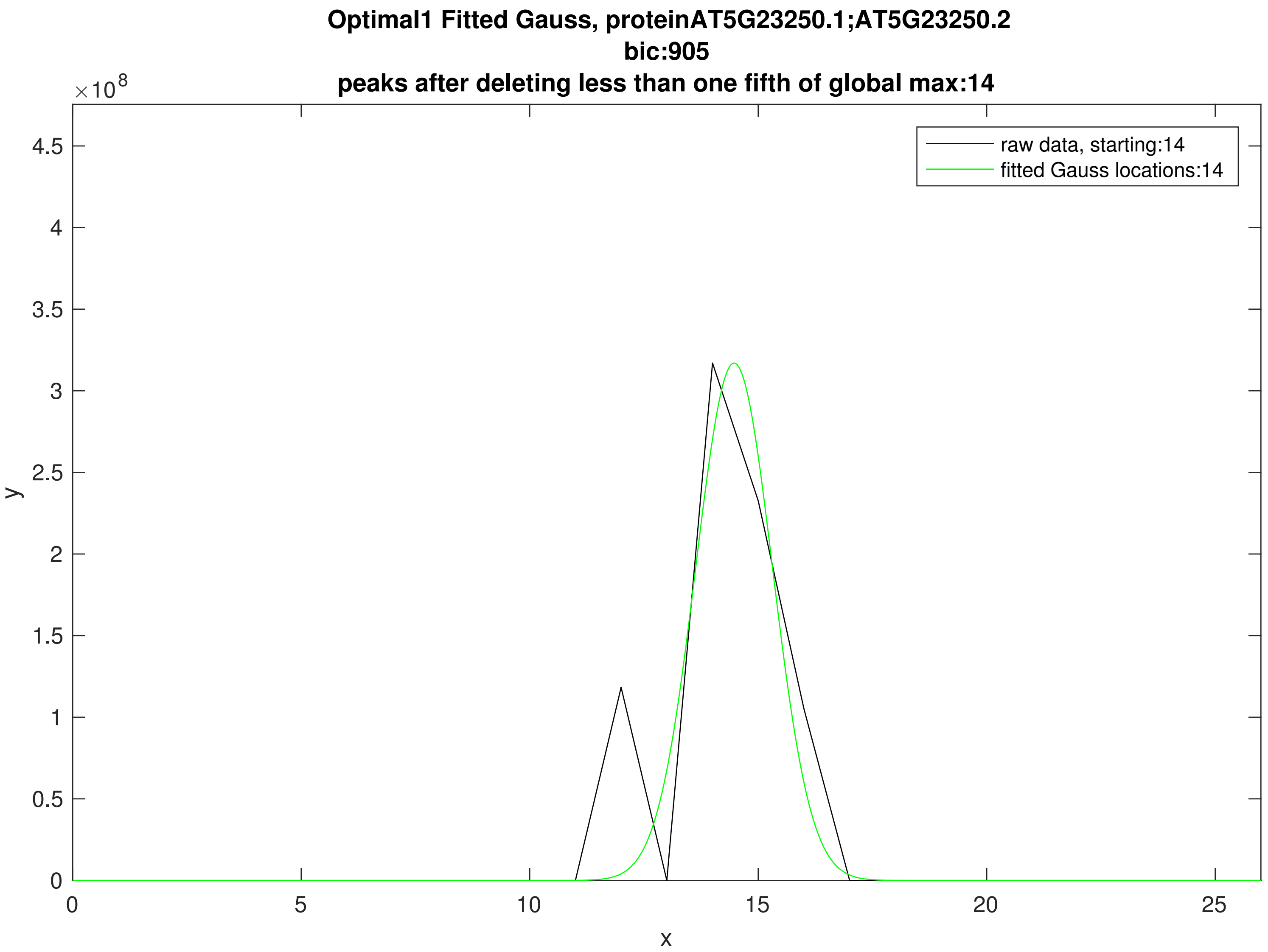
Optimal1 Fitted Gauss, proteinAT5G23250.1;AT5G23250.2
bic:905
peaks after deleting less than one fifth of global max:14
×10^8
raw data, starting:14
fitted Gauss locations:14
y
x
0
0.5
1
1.5
2
2.5
3
3.5
4
4.5
0
5
10
15
20
25

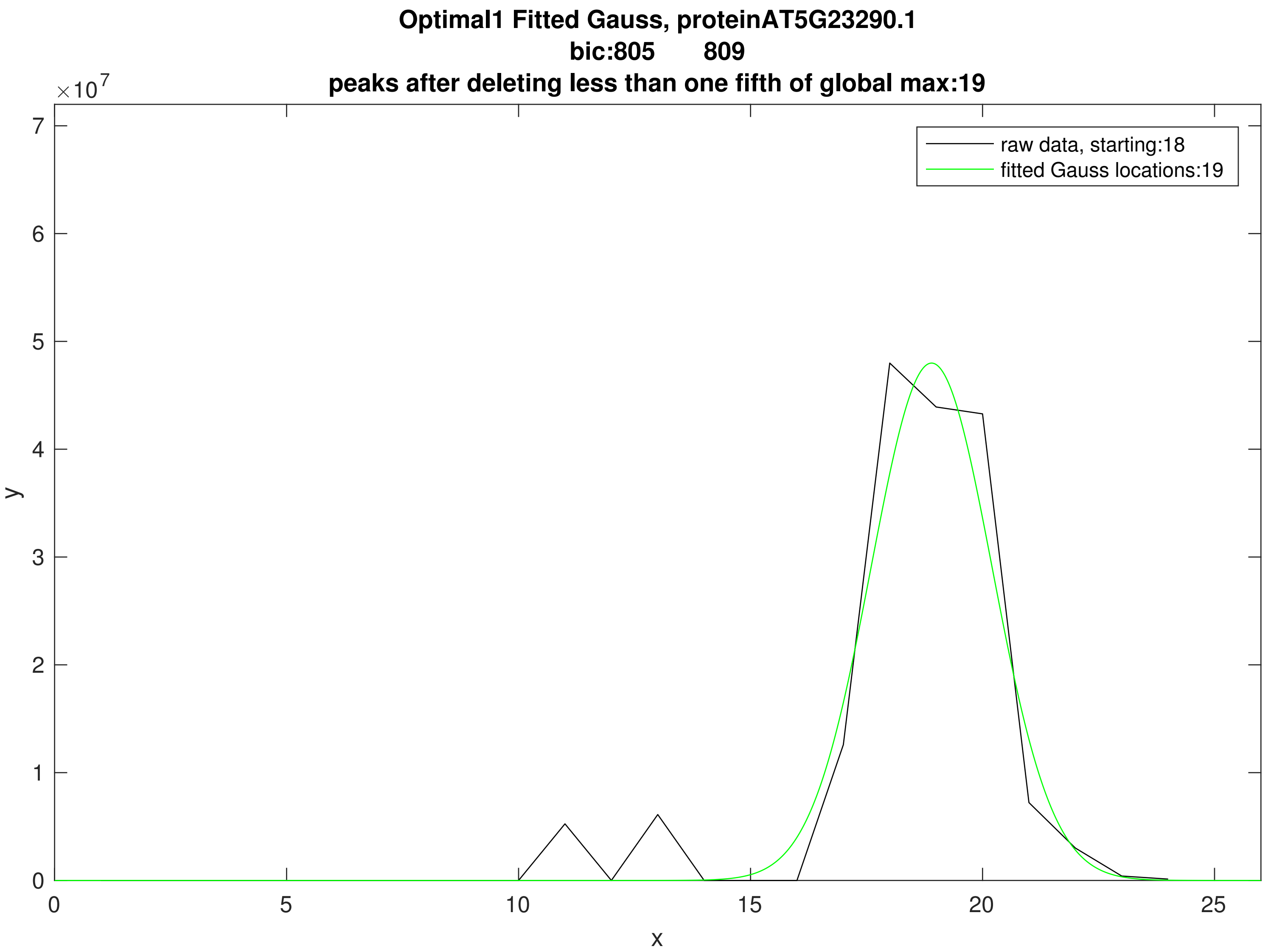

Optimal1 Fitted Gauss, proteinAT5G23290.1
bic:805 809
peaks after deleting less than one fifth of global max:19
×10^7
7
6
5
4
3
2
1
0
y
0
5
10
15
20
25
x
raw data, starting:18
fitted Gauss locations:19

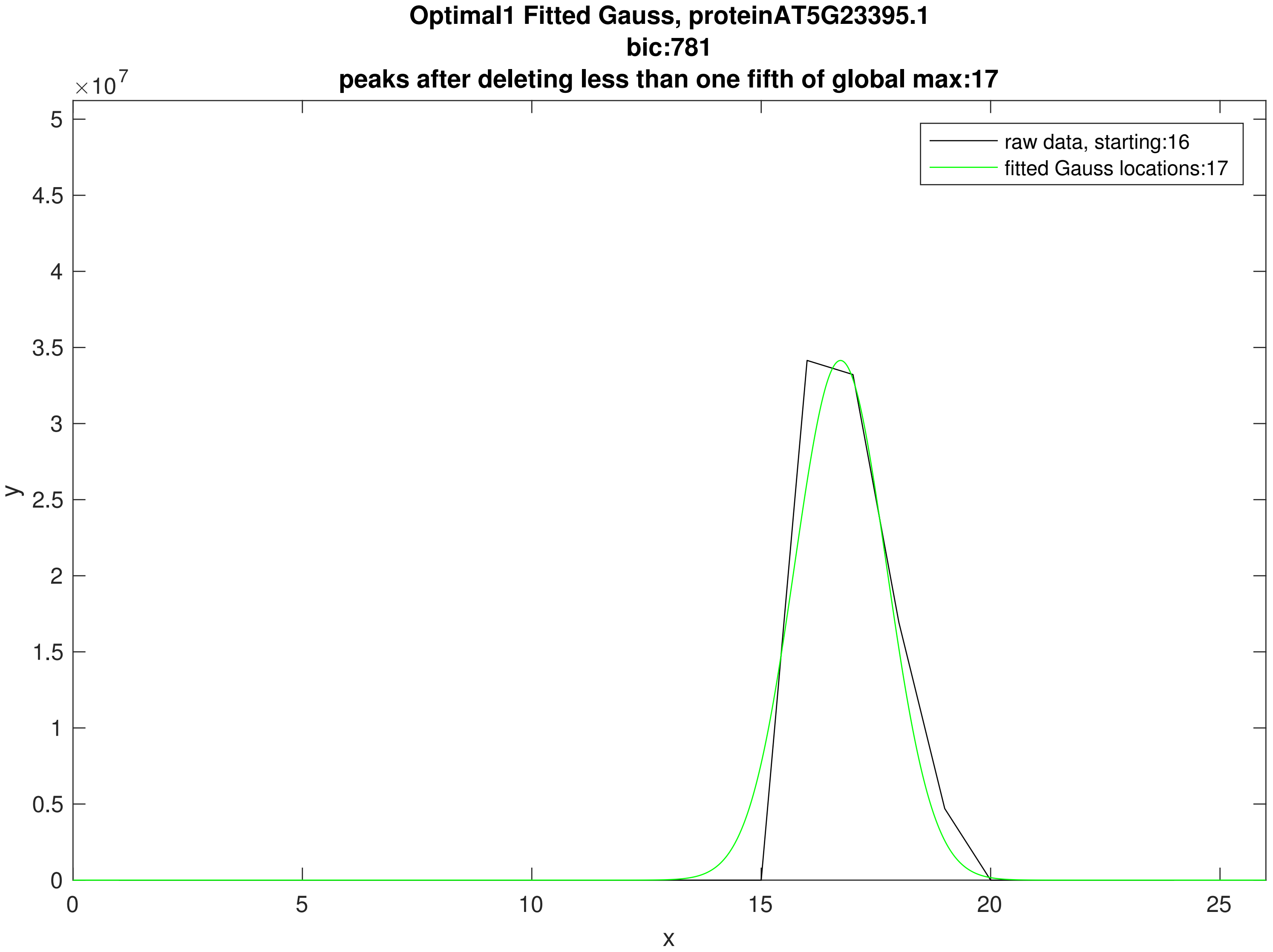
Optimal1 Fitted Gauss, proteinAT5G23395.1
bic:781
peaks after deleting less than one fifth of global max:17
×10^7
y
x
raw data, starting:16
fitted Gauss locations:17
0
0.5
1
1.5
2
2.5
3
3.5
4
4.5
5
0
5
10
15
20
25

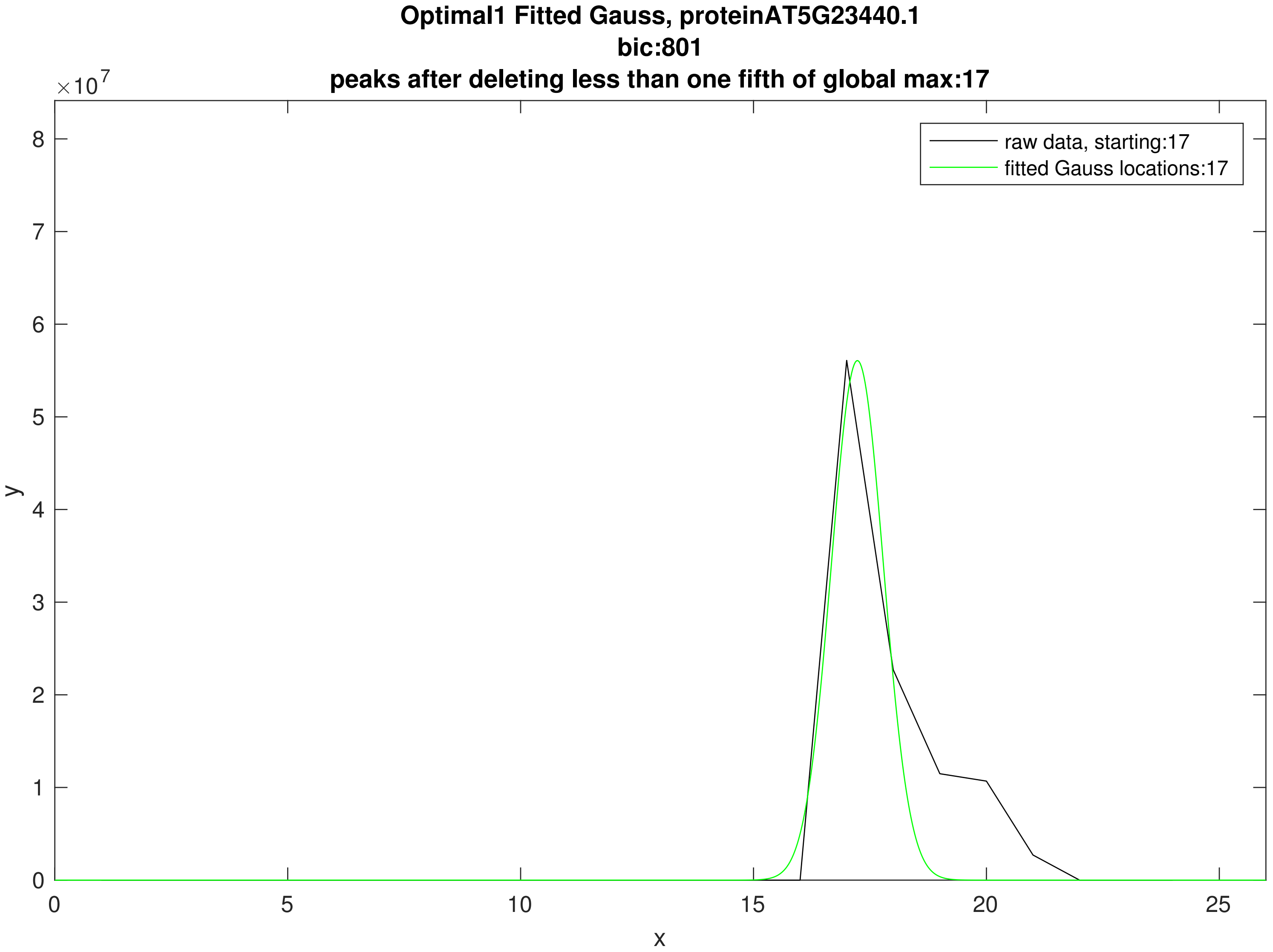

Optimal1 Fitted Gauss, proteinAT5G23440.1
bic:801
peaks after deleting less than one fifth of global max:17
×10^7
raw data, starting:17
fitted Gauss locations:17
y
x
0
1
2
3
4
5
6
7
8
0
5
10
15
20
25

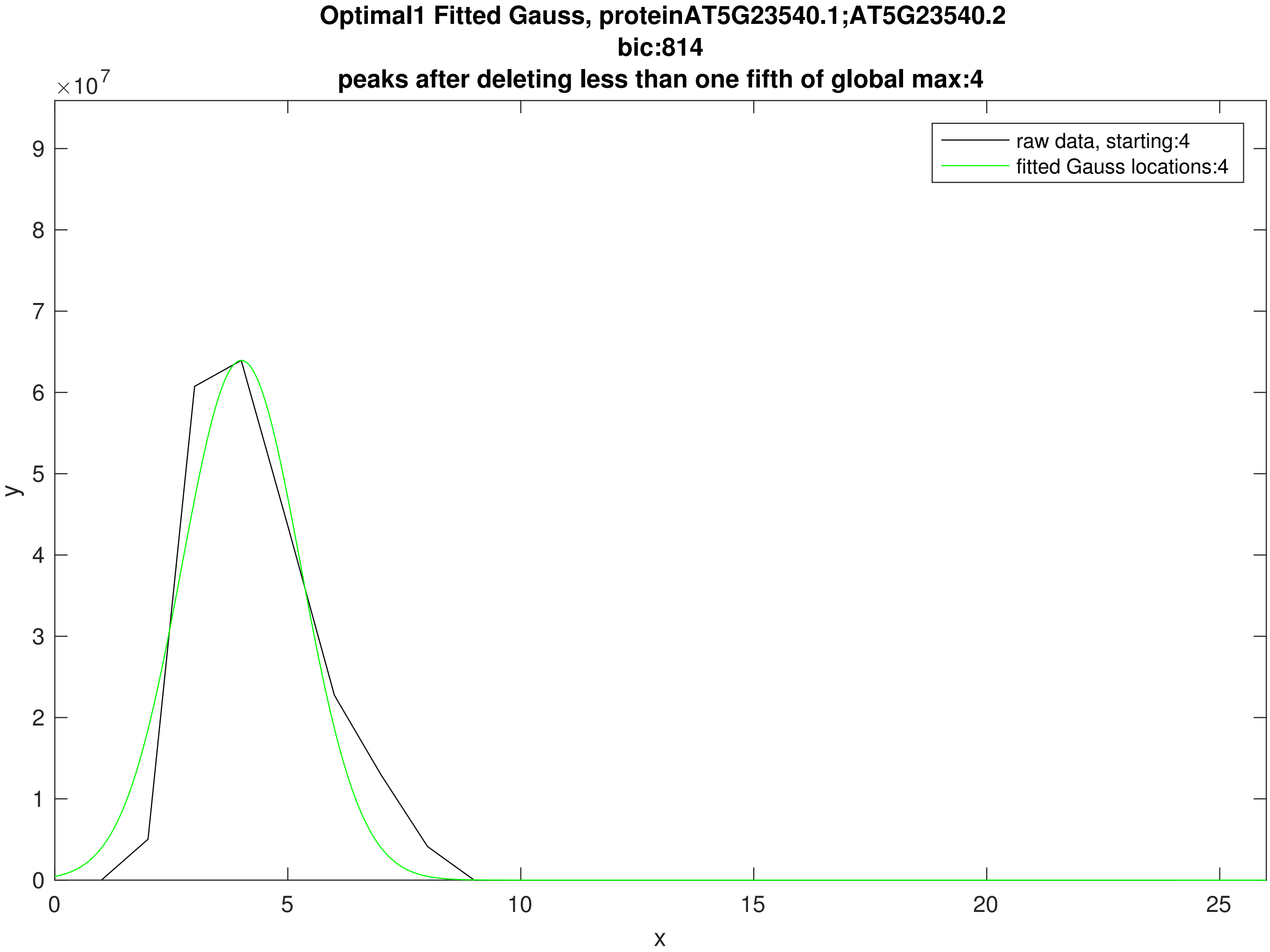
Optimal1 Fitted Gauss, proteinAT5G23540.1;AT5G23540.2
bic:814
peaks after deleting less than one fifth of global max:4
×10^7
raw data, starting:4
fitted Gauss locations:4
y
x
0
1
2
3
4
5
6
7
8
9
0
5
10
15
20
25

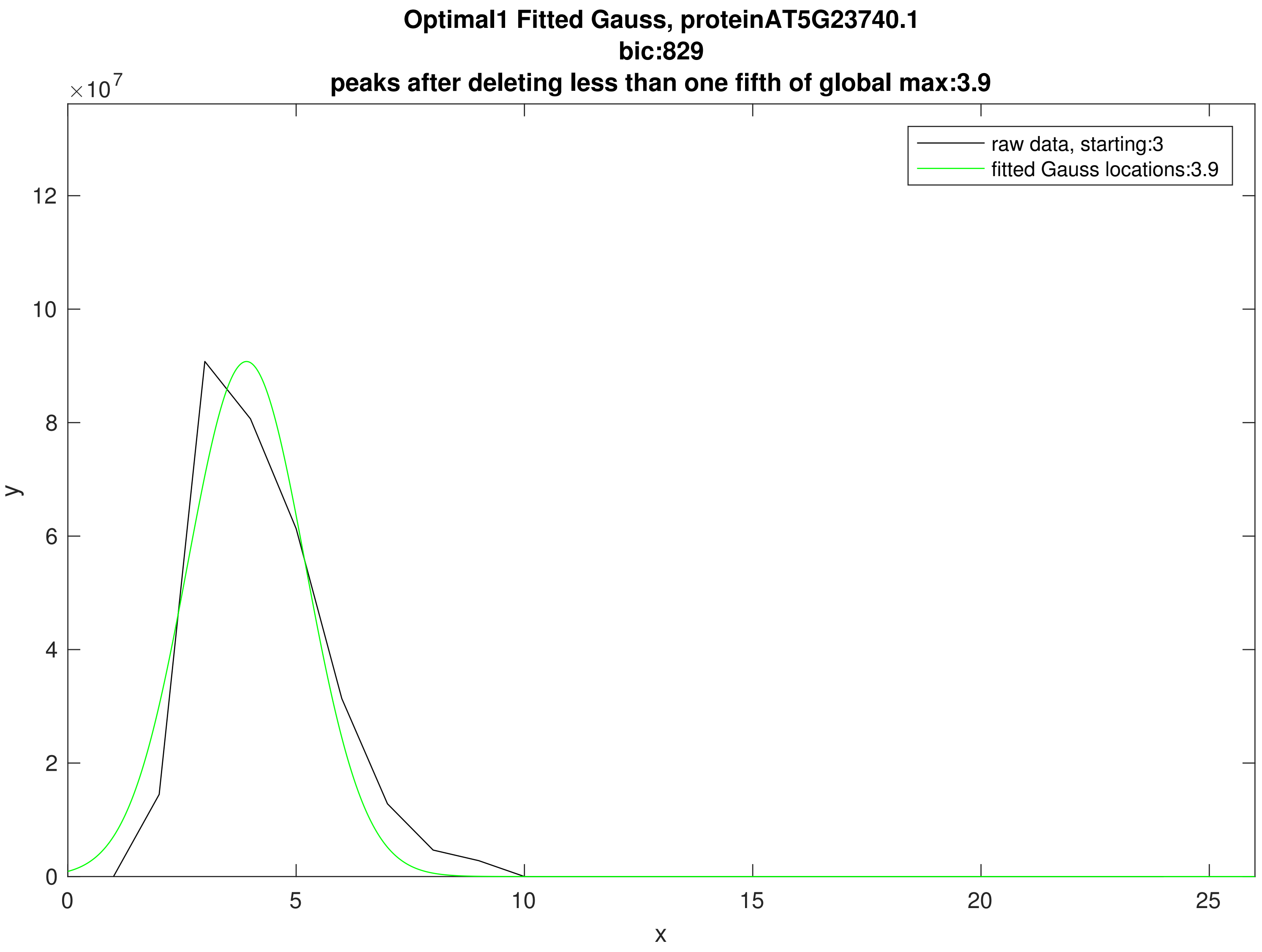

Optimal1 Fitted Gauss, proteinAT5G23740.1
bic:829
peaks after deleting less than one fifth of global max:3.9
×10^7
raw data, starting:3
fitted Gauss locations:3.9
y
x
0
2
4
6
8
10
12
0
5
10
15
20
25

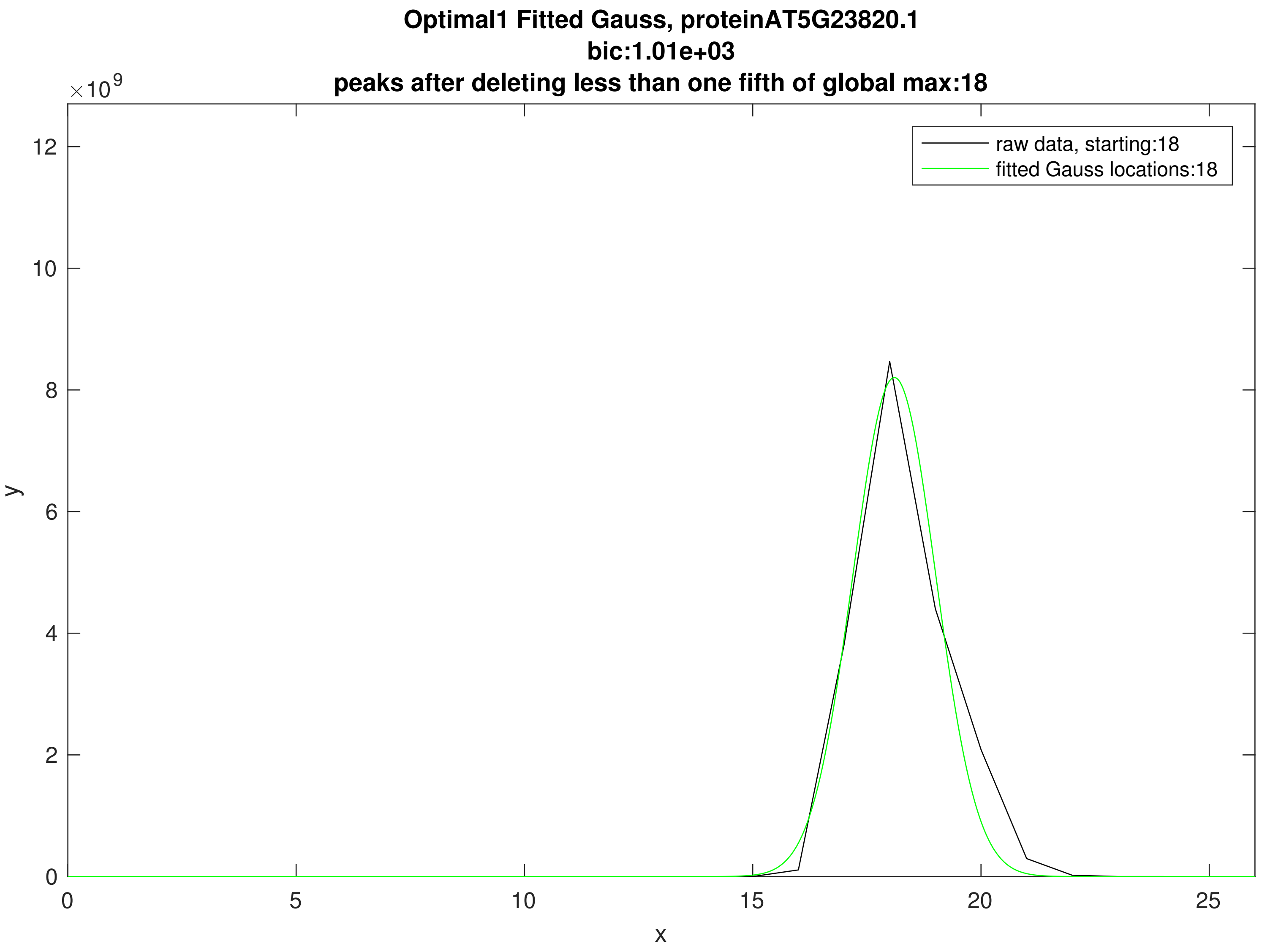

Optimal1 Fitted Gauss, proteinAT5G23820.1
bic:1.01e+03
peaks after deleting less than one fifth of global max:18
×10^9
12
10
8
6
4
2
0
y
0
5
10
15
20
25
x
raw data, starting:18
fitted Gauss locations:18

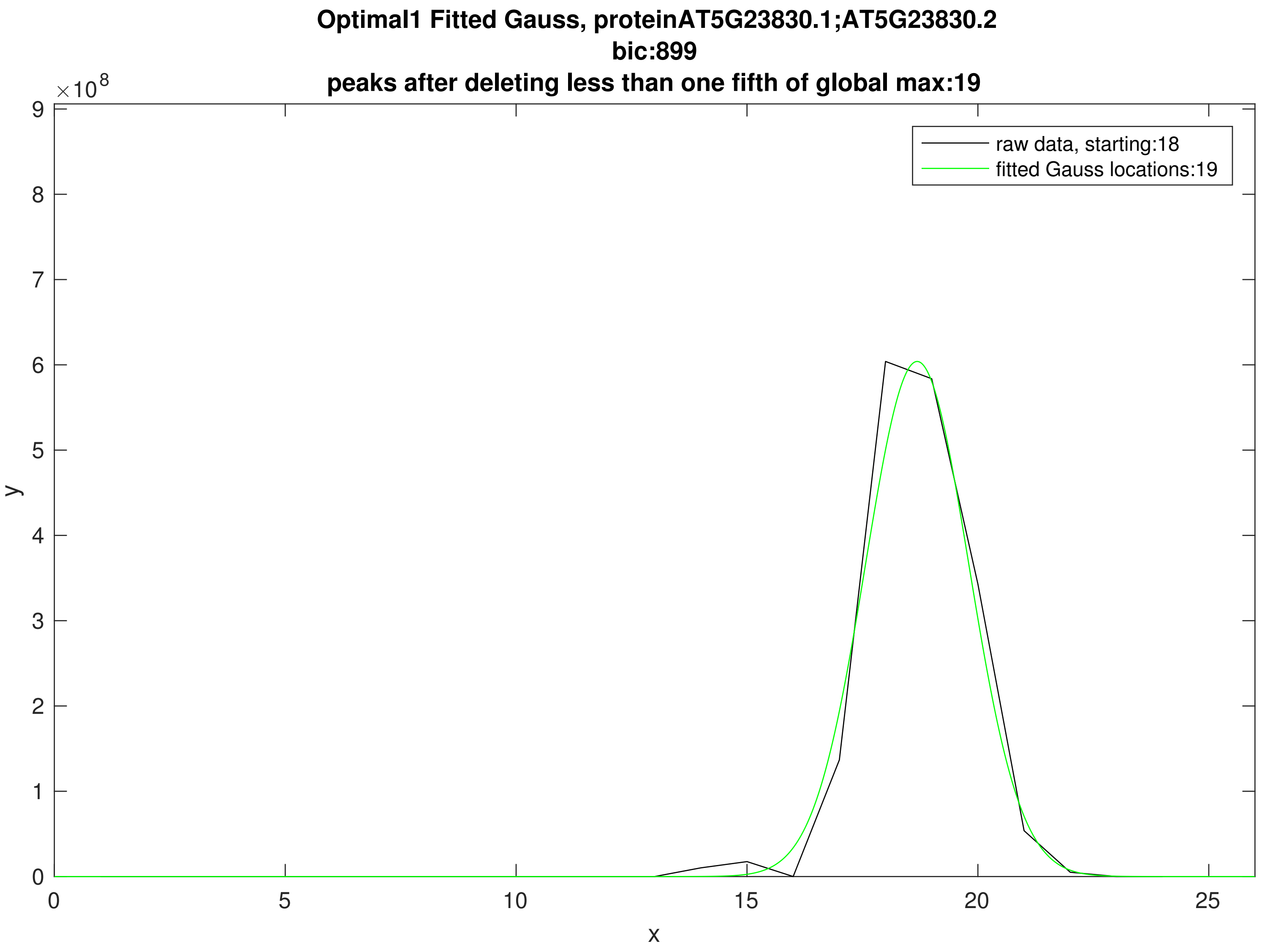

Optimal1 Fitted Gauss, proteinAT5G23830.1;AT5G23830.2
bic:899
peaks after deleting less than one fifth of global max:19
×10^8
0
1
2
3
4
5
6
7
8
9
y
0
5
10
15
20
25
x
raw data, starting:18
fitted Gauss locations:19

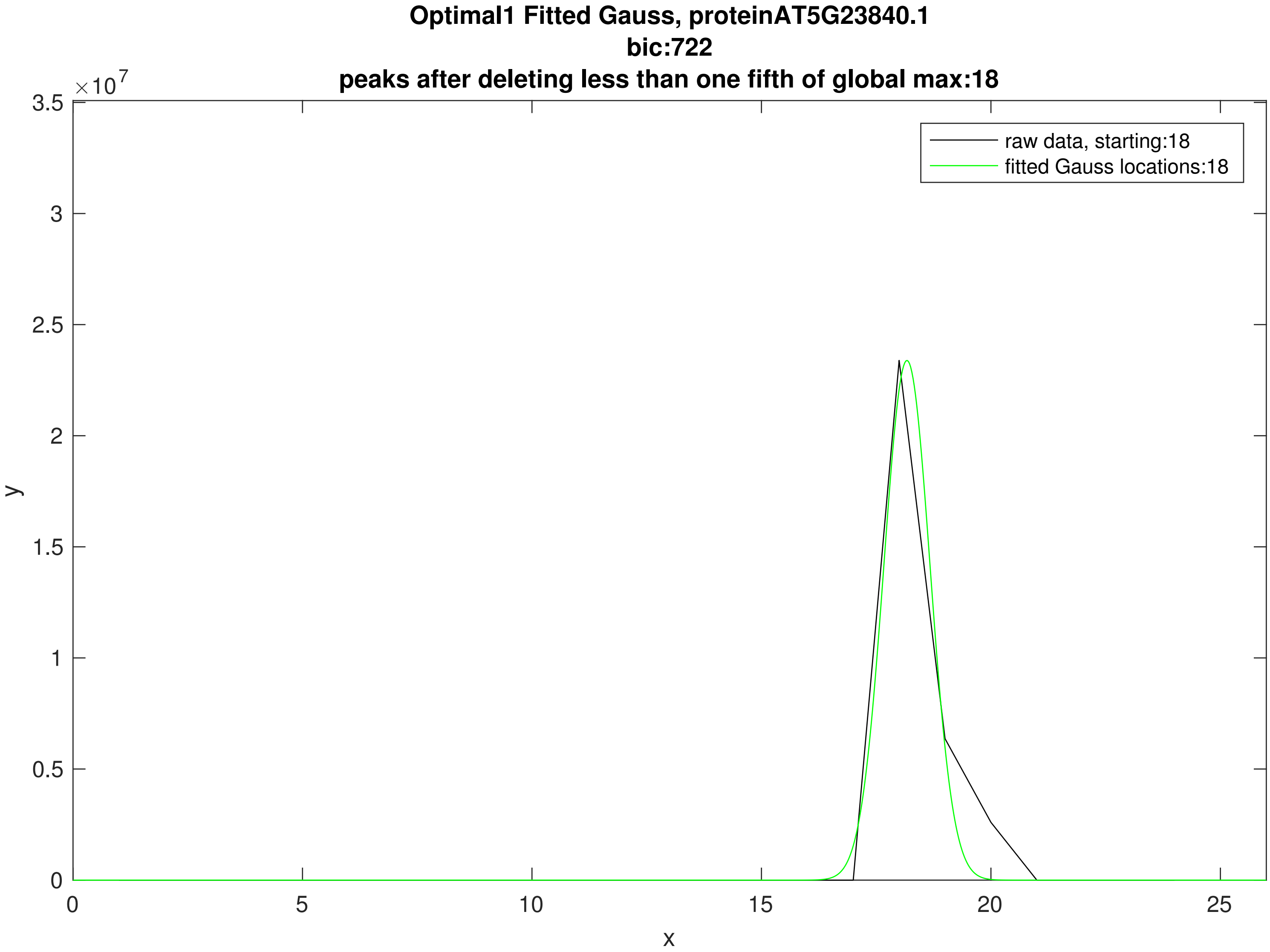
Optimal1 Fitted Gauss, proteinAT5G23840.1
bic:722
peaks after deleting less than one fifth of global max:18
×10^7
3.5
3
2.5
2
1.5
1
0.5
0
y
0
5
10
15
20
25
x
raw data, starting:18
fitted Gauss locations:18

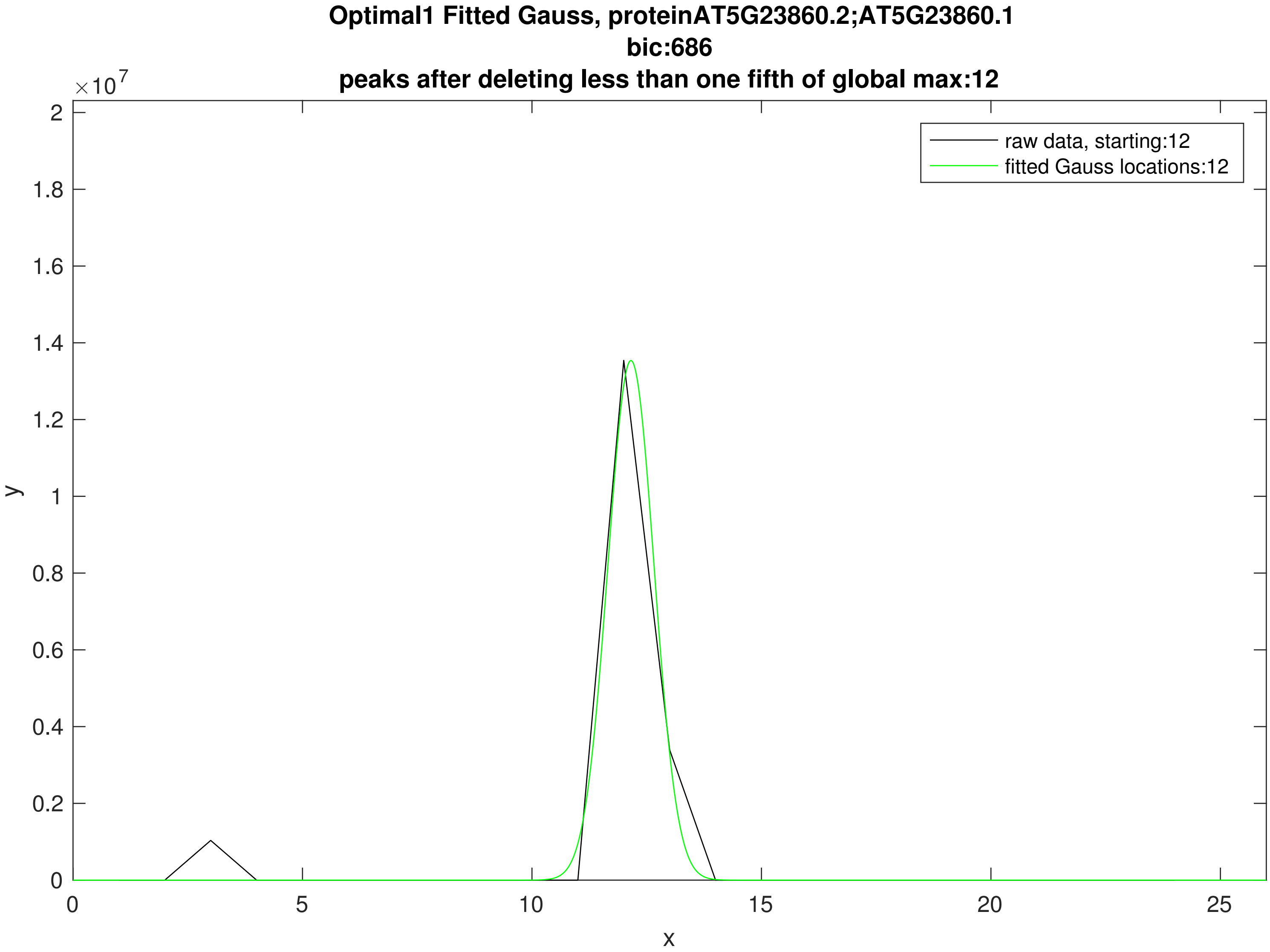

Optimal1 Fitted Gauss, proteinAT5G23860.2;AT5G23860.1
bic:686
peaks after deleting less than one fifth of global max:12
×10^7
2
1.8
1.6
1.4
1.2
1
0.8
0.6
0.4
0.2
0
y
0
5
10
15
20
25
x
raw data, starting:12
fitted Gauss locations:12

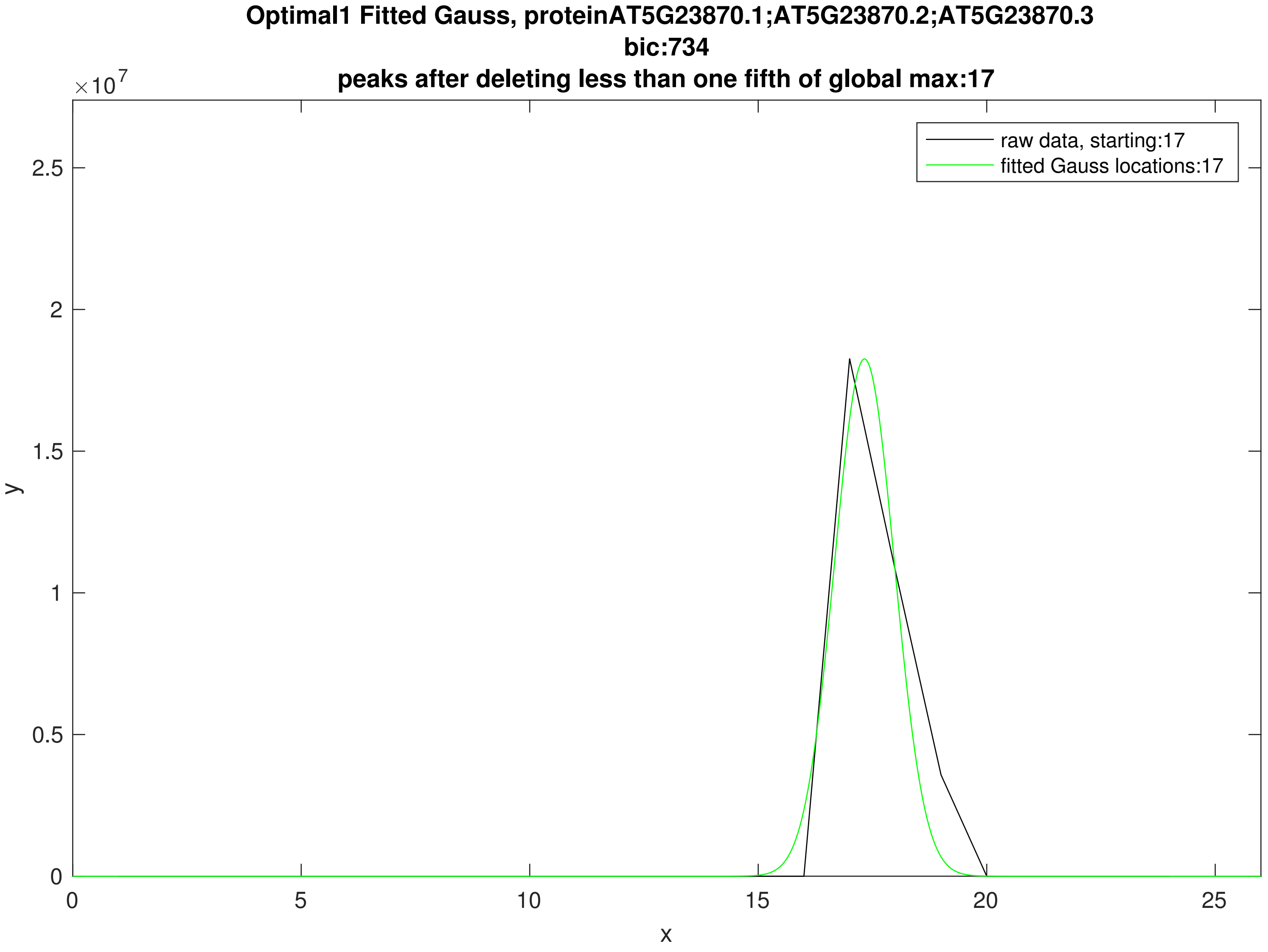
Optimal1 Fitted Gauss, proteinAT5G23870.1;AT5G23870.2;AT5G23870.3
bic:734
peaks after deleting less than one fifth of global max:17
×10^7
raw data, starting:17
fitted Gauss locations:17
y
x
0
0.5
1
1.5
2
2.5
5
10
15
20
25

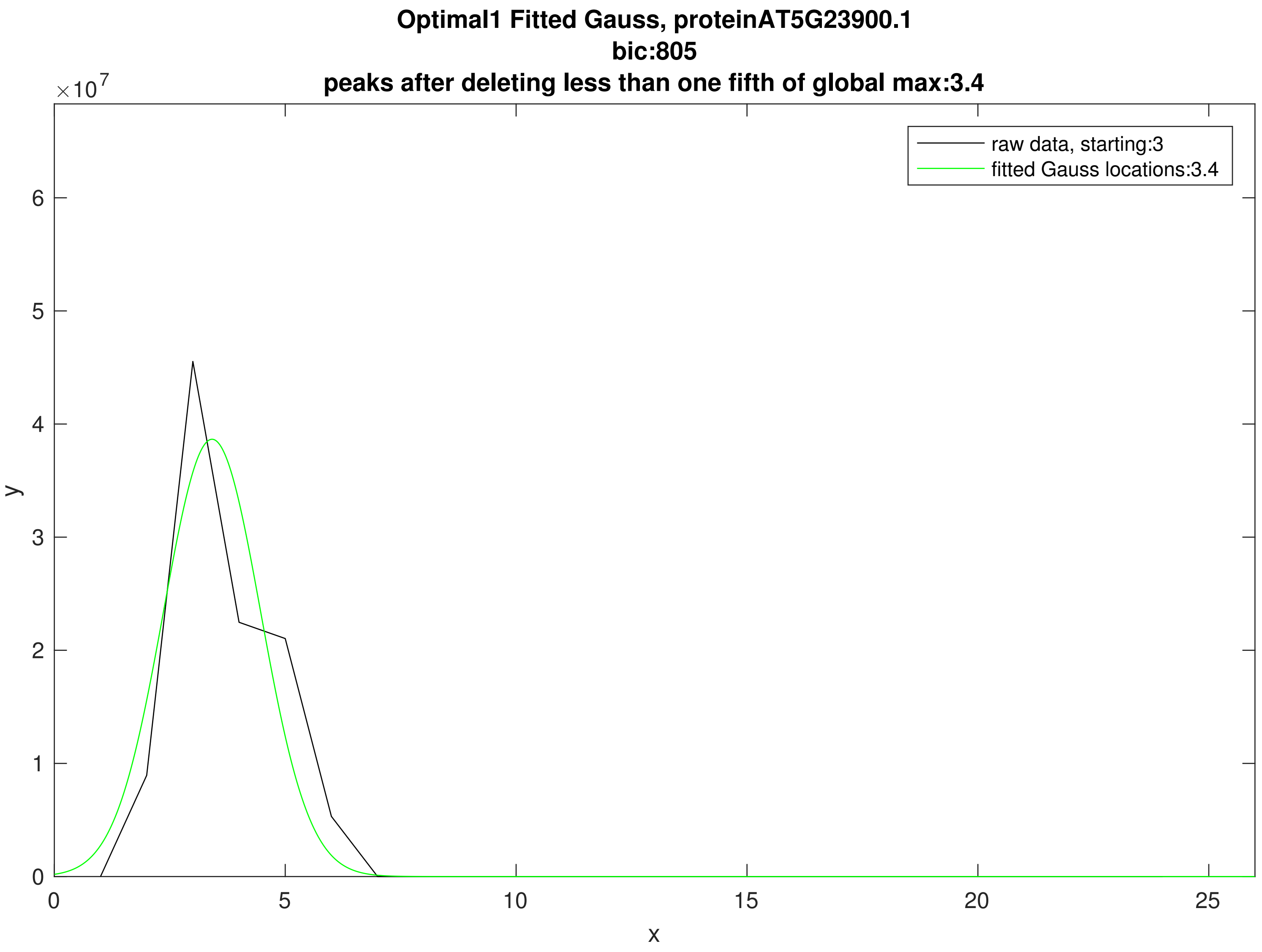

Optimal1 Fitted Gauss, proteinAT5G23900.1
bic:805
peaks after deleting less than one fifth of global max:3.4
×10^7
raw data, starting:3
fitted Gauss locations:3.4
y
x
0
1
2
3
4
5
6
0
5
10
15
20
25

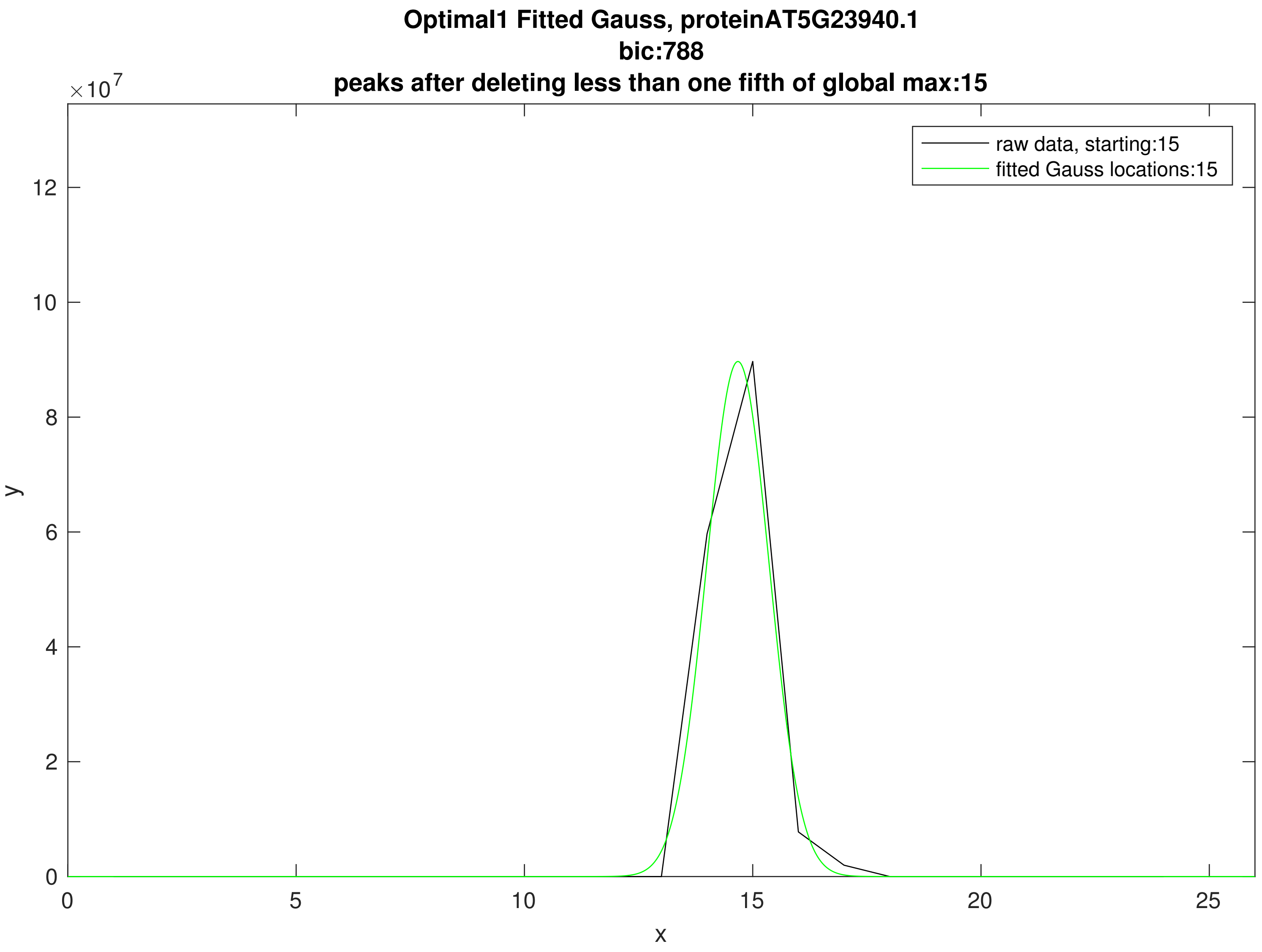

Optimal1 Fitted Gauss, proteinAT5G23940.1
bic:788
peaks after deleting less than one fifth of global max:15
×10^7
raw data, starting:15
fitted Gauss locations:15
12
10
8
6
4
2
0
y
0
5
10
15
20
25
x

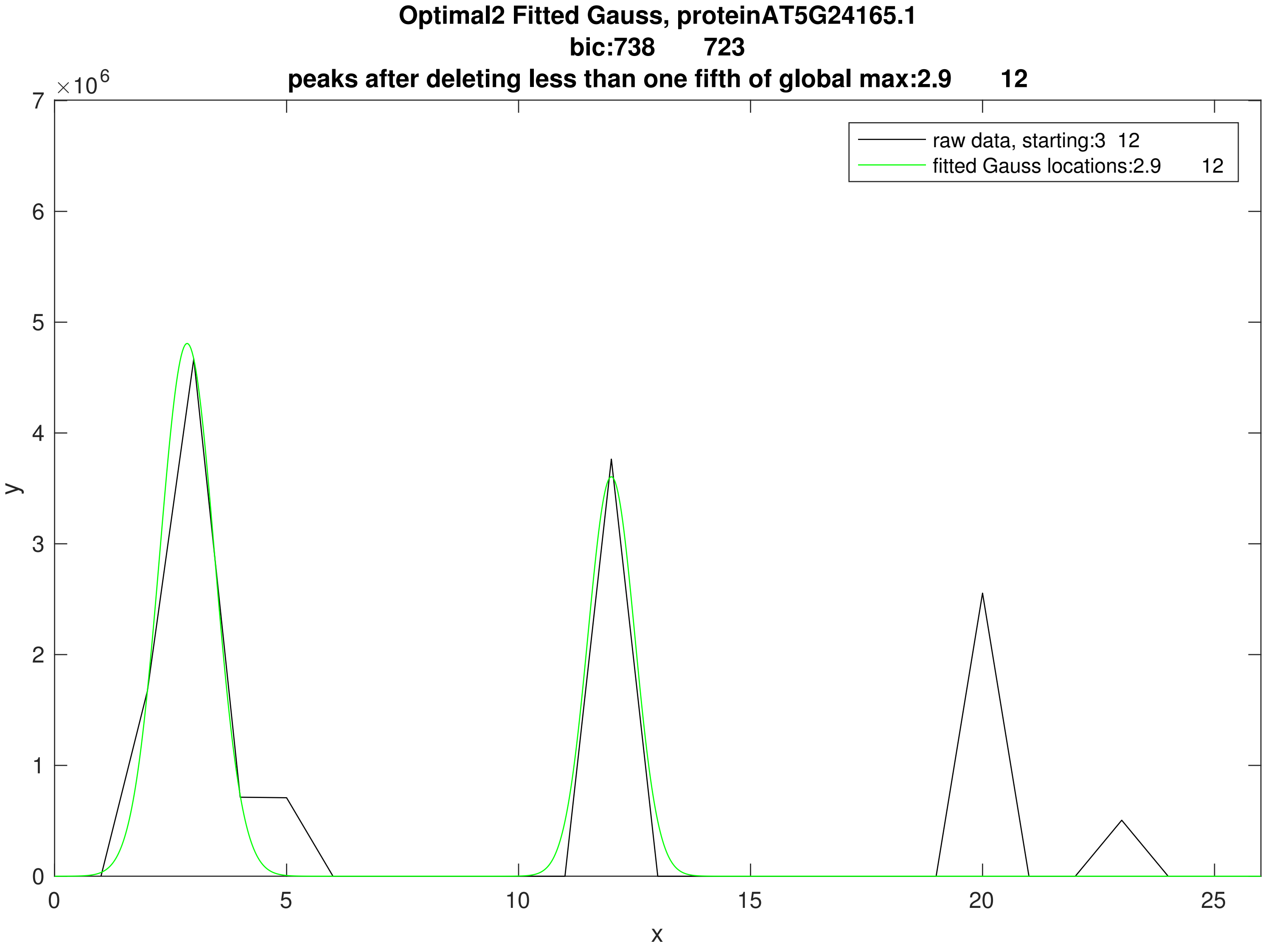

Optimal2 Fitted Gauss, proteinAT5G24165.1
bic:738 723
peaks after deleting less than one fifth of global max:2.9 12
×10^6
raw data, starting:3 12
fitted Gauss locations:2.9 12
y
x
0
1
2
3
4
5
6
7
0
5
10
15
20
25

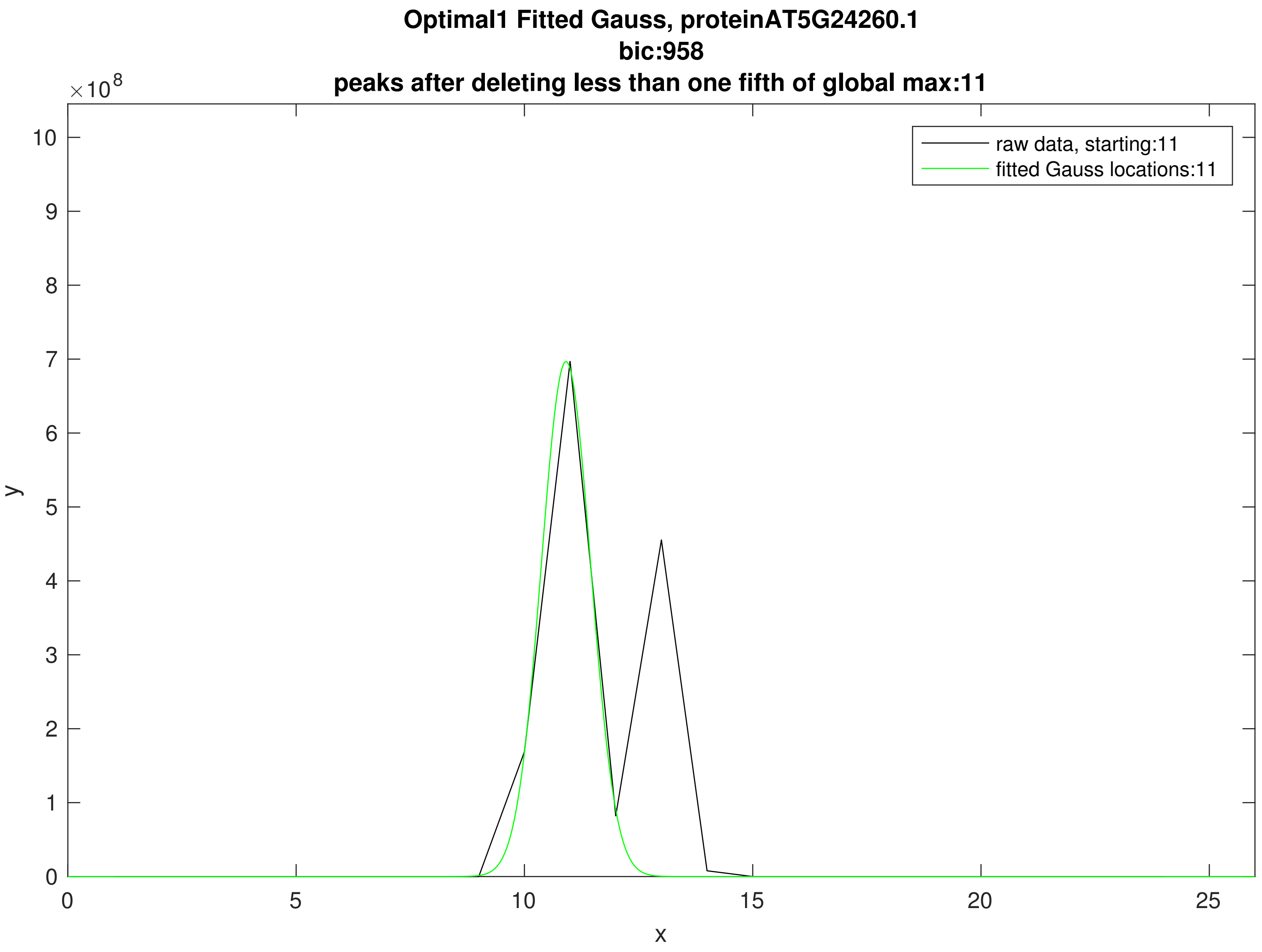
Optimal1 Fitted Gauss, proteinAT5G24260.1
bic:958
peaks after deleting less than one fifth of global max:11
$\times 10^8$
raw data, starting:11
fitted Gauss locations:11
y
x
0
1
2
3
4
5
6
7
8
9
10
0
5
10
15
20
25

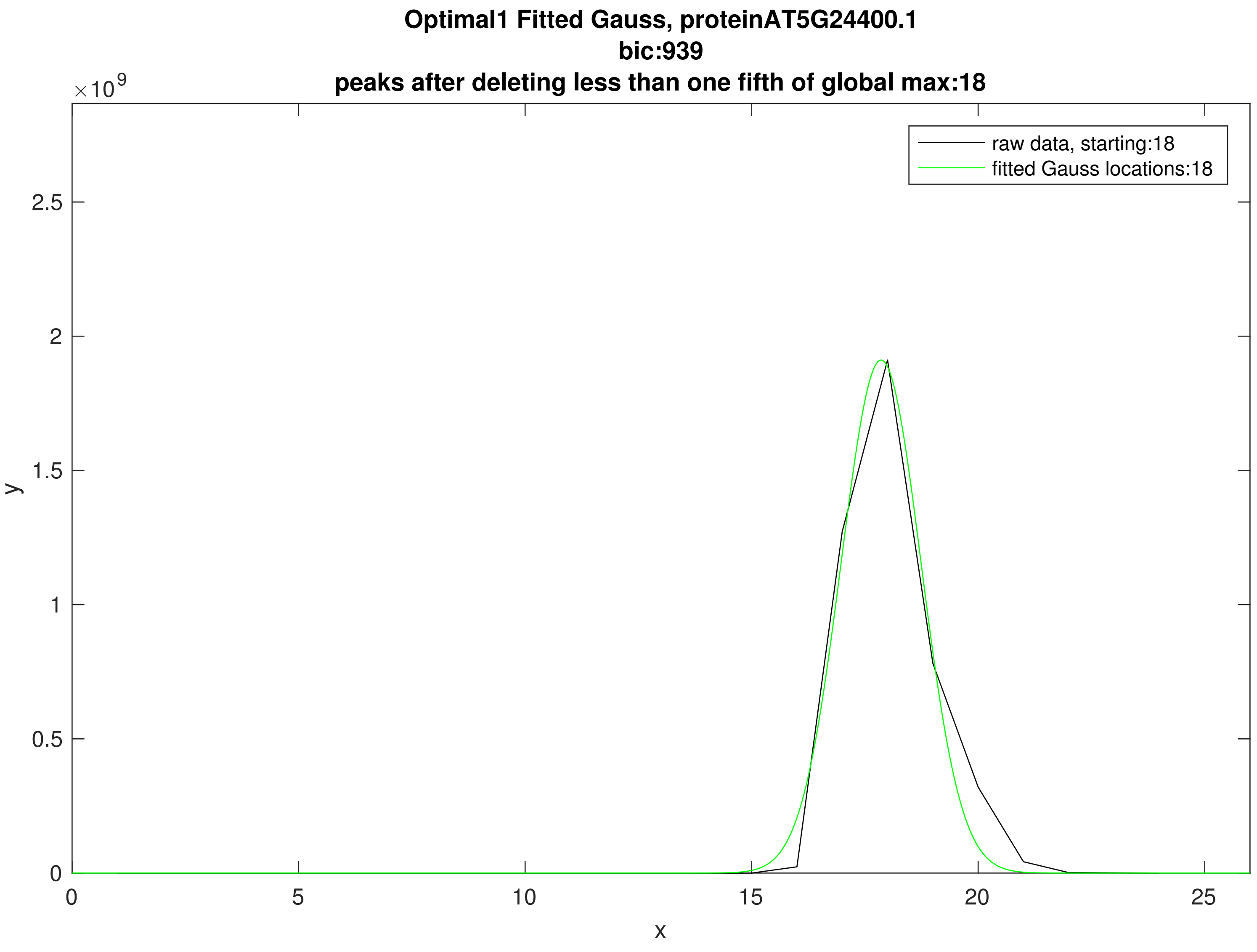

Optimal1 Fitted Gauss, proteinAT5G24400.1
bic:939
peaks after deleting less than one fifth of global max:18
$\times 10^9$
raw data, starting:18
fitted Gauss locations:18
y
x
0
0.5
1
1.5
2
2.5
0
5
10
15
20
25

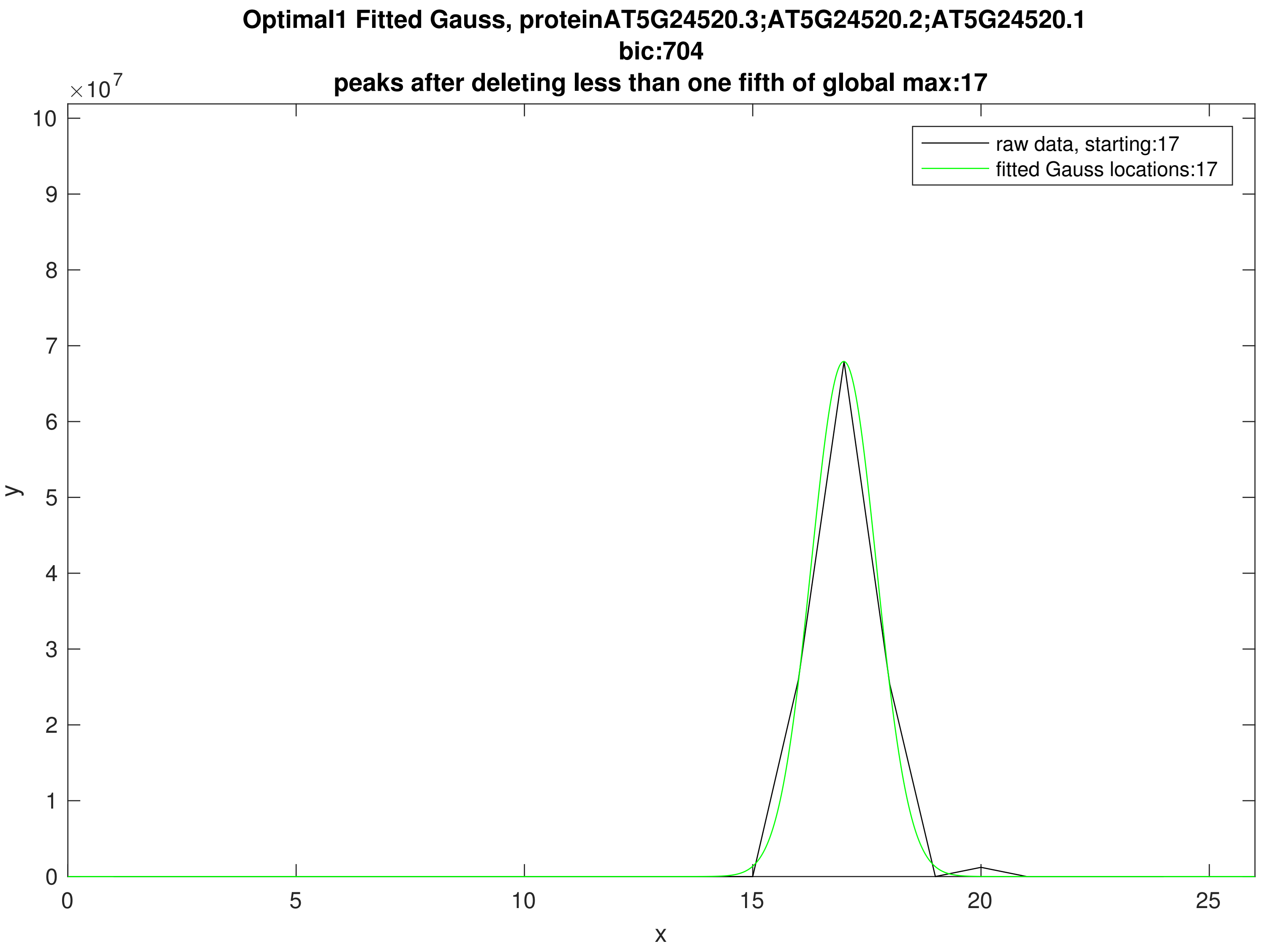
Optimal1 Fitted Gauss, proteinAT5G24520.3;AT5G24520.2;AT5G24520.1
bic:704
peaks after deleting less than one fifth of global max:17
×10^7
raw data, starting:17
fitted Gauss locations:17
y
x
0
1
2
3
4
5
6
7
8
9
10
0
5
10
15
20
25

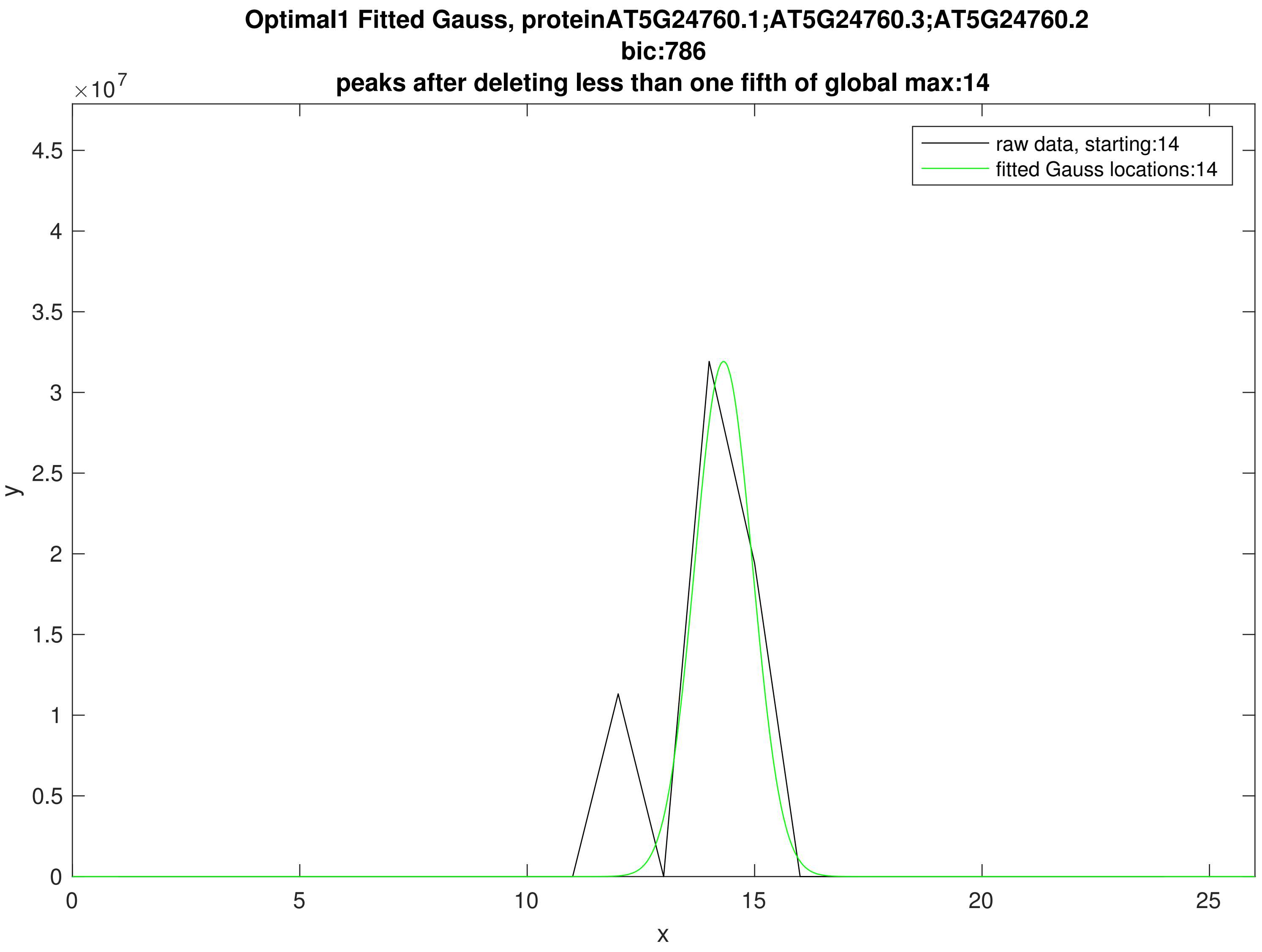

Optimal1 Fitted Gauss, proteinAT5G24760.1;AT5G24760.3;AT5G24760.2
bic:786
peaks after deleting less than one fifth of global max:14
×10^7
raw data, starting:14
fitted Gauss locations:14
y
x
0
0.5
1
1.5
2
2.5
3
3.5
4
4.5
0
5
10
15
20
25

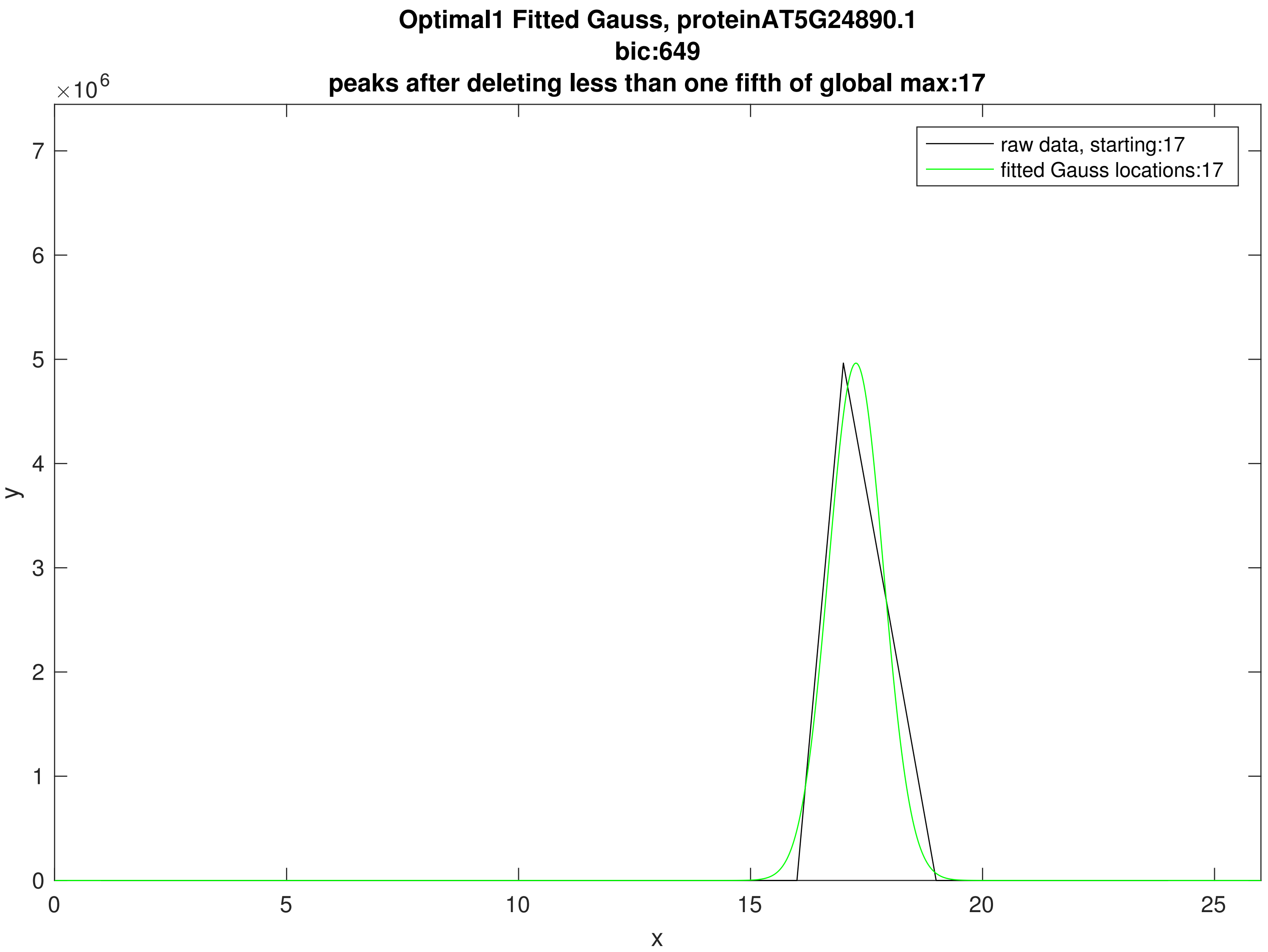
Optimal1 Fitted Gauss, proteinAT5G24890.1
bic:649
peaks after deleting less than one fifth of global max:17
×10^6
raw data, starting:17
fitted Gauss locations:17
y
x
0
1
2
3
4
5
6
7
0
5
10
15
20
25

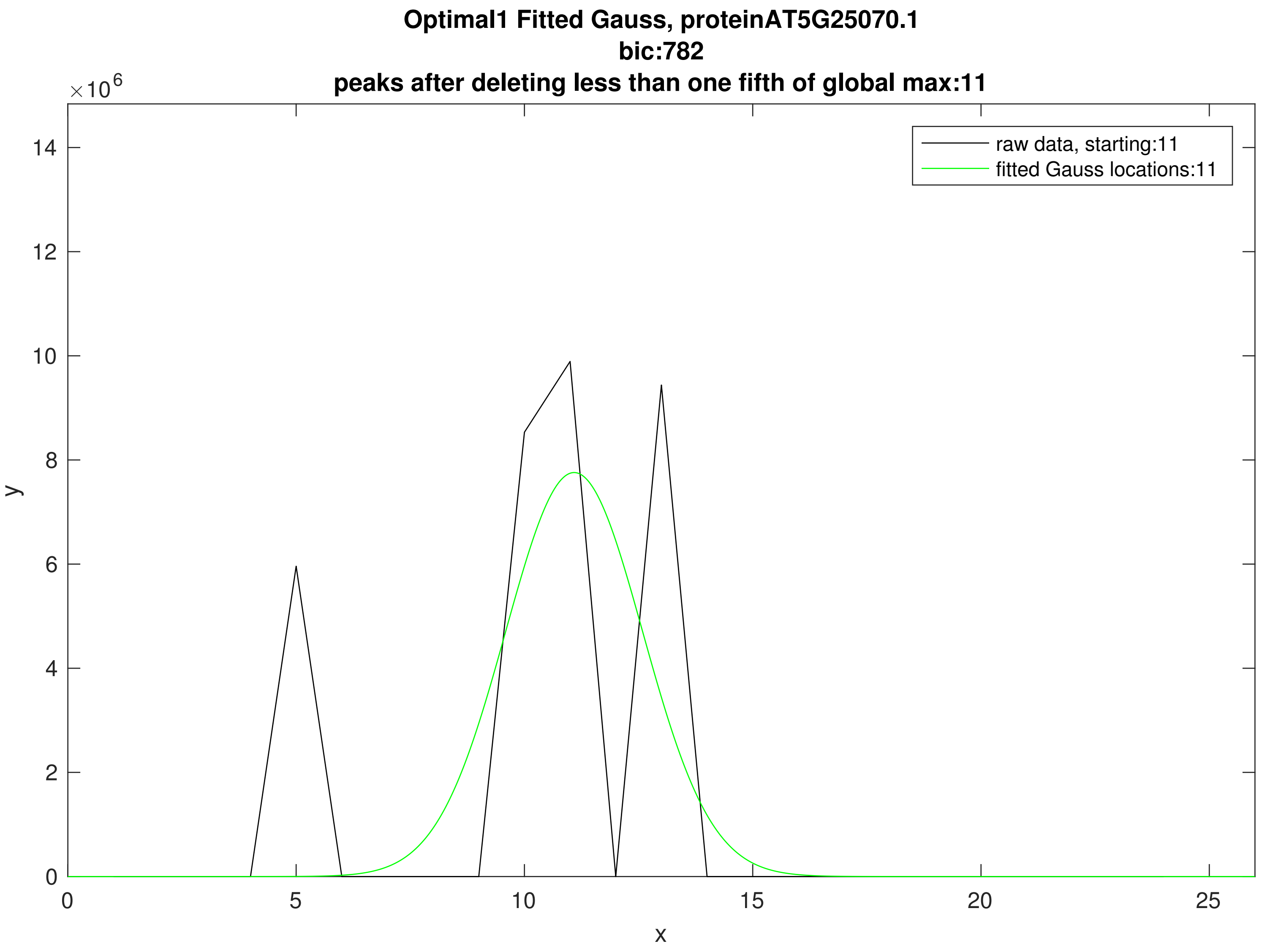

Optimal1 Fitted Gauss, proteinAT5G25070.1
bic:782
peaks after deleting less than one fifth of global max:11
×10^6
14
12
10
8
6
4
2
0
y
0
5
10
15
20
25
x
raw data, starting:11
fitted Gauss locations:11

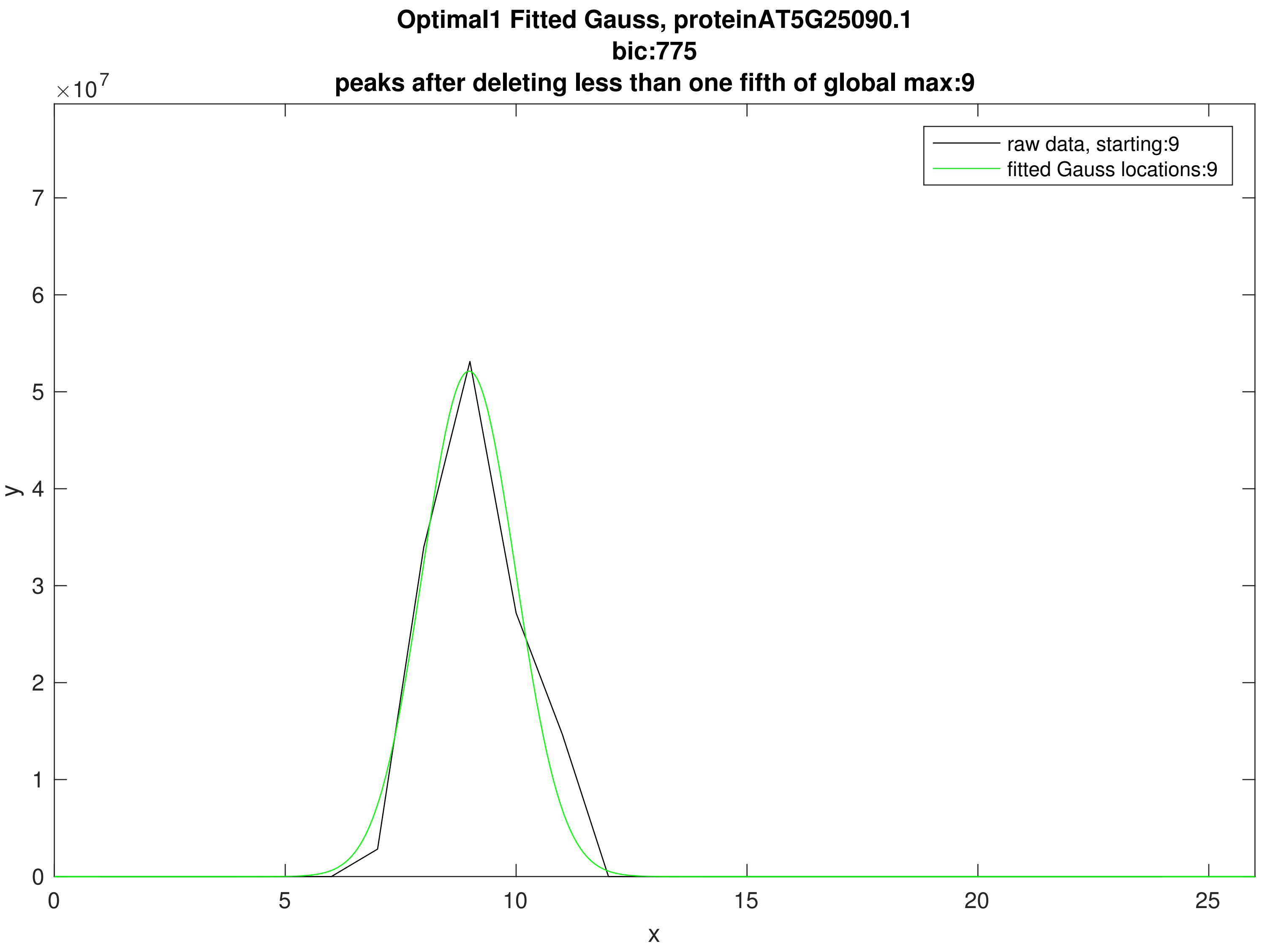

Optimal1 Fitted Gauss, proteinAT5G25090.1
bic:775
peaks after deleting less than one fifth of global max:9
×10^7
raw data, starting:9
fitted Gauss locations:9
y
x
0
1
2
3
4
5
6
7
0
5
10
15
20
25

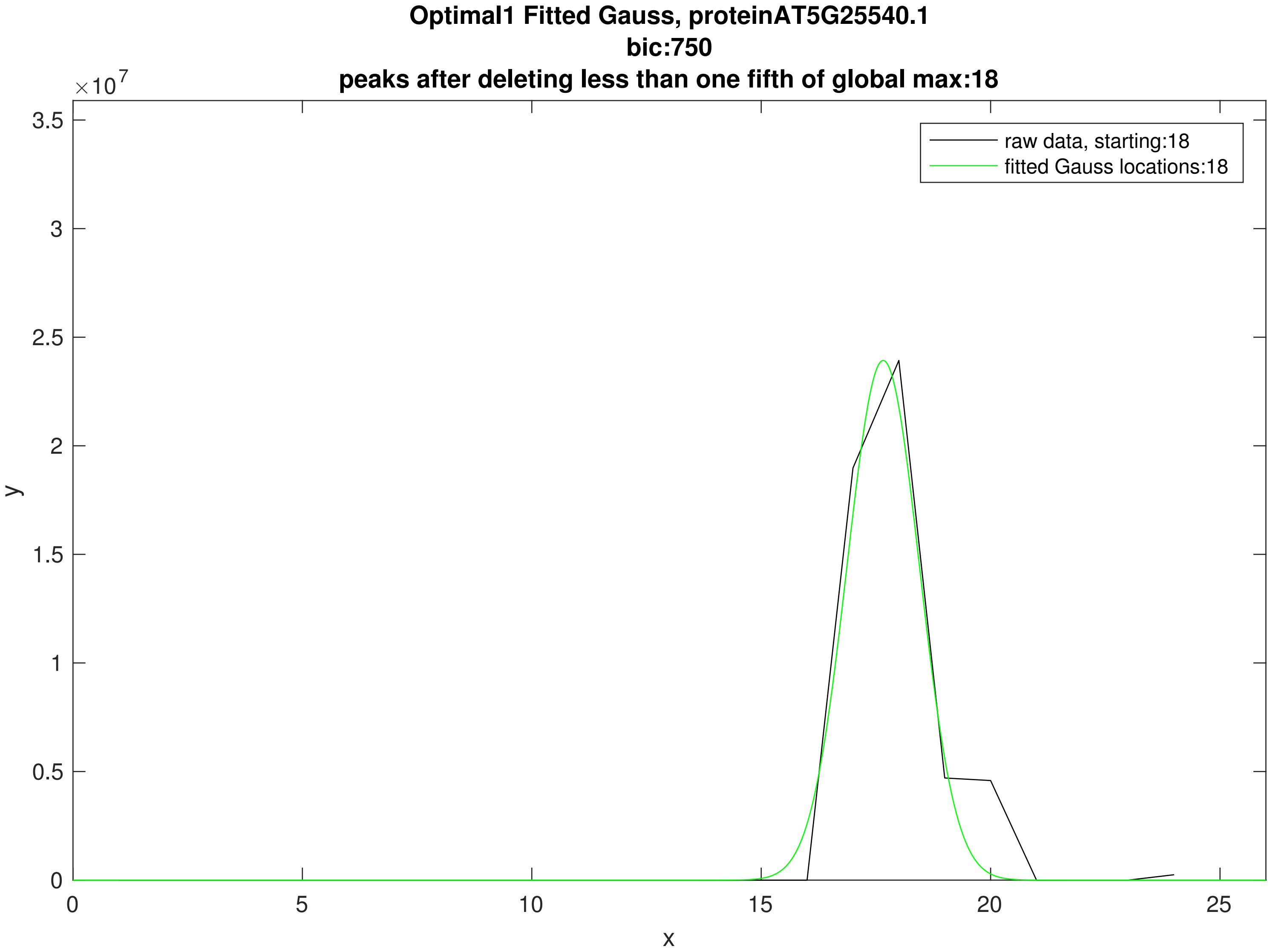
Optimal1 Fitted Gauss, proteinAT5G25540.1
bic:750
peaks after deleting less than one fifth of global max:18
×10^7
3.5
3
2.5
2
1.5
1
0.5
0
y
0
5
10
15
20
25
x
raw data, starting:18
fitted Gauss locations:18

**Optimal1 Fitted Gauss, proteinAT5G25770.1;AT5G25770.2;AT5G25770.3**
**bic:852 860 870**
**peaks after deleting less than one fifth of global max:17**

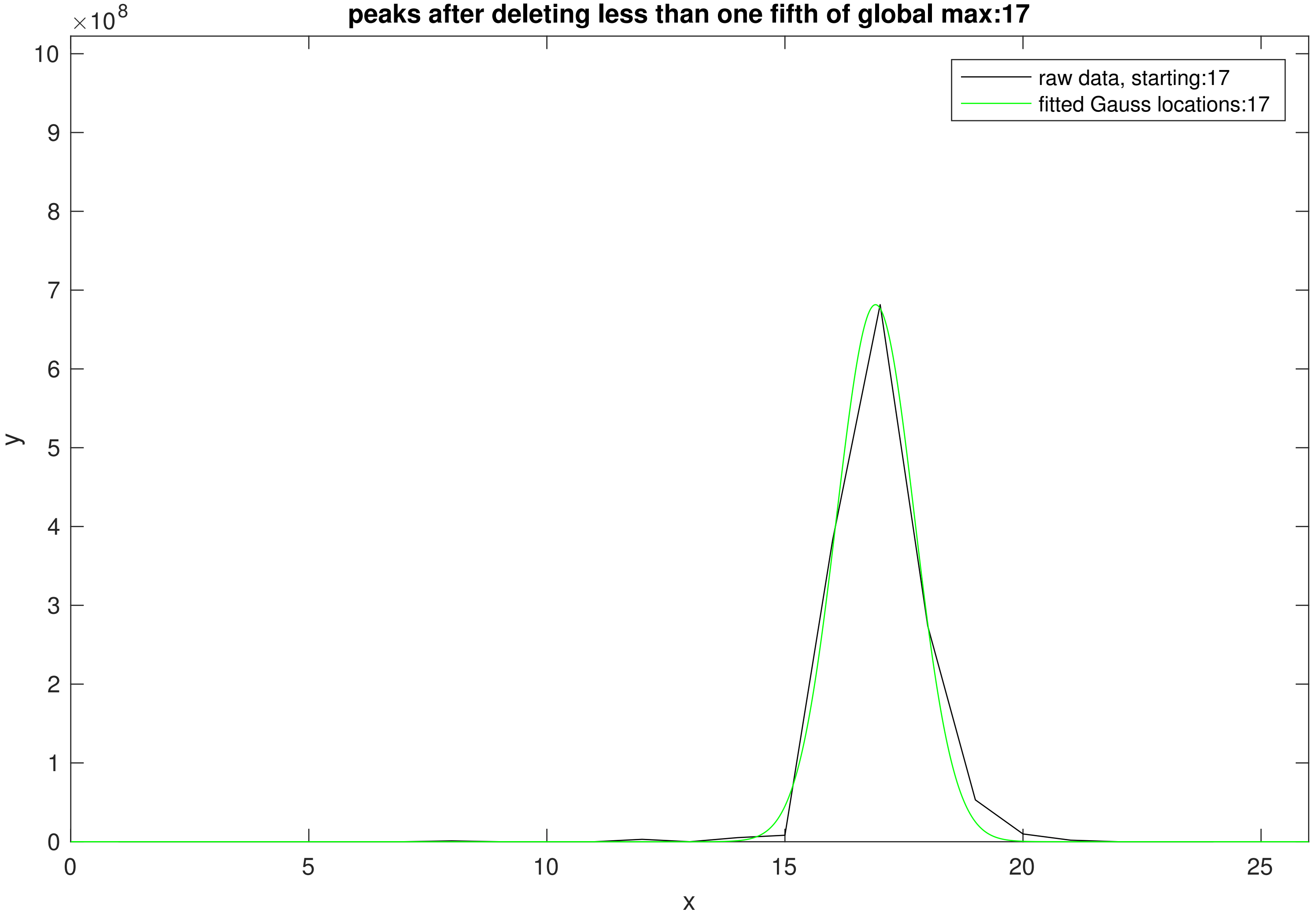

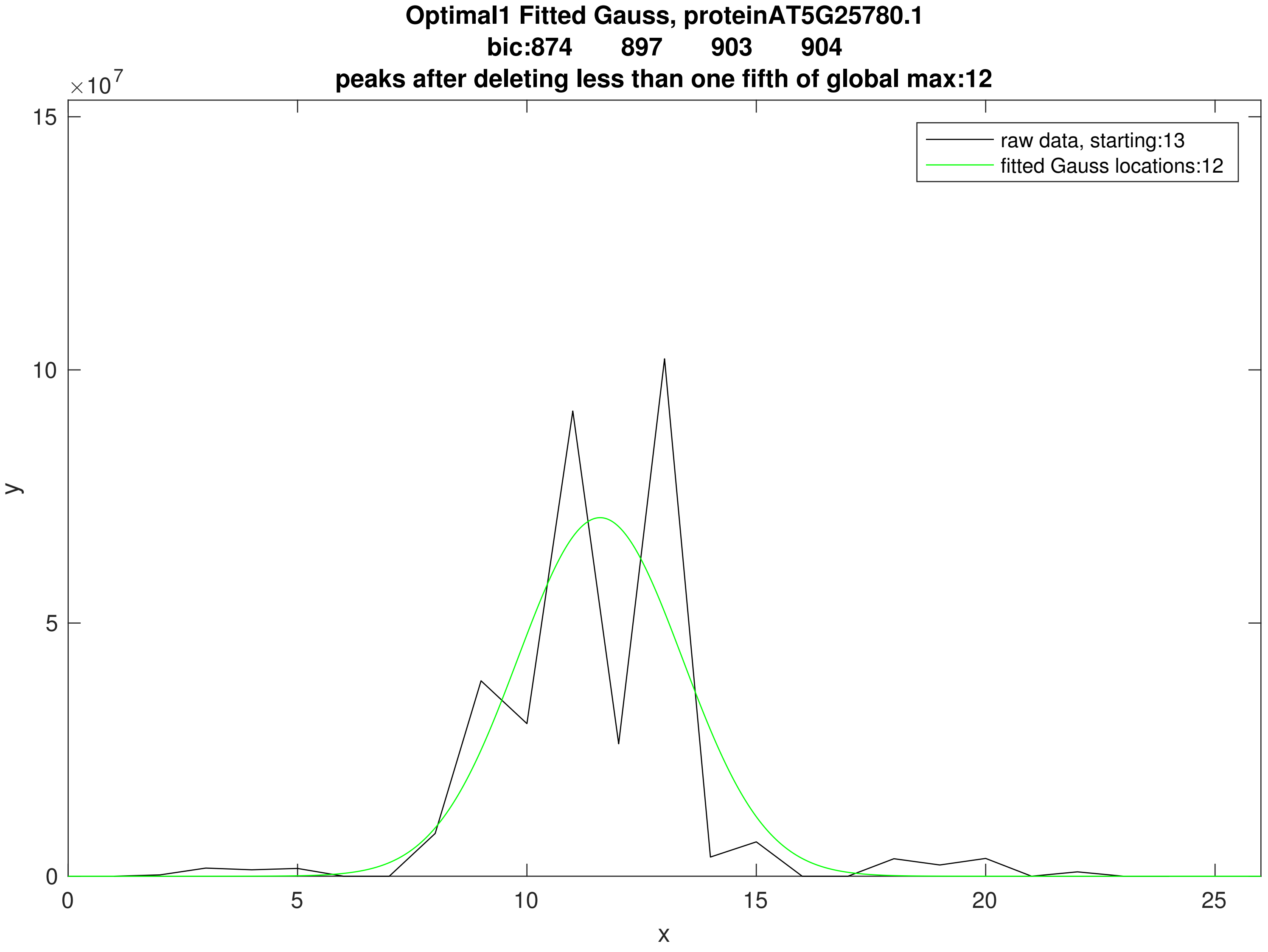

Optimal1 Fitted Gauss, proteinAT5G25780.1
bic:874 897 903 904
peaks after deleting less than one fifth of global max:12
×10^7
raw data, starting:13
fitted Gauss locations:12
y
x
0
5
10
15
20
25

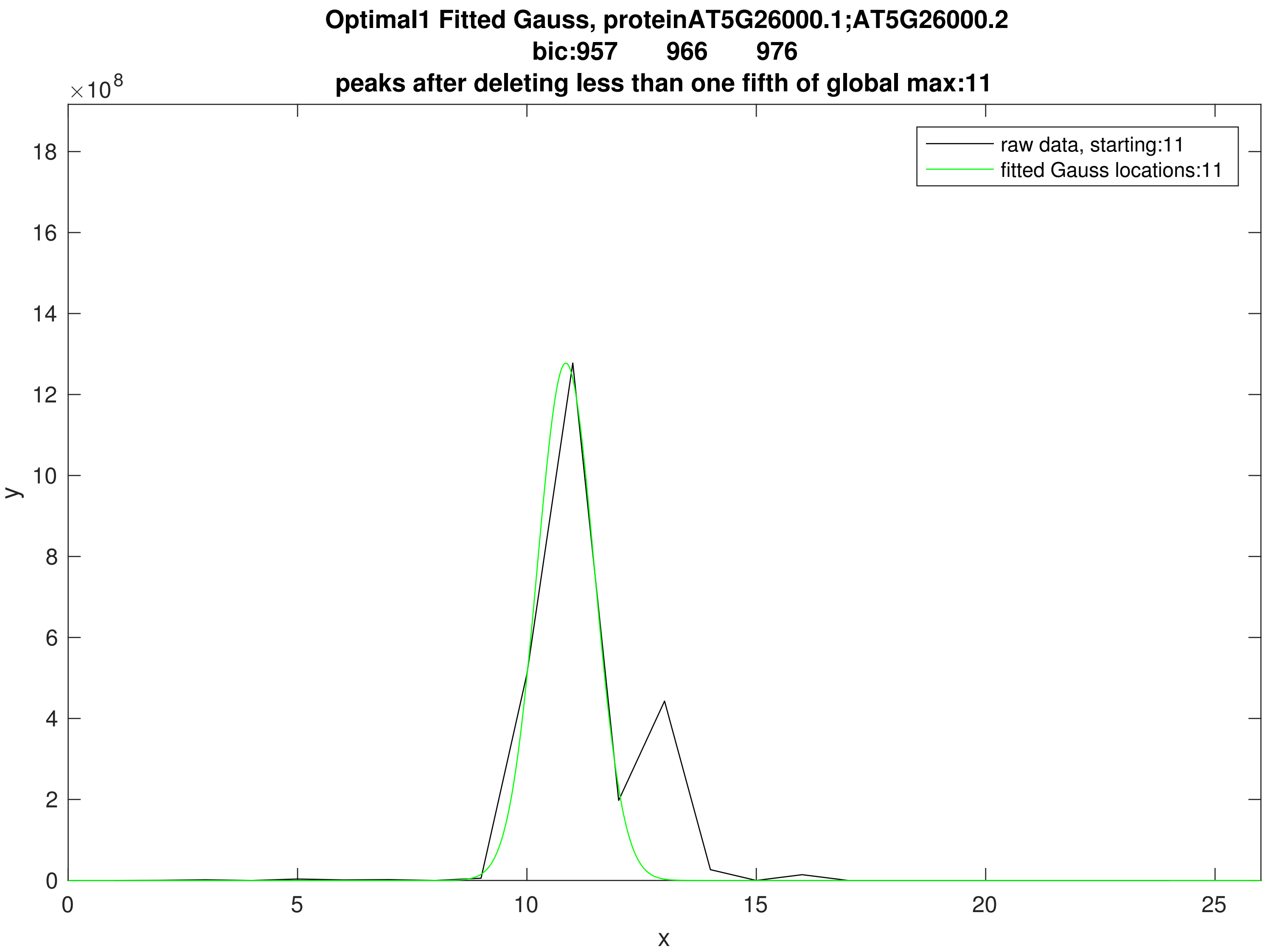
Optimal1 Fitted Gauss, proteinAT5G26000.1;AT5G26000.2
bic:957 966 976
peaks after deleting less than one fifth of global max:11
raw data, starting:11
fitted Gauss locations:11
×10^8
y
x

Optimal1 Fitted Gauss, proteinAT5G26260.1
bic:748
peaks after deleting less than one fifth of global max:3.8

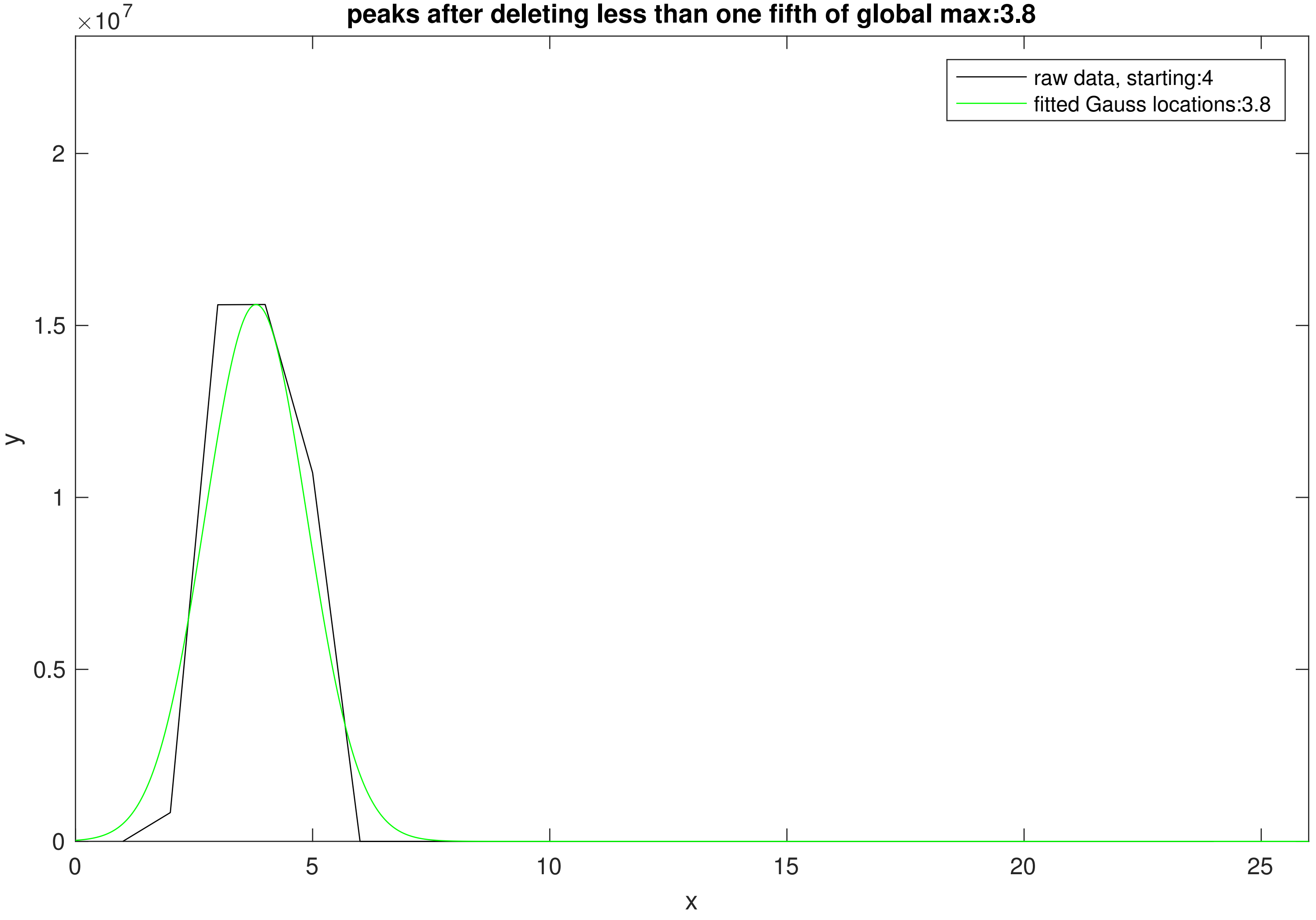

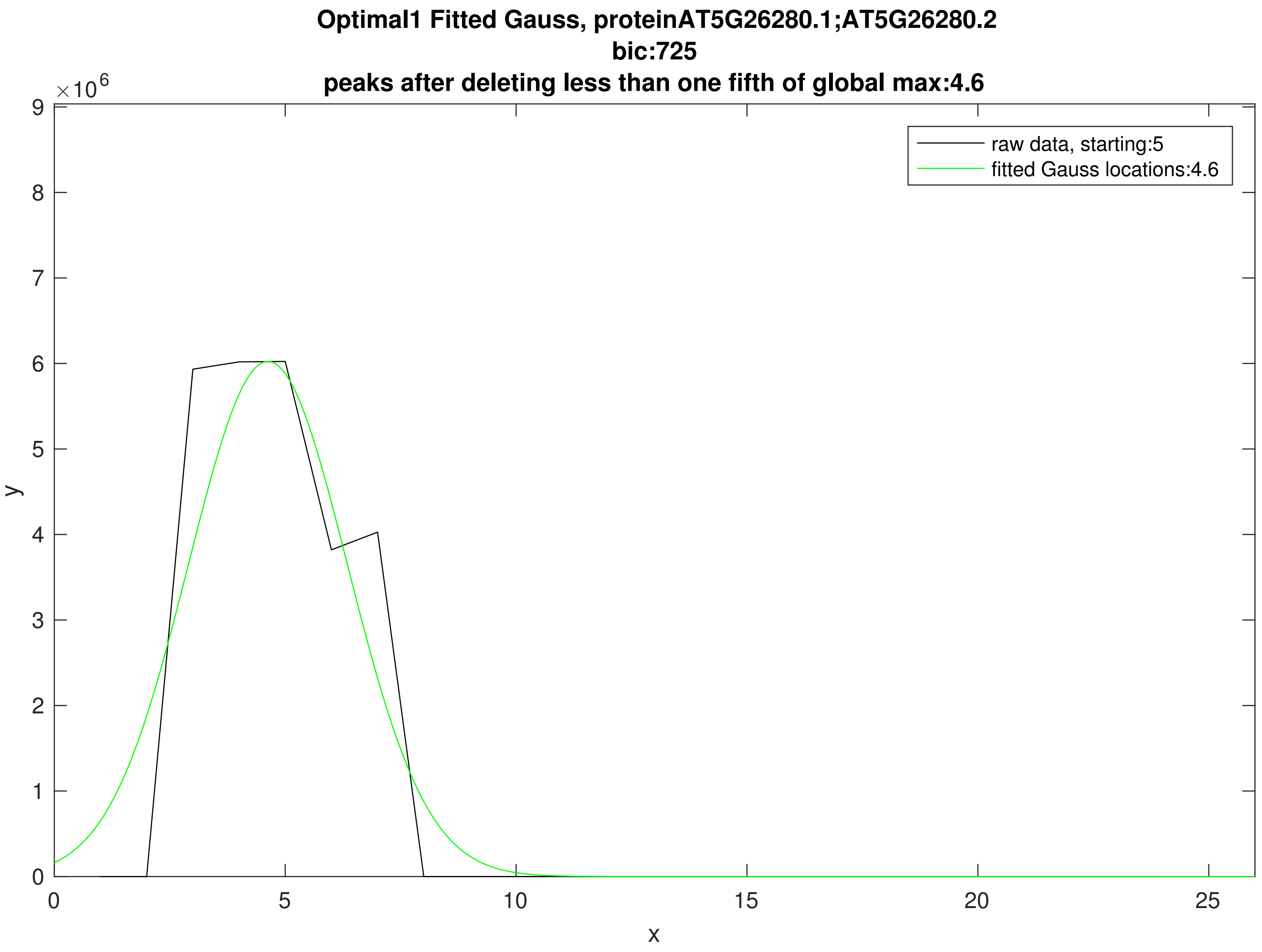
Optimal1 Fitted Gauss, proteinAT5G26280.1;AT5G26280.2
bic:725
peaks after deleting less than one fifth of global max:4.6
×10^6
raw data, starting:5
fitted Gauss locations:4.6
y
x
0
1
2
3
4
5
6
7
8
9
0
5
10
15
20
25

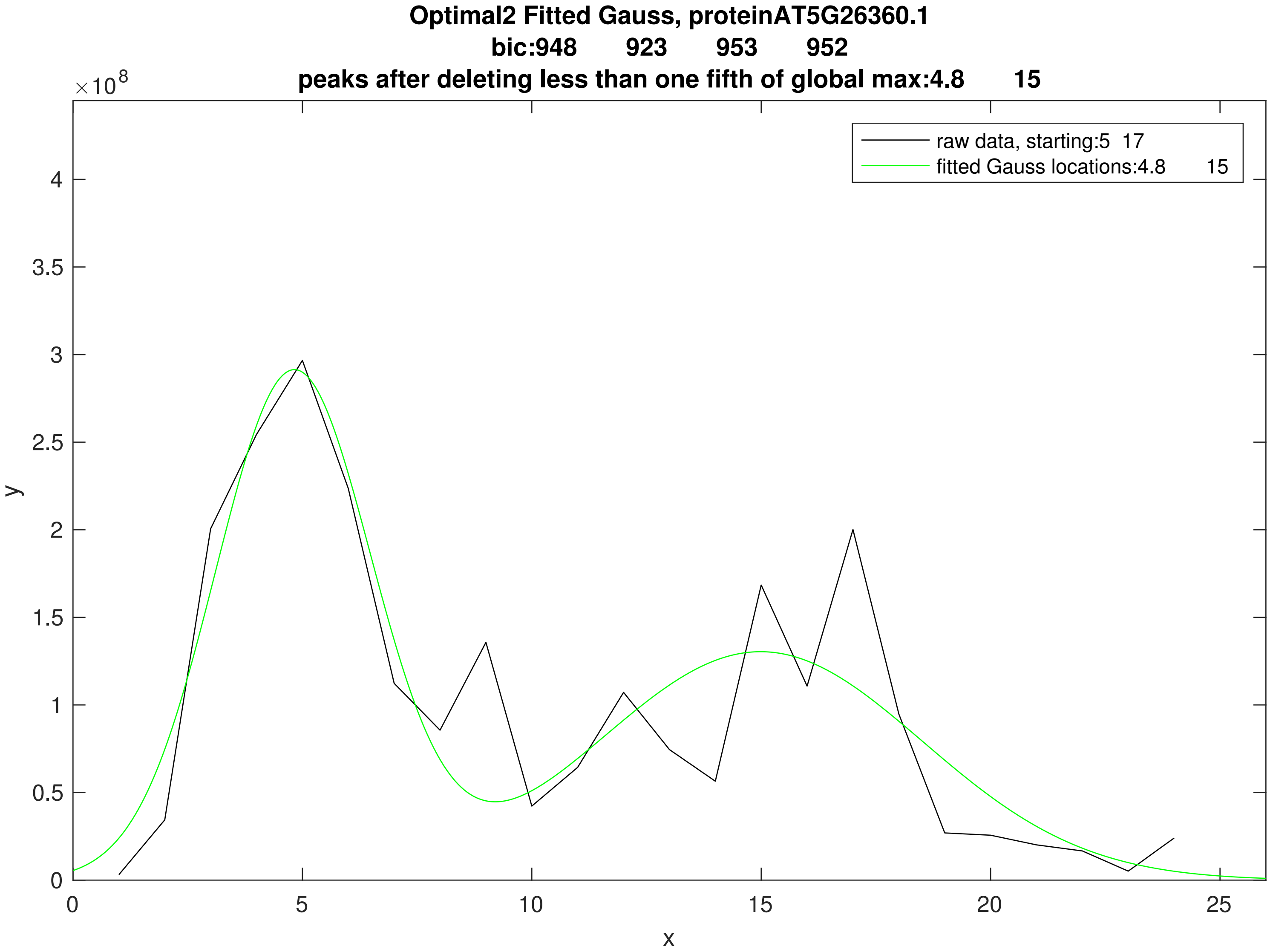
Optimal2 Fitted Gauss, proteinAT5G26360.1
bic:948 923 953 952
peaks after deleting less than one fifth of global max:4.8 15
×10^8
raw data, starting:5 17
fitted Gauss locations:4.8 15
y
x
0
0.5
1
1.5
2
2.5
3
3.5
4
5
10
15
20
25

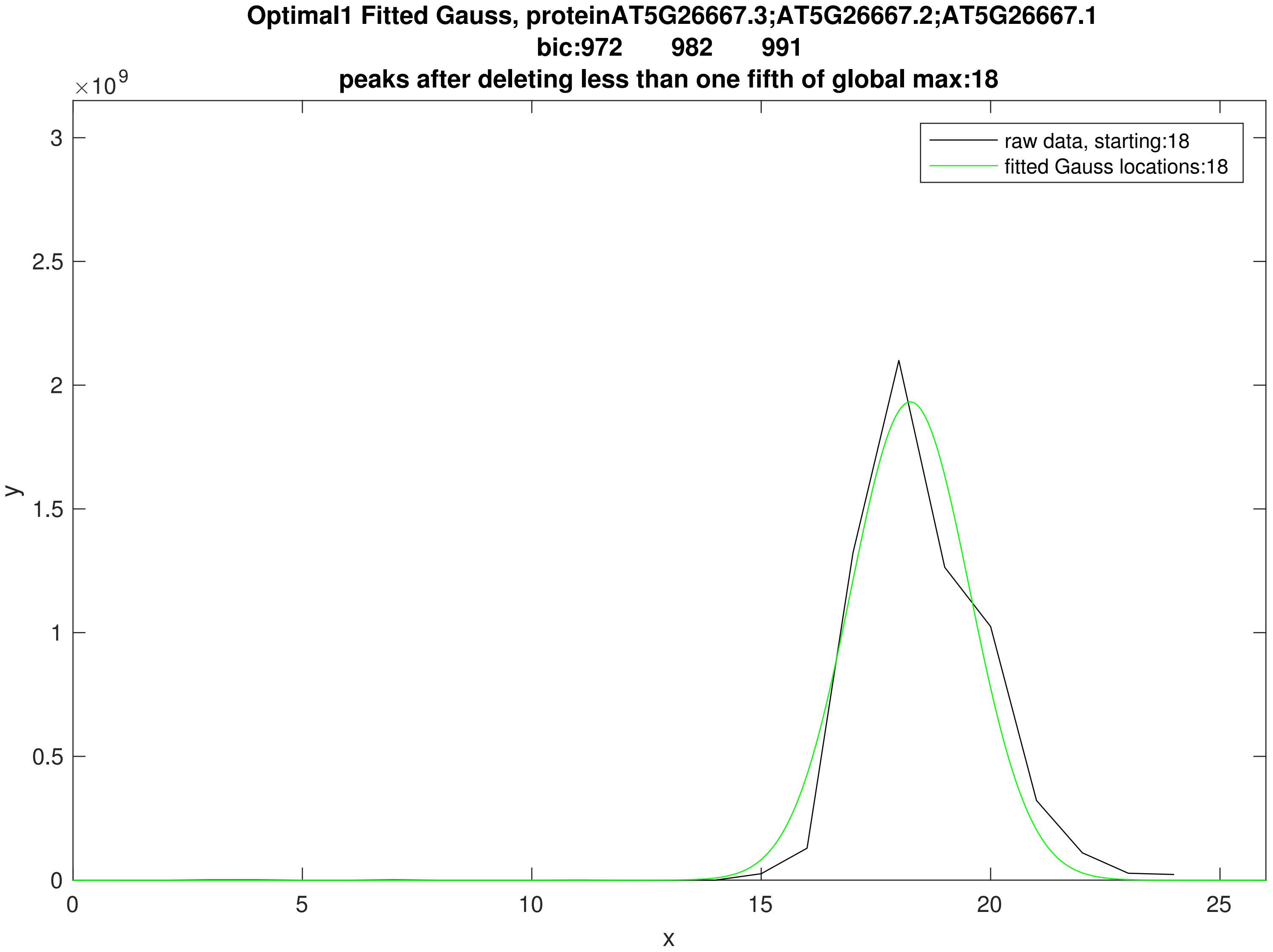
Optimal1 Fitted Gauss, proteinAT5G26667.3;AT5G26667.2;AT5G26667.1
bic:972 982 991
peaks after deleting less than one fifth of global max:18
×10^9
raw data, starting:18
fitted Gauss locations:18
y
x
0
0.5
1
1.5
2
2.5
3
0
5
10
15
20
25

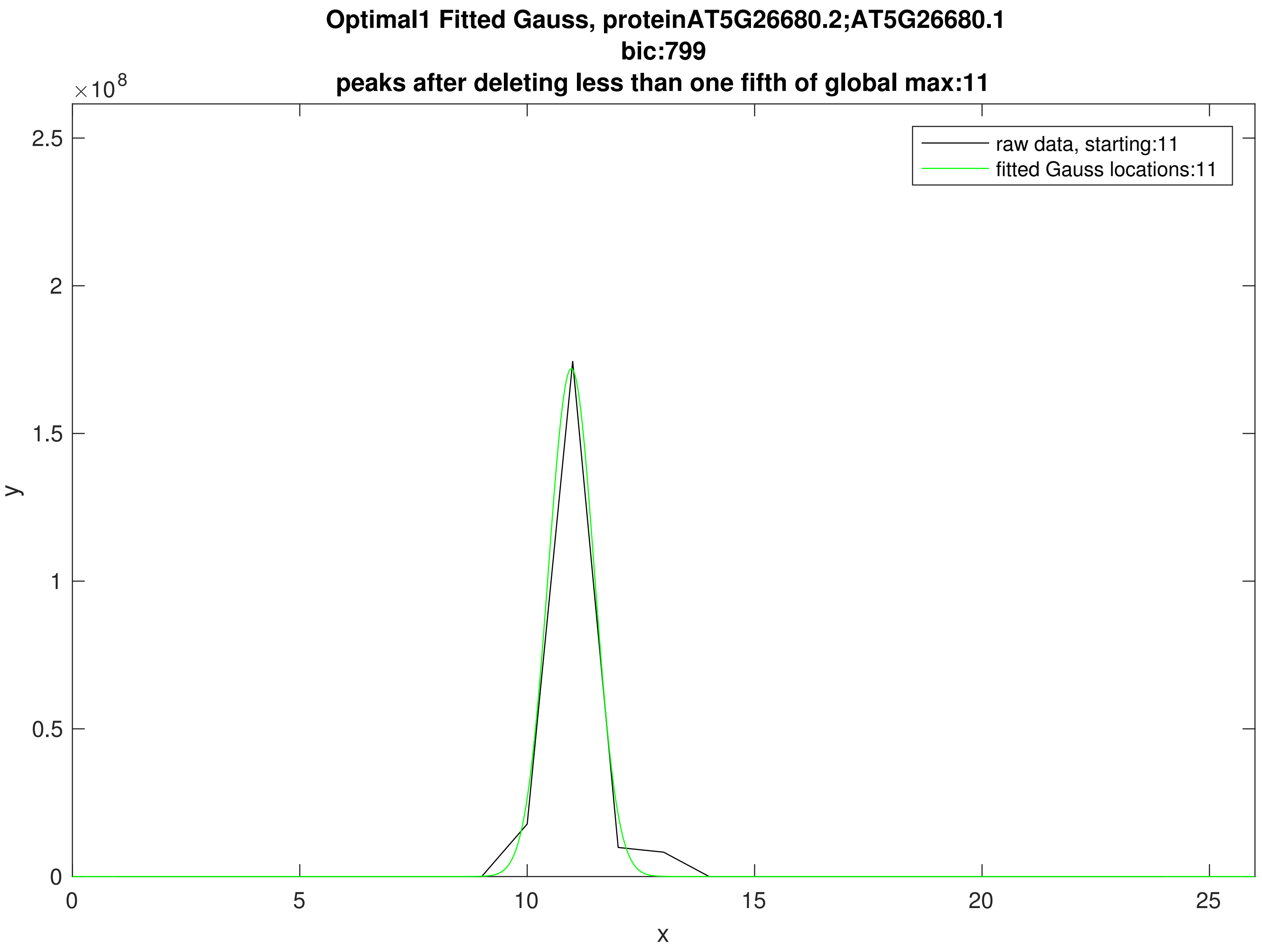

Optimal1 Fitted Gauss, proteinAT5G26680.2;AT5G26680.1
bic:799
peaks after deleting less than one fifth of global max:11
×10^8
2.5
2
1.5
1
0.5
0
y
0
5
10
15
20
25
x
raw data, starting:11
fitted Gauss locations:11

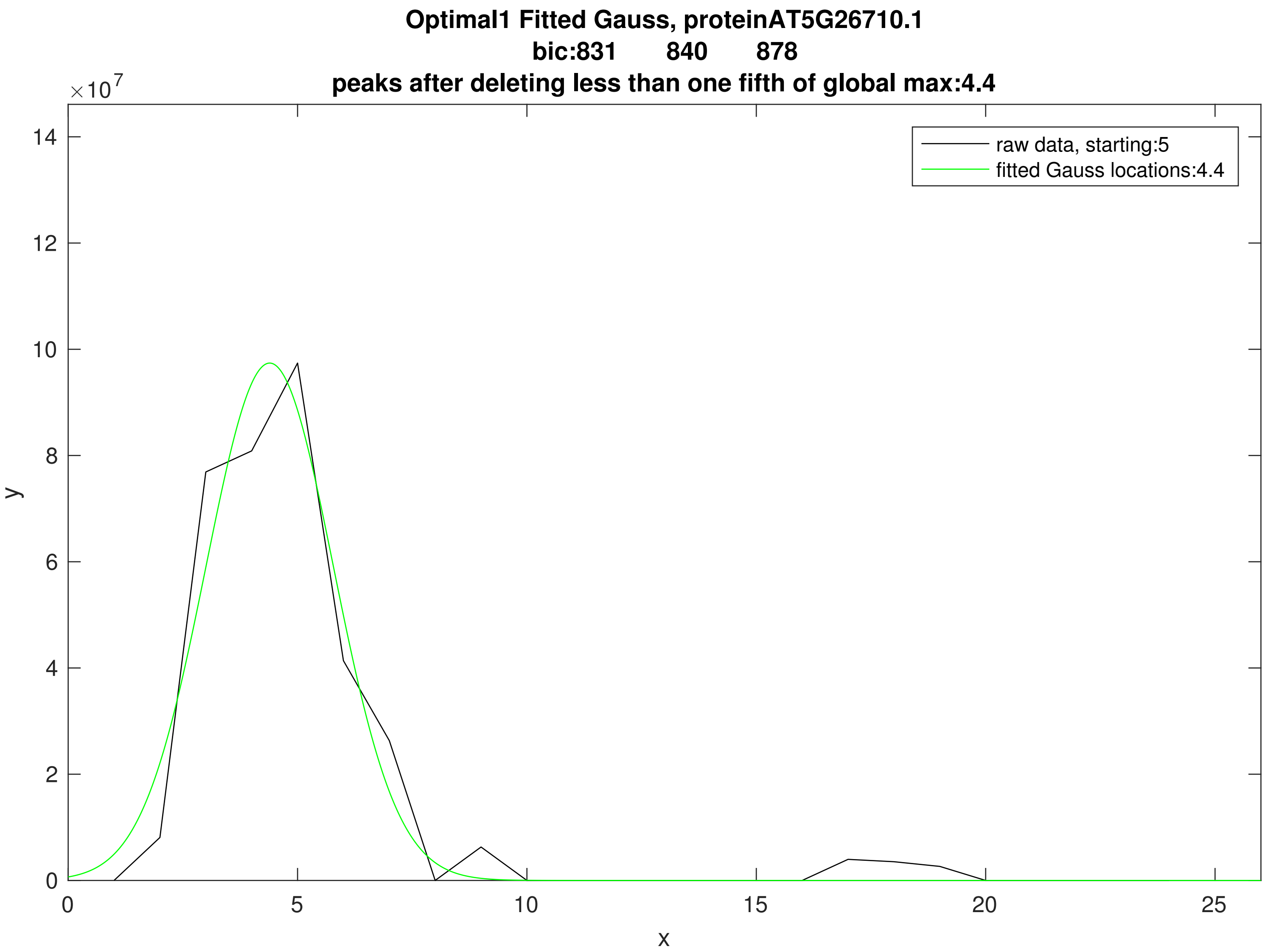

Optimal1 Fitted Gauss, proteinAT5G26710.1
bic:831 840 878
peaks after deleting less than one fifth of global max:4.4
×10^7
raw data, starting:5
fitted Gauss locations:4.4
y
x
0
2
4
6
8
10
12
14
0
5
10
15
20
25

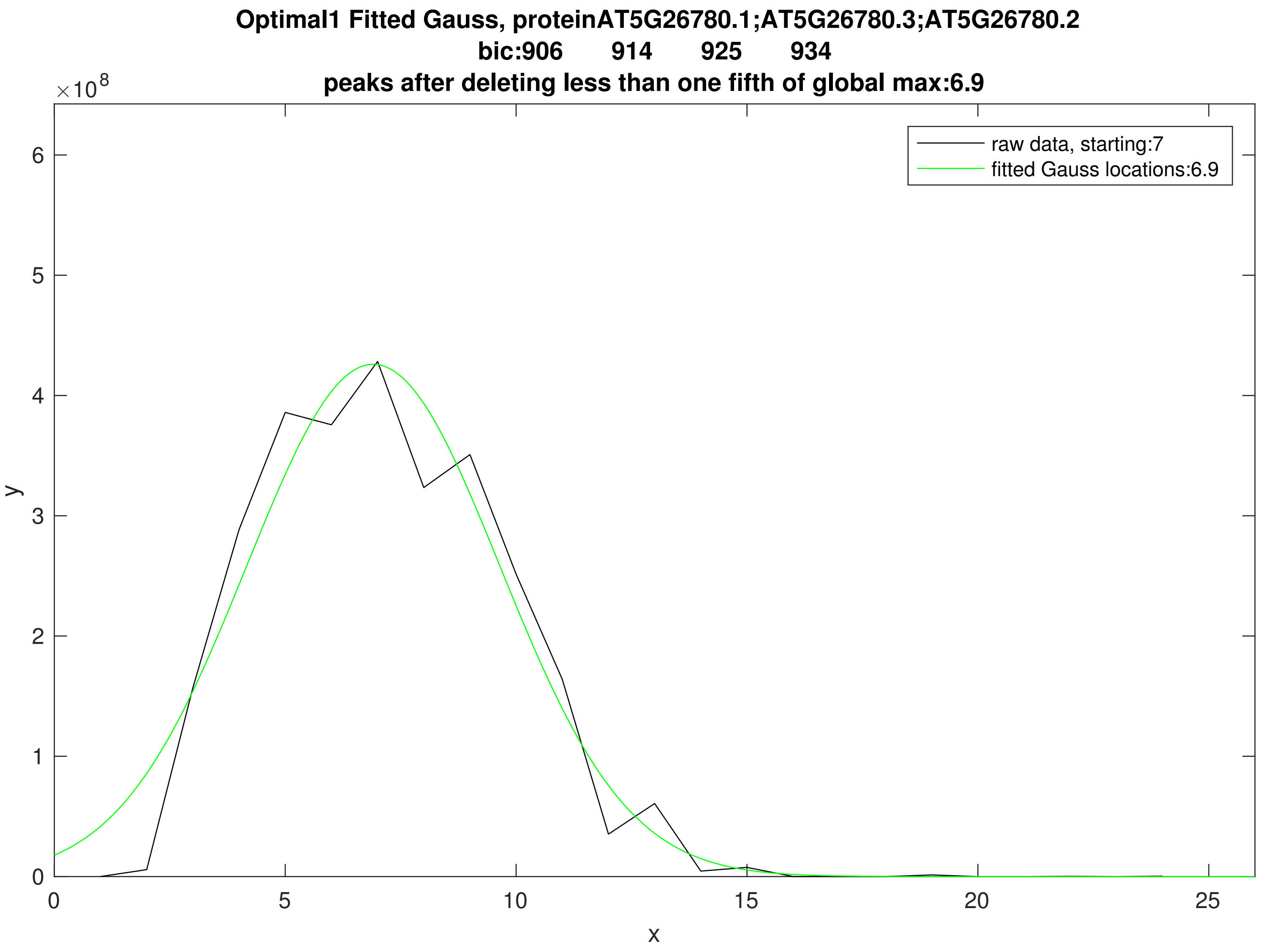

Optimal1 Fitted Gauss, proteinAT5G26780.1;AT5G26780.3;AT5G26780.2
bic:906 914 925 934
peaks after deleting less than one fifth of global max:6.9
×10^8
raw data, starting:7
fitted Gauss locations:6.9
y
x
0
1
2
3
4
5
6
0
5
10
15
20
25

Optimal1 Fitted Gauss, proteinAT5G26830.1
bic:758 770
peaks after deleting less than one fifth of global max:10

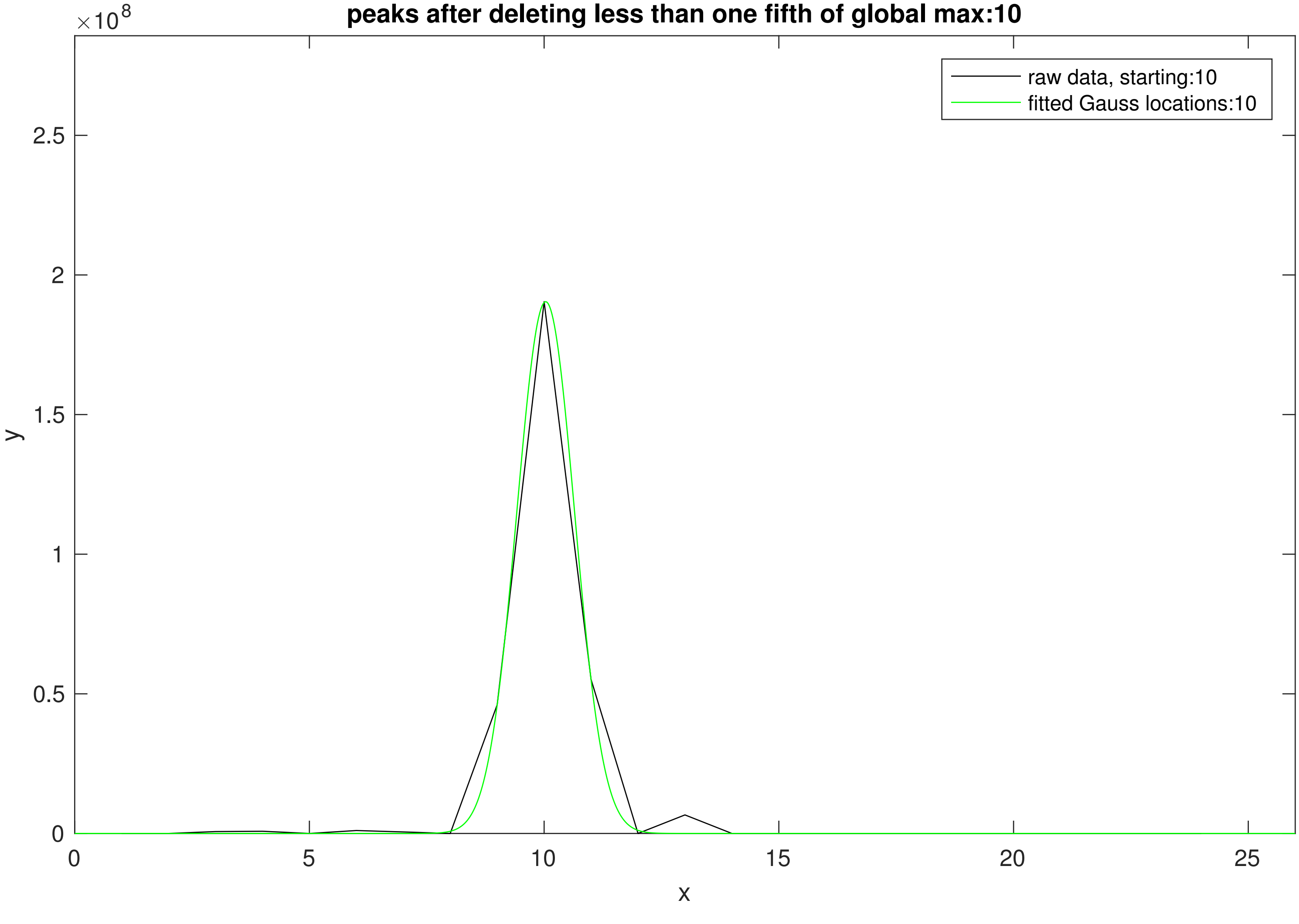

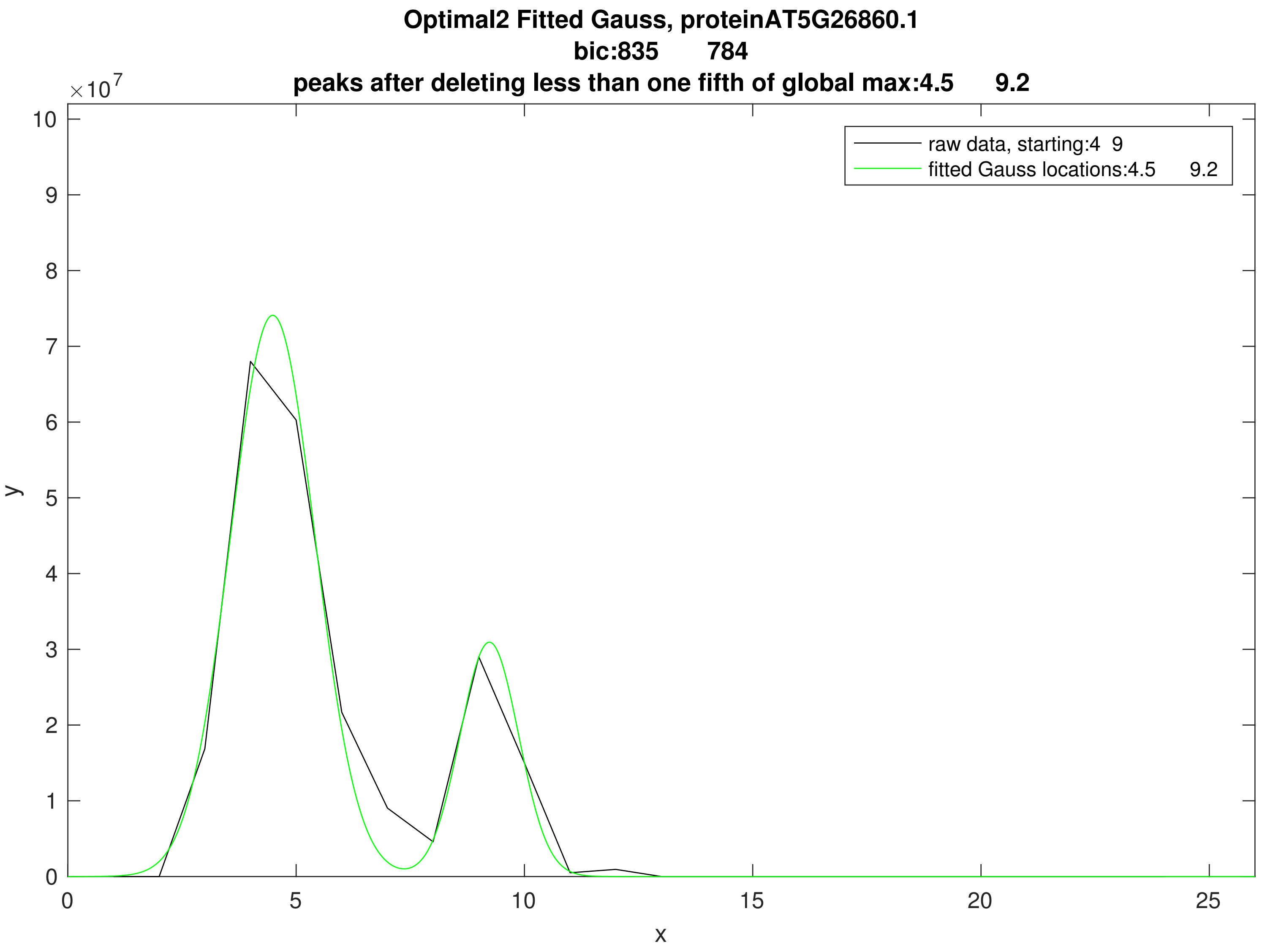

Optimal2 Fitted Gauss, proteinAT5G26860.1
bic:835 784
peaks after deleting less than one fifth of global max:4.5 9.2
×10^7
raw data, starting:4 9
fitted Gauss locations:4.5 9.2
y
x
0
1
2
3
4
5
6
7
8
9
10
0
5
10
15
20
25

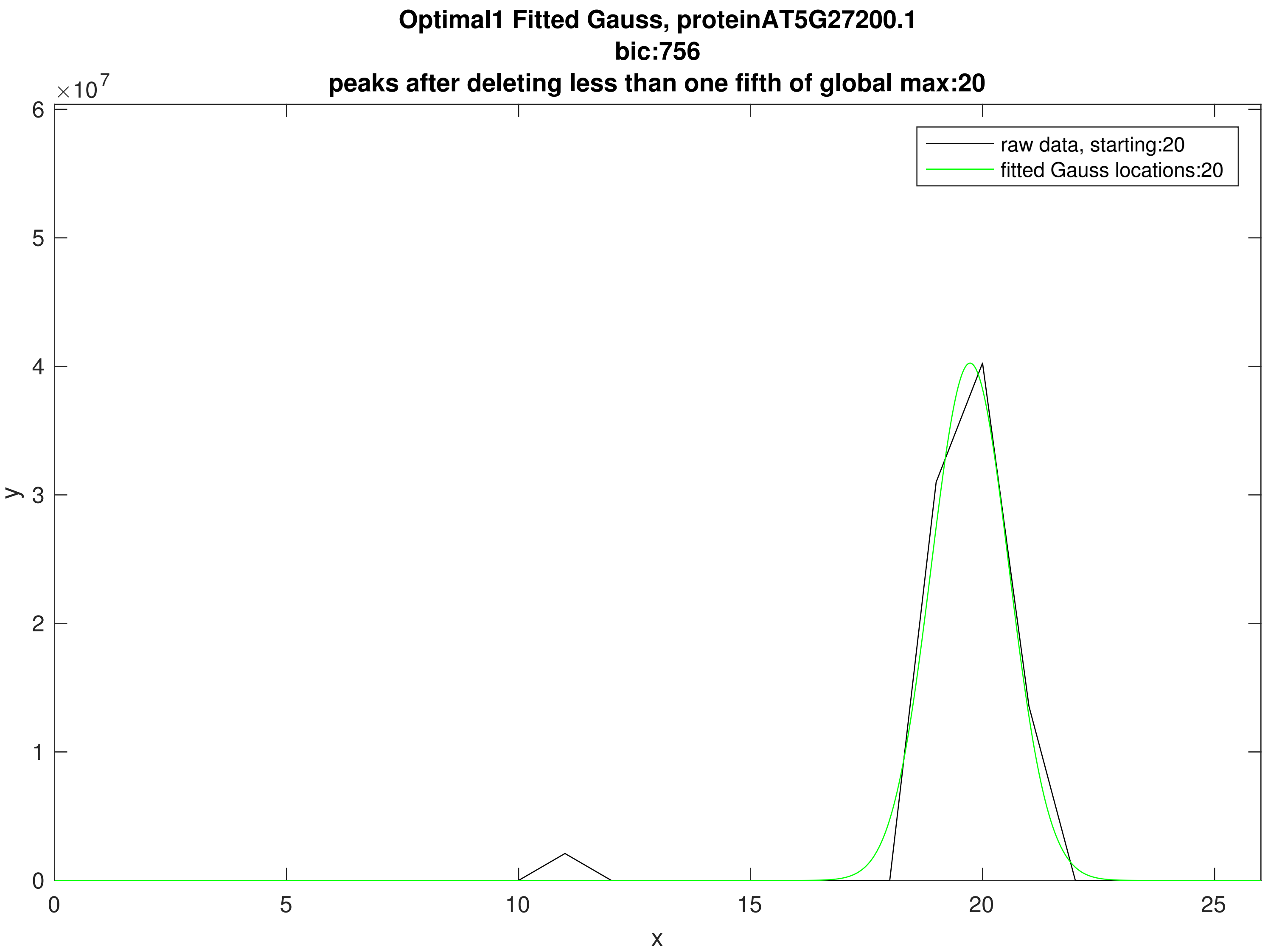
Optimal1 Fitted Gauss, proteinAT5G27200.1
bic:756
peaks after deleting less than one fifth of global max:20
×10^7
raw data, starting:20
fitted Gauss locations:20
y
x
0
1
2
3
4
5
6
0
5
10
15
20
25

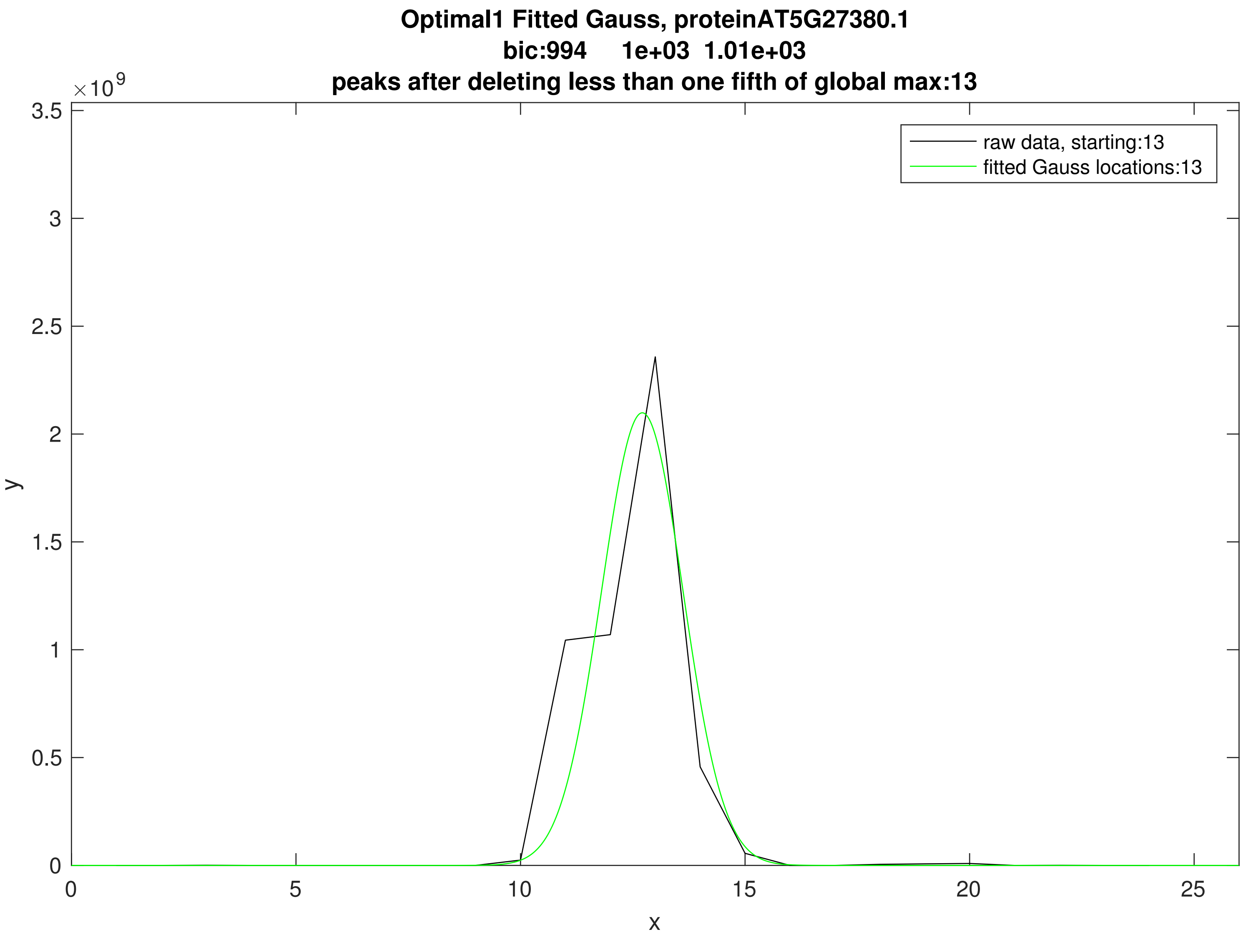

Optimal1 Fitted Gauss, proteinAT5G27380.1
bic:994 1e+03 1.01e+03
peaks after deleting less than one fifth of global max:13
×10^9
3.5
3
2.5
2
1.5
1
0.5
0
y
0
5
10
15
20
25
x
raw data, starting:13
fitted Gauss locations:13

#### Optimal1 Fitted Gauss, proteinAT5G27410.1;AT5G27410.2
#### bic:908 917
#### peaks after deleting less than one fifth of global max:6.9

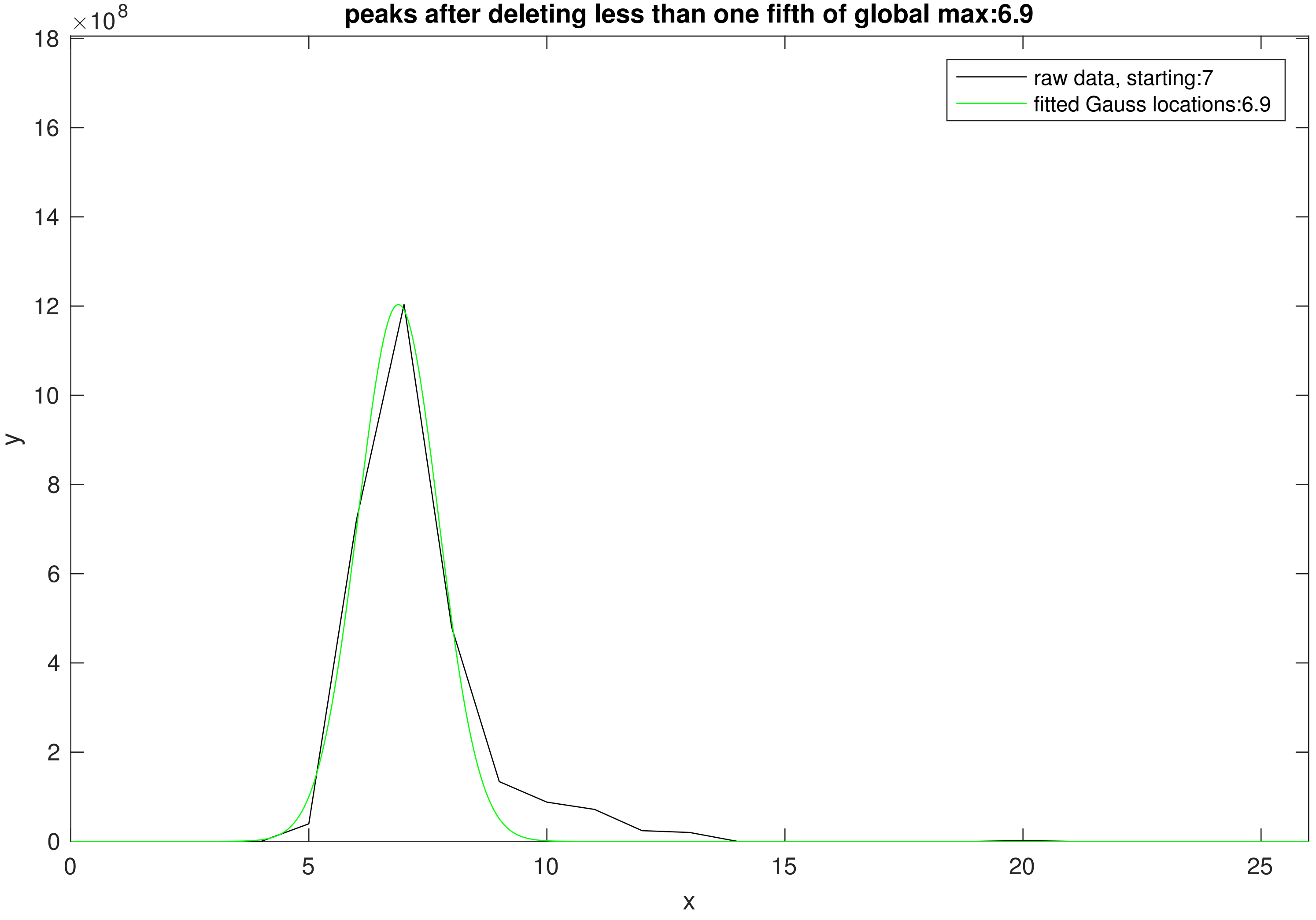

#### Optimal1 Fitted Gauss, proteinAT5G27450.3;AT5G27450.2;AT5G27450.1
#### bic:948 957
#### peaks after deleting less than one fifth of global max:14

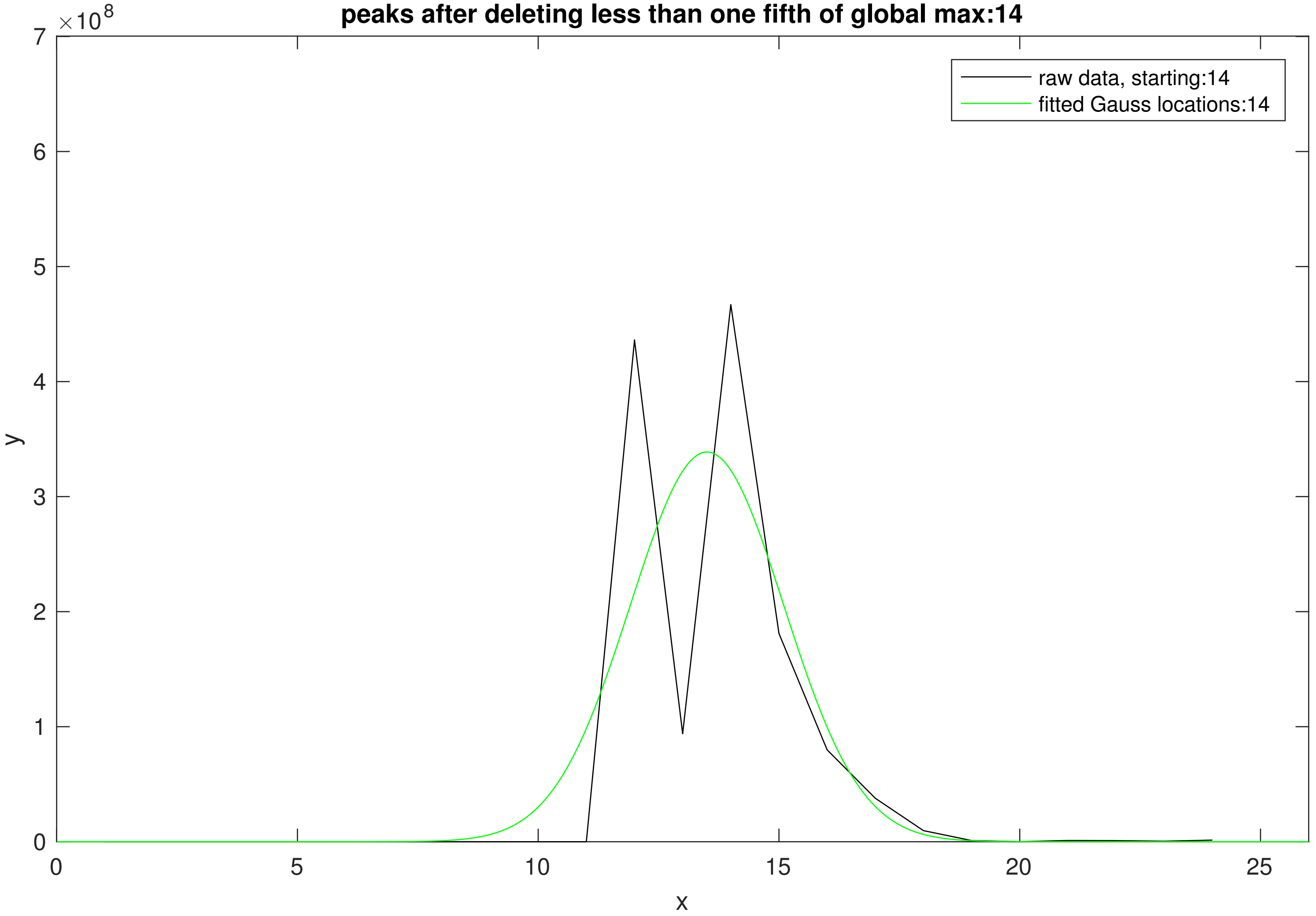

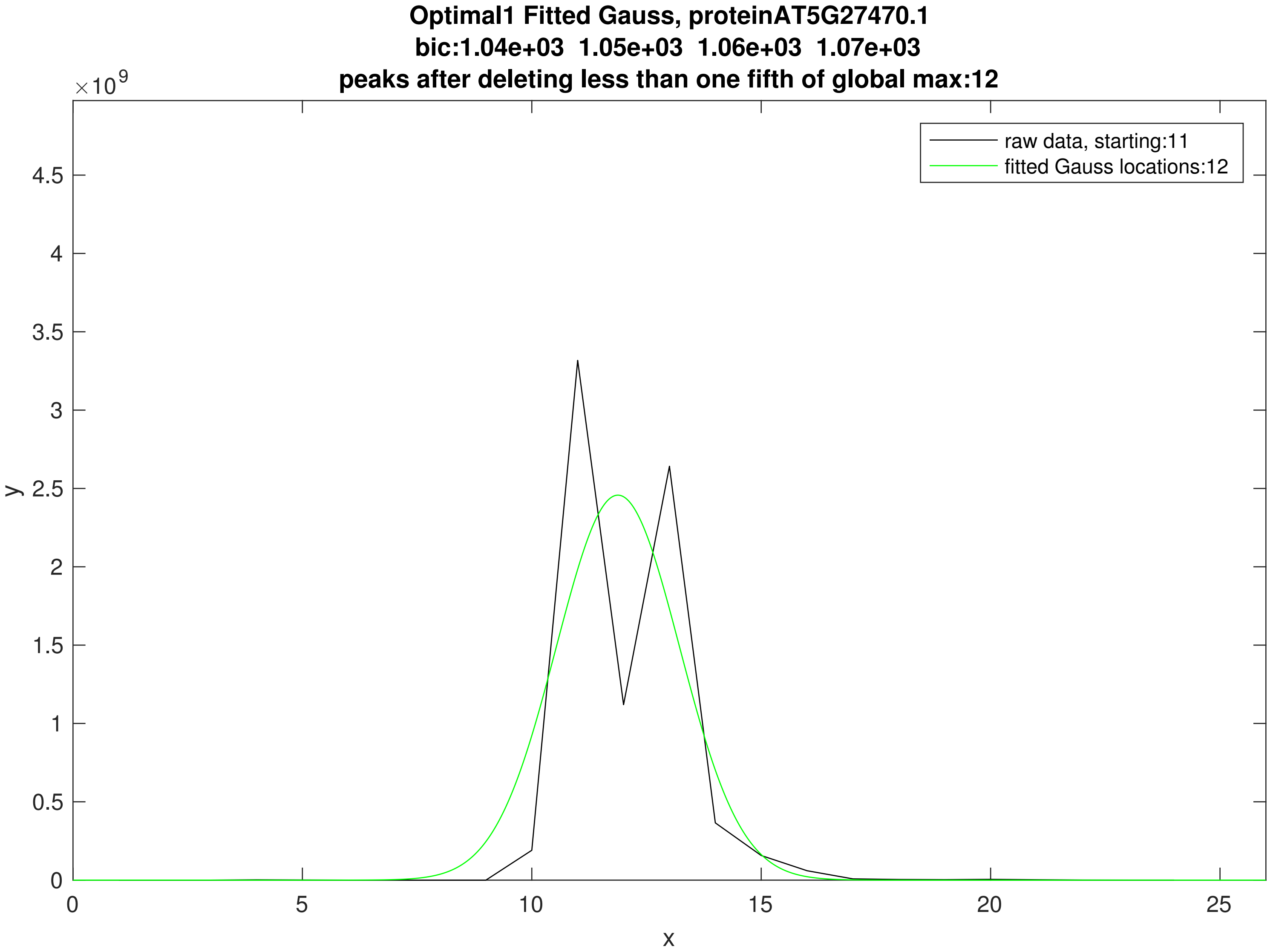

Optimal1 Fitted Gauss, proteinAT5G27470.1
bic:1.04e+03 1.05e+03 1.06e+03 1.07e+03
peaks after deleting less than one fifth of global max:12
×10^9
raw data, starting:11
fitted Gauss locations:12
4.5
4
3.5
3
2.5
2
1.5
1
0.5
0
y
0
5
10
15
20
25
x

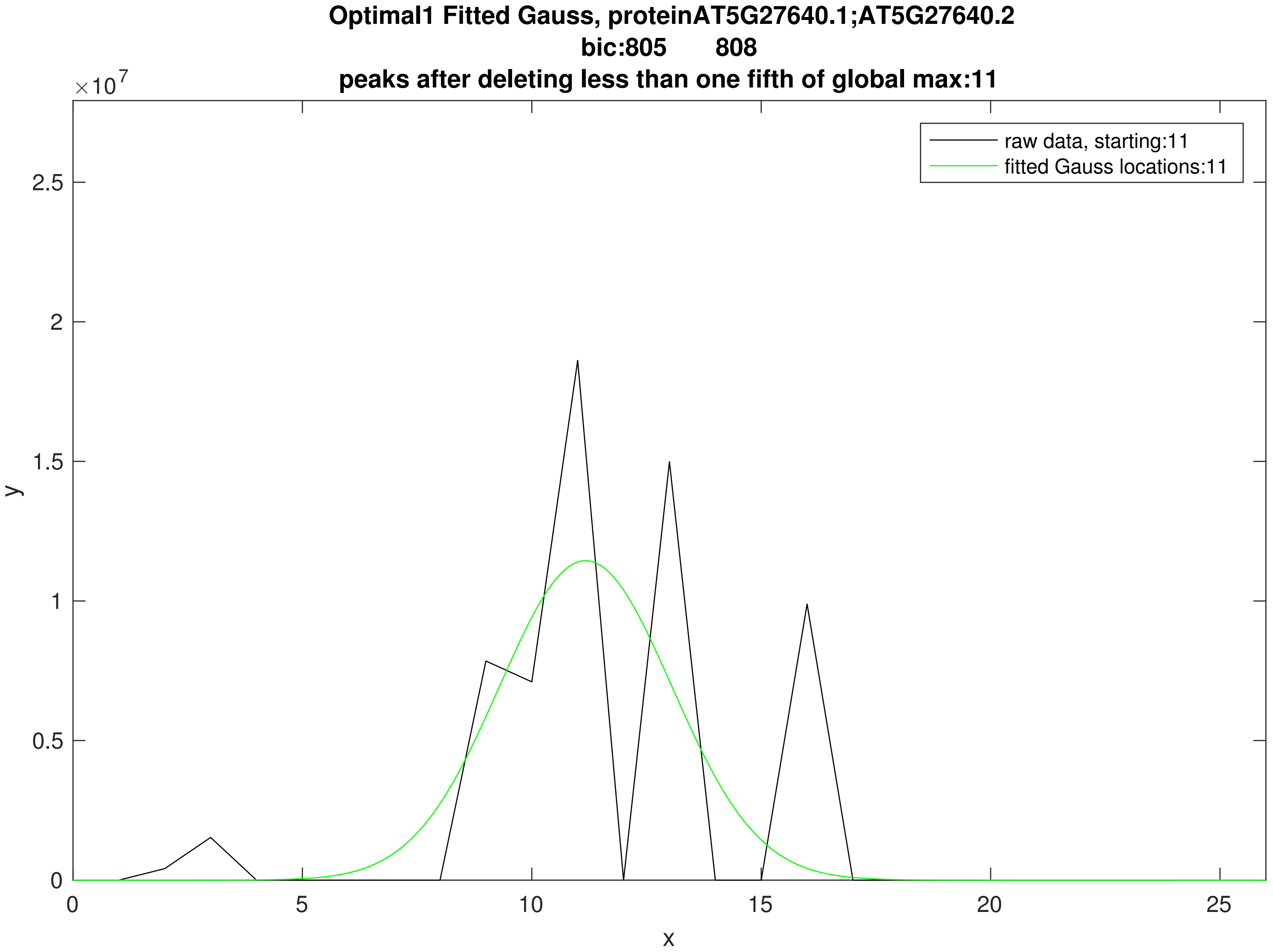

Optimal1 Fitted Gauss, proteinAT5G27640.1;AT5G27640.2
bic:805 808
peaks after deleting less than one fifth of global max:11
×10^7
raw data, starting:11
fitted Gauss locations:11
2.5
2
1.5
1
0.5
0
y
0
5
10
15
20
25
x

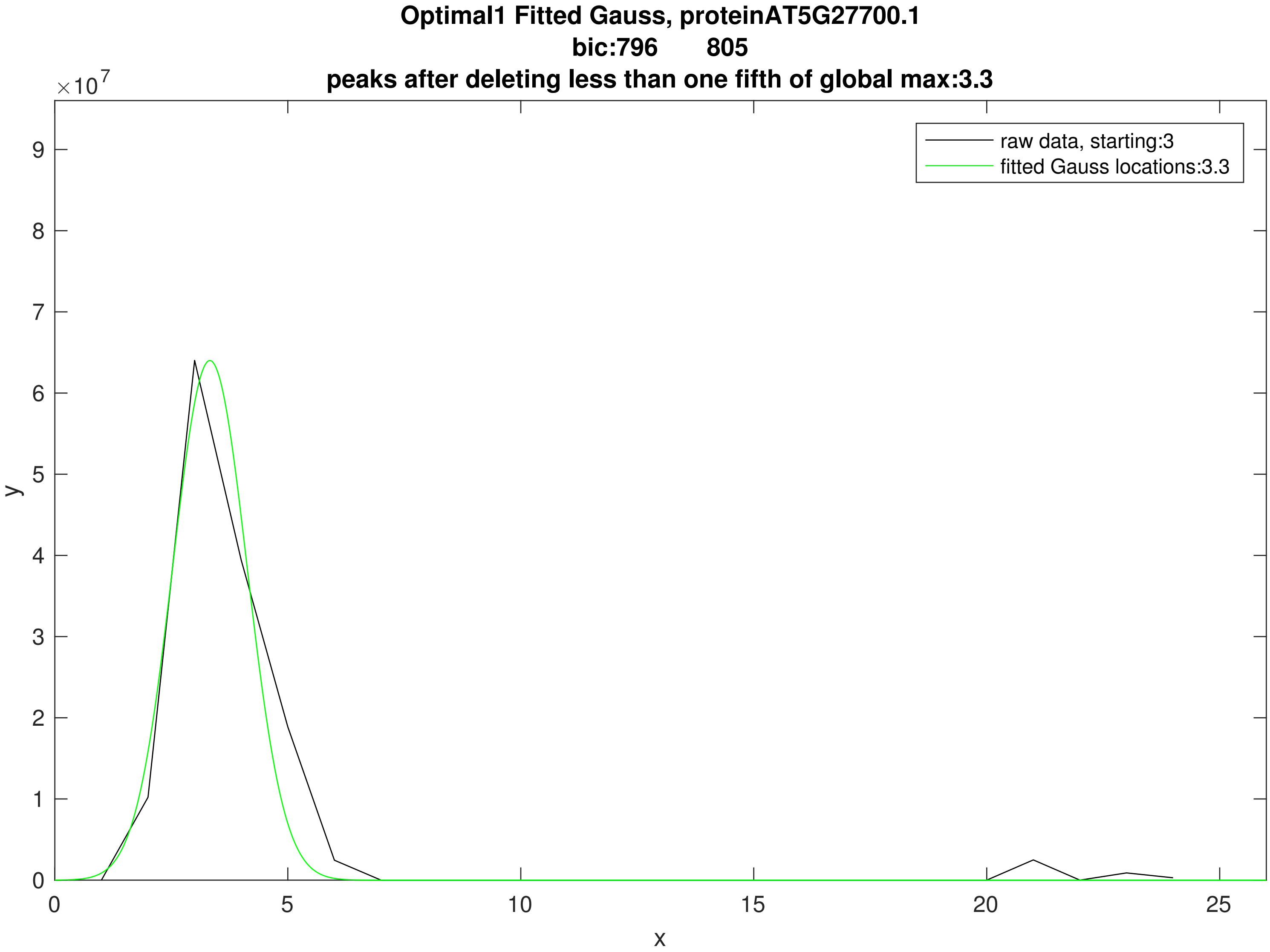

Optimal1 Fitted Gauss, proteinAT5G27700.1
bic:796 805
peaks after deleting less than one fifth of global max:3.3
×10^7
raw data, starting:3
fitted Gauss locations:3.3
y
x
0
1
2
3
4
5
6
7
8
9
0
5
10
15
20
25

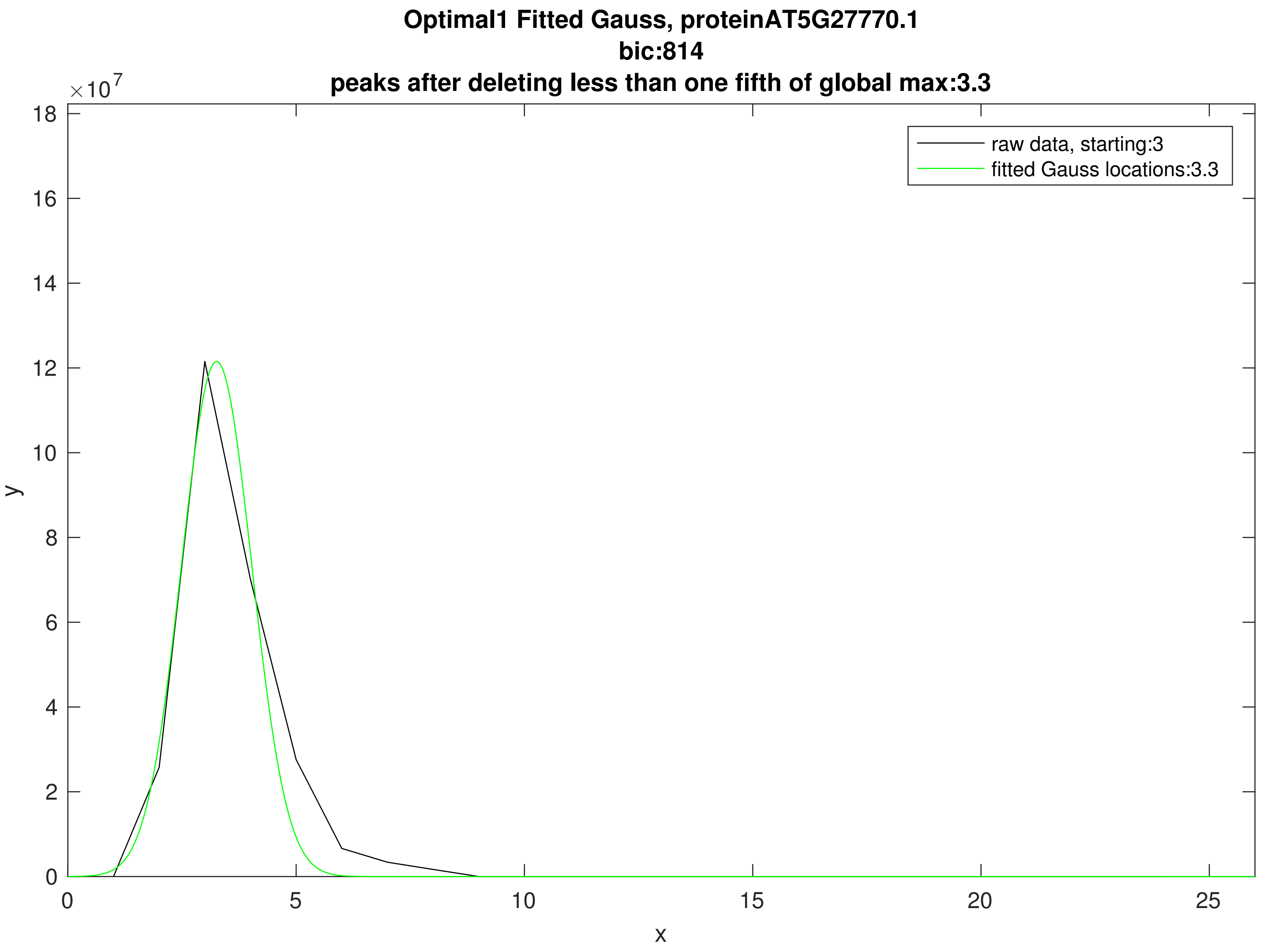

Optimal1 Fitted Gauss, proteinAT5G27770.1
bic:814
peaks after deleting less than one fifth of global max:3.3
×10^7
18
16
14
12
10
8
6
4
2
0
y
0
5
10
15
20
25
x
raw data, starting:3
fitted Gauss locations:3.3

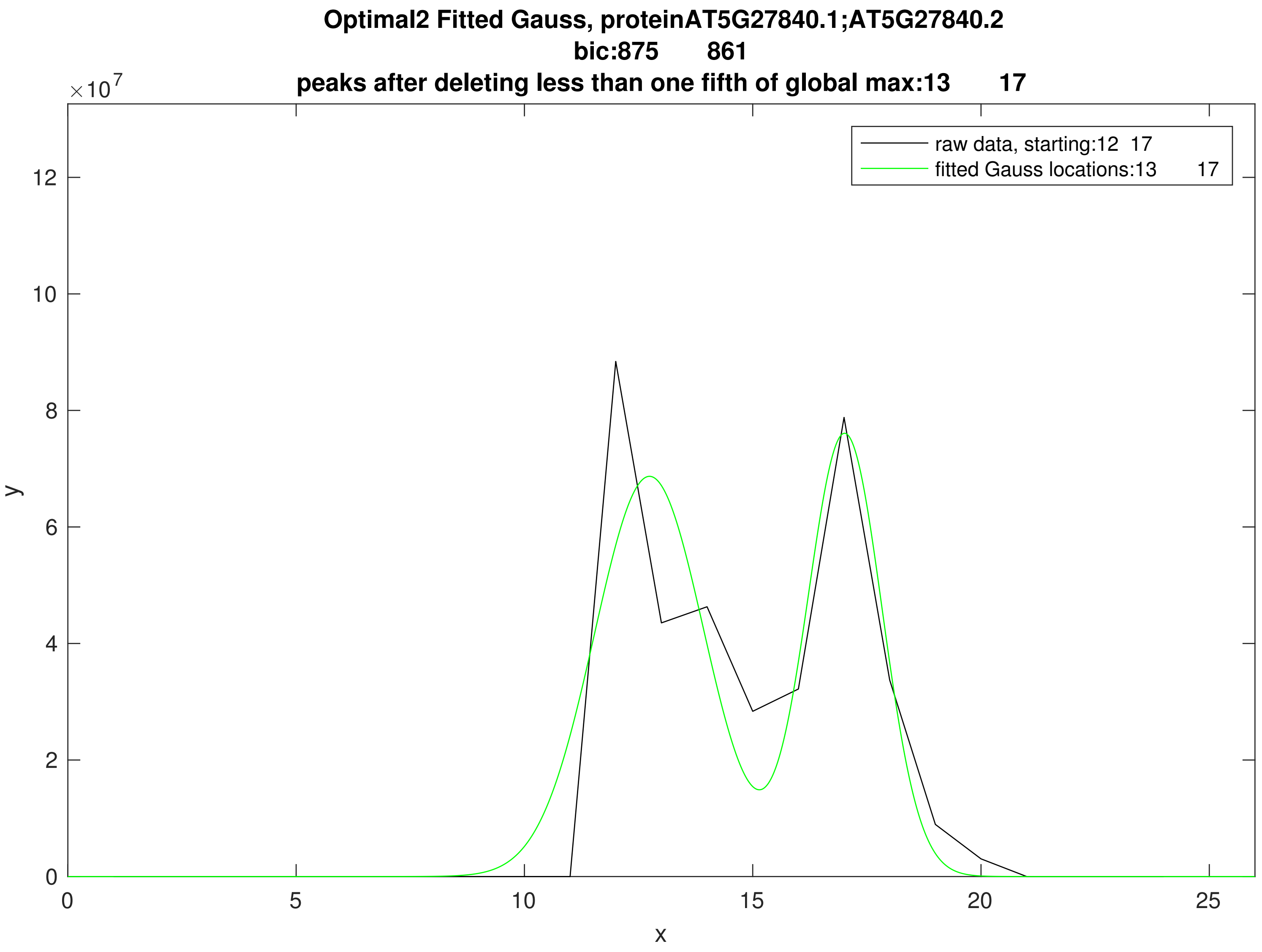

Optimal2 Fitted Gauss, proteinAT5G27840.1;AT5G27840.2
bic:875 861
peaks after deleting less than one fifth of global max:13 17
×10^7
raw data, starting:12 17
fitted Gauss locations:13 17
y
x
0
2
4
6
8
10
12
0
5
10
15
20
25

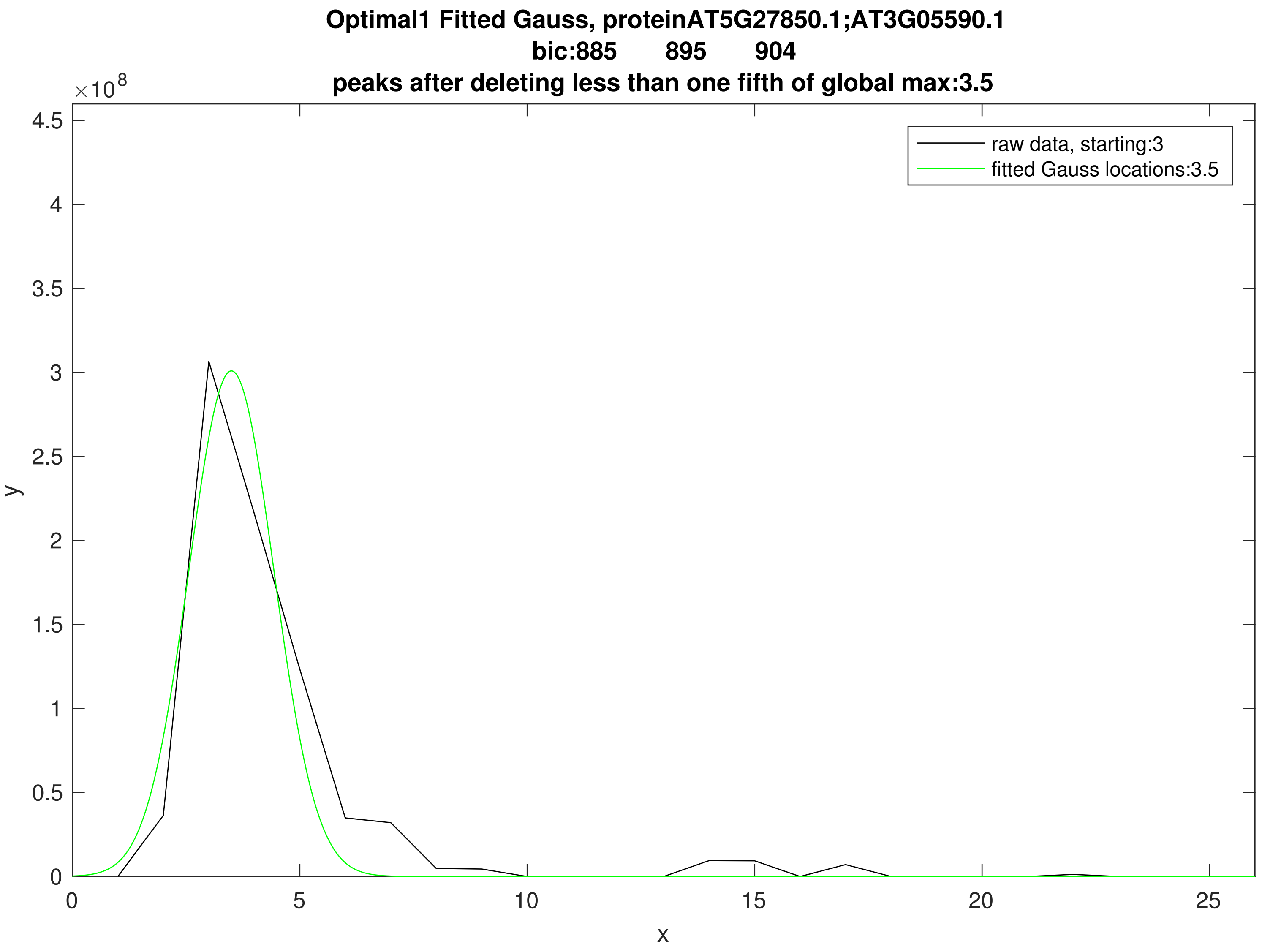
Optimal1 Fitted Gauss, proteinAT5G27850.1;AT3G05590.1
bic:885 895 904
peaks after deleting less than one fifth of global max:3.5
×10^8
4.5
4
3.5
3
2.5
2
1.5
1
0.5
0
y
0
5
10
15
20
25
x
raw data, starting:3
fitted Gauss locations:3.5

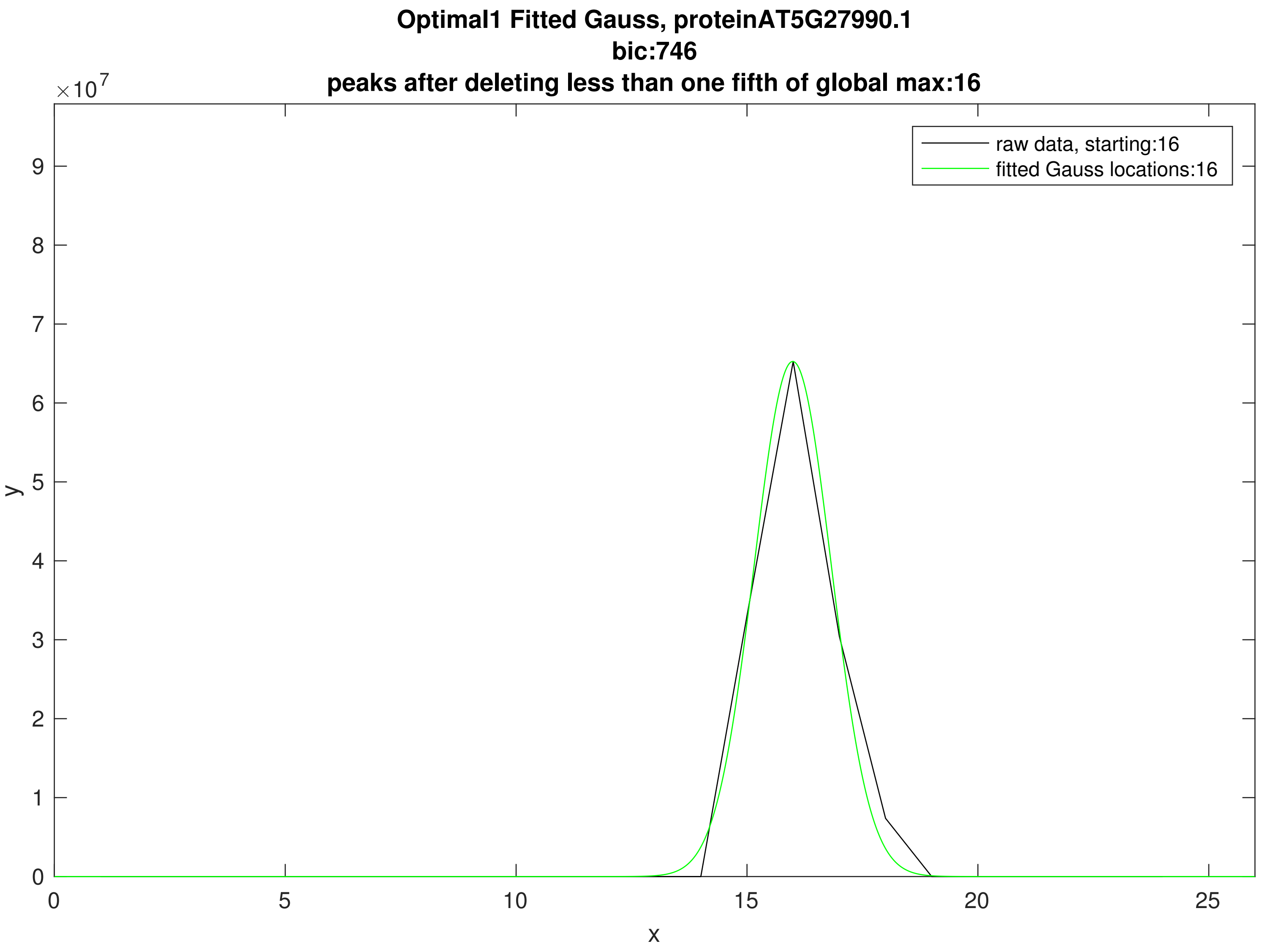
Optimal1 Fitted Gauss, proteinAT5G27990.1
bic:746
peaks after deleting less than one fifth of global max:16
×10^7
raw data, starting:16
fitted Gauss locations:16
y
x
0
1
2
3
4
5
6
7
8
9
0
5
10
15
20
25

**Optimal1 Fitted Gauss, proteinAT5G28020.6;AT5G28020.4;AT5G28020.3;AT5G28020.2;AT5G28020.1;AT5G2802[illegible]**
**bic:928 938 947**
**peaks after deleting less than one fifth of global max:15**

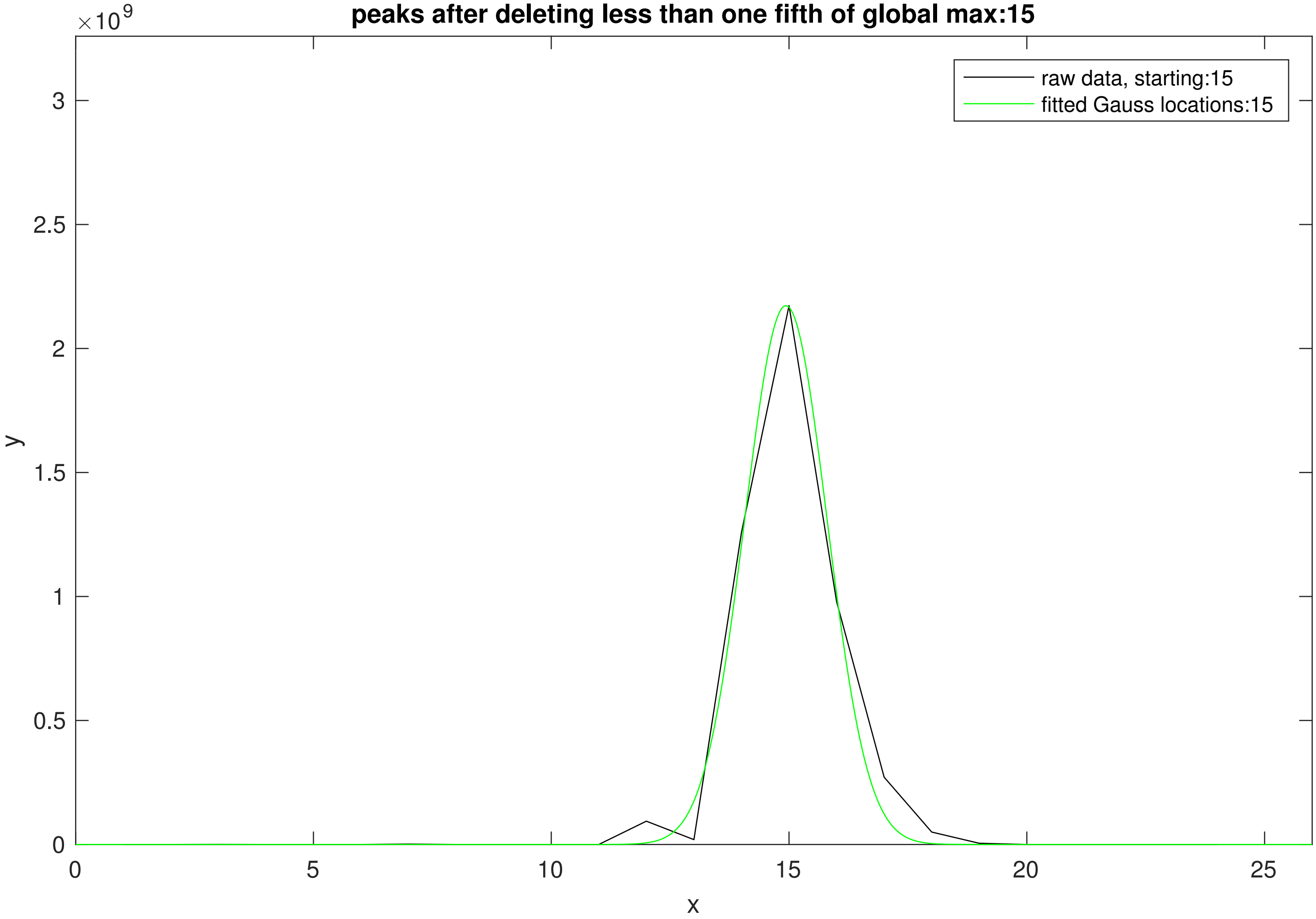

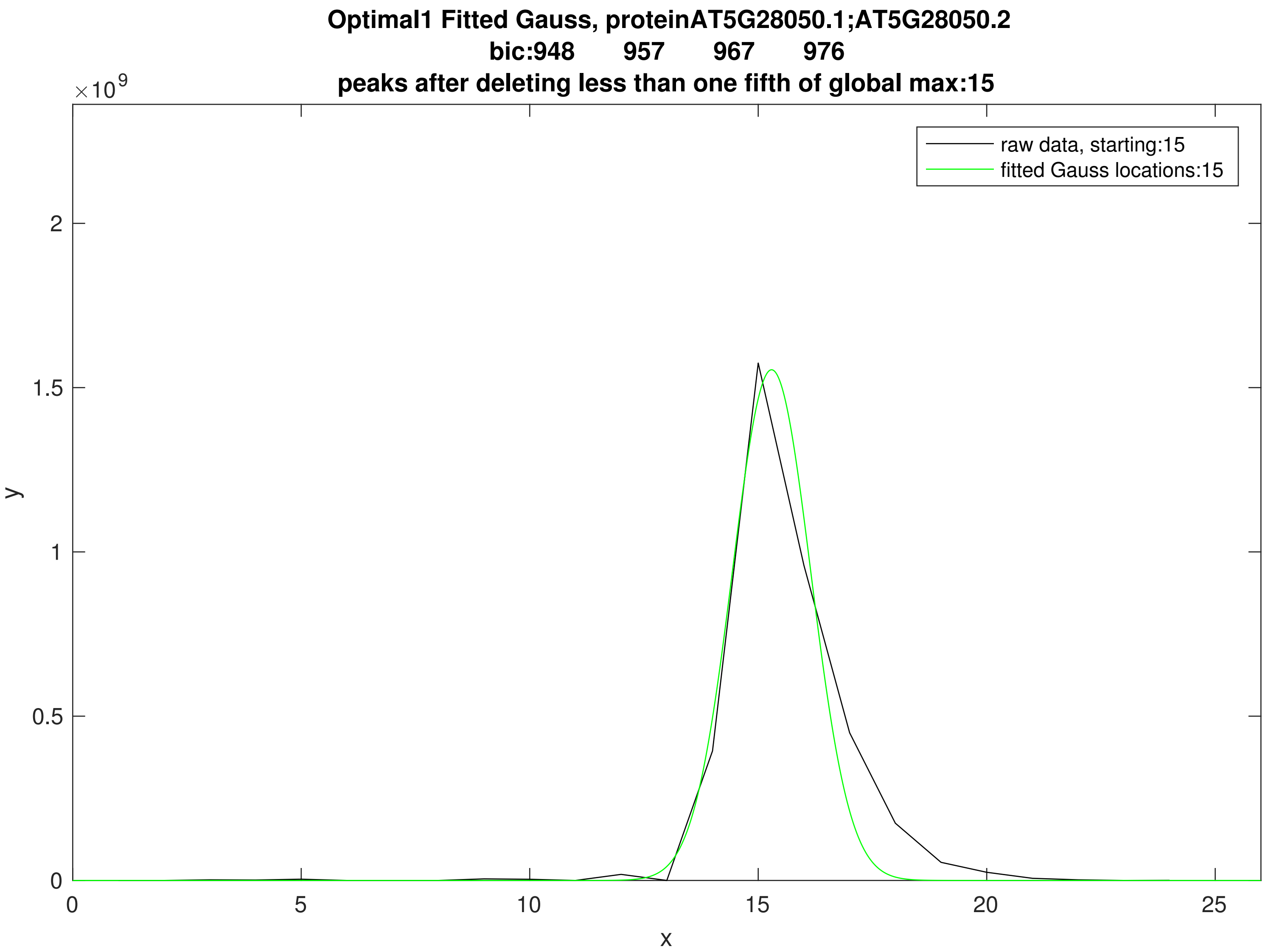
Optimal1 Fitted Gauss, proteinAT5G28050.1;AT5G28050.2
bic:948 957 967 976
peaks after deleting less than one fifth of global max:15
×10^9
raw data, starting:15
fitted Gauss locations:15
y
x
0
0.5
1
1.5
2
0
5
10
15
20
25

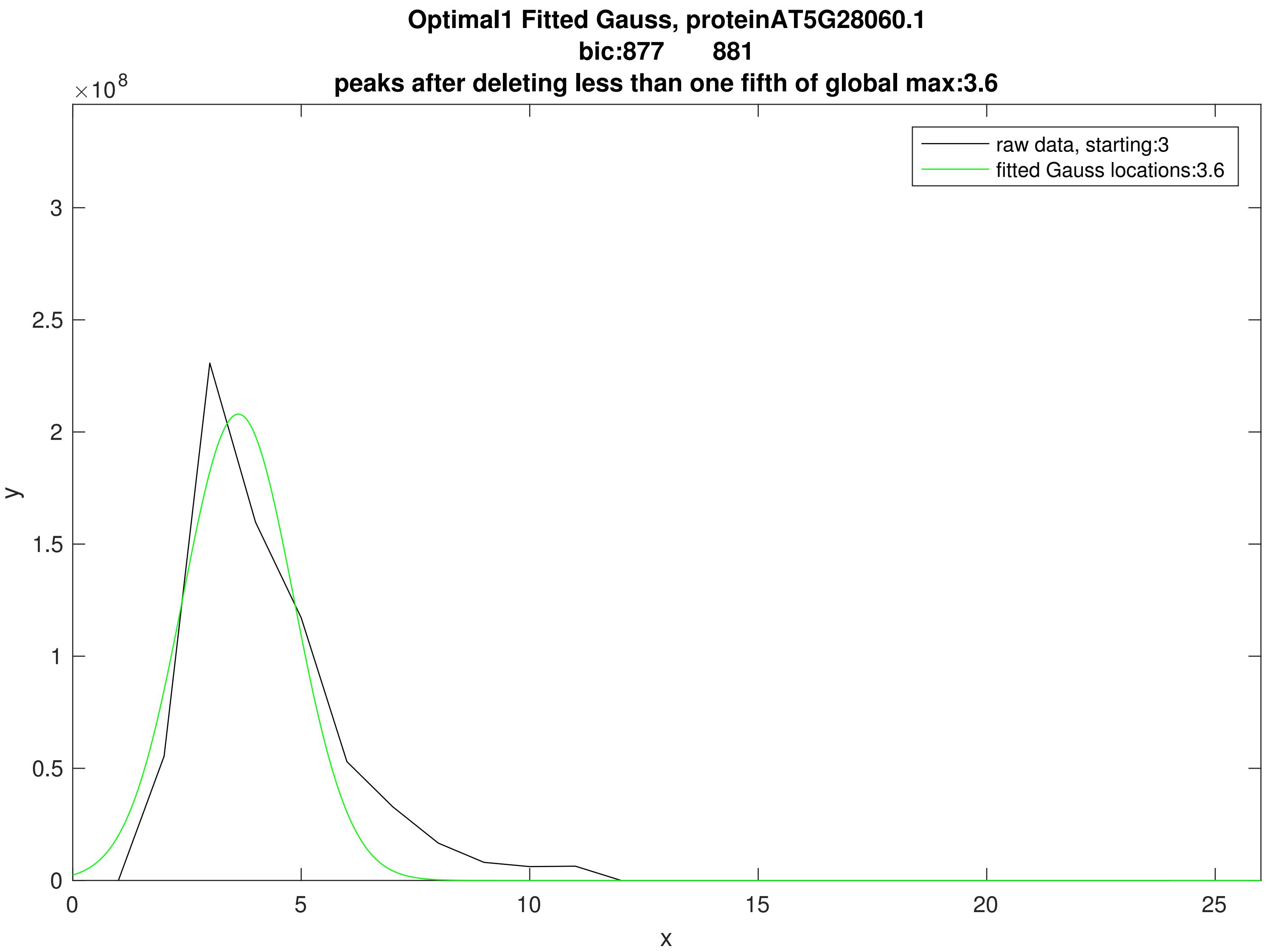

Optimal1 Fitted Gauss, proteinAT5G28060.1
bic:877 881
peaks after deleting less than one fifth of global max:3.6
×10^8
raw data, starting:3
fitted Gauss locations:3.6
y
x
0
0.5
1
1.5
2
2.5
3
0
5
10
15
20
25

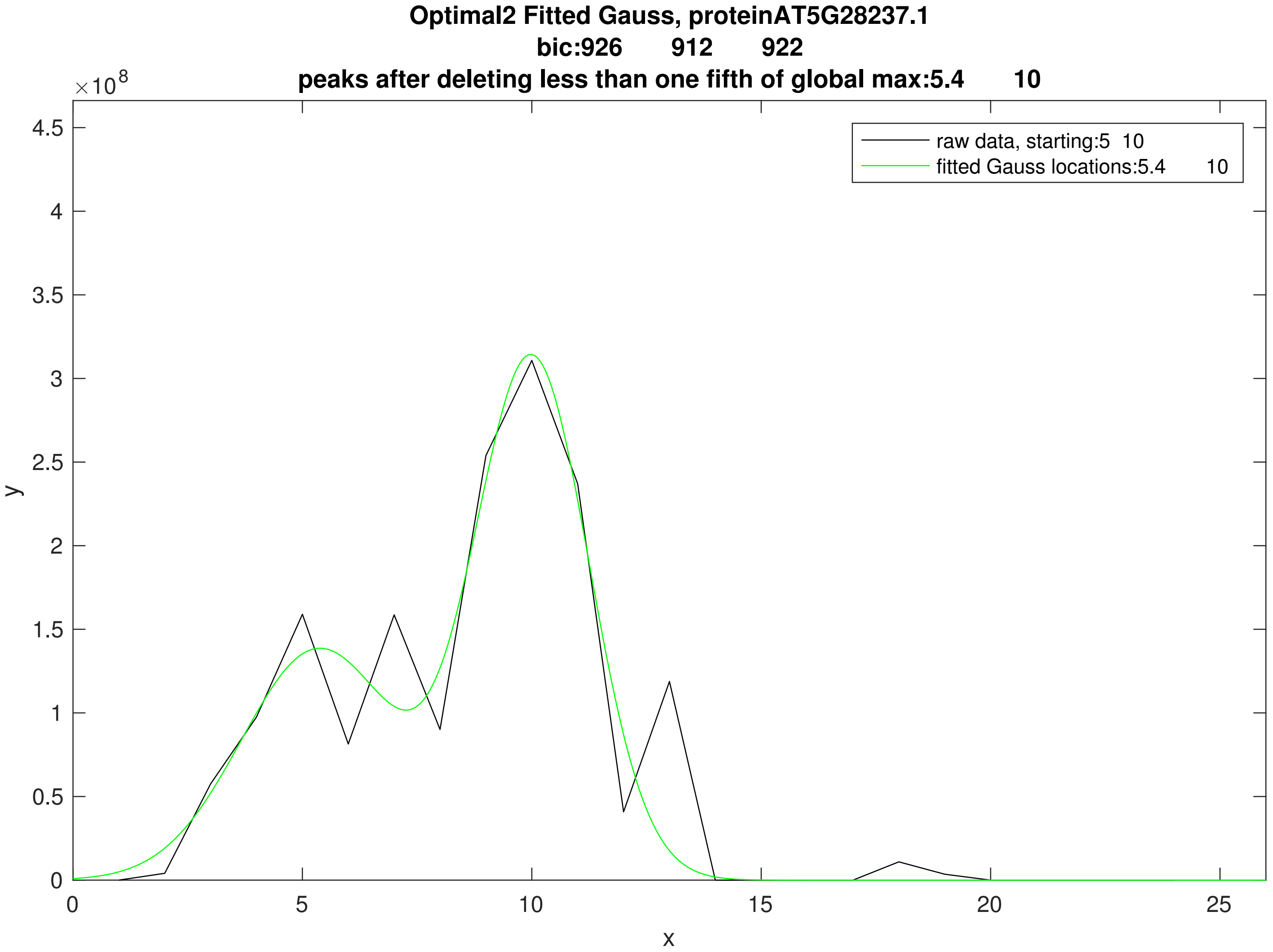
Optimal2 Fitted Gauss, proteinAT5G28237.1
bic:926 912 922
peaks after deleting less than one fifth of global max:5.4 10
×10^8
4.5
4
3.5
3
2.5
2
1.5
1
0.5
0
y
0
5
10
15
20
25
x
raw data, starting:5 10
fitted Gauss locations:5.4 10

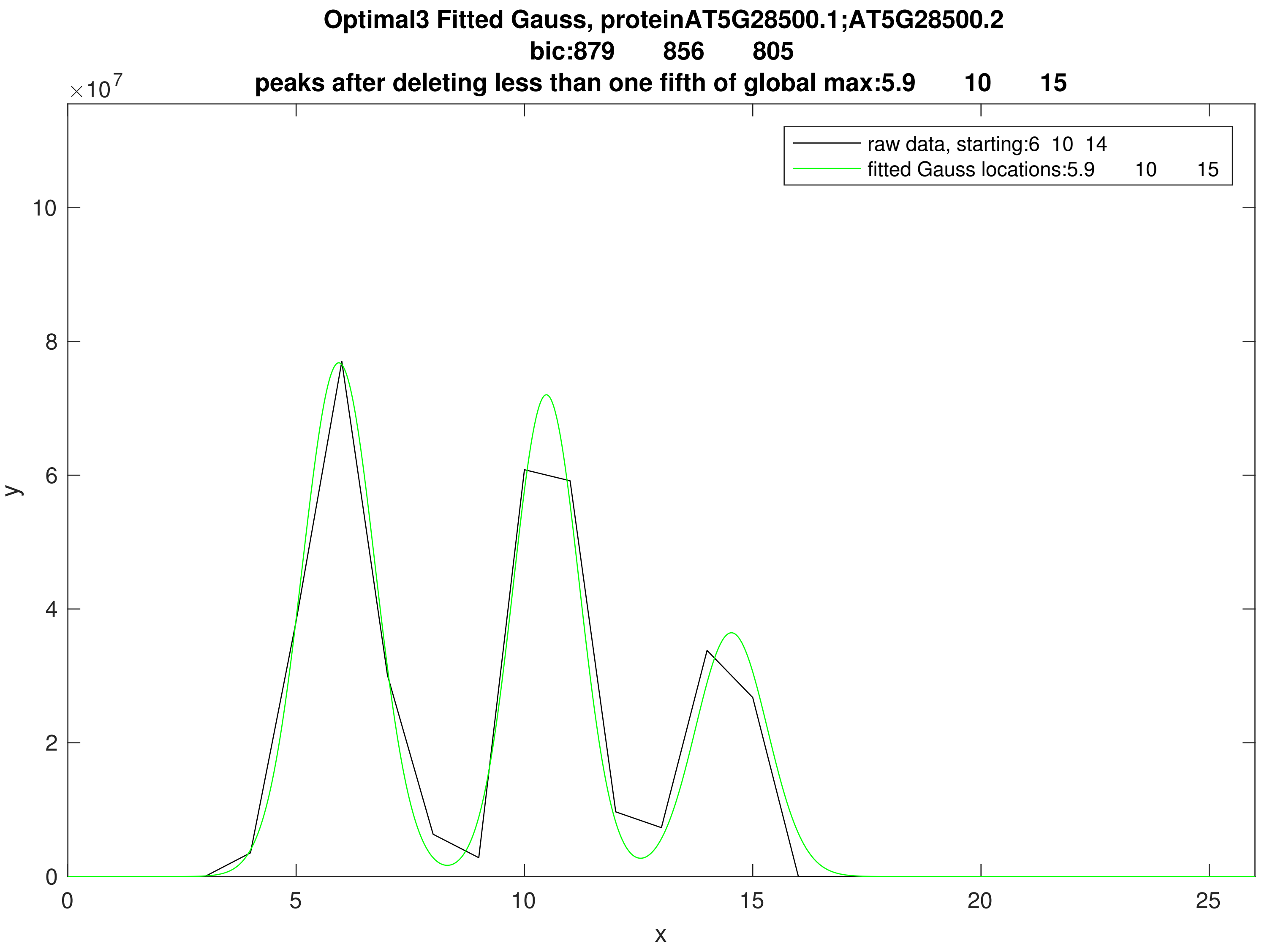
Optimal3 Fitted Gauss, proteinAT5G28500.1;AT5G28500.2
bic:879 856 805
peaks after deleting less than one fifth of global max:5.9 10 15
×10^7
raw data, starting:6 10 14
fitted Gauss locations:5.9 10 15
y
x
0
2
4
6
8
10
5
15
20
25

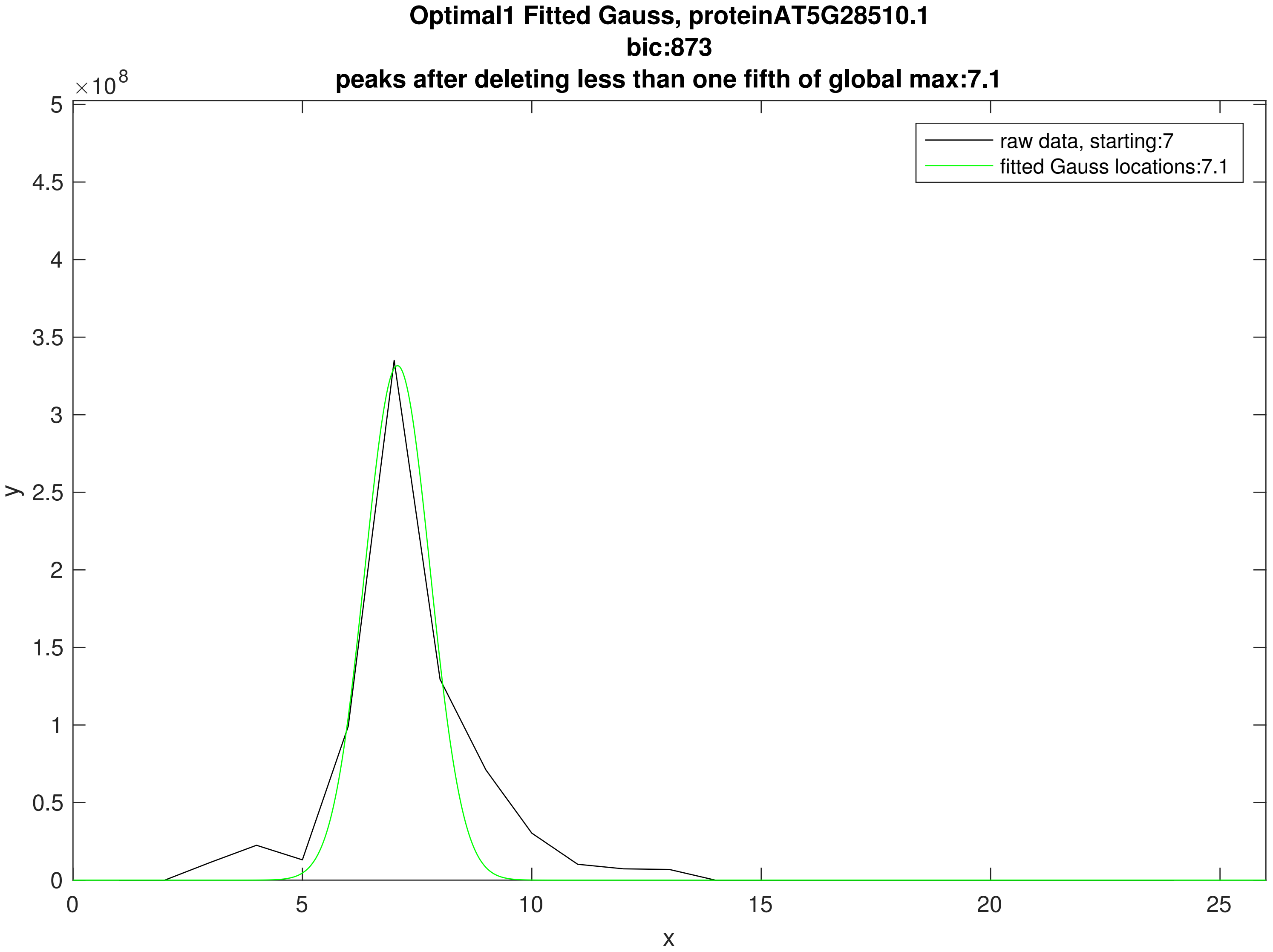
Optimal1 Fitted Gauss, proteinAT5G28510.1
bic:873
peaks after deleting less than one fifth of global max:7.1
×10^8
raw data, starting:7
fitted Gauss locations:7.1
y
x
0
0.5
1
1.5
2
2.5
3
3.5
4
4.5
5
0
5
10
15
20
25

**Optimal3 Fitted Gauss, proteinAT5G28540.1**
**bic:898 901 886 892**
**peaks after deleting less than one fifth of global max:13**

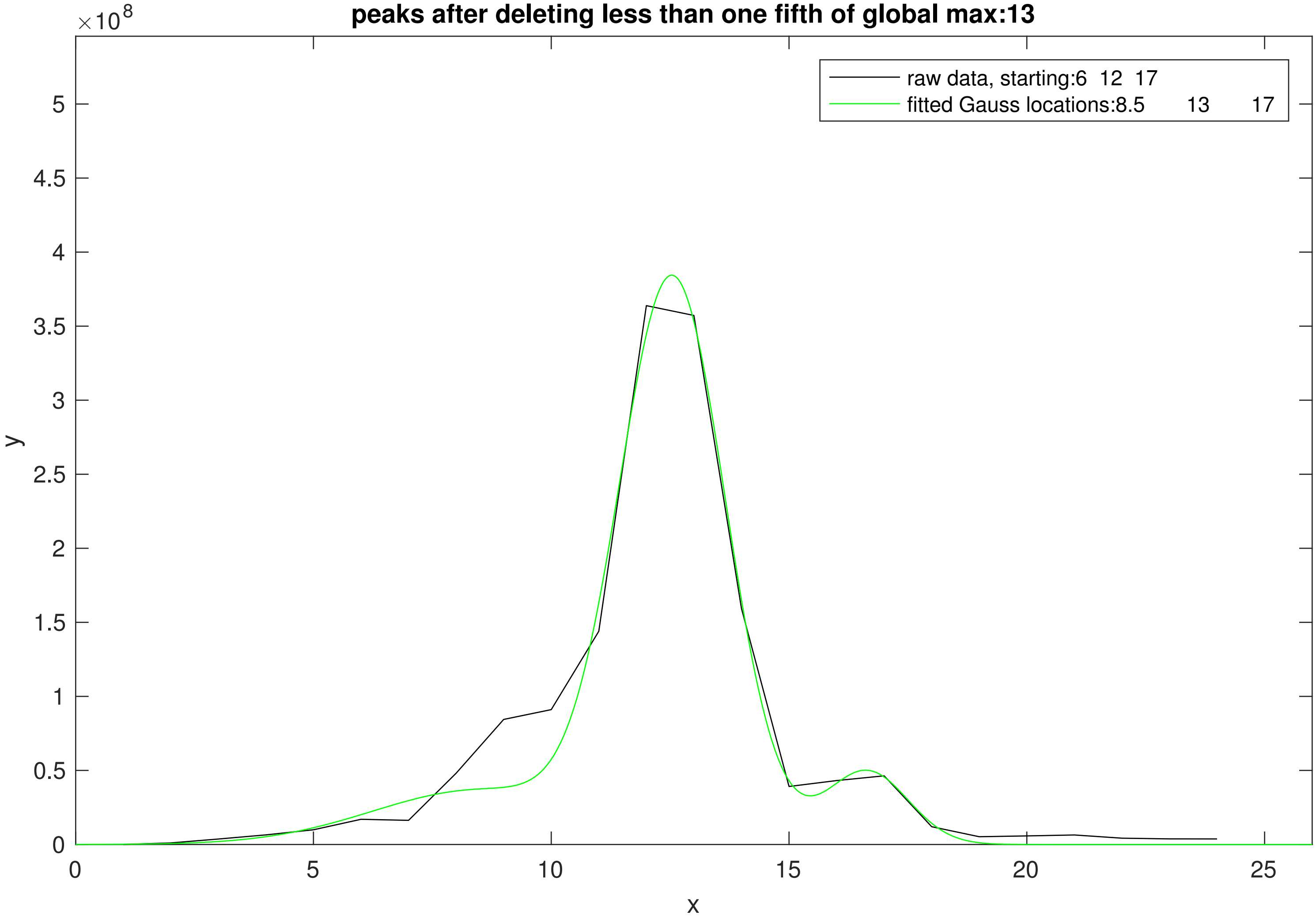

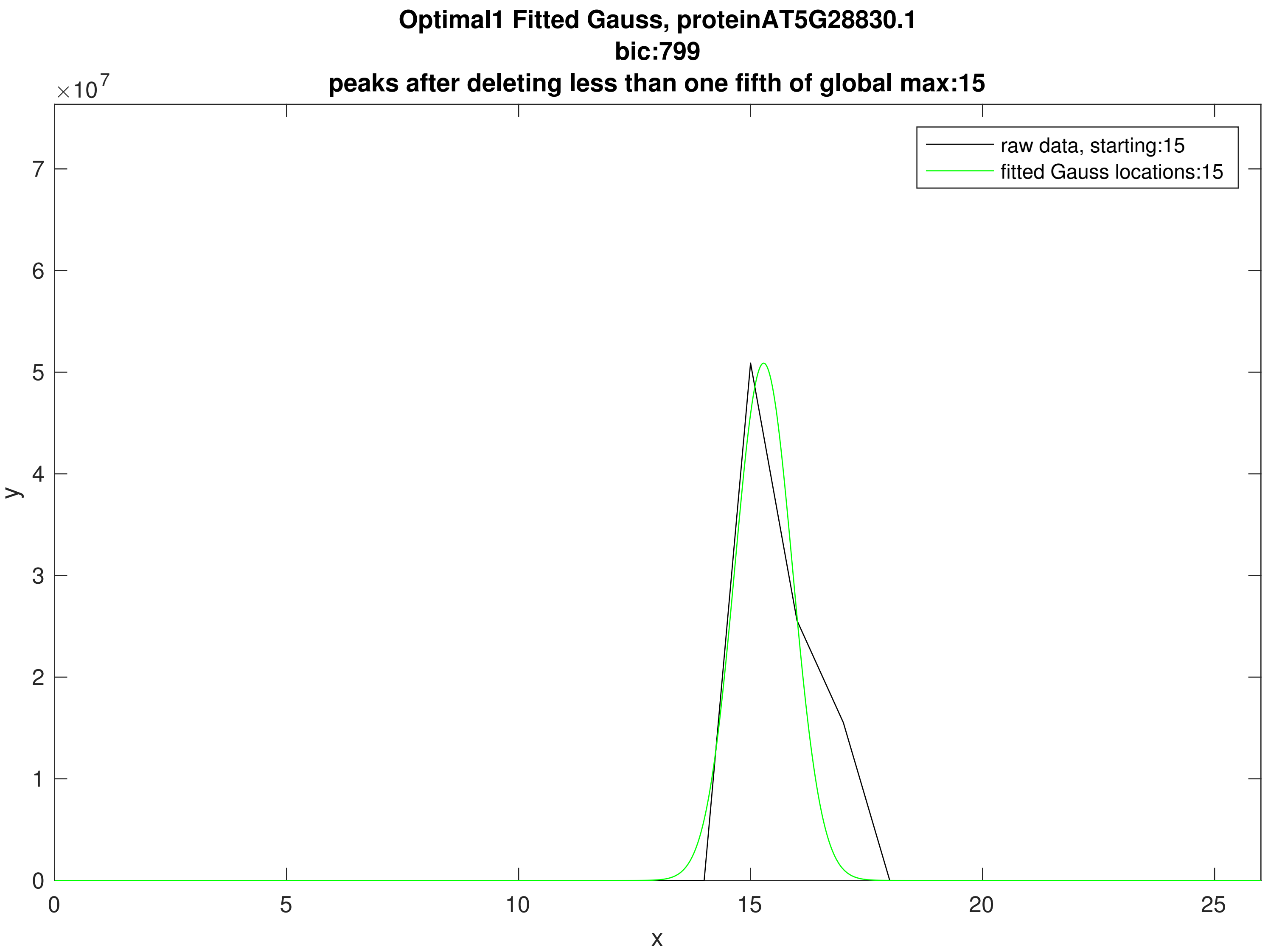

Optimal1 Fitted Gauss, proteinAT5G28830.1
bic:799
peaks after deleting less than one fifth of global max:15
×10^7
raw data, starting:15
fitted Gauss locations:15
y
x
0
1
2
3
4
5
6
7
0
5
10
15
20
25

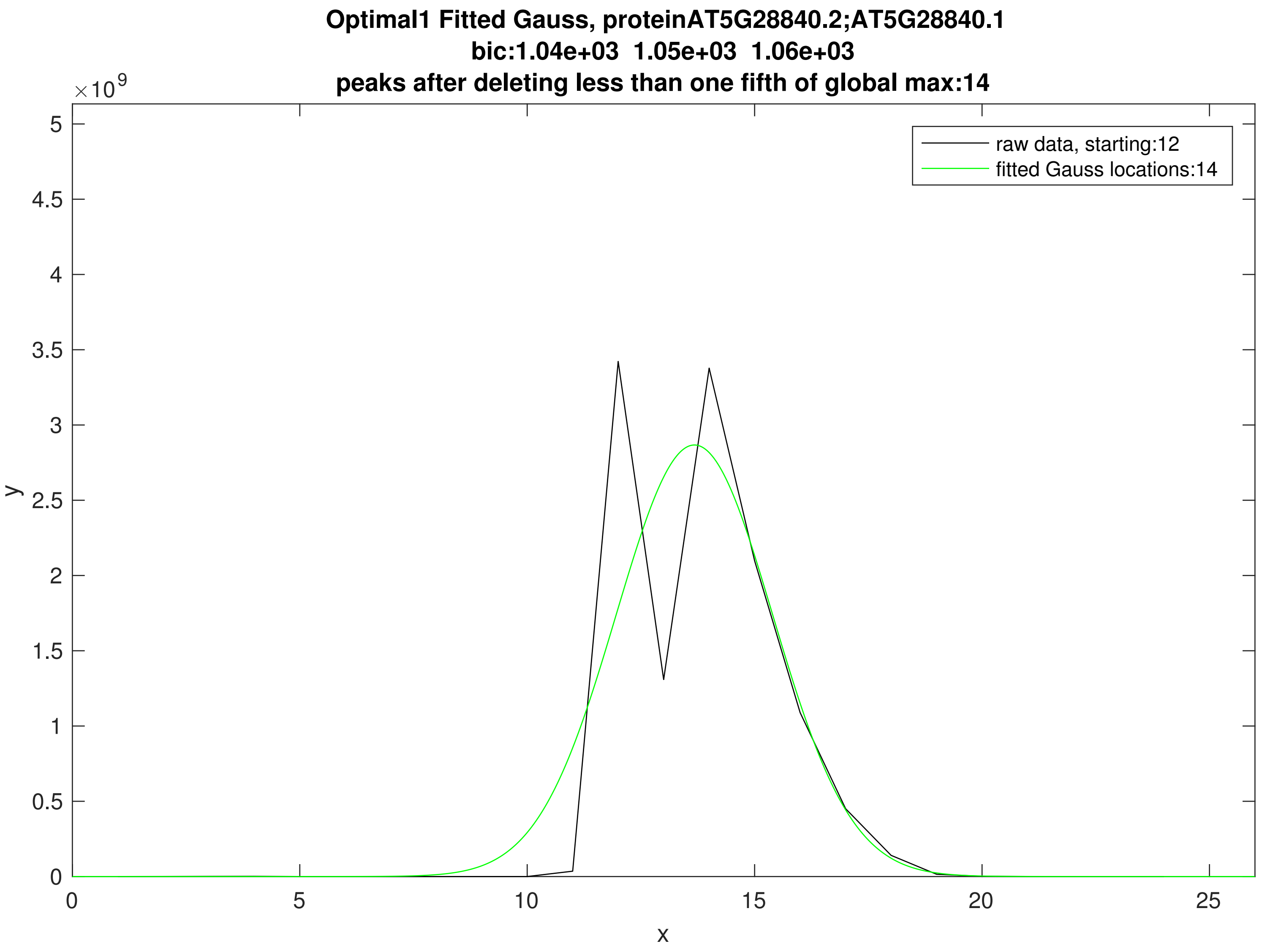
Optimal1 Fitted Gauss, proteinAT5G28840.2;AT5G28840.1
bic:1.04e+03  1.05e+03  1.06e+03
peaks after deleting less than one fifth of global max:14
×10^9
raw data, starting:12
fitted Gauss locations:14
y
x
0
0.5
1
1.5
2
2.5
3
3.5
4
4.5
5
0
5
10
15
20
25

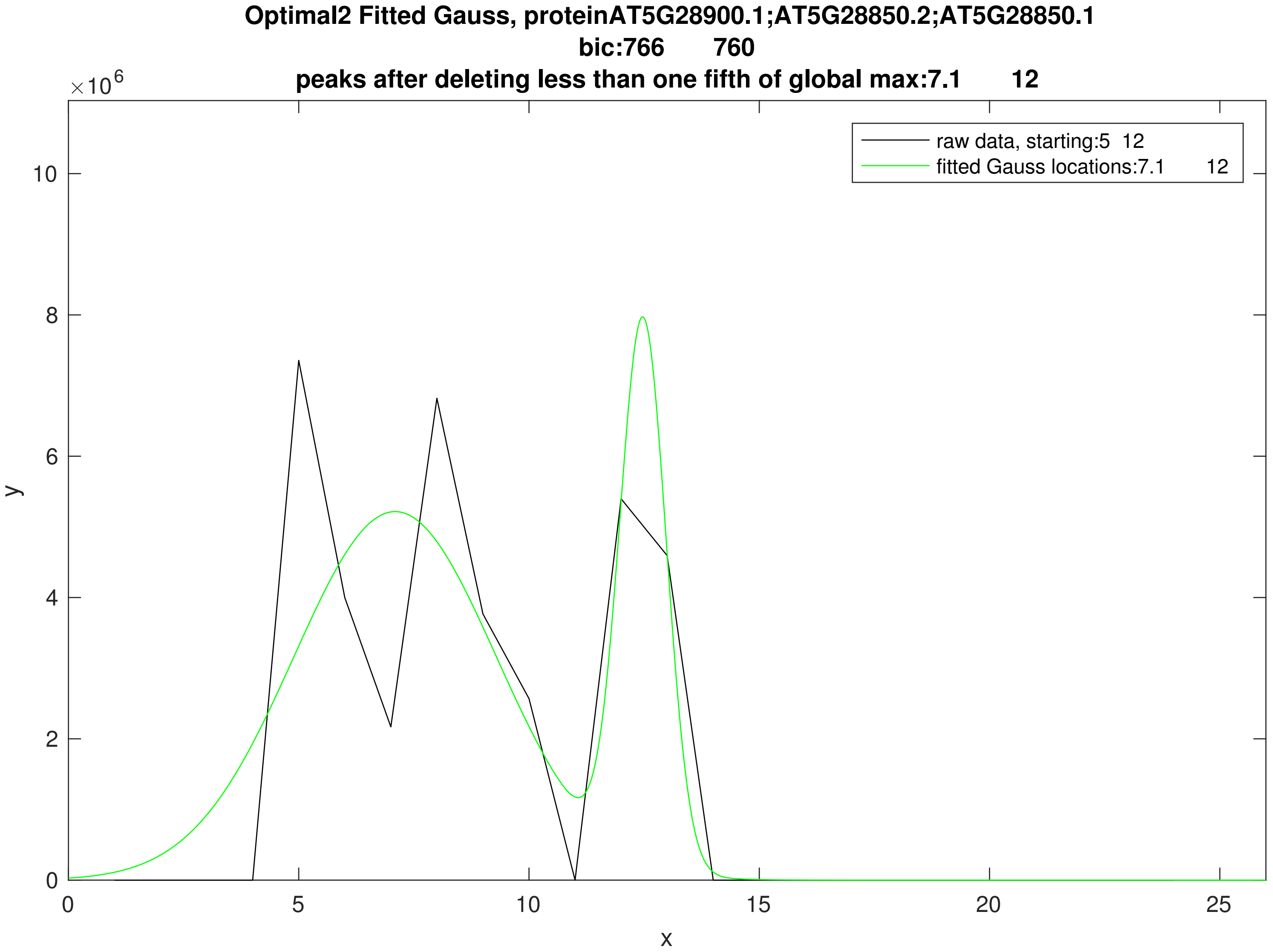

Optimal2 Fitted Gauss, proteinAT5G28900.1;AT5G28850.2;AT5G28850.1
bic:766 760
peaks after deleting less than one fifth of global max:7.1 12
×10^6
raw data, starting:5 12
fitted Gauss locations:7.1 12
y
x
0
2
4
6
8
10
0
5
10
15
20
25

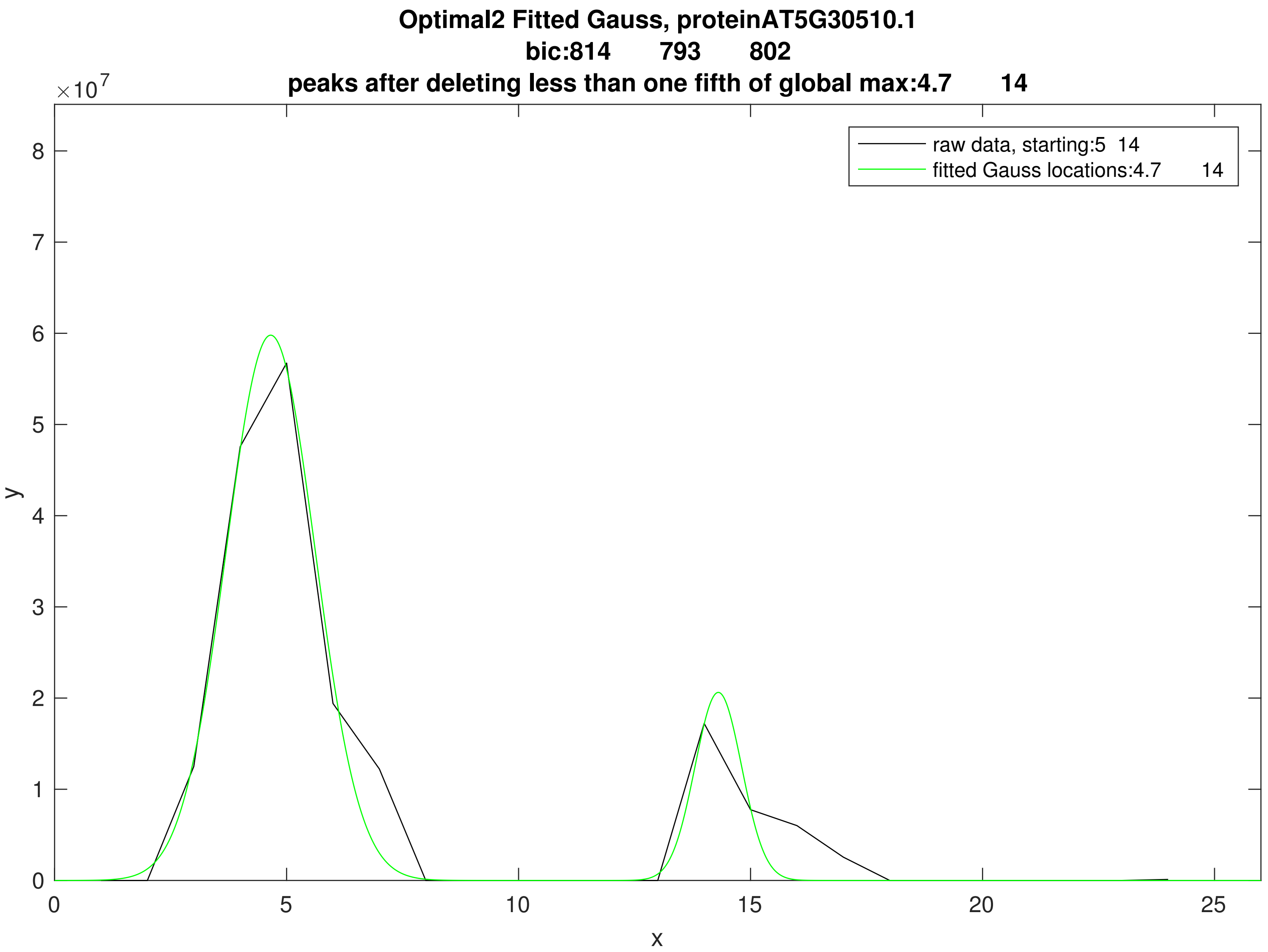
Optimal2 Fitted Gauss, proteinAT5G30510.1
bic:814 793 802
peaks after deleting less than one fifth of global max:4.7 14
×10^7
raw data, starting:5 14
fitted Gauss locations:4.7 14
y
x
0
1
2
3
4
5
6
7
8
0
5
10
15
20
25

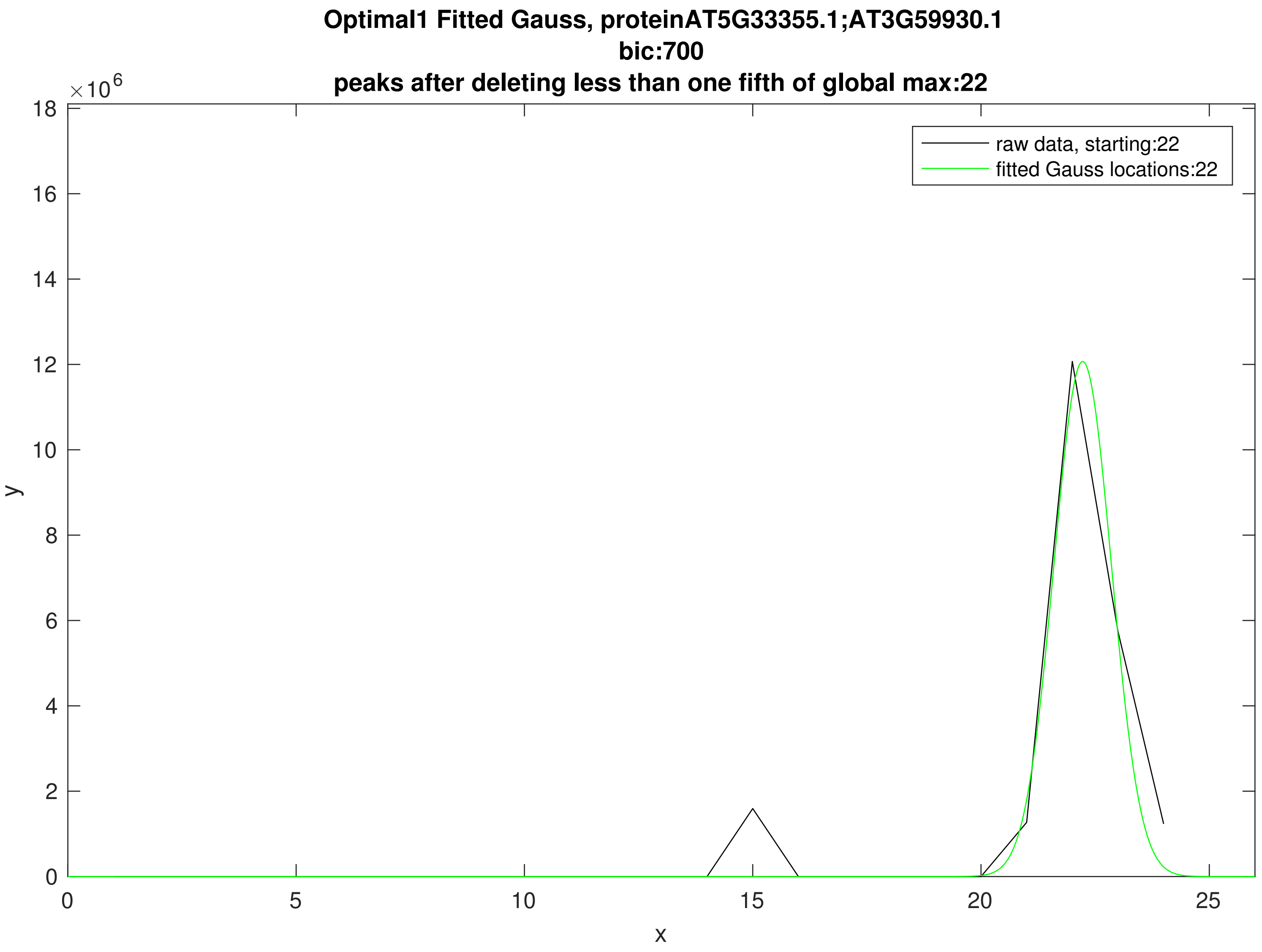
Optimal1 Fitted Gauss, proteinAT5G33355.1;AT3G59930.1
bic:700
peaks after deleting less than one fifth of global max:22
×10^6
raw data, starting:22
fitted Gauss locations:22
y
x
0
2
4
6
8
10
12
14
16
18
0
5
10
15
20
25

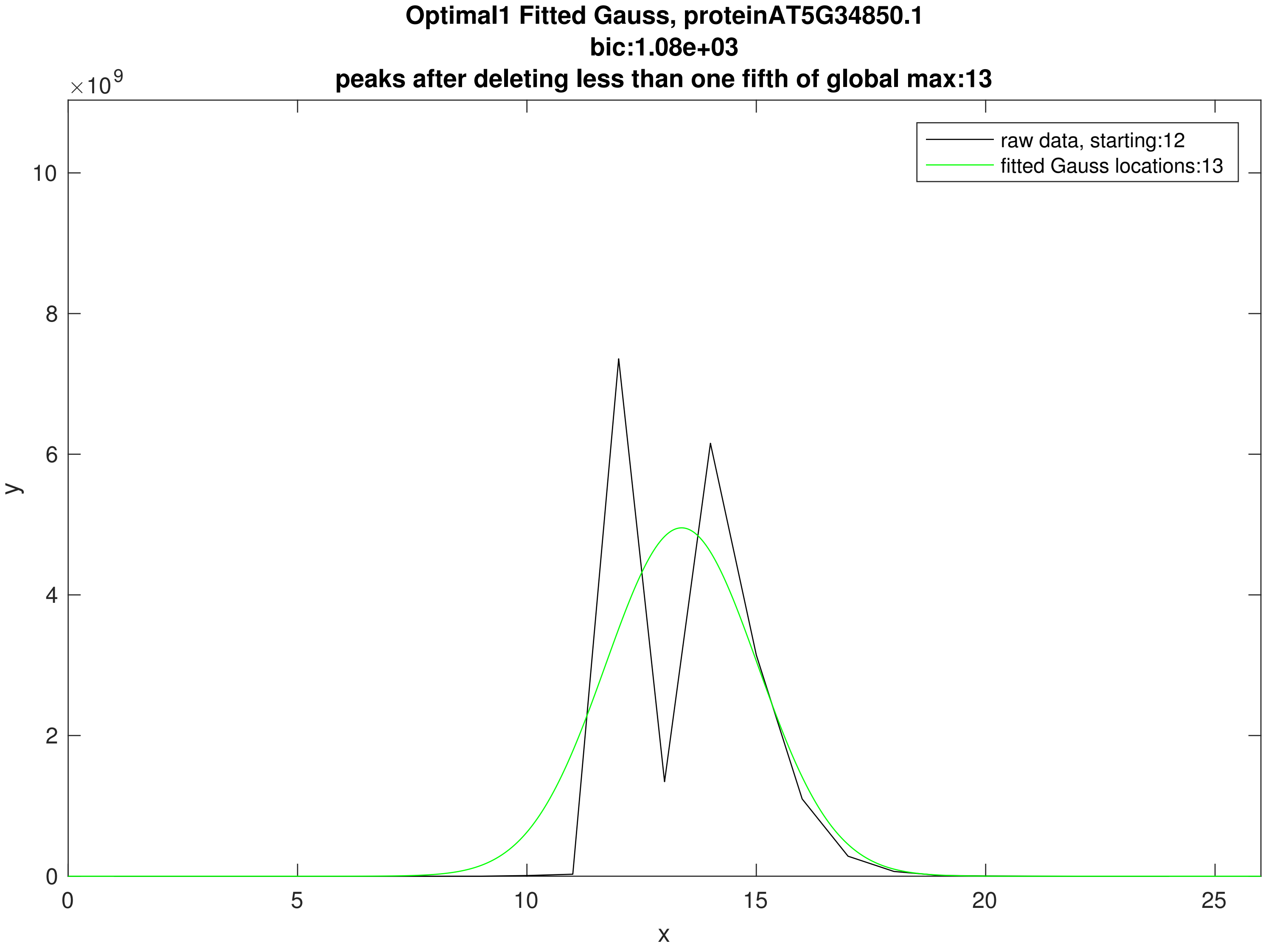
Optimal1 Fitted Gauss, proteinAT5G34850.1
bic:1.08e+03
peaks after deleting less than one fifth of global max:13
×10^9
raw data, starting:12
fitted Gauss locations:13
y
x
0
2
4
6
8
10
0
5
10
15
20
25

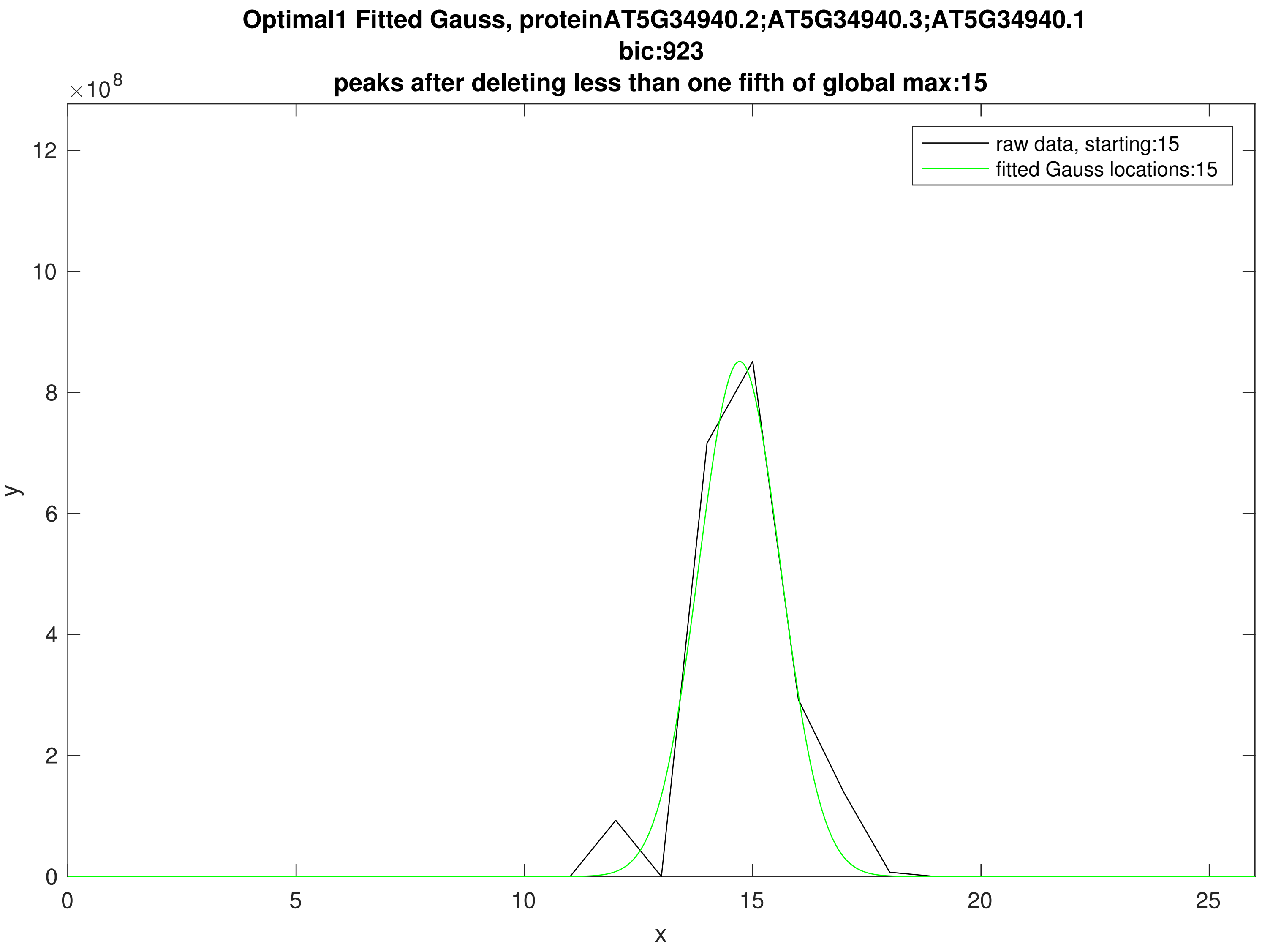

Optimal1 Fitted Gauss, proteinAT5G34940.2;AT5G34940.3;AT5G34940.1
bic:923
peaks after deleting less than one fifth of global max:15
×10^8
12
10
8
6
4
2
0
y
0
5
10
15
20
25
x
raw data, starting:15
fitted Gauss locations:15

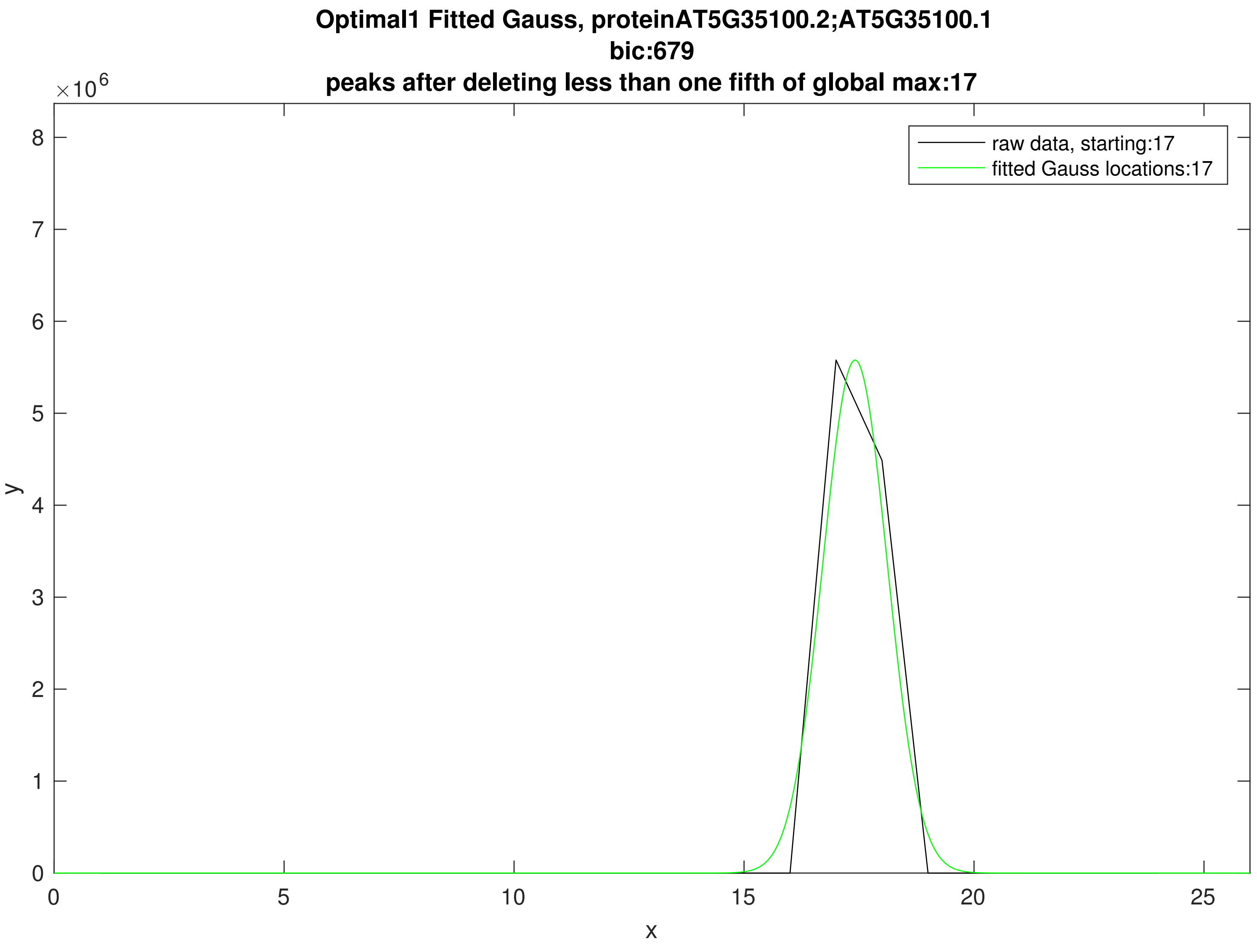
Optimal1 Fitted Gauss, proteinAT5G35100.2;AT5G35100.1
bic:679
peaks after deleting less than one fifth of global max:17
×10^6
raw data, starting:17
fitted Gauss locations:17
y
x
0
1
2
3
4
5
6
7
8
0
5
10
15
20
25

**Optimal1 Fitted Gauss, proteinAT5G35360.3;AT5G35360.1;AT5G35360.2**
**bic:919 925 934**
**peaks after deleting less than one fifth of global max:12**

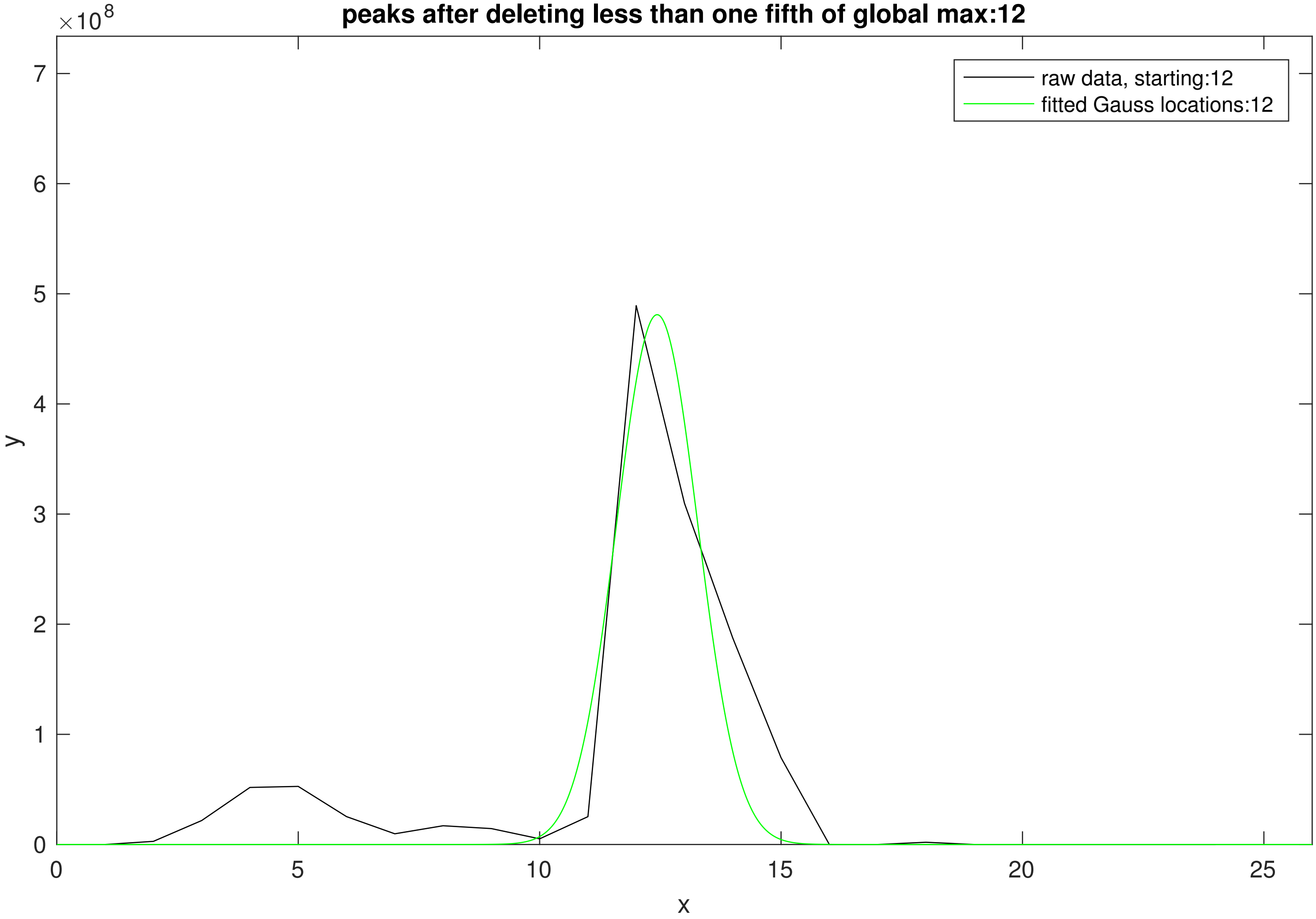

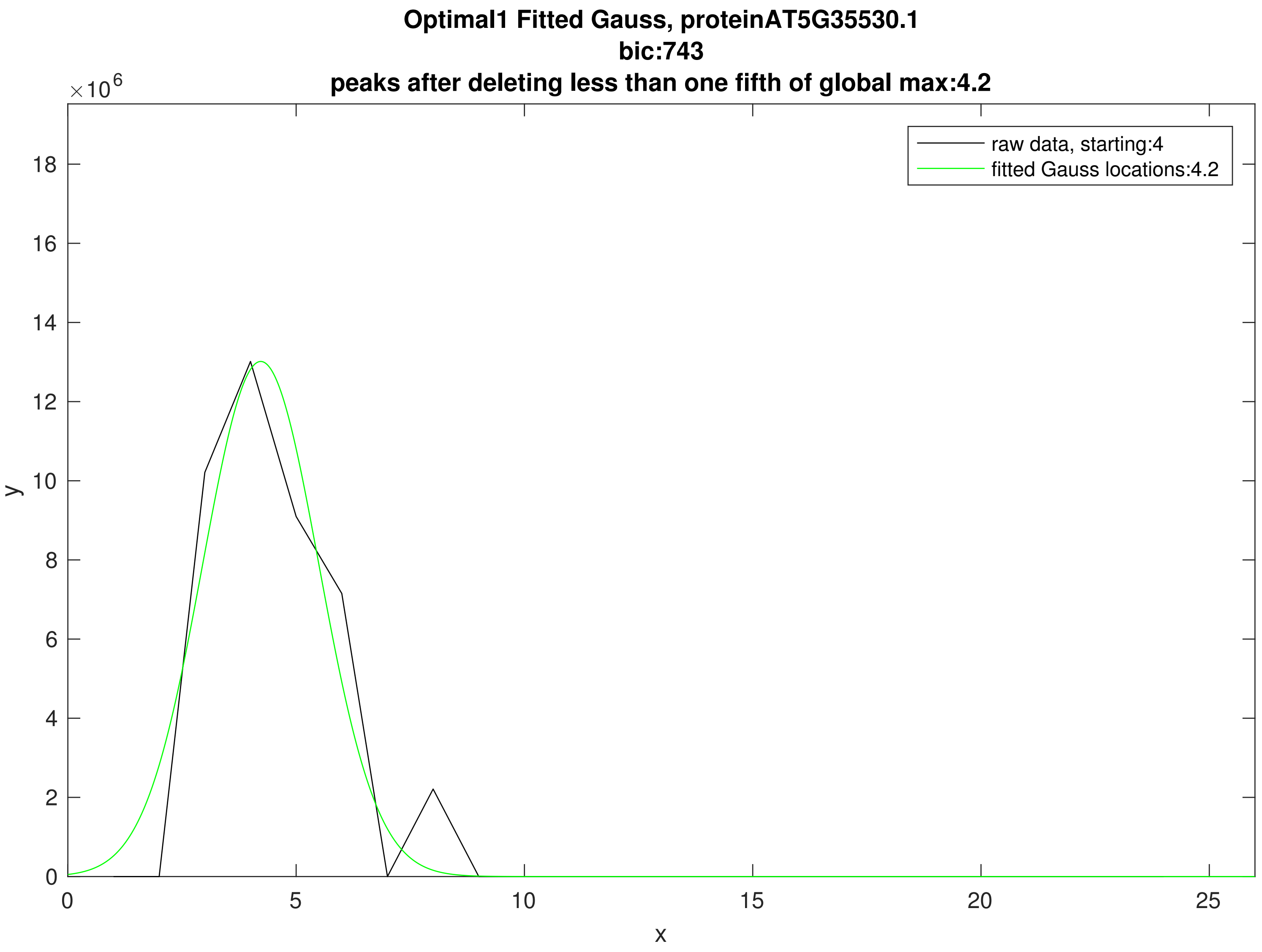

Optimal1 Fitted Gauss, proteinAT5G35530.1
bic:743
peaks after deleting less than one fifth of global max:4.2
×10^6
raw data, starting:4
fitted Gauss locations:4.2
y
0
2
4
6
8
10
12
14
16
18
x
0
5
10
15
20
25

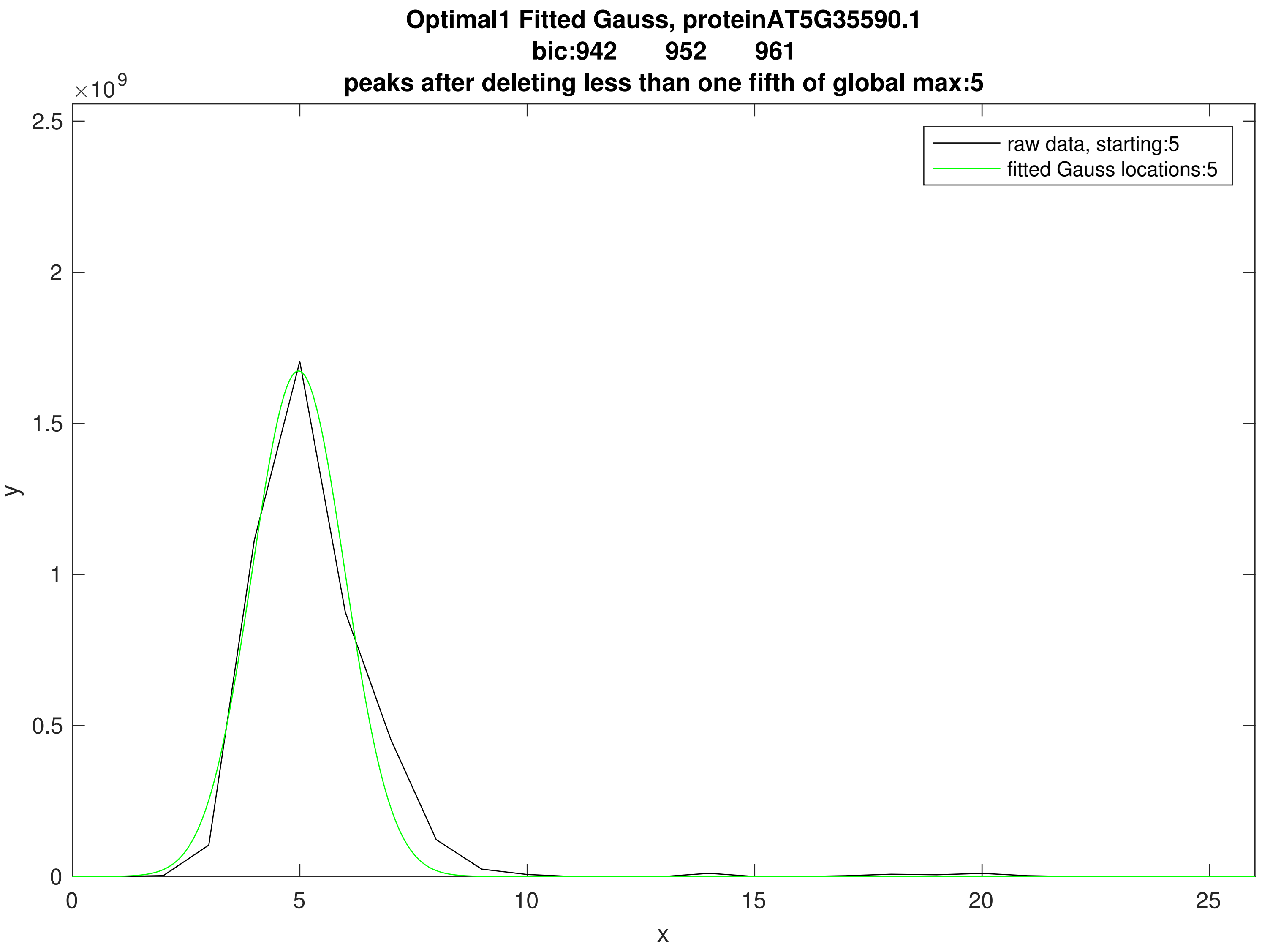

Optimal1 Fitted Gauss, proteinAT5G35590.1
bic:942 952 961
peaks after deleting less than one fifth of global max:5
×10^9
2.5
2
1.5
1
0.5
0
y
0
5
10
15
20
25
x
raw data, starting:5
fitted Gauss locations:5

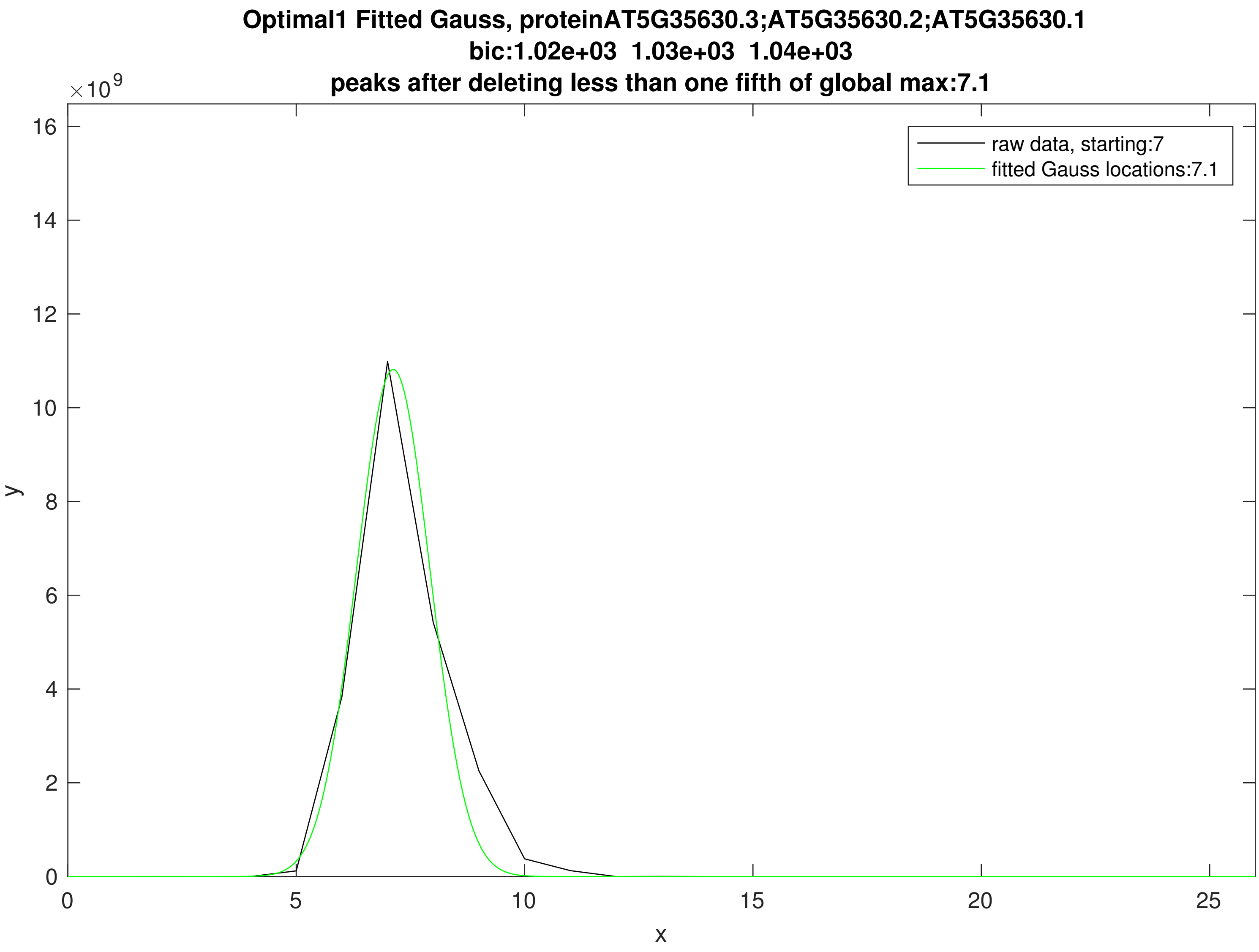

Optimal1 Fitted Gauss, proteinAT5G35630.3;AT5G35630.2;AT5G35630.1
bic:1.02e+03 1.03e+03 1.04e+03
peaks after deleting less than one fifth of global max:7.1
×10^9
raw data, starting:7
fitted Gauss locations:7.1
y
x
0
2
4
6
8
10
12
14
16
0
5
10
15
20
25

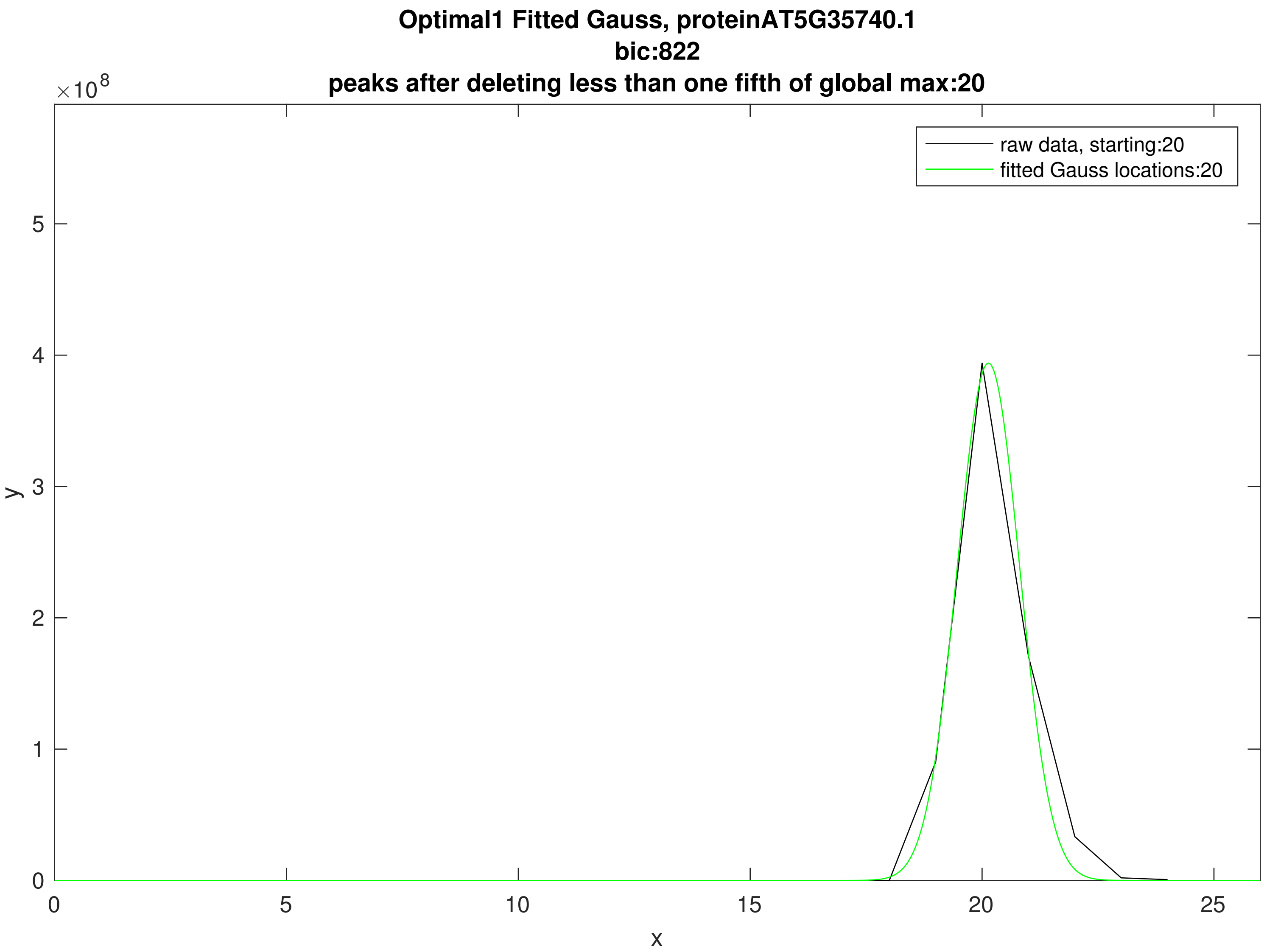

Optimal1 Fitted Gauss, proteinAT5G35740.1
bic:822
peaks after deleting less than one fifth of global max:20
×10^8
raw data, starting:20
fitted Gauss locations:20
y
x
0
1
2
3
4
5
0
5
10
15
20
25

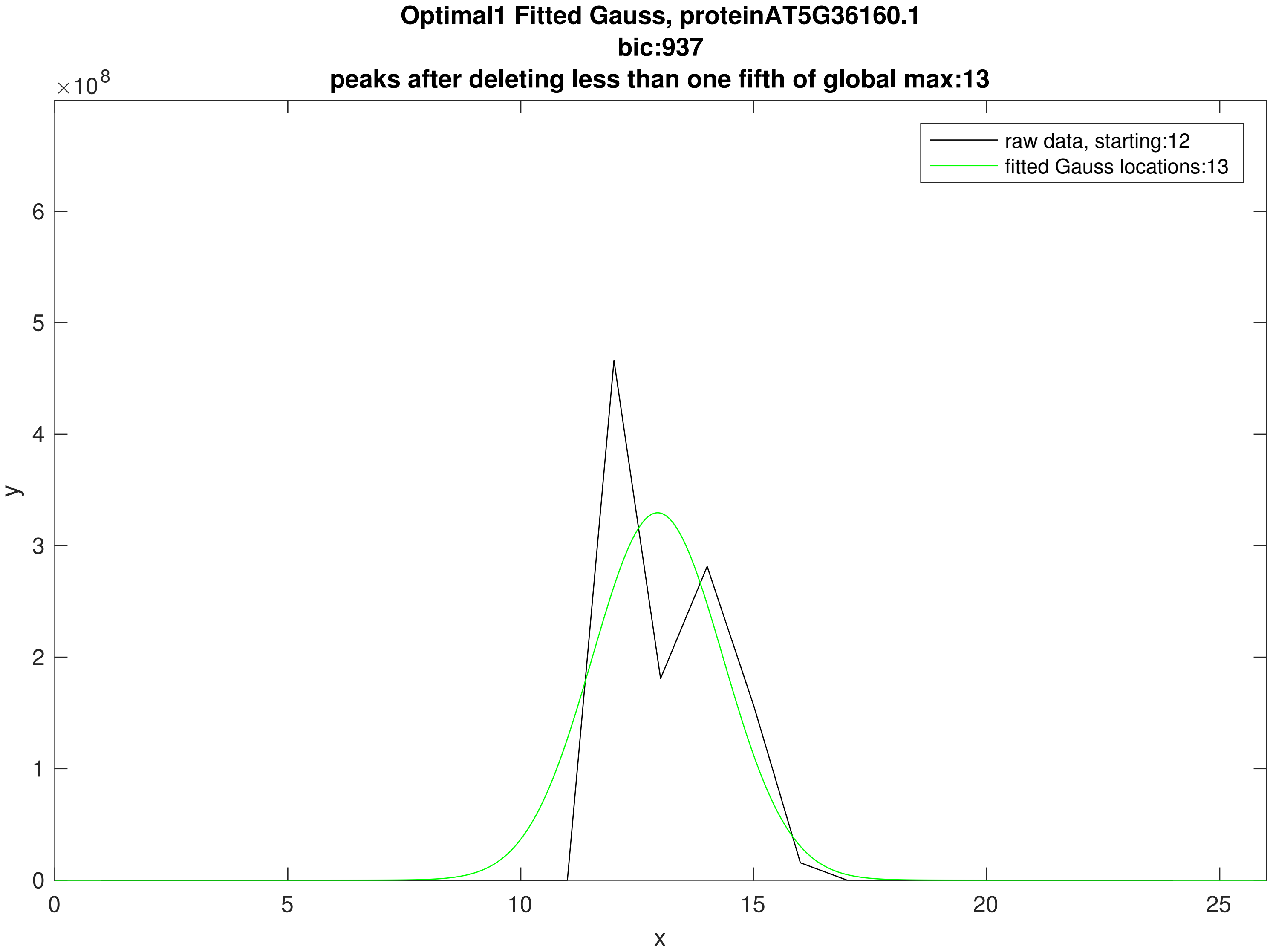
Optimal1 Fitted Gauss, proteinAT5G36160.1
bic:937
peaks after deleting less than one fifth of global max:13
×10^8
raw data, starting:12
fitted Gauss locations:13
y
x
0
1
2
3
4
5
6
0
5
10
15
20
25

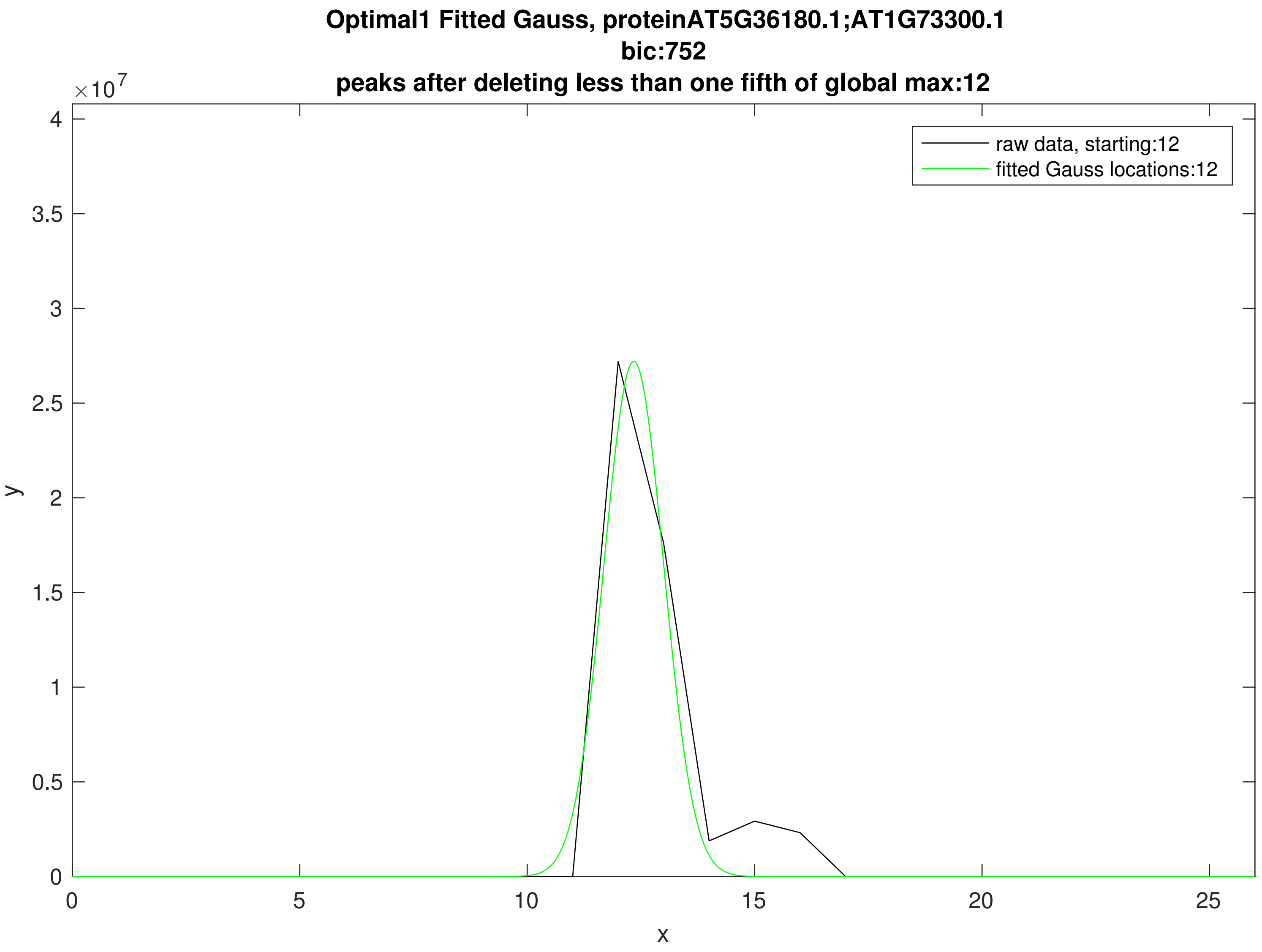
Optimal1 Fitted Gauss, proteinAT5G36180.1;AT1G73300.1
bic:752
peaks after deleting less than one fifth of global max:12
×10^7
raw data, starting:12
fitted Gauss locations:12
y
x
0
0.5
1
1.5
2
2.5
3
3.5
4
0
5
10
15
20
25

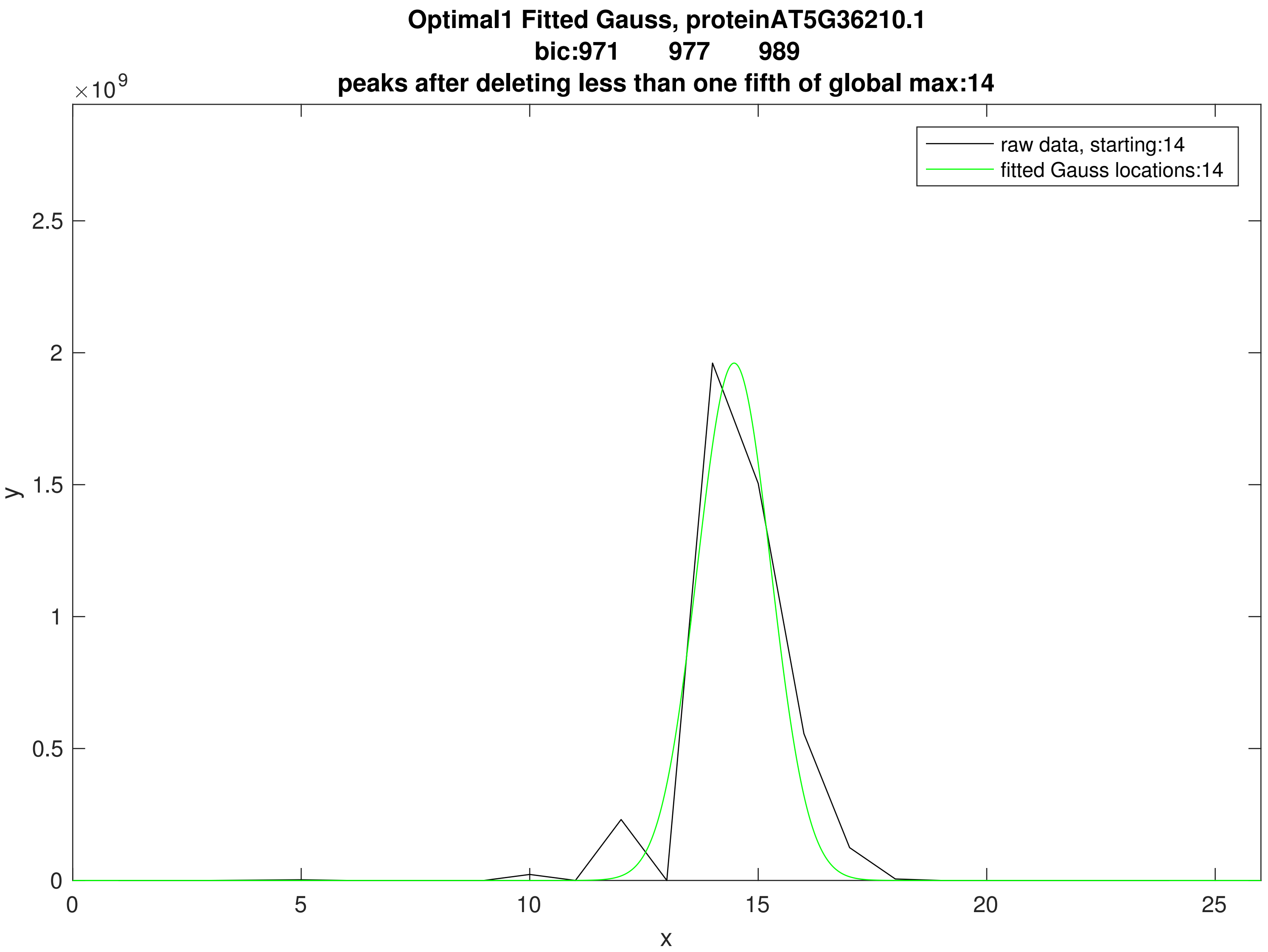
Optimal1 Fitted Gauss, proteinAT5G36210.1
bic:971 977 989
peaks after deleting less than one fifth of global max:14
×10^9
raw data, starting:14
fitted Gauss locations:14
y
x
0
0.5
1
1.5
2
2.5
5
10
15
20
25

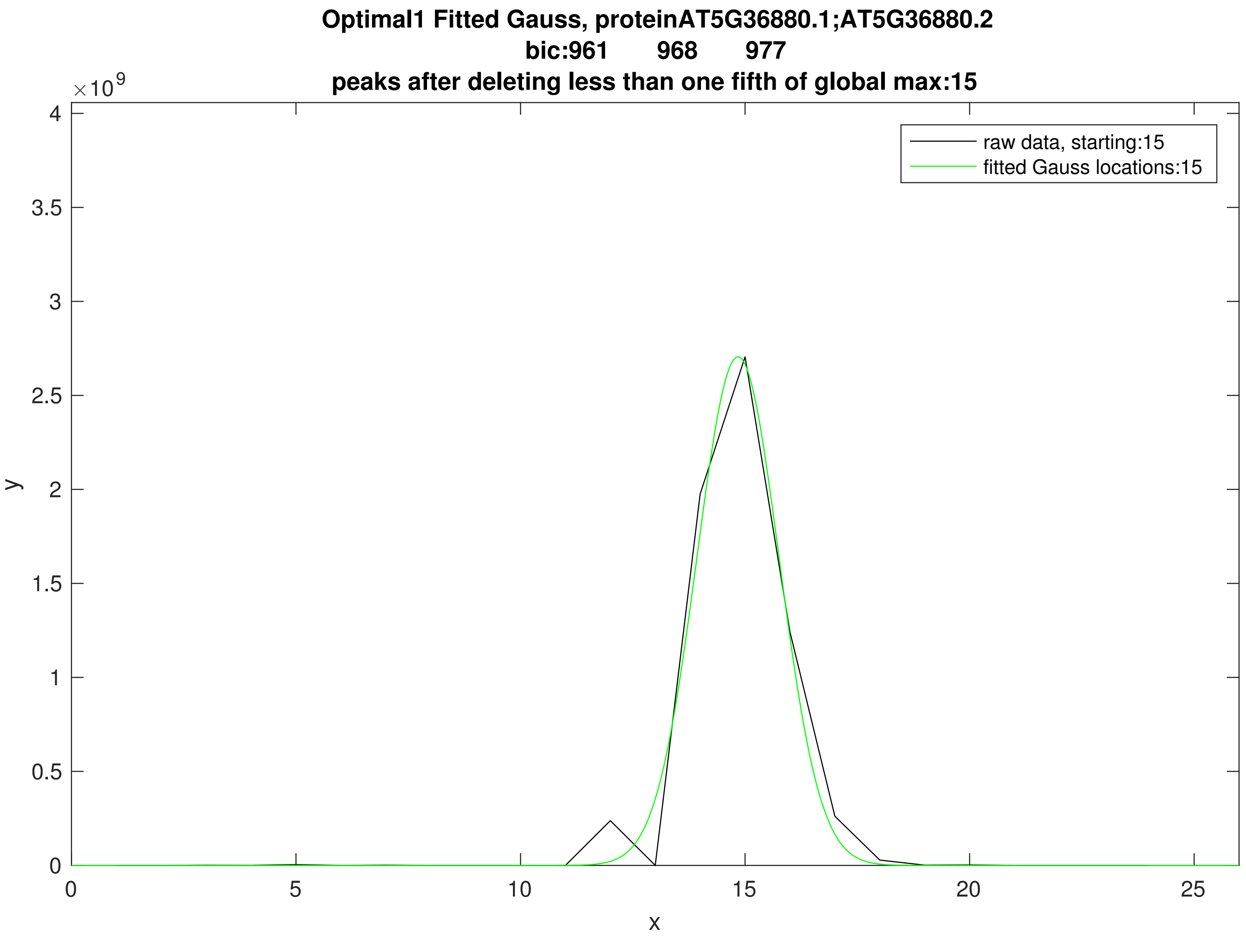

Optimal1 Fitted Gauss, proteinAT5G36880.1;AT5G36880.2
bic:961 968 977
peaks after deleting less than one fifth of global max:15
×10^9
raw data, starting:15
fitted Gauss locations:15
y
x
0
0.5
1
1.5
2
2.5
3
3.5
4
0
5
10
15
20
25

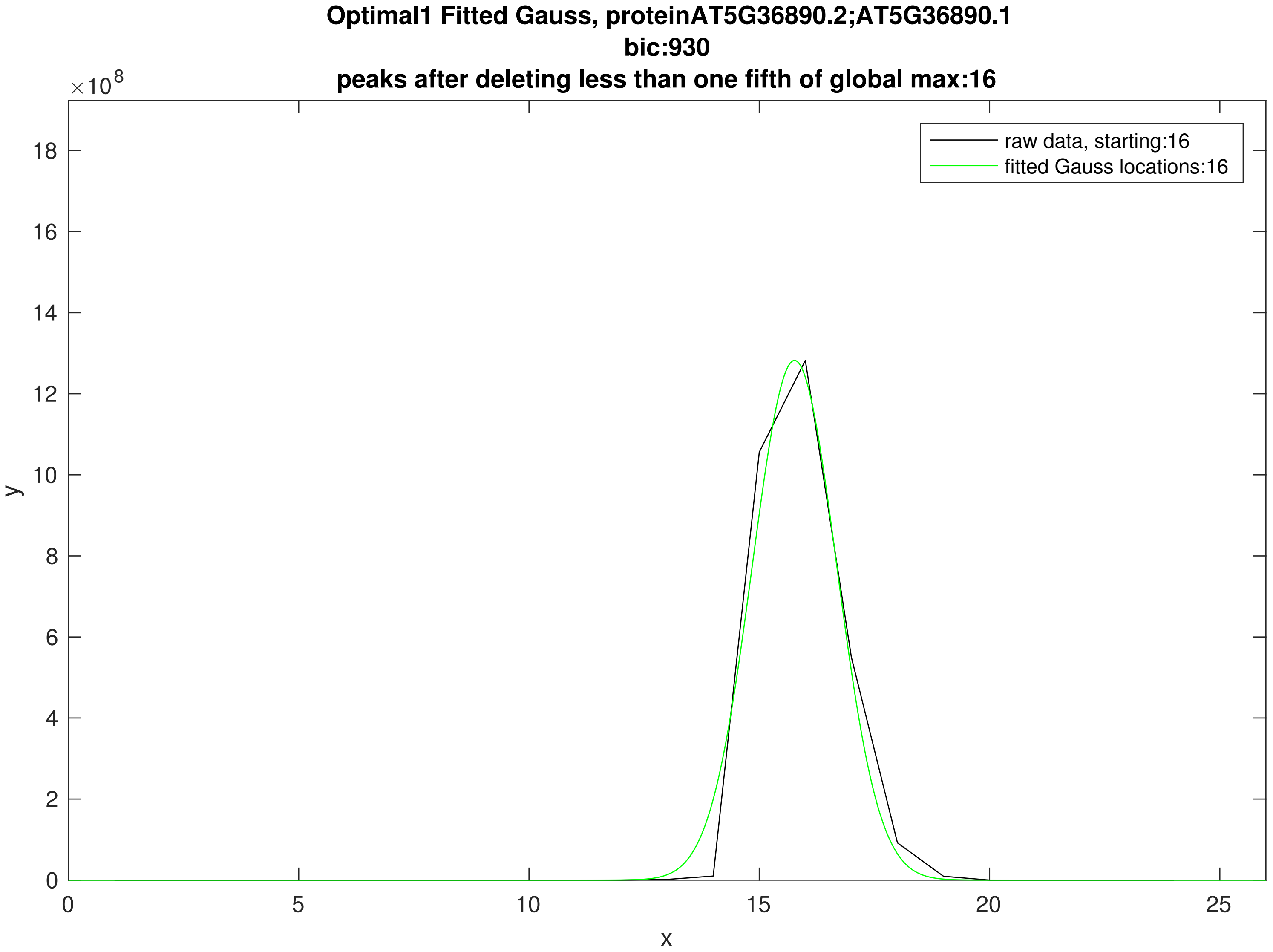

Optimal1 Fitted Gauss, proteinAT5G36890.2;AT5G36890.1
bic:930
peaks after deleting less than one fifth of global max:16
×10^8
raw data, starting:16
fitted Gauss locations:16
y
x
0
2
4
6
8
10
12
14
16
18
5
15
20
25

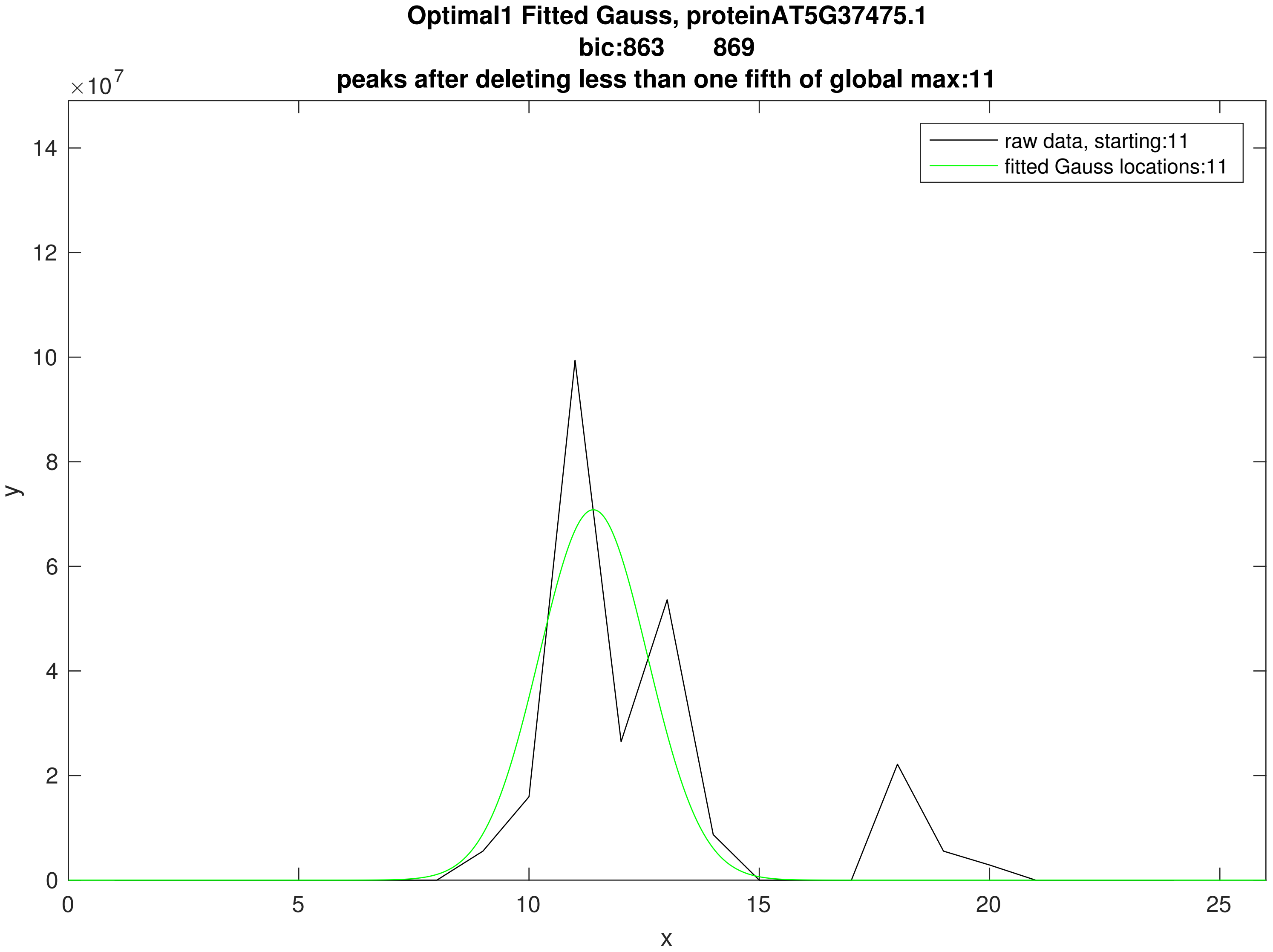
Optimal1 Fitted Gauss, proteinAT5G37475.1
bic:863 869
peaks after deleting less than one fifth of global max:11
×10^7
14
12
10
8
6
4
2
0
y
0
5
10
15
20
25
x
raw data, starting:11
fitted Gauss locations:11

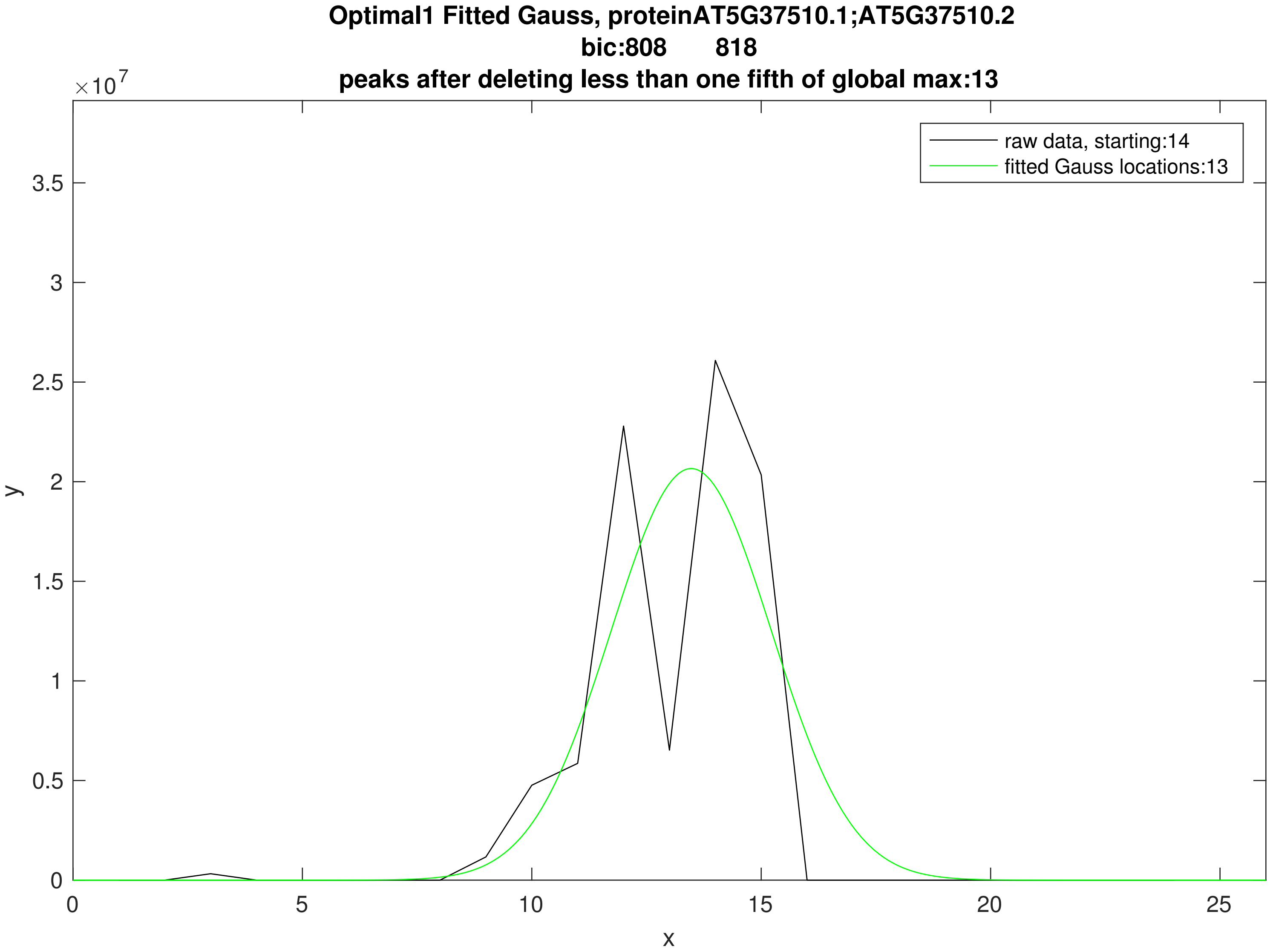

Optimal1 Fitted Gauss, proteinAT5G37510.1;AT5G37510.2
bic:808 818
peaks after deleting less than one fifth of global max:13
×10^7
raw data, starting:14
fitted Gauss locations:13
y
x
0
0.5
1
1.5
2
2.5
3
3.5
0
5
10
15
20
25

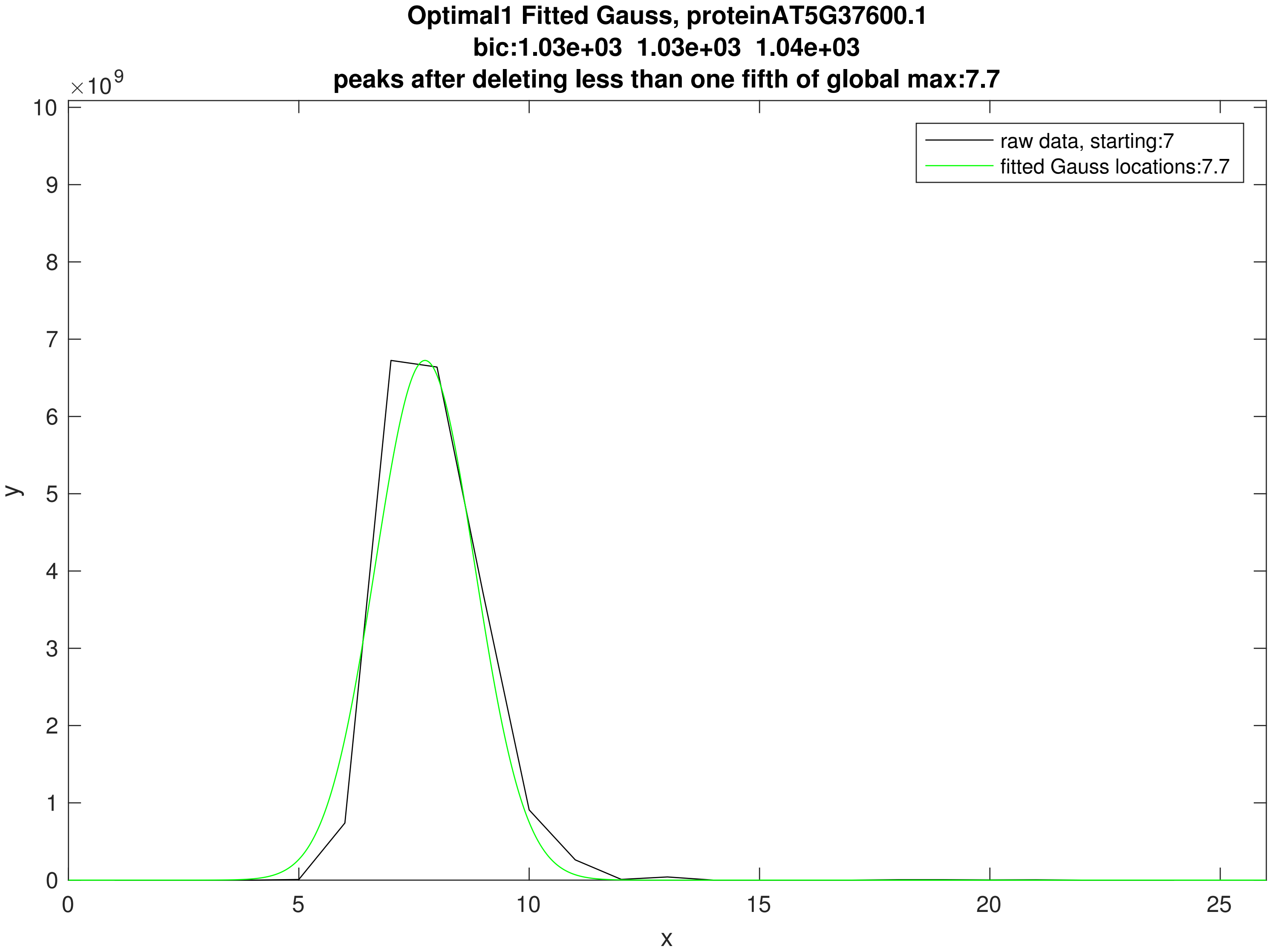
Optimal1 Fitted Gauss, proteinAT5G37600.1
bic:1.03e+03  1.03e+03  1.04e+03
peaks after deleting less than one fifth of global max:7.7
×10^9
raw data, starting:7
fitted Gauss locations:7.7
y
x
0
1
2
3
4
5
6
7
8
9
10
0
5
10
15
20
25

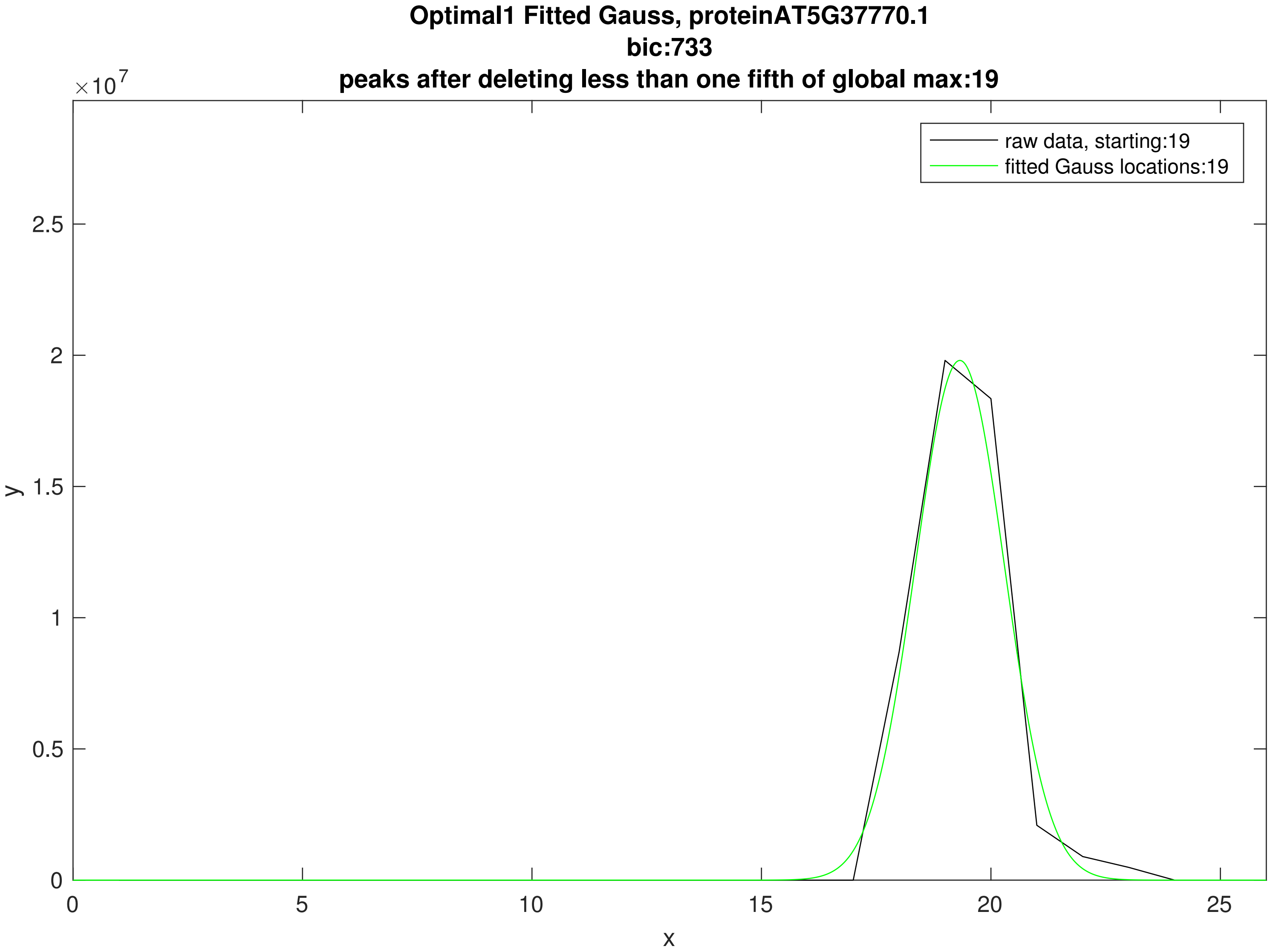
Optimal1 Fitted Gauss, proteinAT5G37770.1
bic:733
peaks after deleting less than one fifth of global max:19
×10^7
raw data, starting:19
fitted Gauss locations:19
y
x
0
0.5
1
1.5
2
2.5
0
5
10
15
20
25

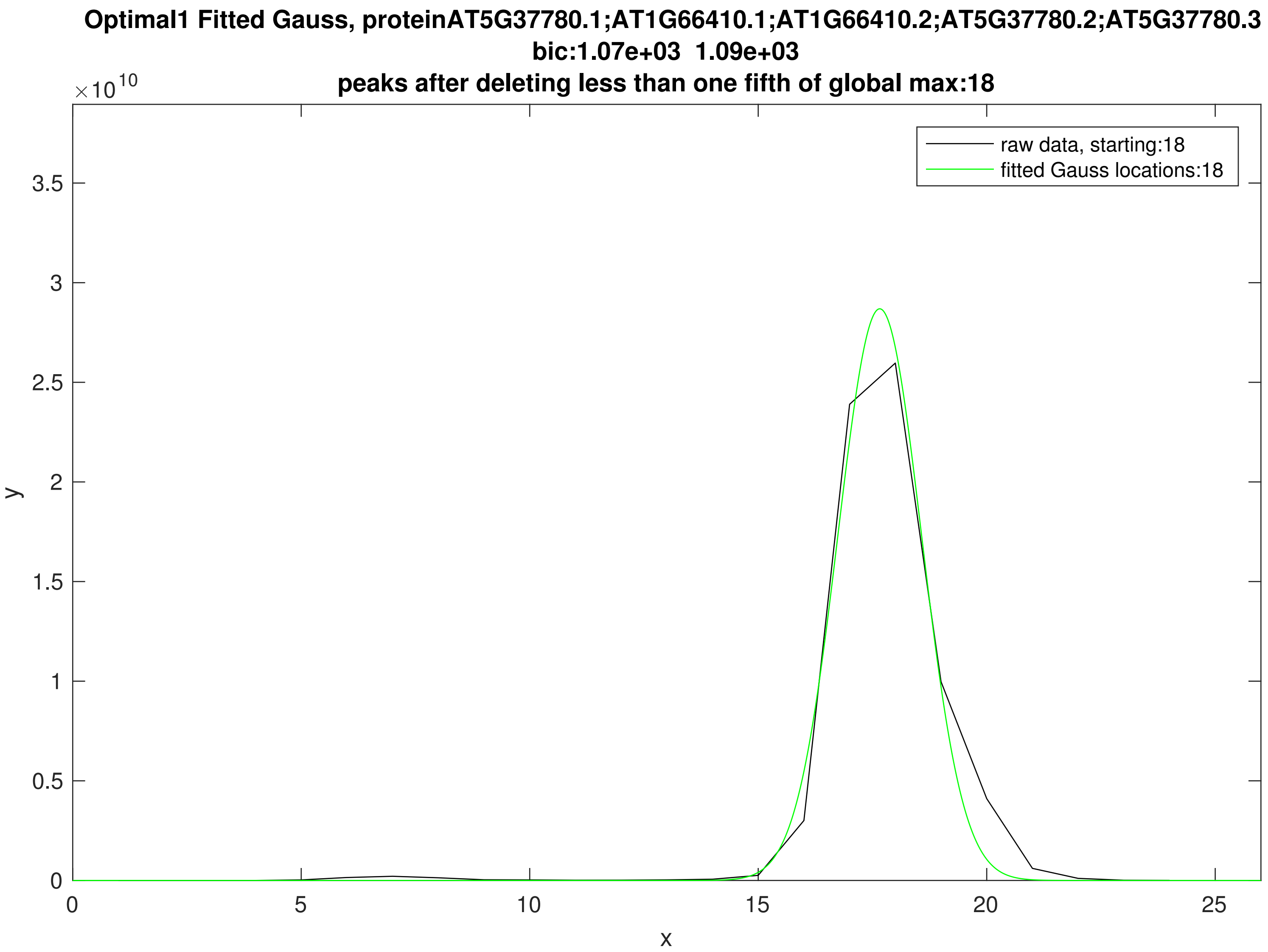
Optimal1 Fitted Gauss, proteinAT5G37780.1;AT1G66410.1;AT1G66410.2;AT5G37780.2;AT5G37780.3
bic:1.07e+03 1.09e+03
peaks after deleting less than one fifth of global max:18
×10^10
3.5
3
2.5
2
1.5
1
0.5
0
y
0
5
10
15
20
25
x
raw data, starting:18
fitted Gauss locations:18

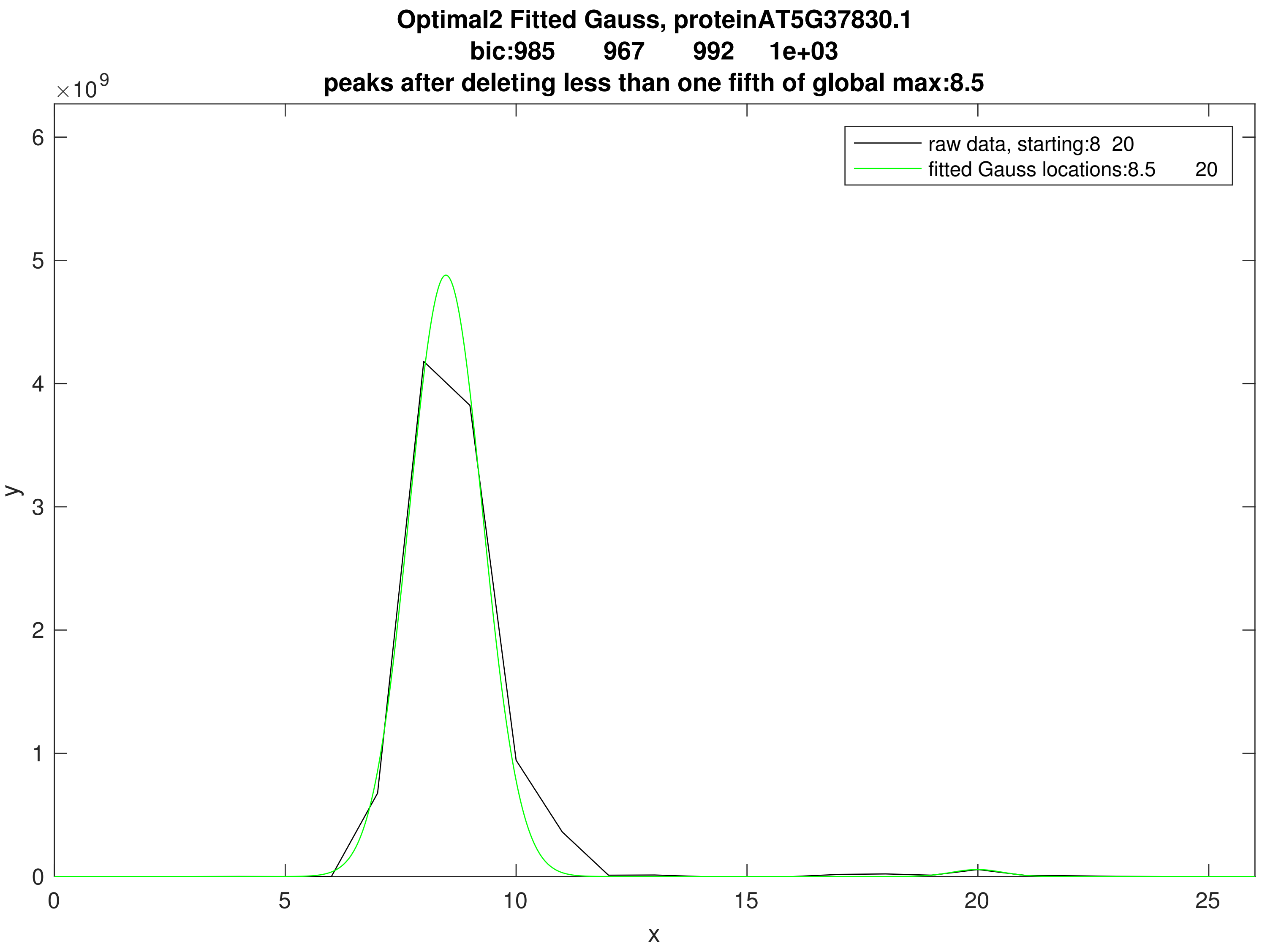

Optimal2 Fitted Gauss, proteinAT5G37830.1
bic:985 967 992 1e+03
peaks after deleting less than one fifth of global max:8.5
×10^9
6
5
4
3
2
1
0
y
0
5
10
15
20
25
x
raw data, starting:8 20
fitted Gauss locations:8.5 20

Optimal1 Fitted Gauss, proteinAT5G37850.2;AT5G37850.1;AT5G37850.3
bic:886
peaks after deleting less than one fifth of global max:15

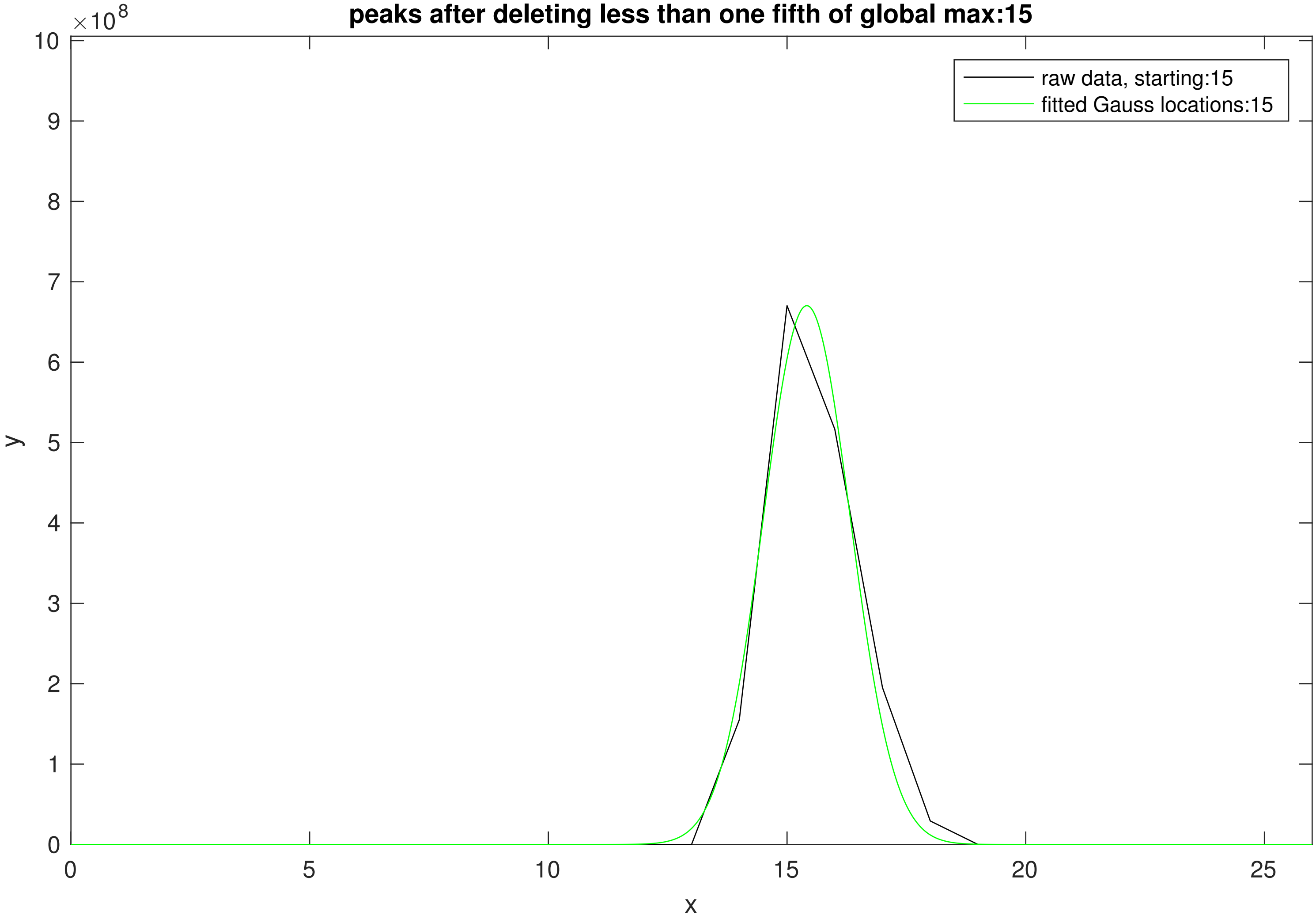

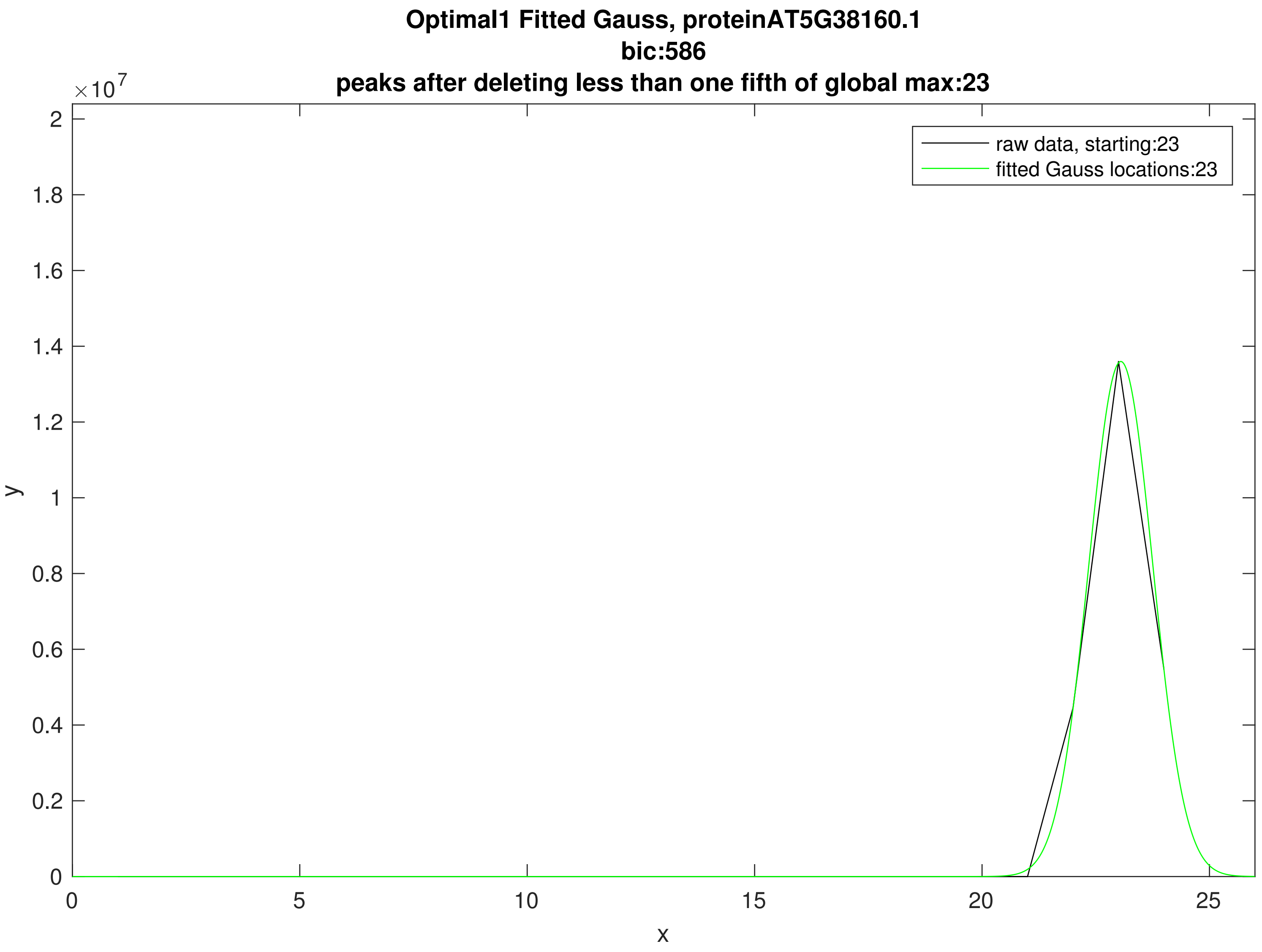

Optimal1 Fitted Gauss, proteinAT5G38160.1
bic:586
peaks after deleting less than one fifth of global max:23
×10^7
raw data, starting:23
fitted Gauss locations:23
y
x
0
5
10
15
20
25
0
0.2
0.4
0.6
0.8
1
1.2
1.4
1.6
1.8
2

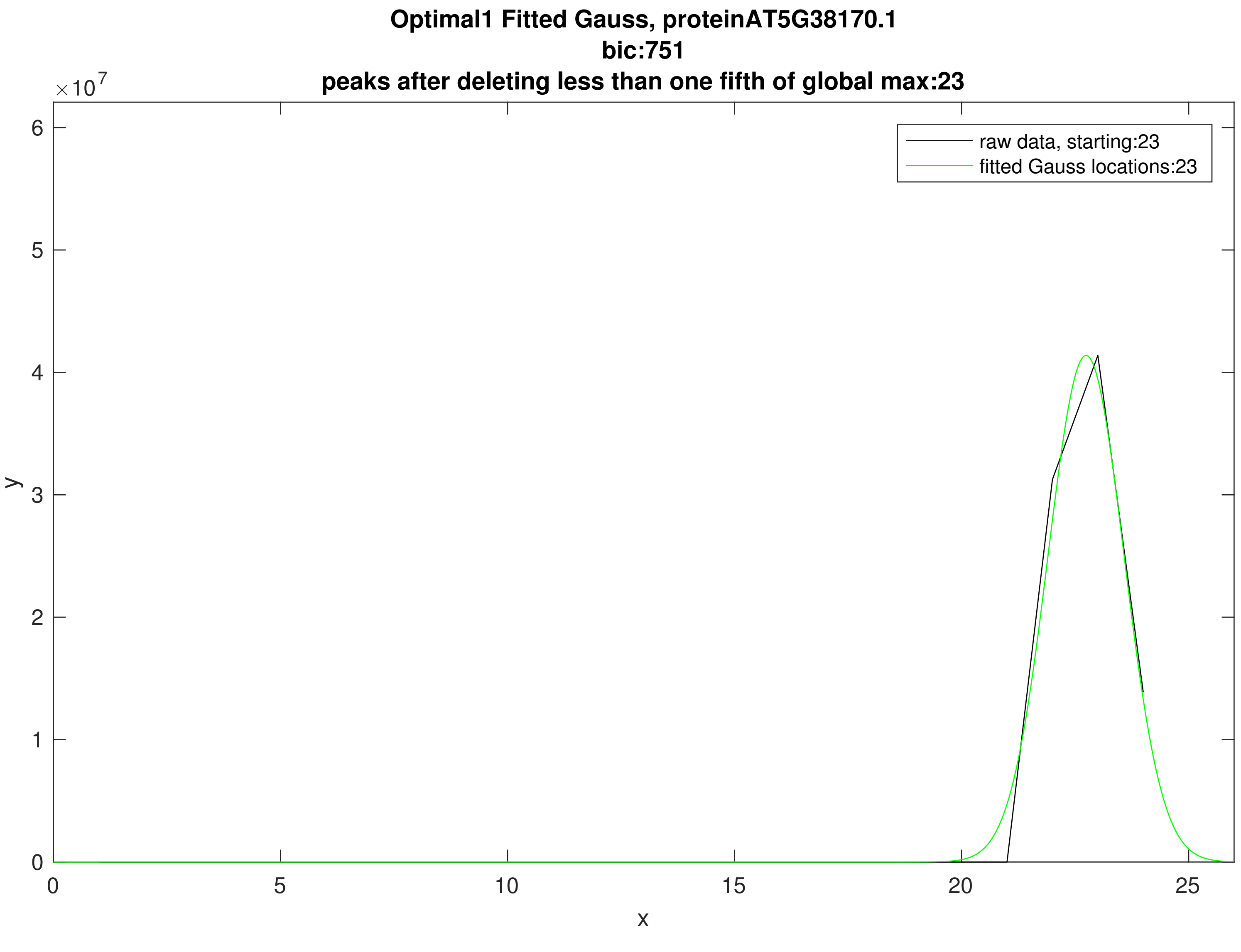
Optimal1 Fitted Gauss, proteinAT5G38170.1
bic:751
peaks after deleting less than one fifth of global max:23
×10^7
raw data, starting:23
fitted Gauss locations:23
y
x
0
1
2
3
4
5
6
0
5
10
15
20
25

Optimal1 Fitted Gauss, proteinAT5G38420.1;AT5G38410.1;AT5G38410.3;AT5G38410.2
bic:1.08e+03 1.09e+03 1.1e+03
peaks after deleting less than one fifth of global max:6.8

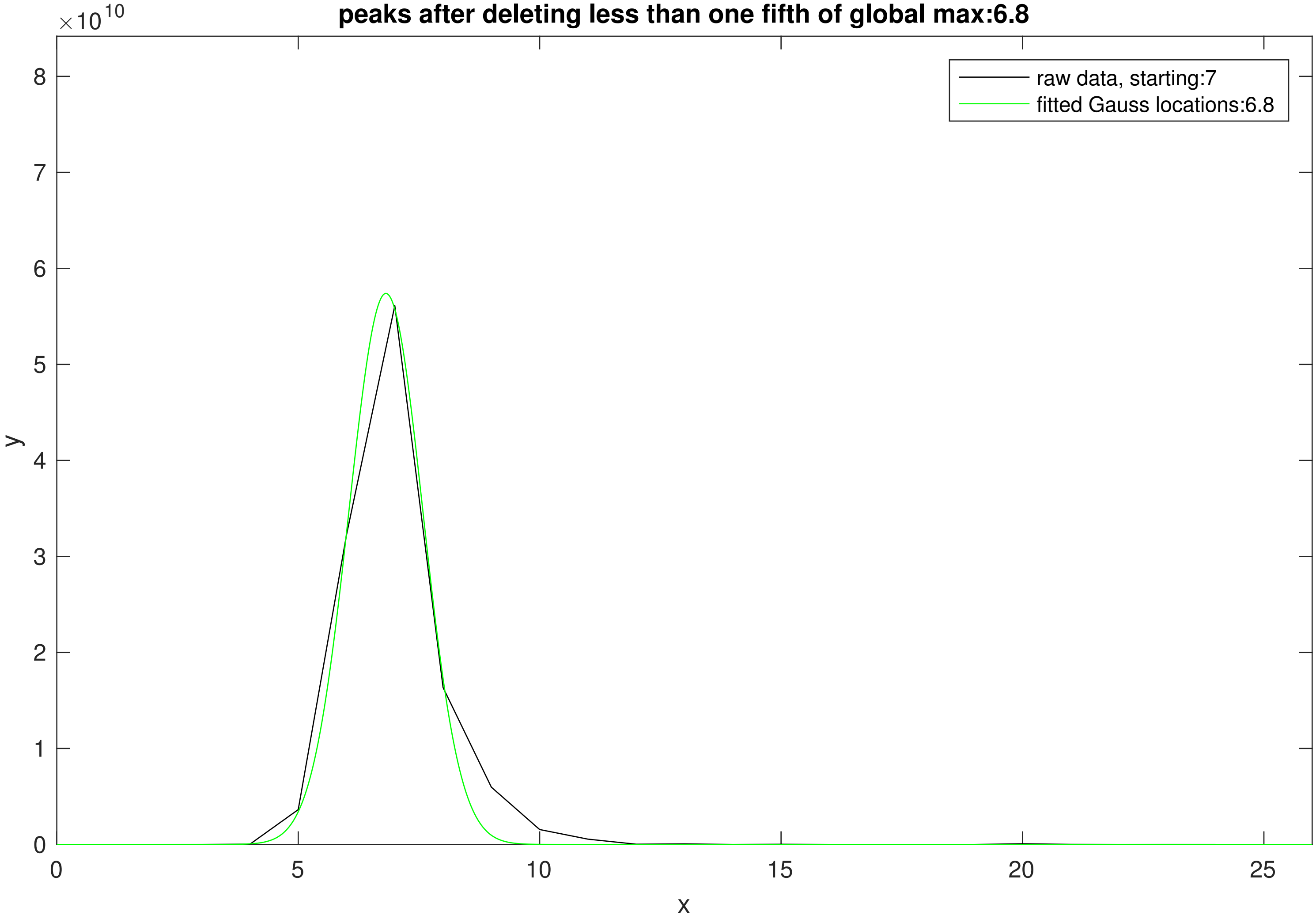

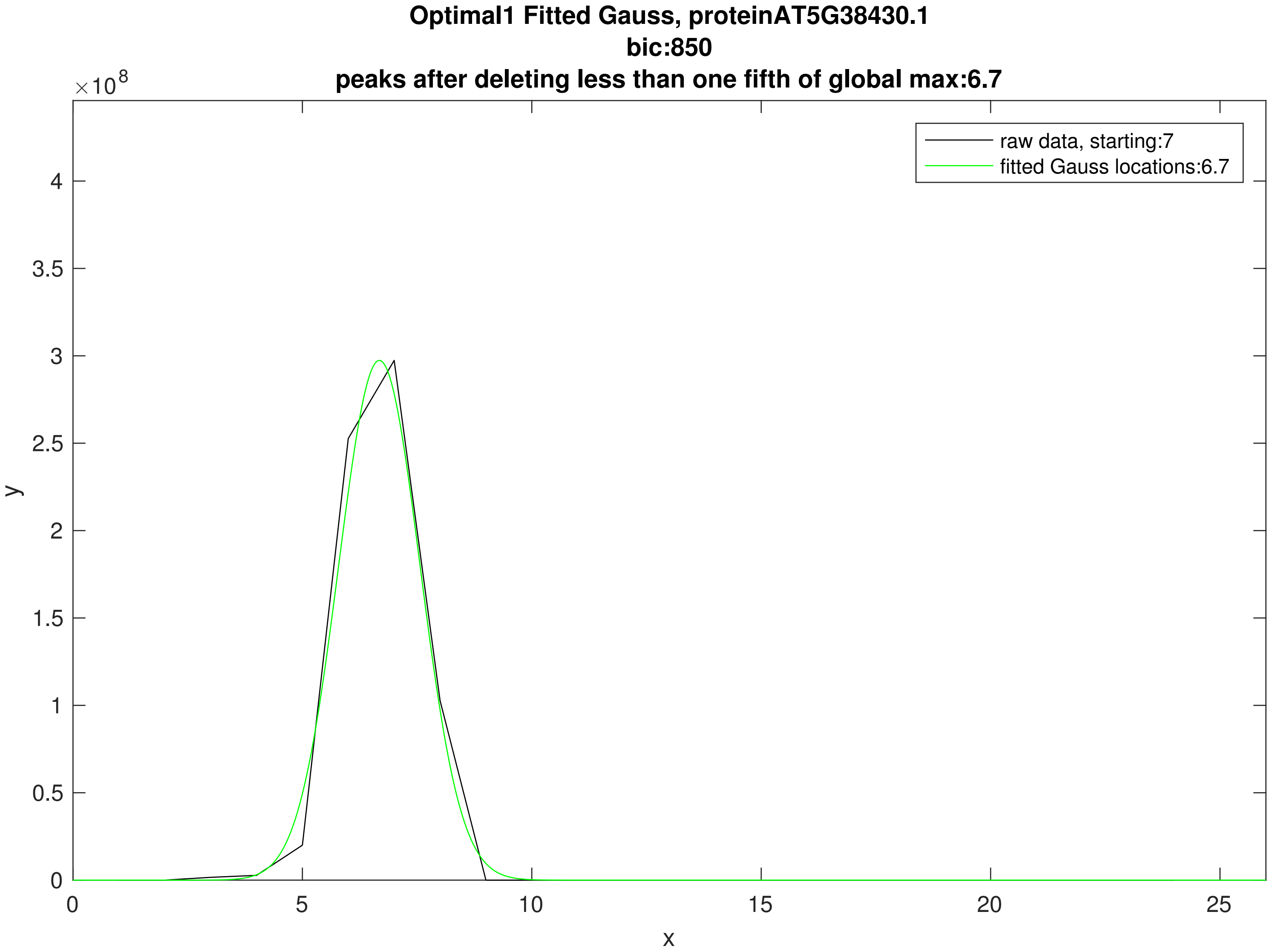
Optimal1 Fitted Gauss, proteinAT5G38430.1
bic:850
peaks after deleting less than one fifth of global max:6.7
×10^8
raw data, starting:7
fitted Gauss locations:6.7
4
3.5
3
2.5
2
1.5
1
0.5
0
y
0
5
10
15
20
25
x

**Optimal2 Fitted Gauss, proteinAT5G38470.1;AT5G38470.2**
**bic:926 916**
**peaks after deleting less than one fifth of global max:13**

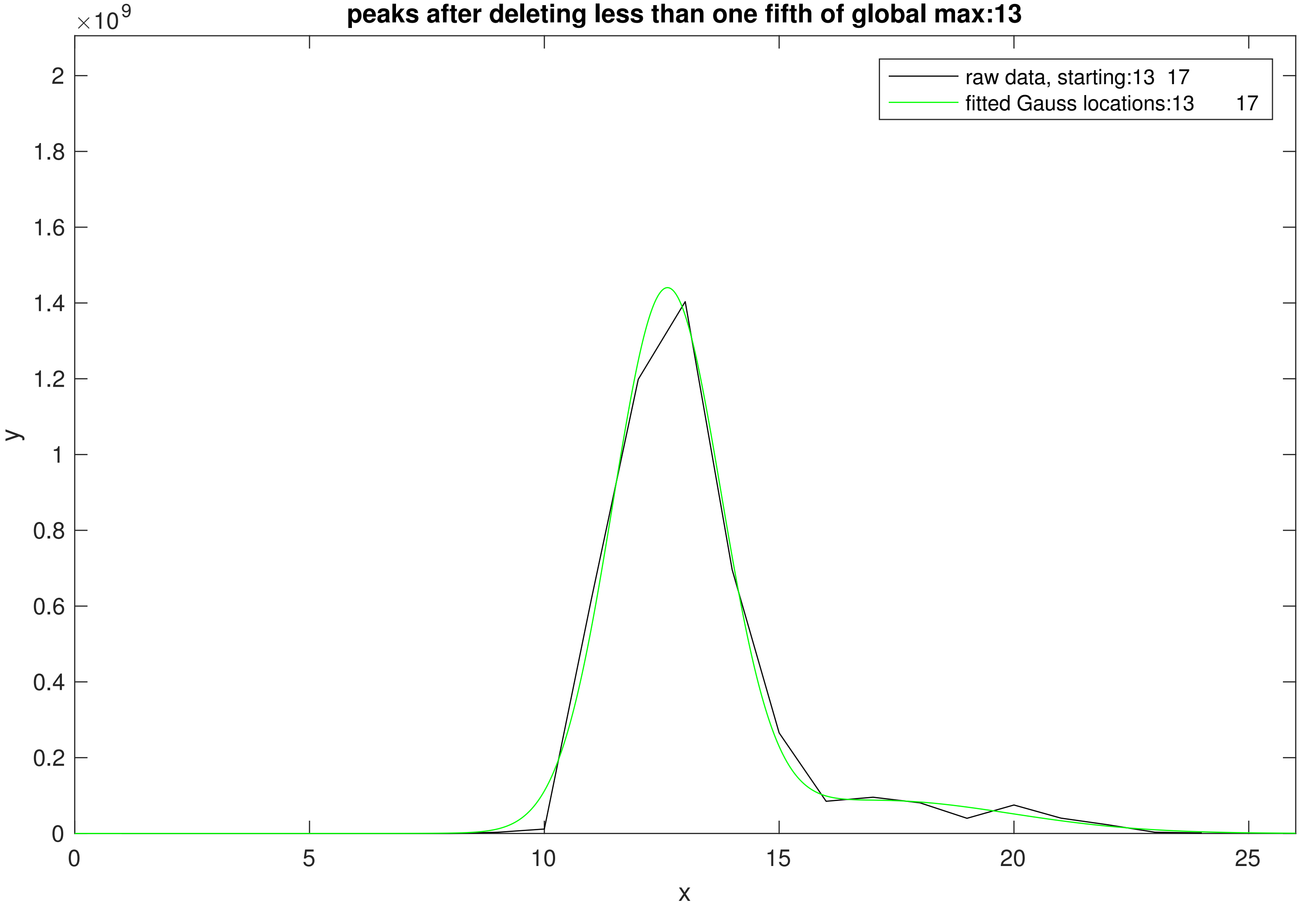

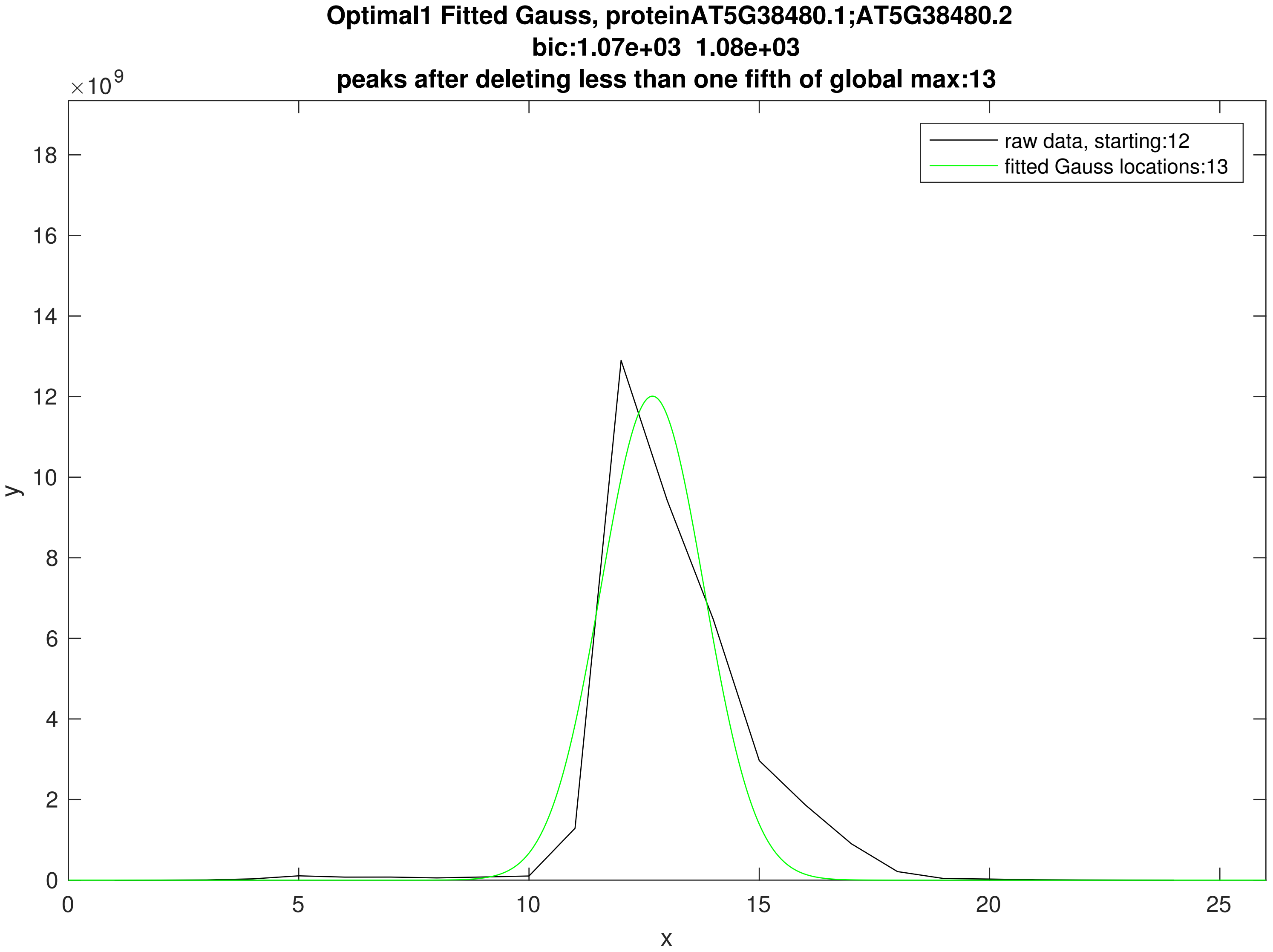
Optimal1 Fitted Gauss, proteinAT5G38480.1;AT5G38480.2
bic:1.07e+03  1.08e+03
peaks after deleting less than one fifth of global max:13
×10^9
raw data, starting:12
fitted Gauss locations:13
y
x
0
2
4
6
8
10
12
14
16
18
0
5
10
15
20
25

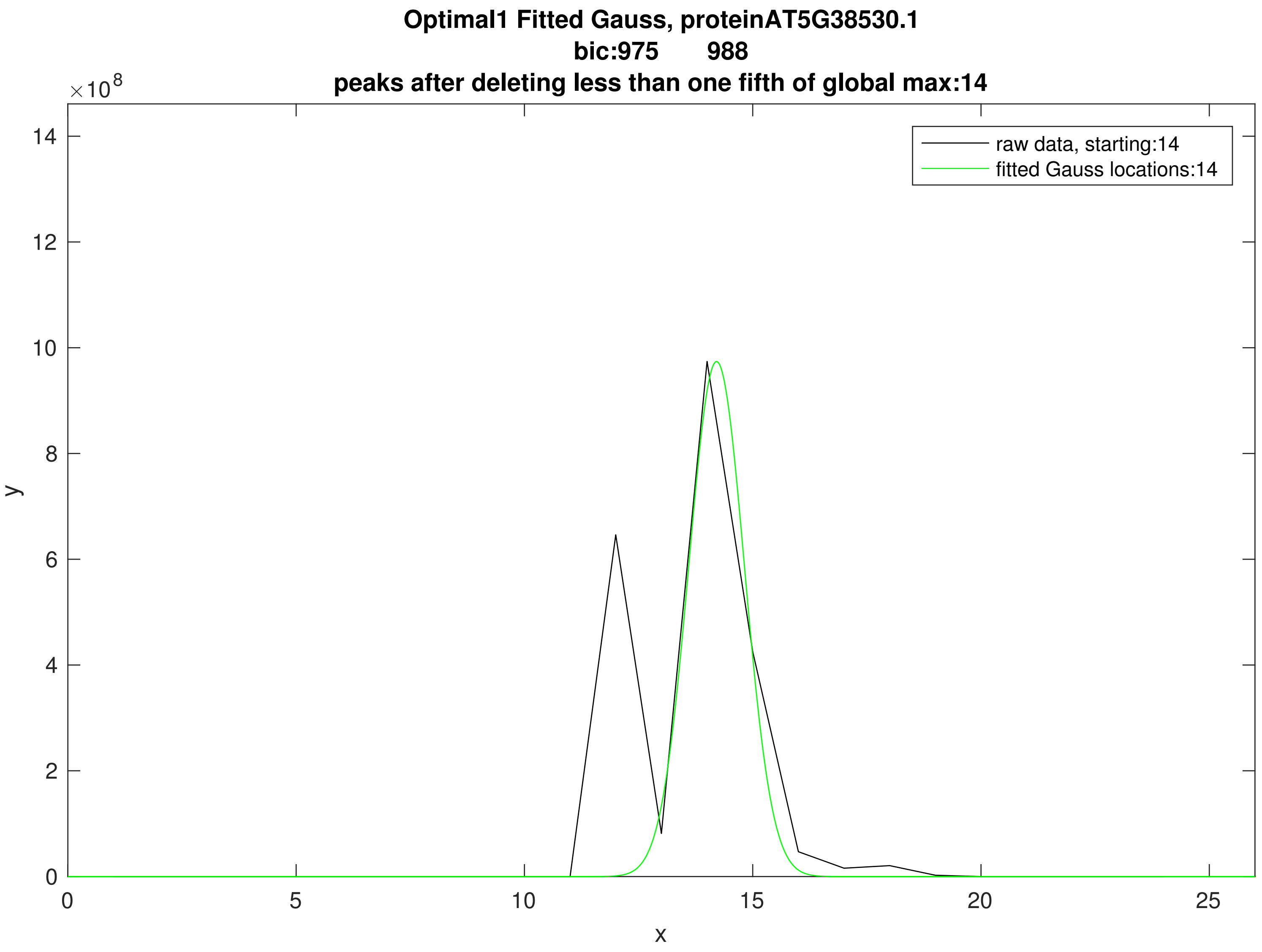
Optimal1 Fitted Gauss, proteinAT5G38530.1
bic:975 988
peaks after deleting less than one fifth of global max:14
×10^8
14
12
10
8
6
4
2
0
y
0
5
10
15
20
25
x
raw data, starting:14
fitted Gauss locations:14

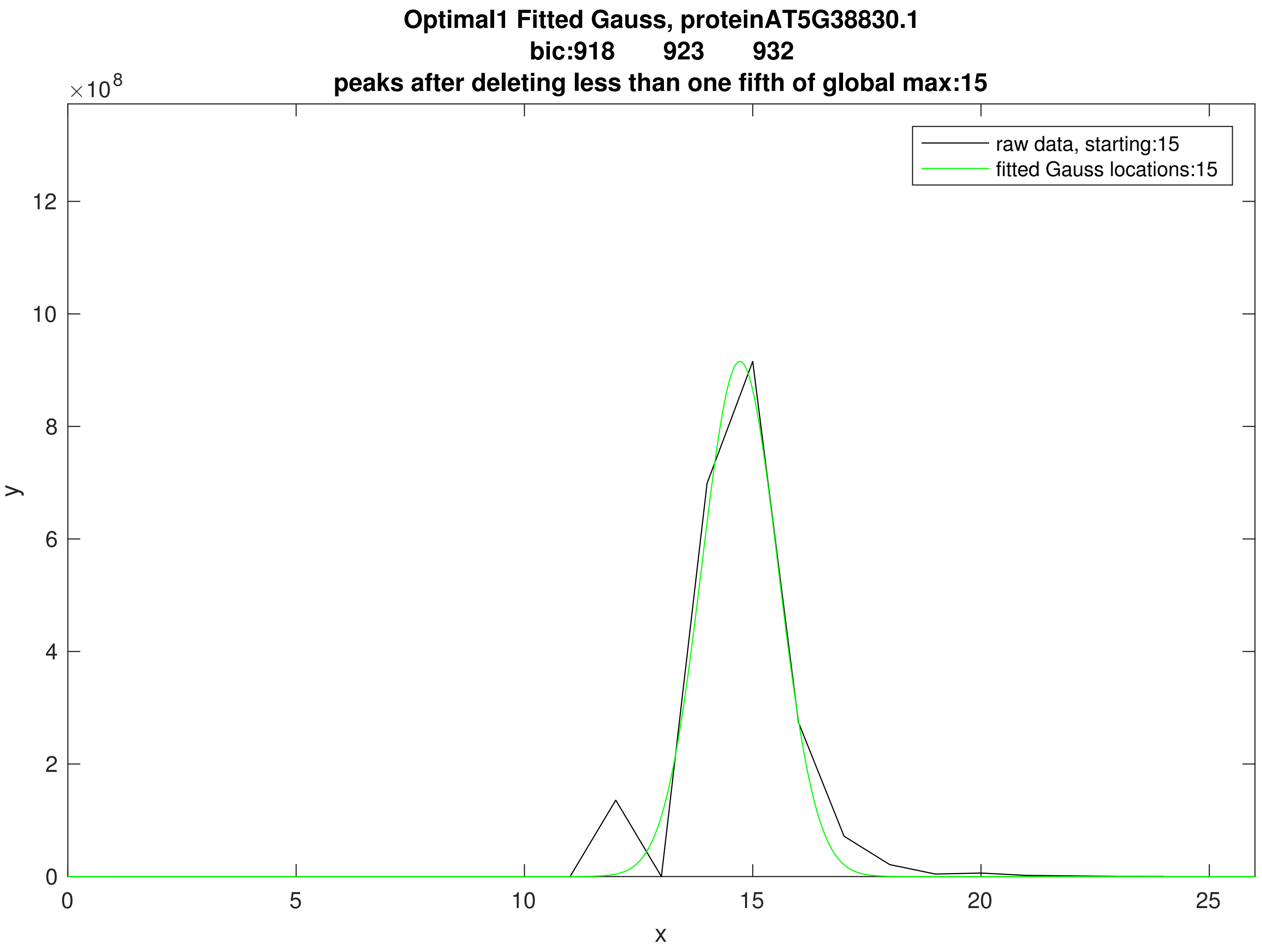
Optimal1 Fitted Gauss, proteinAT5G38830.1
bic:918 923 932
peaks after deleting less than one fifth of global max:15
×10^8
raw data, starting:15
fitted Gauss locations:15
y
x
0
2
4
6
8
10
12
0
5
10
15
20
25

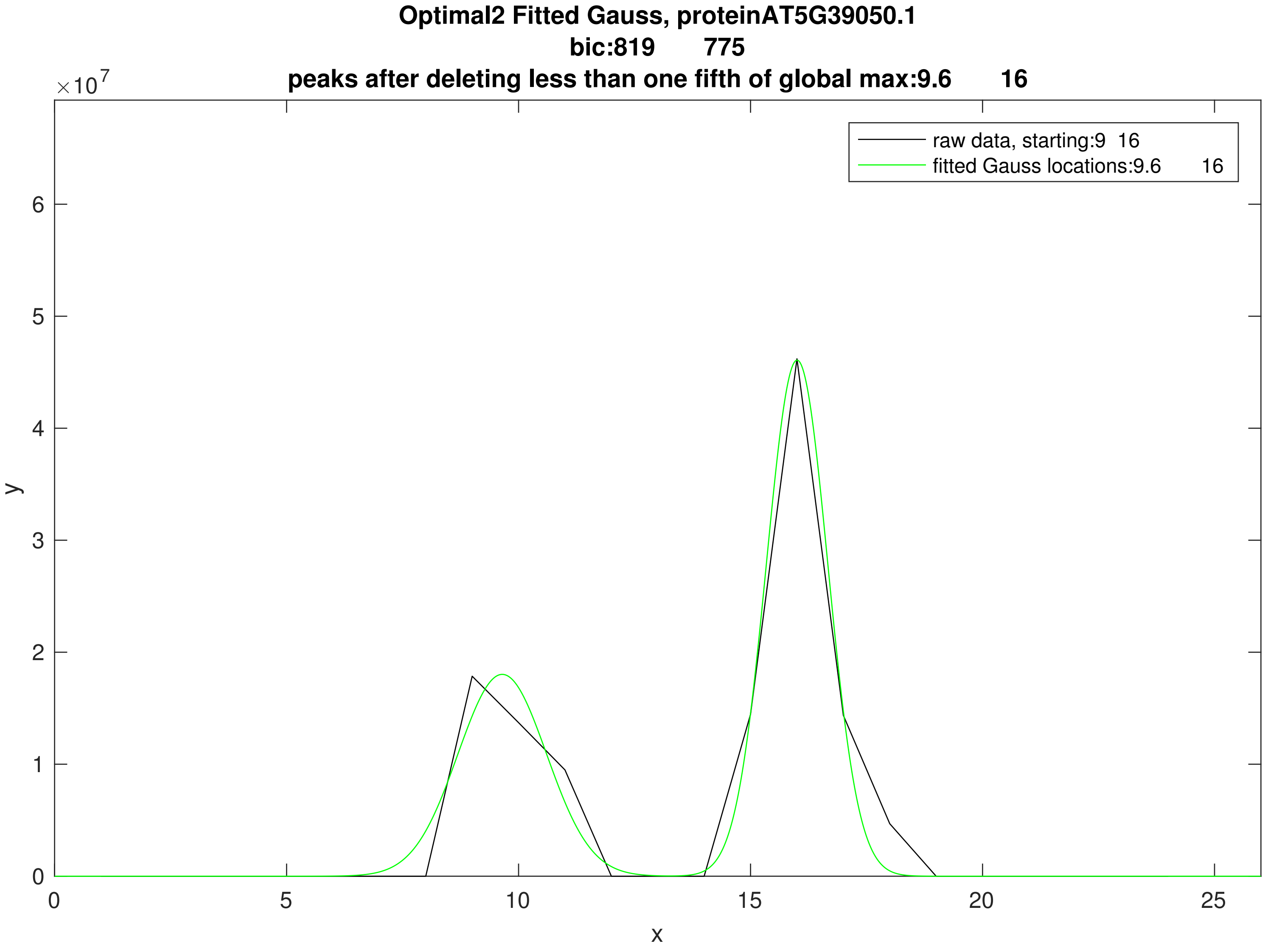

Optimal2 Fitted Gauss, proteinAT5G39050.1
bic:819 775
peaks after deleting less than one fifth of global max:9.6 16
×10^7
raw data, starting:9 16
fitted Gauss locations:9.6 16
0
1
2
3
4
5
6
y
0
5
10
15
20
25
x

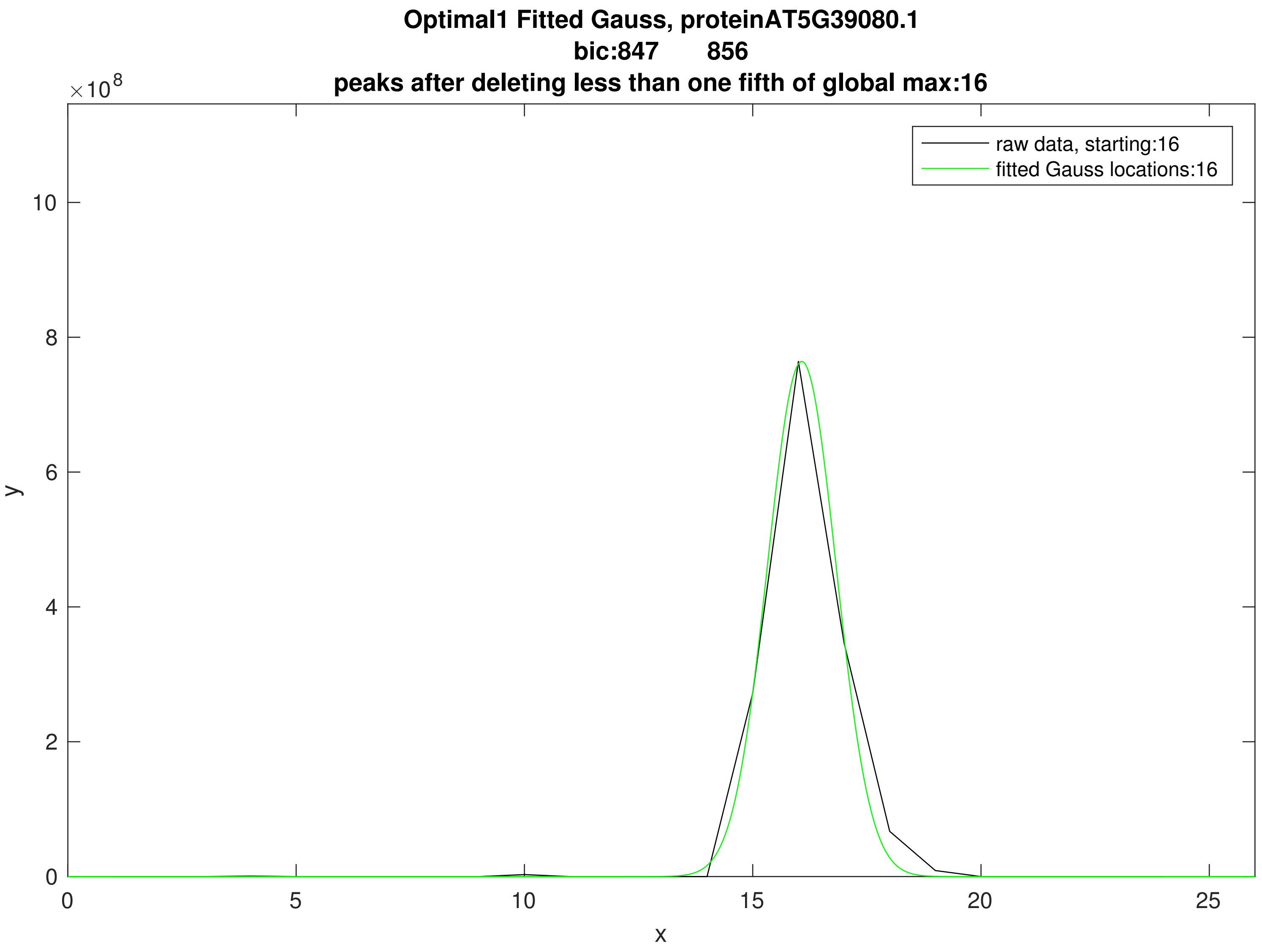

Optimal1 Fitted Gauss, proteinAT5G39080.1
bic:847 856
peaks after deleting less than one fifth of global max:16
×10^8
raw data, starting:16
fitted Gauss locations:16
y
x
0
2
4
6
8
10
5
15
20
25

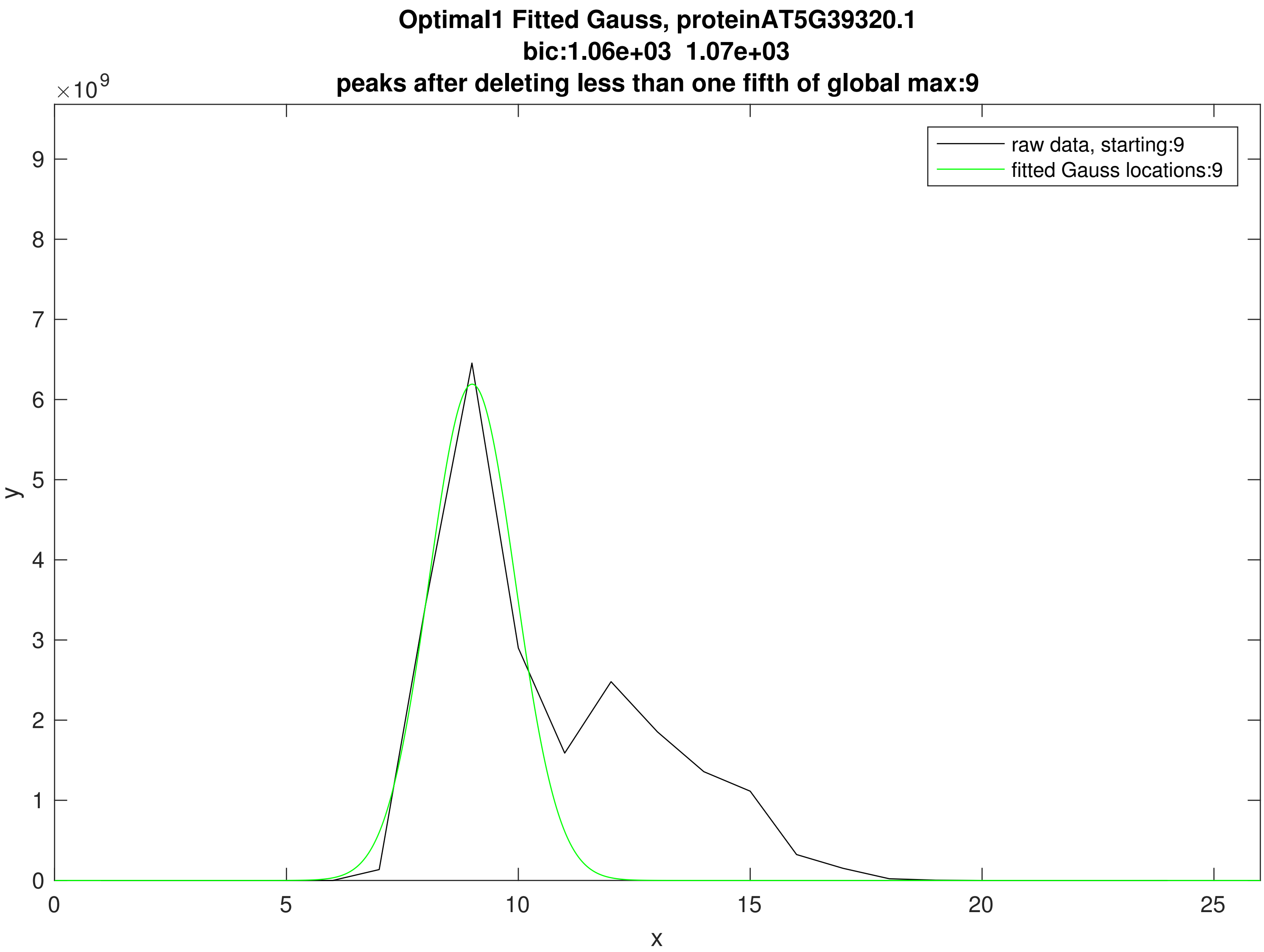
Optimal1 Fitted Gauss, proteinAT5G39320.1
bic:1.06e+03  1.07e+03
peaks after deleting less than one fifth of global max:9
×10^9
raw data, starting:9
fitted Gauss locations:9
y
x
0
1
2
3
4
5
6
7
8
9
0
5
10
15
20
25

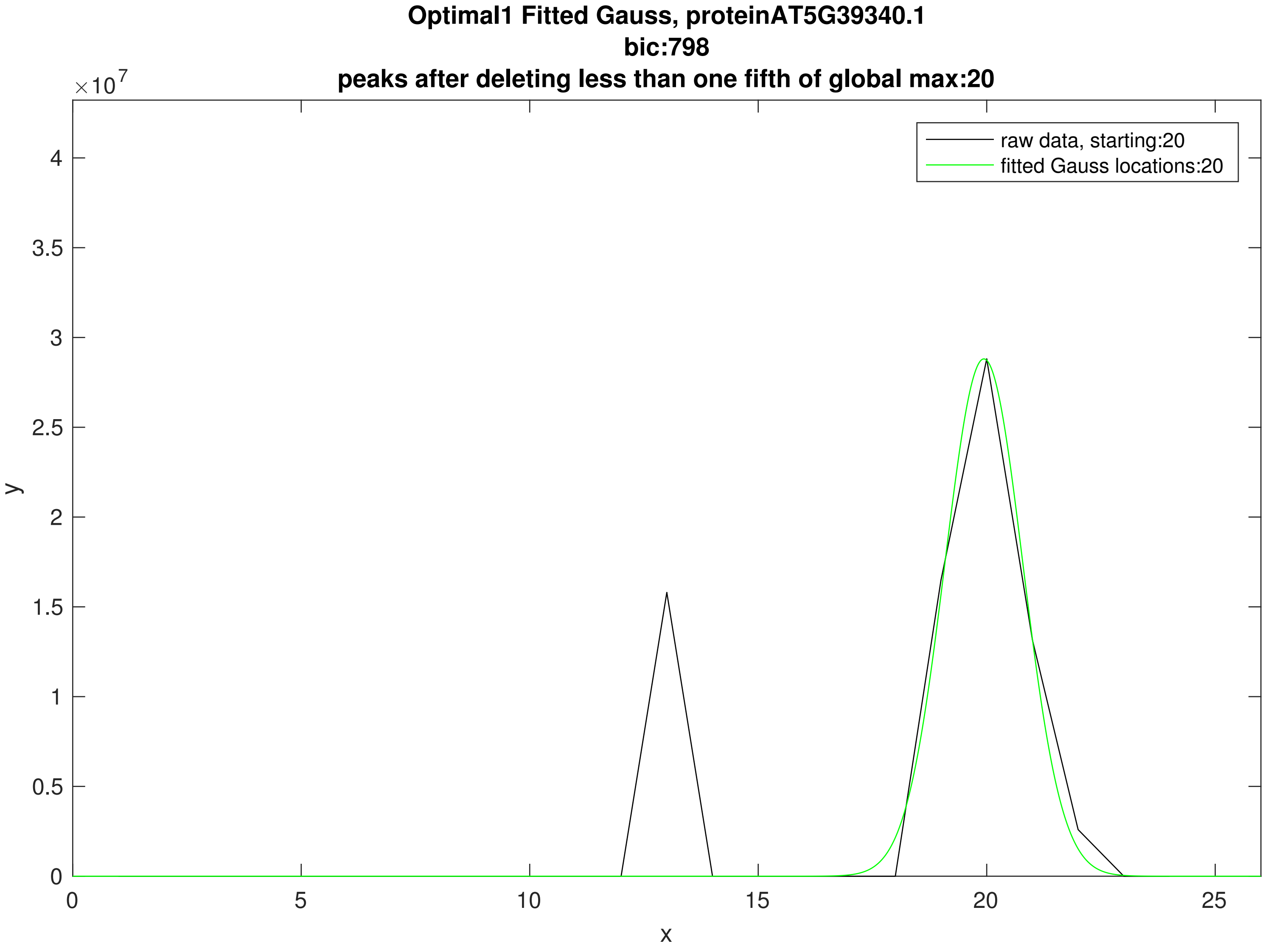

Optimal1 Fitted Gauss, proteinAT5G39340.1
bic:798
peaks after deleting less than one fifth of global max:20
×10^7
raw data, starting:20
fitted Gauss locations:20
y
x
0
0.5
1
1.5
2
2.5
3
3.5
4
0
5
10
15
20
25

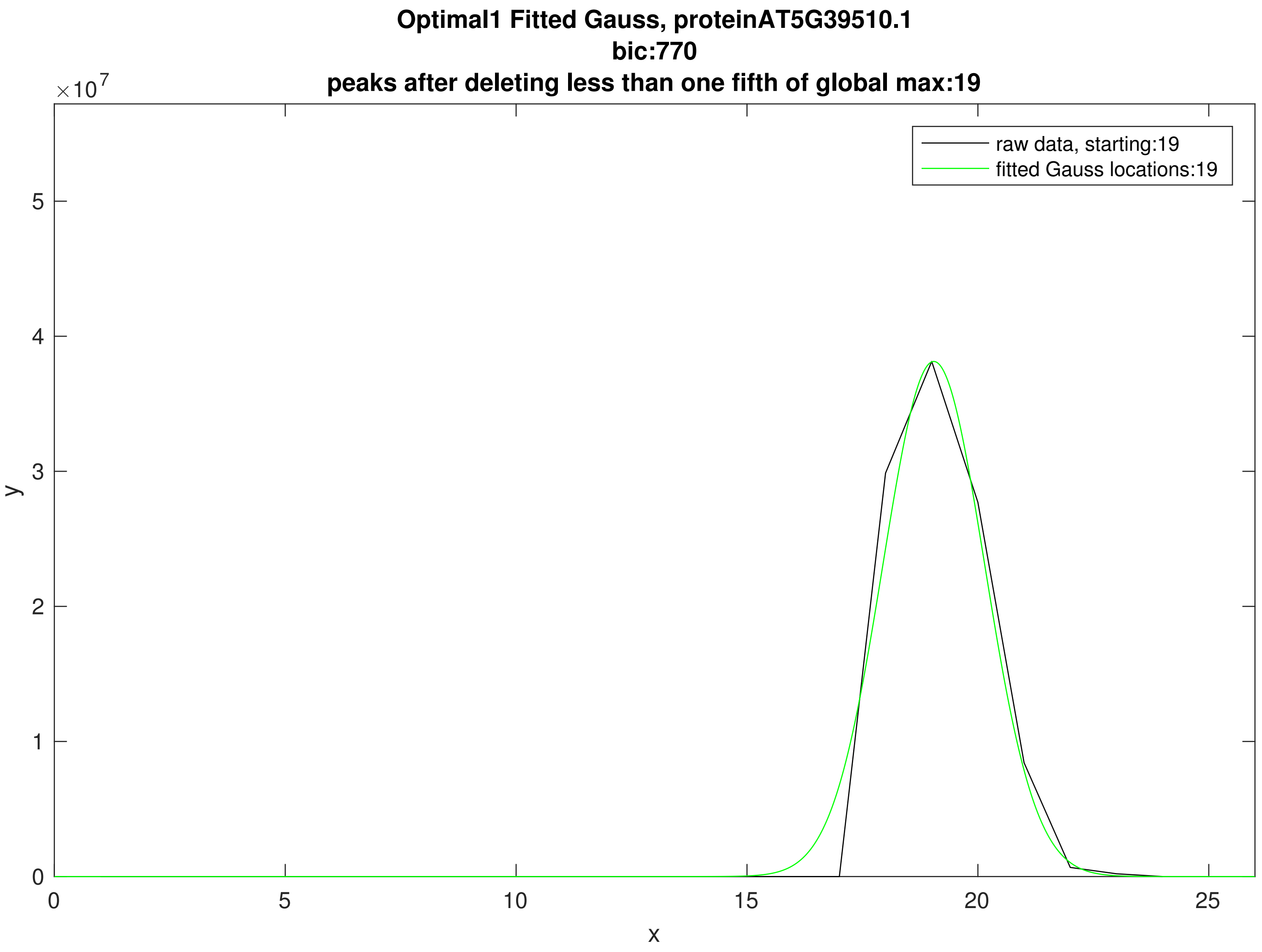
Optimal1 Fitted Gauss, proteinAT5G39510.1
bic:770
peaks after deleting less than one fifth of global max:19
×10^7
raw data, starting:19
fitted Gauss locations:19
y
x
0
1
2
3
4
5
0
5
10
15
20
25

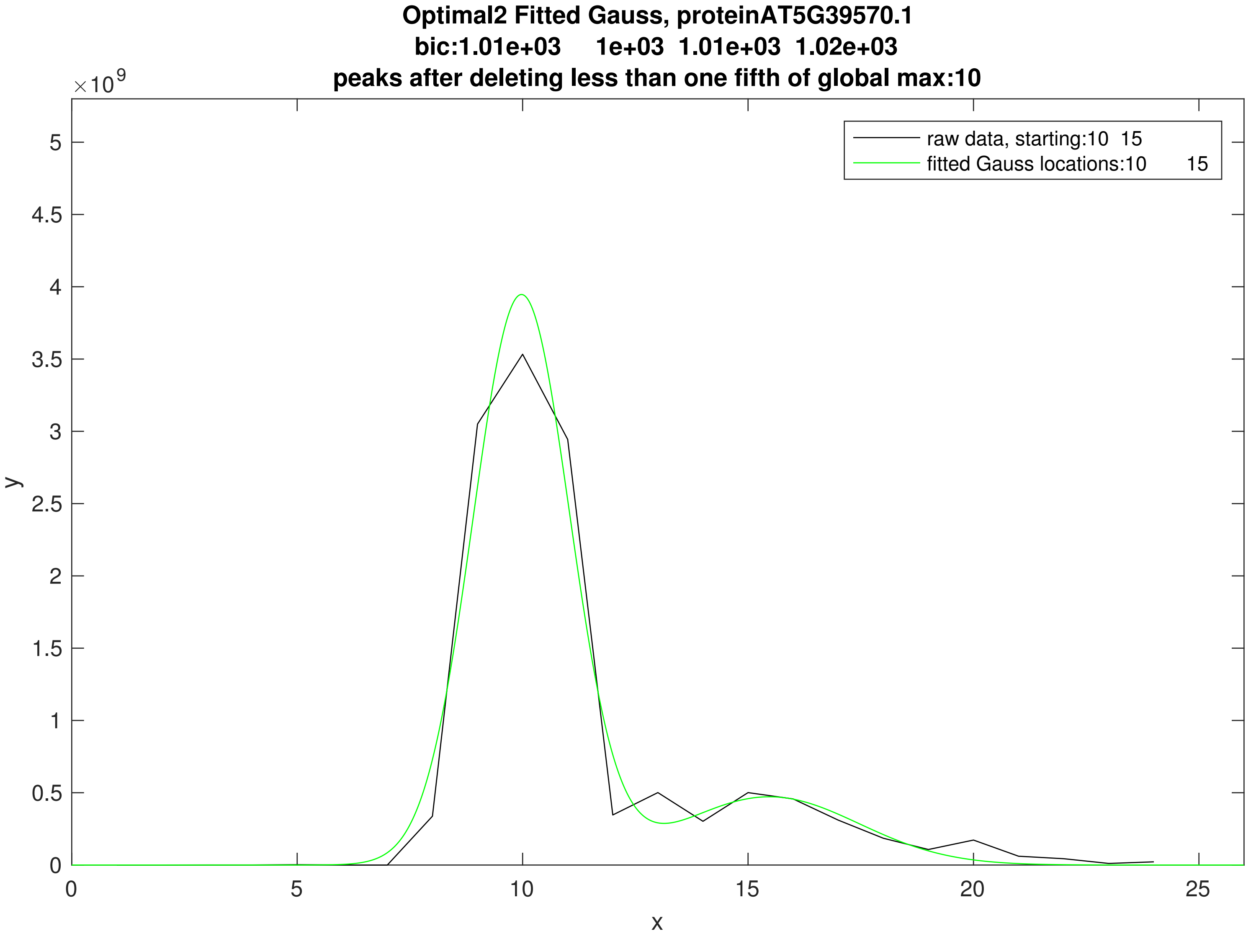
Optimal2 Fitted Gauss, proteinAT5G39570.1
bic:1.01e+03 1e+03 1.01e+03 1.02e+03
peaks after deleting less than one fifth of global max:10
×10^9
raw data, starting:10 15
fitted Gauss locations:10 15
y
x
0
0.5
1
1.5
2
2.5
3
3.5
4
4.5
5
0
5
10
15
20
25

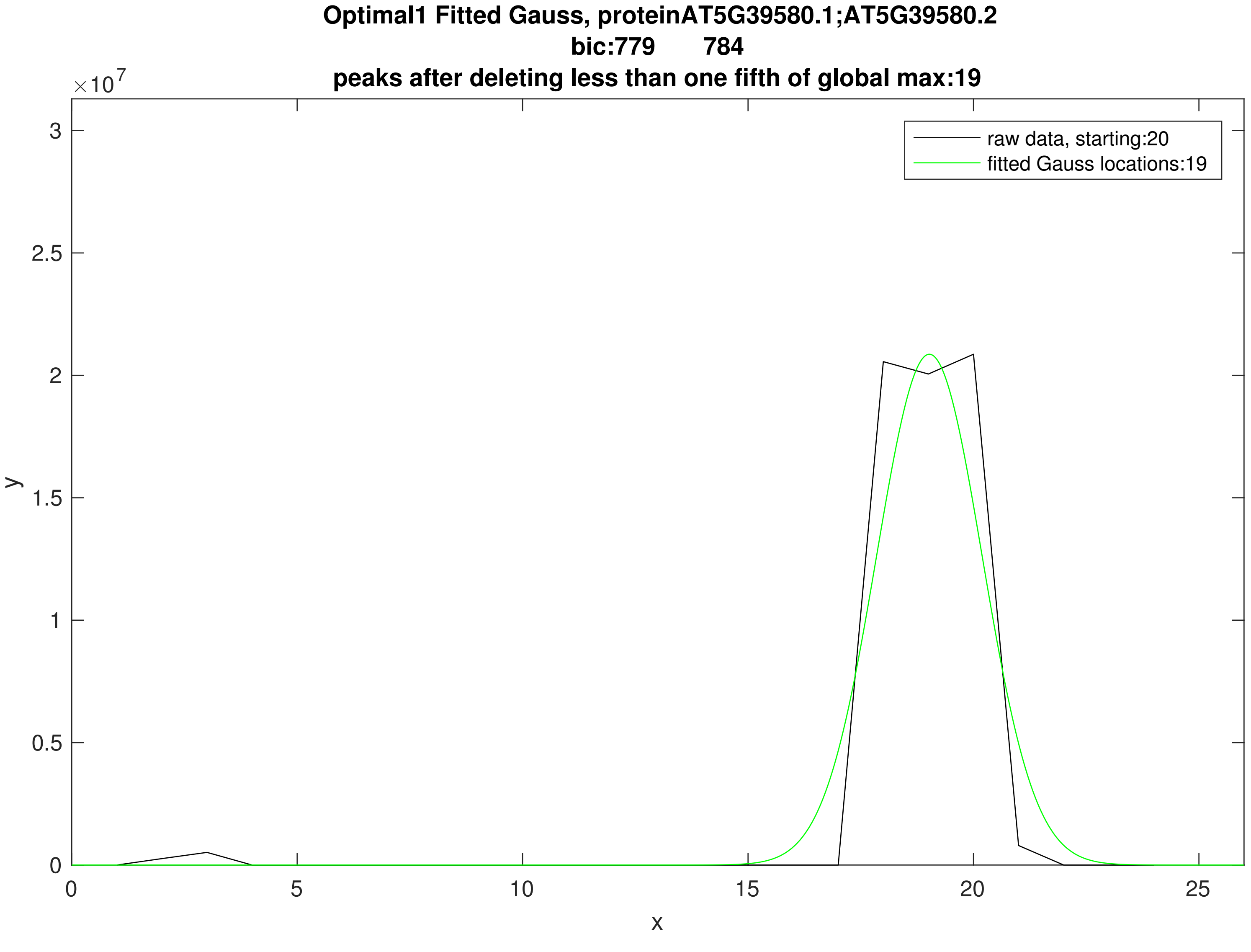
Optimal1 Fitted Gauss, proteinAT5G39580.1;AT5G39580.2
bic:779 784
peaks after deleting less than one fifth of global max:19
×10^7
3
2.5
2
1.5
1
0.5
0
y
0
5
10
15
20
25
x
raw data, starting:20
fitted Gauss locations:19

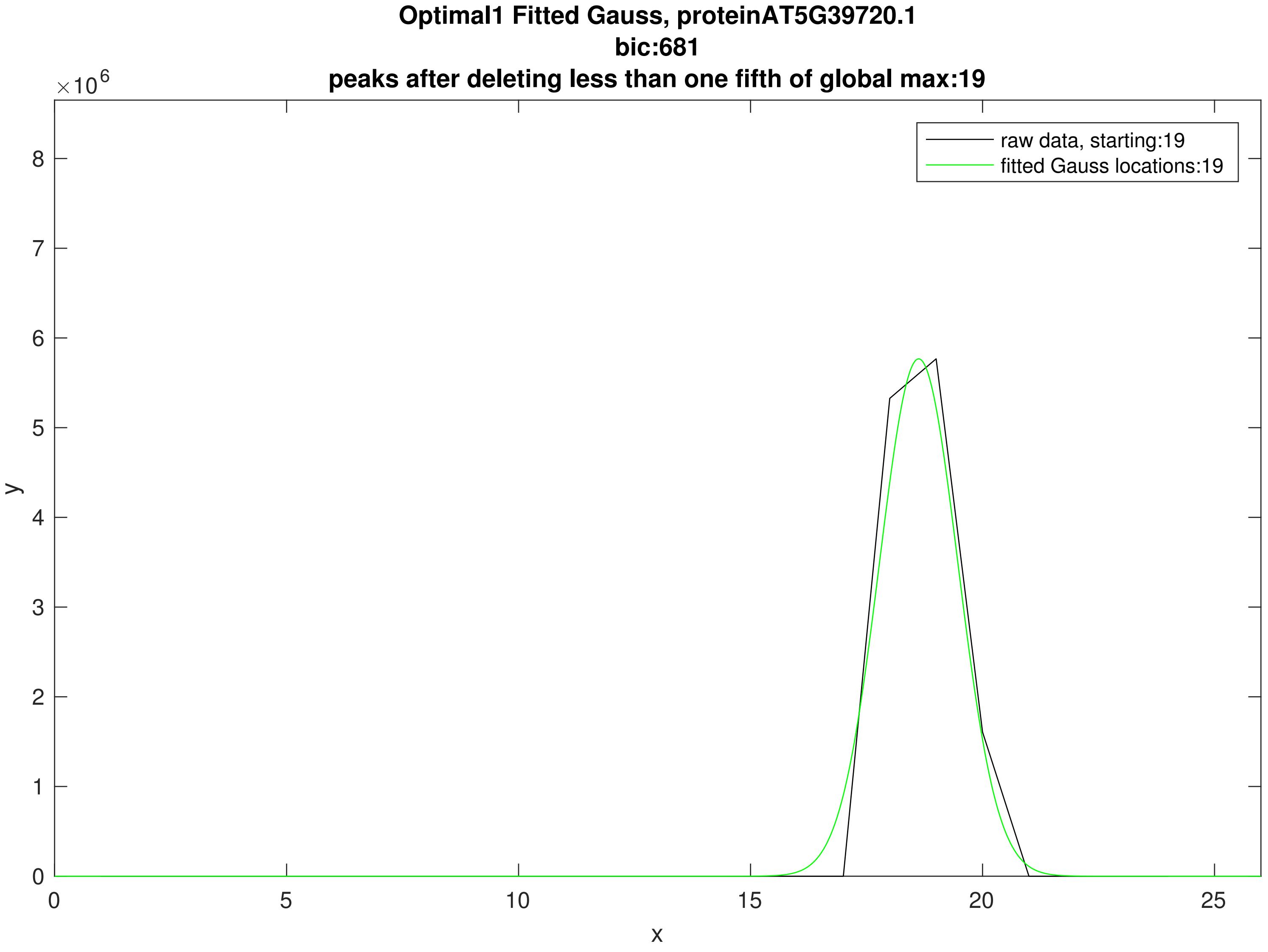
Optimal1 Fitted Gauss, proteinAT5G39720.1
bic:681
peaks after deleting less than one fifth of global max:19
×10^6
raw data, starting:19
fitted Gauss locations:19
y
x
0
1
2
3
4
5
6
7
8
0
5
10
15
20
25

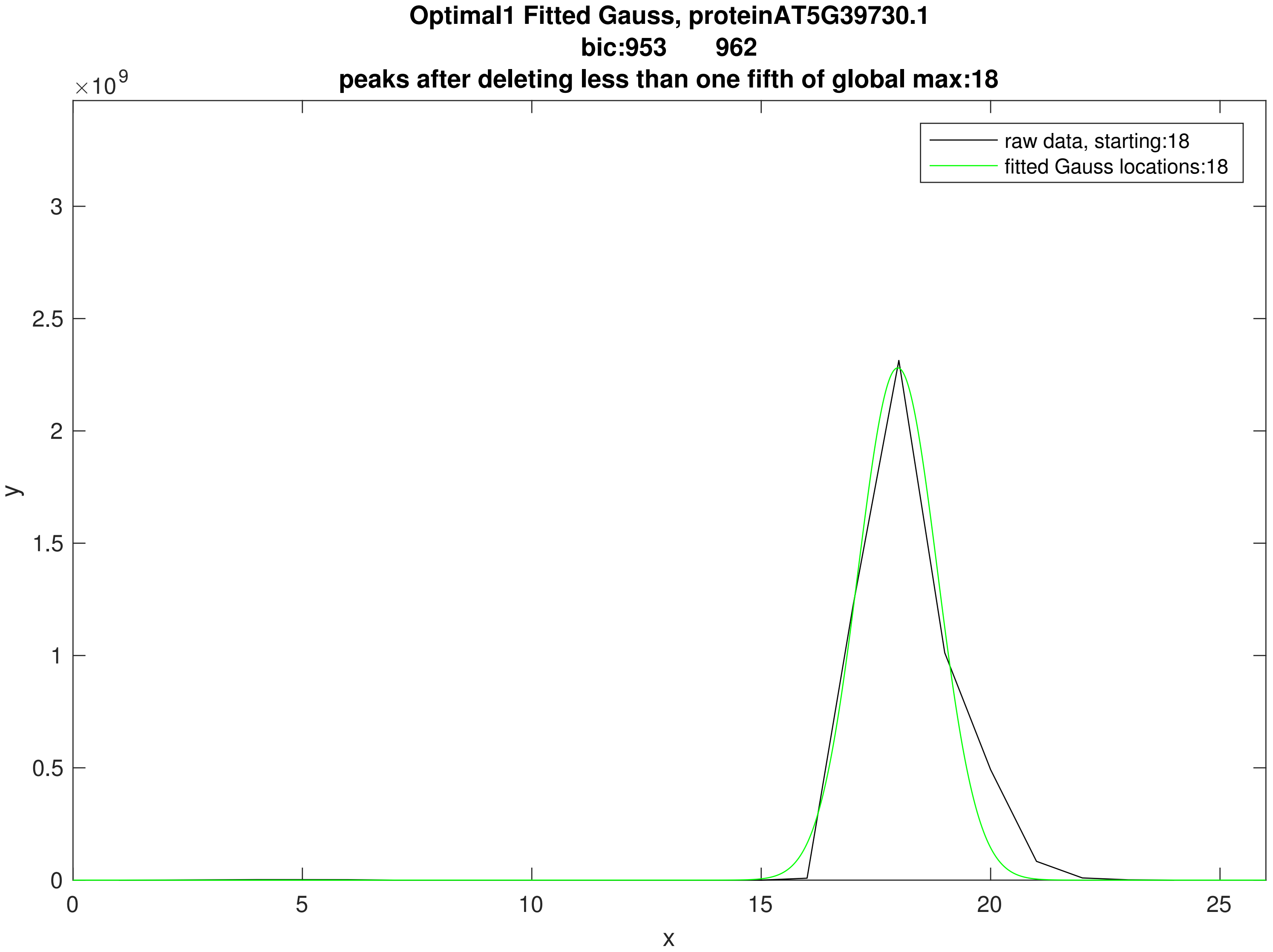

Optimal1 Fitted Gauss, proteinAT5G39730.1
bic:953 962
peaks after deleting less than one fifth of global max:18
×10^9
raw data, starting:18
fitted Gauss locations:18
y
x
0
0.5
1
1.5
2
2.5
3
5
10
15
20
25

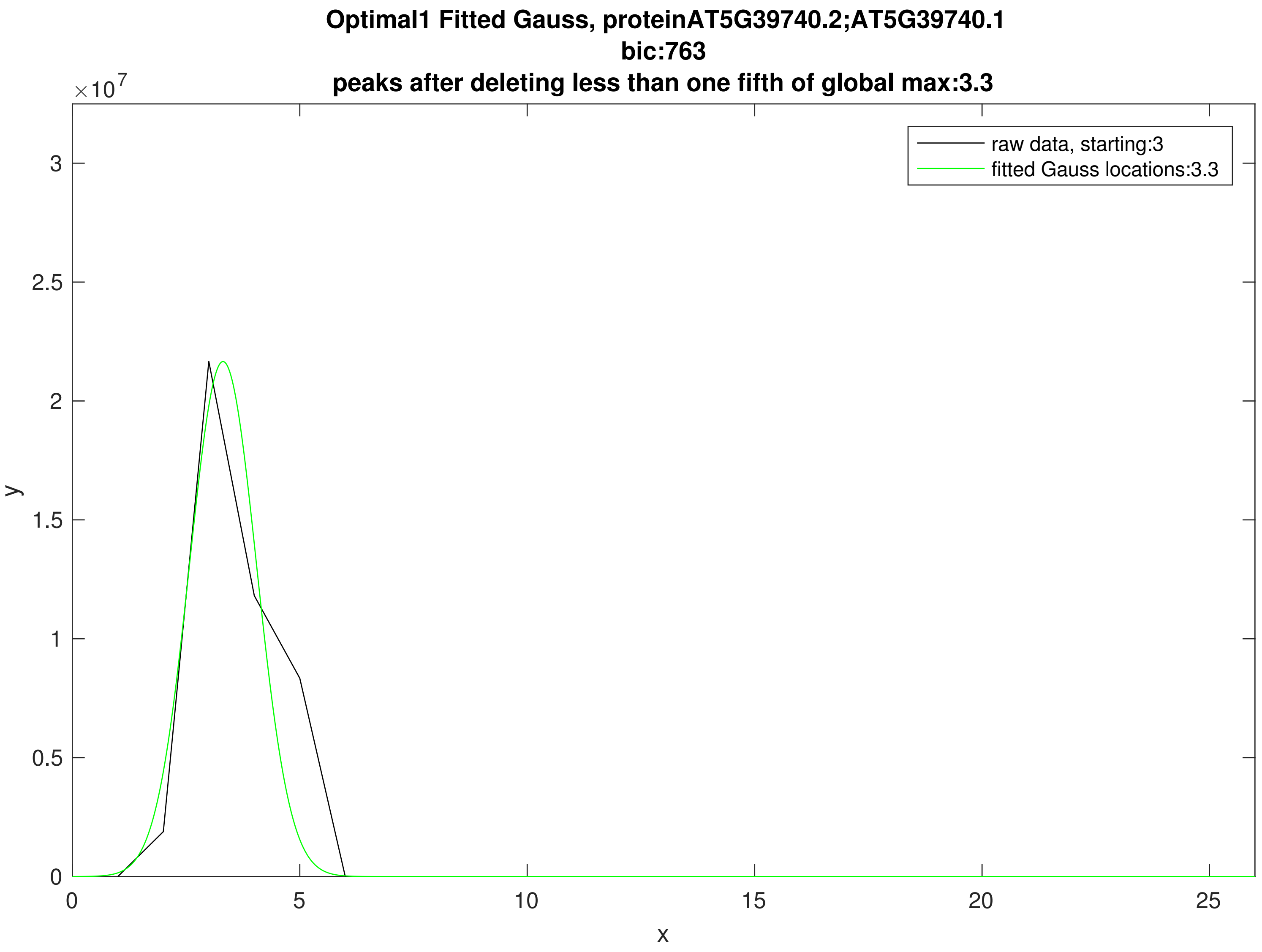
Optimal1 Fitted Gauss, proteinAT5G39740.2;AT5G39740.1
bic:763
peaks after deleting less than one fifth of global max:3.3
×10^7
raw data, starting:3
fitted Gauss locations:3.3
y
x
0
0.5
1
1.5
2
2.5
3
0
5
10
15
20
25

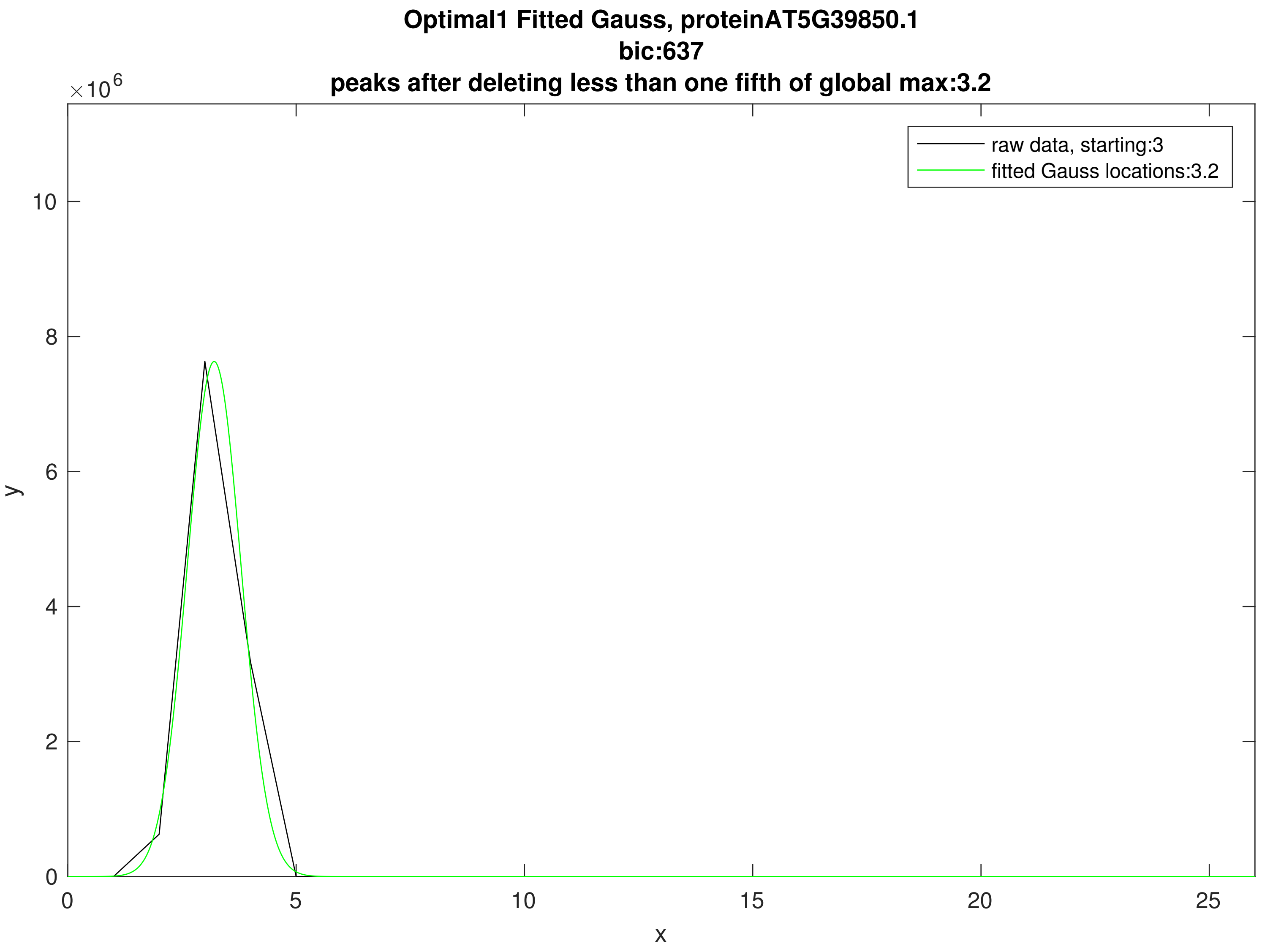

Optimal1 Fitted Gauss, proteinAT5G39850.1
bic:637
peaks after deleting less than one fifth of global max:3.2
×10^6
raw data, starting:3
fitted Gauss locations:3.2
10
8
6
4
2
0
y
0
5
10
15
20
25
x

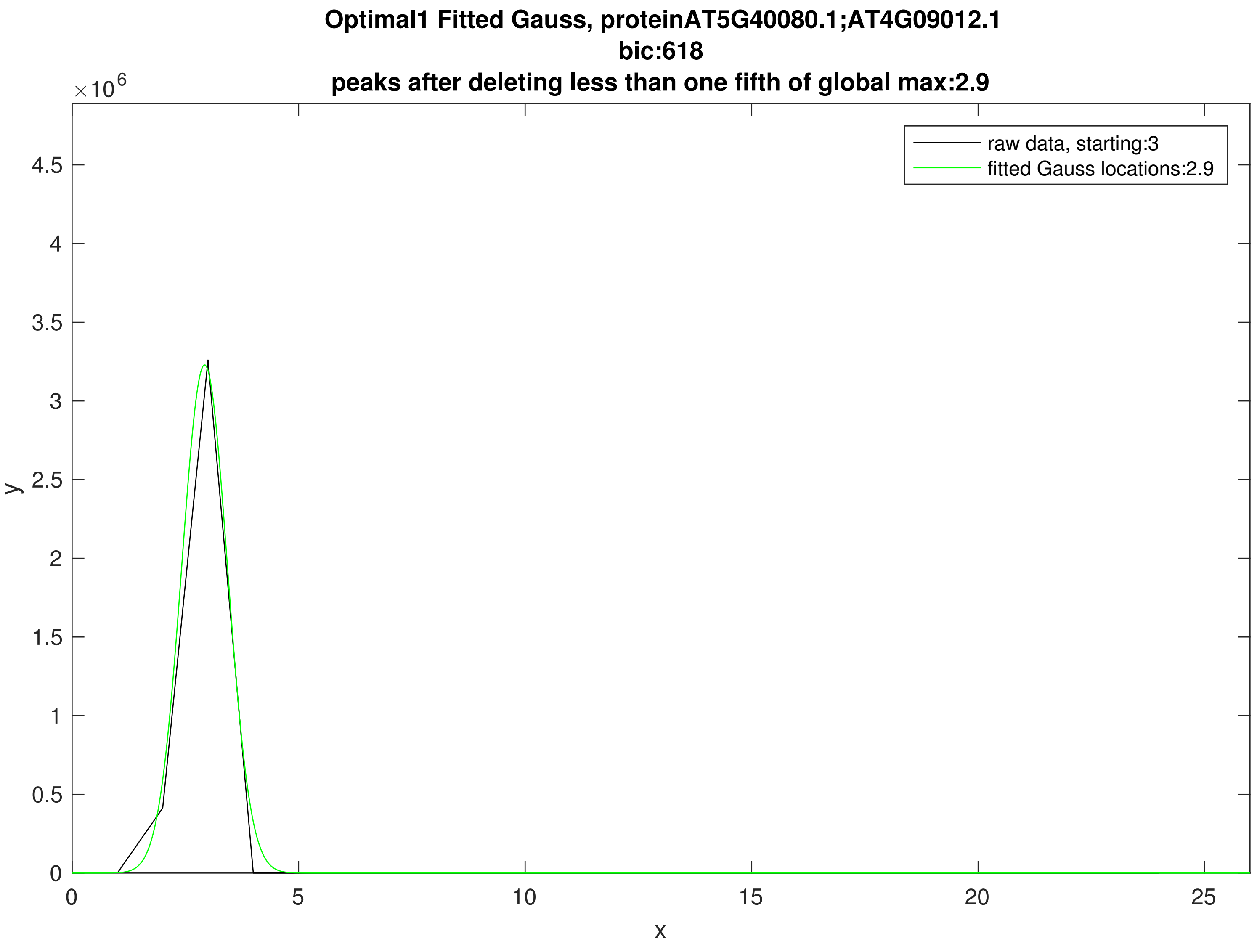
Optimal1 Fitted Gauss, proteinAT5G40080.1;AT4G09012.1
bic:618
peaks after deleting less than one fifth of global max:2.9
×10^6
raw data, starting:3
fitted Gauss locations:2.9
4.5
4
3.5
3
2.5
2
1.5
1
0.5
0
y
0
5
10
15
20
25
x

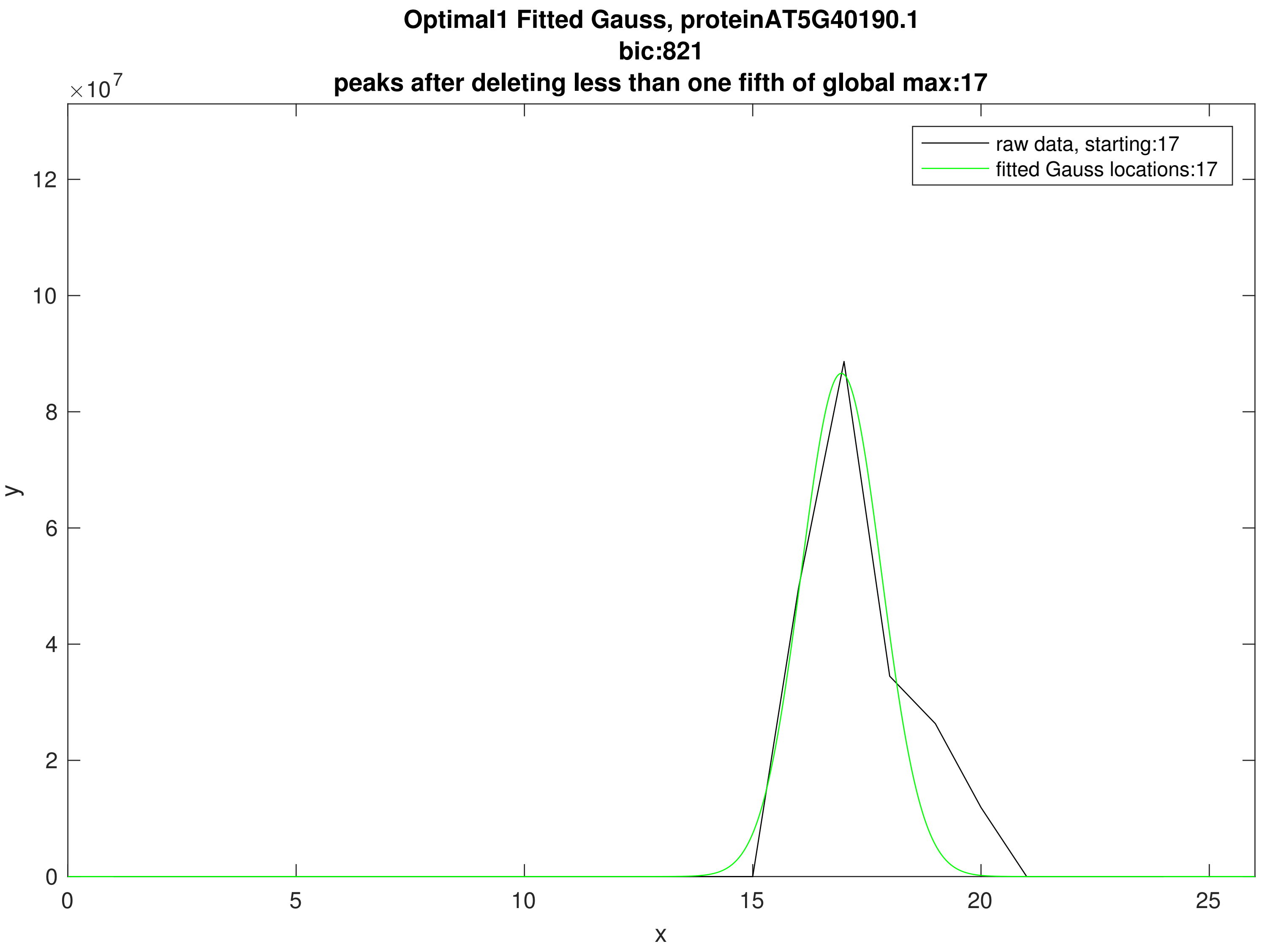
Optimal1 Fitted Gauss, proteinAT5G40190.1
bic:821
peaks after deleting less than one fifth of global max:17
×10^7
y
x
raw data, starting:17
fitted Gauss locations:17
0
2
4
6
8
10
12
0
5
10
15
20
25

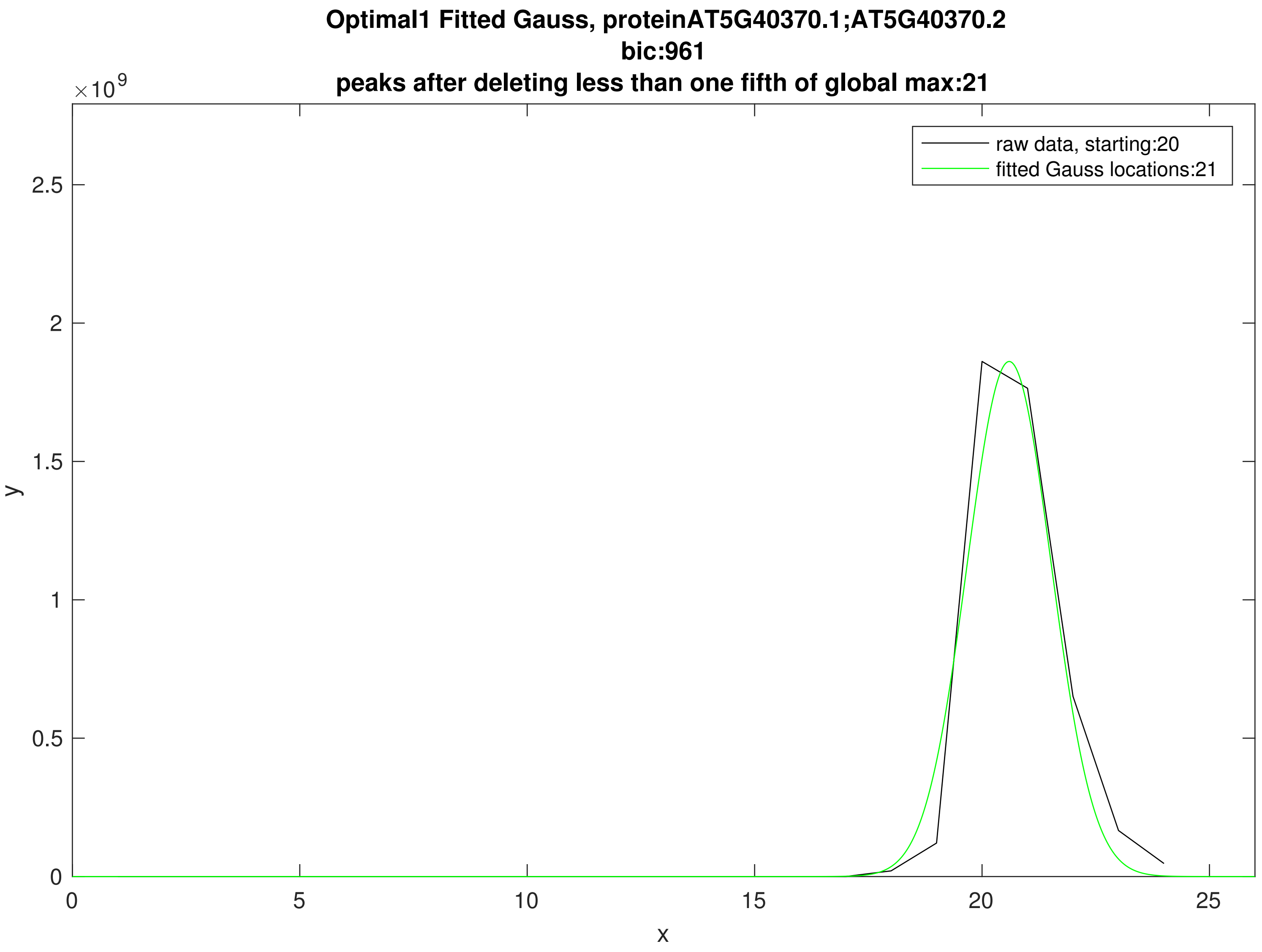
Optimal1 Fitted Gauss, proteinAT5G40370.1;AT5G40370.2
bic:961
peaks after deleting less than one fifth of global max:21
×10^9
raw data, starting:20
fitted Gauss locations:21
y
x
0
0.5
1
1.5
2
2.5
5
10
15
20
25

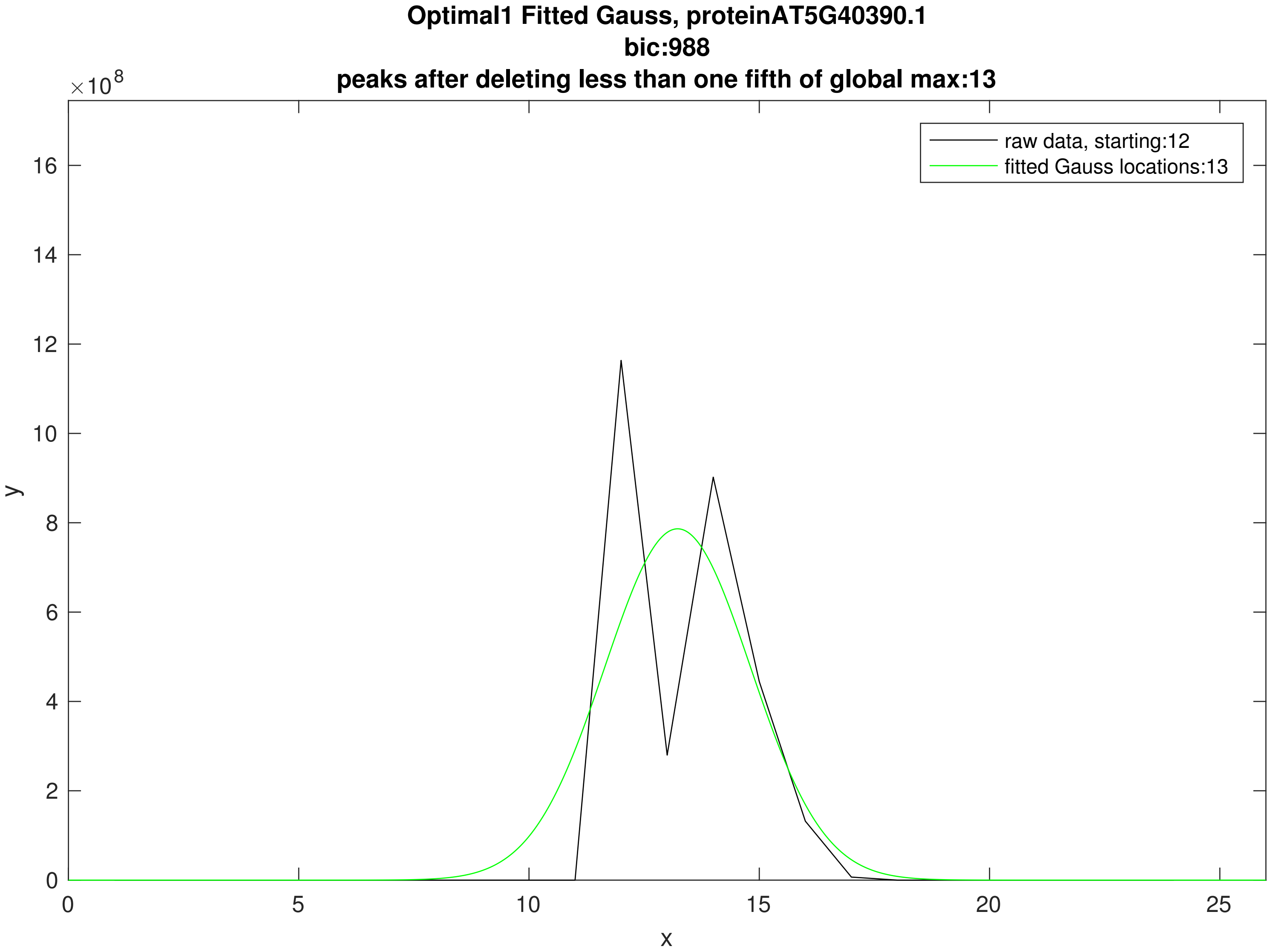
Optimal1 Fitted Gauss, proteinAT5G40390.1
bic:988
peaks after deleting less than one fifth of global max:13
×10^8
raw data, starting:12
fitted Gauss locations:13
y
x
0
2
4
6
8
10
12
14
16
0
5
10
15
20
25

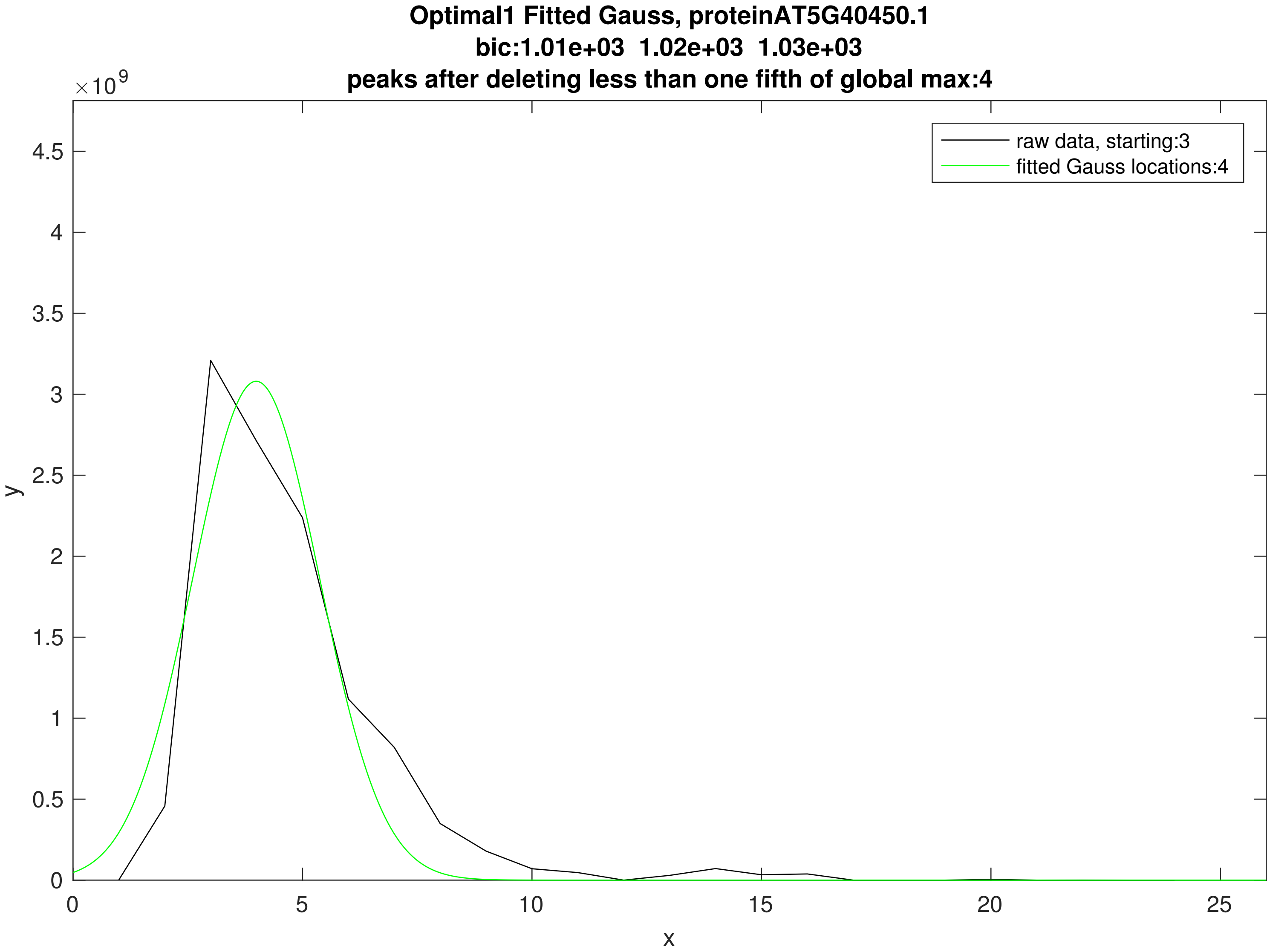

Optimal1 Fitted Gauss, proteinAT5G40450.1
bic:1.01e+03 1.02e+03 1.03e+03
peaks after deleting less than one fifth of global max:4
×10^9
raw data, starting:3
fitted Gauss locations:4
4.5
4
3.5
3
2.5
2
1.5
1
0.5
0
y
0
5
10
15
20
25
x

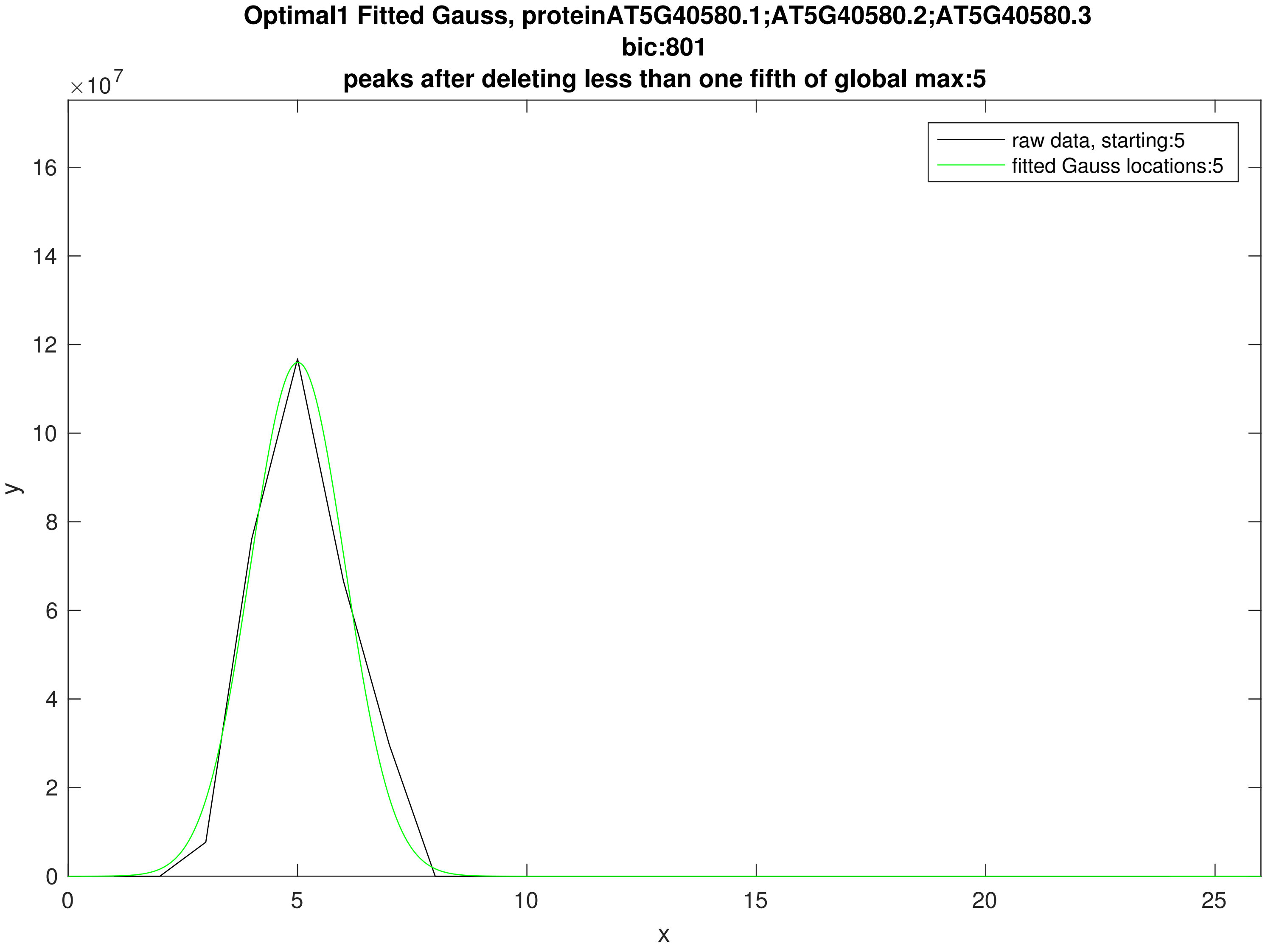

Optimal1 Fitted Gauss, proteinAT5G40580.1;AT5G40580.2;AT5G40580.3
bic:801
peaks after deleting less than one fifth of global max:5
×10^7
raw data, starting:5
fitted Gauss locations:5
y
x
0
2
4
6
8
10
12
14
16
0
5
10
15
20
25

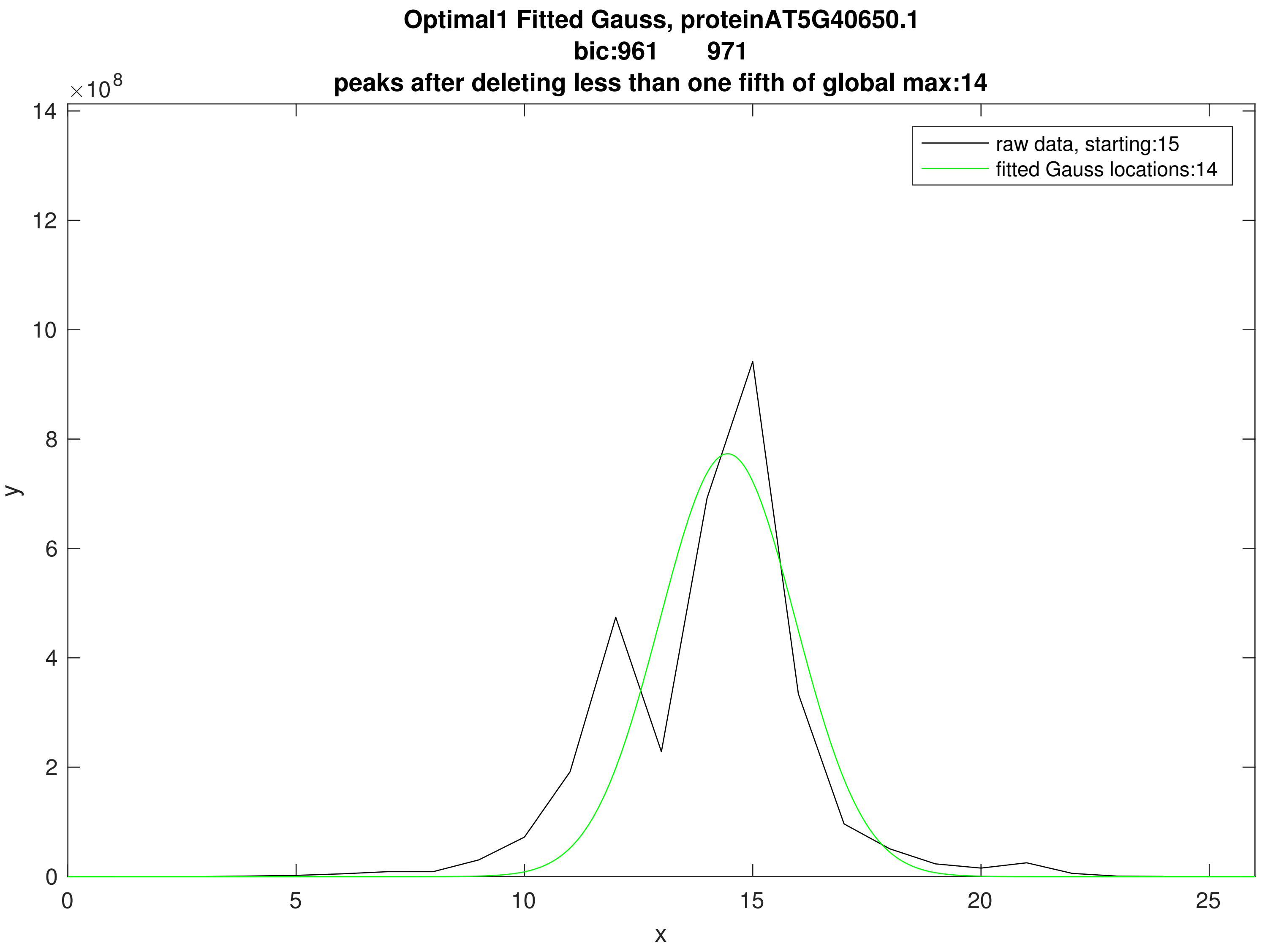

Optimal1 Fitted Gauss, proteinAT5G40650.1
bic:961 971
peaks after deleting less than one fifth of global max:14
×10^8
14
12
10
8
6
4
2
0
0
5
10
15
20
25
x
y
raw data, starting:15
fitted Gauss locations:14

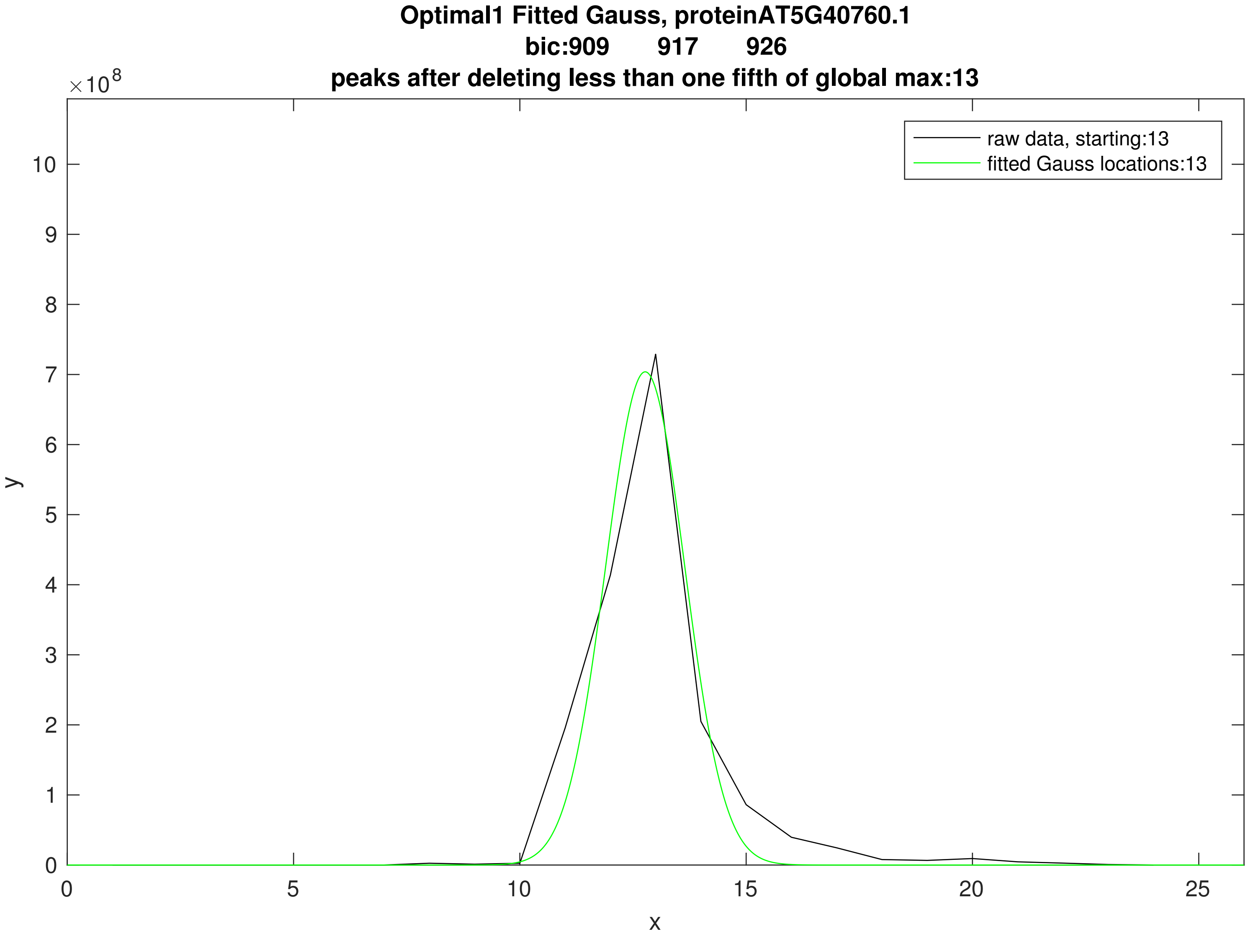

Optimal1 Fitted Gauss, proteinAT5G40760.1
bic:909 917 926
peaks after deleting less than one fifth of global max:13
×10^8
raw data, starting:13
fitted Gauss locations:13
y
x
0
1
2
3
4
5
6
7
8
9
10
0
5
10
15
20
25

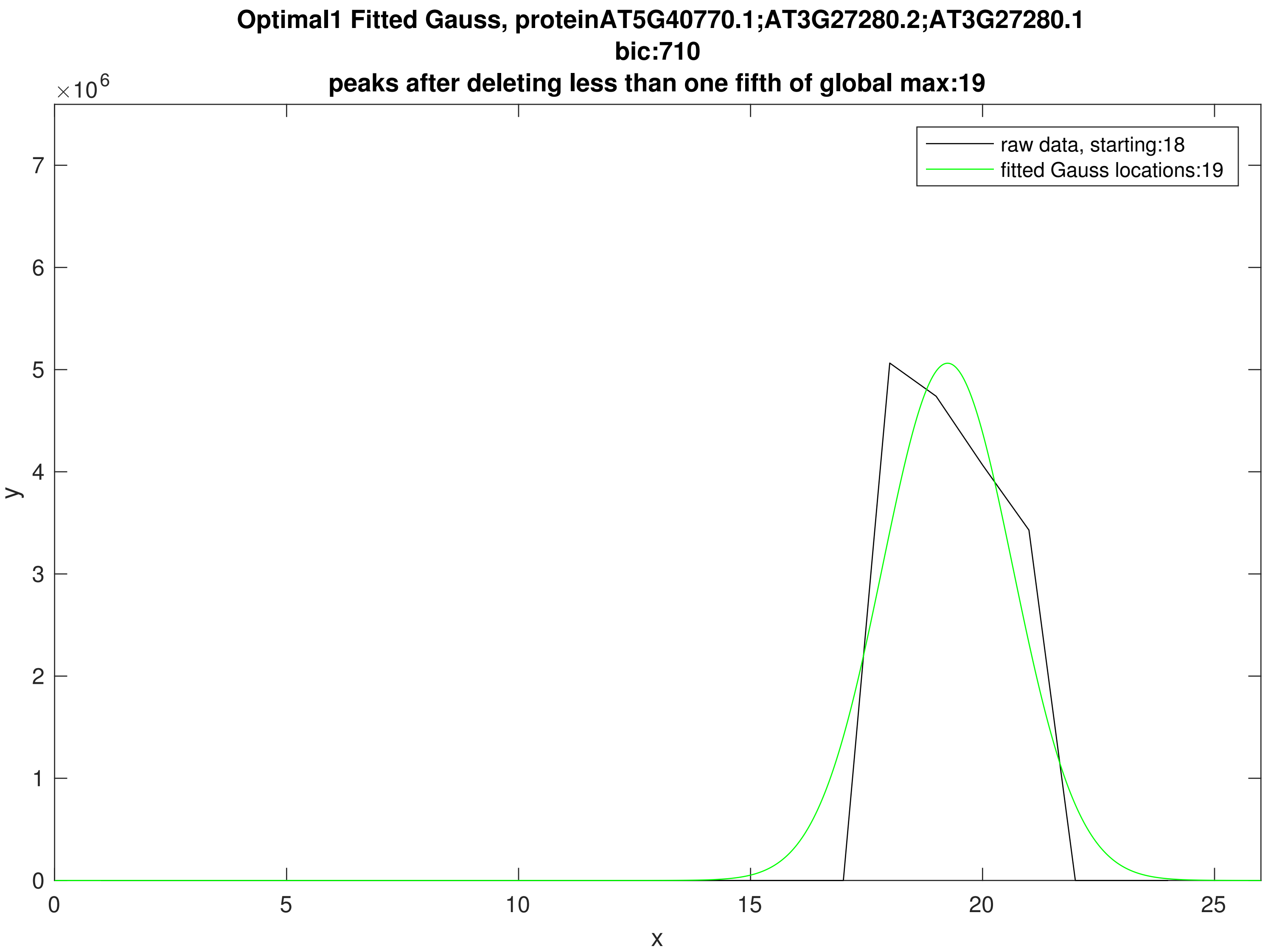

Optimal1 Fitted Gauss, proteinAT5G40770.1;AT3G27280.2;AT3G27280.1
bic:710
peaks after deleting less than one fifth of global max:19
×10^6
raw data, starting:18
fitted Gauss locations:19
y
x
0
1
2
3
4
5
6
7
0
5
10
15
20
25

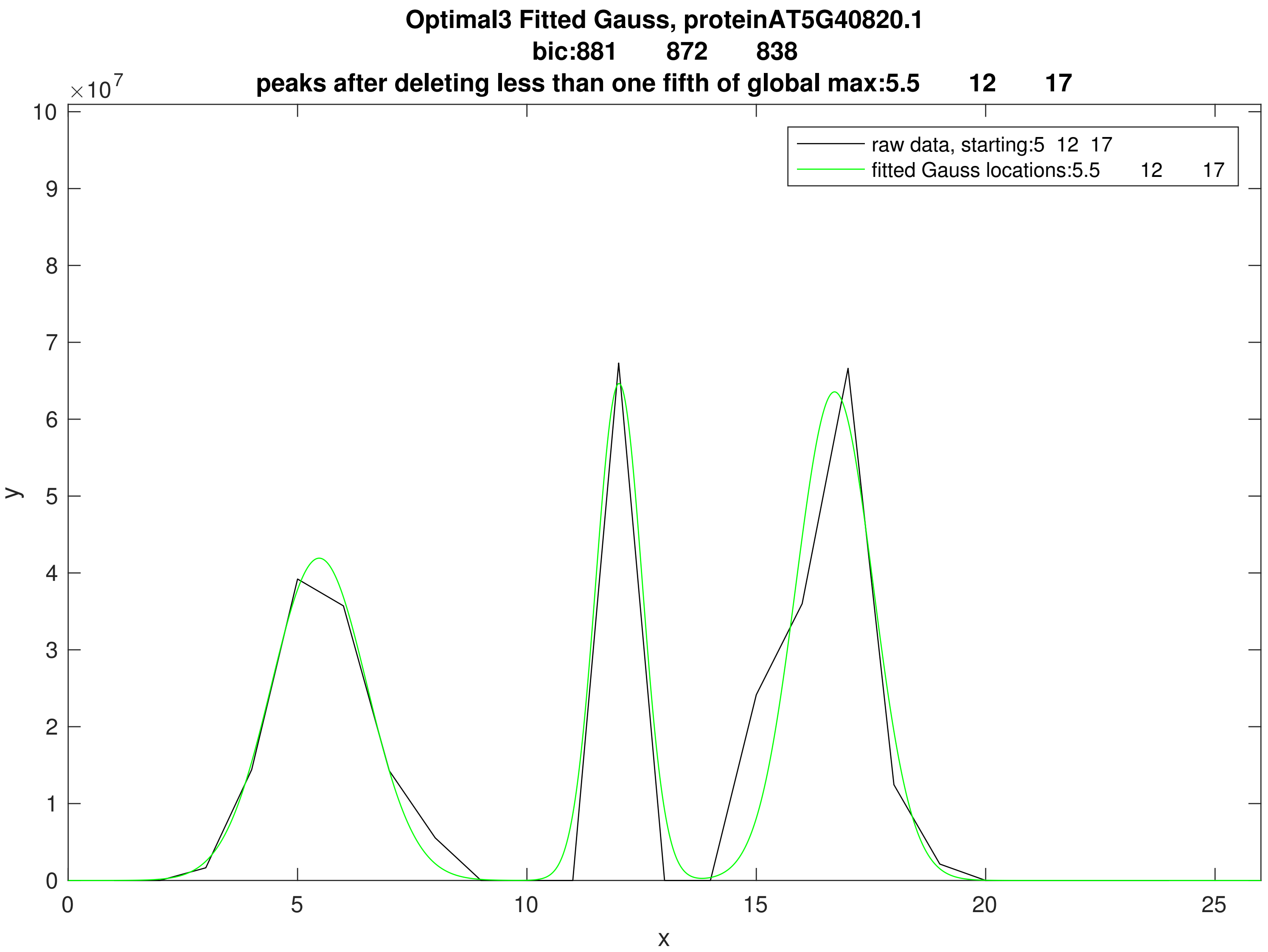

Optimal3 Fitted Gauss, proteinAT5G40820.1
bic:881 872 838
peaks after deleting less than one fifth of global max:5.5 12 17
×10^7
raw data, starting:5 12 17
fitted Gauss locations:5.5 12 17
y
x

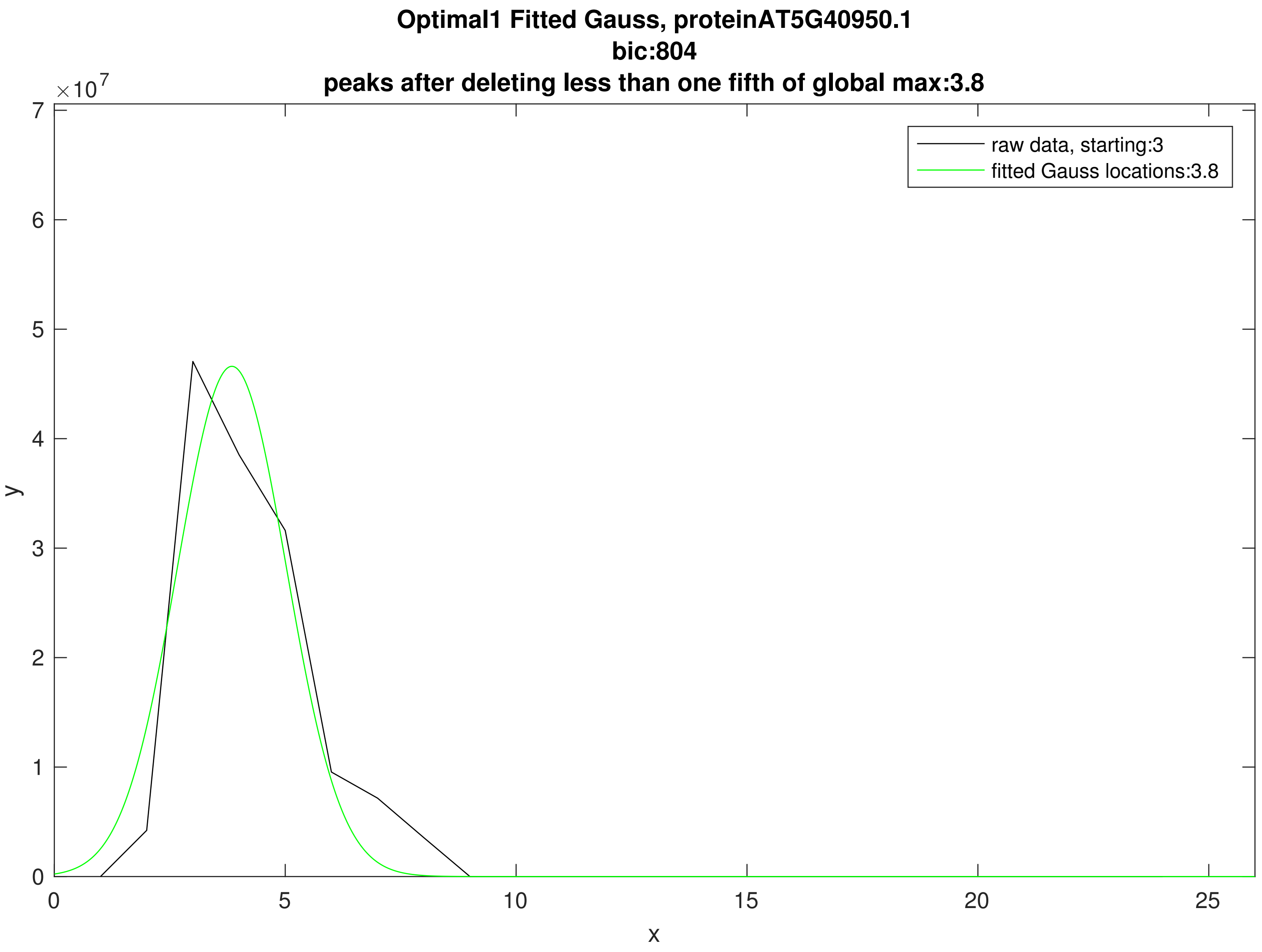
Optimal1 Fitted Gauss, proteinAT5G40950.1
bic:804
peaks after deleting less than one fifth of global max:3.8
×10^7
raw data, starting:3
fitted Gauss locations:3.8
y
x
0
1
2
3
4
5
6
7
0
5
10
15
20
25

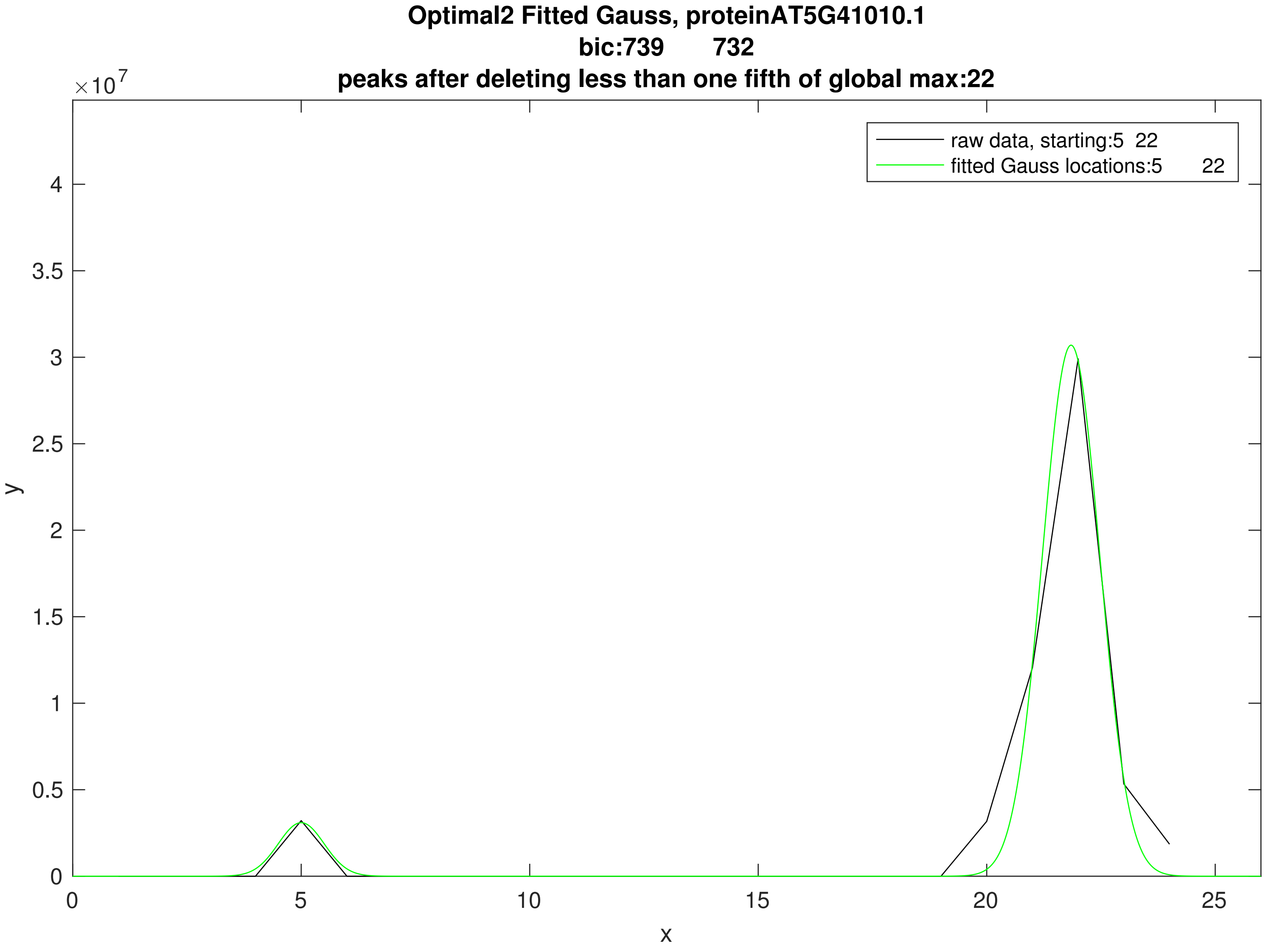
Optimal2 Fitted Gauss, proteinAT5G41010.1
bic:739 732
peaks after deleting less than one fifth of global max:22
×10^7
raw data, starting:5 22
fitted Gauss locations:5 22
y
x
0
0.5
1
1.5
2
2.5
3
3.5
4
5
10
15
20
25

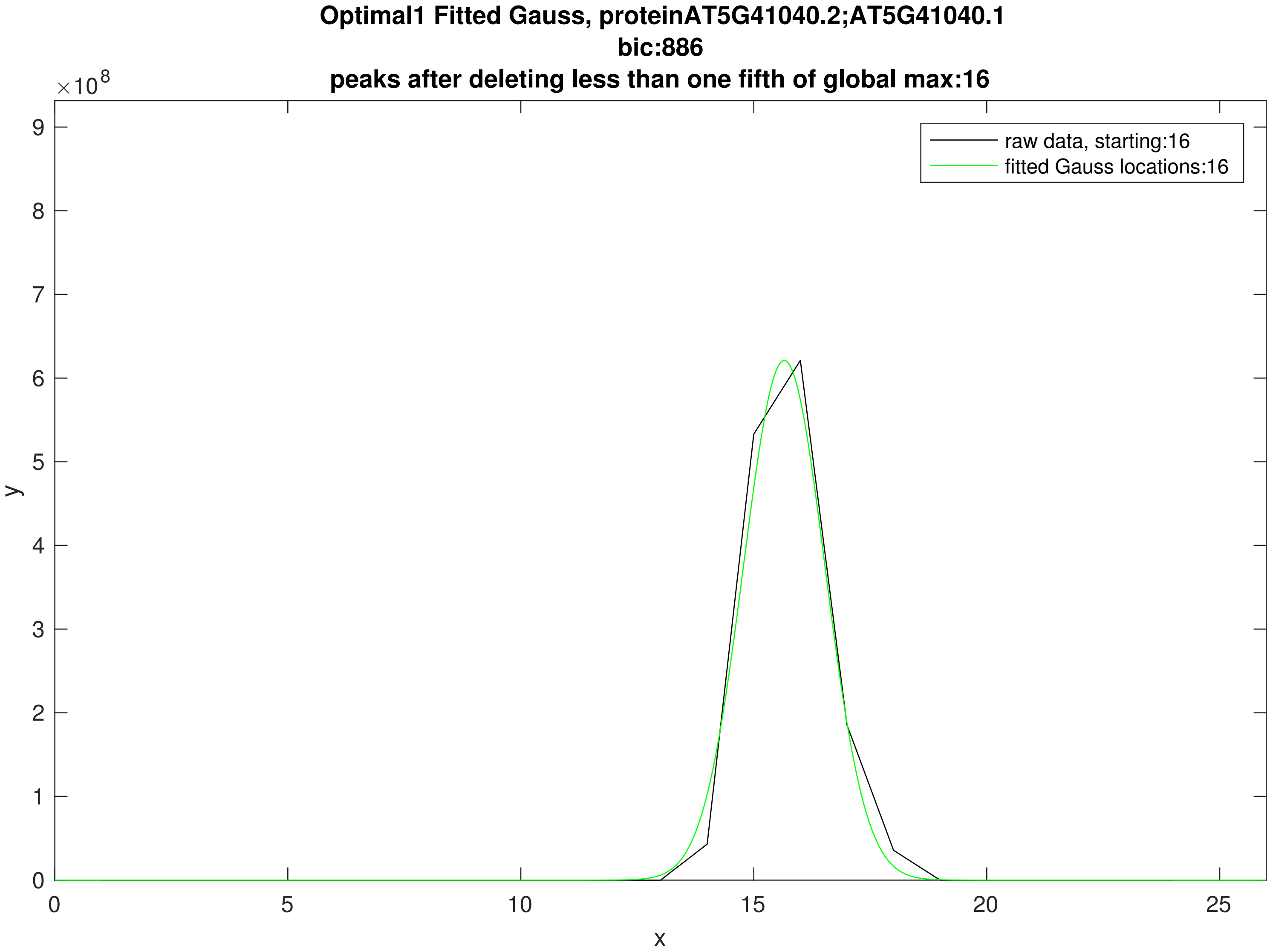

Optimal1 Fitted Gauss, proteinAT5G41040.2;AT5G41040.1
bic:886
peaks after deleting less than one fifth of global max:16
×10^8
raw data, starting:16
fitted Gauss locations:16
y
x
0
1
2
3
4
5
6
7
8
9
0
5
10
15
20
25

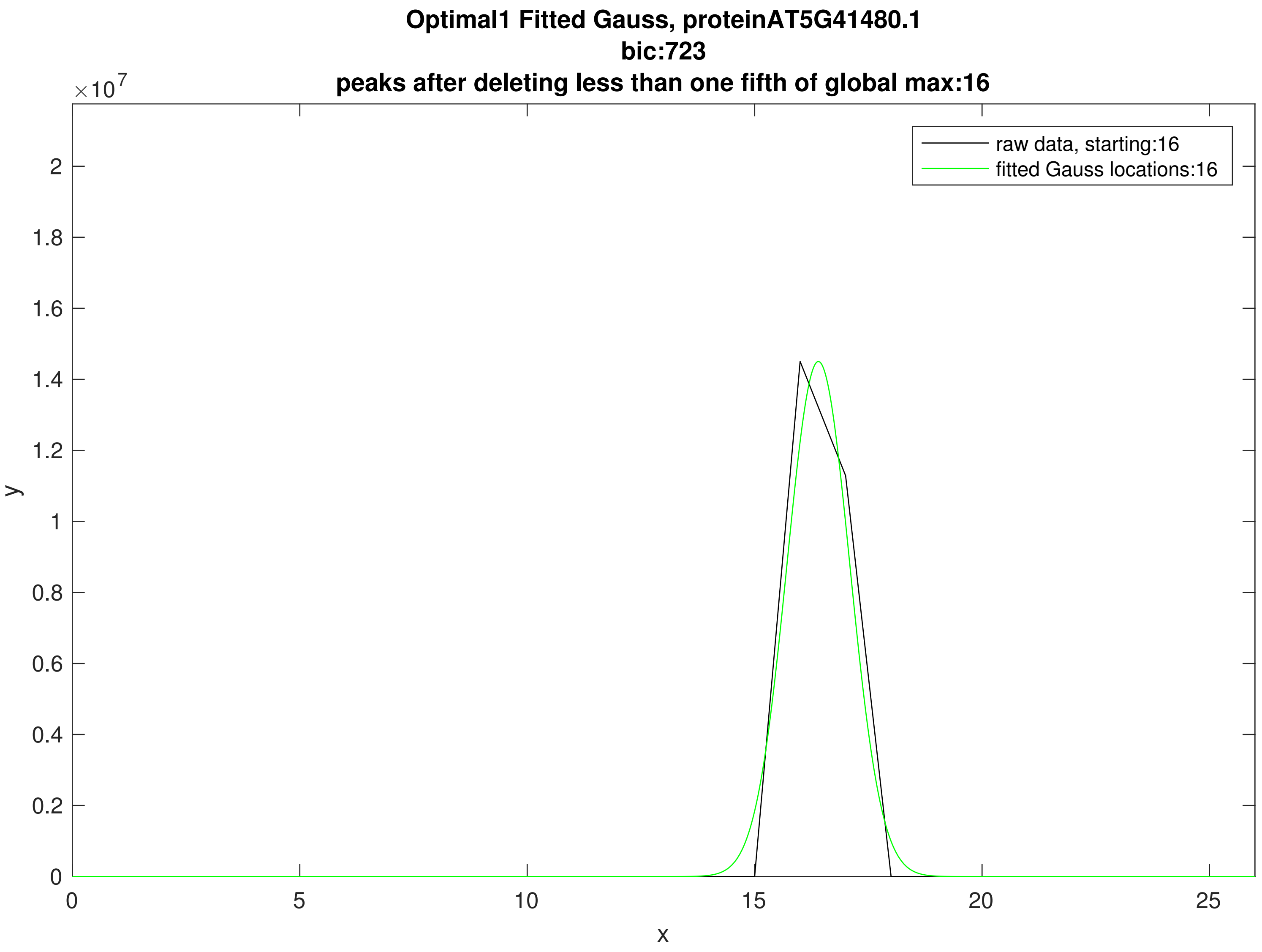
Optimal1 Fitted Gauss, proteinAT5G41480.1
bic:723
peaks after deleting less than one fifth of global max:16
×10^7
raw data, starting:16
fitted Gauss locations:16
y
x
0
0.2
0.4
0.6
0.8
1
1.2
1.4
1.6
1.8
2
0
5
10
15
20
25

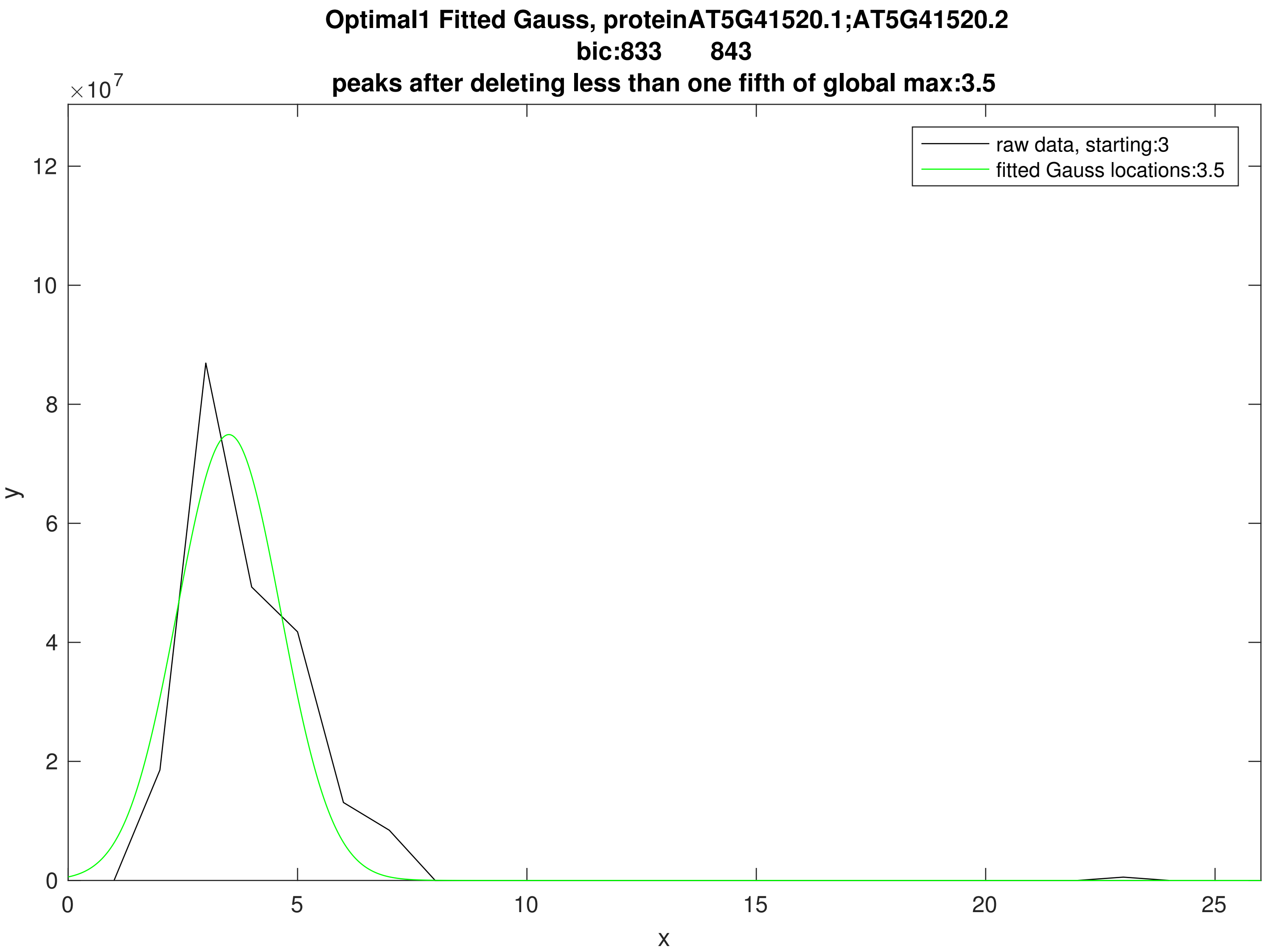
Optimal1 Fitted Gauss, proteinAT5G41520.1;AT5G41520.2
bic:833 843
peaks after deleting less than one fifth of global max:3.5
×10^7
raw data, starting:3
fitted Gauss locations:3.5
y
x
0
2
4
6
8
10
12
0
5
10
15
20
25

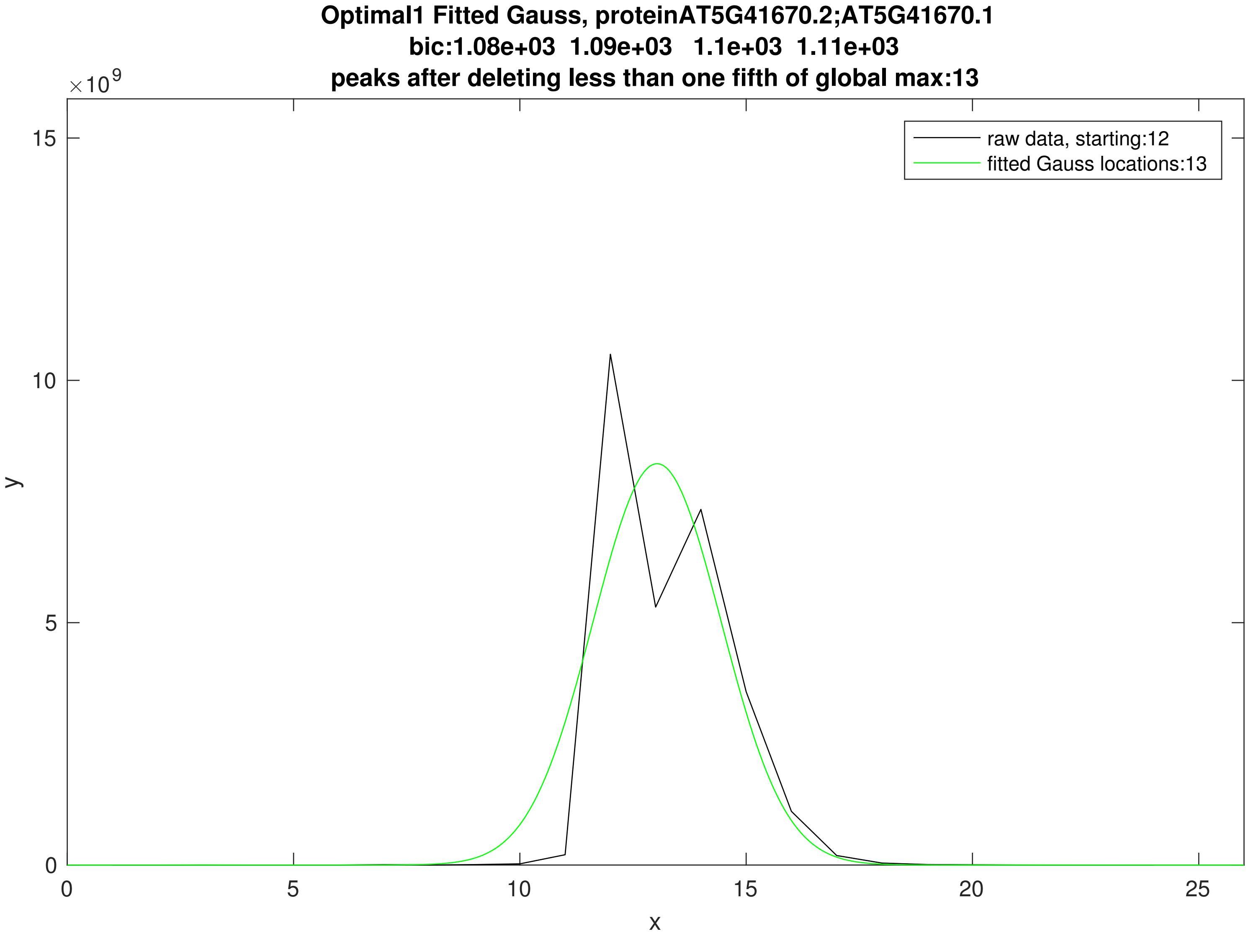

Optimal1 Fitted Gauss, proteinAT5G41670.2;AT5G41670.1
bic:1.08e+03 1.09e+03 1.1e+03 1.11e+03
peaks after deleting less than one fifth of global max:13
×10^9
raw data, starting:12
fitted Gauss locations:13
15
10
5
0
y
0
5
10
15
20
25
x

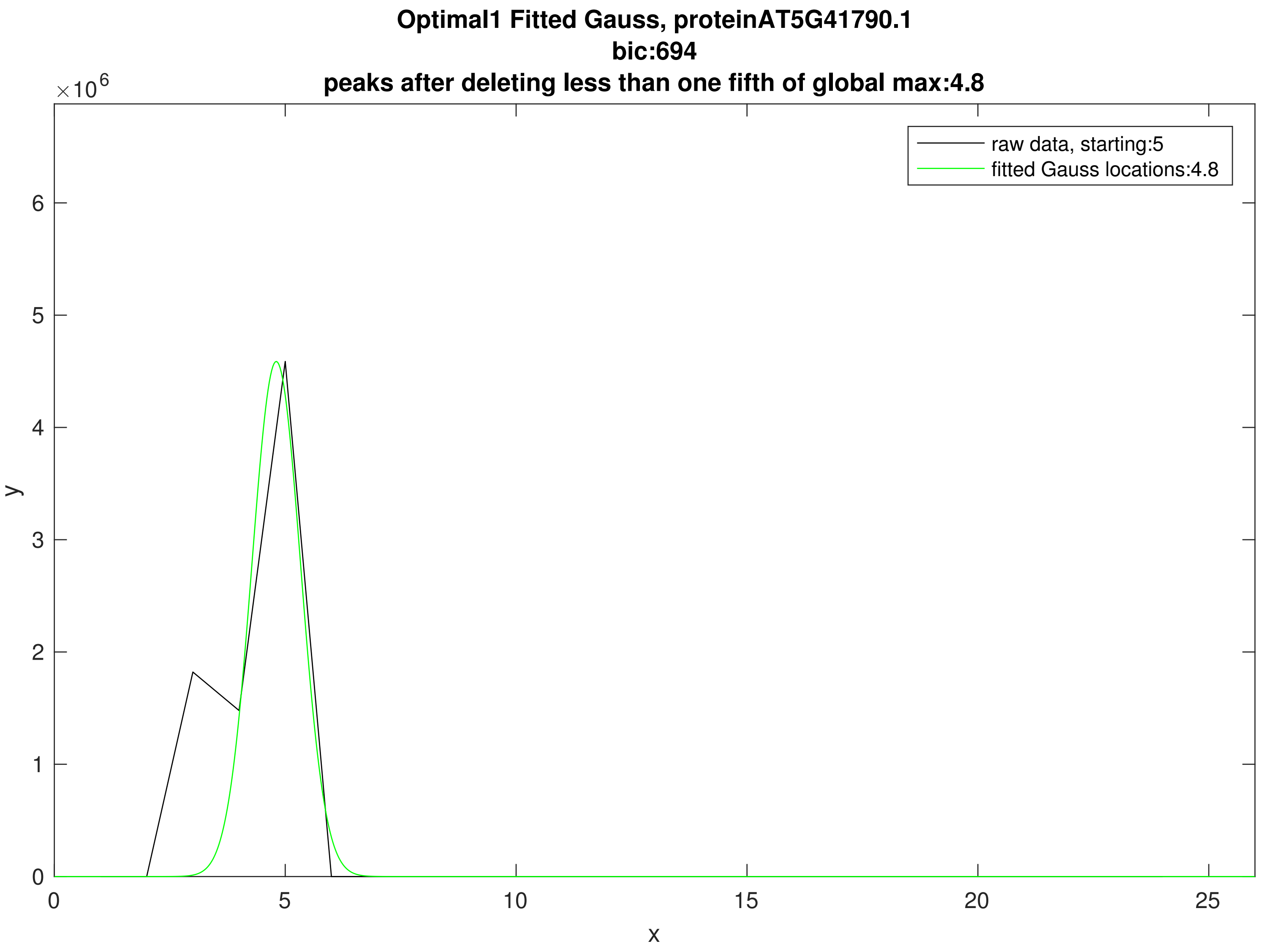

Optimal1 Fitted Gauss, proteinAT5G41790.1
bic:694
peaks after deleting less than one fifth of global max:4.8
×10^6
raw data, starting:5
fitted Gauss locations:4.8
y
x
0
1
2
3
4
5
6
0
5
10
15
20
25

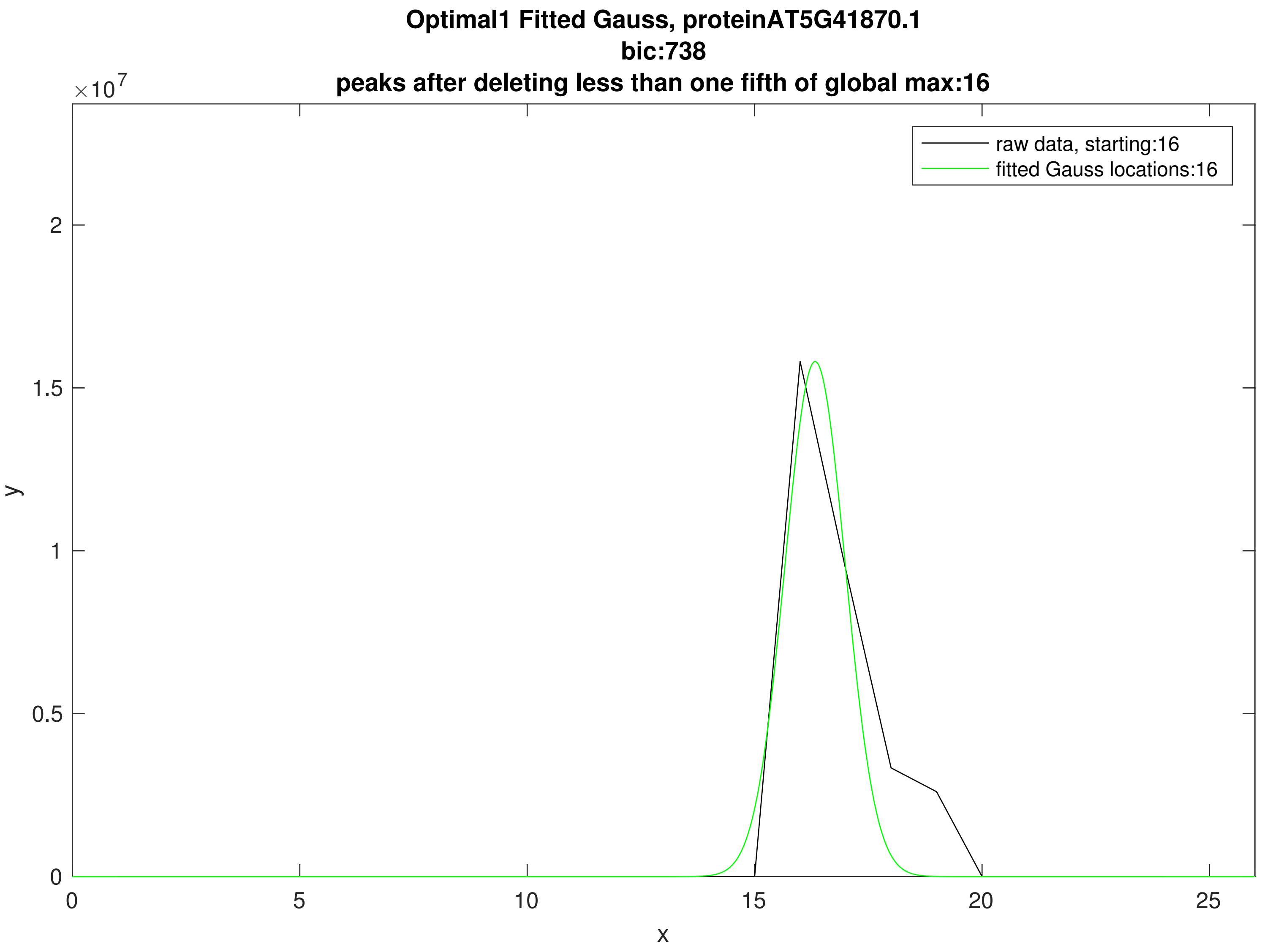
Optimal1 Fitted Gauss, proteinAT5G41870.1
bic:738
peaks after deleting less than one fifth of global max:16
×10^7
raw data, starting:16
fitted Gauss locations:16
y
x
0
0.5
1
1.5
2
0
5
10
15
20
25

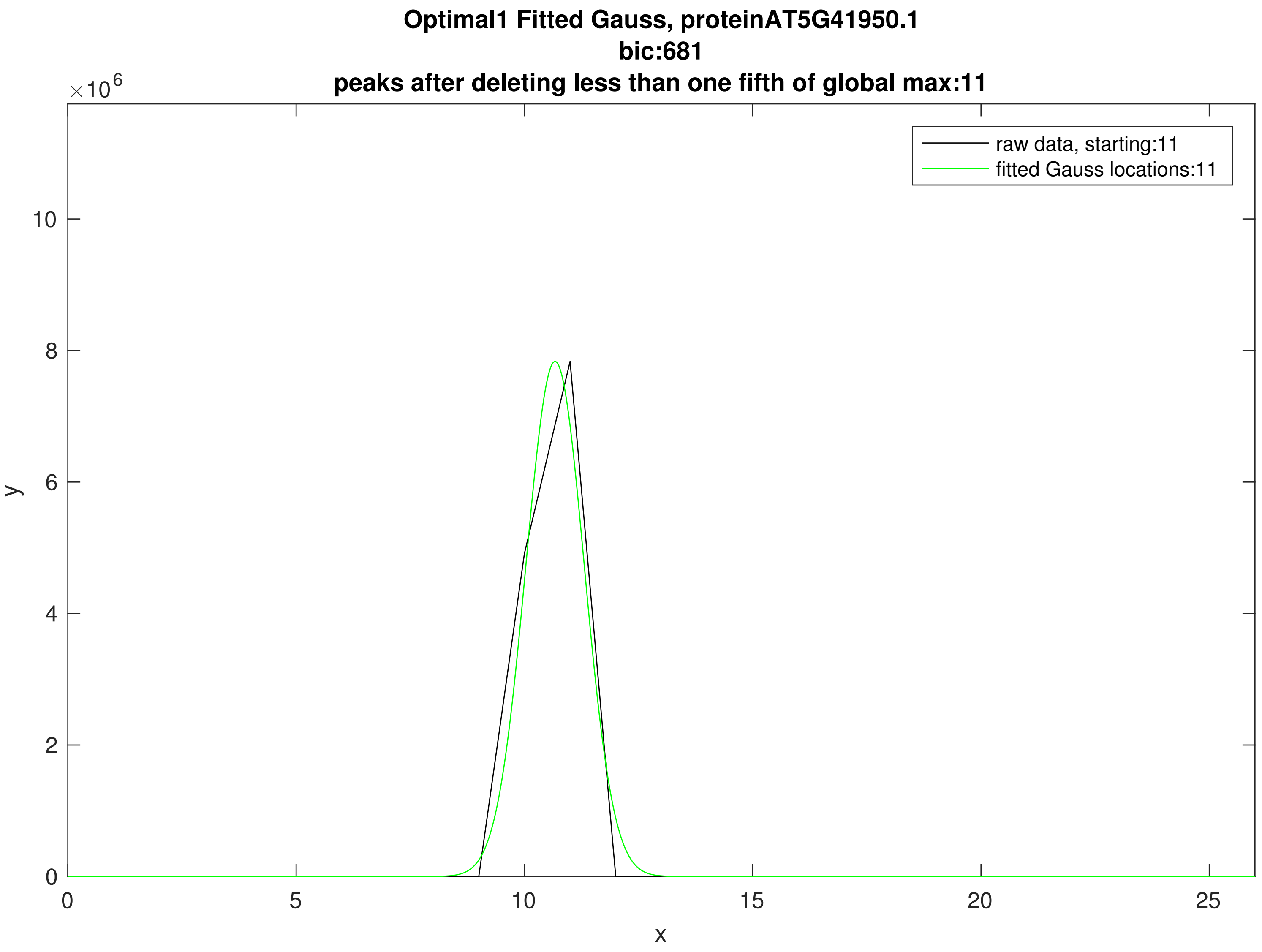
Optimal1 Fitted Gauss, proteinAT5G41950.1
bic:681
peaks after deleting less than one fifth of global max:11
×10^6
raw data, starting:11
fitted Gauss locations:11
y
x
0
2
4
6
8
10
0
5
10
15
20
25

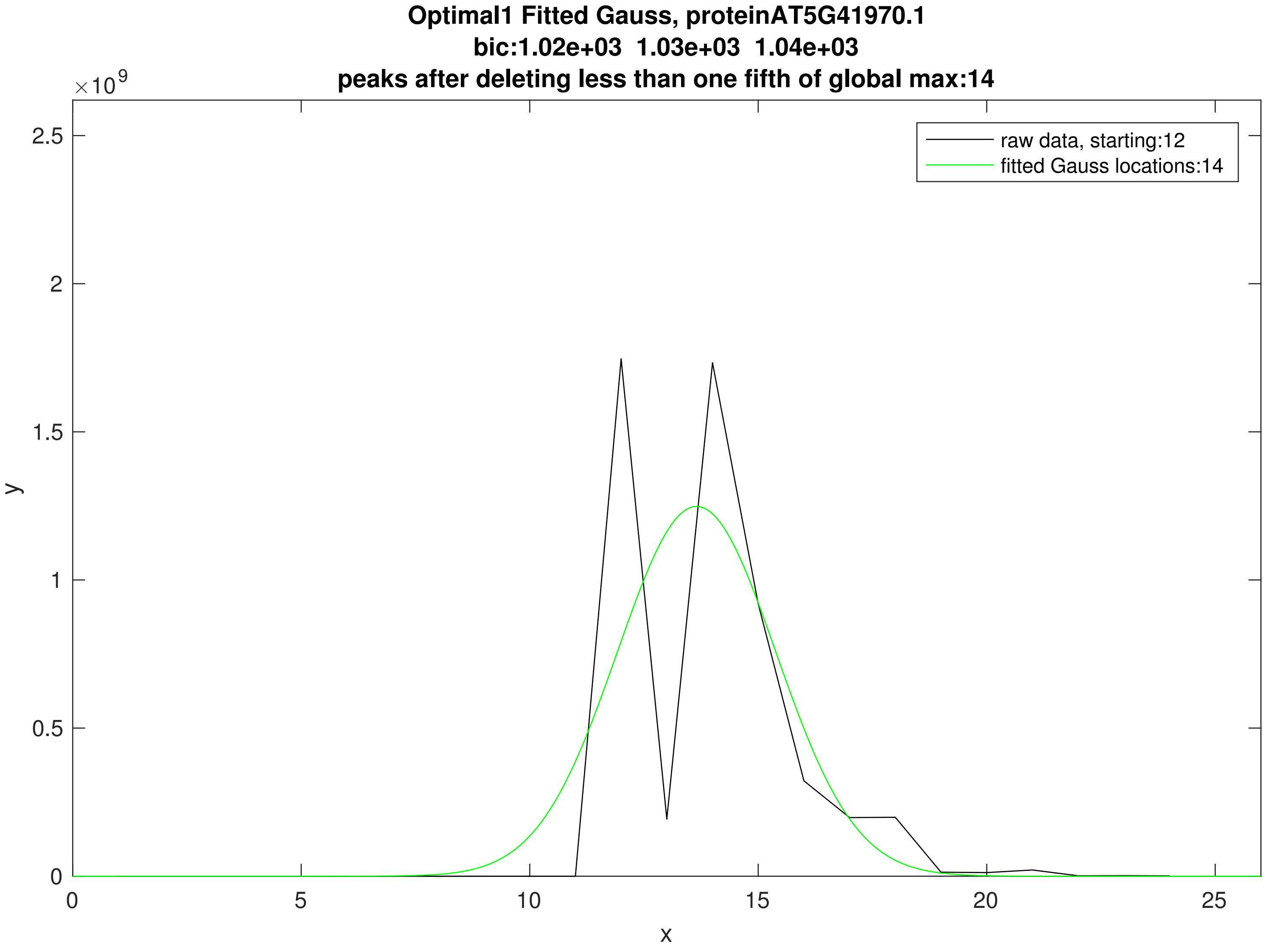

Optimal1 Fitted Gauss, proteinAT5G41970.1
bic:1.02e+03 1.03e+03 1.04e+03
peaks after deleting less than one fifth of global max:14
×10^9
2.5
2
1.5
1
0.5
0
y
0
5
10
15
20
25
x
raw data, starting:12
fitted Gauss locations:14

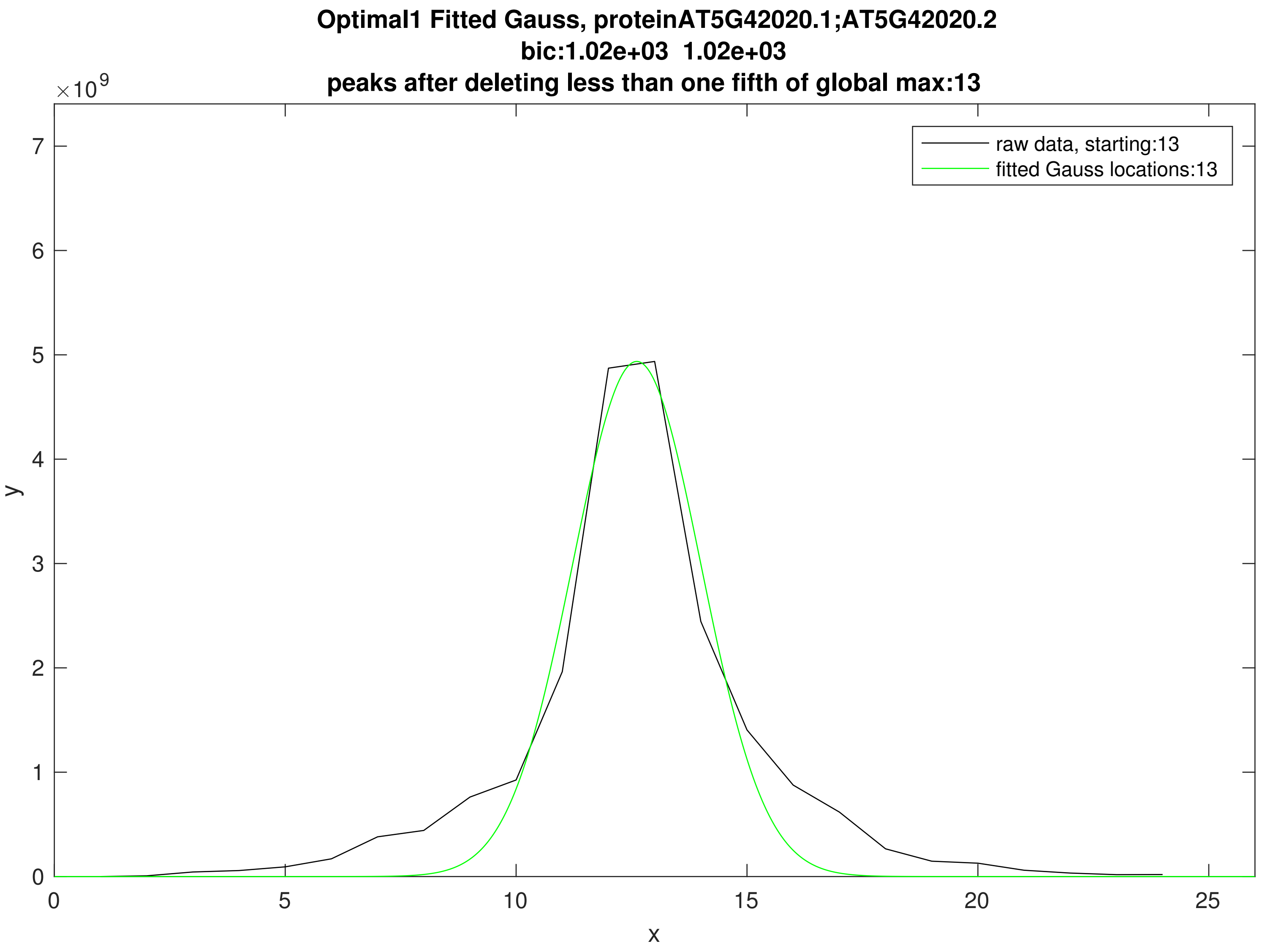

Optimal1 Fitted Gauss, proteinAT5G42020.1;AT5G42020.2
bic:1.02e+03  1.02e+03
peaks after deleting less than one fifth of global max:13
×10^9
raw data, starting:13
fitted Gauss locations:13
y
x
0
1
2
3
4
5
6
7
0
5
10
15
20
25

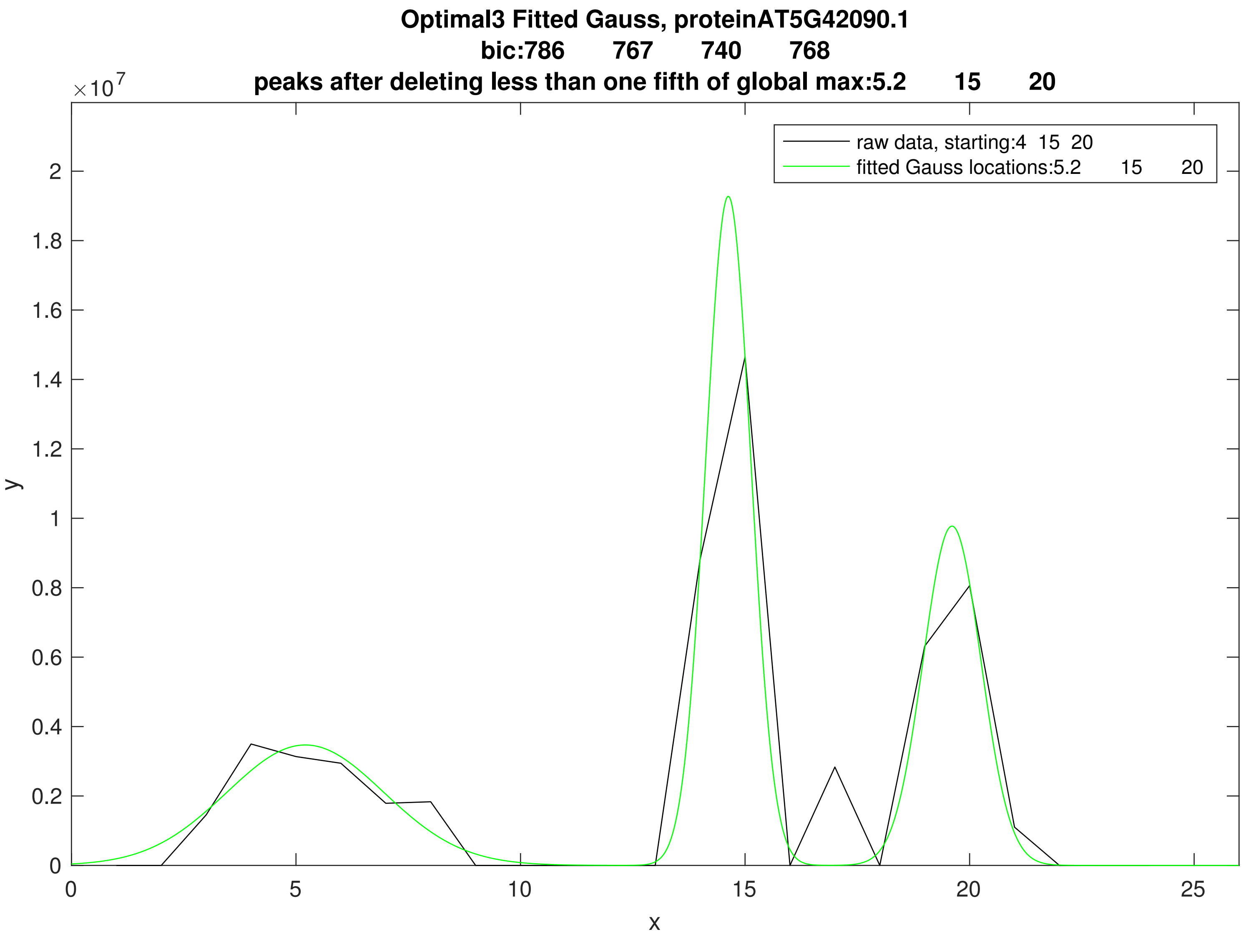

Optimal3 Fitted Gauss, proteinAT5G42090.1
bic:786 767 740 768
peaks after deleting less than one fifth of global max:5.2 15 20
×10^7
raw data, starting:4 15 20
fitted Gauss locations:5.2 15 20
y
x
0
0.2
0.4
0.6
0.8
1
1.2
1.4
1.6
1.8
2
5
10
15
20
25

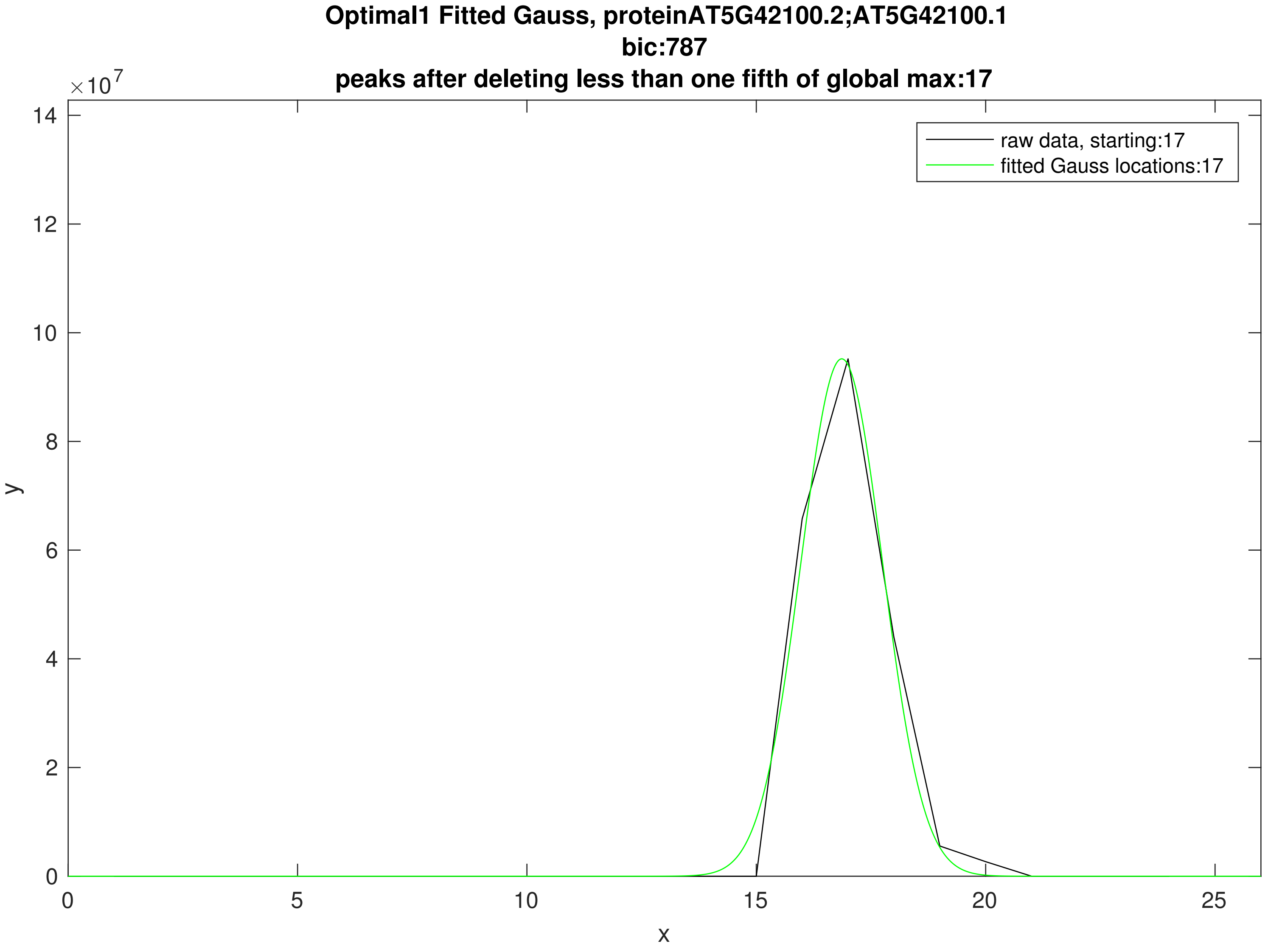
Optimal1 Fitted Gauss, proteinAT5G42100.2;AT5G42100.1
bic:787
peaks after deleting less than one fifth of global max:17
×10^7
raw data, starting:17
fitted Gauss locations:17
y
x
0
2
4
6
8
10
12
14
0
5
10
15
20
25

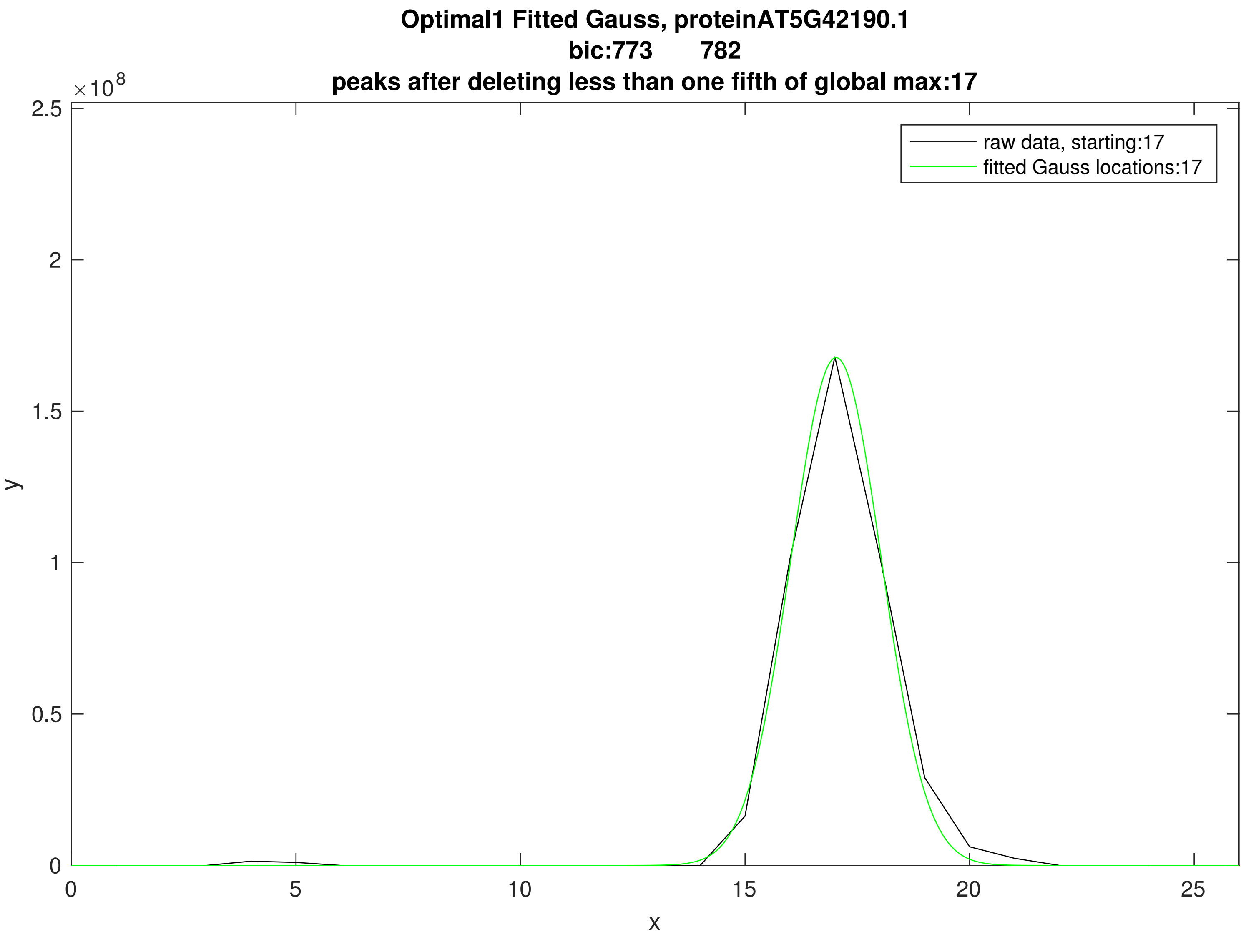

Optimal1 Fitted Gauss, proteinAT5G42190.1
bic:773 782
peaks after deleting less than one fifth of global max:17
×10^8
2.5
2
1.5
1
0.5
0
y
0
5
10
15
20
25
x
raw data, starting:17
fitted Gauss locations:17

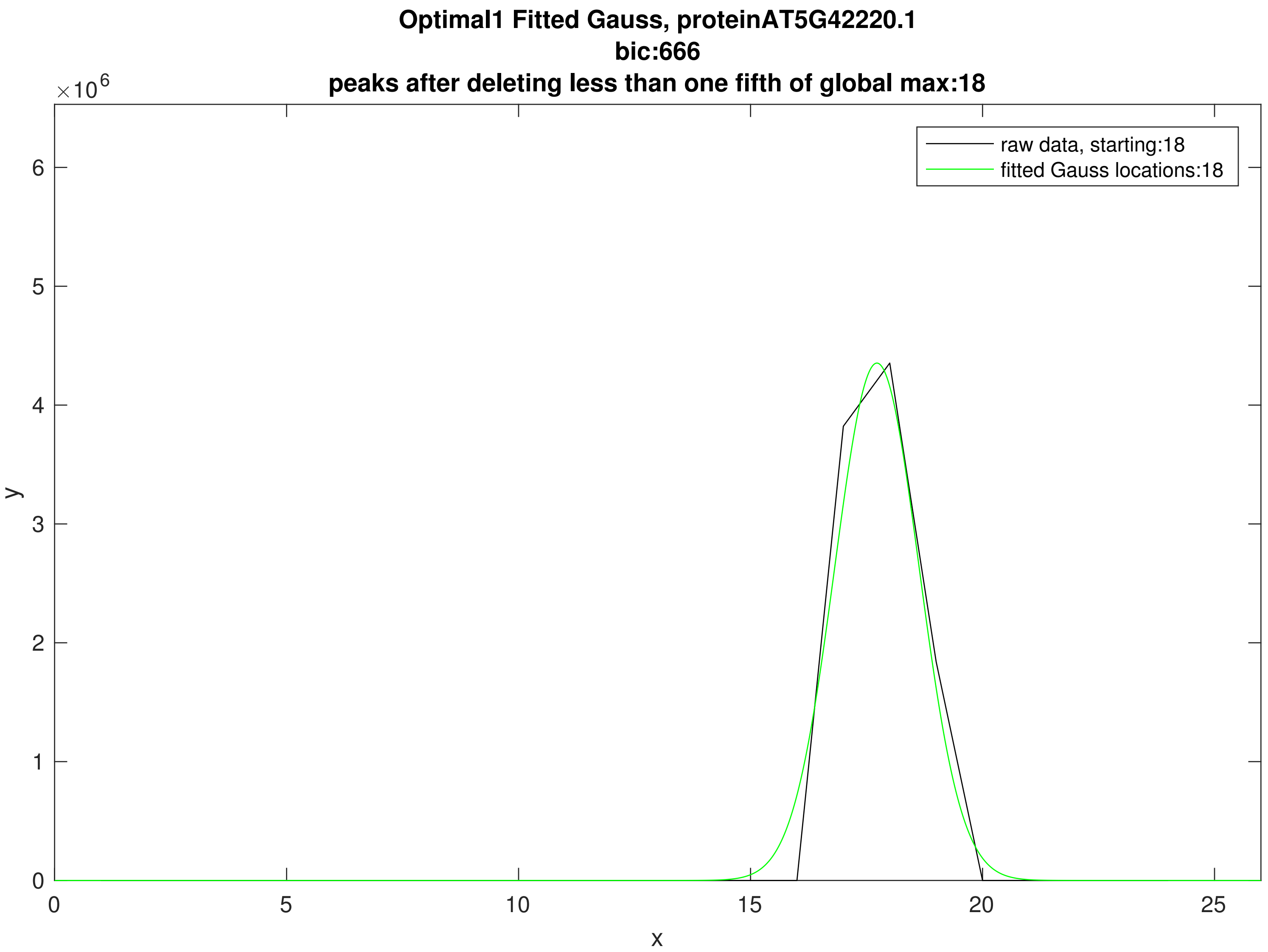
Optimal1 Fitted Gauss, proteinAT5G42220.1
bic:666
peaks after deleting less than one fifth of global max:18
×10^6
6
5
4
3
2
1
0
y
0
5
10
15
20
25
x
raw data, starting:18
fitted Gauss locations:18

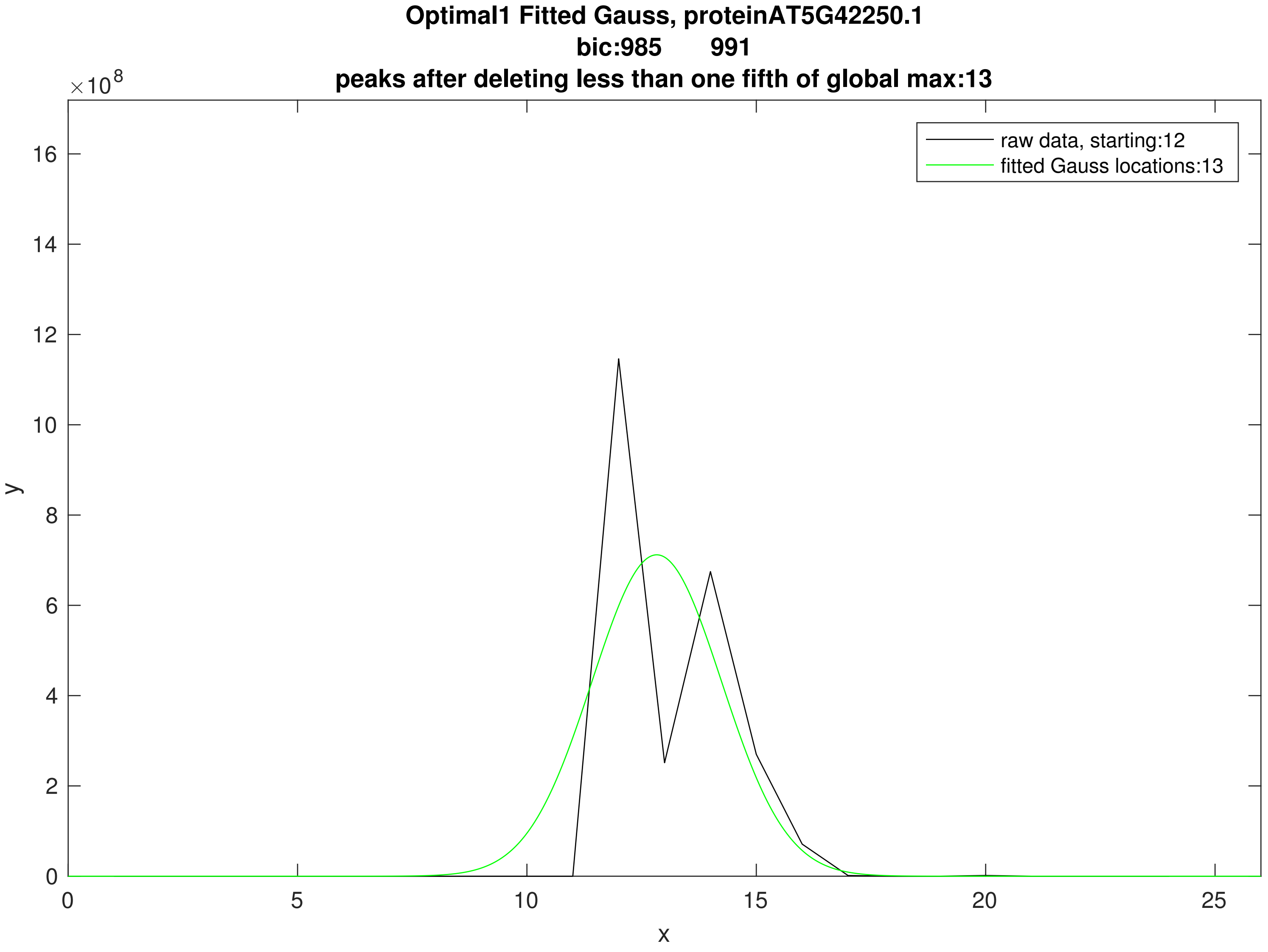

Optimal1 Fitted Gauss, proteinAT5G42250.1
bic:985 991
peaks after deleting less than one fifth of global max:13
×10^8
raw data, starting:12
fitted Gauss locations:13
16
14
12
10
8
6
4
2
0
y
0
5
10
15
20
25
x

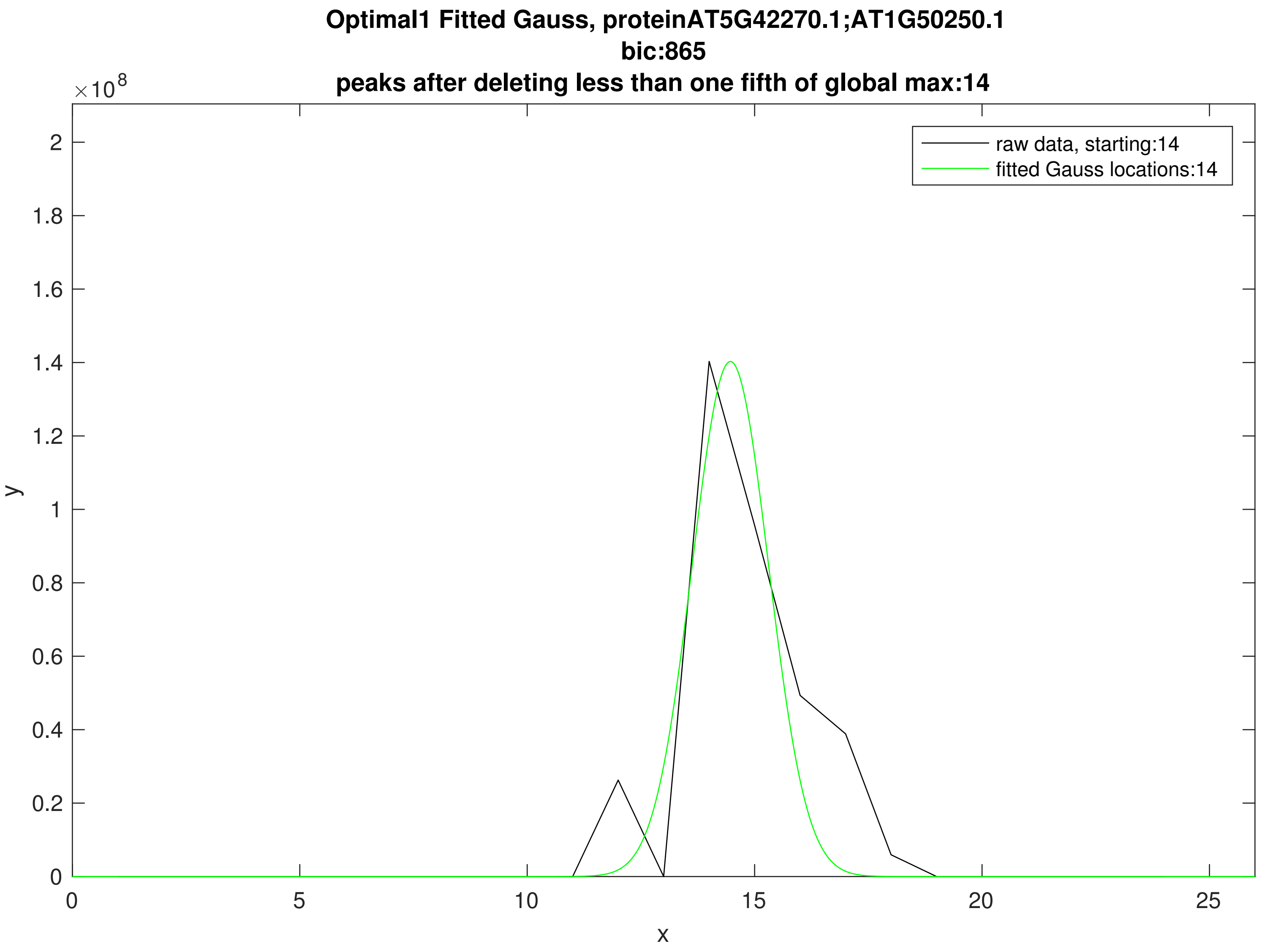
Optimal1 Fitted Gauss, proteinAT5G42270.1;AT1G50250.1
bic:865
peaks after deleting less than one fifth of global max:14
×10^8
raw data, starting:14
fitted Gauss locations:14
y
x
0
0.2
0.4
0.6
0.8
1
1.2
1.4
1.6
1.8
2
0
5
10
15
20
25

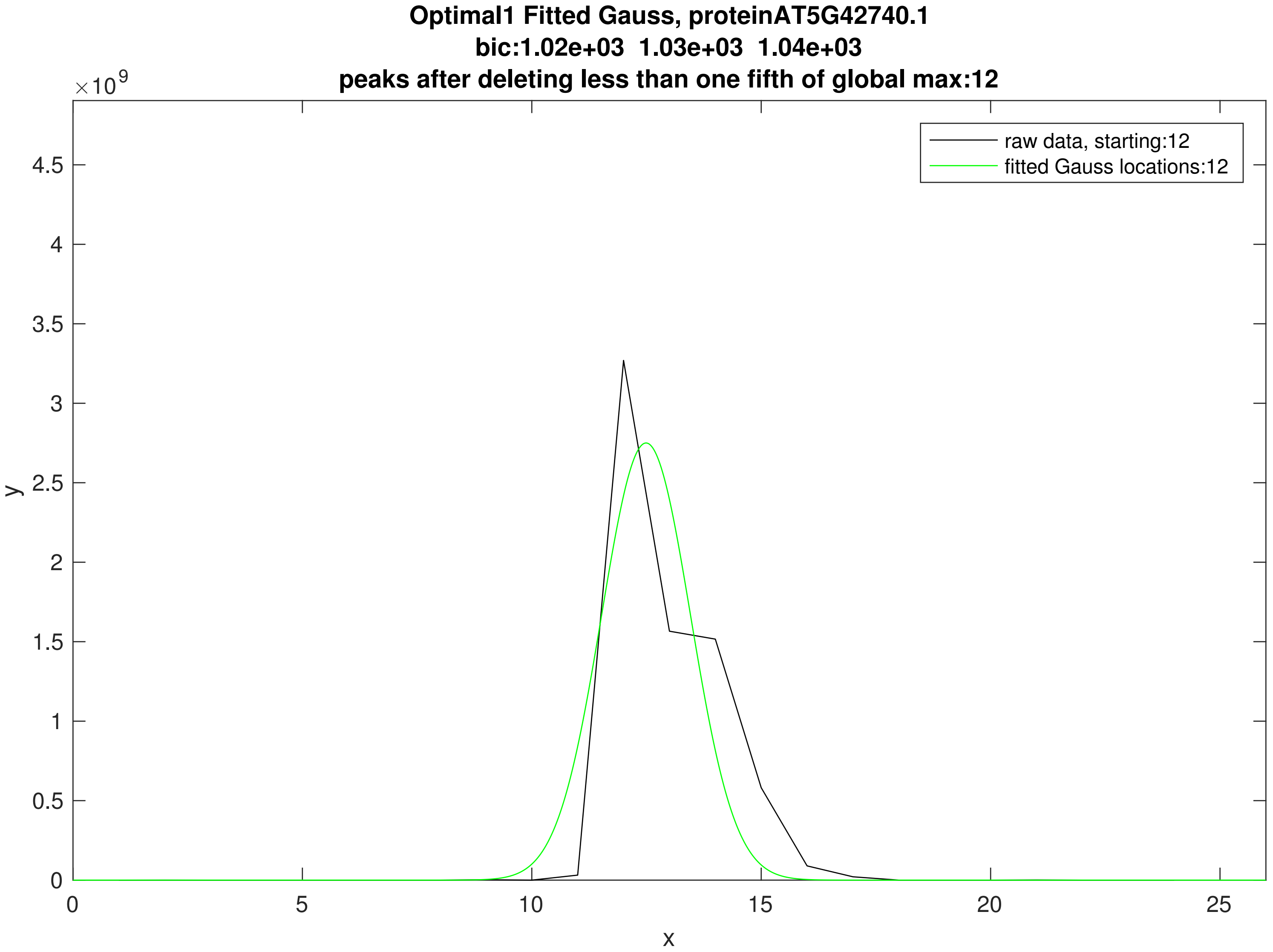

Optimal1 Fitted Gauss, proteinAT5G42740.1
bic:1.02e+03 1.03e+03 1.04e+03
peaks after deleting less than one fifth of global max:12
×10^9
4.5
4
3.5
3
2.5
2
1.5
1
0.5
0
y
0
5
10
15
20
25
x
raw data, starting:12
fitted Gauss locations:12

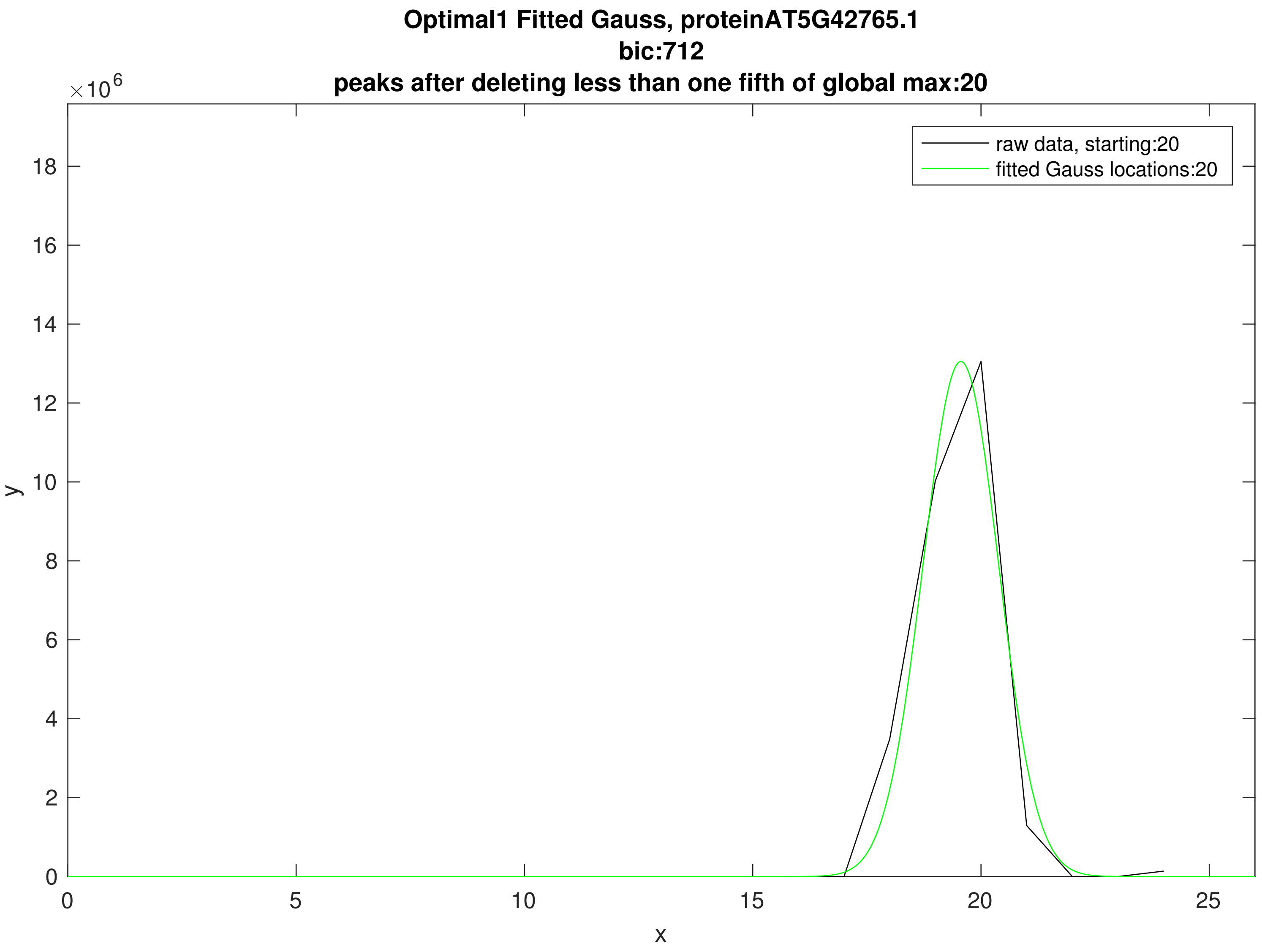
Optimal1 Fitted Gauss, proteinAT5G42765.1
bic:712
peaks after deleting less than one fifth of global max:20
×10^6
raw data, starting:20
fitted Gauss locations:20
y
x
0
2
4
6
8
10
12
14
16
18
0
5
10
15
20
25

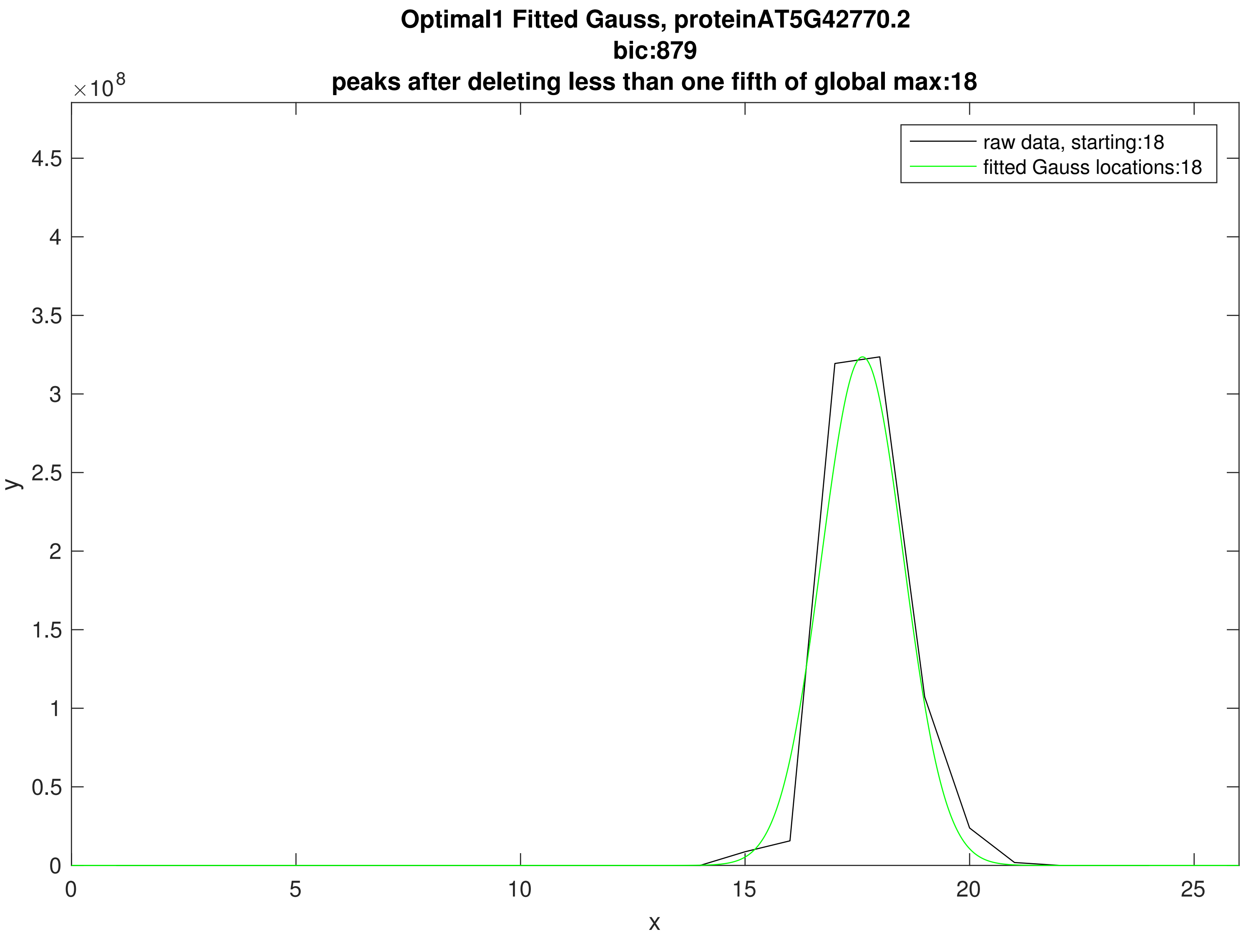

Optimal1 Fitted Gauss, proteinAT5G42770.2
bic:879
peaks after deleting less than one fifth of global max:18
×10^8
4.5
4
3.5
3
2.5
2
1.5
1
0.5
0
y
0
5
10
15
20
25
x
raw data, starting:18
fitted Gauss locations:18

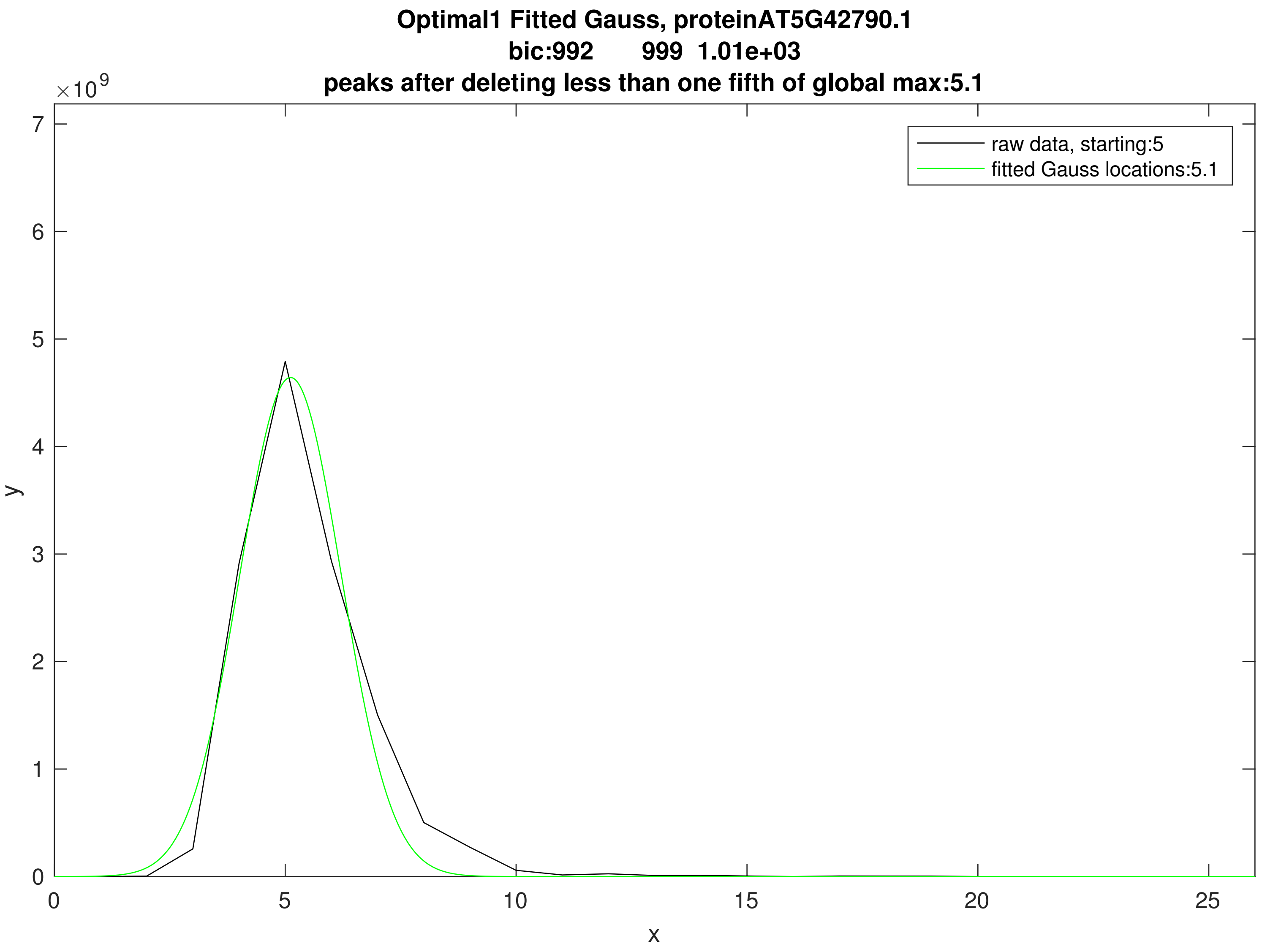

Optimal1 Fitted Gauss, proteinAT5G42790.1
bic:992 999 1.01e+03
peaks after deleting less than one fifth of global max:5.1
×10^9
raw data, starting:5
fitted Gauss locations:5.1
y
x
0
1
2
3
4
5
6
7
10
15
20
25

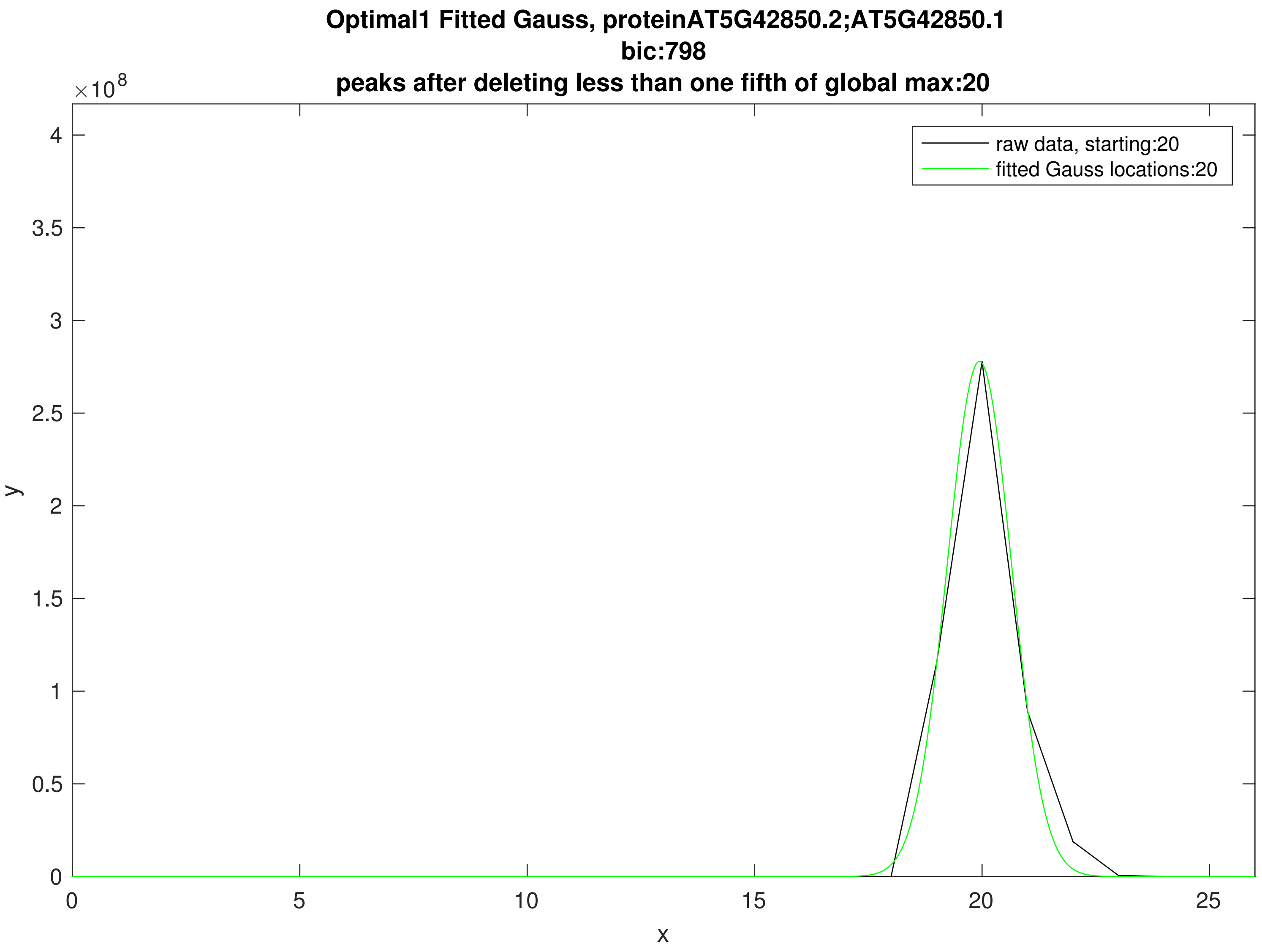
Optimal1 Fitted Gauss, proteinAT5G42850.2;AT5G42850.1
bic:798
peaks after deleting less than one fifth of global max:20
×10^8
4
3.5
3
2.5
2
1.5
1
0.5
0
y
0
5
10
15
20
25
x
raw data, starting:20
fitted Gauss locations:20

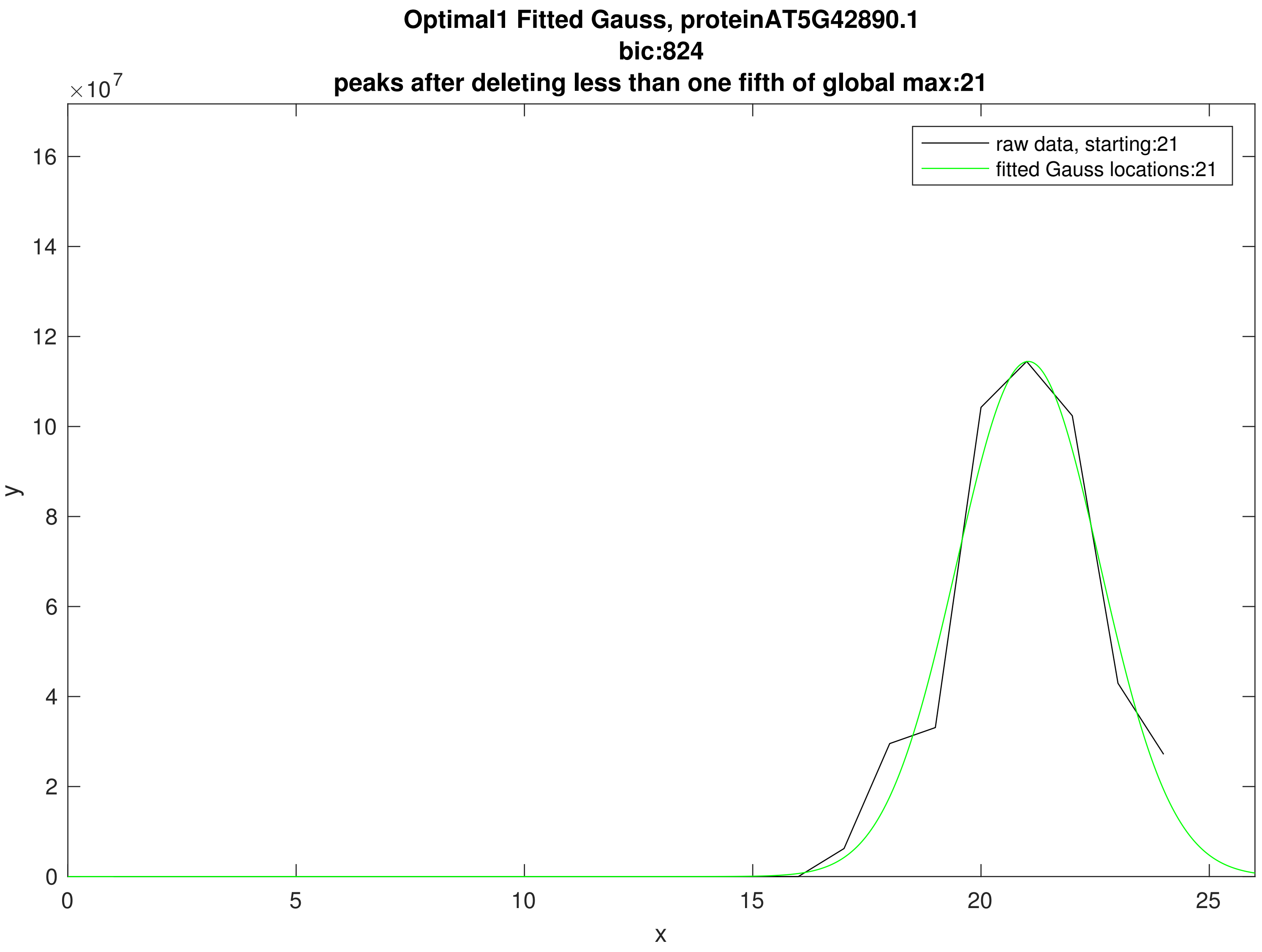
Optimal1 Fitted Gauss, proteinAT5G42890.1
bic:824
peaks after deleting less than one fifth of global max:21
×10^7
16
14
12
10
8
6
4
2
0
y
0
5
10
15
20
25
x
raw data, starting:21
fitted Gauss locations:21

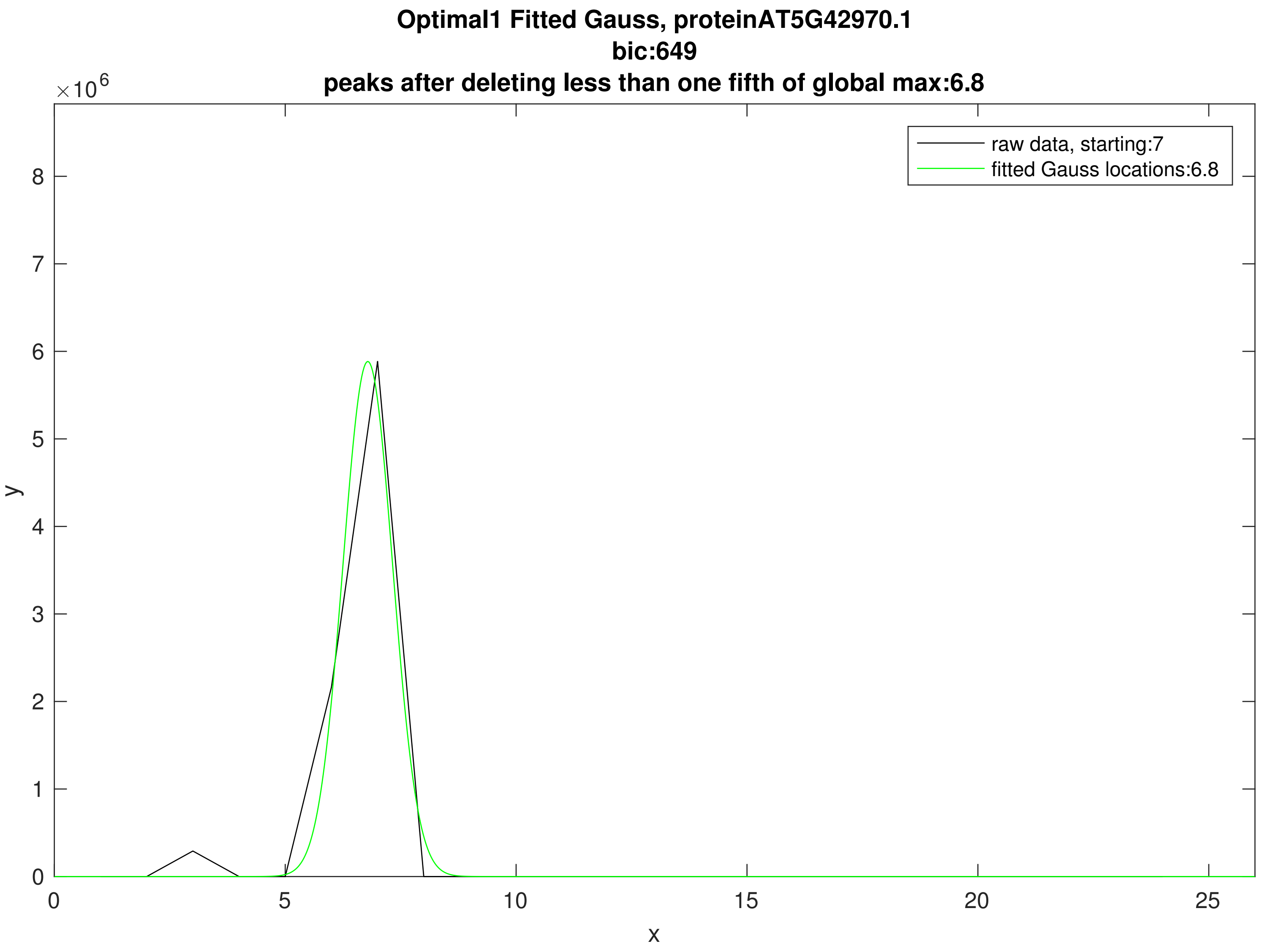

Optimal1 Fitted Gauss, proteinAT5G42970.1
bic:649
peaks after deleting less than one fifth of global max:6.8
×10^6
raw data, starting:7
fitted Gauss locations:6.8
y
x
0
1
2
3
4
5
6
7
8
0
5
10
15
20
25

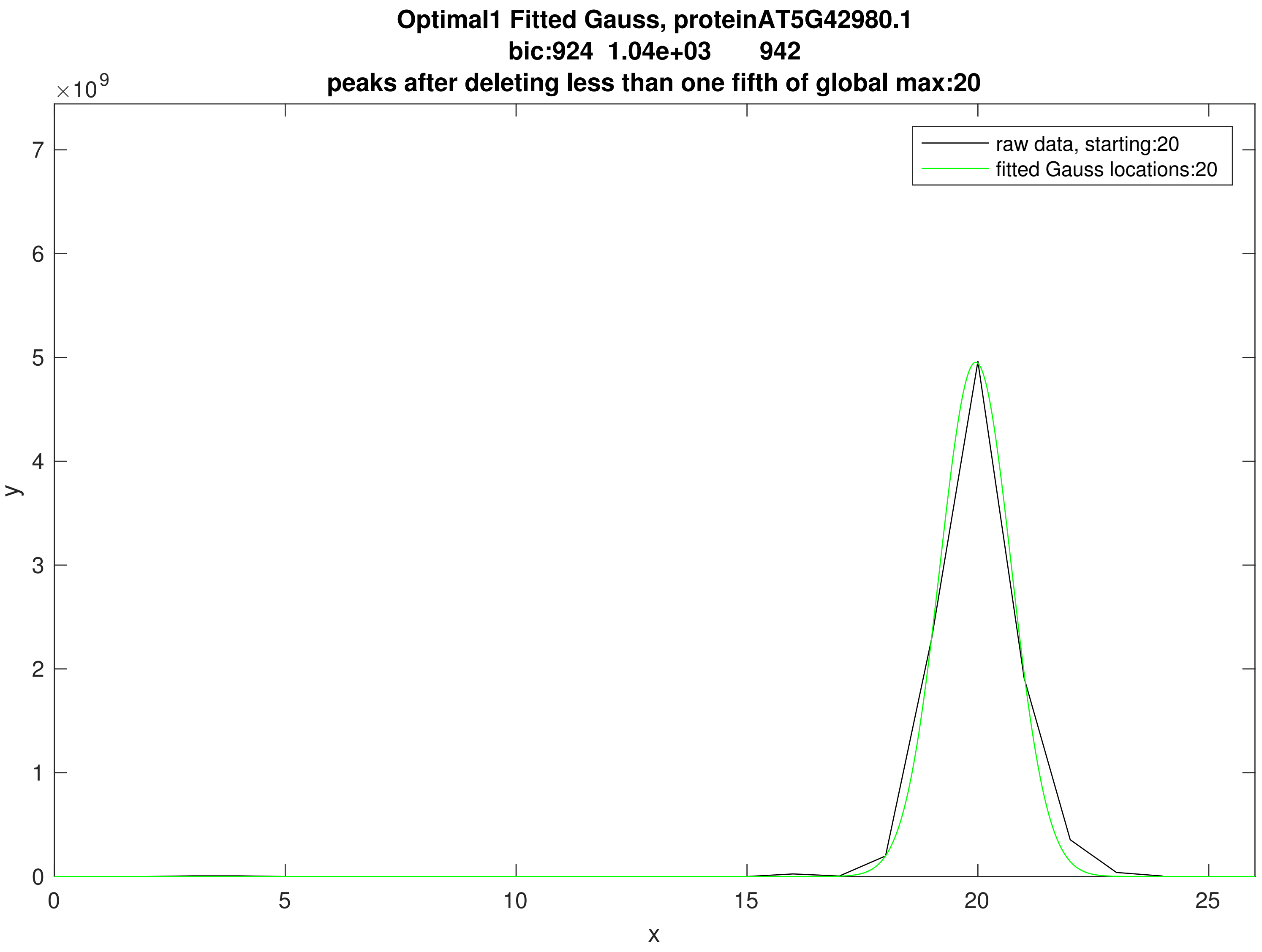

Optimal1 Fitted Gauss, proteinAT5G42980.1
bic:924 1.04e+03 942
peaks after deleting less than one fifth of global max:20
×10^9
raw data, starting:20
fitted Gauss locations:20
y
x

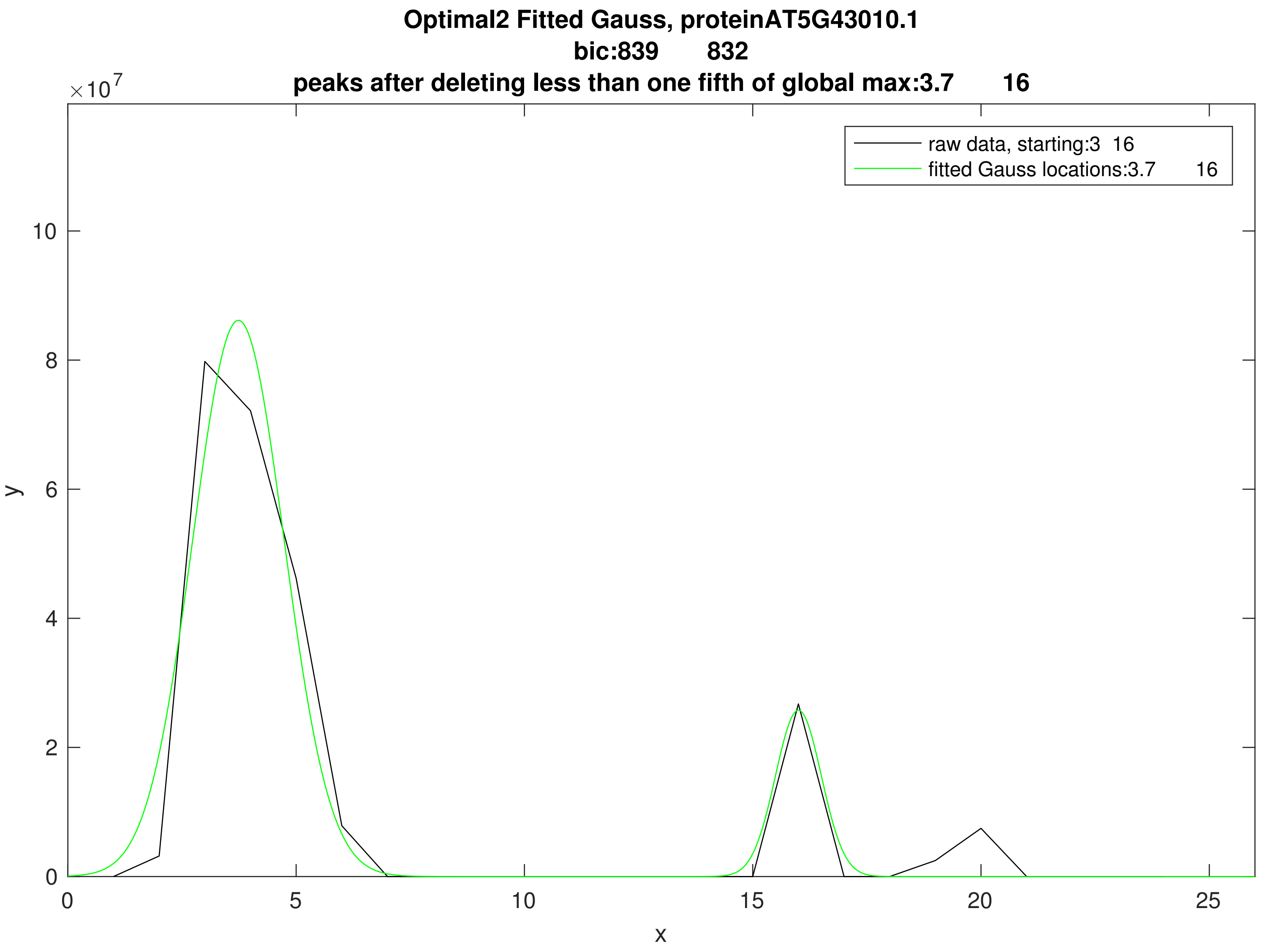

Optimal2 Fitted Gauss, proteinAT5G43010.1
bic:839 832
peaks after deleting less than one fifth of global max:3.7 16
×10^7
raw data, starting:3 16
fitted Gauss locations:3.7 16
y
x
0
2
4
6
8
10
0
5
10
15
20
25

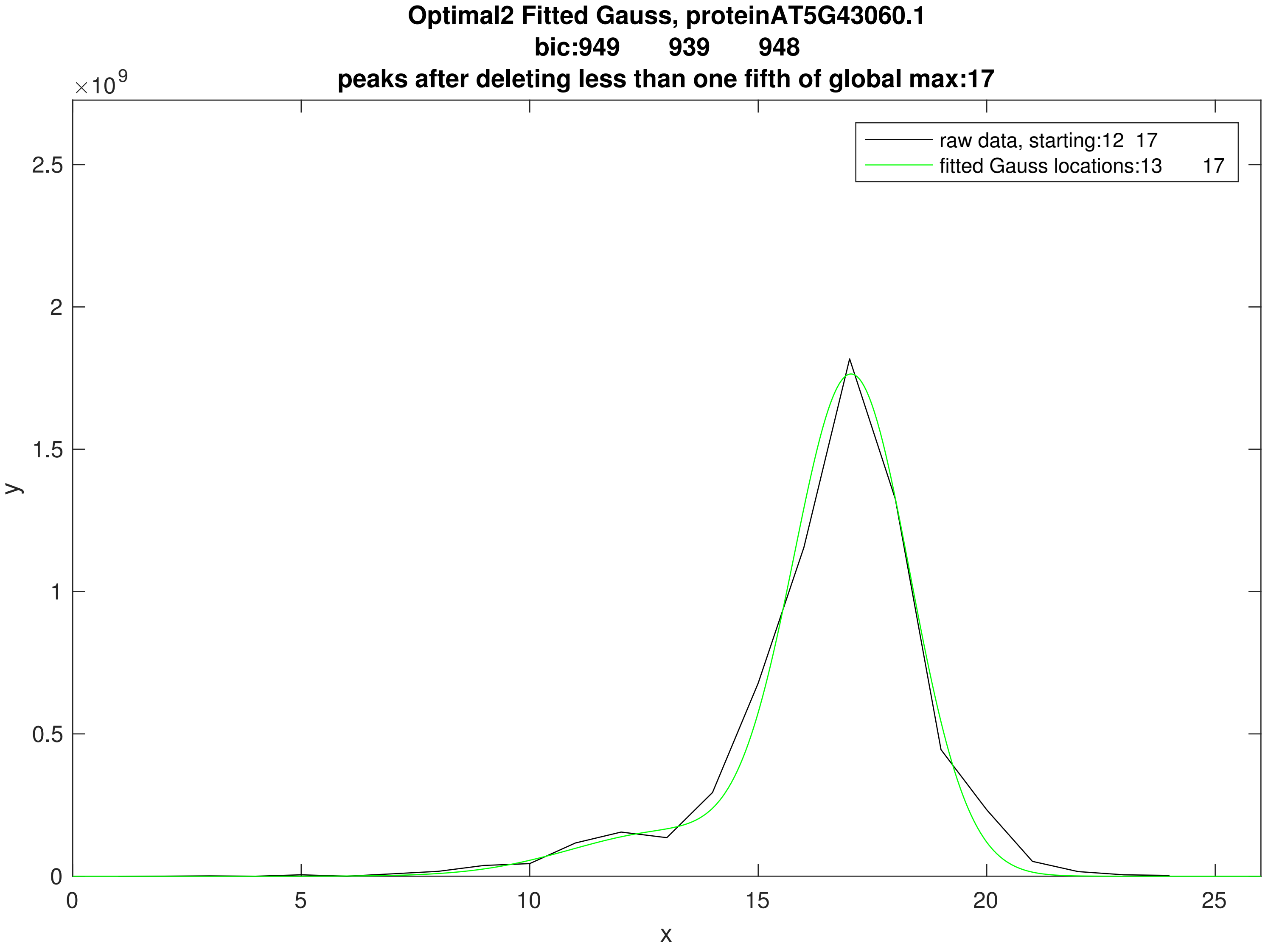
Optimal2 Fitted Gauss, proteinAT5G43060.1
bic:949 939 948
peaks after deleting less than one fifth of global max:17
×10^9
raw data, starting:12 17
fitted Gauss locations:13 17
y
x
0
0.5
1
1.5
2
2.5
5
10
15
20
25

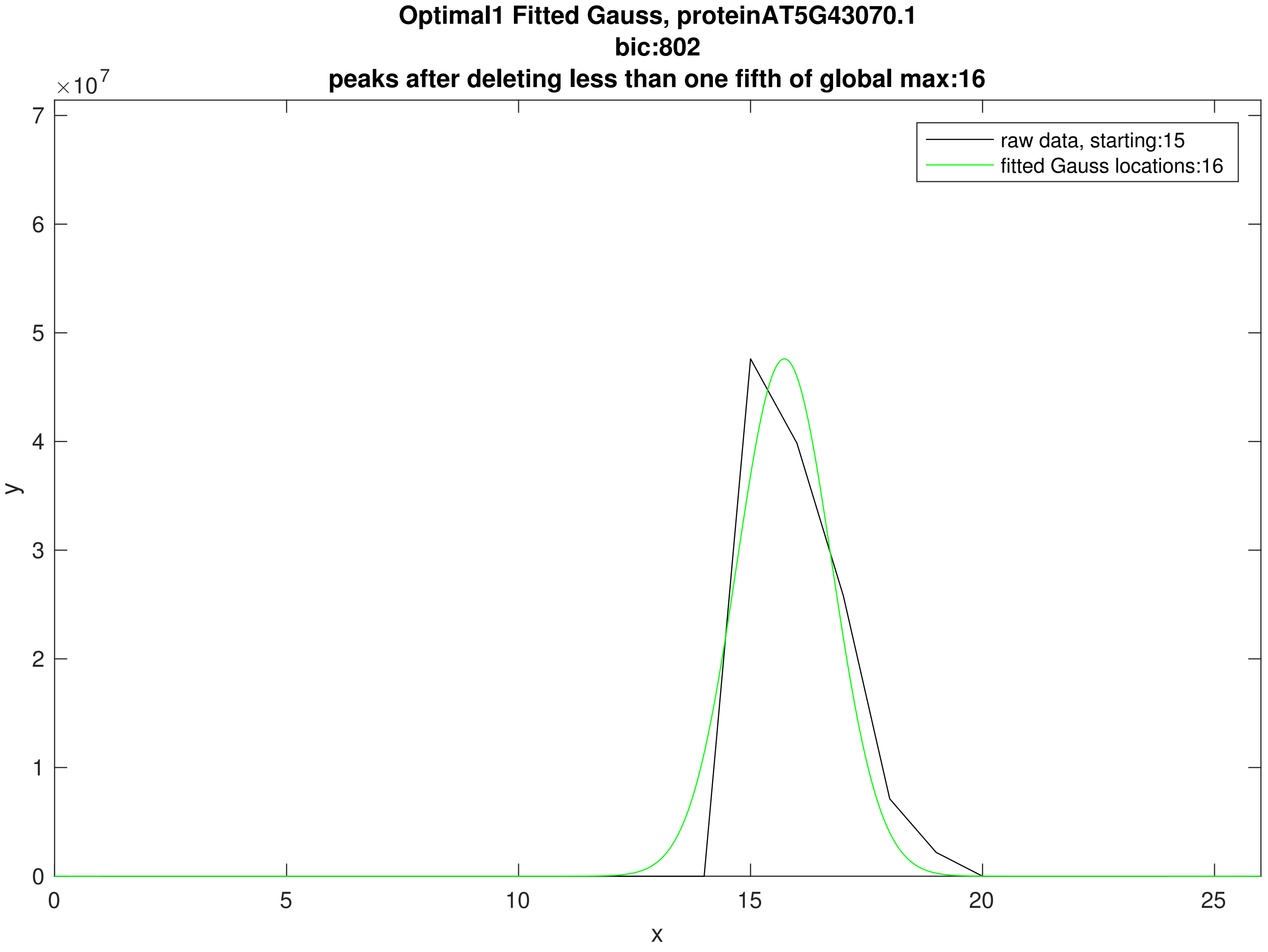

Optimal1 Fitted Gauss, proteinAT5G43070.1
bic:802
peaks after deleting less than one fifth of global max:16
×10^7
raw data, starting:15
fitted Gauss locations:16
y
x
0
1
2
3
4
5
6
7
0
5
10
15
20
25

**Optimal1 Fitted Gauss, proteinAT5G43100.1**
**bic:722**
**peaks after deleting less than one fifth of global max:16**

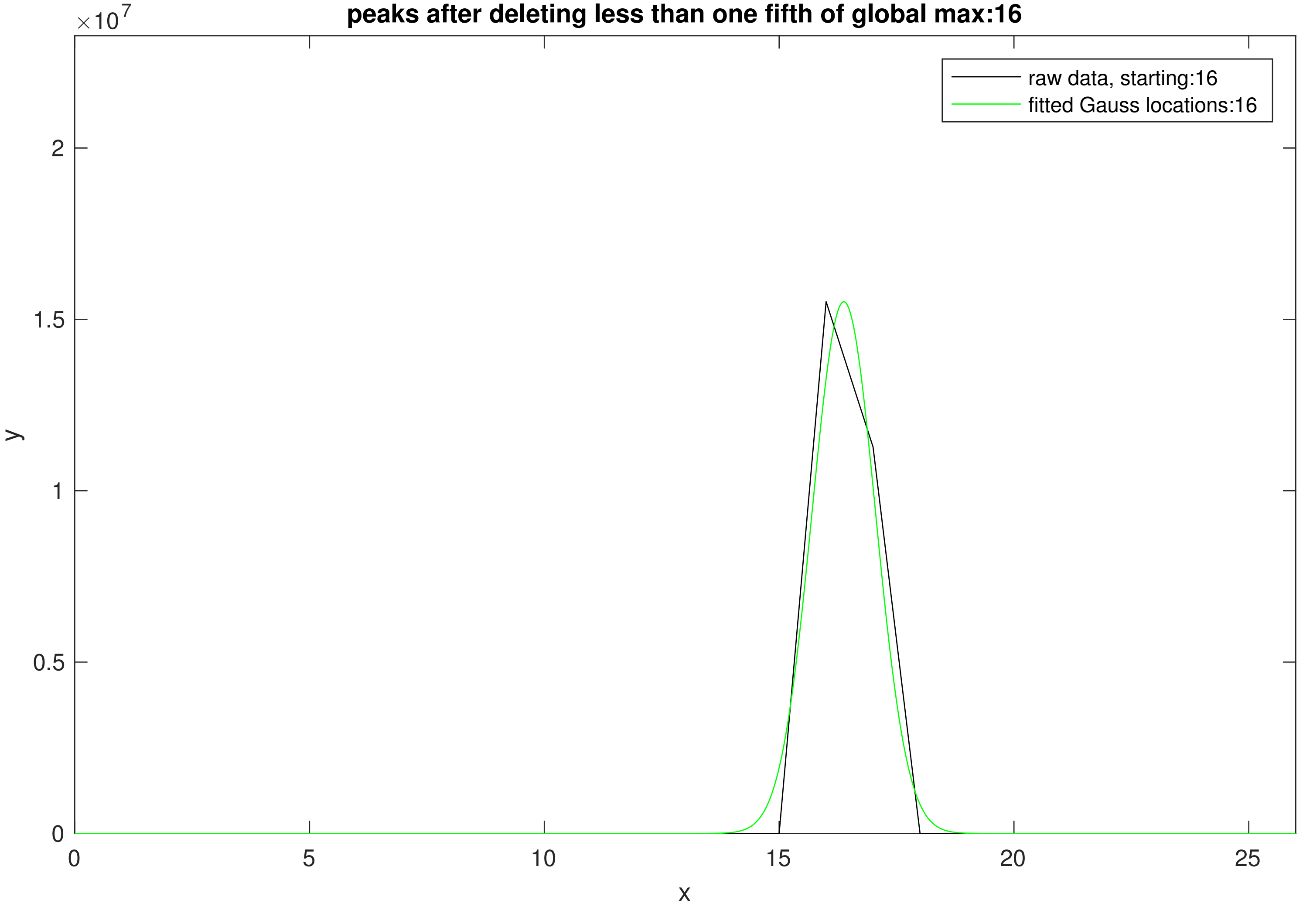

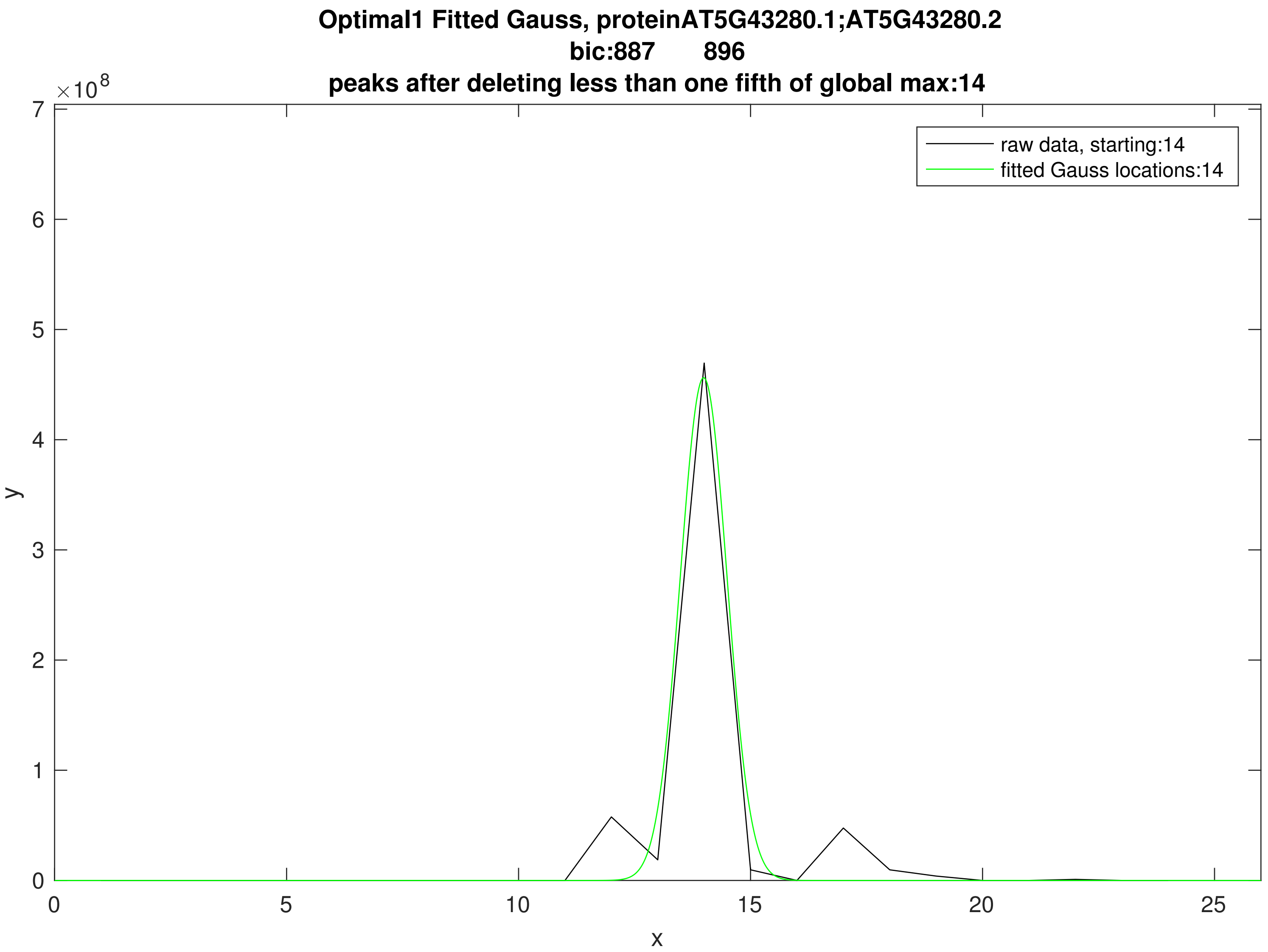
Optimal1 Fitted Gauss, proteinAT5G43280.1;AT5G43280.2
bic:887 896
peaks after deleting less than one fifth of global max:14
raw data, starting:14
fitted Gauss locations:14
×10^8
y
x

**Optimal1 Fitted Gauss, proteinAT5G43330.1**
**bic:968 978**
**peaks after deleting less than one fifth of global max:15**

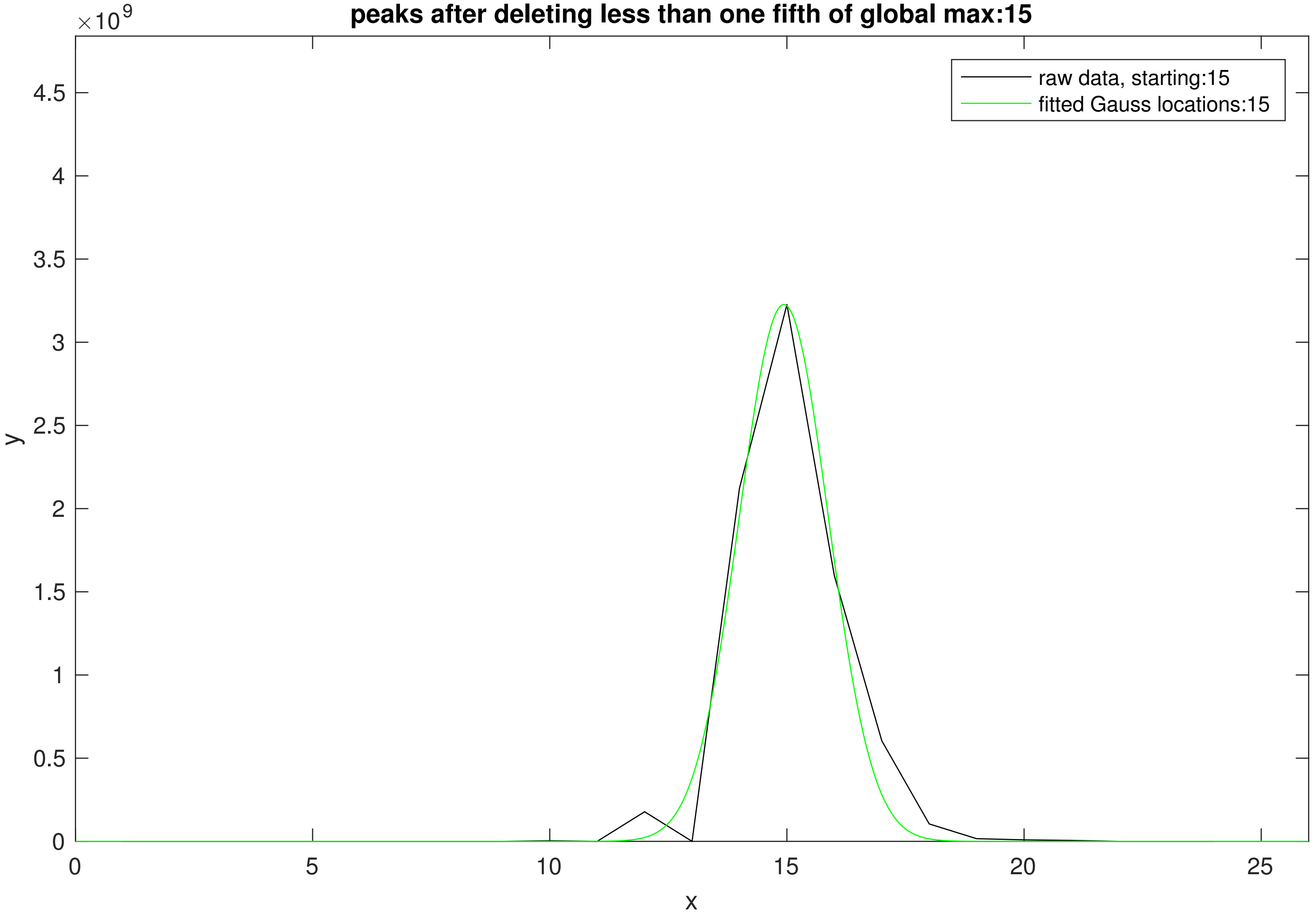

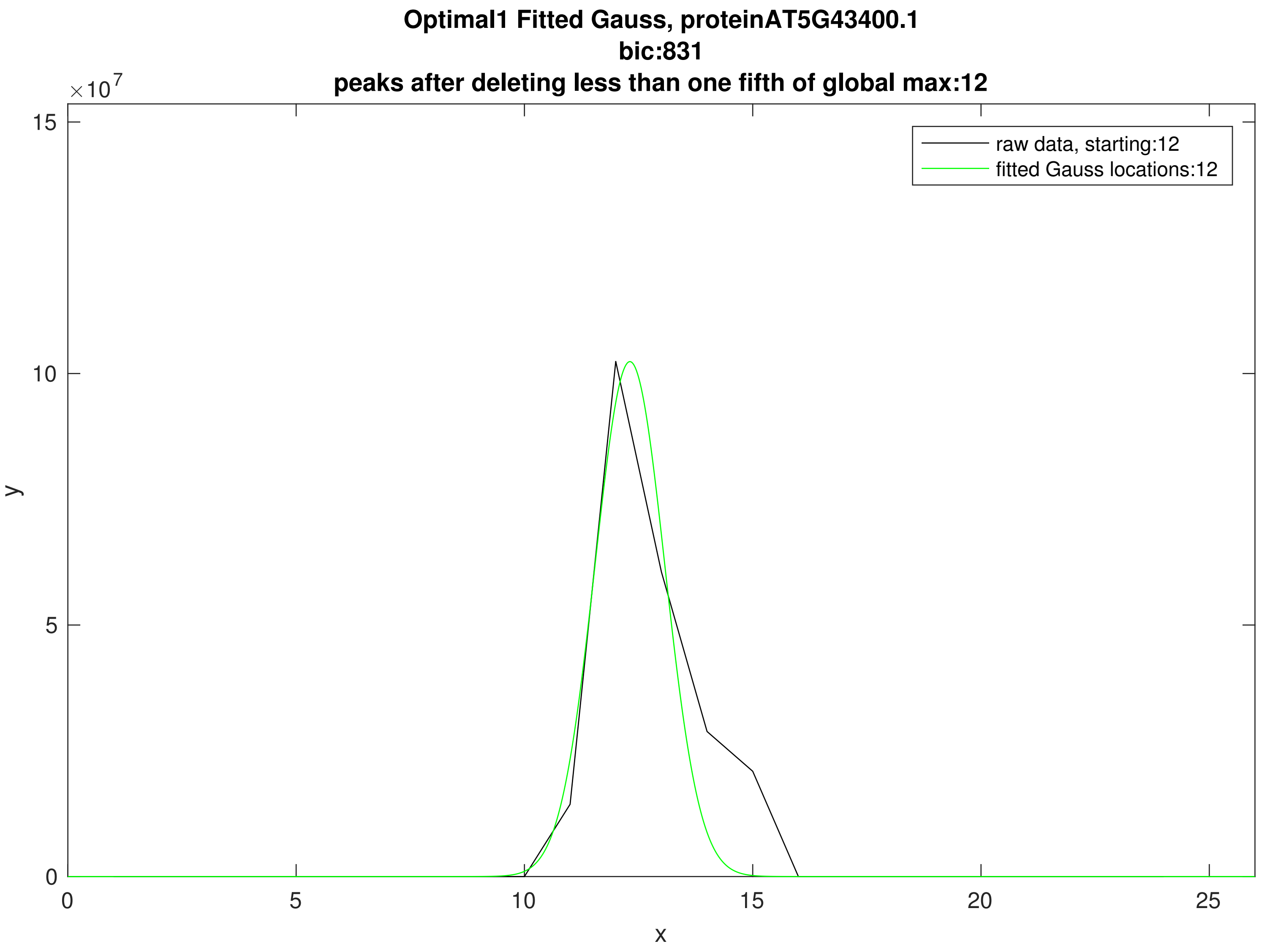

Optimal1 Fitted Gauss, proteinAT5G43400.1
bic:831
peaks after deleting less than one fifth of global max:12
×10^7
15
10
5
0
y
raw data, starting:12
fitted Gauss locations:12
0
5
10
15
20
25
x

**Optimal1 Fitted Gauss, proteinAT5G43430.1;AT5G43430.3;AT5G43430.2**
**bic:781**
**peaks after deleting less than one fifth of global max:15**

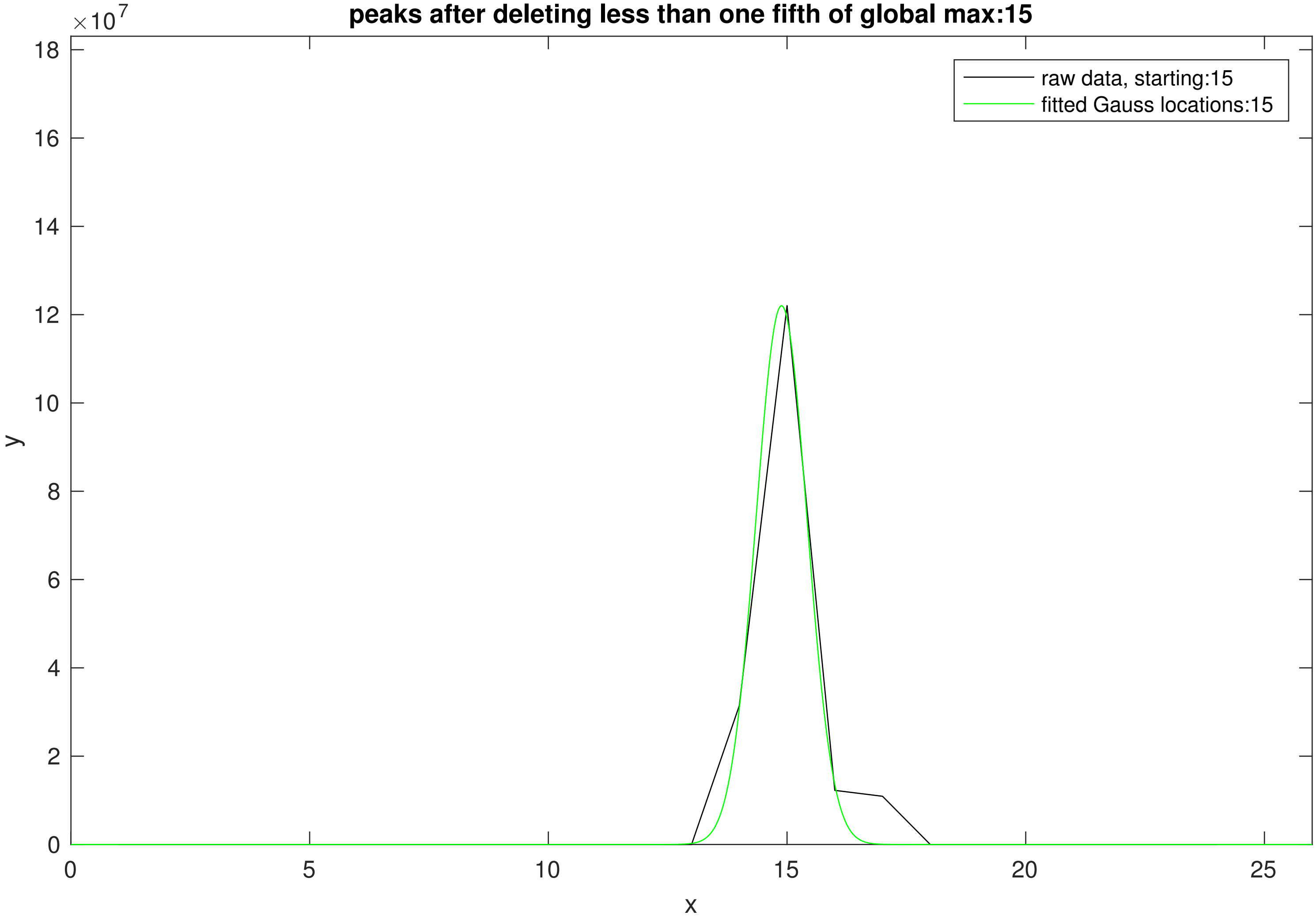

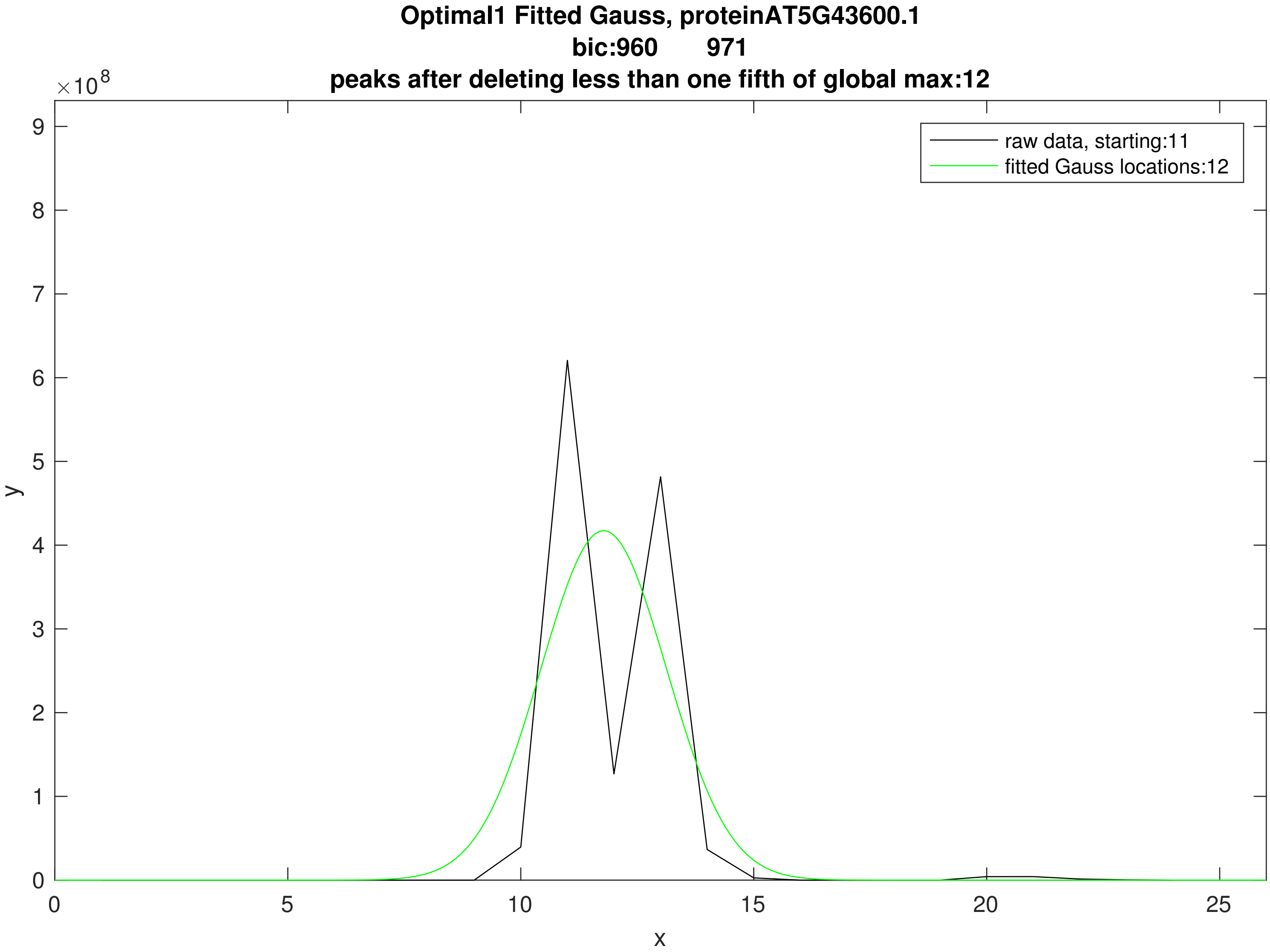

Optimal1 Fitted Gauss, proteinAT5G43600.1
bic:960 971
peaks after deleting less than one fifth of global max:12
×10^8
9
8
7
6
5
4
3
2
1
0
y
0
5
10
15
20
25
x
raw data, starting:11
fitted Gauss locations:12

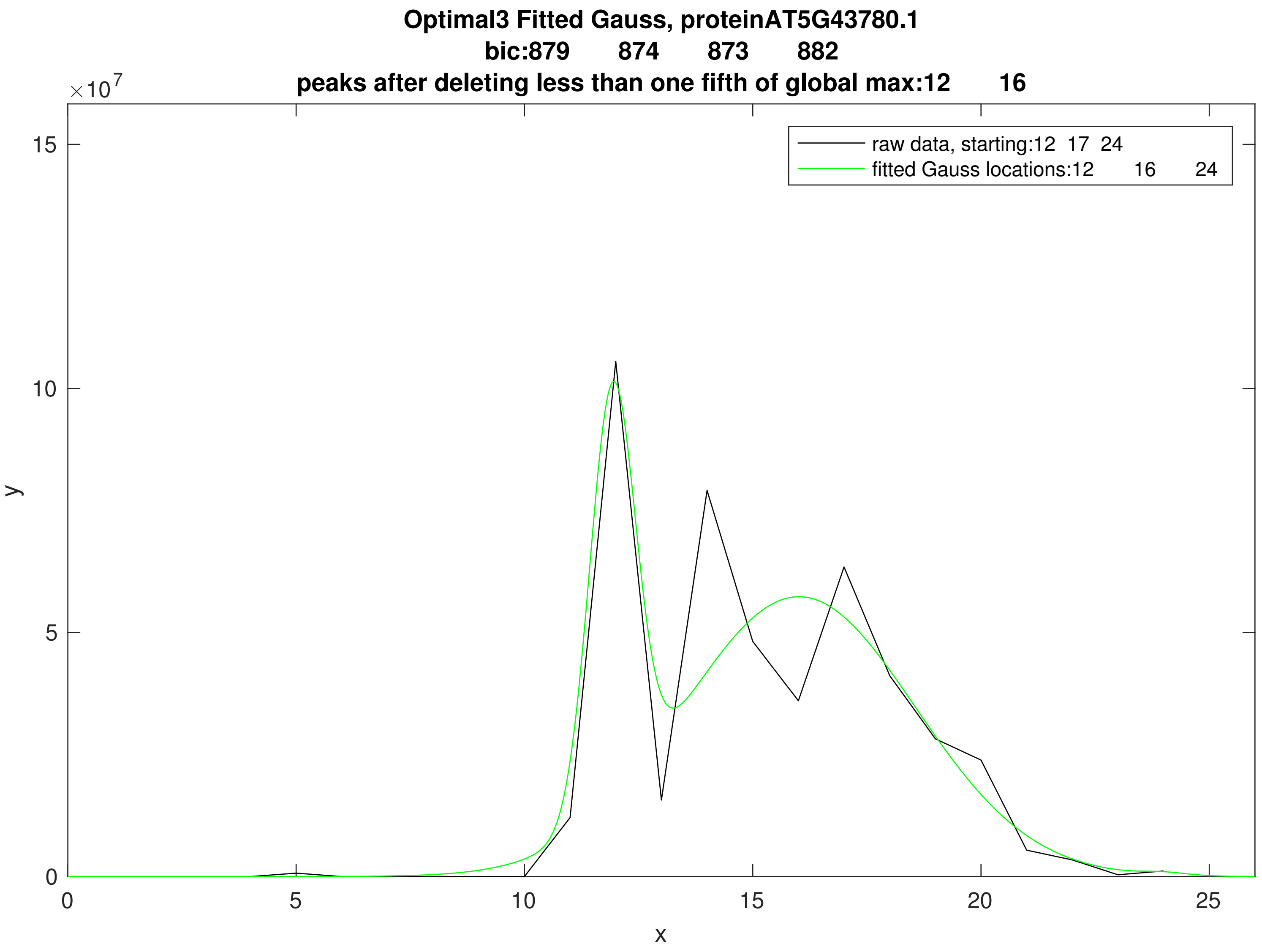

Optimal3 Fitted Gauss, proteinAT5G43780.1
bic:879 874 873 882
peaks after deleting less than one fifth of global max:12 16
×10^7
15
10
5
0
y
0
5
10
15
20
25
x
raw data, starting:12 17 24
fitted Gauss locations:12 16 24

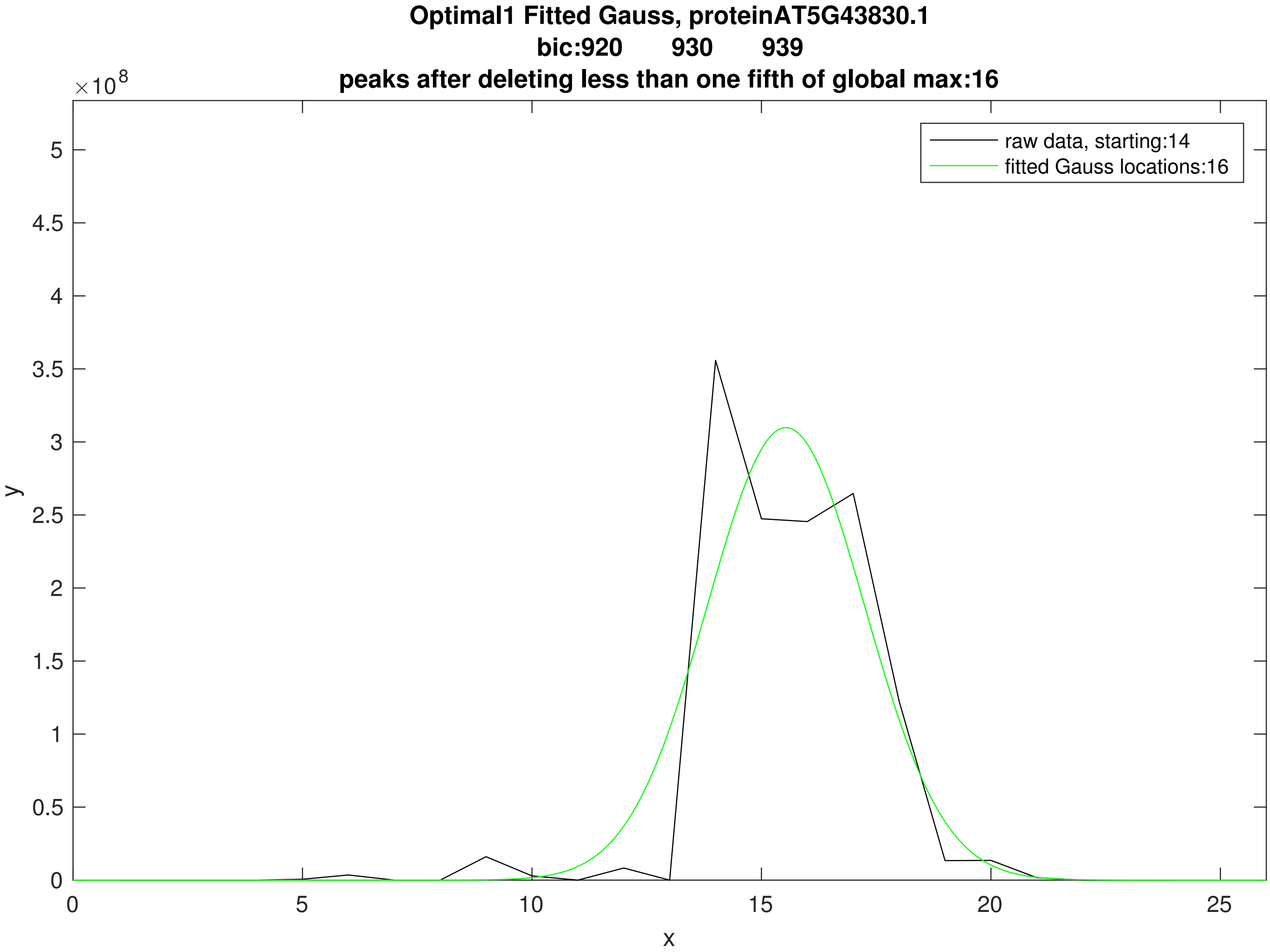
Optimal1 Fitted Gauss, proteinAT5G43830.1
bic:920 930 939
peaks after deleting less than one fifth of global max:16
×10^8
raw data, starting:14
fitted Gauss locations:16
y
x
0
0.5
1
1.5
2
2.5
3
3.5
4
4.5
5
5
10
15
20
25

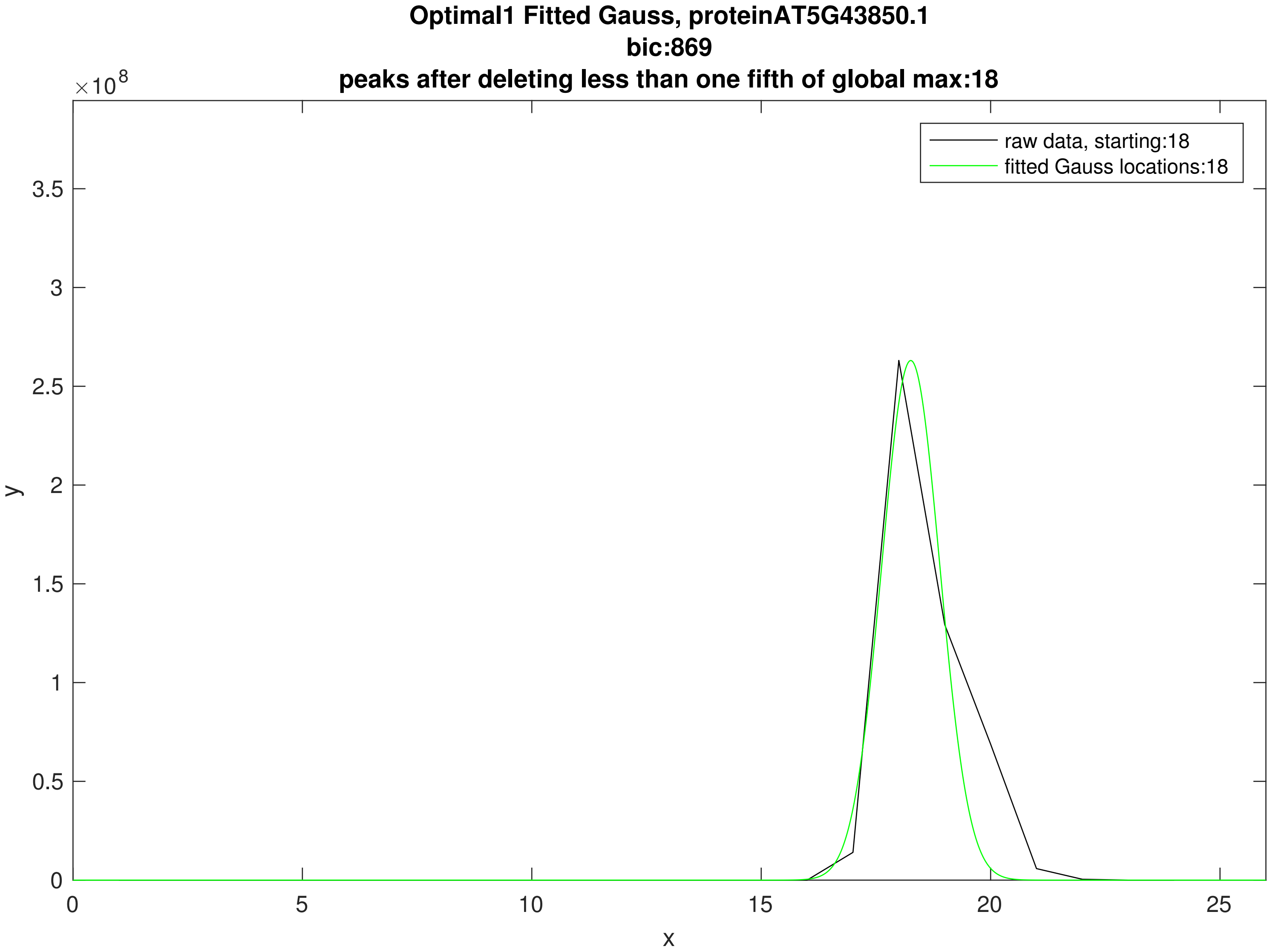

Optimal1 Fitted Gauss, proteinAT5G43850.1
bic:869
peaks after deleting less than one fifth of global max:18
×10^8
raw data, starting:18
fitted Gauss locations:18
3.5
3
2.5
2
1.5
1
0.5
0
y
0
5
10
15
20
25
x

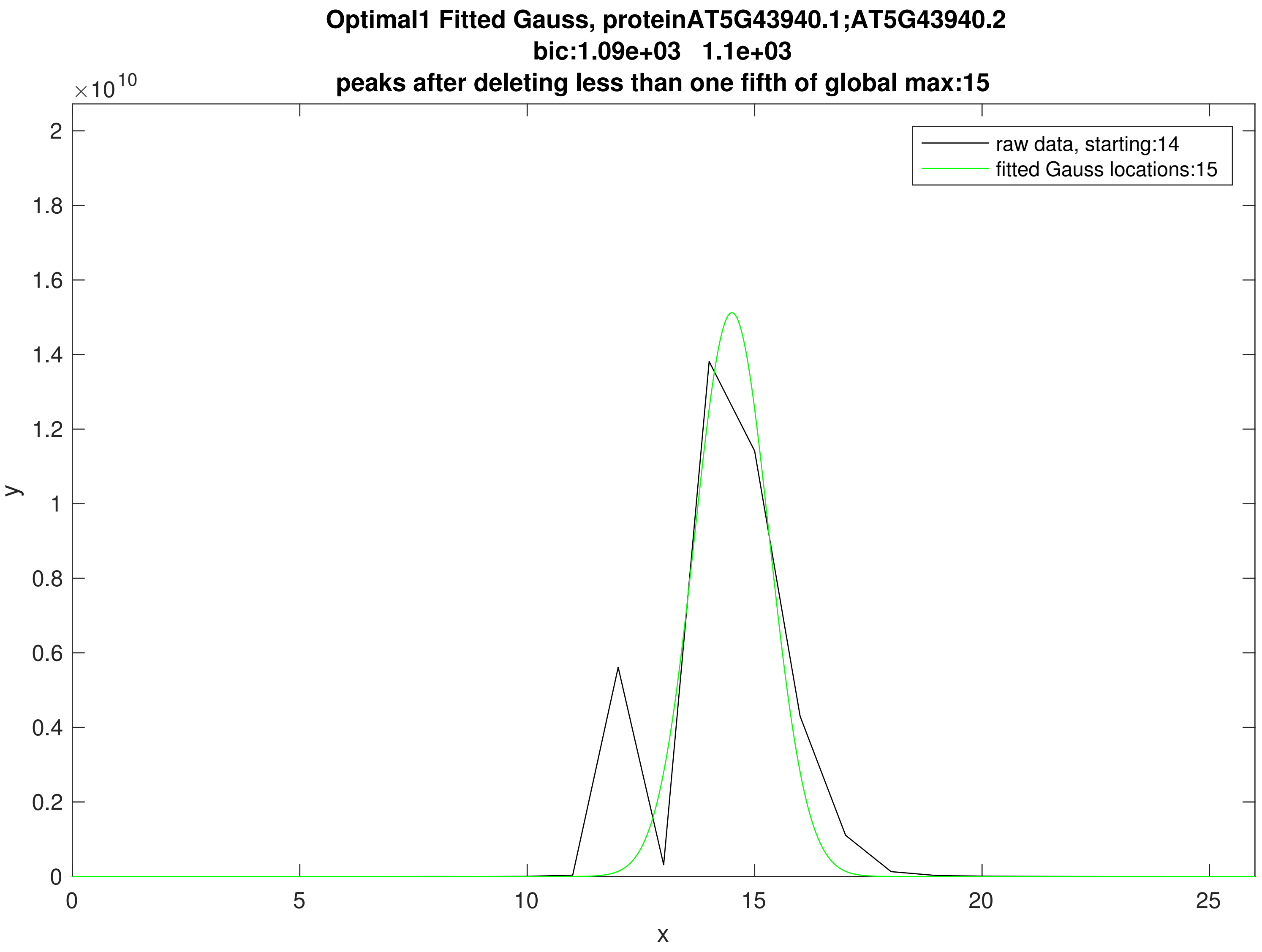
Optimal1 Fitted Gauss, proteinAT5G43940.1;AT5G43940.2
bic:1.09e+03 1.1e+03
peaks after deleting less than one fifth of global max:15
×10^10
raw data, starting:14
fitted Gauss locations:15
y
x
0
0.2
0.4
0.6
0.8
1
1.2
1.4
1.6
1.8
2
0
5
10
15
20
25

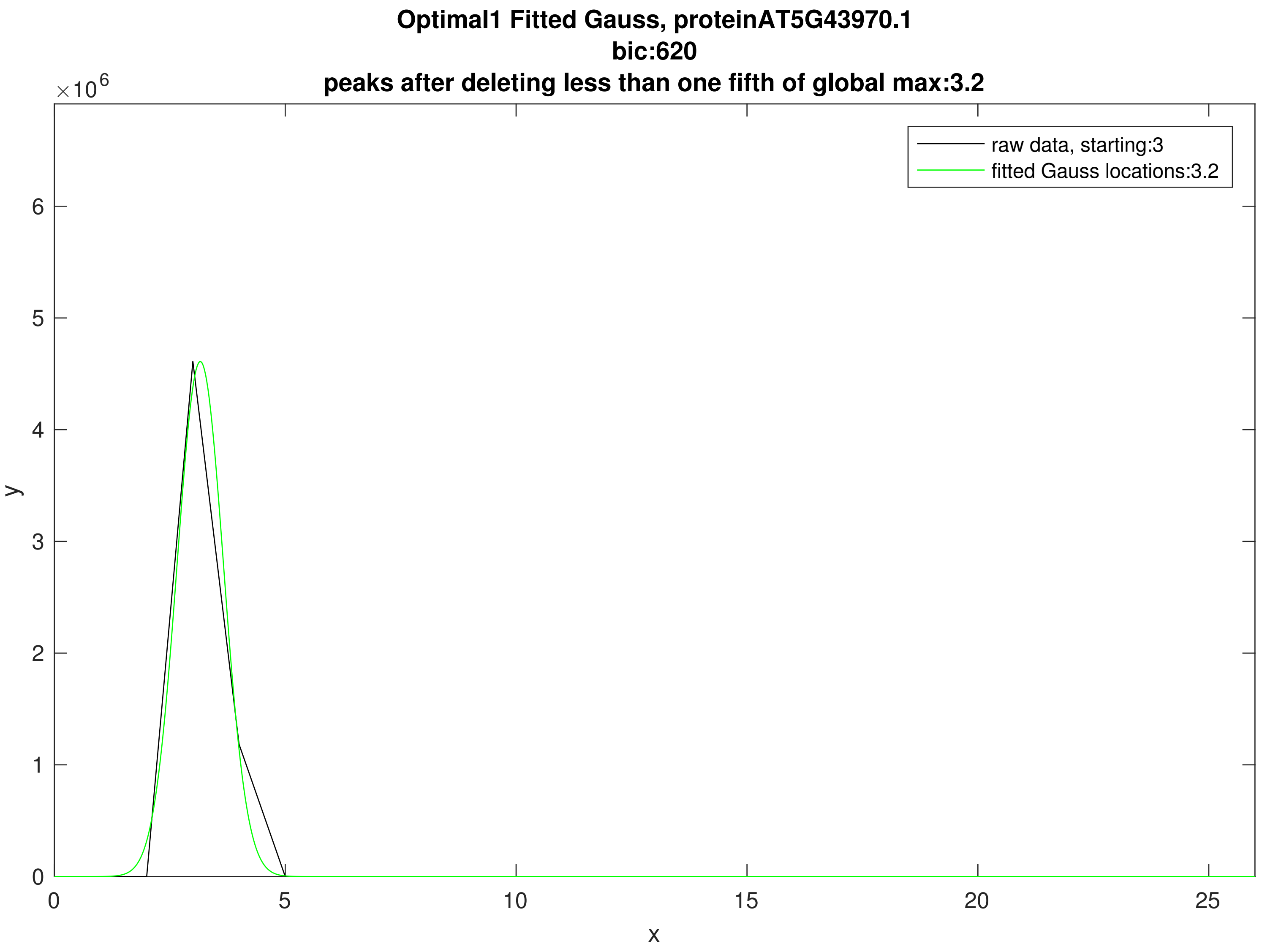

Optimal1 Fitted Gauss, proteinAT5G43970.1
bic:620
peaks after deleting less than one fifth of global max:3.2
×10^6
raw data, starting:3
fitted Gauss locations:3.2
y
x
0
1
2
3
4
5
6
0
5
10
15
20
25

Optimal1 Fitted Gauss, proteinAT5G44020.1
bic:965 966 976
peaks after deleting less than one fifth of global max:9.6

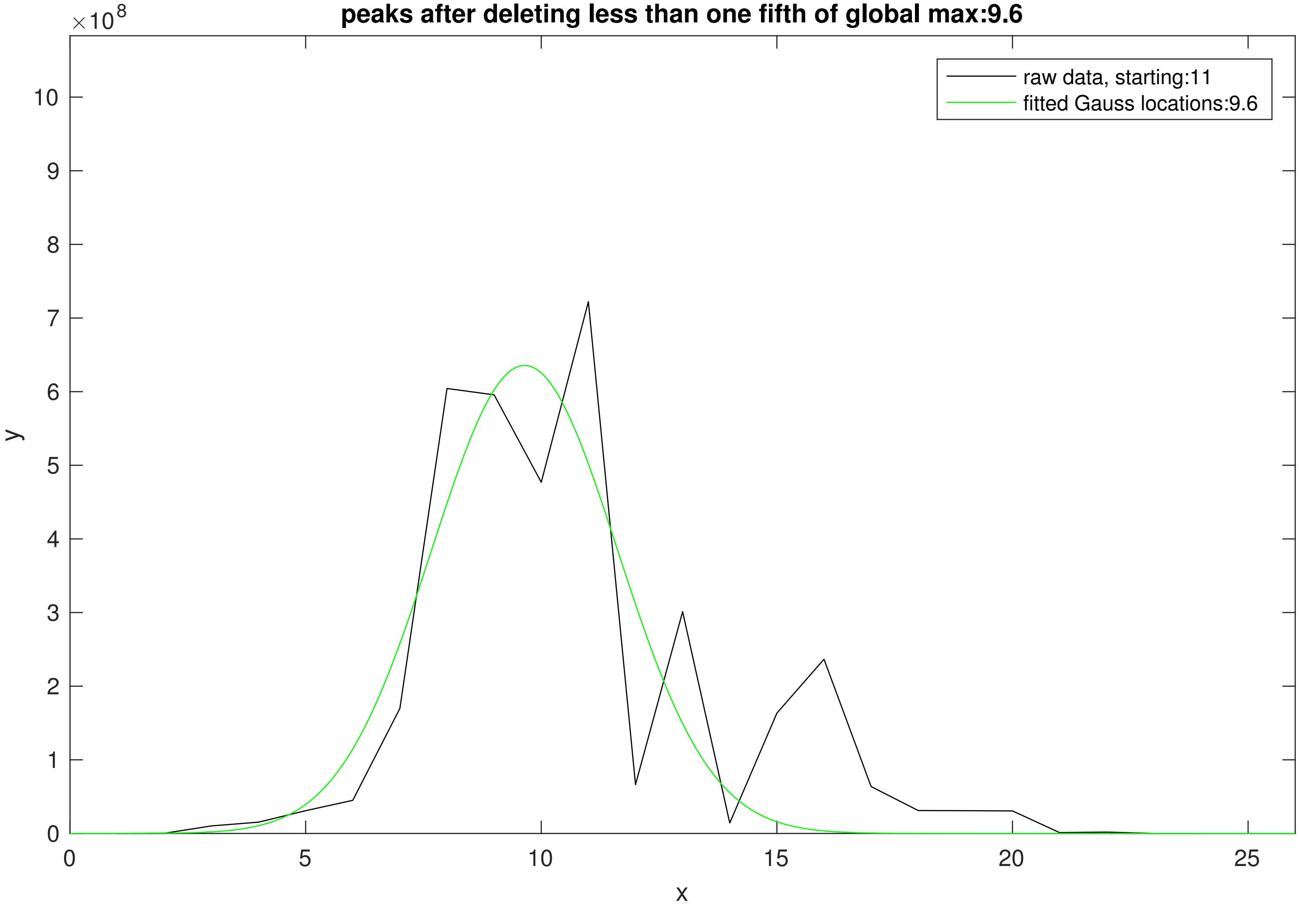

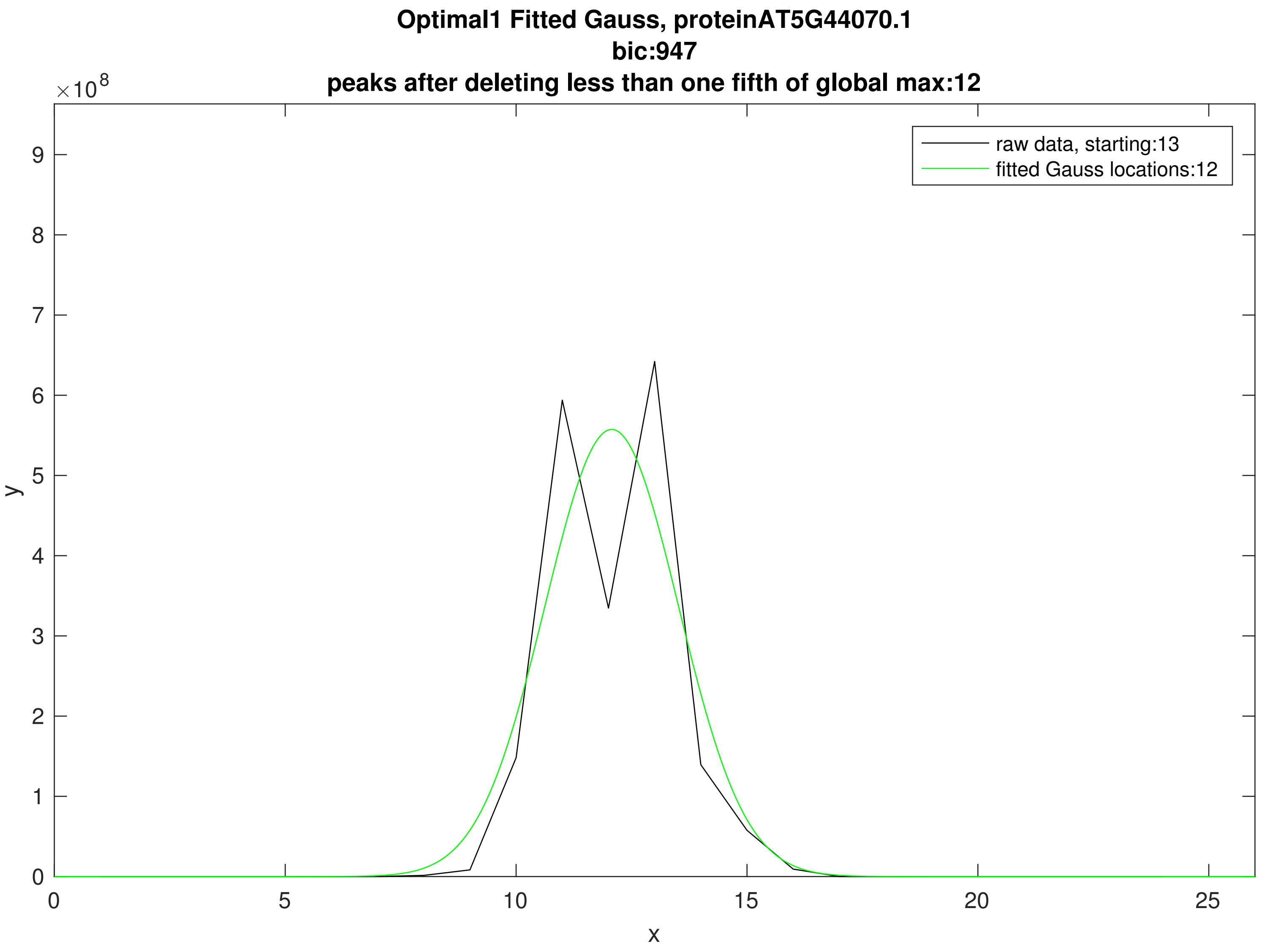
Optimal1 Fitted Gauss, proteinAT5G44070.1
bic:947
peaks after deleting less than one fifth of global max:12
×10^8
raw data, starting:13
fitted Gauss locations:12
y
x
0
1
2
3
4
5
6
7
8
9
10
15
20
25

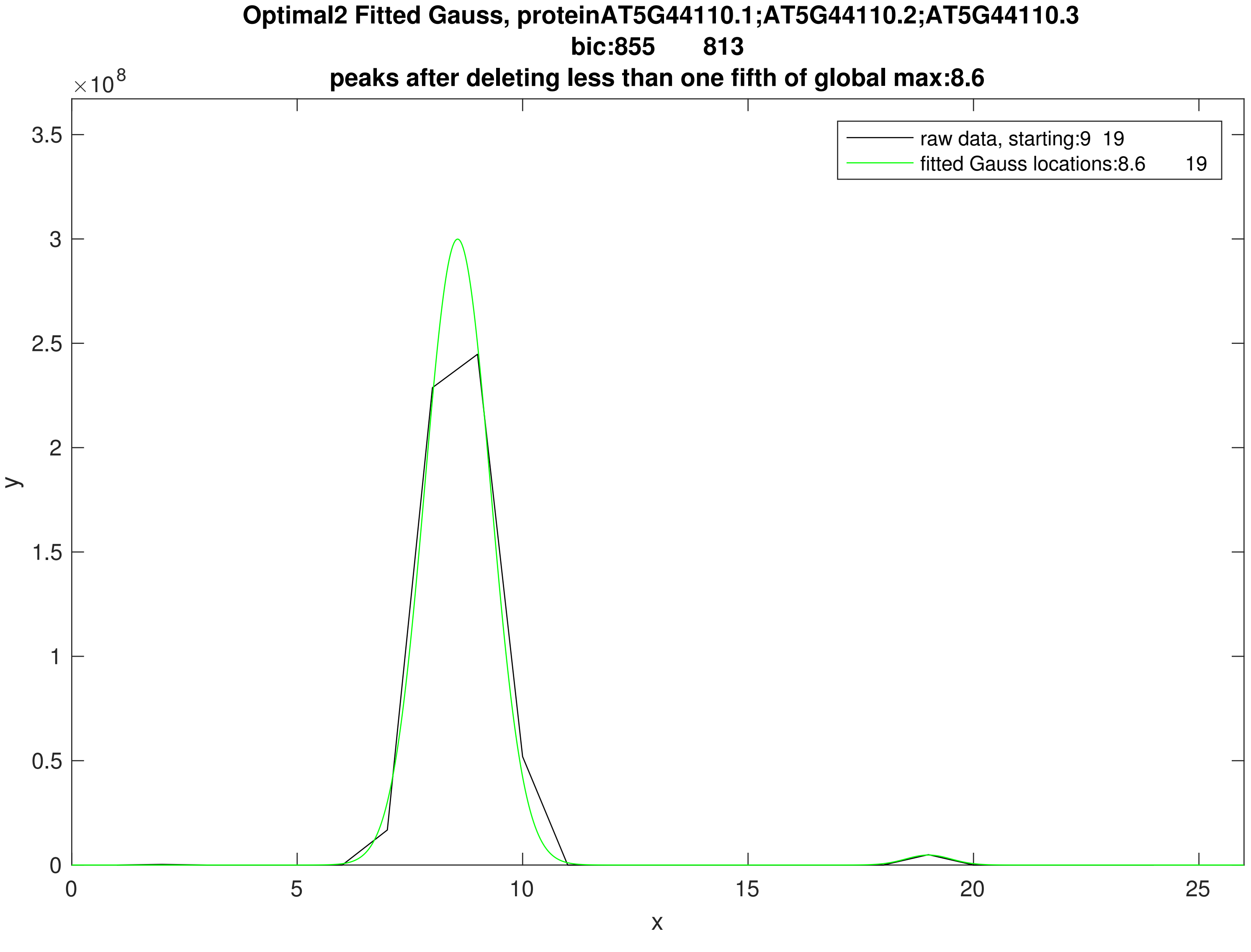

Optimal2 Fitted Gauss, proteinAT5G44110.1;AT5G44110.2;AT5G44110.3
bic:855 813
peaks after deleting less than one fifth of global max:8.6
×10^8
raw data, starting:9 19
fitted Gauss locations:8.6 19
3.5
3
2.5
2
1.5
1
0.5
0
y
0
5
10
15
20
25
x

**Optimal1 Fitted Gauss, proteinAT5G44120.3;AT5G44120.2;AT5G44120.1**
**bic:1.03e+03  1.04e+03  1.05e+03**
**peaks after deleting less than one fifth of global max:11**

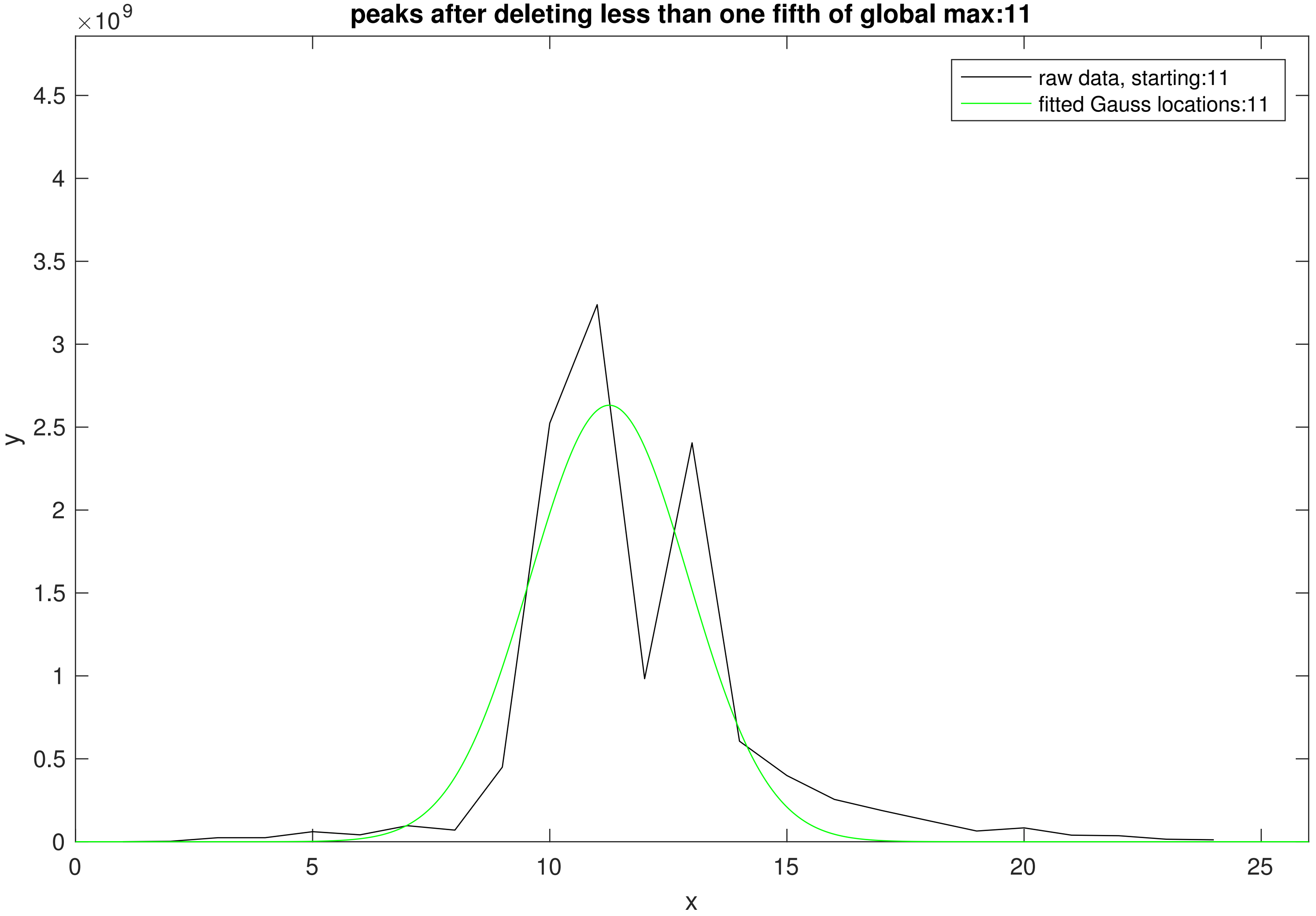

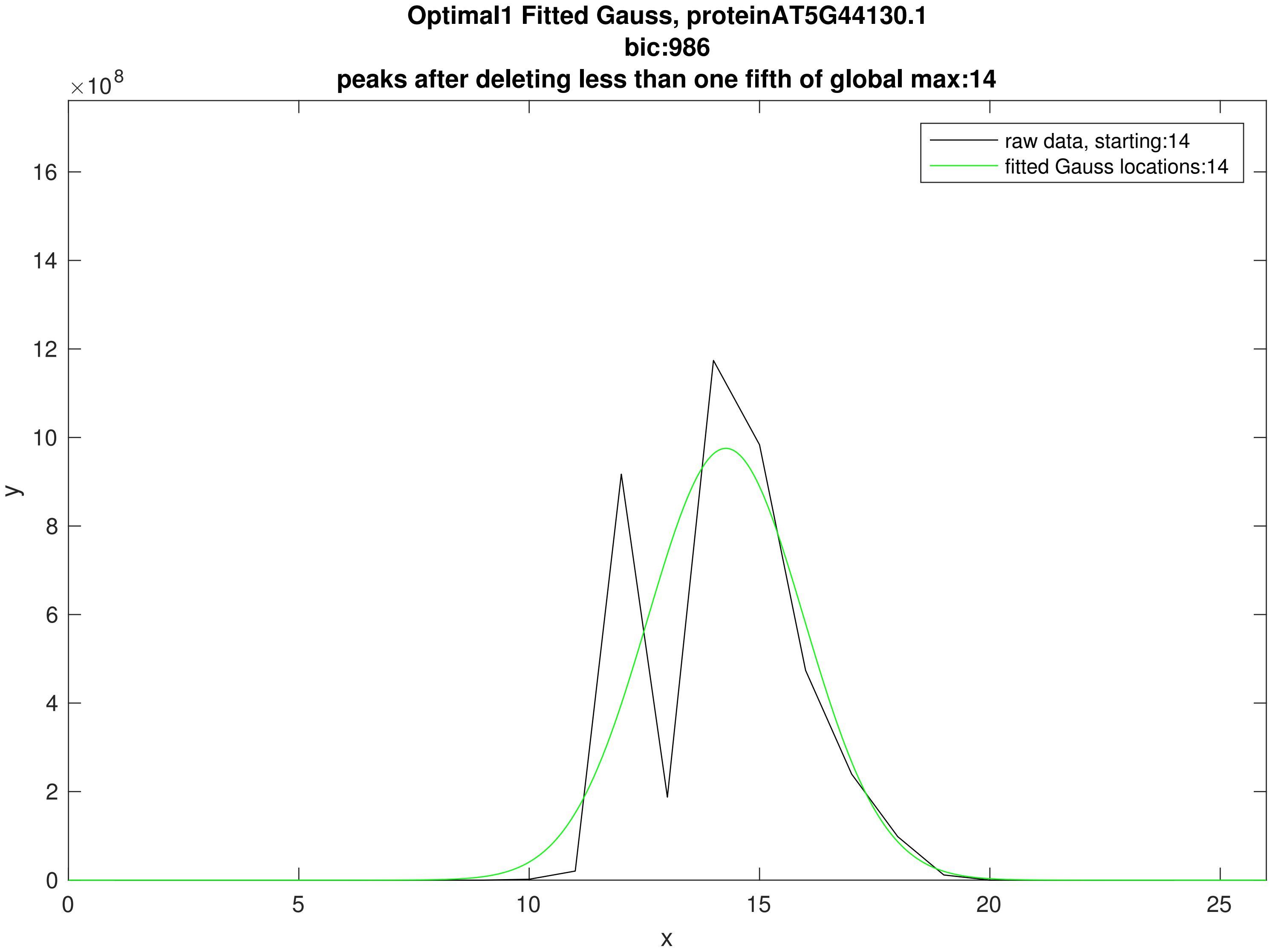
Optimal1 Fitted Gauss, proteinAT5G44130.1
bic:986
peaks after deleting less than one fifth of global max:14
×10^8
16
14
12
10
8
6
4
2
0
y
0
5
10
15
20
25
x
raw data, starting:14
fitted Gauss locations:14

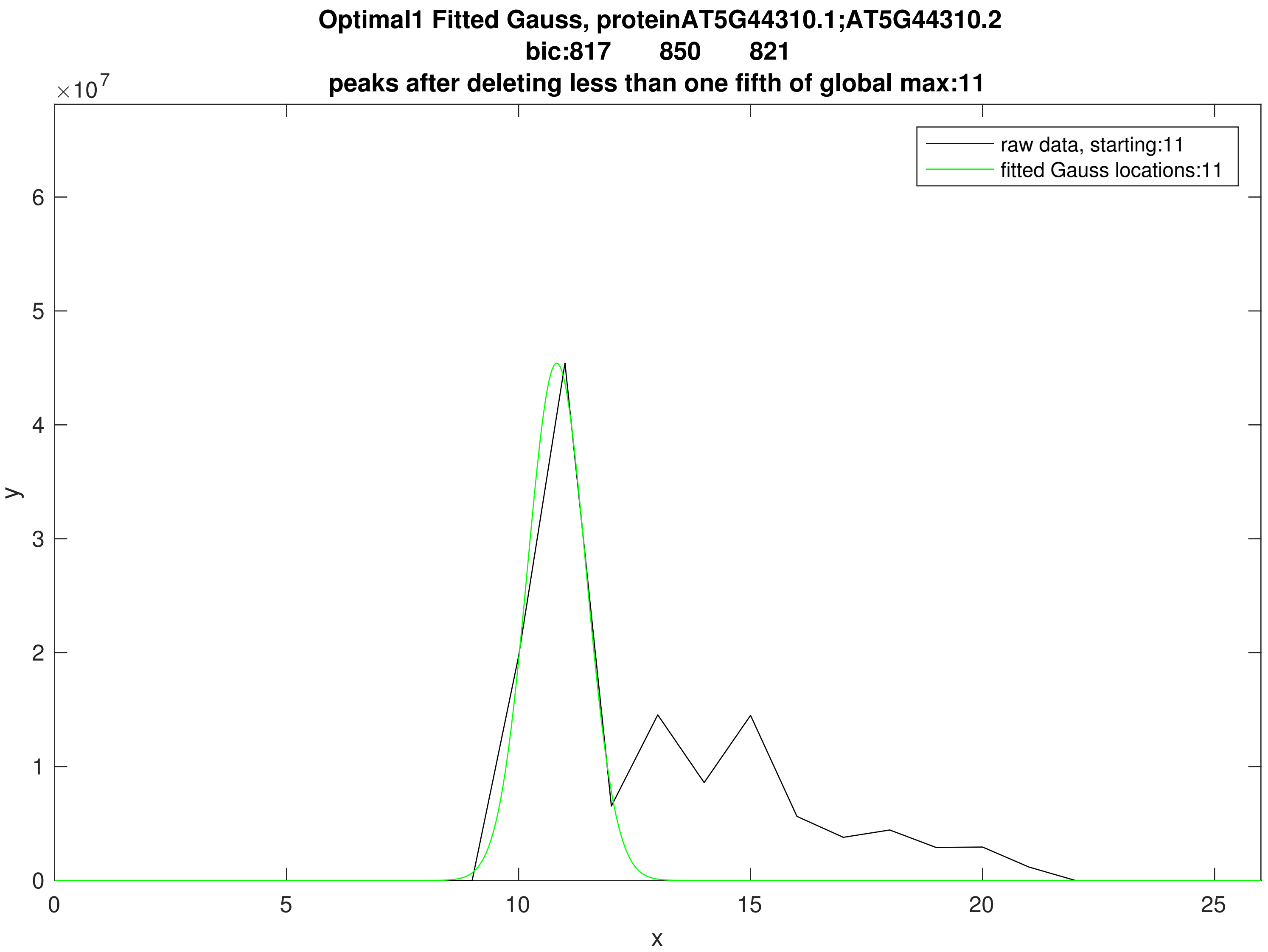

Optimal1 Fitted Gauss, proteinAT5G44310.1;AT5G44310.2
bic:817 850 821
peaks after deleting less than one fifth of global max:11
×10^7
raw data, starting:11
fitted Gauss locations:11
y
x
0
1
2
3
4
5
6
0
5
10
15
20
25

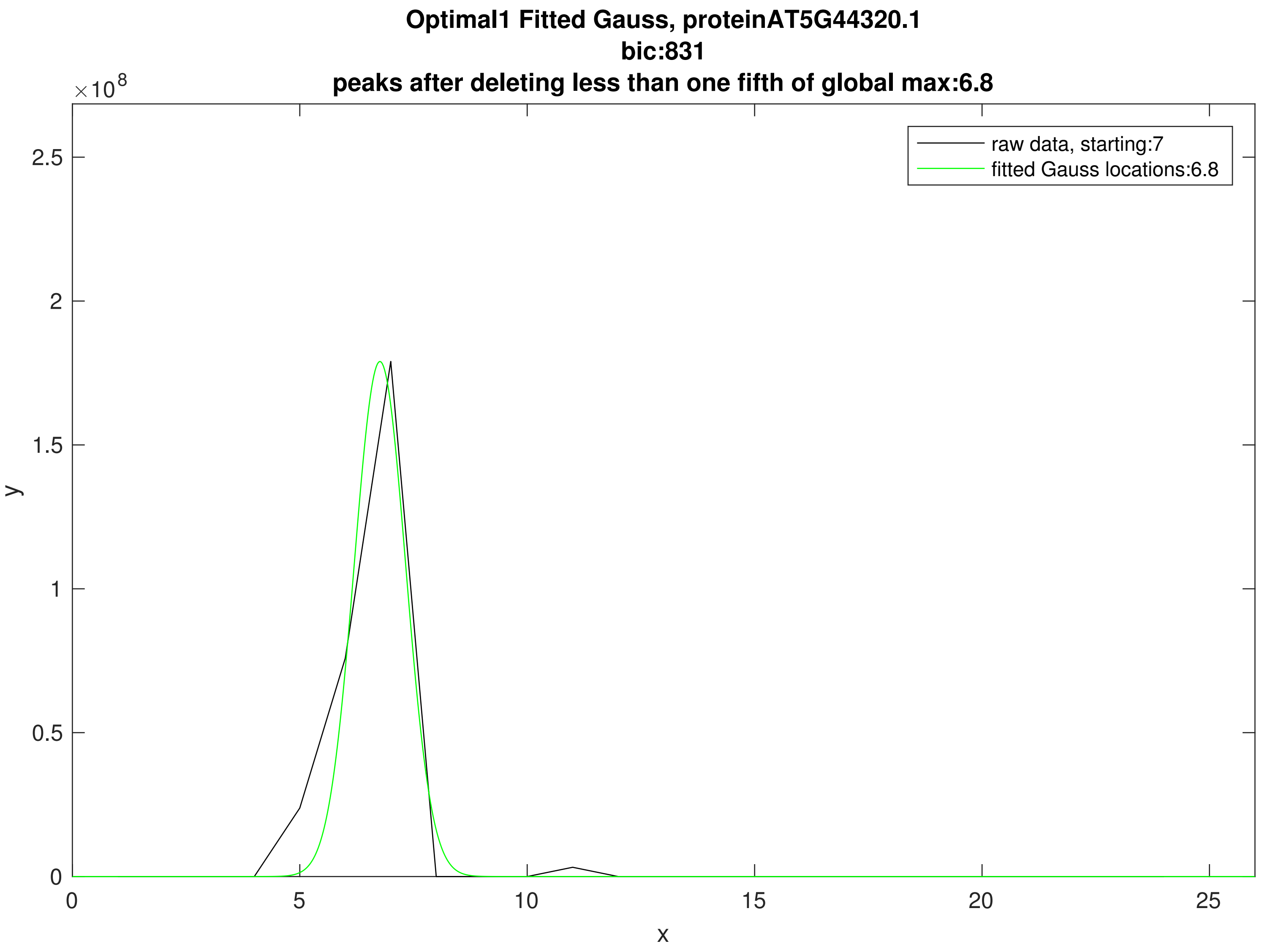

Optimal1 Fitted Gauss, proteinAT5G44320.1
bic:831
peaks after deleting less than one fifth of global max:6.8
×10^8
2.5
2
1.5
1
0.5
0
y
0
5
10
15
20
25
x
raw data, starting:7
fitted Gauss locations:6.8

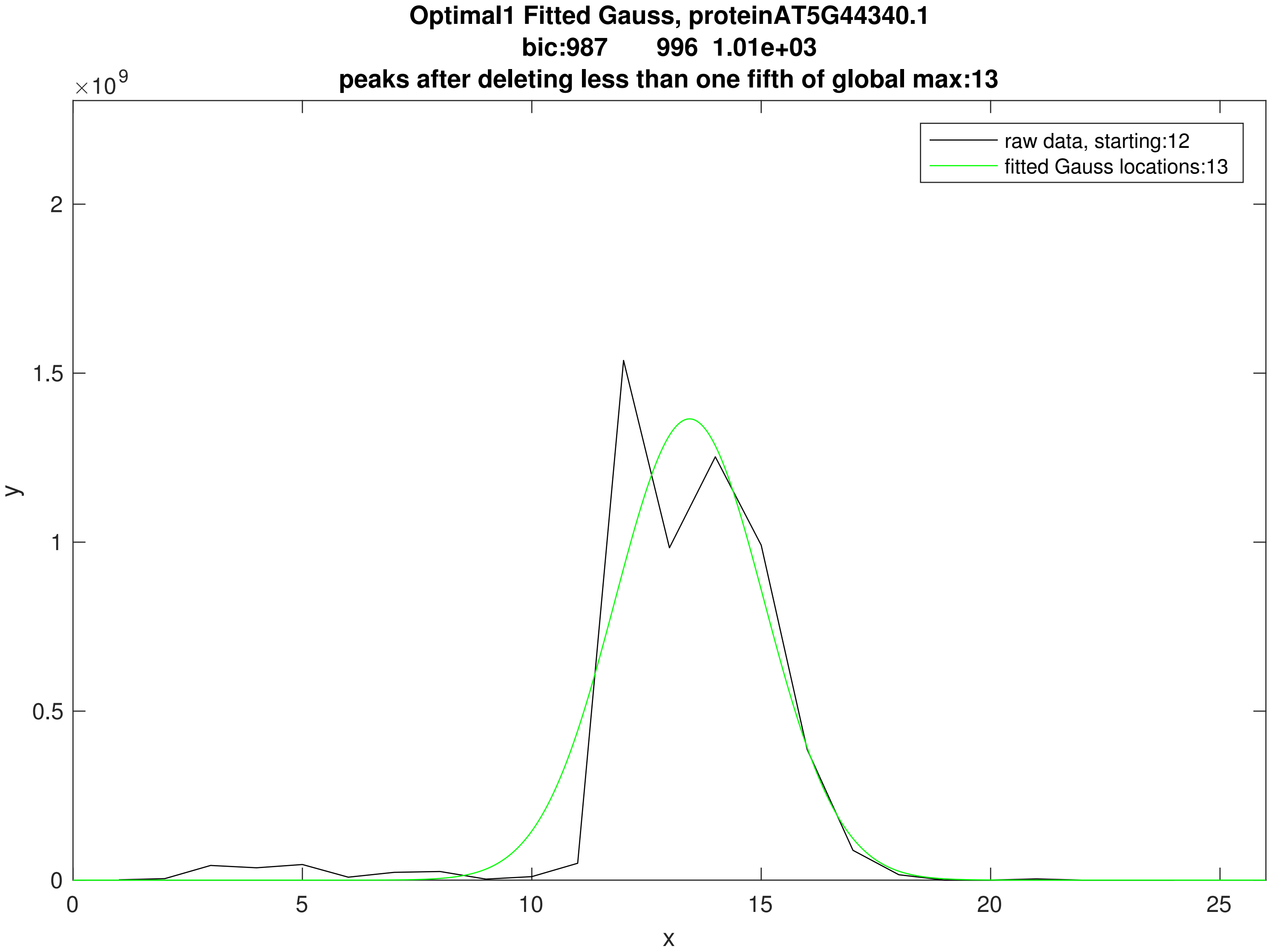
Optimal1 Fitted Gauss, proteinAT5G44340.1
bic:987 996 1.01e+03
peaks after deleting less than one fifth of global max:13
×10^9
raw data, starting:12
fitted Gauss locations:13
y
x
0
0.5
1
1.5
2
5
10
15
20
25

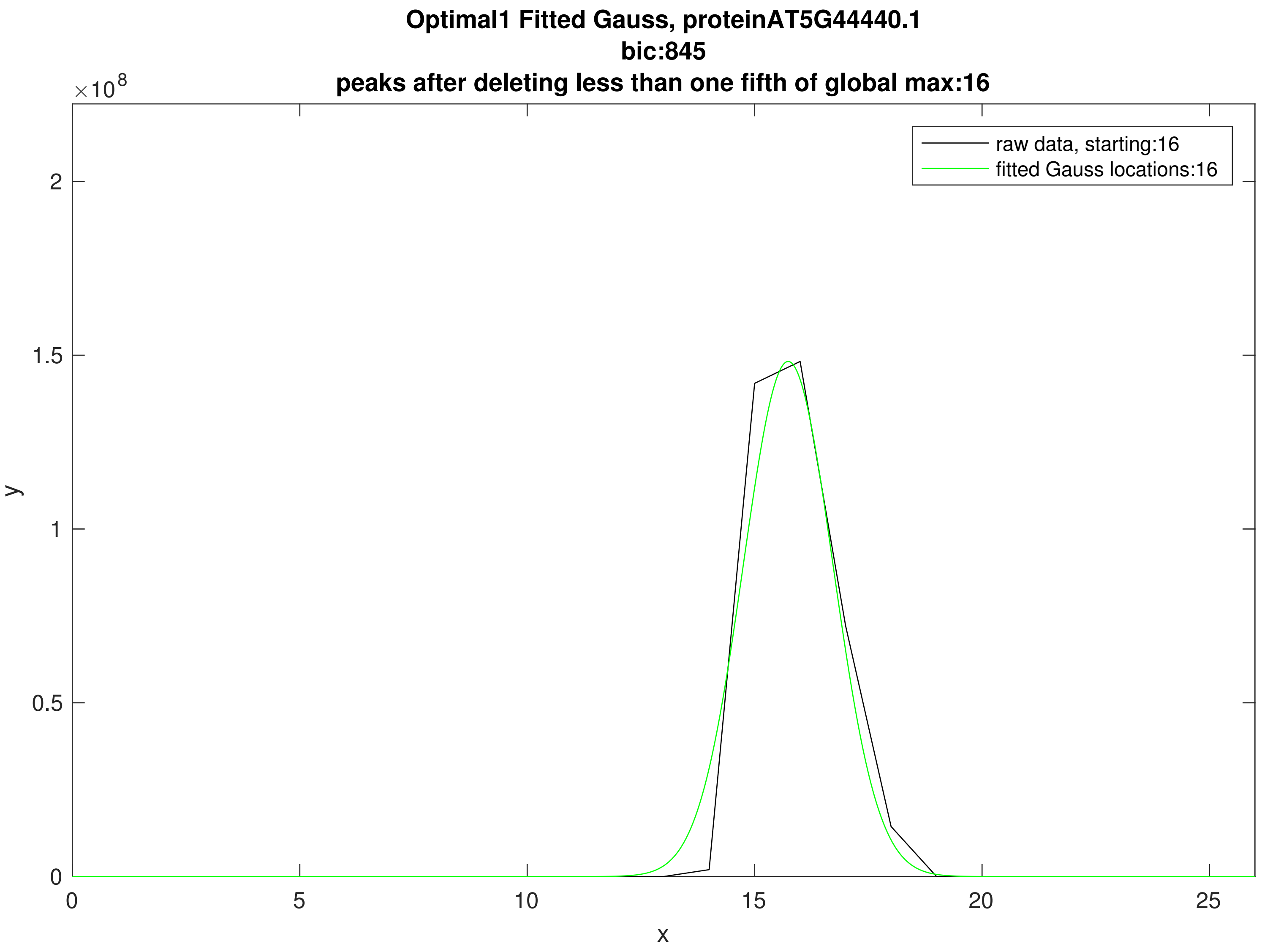
Optimal1 Fitted Gauss, proteinAT5G44440.1
bic:845
peaks after deleting less than one fifth of global max:16
×10^8
raw data, starting:16
fitted Gauss locations:16
y
x
0
0.5
1
1.5
2
5
10
15
20
25

**Optimal1 Fitted Gauss, proteinAT5G44460.1**
**bic:852**
**peaks after deleting less than one fifth of global max:18**

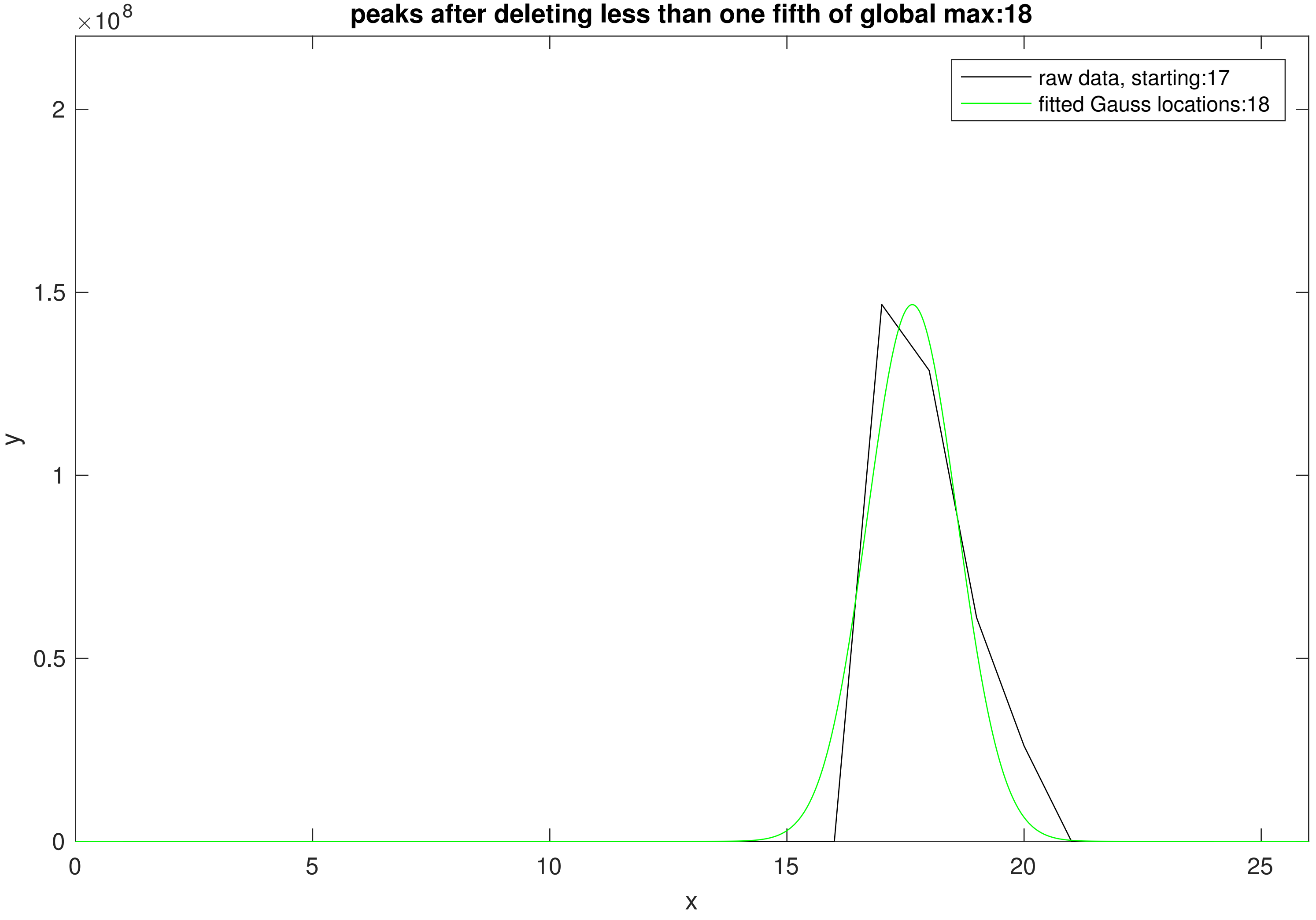

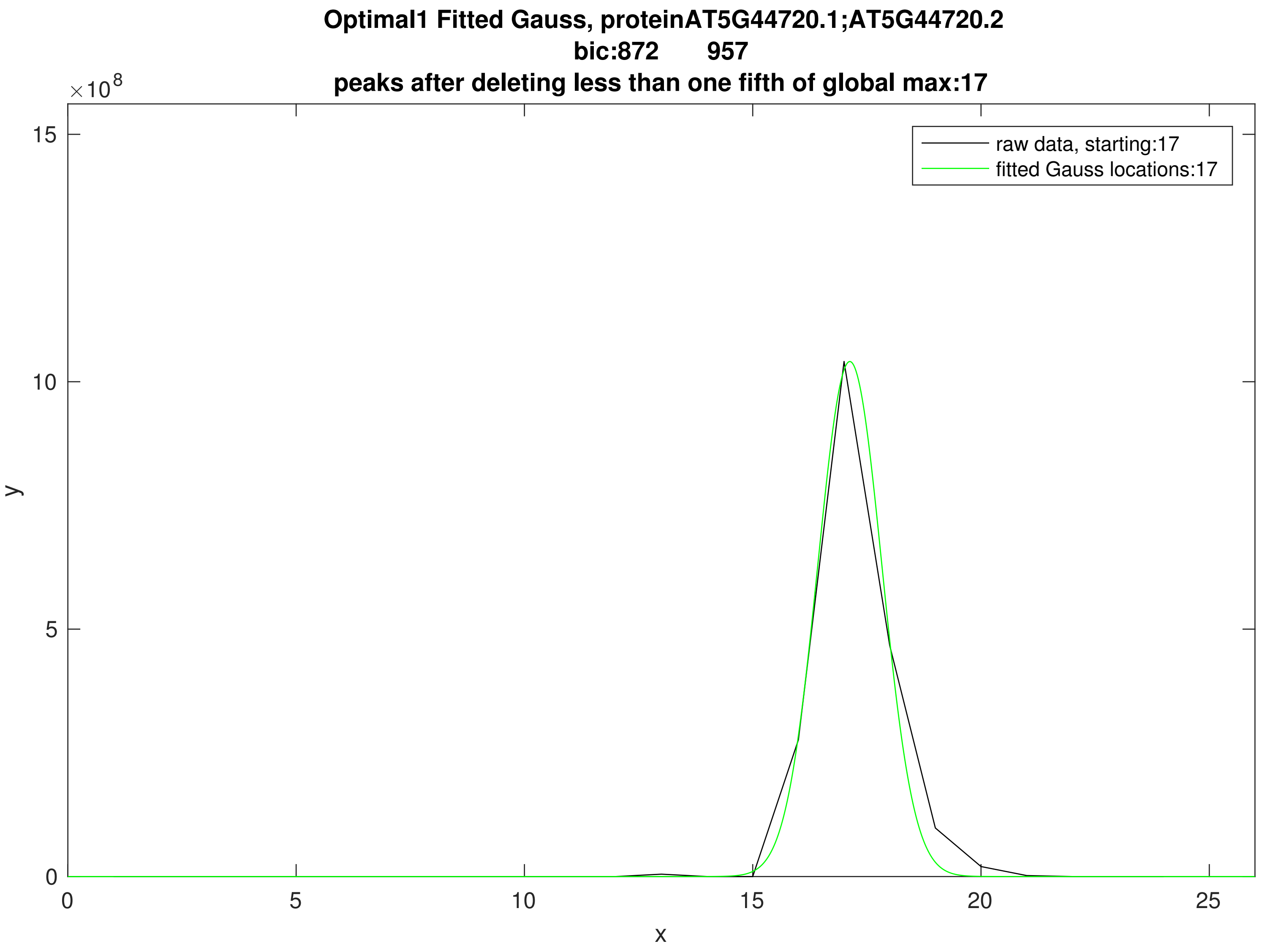
Optimal1 Fitted Gauss, proteinAT5G44720.1;AT5G44720.2
bic:872 957
peaks after deleting less than one fifth of global max:17
×10^8
15
10
5
0
y
0
5
10
15
20
25
x
raw data, starting:17
fitted Gauss locations:17

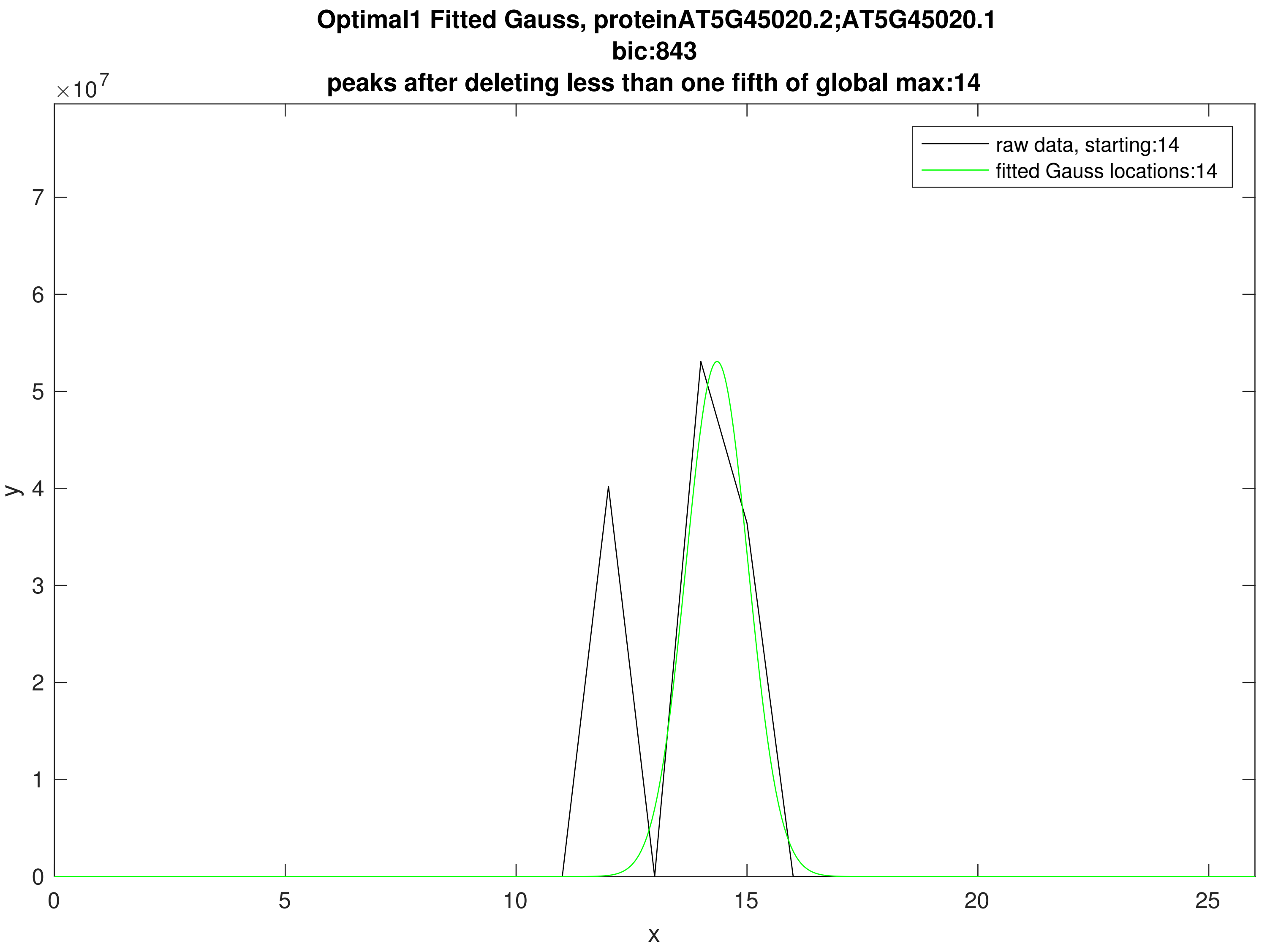
Optimal1 Fitted Gauss, proteinAT5G45020.2;AT5G45020.1
bic:843
peaks after deleting less than one fifth of global max:14
×10^7
raw data, starting:14
fitted Gauss locations:14
y
x
0
1
2
3
4
5
6
7
0
5
10
15
20
25

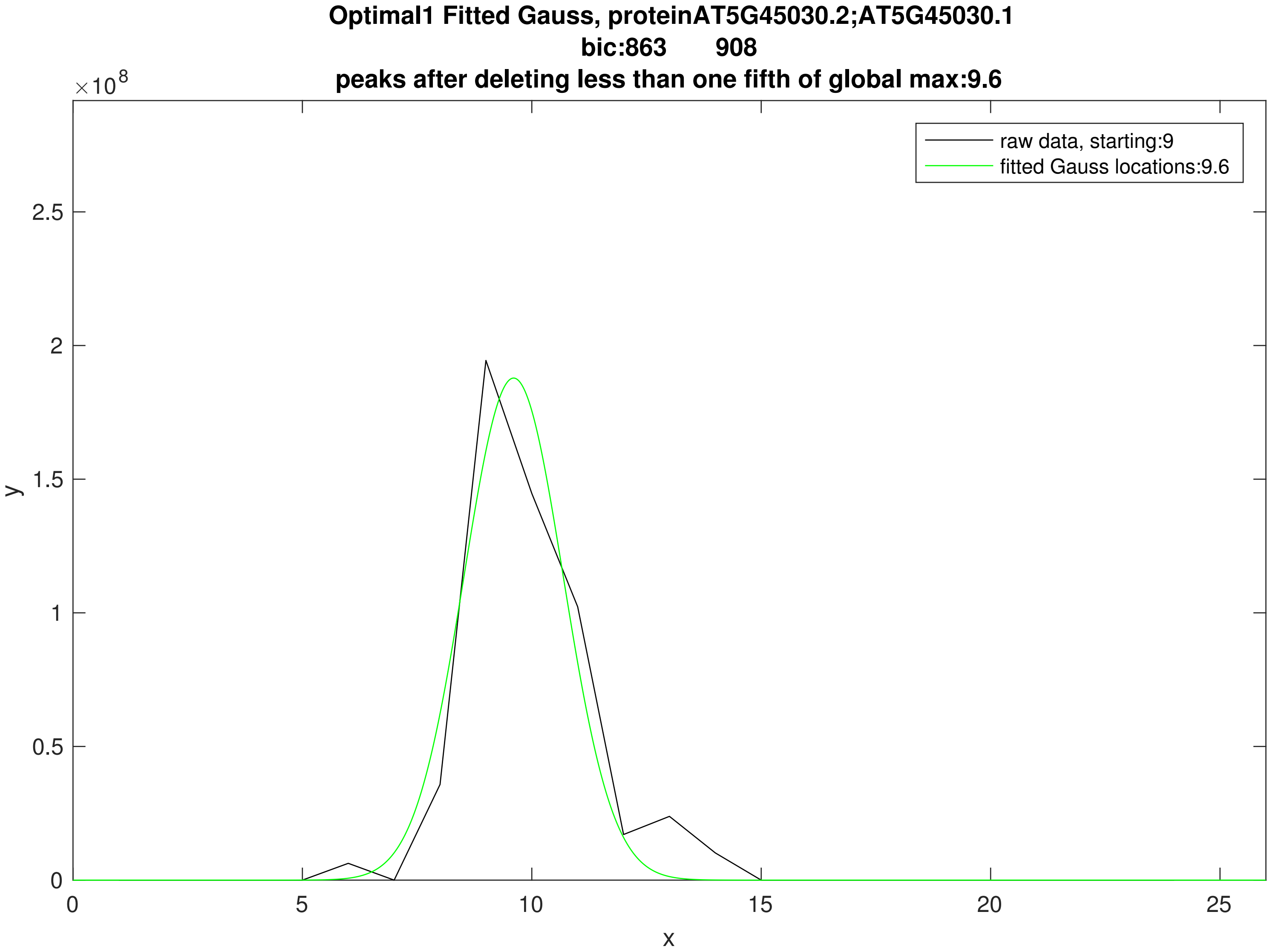
Optimal1 Fitted Gauss, proteinAT5G45030.2;AT5G45030.1
bic:863 908
peaks after deleting less than one fifth of global max:9.6
×10^8
raw data, starting:9
fitted Gauss locations:9.6
y
x
0
0.5
1
1.5
2
2.5
0
5
10
15
20
25

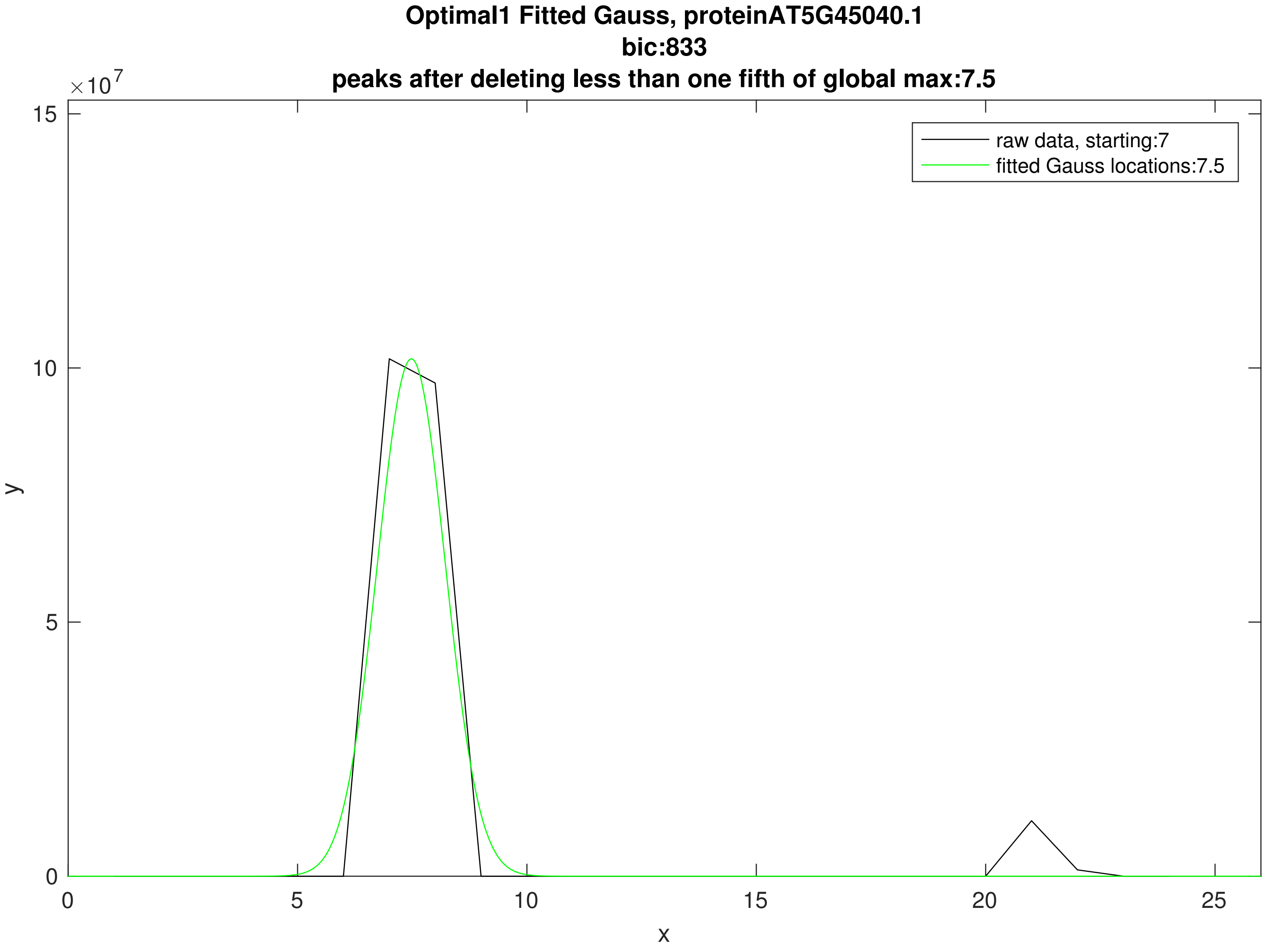

Optimal1 Fitted Gauss, proteinAT5G45040.1
bic:833
peaks after deleting less than one fifth of global max:7.5
×10^7
15
10
5
0
y
0
5
10
15
20
25
x
raw data, starting:7
fitted Gauss locations:7.5

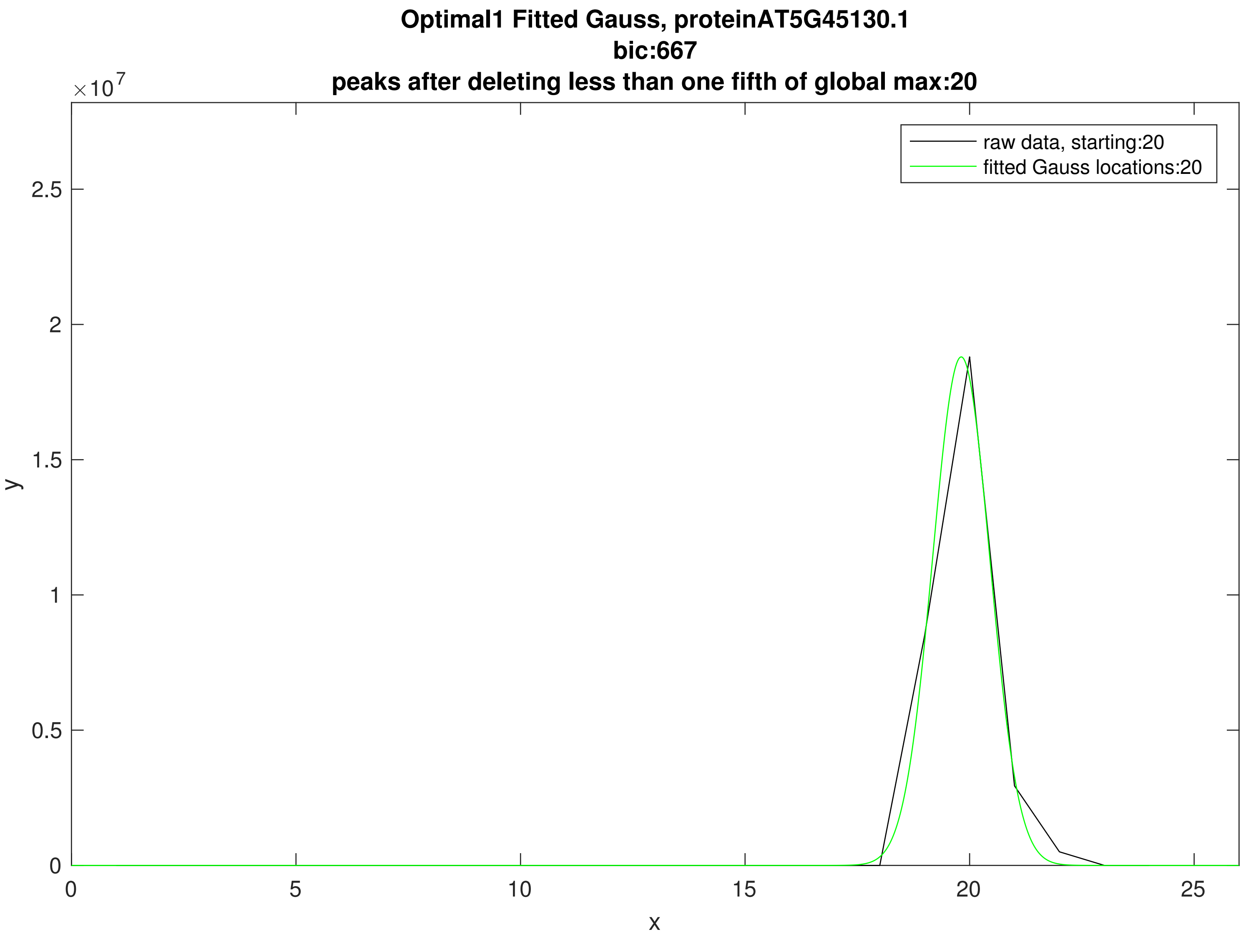

Optimal1 Fitted Gauss, proteinAT5G45130.1
bic:667
peaks after deleting less than one fifth of global max:20
×10^7
raw data, starting:20
fitted Gauss locations:20
2.5
2
1.5
1
0.5
0
y
0
5
10
15
20
25
x

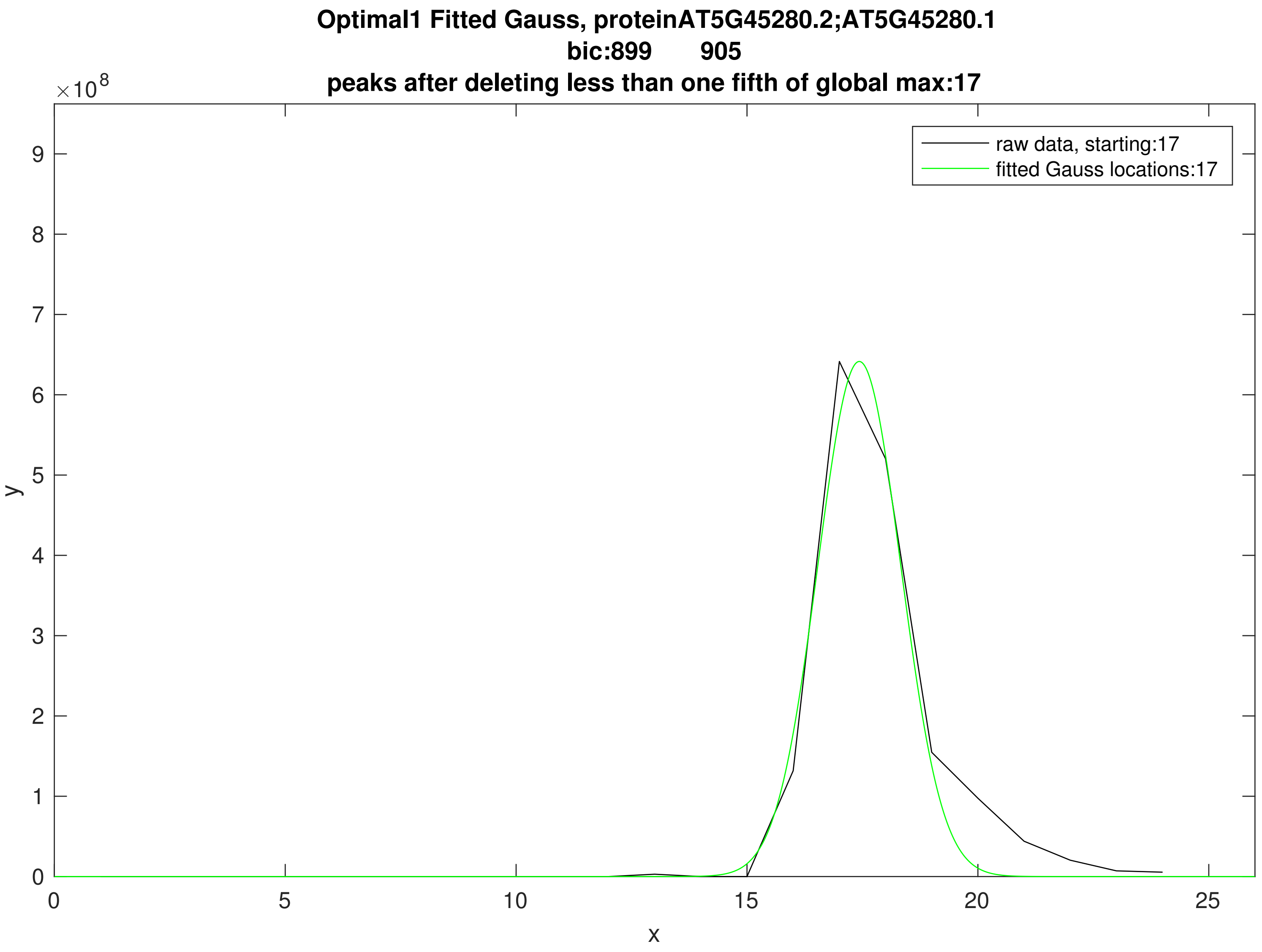

Optimal1 Fitted Gauss, proteinAT5G45280.2;AT5G45280.1
bic:899 905
peaks after deleting less than one fifth of global max:17
×10^8
raw data, starting:17
fitted Gauss locations:17
y
x
0
1
2
3
4
5
6
7
8
9
0
5
10
15
20
25

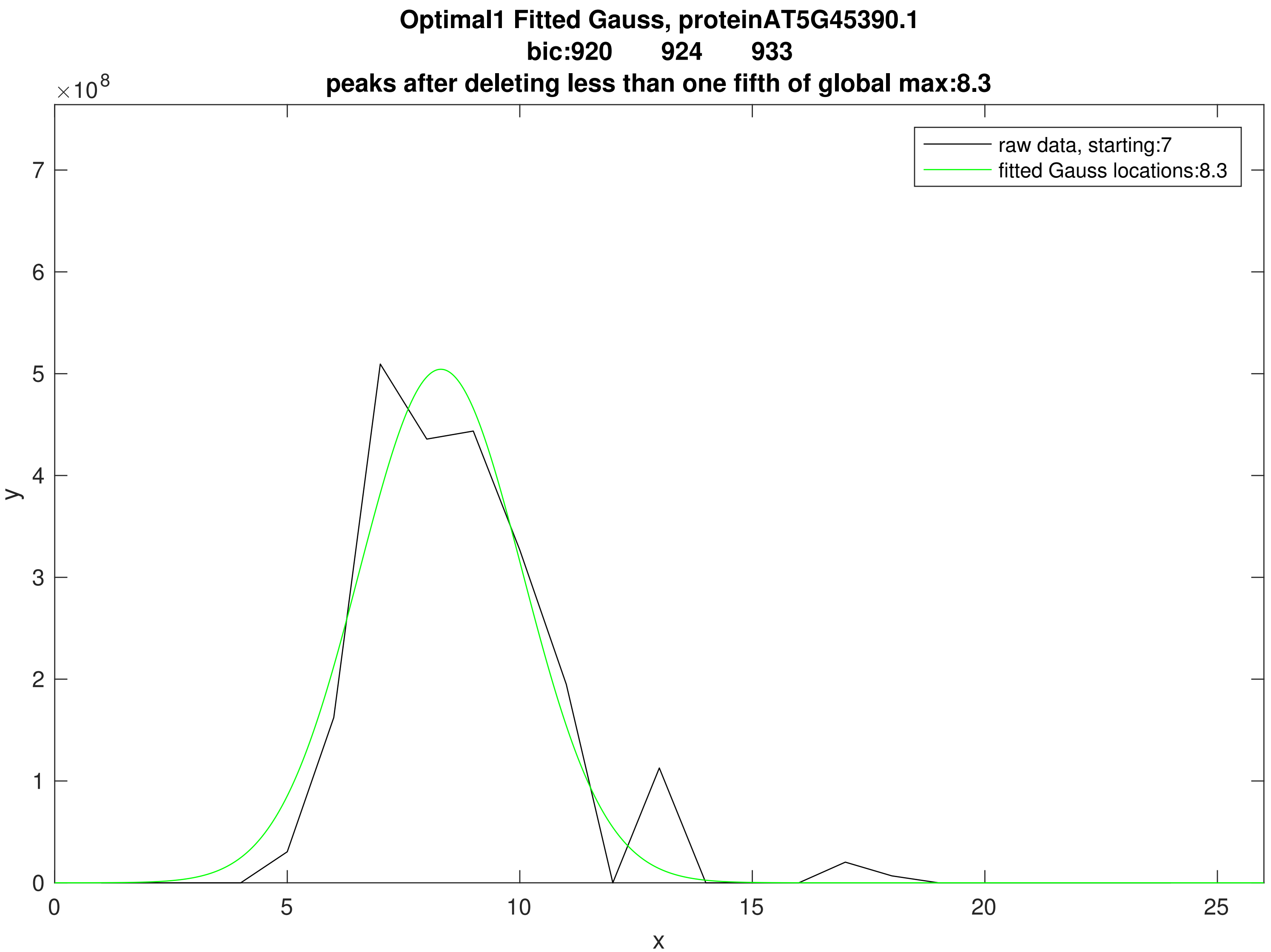
Optimal1 Fitted Gauss, proteinAT5G45390.1
bic:920 924 933
peaks after deleting less than one fifth of global max:8.3
×10^8
raw data, starting:7
fitted Gauss locations:8.3
y
x
0
1
2
3
4
5
6
7
0
5
10
15
20
25

Optimal1 Fitted Gauss, proteinAT5G45620.2;AT5G45620.1
bic:747
peaks after deleting less than one fifth of global max:4

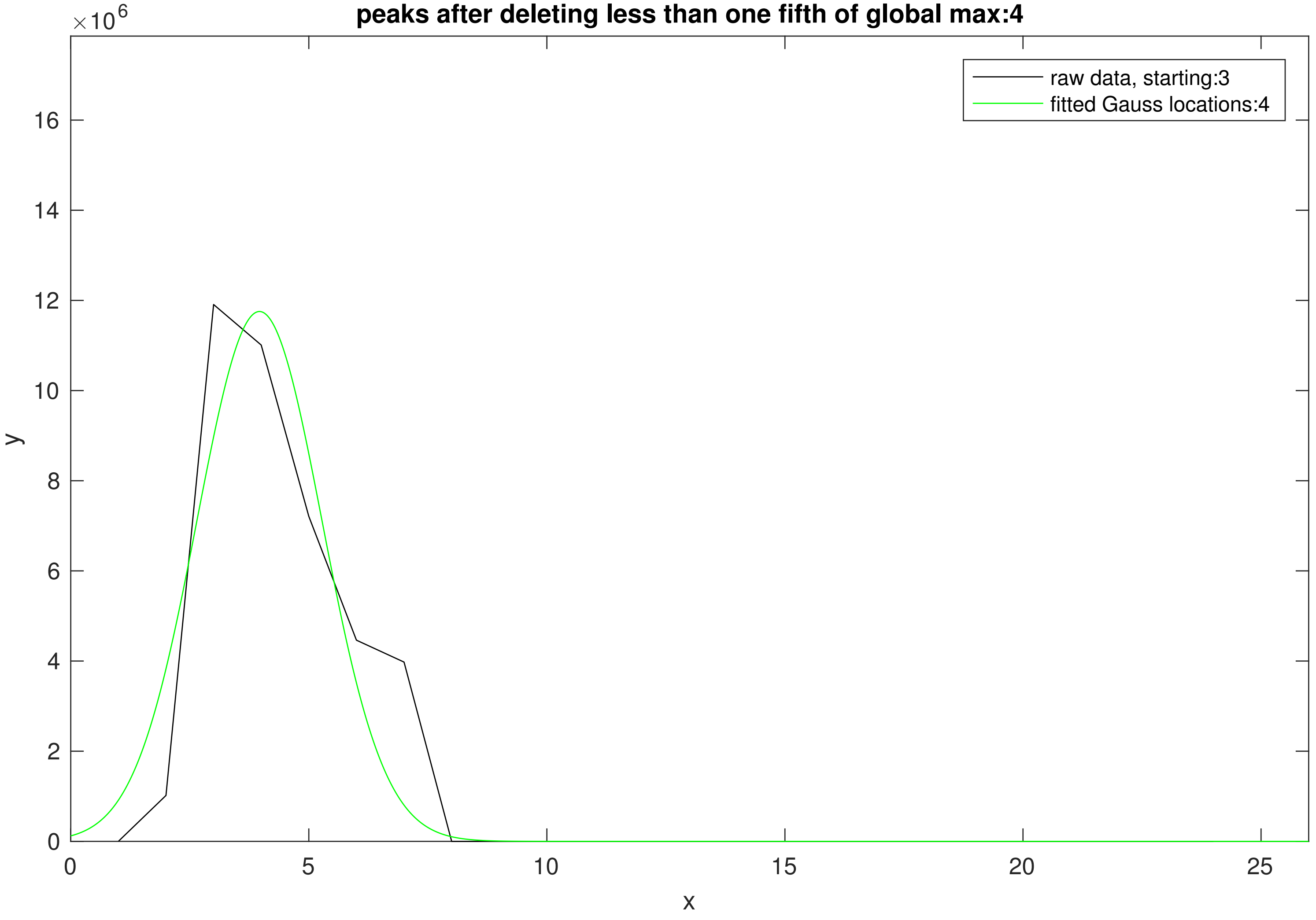

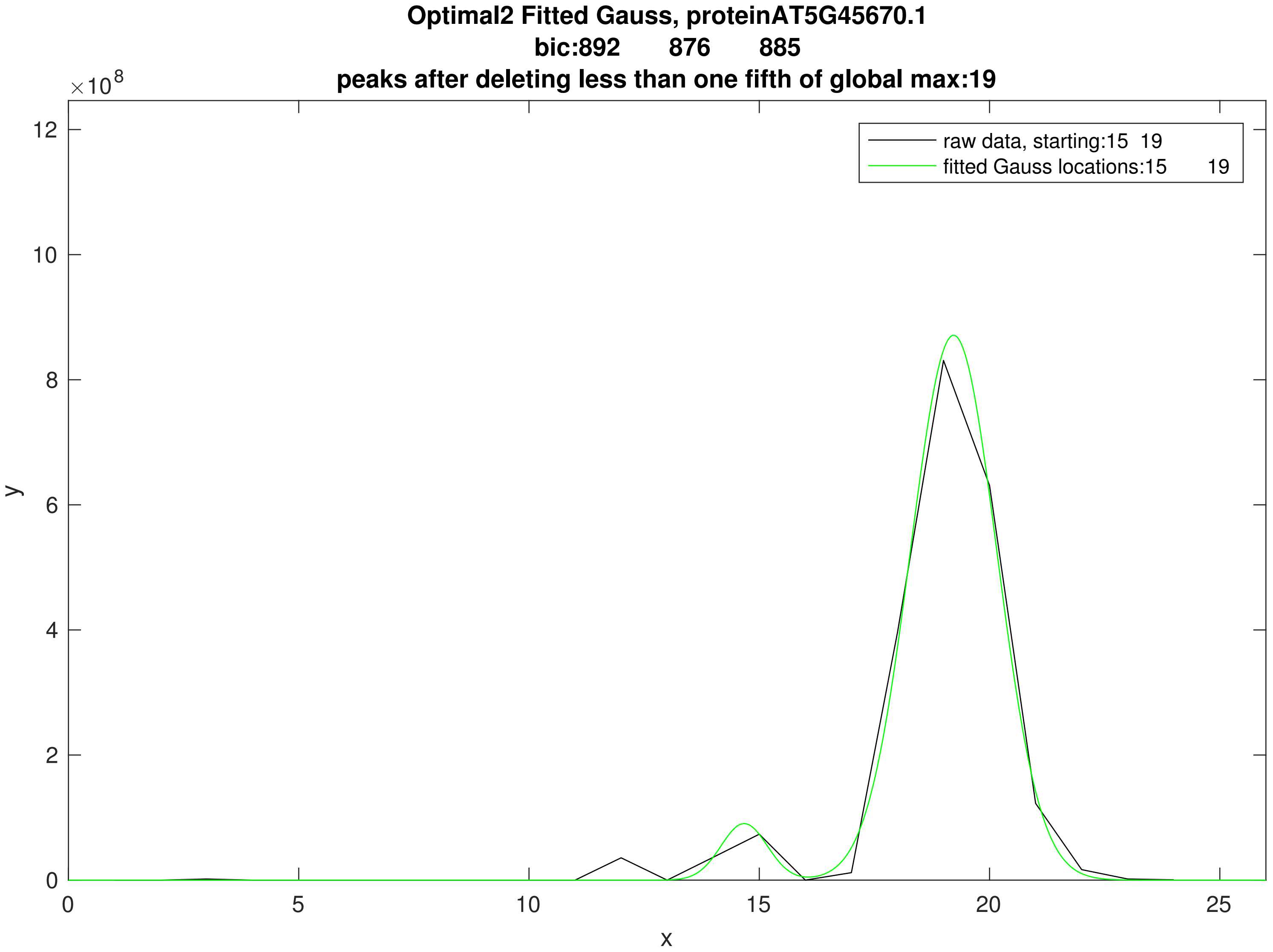
Optimal2 Fitted Gauss, proteinAT5G45670.1
bic:892 876 885
peaks after deleting less than one fifth of global max:19
×10^8
raw data, starting:15 19
fitted Gauss locations:15 19
y
x
0
2
4
6
8
10
12
0
5
10
15
20
25

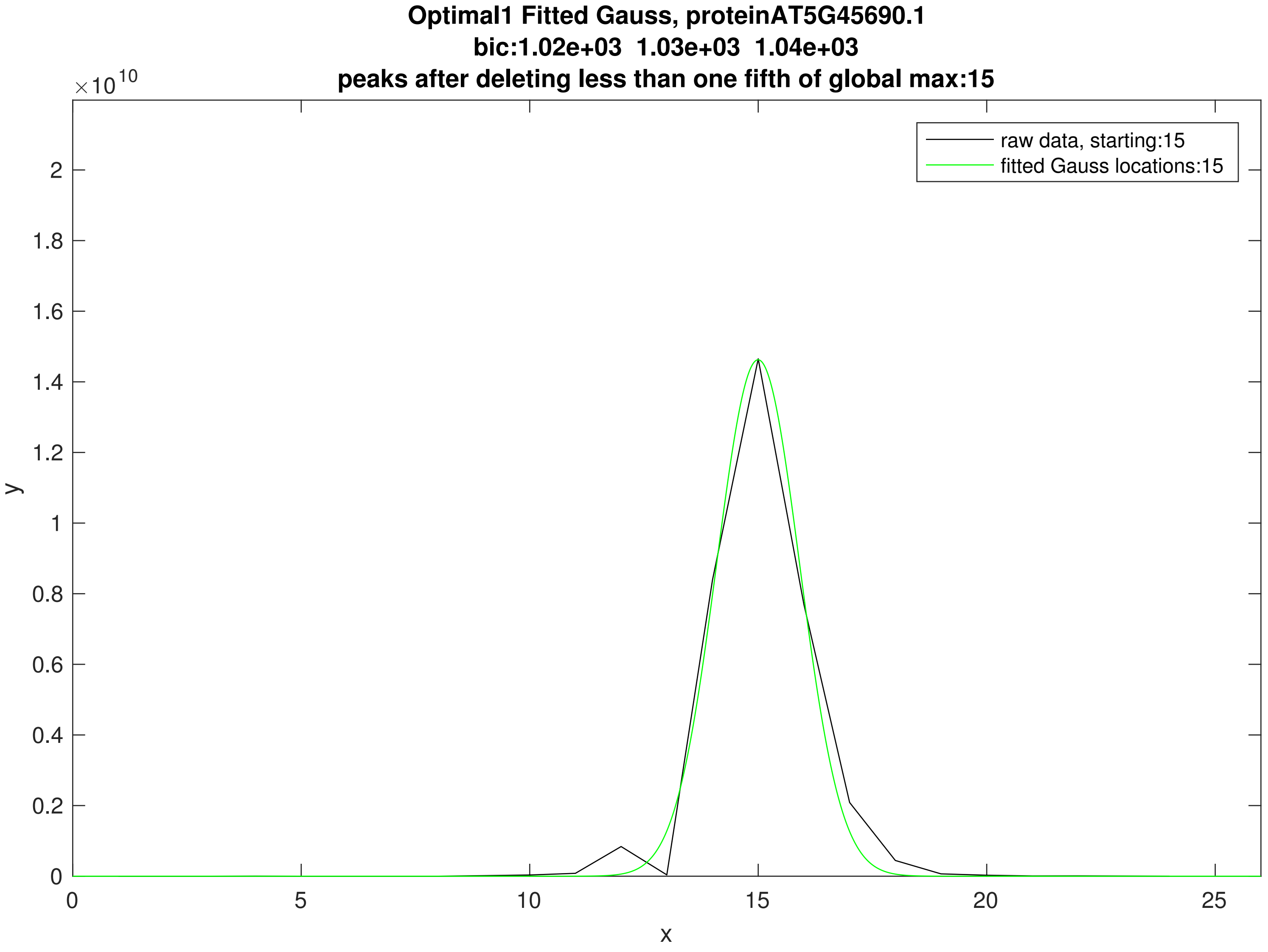
Optimal1 Fitted Gauss, proteinAT5G45690.1
bic:1.02e+03 1.03e+03 1.04e+03
peaks after deleting less than one fifth of global max:15
×10^10
raw data, starting:15
fitted Gauss locations:15
2
1.8
1.6
1.4
1.2
1
0.8
0.6
0.4
0.2
0
y
0
5
10
15
20
25
x

**Optimal1 Fitted Gauss, proteinAT5G45775.2;AT4G18730.1;AT3G58700.1;AT2G42740.1;AT5G45775.1**
**bic:876 882**
**peaks after deleting less than one fifth of global max:3.4**

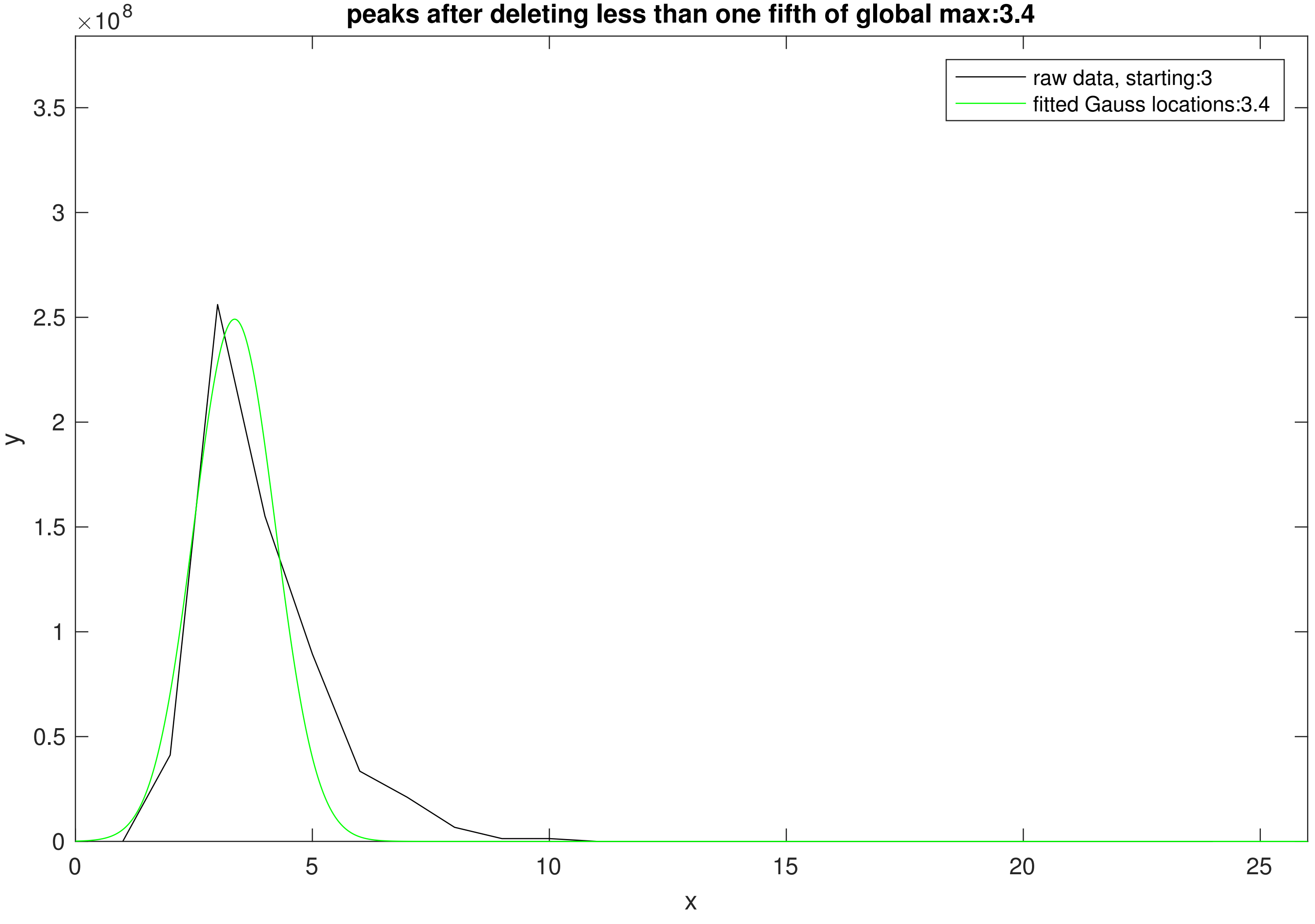

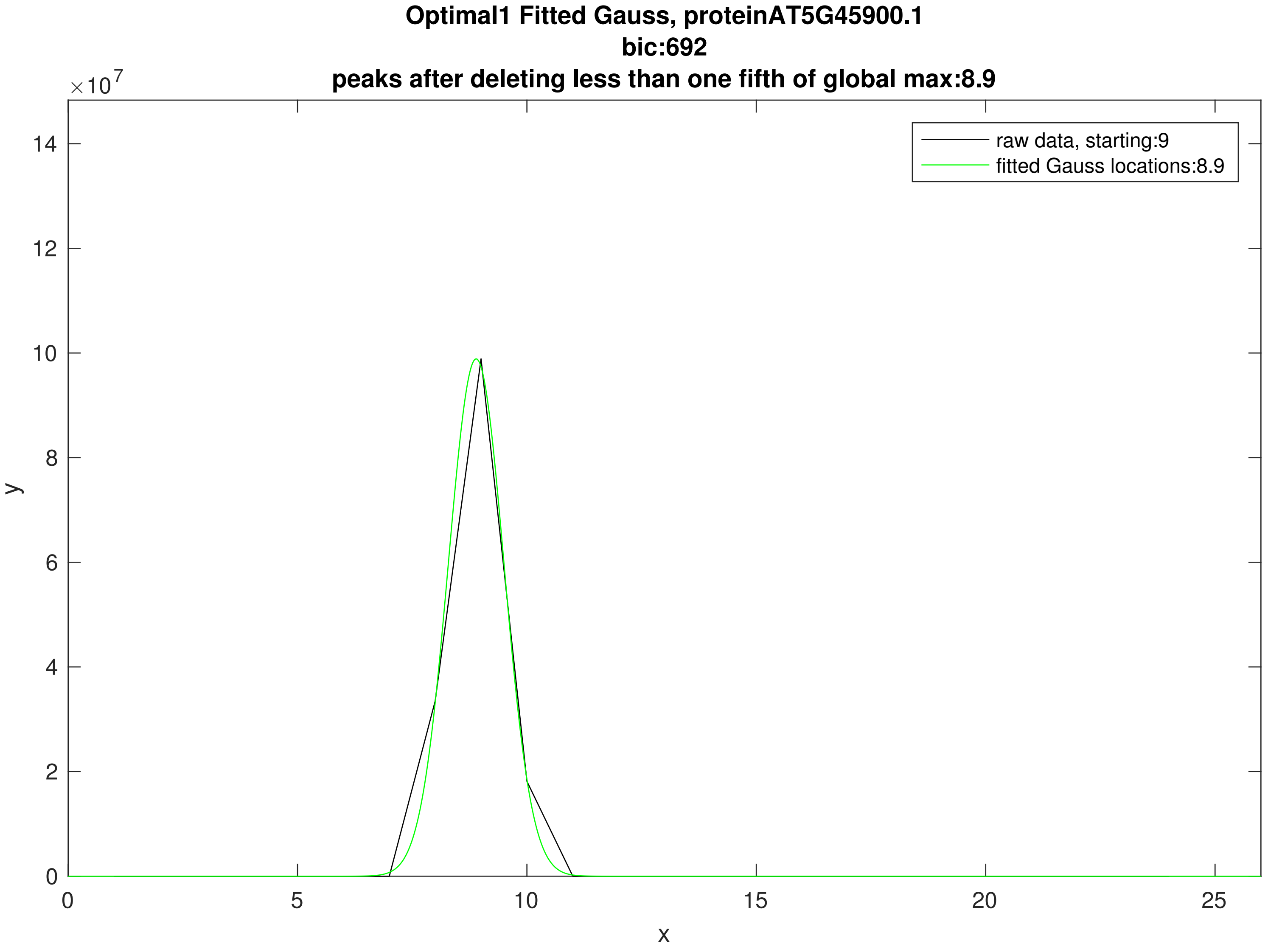
Optimal1 Fitted Gauss, proteinAT5G45900.1
bic:692
peaks after deleting less than one fifth of global max:8.9
×10^7
raw data, starting:9
fitted Gauss locations:8.9
14
12
10
8
6
4
2
0
y
0
5
10
15
20
25
x

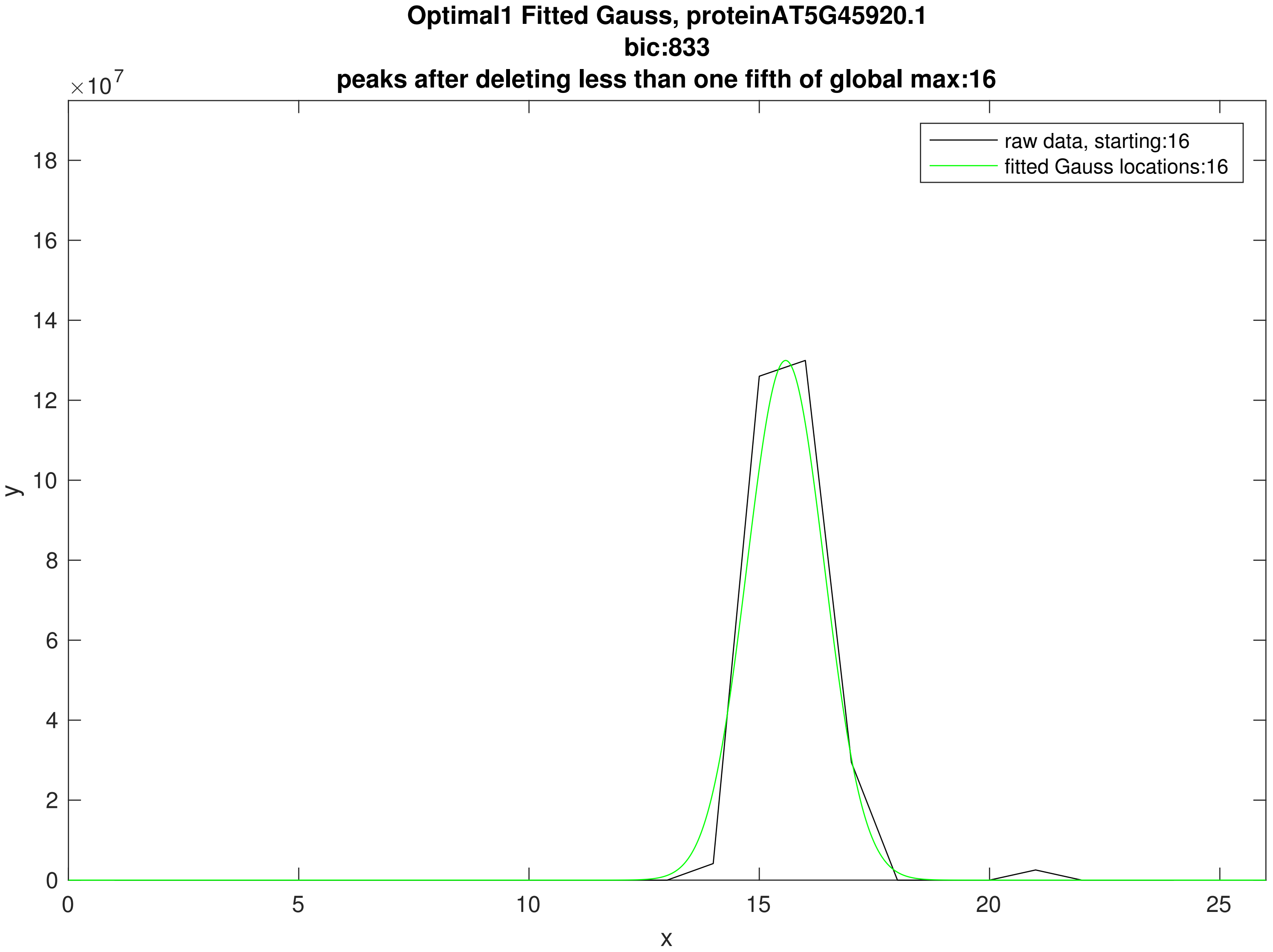
Optimal1 Fitted Gauss, proteinAT5G45920.1
bic:833
peaks after deleting less than one fifth of global max:16
×10^7
raw data, starting:16
fitted Gauss locations:16
y
x
0
2
4
6
8
10
12
14
16
18
5
15
20
25

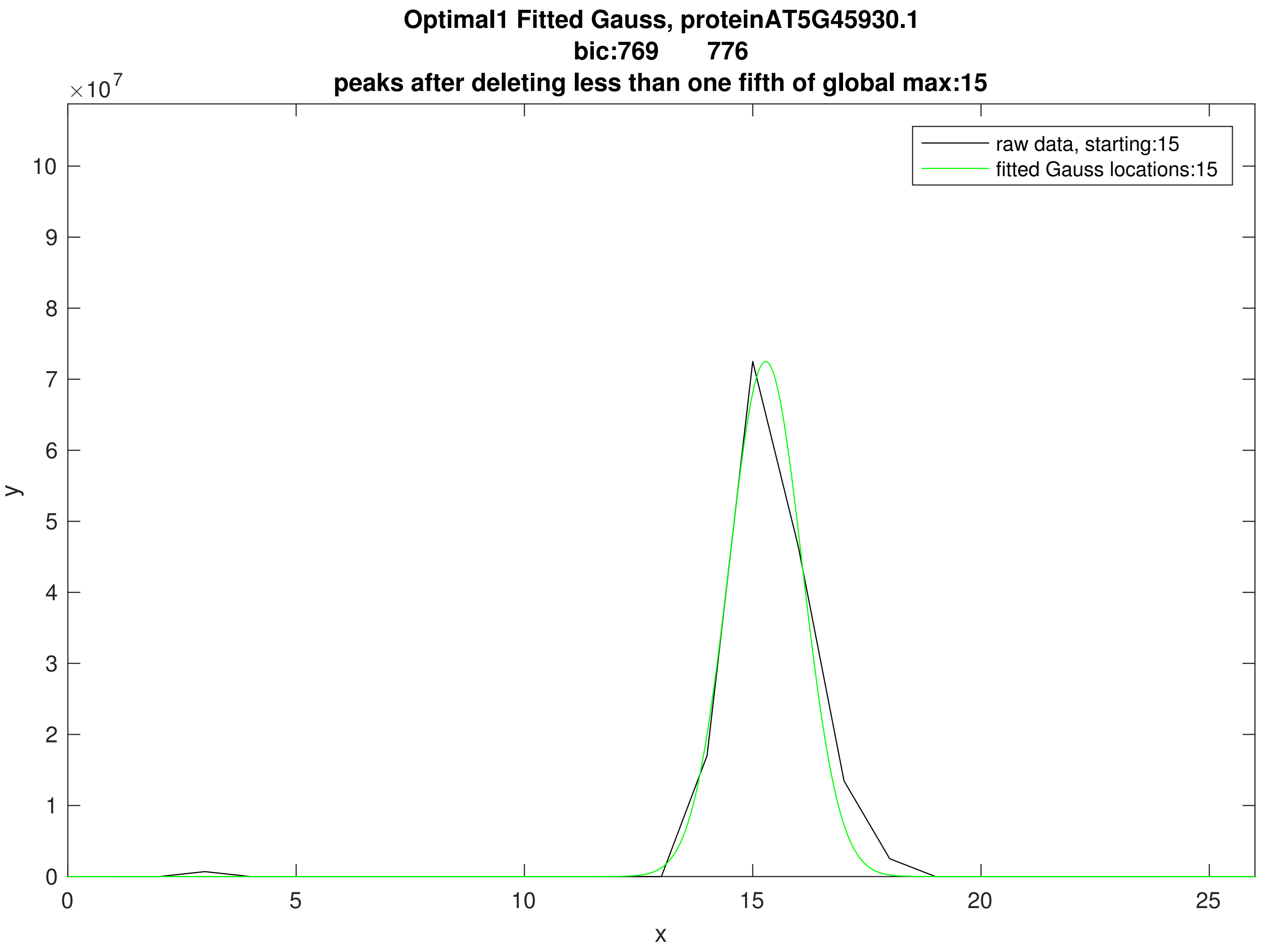
Optimal1 Fitted Gauss, proteinAT5G45930.1
bic:769 776
peaks after deleting less than one fifth of global max:15
×10^7
raw data, starting:15
fitted Gauss locations:15
y
x
0
1
2
3
4
5
6
7
8
9
10
0
5
10
15
20
25

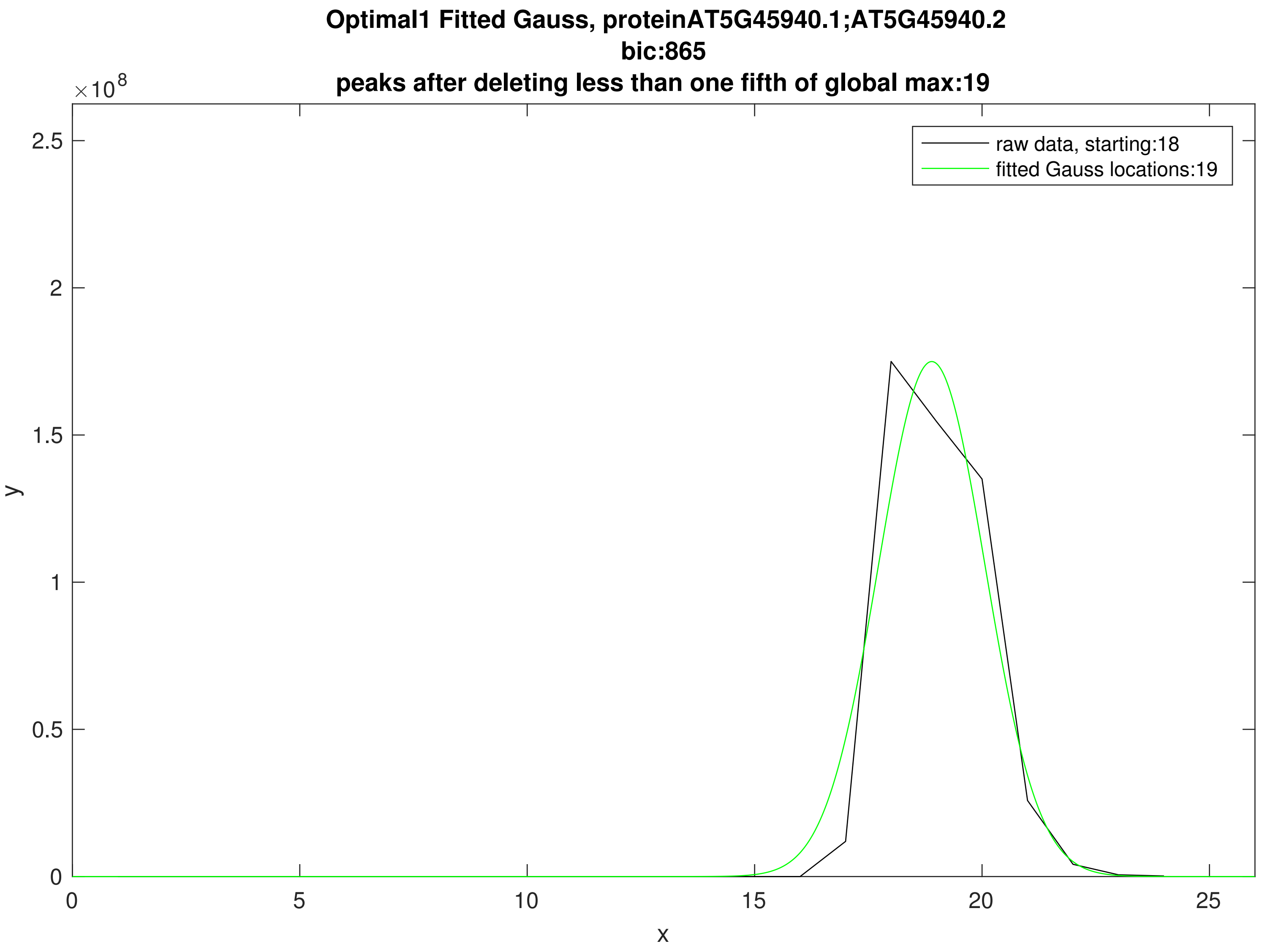

Optimal1 Fitted Gauss, proteinAT5G45940.1;AT5G45940.2
bic:865
peaks after deleting less than one fifth of global max:19
×10^8
raw data, starting:18
fitted Gauss locations:19
y
x
0
0.5
1
1.5
2
2.5
5
10
15
20
25

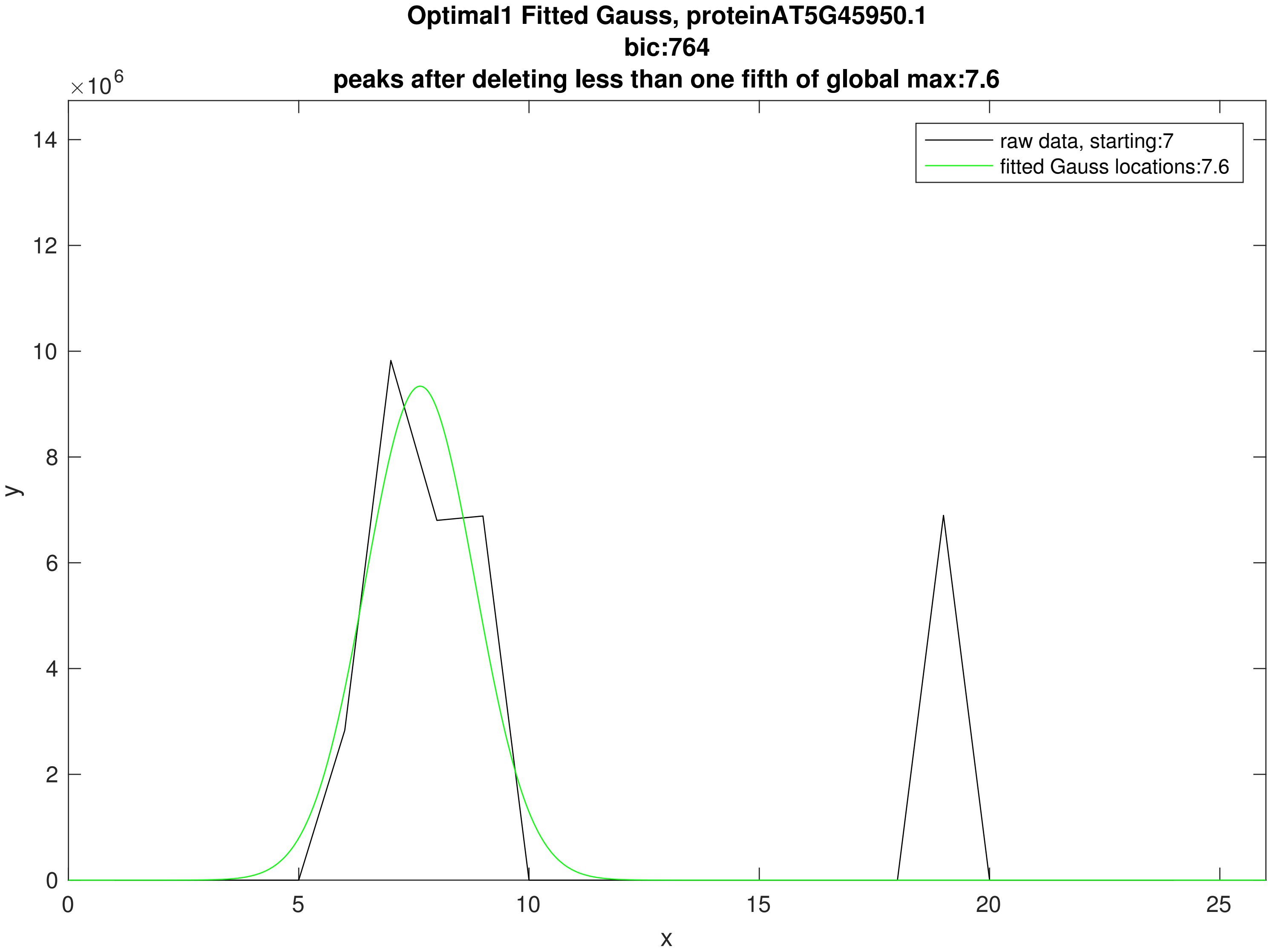

Optimal1 Fitted Gauss, proteinAT5G45950.1
bic:764
peaks after deleting less than one fifth of global max:7.6
×10^6
raw data, starting:7
fitted Gauss locations:7.6
y
x
0
2
4
6
8
10
12
14
0
5
10
15
20
25

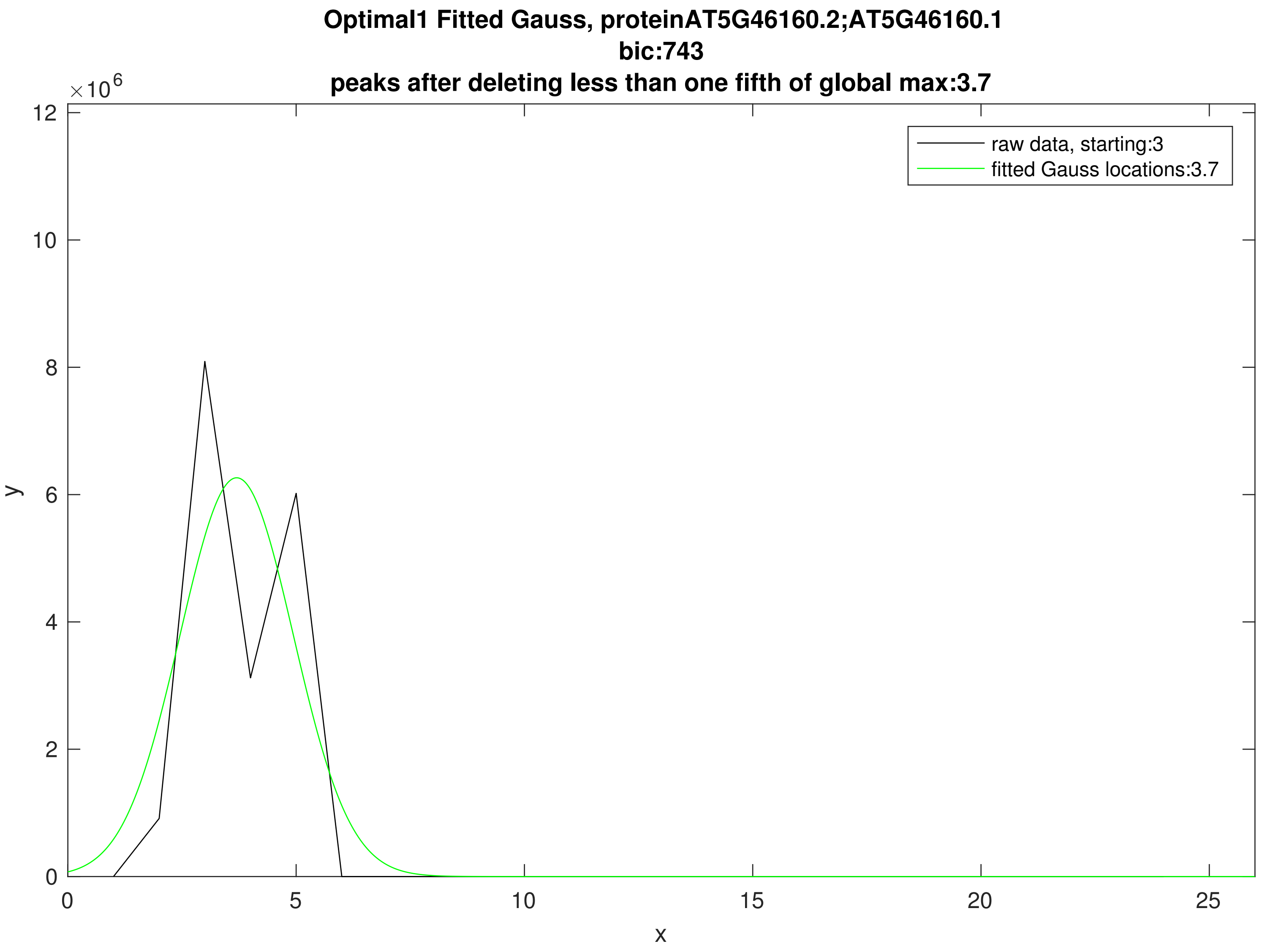
Optimal1 Fitted Gauss, proteinAT5G46160.2;AT5G46160.1
bic:743
peaks after deleting less than one fifth of global max:3.7
raw data, starting:3
fitted Gauss locations:3.7
y
x

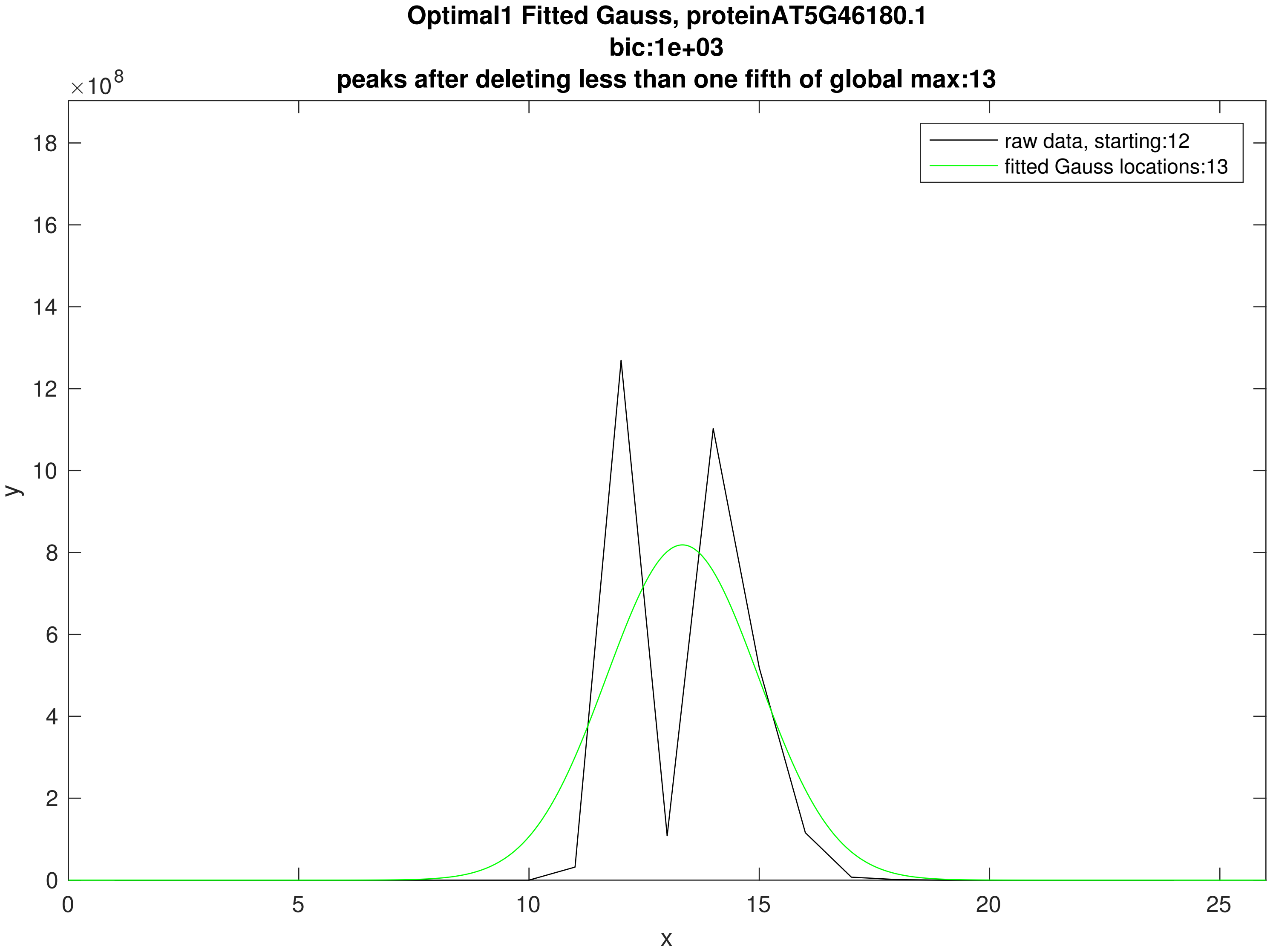
Optimal1 Fitted Gauss, proteinAT5G46180.1
bic:1e+03
peaks after deleting less than one fifth of global max:13
×10^8
raw data, starting:12
fitted Gauss locations:13
y
x
0
2
4
6
8
10
12
14
16
18
0
5
10
15
20
25

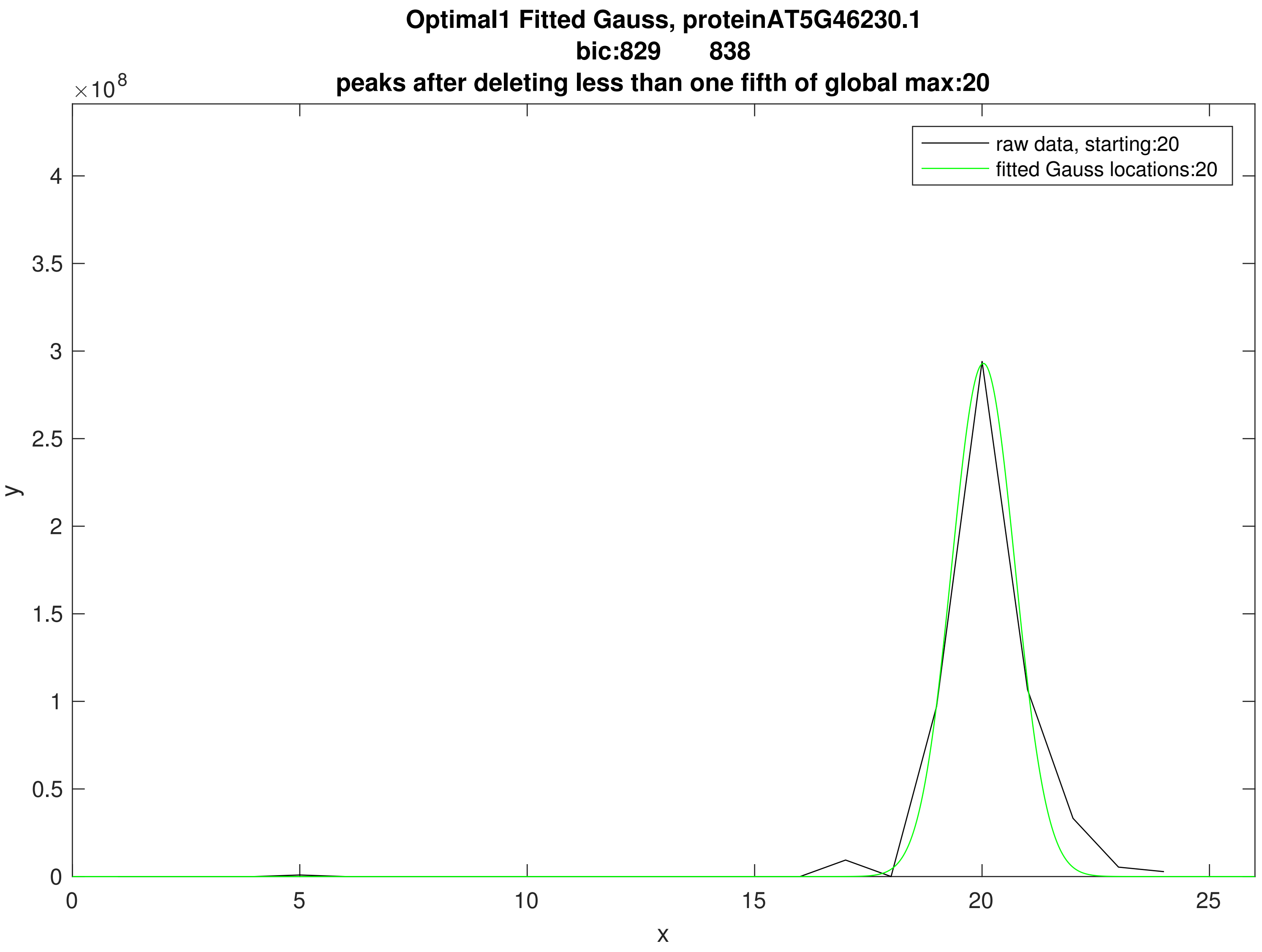
Optimal1 Fitted Gauss, proteinAT5G46230.1
bic:829 838
peaks after deleting less than one fifth of global max:20
×10^8
raw data, starting:20
fitted Gauss locations:20
y
x
0
0.5
1
1.5
2
2.5
3
3.5
4
5
10
15
20
25

Optimal2 Fitted Gauss, proteinAT5G46260.1
bic:869 747
peaks after deleting less than one fifth of global max:8.8 16

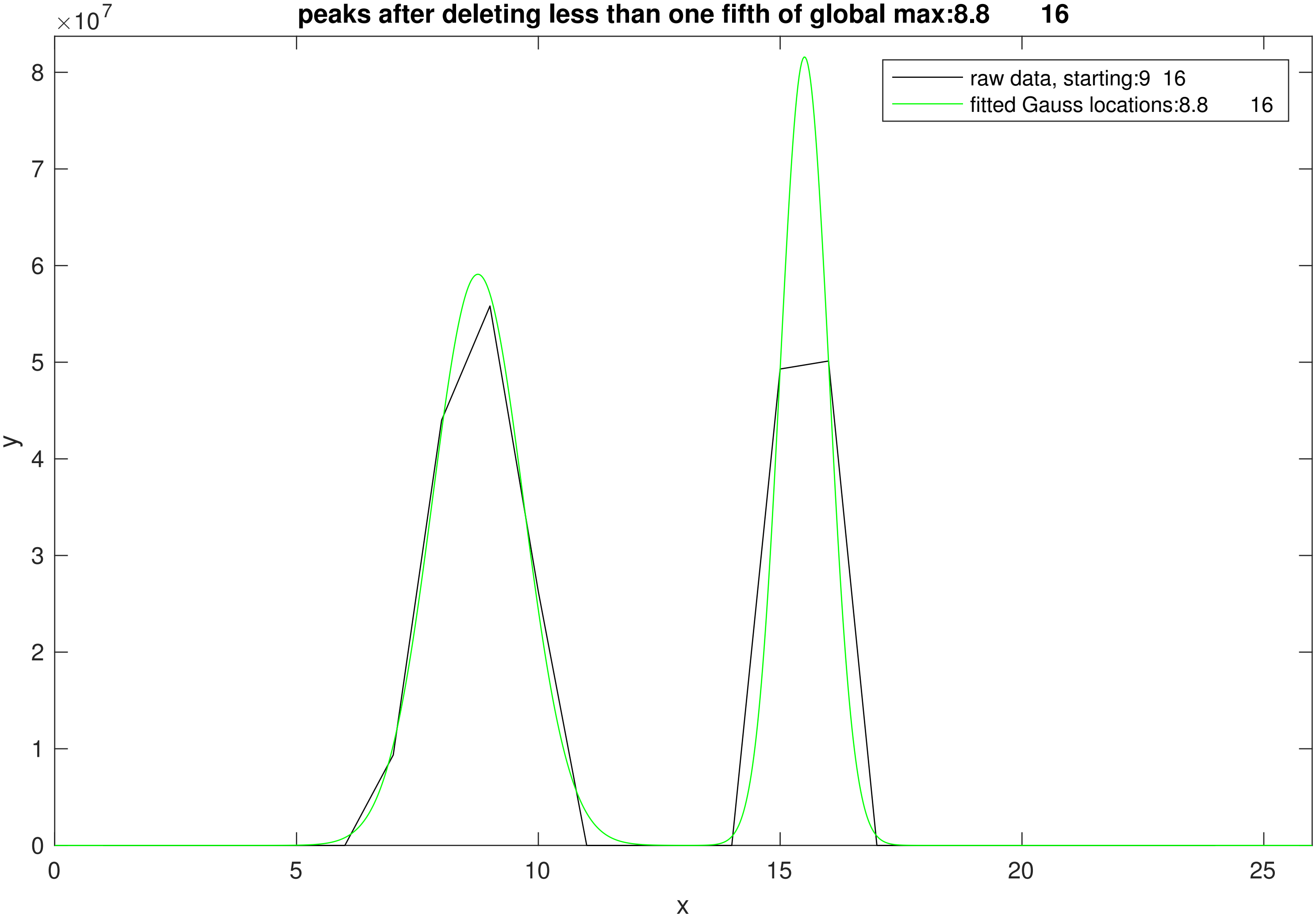

#### Optimal1 Fitted Gauss, proteinAT5G46290.1;AT5G46290.2;AT5G46290.3
#### bic:1.01e+03  1.02e+03  1.03e+03
#### peaks after deleting less than one fifth of global max:15

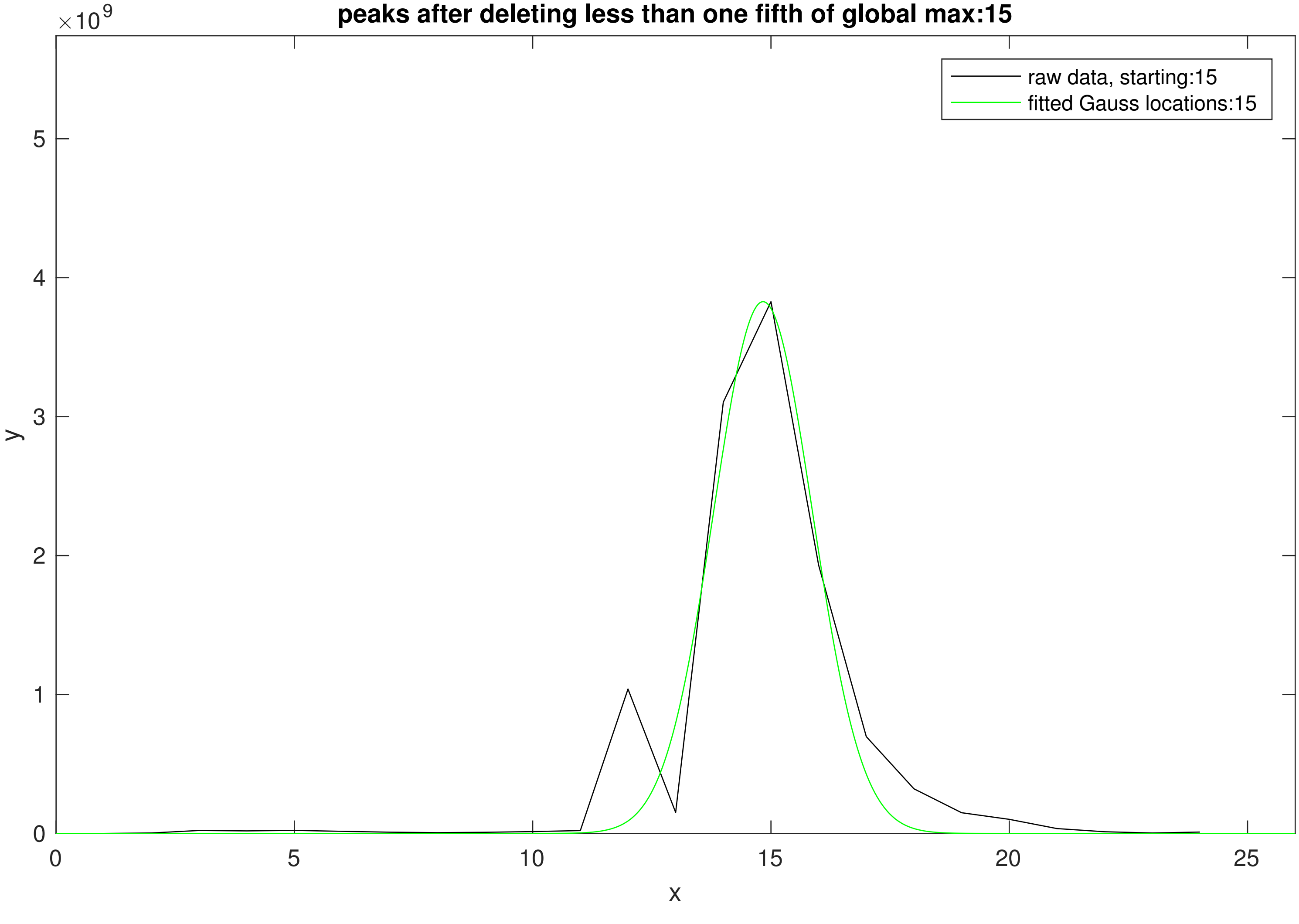

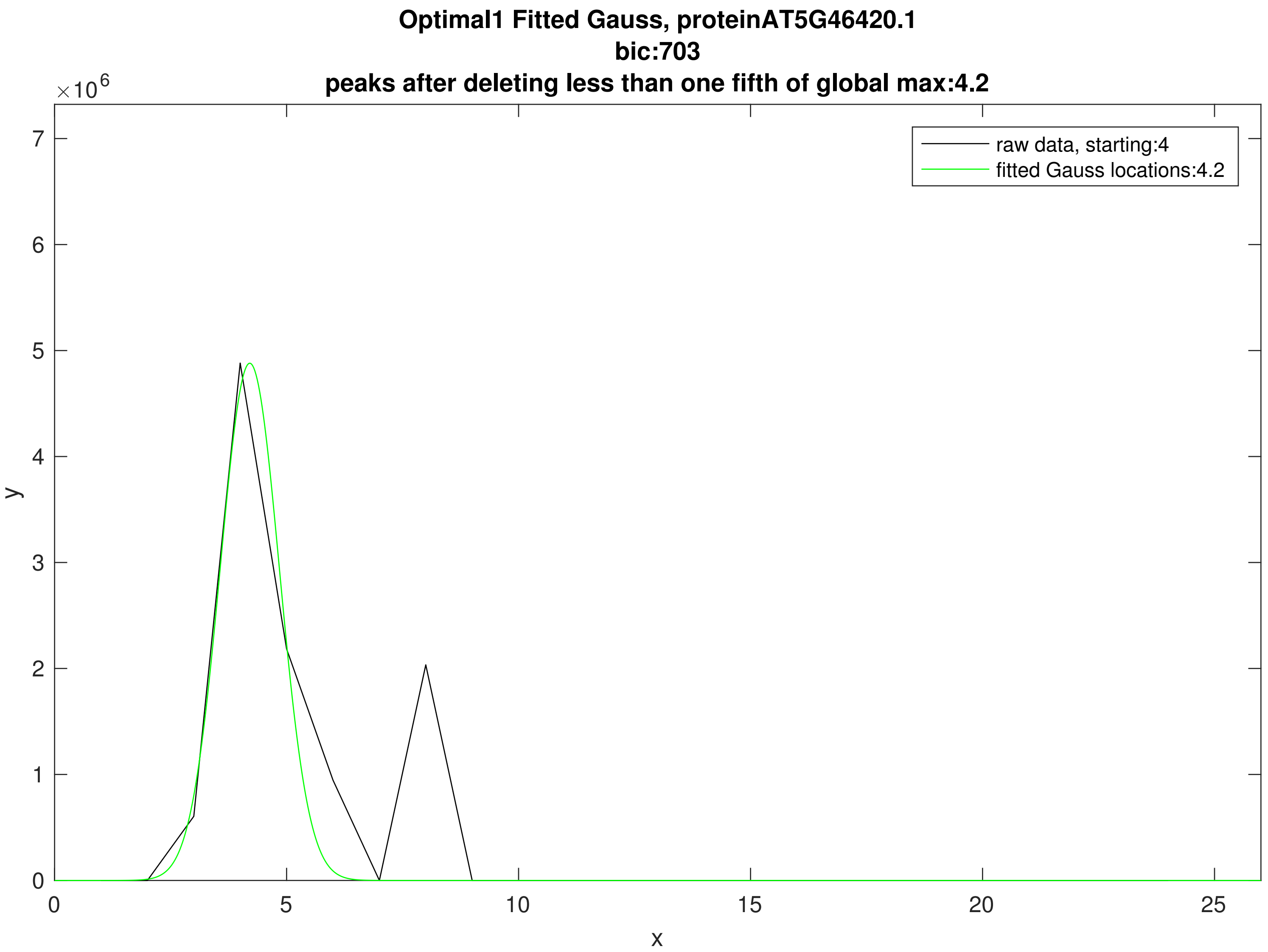

Optimal1 Fitted Gauss, proteinAT5G46420.1
bic:703
peaks after deleting less than one fifth of global max:4.2
×10^6
raw data, starting:4
fitted Gauss locations:4.2
y
x
0
1
2
3
4
5
6
7
0
5
10
15
20
25

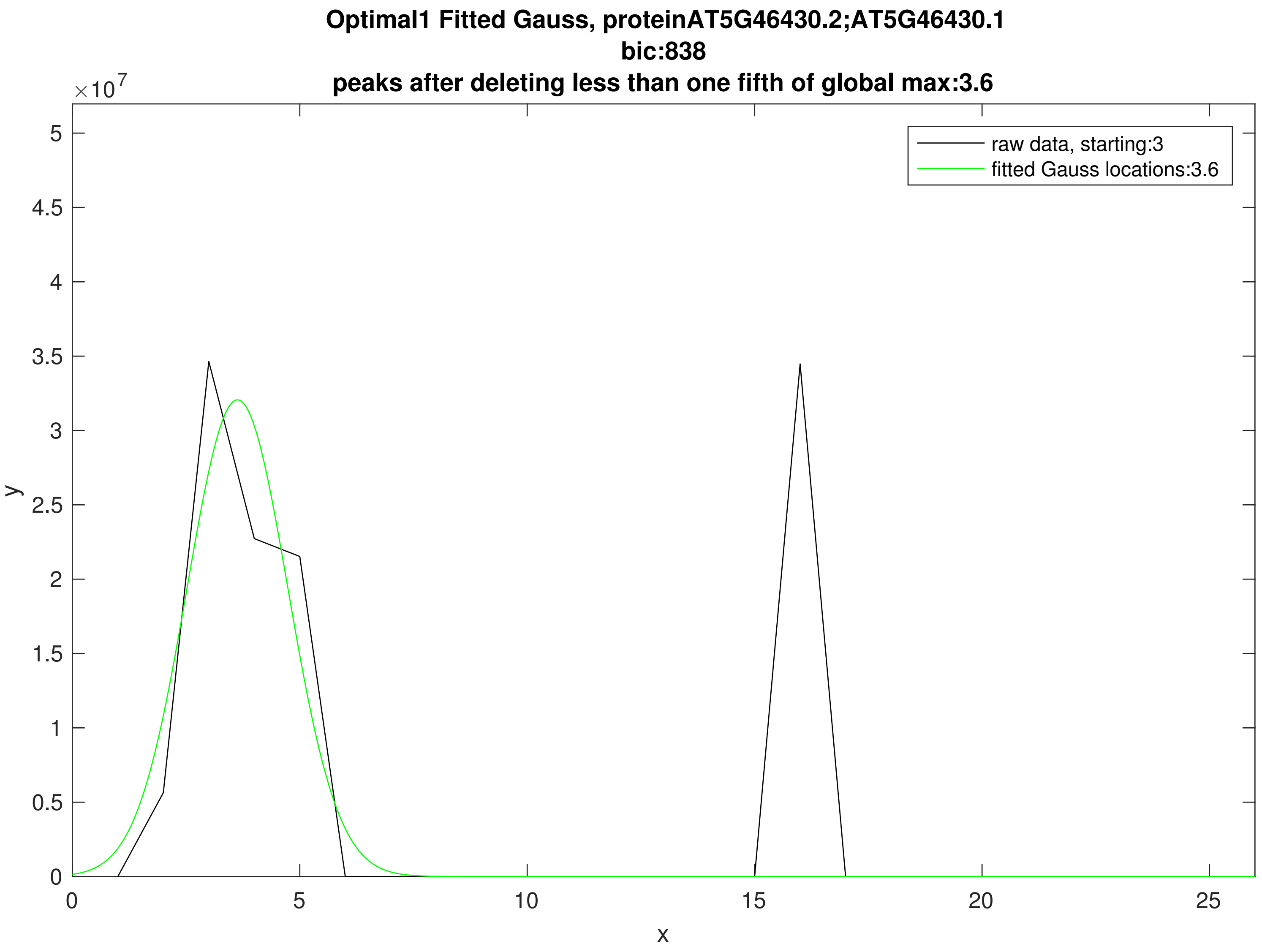

Optimal1 Fitted Gauss, proteinAT5G46430.2;AT5G46430.1
bic:838
peaks after deleting less than one fifth of global max:3.6
×10^7
raw data, starting:3
fitted Gauss locations:3.6
y
x
0
0.5
1
1.5
2
2.5
3
3.5
4
4.5
5
0
5
10
15
20
25

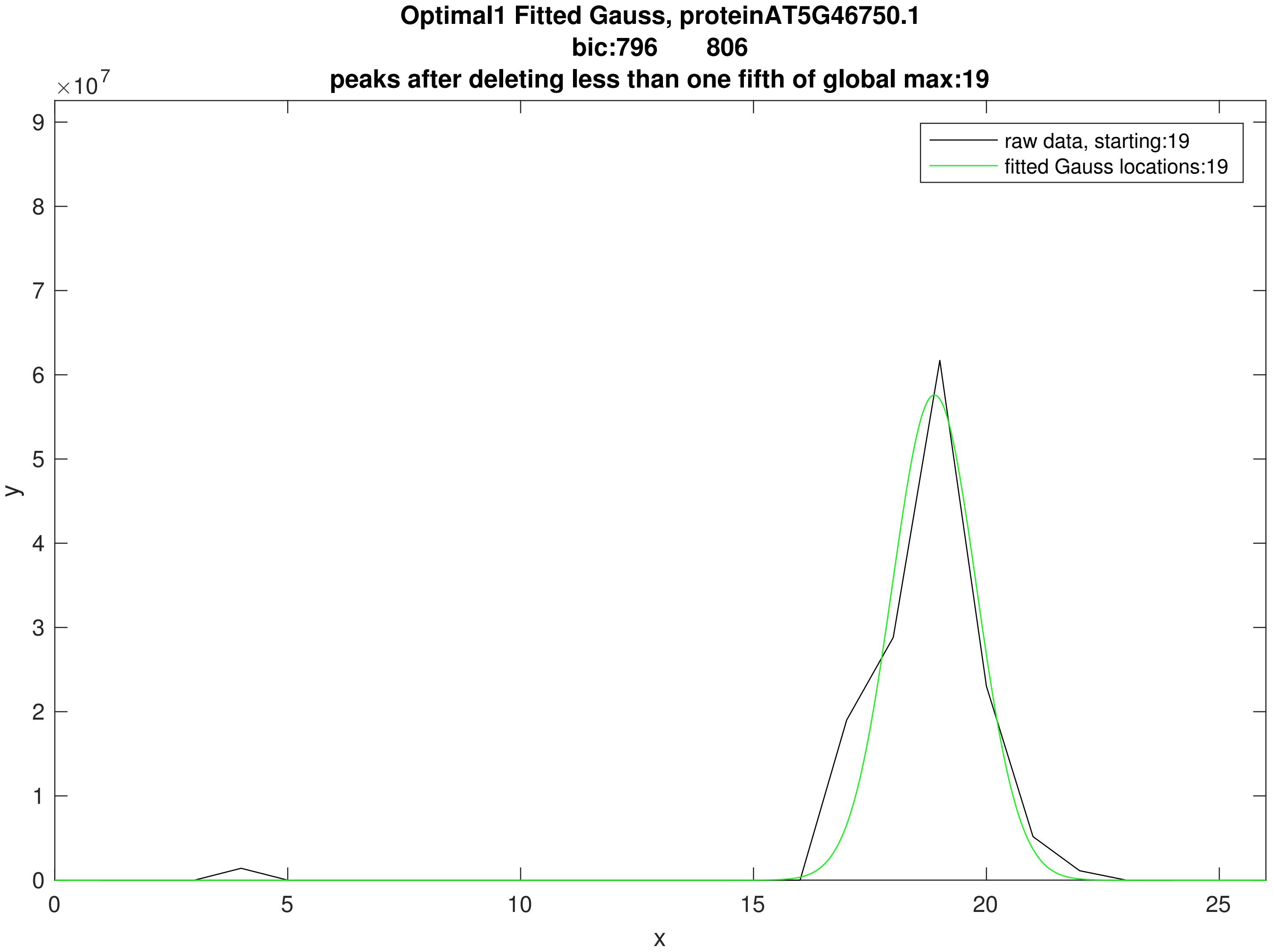
Optimal1 Fitted Gauss, proteinAT5G46750.1
bic:796 806
peaks after deleting less than one fifth of global max:19
×10^7
raw data, starting:19
fitted Gauss locations:19
y
x
0
1
2
3
4
5
6
7
8
9
0
5
10
15
20
25

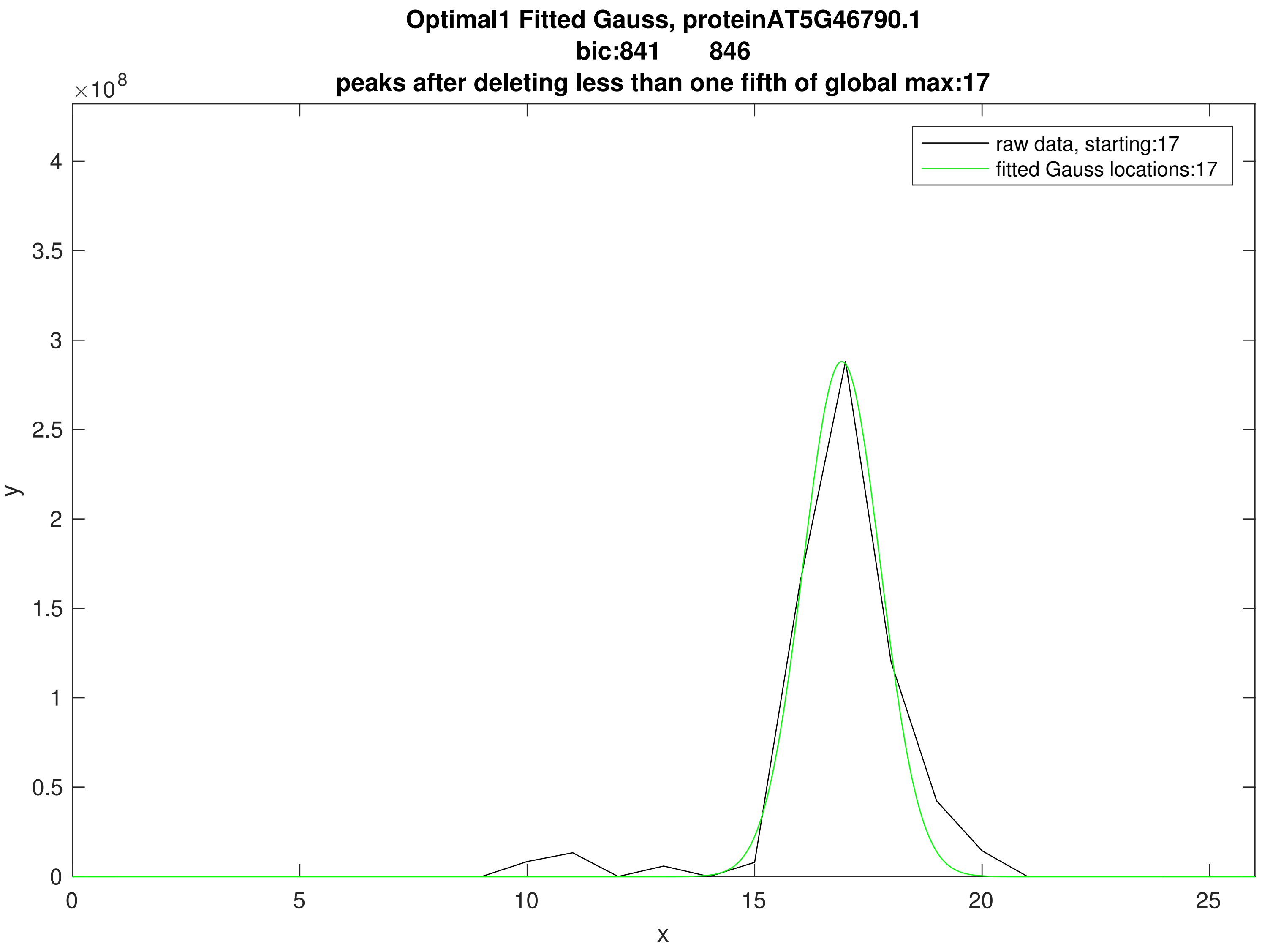
Optimal1 Fitted Gauss, proteinAT5G46790.1
bic:841 846
peaks after deleting less than one fifth of global max:17
×10^8
raw data, starting:17
fitted Gauss locations:17
y
x
0
0.5
1
1.5
2
2.5
3
3.5
4
0
5
10
15
20
25

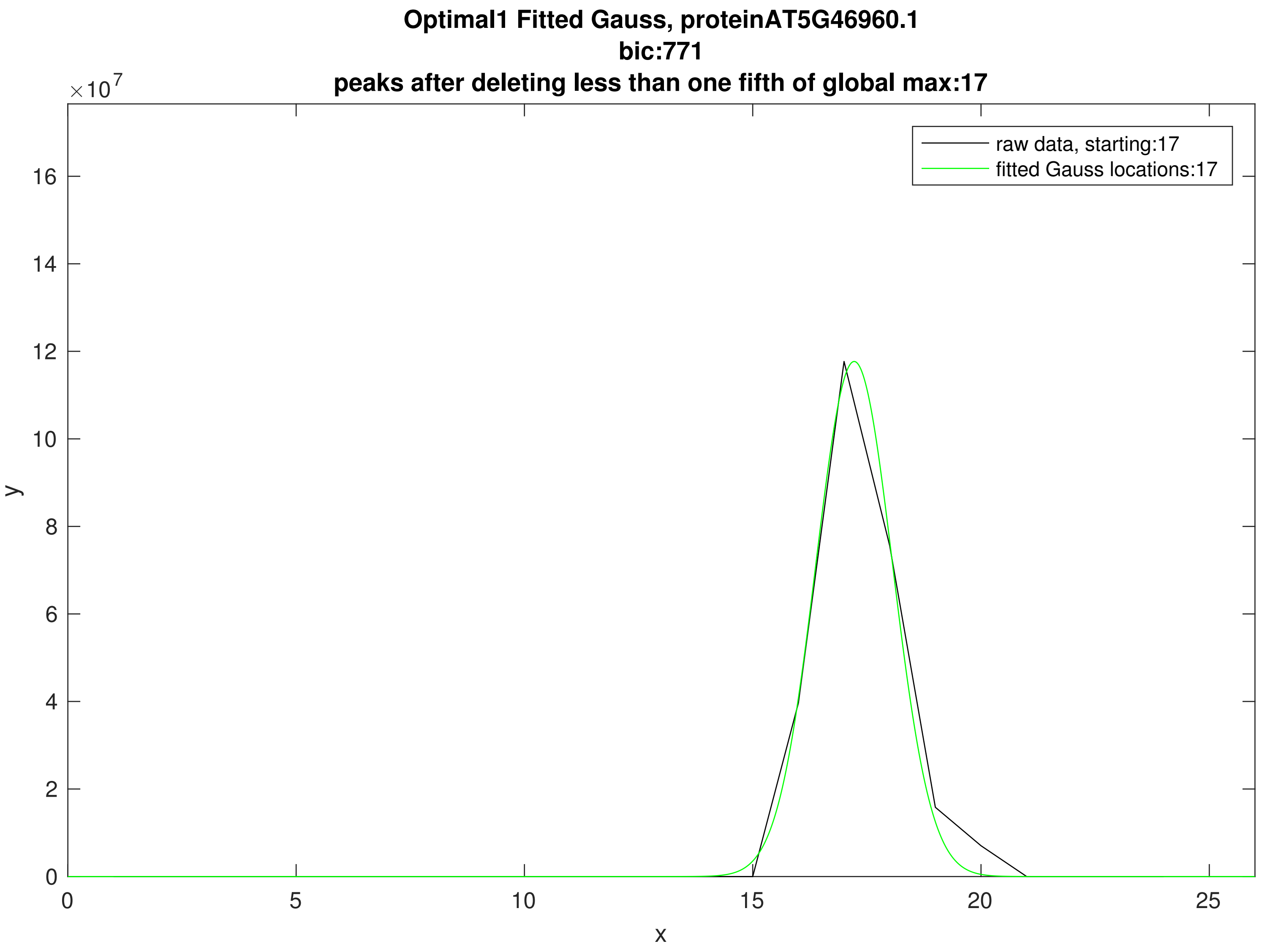
Optimal1 Fitted Gauss, proteinAT5G46960.1
bic:771
peaks after deleting less than one fifth of global max:17
×10^7
raw data, starting:17
fitted Gauss locations:17
y
x
0
2
4
6
8
10
12
14
16
0
5
10
15
20
25

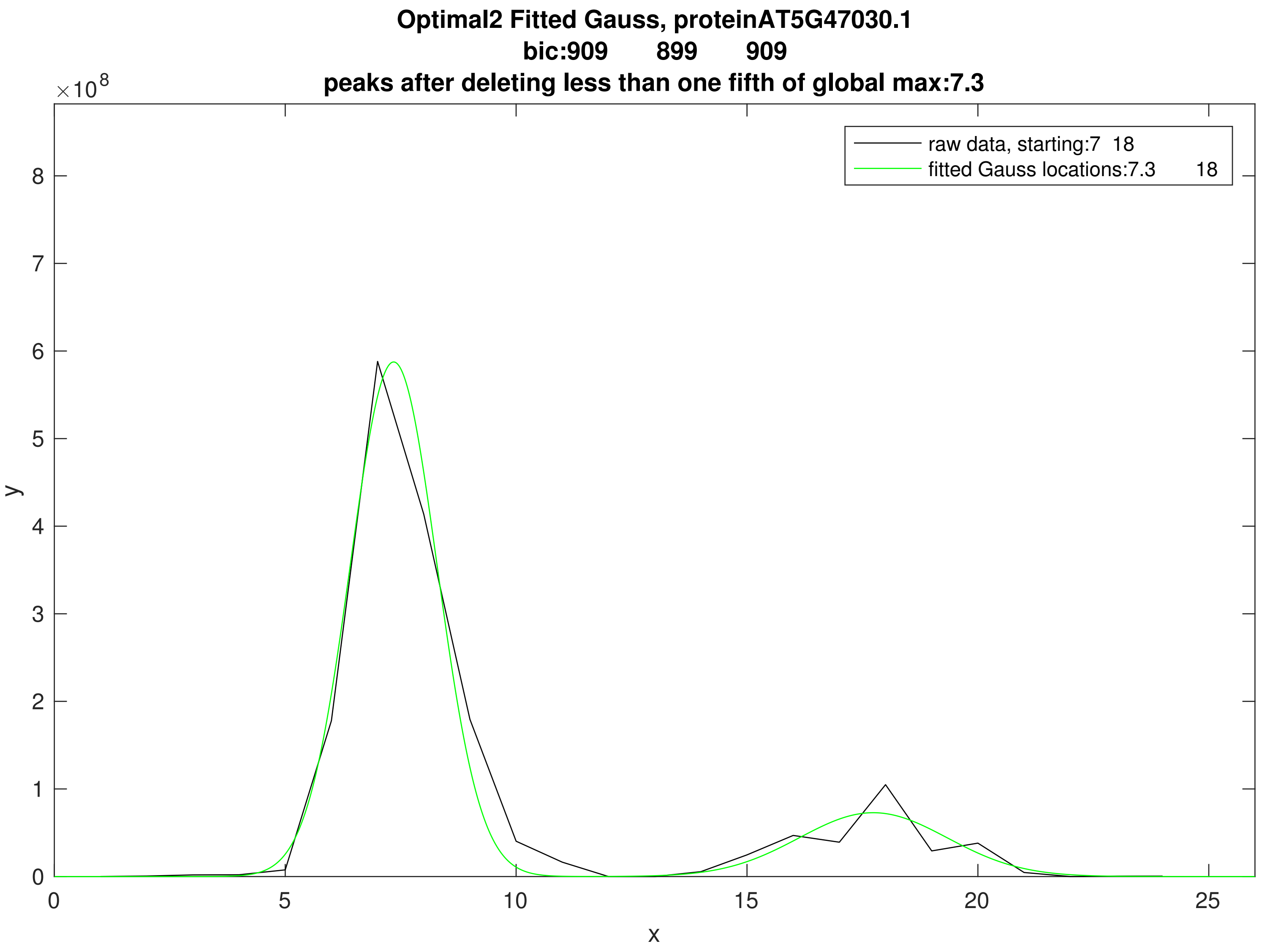
Optimal2 Fitted Gauss, proteinAT5G47030.1
bic:909 899 909
peaks after deleting less than one fifth of global max:7.3
×10^8
raw data, starting:7 18
fitted Gauss locations:7.3 18
y
x
0
1
2
3
4
5
6
7
8
0
5
10
15
20
25

**Optimal2 Fitted Gauss, proteinAT5G47190.1**
**bic:833 810**
**peaks after deleting less than one fifth of global max:4.8 17**

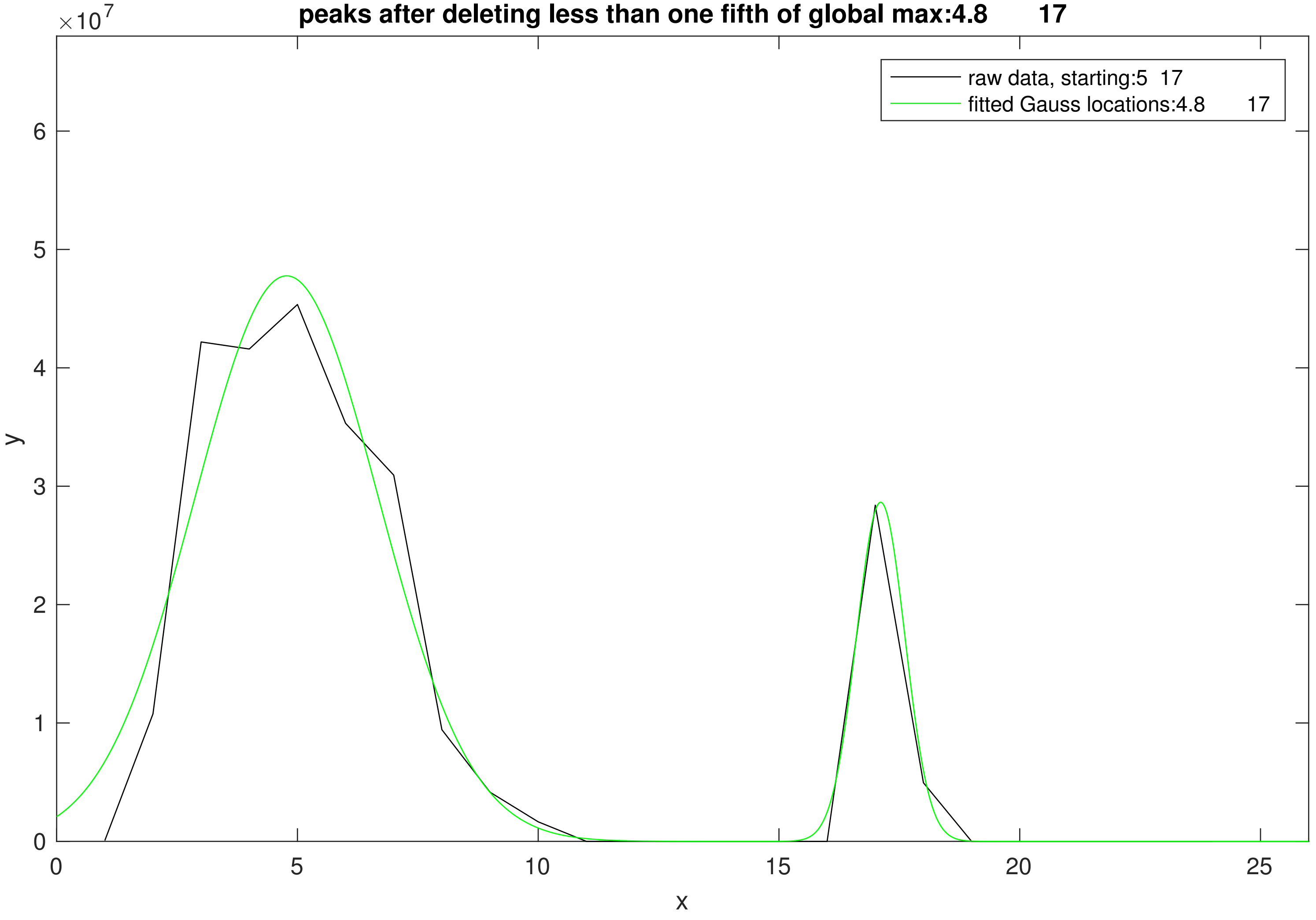

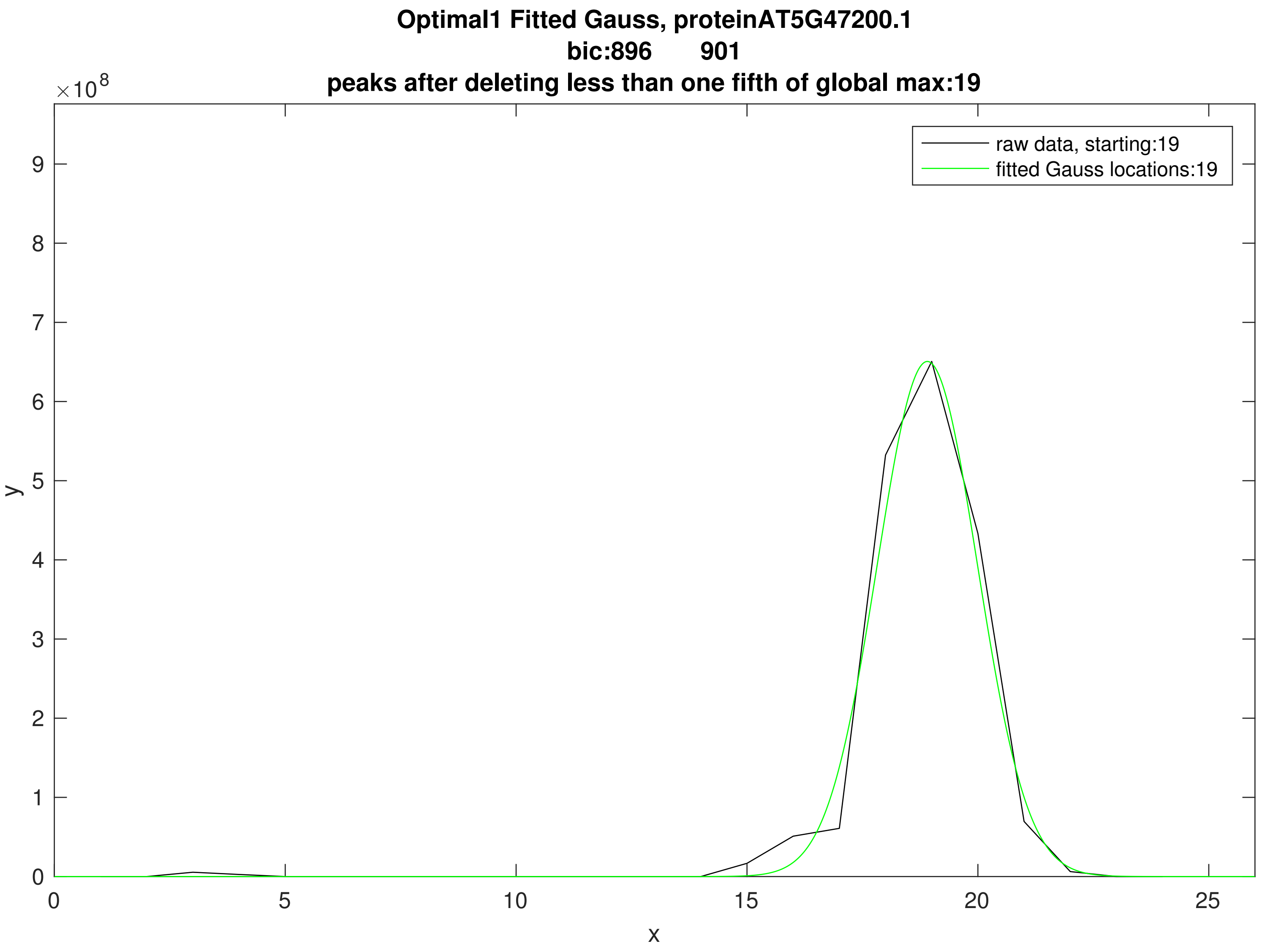
Optimal1 Fitted Gauss, proteinAT5G47200.1
bic:896 901
peaks after deleting less than one fifth of global max:19
×10^8
raw data, starting:19
fitted Gauss locations:19
y
x
0
1
2
3
4
5
6
7
8
9
0
5
10
15
20
25

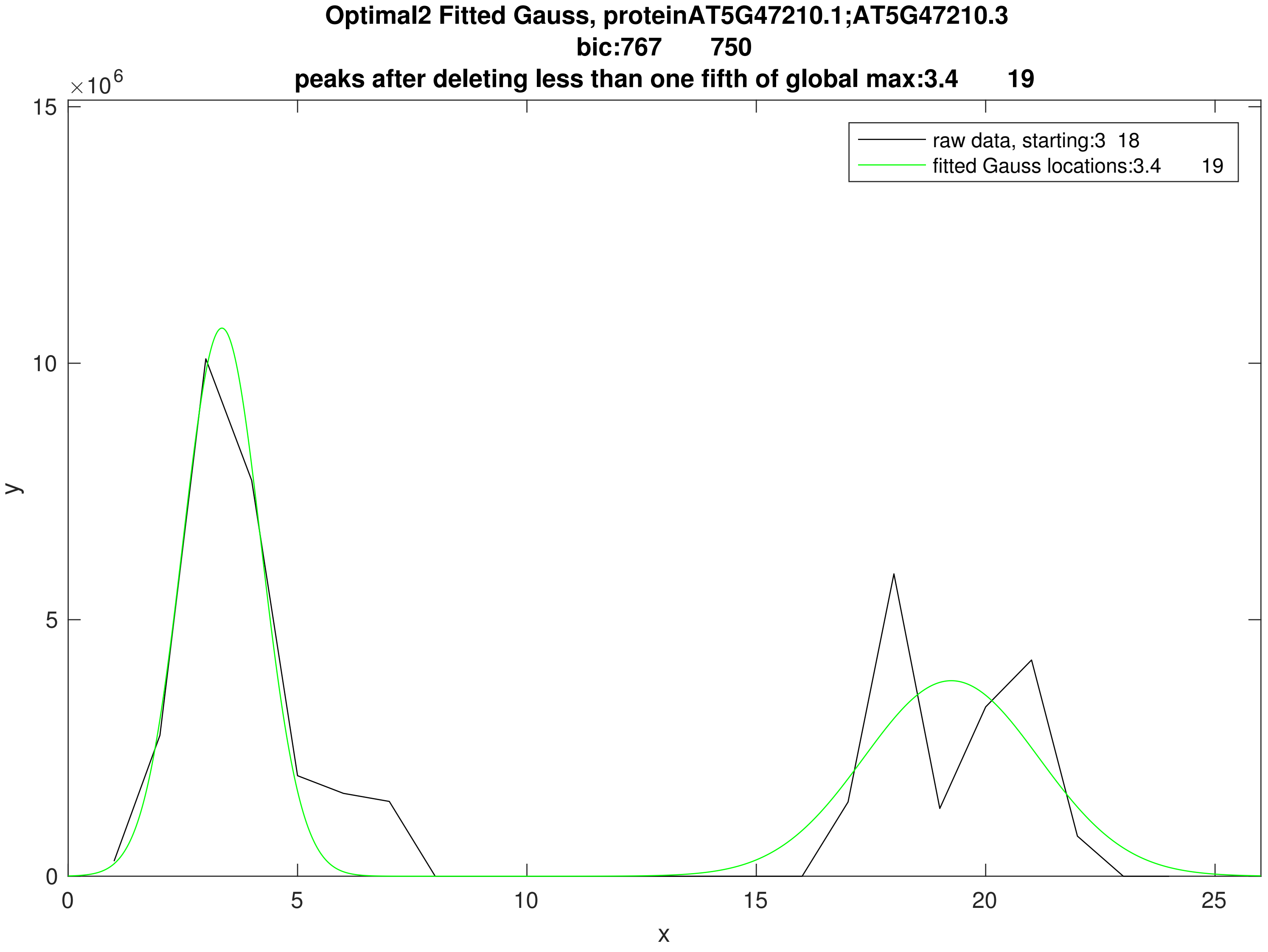

Optimal2 Fitted Gauss, proteinAT5G47210.1;AT5G47210.3
bic:767 750
peaks after deleting less than one fifth of global max:3.4 19
×10^6
raw data, starting:3 18
fitted Gauss locations:3.4 19
y
x
0
5
10
15
20
25

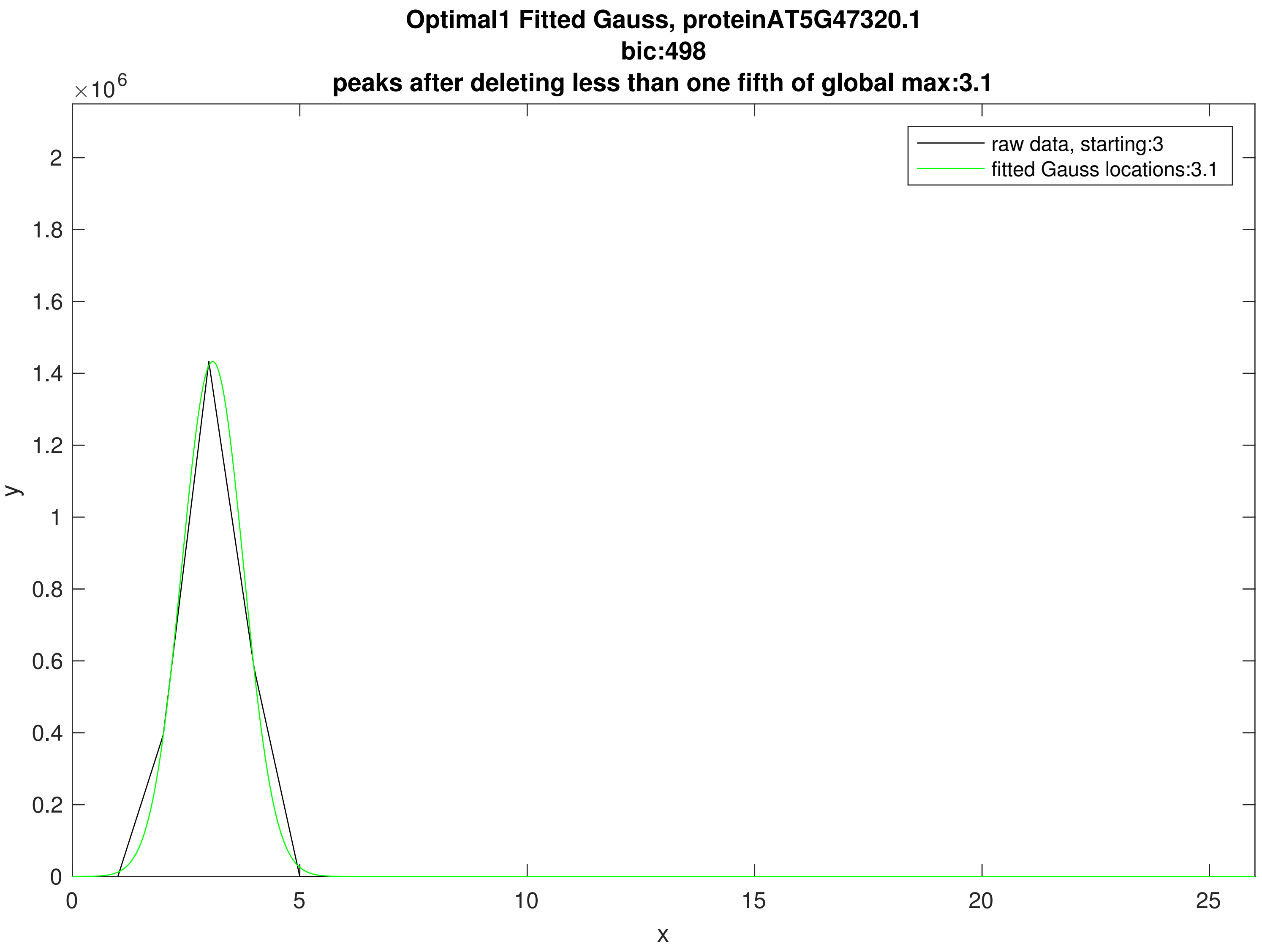
Optimal1 Fitted Gauss, proteinAT5G47320.1
bic:498
peaks after deleting less than one fifth of global max:3.1
×10^6
raw data, starting:3
fitted Gauss locations:3.1
y
x
0
0.2
0.4
0.6
0.8
1
1.2
1.4
1.6
1.8
2
0
5
10
15
20
25

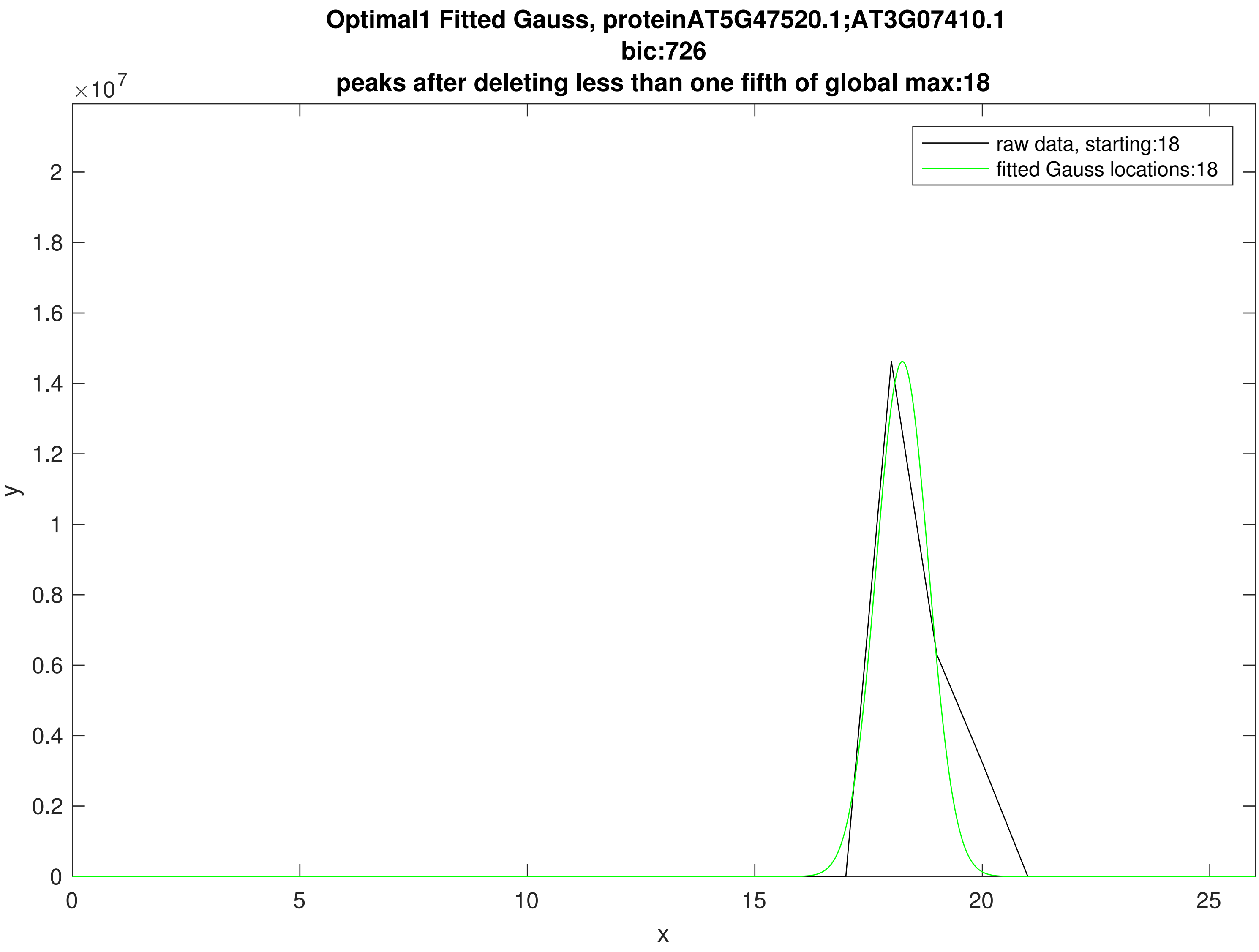

Optimal1 Fitted Gauss, proteinAT5G47520.1;AT3G07410.1
bic:726
peaks after deleting less than one fifth of global max:18
×10^7
2
1.8
1.6
1.4
1.2
1
0.8
0.6
0.4
0.2
0
y
0
5
10
15
20
25
x
raw data, starting:18
fitted Gauss locations:18

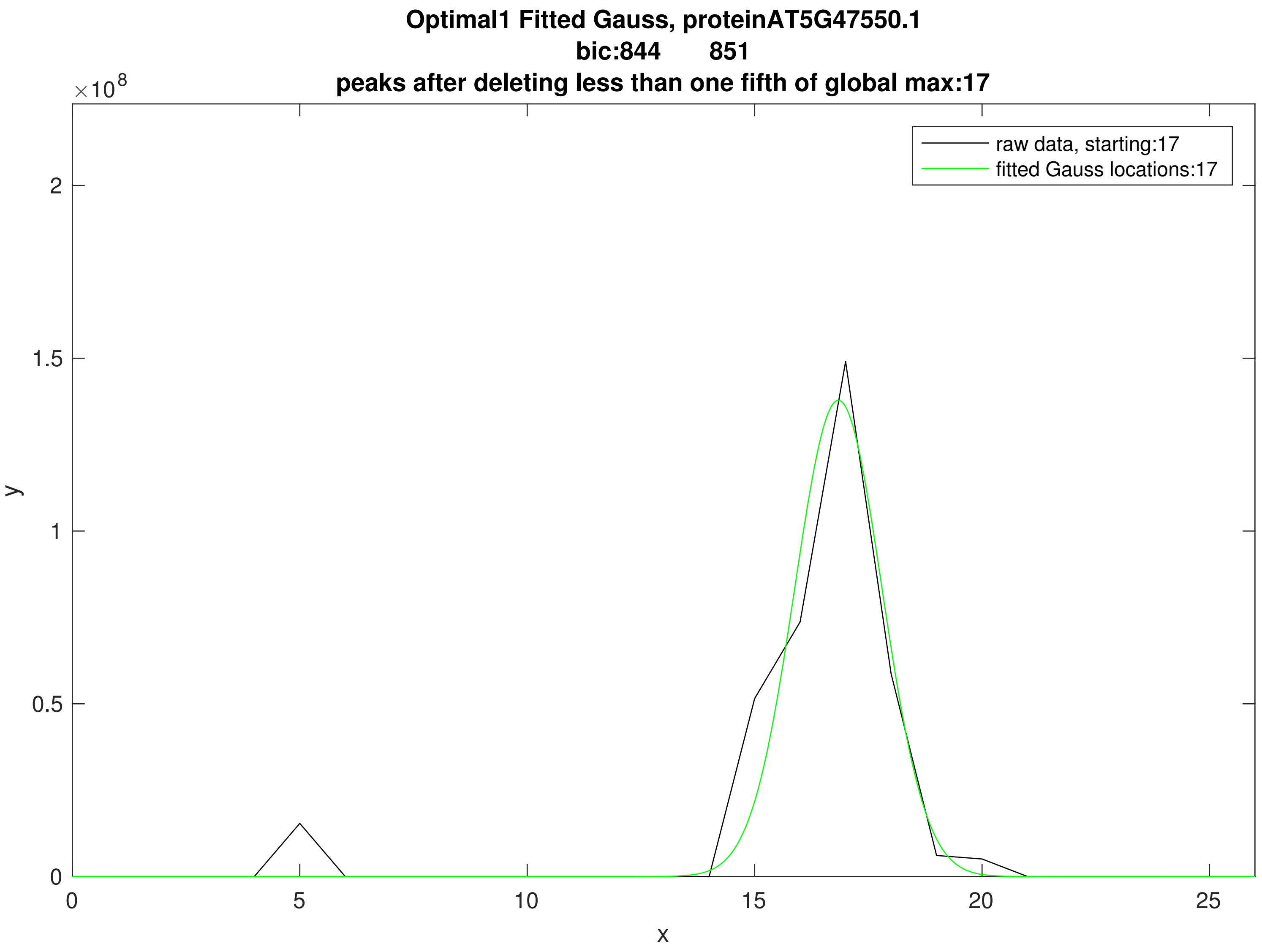
Optimal1 Fitted Gauss, proteinAT5G47550.1
bic:844 851
peaks after deleting less than one fifth of global max:17
×10^8
raw data, starting:17
fitted Gauss locations:17
y
x
0
0.5
1
1.5
2
0
5
10
15
20
25

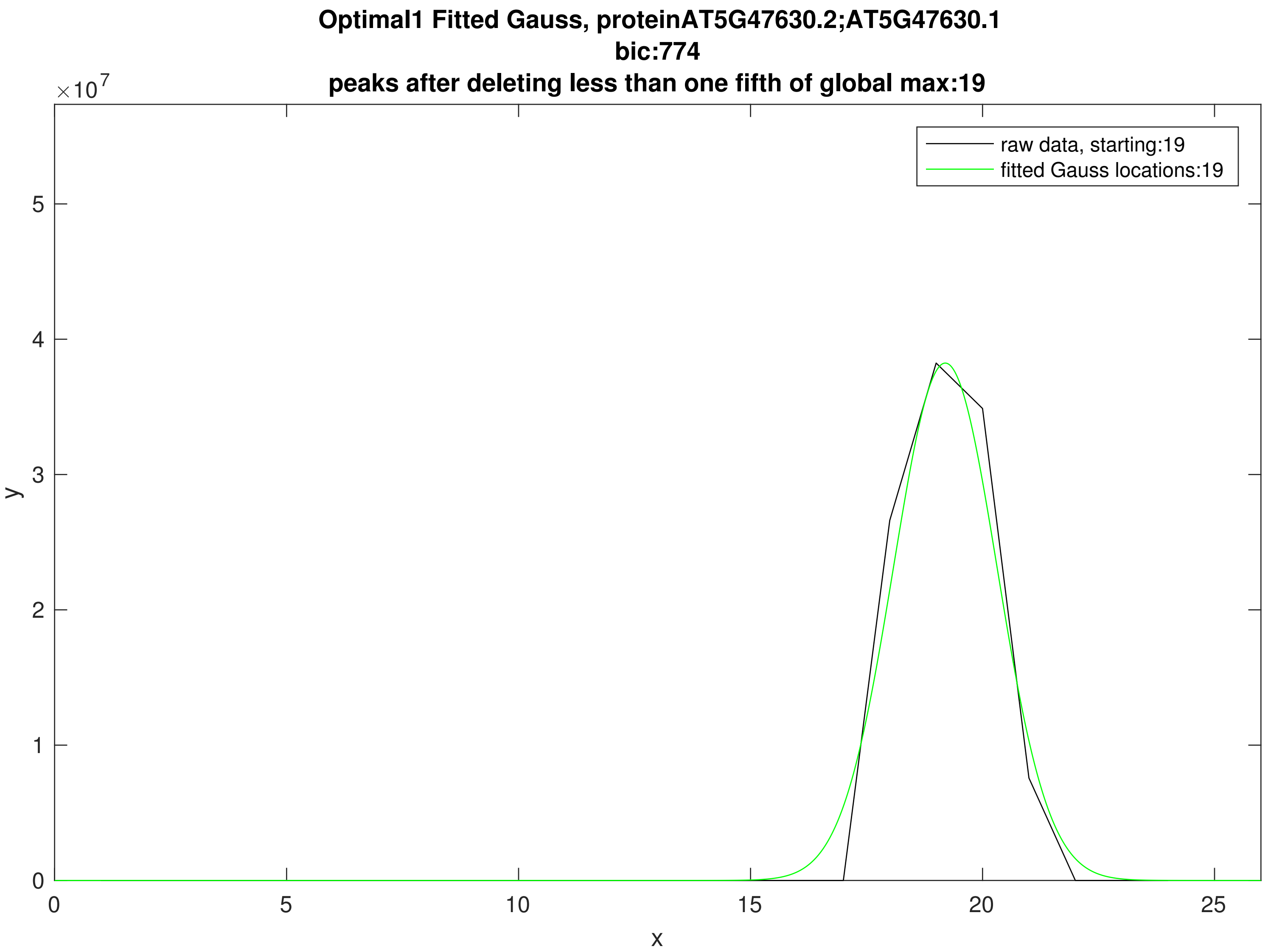
Optimal1 Fitted Gauss, proteinAT5G47630.2;AT5G47630.1
bic:774
peaks after deleting less than one fifth of global max:19
×10^7
raw data, starting:19
fitted Gauss locations:19
y
x
0
1
2
3
4
5
0
5
10
15
20
25

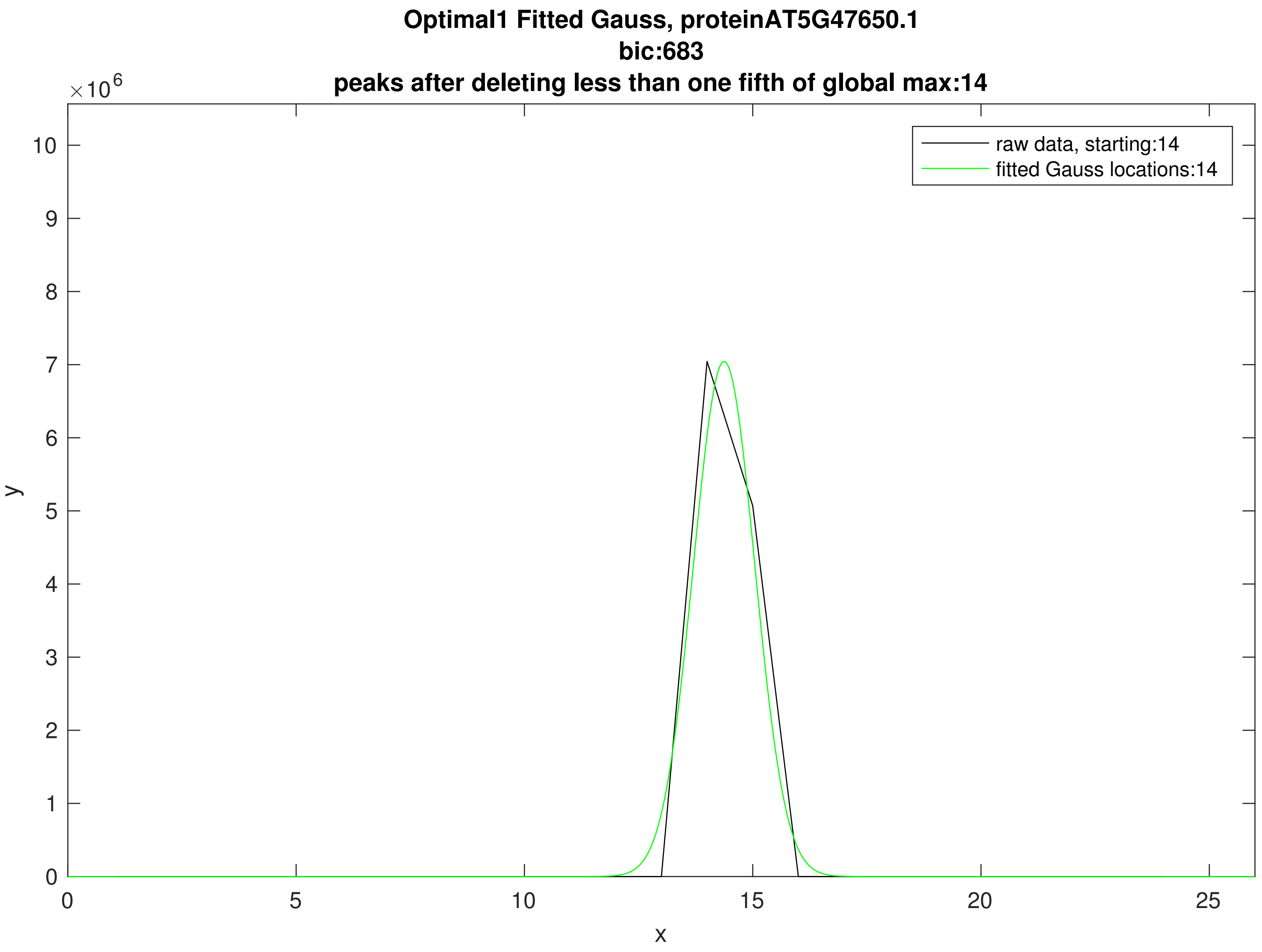

Optimal1 Fitted Gauss, proteinAT5G47650.1
bic:683
peaks after deleting less than one fifth of global max:14
×10^6
raw data, starting:14
fitted Gauss locations:14
y
x
0
1
2
3
4
5
6
7
8
9
10
0
5
10
15
20
25

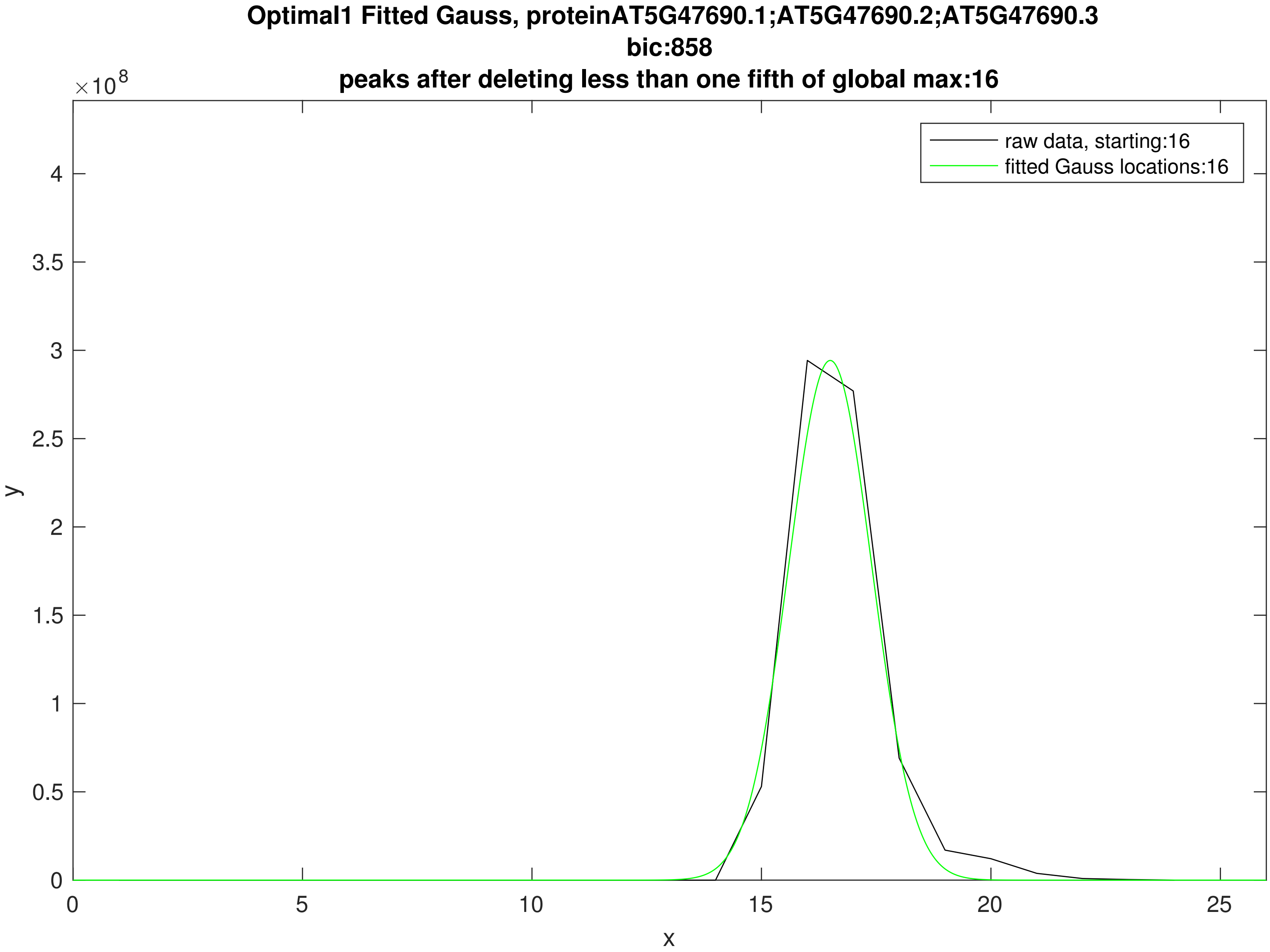
Optimal1 Fitted Gauss, proteinAT5G47690.1;AT5G47690.2;AT5G47690.3
bic:858
peaks after deleting less than one fifth of global max:16
×10^8
raw data, starting:16
fitted Gauss locations:16
0
0.5
1
1.5
2
2.5
3
3.5
4
y
0
5
10
15
20
25
x

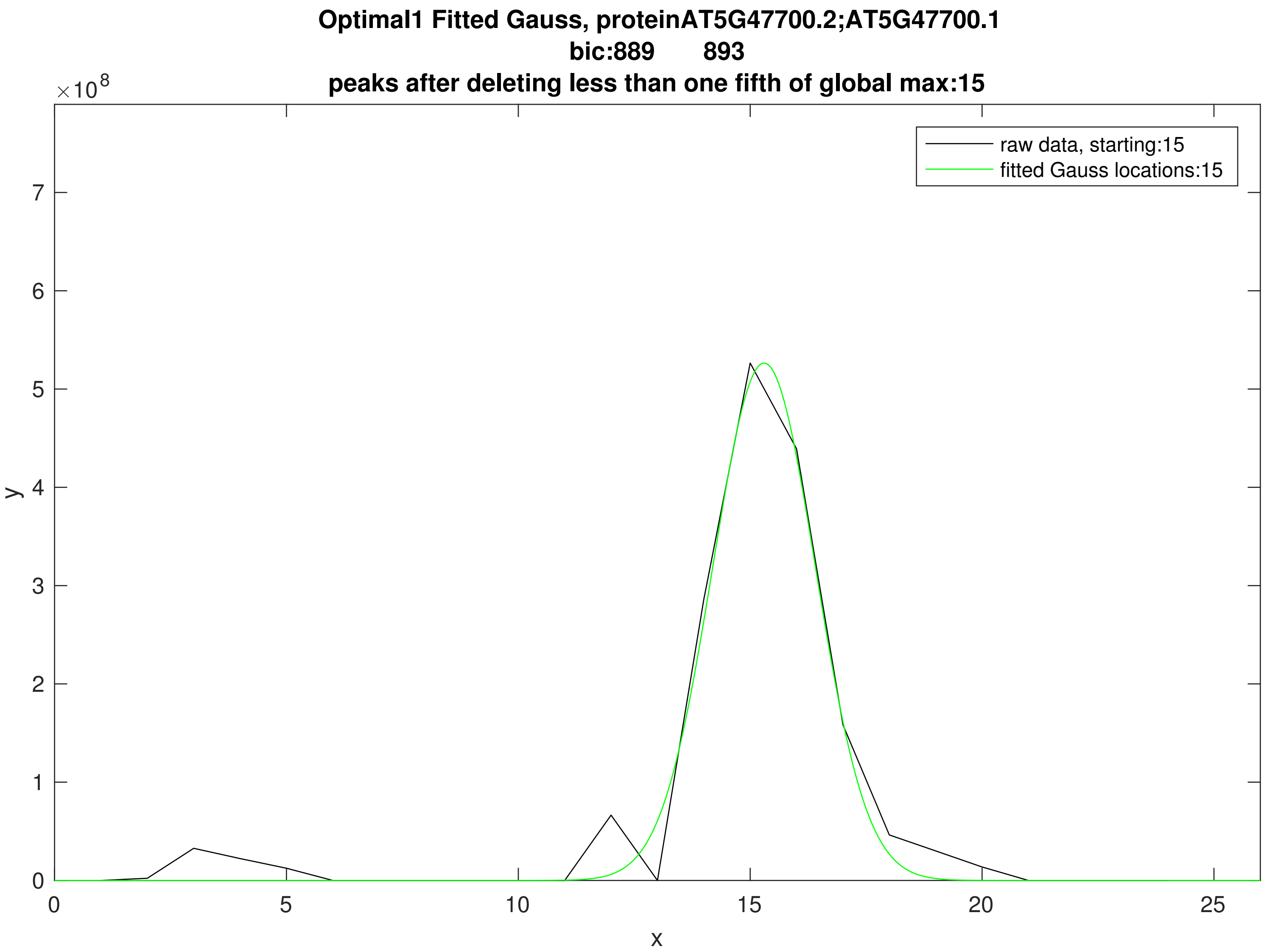

Optimal1 Fitted Gauss, proteinAT5G47700.2;AT5G47700.1
bic:889 893
peaks after deleting less than one fifth of global max:15
×10^8
raw data, starting:15
fitted Gauss locations:15
y
x
0
1
2
3
4
5
6
7
0
5
10
15
20
25

**Optimal1 Fitted Gauss, proteinAT5G47720.3;AT5G47720.1;AT5G47720.4;AT5G47720.2;AT5G47720.5**
**bic:940**
**peaks after deleting less than one fifth of global max:10**

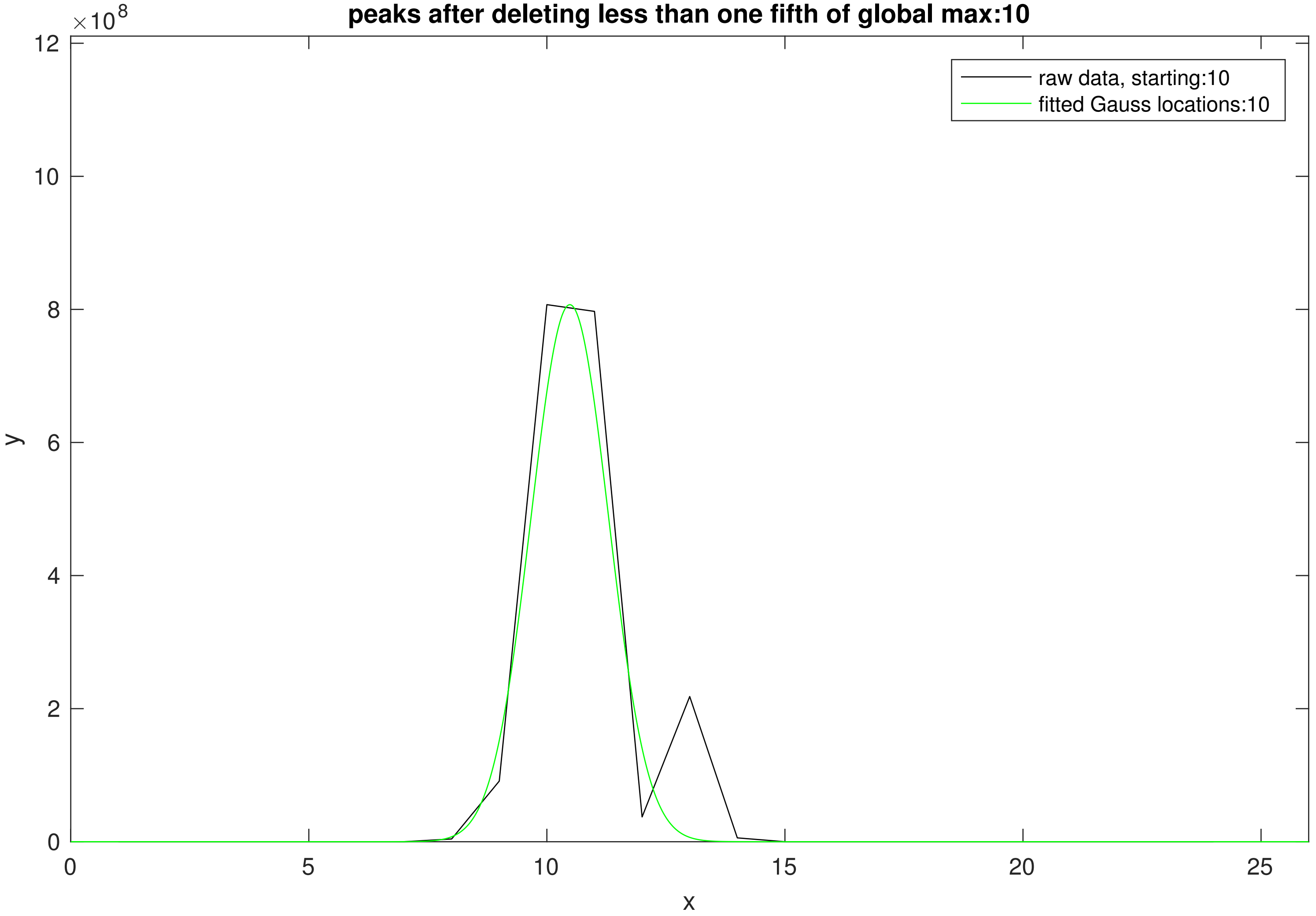

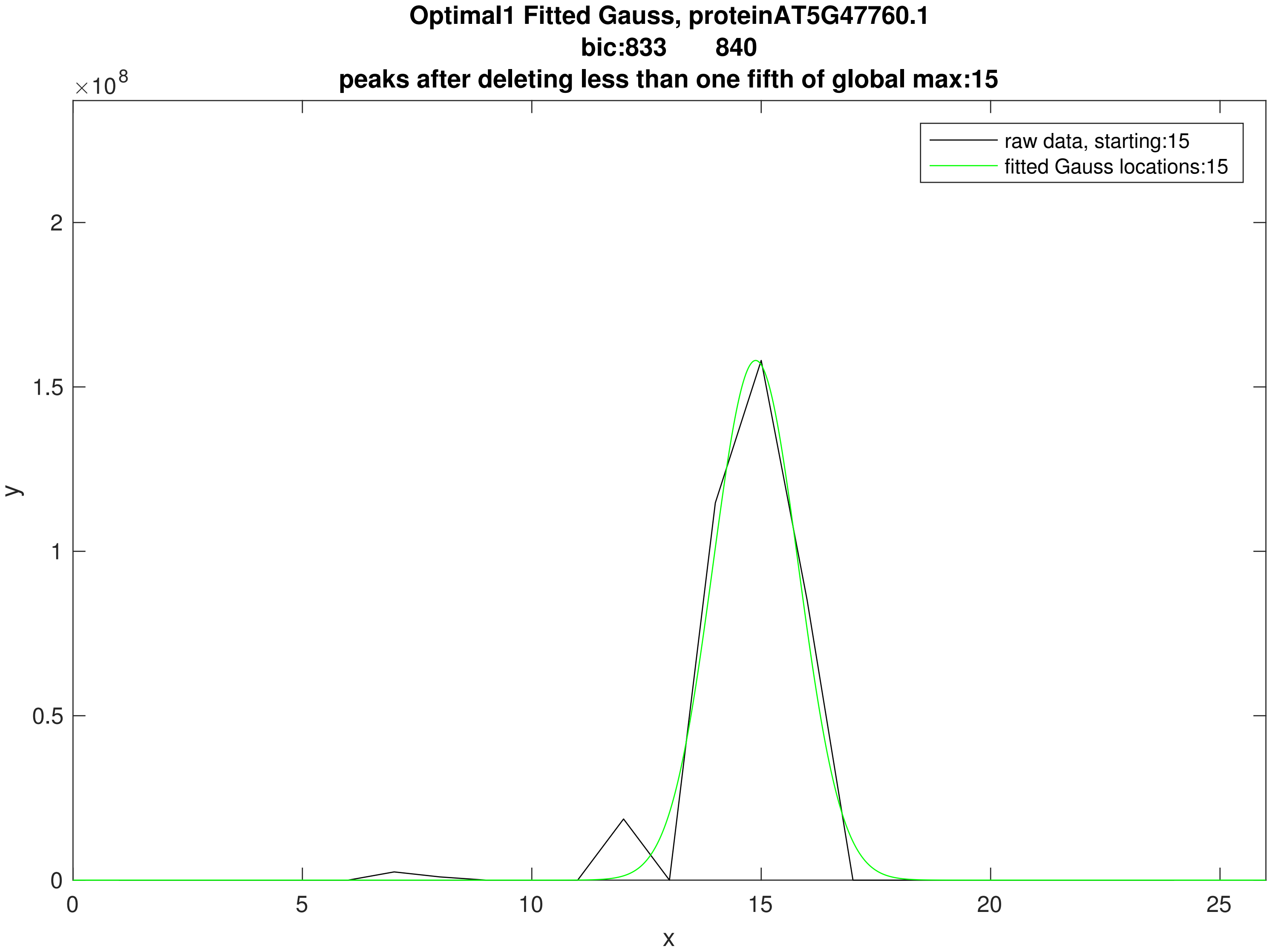
Optimal1 Fitted Gauss, proteinAT5G47760.1
bic:833 840
peaks after deleting less than one fifth of global max:15
×10^8
raw data, starting:15
fitted Gauss locations:15
y
x
0
0.5
1
1.5
2
0
5
10
15
20
25

**Optimal1 Fitted Gauss, proteinAT5G47770.1**
**bic:1.07e+03  1.08e+03  1.08e+03**
**peaks after deleting less than one fifth of global max:14**

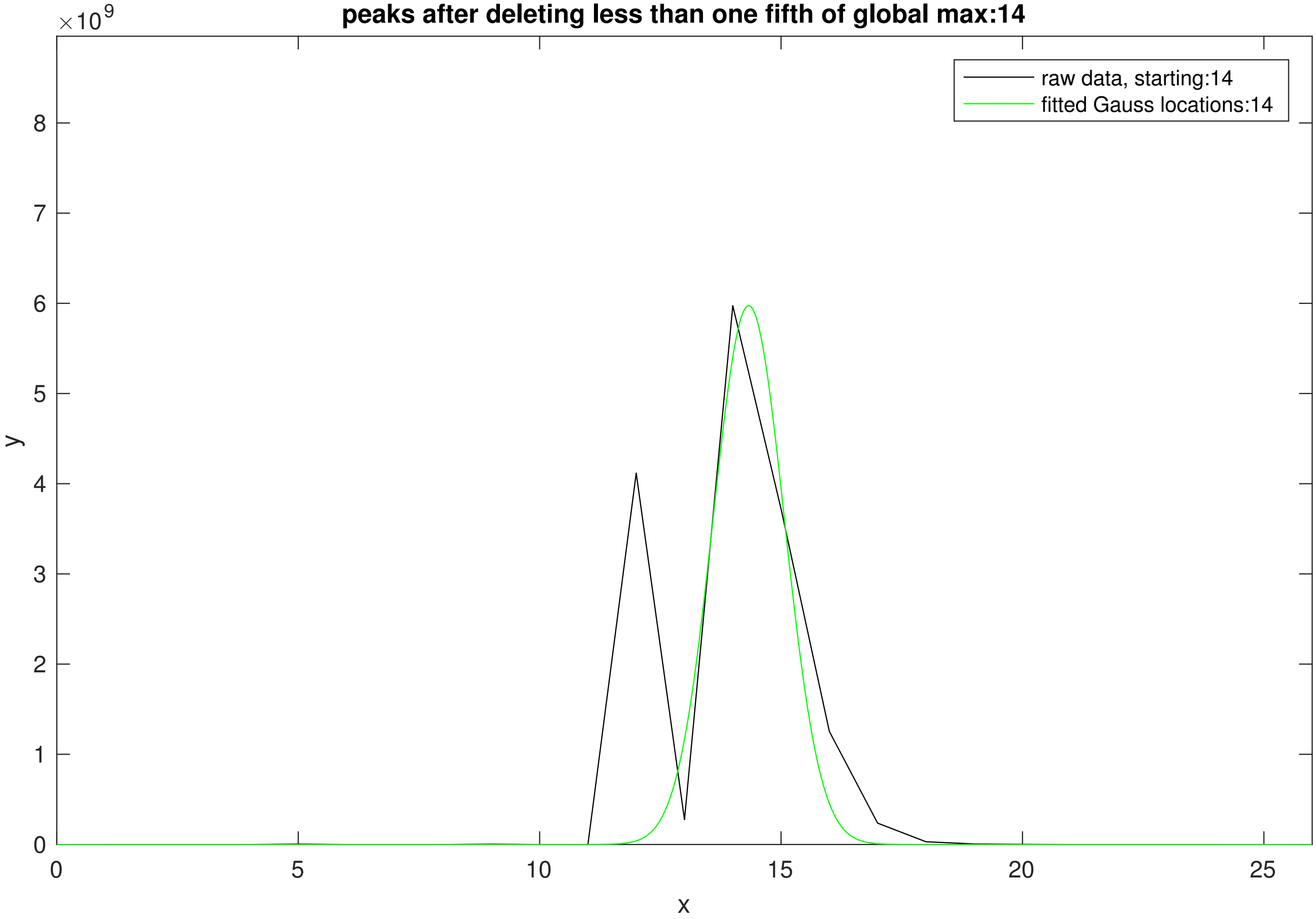

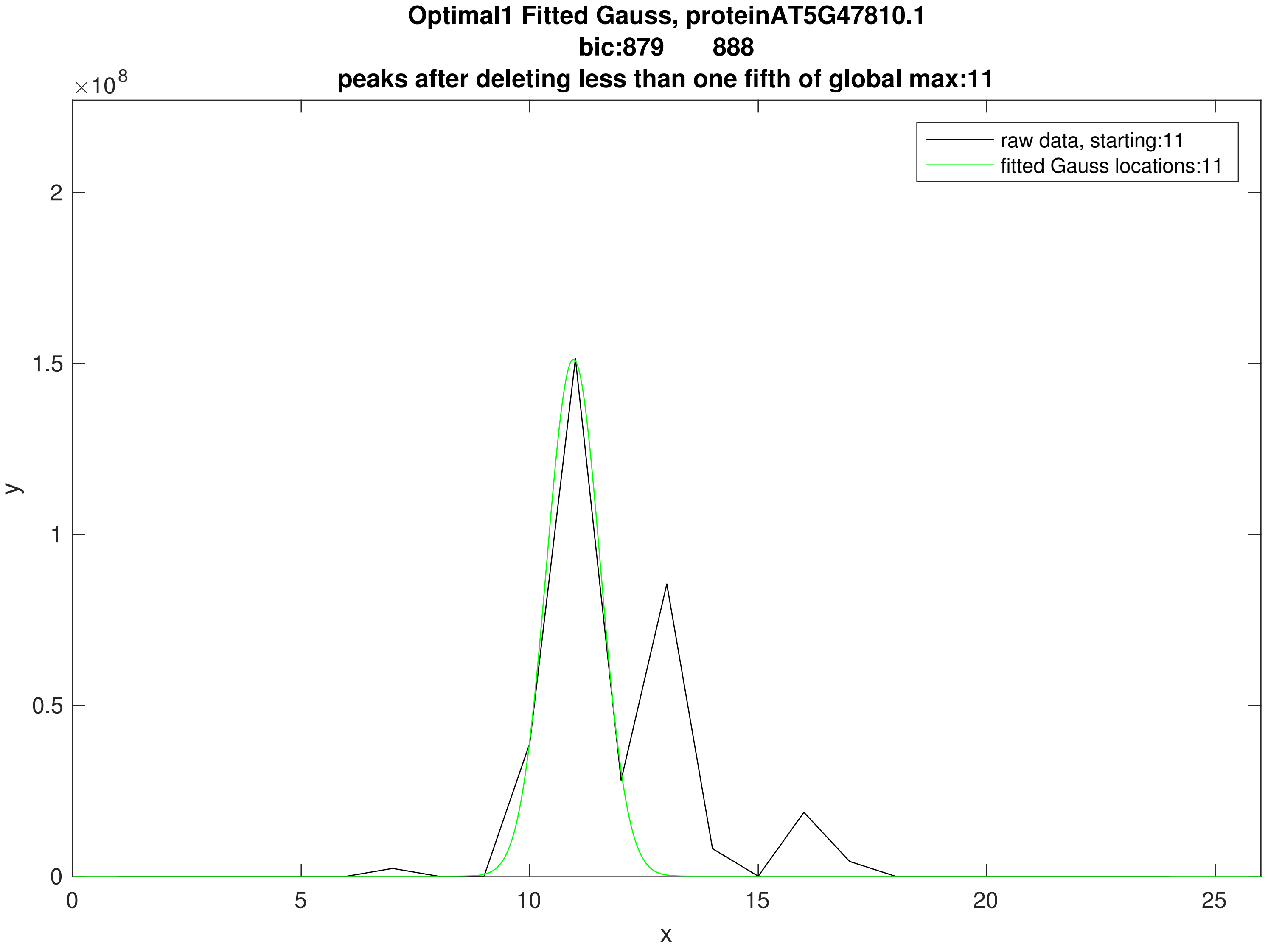

Optimal1 Fitted Gauss, proteinAT5G47810.1
bic:879 888
peaks after deleting less than one fifth of global max:11
×10^8
raw data, starting:11
fitted Gauss locations:11
y
x
0
0.5
1
1.5
2
0
5
10
15
20
25

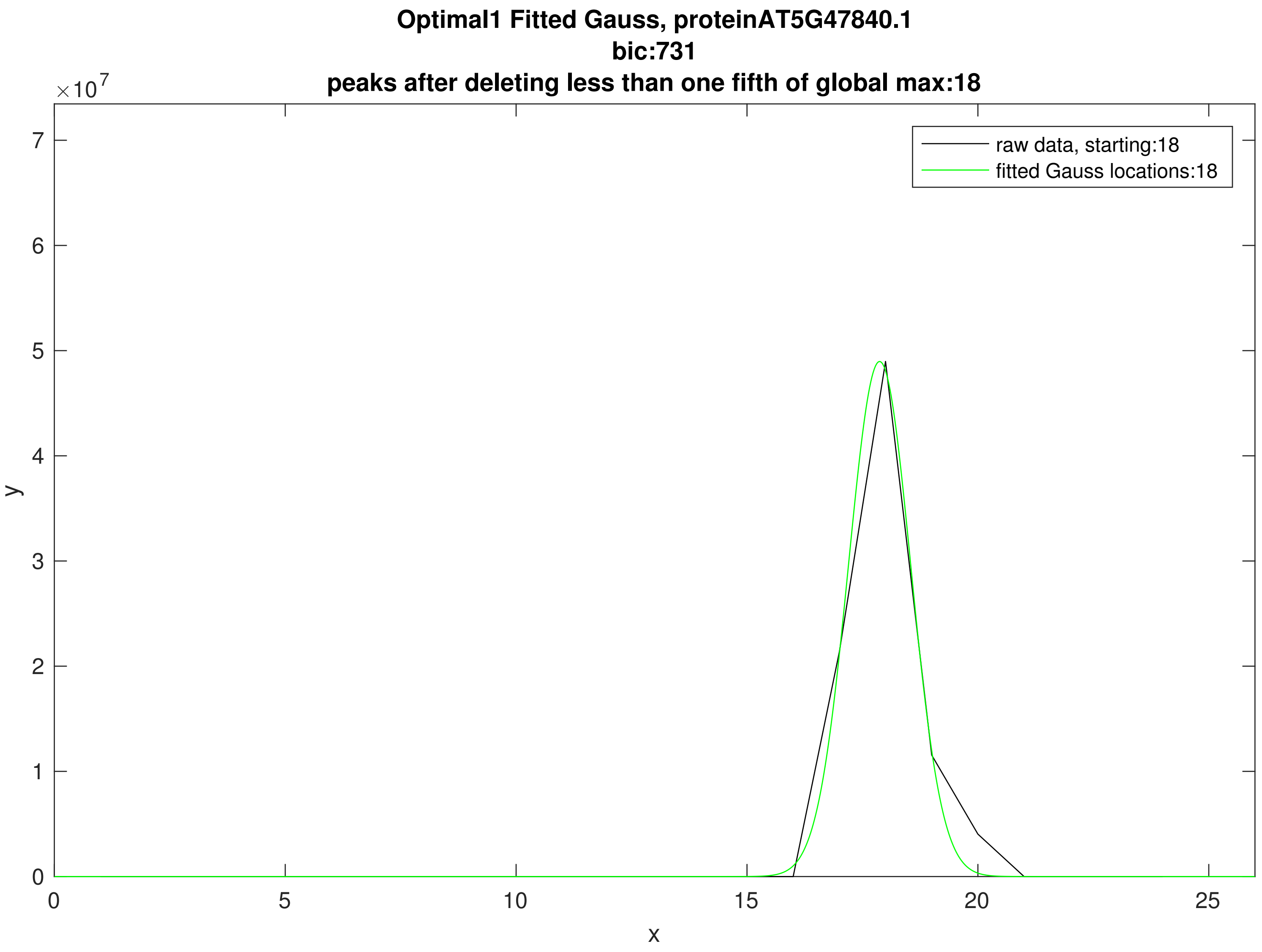
Optimal1 Fitted Gauss, proteinAT5G47840.1
bic:731
peaks after deleting less than one fifth of global max:18
×10^7
raw data, starting:18
fitted Gauss locations:18
y
x
0
1
2
3
4
5
6
7
0
5
10
15
20
25

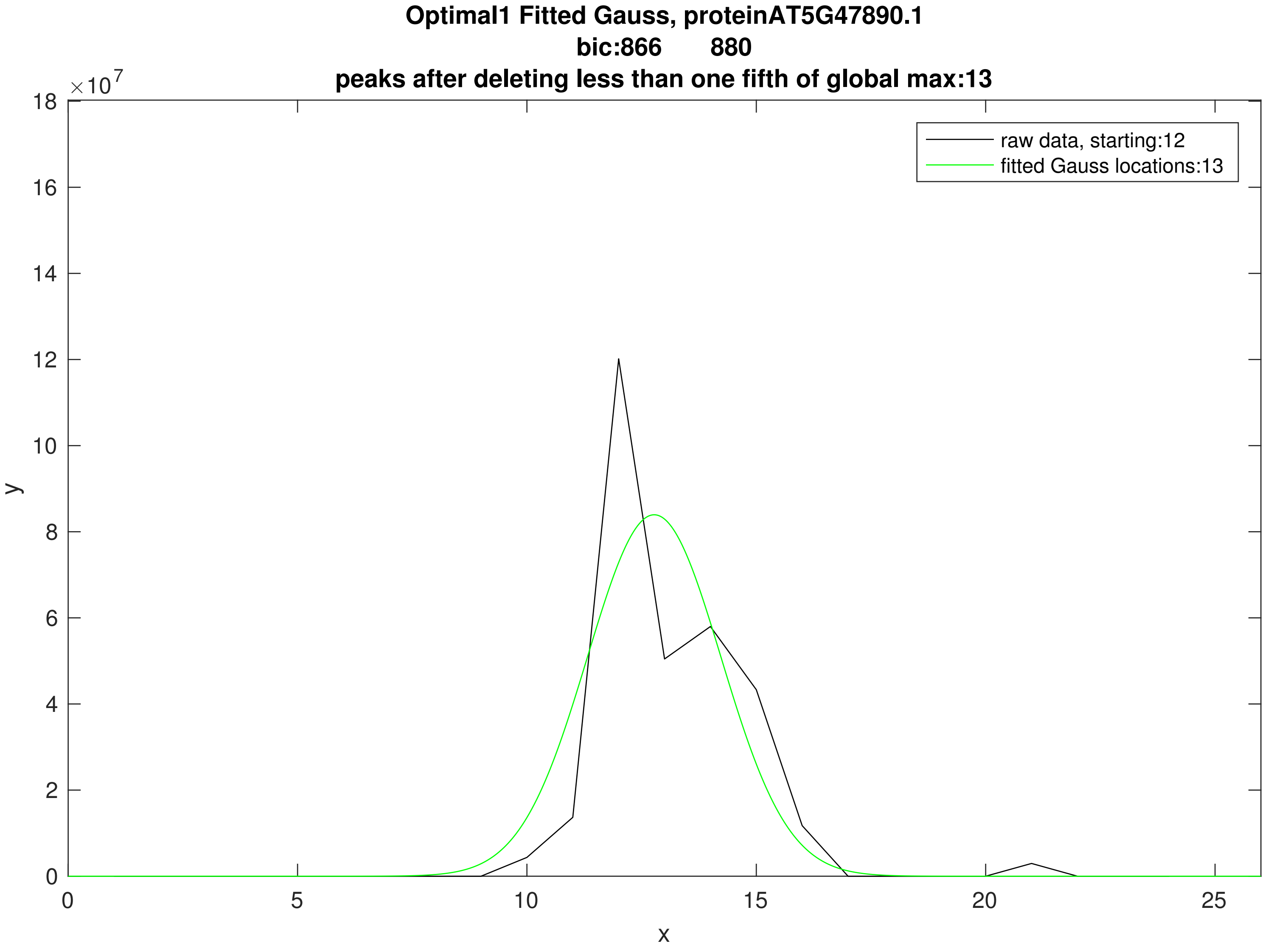
Optimal1 Fitted Gauss, proteinAT5G47890.1
bic:866 880
peaks after deleting less than one fifth of global max:13
raw data, starting:12
fitted Gauss locations:13
y
x

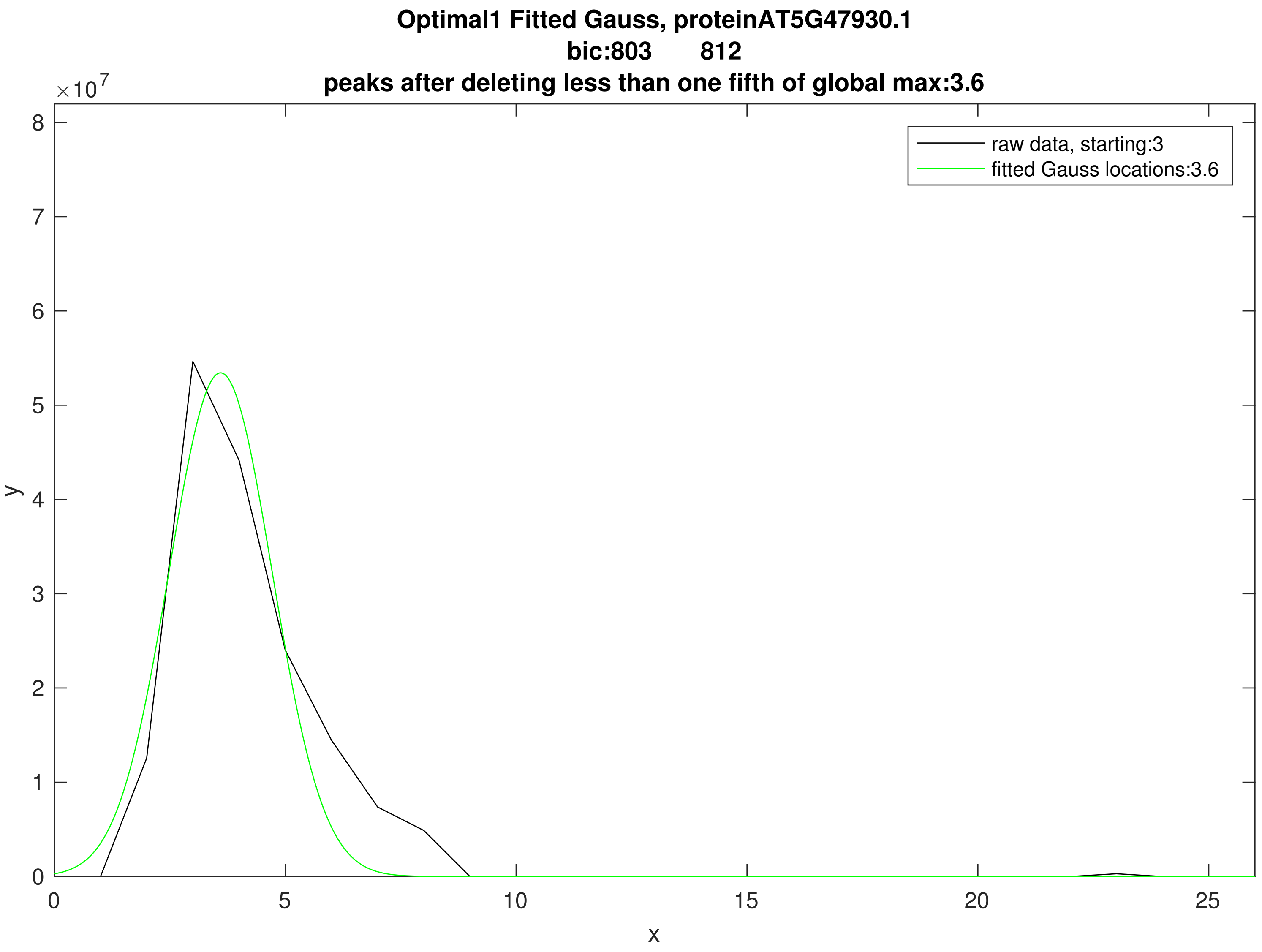
Optimal1 Fitted Gauss, proteinAT5G47930.1
bic:803 812
peaks after deleting less than one fifth of global max:3.6
×10^7
raw data, starting:3
fitted Gauss locations:3.6
y
x
0
1
2
3
4
5
6
7
8
0
5
10
15
20
25

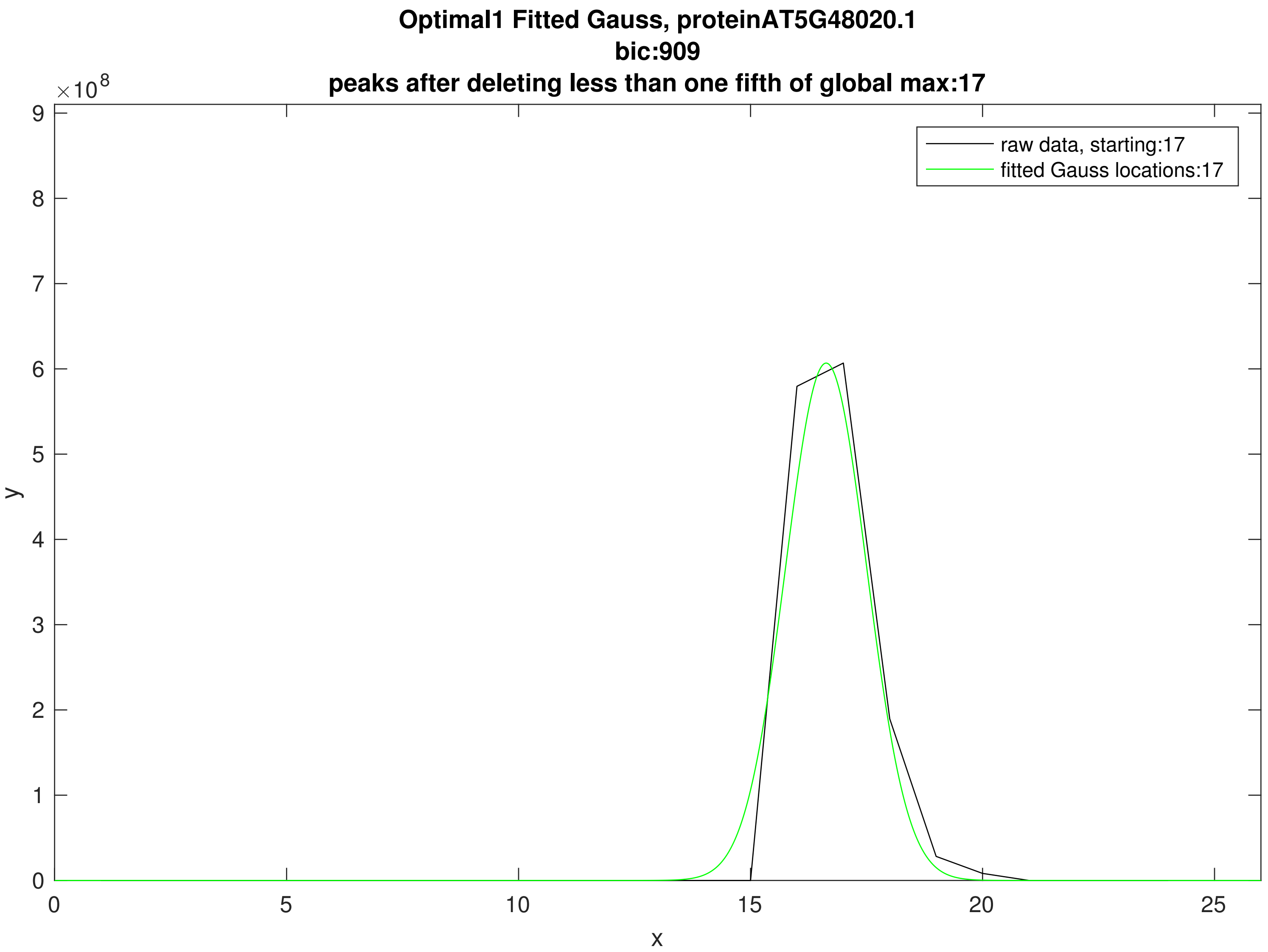

Optimal1 Fitted Gauss, proteinAT5G48020.1
bic:909
peaks after deleting less than one fifth of global max:17
×10^8
9
8
7
6
5
4
3
2
1
0
y
0
5
10
15
20
25
x
raw data, starting:17
fitted Gauss locations:17

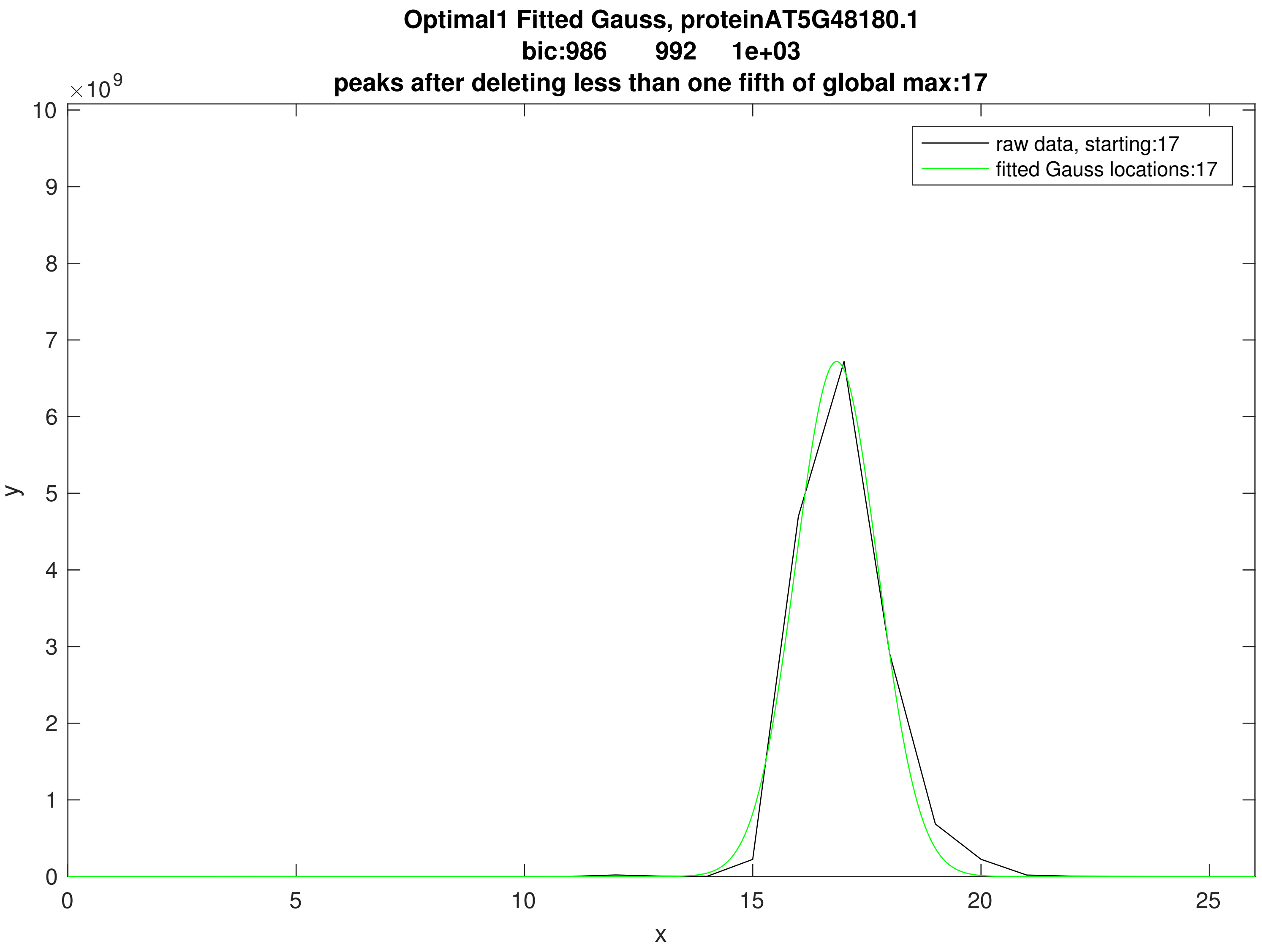

Optimal1 Fitted Gauss, proteinAT5G48180.1
bic:986 992 1e+03
peaks after deleting less than one fifth of global max:17
×10^9
10
9
8
7
6
5
4
3
2
1
0
y
0
5
10
15
20
25
x
raw data, starting:17
fitted Gauss locations:17

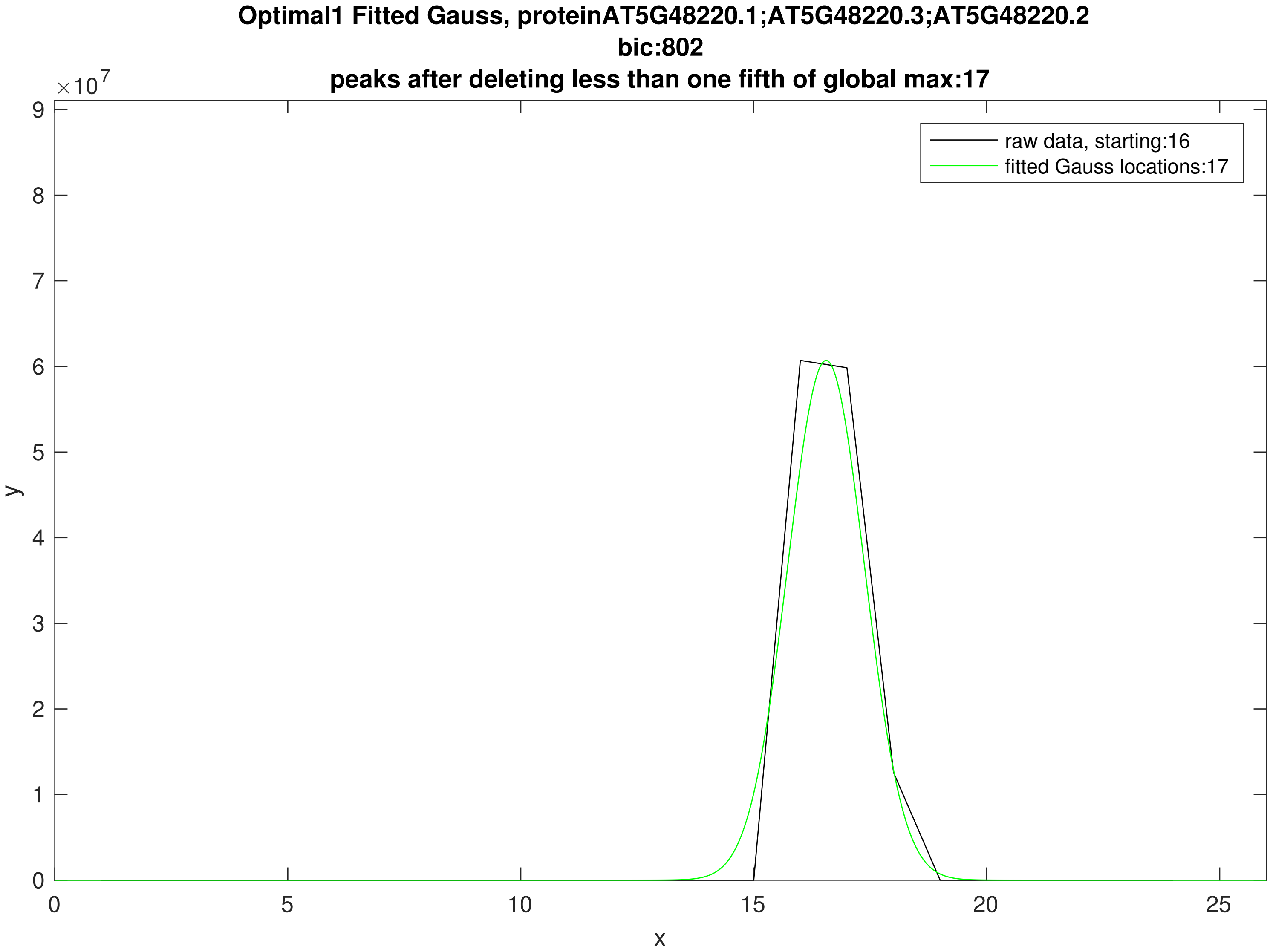
Optimal1 Fitted Gauss, proteinAT5G48220.1;AT5G48220.3;AT5G48220.2
bic:802
peaks after deleting less than one fifth of global max:17
×10^7
raw data, starting:16
fitted Gauss locations:17
y
x
0
1
2
3
4
5
6
7
8
9
0
5
10
15
20
25

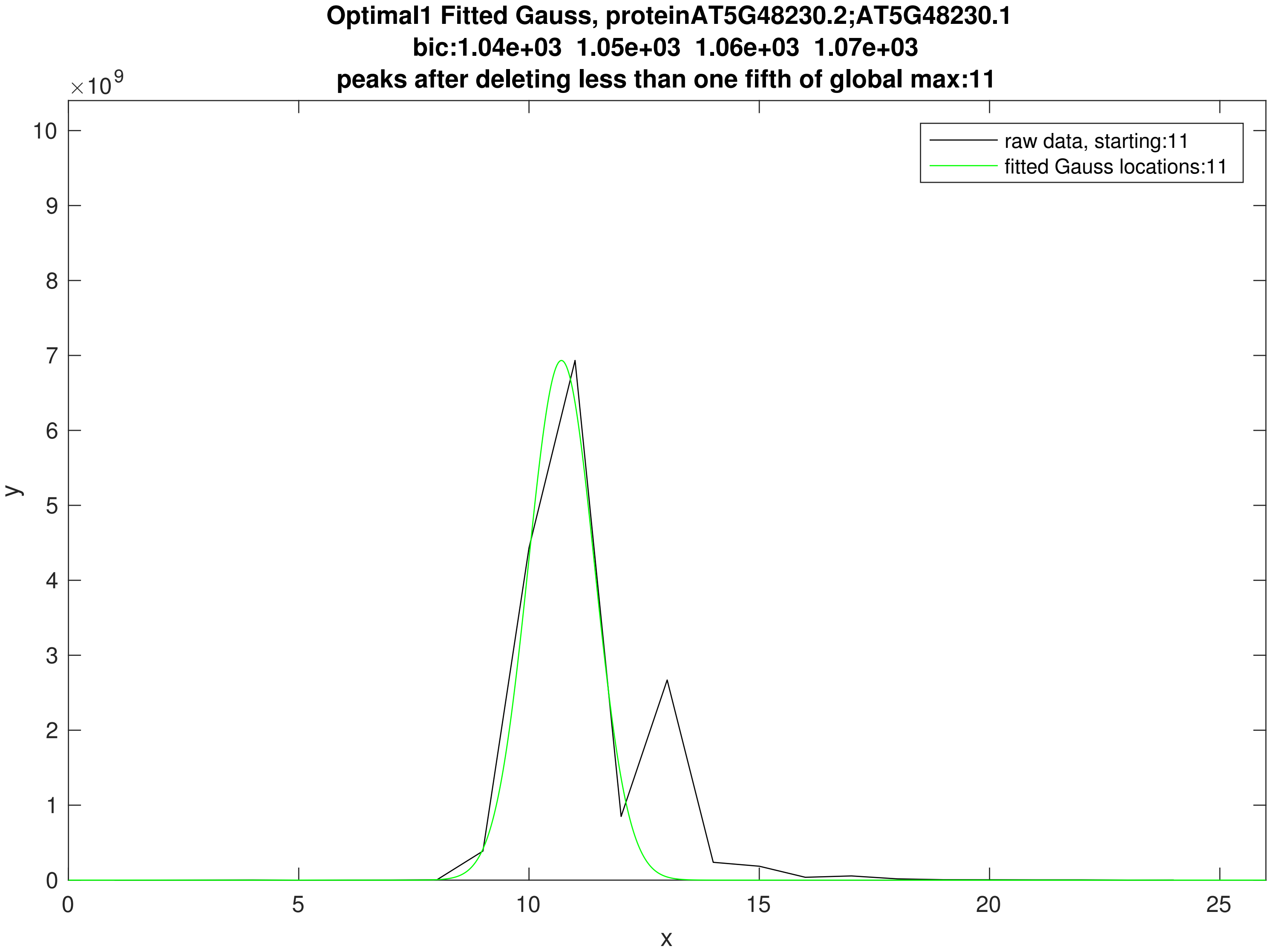

Optimal1 Fitted Gauss, proteinAT5G48230.2;AT5G48230.1
bic:1.04e+03 1.05e+03 1.06e+03 1.07e+03
peaks after deleting less than one fifth of global max:11
×10^9
raw data, starting:11
fitted Gauss locations:11
y
x
0
1
2
3
4
5
6
7
8
9
10
0
5
10
15
20
25

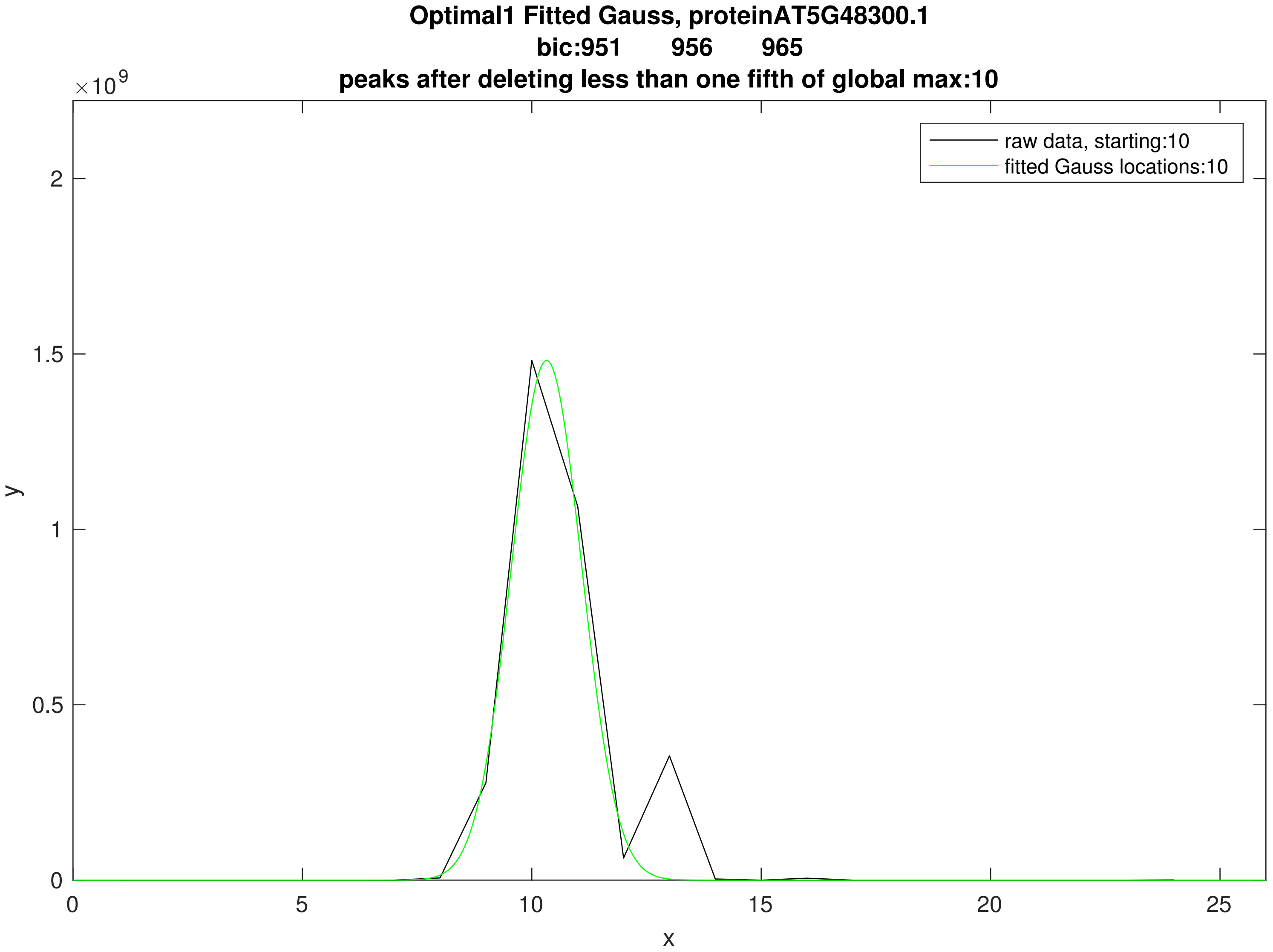
Optimal1 Fitted Gauss, proteinAT5G48300.1
bic:951 956 965
peaks after deleting less than one fifth of global max:10
×10^9
raw data, starting:10
fitted Gauss locations:10
y
x
0
0.5
1
1.5
2
0
5
10
15
20
25

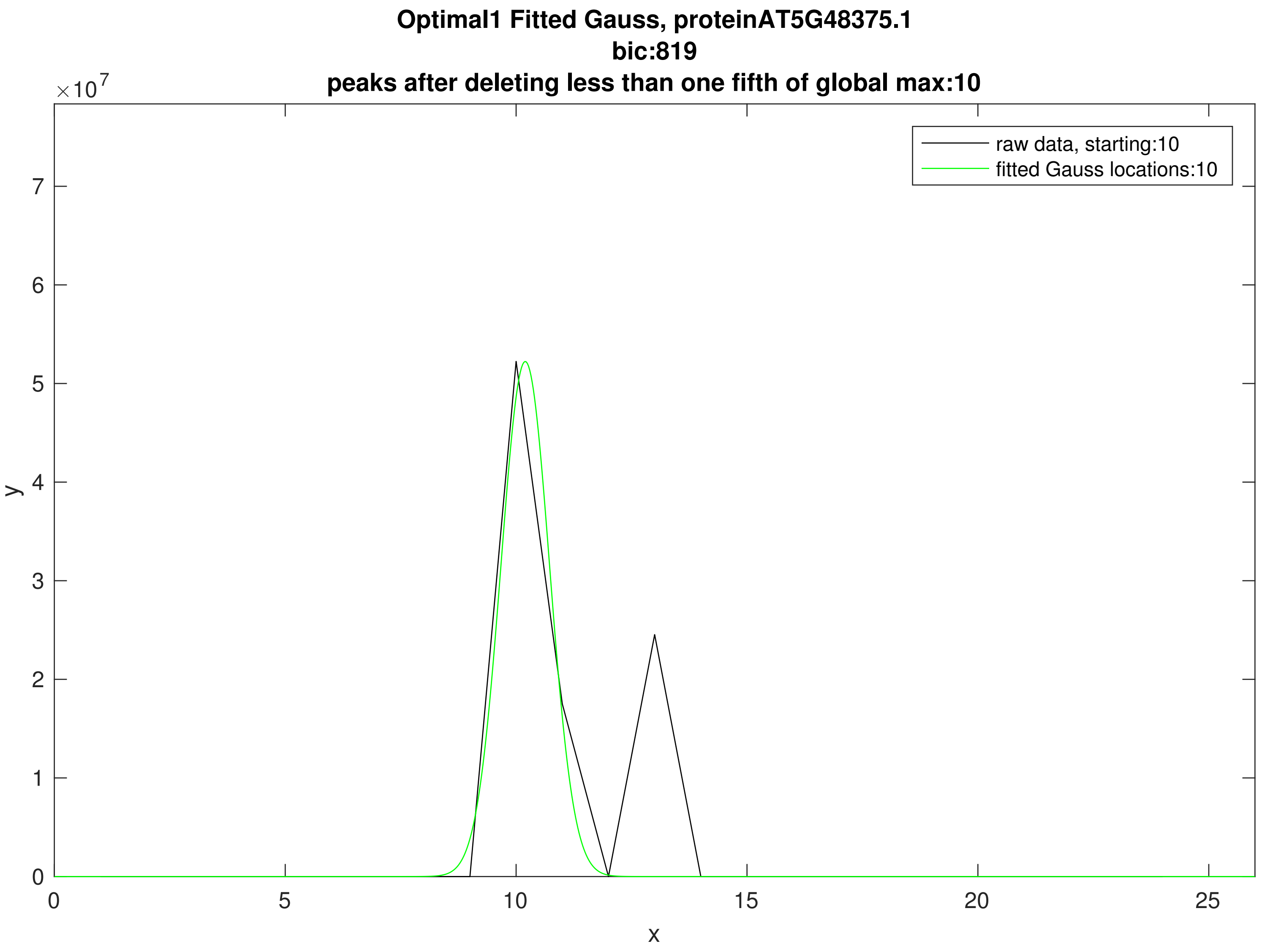

Optimal1 Fitted Gauss, proteinAT5G48375.1
bic:819
peaks after deleting less than one fifth of global max:10
×10^7
raw data, starting:10
fitted Gauss locations:10
y
x
0
1
2
3
4
5
6
7
0
5
10
15
20
25

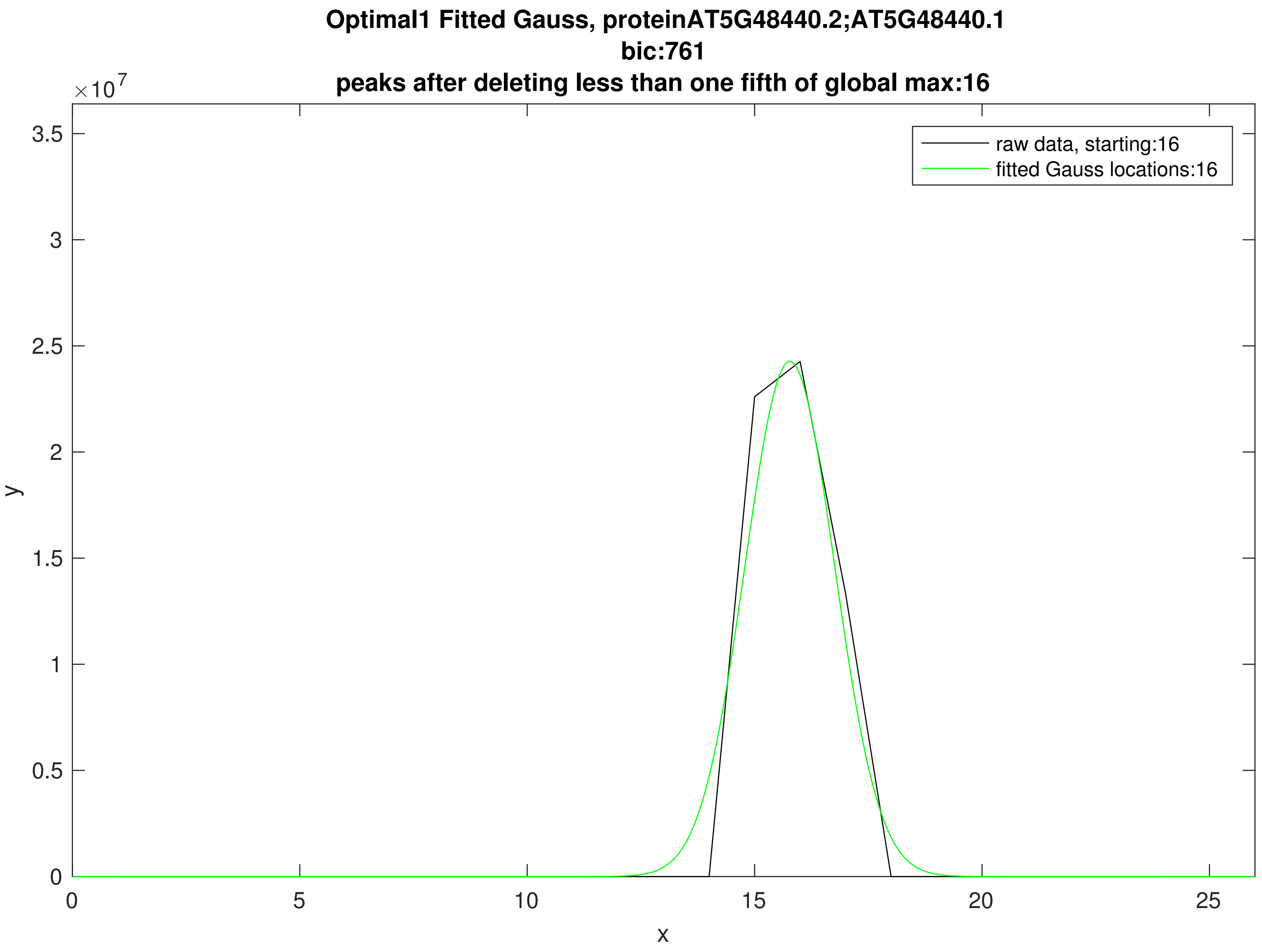

Optimal1 Fitted Gauss, proteinAT5G48440.2;AT5G48440.1
bic:761
peaks after deleting less than one fifth of global max:16
×10^7
3.5
3
2.5
2
1.5
1
0.5
0
y
0
5
10
15
20
25
x
raw data, starting:16
fitted Gauss locations:16

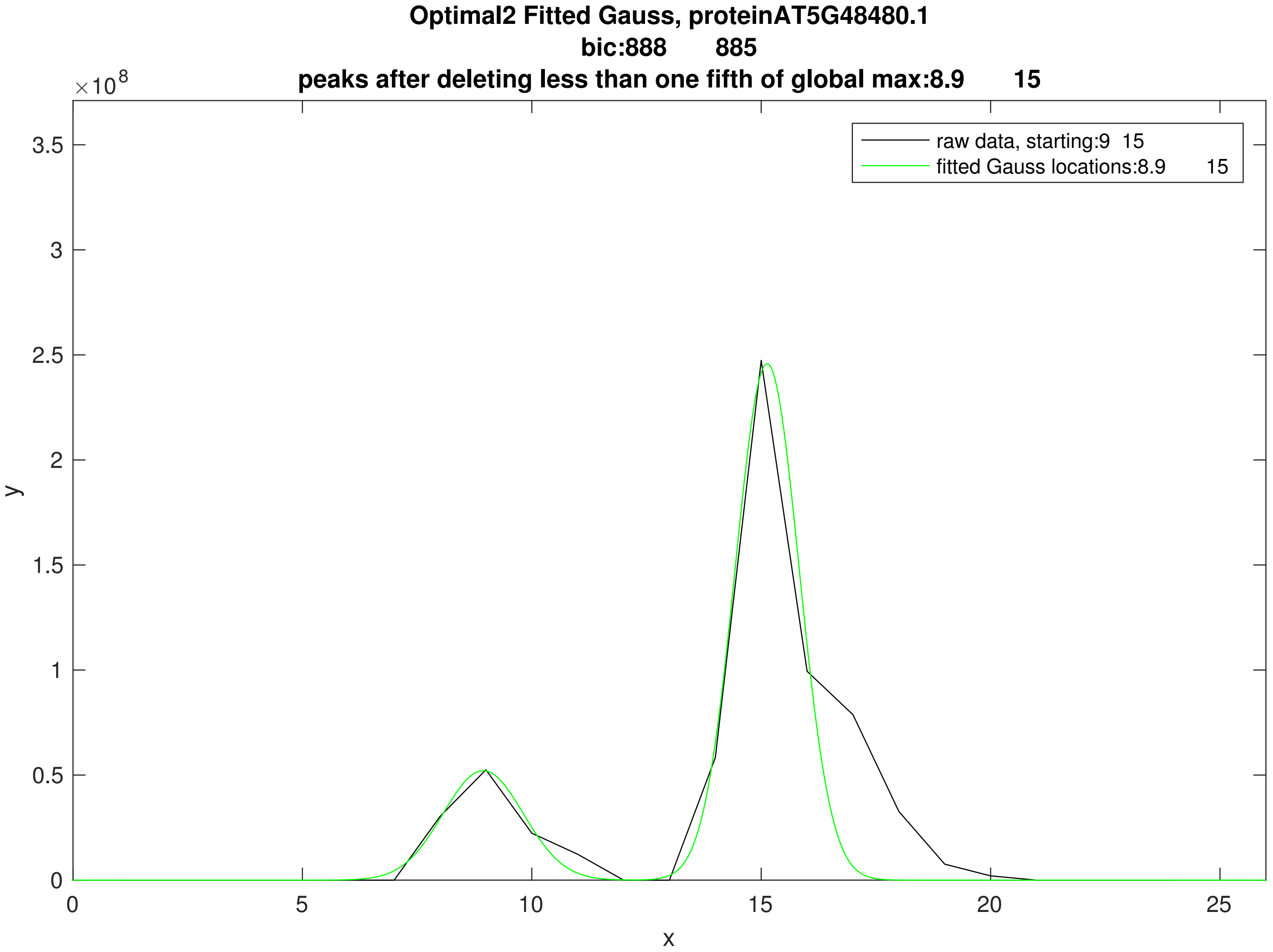

Optimal2 Fitted Gauss, proteinAT5G48480.1
bic:888 885
peaks after deleting less than one fifth of global max:8.9 15
×10^8
raw data, starting:9 15
fitted Gauss locations:8.9 15
3.5
3
2.5
2
1.5
1
0.5
0
y
0
5
10
15
20
25
x

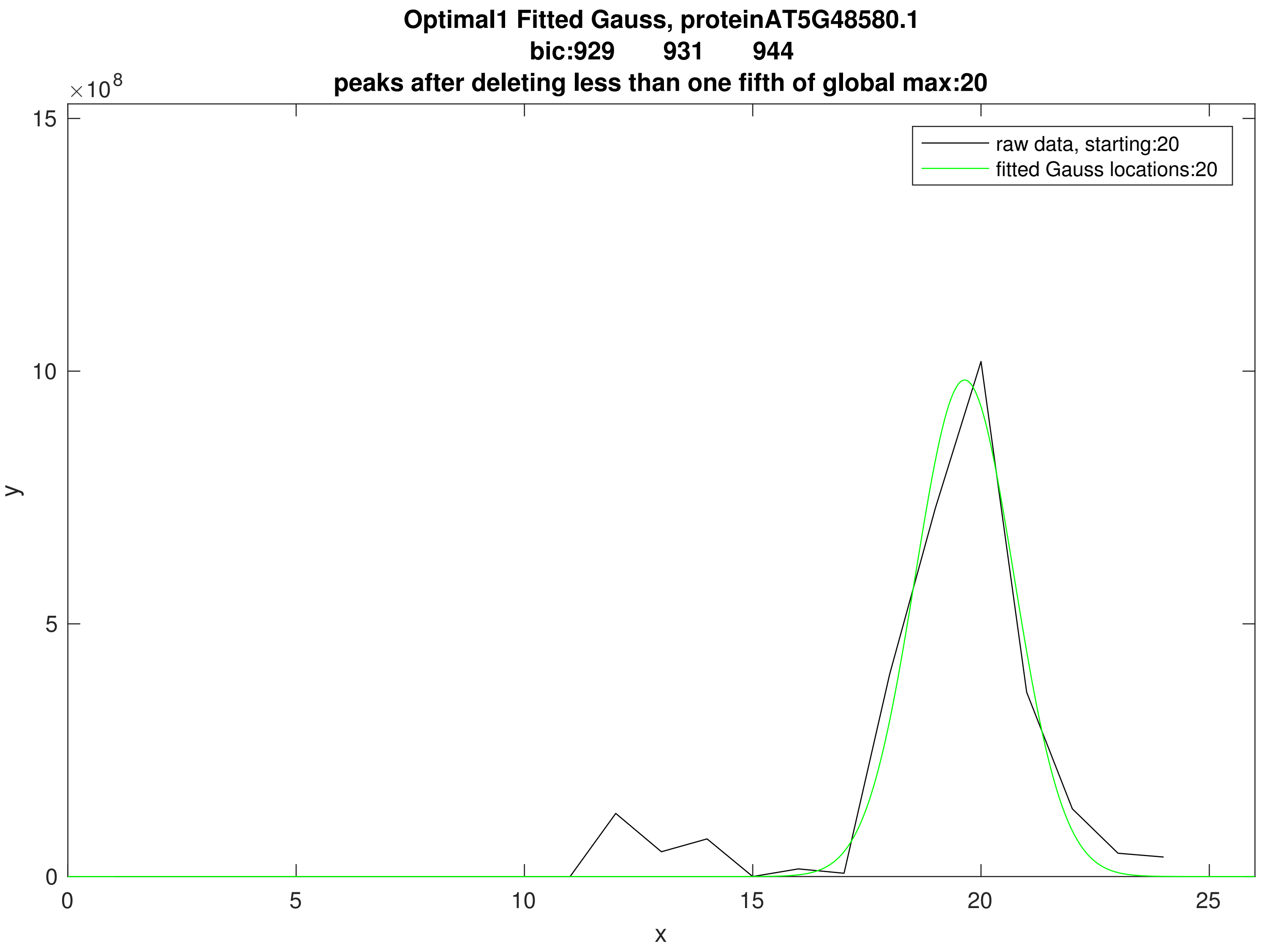
Optimal1 Fitted Gauss, proteinAT5G48580.1
bic:929 931 944
peaks after deleting less than one fifth of global max:20
×10^8
15
10
5
0
y
0
5
10
15
20
25
x
raw data, starting:20
fitted Gauss locations:20

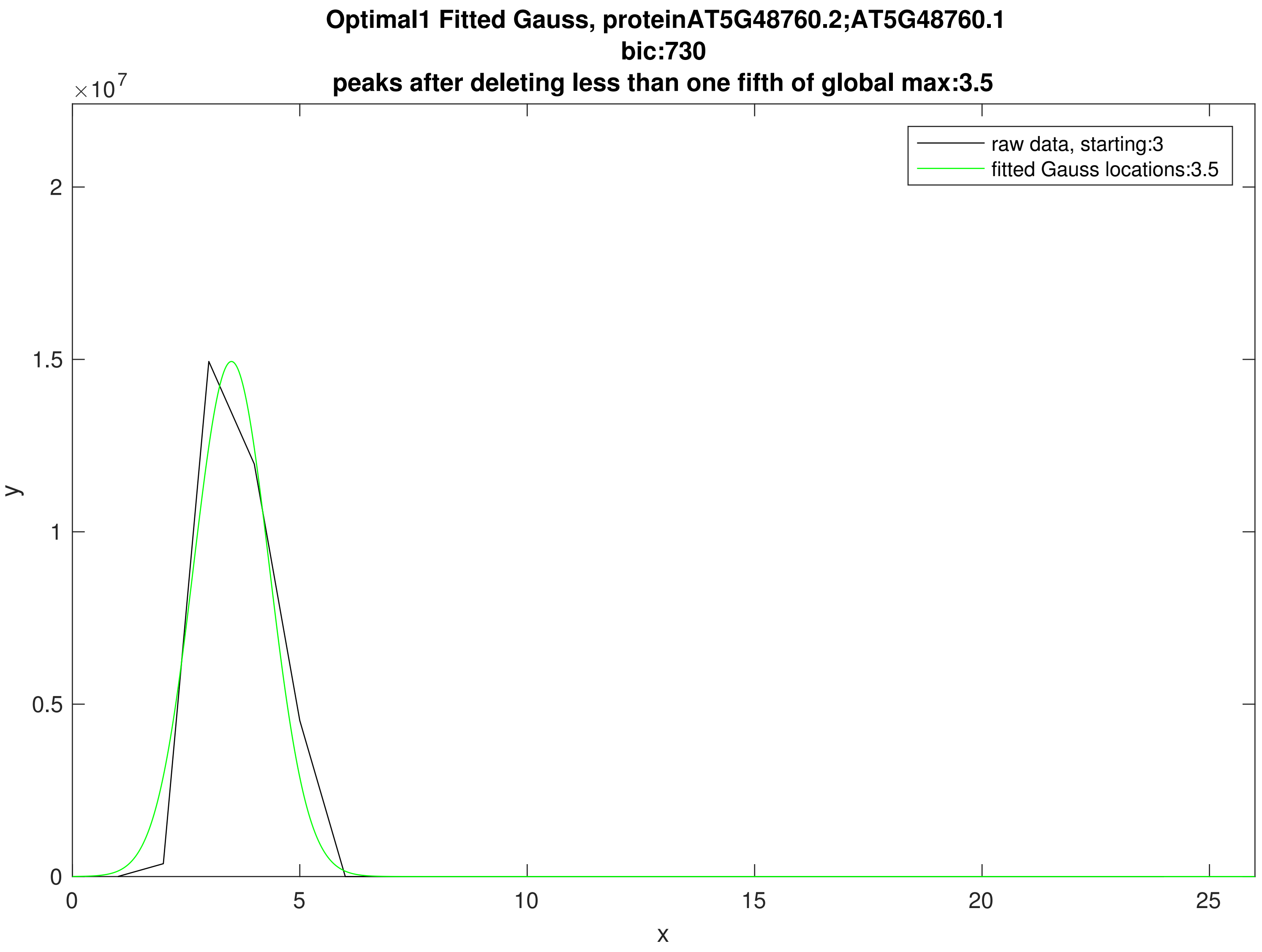
Optimal1 Fitted Gauss, proteinAT5G48760.2;AT5G48760.1
bic:730
peaks after deleting less than one fifth of global max:3.5
×10^7
raw data, starting:3
fitted Gauss locations:3.5
y
x
0
0.5
1
1.5
2
5
10
15
20
25

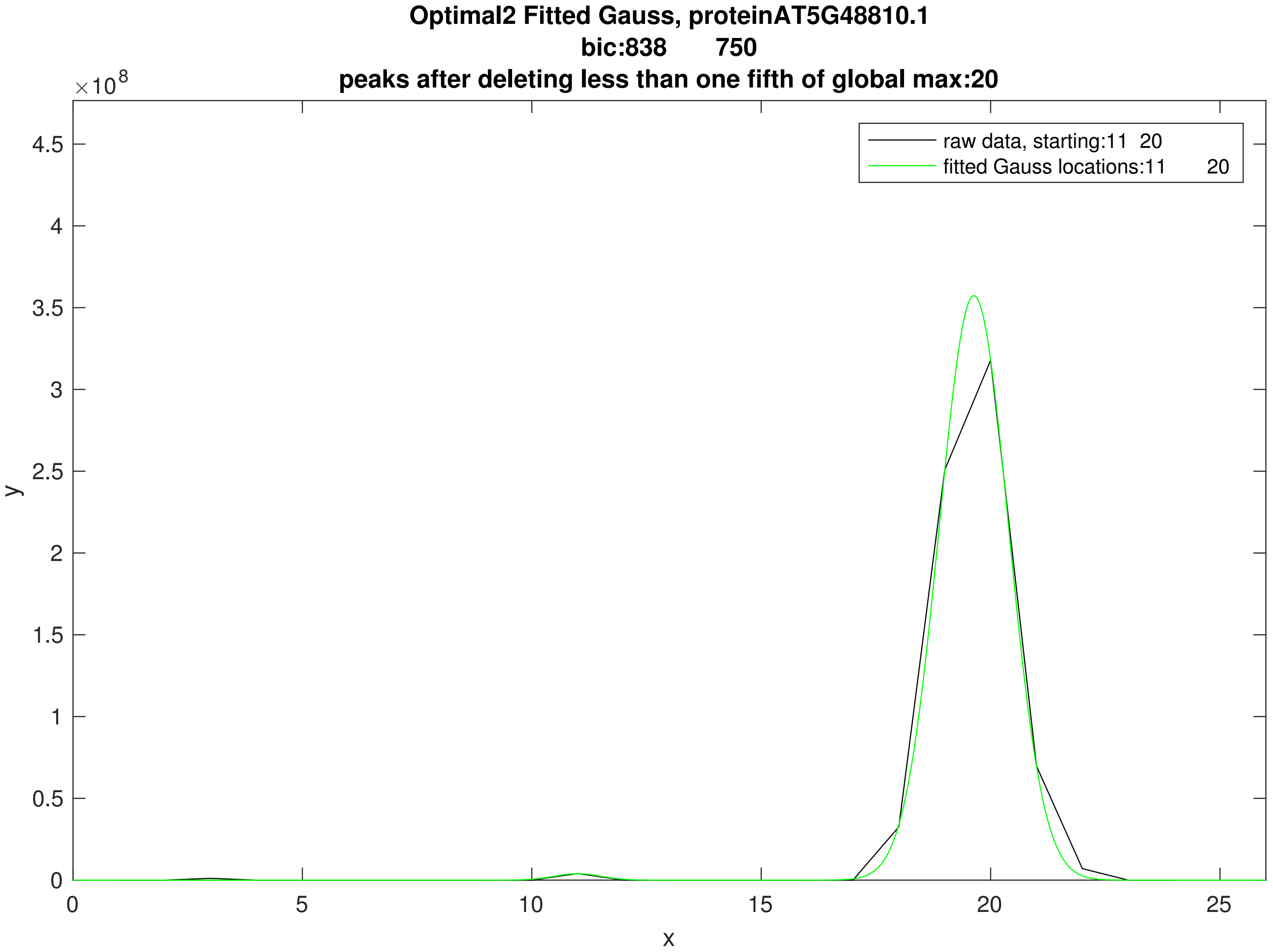
Optimal2 Fitted Gauss, proteinAT5G48810.1
bic:838 750
peaks after deleting less than one fifth of global max:20
×10^8
raw data, starting:11 20
fitted Gauss locations:11 20
y
x
0
0.5
1
1.5
2
2.5
3
3.5
4
4.5
5
10
15
20
25

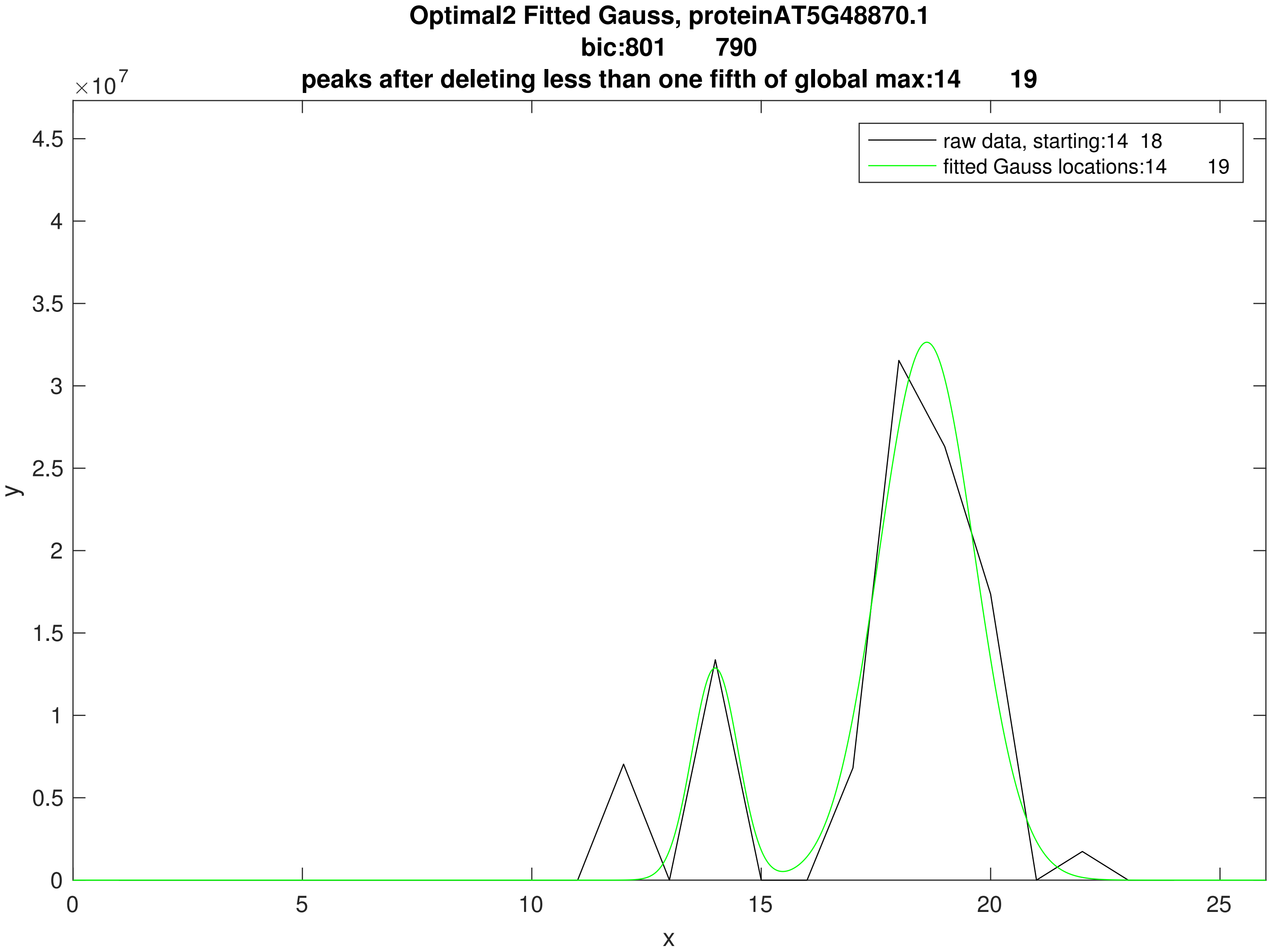
Optimal2 Fitted Gauss, proteinAT5G48870.1
bic:801 790
peaks after deleting less than one fifth of global max:14 19
×10^7
raw data, starting:14 18
fitted Gauss locations:14 19
4.5
4
3.5
3
2.5
2
1.5
1
0.5
0
y
0
5
10
15
20
25
x

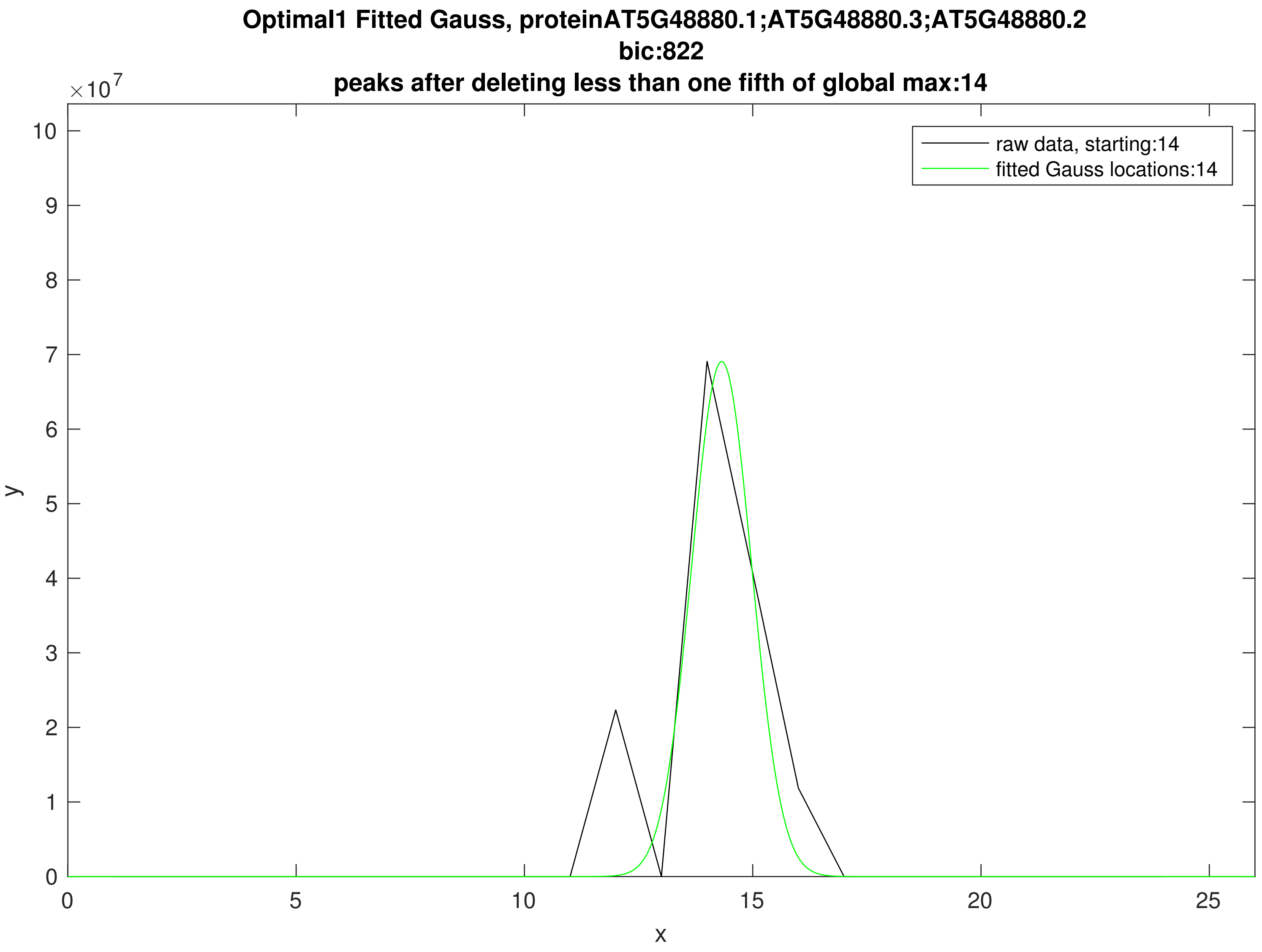

Optimal1 Fitted Gauss, proteinAT5G48880.1;AT5G48880.3;AT5G48880.2
bic:822
peaks after deleting less than one fifth of global max:14
×10^7
raw data, starting:14
fitted Gauss locations:14
y
x
0
1
2
3
4
5
6
7
8
9
10
0
5
10
15
20
25

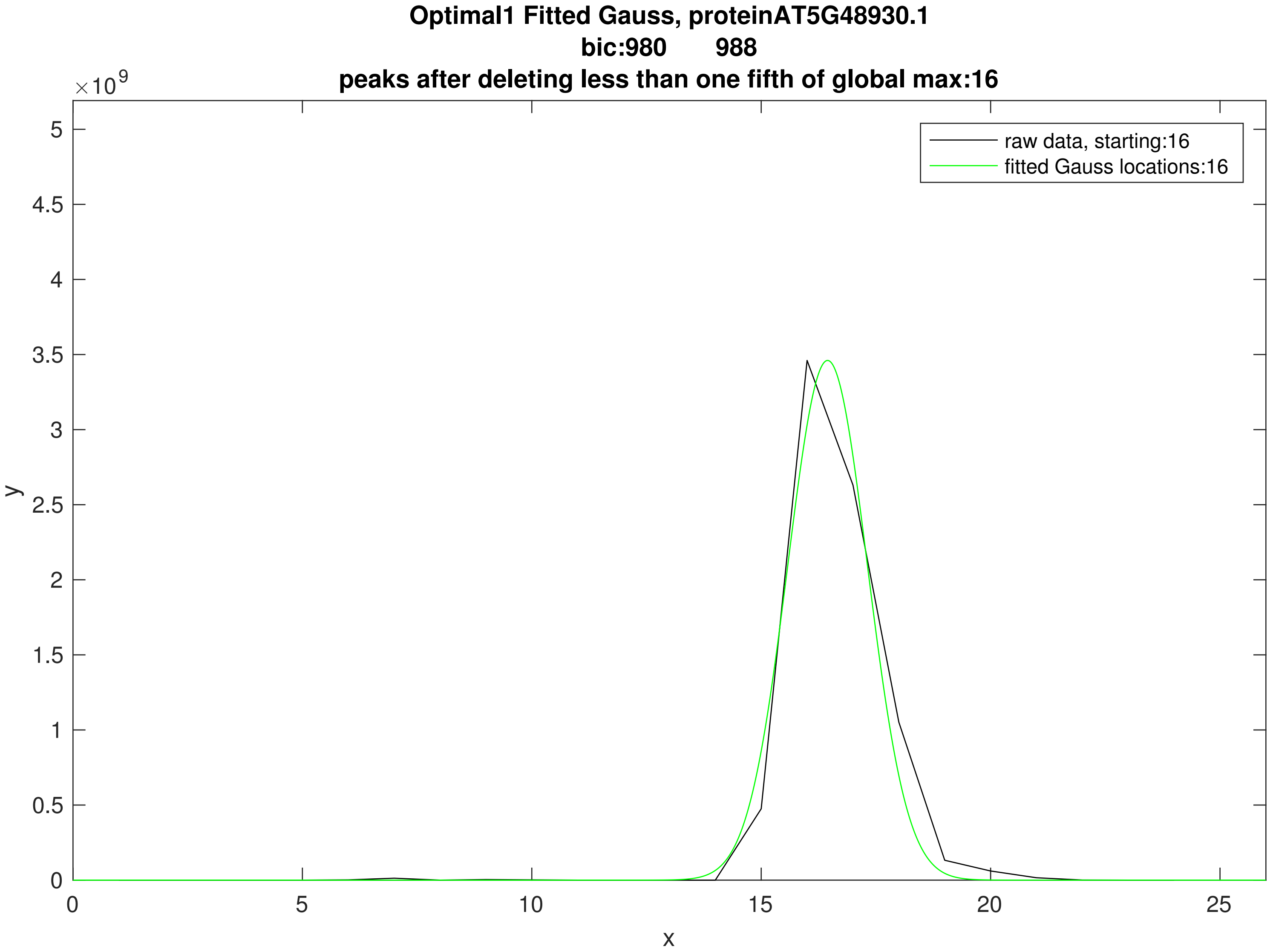
Optimal1 Fitted Gauss, proteinAT5G48930.1
bic:980 988
peaks after deleting less than one fifth of global max:16
×10^9
raw data, starting:16
fitted Gauss locations:16
y
x
0
0.5
1
1.5
2
2.5
3
3.5
4
4.5
5
0
5
10
15
20
25

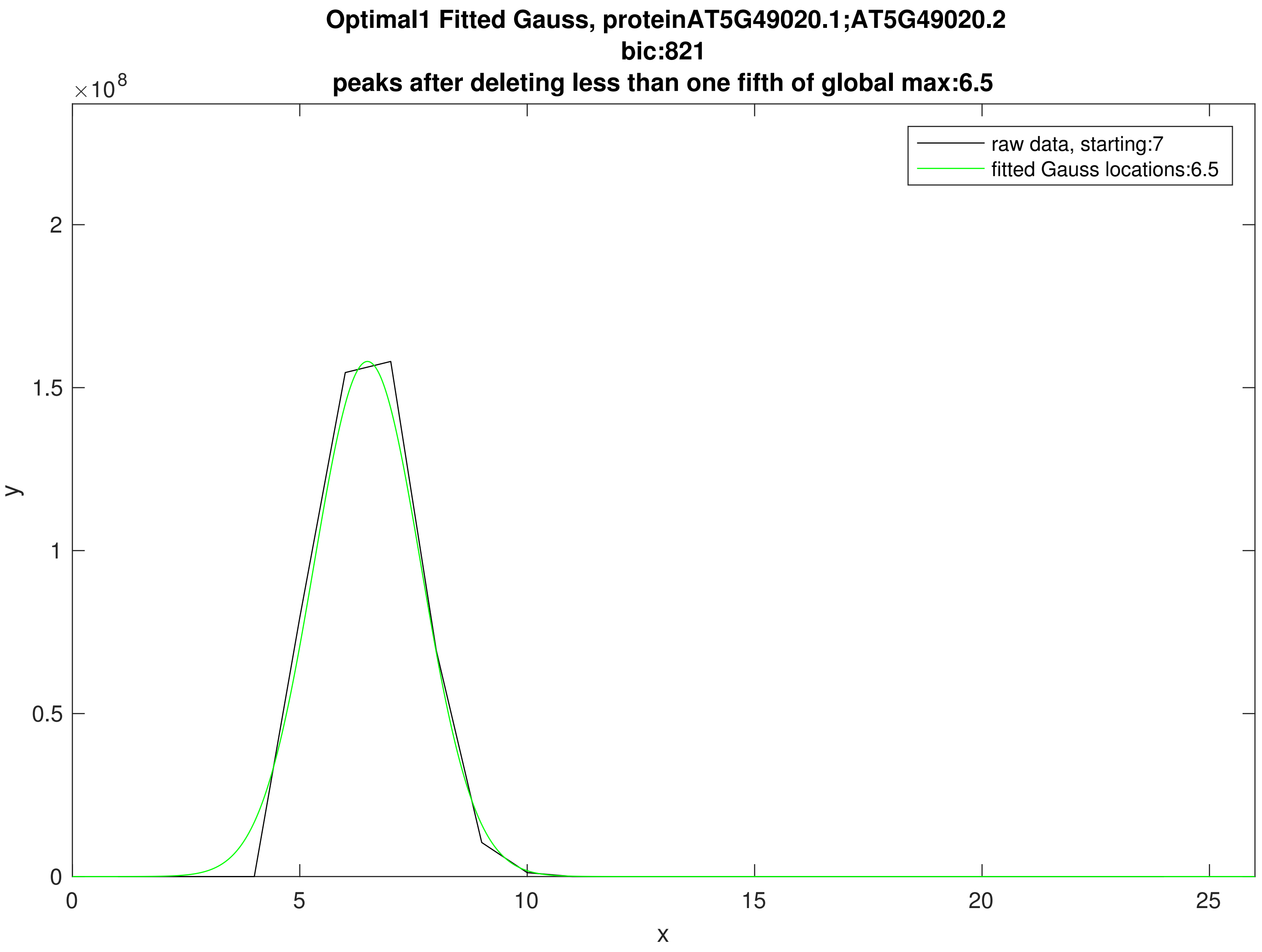

Optimal1 Fitted Gauss, proteinAT5G49020.1;AT5G49020.2
bic:821
peaks after deleting less than one fifth of global max:6.5
×10^8
raw data, starting:7
fitted Gauss locations:6.5
y
x
0
0.5
1
1.5
2
0
5
10
15
20
25

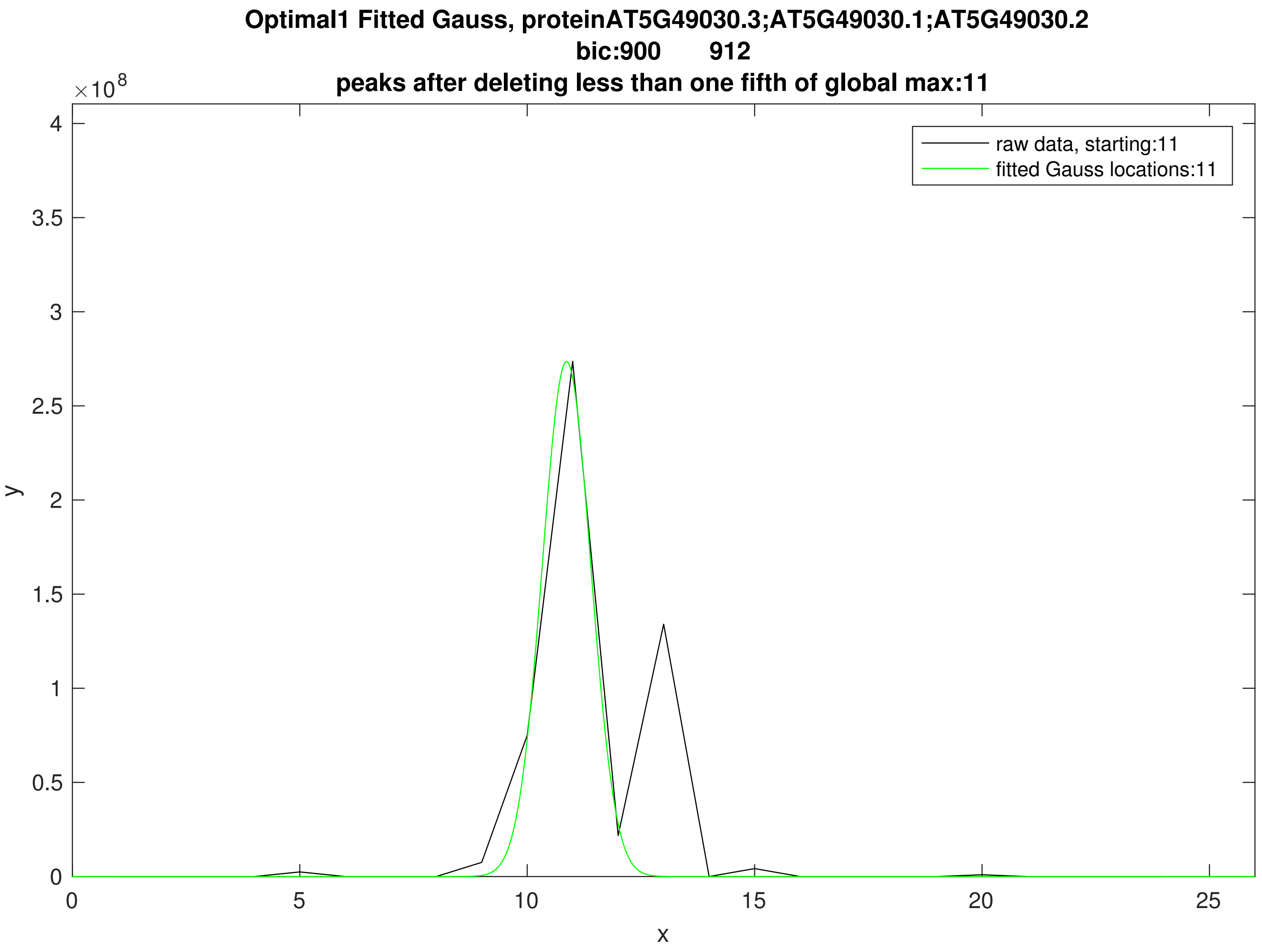

Optimal1 Fitted Gauss, proteinAT5G49030.3;AT5G49030.1;AT5G49030.2
bic:900 912
peaks after deleting less than one fifth of global max:11
×10^8
raw data, starting:11
fitted Gauss locations:11
y
x
0
0.5
1
1.5
2
2.5
3
3.5
4
5
10
15
20
25

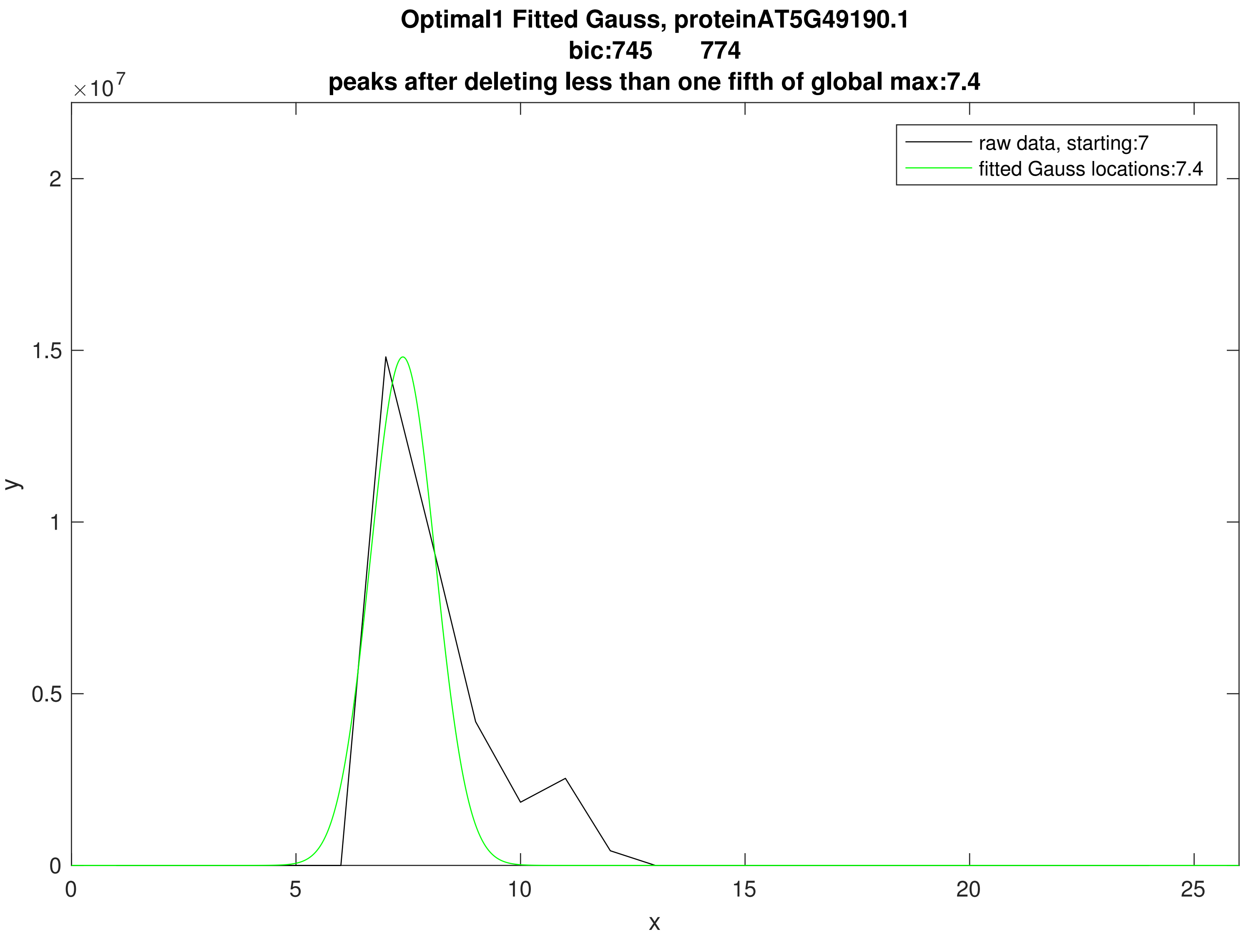

Optimal1 Fitted Gauss, proteinAT5G49190.1
bic:745 774
peaks after deleting less than one fifth of global max:7.4
×10^7
raw data, starting:7
fitted Gauss locations:7.4
y
x
0
0.5
1
1.5
2
5
10
15
20
25

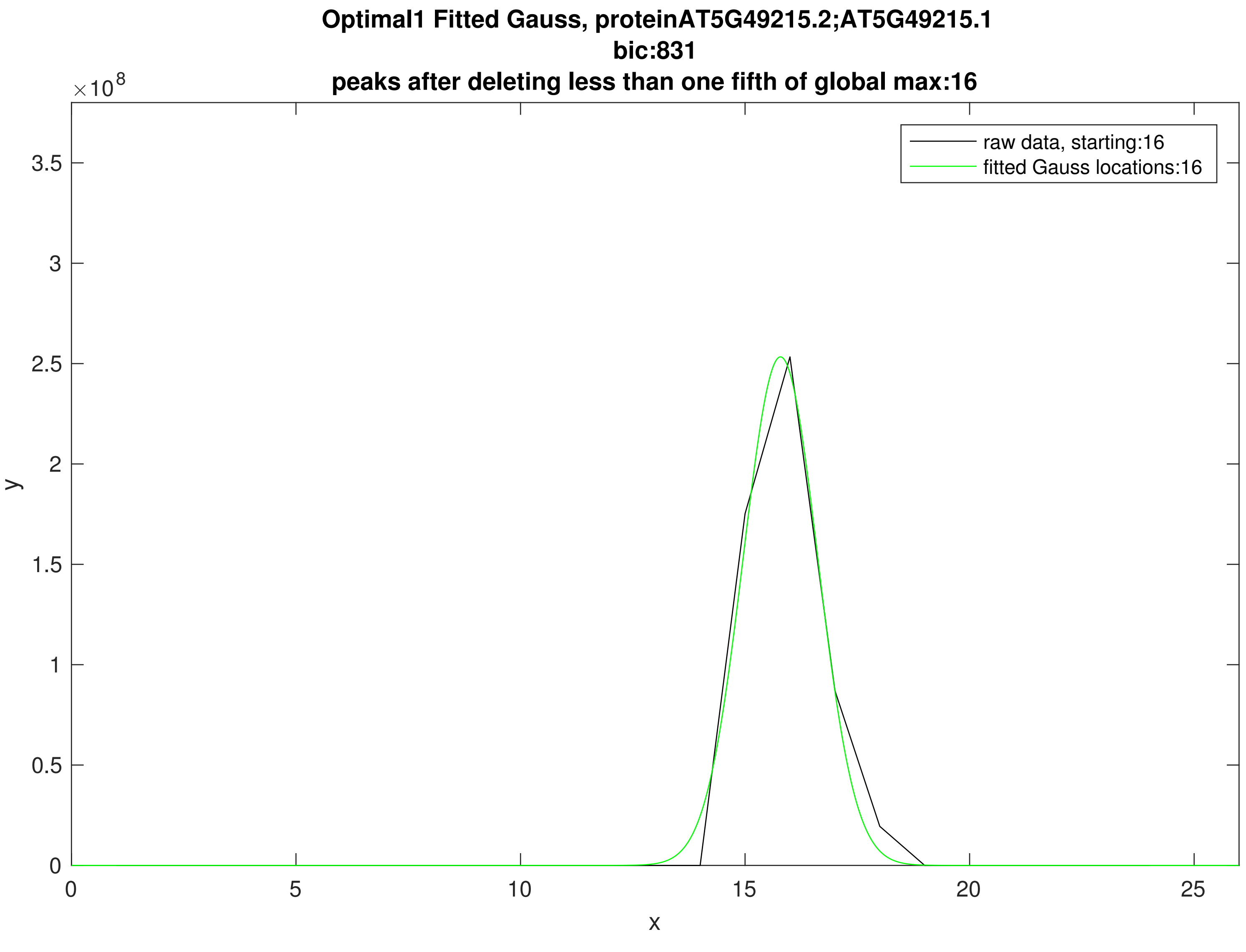
Optimal1 Fitted Gauss, proteinAT5G49215.2;AT5G49215.1
bic:831
peaks after deleting less than one fifth of global max:16
×10^8
raw data, starting:16
fitted Gauss locations:16
y
x
0
0.5
1
1.5
2
2.5
3
3.5
0
5
10
15
20
25

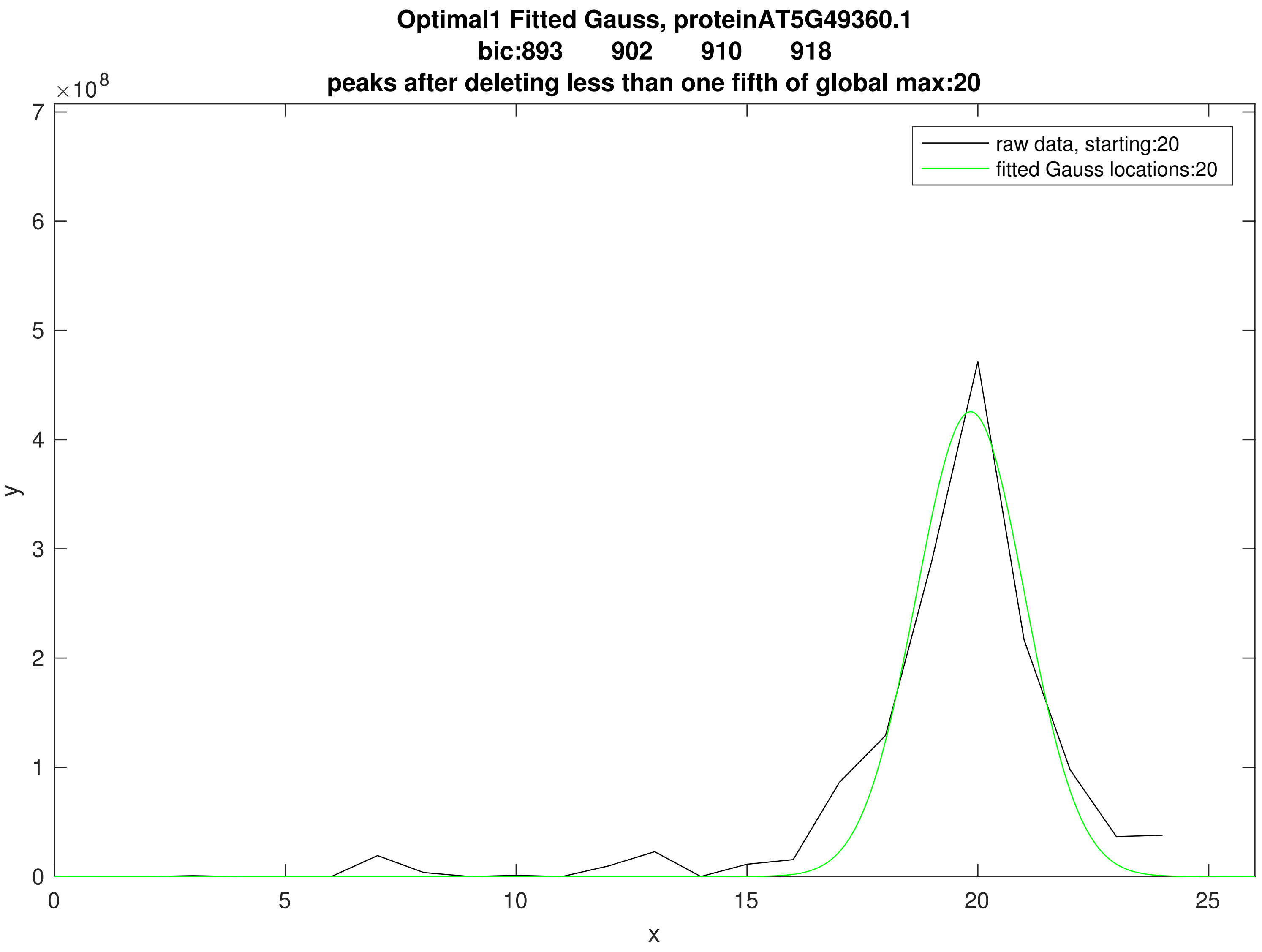
Optimal1 Fitted Gauss, proteinAT5G49360.1
bic:893 902 910 918
peaks after deleting less than one fifth of global max:20
×10^8
raw data, starting:20
fitted Gauss locations:20
y
x
0
1
2
3
4
5
6
7
0
5
10
15
20
25

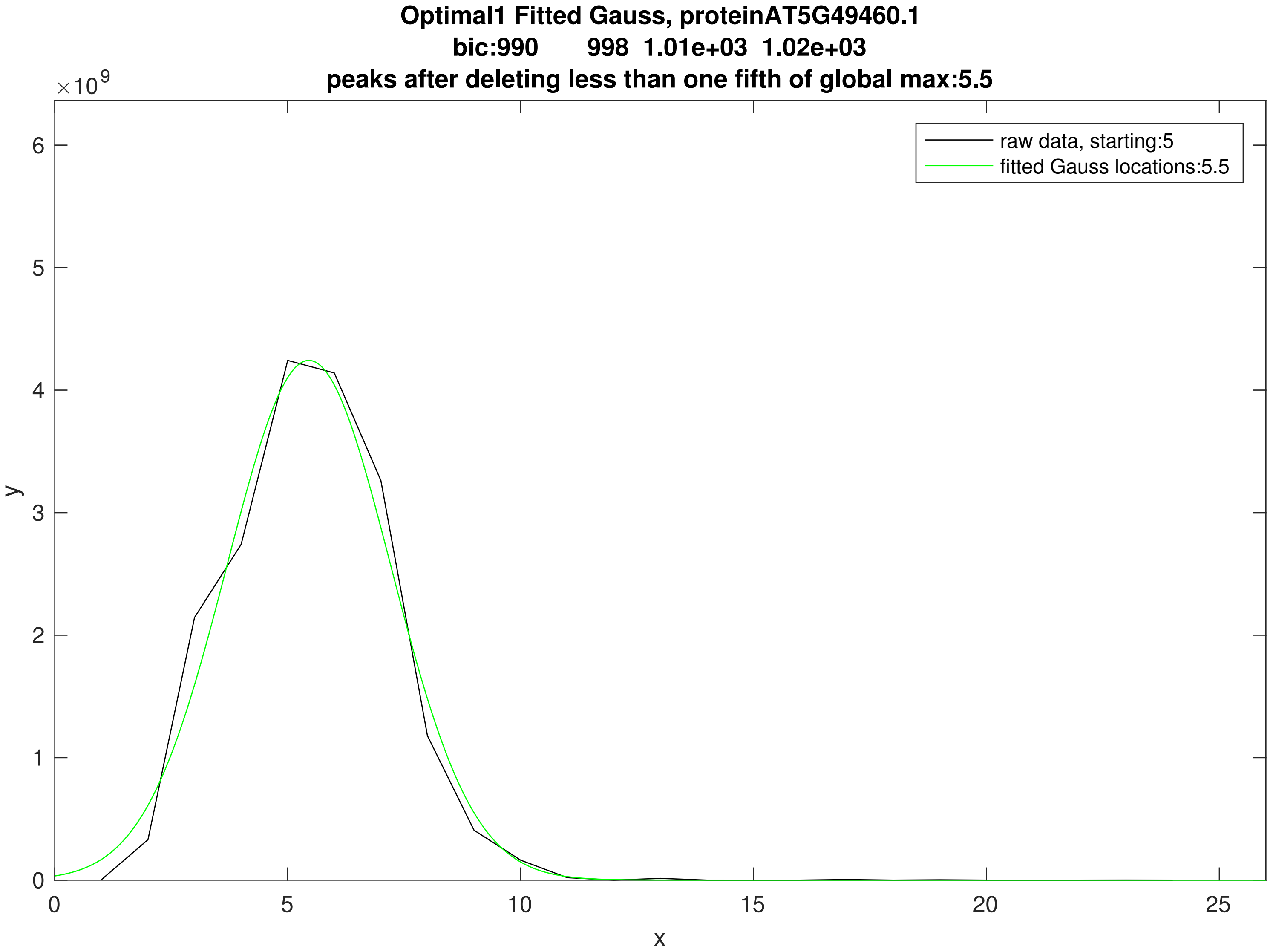

Optimal1 Fitted Gauss, proteinAT5G49460.1
bic:990 998 1.01e+03 1.02e+03
peaks after deleting less than one fifth of global max:5.5
×10^9
raw data, starting:5
fitted Gauss locations:5.5
y
x
0
1
2
3
4
5
6
10
15
20
25

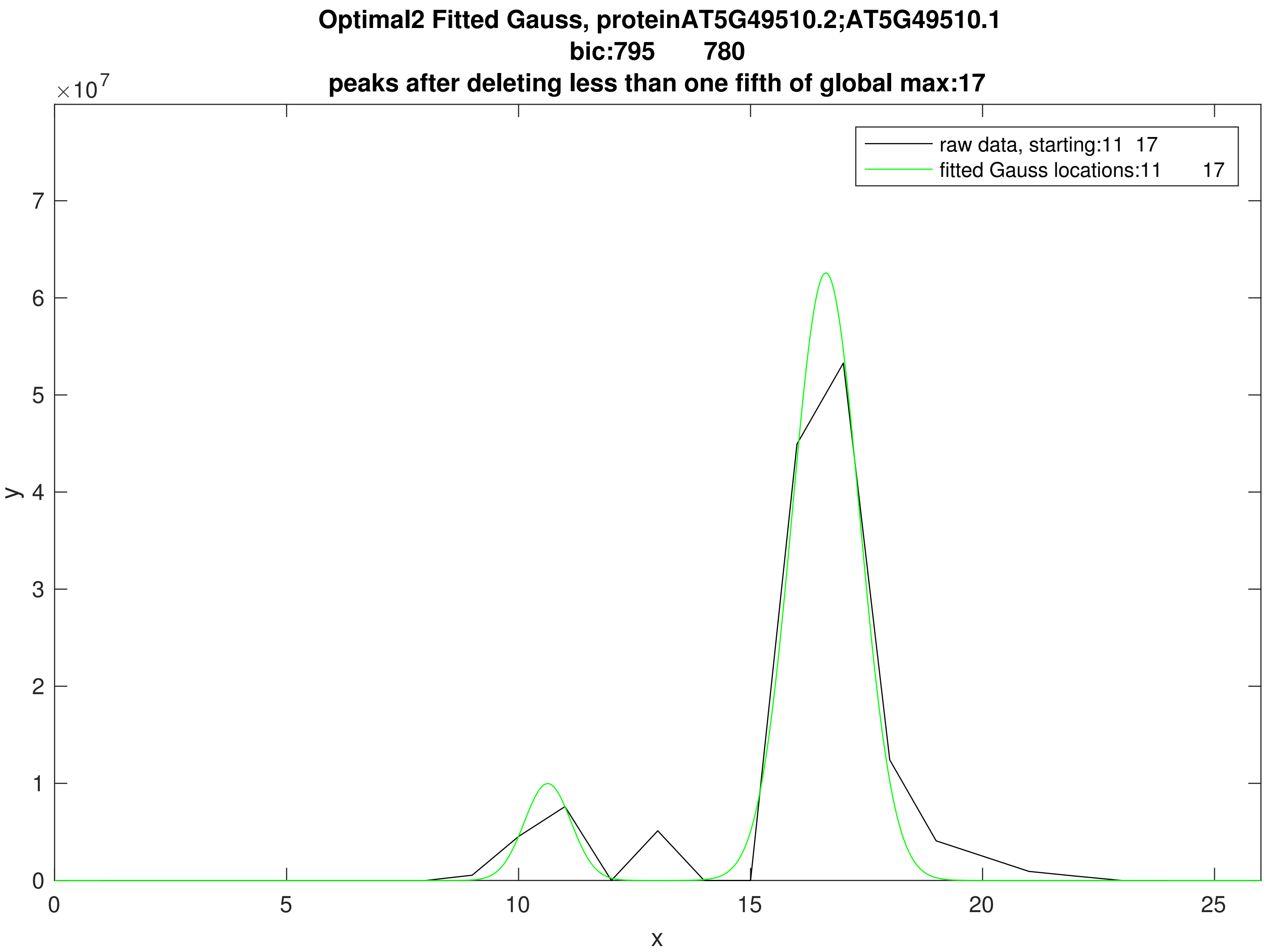

Optimal2 Fitted Gauss, proteinAT5G49510.2;AT5G49510.1
bic:795 780
peaks after deleting less than one fifth of global max:17
×10^7
raw data, starting:11 17
fitted Gauss locations:11 17
y
x
0
1
2
3
4
5
6
7
0
5
10
15
20
25

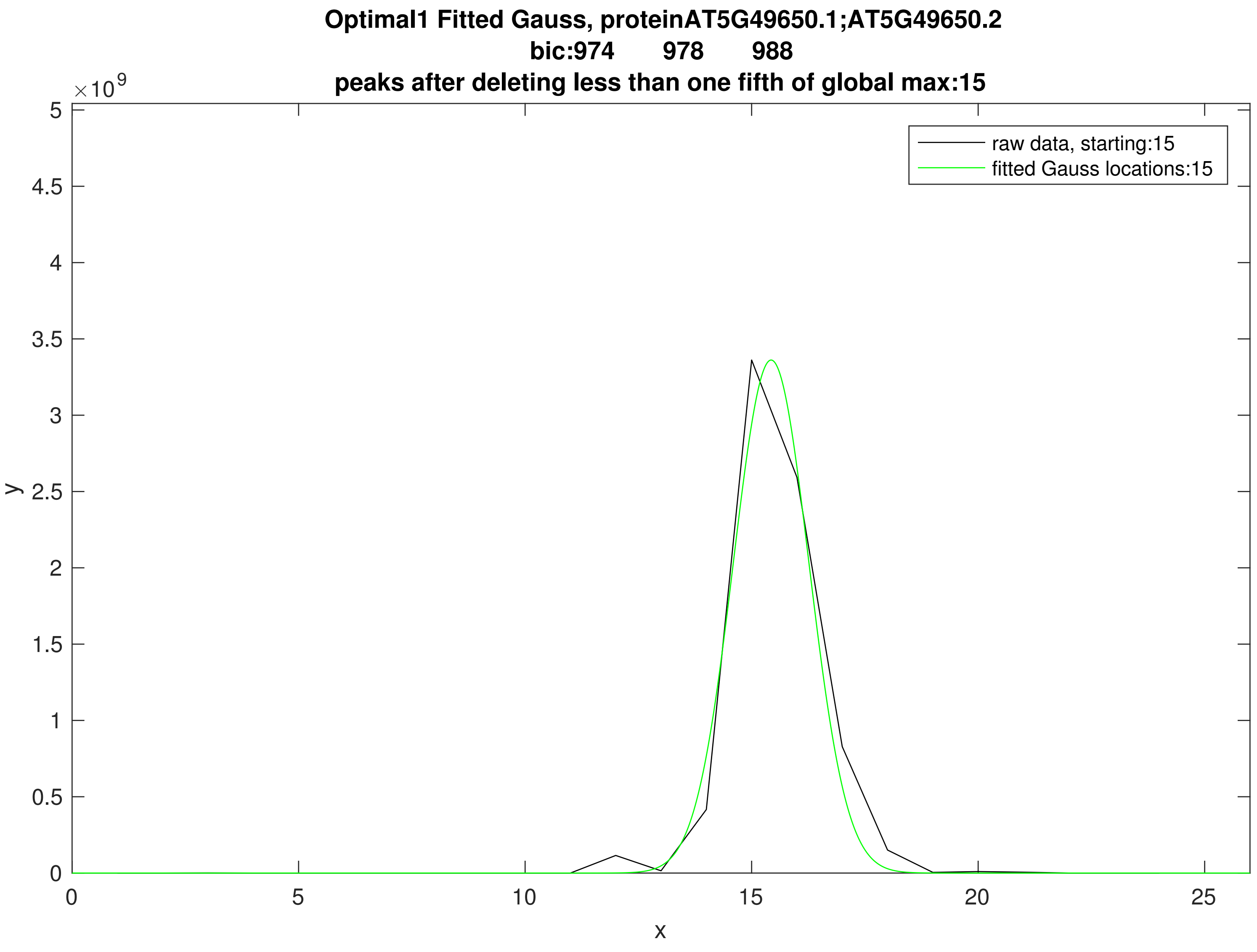

Optimal1 Fitted Gauss, proteinAT5G49650.1;AT5G49650.2
bic:974 978 988
peaks after deleting less than one fifth of global max:15
×10^9
raw data, starting:15
fitted Gauss locations:15
y
x
0
0.5
1
1.5
2
2.5
3
3.5
4
4.5
5
0
5
10
15
20
25

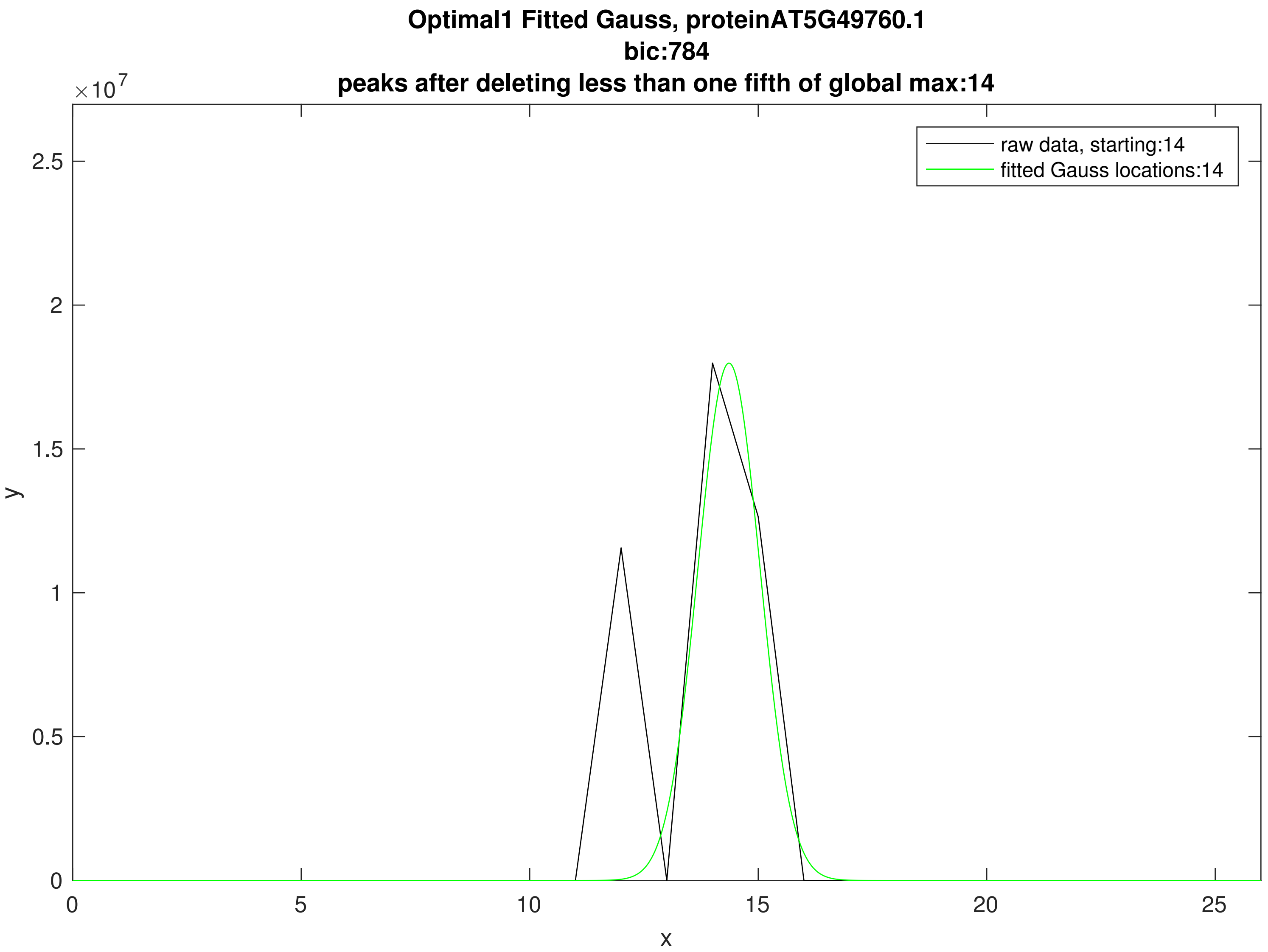
Optimal1 Fitted Gauss, proteinAT5G49760.1
bic:784
peaks after deleting less than one fifth of global max:14
×10^7
raw data, starting:14
fitted Gauss locations:14
y
x
0
0.5
1
1.5
2
2.5
0
5
10
15
20
25

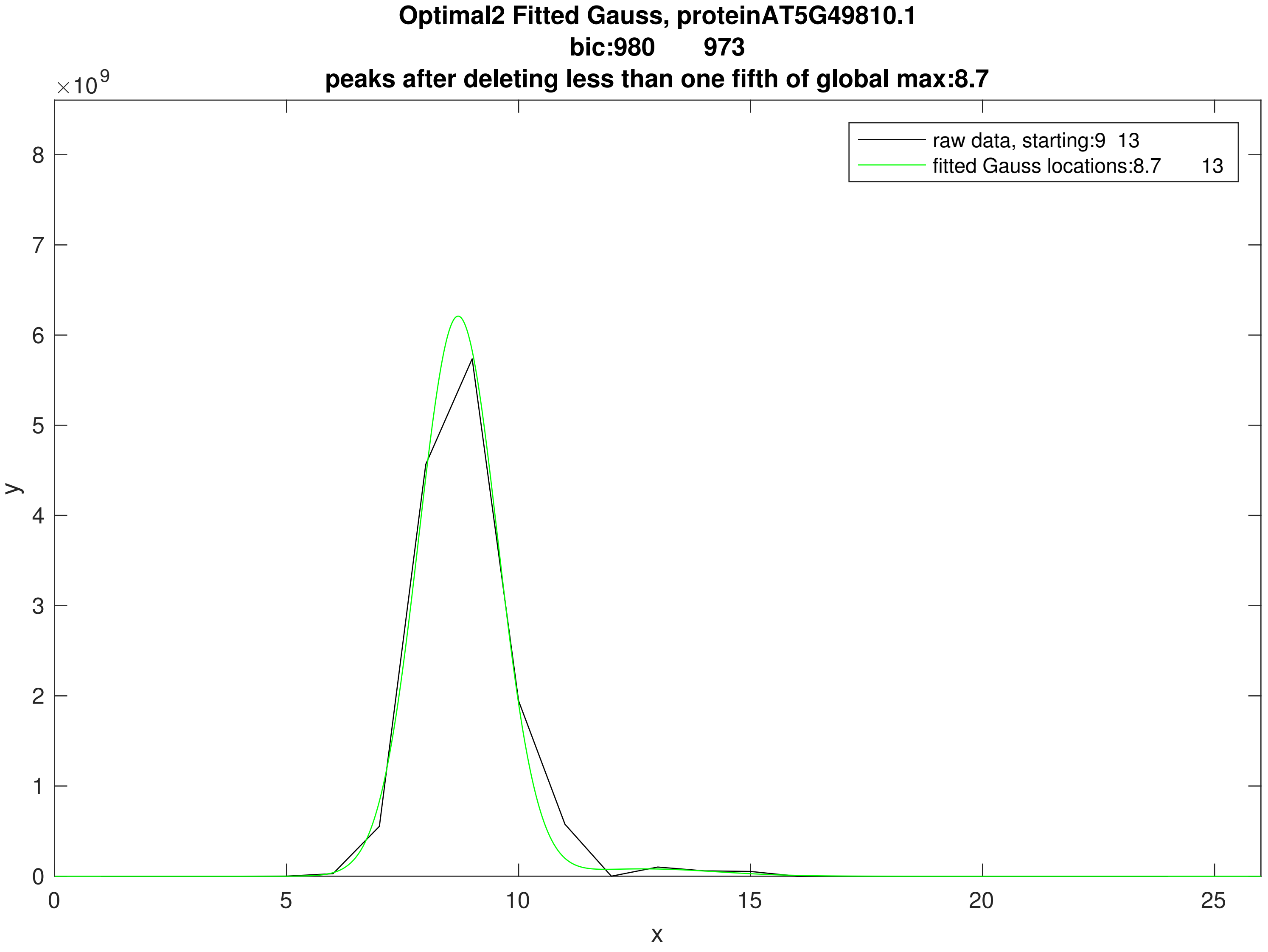

Optimal2 Fitted Gauss, proteinAT5G49810.1
bic:980 973
peaks after deleting less than one fifth of global max:8.7
×10^9
raw data, starting:9 13
fitted Gauss locations:8.7 13
y
x
0
1
2
3
4
5
6
7
8
0
5
10
15
20
25

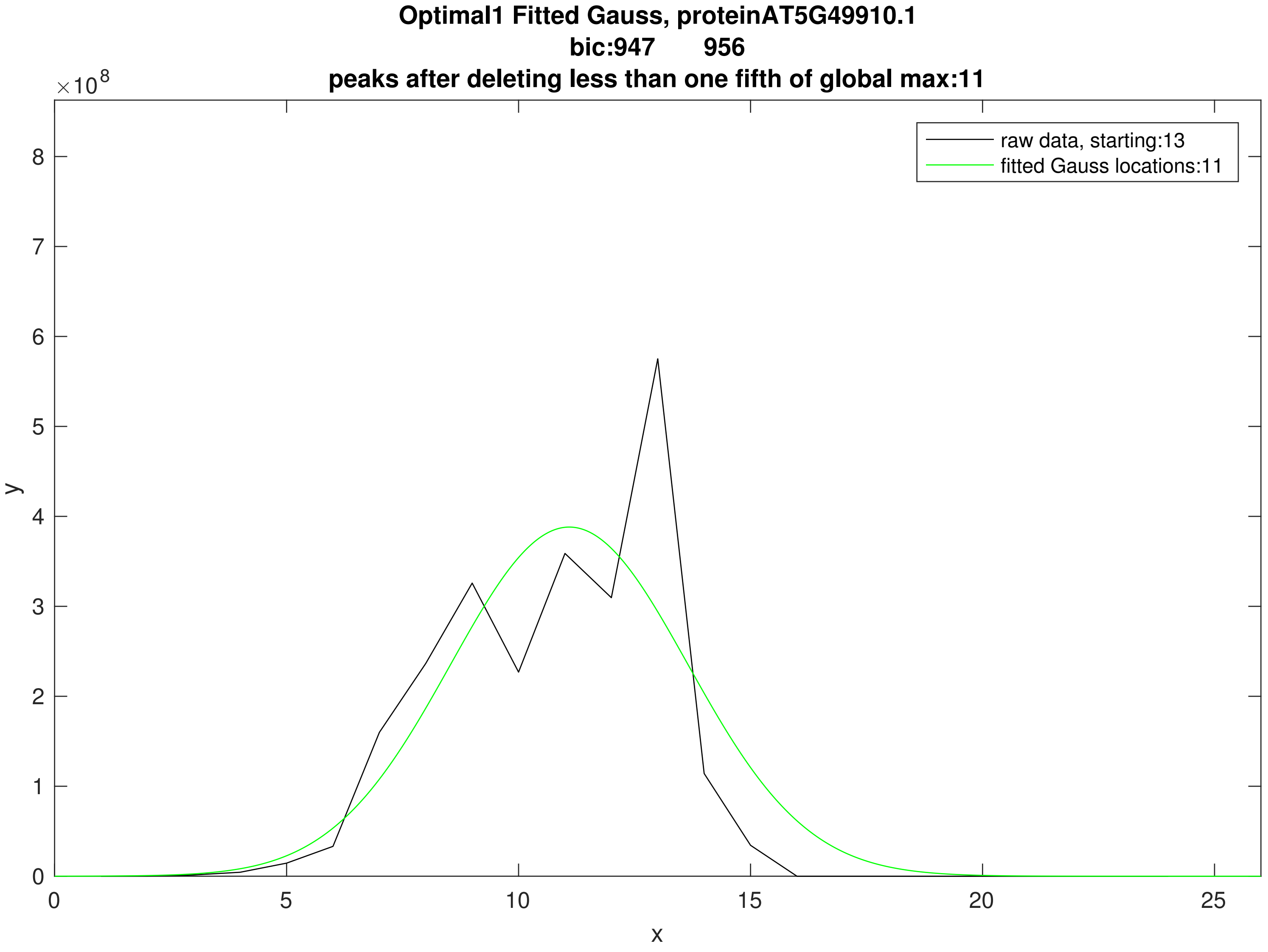
Optimal1 Fitted Gauss, proteinAT5G49910.1
bic:947 956
peaks after deleting less than one fifth of global max:11
×10^8
raw data, starting:13
fitted Gauss locations:11
y
x
0
1
2
3
4
5
6
7
8
0
5
10
15
20
25

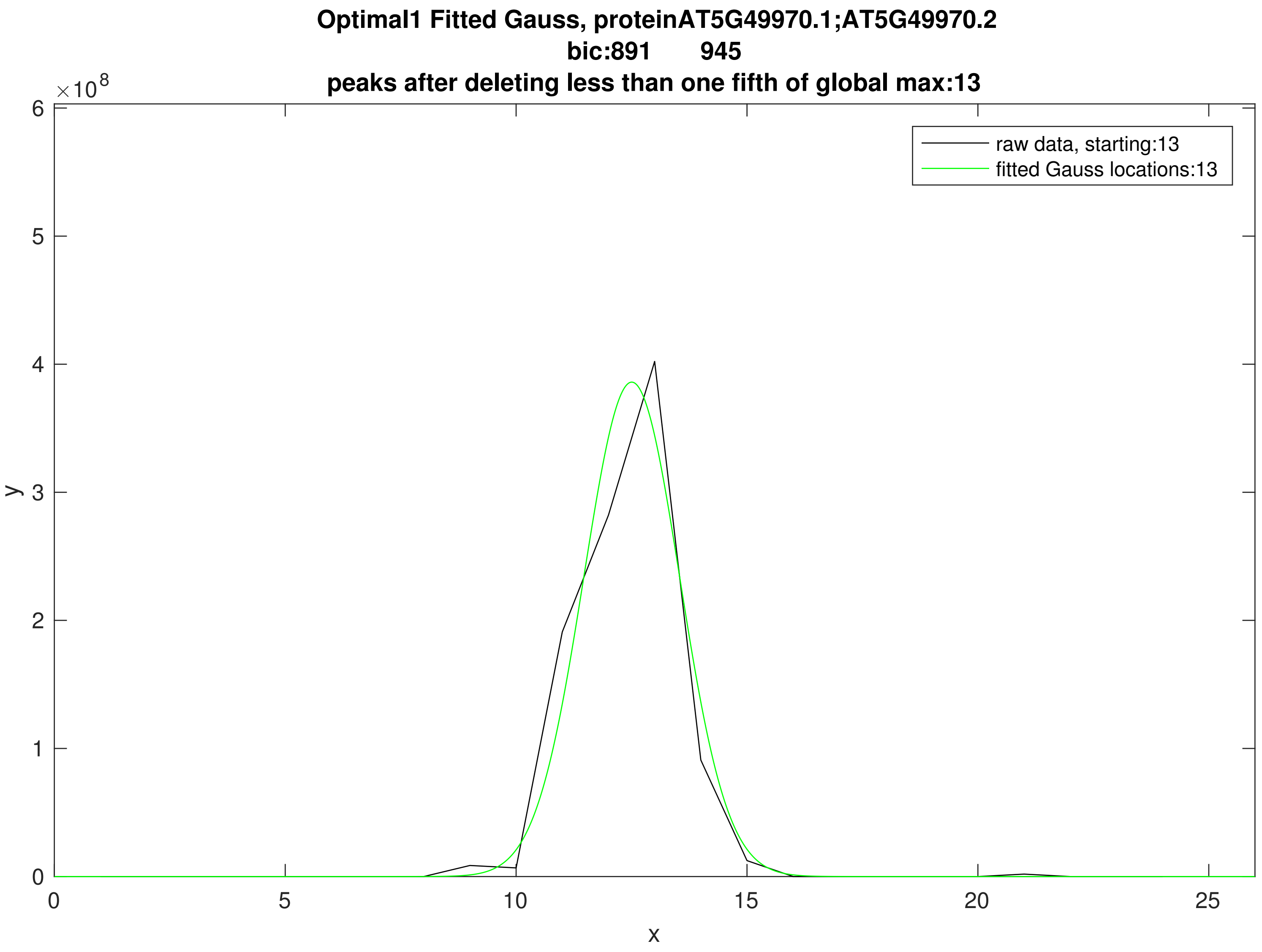

Optimal1 Fitted Gauss, proteinAT5G49970.1;AT5G49970.2
bic:891 945
peaks after deleting less than one fifth of global max:13
×10^8
6
5
4
3
2
1
0
y
0
5
10
15
20
25
x
raw data, starting:13
fitted Gauss locations:13

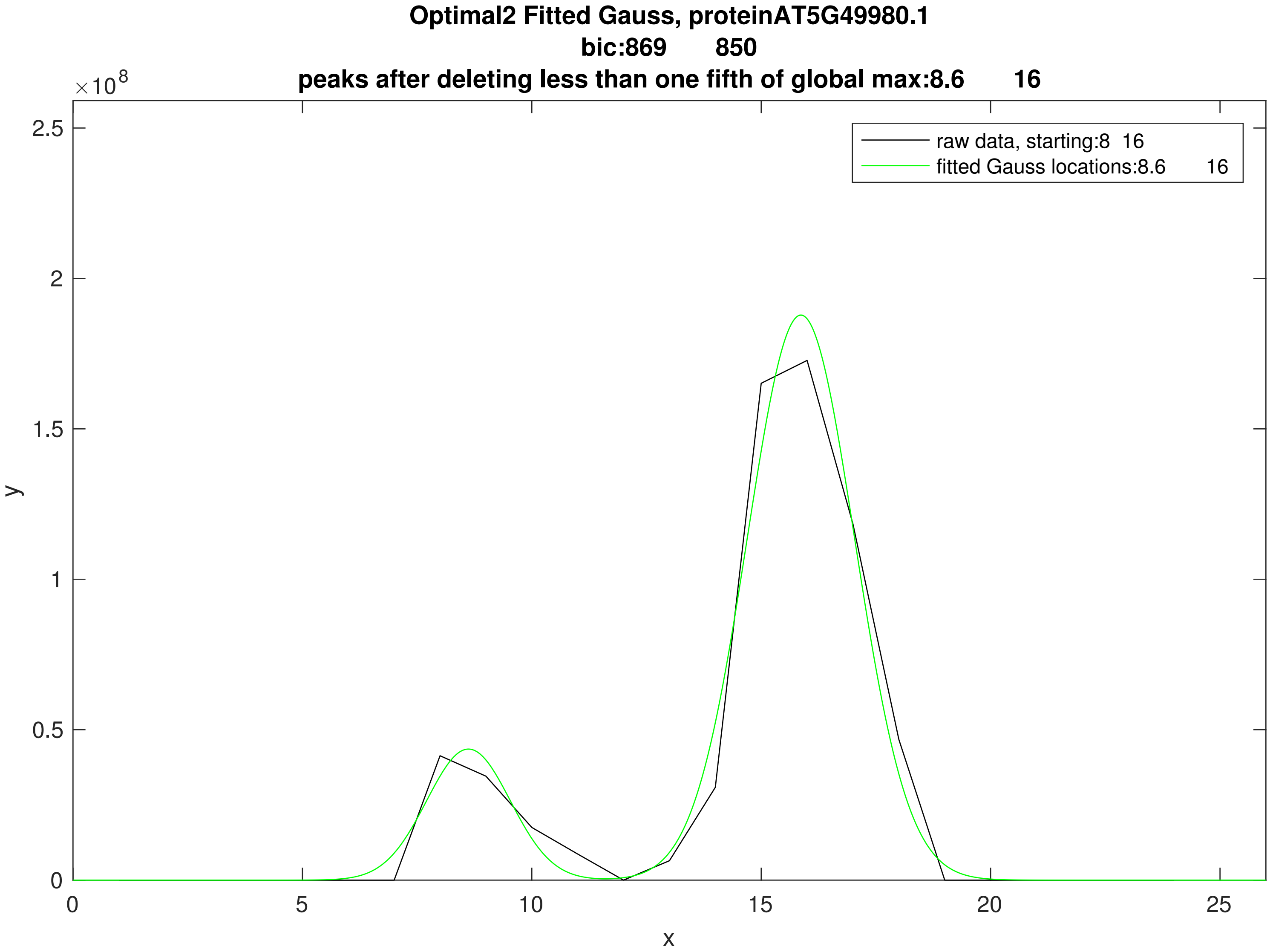

Optimal2 Fitted Gauss, proteinAT5G49980.1
bic:869 850
peaks after deleting less than one fifth of global max:8.6 16
×10^8
raw data, starting:8 16
fitted Gauss locations:8.6 16
2.5
2
1.5
1
0.5
0
y
0
5
10
15
20
25
x

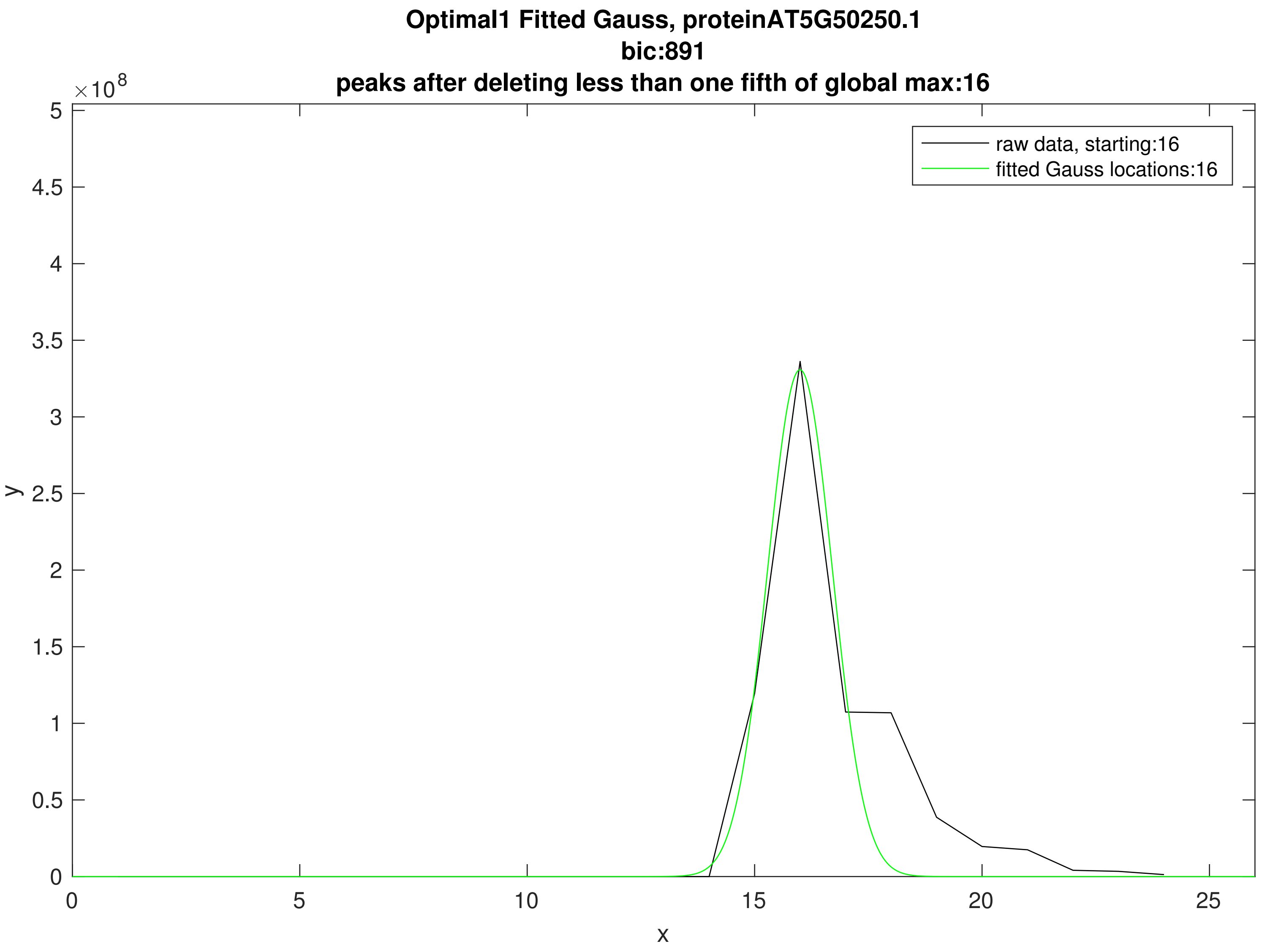
Optimal1 Fitted Gauss, proteinAT5G50250.1
bic:891
peaks after deleting less than one fifth of global max:16
×10^8
raw data, starting:16
fitted Gauss locations:16
y
x
0
0.5
1
1.5
2
2.5
3
3.5
4
4.5
5
0
5
10
15
20
25

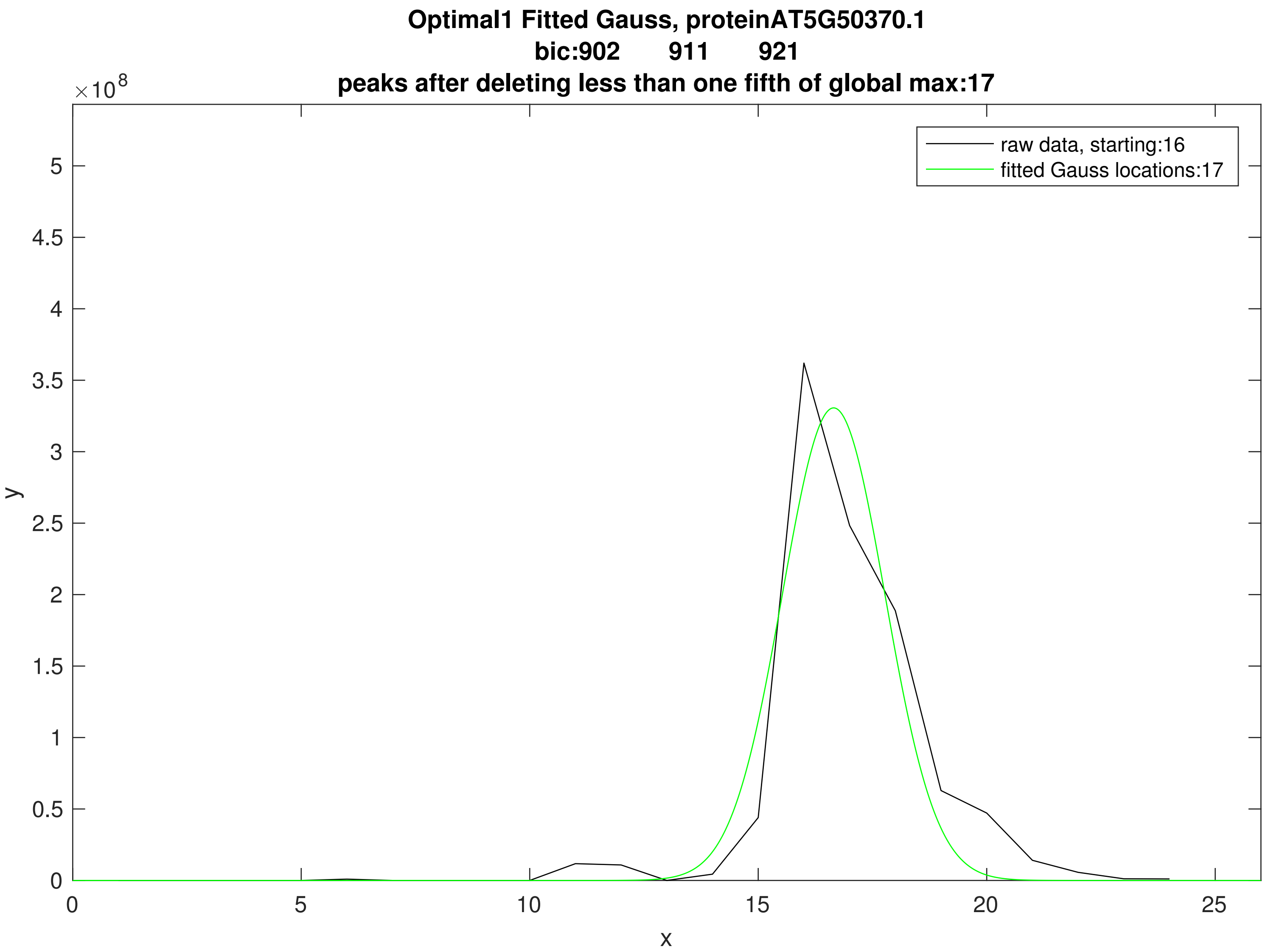

Optimal1 Fitted Gauss, proteinAT5G50370.1
bic:902 911 921
peaks after deleting less than one fifth of global max:17
×10^8
raw data, starting:16
fitted Gauss locations:17
y
x
0
0.5
1
1.5
2
2.5
3
3.5
4
4.5
5
0
5
10
15
20
25

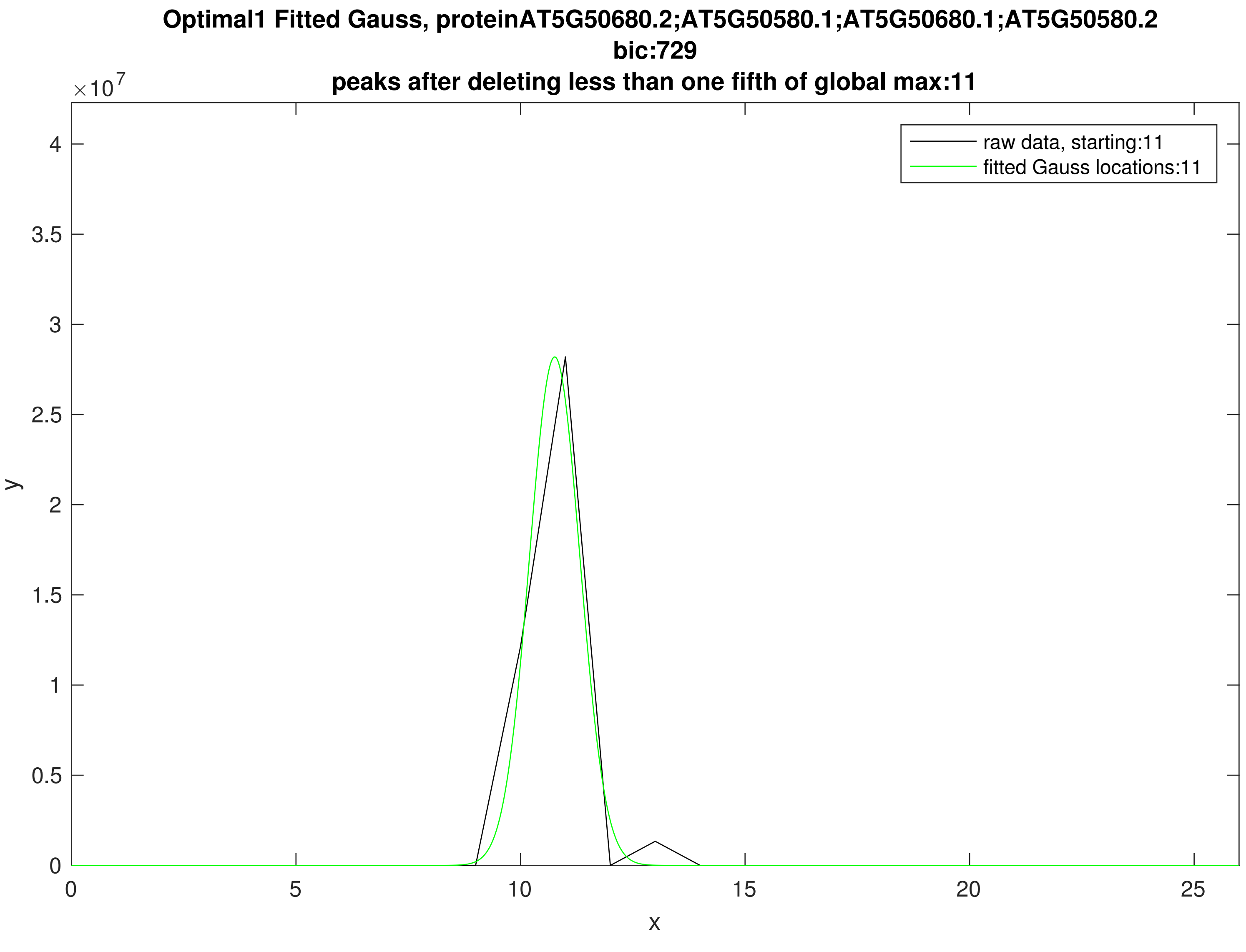

Optimal1 Fitted Gauss, proteinAT5G50680.2;AT5G50580.1;AT5G50680.1;AT5G50580.2
bic:729
peaks after deleting less than one fifth of global max:11
×10^7
4
3.5
3
2.5
2
1.5
1
0.5
0
y
0
5
10
15
20
25
x
raw data, starting:11
fitted Gauss locations:11

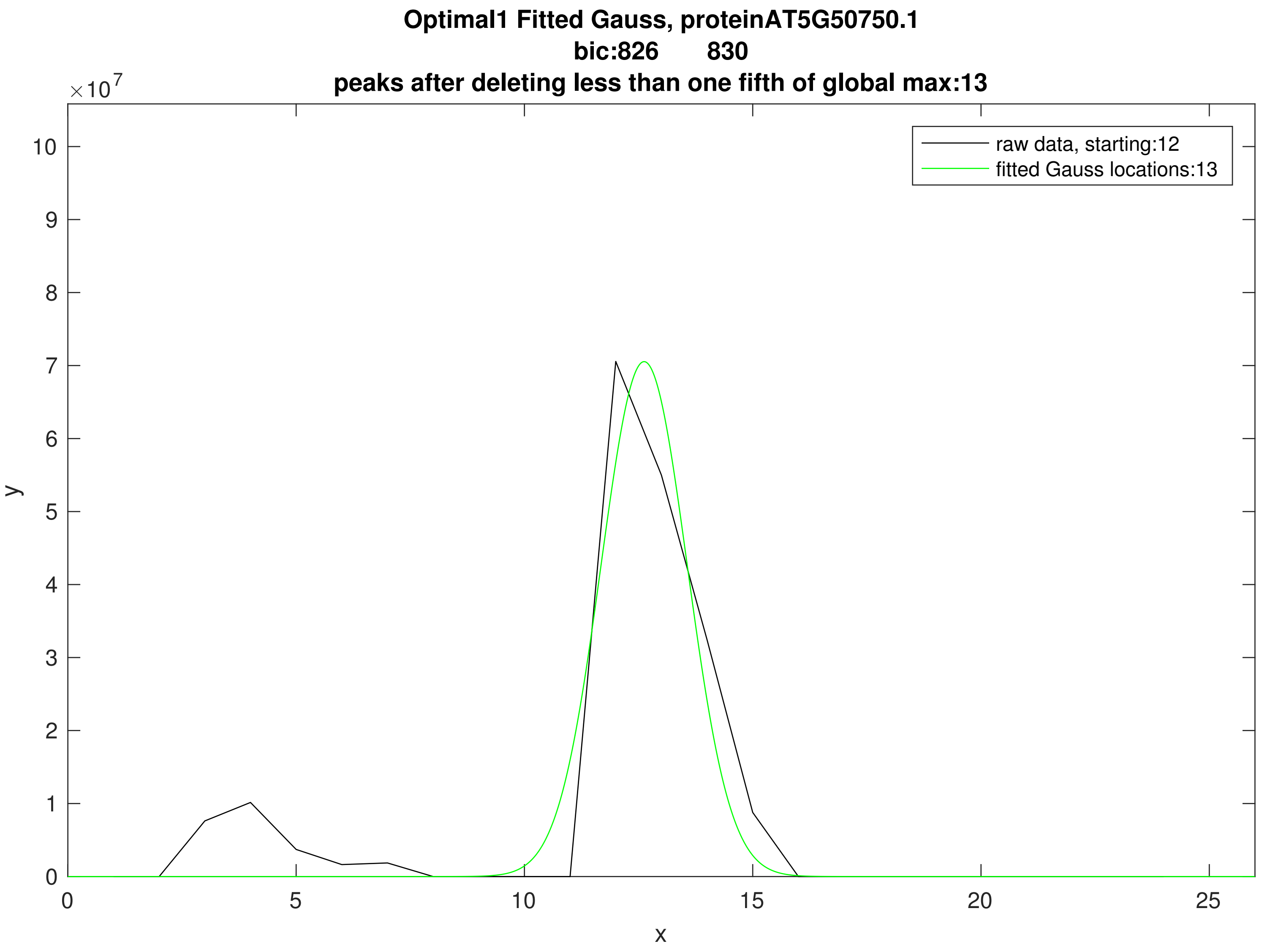
Optimal1 Fitted Gauss, proteinAT5G50750.1
bic:826 830
peaks after deleting less than one fifth of global max:13
×10^7
raw data, starting:12
fitted Gauss locations:13
y
x
0
1
2
3
4
5
6
7
8
9
10
0
5
10
15
20
25

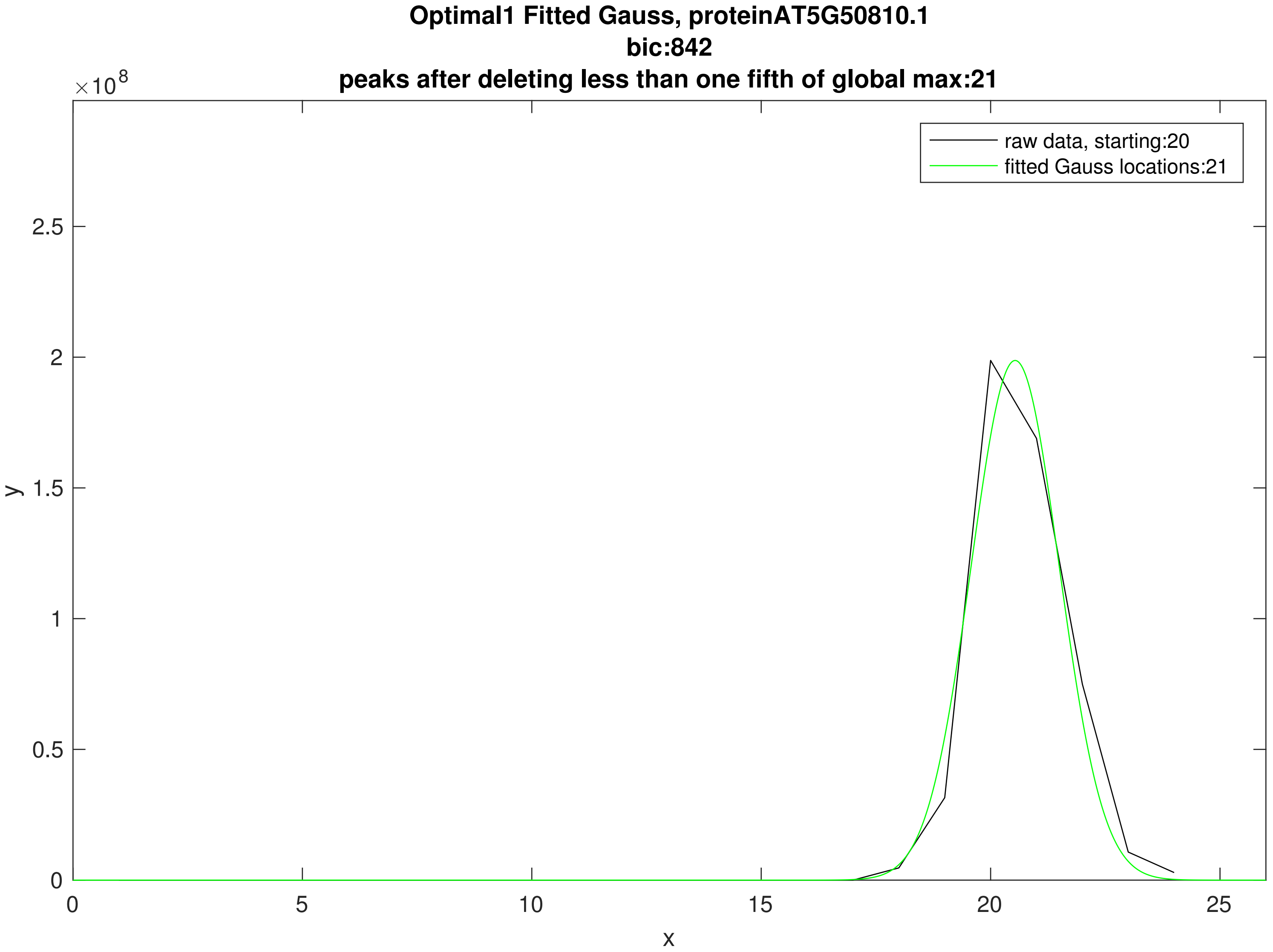
Optimal1 Fitted Gauss, proteinAT5G50810.1
bic:842
peaks after deleting less than one fifth of global max:21
×10^8
raw data, starting:20
fitted Gauss locations:21
2.5
2
1.5
1
0.5
0
y
0
5
10
15
20
25
x

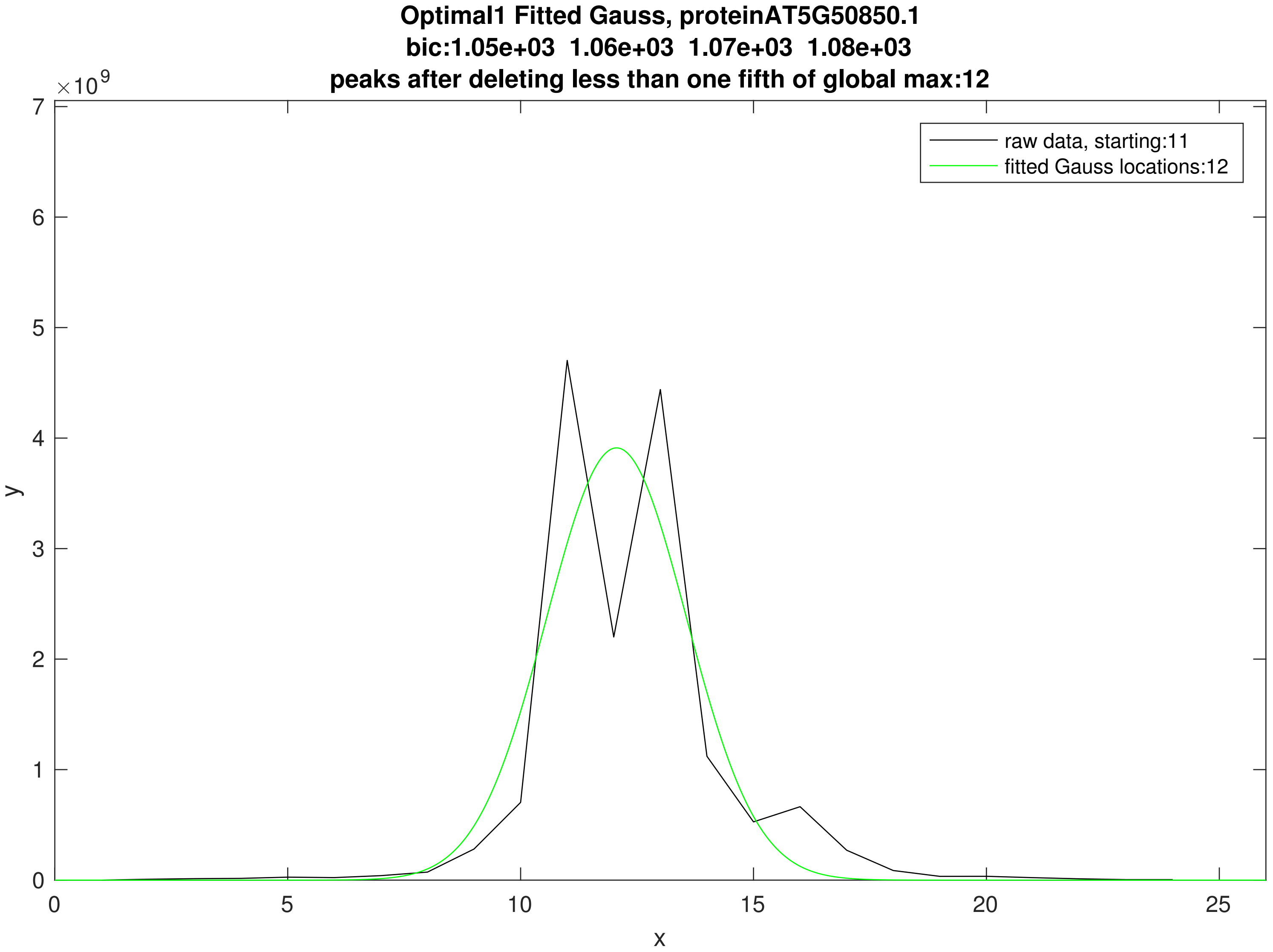
Optimal1 Fitted Gauss, proteinAT5G50850.1
bic:1.05e+03 1.06e+03 1.07e+03 1.08e+03
peaks after deleting less than one fifth of global max:12
×10^9
raw data, starting:11
fitted Gauss locations:12
y
x
0
1
2
3
4
5
6
7
0
5
10
15
20
25

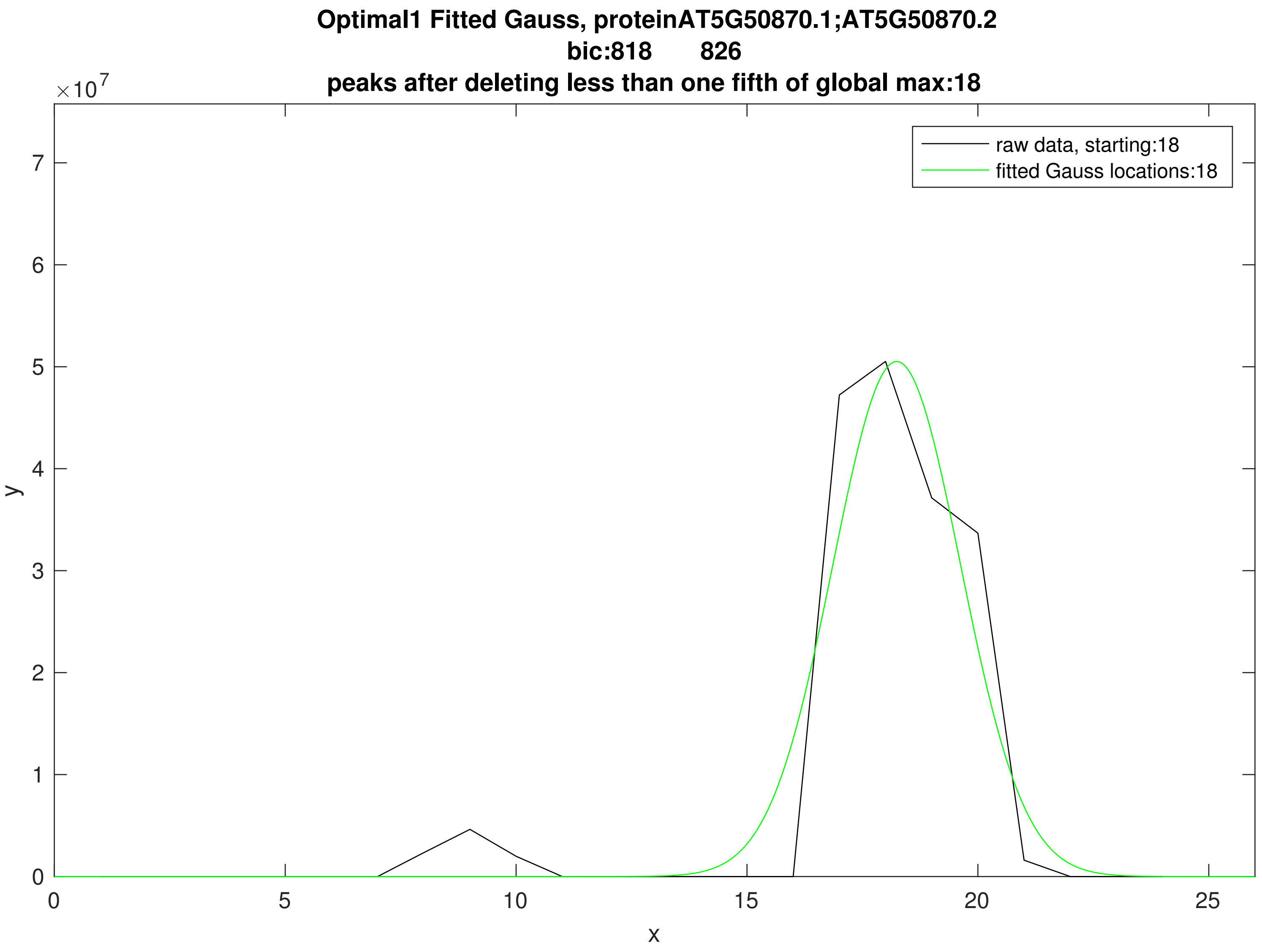
Optimal1 Fitted Gauss, proteinAT5G50870.1;AT5G50870.2
bic:818 826
peaks after deleting less than one fifth of global max:18
×10^7
raw data, starting:18
fitted Gauss locations:18
y
x
0
1
2
3
4
5
6
7
0
5
10
15
20
25

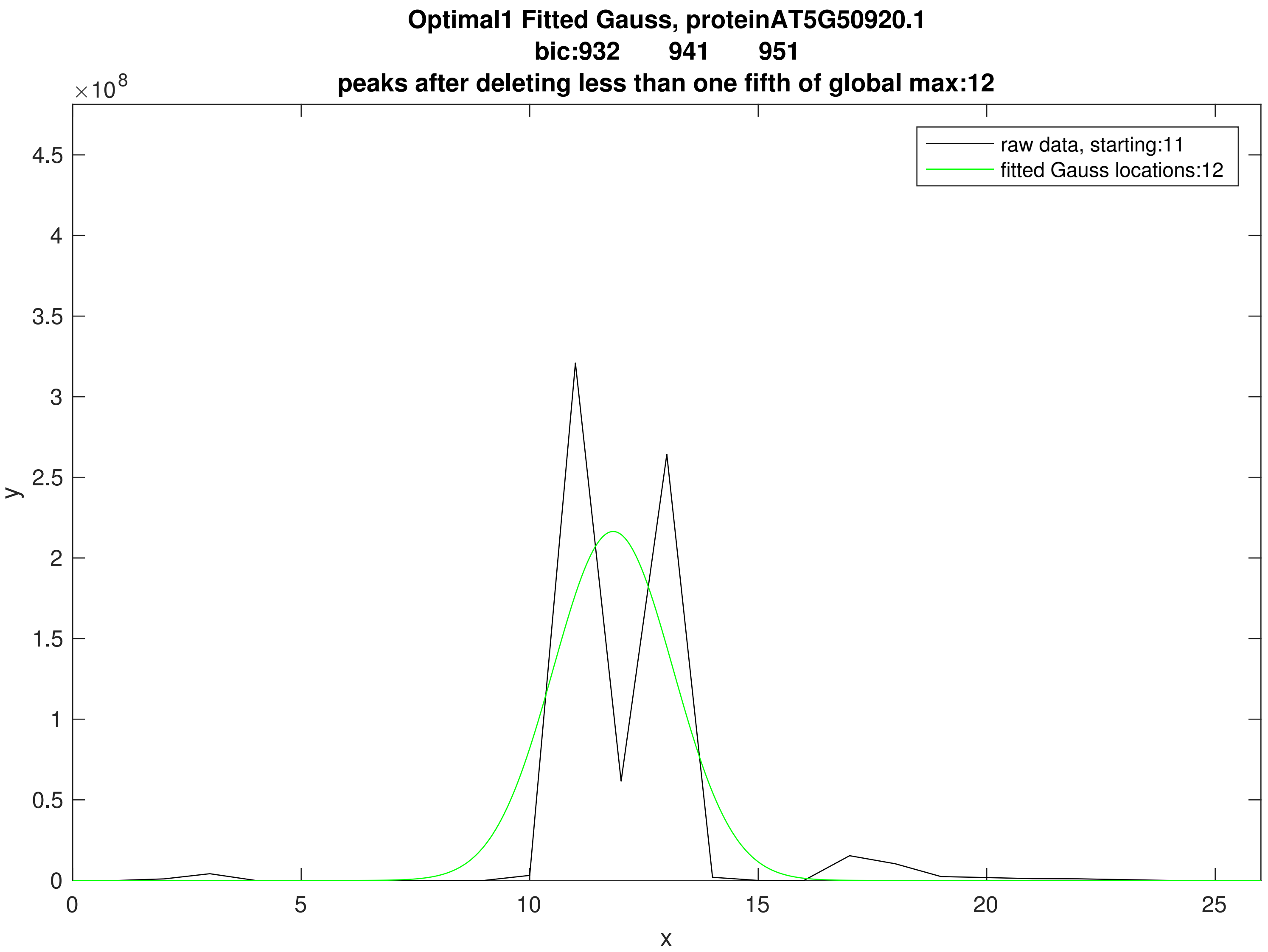
Optimal1 Fitted Gauss, proteinAT5G50920.1
bic:932 941 951
peaks after deleting less than one fifth of global max:12
×10^8
raw data, starting:11
fitted Gauss locations:12
y
x
0
0.5
1
1.5
2
2.5
3
3.5
4
4.5
0
5
10
15
20
25

#### Optimal1 Fitted Gauss, proteinAT5G50960.1
#### bic:782
#### peaks after deleting less than one fifth of global max:13

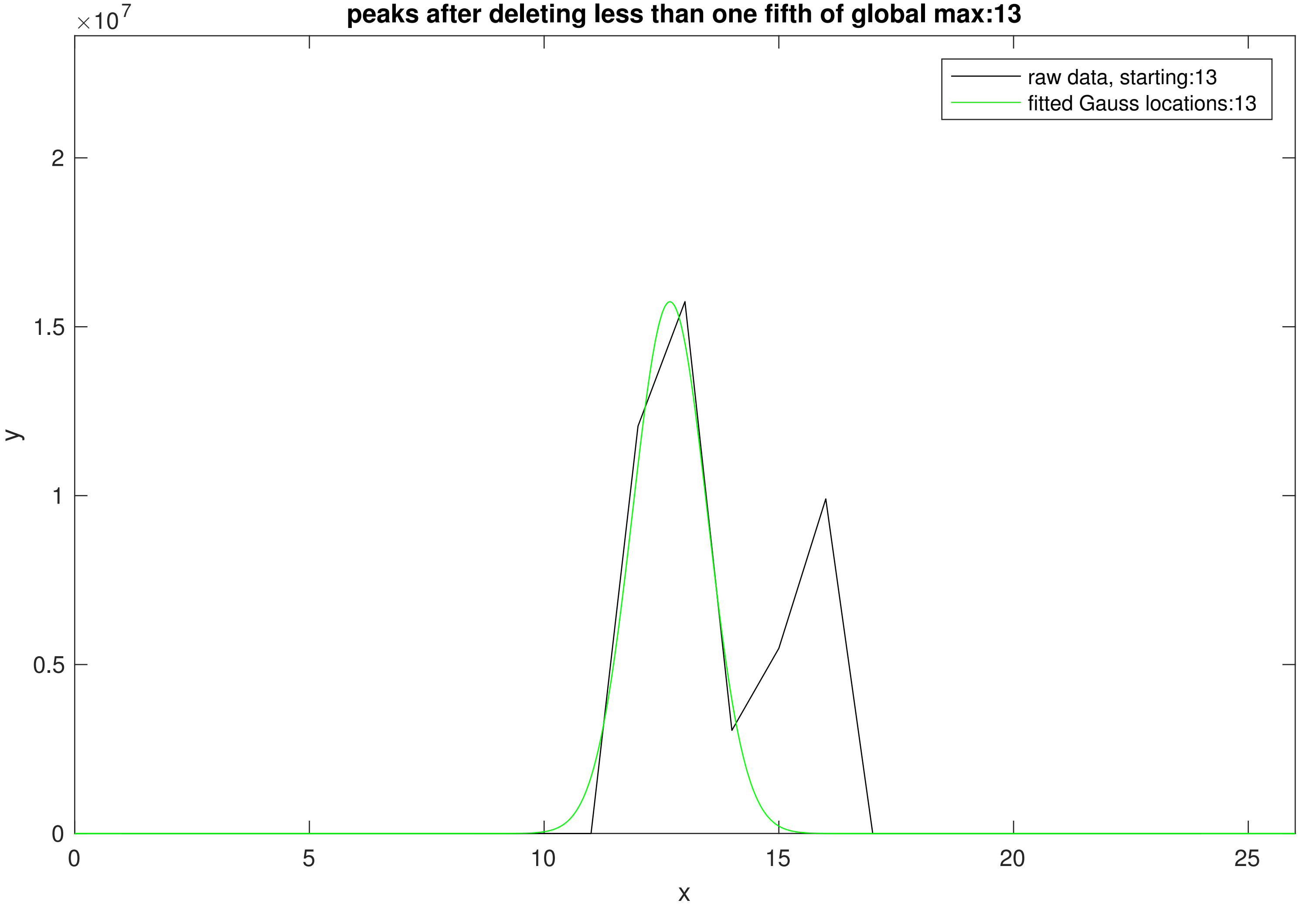

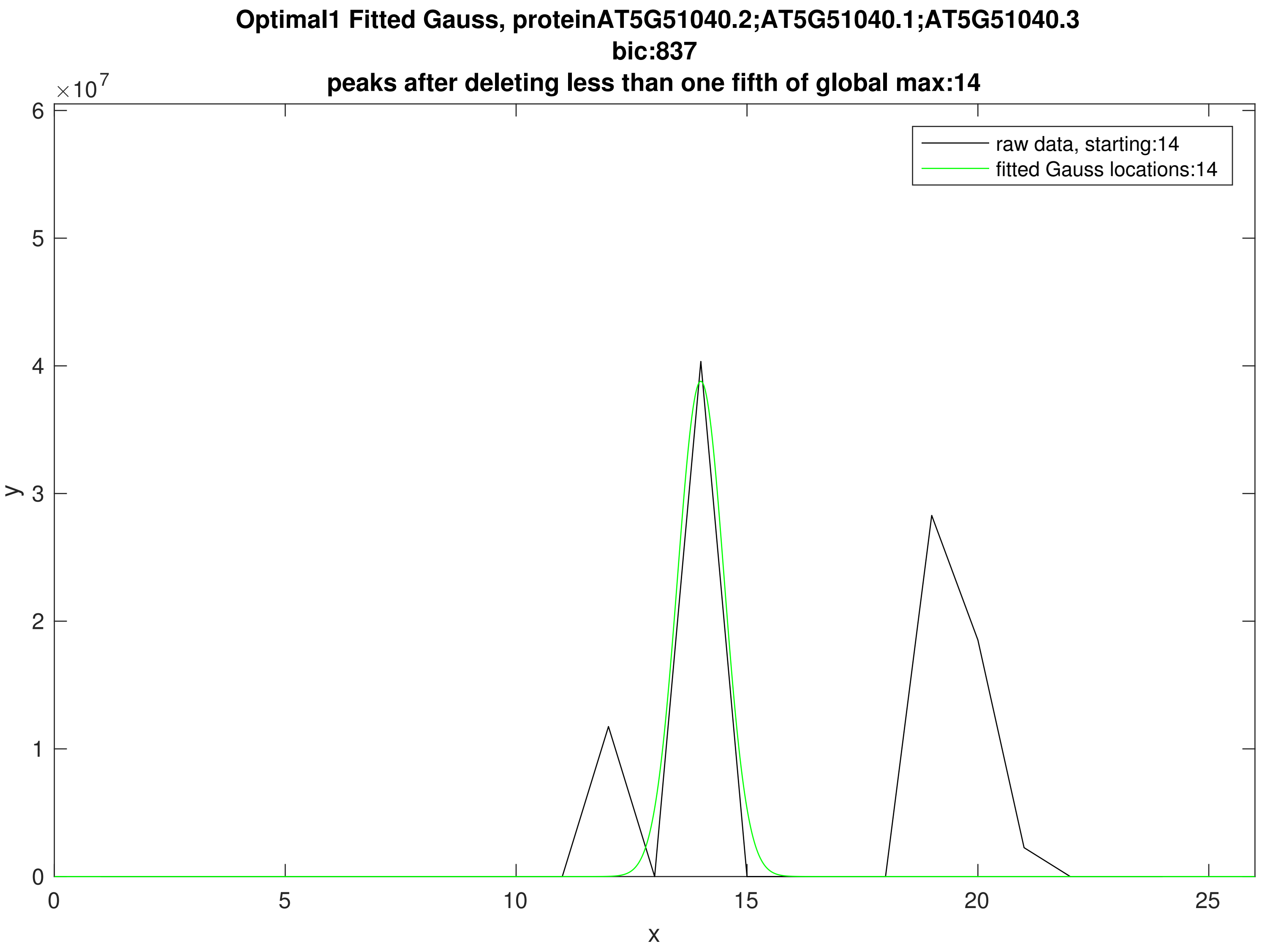
Optimal1 Fitted Gauss, proteinAT5G51040.2;AT5G51040.1;AT5G51040.3
bic:837
peaks after deleting less than one fifth of global max:14
×10^7
6
5
4
3
2
1
0
y
0
5
10
15
20
25
x
raw data, starting:14
fitted Gauss locations:14

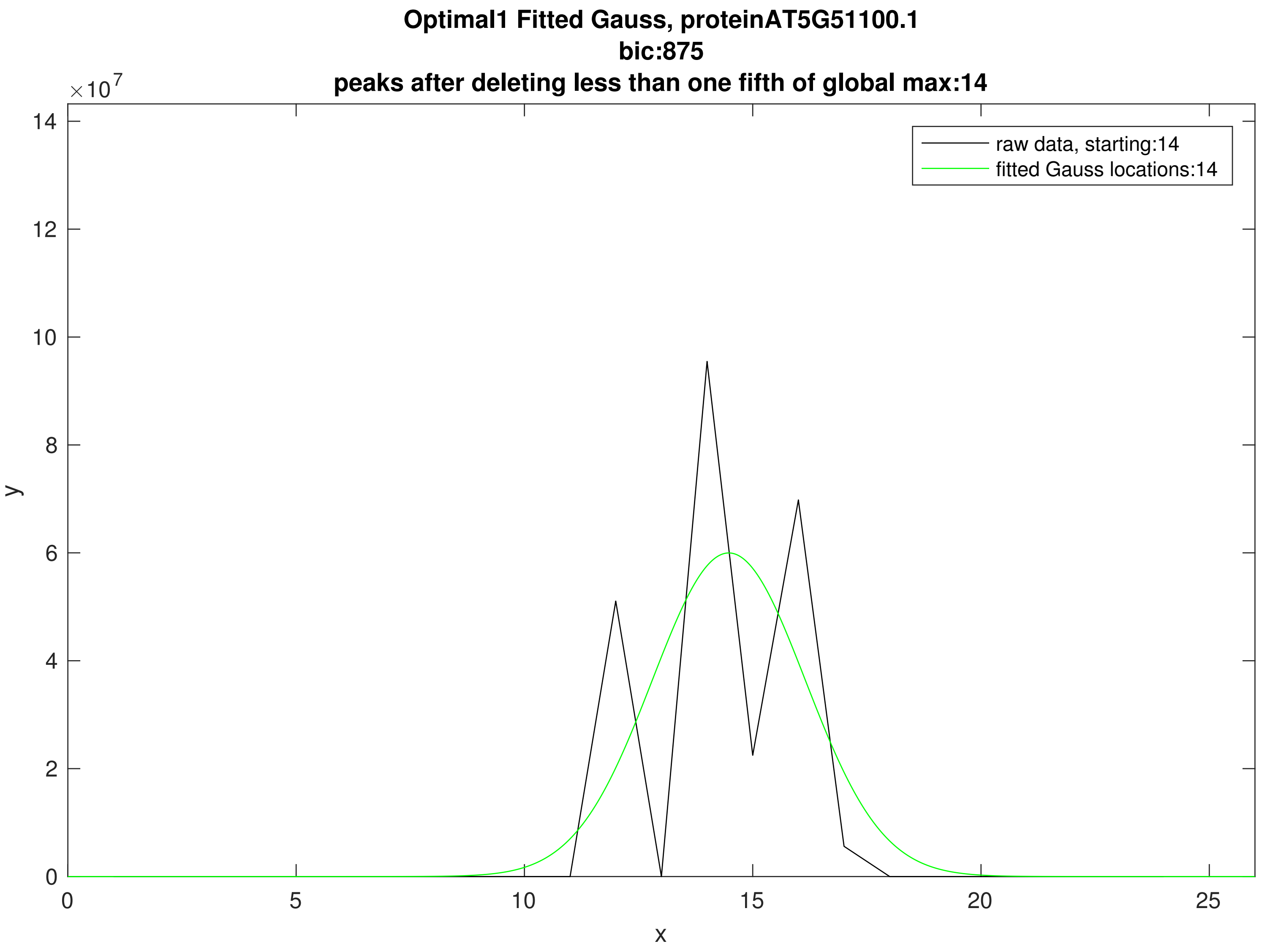
Optimal1 Fitted Gauss, proteinAT5G51100.1
bic:875
peaks after deleting less than one fifth of global max:14
raw data, starting:14
fitted Gauss locations:14
x
y

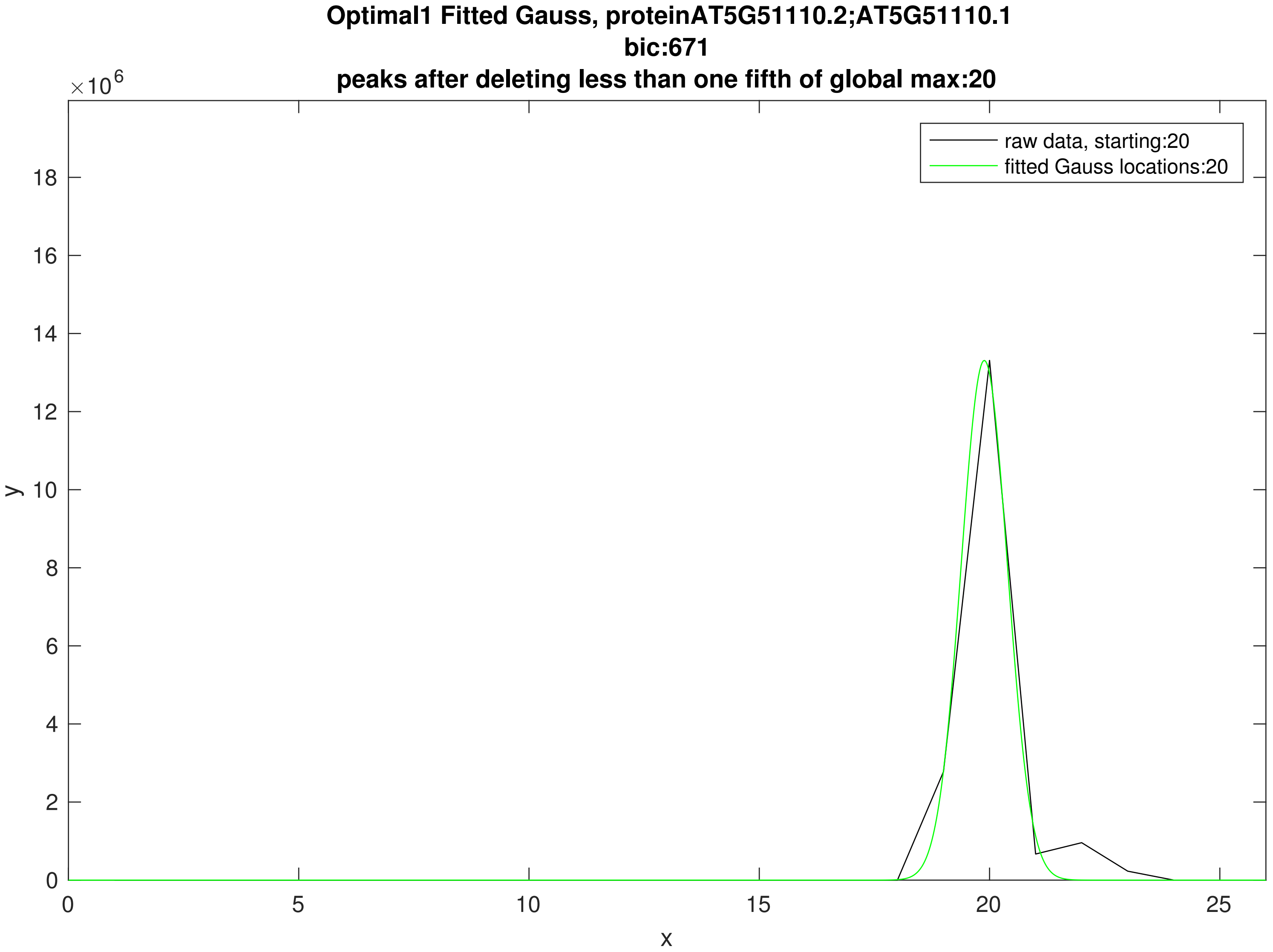

Optimal1 Fitted Gauss, proteinAT5G51110.2;AT5G51110.1
bic:671
peaks after deleting less than one fifth of global max:20
×10^6
raw data, starting:20
fitted Gauss locations:20
y
x
0
2
4
6
8
10
12
14
16
18
0
5
10
15
20
25

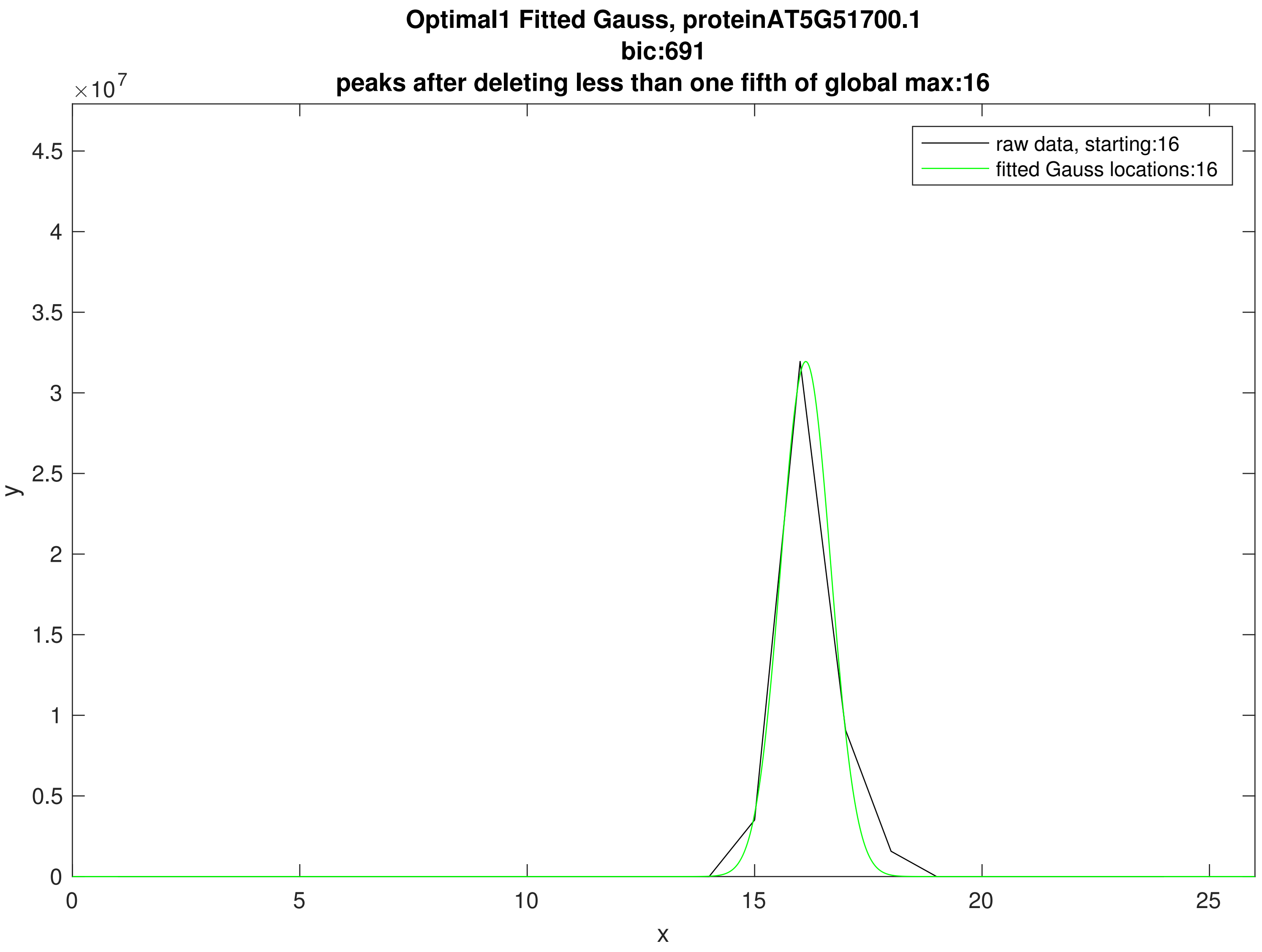
Optimal1 Fitted Gauss, proteinAT5G51700.1
bic:691
peaks after deleting less than one fifth of global max:16
×10^7
raw data, starting:16
fitted Gauss locations:16
y
x
0
0.5
1
1.5
2
2.5
3
3.5
4
4.5
5
10
15
20
25

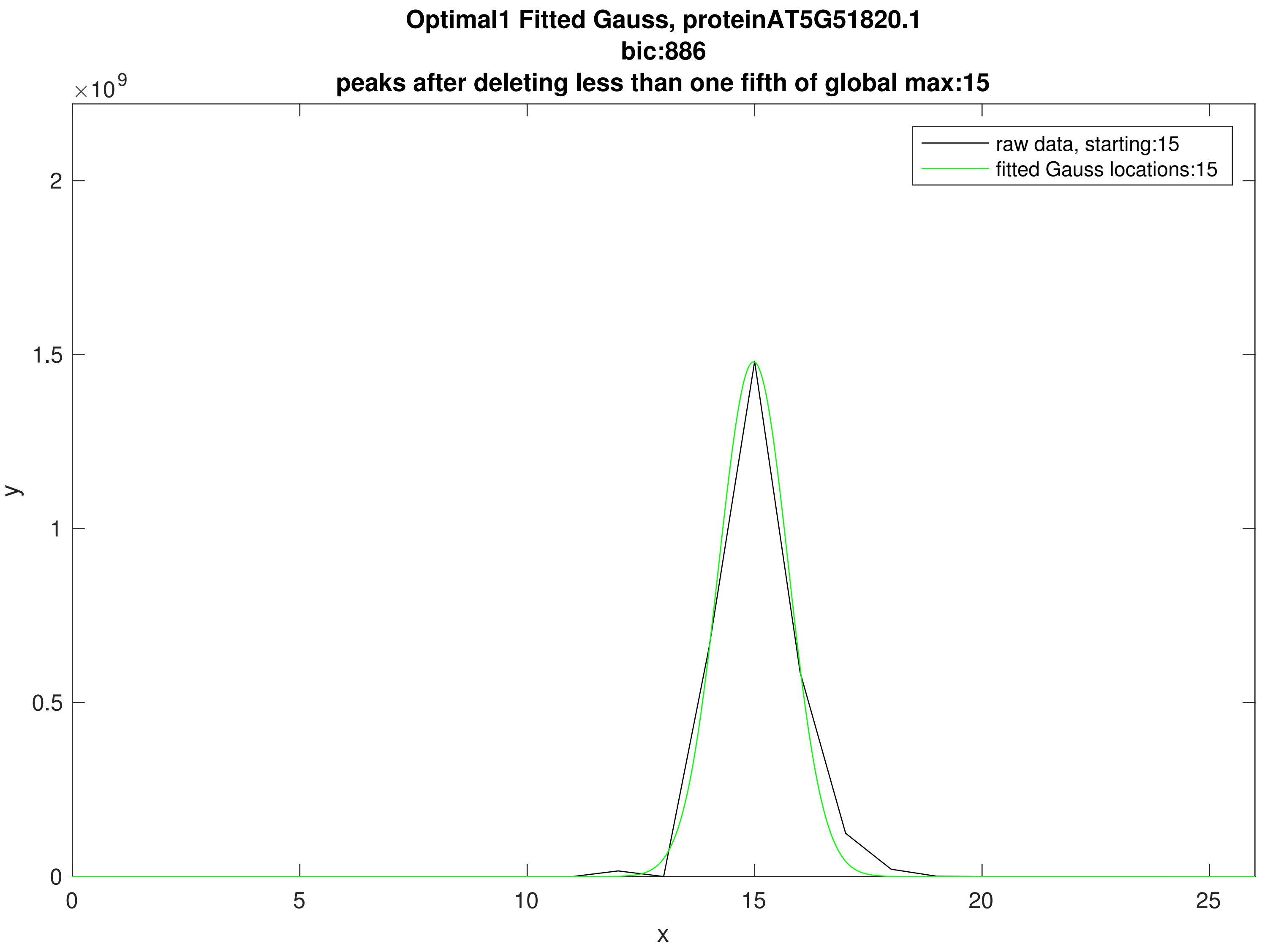
Optimal1 Fitted Gauss, proteinAT5G51820.1
bic:886
peaks after deleting less than one fifth of global max:15
×10^9
2
1.5
1
0.5
0
y
0
5
10
15
20
25
x
raw data, starting:15
fitted Gauss locations:15

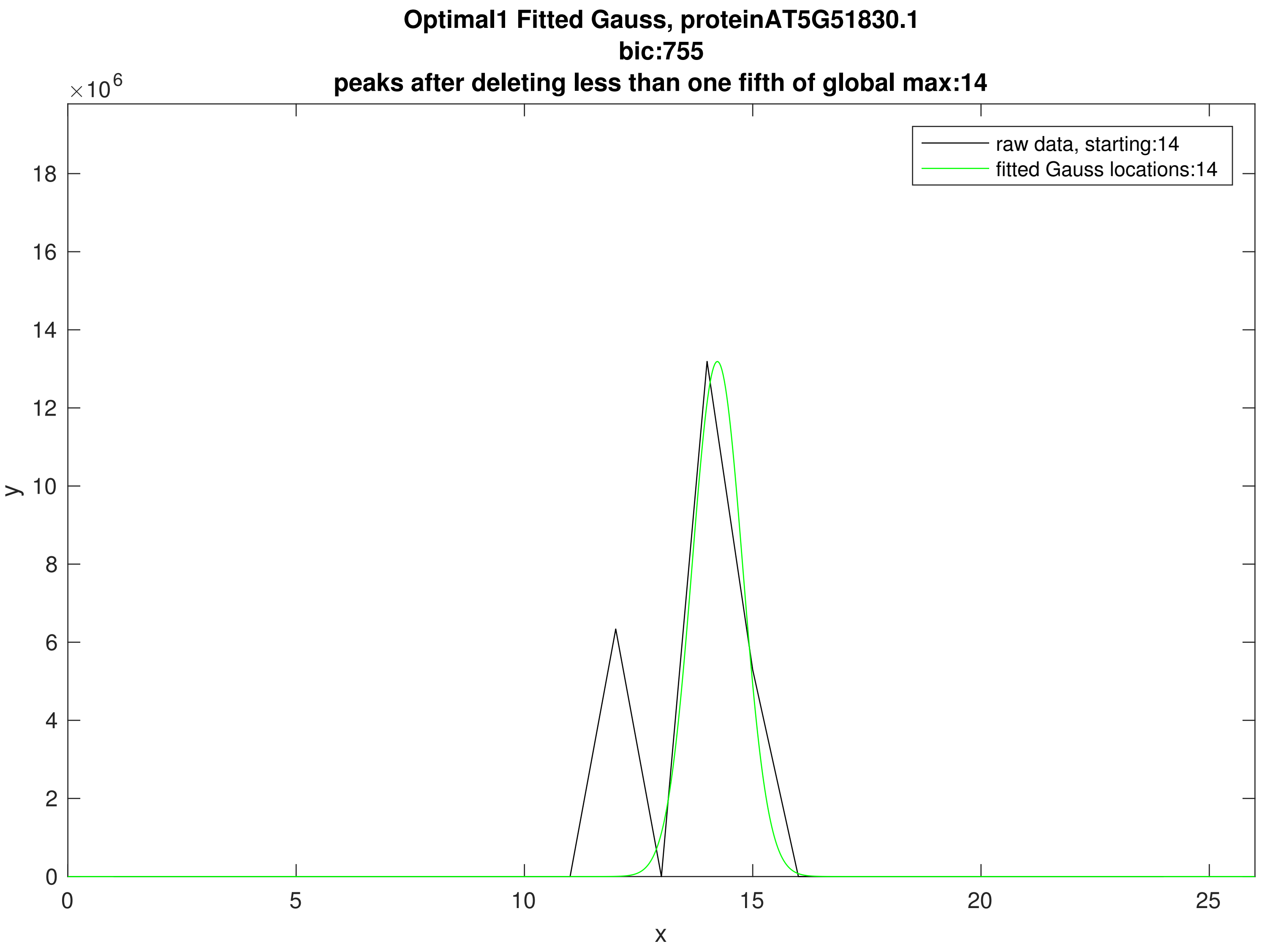
Optimal1 Fitted Gauss, proteinAT5G51830.1
bic:755
peaks after deleting less than one fifth of global max:14
×10^6
raw data, starting:14
fitted Gauss locations:14
y
x
0
2
4
6
8
10
12
14
16
18
0
5
10
15
20
25

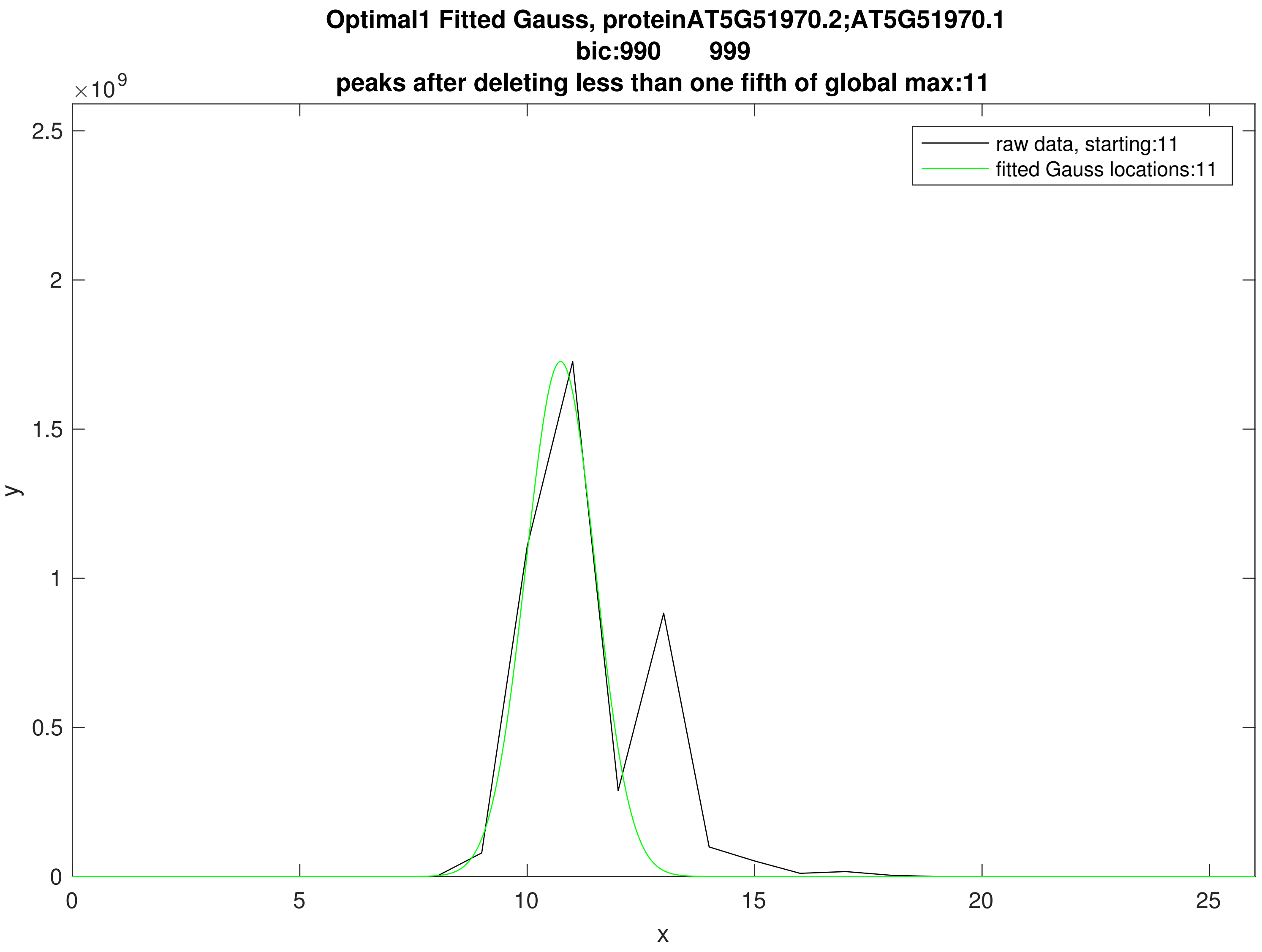
Optimal1 Fitted Gauss, proteinAT5G51970.2;AT5G51970.1
bic:990 999
peaks after deleting less than one fifth of global max:11
×10^9
raw data, starting:11
fitted Gauss locations:11
y
x
0
0.5
1
1.5
2
2.5
5
10
15
20
25

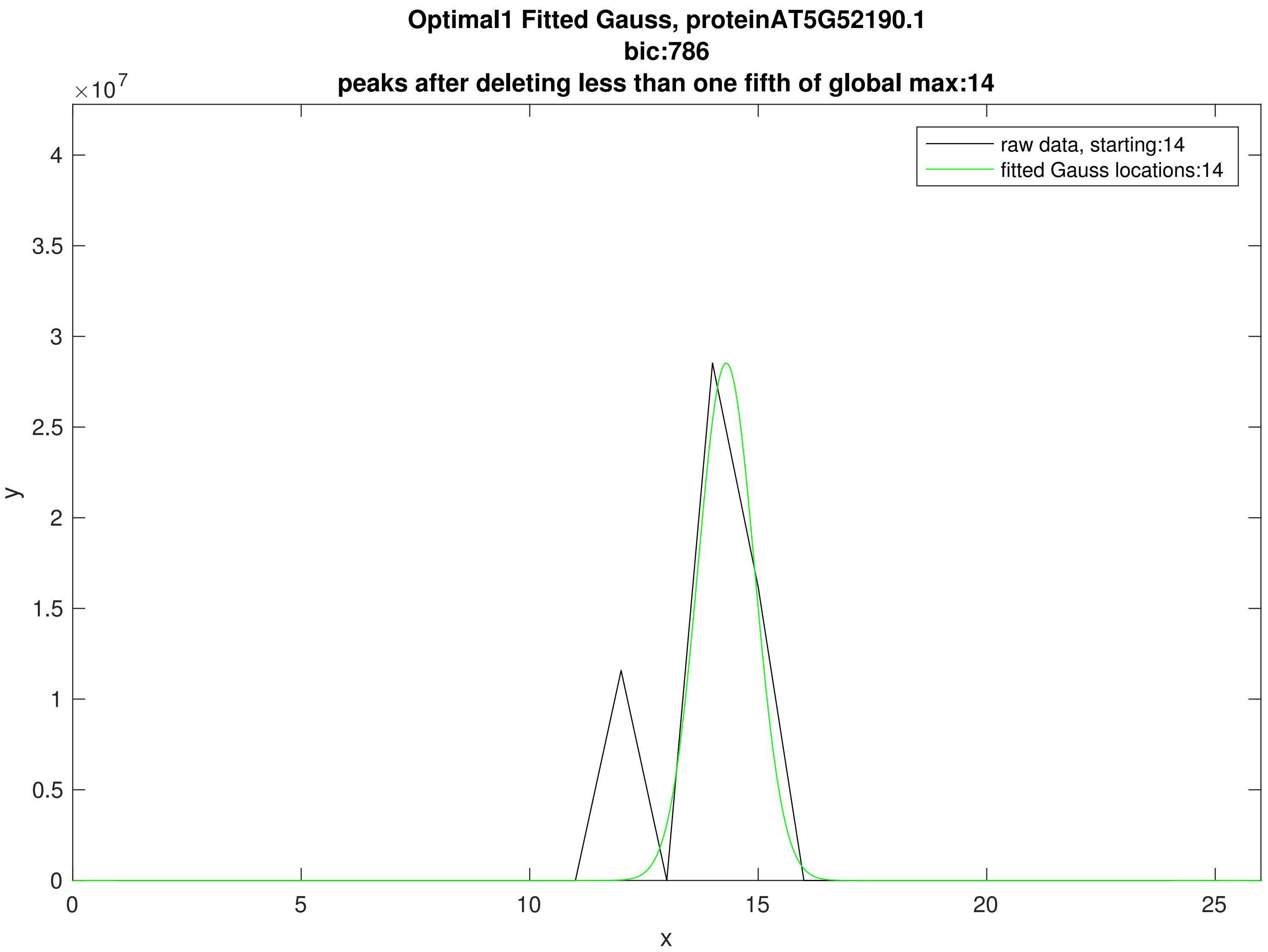

Optimal1 Fitted Gauss, proteinAT5G52190.1
bic:786
peaks after deleting less than one fifth of global max:14
×10^7
raw data, starting:14
fitted Gauss locations:14
y
x
0
0.5
1
1.5
2
2.5
3
3.5
4
0
5
10
15
20
25

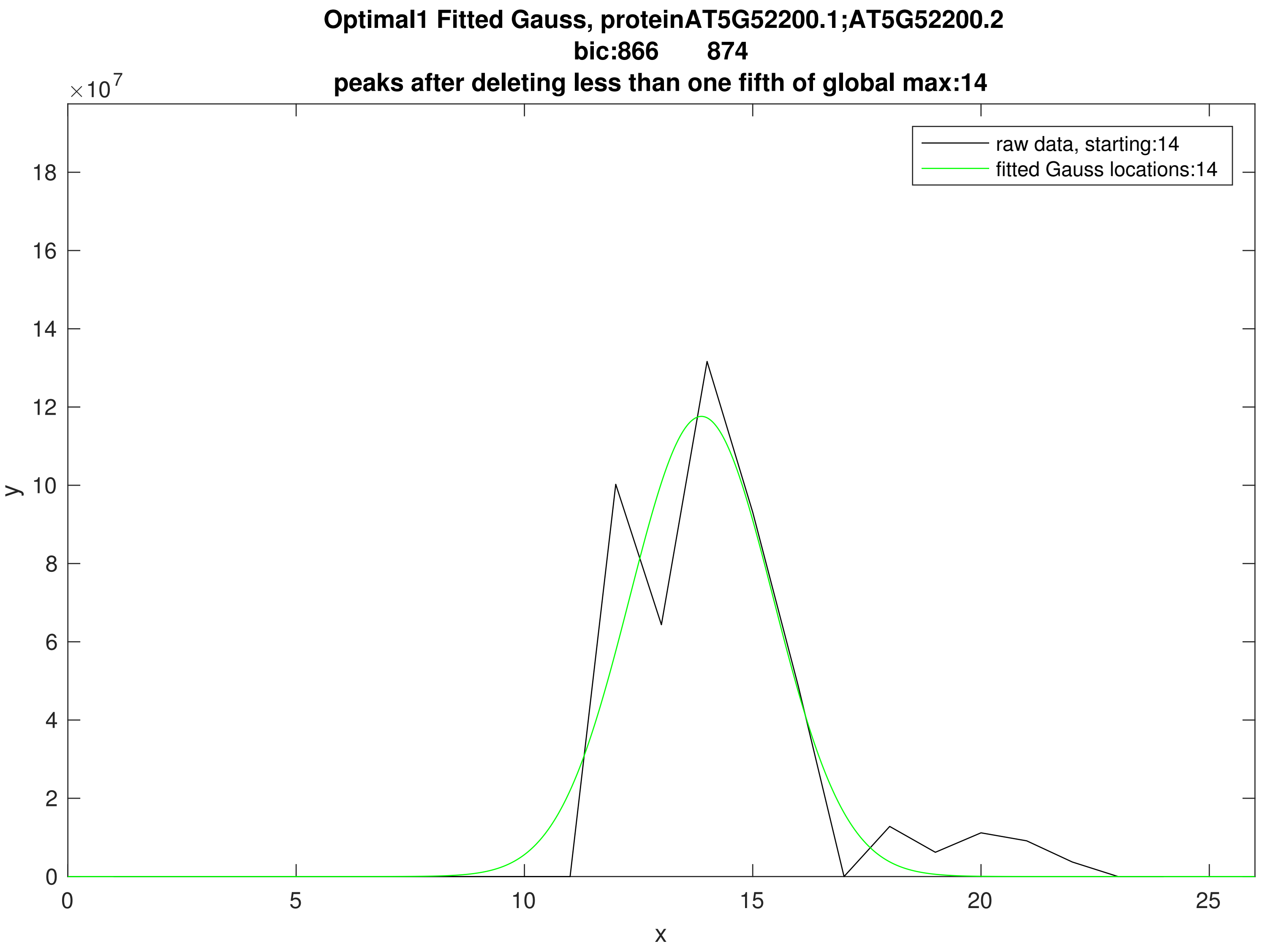

Optimal1 Fitted Gauss, proteinAT5G52200.1;AT5G52200.2
bic:866 874
peaks after deleting less than one fifth of global max:14
×10^7
raw data, starting:14
fitted Gauss locations:14
y
x
0
2
4
6
8
10
12
14
16
18
0
5
10
15
20
25

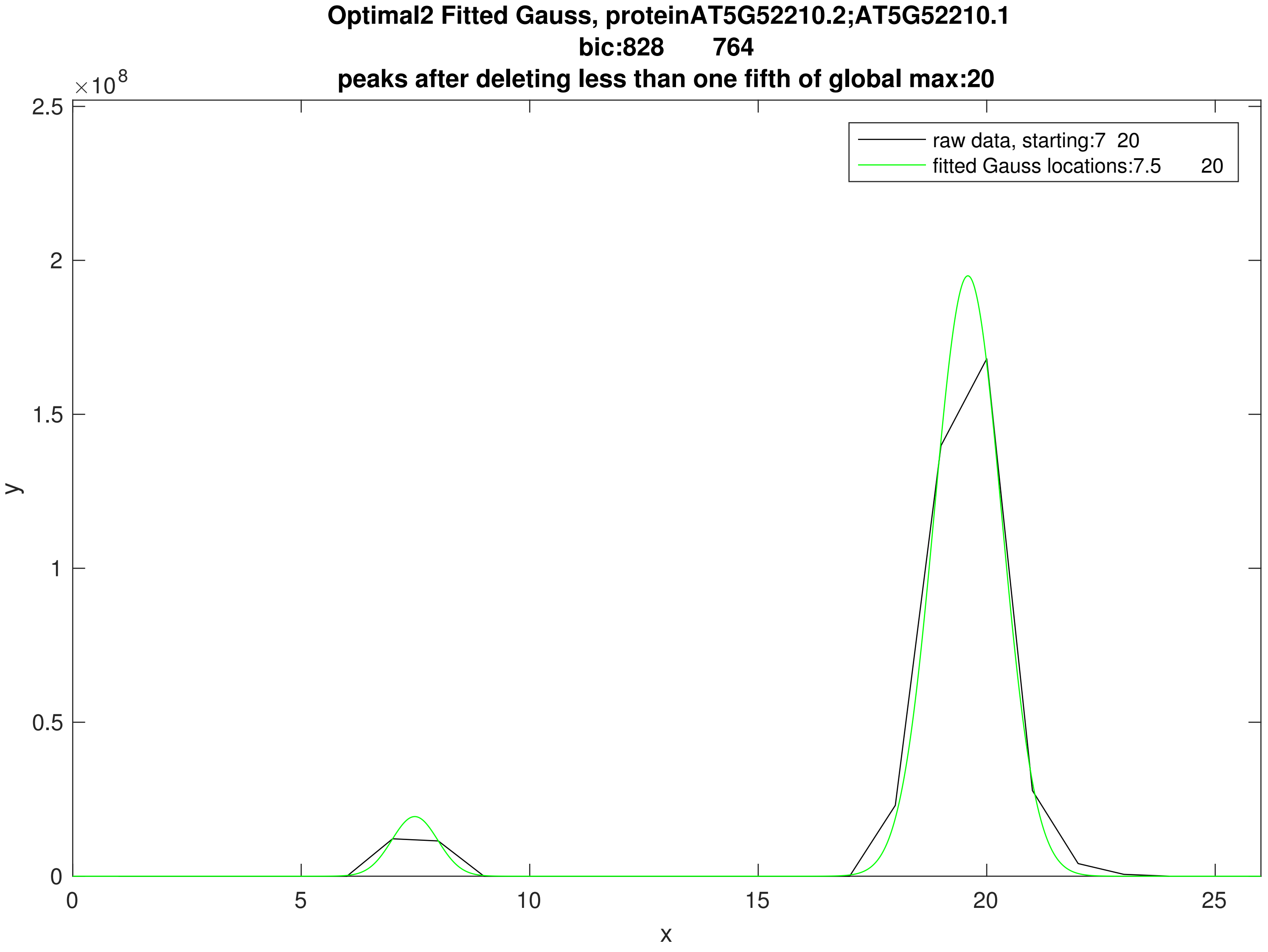

Optimal2 Fitted Gauss, proteinAT5G52210.2;AT5G52210.1
bic:828 764
peaks after deleting less than one fifth of global max:20
×10^8
2.5
2
1.5
1
0.5
0
y
0
5
10
15
20
25
x
raw data, starting:7 20
fitted Gauss locations:7.5 20

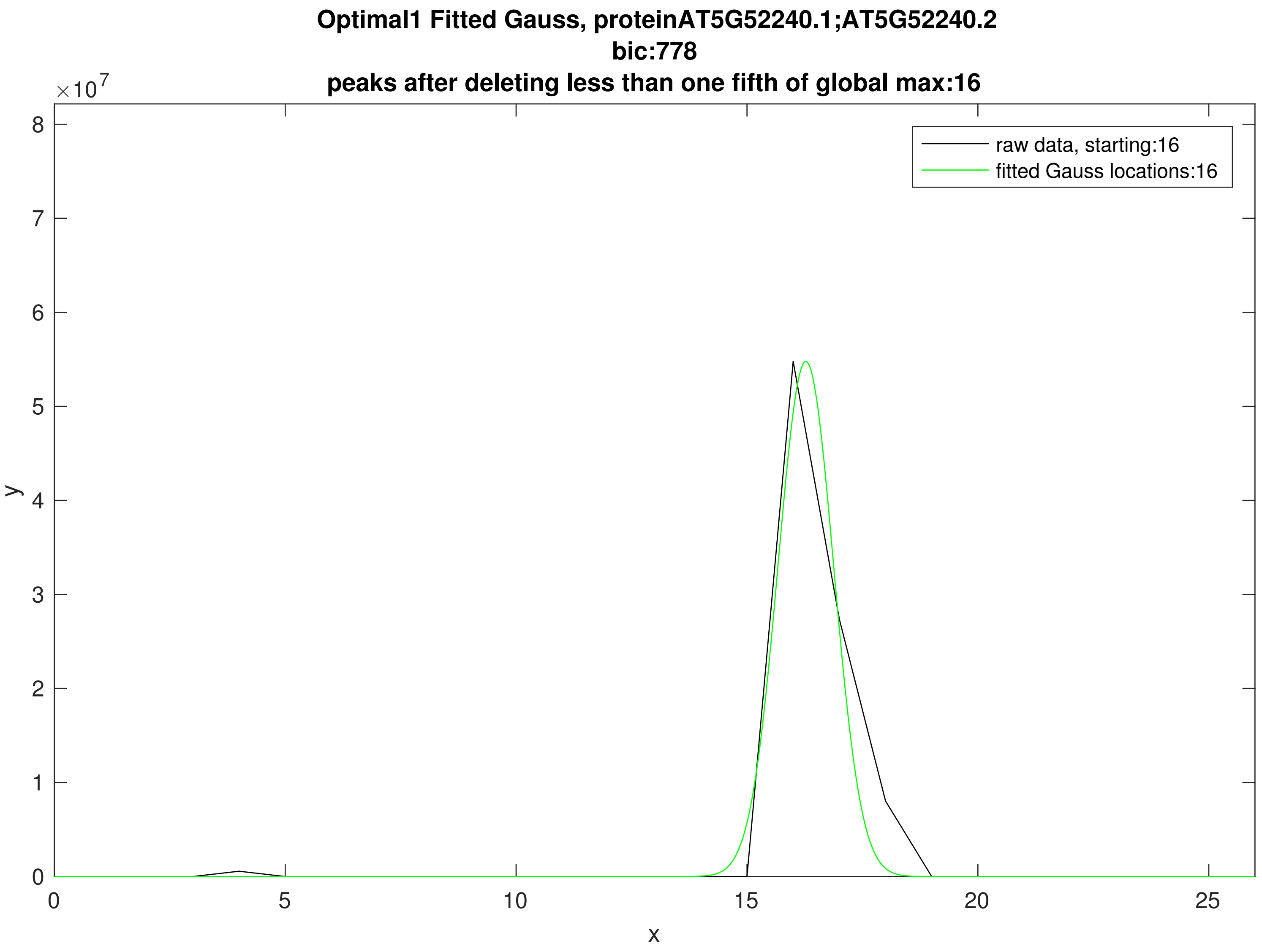
Optimal1 Fitted Gauss, proteinAT5G52240.1;AT5G52240.2
bic:778
peaks after deleting less than one fifth of global max:16
×10^7
raw data, starting:16
fitted Gauss locations:16
y
x
0
1
2
3
4
5
6
7
8
0
5
10
15
20
25

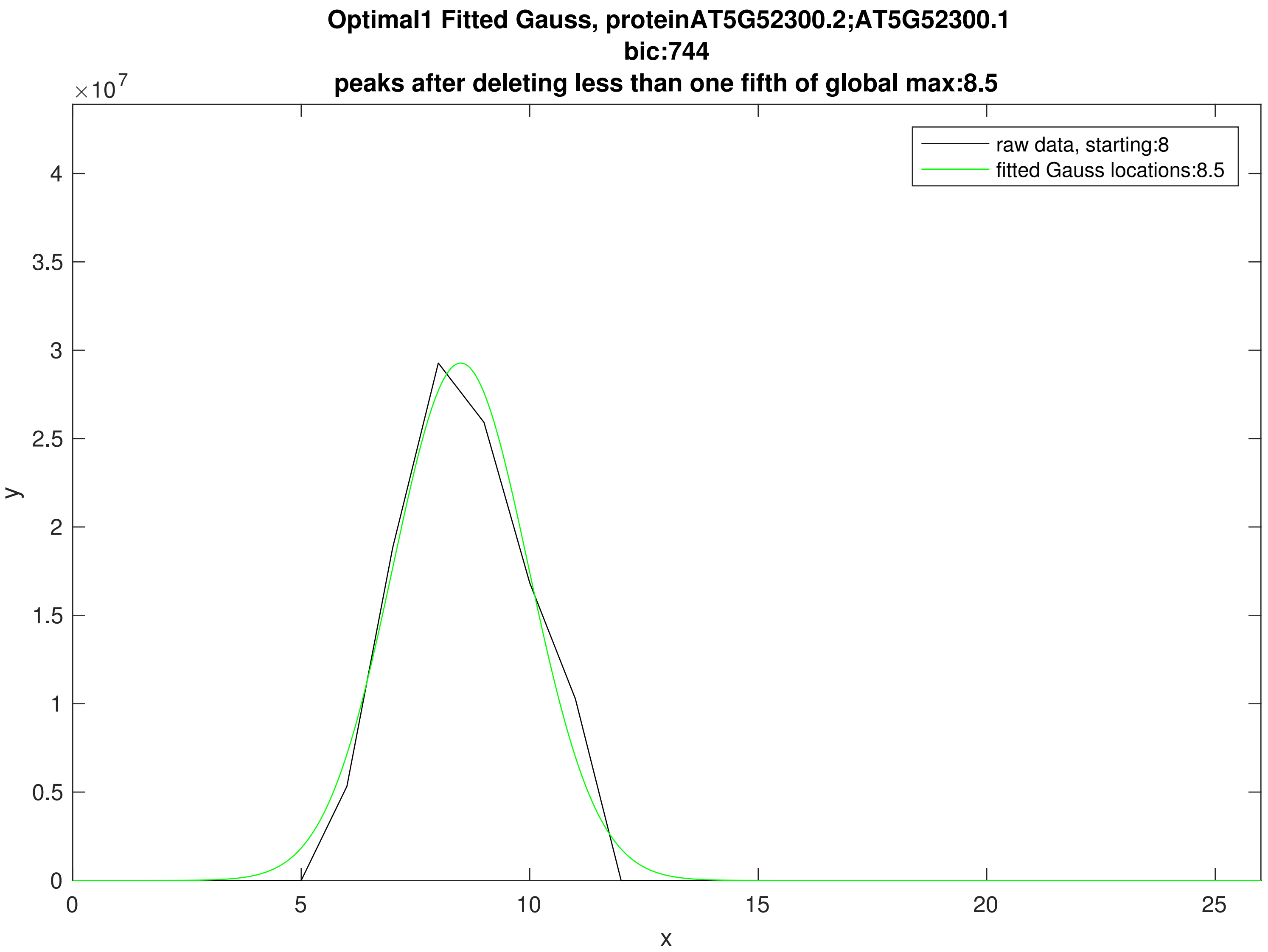
Optimal1 Fitted Gauss, proteinAT5G52300.2;AT5G52300.1
bic:744
peaks after deleting less than one fifth of global max:8.5
×10^7
raw data, starting:8
fitted Gauss locations:8.5
y
x
0
0.5
1
1.5
2
2.5
3
3.5
4
0
5
10
15
20
25

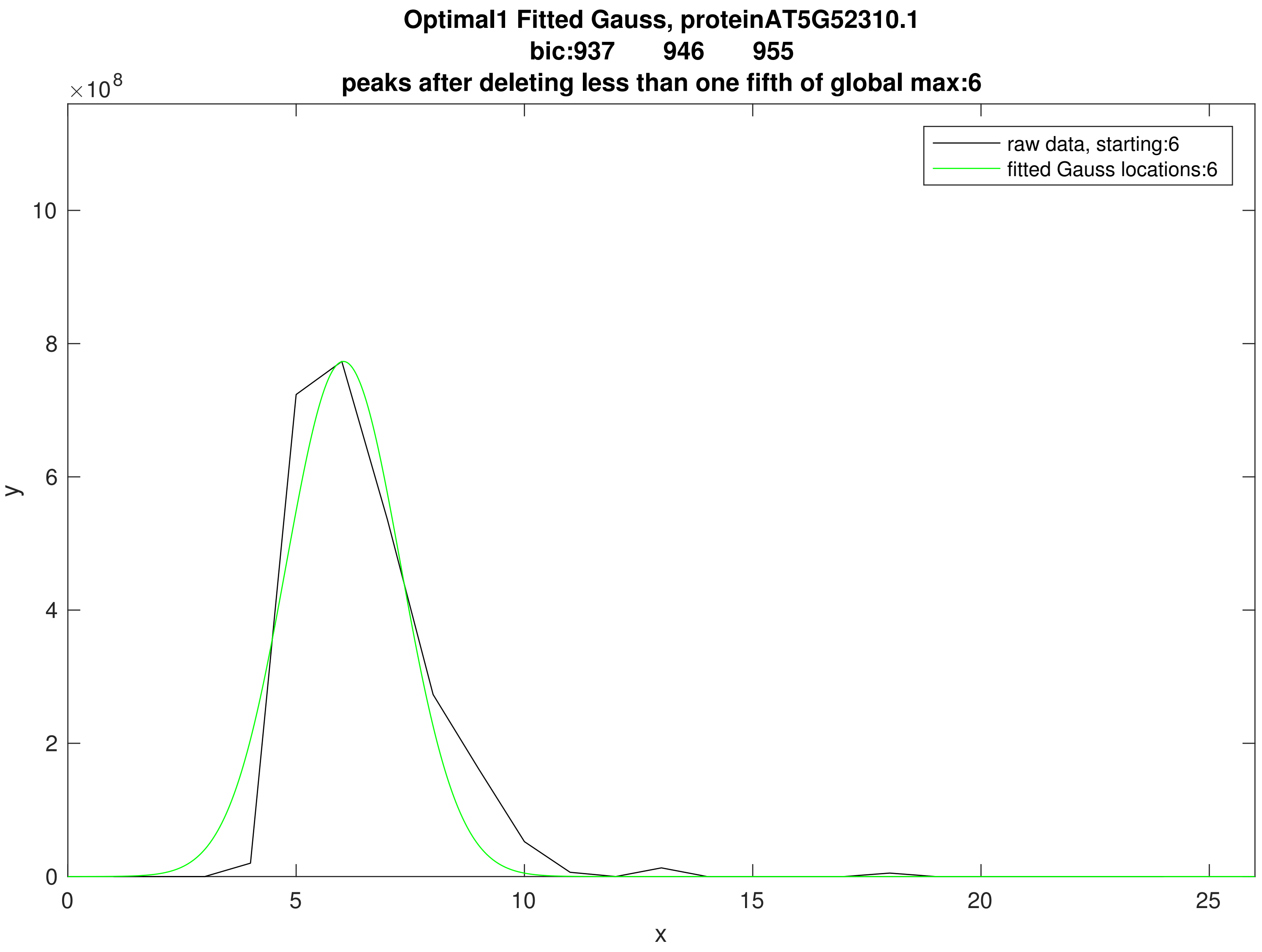
Optimal1 Fitted Gauss, proteinAT5G52310.1
bic:937 946 955
peaks after deleting less than one fifth of global max:6
×10^8
raw data, starting:6
fitted Gauss locations:6
y
x
0
2
4
6
8
10
0
5
10
15
20
25

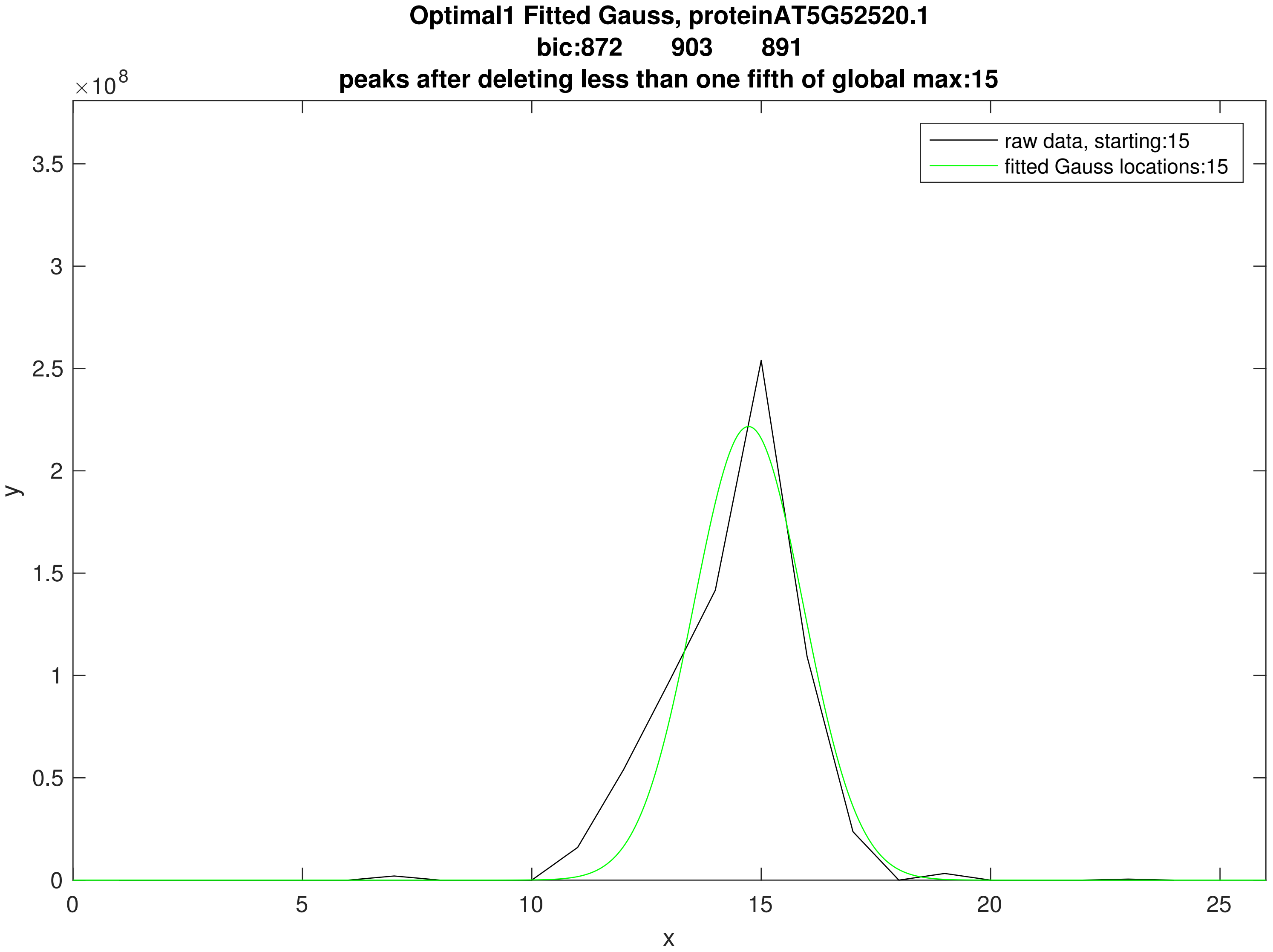

Optimal1 Fitted Gauss, proteinAT5G52520.1
bic:872 903 891
peaks after deleting less than one fifth of global max:15
×10^8
3.5
3
2.5
2
1.5
1
0.5
0
y
0
5
10
15
20
25
x
raw data, starting:15
fitted Gauss locations:15

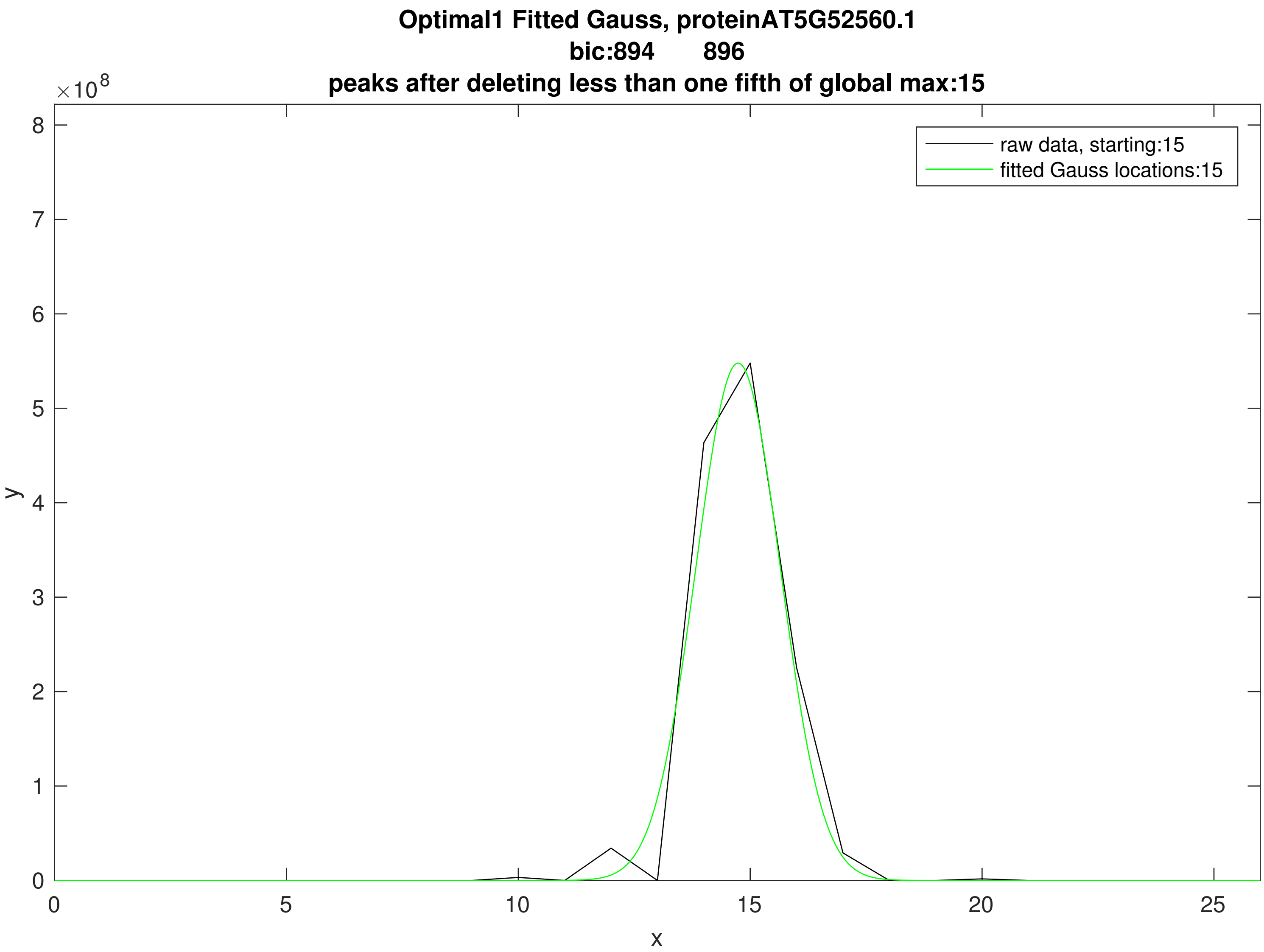

Optimal1 Fitted Gauss, proteinAT5G52560.1
bic:894 896
peaks after deleting less than one fifth of global max:15
×10^8
raw data, starting:15
fitted Gauss locations:15
y
x
0
1
2
3
4
5
6
7
8
0
5
10
15
20
25

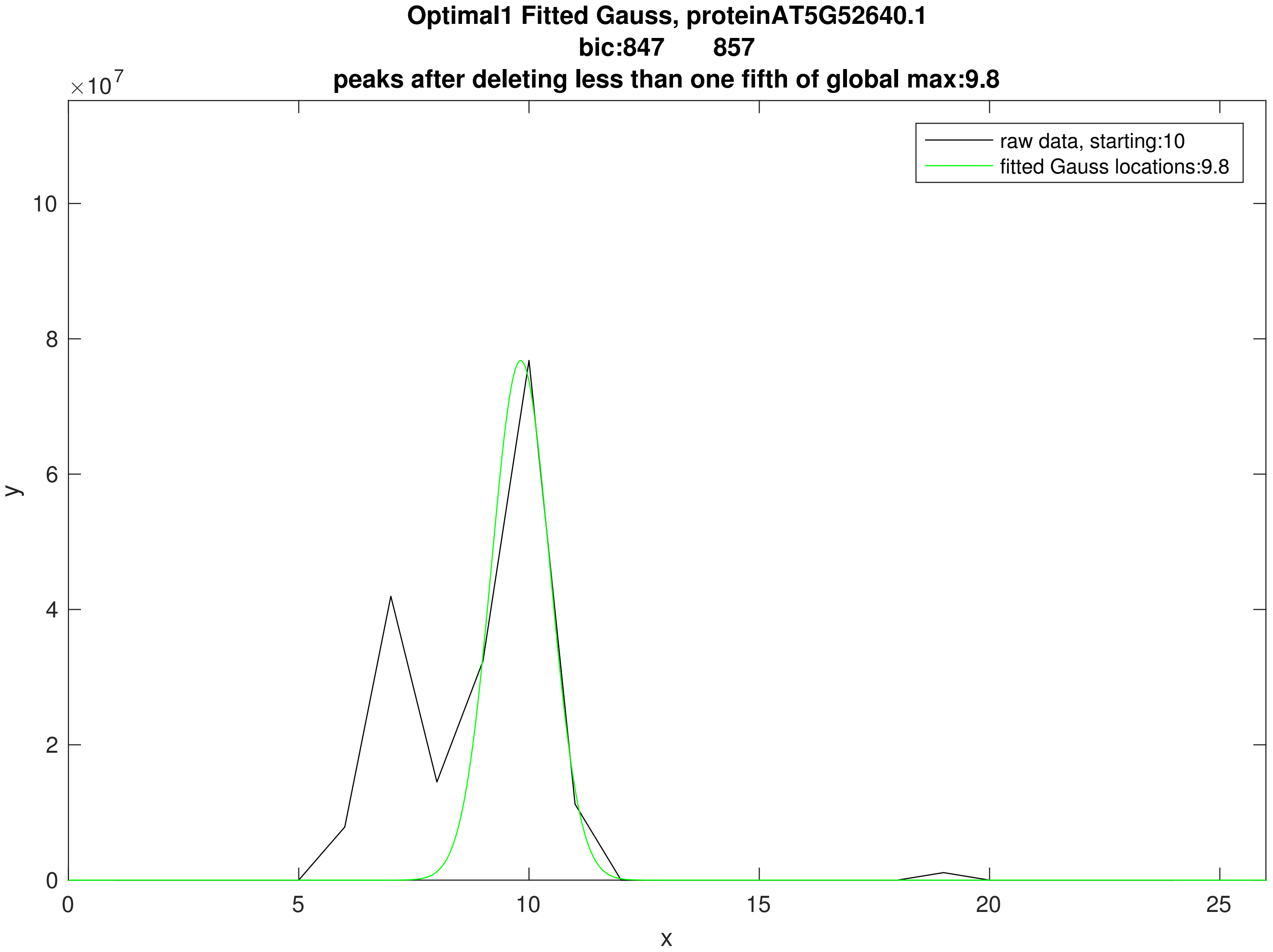
Optimal1 Fitted Gauss, proteinAT5G52640.1
bic:847 857
peaks after deleting less than one fifth of global max:9.8
×10^7
raw data, starting:10
fitted Gauss locations:9.8
y
x
0
2
4
6
8
10
0
5
10
15
20
25

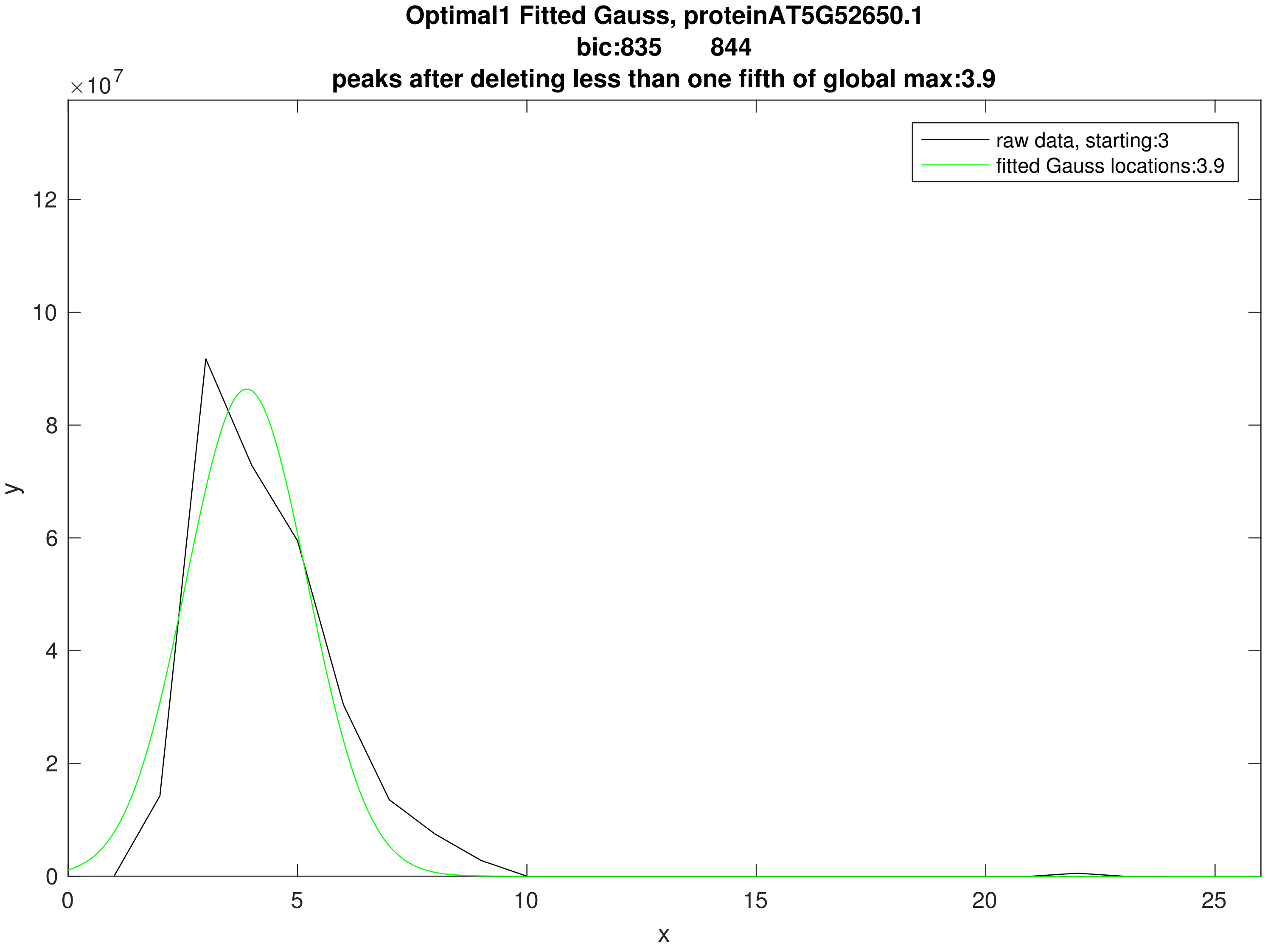

Optimal1 Fitted Gauss, proteinAT5G52650.1
bic:835 844
peaks after deleting less than one fifth of global max:3.9
×10^7
raw data, starting:3
fitted Gauss locations:3.9
y
x
0
2
4
6
8
10
12
0
5
10
15
20
25

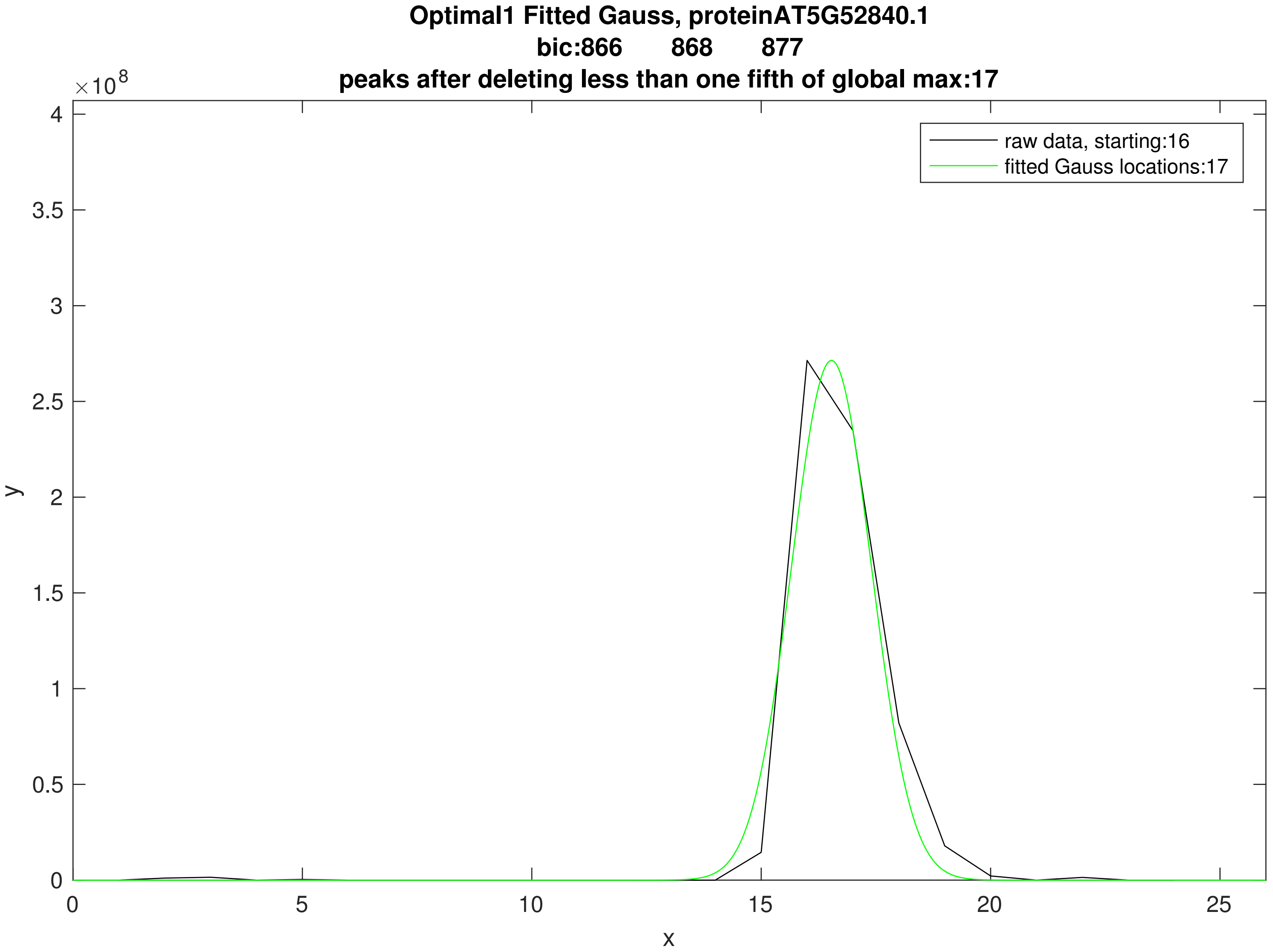
Optimal1 Fitted Gauss, proteinAT5G52840.1
bic:866 868 877
peaks after deleting less than one fifth of global max:17
×10^8
4
3.5
3
2.5
2
1.5
1
0.5
0
y
0
5
10
15
20
25
x
raw data, starting:16
fitted Gauss locations:17

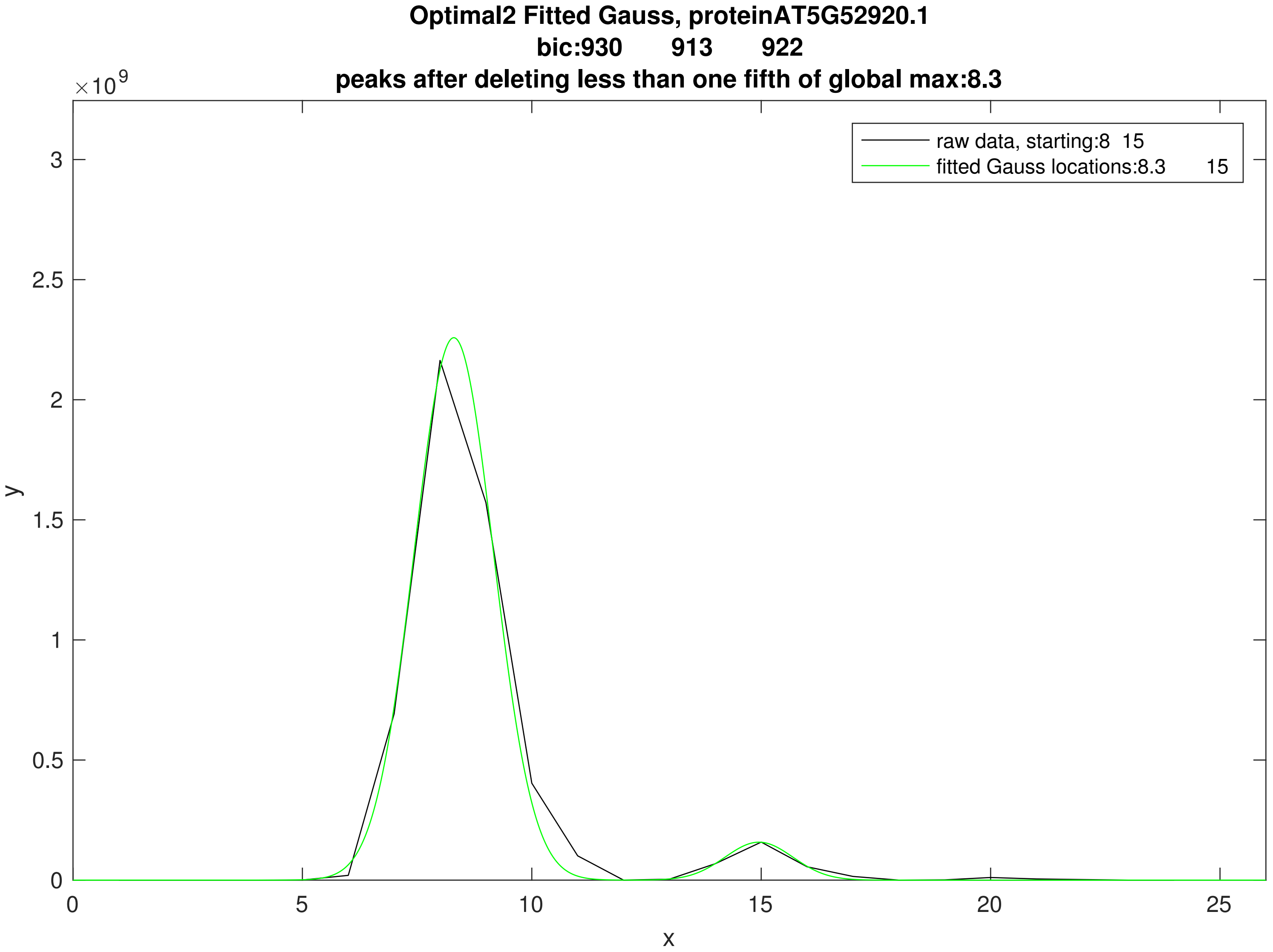

Optimal2 Fitted Gauss, proteinAT5G52920.1
bic:930 913 922
peaks after deleting less than one fifth of global max:8.3
×10^9
raw data, starting:8 15
fitted Gauss locations:8.3 15
y
x
0
0.5
1
1.5
2
2.5
3
5
10
15
20
25

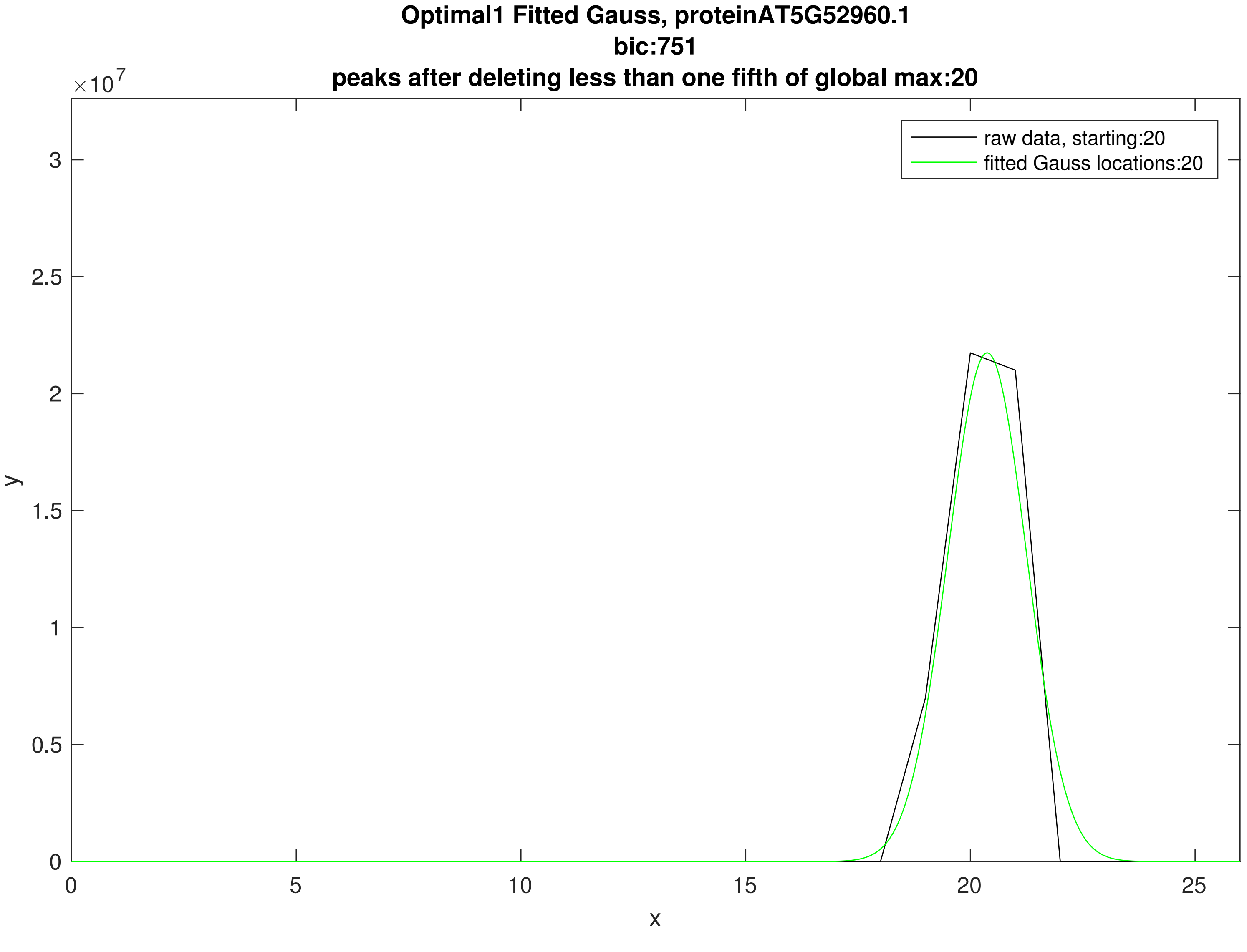
Optimal1 Fitted Gauss, proteinAT5G52960.1
bic:751
peaks after deleting less than one fifth of global max:20
$\times 10^7$
raw data, starting:20
fitted Gauss locations:20
3
2.5
2
1.5
1
0.5
0
y
0
5
10
15
20
25
x

Optimal1 Fitted Gauss, proteinAT5G53070.1
bic:632
peaks after deleting less than one fifth of global max:3.3

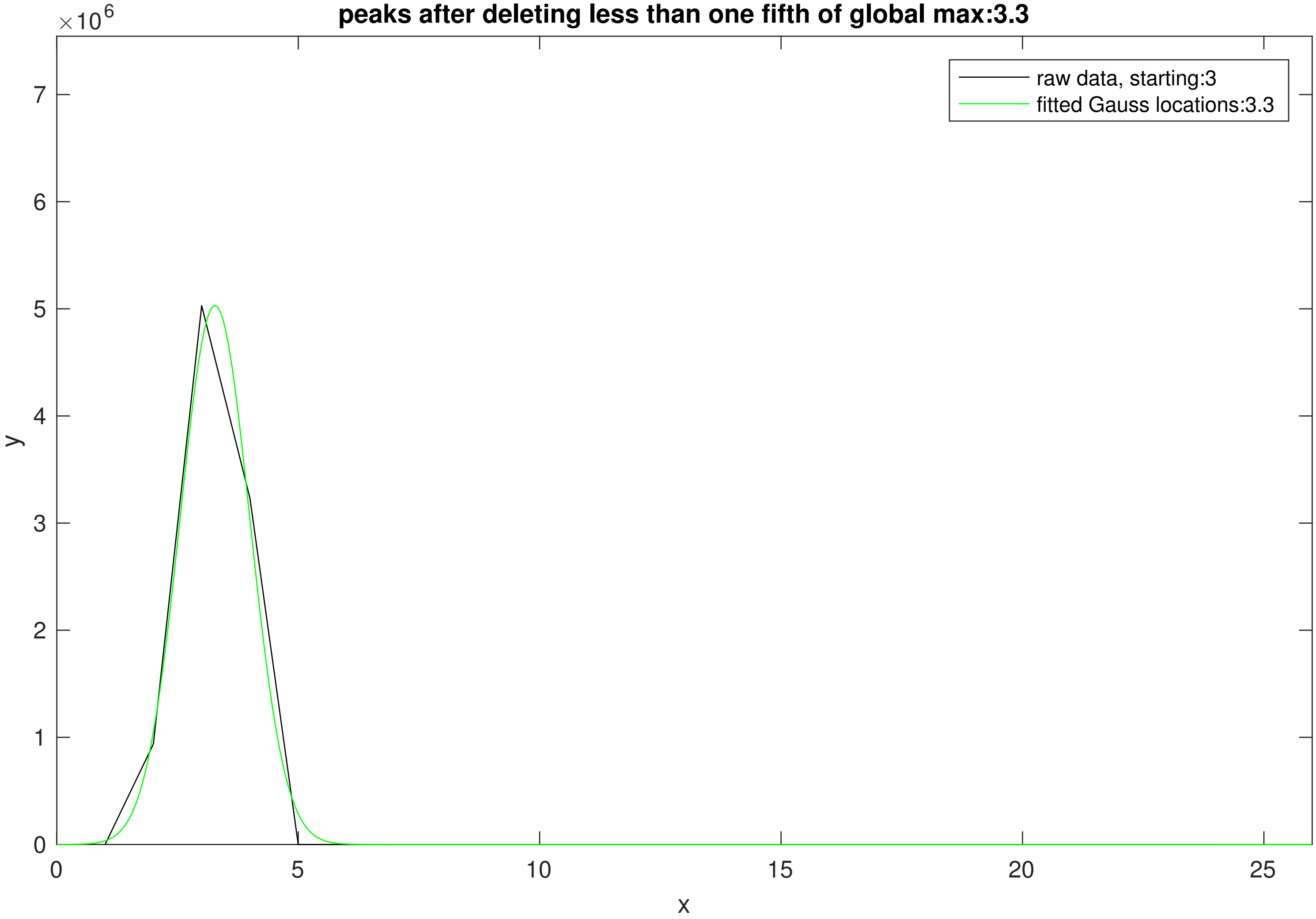

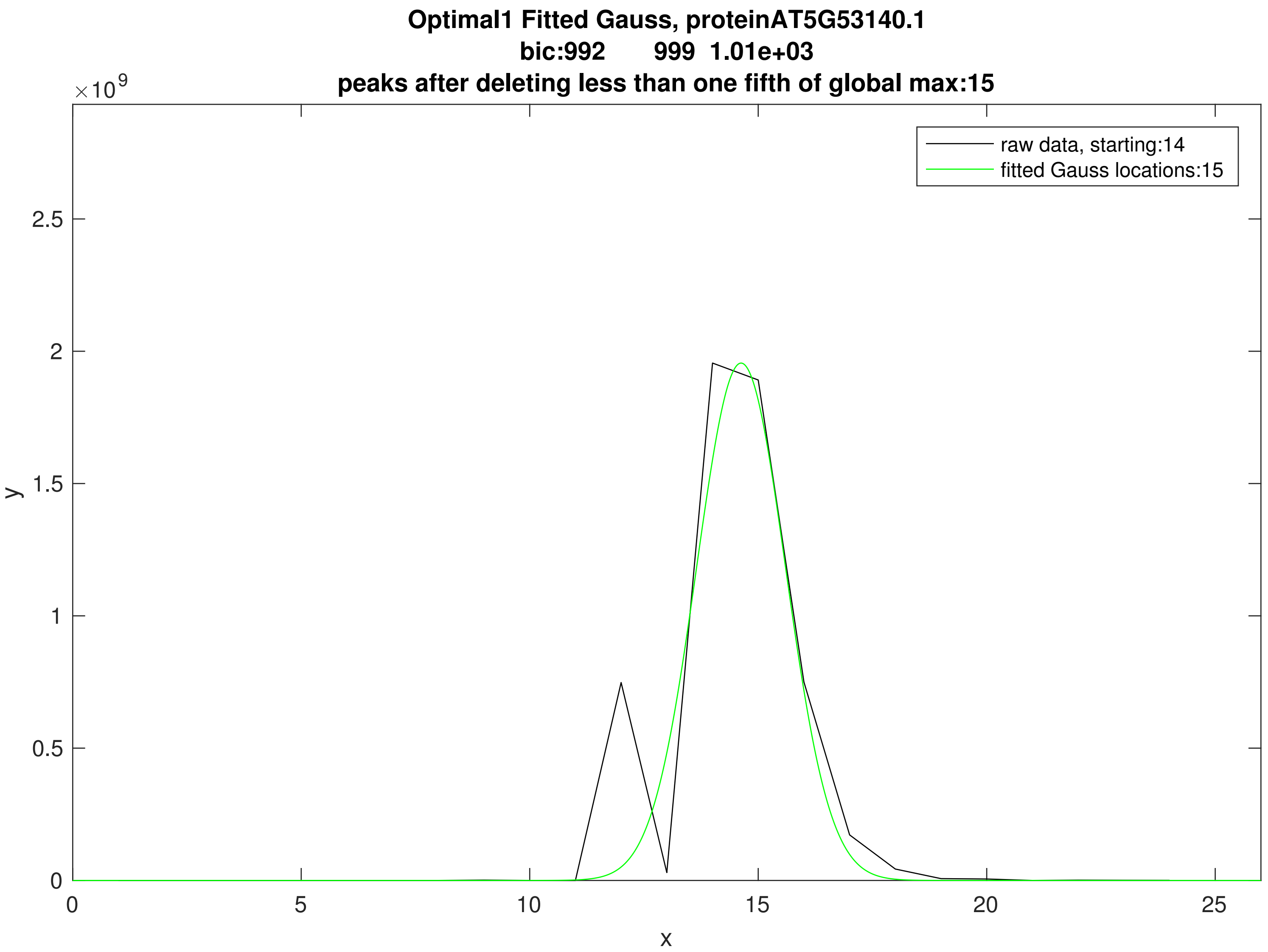

Optimal1 Fitted Gauss, proteinAT5G53140.1
bic:992 999 1.01e+03
peaks after deleting less than one fifth of global max:15
×10^9
raw data, starting:14
fitted Gauss locations:15
y
x
0
0.5
1
1.5
2
2.5
5
10
15
20
25

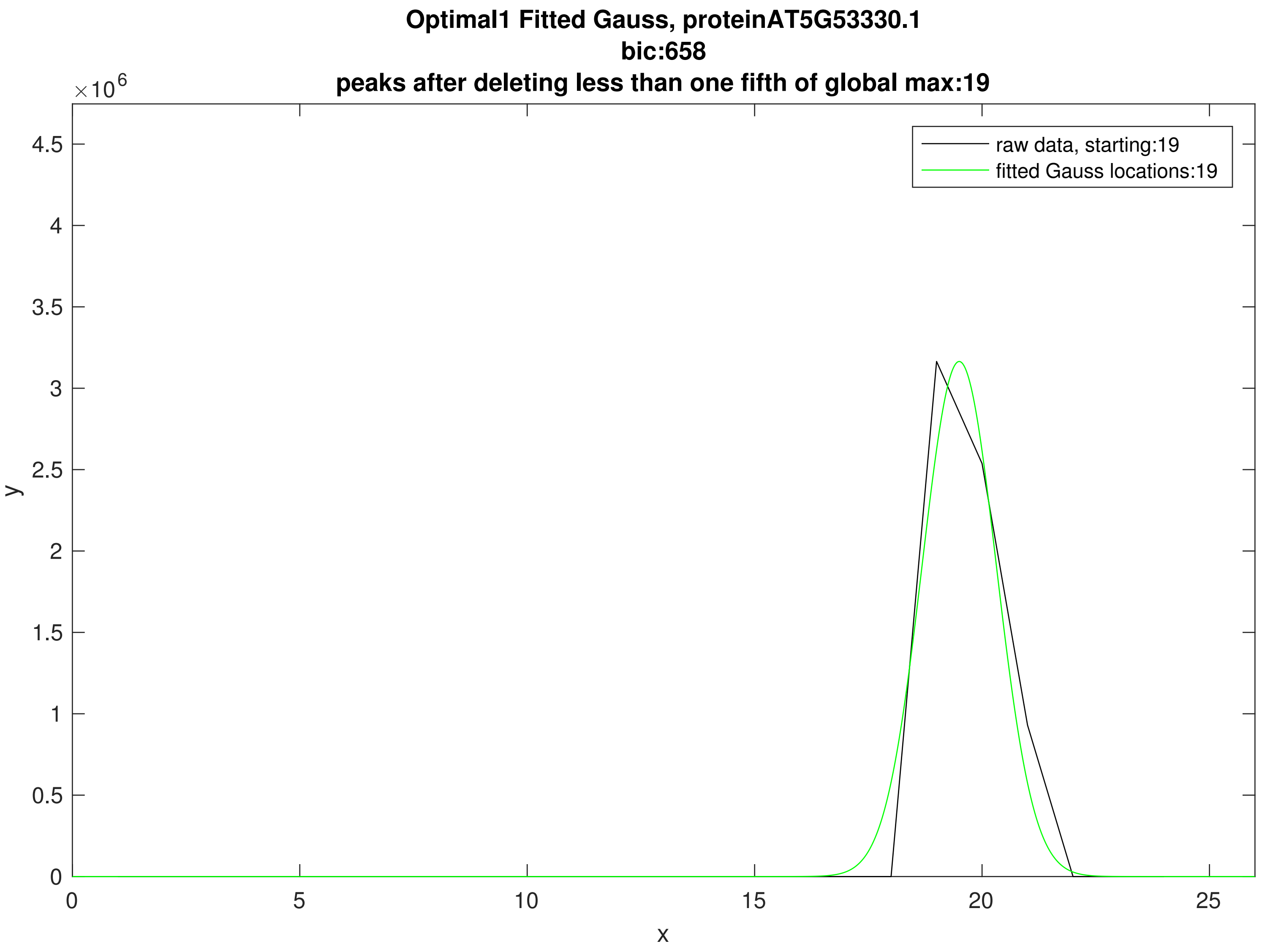
Optimal1 Fitted Gauss, proteinAT5G53330.1
bic:658
peaks after deleting less than one fifth of global max:19
×10^6
raw data, starting:19
fitted Gauss locations:19
y
x
4.5
4
3.5
3
2.5
2
1.5
1
0.5
0
0
5
10
15
20
25

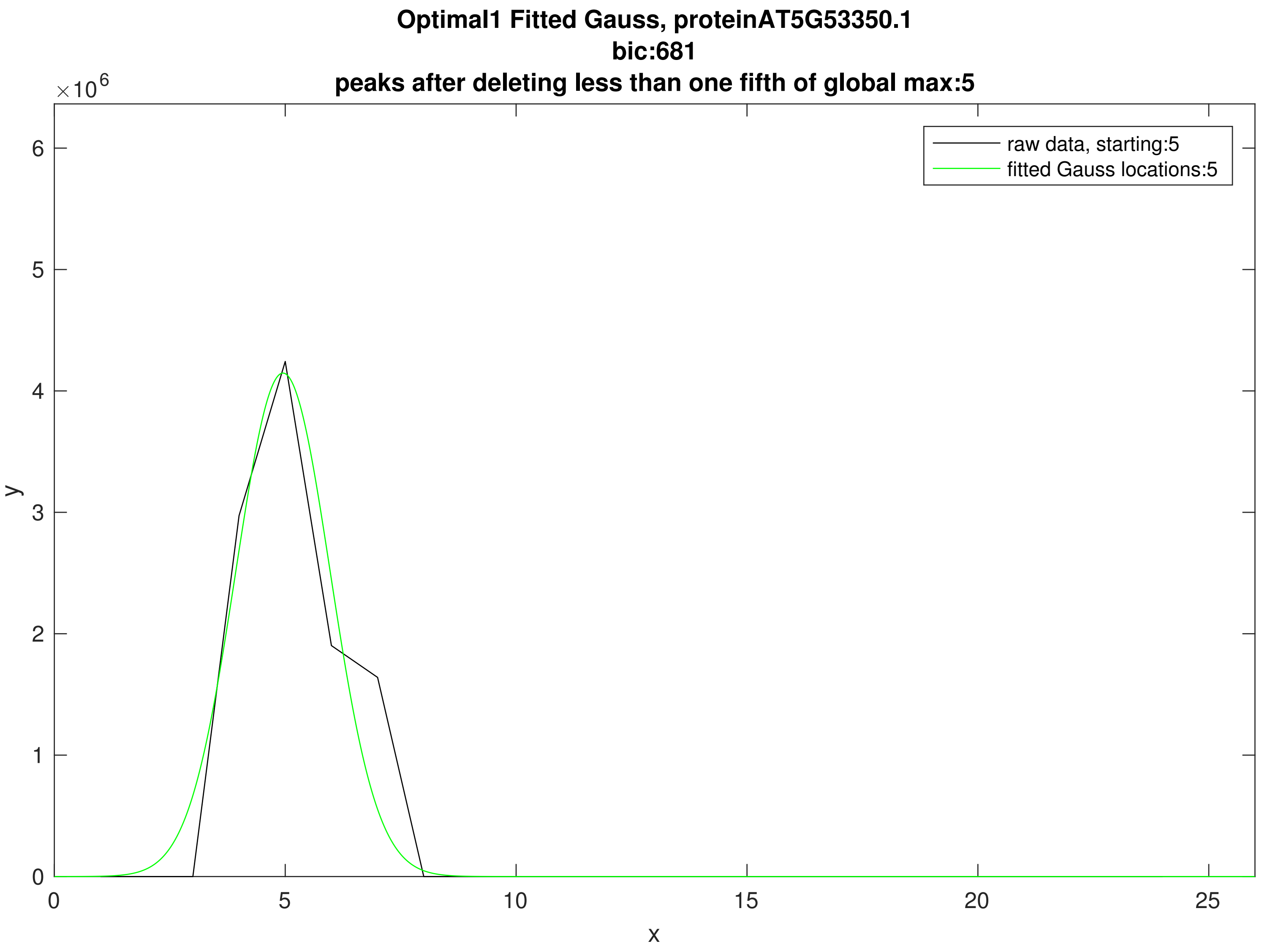
Optimal1 Fitted Gauss, proteinAT5G53350.1
bic:681
peaks after deleting less than one fifth of global max:5
×10^6
raw data, starting:5
fitted Gauss locations:5
y
x
0
1
2
3
4
5
6
0
5
10
15
20
25

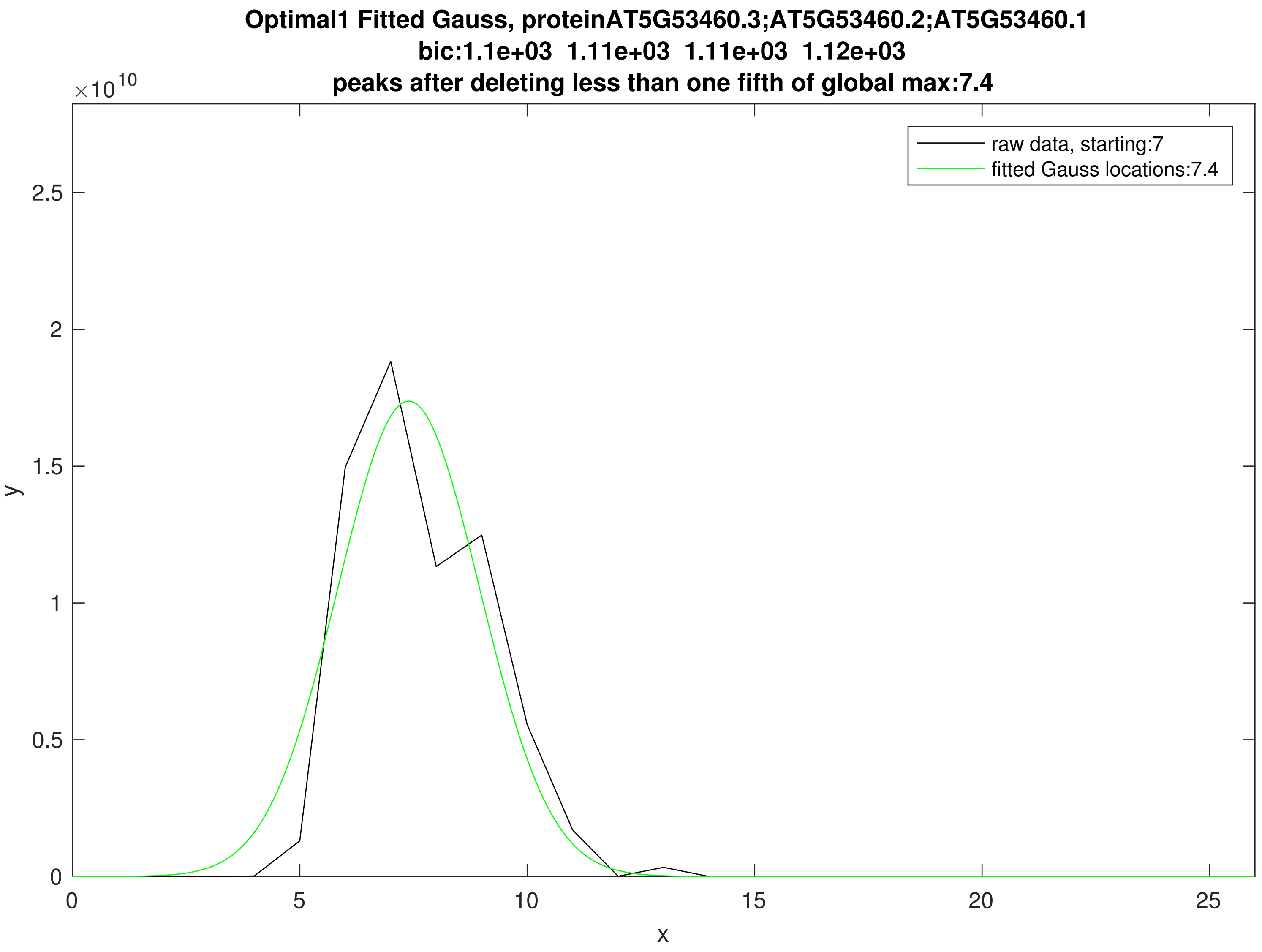

Optimal1 Fitted Gauss, proteinAT5G53460.3;AT5G53460.2;AT5G53460.1
bic:1.1e+03 1.11e+03 1.11e+03 1.12e+03
peaks after deleting less than one fifth of global max:7.4
×10^10
raw data, starting:7
fitted Gauss locations:7.4
y
x
0
0.5
1
1.5
2
2.5
5
10
15
20
25

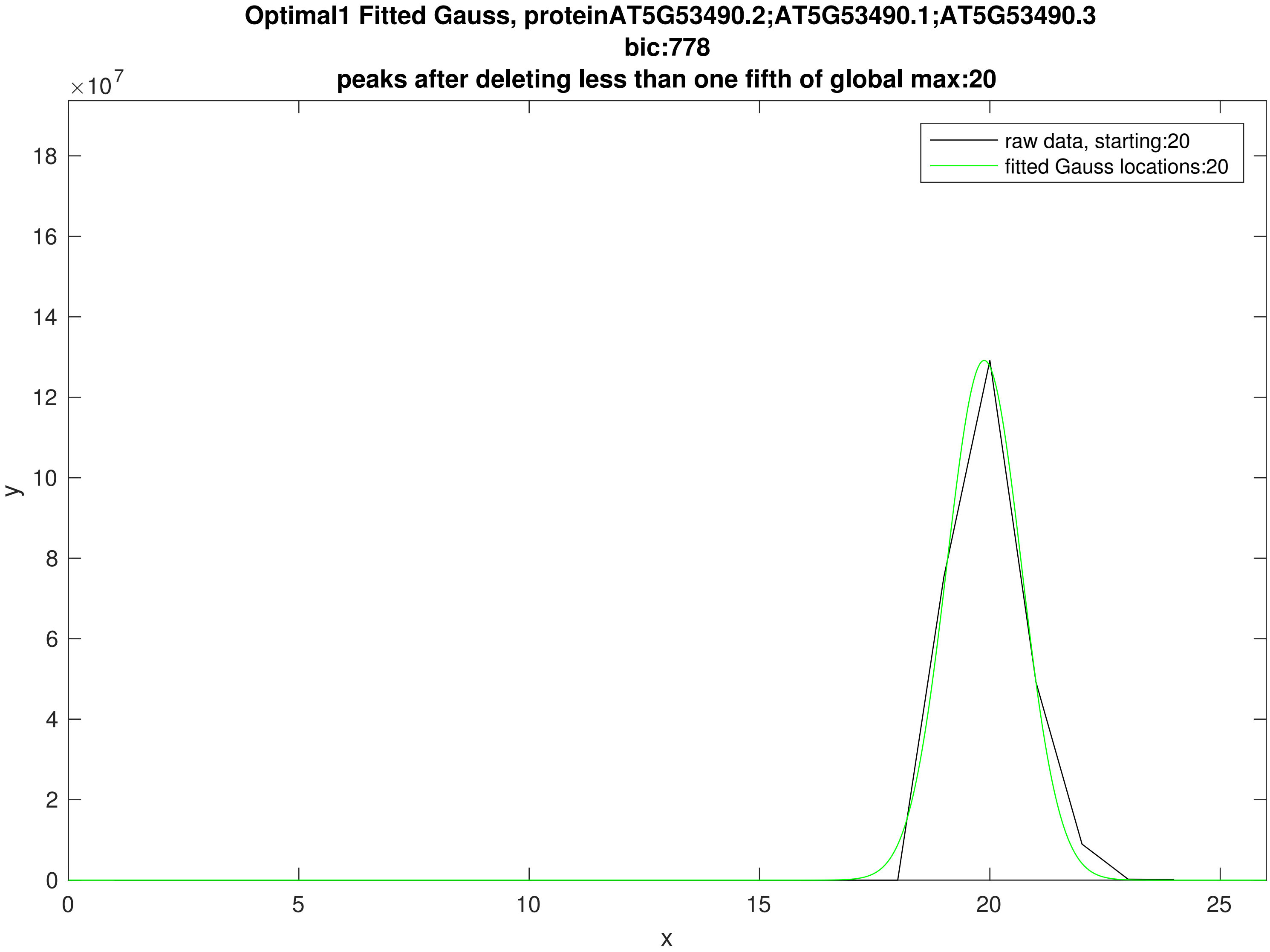
Optimal1 Fitted Gauss, proteinAT5G53490.2;AT5G53490.1;AT5G53490.3
bic:778
peaks after deleting less than one fifth of global max:20
×10^7
18
16
14
12
10
8
6
4
2
0
y
0
5
10
15
20
25
x
raw data, starting:20
fitted Gauss locations:20

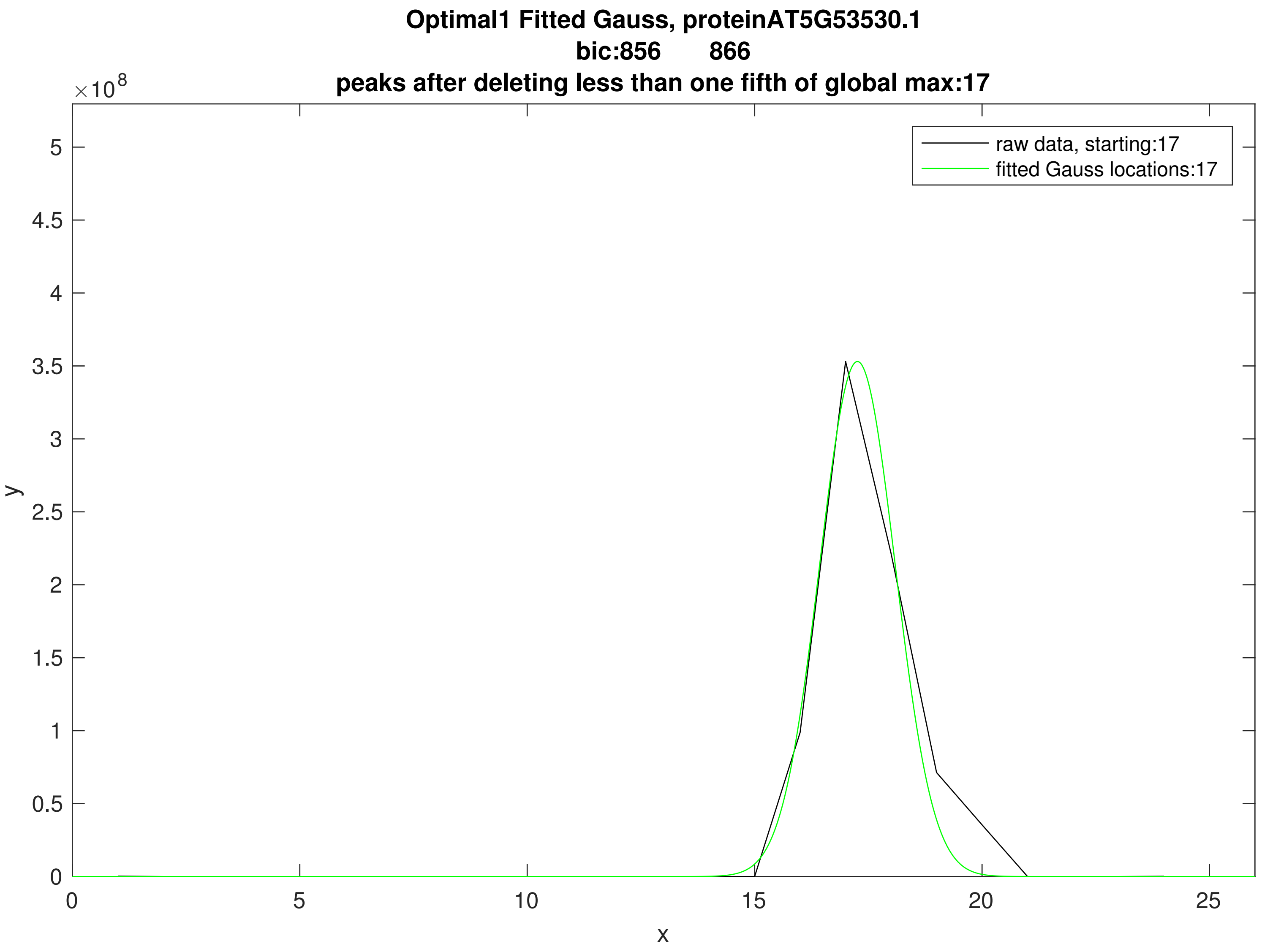

Optimal1 Fitted Gauss, proteinAT5G53530.1
bic:856 866
peaks after deleting less than one fifth of global max:17
×10^8
5
4.5
4
3.5
3
2.5
2
1.5
1
0.5
0
y
0
5
10
15
20
25
x
raw data, starting:17
fitted Gauss locations:17

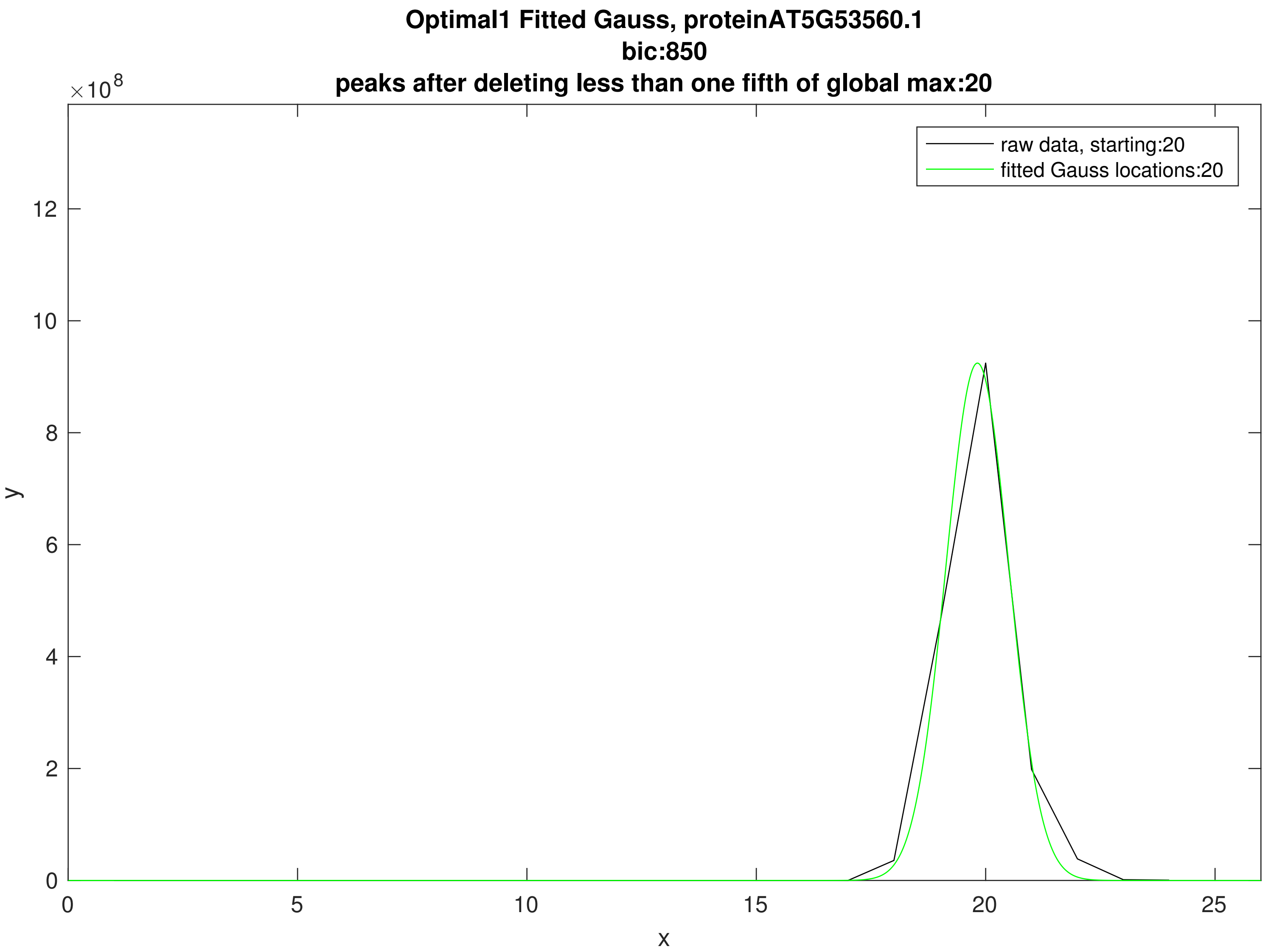
Optimal1 Fitted Gauss, proteinAT5G53560.1
bic:850
peaks after deleting less than one fifth of global max:20
×10^8
raw data, starting:20
fitted Gauss locations:20
12
10
8
6
4
2
0
y
0
5
10
15
20
25
x

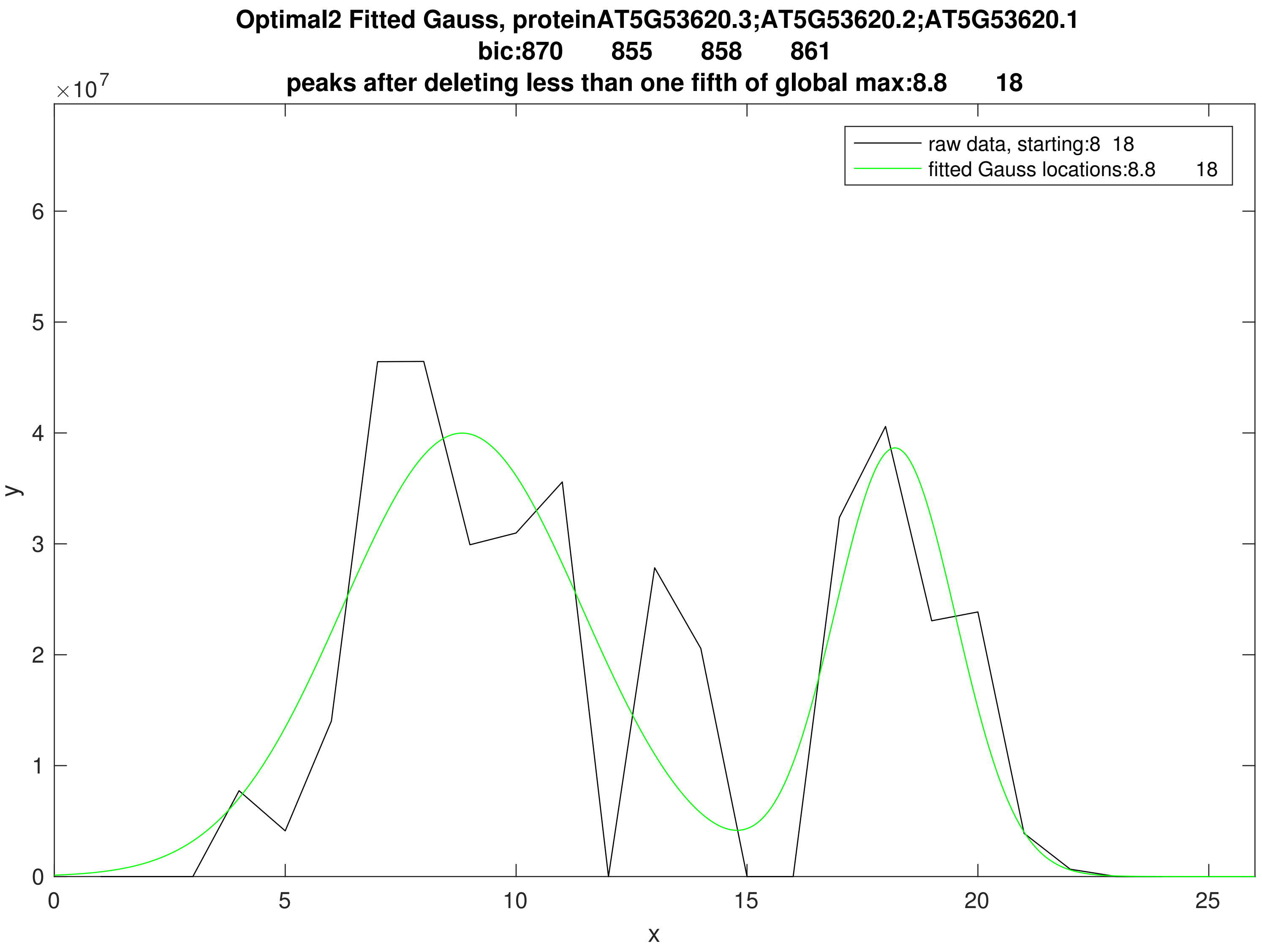

Optimal2 Fitted Gauss, proteinAT5G53620.3;AT5G53620.2;AT5G53620.1
bic:870 855 858 861
peaks after deleting less than one fifth of global max:8.8 18
×10^7
raw data, starting:8 18
fitted Gauss locations:8.8 18
y
x
0
1
2
3
4
5
6
0
5
10
15
20
25

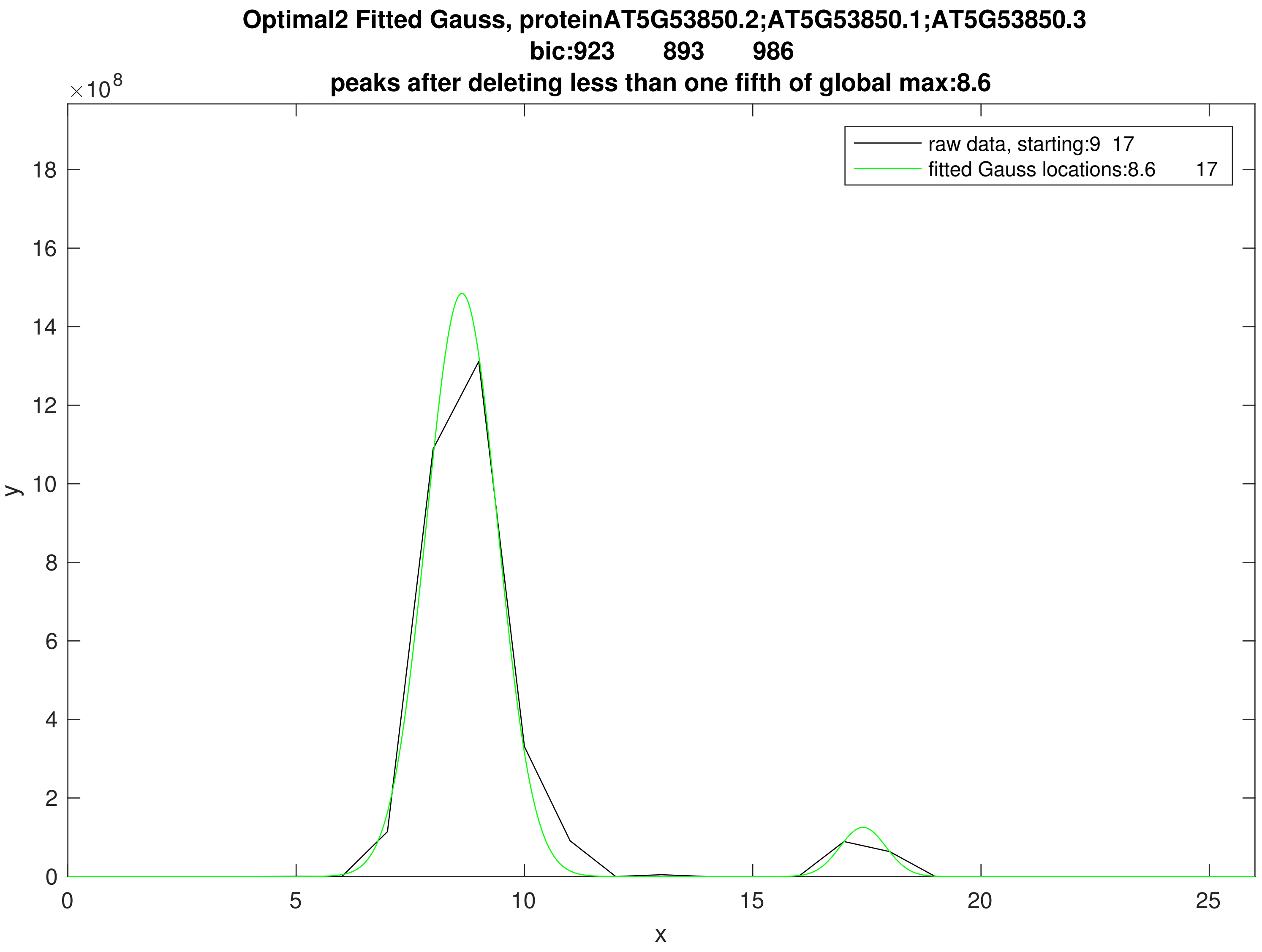

Optimal2 Fitted Gauss, proteinAT5G53850.2;AT5G53850.1;AT5G53850.3
bic:923 893 986
peaks after deleting less than one fifth of global max:8.6
×10^8
raw data, starting:9 17
fitted Gauss locations:8.6 17
y
x
0 2 4 6 8 10 12 14 16 18
0 5 10 15 20 25

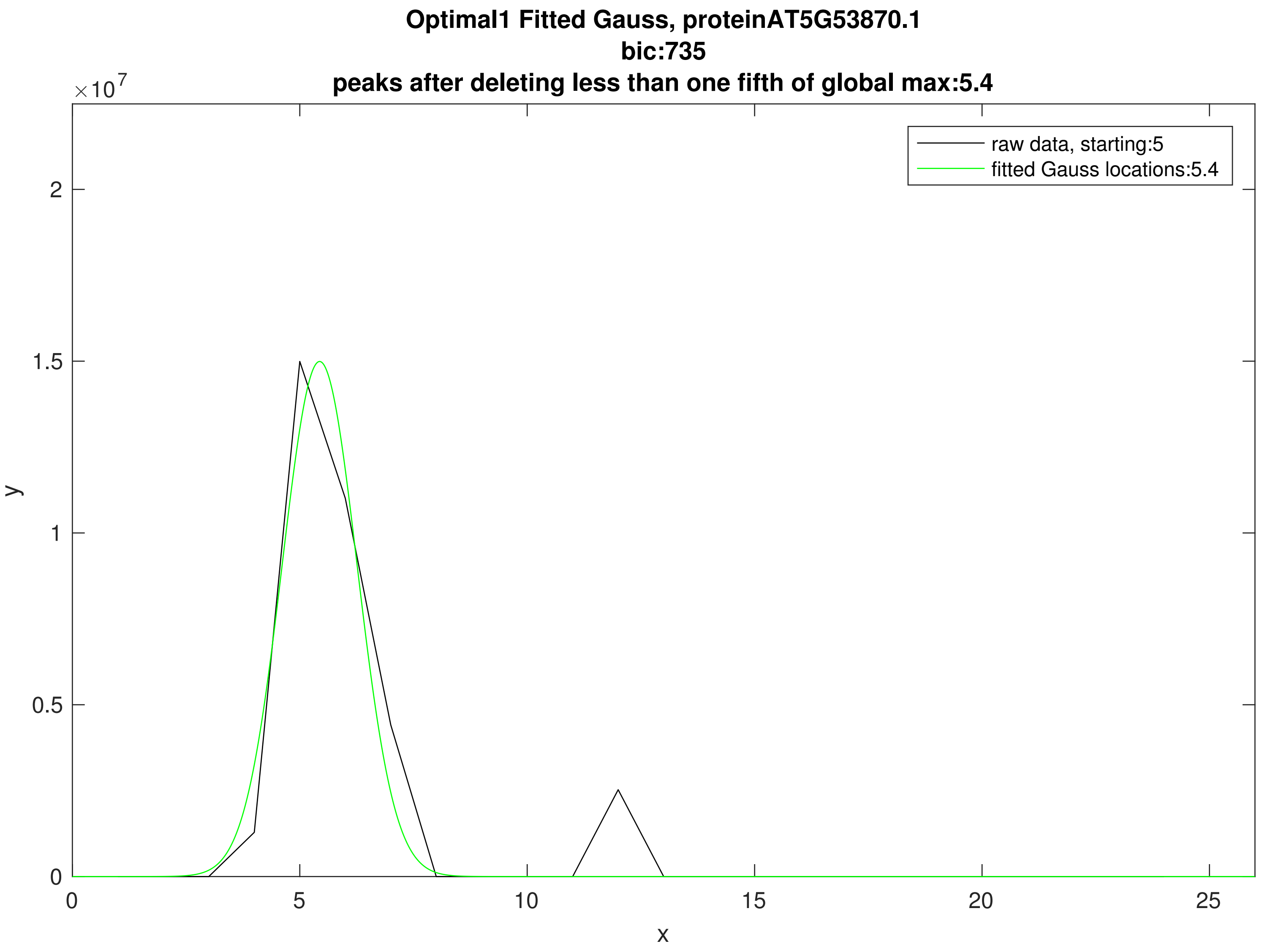
Optimal1 Fitted Gauss, proteinAT5G53870.1
bic:735
peaks after deleting less than one fifth of global max:5.4
×10^7
raw data, starting:5
fitted Gauss locations:5.4
y
x
0
0.5
1
1.5
2
0
5
10
15
20
25

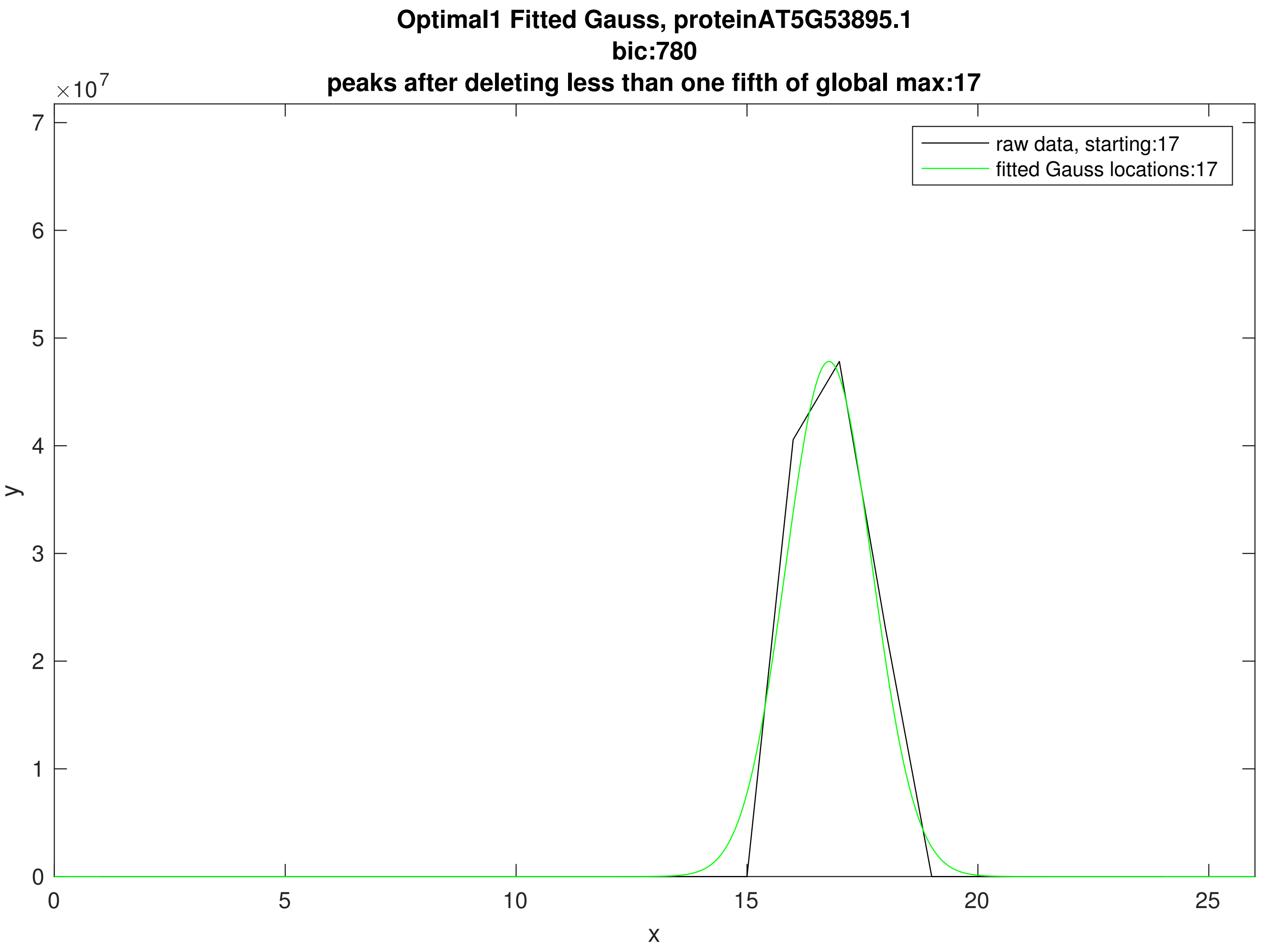
Optimal1 Fitted Gauss, proteinAT5G53895.1
bic:780
peaks after deleting less than one fifth of global max:17
×10^7
raw data, starting:17
fitted Gauss locations:17
y
x
0
1
2
3
4
5
6
7
0
5
10
15
20
25

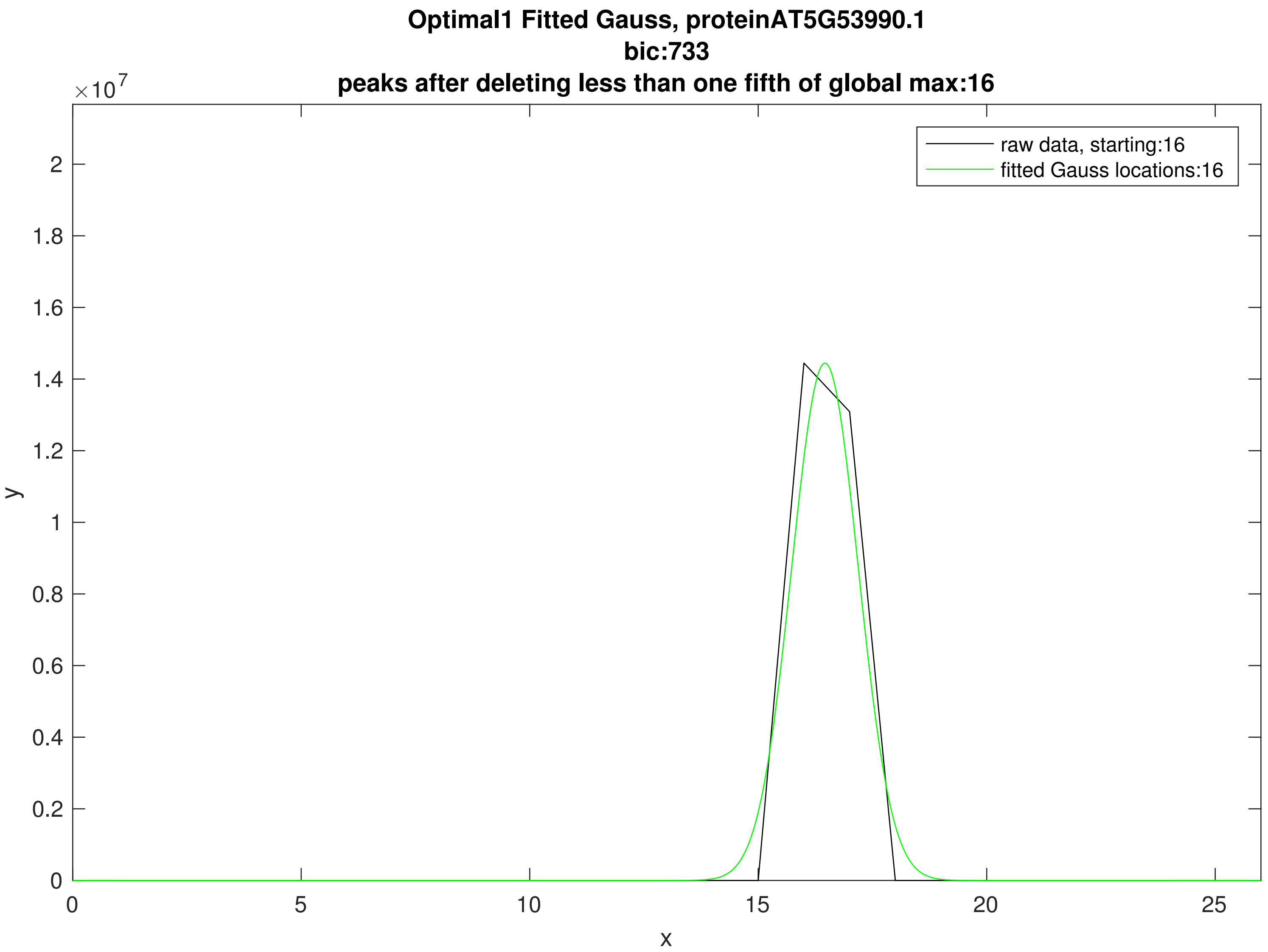
Optimal1 Fitted Gauss, proteinAT5G53990.1
bic:733
peaks after deleting less than one fifth of global max:16
×10^7
raw data, starting:16
fitted Gauss locations:16
y
x
0
0.2
0.4
0.6
0.8
1
1.2
1.4
1.6
1.8
2
0
5
10
15
20
25

#### Optimal1 Fitted Gauss, proteinAT5G54080.2;AT5G54080.1
#### bic:876
#### peaks after deleting less than one fifth of global max:8.6

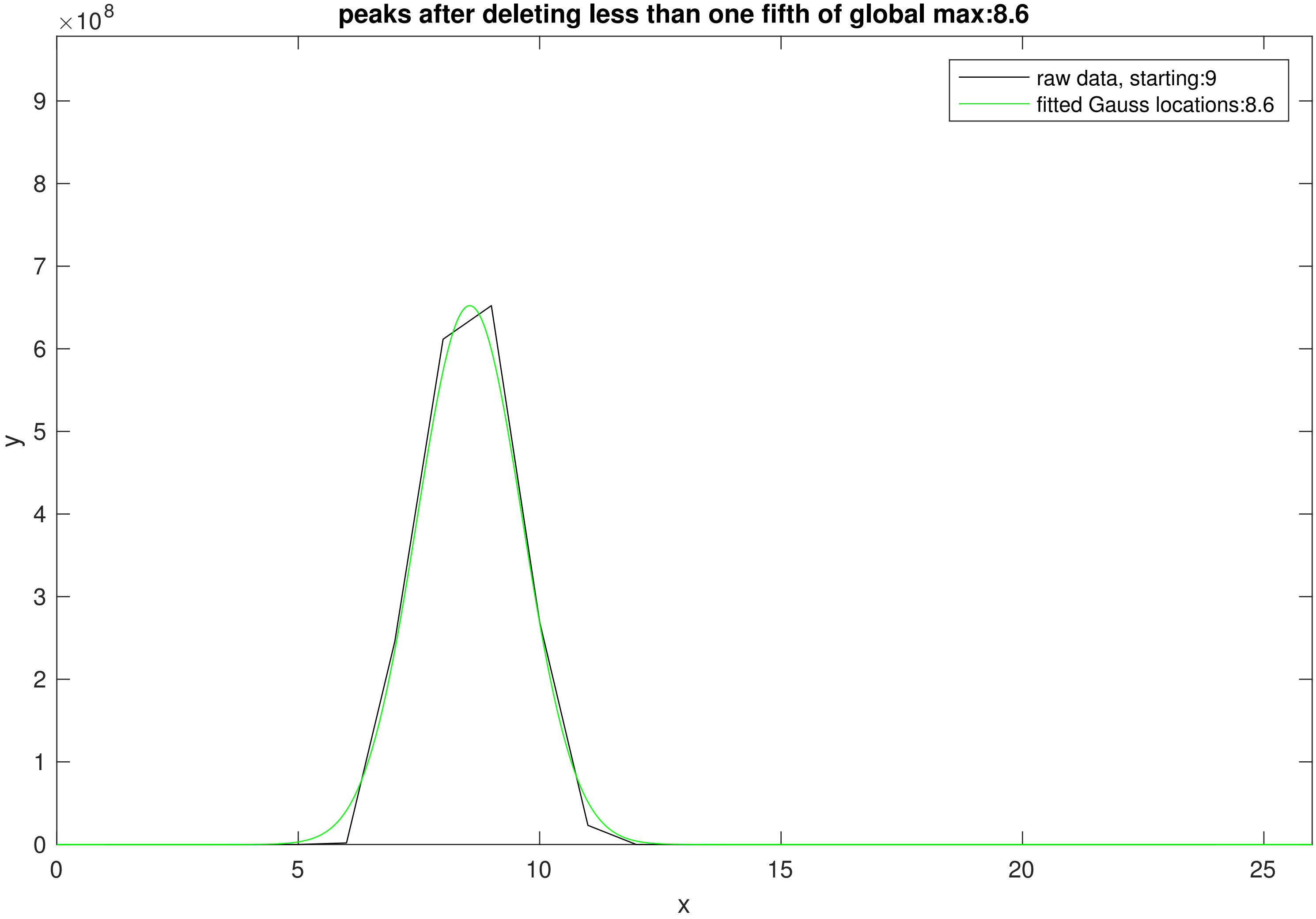

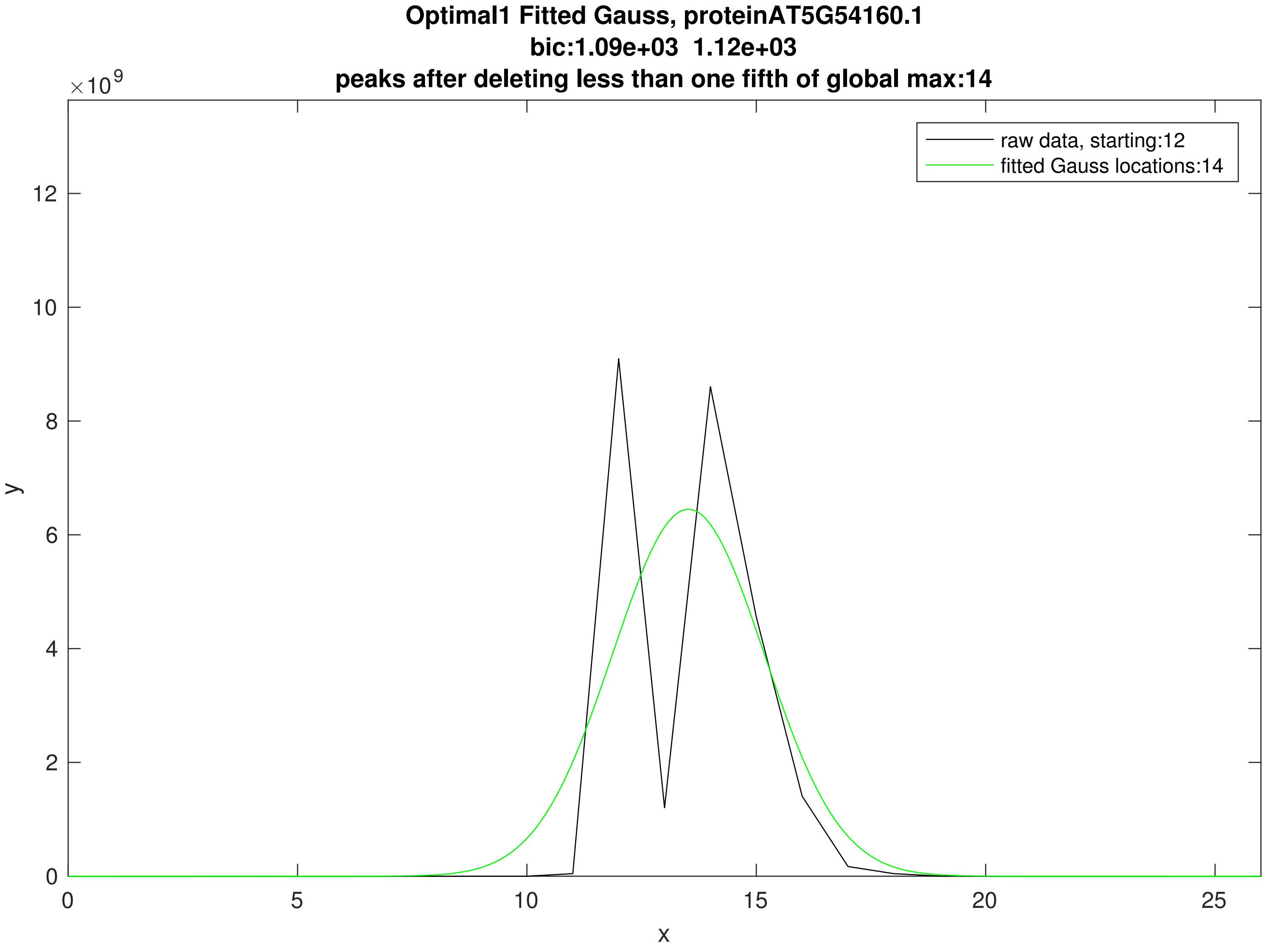
Optimal1 Fitted Gauss, proteinAT5G54160.1
bic:1.09e+03  1.12e+03
peaks after deleting less than one fifth of global max:14
×10^9
raw data, starting:12
fitted Gauss locations:14
y
x
0
2
4
6
8
10
12
0
5
10
15
20
25

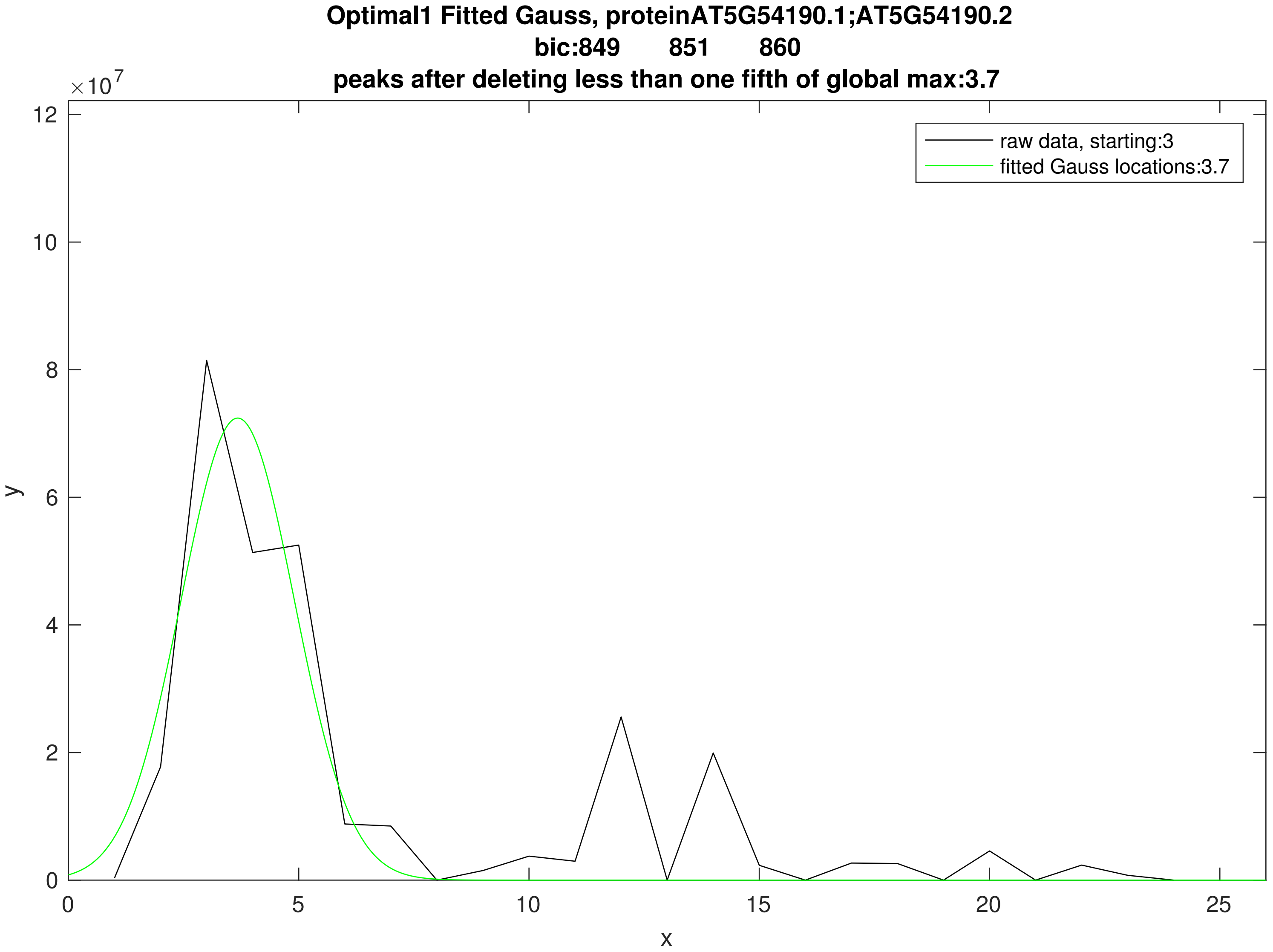
Optimal1 Fitted Gauss, proteinAT5G54190.1;AT5G54190.2
bic:849 851 860
peaks after deleting less than one fifth of global max:3.7
×10^7
raw data, starting:3
fitted Gauss locations:3.7
y
x
0
2
4
6
8
10
12
0
5
10
15
20
25

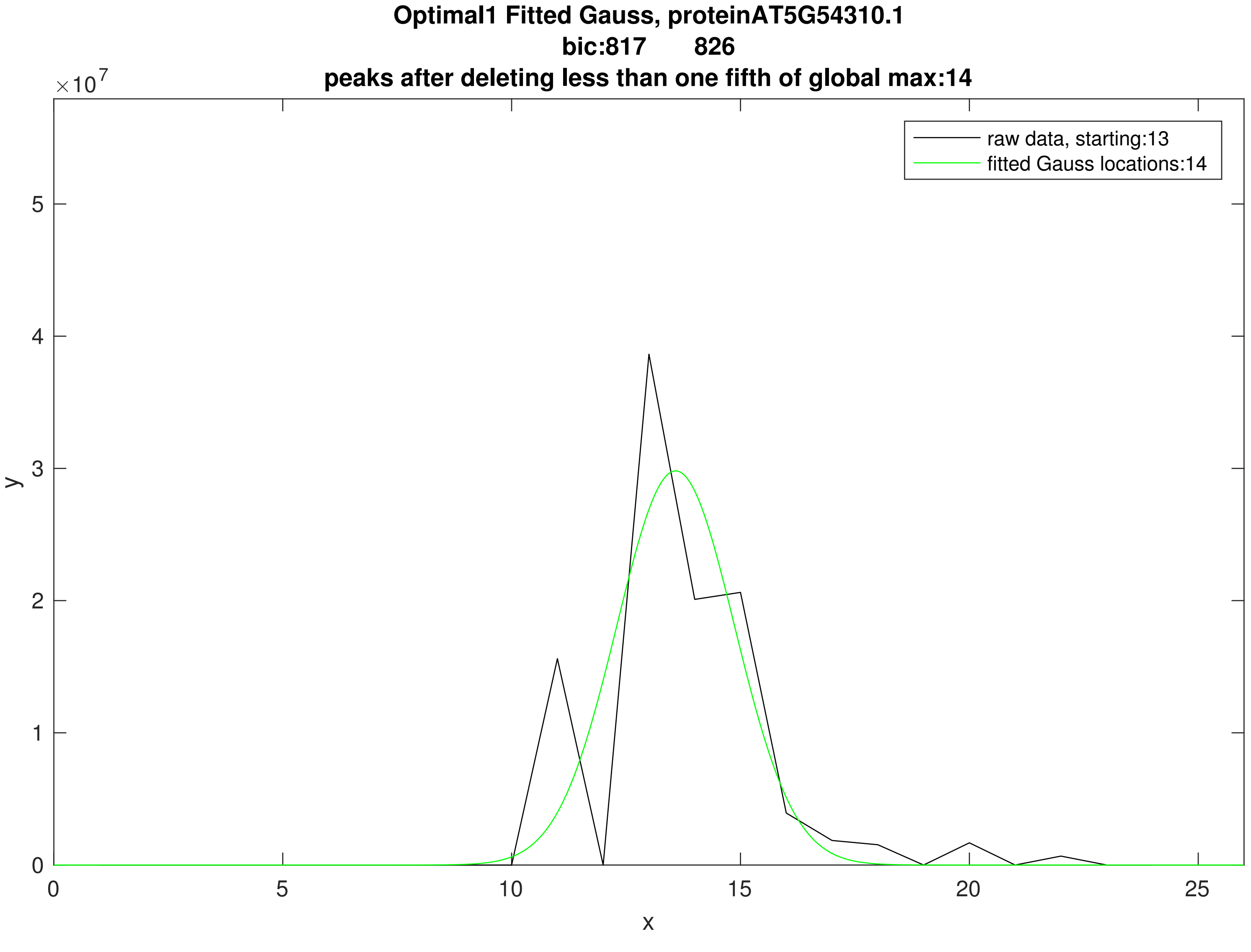

Optimal1 Fitted Gauss, proteinAT5G54310.1
bic:817 826
peaks after deleting less than one fifth of global max:14
×10^7
raw data, starting:13
fitted Gauss locations:14
y
x
0
1
2
3
4
5
0
5
10
15
20
25

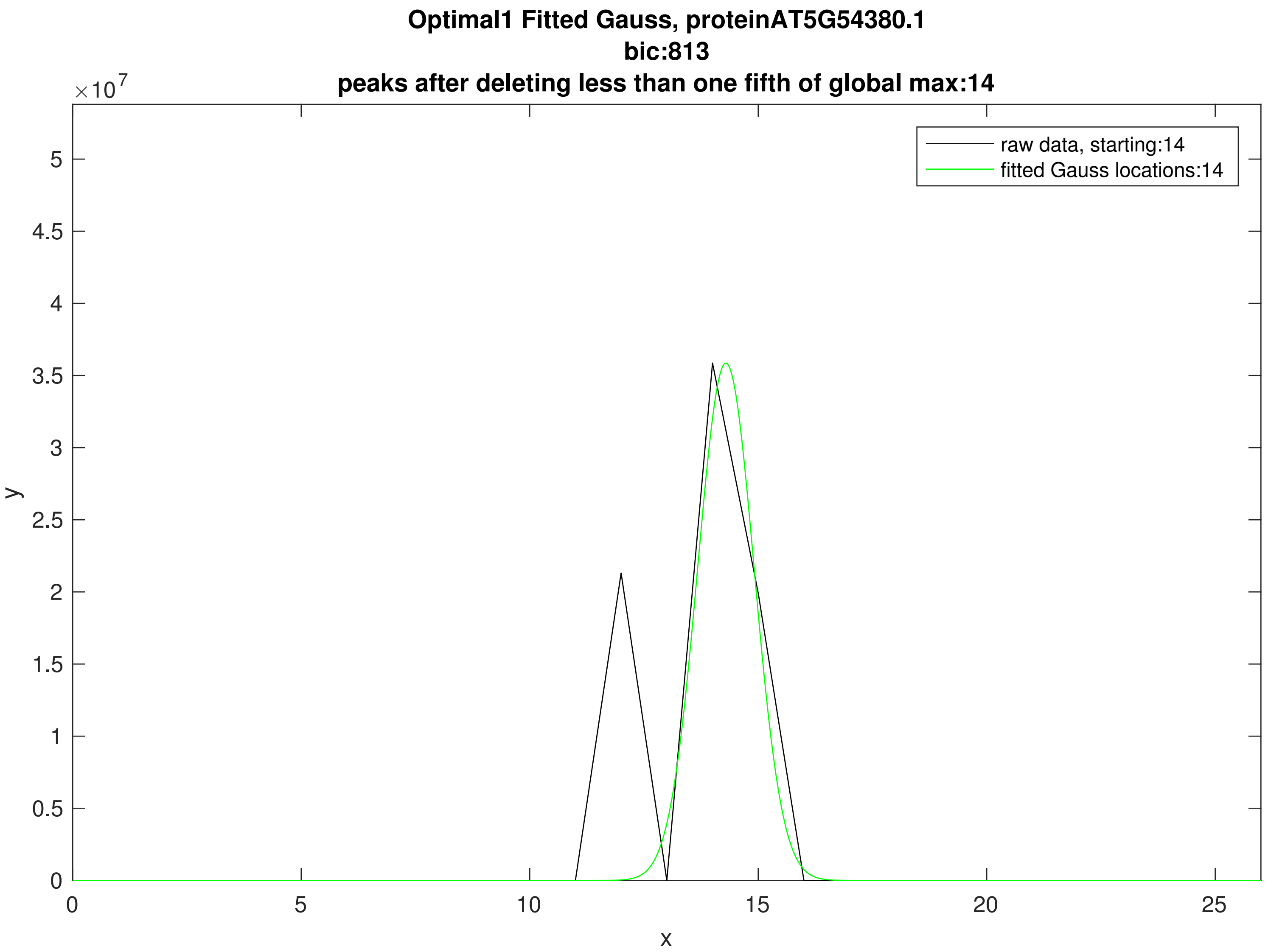
Optimal1 Fitted Gauss, proteinAT5G54380.1
bic:813
peaks after deleting less than one fifth of global max:14
×10^7
raw data, starting:14
fitted Gauss locations:14
y
x
0
0.5
1
1.5
2
2.5
3
3.5
4
4.5
5
0
5
10
15
20
25

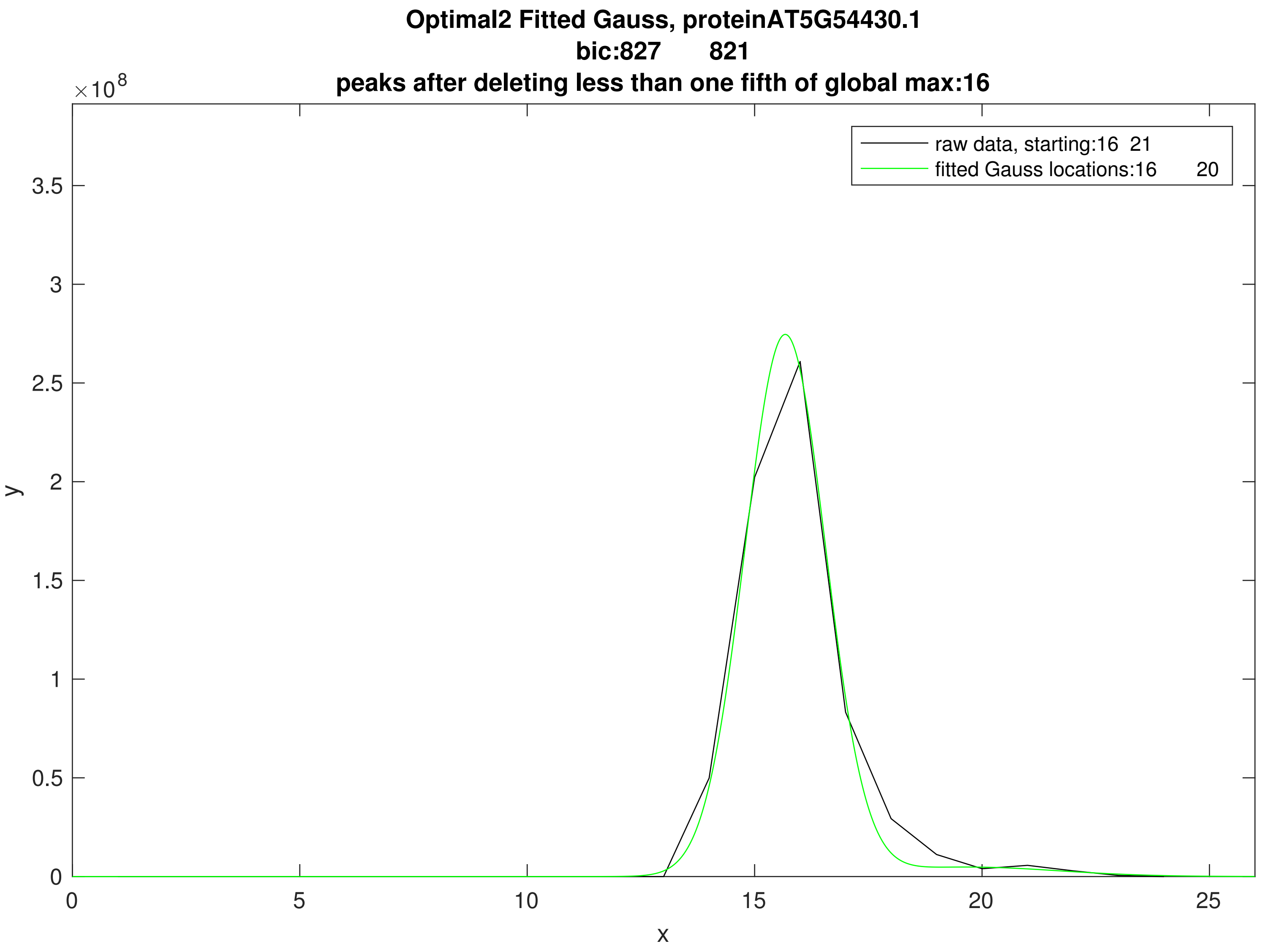
Optimal2 Fitted Gauss, proteinAT5G54430.1
bic:827 821
peaks after deleting less than one fifth of global max:16
×10^8
raw data, starting:16 21
fitted Gauss locations:16 20
y
x
0
0.5
1
1.5
2
2.5
3
3.5
0
5
10
15
20
25

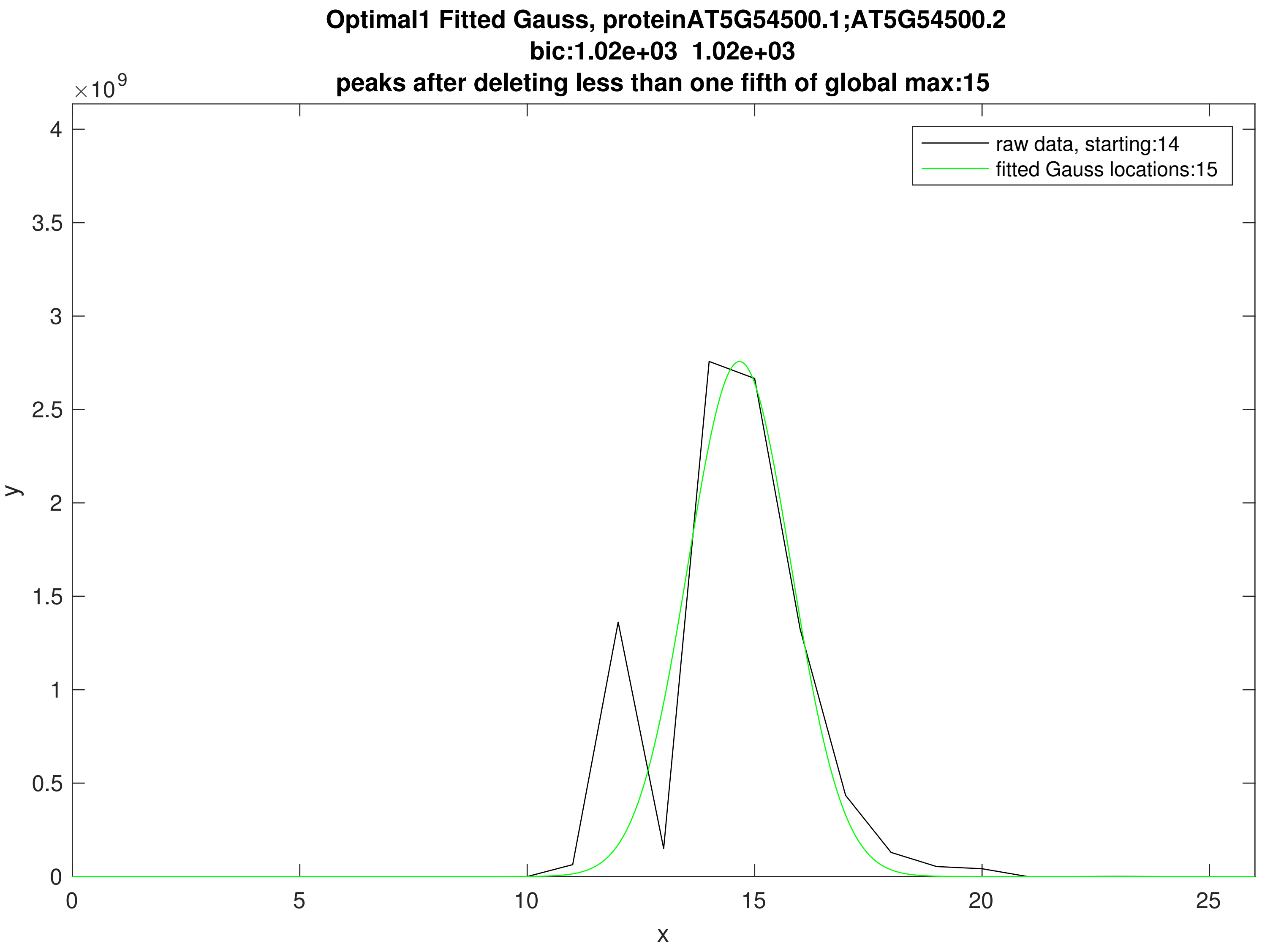

Optimal1 Fitted Gauss, proteinAT5G54500.1;AT5G54500.2
bic:1.02e+03  1.02e+03
peaks after deleting less than one fifth of global max:15
×10^9
raw data, starting:14
fitted Gauss locations:15
y
x
0
0.5
1
1.5
2
2.5
3
3.5
4
5
10
15
20
25

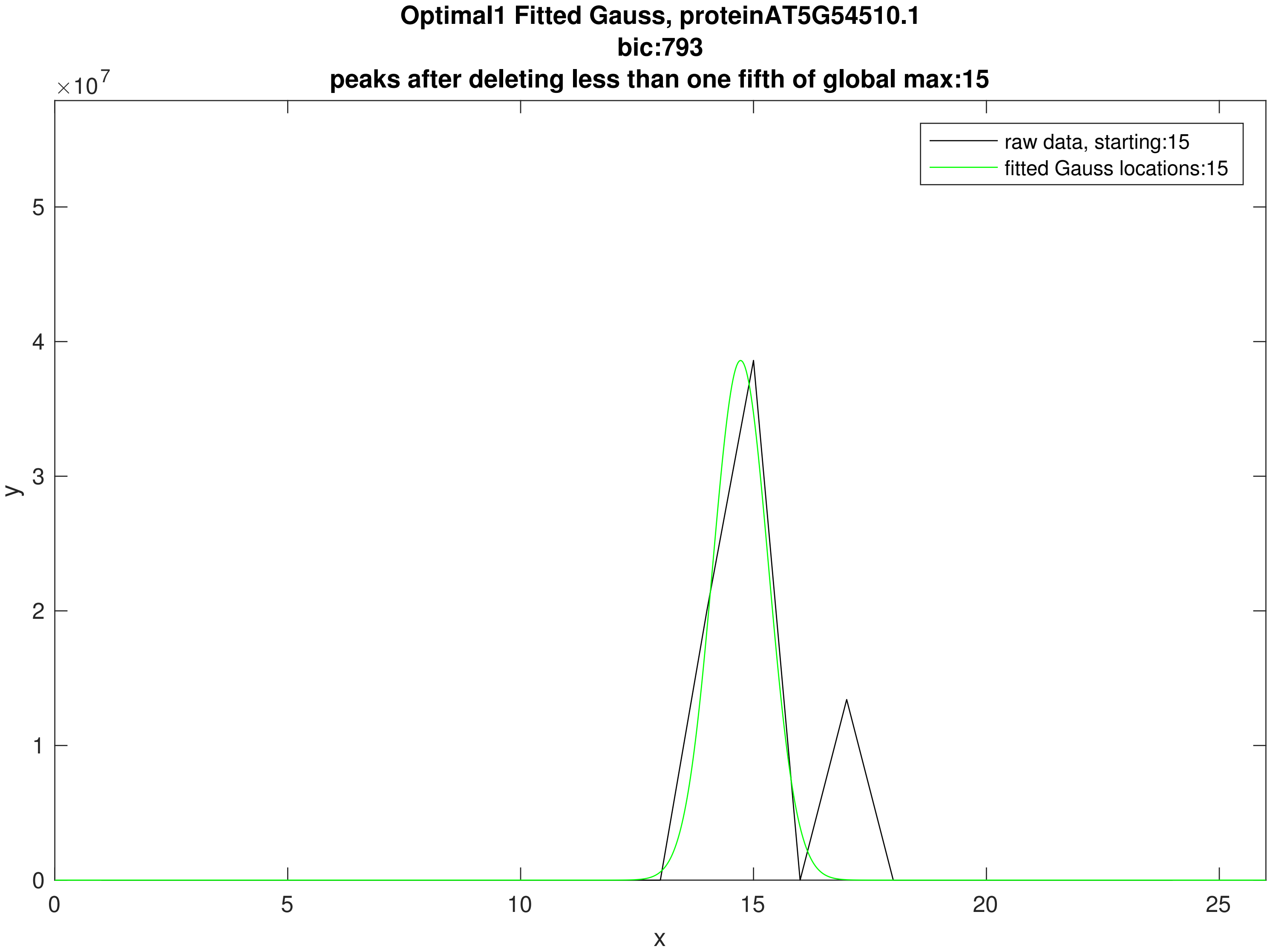
Optimal1 Fitted Gauss, proteinAT5G54510.1
bic:793
peaks after deleting less than one fifth of global max:15
×10^7
raw data, starting:15
fitted Gauss locations:15
5
4
3
2
1
0
y
0
5
10
15
20
25
x

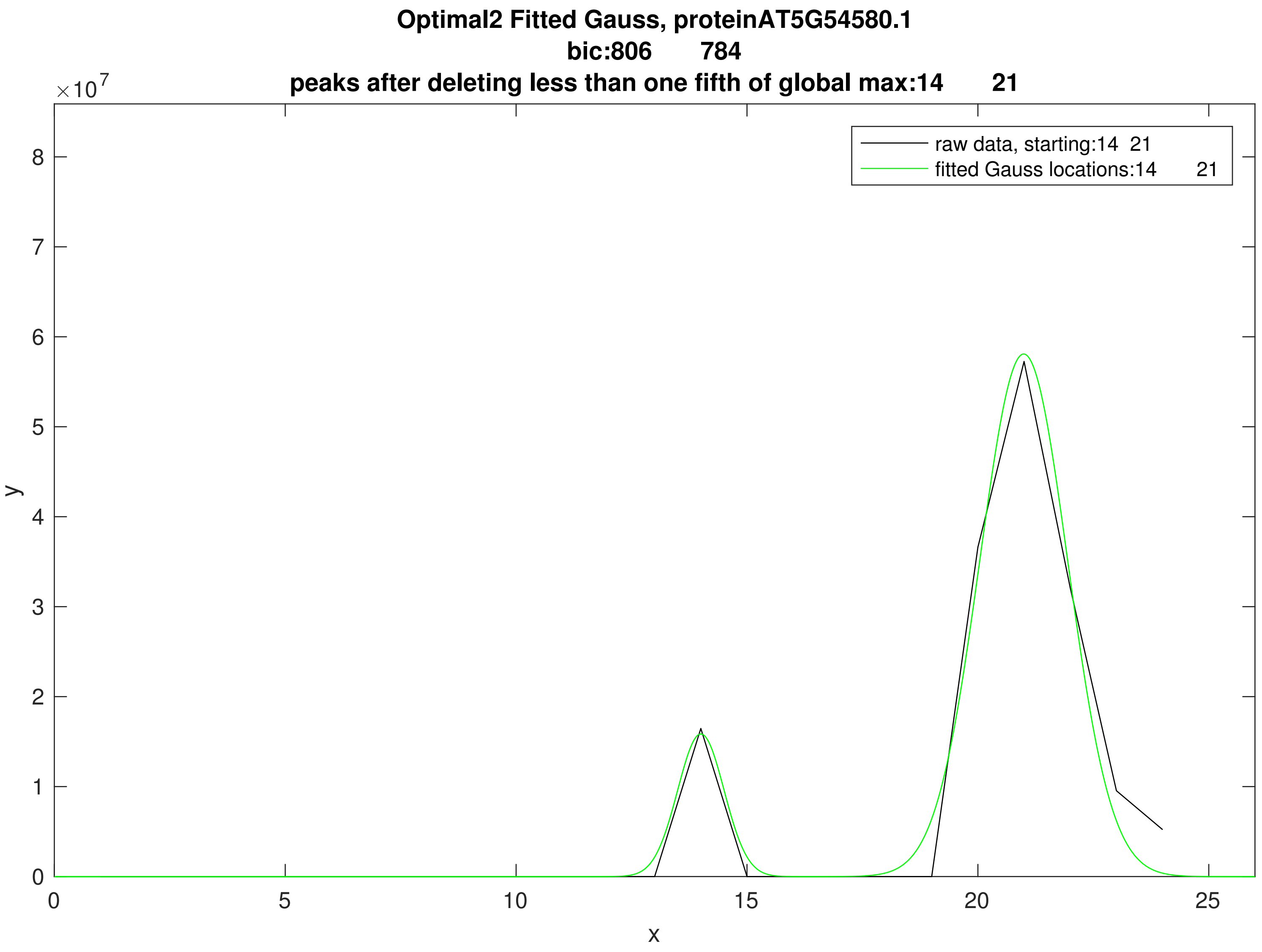

Optimal2 Fitted Gauss, proteinAT5G54580.1
bic:806 784
peaks after deleting less than one fifth of global max:14 21
×10^7
raw data, starting:14 21
fitted Gauss locations:14 21
y
x
0
1
2
3
4
5
6
7
8
0
5
10
15
20
25

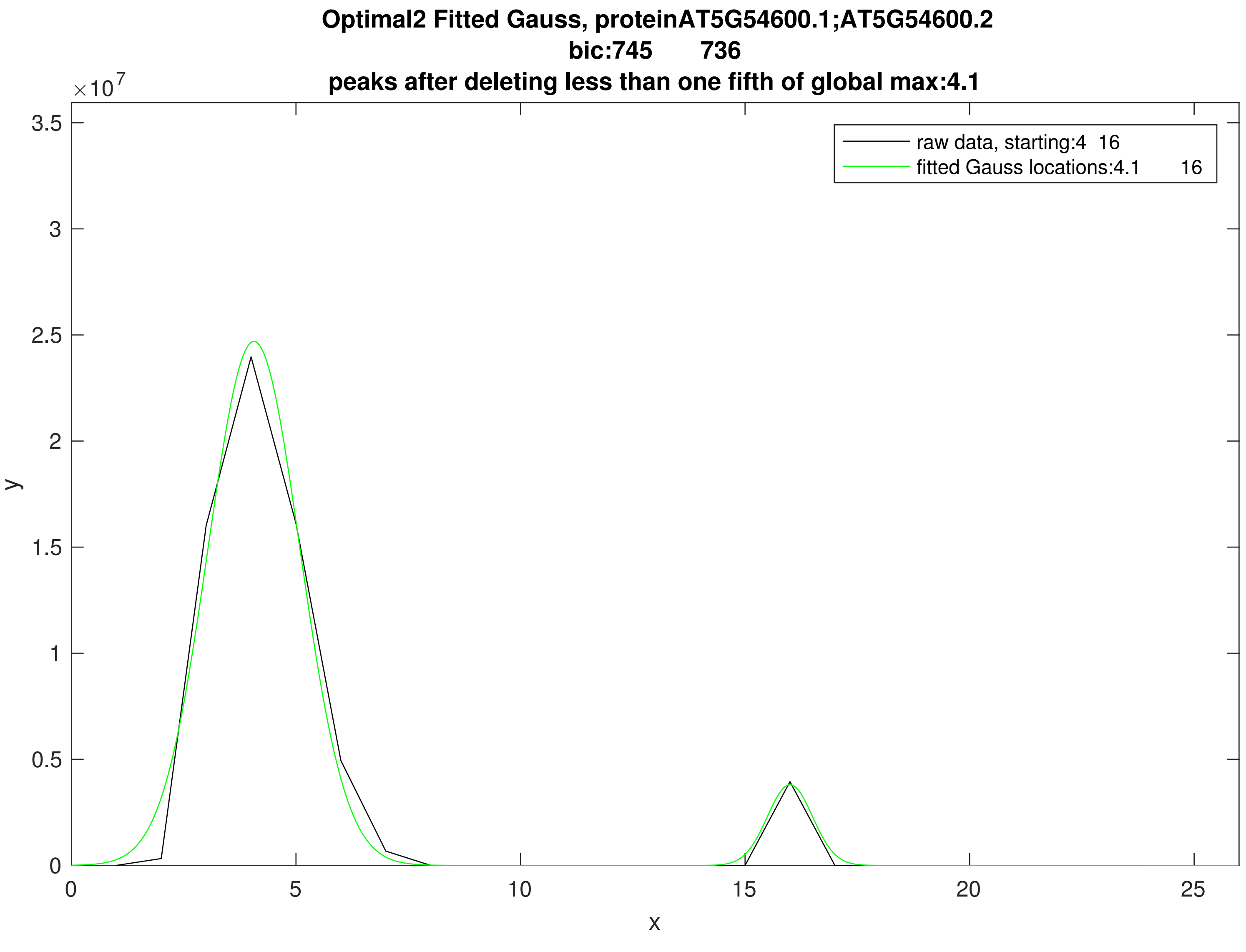

Optimal2 Fitted Gauss, proteinAT5G54600.1;AT5G54600.2
bic:745 736
peaks after deleting less than one fifth of global max:4.1
×10^7
3.5
3
2.5
2
1.5
1
0.5
0
y
raw data, starting:4 16
fitted Gauss locations:4.1 16
0
5
10
15
20
25
x

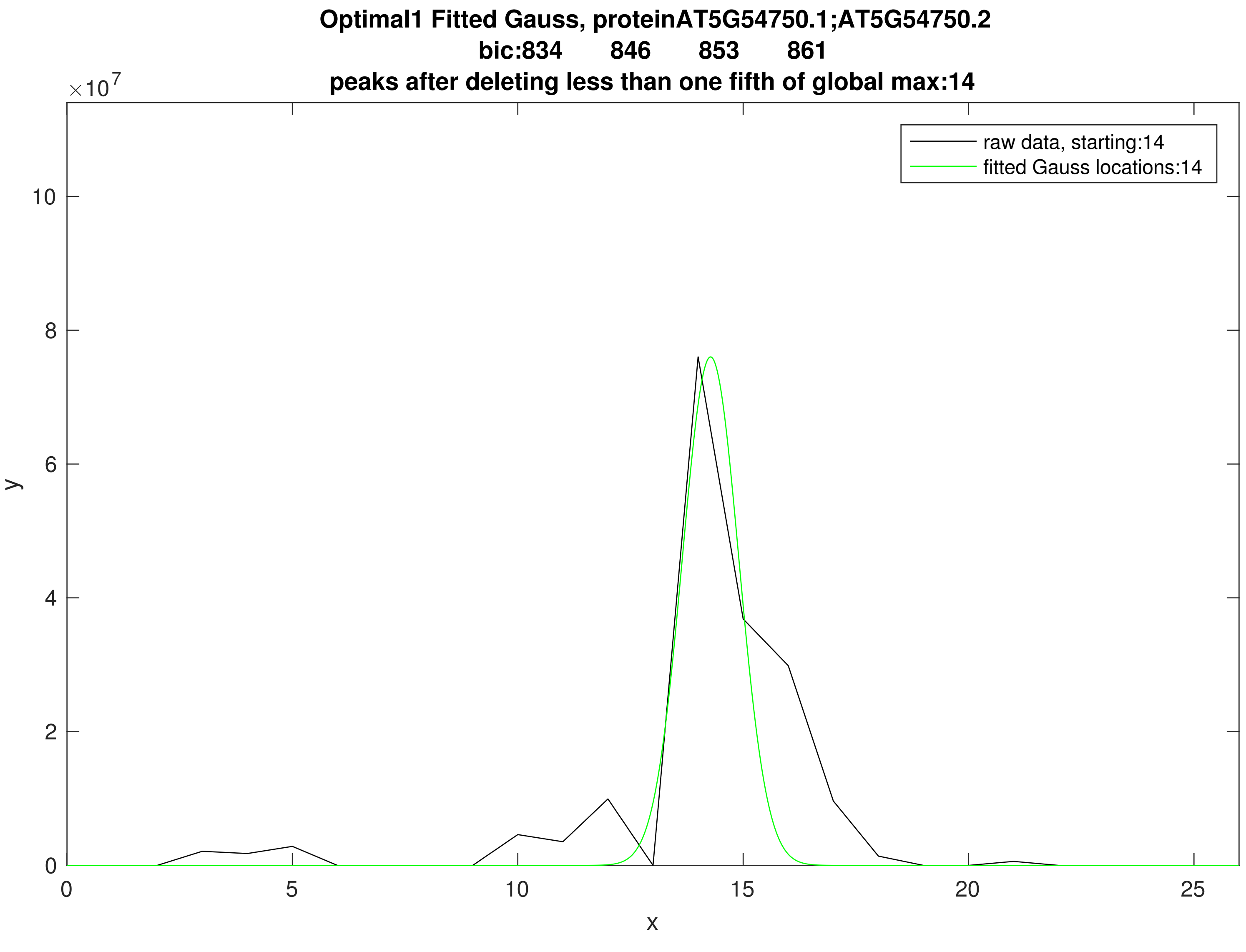

Optimal1 Fitted Gauss, proteinAT5G54750.1;AT5G54750.2
bic:834 846 853 861
peaks after deleting less than one fifth of global max:14
×10^7
raw data, starting:14
fitted Gauss locations:14
y
x
0
2
4
6
8
10
0
5
10
15
20
25

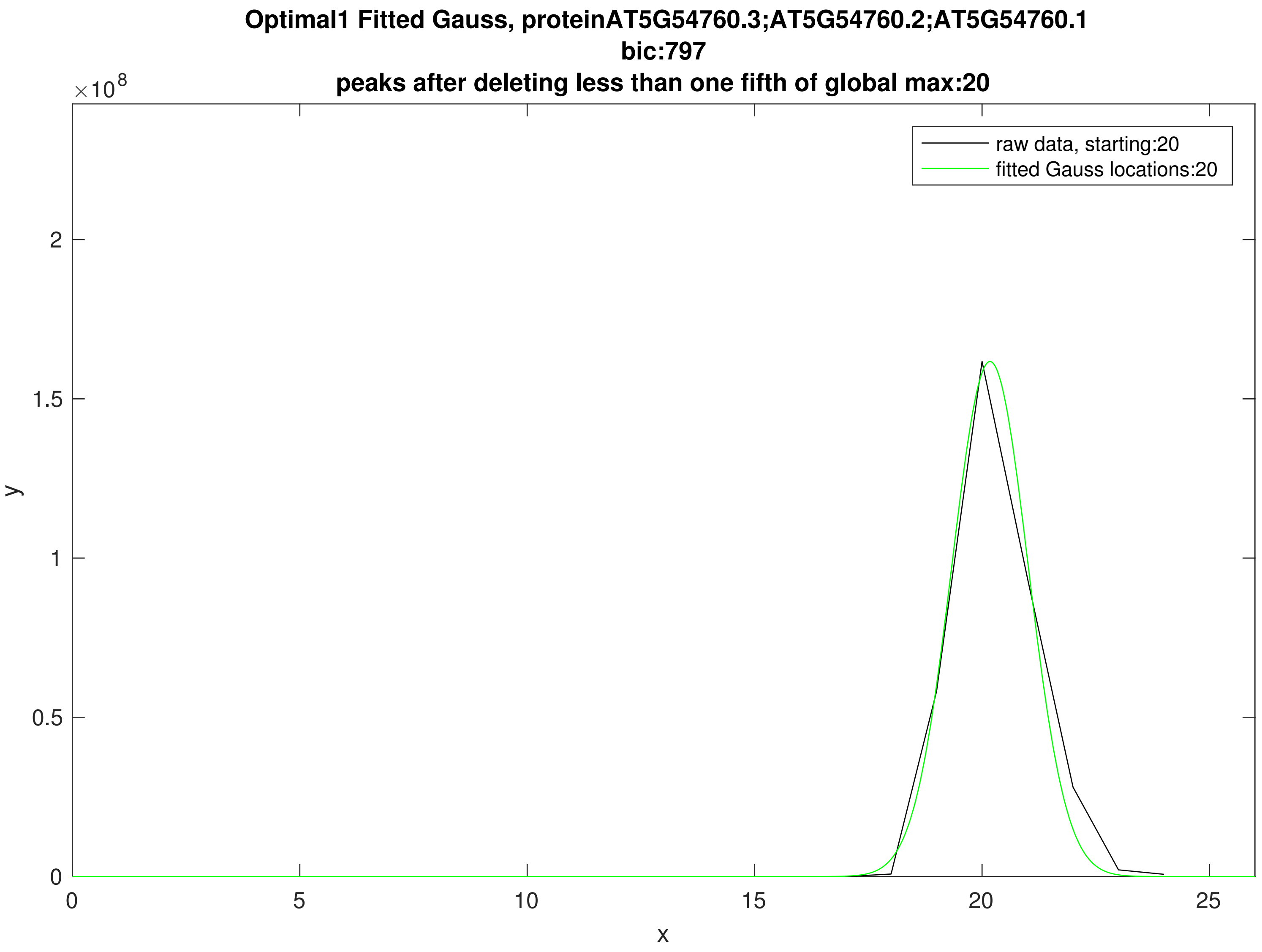

Optimal1 Fitted Gauss, proteinAT5G54760.3;AT5G54760.2;AT5G54760.1
bic:797
peaks after deleting less than one fifth of global max:20
×10^8
raw data, starting:20
fitted Gauss locations:20
2
1.5
1
0.5
0
y
0
5
10
15
20
25
x

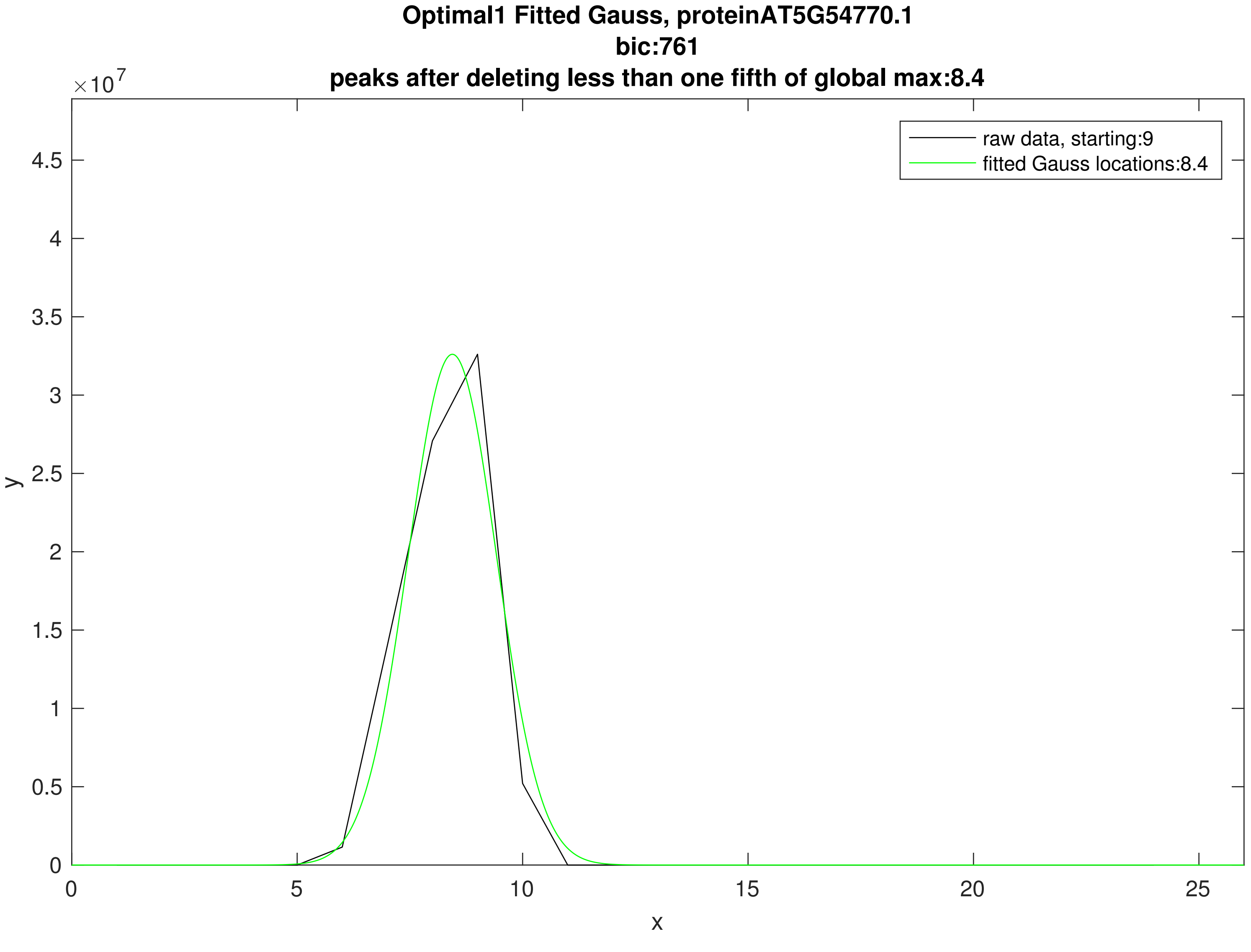
Optimal1 Fitted Gauss, proteinAT5G54770.1
bic:761
peaks after deleting less than one fifth of global max:8.4
×10^7
4.5
4
3.5
3
2.5
2
1.5
1
0.5
0
y
0
5
10
15
20
25
x
raw data, starting:9
fitted Gauss locations:8.4

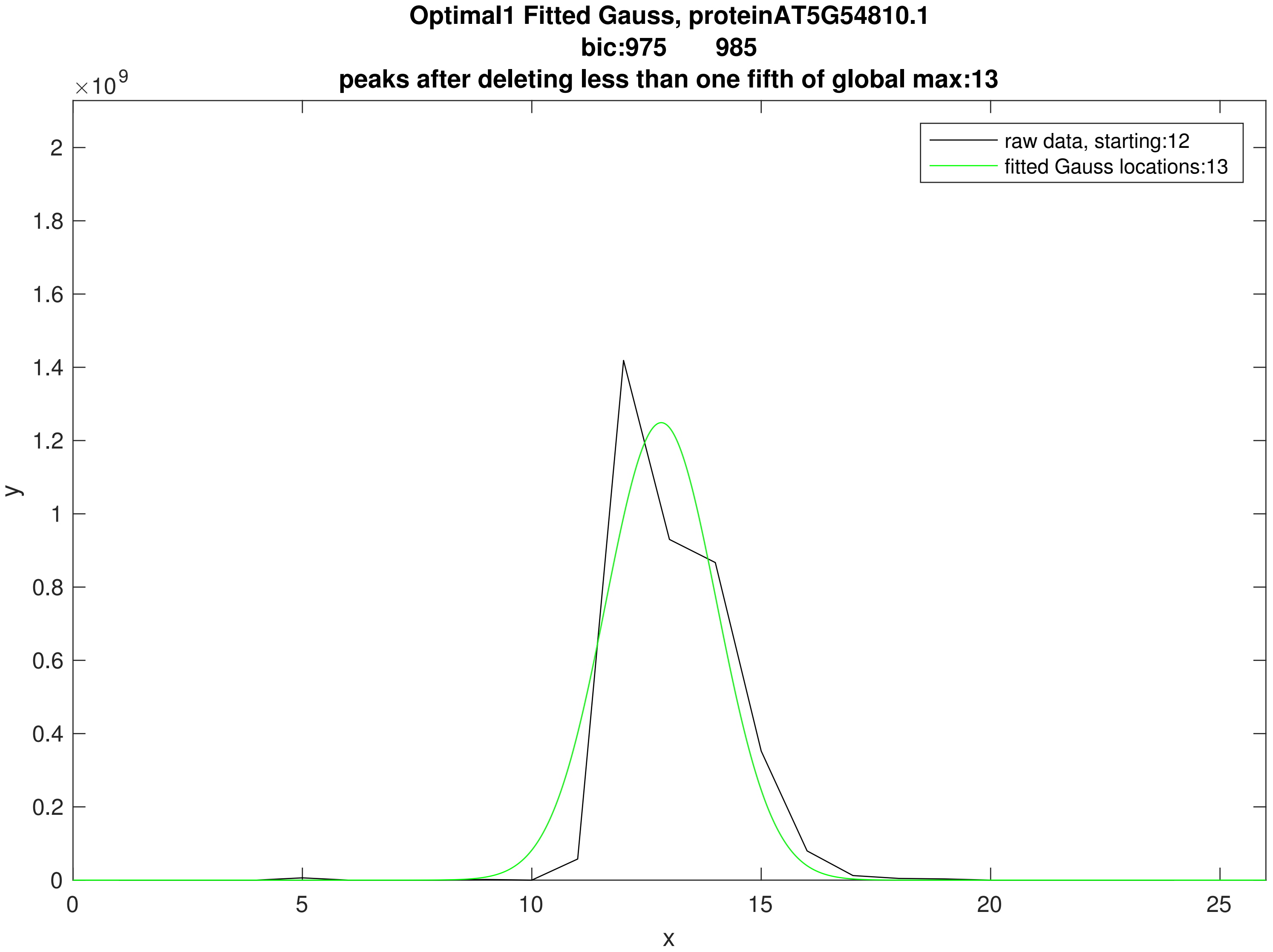
Optimal1 Fitted Gauss, proteinAT5G54810.1
bic:975 985
peaks after deleting less than one fifth of global max:13
×10^9
raw data, starting:12
fitted Gauss locations:13
y
x
0
0.2
0.4
0.6
0.8
1
1.2
1.4
1.6
1.8
2
5
10
15
20
25

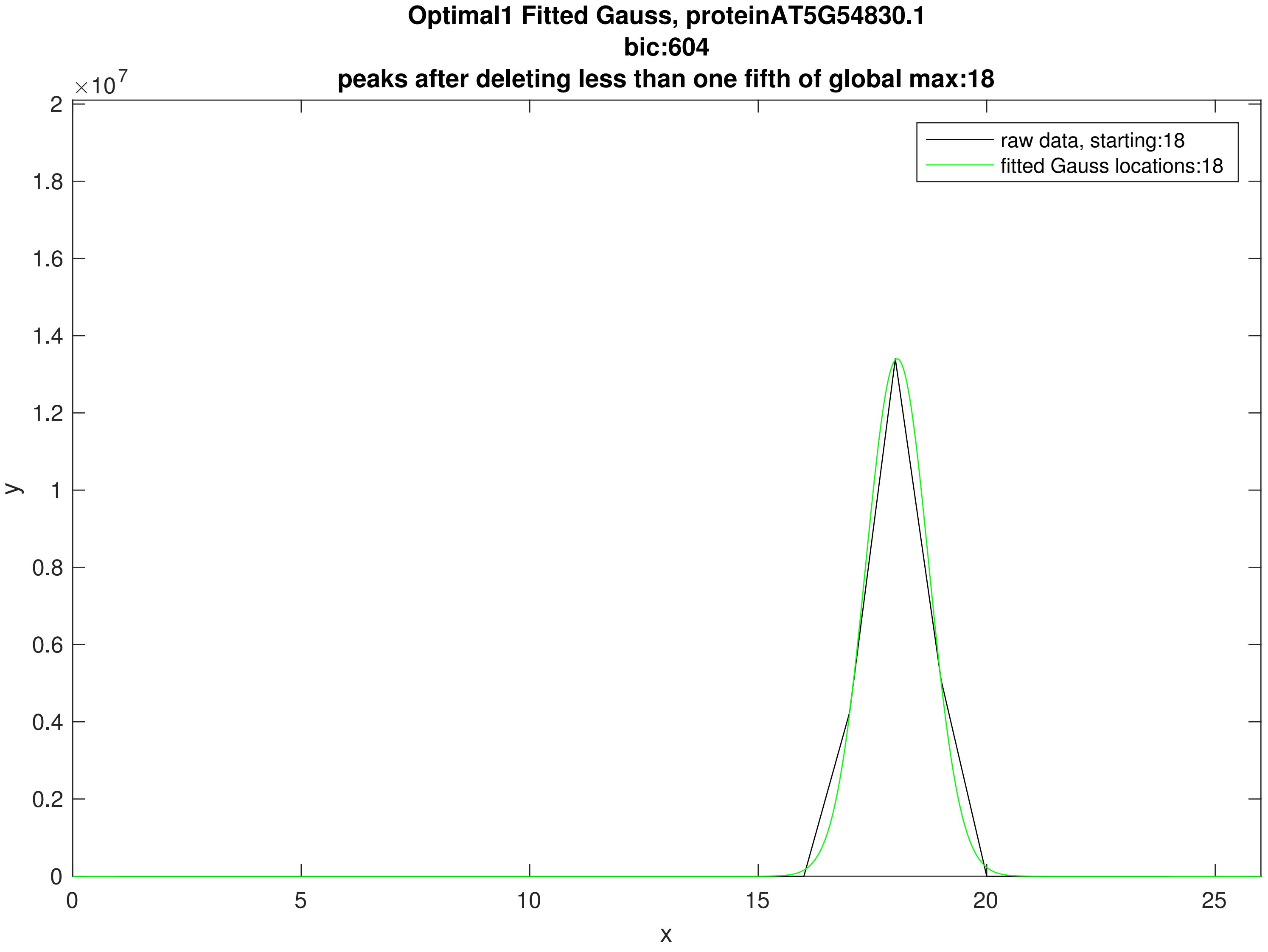

Optimal1 Fitted Gauss, proteinAT5G54830.1
bic:604
peaks after deleting less than one fifth of global max:18
×10^7
raw data, starting:18
fitted Gauss locations:18
y
x
0
0.2
0.4
0.6
0.8
1
1.2
1.4
1.6
1.8
2
0
5
10
15
20
25

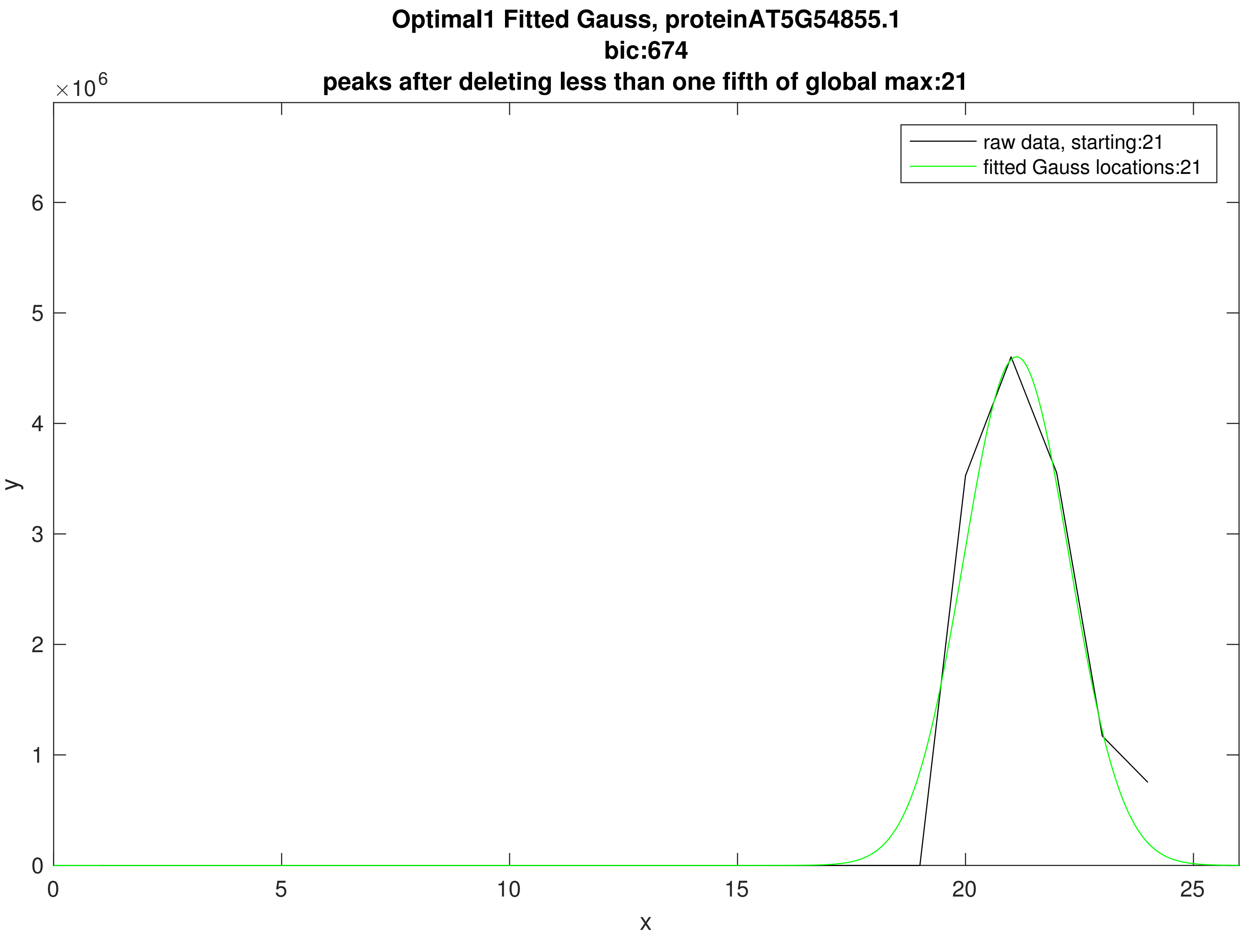
Optimal1 Fitted Gauss, proteinAT5G54855.1
bic:674
peaks after deleting less than one fifth of global max:21
×10^6
raw data, starting:21
fitted Gauss locations:21
y
x
0
1
2
3
4
5
6
0
5
10
15
20
25

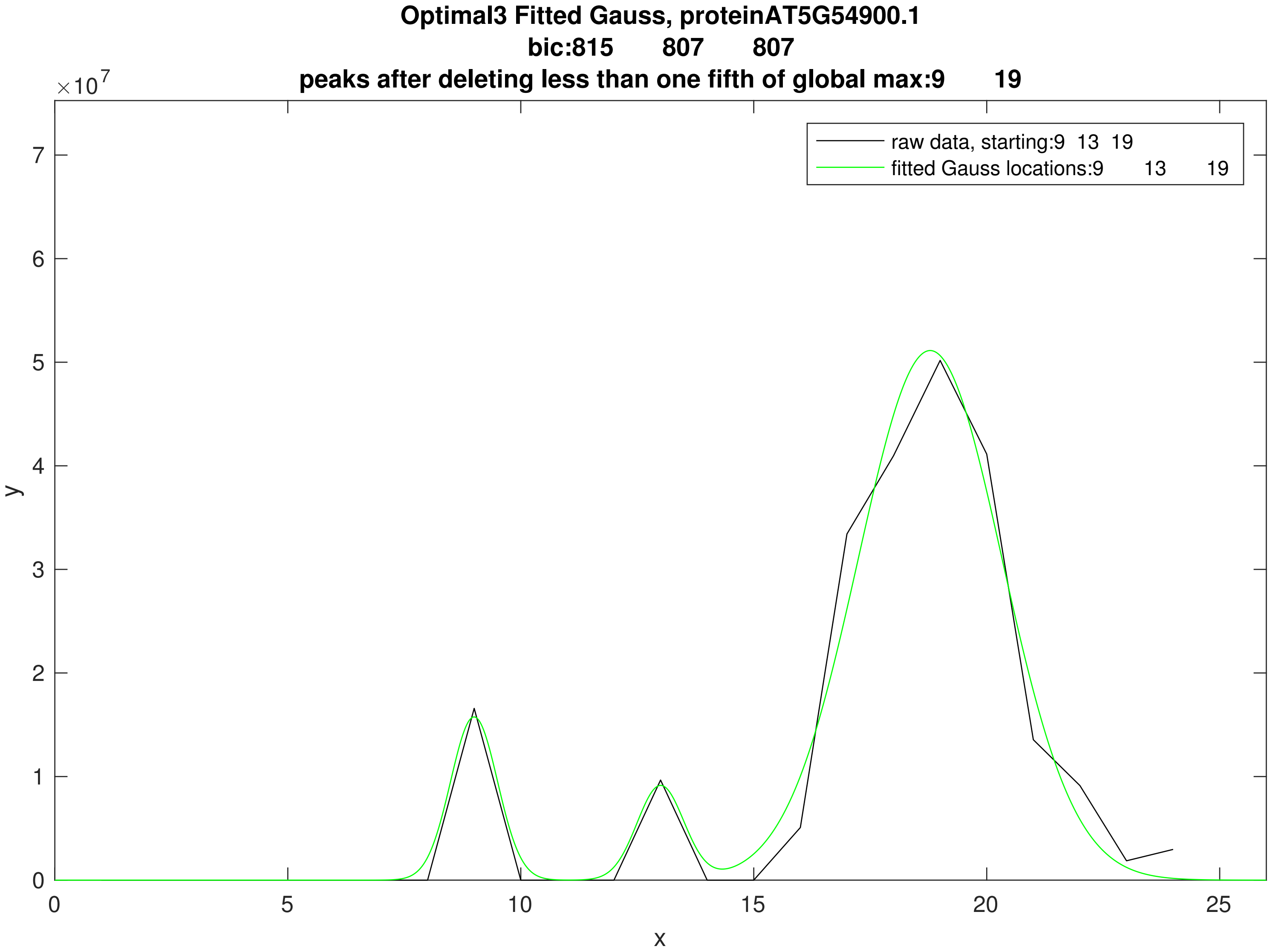
Optimal3 Fitted Gauss, proteinAT5G54900.1
bic:815 807 807
peaks after deleting less than one fifth of global max:9 19
×10^7
raw data, starting:9 13 19
fitted Gauss locations:9 13 19
y
x
0
1
2
3
4
5
6
7
0
5
10
15
20
25

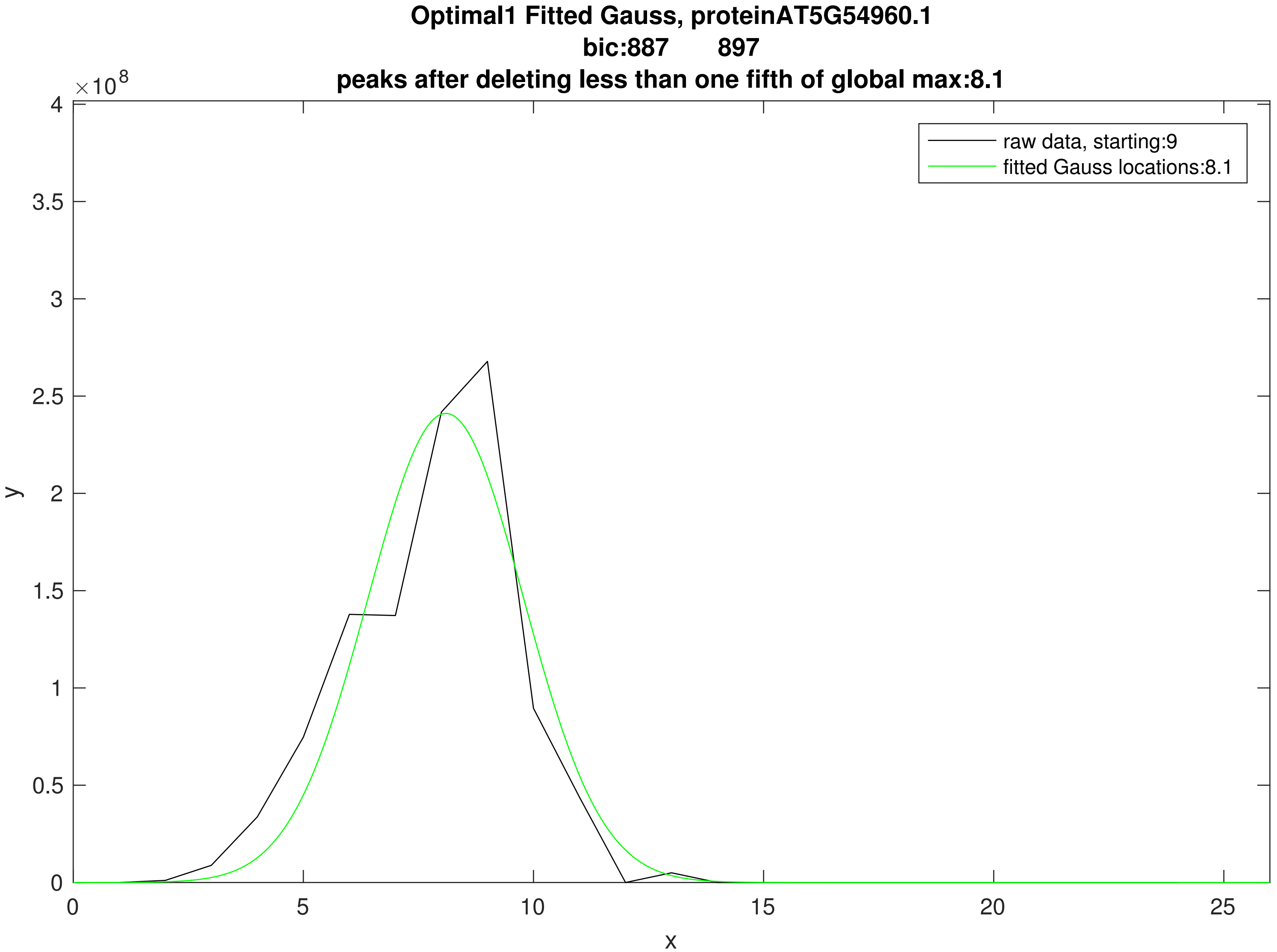

Optimal1 Fitted Gauss, proteinAT5G54960.1
bic:887 897
peaks after deleting less than one fifth of global max:8.1
×10^8
raw data, starting:9
fitted Gauss locations:8.1
y
x
0
0.5
1
1.5
2
2.5
3
3.5
4
5
10
15
20
25

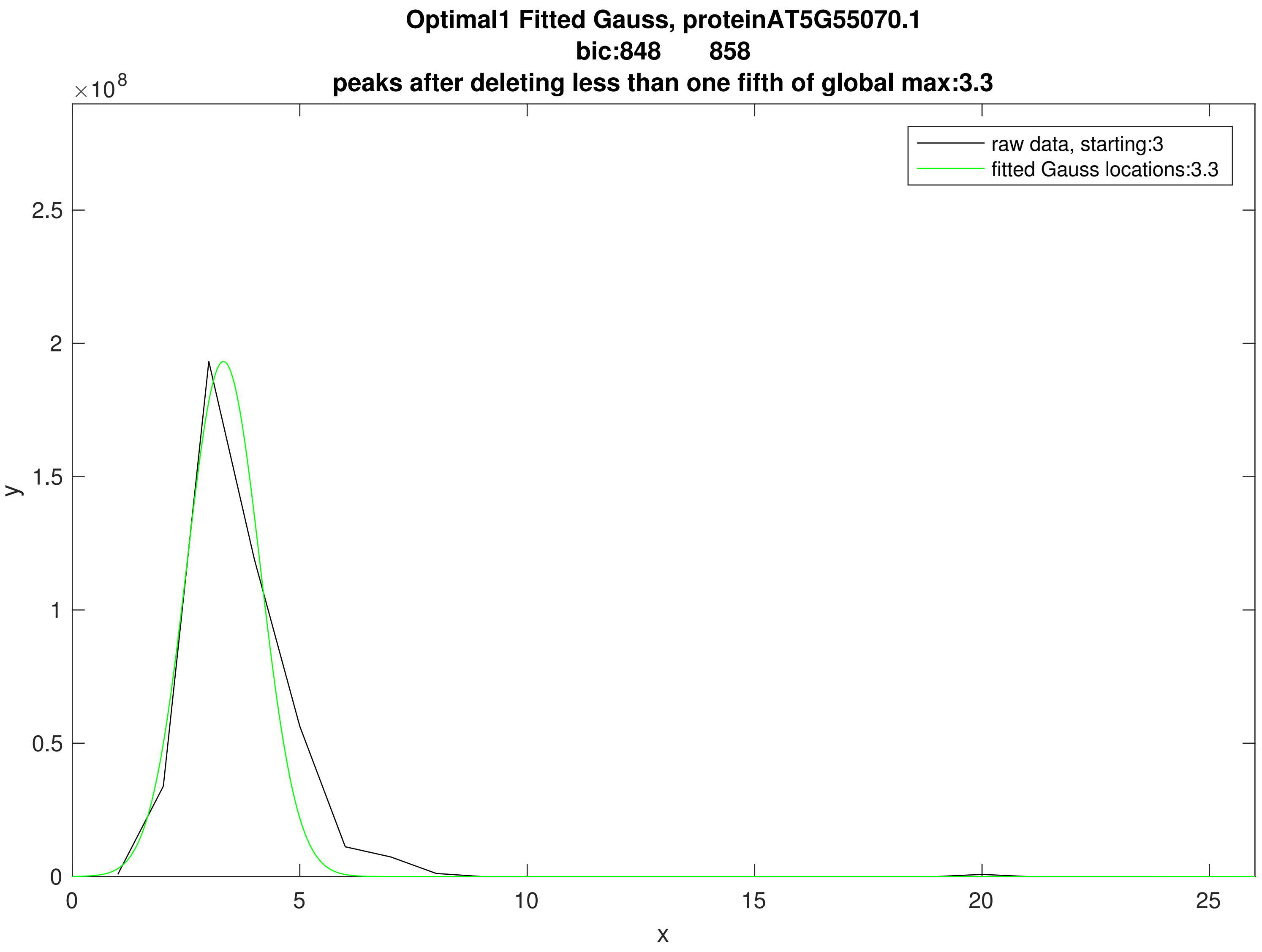
Optimal1 Fitted Gauss, proteinAT5G55070.1
bic:848 858
peaks after deleting less than one fifth of global max:3.3
×10^8
raw data, starting:3
fitted Gauss locations:3.3
y
x
0
0.5
1
1.5
2
2.5
5
10
15
20
25

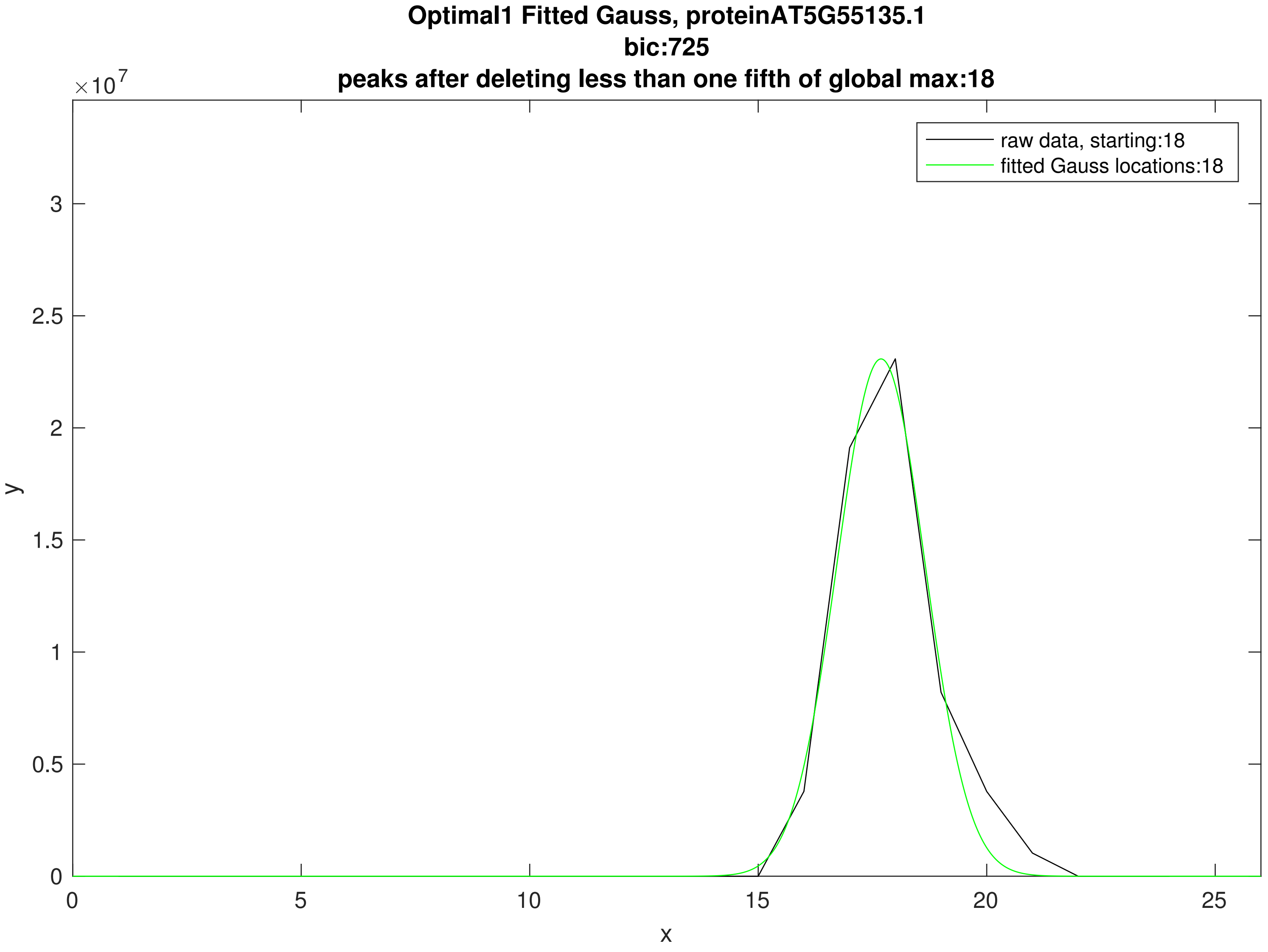
Optimal1 Fitted Gauss, proteinAT5G55135.1
bic:725
peaks after deleting less than one fifth of global max:18
×10^7
raw data, starting:18
fitted Gauss locations:18
3
2.5
2
1.5
1
0.5
0
y
0
5
10
15
20
25
x

#### Optimal1 Fitted Gauss, proteinAT5G55160.1;AT5G55160.2
#### bic:914 923 932
#### peaks after deleting less than one fifth of global max:20

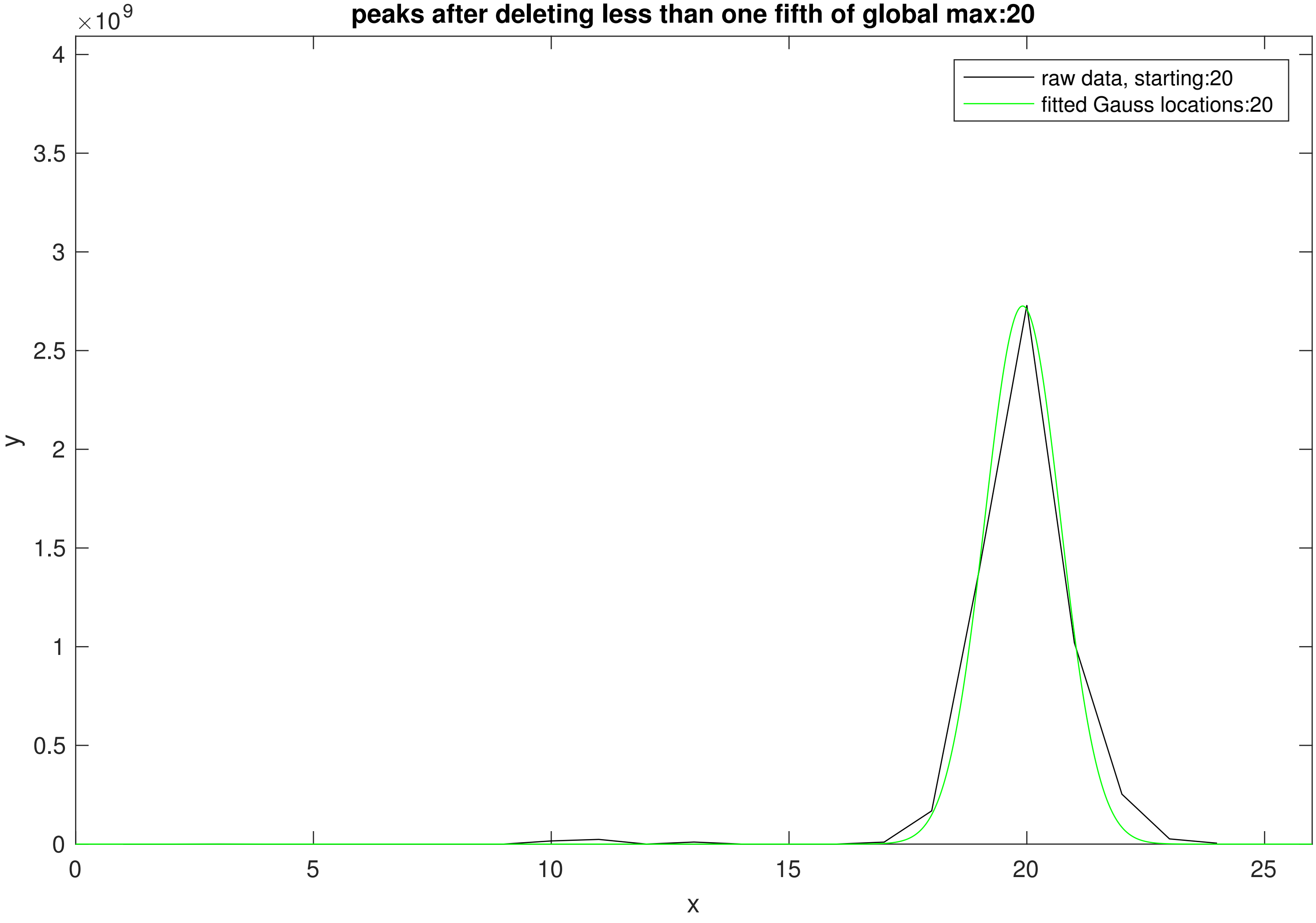

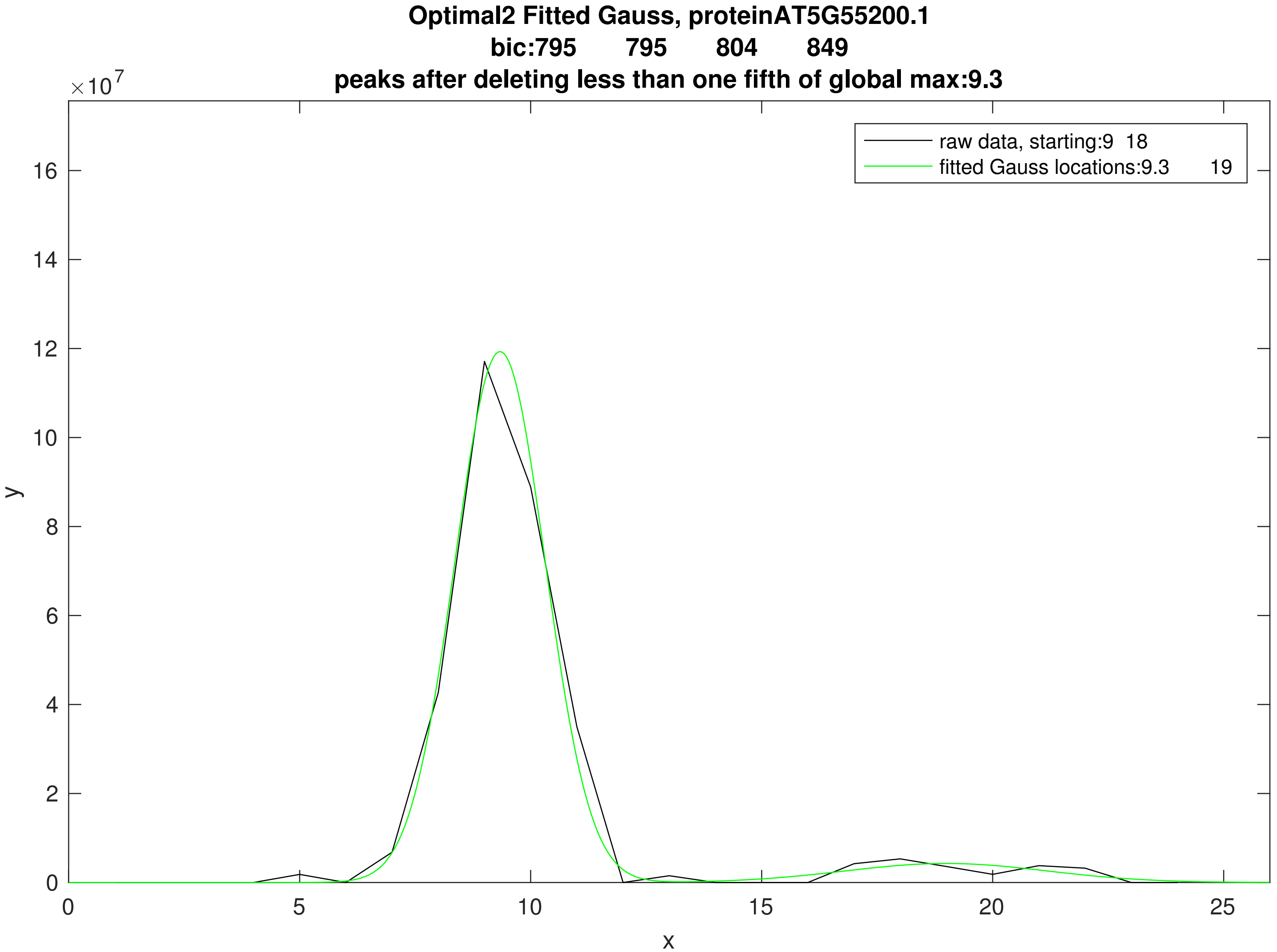
Optimal2 Fitted Gauss, proteinAT5G55200.1
bic:795 795 804 849
peaks after deleting less than one fifth of global max:9.3
×10^7
raw data, starting:9 18
fitted Gauss locations:9.3 19
16
14
12
10
8
6
4
2
0
y
0
5
10
15
20
25
x

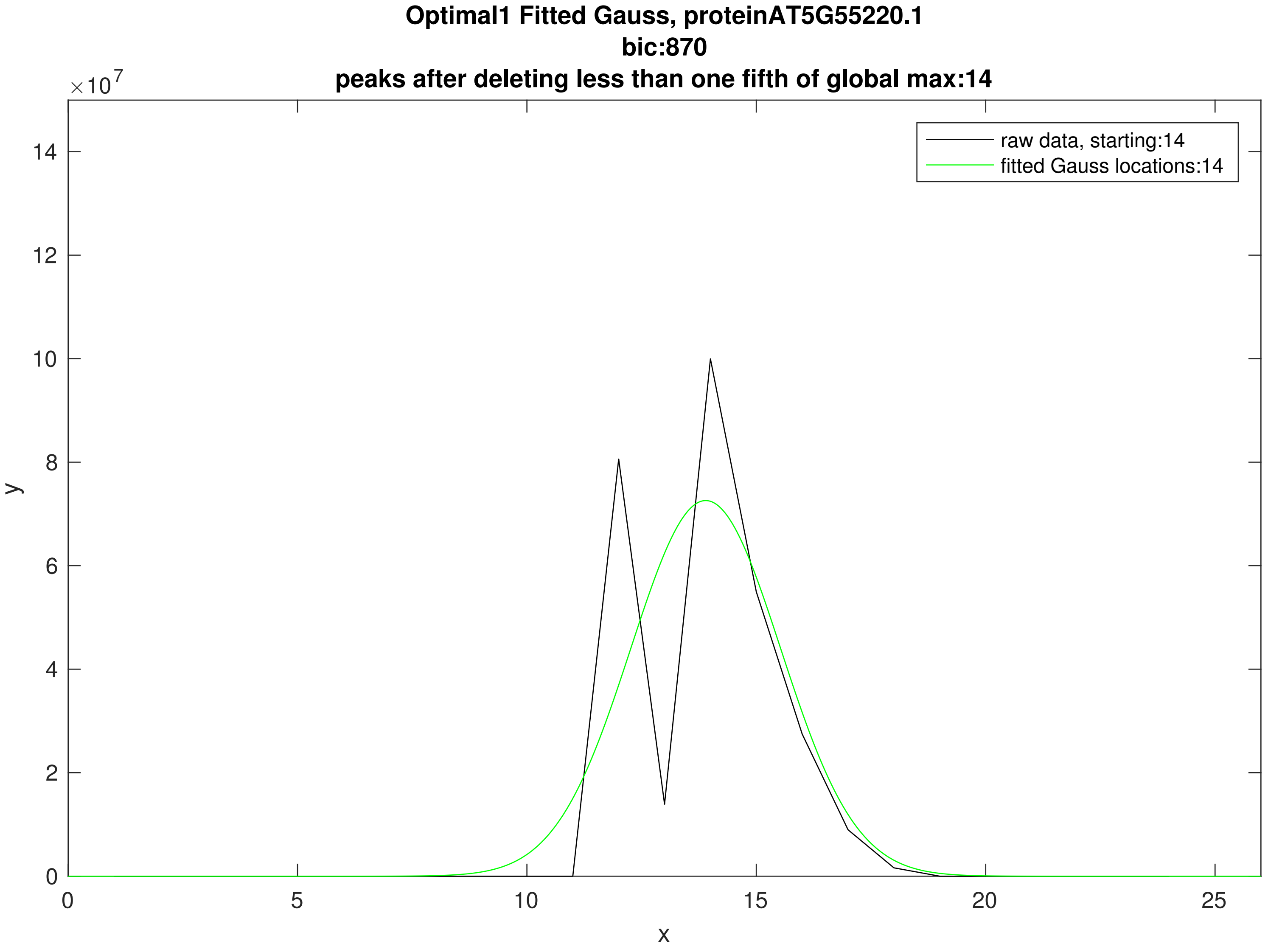
Optimal1 Fitted Gauss, proteinAT5G55220.1
bic:870
peaks after deleting less than one fifth of global max:14
×10^7
raw data, starting:14
fitted Gauss locations:14
y
x
0
2
4
6
8
10
12
14
0
5
10
15
20
25

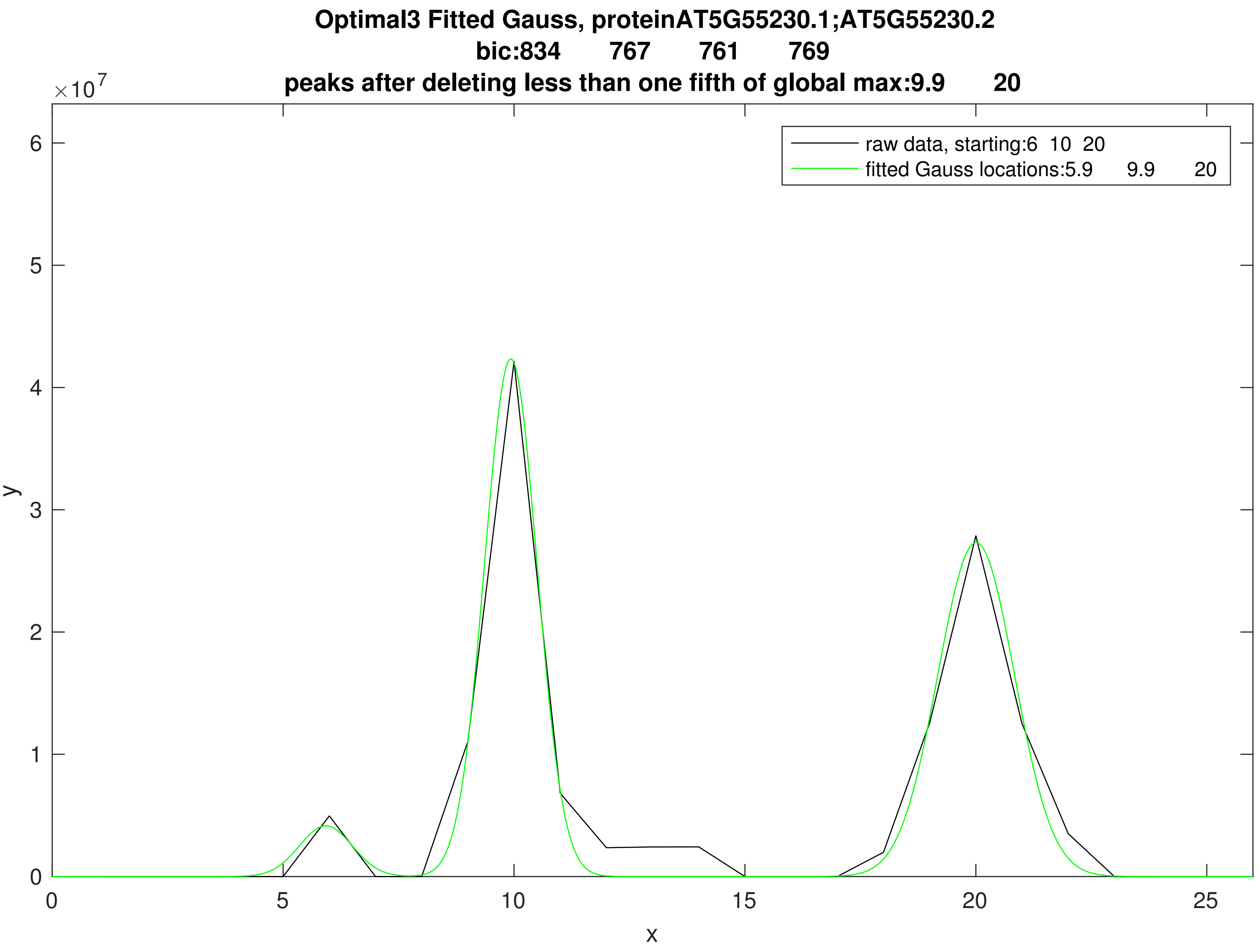

Optimal3 Fitted Gauss, proteinAT5G55230.1;AT5G55230.2
bic:834 767 761 769
peaks after deleting less than one fifth of global max:9.9 20
raw data, starting:6 10 20
fitted Gauss locations:5.9 9.9 20
×10^7
y
x

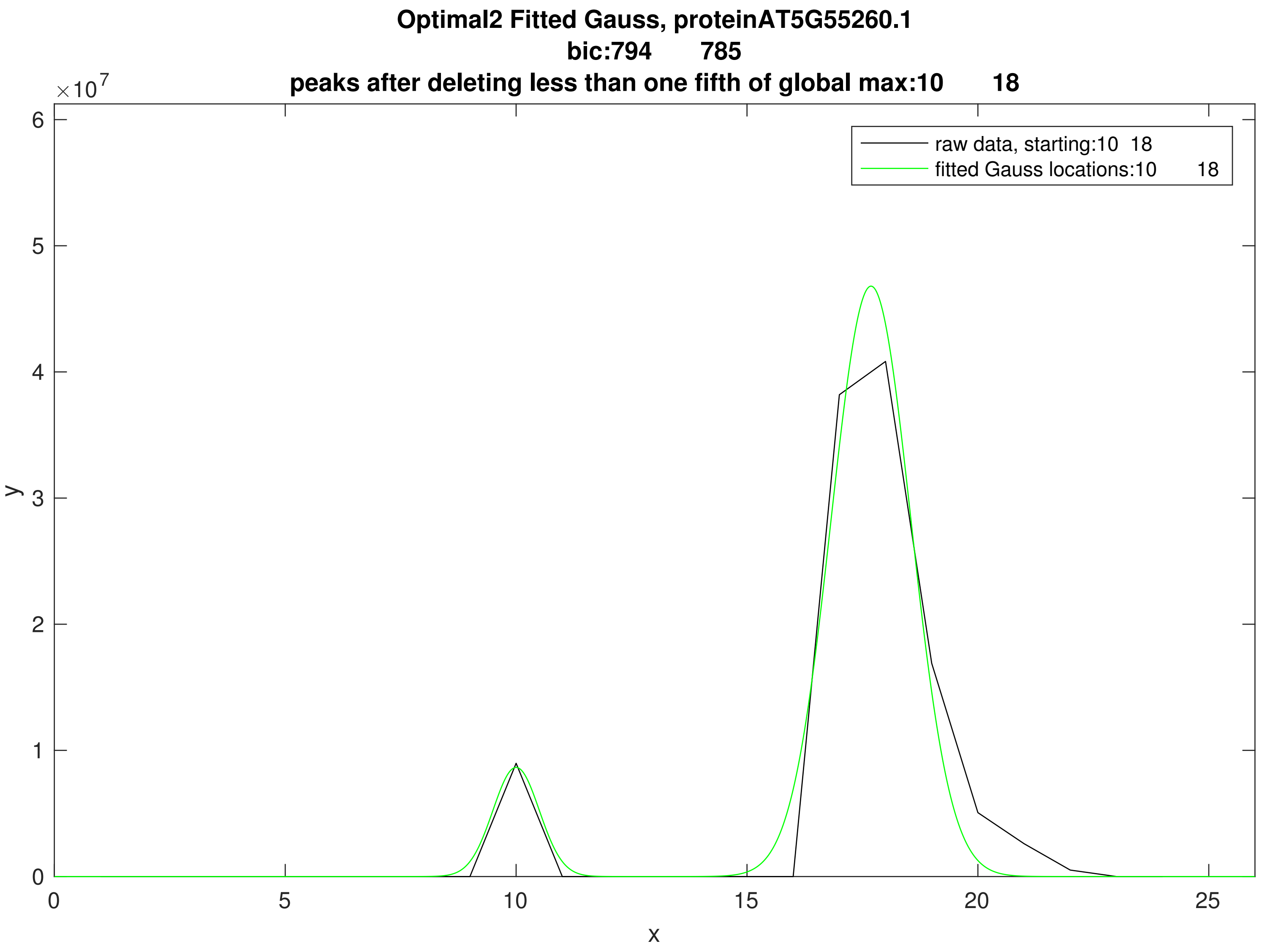

Optimal2 Fitted Gauss, proteinAT5G55260.1
bic:794 785
peaks after deleting less than one fifth of global max:10 18
×10^7
raw data, starting:10 18
fitted Gauss locations:10 18
y
x
0
1
2
3
4
5
6
0
5
10
15
20
25

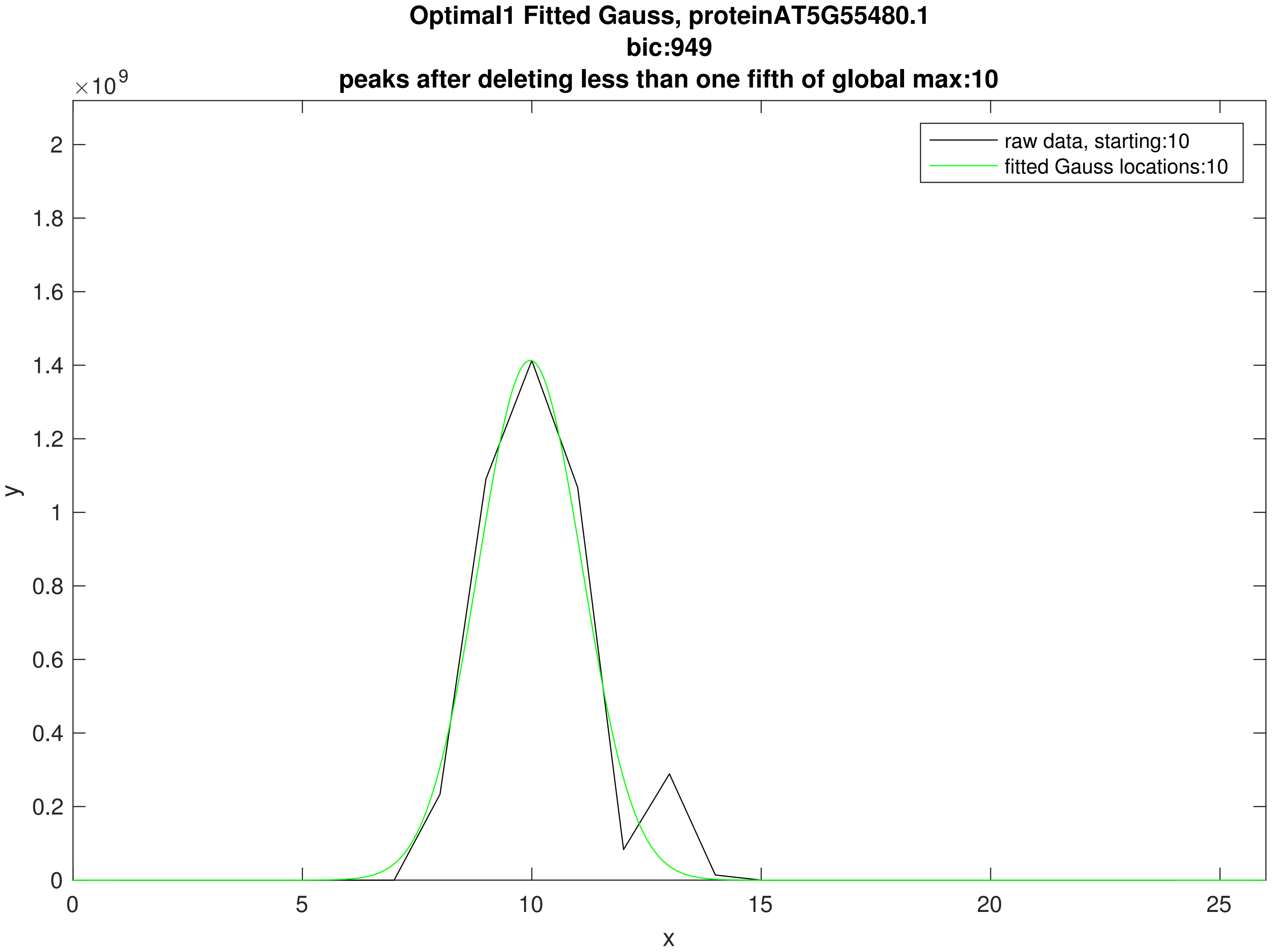
Optimal1 Fitted Gauss, proteinAT5G55480.1
bic:949
peaks after deleting less than one fifth of global max:10
×10^9
2
1.8
1.6
1.4
1.2
1
0.8
0.6
0.4
0.2
0
y
0
5
10
15
20
25
x
raw data, starting:10
fitted Gauss locations:10

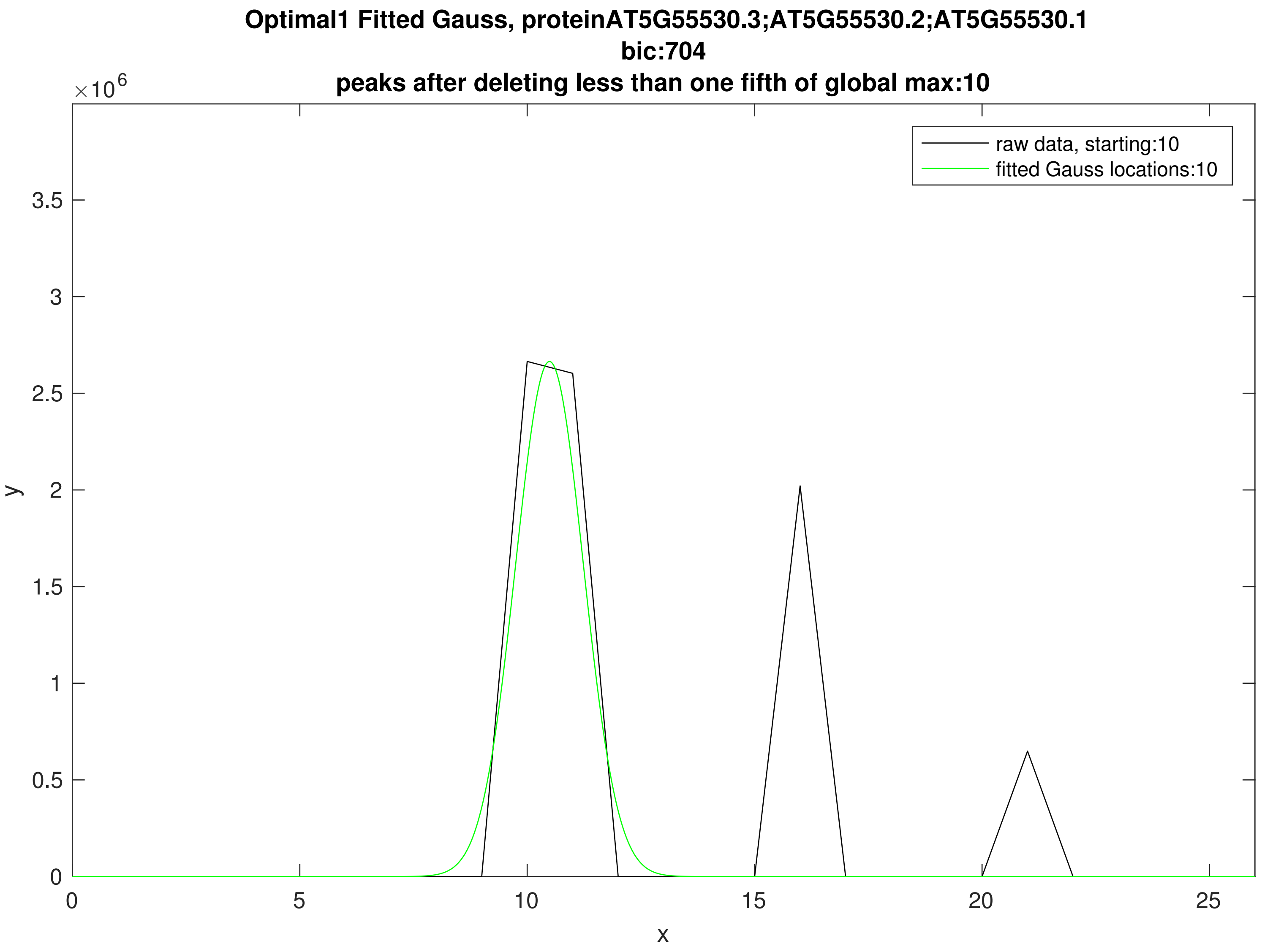
Optimal1 Fitted Gauss, proteinAT5G55530.3;AT5G55530.2;AT5G55530.1
bic:704
peaks after deleting less than one fifth of global max:10
×10^6
raw data, starting:10
fitted Gauss locations:10
3.5
3
2.5
2
1.5
1
0.5
0
y
0
5
10
15
20
25
x

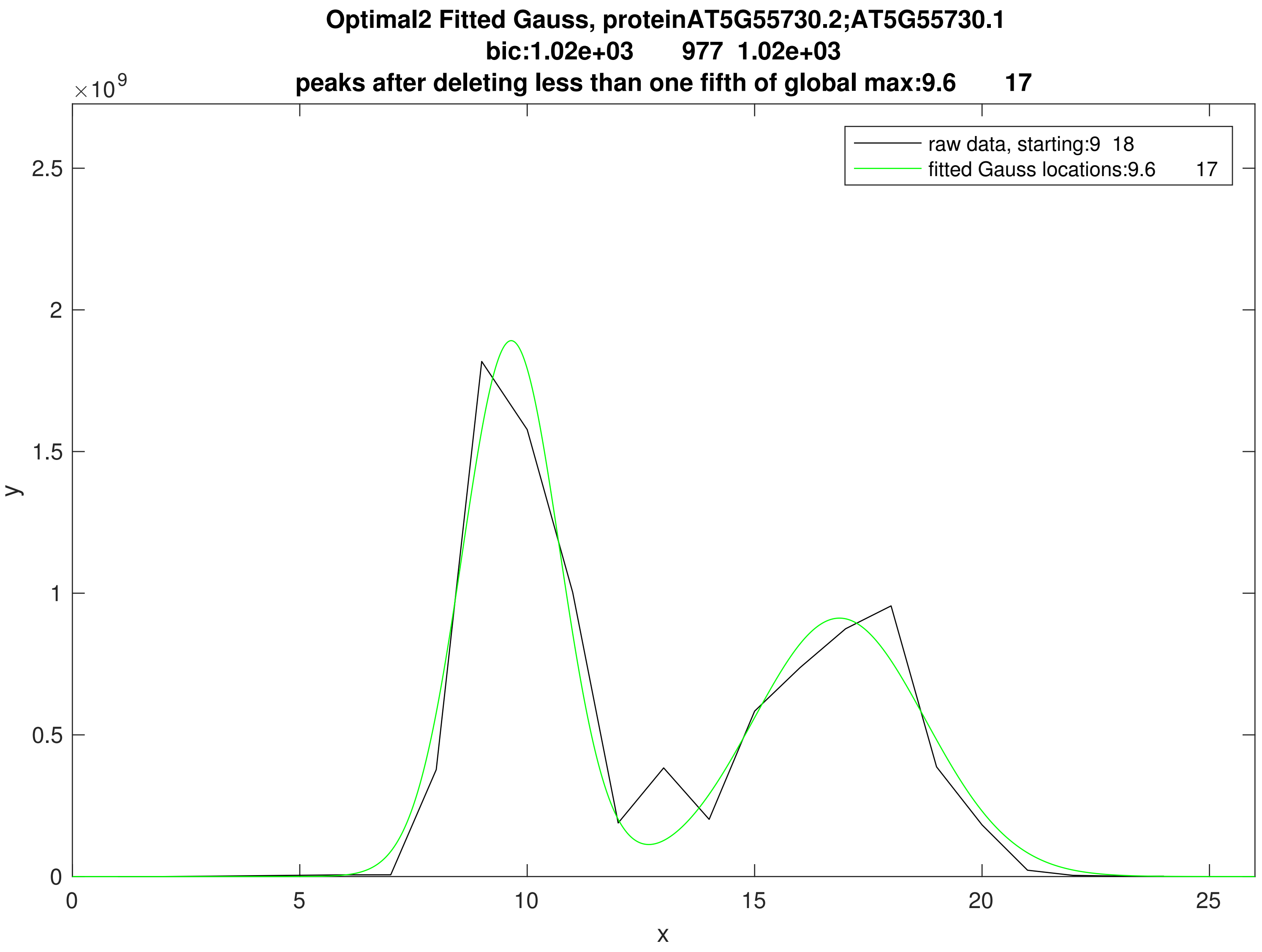

Optimal2 Fitted Gauss, proteinAT5G55730.2;AT5G55730.1
bic:1.02e+03 977 1.02e+03
peaks after deleting less than one fifth of global max:9.6 17
×10^9
raw data, starting:9 18
fitted Gauss locations:9.6 17
y
x
0
0.5
1
1.5
2
2.5
5
10
15
20
25

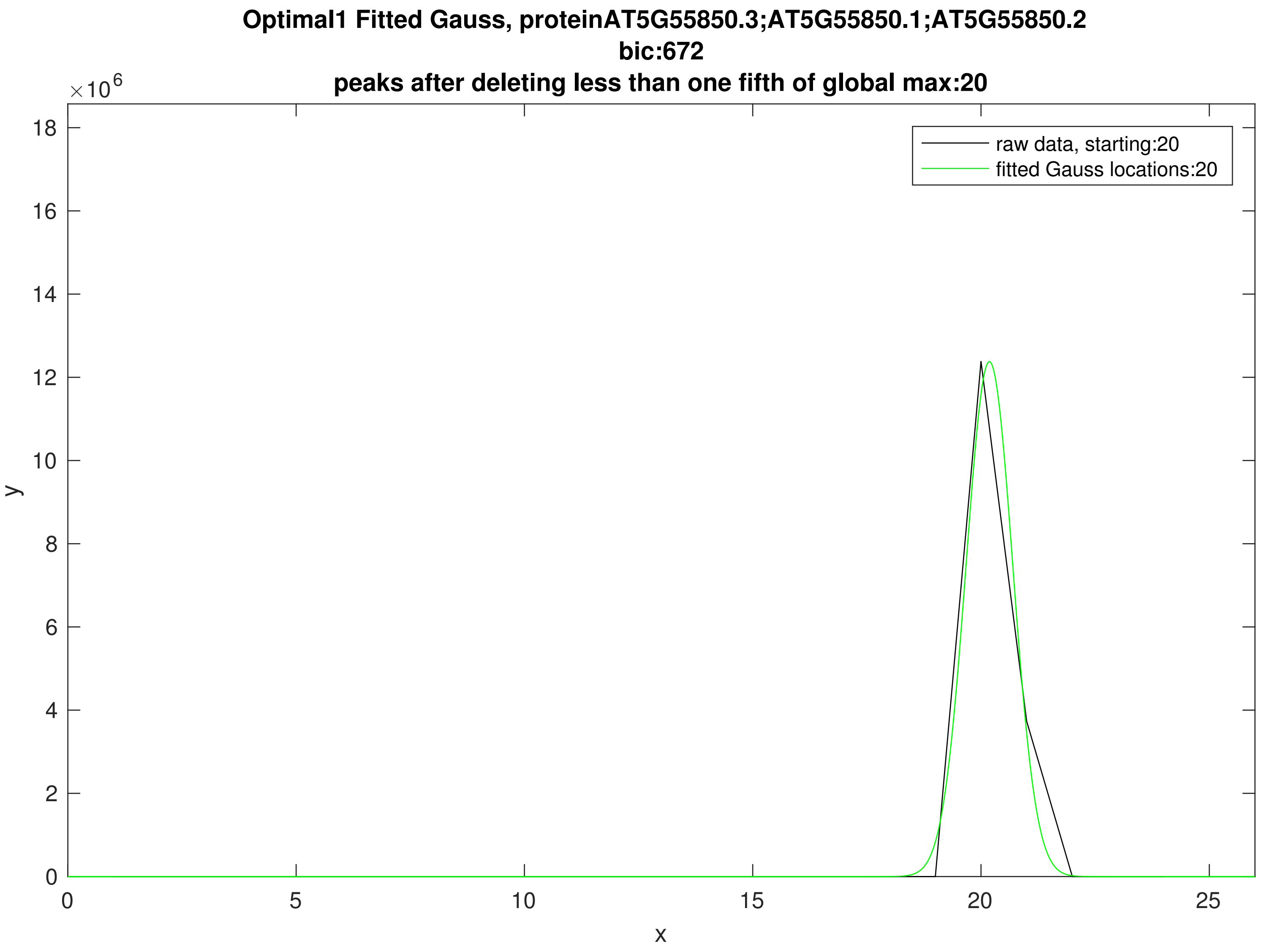

Optimal1 Fitted Gauss, proteinAT5G55850.3;AT5G55850.1;AT5G55850.2
bic:672
peaks after deleting less than one fifth of global max:20
×10^6
raw data, starting:20
fitted Gauss locations:20
y
x
0
2
4
6
8
10
12
14
16
18
0
5
10
15
20
25

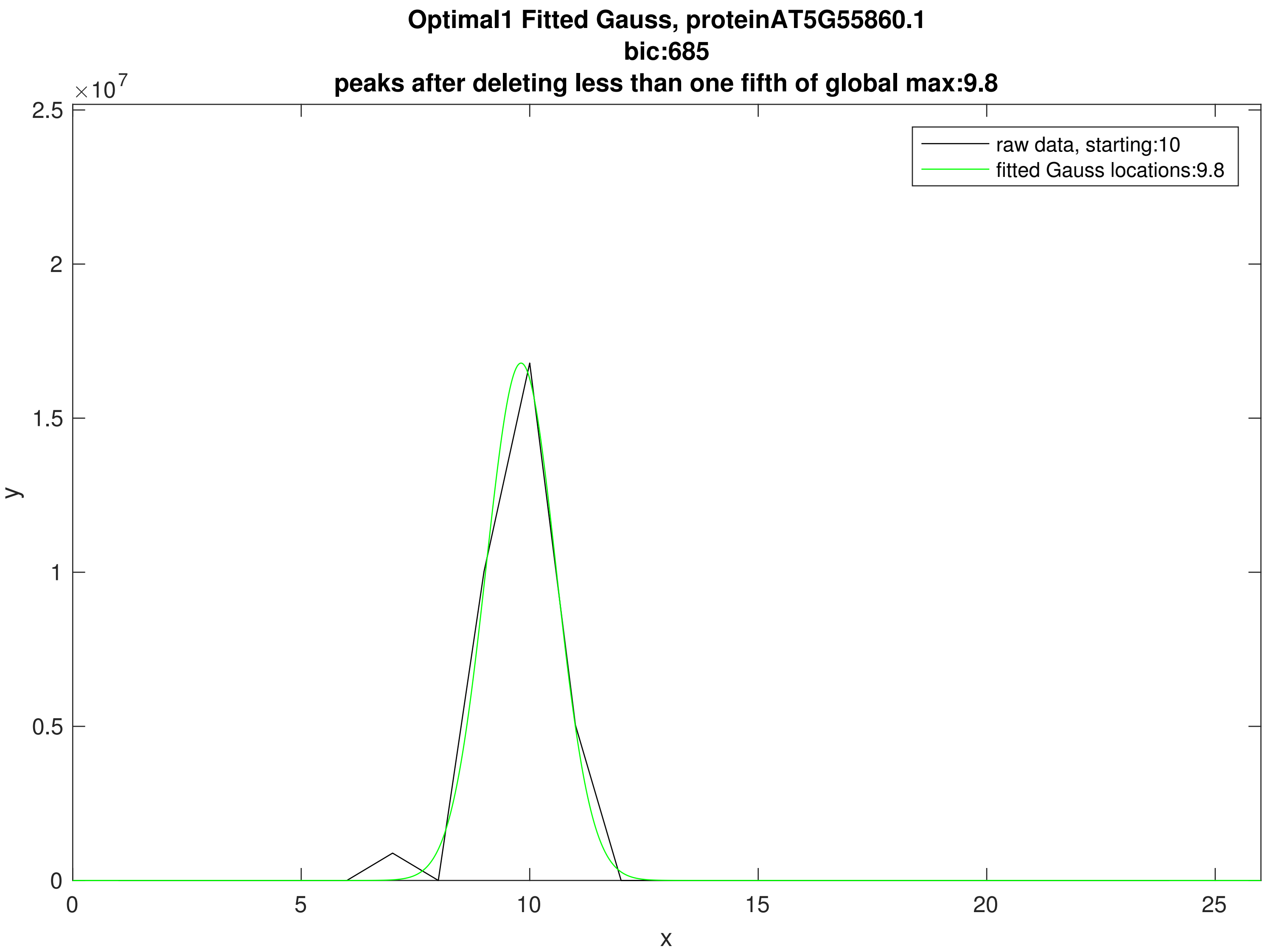
Optimal1 Fitted Gauss, proteinAT5G55860.1
bic:685
peaks after deleting less than one fifth of global max:9.8
$\times 10^7$
2.5
2
1.5
1
0.5
0
y
0
5
10
15
20
25
x
raw data, starting:10
fitted Gauss locations:9.8

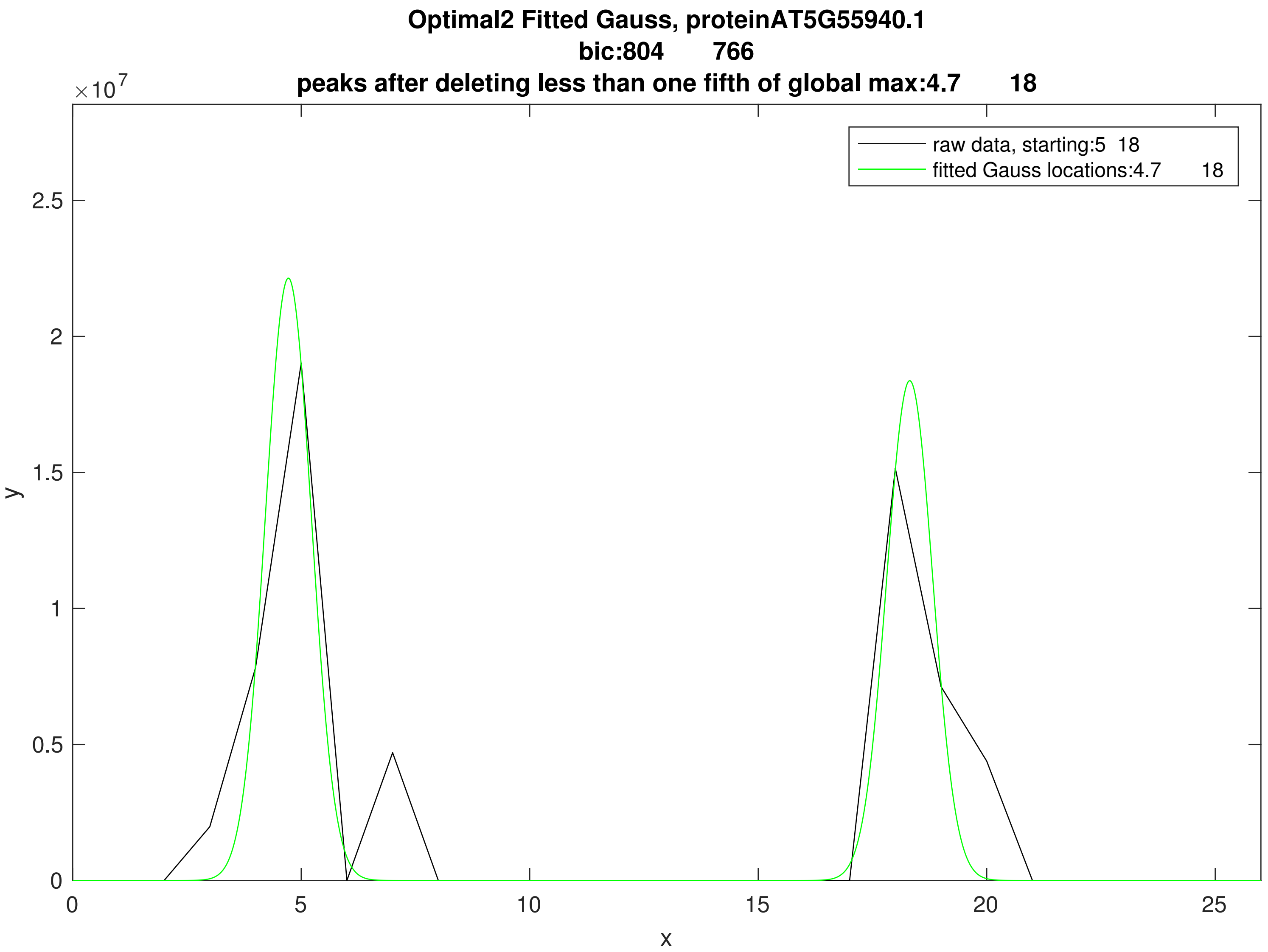

Optimal2 Fitted Gauss, proteinAT5G55940.1
bic:804 766
peaks after deleting less than one fifth of global max:4.7 18
×10^7
raw data, starting:5 18
fitted Gauss locations:4.7 18
y
x
0
0.5
1
1.5
2
2.5
5
10
15
20
25

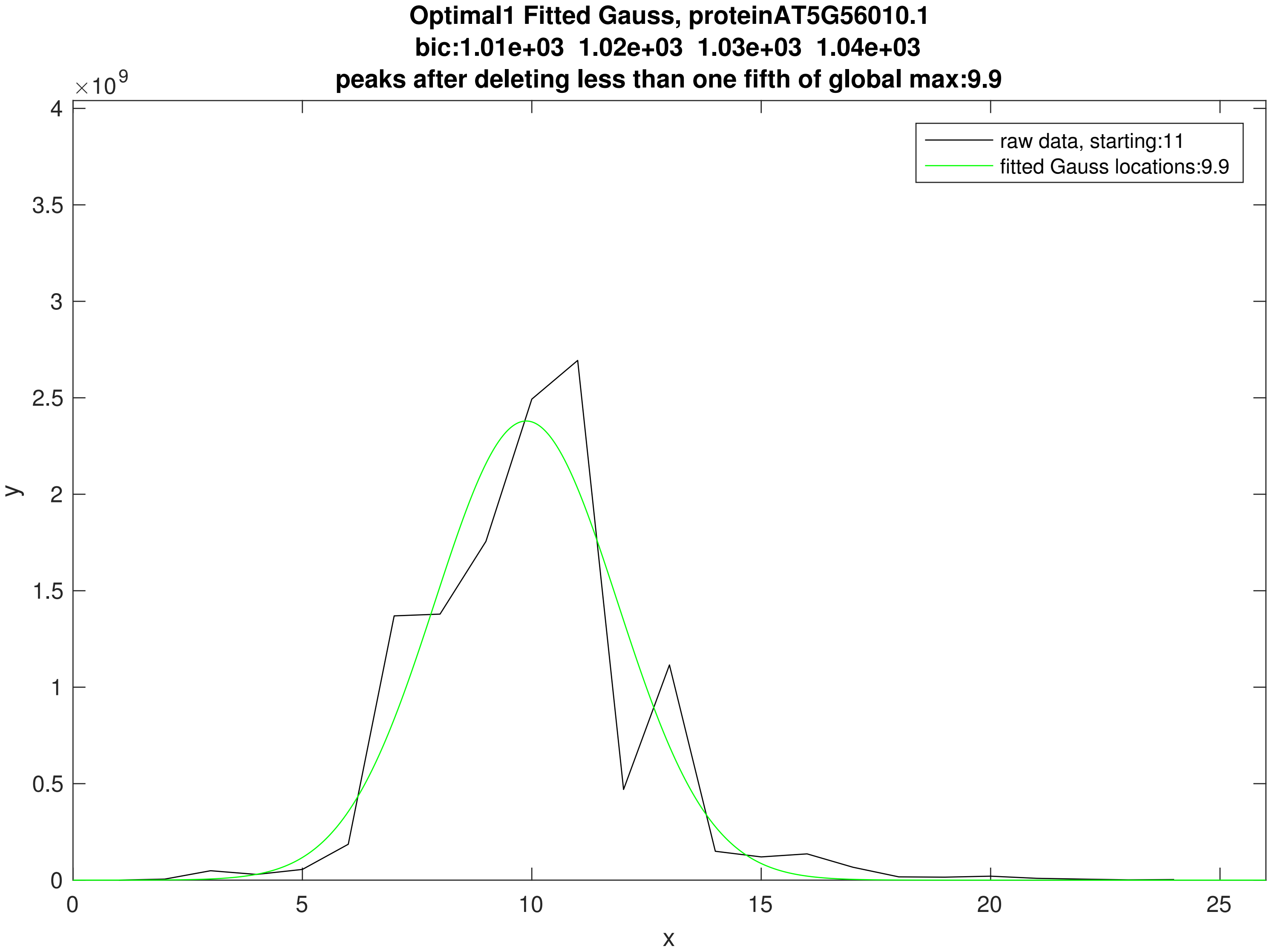

Optimal1 Fitted Gauss, proteinAT5G56010.1
bic:1.01e+03 1.02e+03 1.03e+03 1.04e+03
peaks after deleting less than one fifth of global max:9.9
×10^9
raw data, starting:11
fitted Gauss locations:9.9
y
x
0
0.5
1
1.5
2
2.5
3
3.5
4
0
5
10
15
20
25

**Optimal1 Fitted Gauss, proteinAT5G56030.1;AT5G56030.2**
**bic:833 842**
**peaks after deleting less than one fifth of global max:9.3**

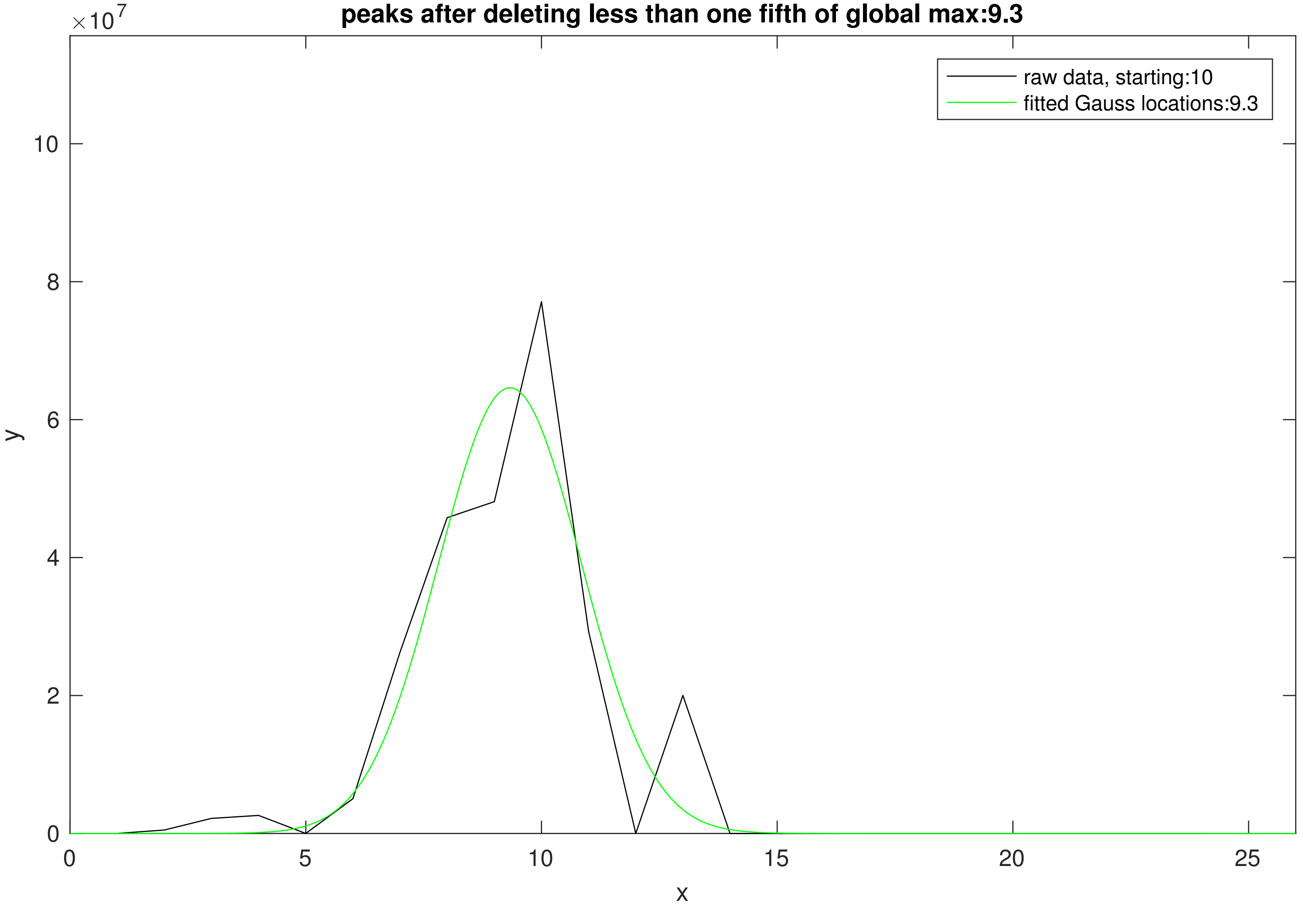

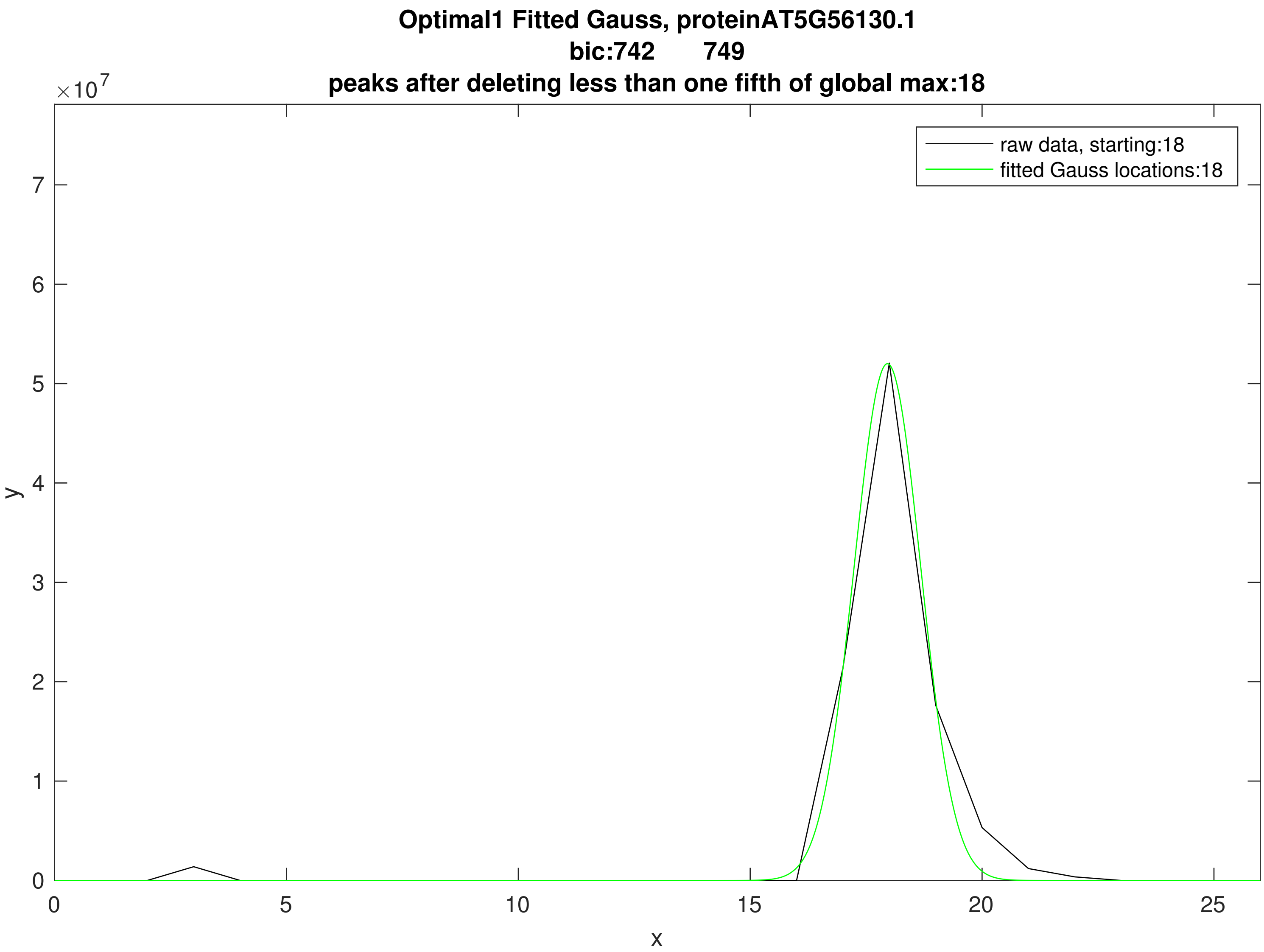
Optimal1 Fitted Gauss, proteinAT5G56130.1
bic:742 749
peaks after deleting less than one fifth of global max:18
×10^7
raw data, starting:18
fitted Gauss locations:18
y
x
0
1
2
3
4
5
6
7
0
5
10
15
20
25

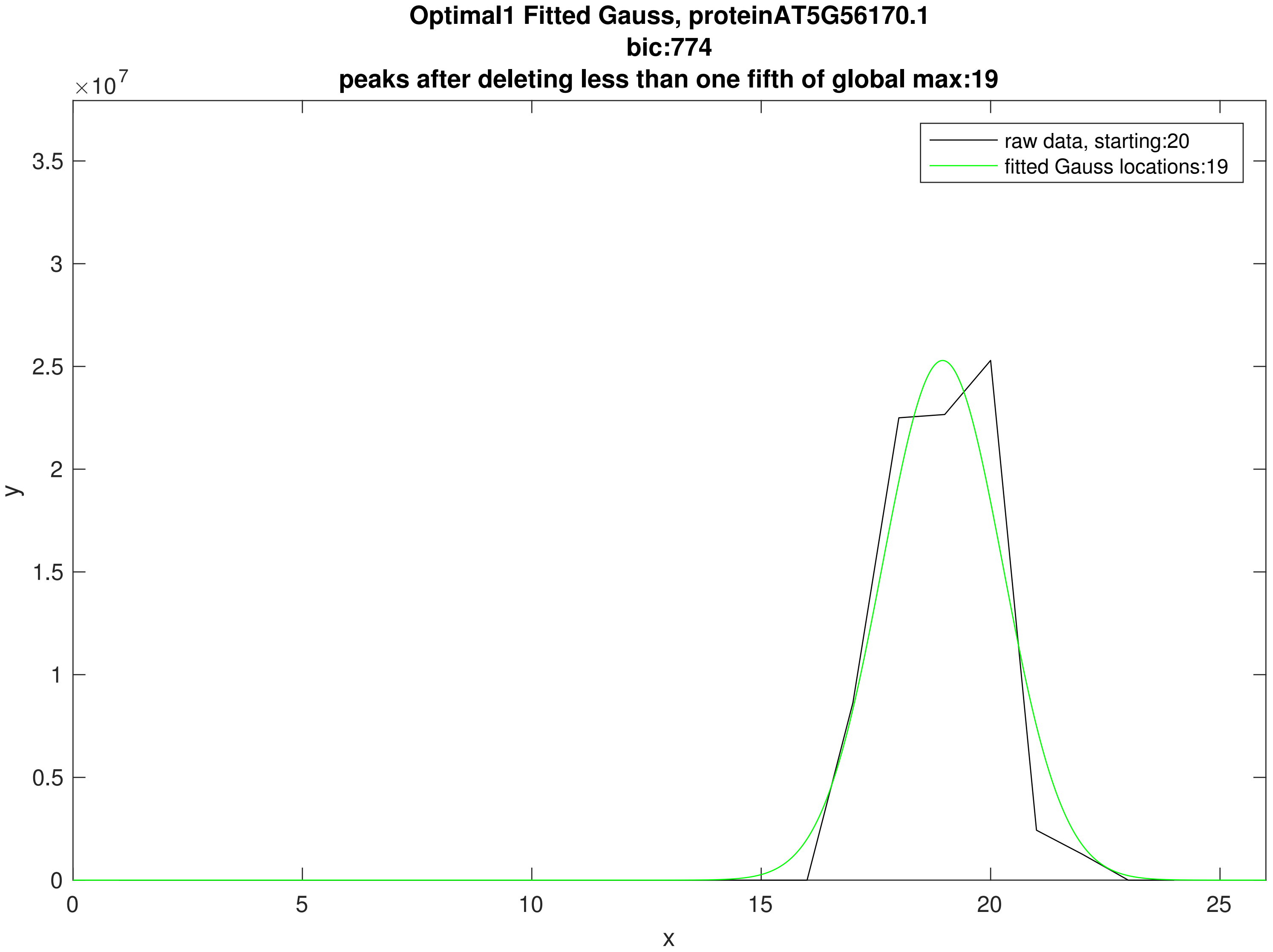
Optimal1 Fitted Gauss, proteinAT5G56170.1
bic:774
peaks after deleting less than one fifth of global max:19
×10^7
raw data, starting:20
fitted Gauss locations:19
3.5
3
2.5
2
1.5
1
0.5
0
y
0
5
10
15
20
25
x

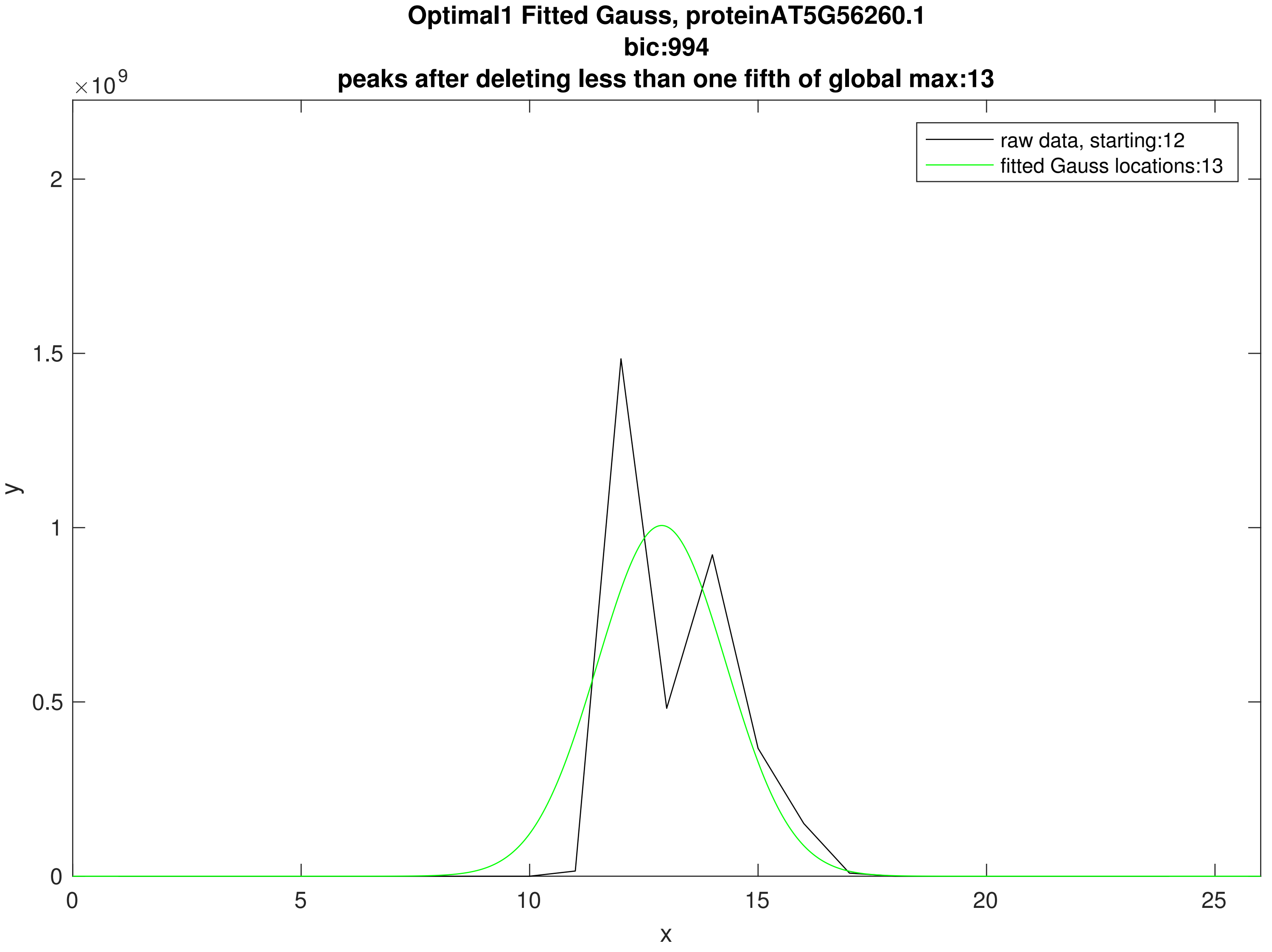

Optimal1 Fitted Gauss, proteinAT5G56260.1
bic:994
peaks after deleting less than one fifth of global max:13
×10^9
raw data, starting:12
fitted Gauss locations:13
y
x
0
0.5
1
1.5
2
0
5
10
15
20
25

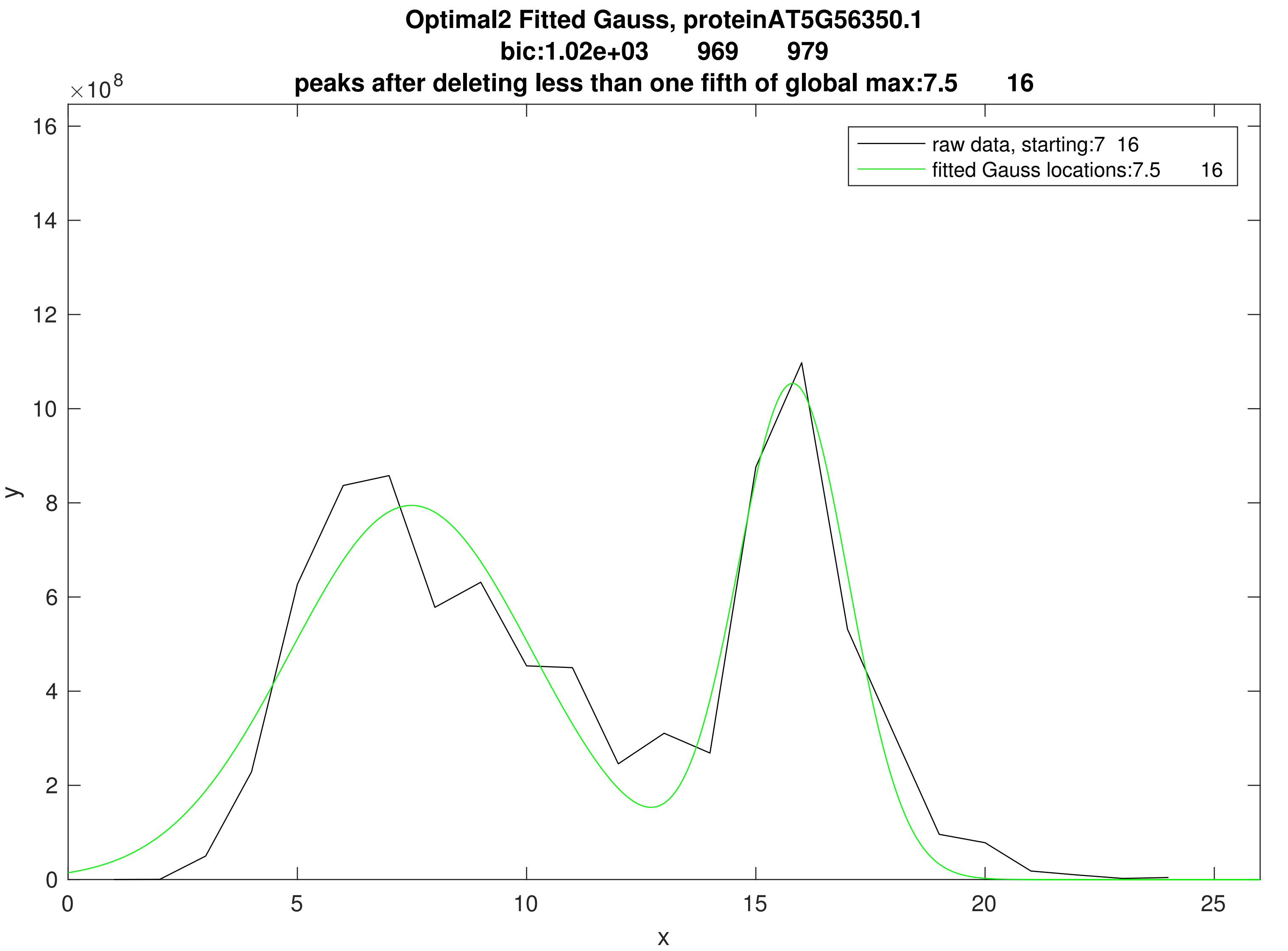
Optimal2 Fitted Gauss, proteinAT5G56350.1
bic:1.02e+03 969 979
peaks after deleting less than one fifth of global max:7.5 16
×10^8
raw data, starting:7 16
fitted Gauss locations:7.5 16
y
x
0
2
4
6
8
10
12
14
16
0
5
10
15
20
25

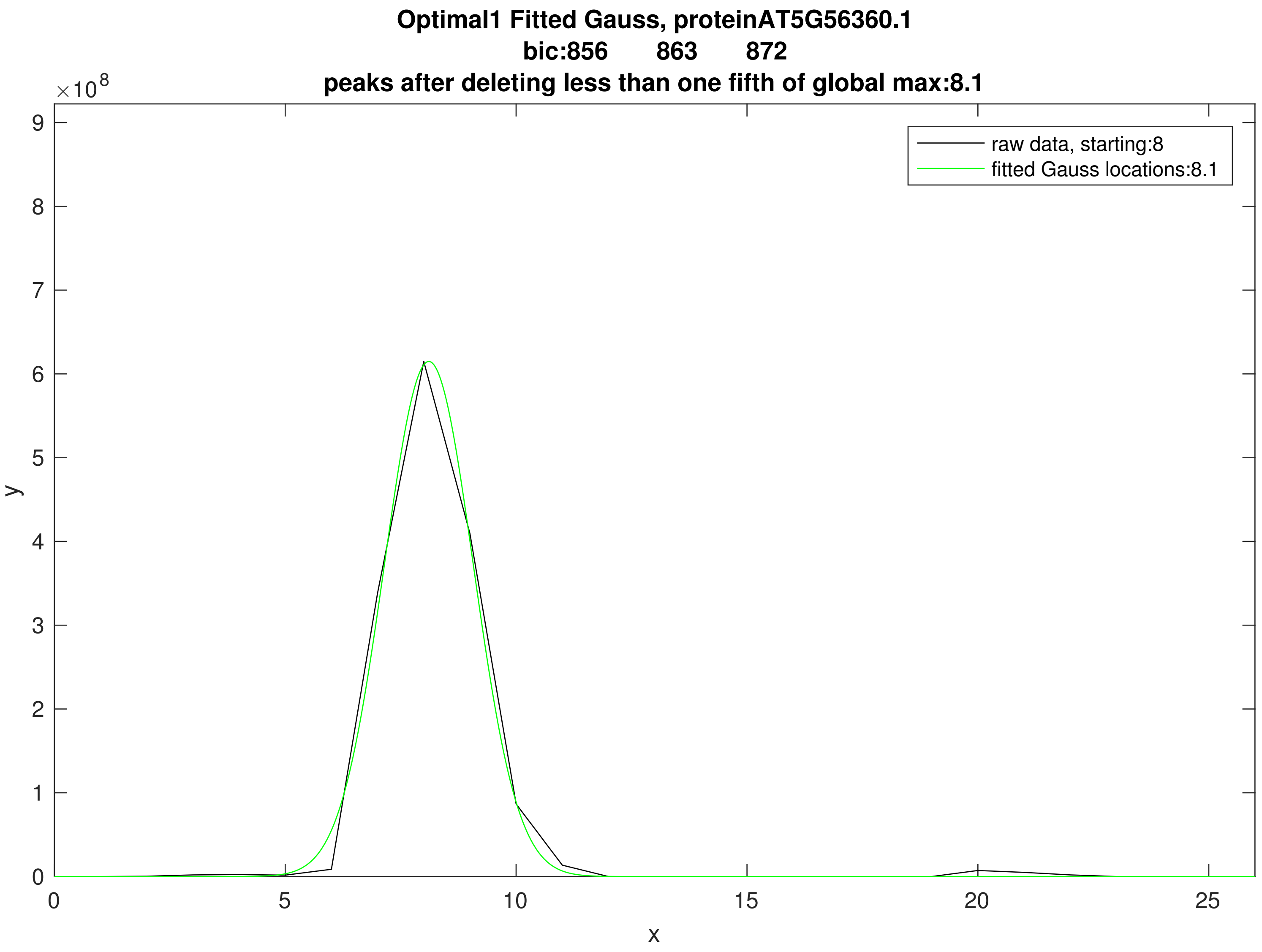
Optimal1 Fitted Gauss, proteinAT5G56360.1
bic:856 863 872
peaks after deleting less than one fifth of global max:8.1
×10^8
raw data, starting:8
fitted Gauss locations:8.1
y
x
0
1
2
3
4
5
6
7
8
9
10
15
20
25

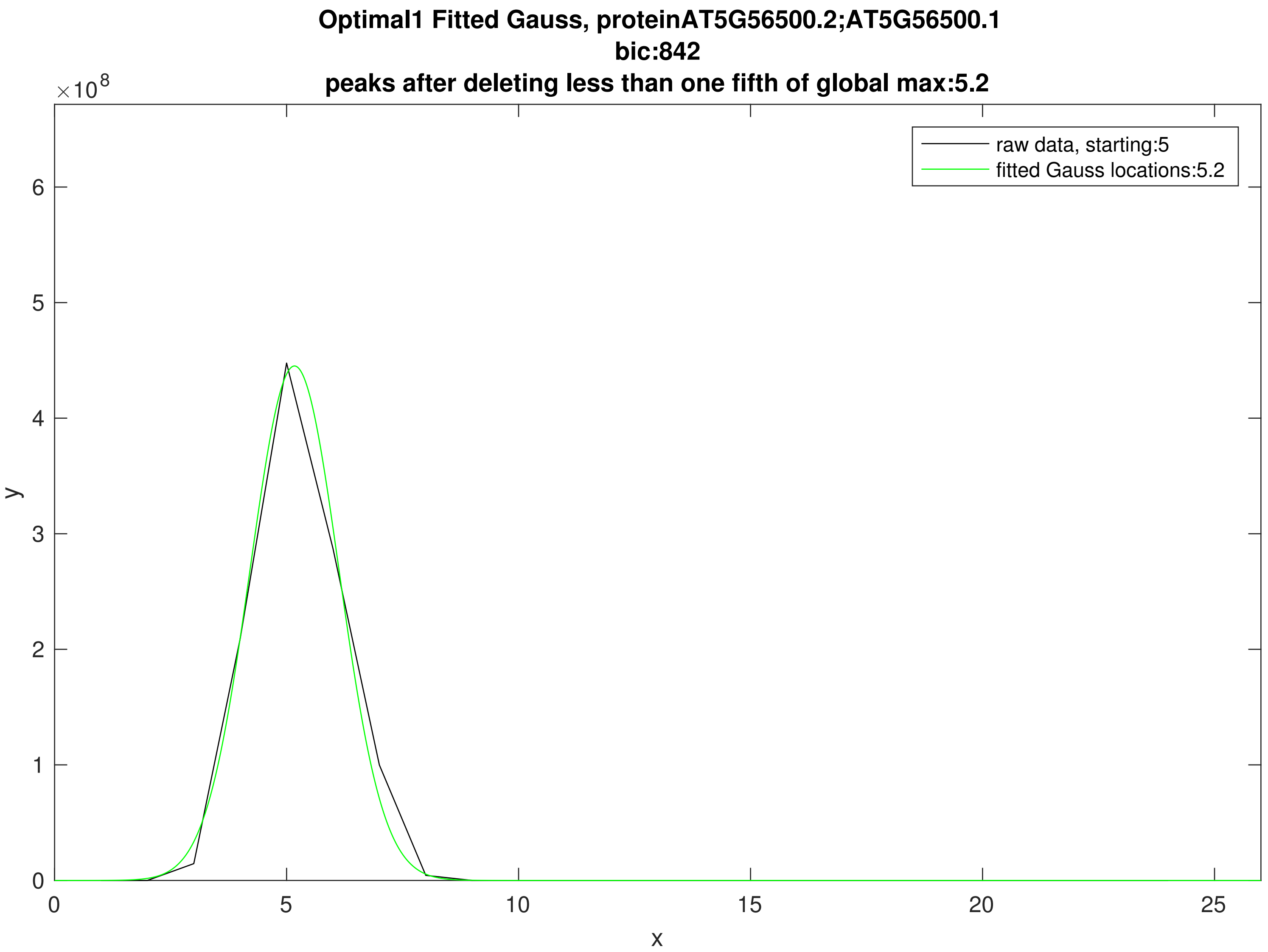
Optimal1 Fitted Gauss, proteinAT5G56500.2;AT5G56500.1
bic:842
peaks after deleting less than one fifth of global max:5.2
×10^8
raw data, starting:5
fitted Gauss locations:5.2
y
x
0
1
2
3
4
5
6
0
5
10
15
20
25

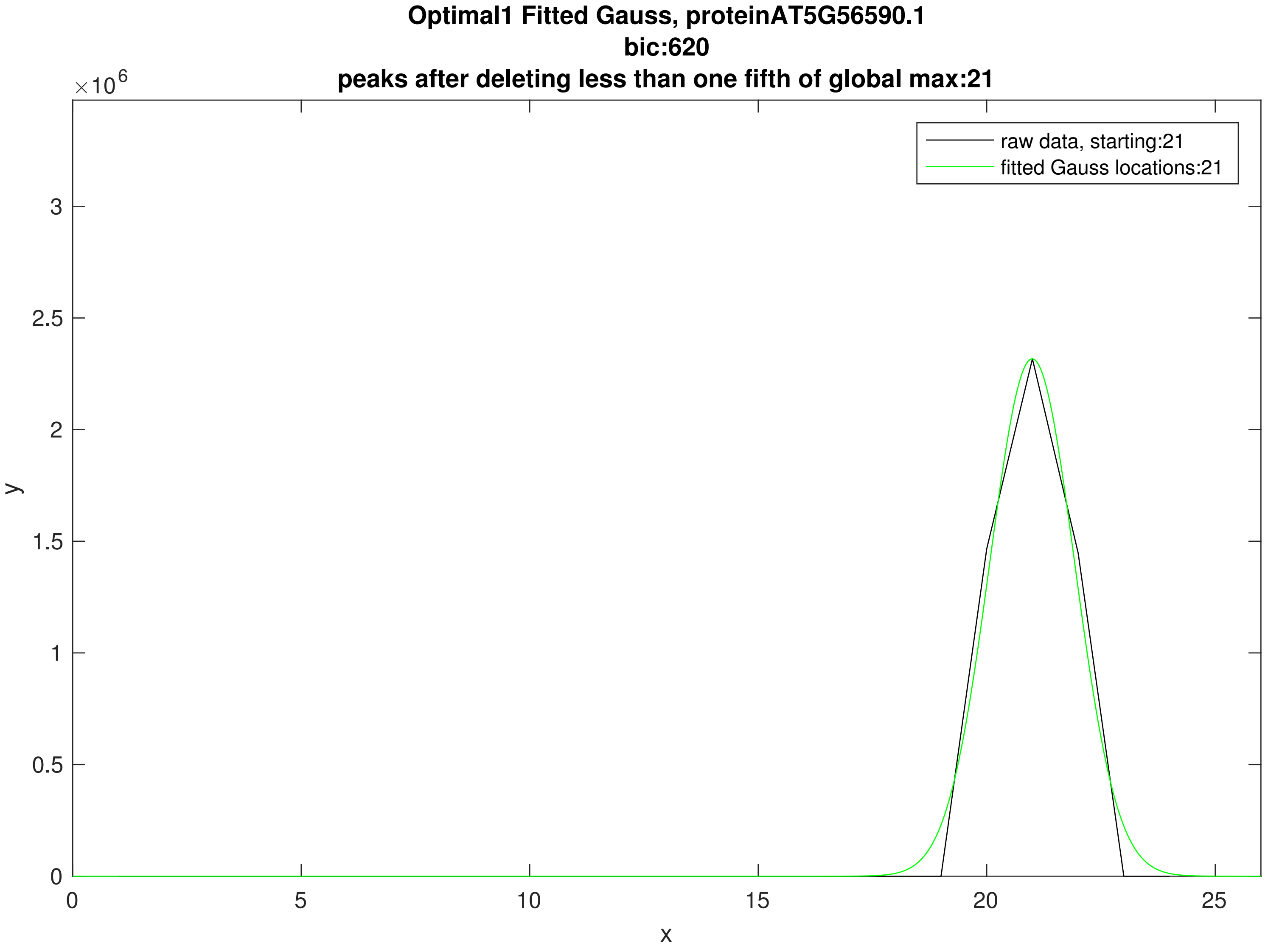

Optimal1 Fitted Gauss, proteinAT5G56590.1
bic:620
peaks after deleting less than one fifth of global max:21
×10^6
raw data, starting:21
fitted Gauss locations:21
3
2.5
2
1.5
1
0.5
0
y
0
5
10
15
20
25
x

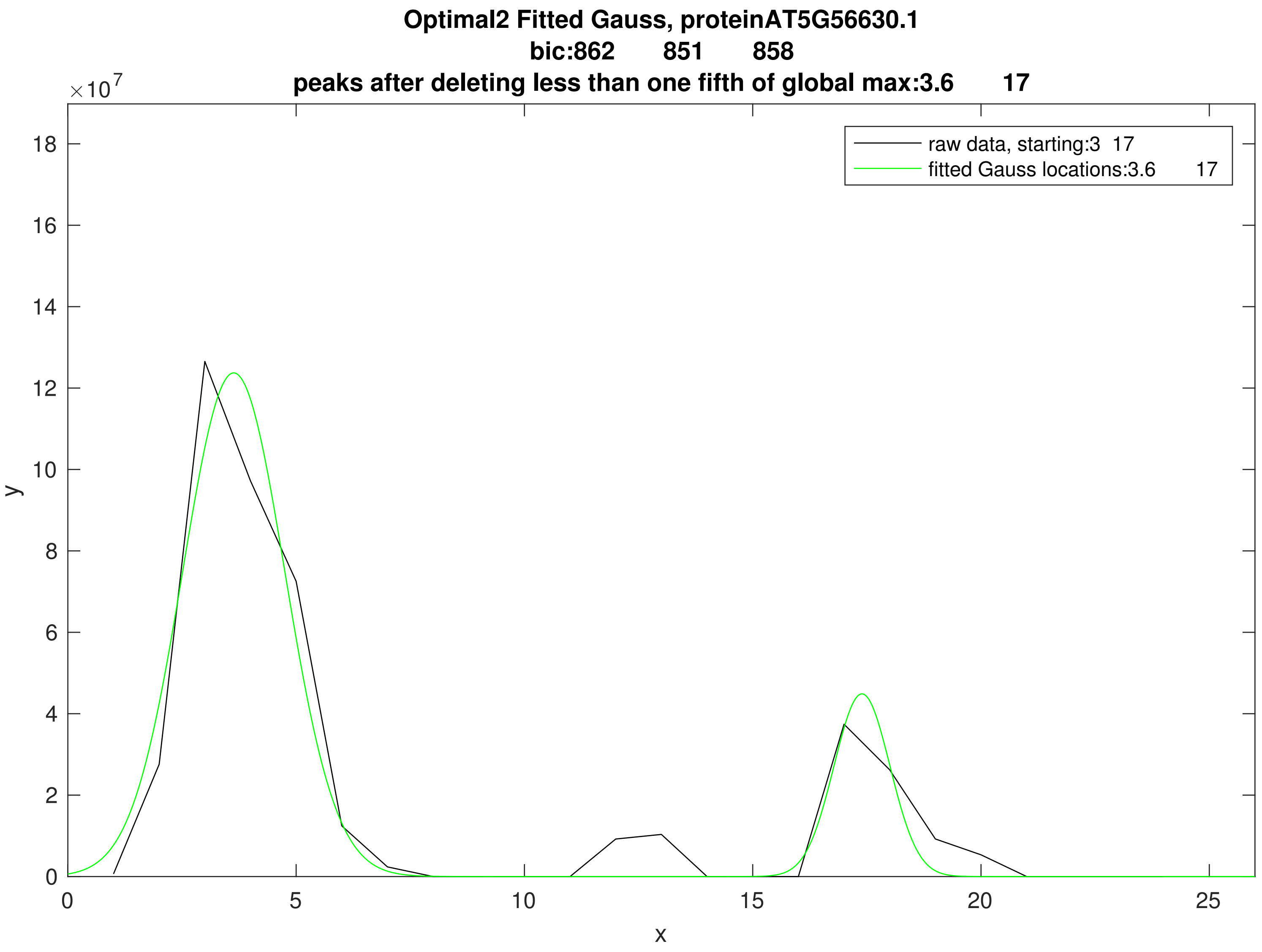

Optimal2 Fitted Gauss, proteinAT5G56630.1
bic:862 851 858
peaks after deleting less than one fifth of global max:3.6 17
×10^7
raw data, starting:3 17
fitted Gauss locations:3.6 17
y
x
0
2
4
6
8
10
12
14
16
18
0
5
10
15
20
25

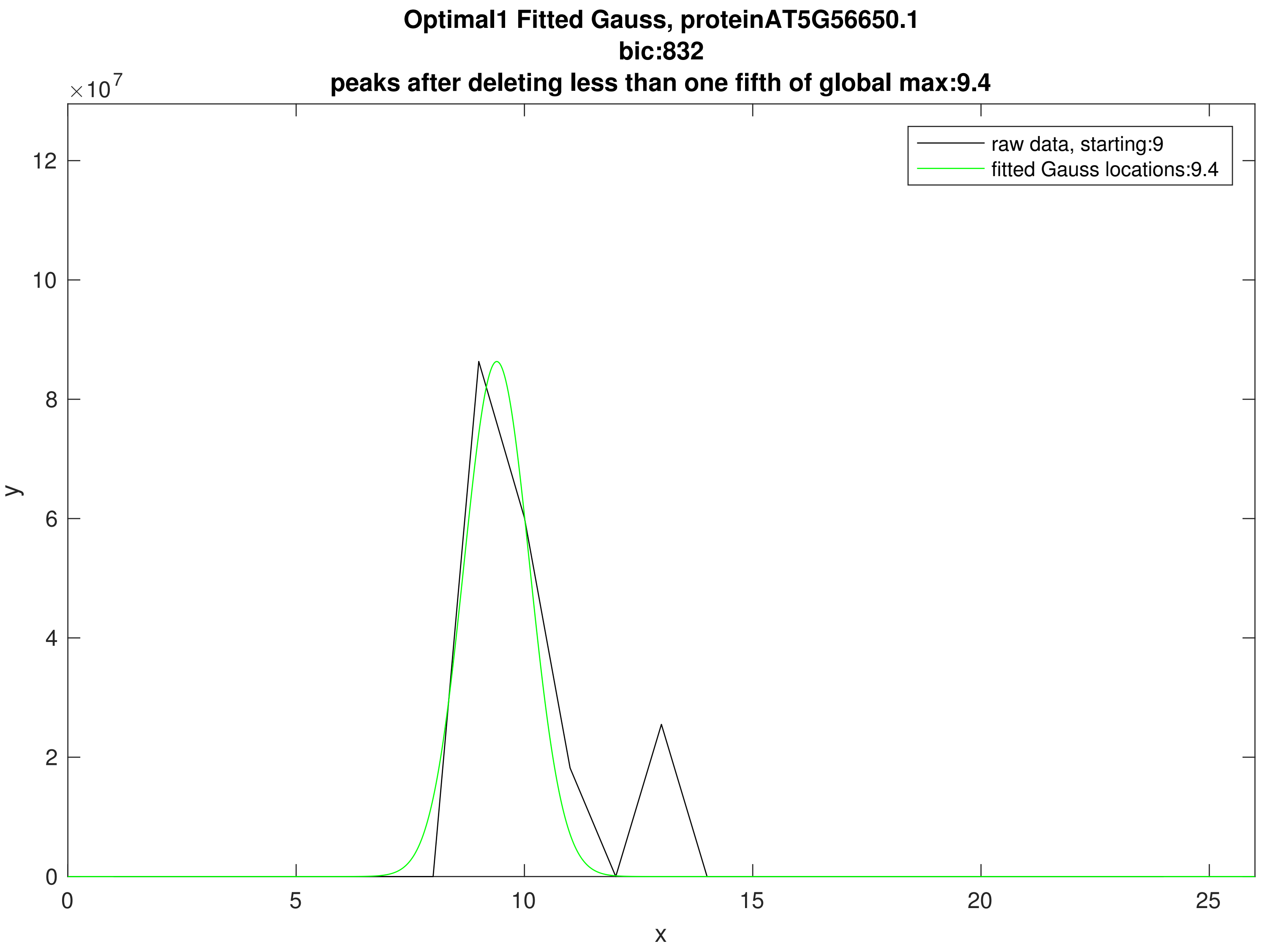

Optimal1 Fitted Gauss, proteinAT5G56650.1
bic:832
peaks after deleting less than one fifth of global max:9.4
×10^7
12
10
8
6
4
2
0
y
0
5
10
15
20
25
x
raw data, starting:9
fitted Gauss locations:9.4

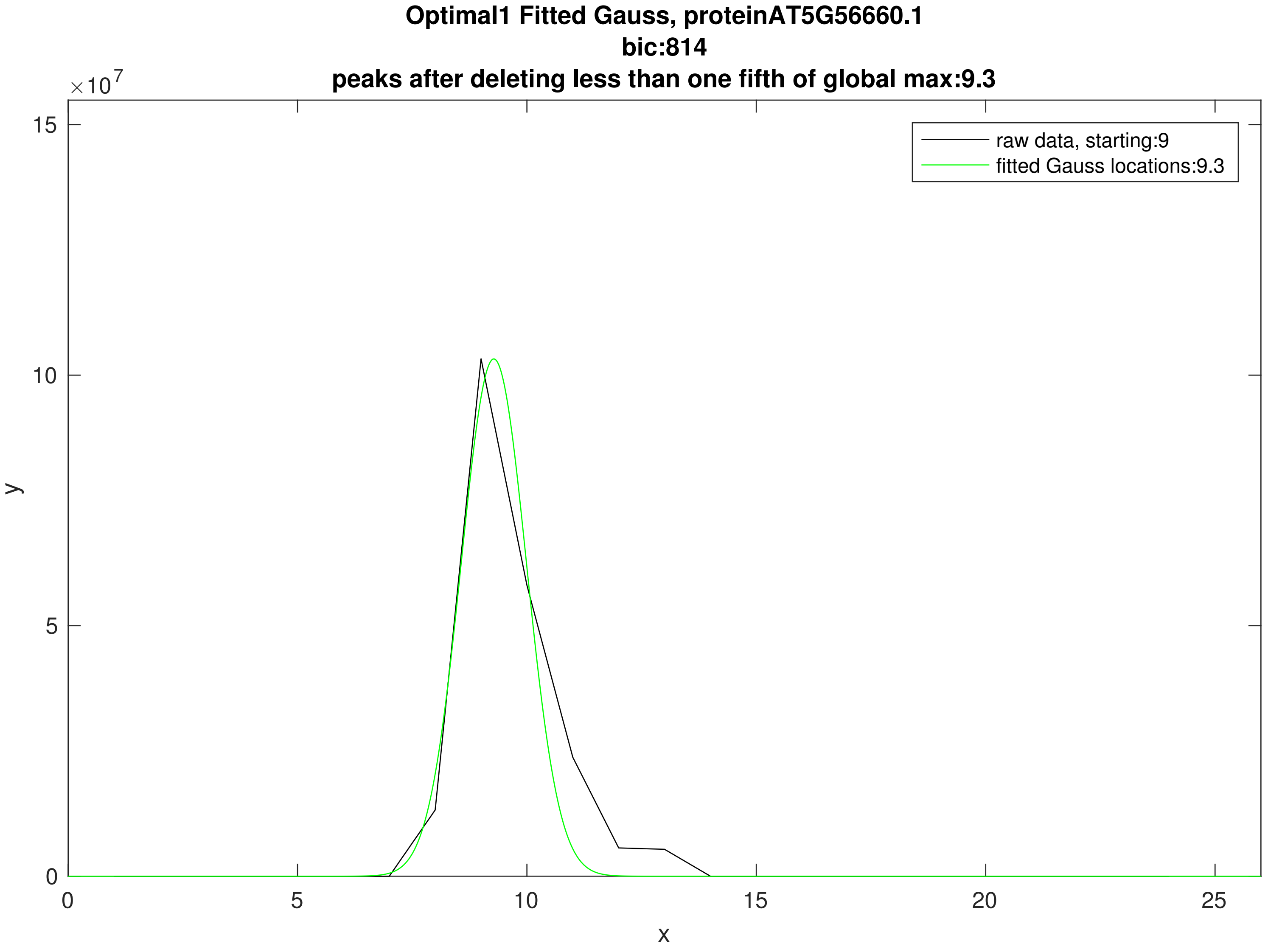

Optimal1 Fitted Gauss, proteinAT5G56660.1
bic:814
peaks after deleting less than one fifth of global max:9.3
×10^7
15
10
5
0
y
0
5
10
15
20
25
x
raw data, starting:9
fitted Gauss locations:9.3

Optimal1 Fitted Gauss, proteinAT5G56670.1;AT4G29390.1;AT2G19750.1
bic:818 826
peaks after deleting less than one fifth of global max:3.7

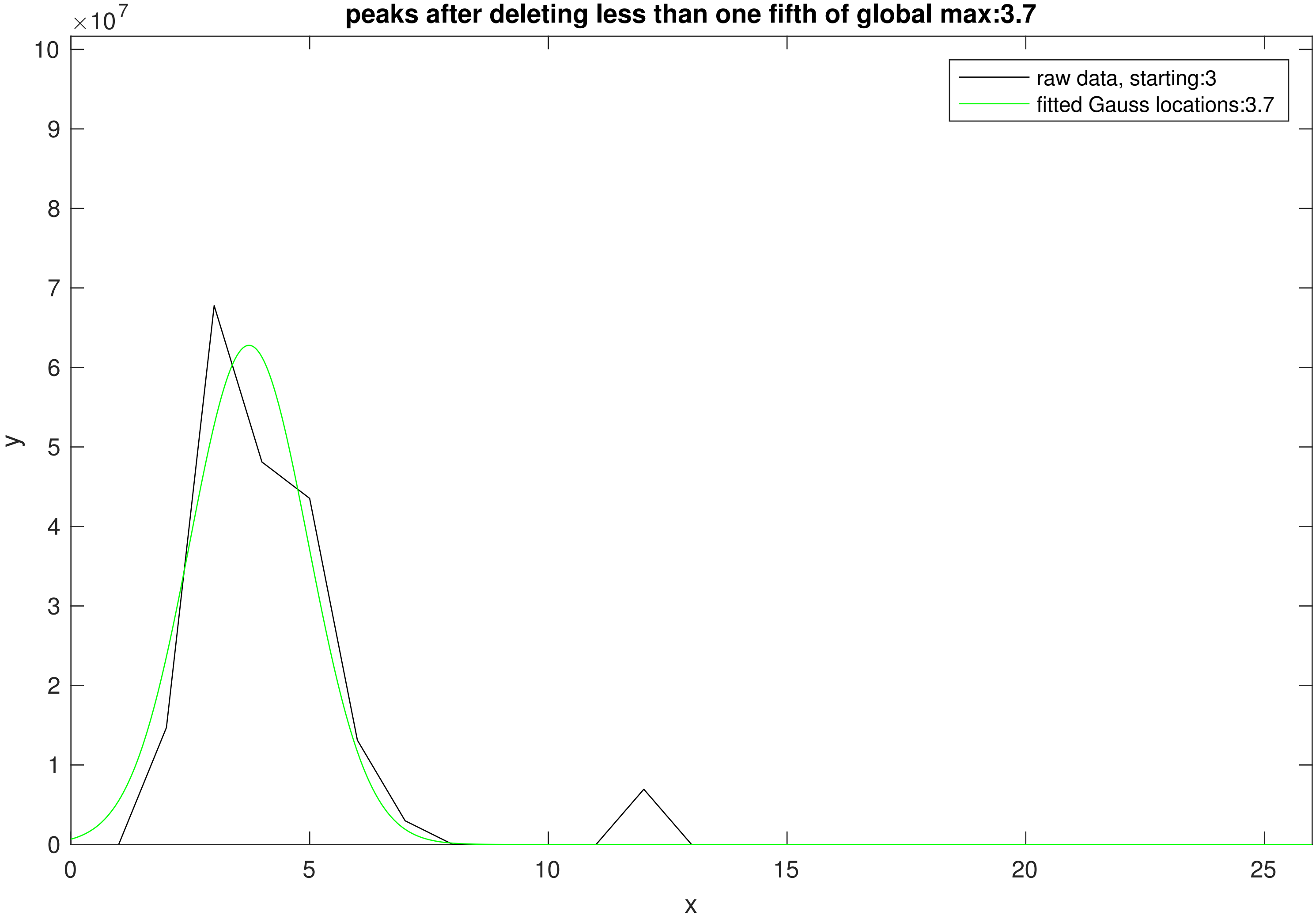

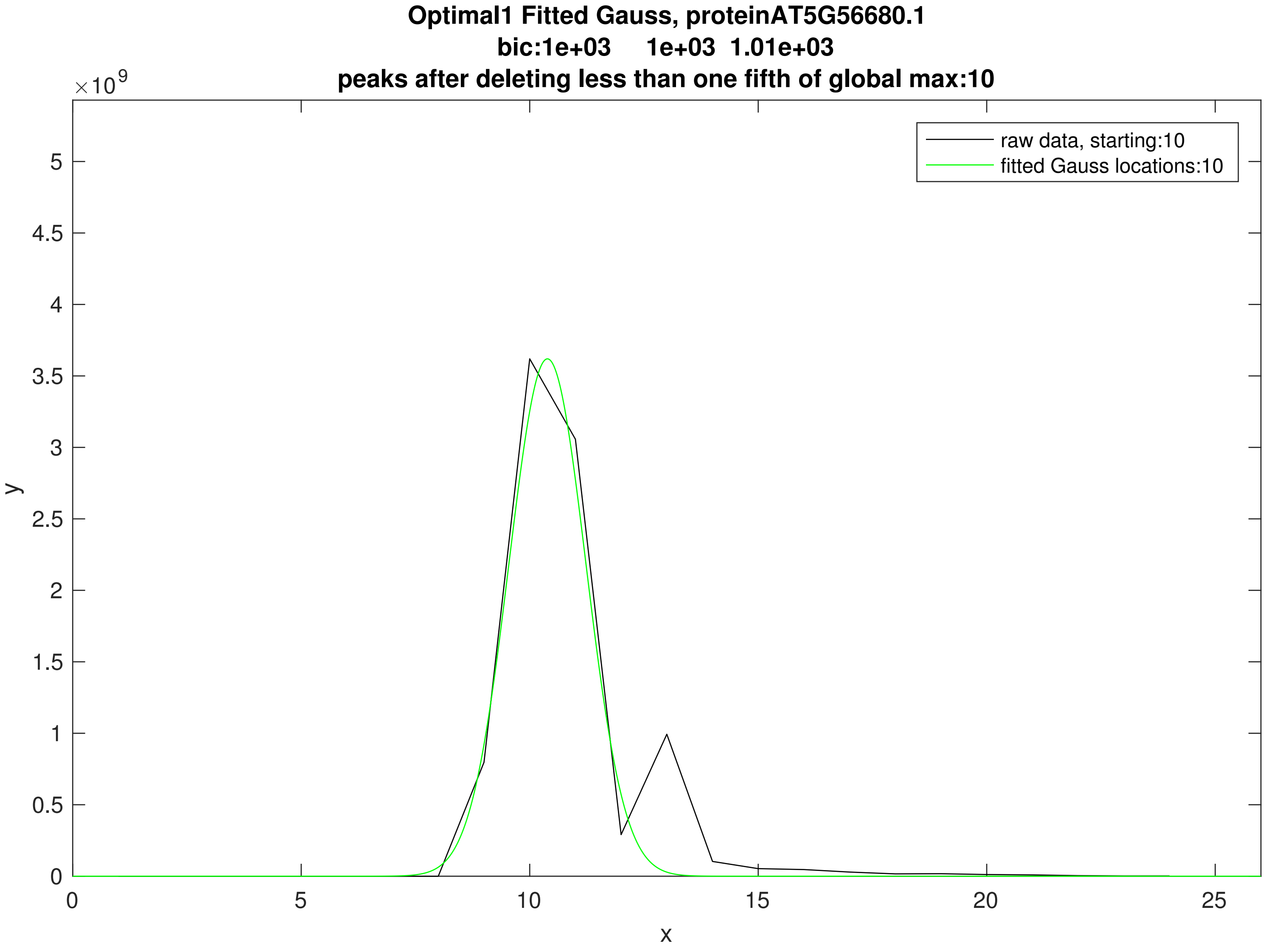

Optimal1 Fitted Gauss, proteinAT5G56680.1
bic:1e+03 1e+03 1.01e+03
peaks after deleting less than one fifth of global max:10
×10^9
raw data, starting:10
fitted Gauss locations:10
5
4.5
4
3.5
3
2.5
2
1.5
1
0.5
0
y
0
5
10
15
20
25
x

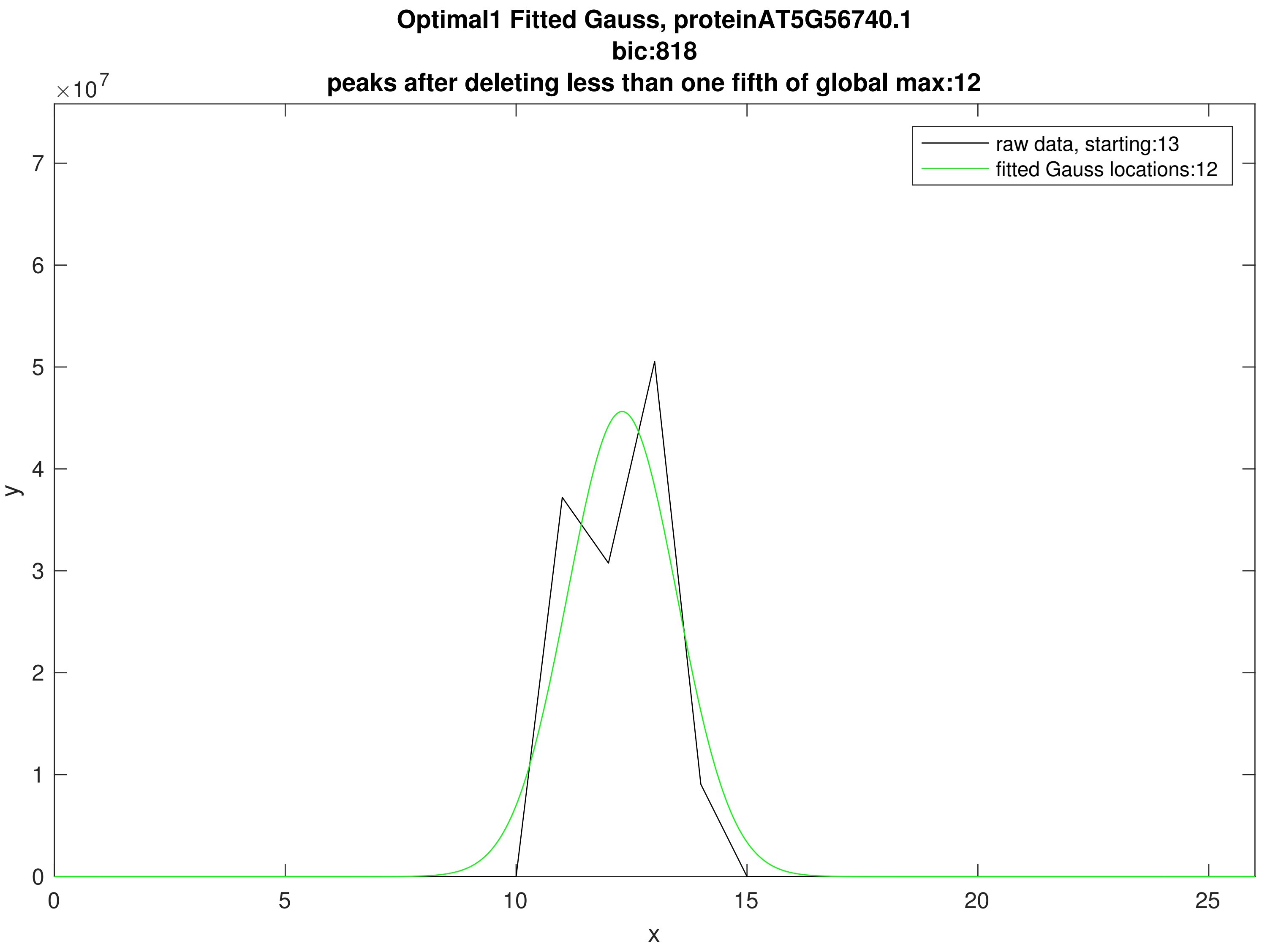

Optimal1 Fitted Gauss, proteinAT5G56740.1
bic:818
peaks after deleting less than one fifth of global max:12
×10^7
raw data, starting:13
fitted Gauss locations:12
y
x
0
1
2
3
4
5
6
7
0
5
10
15
20
25

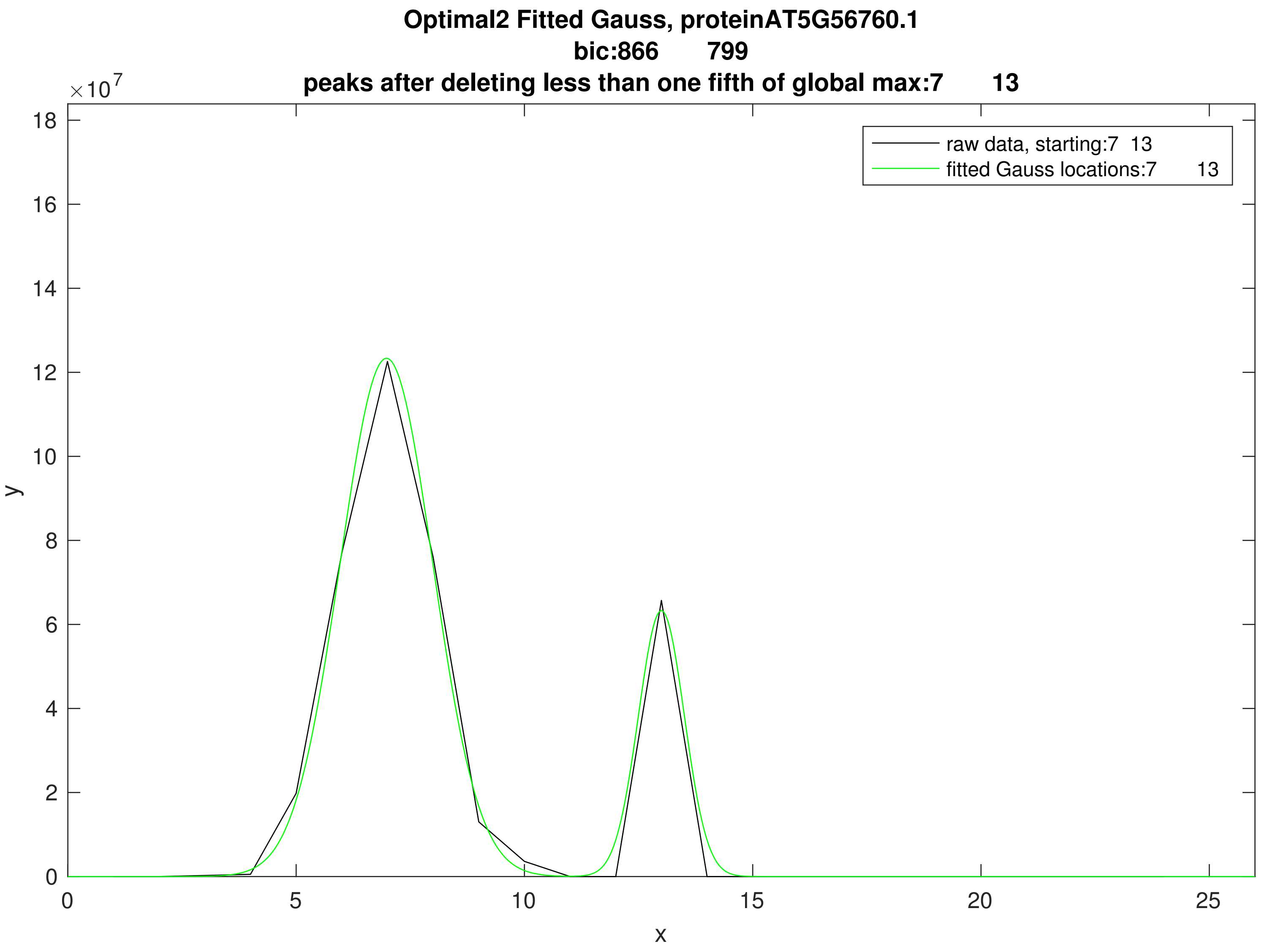

Optimal2 Fitted Gauss, proteinAT5G56760.1
bic:866 799
peaks after deleting less than one fifth of global max:7 13
×10^7
18
16
14
12
10
8
6
4
2
0
y
0
5
10
15
20
25
x
raw data, starting:7 13
fitted Gauss locations:7 13

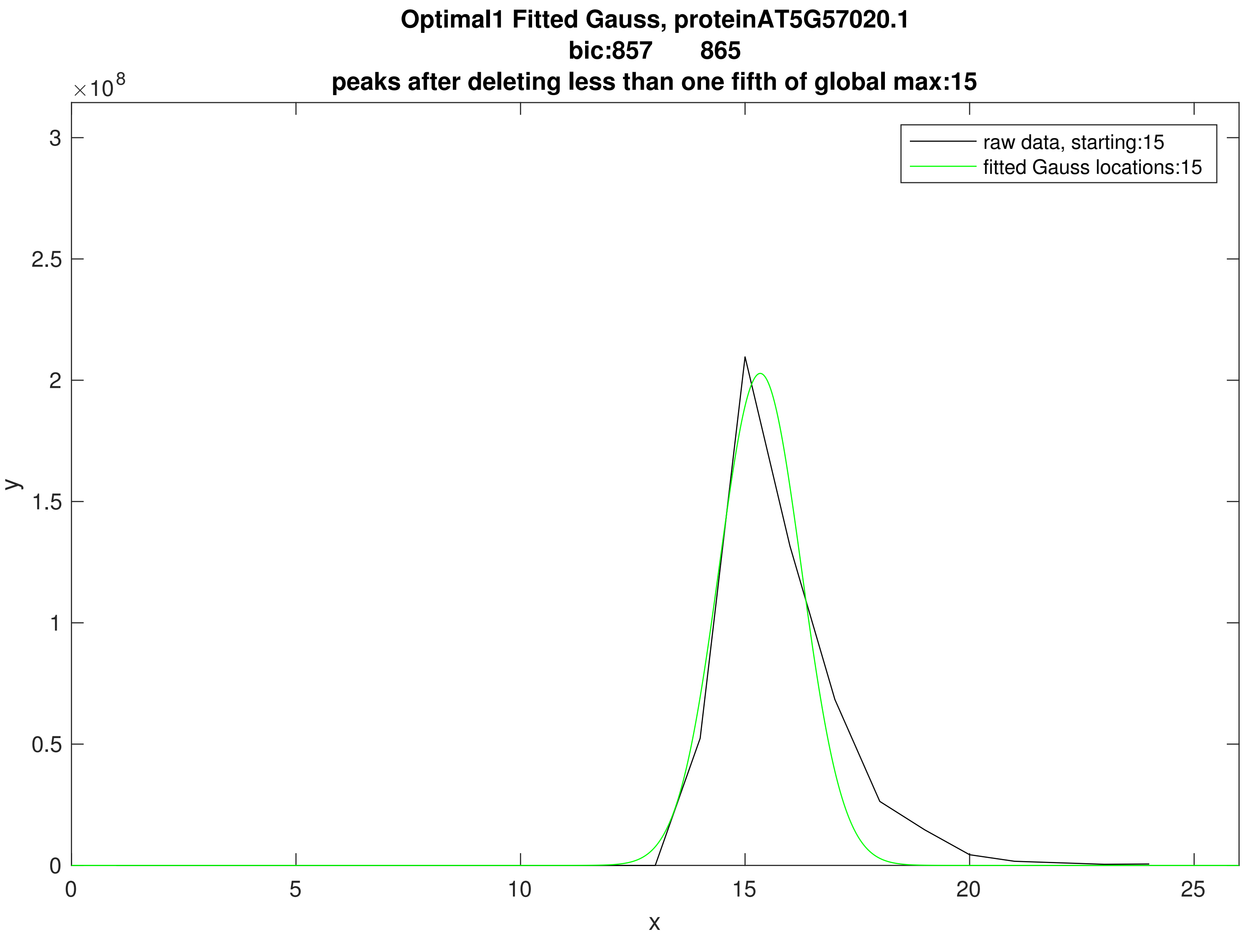
Optimal1 Fitted Gauss, proteinAT5G57020.1
bic:857 865
peaks after deleting less than one fifth of global max:15
×10^8
raw data, starting:15
fitted Gauss locations:15
y
x
0
0.5
1
1.5
2
2.5
3
0
5
10
15
20
25

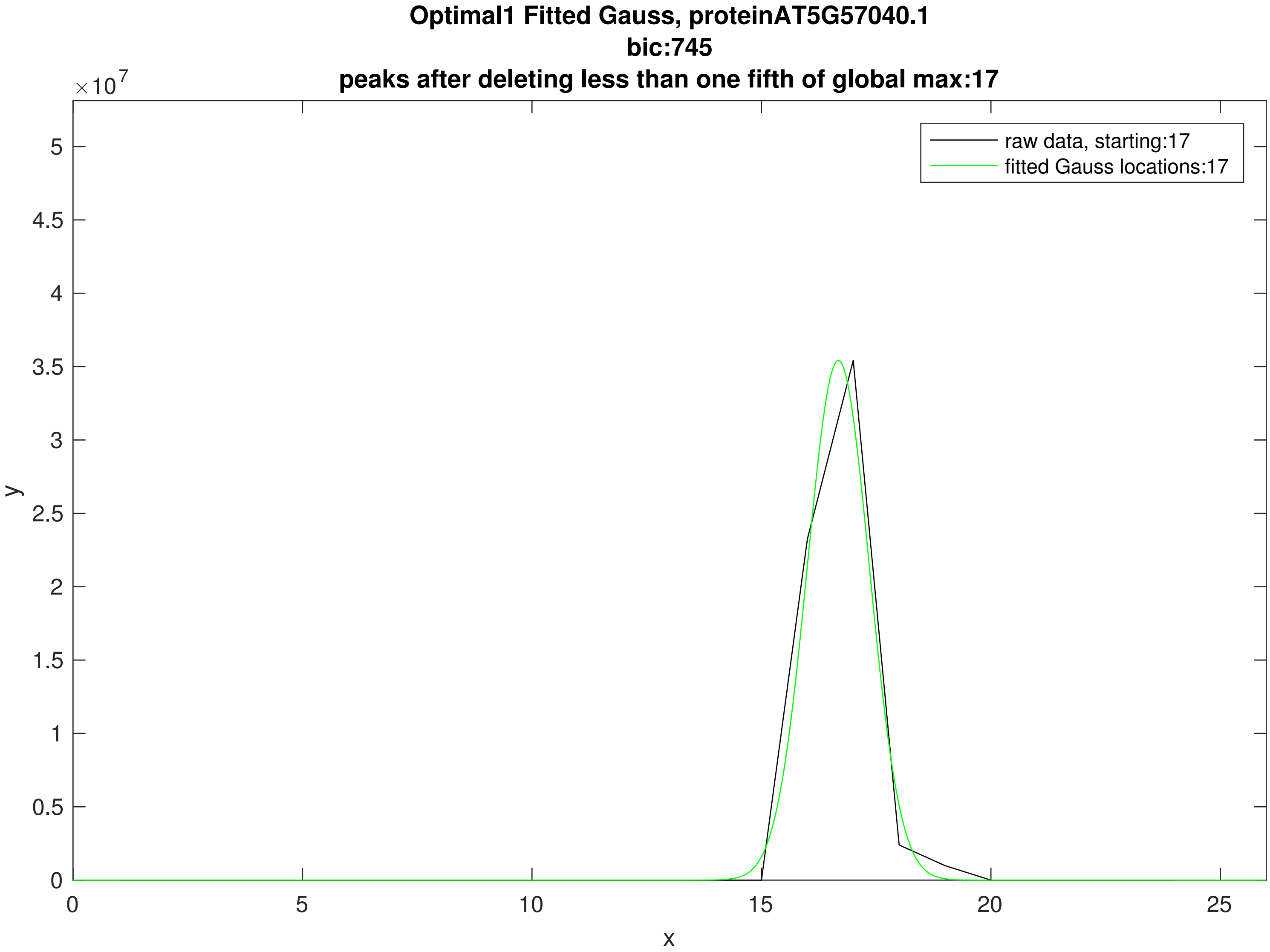

Optimal1 Fitted Gauss, proteinAT5G57040.1
bic:745
peaks after deleting less than one fifth of global max:17
×10^7
raw data, starting:17
fitted Gauss locations:17
y
x
0
0.5
1
1.5
2
2.5
3
3.5
4
4.5
5
0
5
10
15
20
25

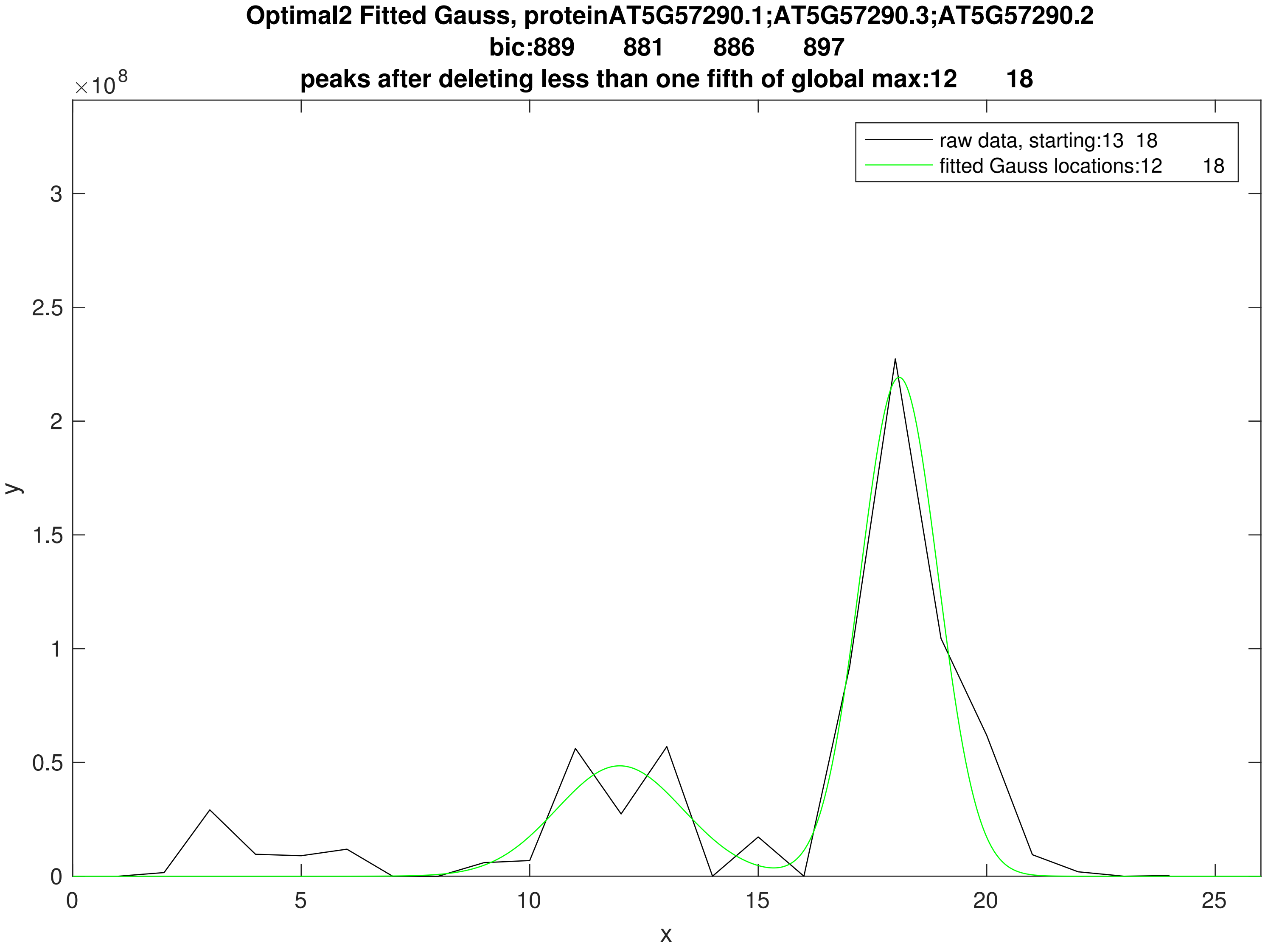

Optimal2 Fitted Gauss, proteinAT5G57290.1;AT5G57290.3;AT5G57290.2
bic:889 881 886 897
peaks after deleting less than one fifth of global max:12 18
×10^8
raw data, starting:13 18
fitted Gauss locations:12 18
y
x
0
0.5
1
1.5
2
2.5
3
5
10
15
20
25

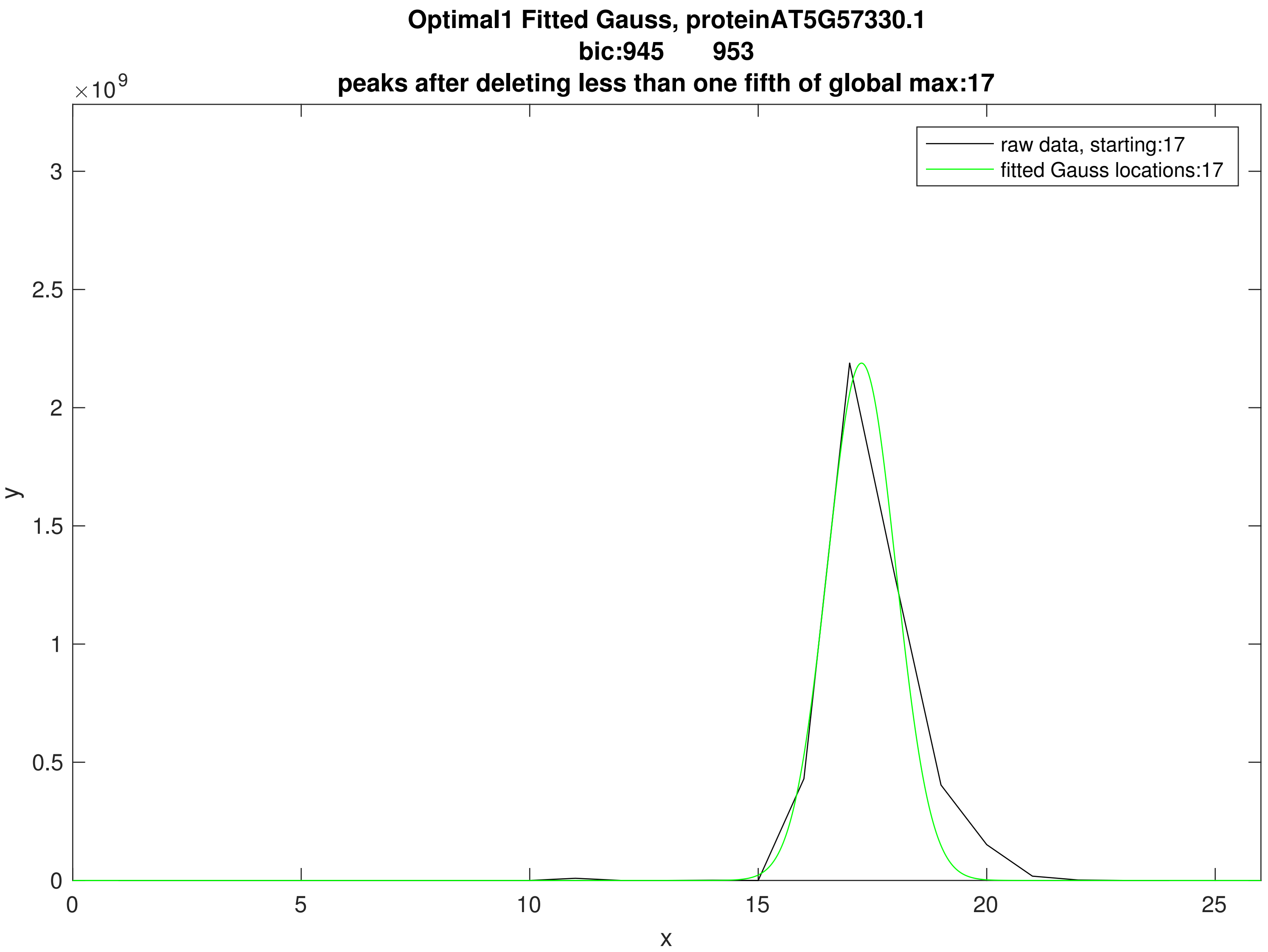

Optimal1 Fitted Gauss, proteinAT5G57330.1
bic:945 953
peaks after deleting less than one fifth of global max:17
×10^9
3
2.5
2
1.5
1
0.5
0
y
0
5
10
15
20
25
x
raw data, starting:17
fitted Gauss locations:17

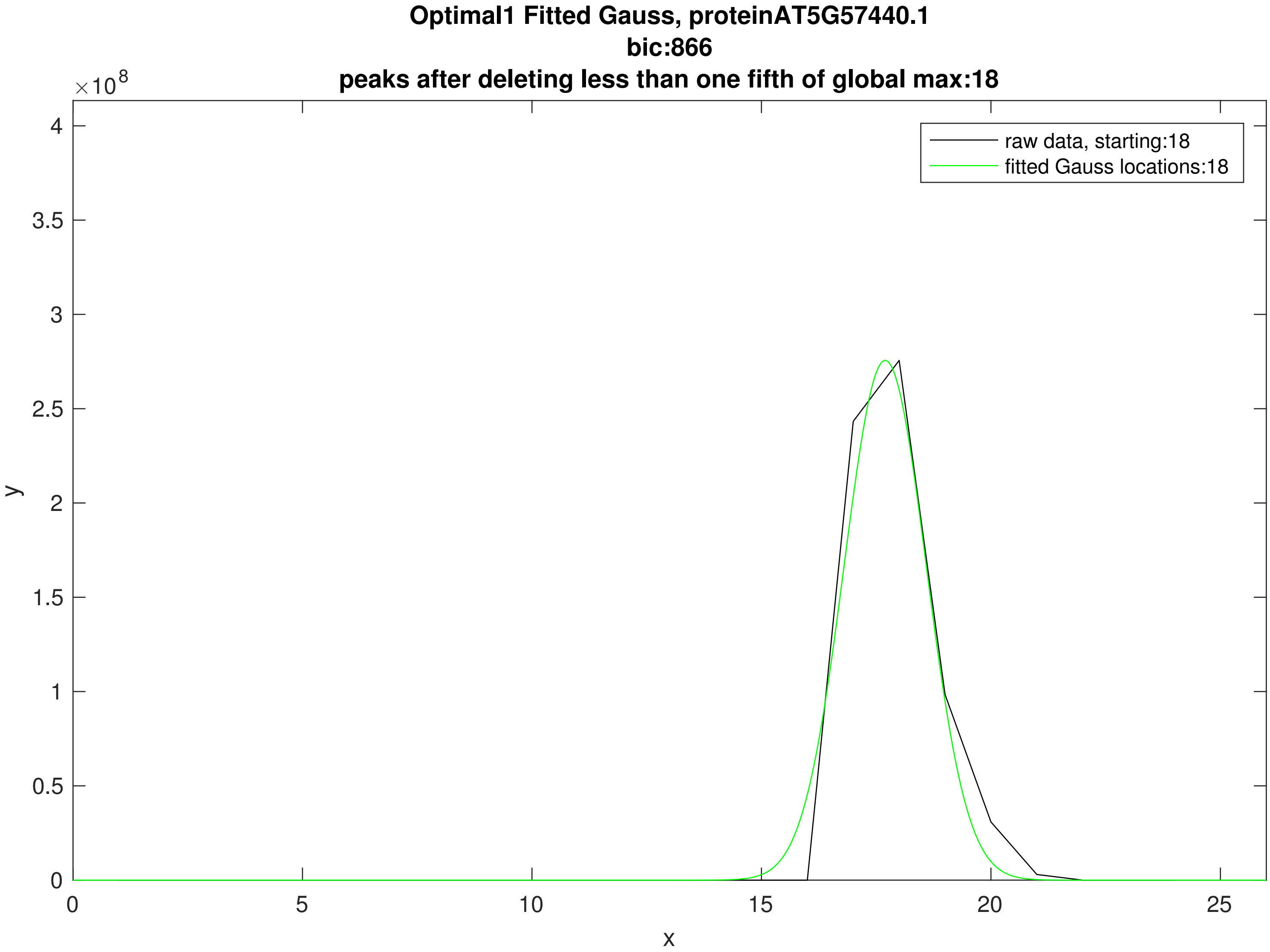
Optimal1 Fitted Gauss, proteinAT5G57440.1
bic:866
peaks after deleting less than one fifth of global max:18
×10^8
4
3.5
3
2.5
2
1.5
1
0.5
0
y
0
5
10
15
20
25
x
raw data, starting:18
fitted Gauss locations:18

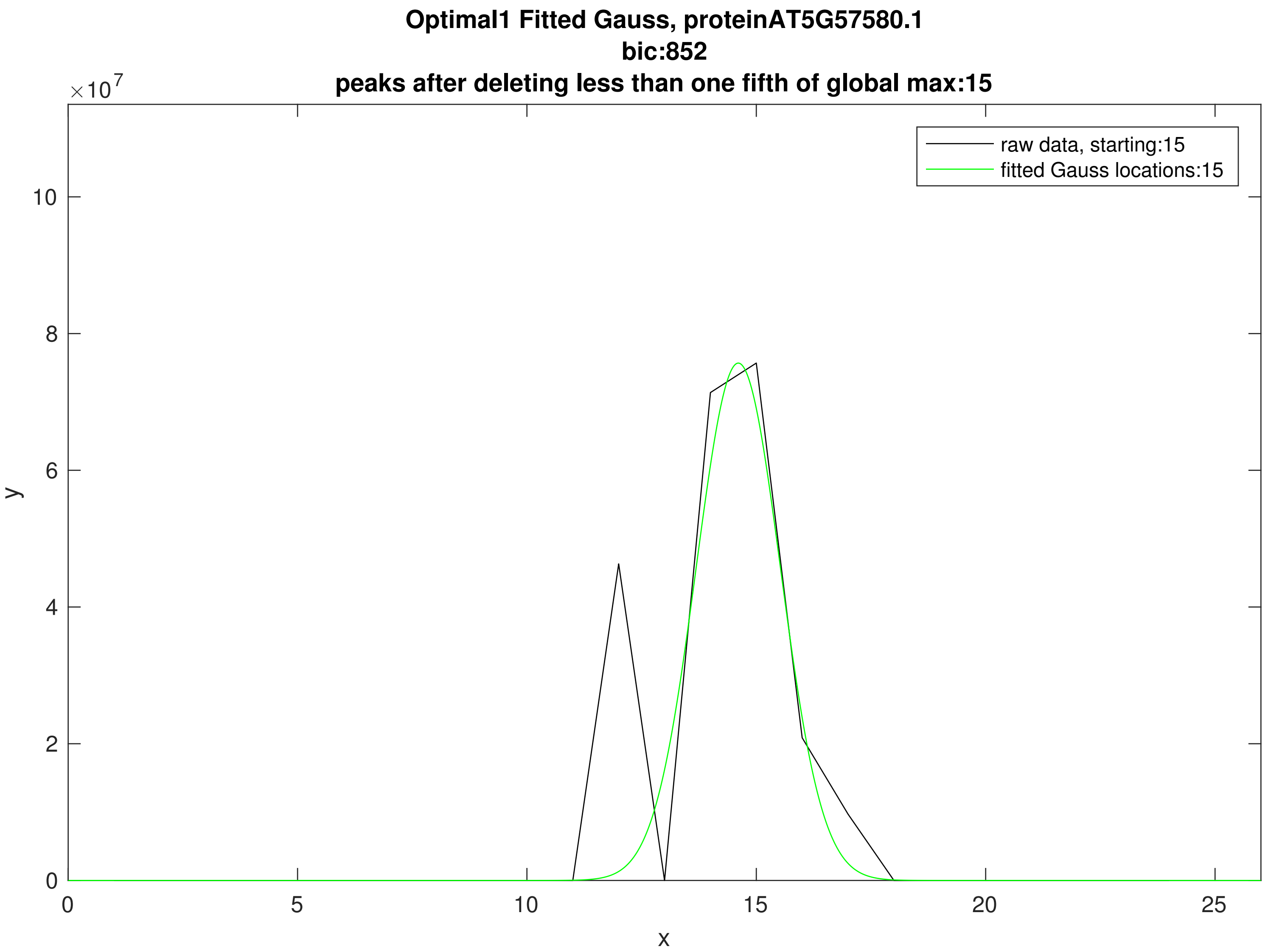

Optimal1 Fitted Gauss, proteinAT5G57580.1
bic:852
peaks after deleting less than one fifth of global max:15
×10^7
raw data, starting:15
fitted Gauss locations:15
10
8
6
4
2
0
y
0
5
10
15
20
25
x

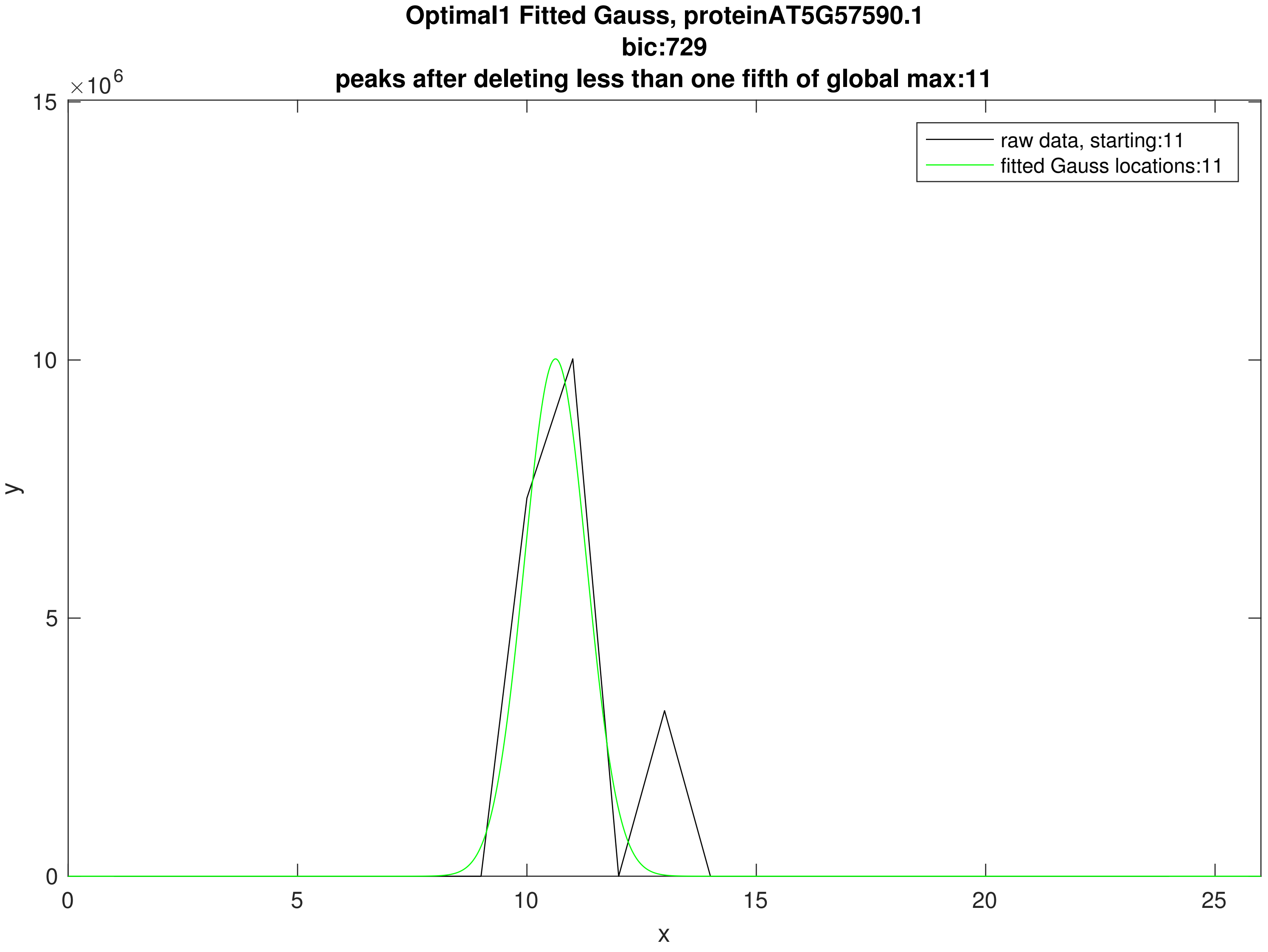
Optimal1 Fitted Gauss, proteinAT5G57590.1
bic:729
peaks after deleting less than one fifth of global max:11
raw data, starting:11
fitted Gauss locations:11
×10^6
15
10
5
0
y
0
5
10
15
20
25
x

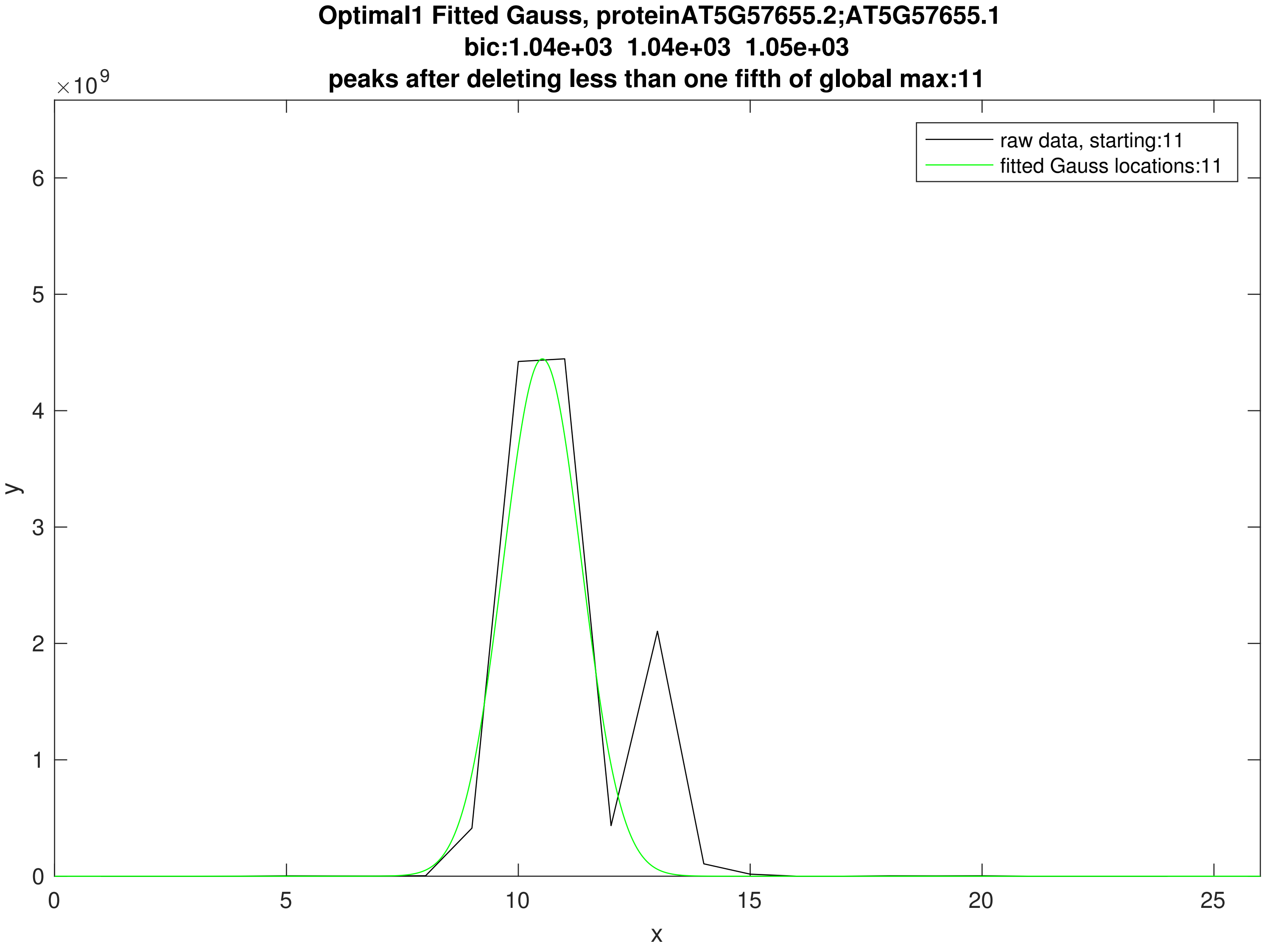

Optimal1 Fitted Gauss, proteinAT5G57655.2;AT5G57655.1
bic:1.04e+03 1.04e+03 1.05e+03
peaks after deleting less than one fifth of global max:11
×10^9
raw data, starting:11
fitted Gauss locations:11
y
x
0
1
2
3
4
5
6
0
5
10
15
20
25

Optimal1 Fitted Gauss, proteinAT5G57700.2;AT5G57700.4;AT5G57700.1;AT5G57700.3
bic:919 929
peaks after deleting less than one fifth of global max:13

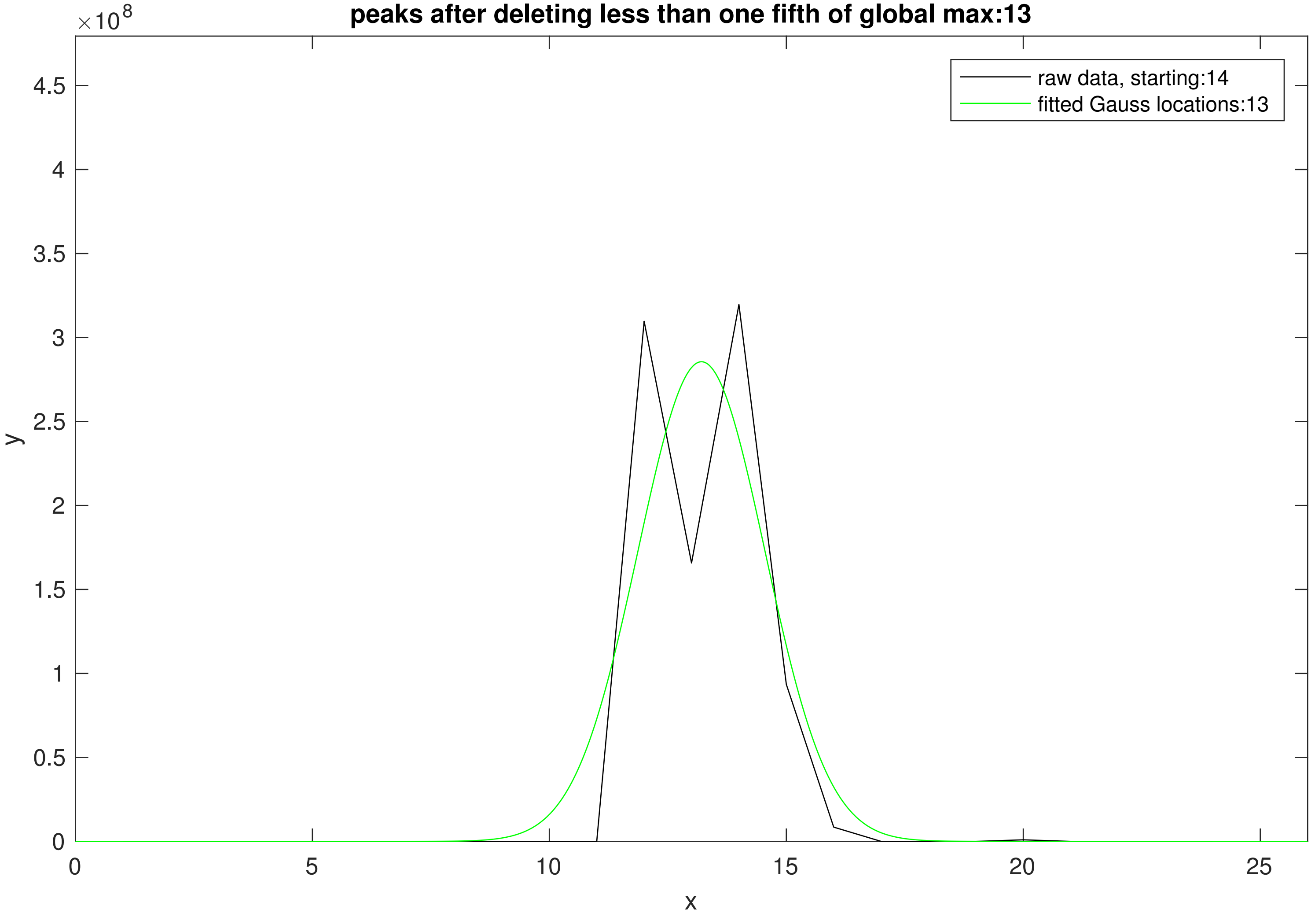

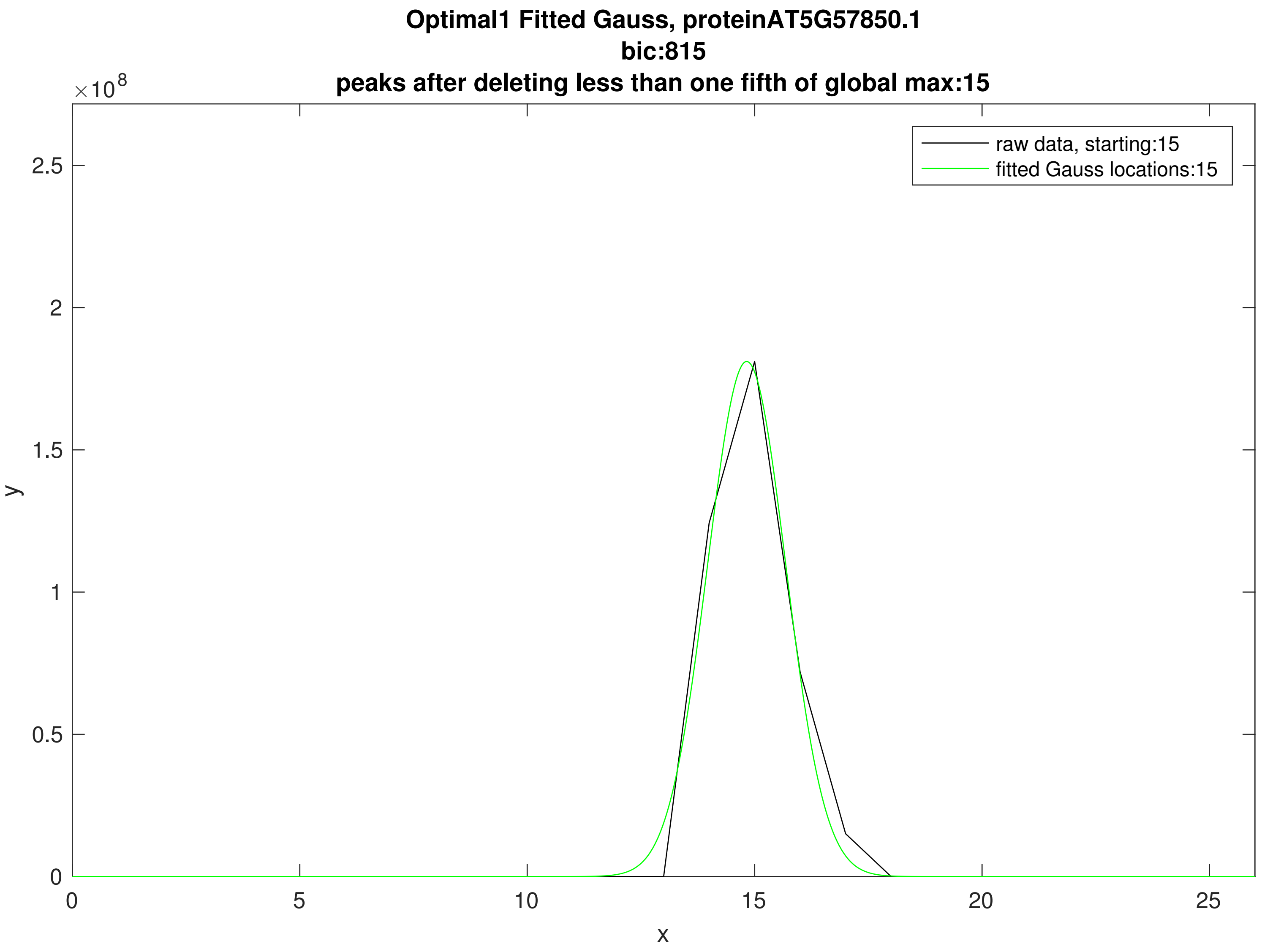

Optimal1 Fitted Gauss, proteinAT5G57850.1
bic:815
peaks after deleting less than one fifth of global max:15
×10^8
raw data, starting:15
fitted Gauss locations:15
2.5
2
1.5
1
0.5
0
y
0
5
10
15
20
25
x

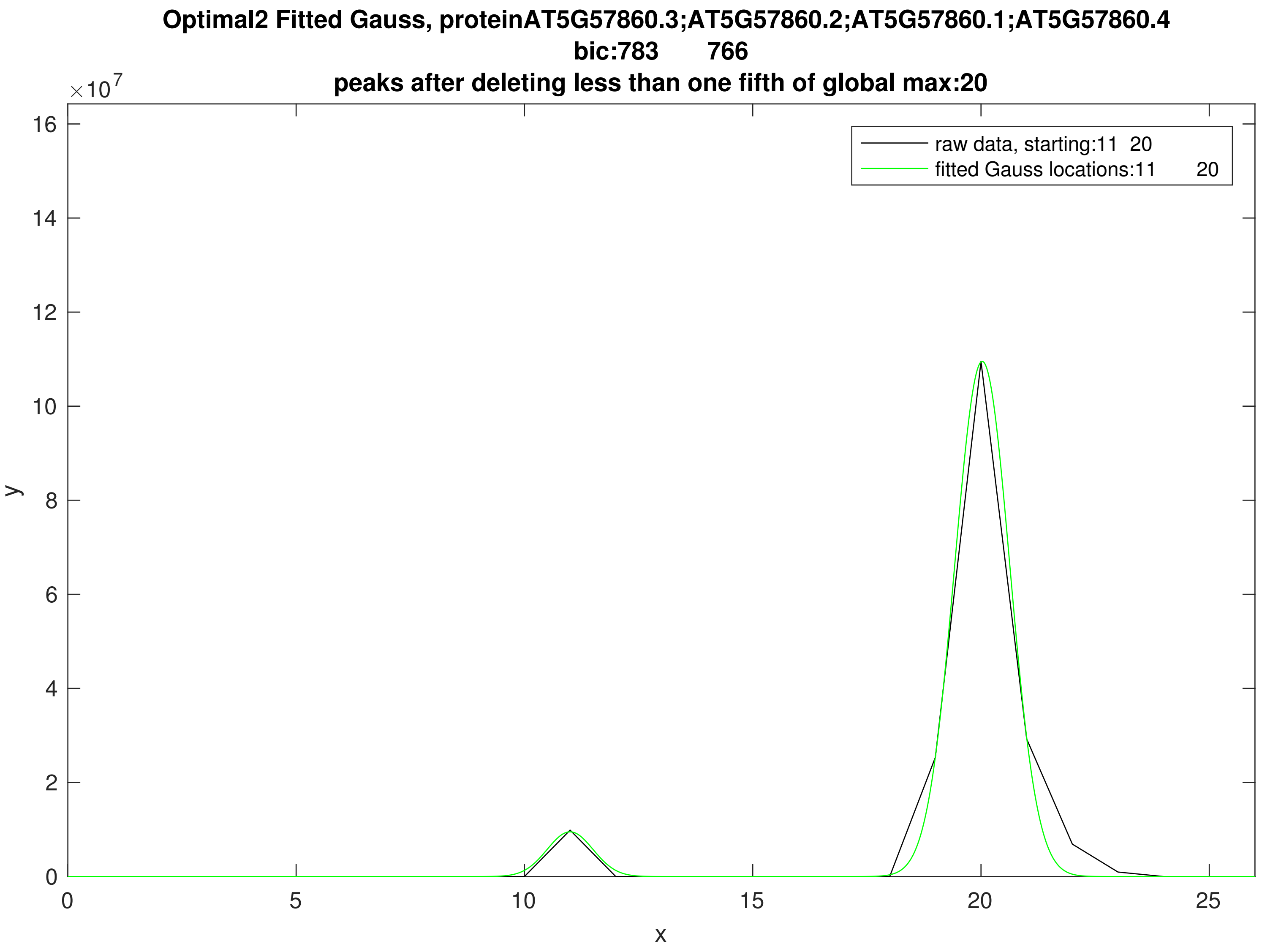
Optimal2 Fitted Gauss, proteinAT5G57860.3;AT5G57860.2;AT5G57860.1;AT5G57860.4
bic:783 766
peaks after deleting less than one fifth of global max:20
×10^7
raw data, starting:11 20
fitted Gauss locations:11 20
y
x
0
2
4
6
8
10
12
14
16
0
5
10
15
20
25

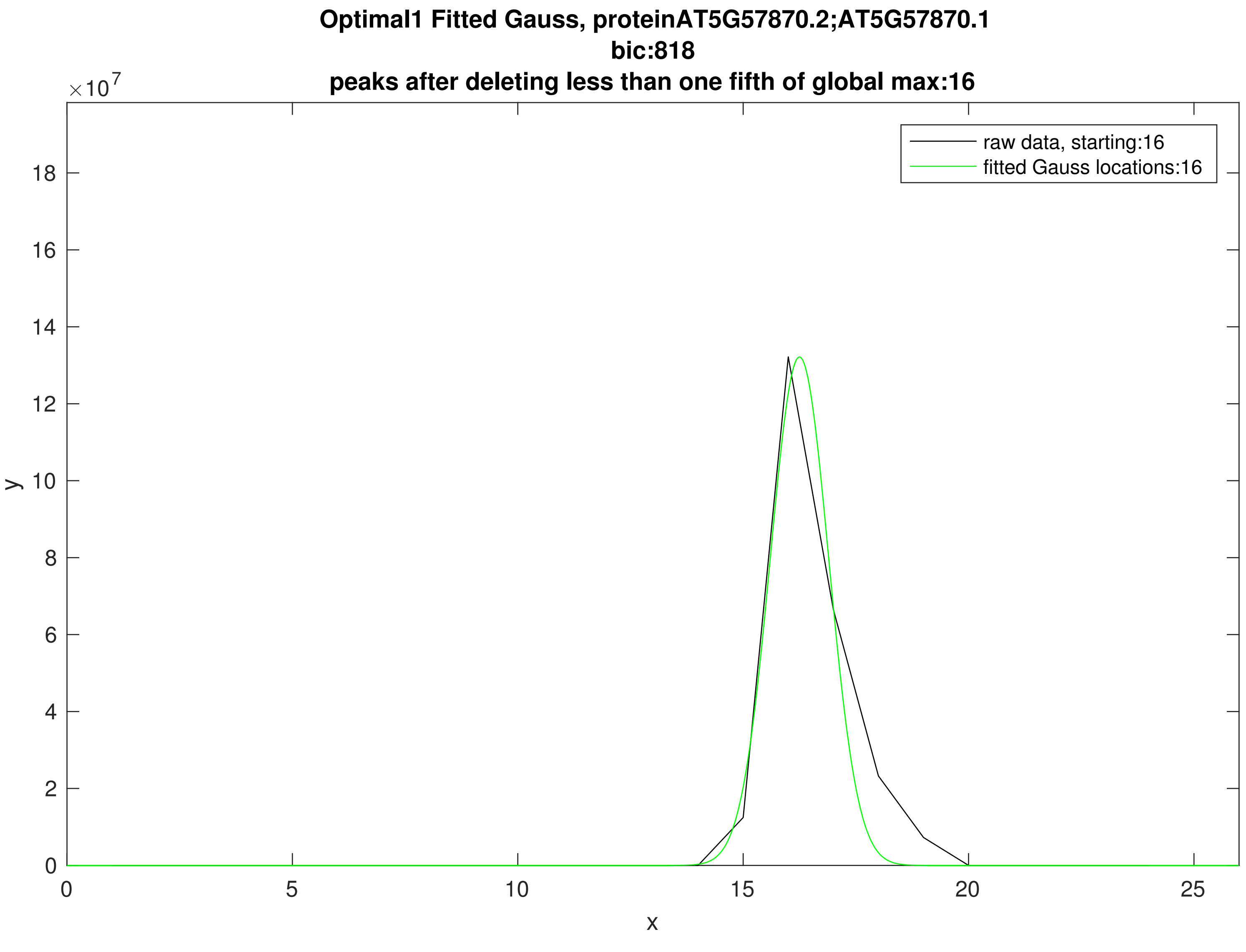

Optimal1 Fitted Gauss, proteinAT5G57870.2;AT5G57870.1
bic:818
peaks after deleting less than one fifth of global max:16
×10^7
raw data, starting:16
fitted Gauss locations:16
y
x
0
2
4
6
8
10
12
14
16
18
0
5
10
15
20
25

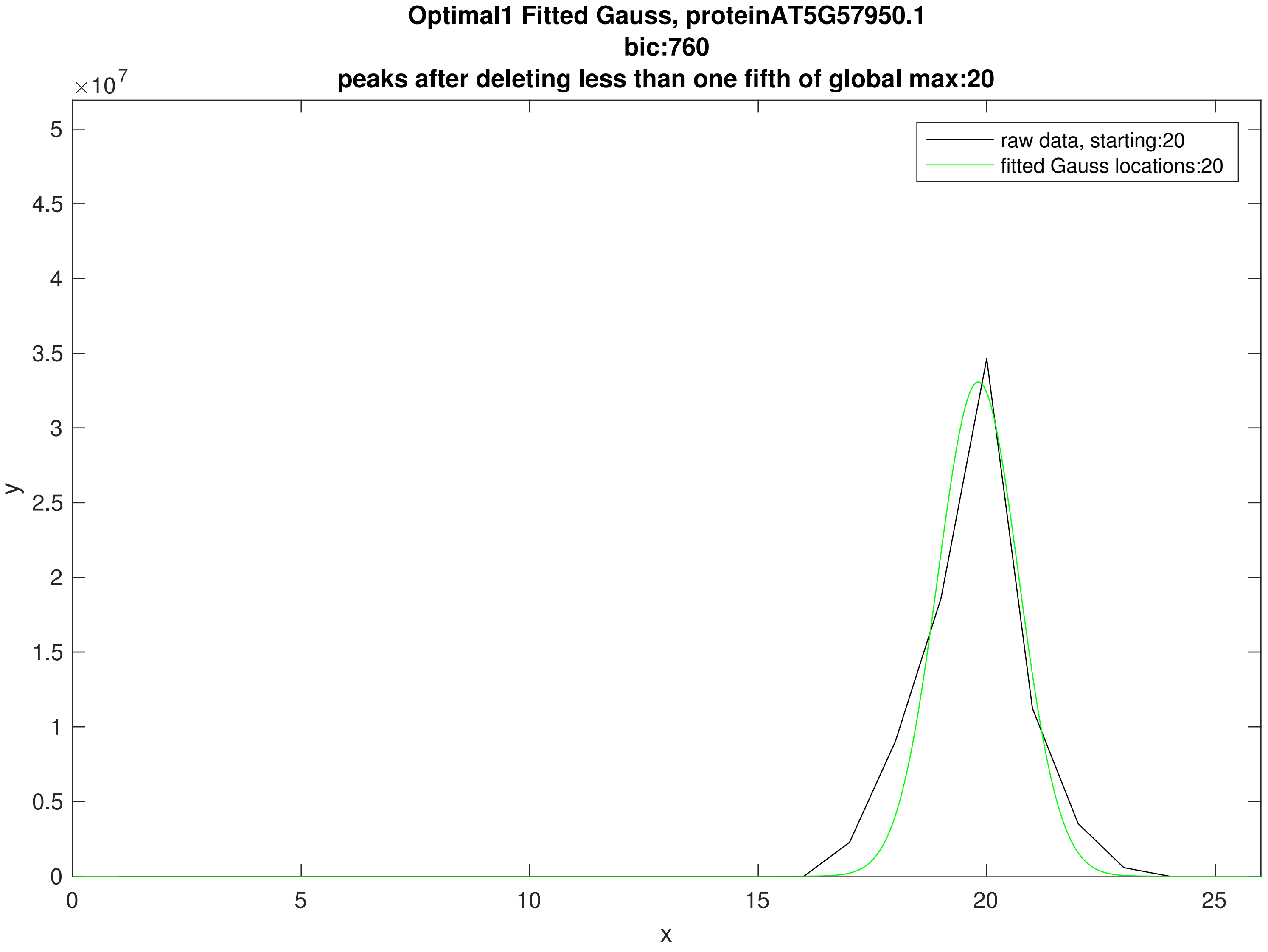

Optimal1 Fitted Gauss, proteinAT5G57950.1
bic:760
peaks after deleting less than one fifth of global max:20
×10^7
raw data, starting:20
fitted Gauss locations:20
y
x
0
0.5
1
1.5
2
2.5
3
3.5
4
4.5
5
0
5
10
15
20
25

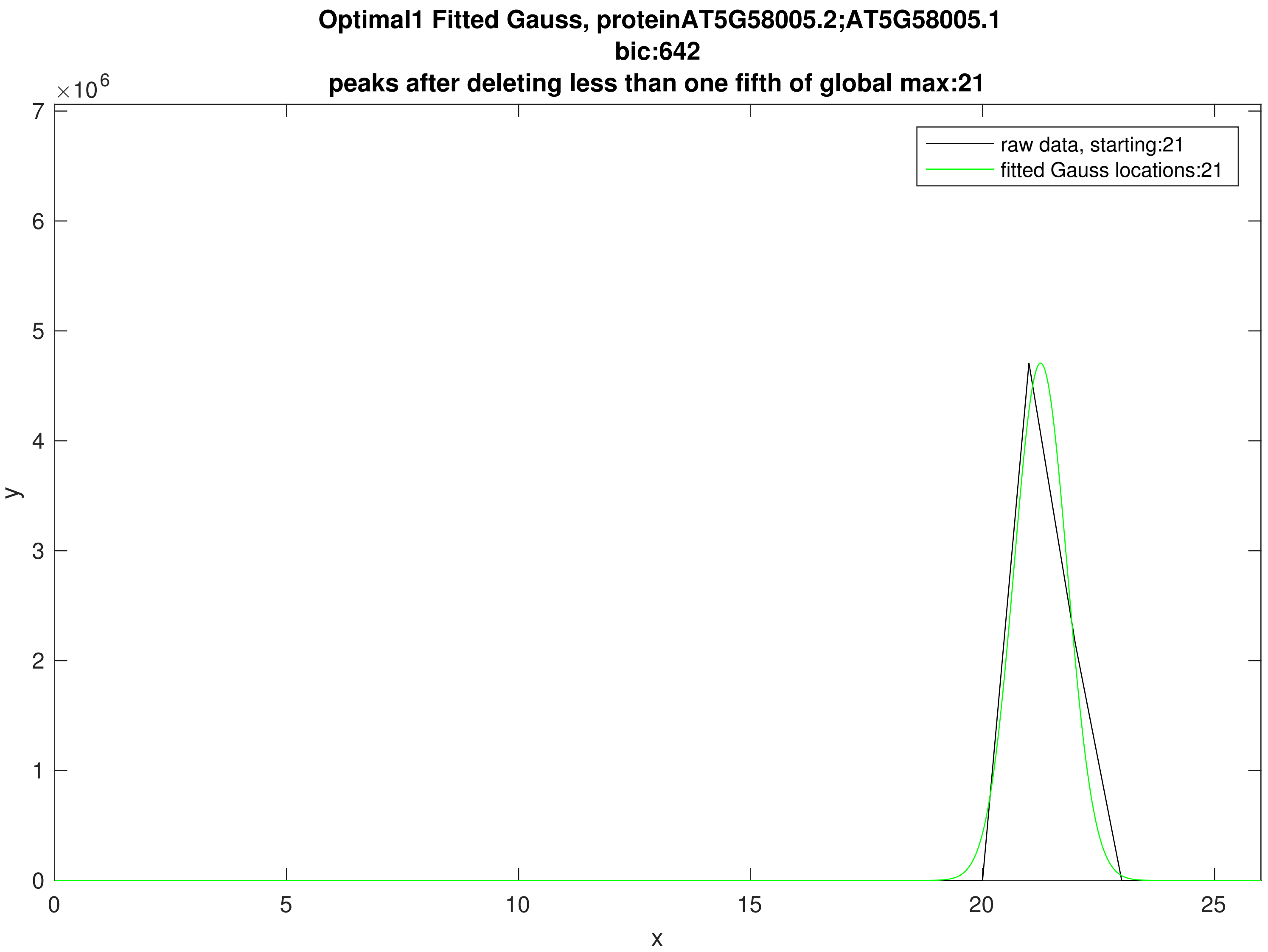

Optimal1 Fitted Gauss, proteinAT5G58005.2;AT5G58005.1
bic:642
peaks after deleting less than one fifth of global max:21
×10^6
raw data, starting:21
fitted Gauss locations:21
y
x
0
1
2
3
4
5
6
7
0
5
10
15
20
25

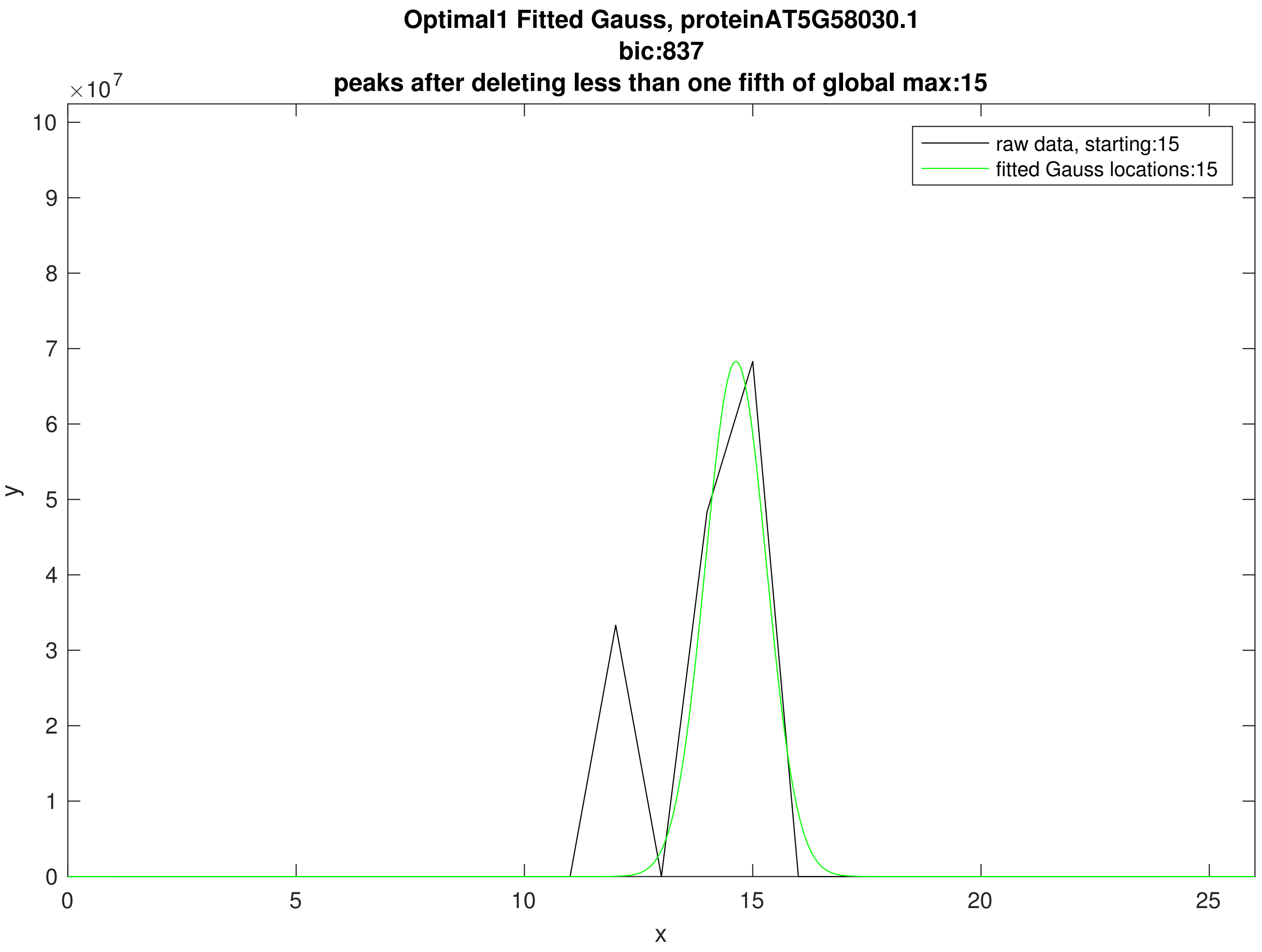
Optimal1 Fitted Gauss, proteinAT5G58030.1
bic:837
peaks after deleting less than one fifth of global max:15
×10^7
raw data, starting:15
fitted Gauss locations:15
y
x
0
1
2
3
4
5
6
7
8
9
10
0
5
10
15
20
25

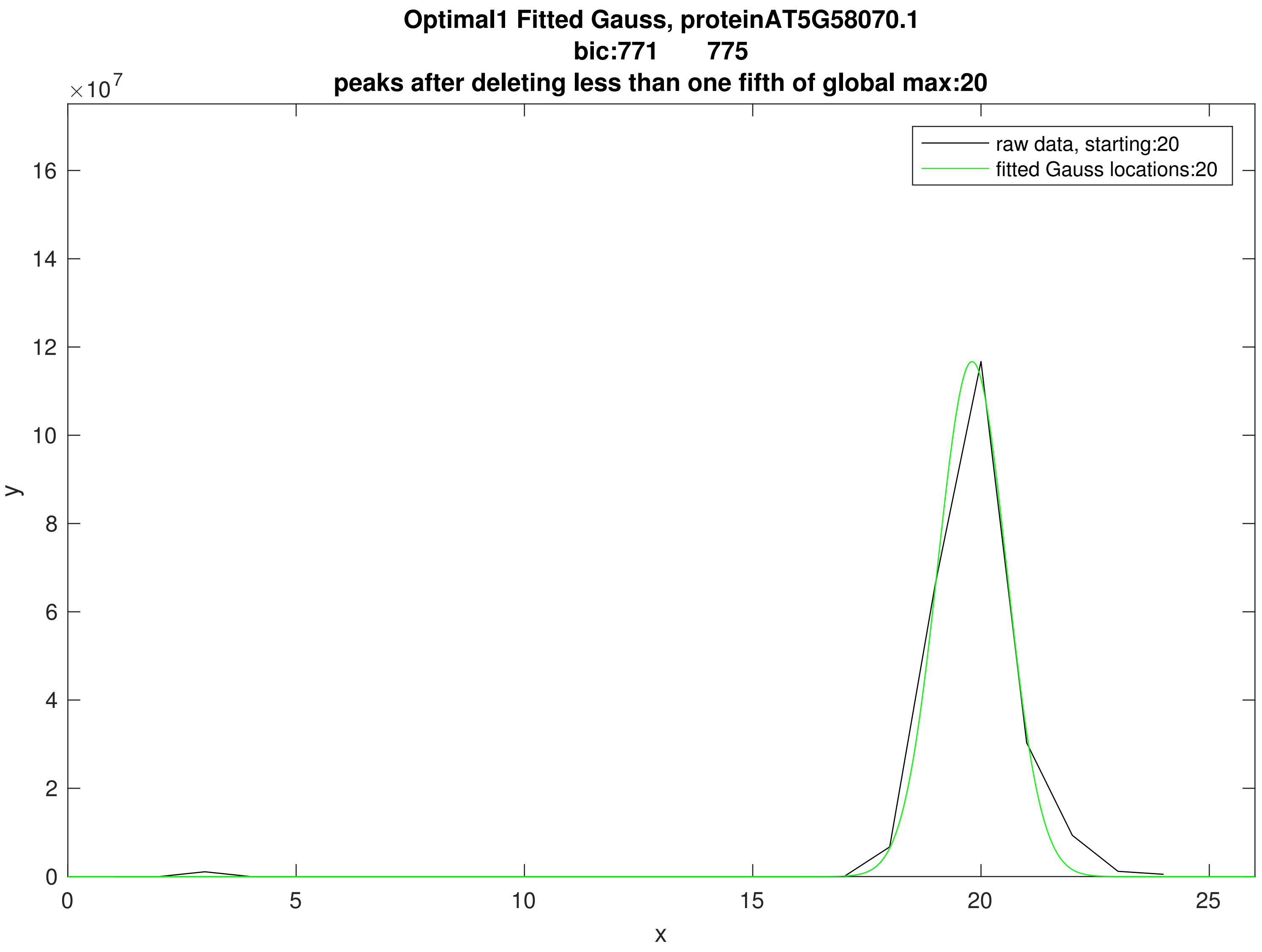
Optimal1 Fitted Gauss, proteinAT5G58070.1
bic:771 775
peaks after deleting less than one fifth of global max:20
×10^7
raw data, starting:20
fitted Gauss locations:20
16
14
12
10
8
6
4
2
0
y
0
5
10
15
20
25
x

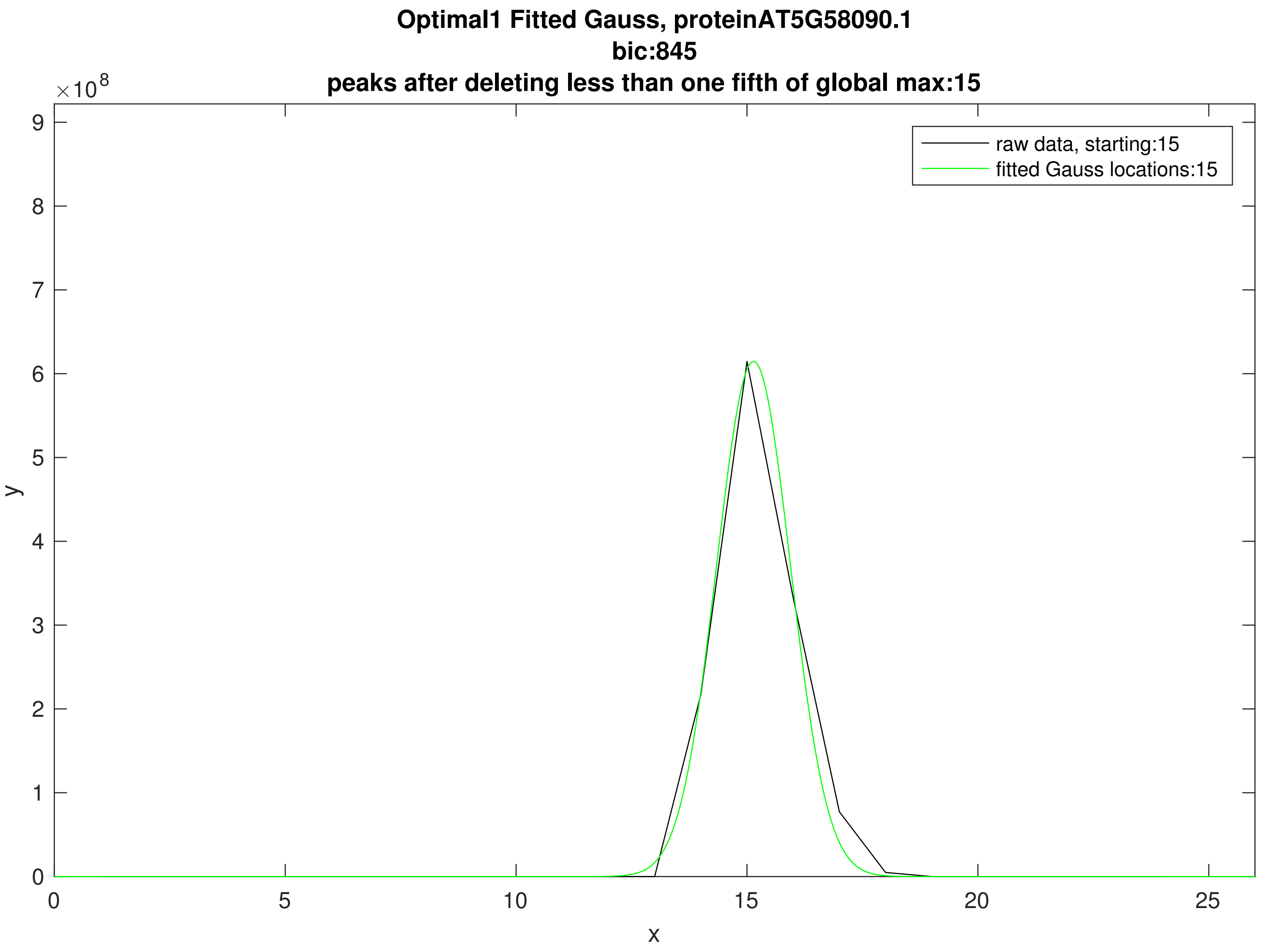
Optimal1 Fitted Gauss, proteinAT5G58090.1
bic:845
peaks after deleting less than one fifth of global max:15
×10^8
raw data, starting:15
fitted Gauss locations:15
y
x
0
1
2
3
4
5
6
7
8
9
0
5
10
15
20
25

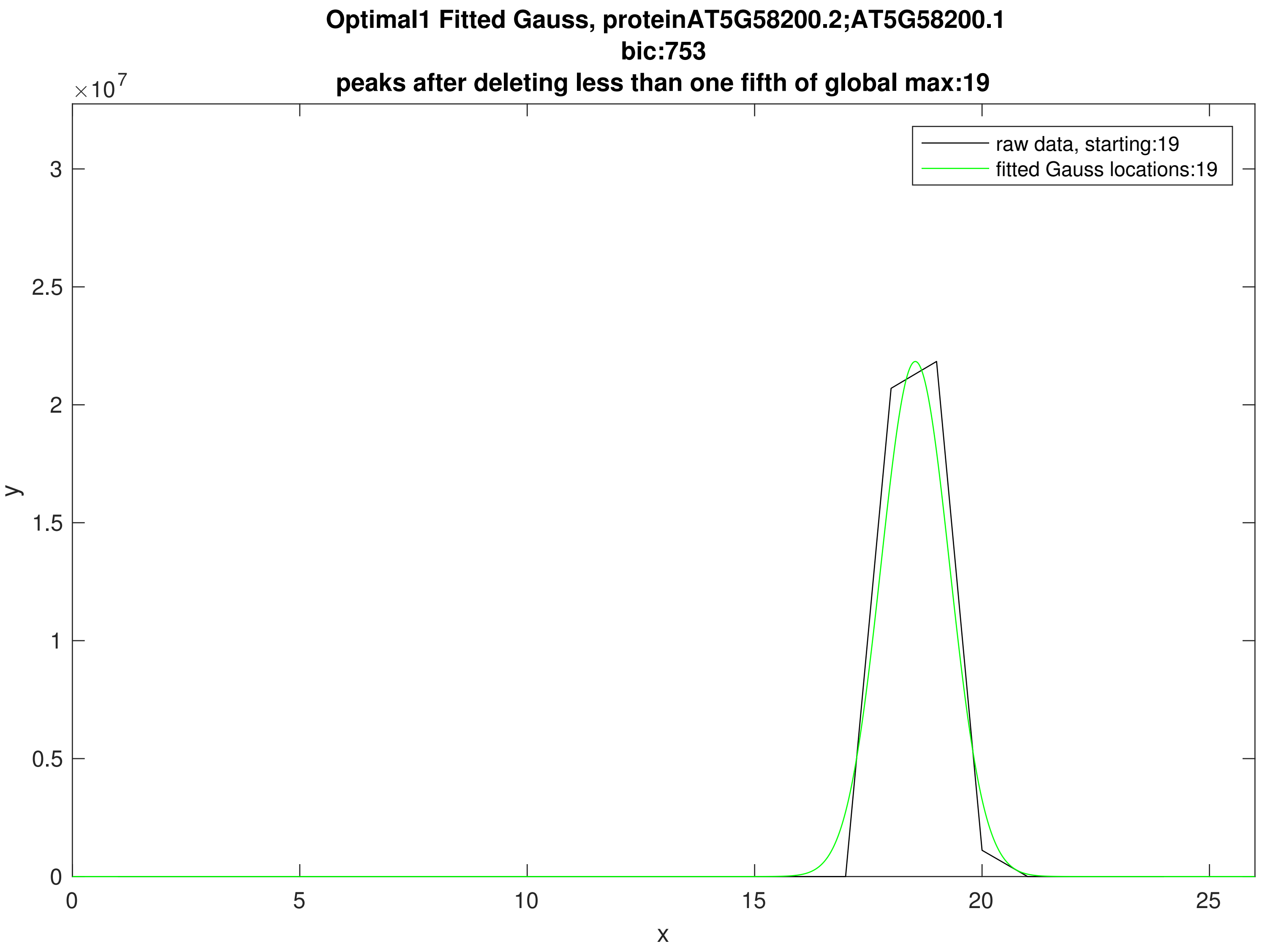

Optimal1 Fitted Gauss, proteinAT5G58200.2;AT5G58200.1
bic:753
peaks after deleting less than one fifth of global max:19
×10^7
3
2.5
2
1.5
1
0.5
0
y
0
5
10
15
20
25
x
raw data, starting:19
fitted Gauss locations:19

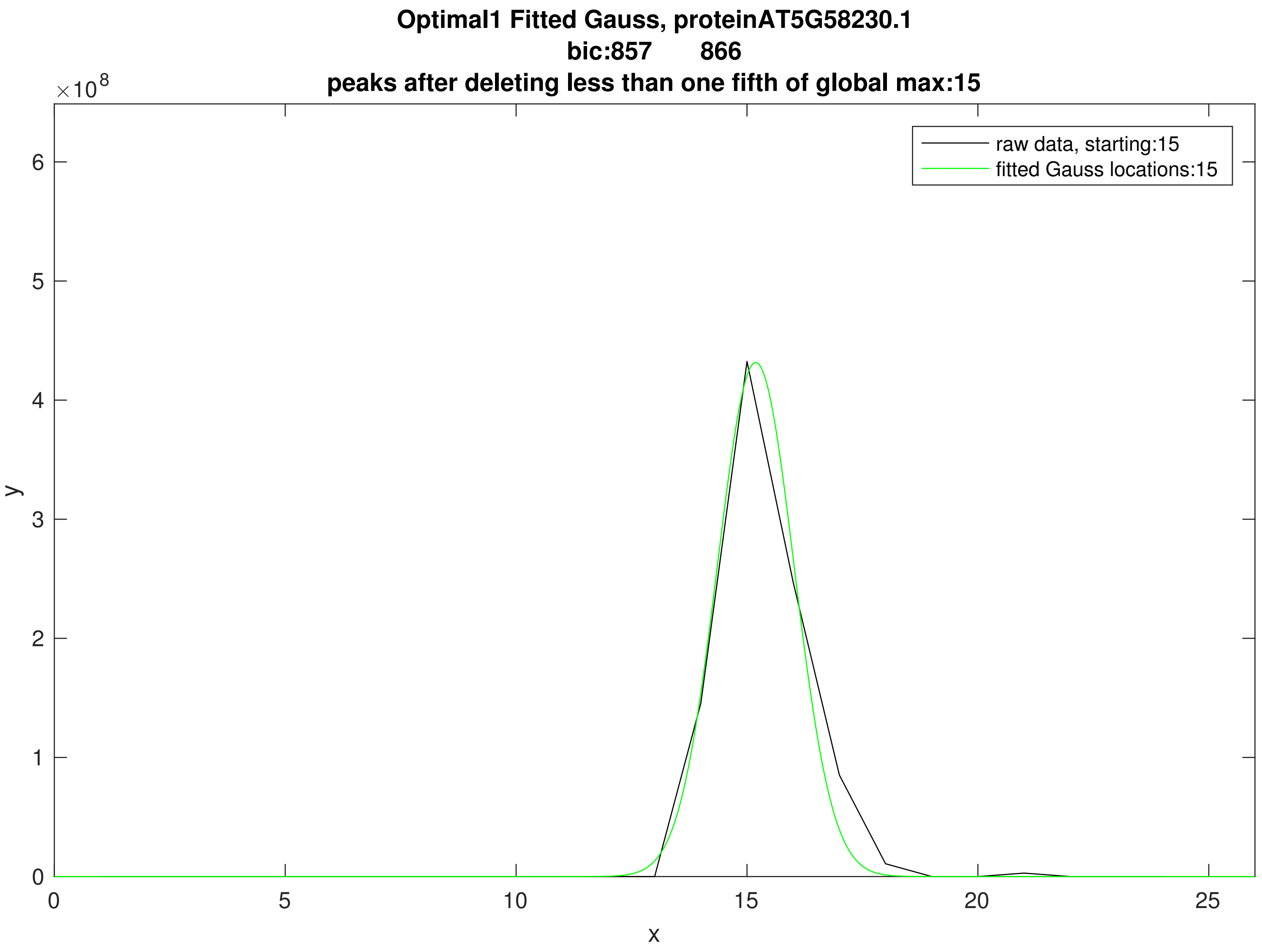

Optimal1 Fitted Gauss, proteinAT5G58230.1
bic:857 866
peaks after deleting less than one fifth of global max:15
×10^8
raw data, starting:15
fitted Gauss locations:15
y
x
0
1
2
3
4
5
6
0
5
10
15
20
25

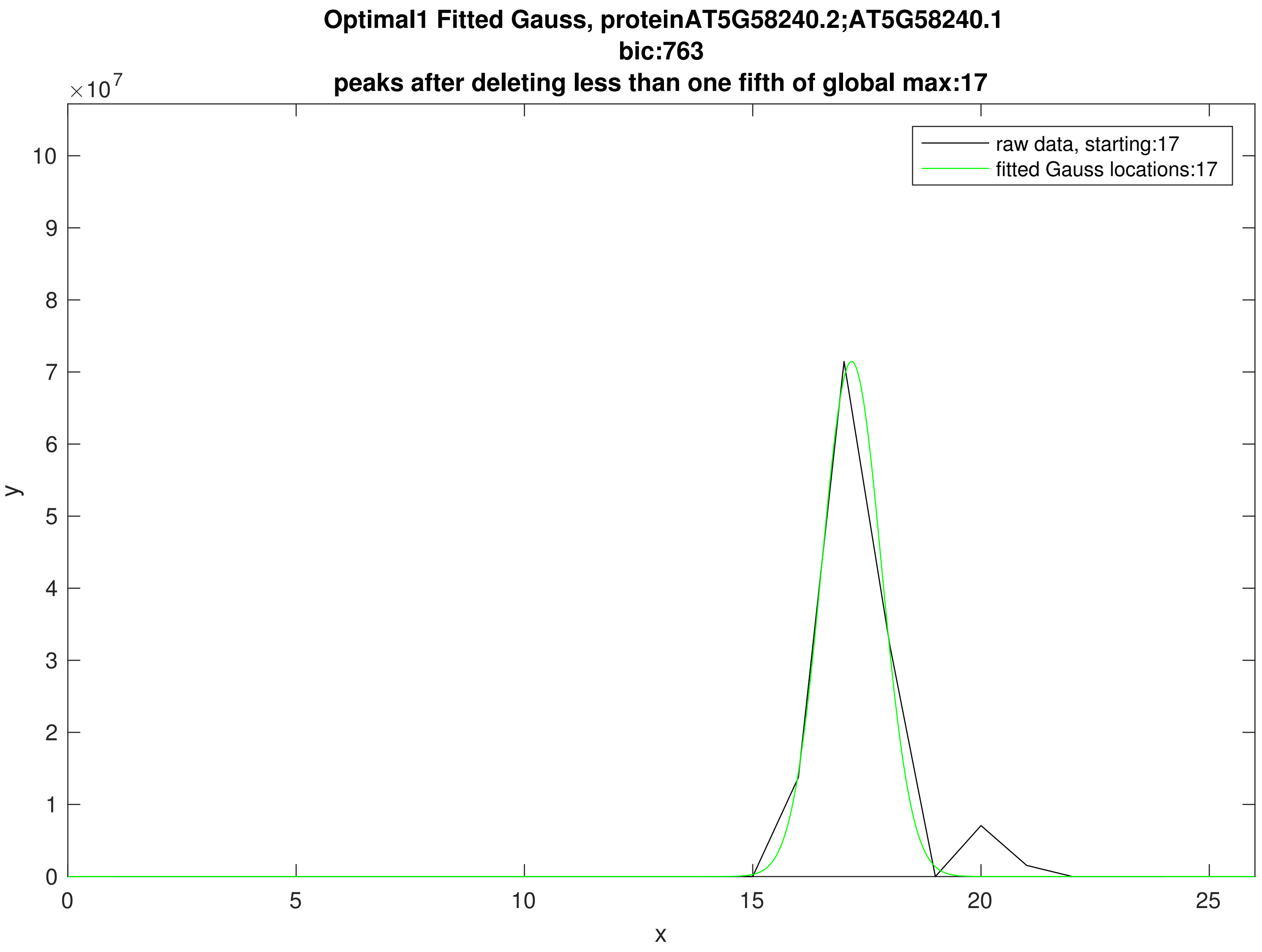
Optimal1 Fitted Gauss, proteinAT5G58240.2;AT5G58240.1
bic:763
peaks after deleting less than one fifth of global max:17
×10^7
raw data, starting:17
fitted Gauss locations:17
y
x
0
1
2
3
4
5
6
7
8
9
10
0
5
10
15
20
25

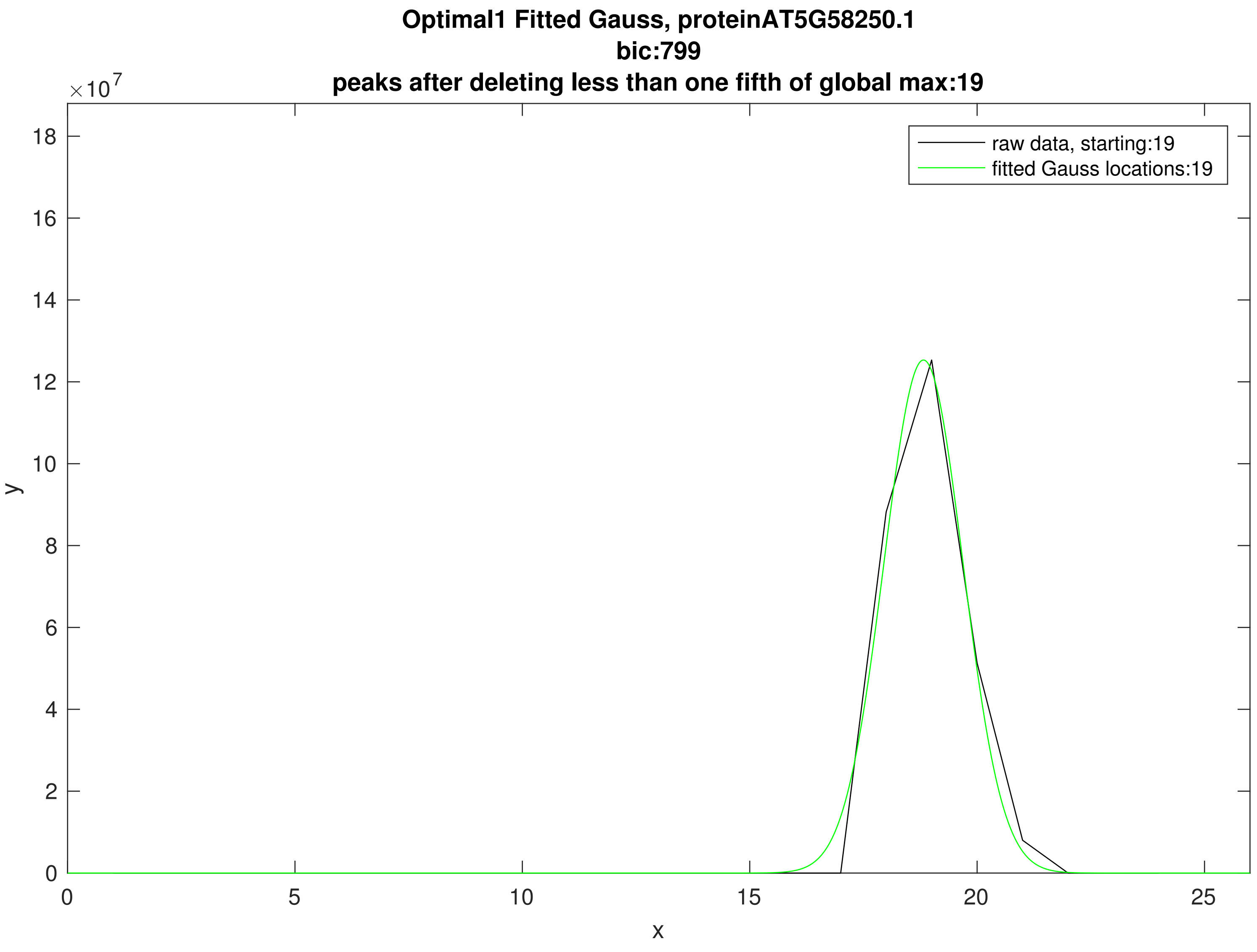
Optimal1 Fitted Gauss, proteinAT5G58250.1
bic:799
peaks after deleting less than one fifth of global max:19
×10^7
raw data, starting:19
fitted Gauss locations:19
y
x
0
2
4
6
8
10
12
14
16
18
0
5
10
15
20
25

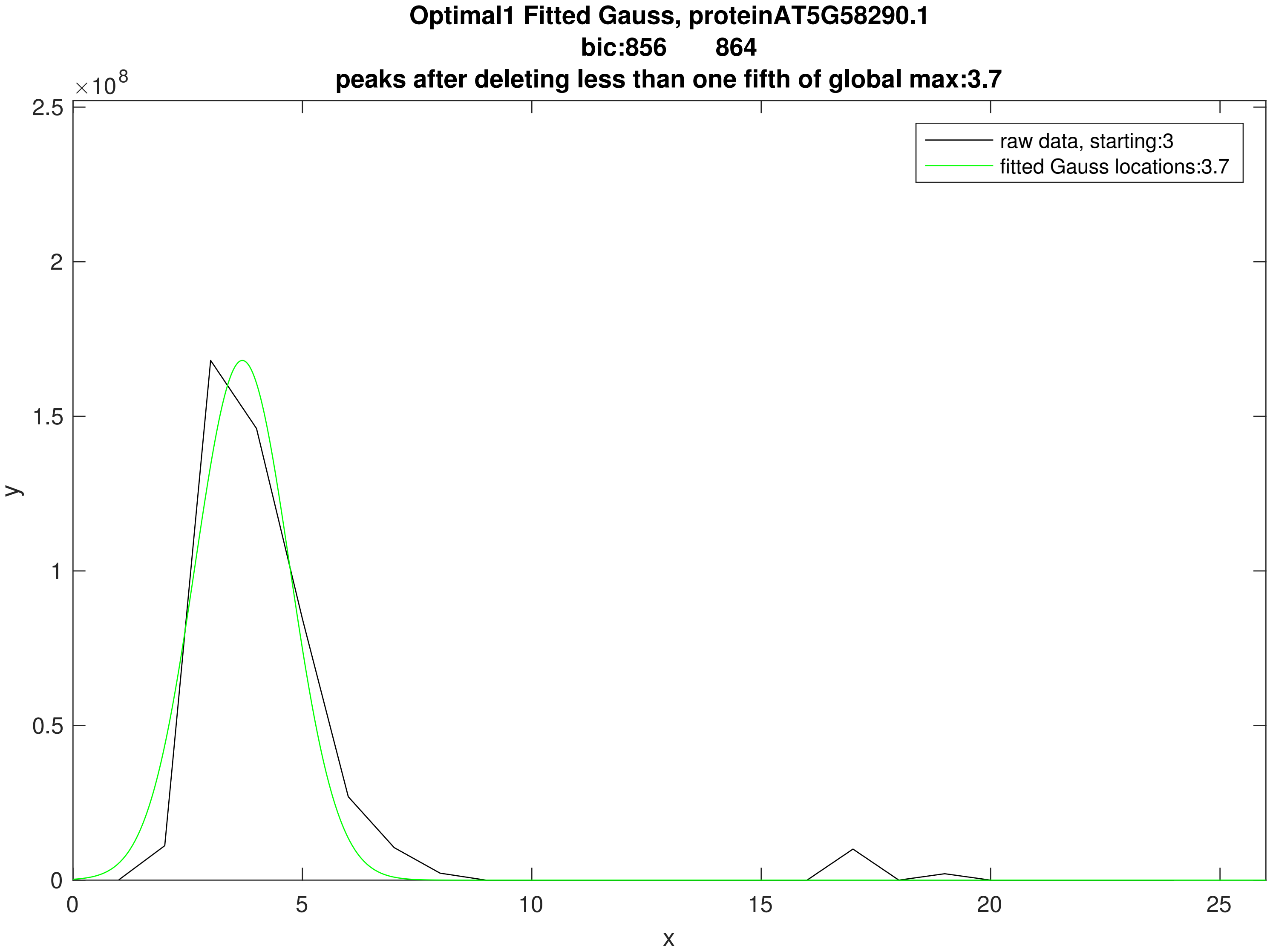
Optimal1 Fitted Gauss, proteinAT5G58290.1
bic:856 864
peaks after deleting less than one fifth of global max:3.7
raw data, starting:3
fitted Gauss locations:3.7
$\times 10^8$
2.5
2
1.5
1
0.5
0
y
0
5
10
15
20
25
x

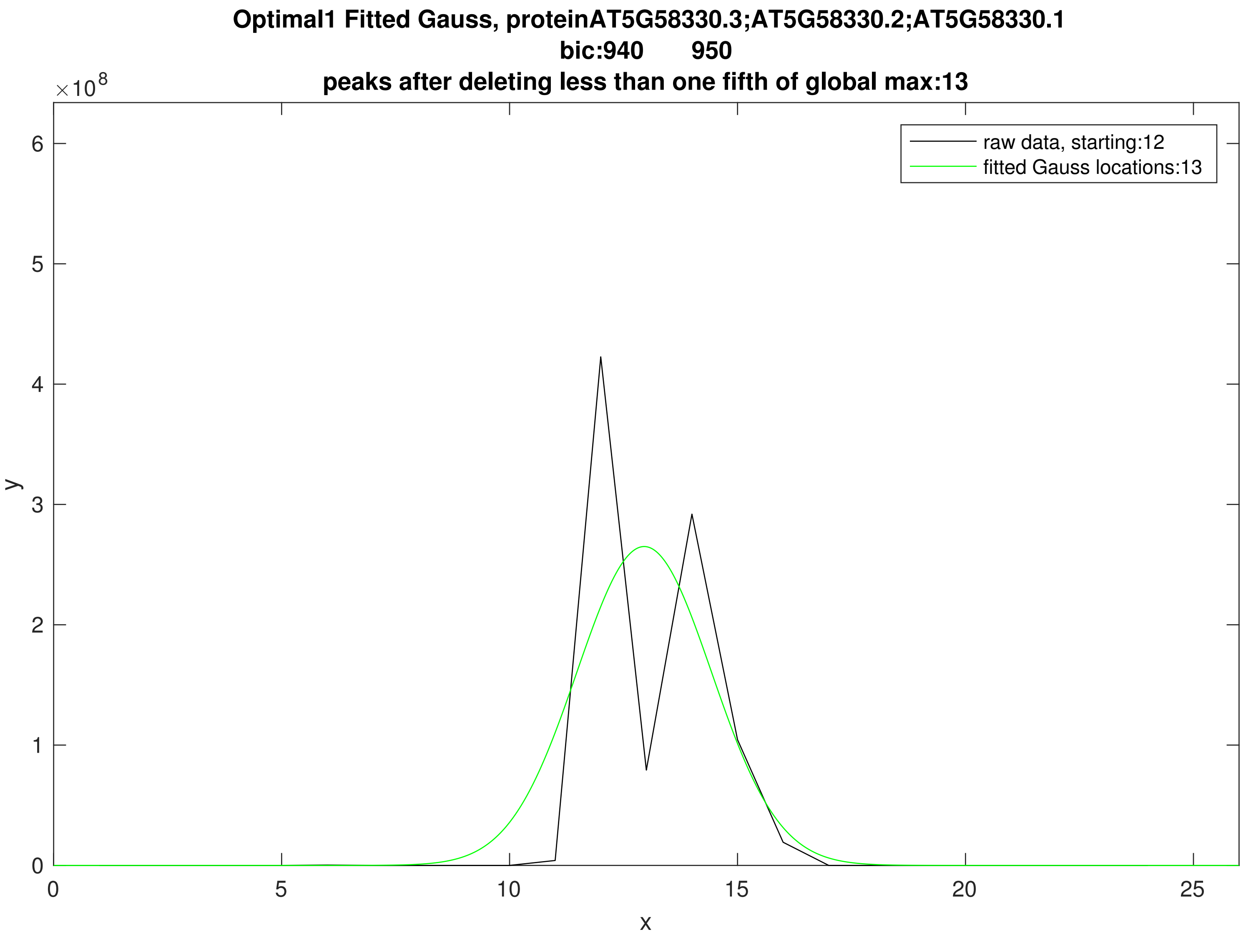
Optimal1 Fitted Gauss, proteinAT5G58330.3;AT5G58330.2;AT5G58330.1
bic:940 950
peaks after deleting less than one fifth of global max:13
×10^8
raw data, starting:12
fitted Gauss locations:13
y
x
0
1
2
3
4
5
6
0
5
10
15
20
25

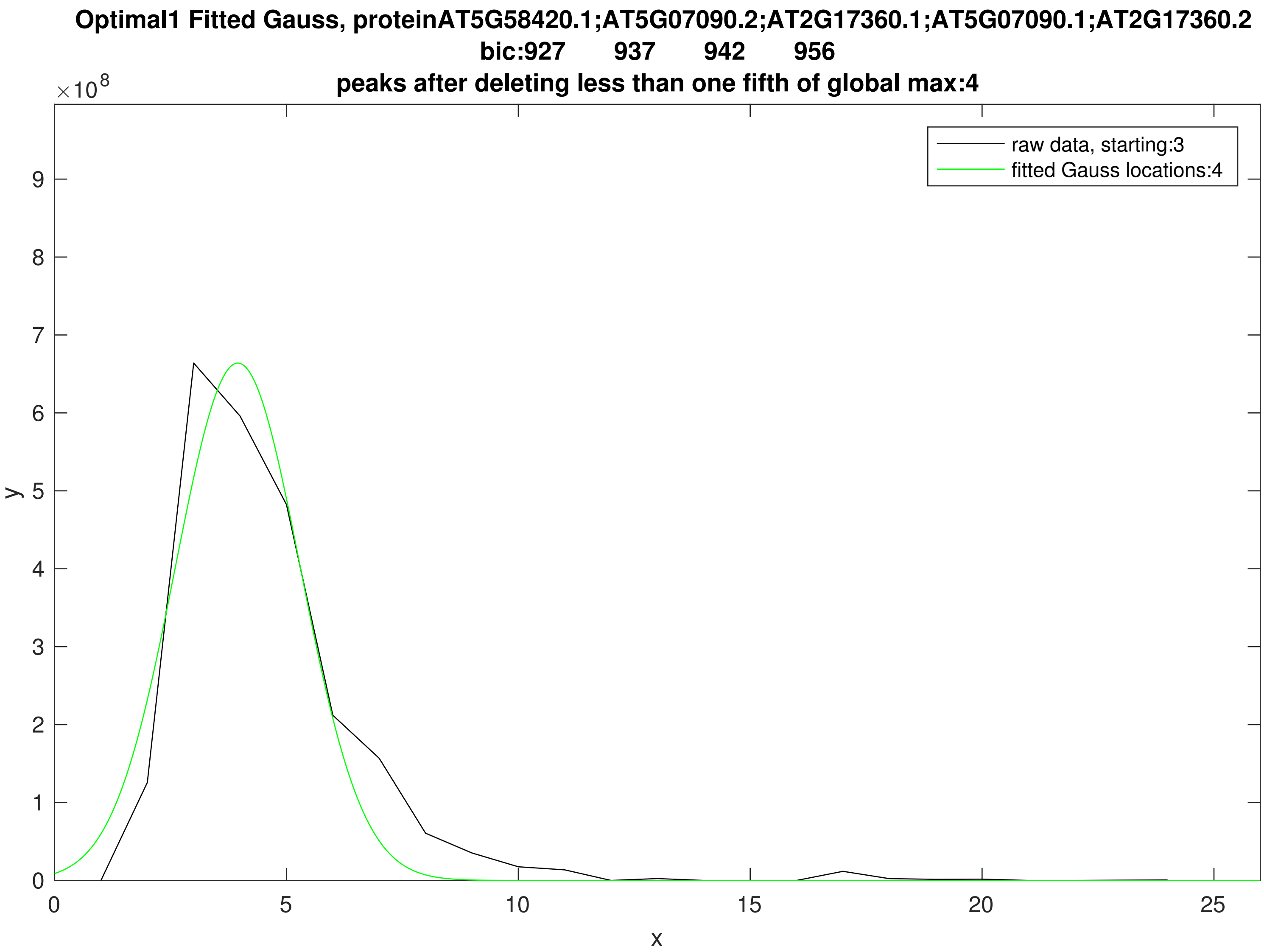

Optimal1 Fitted Gauss, proteinAT5G58420.1;AT5G07090.2;AT2G17360.1;AT5G07090.1;AT2G17360.2
bic:927 937 942 956
peaks after deleting less than one fifth of global max:4
×10^8
raw data, starting:3
fitted Gauss locations:4
y
x
0
1
2
3
4
5
6
7
8
9
0
5
10
15
20
25

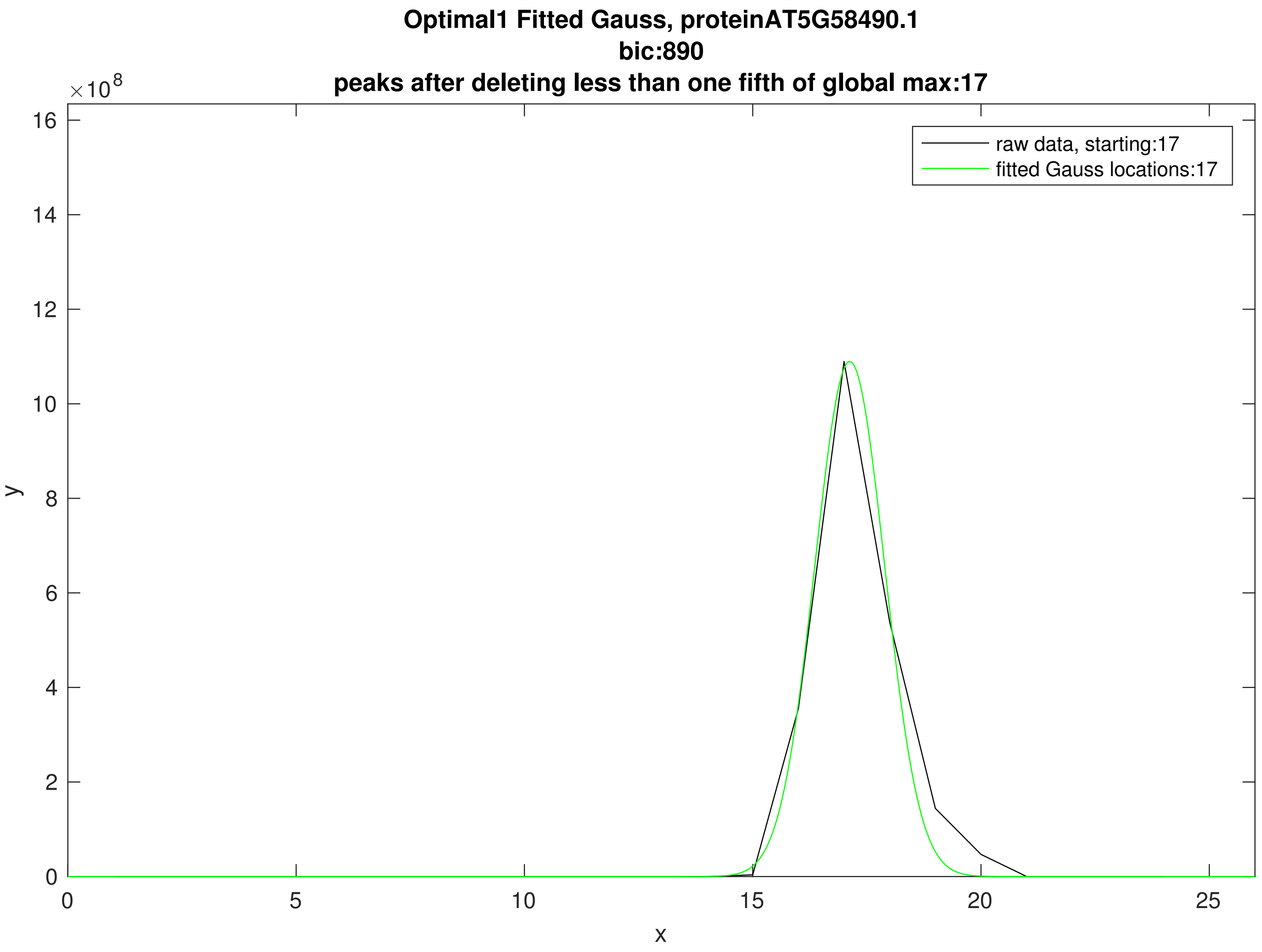

Optimal1 Fitted Gauss, proteinAT5G58490.1
bic:890
peaks after deleting less than one fifth of global max:17
×10^8
raw data, starting:17
fitted Gauss locations:17
y
x
0
2
4
6
8
10
12
14
16
0
5
10
15
20
25

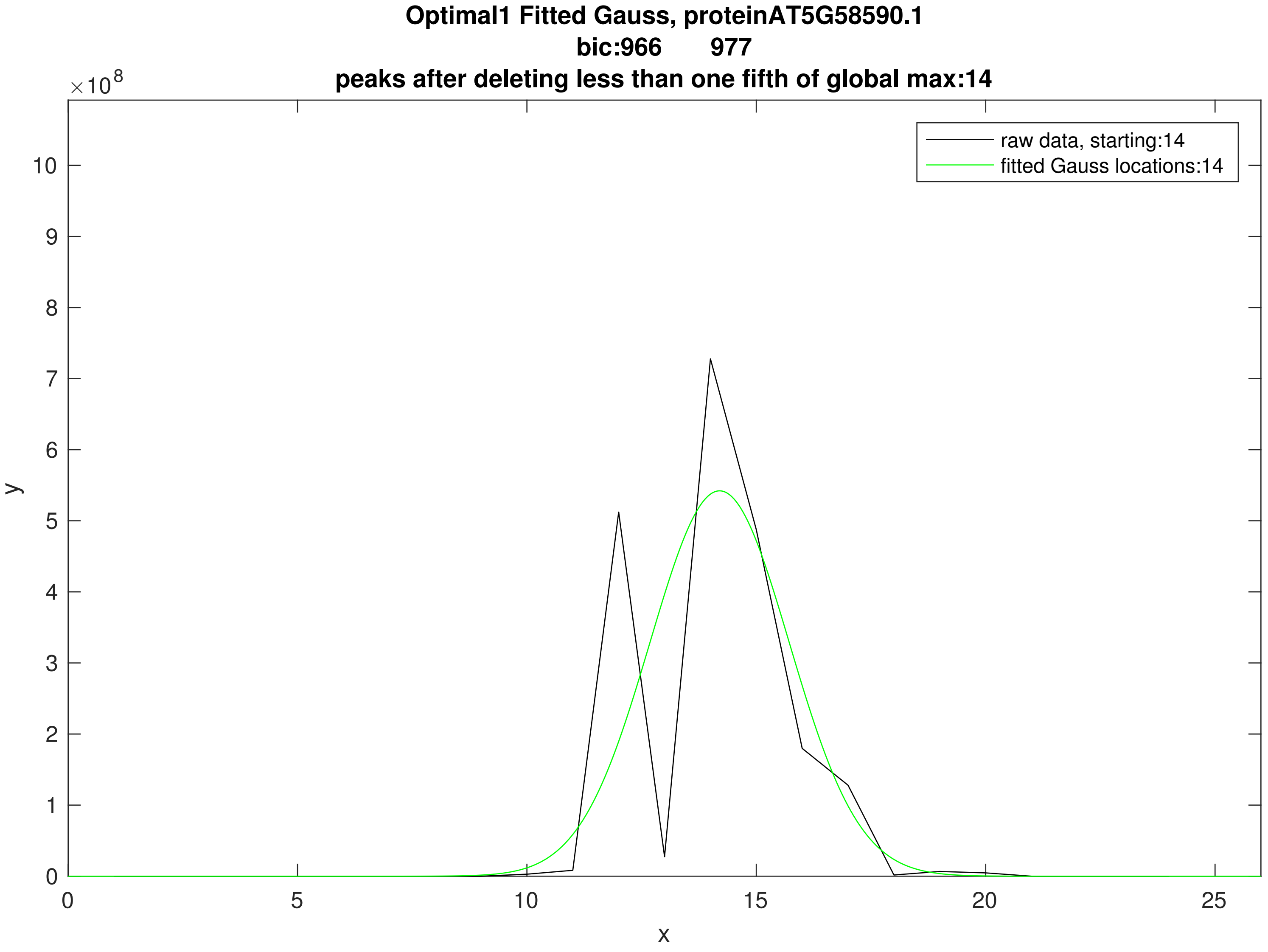

Optimal1 Fitted Gauss, proteinAT5G58590.1
bic:966 977
peaks after deleting less than one fifth of global max:14
×10^8
raw data, starting:14
fitted Gauss locations:14
y
x
0
1
2
3
4
5
6
7
8
9
10
0
5
10
15
20
25

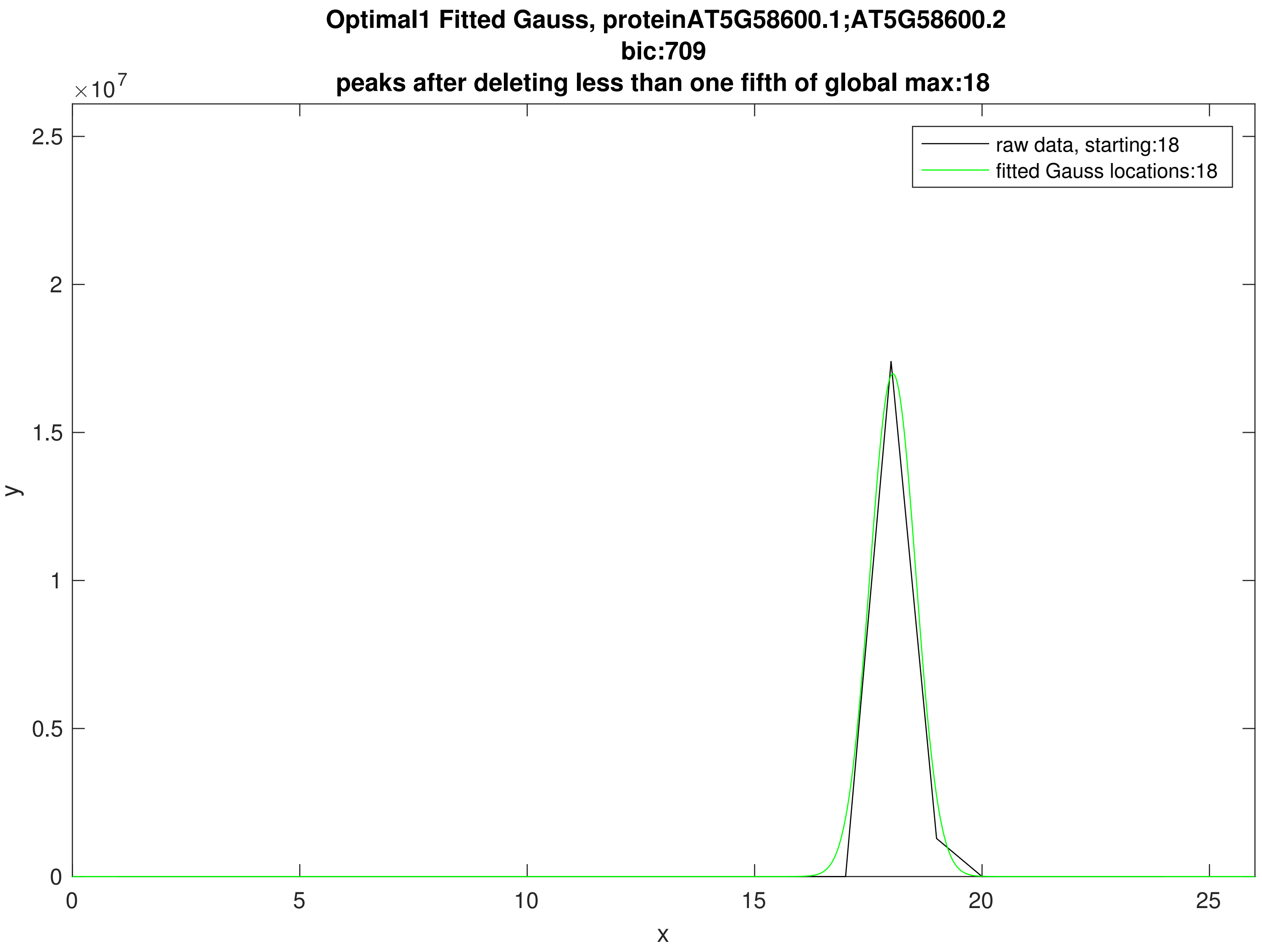
Optimal1 Fitted Gauss, proteinAT5G58600.1;AT5G58600.2
bic:709
peaks after deleting less than one fifth of global max:18
×10^7
2.5
2
1.5
1
0.5
0
y
0
5
10
15
20
25
x
raw data, starting:18
fitted Gauss locations:18

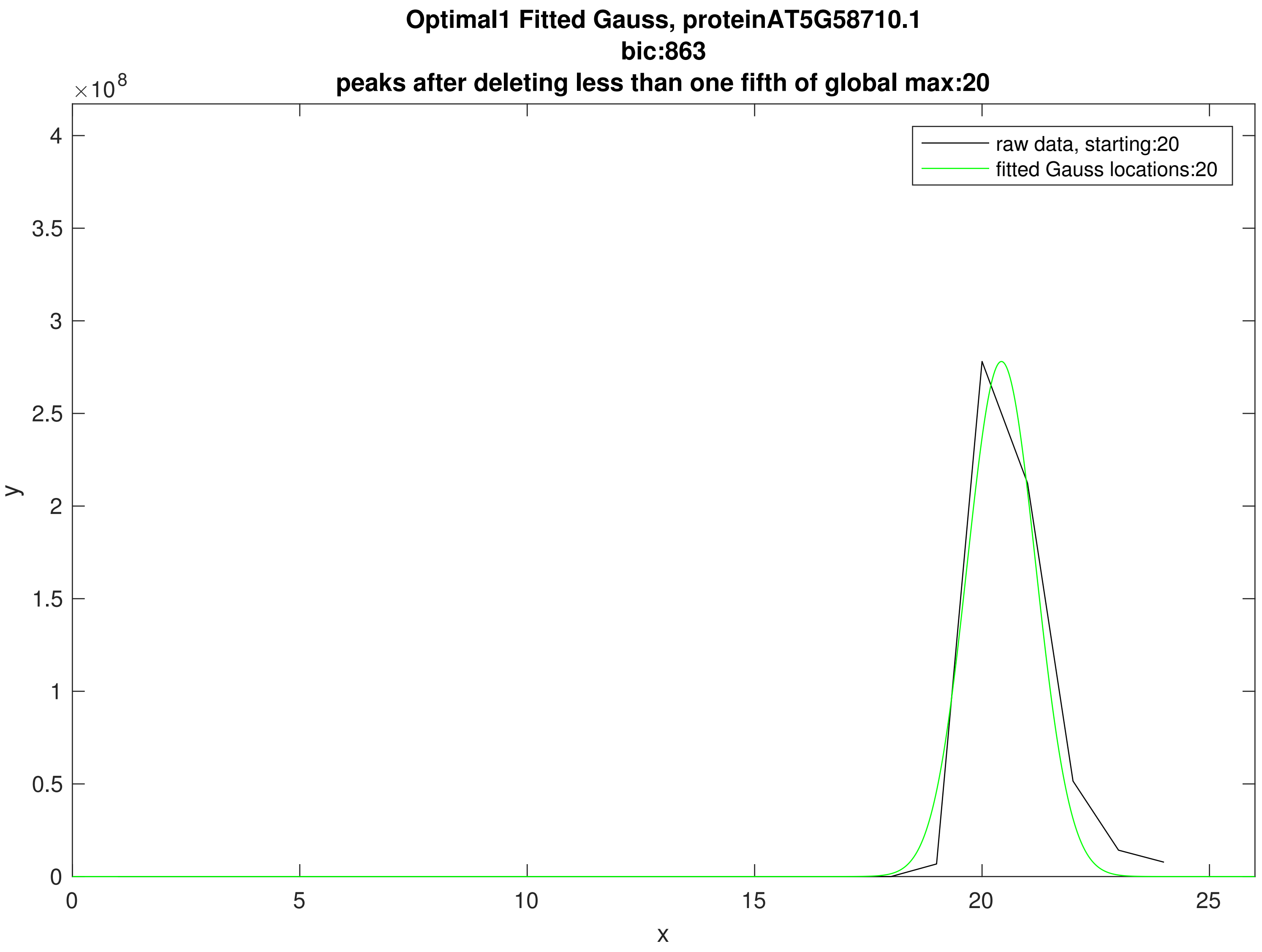
Optimal1 Fitted Gauss, proteinAT5G58710.1
bic:863
peaks after deleting less than one fifth of global max:20
×10^8
raw data, starting:20
fitted Gauss locations:20
y
x
0
0.5
1
1.5
2
2.5
3
3.5
4
5
10
15
20
25

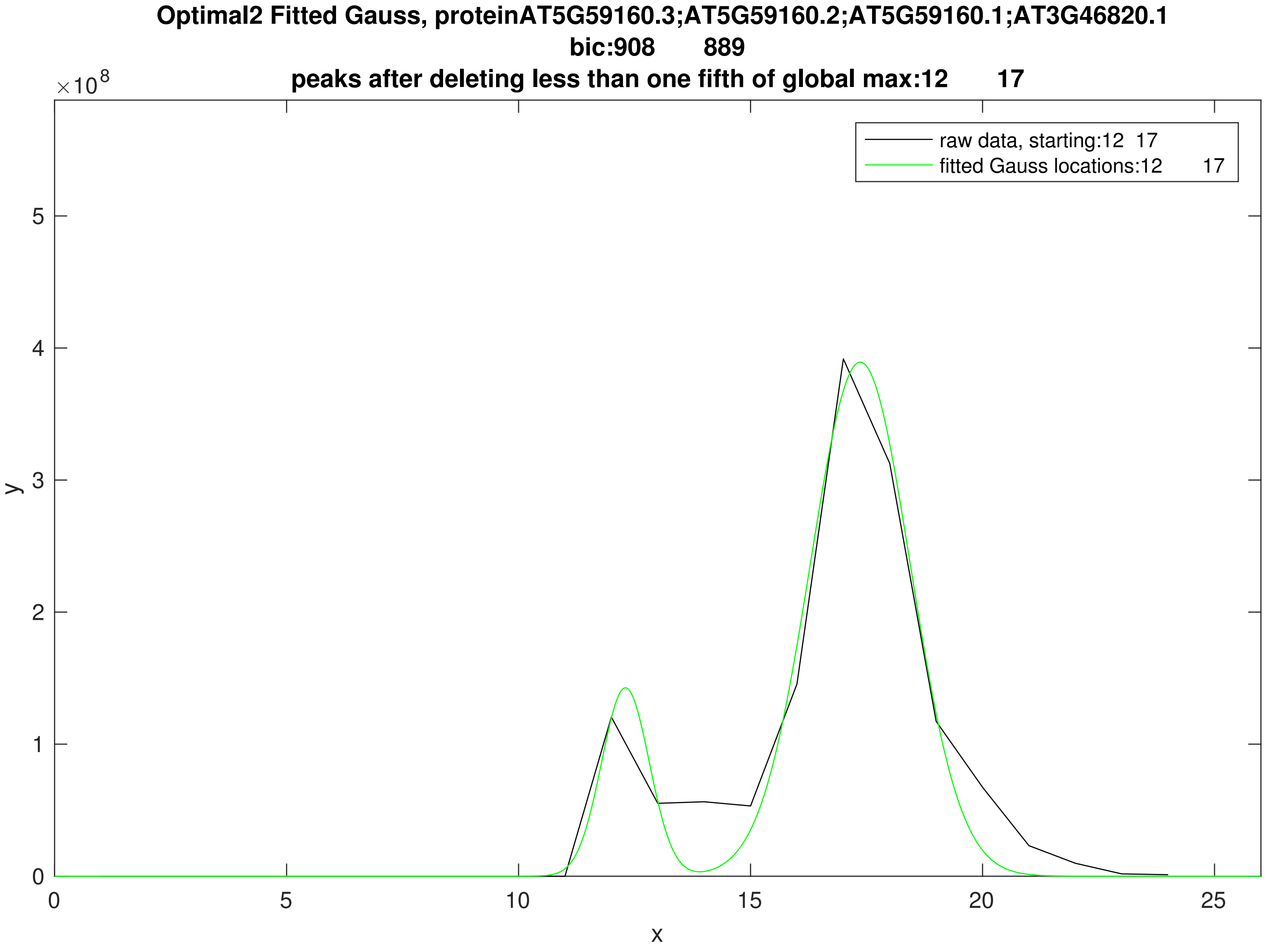

Optimal2 Fitted Gauss, proteinAT5G59160.3;AT5G59160.2;AT5G59160.1;AT3G46820.1
bic:908 889
peaks after deleting less than one fifth of global max:12 17
×10^8
raw data, starting:12 17
fitted Gauss locations:12 17
y
x
0
1
2
3
4
5
0
5
10
15
20
25

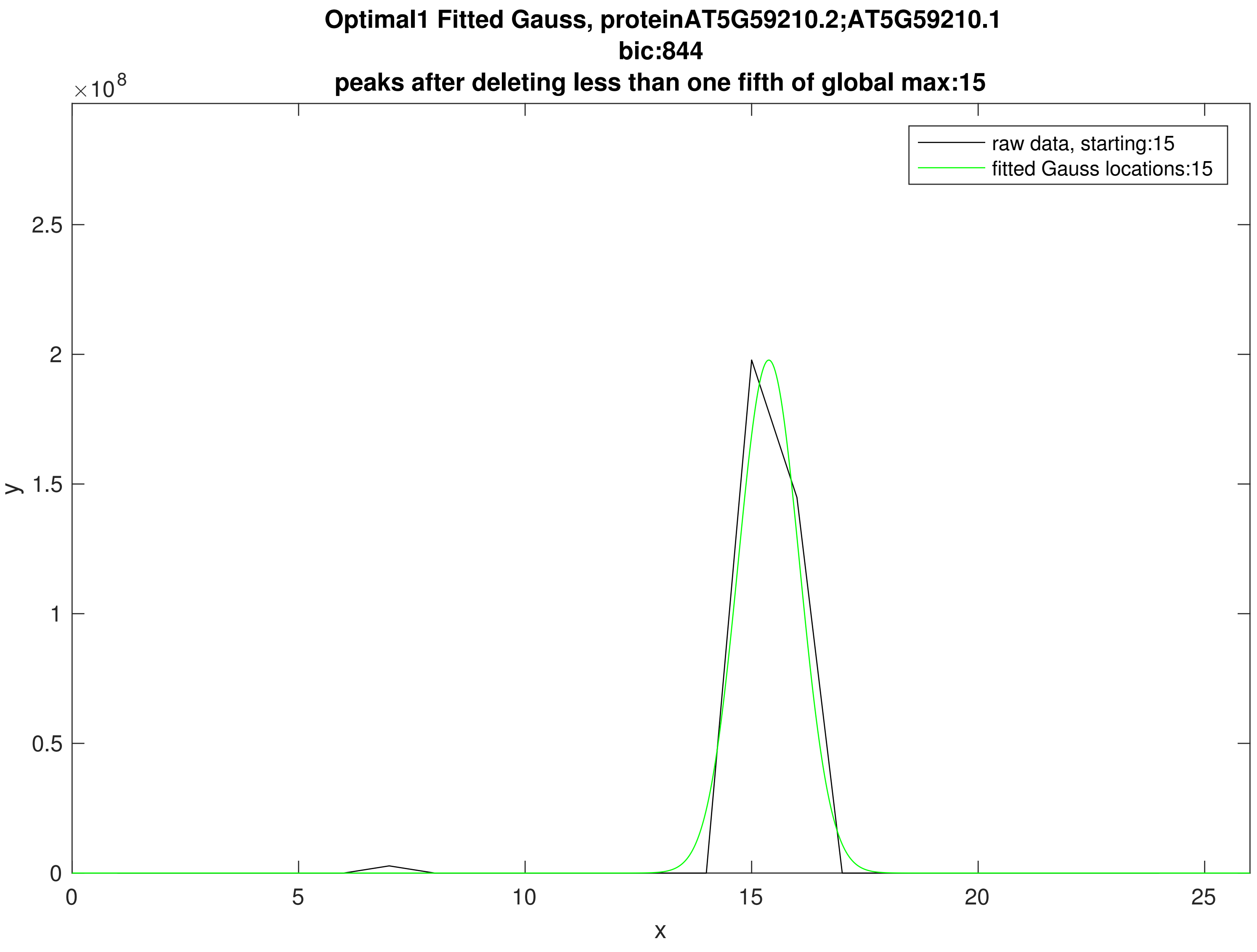
Optimal1 Fitted Gauss, proteinAT5G59210.2;AT5G59210.1
bic:844
peaks after deleting less than one fifth of global max:15
×10^8
raw data, starting:15
fitted Gauss locations:15
y
x
0
0.5
1
1.5
2
2.5
0
5
10
15
20
25

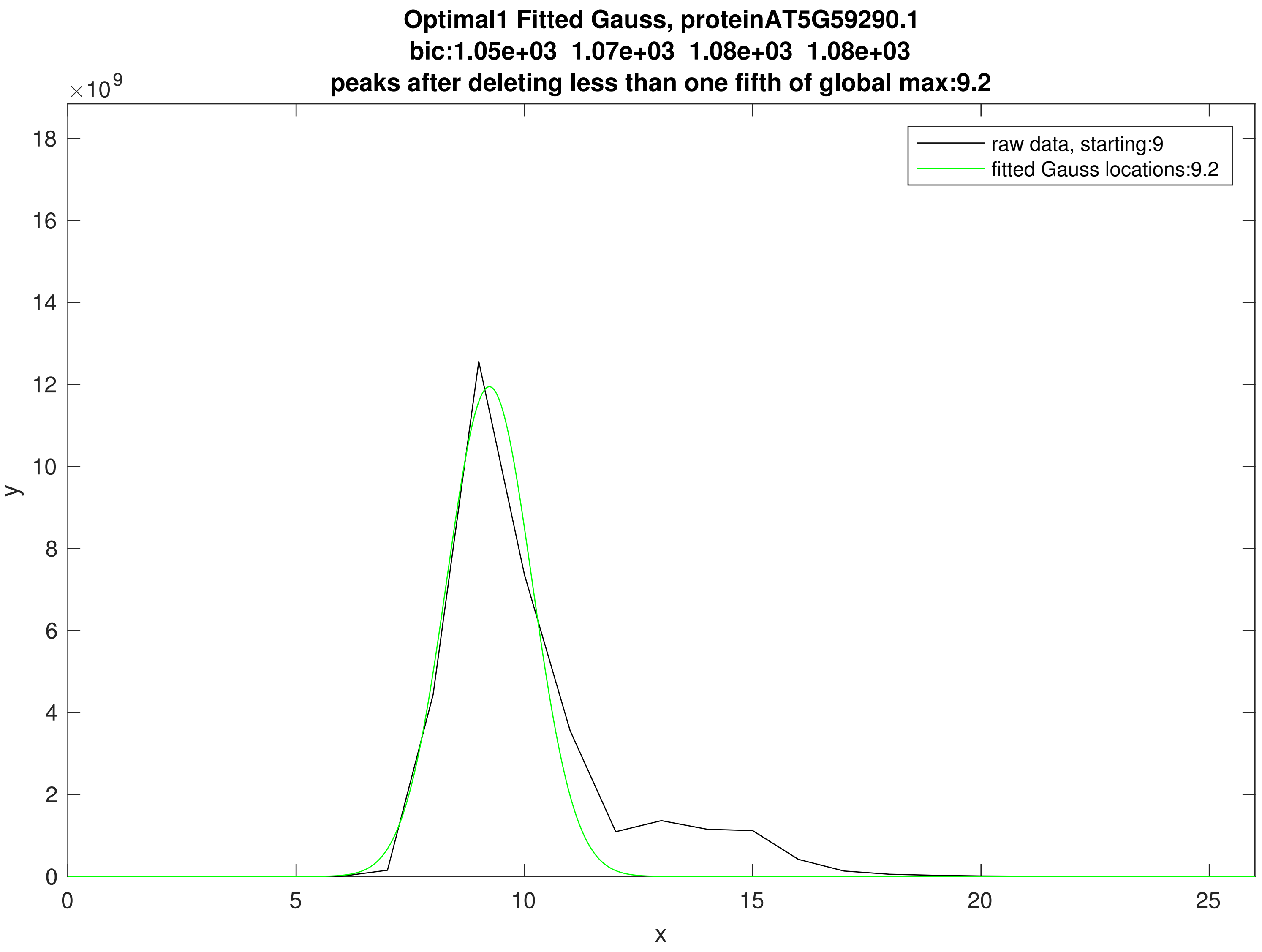
Optimal1 Fitted Gauss, proteinAT5G59290.1
bic:1.05e+03  1.07e+03  1.08e+03  1.08e+03
peaks after deleting less than one fifth of global max:9.2
×10^9
raw data, starting:9
fitted Gauss locations:9.2
y
x
0
2
4
6
8
10
12
14
16
18
0
5
10
15
20
25

Optimal1 Fitted Gauss, proteinAT5G59300.1
bic:743
peaks after deleting less than one fifth of global max:18

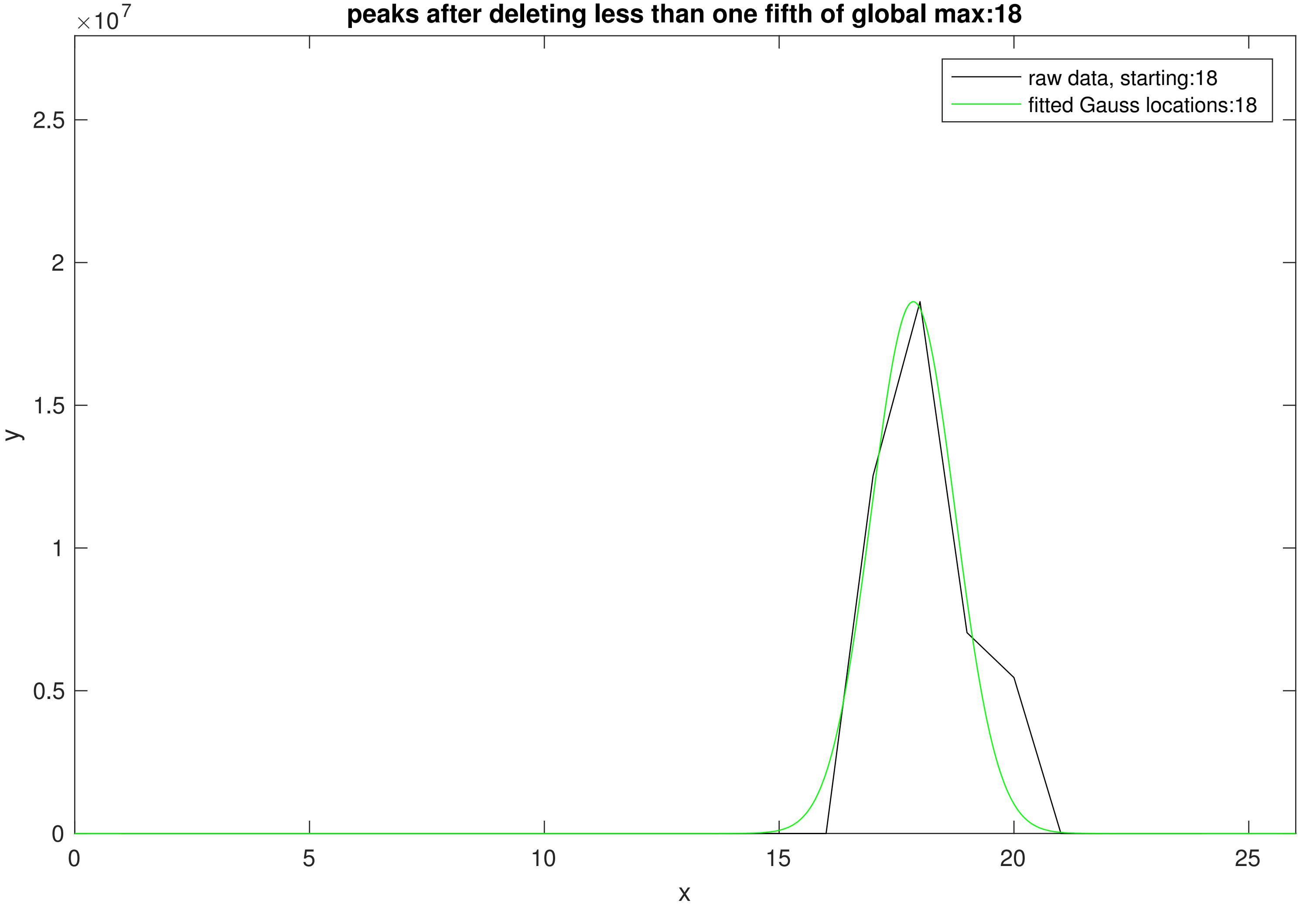

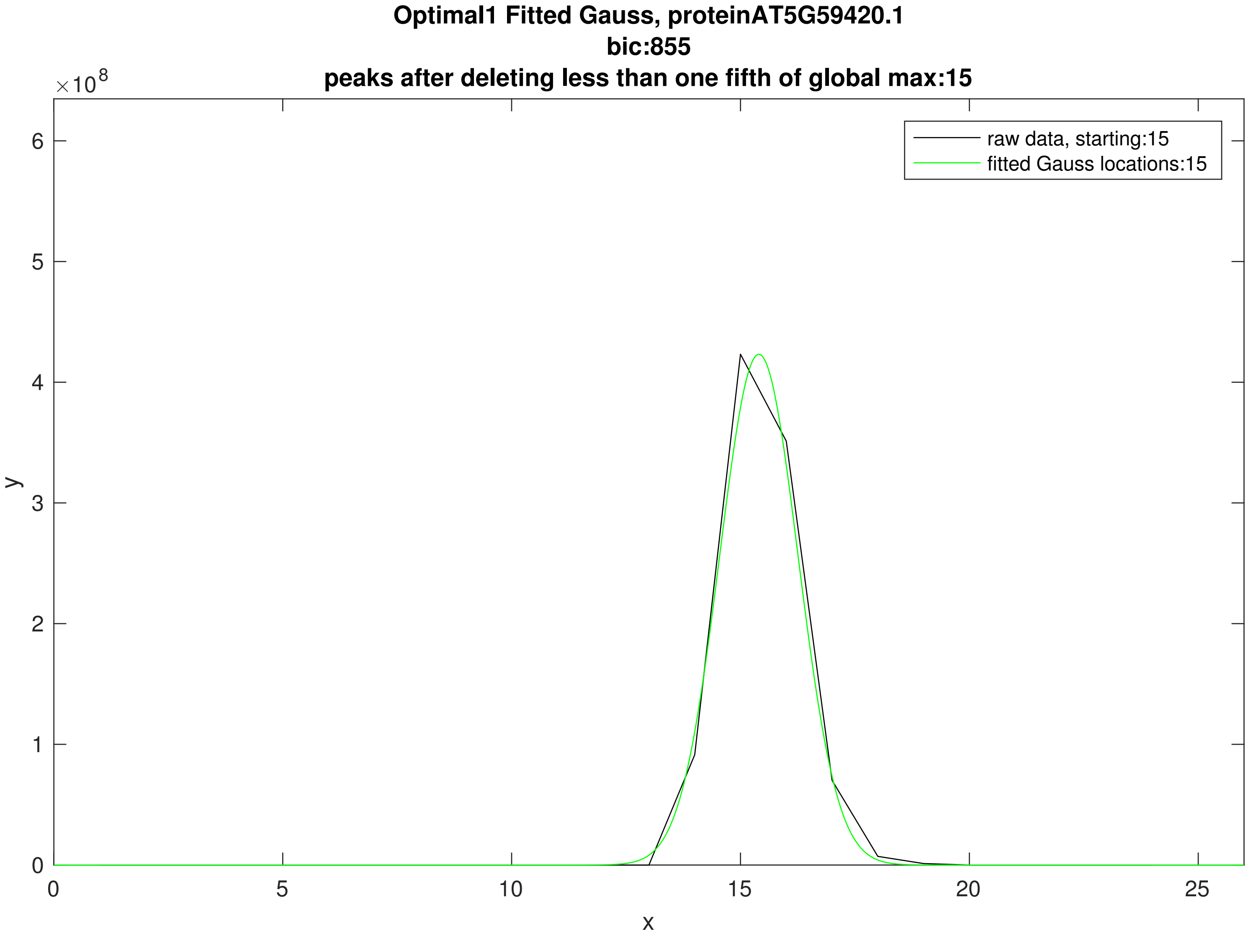
Optimal1 Fitted Gauss, proteinAT5G59420.1
bic:855
peaks after deleting less than one fifth of global max:15
×10^8
raw data, starting:15
fitted Gauss locations:15
0
1
2
3
4
5
6
y
0
5
10
15
20
25
x

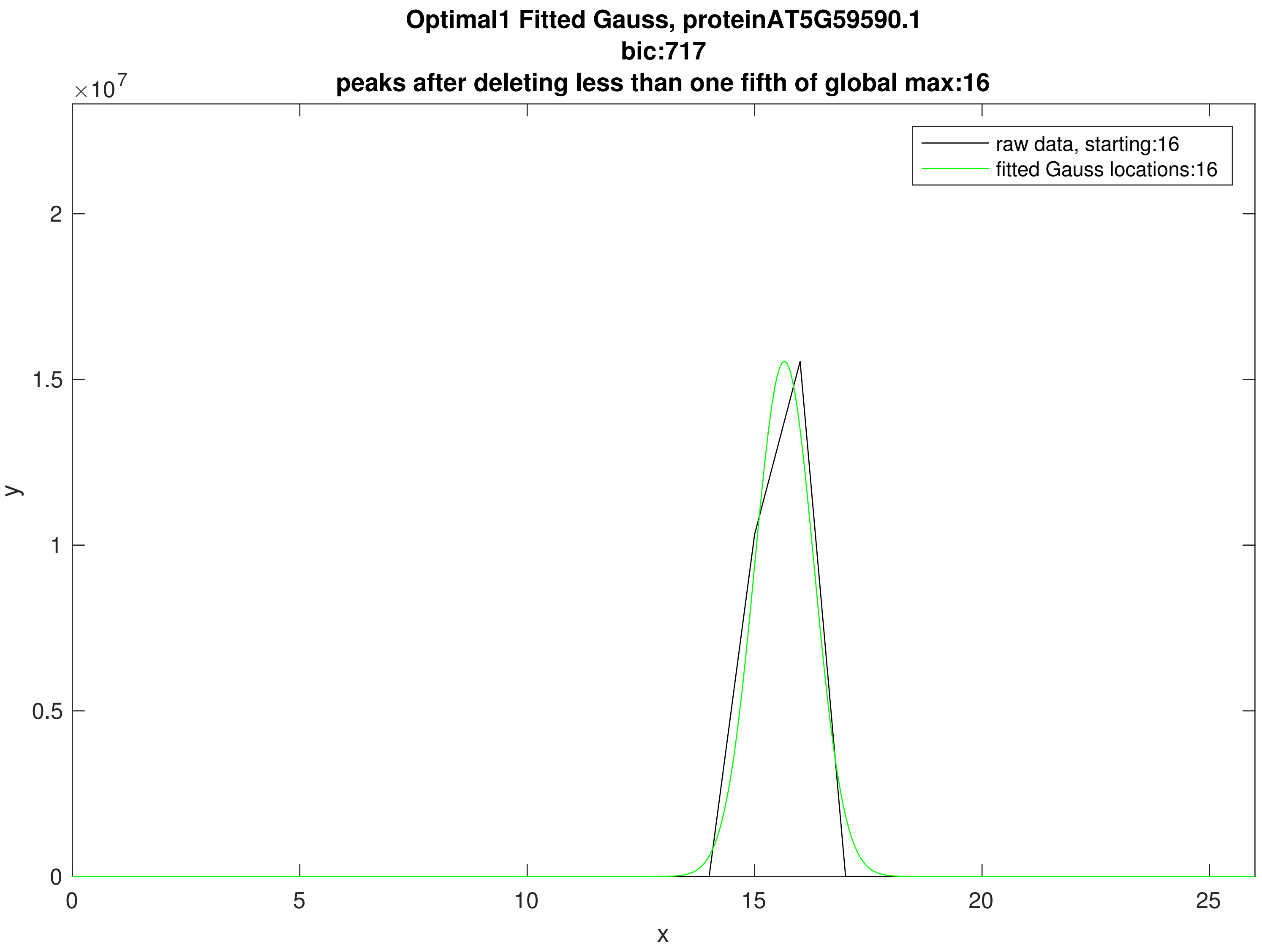

Optimal1 Fitted Gauss, proteinAT5G59590.1
bic:717
peaks after deleting less than one fifth of global max:16
×10^7
raw data, starting:16
fitted Gauss locations:16
y
x
0
0.5
1
1.5
2
0
5
10
15
20
25

#### Optimal1 Fitted Gauss, proteinAT5G59850.1;AT1G07770.2;AT1G07770.1
#### bic:863 872
#### peaks after deleting less than one fifth of global max:3.7

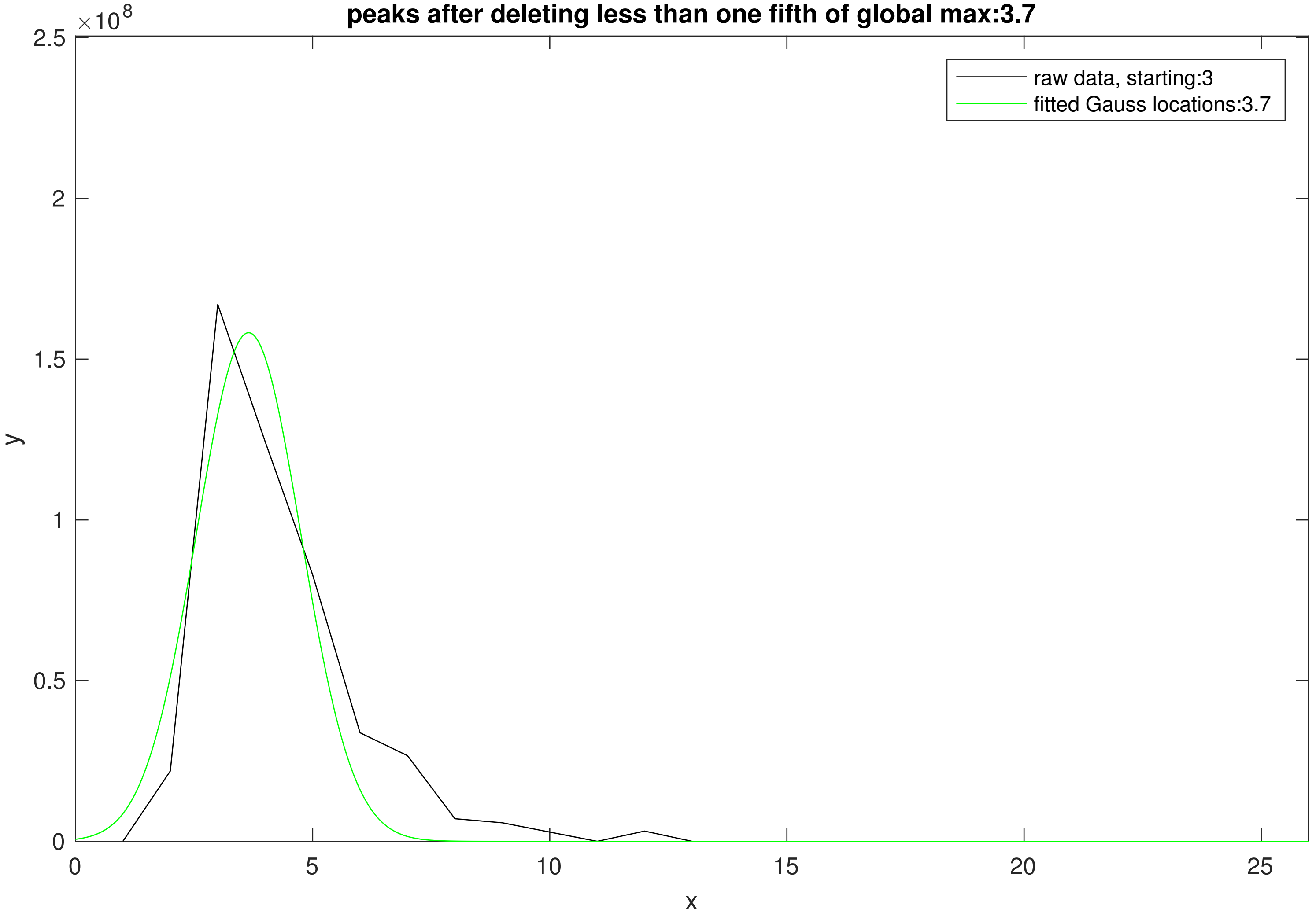

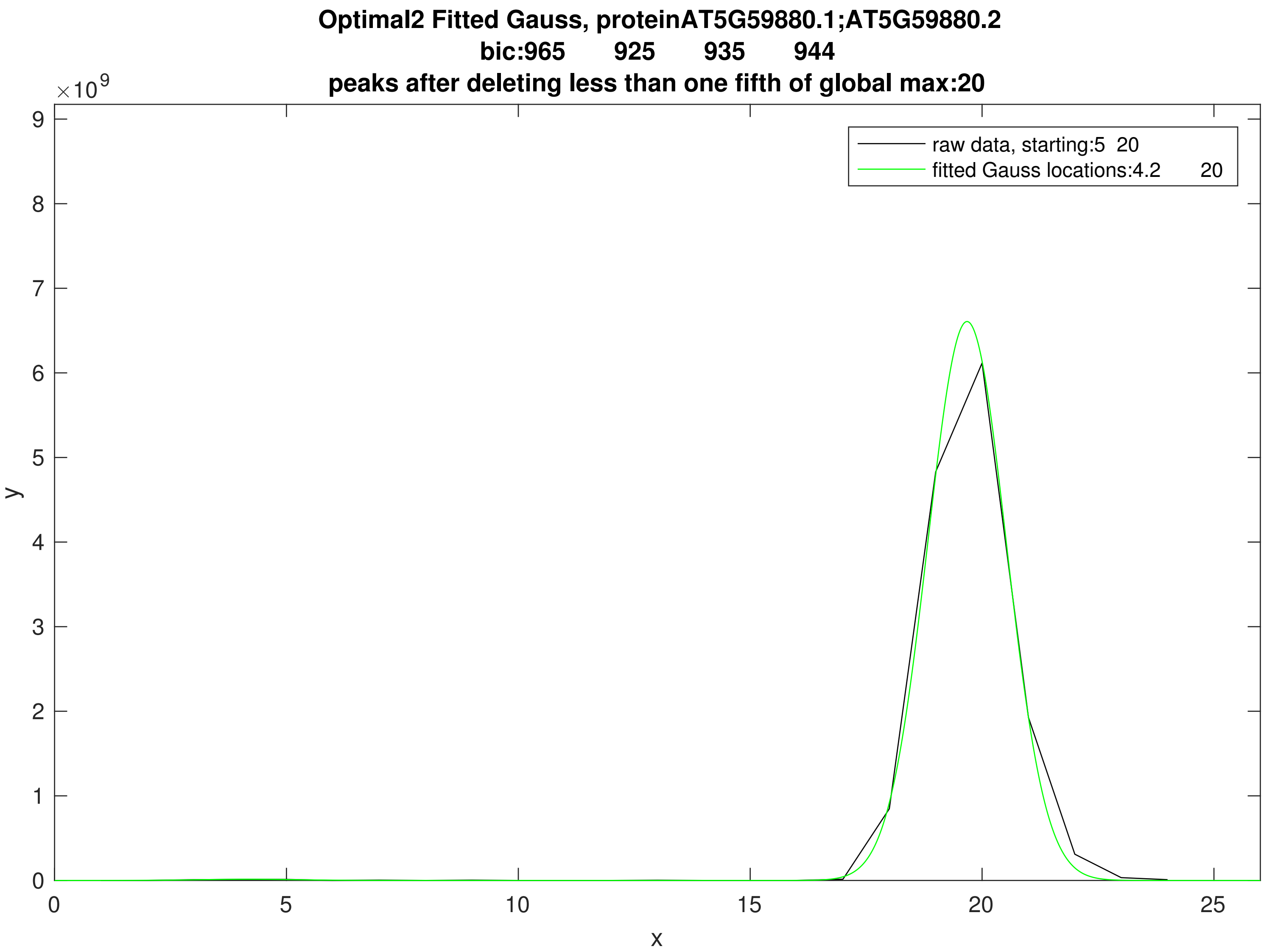
Optimal2 Fitted Gauss, proteinAT5G59880.1;AT5G59880.2
bic:965 925 935 944
peaks after deleting less than one fifth of global max:20
raw data, starting:5 20
fitted Gauss locations:4.2 20
×10^9
y
x
0
1
2
3
4
5
6
7
8
9
0
5
10
15
20
25

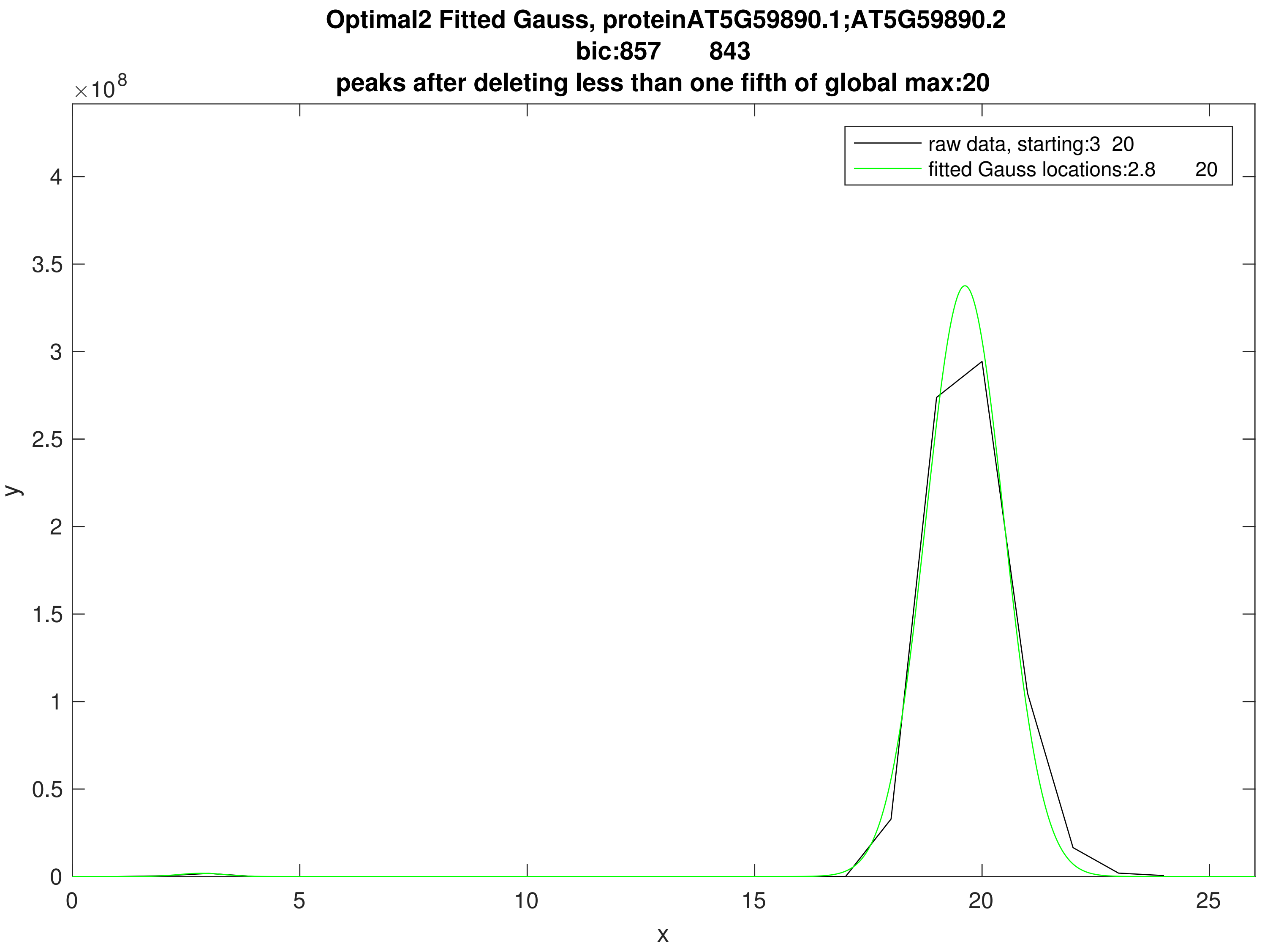

Optimal2 Fitted Gauss, proteinAT5G59890.1;AT5G59890.2
bic:857 843
peaks after deleting less than one fifth of global max:20
×10^8
raw data, starting:3 20
fitted Gauss locations:2.8 20
y
x
0
0.5
1
1.5
2
2.5
3
3.5
4
5
10
15
20
25

s, proteinAT5G59970.1;AT5G59690.1;AT3G53730.1;AT3G46320.1;AT3G45930.1;AT2G28740.1;AT1G07820.2;AT1G0782
bic:681 710
peaks after deleting less than one fifth of global max:2.9

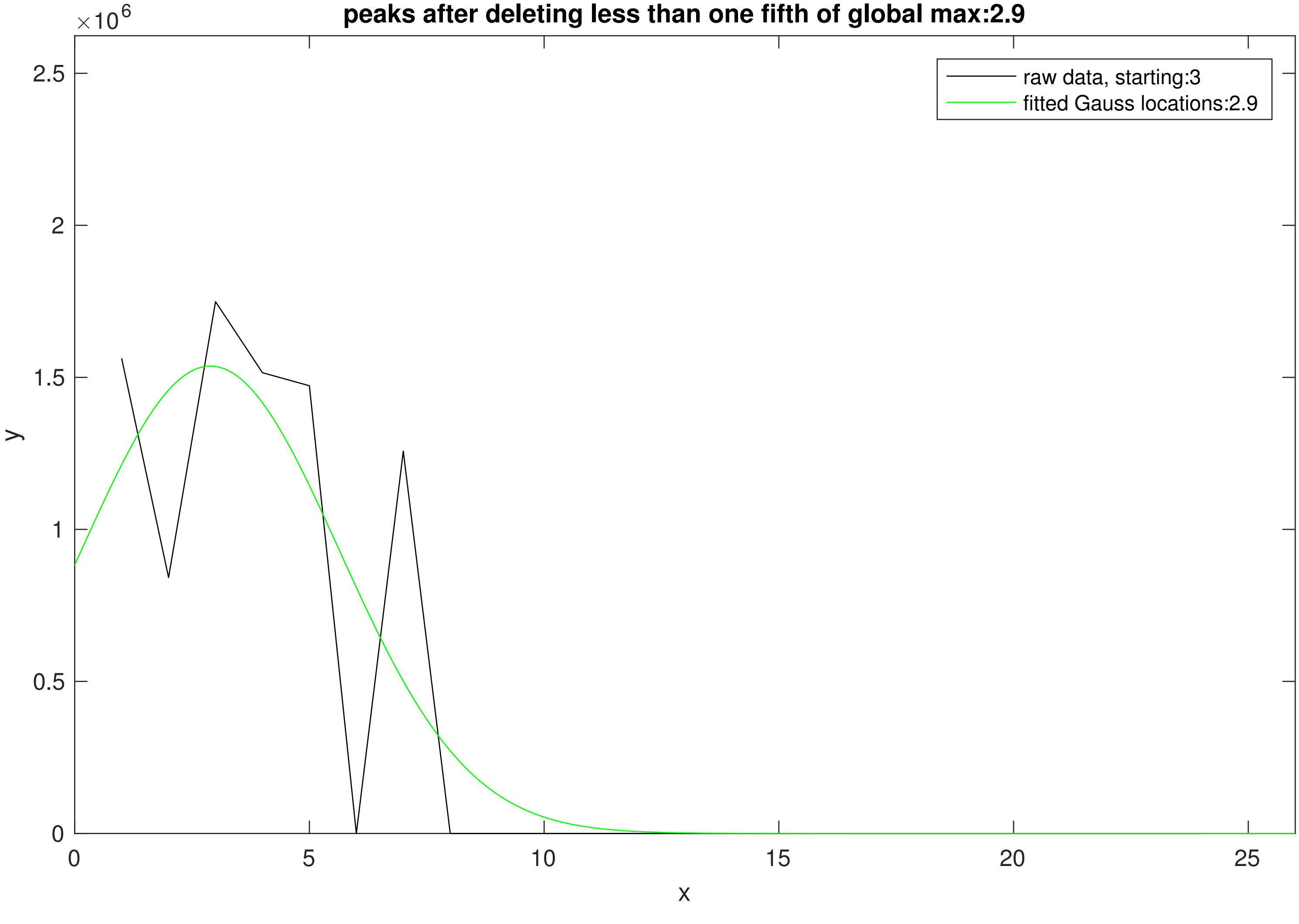

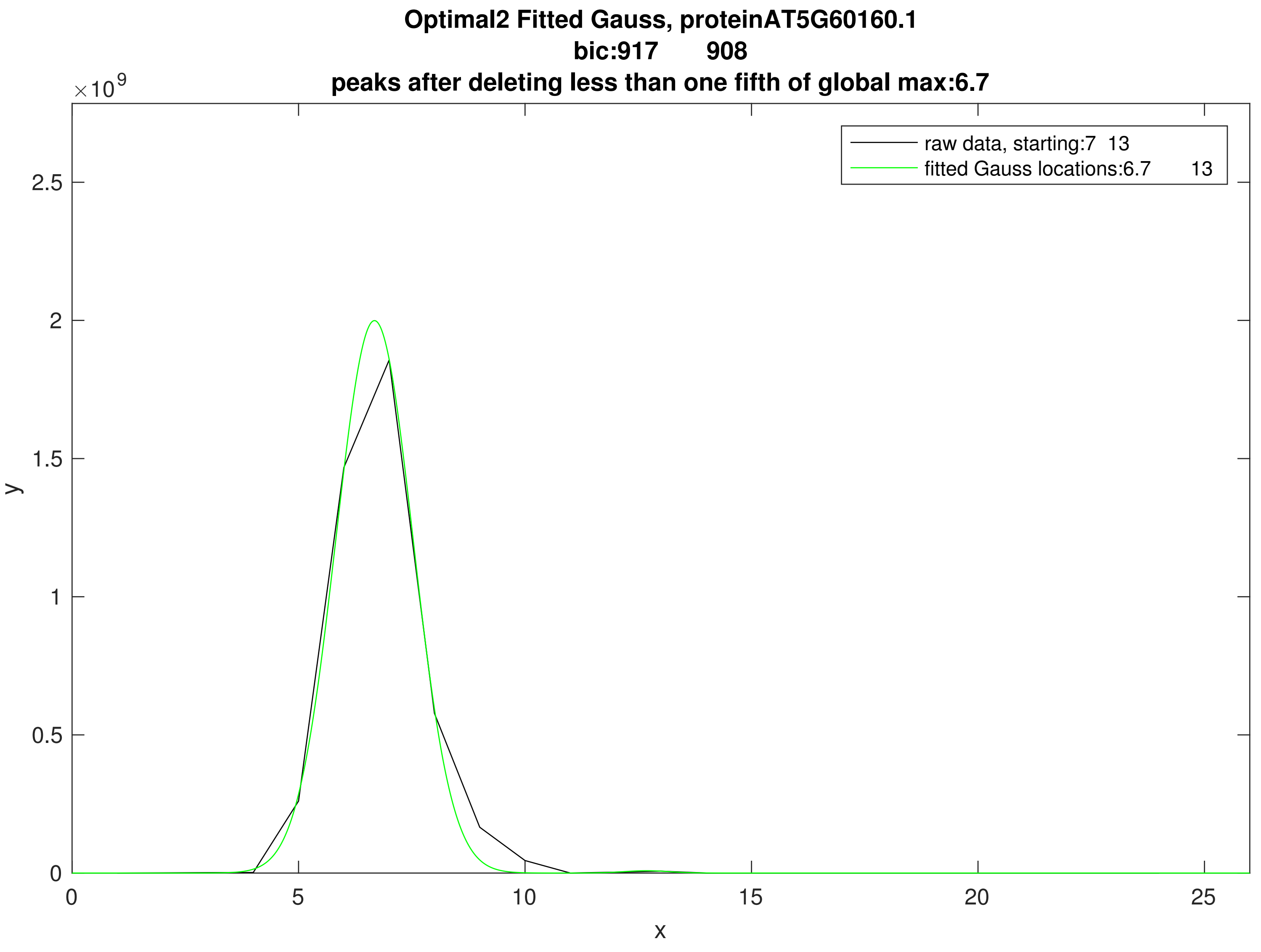

Optimal2 Fitted Gauss, proteinAT5G60160.1
bic:917 908
peaks after deleting less than one fifth of global max:6.7
×10^9
raw data, starting:7 13
fitted Gauss locations:6.7 13
y
x
0
0.5
1
1.5
2
2.5
5
10
15
20
25

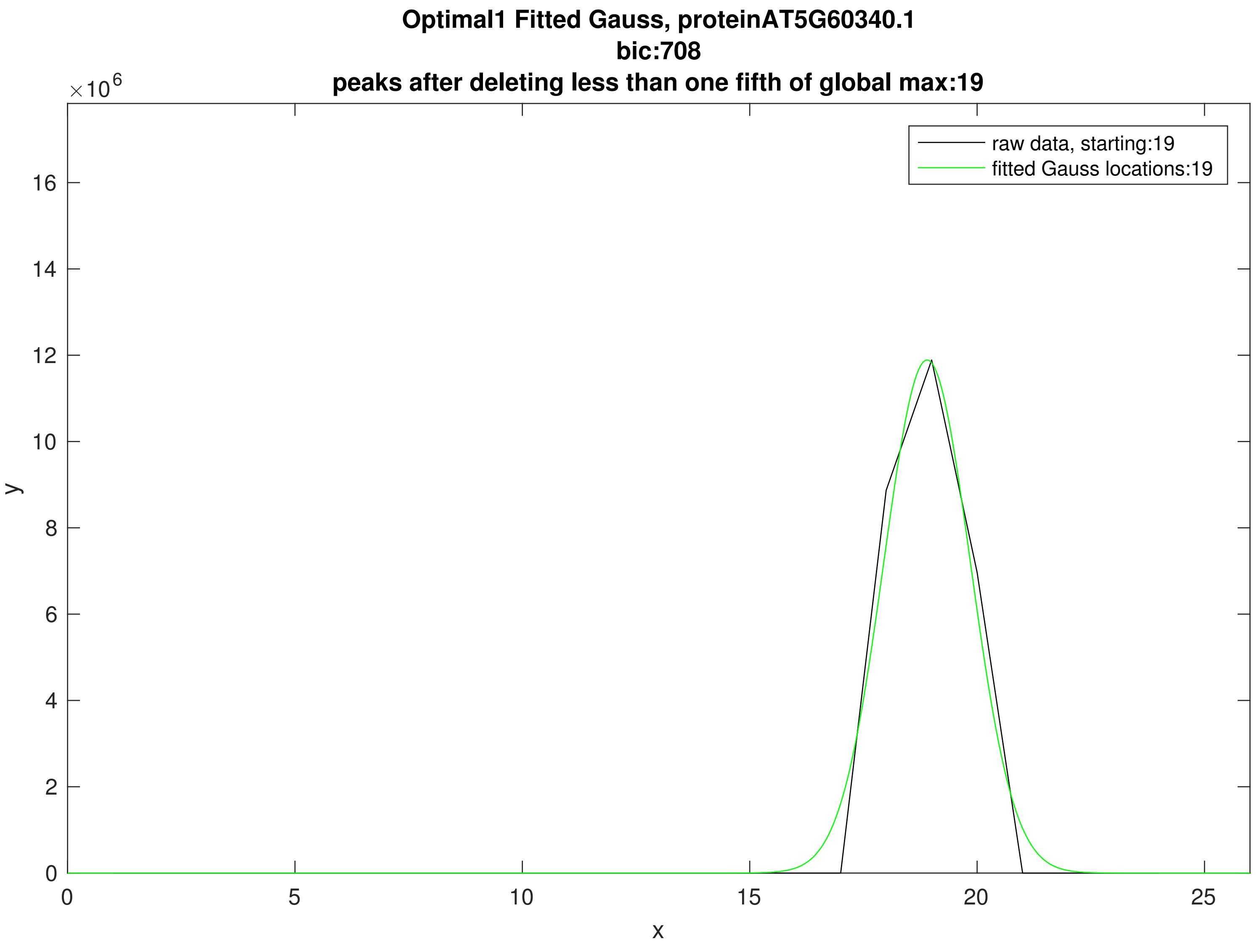
Optimal1 Fitted Gauss, proteinAT5G60340.1
bic:708
peaks after deleting less than one fifth of global max:19
×10^6
raw data, starting:19
fitted Gauss locations:19
y
x
0
2
4
6
8
10
12
14
16
0
5
10
15
20
25

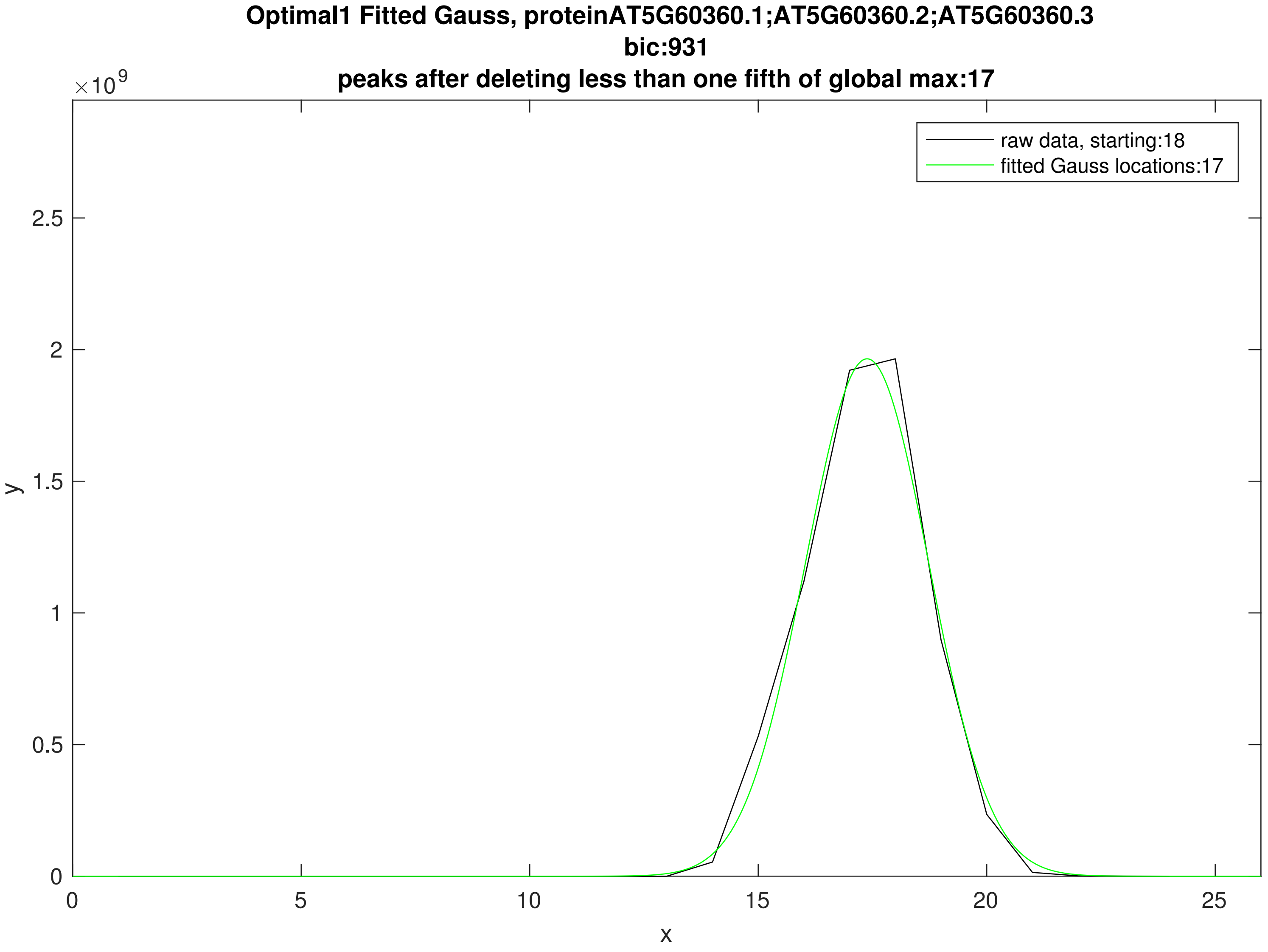
Optimal1 Fitted Gauss, proteinAT5G60360.1;AT5G60360.2;AT5G60360.3
bic:931
peaks after deleting less than one fifth of global max:17
×10^9
raw data, starting:18
fitted Gauss locations:17
y
x
0
0.5
1
1.5
2
2.5
0
5
10
15
20
25

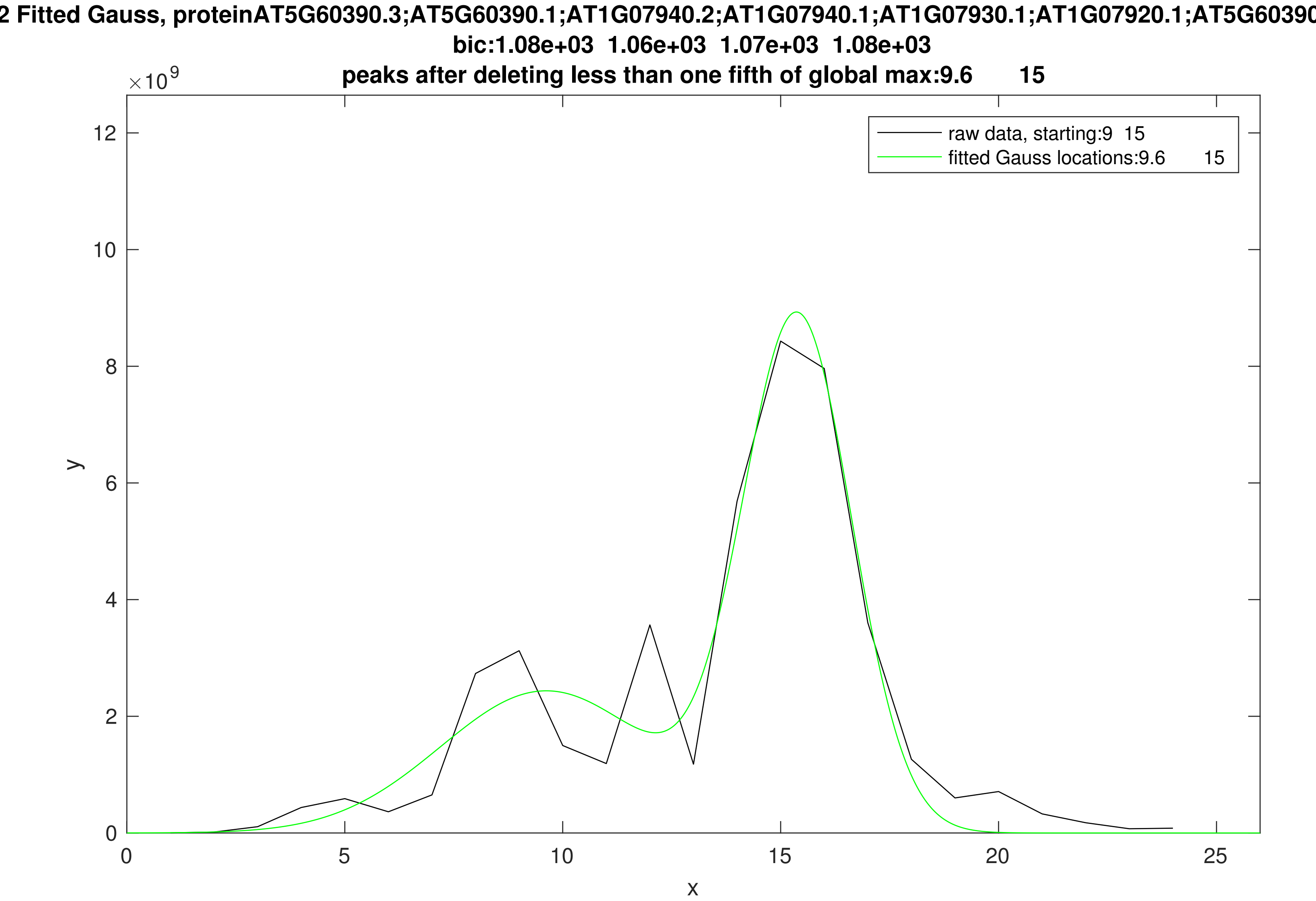
2 Fitted Gauss, proteinAT5G60390.3;AT5G60390.1;AT1G07940.2;AT1G07940.1;AT1G07930.1;AT1G07920.1;AT5G60390
bic:1.08e+03 1.06e+03 1.07e+03 1.08e+03
peaks after deleting less than one fifth of global max:9.6 15
×10^9
raw data, starting:9 15
fitted Gauss locations:9.6 15
y
x
0
2
4
6
8
10
12
0
5
10
15
20
25

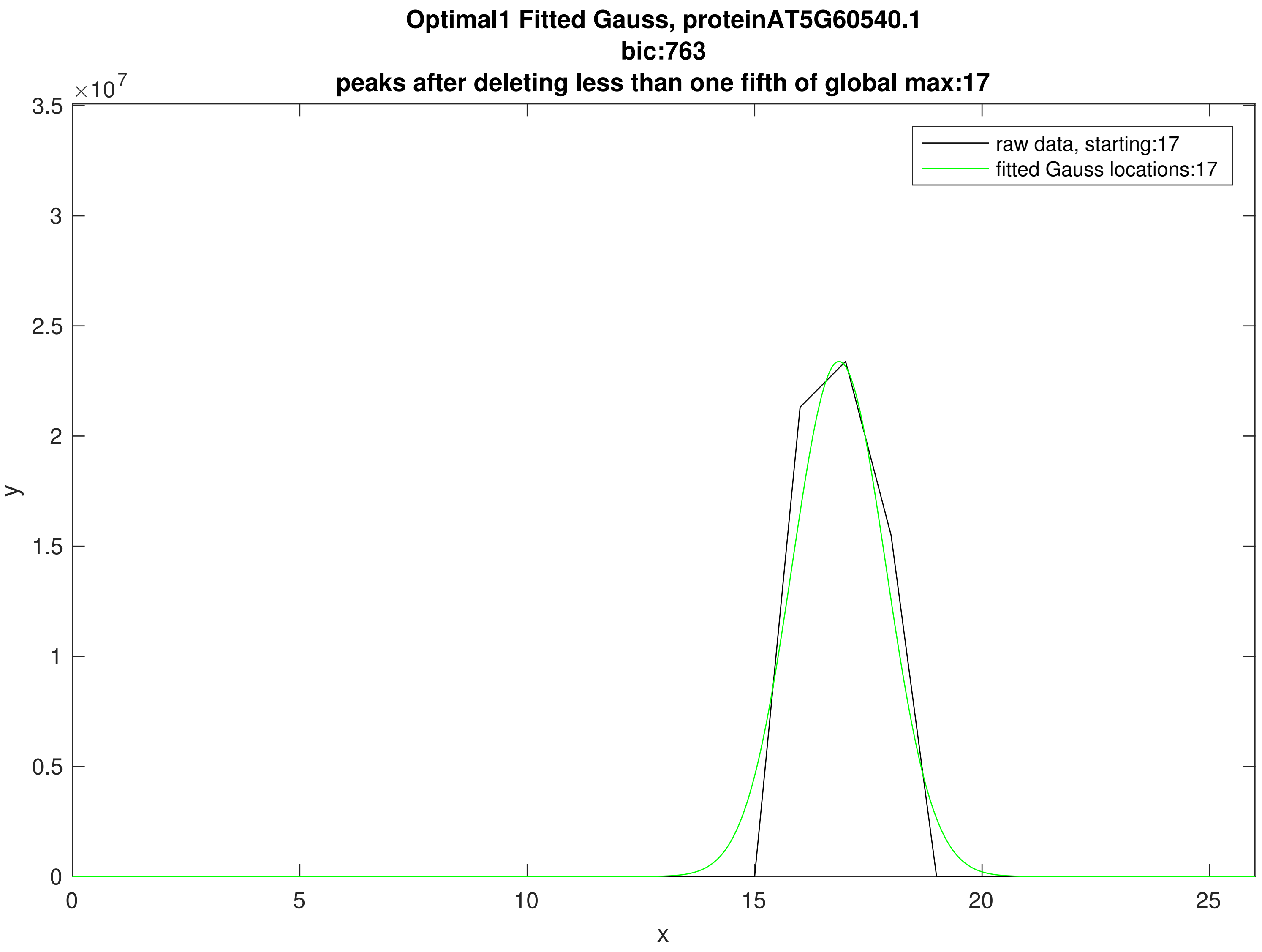

Optimal1 Fitted Gauss, proteinAT5G60540.1
bic:763
peaks after deleting less than one fifth of global max:17
×10^7
3.5
3
2.5
2
1.5
1
0.5
0
y
0
5
10
15
20
25
x
raw data, starting:17
fitted Gauss locations:17

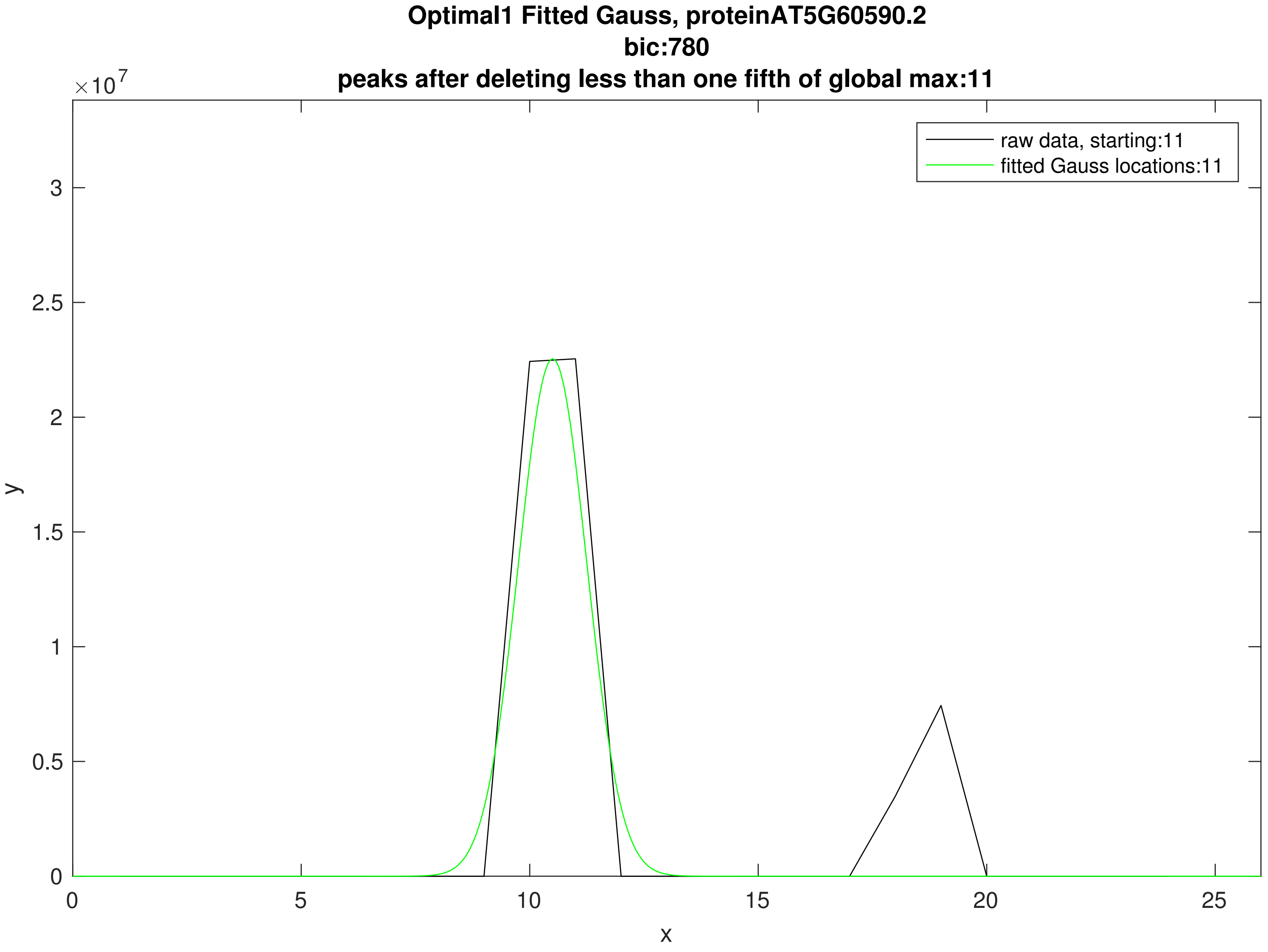

Optimal1 Fitted Gauss, proteinAT5G60590.2
bic:780
peaks after deleting less than one fifth of global max:11
×10^7
raw data, starting:11
fitted Gauss locations:11
y
x
0
0.5
1
1.5
2
2.5
3
5
10
15
20
25

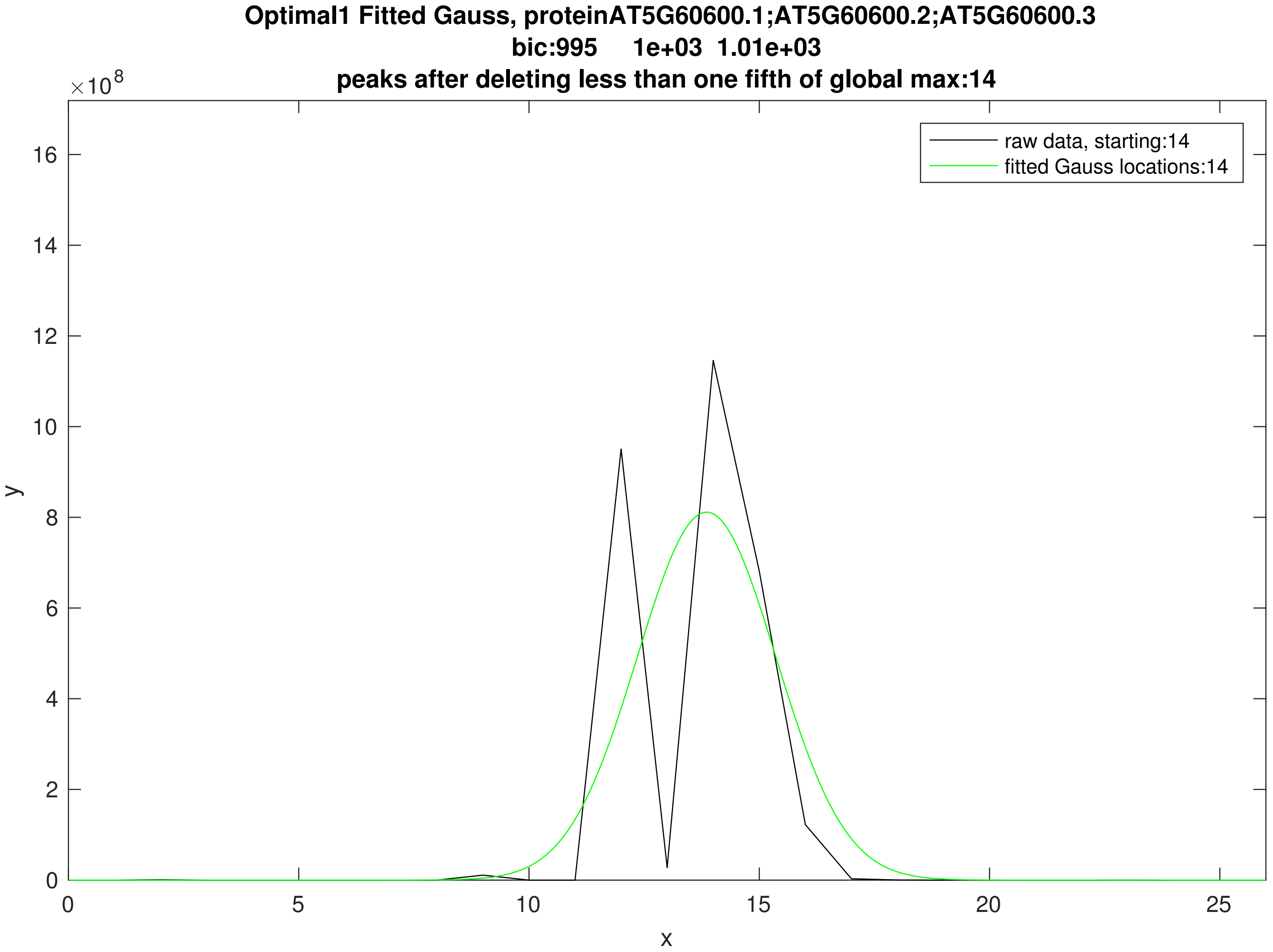

Optimal1 Fitted Gauss, proteinAT5G60600.1;AT5G60600.2;AT5G60600.3
bic:995 1e+03 1.01e+03
peaks after deleting less than one fifth of global max:14
×10^8
raw data, starting:14
fitted Gauss locations:14
y
x
0
2
4
6
8
10
12
14
16
0
5
10
15
20
25

**Optimal2 Fitted Gauss, proteinAT5G60640.1;AT5G60640.3;AT5G60640.2**
**bic:1.02e+03  1.02e+03  1.03e+03**
**peaks after deleting less than one fifth of global max:11       17**

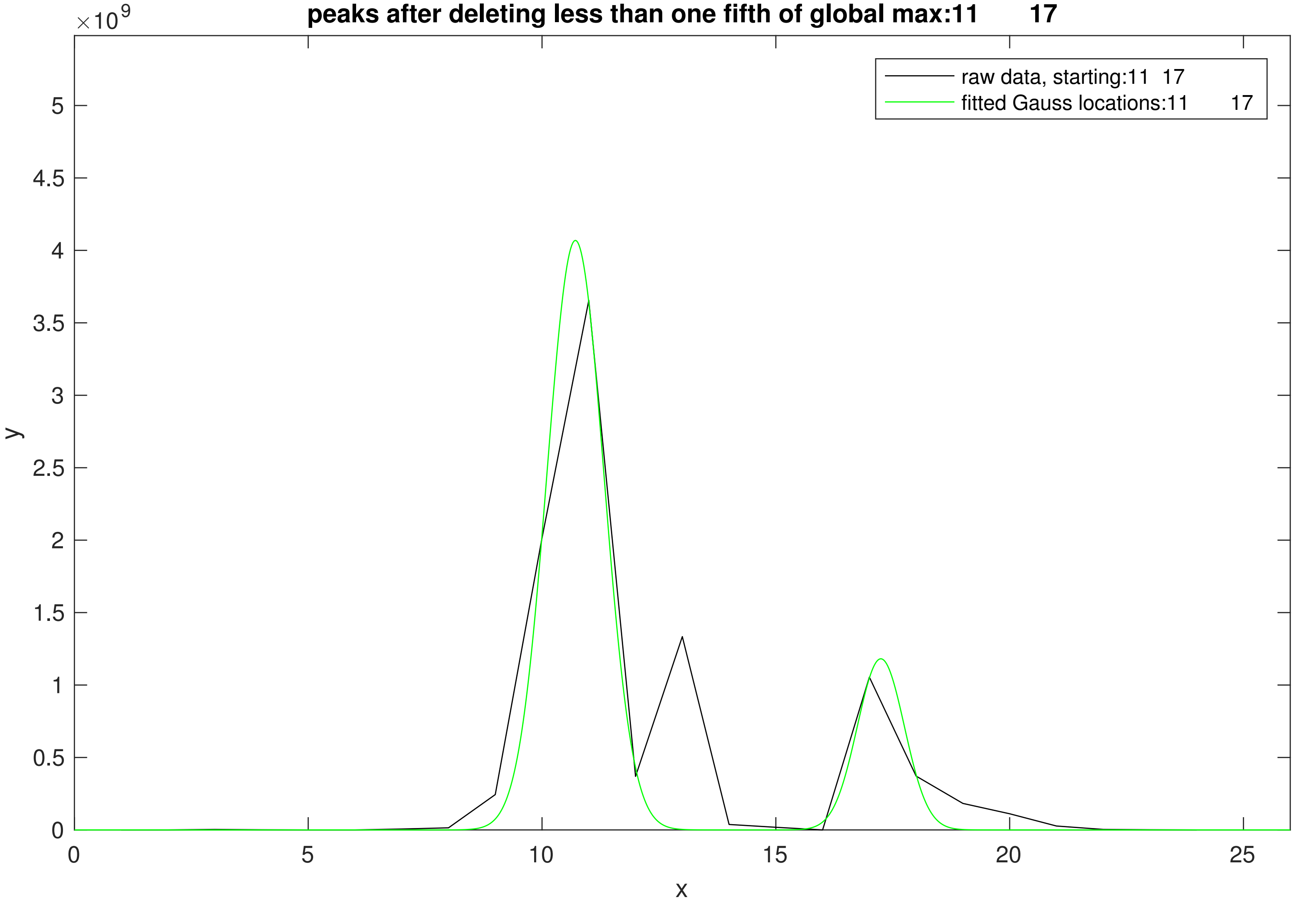

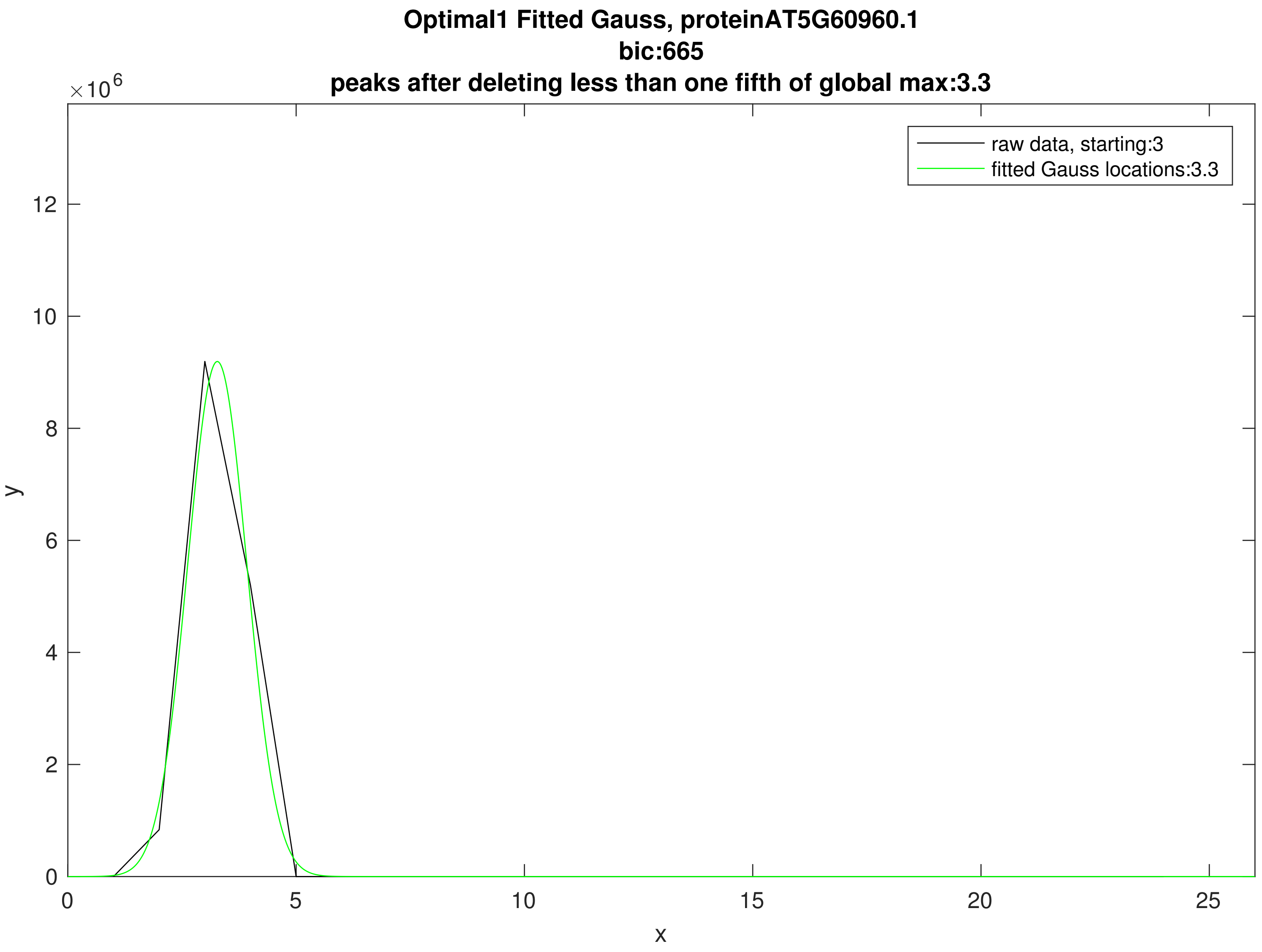
Optimal1 Fitted Gauss, proteinAT5G60960.1
bic:665
peaks after deleting less than one fifth of global max:3.3
×10^6
raw data, starting:3
fitted Gauss locations:3.3
y
x
0
2
4
6
8
10
12
0
5
10
15
20
25

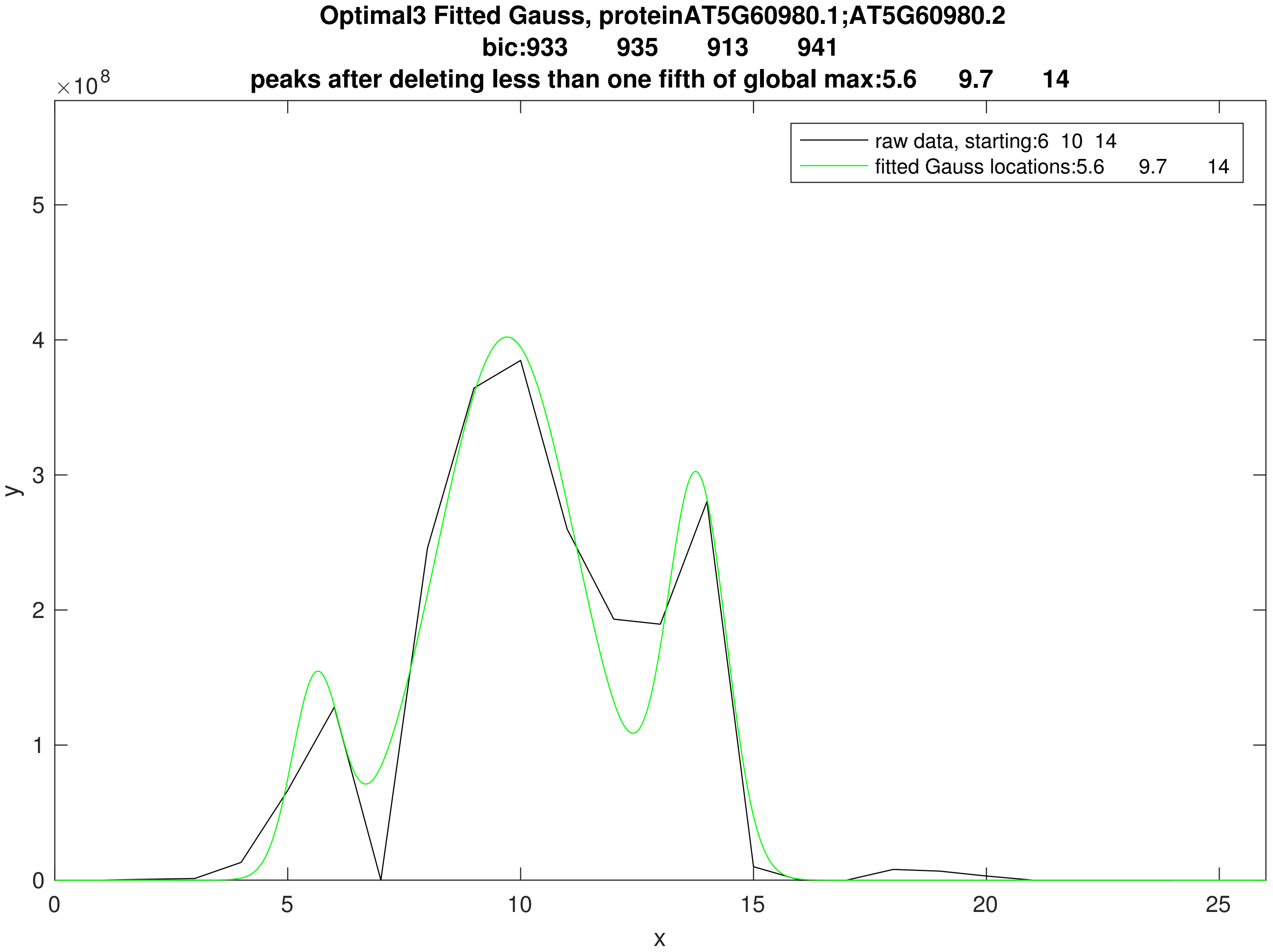

Optimal3 Fitted Gauss, proteinAT5G60980.1;AT5G60980.2
bic:933 935 913 941
peaks after deleting less than one fifth of global max:5.6 9.7 14
×10^8
raw data, starting:6 10 14
fitted Gauss locations:5.6 9.7 14
y
x
0
1
2
3
4
5
0
5
10
15
20
25

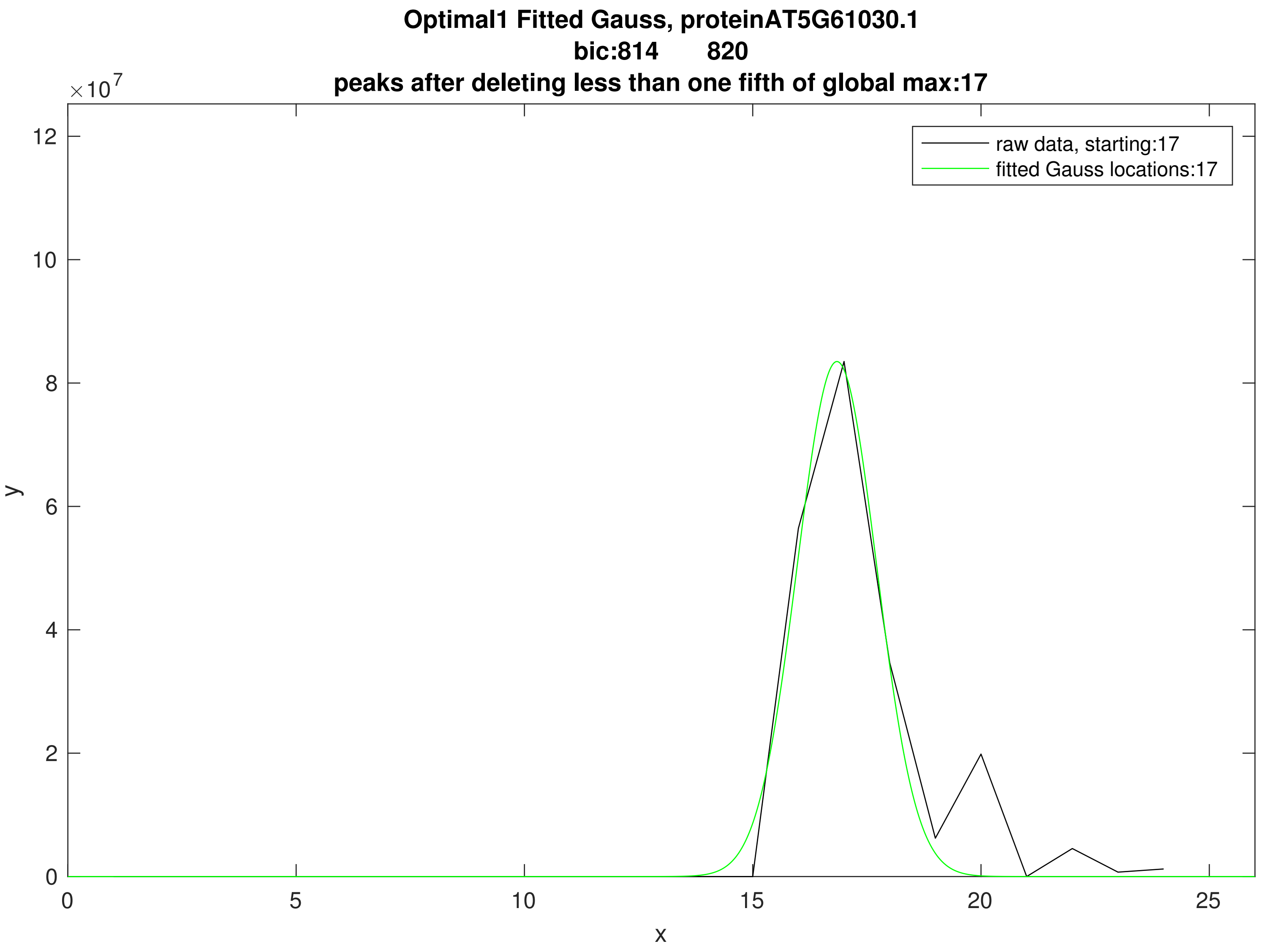

Optimal1 Fitted Gauss, proteinAT5G61030.1
bic:814 820
peaks after deleting less than one fifth of global max:17
×10^7
12
10
8
6
4
2
0
y
0
5
10
15
20
25
x
raw data, starting:17
fitted Gauss locations:17

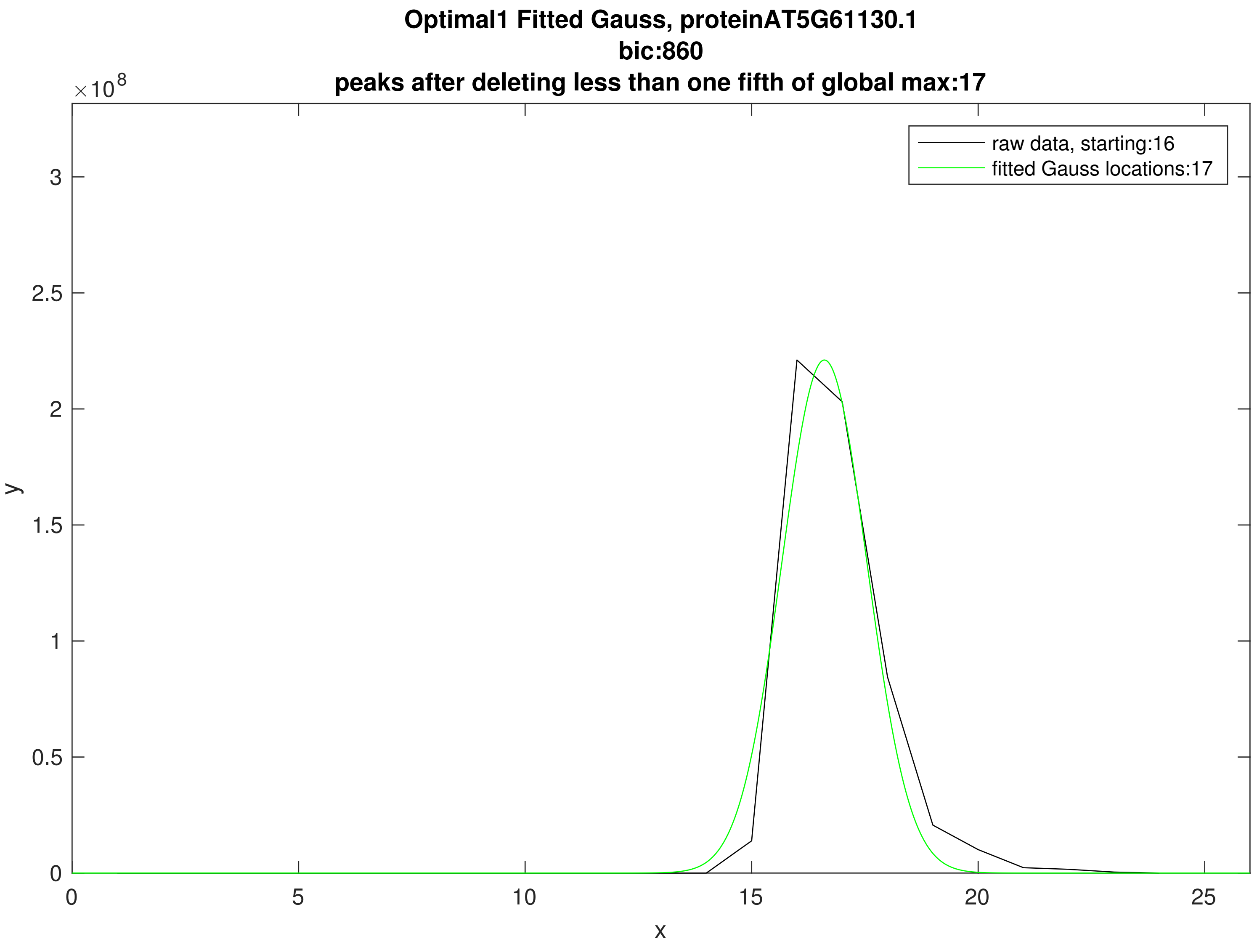
Optimal1 Fitted Gauss, proteinAT5G61130.1
bic:860
peaks after deleting less than one fifth of global max:17
×10^8
raw data, starting:16
fitted Gauss locations:17
y
x
0
0.5
1
1.5
2
2.5
3
0
5
10
15
20
25

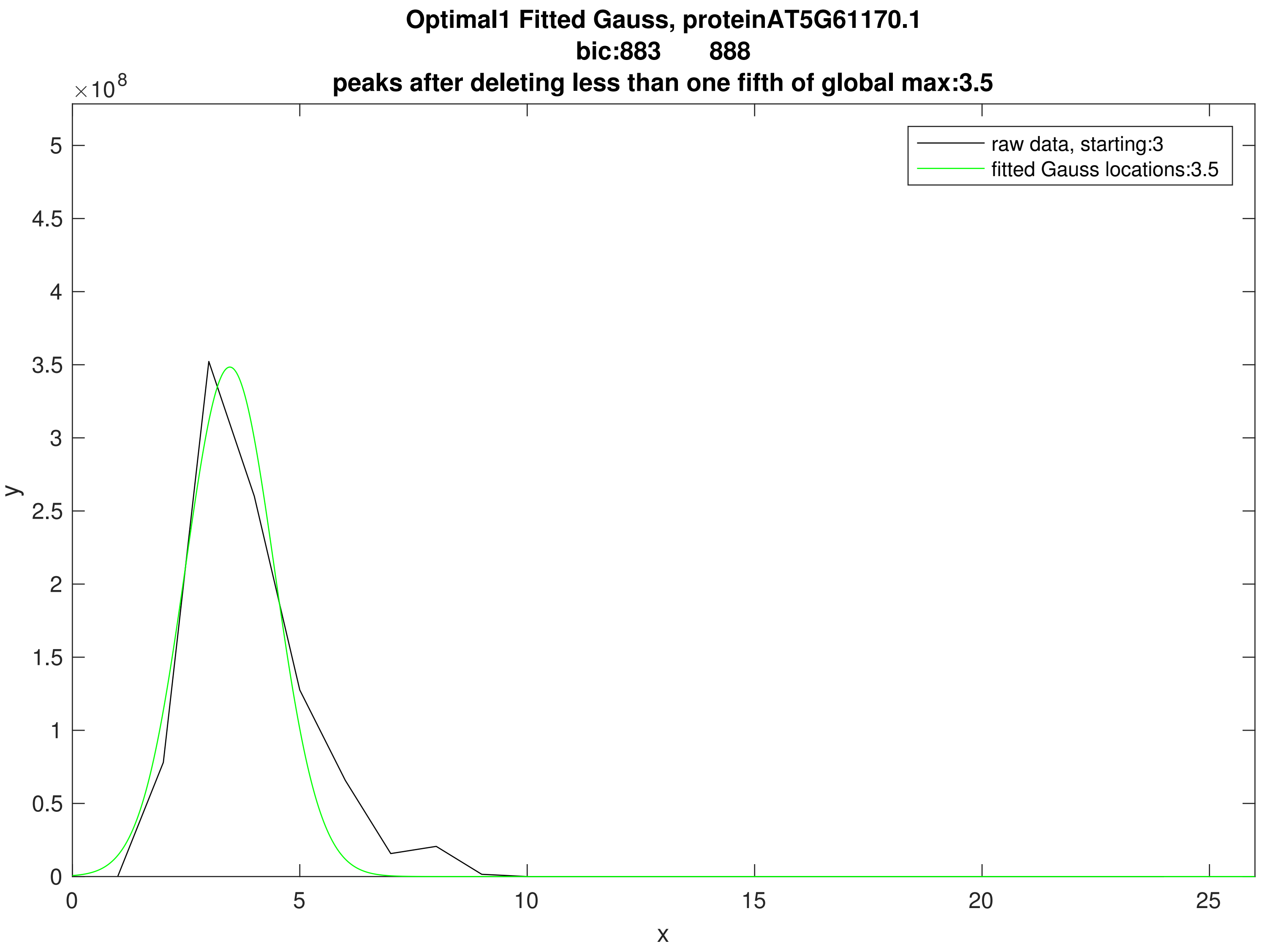
Optimal1 Fitted Gauss, proteinAT5G61170.1
bic:883 888
peaks after deleting less than one fifth of global max:3.5
×10^8
raw data, starting:3
fitted Gauss locations:3.5
y
x
0
0.5
1
1.5
2
2.5
3
3.5
4
4.5
5
0
5
10
15
20
25

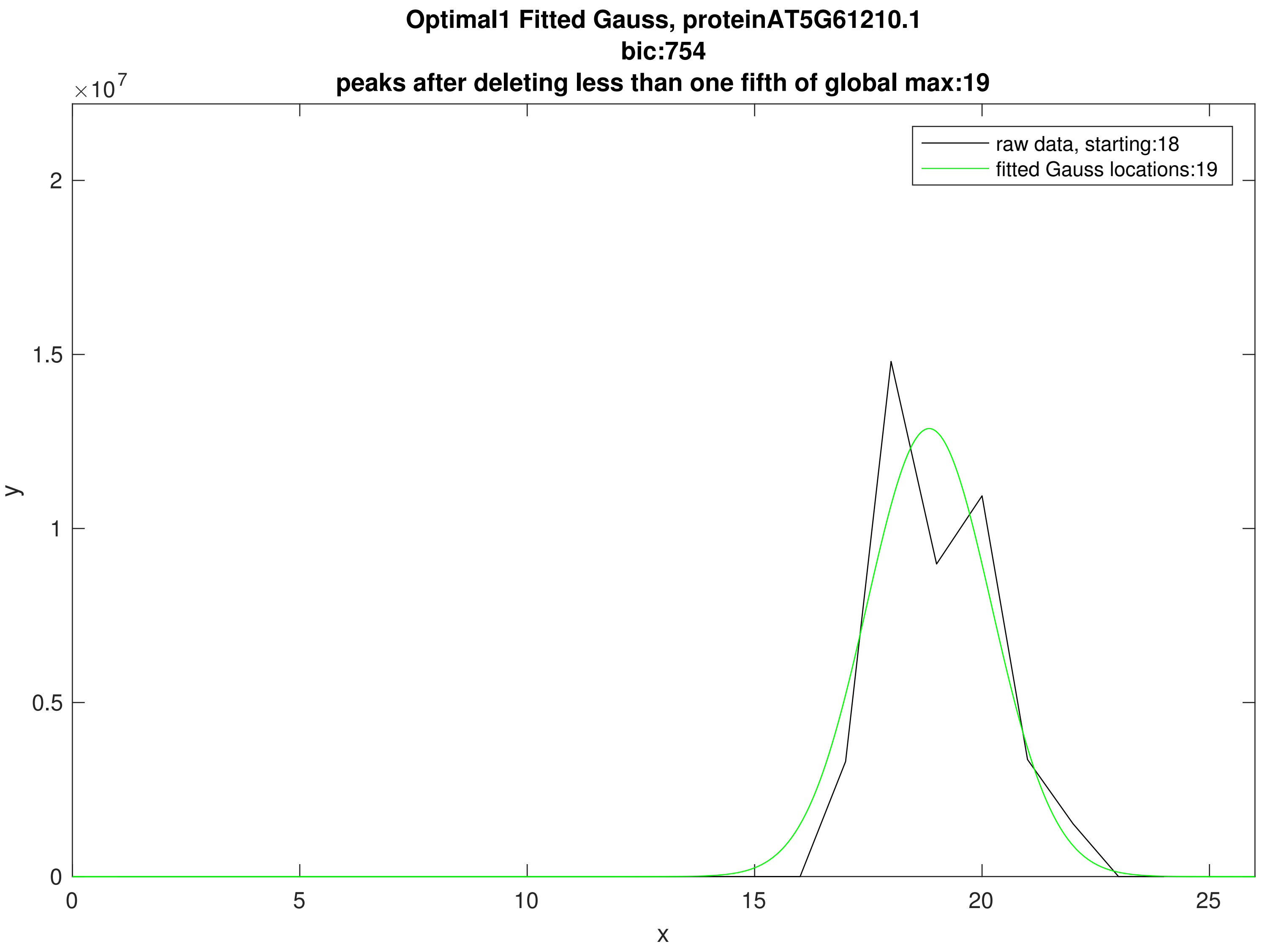

Optimal1 Fitted Gauss, proteinAT5G61210.1
bic:754
peaks after deleting less than one fifth of global max:19
×10^7
raw data, starting:18
fitted Gauss locations:19
y
x
0
0.5
1
1.5
2
0
5
10
15
20
25

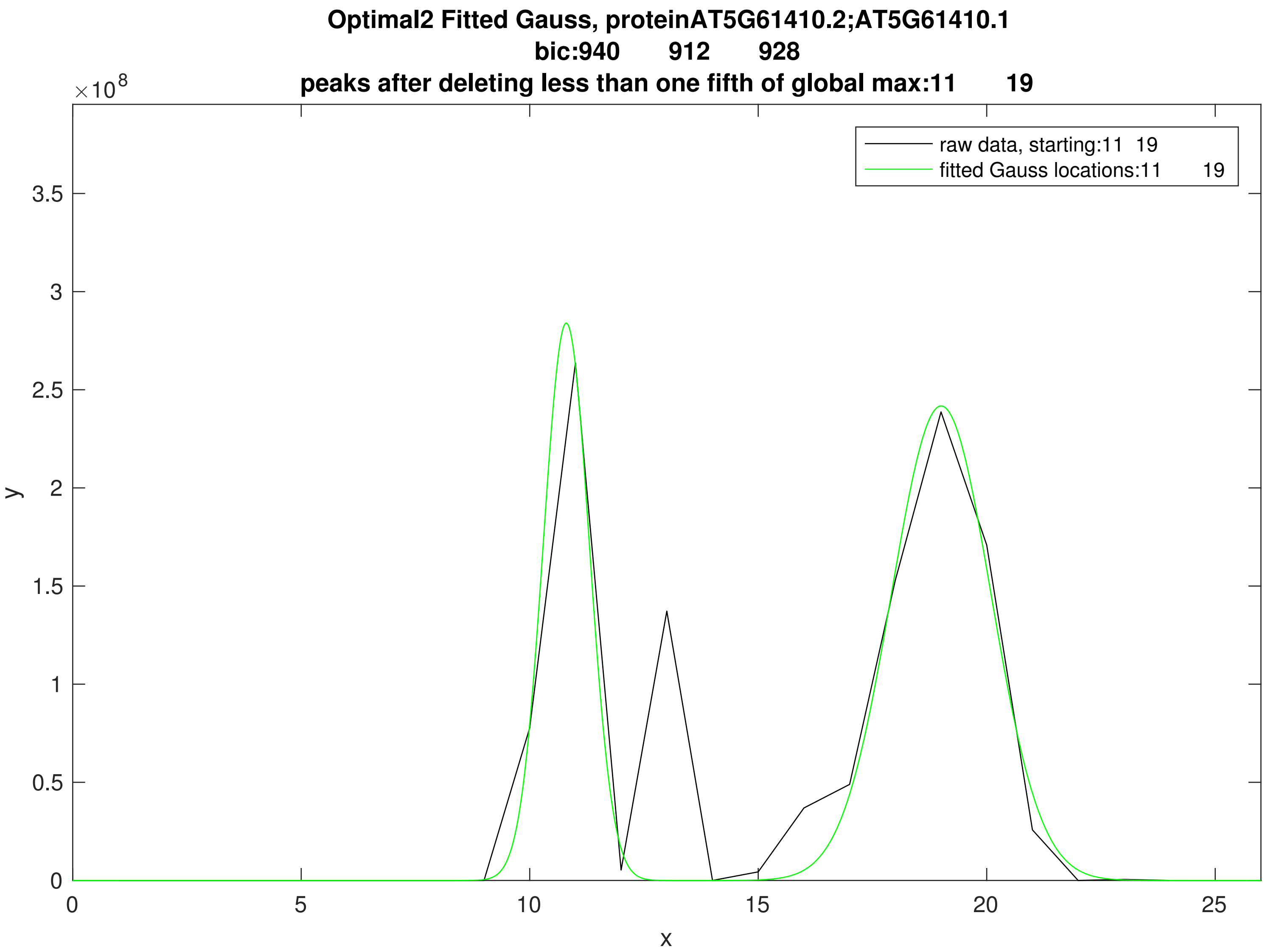

Optimal2 Fitted Gauss, proteinAT5G61410.2;AT5G61410.1
bic:940 912 928
peaks after deleting less than one fifth of global max:11 19
×10^8
raw data, starting:11 19
fitted Gauss locations:11 19
3.5
3
2.5
2
1.5
1
0.5
0
y
0
5
10
15
20
25
x

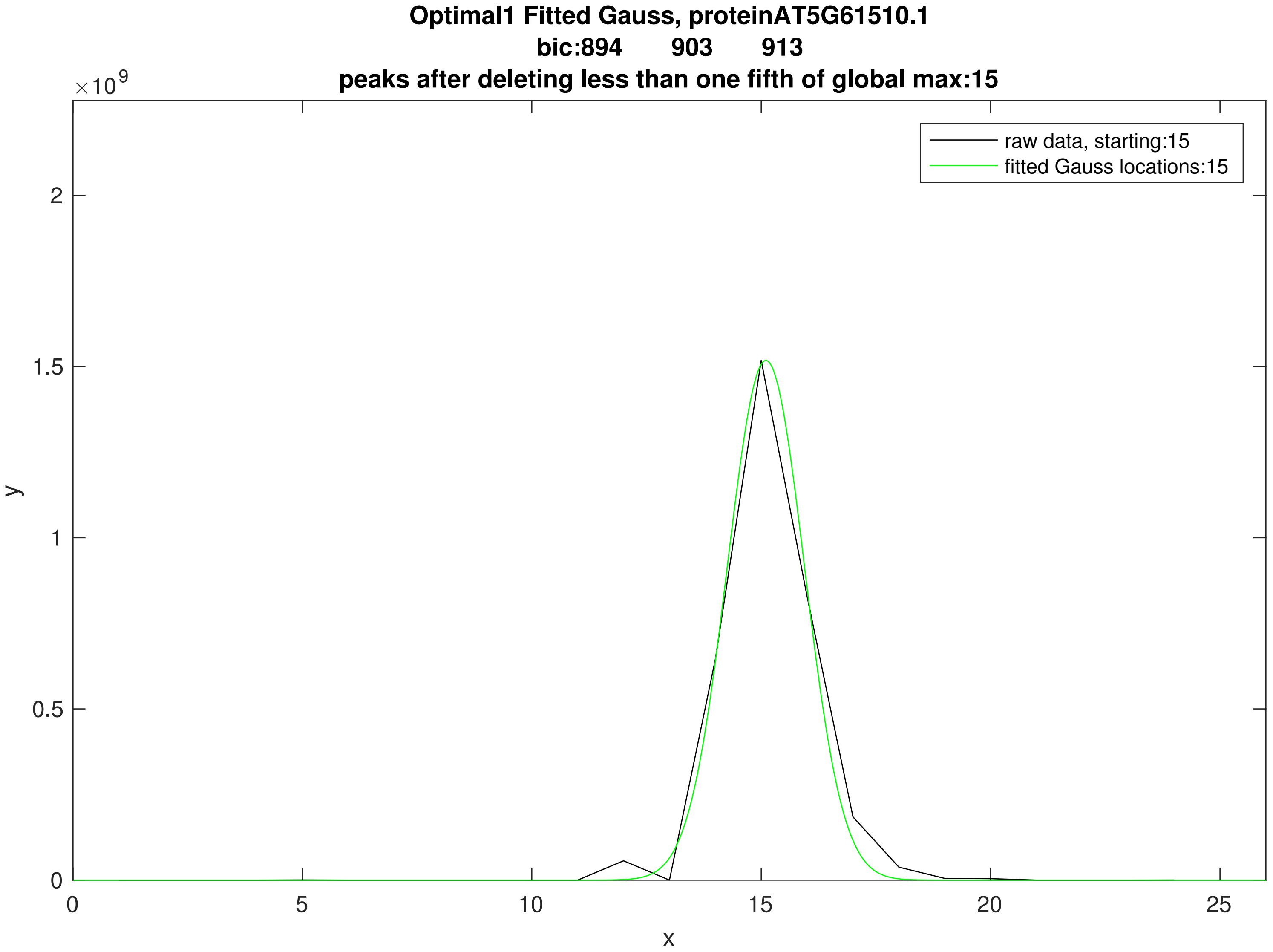
Optimal1 Fitted Gauss, proteinAT5G61510.1
bic:894 903 913
peaks after deleting less than one fifth of global max:15
×10^9
raw data, starting:15
fitted Gauss locations:15
y
x
0
0.5
1
1.5
2
0
5
10
15
20
25

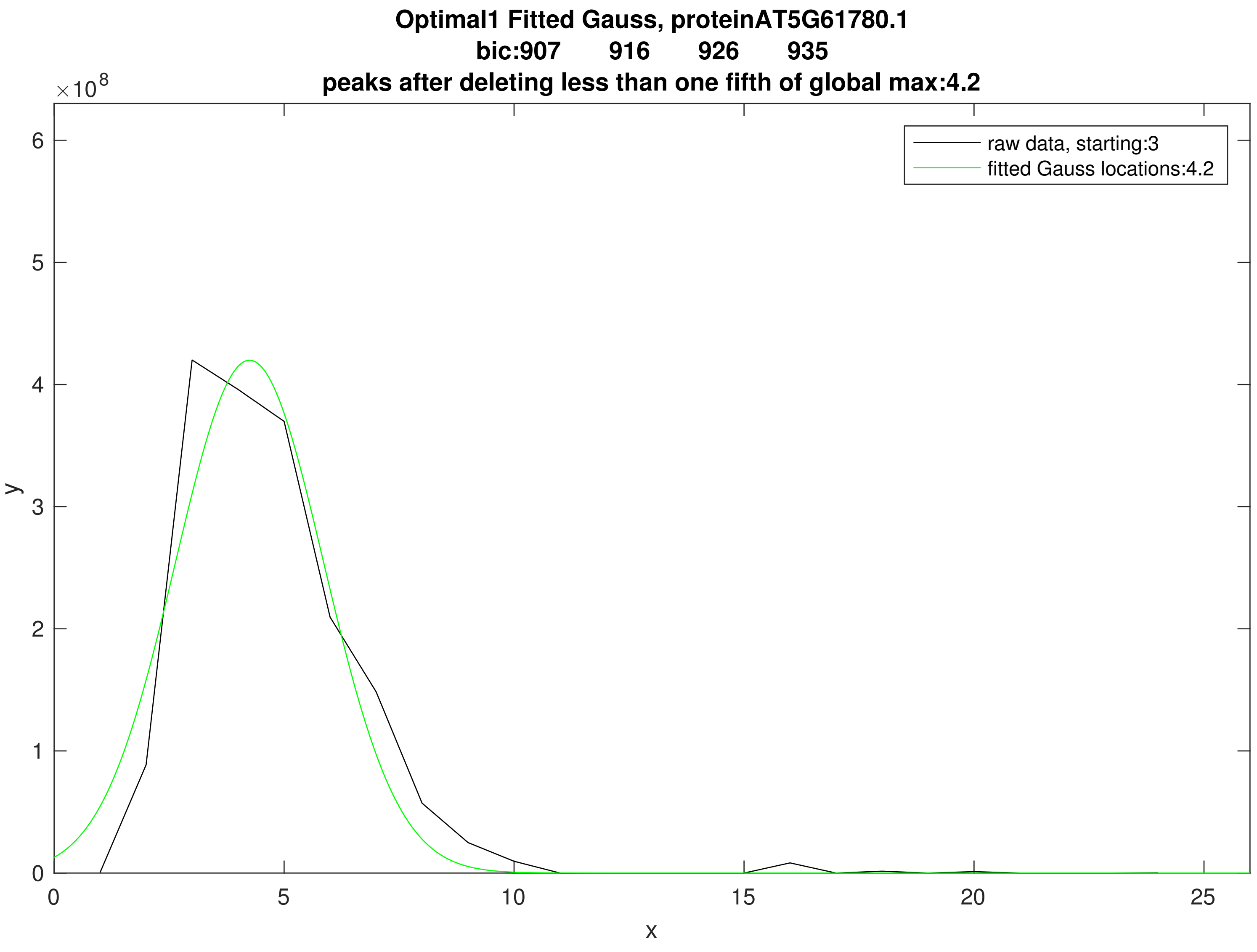
Optimal1 Fitted Gauss, proteinAT5G61780.1
bic:907 916 926 935
peaks after deleting less than one fifth of global max:4.2
×10^8
raw data, starting:3
fitted Gauss locations:4.2
y
x
0
1
2
3
4
5
6
0
5
10
15
20
25

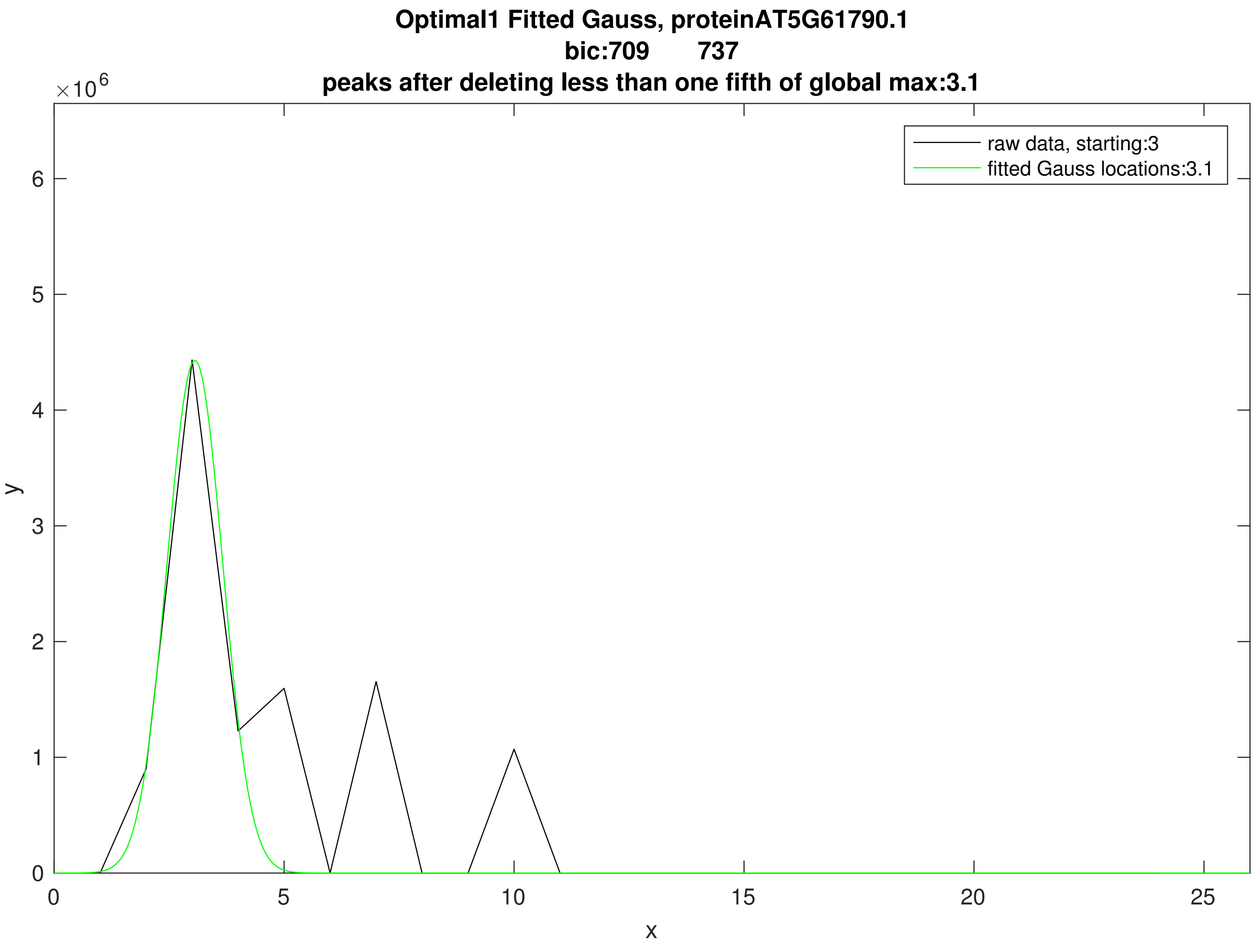

Optimal1 Fitted Gauss, proteinAT5G61790.1
bic:709 737
peaks after deleting less than one fifth of global max:3.1
×10^6
raw data, starting:3
fitted Gauss locations:3.1
y
x
0
1
2
3
4
5
6
0
5
10
15
20
25

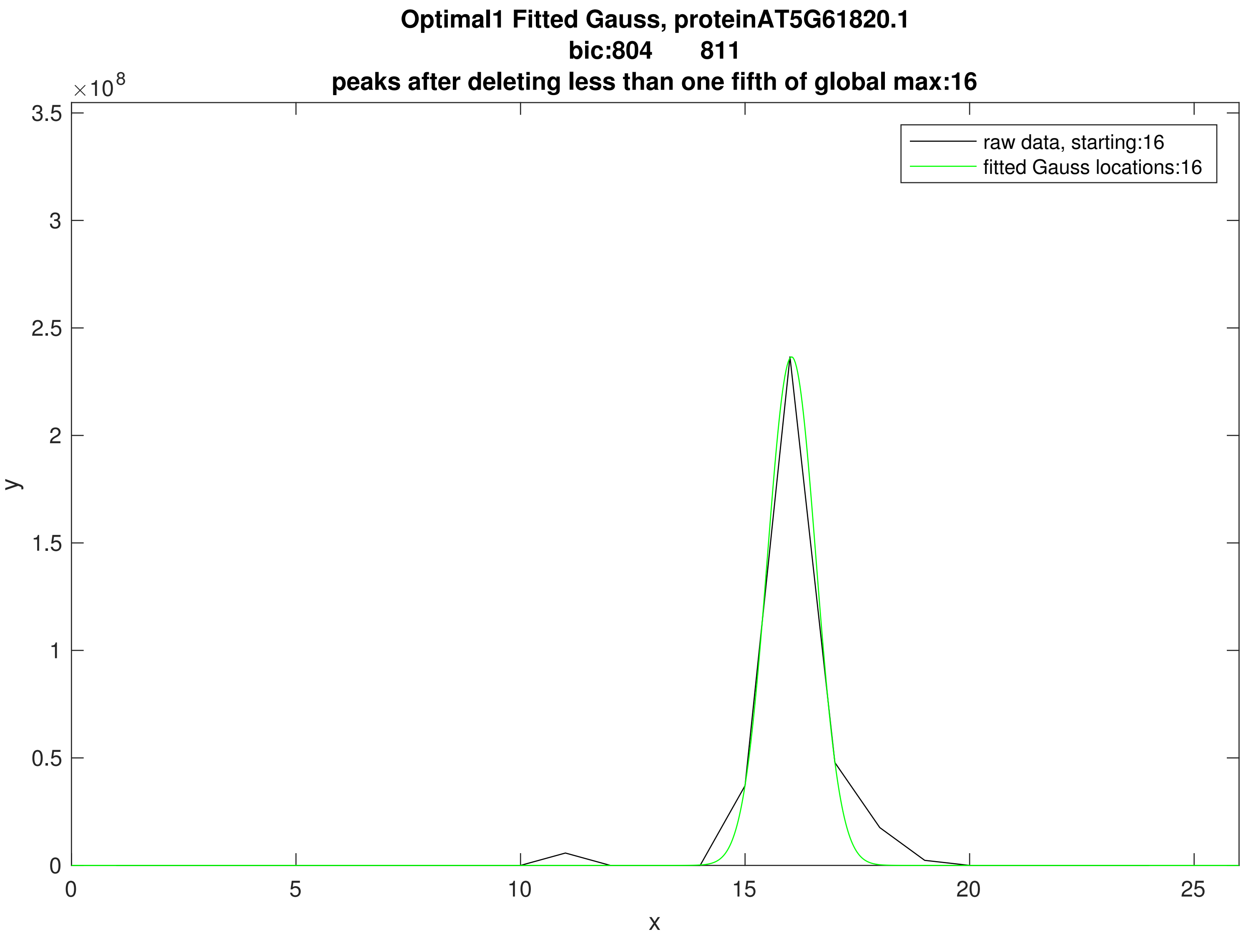
Optimal1 Fitted Gauss, proteinAT5G61820.1
bic:804 811
peaks after deleting less than one fifth of global max:16
×10^8
3.5
3
2.5
2
1.5
1
0.5
0
y
0
5
10
15
20
25
x
raw data, starting:16
fitted Gauss locations:16

#### Optimal1 Fitted Gauss, proteinAT5G62180.1
#### bic:822 826
#### peaks after deleting less than one fifth of global max:17

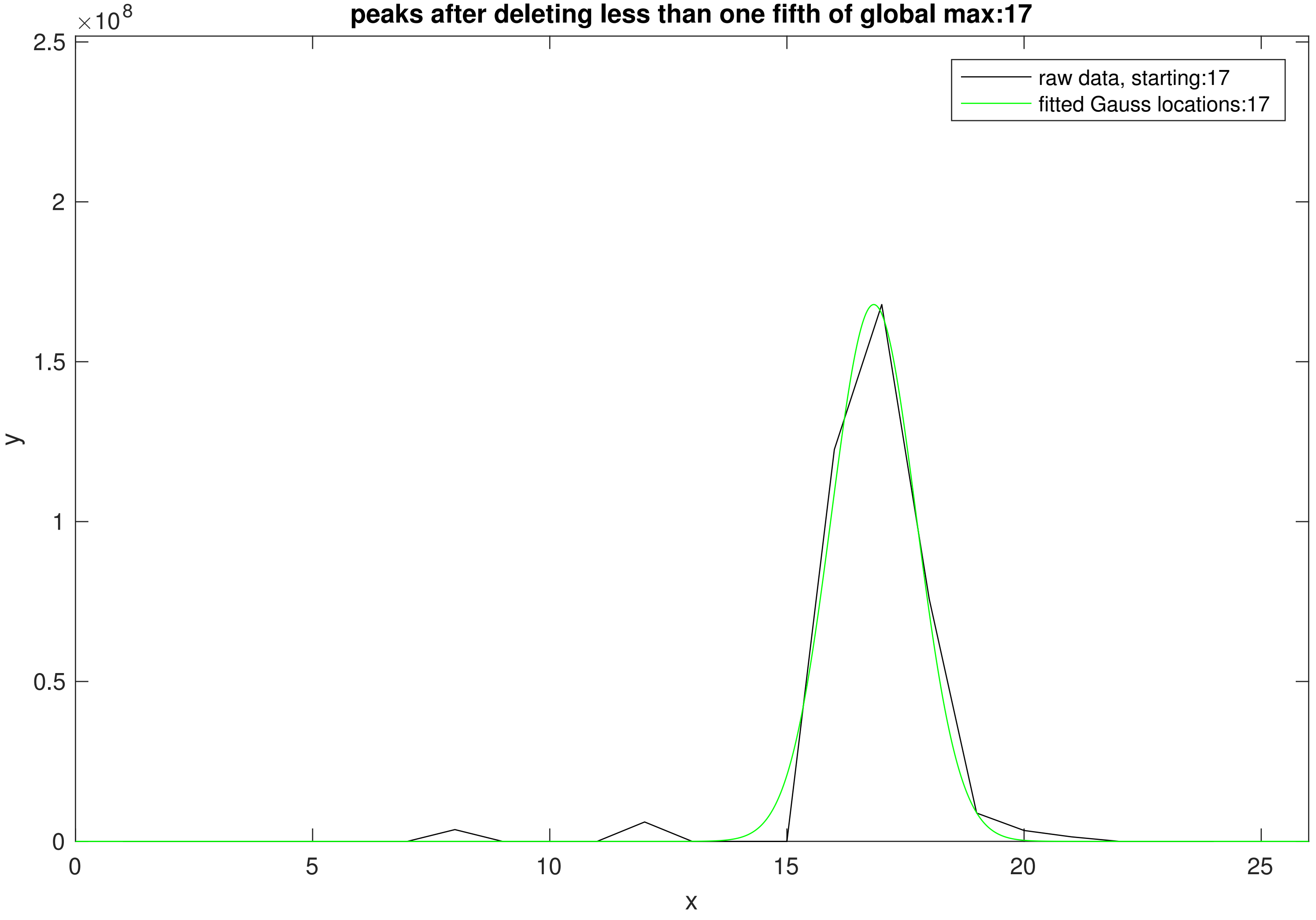

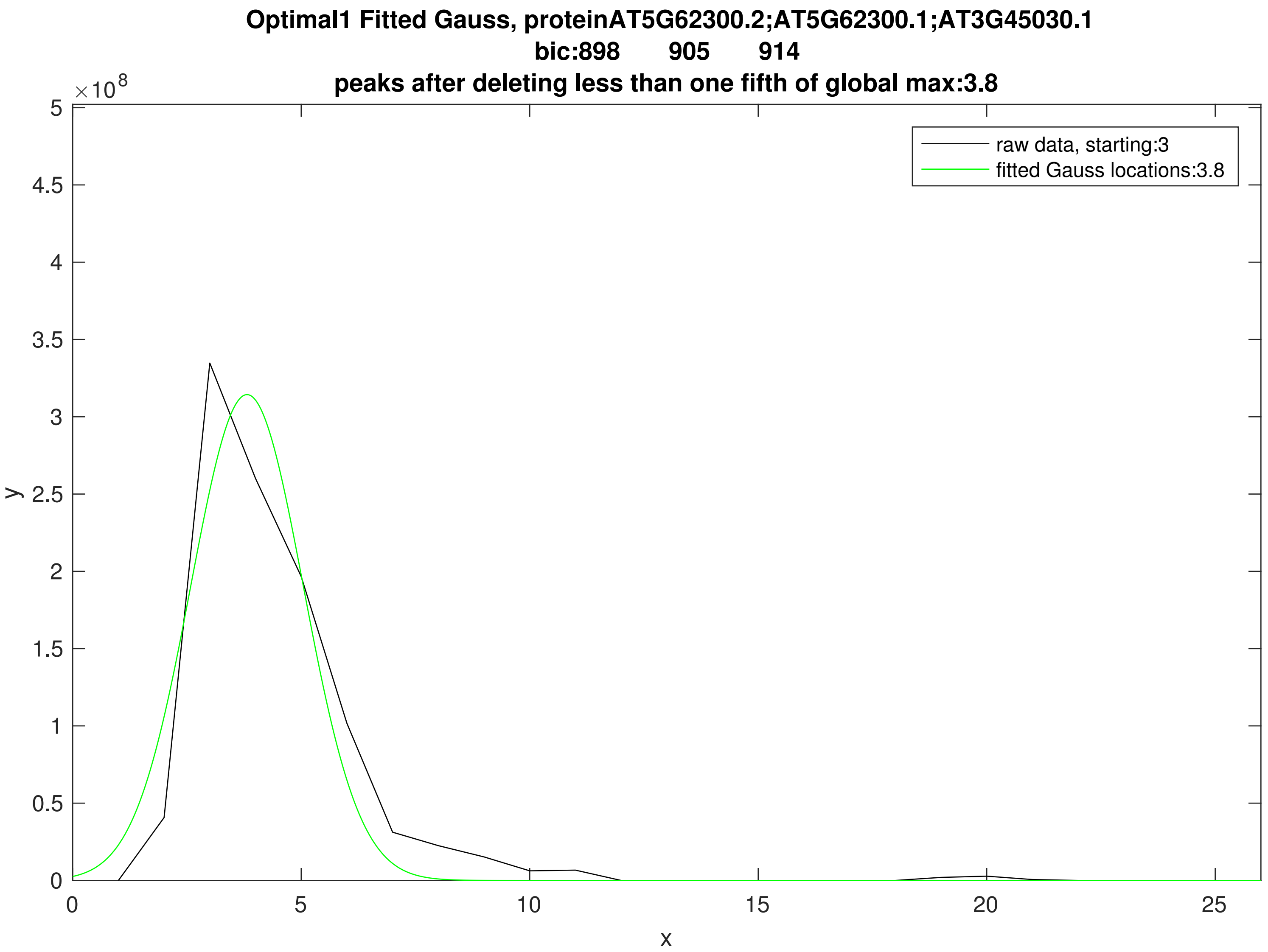

Optimal1 Fitted Gauss, proteinAT5G62300.2;AT5G62300.1;AT3G45030.1
bic:898 905 914
peaks after deleting less than one fifth of global max:3.8
×10^8
raw data, starting:3
fitted Gauss locations:3.8
y
x

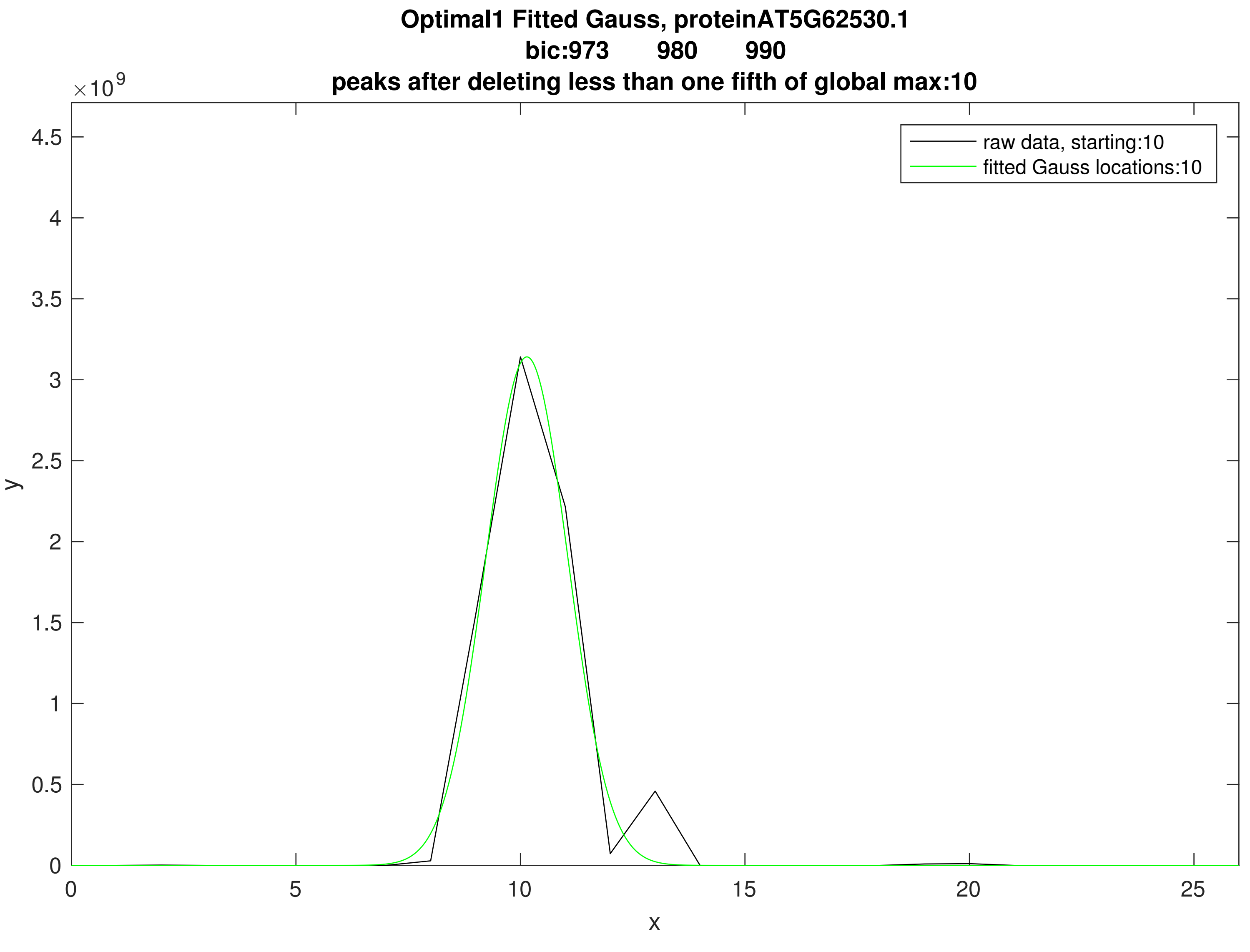

Optimal1 Fitted Gauss, proteinAT5G62530.1
bic:973 980 990
peaks after deleting less than one fifth of global max:10
×10^9
raw data, starting:10
fitted Gauss locations:10
y
x

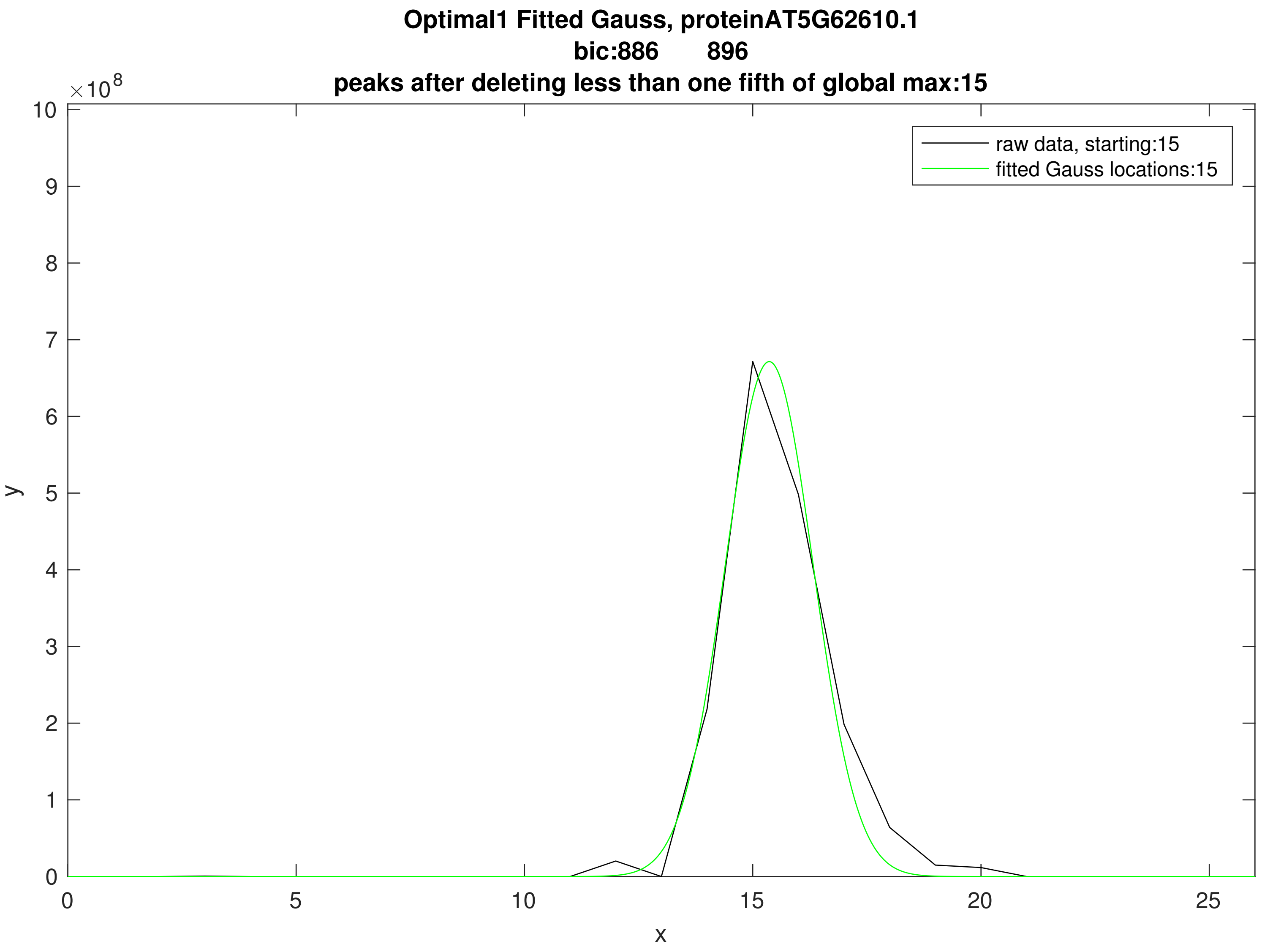

Optimal1 Fitted Gauss, proteinAT5G62610.1
bic:886 896
peaks after deleting less than one fifth of global max:15
×10^8
raw data, starting:15
fitted Gauss locations:15
y
x
0
1
2
3
4
5
6
7
8
9
10
0
5
10
15
20
25

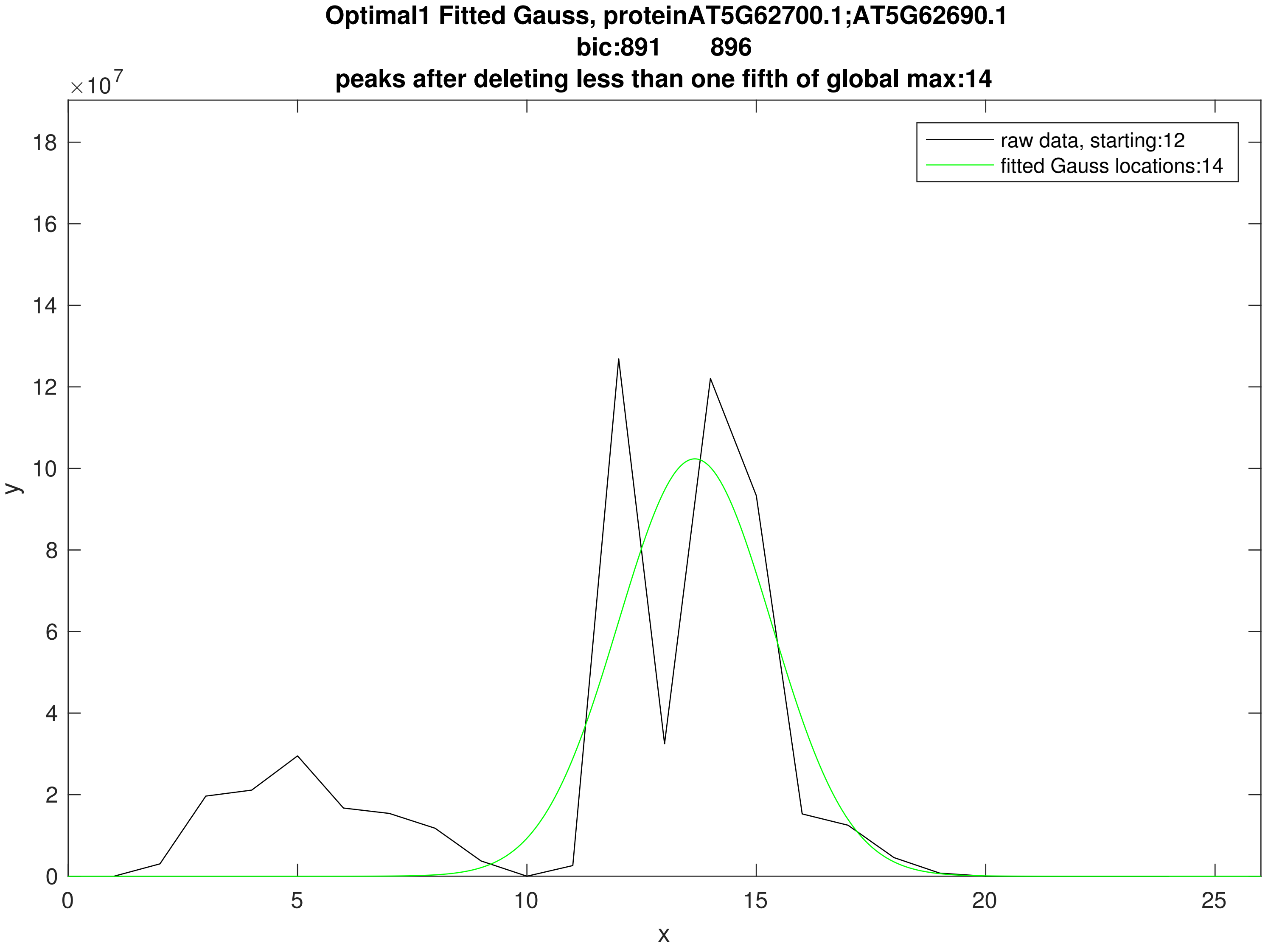

Optimal1 Fitted Gauss, proteinAT5G62700.1;AT5G62690.1
bic:891 896
peaks after deleting less than one fifth of global max:14
×10^7
raw data, starting:12
fitted Gauss locations:14
y
x
0
2
4
6
8
10
12
14
16
18
0
5
10
15
20
25

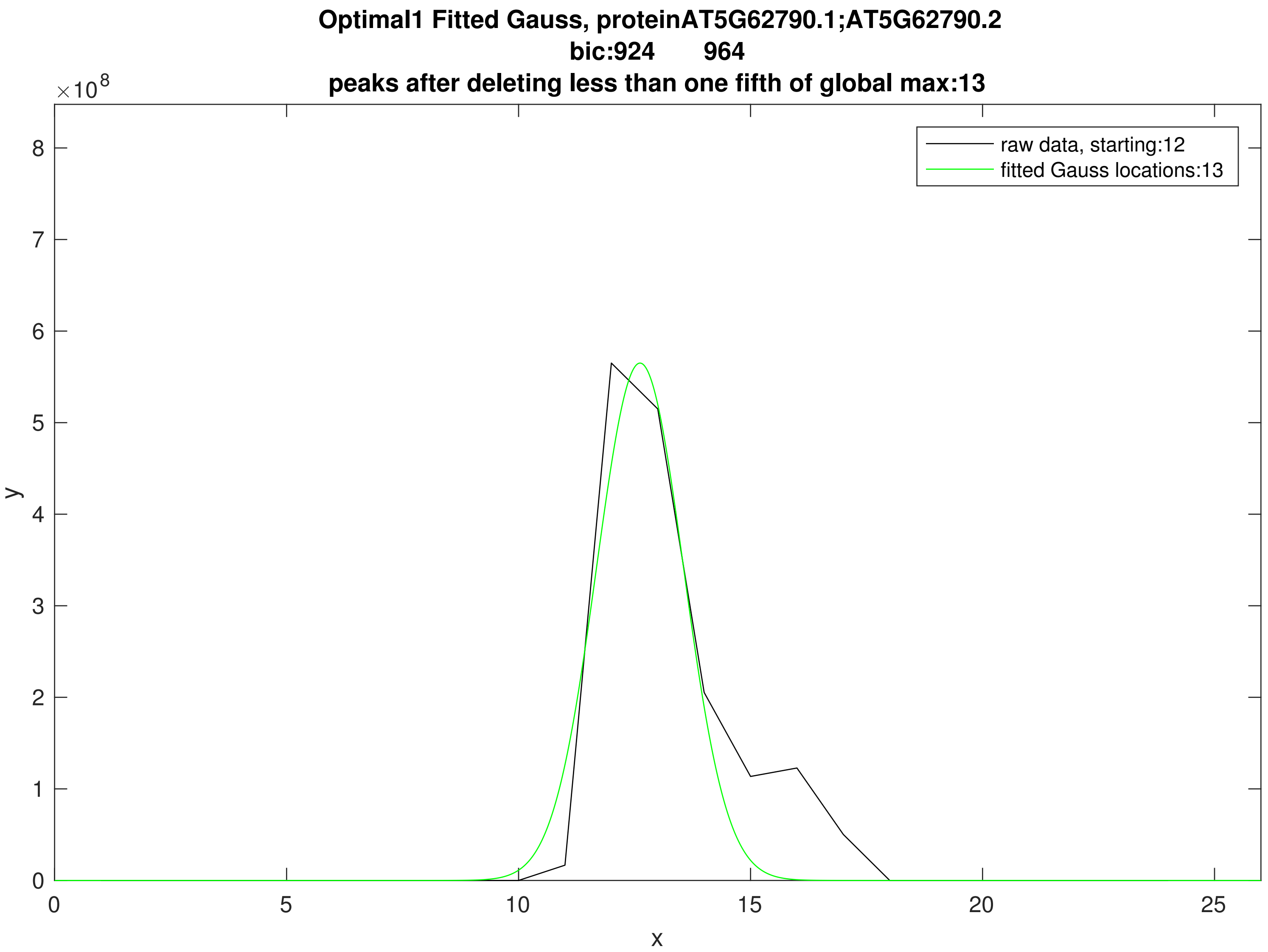

Optimal1 Fitted Gauss, proteinAT5G62790.1;AT5G62790.2
bic:924 964
peaks after deleting less than one fifth of global max:13
×10^8
raw data, starting:12
fitted Gauss locations:13
y
x
0
1
2
3
4
5
6
7
8
0
5
10
15
20
25

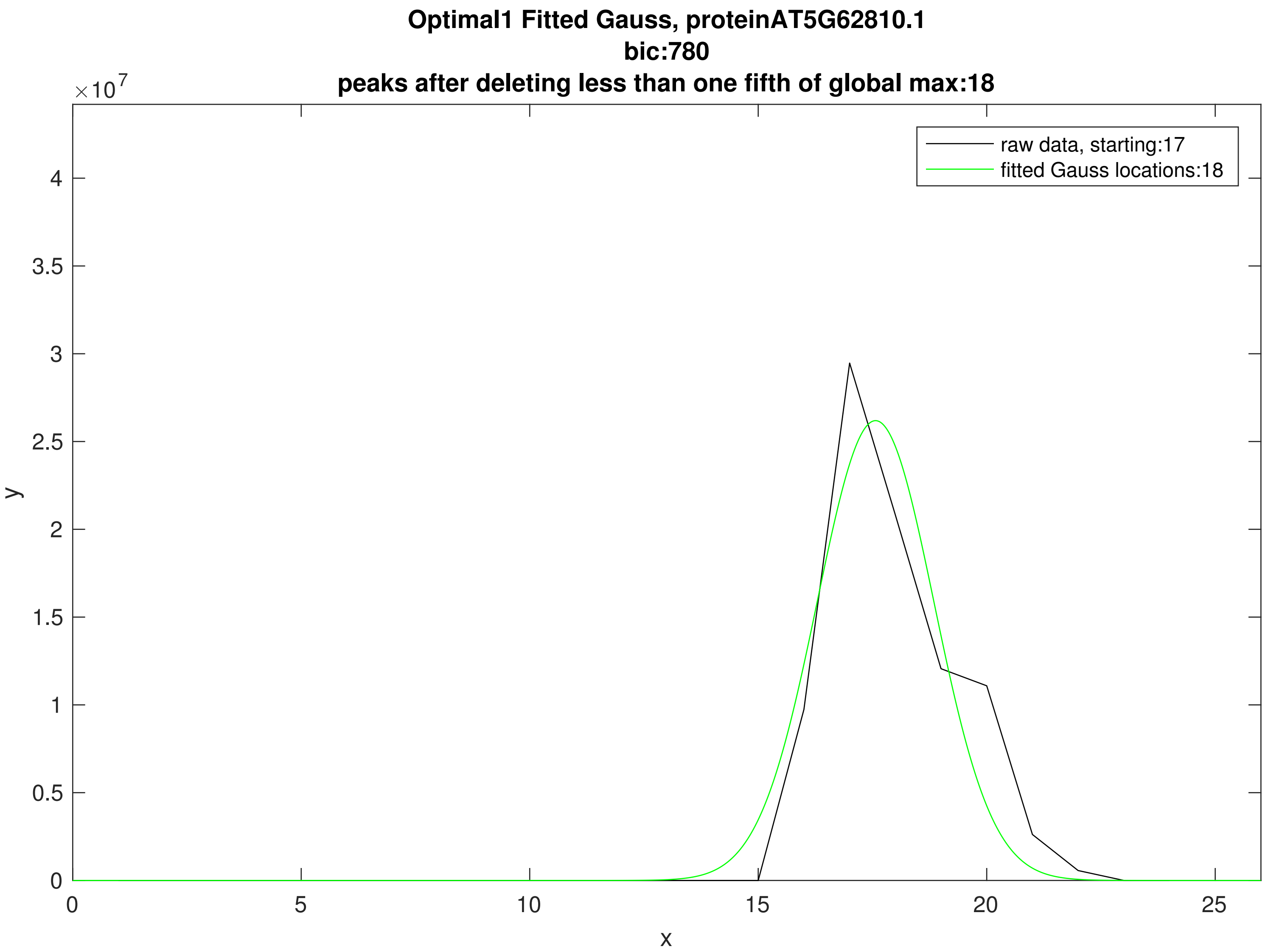
Optimal1 Fitted Gauss, proteinAT5G62810.1
bic:780
peaks after deleting less than one fifth of global max:18
×10^7
raw data, starting:17
fitted Gauss locations:18
y
x
0
0.5
1
1.5
2
2.5
3
3.5
4
5
10
15
20
25

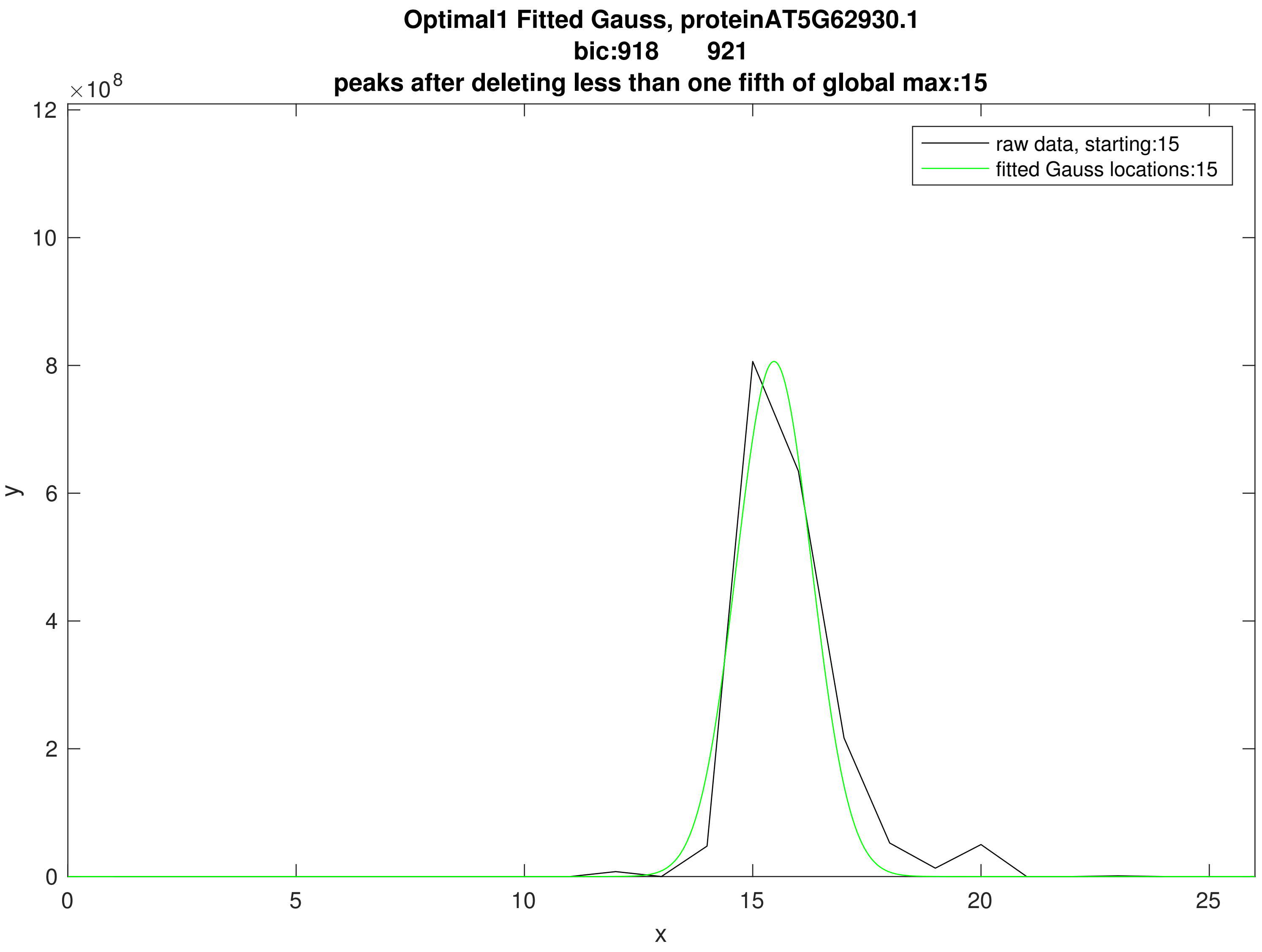

Optimal1 Fitted Gauss, proteinAT5G62930.1
bic:918 921
peaks after deleting less than one fifth of global max:15
×10^8
12
10
8
6
4
2
0
y
0
5
10
15
20
25
x
raw data, starting:15
fitted Gauss locations:15

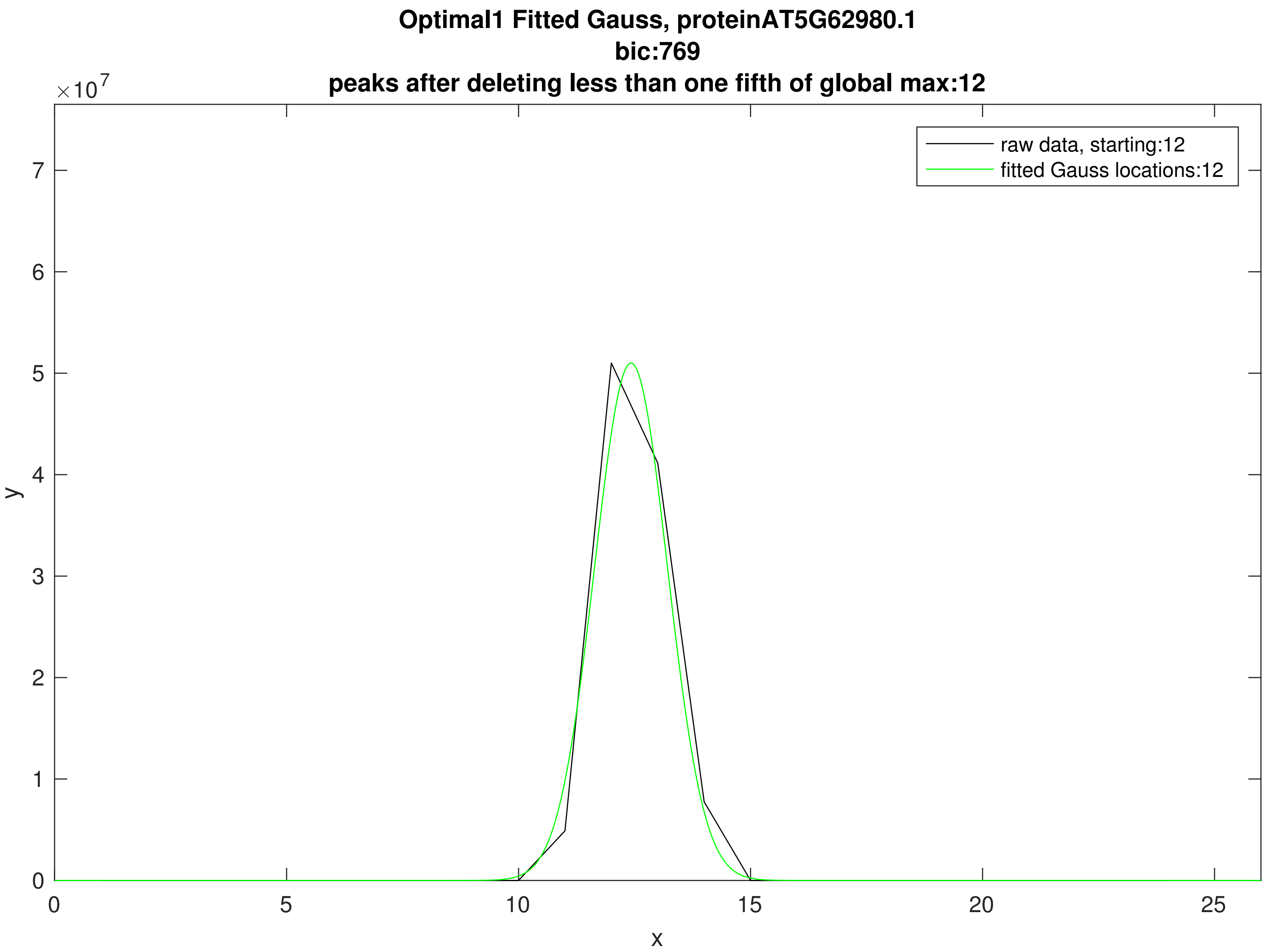

Optimal1 Fitted Gauss, proteinAT5G62980.1
bic:769
peaks after deleting less than one fifth of global max:12
×10^7
raw data, starting:12
fitted Gauss locations:12
y
x
0
1
2
3
4
5
6
7
0
5
10
15
20
25

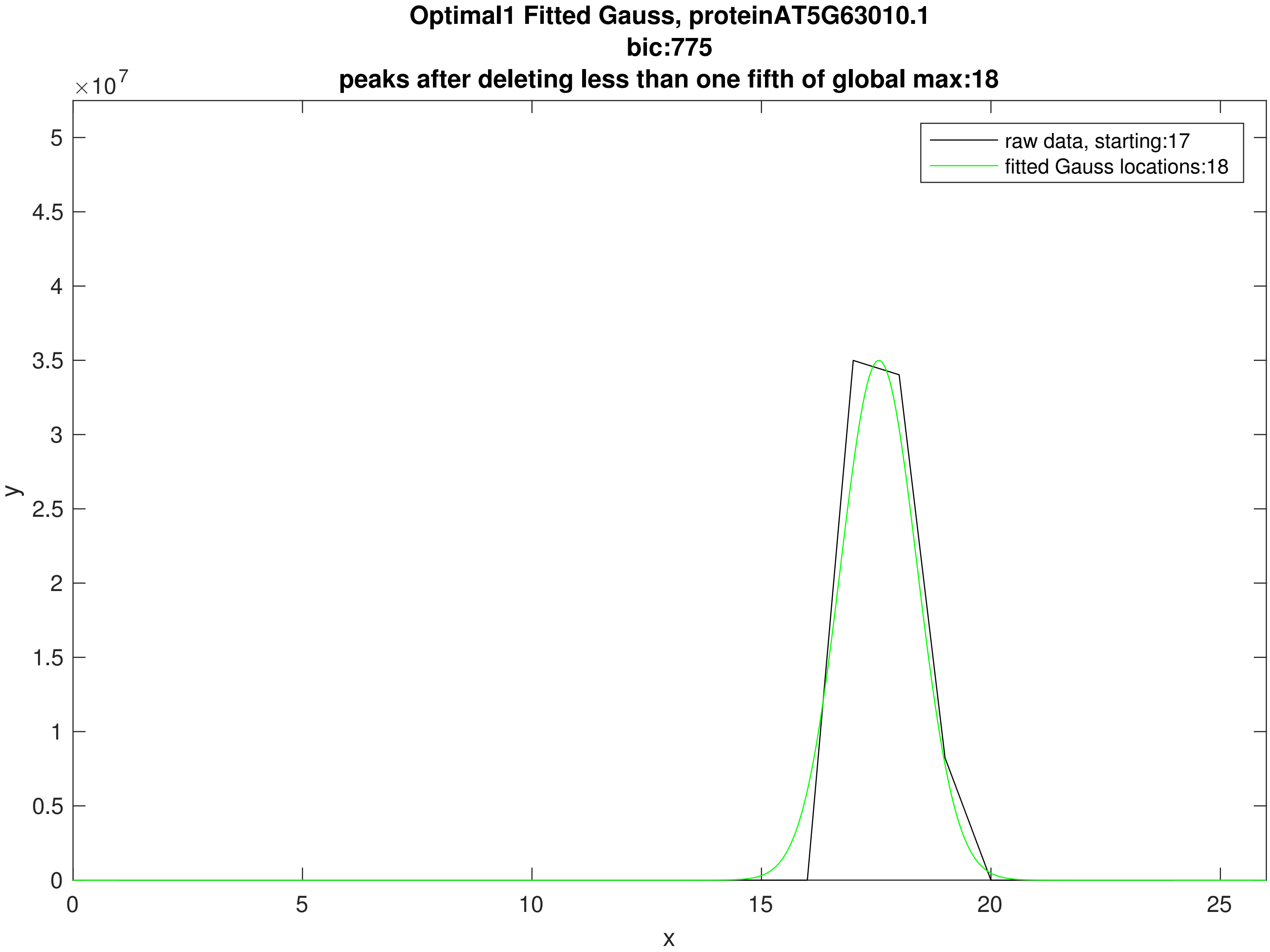
Optimal1 Fitted Gauss, proteinAT5G63010.1
bic:775
peaks after deleting less than one fifth of global max:18
×10^7
raw data, starting:17
fitted Gauss locations:18
y
x
0
0.5
1
1.5
2
2.5
3
3.5
4
4.5
5
0
5
10
15
20
25

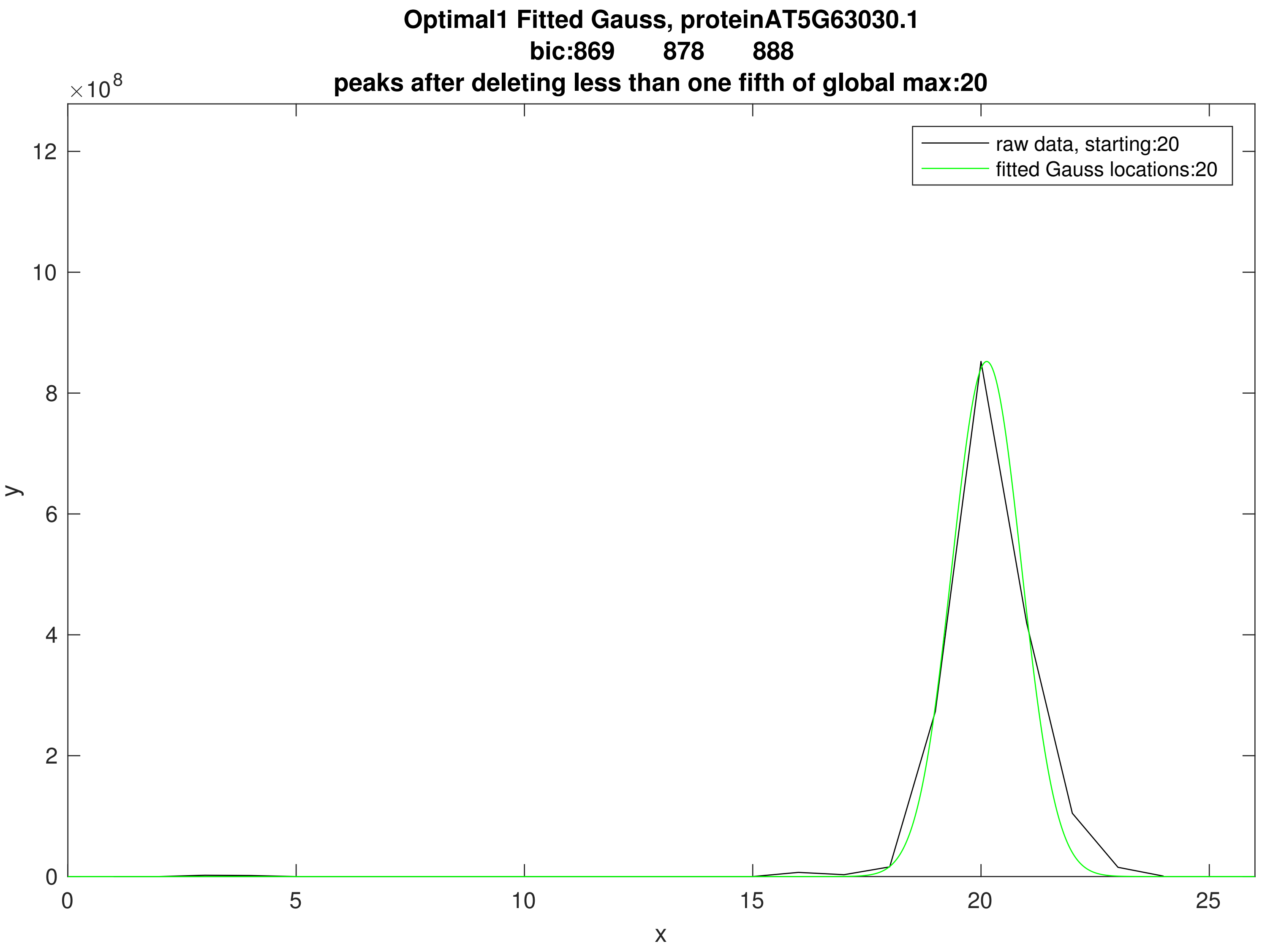
Optimal1 Fitted Gauss, proteinAT5G63030.1
bic:869 878 888
peaks after deleting less than one fifth of global max:20
×10^8
12
10
8
6
4
2
0
y
0
5
10
15
20
25
x
raw data, starting:20
fitted Gauss locations:20

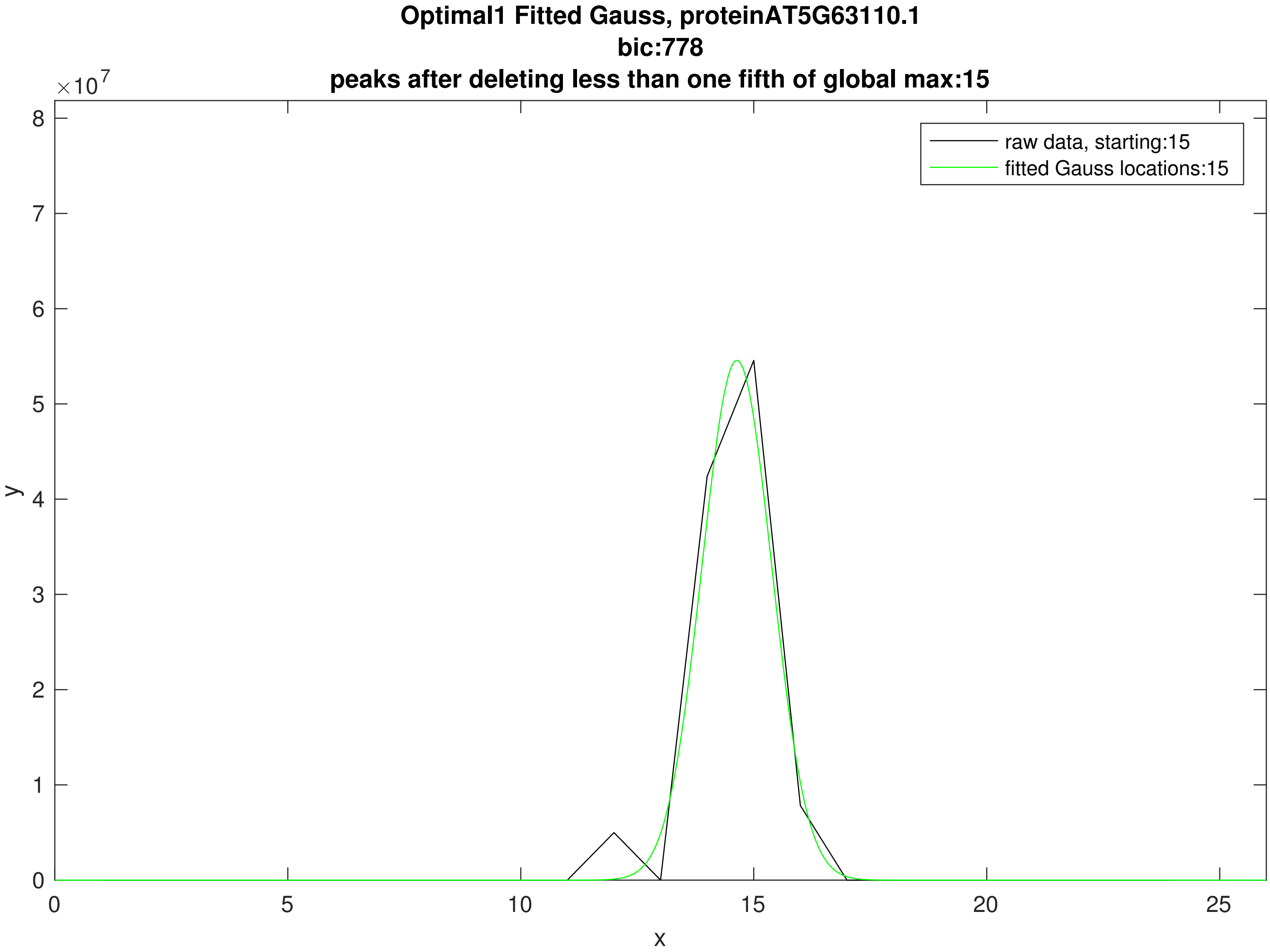

Optimal1 Fitted Gauss, proteinAT5G63110.1
bic:778
peaks after deleting less than one fifth of global max:15
×10^7
8
7
6
5
4
3
2
1
0
y
0
5
10
15
20
25
x
raw data, starting:15
fitted Gauss locations:15

**Optimal1 Fitted Gauss, proteinAT5G63310.1**
**bic:806**
**peaks after deleting less than one fifth of global max:4.6**

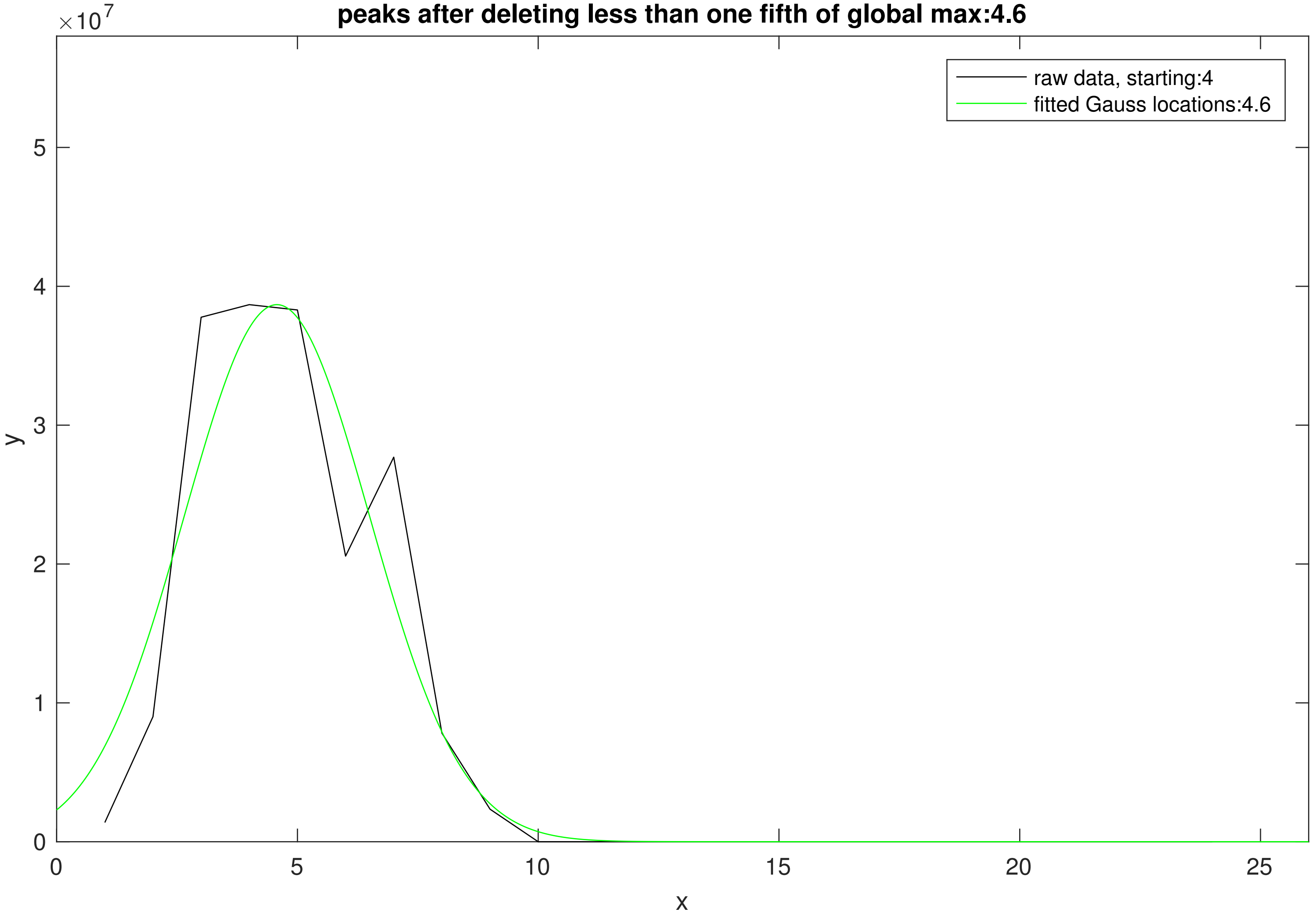

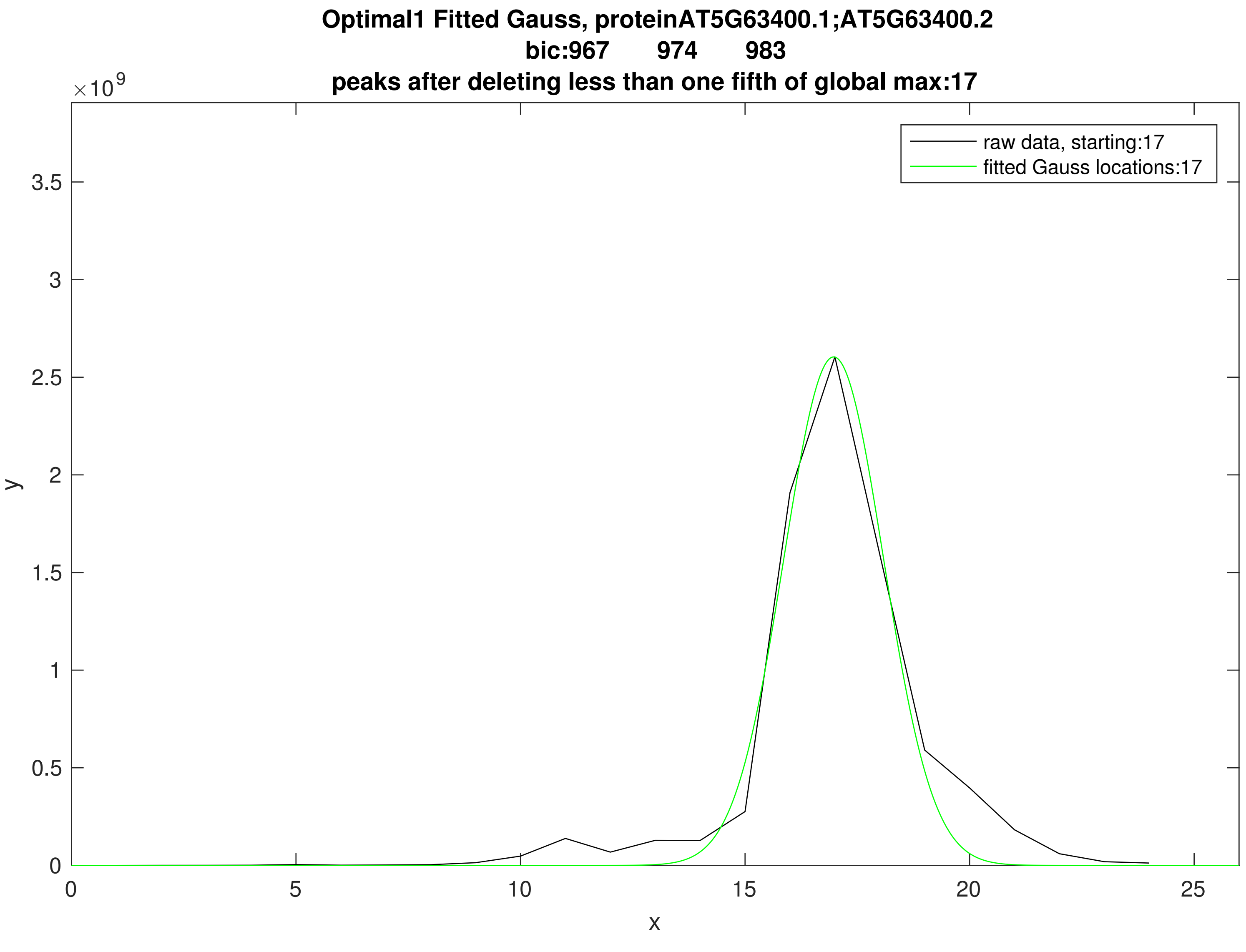
Optimal1 Fitted Gauss, proteinAT5G63400.1;AT5G63400.2
bic:967 974 983
peaks after deleting less than one fifth of global max:17
×10^9
raw data, starting:17
fitted Gauss locations:17
y
x
0
0.5
1
1.5
2
2.5
3
3.5
5
10
15
20
25

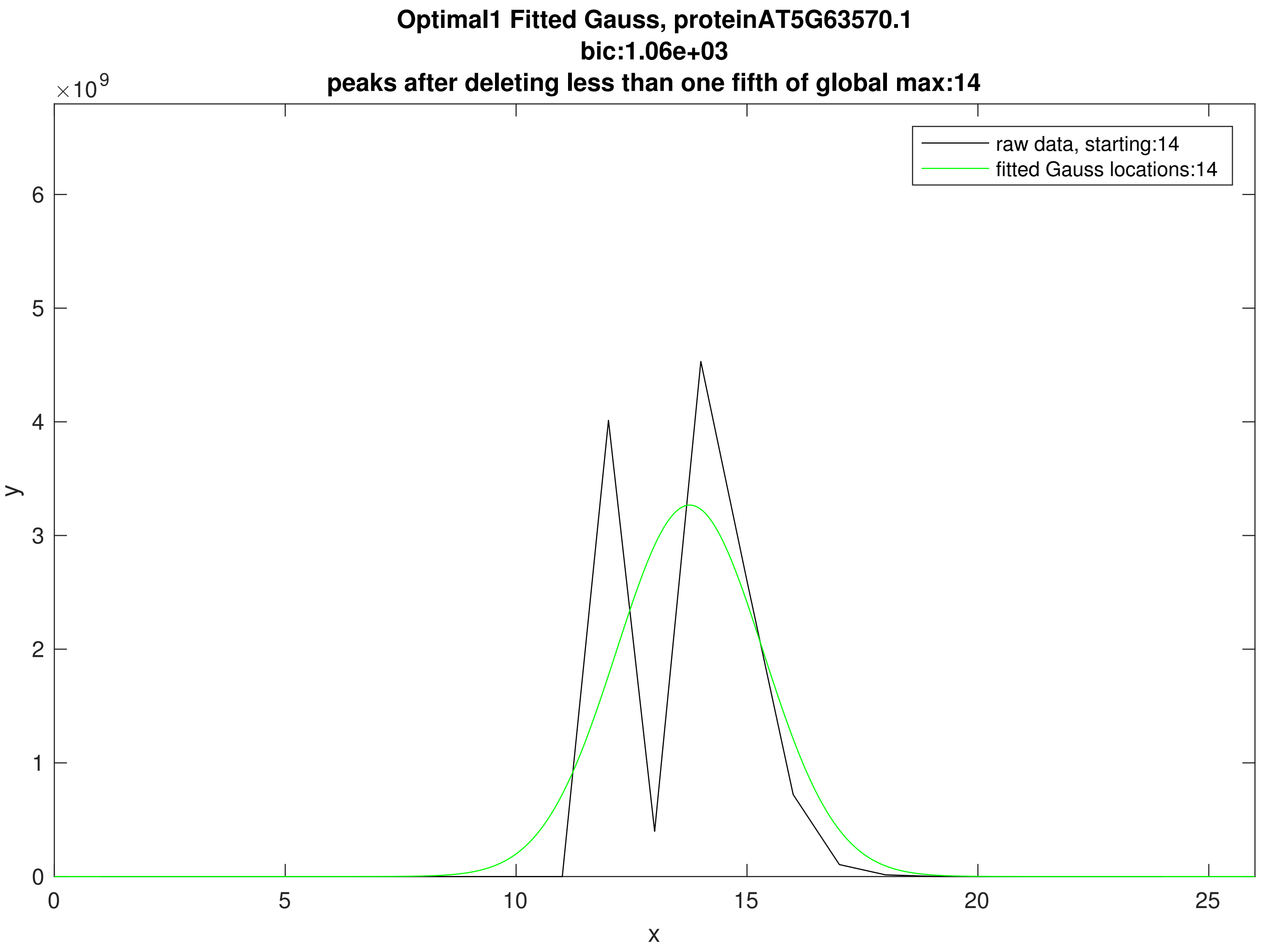

Optimal1 Fitted Gauss, proteinAT5G63570.1
bic:1.06e+03
peaks after deleting less than one fifth of global max:14
×10^9
raw data, starting:14
fitted Gauss locations:14
y
x
0
1
2
3
4
5
6
0
5
10
15
20
25

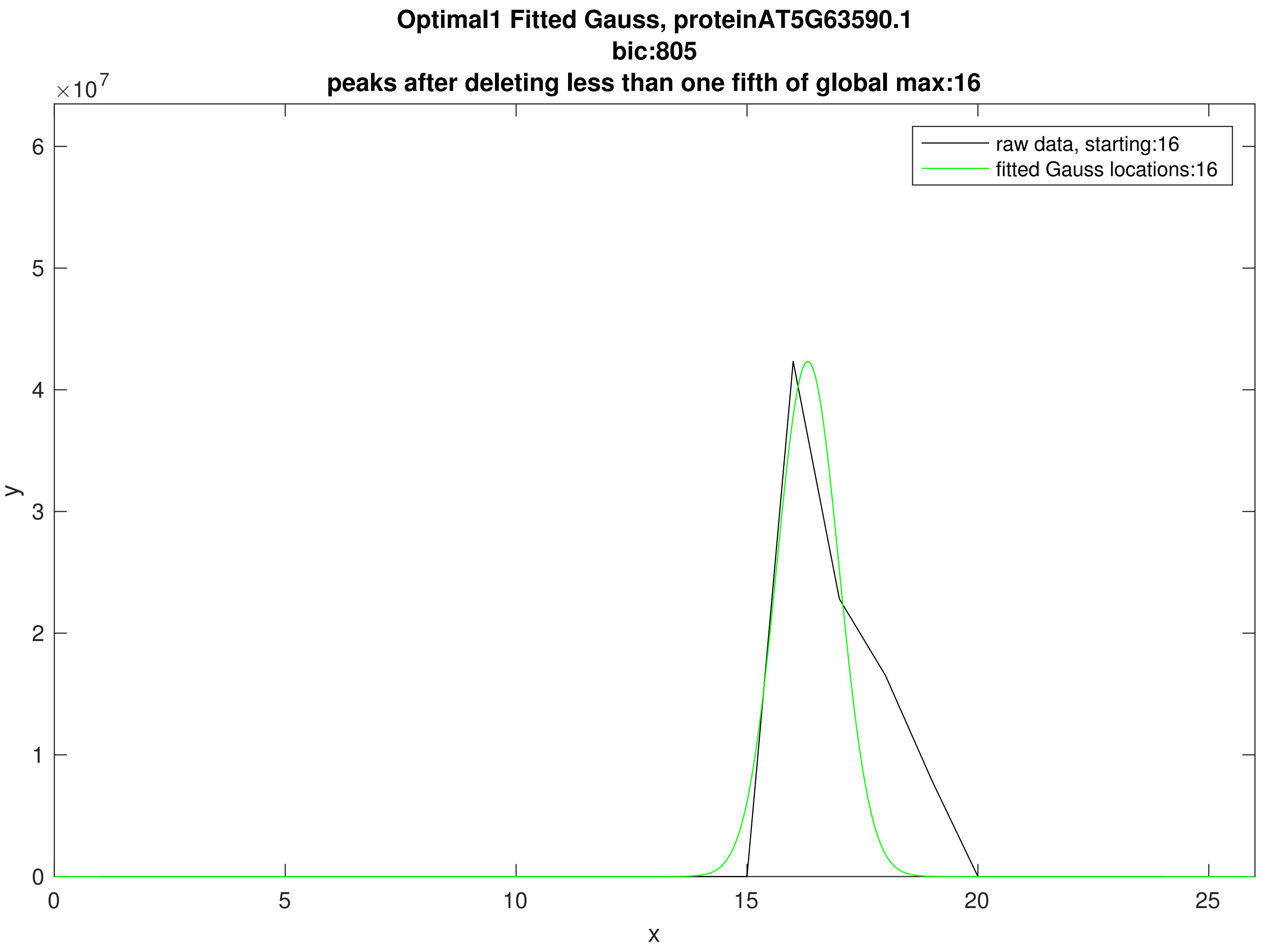
Optimal1 Fitted Gauss, proteinAT5G63590.1
bic:805
peaks after deleting less than one fifth of global max:16
×10^7
6
5
4
3
2
1
0
y
0
5
10
15
20
25
x
raw data, starting:16
fitted Gauss locations:16

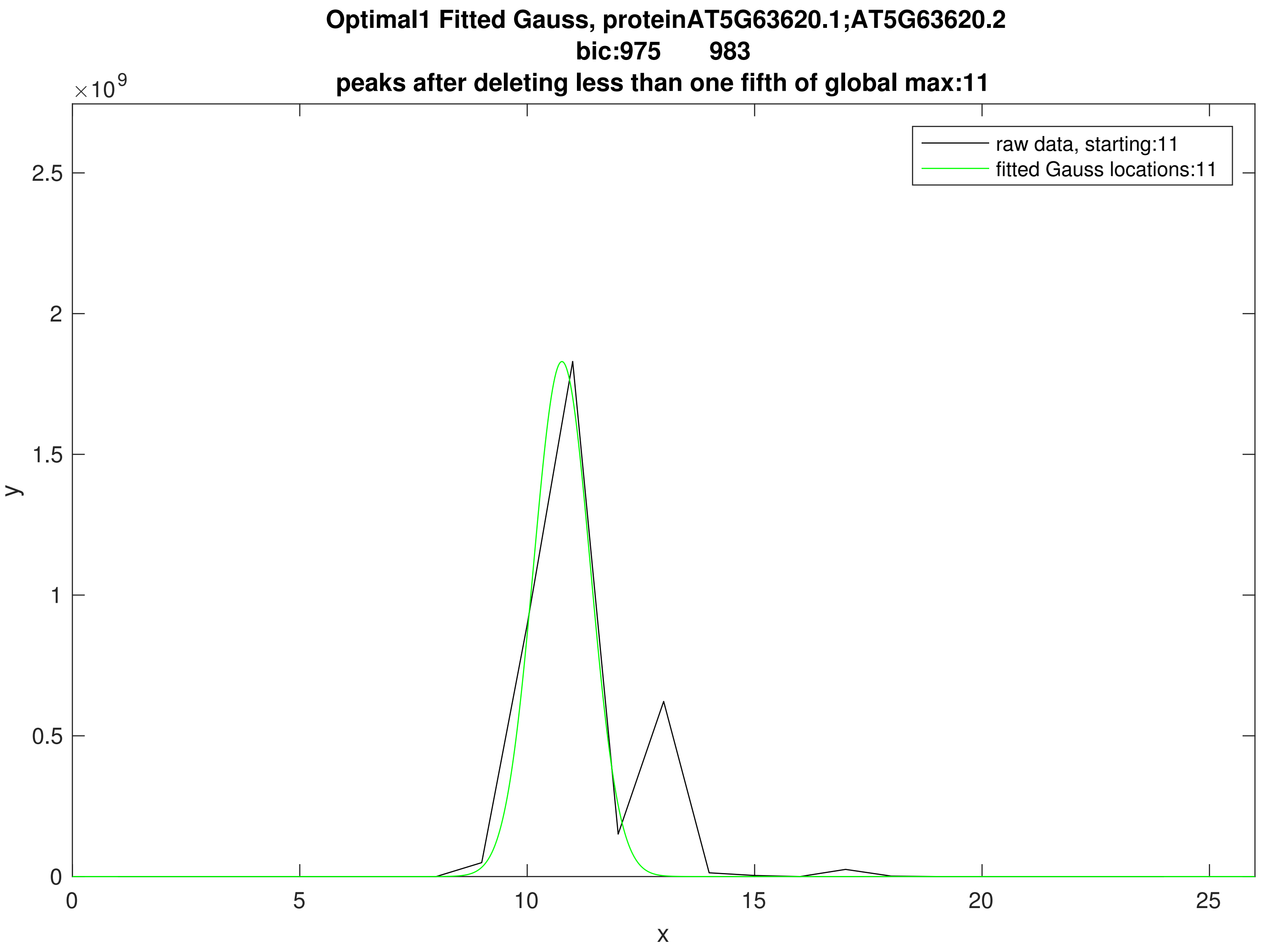
Optimal1 Fitted Gauss, proteinAT5G63620.1;AT5G63620.2
bic:975 983
peaks after deleting less than one fifth of global max:11
×10^9
raw data, starting:11
fitted Gauss locations:11
y
x
0
0.5
1
1.5
2
2.5
0
5
10
15
20
25

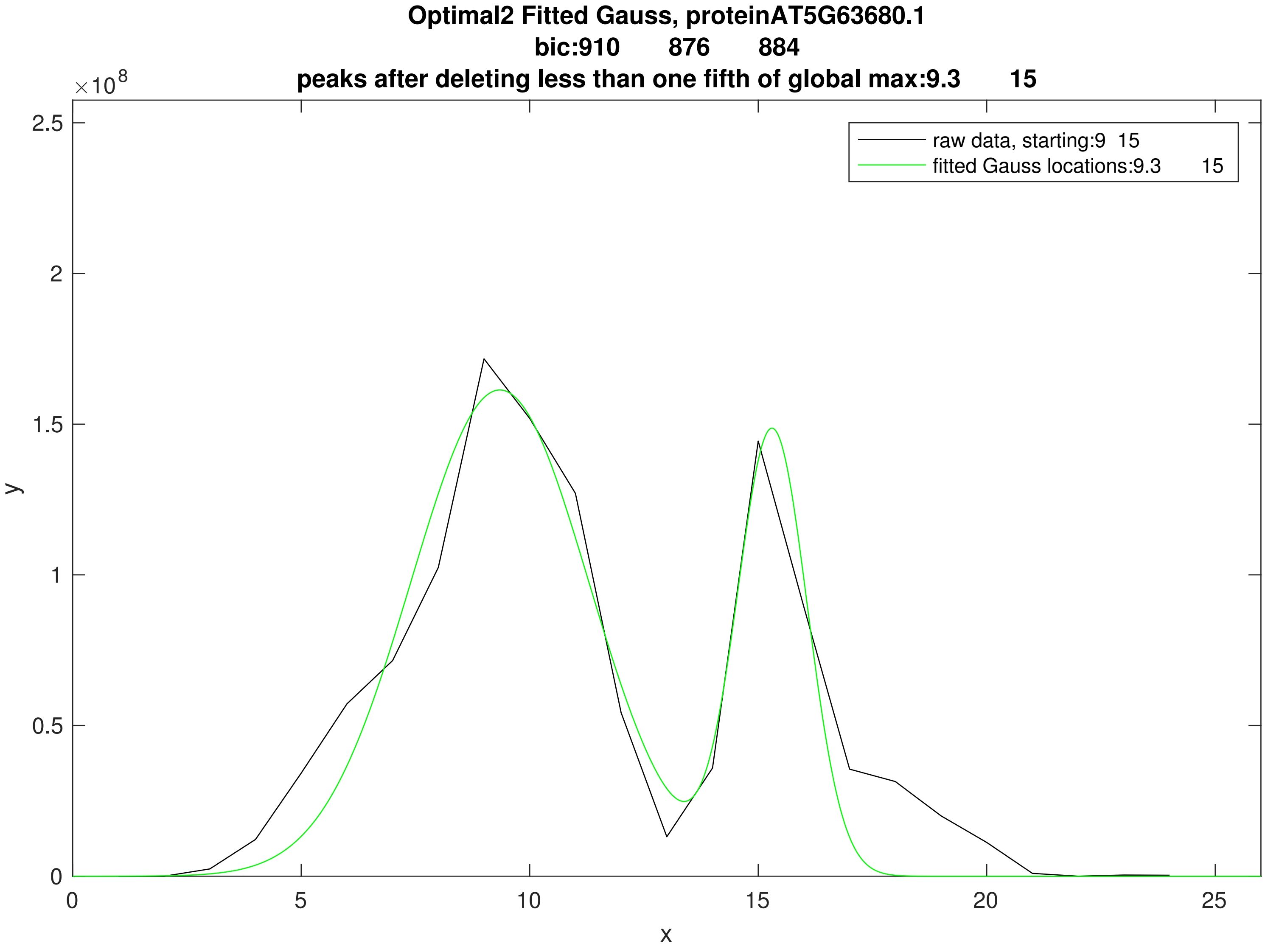

Optimal2 Fitted Gauss, proteinAT5G63680.1
bic:910 876 884
peaks after deleting less than one fifth of global max:9.3 15
×10^8
raw data, starting:9 15
fitted Gauss locations:9.3 15
y
x
0
0.5
1
1.5
2
2.5
5
10
15
20
25

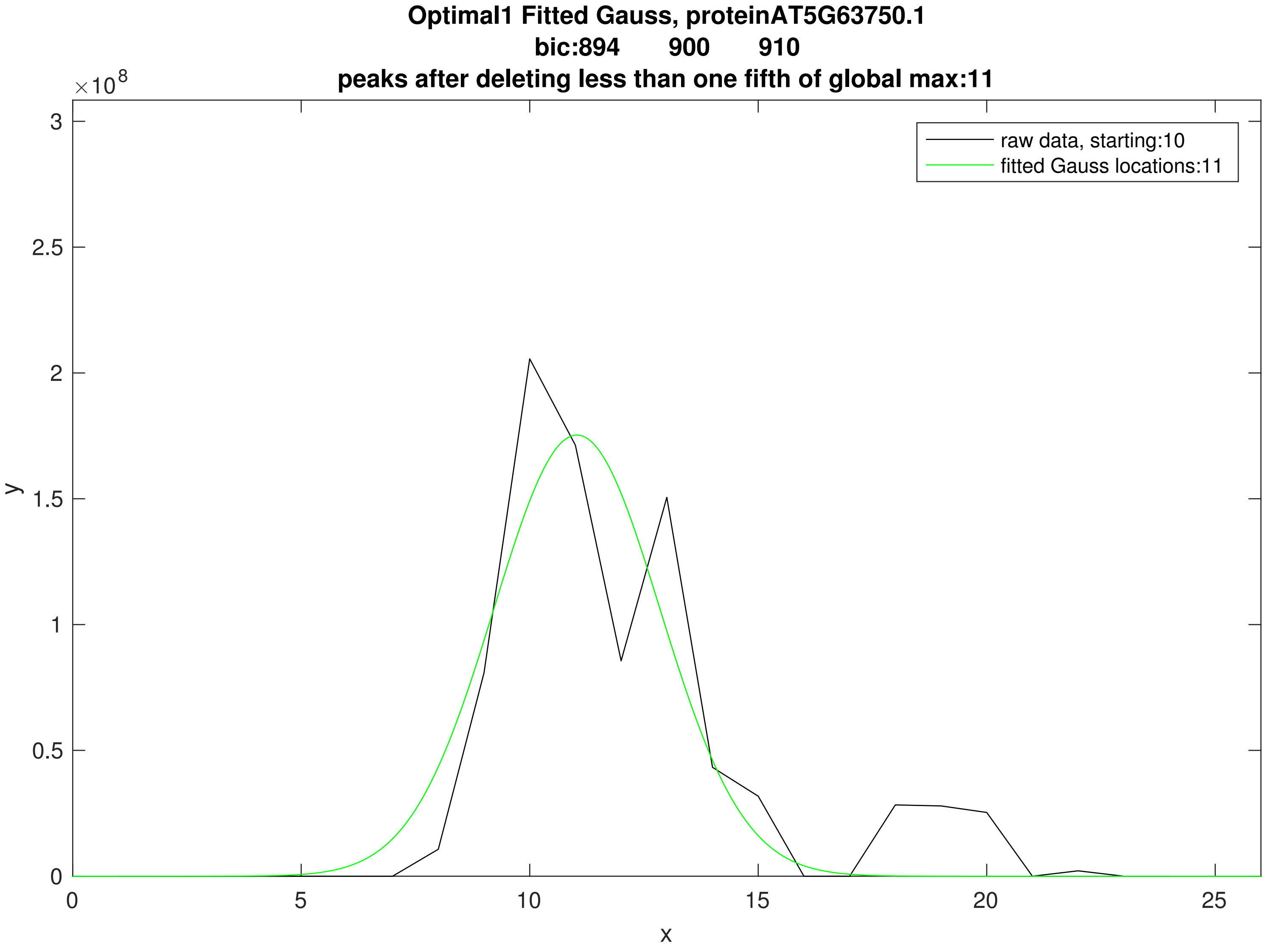
Optimal1 Fitted Gauss, proteinAT5G63750.1
bic:894 900 910
peaks after deleting less than one fifth of global max:11
×10^8
raw data, starting:10
fitted Gauss locations:11
y
x
0
0.5
1
1.5
2
2.5
3
0
5
10
15
20
25

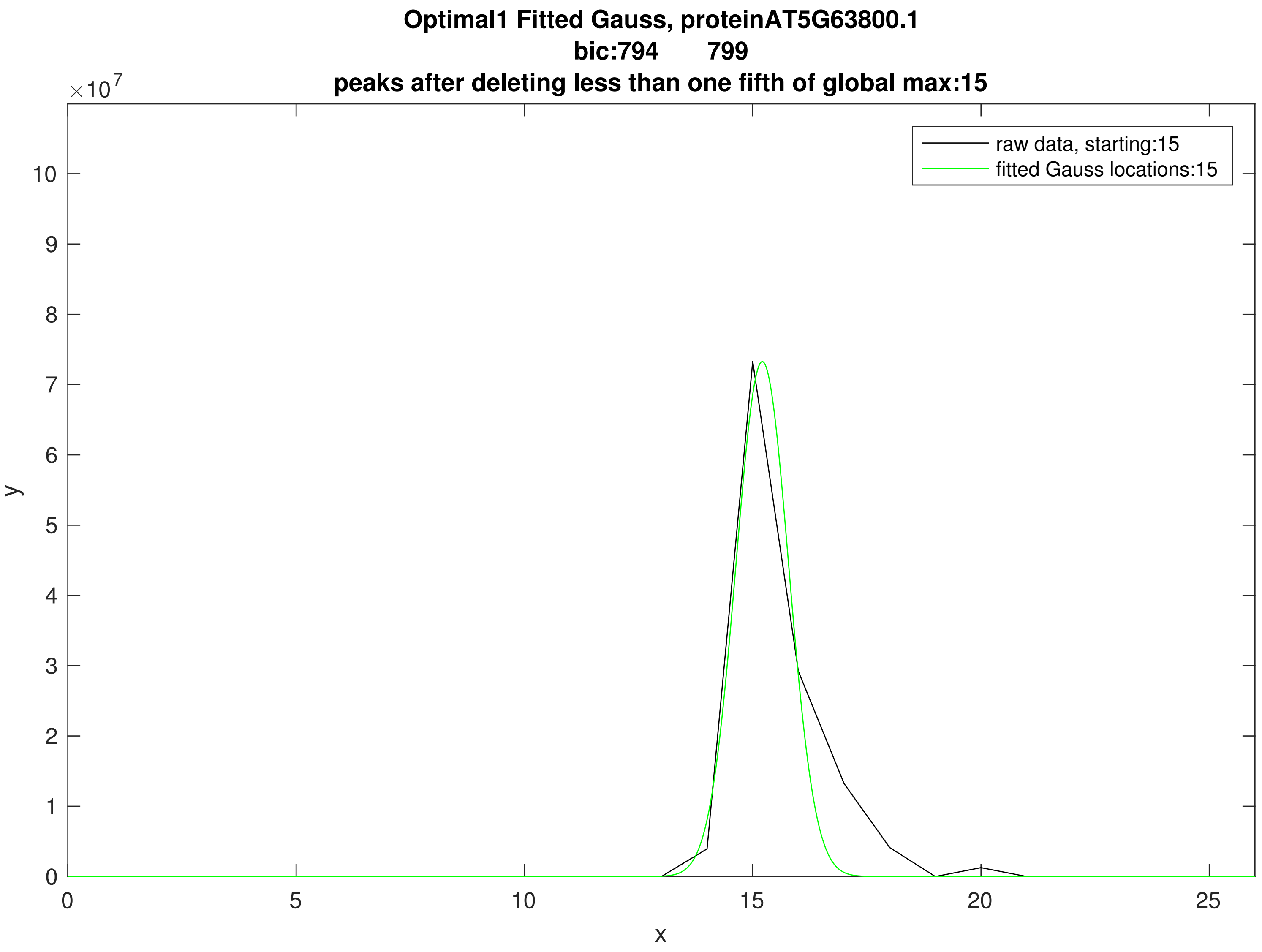
Optimal1 Fitted Gauss, proteinAT5G63800.1
bic:794 799
peaks after deleting less than one fifth of global max:15
×10^7
raw data, starting:15
fitted Gauss locations:15
y
x
0
1
2
3
4
5
6
7
8
9
10
0
5
10
15
20
25

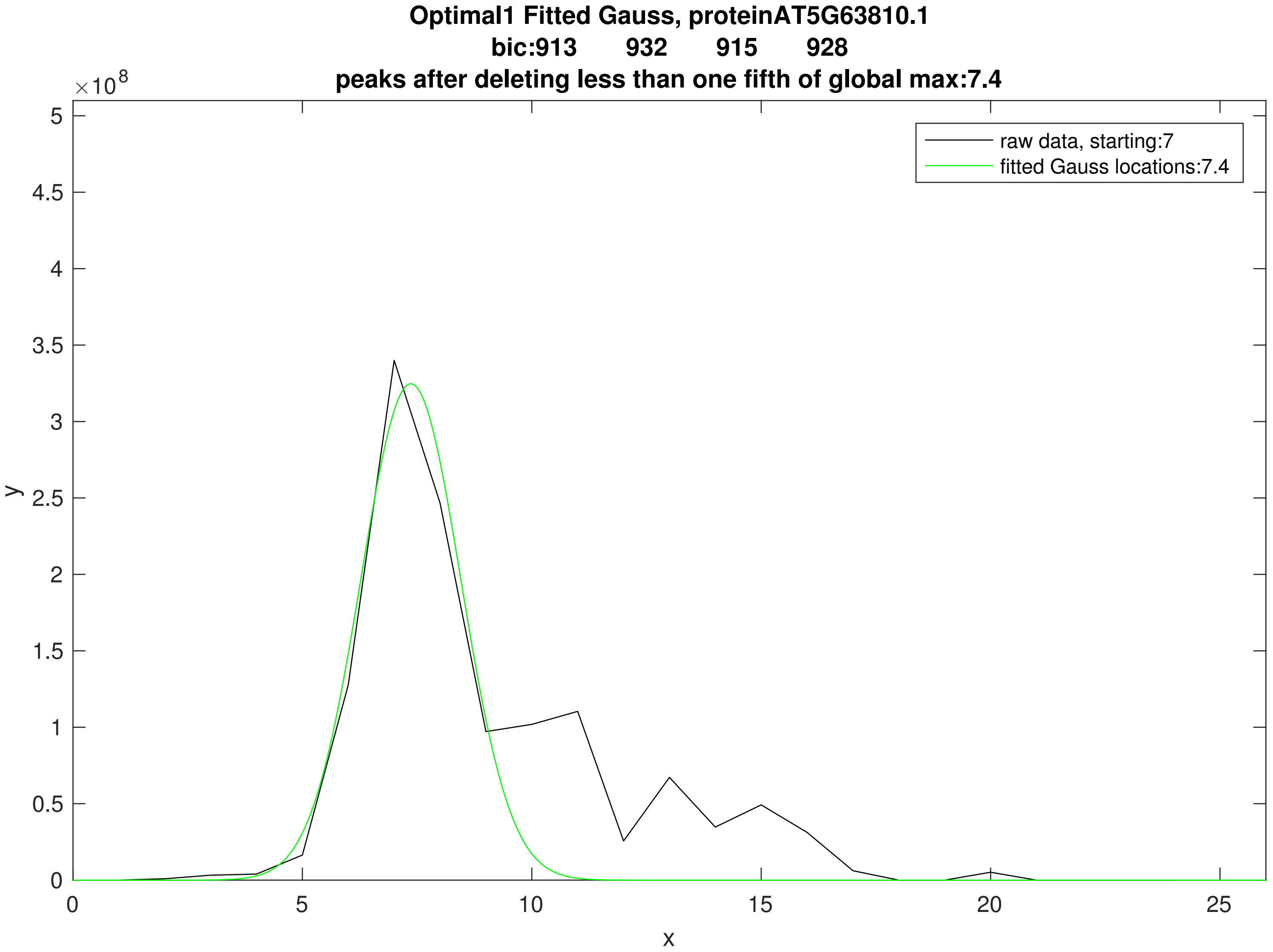
Optimal1 Fitted Gauss, proteinAT5G63810.1
bic:913 932 915 928
peaks after deleting less than one fifth of global max:7.4
×10^8
raw data, starting:7
fitted Gauss locations:7.4
y
x
0
0.5
1
1.5
2
2.5
3
3.5
4
4.5
5
0
5
10
15
20
25

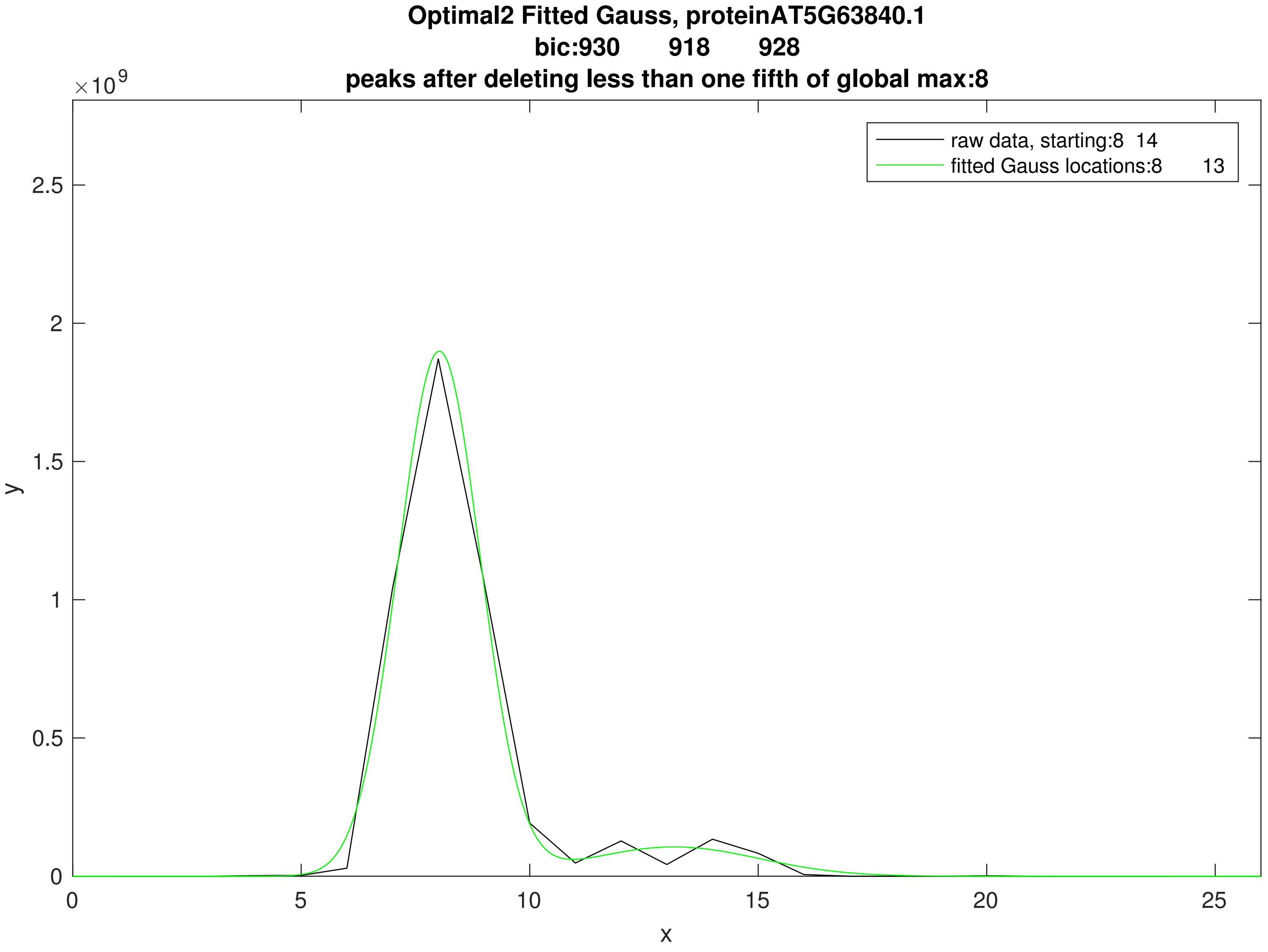

Optimal2 Fitted Gauss, proteinAT5G63840.1
bic:930 918 928
peaks after deleting less than one fifth of global max:8
×10^9
raw data, starting:8 14
fitted Gauss locations:8 13
y
x
0
0.5
1
1.5
2
2.5
5
10
15
20
25

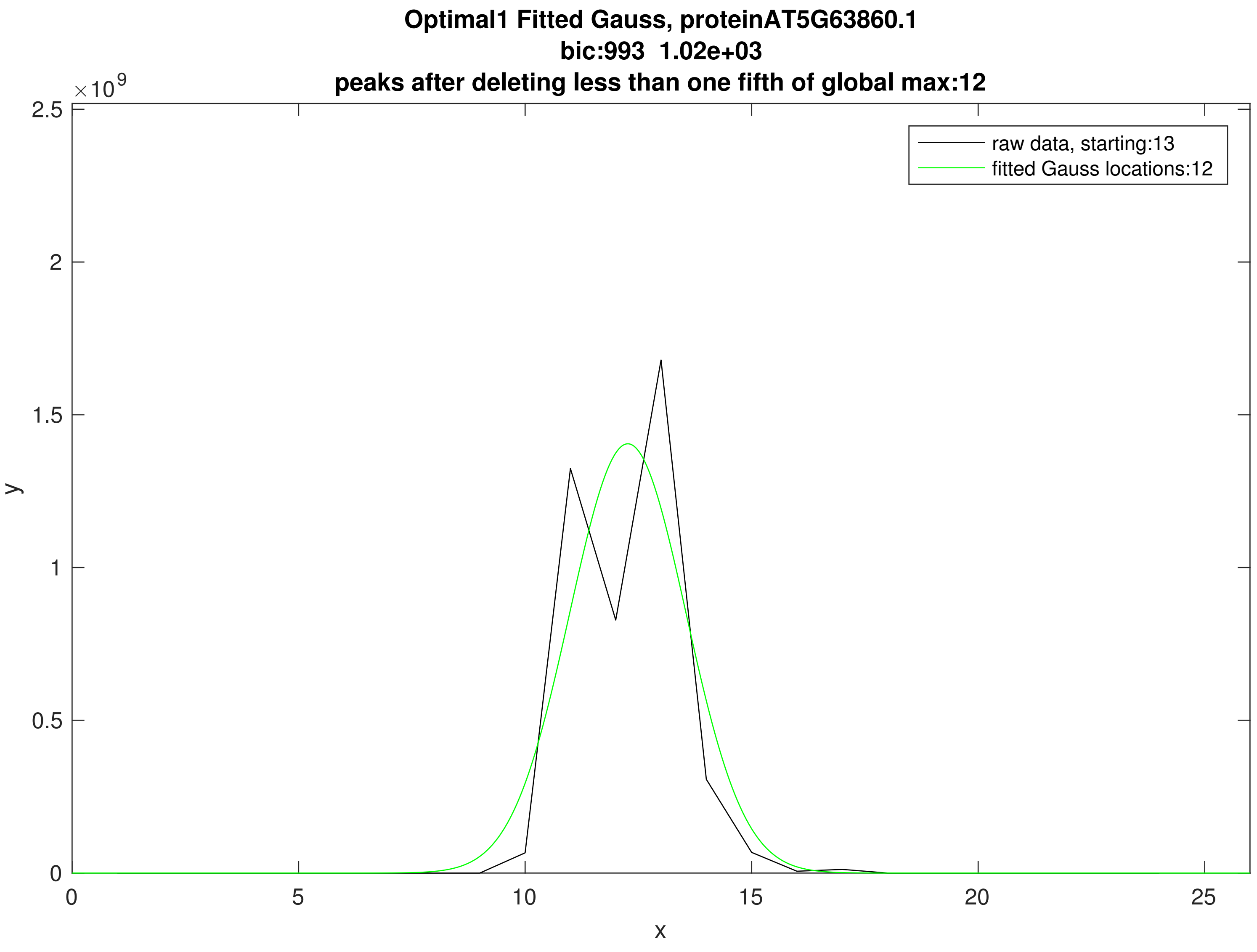
Optimal1 Fitted Gauss, proteinAT5G63860.1
bic:993 1.02e+03
peaks after deleting less than one fifth of global max:12
×10^9
2.5
2
1.5
1
0.5
0
y
0
5
10
15
20
25
x
raw data, starting:13
fitted Gauss locations:12

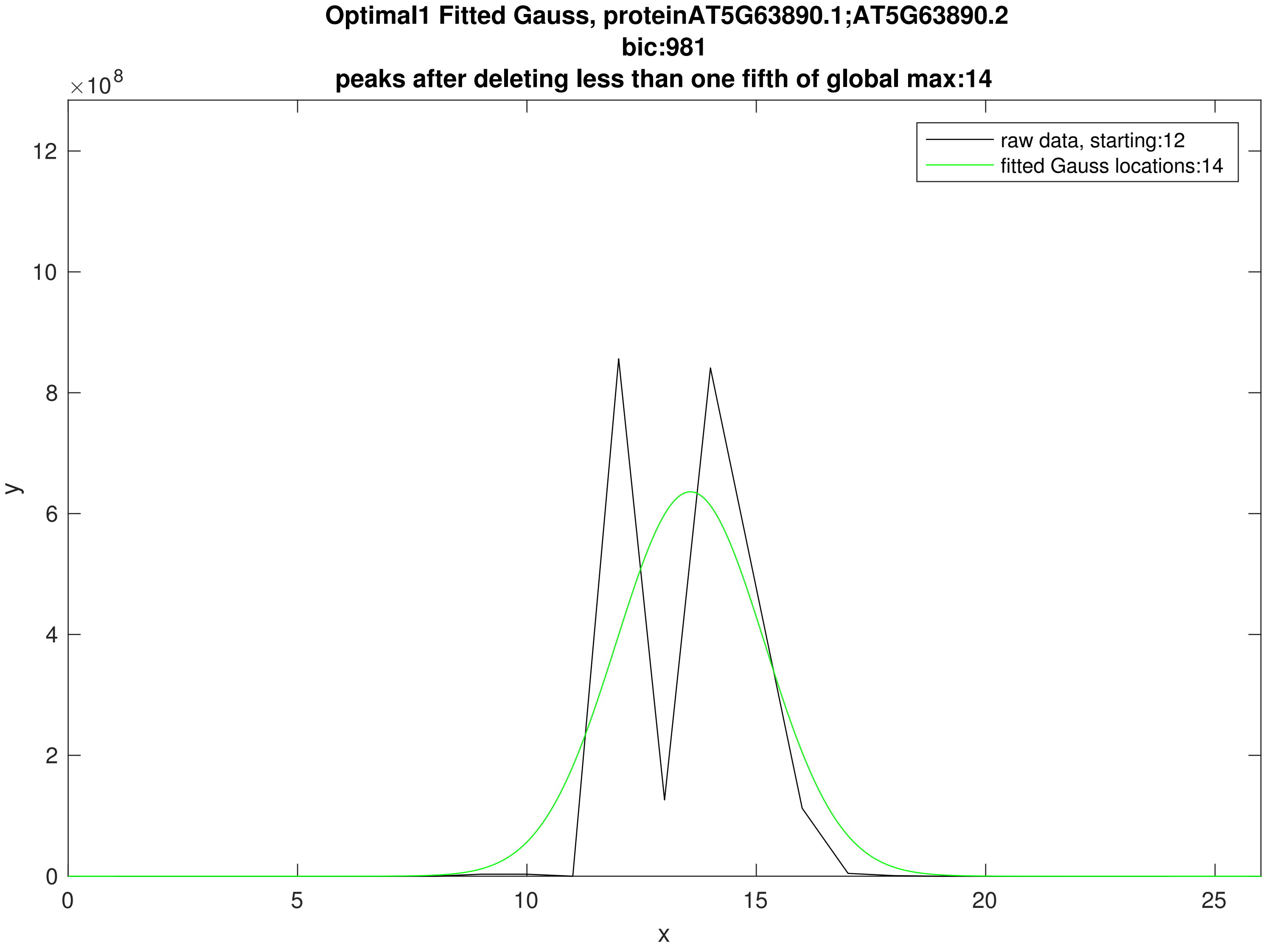
Optimal1 Fitted Gauss, proteinAT5G63890.1;AT5G63890.2
bic:981
peaks after deleting less than one fifth of global max:14
×10^8
raw data, starting:12
fitted Gauss locations:14
y
x
0
2
4
6
8
10
12
0
5
10
15
20
25

#### Optimal2 Fitted Gauss, proteinAT5G63980.1
#### bic:987 969
#### peaks after deleting less than one fifth of global max:16

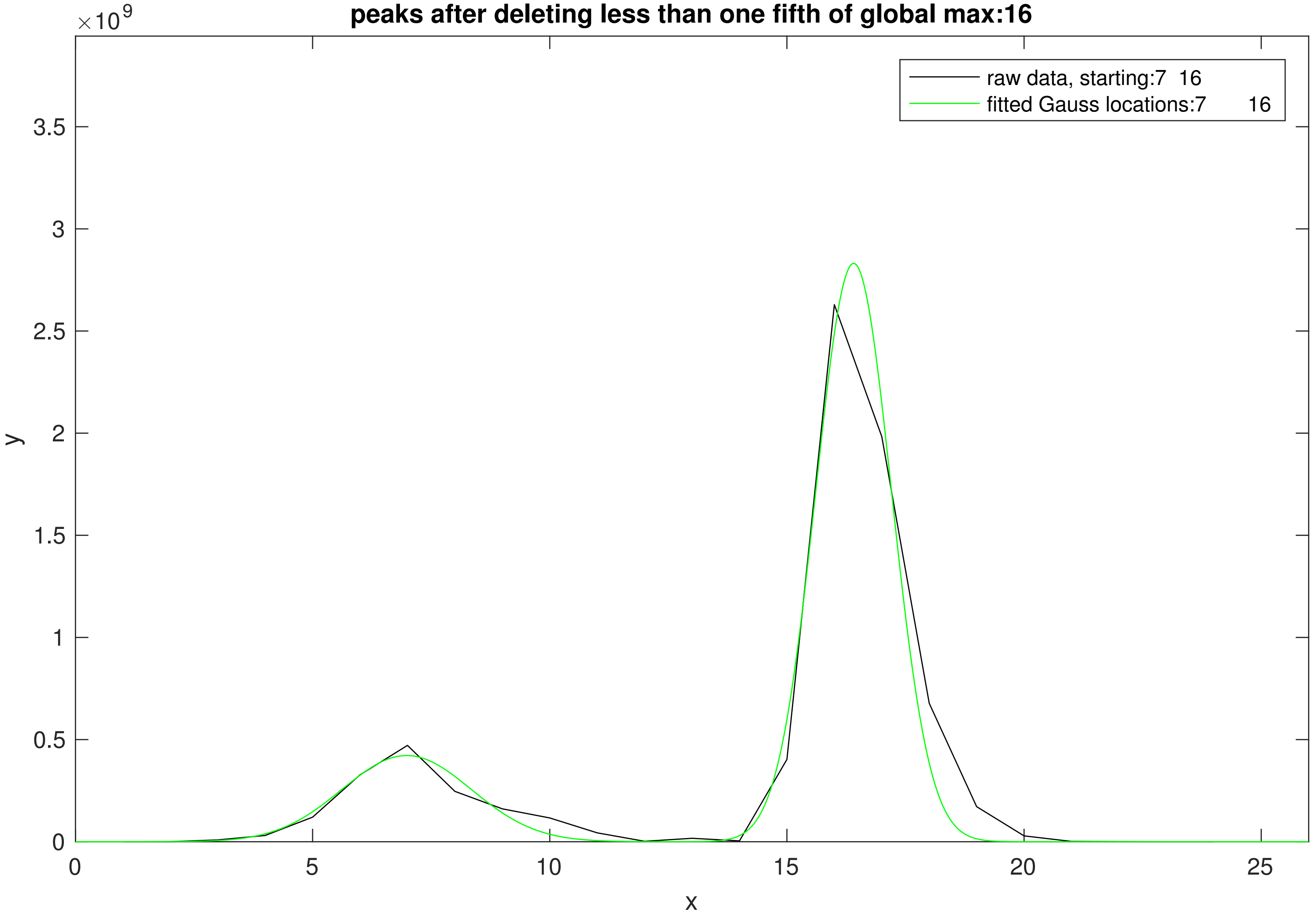

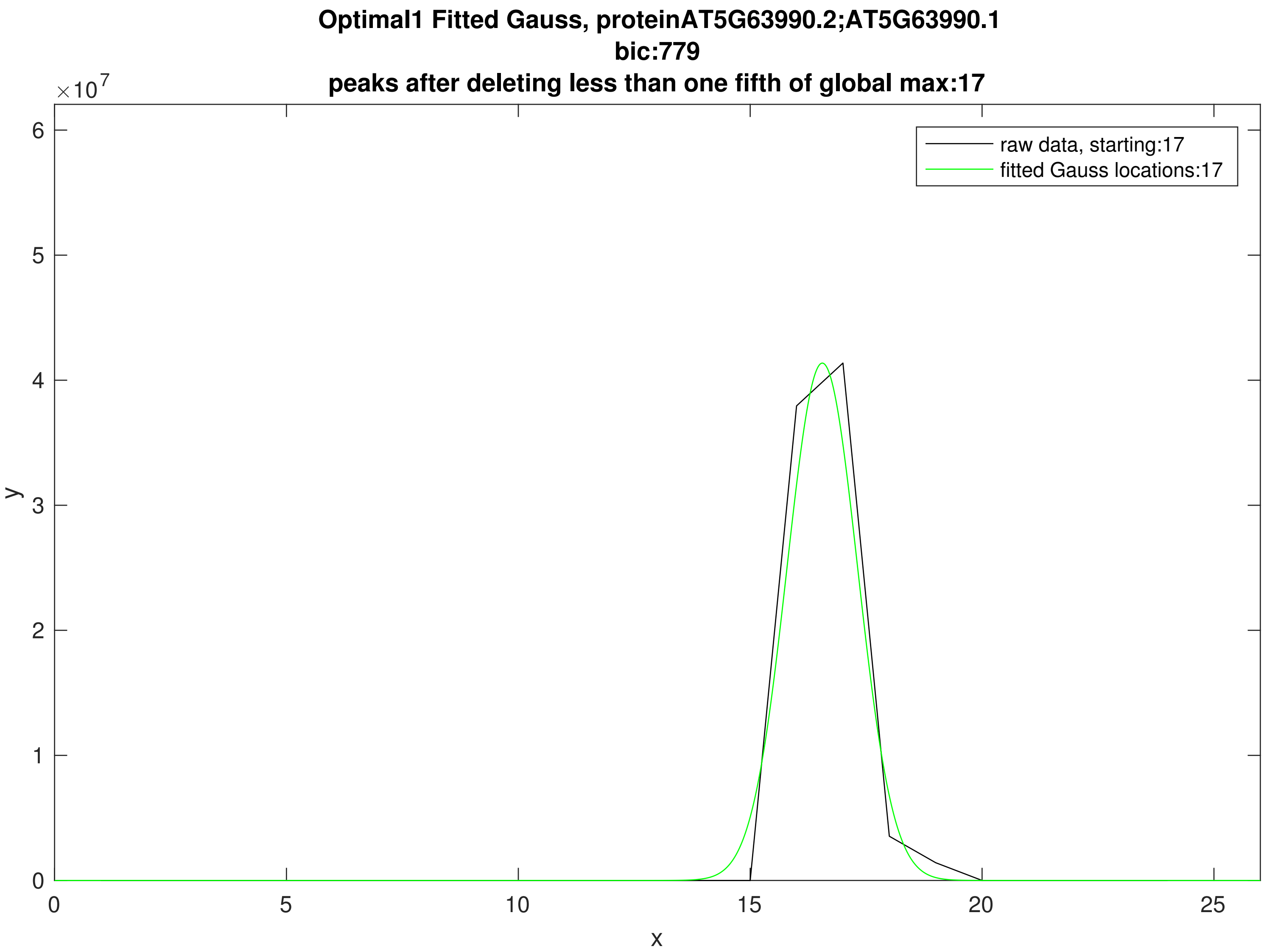
Optimal1 Fitted Gauss, proteinAT5G63990.2;AT5G63990.1
bic:779
peaks after deleting less than one fifth of global max:17
×10^7
raw data, starting:17
fitted Gauss locations:17
y
x
0
1
2
3
4
5
6
0
5
10
15
20
25

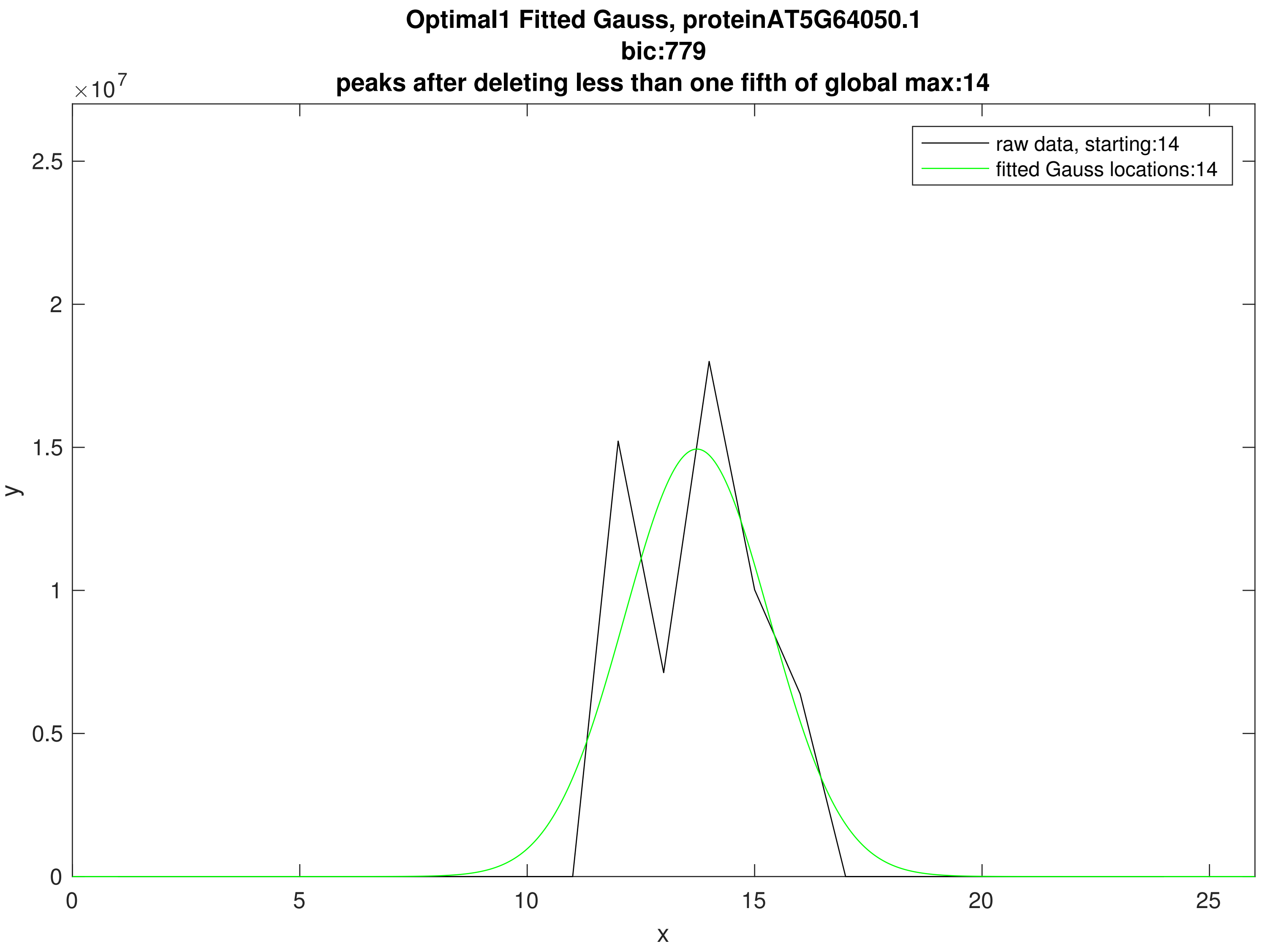

Optimal1 Fitted Gauss, proteinAT5G64050.1
bic:779
peaks after deleting less than one fifth of global max:14
×10^7
2.5
2
1.5
1
0.5
0
y
0
5
10
15
20
25
x
raw data, starting:14
fitted Gauss locations:14

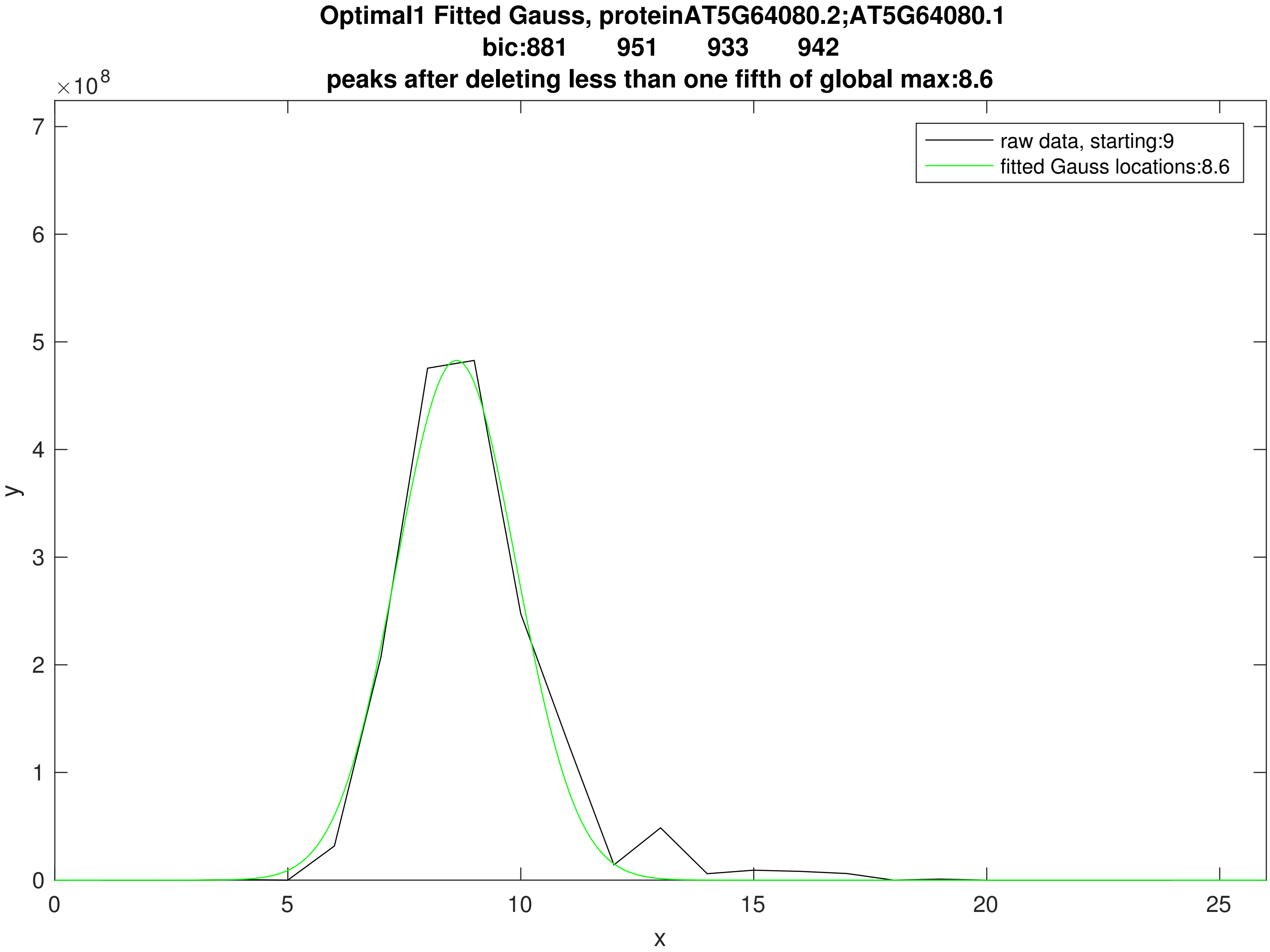

Optimal1 Fitted Gauss, proteinAT5G64080.2;AT5G64080.1
bic:881 951 933 942
peaks after deleting less than one fifth of global max:8.6
×10^8
raw data, starting:9
fitted Gauss locations:8.6
y
x
0
1
2
3
4
5
6
7
10
15
20
25

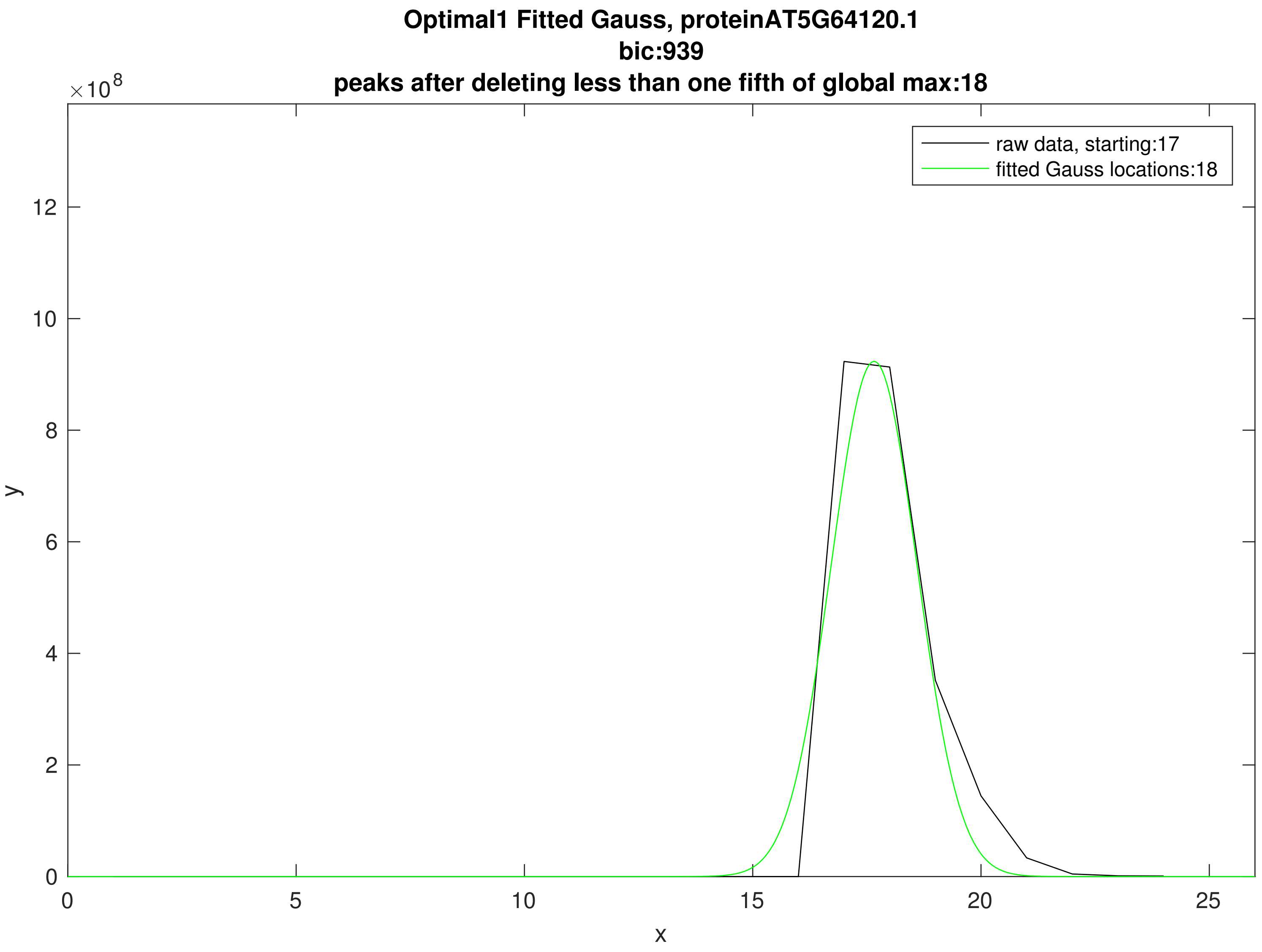

Optimal1 Fitted Gauss, proteinAT5G64120.1
bic:939
peaks after deleting less than one fifth of global max:18
×10^8
raw data, starting:17
fitted Gauss locations:18
y
x
0
2
4
6
8
10
12
0
5
10
15
20
25

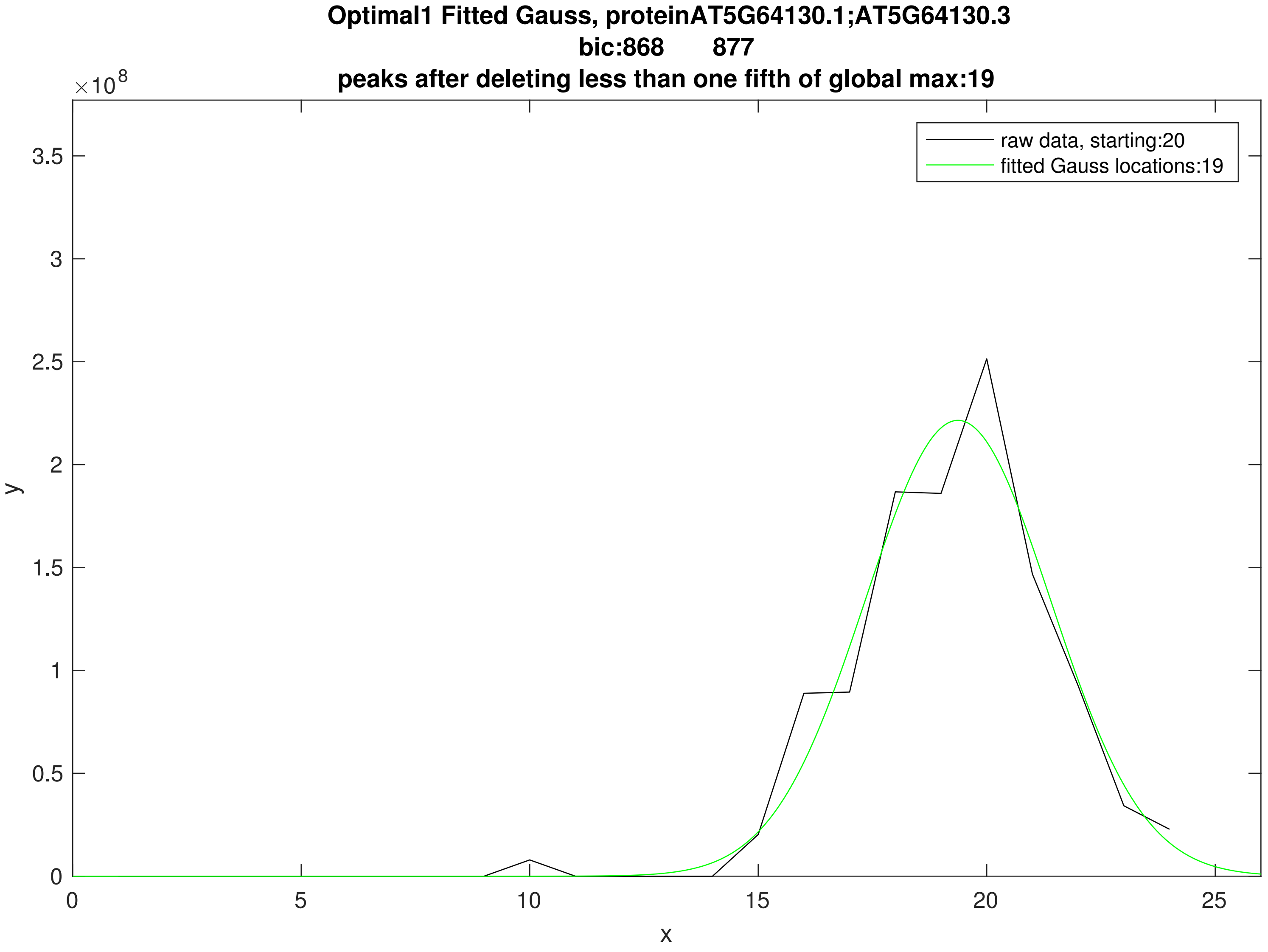
Optimal1 Fitted Gauss, proteinAT5G64130.1;AT5G64130.3
bic:868 877
peaks after deleting less than one fifth of global max:19
×10^8
3.5
3
2.5
2
1.5
1
0.5
0
y
0
5
10
15
20
25
x
raw data, starting:20
fitted Gauss locations:19

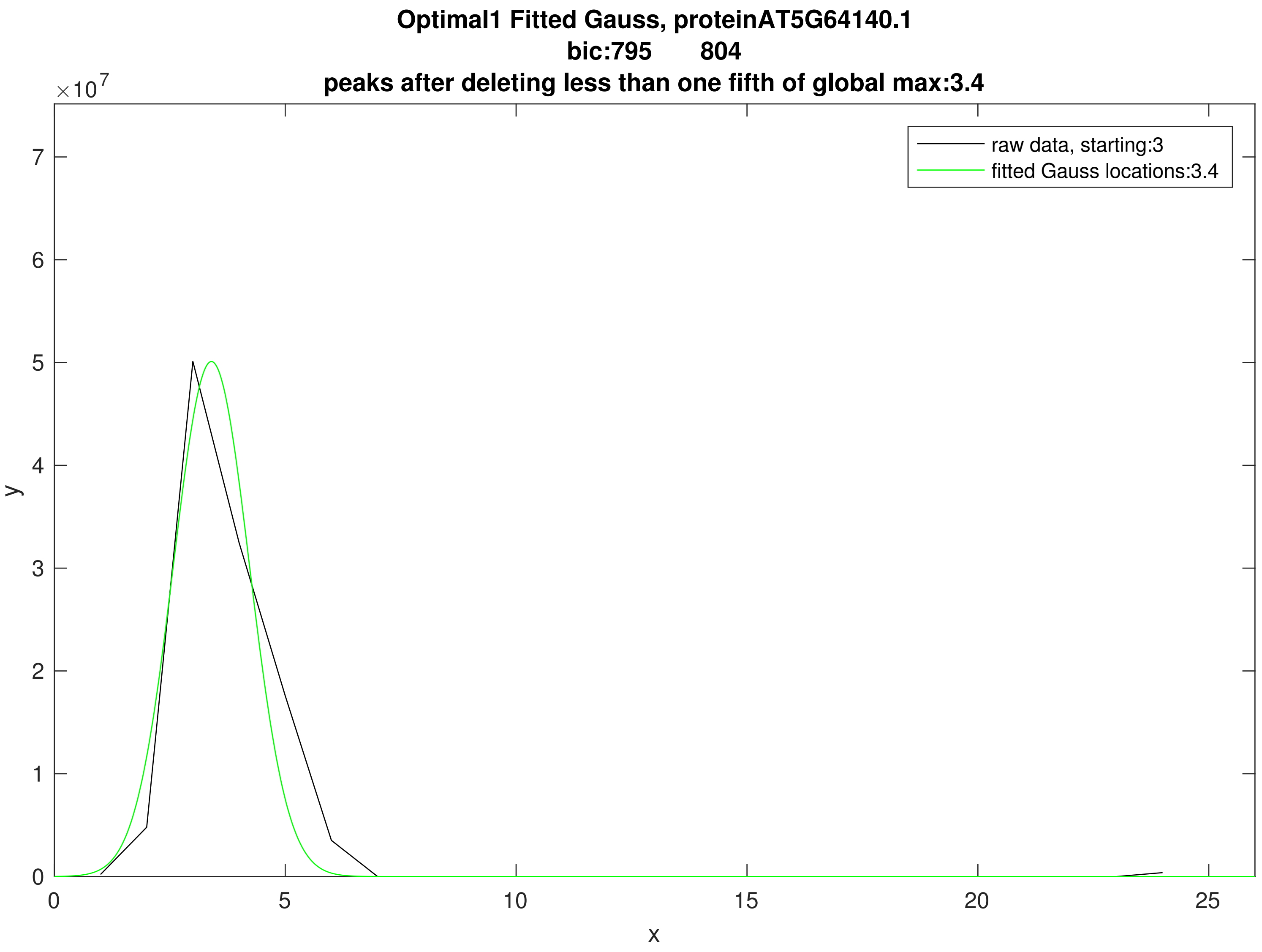
Optimal1 Fitted Gauss, proteinAT5G64140.1
bic:795 804
peaks after deleting less than one fifth of global max:3.4
×10^7
raw data, starting:3
fitted Gauss locations:3.4
y
x
0
1
2
3
4
5
6
7
0
5
10
15
20
25

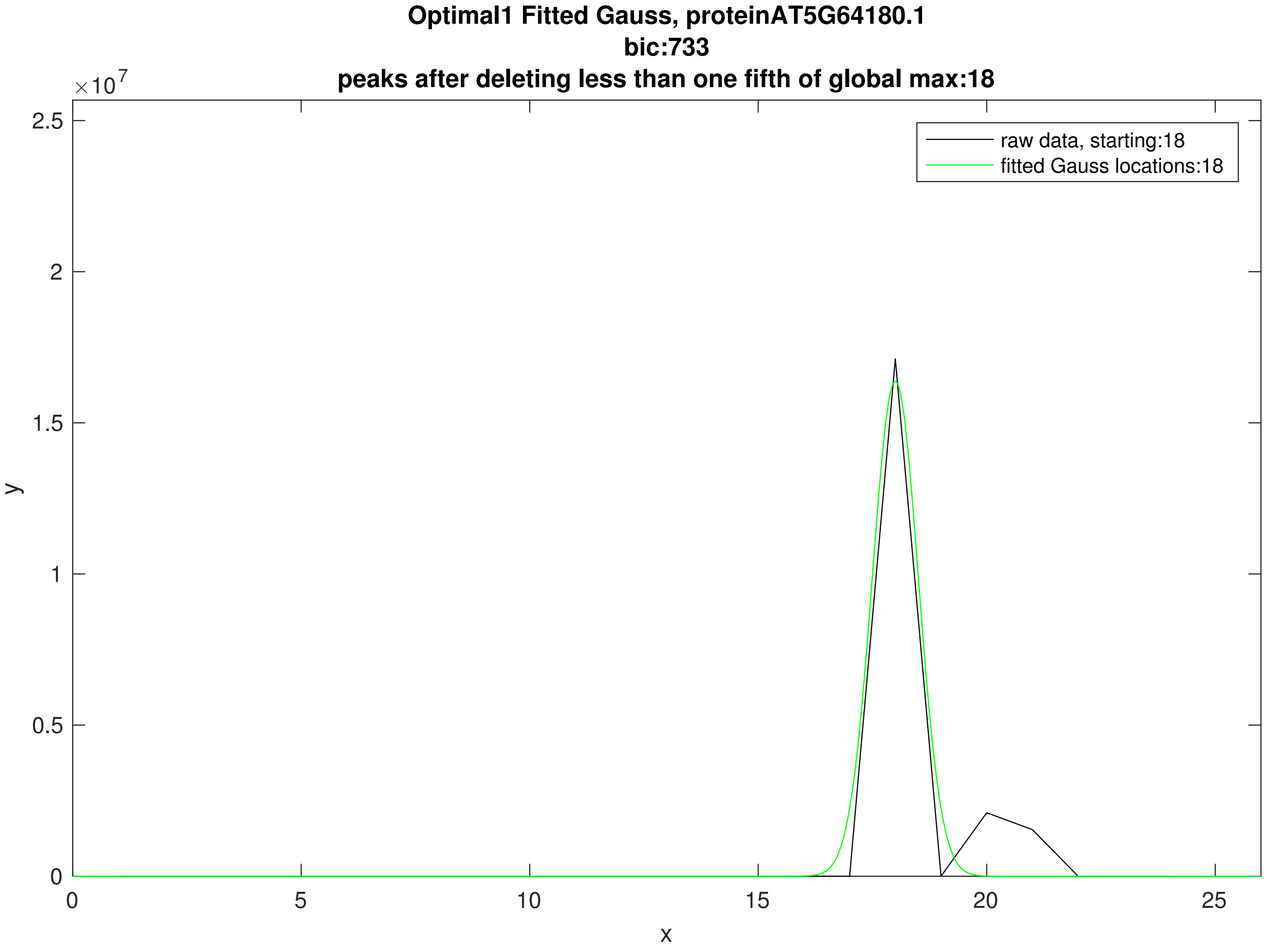
Optimal1 Fitted Gauss, proteinAT5G64180.1
bic:733
peaks after deleting less than one fifth of global max:18
×10^7
2.5
2
1.5
1
0.5
0
y
0
5
10
15
20
25
x
raw data, starting:18
fitted Gauss locations:18

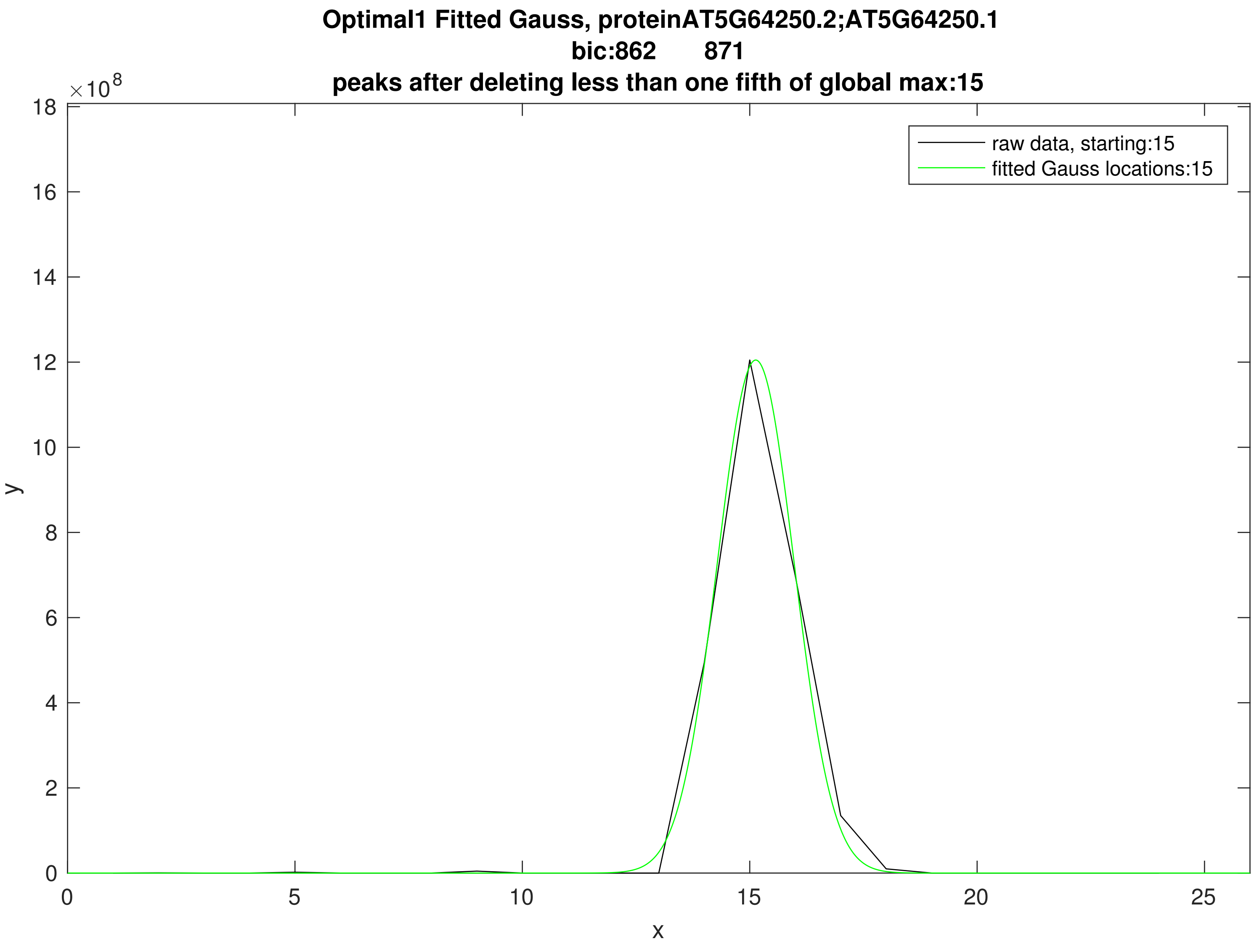

Optimal1 Fitted Gauss, proteinAT5G64250.2;AT5G64250.1
bic:862 871
peaks after deleting less than one fifth of global max:15
×10^8
18
16
14
12
10
8
6
4
2
0
y
0
5
10
15
20
25
x
raw data, starting:15
fitted Gauss locations:15

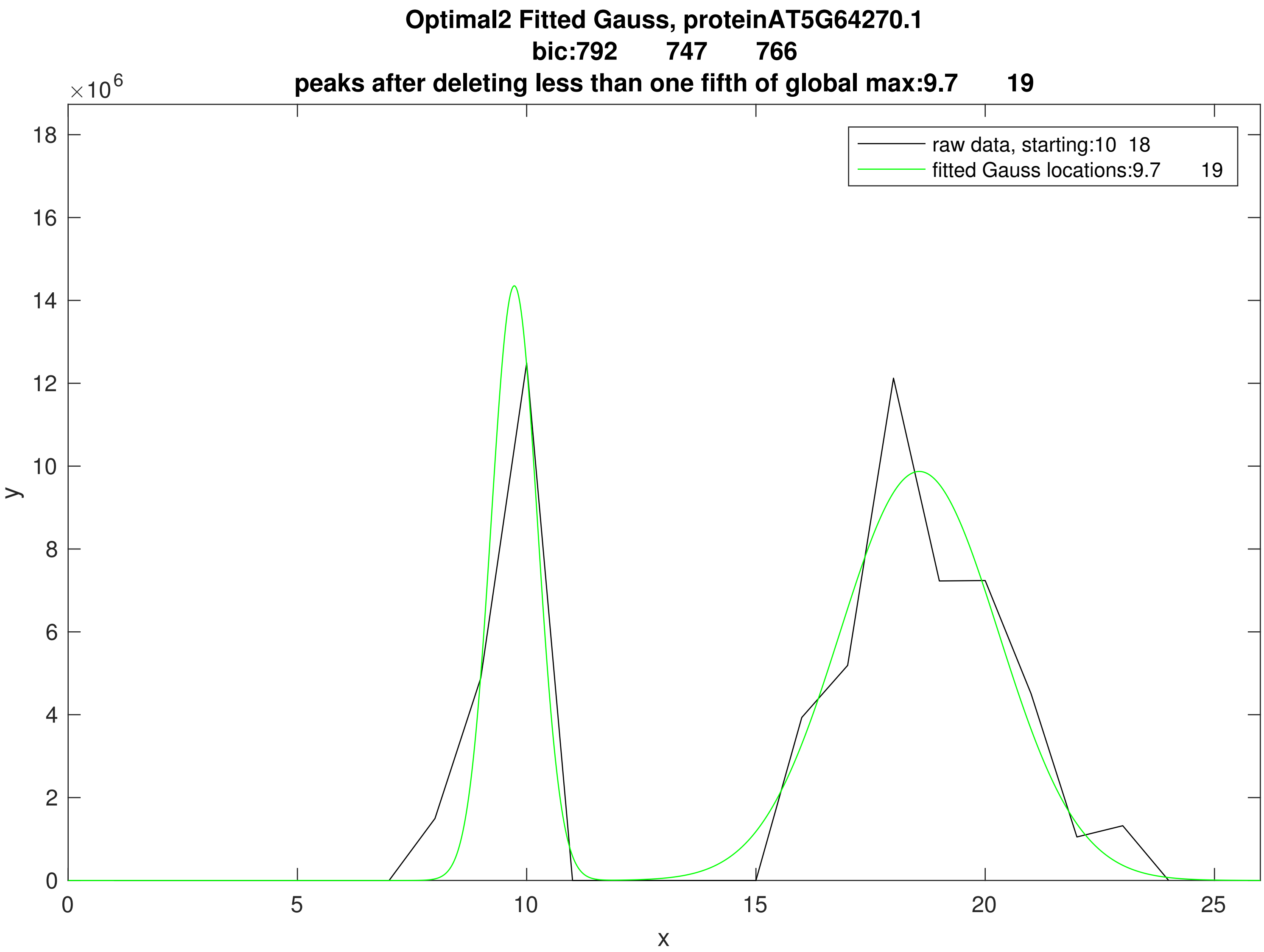
Optimal2 Fitted Gauss, proteinAT5G64270.1
bic:792 747 766
peaks after deleting less than one fifth of global max:9.7 19
×10^6
raw data, starting:10 18
fitted Gauss locations:9.7 19
y
x
0
2
4
6
8
10
12
14
16
18
0
5
10
15
20
25

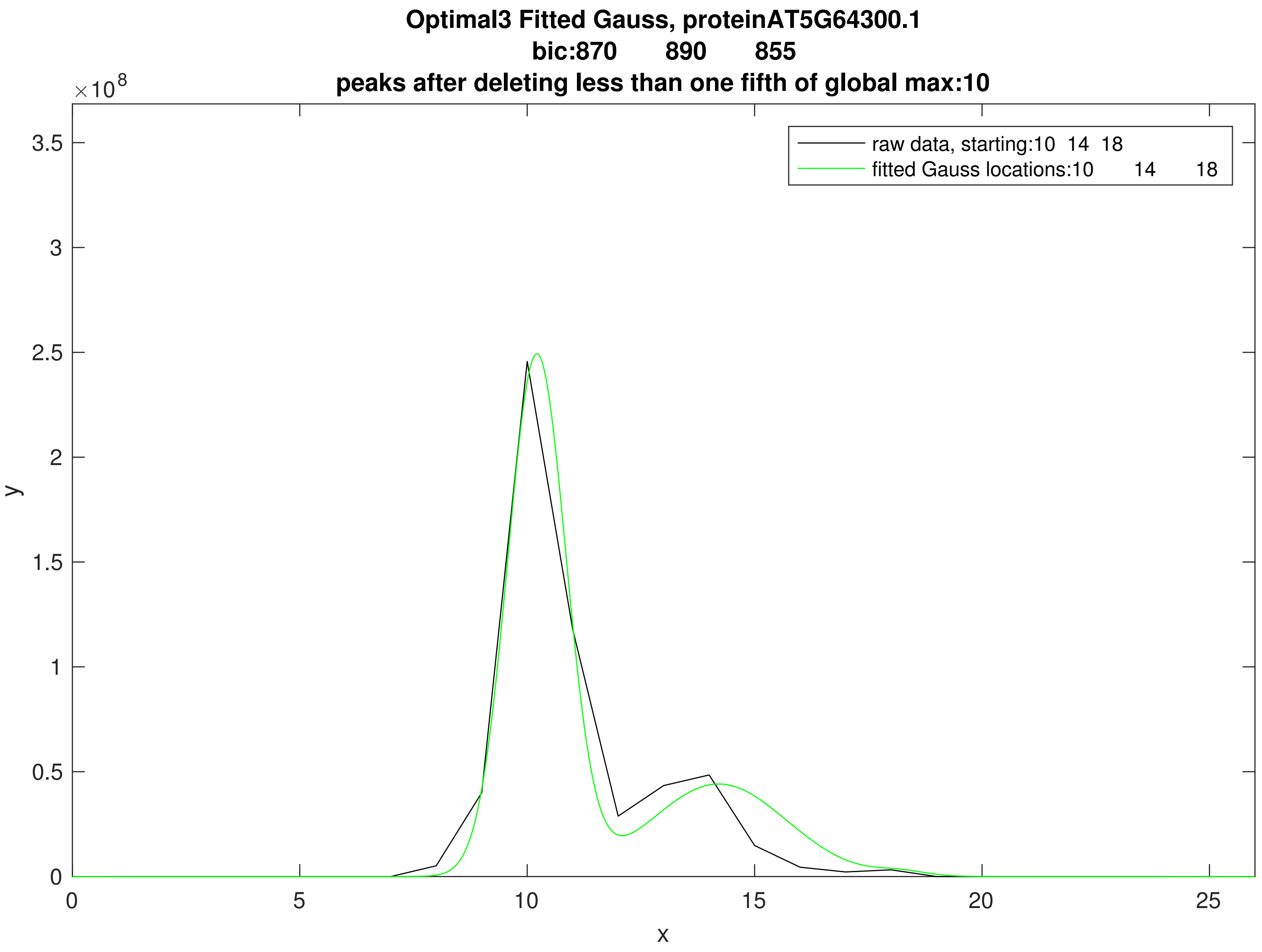
Optimal3 Fitted Gauss, proteinAT5G64300.1
bic:870 890 855
peaks after deleting less than one fifth of global max:10
×10^8
raw data, starting:10 14 18
fitted Gauss locations:10 14 18
3.5
3
2.5
2
1.5
1
0.5
0
y
0
5
10
15
20
25
x

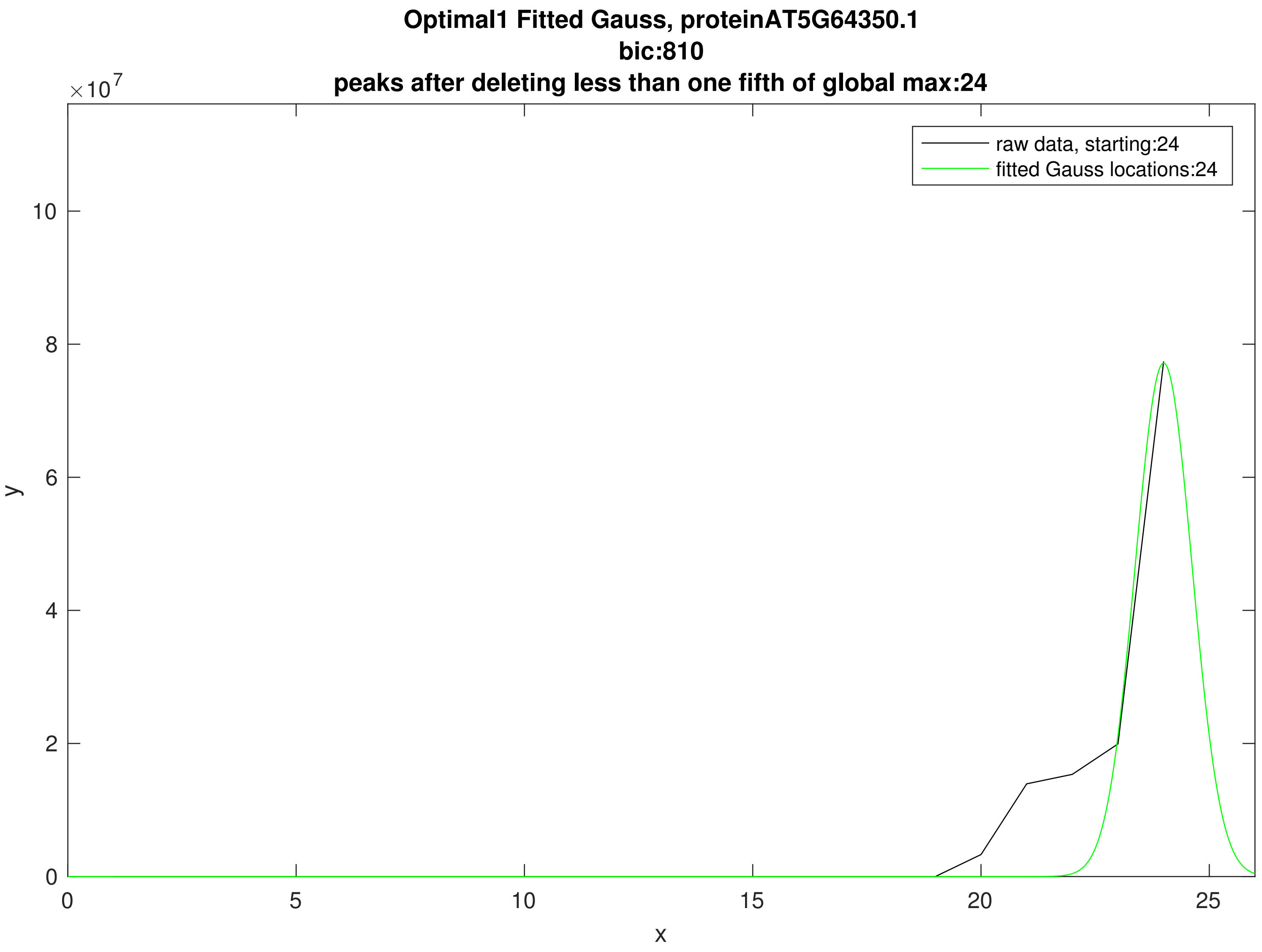
Optimal1 Fitted Gauss, proteinAT5G64350.1
bic:810
peaks after deleting less than one fifth of global max:24
×10^7
raw data, starting:24
fitted Gauss locations:24
y
x
0
2
4
6
8
10
0
5
10
15
20
25

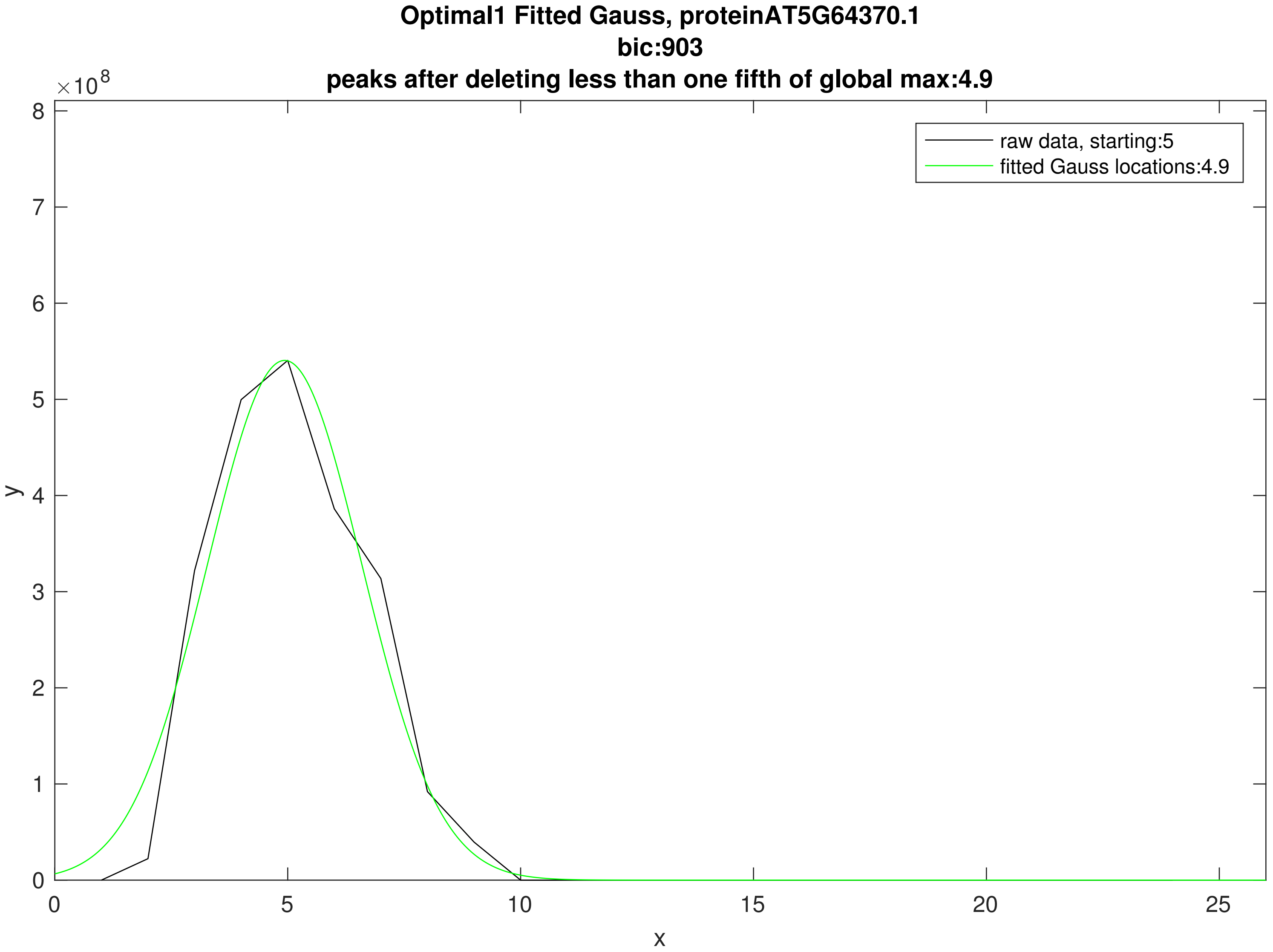
Optimal1 Fitted Gauss, proteinAT5G64370.1
bic:903
peaks after deleting less than one fifth of global max:4.9
×10^8
8
7
6
5
4
3
2
1
0
y
0
5
10
15
20
25
x
raw data, starting:5
fitted Gauss locations:4.9

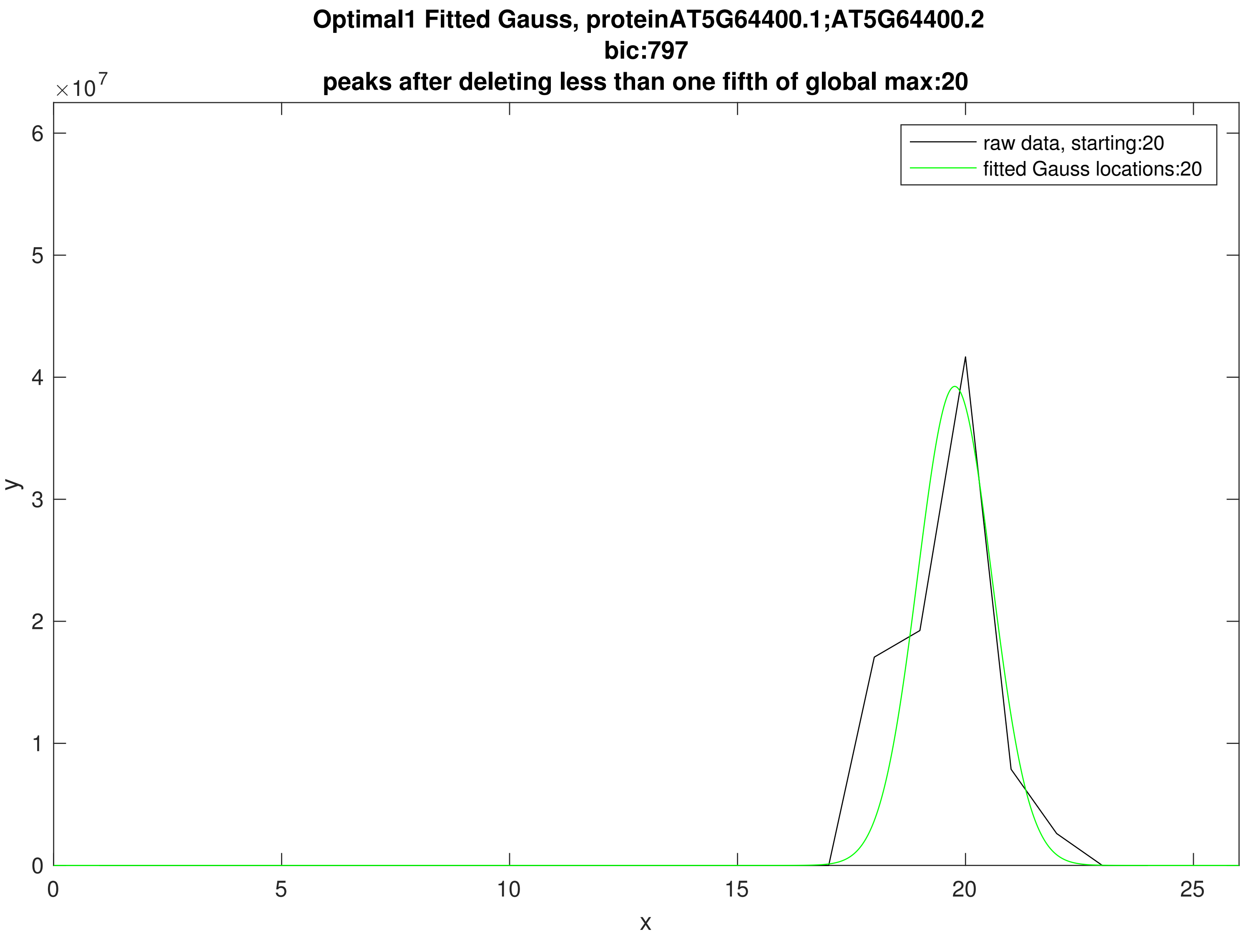

Optimal1 Fitted Gauss, proteinAT5G64400.1;AT5G64400.2
bic:797
peaks after deleting less than one fifth of global max:20
×10^7
raw data, starting:20
fitted Gauss locations:20
y
x
0
1
2
3
4
5
6
0
5
10
15
20
25

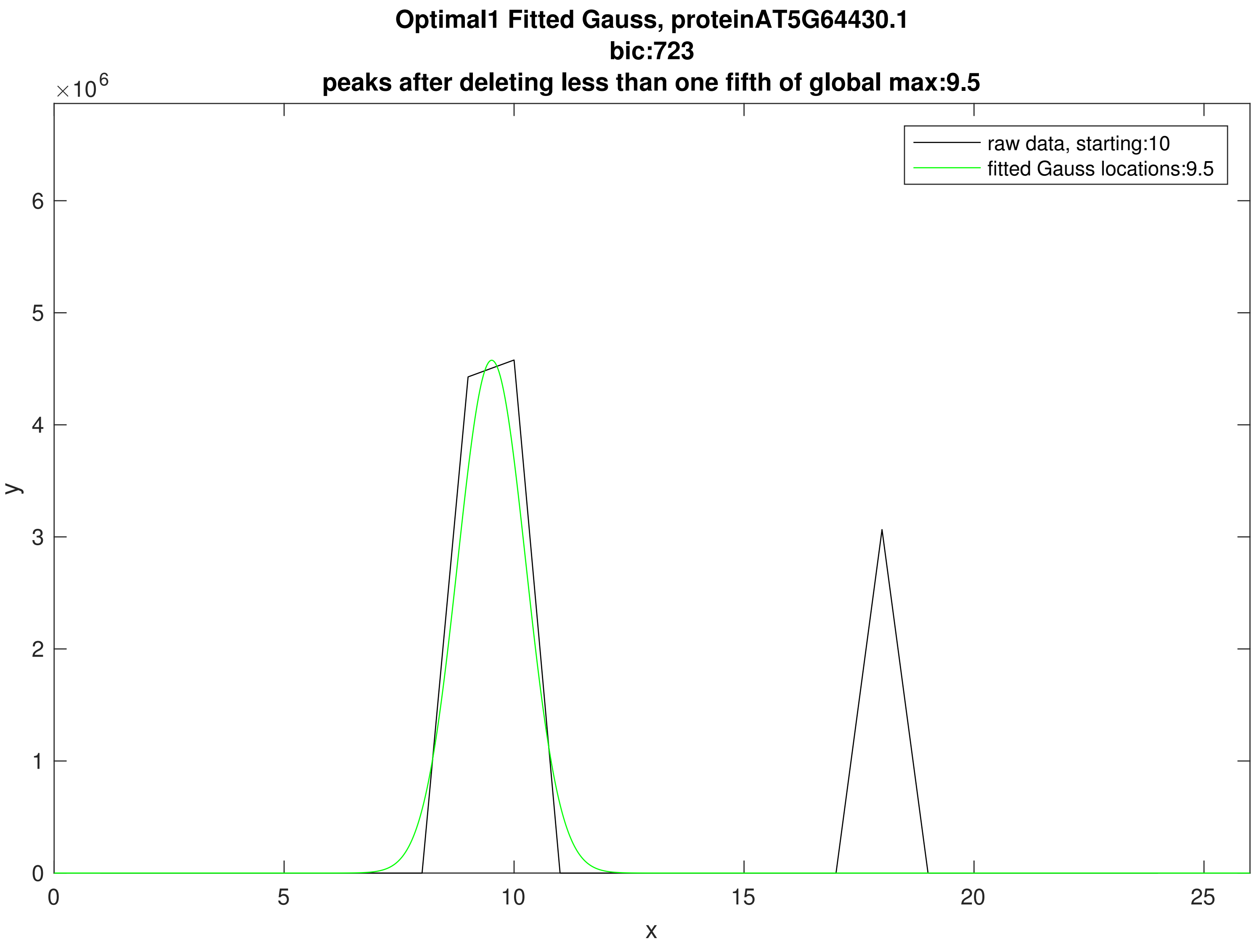

Optimal1 Fitted Gauss, proteinAT5G64430.1
bic:723
peaks after deleting less than one fifth of global max:9.5
×10^6
raw data, starting:10
fitted Gauss locations:9.5
y
x
0
1
2
3
4
5
6
0
5
10
15
20
25

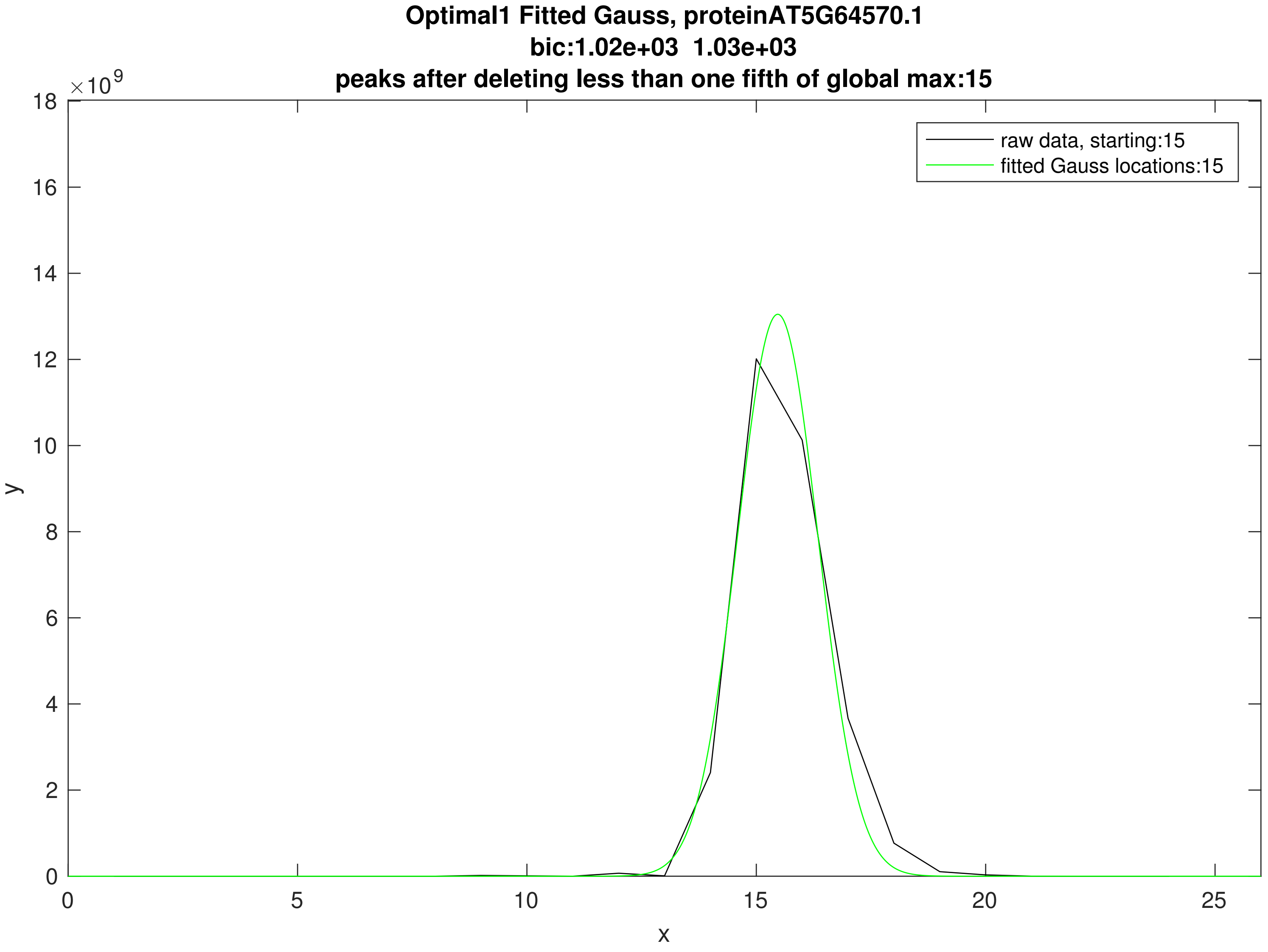
Optimal1 Fitted Gauss, proteinAT5G64570.1
bic:1.02e+03 1.03e+03
peaks after deleting less than one fifth of global max:15
×10^9
18
16
14
12
10
8
6
4
2
0
y
0
5
10
15
20
25
x
raw data, starting:15
fitted Gauss locations:15

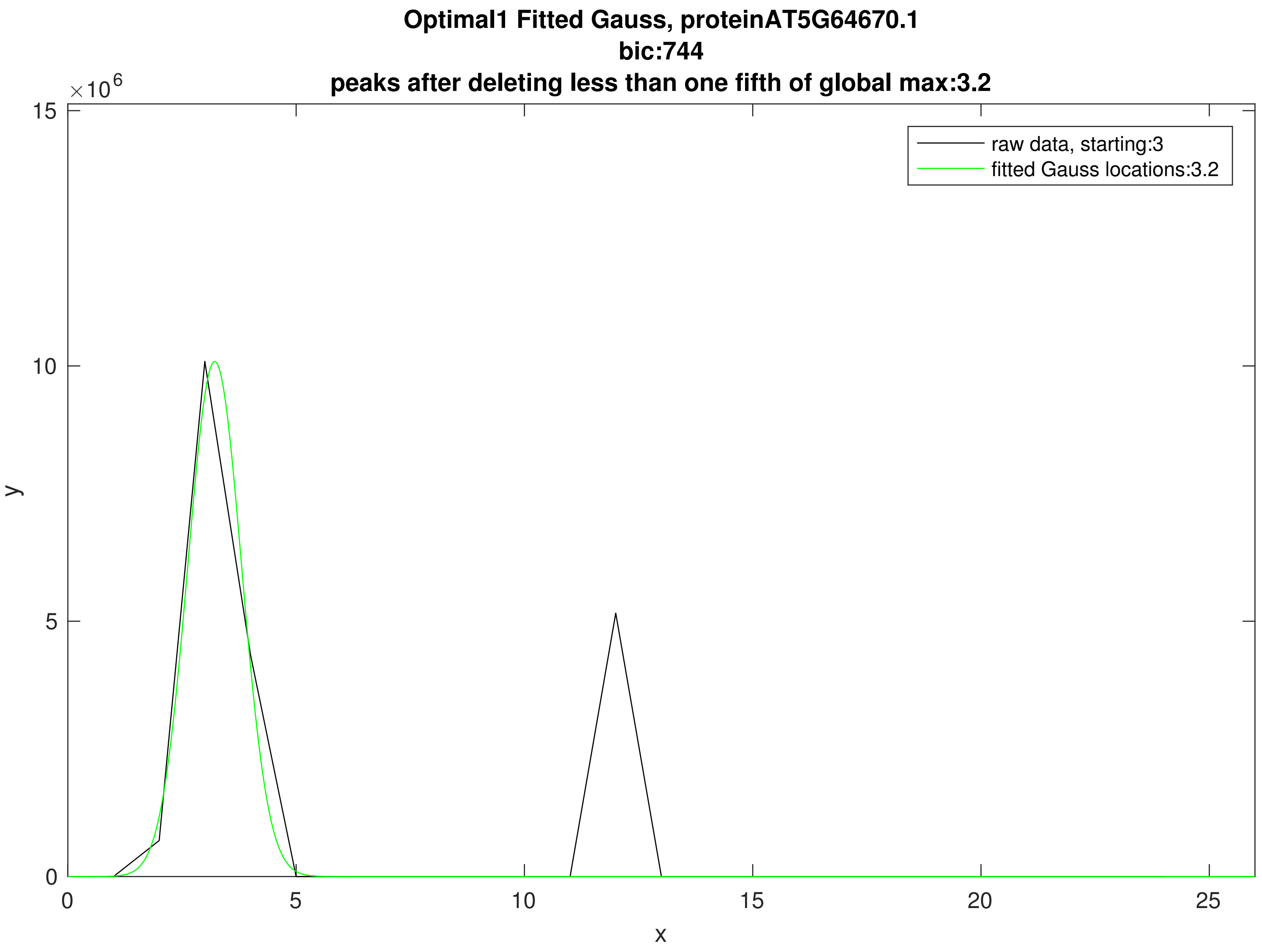
Optimal1 Fitted Gauss, proteinAT5G64670.1
bic:744
peaks after deleting less than one fifth of global max:3.2
×10^6
15
10
5
0
y
0
5
10
15
20
25
x
raw data, starting:3
fitted Gauss locations:3.2

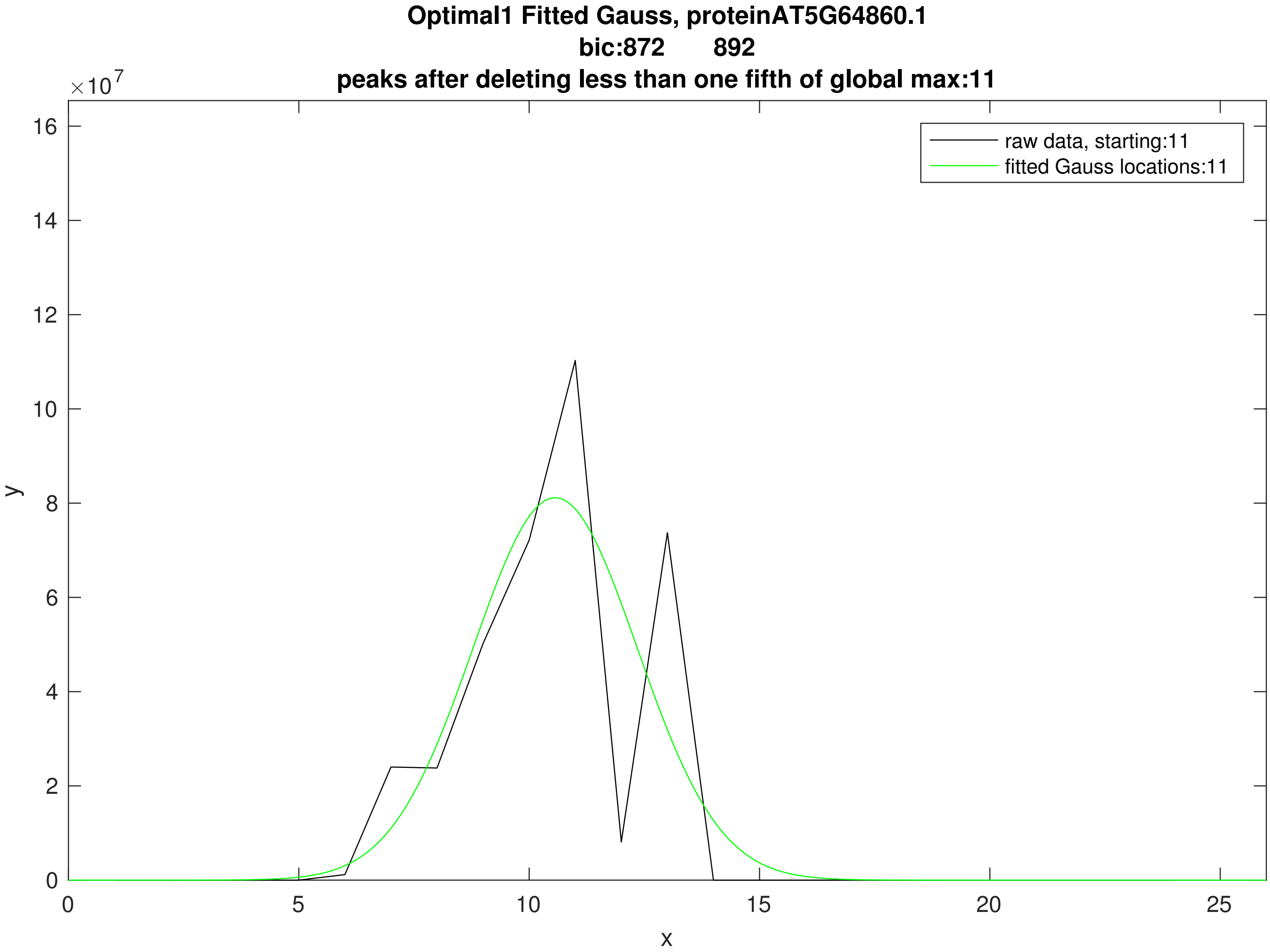
Optimal1 Fitted Gauss, proteinAT5G64860.1
bic:872 892
peaks after deleting less than one fifth of global max:11
×10^7
16
14
12
10
8
6
4
2
0
y
0
5
10
15
20
25
x
raw data, starting:11
fitted Gauss locations:11

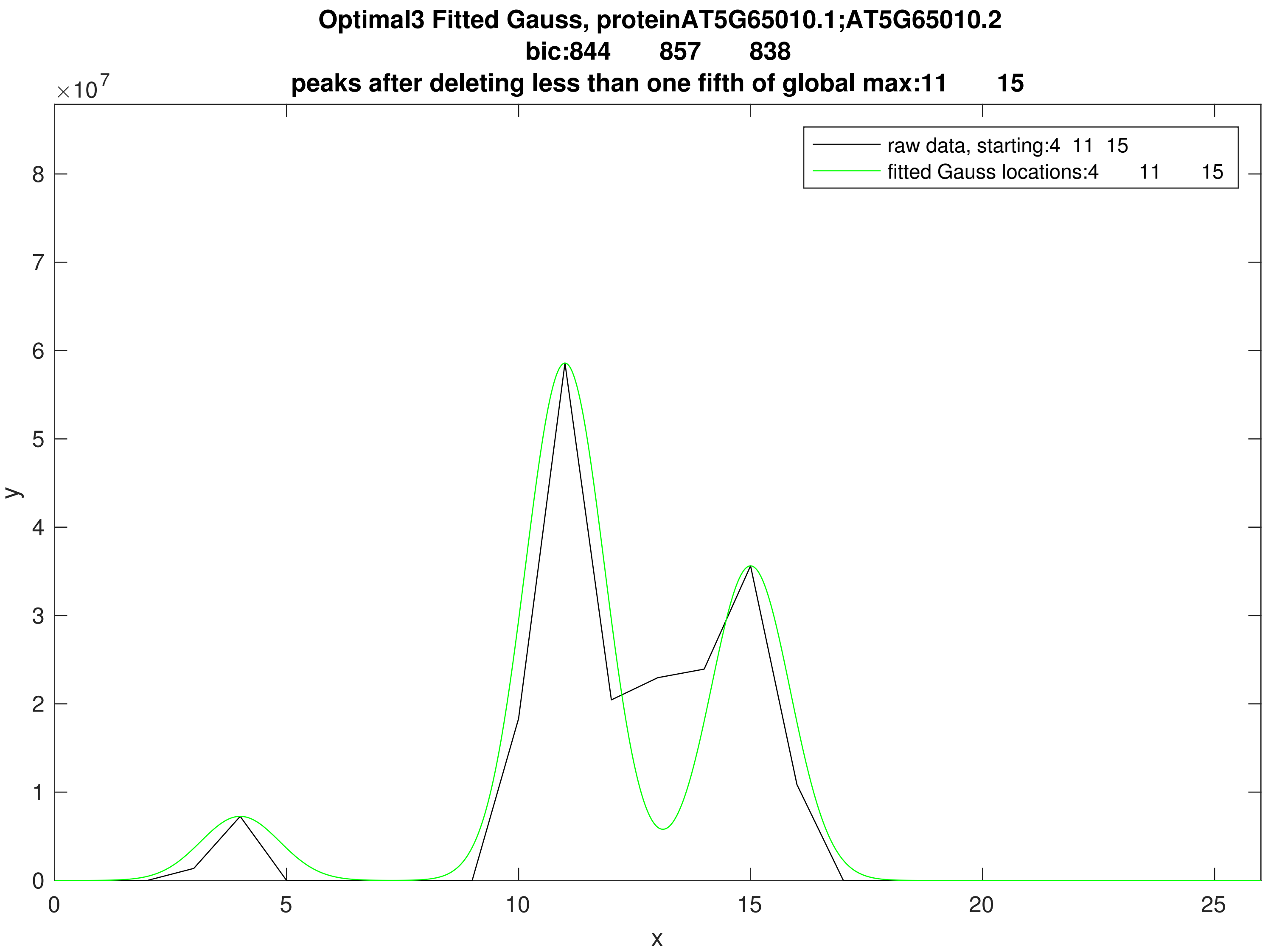
Optimal3 Fitted Gauss, proteinAT5G65010.1;AT5G65010.2
bic:844 857 838
peaks after deleting less than one fifth of global max:11 15
×10^7
raw data, starting:4 11 15
fitted Gauss locations:4 11 15
y
x
0
1
2
3
4
5
6
7
8
0
5
10
15
20
25

Optimal1 Fitted Gauss, proteinAT5G65020.1;AT5G65020.2
bic:1.09e+03 1.1e+03 1.11e+03
peaks after deleting less than one fifth of global max:15

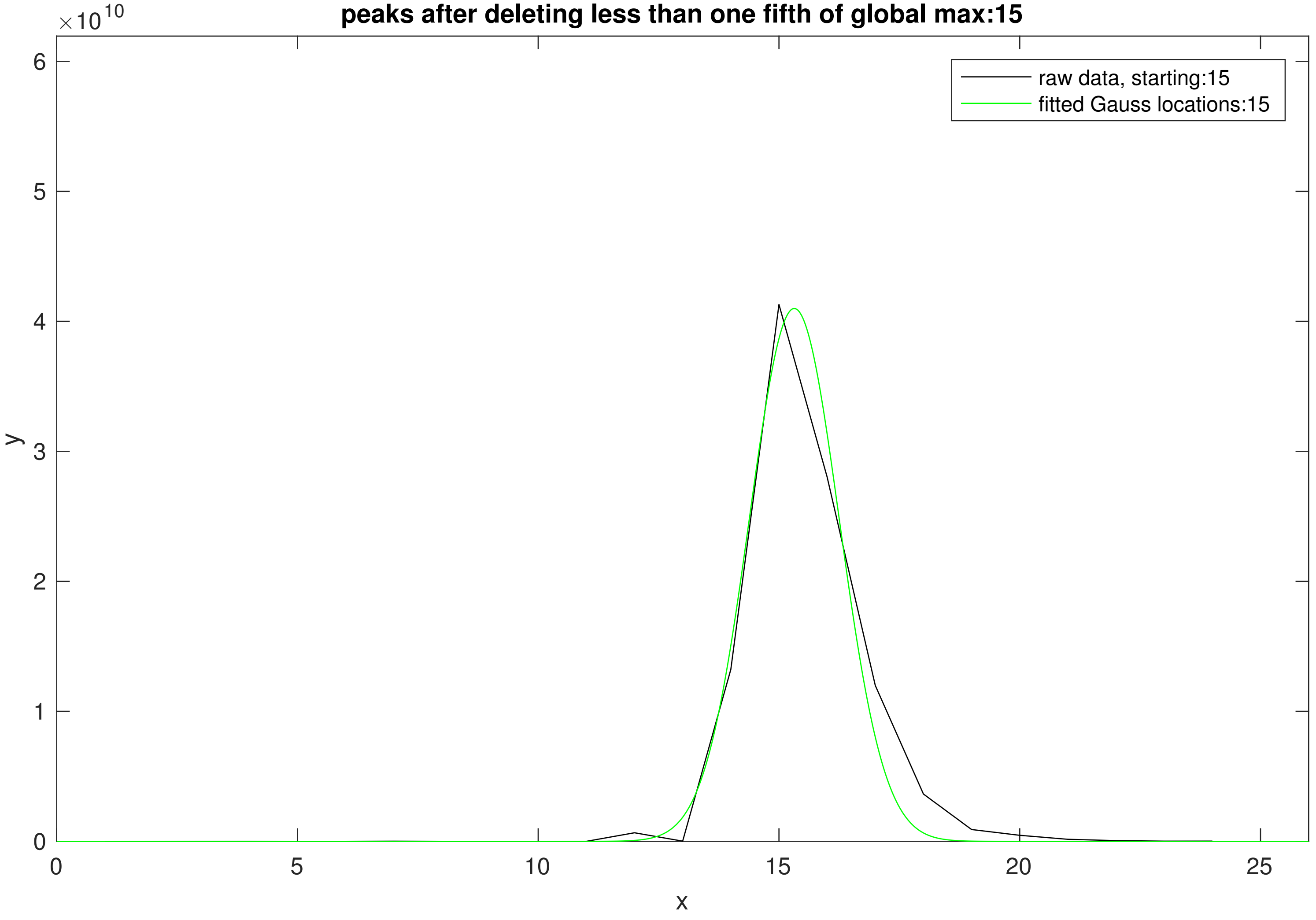

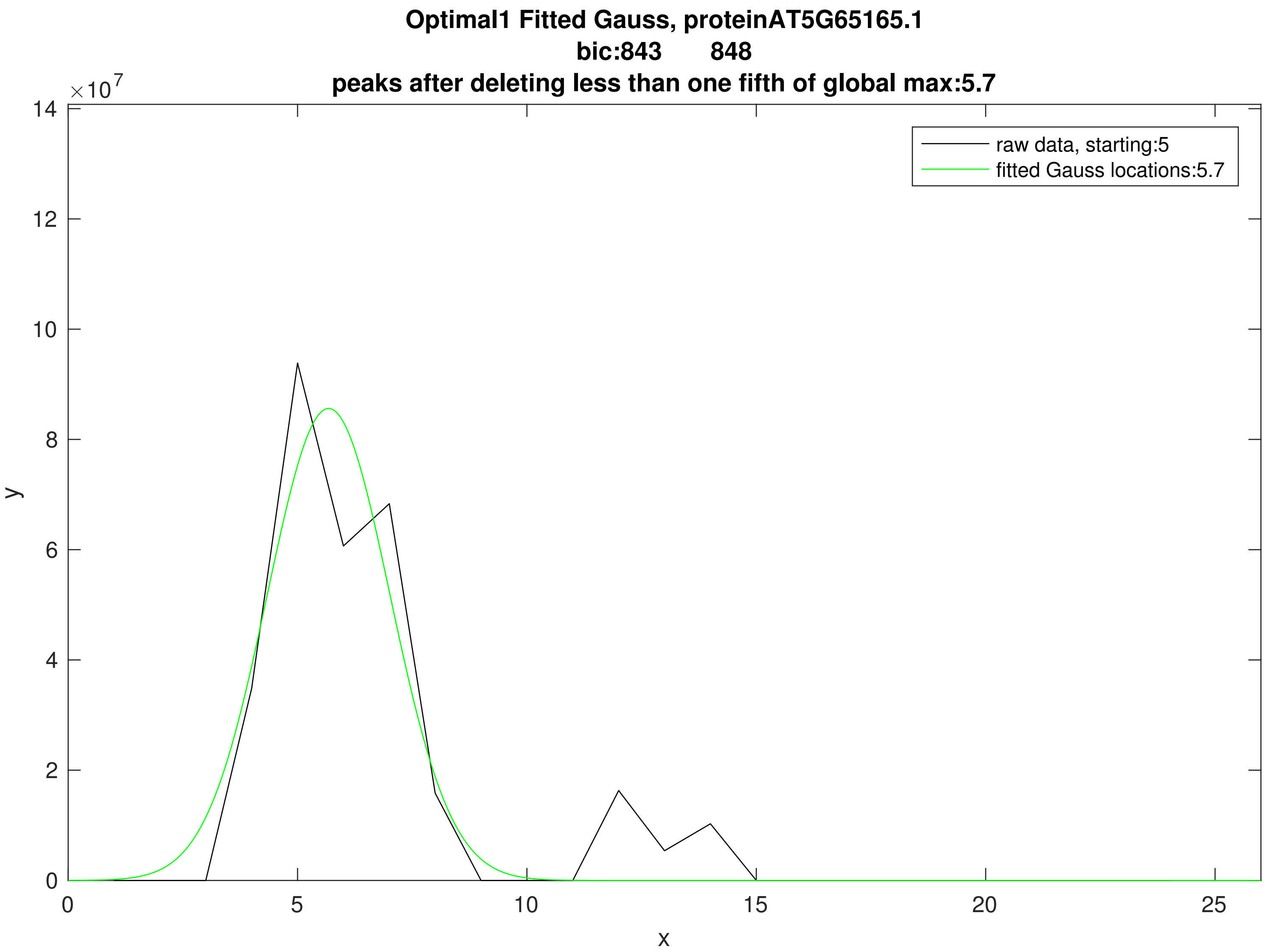
Optimal1 Fitted Gauss, proteinAT5G65165.1
bic:843 848
peaks after deleting less than one fifth of global max:5.7
×10^7
raw data, starting:5
fitted Gauss locations:5.7
y
x
0
2
4
6
8
10
12
14
5
15
20
25

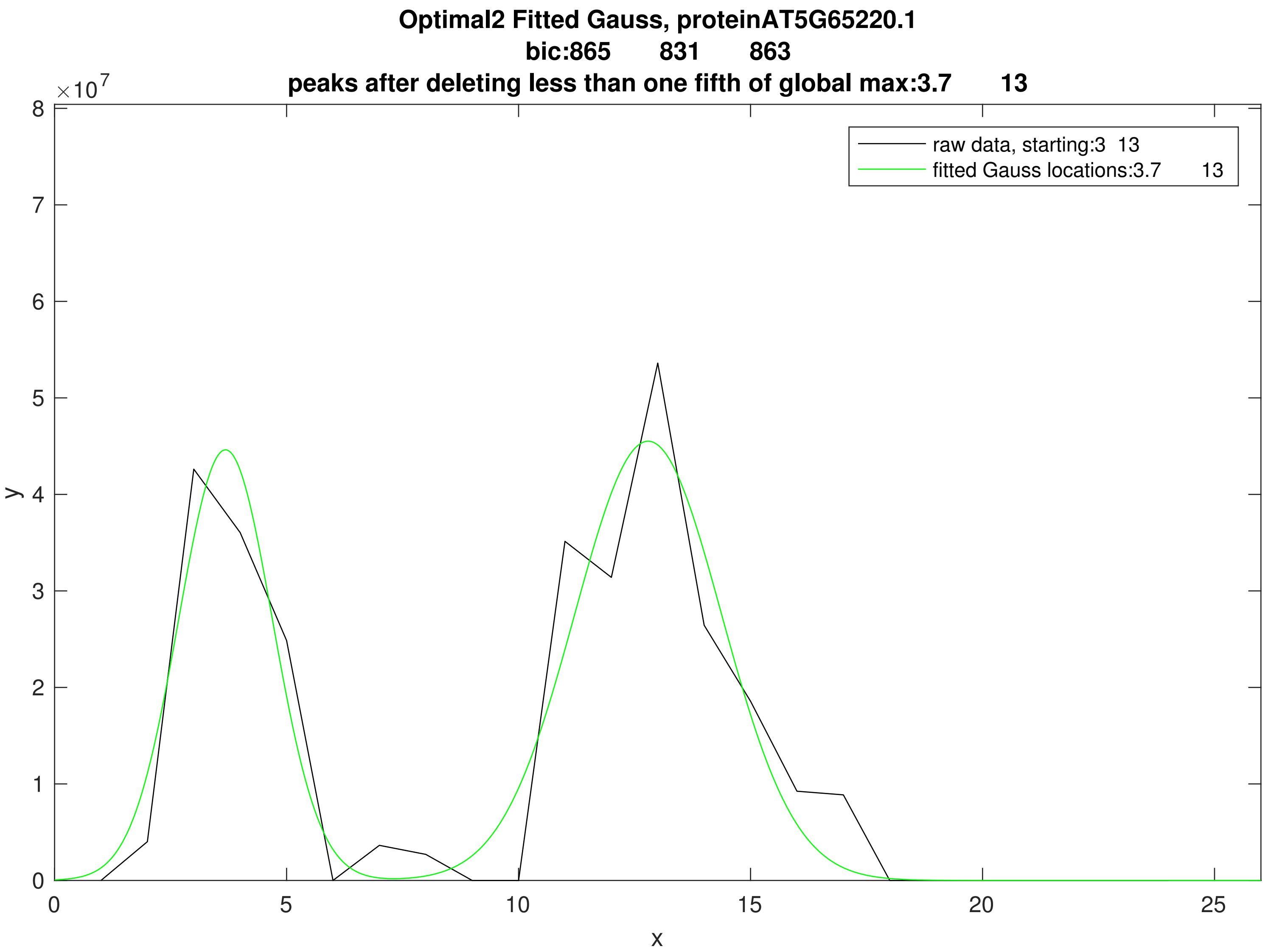
Optimal2 Fitted Gauss, proteinAT5G65220.1
bic:865 831 863
peaks after deleting less than one fifth of global max:3.7 13
×10^7
raw data, starting:3 13
fitted Gauss locations:3.7 13
y
x
0
1
2
3
4
5
6
7
8
0
5
10
15
20
25

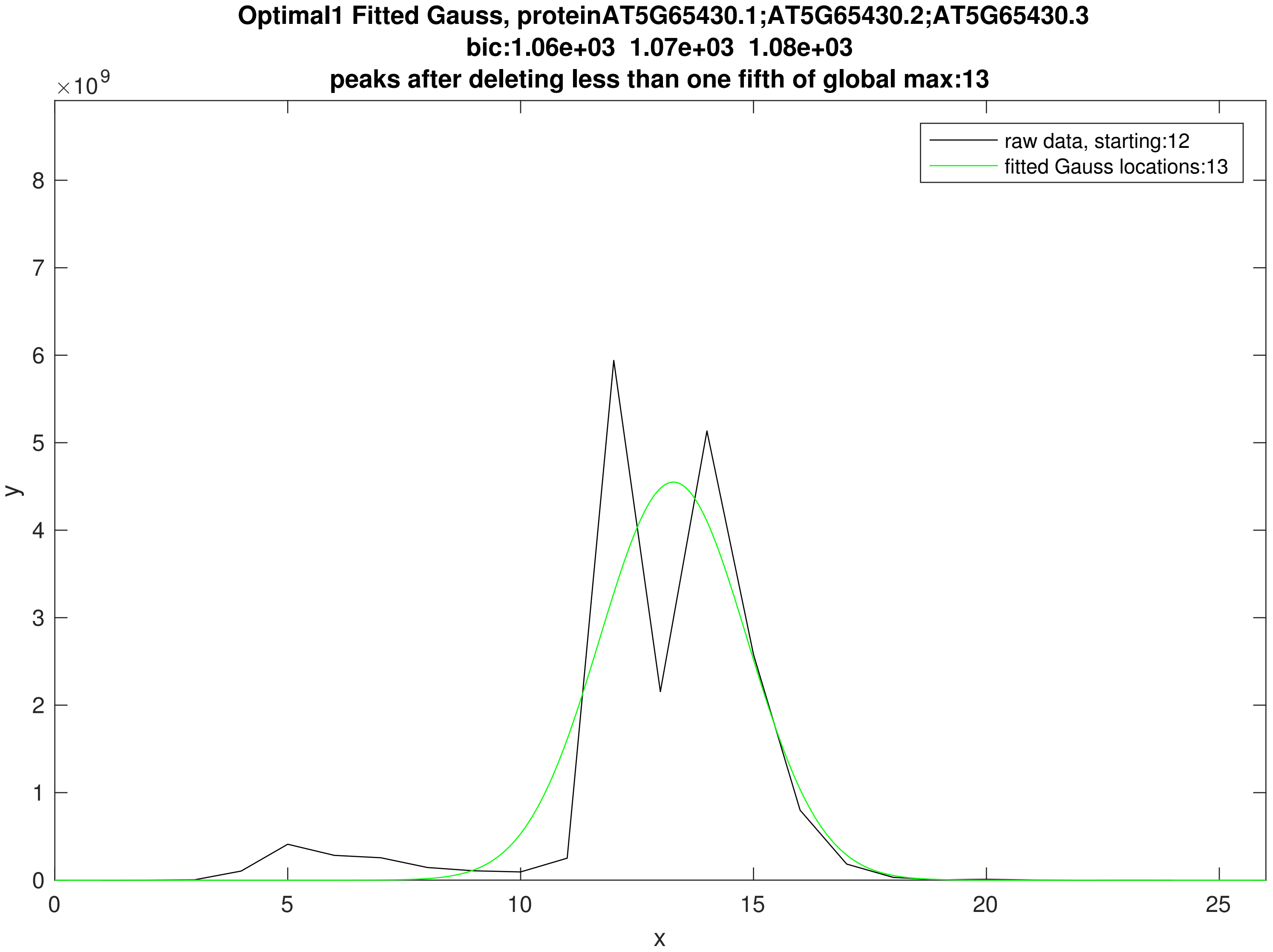

Optimal1 Fitted Gauss, proteinAT5G65430.1;AT5G65430.2;AT5G65430.3
bic:1.06e+03 1.07e+03 1.08e+03
peaks after deleting less than one fifth of global max:13
×10^9
raw data, starting:12
fitted Gauss locations:13
y
x
0
1
2
3
4
5
6
7
8
10
15
20
25

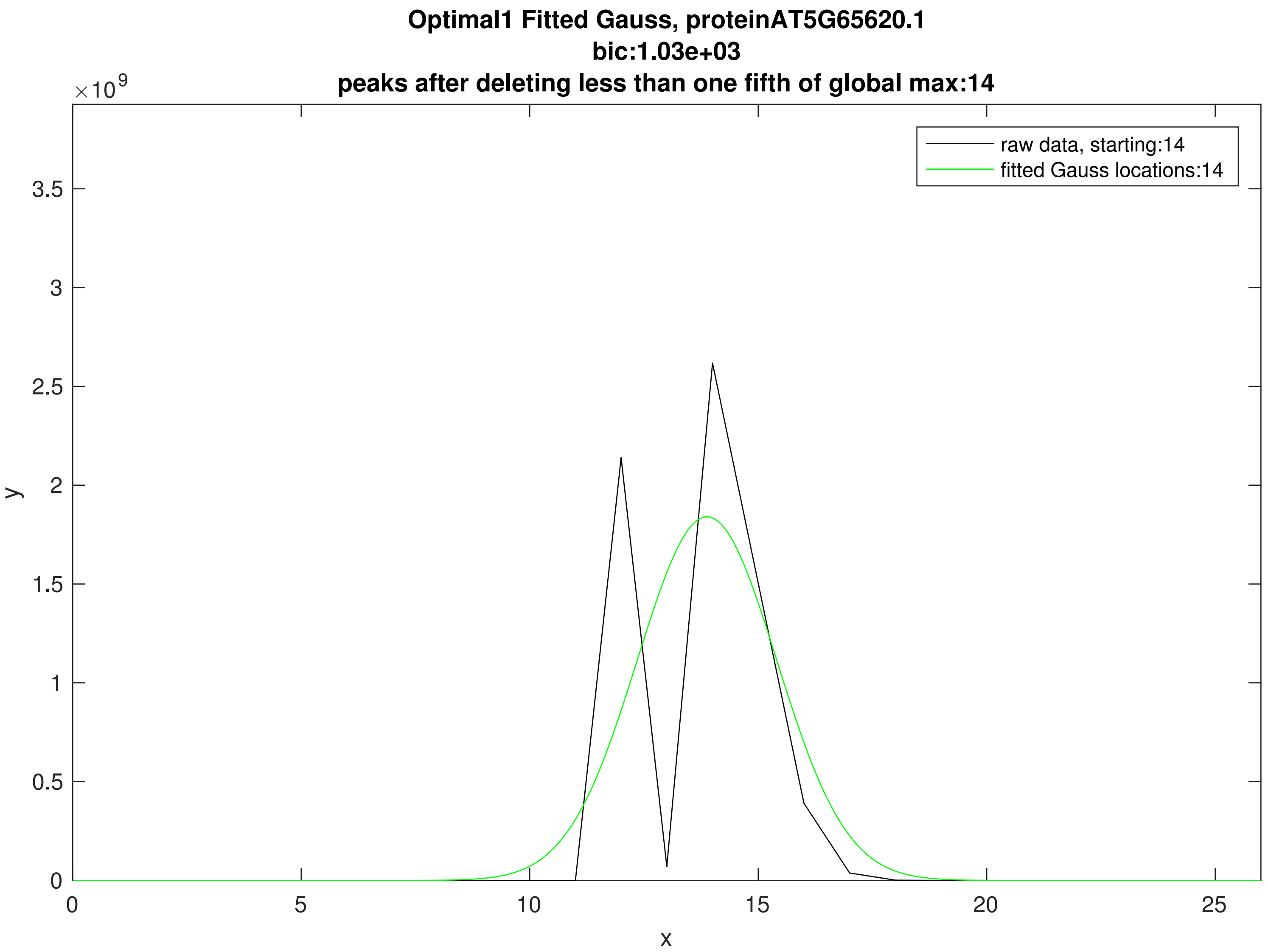

Optimal1 Fitted Gauss, proteinAT5G65620.1
bic:1.03e+03
peaks after deleting less than one fifth of global max:14
×10^9
raw data, starting:14
fitted Gauss locations:14
y
x
0
0.5
1
1.5
2
2.5
3
3.5
0
5
10
15
20
25

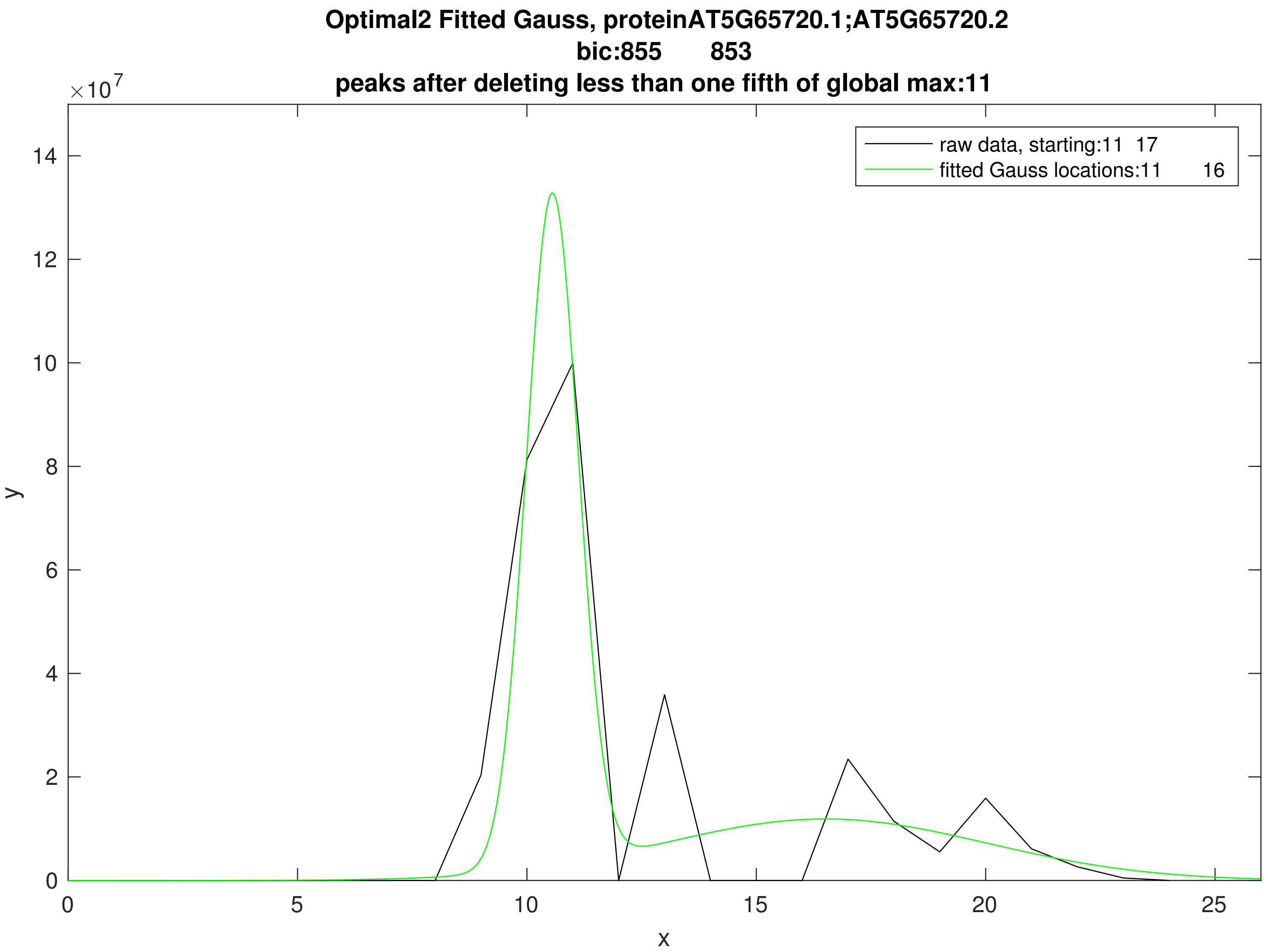
Optimal2 Fitted Gauss, proteinAT5G65720.1;AT5G65720.2
bic:855 853
peaks after deleting less than one fifth of global max:11
×10^7
raw data, starting:11 17
fitted Gauss locations:11 16
y
x
0
2
4
6
8
10
12
14
5
15
20
25

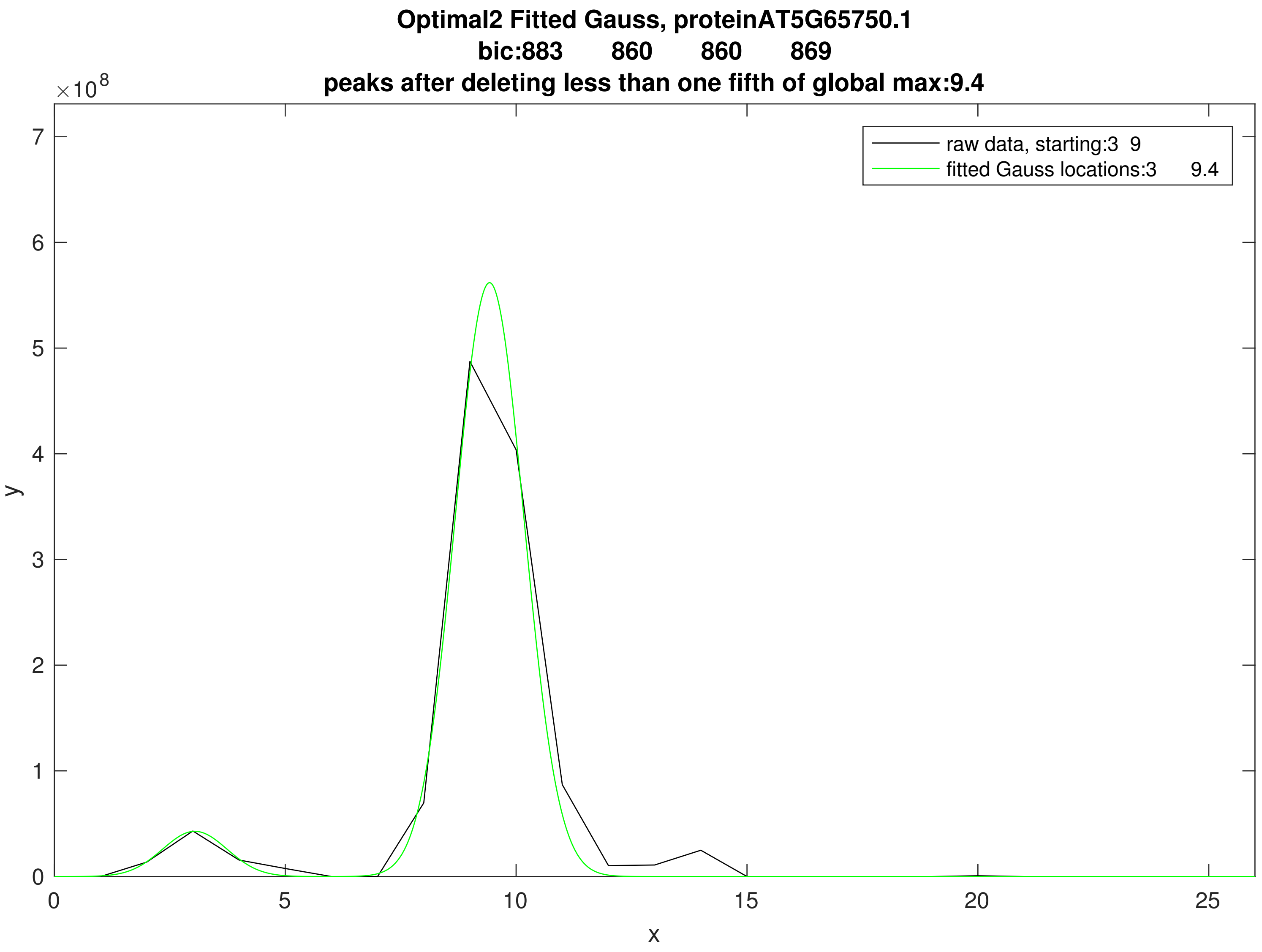
Optimal2 Fitted Gauss, proteinAT5G65750.1
bic:883 860 860 869
peaks after deleting less than one fifth of global max:9.4
×10^8
raw data, starting:3 9
fitted Gauss locations:3 9.4
y
x
0
1
2
3
4
5
6
7
0
5
10
15
20
25

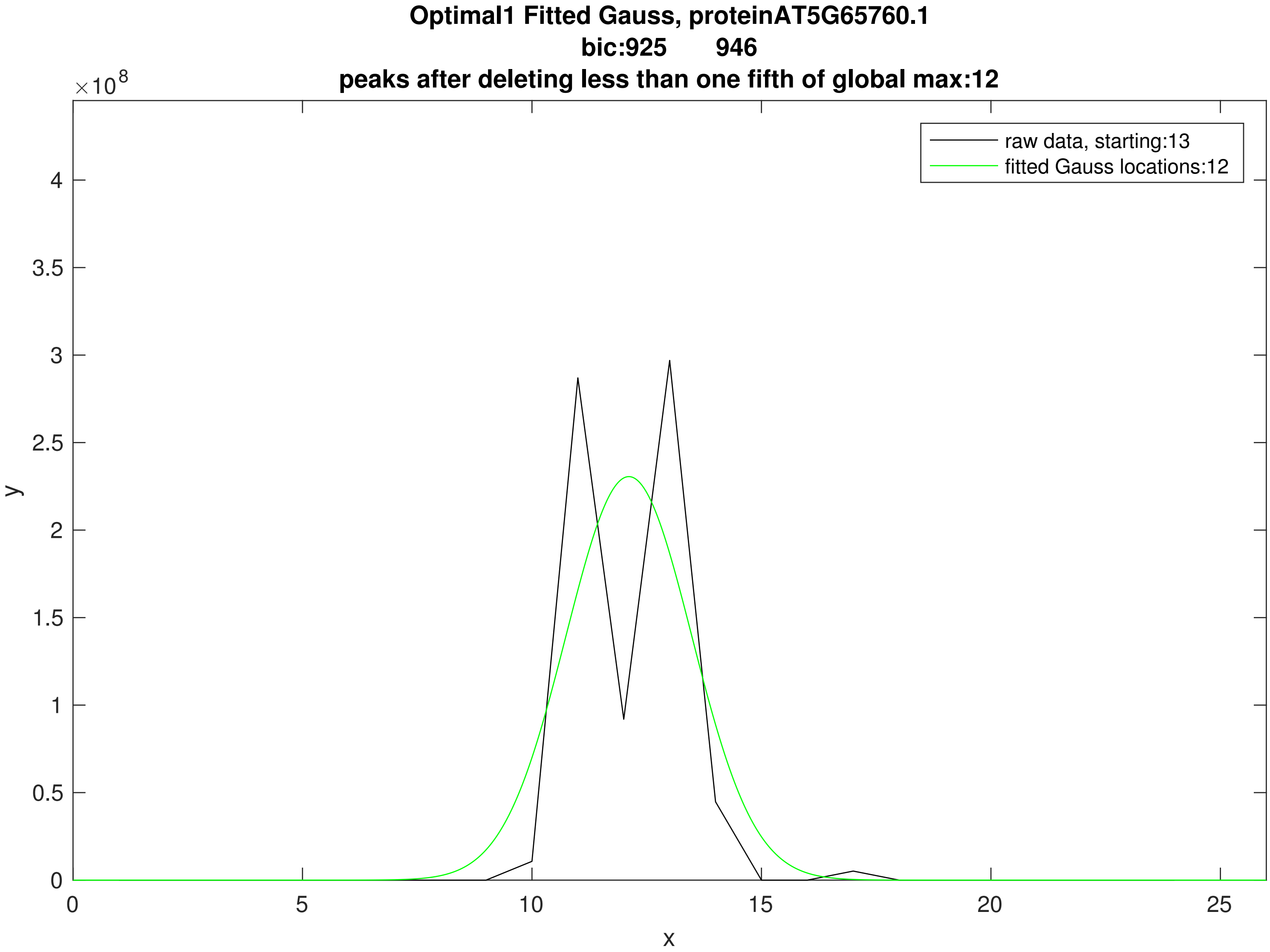
Optimal1 Fitted Gauss, proteinAT5G65760.1
bic:925 946
peaks after deleting less than one fifth of global max:12
×10^8
raw data, starting:13
fitted Gauss locations:12
y
x
0
0.5
1
1.5
2
2.5
3
3.5
4
5
10
15
20
25

**Optimal1 Fitted Gauss, proteinAT5G65770.3;AT5G65770.1;AT5G65780.2;AT5G65770.2**
**bic:717 720**
**peaks after deleting less than one fifth of global max:7.2**

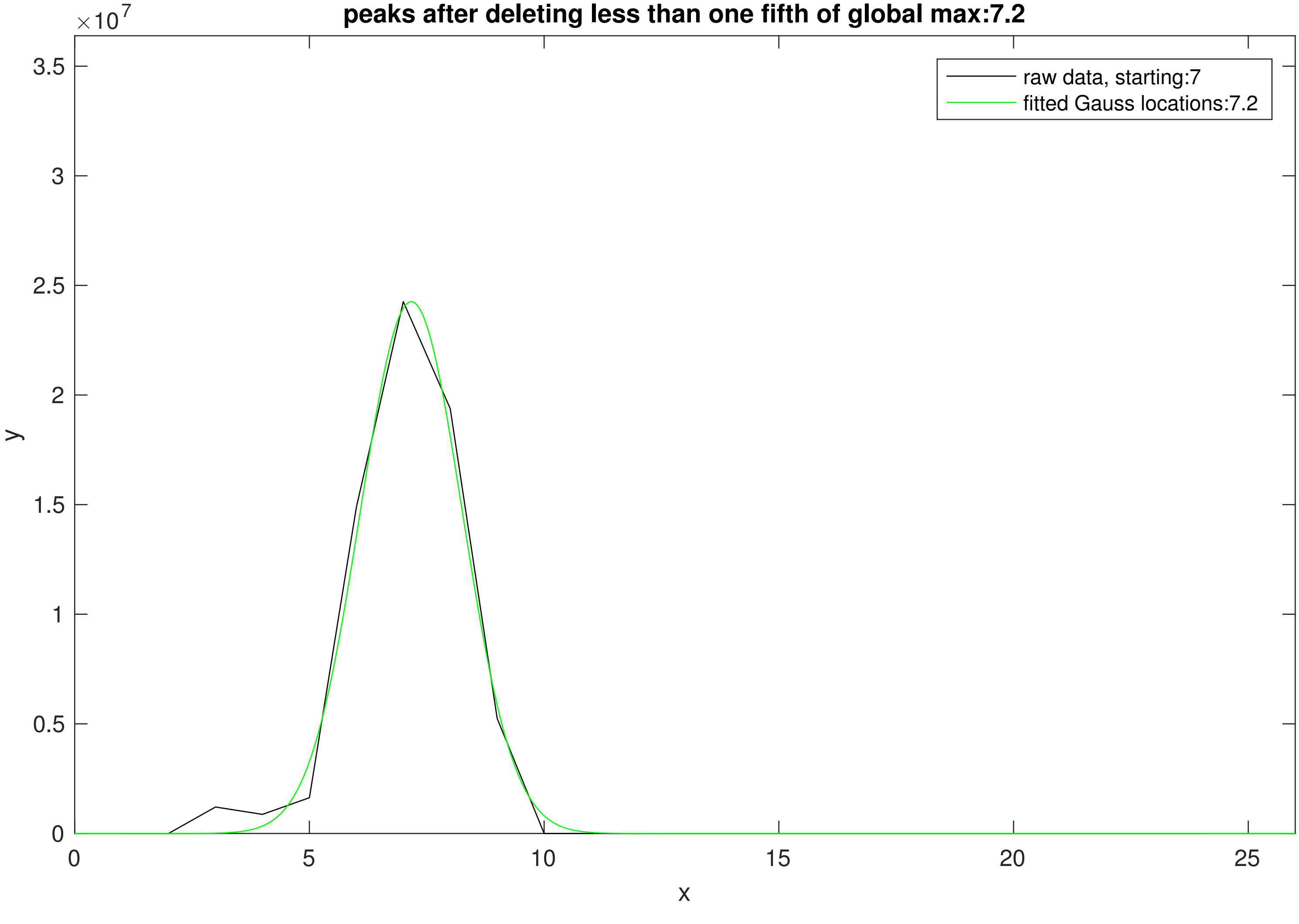

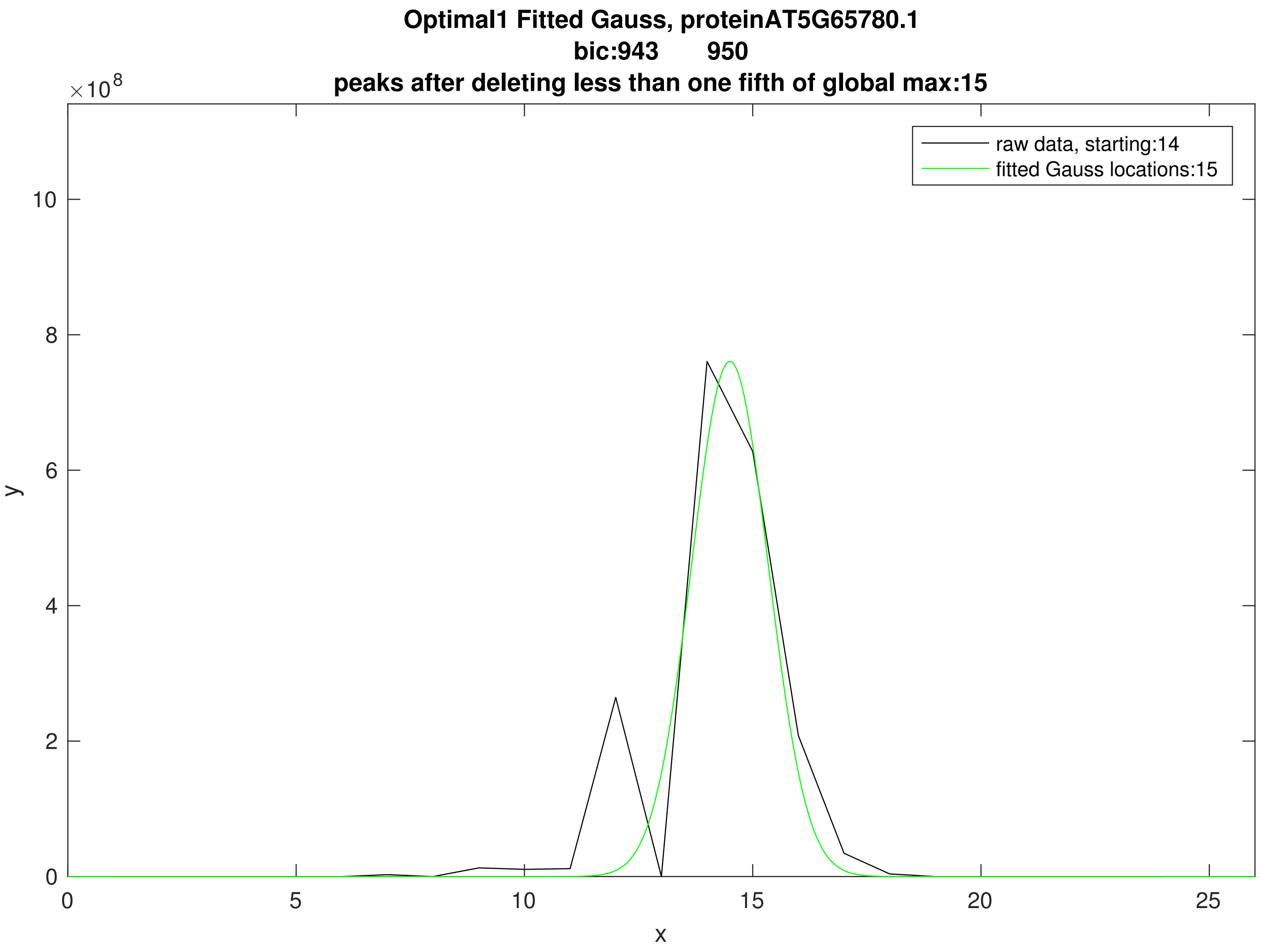

Optimal1 Fitted Gauss, proteinAT5G65780.1
bic:943 950
peaks after deleting less than one fifth of global max:15
×10^8
raw data, starting:14
fitted Gauss locations:15
y
x
0
2
4
6
8
10
0
5
10
15
20
25

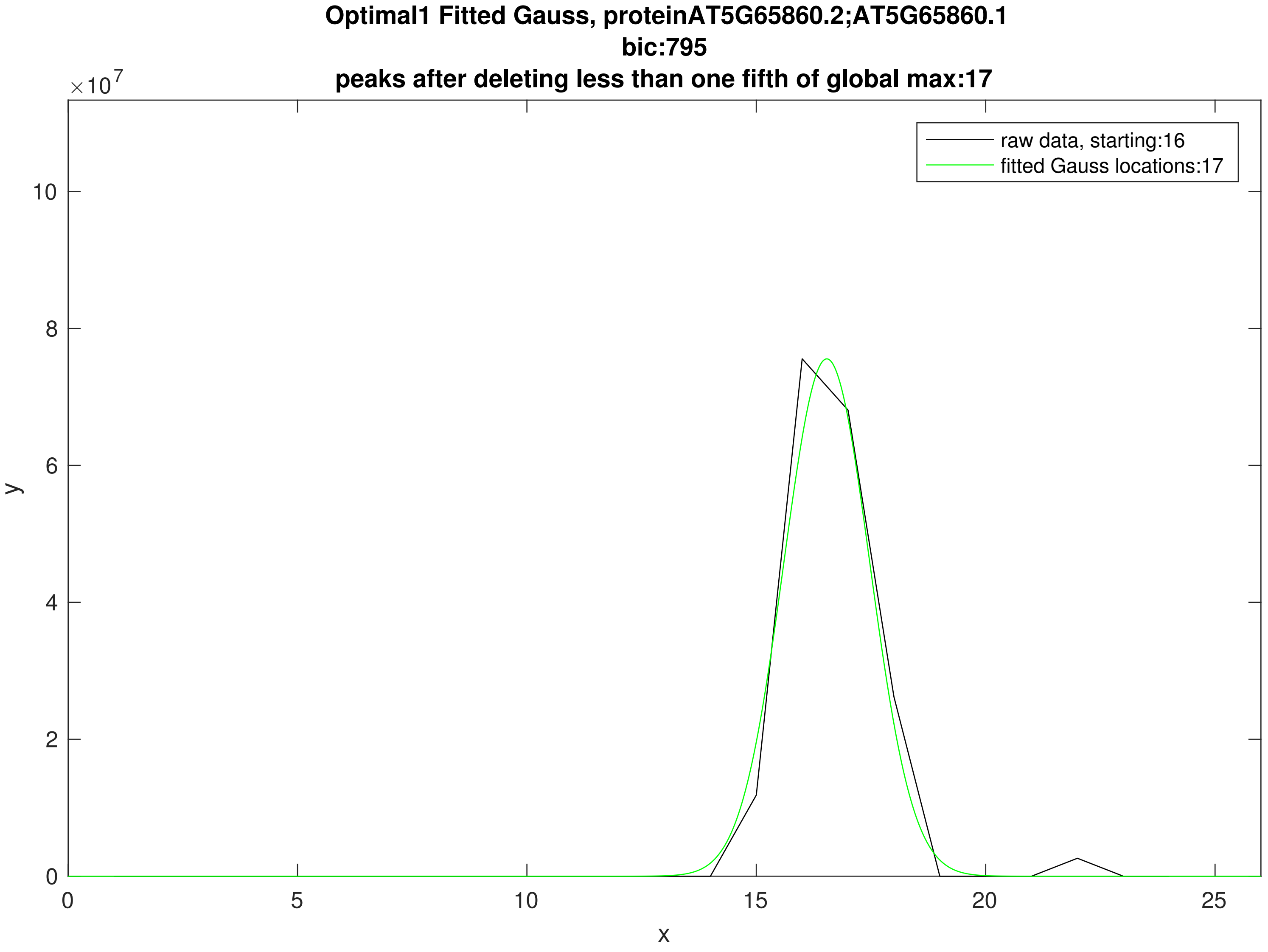

Optimal1 Fitted Gauss, proteinAT5G65860.2;AT5G65860.1
bic:795
peaks after deleting less than one fifth of global max:17
×10^7
raw data, starting:16
fitted Gauss locations:17
y
x
0
2
4
6
8
10
0
5
10
15
20
25

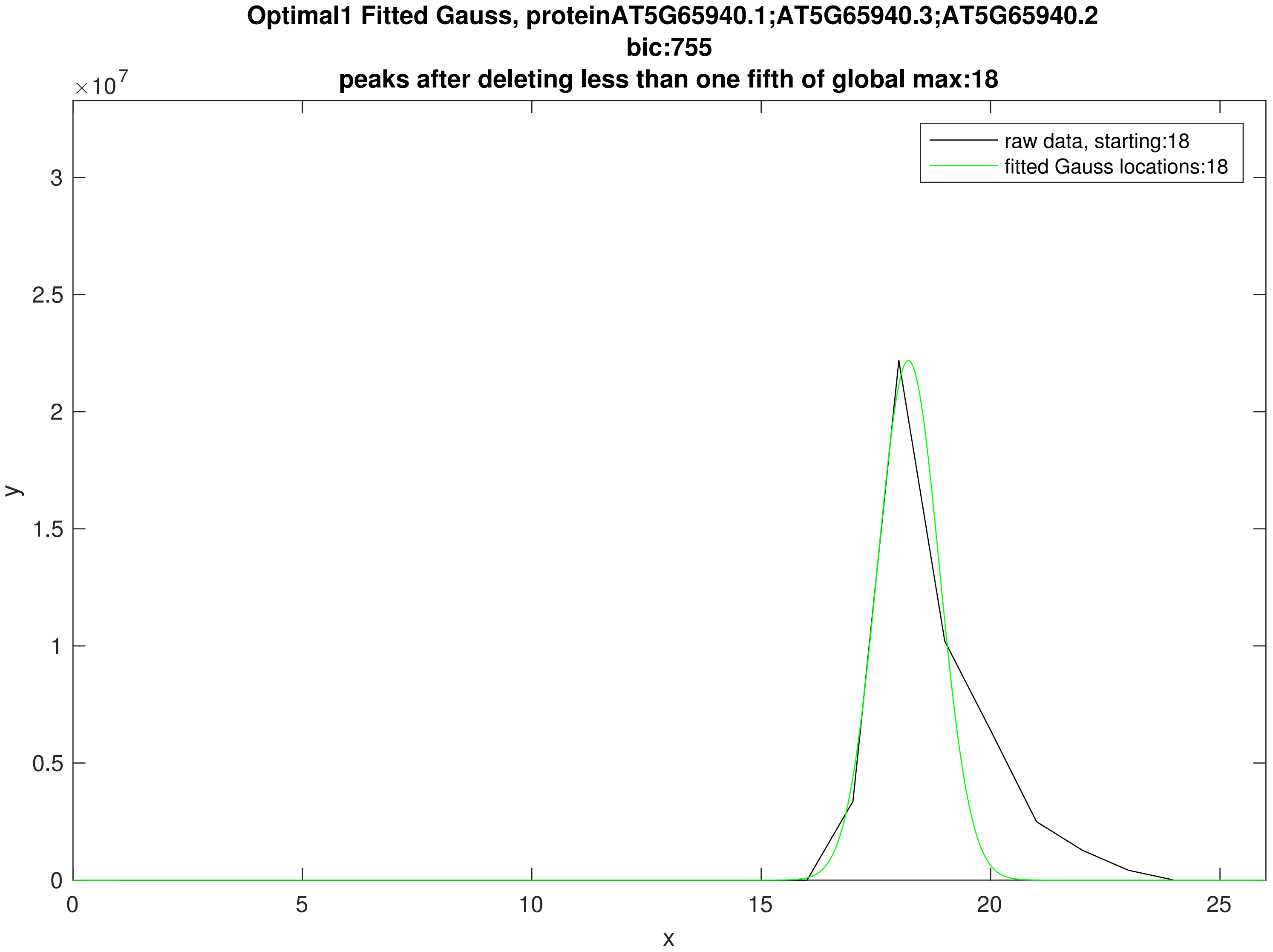
Optimal1 Fitted Gauss, proteinAT5G65940.1;AT5G65940.3;AT5G65940.2
bic:755
peaks after deleting less than one fifth of global max:18
×10^7
raw data, starting:18
fitted Gauss locations:18
y
x
0
0.5
1
1.5
2
2.5
3
0
5
10
15
20
25

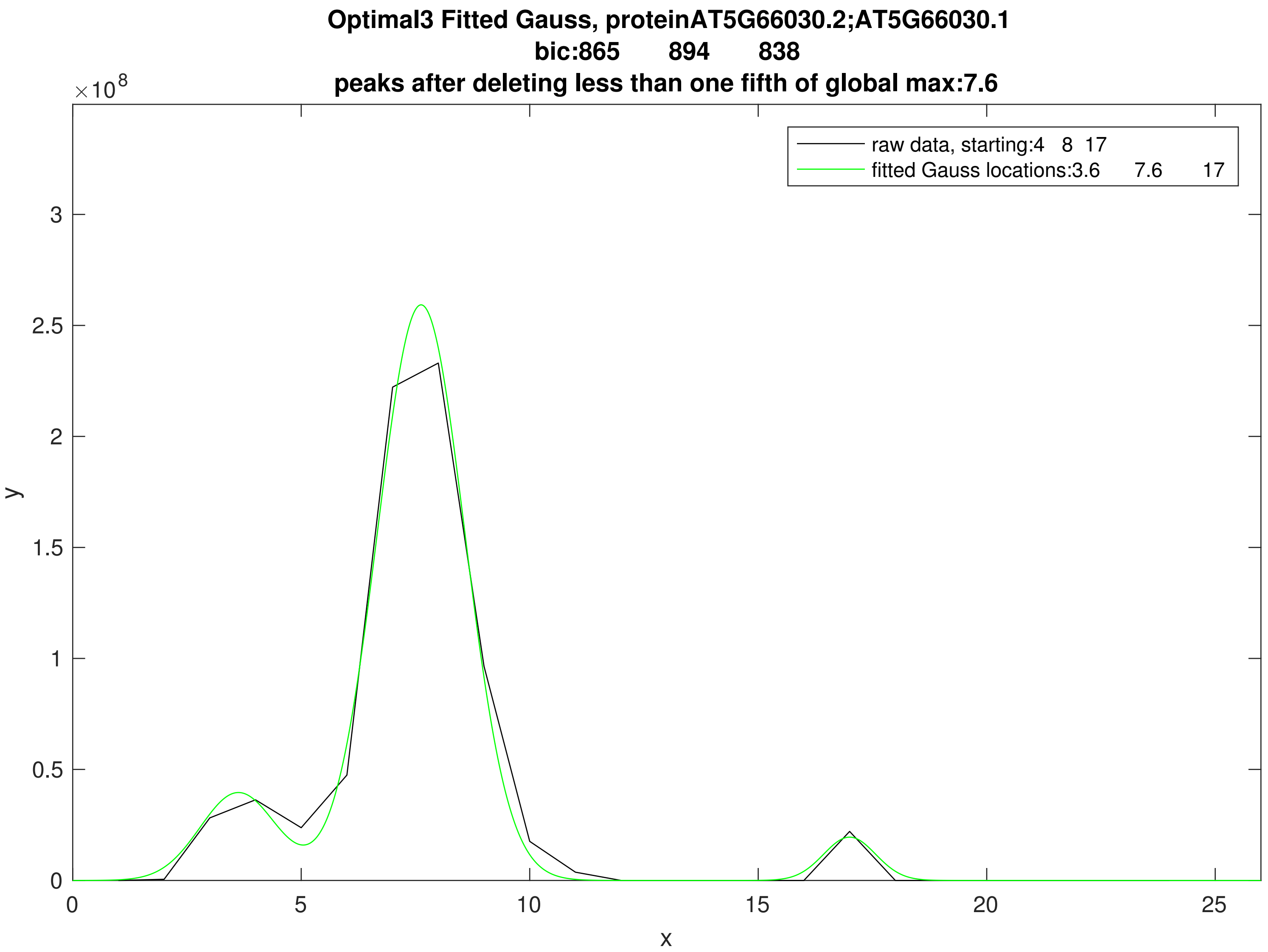

Optimal3 Fitted Gauss, proteinAT5G66030.2;AT5G66030.1
bic:865 894 838
peaks after deleting less than one fifth of global max:7.6
×10^8
raw data, starting:4 8 17
fitted Gauss locations:3.6 7.6 17
3
2.5
2
1.5
1
0.5
0
y
0
5
10
15
20
25
x

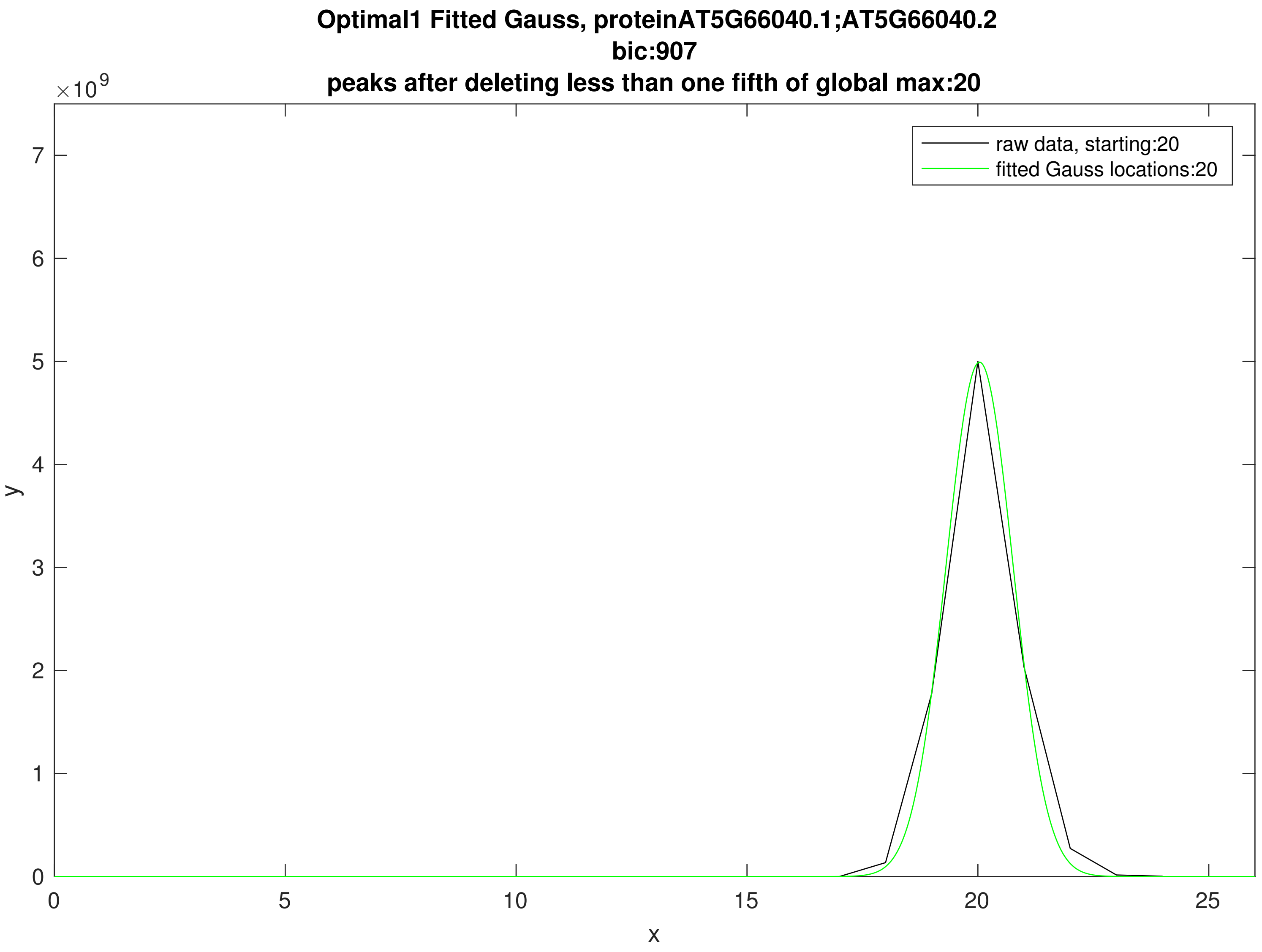

Optimal1 Fitted Gauss, proteinAT5G66040.1;AT5G66040.2
bic:907
peaks after deleting less than one fifth of global max:20
×10^9
raw data, starting:20
fitted Gauss locations:20
y
x
0
1
2
3
4
5
6
7
0
5
10
15
20
25

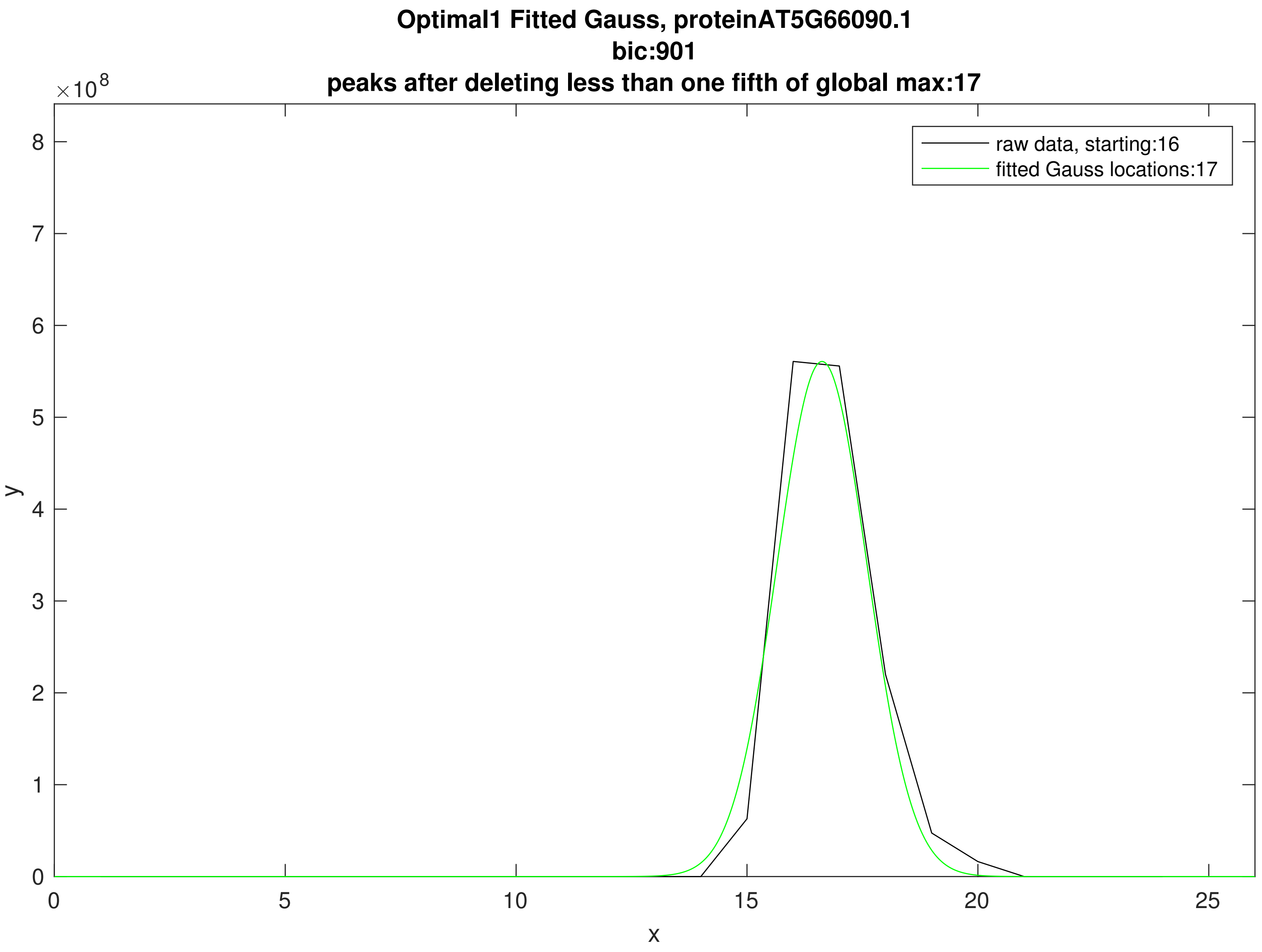
Optimal1 Fitted Gauss, proteinAT5G66090.1
bic:901
peaks after deleting less than one fifth of global max:17
×10^8
raw data, starting:16
fitted Gauss locations:17
y
x
0
1
2
3
4
5
6
7
8
0
5
10
15
20
25

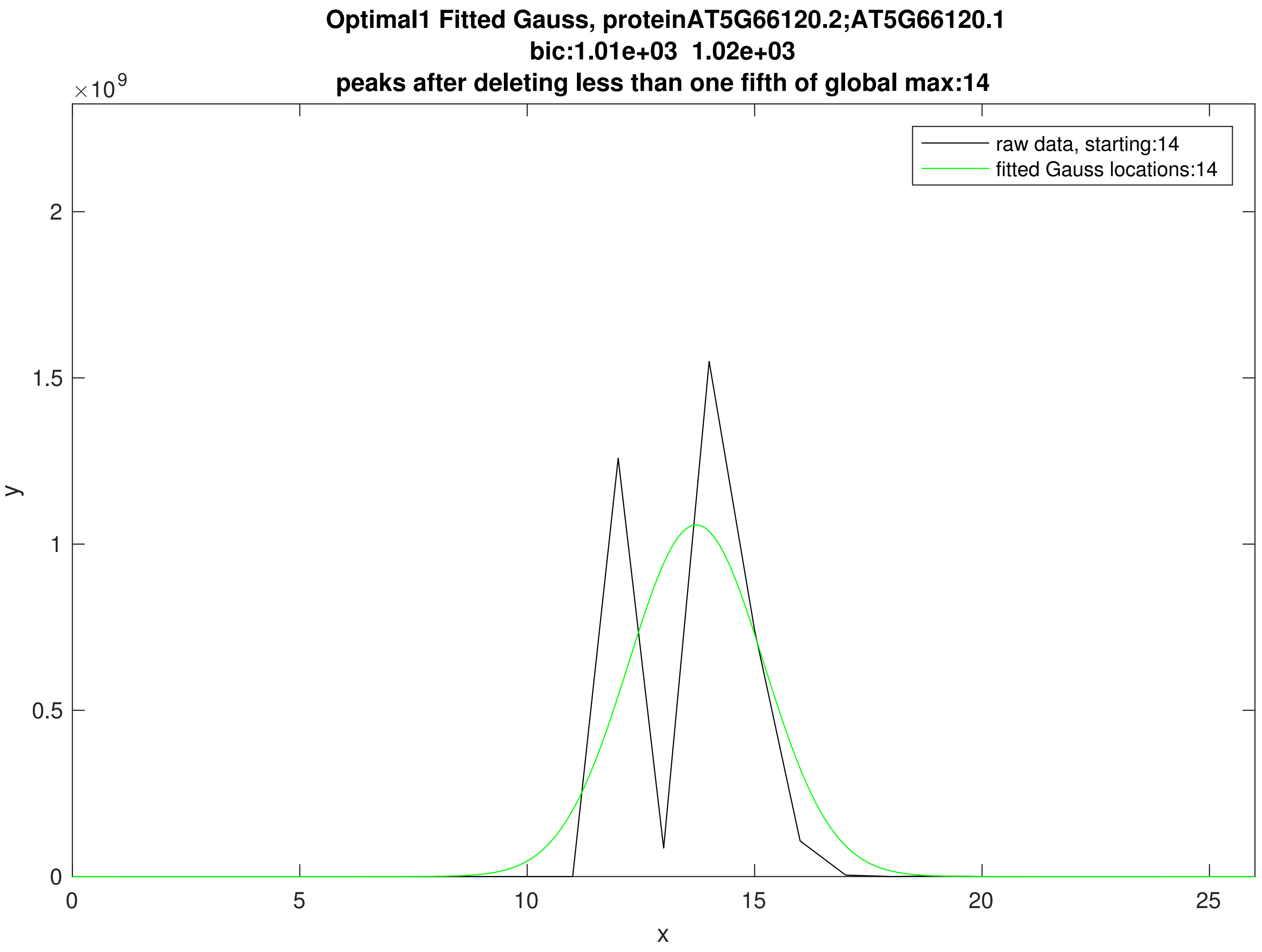
Optimal1 Fitted Gauss, proteinAT5G66120.2;AT5G66120.1
bic:1.01e+03  1.02e+03
peaks after deleting less than one fifth of global max:14
×10^9
raw data, starting:14
fitted Gauss locations:14
y
x
0
0.5
1
1.5
2
0
5
10
15
20
25

**Optimal1 Fitted Gauss, proteinAT5G66140.1**
**bic:845**
**peaks after deleting less than one fifth of global max:5.1**

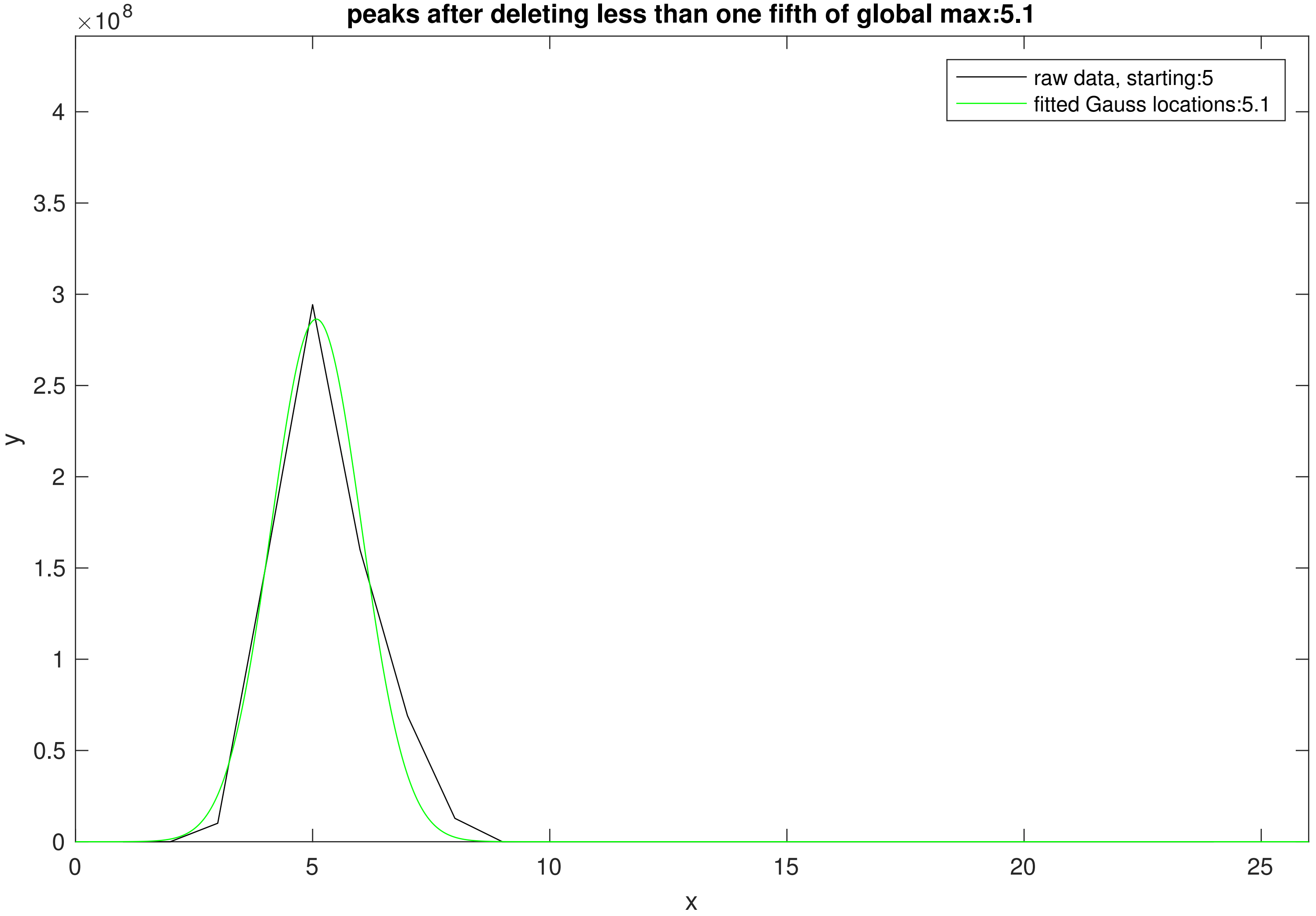

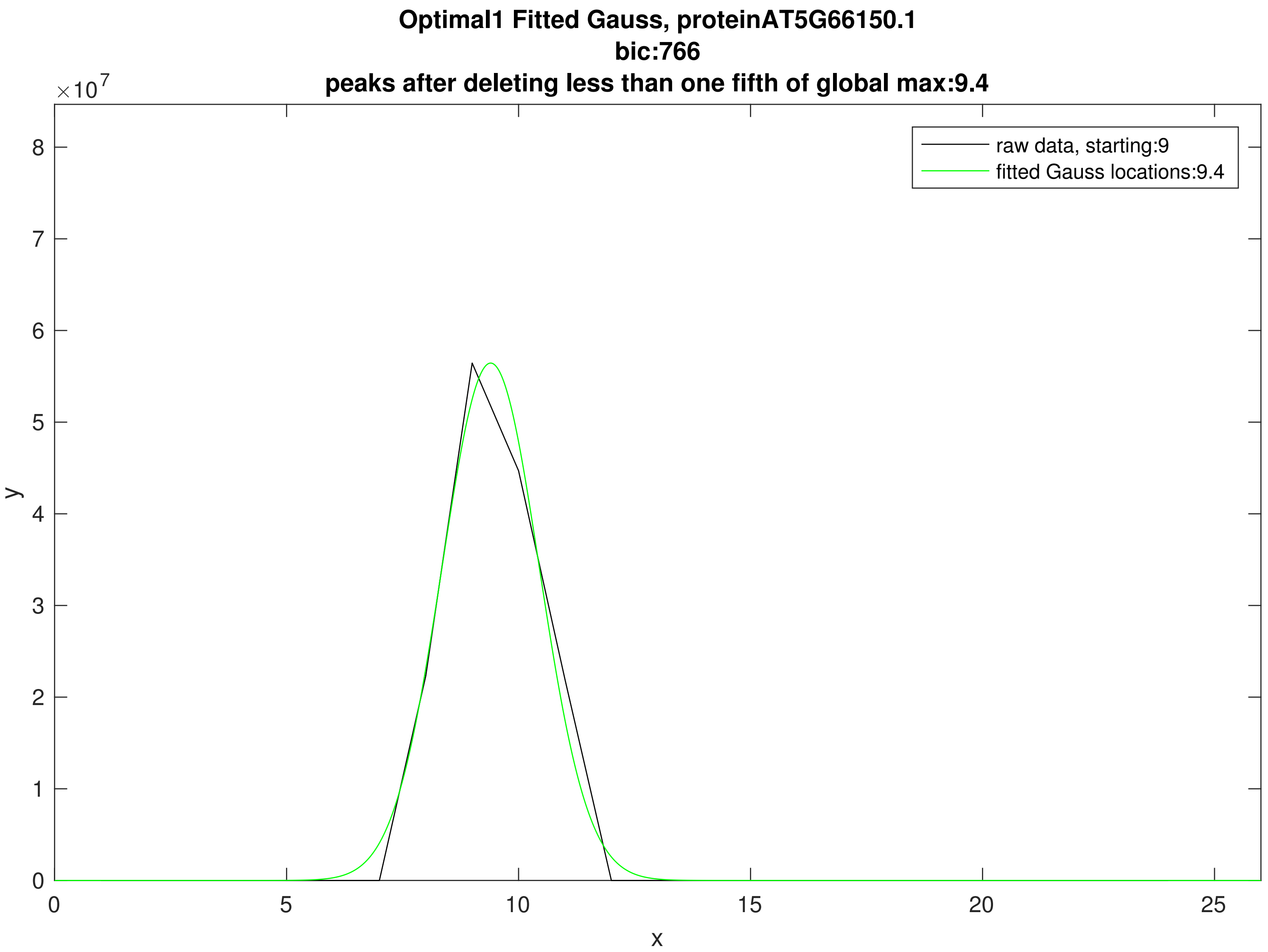
Optimal1 Fitted Gauss, proteinAT5G66150.1
bic:766
peaks after deleting less than one fifth of global max:9.4
×10^7
raw data, starting:9
fitted Gauss locations:9.4
y
x
0
1
2
3
4
5
6
7
8
0
5
10
15
20
25

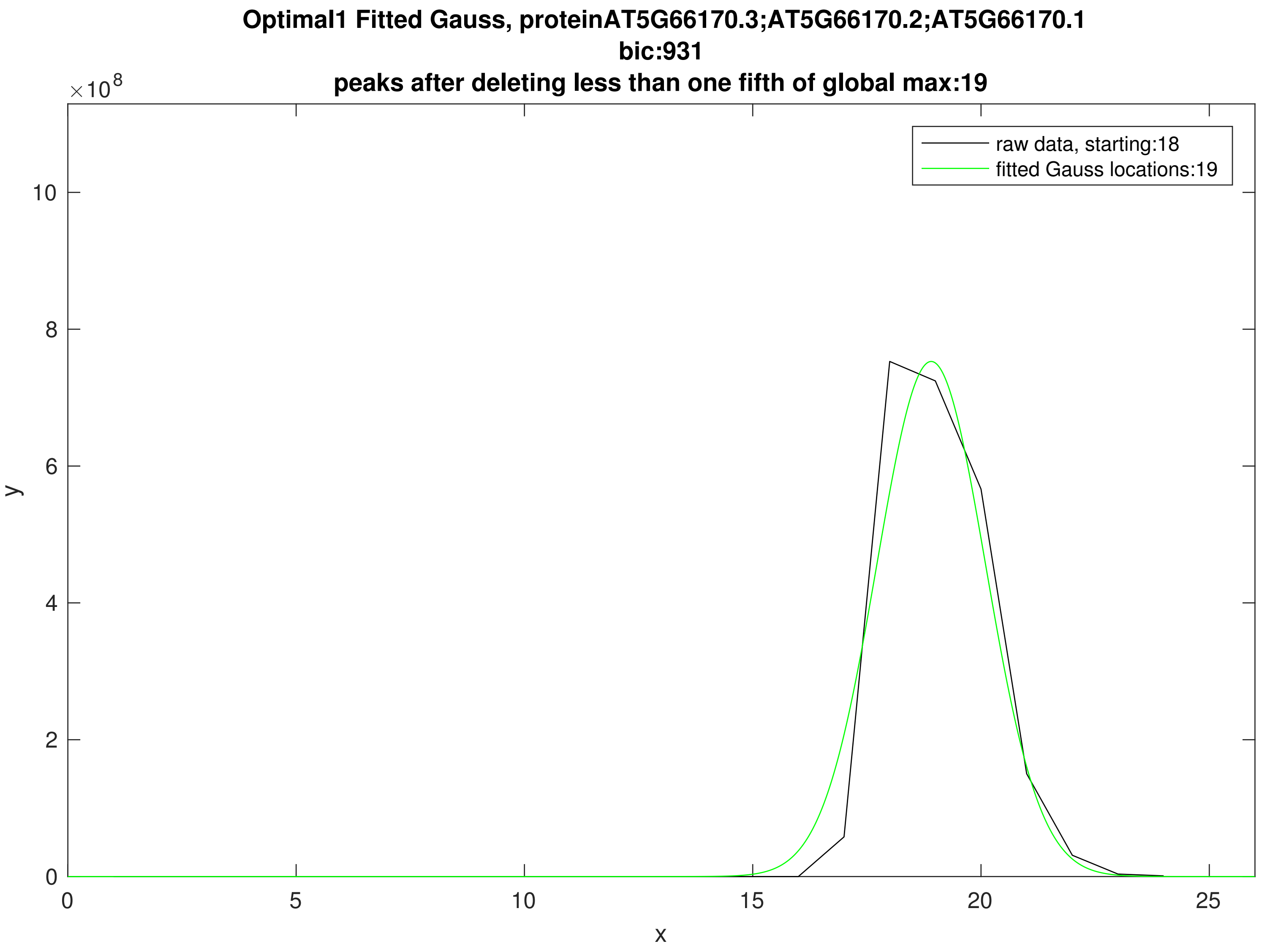
Optimal1 Fitted Gauss, proteinAT5G66170.3;AT5G66170.2;AT5G66170.1
bic:931
peaks after deleting less than one fifth of global max:19
×10^8
raw data, starting:18
fitted Gauss locations:19
y
x
0
2
4
6
8
10
0
5
10
15
20
25

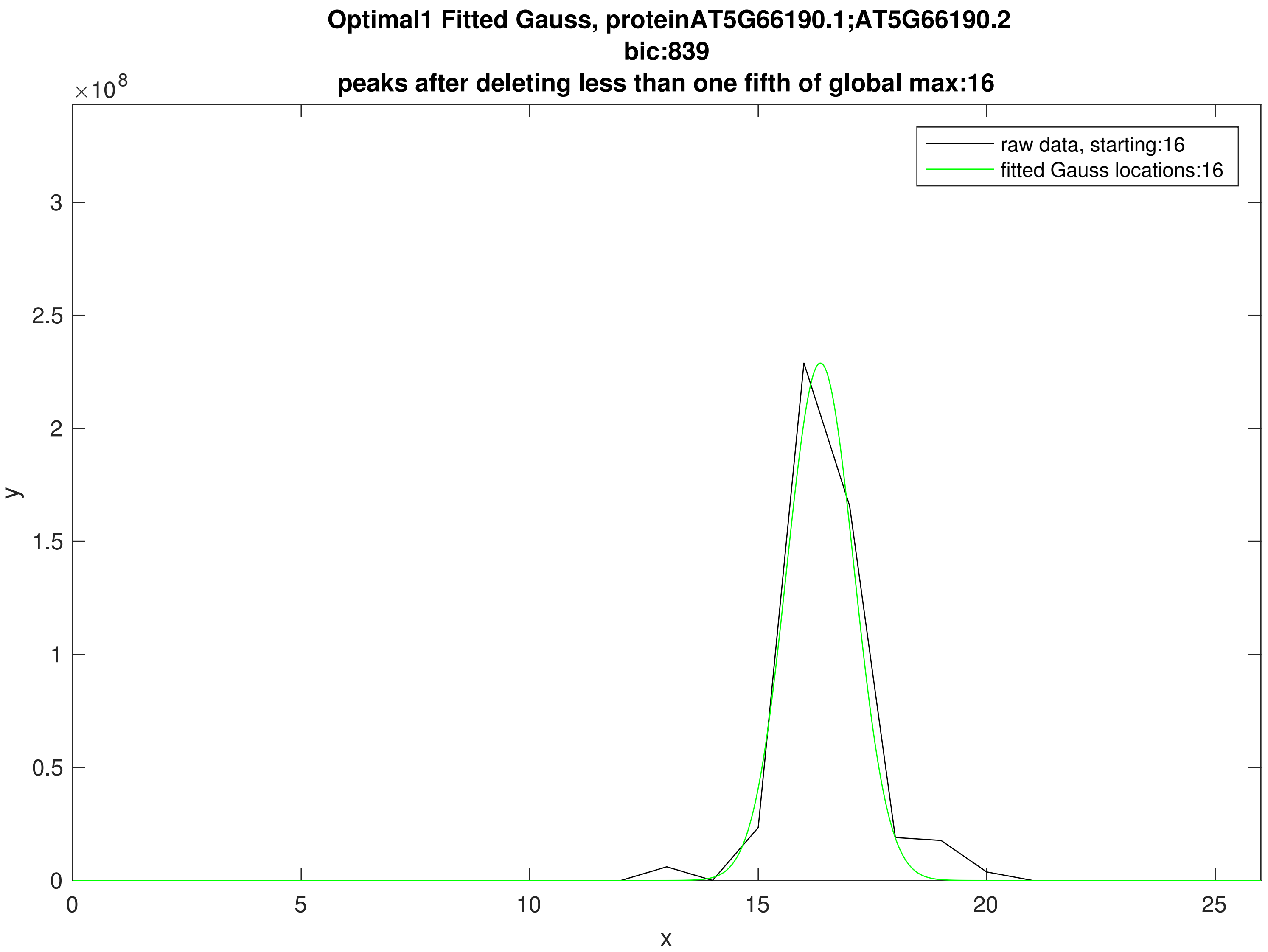
Optimal1 Fitted Gauss, proteinAT5G66190.1;AT5G66190.2
bic:839
peaks after deleting less than one fifth of global max:16
×10^8
raw data, starting:16
fitted Gauss locations:16
y
x
0
0.5
1
1.5
2
2.5
3
0
5
10
15
20
25

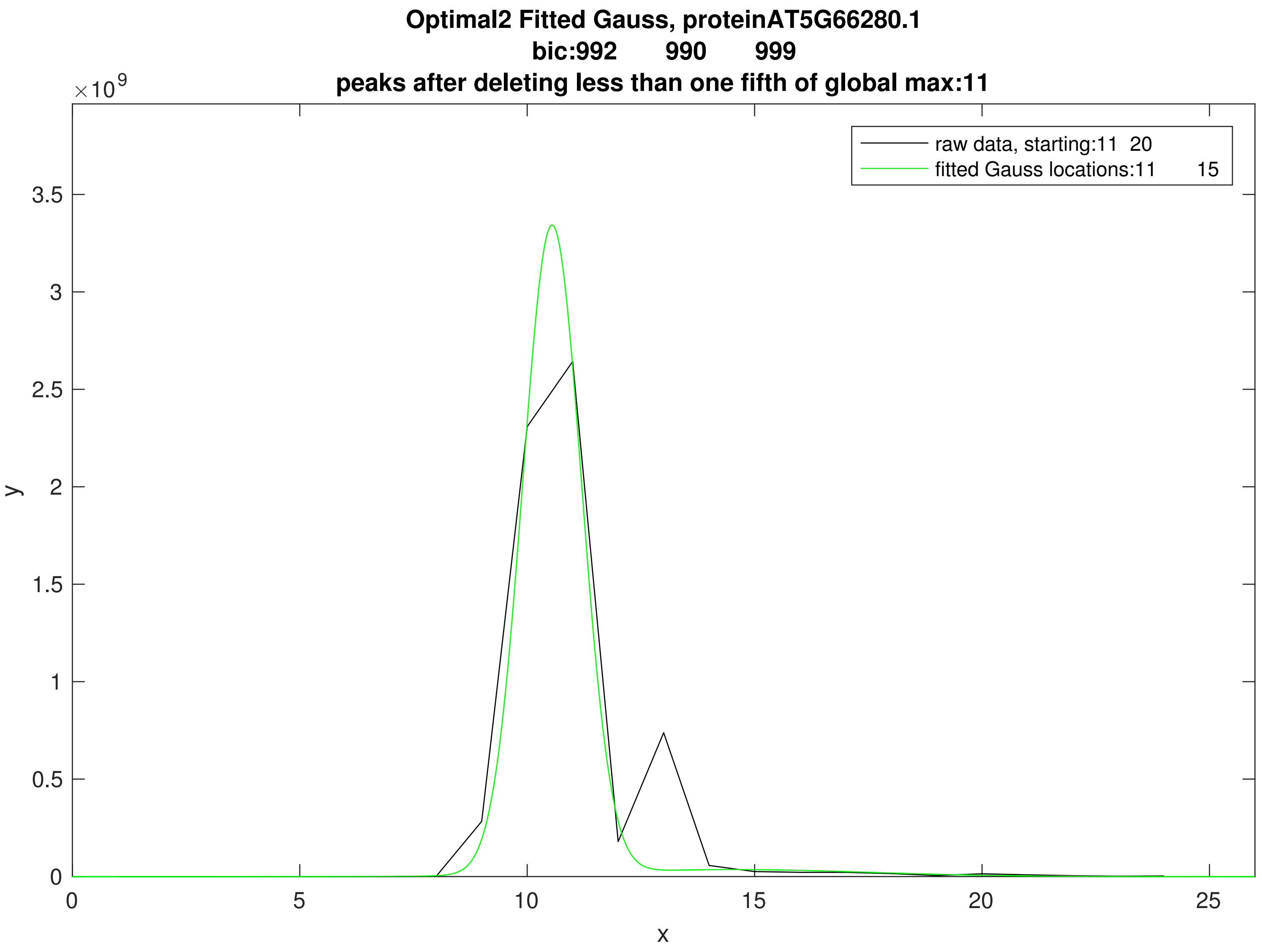

Optimal2 Fitted Gauss, proteinAT5G66280.1
bic:992 990 999
peaks after deleting less than one fifth of global max:11
×10^9
raw data, starting:11 20
fitted Gauss locations:11 15
y
x
0
0.5
1
1.5
2
2.5
3
3.5
0
5
10
15
20
25

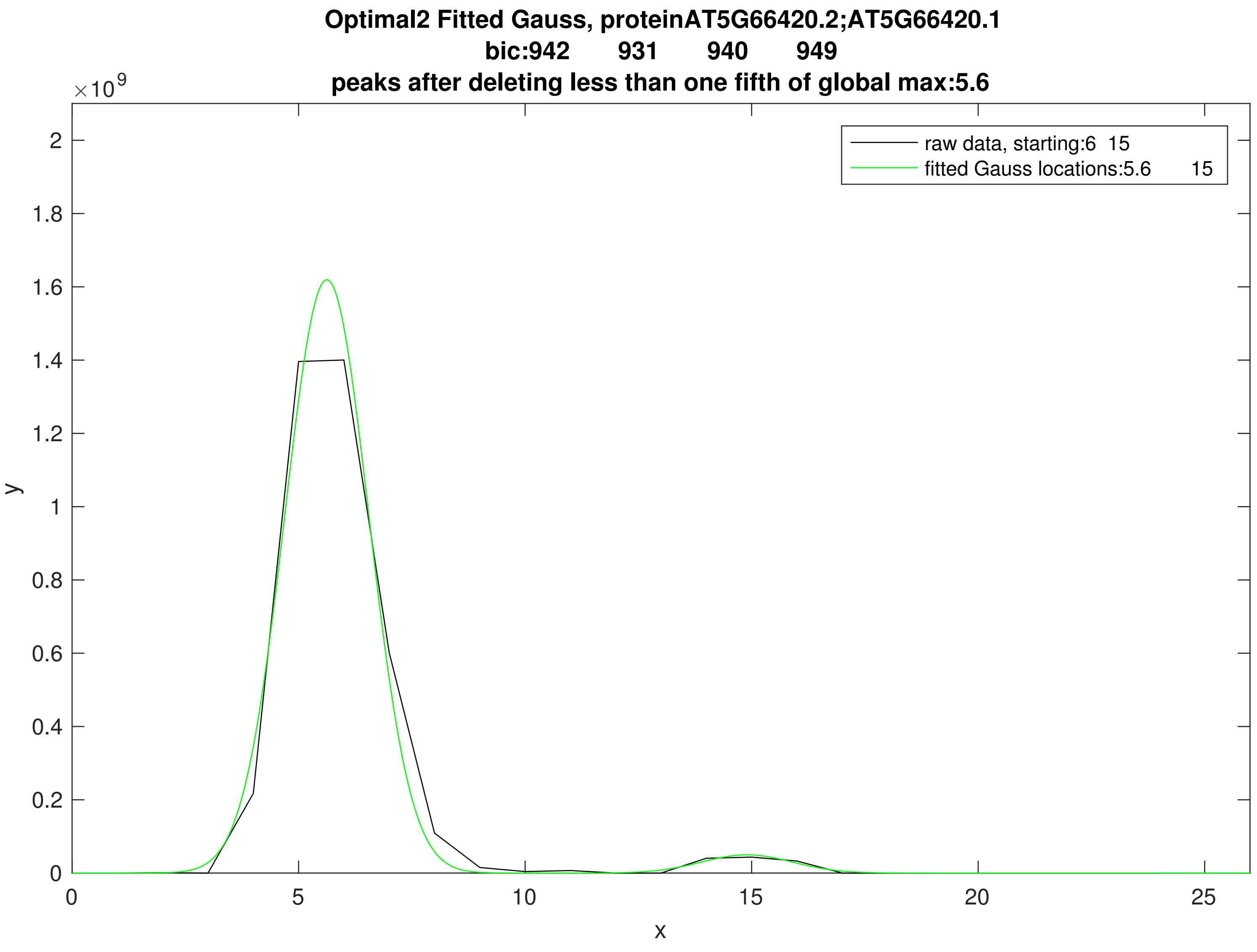
Optimal2 Fitted Gauss, proteinAT5G66420.2;AT5G66420.1
bic:942 931 940 949
peaks after deleting less than one fifth of global max:5.6
×10^9
raw data, starting:6 15
fitted Gauss locations:5.6 15
y
x
0
0.2
0.4
0.6
0.8
1
1.2
1.4
1.6
1.8
2
0
5
10
15
20
25

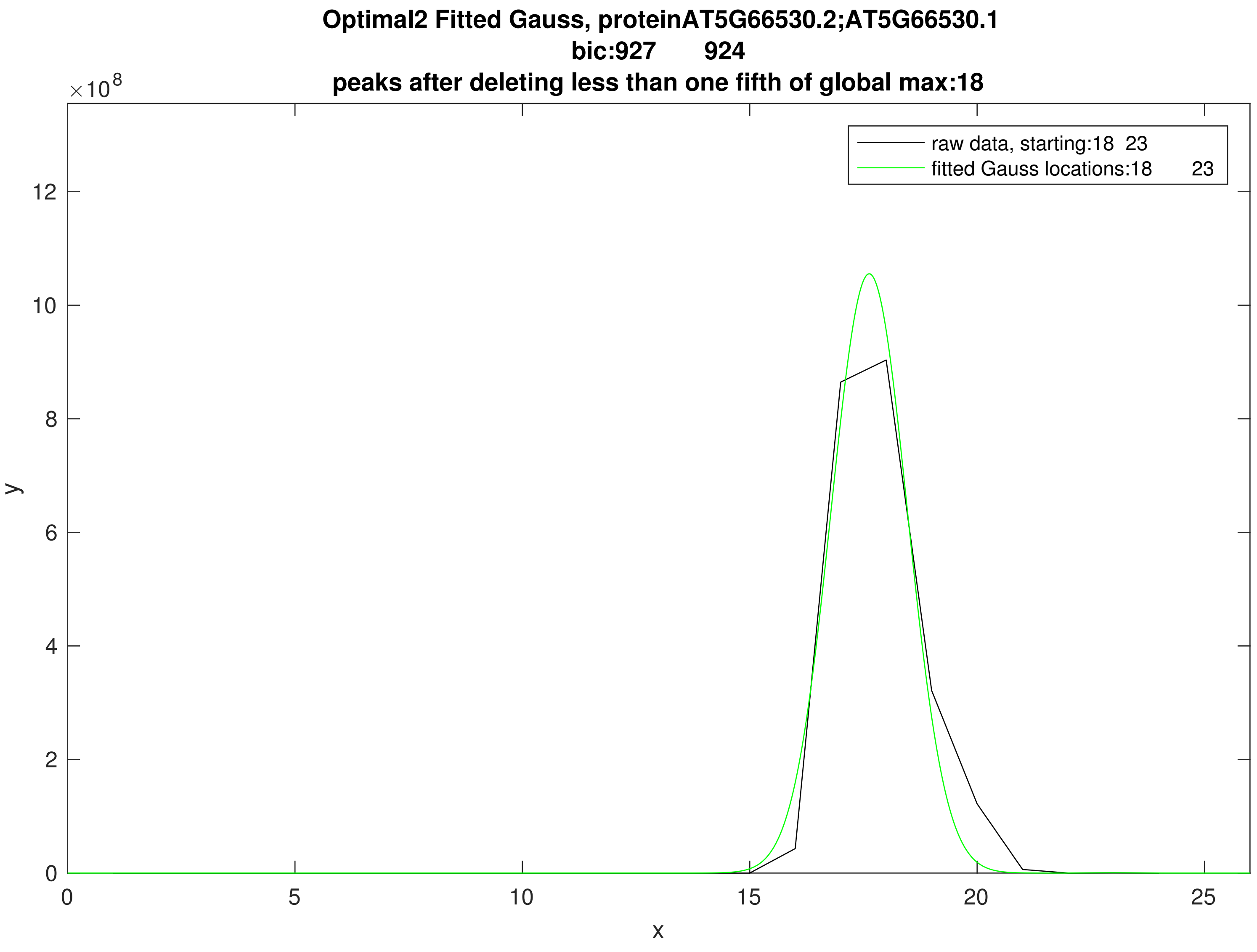

Optimal2 Fitted Gauss, proteinAT5G66530.2;AT5G66530.1
bic:927 924
peaks after deleting less than one fifth of global max:18
×10^8
raw data, starting:18 23
fitted Gauss locations:18 23
y
x
0
2
4
6
8
10
12
0
5
10
15
20
25

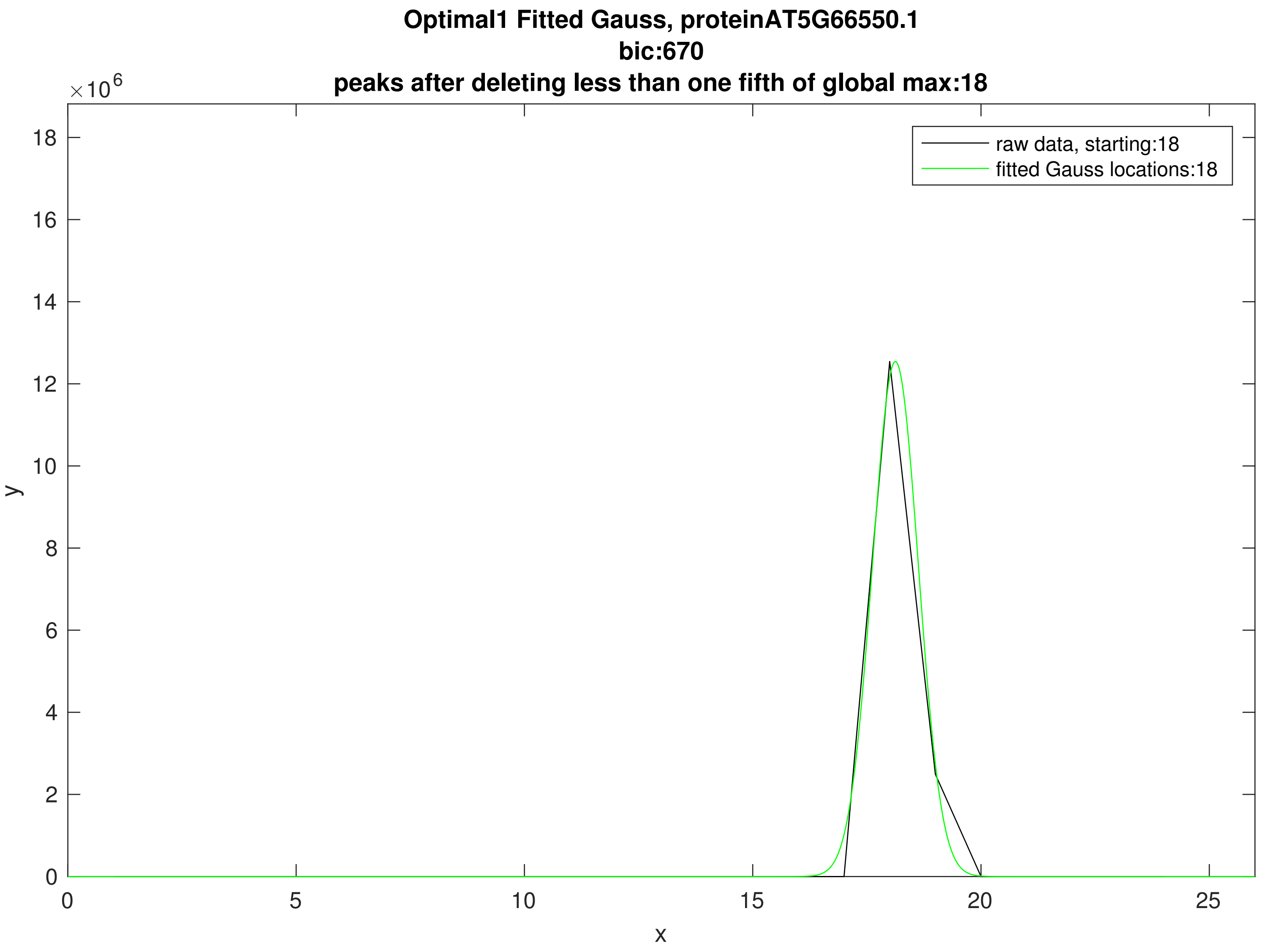
Optimal1 Fitted Gauss, proteinAT5G66550.1
bic:670
peaks after deleting less than one fifth of global max:18
×10^6
raw data, starting:18
fitted Gauss locations:18
y
x
0
2
4
6
8
10
12
14
16
18
0
5
10
15
20
25

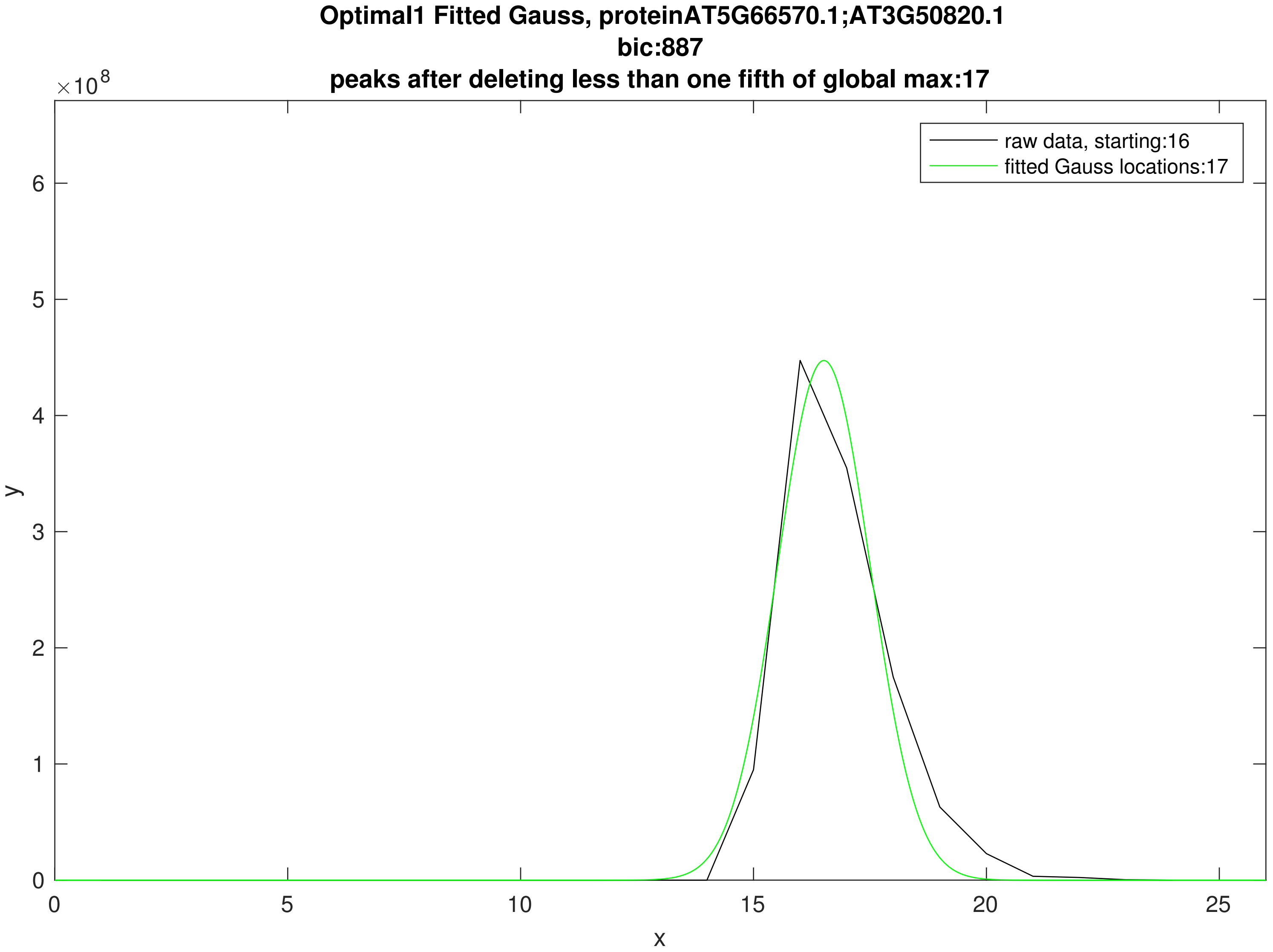

Optimal1 Fitted Gauss, proteinAT5G66570.1;AT3G50820.1
bic:887
peaks after deleting less than one fifth of global max:17
×10^8
raw data, starting:16
fitted Gauss locations:17
y
x
0
1
2
3
4
5
6
0
5
10
15
20
25

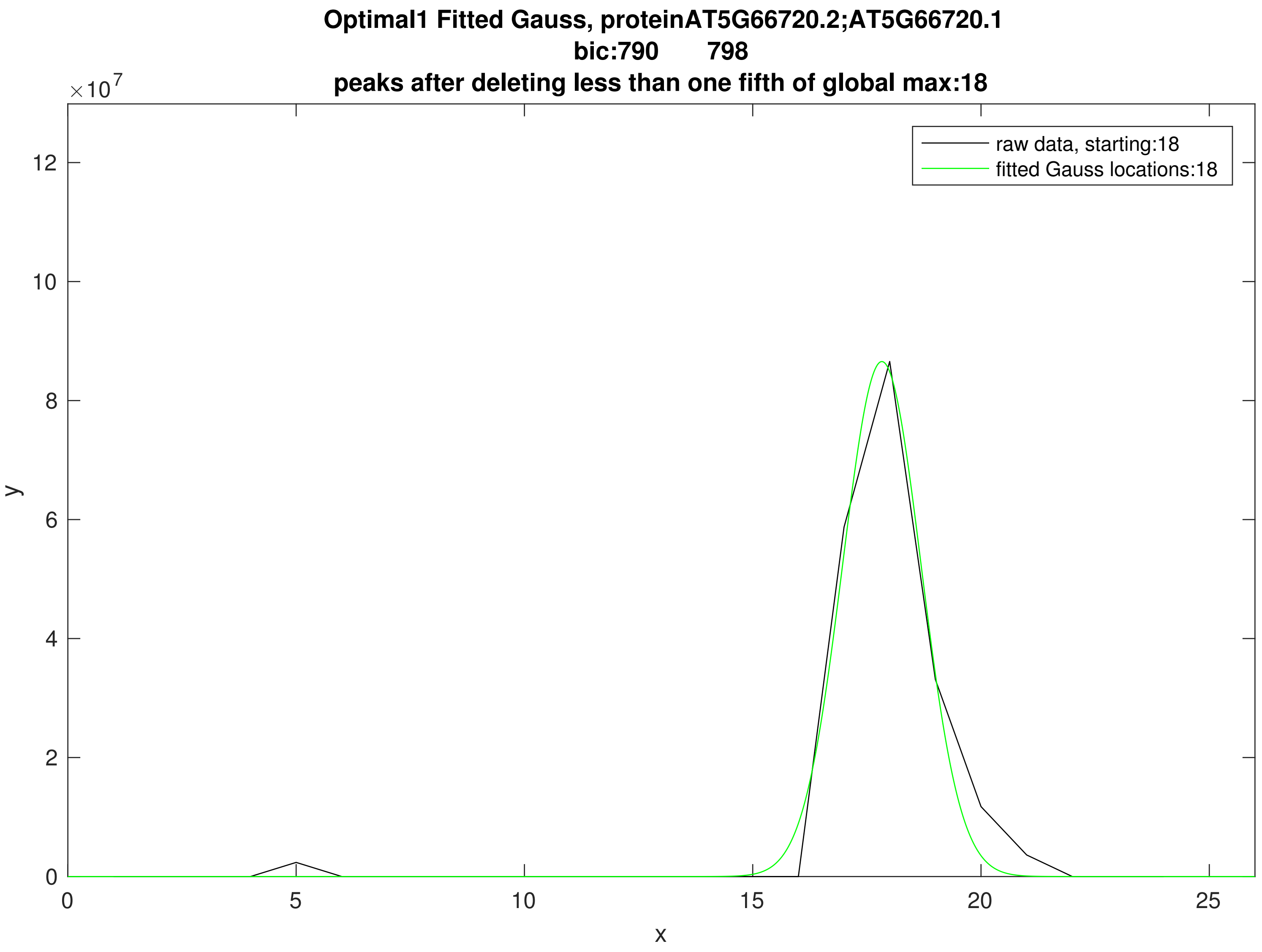

Optimal1 Fitted Gauss, proteinAT5G66720.2;AT5G66720.1
bic:790 798
peaks after deleting less than one fifth of global max:18
×10^7
raw data, starting:18
fitted Gauss locations:18
y
x
0
2
4
6
8
10
12
0
5
10
15
20
25

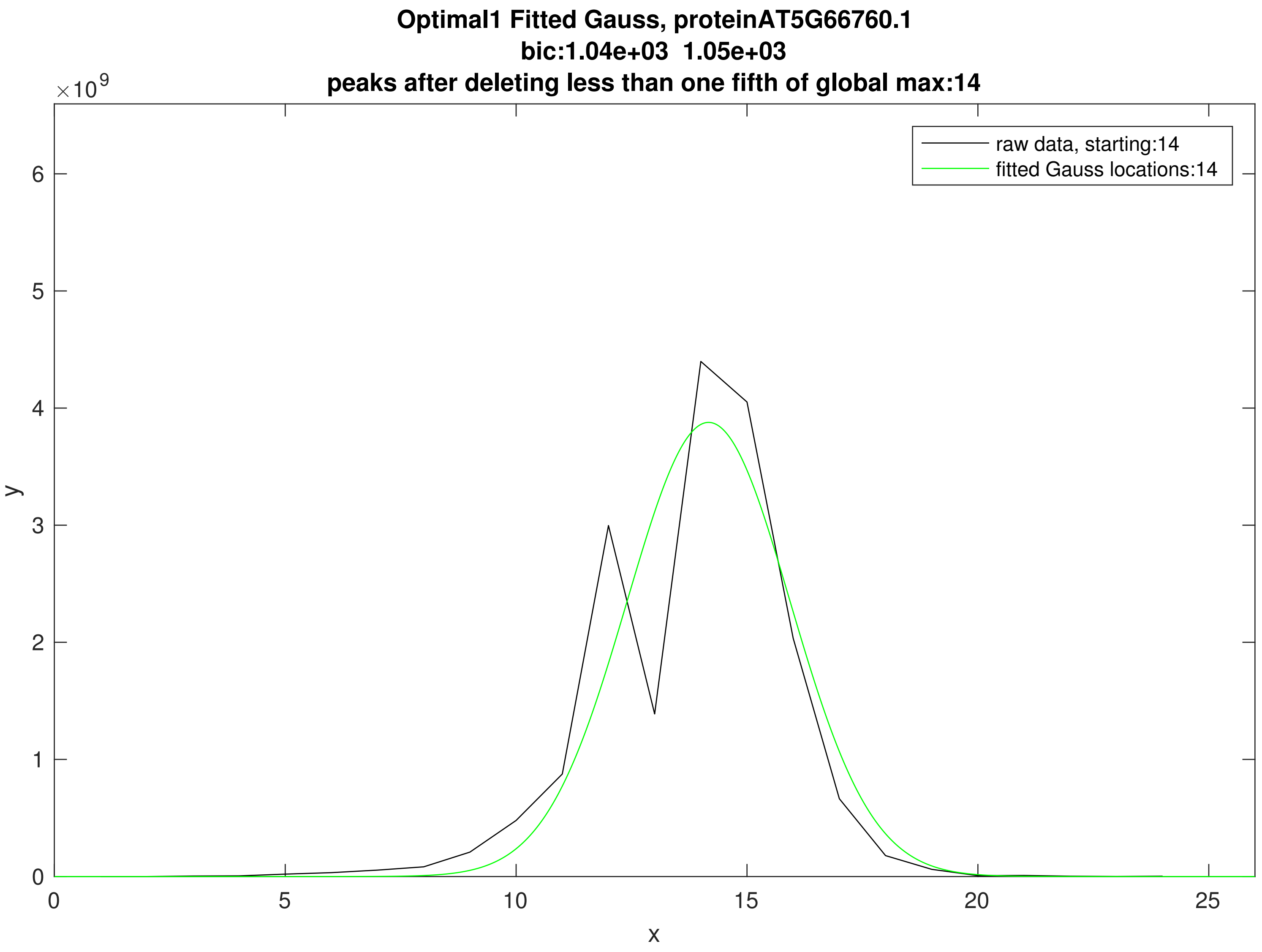
Optimal1 Fitted Gauss, proteinAT5G66760.1
bic:1.04e+03  1.05e+03
peaks after deleting less than one fifth of global max:14
×10^9
raw data, starting:14
fitted Gauss locations:14
y
x
0
1
2
3
4
5
6
0
5
10
15
20
25

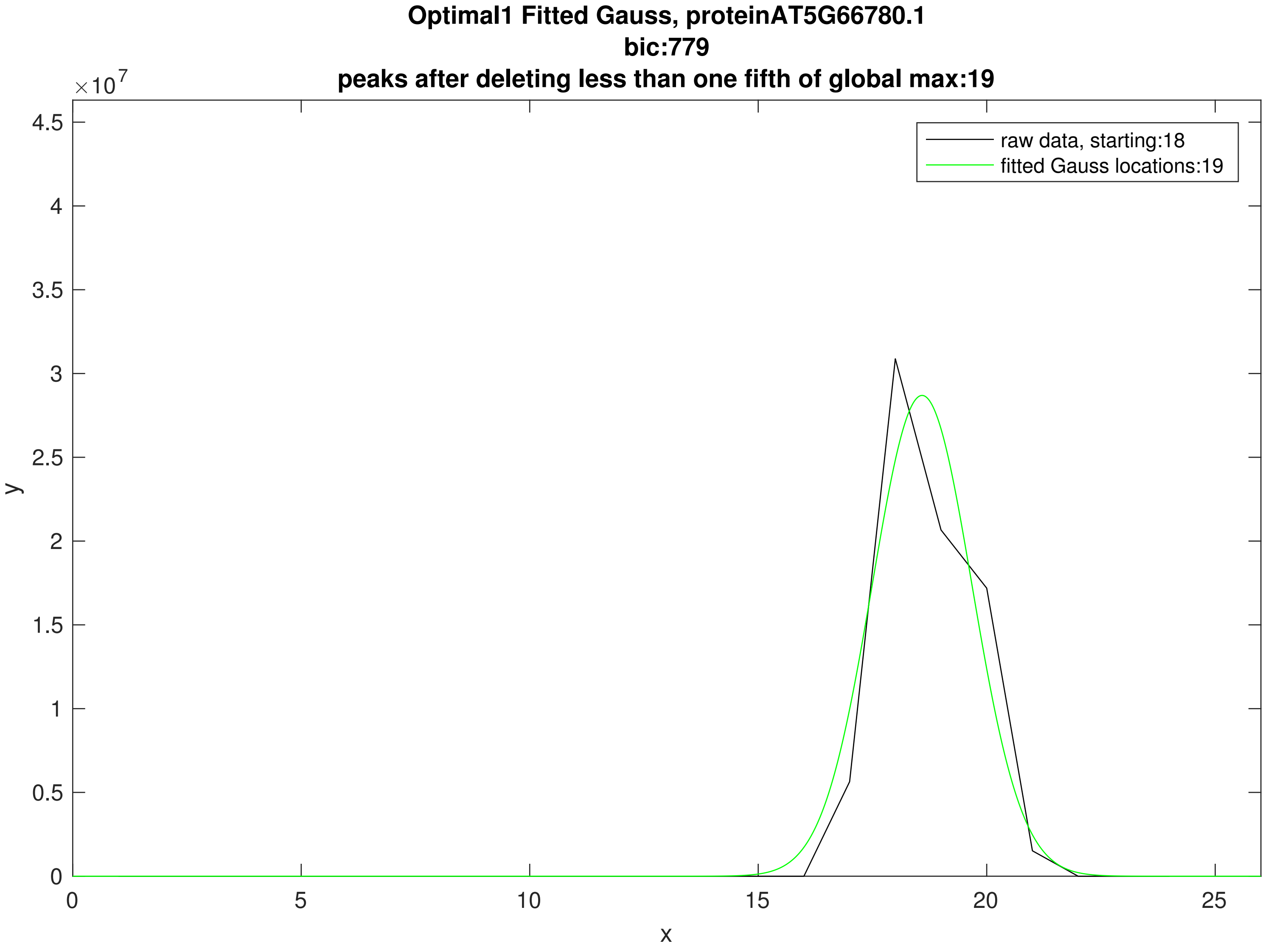
Optimal1 Fitted Gauss, proteinAT5G66780.1
bic:779
peaks after deleting less than one fifth of global max:19
×10^7
raw data, starting:18
fitted Gauss locations:19
y
x
0
0.5
1
1.5
2
2.5
3
3.5
4
4.5
5
10
15
20
25

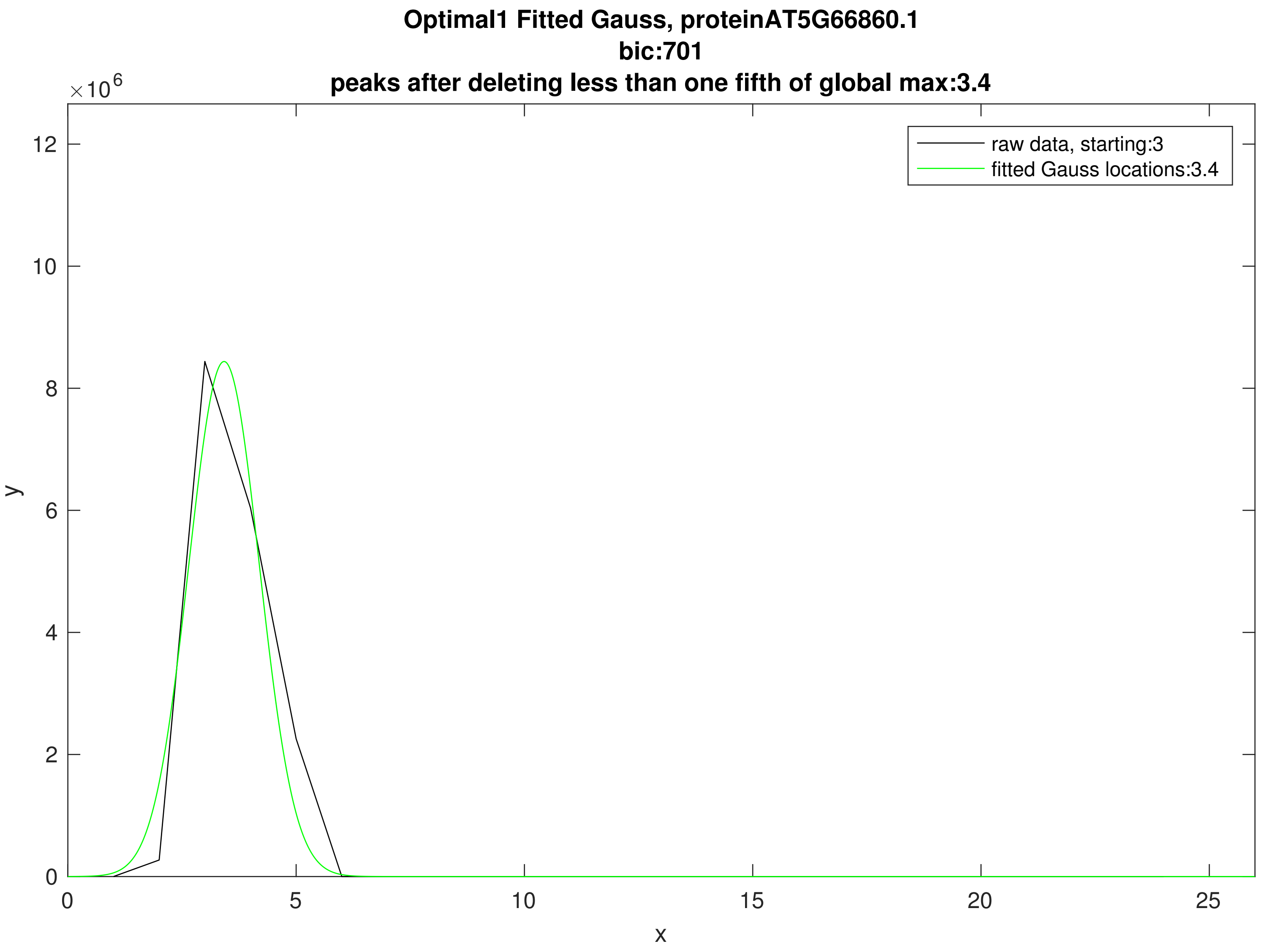

Optimal1 Fitted Gauss, proteinAT5G66860.1
bic:701
peaks after deleting less than one fifth of global max:3.4
×10^6
raw data, starting:3
fitted Gauss locations:3.4
y
x
0
2
4
6
8
10
12
0
5
10
15
20
25

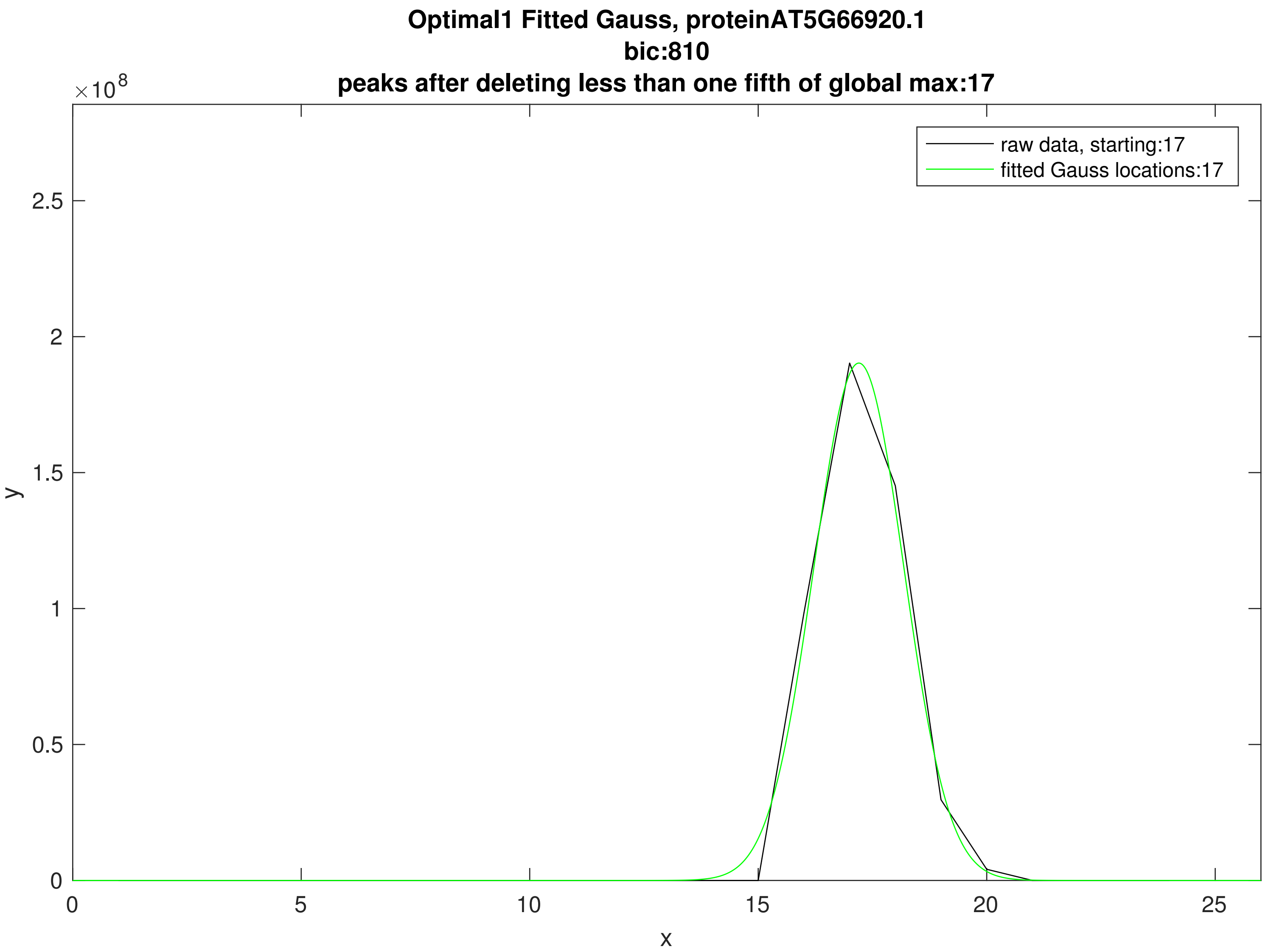
Optimal1 Fitted Gauss, proteinAT5G66920.1
bic:810
peaks after deleting less than one fifth of global max:17
×10^8
raw data, starting:17
fitted Gauss locations:17
y
x
2.5
2
1.5
1
0.5
0
0
5
10
15
20
25

#### Optimal1 Fitted Gauss, proteinAT5G67130.1
#### bic:843
#### peaks after deleting less than one fifth of global max:17

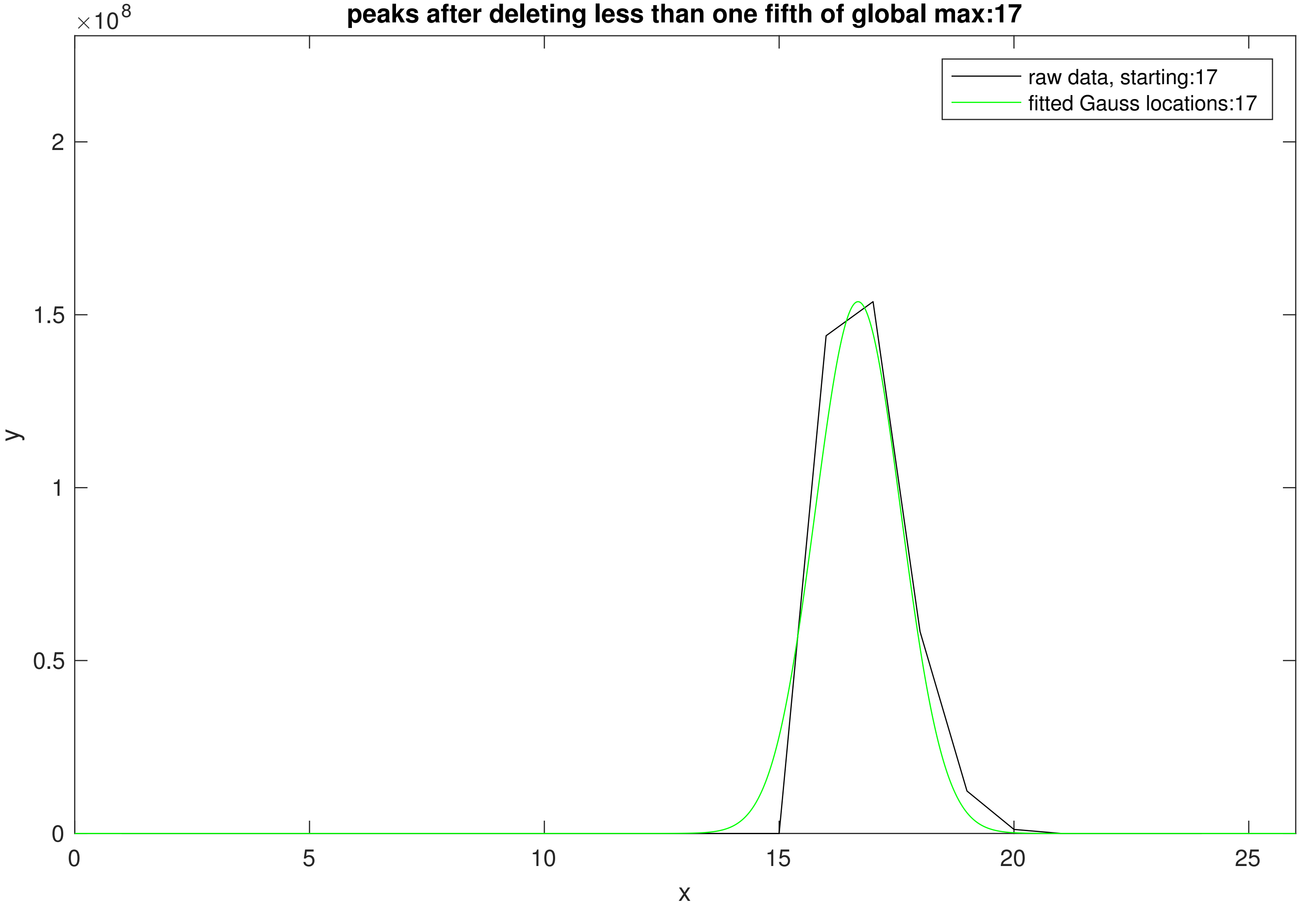

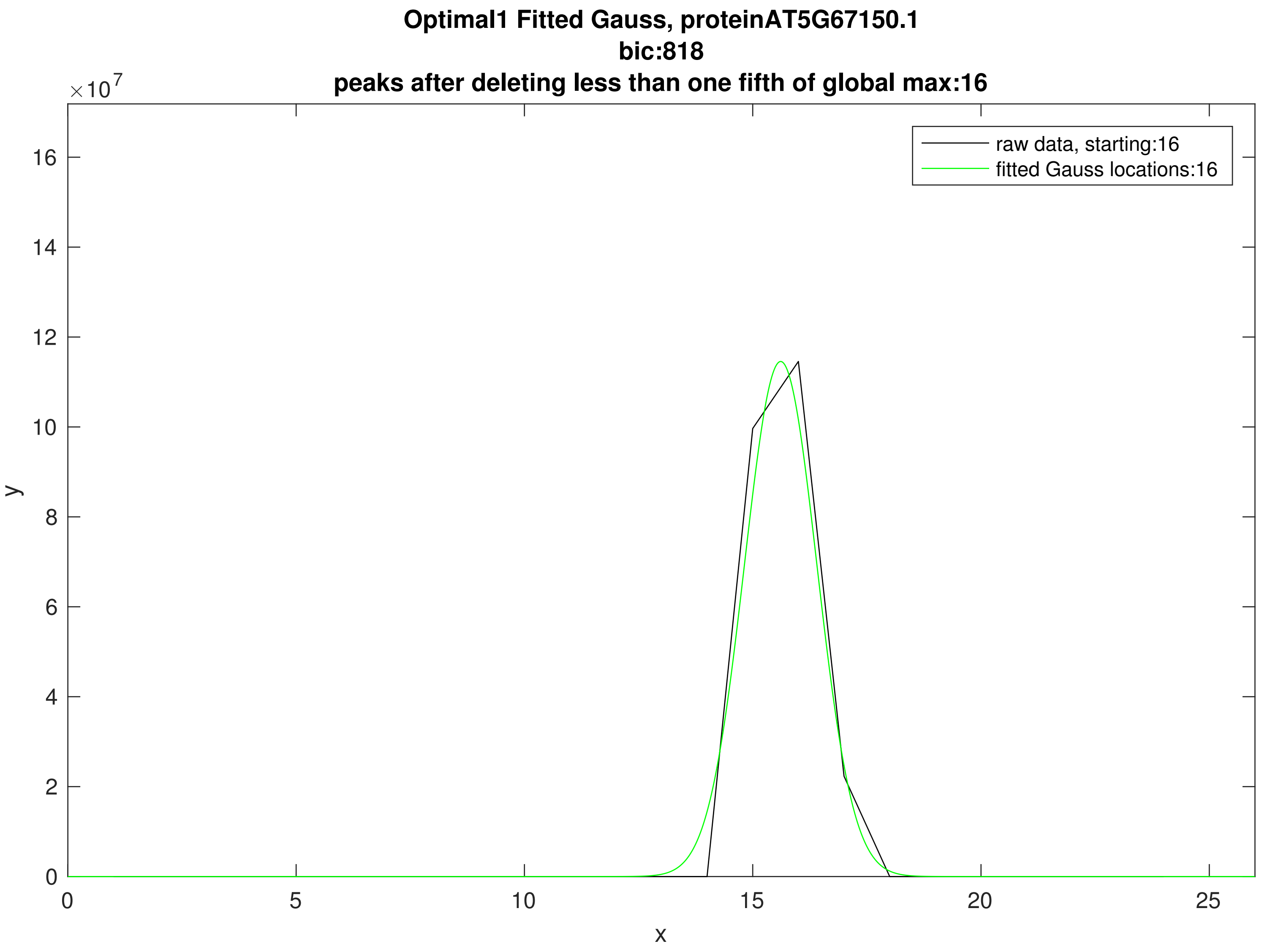

Optimal1 Fitted Gauss, proteinAT5G67150.1
bic:818
peaks after deleting less than one fifth of global max:16
×10^7
raw data, starting:16
fitted Gauss locations:16
y
x
0
2
4
6
8
10
12
14
16
0
5
10
15
20
25

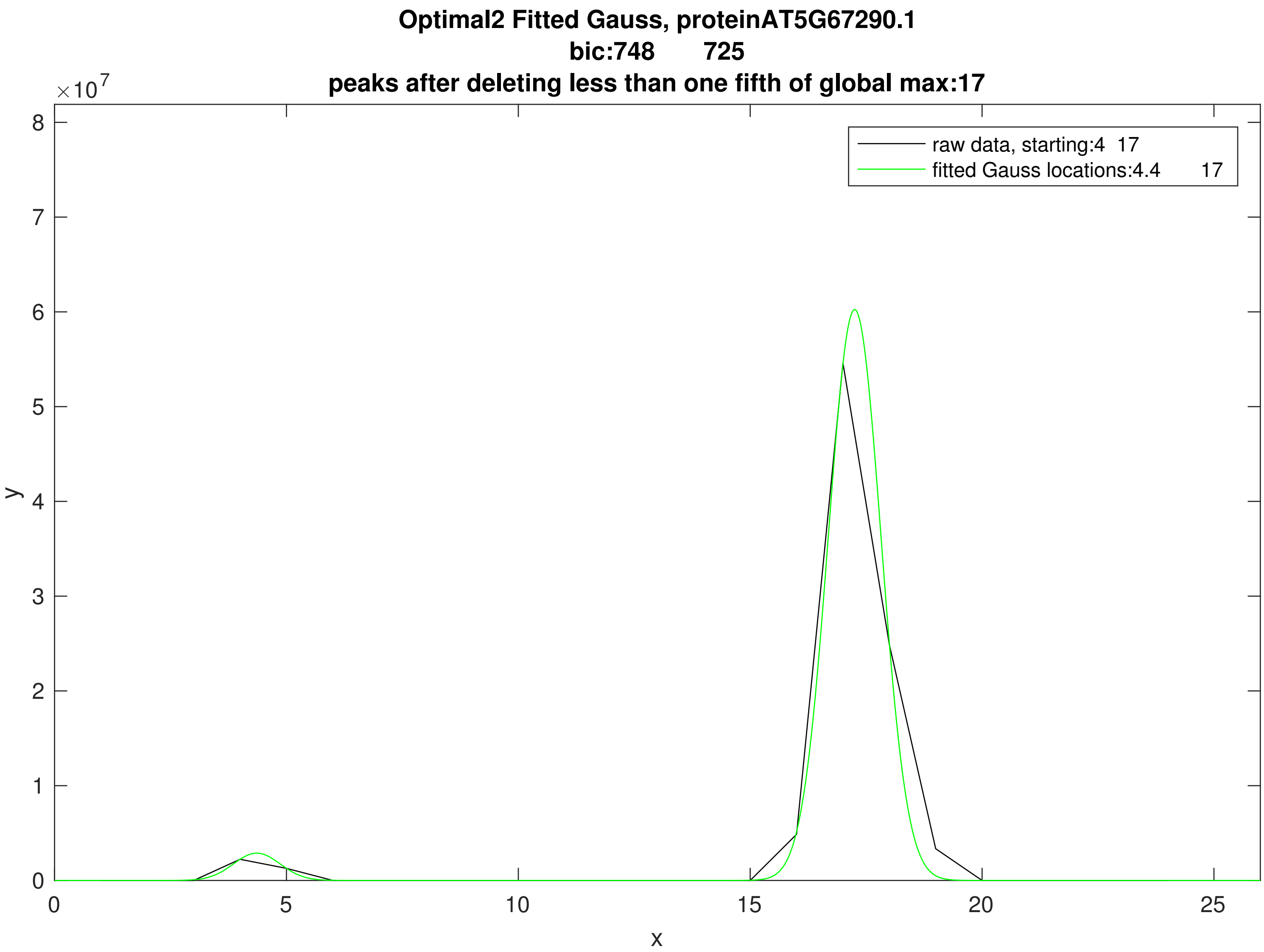

Optimal2 Fitted Gauss, proteinAT5G67290.1
bic:748 725
peaks after deleting less than one fifth of global max:17
×10^7
raw data, starting:4 17
fitted Gauss locations:4.4 17
y
x
0
1
2
3
4
5
6
7
8
0
5
10
15
20
25

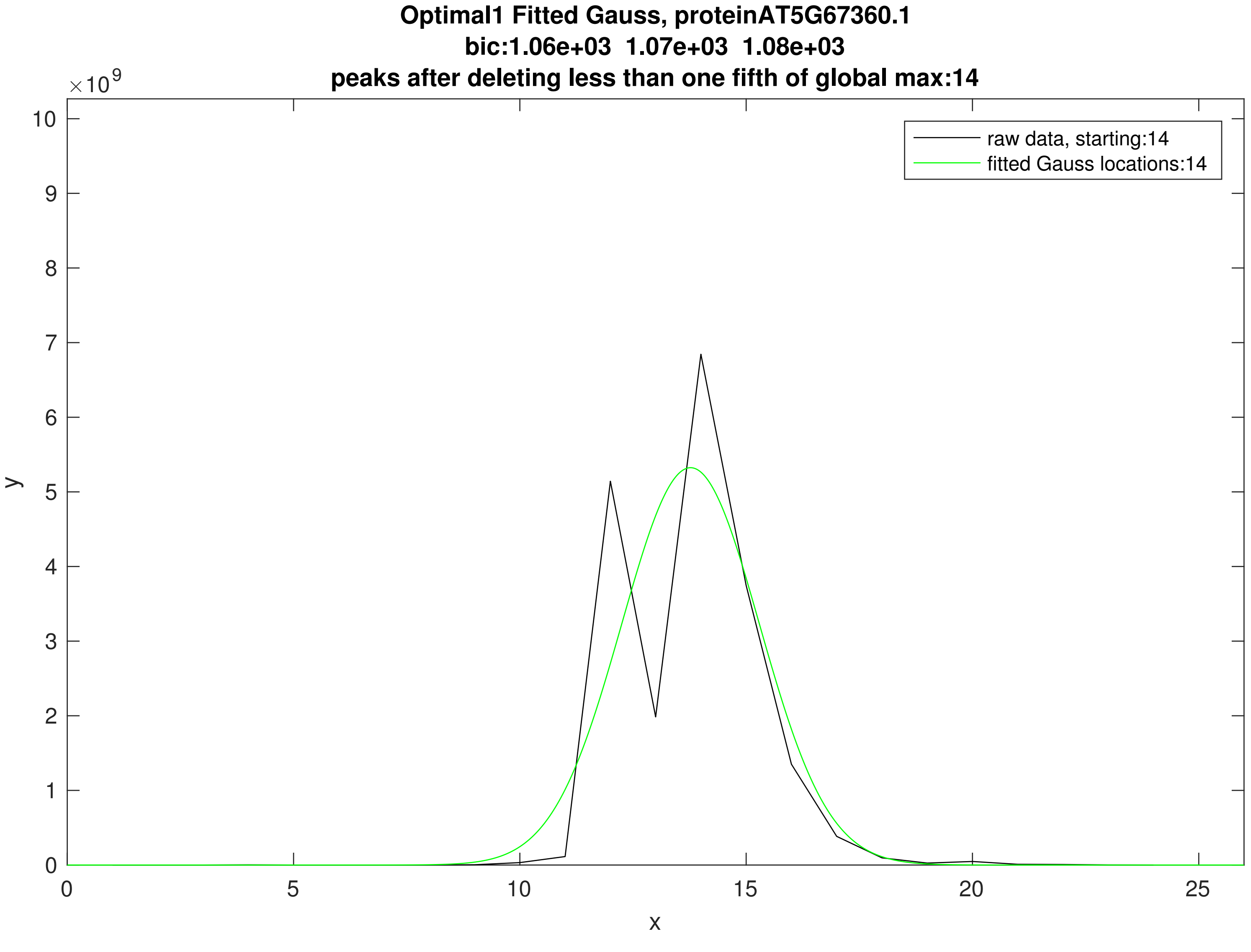

Optimal1 Fitted Gauss, proteinAT5G67360.1
bic:1.06e+03 1.07e+03 1.08e+03
peaks after deleting less than one fifth of global max:14
×10^9
raw data, starting:14
fitted Gauss locations:14
y
x
0
1
2
3
4
5
6
7
8
9
10
0
5
10
15
20
25

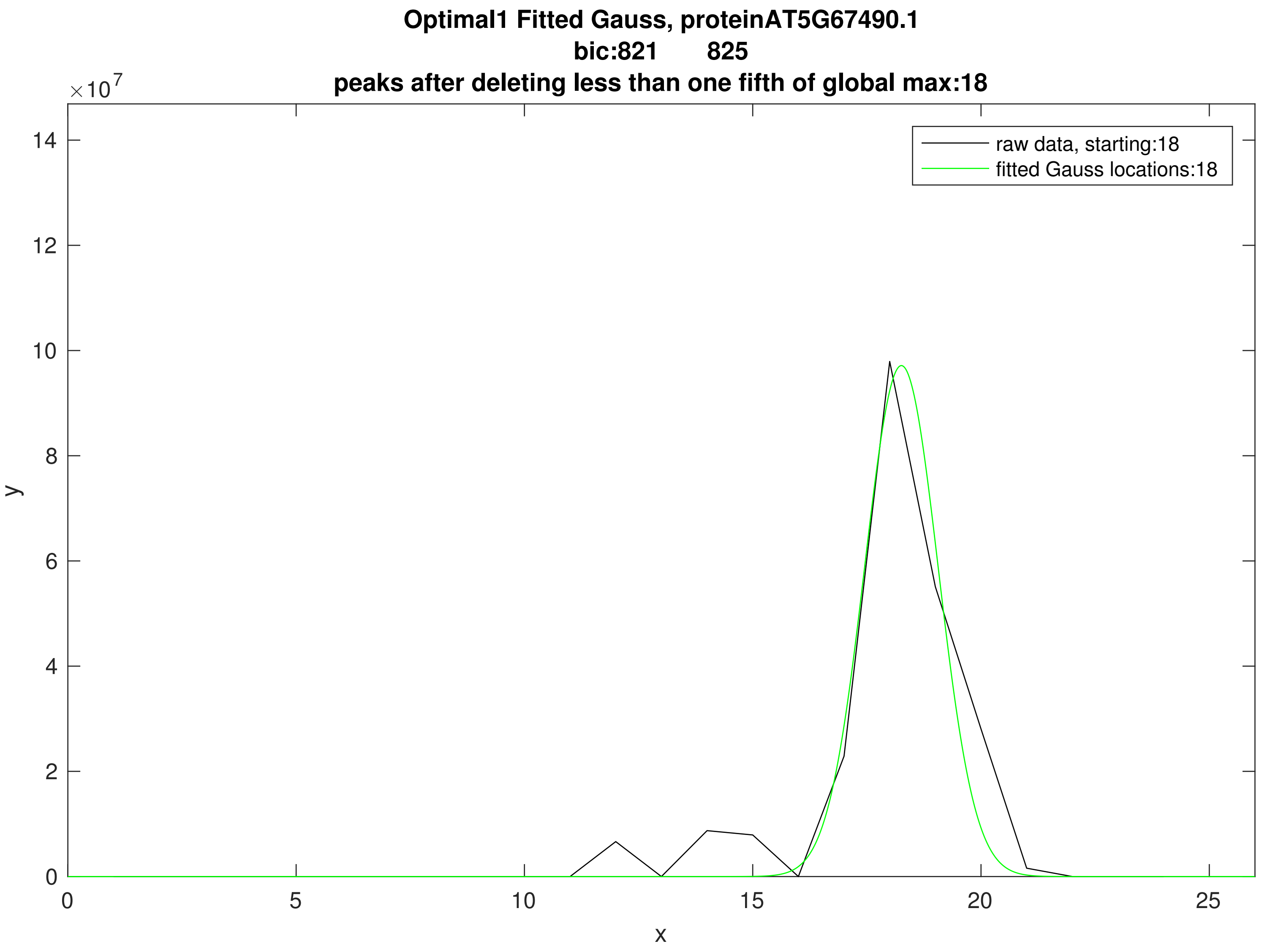
Optimal1 Fitted Gauss, proteinAT5G67490.1
bic:821 825
peaks after deleting less than one fifth of global max:18
×10^7
raw data, starting:18
fitted Gauss locations:18
y
x
0
2
4
6
8
10
12
14
0
5
10
15
20
25

**Optimal1 Fitted Gauss, proteinAT5G67510.1**
**bic:536**
**peaks after deleting less than one fifth of global max:3.1**

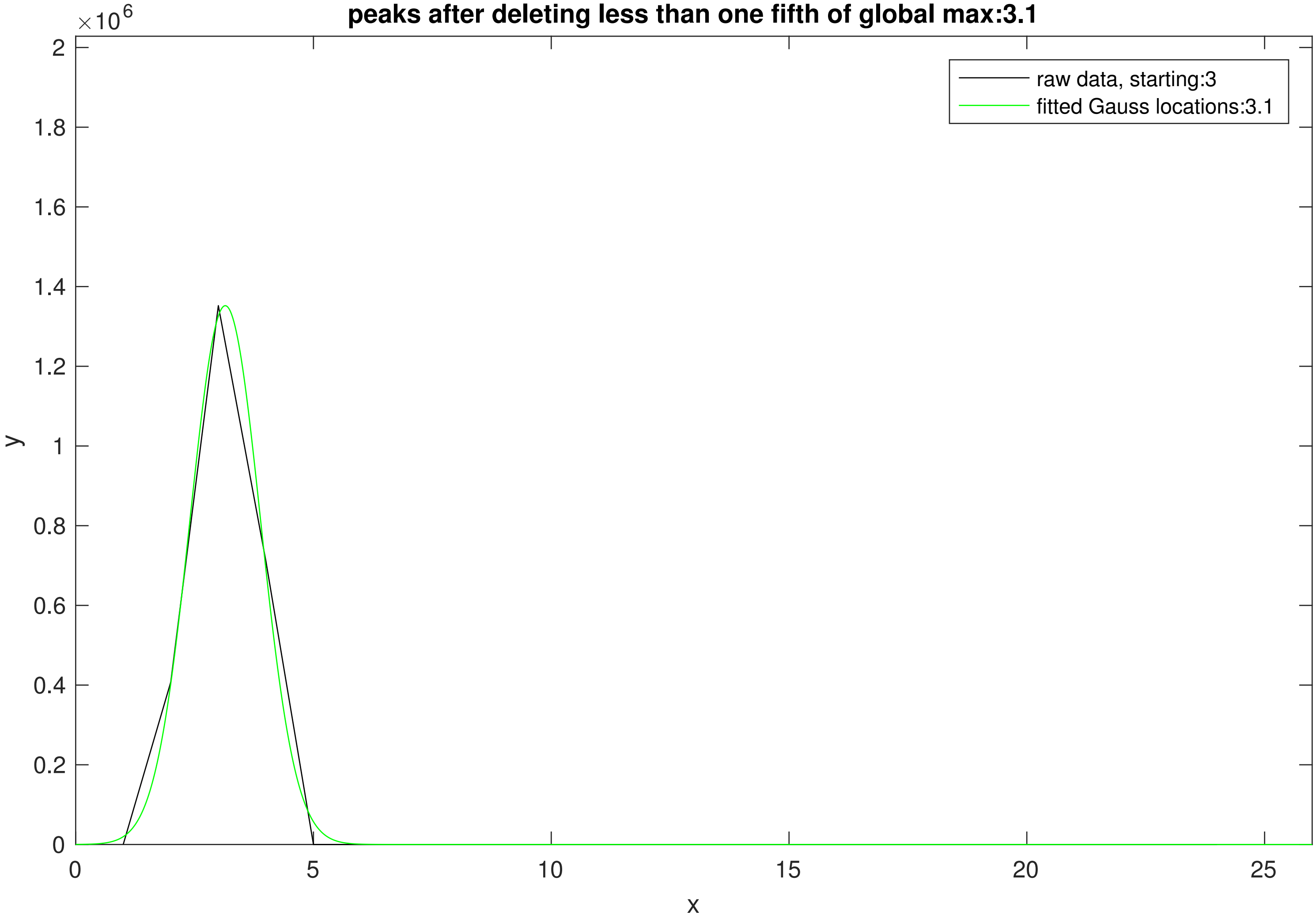

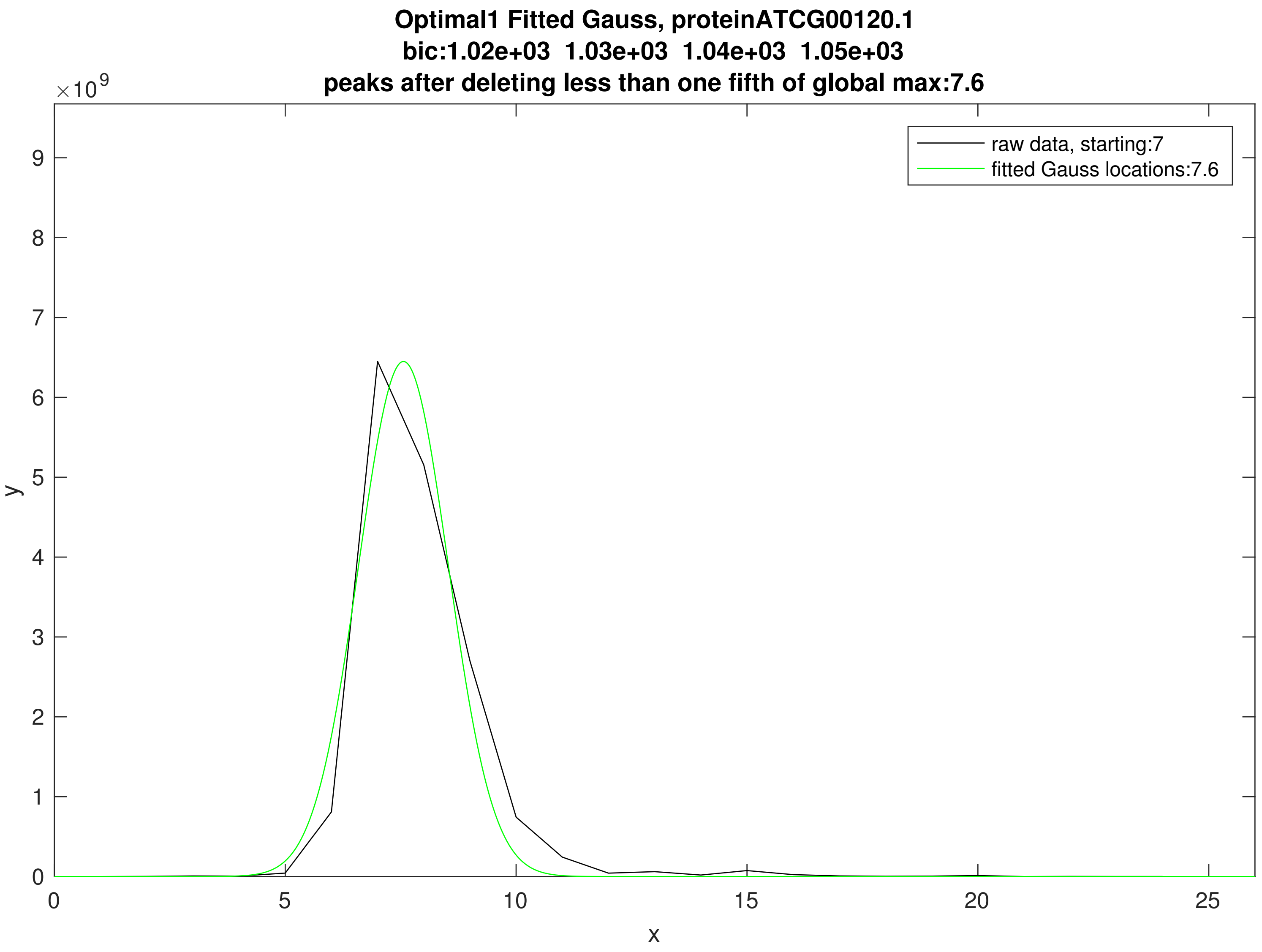

Optimal1 Fitted Gauss, proteinATCG00120.1
bic:1.02e+03 1.03e+03 1.04e+03 1.05e+03
peaks after deleting less than one fifth of global max:7.6
×10^9
raw data, starting:7
fitted Gauss locations:7.6
y
x
0
1
2
3
4
5
6
7
8
9
10
15
20
25

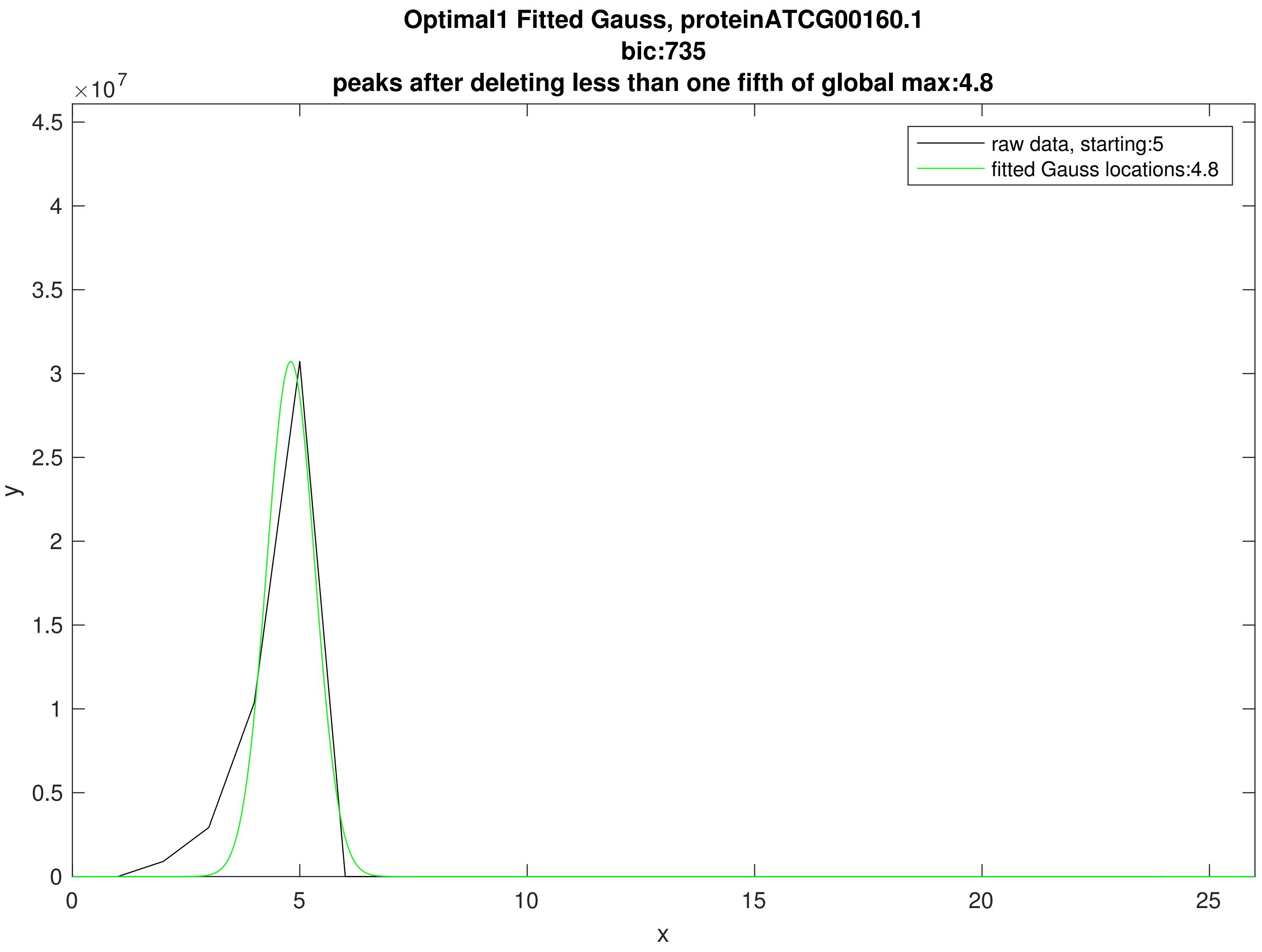
Optimal1 Fitted Gauss, proteinATCG00160.1
bic:735
peaks after deleting less than one fifth of global max:4.8
×10^7
4.5
4
3.5
3
2.5
2
1.5
1
0.5
0
y
0
5
10
15
20
25
x
raw data, starting:5
fitted Gauss locations:4.8

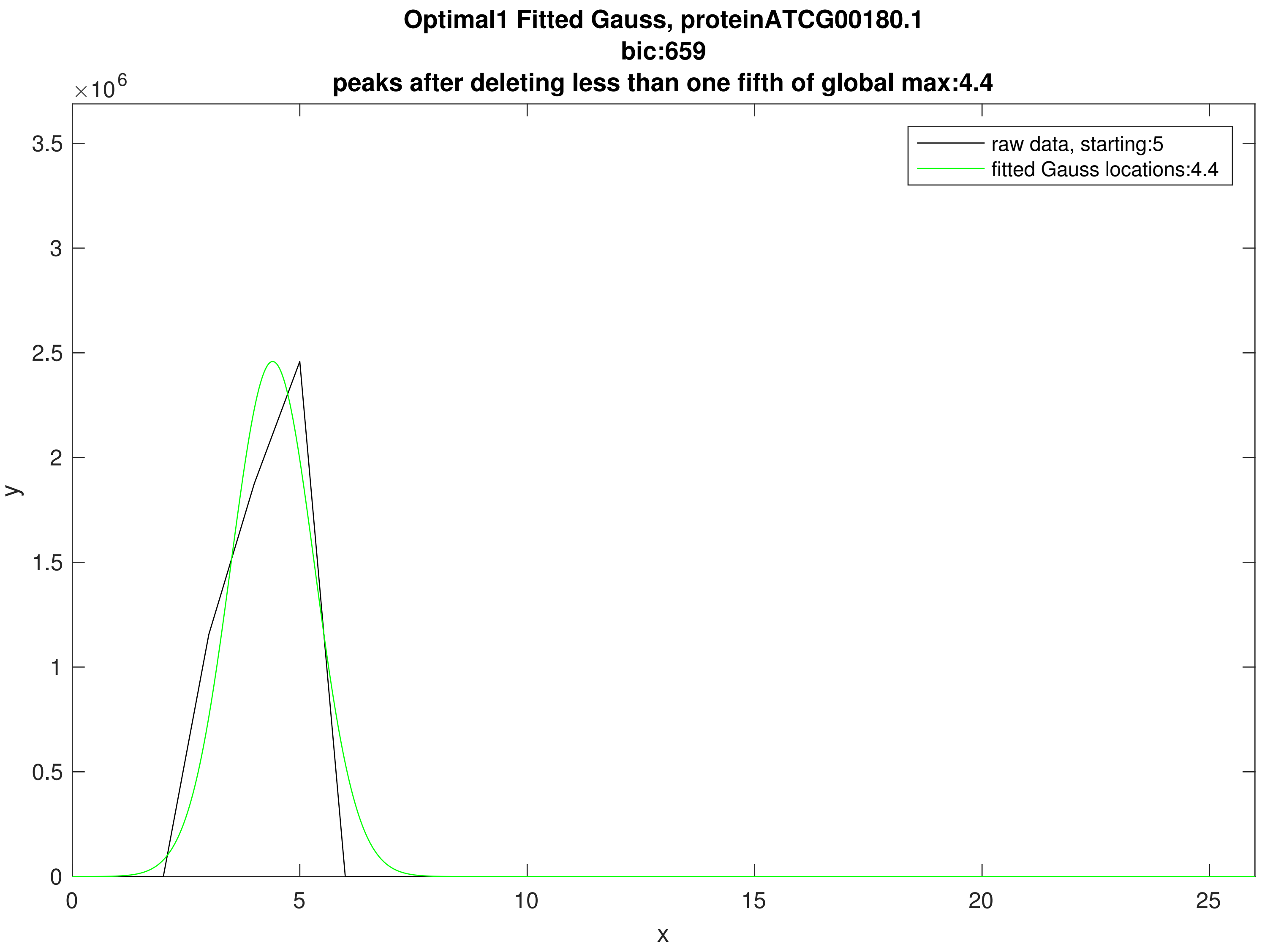

Optimal1 Fitted Gauss, proteinATCG00180.1
bic:659
peaks after deleting less than one fifth of global max:4.4
×10^6
raw data, starting:5
fitted Gauss locations:4.4
3.5
3
2.5
2
1.5
1
0.5
0
y
0
5
10
15
20
25
x

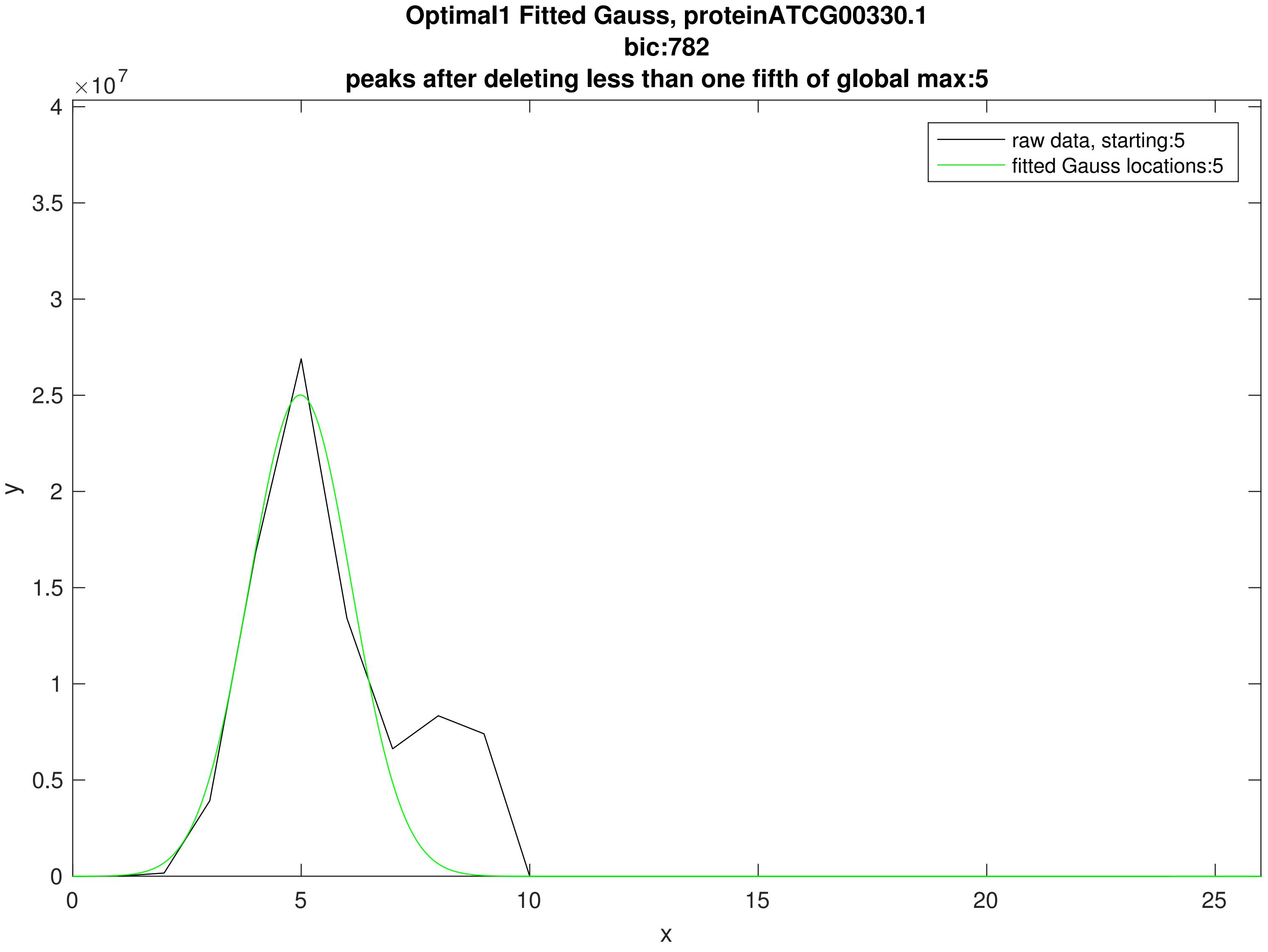
Optimal1 Fitted Gauss, proteinATCG00330.1
bic:782
peaks after deleting less than one fifth of global max:5
×10^7
raw data, starting:5
fitted Gauss locations:5
y
x
0
0.5
1
1.5
2
2.5
3
3.5
4
5
10
15
20
25

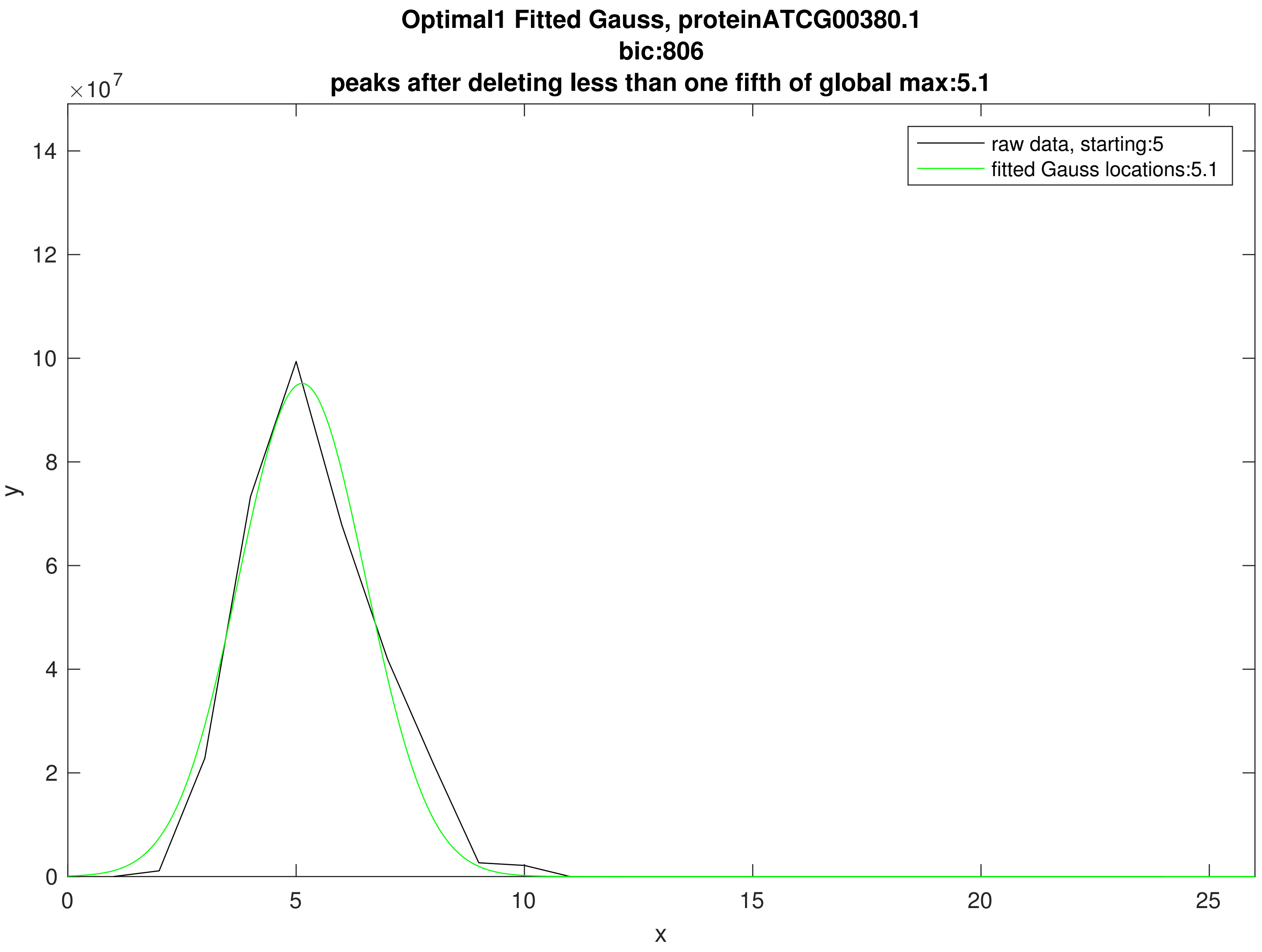
Optimal1 Fitted Gauss, proteinATCG00380.1
bic:806
peaks after deleting less than one fifth of global max:5.1
×10^7
14
12
10
8
6
4
2
0
y
0
5
10
15
20
25
x
raw data, starting:5
fitted Gauss locations:5.1

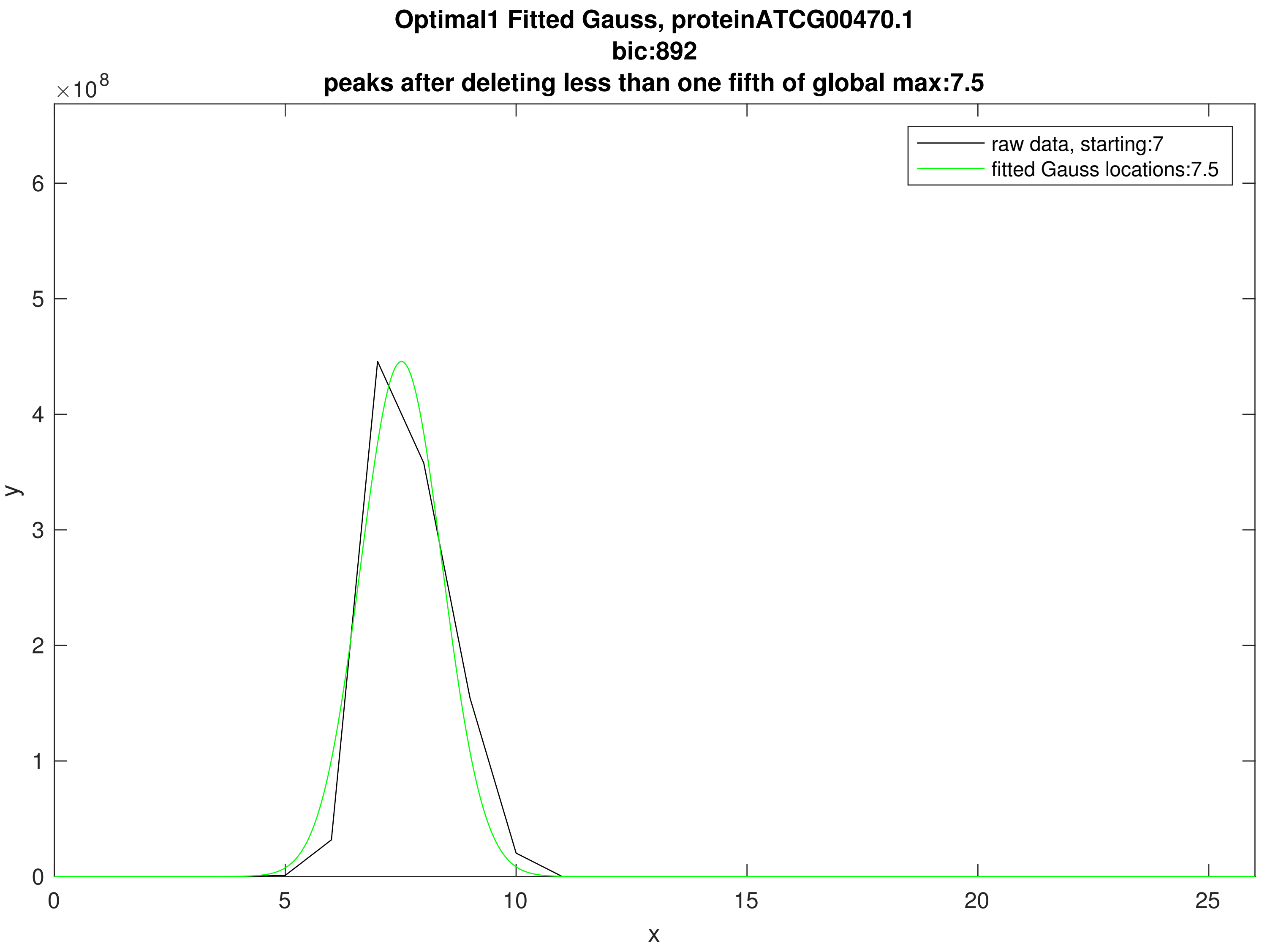

Optimal1 Fitted Gauss, proteinATCG00470.1
bic:892
peaks after deleting less than one fifth of global max:7.5
×10^8
raw data, starting:7
fitted Gauss locations:7.5
y
x
0
1
2
3
4
5
6
0
5
10
15
20
25

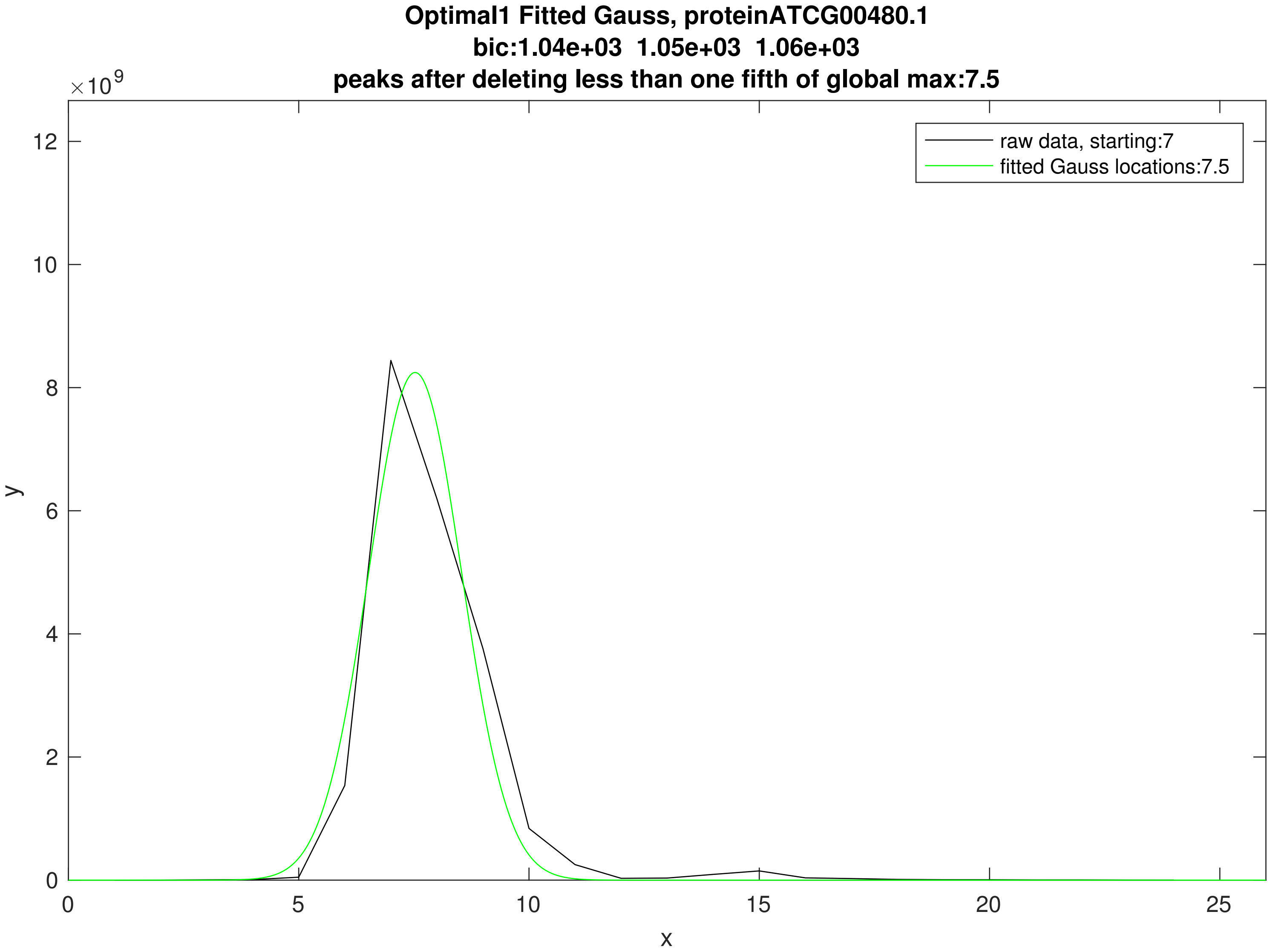
Optimal1 Fitted Gauss, proteinATCG00480.1
bic:1.04e+03  1.05e+03  1.06e+03
peaks after deleting less than one fifth of global max:7.5
×10^9
raw data, starting:7
fitted Gauss locations:7.5
y
x
0
2
4
6
8
10
12
5
15
20
25

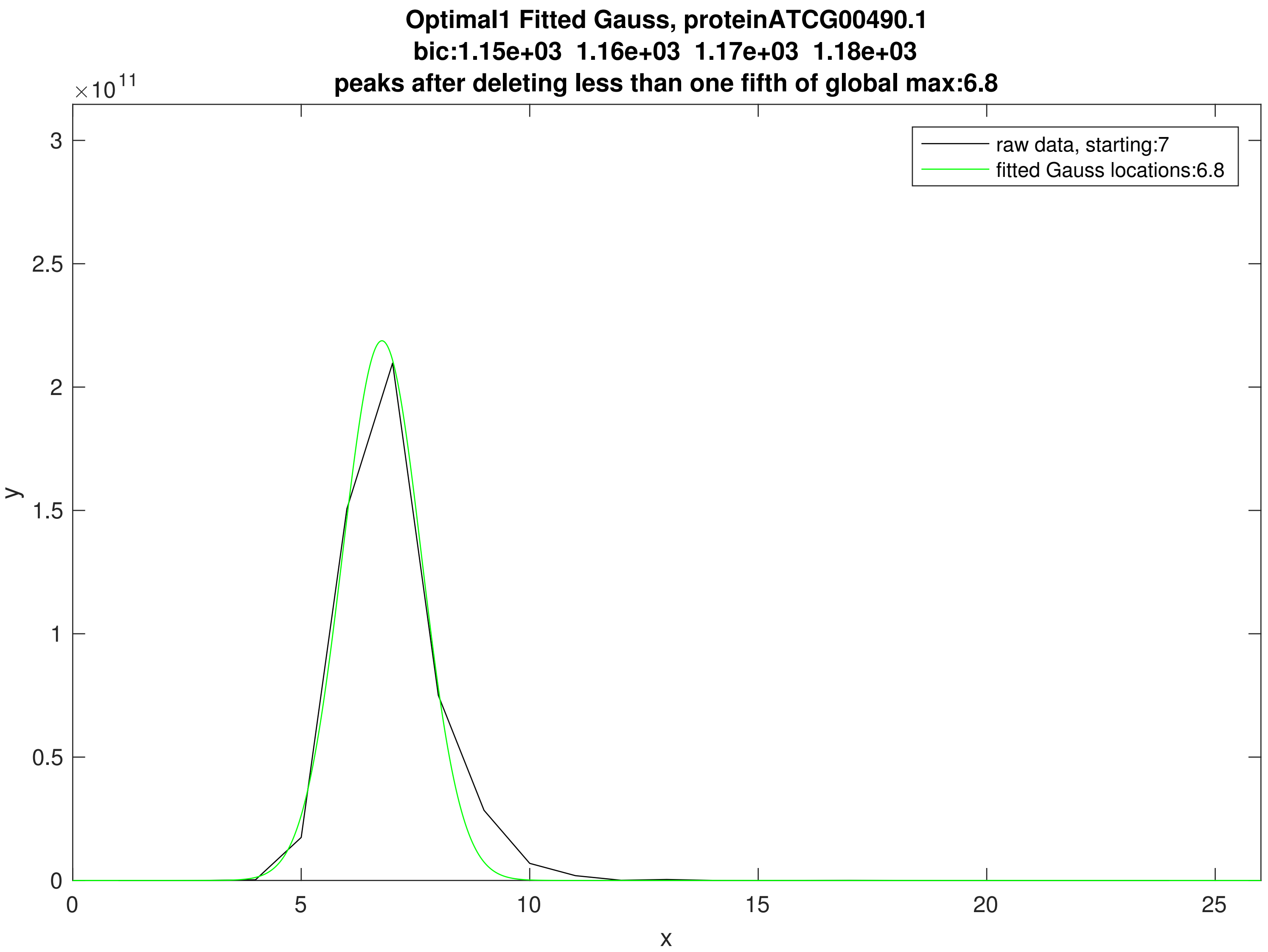
Optimal1 Fitted Gauss, proteinATCG00490.1
bic:1.15e+03 1.16e+03 1.17e+03 1.18e+03
peaks after deleting less than one fifth of global max:6.8
×10^11
raw data, starting:7
fitted Gauss locations:6.8
3
2.5
2
1.5
1
0.5
0
y
0
5
10
15
20
25
x

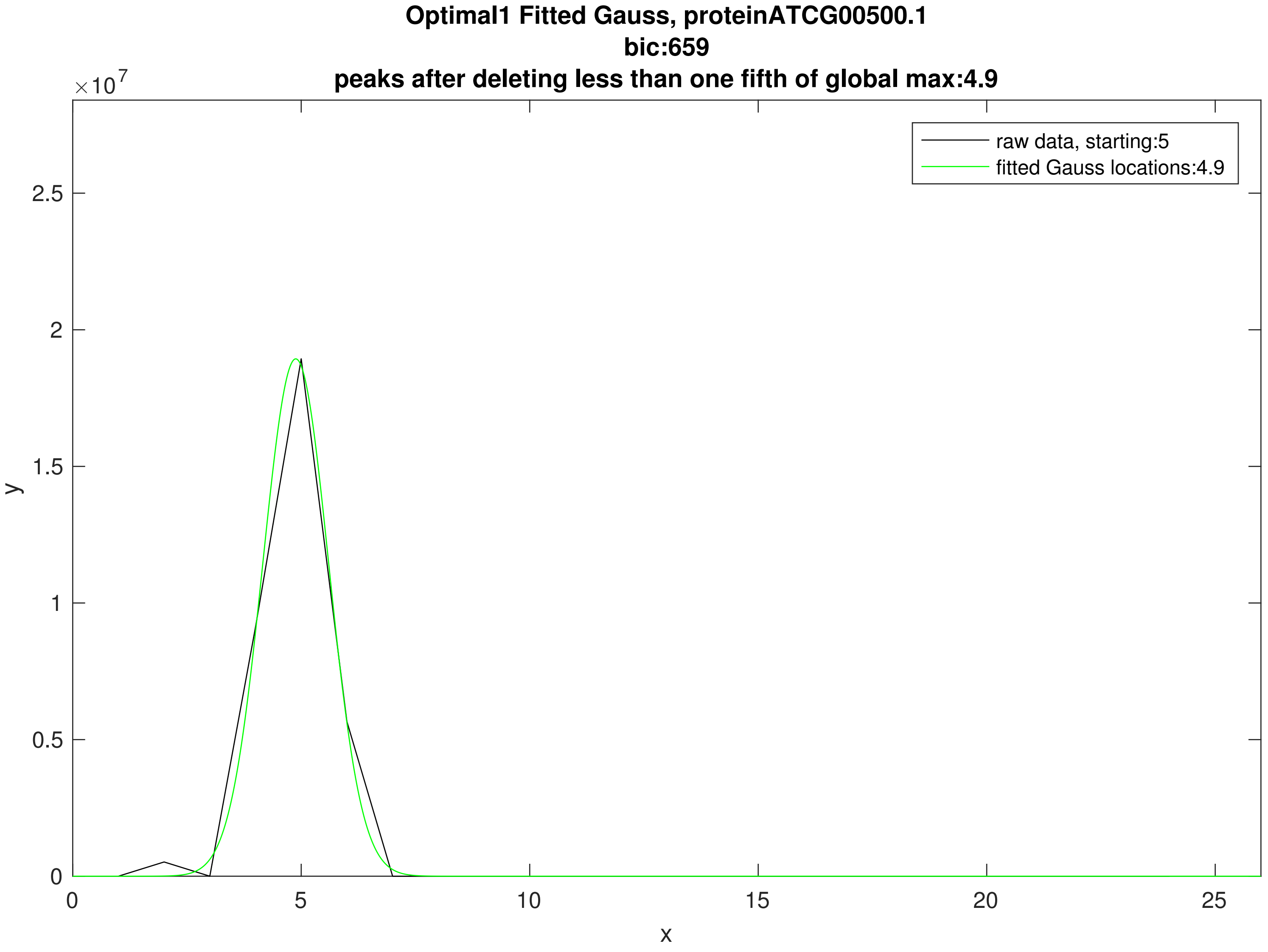
Optimal1 Fitted Gauss, proteinATCG00500.1
bic:659
peaks after deleting less than one fifth of global max:4.9
×10^7
raw data, starting:5
fitted Gauss locations:4.9
y
x
0
0.5
1
1.5
2
2.5
5
10
15
20
25

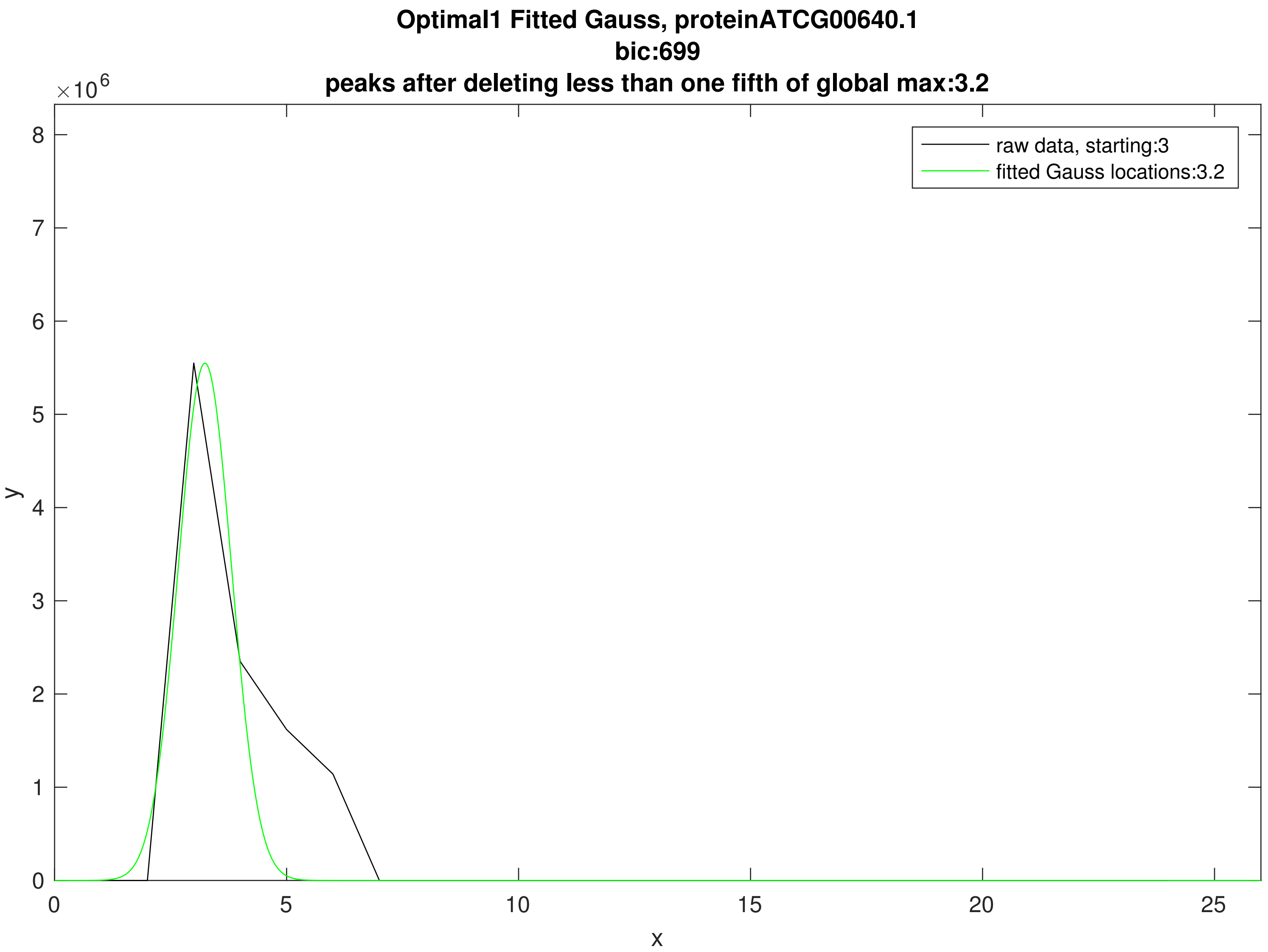
Optimal1 Fitted Gauss, proteinATCG00640.1
bic:699
peaks after deleting less than one fifth of global max:3.2
×10^6
raw data, starting:3
fitted Gauss locations:3.2
y
x
0
1
2
3
4
5
6
7
8
0
5
10
15
20
25

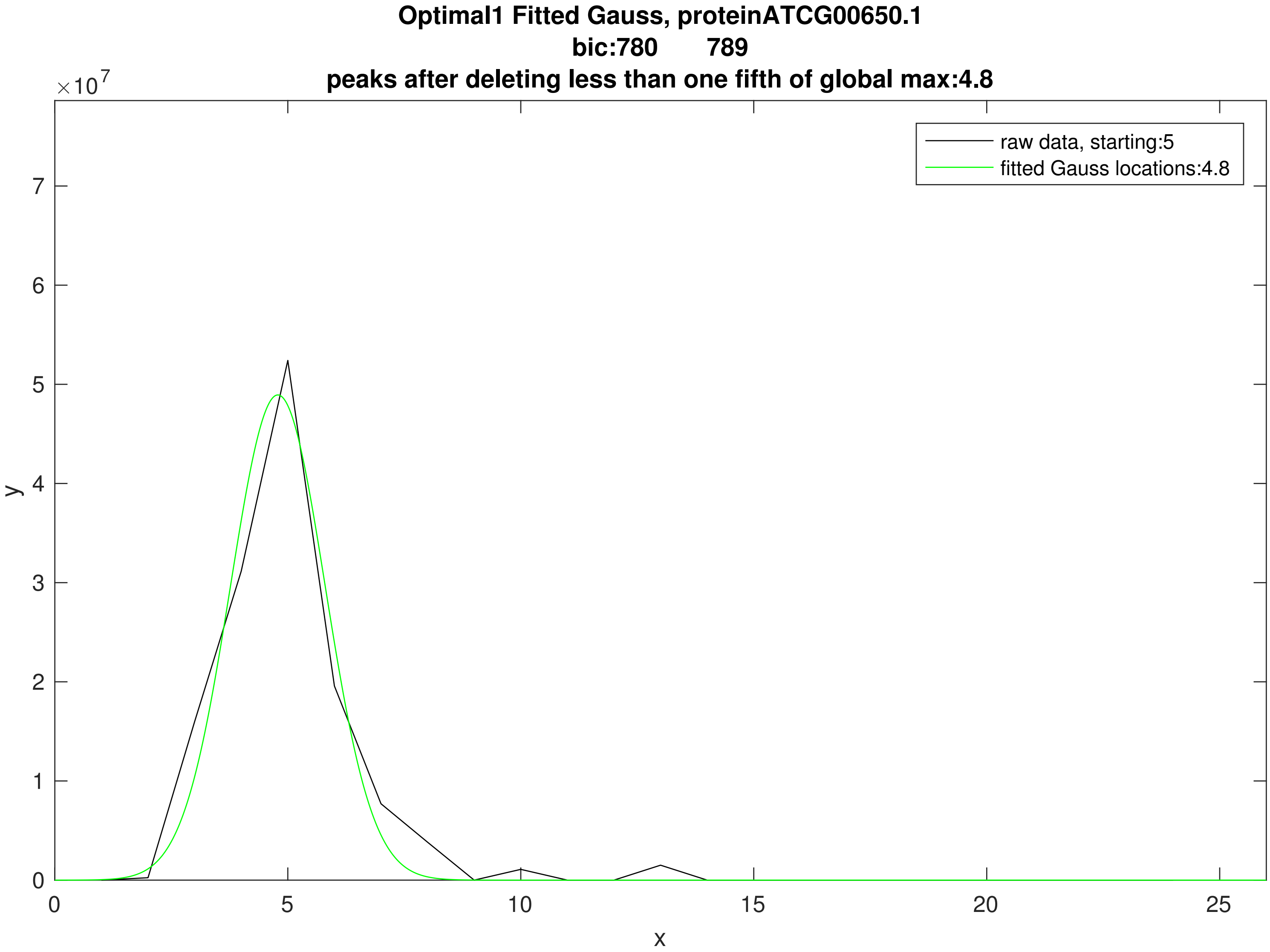

Optimal1 Fitted Gauss, proteinATCG00650.1
bic:780 789
peaks after deleting less than one fifth of global max:4.8
×10^7
raw data, starting:5
fitted Gauss locations:4.8
y
x
0
1
2
3
4
5
6
7
0
5
10
15
20
25

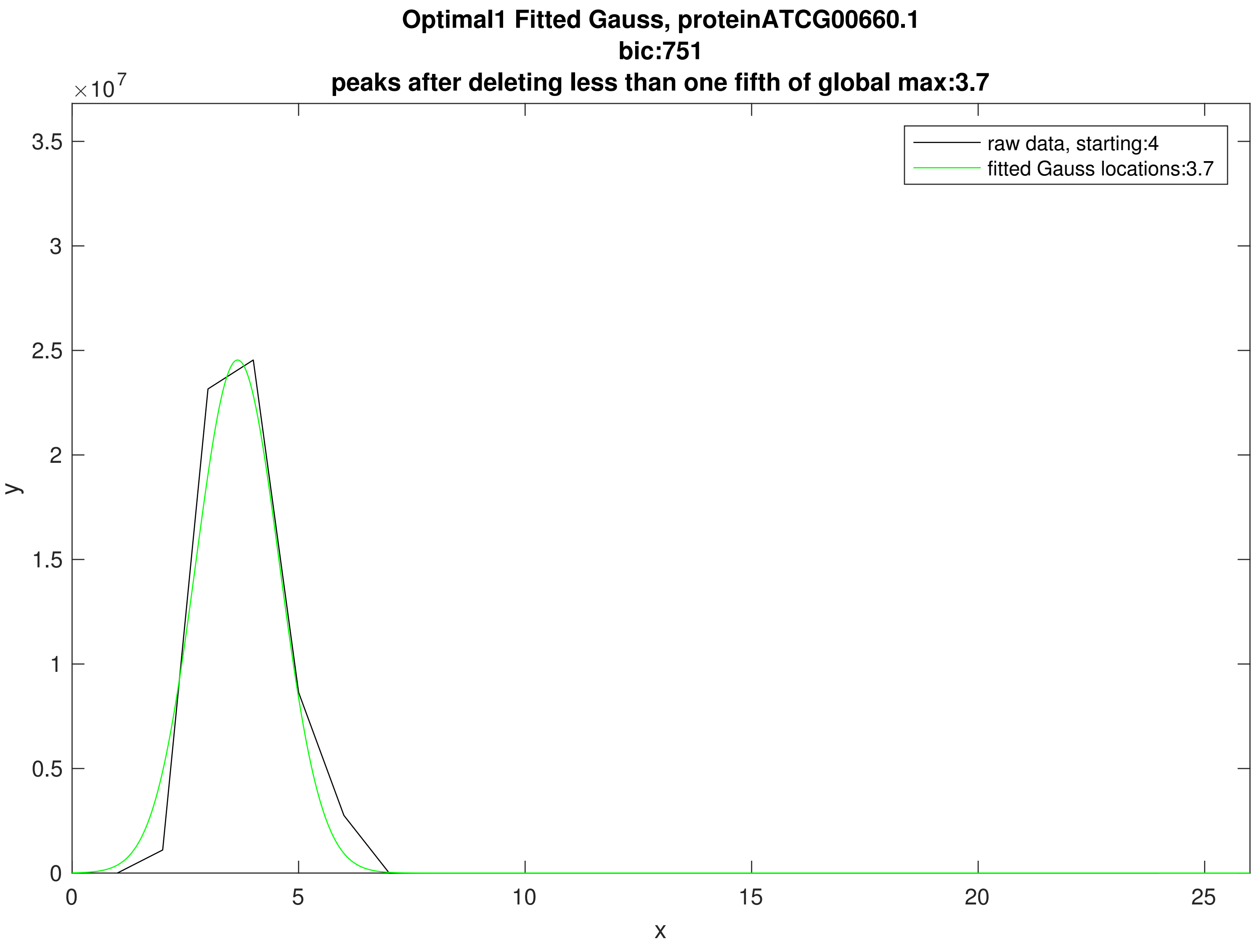

Optimal1 Fitted Gauss, proteinATCG00660.1
bic:751
peaks after deleting less than one fifth of global max:3.7
×10^7
raw data, starting:4
fitted Gauss locations:3.7
3.5
3
2.5
2
1.5
1
0.5
0
y
0
5
10
15
20
25
x

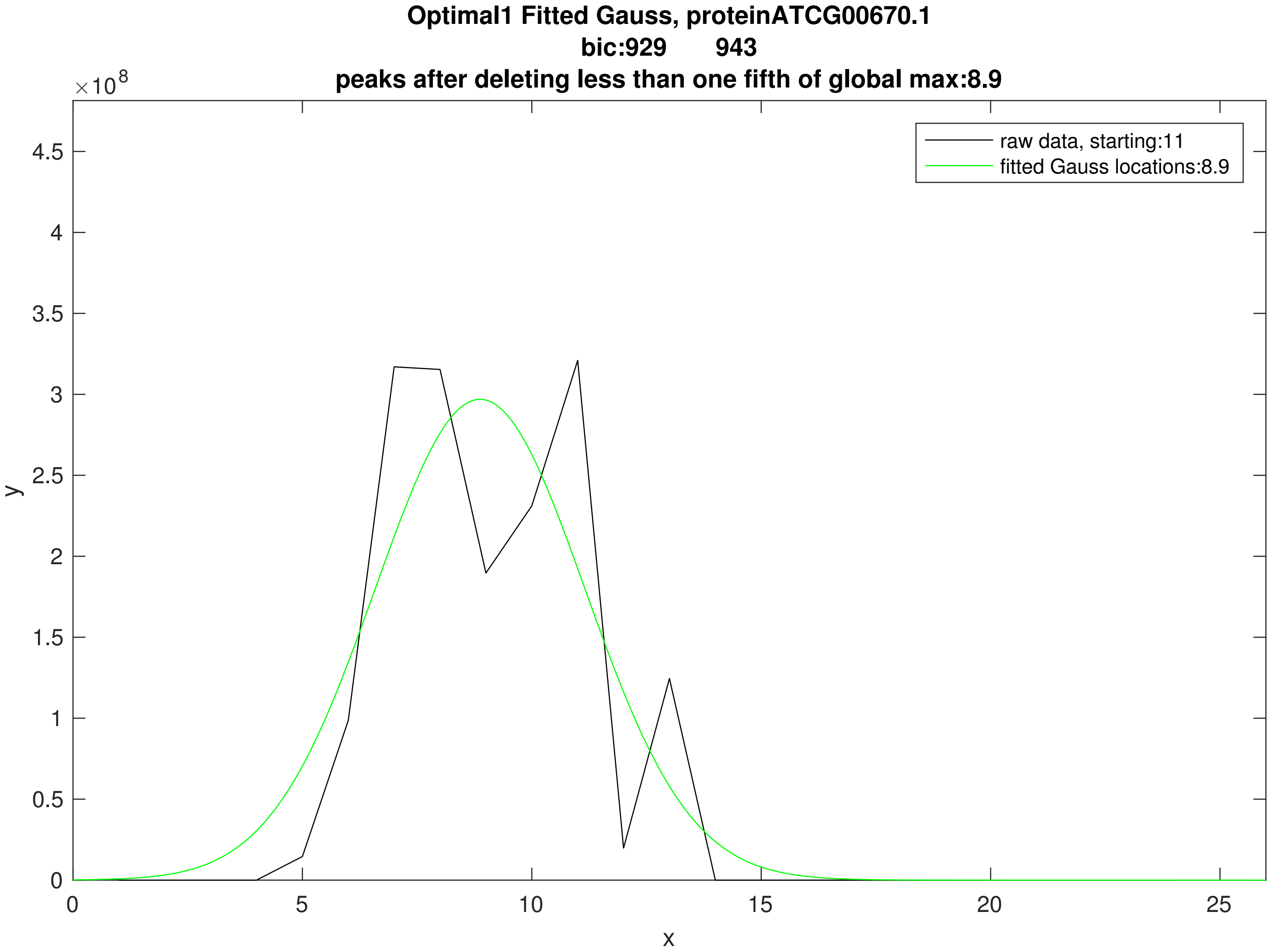
Optimal1 Fitted Gauss, proteinATCG00670.1
bic:929 943
peaks after deleting less than one fifth of global max:8.9
×10^8
raw data, starting:11
fitted Gauss locations:8.9
y
x
0
0.5
1
1.5
2
2.5
3
3.5
4
4.5
0
5
10
15
20
25

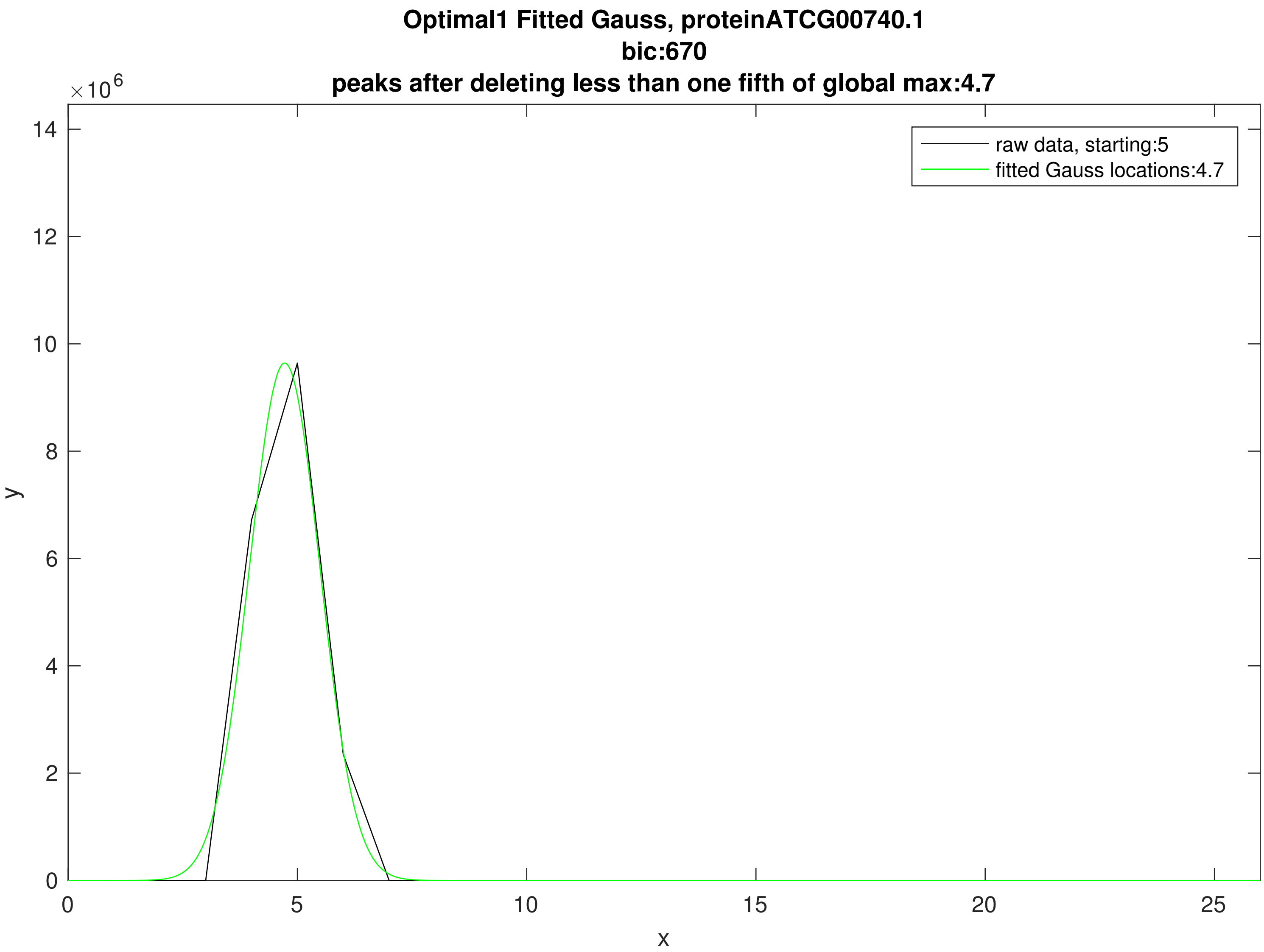

Optimal1 Fitted Gauss, proteinATCG00740.1
bic:670
peaks after deleting less than one fifth of global max:4.7
×10^6
raw data, starting:5
fitted Gauss locations:4.7
y
x

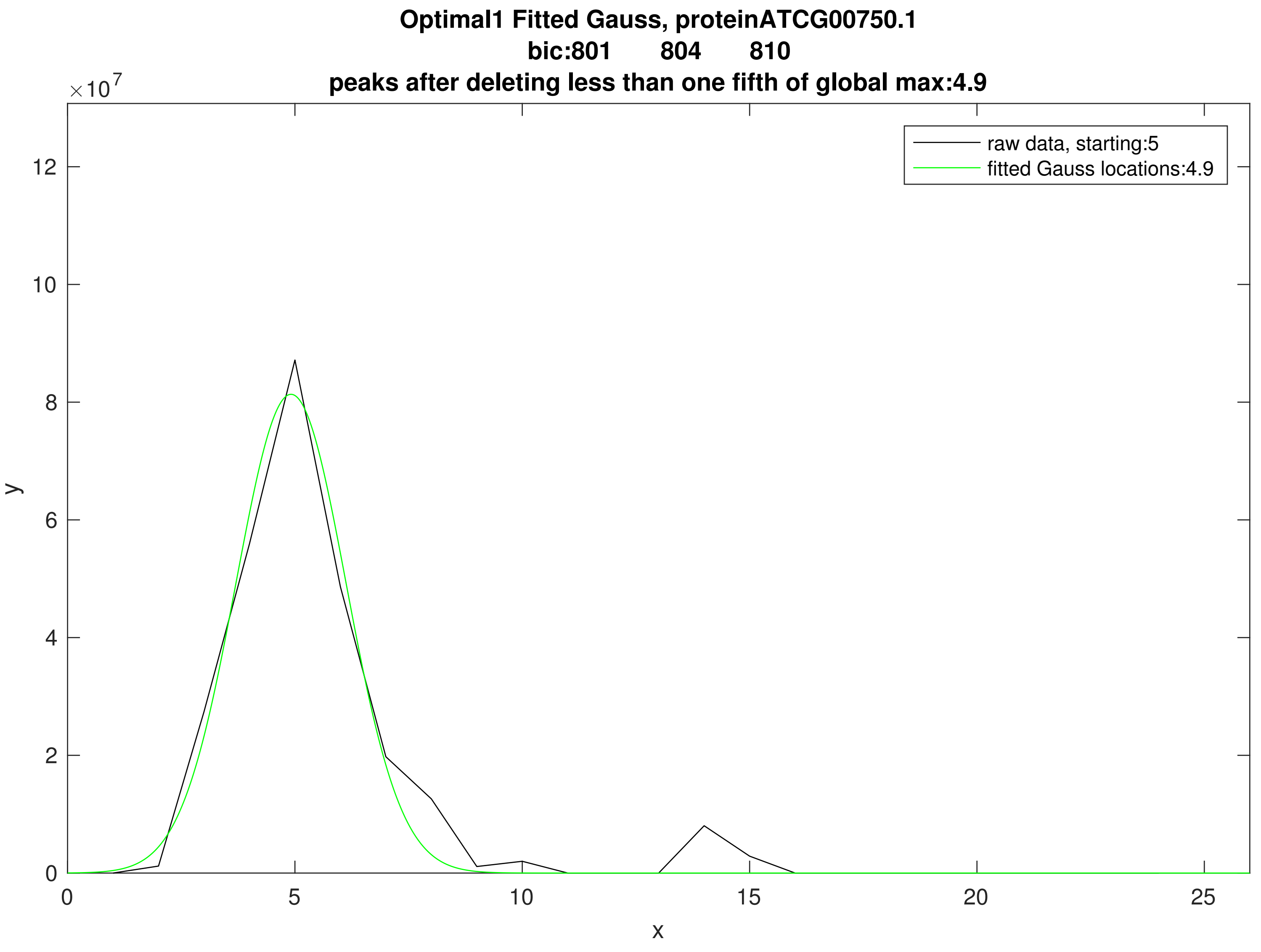

Optimal1 Fitted Gauss, proteinATCG00750.1
bic:801 804 810
peaks after deleting less than one fifth of global max:4.9
×10^7
raw data, starting:5
fitted Gauss locations:4.9
y
x
0
2
4
6
8
10
12
0
5
10
15
20
25

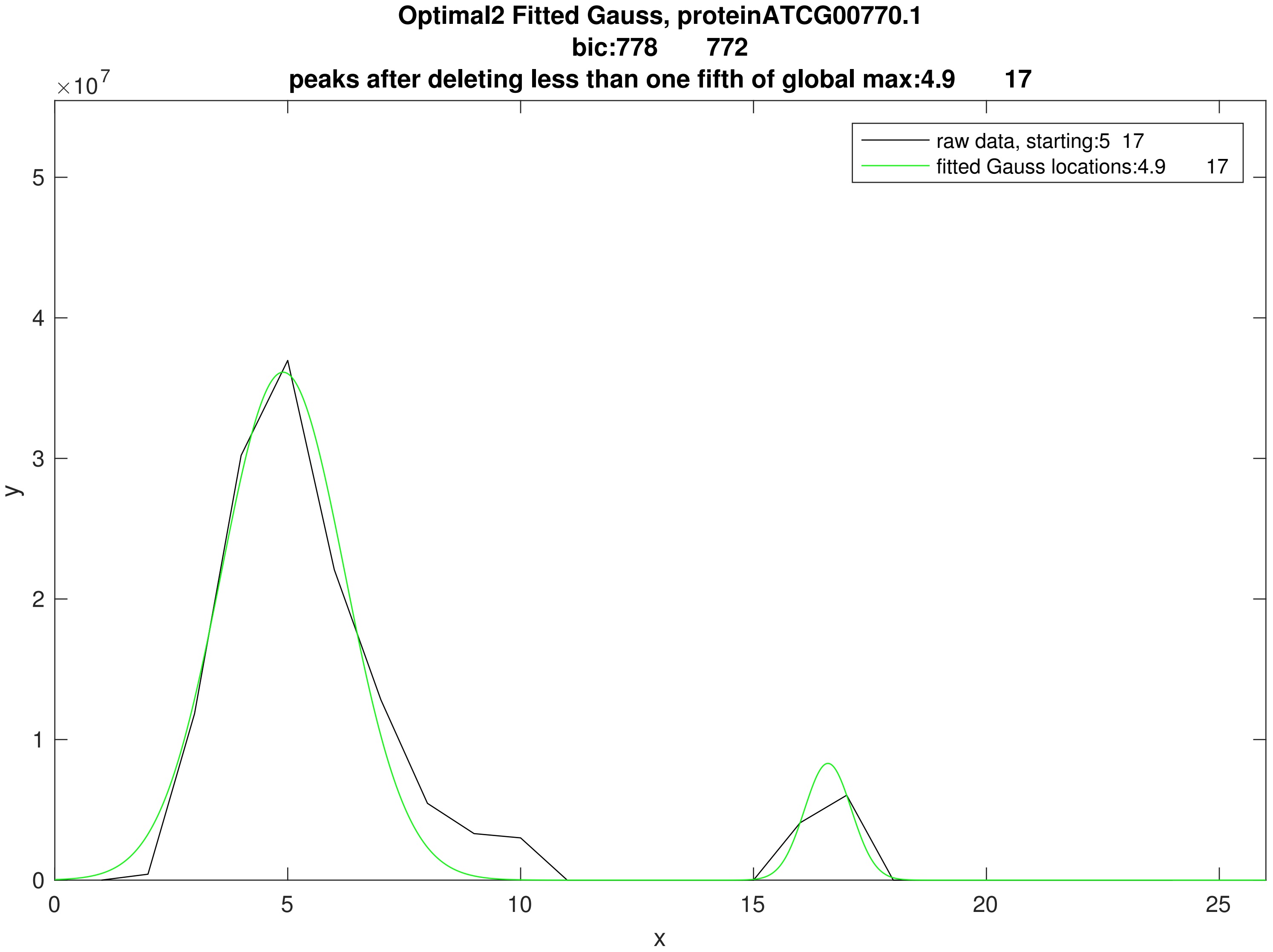
Optimal2 Fitted Gauss, proteinATCG00770.1
bic:778 772
peaks after deleting less than one fifth of global max:4.9 17
×10^7
raw data, starting:5 17
fitted Gauss locations:4.9 17
y
x
0
1
2
3
4
5
0
5
10
15
20
25

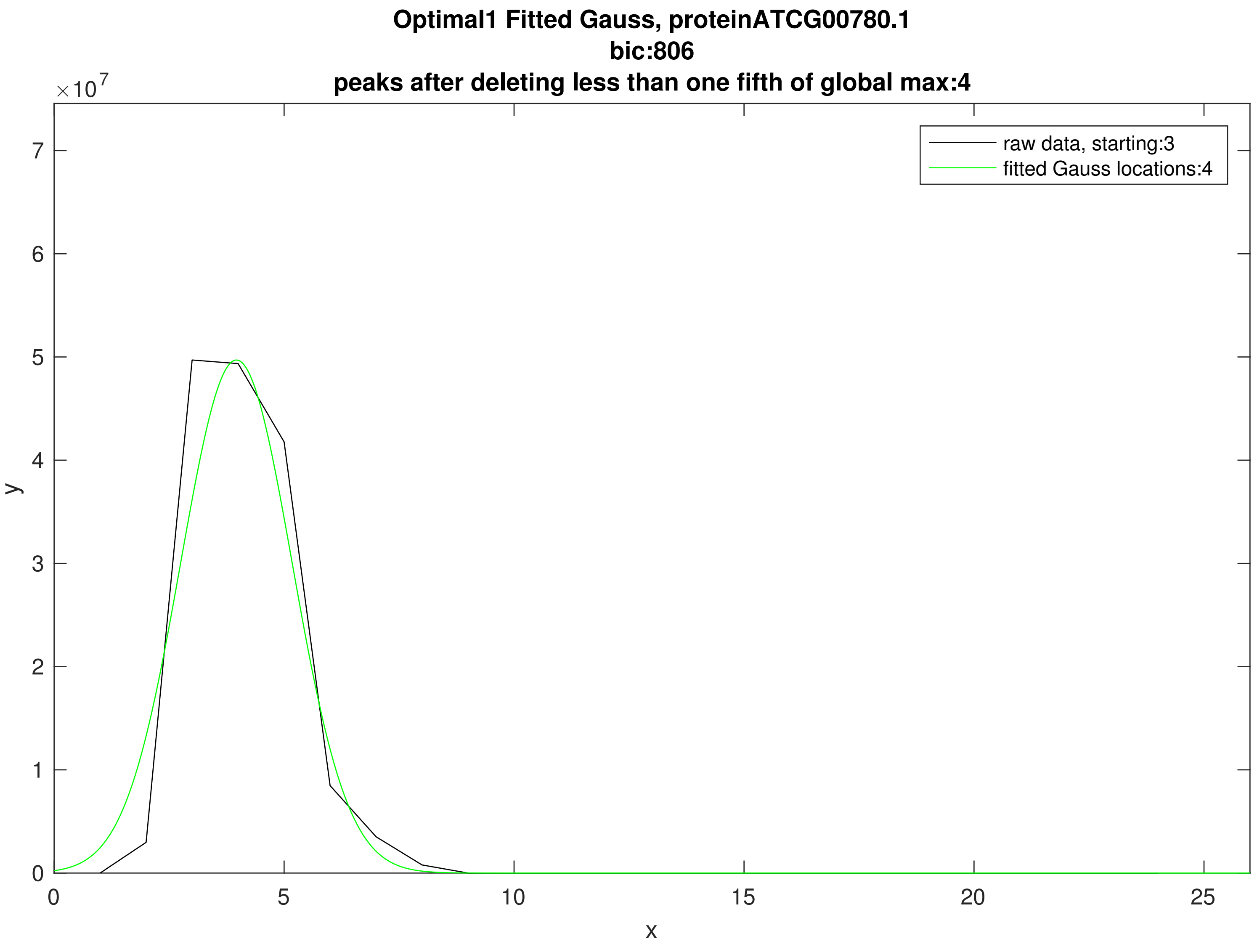
Optimal1 Fitted Gauss, proteinATCG00780.1
bic:806
peaks after deleting less than one fifth of global max:4
×10^7
raw data, starting:3
fitted Gauss locations:4
y
x
0
1
2
3
4
5
6
7
0
5
10
15
20
25

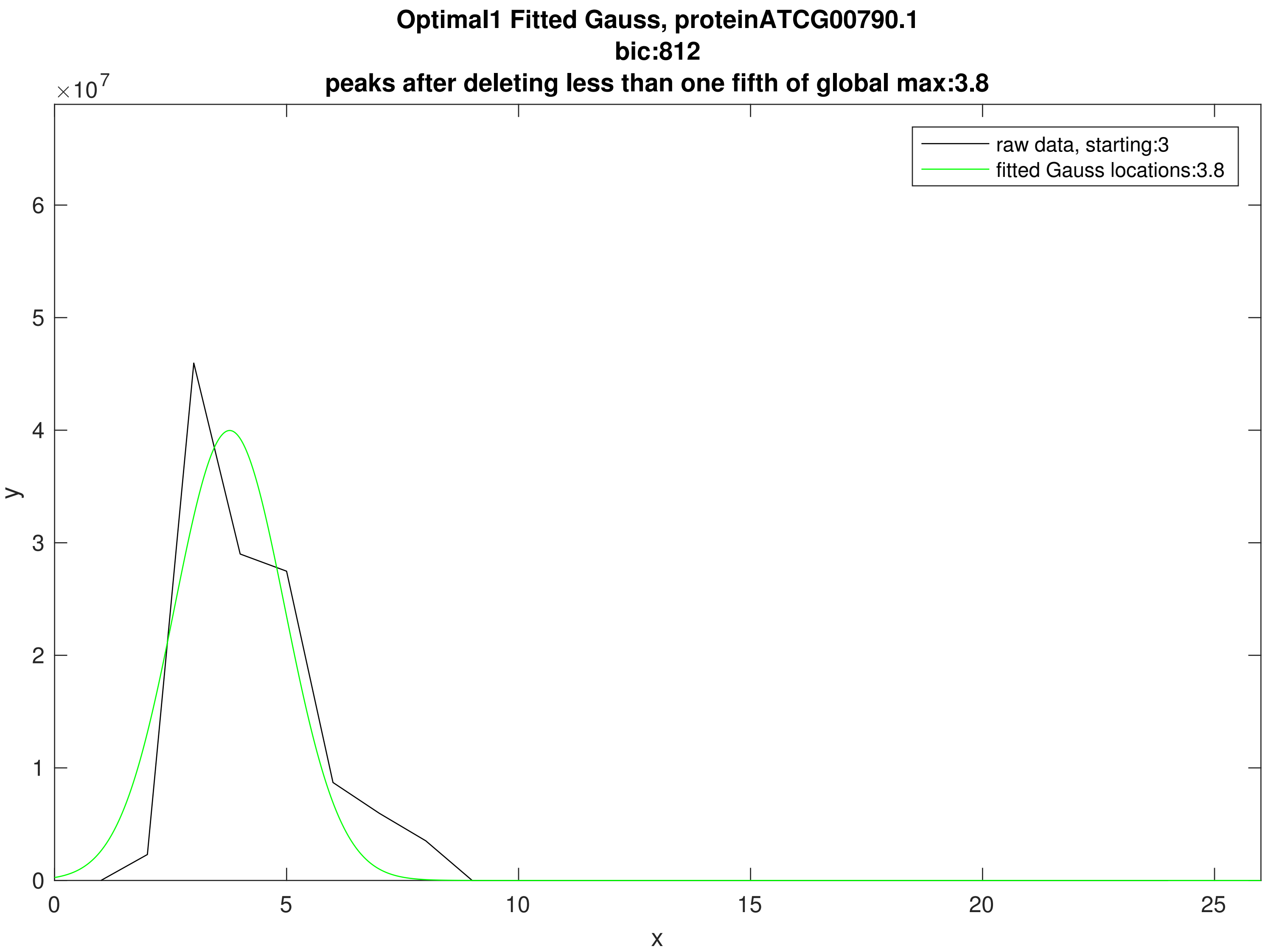
Optimal1 Fitted Gauss, proteinATCG00790.1
bic:812
peaks after deleting less than one fifth of global max:3.8
×10^7
raw data, starting:3
fitted Gauss locations:3.8
0
1
2
3
4
5
6
y
0
5
10
15
20
25
x

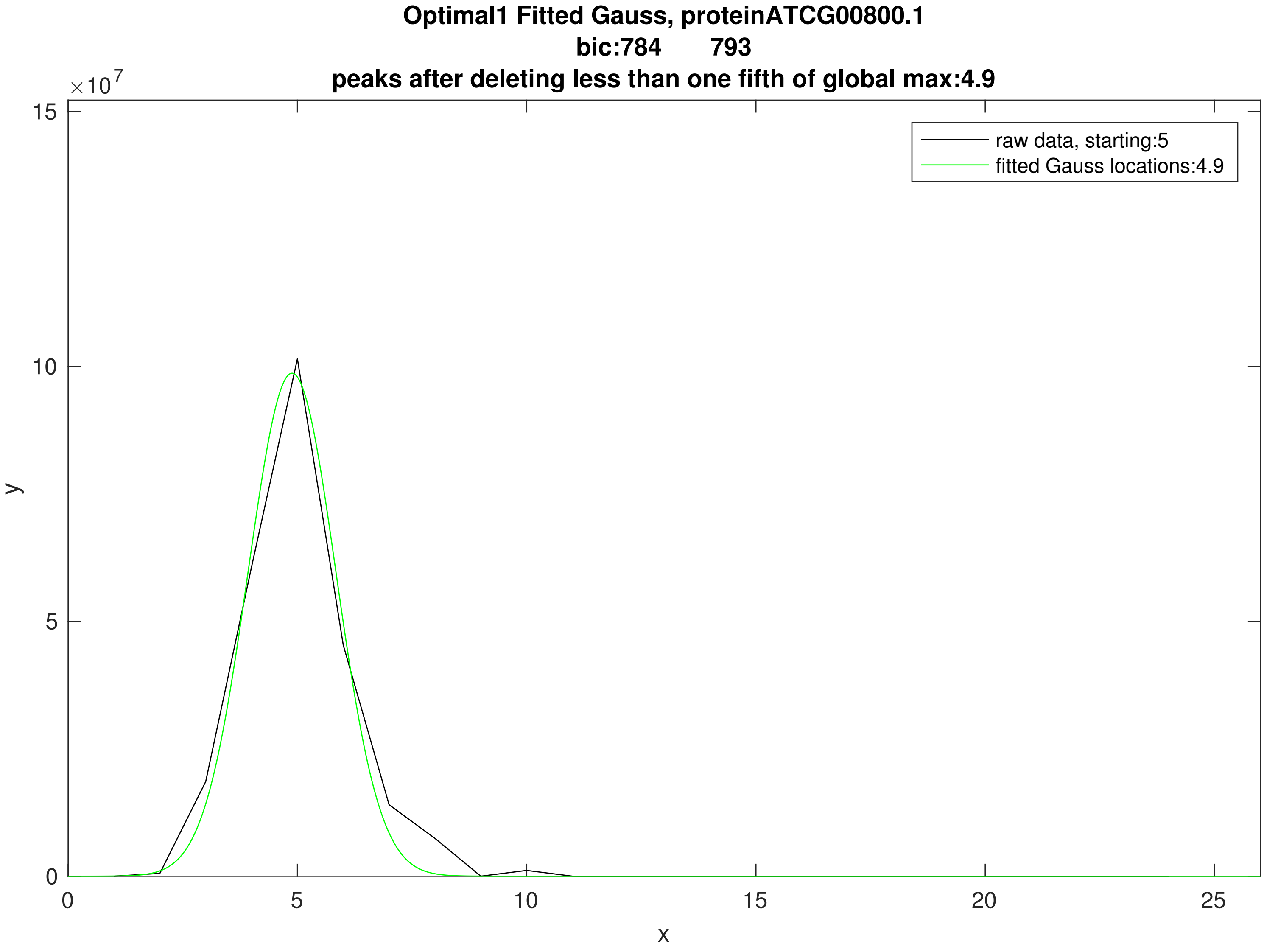
Optimal1 Fitted Gauss, proteinATCG00800.1
bic:784 793
peaks after deleting less than one fifth of global max:4.9
×10^7
raw data, starting:5
fitted Gauss locations:4.9
y
x
0
5
10
15
20
25

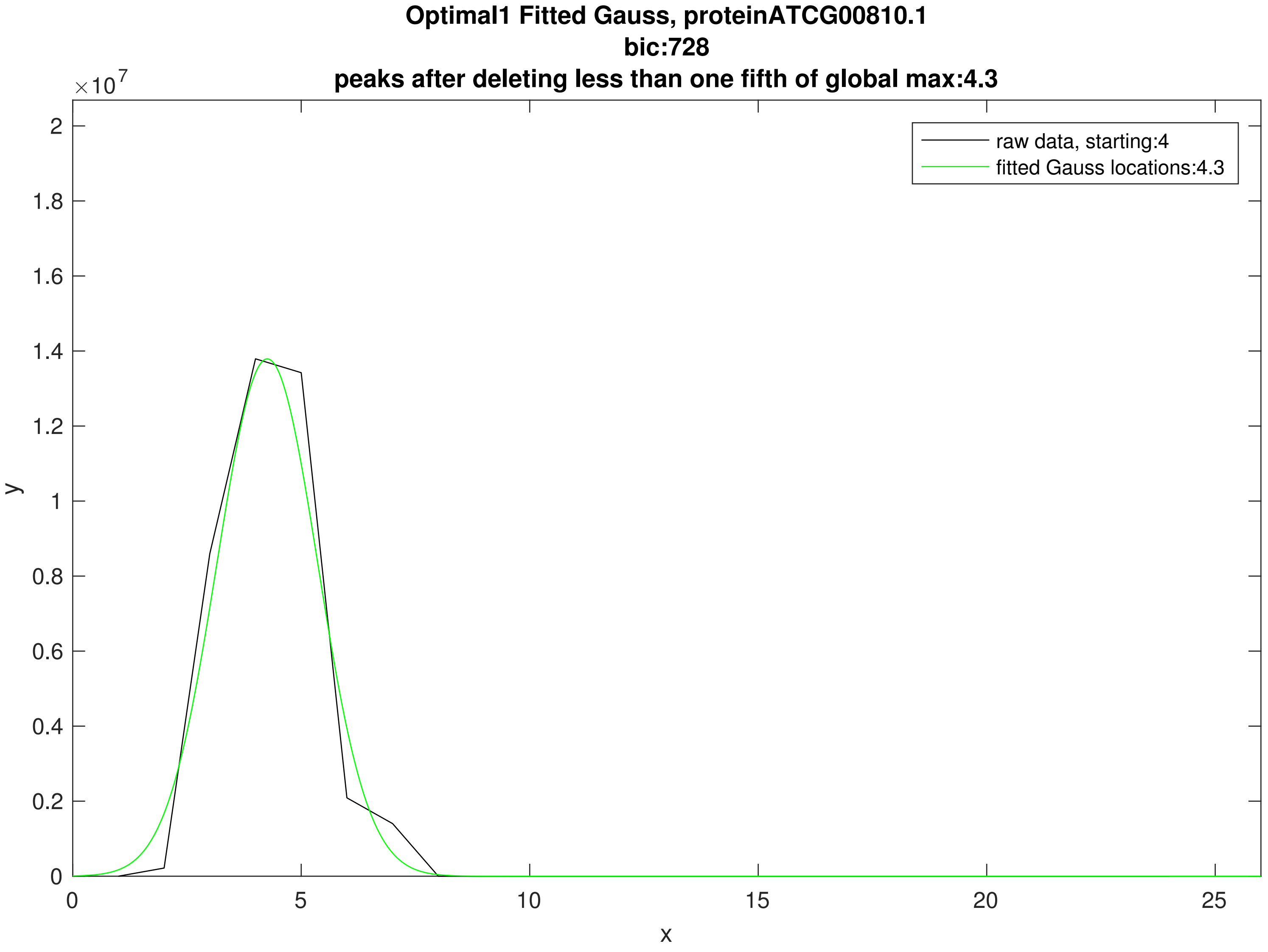

Optimal1 Fitted Gauss, proteinATCG00810.1
bic:728
peaks after deleting less than one fifth of global max:4.3
×10^7
raw data, starting:4
fitted Gauss locations:4.3
y
x

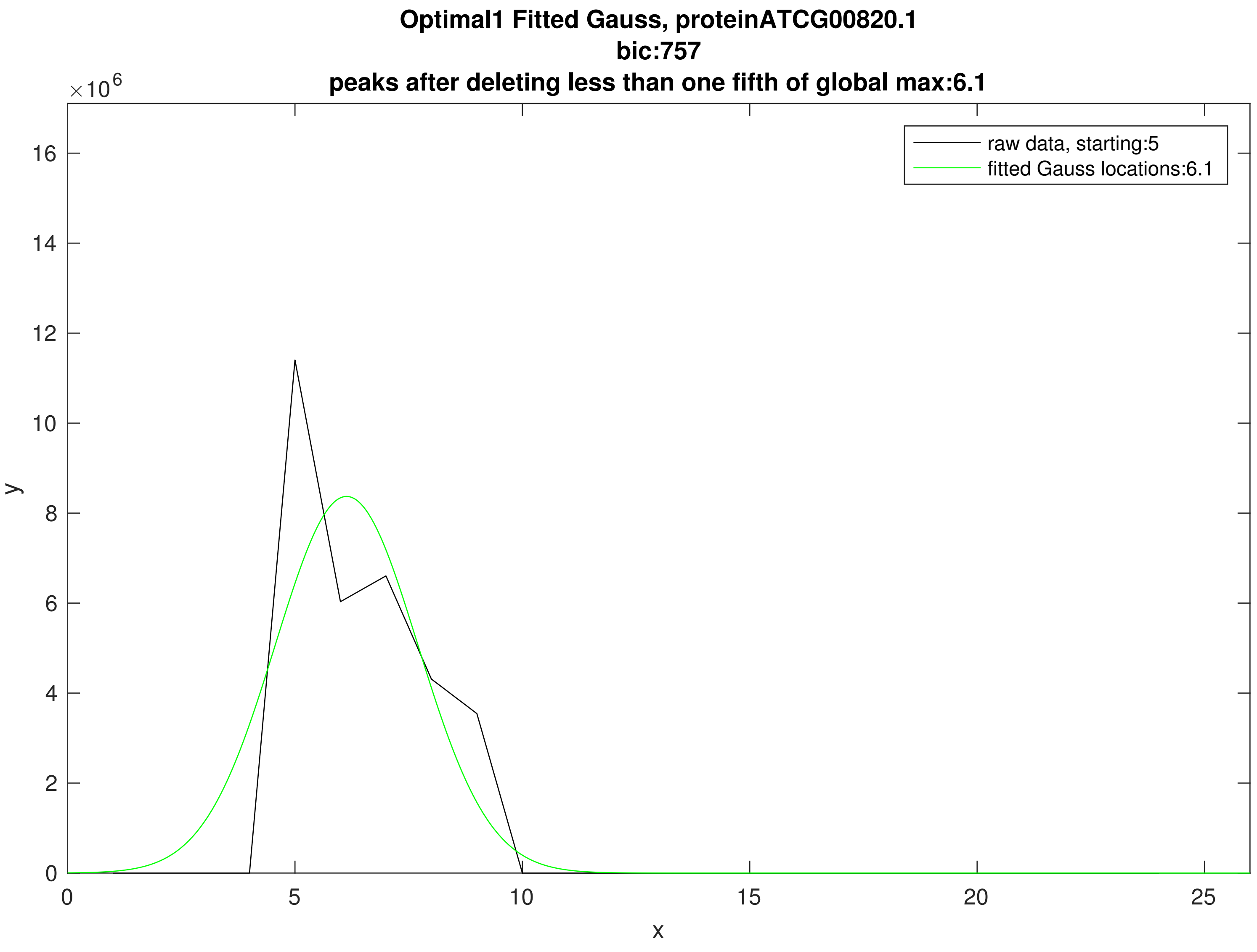
Optimal1 Fitted Gauss, proteinATCG00820.1
bic:757
peaks after deleting less than one fifth of global max:6.1
×10^6
raw data, starting:5
fitted Gauss locations:6.1
y
x
0
2
4
6
8
10
12
14
16
0
5
10
15
20
25

**Optimal1 Fitted Gauss, proteinATCG01060.1**
**bic:753**
**peaks after deleting less than one fifth of global max:21**

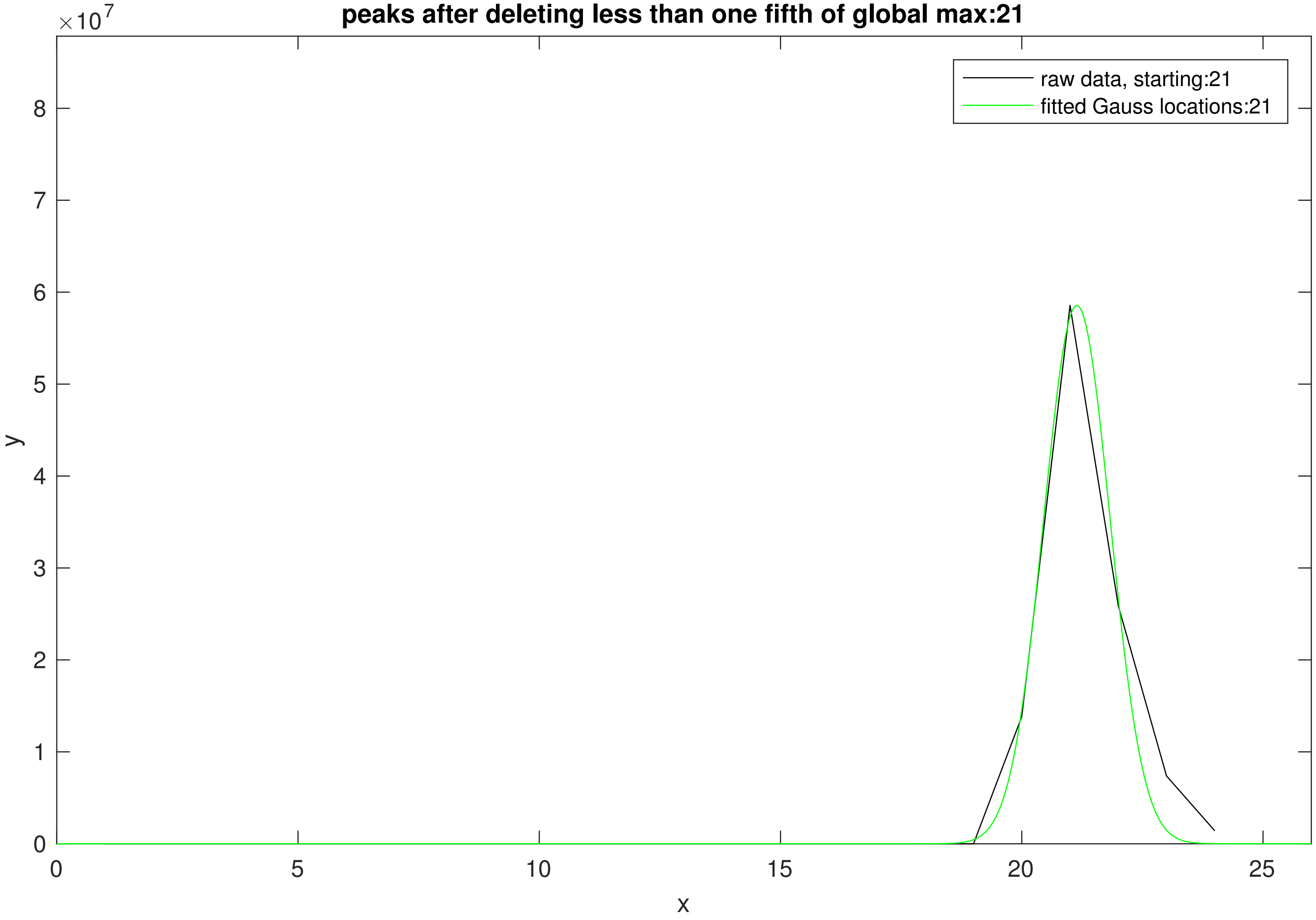

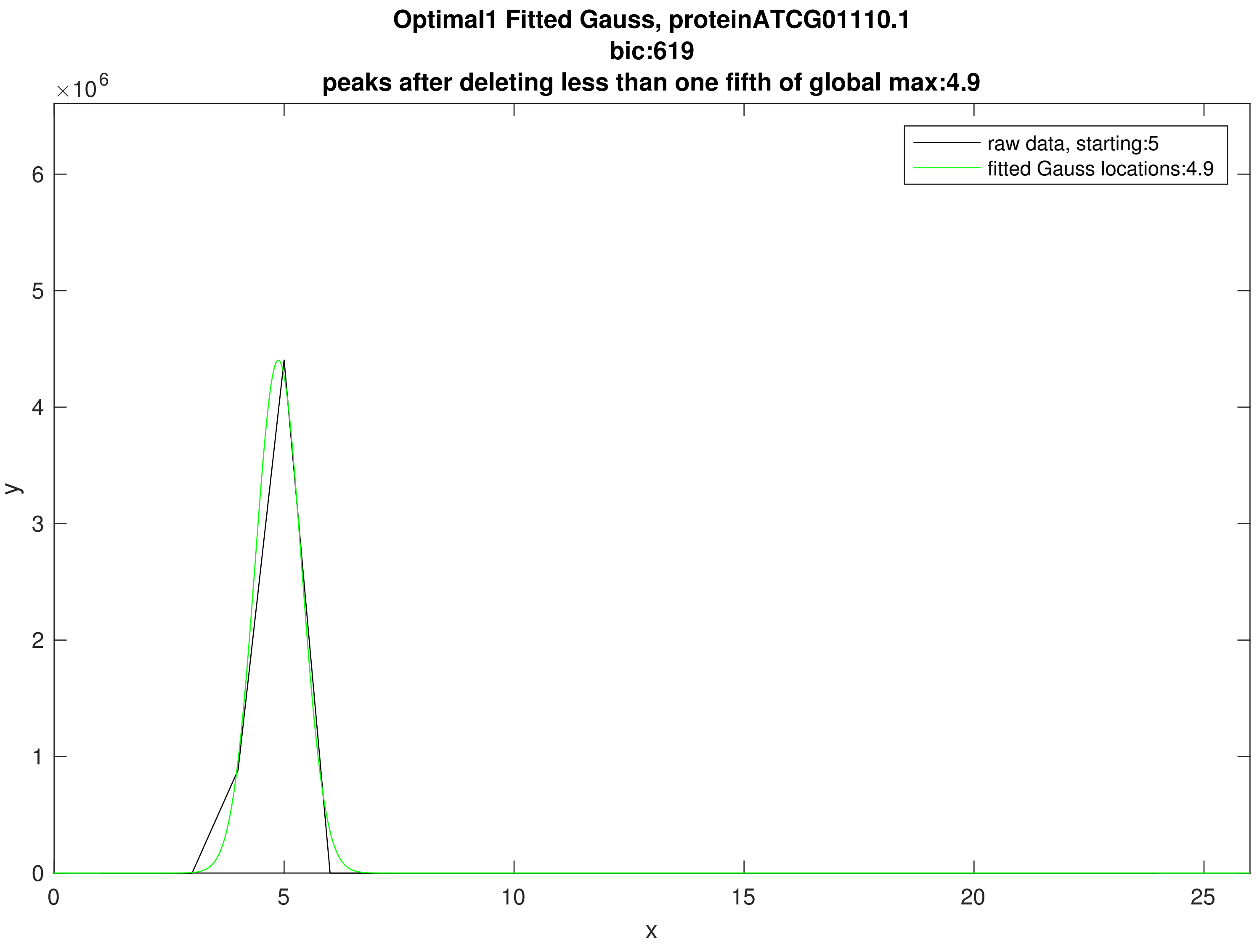
Optimal1 Fitted Gauss, proteinATCG01110.1
bic:619
peaks after deleting less than one fifth of global max:4.9
×10^6
raw data, starting:5
fitted Gauss locations:4.9
0
1
2
3
4
5
6
y
0
5
10
15
20
25
x

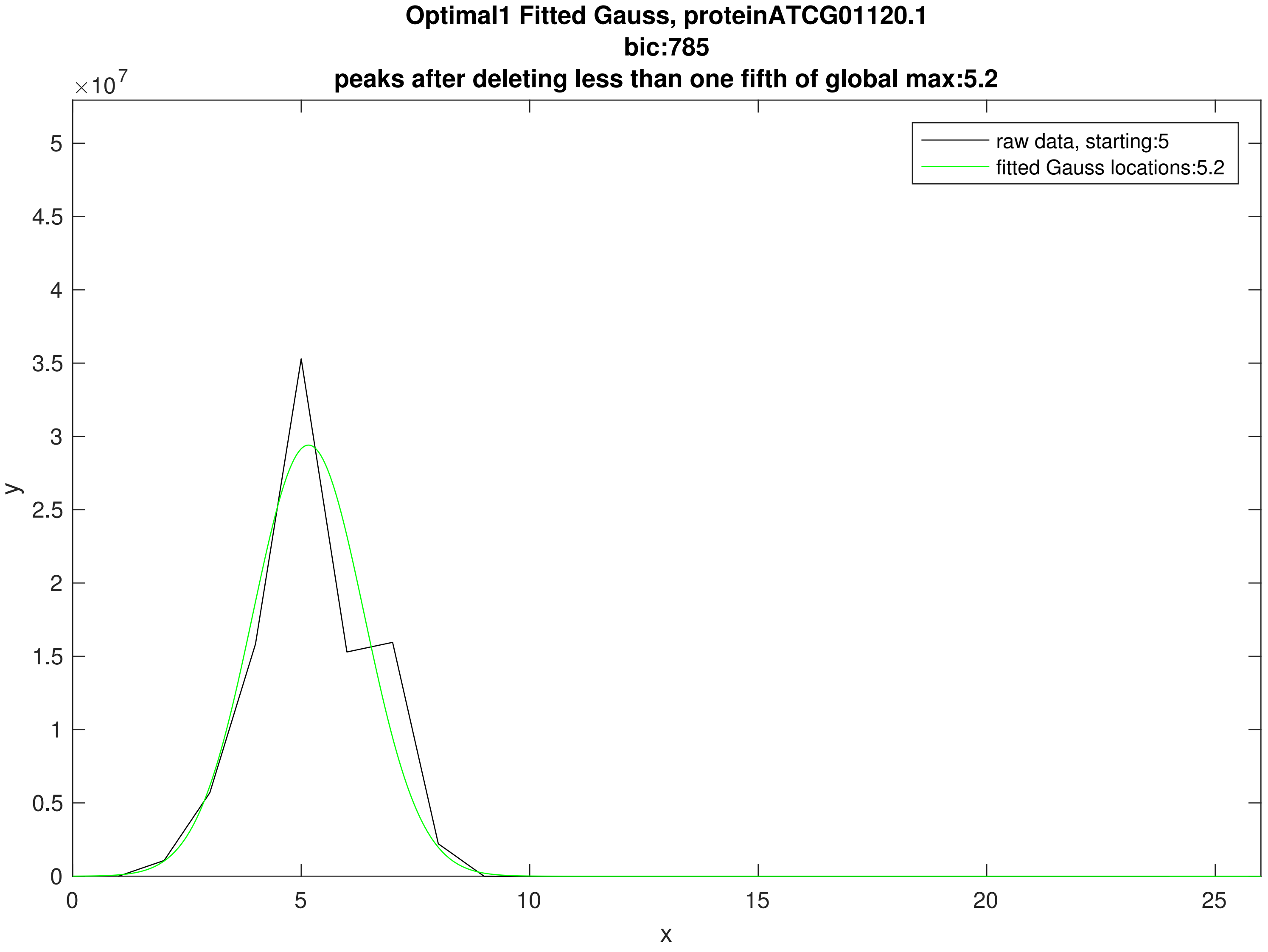
Optimal1 Fitted Gauss, proteinATCG01120.1
bic:785
peaks after deleting less than one fifth of global max:5.2
×10^7
raw data, starting:5
fitted Gauss locations:5.2
y
x
0
0.5
1
1.5
2
2.5
3
3.5
4
4.5
5
0
5
10
15
20
25

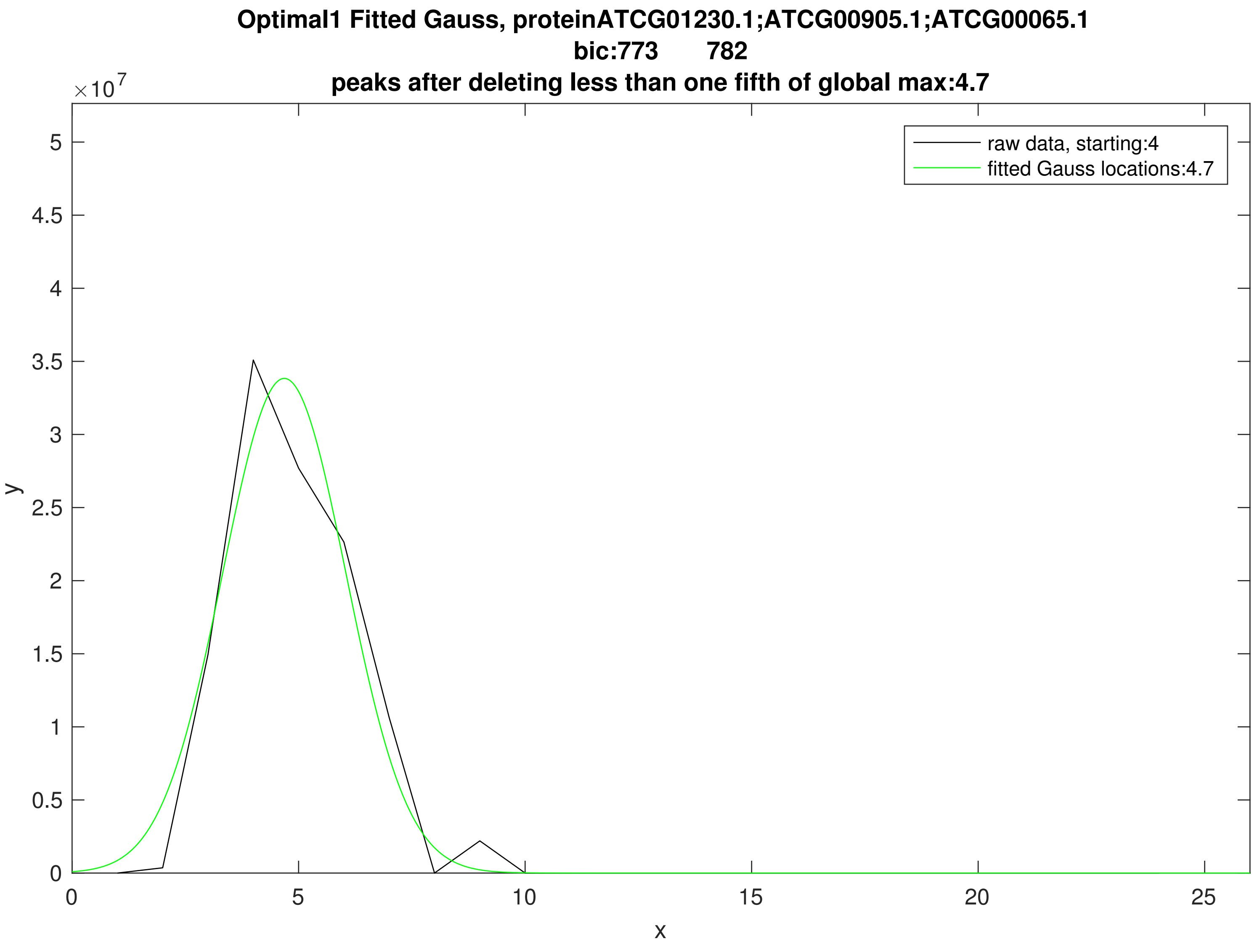
Optimal1 Fitted Gauss, proteinATCG01230.1;ATCG00905.1;ATCG00065.1
bic:773 782
peaks after deleting less than one fifth of global max:4.7
×10^7
raw data, starting:4
fitted Gauss locations:4.7
y
x
0
0.5
1
1.5
2
2.5
3
3.5
4
4.5
5
0
5
10
15
20
25

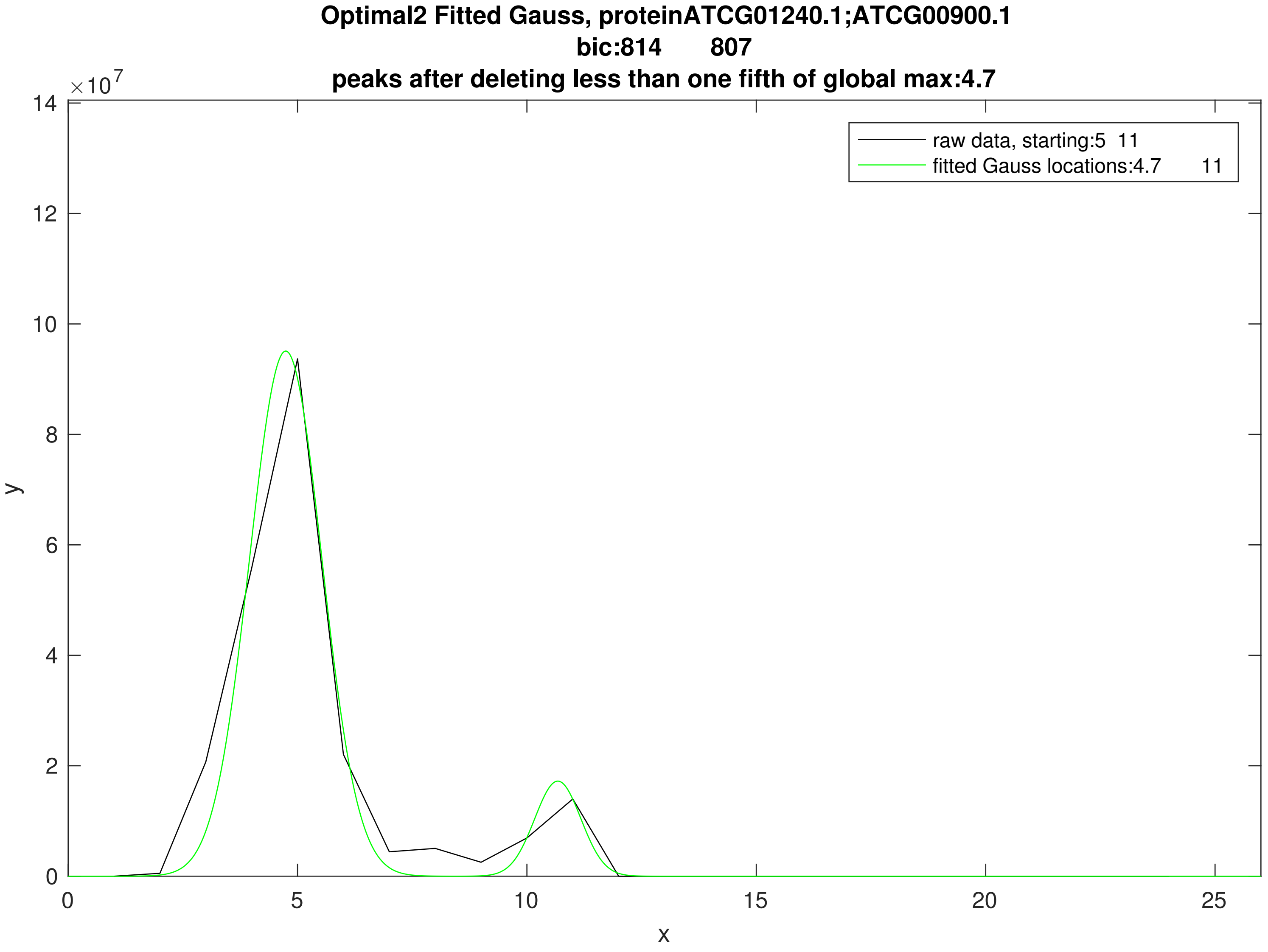

Optimal2 Fitted Gauss, proteinATCG01240.1;ATCG00900.1
bic:814 807
peaks after deleting less than one fifth of global max:4.7
×10^7
14
12
10
8
6
4
2
0
y
0
5
10
15
20
25
x
raw data, starting:5 11
fitted Gauss locations:4.7 11

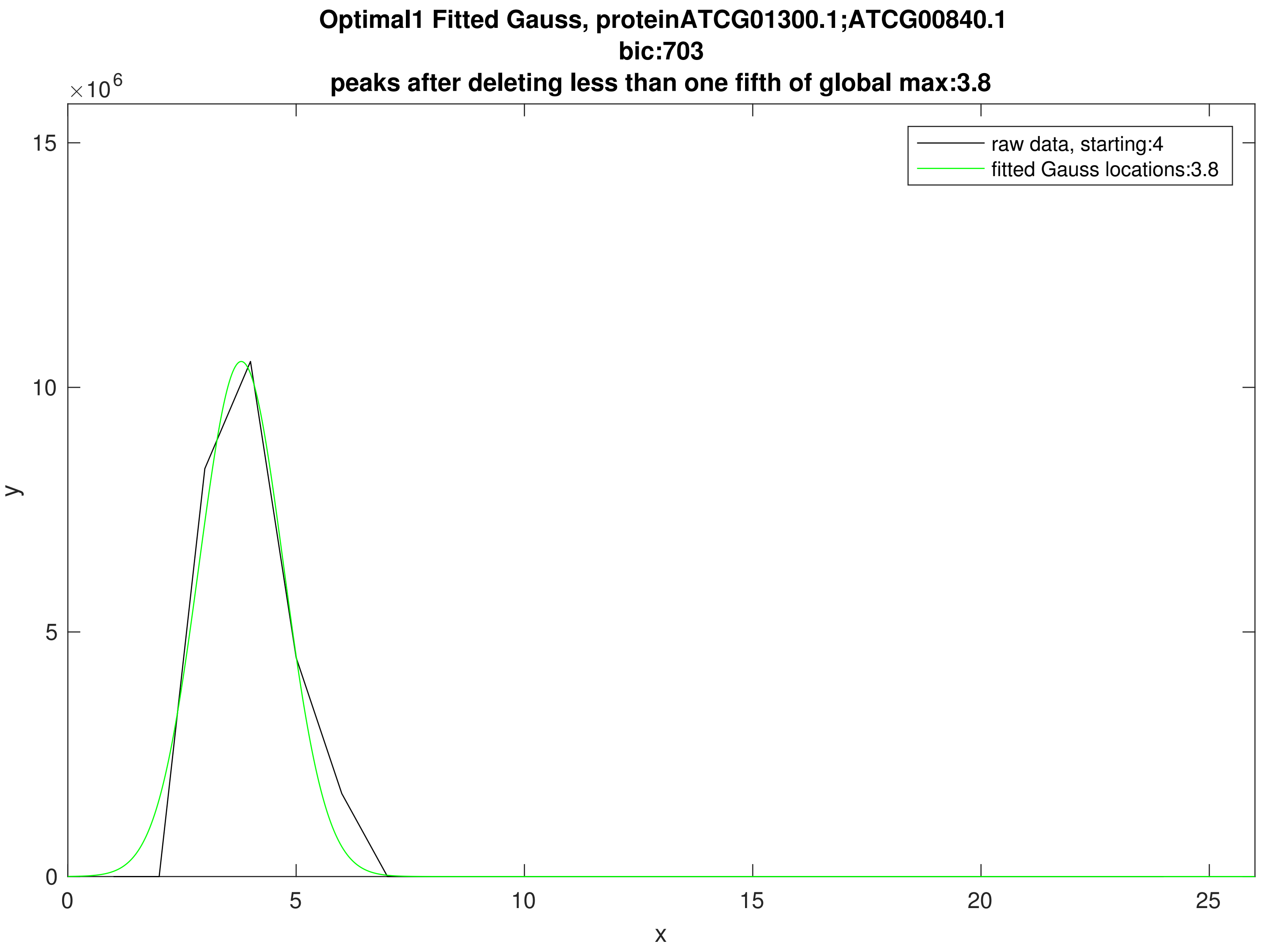

Optimal1 Fitted Gauss, proteinATCG01300.1;ATCG00840.1
bic:703
peaks after deleting less than one fifth of global max:3.8
×10^6
15
10
5
0
y
0
5
10
15
20
25
x
raw data, starting:4
fitted Gauss locations:3.8

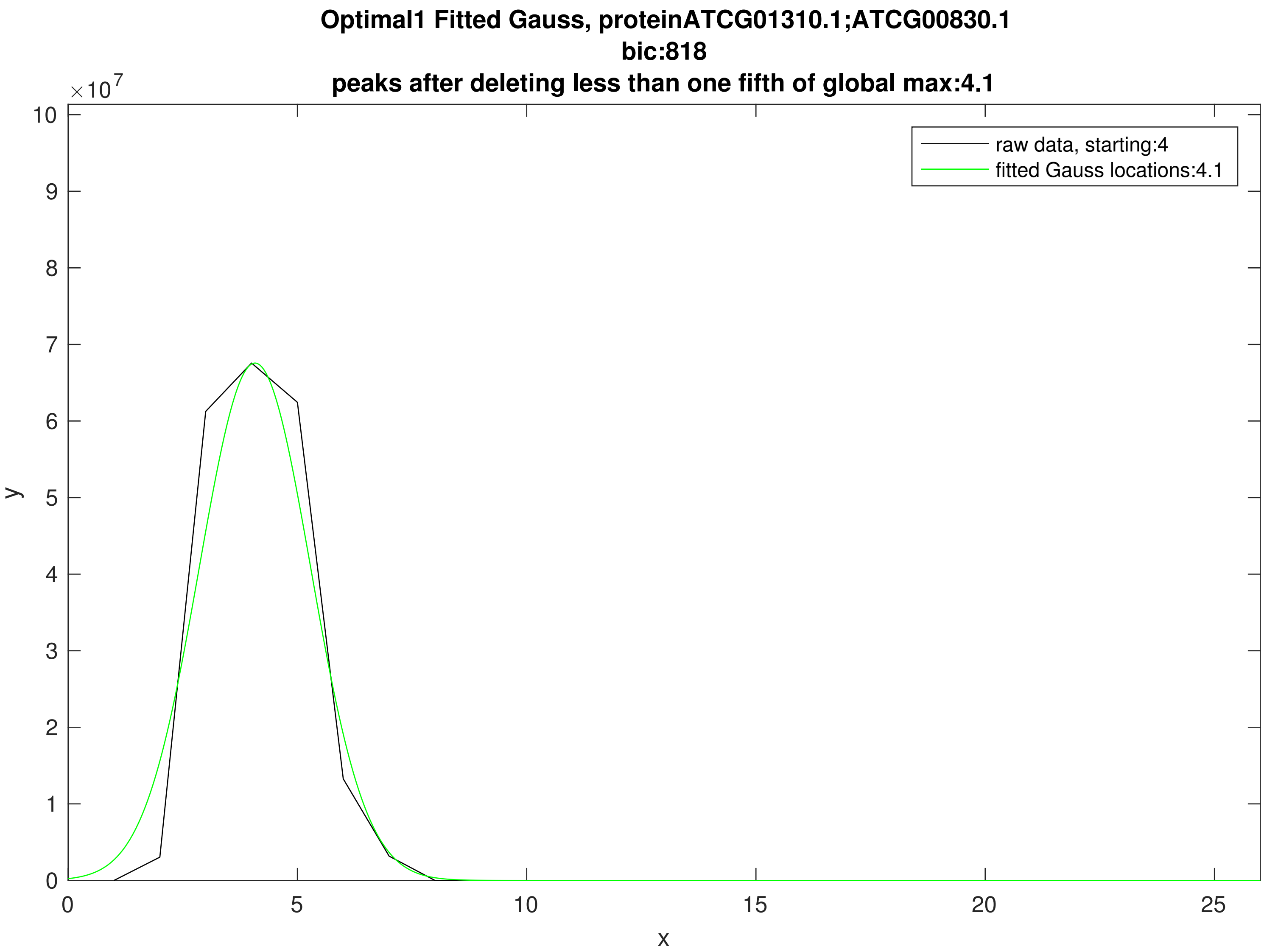
Optimal1 Fitted Gauss, proteinATCG01310.1;ATCG00830.1
bic:818
peaks after deleting less than one fifth of global max:4.1
×10^7
10
9
8
7
6
5
4
3
2
1
0
y
0
5
10
15
20
25
x
raw data, starting:4
fitted Gauss locations:4.1

**Optimal1 Fitted Gauss, proteinATMG00510.1**
**bic:701**
**peaks after deleting less than one fifth of global max:5**

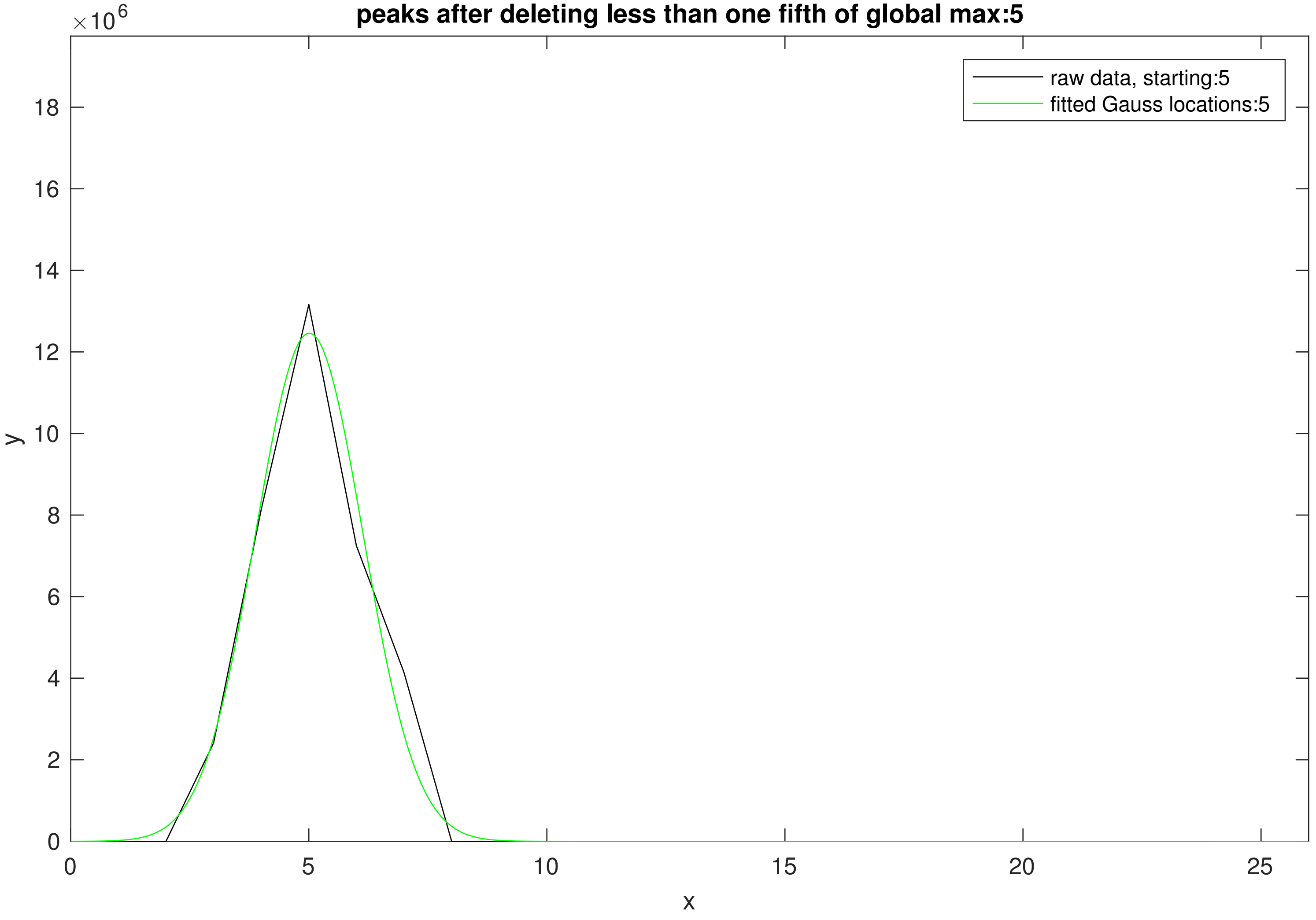

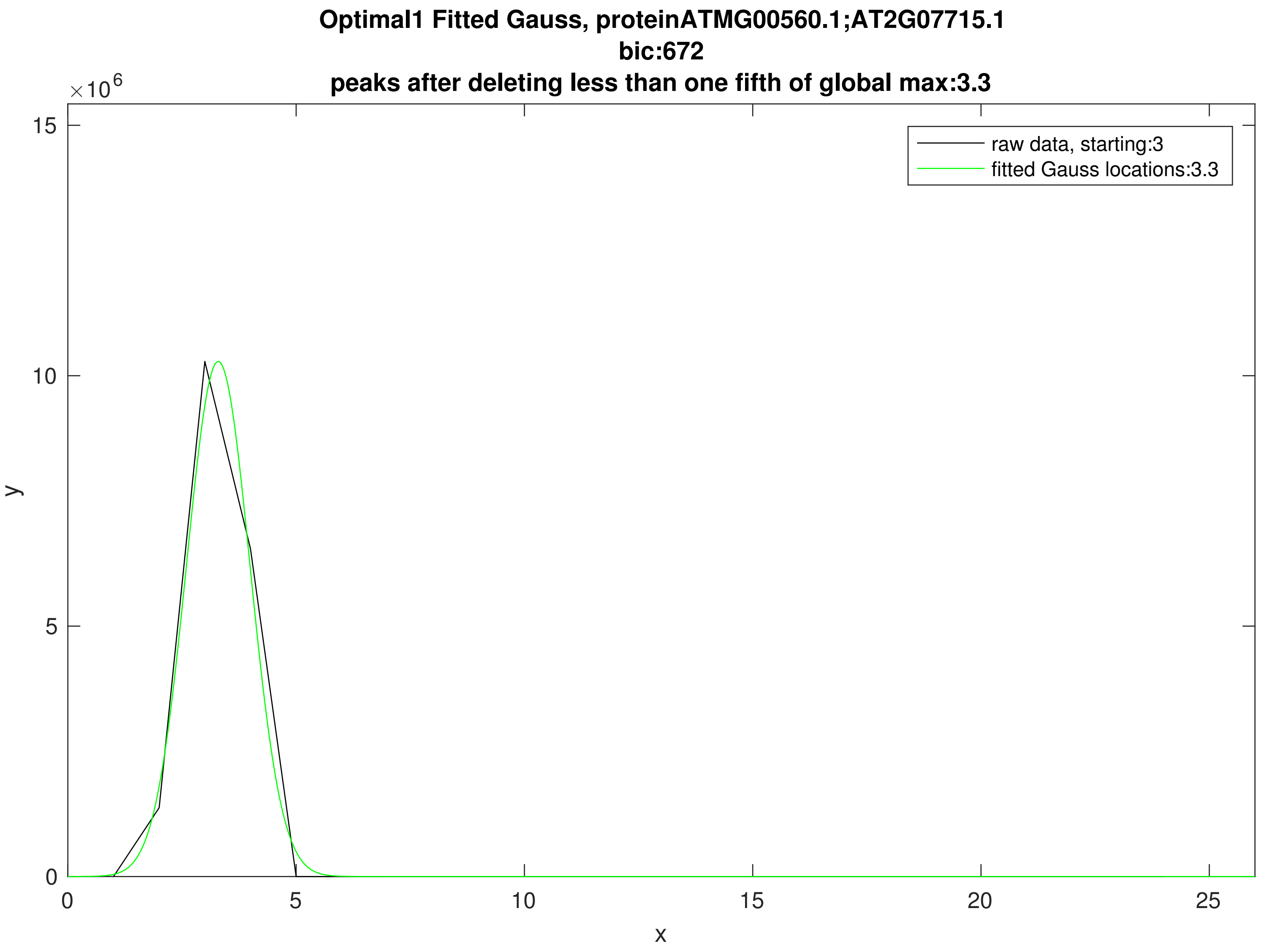

Optimal1 Fitted Gauss, proteinATMG00560.1;AT2G07715.1
bic:672
peaks after deleting less than one fifth of global max:3.3
×10^6
15
10
5
0
y
0
5
10
15
20
25
x
raw data, starting:3
fitted Gauss locations:3.3

### $SEC_{Untreated}$ replicate 2 elution profiles

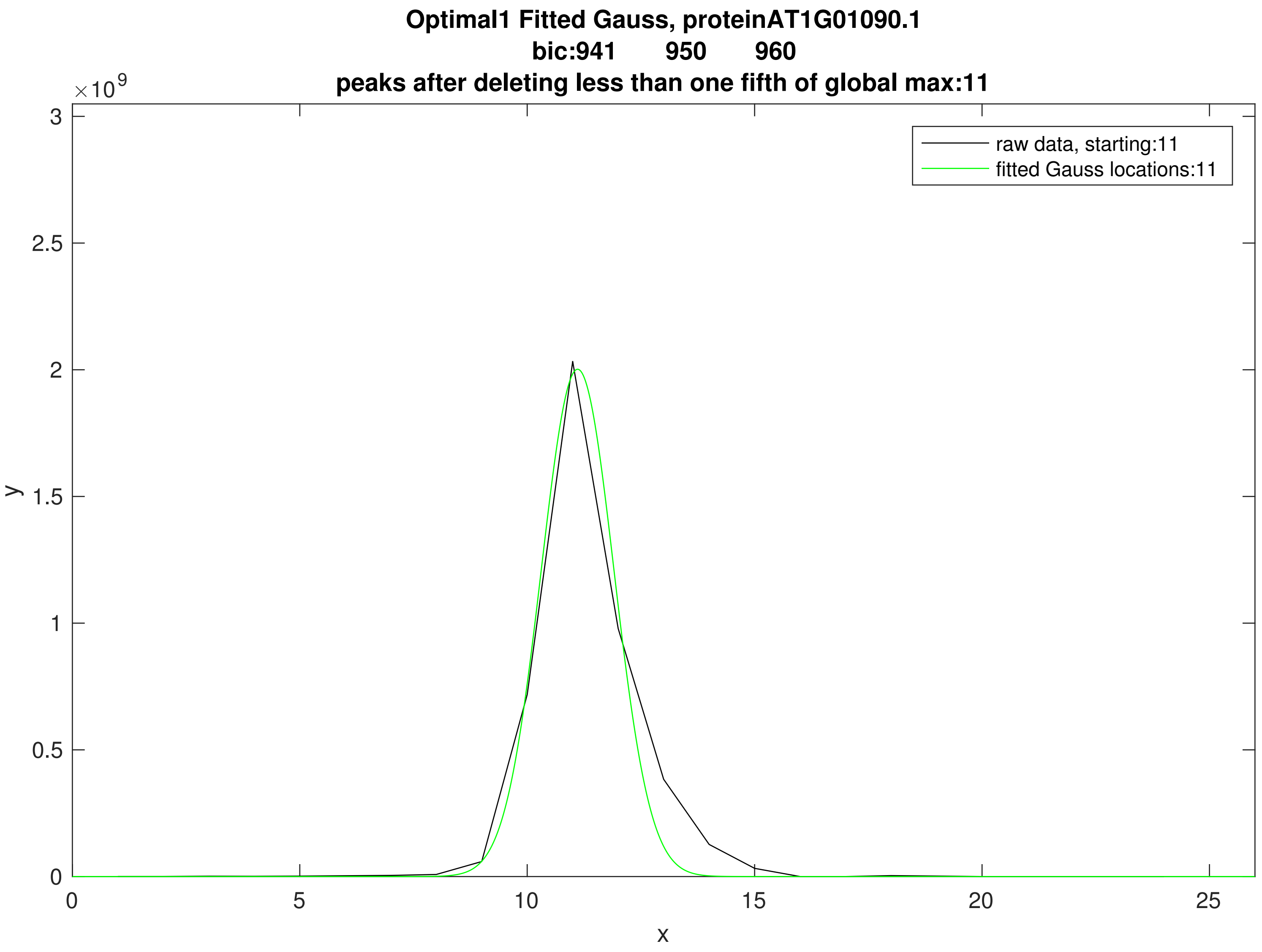

Optimal1 Fitted Gauss, proteinAT1G01090.1
bic:941 950 960
peaks after deleting less than one fifth of global max:11
×10^9
raw data, starting:11
fitted Gauss locations:11
y
x
0
0.5
1
1.5
2
2.5
3
0
5
10
15
20
25

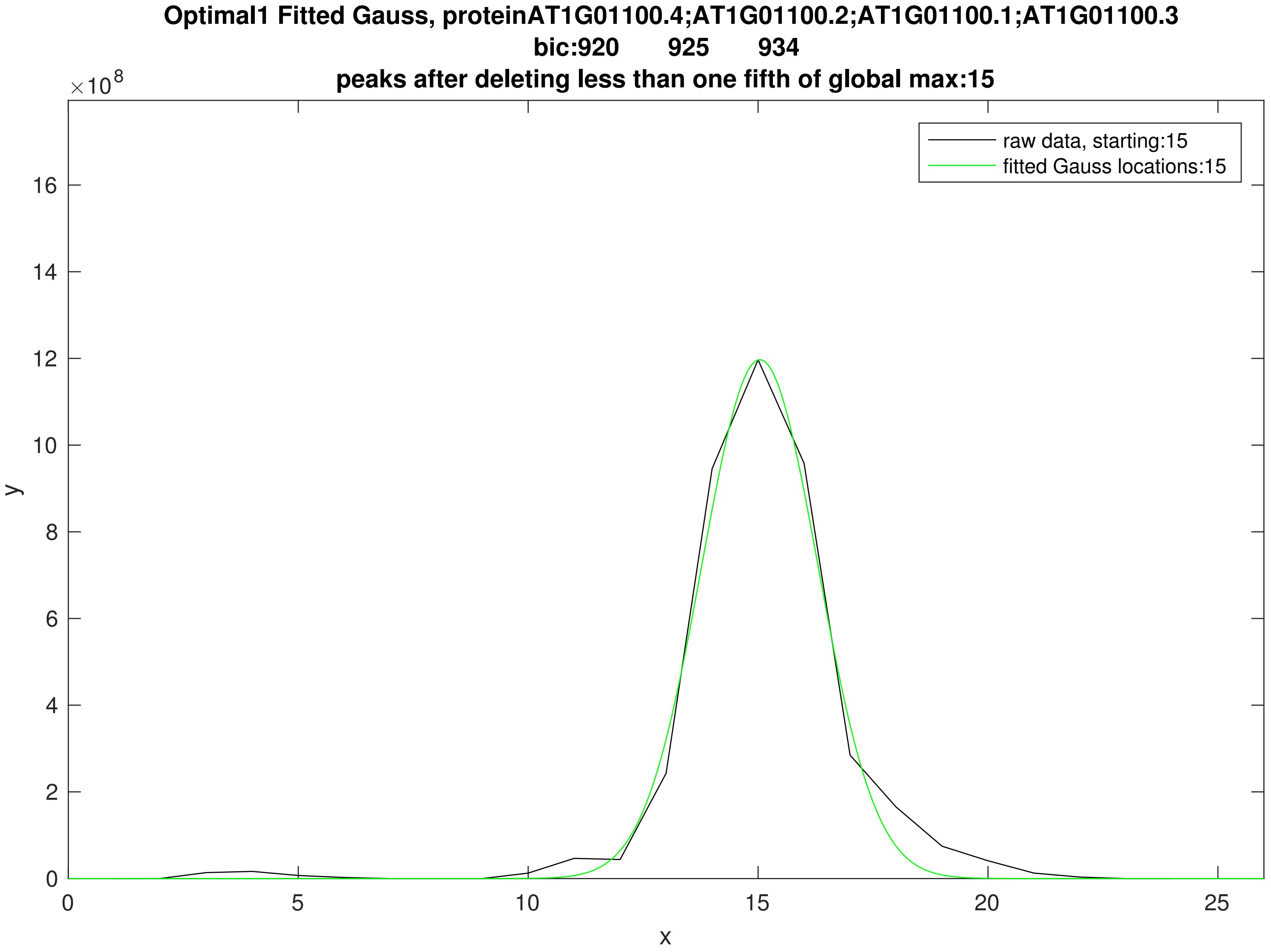

Optimal1 Fitted Gauss, proteinAT1G01100.4;AT1G01100.2;AT1G01100.1;AT1G01100.3
bic:920 925 934
peaks after deleting less than one fifth of global max:15
×10^8
raw data, starting:15
fitted Gauss locations:15
16
14
12
10
8
6
4
2
0
y
0
5
10
15
20
25
x

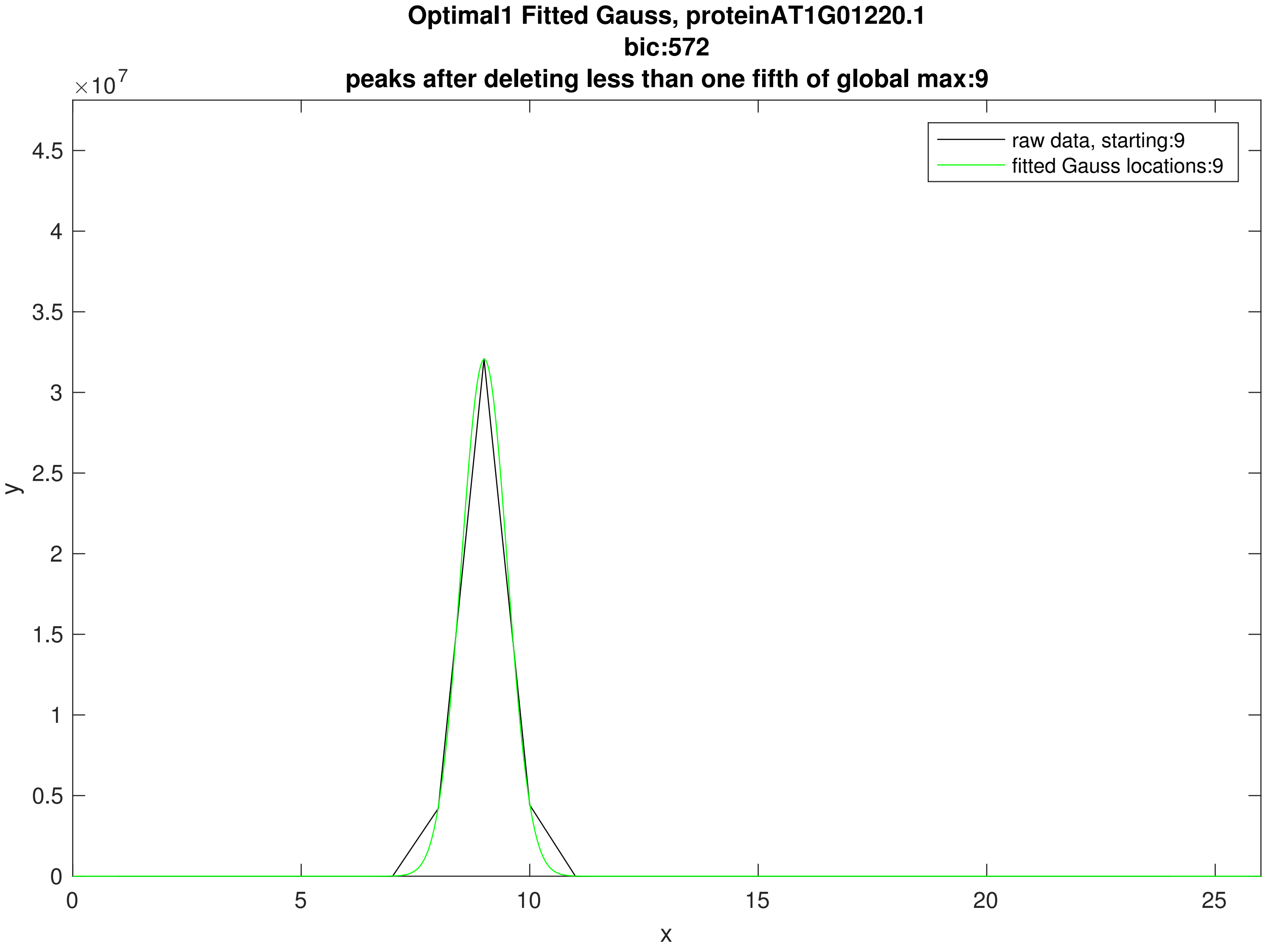
Optimal1 Fitted Gauss, proteinAT1G01220.1
bic:572
peaks after deleting less than one fifth of global max:9
×10^7
raw data, starting:9
fitted Gauss locations:9
y
x
0
0.5
1
1.5
2
2.5
3
3.5
4
4.5
0
5
10
15
20
25

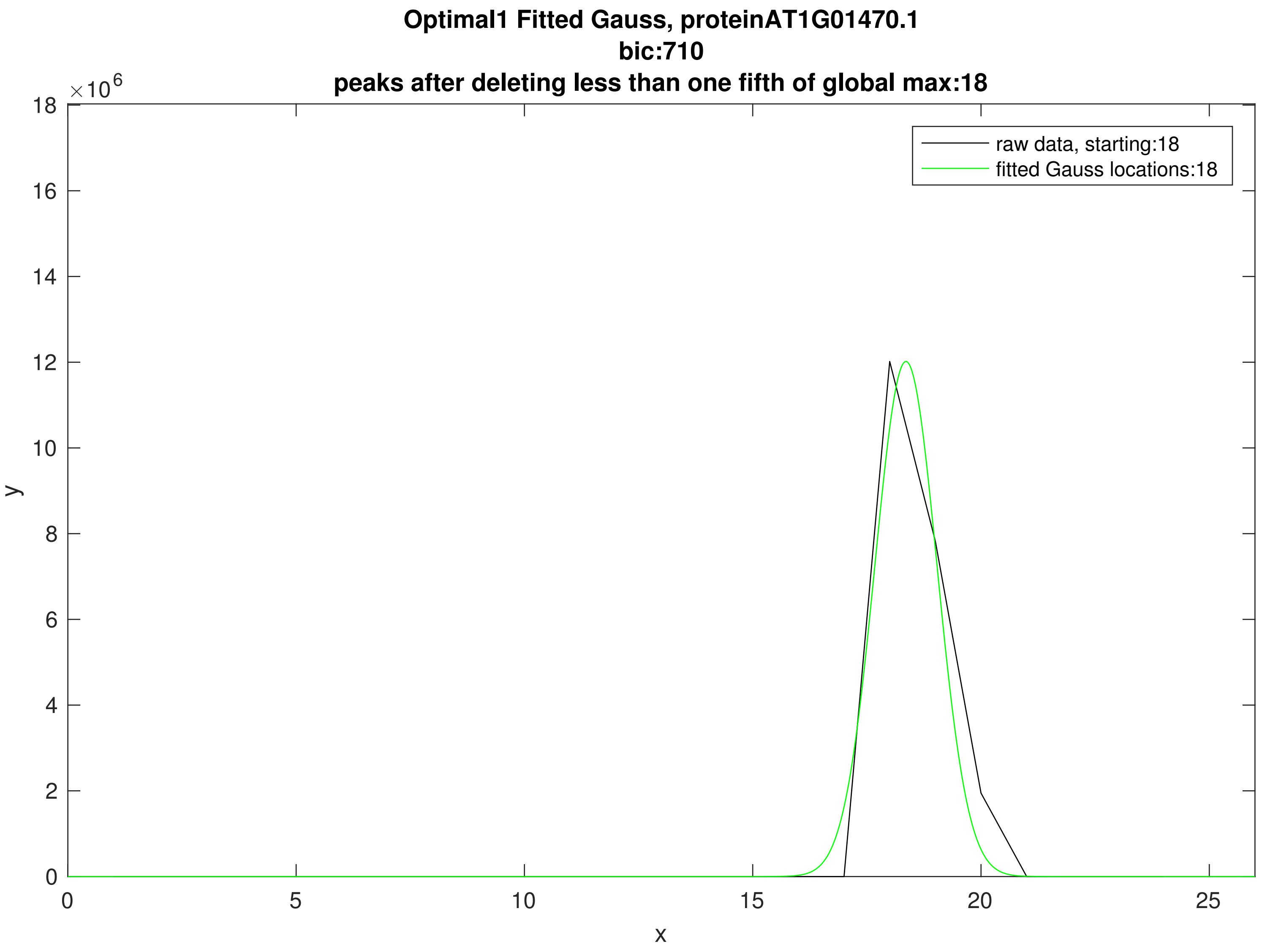

Optimal1 Fitted Gauss, proteinAT1G01470.1
bic:710
peaks after deleting less than one fifth of global max:18
×10^6
18
16
14
12
10
8
6
4
2
0
y
0
5
10
15
20
25
x
raw data, starting:18
fitted Gauss locations:18

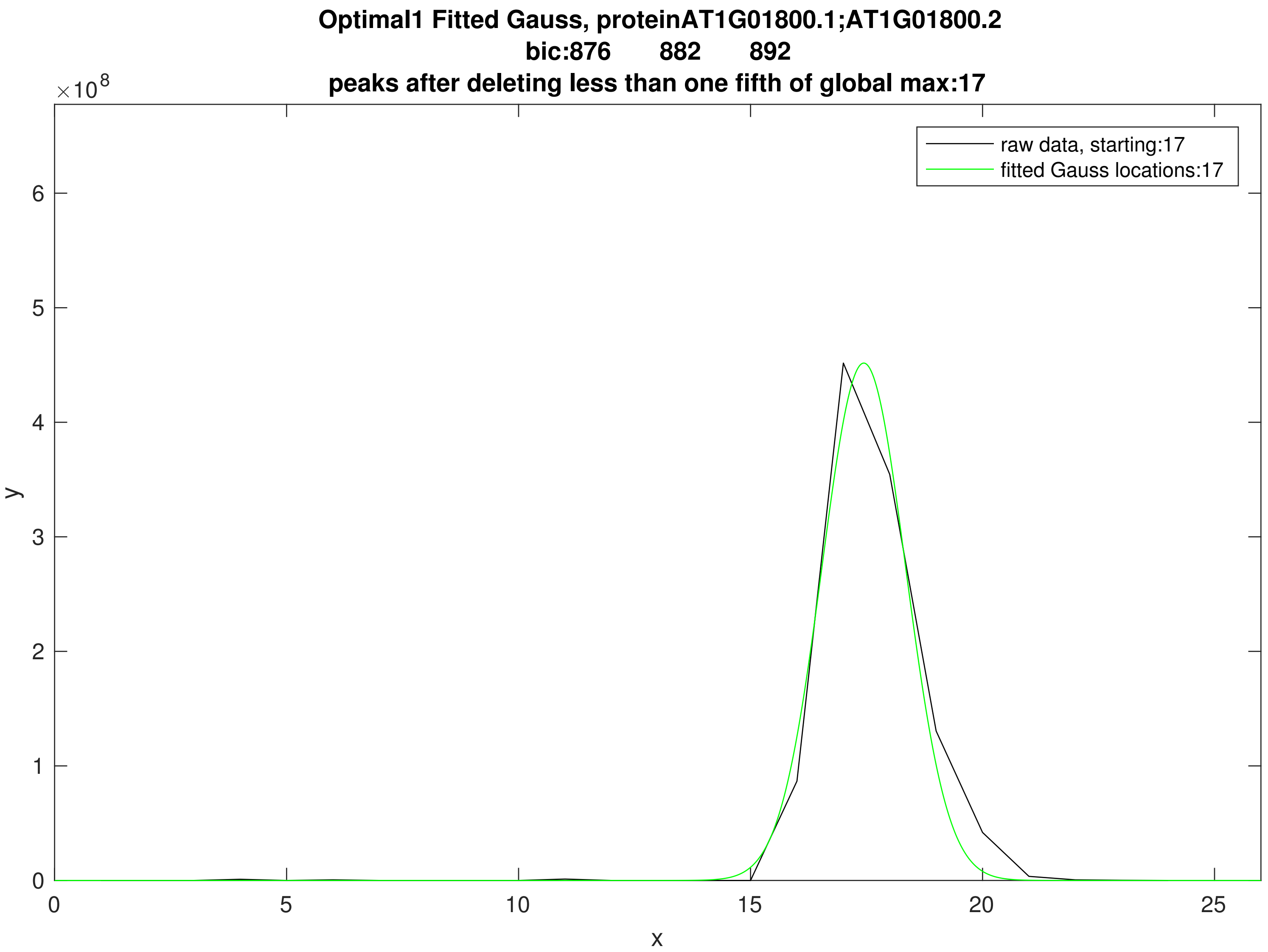
Optimal1 Fitted Gauss, proteinAT1G01800.1;AT1G01800.2
bic:876 882 892
peaks after deleting less than one fifth of global max:17
×10^8
raw data, starting:17
fitted Gauss locations:17
y
x
0
1
2
3
4
5
6
0
5
10
15
20
25

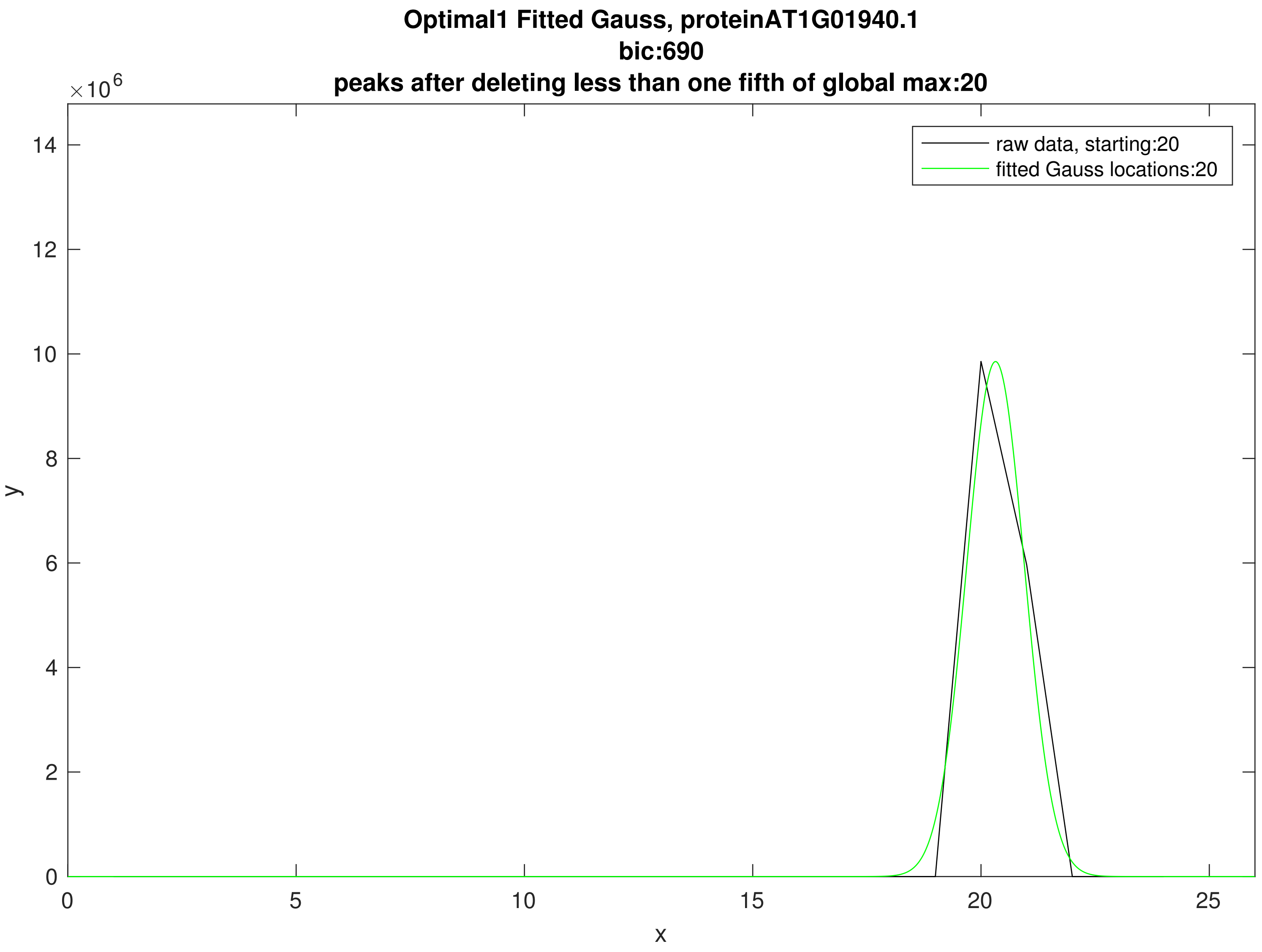
Optimal1 Fitted Gauss, proteinAT1G01940.1
bic:690
peaks after deleting less than one fifth of global max:20
×10^6
14
12
10
8
6
4
2
0
y
0
5
10
15
20
25
x
raw data, starting:20
fitted Gauss locations:20

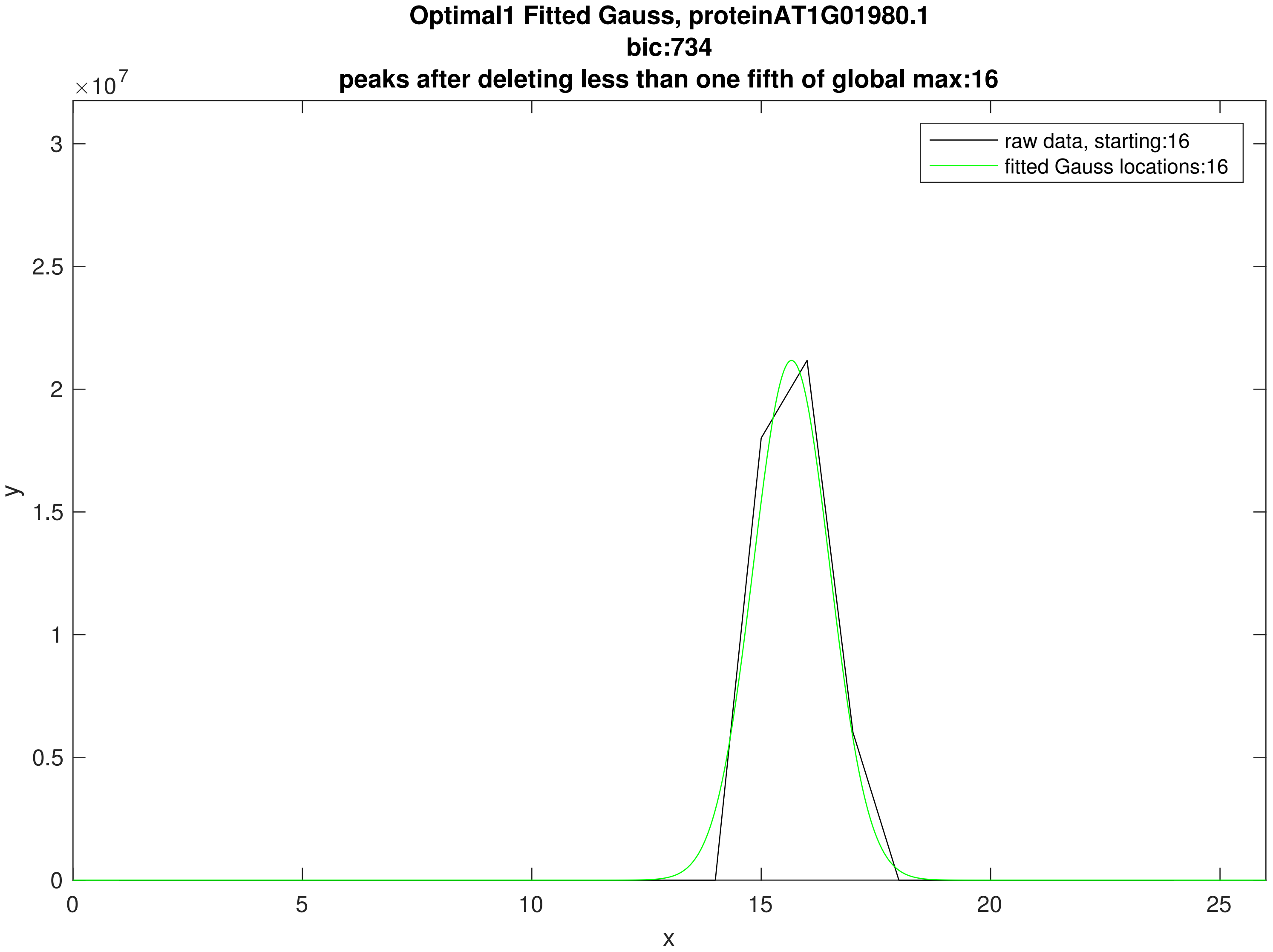
Optimal1 Fitted Gauss, proteinAT1G01980.1
bic:734
peaks after deleting less than one fifth of global max:16
×10^7
3
2.5
2
1.5
1
0.5
0
y
0
5
10
15
20
25
x
raw data, starting:16
fitted Gauss locations:16

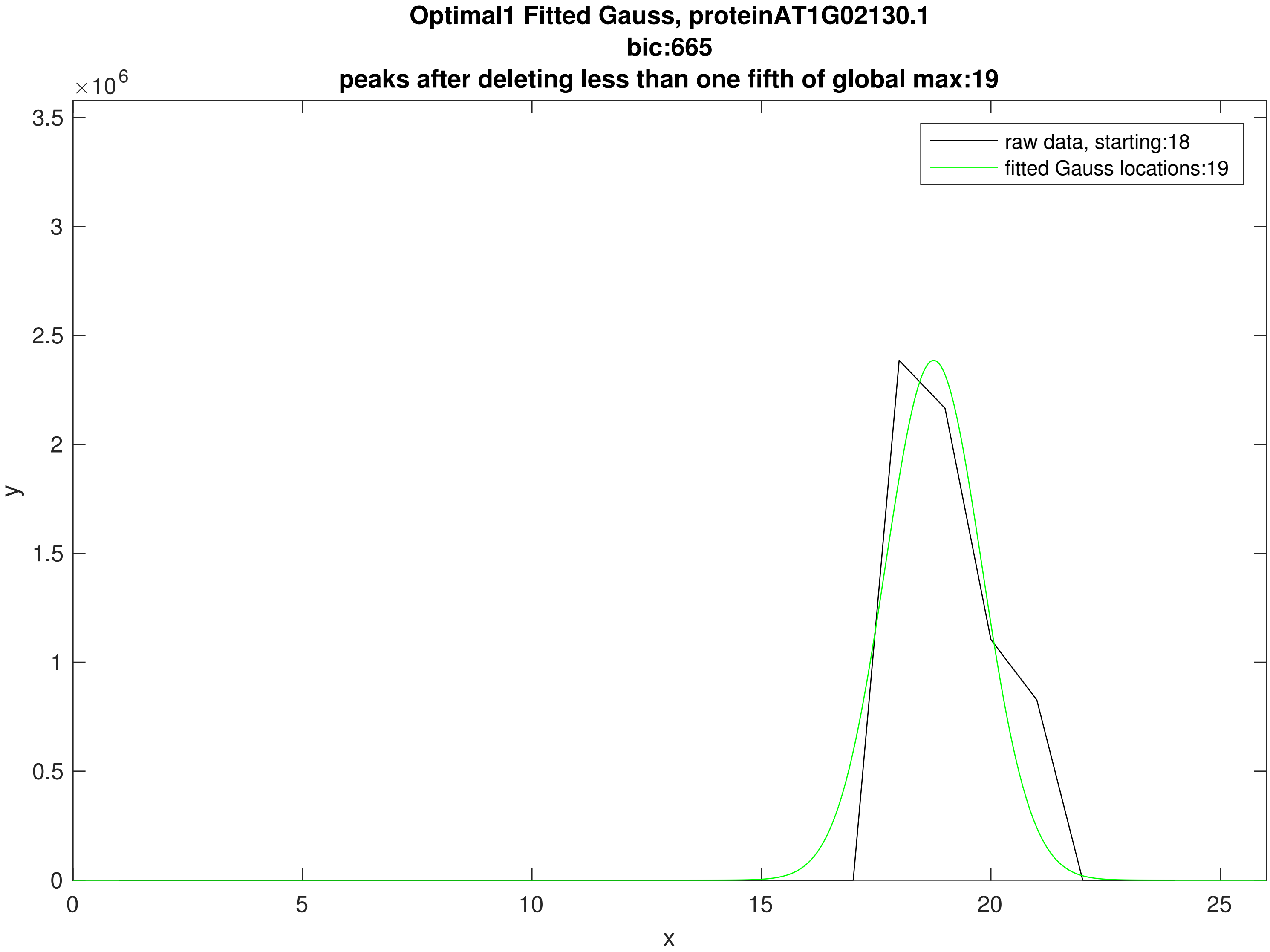

Optimal1 Fitted Gauss, proteinAT1G02130.1
bic:665
peaks after deleting less than one fifth of global max:19
×10^6
3.5
3
2.5
2
1.5
1
0.5
0
y
0
5
10
15
20
25
x
raw data, starting:18
fitted Gauss locations:19

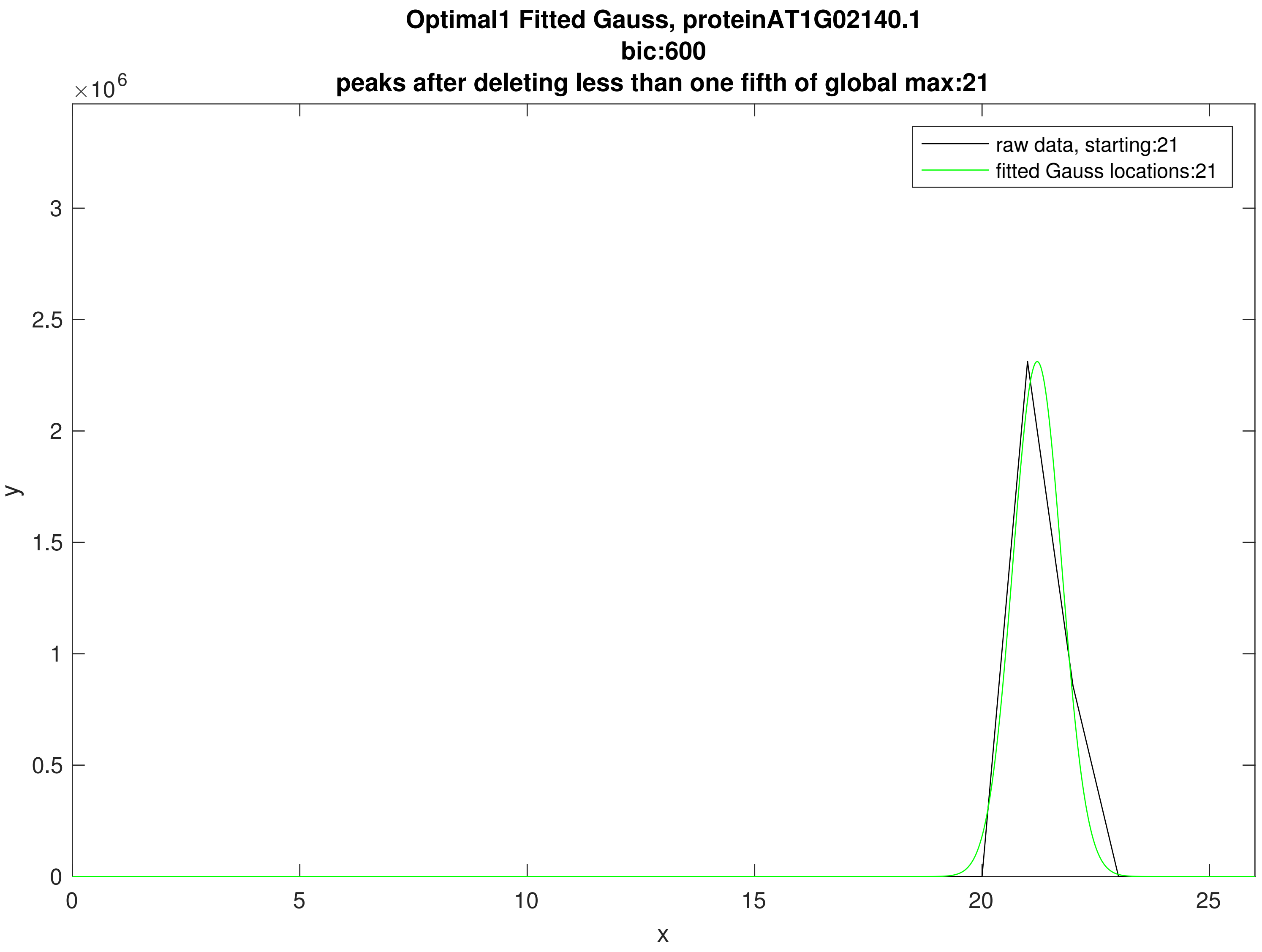
Optimal1 Fitted Gauss, proteinAT1G02140.1
bic:600
peaks after deleting less than one fifth of global max:21
×10^6
raw data, starting:21
fitted Gauss locations:21
y
x
0
0.5
1
1.5
2
2.5
3
0
5
10
15
20
25

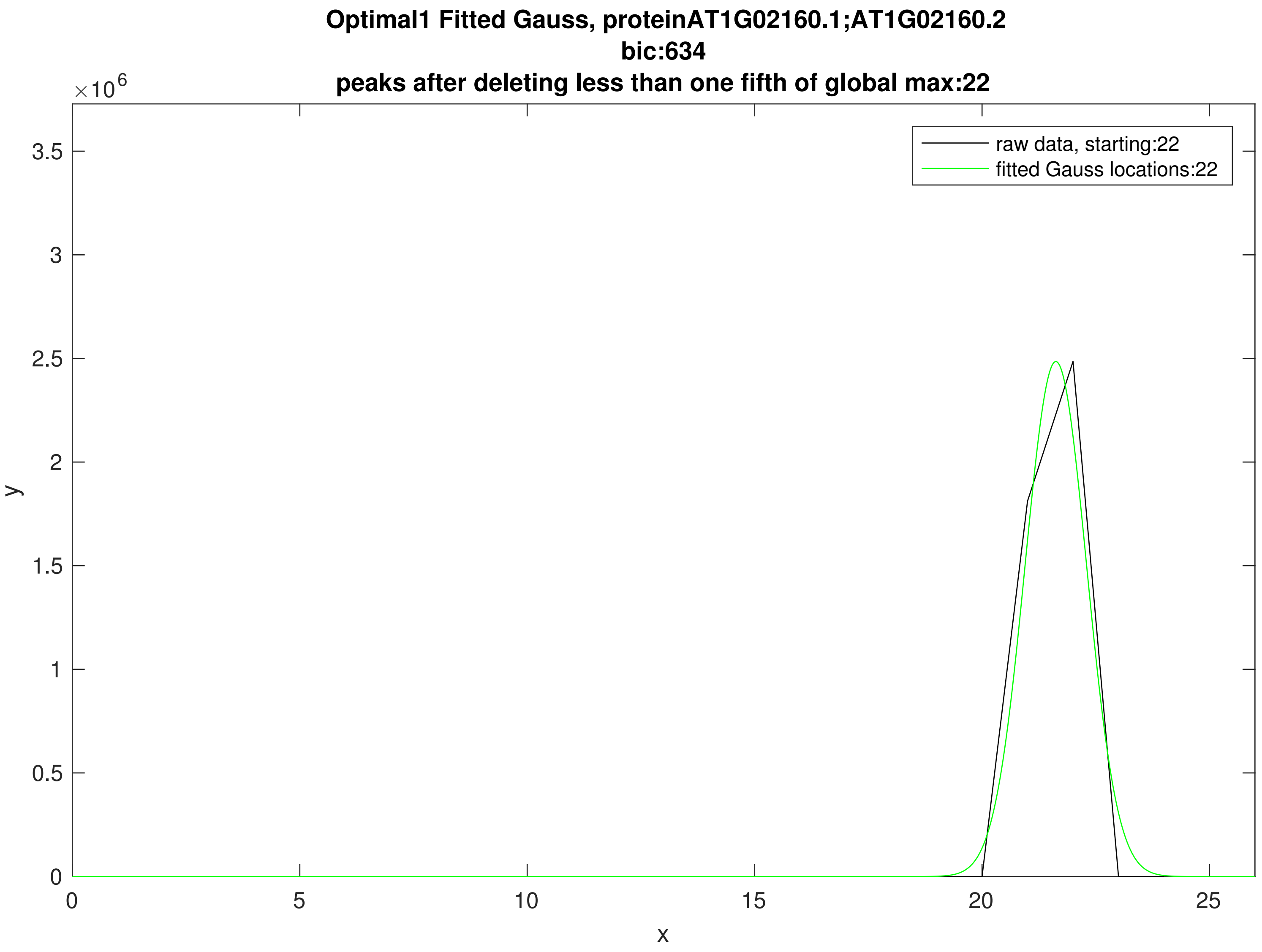

Optimal1 Fitted Gauss, proteinAT1G02160.1;AT1G02160.2
bic:634
peaks after deleting less than one fifth of global max:22
×10^6
3.5
3
2.5
2
1.5
1
0.5
0
y
0
5
10
15
20
25
x
raw data, starting:22
fitted Gauss locations:22

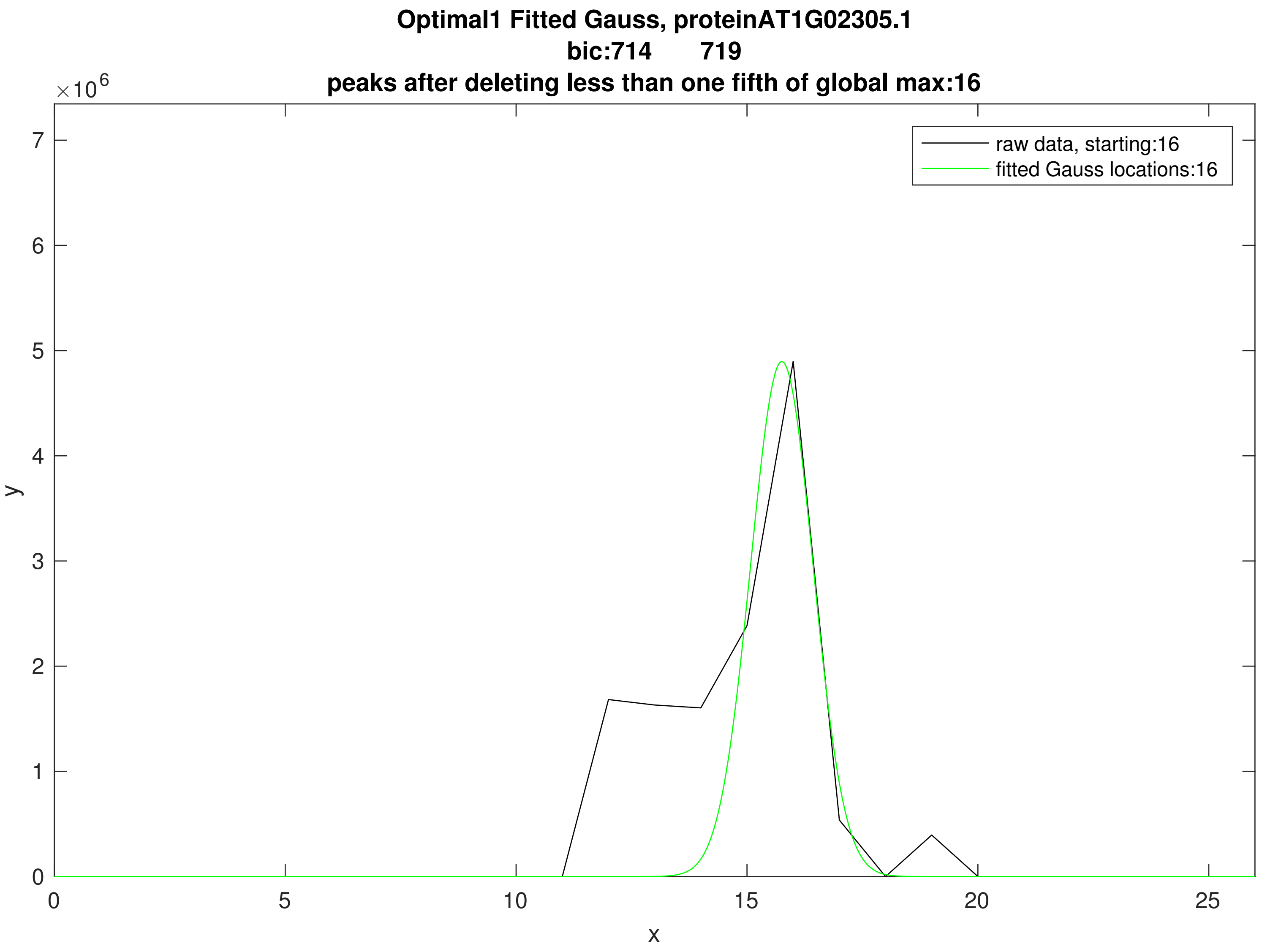

Optimal1 Fitted Gauss, proteinAT1G02305.1
bic:714 719
peaks after deleting less than one fifth of global max:16
×10^6
raw data, starting:16
fitted Gauss locations:16
y
x
0
1
2
3
4
5
6
7
0
5
10
15
20
25

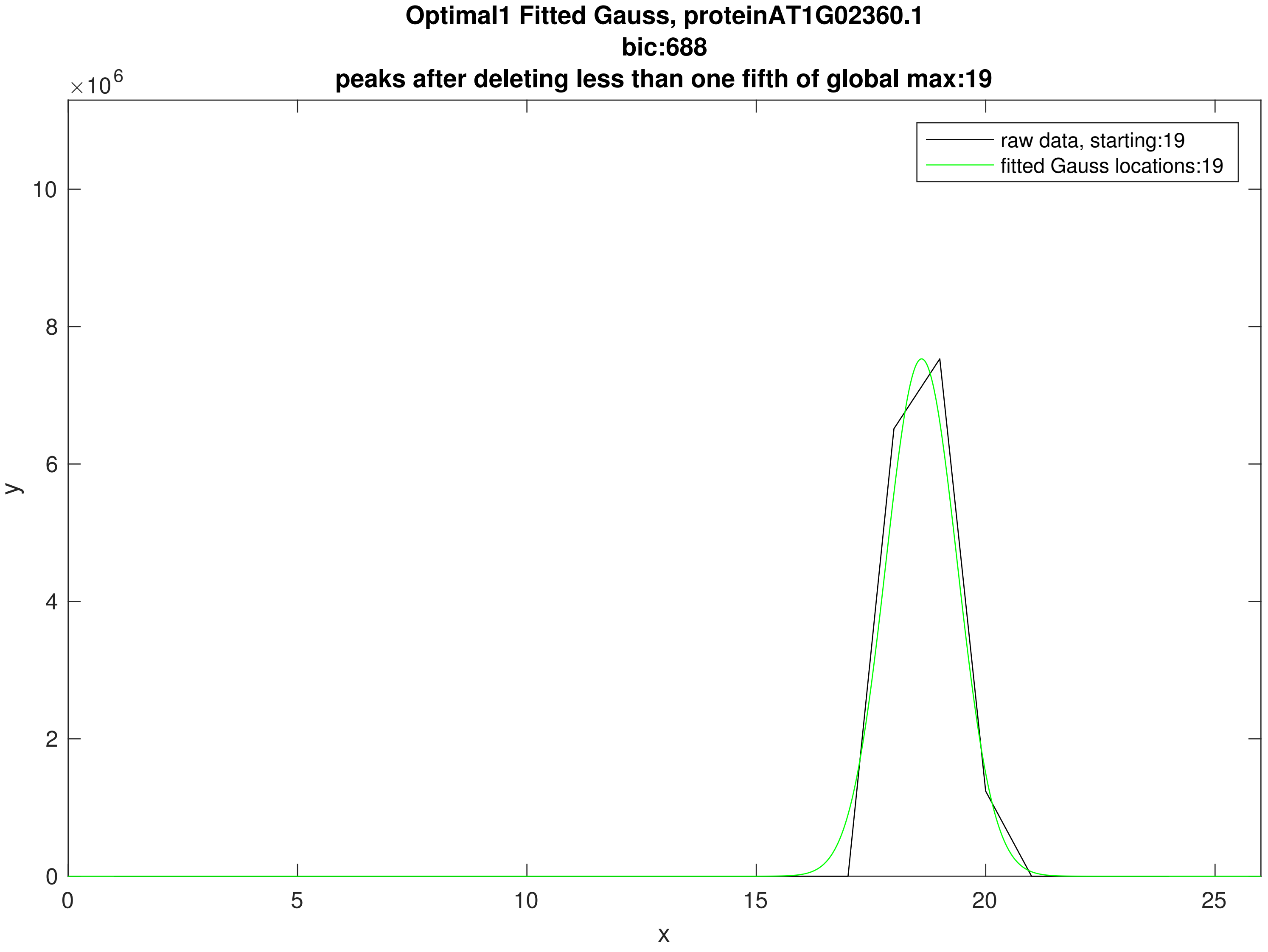
Optimal1 Fitted Gauss, proteinAT1G02360.1
bic:688
peaks after deleting less than one fifth of global max:19
×10^6
raw data, starting:19
fitted Gauss locations:19
10
8
6
4
2
0
y
0
5
10
15
20
25
x

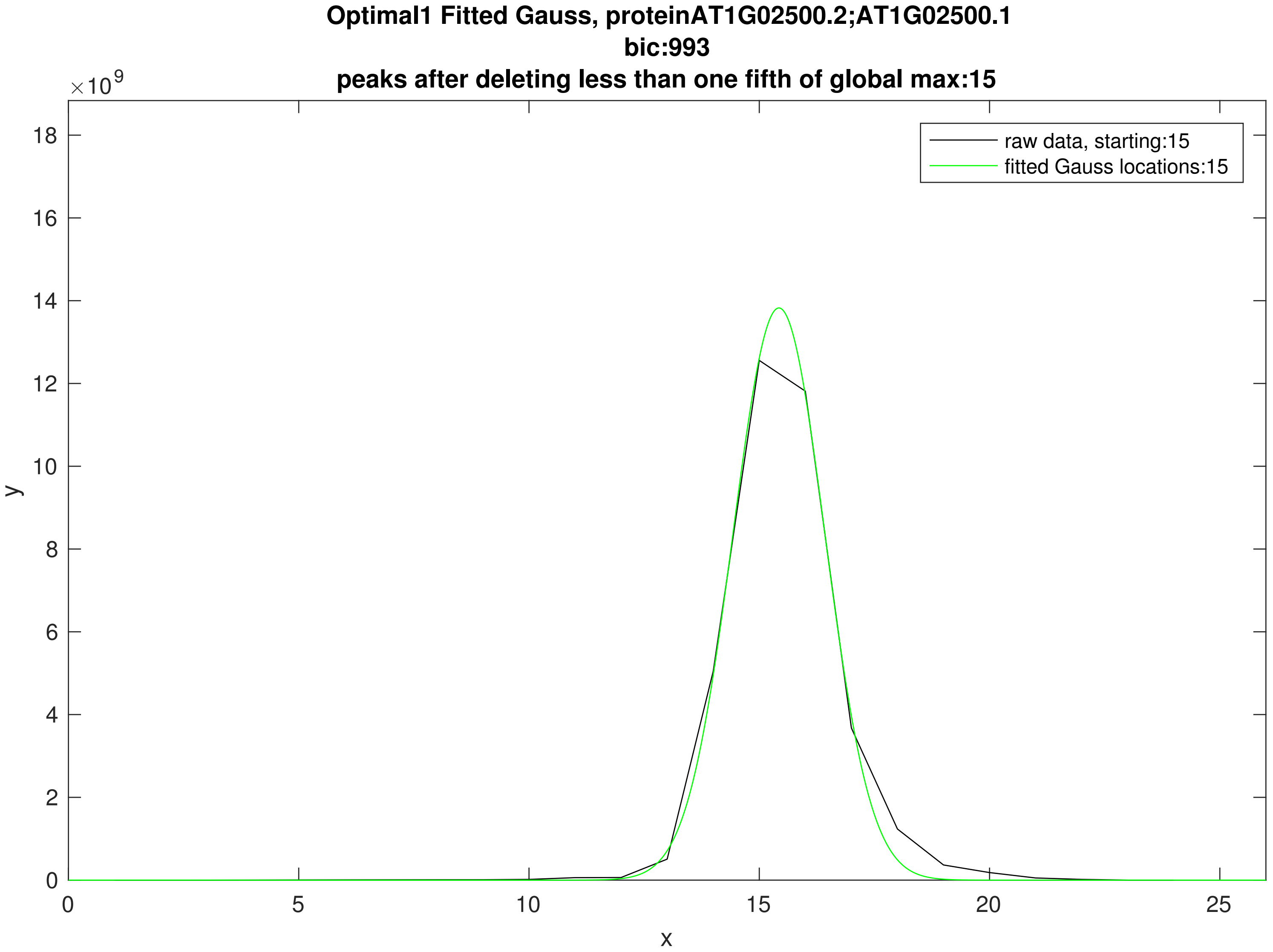
Optimal1 Fitted Gauss, proteinAT1G02500.2;AT1G02500.1
bic:993
peaks after deleting less than one fifth of global max:15
×10^9
raw data, starting:15
fitted Gauss locations:15
y
x
0
2
4
6
8
10
12
14
16
18
0
5
10
15
20
25

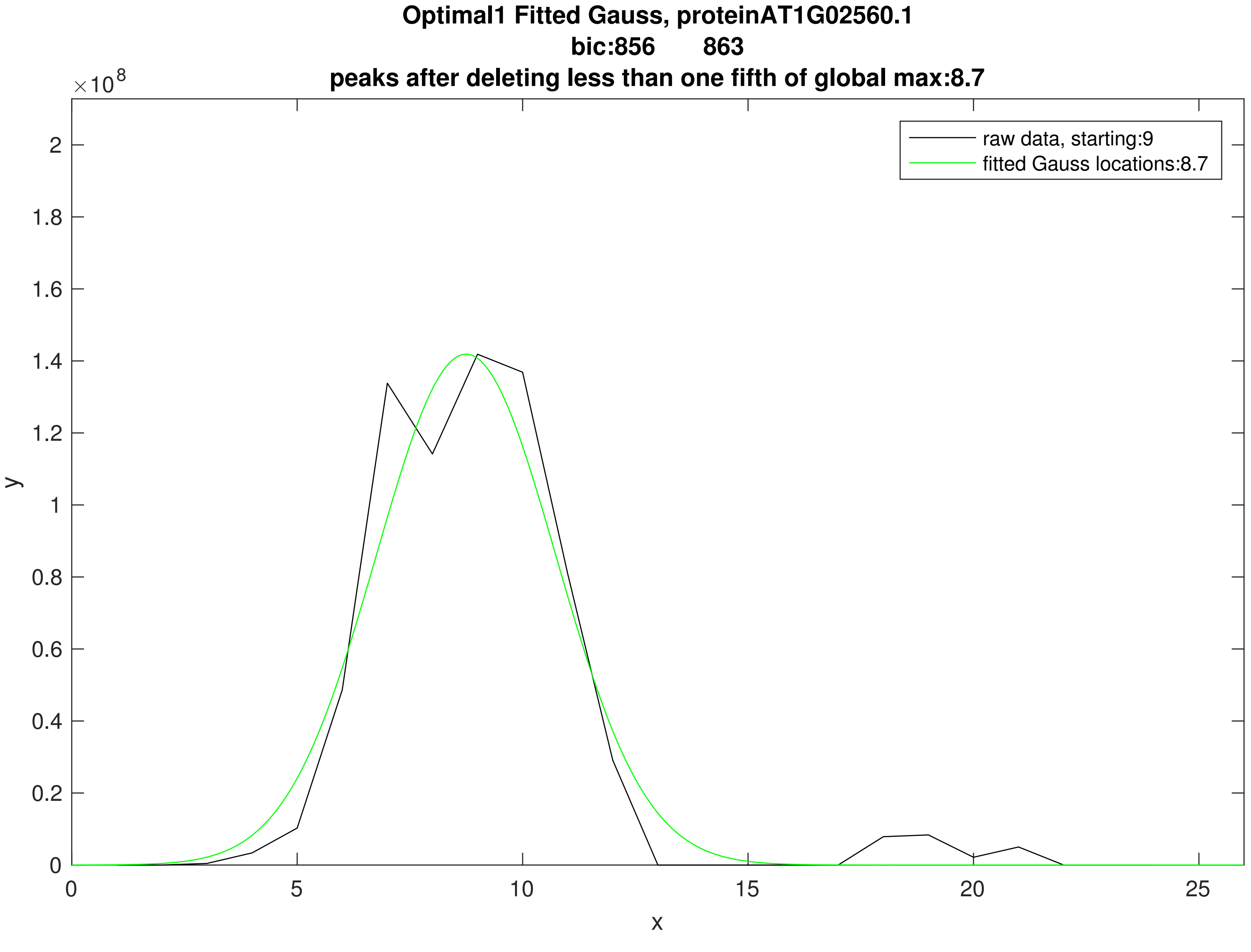

Optimal1 Fitted Gauss, proteinAT1G02560.1
bic:856 863
peaks after deleting less than one fifth of global max:8.7
×10^8
raw data, starting:9
fitted Gauss locations:8.7
y
x
0
0.2
0.4
0.6
0.8
1
1.2
1.4
1.6
1.8
2
5
10
15
20
25

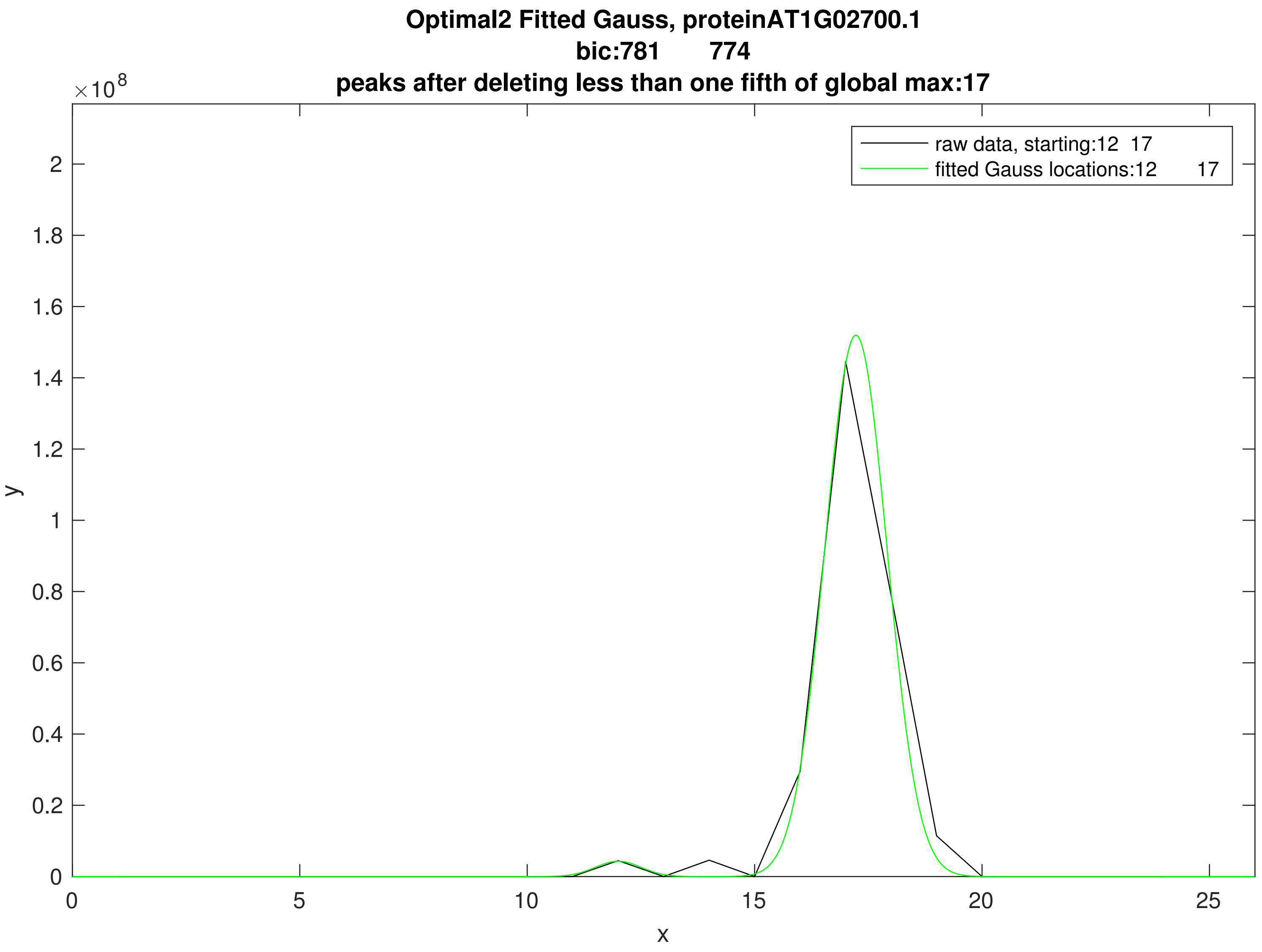

Optimal2 Fitted Gauss, proteinAT1G02700.1
bic:781 774
peaks after deleting less than one fifth of global max:17
×10^8
raw data, starting:12 17
fitted Gauss locations:12 17
y
x
0
0.2
0.4
0.6
0.8
1
1.2
1.4
1.6
1.8
2
5
10
15
20
25

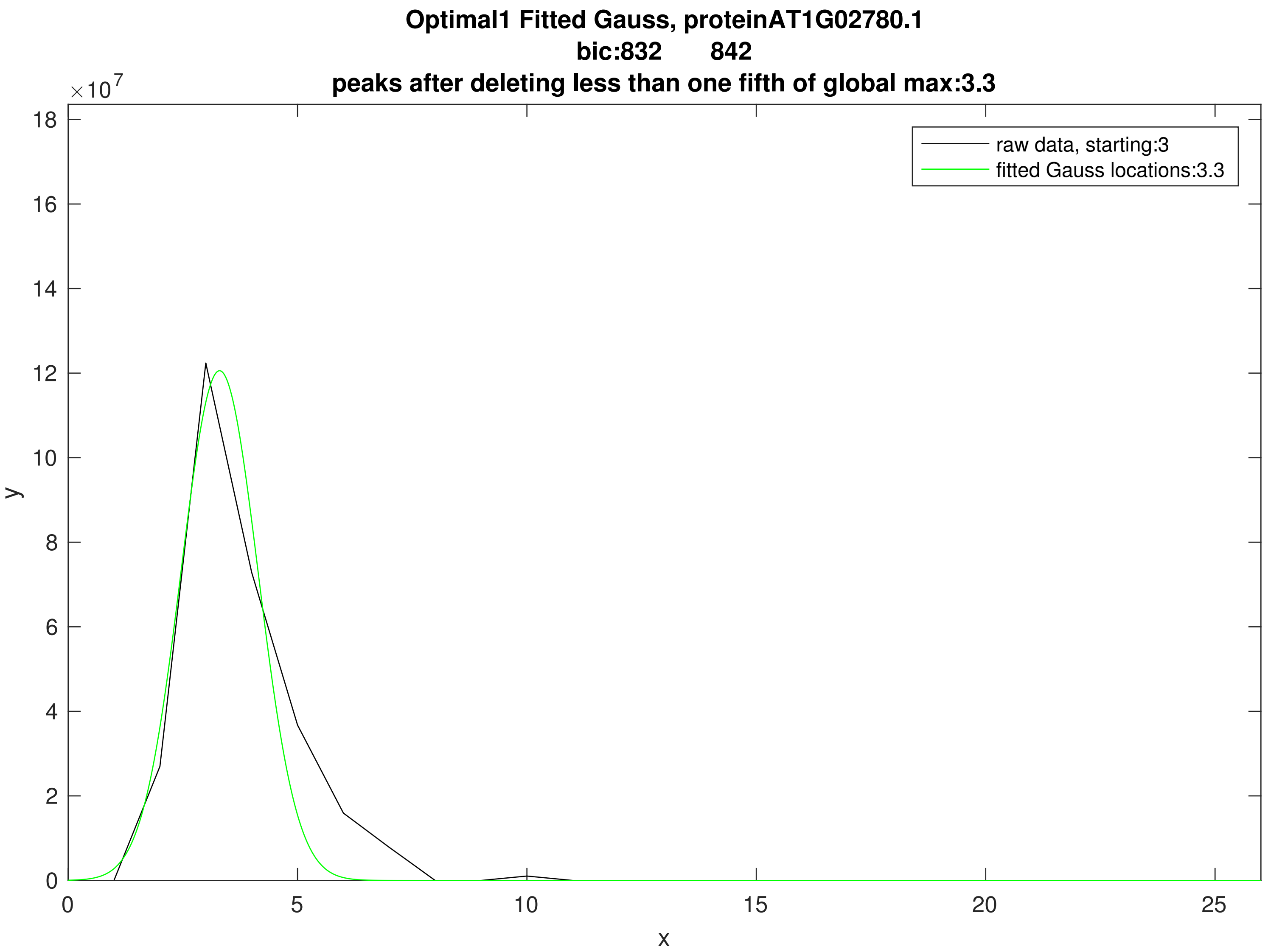

Optimal1 Fitted Gauss, proteinAT1G02780.1
bic:832 842
peaks after deleting less than one fifth of global max:3.3
×10^7
raw data, starting:3
fitted Gauss locations:3.3
y
x
0
2
4
6
8
10
12
14
16
18
0
5
10
15
20
25

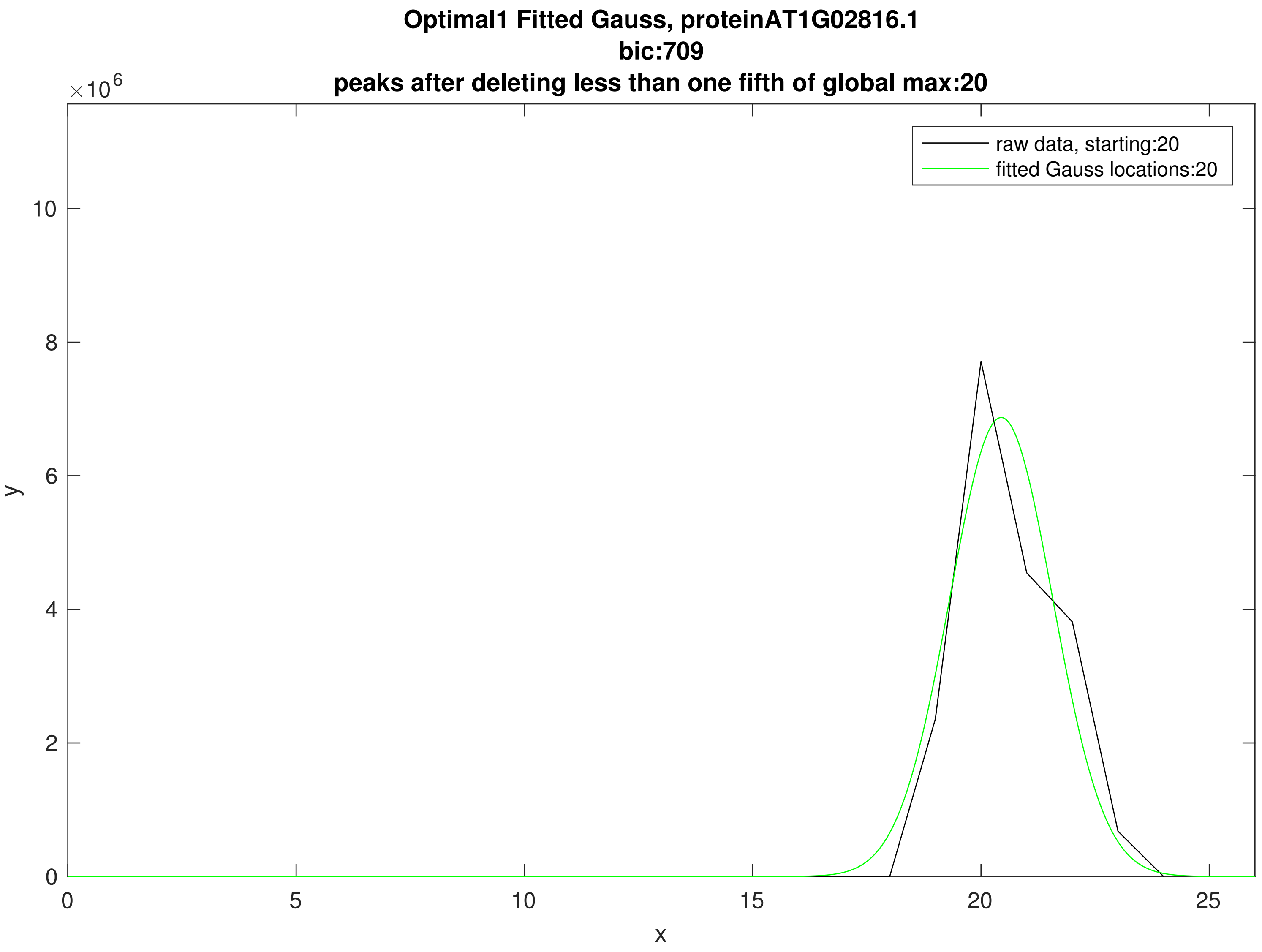

Optimal1 Fitted Gauss, proteinAT1G02816.1
bic:709
peaks after deleting less than one fifth of global max:20
×10^6
raw data, starting:20
fitted Gauss locations:20
10
8
6
4
2
0
y
0
5
10
15
20
25
x

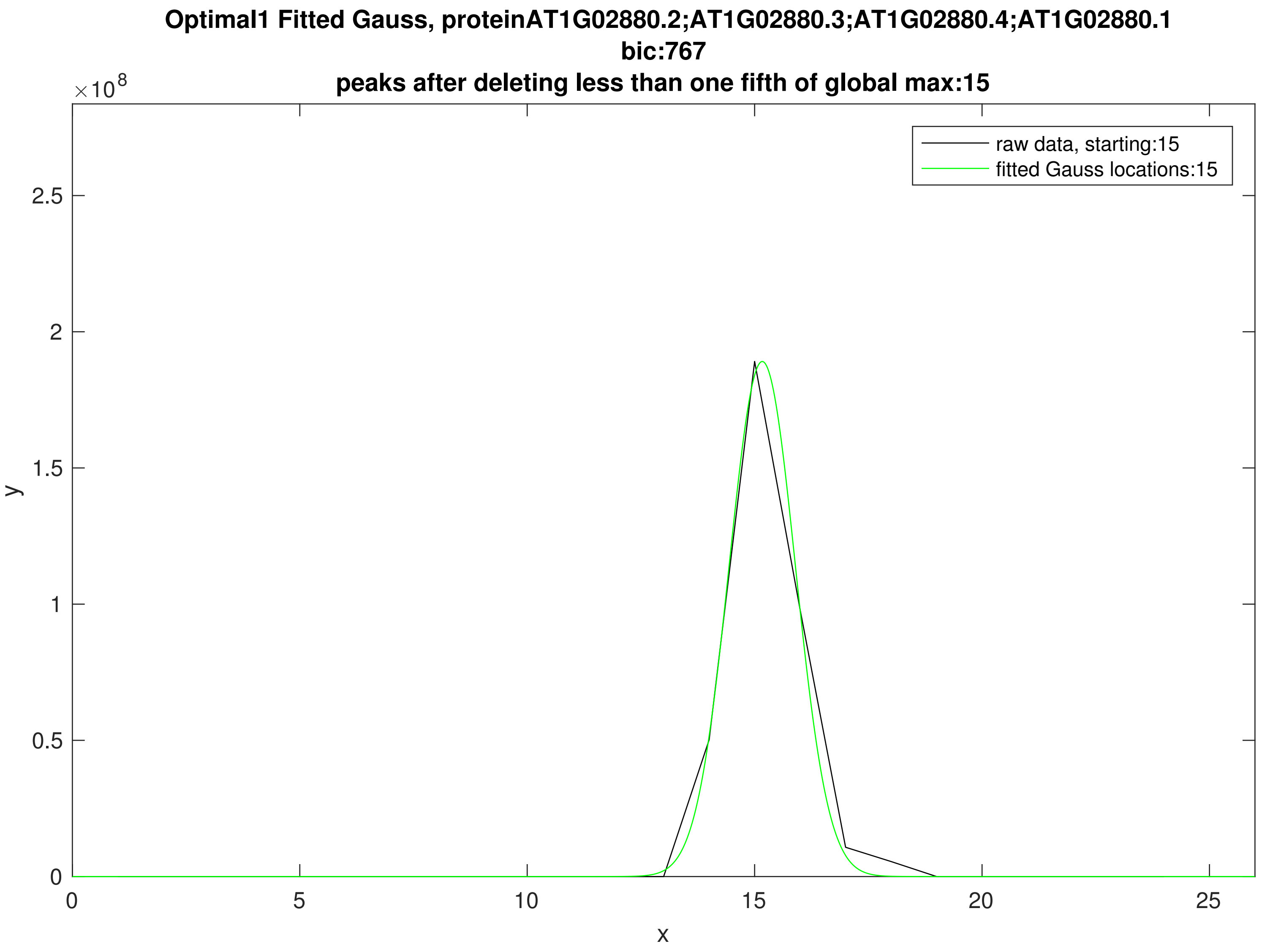
Optimal1 Fitted Gauss, proteinAT1G02880.2;AT1G02880.3;AT1G02880.4;AT1G02880.1
bic:767
peaks after deleting less than one fifth of global max:15
×10^8
raw data, starting:15
fitted Gauss locations:15
y
x
0
0.5
1
1.5
2
2.5
0
5
10
15
20
25

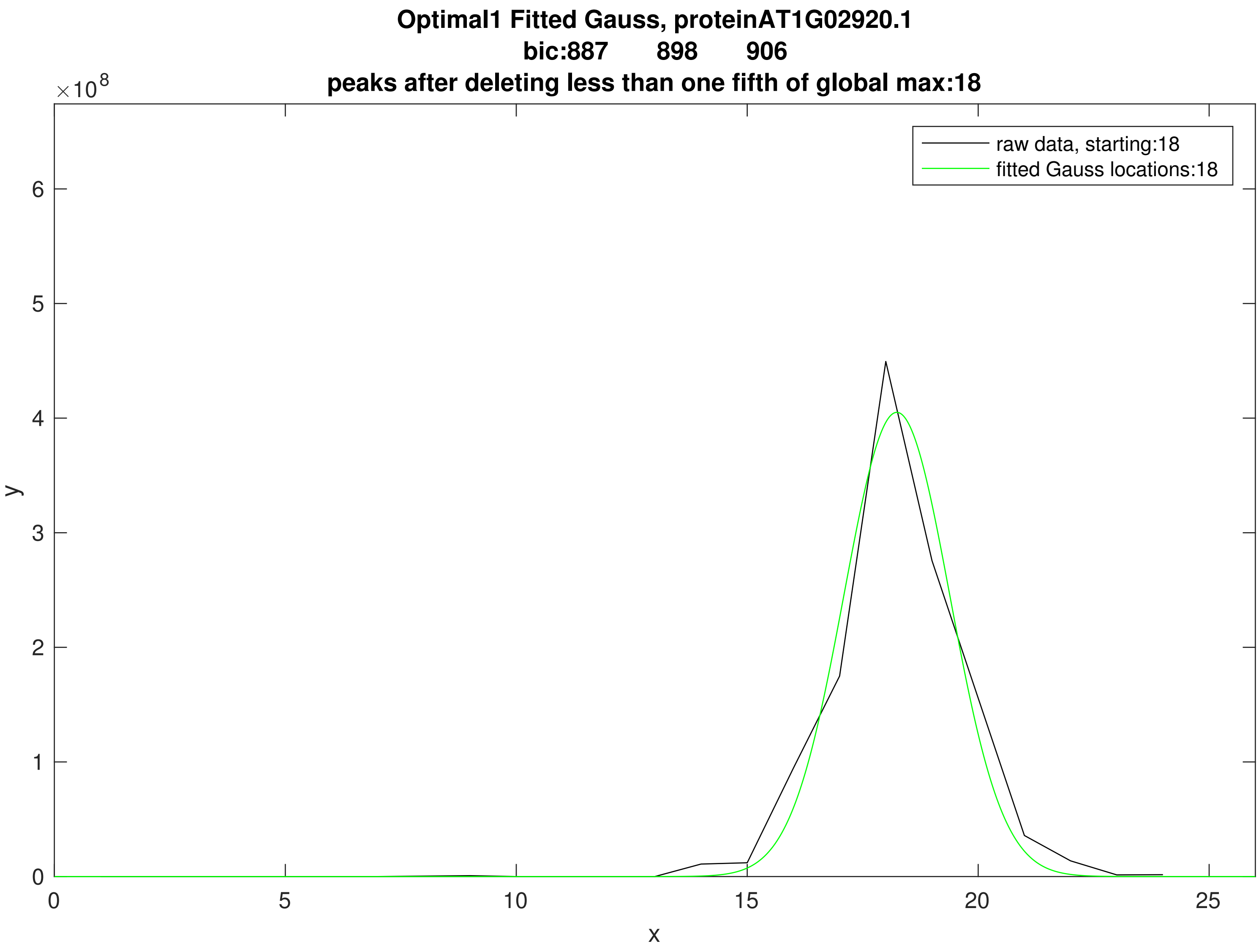
Optimal1 Fitted Gauss, proteinAT1G02920.1
bic:887 898 906
peaks after deleting less than one fifth of global max:18
×10^8
raw data, starting:18
fitted Gauss locations:18
y
x
0
1
2
3
4
5
6
0
5
10
15
20
25

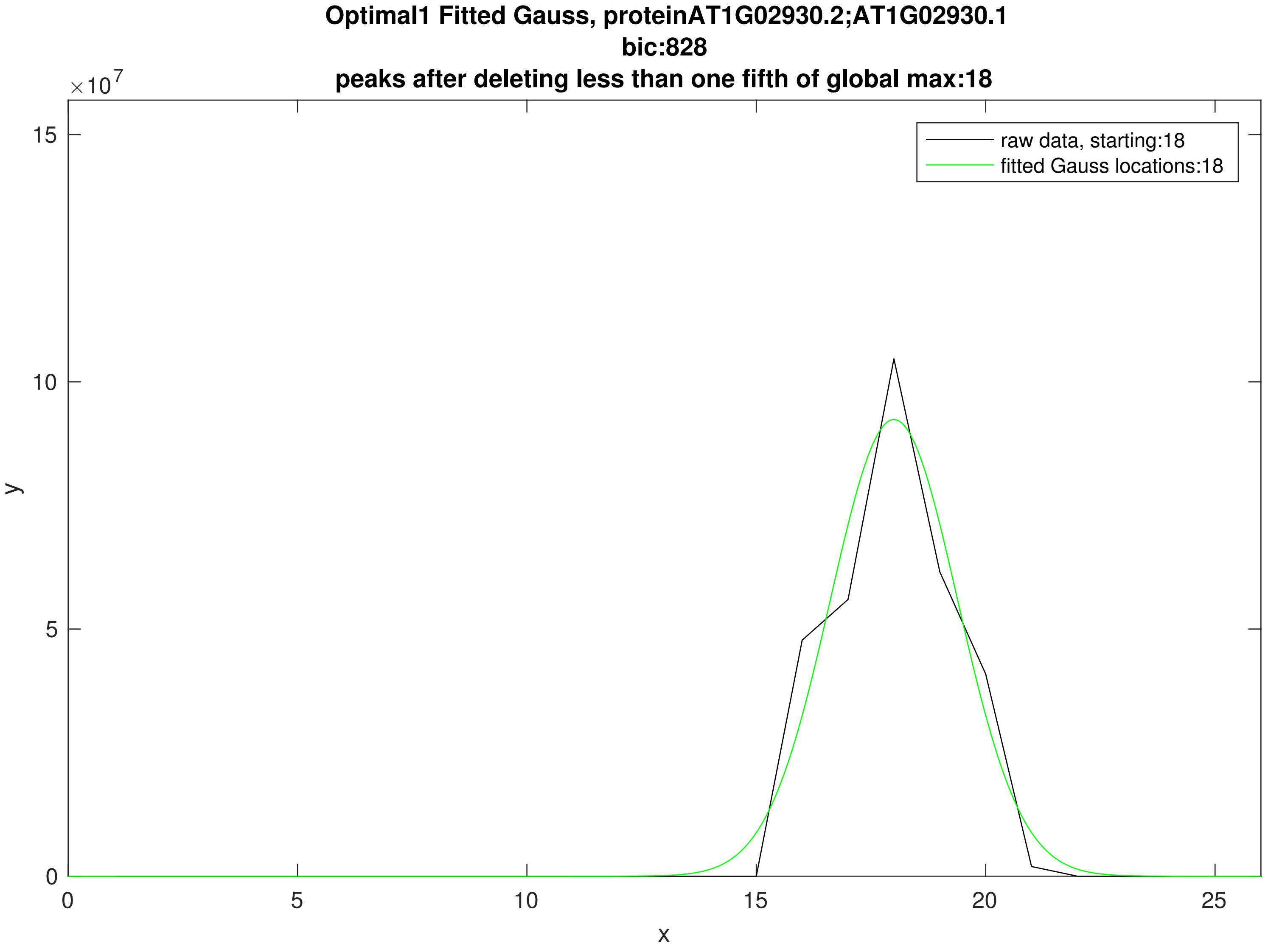

Optimal1 Fitted Gauss, proteinAT1G02930.2;AT1G02930.1
bic:828
peaks after deleting less than one fifth of global max:18
×10^7
15
10
5
0
y
0
5
10
15
20
25
x
raw data, starting:18
fitted Gauss locations:18

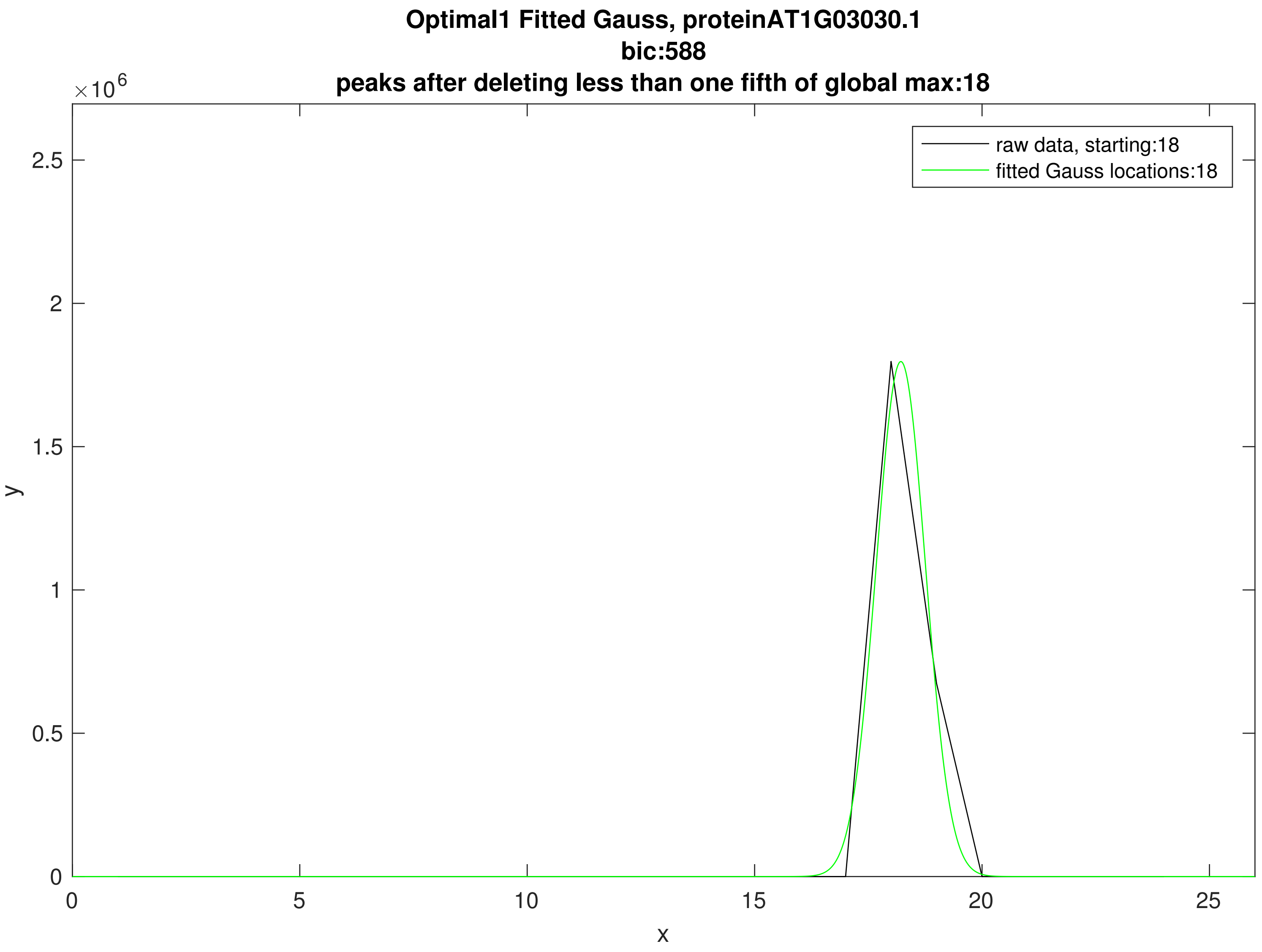

Optimal1 Fitted Gauss, proteinAT1G03030.1
bic:588
peaks after deleting less than one fifth of global max:18
×10^6
2.5
2
1.5
1
0.5
0
y
0
5
10
15
20
25
x
raw data, starting:18
fitted Gauss locations:18

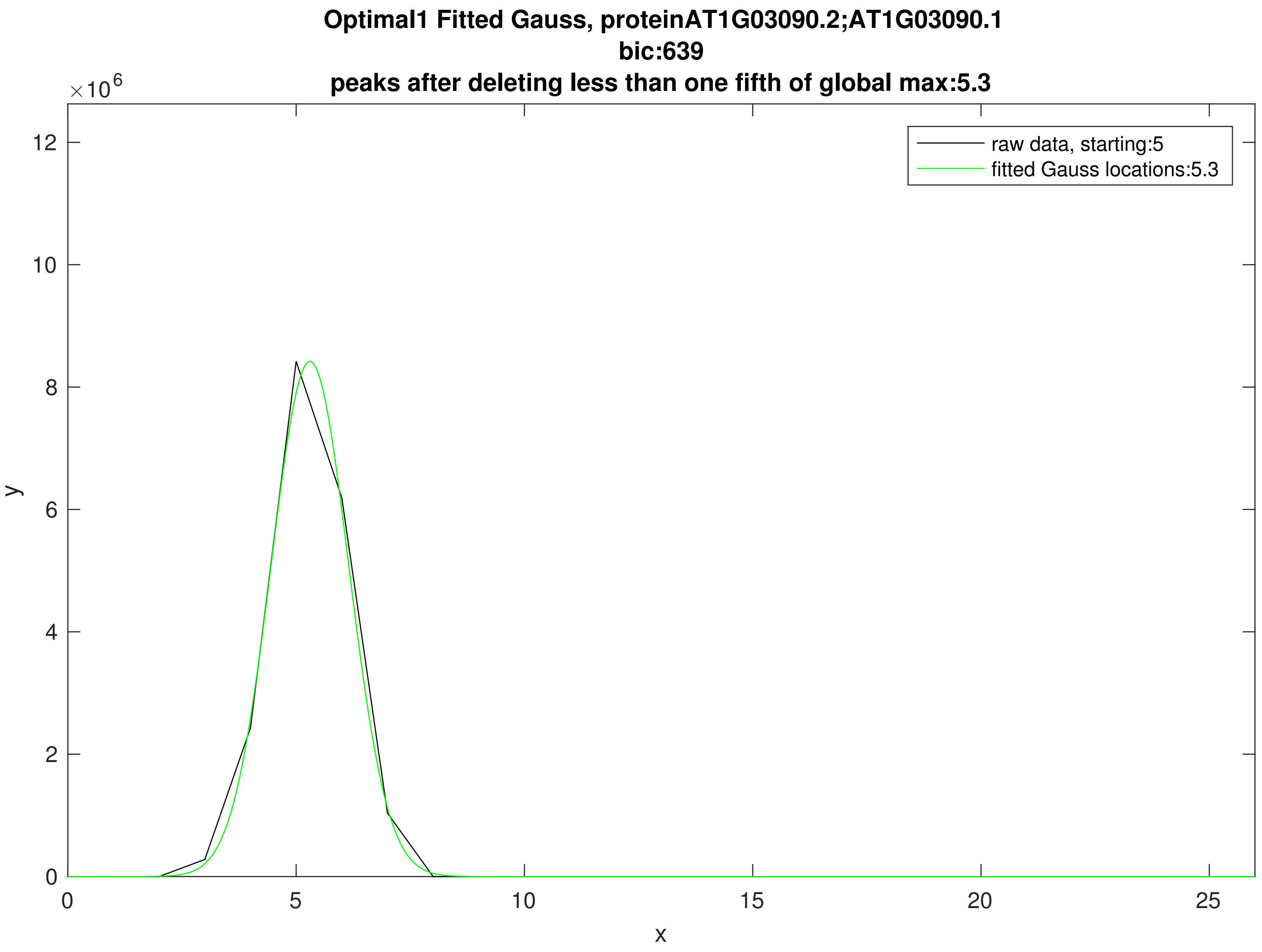
Optimal1 Fitted Gauss, proteinAT1G03090.2;AT1G03090.1
bic:639
peaks after deleting less than one fifth of global max:5.3
×10^6
raw data, starting:5
fitted Gauss locations:5.3
y
x
0
2
4
6
8
10
12
0
5
10
15
20
25

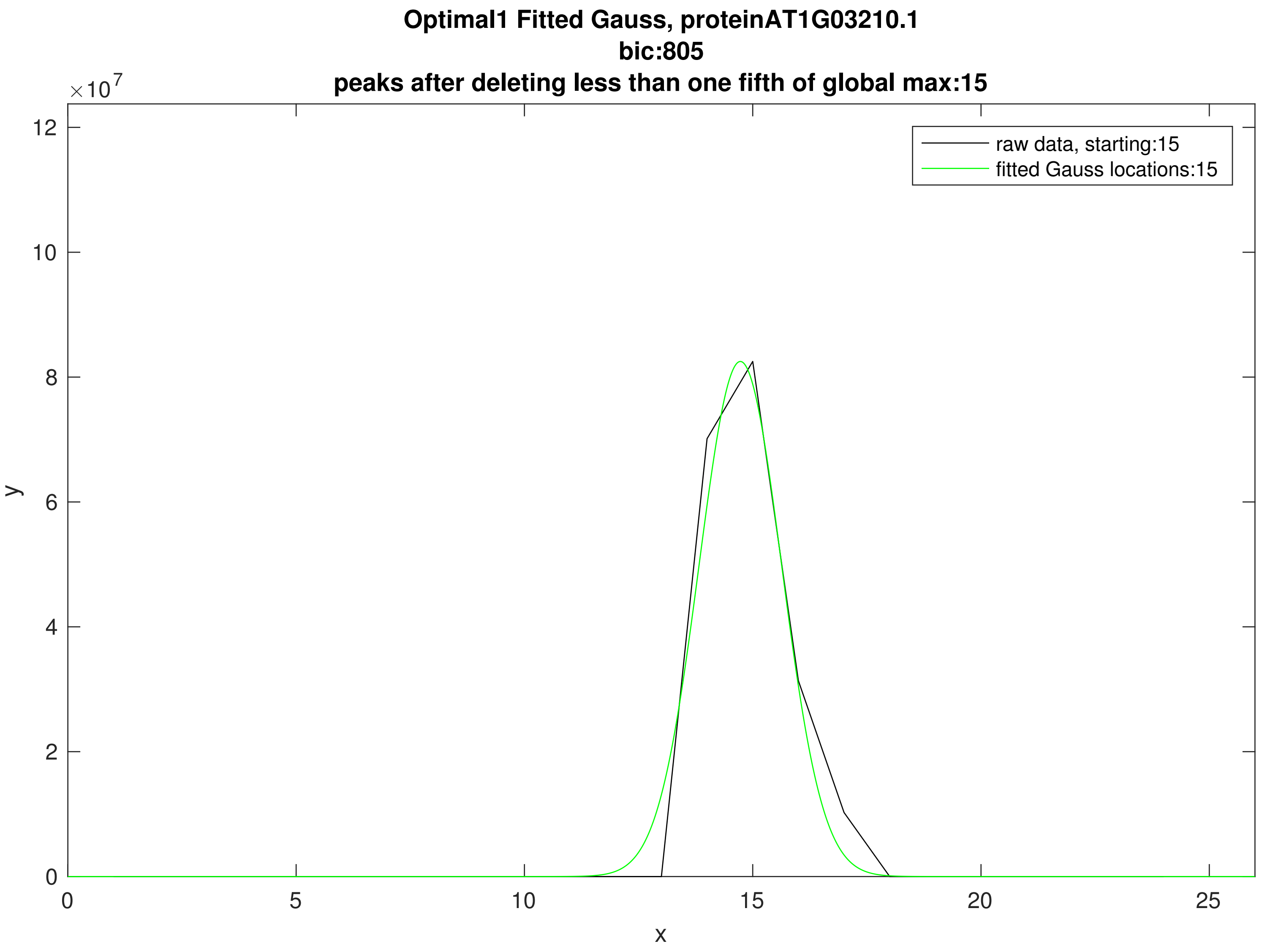
Optimal1 Fitted Gauss, proteinAT1G03210.1
bic:805
peaks after deleting less than one fifth of global max:15
×10^7
12
10
8
6
4
2
0
y
0
5
10
15
20
25
x
raw data, starting:15
fitted Gauss locations:15

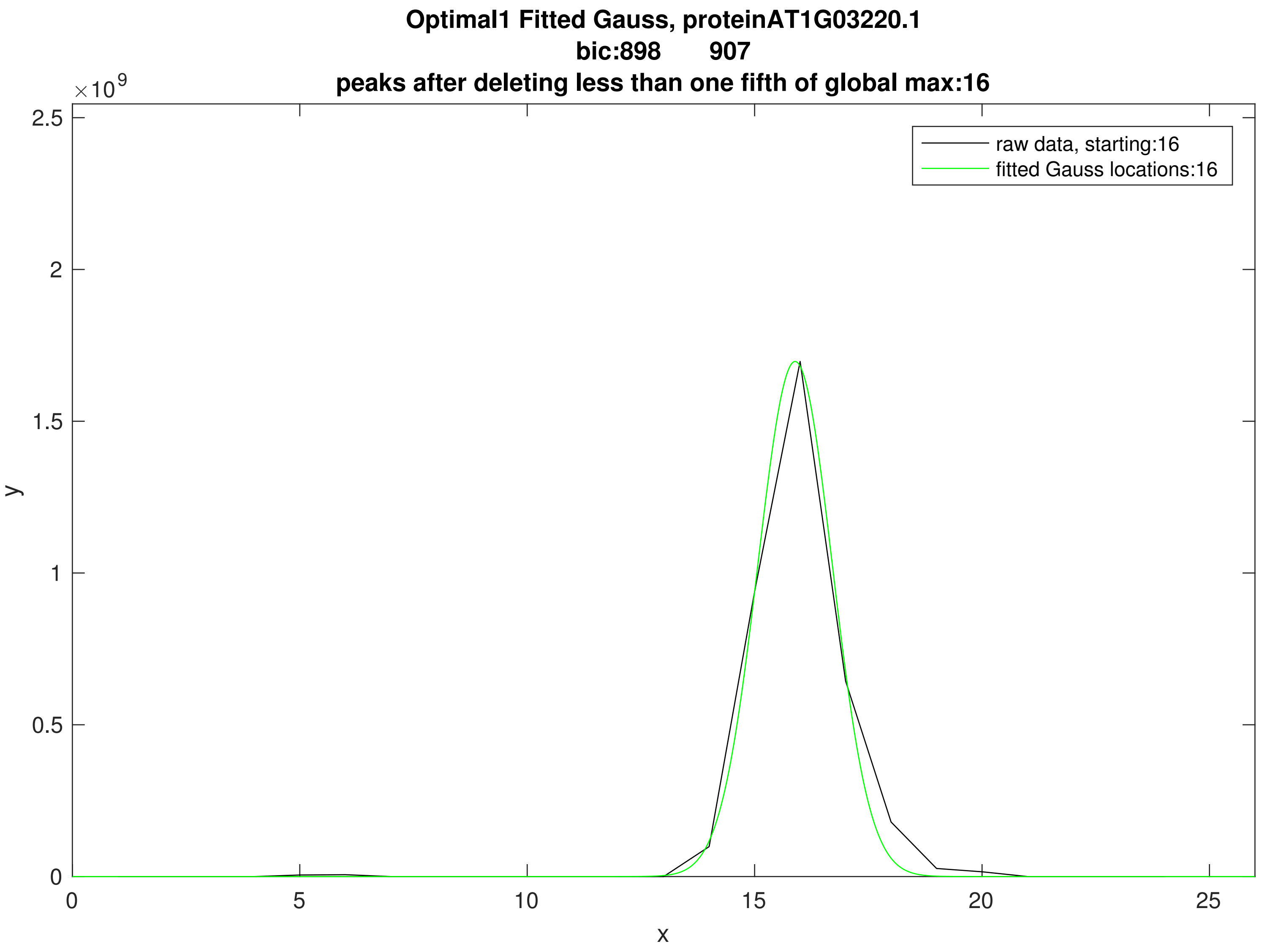
Optimal1 Fitted Gauss, proteinAT1G03220.1
bic:898 907
peaks after deleting less than one fifth of global max:16
×10^9
2.5
2
1.5
1
0.5
0
y
0
5
10
15
20
25
x
raw data, starting:16
fitted Gauss locations:16

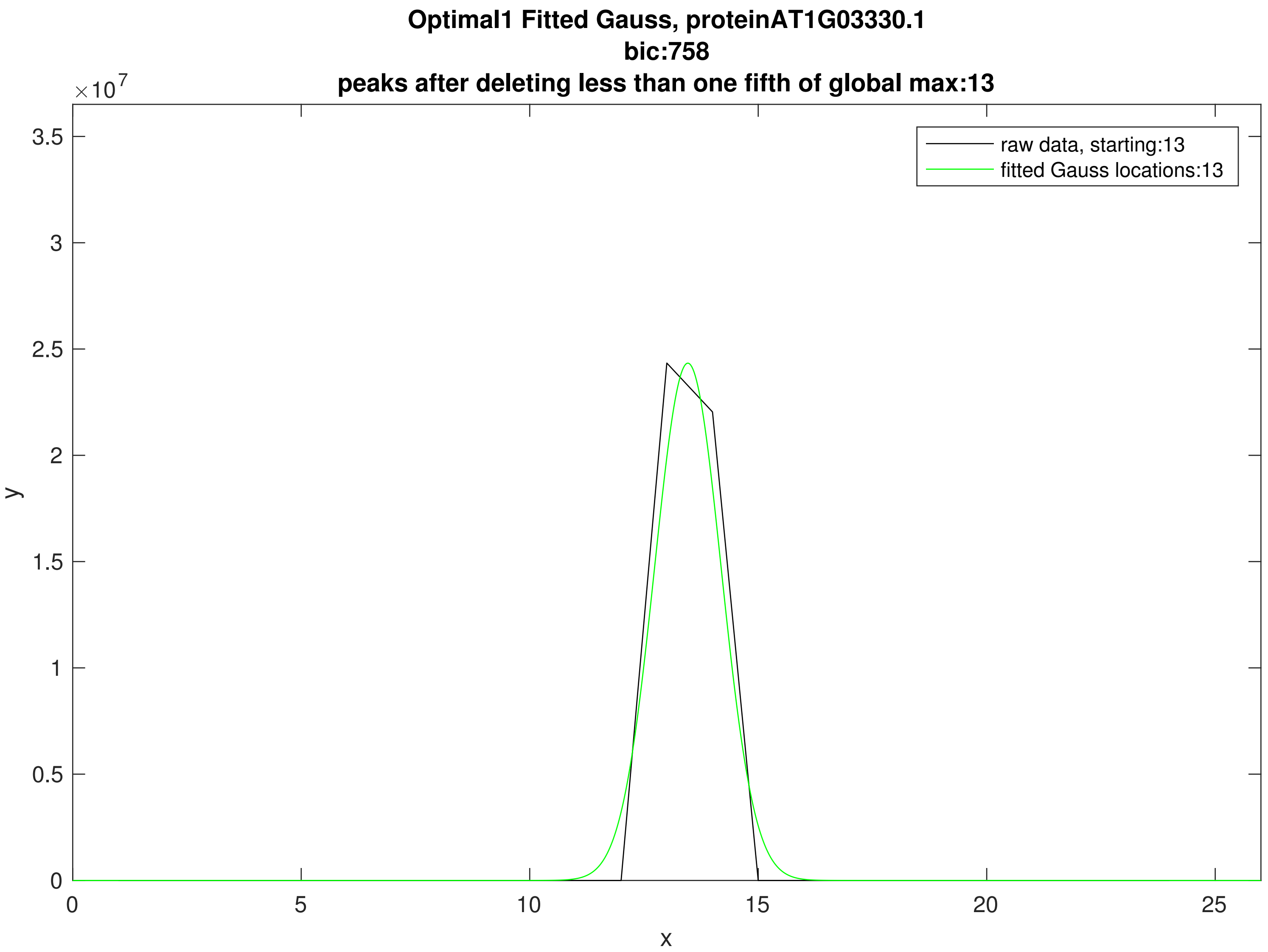
Optimal1 Fitted Gauss, proteinAT1G03330.1
bic:758
peaks after deleting less than one fifth of global max:13
×10^7
raw data, starting:13
fitted Gauss locations:13
3.5
3
2.5
2
1.5
1
0.5
0
y
0
5
10
15
20
25
x

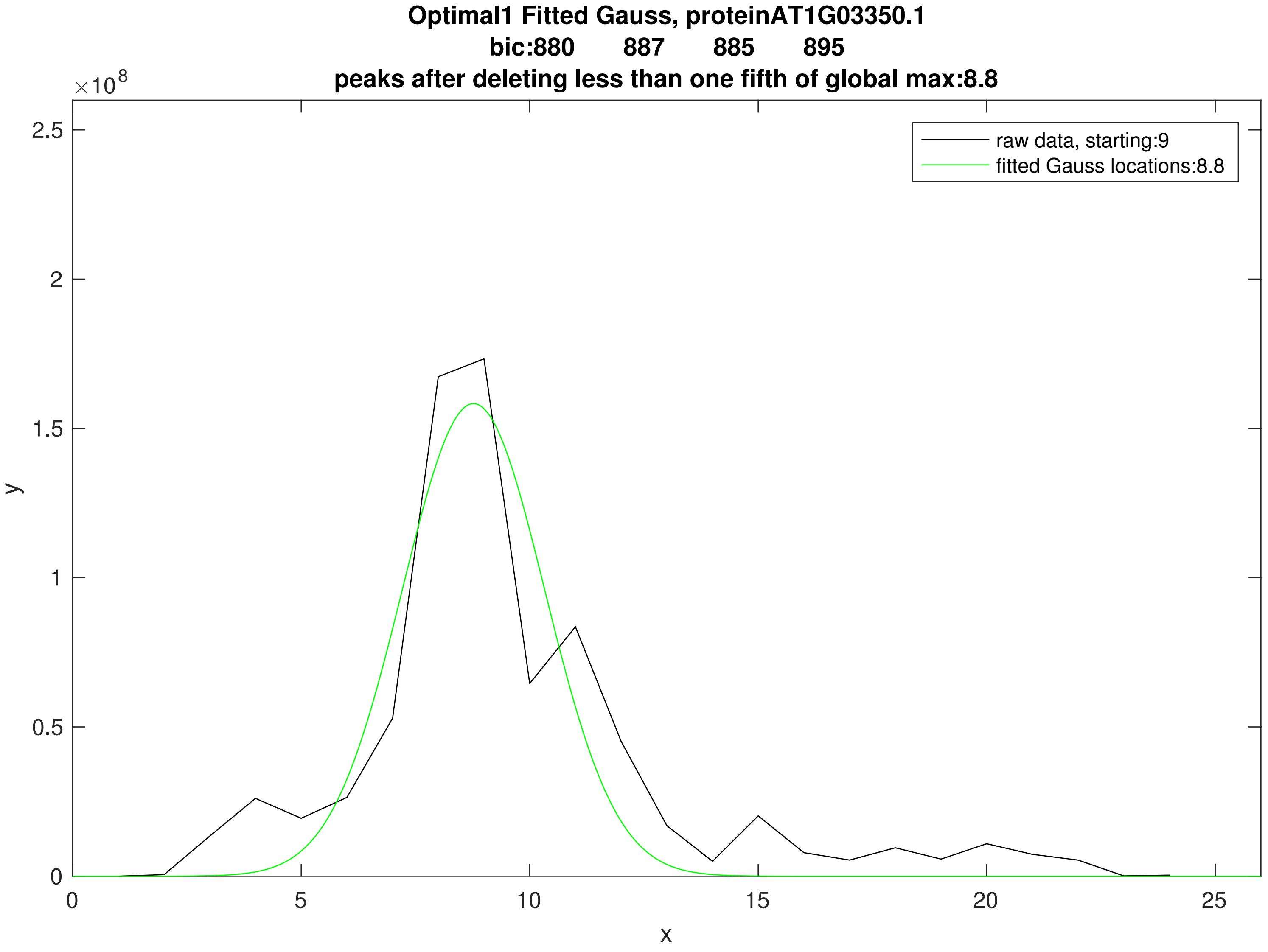

Optimal1 Fitted Gauss, proteinAT1G03350.1
bic:880 887 885 895
peaks after deleting less than one fifth of global max:8.8
×10^8
2.5
2
1.5
1
0.5
0
y
0
5
10
15
20
25
x
raw data, starting:9
fitted Gauss locations:8.8

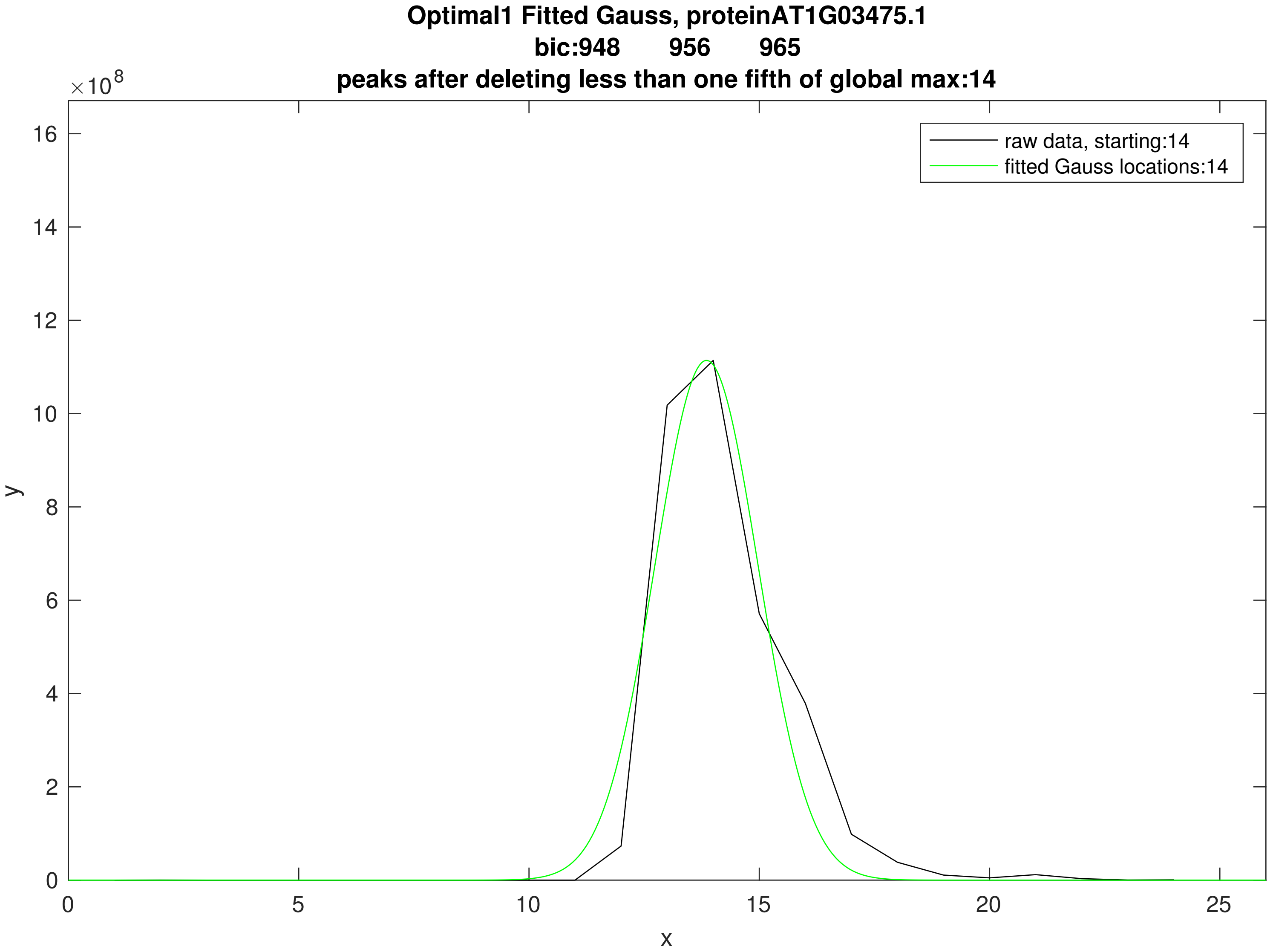

Optimal1 Fitted Gauss, proteinAT1G03475.1
bic:948 956 965
peaks after deleting less than one fifth of global max:14
×10^8
16
14
12
10
8
6
4
2
0
y
0
5
10
15
20
25
x
raw data, starting:14
fitted Gauss locations:14

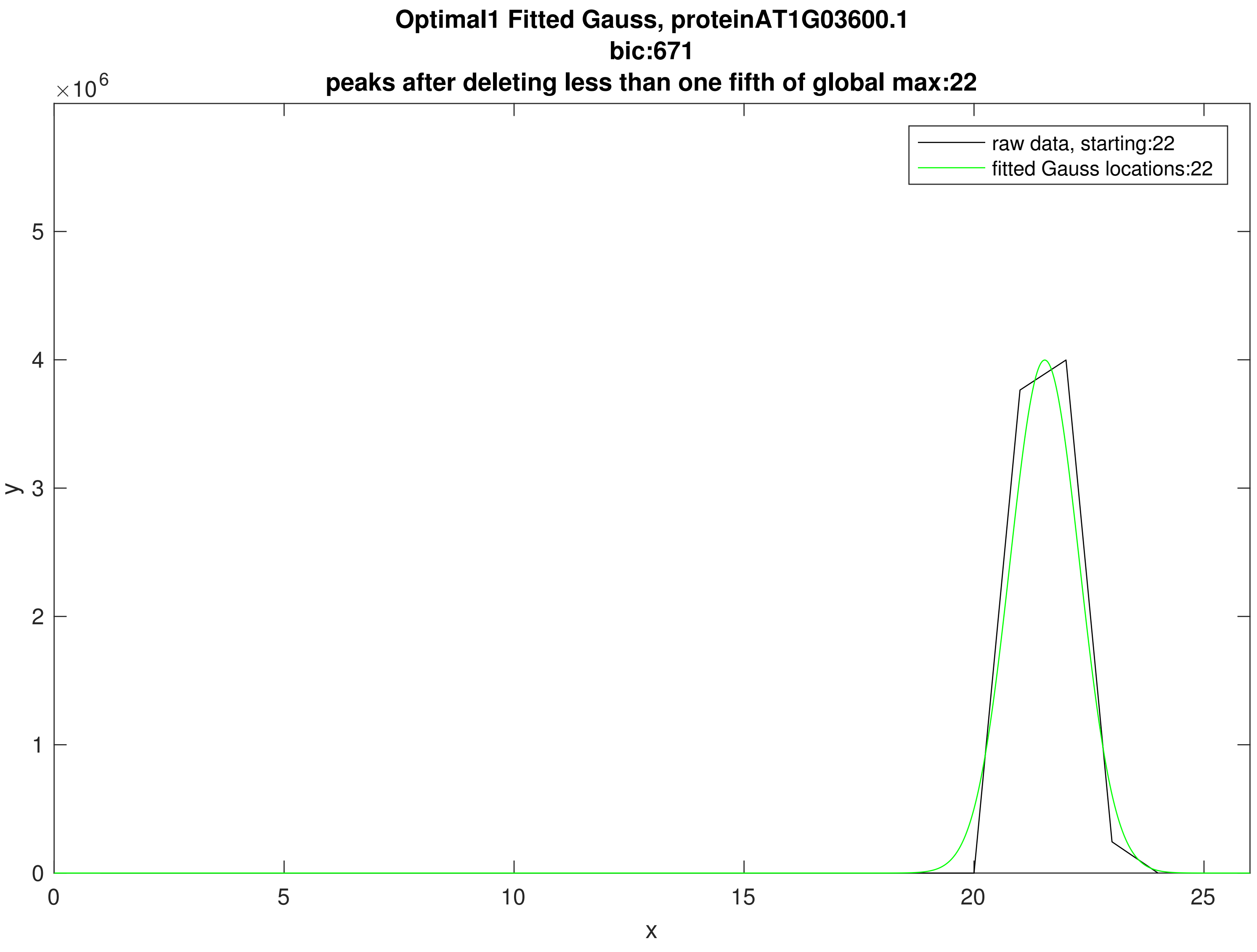
Optimal1 Fitted Gauss, proteinAT1G03600.1
bic:671
peaks after deleting less than one fifth of global max:22
×10^6
raw data, starting:22
fitted Gauss locations:22
y
x
0
1
2
3
4
5
0
5
10
15
20
25

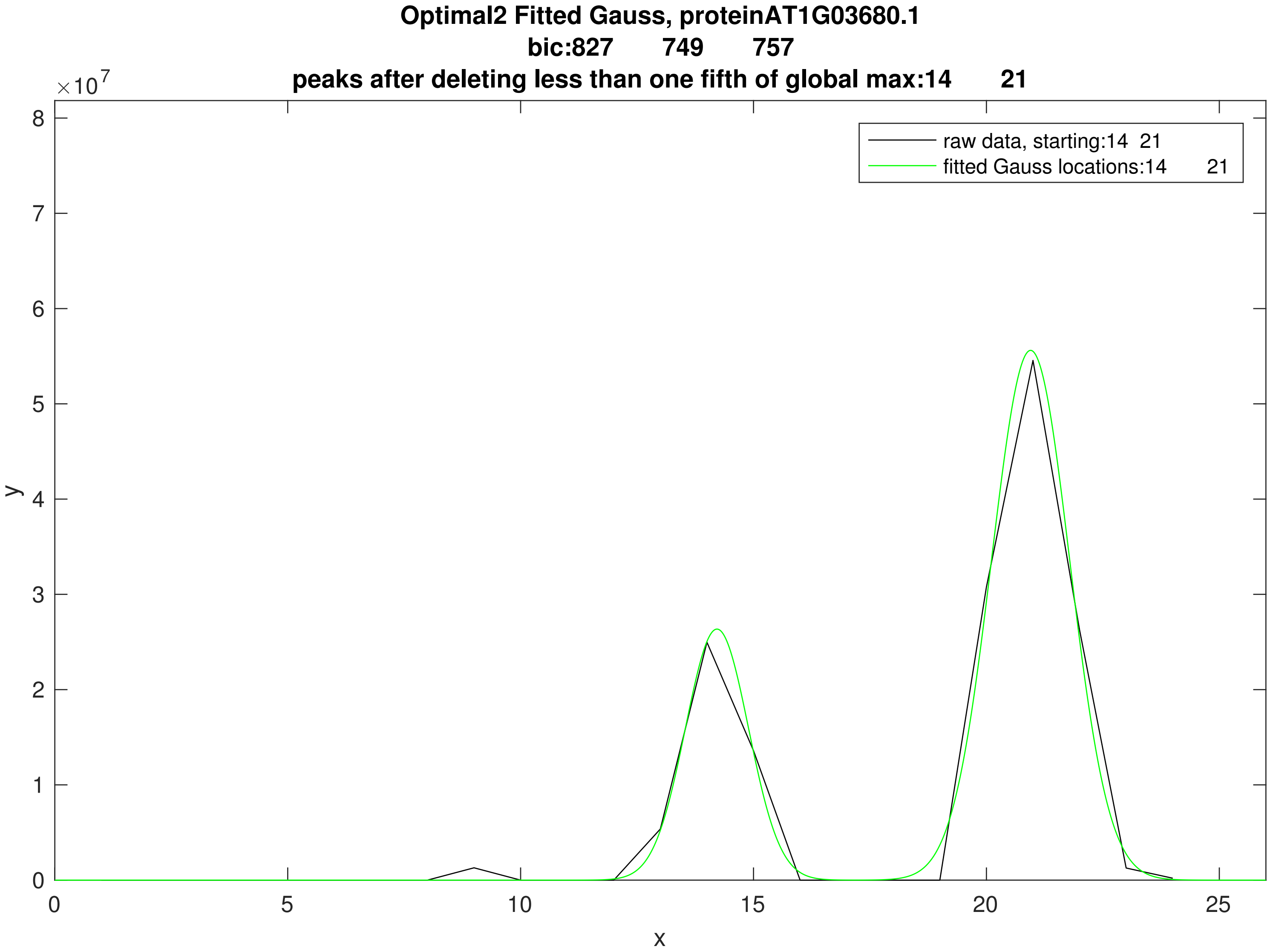
Optimal2 Fitted Gauss, proteinAT1G03680.1
bic:827 749 757
peaks after deleting less than one fifth of global max:14 21
raw data, starting:14 21
fitted Gauss locations:14 21
×10^7
y
x

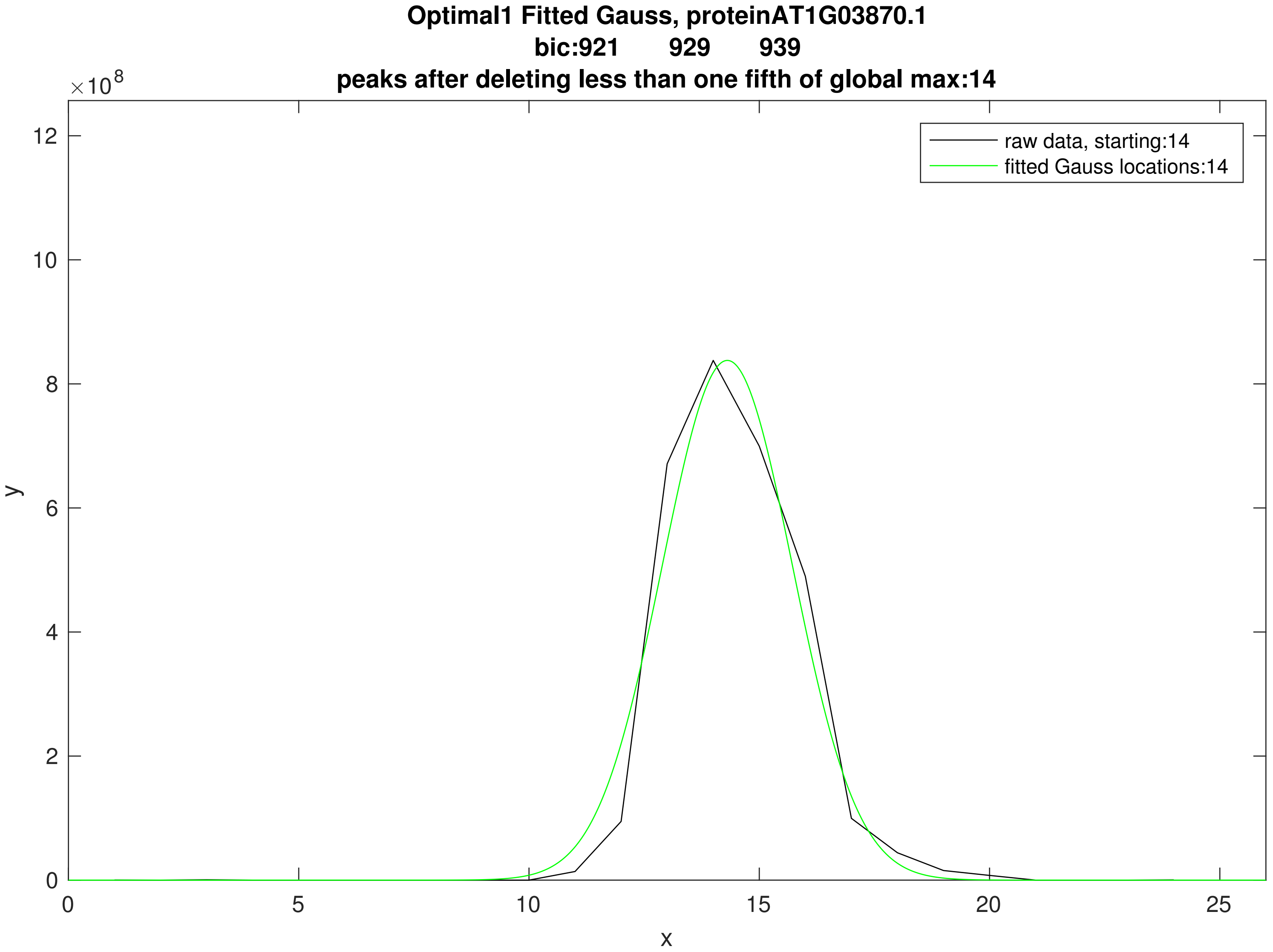
Optimal1 Fitted Gauss, proteinAT1G03870.1
bic:921 929 939
peaks after deleting less than one fifth of global max:14
×10^8
12
10
8
6
4
2
0
y
0
5
10
15
20
25
x
raw data, starting:14
fitted Gauss locations:14

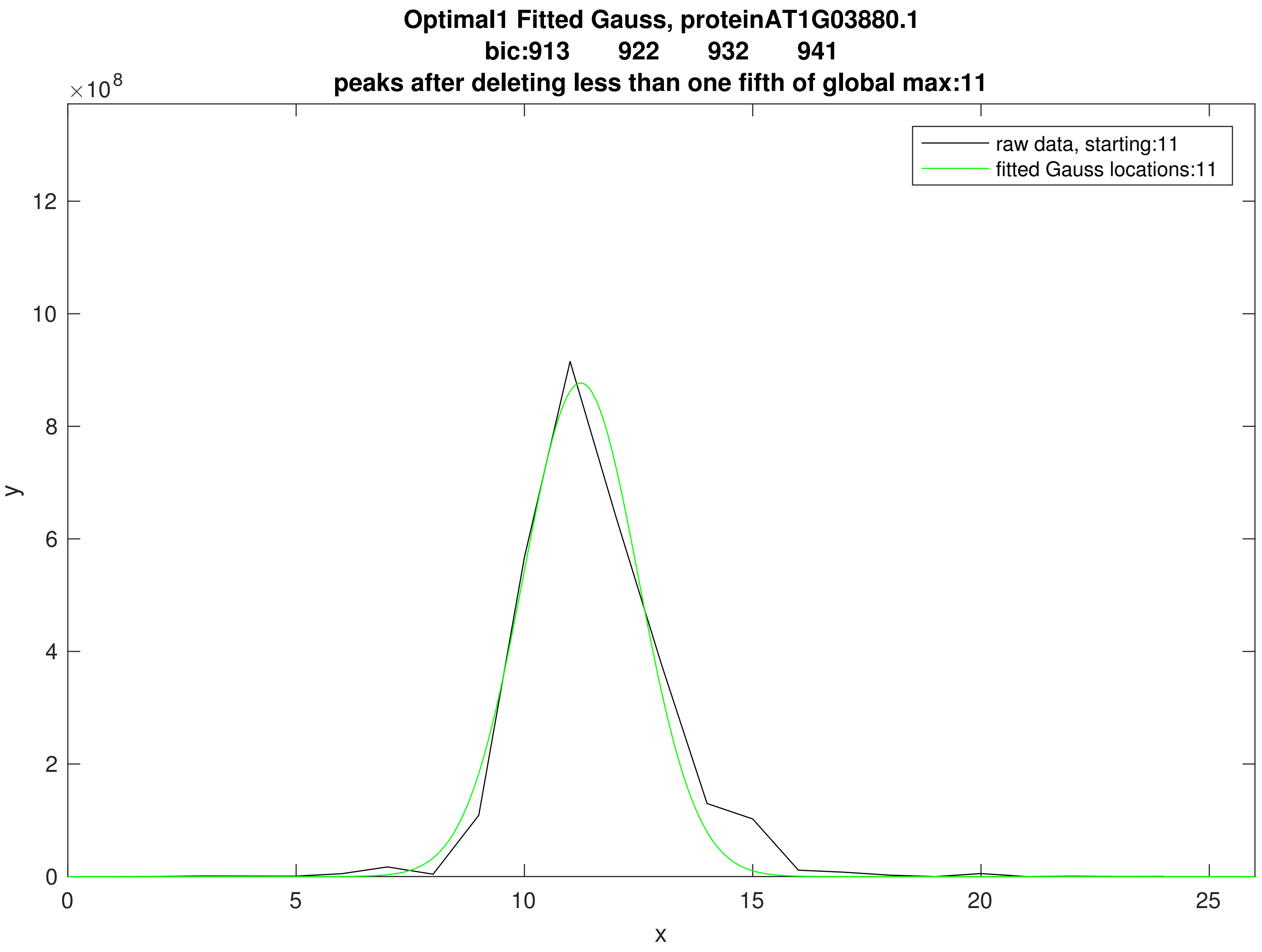
Optimal1 Fitted Gauss, proteinAT1G03880.1
bic:913 922 932 941
peaks after deleting less than one fifth of global max:11
×10^8
raw data, starting:11
fitted Gauss locations:11
12
10
8
6
4
2
0
y
0
5
10
15
20
25
x

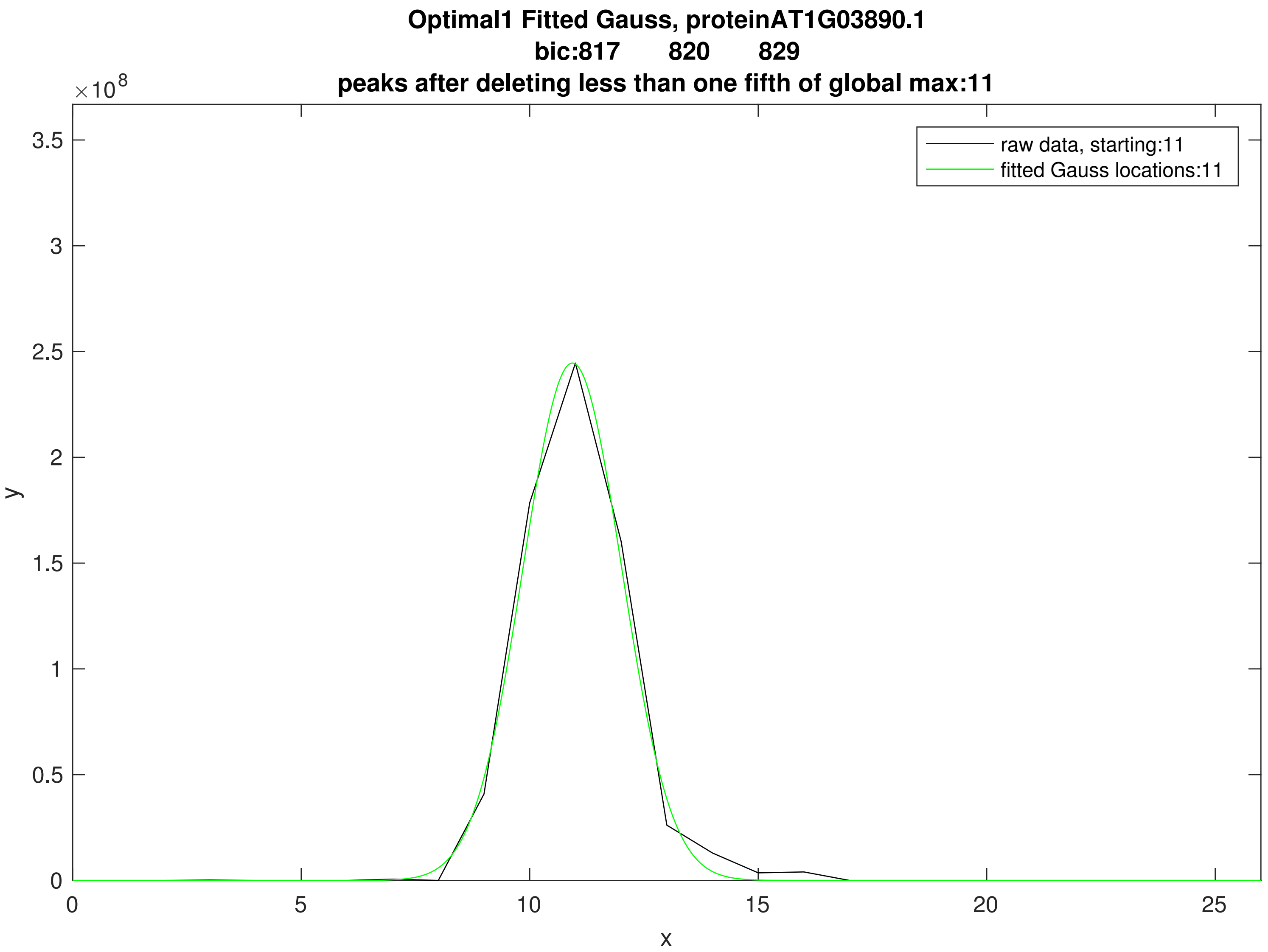

Optimal1 Fitted Gauss, proteinAT1G03890.1
bic:817 820 829
peaks after deleting less than one fifth of global max:11
×10^8
3.5
3
2.5
2
1.5
1
0.5
0
y
0
5
10
15
20
25
x
raw data, starting:11
fitted Gauss locations:11

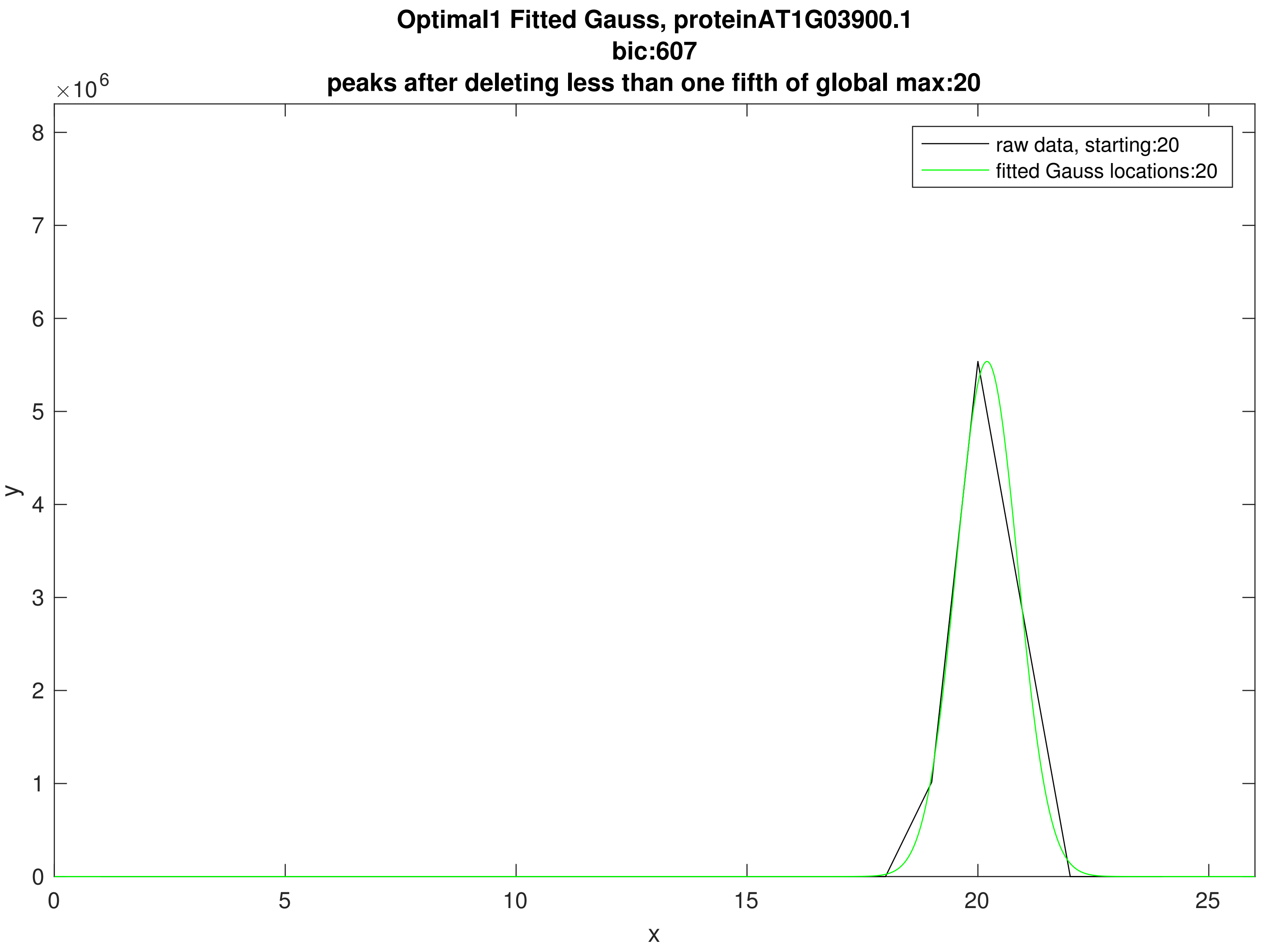
Optimal1 Fitted Gauss, proteinAT1G03900.1
bic:607
peaks after deleting less than one fifth of global max:20
×10^6
raw data, starting:20
fitted Gauss locations:20
y
x
0
1
2
3
4
5
6
7
8
0
5
10
15
20
25

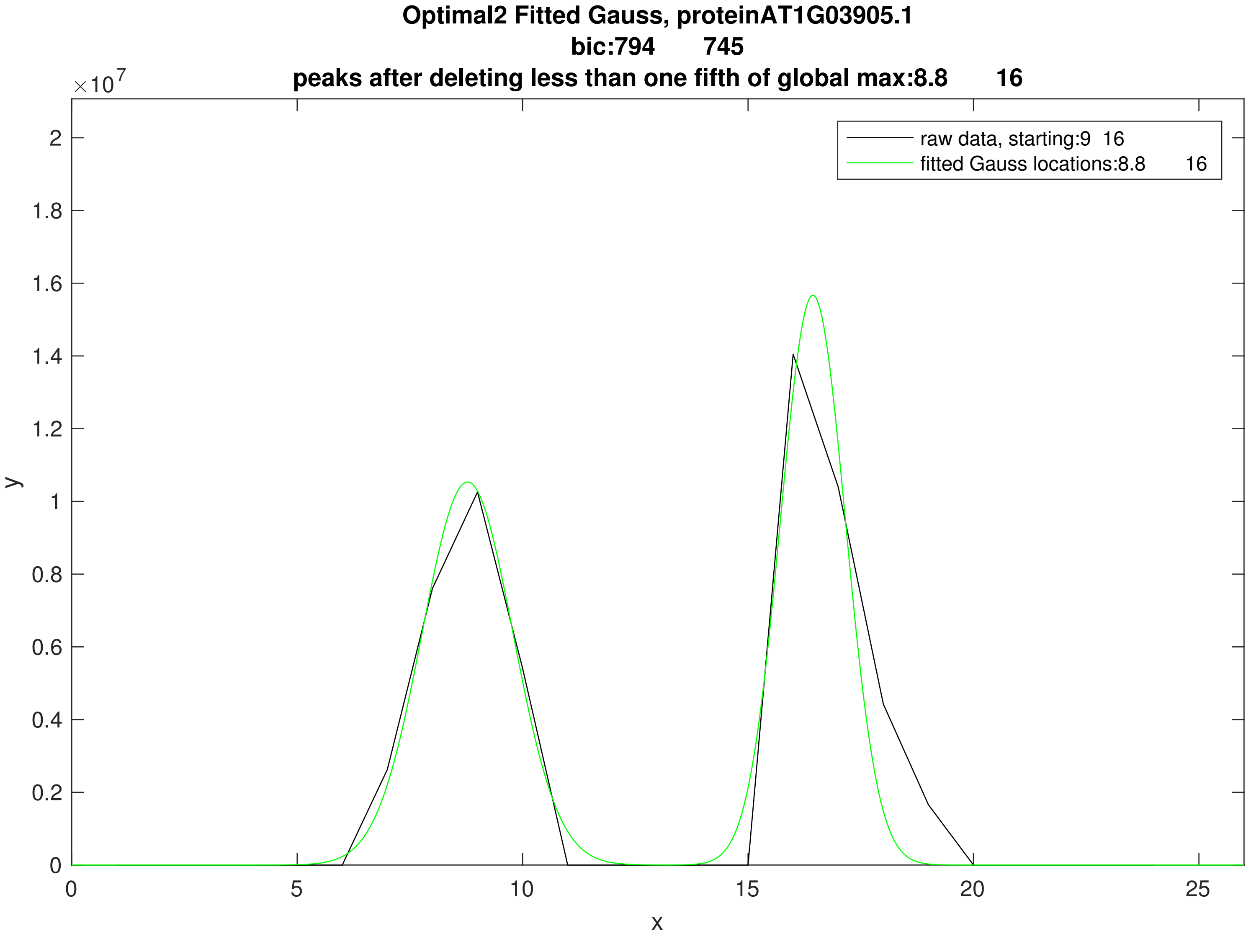

Optimal2 Fitted Gauss, proteinAT1G03905.1
bic:794 745
peaks after deleting less than one fifth of global max:8.8 16
×10^7
raw data, starting:9 16
fitted Gauss locations:8.8 16
2
1.8
1.6
1.4
1.2
1
0.8
0.6
0.4
0.2
0
y
0
5
10
15
20
25
x

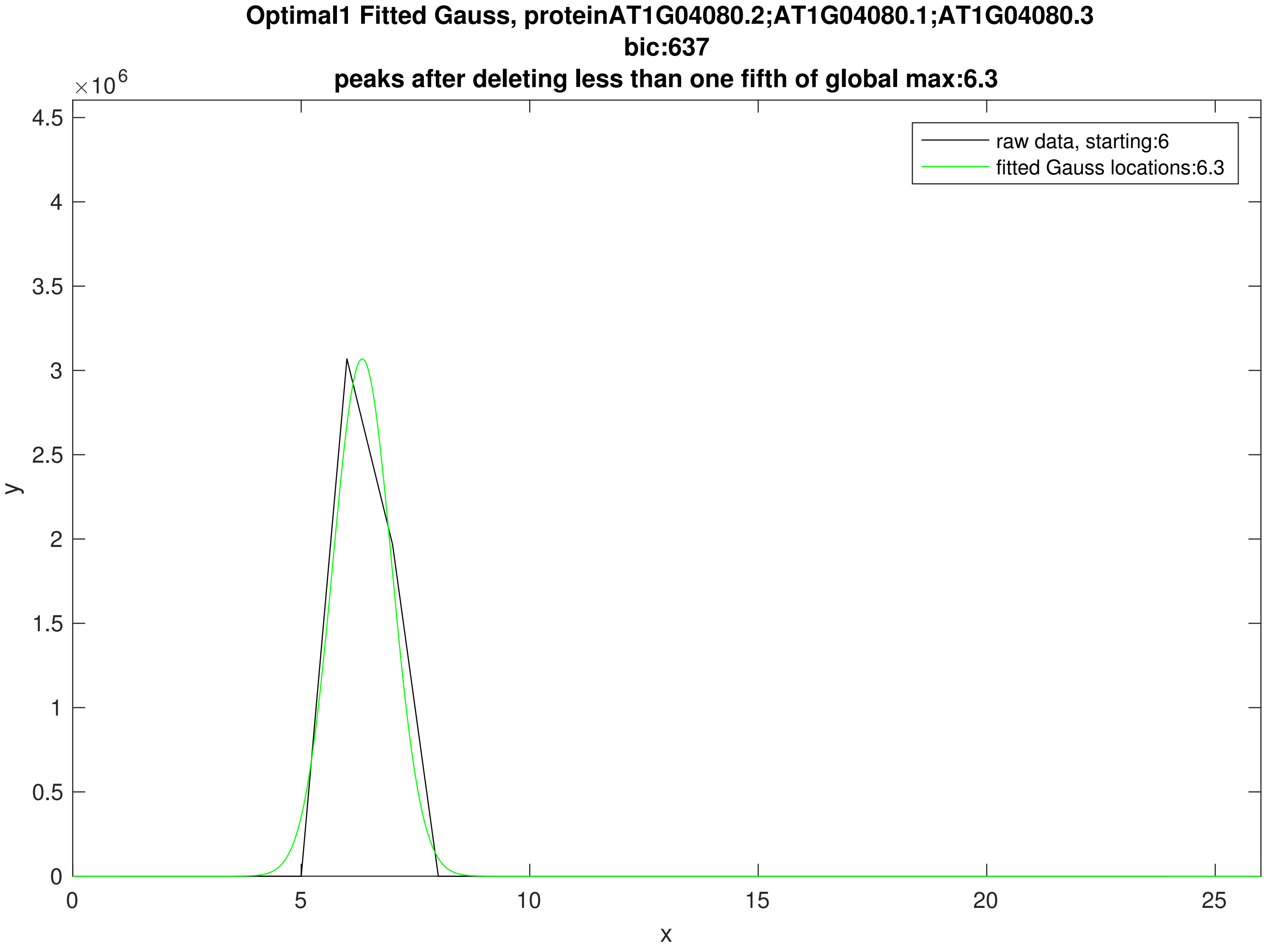
Optimal1 Fitted Gauss, proteinAT1G04080.2;AT1G04080.1;AT1G04080.3
bic:637
peaks after deleting less than one fifth of global max:6.3
×10^6
raw data, starting:6
fitted Gauss locations:6.3
y
x
0
0.5
1
1.5
2
2.5
3
3.5
4
4.5
5
10
15
20
25

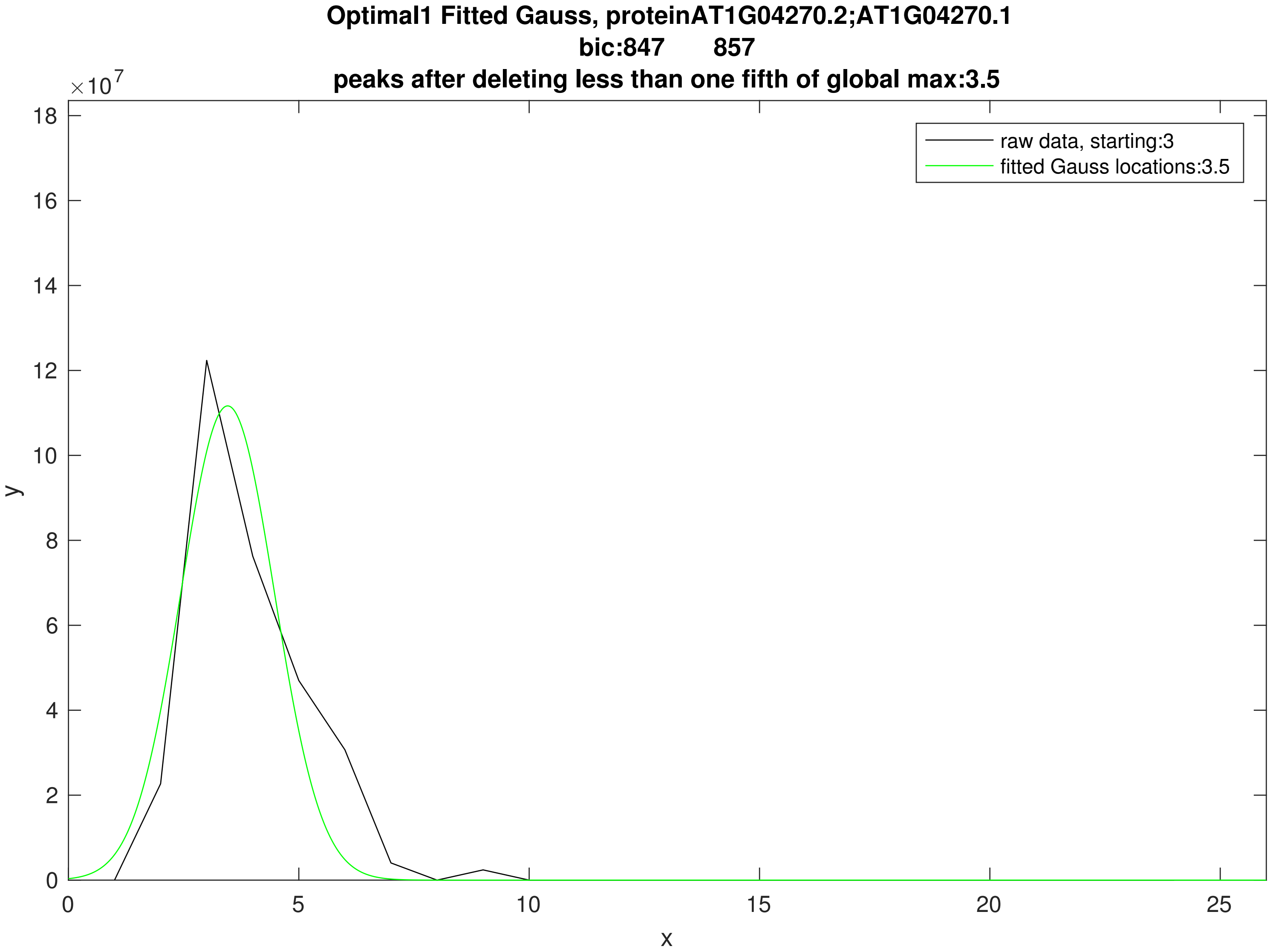

Optimal1 Fitted Gauss, proteinAT1G04270.2;AT1G04270.1
bic:847 857
peaks after deleting less than one fifth of global max:3.5
×10^7
18
16
14
12
10
8
6
4
2
0
y
0
5
10
15
20
25
x
raw data, starting:3
fitted Gauss locations:3.5

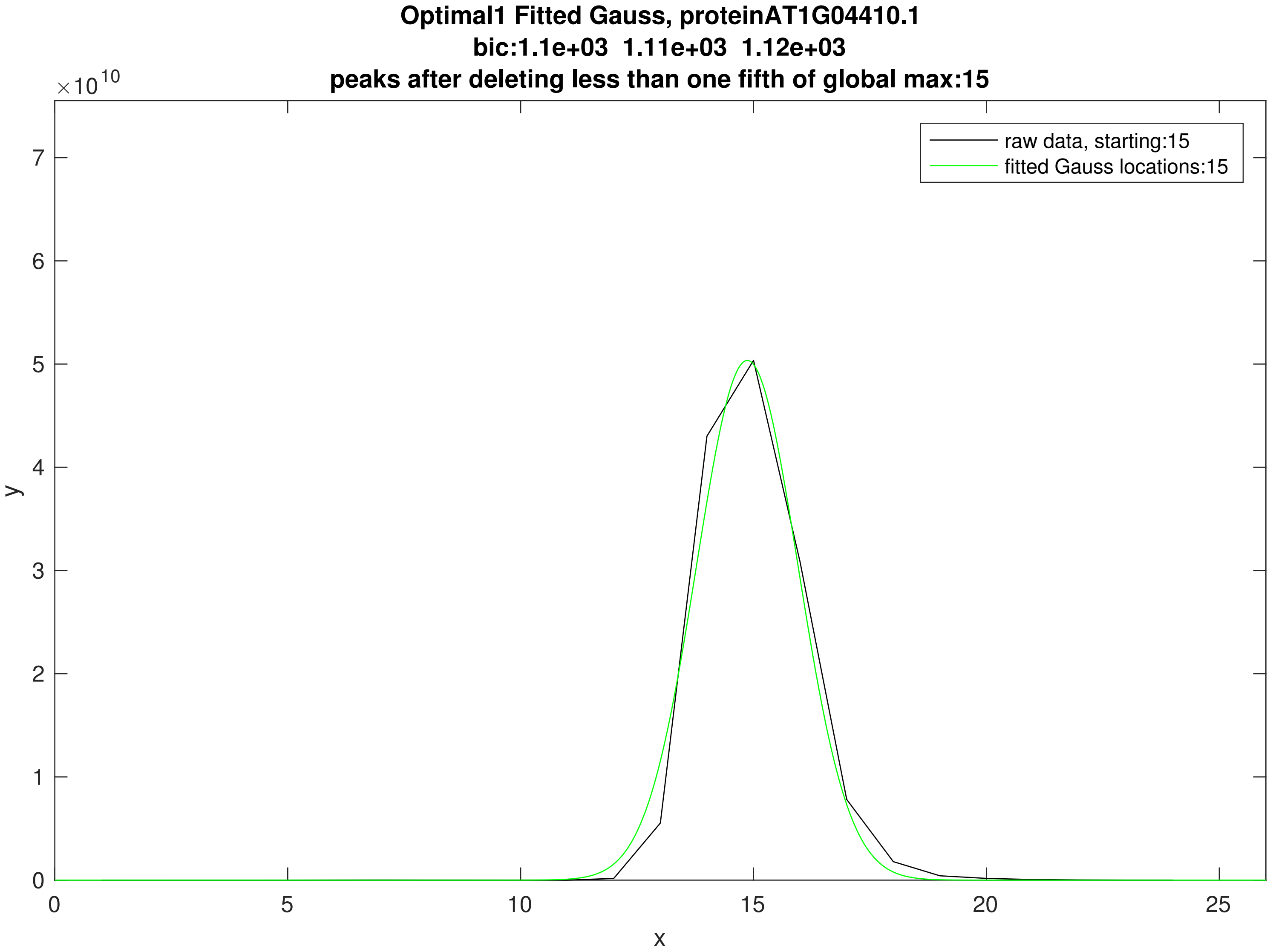
Optimal1 Fitted Gauss, proteinAT1G04410.1
bic:1.1e+03 1.11e+03 1.12e+03
peaks after deleting less than one fifth of global max:15
×10^10
raw data, starting:15
fitted Gauss locations:15
y
x
0
1
2
3
4
5
6
7
0
5
10
15
20
25

#### Optimal1 Fitted Gauss, proteinAT1G04420.1
#### bic:740
#### peaks after deleting less than one fifth of global max:14

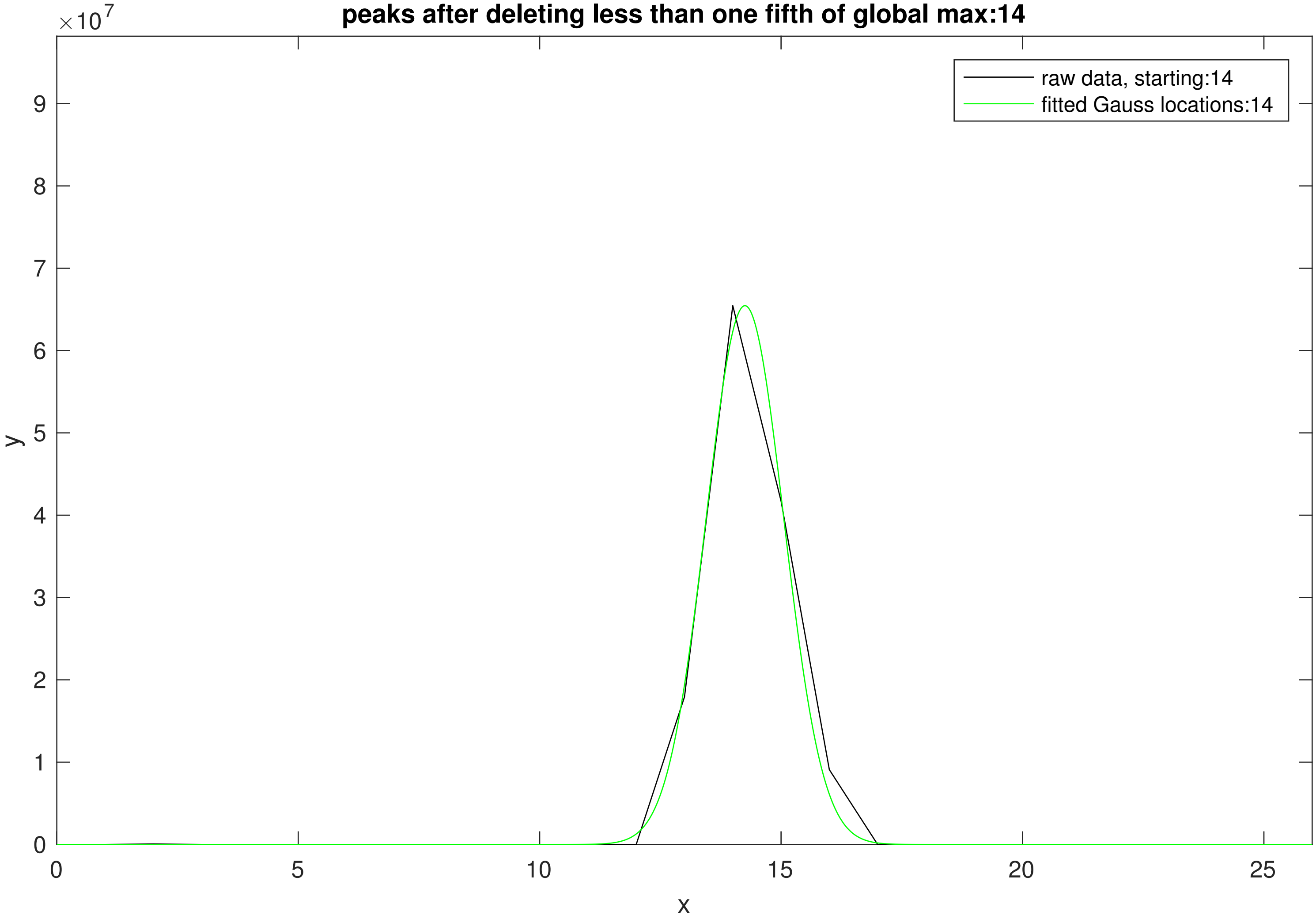

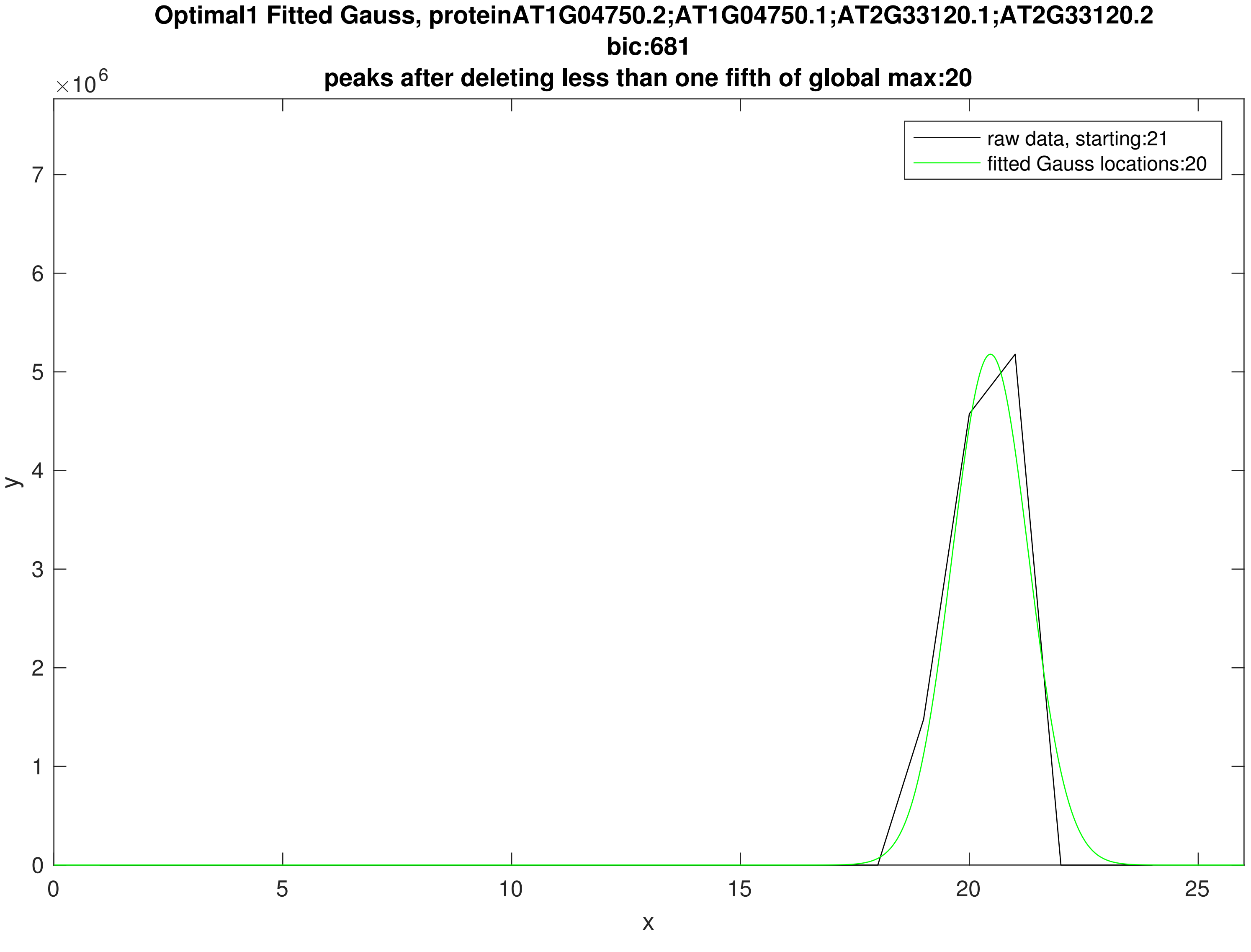

Optimal1 Fitted Gauss, proteinAT1G04750.2;AT1G04750.1;AT2G33120.1;AT2G33120.2
bic:681
peaks after deleting less than one fifth of global max:20
×10^6
raw data, starting:21
fitted Gauss locations:20
y
x
0
1
2
3
4
5
6
7
0
5
10
15
20
25

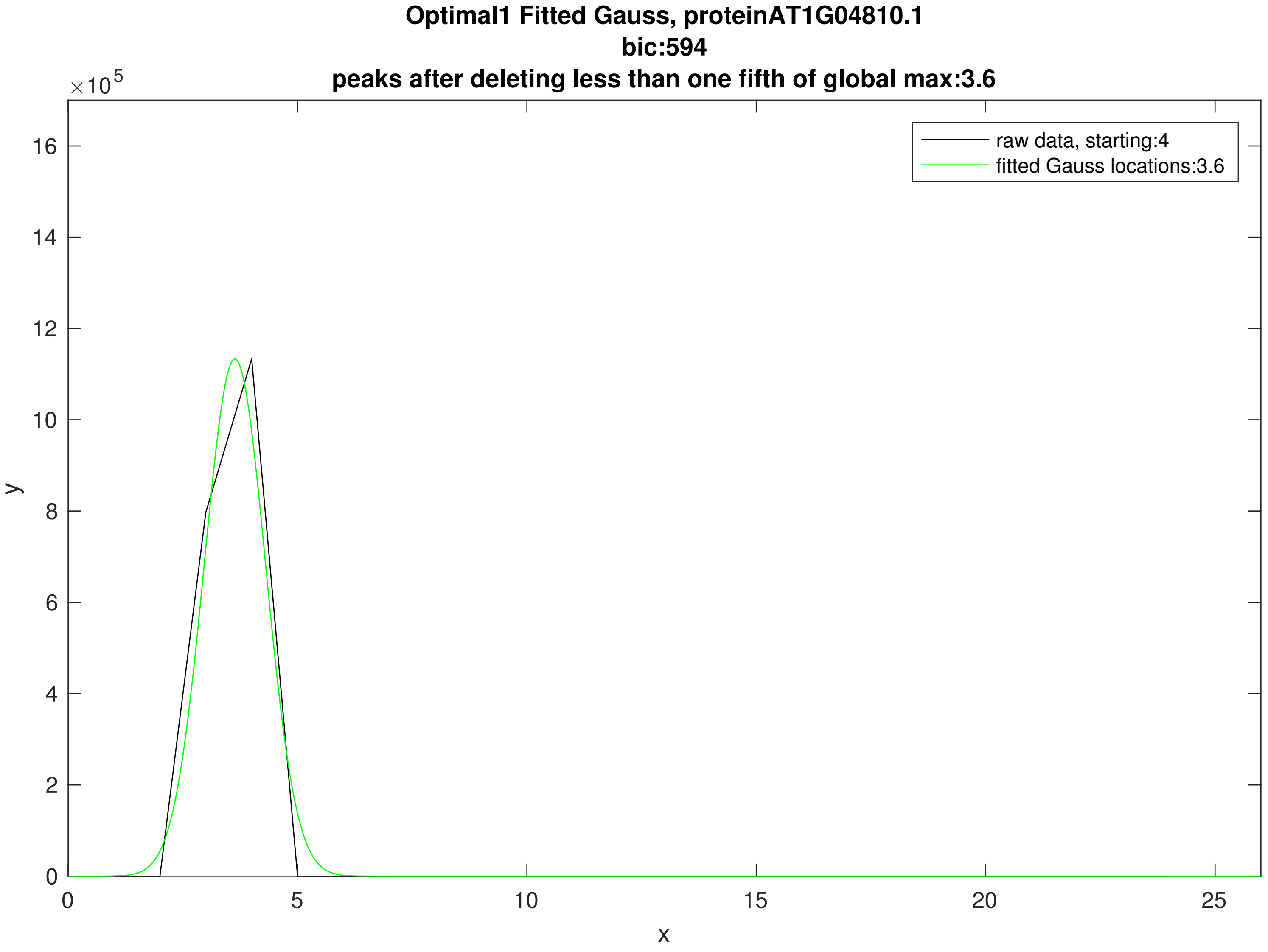
Optimal1 Fitted Gauss, proteinAT1G04810.1
bic:594
peaks after deleting less than one fifth of global max:3.6
×10^5
raw data, starting:4
fitted Gauss locations:3.6
y
x
0
2
4
6
8
10
12
14
16
0
5
10
15
20
25

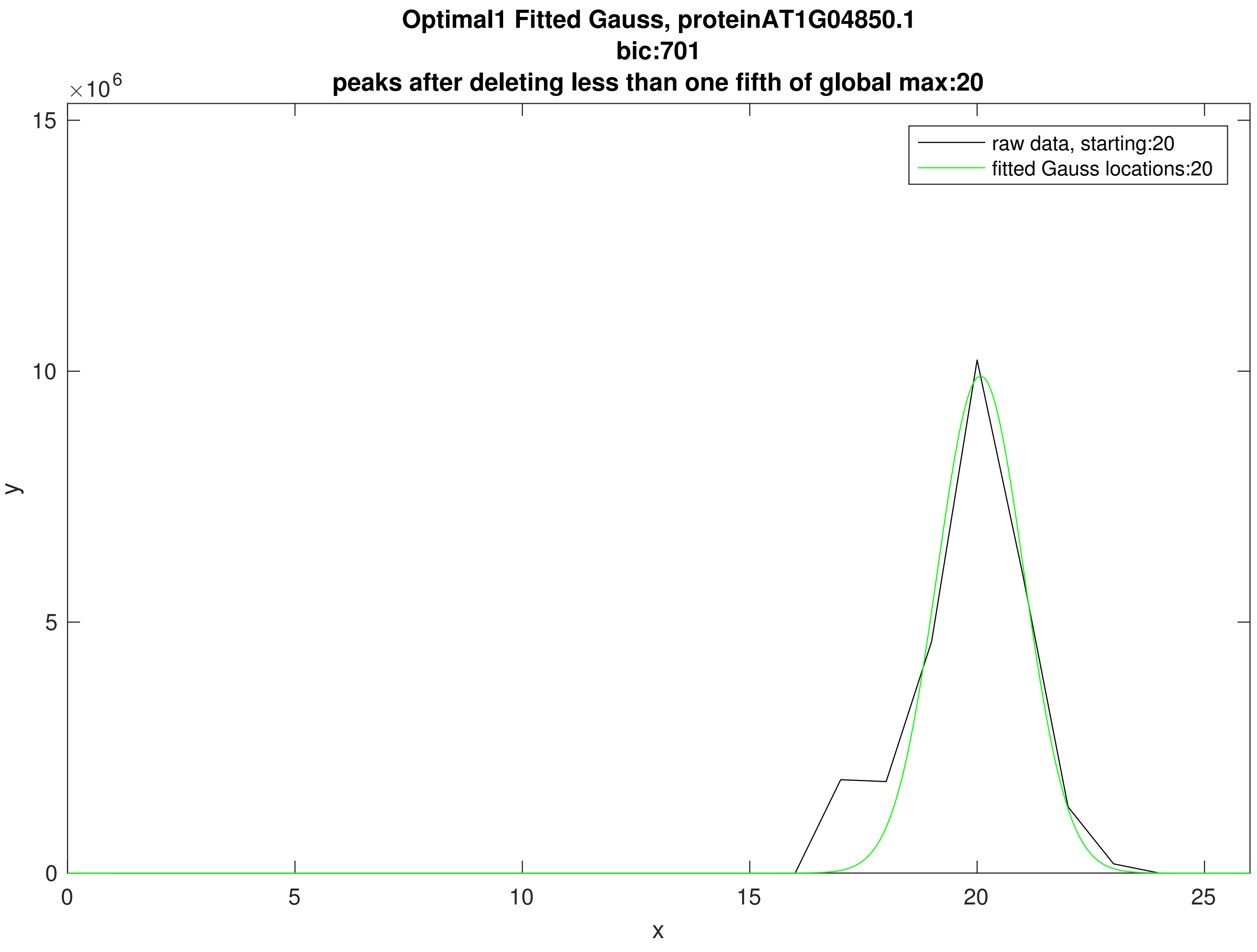

Optimal1 Fitted Gauss, proteinAT1G04850.1
bic:701
peaks after deleting less than one fifth of global max:20
×10^6
15
10
5
0
y
0
5
10
15
20
25
x
raw data, starting:20
fitted Gauss locations:20

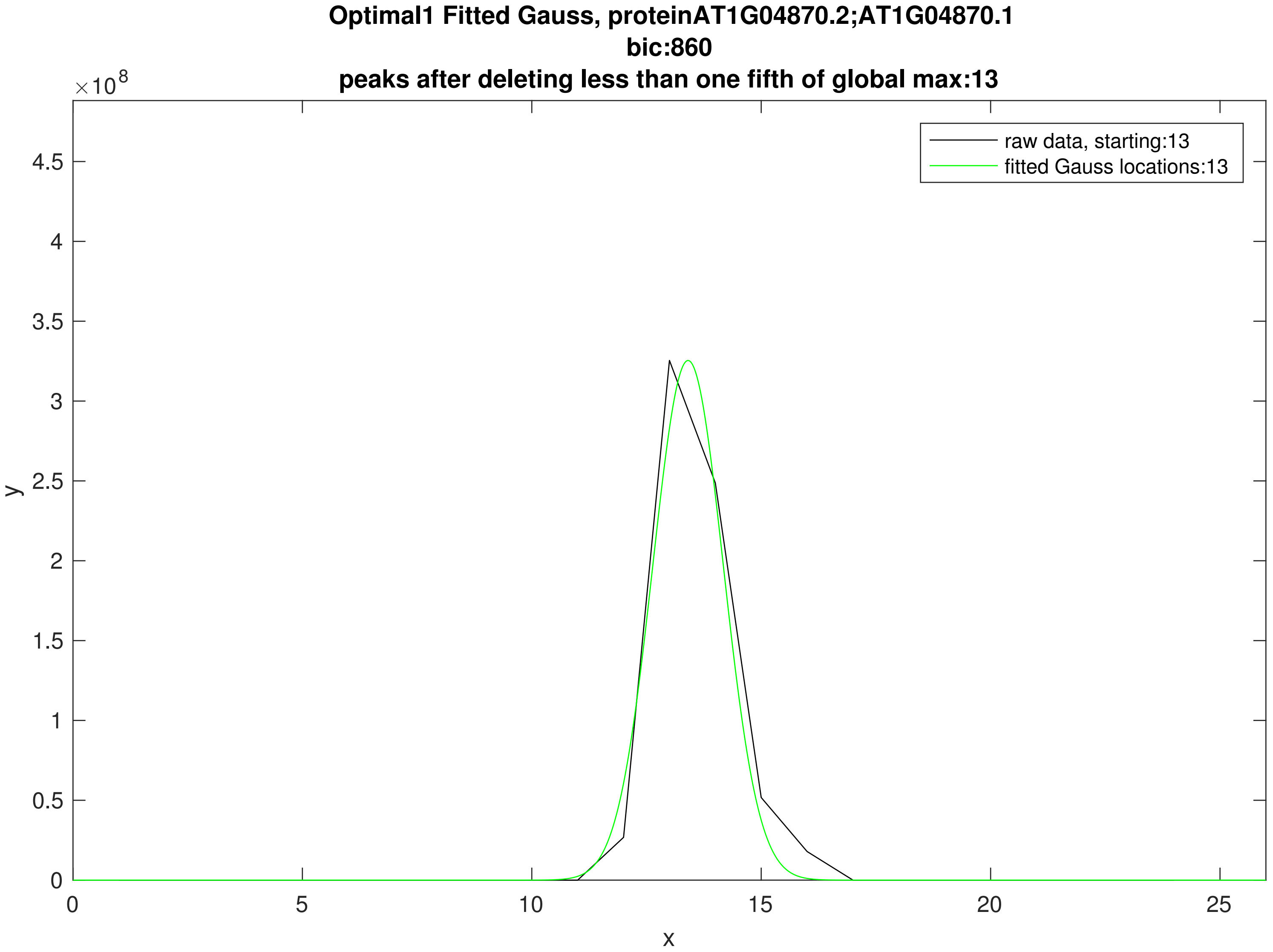
Optimal1 Fitted Gauss, proteinAT1G04870.2;AT1G04870.1
bic:860
peaks after deleting less than one fifth of global max:13
×10^8
raw data, starting:13
fitted Gauss locations:13
y
x
0
0.5
1
1.5
2
2.5
3
3.5
4
4.5
0
5
10
15
20
25

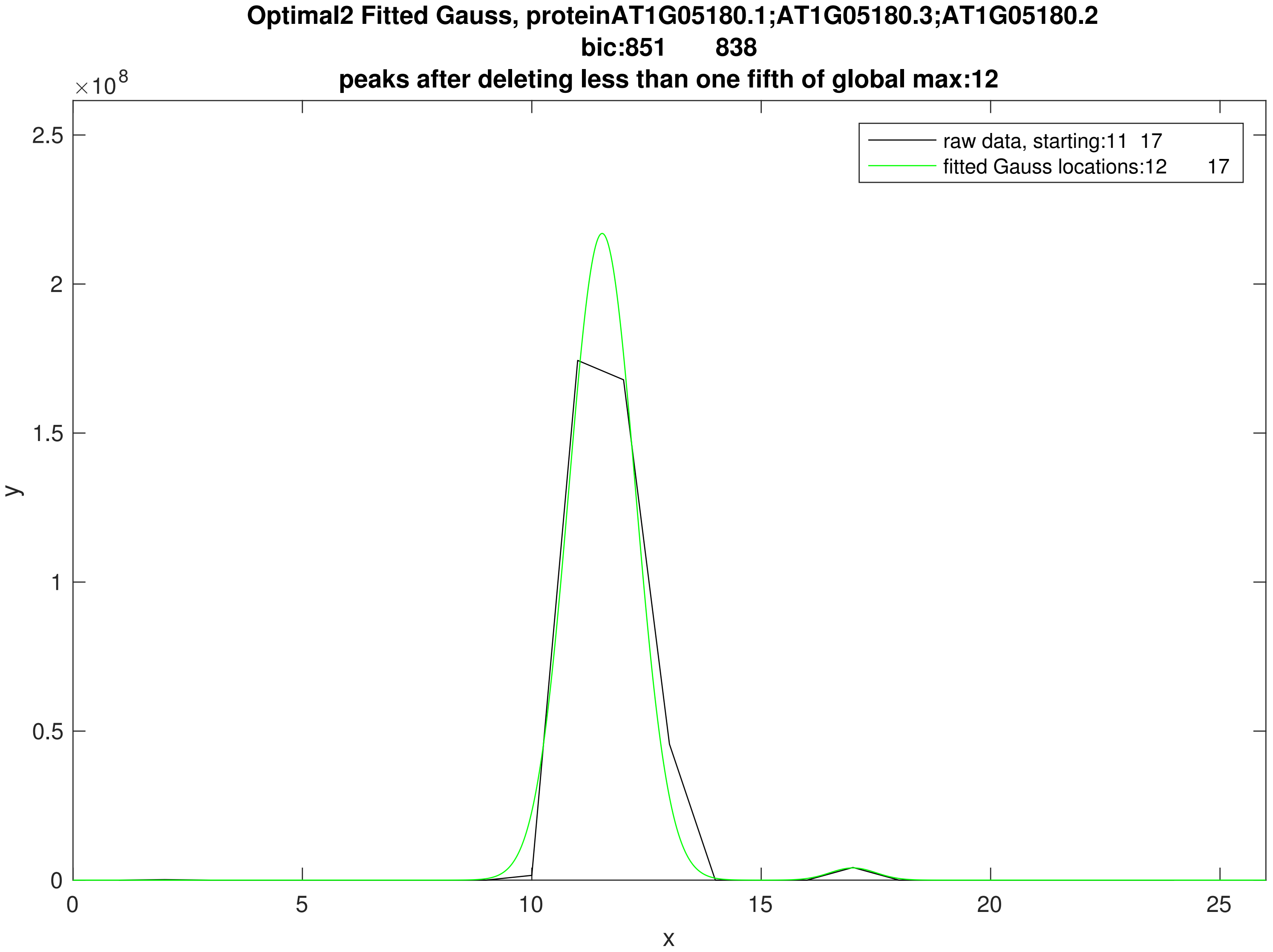
Optimal2 Fitted Gauss, proteinAT1G05180.1;AT1G05180.3;AT1G05180.2
bic:851 838
peaks after deleting less than one fifth of global max:12
×10^8
raw data, starting:11 17
fitted Gauss locations:12 17
y
x
0
0.5
1
1.5
2
2.5
5
10
15
20
25

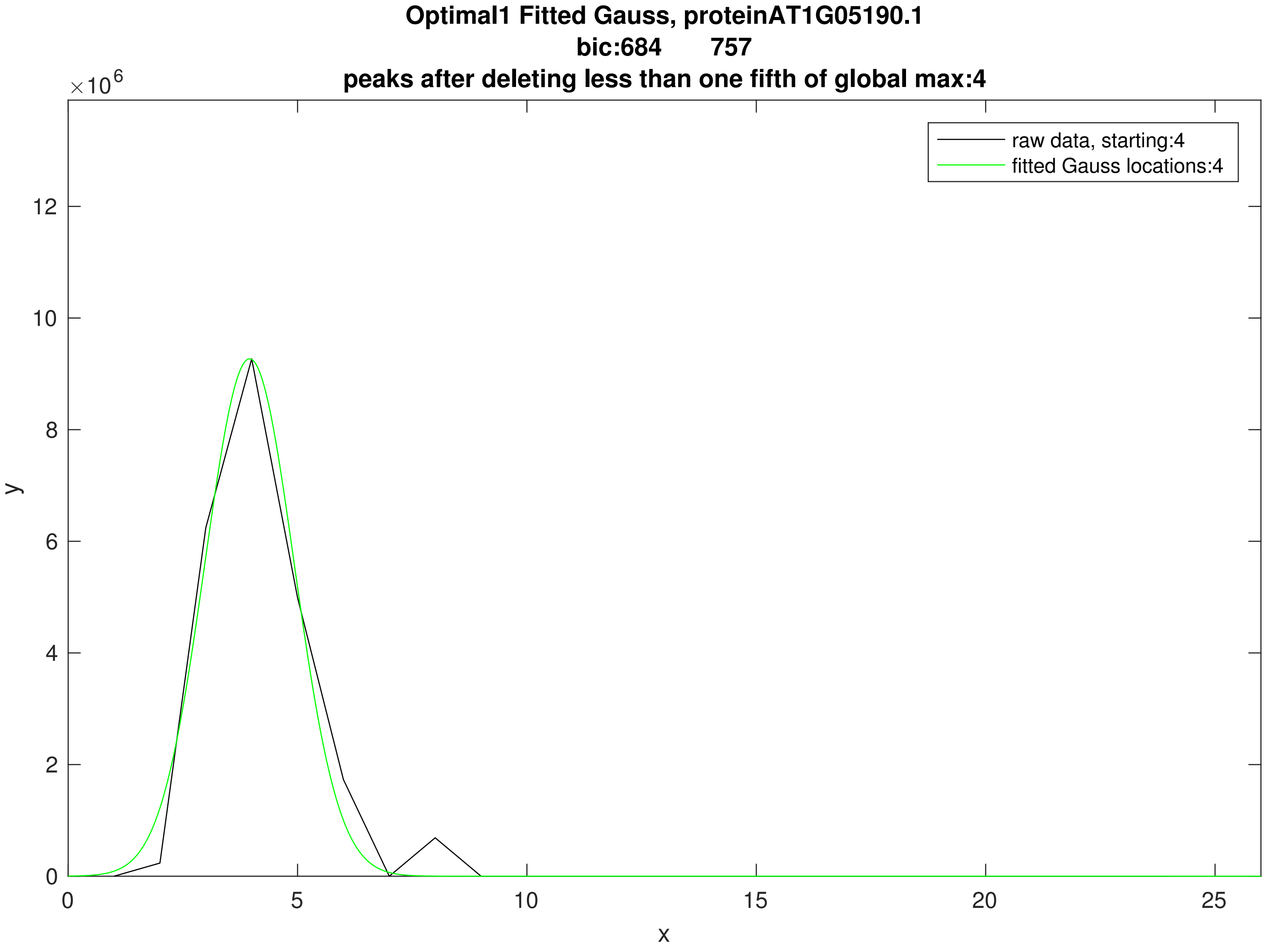
Optimal1 Fitted Gauss, proteinAT1G05190.1
bic:684 757
peaks after deleting less than one fifth of global max:4
×10^6
raw data, starting:4
fitted Gauss locations:4
y
x
0
2
4
6
8
10
12
0
5
10
15
20
25

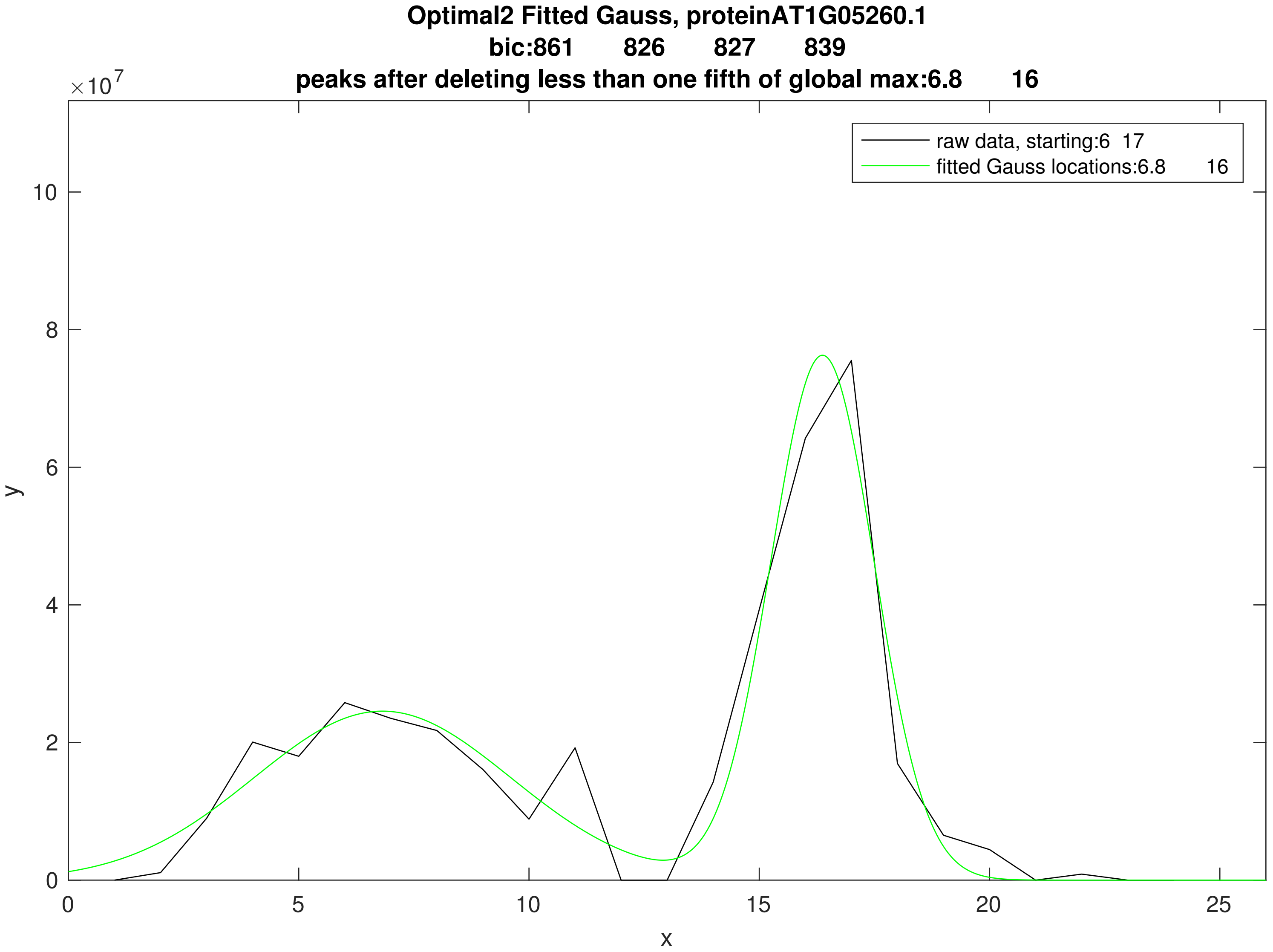

Optimal2 Fitted Gauss, proteinAT1G05260.1
bic:861 826 827 839
peaks after deleting less than one fifth of global max:6.8 16
×10^7
raw data, starting:6 17
fitted Gauss locations:6.8 16
10
8
6
4
2
0
y
0
5
10
15
20
25
x

Optimal1 Fitted Gauss, proteinAT1G05320.3;AT1G05320.2;AT1G05320.1
bic:677
peaks after deleting less than one fifth of global max:11

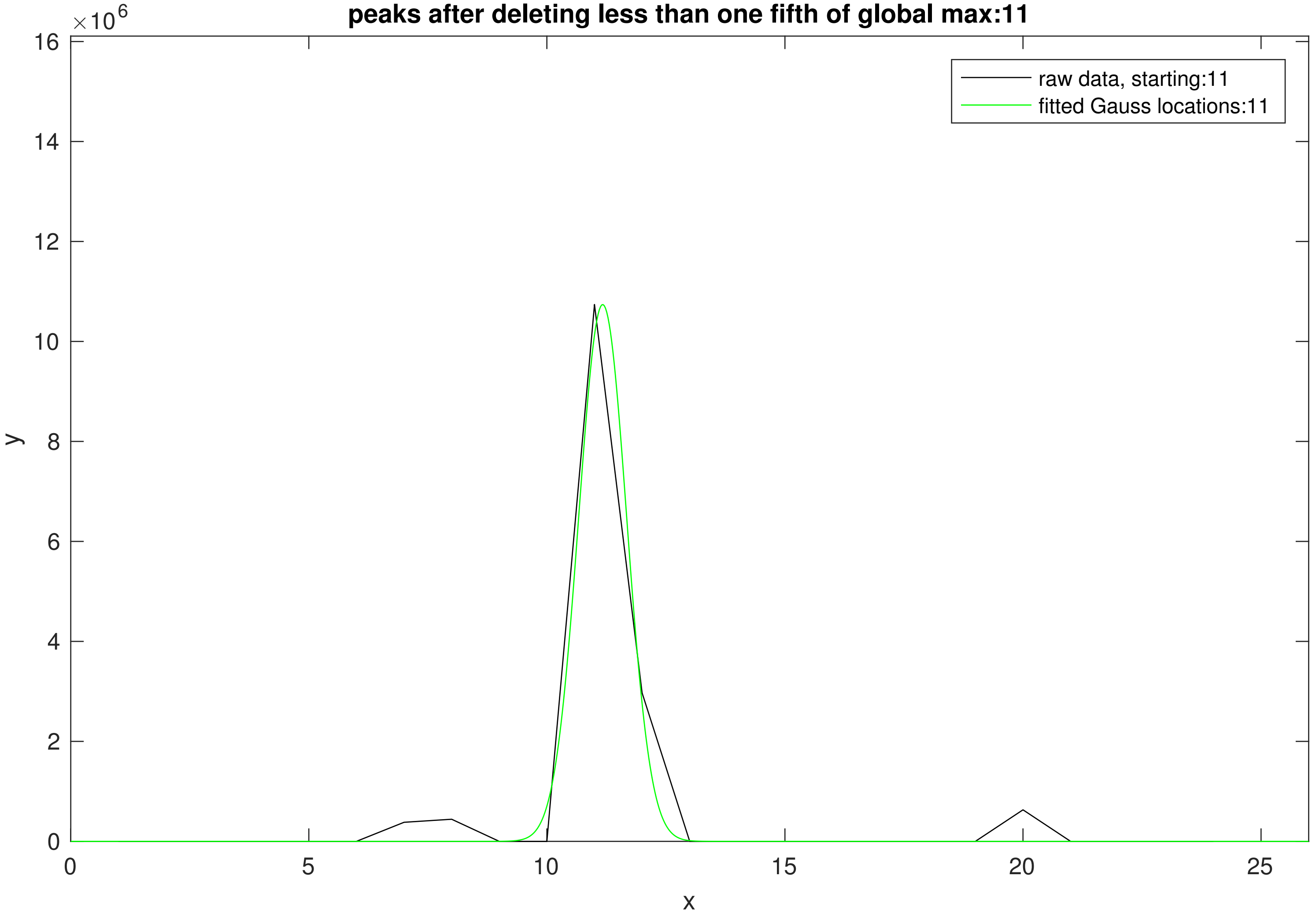

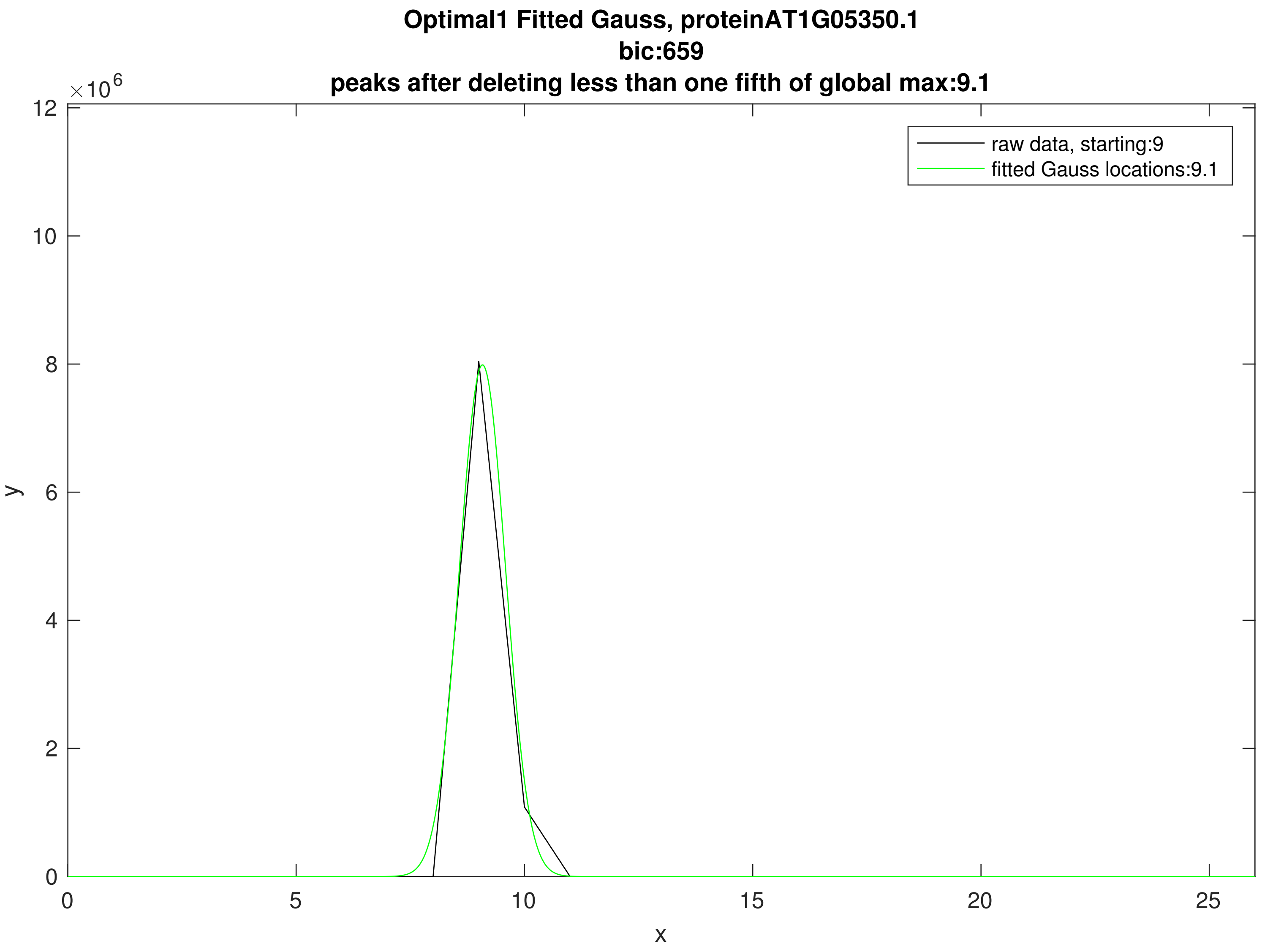

Optimal1 Fitted Gauss, proteinAT1G05350.1
bic:659
peaks after deleting less than one fifth of global max:9.1
×10^6
12
10
8
6
4
2
0
y
0
5
10
15
20
25
x
raw data, starting:9
fitted Gauss locations:9.1

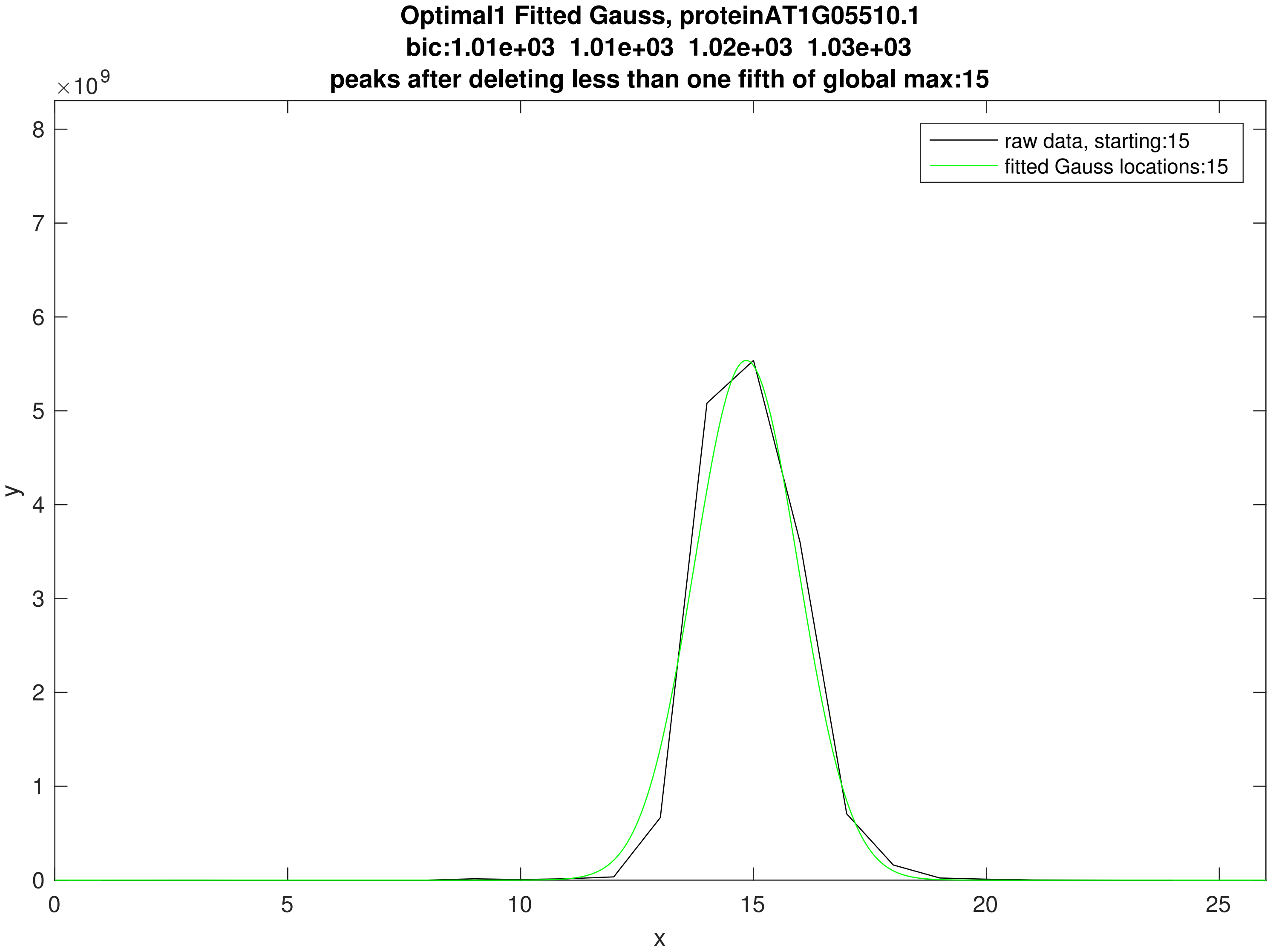

Optimal1 Fitted Gauss, proteinAT1G05510.1
bic:1.01e+03 1.01e+03 1.02e+03 1.03e+03
peaks after deleting less than one fifth of global max:15
×10^9
raw data, starting:15
fitted Gauss locations:15
y
x
0
1
2
3
4
5
6
7
8
0
5
10
15
20
25

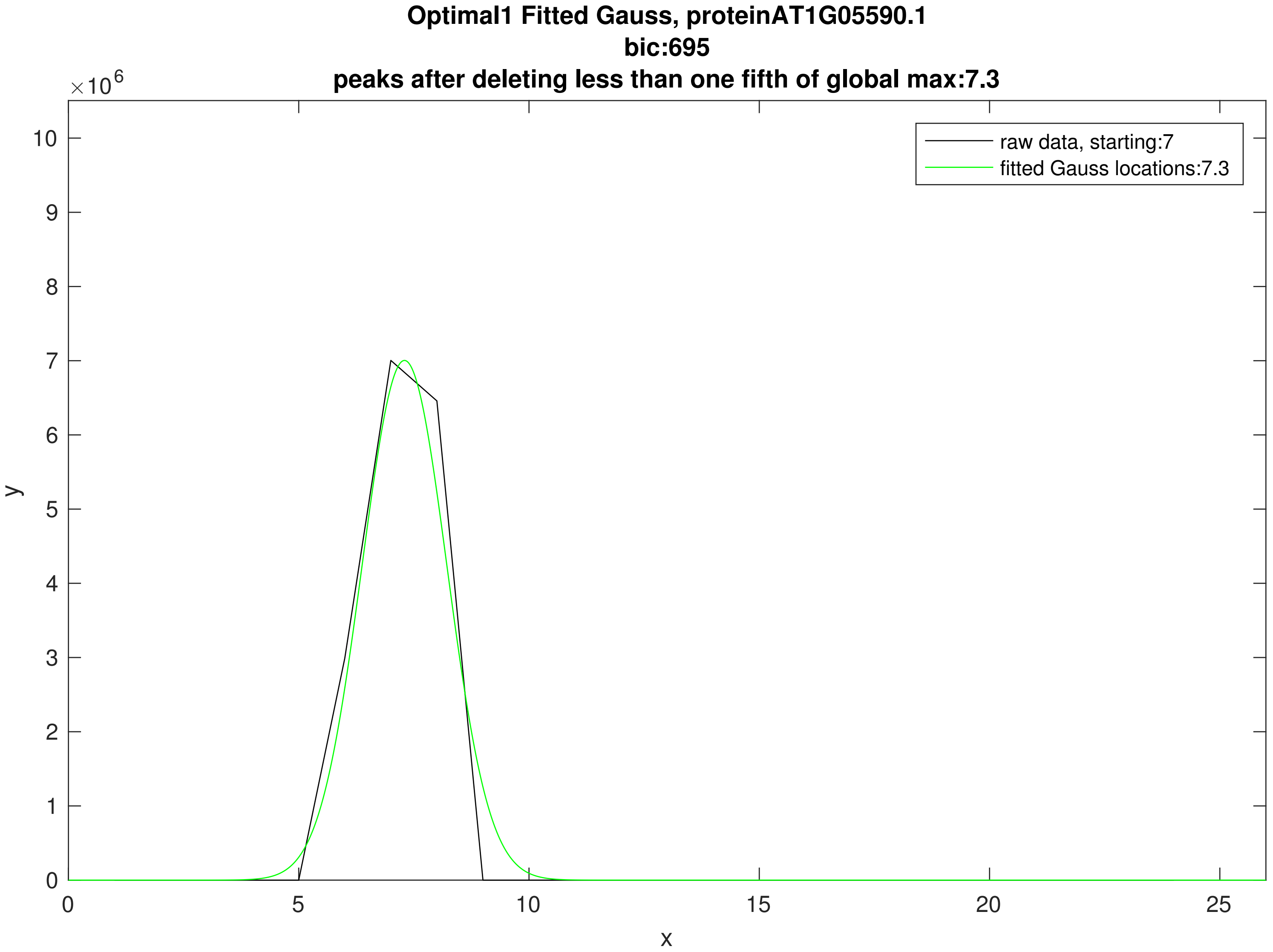
Optimal1 Fitted Gauss, proteinAT1G05590.1
bic:695
peaks after deleting less than one fifth of global max:7.3
×10^6
raw data, starting:7
fitted Gauss locations:7.3
y
x
0
1
2
3
4
5
6
7
8
9
10
0
5
10
15
20
25

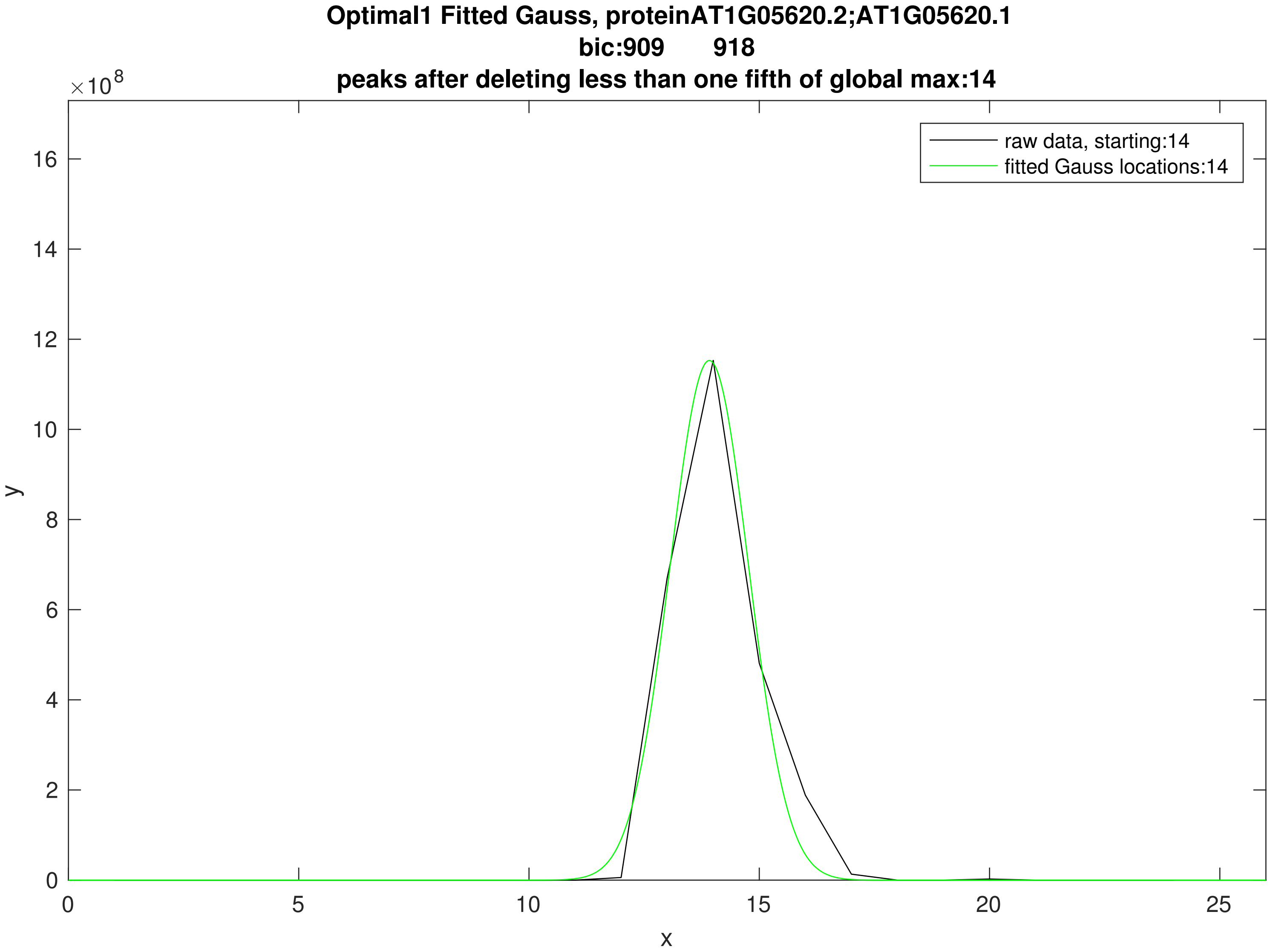
Optimal1 Fitted Gauss, proteinAT1G05620.2;AT1G05620.1
bic:909 918
peaks after deleting less than one fifth of global max:14
×10^8
16
14
12
10
8
6
4
2
0
y
0
5
10
15
20
25
x
raw data, starting:14
fitted Gauss locations:14

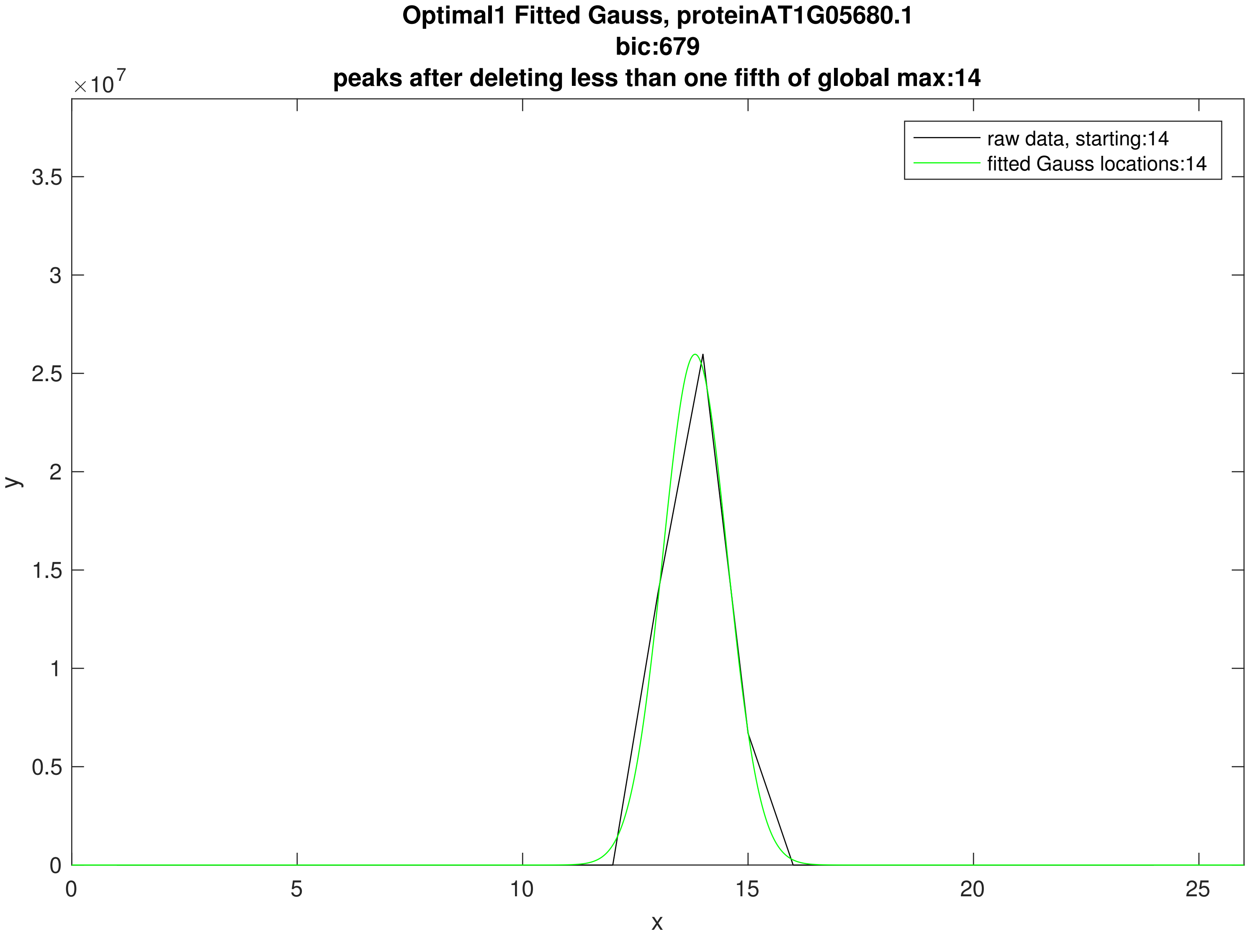
Optimal1 Fitted Gauss, proteinAT1G05680.1
bic:679
peaks after deleting less than one fifth of global max:14
×10^7
3.5
3
2.5
2
1.5
1
0.5
0
y
0
5
10
15
20
25
x
raw data, starting:14
fitted Gauss locations:14

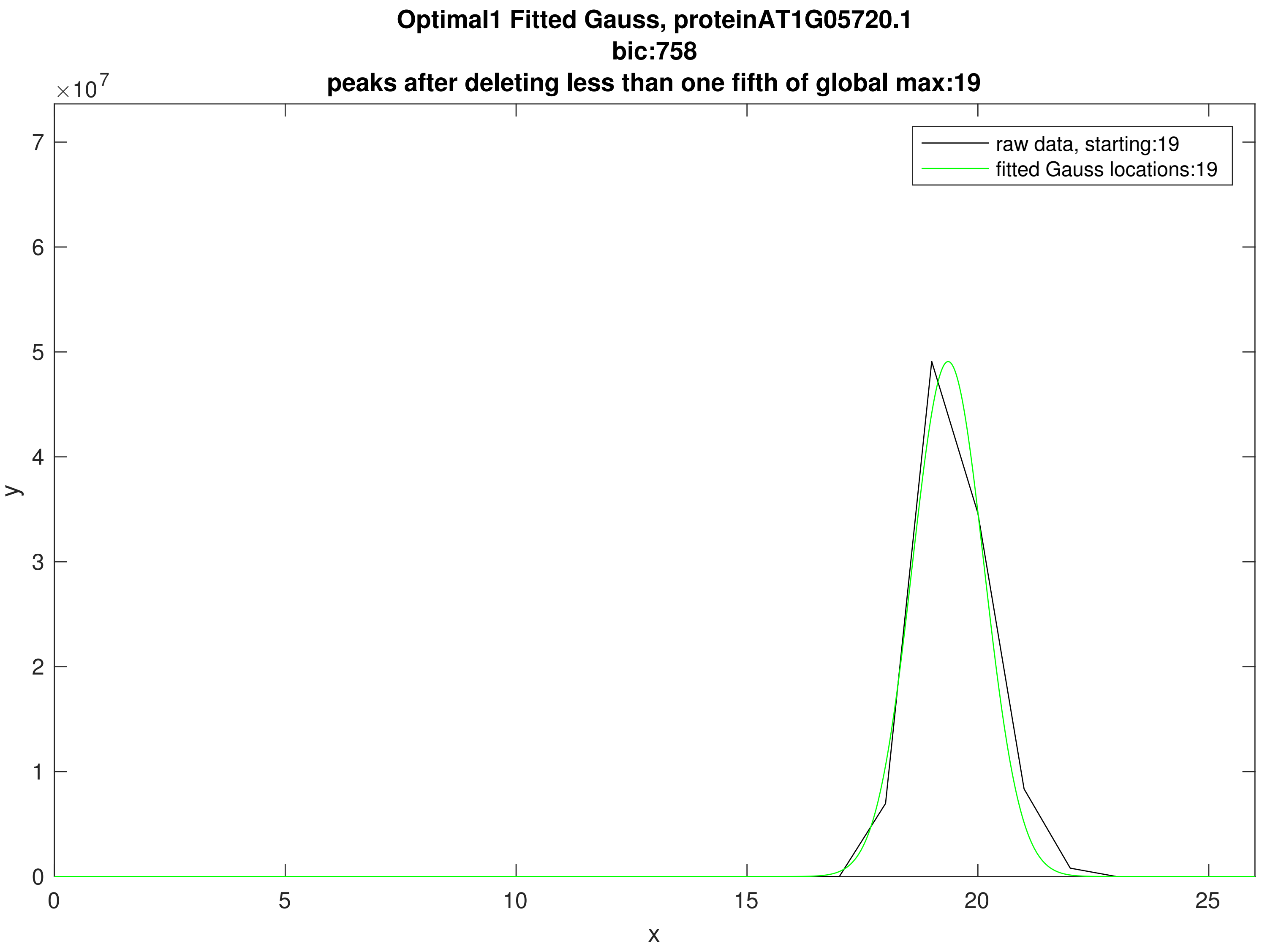

Optimal1 Fitted Gauss, proteinAT1G05720.1
bic:758
peaks after deleting less than one fifth of global max:19
×10^7
raw data, starting:19
fitted Gauss locations:19
y
x
0
1
2
3
4
5
6
7
0
5
10
15
20
25

#### Optimal1 Fitted Gauss, proteinAT1G06000.1
#### bic:900 909
#### peaks after deleting less than one fifth of global max:16

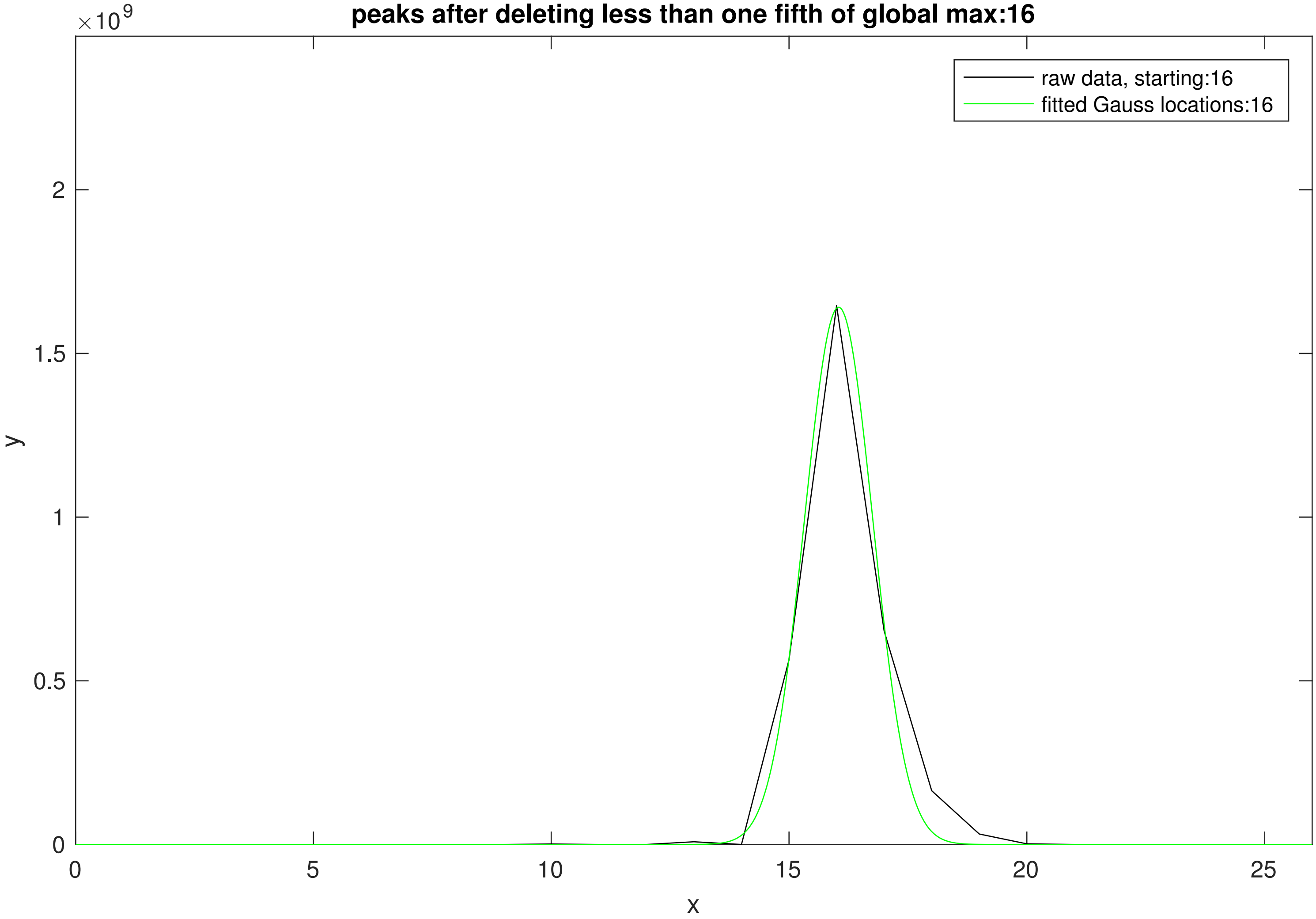

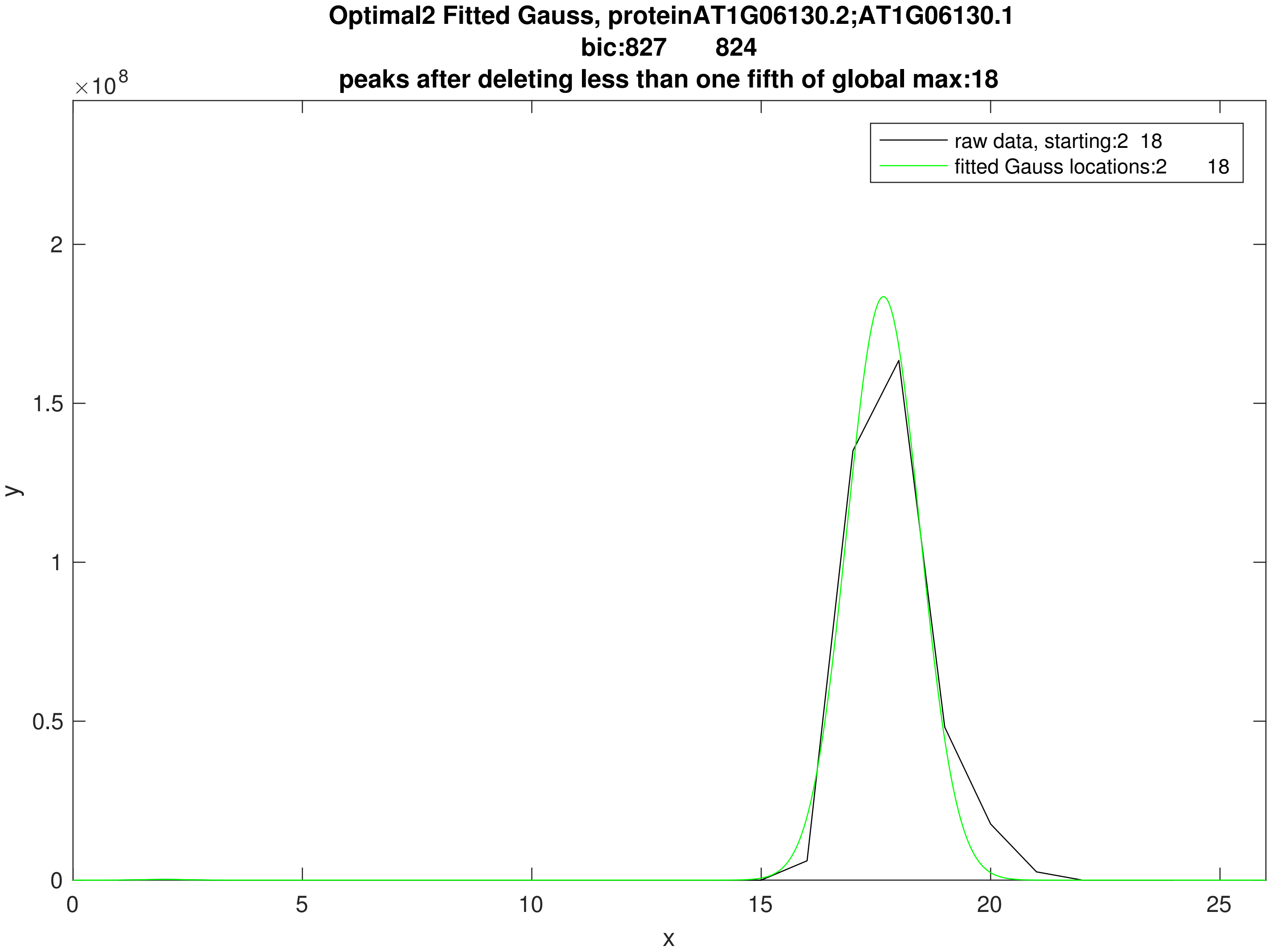
Optimal2 Fitted Gauss, proteinAT1G06130.2;AT1G06130.1
bic:827 824
peaks after deleting less than one fifth of global max:18
×10^8
raw data, starting:2 18
fitted Gauss locations:2 18
y
x
0
0.5
1
1.5
2
0
5
10
15
20
25

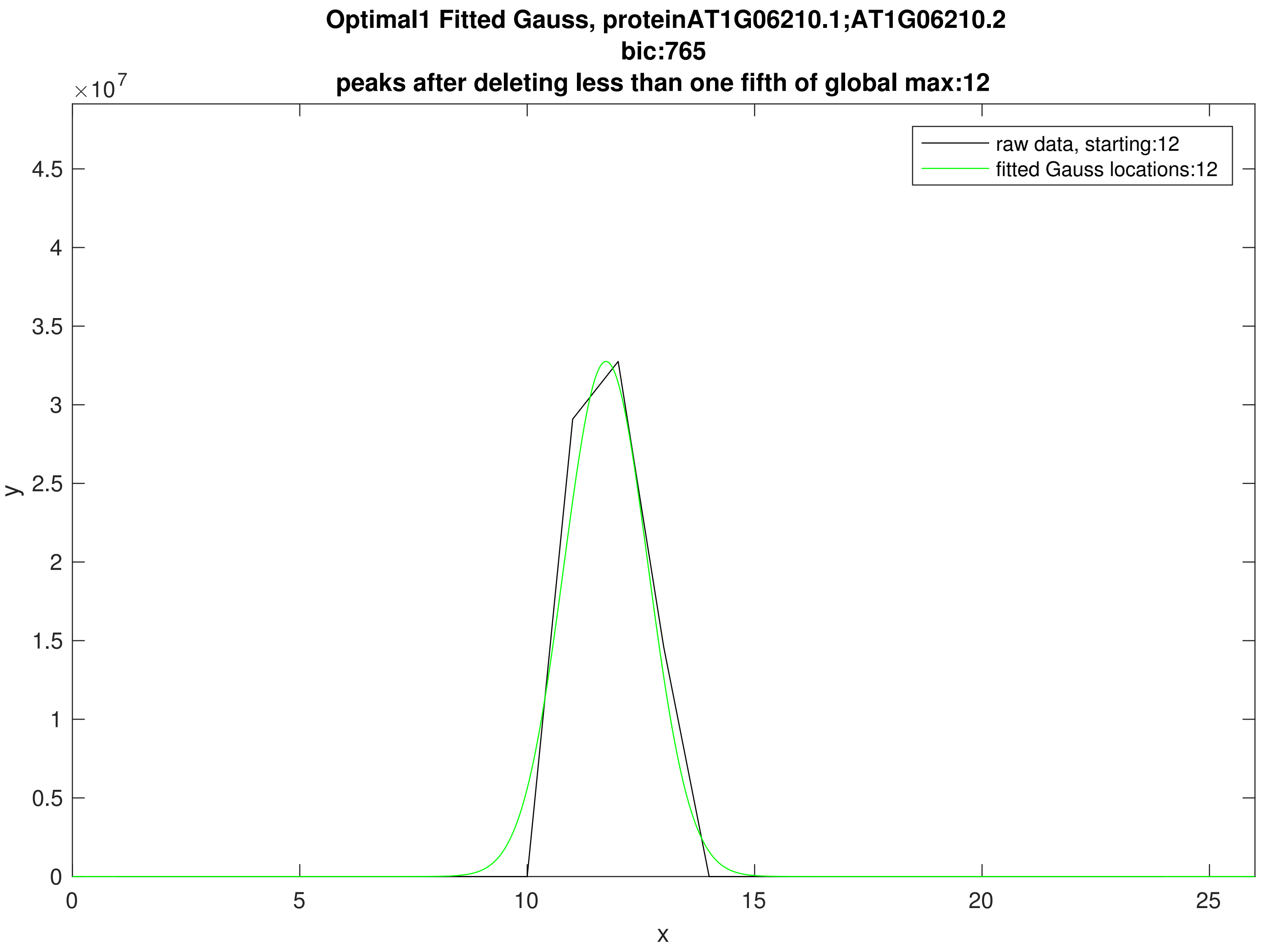

Optimal1 Fitted Gauss, proteinAT1G06210.1;AT1G06210.2
bic:765
peaks after deleting less than one fifth of global max:12
×10^7
4.5
4
3.5
3
2.5
2
1.5
1
0.5
0
y
0
5
10
15
20
25
x
raw data, starting:12
fitted Gauss locations:12

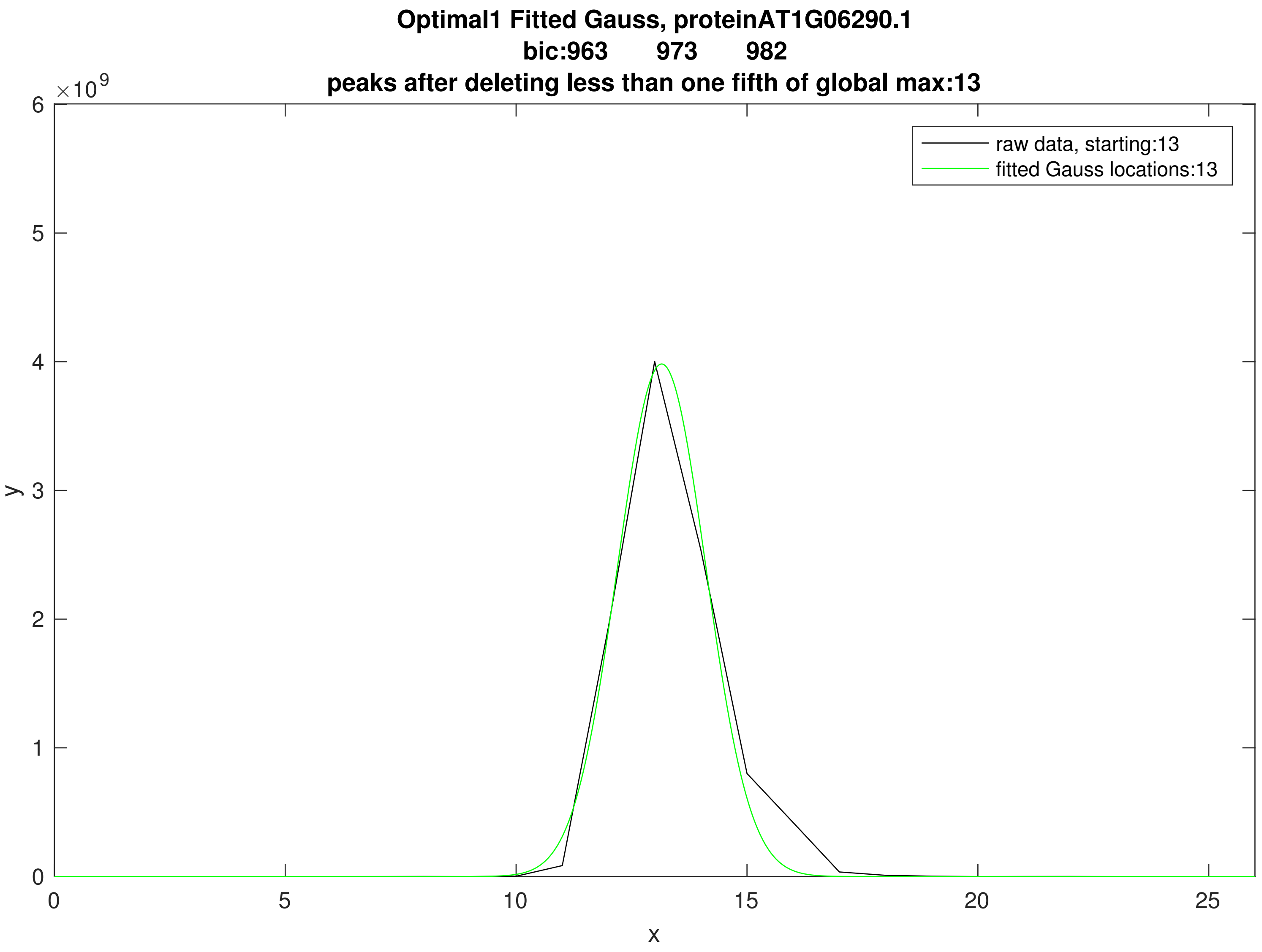

Optimal1 Fitted Gauss, proteinAT1G06290.1
bic:963 973 982
peaks after deleting less than one fifth of global max:13
×10^9
6
5
4
3
2
1
0
y
0
5
10
15
20
25
x
raw data, starting:13
fitted Gauss locations:13

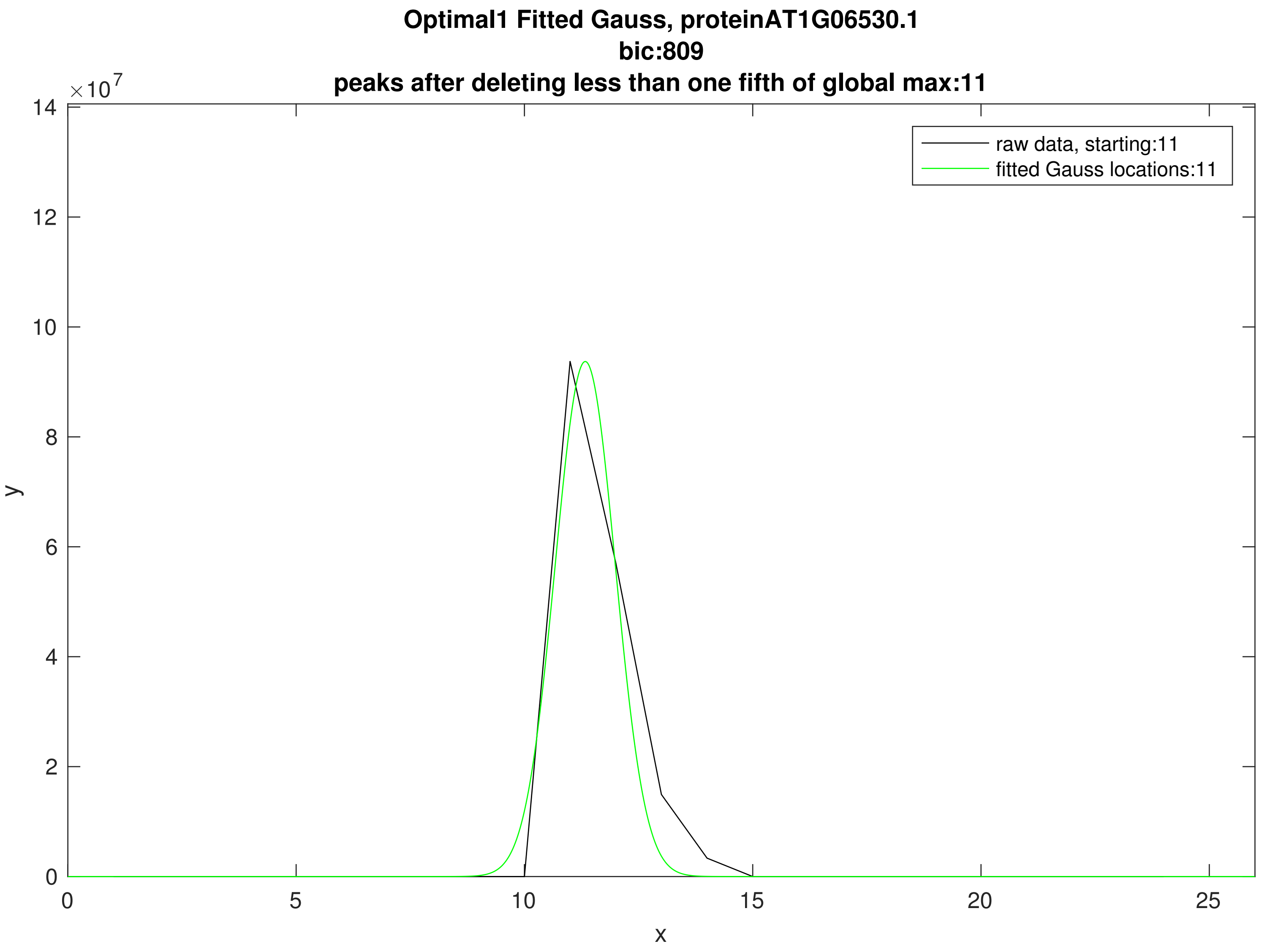
Optimal1 Fitted Gauss, proteinAT1G06530.1
bic:809
peaks after deleting less than one fifth of global max:11
×10^7
14
12
10
8
6
4
2
0
y
raw data, starting:11
fitted Gauss locations:11
0
5
10
15
20
25
x

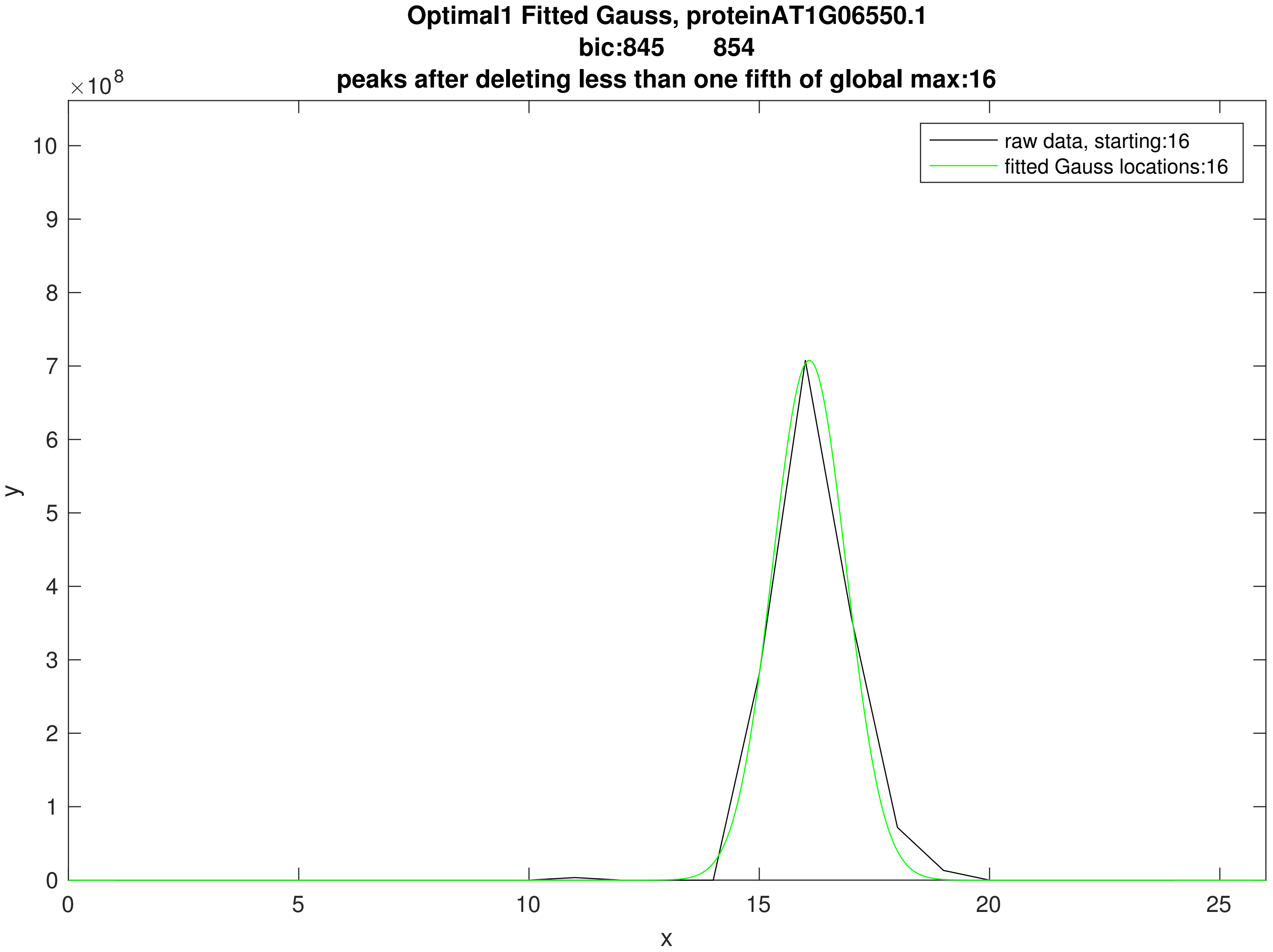
Optimal1 Fitted Gauss, proteinAT1G06550.1
bic:845 854
peaks after deleting less than one fifth of global max:16
raw data, starting:16
fitted Gauss locations:16
$\times 10^8$
y
x
0
1
2
3
4
5
6
7
8
9
10
0
5
10
15
20
25

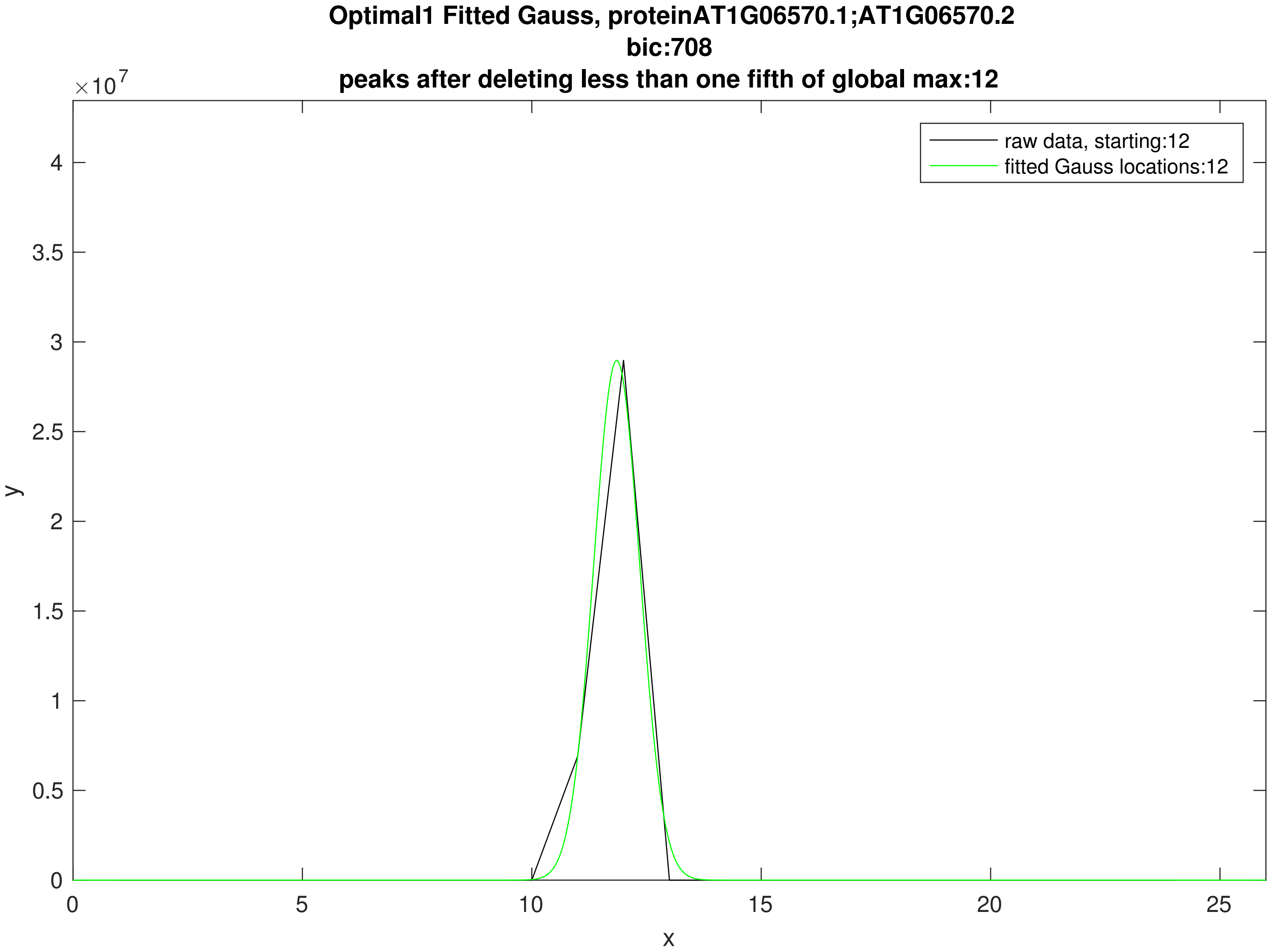
Optimal1 Fitted Gauss, proteinAT1G06570.1;AT1G06570.2
bic:708
peaks after deleting less than one fifth of global max:12
×10^7
4
3.5
3
2.5
2
1.5
1
0.5
0
y
0
5
10
15
20
25
x
raw data, starting:12
fitted Gauss locations:12

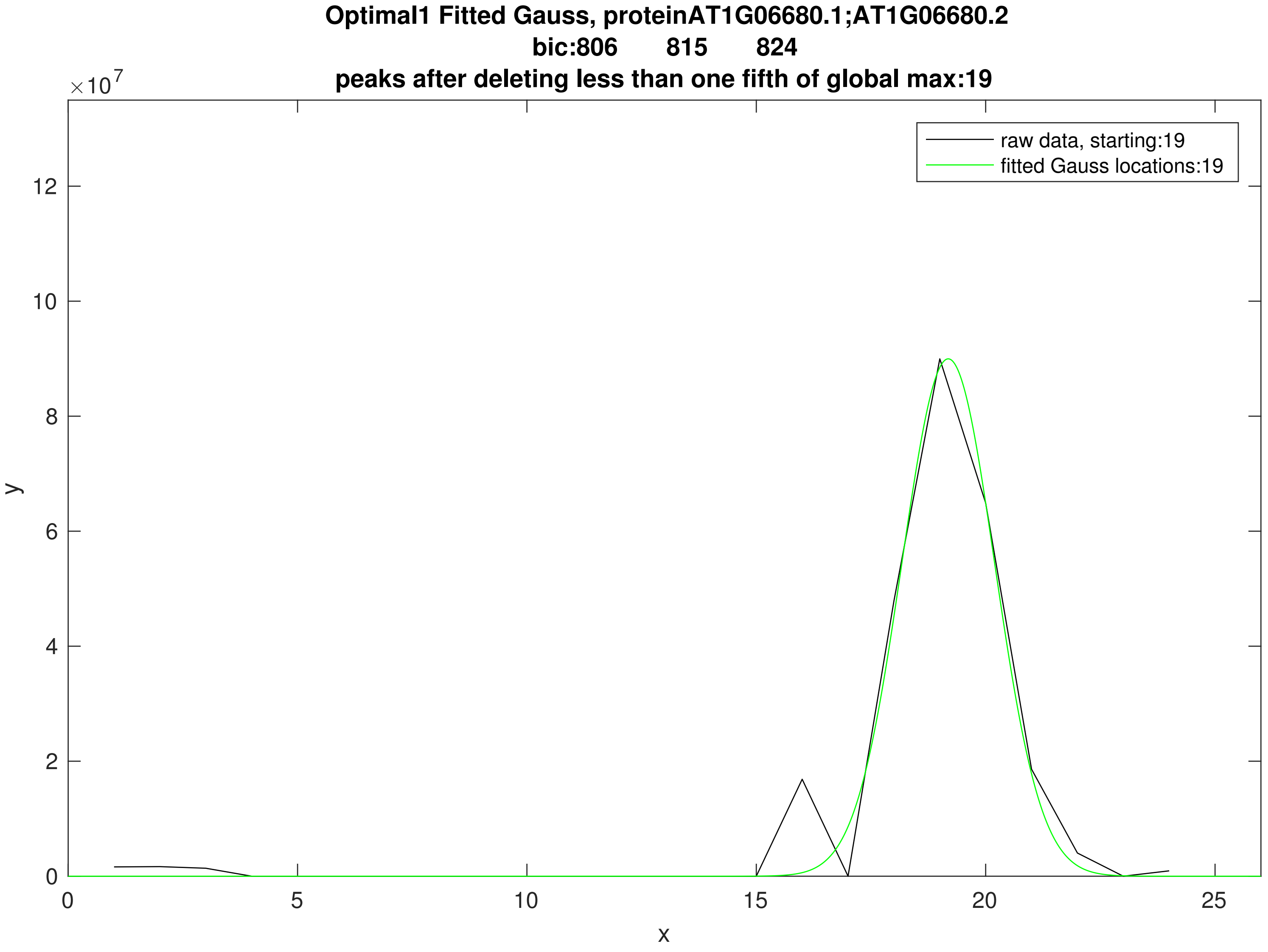
Optimal1 Fitted Gauss, proteinAT1G06680.1;AT1G06680.2
bic:806 815 824
peaks after deleting less than one fifth of global max:19
×10^7
raw data, starting:19
fitted Gauss locations:19
12
10
8
6
4
2
0
y
0
5
10
15
20
25
x

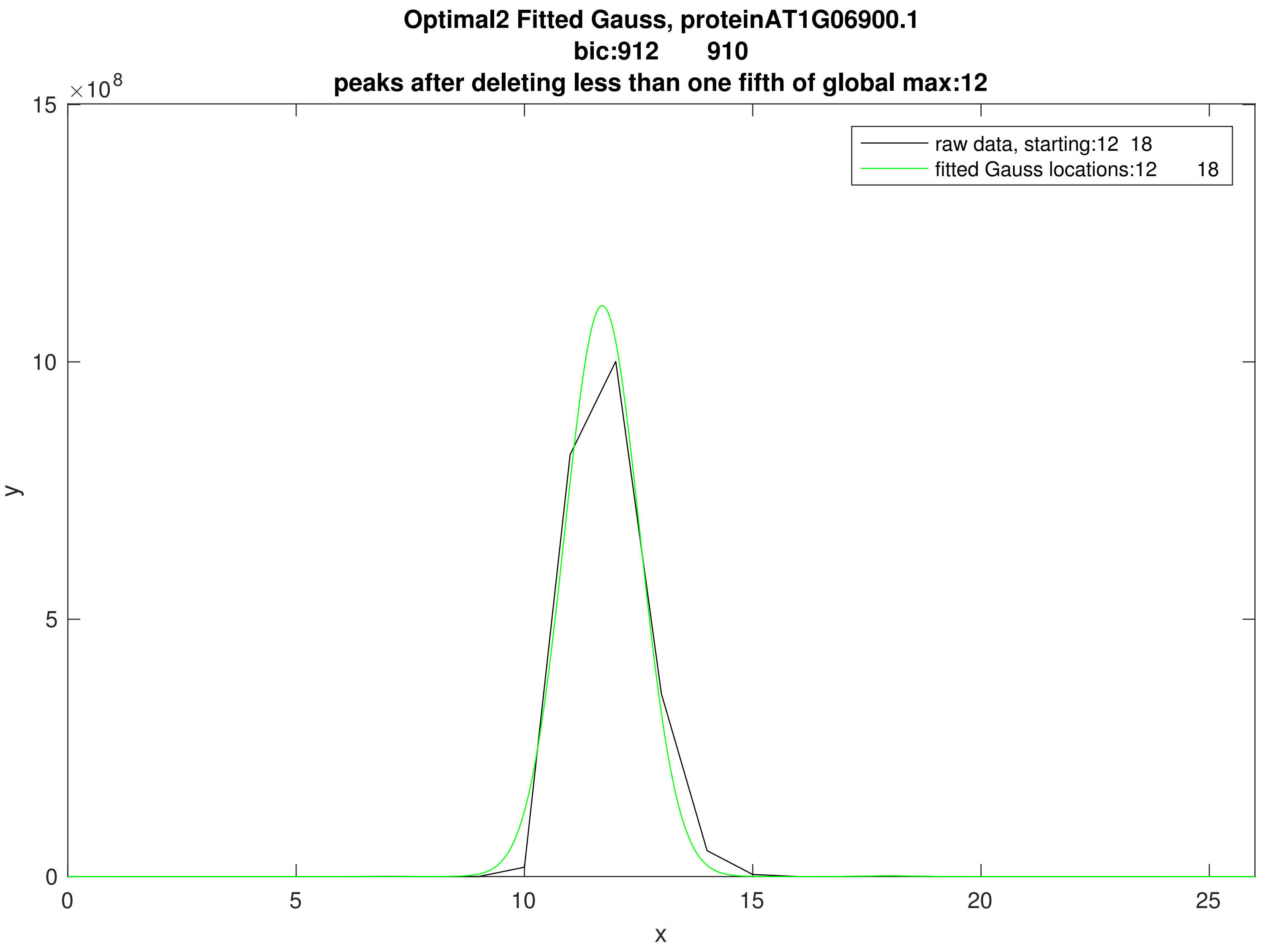
Optimal2 Fitted Gauss, proteinAT1G06900.1
bic:912 910
peaks after deleting less than one fifth of global max:12
×10^8
15
10
5
0
y
0
5
10
15
20
25
x
raw data, starting:12 18
fitted Gauss locations:12 18

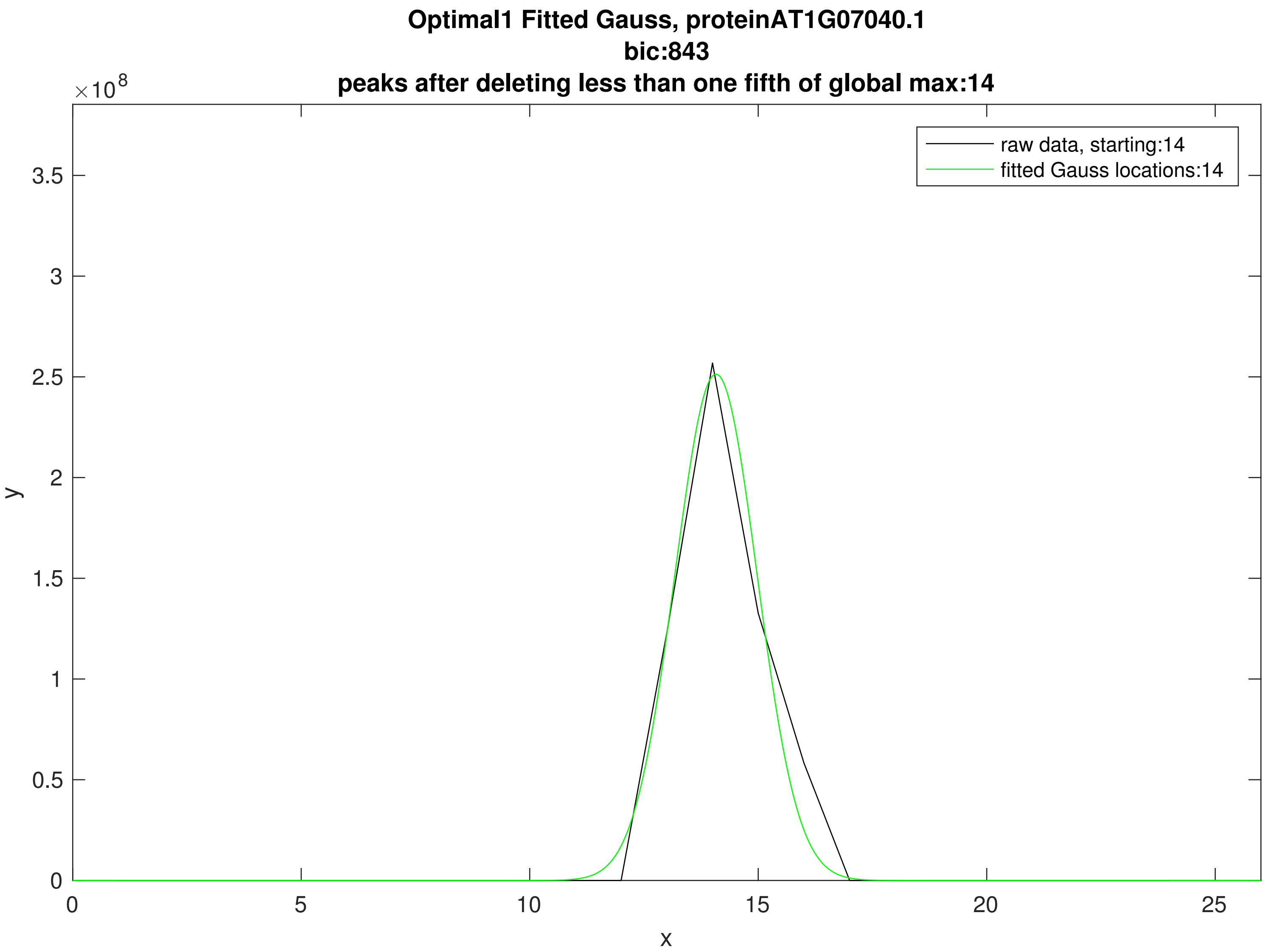
Optimal1 Fitted Gauss, proteinAT1G07040.1
bic:843
peaks after deleting less than one fifth of global max:14
×10^8
3.5
3
2.5
2
1.5
1
0.5
0
y
0
5
10
15
20
25
x
raw data, starting:14
fitted Gauss locations:14

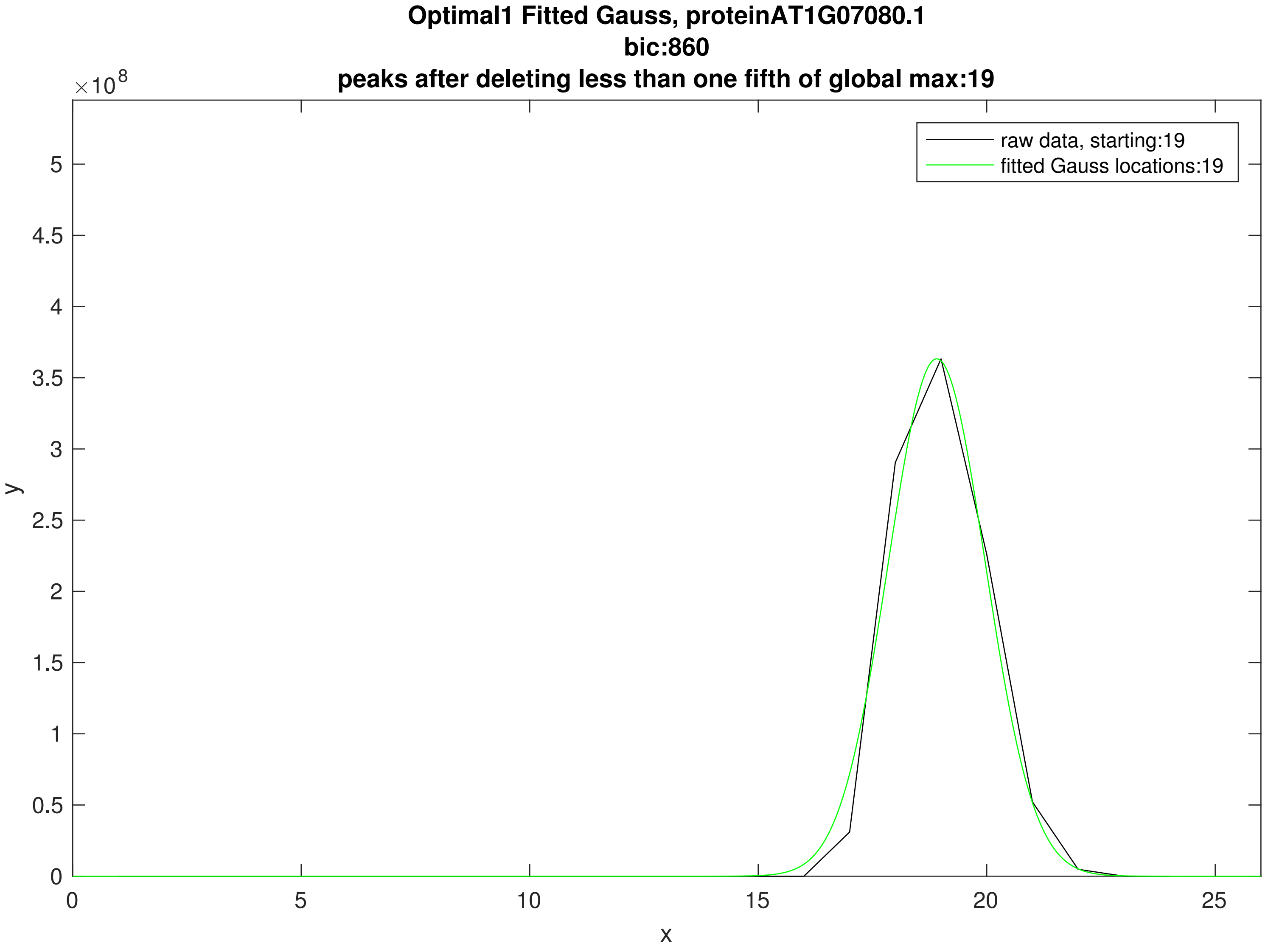

Optimal1 Fitted Gauss, proteinAT1G07080.1
bic:860
peaks after deleting less than one fifth of global max:19
×10^8
raw data, starting:19
fitted Gauss locations:19
y
x
0
0.5
1
1.5
2
2.5
3
3.5
4
4.5
5
0
5
10
15
20
25

**Optimal1 Fitted Gauss, proteinAT1G07140.1**
**bic:1.01e+03  1.02e+03  1.03e+03  1.04e+03**
**peaks after deleting less than one fifth of global max:14**

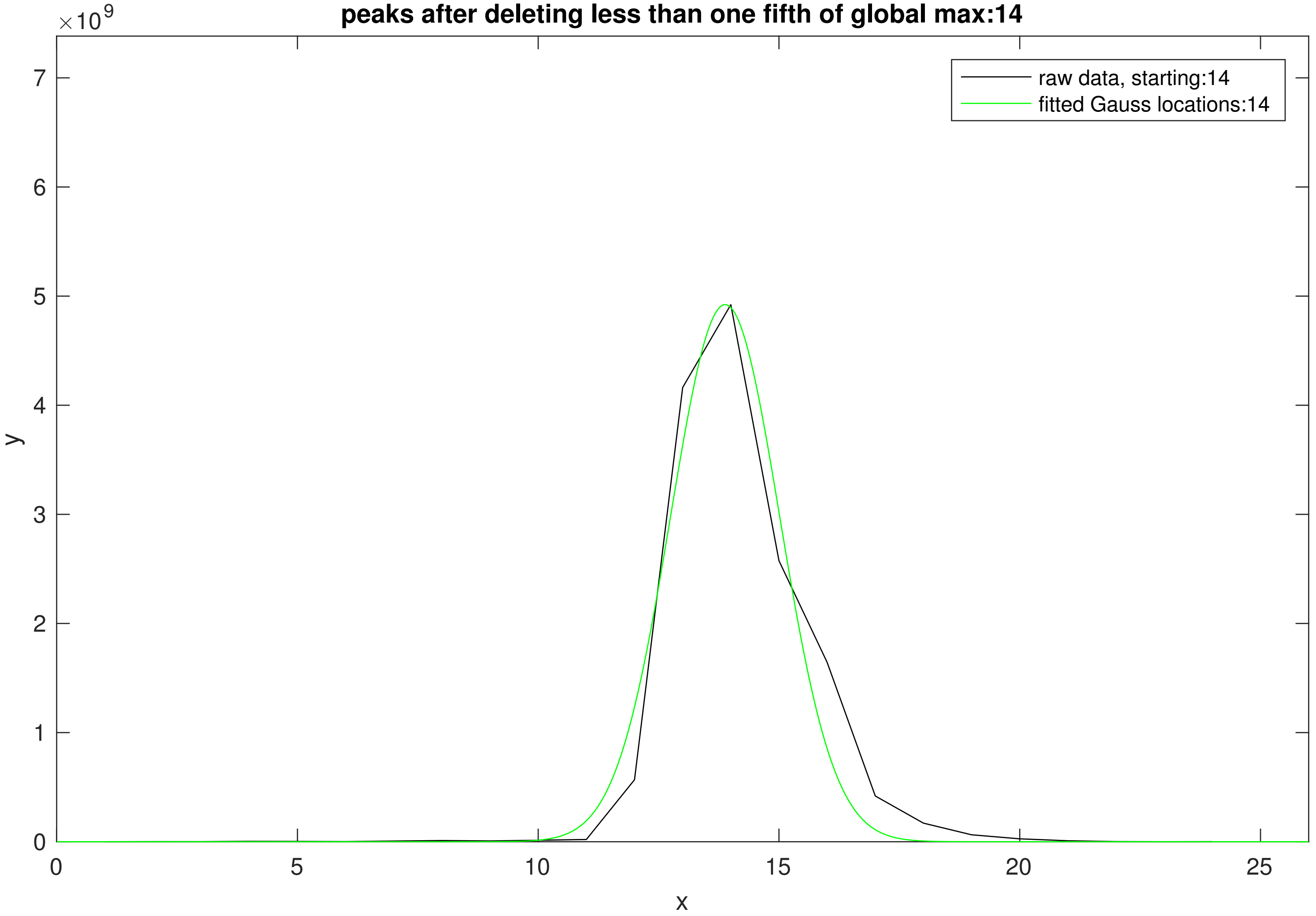

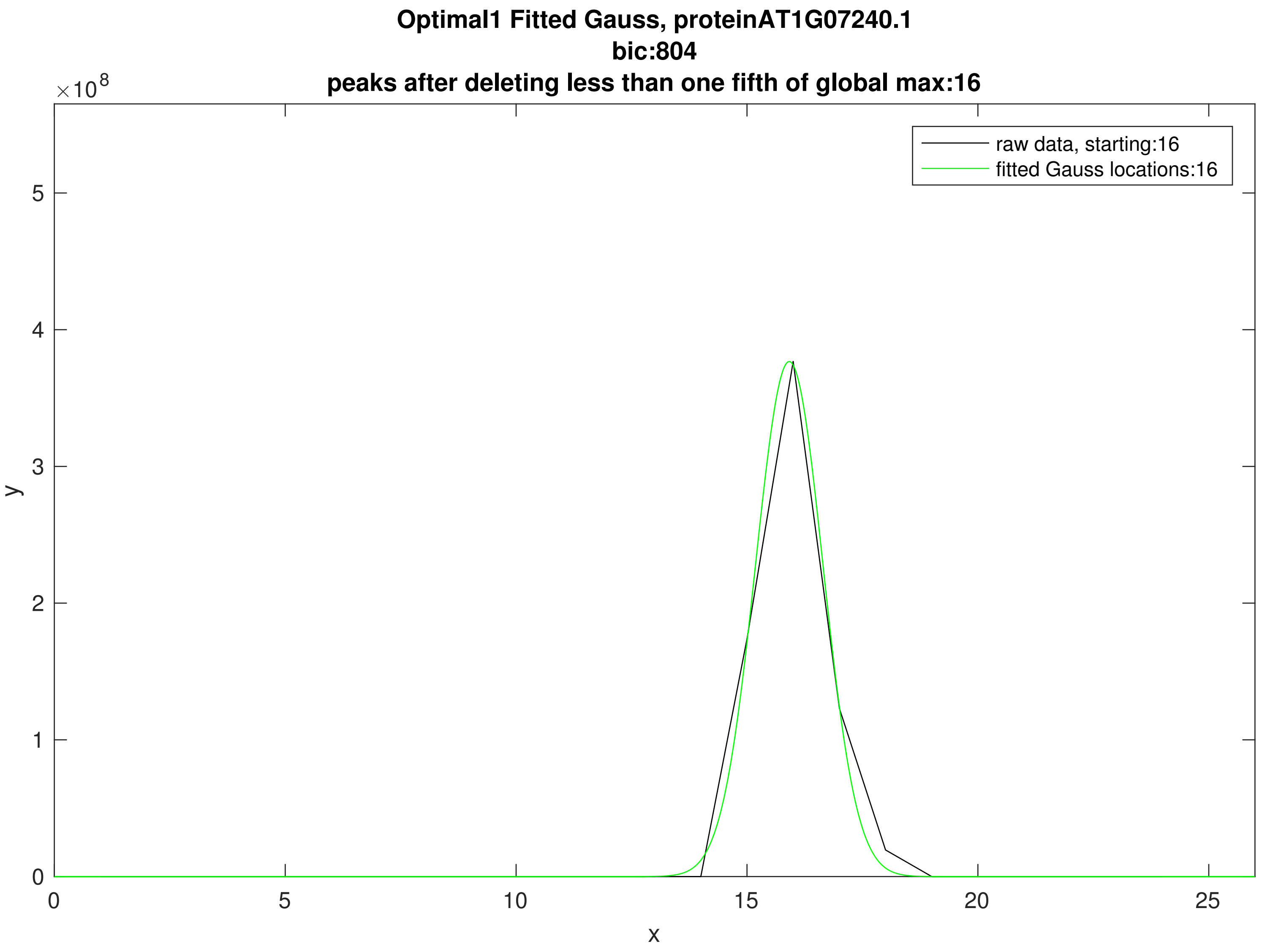
Optimal1 Fitted Gauss, proteinAT1G07240.1
bic:804
peaks after deleting less than one fifth of global max:16
×10^8
raw data, starting:16
fitted Gauss locations:16
y
x
0
1
2
3
4
5
0
5
10
15
20
25

#### Optimal2 Fitted Gauss, proteinAT1G07250.1
#### bic:814 813
#### peaks after deleting less than one fifth of global max:16

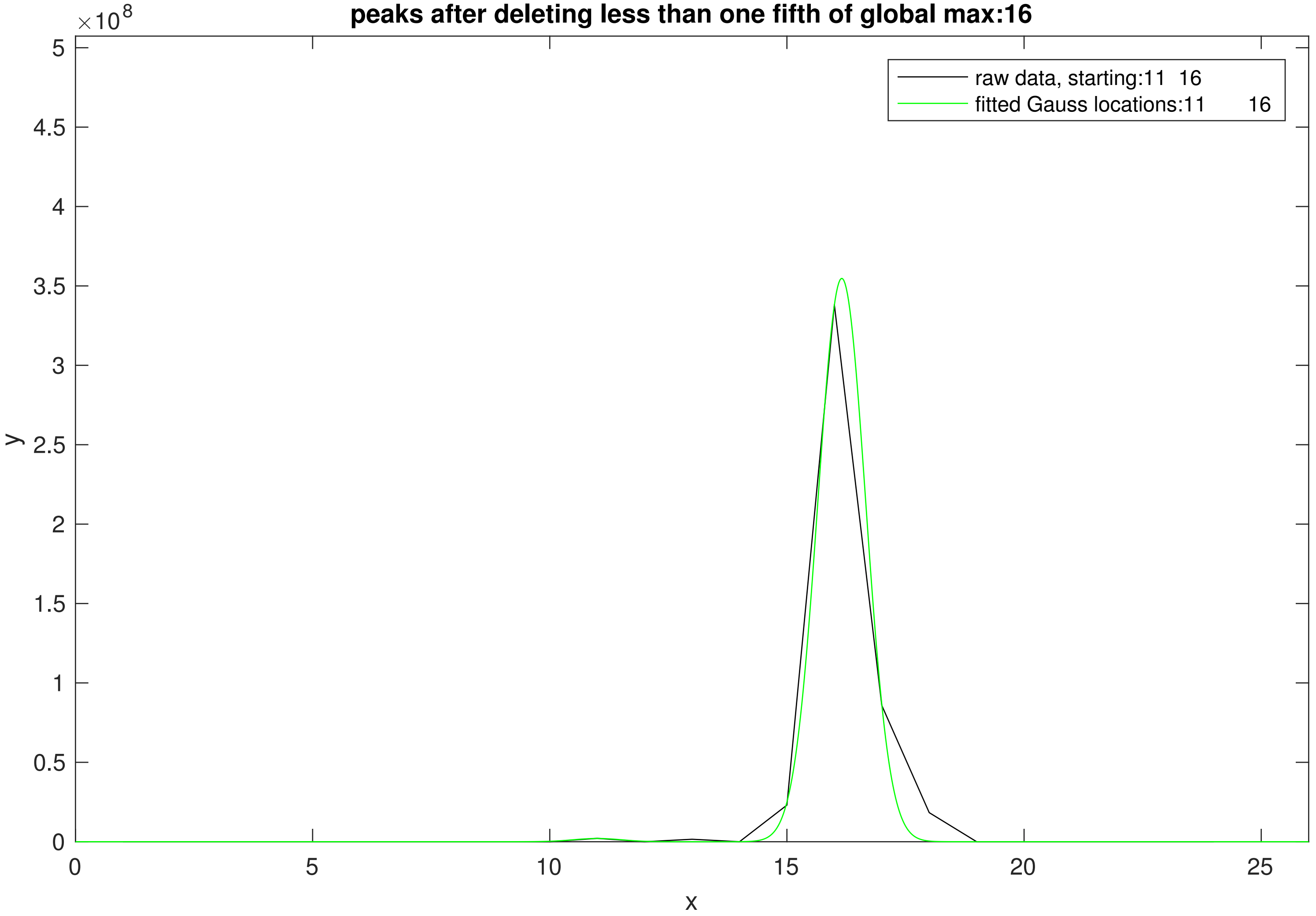

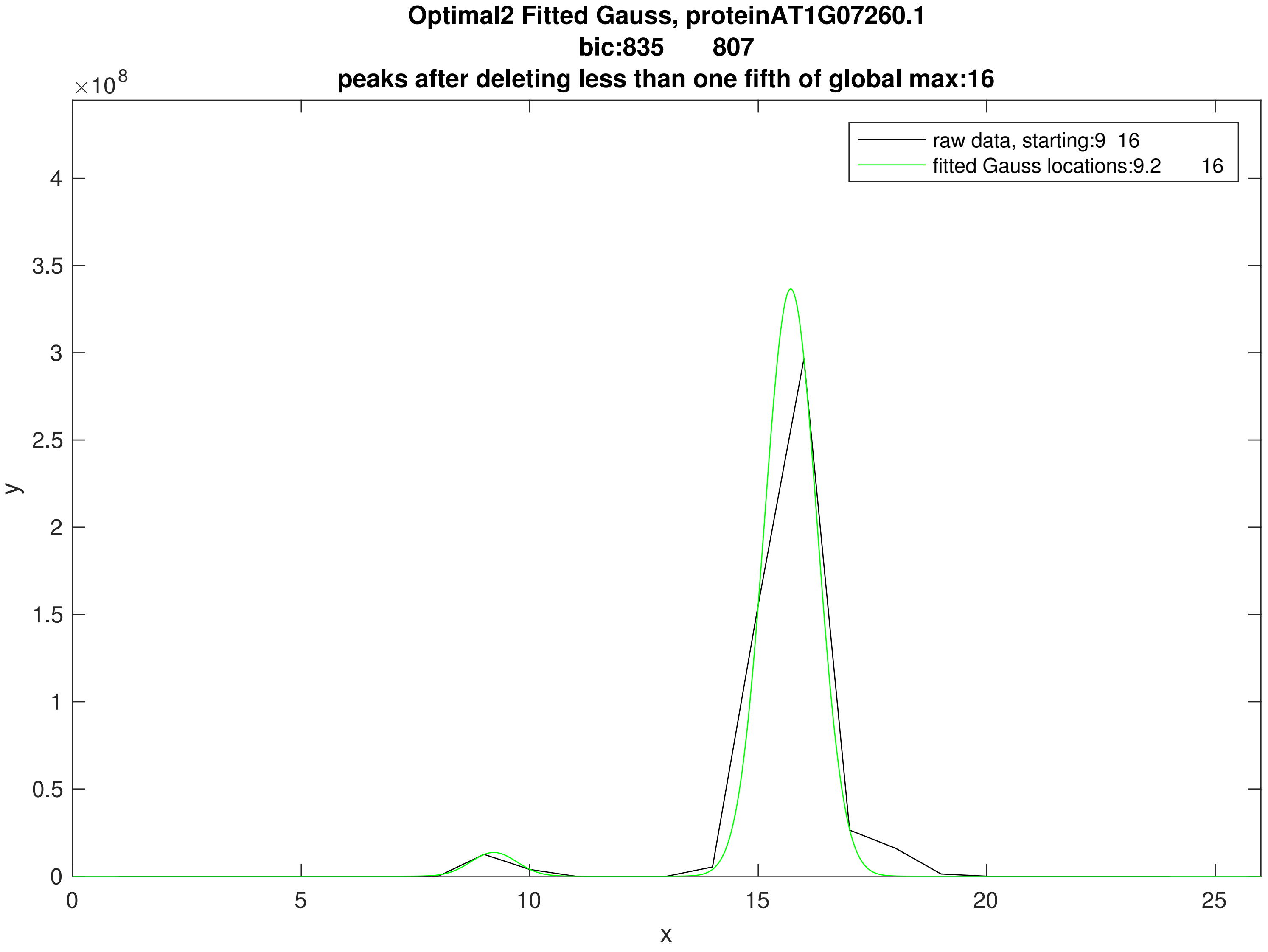
Optimal2 Fitted Gauss, proteinAT1G07260.1
bic:835 807
peaks after deleting less than one fifth of global max:16
raw data, starting:9 16
fitted Gauss locations:9.2 16
×10^8
y
x
0
0.5
1
1.5
2
2.5
3
3.5
4
5
10
15
20
25

Optimal1 Fitted Gauss, proteinAT1G07320.4;AT1G07320.3;AT1G07320.2;AT1G07320.1
bic:672
peaks after deleting less than one fifth of global max:4.1

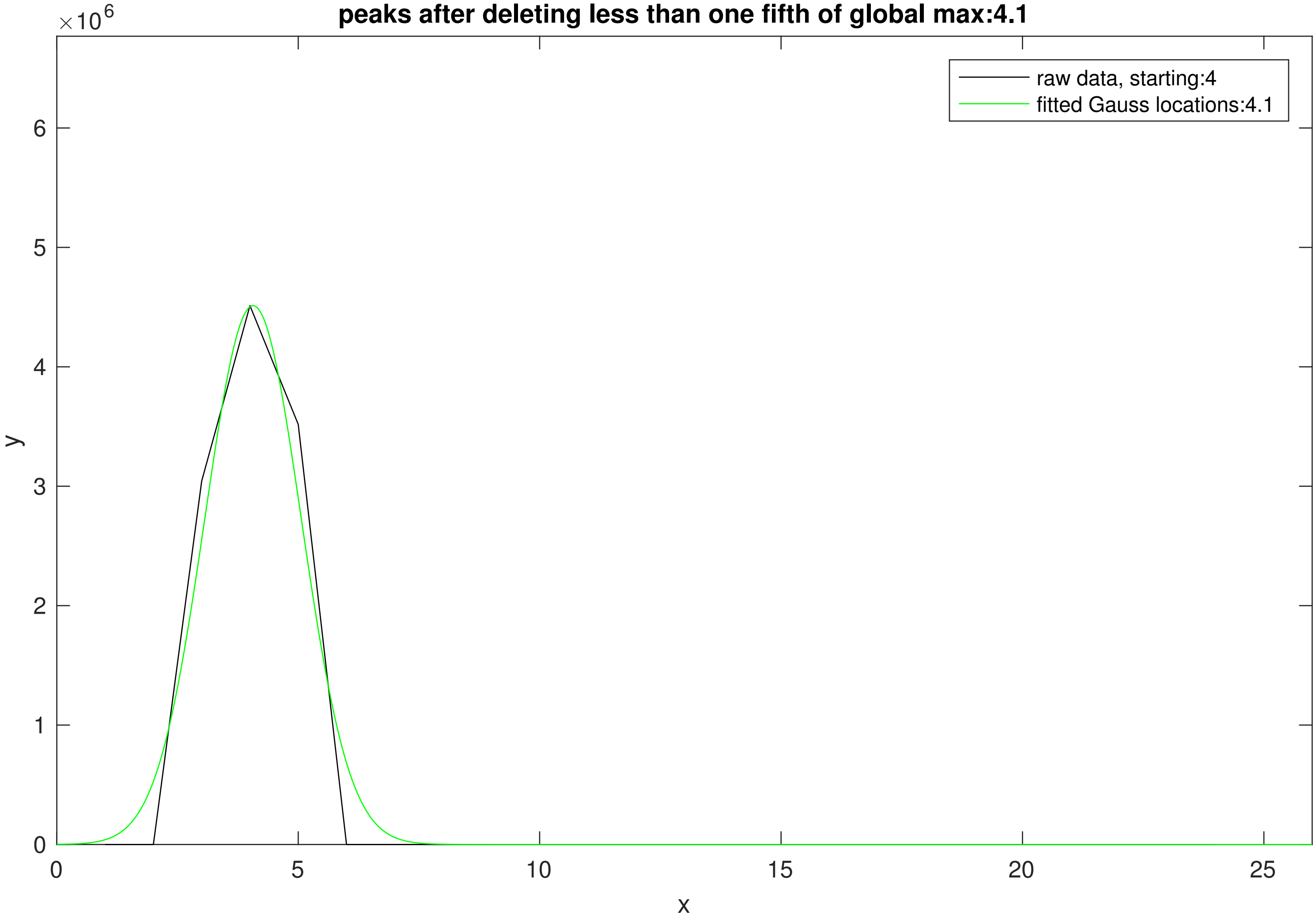

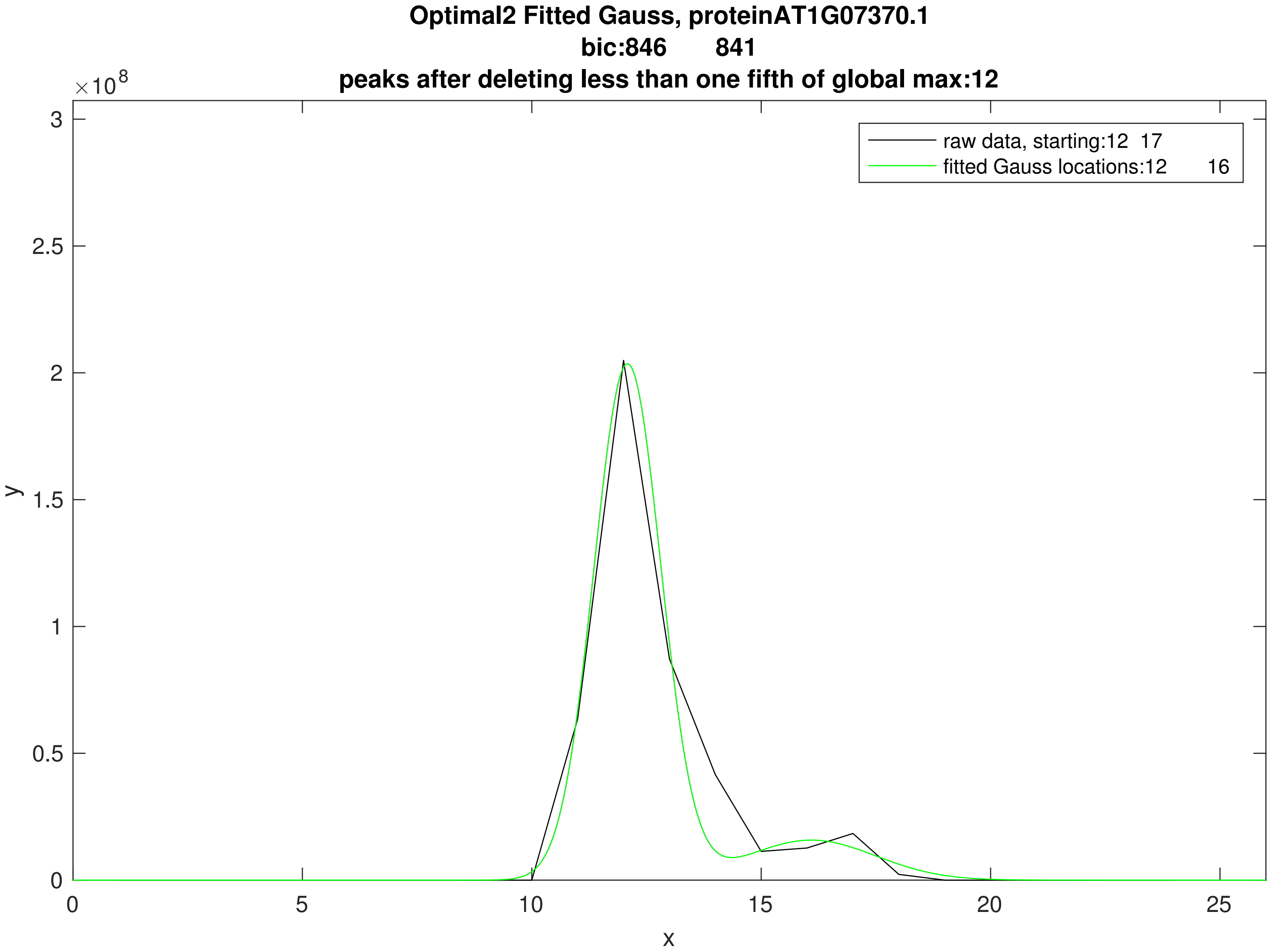
Optimal2 Fitted Gauss, proteinAT1G07370.1
bic:846 841
peaks after deleting less than one fifth of global max:12
×10^8
raw data, starting:12 17
fitted Gauss locations:12 16
y
x
0
0.5
1
1.5
2
2.5
3
0
5
10
15
20
25

#### Optimal1 Fitted Gauss, proteinAT1G07440.2;AT1G07440.1
#### bic:694
#### peaks after deleting less than one fifth of global max:14

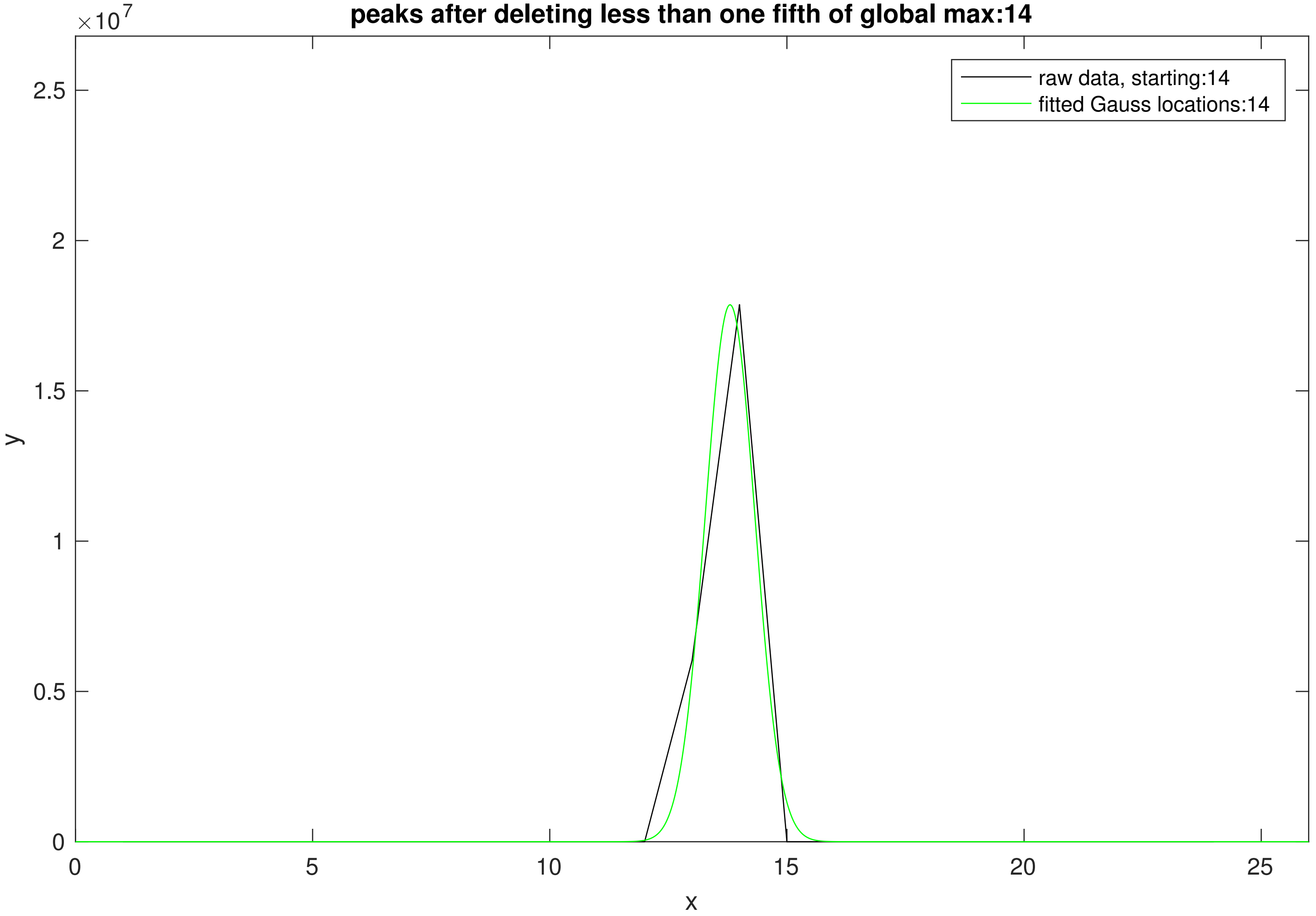

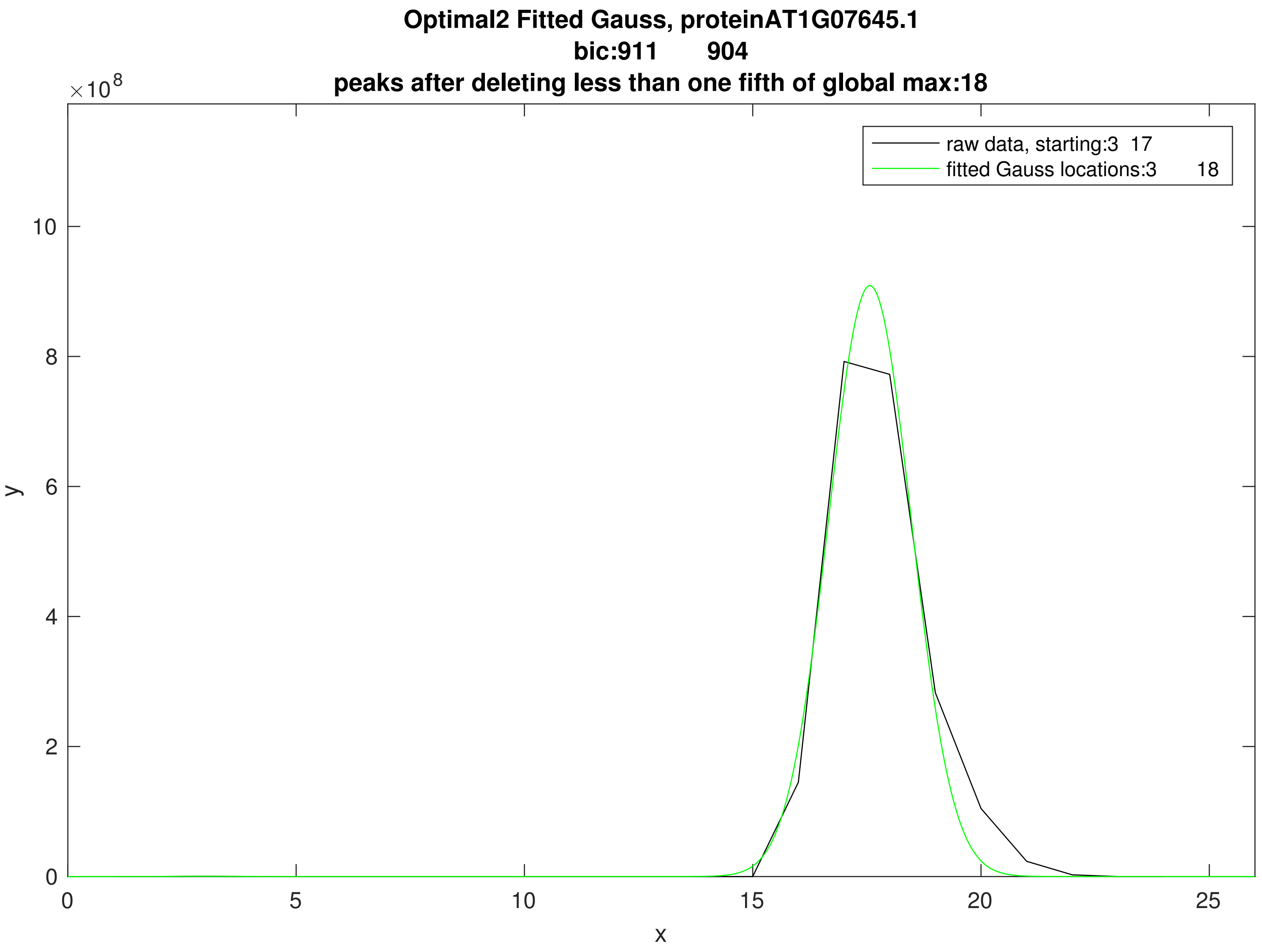
Optimal2 Fitted Gauss, proteinAT1G07645.1
bic:911 904
peaks after deleting less than one fifth of global max:18
×10^8
raw data, starting:3 17
fitted Gauss locations:3 18
y
x
0
2
4
6
8
10
0
5
10
15
20
25

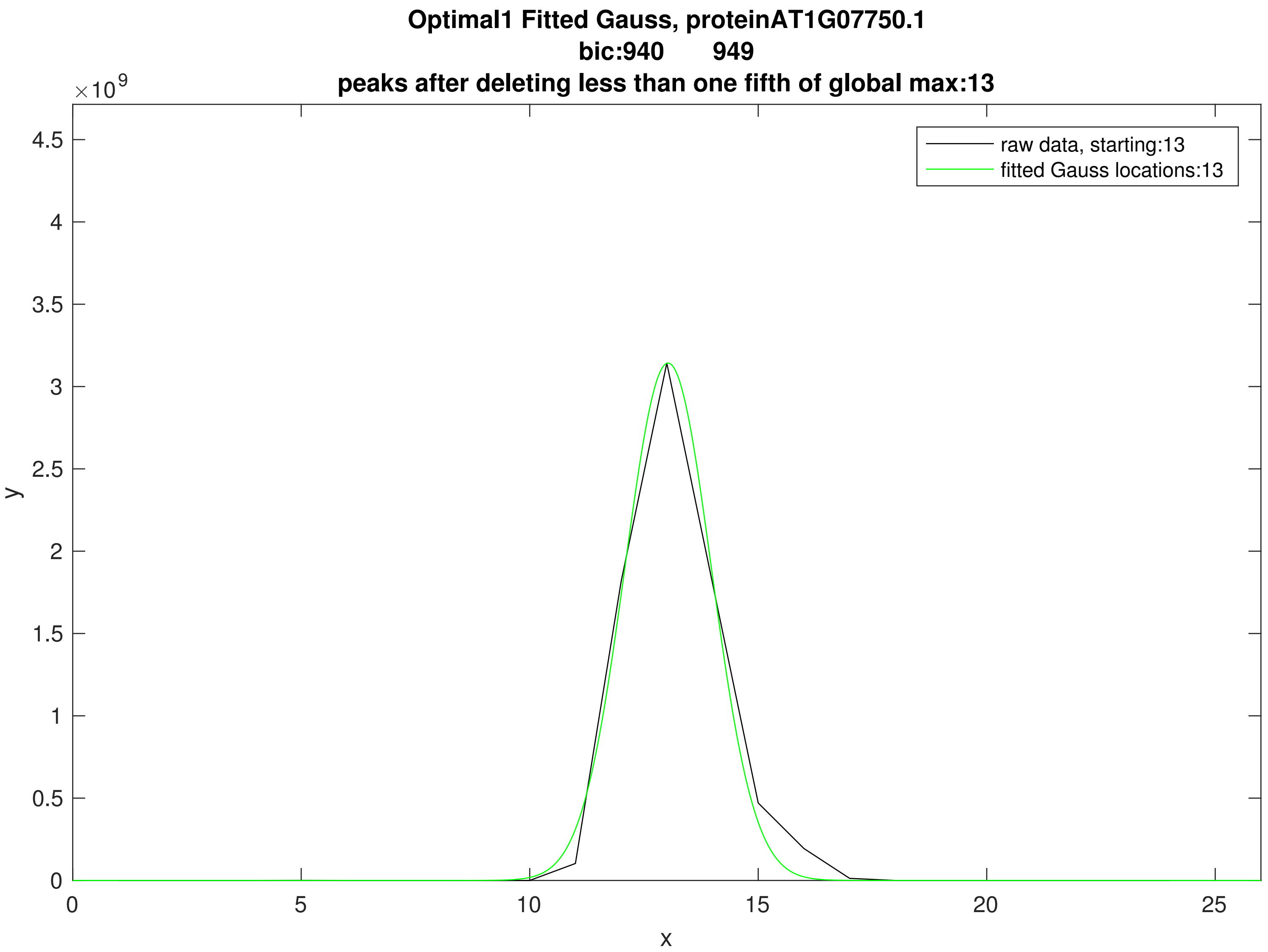
Optimal1 Fitted Gauss, proteinAT1G07750.1
bic:940 949
peaks after deleting less than one fifth of global max:13
×10^9
raw data, starting:13
fitted Gauss locations:13
y
x
0
0.5
1
1.5
2
2.5
3
3.5
4
4.5
5
10
15
20
25

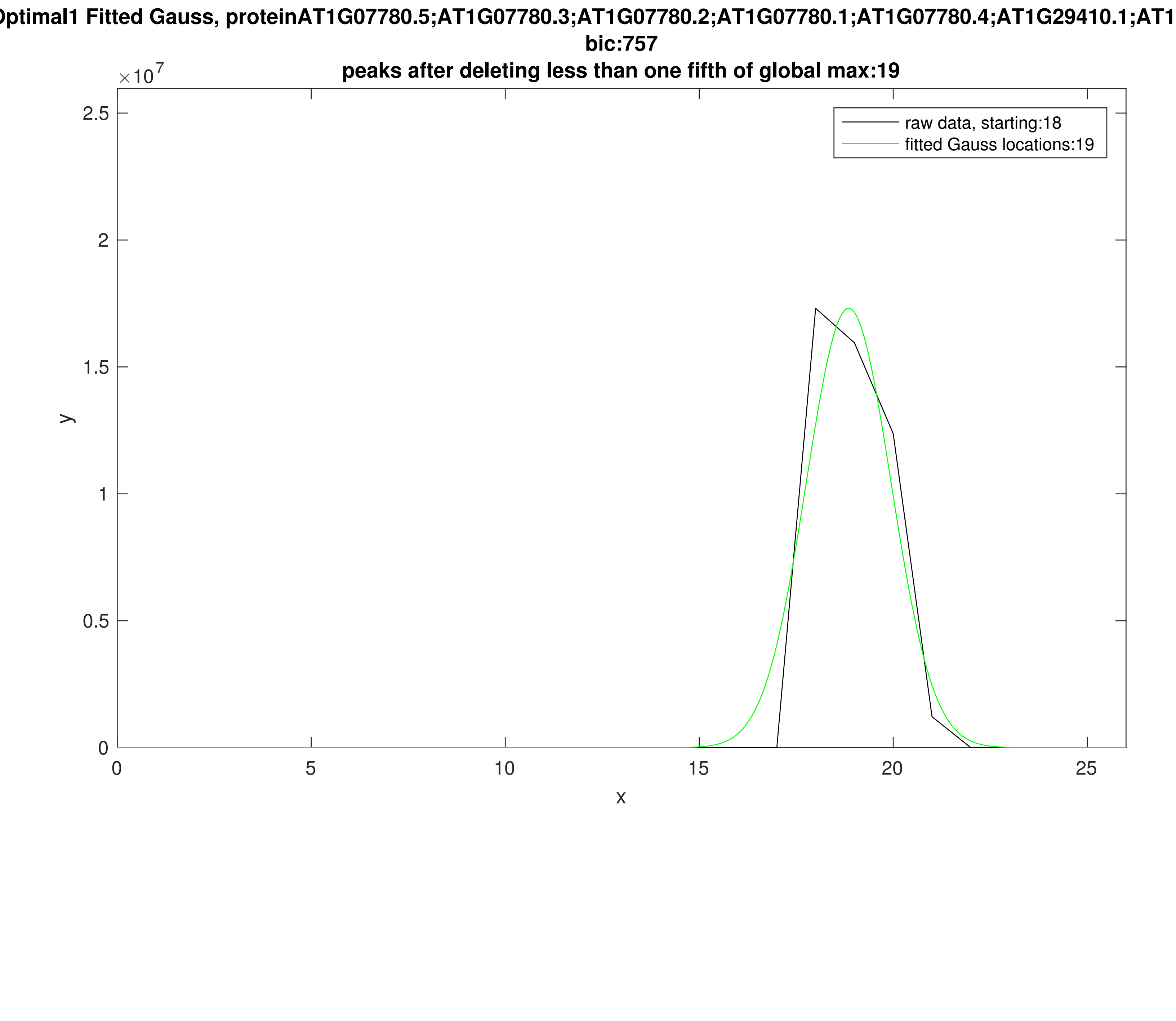

Optimal1 Fitted Gauss, proteinAT1G07780.5;AT1G07780.3;AT1G07780.2;AT1G07780.1;AT1G07780.4;AT1G29410.1;AT1
bic:757
peaks after deleting less than one fifth of global max:19
×10^7
raw data, starting:18
fitted Gauss locations:19
y
x
0
0.5
1
1.5
2
2.5
0
5
10
15
20
25

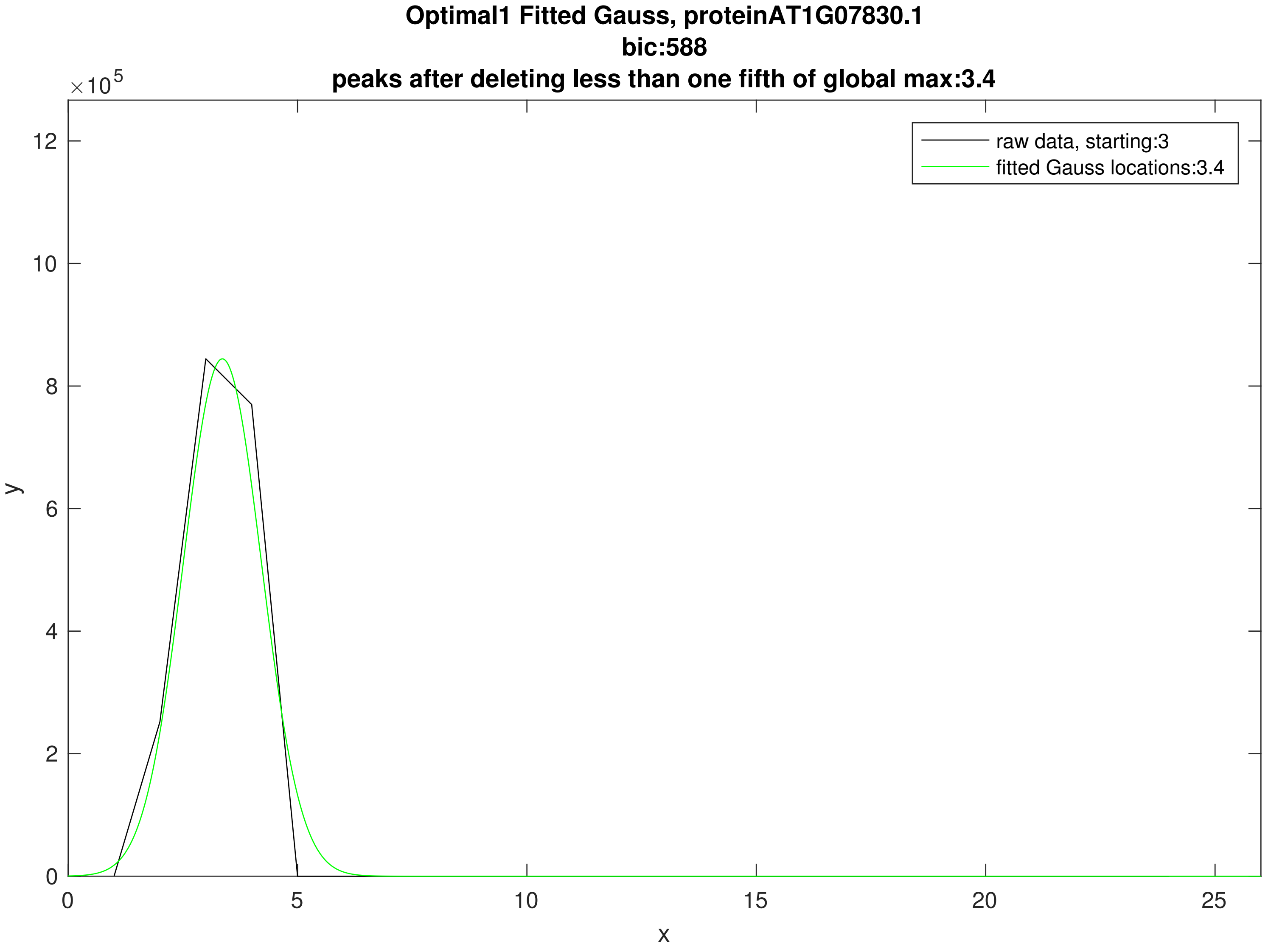
Optimal1 Fitted Gauss, proteinAT1G07830.1
bic:588
peaks after deleting less than one fifth of global max:3.4
×10^5
raw data, starting:3
fitted Gauss locations:3.4
y
x
0
2
4
6
8
10
12
0
5
10
15
20
25

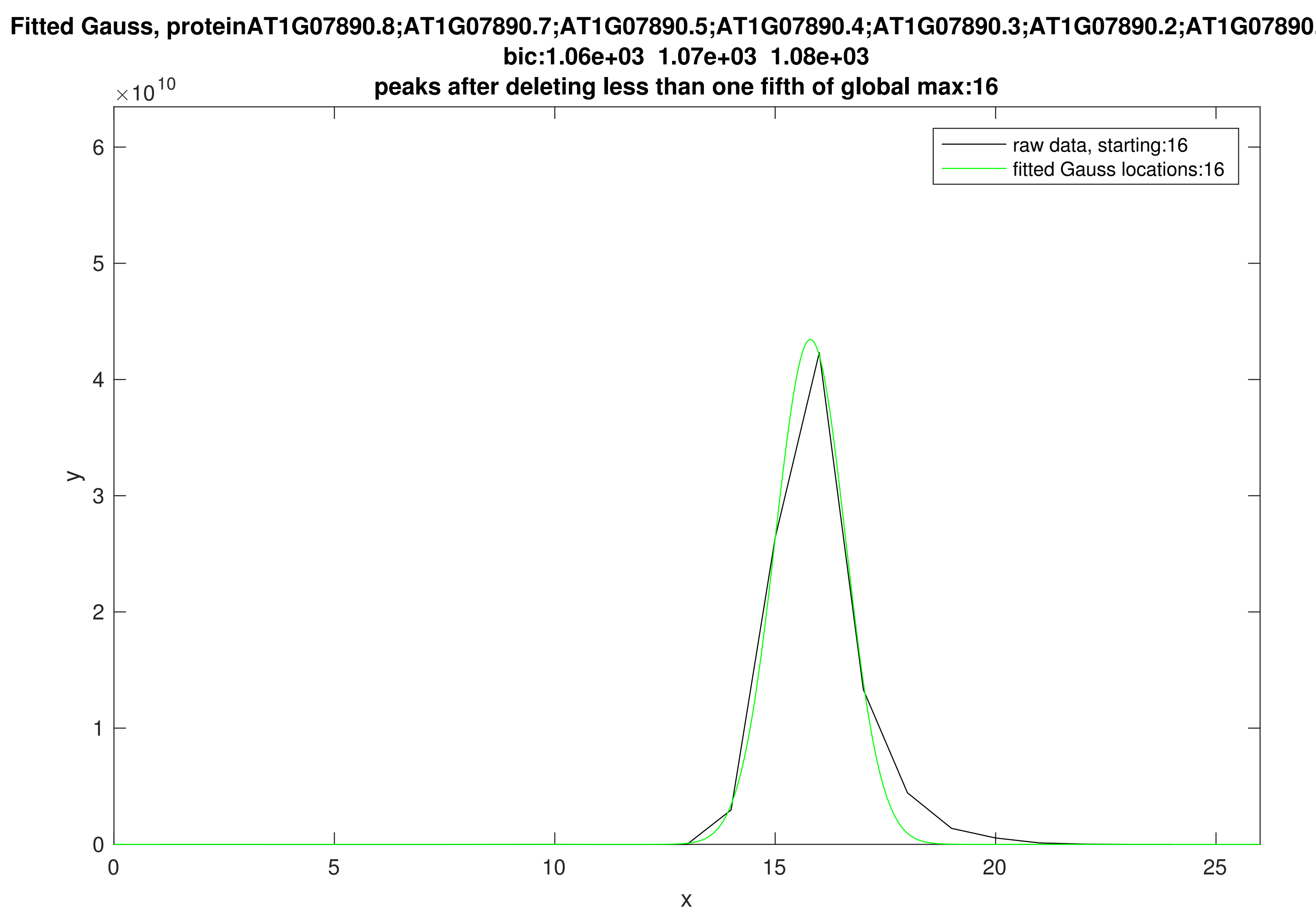
Fitted Gauss, proteinAT1G07890.8;AT1G07890.7;AT1G07890.5;AT1G07890.4;AT1G07890.3;AT1G07890.2;AT1G07890.
bic:1.06e+03 1.07e+03 1.08e+03
peaks after deleting less than one fifth of global max:16
×10^10
raw data, starting:16
fitted Gauss locations:16
y
x
0
1
2
3
4
5
6
0
5
10
15
20
25

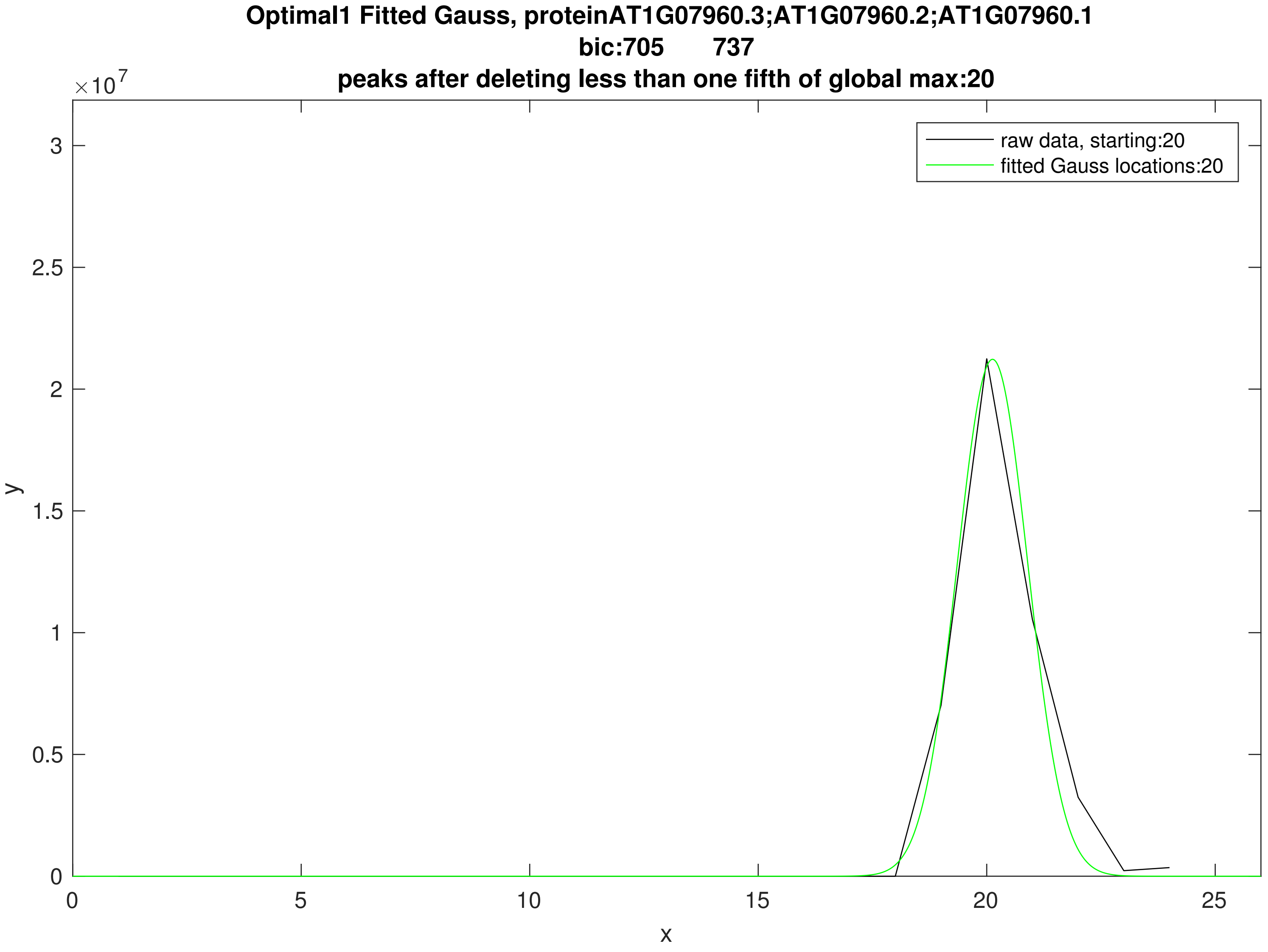
Optimal1 Fitted Gauss, proteinAT1G07960.3;AT1G07960.2;AT1G07960.1
bic:705 737
peaks after deleting less than one fifth of global max:20
×10^7
raw data, starting:20
fitted Gauss locations:20
y
x
0
0.5
1
1.5
2
2.5
3
0
5
10
15
20
25

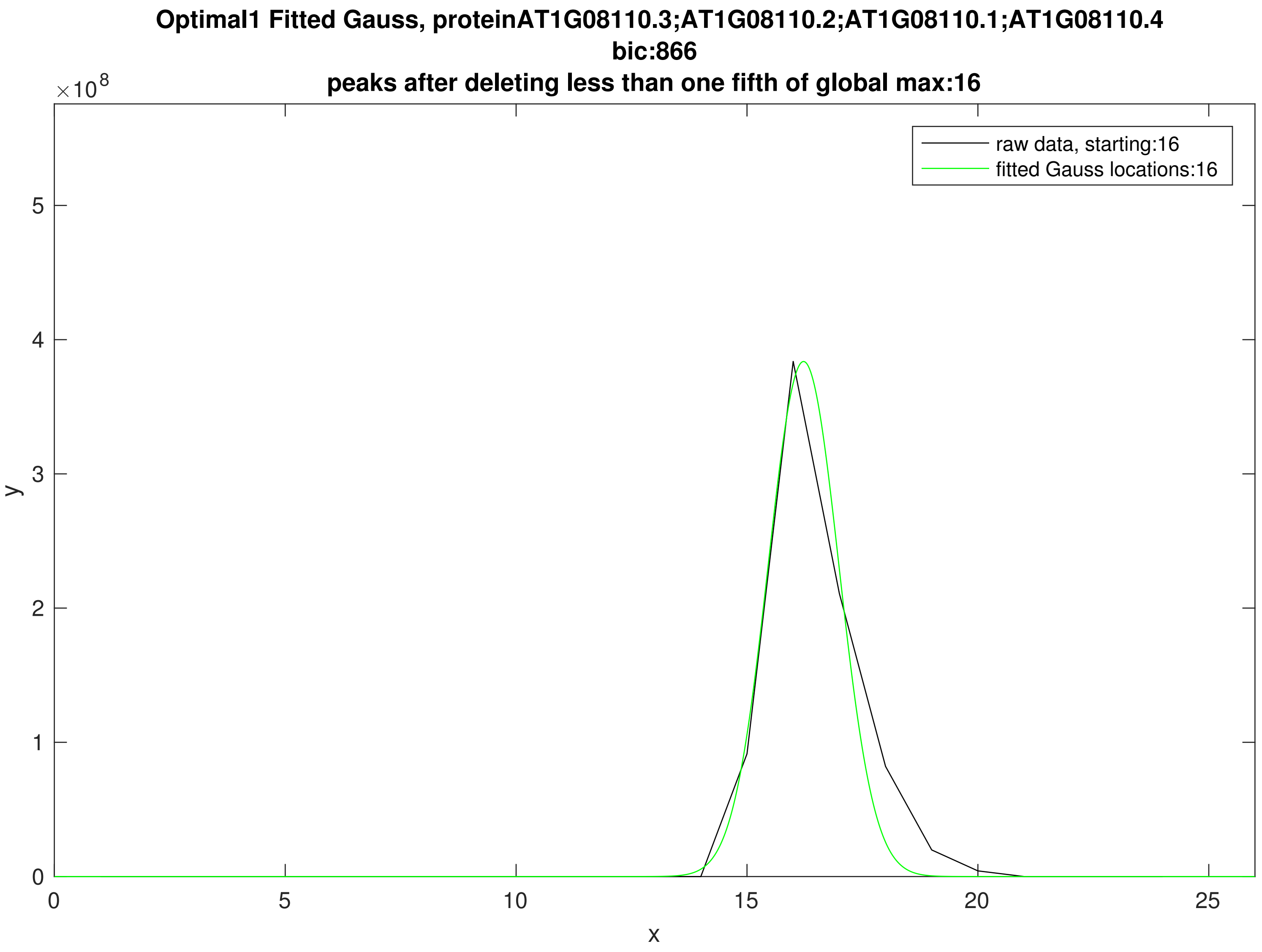
Optimal1 Fitted Gauss, proteinAT1G08110.3;AT1G08110.2;AT1G08110.1;AT1G08110.4
bic:866
peaks after deleting less than one fifth of global max:16
×10^8
raw data, starting:16
fitted Gauss locations:16
y
x
0
1
2
3
4
5
0
5
10
15
20
25

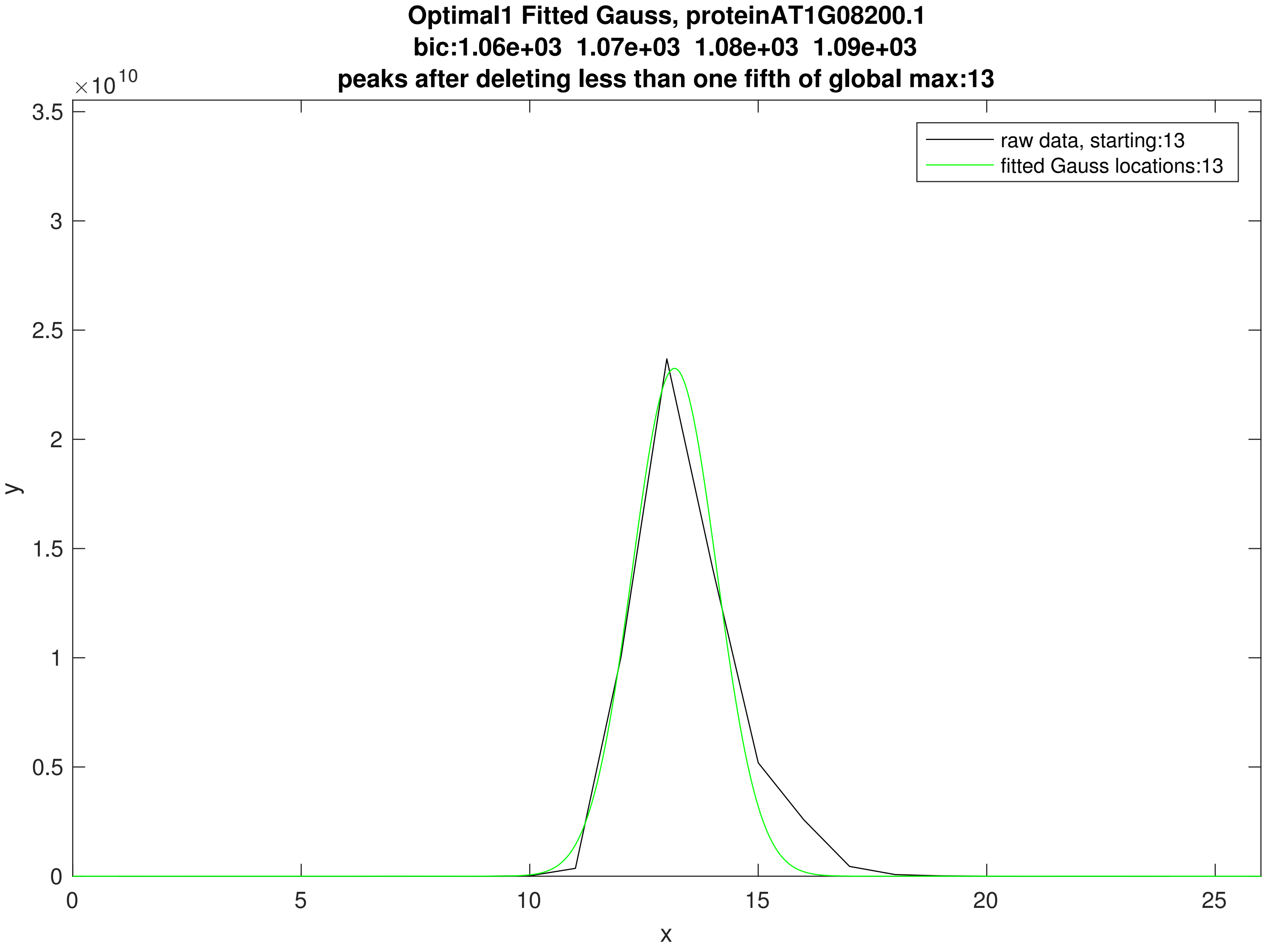
Optimal1 Fitted Gauss, proteinAT1G08200.1
bic:1.06e+03 1.07e+03 1.08e+03 1.09e+03
peaks after deleting less than one fifth of global max:13
×10^10
3.5
3
2.5
2
1.5
1
0.5
0
y
0
5
10
15
20
25
x
raw data, starting:13
fitted Gauss locations:13

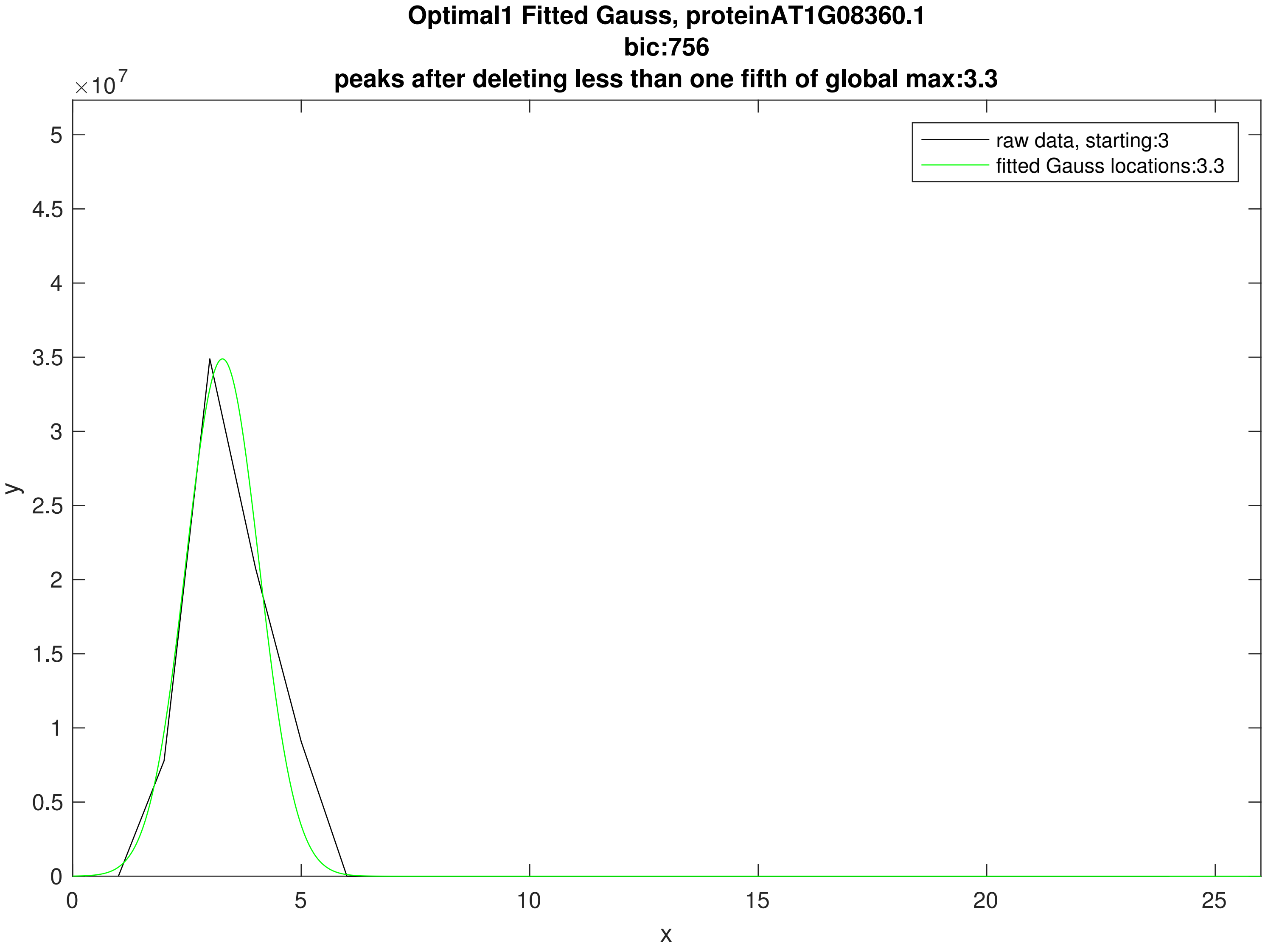
Optimal1 Fitted Gauss, proteinAT1G08360.1
bic:756
peaks after deleting less than one fifth of global max:3.3
×10^7
raw data, starting:3
fitted Gauss locations:3.3
y
x
0
0.5
1
1.5
2
2.5
3
3.5
4
4.5
5
0
5
10
15
20
25

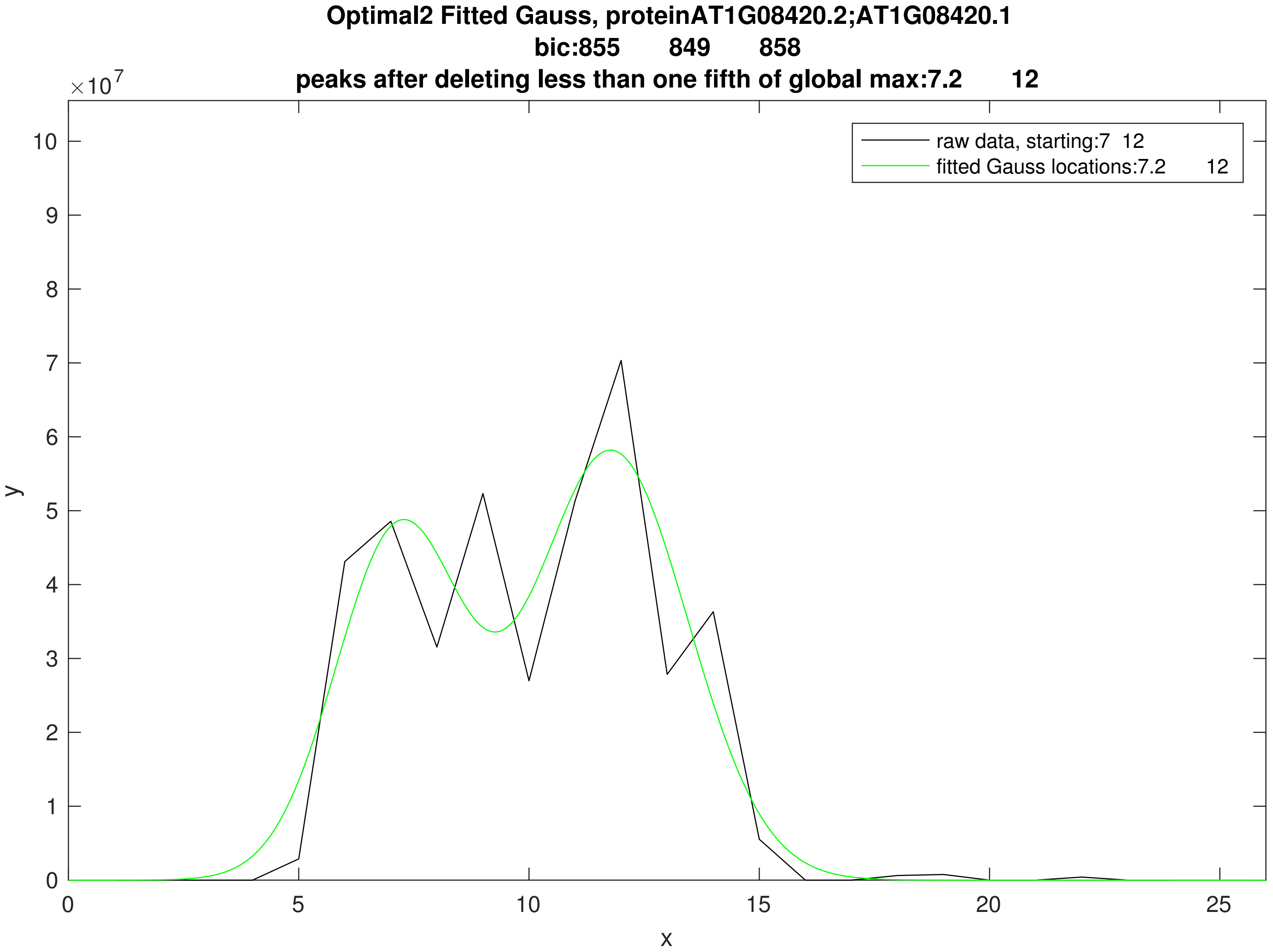
Optimal2 Fitted Gauss, proteinAT1G08420.2;AT1G08420.1
bic:855 849 858
peaks after deleting less than one fifth of global max:7.2 12
×10^7
raw data, starting:7 12
fitted Gauss locations:7.2 12
y
x
0
5
10
15
20
25
1
2
3
4
6
7
8
9

**Optimal2 Fitted Gauss, proteinAT1G08490.1**
**bic:805 800**
**peaks after deleting less than one fifth of global max:13**

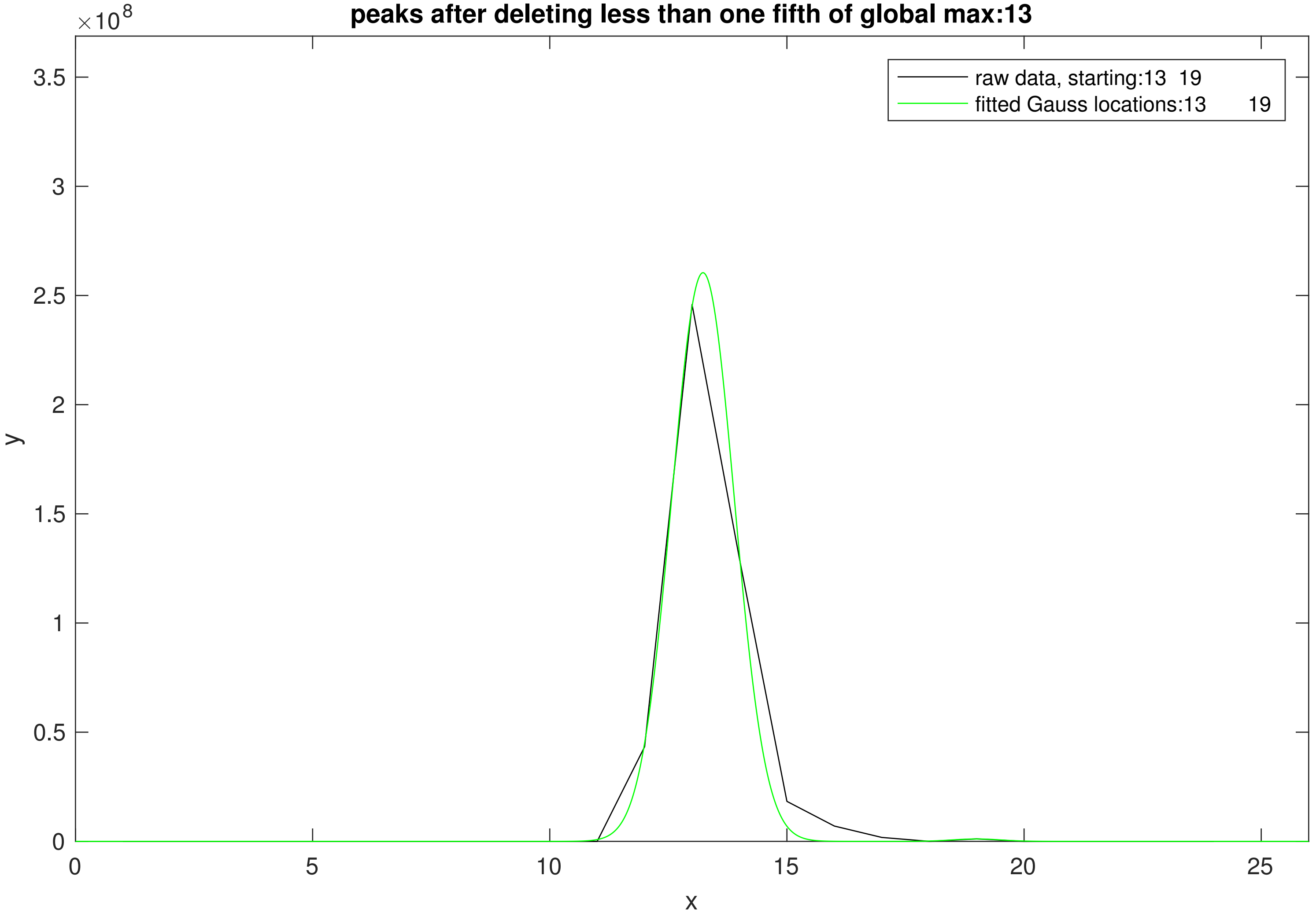

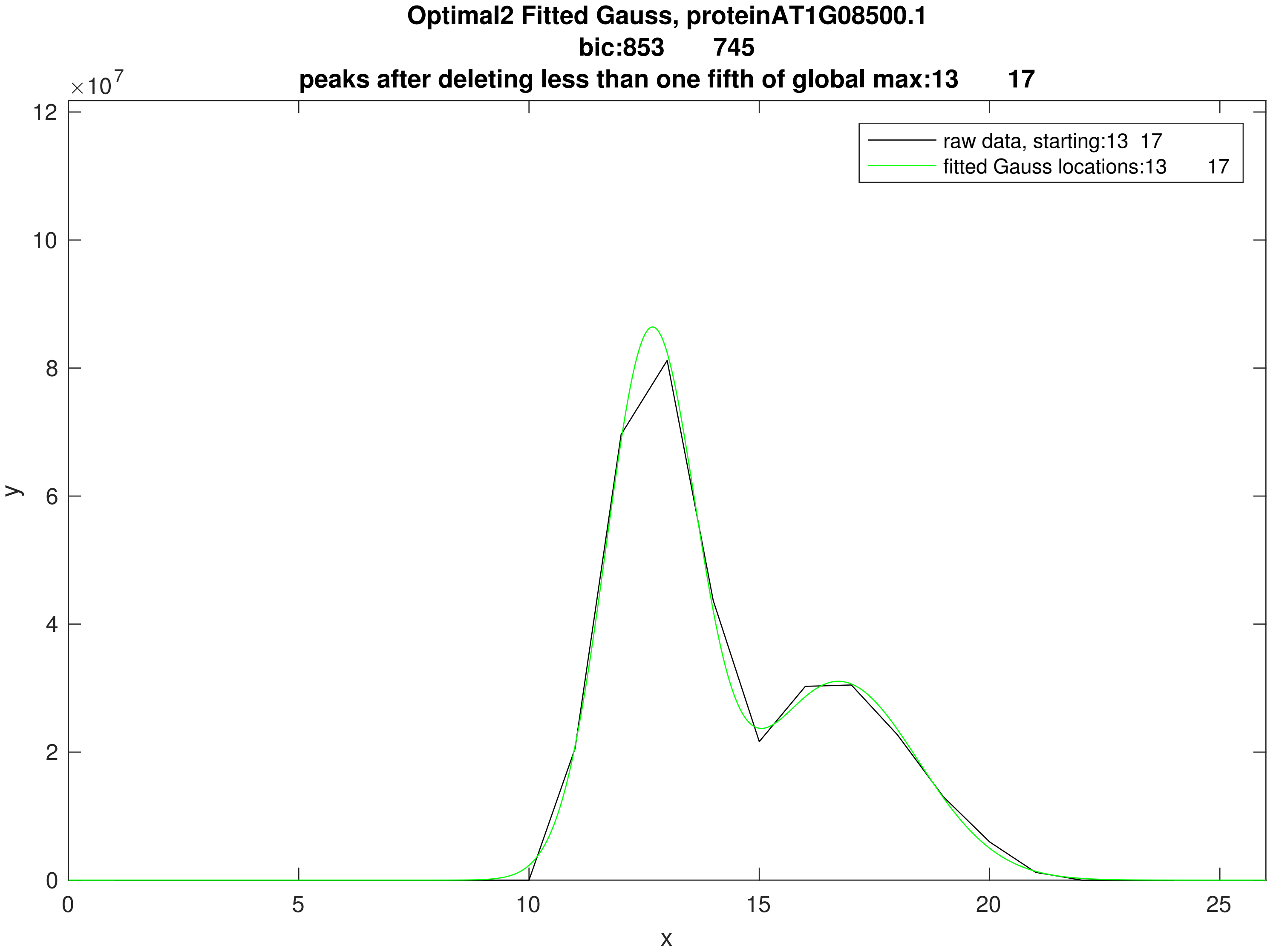

Optimal2 Fitted Gauss, proteinAT1G08500.1
bic:853 745
peaks after deleting less than one fifth of global max:13 17
×10^7
12
10
8
6
4
2
0
y
0
5
10
15
20
25
x
raw data, starting:13 17
fitted Gauss locations:13 17

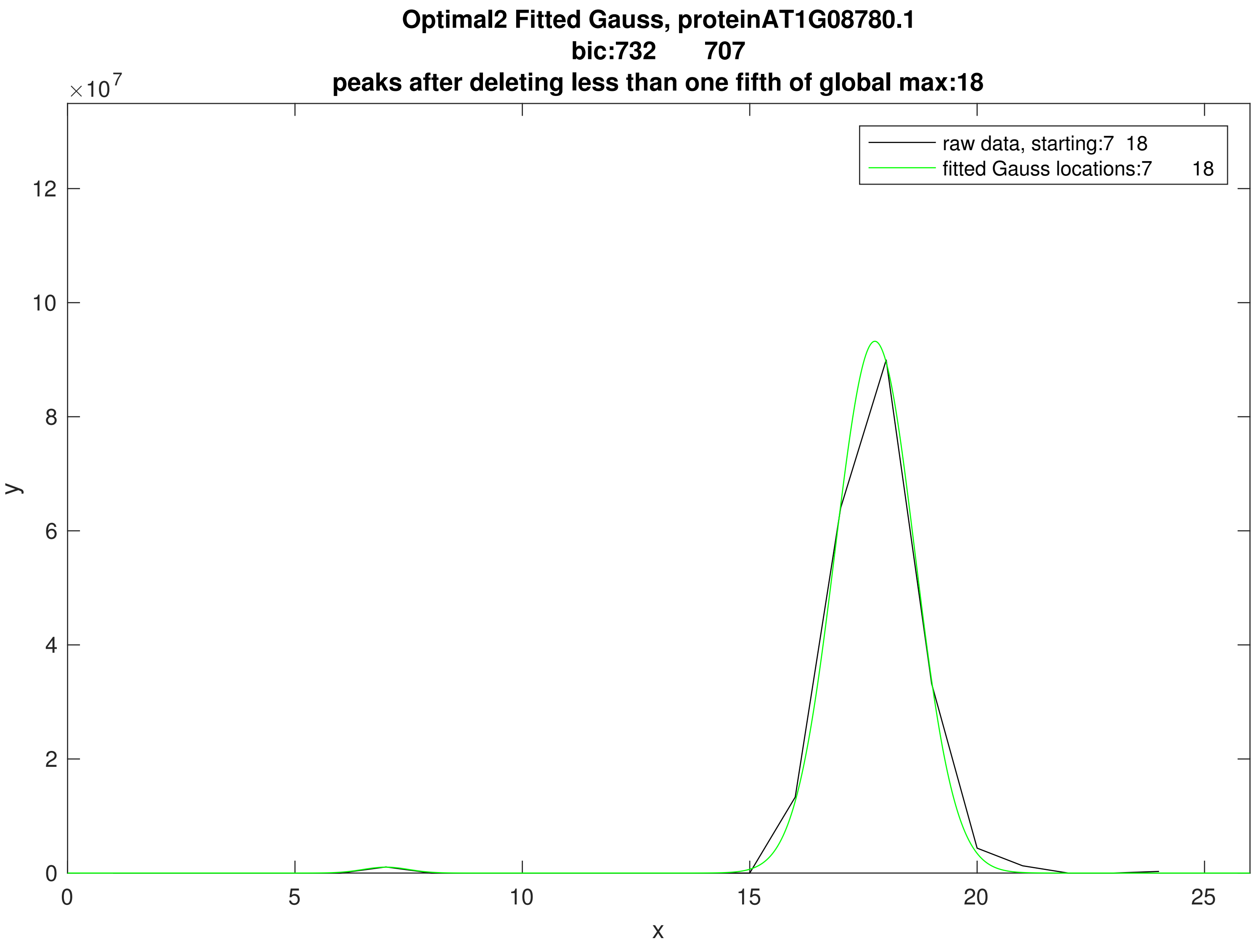

Optimal2 Fitted Gauss, proteinAT1G08780.1
bic:732 707
peaks after deleting less than one fifth of global max:18
×10^7
raw data, starting:7 18
fitted Gauss locations:7 18
y
x
0
2
4
6
8
10
12
0
5
10
15
20
25

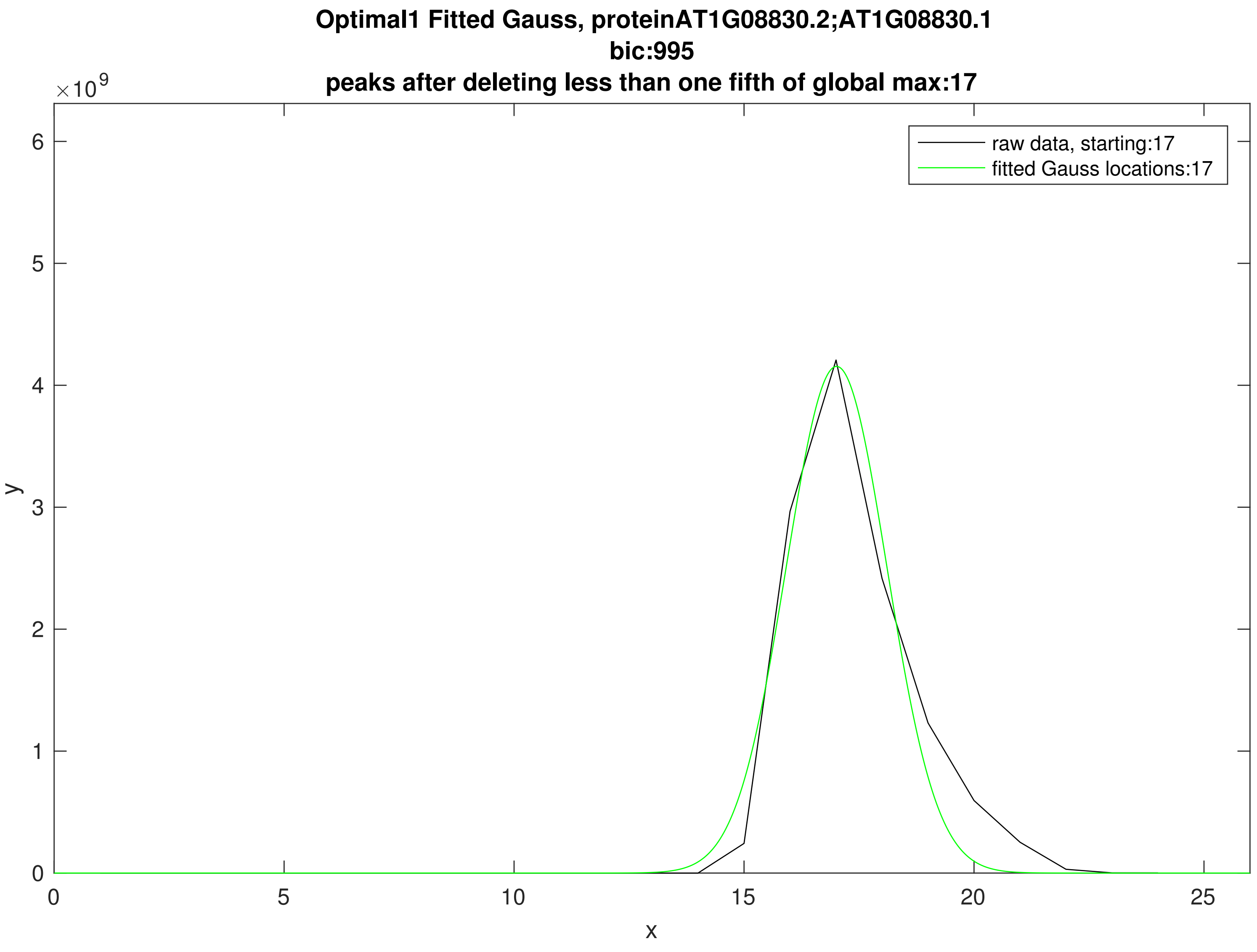
Optimal1 Fitted Gauss, proteinAT1G08830.2;AT1G08830.1
bic:995
peaks after deleting less than one fifth of global max:17
×10^9
raw data, starting:17
fitted Gauss locations:17
y
x
0
1
2
3
4
5
6
0
5
10
15
20
25

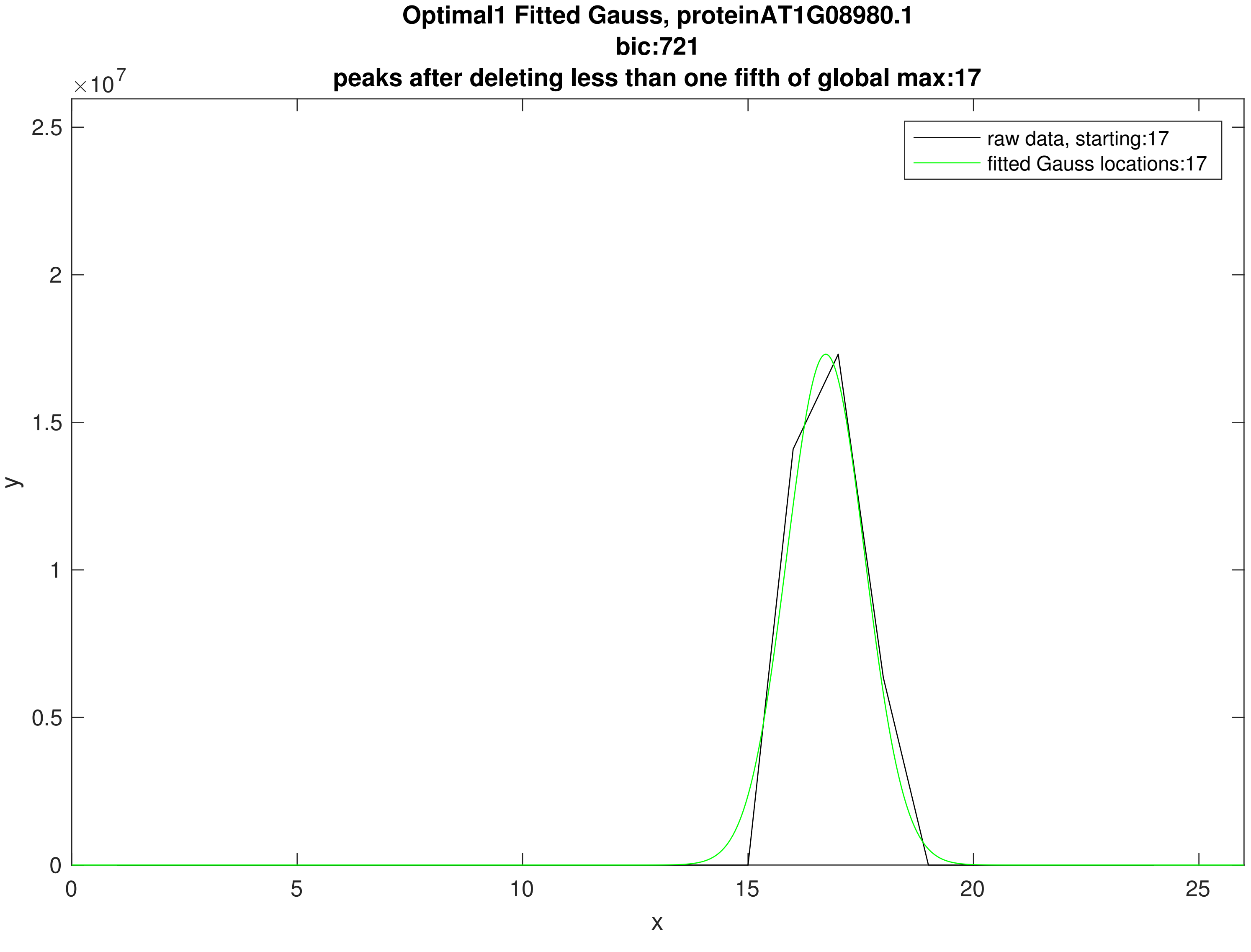
Optimal1 Fitted Gauss, proteinAT1G08980.1
bic:721
peaks after deleting less than one fifth of global max:17
×10^7
2.5
2
1.5
1
0.5
0
y
0
5
10
15
20
25
x
raw data, starting:17
fitted Gauss locations:17

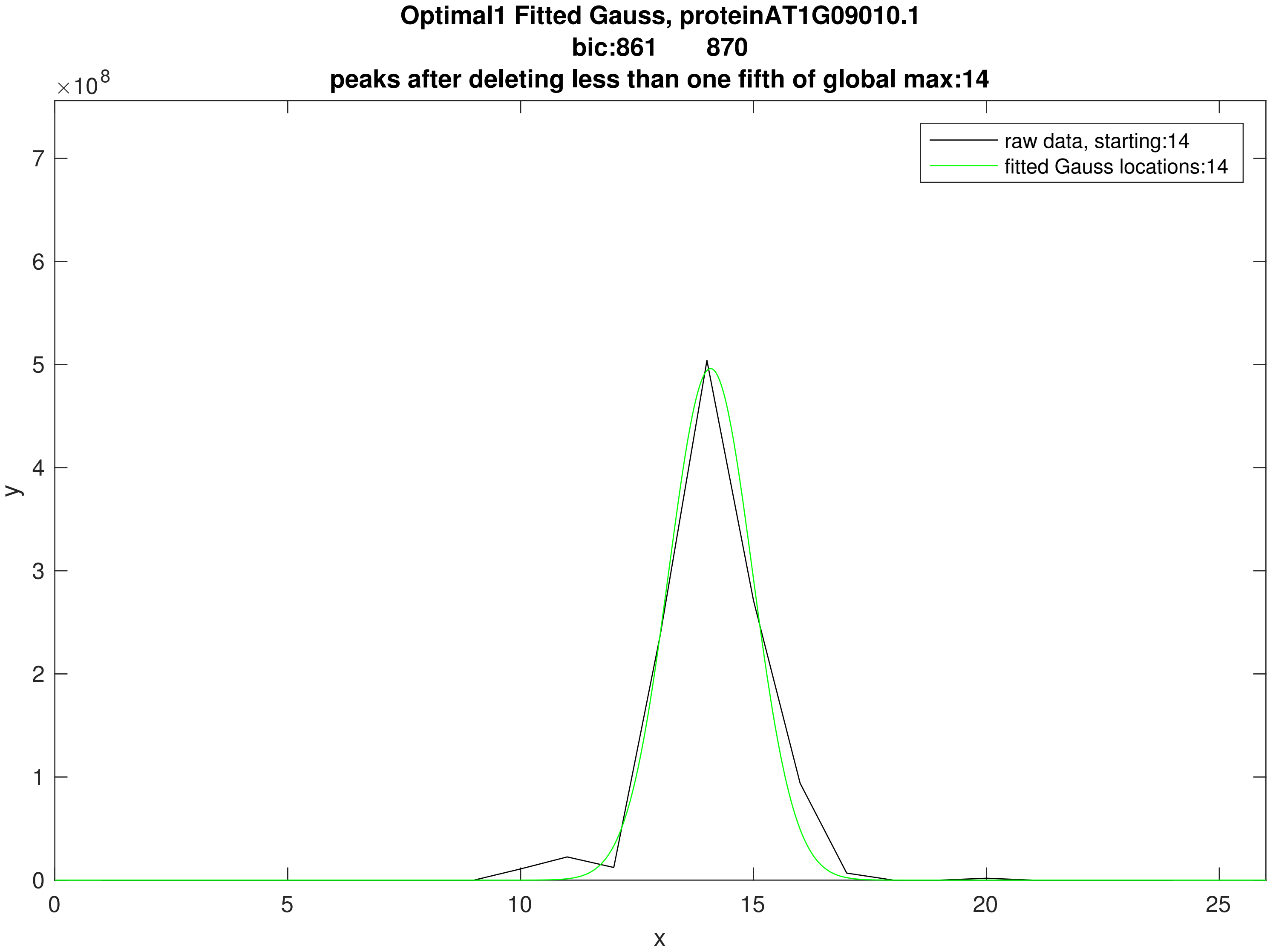
Optimal1 Fitted Gauss, proteinAT1G09010.1
bic:861 870
peaks after deleting less than one fifth of global max:14
×10^8
raw data, starting:14
fitted Gauss locations:14
y
x
0
1
2
3
4
5
6
7
0
5
10
15
20
25

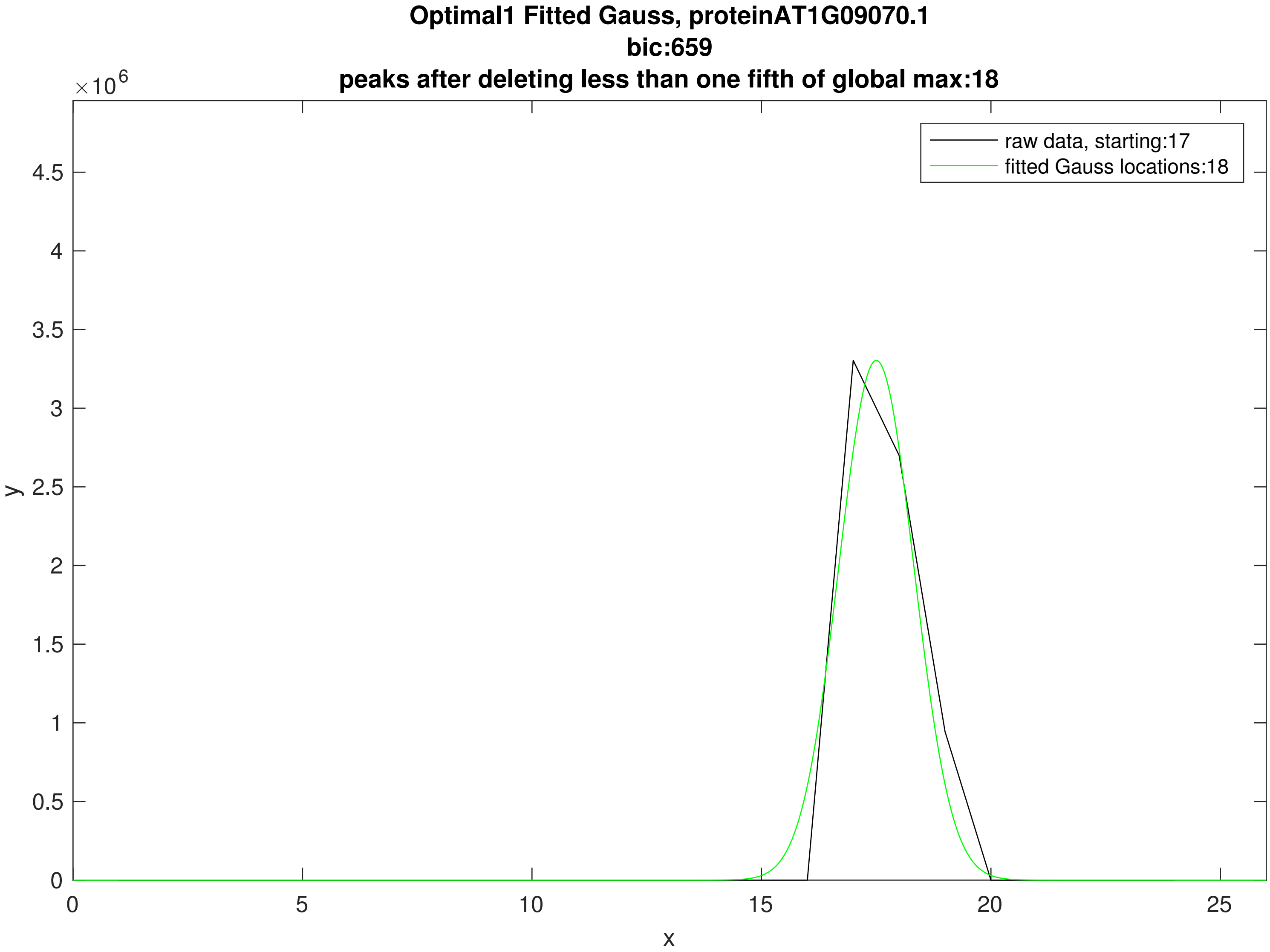
Optimal1 Fitted Gauss, proteinAT1G09070.1
bic:659
peaks after deleting less than one fifth of global max:18
×10^6
4.5
4
3.5
3
2.5
2
1.5
1
0.5
0
y
0
5
10
15
20
25
x
raw data, starting:17
fitted Gauss locations:18

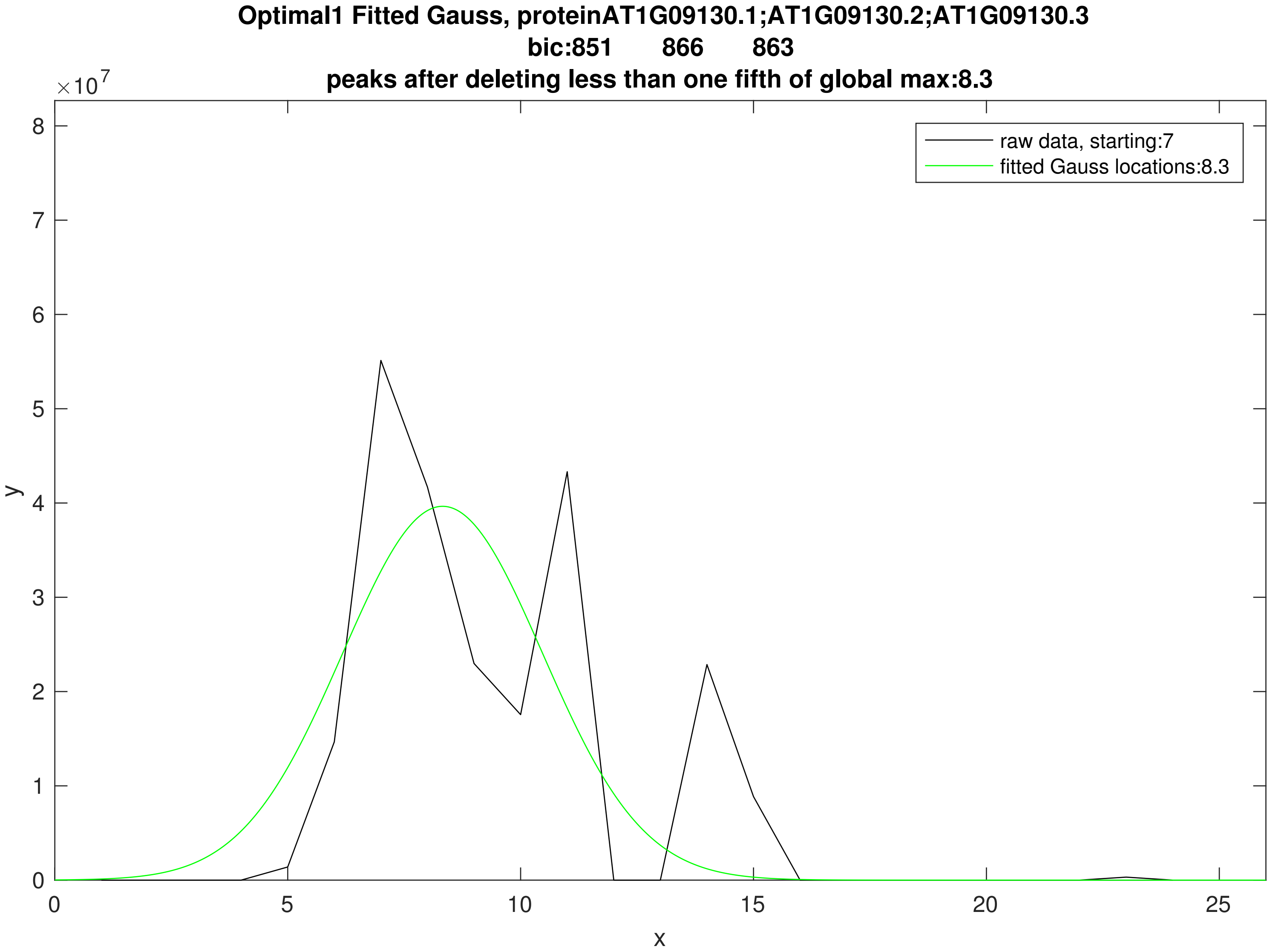
Optimal1 Fitted Gauss, proteinAT1G09130.1;AT1G09130.2;AT1G09130.3
bic:851 866 863
peaks after deleting less than one fifth of global max:8.3
×10^7
raw data, starting:7
fitted Gauss locations:8.3
y
x
0
1
2
3
4
5
6
7
8
0
5
10
15
20
25

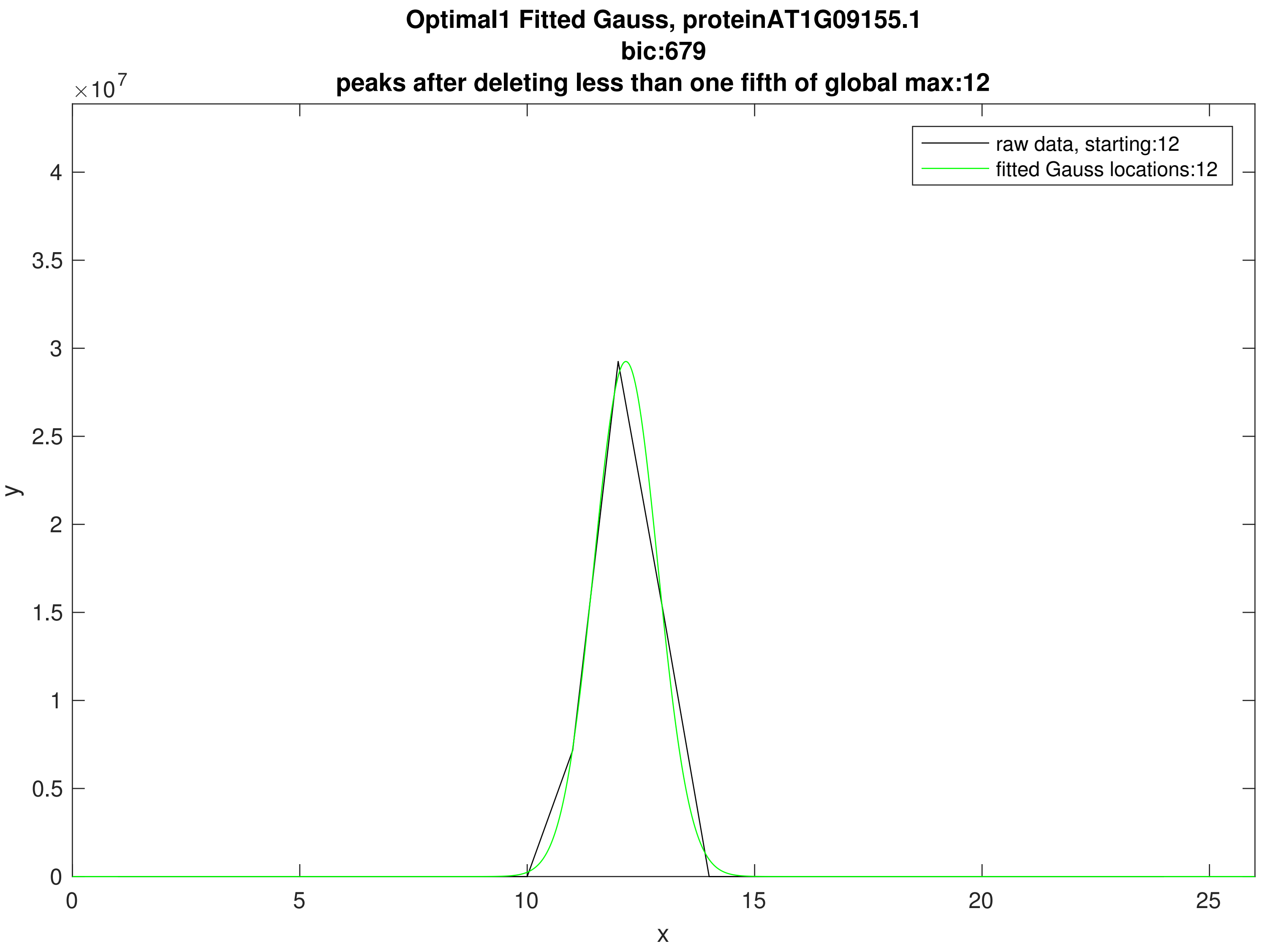

Optimal1 Fitted Gauss, proteinAT1G09155.1
bic:679
peaks after deleting less than one fifth of global max:12
×10^7
raw data, starting:12
fitted Gauss locations:12
y
x
0
0.5
1
1.5
2
2.5
3
3.5
4
5
10
15
20
25

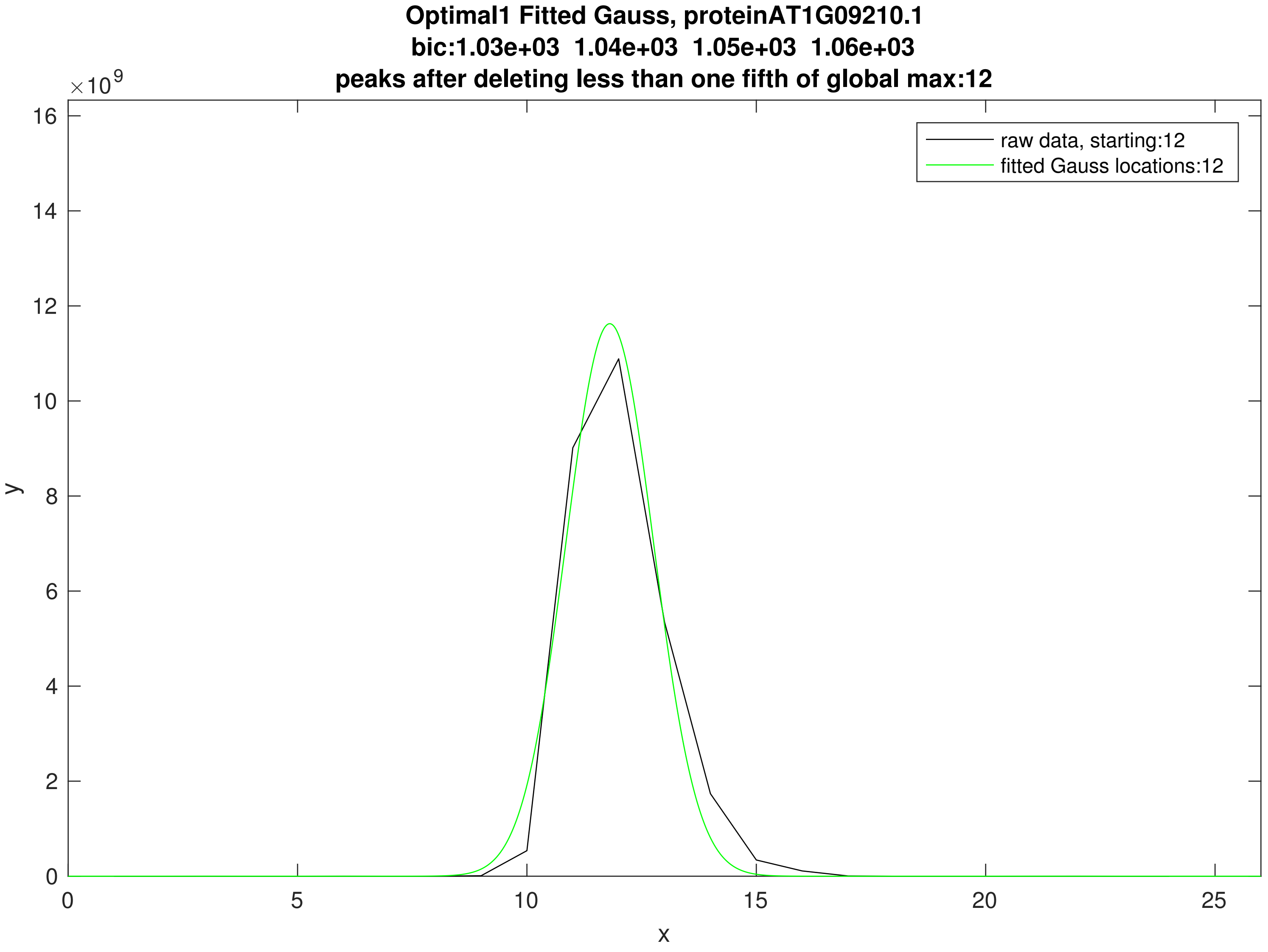

Optimal1 Fitted Gauss, proteinAT1G09210.1
bic:1.03e+03  1.04e+03  1.05e+03  1.06e+03
peaks after deleting less than one fifth of global max:12
×10^9
16
14
12
10
8
6
4
2
0
y
0
5
10
15
20
25
x
raw data, starting:12
fitted Gauss locations:12

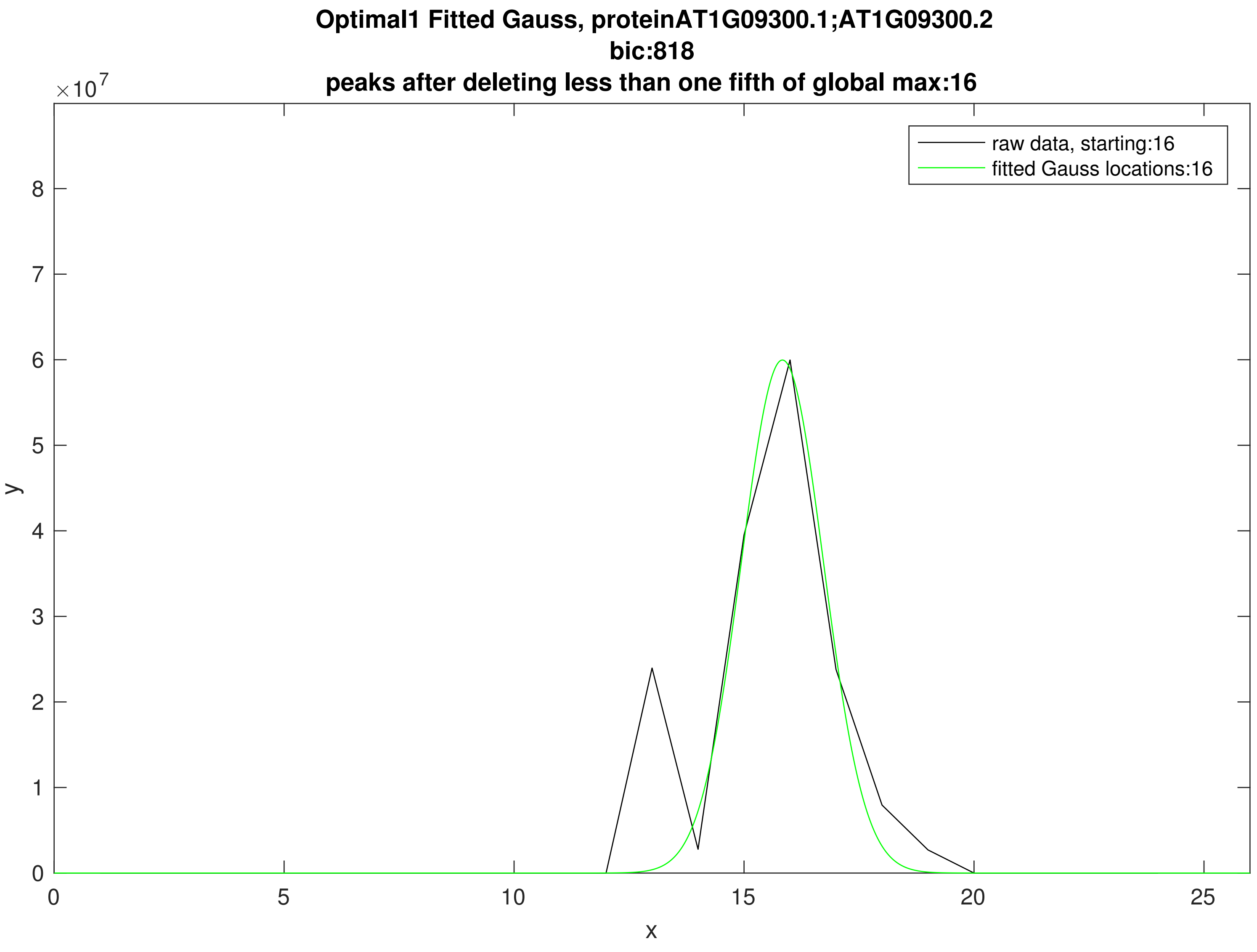
Optimal1 Fitted Gauss, proteinAT1G09300.1;AT1G09300.2
bic:818
peaks after deleting less than one fifth of global max:16
×10^7
raw data, starting:16
fitted Gauss locations:16
y
x
0
1
2
3
4
5
6
7
8
0
5
10
15
20
25

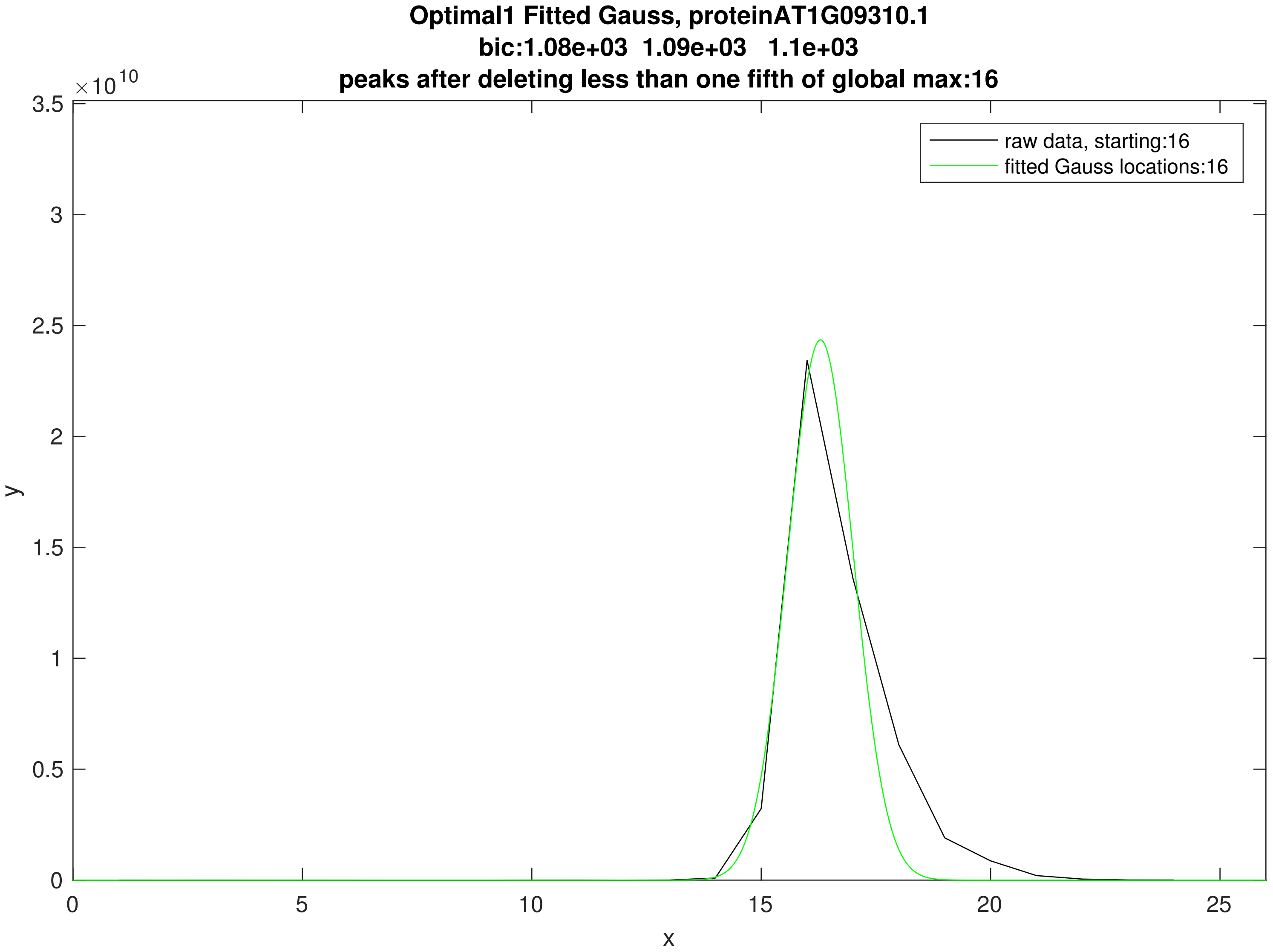

Optimal1 Fitted Gauss, proteinAT1G09310.1
bic:1.08e+03 1.09e+03 1.1e+03
peaks after deleting less than one fifth of global max:16
×10^10
raw data, starting:16
fitted Gauss locations:16
y
x
0
0.5
1
1.5
2
2.5
3
3.5
0
5
10
15
20
25

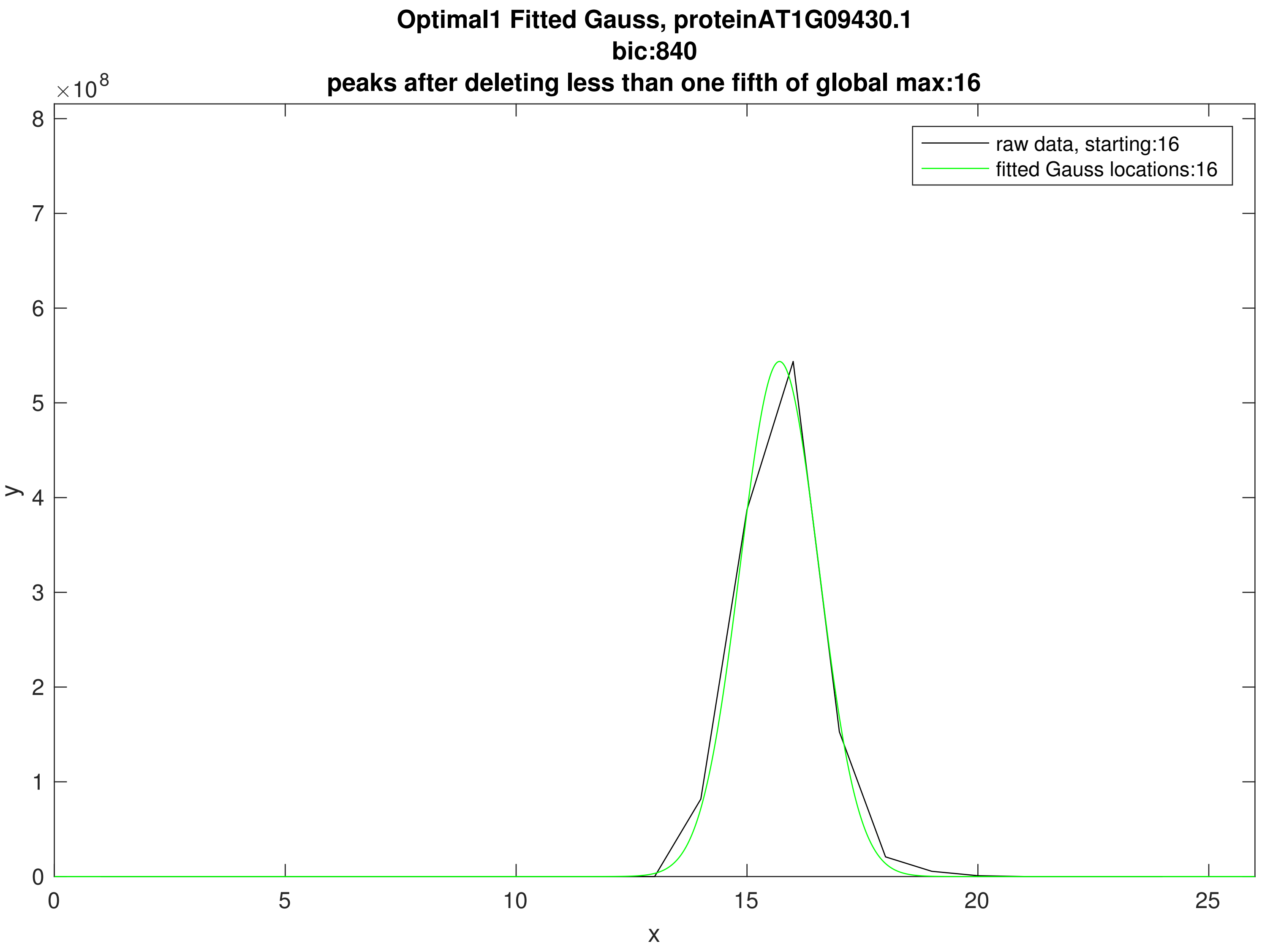
Optimal1 Fitted Gauss, proteinAT1G09430.1
bic:840
peaks after deleting less than one fifth of global max:16
×10^8
raw data, starting:16
fitted Gauss locations:16
y
x
0
1
2
3
4
5
6
7
8
0
5
10
15
20
25

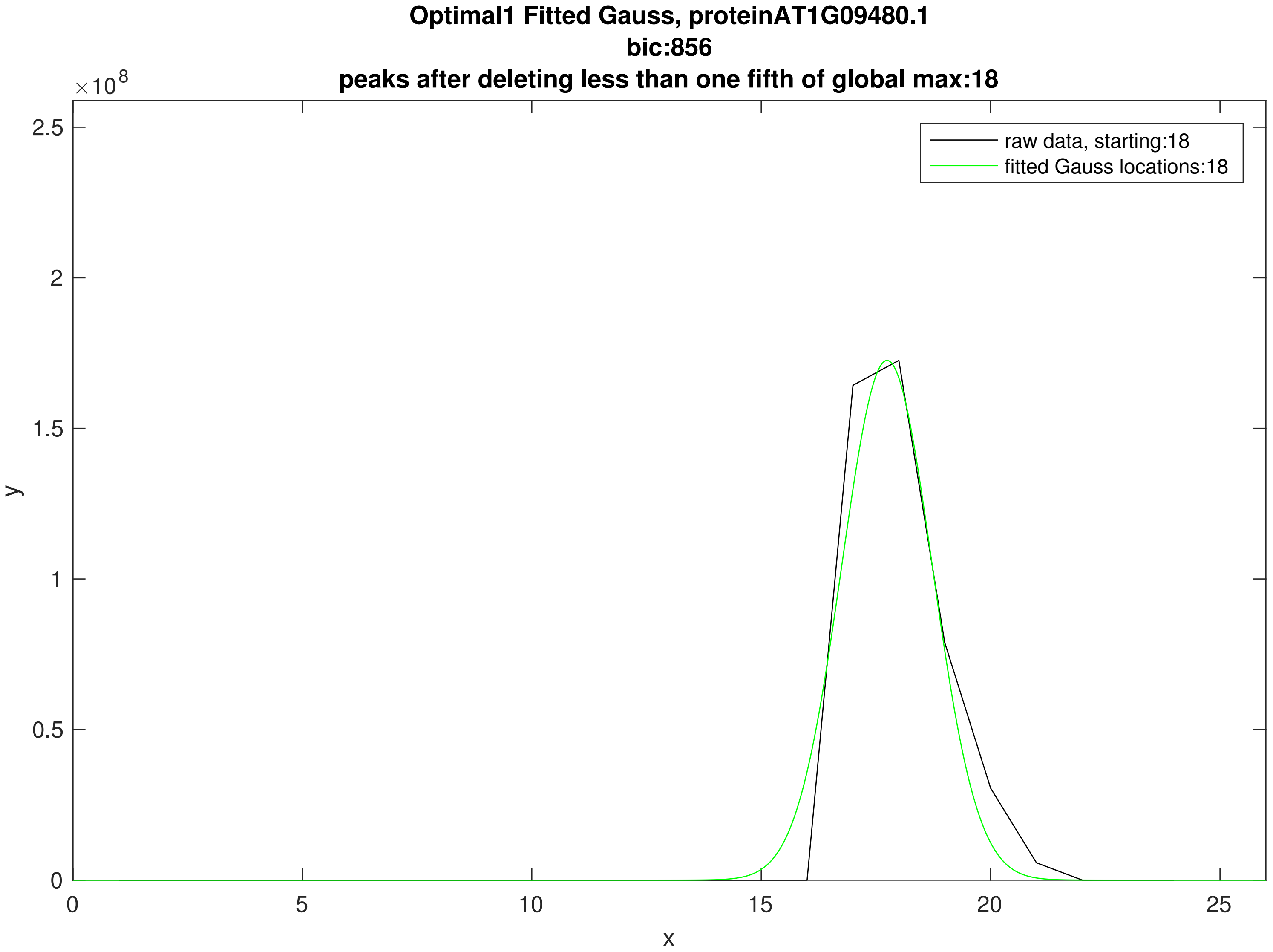
Optimal1 Fitted Gauss, proteinAT1G09480.1
bic:856
peaks after deleting less than one fifth of global max:18
×10^8
2.5
2
1.5
1
0.5
0
y
0
5
10
15
20
25
x
raw data, starting:18
fitted Gauss locations:18

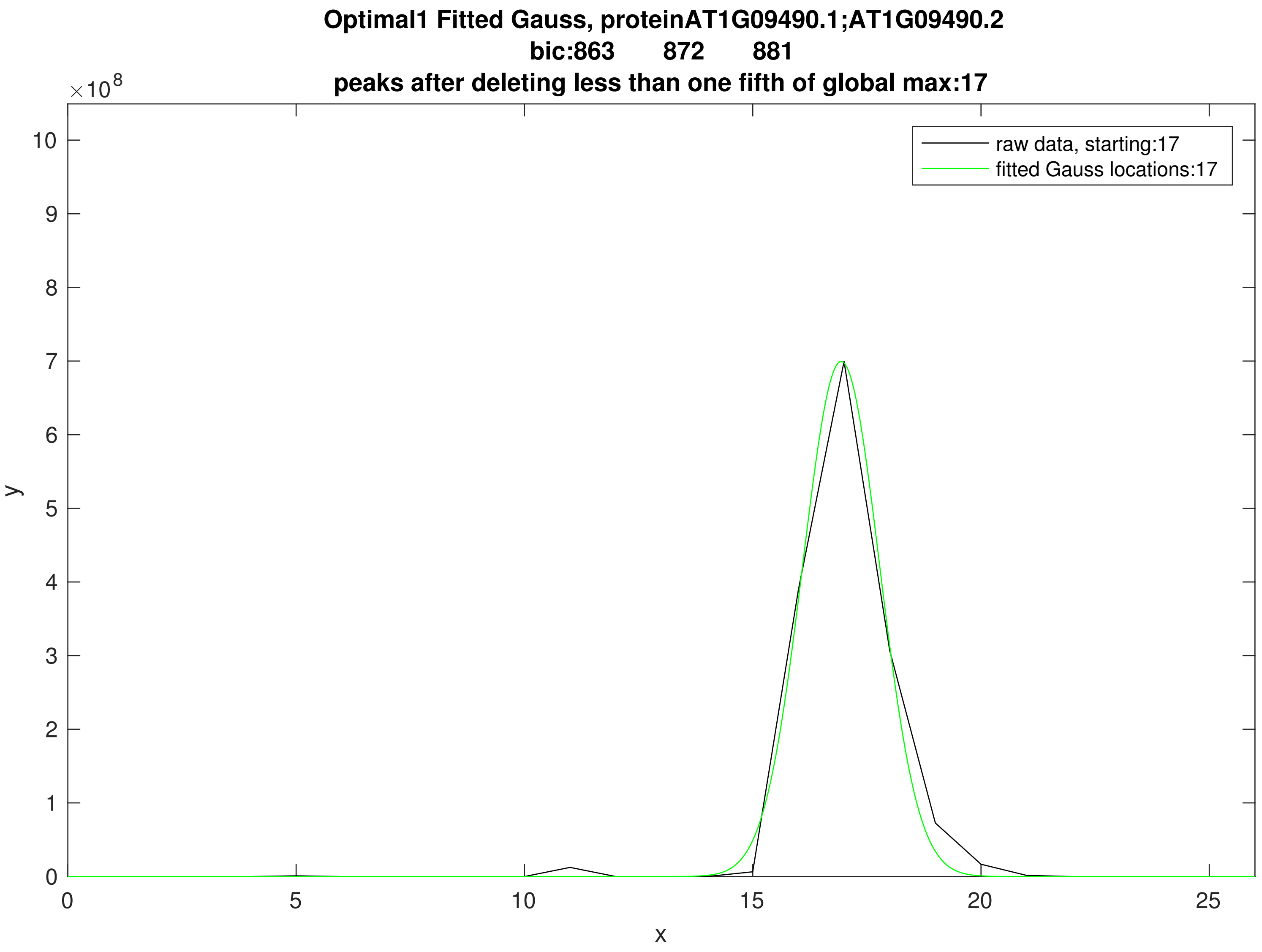

Optimal1 Fitted Gauss, proteinAT1G09490.1;AT1G09490.2
bic:863 872 881
peaks after deleting less than one fifth of global max:17
×10^8
raw data, starting:17
fitted Gauss locations:17
y
x
0
1
2
3
4
5
6
7
8
9
10
0
5
10
15
20
25

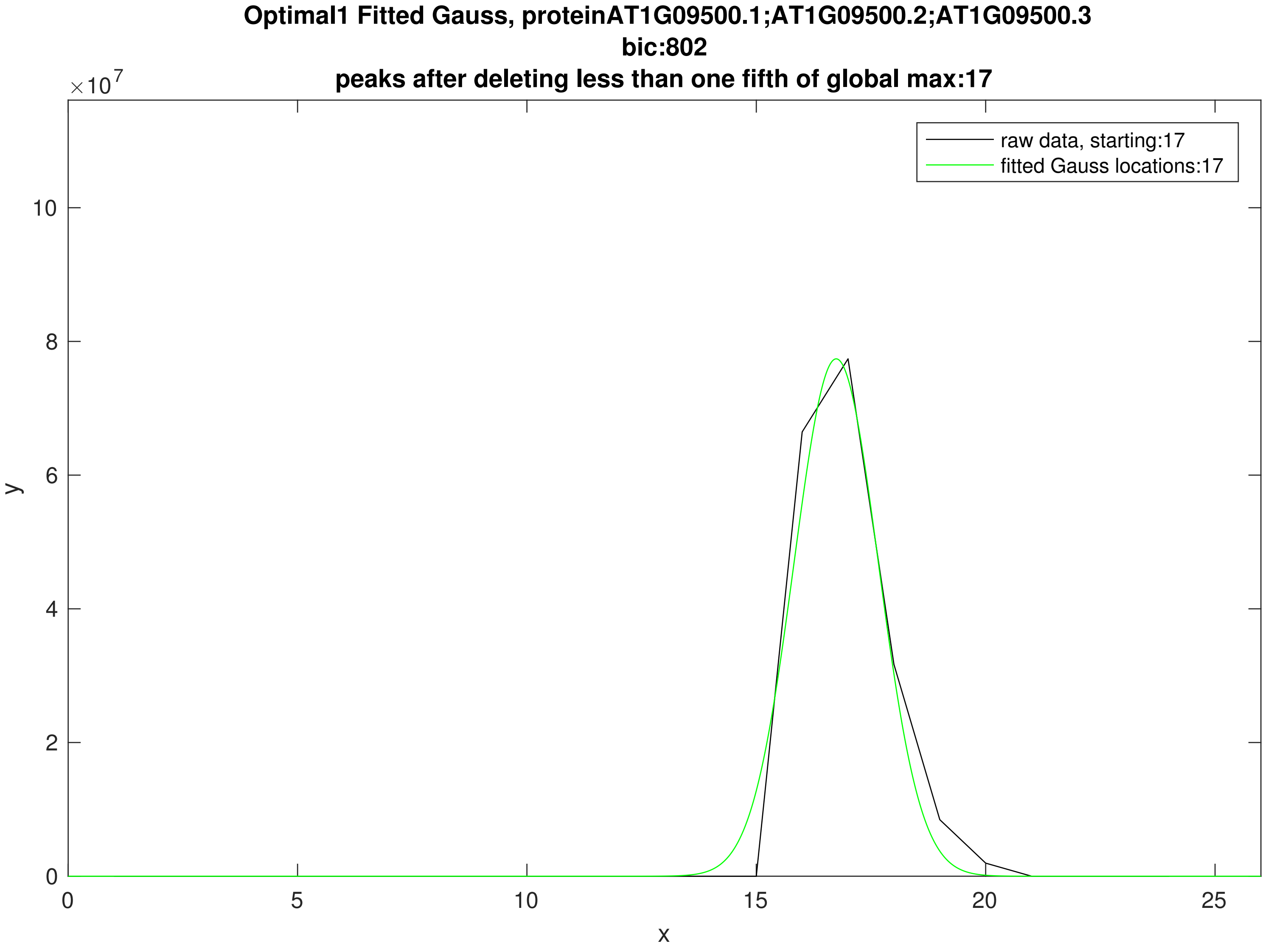
Optimal1 Fitted Gauss, proteinAT1G09500.1;AT1G09500.2;AT1G09500.3
bic:802
peaks after deleting less than one fifth of global max:17
×10^7
raw data, starting:17
fitted Gauss locations:17
y
x
0
2
4
6
8
10
0
5
10
15
20
25

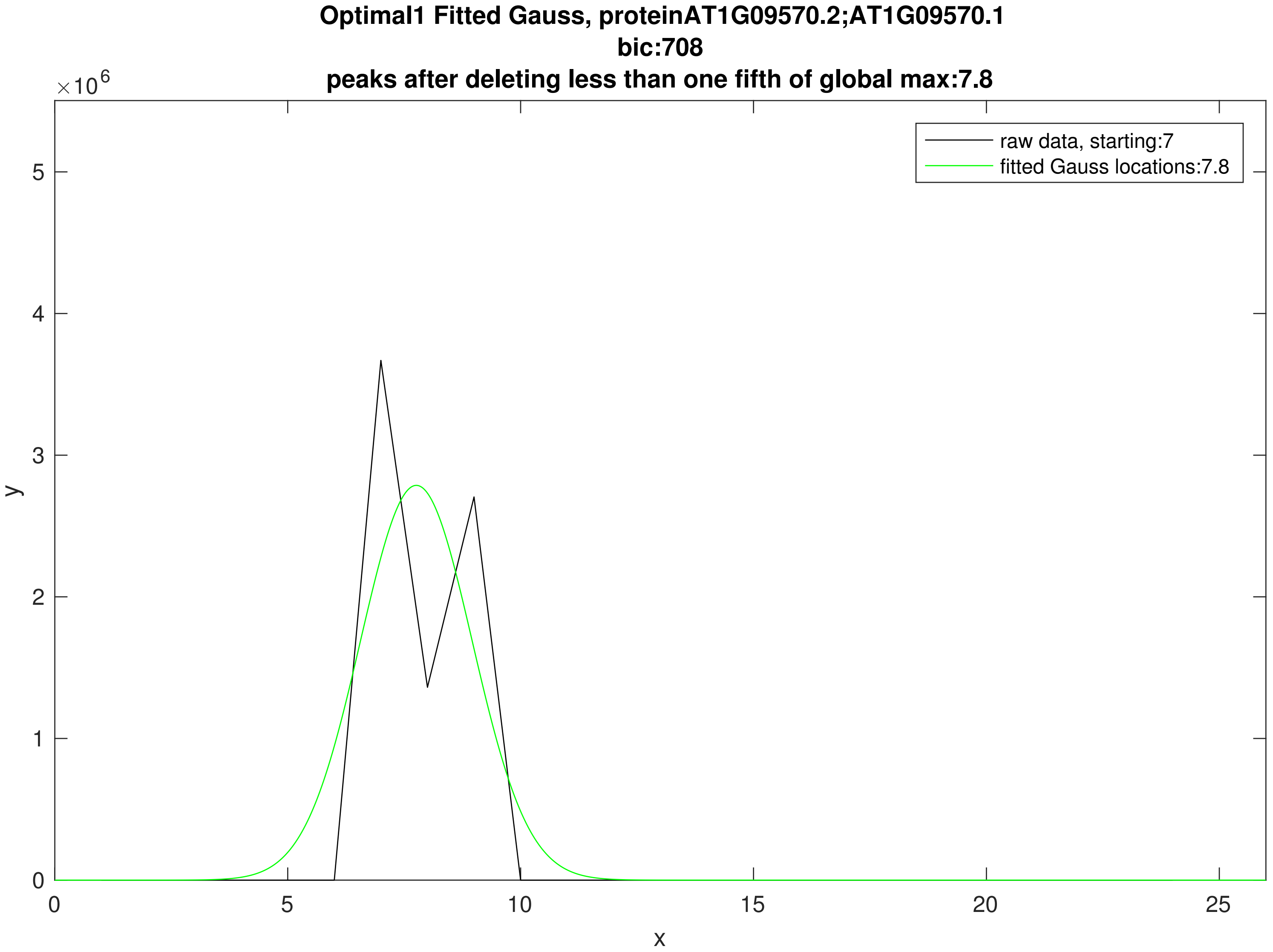
Optimal1 Fitted Gauss, proteinAT1G09570.2;AT1G09570.1
bic:708
peaks after deleting less than one fifth of global max:7.8
×10^6
raw data, starting:7
fitted Gauss locations:7.8
y
x
0
1
2
3
4
5
0
5
10
15
20
25

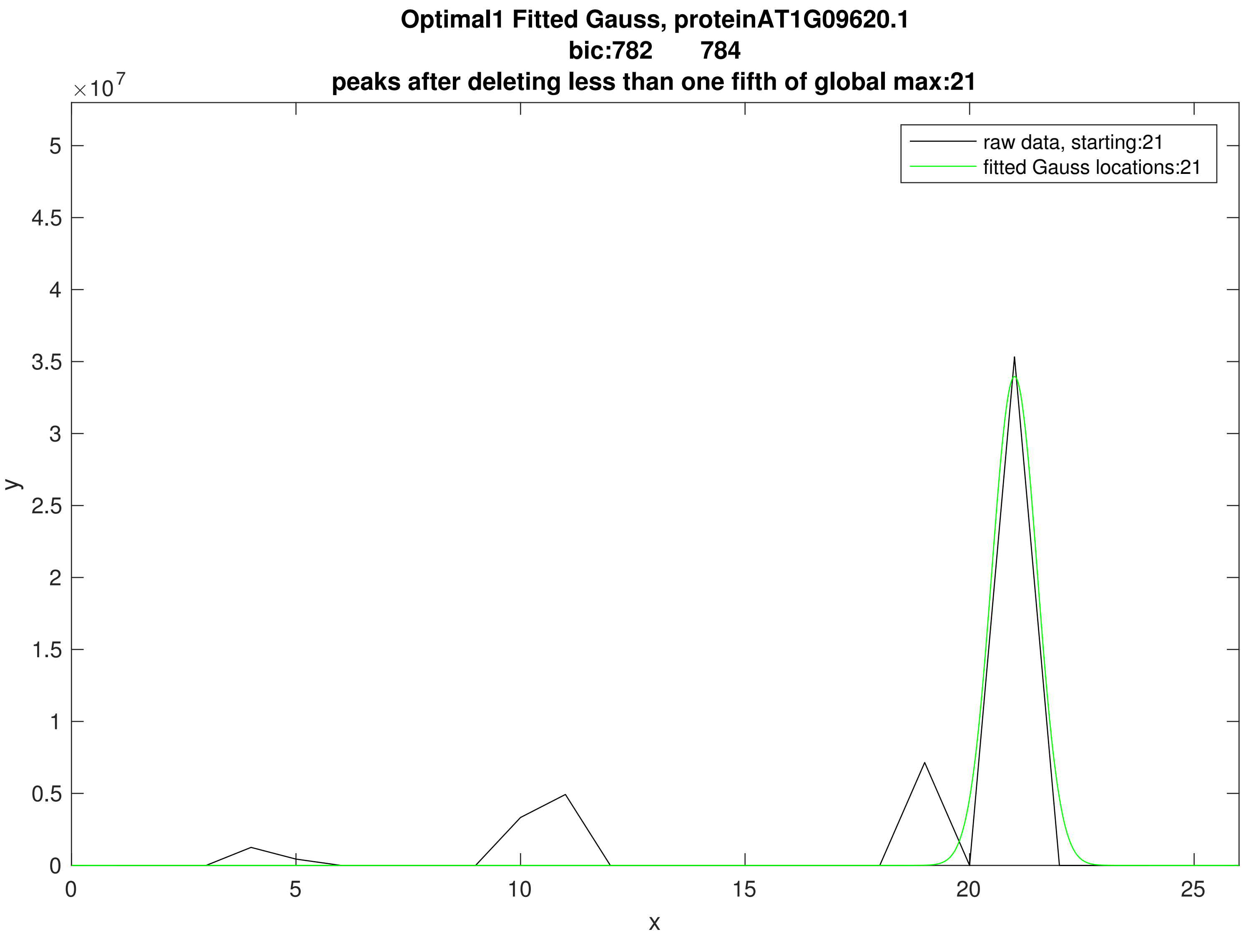
Optimal1 Fitted Gauss, proteinAT1G09620.1
bic:782 784
peaks after deleting less than one fifth of global max:21
×10^7
5
4.5
4
3.5
3
2.5
2
1.5
1
0.5
0
y
0
5
10
15
20
25
x
raw data, starting:21
fitted Gauss locations:21

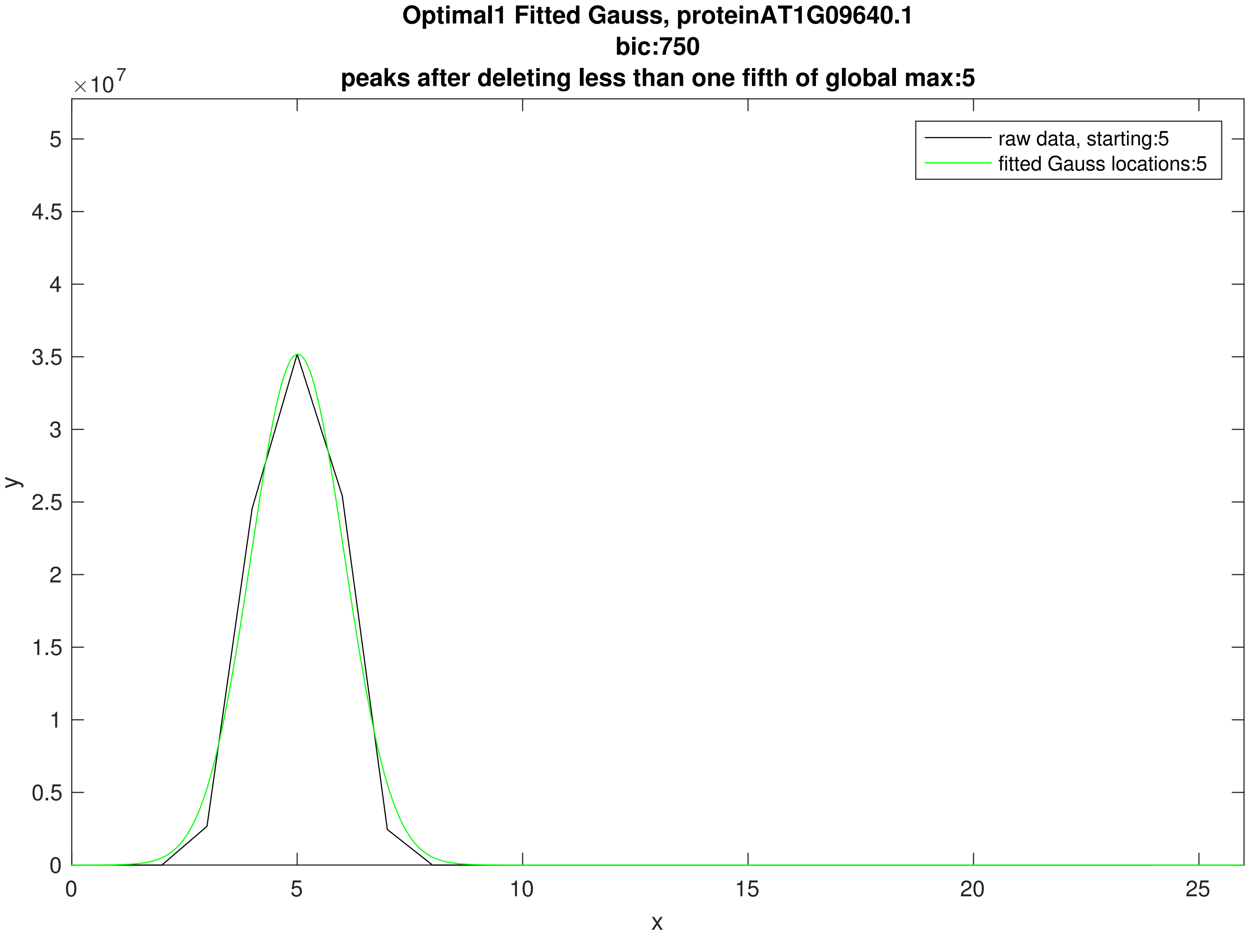

Optimal1 Fitted Gauss, proteinAT1G09640.1
bic:750
peaks after deleting less than one fifth of global max:5
×10^7
raw data, starting:5
fitted Gauss locations:5
y
x
0
0.5
1
1.5
2
2.5
3
3.5
4
4.5
5
0
5
10
15
20
25

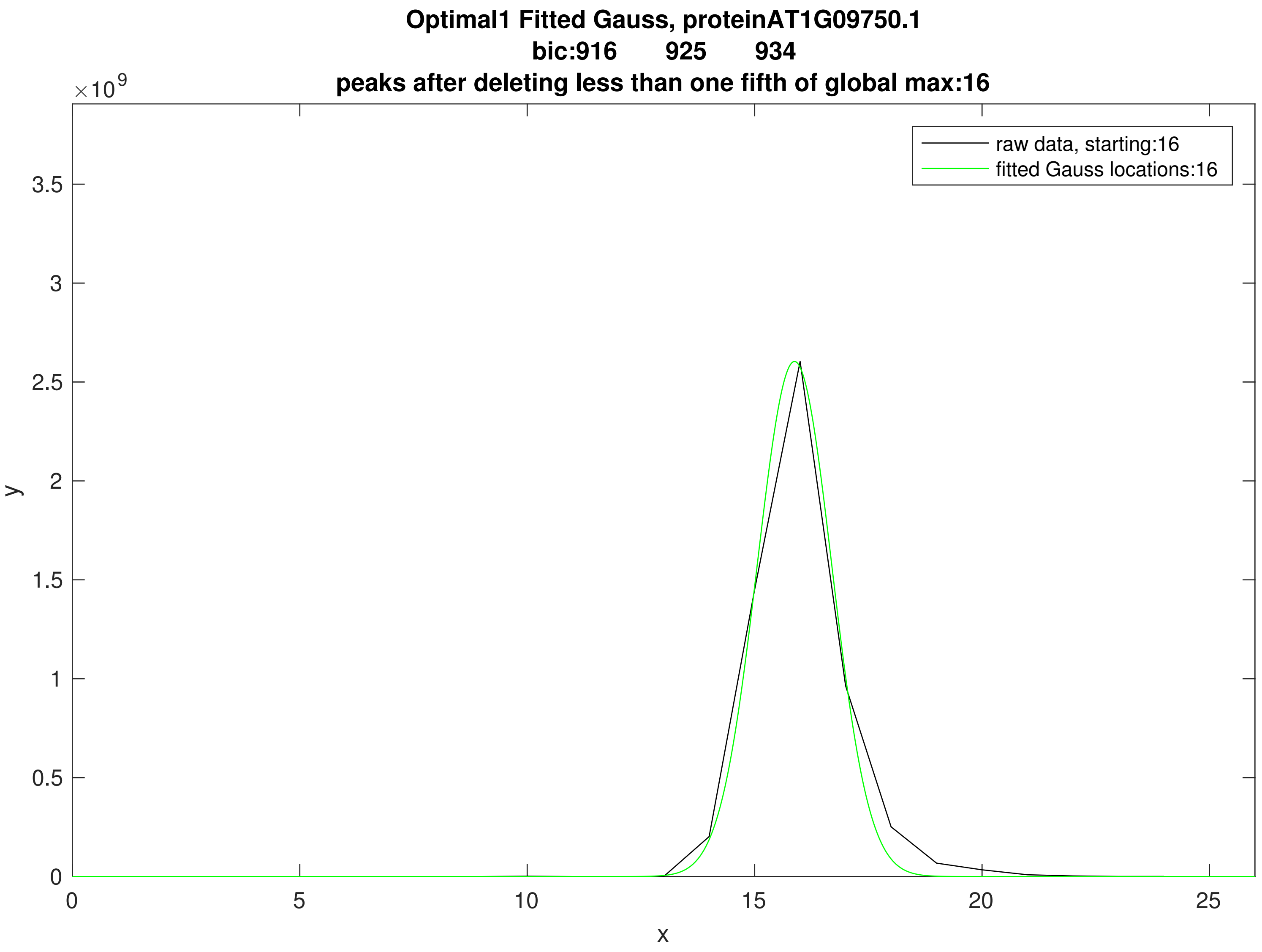

Optimal1 Fitted Gauss, proteinAT1G09750.1
bic:916 925 934
peaks after deleting less than one fifth of global max:16
×10^9
raw data, starting:16
fitted Gauss locations:16
y
x
0
0.5
1
1.5
2
2.5
3
3.5
5
10
15
20
25

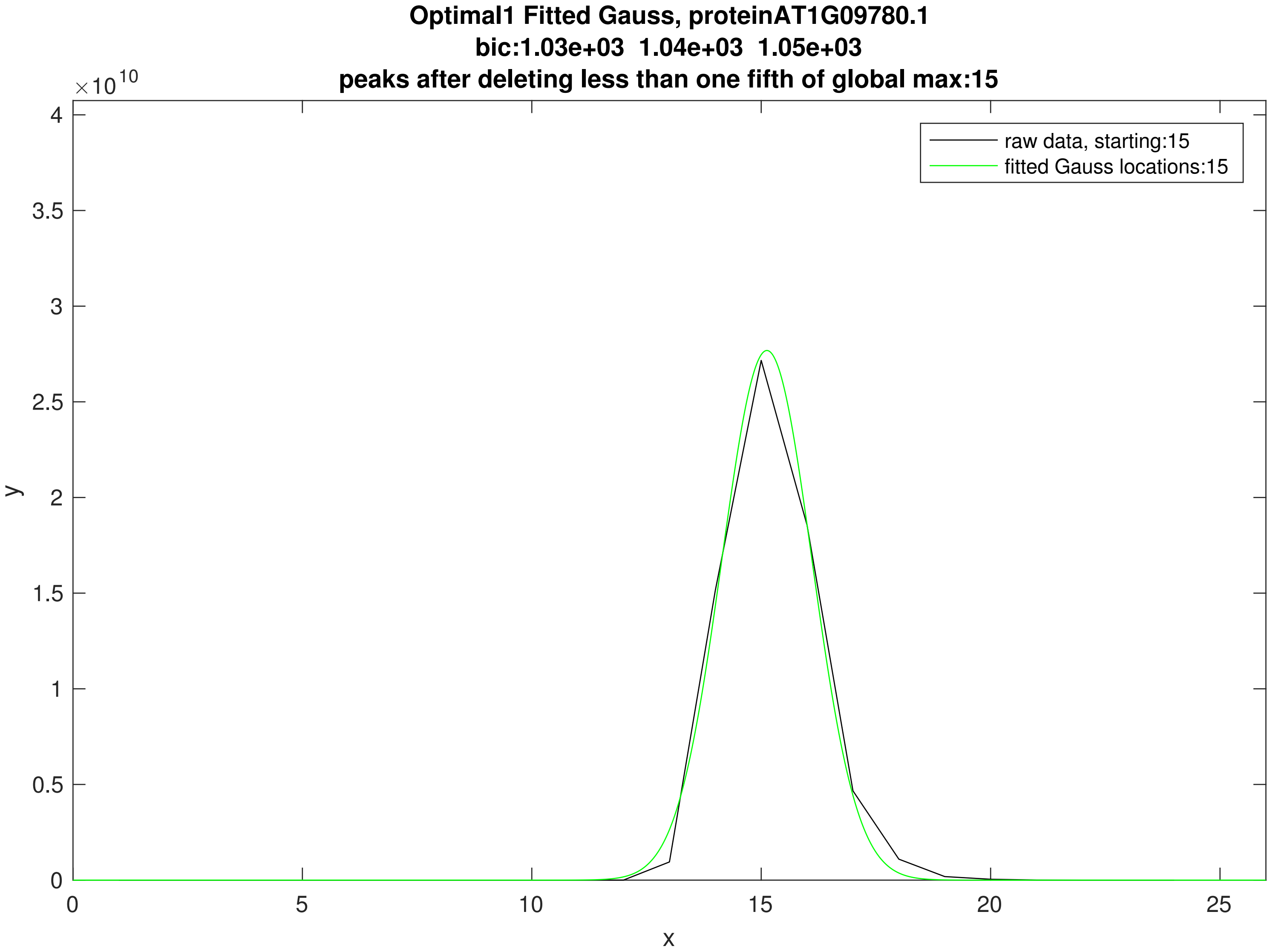
Optimal1 Fitted Gauss, proteinAT1G09780.1
bic:1.03e+03 1.04e+03 1.05e+03
peaks after deleting less than one fifth of global max:15
×10^10
raw data, starting:15
fitted Gauss locations:15
y
x
0
0.5
1
1.5
2
2.5
3
3.5
4
5
10
15
20
25

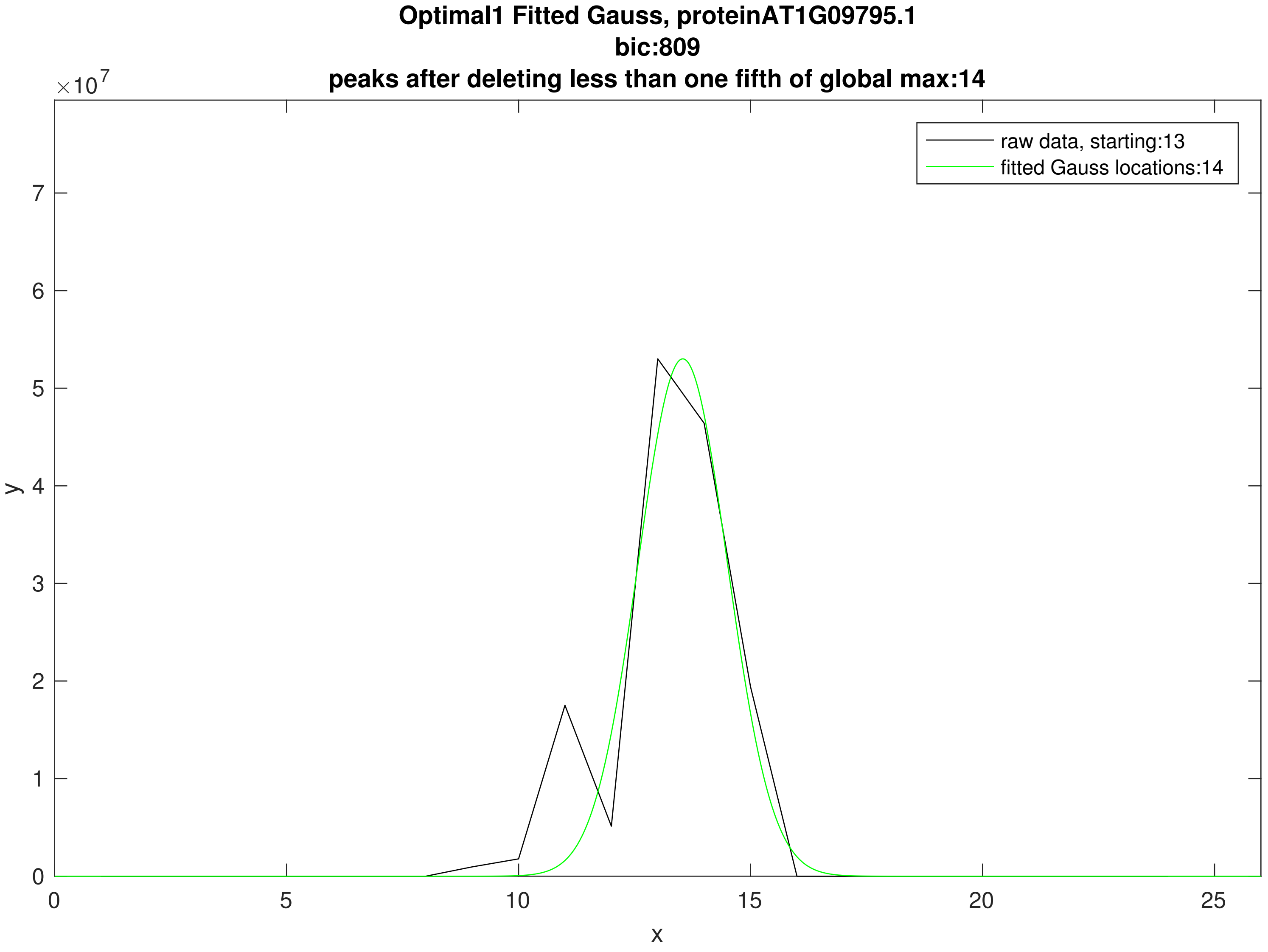
Optimal1 Fitted Gauss, proteinAT1G09795.1
bic:809
peaks after deleting less than one fifth of global max:14
×10^7
raw data, starting:13
fitted Gauss locations:14
y
x
0
1
2
3
4
5
6
7
0
5
10
15
20
25

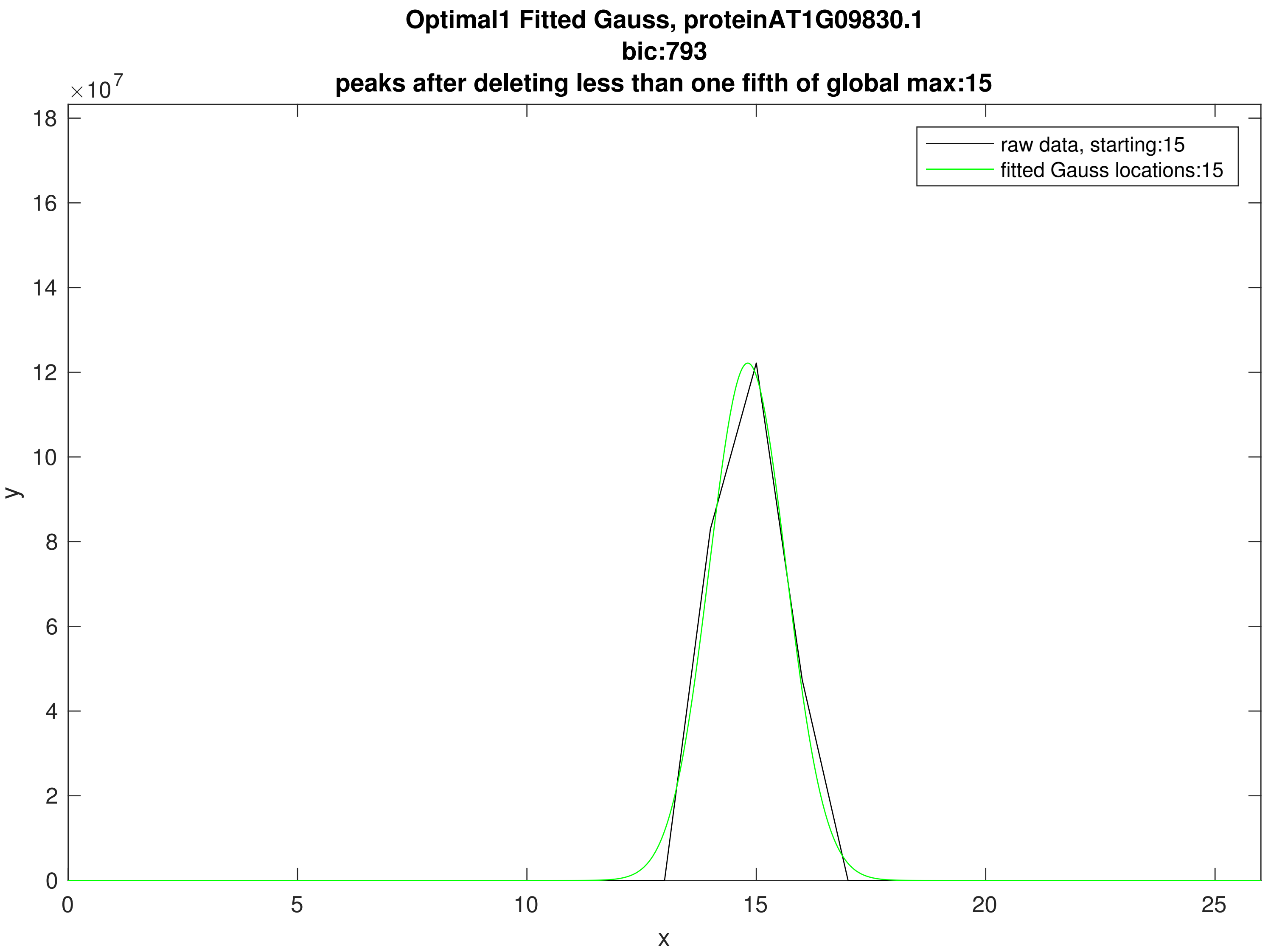

Optimal1 Fitted Gauss, proteinAT1G09830.1
bic:793
peaks after deleting less than one fifth of global max:15
×10^7
18
16
14
12
10
8
6
4
2
0
y
0
5
10
15
20
25
x
raw data, starting:15
fitted Gauss locations:15

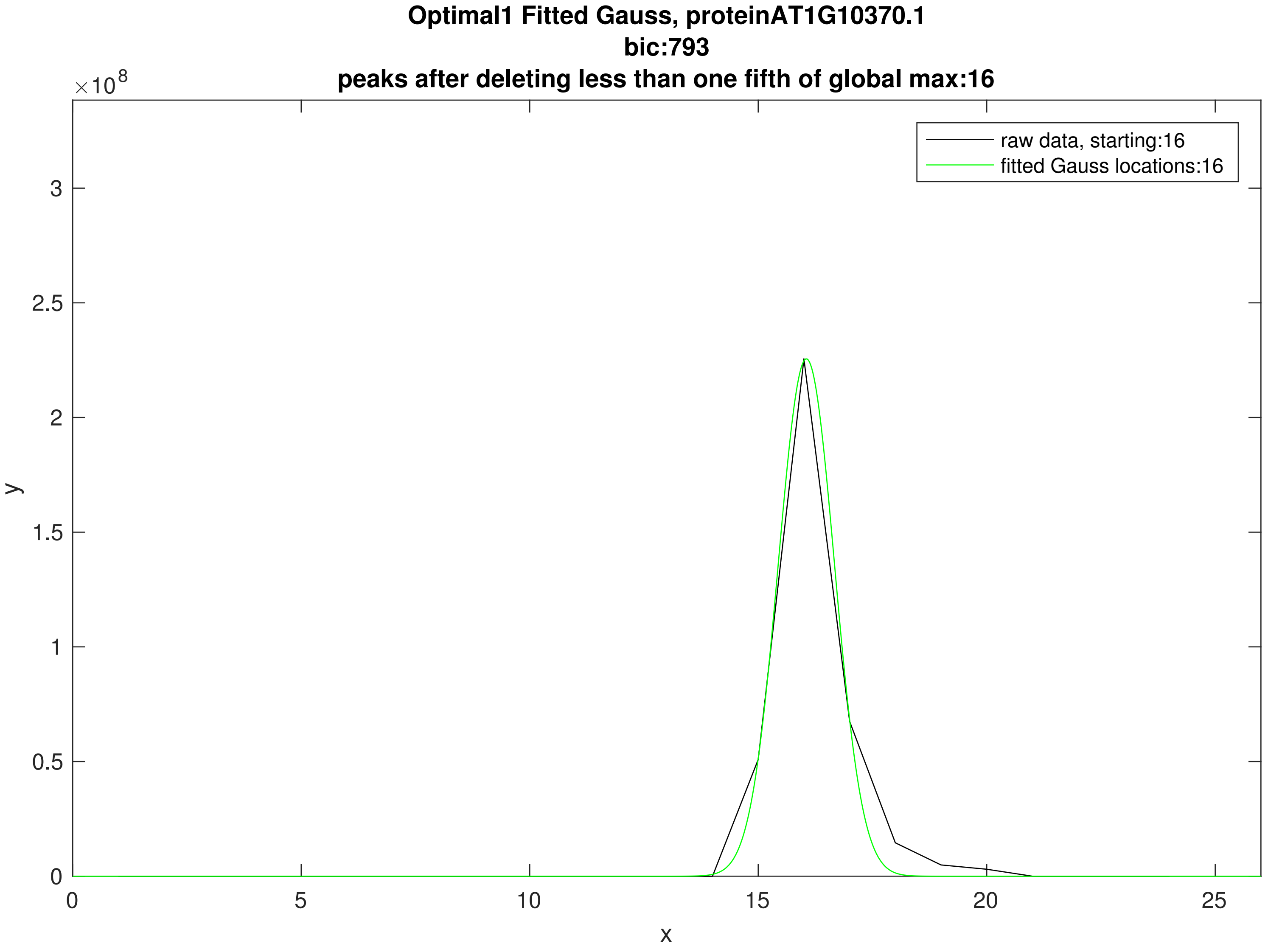
Optimal1 Fitted Gauss, proteinAT1G10370.1
bic:793
peaks after deleting less than one fifth of global max:16
×10^8
raw data, starting:16
fitted Gauss locations:16
y
x
0
0.5
1
1.5
2
2.5
3
0
5
10
15
20
25

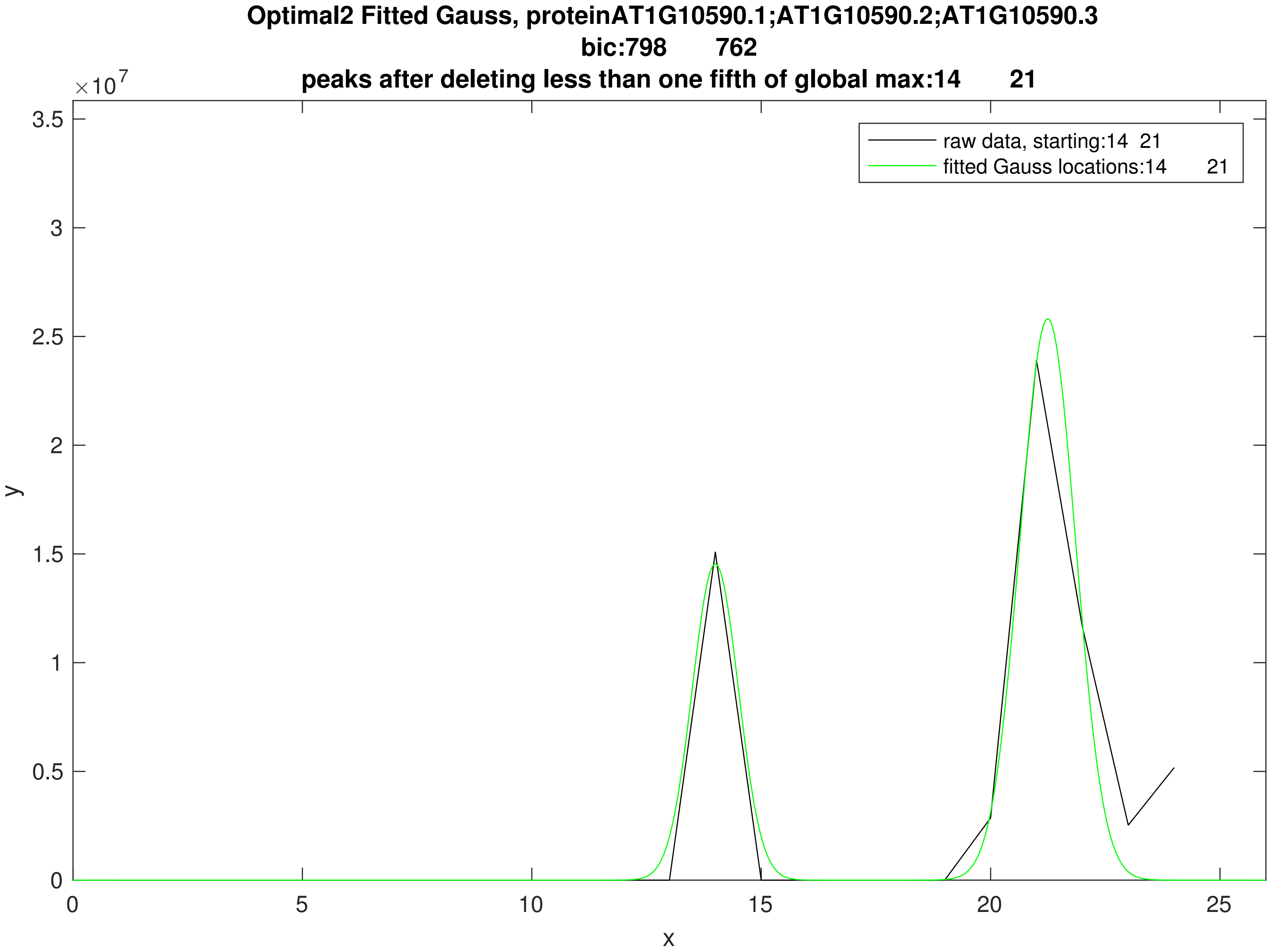
Optimal2 Fitted Gauss, proteinAT1G10590.1;AT1G10590.2;AT1G10590.3
bic:798 762
peaks after deleting less than one fifth of global max:14 21
×10^7
3.5
3
2.5
2
1.5
1
0.5
0
y
0
5
10
15
20
25
x
raw data, starting:14 21
fitted Gauss locations:14 21

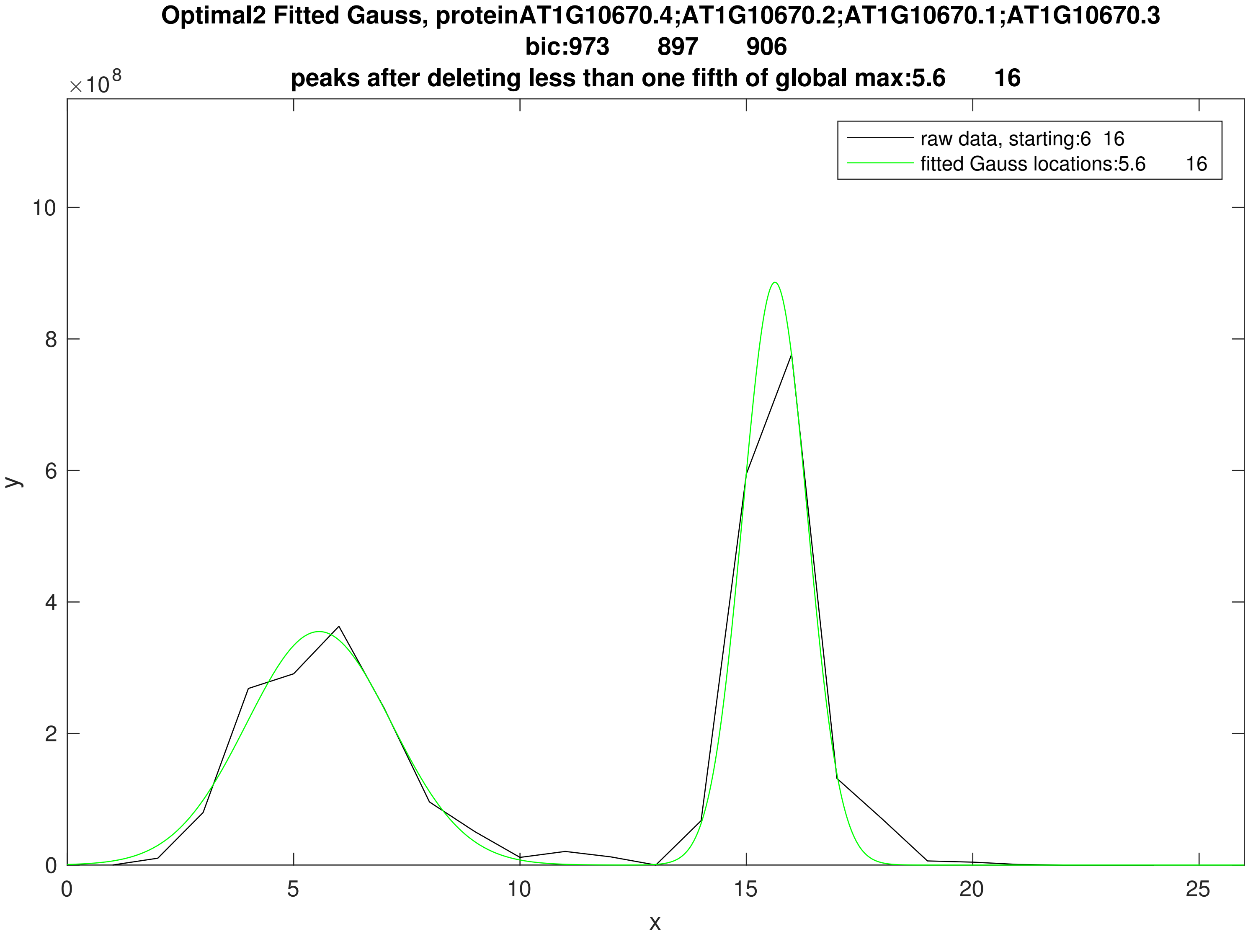
Optimal2 Fitted Gauss, proteinAT1G10670.4;AT1G10670.2;AT1G10670.1;AT1G10670.3
bic:973 897 906
peaks after deleting less than one fifth of global max:5.6 16
×10^8
raw data, starting:6 16
fitted Gauss locations:5.6 16
y
x
0
2
4
6
8
10
0
5
10
15
20
25

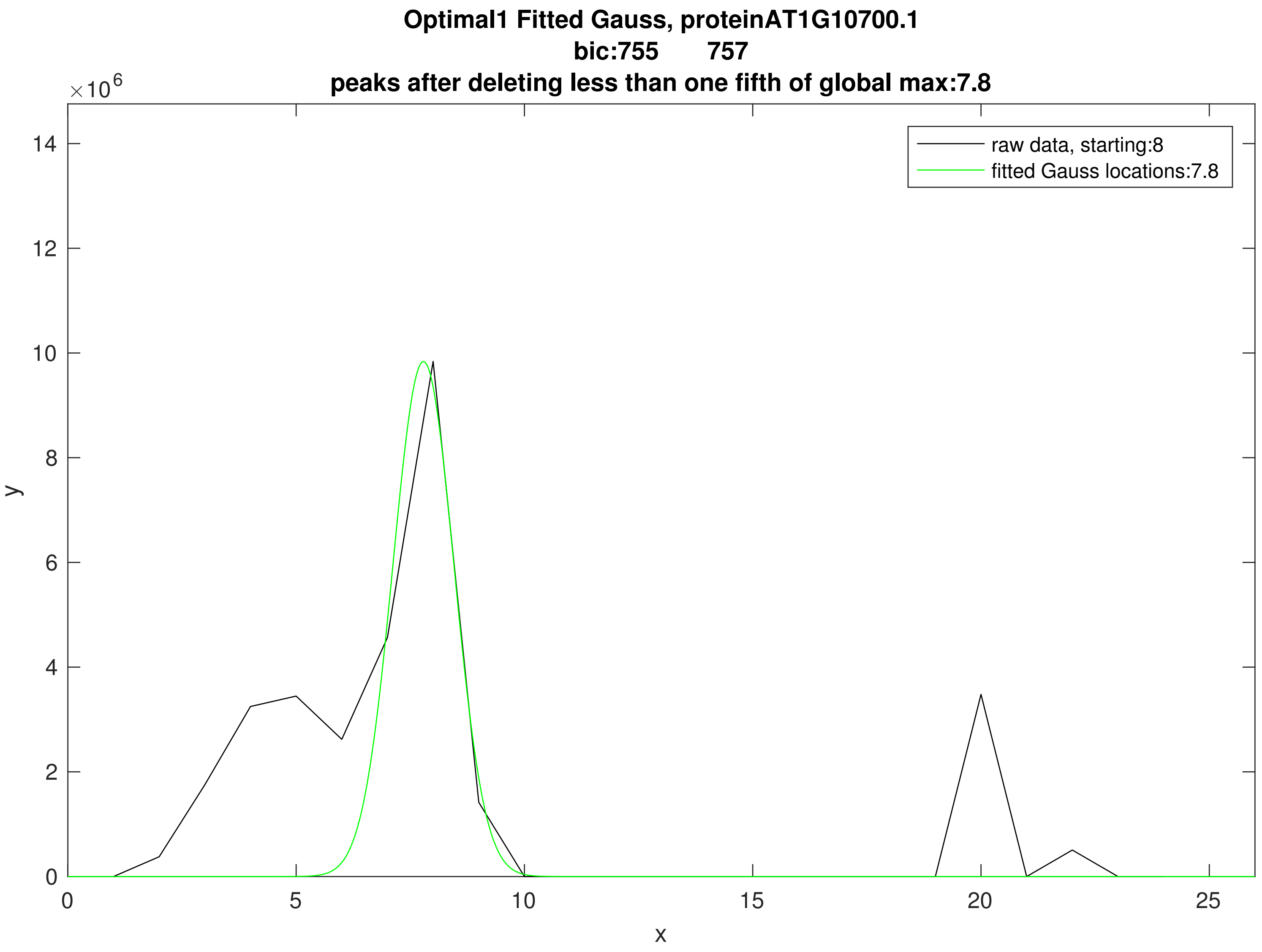
Optimal1 Fitted Gauss, proteinAT1G10700.1
bic:755 757
peaks after deleting less than one fifth of global max:7.8
×10^6
14
12
10
8
6
4
2
0
y
0
5
10
15
20
25
x
raw data, starting:8
fitted Gauss locations:7.8

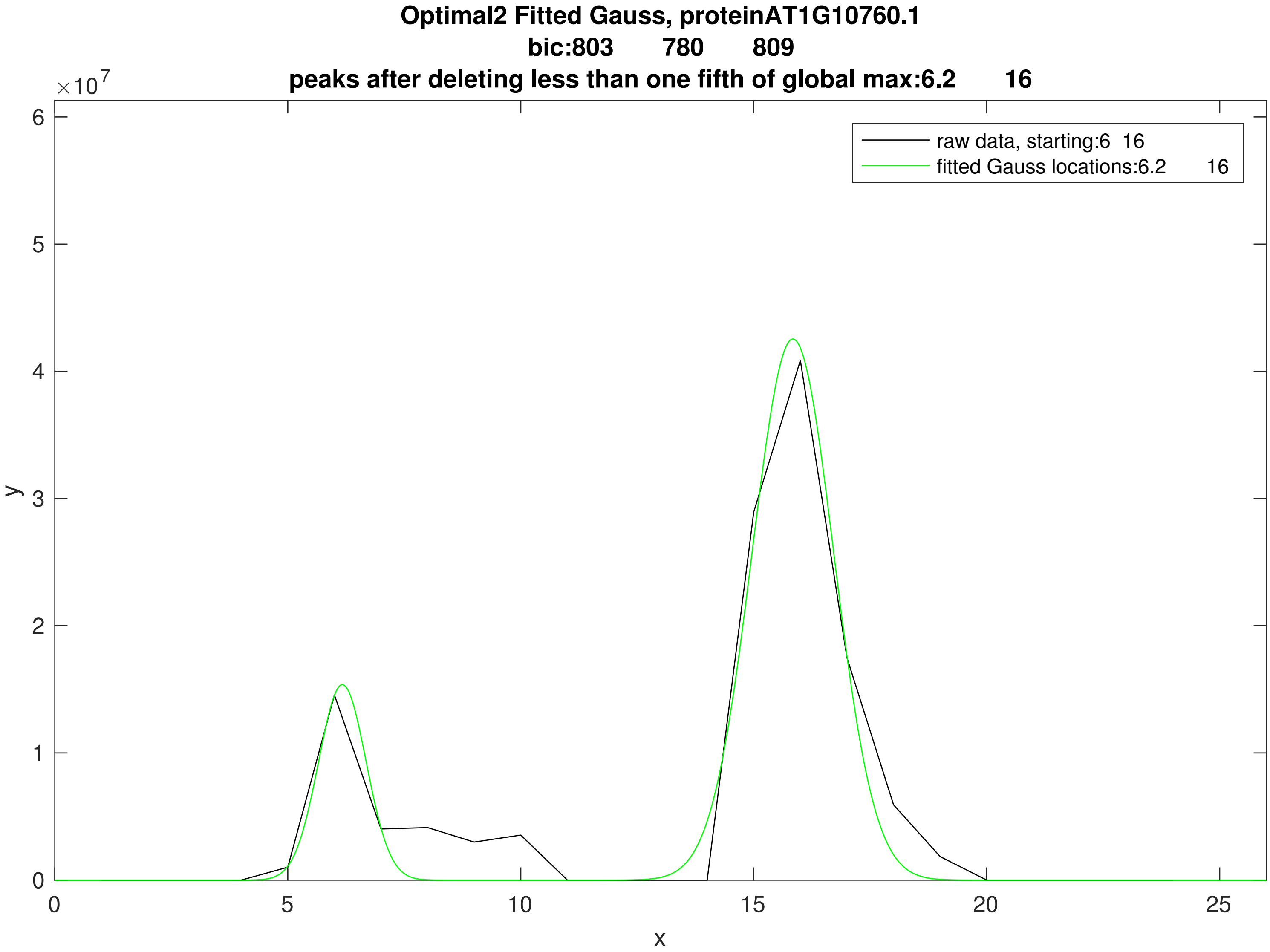

Optimal2 Fitted Gauss, proteinAT1G10760.1
bic:803 780 809
peaks after deleting less than one fifth of global max:6.2 16
×10^7
raw data, starting:6 16
fitted Gauss locations:6.2 16
y
x
0
1
2
3
4
5
6
0
5
10
15
20
25

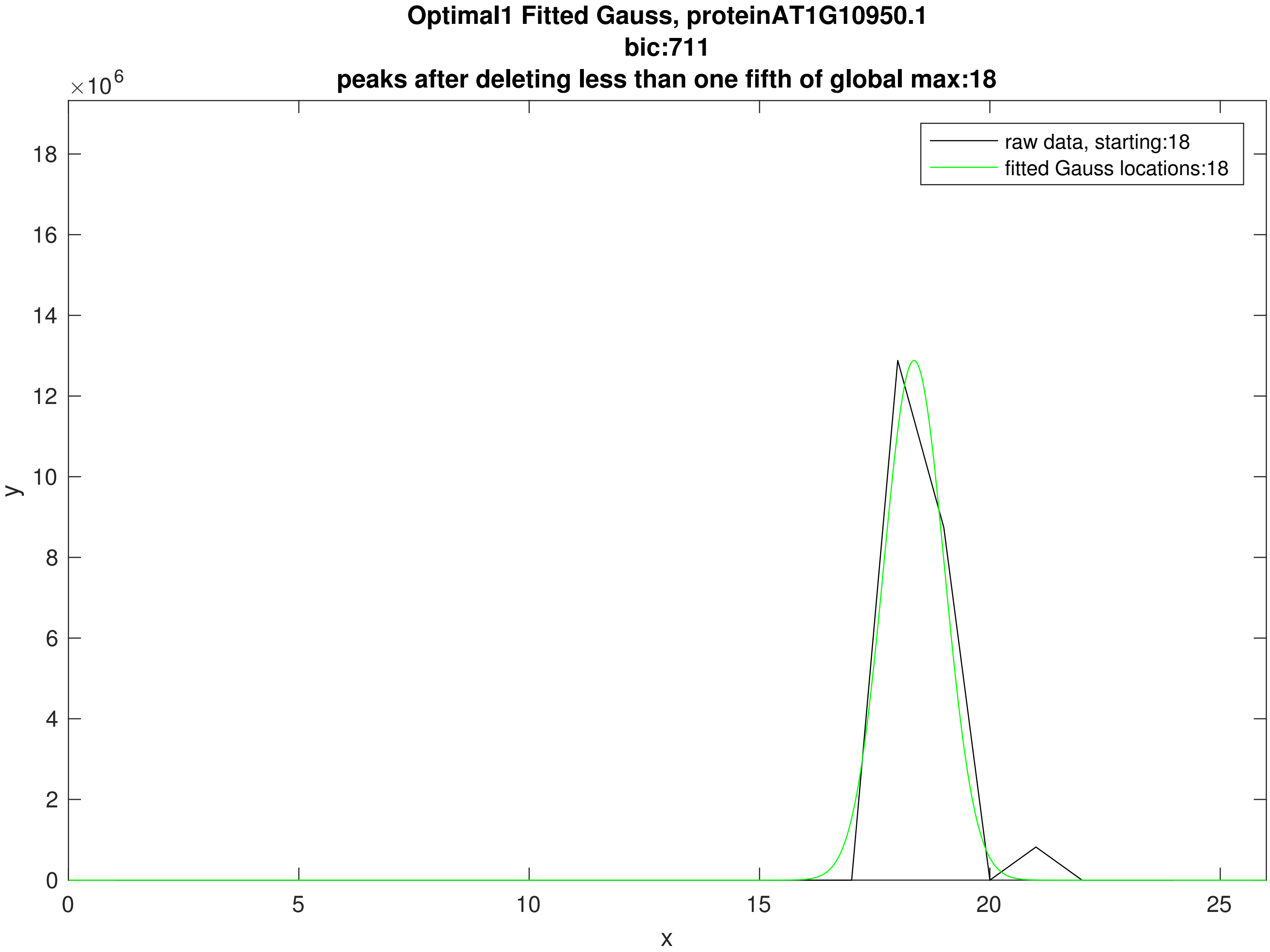
Optimal1 Fitted Gauss, proteinAT1G10950.1
bic:711
peaks after deleting less than one fifth of global max:18
×10^6
raw data, starting:18
fitted Gauss locations:18
y
x
0
2
4
6
8
10
12
14
16
18
0
5
10
15
20
25

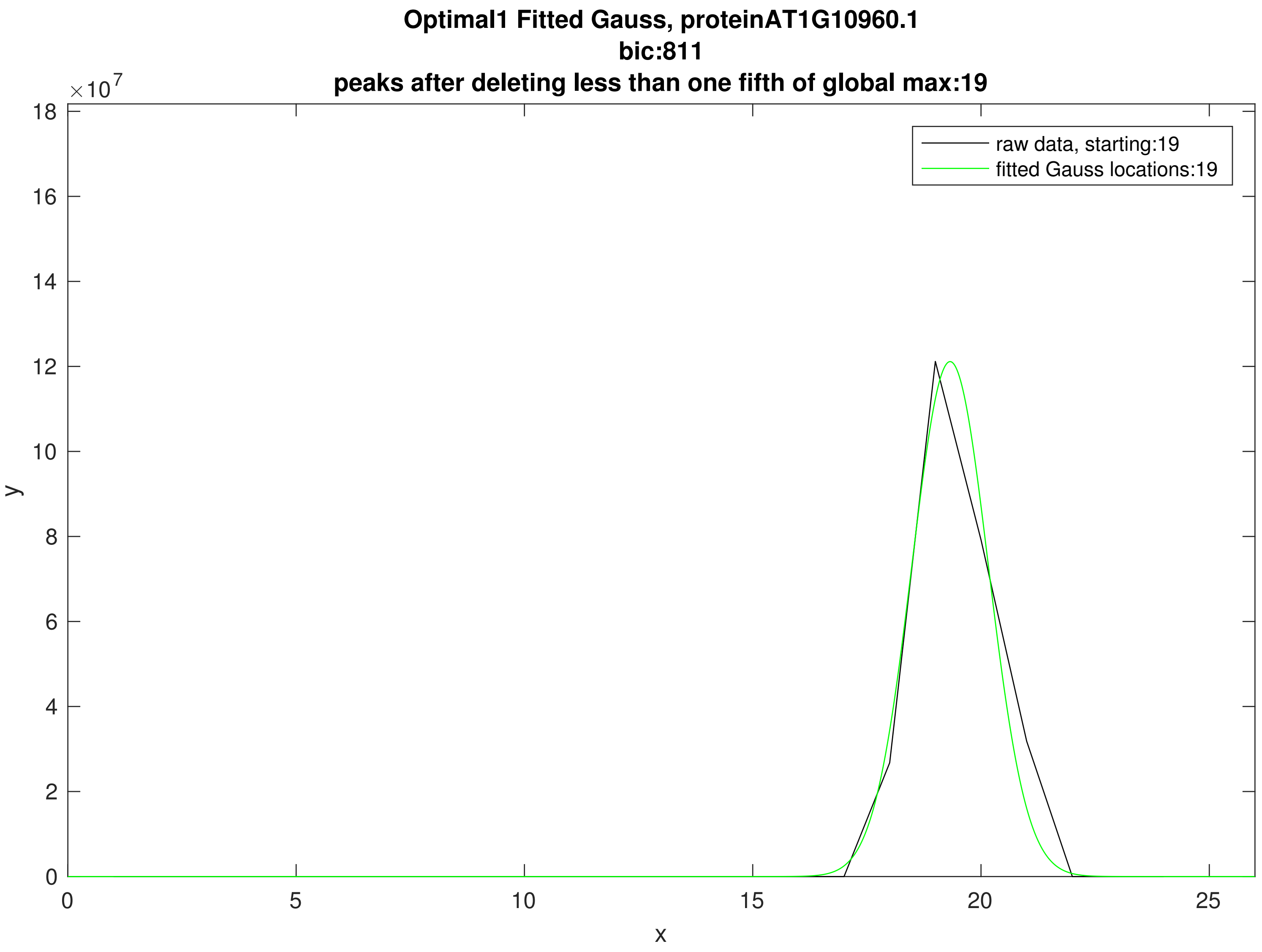
Optimal1 Fitted Gauss, proteinAT1G10960.1
bic:811
peaks after deleting less than one fifth of global max:19
×10^7
18
16
14
12
10
8
6
4
2
0
y
0
5
10
15
20
25
x
raw data, starting:19
fitted Gauss locations:19

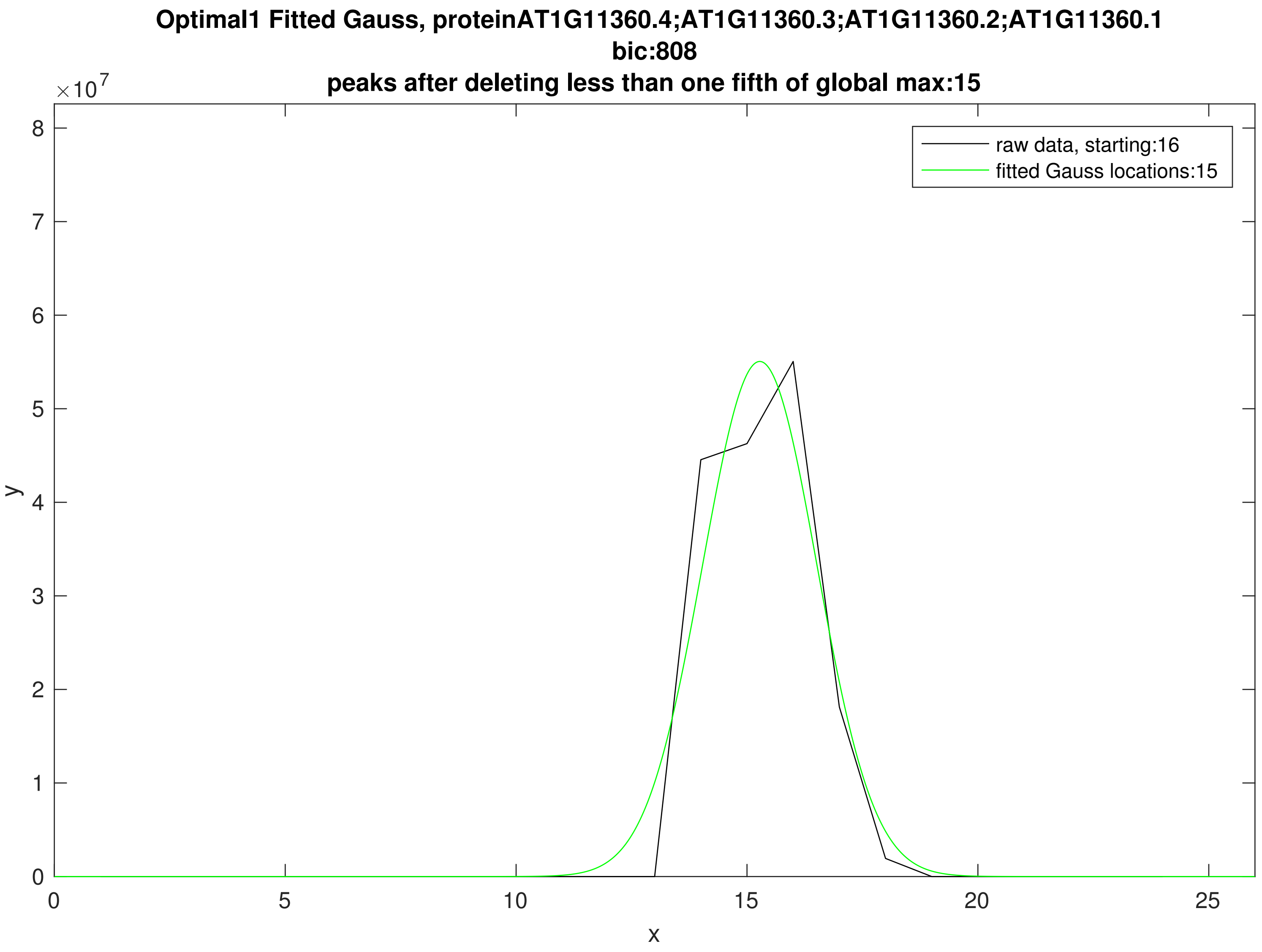

Optimal1 Fitted Gauss, proteinAT1G11360.4;AT1G11360.3;AT1G11360.2;AT1G11360.1
bic:808
peaks after deleting less than one fifth of global max:15
×10^7
raw data, starting:16
fitted Gauss locations:15
y
x
0
1
2
3
4
5
6
7
8
0
5
10
15
20
25

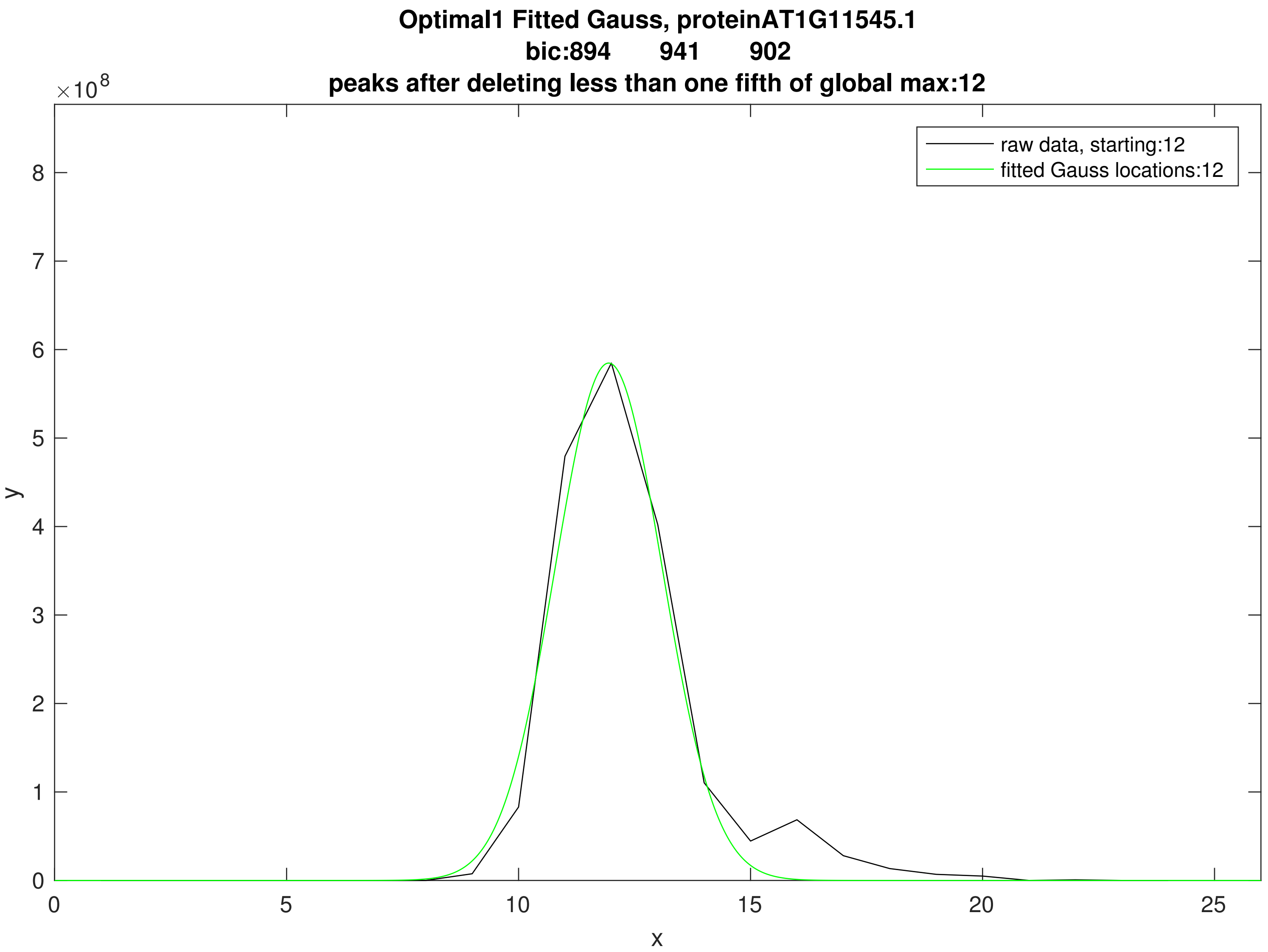

Optimal1 Fitted Gauss, proteinAT1G11545.1
bic:894 941 902
peaks after deleting less than one fifth of global max:12
×10^8
raw data, starting:12
fitted Gauss locations:12
y
x
0
1
2
3
4
5
6
7
8
0
5
10
15
20
25

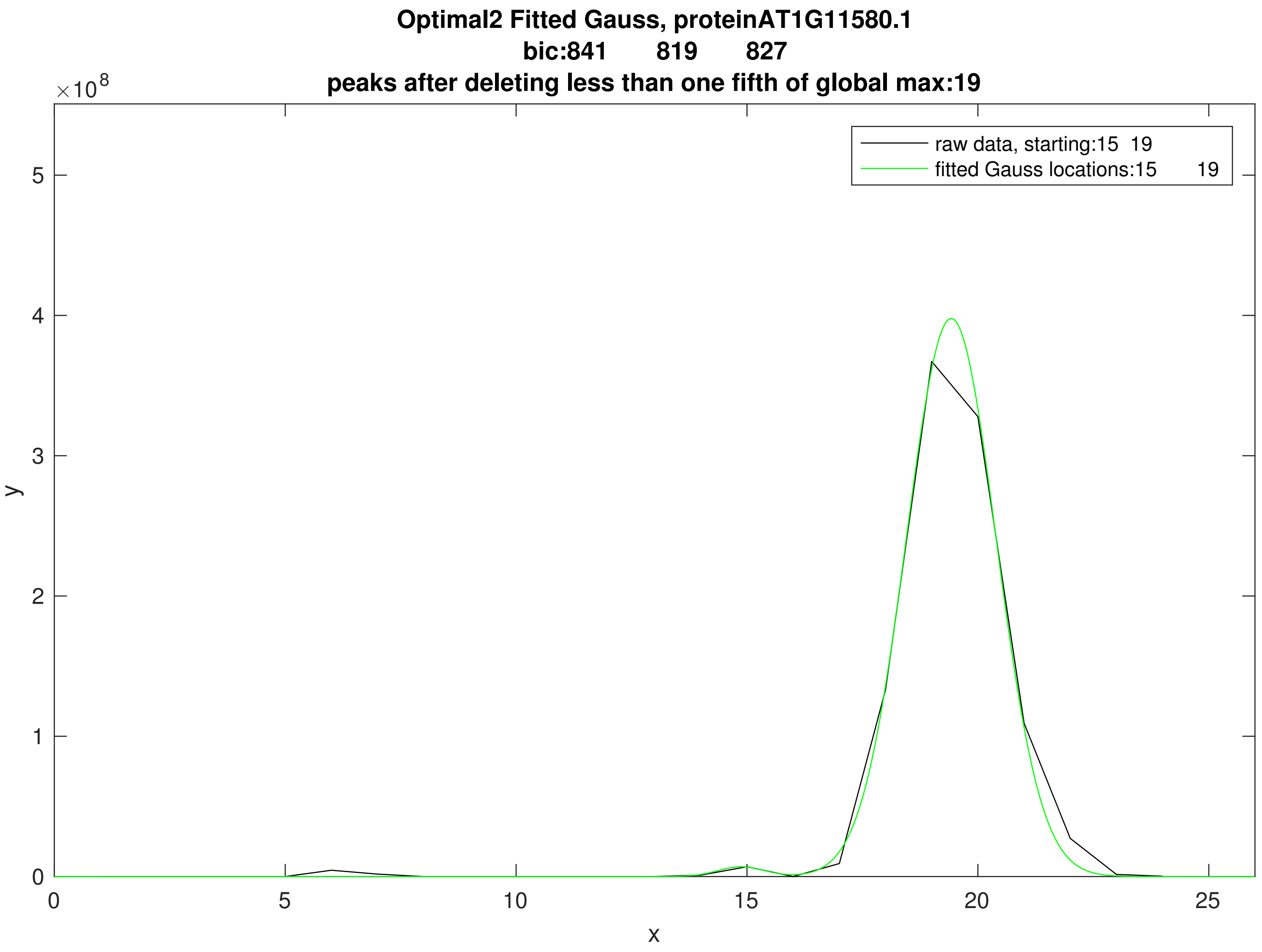
Optimal2 Fitted Gauss, proteinAT1G11580.1
bic:841 819 827
peaks after deleting less than one fifth of global max:19
×10^8
raw data, starting:15 19
fitted Gauss locations:15 19
y
x
0
1
2
3
4
5
0
5
10
15
20
25

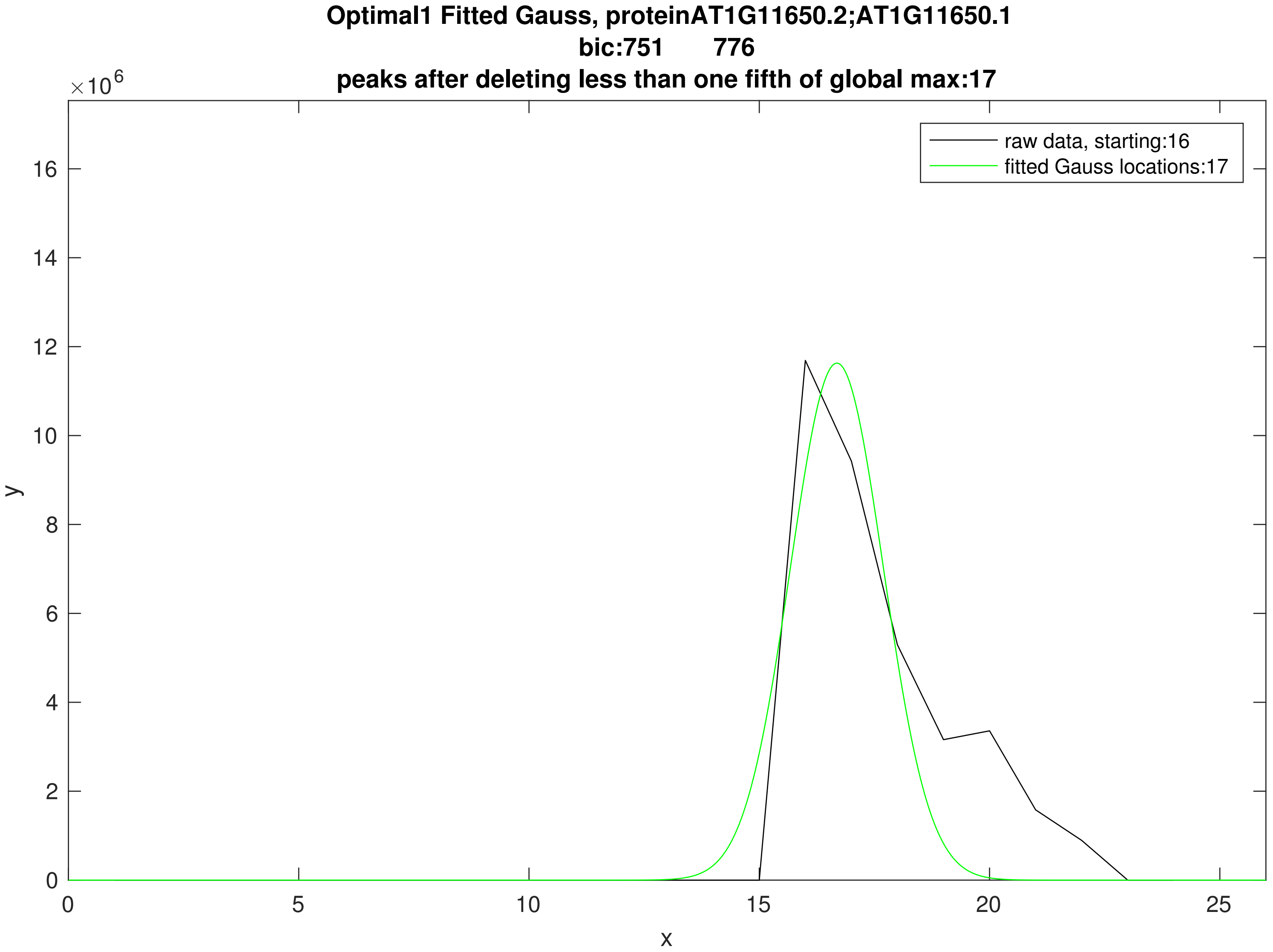

Optimal1 Fitted Gauss, proteinAT1G11650.2;AT1G11650.1
bic:751 776
peaks after deleting less than one fifth of global max:17
×10^6
raw data, starting:16
fitted Gauss locations:17
y
x
0
2
4
6
8
10
12
14
16
0
5
10
15
20
25

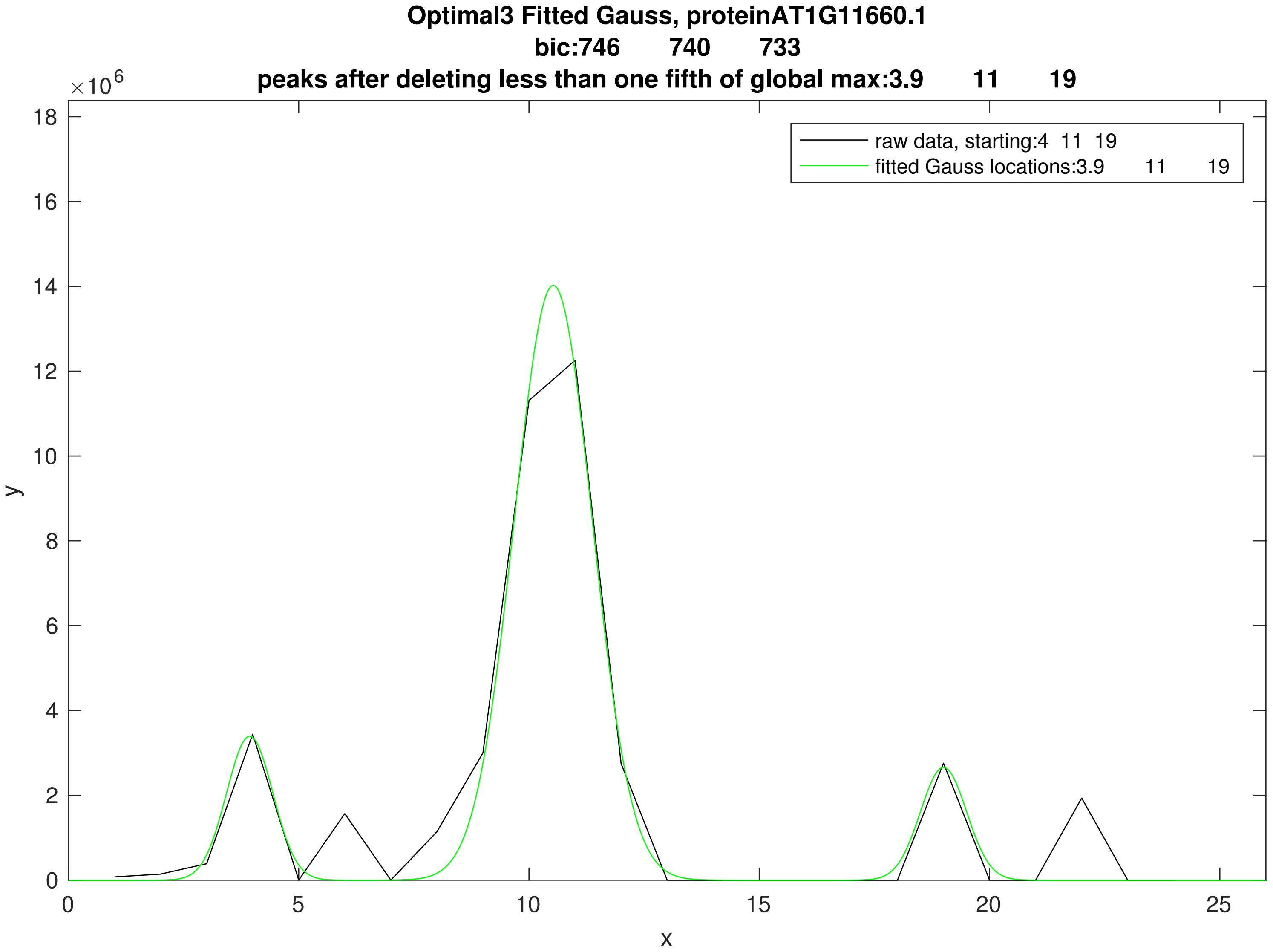
Optimal3 Fitted Gauss, proteinAT1G11660.1
bic:746 740 733
peaks after deleting less than one fifth of global max:3.9 11 19
×10^6
raw data, starting:4 11 19
fitted Gauss locations:3.9 11 19
y
x
0
2
4
6
8
10
12
14
16
18
0
5
10
15
20
25

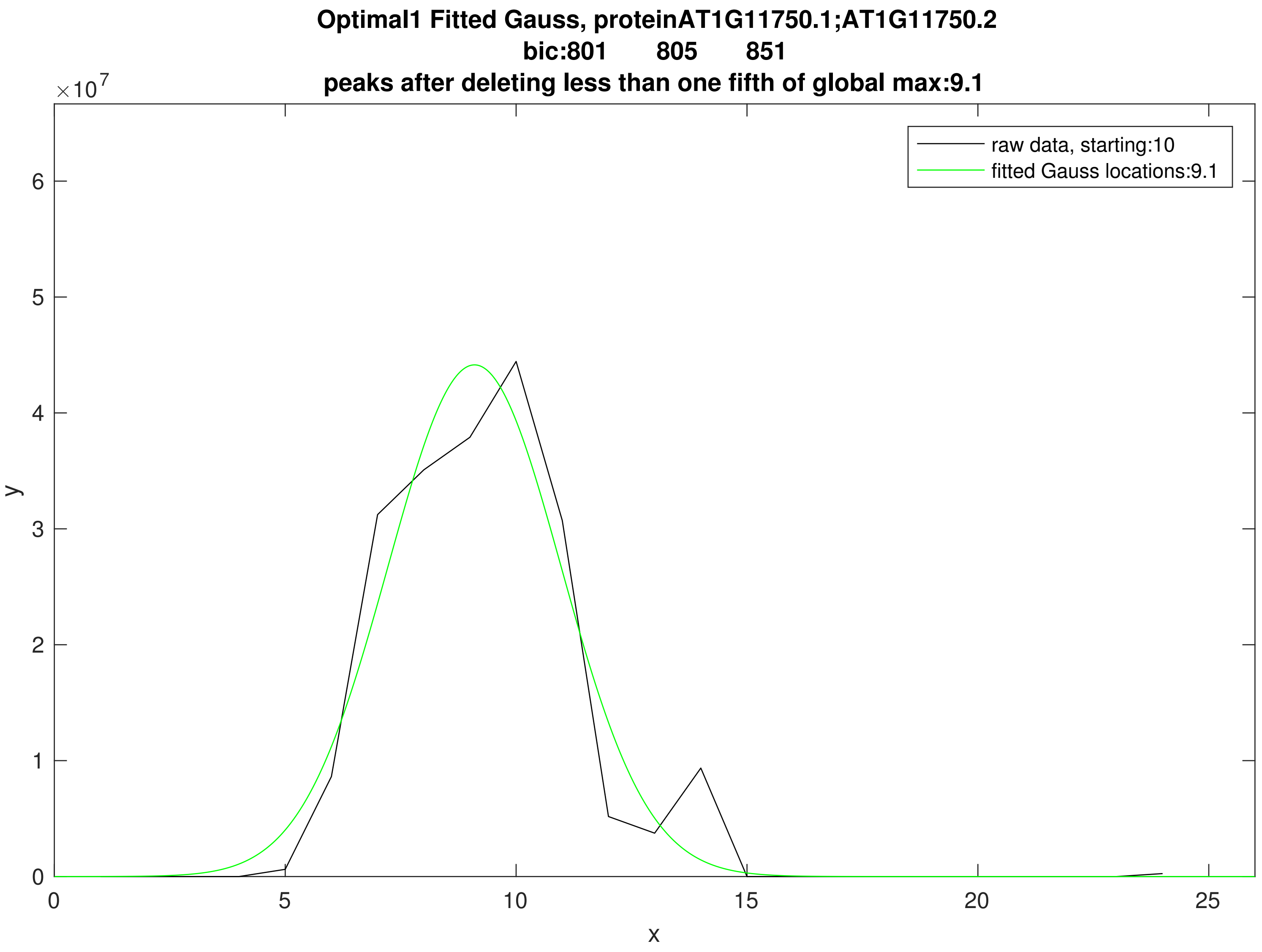
Optimal1 Fitted Gauss, proteinAT1G11750.1;AT1G11750.2
bic:801 805 851
peaks after deleting less than one fifth of global max:9.1
×10^7
raw data, starting:10
fitted Gauss locations:9.1
y
x
0
1
2
3
4
5
6
0
5
10
15
20
25

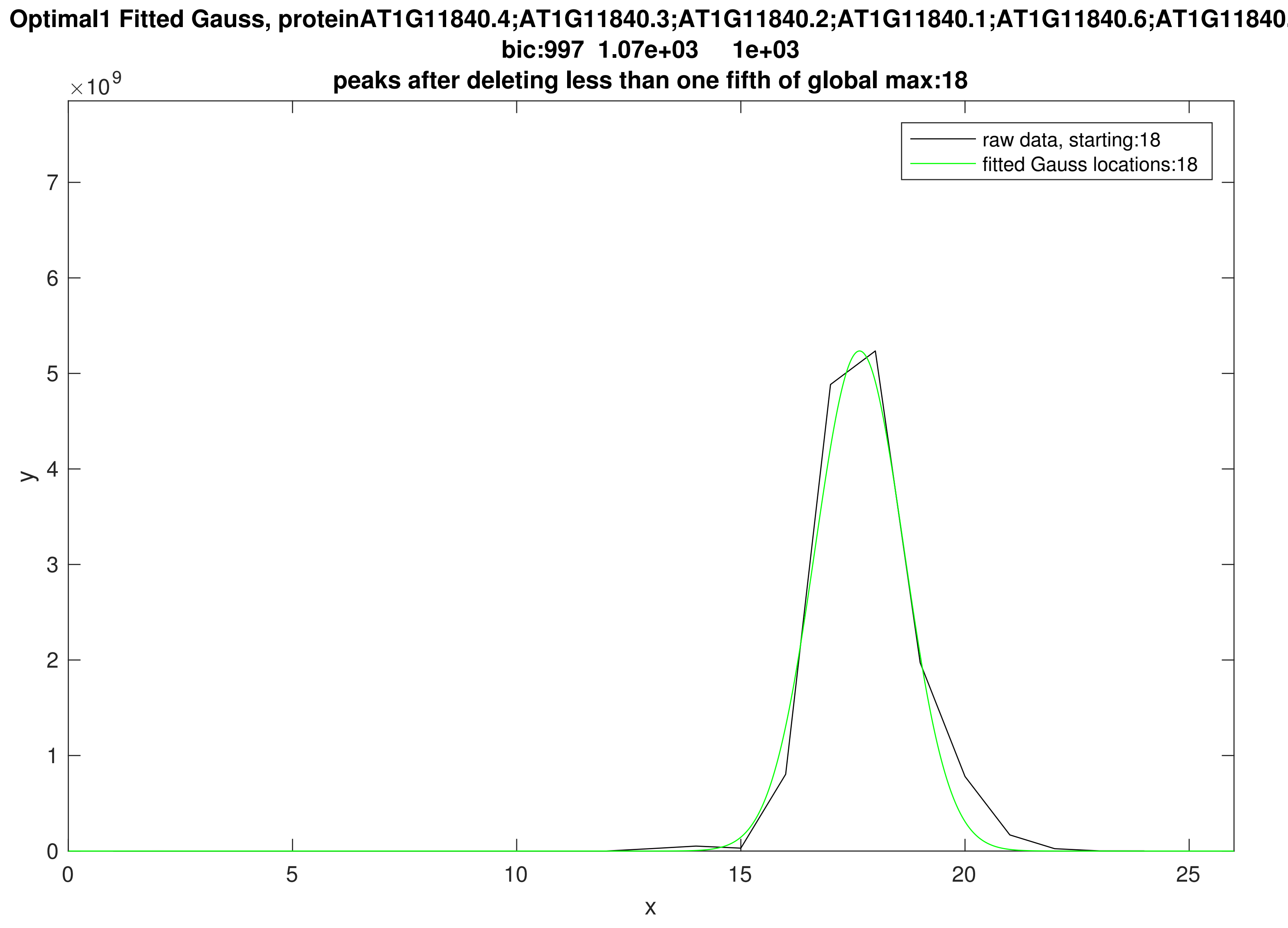

Optimal1 Fitted Gauss, proteinAT1G11840.4;AT1G11840.3;AT1G11840.2;AT1G11840.1;AT1G11840.6;AT1G11840.
bic:997 1.07e+03 1e+03
peaks after deleting less than one fifth of global max:18
×10^9
raw data, starting:18
fitted Gauss locations:18
y
x
0
1
2
3
4
5
6
7
0
5
10
15
20
25

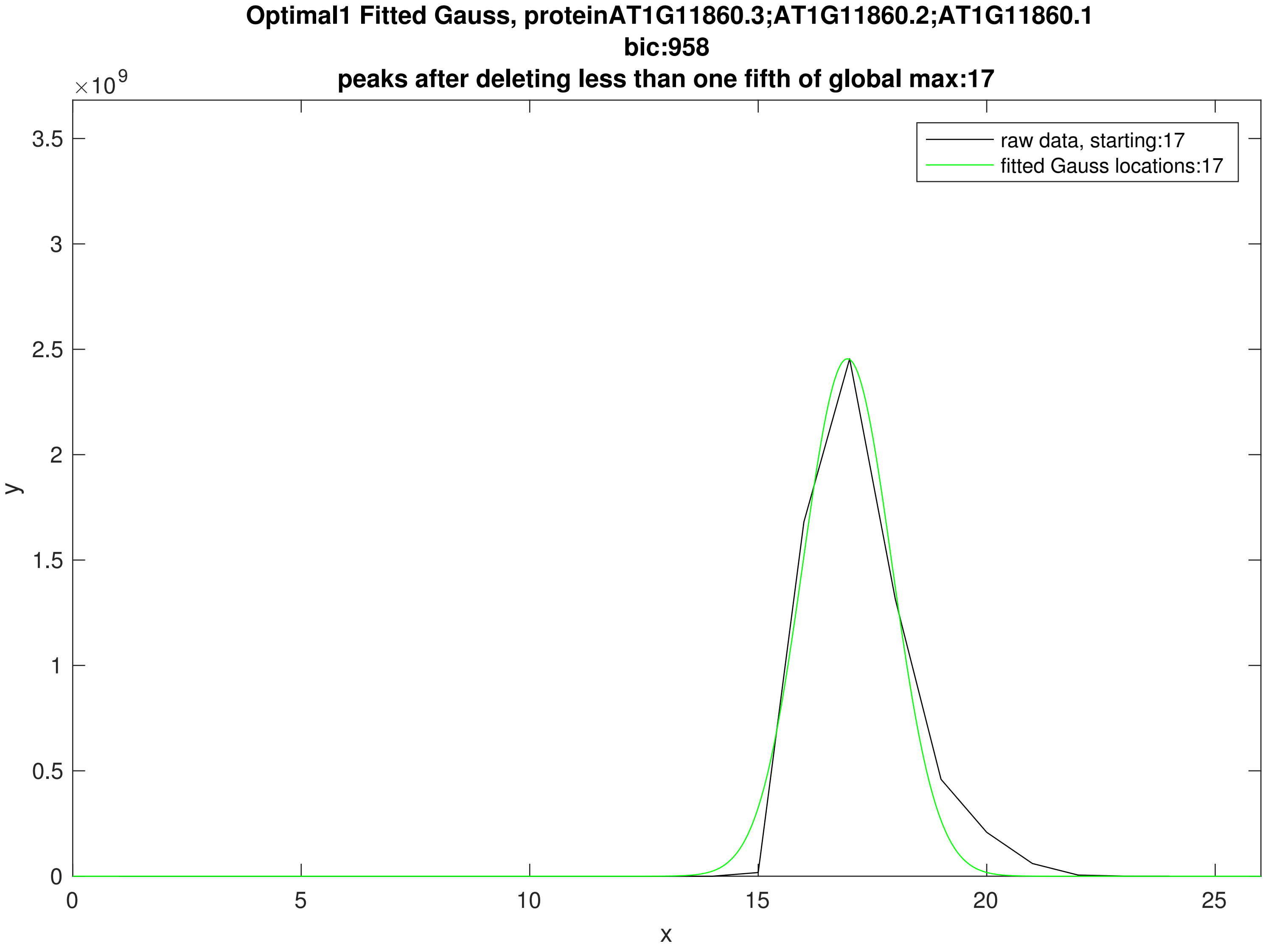

Optimal1 Fitted Gauss, proteinAT1G11860.3;AT1G11860.2;AT1G11860.1
bic:958
peaks after deleting less than one fifth of global max:17
×10^9
3.5
3
2.5
2
1.5
1
0.5
0
y
0
5
10
15
20
25
x
raw data, starting:17
fitted Gauss locations:17

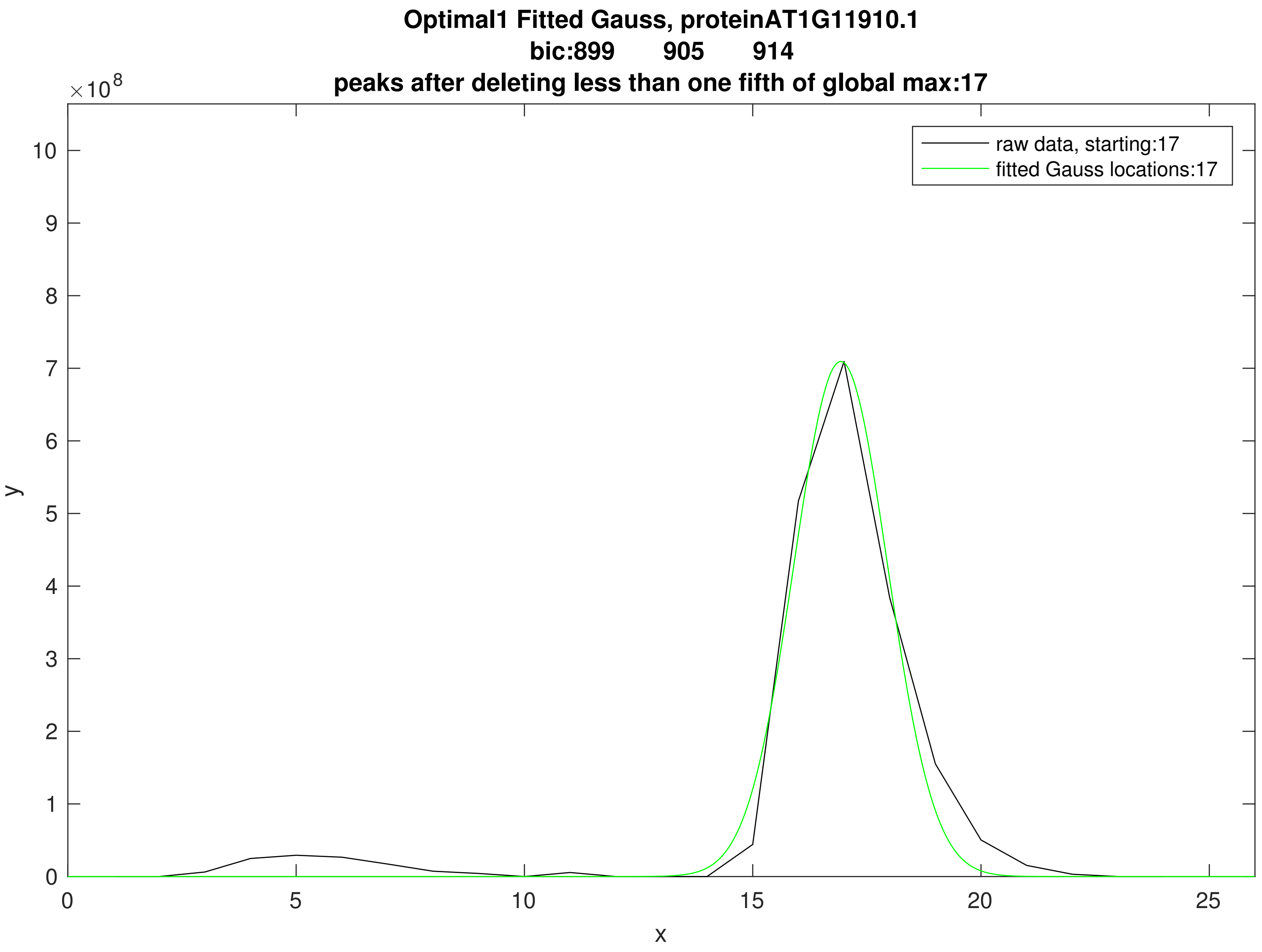
Optimal1 Fitted Gauss, proteinAT1G11910.1
bic:899 905 914
peaks after deleting less than one fifth of global max:17
×10^8
raw data, starting:17
fitted Gauss locations:17
y
x
0
1
2
3
4
5
6
7
8
9
10
0
5
10
15
20
25

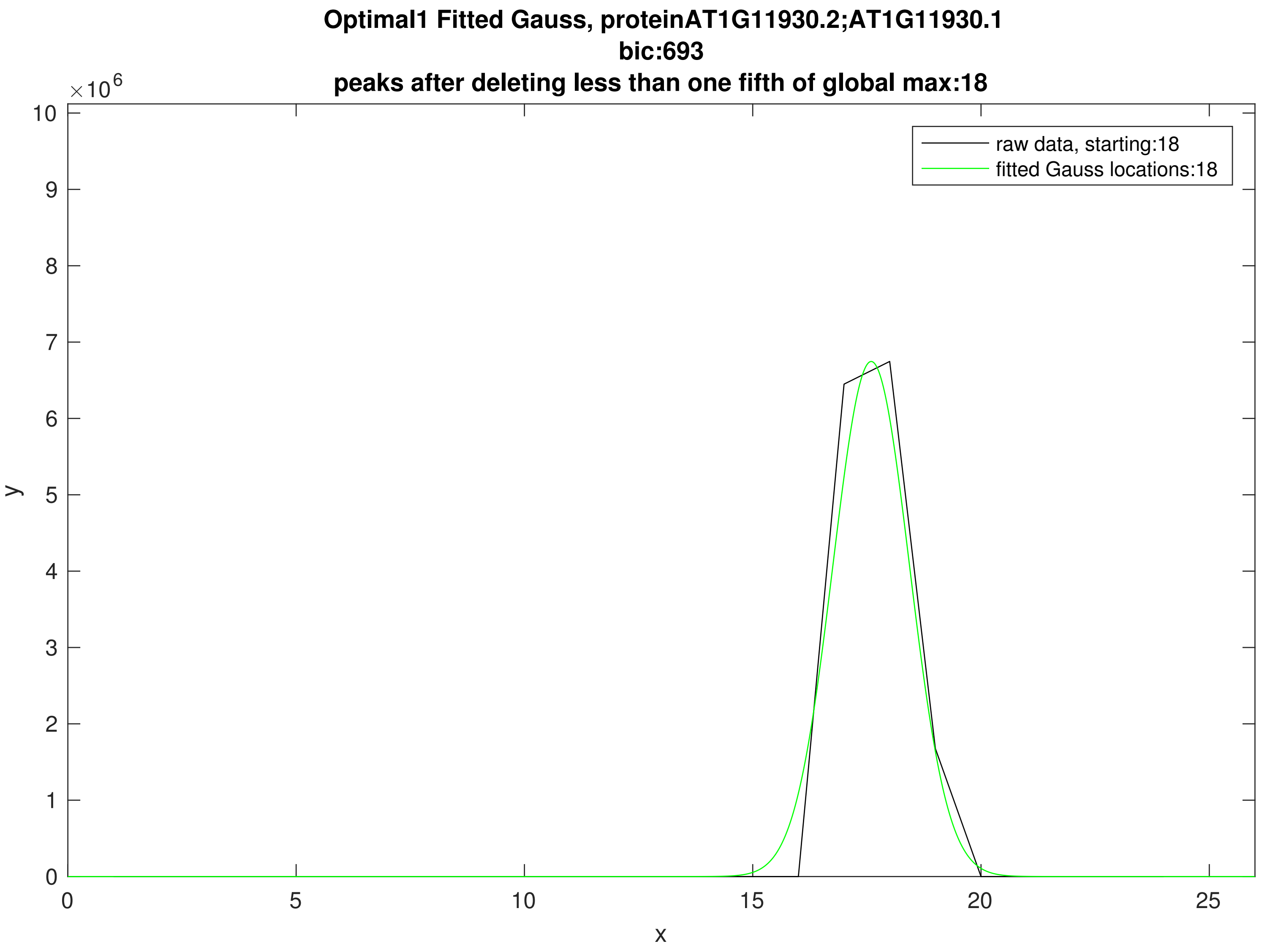

Optimal1 Fitted Gauss, proteinAT1G11930.2;AT1G11930.1
bic:693
peaks after deleting less than one fifth of global max:18
×10^6
raw data, starting:18
fitted Gauss locations:18
y
x
0
1
2
3
4
5
6
7
8
9
10
0
5
10
15
20
25

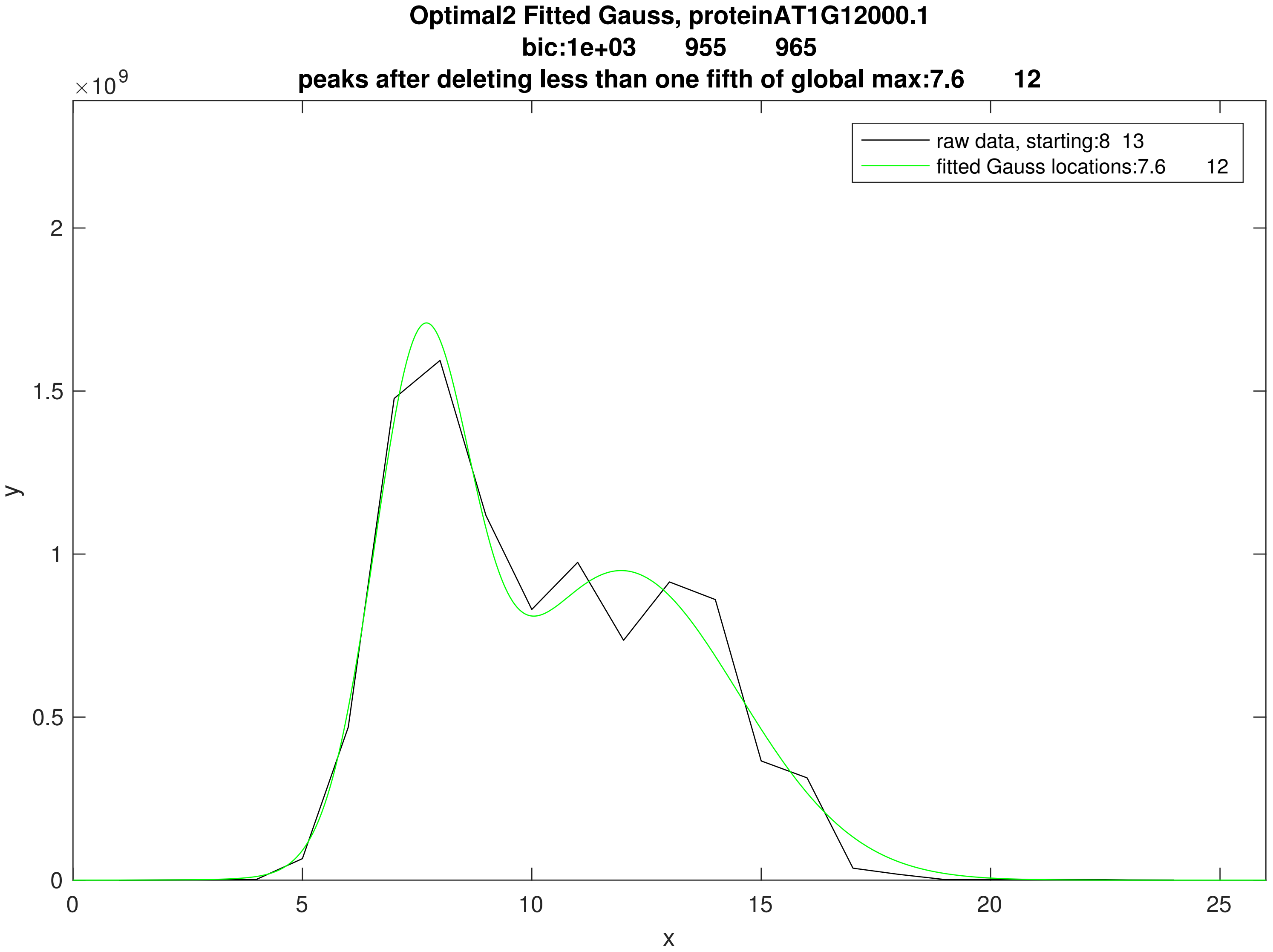
Optimal2 Fitted Gauss, proteinAT1G12000.1
bic:1e+03 955 965
peaks after deleting less than one fifth of global max:7.6 12
×10^9
raw data, starting:8 13
fitted Gauss locations:7.6 12
y
x
0
0.5
1
1.5
2
5
10
15
20
25

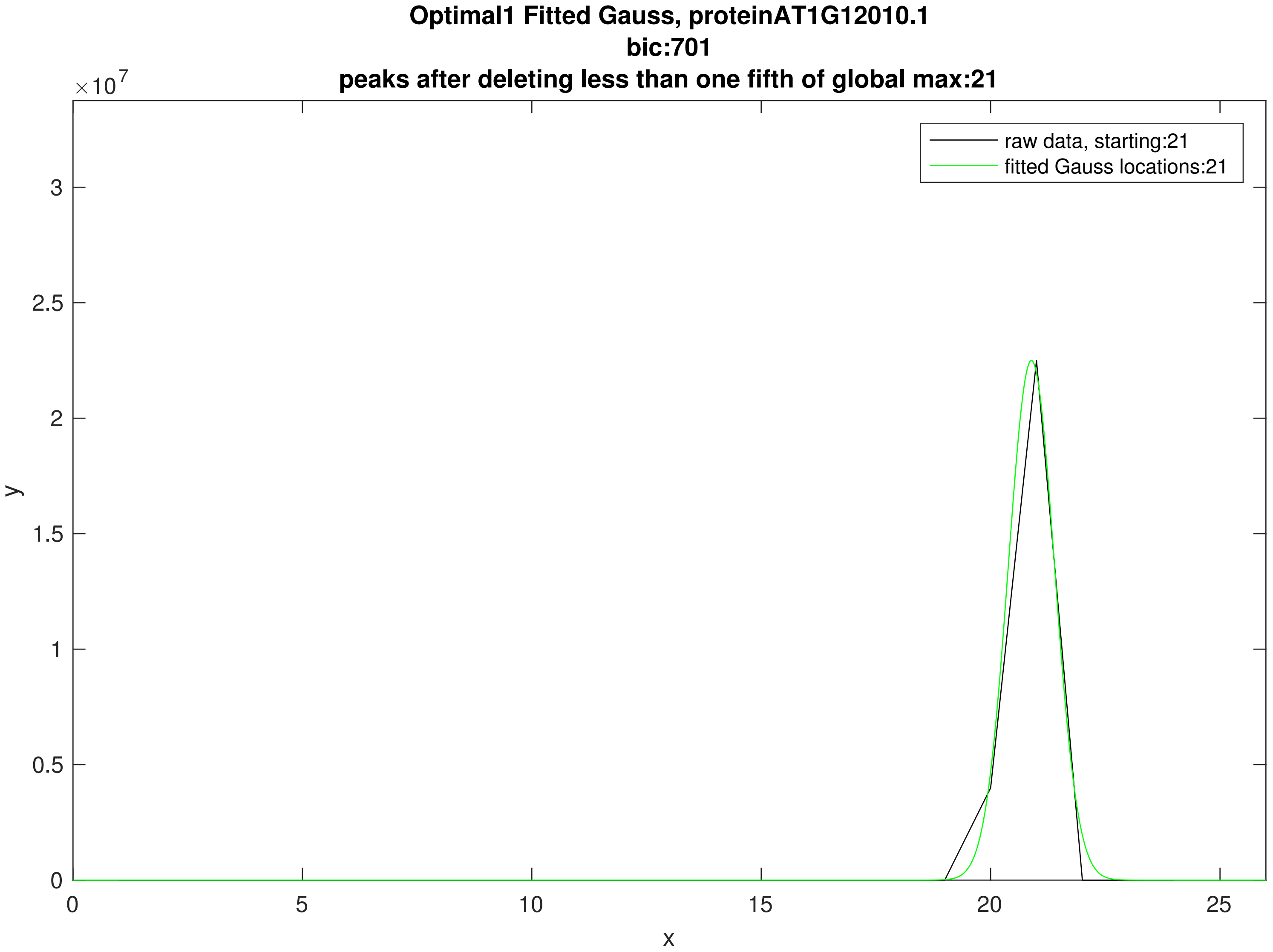
Optimal1 Fitted Gauss, proteinAT1G12010.1
bic:701
peaks after deleting less than one fifth of global max:21
×10^7
raw data, starting:21
fitted Gauss locations:21
y
x
0
0.5
1
1.5
2
2.5
3
0
5
10
15
20
25

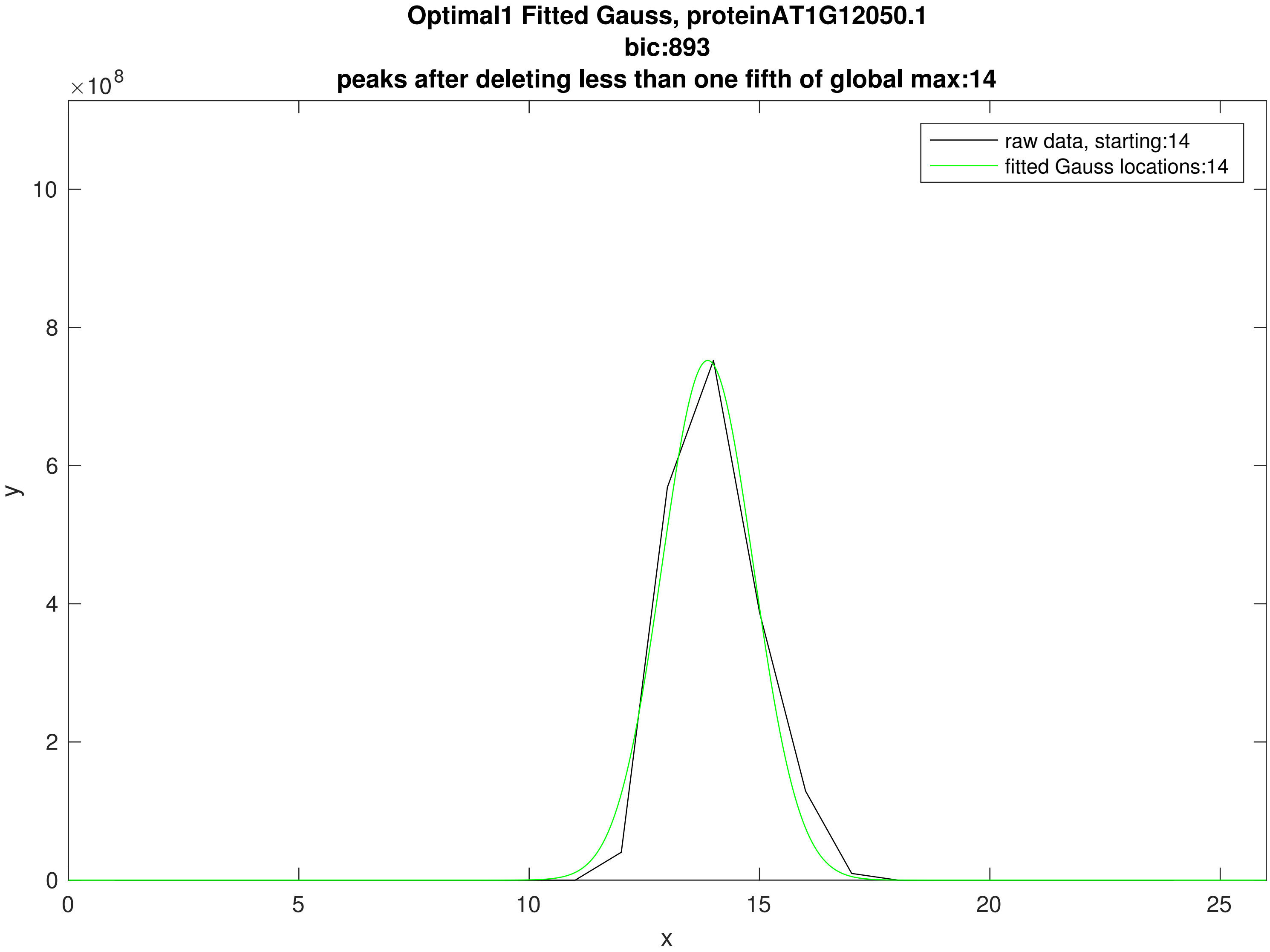
Optimal1 Fitted Gauss, proteinAT1G12050.1
bic:893
peaks after deleting less than one fifth of global max:14
×10^8
raw data, starting:14
fitted Gauss locations:14
y
x
0
2
4
6
8
10
0
5
10
15
20
25

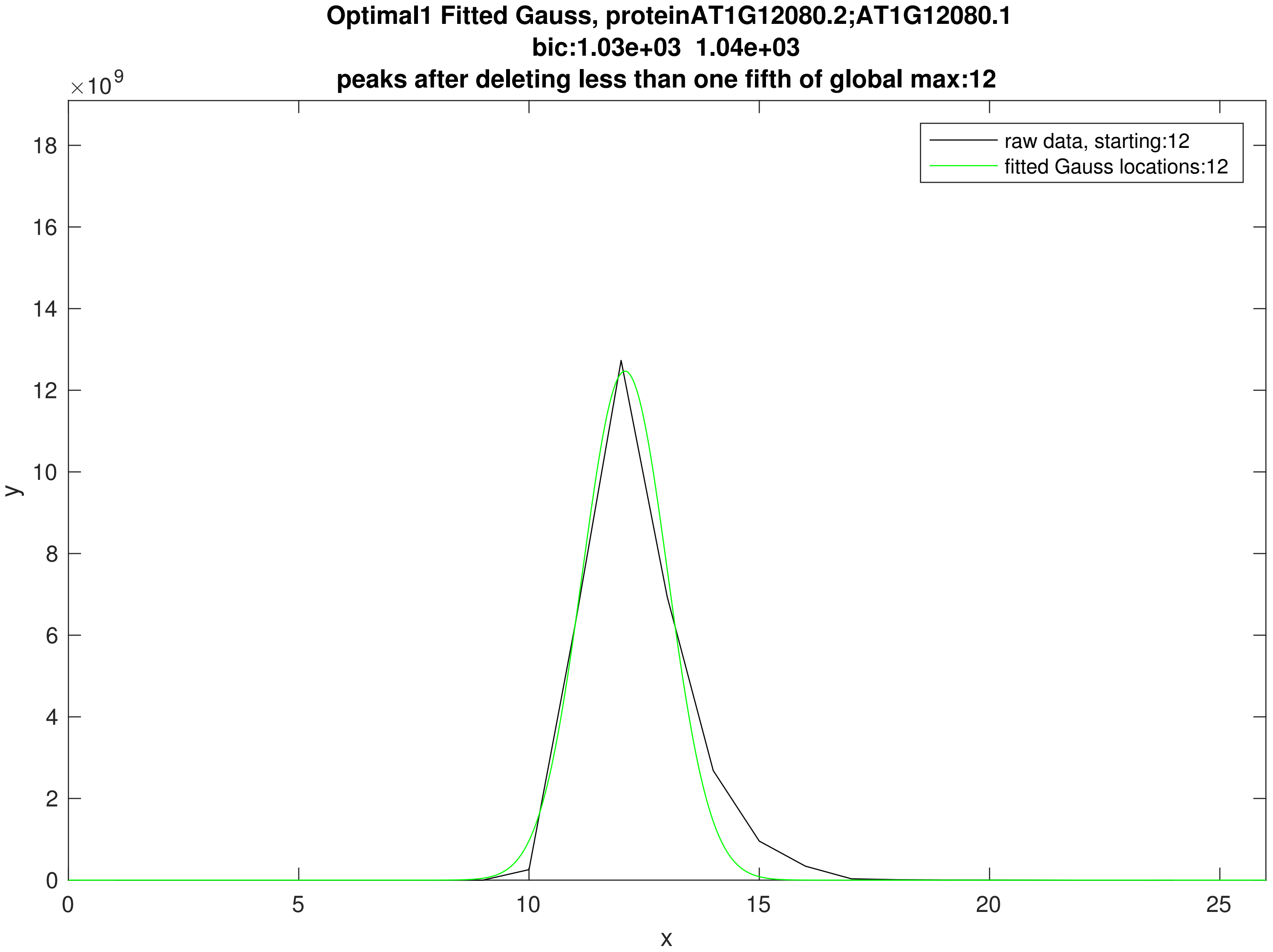
Optimal1 Fitted Gauss, proteinAT1G12080.2;AT1G12080.1
bic:1.03e+03  1.04e+03
peaks after deleting less than one fifth of global max:12
×10^9
raw data, starting:12
fitted Gauss locations:12
y
x
0
2
4
6
8
10
12
14
16
18
0
5
10
15
20
25

Optimal1 Fitted Gauss, proteinAT1G12140.1;AT1G12140.2
bic:730
peaks after deleting less than one fifth of global max:16

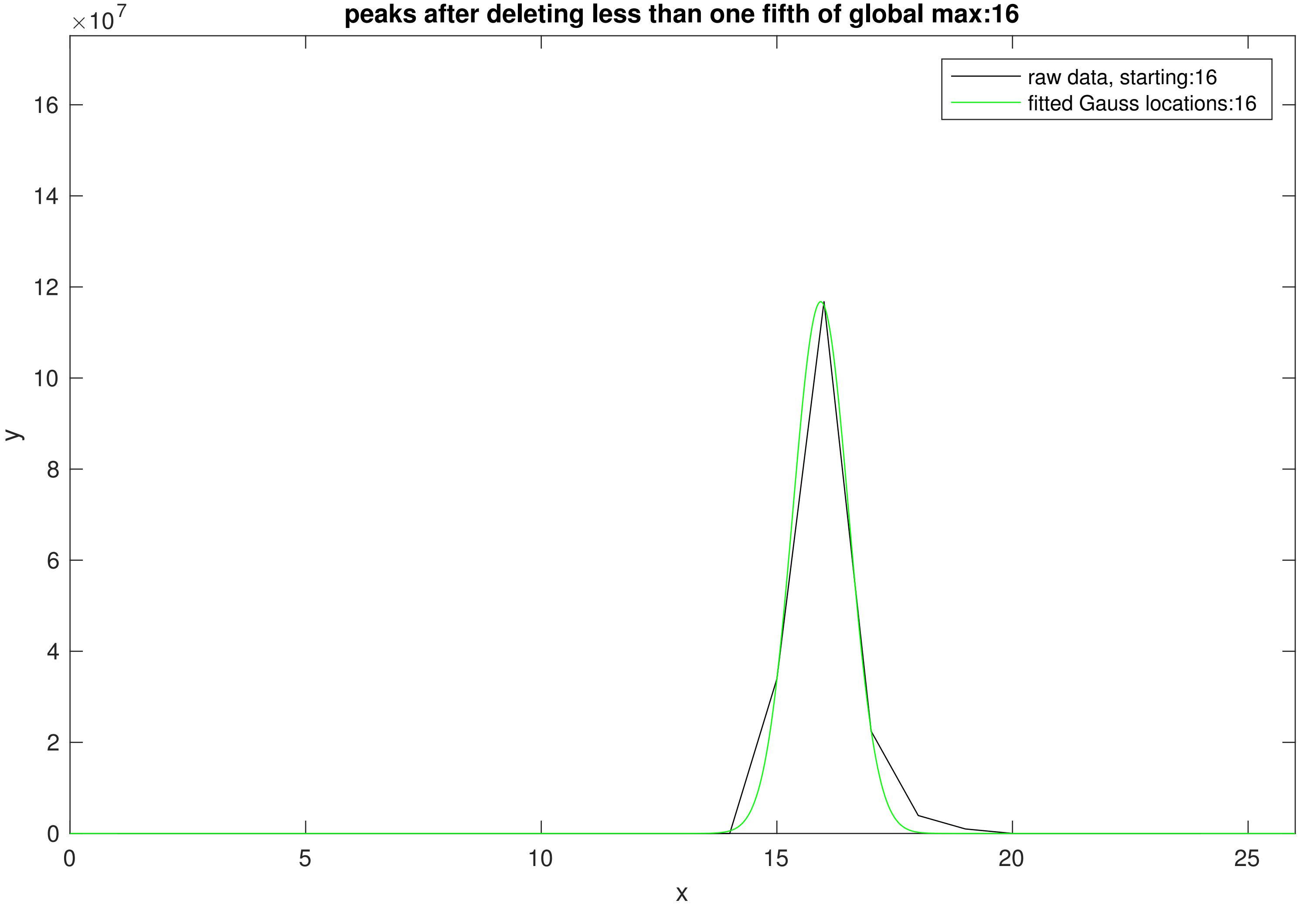

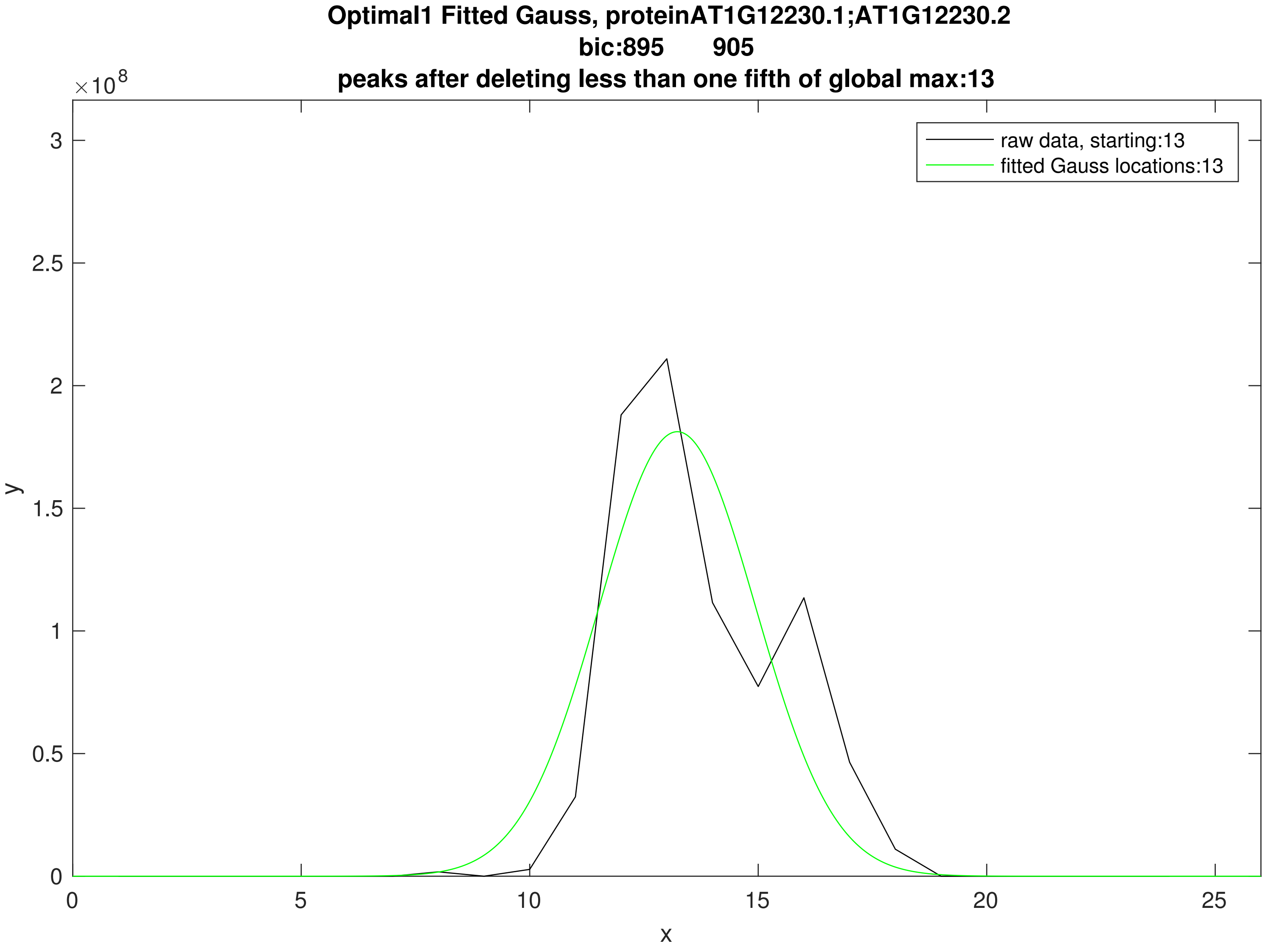
Optimal1 Fitted Gauss, proteinAT1G12230.1;AT1G12230.2
bic:895 905
peaks after deleting less than one fifth of global max:13
×10^8
3
2.5
2
1.5
1
0.5
0
y
0
5
10
15
20
25
x
raw data, starting:13
fitted Gauss locations:13

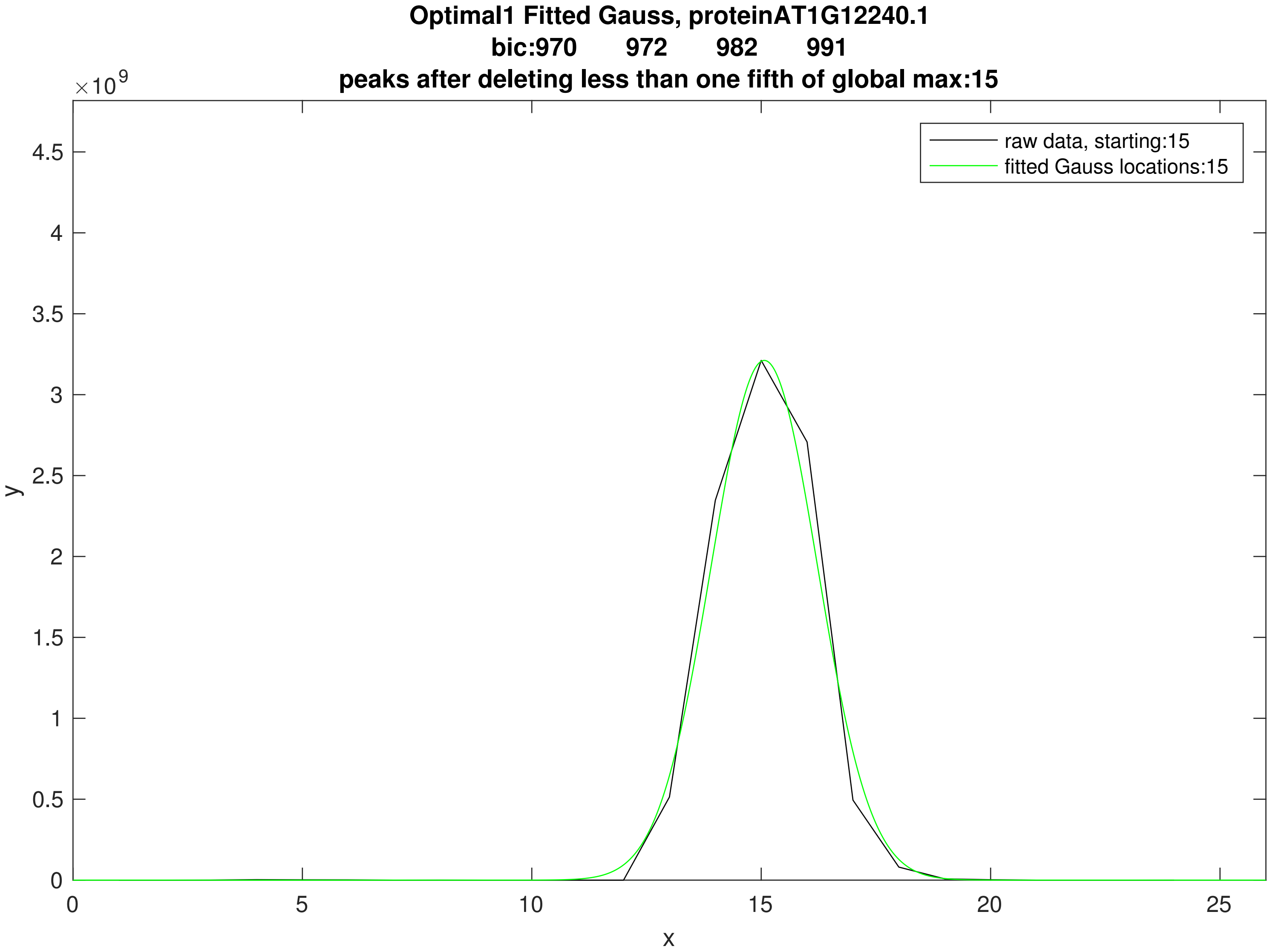
Optimal1 Fitted Gauss, proteinAT1G12240.1
bic:970 972 982 991
peaks after deleting less than one fifth of global max:15
×10^9
raw data, starting:15
fitted Gauss locations:15
4.5
4
3.5
3
2.5
2
1.5
1
0.5
0
y
0
5
10
15
20
25
x

#### Optimal1 Fitted Gauss, proteinAT1G12270.1
#### bic:706
#### peaks after deleting less than one fifth of global max:11

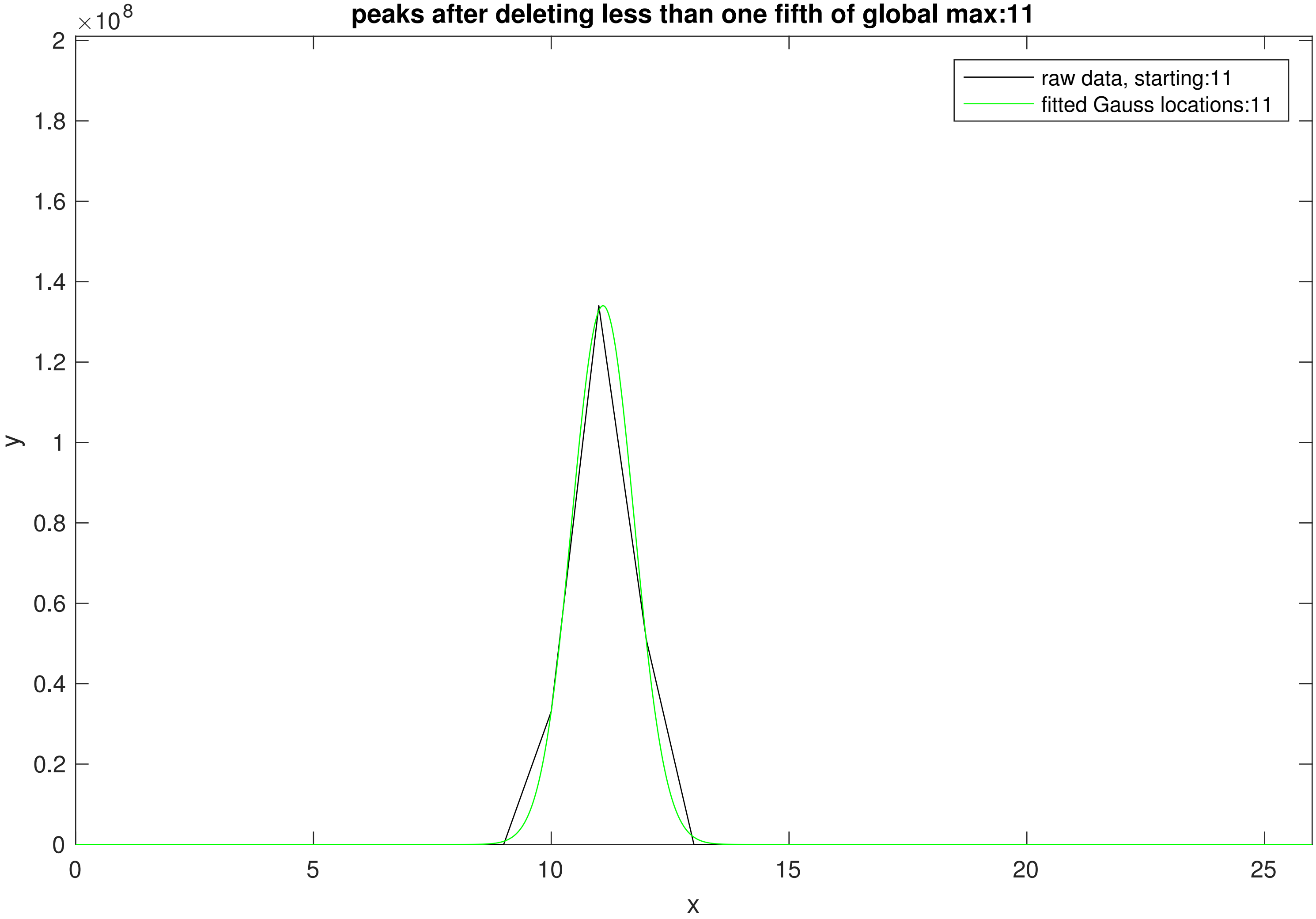

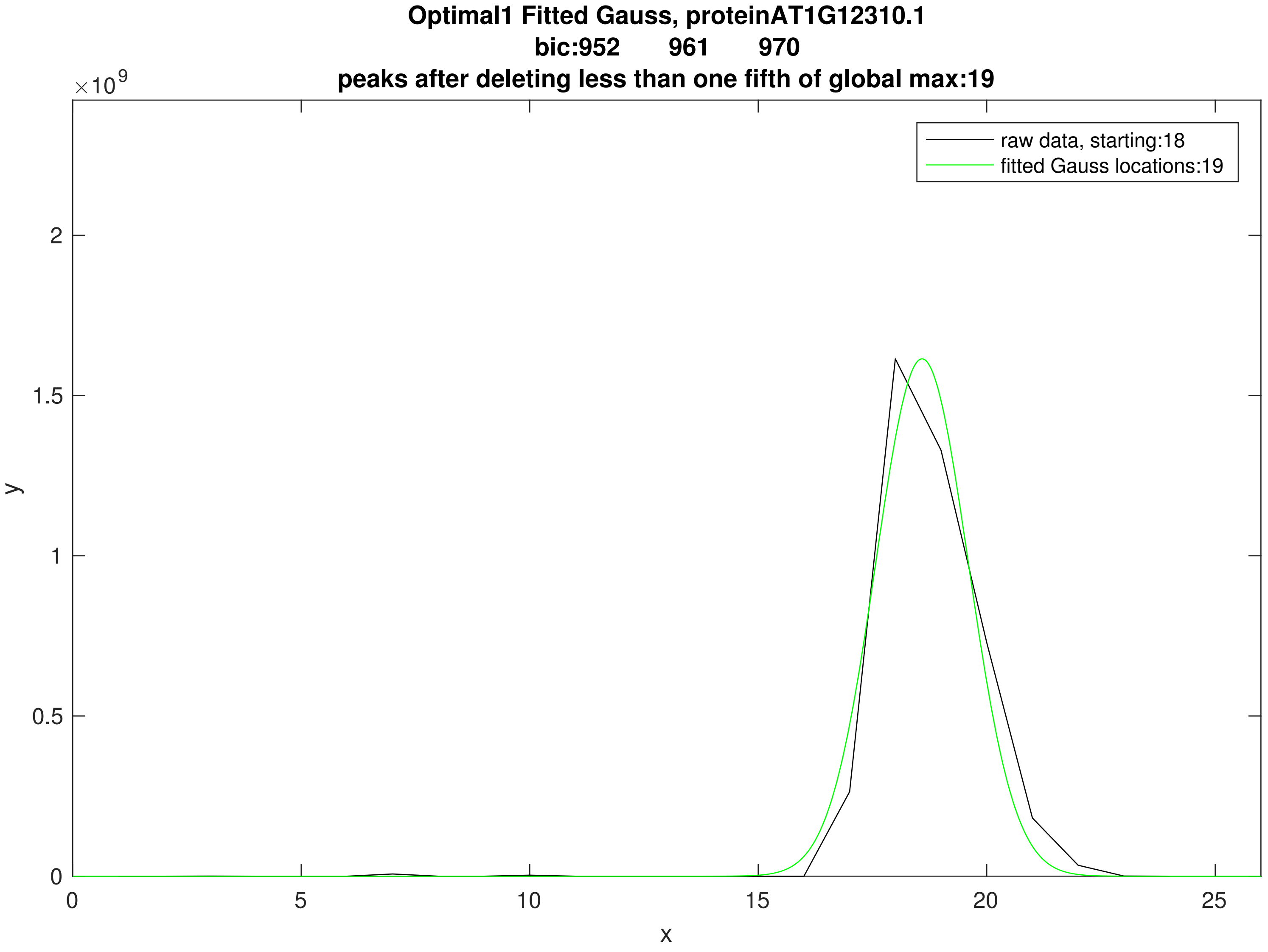

Optimal1 Fitted Gauss, proteinAT1G12310.1
bic:952 961 970
peaks after deleting less than one fifth of global max:19
×10^9
raw data, starting:18
fitted Gauss locations:19
y
x
0
0.5
1
1.5
2
0
5
10
15
20
25

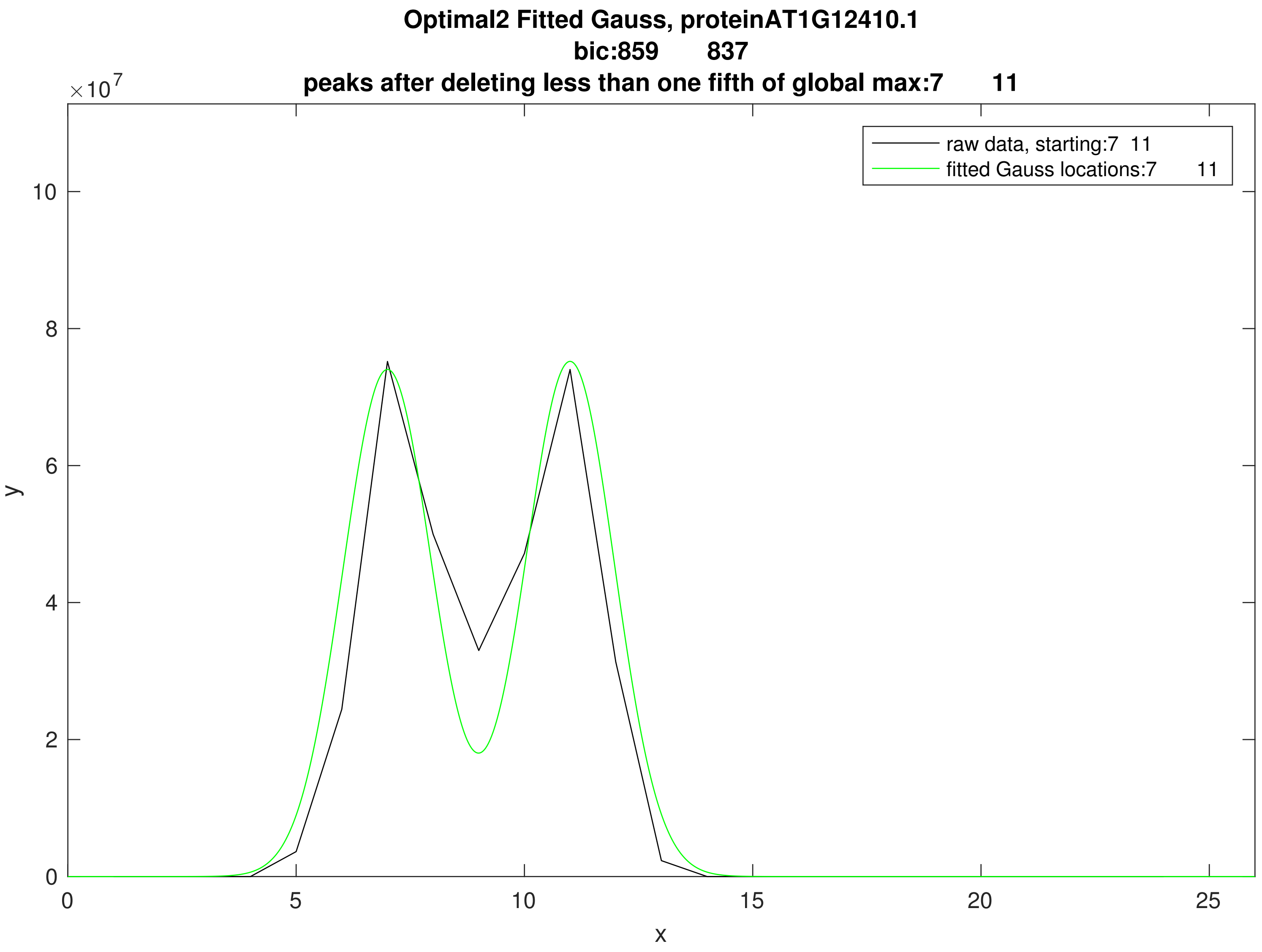
Optimal2 Fitted Gauss, proteinAT1G12410.1
bic:859 837
peaks after deleting less than one fifth of global max:7 11
×10^7
raw data, starting:7 11
fitted Gauss locations:7 11
y
x
0
2
4
6
8
10
0
5
10
15
20
25

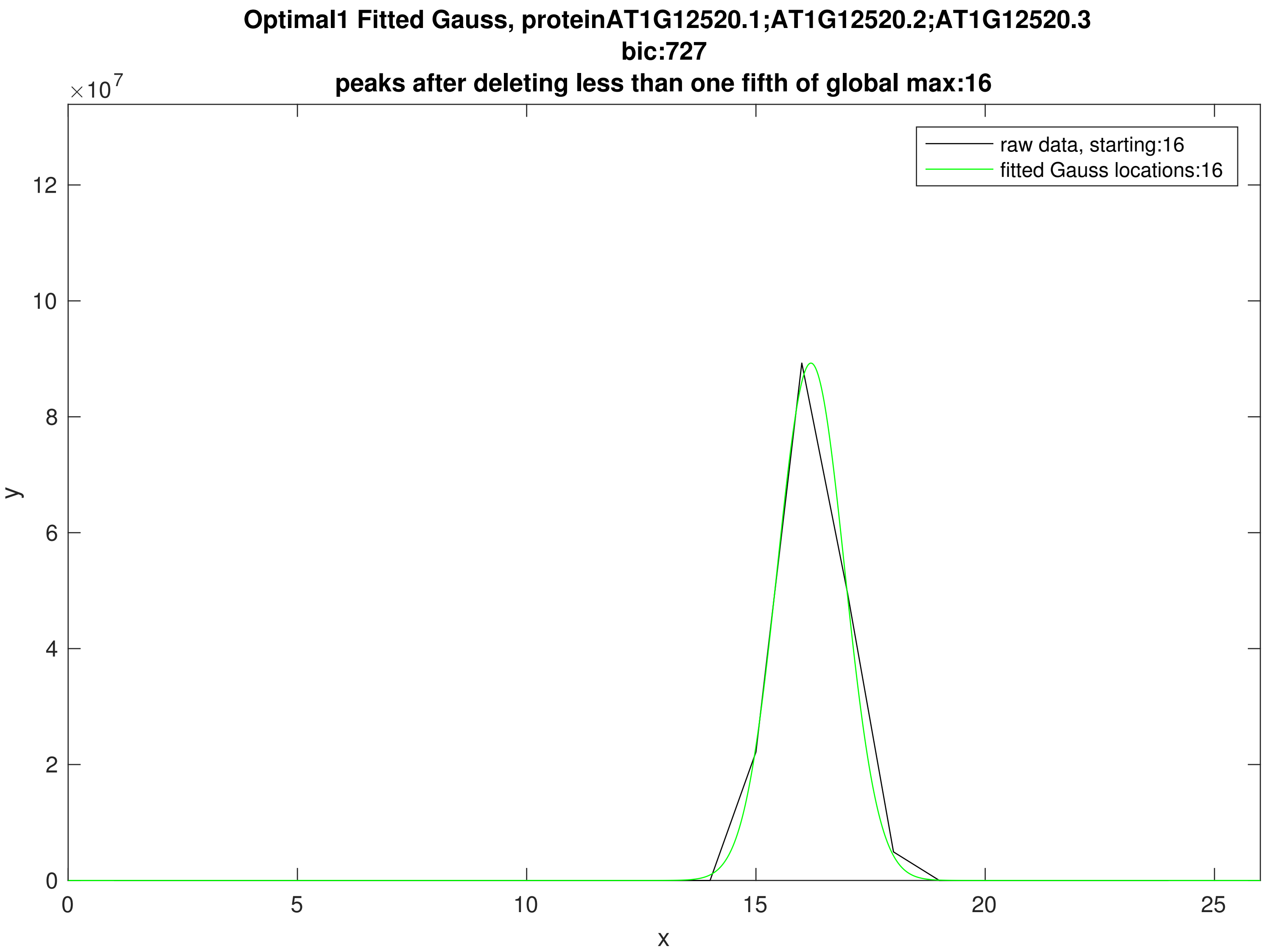
Optimal1 Fitted Gauss, proteinAT1G12520.1;AT1G12520.2;AT1G12520.3
bic:727
peaks after deleting less than one fifth of global max:16
×10^7
raw data, starting:16
fitted Gauss locations:16
y
x
0
2
4
6
8
10
12
0
5
10
15
20
25

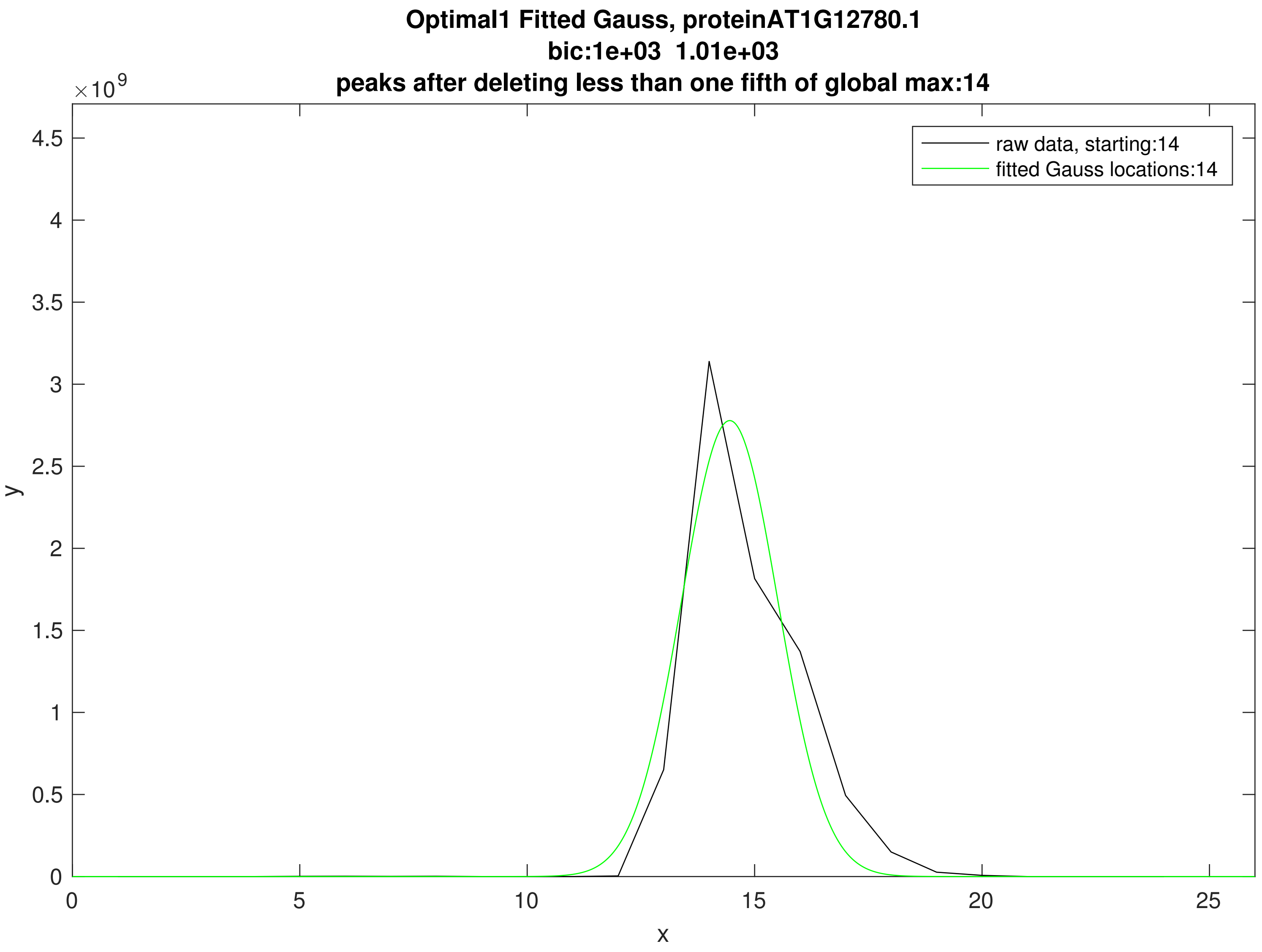
Optimal1 Fitted Gauss, proteinAT1G12780.1
bic:1e+03 1.01e+03
peaks after deleting less than one fifth of global max:14
×10^9
4.5
4
3.5
3
2.5
2
1.5
1
0.5
0
y
0
5
10
15
20
25
x
raw data, starting:14
fitted Gauss locations:14

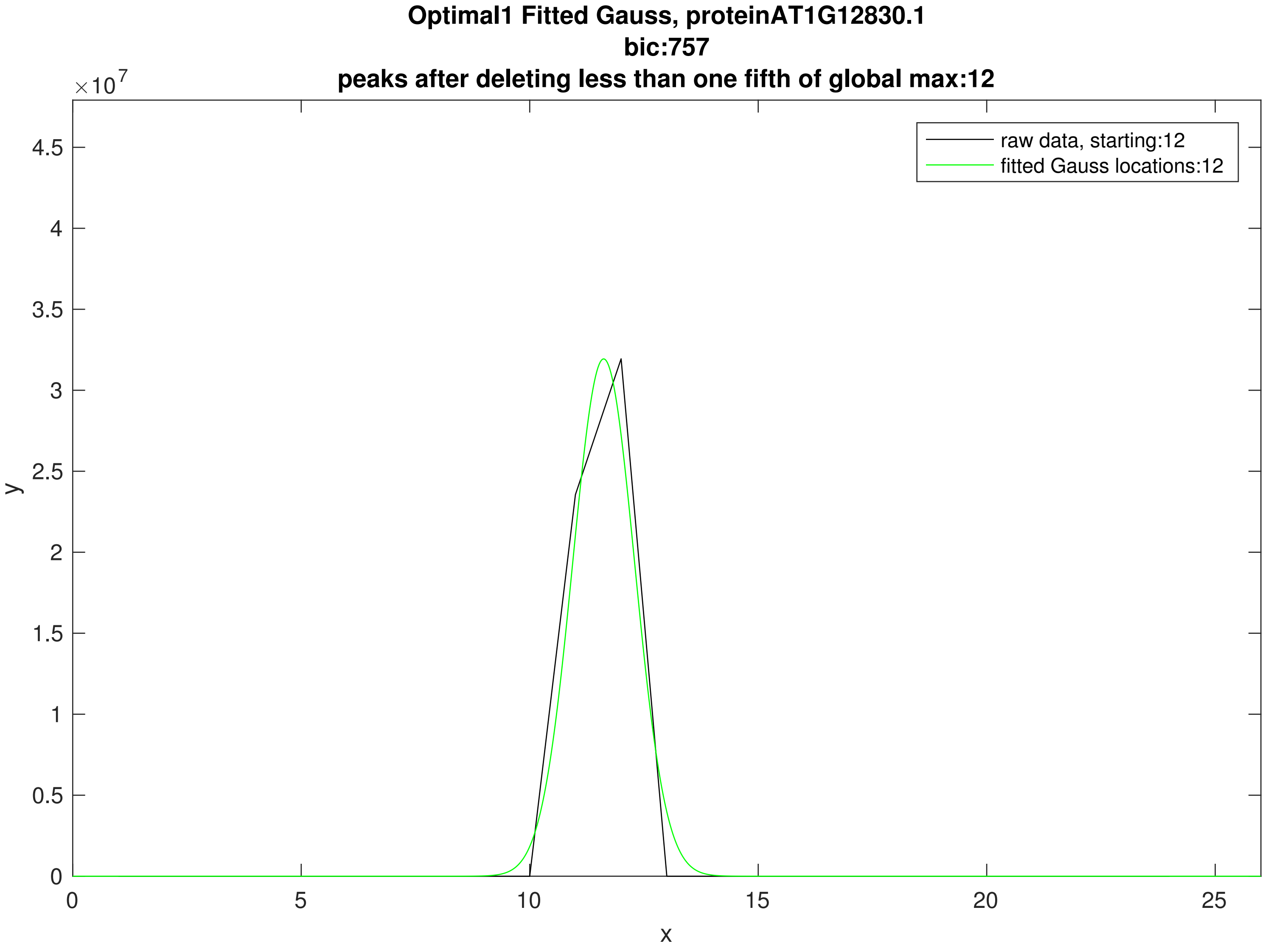
Optimal1 Fitted Gauss, proteinAT1G12830.1
bic:757
peaks after deleting less than one fifth of global max:12
×10^7
raw data, starting:12
fitted Gauss locations:12
y
x
0
0.5
1
1.5
2
2.5
3
3.5
4
4.5
0
5
10
15
20
25

Optimal1 Fitted Gauss, proteinAT1G12840.1
bic:800
peaks after deleting less than one fifth of global max:16

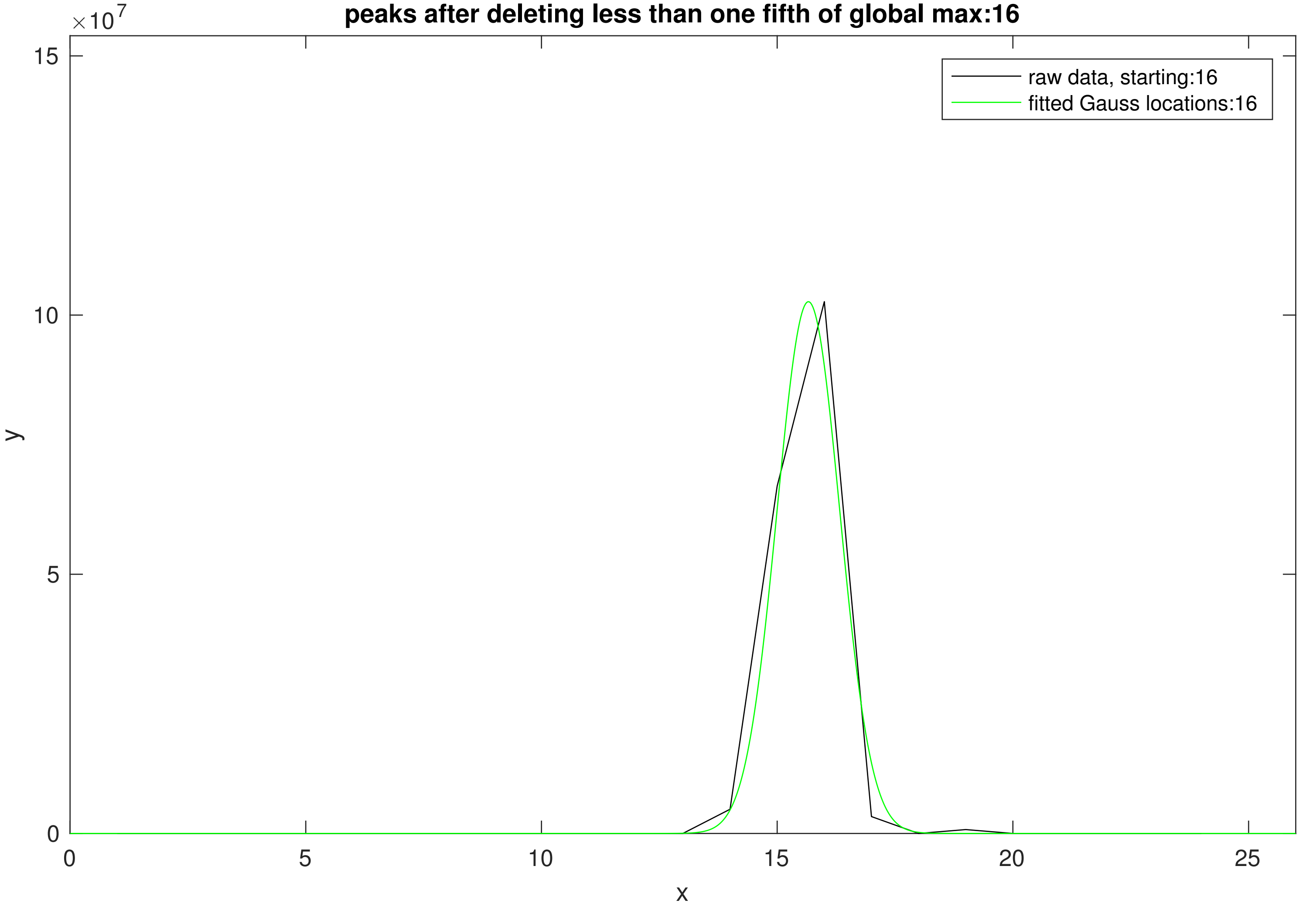

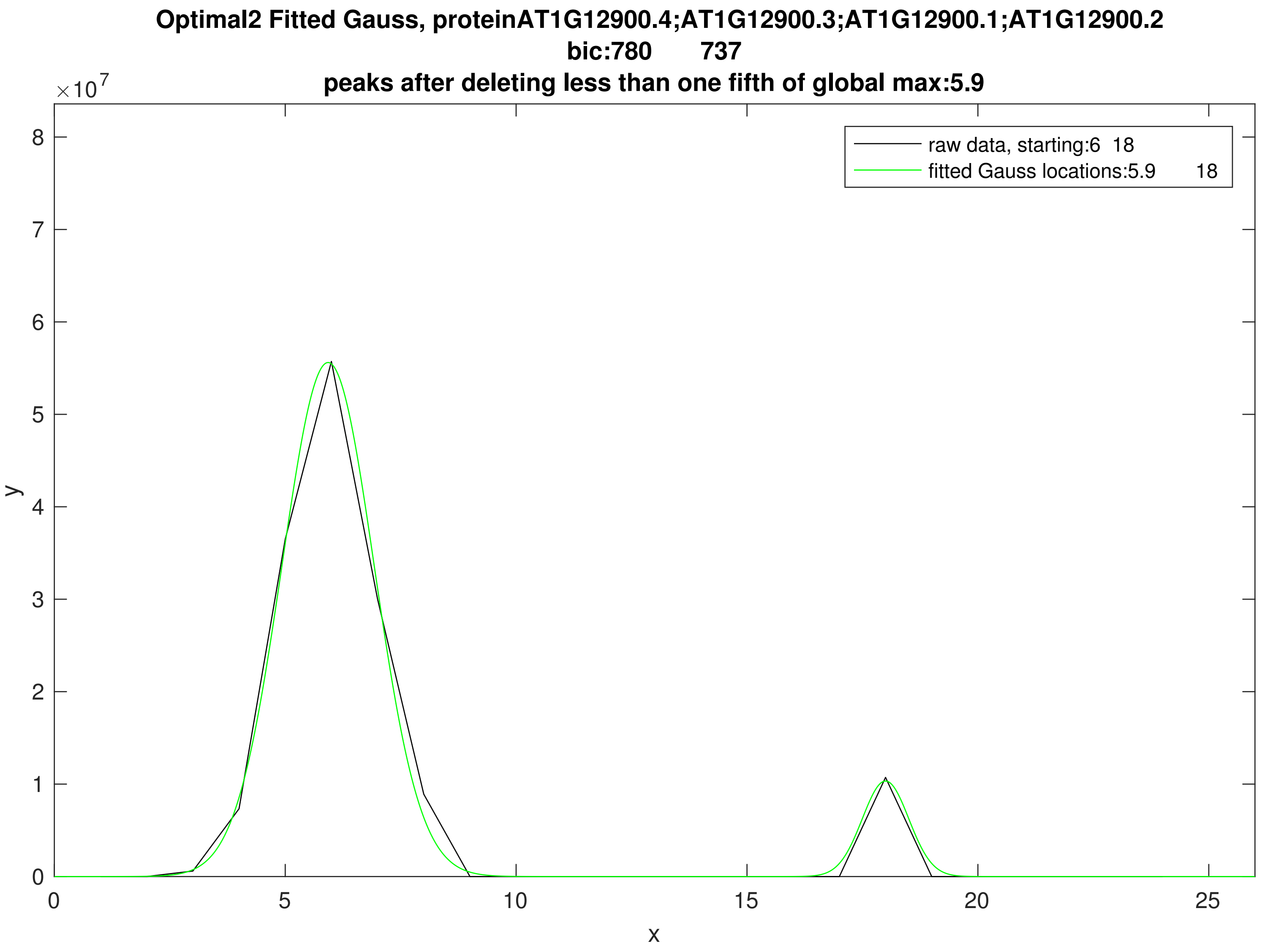

Optimal2 Fitted Gauss, proteinAT1G12900.4;AT1G12900.3;AT1G12900.1;AT1G12900.2
bic:780 737
peaks after deleting less than one fifth of global max:5.9
×10^7
8
7
6
5
4
3
2
1
0
y
0
5
10
15
20
25
x
raw data, starting:6 18
fitted Gauss locations:5.9 18

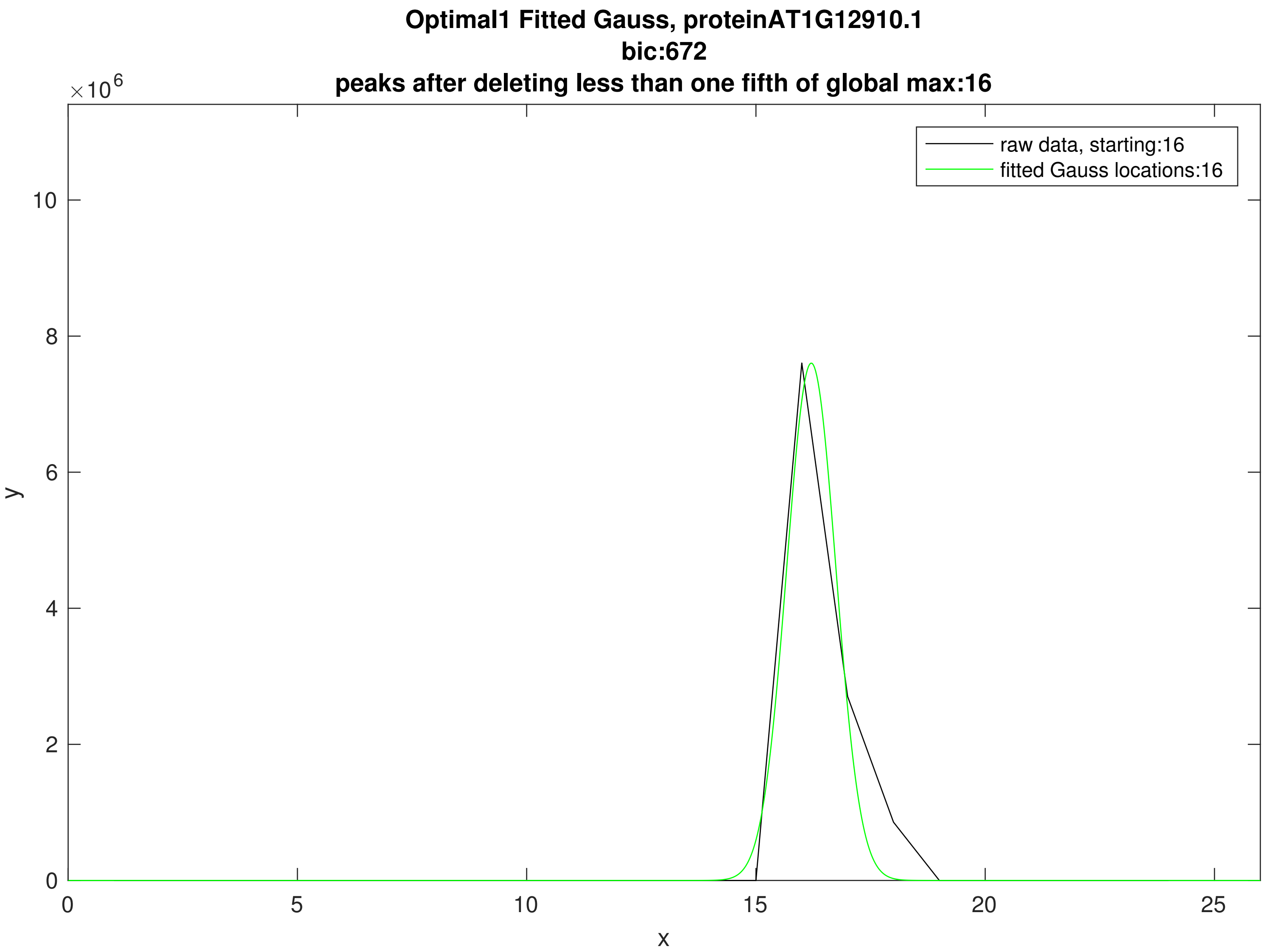

Optimal1 Fitted Gauss, proteinAT1G12910.1
bic:672
peaks after deleting less than one fifth of global max:16
×10^6
raw data, starting:16
fitted Gauss locations:16
10
8
6
4
2
0
y
0
5
10
15
20
25
x

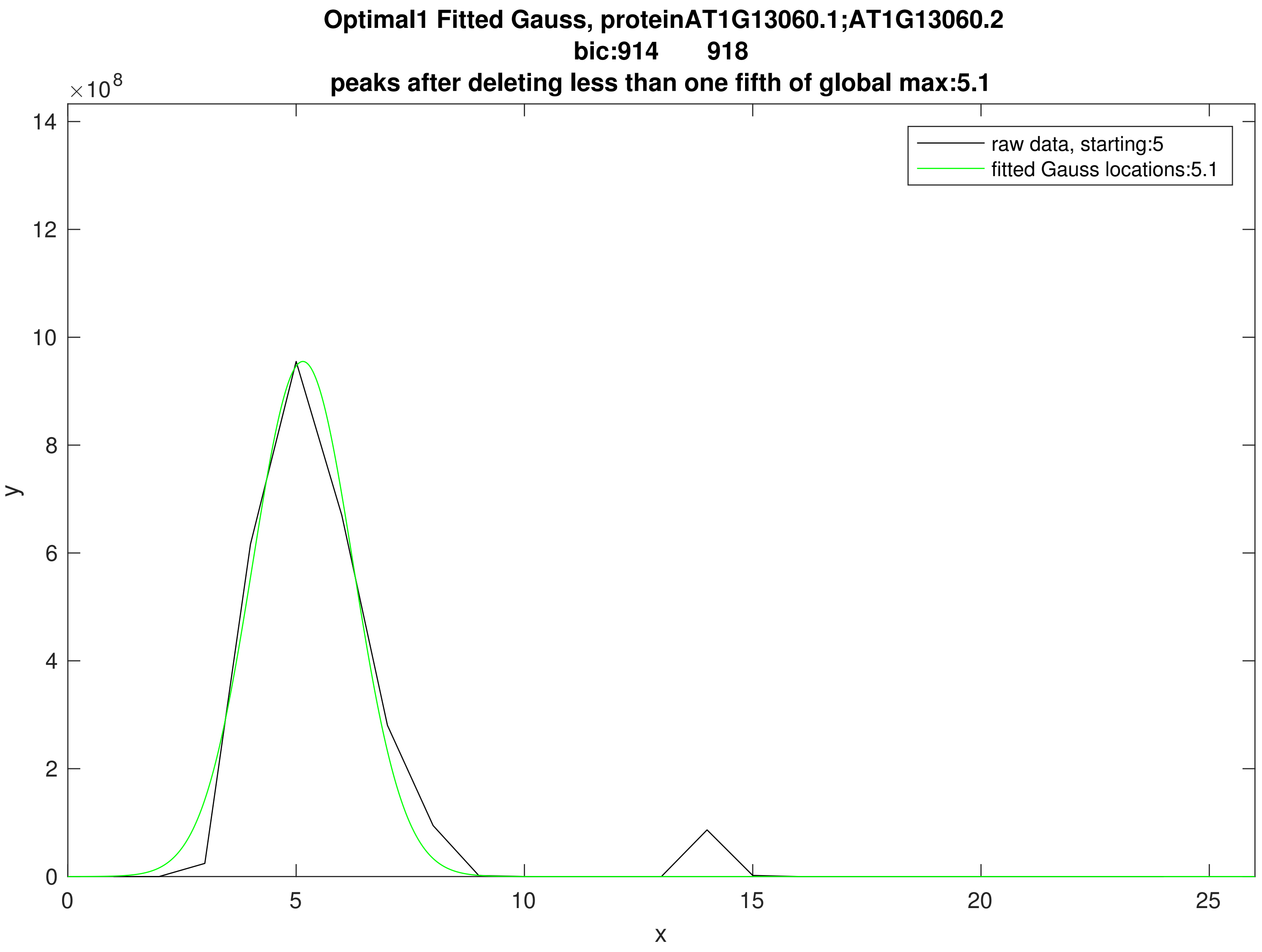

Optimal1 Fitted Gauss, proteinAT1G13060.1;AT1G13060.2
bic:914 918
peaks after deleting less than one fifth of global max:5.1
×10^8
14
12
10
8
6
4
2
0
y
0
5
10
15
20
25
x
raw data, starting:5
fitted Gauss locations:5.1

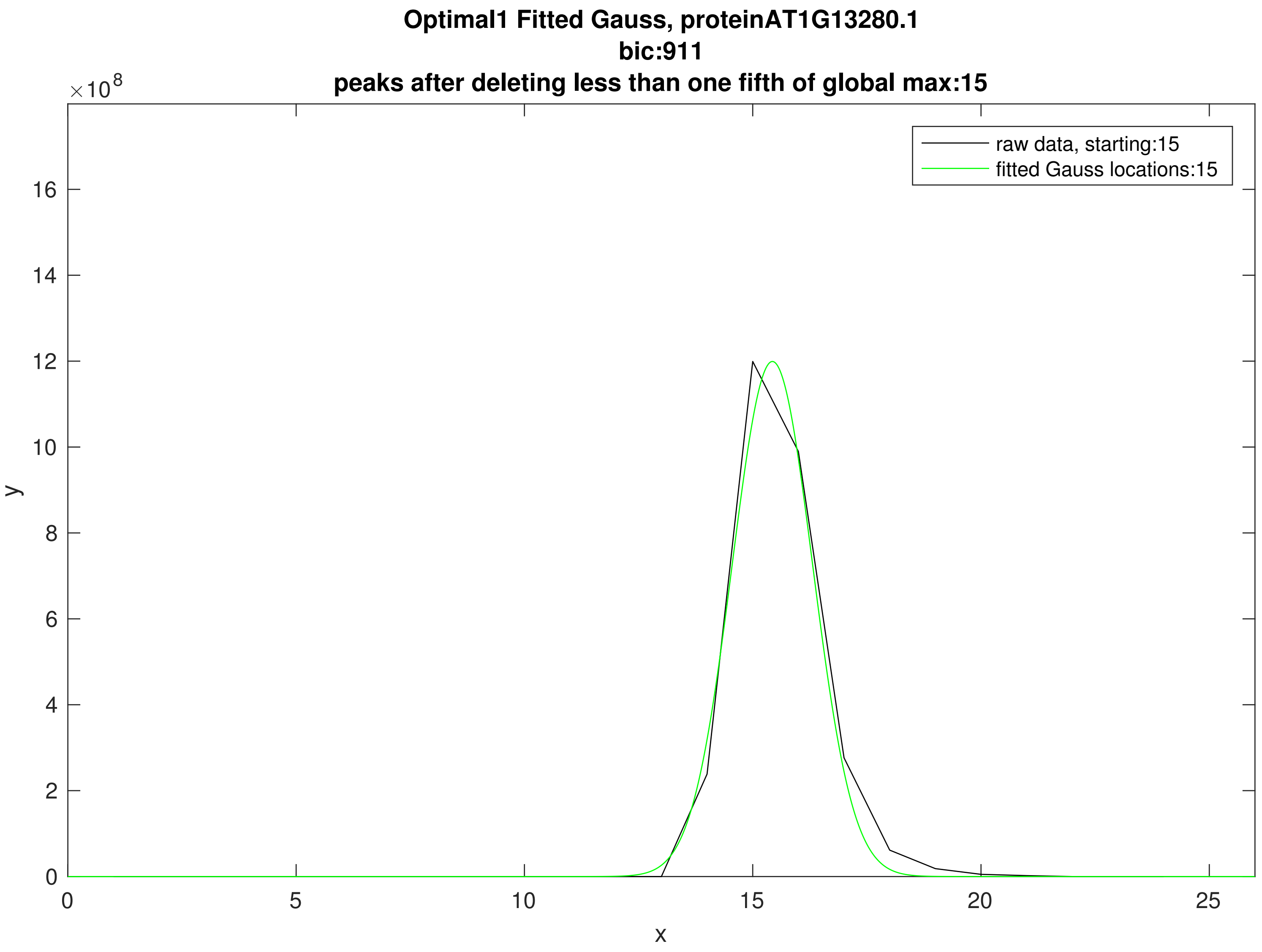
Optimal1 Fitted Gauss, proteinAT1G13280.1
bic:911
peaks after deleting less than one fifth of global max:15
×10^8
raw data, starting:15
fitted Gauss locations:15
16
14
12
10
8
6
4
2
0
y
0
5
10
15
20
25
x

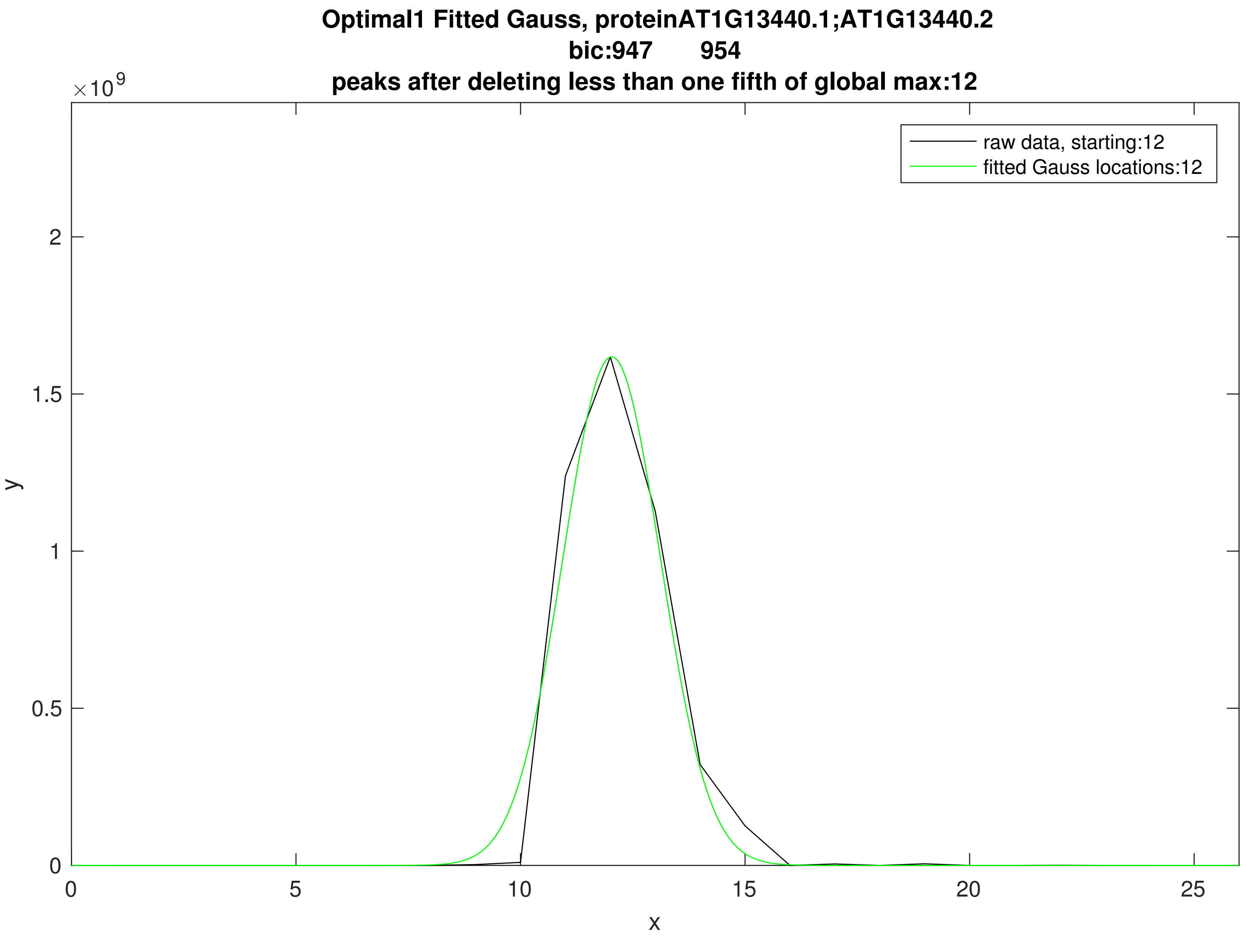
Optimal1 Fitted Gauss, proteinAT1G13440.1;AT1G13440.2
bic:947 954
peaks after deleting less than one fifth of global max:12
×10^9
raw data, starting:12
fitted Gauss locations:12
y
x
0
0.5
1
1.5
2
0
5
10
15
20
25

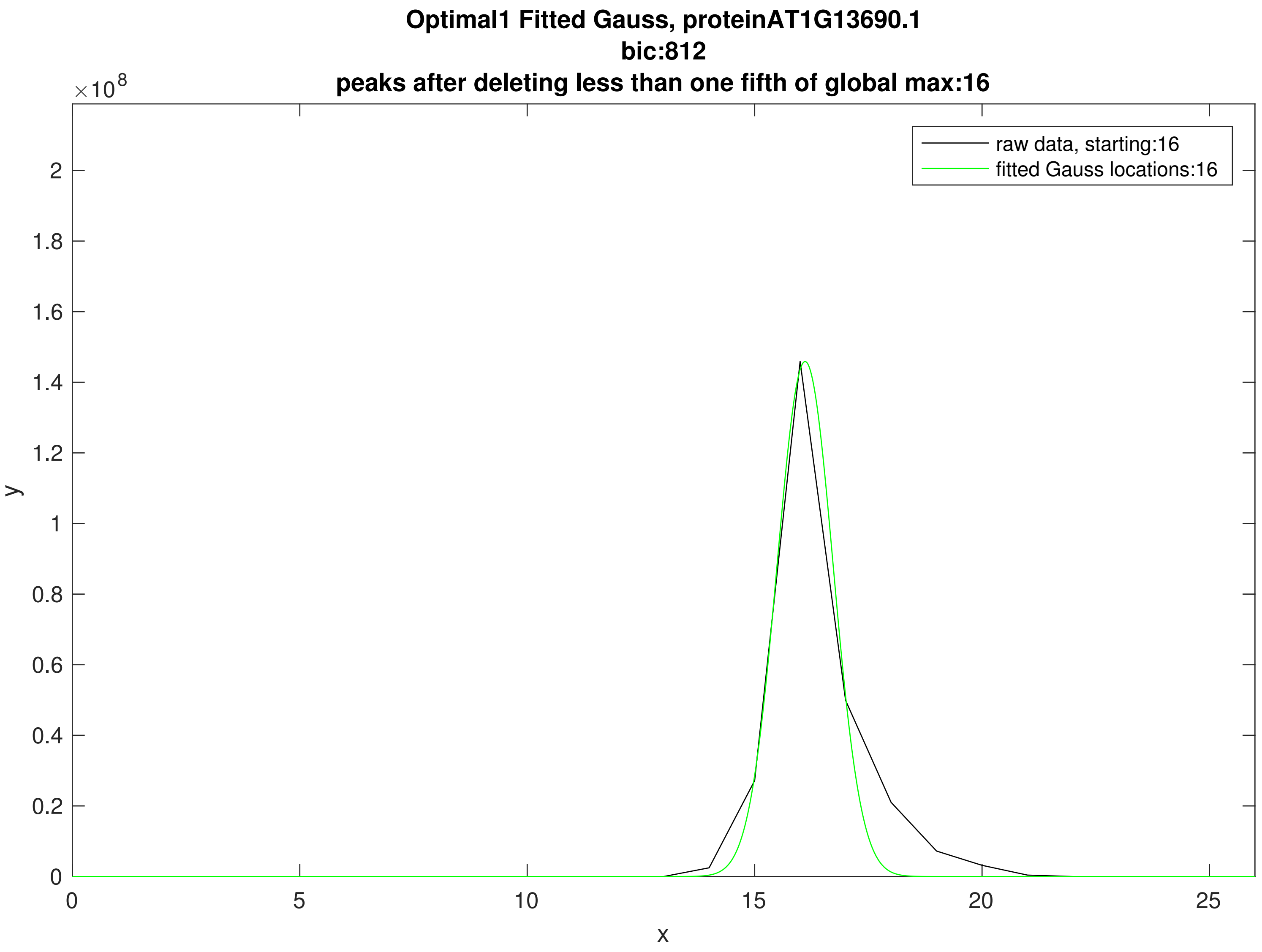
Optimal1 Fitted Gauss, proteinAT1G13690.1
bic:812
peaks after deleting less than one fifth of global max:16
×10^8
raw data, starting:16
fitted Gauss locations:16
2
1.8
1.6
1.4
1.2
1
0.8
0.6
0.4
0.2
0
y
0
5
10
15
20
25
x

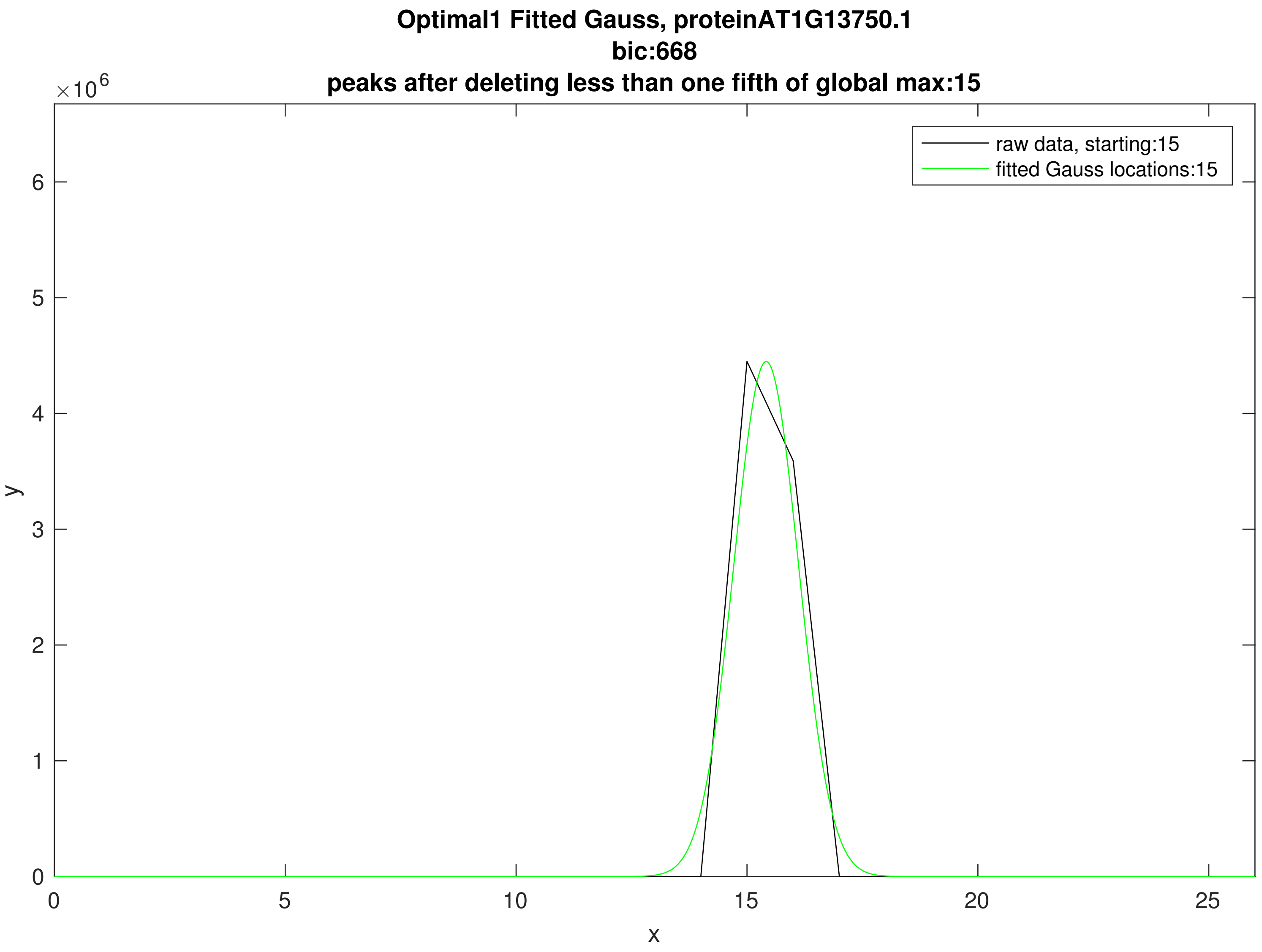
Optimal1 Fitted Gauss, proteinAT1G13750.1
bic:668
peaks after deleting less than one fifth of global max:15
×10^6
6
5
4
3
2
1
0
y
0
5
10
15
20
25
x
raw data, starting:15
fitted Gauss locations:15

**Optimal2 Fitted Gauss, proteinAT1G13900.1**
**bic:885 870 879**
**peaks after deleting less than one fifth of global max:11**

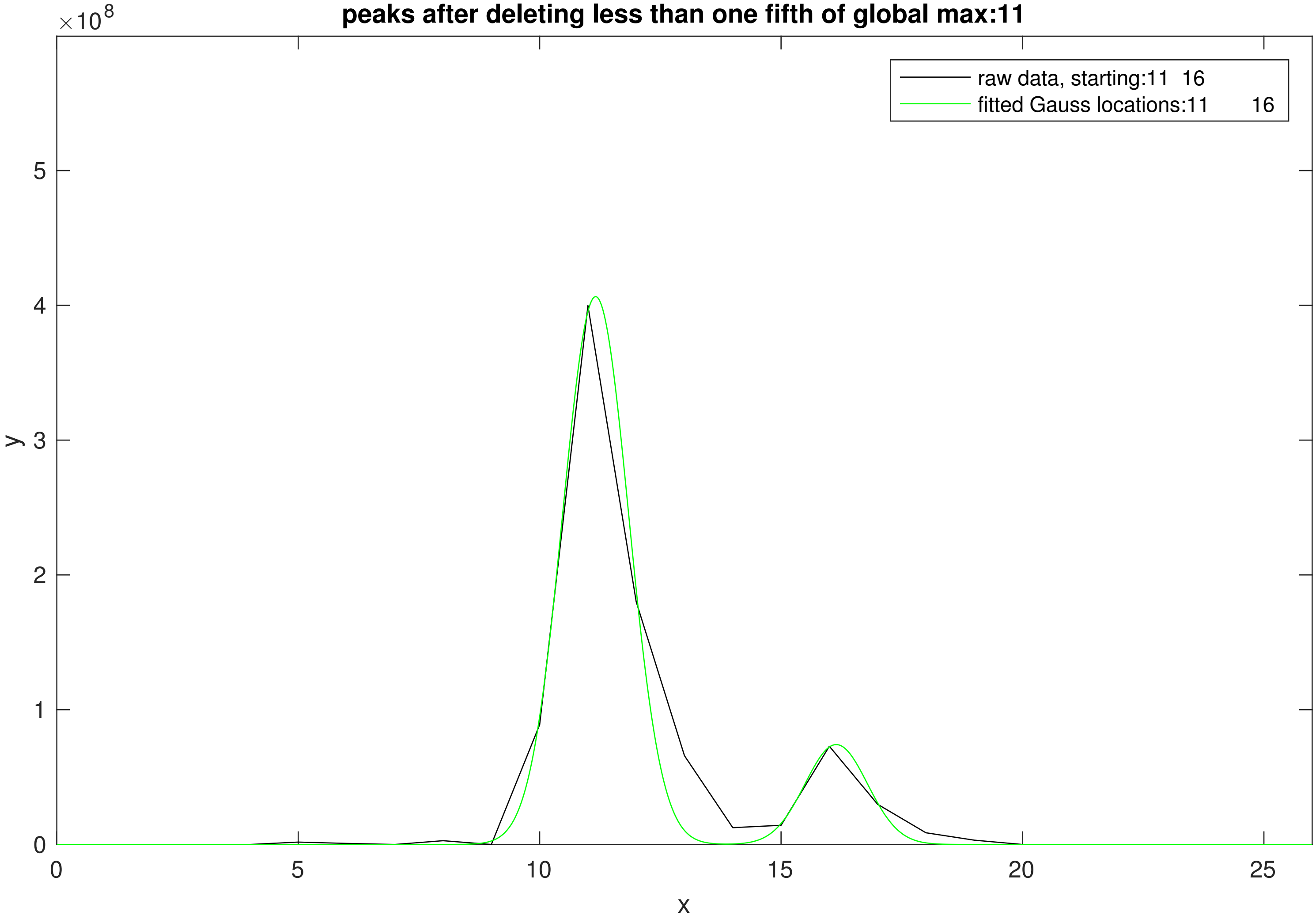

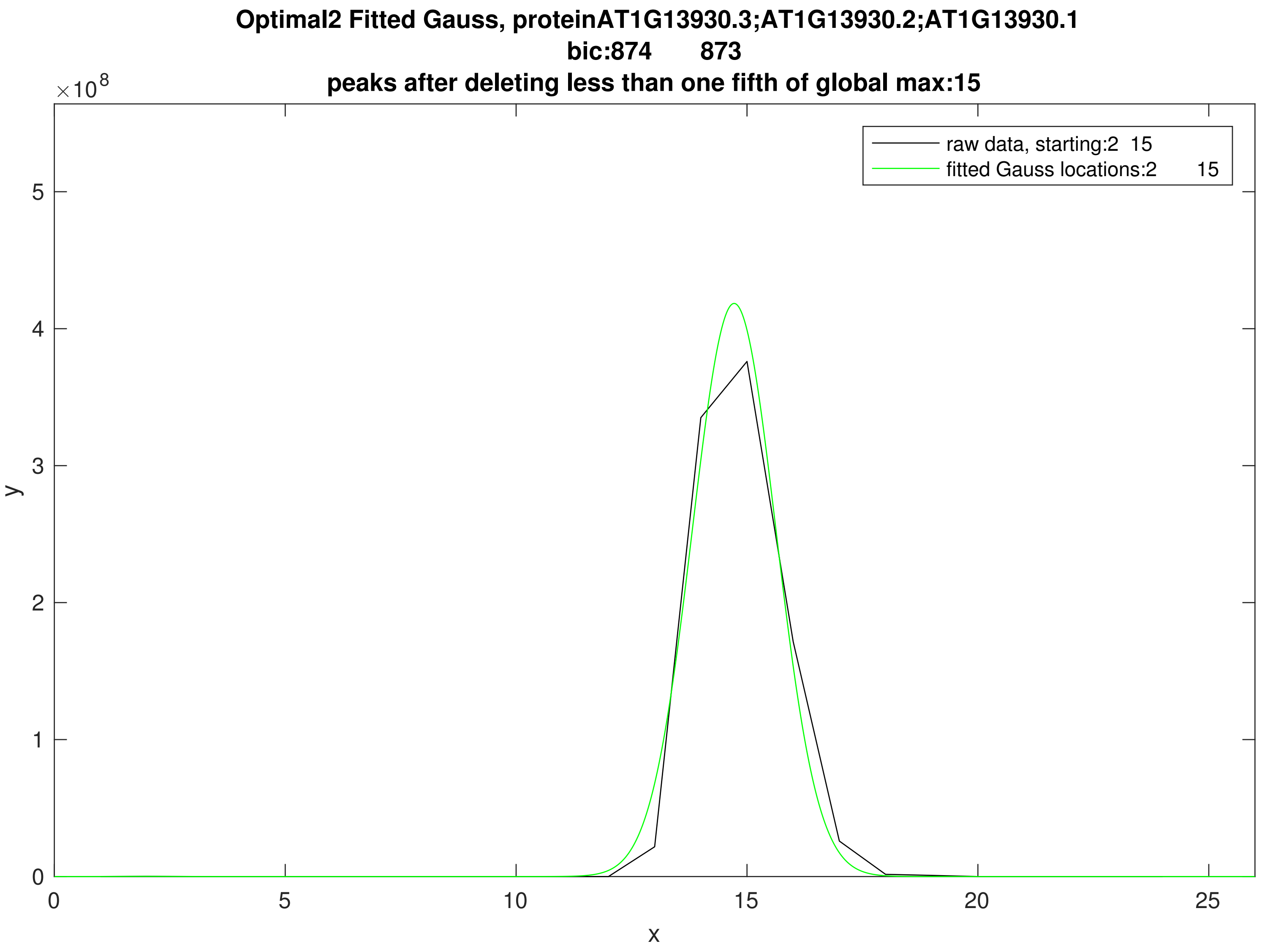
Optimal2 Fitted Gauss, proteinAT1G13930.3;AT1G13930.2;AT1G13930.1
bic:874 873
peaks after deleting less than one fifth of global max:15
×10^8
raw data, starting:2 15
fitted Gauss locations:2 15
y
x
0
1
2
3
4
5
0
5
10
15
20
25

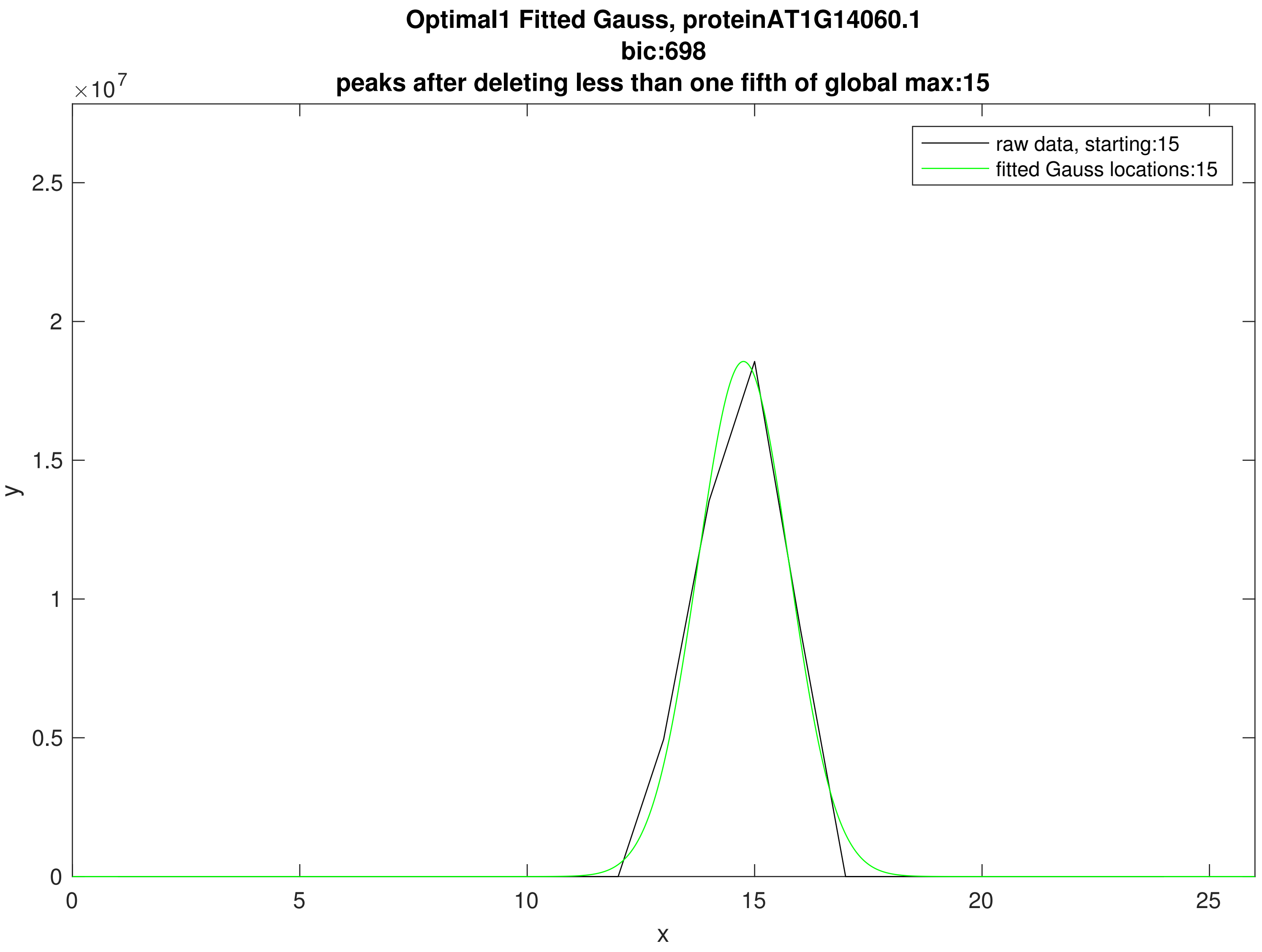

Optimal1 Fitted Gauss, proteinAT1G14060.1
bic:698
peaks after deleting less than one fifth of global max:15
×10^7
2.5
2
1.5
1
0.5
0
y
0
5
10
15
20
25
x
raw data, starting:15
fitted Gauss locations:15

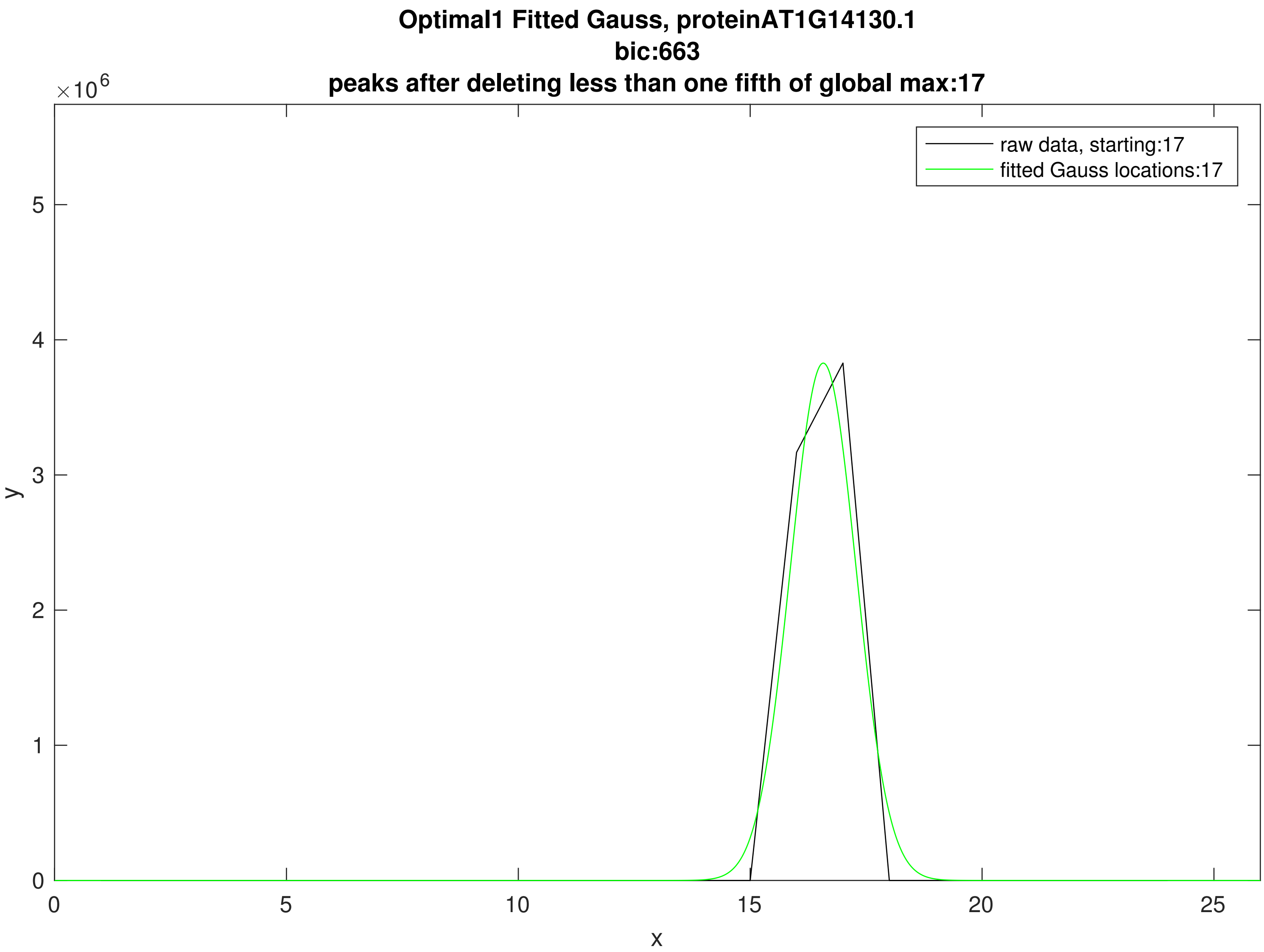
Optimal1 Fitted Gauss, proteinAT1G14130.1
bic:663
peaks after deleting less than one fifth of global max:17
×10^6
raw data, starting:17
fitted Gauss locations:17
y
x
0
1
2
3
4
5
0
5
10
15
20
25

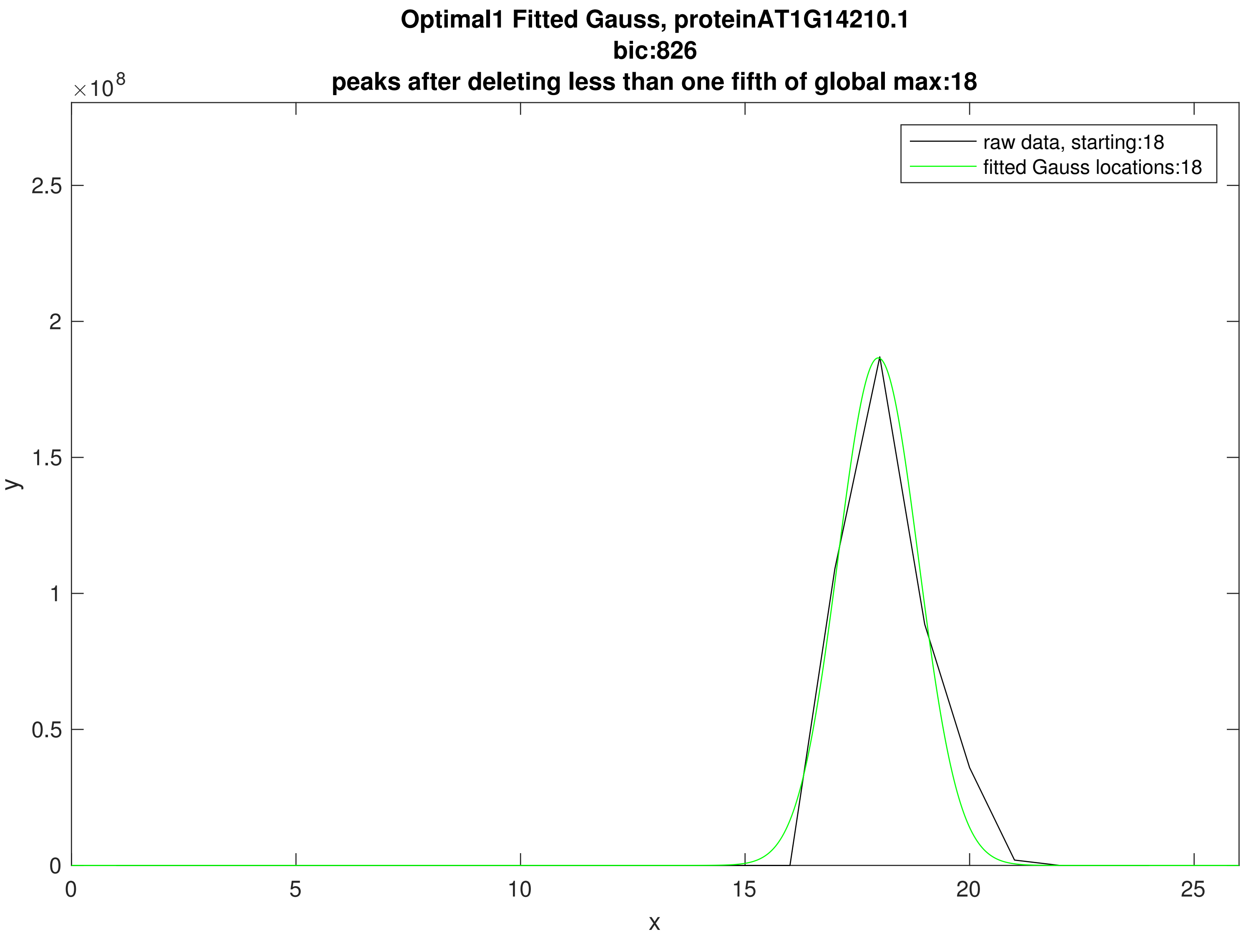
Optimal1 Fitted Gauss, proteinAT1G14210.1
bic:826
peaks after deleting less than one fifth of global max:18
×10^8
raw data, starting:18
fitted Gauss locations:18
y
x
0
0.5
1
1.5
2
2.5
0
5
10
15
20
25

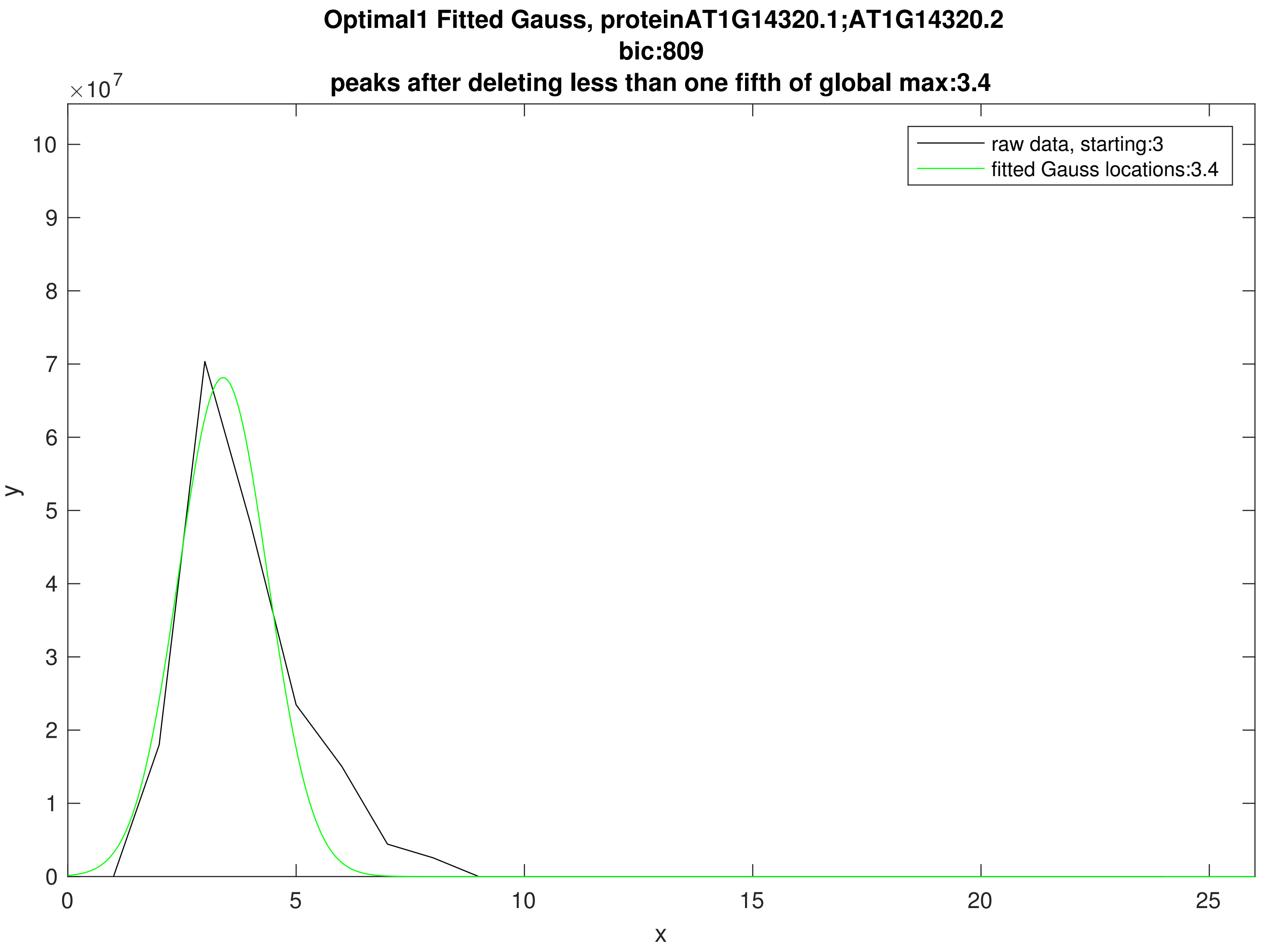
Optimal1 Fitted Gauss, proteinAT1G14320.1;AT1G14320.2
bic:809
peaks after deleting less than one fifth of global max:3.4
×10^7
raw data, starting:3
fitted Gauss locations:3.4
y
x
0
1
2
3
4
5
6
7
8
9
10
0
5
10
15
20
25

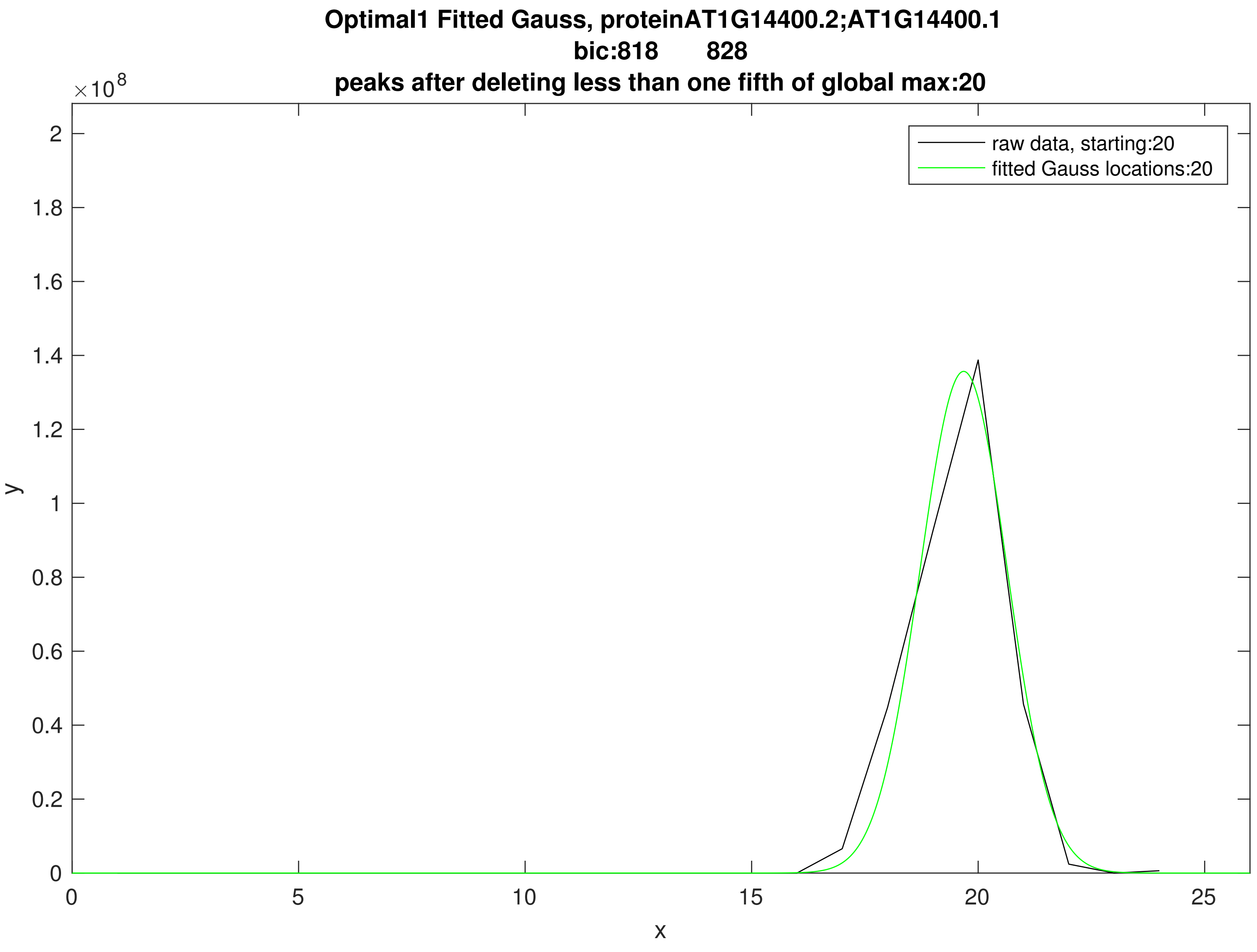

Optimal1 Fitted Gauss, proteinAT1G14400.2;AT1G14400.1
bic:818 828
peaks after deleting less than one fifth of global max:20
×10^8
raw data, starting:20
fitted Gauss locations:20
y
x
0
0.2
0.4
0.6
0.8
1
1.2
1.4
1.6
1.8
2
0
5
10
15
20
25

Optimal1 Fitted Gauss, proteinAT1G14540.1
bic:704
peaks after deleting less than one fifth of global max:18

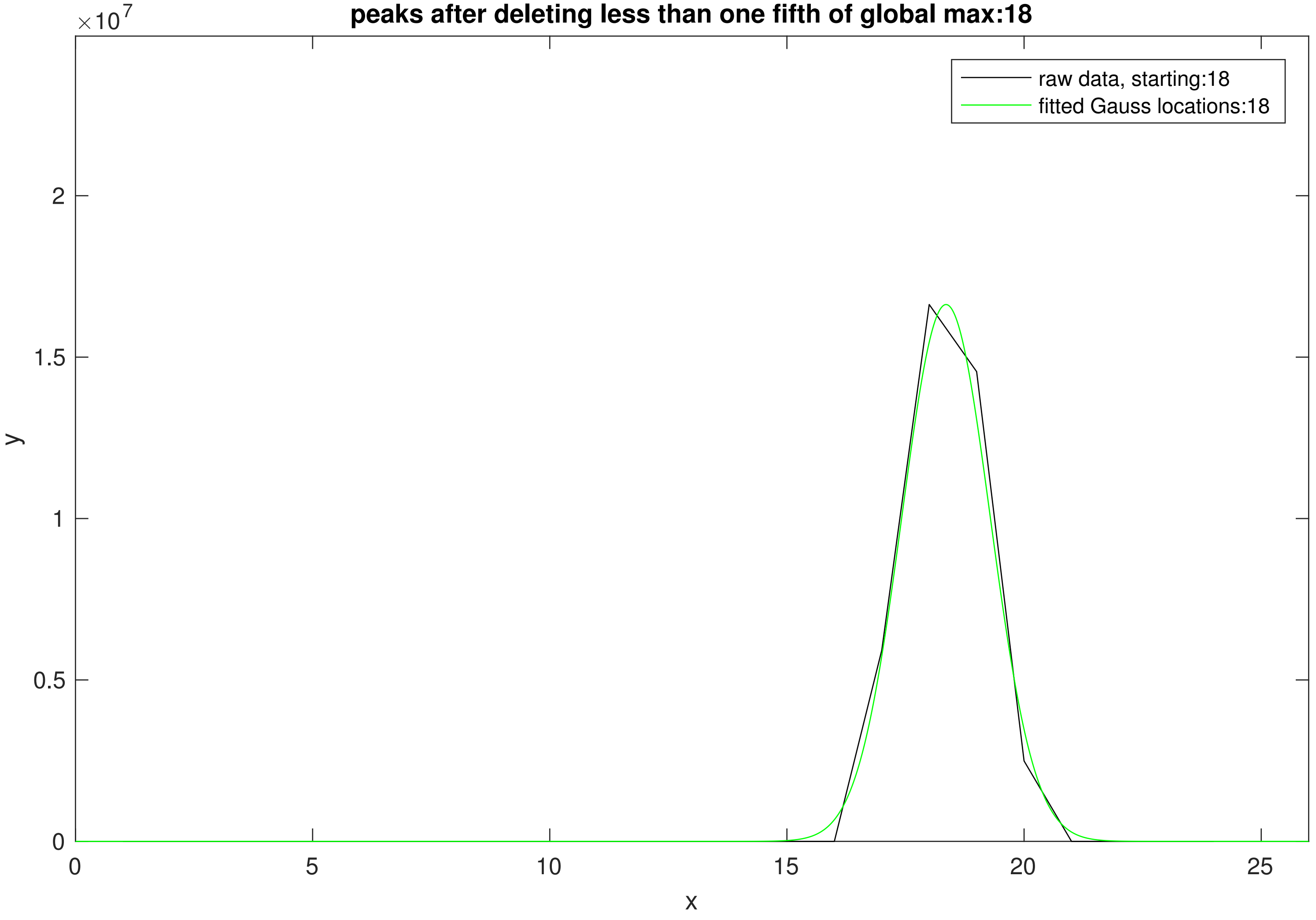

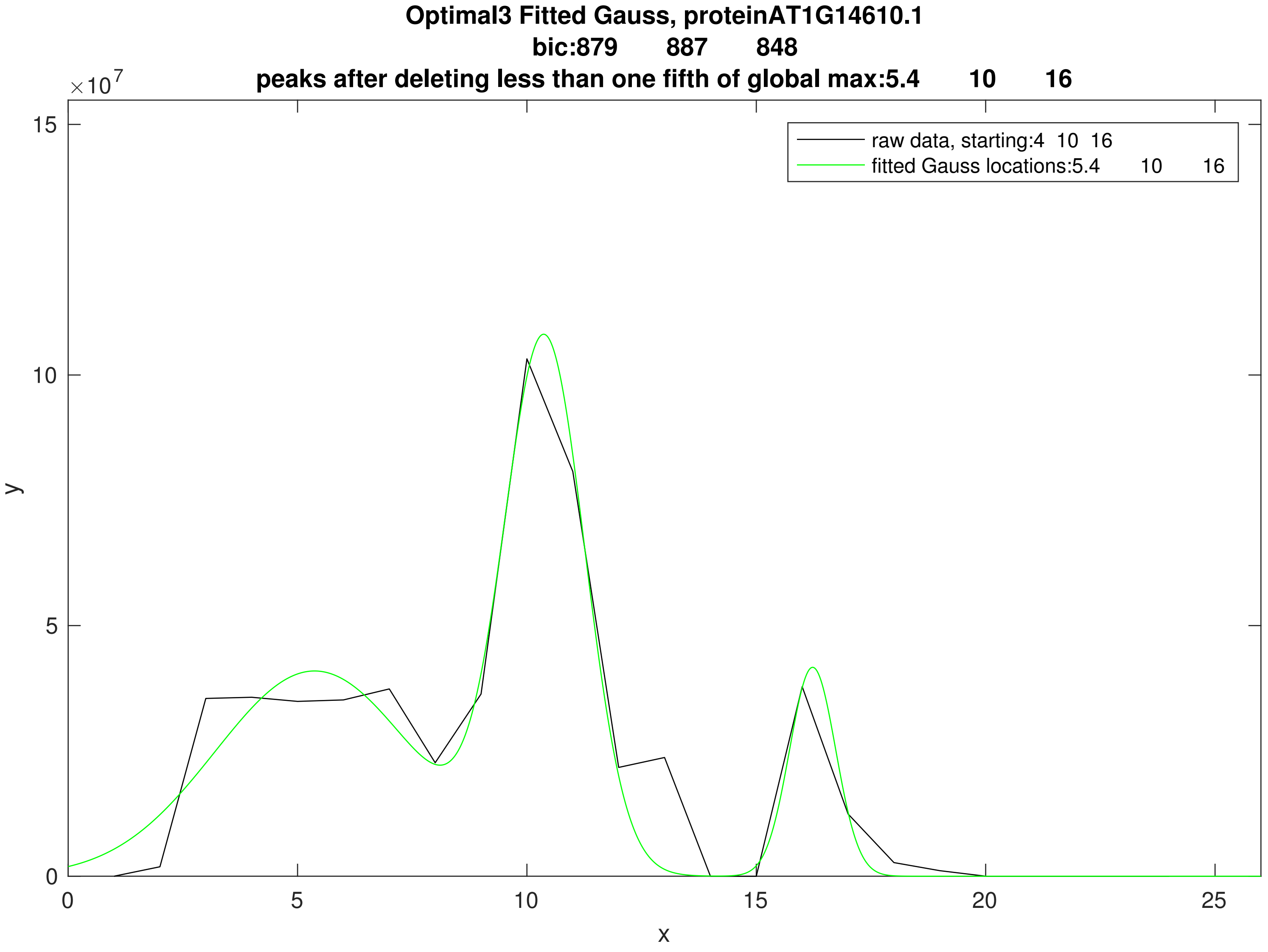
Optimal3 Fitted Gauss, proteinAT1G14610.1
bic:879 887 848
peaks after deleting less than one fifth of global max:5.4 10 16
×10^7
raw data, starting:4 10 16
fitted Gauss locations:5.4 10 16
y
x
0
5
10
15
20
25

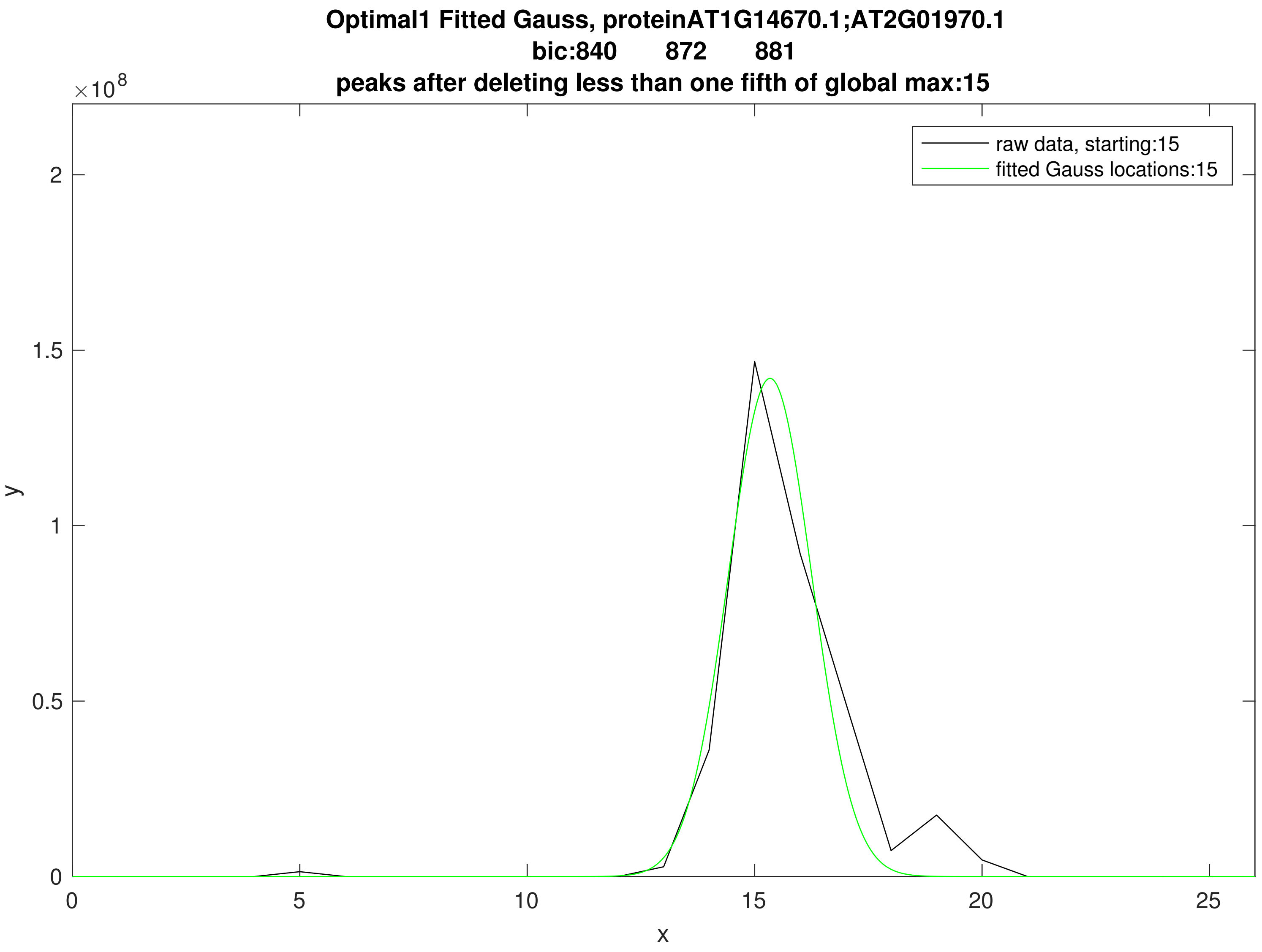

Optimal1 Fitted Gauss, proteinAT1G14670.1;AT2G01970.1
bic:840 872 881
peaks after deleting less than one fifth of global max:15
×10^8
raw data, starting:15
fitted Gauss locations:15
y
x
0
0.5
1
1.5
2
0
5
10
15
20
25

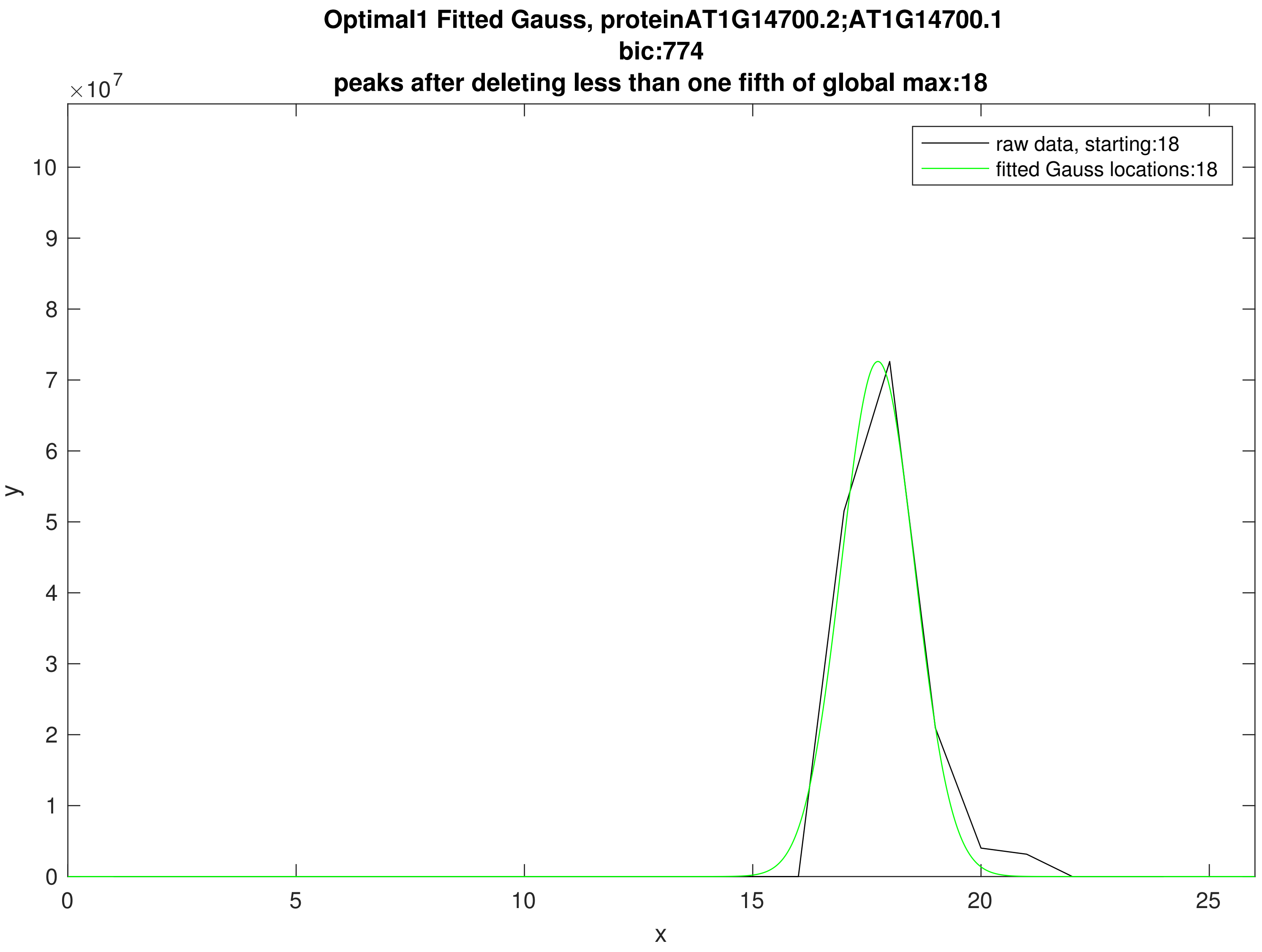

Optimal1 Fitted Gauss, proteinAT1G14700.2;AT1G14700.1
bic:774
peaks after deleting less than one fifth of global max:18
×10^7
raw data, starting:18
fitted Gauss locations:18
y
x
0
1
2
3
4
5
6
7
8
9
10
0
5
10
15
20
25

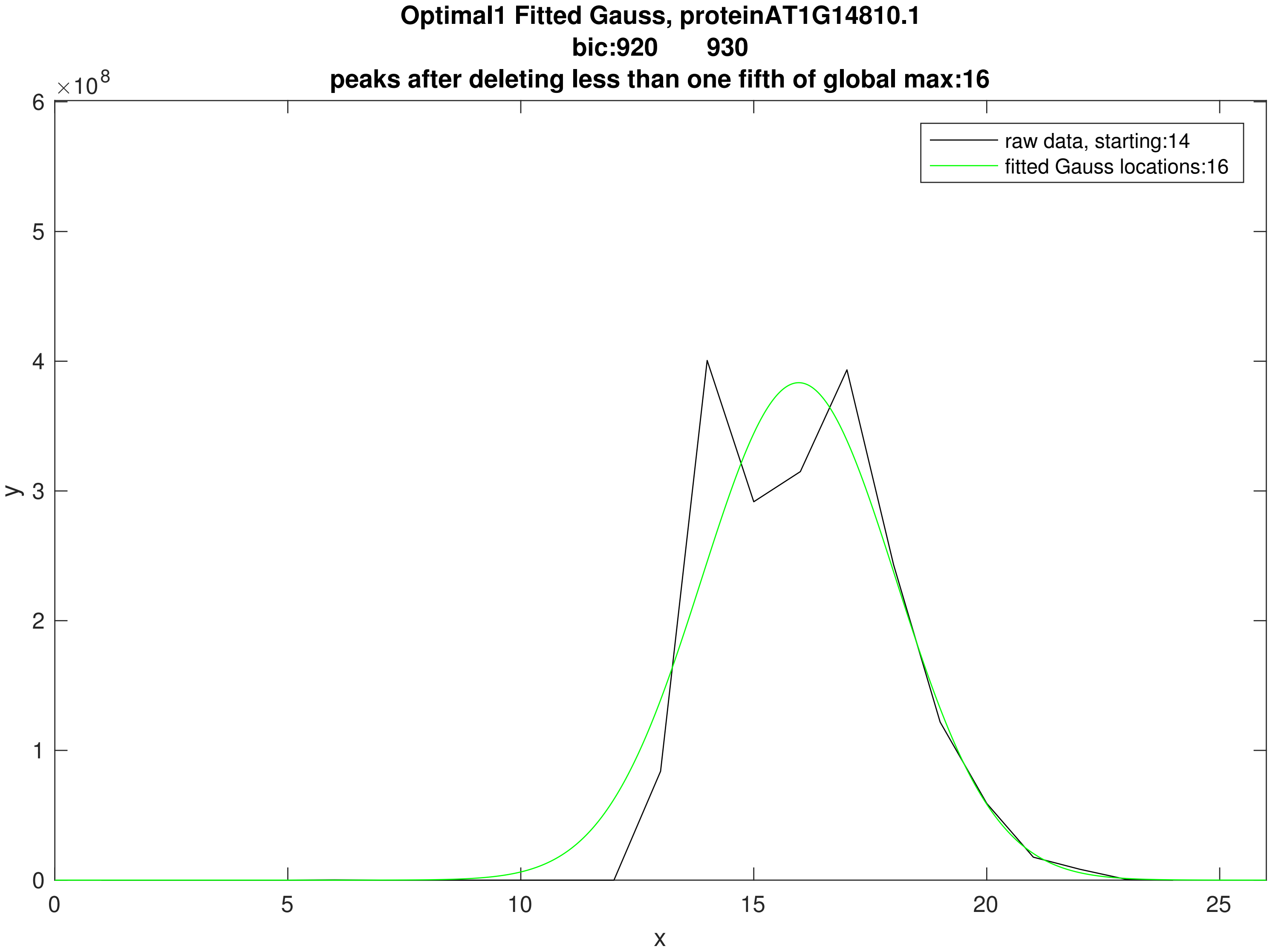
Optimal1 Fitted Gauss, proteinAT1G14810.1
bic:920 930
peaks after deleting less than one fifth of global max:16
×10^8
6
5
4
3
2
1
0
y
0
5
10
15
20
25
x
raw data, starting:14
fitted Gauss locations:16

#### Optimal1 Fitted Gauss, proteinAT1G14930.1
#### bic:841 929
#### peaks after deleting less than one fifth of global max:20

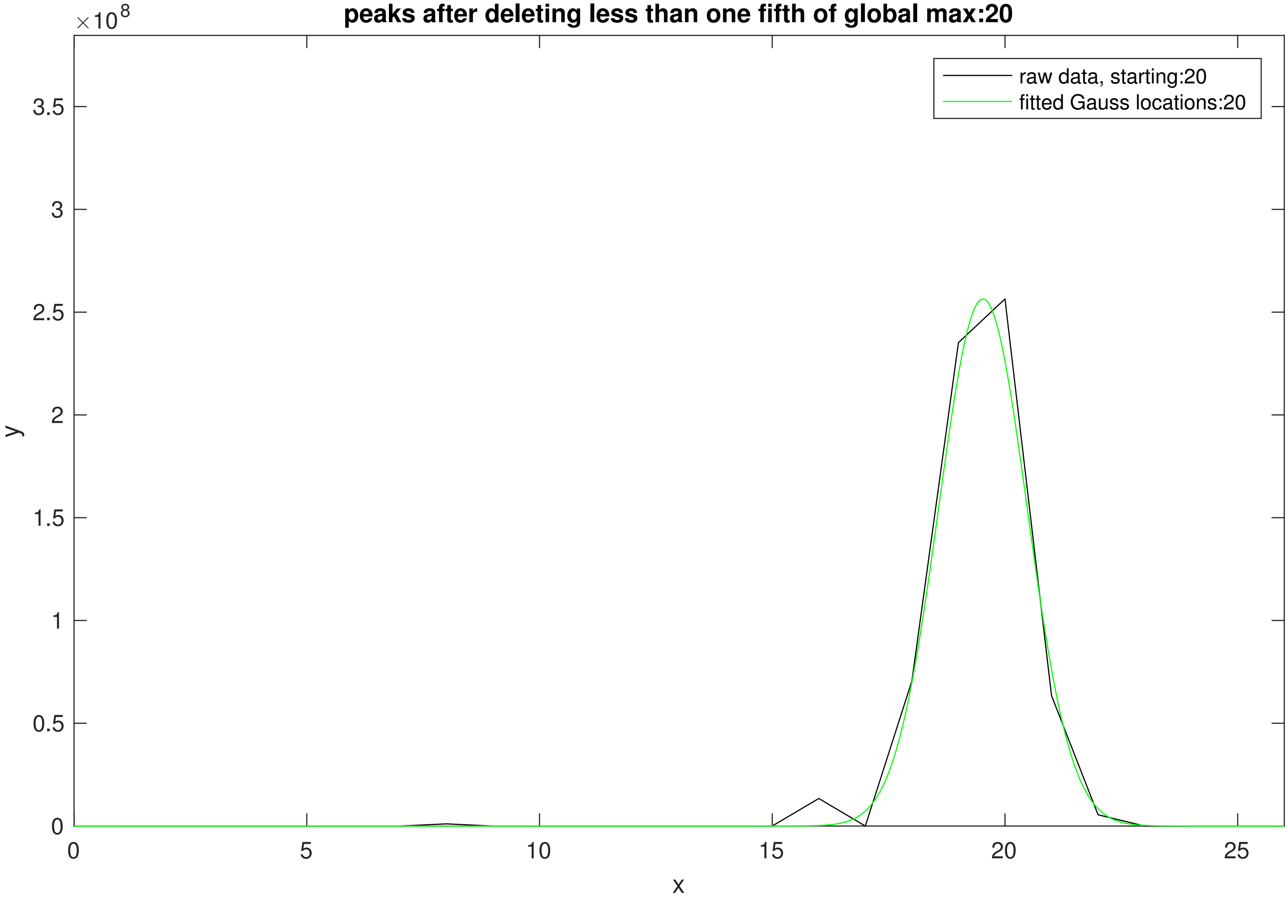

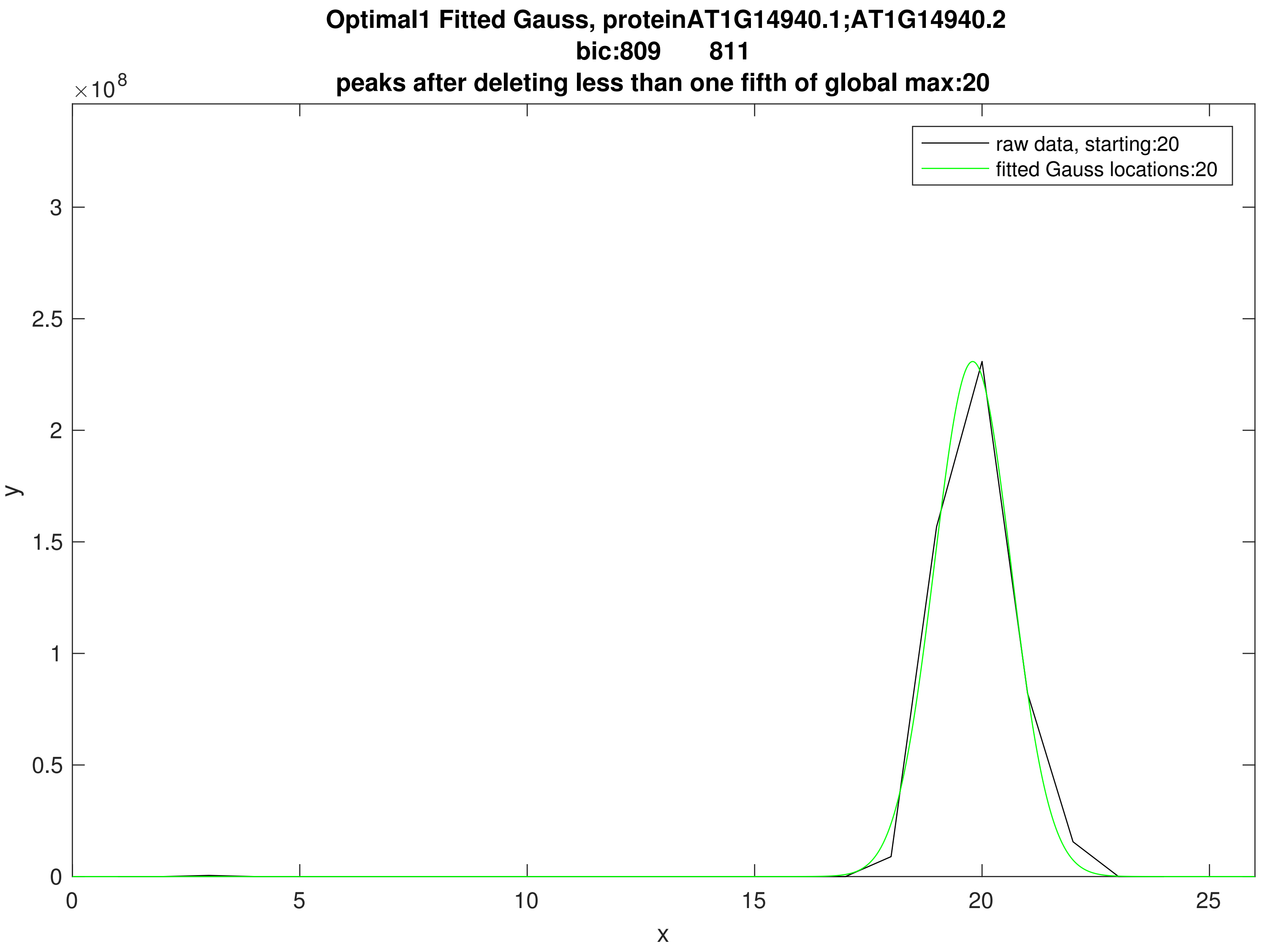
Optimal1 Fitted Gauss, proteinAT1G14940.1;AT1G14940.2
bic:809 811
peaks after deleting less than one fifth of global max:20
×10^8
raw data, starting:20
fitted Gauss locations:20
y
x
0
0.5
1
1.5
2
2.5
3
5
10
15
20
25

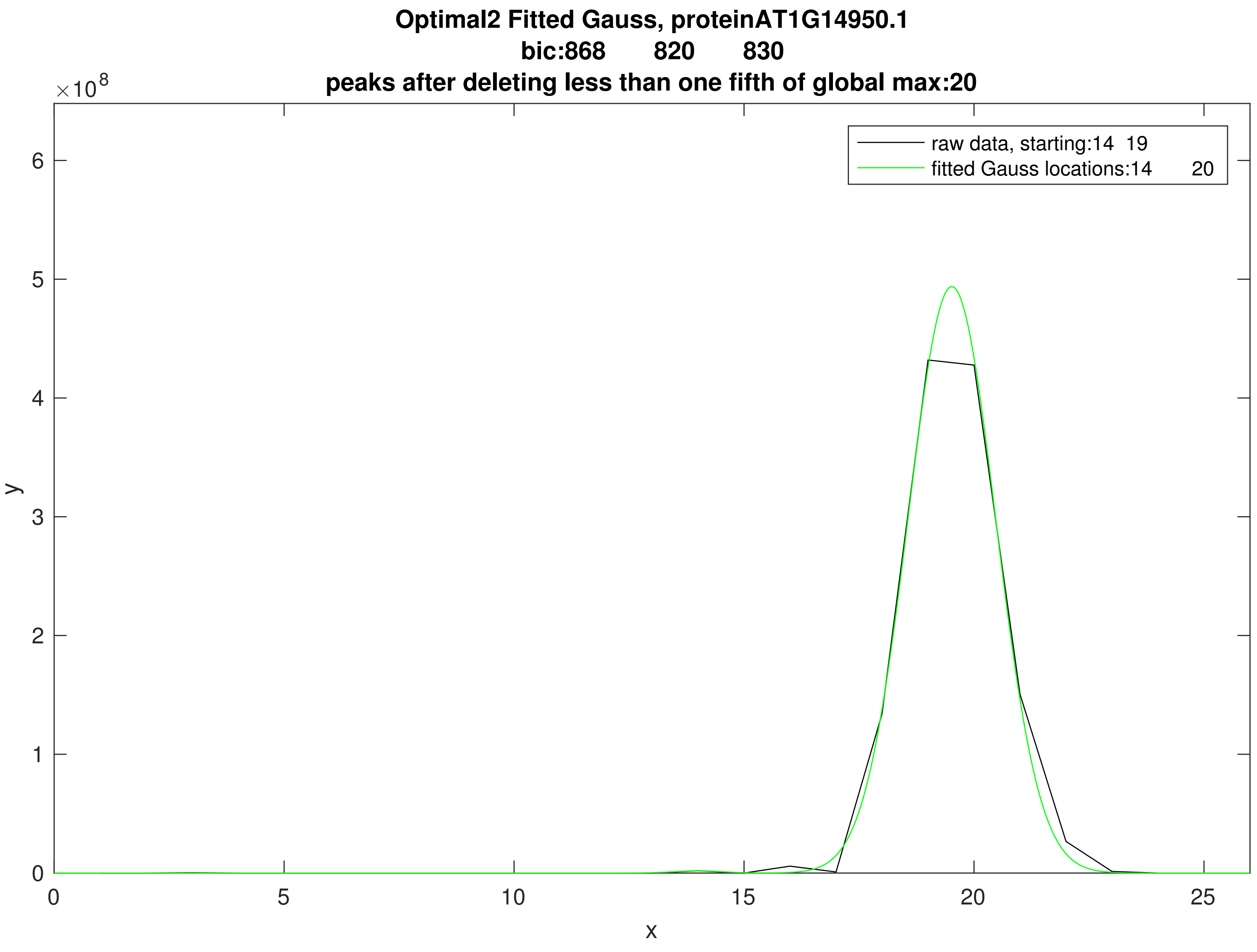
Optimal2 Fitted Gauss, proteinAT1G14950.1
bic:868 820 830
peaks after deleting less than one fifth of global max:20
×10^8
raw data, starting:14 19
fitted Gauss locations:14 20
y
x
0
1
2
3
4
5
6
0
5
10
15
20
25

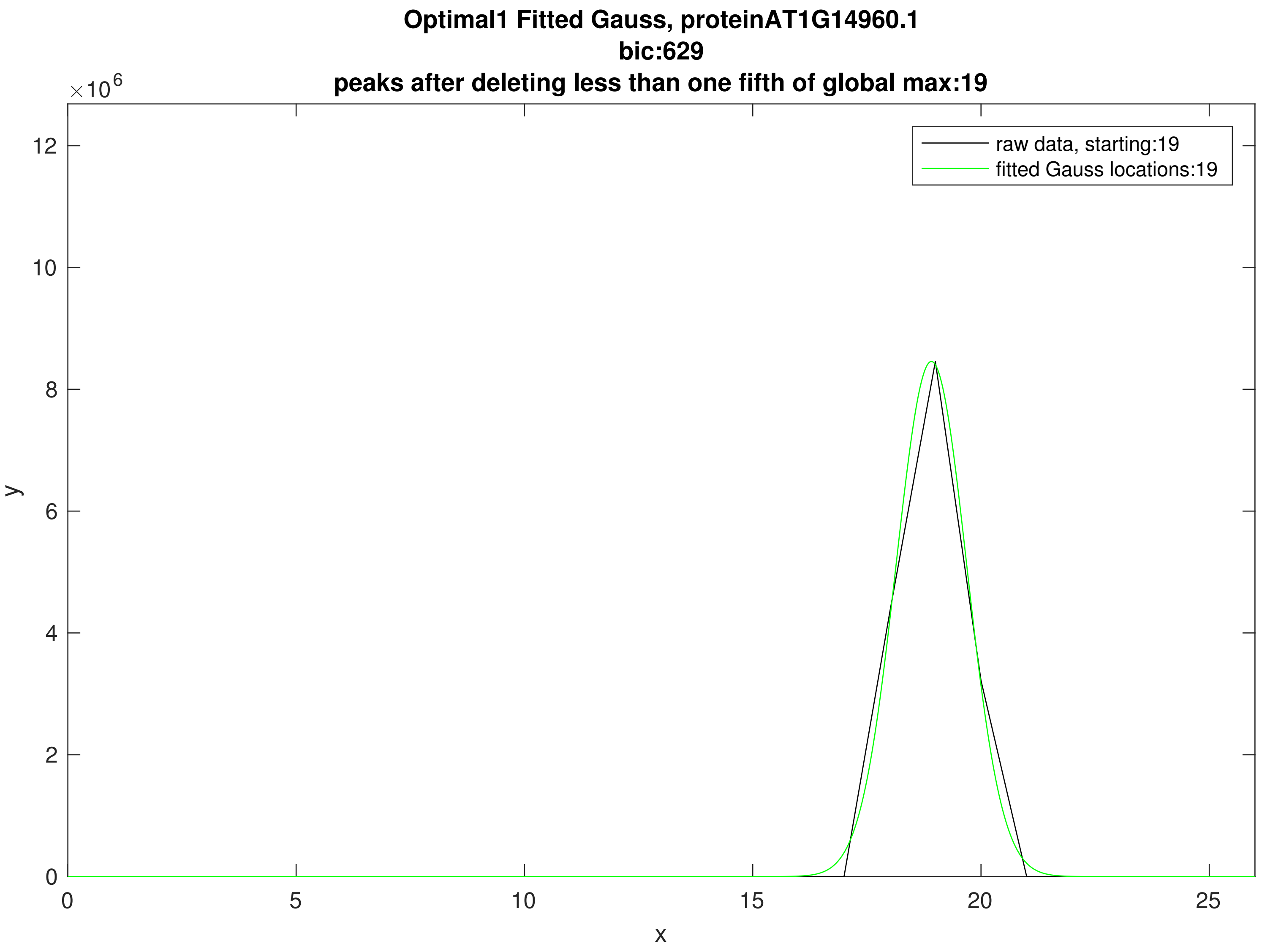
Optimal1 Fitted Gauss, proteinAT1G14960.1
bic:629
peaks after deleting less than one fifth of global max:19
×10^6
raw data, starting:19
fitted Gauss locations:19
y
x
0
2
4
6
8
10
12
0
5
10
15
20
25

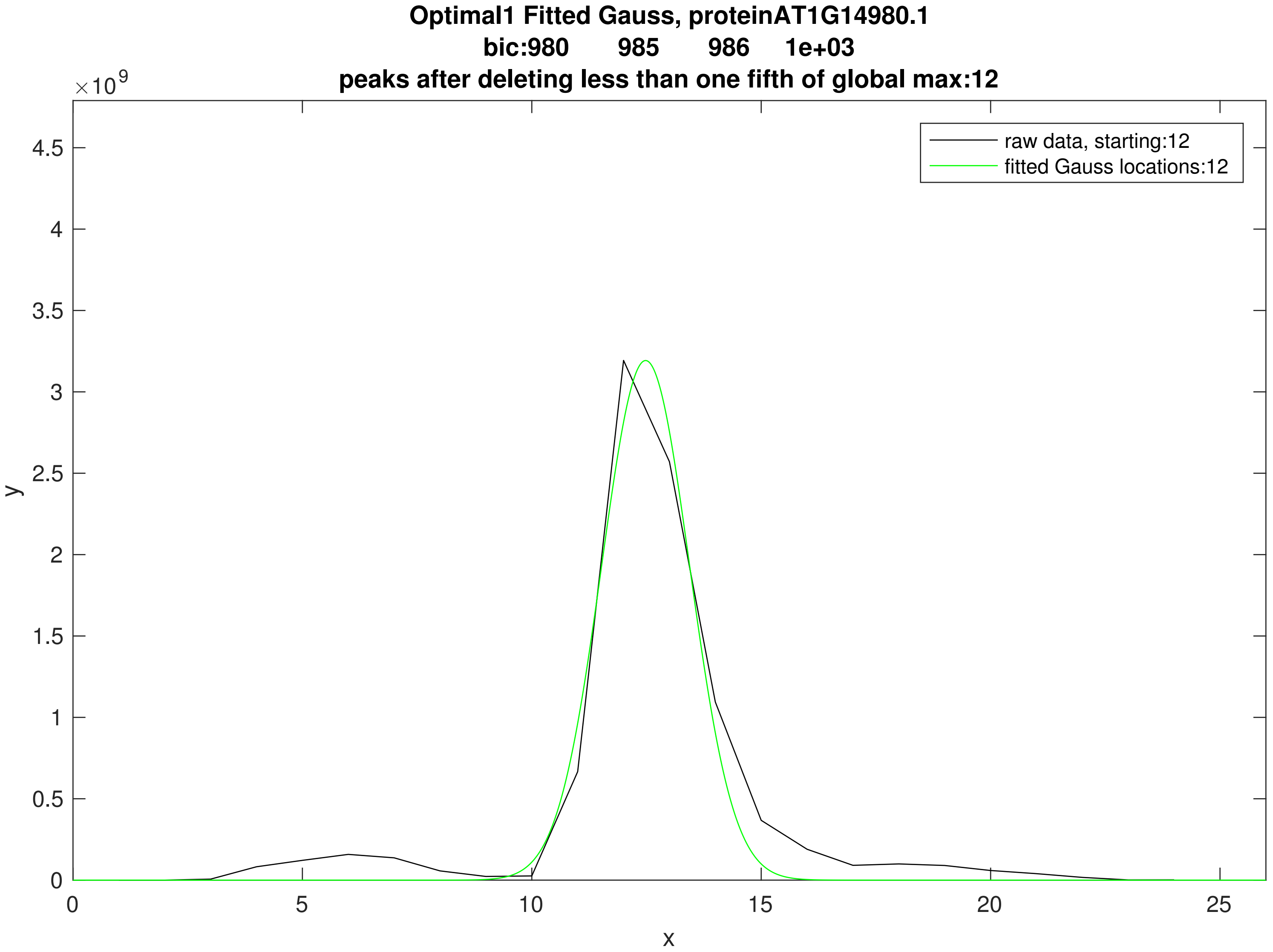

Optimal1 Fitted Gauss, proteinAT1G14980.1
bic:980 985 986 1e+03
peaks after deleting less than one fifth of global max:12
×10^9
4.5
4
3.5
3
2.5
2
1.5
1
0.5
0
y
0
5
10
15
20
25
x
raw data, starting:12
fitted Gauss locations:12

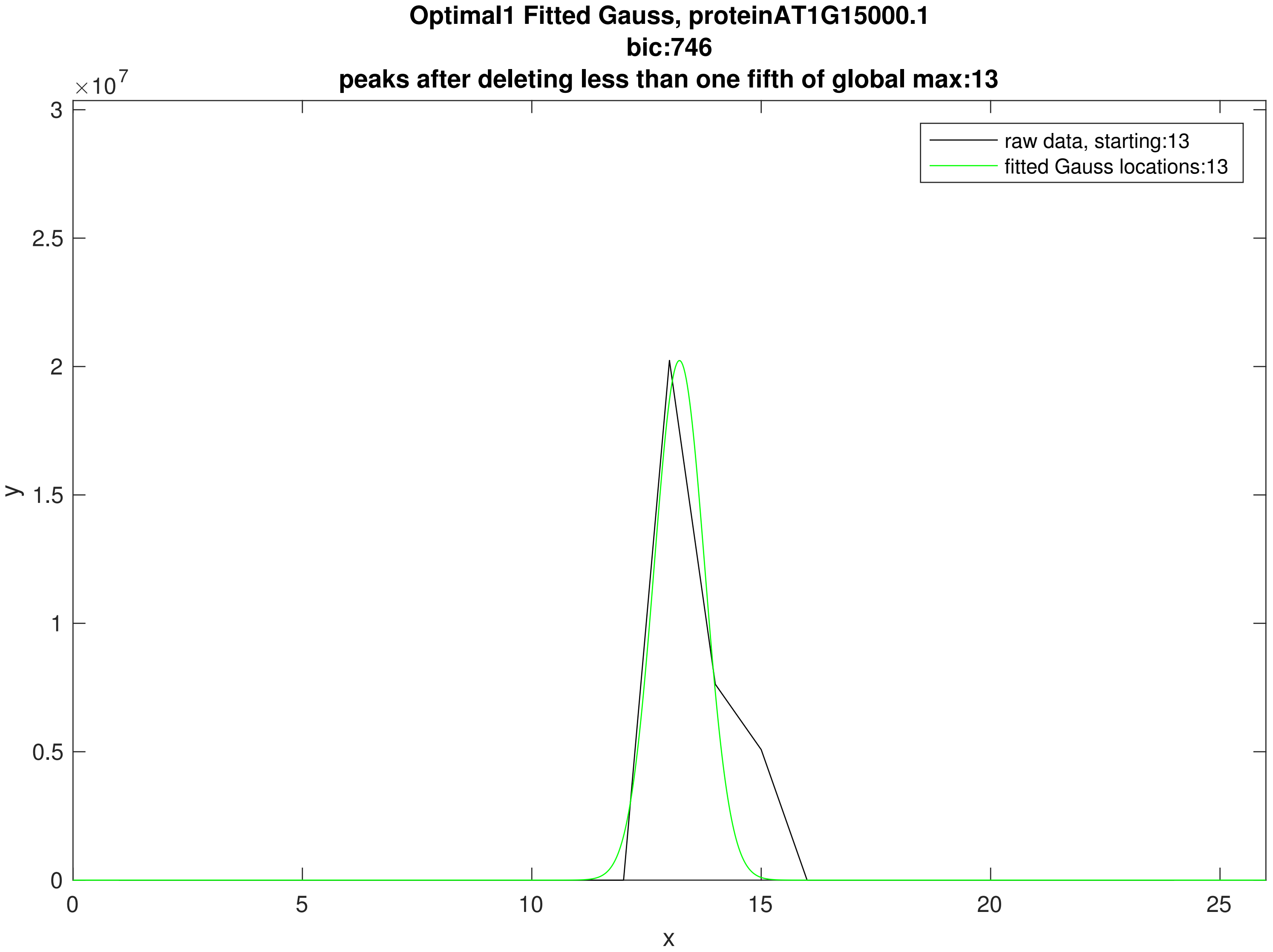

Optimal1 Fitted Gauss, proteinAT1G15000.1
bic:746
peaks after deleting less than one fifth of global max:13
×10^7
3
2.5
2
1.5
1
0.5
0
y
0
5
10
15
20
25
x
raw data, starting:13
fitted Gauss locations:13

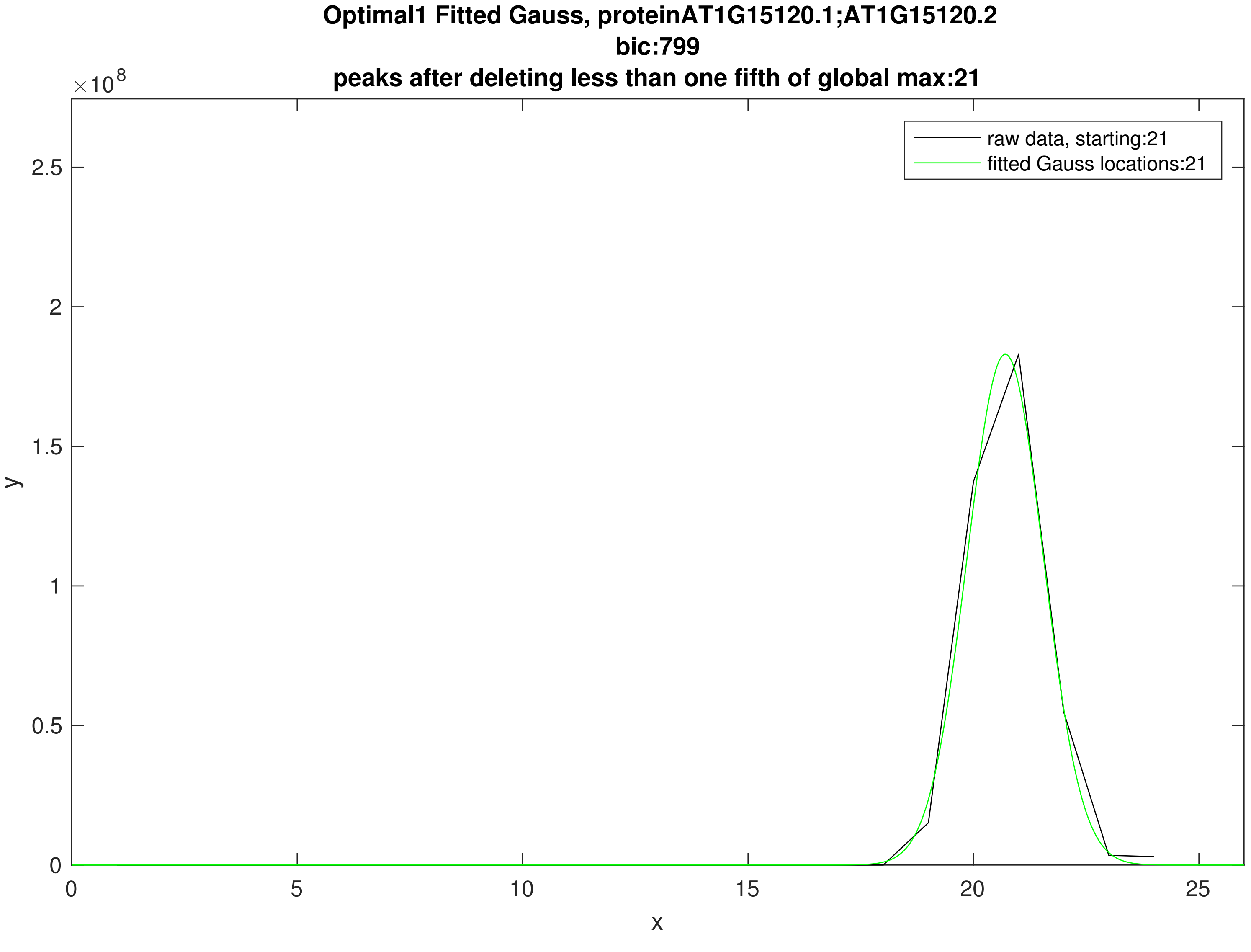
Optimal1 Fitted Gauss, proteinAT1G15120.1;AT1G15120.2
bic:799
peaks after deleting less than one fifth of global max:21
×10^8
2.5
2
1.5
1
0.5
0
y
0
5
10
15
20
25
x
raw data, starting:21
fitted Gauss locations:21

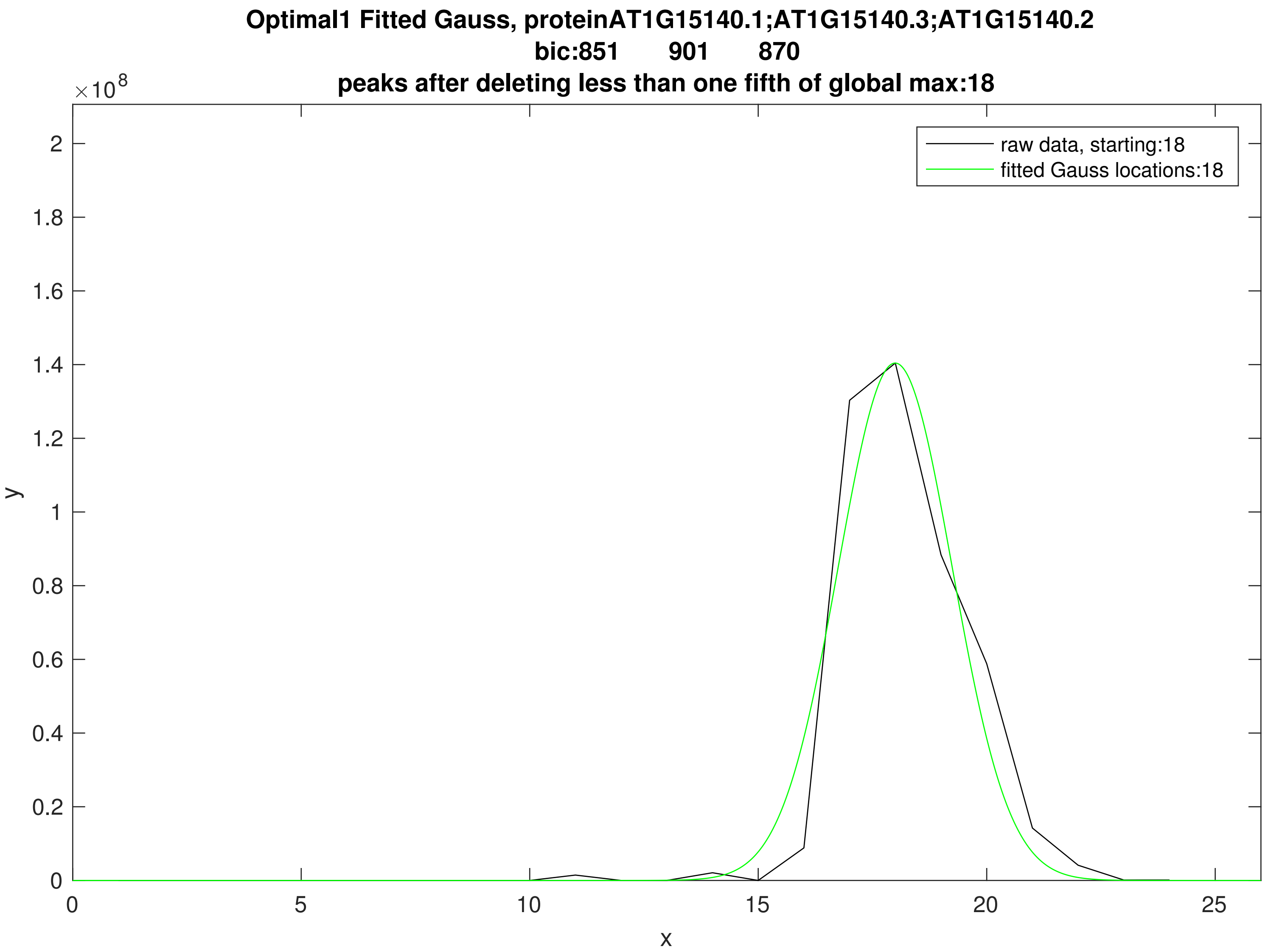

Optimal1 Fitted Gauss, proteinAT1G15140.1;AT1G15140.3;AT1G15140.2
bic:851 901 870
peaks after deleting less than one fifth of global max:18
×10^8
raw data, starting:18
fitted Gauss locations:18
y
x
0
0.2
0.4
0.6
0.8
1
1.2
1.4
1.6
1.8
2
0
5
10
15
20
25

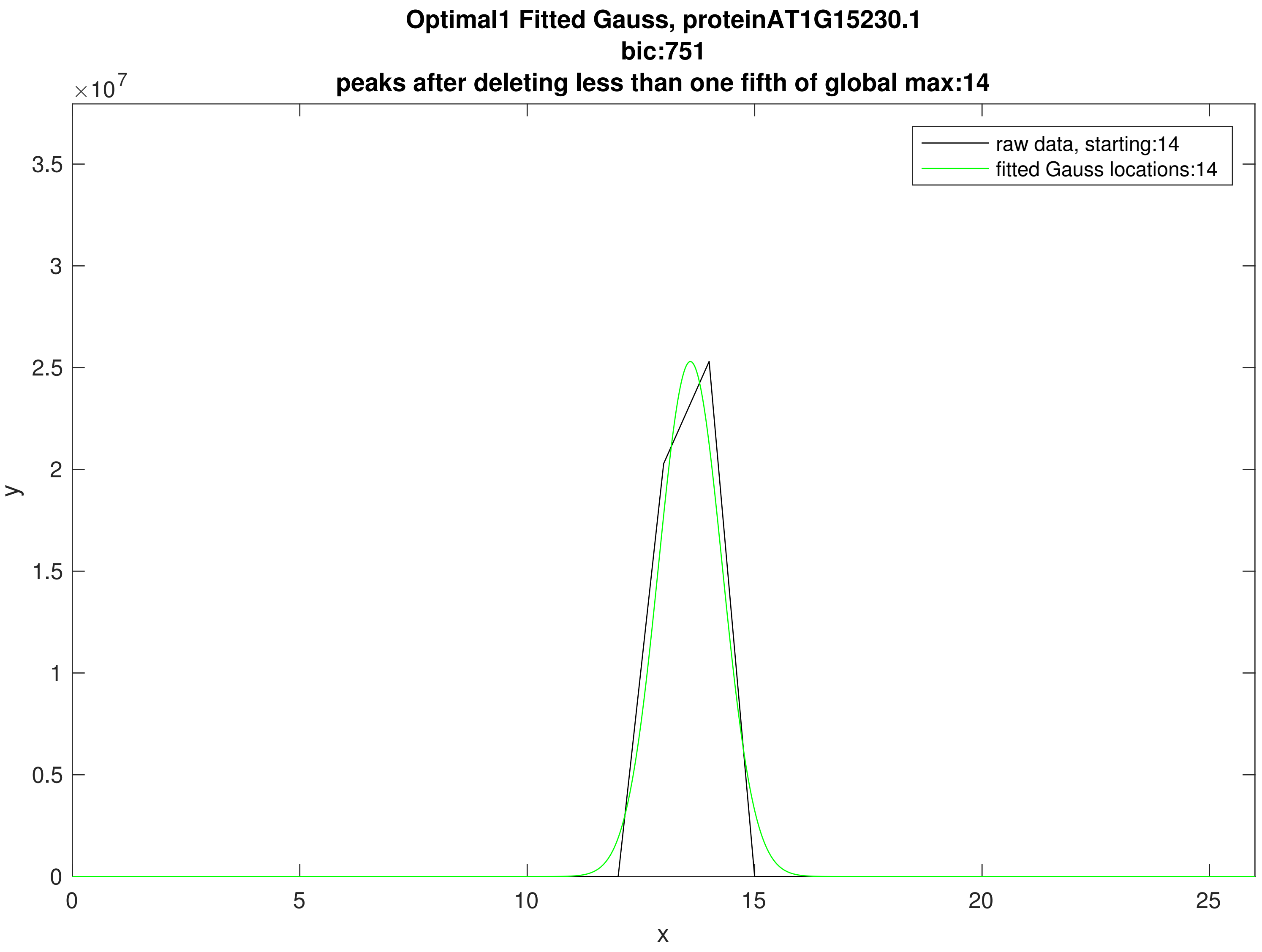
Optimal1 Fitted Gauss, proteinAT1G15230.1
bic:751
peaks after deleting less than one fifth of global max:14
×10^7
raw data, starting:14
fitted Gauss locations:14
3.5
3
2.5
2
1.5
1
0.5
0
y
0
5
10
15
20
25
x

**Optimal1 Fitted Gauss, proteinAT1G15270.1**
**bic:671**
**peaks after deleting less than one fifth of global max:20**

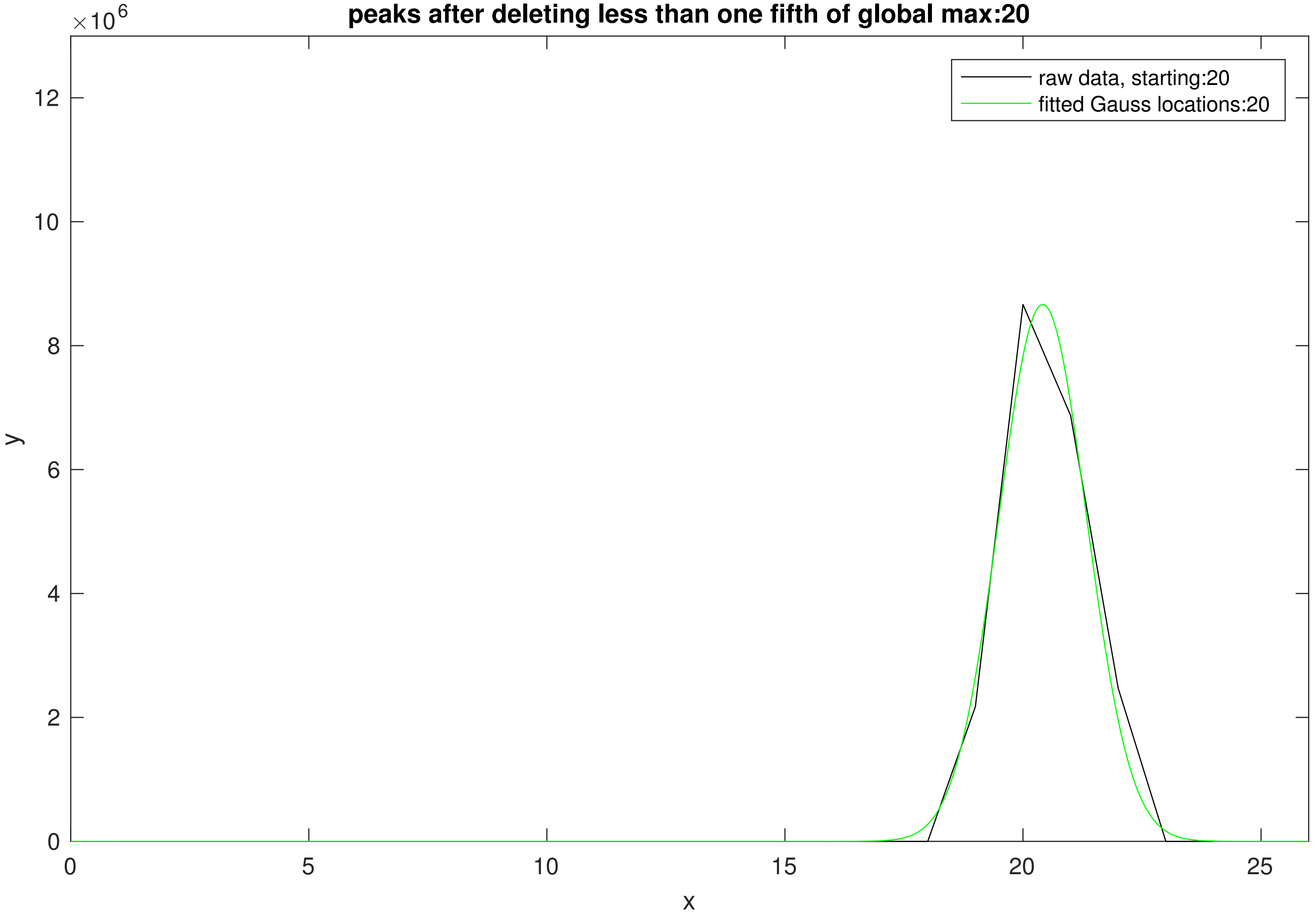

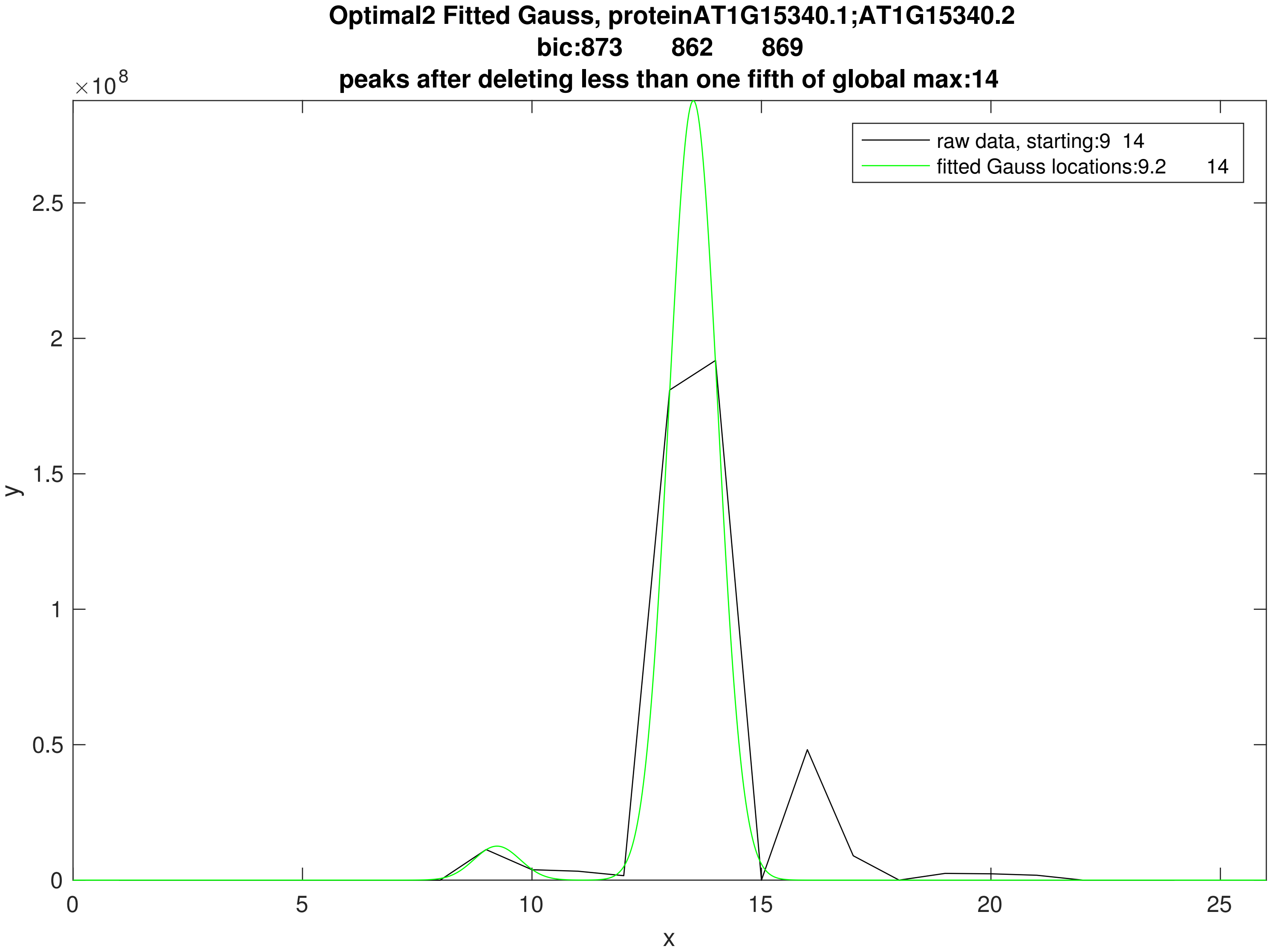

Optimal2 Fitted Gauss, proteinAT1G15340.1;AT1G15340.2
bic:873 862 869
peaks after deleting less than one fifth of global max:14
×10^8
raw data, starting:9 14
fitted Gauss locations:9.2 14
y
x
0
0.5
1
1.5
2
2.5
0
5
10
15
20
25

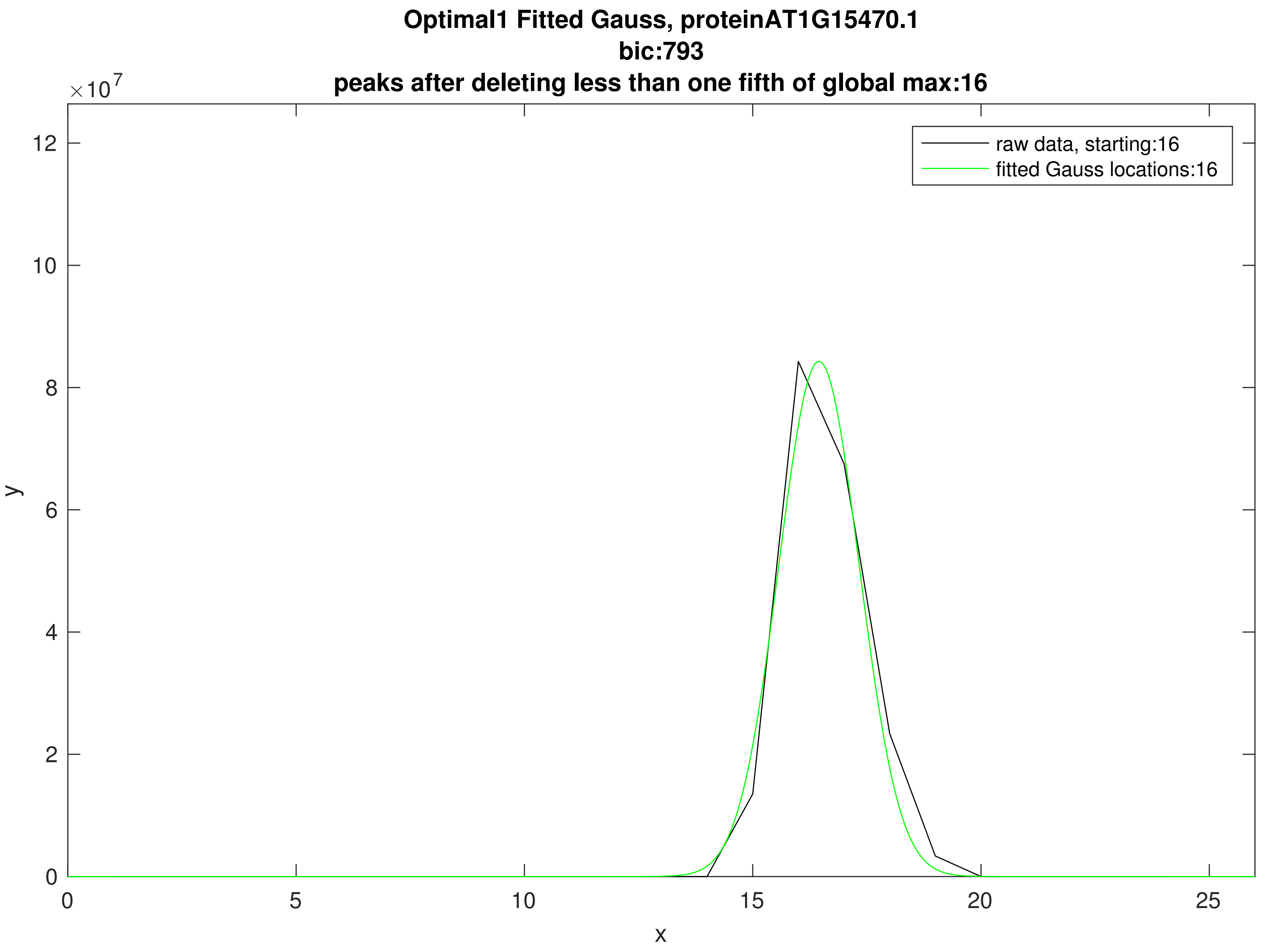

Optimal1 Fitted Gauss, proteinAT1G15470.1
bic:793
peaks after deleting less than one fifth of global max:16
×10^7
12
10
8
6
4
2
0
y
0
5
10
15
20
25
x
raw data, starting:16
fitted Gauss locations:16

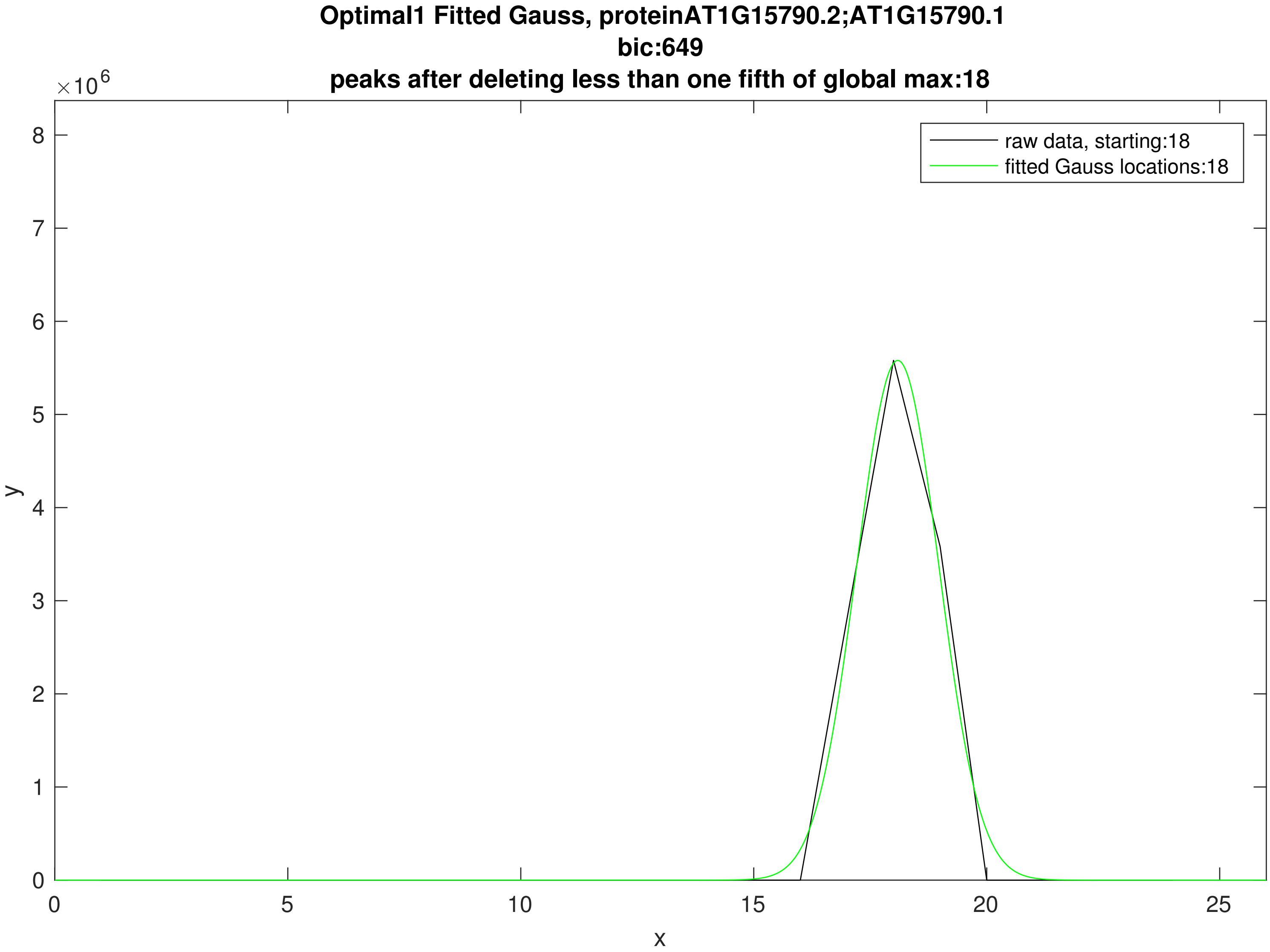

Optimal1 Fitted Gauss, proteinAT1G15790.2;AT1G15790.1
bic:649
peaks after deleting less than one fifth of global max:18
×10^6
raw data, starting:18
fitted Gauss locations:18
y
x
0
1
2
3
4
5
6
7
8
0
5
10
15
20
25

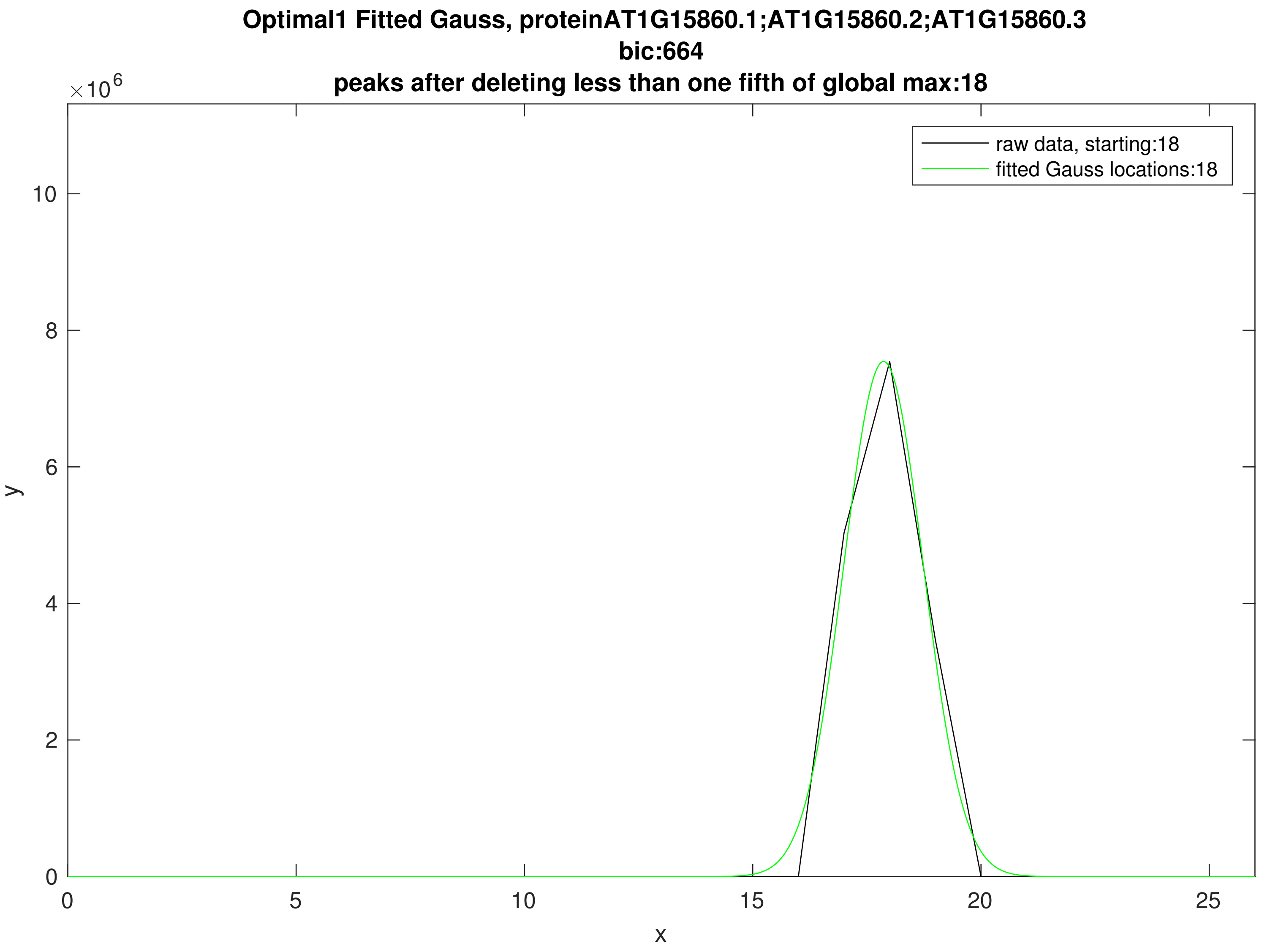
Optimal1 Fitted Gauss, proteinAT1G15860.1;AT1G15860.2;AT1G15860.3
bic:664
peaks after deleting less than one fifth of global max:18
×10^6
raw data, starting:18
fitted Gauss locations:18
y
x
0
2
4
6
8
10
0
5
10
15
20
25

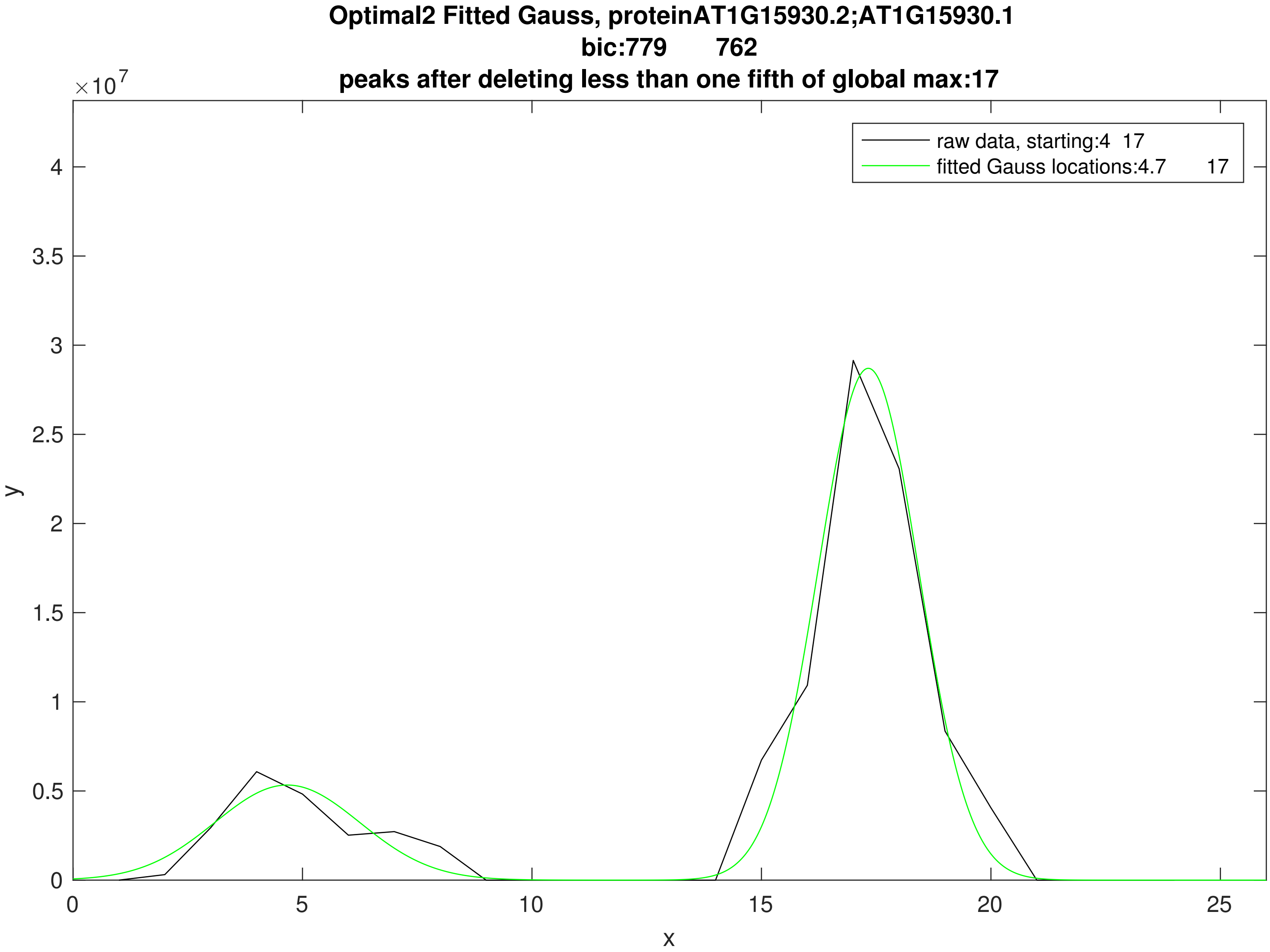
Optimal2 Fitted Gauss, proteinAT1G15930.2;AT1G15930.1
bic:779 762
peaks after deleting less than one fifth of global max:17
×10^7
raw data, starting:4 17
fitted Gauss locations:4.7 17
y
x
0
0.5
1
1.5
2
2.5
3
3.5
4
5
10
15
20
25

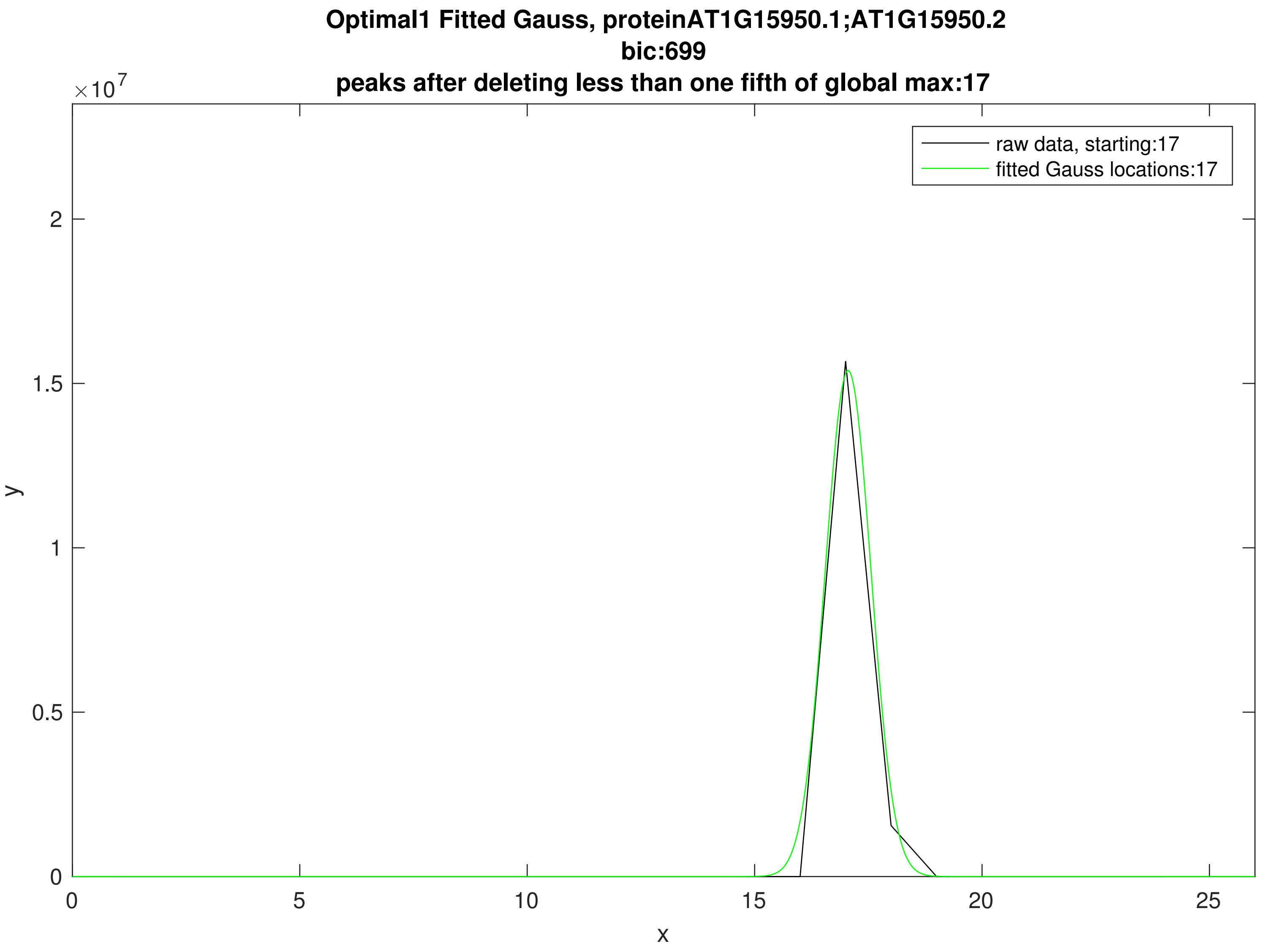
Optimal1 Fitted Gauss, proteinAT1G15950.1;AT1G15950.2
bic:699
peaks after deleting less than one fifth of global max:17
×10^7
raw data, starting:17
fitted Gauss locations:17
y
x
0
0.5
1
1.5
2
5
10
15
20
25

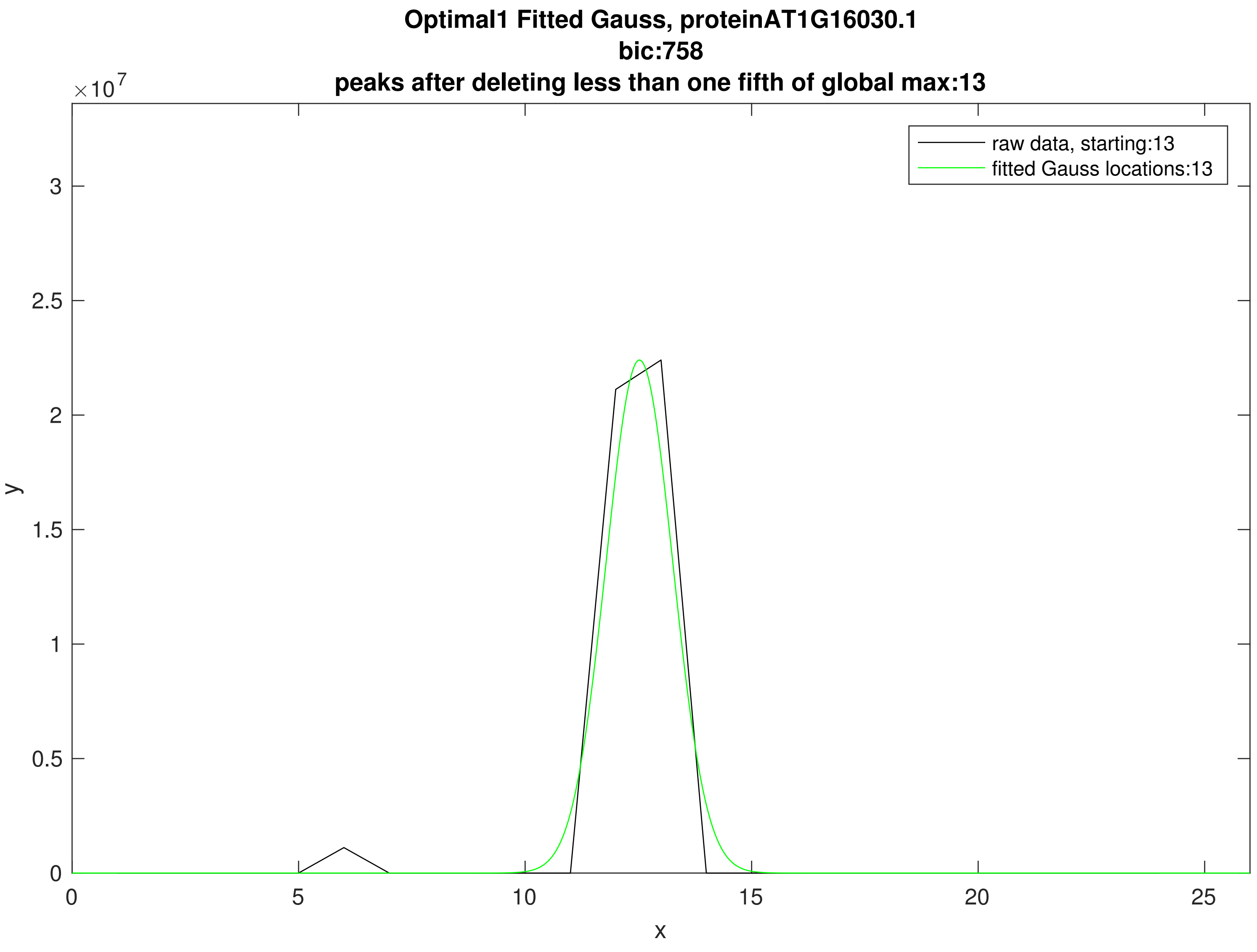

Optimal1 Fitted Gauss, proteinAT1G16030.1
bic:758
peaks after deleting less than one fifth of global max:13
×10^7
raw data, starting:13
fitted Gauss locations:13
y
x
0
0.5
1
1.5
2
2.5
3
0
5
10
15
20
25

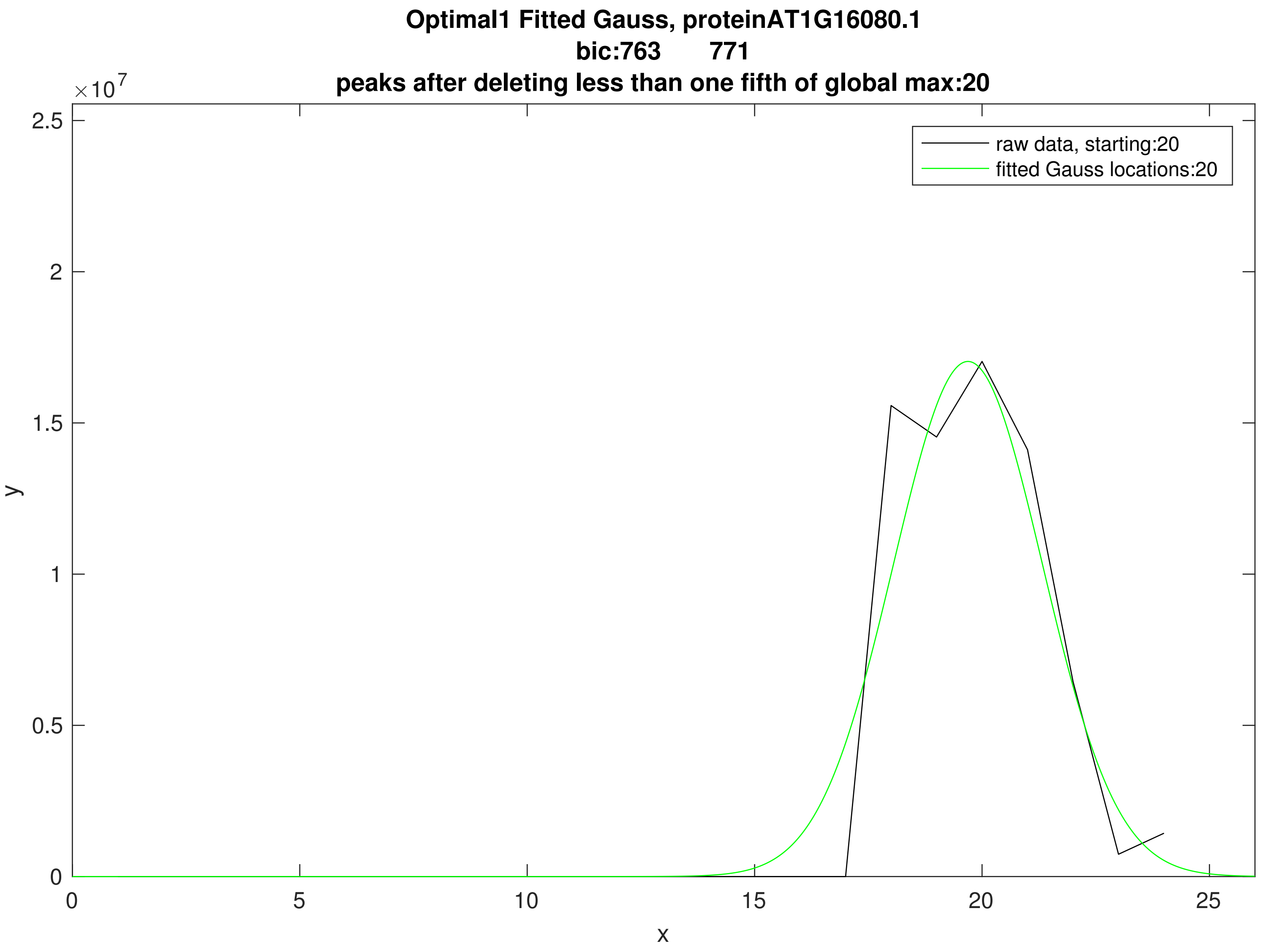

Optimal1 Fitted Gauss, proteinAT1G16080.1
bic:763 771
peaks after deleting less than one fifth of global max:20
×10^7
2.5
2
1.5
1
0.5
0
y
0
5
10
15
20
25
x
raw data, starting:20
fitted Gauss locations:20

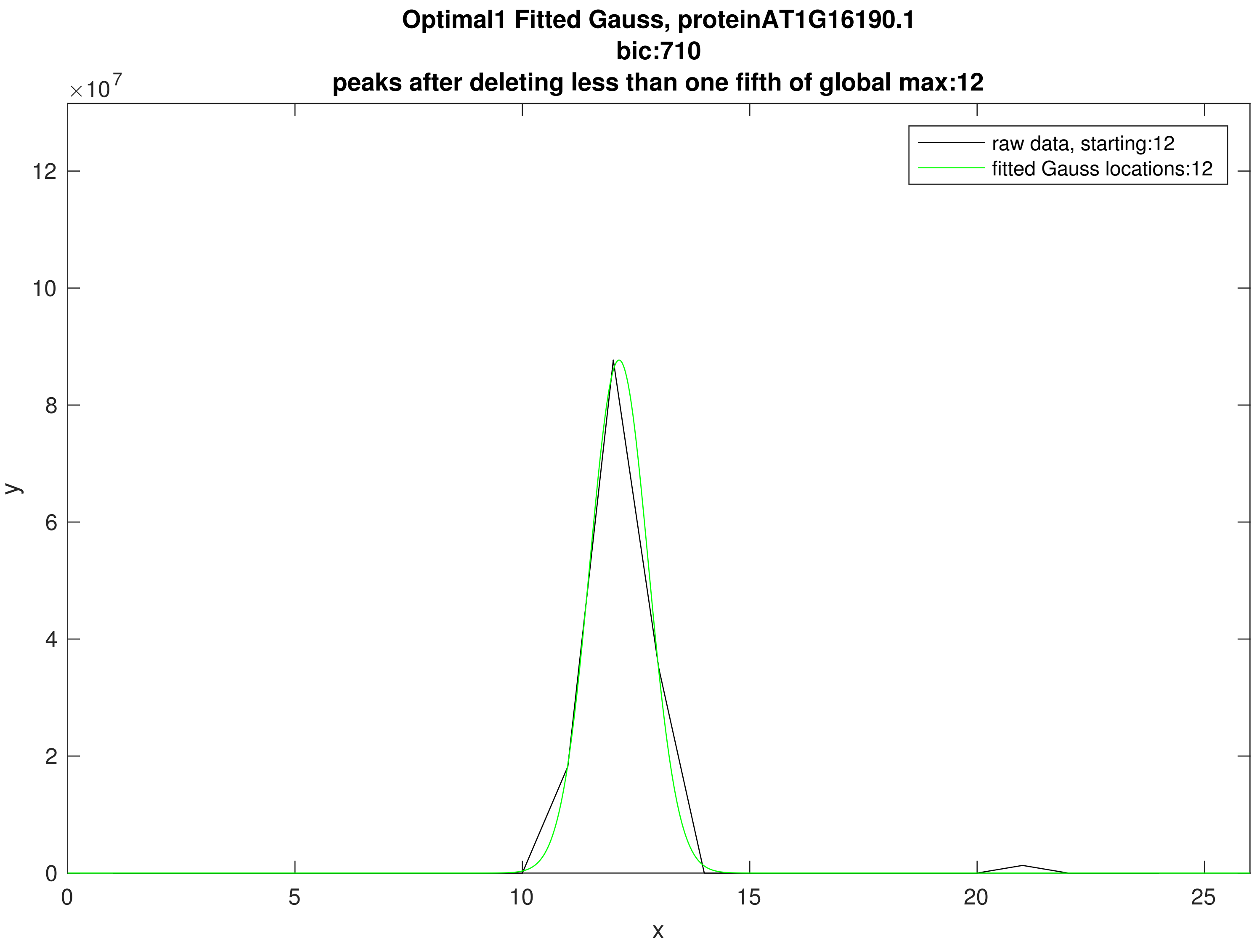
Optimal1 Fitted Gauss, proteinAT1G16190.1
bic:710
peaks after deleting less than one fifth of global max:12
×10^7
raw data, starting:12
fitted Gauss locations:12
y
x
0
2
4
6
8
10
12
0
5
10
15
20
25

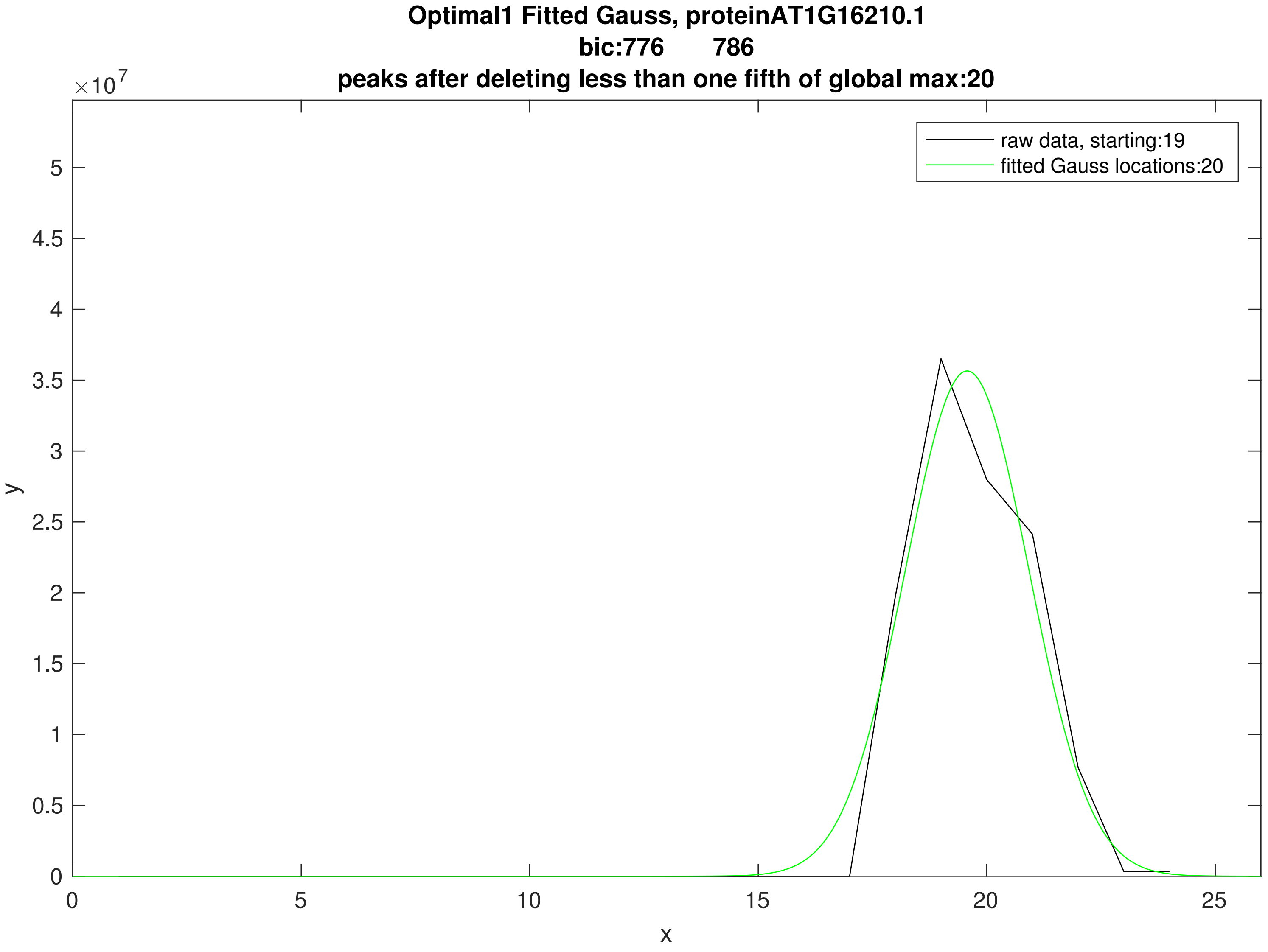
Optimal1 Fitted Gauss, proteinAT1G16210.1
bic:776 786
peaks after deleting less than one fifth of global max:20
×10^7
raw data, starting:19
fitted Gauss locations:20
y
x
0
5
10
15
20
25
0.5
1
1.5
2
2.5
3
3.5
4
4.5

Optimal1 Fitted Gauss, proteinAT1G16300.1
bic:791
peaks after deleting less than one fifth of global max:11

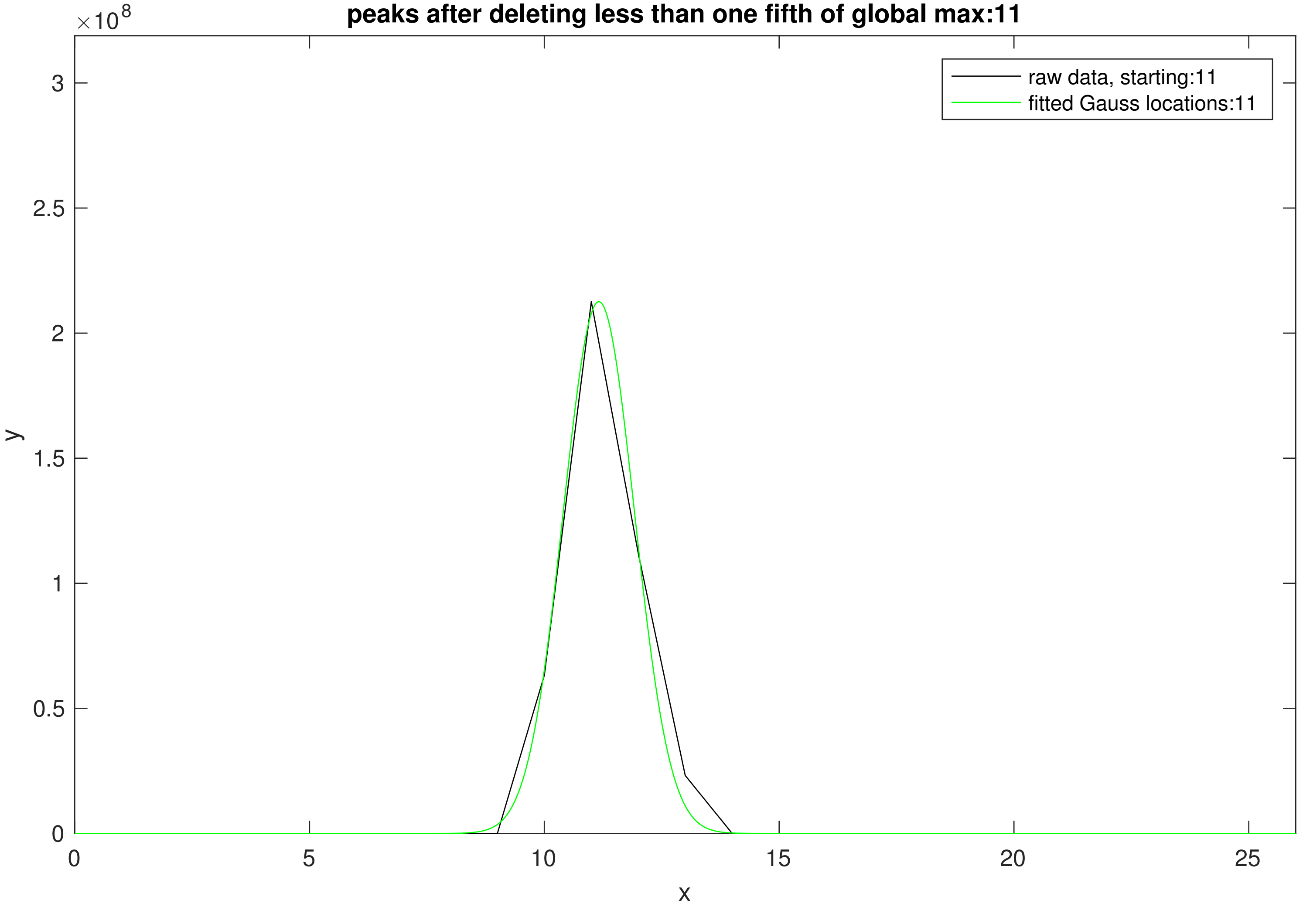

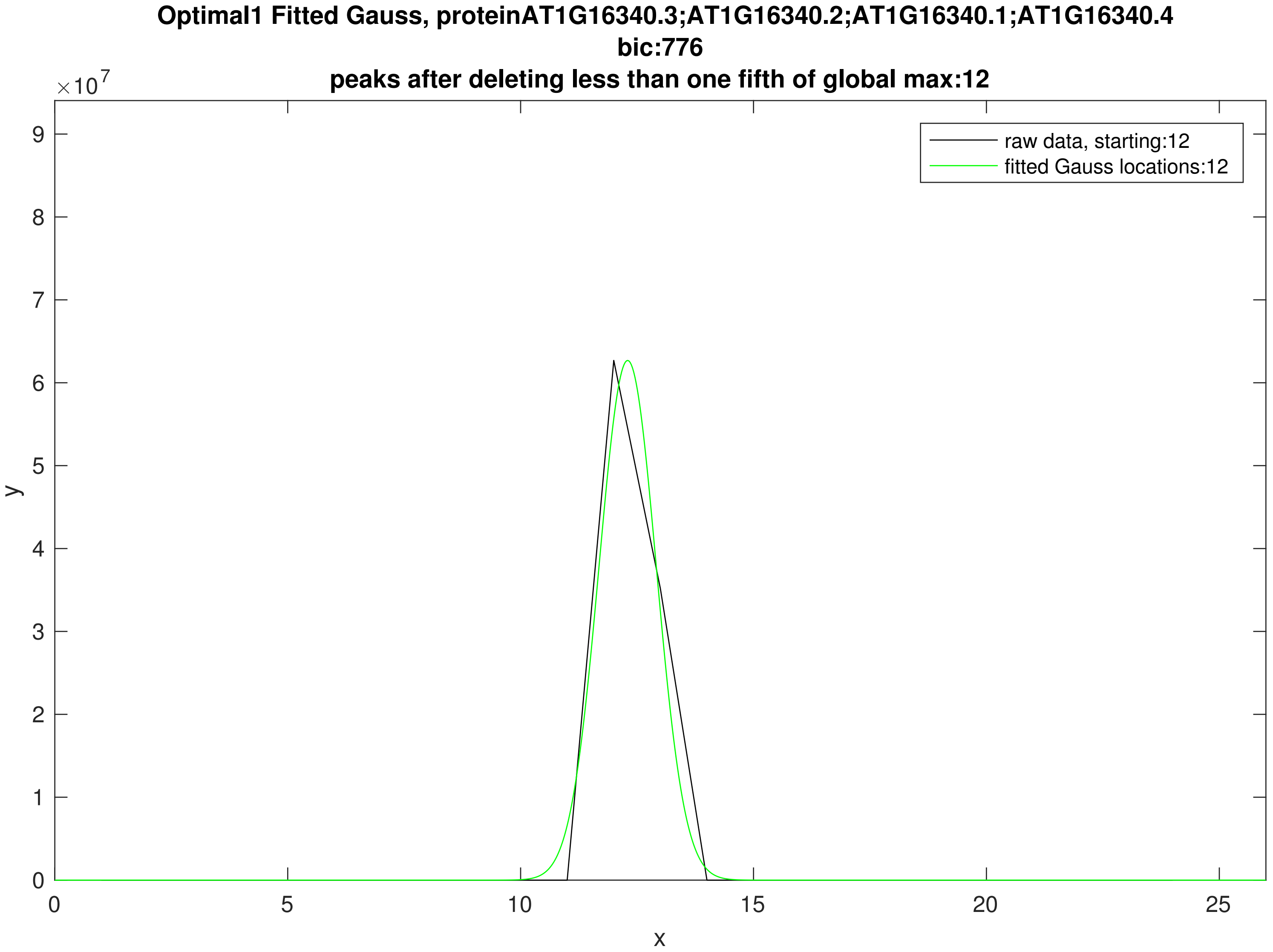
Optimal1 Fitted Gauss, proteinAT1G16340.3;AT1G16340.2;AT1G16340.1;AT1G16340.4
bic:776
peaks after deleting less than one fifth of global max:12
×10^7
raw data, starting:12
fitted Gauss locations:12
y
x
0
1
2
3
4
5
6
7
8
9
0
5
10
15
20
25

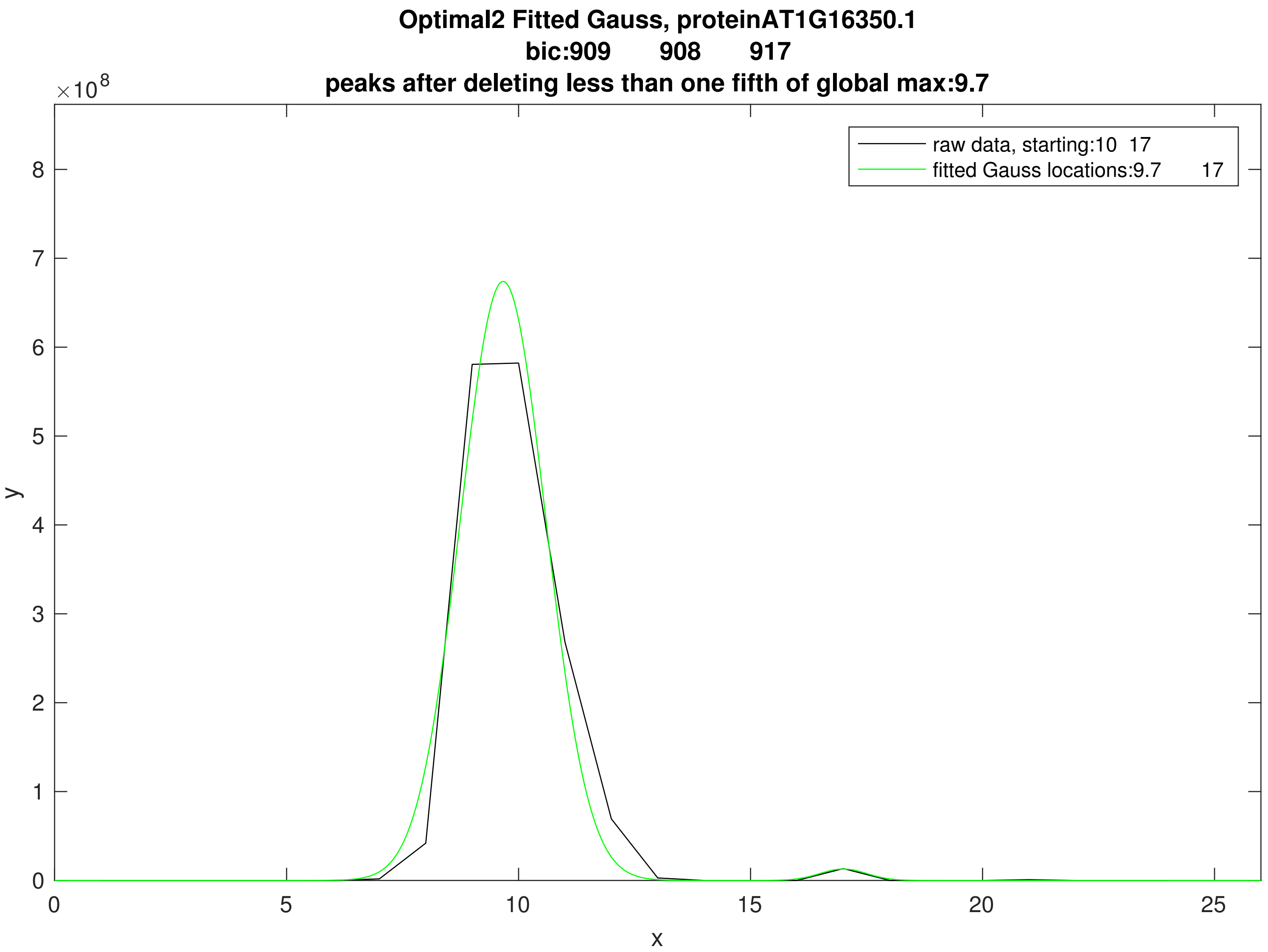

Optimal2 Fitted Gauss, proteinAT1G16350.1
bic:909 908 917
peaks after deleting less than one fifth of global max:9.7
×10^8
raw data, starting:10 17
fitted Gauss locations:9.7 17
y
x
0
1
2
3
4
5
6
7
8
0
5
10
15
20
25

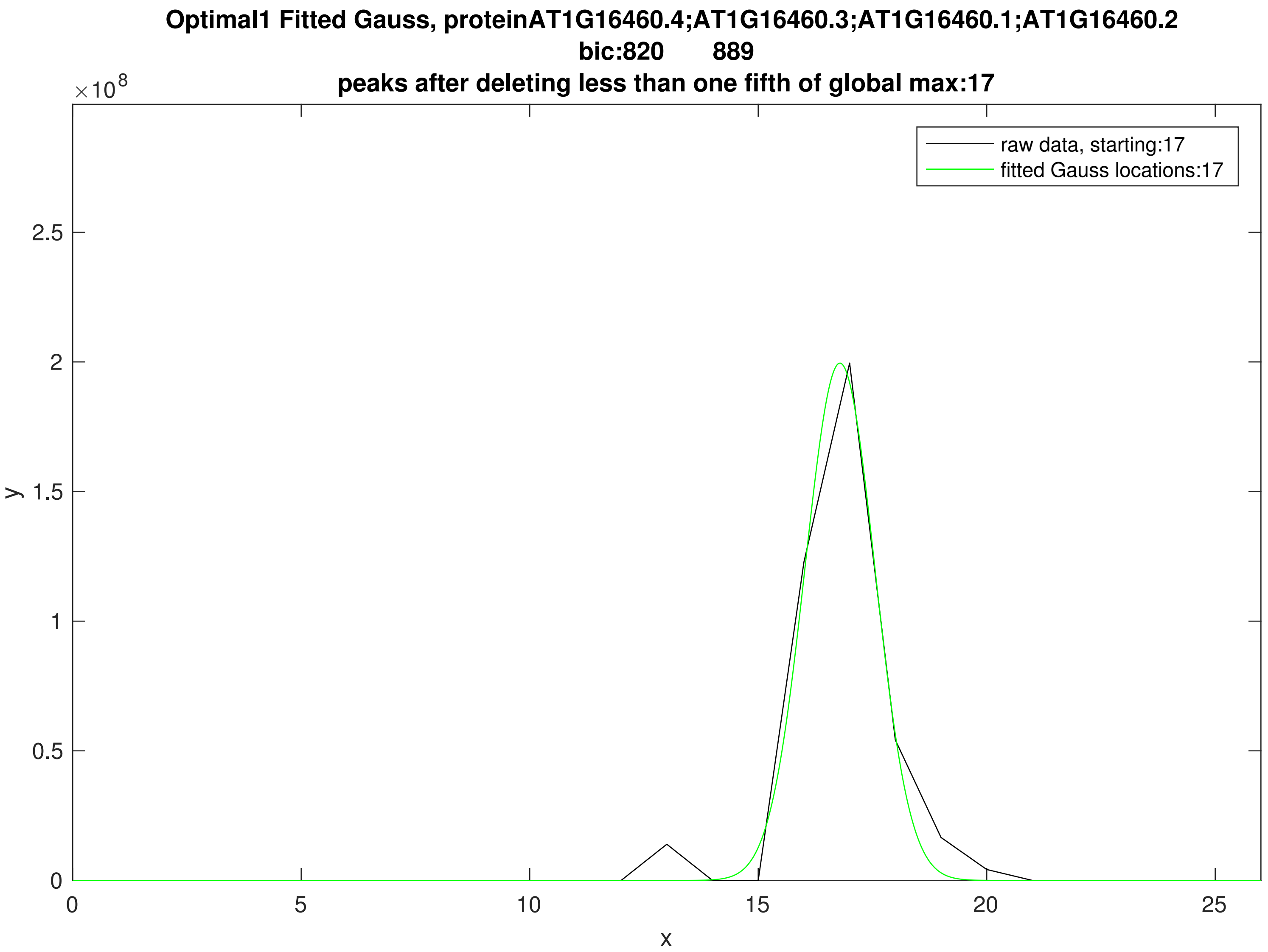
Optimal1 Fitted Gauss, proteinAT1G16460.4;AT1G16460.3;AT1G16460.1;AT1G16460.2
bic:820 889
peaks after deleting less than one fifth of global max:17
×10^8
raw data, starting:17
fitted Gauss locations:17
y
x
0
0.5
1
1.5
2
2.5
5
10
15
20
25

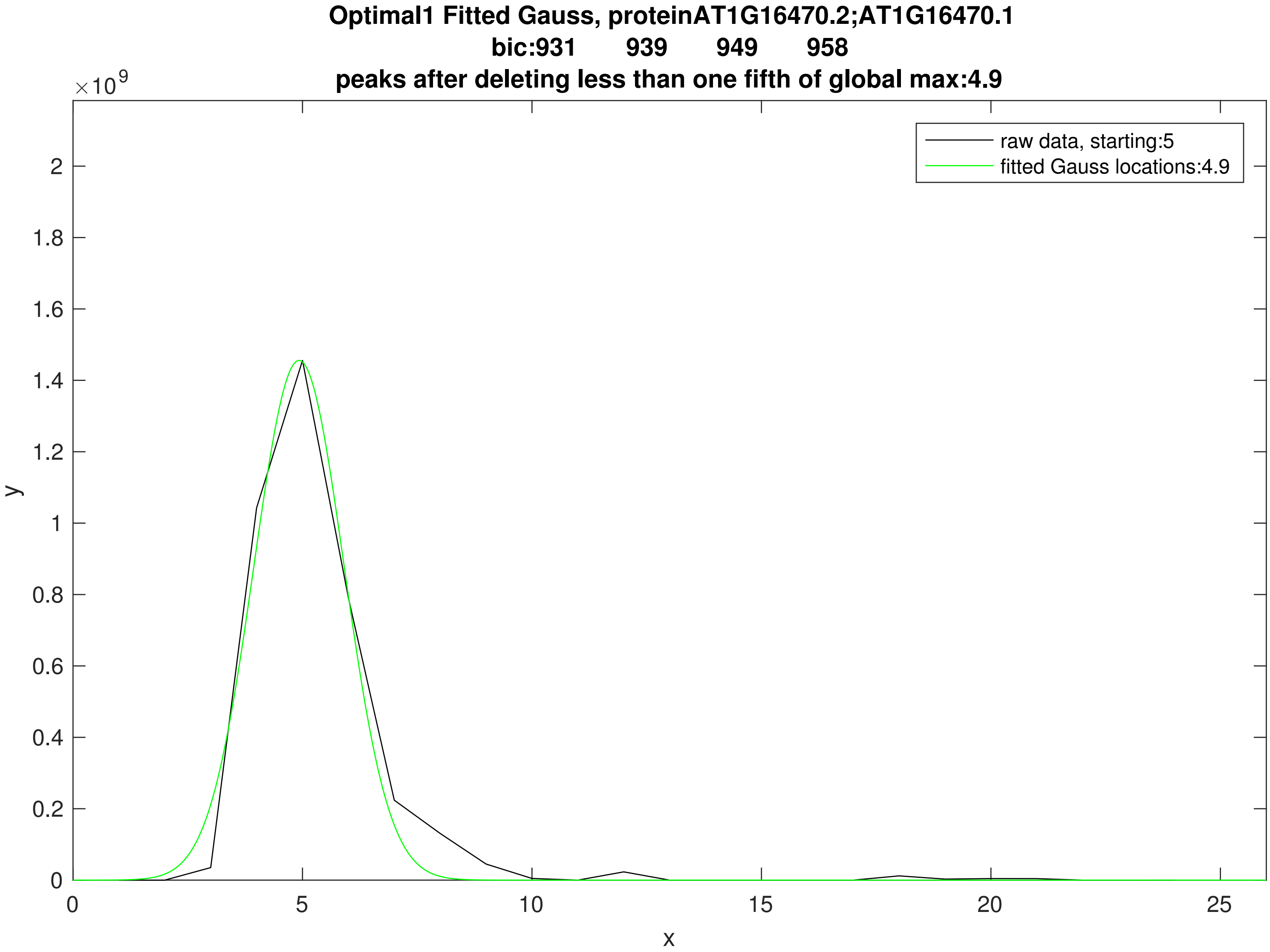

Optimal1 Fitted Gauss, proteinAT1G16470.2;AT1G16470.1
bic:931 939 949 958
peaks after deleting less than one fifth of global max:4.9
×10^9
raw data, starting:5
fitted Gauss locations:4.9
2
1.8
1.6
1.4
1.2
1
0.8
0.6
0.4
0.2
0
y
0
5
10
15
20
25
x

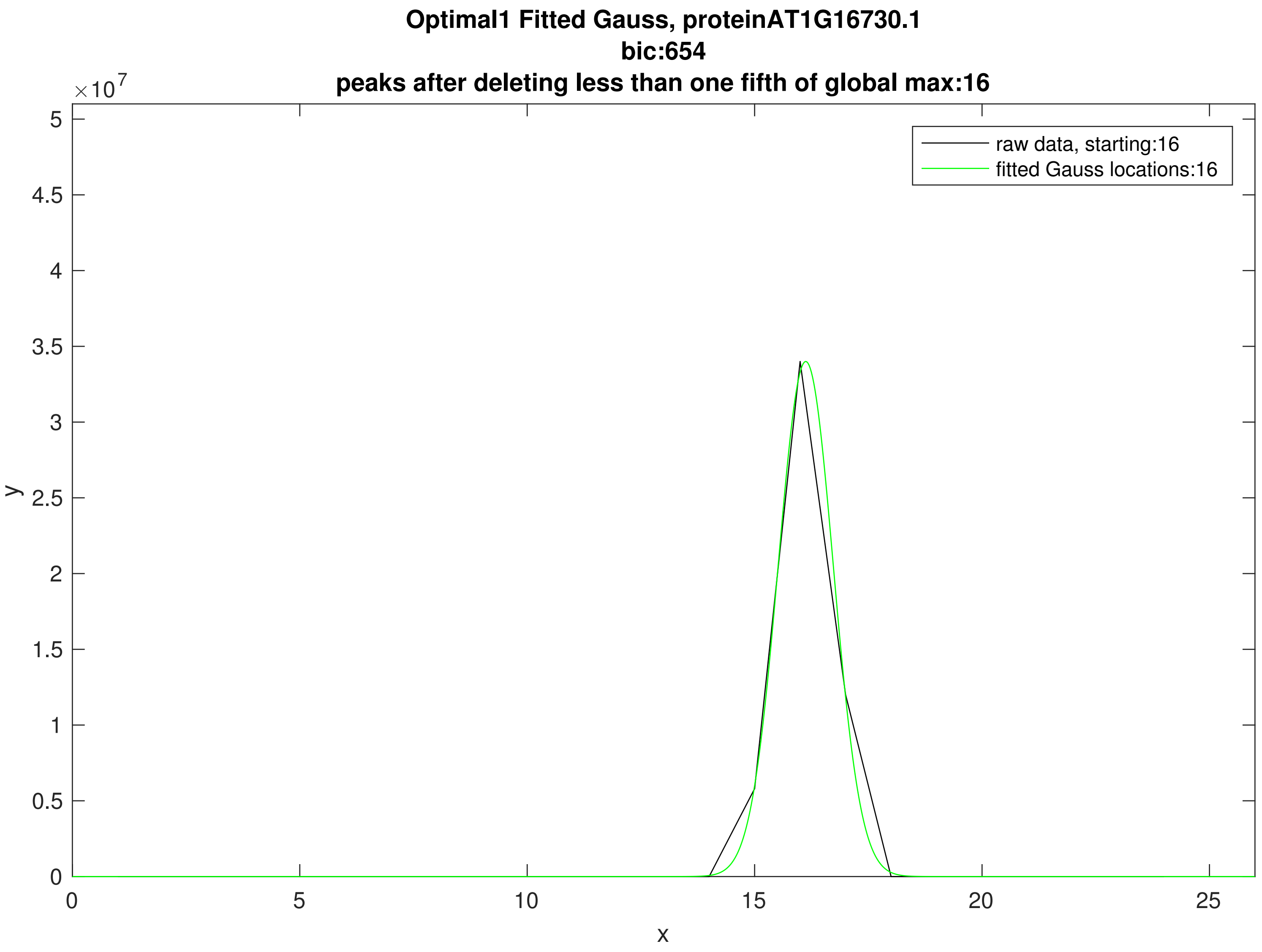

Optimal1 Fitted Gauss, proteinAT1G16730.1
bic:654
peaks after deleting less than one fifth of global max:16
×10^7
raw data, starting:16
fitted Gauss locations:16
y
x
0
0.5
1
1.5
2
2.5
3
3.5
4
4.5
5
0
5
10
15
20
25

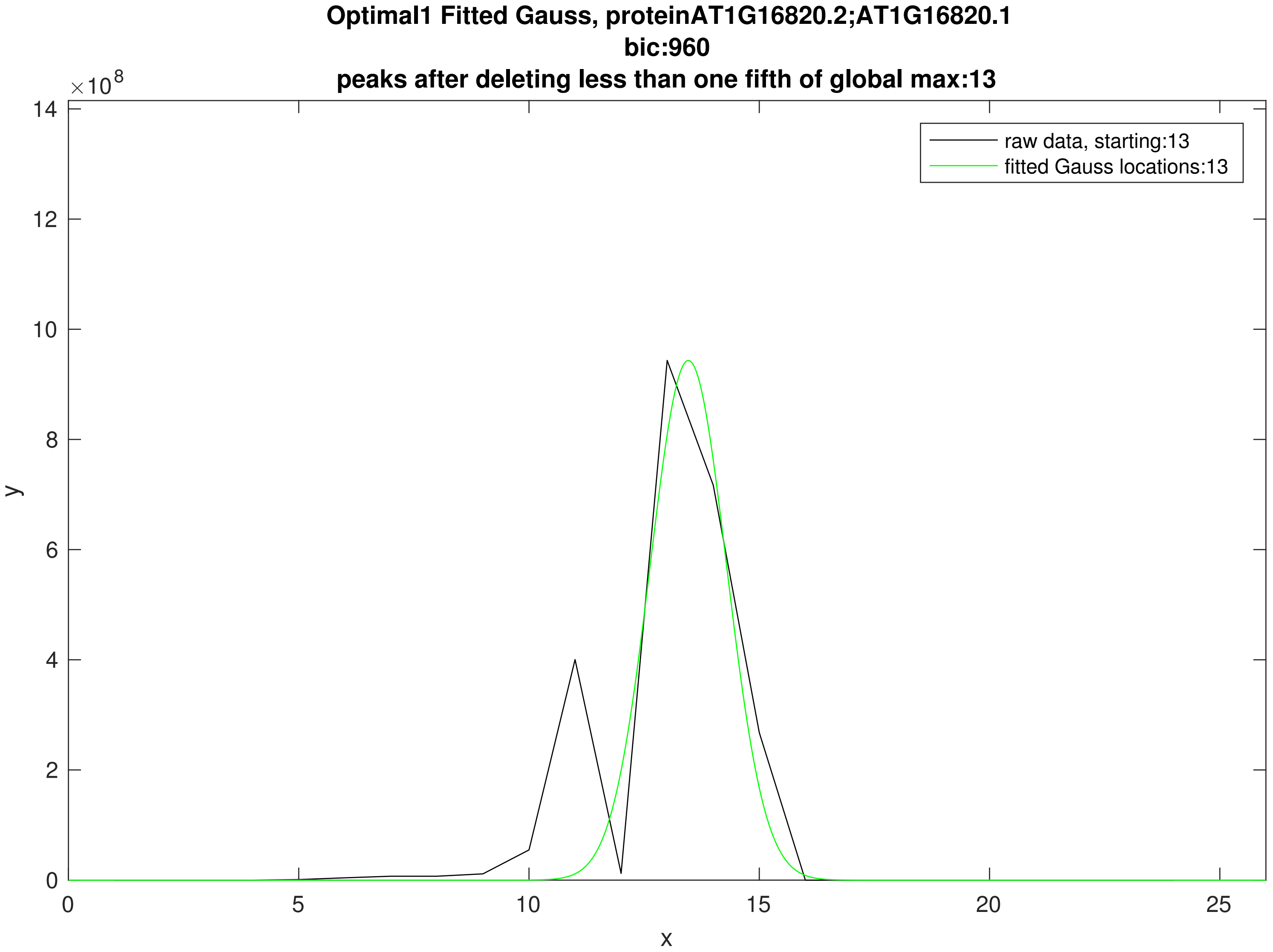
Optimal1 Fitted Gauss, proteinAT1G16820.2;AT1G16820.1
bic:960
peaks after deleting less than one fifth of global max:13
×10^8
14
12
10
8
6
4
2
0
y
0
5
10
15
20
25
x
raw data, starting:13
fitted Gauss locations:13

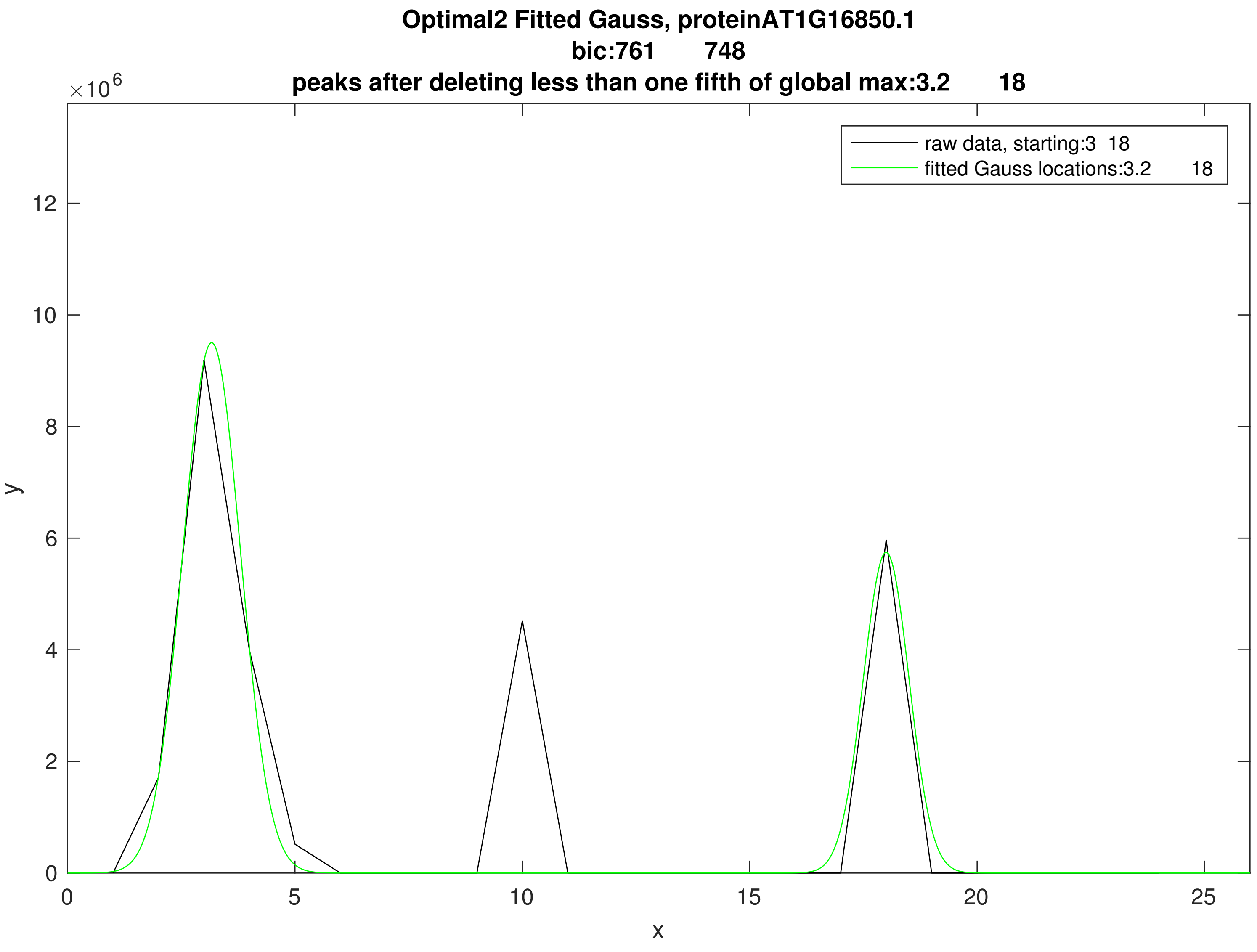
Optimal2 Fitted Gauss, proteinAT1G16850.1
bic:761 748
peaks after deleting less than one fifth of global max:3.2 18
×10^6
raw data, starting:3 18
fitted Gauss locations:3.2 18
y
x
0
2
4
6
8
10
12
0
5
10
15
20
25

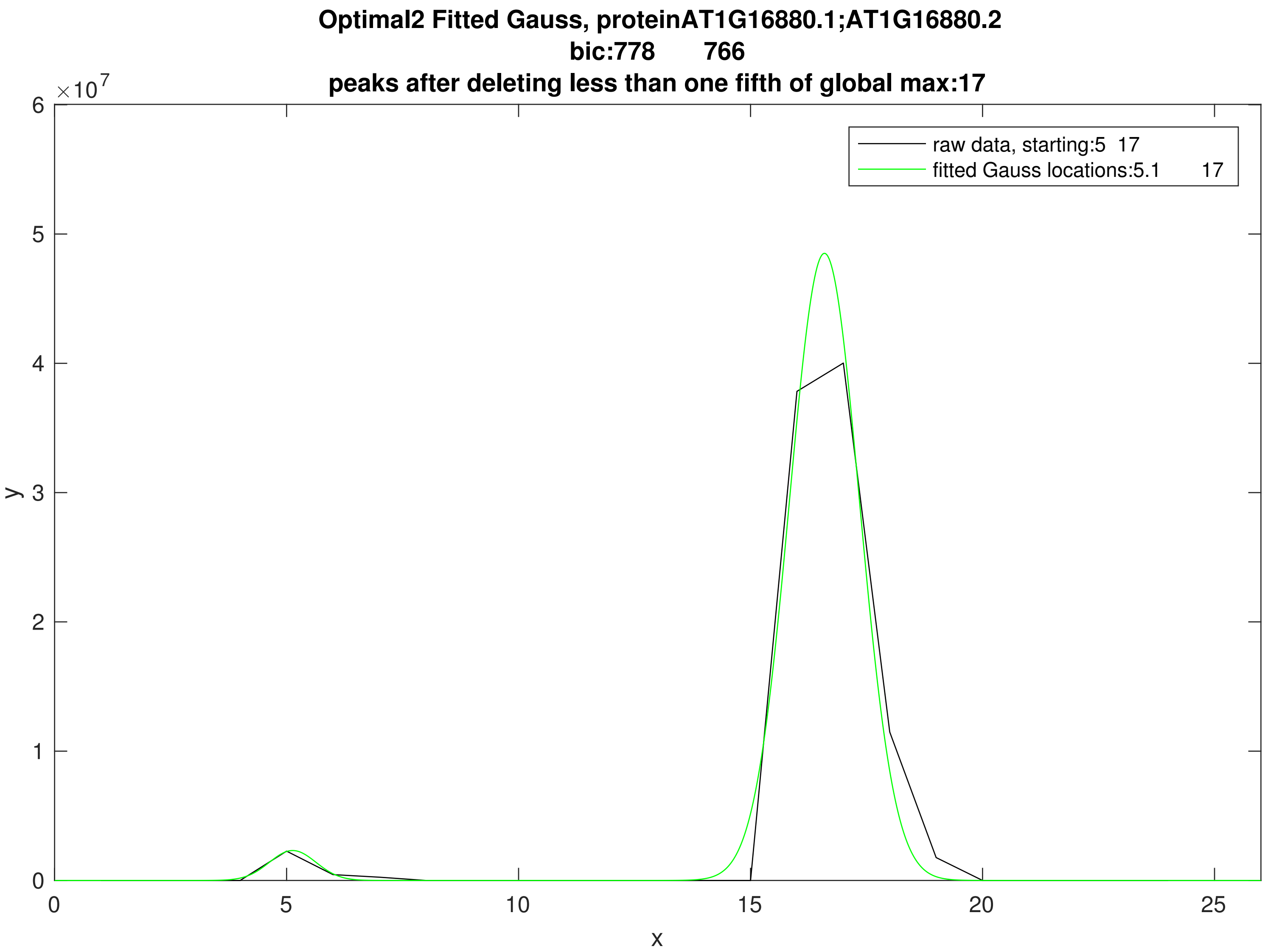
Optimal2 Fitted Gauss, proteinAT1G16880.1;AT1G16880.2
bic:778 766
peaks after deleting less than one fifth of global max:17
×10^7
raw data, starting:5 17
fitted Gauss locations:5.1 17
y
x
0
1
2
3
4
5
6
0
5
10
15
20
25

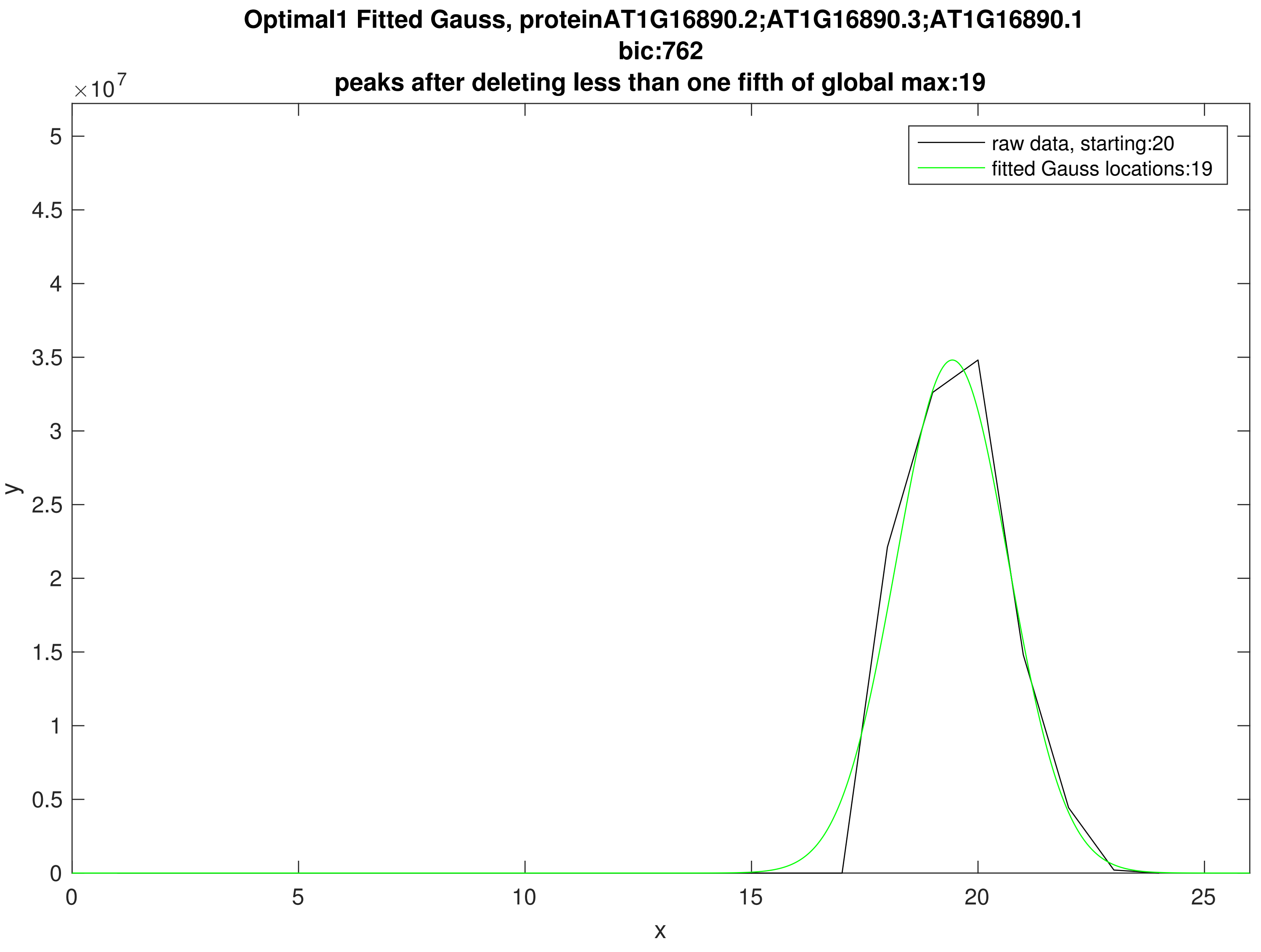
Optimal1 Fitted Gauss, proteinAT1G16890.2;AT1G16890.3;AT1G16890.1
bic:762
peaks after deleting less than one fifth of global max:19
×10^7
raw data, starting:20
fitted Gauss locations:19
y
x
0
0.5
1
1.5
2
2.5
3
3.5
4
4.5
5
0
5
10
15
20
25

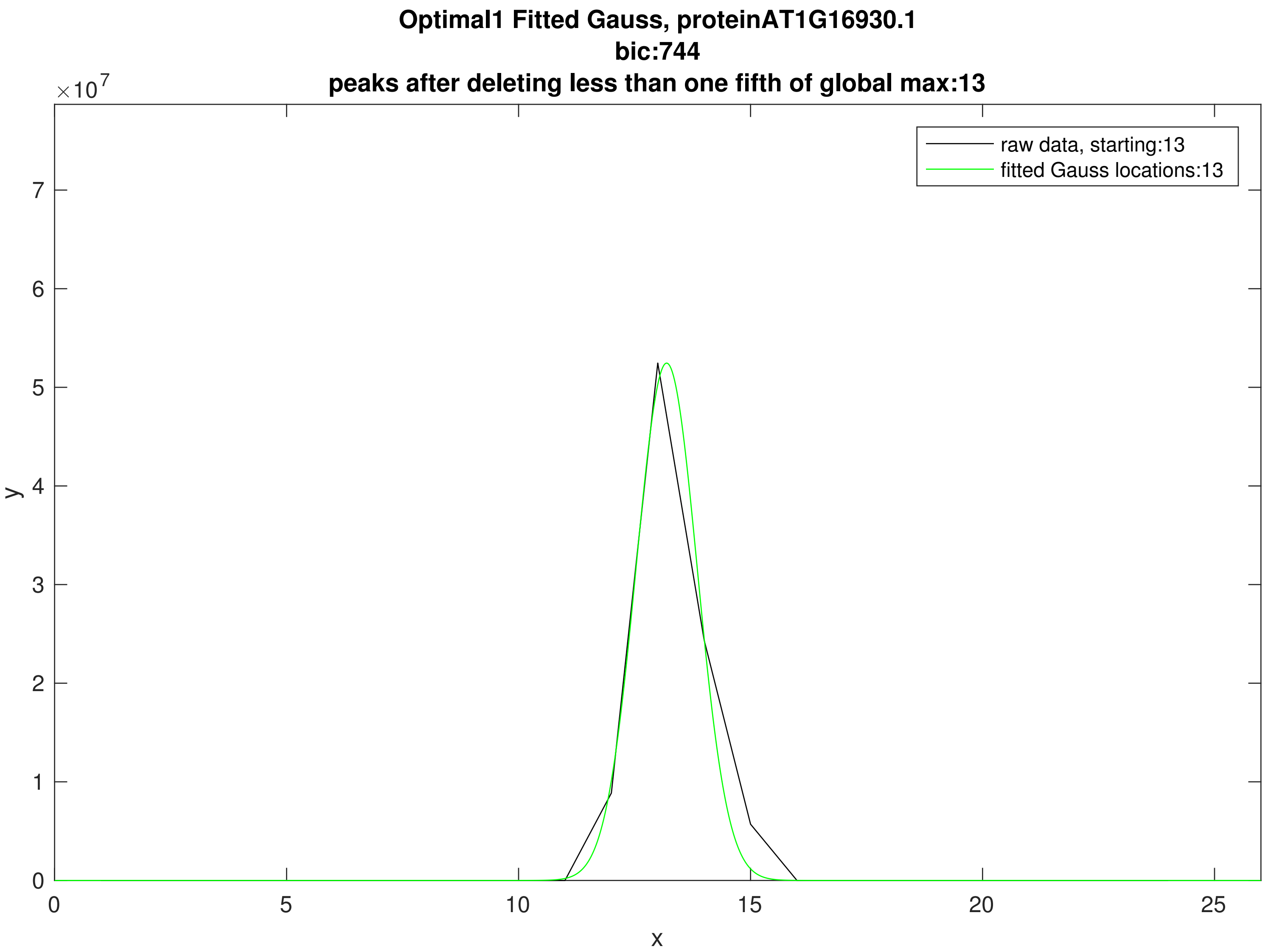
Optimal1 Fitted Gauss, proteinAT1G16930.1
bic:744
peaks after deleting less than one fifth of global max:13
×10^7
raw data, starting:13
fitted Gauss locations:13
y
x
0
1
2
3
4
5
6
7
0
5
10
15
20
25

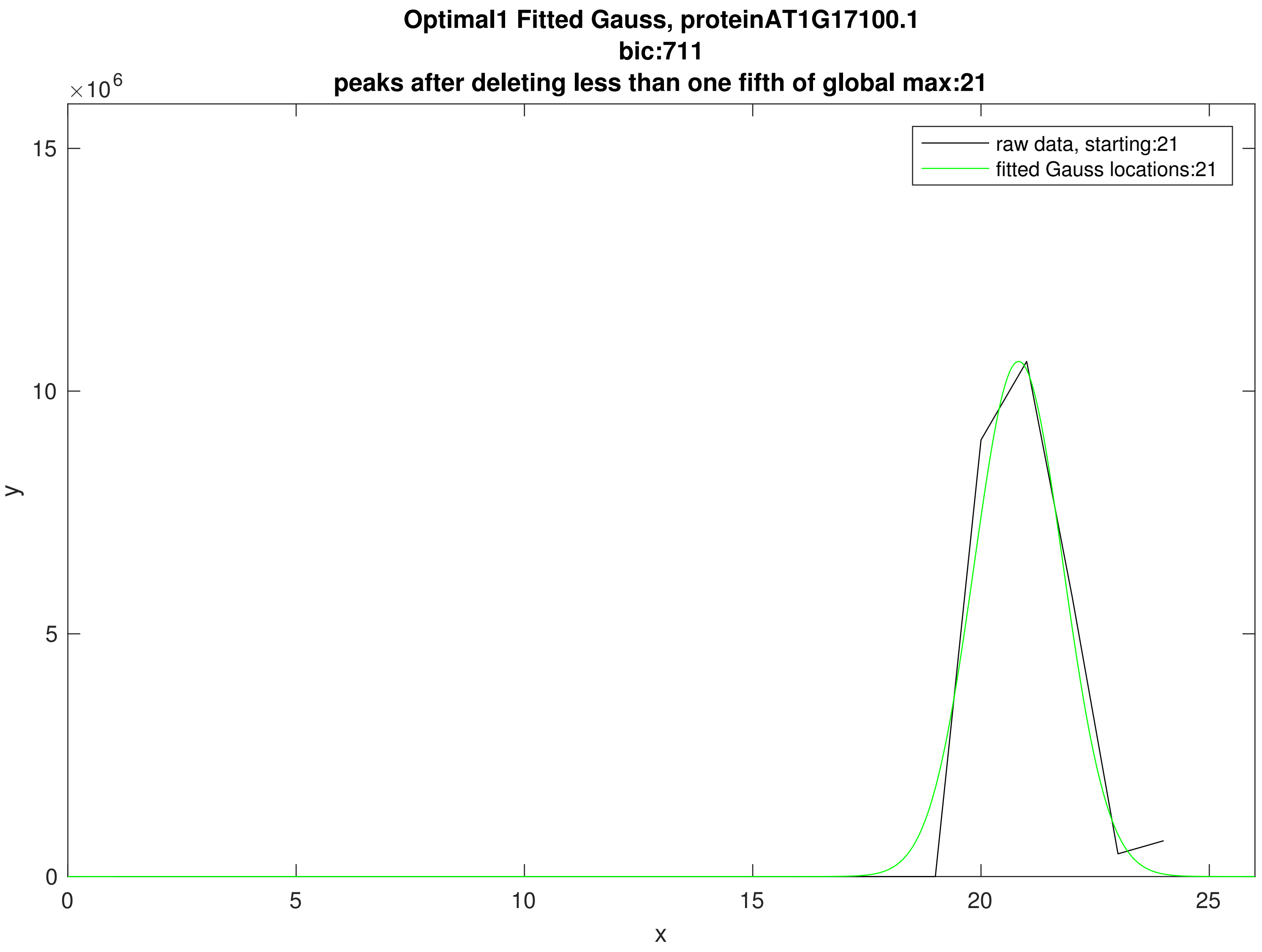
Optimal1 Fitted Gauss, proteinAT1G17100.1
bic:711
peaks after deleting less than one fifth of global max:21
×10^6
15
10
5
0
y
0
5
10
15
20
25
x
raw data, starting:21
fitted Gauss locations:21

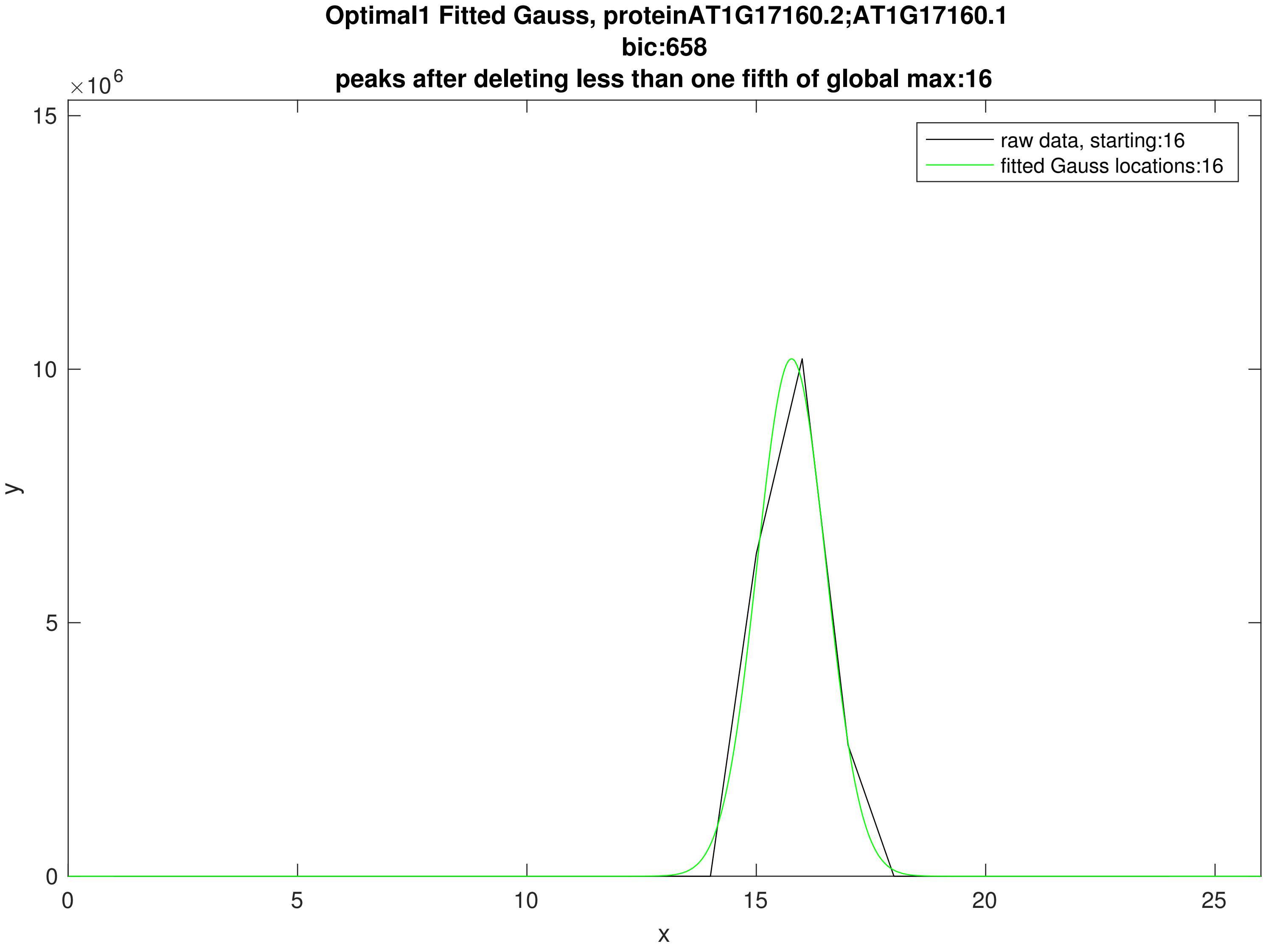
Optimal1 Fitted Gauss, proteinAT1G17160.2;AT1G17160.1
bic:658
peaks after deleting less than one fifth of global max:16
×10^6
raw data, starting:16
fitted Gauss locations:16
y
x
0
5
10
15
20
25

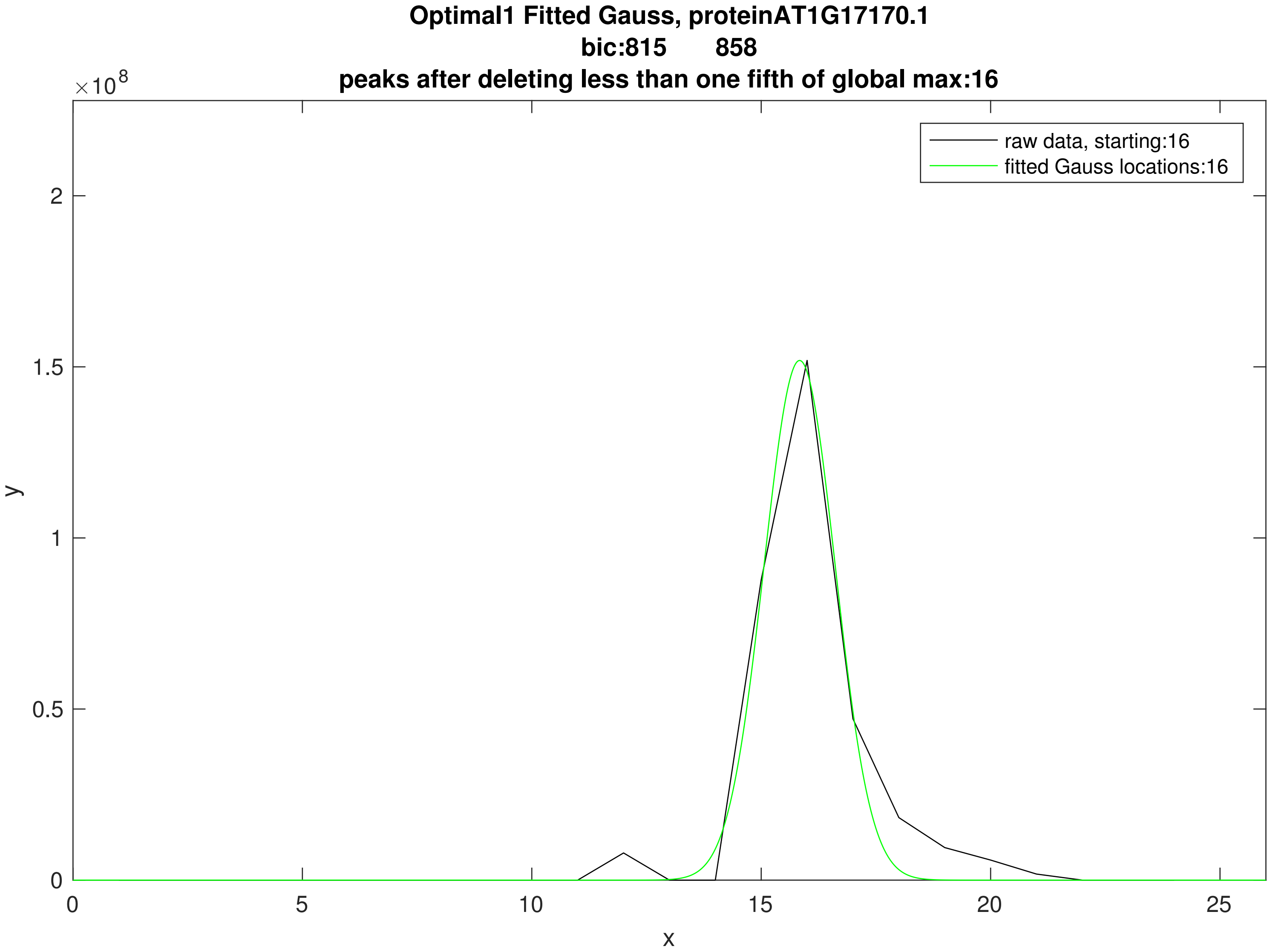
Optimal1 Fitted Gauss, proteinAT1G17170.1
bic:815 858
peaks after deleting less than one fifth of global max:16
×10^8
raw data, starting:16
fitted Gauss locations:16
y
x
0
0.5
1
1.5
2
0
5
10
15
20
25

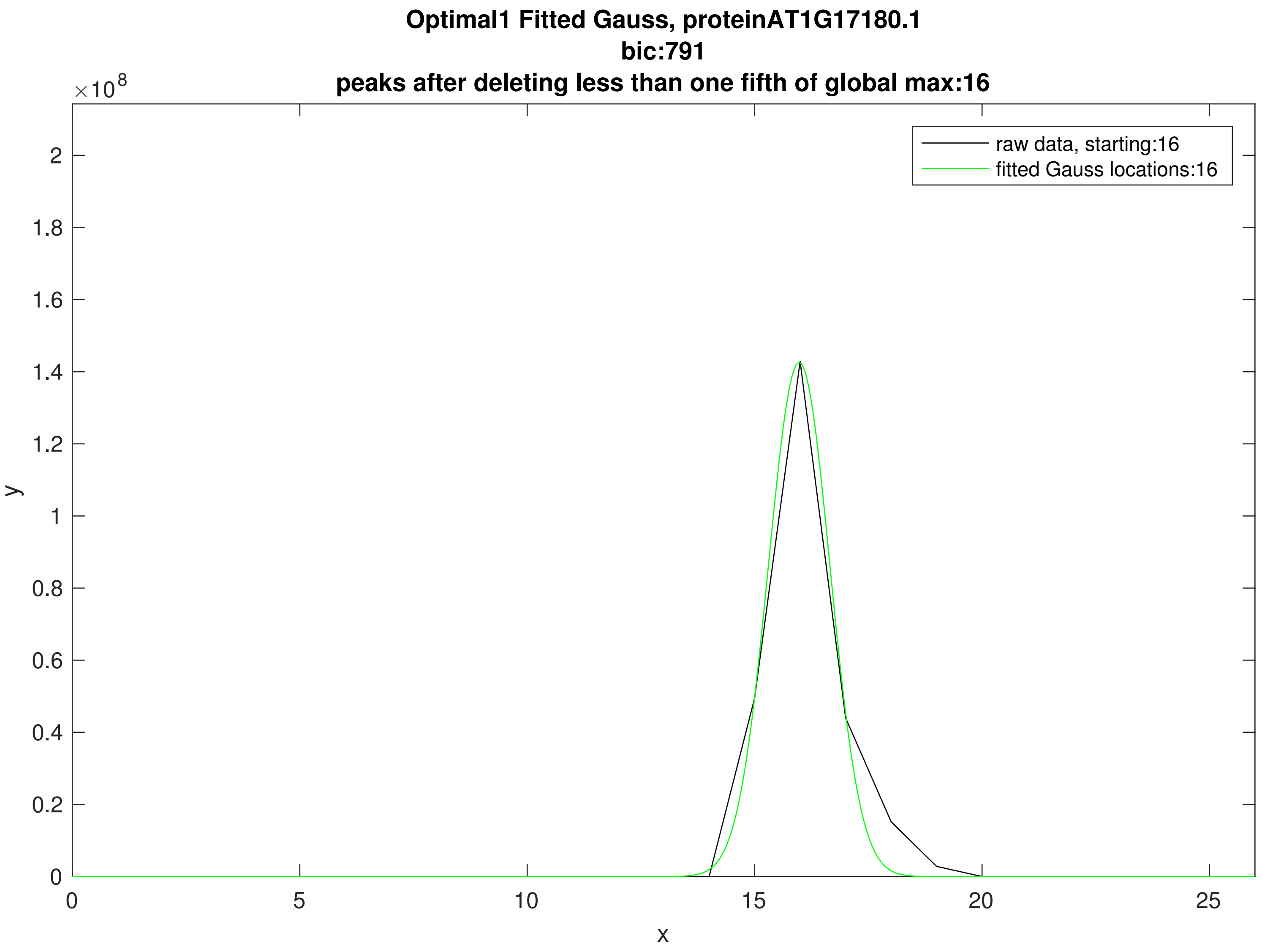
Optimal1 Fitted Gauss, proteinAT1G17180.1
bic:791
peaks after deleting less than one fifth of global max:16
×10^8
raw data, starting:16
fitted Gauss locations:16
y
x

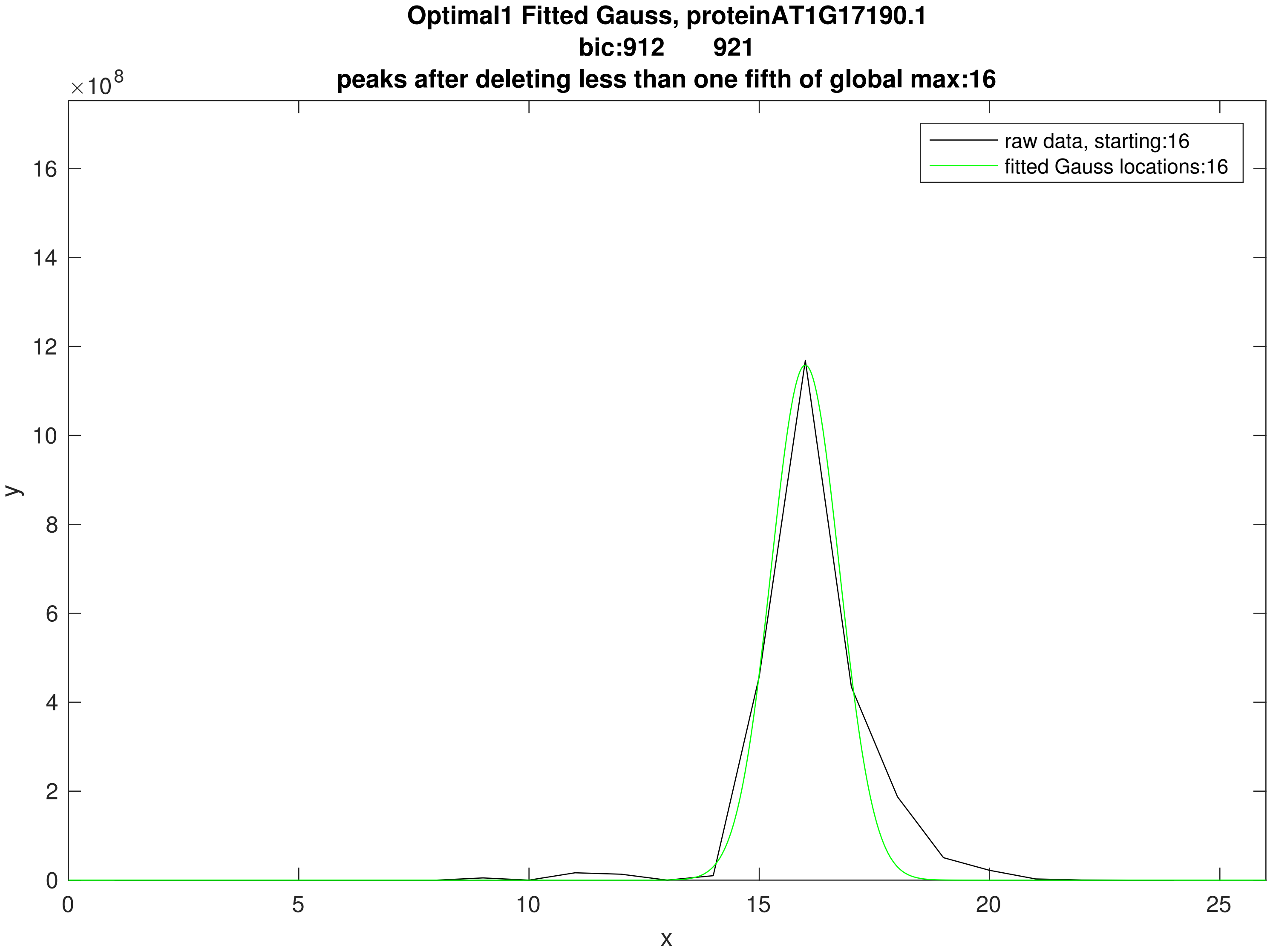

Optimal1 Fitted Gauss, proteinAT1G17190.1
bic:912 921
peaks after deleting less than one fifth of global max:16
×10^8
raw data, starting:16
fitted Gauss locations:16
y
x
0
2
4
6
8
10
12
14
16
0
5
10
15
20
25

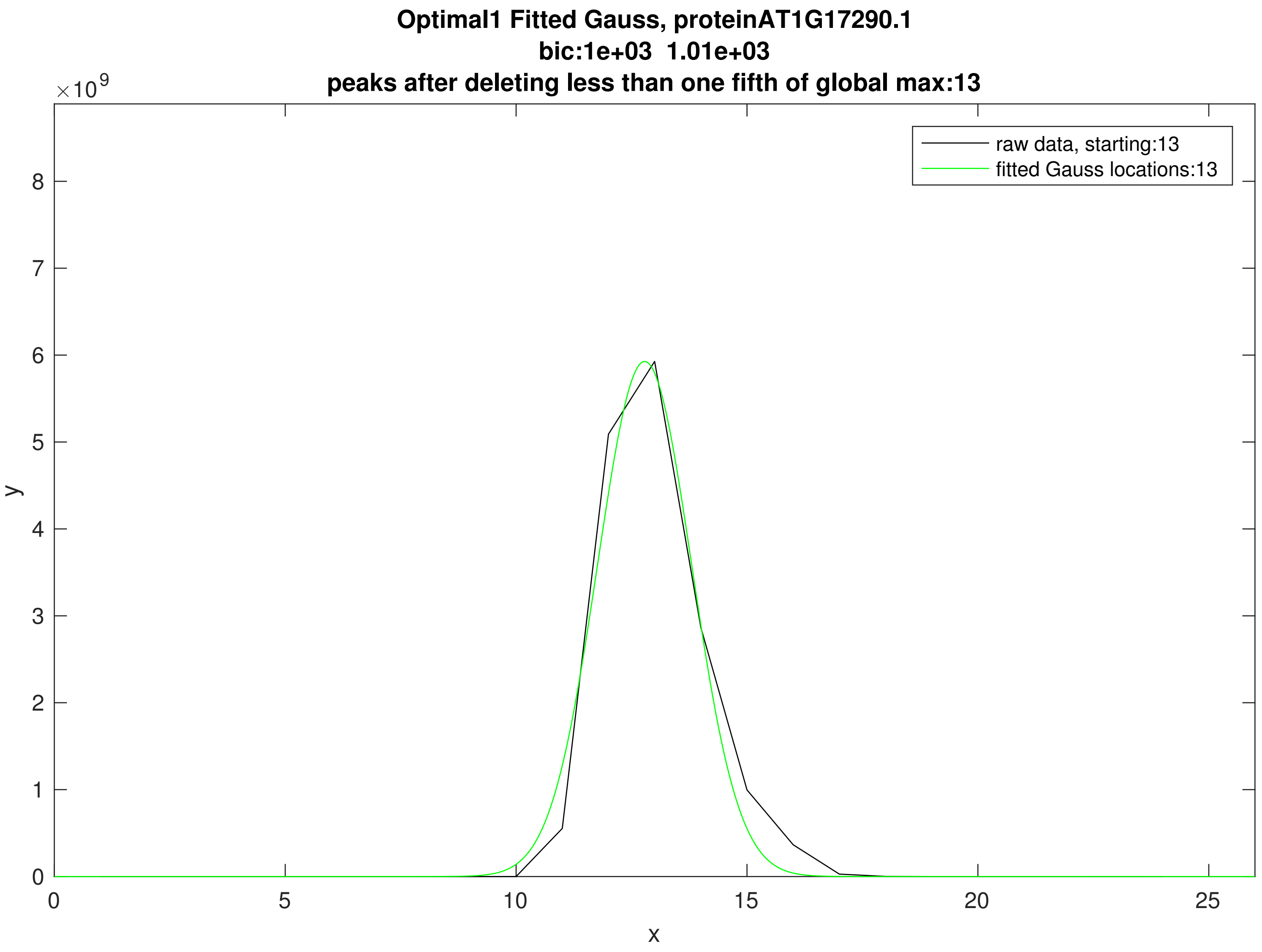
Optimal1 Fitted Gauss, proteinAT1G17290.1
bic:1e+03  1.01e+03
peaks after deleting less than one fifth of global max:13
×10^9
raw data, starting:13
fitted Gauss locations:13
y
x
0
1
2
3
4
5
6
7
8
0
5
10
15
20
25

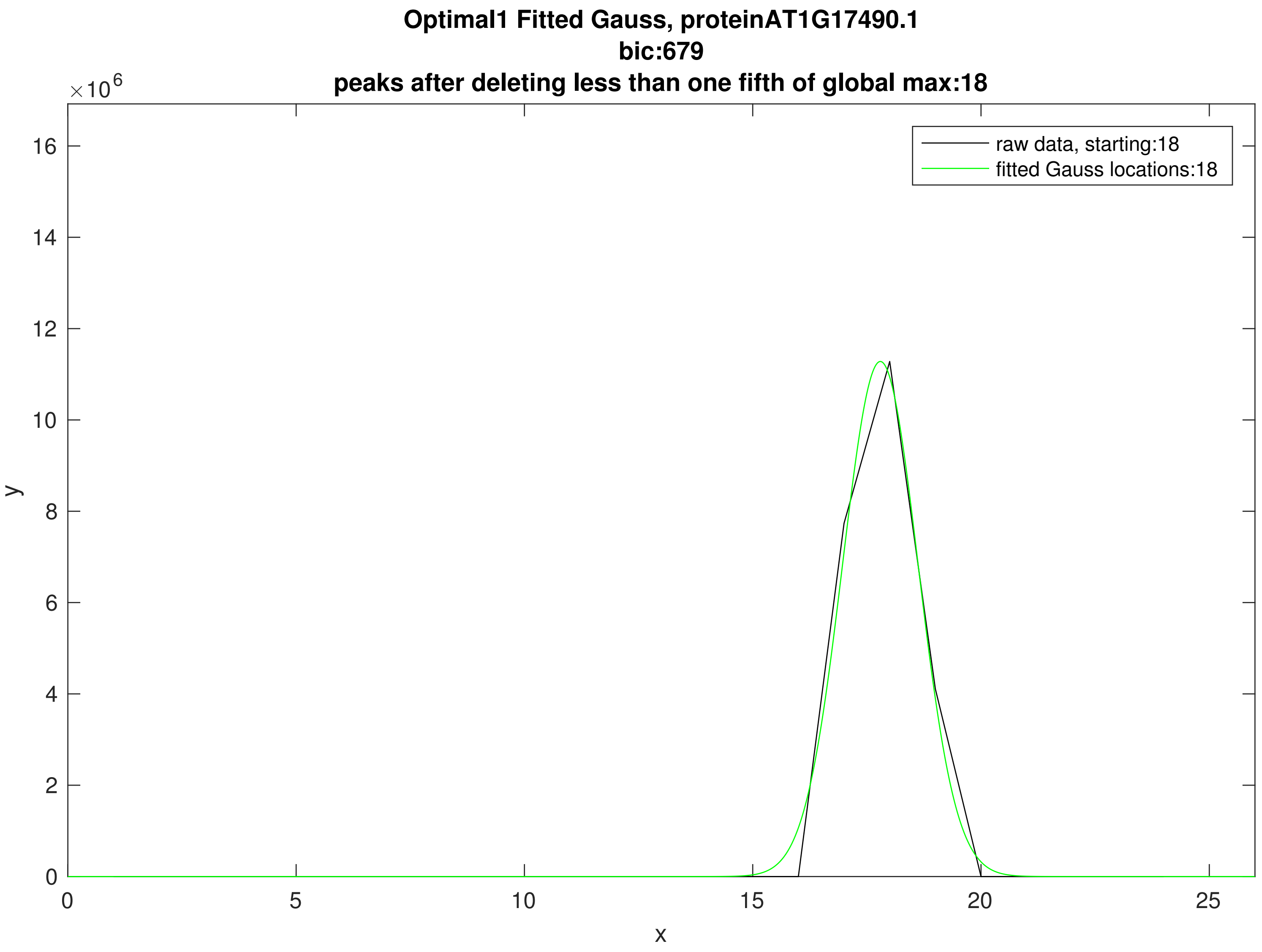

Optimal1 Fitted Gauss, proteinAT1G17490.1
bic:679
peaks after deleting less than one fifth of global max:18
×10^6
raw data, starting:18
fitted Gauss locations:18
y
x
0
2
4
6
8
10
12
14
16
0
5
10
15
20
25

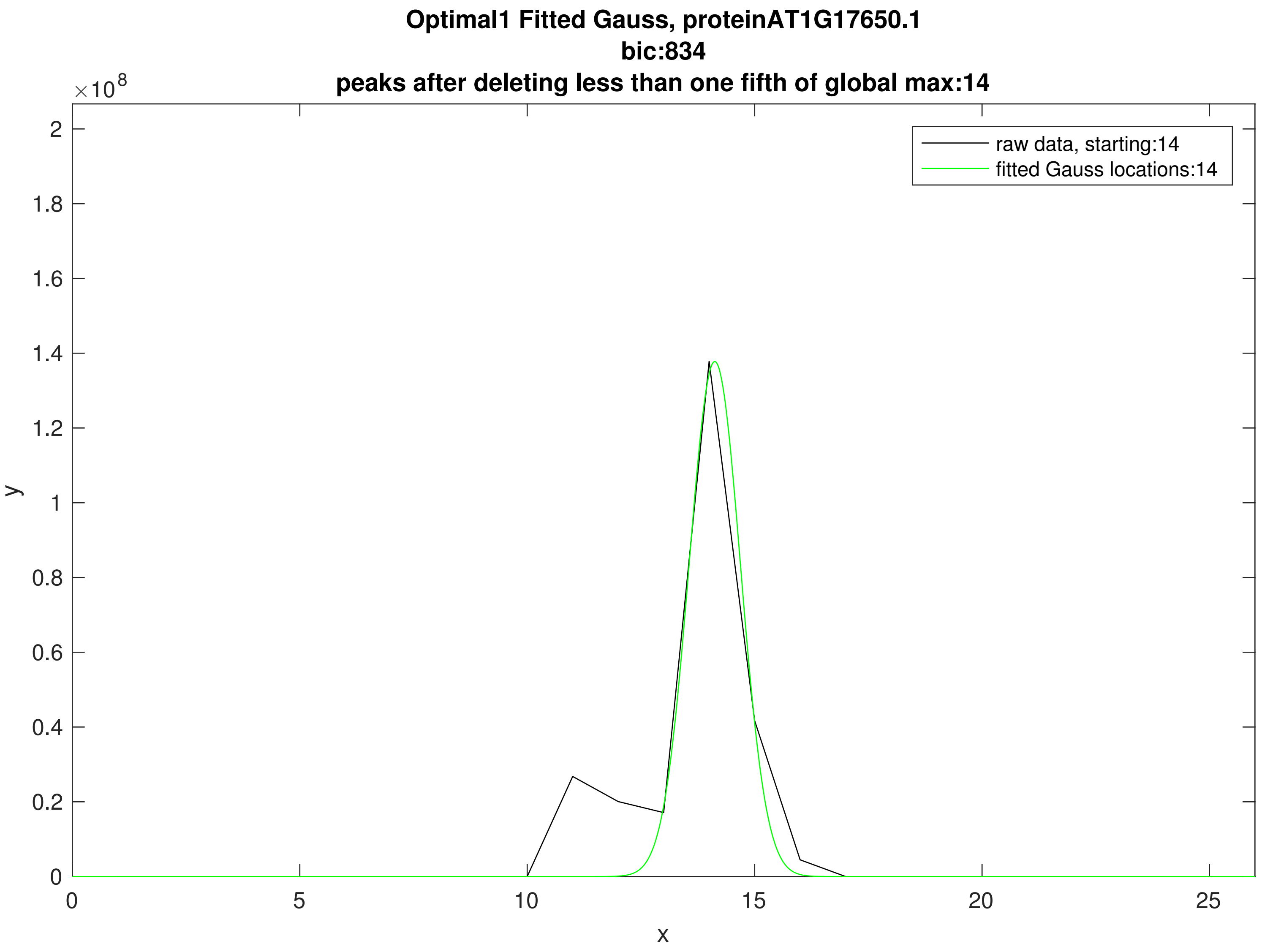

Optimal1 Fitted Gauss, proteinAT1G17650.1
bic:834
peaks after deleting less than one fifth of global max:14
×10^8
2
1.8
1.6
1.4
1.2
1
0.8
0.6
0.4
0.2
0
y
0
5
10
15
20
25
x
raw data, starting:14
fitted Gauss locations:14

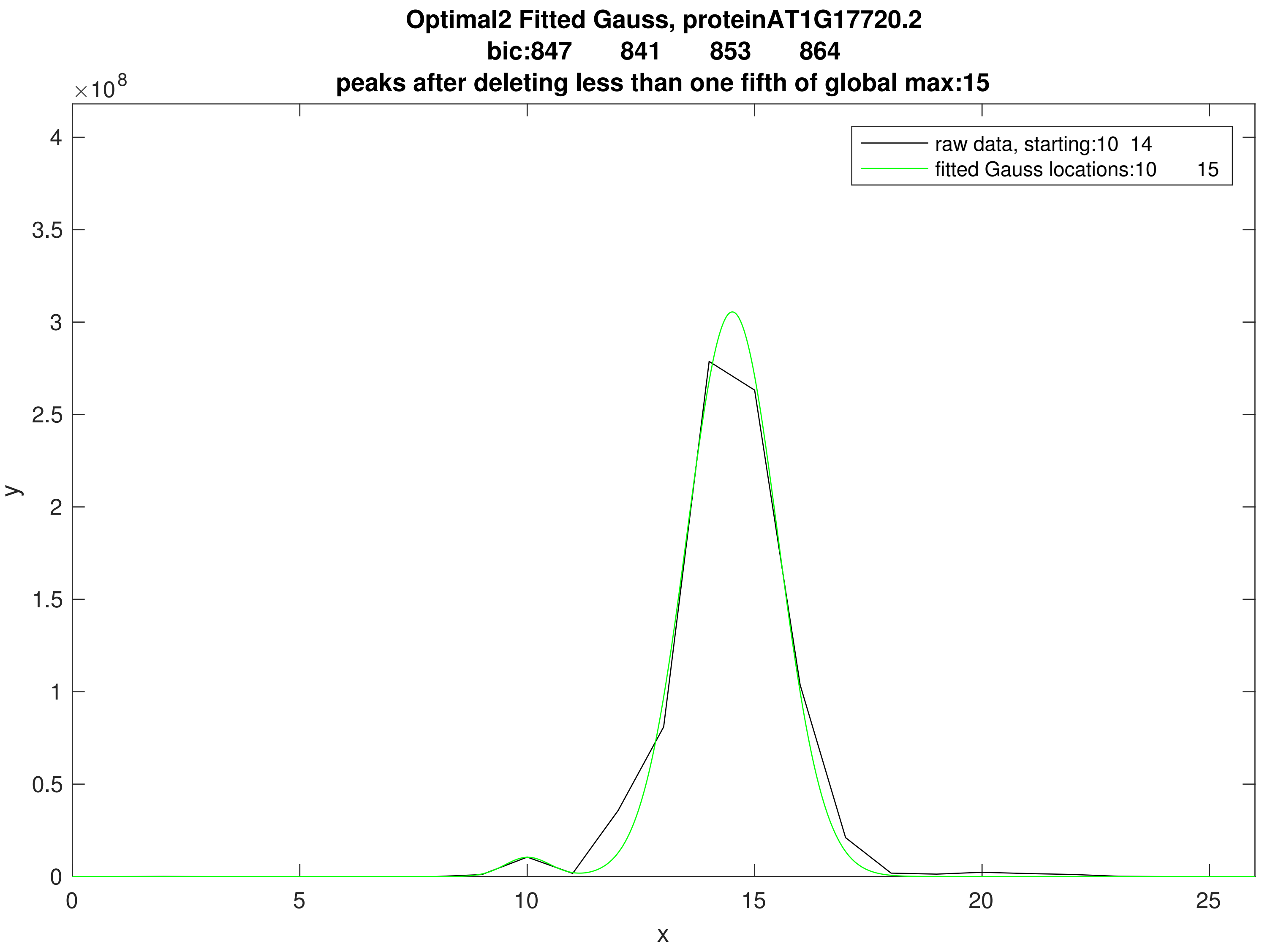

Optimal2 Fitted Gauss, proteinAT1G17720.2
bic:847 841 853 864
peaks after deleting less than one fifth of global max:15
raw data, starting:10 14
fitted Gauss locations:10 15
×10^8
y
x

**Optimal1 Fitted Gauss, proteinAT1G17745.1;AT1G17745.2**
**bic:837**
**peaks after deleting less than one fifth of global max:10**

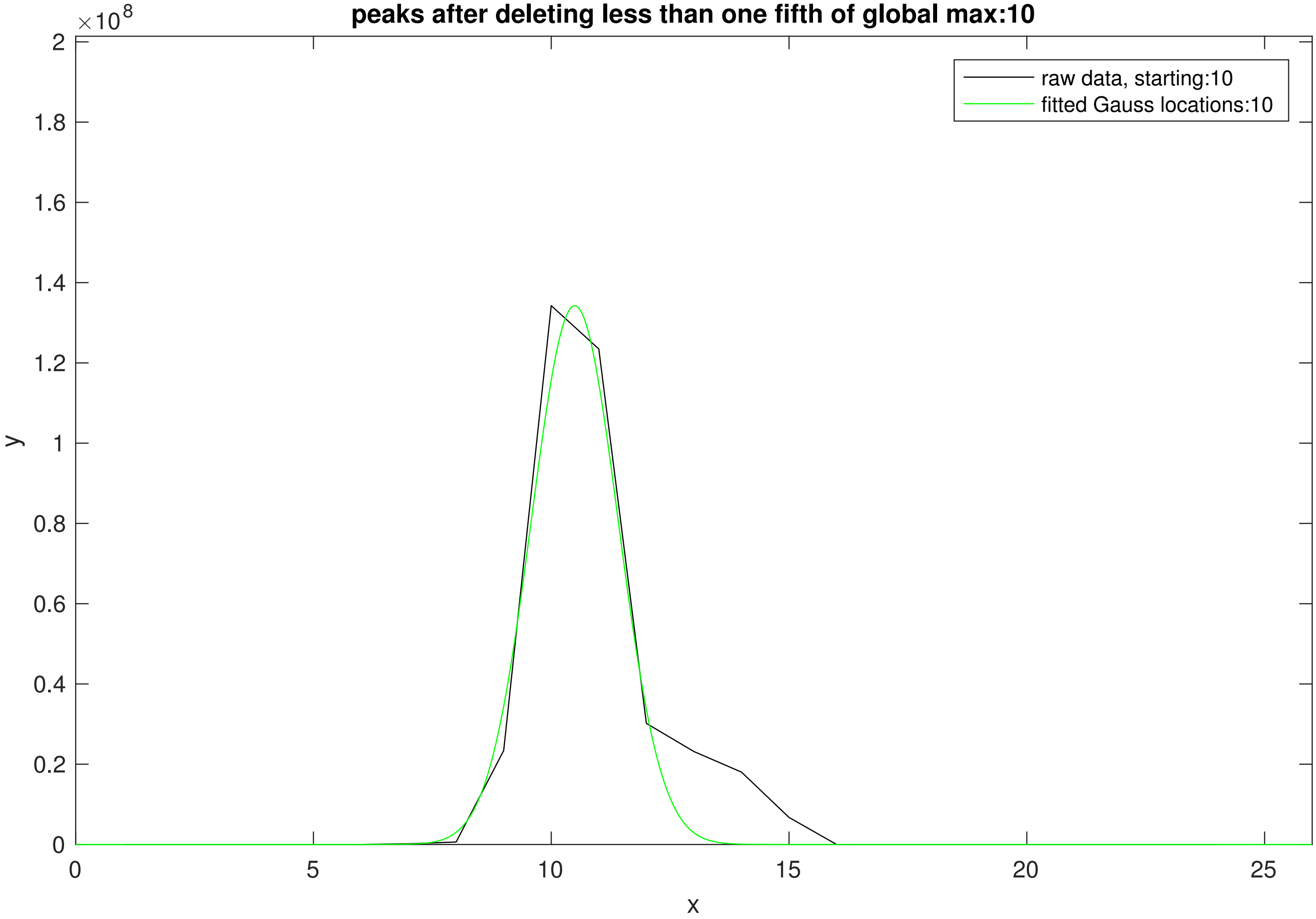

**Optimal2 Fitted Gauss, proteinAT1G17860.1**
**bic:675 656**
**peaks after deleting less than one fifth of global max:21**

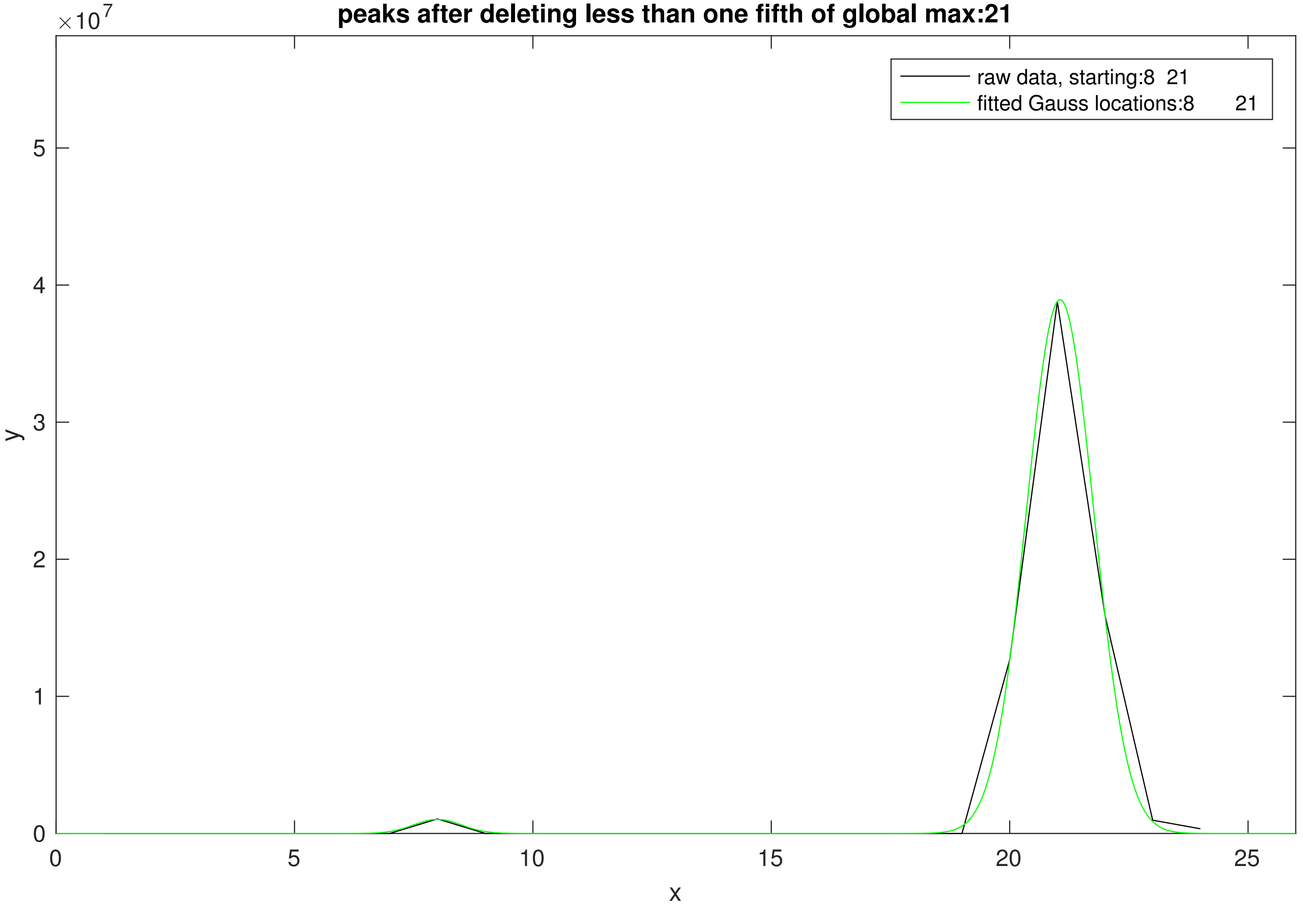

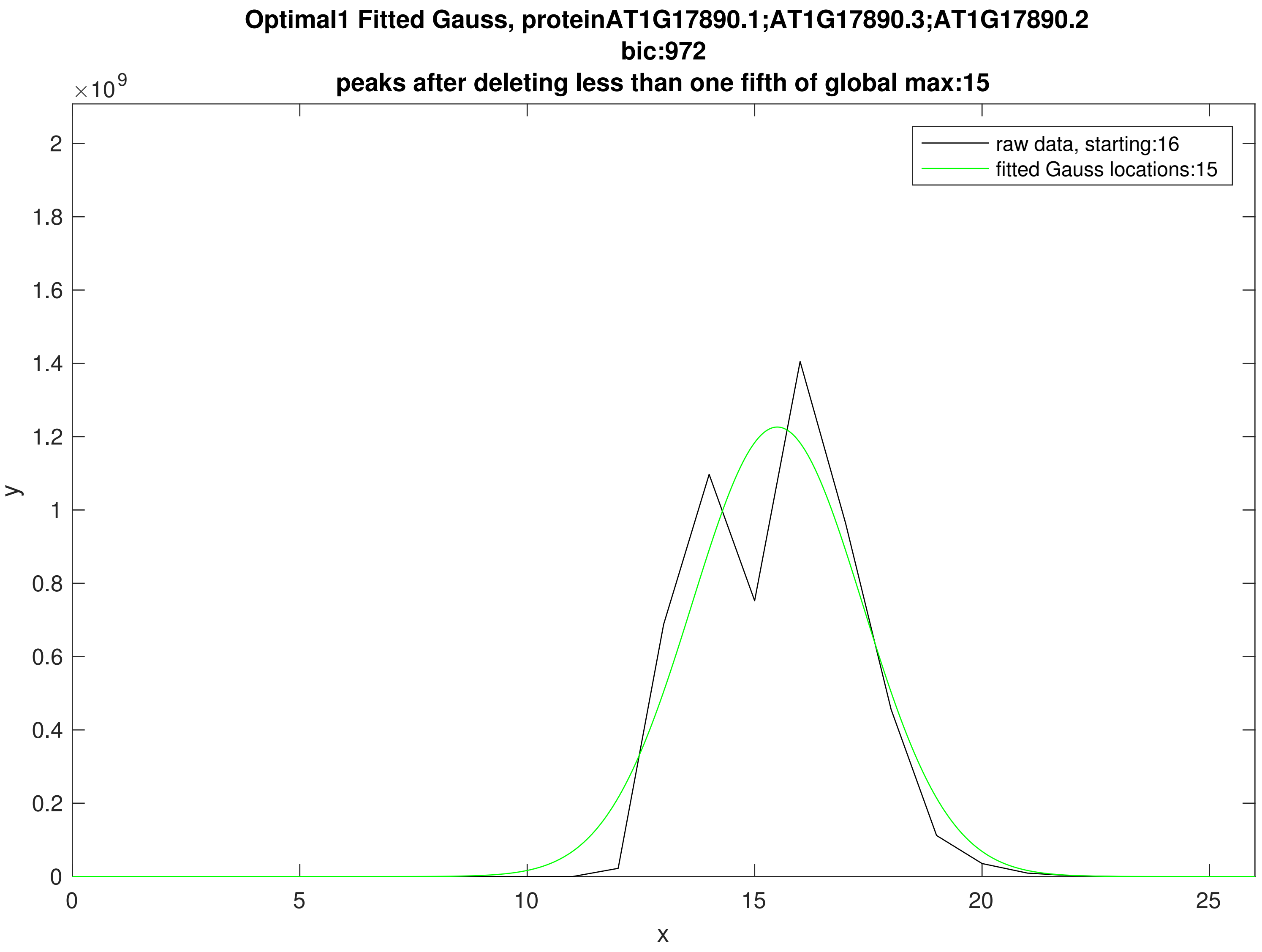
Optimal1 Fitted Gauss, proteinAT1G17890.1;AT1G17890.3;AT1G17890.2
bic:972
peaks after deleting less than one fifth of global max:15
×10^9
2
1.8
1.6
1.4
1.2
1
0.8
0.6
0.4
0.2
0
y
0
5
10
15
20
25
x
raw data, starting:16
fitted Gauss locations:15

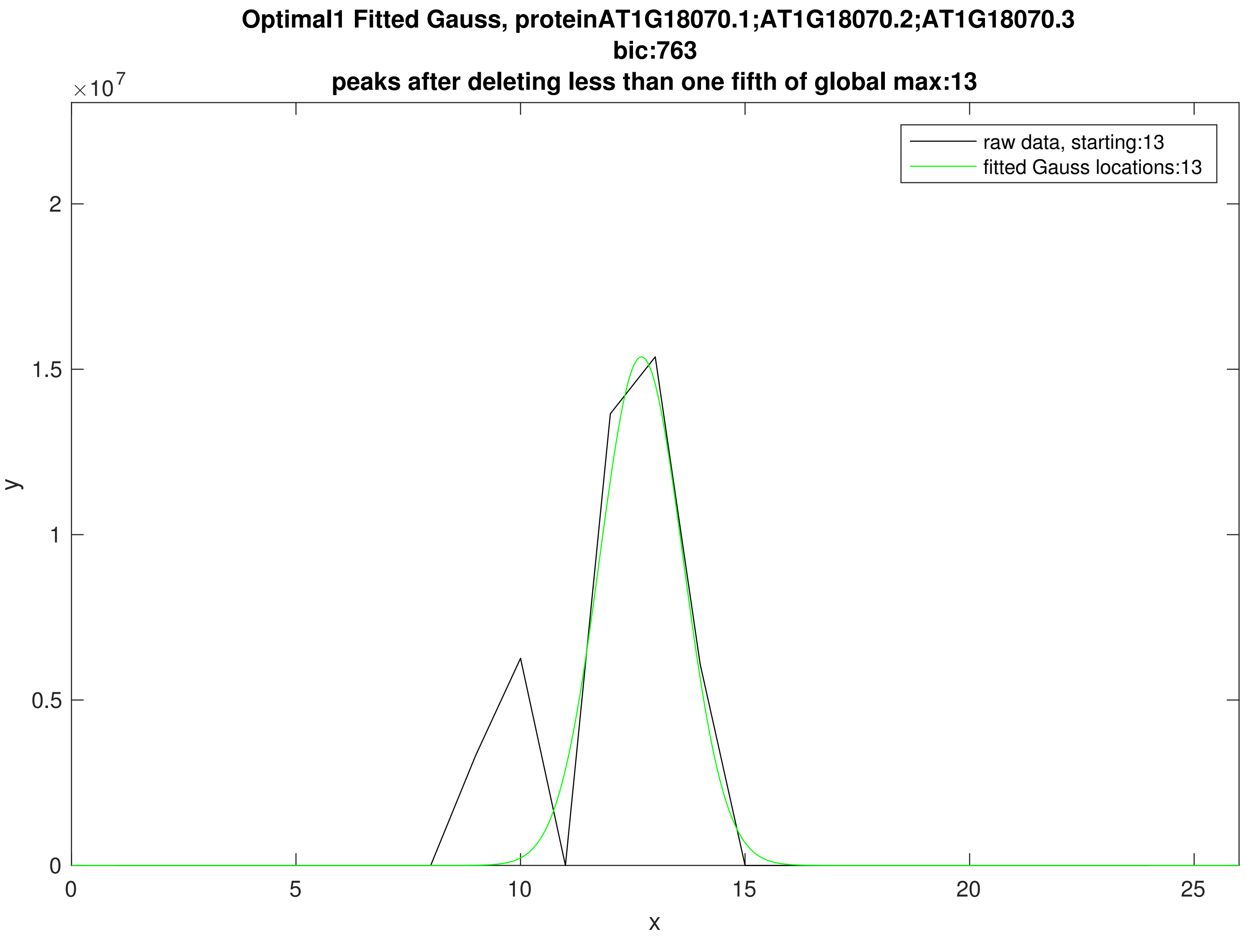
Optimal1 Fitted Gauss, proteinAT1G18070.1;AT1G18070.2;AT1G18070.3
bic:763
peaks after deleting less than one fifth of global max:13
×10^7
raw data, starting:13
fitted Gauss locations:13
y
x
0
0.5
1
1.5
2
0
5
10
15
20
25

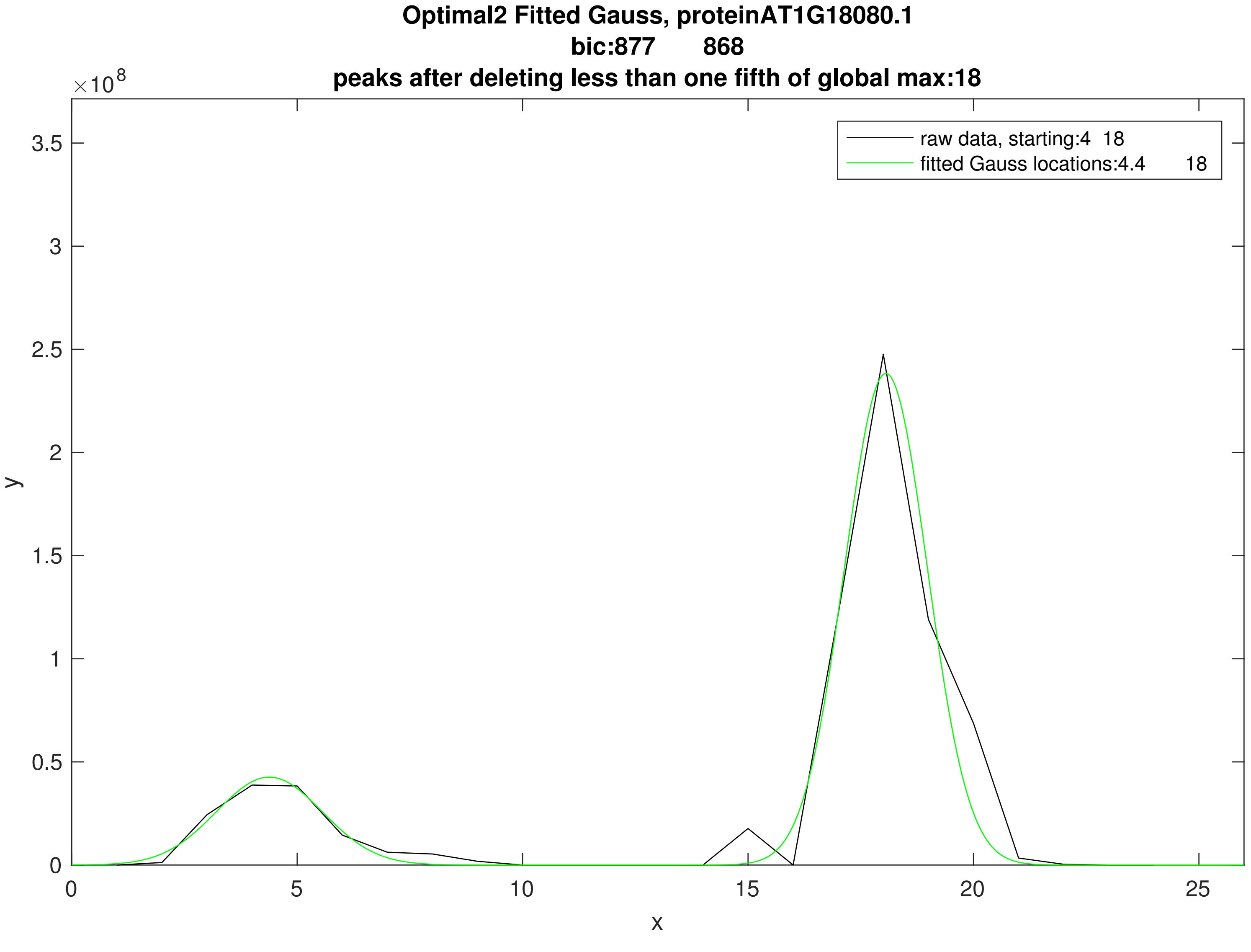
Optimal2 Fitted Gauss, proteinAT1G18080.1
bic:877 868
peaks after deleting less than one fifth of global max:18
×10^8
raw data, starting:4 18
fitted Gauss locations:4.4 18
y
x
0
0.5
1
1.5
2
2.5
3
3.5
0
5
10
15
20
25

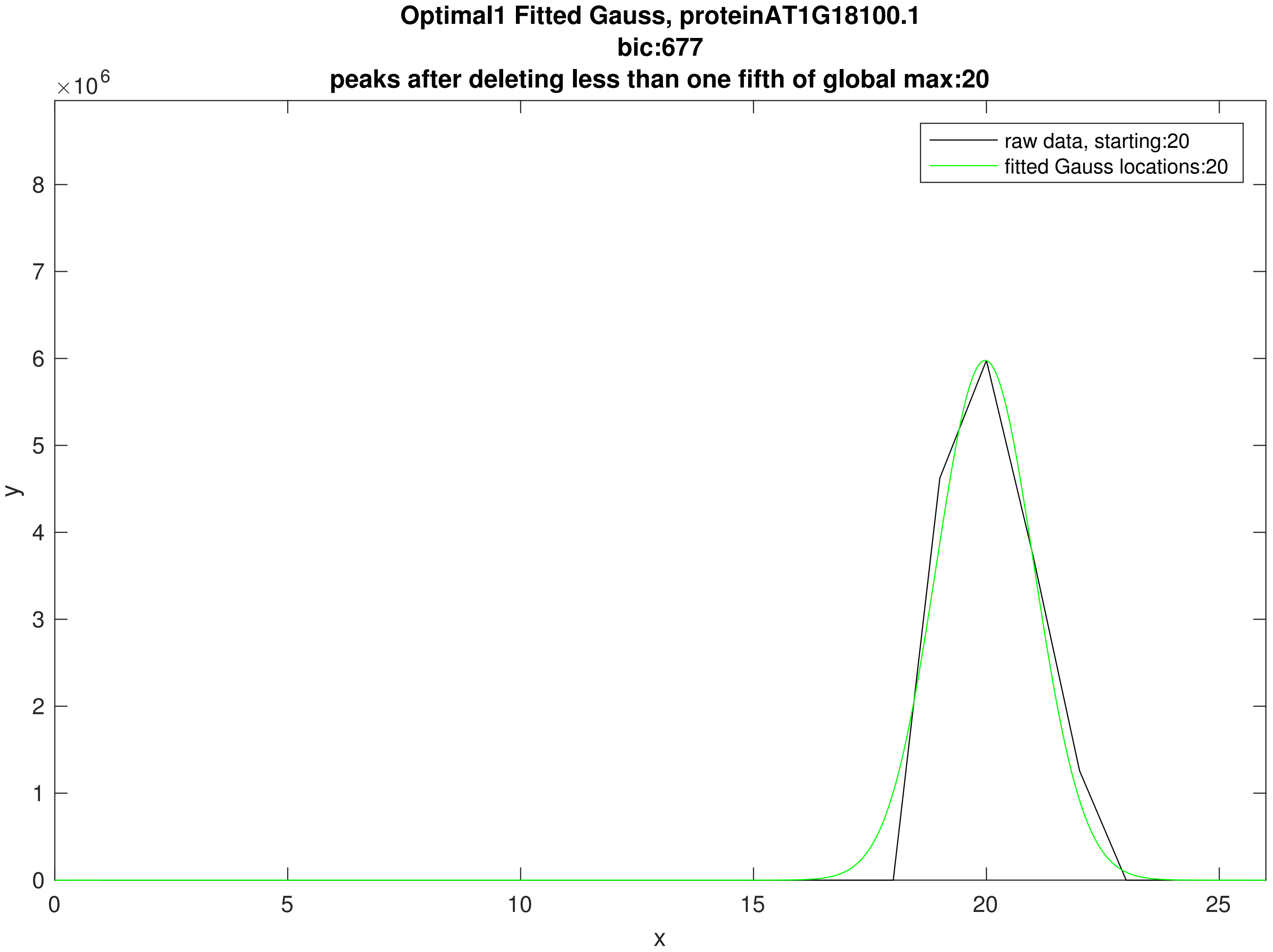
Optimal1 Fitted Gauss, proteinAT1G18100.1
bic:677
peaks after deleting less than one fifth of global max:20
×10^6
raw data, starting:20
fitted Gauss locations:20
y
x
0
1
2
3
4
5
6
7
8
0
5
10
15
20
25

#### Optimal1 Fitted Gauss, proteinAT1G18210.2;AT1G18210.1
#### bic:910 914
#### peaks after deleting less than one fifth of global max:18

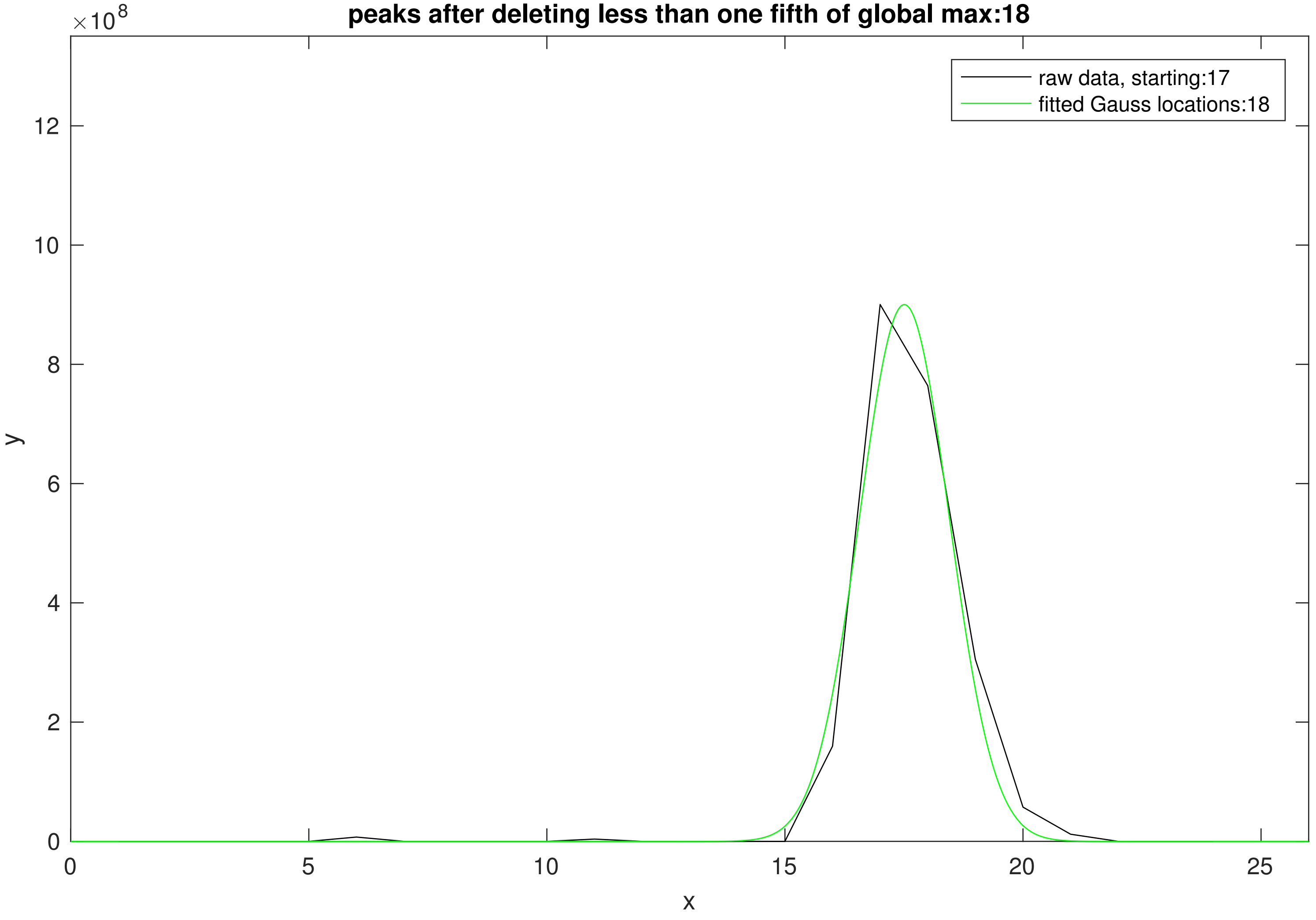

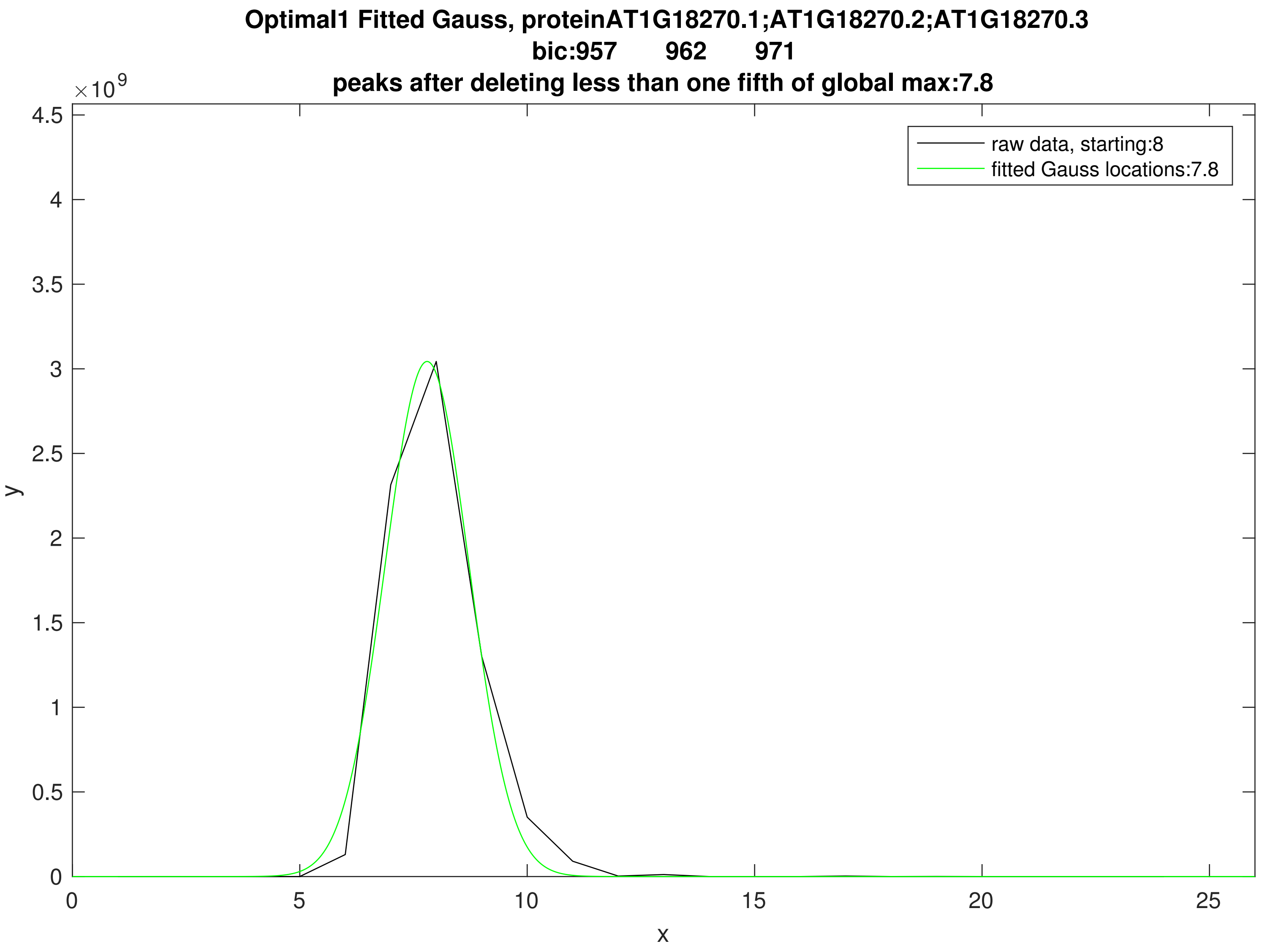
Optimal1 Fitted Gauss, proteinAT1G18270.1;AT1G18270.2;AT1G18270.3
bic:957 962 971
peaks after deleting less than one fifth of global max:7.8
×10^9
4.5
4
3.5
3
2.5
2
1.5
1
0.5
0
y
0
5
10
15
20
25
x
raw data, starting:8
fitted Gauss locations:7.8

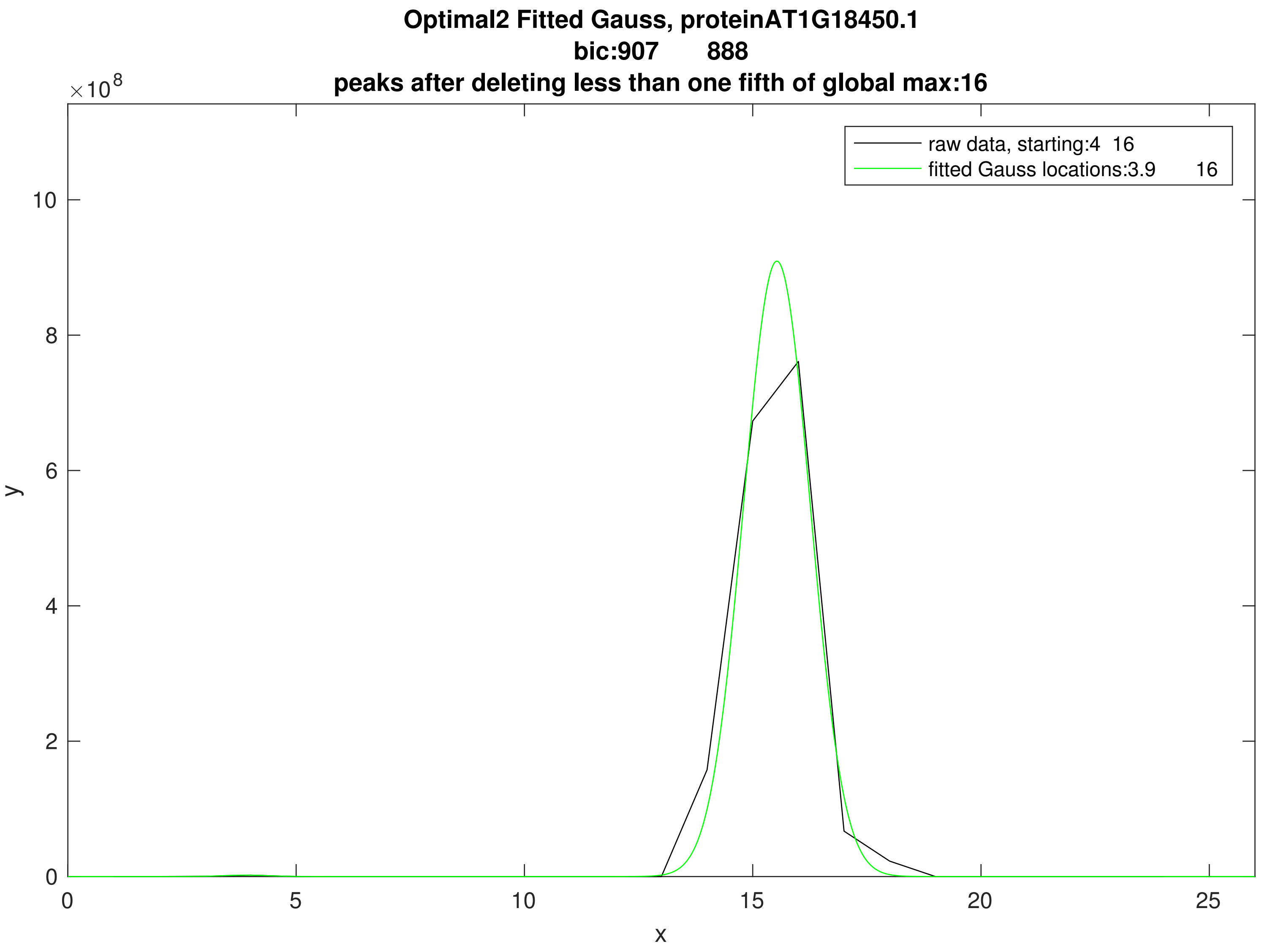
Optimal2 Fitted Gauss, proteinAT1G18450.1
bic:907 888
peaks after deleting less than one fifth of global max:16
×10^8
raw data, starting:4 16
fitted Gauss locations:3.9 16
10
8
6
4
2
0
y
0
5
10
15
20
25
x

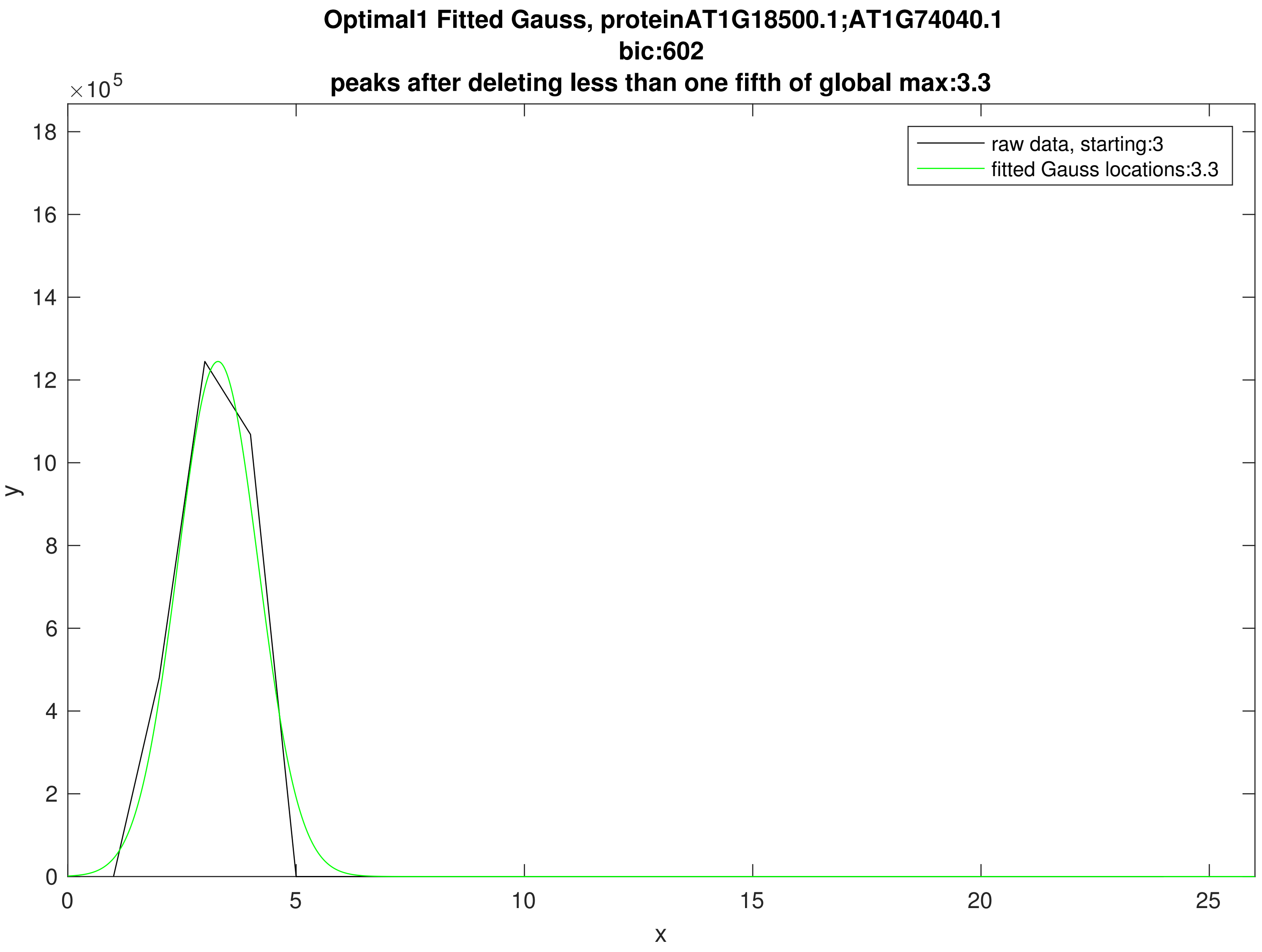
Optimal1 Fitted Gauss, proteinAT1G18500.1;AT1G74040.1
bic:602
peaks after deleting less than one fifth of global max:3.3
×10^5
raw data, starting:3
fitted Gauss locations:3.3
y
x
0
2
4
6
8
10
12
14
16
18
0
5
10
15
20
25

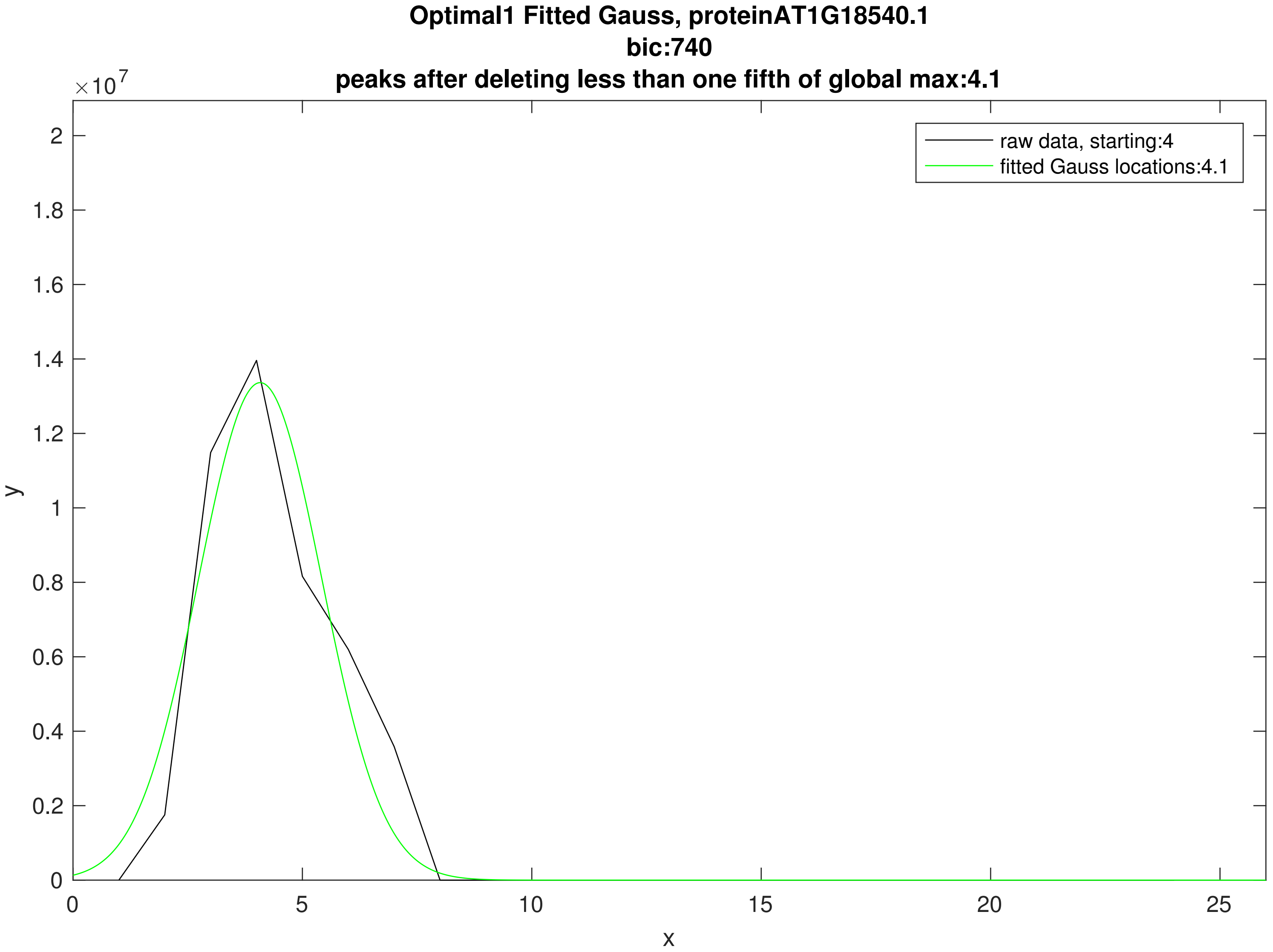

Optimal1 Fitted Gauss, proteinAT1G18540.1
bic:740
peaks after deleting less than one fifth of global max:4.1
×10^7
raw data, starting:4
fitted Gauss locations:4.1
y
x

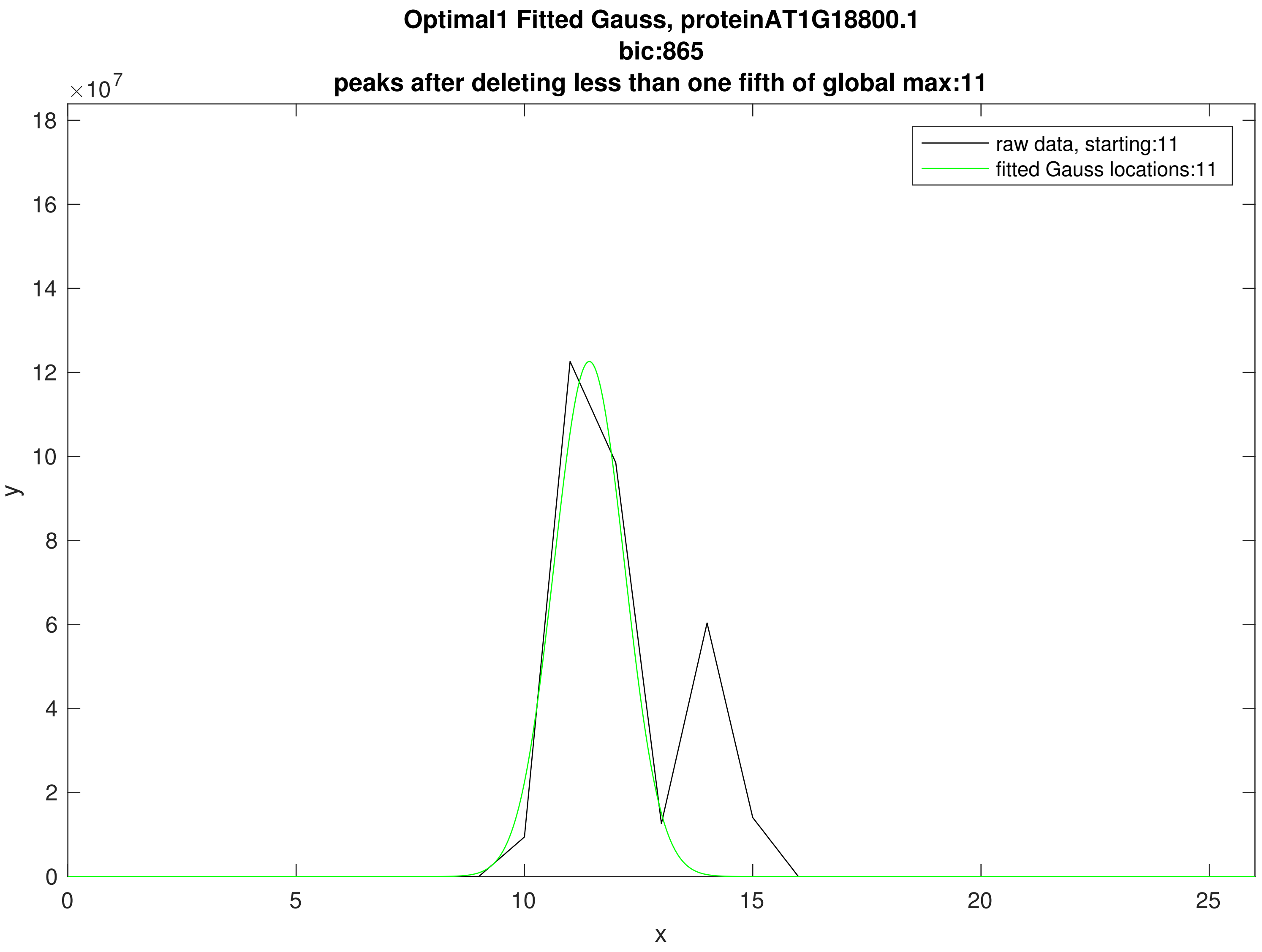
Optimal1 Fitted Gauss, proteinAT1G18800.1
bic:865
peaks after deleting less than one fifth of global max:11
×10^7
raw data, starting:11
fitted Gauss locations:11
y
x
0
2
4
6
8
10
12
14
16
18
0
5
10
15
20
25

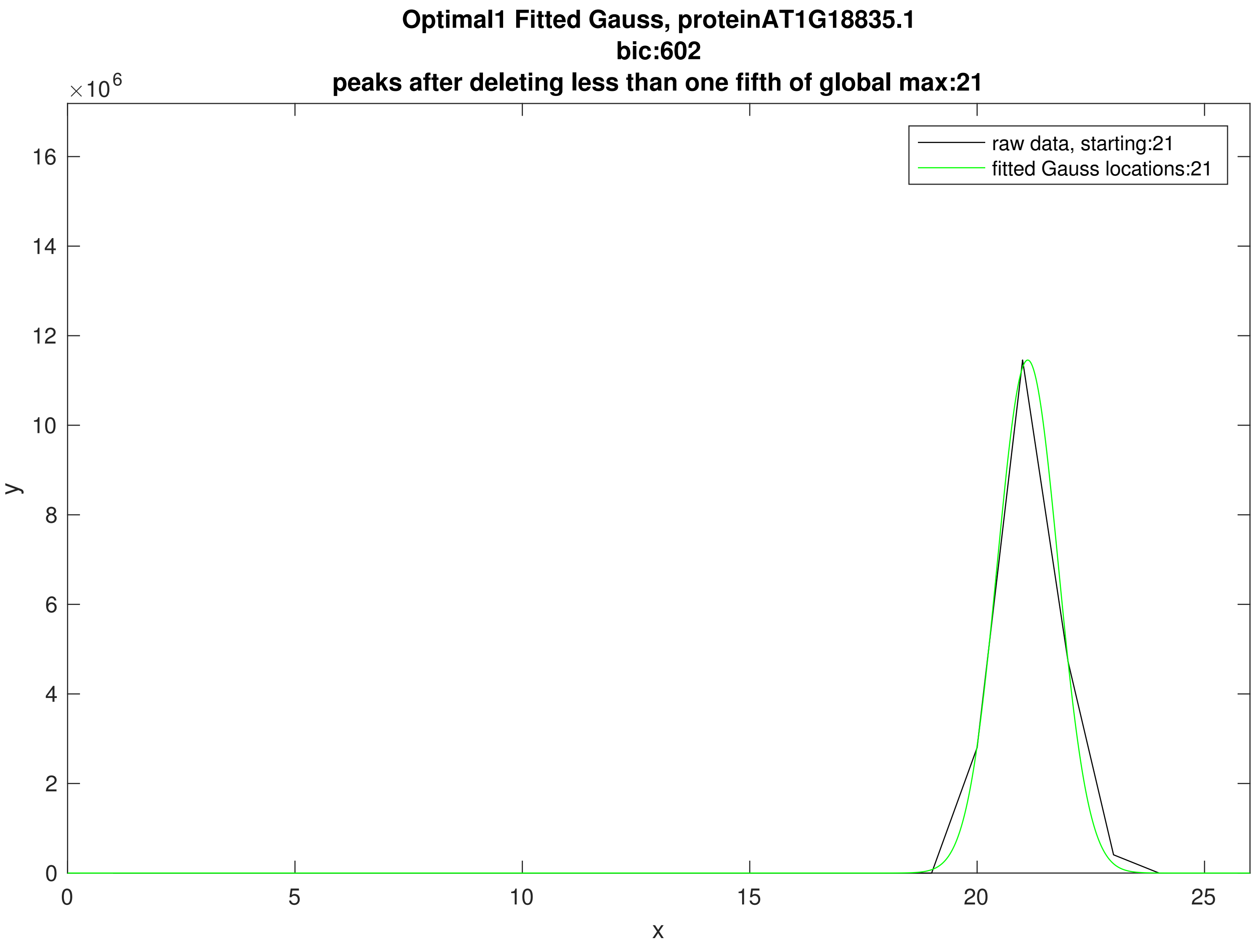

Optimal1 Fitted Gauss, proteinAT1G18835.1
bic:602
peaks after deleting less than one fifth of global max:21
×10^6
raw data, starting:21
fitted Gauss locations:21
16
14
12
10
8
6
4
2
0
y
0
5
10
15
20
25
x

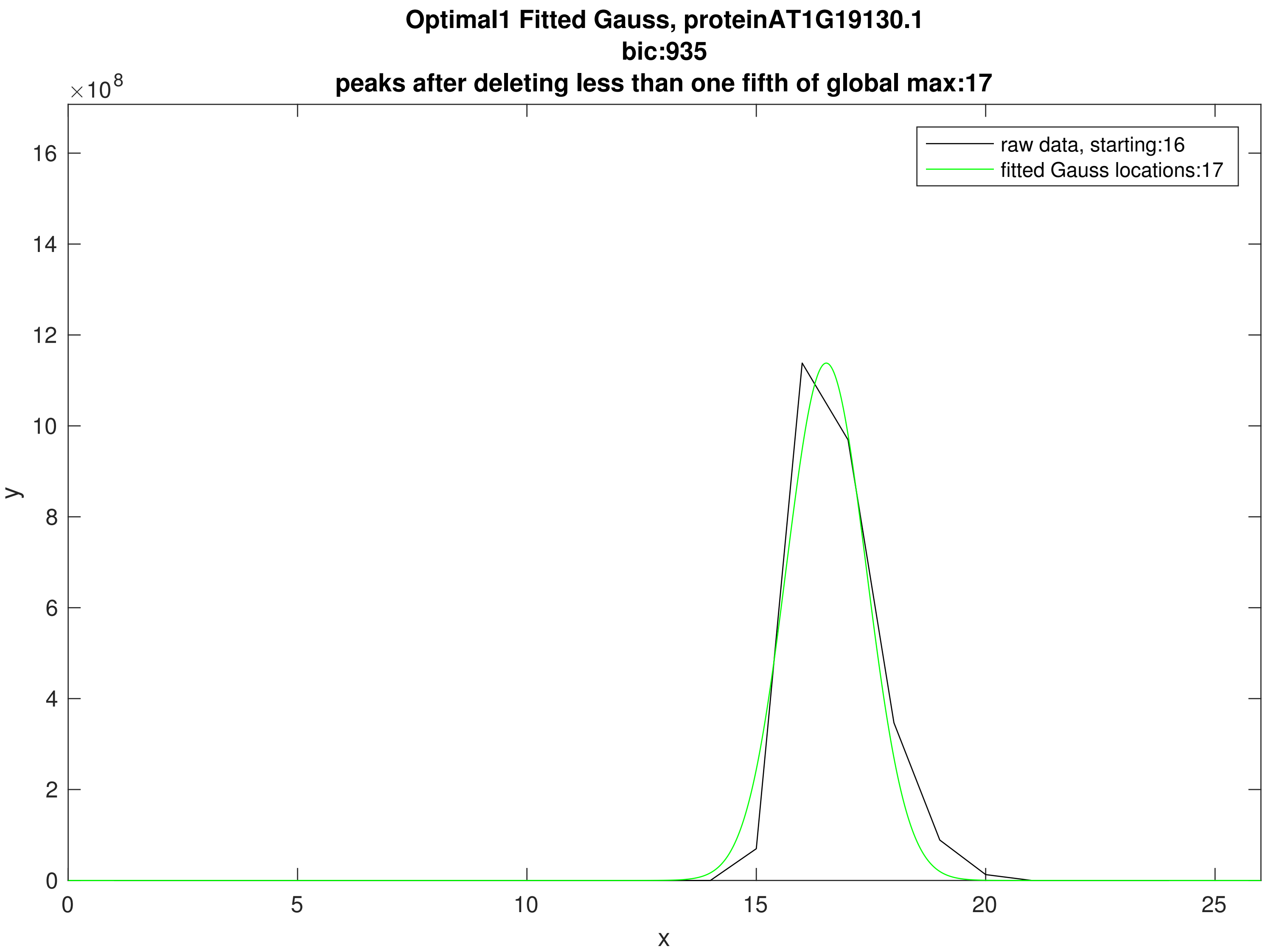
Optimal1 Fitted Gauss, proteinAT1G19130.1
bic:935
peaks after deleting less than one fifth of global max:17
×10^8
16
14
12
10
8
6
4
2
0
y
0
5
10
15
20
25
x
raw data, starting:16
fitted Gauss locations:17

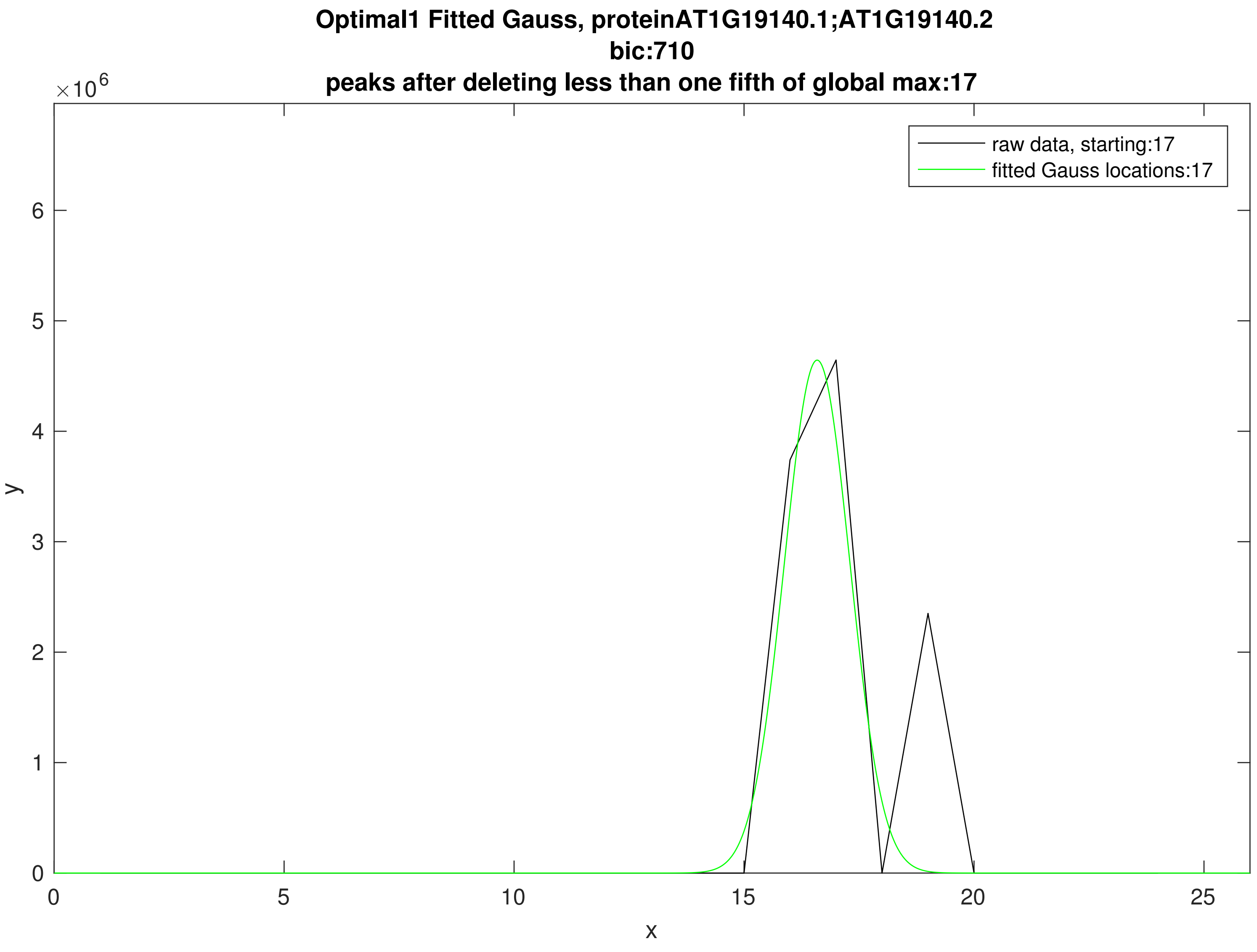

Optimal1 Fitted Gauss, proteinAT1G19140.1;AT1G19140.2
bic:710
peaks after deleting less than one fifth of global max:17
×10^6
raw data, starting:17
fitted Gauss locations:17
y
x
0
1
2
3
4
5
6
0
5
10
15
20
25

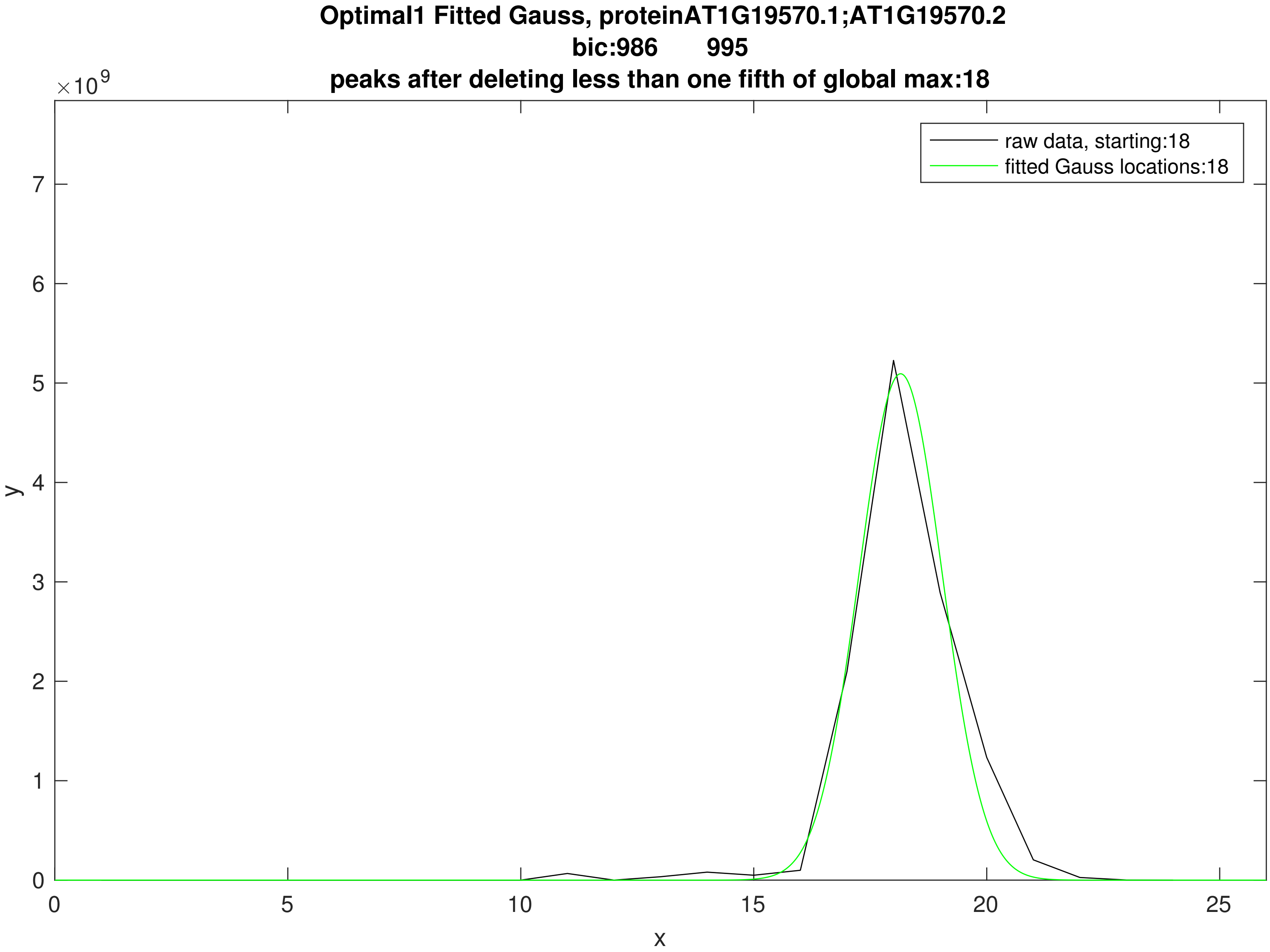
Optimal1 Fitted Gauss, proteinAT1G19570.1;AT1G19570.2
bic:986 995
peaks after deleting less than one fifth of global max:18
×10^9
raw data, starting:18
fitted Gauss locations:18
y
x
0
1
2
3
4
5
6
7
0
5
10
15
20
25

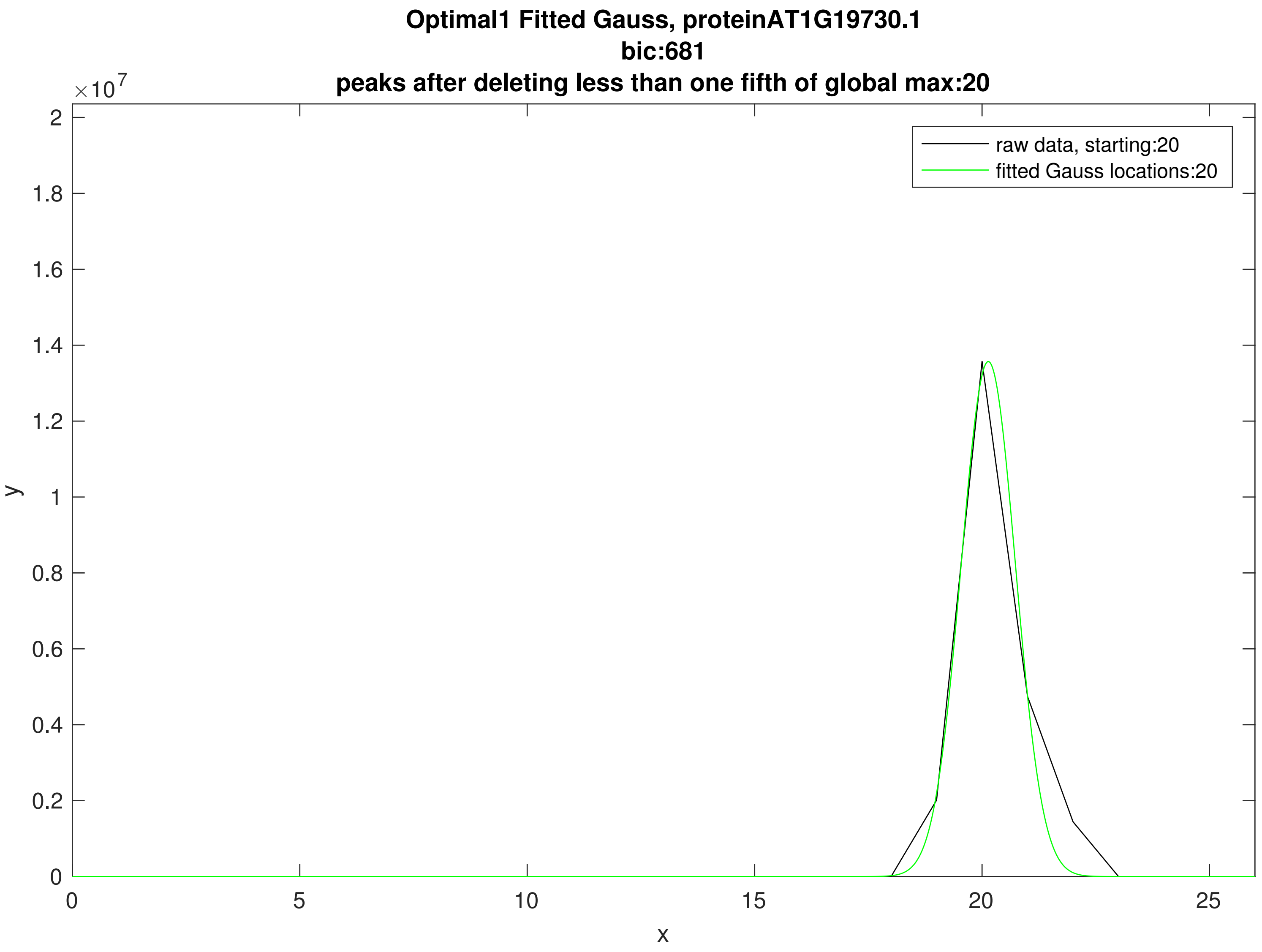
Optimal1 Fitted Gauss, proteinAT1G19730.1
bic:681
peaks after deleting less than one fifth of global max:20
×10^7
raw data, starting:20
fitted Gauss locations:20
y
x
0
0.2
0.4
0.6
0.8
1
1.2
1.4
1.6
1.8
2
5
10
15
20
25

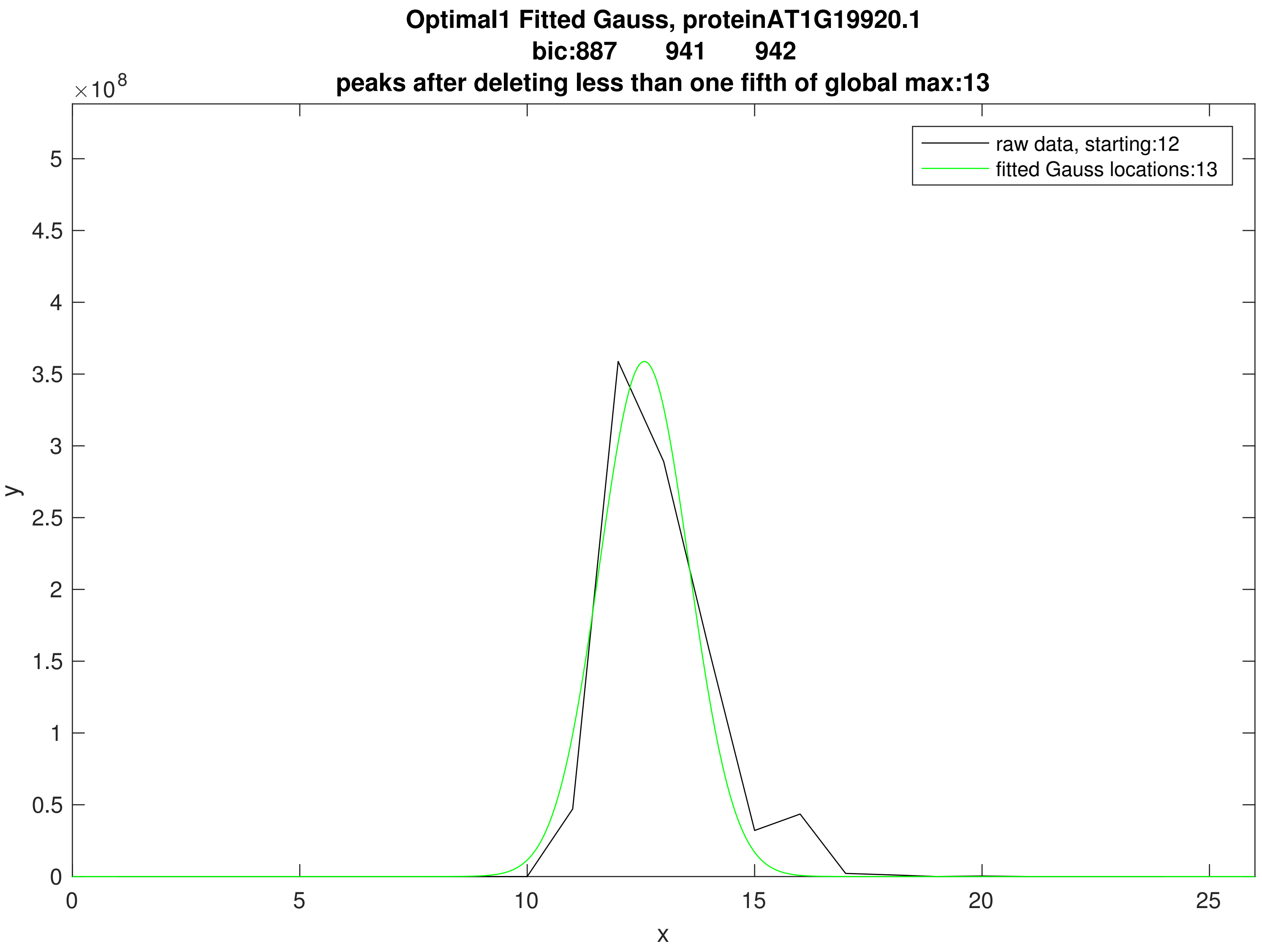
Optimal1 Fitted Gauss, proteinAT1G19920.1
bic:887 941 942
peaks after deleting less than one fifth of global max:13
×10^8
raw data, starting:12
fitted Gauss locations:13
y
x
0
0.5
1
1.5
2
2.5
3
3.5
4
4.5
5
0
5
10
15
20
25

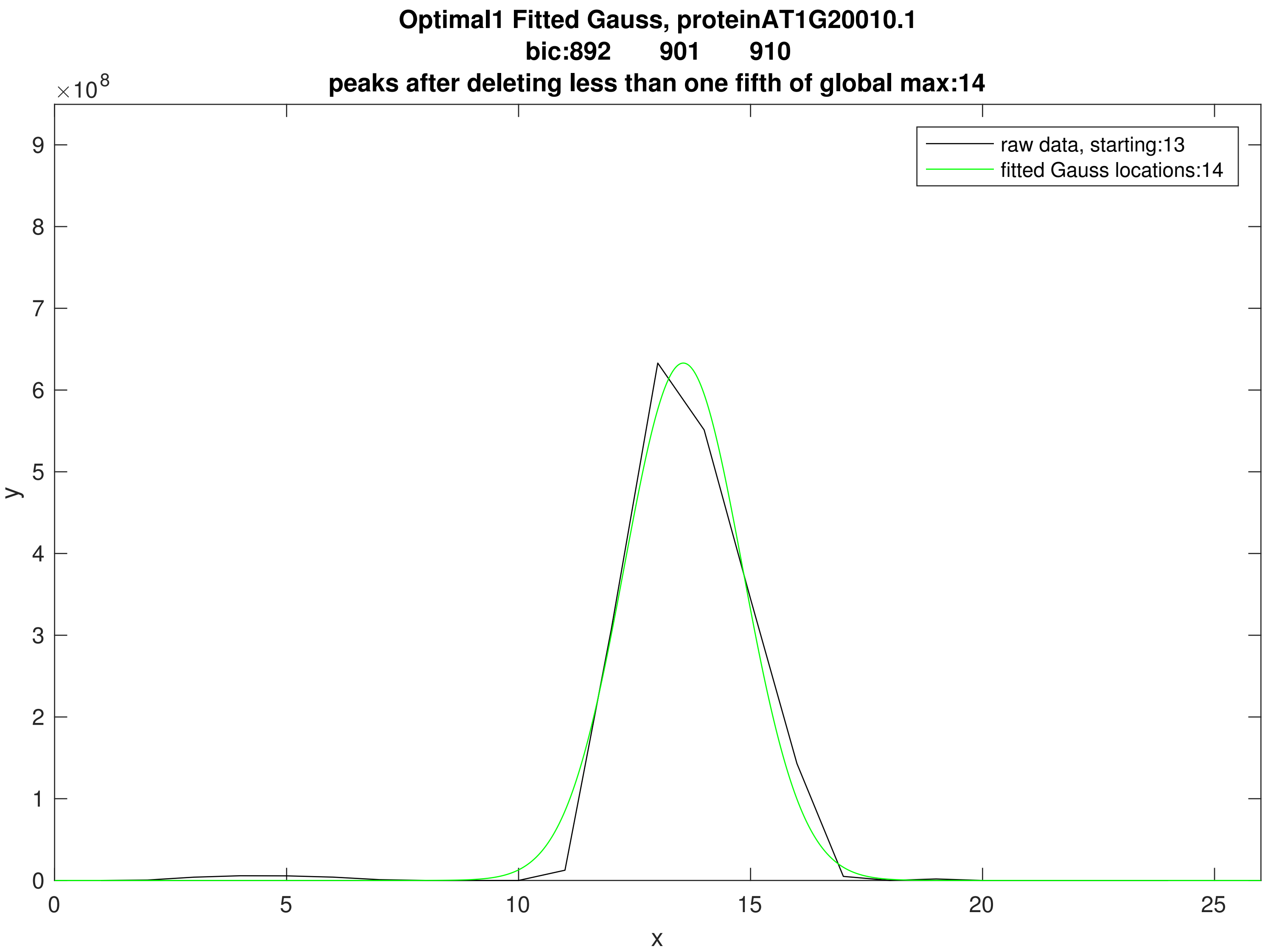
Optimal1 Fitted Gauss, proteinAT1G20010.1
bic:892 901 910
peaks after deleting less than one fifth of global max:14
×10^8
raw data, starting:13
fitted Gauss locations:14
y
x
0
1
2
3
4
5
6
7
8
9
0
5
10
15
20
25

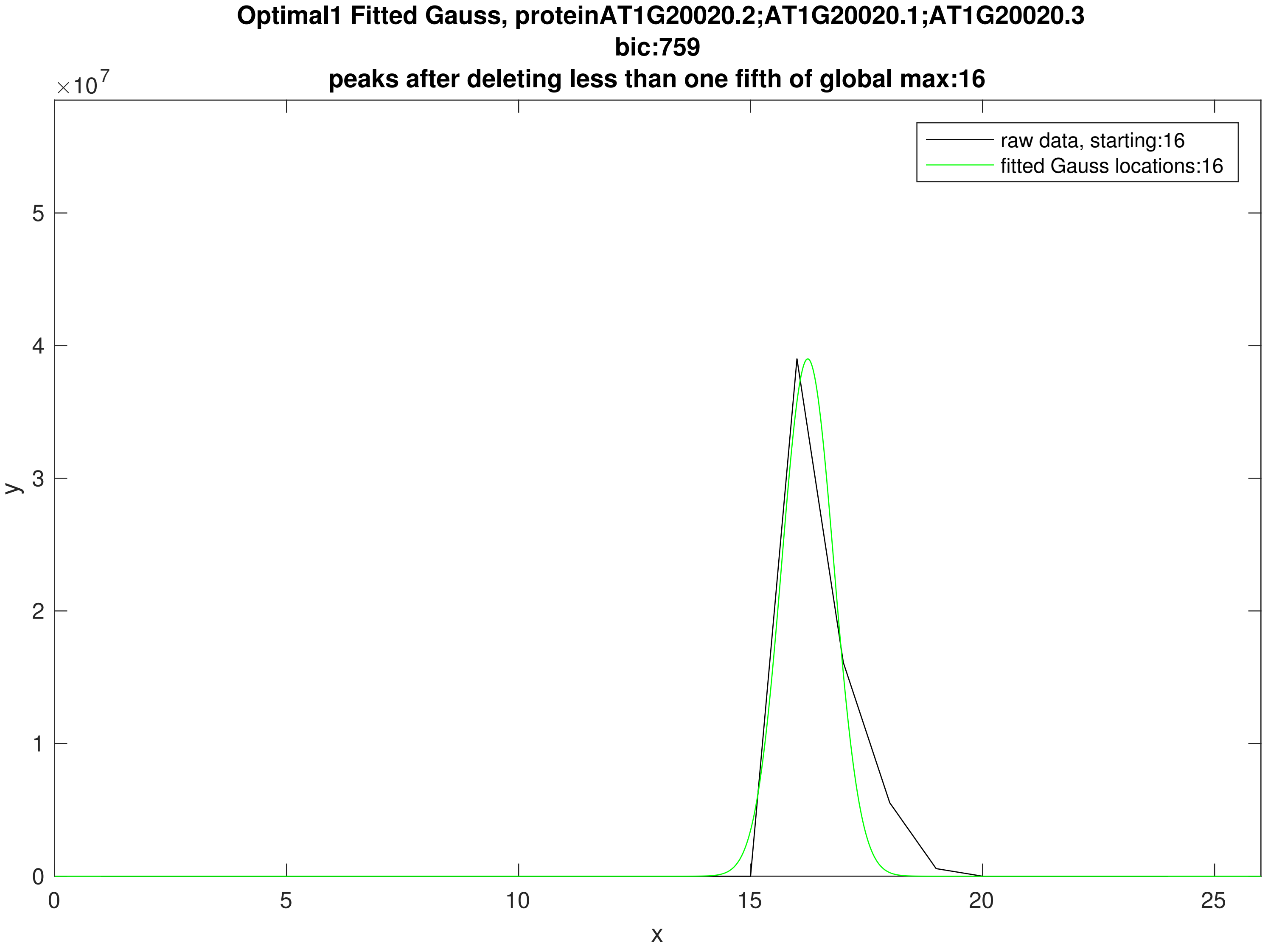
Optimal1 Fitted Gauss, proteinAT1G20020.2;AT1G20020.1;AT1G20020.3
bic:759
peaks after deleting less than one fifth of global max:16
×10^7
raw data, starting:16
fitted Gauss locations:16
y
x
0
1
2
3
4
5
0
5
10
15
20
25

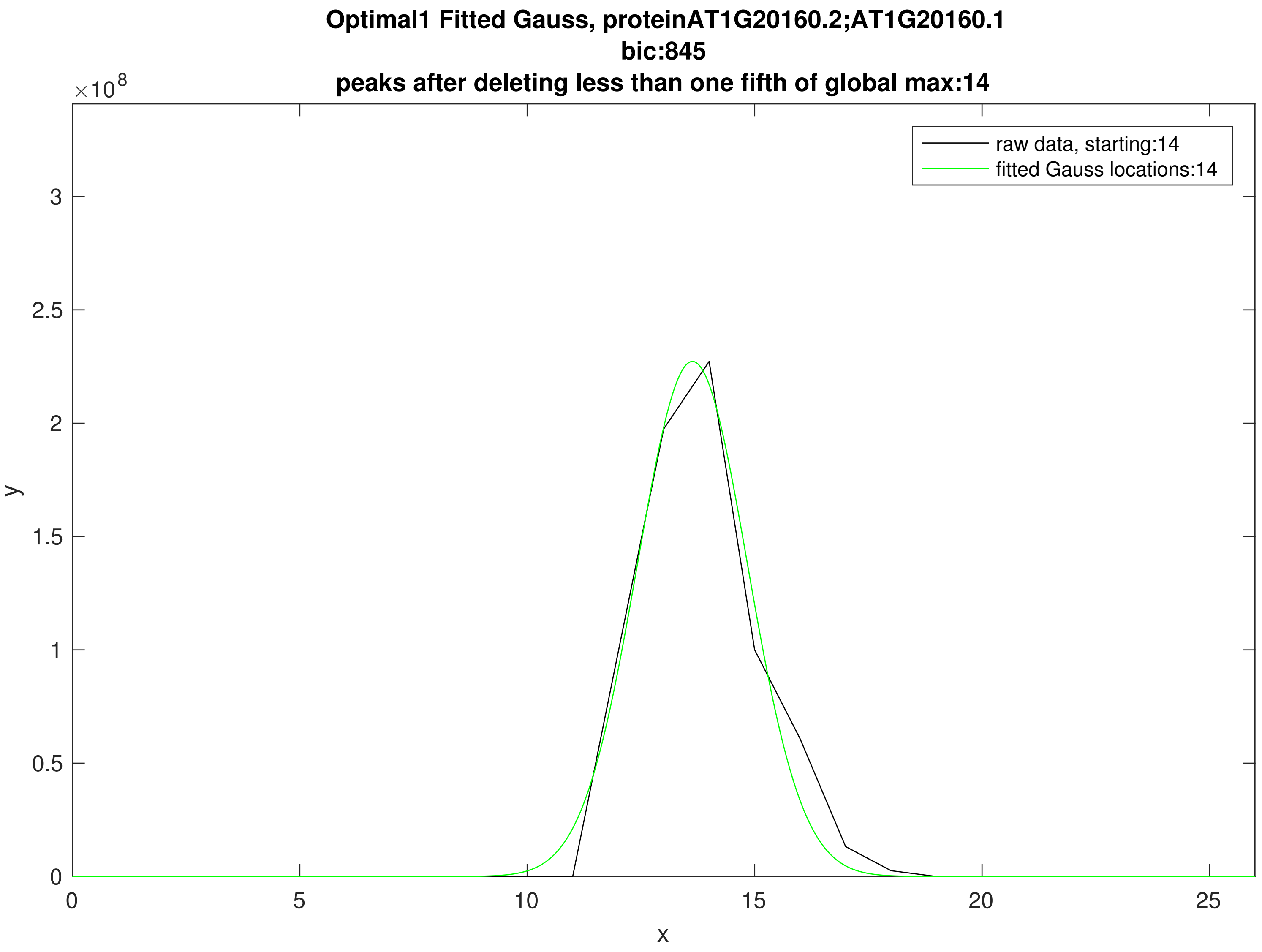

Optimal1 Fitted Gauss, proteinAT1G20160.2;AT1G20160.1
bic:845
peaks after deleting less than one fifth of global max:14
×10^8
raw data, starting:14
fitted Gauss locations:14
y
x
0
0.5
1
1.5
2
2.5
3
0
5
10
15
20
25

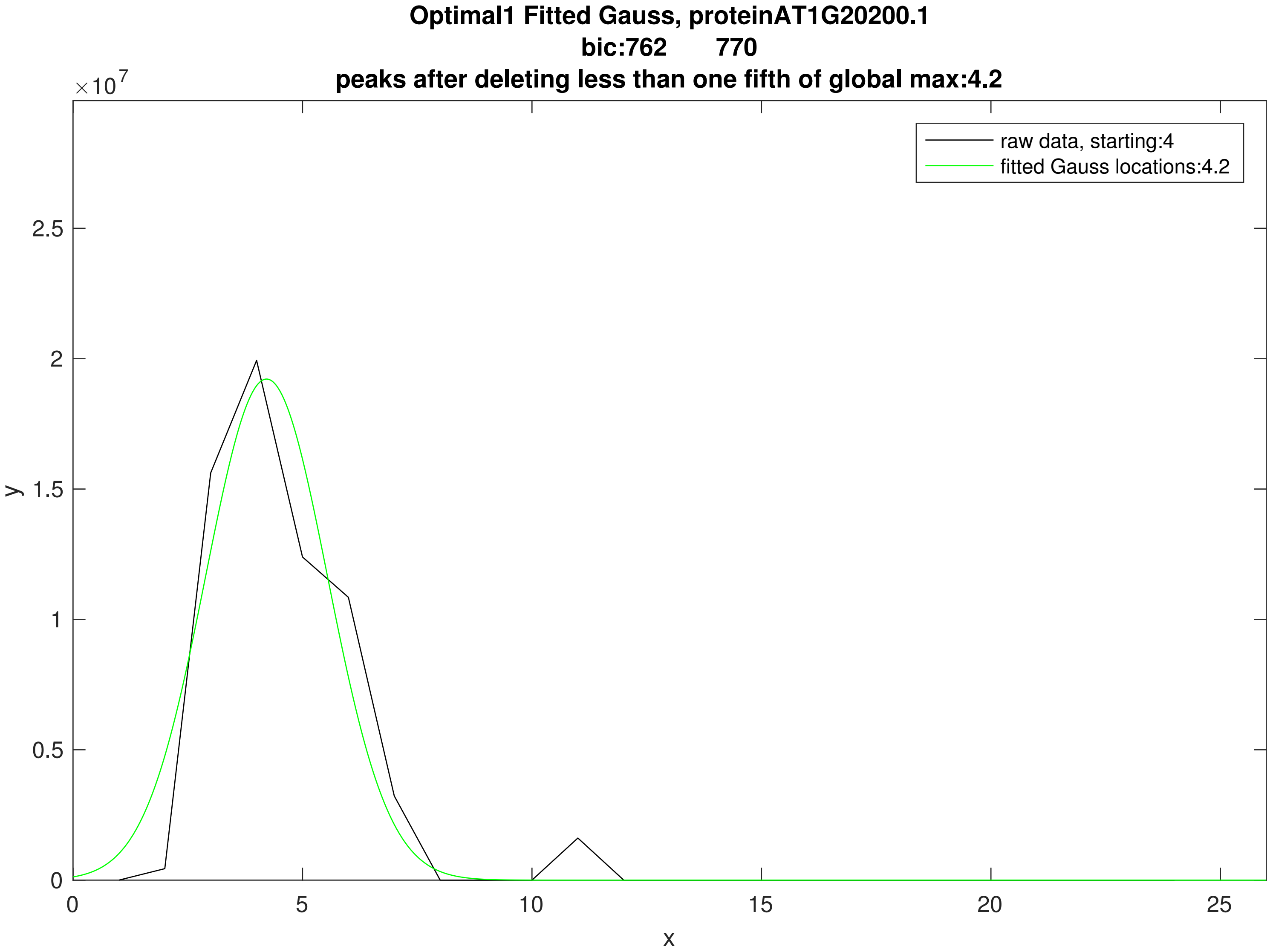
Optimal1 Fitted Gauss, proteinAT1G20200.1
bic:762 770
peaks after deleting less than one fifth of global max:4.2
×10^7
raw data, starting:4
fitted Gauss locations:4.2
2.5
2
1.5
1
0.5
0
y
0
5
10
15
20
25
x

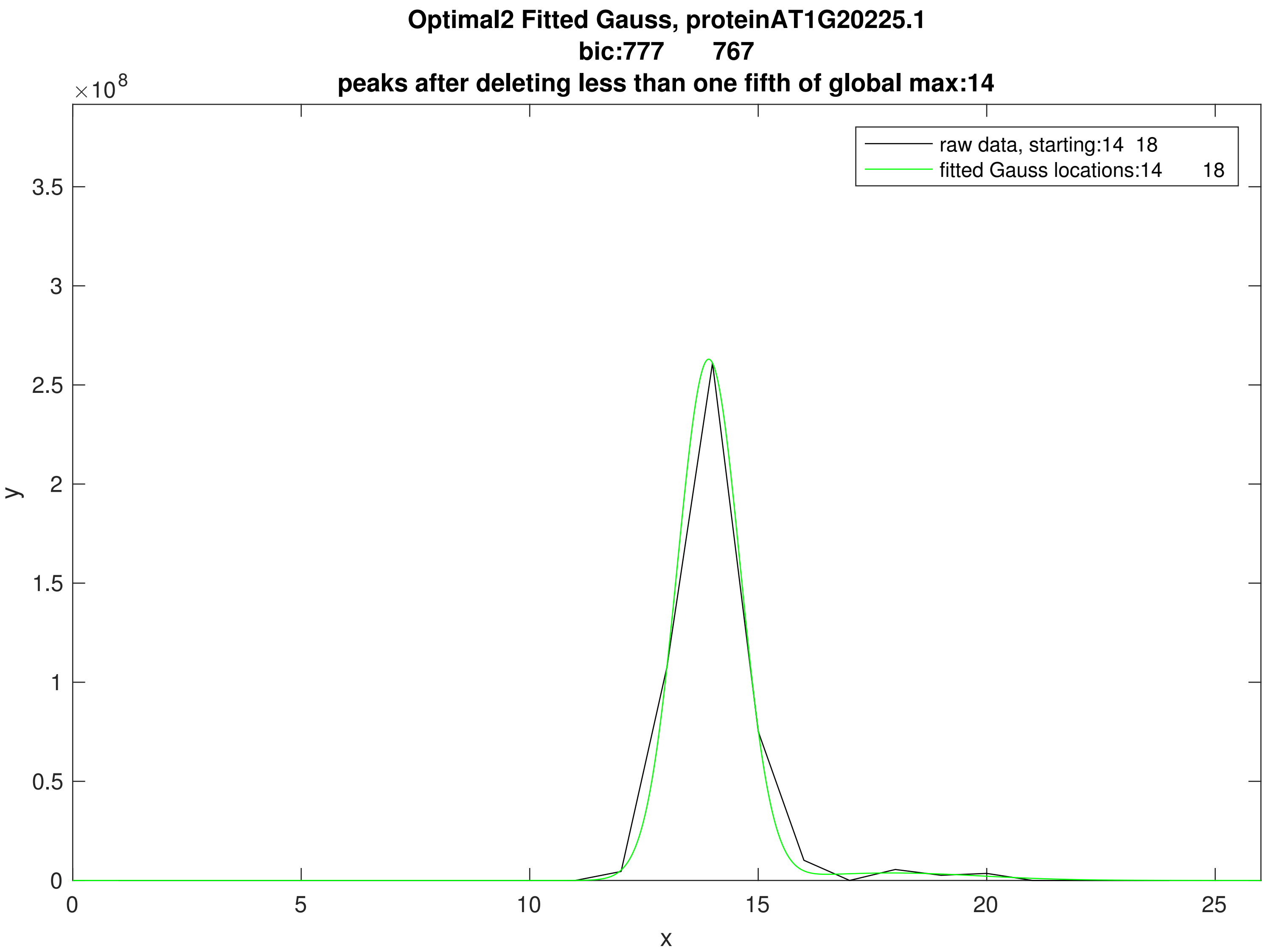

Optimal2 Fitted Gauss, proteinAT1G20225.1
bic:777 767
peaks after deleting less than one fifth of global max:14
×10^8
raw data, starting:14 18
fitted Gauss locations:14 18
3.5
3
2.5
2
1.5
1
0.5
0
y
0
5
10
15
20
25
x

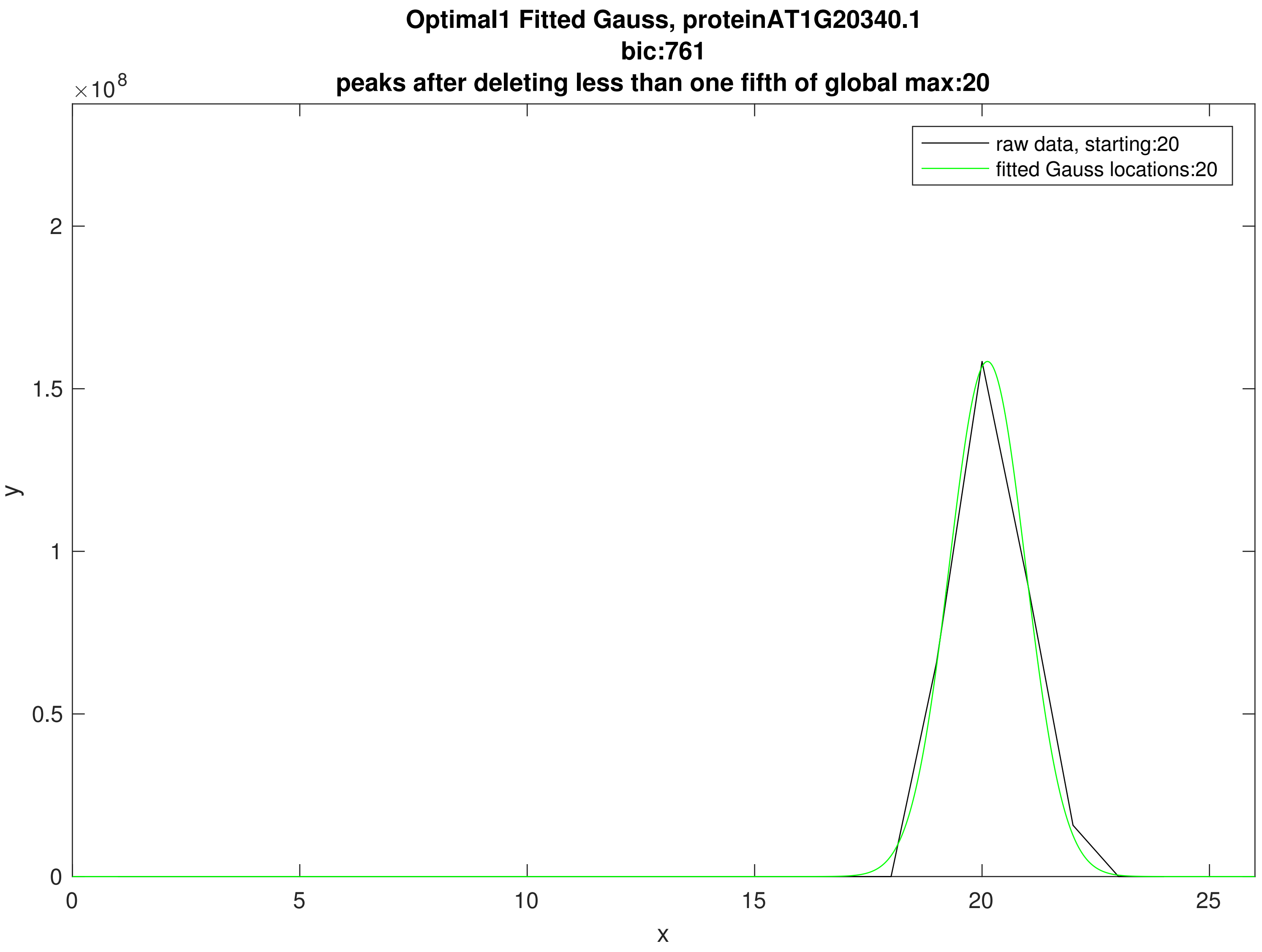
Optimal1 Fitted Gauss, proteinAT1G20340.1
bic:761
peaks after deleting less than one fifth of global max:20
×10^8
raw data, starting:20
fitted Gauss locations:20
y
x
0
0.5
1
1.5
2
0
5
10
15
20
25

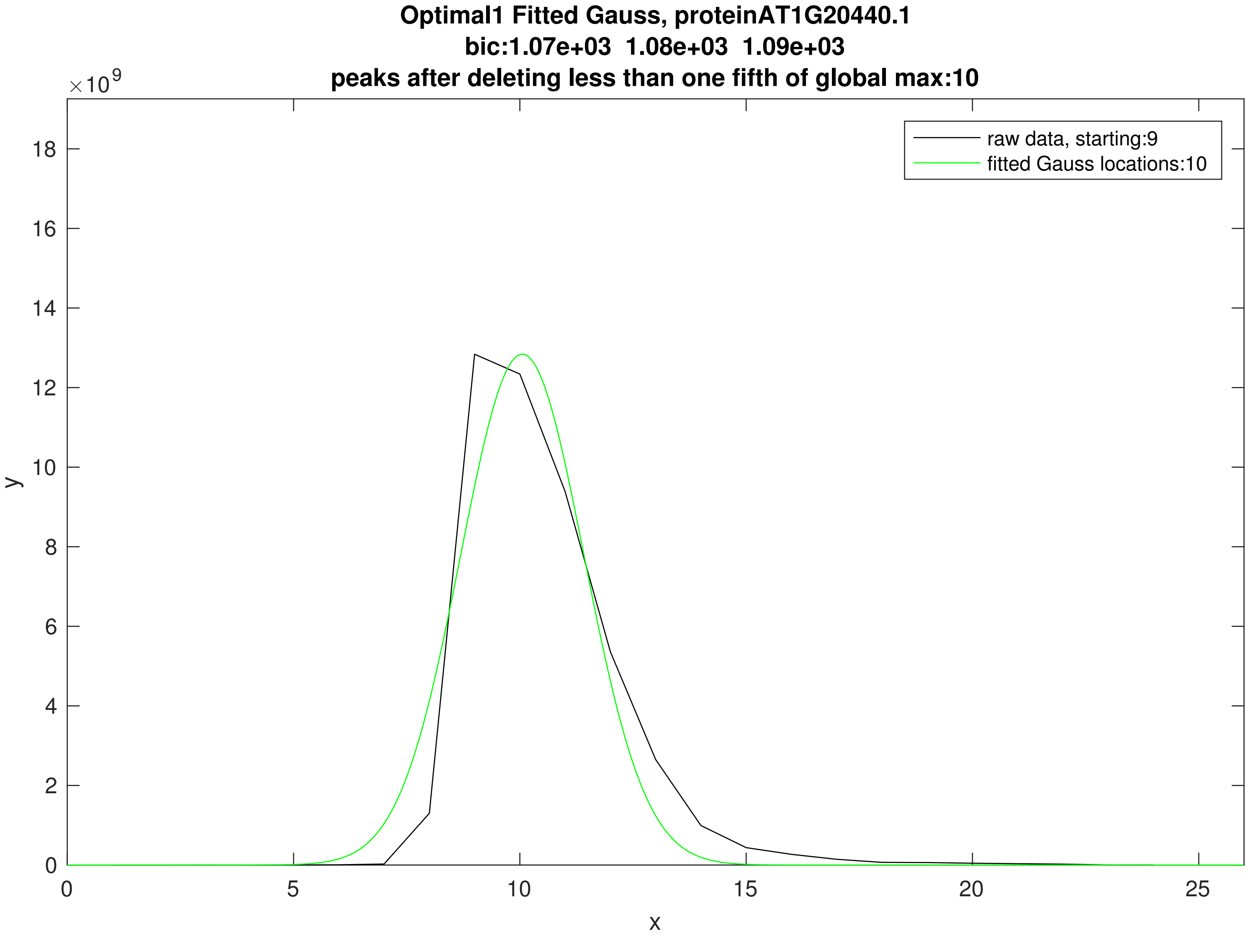

Optimal1 Fitted Gauss, proteinAT1G20440.1
bic:1.07e+03  1.08e+03  1.09e+03
peaks after deleting less than one fifth of global max:10
×10^9
raw data, starting:9
fitted Gauss locations:10
y
x
0
2
4
6
8
10
12
14
16
18
0
5
10
15
20
25

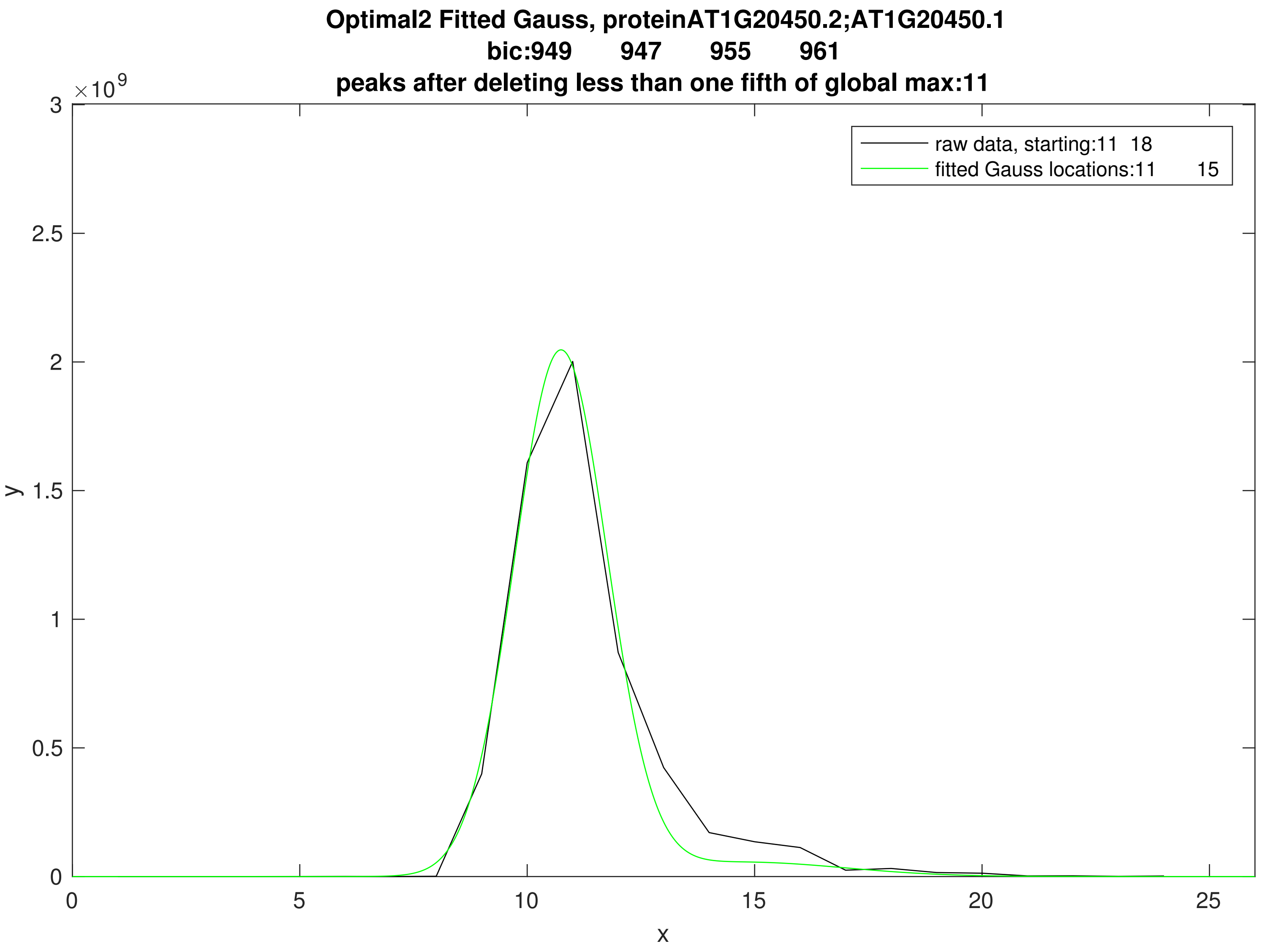
Optimal2 Fitted Gauss, proteinAT1G20450.2;AT1G20450.1
bic:949 947 955 961
peaks after deleting less than one fifth of global max:11
raw data, starting:11 18
fitted Gauss locations:11 15
$\times 10^9$
y
x
0
0.5
1
1.5
2
2.5
3
5
10
15
20
25

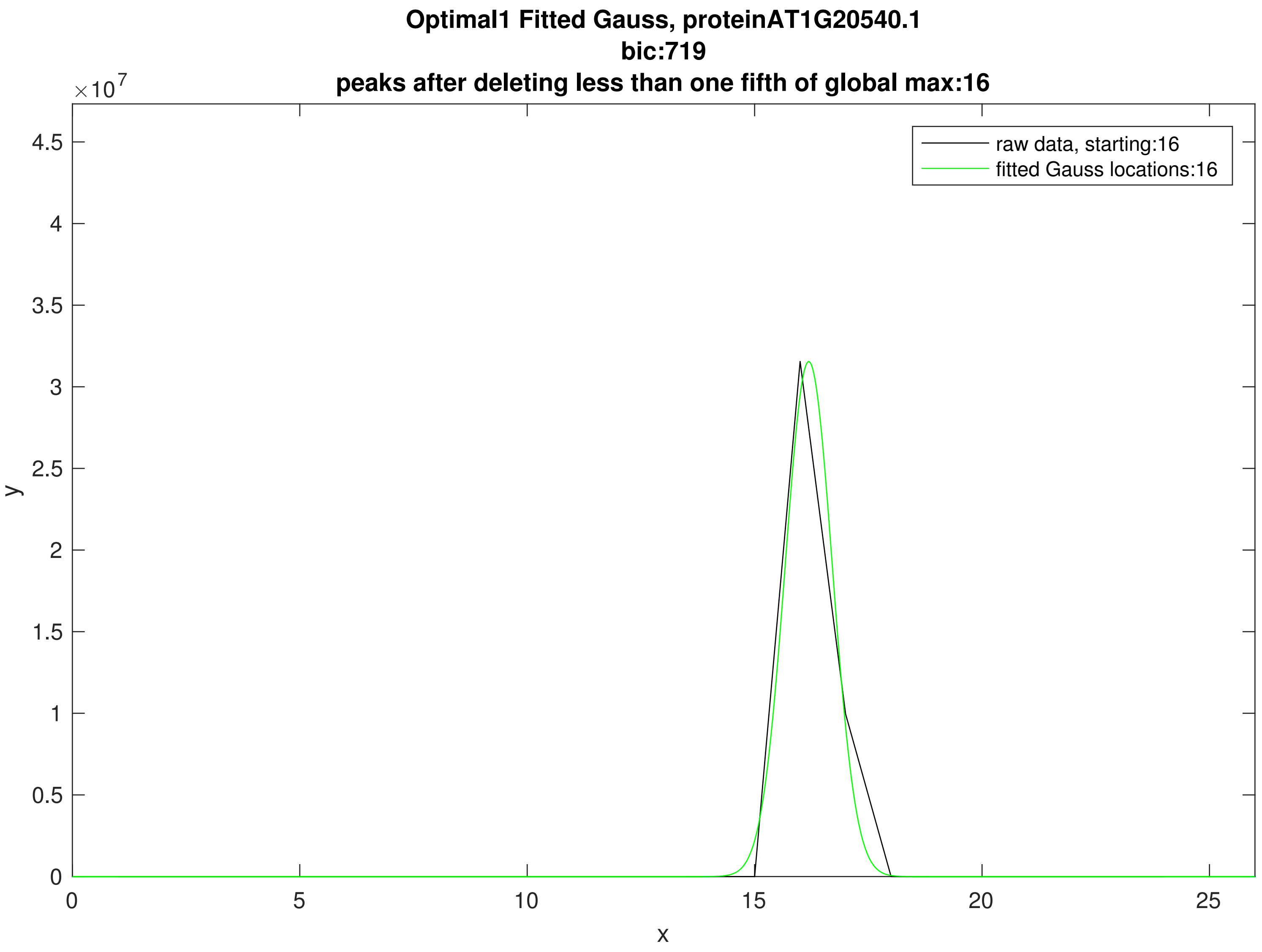
Optimal1 Fitted Gauss, proteinAT1G20540.1
bic:719
peaks after deleting less than one fifth of global max:16
×10^7
4.5
4
3.5
3
2.5
2
1.5
1
0.5
0
y
0
5
10
15
20
25
x
raw data, starting:16
fitted Gauss locations:16

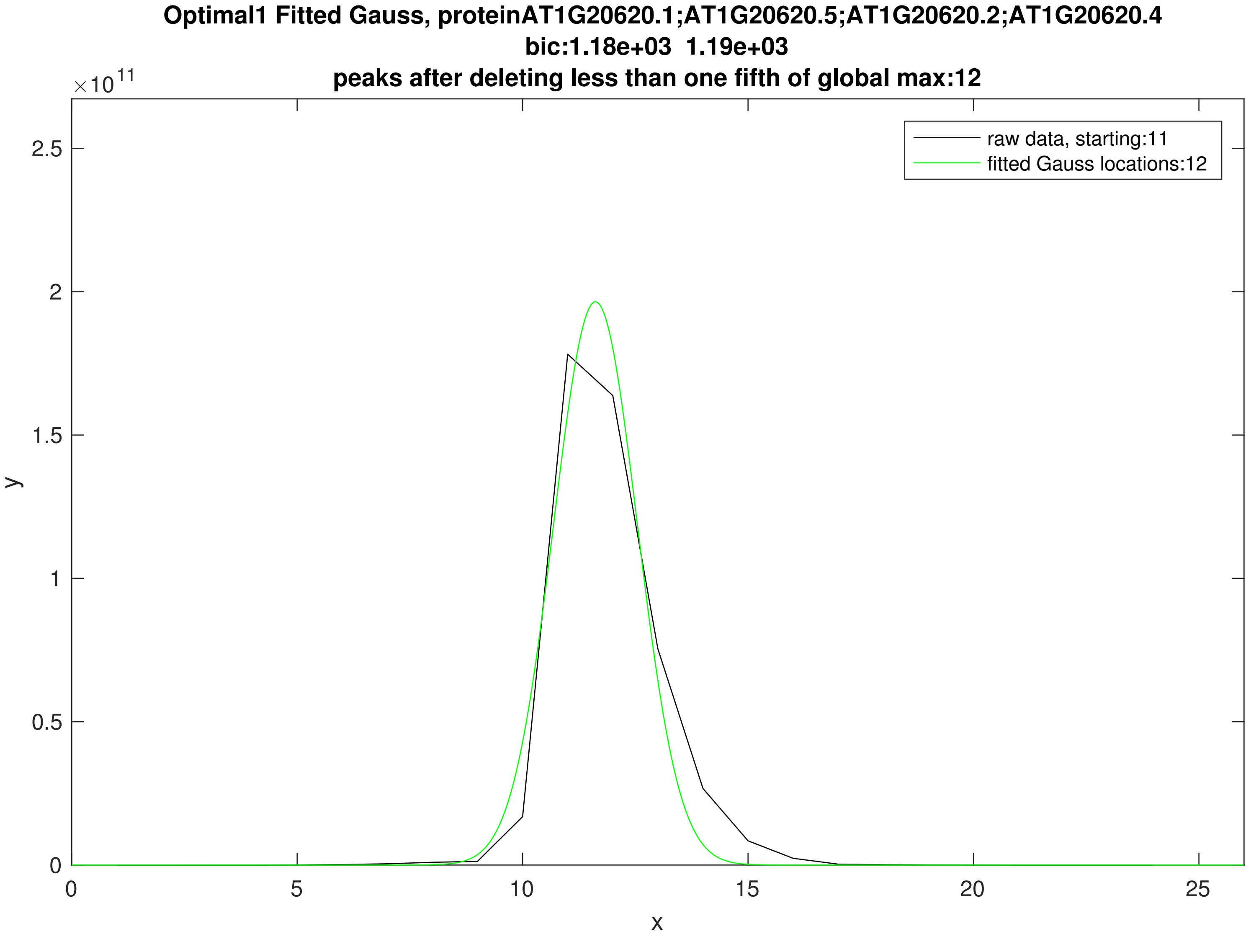

Optimal1 Fitted Gauss, proteinAT1G20620.1;AT1G20620.5;AT1G20620.2;AT1G20620.4
bic:1.18e+03  1.19e+03
peaks after deleting less than one fifth of global max:12
×10^11
2.5
2
1.5
1
0.5
0
y
0
5
10
15
20
25
x
raw data, starting:11
fitted Gauss locations:12

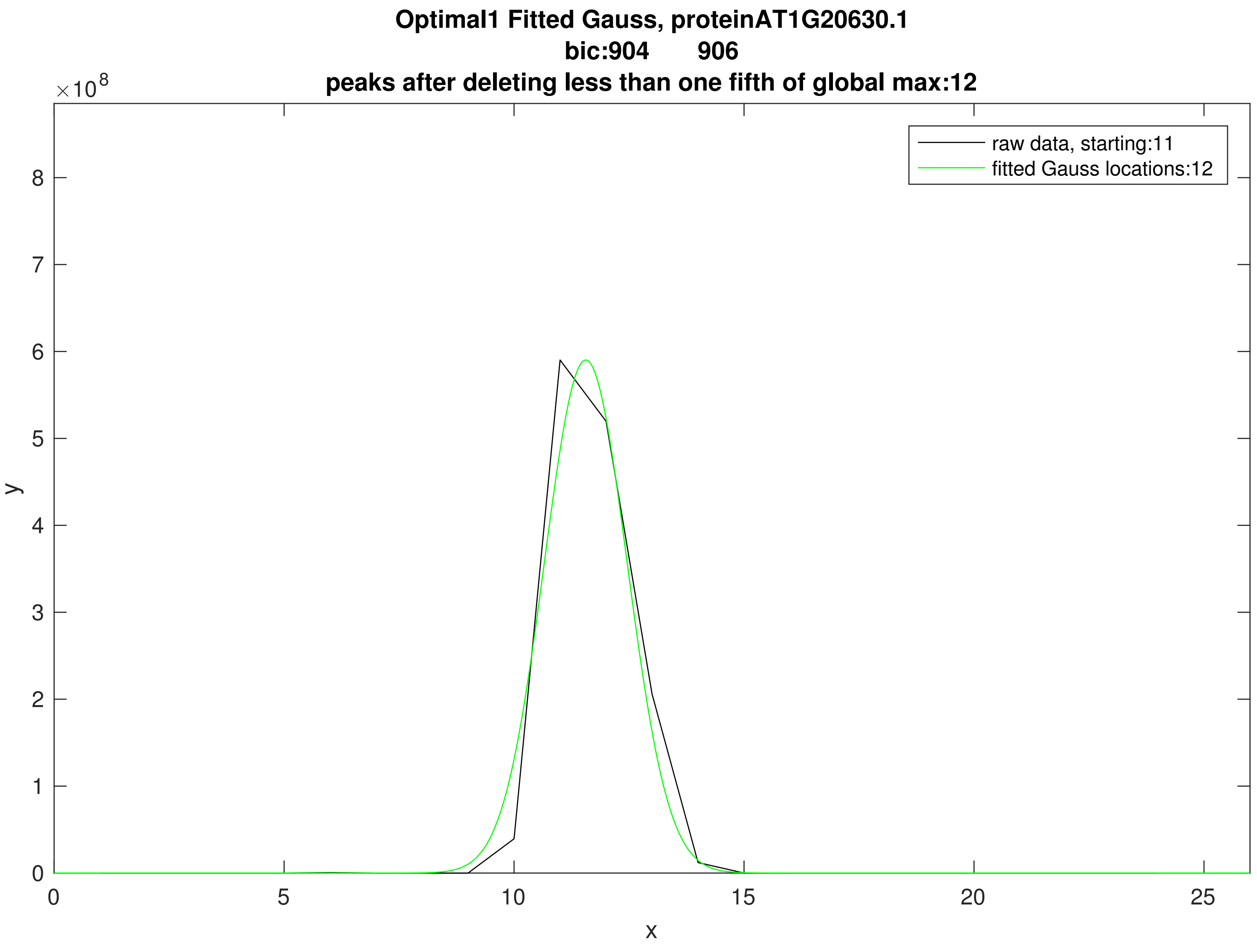

Optimal1 Fitted Gauss, proteinAT1G20630.1
bic:904 906
peaks after deleting less than one fifth of global max:12
×10^8
raw data, starting:11
fitted Gauss locations:12
y
x
0
1
2
3
4
5
6
7
8
0
5
10
15
20
25

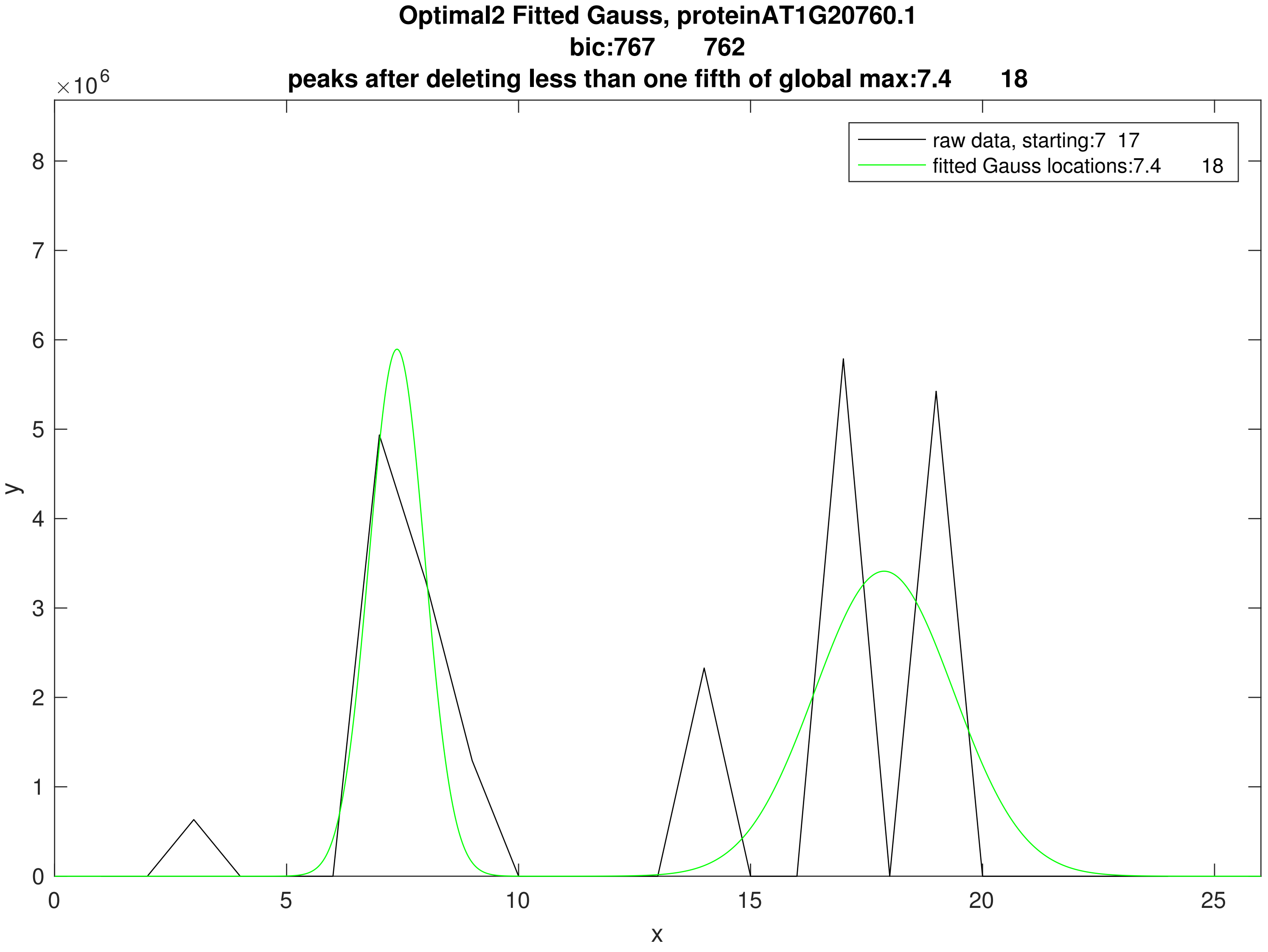
Optimal2 Fitted Gauss, proteinAT1G20760.1
bic:767 762
peaks after deleting less than one fifth of global max:7.4 18
×10^6
raw data, starting:7 17
fitted Gauss locations:7.4 18
y
x
0
1
2
3
4
5
6
7
8
0
5
10
15
20
25

**Optimal1 Fitted Gauss, proteinAT1G20950.1**
**bic:987 996**
**peaks after deleting less than one fifth of global max:8.6**

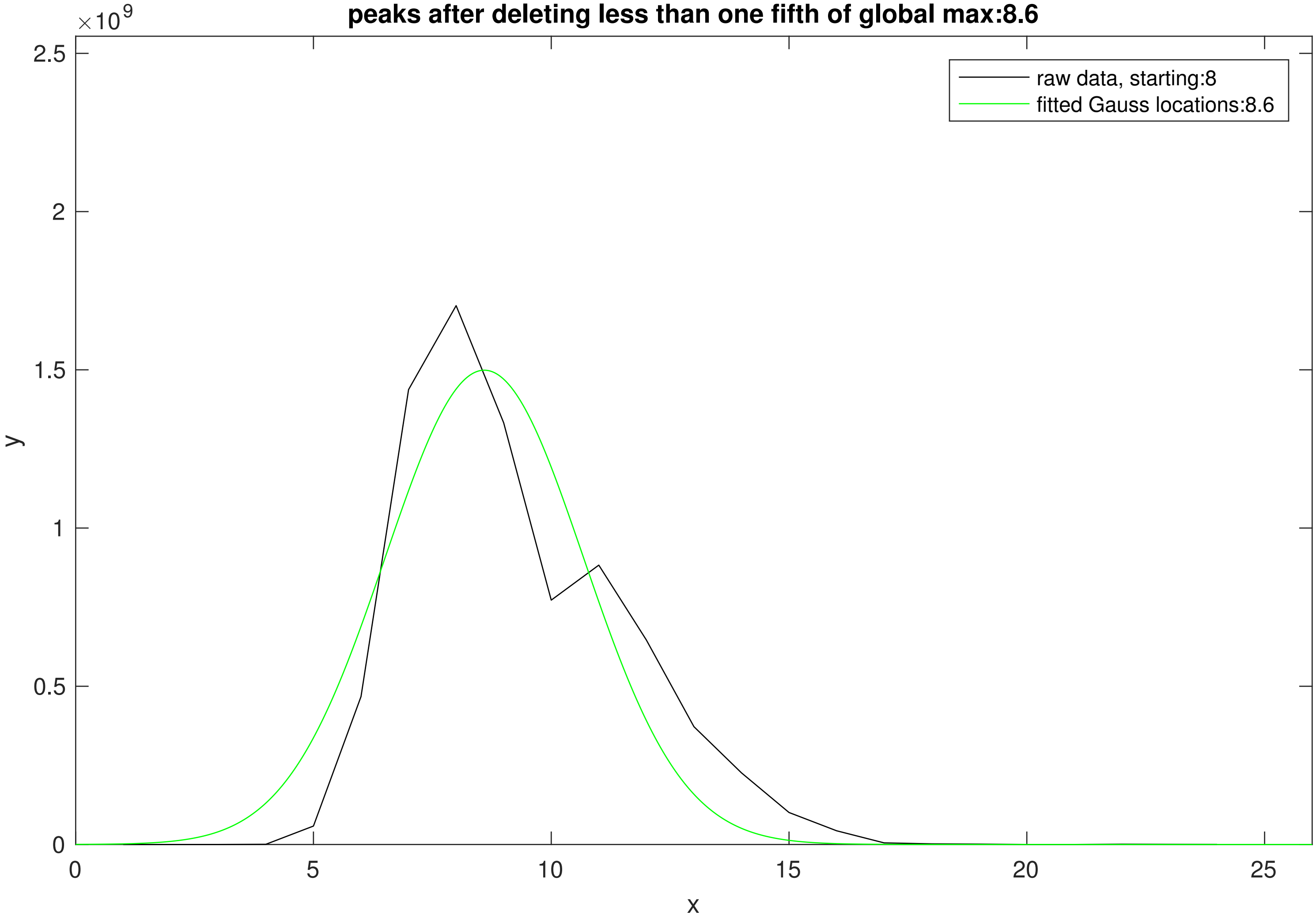

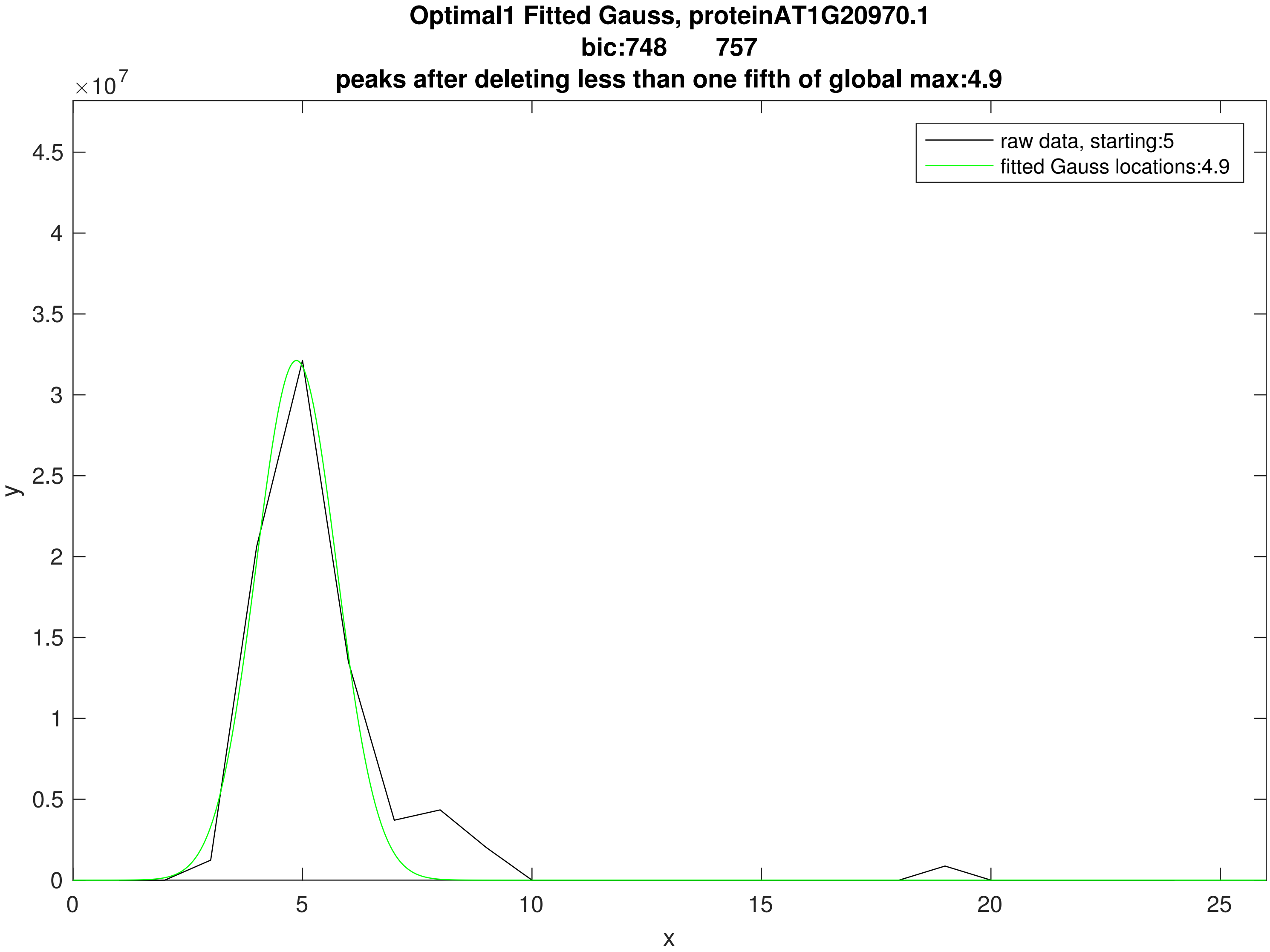

Optimal1 Fitted Gauss, proteinAT1G20970.1
bic:748 757
peaks after deleting less than one fifth of global max:4.9
×10^7
4.5
4
3.5
3
2.5
2
1.5
1
0.5
0
y
0
5
10
15
20
25
x
raw data, starting:5
fitted Gauss locations:4.9

**Optimal1 Fitted Gauss, proteinAT1G21065.1**
**bic:776**
**peaks after deleting less than one fifth of global max:17**

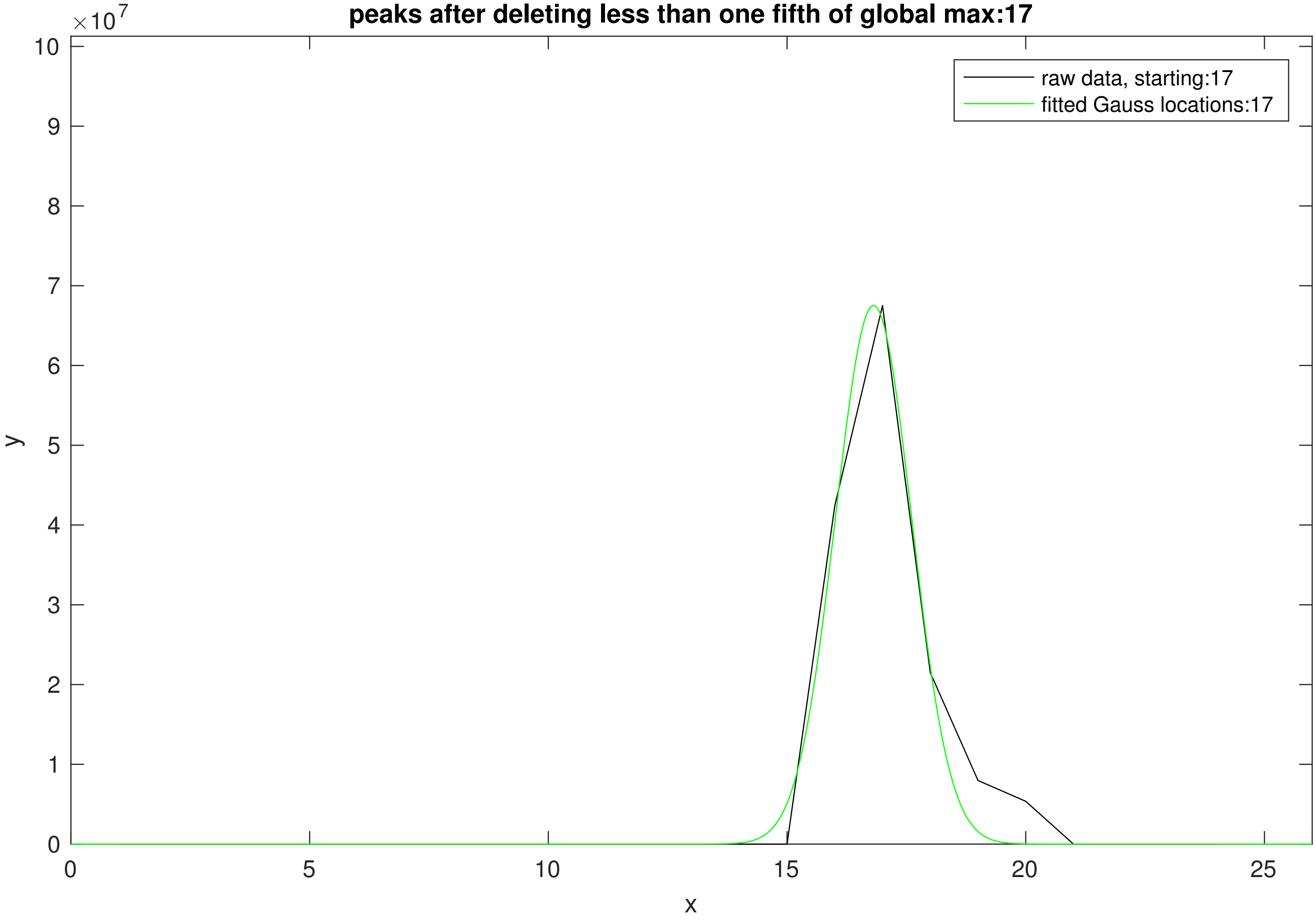

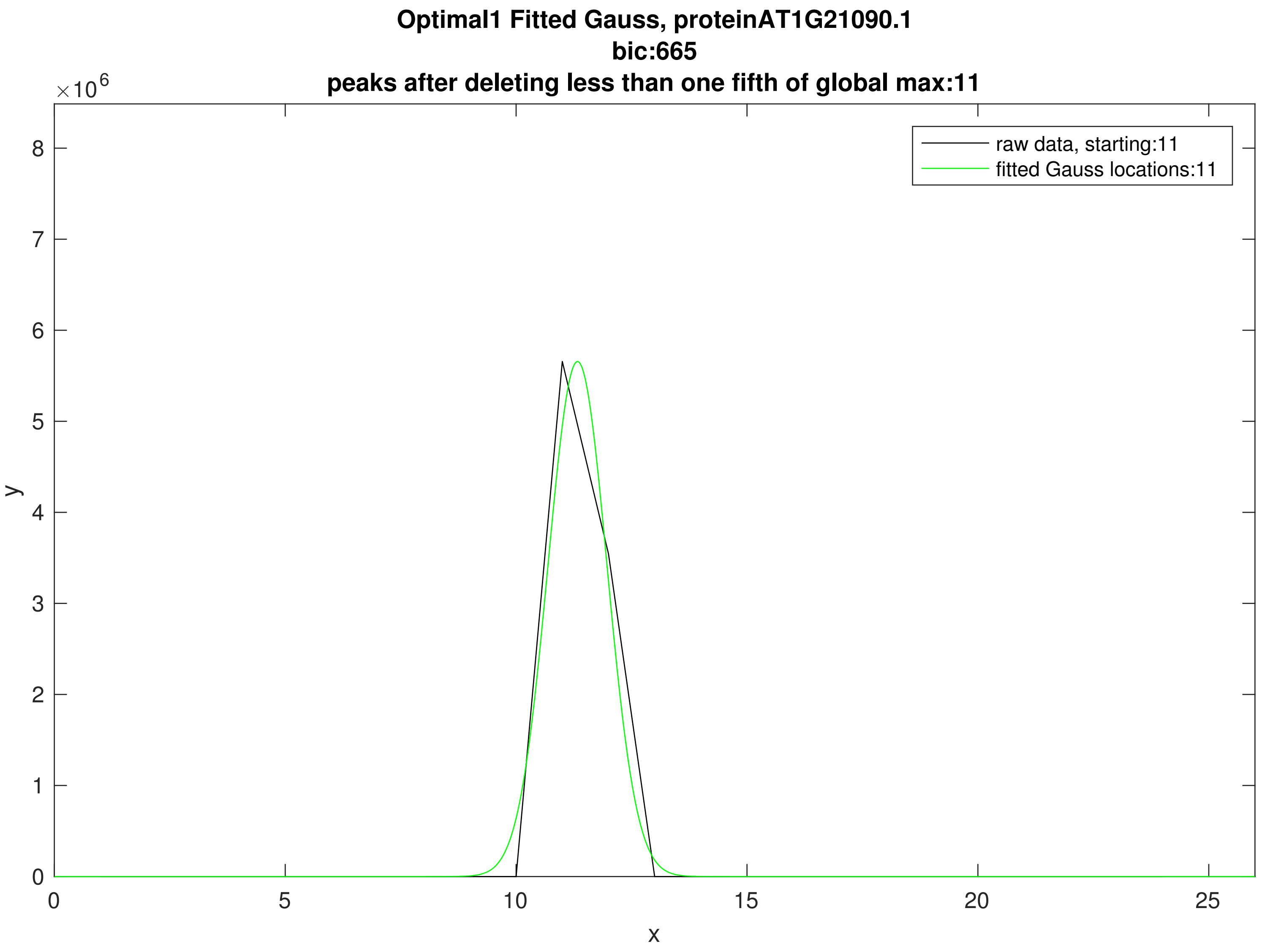
Optimal1 Fitted Gauss, proteinAT1G21090.1
bic:665
peaks after deleting less than one fifth of global max:11
×10^6
raw data, starting:11
fitted Gauss locations:11
y
x
0
1
2
3
4
5
6
7
8
0
5
10
15
20
25

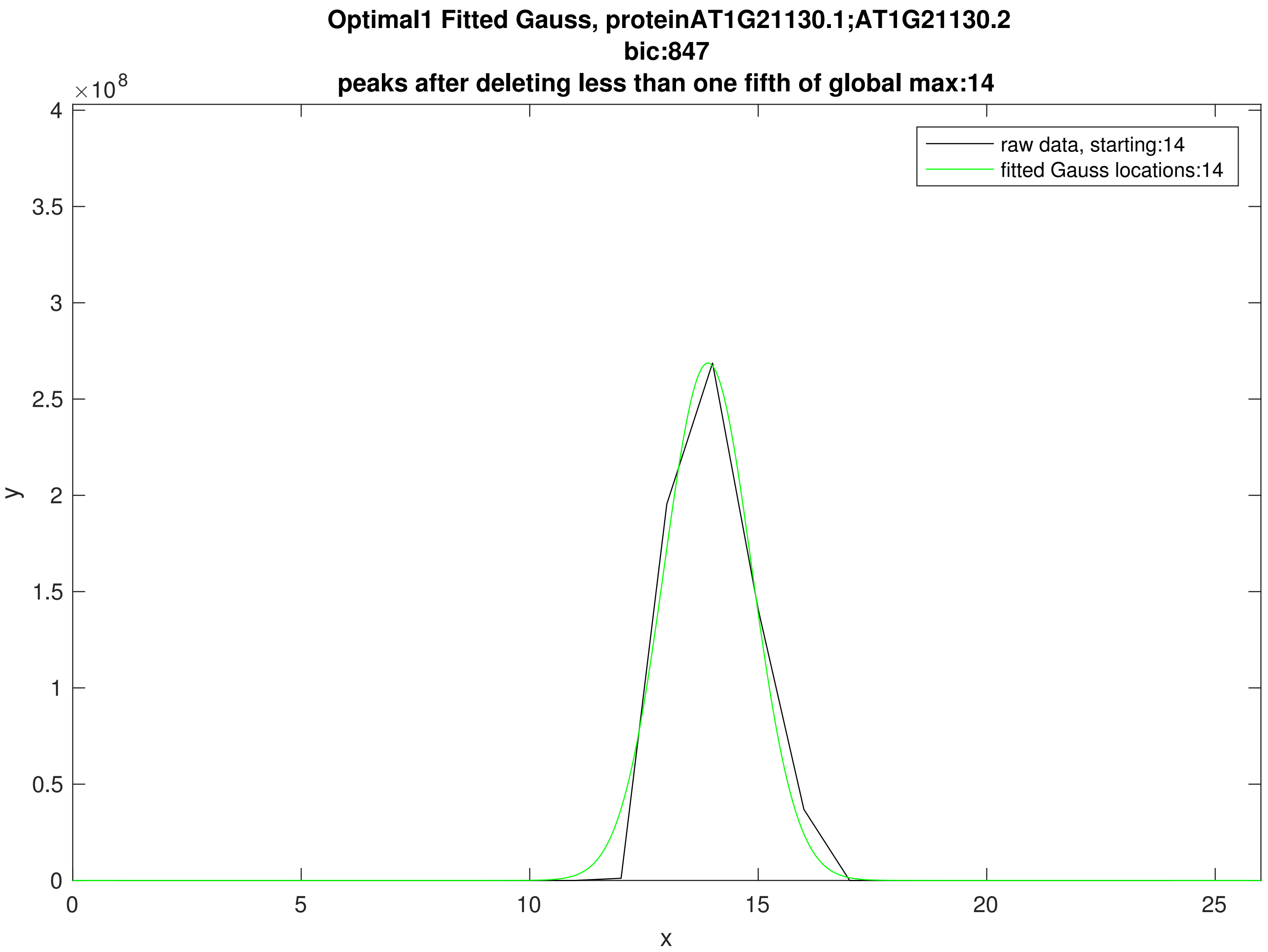
Optimal1 Fitted Gauss, proteinAT1G21130.1;AT1G21130.2
bic:847
peaks after deleting less than one fifth of global max:14
×10^8
4
3.5
3
2.5
2
1.5
1
0.5
0
y
0
5
10
15
20
25
x
raw data, starting:14
fitted Gauss locations:14

Optimal1 Fitted Gauss, proteinAT1G21350.1;AT1G21350.2;AT1G21350.3
bic:706
peaks after deleting less than one fifth of global max:19

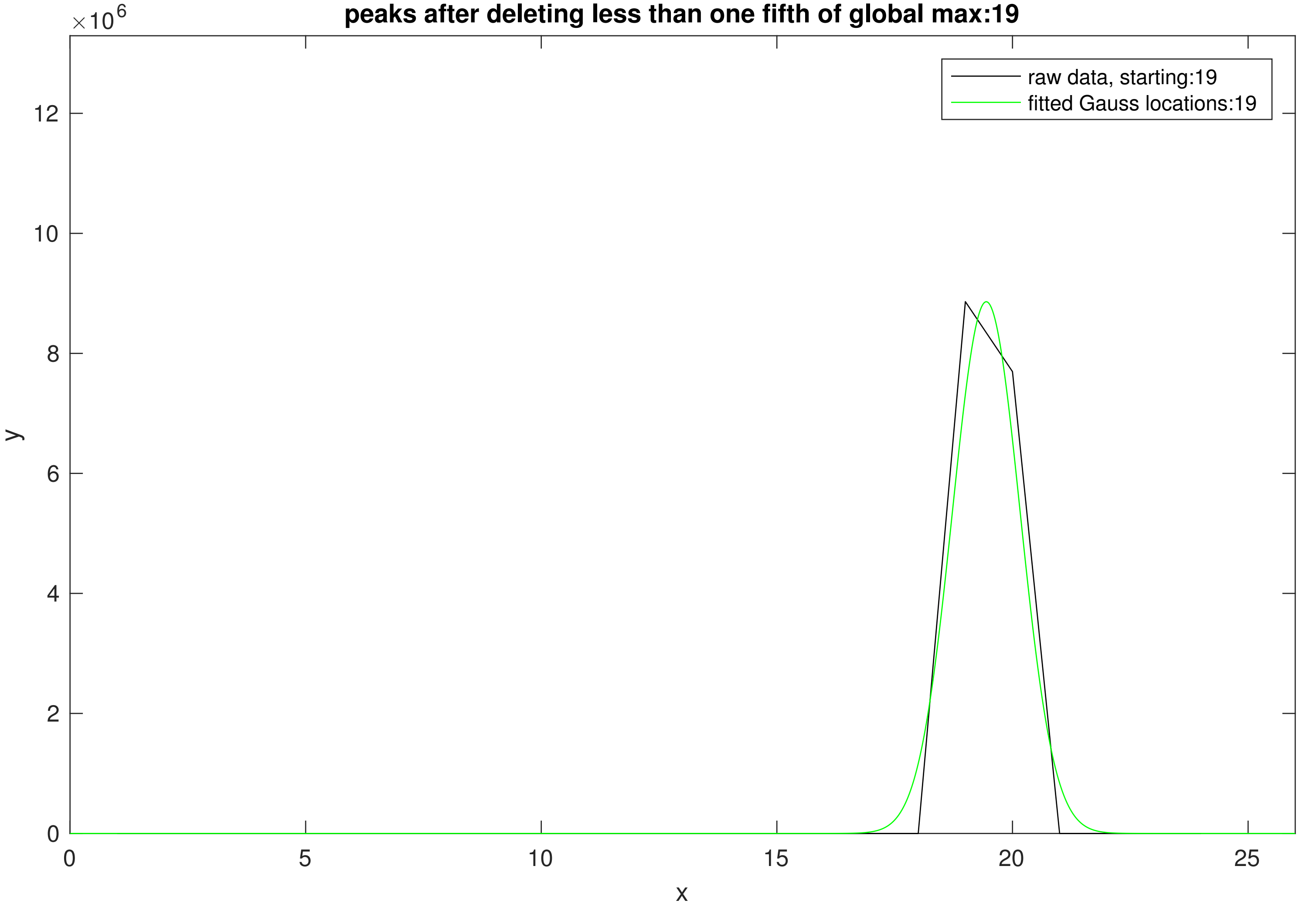

#### Optimal1 Fitted Gauss, proteinAT1G21400.1
#### bic:704
#### peaks after deleting less than one fifth of global max:12

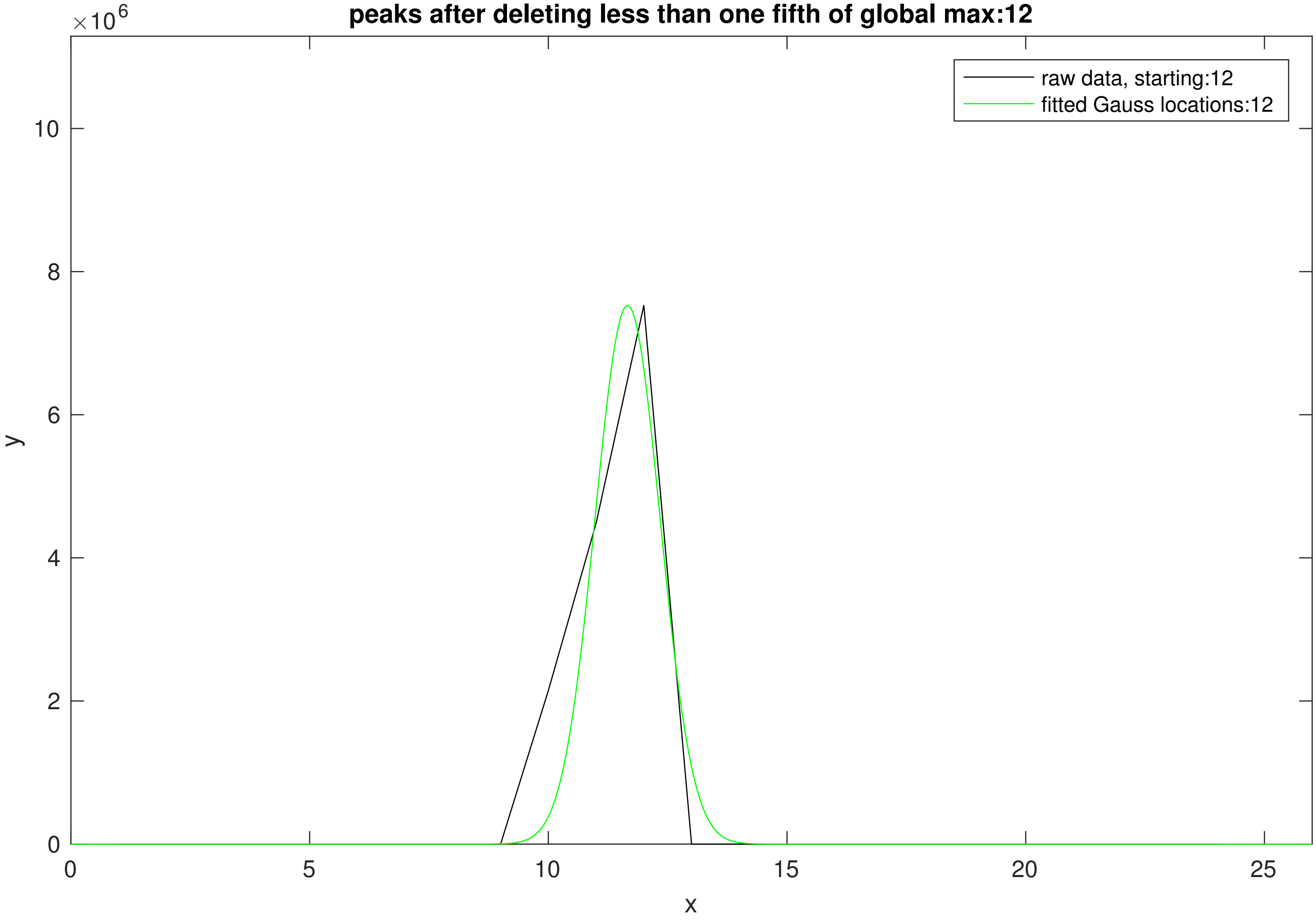

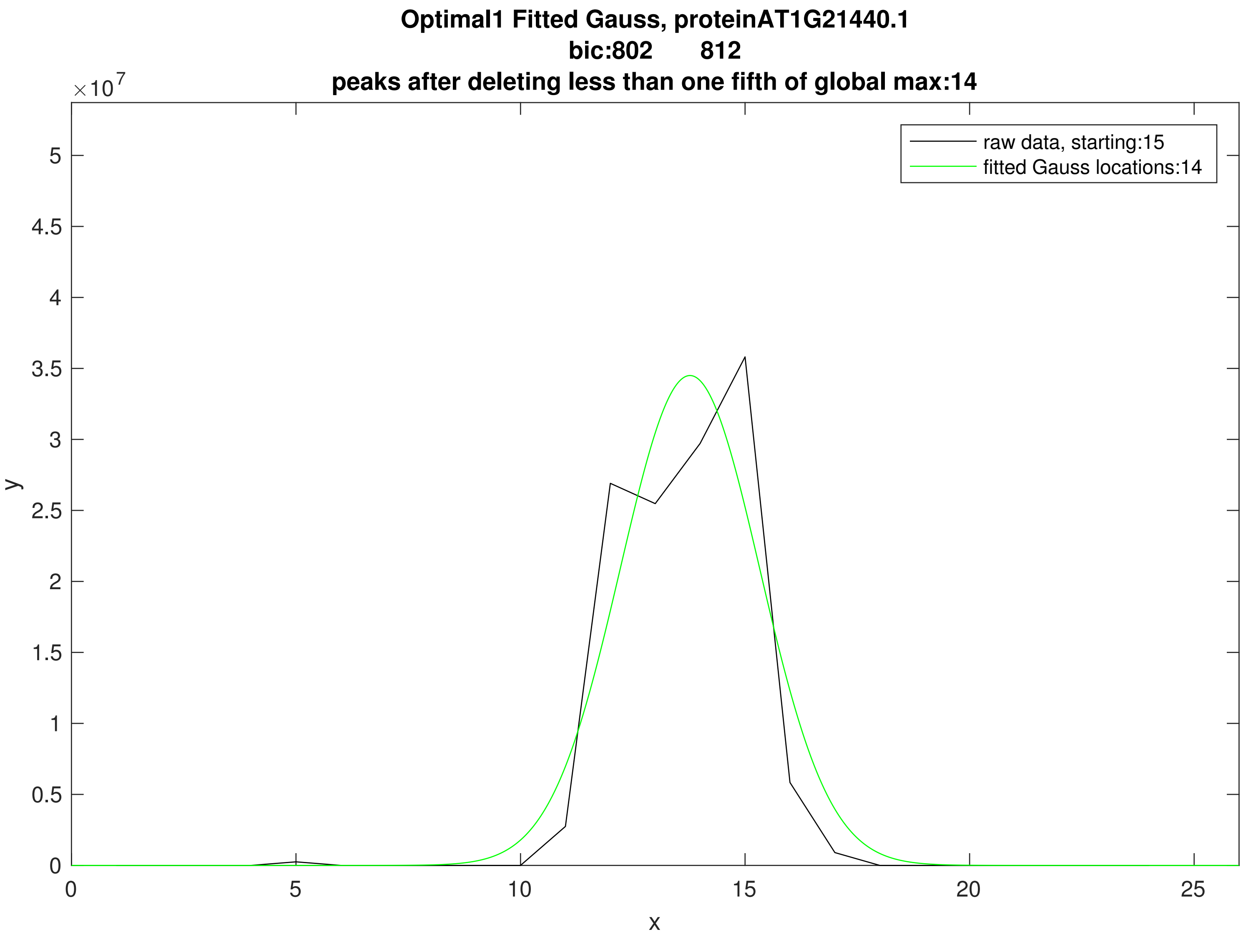

Optimal1 Fitted Gauss, proteinAT1G21440.1
bic:802 812
peaks after deleting less than one fifth of global max:14
×10^7
raw data, starting:15
fitted Gauss locations:14
y
x
0
0.5
1
1.5
2
2.5
3
3.5
4
4.5
5
0
5
10
15
20
25

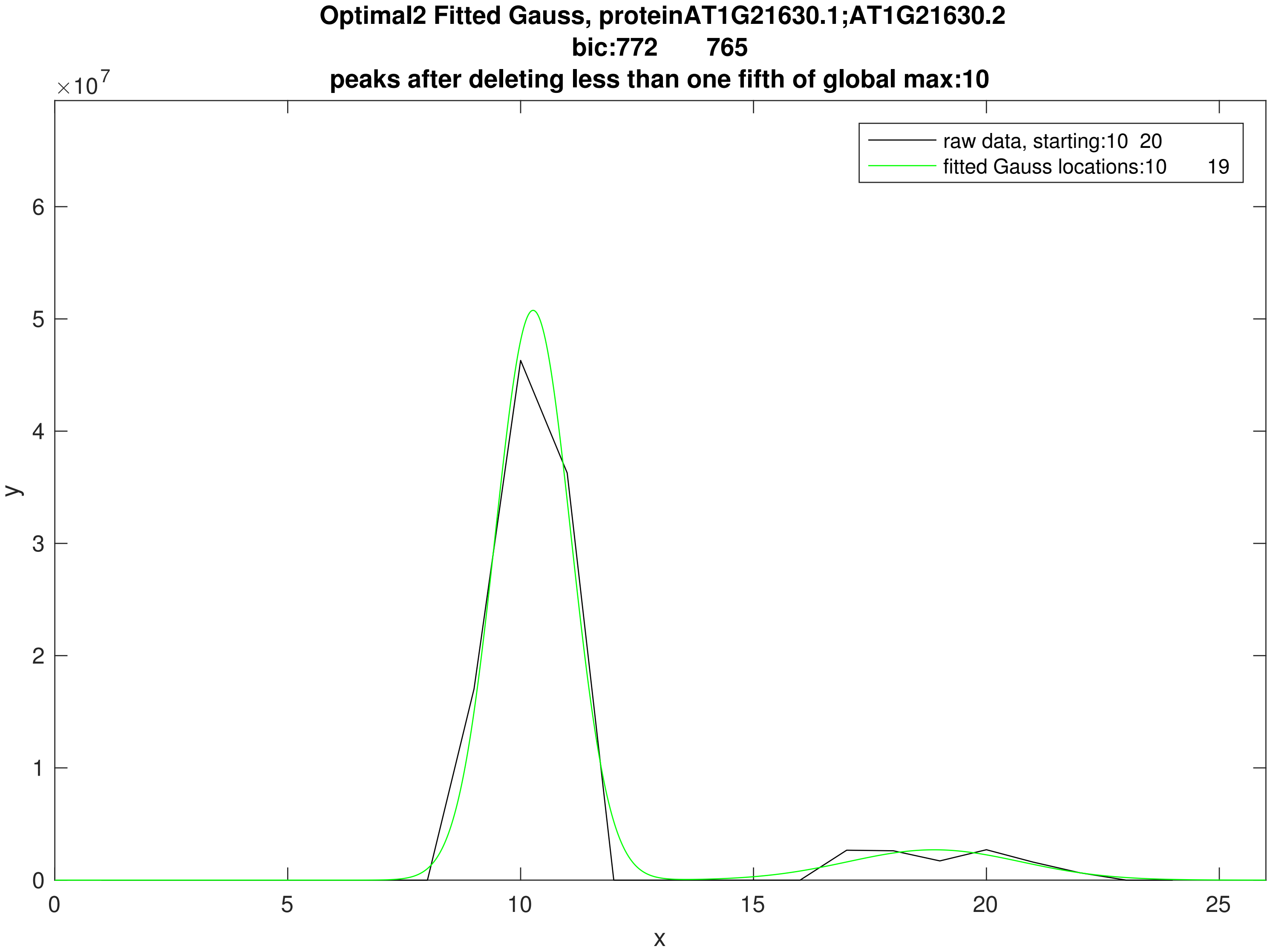

Optimal2 Fitted Gauss, proteinAT1G21630.1;AT1G21630.2
bic:772 765
peaks after deleting less than one fifth of global max:10
×10^7
raw data, starting:10 20
fitted Gauss locations:10 19
y
x
0
1
2
3
4
5
6
0
5
10
15
20
25

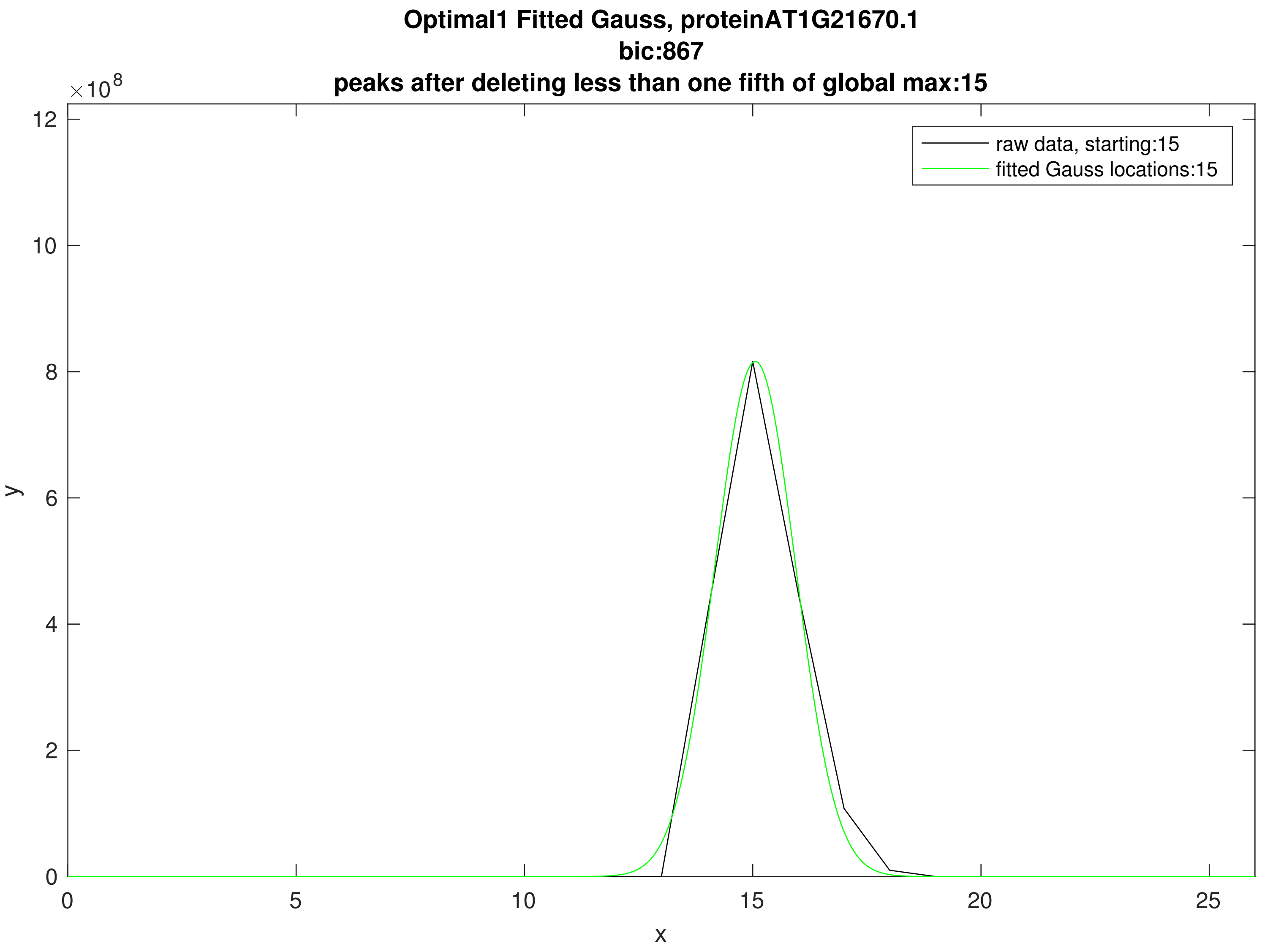
Optimal1 Fitted Gauss, proteinAT1G21670.1
bic:867
peaks after deleting less than one fifth of global max:15
×10^8
12
10
8
6
4
2
0
y
0
5
10
15
20
25
x
raw data, starting:15
fitted Gauss locations:15

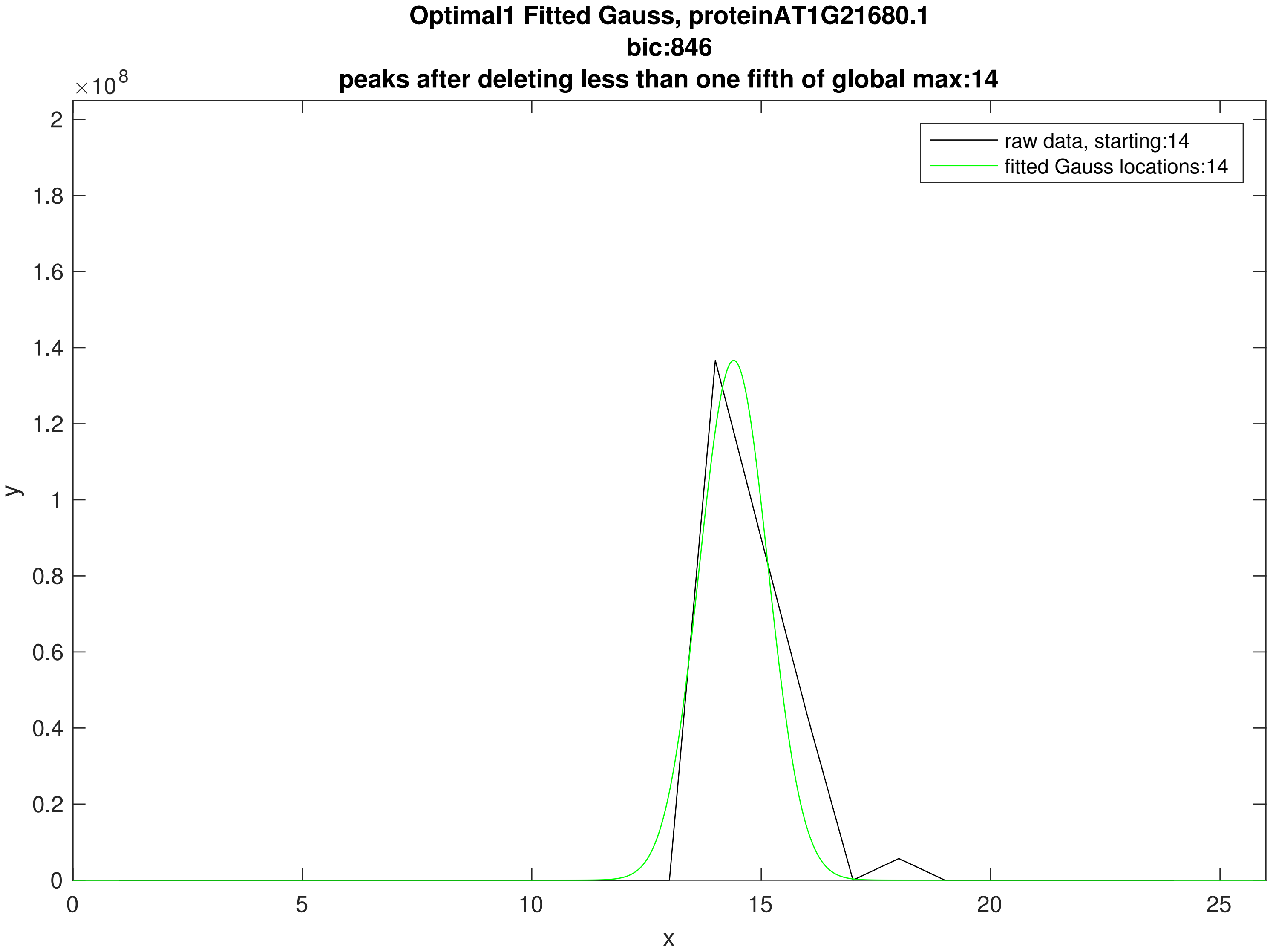

Optimal1 Fitted Gauss, proteinAT1G21680.1
bic:846
peaks after deleting less than one fifth of global max:14
×10^8
raw data, starting:14
fitted Gauss locations:14
y
x
0
0.2
0.4
0.6
0.8
1
1.2
1.4
1.6
1.8
2
0
5
10
15
20
25

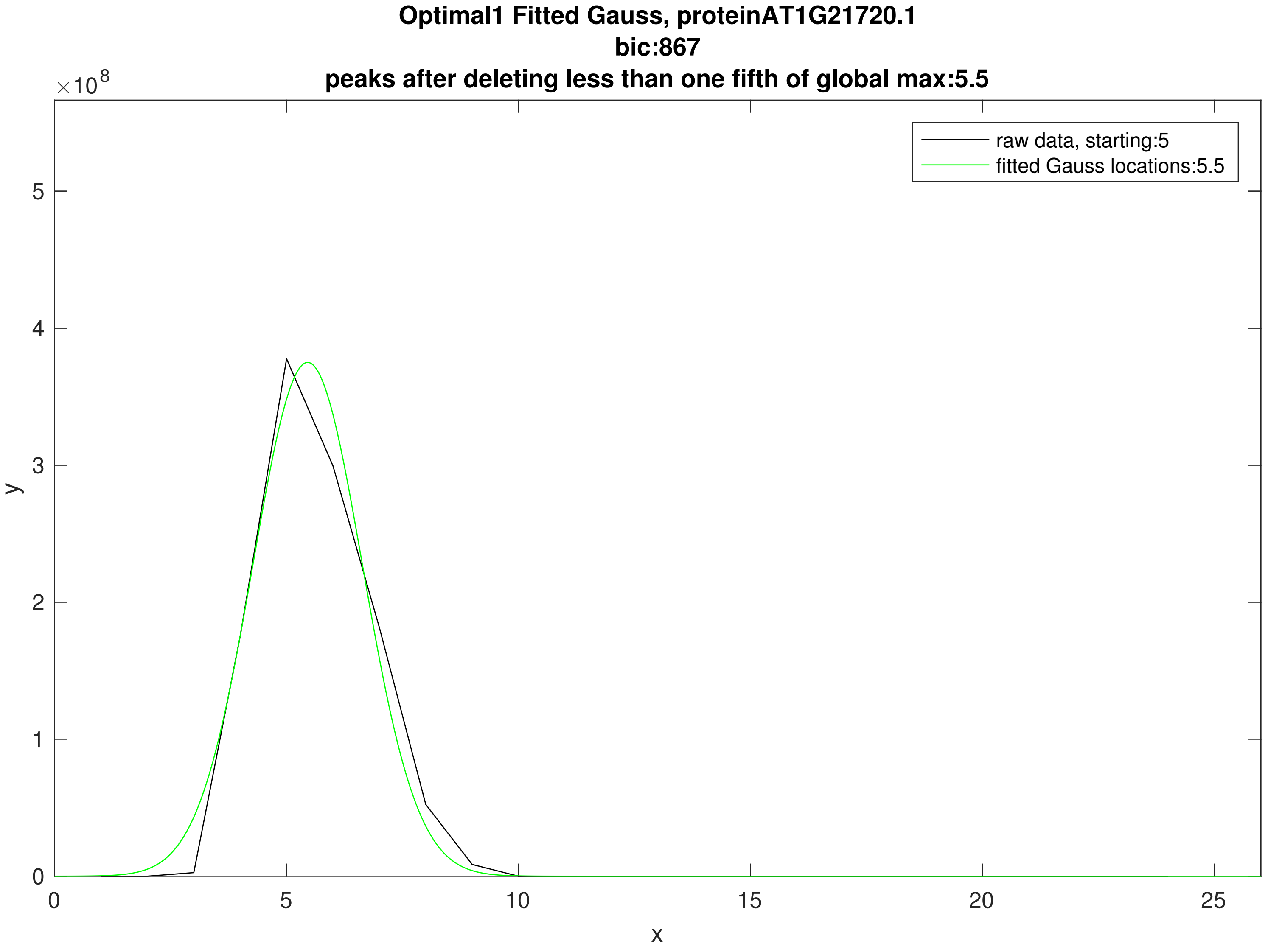

Optimal1 Fitted Gauss, proteinAT1G21720.1
bic:867
peaks after deleting less than one fifth of global max:5.5
×10^8
raw data, starting:5
fitted Gauss locations:5.5
y
x
0
1
2
3
4
5
0
5
10
15
20
25

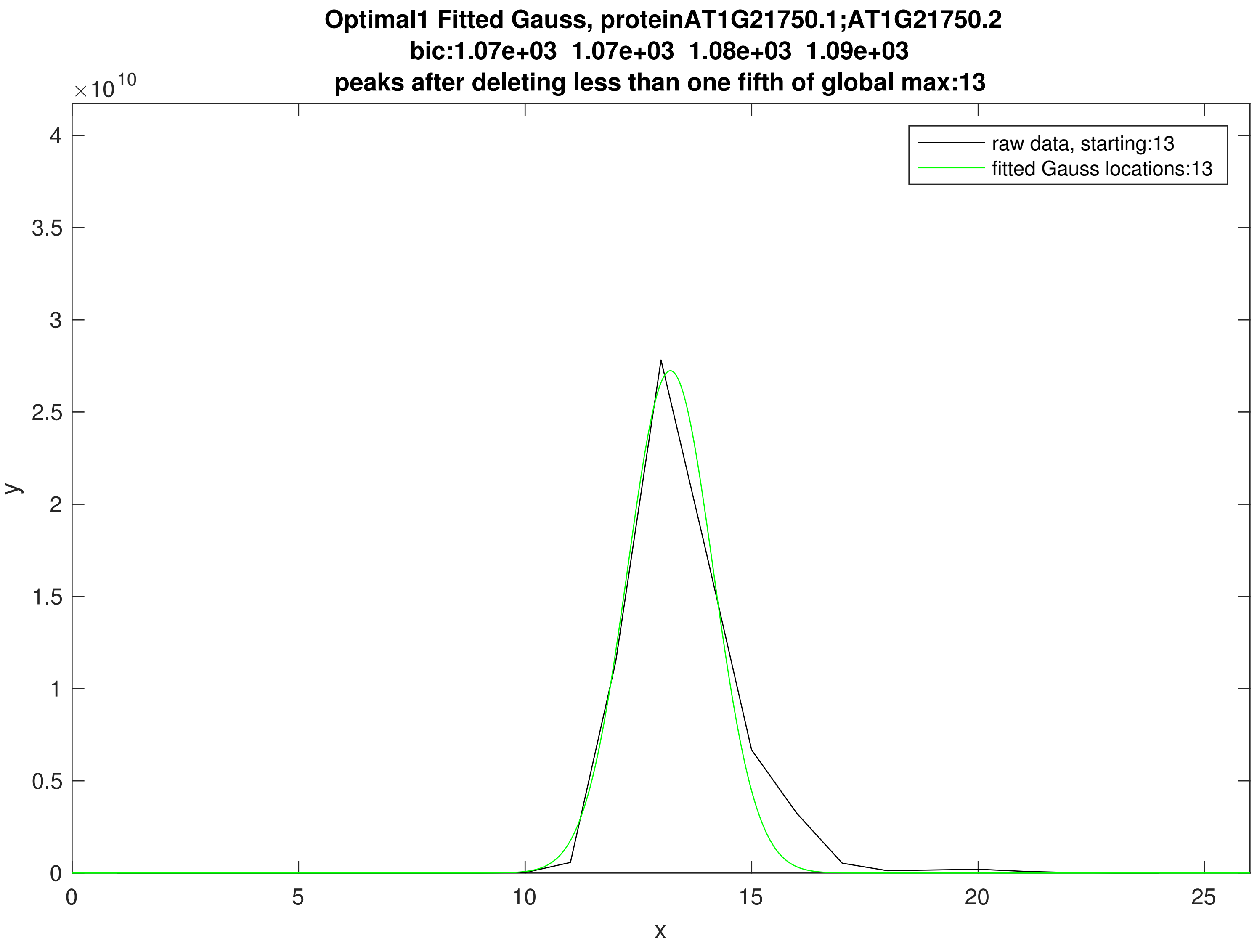
Optimal1 Fitted Gauss, proteinAT1G21750.1;AT1G21750.2
bic:1.07e+03  1.07e+03  1.08e+03  1.09e+03
peaks after deleting less than one fifth of global max:13
×10^10
raw data, starting:13
fitted Gauss locations:13
y
x
0
0.5
1
1.5
2
2.5
3
3.5
4
0
5
10
15
20
25

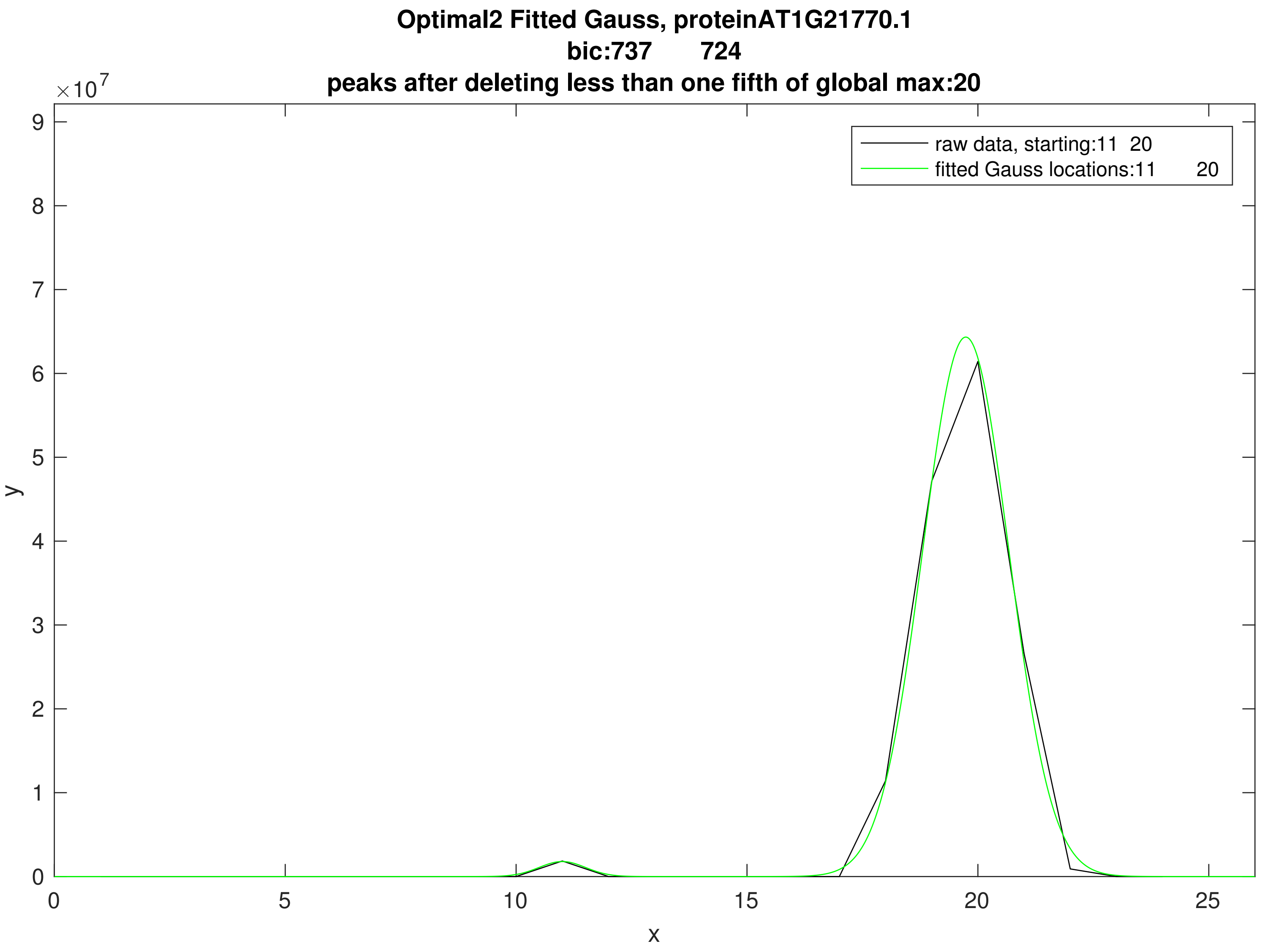

Optimal2 Fitted Gauss, proteinAT1G21770.1
bic:737 724
peaks after deleting less than one fifth of global max:20
×10^7
raw data, starting:11 20
fitted Gauss locations:11 20
y
x
0
1
2
3
4
5
6
7
8
9
0
5
10
15
20
25

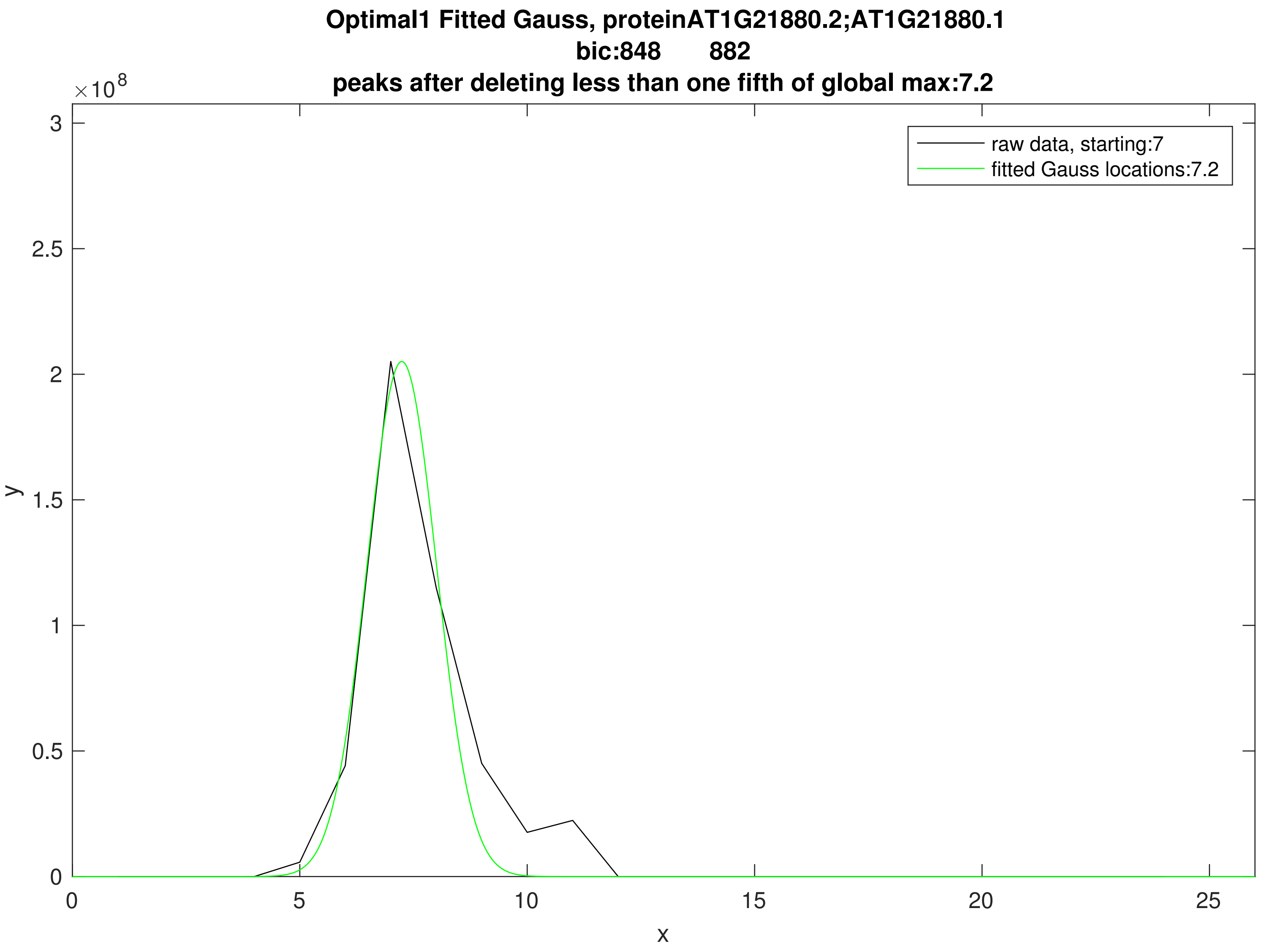

Optimal1 Fitted Gauss, proteinAT1G21880.2;AT1G21880.1
bic:848 882
peaks after deleting less than one fifth of global max:7.2
×10^8
raw data, starting:7
fitted Gauss locations:7.2
y
x
0
0.5
1
1.5
2
2.5
3
0
5
10
15
20
25

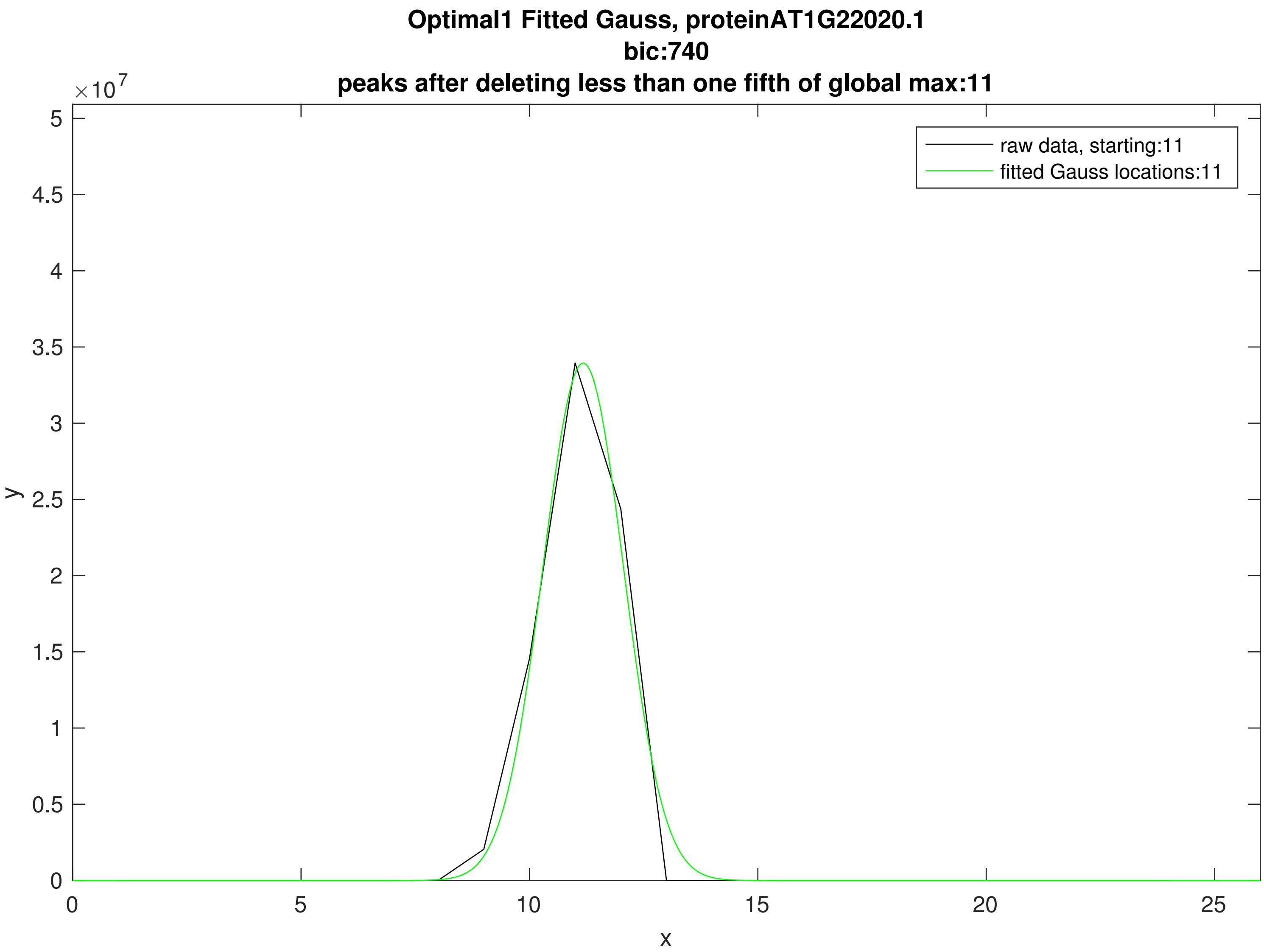
Optimal1 Fitted Gauss, proteinAT1G22020.1
bic:740
peaks after deleting less than one fifth of global max:11
×10^7
5
4.5
4
3.5
3
2.5
2
1.5
1
0.5
0
y
0
5
10
15
20
25
x
raw data, starting:11
fitted Gauss locations:11

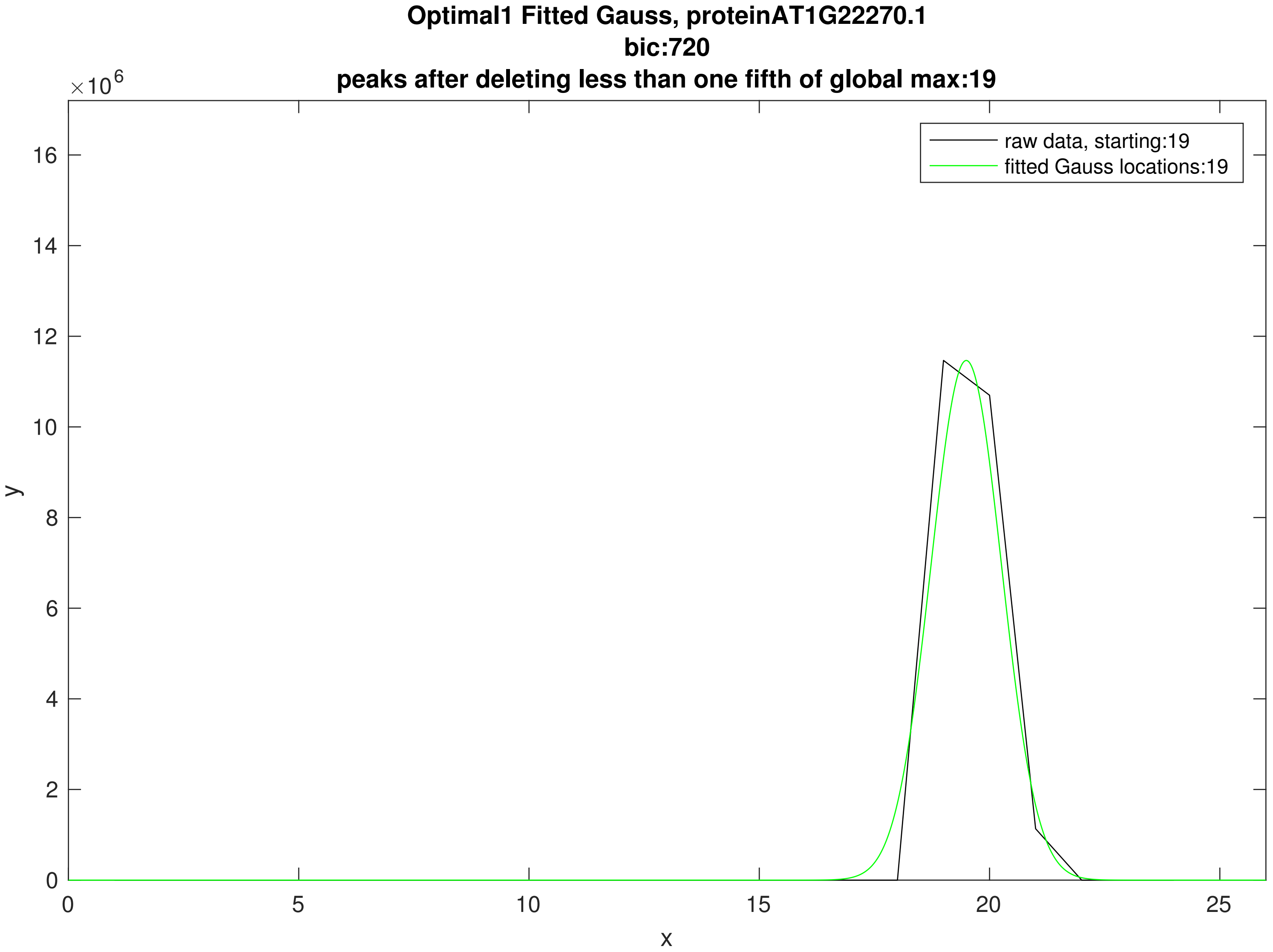
Optimal1 Fitted Gauss, proteinAT1G22270.1
bic:720
peaks after deleting less than one fifth of global max:19
×10^6
raw data, starting:19
fitted Gauss locations:19
y
x
0
2
4
6
8
10
12
14
16
0
5
10
15
20
25

Optimal1 Fitted Gauss, proteinAT1G22300.1;AT1G22300.2;AT1G22300.3
bic:1.02e+03  1.03e+03  1.04e+03  1.04e+03
peaks after deleting less than one fifth of global max:13

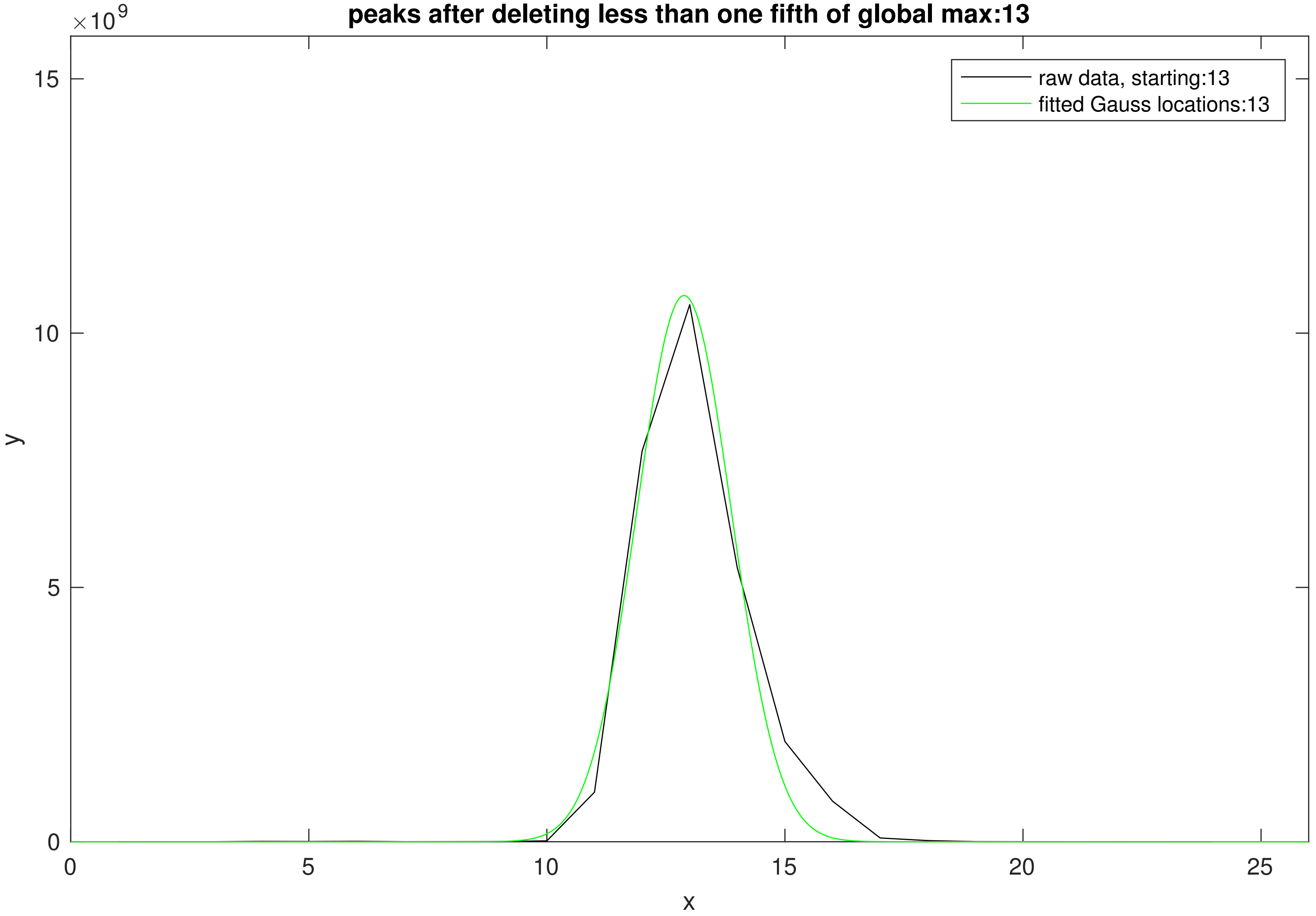

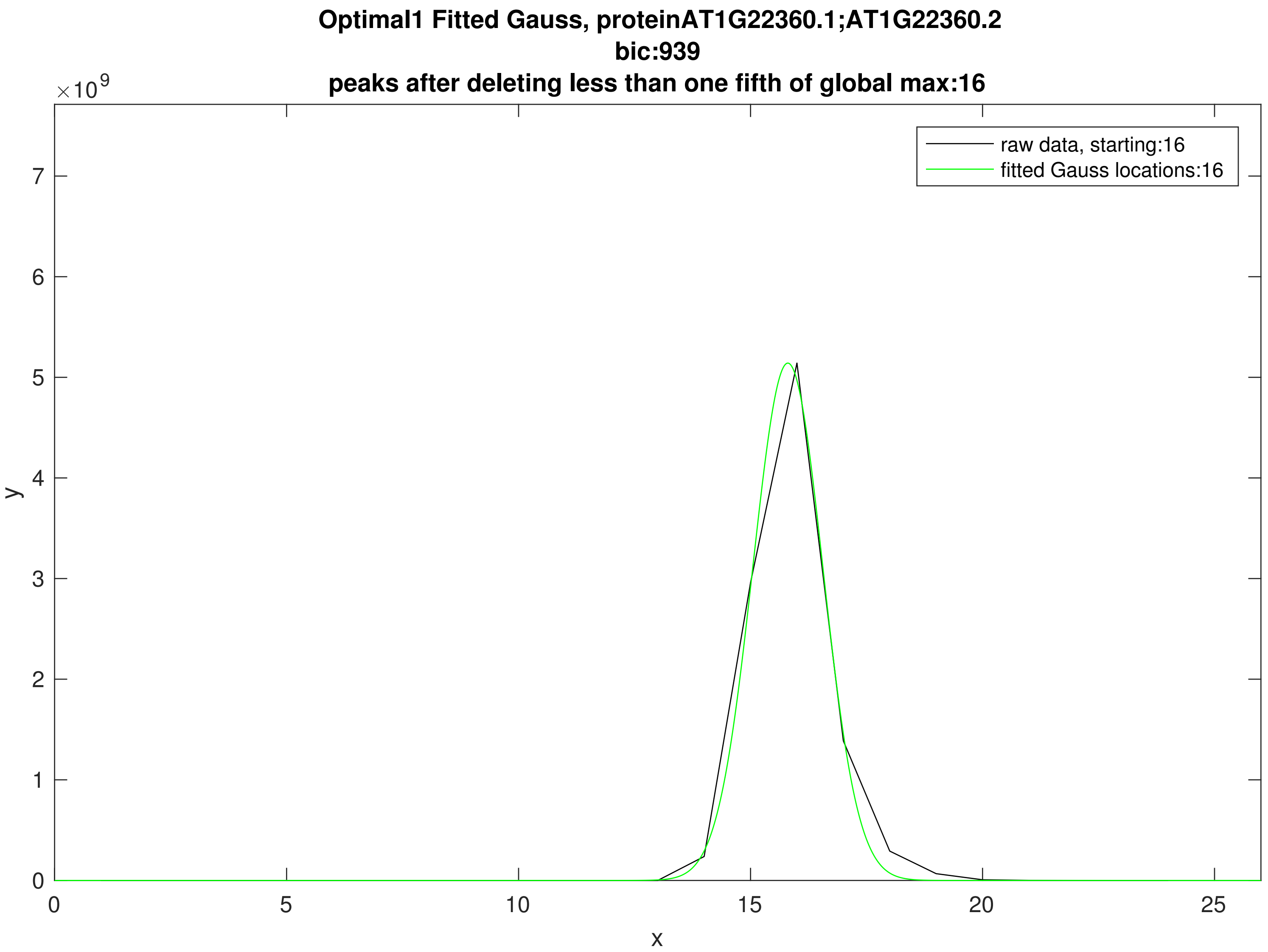
Optimal1 Fitted Gauss, proteinAT1G22360.1;AT1G22360.2
bic:939
peaks after deleting less than one fifth of global max:16
×10^9
raw data, starting:16
fitted Gauss locations:16
y
x
0
1
2
3
4
5
6
7
0
5
10
15
20
25

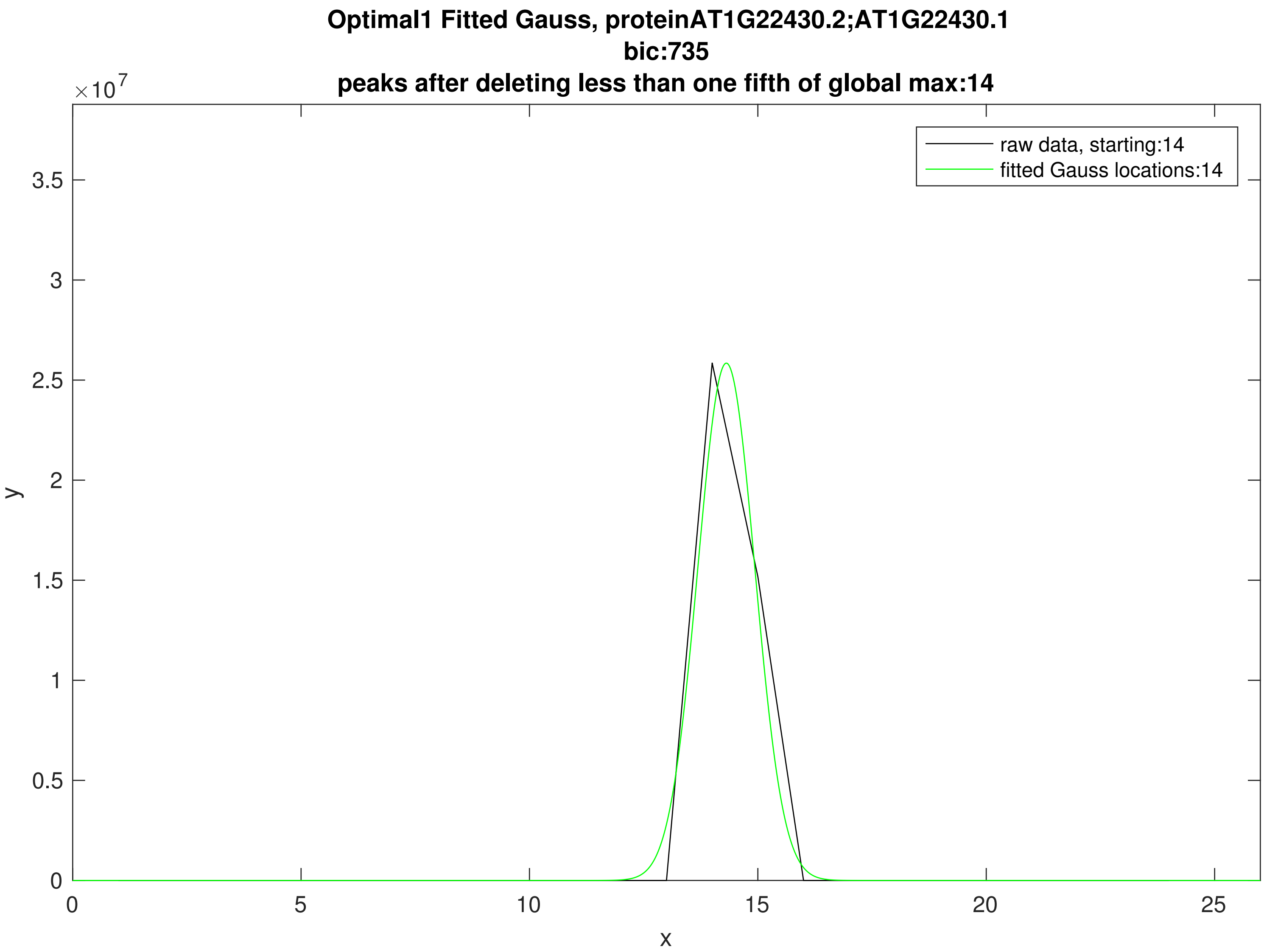
Optimal1 Fitted Gauss, proteinAT1G22430.2;AT1G22430.1
bic:735
peaks after deleting less than one fifth of global max:14
×10^7
3.5
3
2.5
2
1.5
1
0.5
0
y
0
5
10
15
20
25
x
raw data, starting:14
fitted Gauss locations:14

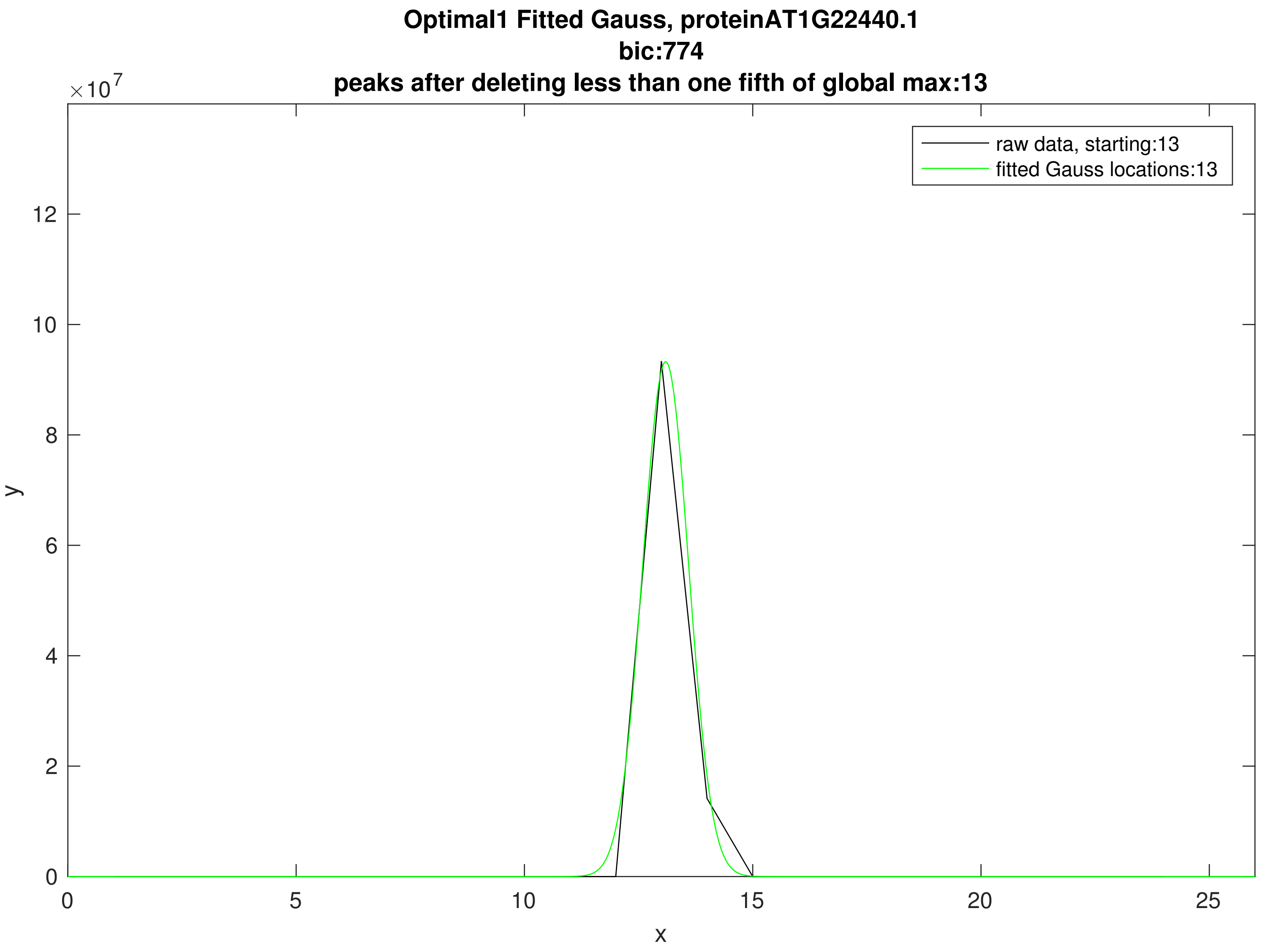
Optimal1 Fitted Gauss, proteinAT1G22440.1
bic:774
peaks after deleting less than one fifth of global max:13
×10^7
raw data, starting:13
fitted Gauss locations:13
y
x
0
2
4
6
8
10
12
0
5
10
15
20
25

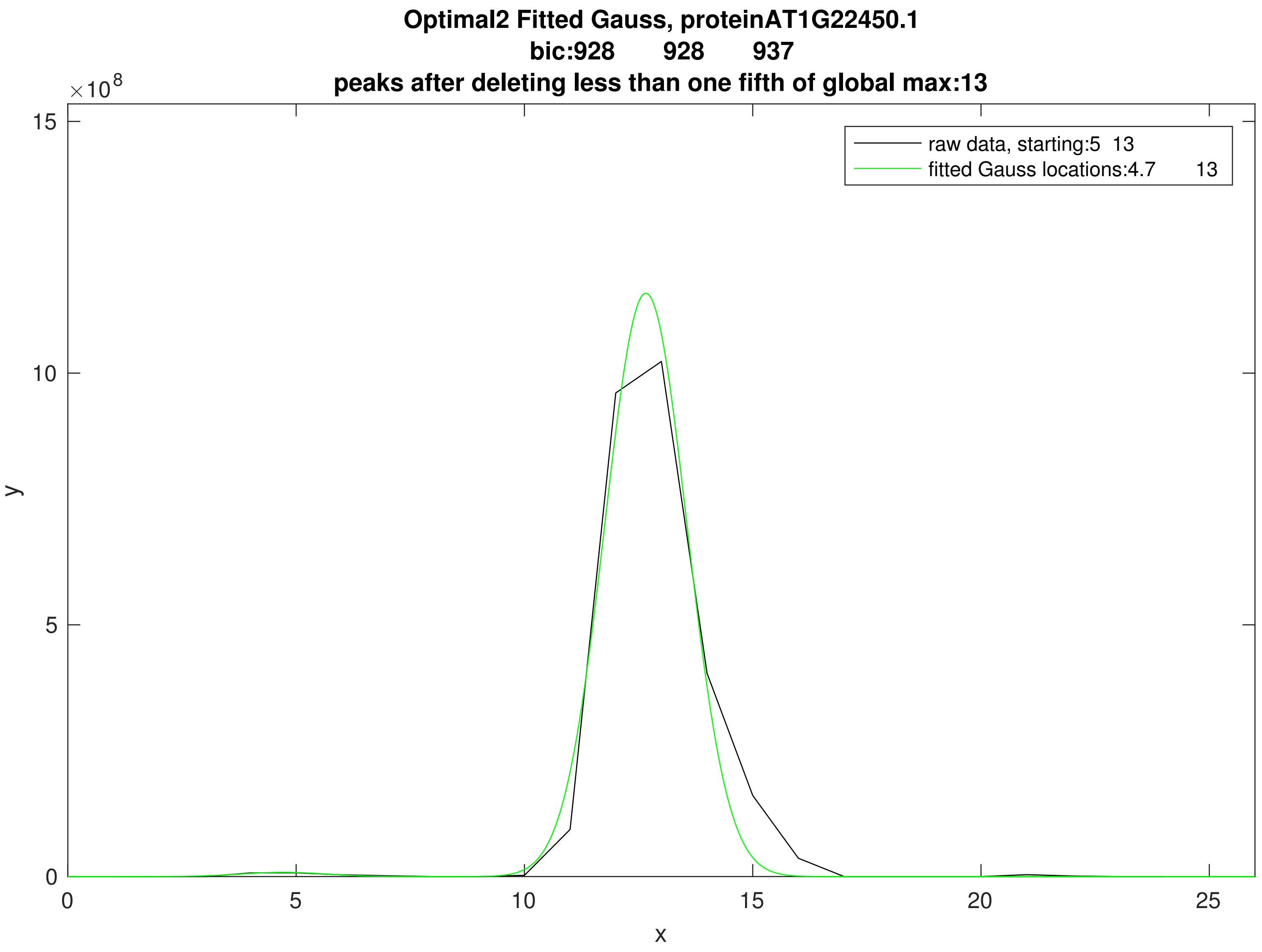

Optimal2 Fitted Gauss, proteinAT1G22450.1
bic:928 928 937
peaks after deleting less than one fifth of global max:13
×10^8
15
10
5
0
y
0
5
10
15
20
25
x
raw data, starting:5 13
fitted Gauss locations:4.7 13

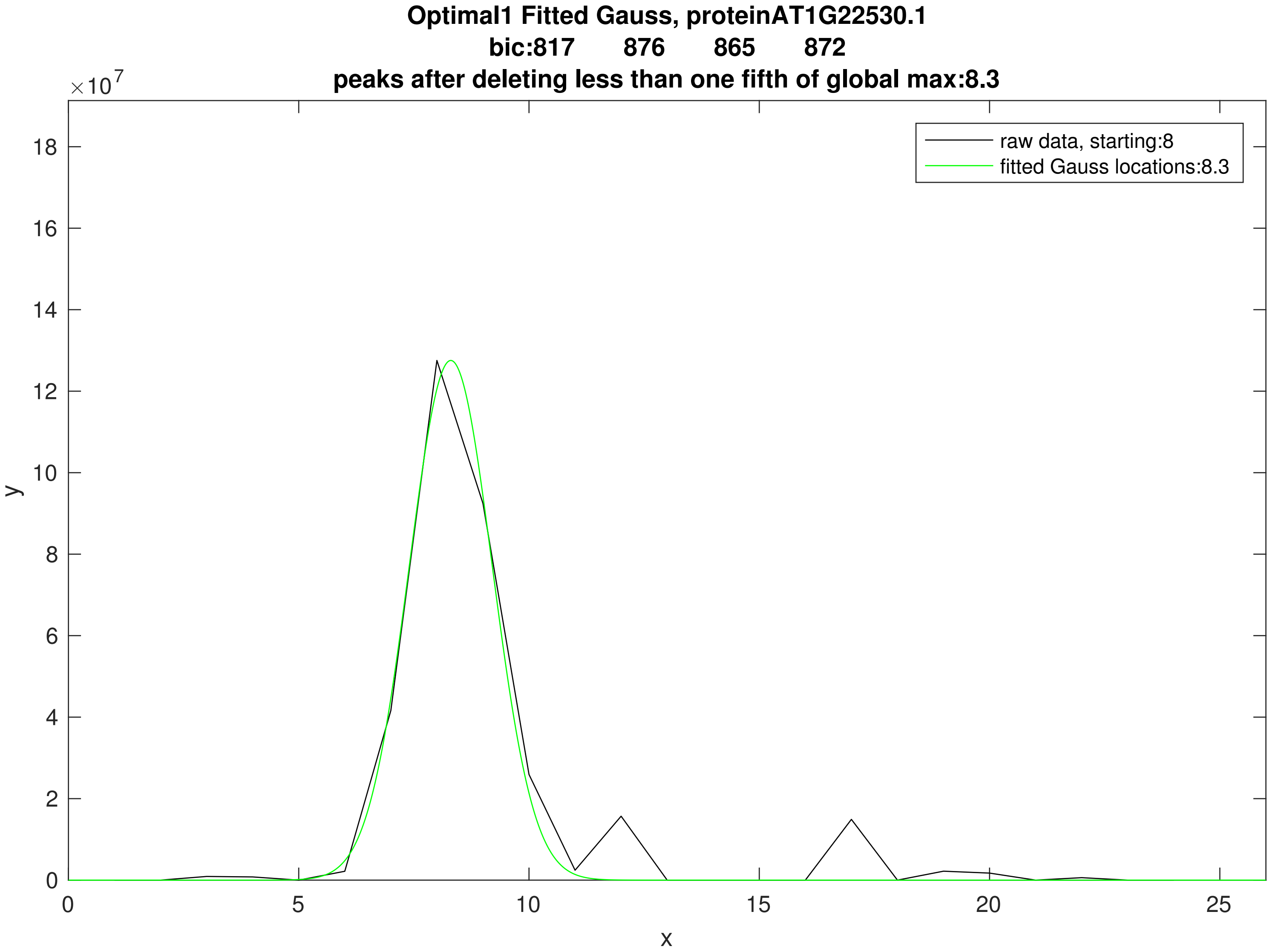

Optimal1 Fitted Gauss, proteinAT1G22530.1
bic:817 876 865 872
peaks after deleting less than one fifth of global max:8.3
×10^7
y
x
raw data, starting:8
fitted Gauss locations:8.3

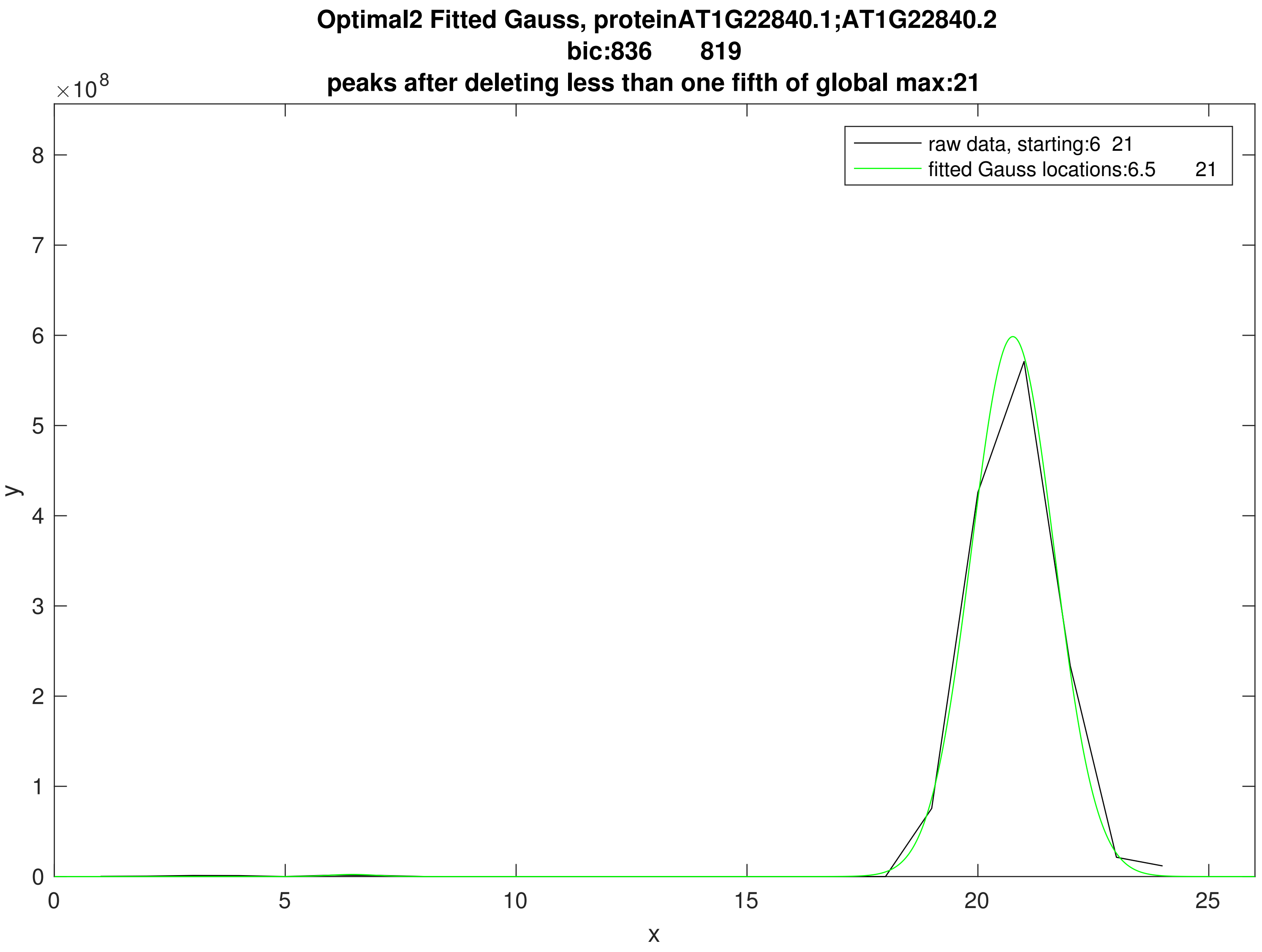

Optimal2 Fitted Gauss, proteinAT1G22840.1;AT1G22840.2
bic:836 819
peaks after deleting less than one fifth of global max:21
×10^8
raw data, starting:6 21
fitted Gauss locations:6.5 21
y
x
0
1
2
3
4
5
6
7
8
0
5
10
15
20
25

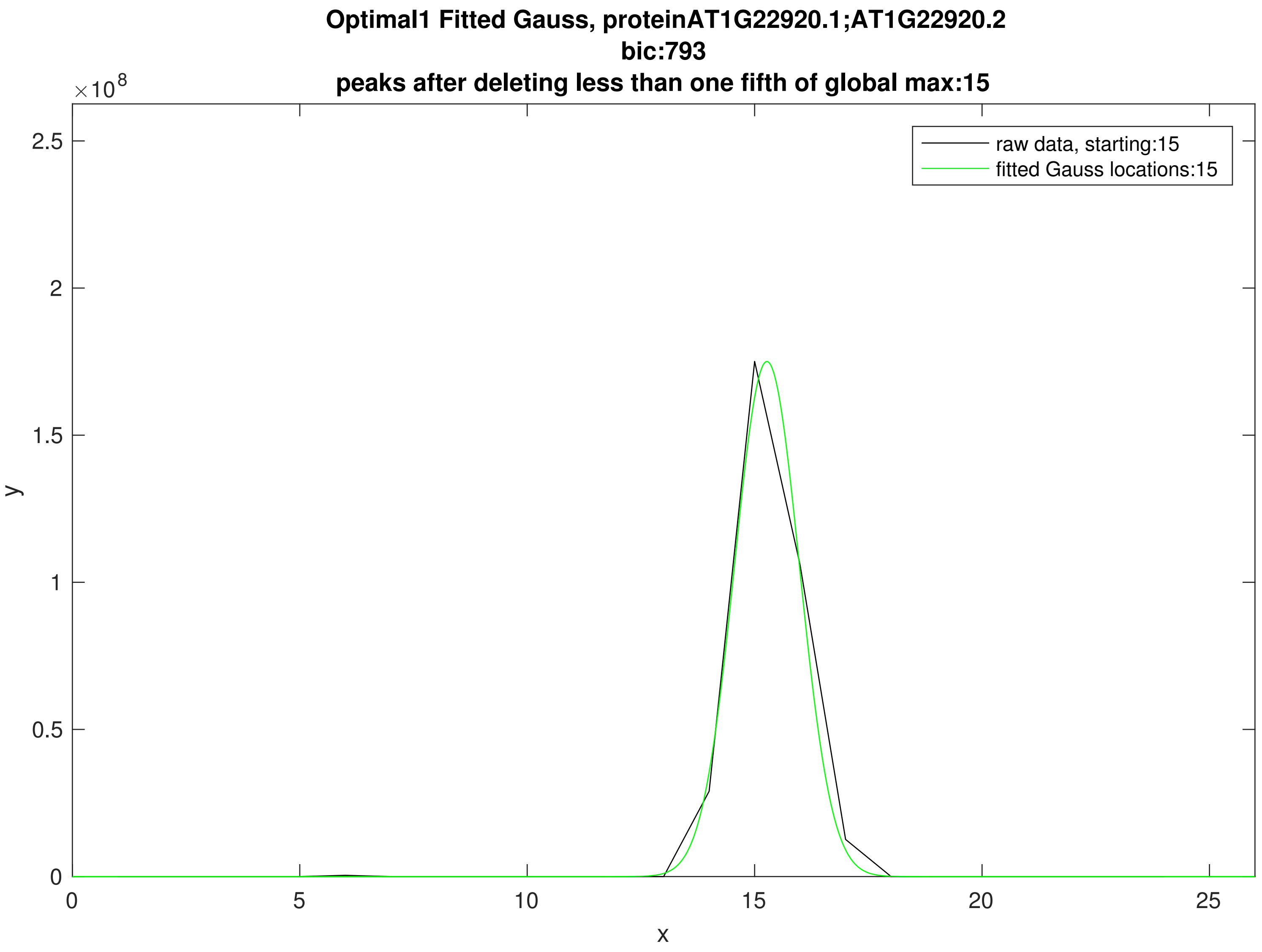

Optimal1 Fitted Gauss, proteinAT1G22920.1;AT1G22920.2
bic:793
peaks after deleting less than one fifth of global max:15
×10^8
2.5
2
1.5
1
0.5
0
y
0
5
10
15
20
25
x
raw data, starting:15
fitted Gauss locations:15

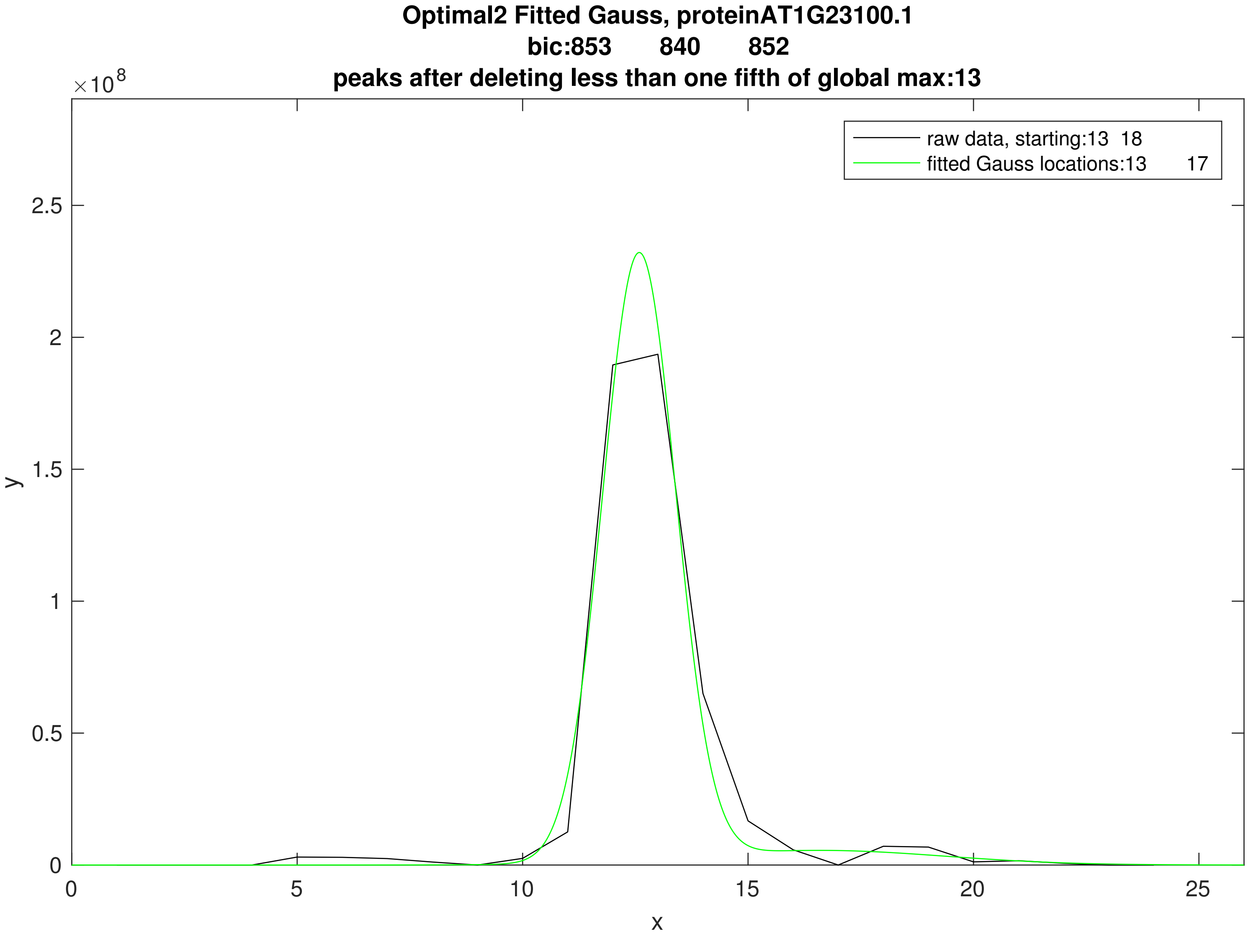

Optimal2 Fitted Gauss, proteinAT1G23100.1
bic:853 840 852
peaks after deleting less than one fifth of global max:13
×10^8
raw data, starting:13 18
fitted Gauss locations:13 17
y
x
0
0.5
1
1.5
2
2.5
0
5
10
15
20
25

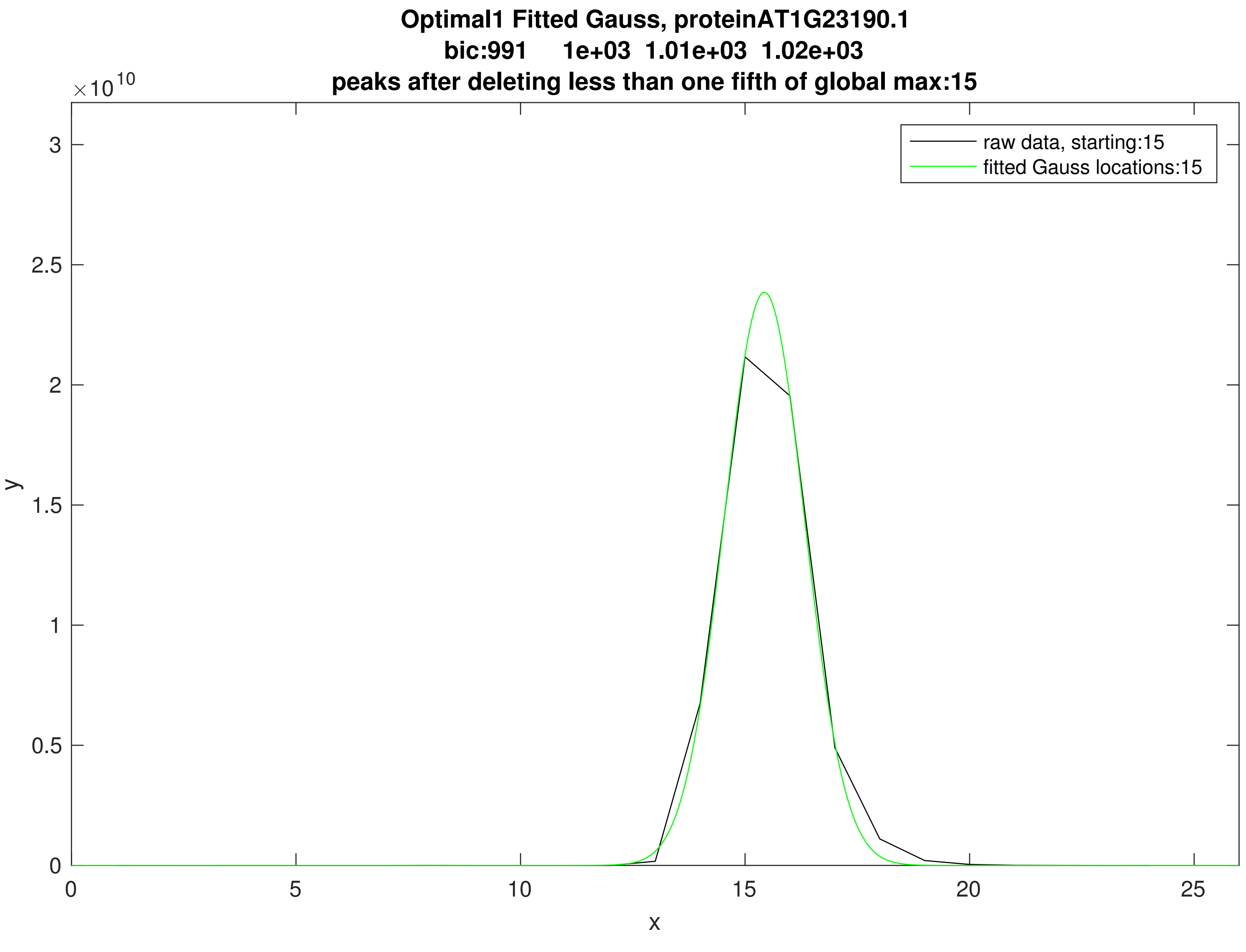
Optimal1 Fitted Gauss, proteinAT1G23190.1
bic:991 1e+03 1.01e+03 1.02e+03
peaks after deleting less than one fifth of global max:15
×10^10
raw data, starting:15
fitted Gauss locations:15
y
x
0
0.5
1
1.5
2
2.5
3
0
5
10
15
20
25

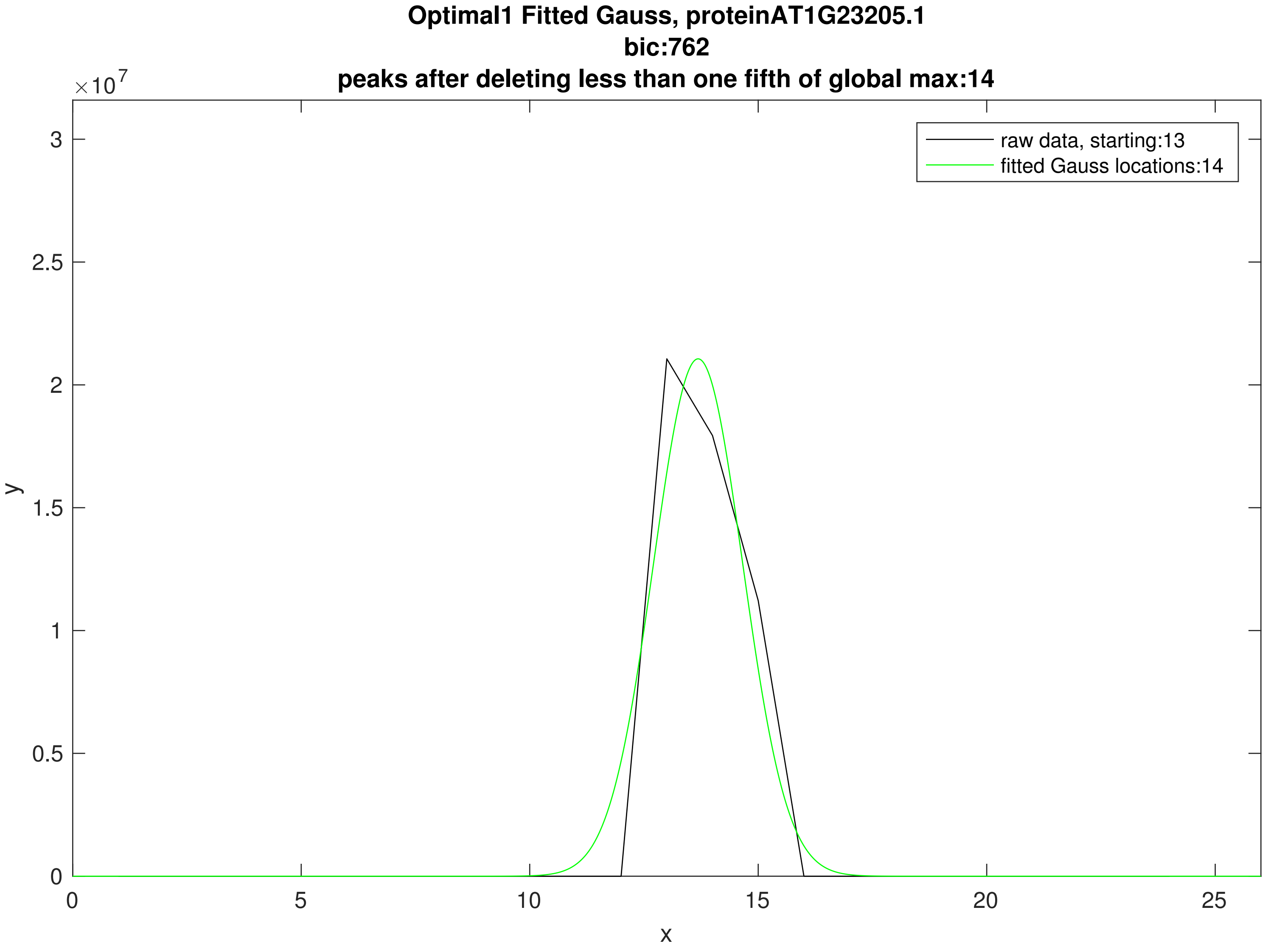
Optimal1 Fitted Gauss, proteinAT1G23205.1
bic:762
peaks after deleting less than one fifth of global max:14
×10^7
raw data, starting:13
fitted Gauss locations:14
3
2.5
2
1.5
1
0.5
0
y
0
5
10
15
20
25
x

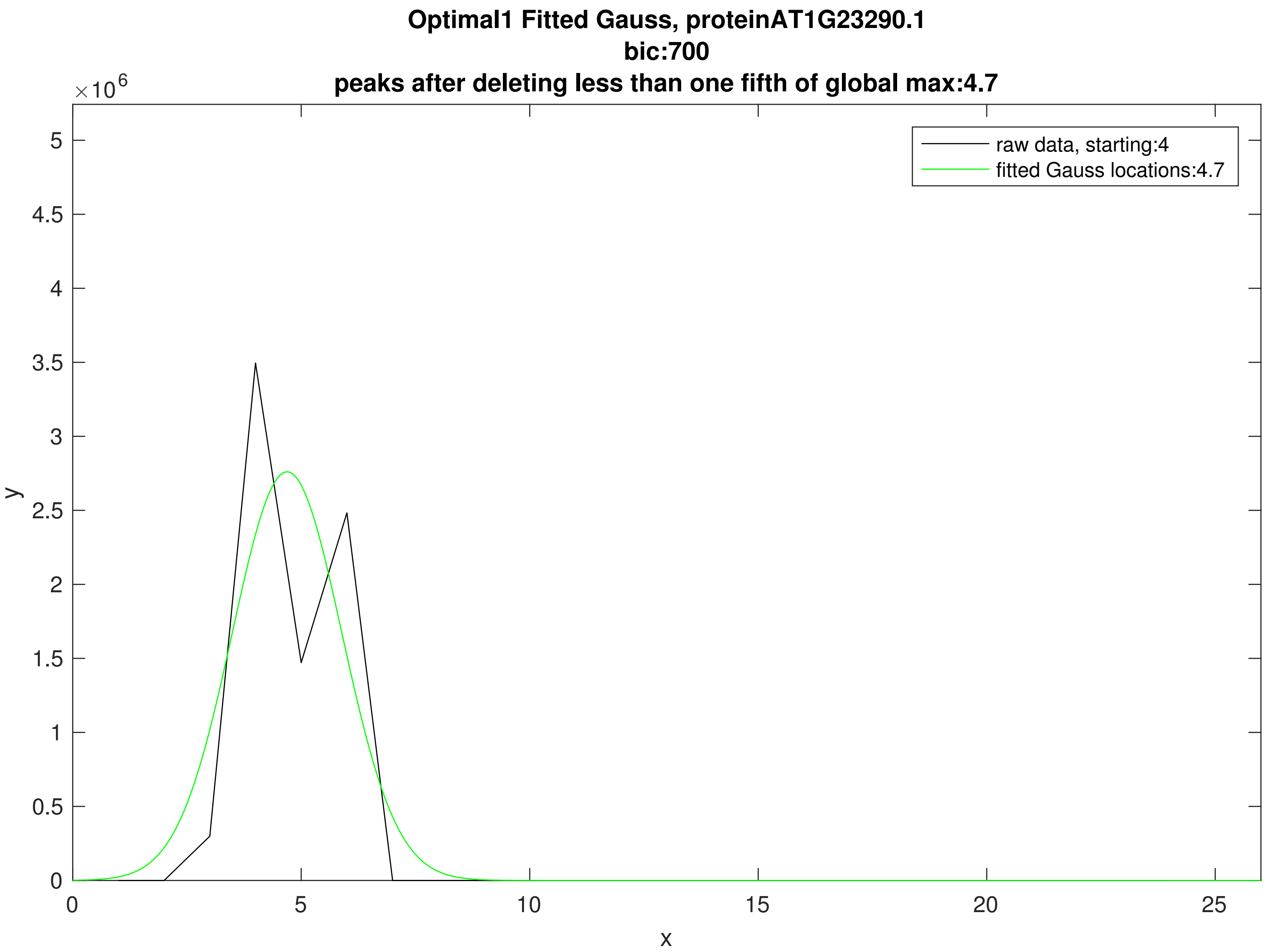
Optimal1 Fitted Gauss, proteinAT1G23290.1
bic:700
peaks after deleting less than one fifth of global max:4.7
raw data, starting:4
fitted Gauss locations:4.7
×10^6
y
x
0
0.5
1
1.5
2
2.5
3
3.5
4
4.5
5
0
5
10
15
20
25

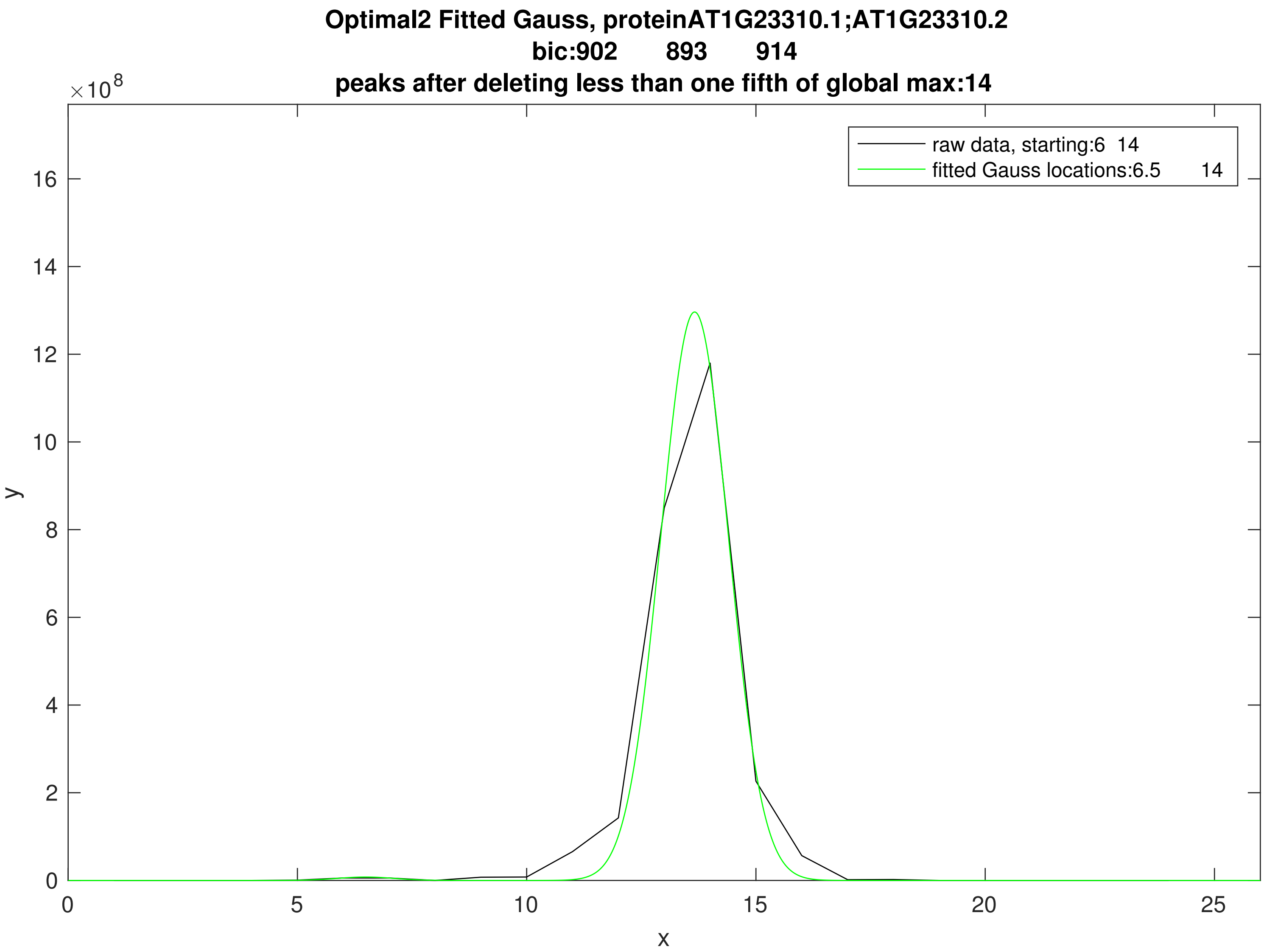

Optimal2 Fitted Gauss, proteinAT1G23310.1;AT1G23310.2
bic:902 893 914
peaks after deleting less than one fifth of global max:14
×10^8
raw data, starting:6 14
fitted Gauss locations:6.5 14
y
x
0
2
4
6
8
10
12
14
16
5
15
20
25

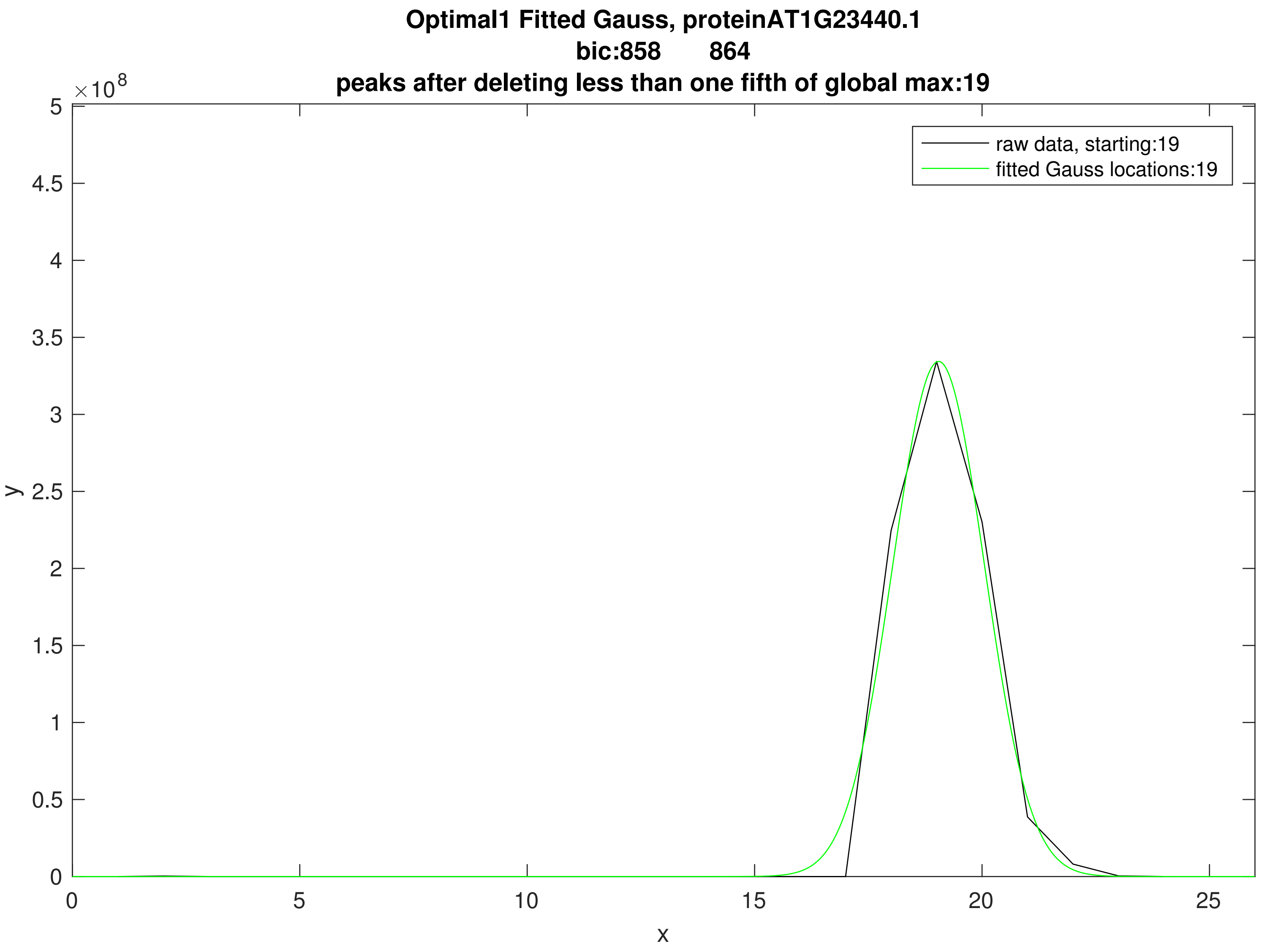
Optimal1 Fitted Gauss, proteinAT1G23440.1
bic:858 864
peaks after deleting less than one fifth of global max:19
×10^8
5
4.5
4
3.5
3
2.5
2
1.5
1
0.5
0
y
0
5
10
15
20
25
x
raw data, starting:19
fitted Gauss locations:19

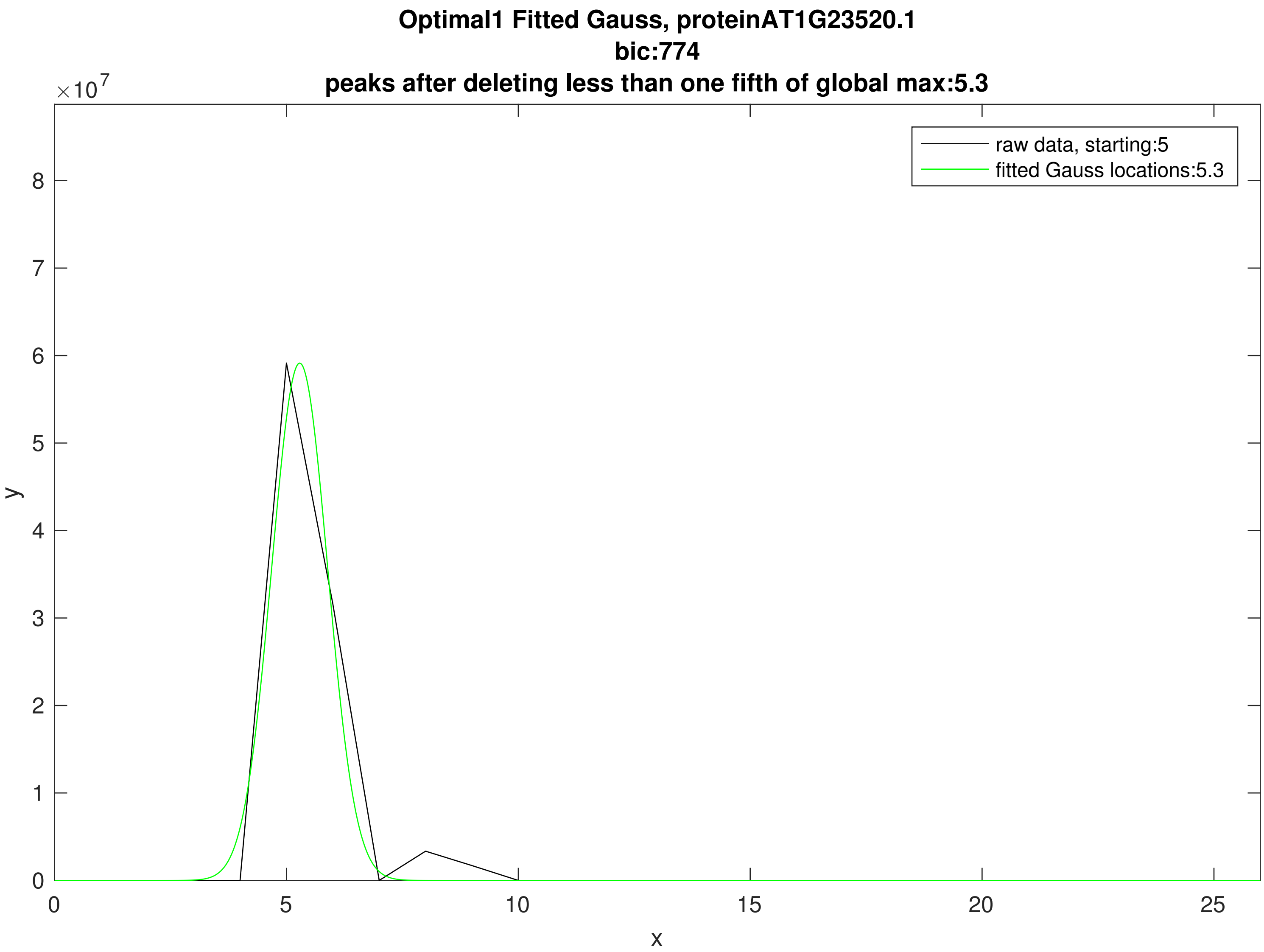

Optimal1 Fitted Gauss, proteinAT1G23520.1
bic:774
peaks after deleting less than one fifth of global max:5.3
×10^7
8
7
6
5
4
3
2
1
0
y
0
5
10
15
20
25
x
raw data, starting:5
fitted Gauss locations:5.3

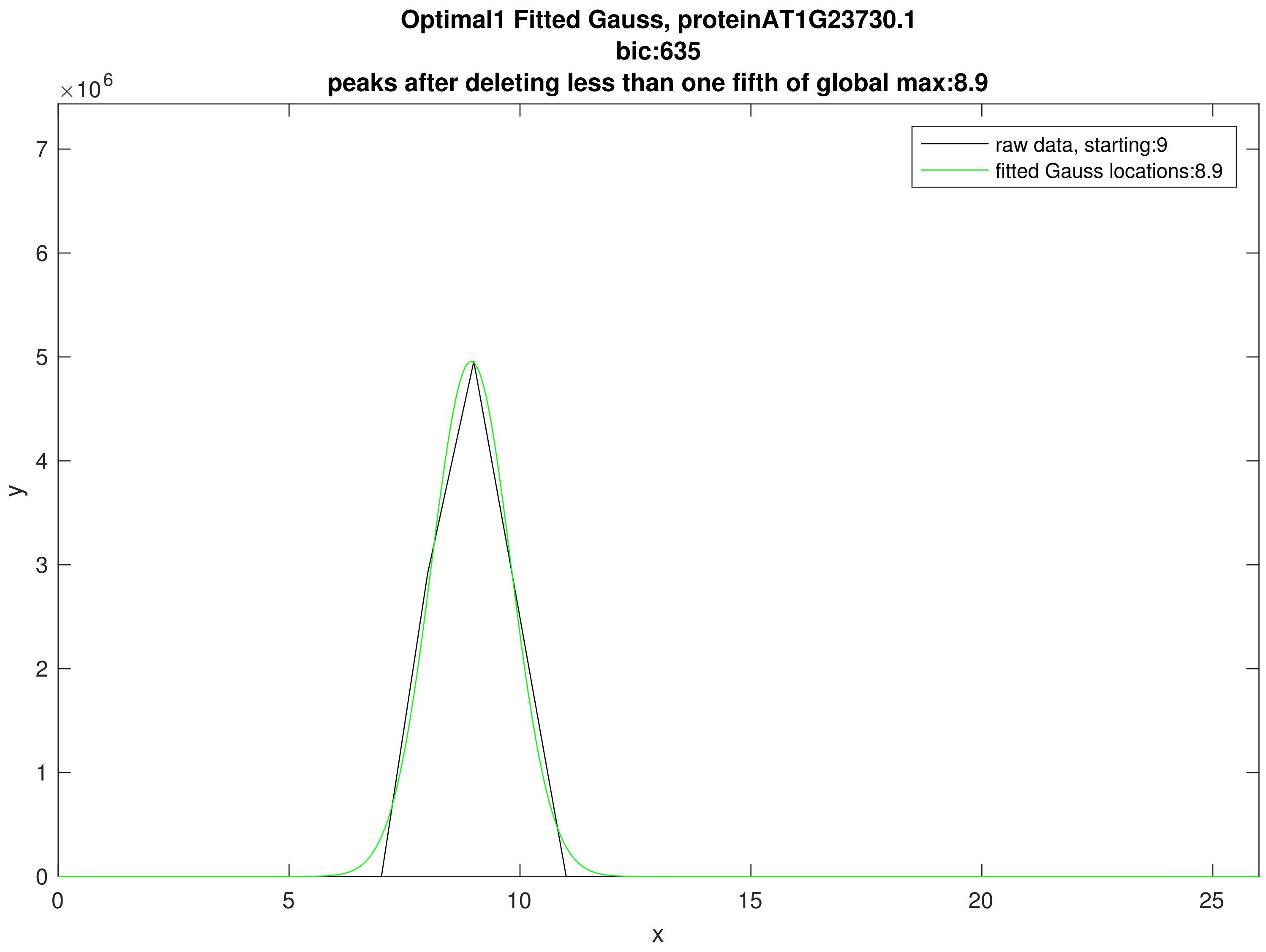

Optimal1 Fitted Gauss, proteinAT1G23730.1
bic:635
peaks after deleting less than one fifth of global max:8.9
×10^6
raw data, starting:9
fitted Gauss locations:8.9
y
x
0
1
2
3
4
5
6
7
0
5
10
15
20
25

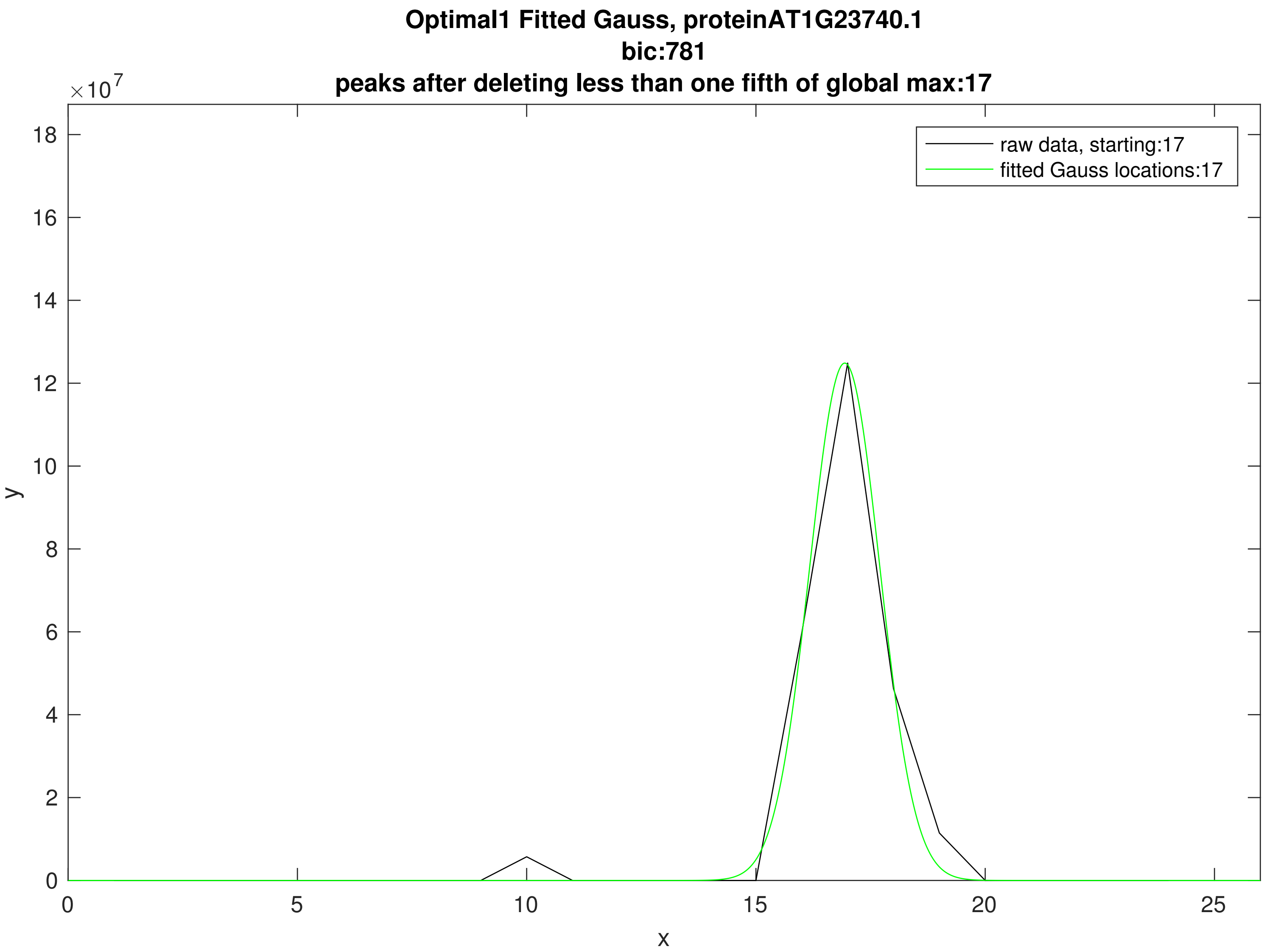

Optimal1 Fitted Gauss, proteinAT1G23740.1
bic:781
peaks after deleting less than one fifth of global max:17
×10^7
18
16
14
12
10
8
6
4
2
0
y
0
5
10
15
20
25
x
raw data, starting:17
fitted Gauss locations:17

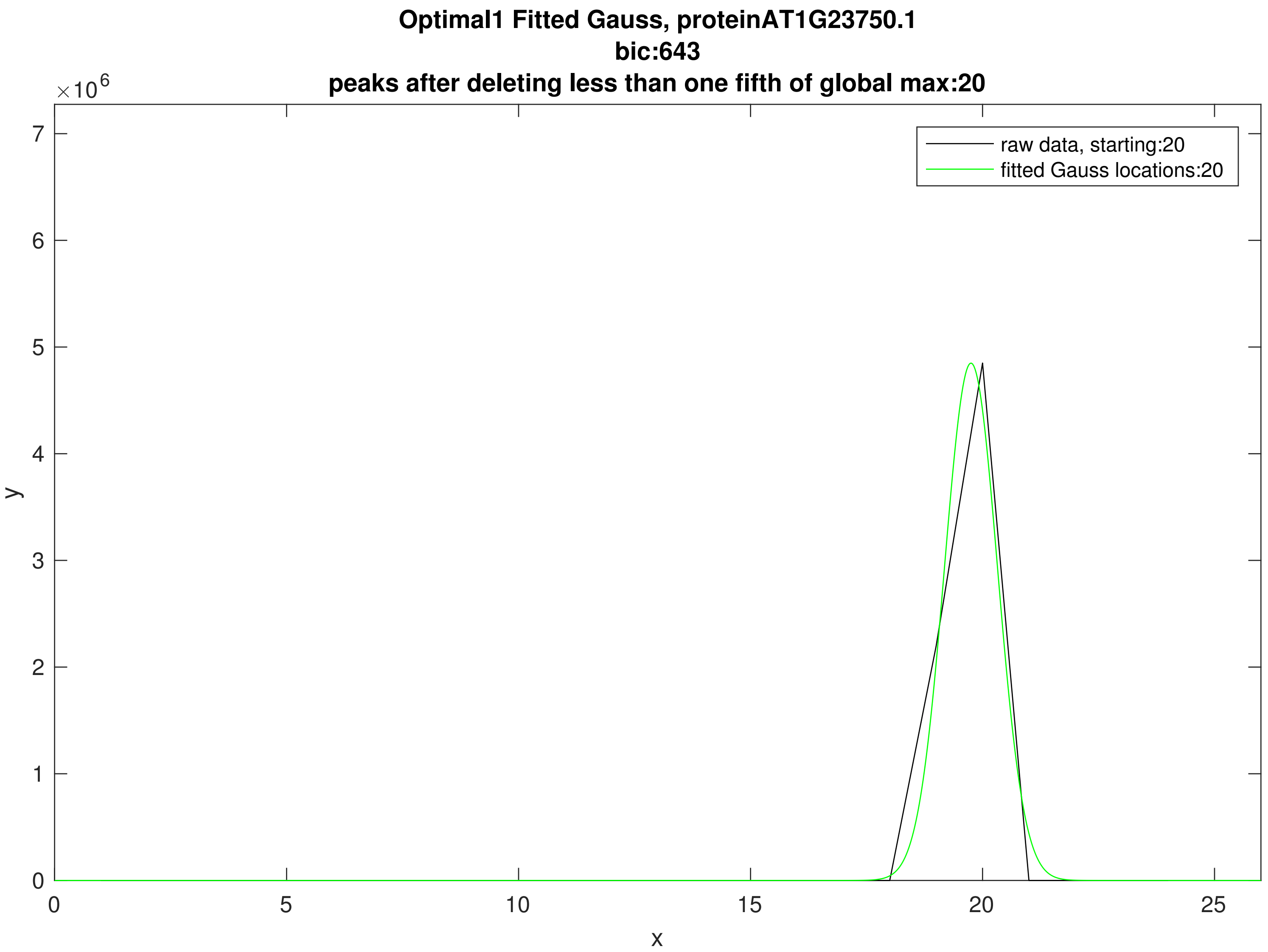
Optimal1 Fitted Gauss, proteinAT1G23750.1
bic:643
peaks after deleting less than one fifth of global max:20
×10^6
raw data, starting:20
fitted Gauss locations:20
y
x
0
1
2
3
4
5
6
7
0
5
10
15
20
25

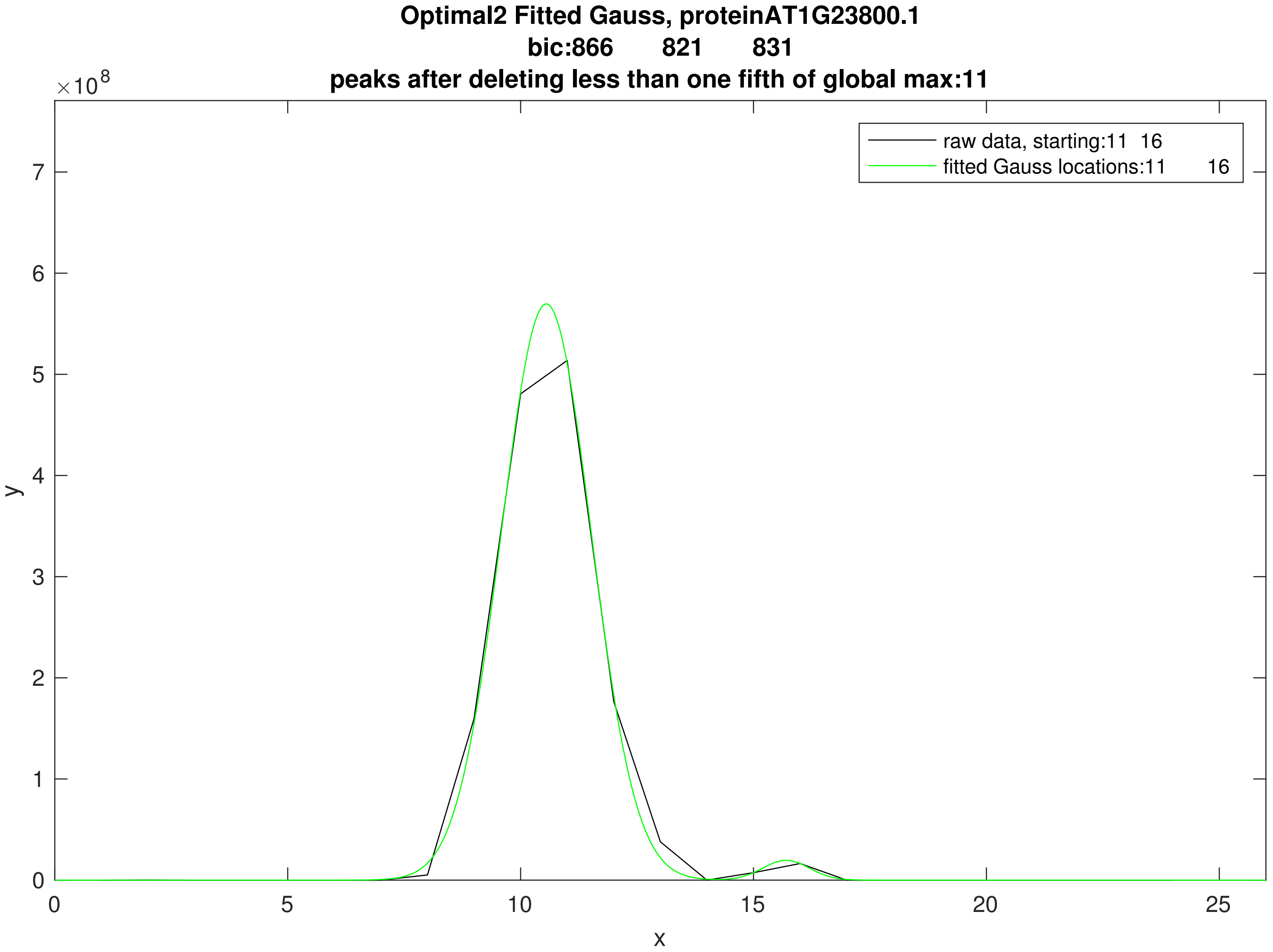

Optimal2 Fitted Gauss, proteinAT1G23800.1
bic:866 821 831
peaks after deleting less than one fifth of global max:11
×10^8
raw data, starting:11 16
fitted Gauss locations:11 16
y
x
0
1
2
3
4
5
6
7
0
5
10
15
20
25

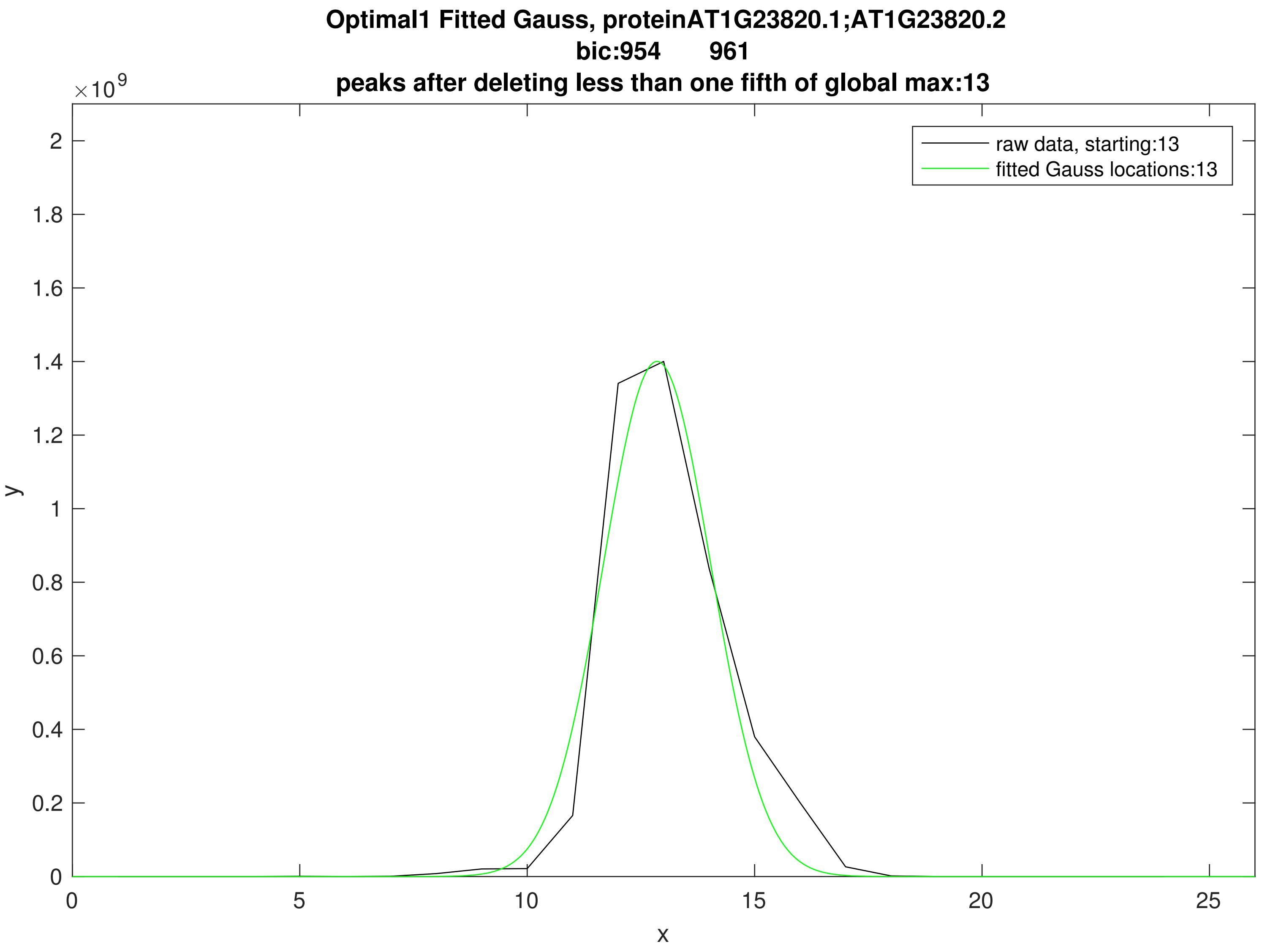

Optimal1 Fitted Gauss, proteinAT1G23820.1;AT1G23820.2
bic:954 961
peaks after deleting less than one fifth of global max:13
×10^9
2
1.8
1.6
1.4
1.2
1
0.8
0.6
0.4
0.2
0
y
0
5
10
15
20
25
x
raw data, starting:13
fitted Gauss locations:13

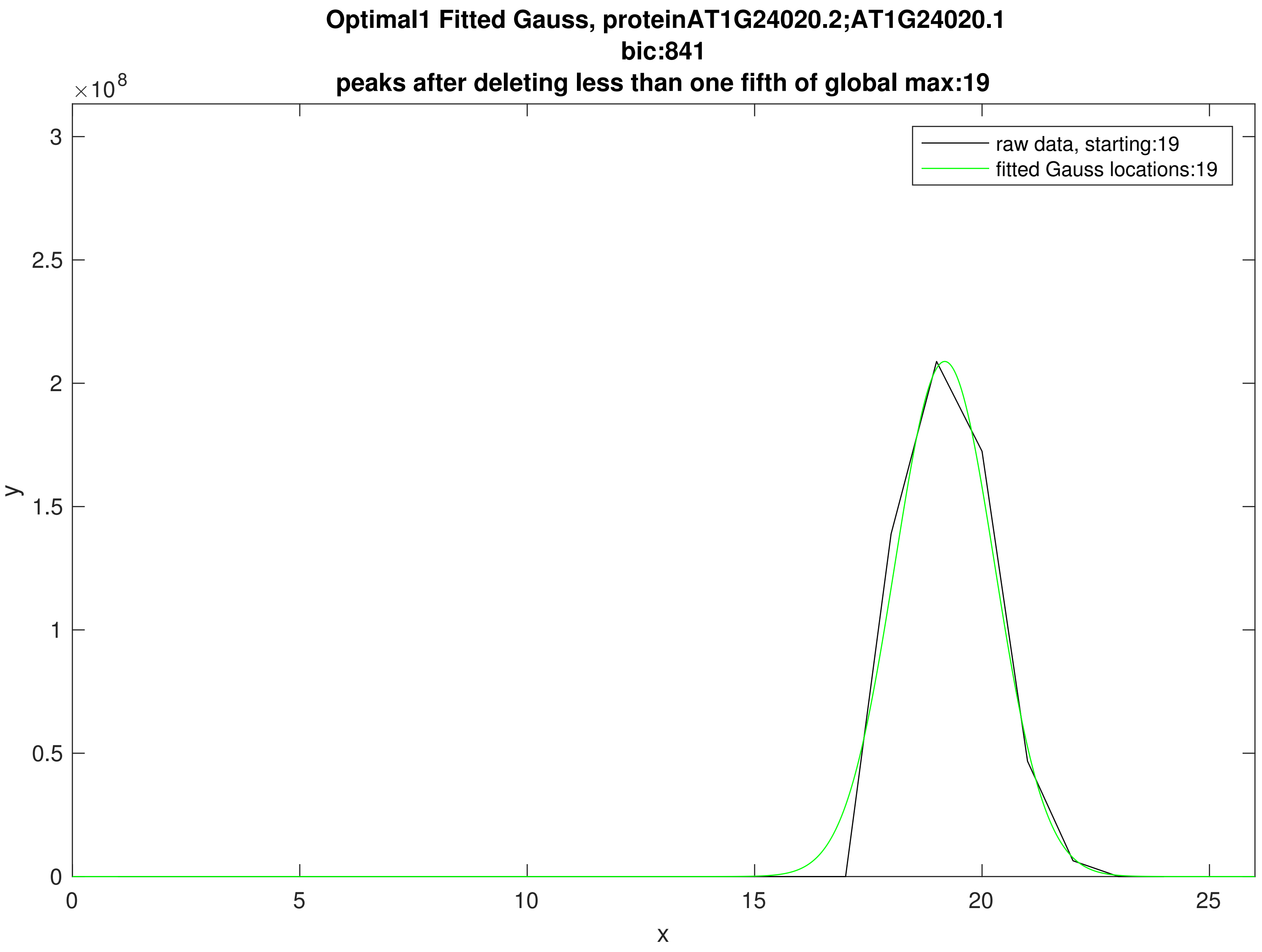

Optimal1 Fitted Gauss, proteinAT1G24020.2;AT1G24020.1
bic:841
peaks after deleting less than one fifth of global max:19
×10^8
3
2.5
2
1.5
1
0.5
0
y
0
5
10
15
20
25
x
raw data, starting:19
fitted Gauss locations:19

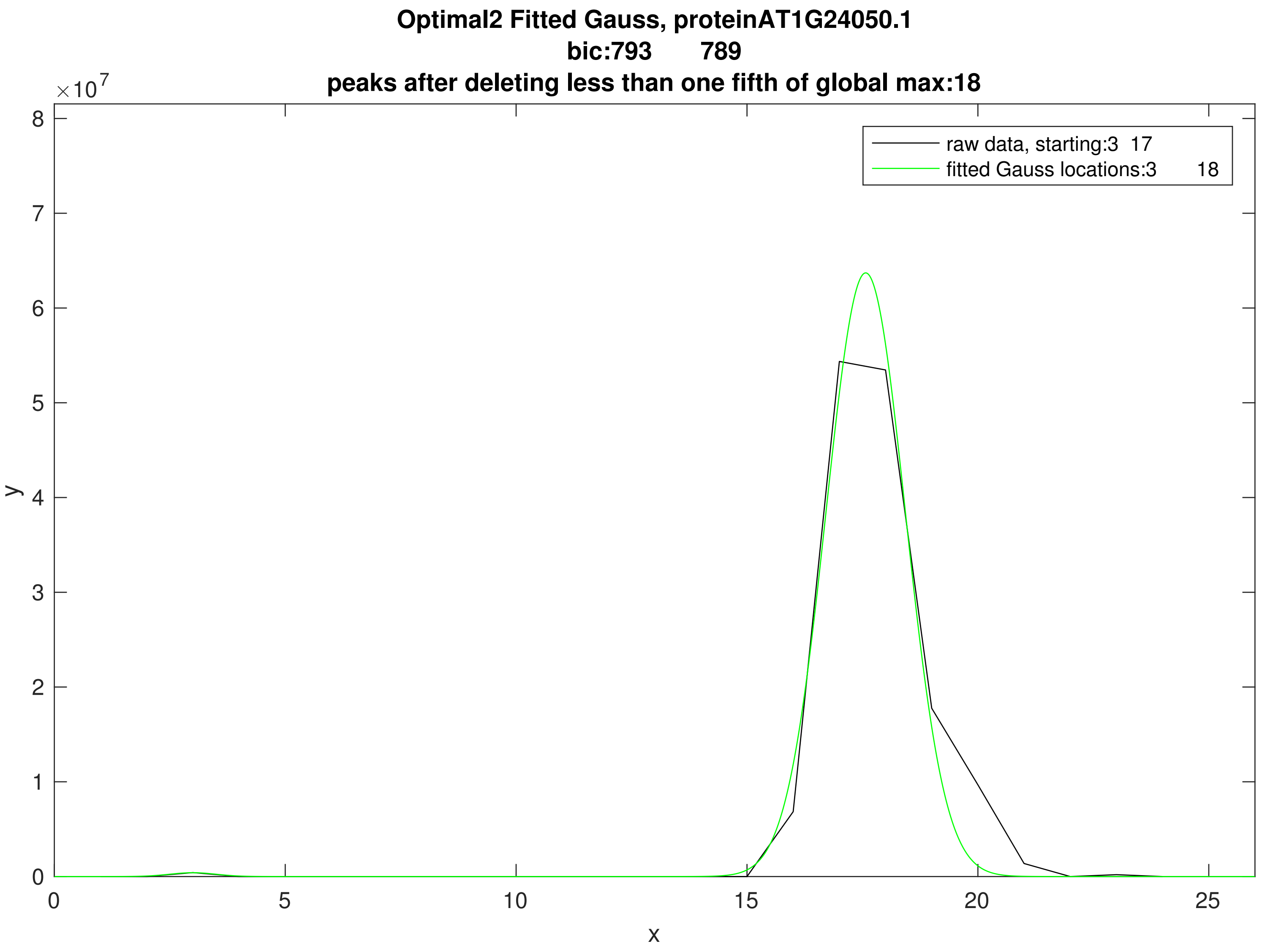

Optimal2 Fitted Gauss, proteinAT1G24050.1
bic:793 789
peaks after deleting less than one fifth of global max:18
×10^7
raw data, starting:3 17
fitted Gauss locations:3 18
y
x
0
1
2
3
4
5
6
7
8
0
5
10
15
20
25

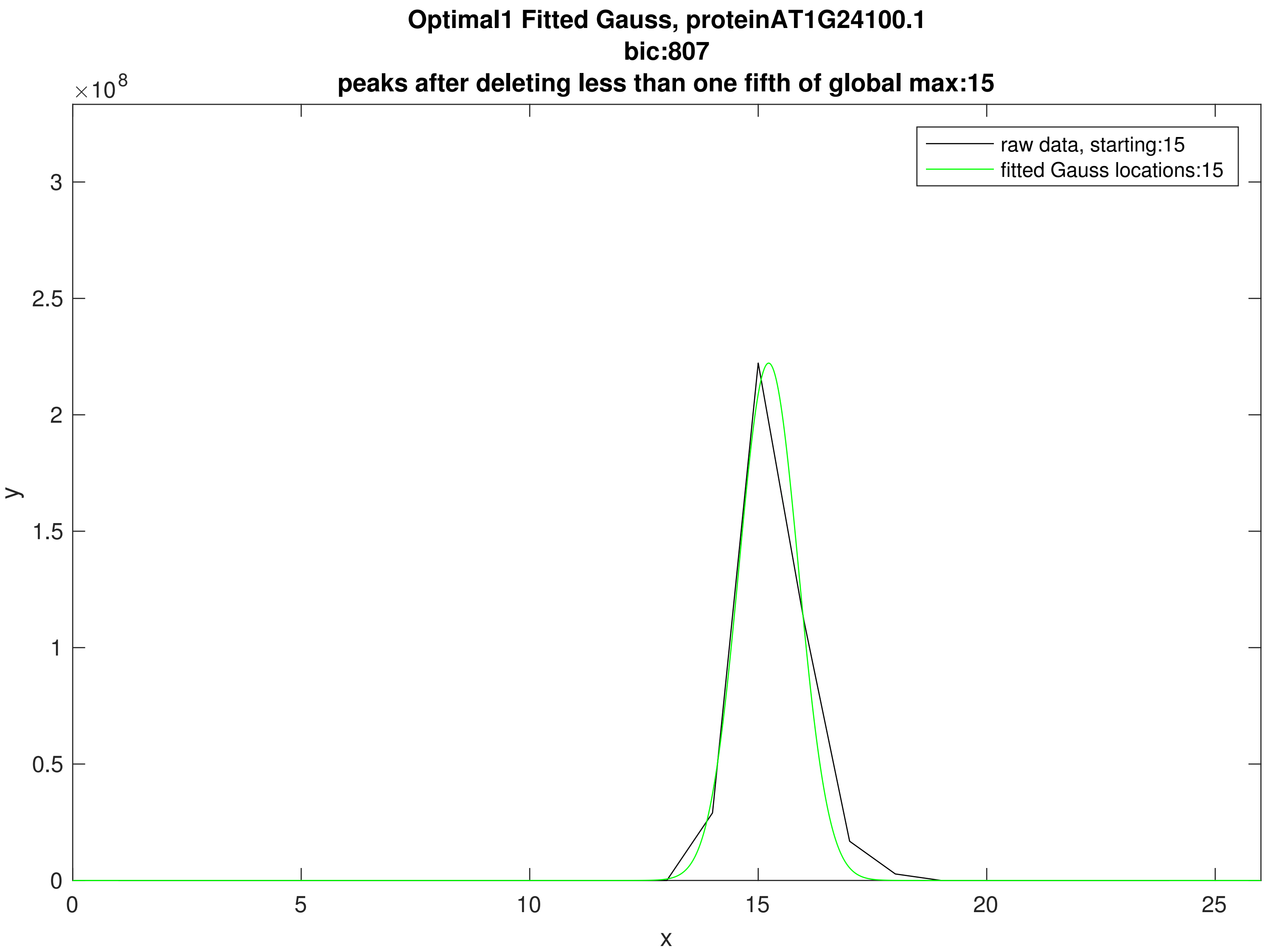
Optimal1 Fitted Gauss, proteinAT1G24100.1
bic:807
peaks after deleting less than one fifth of global max:15
×10^8
raw data, starting:15
fitted Gauss locations:15
y
x
0
0.5
1
1.5
2
2.5
3
0
5
10
15
20
25

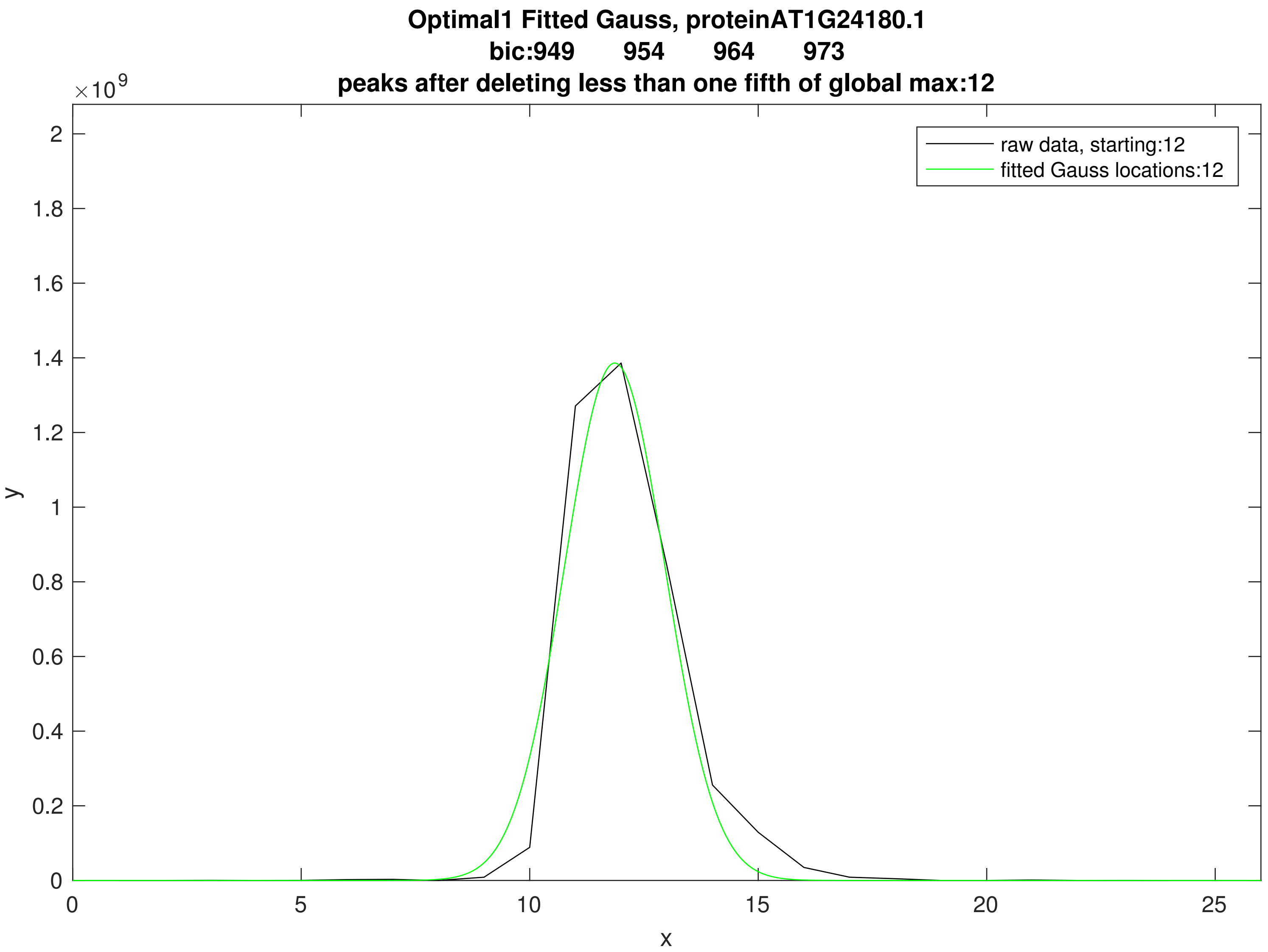
Optimal1 Fitted Gauss, proteinAT1G24180.1
bic:949 954 964 973
peaks after deleting less than one fifth of global max:12
×10^9
raw data, starting:12
fitted Gauss locations:12
y
x
0
0.2
0.4
0.6
0.8
1
1.2
1.4
1.6
1.8
2
5
10
15
20
25

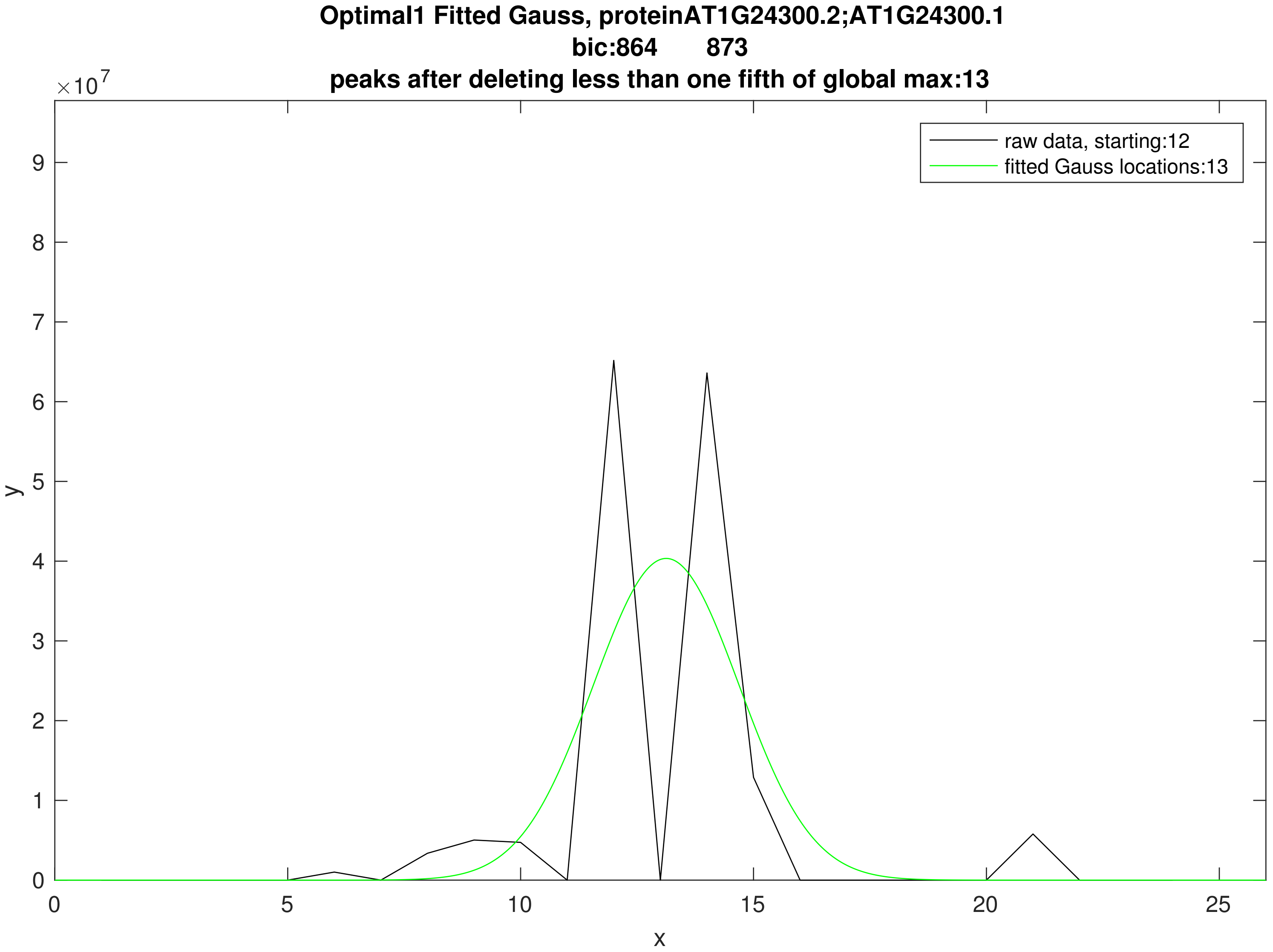
Optimal1 Fitted Gauss, proteinAT1G24300.2;AT1G24300.1
bic:864 873
peaks after deleting less than one fifth of global max:13
×10^7
raw data, starting:12
fitted Gauss locations:13
y
x
0
1
2
3
4
5
6
7
8
9
0
5
10
15
20
25

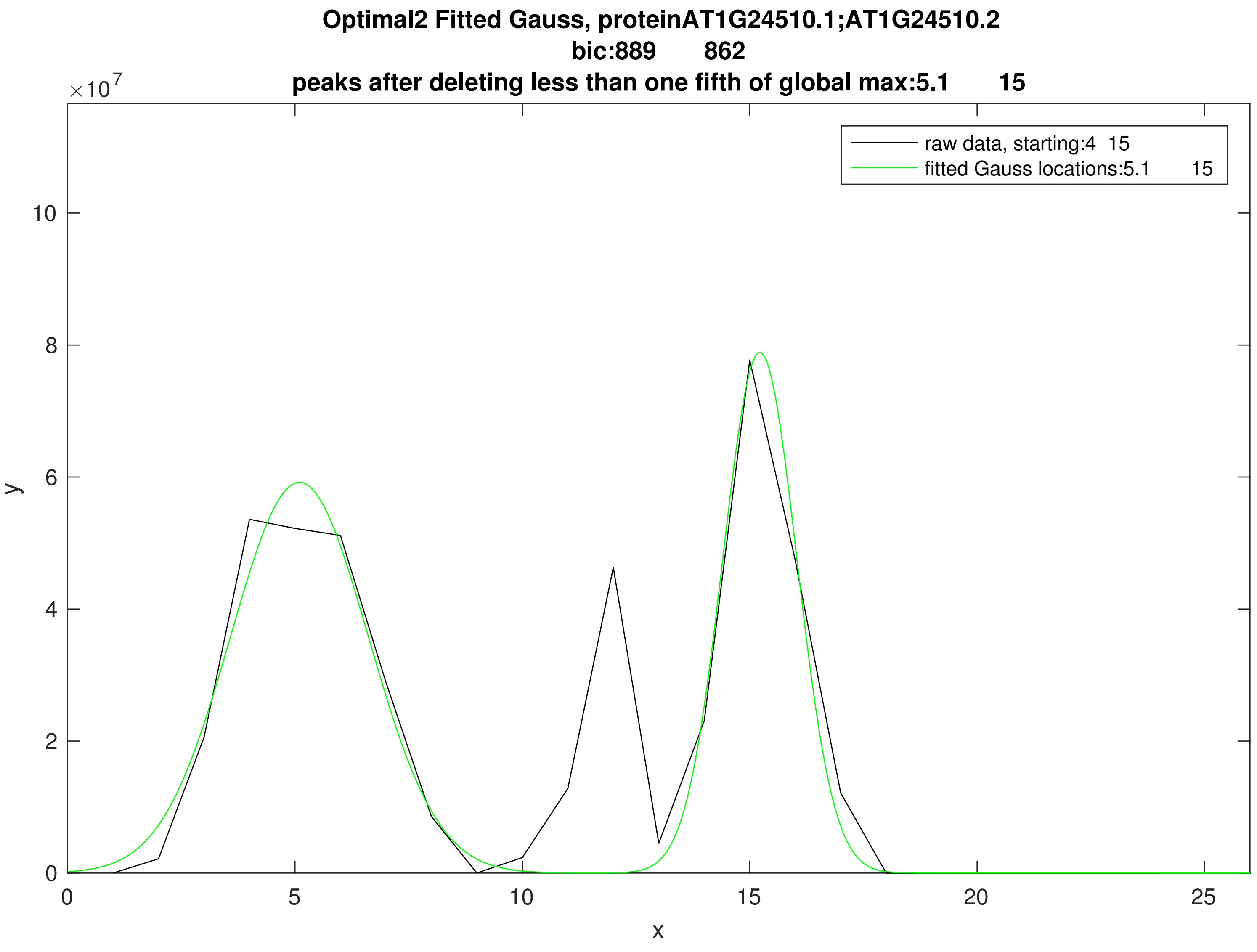

Optimal2 Fitted Gauss, proteinAT1G24510.1;AT1G24510.2
bic:889 862
peaks after deleting less than one fifth of global max:5.1 15
×10^7
raw data, starting:4 15
fitted Gauss locations:5.1 15
10
8
6
4
2
0
y
0
5
10
15
20
25
x

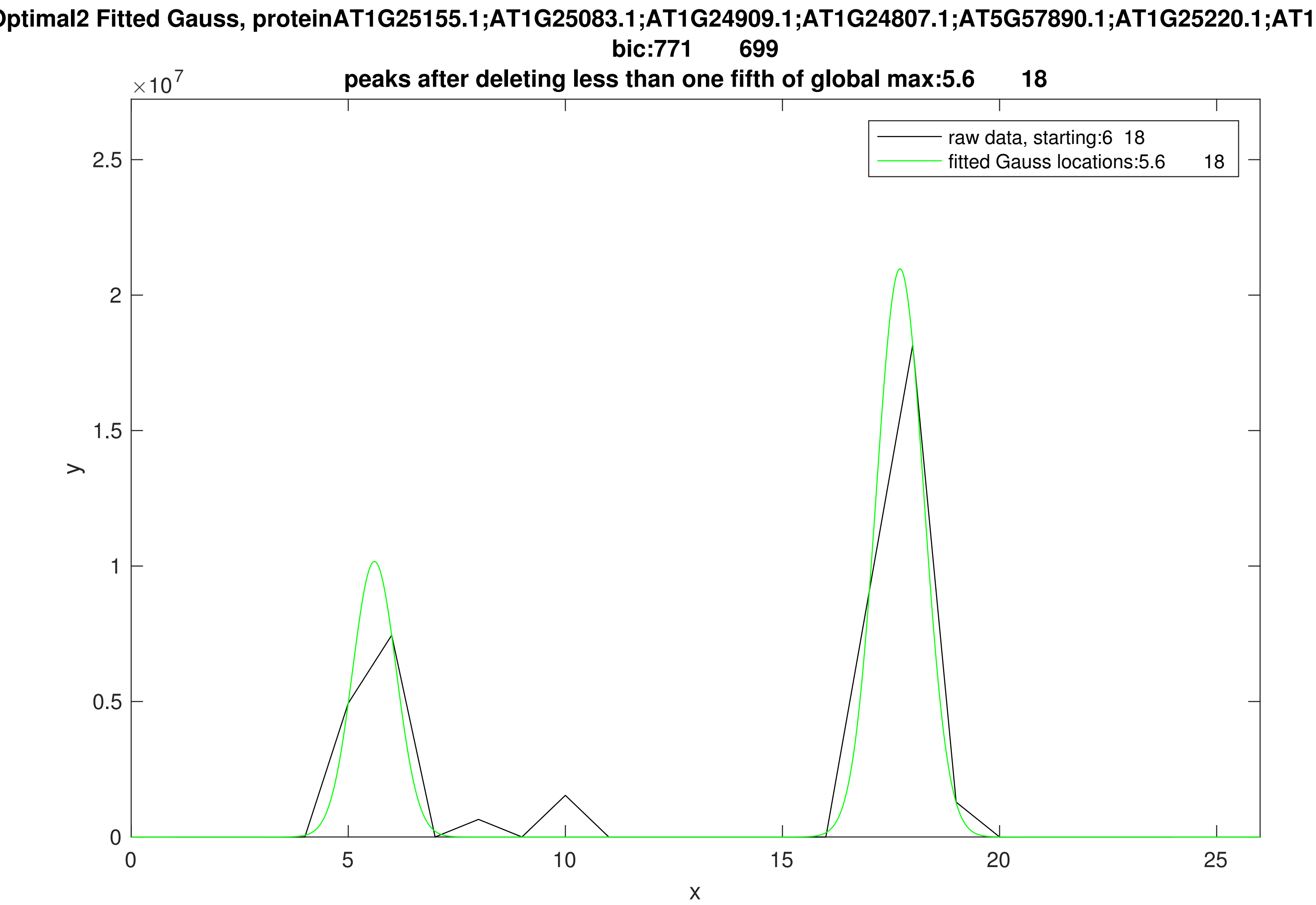
Optimal2 Fitted Gauss, proteinAT1G25155.1;AT1G25083.1;AT1G24909.1;AT1G24807.1;AT5G57890.1;AT1G25220.1;AT1
bic:771 699
peaks after deleting less than one fifth of global max:5.6 18
×10^7
raw data, starting:6 18
fitted Gauss locations:5.6 18
y
x
0
0.5
1
1.5
2
2.5
5
10
15
20
25

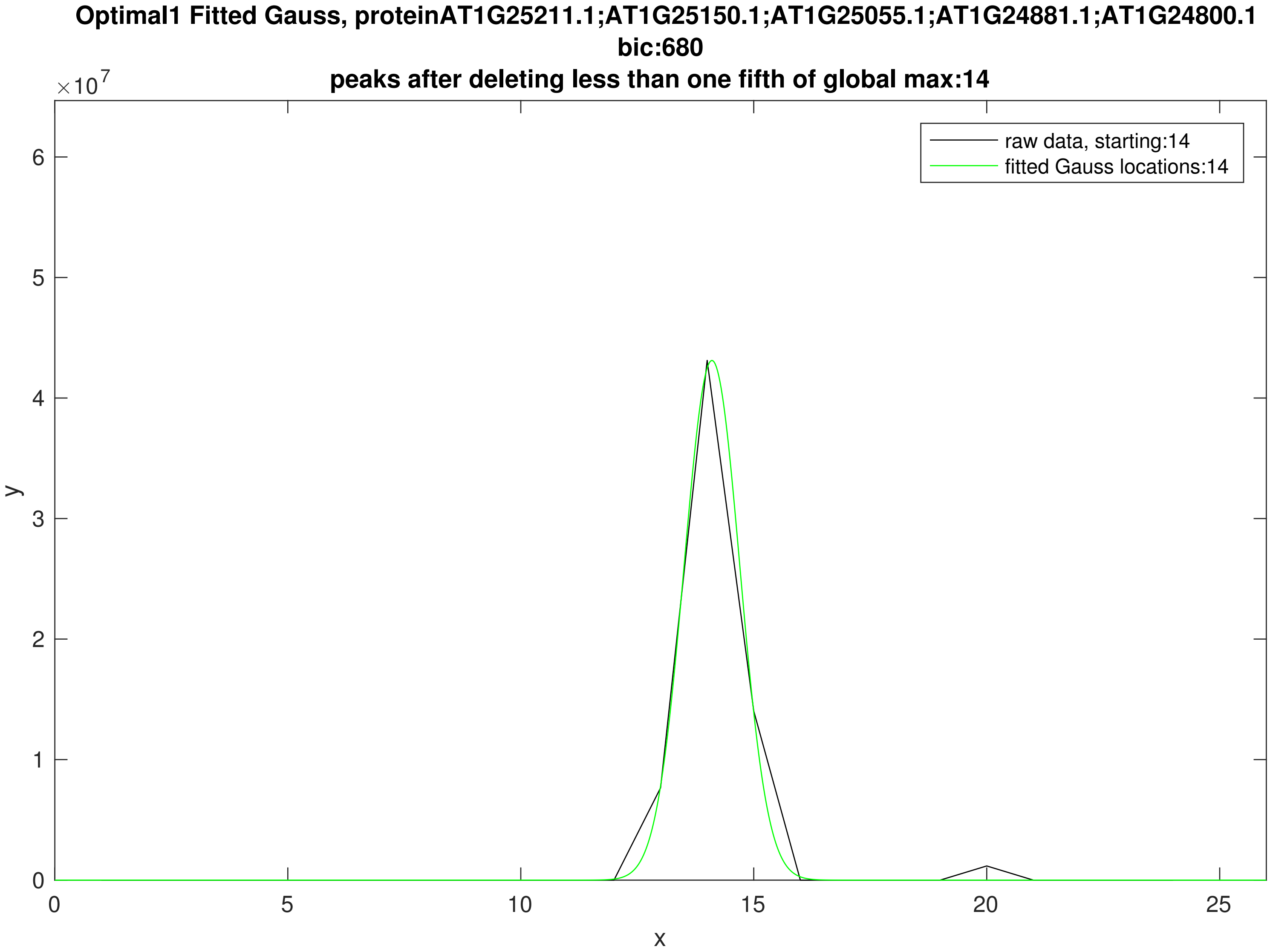
Optimal1 Fitted Gauss, proteinAT1G25211.1;AT1G25150.1;AT1G25055.1;AT1G24881.1;AT1G24800.1
bic:680
peaks after deleting less than one fifth of global max:14
×10^7
raw data, starting:14
fitted Gauss locations:14
y
x
0
1
2
3
4
5
6
0
5
10
15
20
25

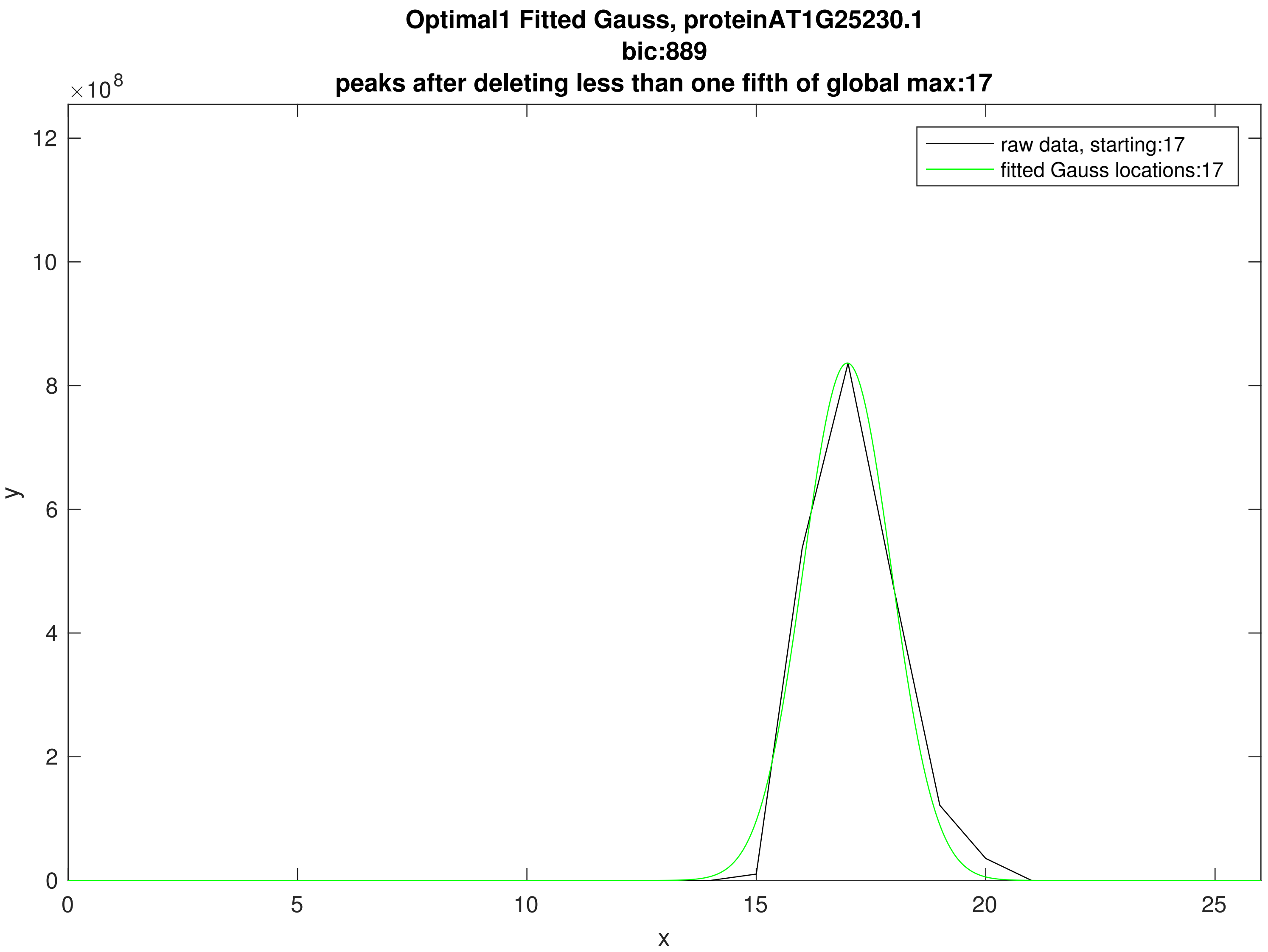
Optimal1 Fitted Gauss, proteinAT1G25230.1
bic:889
peaks after deleting less than one fifth of global max:17
×10^8
12
10
8
6
4
2
0
y
0
5
10
15
20
25
x
raw data, starting:17
fitted Gauss locations:17

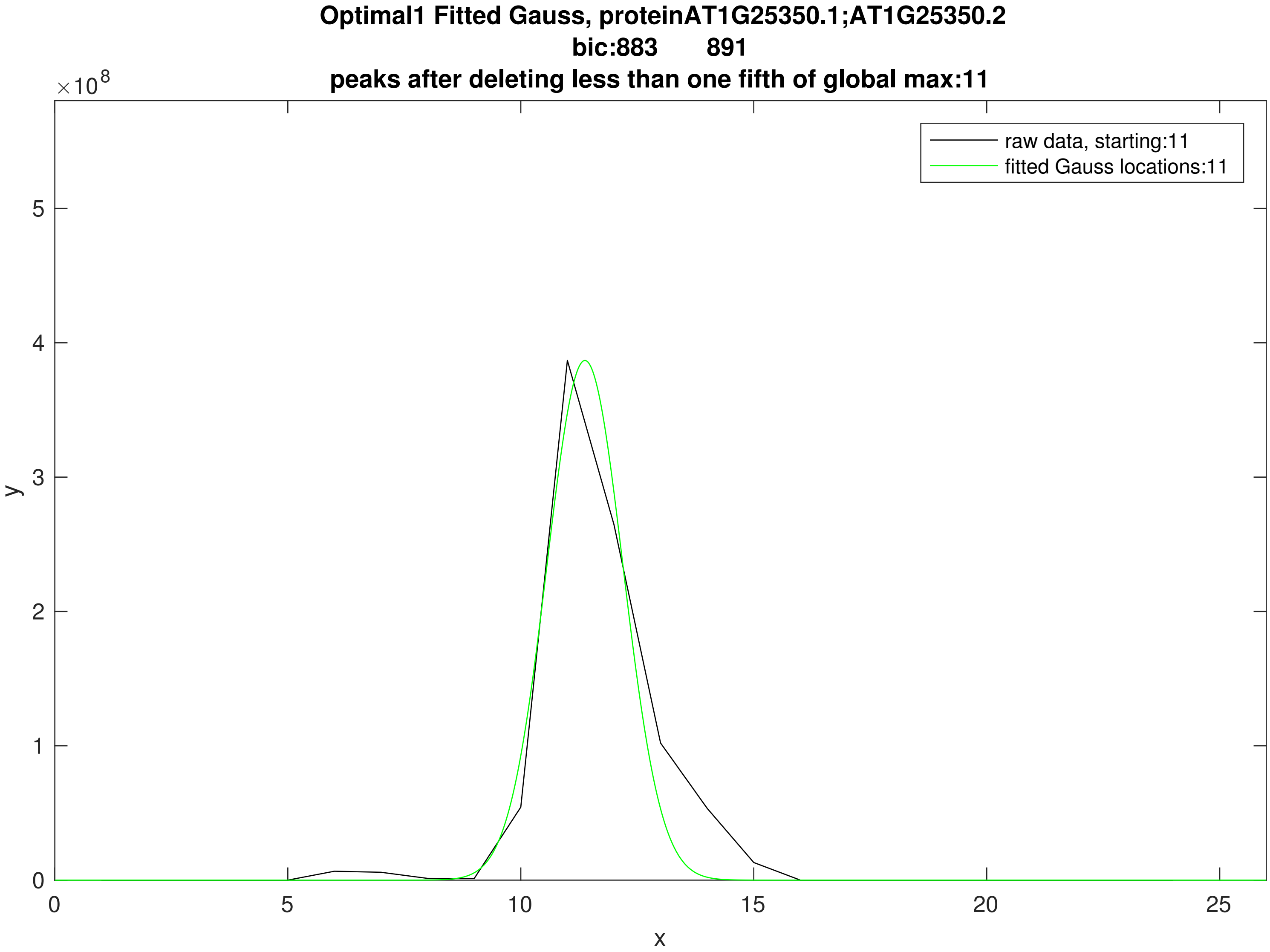
Optimal1 Fitted Gauss, proteinAT1G25350.1;AT1G25350.2
bic:883 891
peaks after deleting less than one fifth of global max:11
×10^8
raw data, starting:11
fitted Gauss locations:11
y
x
0
1
2
3
4
5
0
5
10
15
20
25

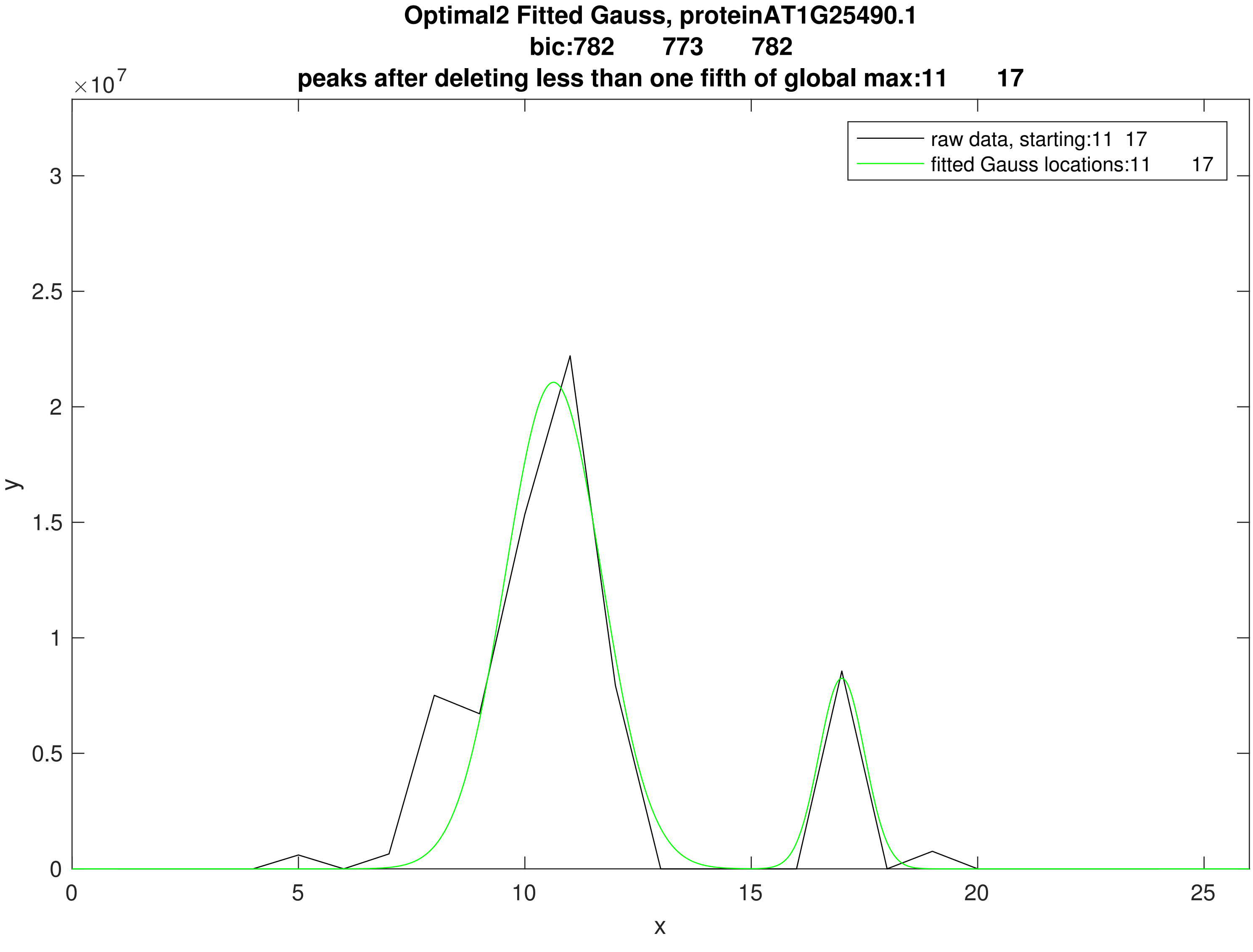
Optimal2 Fitted Gauss, proteinAT1G25490.1
bic:782 773 782
peaks after deleting less than one fifth of global max:11 17
×10^7
raw data, starting:11 17
fitted Gauss locations:11 17
y
x
0
0.5
1
1.5
2
2.5
3
0
5
10
15
20
25

Optimal1 Fitted Gauss, proteinAT1G25570.1
bic:721
peaks after deleting less than one fifth of global max:15

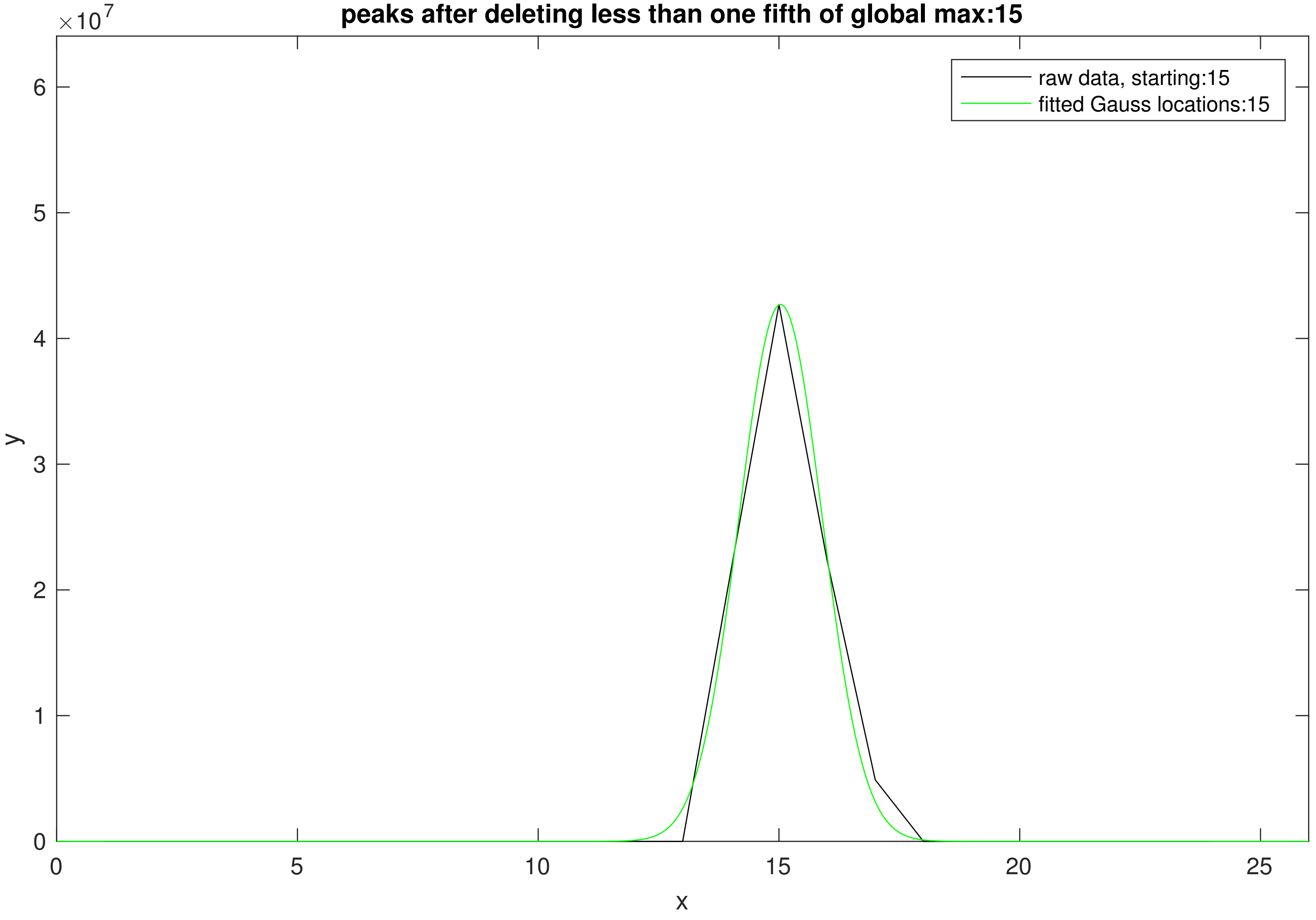

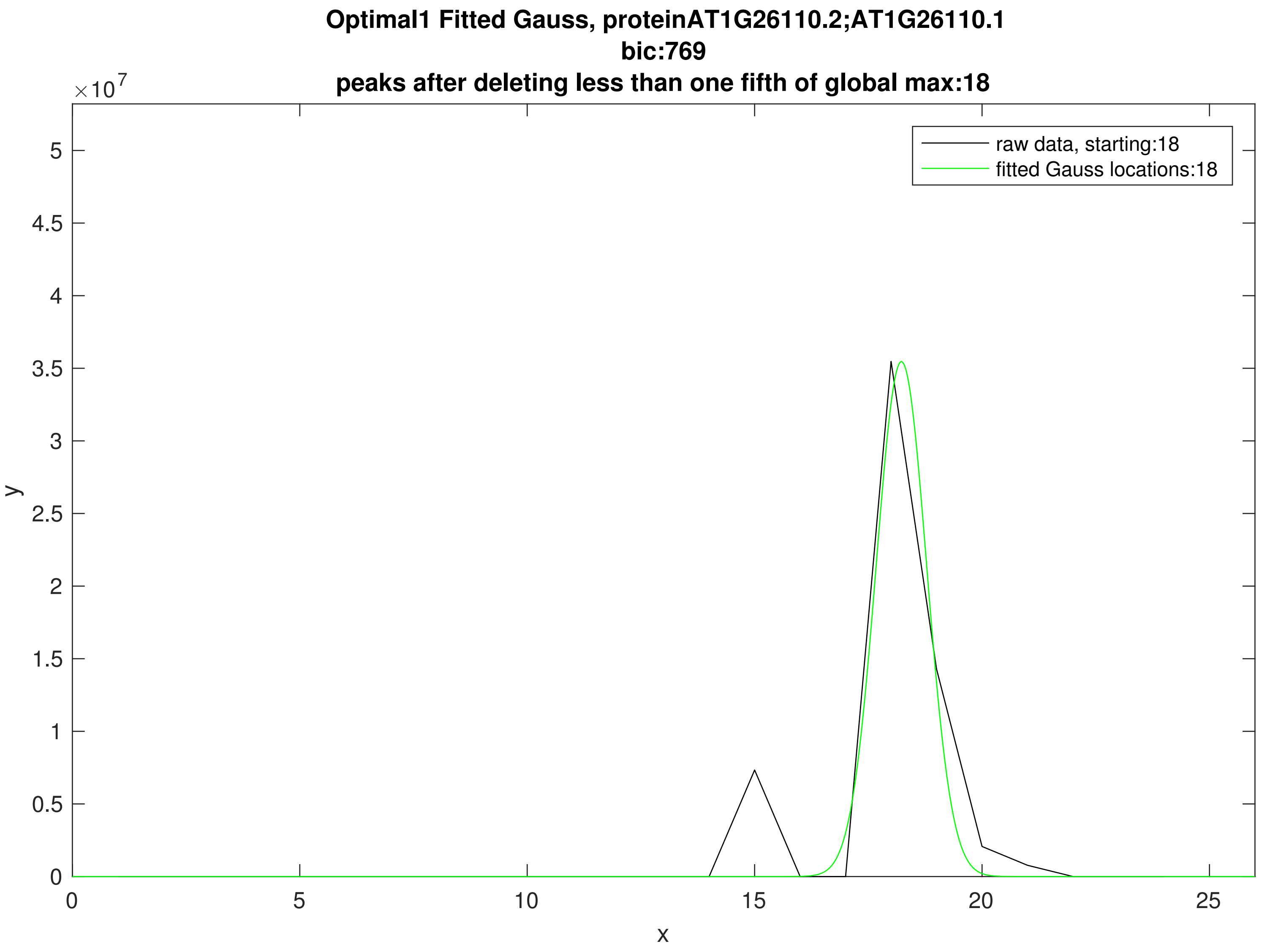
Optimal1 Fitted Gauss, proteinAT1G26110.2;AT1G26110.1
bic:769
peaks after deleting less than one fifth of global max:18
×10^7
raw data, starting:18
fitted Gauss locations:18
y
x
0
0.5
1
1.5
2
2.5
3
3.5
4
4.5
5
0
5
10
15
20
25

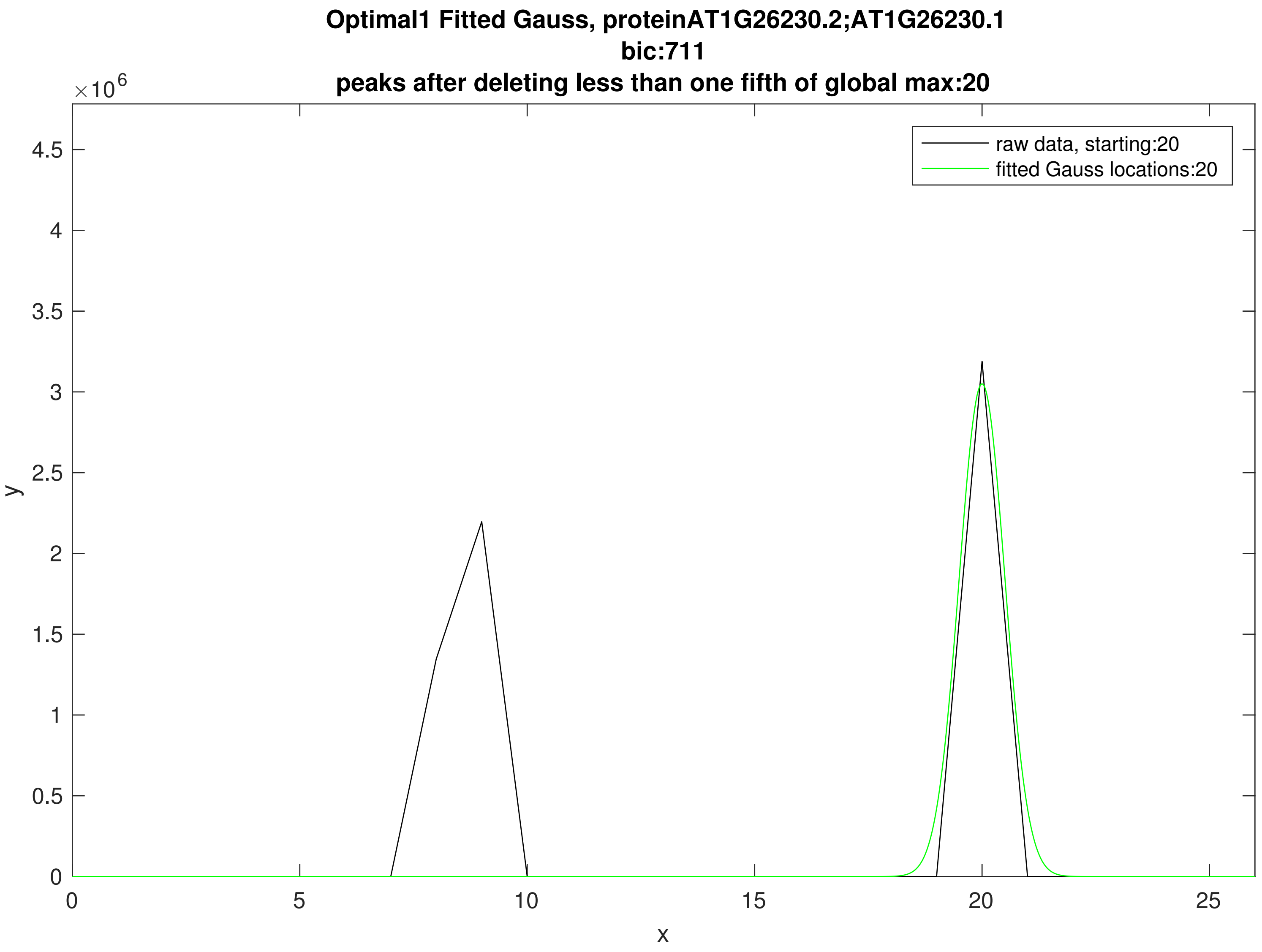
Optimal1 Fitted Gauss, proteinAT1G26230.2;AT1G26230.1
bic:711
peaks after deleting less than one fifth of global max:20
×10^6
4.5
4
3.5
3
2.5
2
1.5
1
0.5
0
y
0
5
10
15
20
25
x
raw data, starting:20
fitted Gauss locations:20

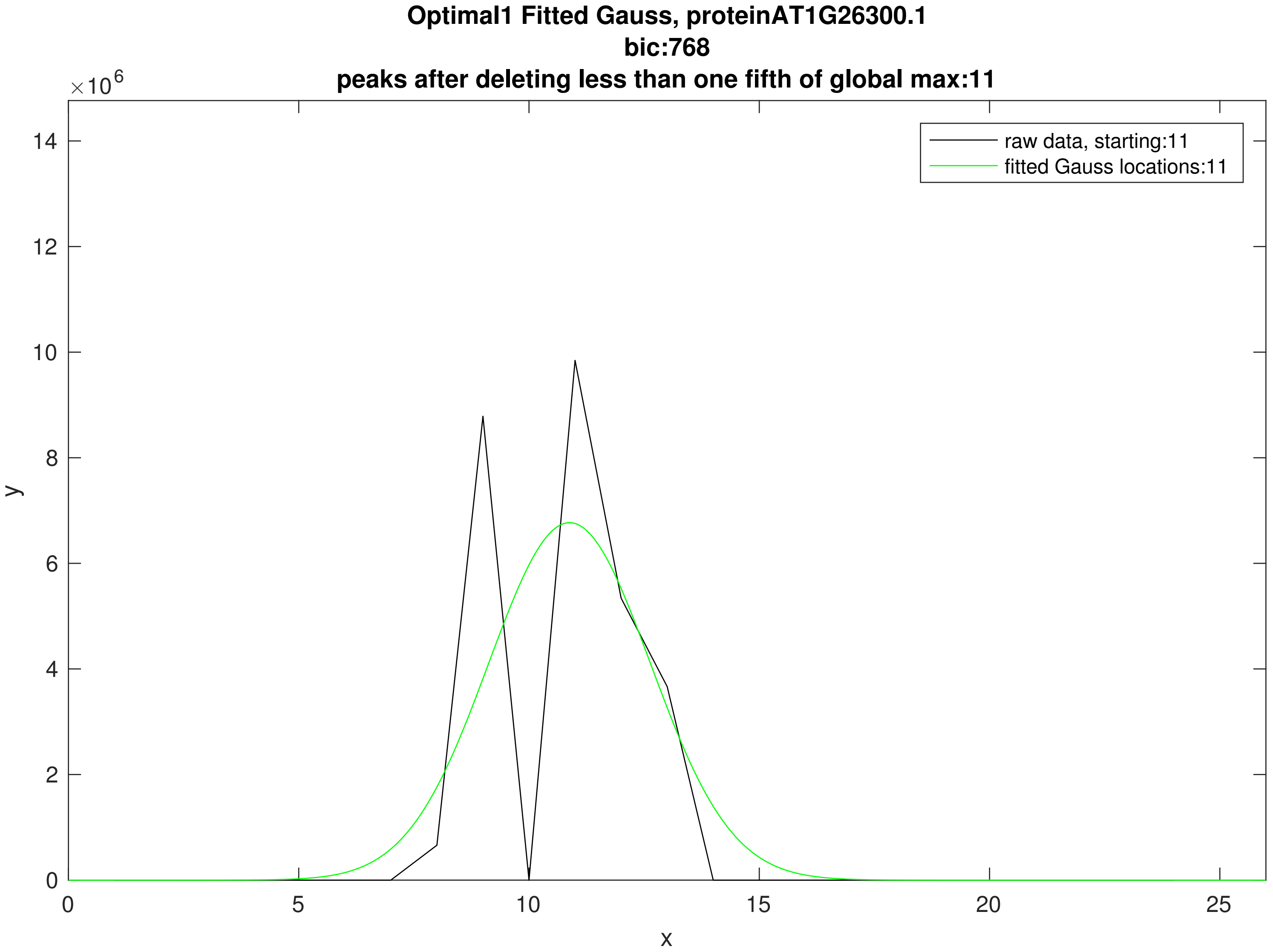

Optimal1 Fitted Gauss, proteinAT1G26300.1
bic:768
peaks after deleting less than one fifth of global max:11
×10^6
14
12
10
8
6
4
2
0
y
0
5
10
15
20
25
x
raw data, starting:11
fitted Gauss locations:11

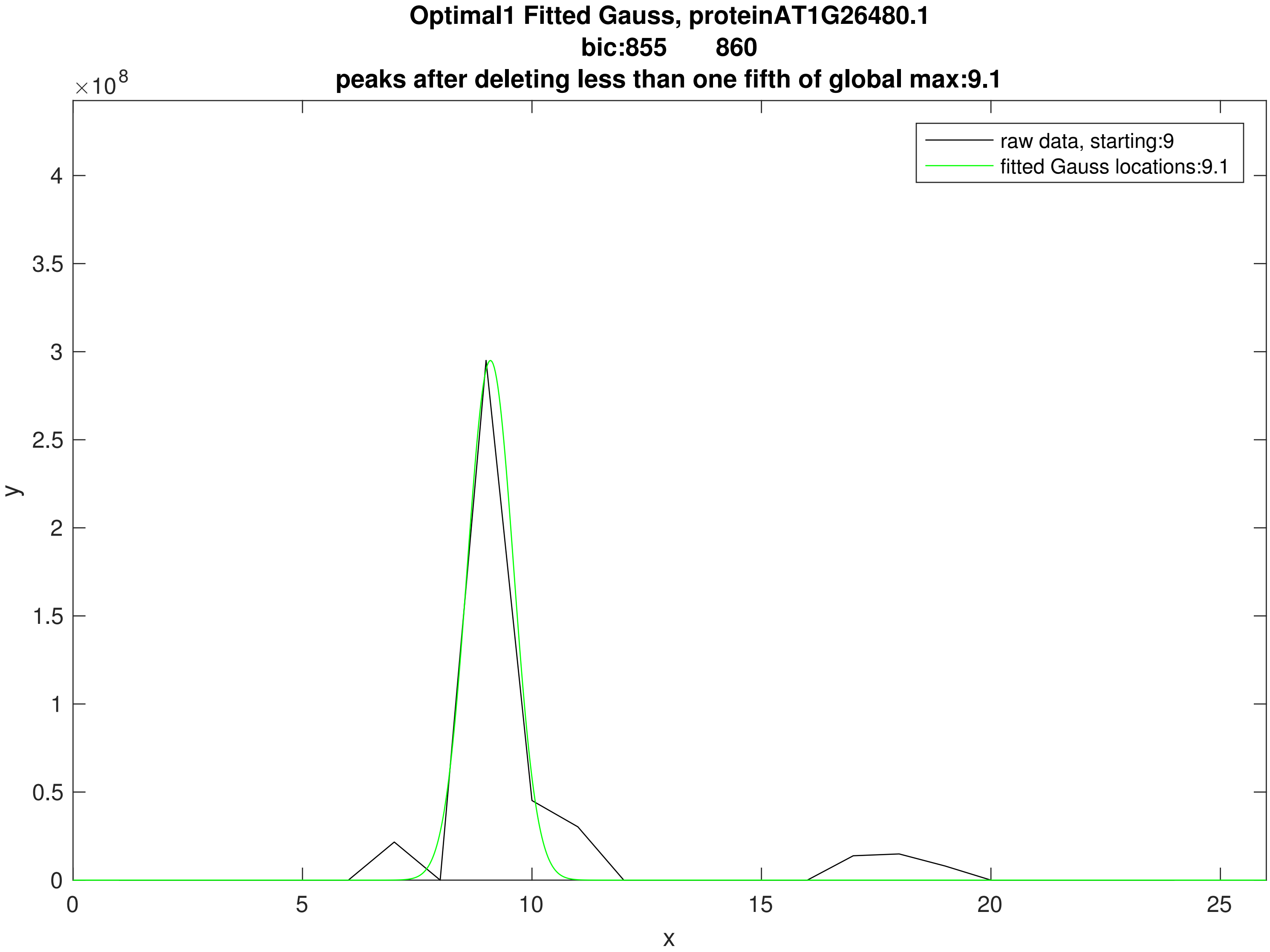
Optimal1 Fitted Gauss, proteinAT1G26480.1
bic:855 860
peaks after deleting less than one fifth of global max:9.1
×10^8
raw data, starting:9
fitted Gauss locations:9.1
y
x
0
0.5
1
1.5
2
2.5
3
3.5
4
5
10
15
20
25

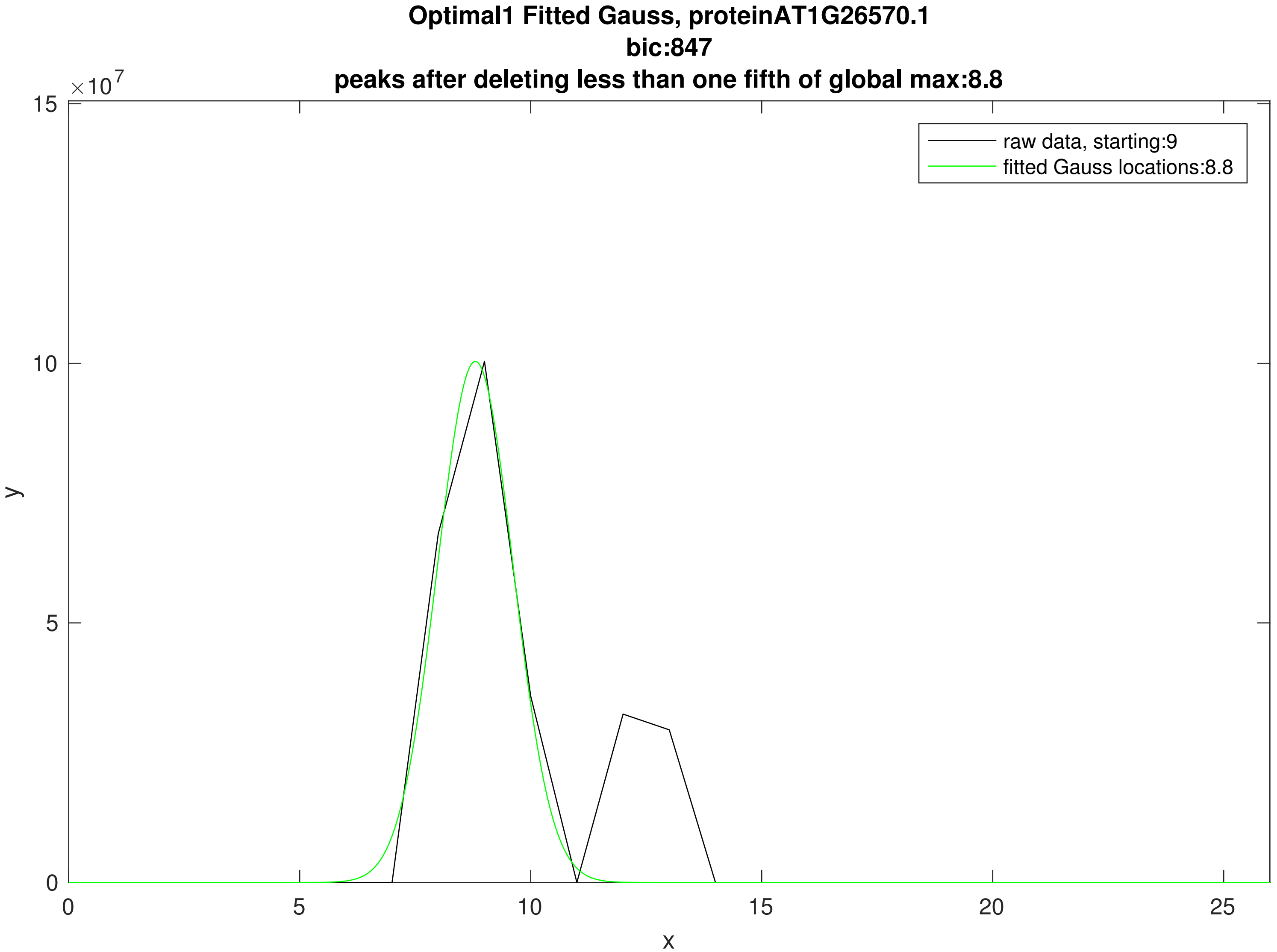
Optimal1 Fitted Gauss, proteinAT1G26570.1
bic:847
peaks after deleting less than one fifth of global max:8.8
×10^7
15
10
5
0
y
0
5
10
15
20
25
x
raw data, starting:9
fitted Gauss locations:8.8

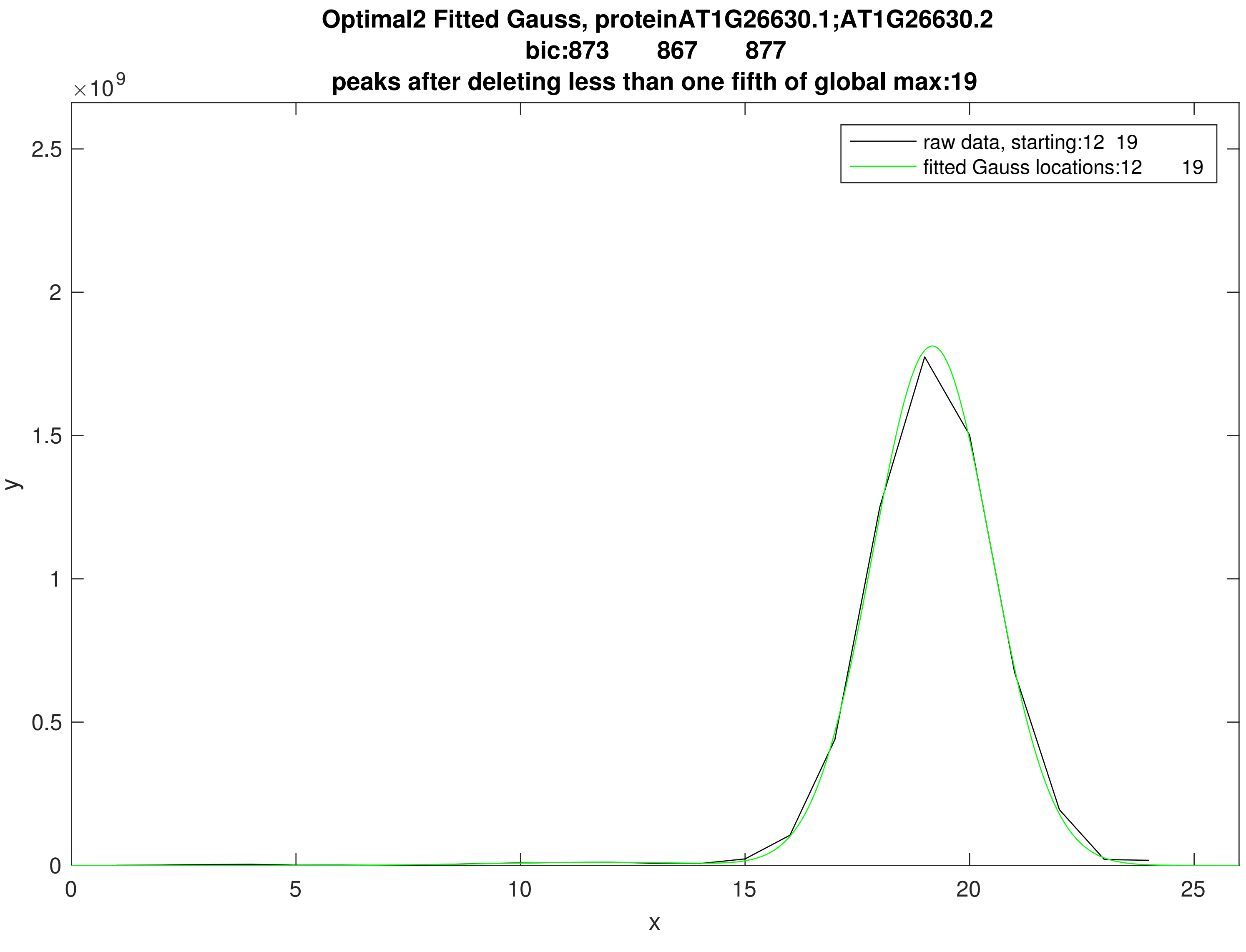
Optimal2 Fitted Gauss, proteinAT1G26630.1;AT1G26630.2
bic:873 867 877
peaks after deleting less than one fifth of global max:19
×10^9
2.5
2
1.5
1
0.5
0
y
0
5
10
15
20
25
x
raw data, starting:12 19
fitted Gauss locations:12 19

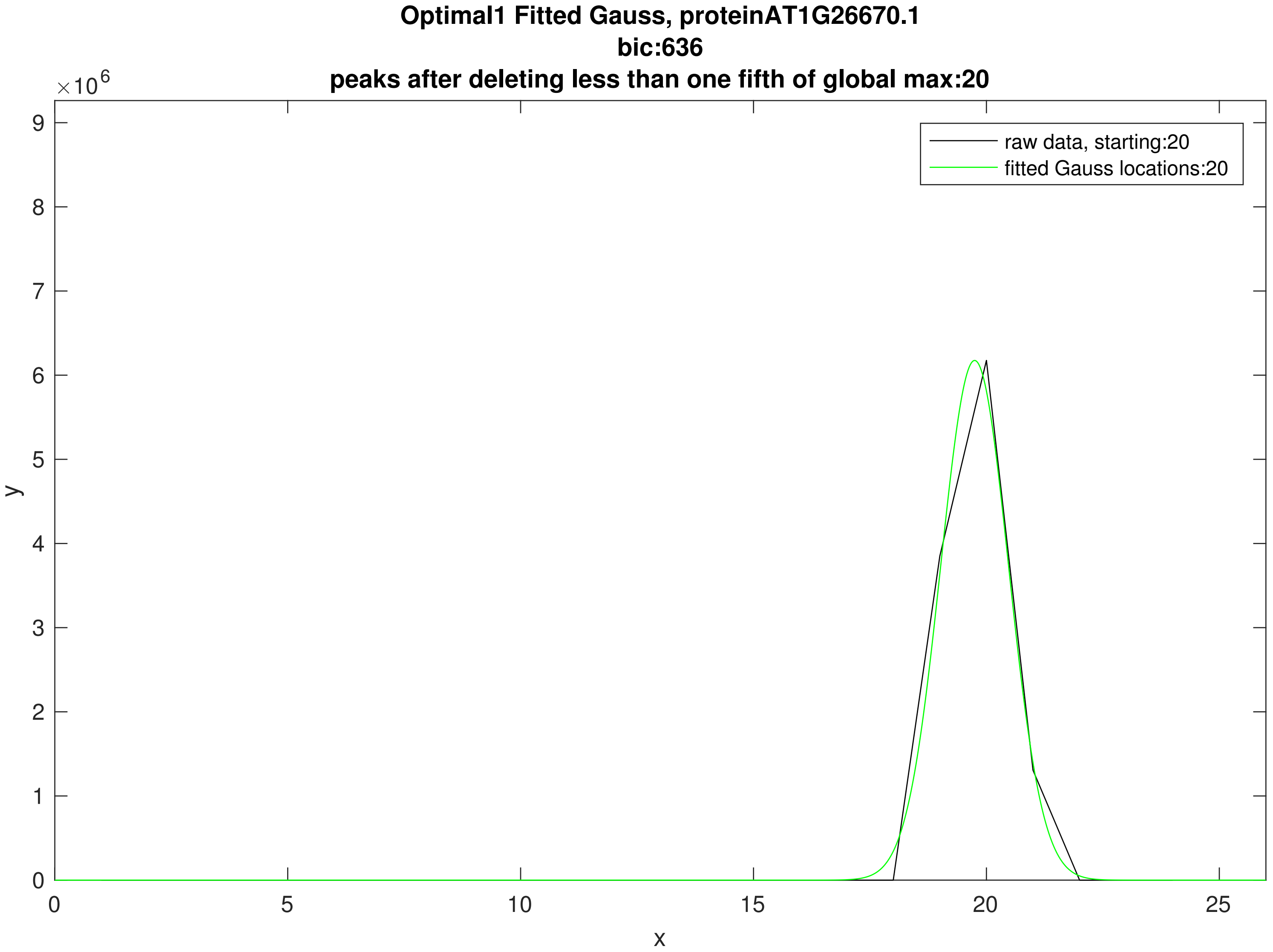
Optimal1 Fitted Gauss, proteinAT1G26670.1
bic:636
peaks after deleting less than one fifth of global max:20
×10^6
raw data, starting:20
fitted Gauss locations:20
y
x
0
1
2
3
4
5
6
7
8
9
0
5
10
15
20
25

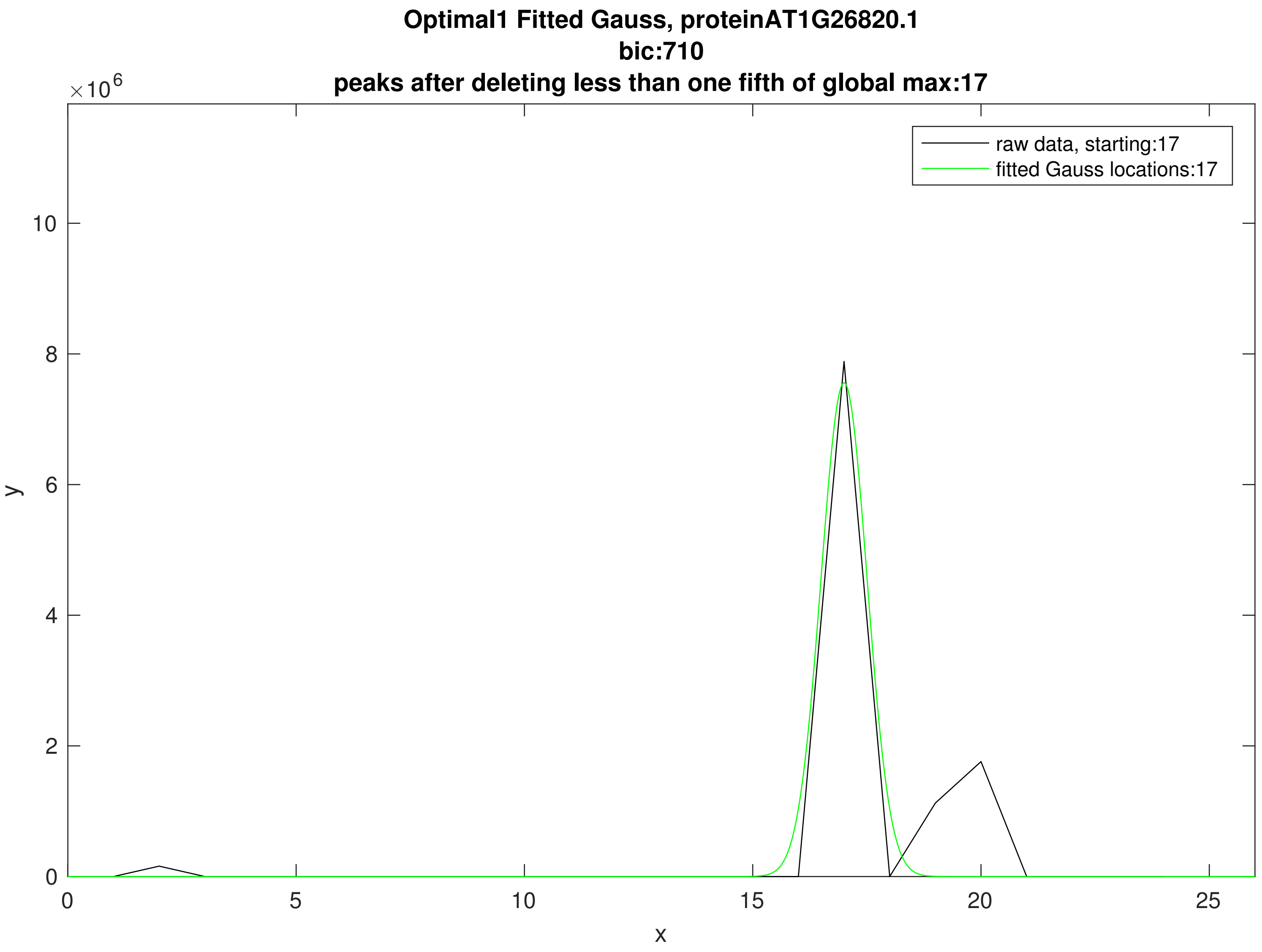

Optimal1 Fitted Gauss, proteinAT1G26820.1
bic:710
peaks after deleting less than one fifth of global max:17
×10^6
raw data, starting:17
fitted Gauss locations:17
10
8
6
4
2
0
y
0
5
10
15
20
25
x

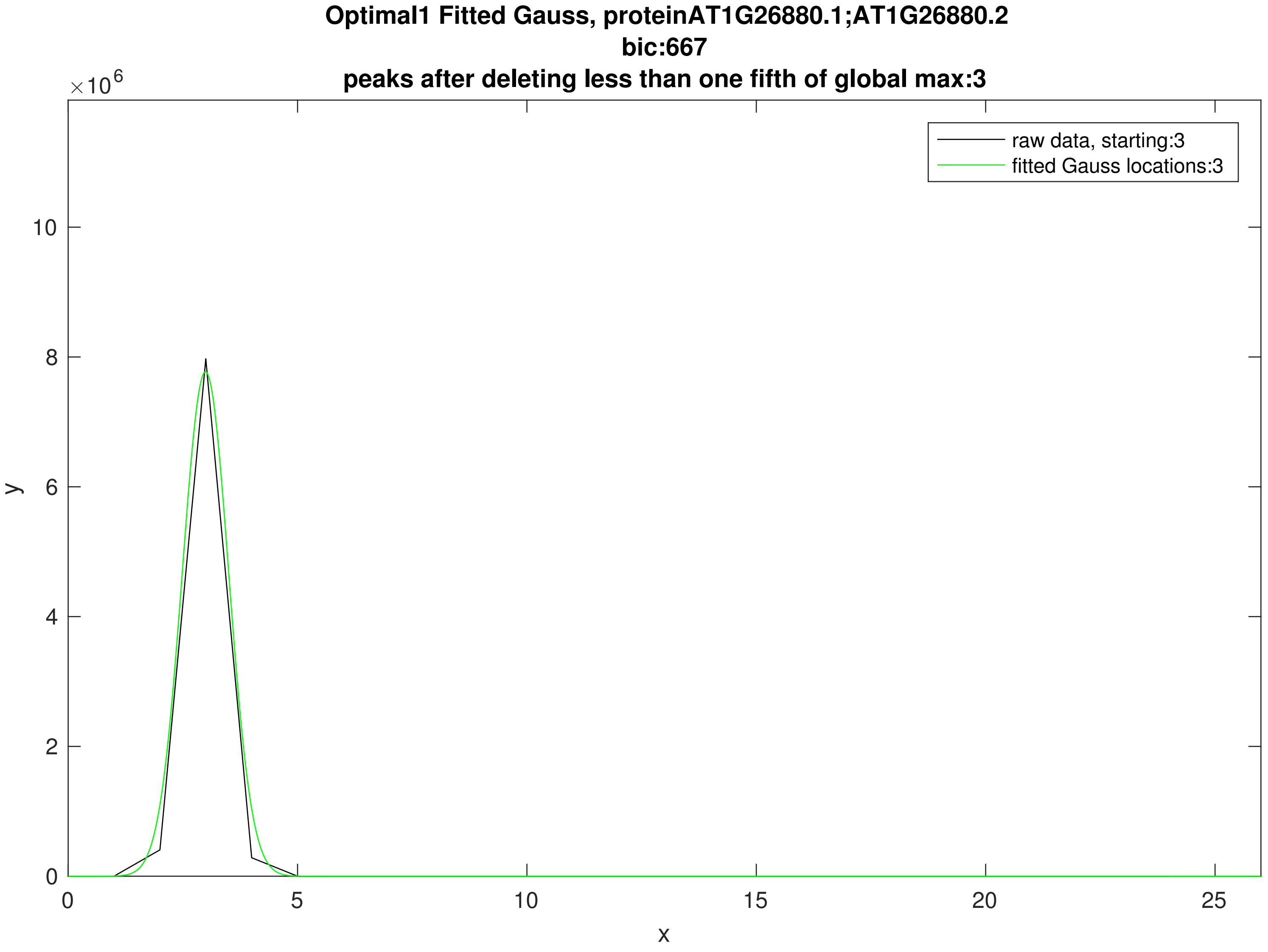
Optimal1 Fitted Gauss, proteinAT1G26880.1;AT1G26880.2
bic:667
peaks after deleting less than one fifth of global max:3
×10^6
raw data, starting:3
fitted Gauss locations:3
y
x
0
2
4
6
8
10
0
5
10
15
20
25

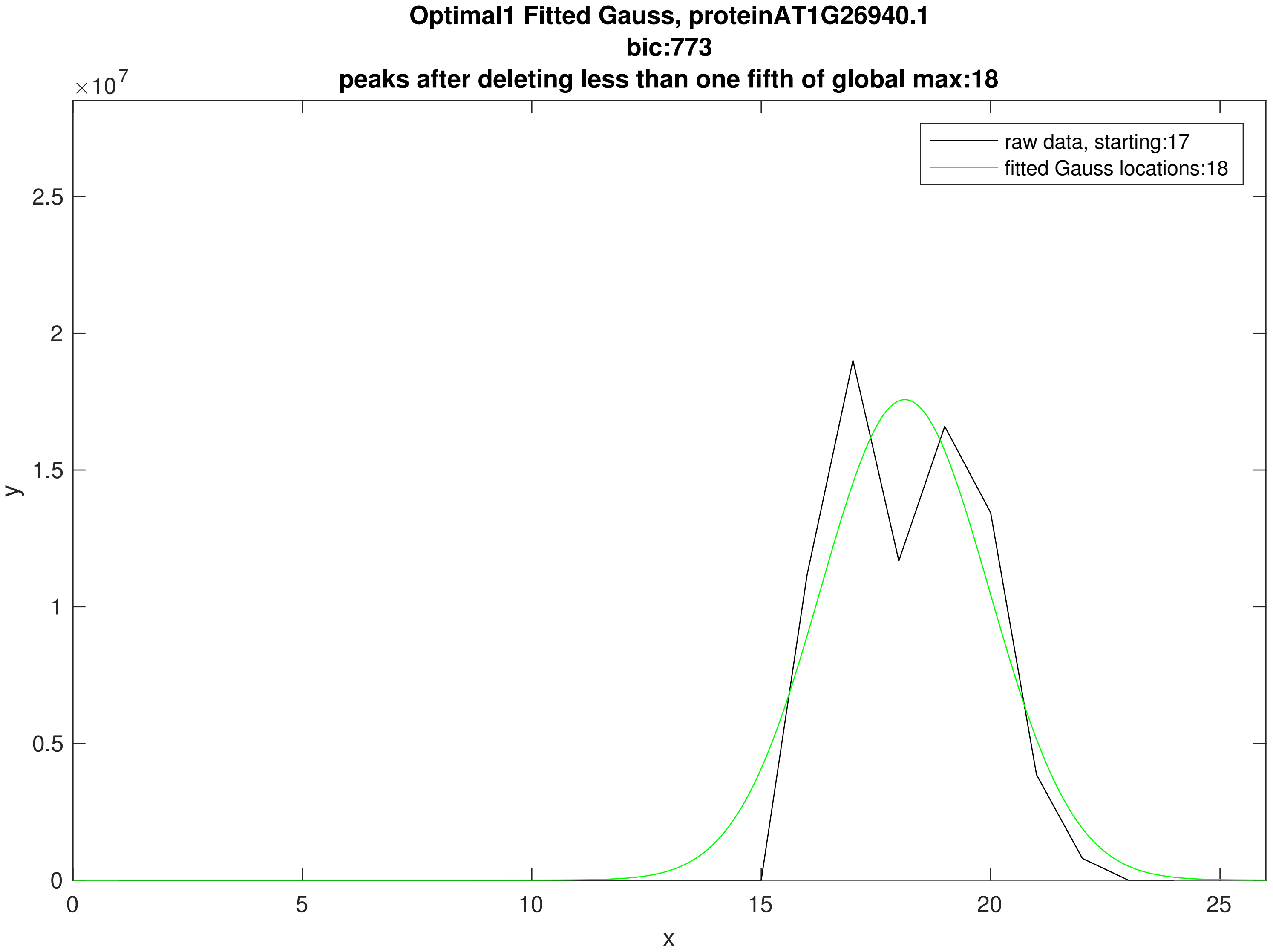
Optimal1 Fitted Gauss, proteinAT1G26940.1
bic:773
peaks after deleting less than one fifth of global max:18
×10^7
raw data, starting:17
fitted Gauss locations:18
2.5
2
1.5
1
0.5
0
y
0
5
10
15
20
25
x

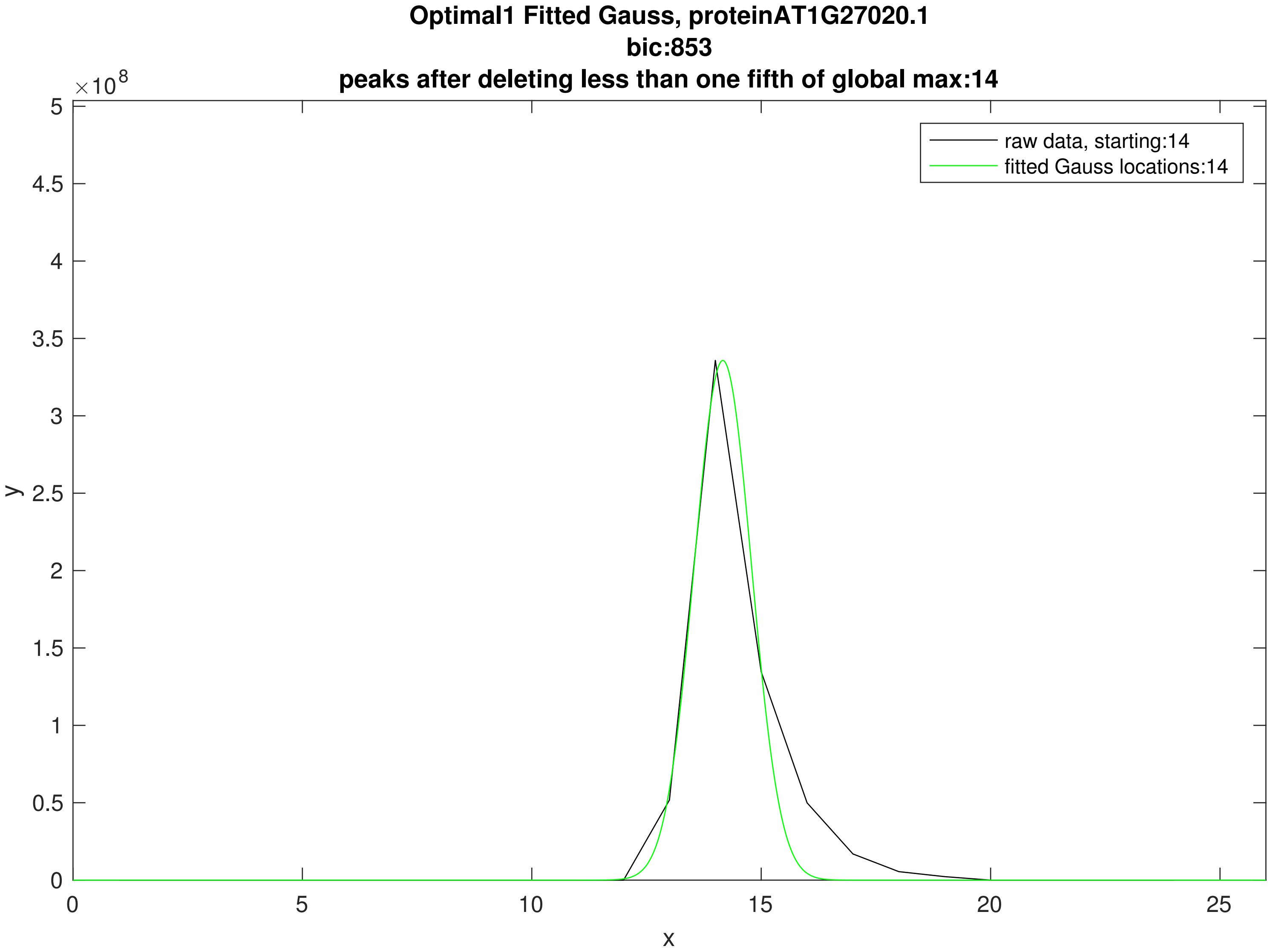

Optimal1 Fitted Gauss, proteinAT1G27020.1
bic:853
peaks after deleting less than one fifth of global max:14
×10^8
5
4.5
4
3.5
3
2.5
2
1.5
1
0.5
0
y
0
5
10
15
20
25
x
raw data, starting:14
fitted Gauss locations:14

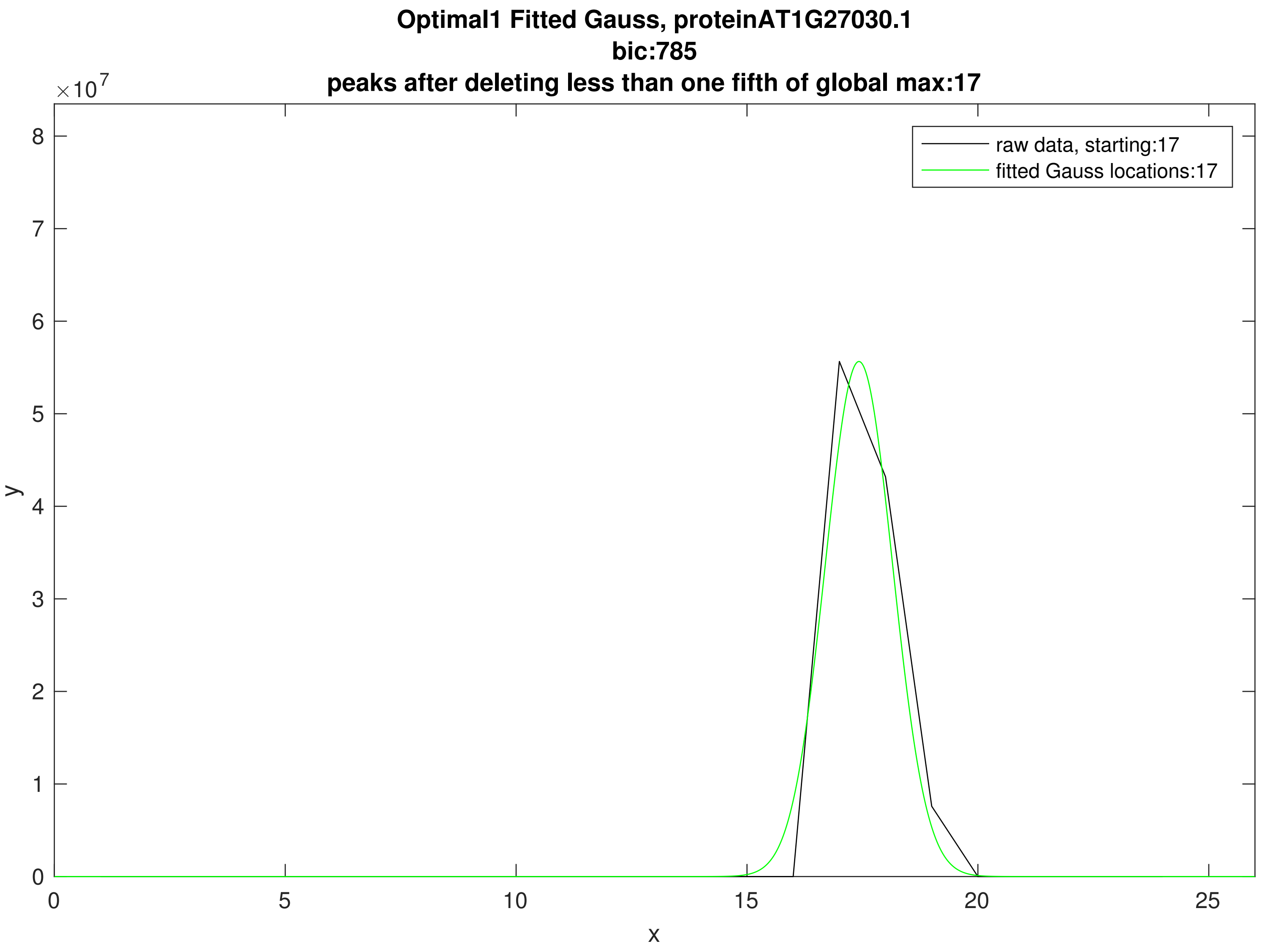

Optimal1 Fitted Gauss, proteinAT1G27030.1
bic:785
peaks after deleting less than one fifth of global max:17
×10^7
raw data, starting:17
fitted Gauss locations:17
y
x
0
1
2
3
4
5
6
7
8
0
5
10
15
20
25

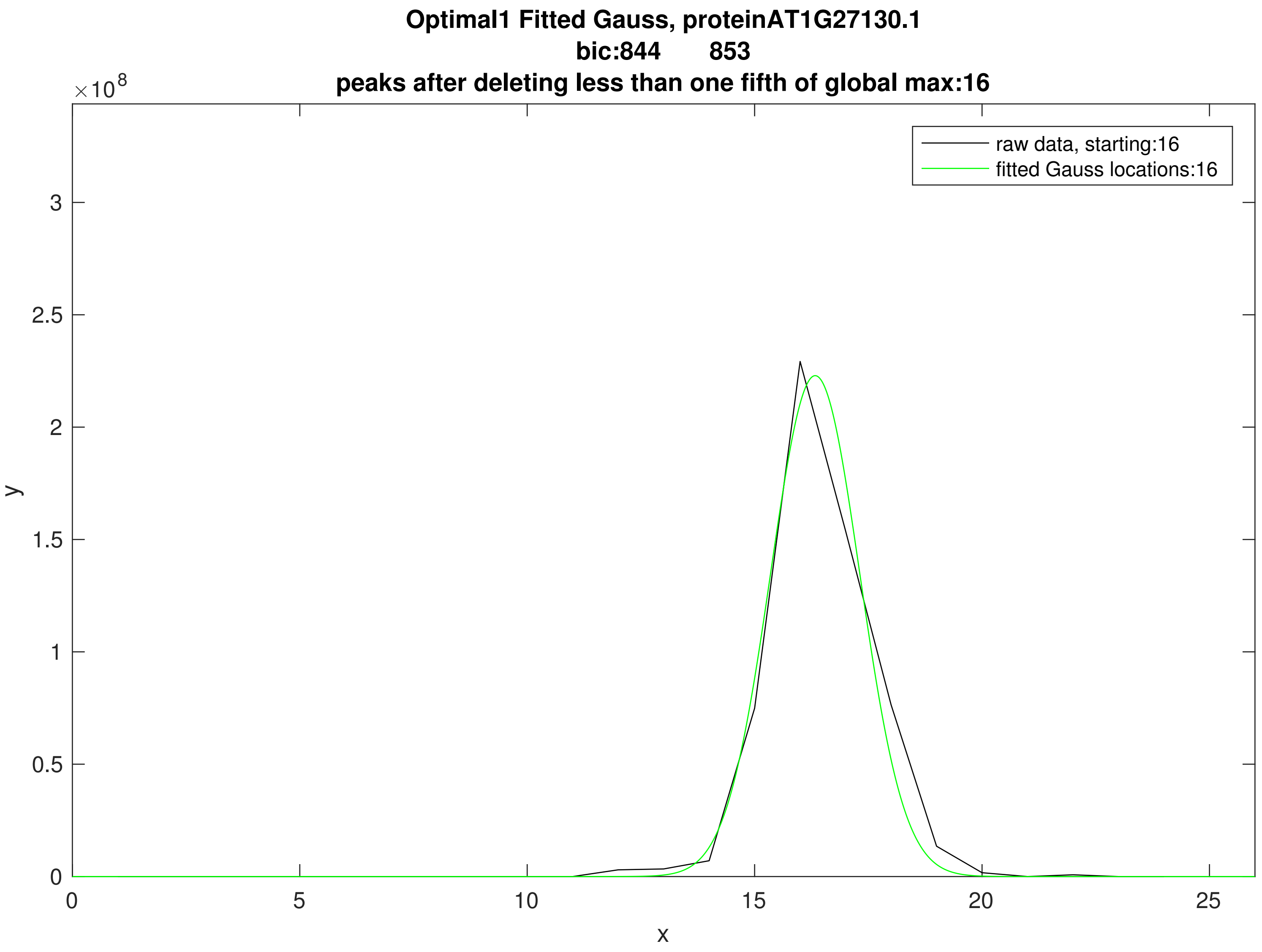

Optimal1 Fitted Gauss, proteinAT1G27130.1
bic:844 853
peaks after deleting less than one fifth of global max:16
×10^8
raw data, starting:16
fitted Gauss locations:16
y
x
0
0.5
1
1.5
2
2.5
3
0
5
10
15
20
25

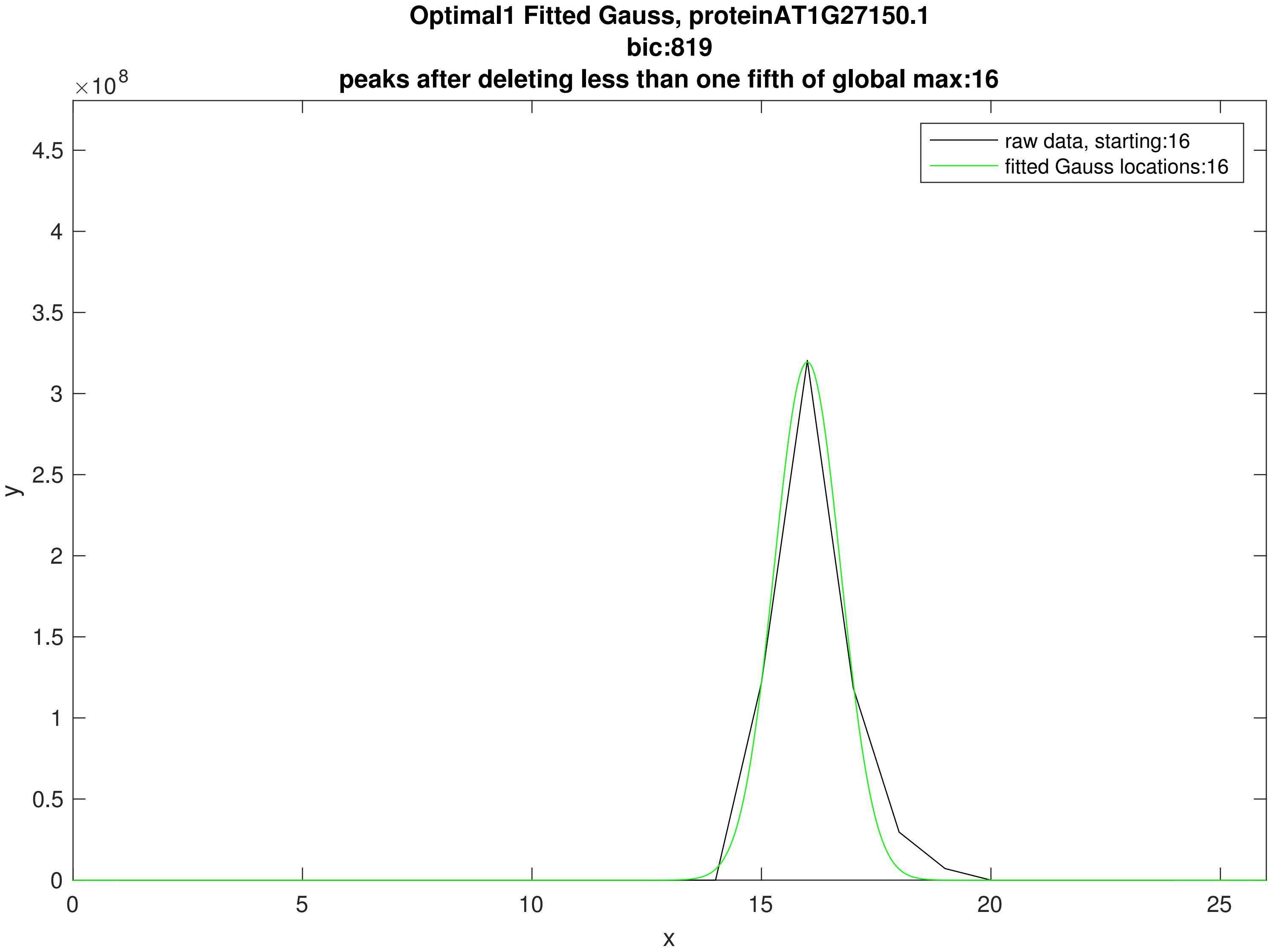
Optimal1 Fitted Gauss, proteinAT1G27150.1
bic:819
peaks after deleting less than one fifth of global max:16
×10^8
raw data, starting:16
fitted Gauss locations:16
y
x
0
0.5
1
1.5
2
2.5
3
3.5
4
4.5
0
5
10
15
20
25

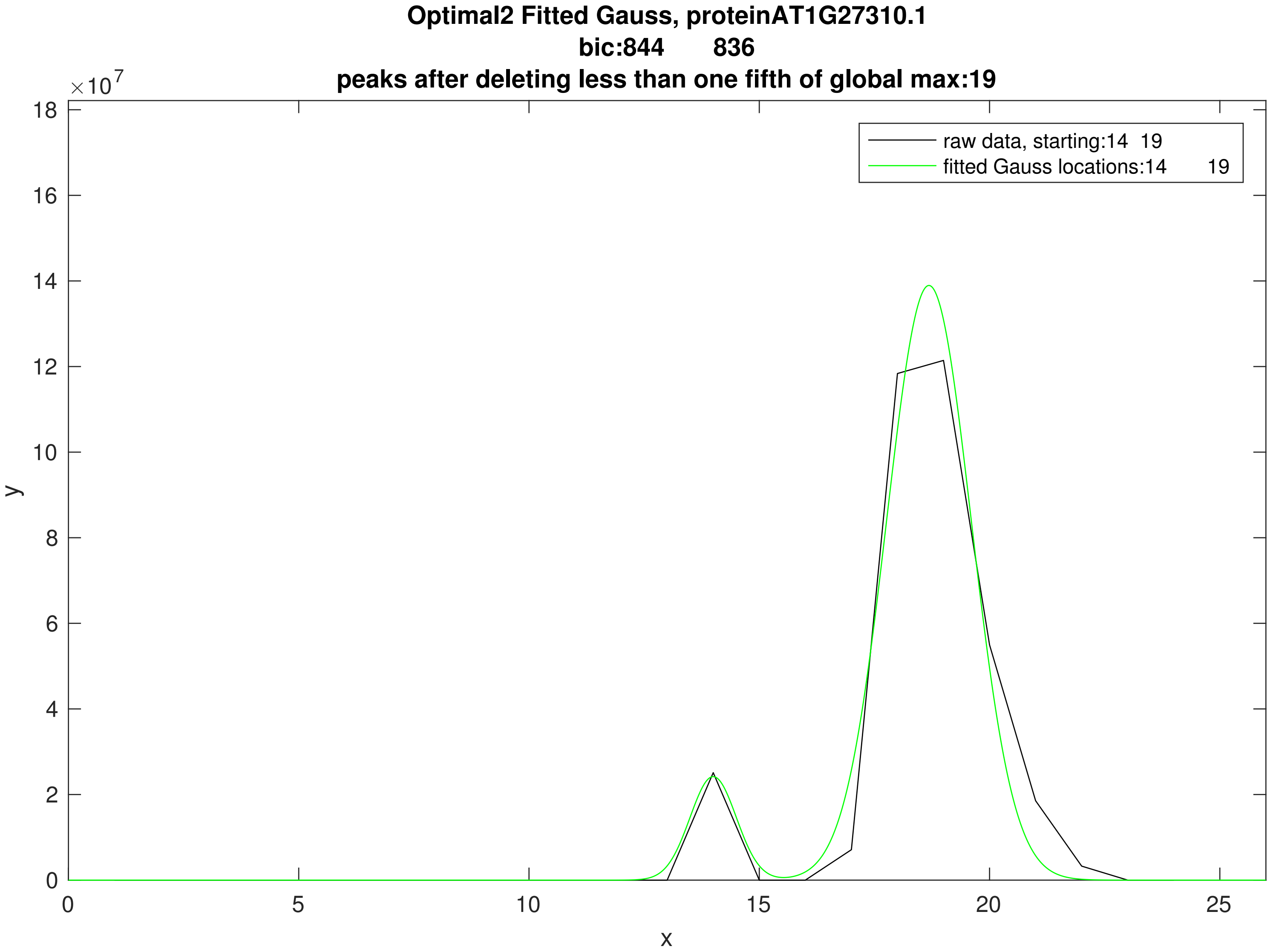
Optimal2 Fitted Gauss, proteinAT1G27310.1
bic:844 836
peaks after deleting less than one fifth of global max:19
raw data, starting:14 19
fitted Gauss locations:14 19
×10^7
y
x
0
2
4
6
8
10
12
14
16
18
0
5
10
15
20
25

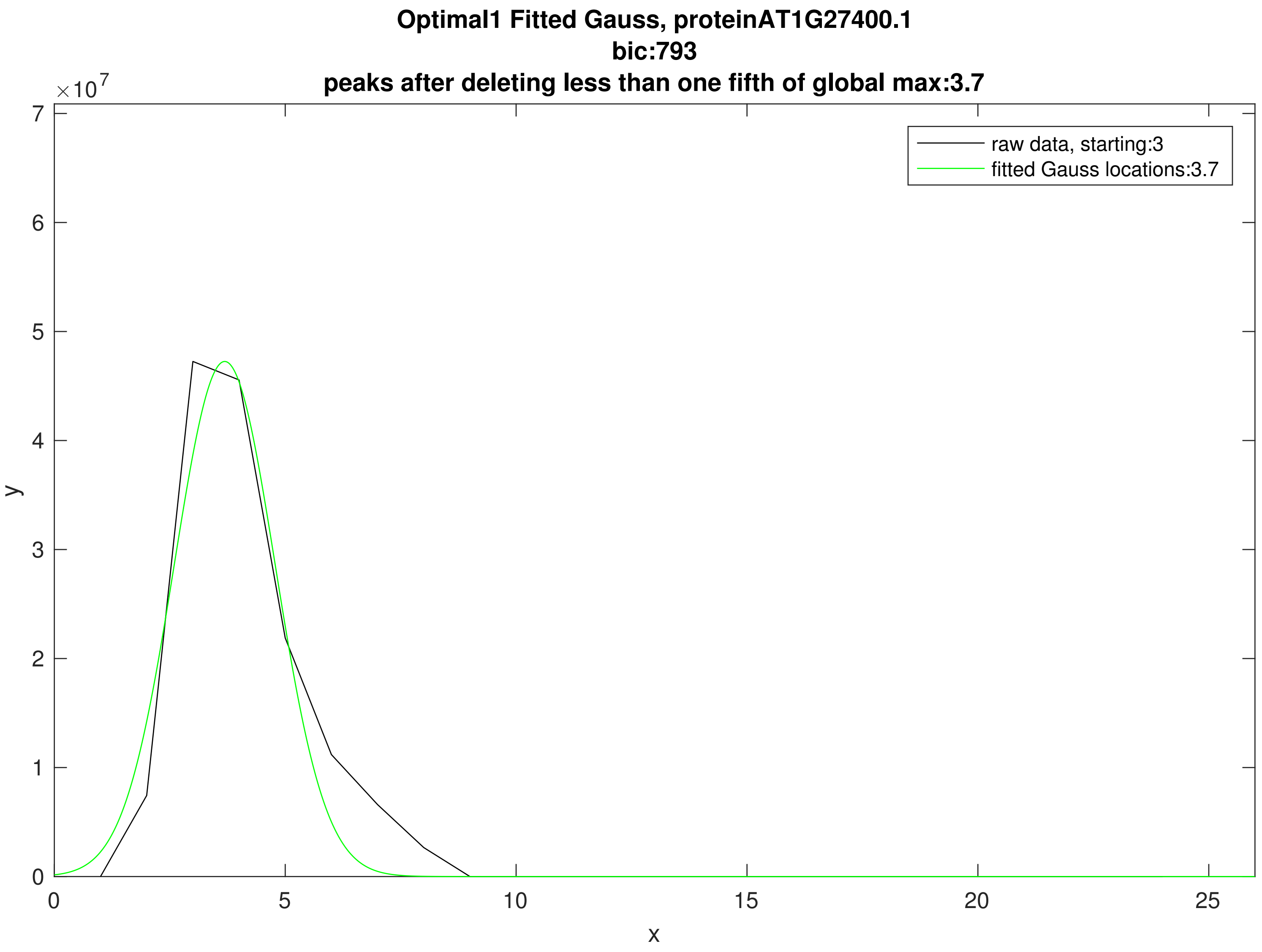
Optimal1 Fitted Gauss, proteinAT1G27400.1
bic:793
peaks after deleting less than one fifth of global max:3.7
×10^7
raw data, starting:3
fitted Gauss locations:3.7
y
x
0
1
2
3
4
5
6
7
0
5
10
15
20
25

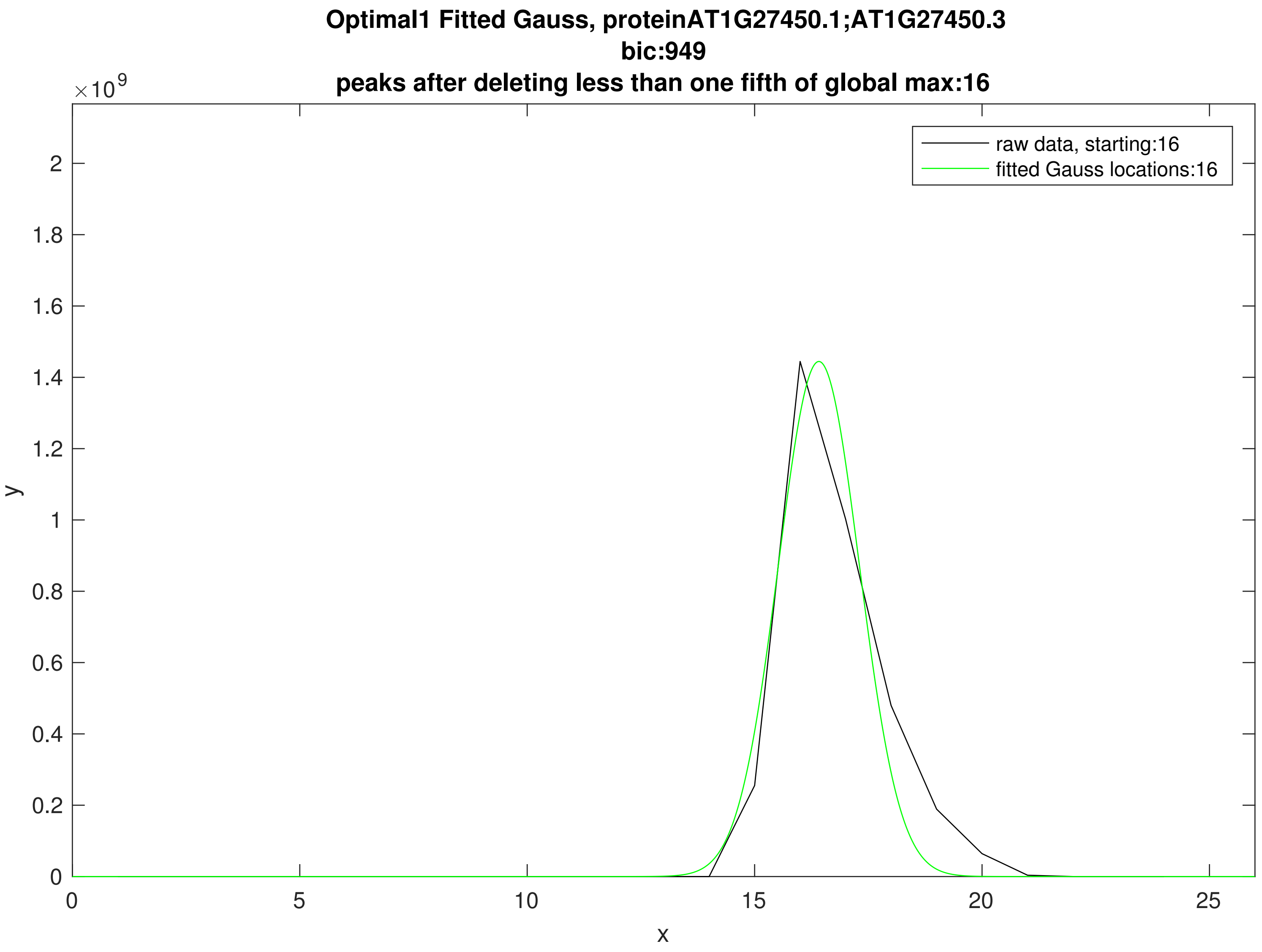

Optimal1 Fitted Gauss, proteinAT1G27450.1;AT1G27450.3
bic:949
peaks after deleting less than one fifth of global max:16
×10^9
raw data, starting:16
fitted Gauss locations:16
y
x
0
0.2
0.4
0.6
0.8
1
1.2
1.4
1.6
1.8
2
0
5
10
15
20
25

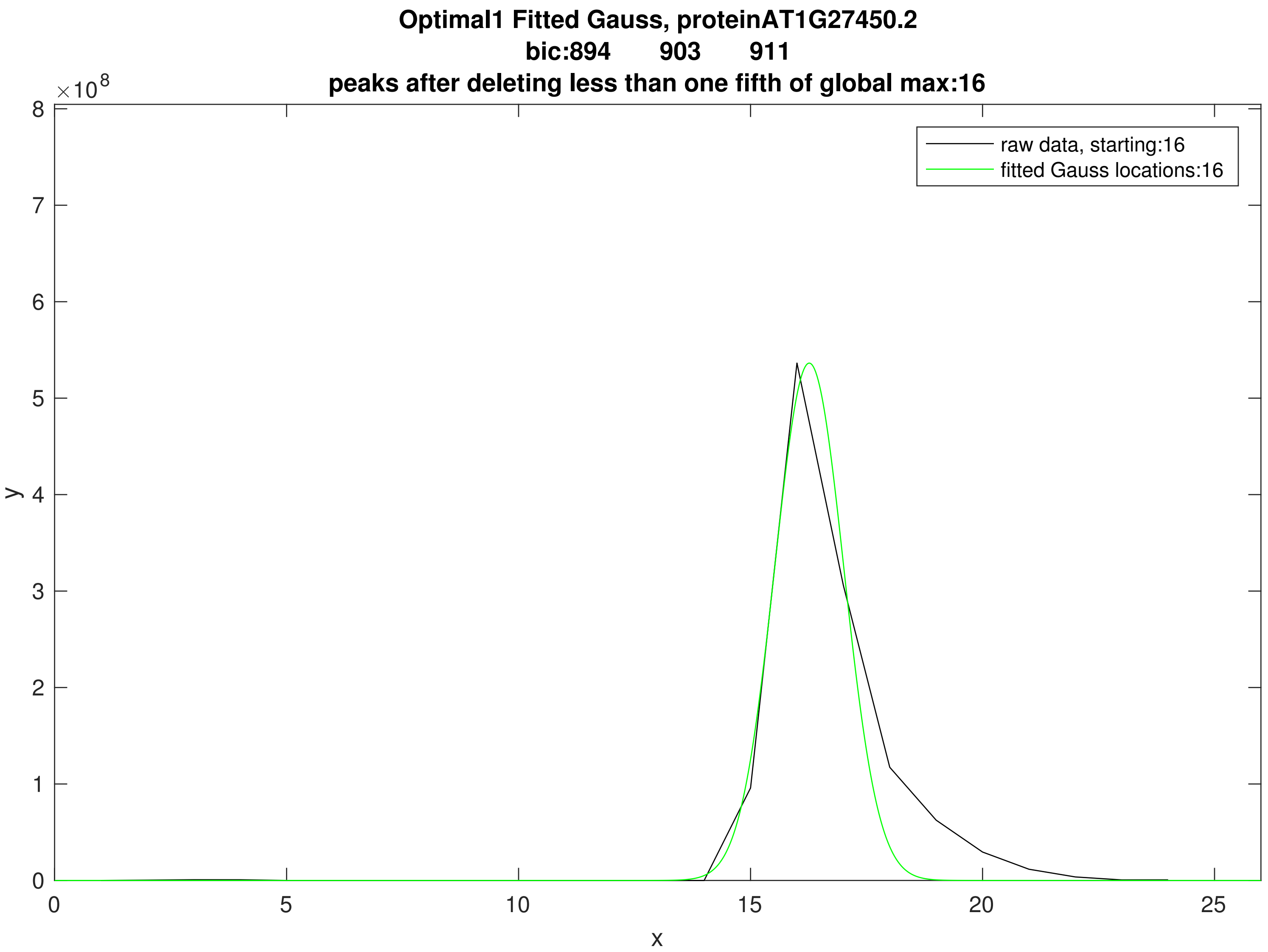
Optimal1 Fitted Gauss, proteinAT1G27450.2
bic:894 903 911
peaks after deleting less than one fifth of global max:16
×10^8
raw data, starting:16
fitted Gauss locations:16
y
x
0
1
2
3
4
5
6
7
8
10
15
20
25

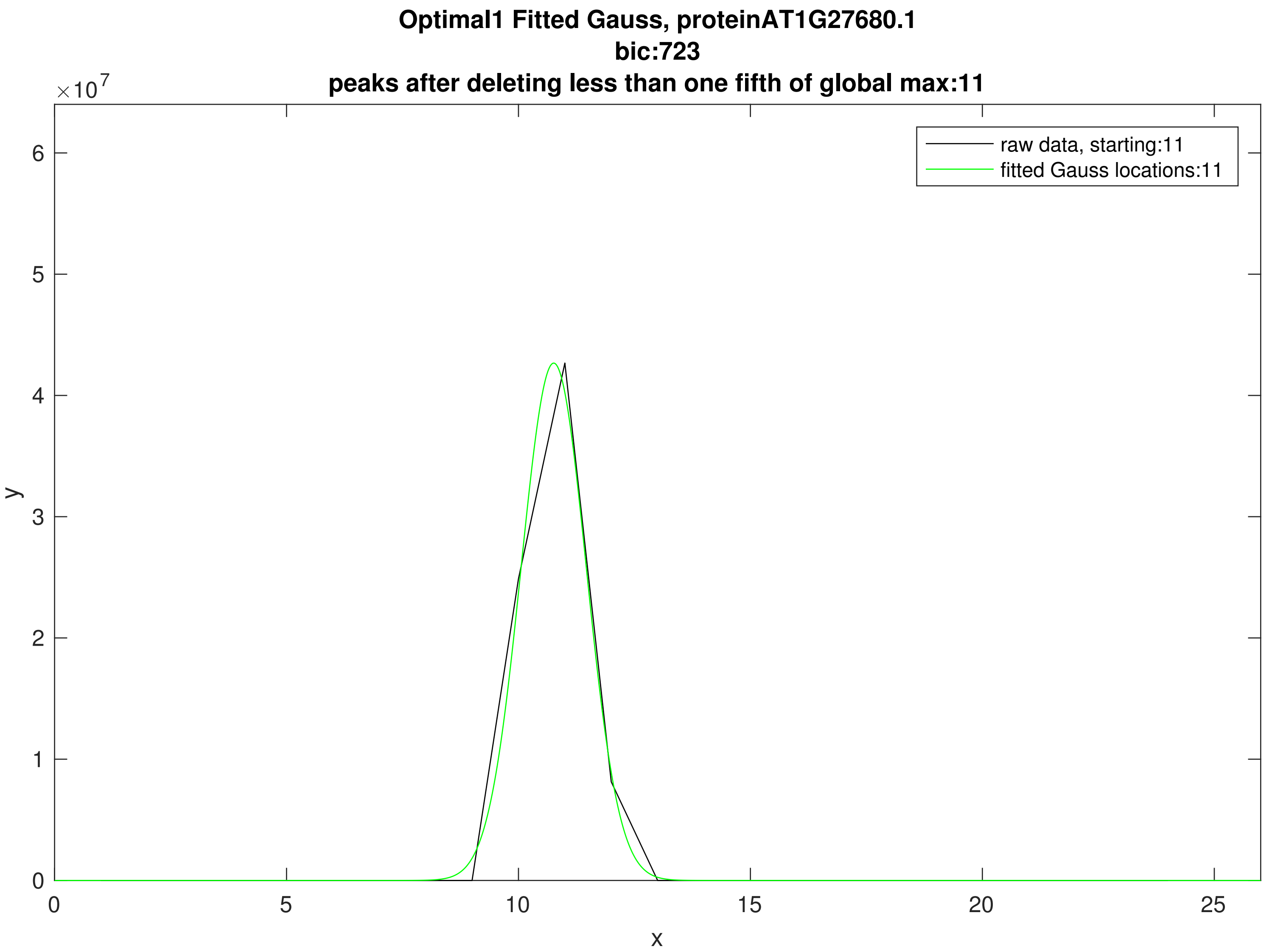
Optimal1 Fitted Gauss, proteinAT1G27680.1
bic:723
peaks after deleting less than one fifth of global max:11
×10^7
y
x
raw data, starting:11
fitted Gauss locations:11
0
1
2
3
4
5
6
0
5
10
15
20
25

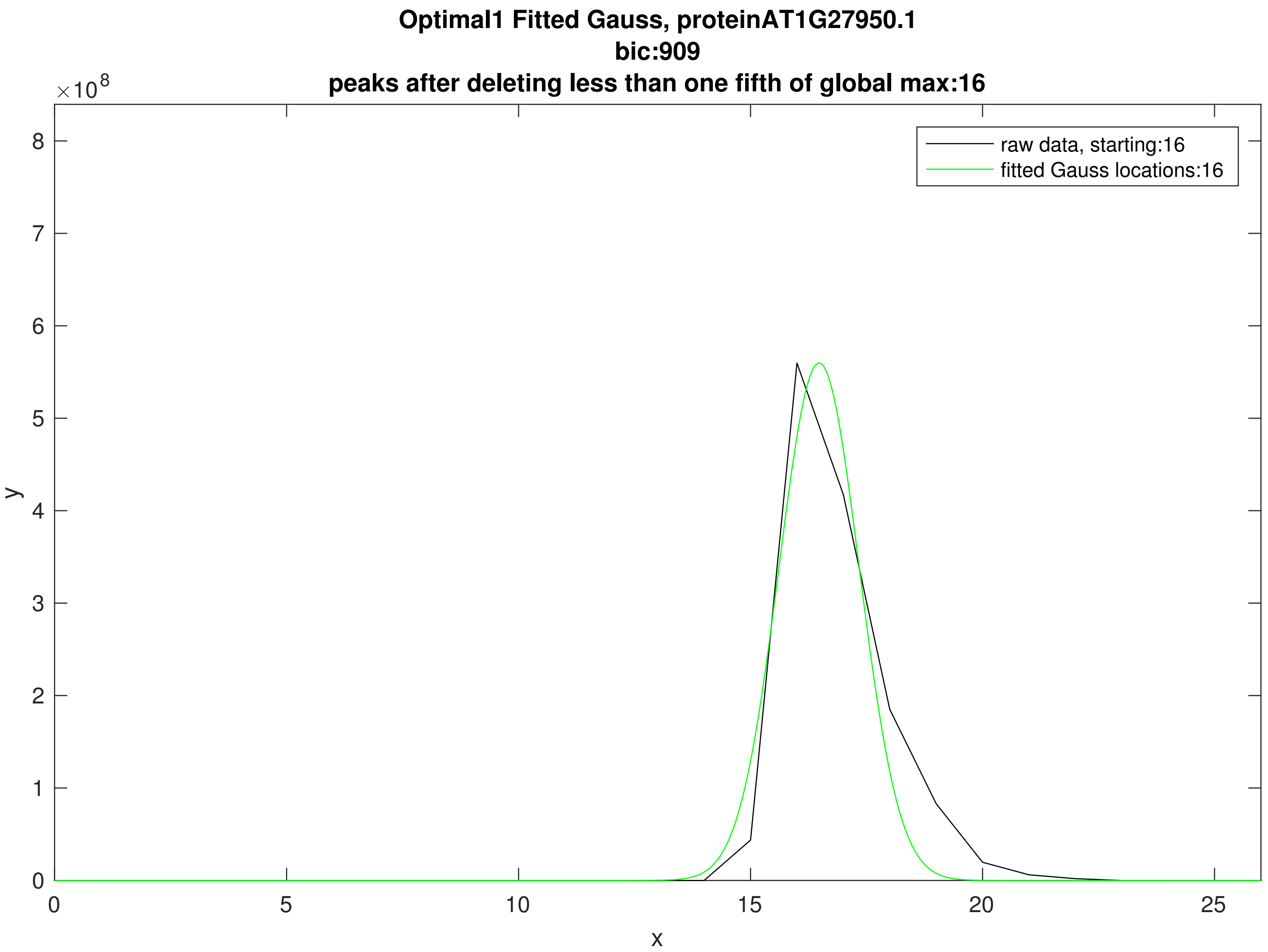
Optimal1 Fitted Gauss, proteinAT1G27950.1
bic:909
peaks after deleting less than one fifth of global max:16
×10^8
raw data, starting:16
fitted Gauss locations:16
y
x
0
1
2
3
4
5
6
7
8
0
5
10
15
20
25

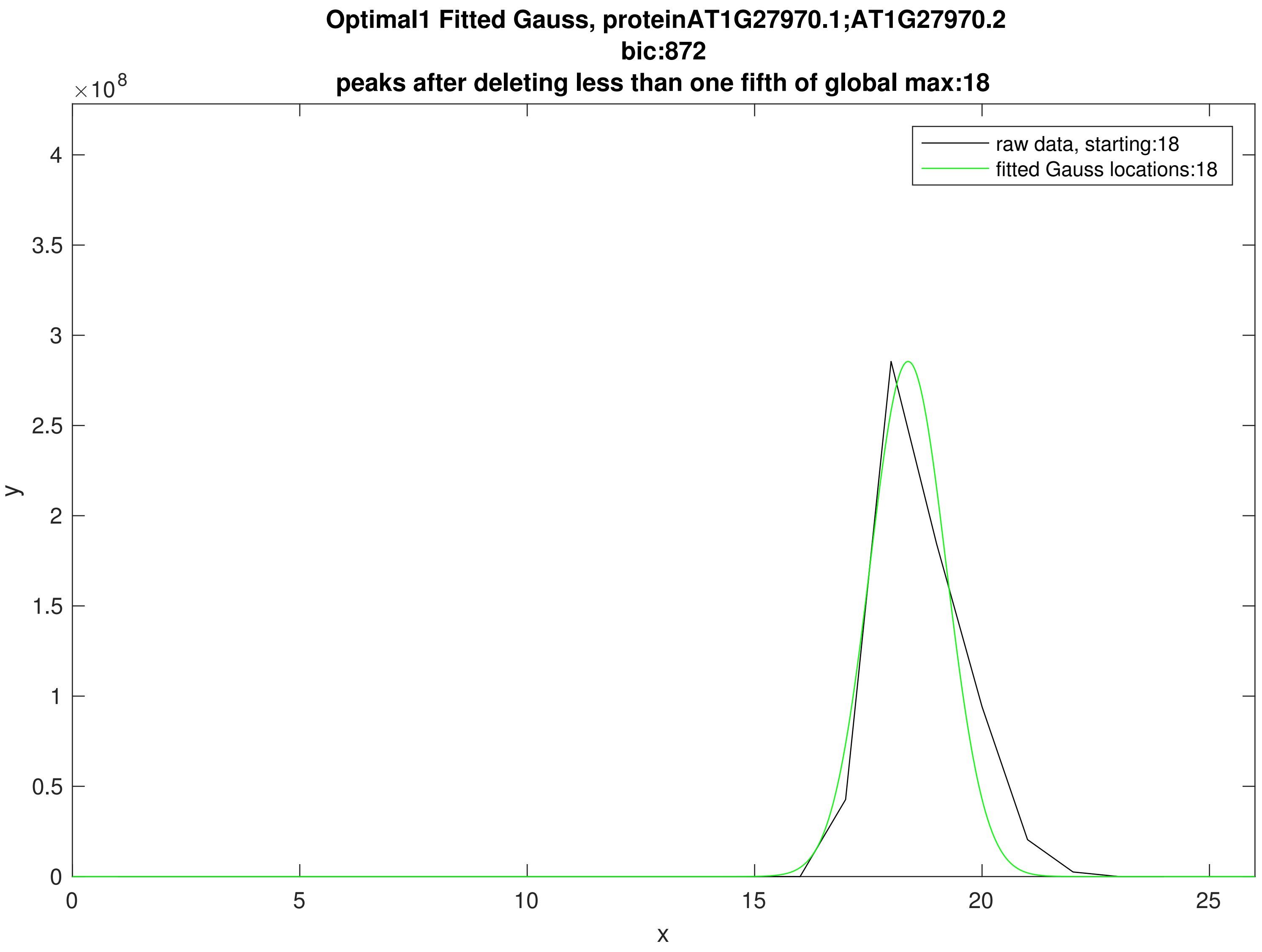

Optimal1 Fitted Gauss, proteinAT1G27970.1;AT1G27970.2
bic:872
peaks after deleting less than one fifth of global max:18
×10^8
raw data, starting:18
fitted Gauss locations:18
y
x
0
0.5
1
1.5
2
2.5
3
3.5
4
0
5
10
15
20
25

**Optimal1 Fitted Gauss, proteinAT1G28130.1;AT1G28130.2**
**bic:950 959**
**peaks after deleting less than one fifth of global max:15**

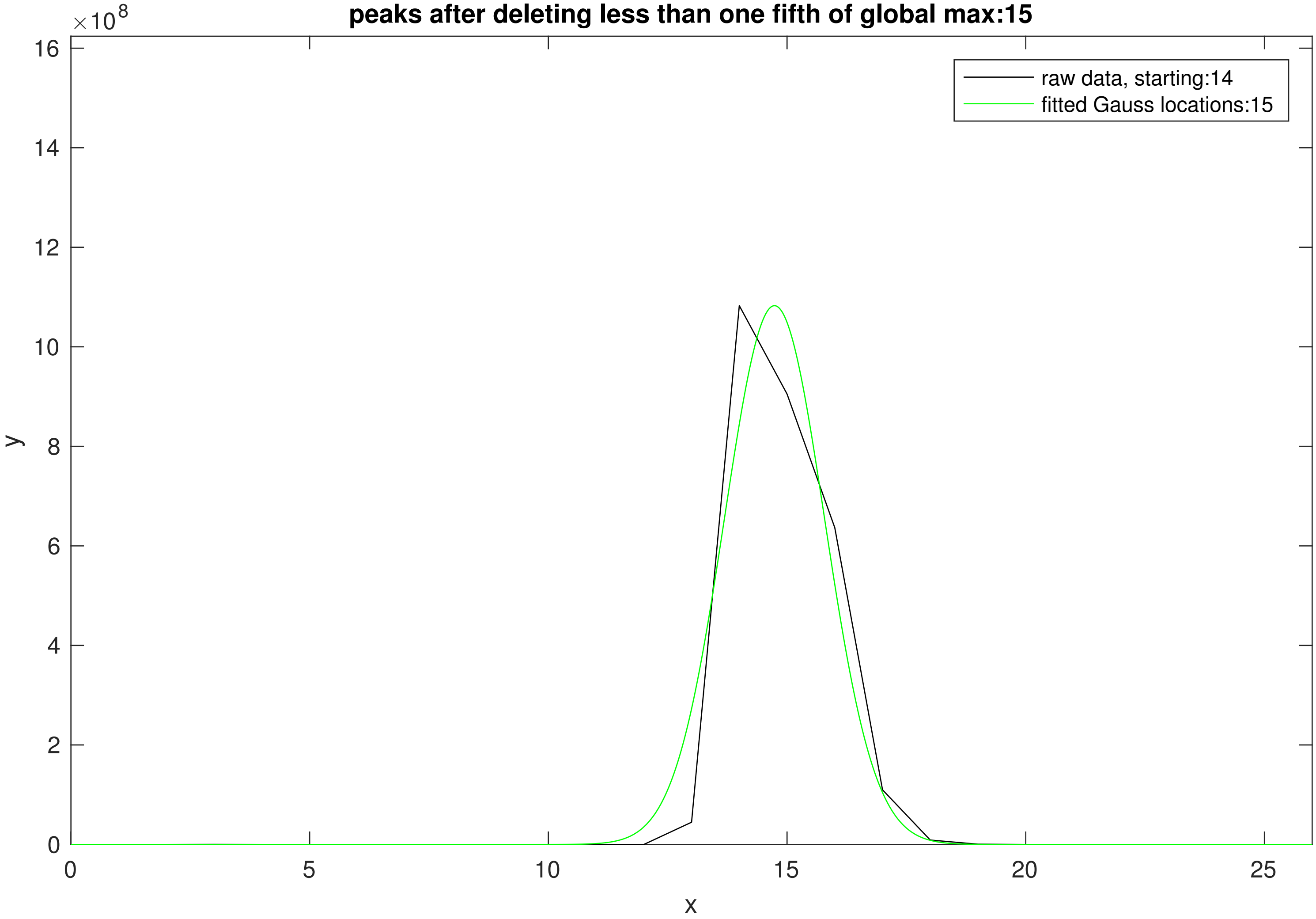

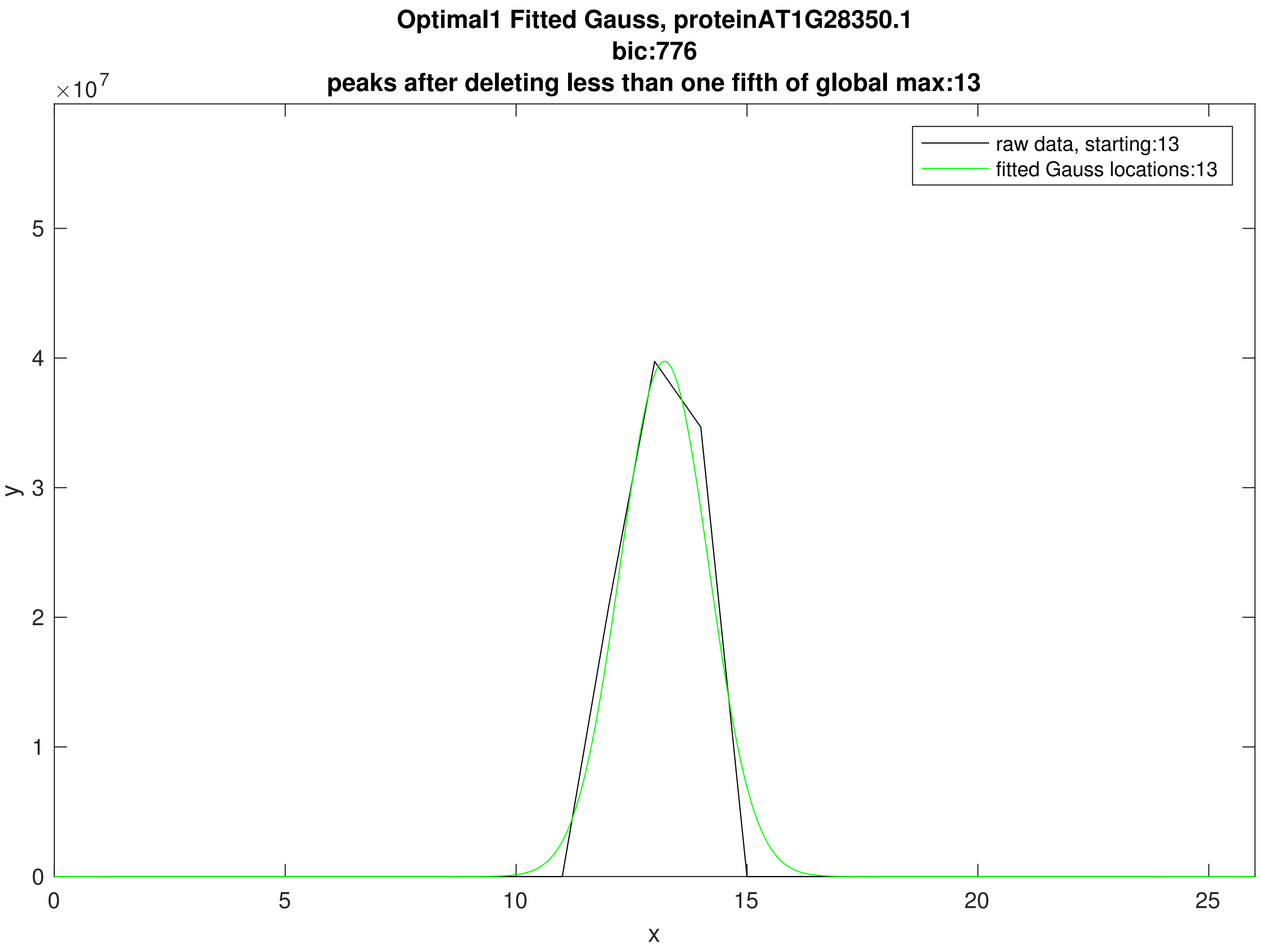

Optimal1 Fitted Gauss, proteinAT1G28350.1
bic:776
peaks after deleting less than one fifth of global max:13
×10^7
raw data, starting:13
fitted Gauss locations:13
0
1
2
3
4
5
y
0
5
10
15
20
25
x

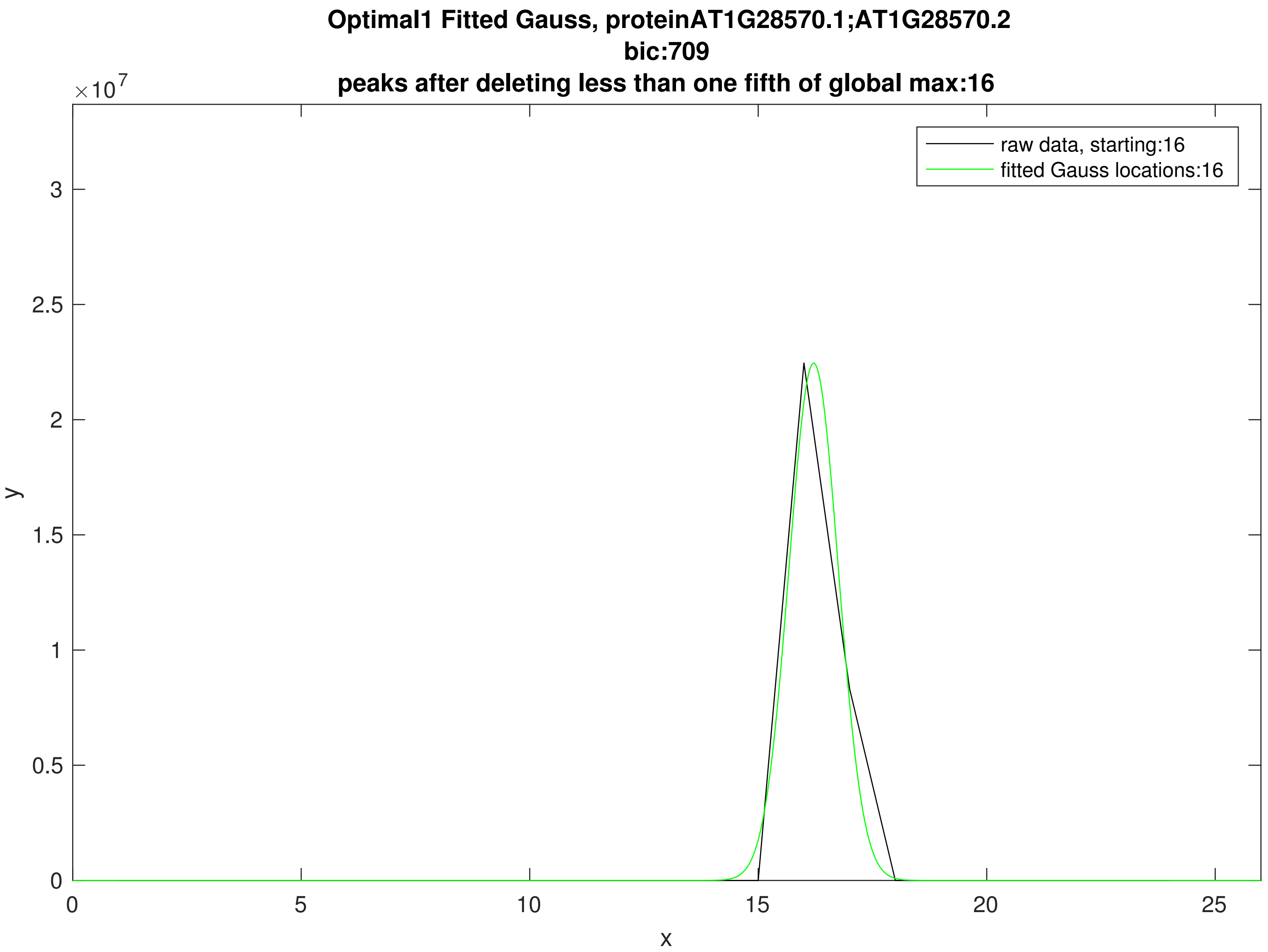
Optimal1 Fitted Gauss, proteinAT1G28570.1;AT1G28570.2
bic:709
peaks after deleting less than one fifth of global max:16
×10^7
raw data, starting:16
fitted Gauss locations:16
y
x
0
0.5
1
1.5
2
2.5
3
0
5
10
15
20
25

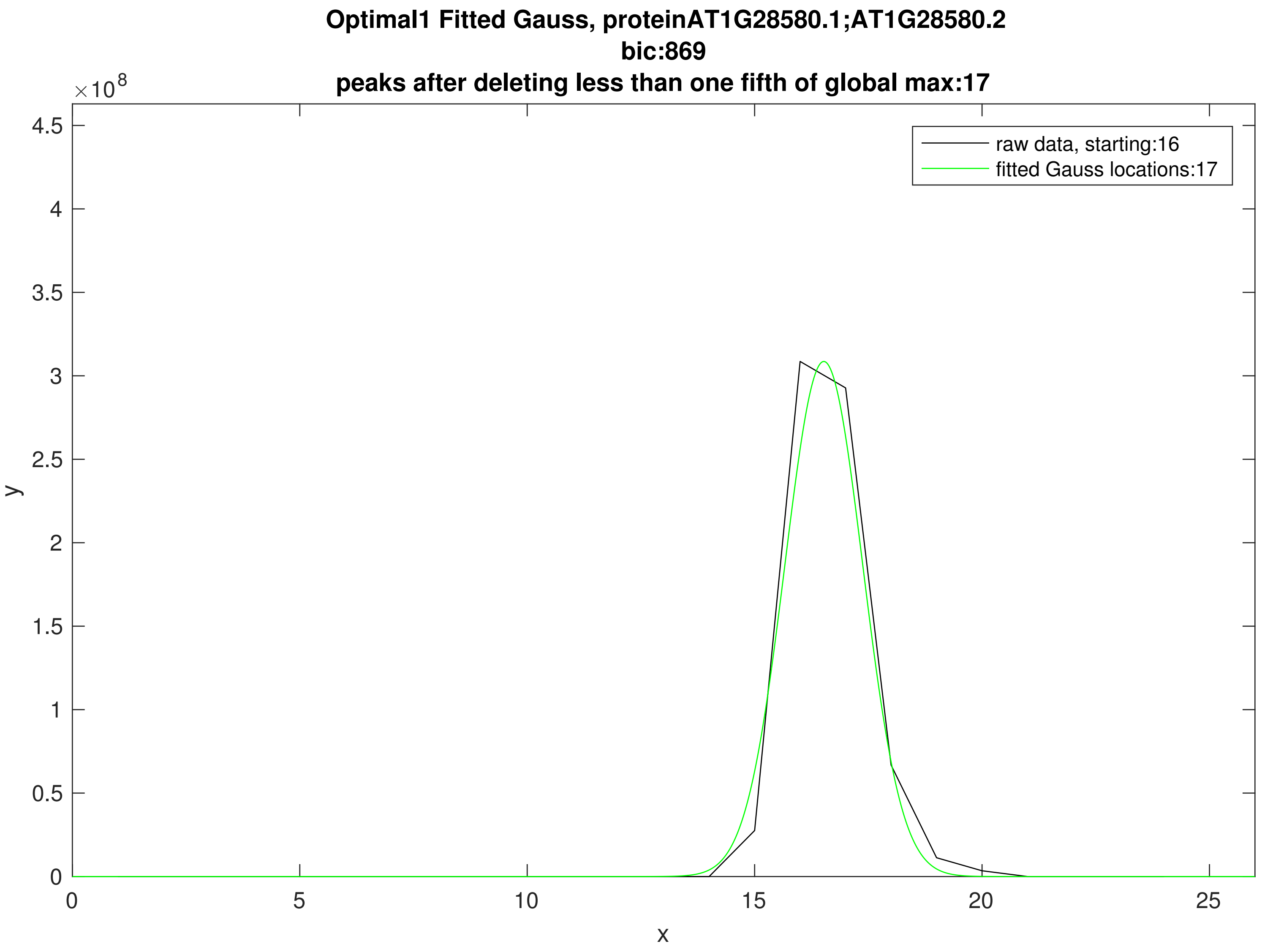
Optimal1 Fitted Gauss, proteinAT1G28580.1;AT1G28580.2
bic:869
peaks after deleting less than one fifth of global max:17
×10^8
raw data, starting:16
fitted Gauss locations:17
4.5
4
3.5
3
2.5
2
1.5
1
0.5
0
y
0
5
10
15
20
25
x

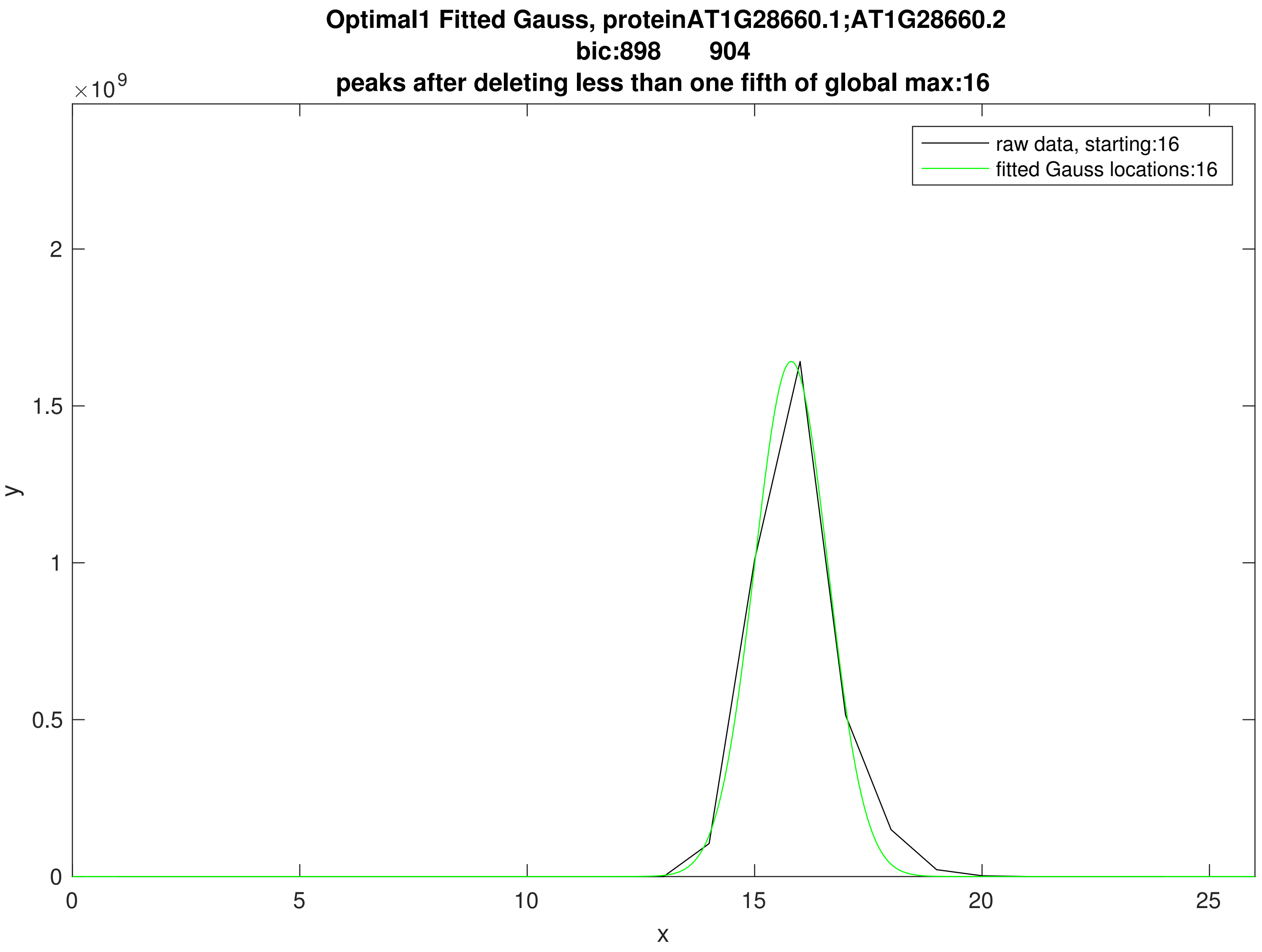

Optimal1 Fitted Gauss, proteinAT1G28660.1;AT1G28660.2
bic:898 904
peaks after deleting less than one fifth of global max:16
×10^9
raw data, starting:16
fitted Gauss locations:16
y
x
0
0.5
1
1.5
2
5
10
15
20
25

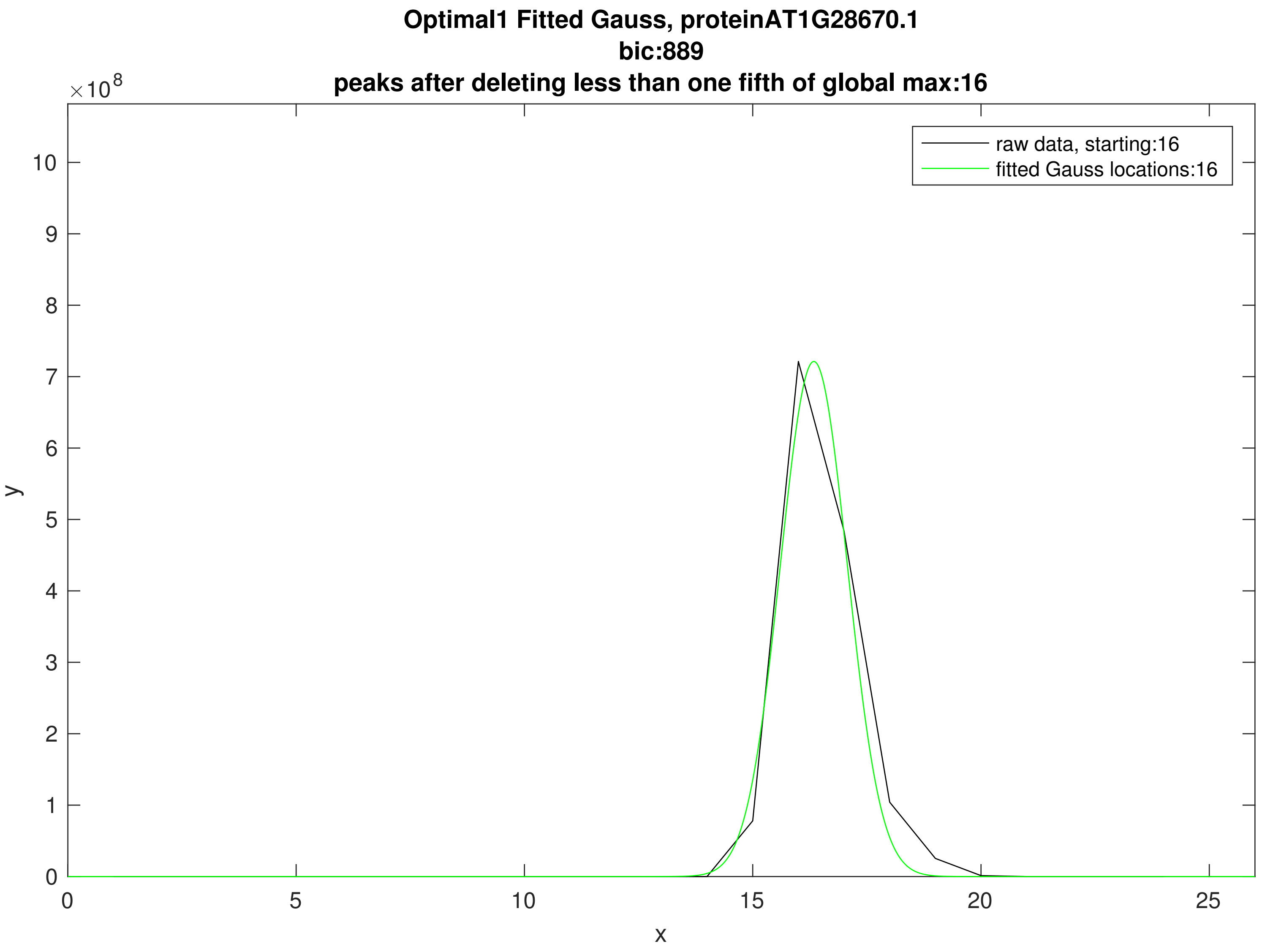
Optimal1 Fitted Gauss, proteinAT1G28670.1
bic:889
peaks after deleting less than one fifth of global max:16
raw data, starting:16
fitted Gauss locations:16
×10^8
y
x

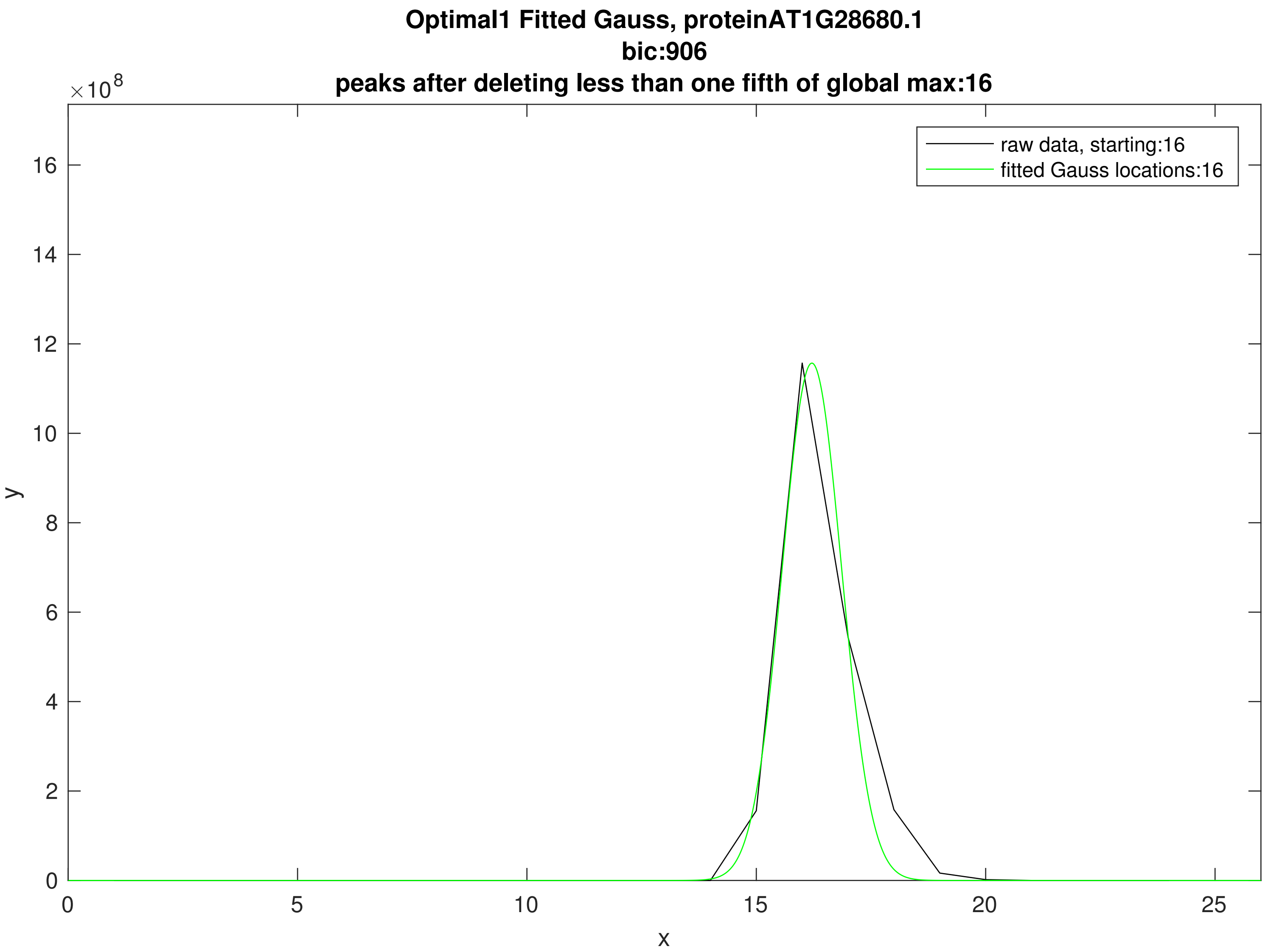

Optimal1 Fitted Gauss, proteinAT1G28680.1
bic:906
peaks after deleting less than one fifth of global max:16
×10^8
raw data, starting:16
fitted Gauss locations:16
16
14
12
10
8
6
4
2
0
y
0
5
10
15
20
25
x

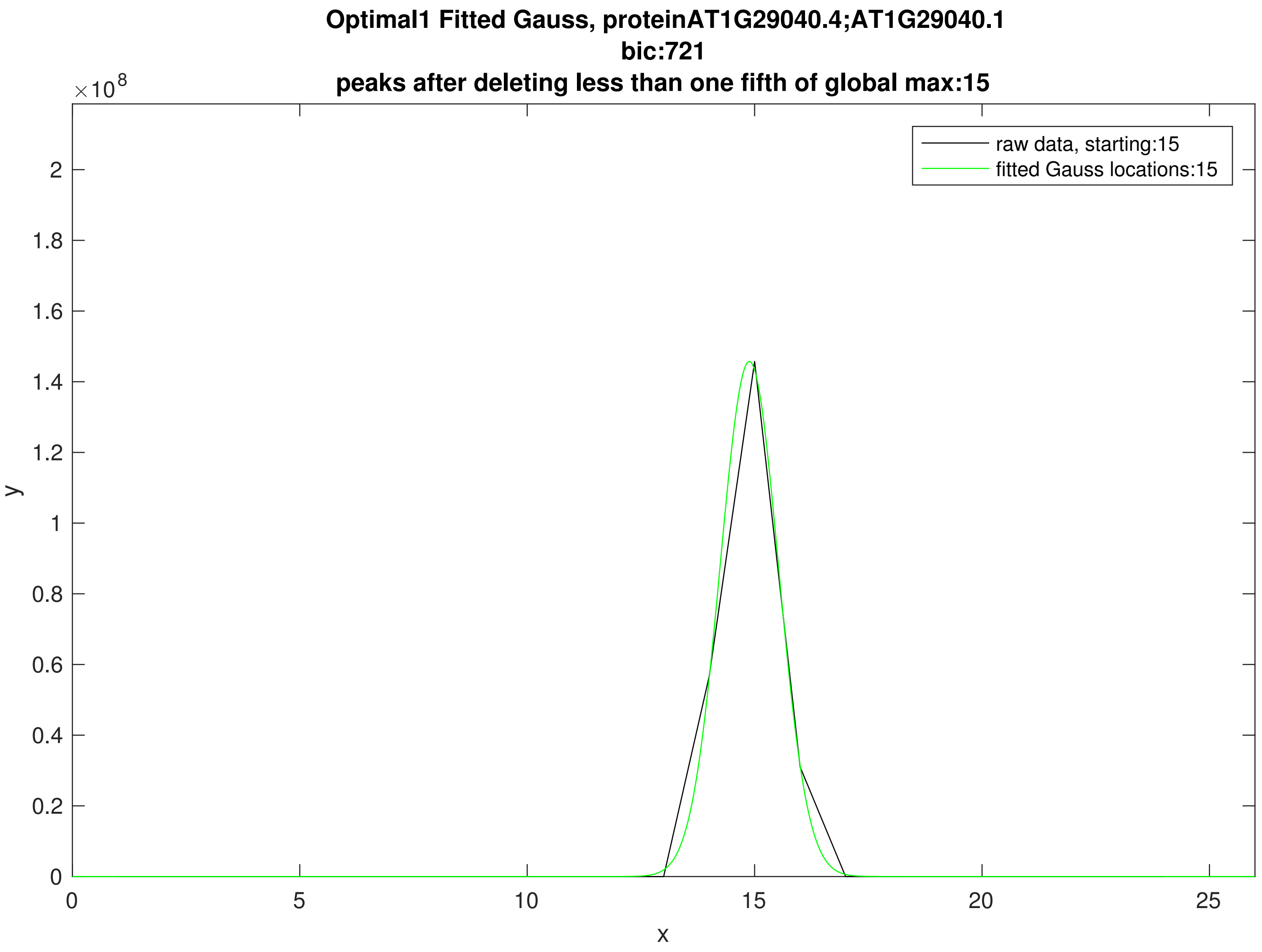
Optimal1 Fitted Gauss, proteinAT1G29040.4;AT1G29040.1
bic:721
peaks after deleting less than one fifth of global max:15
×10^8
raw data, starting:15
fitted Gauss locations:15
y
x
0
5
10
15
20
25
0
0.2
0.4
0.6
0.8
1
1.2
1.4
1.6
1.8
2

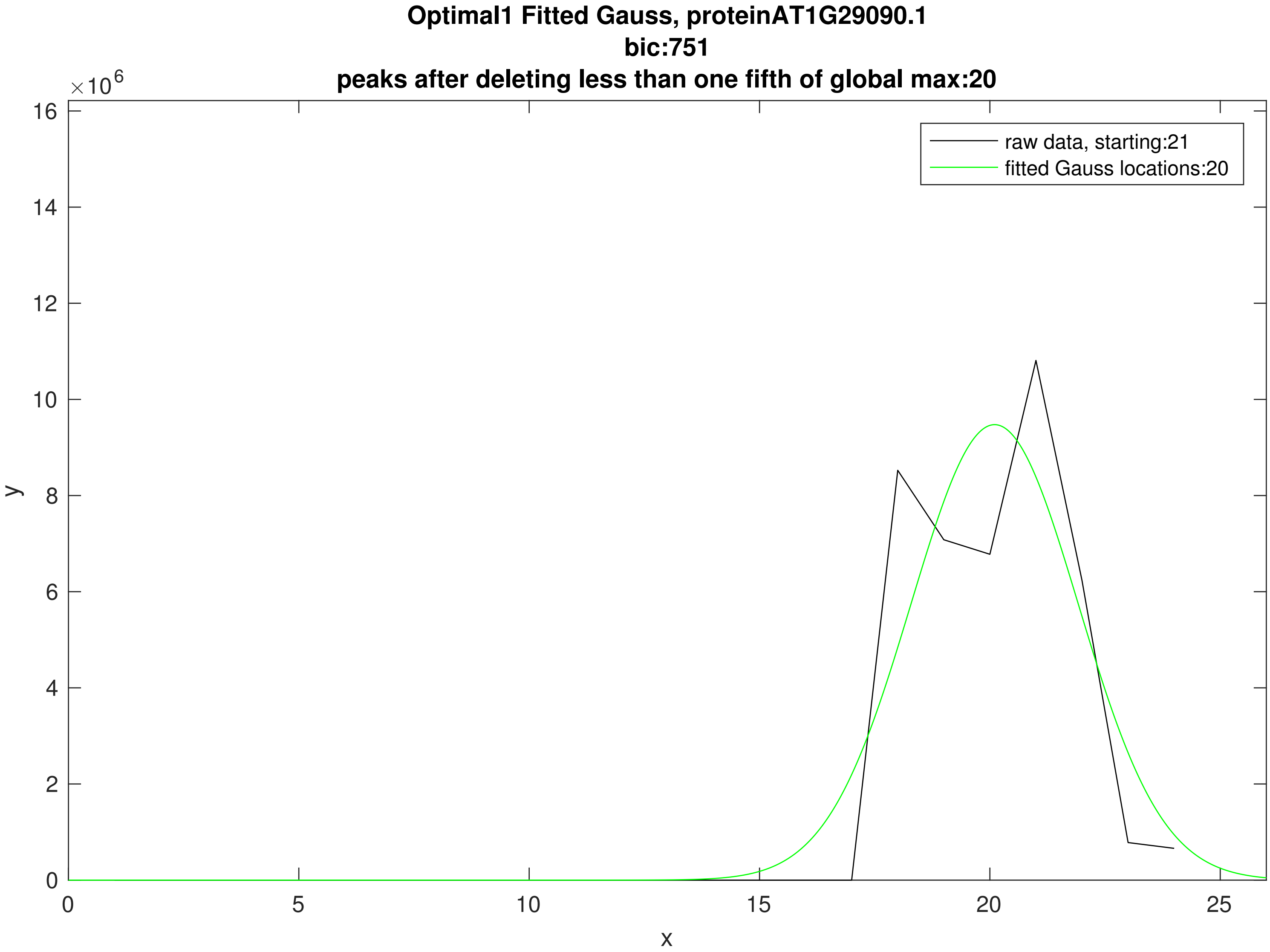

Optimal1 Fitted Gauss, proteinAT1G29090.1
bic:751
peaks after deleting less than one fifth of global max:20
×10^6
16
14
12
10
8
6
4
2
0
y
0
5
10
15
20
25
x
raw data, starting:21
fitted Gauss locations:20

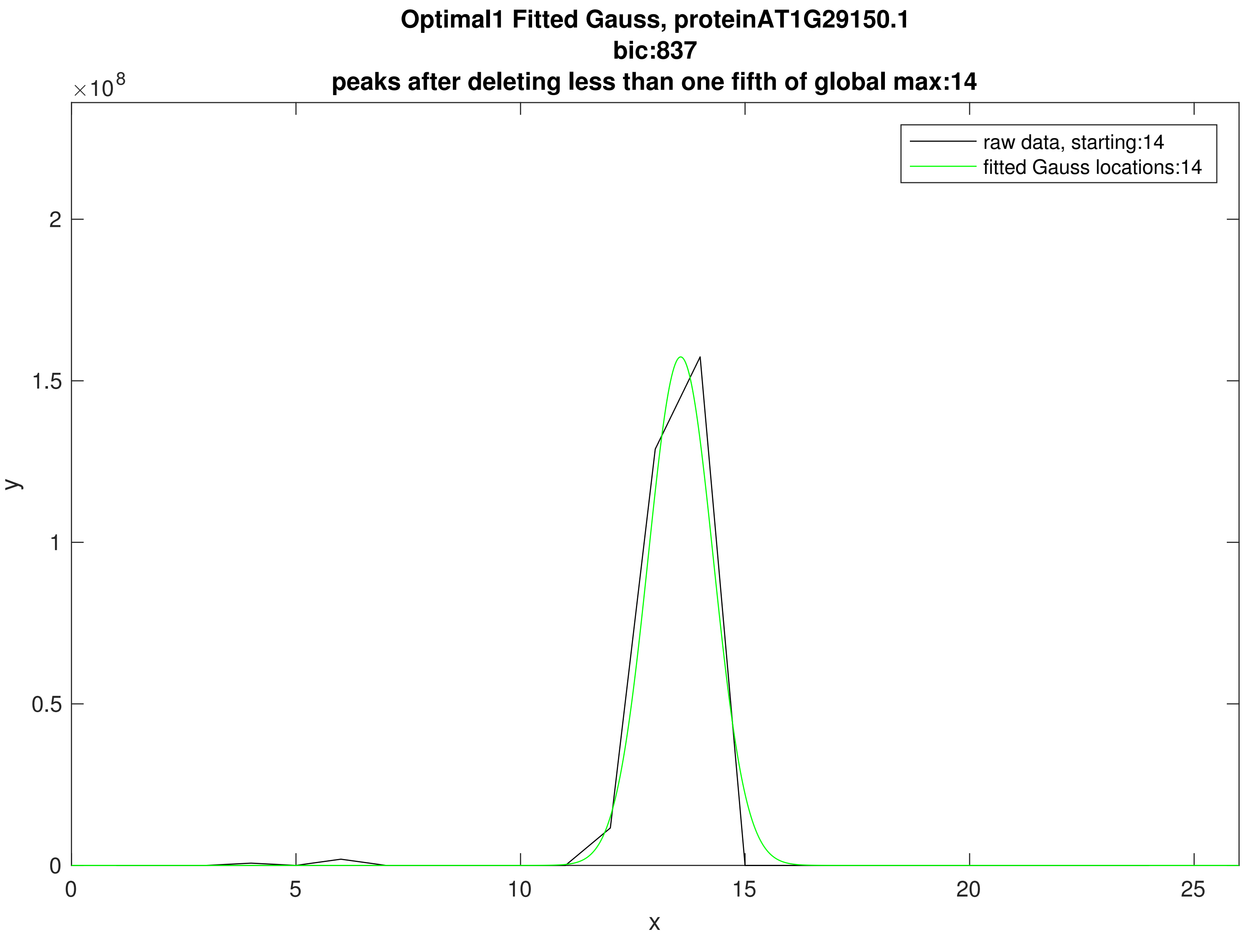

Optimal1 Fitted Gauss, proteinAT1G29150.1
bic:837
peaks after deleting less than one fifth of global max:14
×10^8
raw data, starting:14
fitted Gauss locations:14
y
x
0
0.5
1
1.5
2
0
5
10
15
20
25

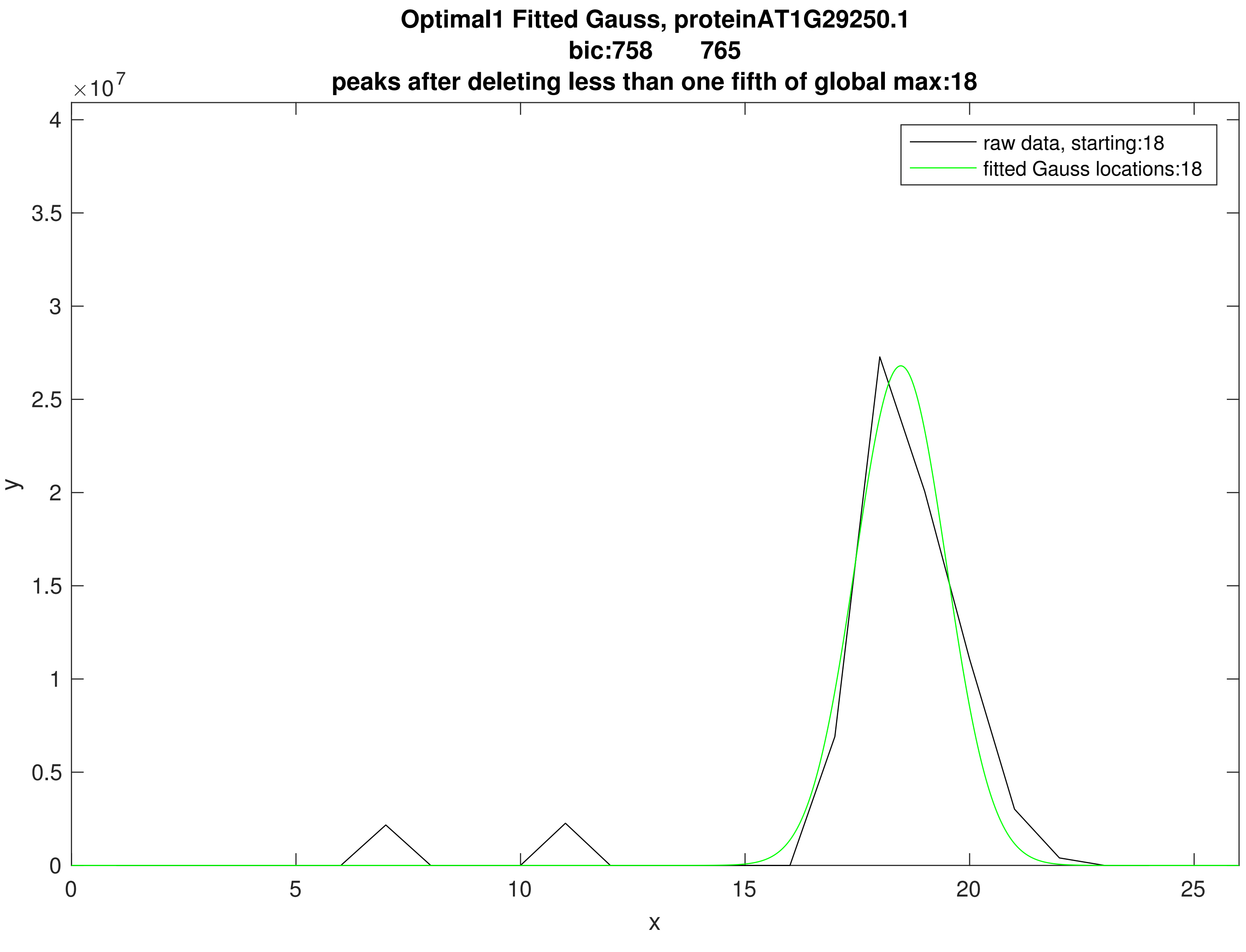
Optimal1 Fitted Gauss, proteinAT1G29250.1
bic:758 765
peaks after deleting less than one fifth of global max:18
×10^7
4
3.5
3
2.5
2
1.5
1
0.5
0
y
0
5
10
15
20
25
x
raw data, starting:18
fitted Gauss locations:18

**Optimal1 Fitted Gauss, proteinAT1G29260.1**
**bic:841 850**
**peaks after deleting less than one fifth of global max:18**

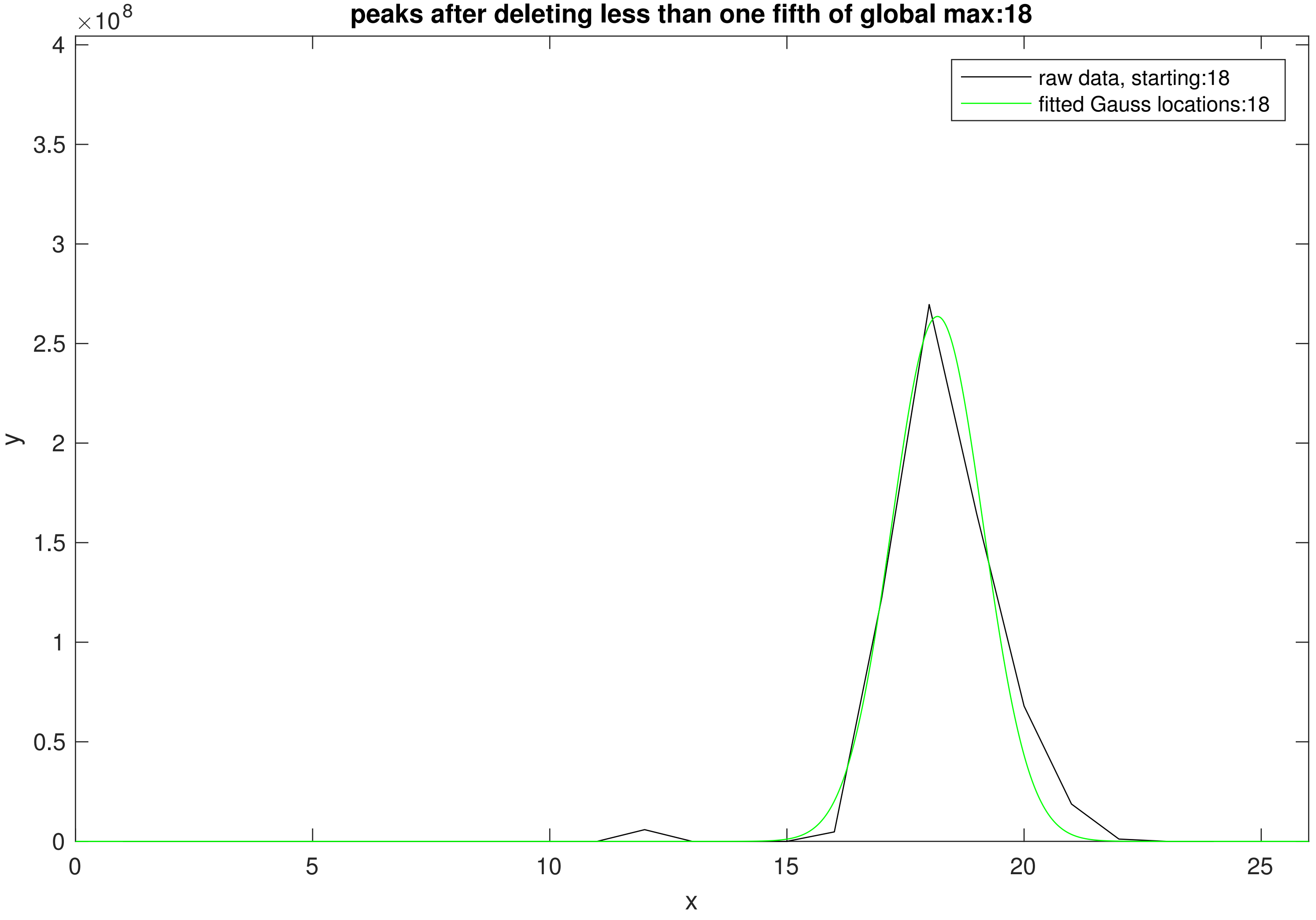

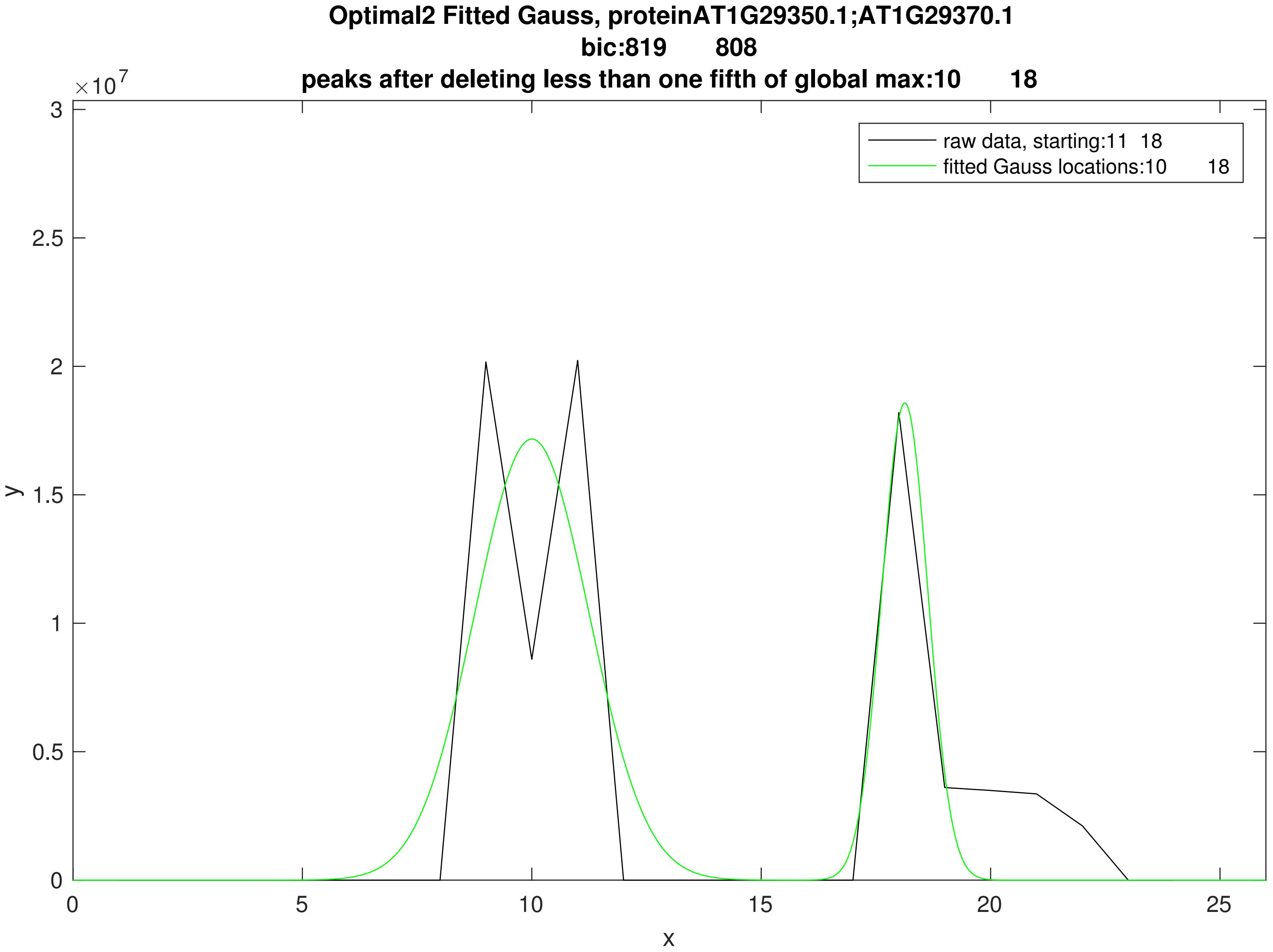
Optimal2 Fitted Gauss, proteinAT1G29350.1;AT1G29370.1
bic:819 808
peaks after deleting less than one fifth of global max:10 18
×10^7
3
2.5
2
1.5
1
0.5
0
y
0
5
10
15
20
25
x
raw data, starting:11 18
fitted Gauss locations:10 18

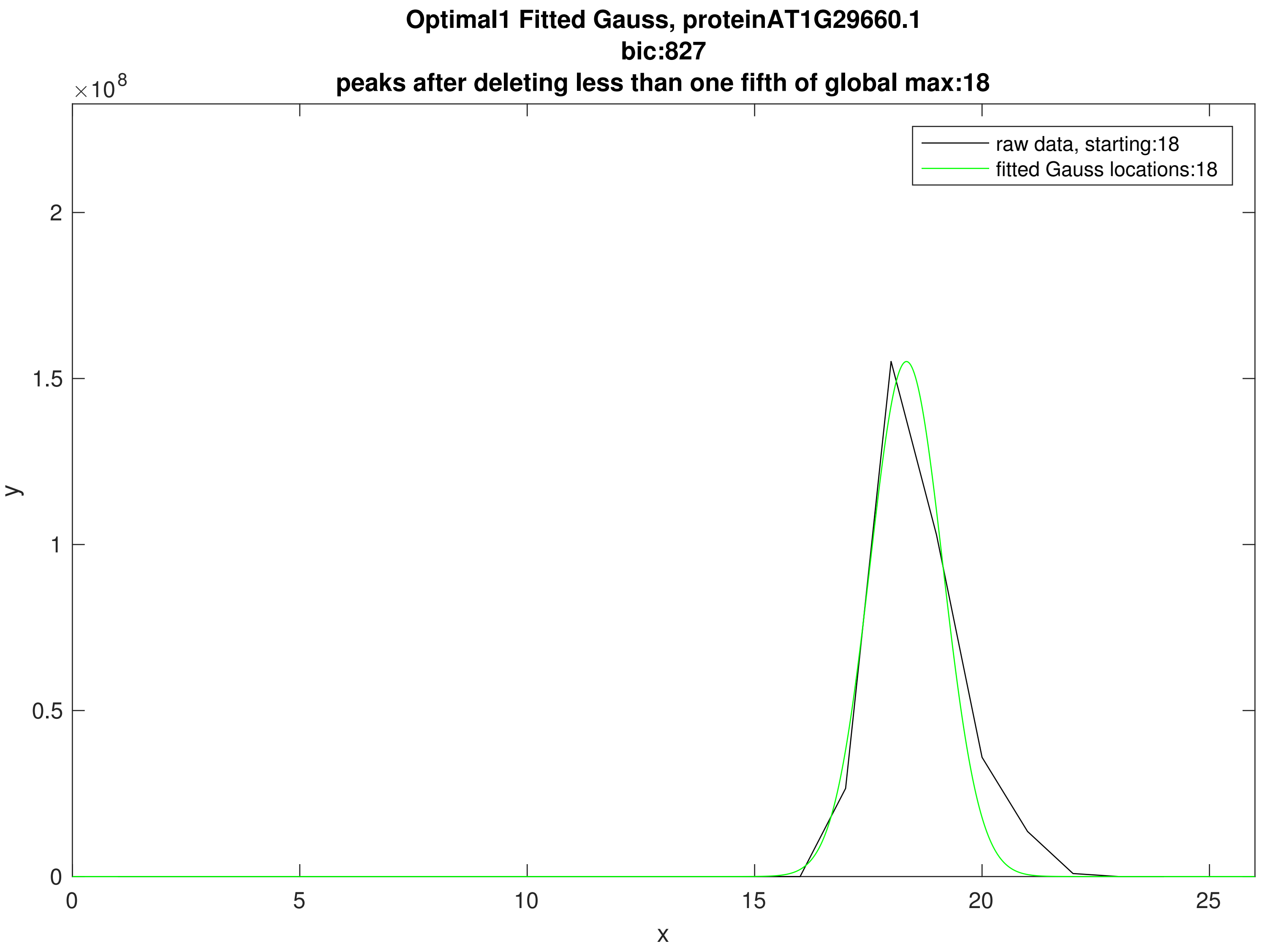
Optimal1 Fitted Gauss, proteinAT1G29660.1
bic:827
peaks after deleting less than one fifth of global max:18
×10^8
raw data, starting:18
fitted Gauss locations:18
y
x
0
0.5
1
1.5
2
0
5
10
15
20
25

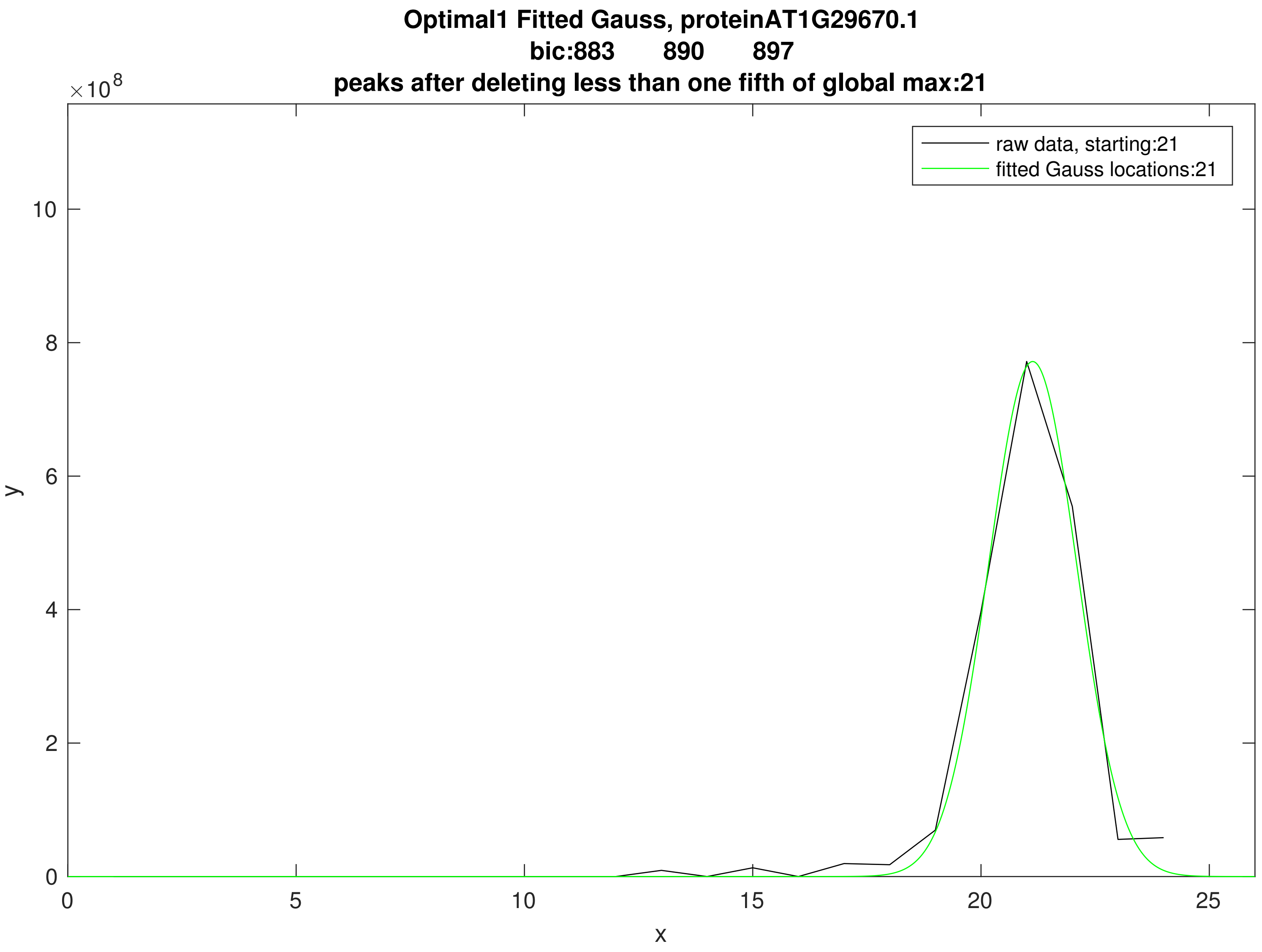
Optimal1 Fitted Gauss, proteinAT1G29670.1
bic:883 890 897
peaks after deleting less than one fifth of global max:21
×10^8
raw data, starting:21
fitted Gauss locations:21
y
x
0
2
4
6
8
10
0
5
10
15
20
25

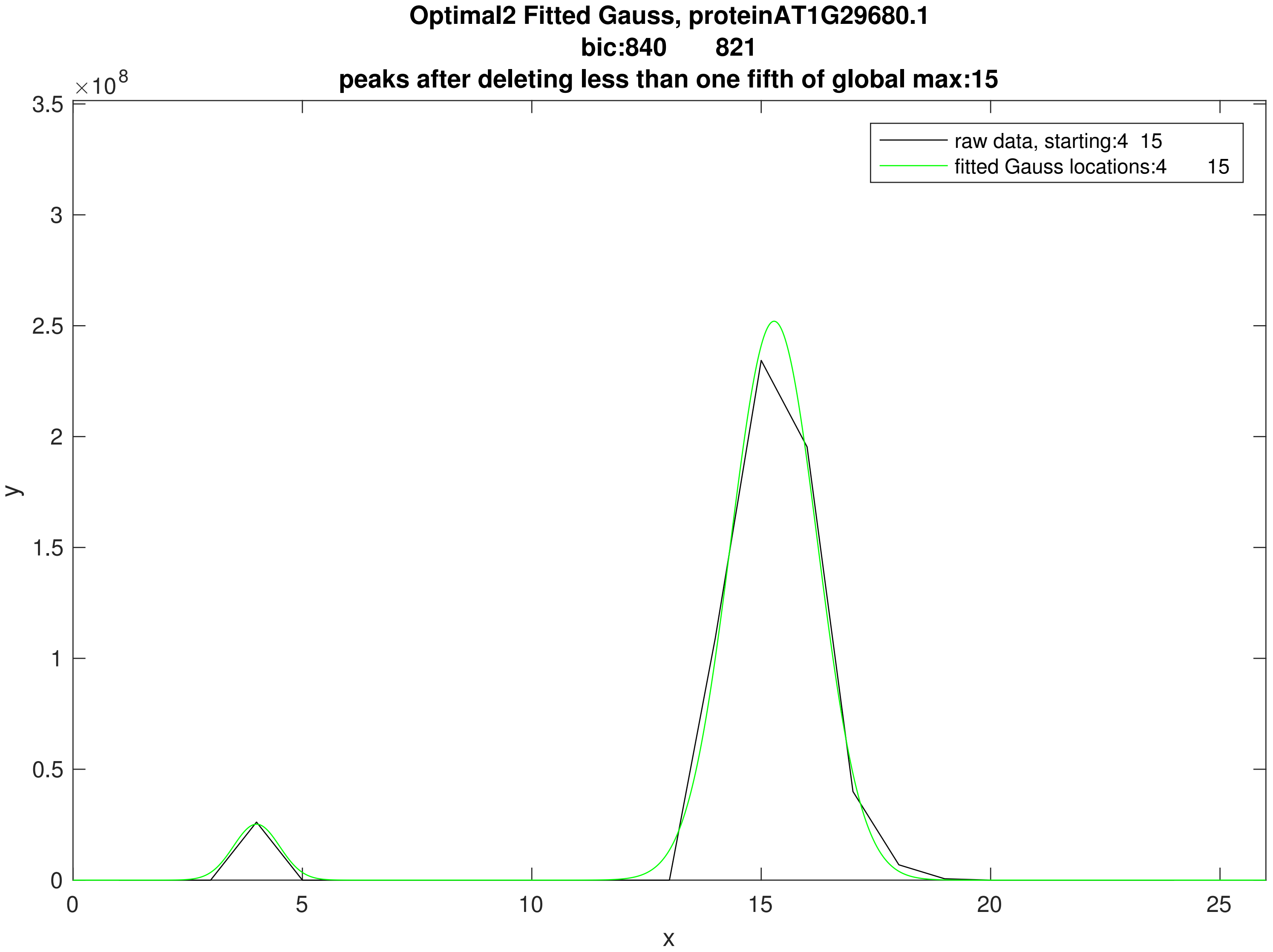

Optimal2 Fitted Gauss, proteinAT1G29680.1
bic:840 821
peaks after deleting less than one fifth of global max:15
×10^8
3.5
3
2.5
2
1.5
1
0.5
0
y
0
5
10
15
20
25
x
raw data, starting:4 15
fitted Gauss locations:4 15

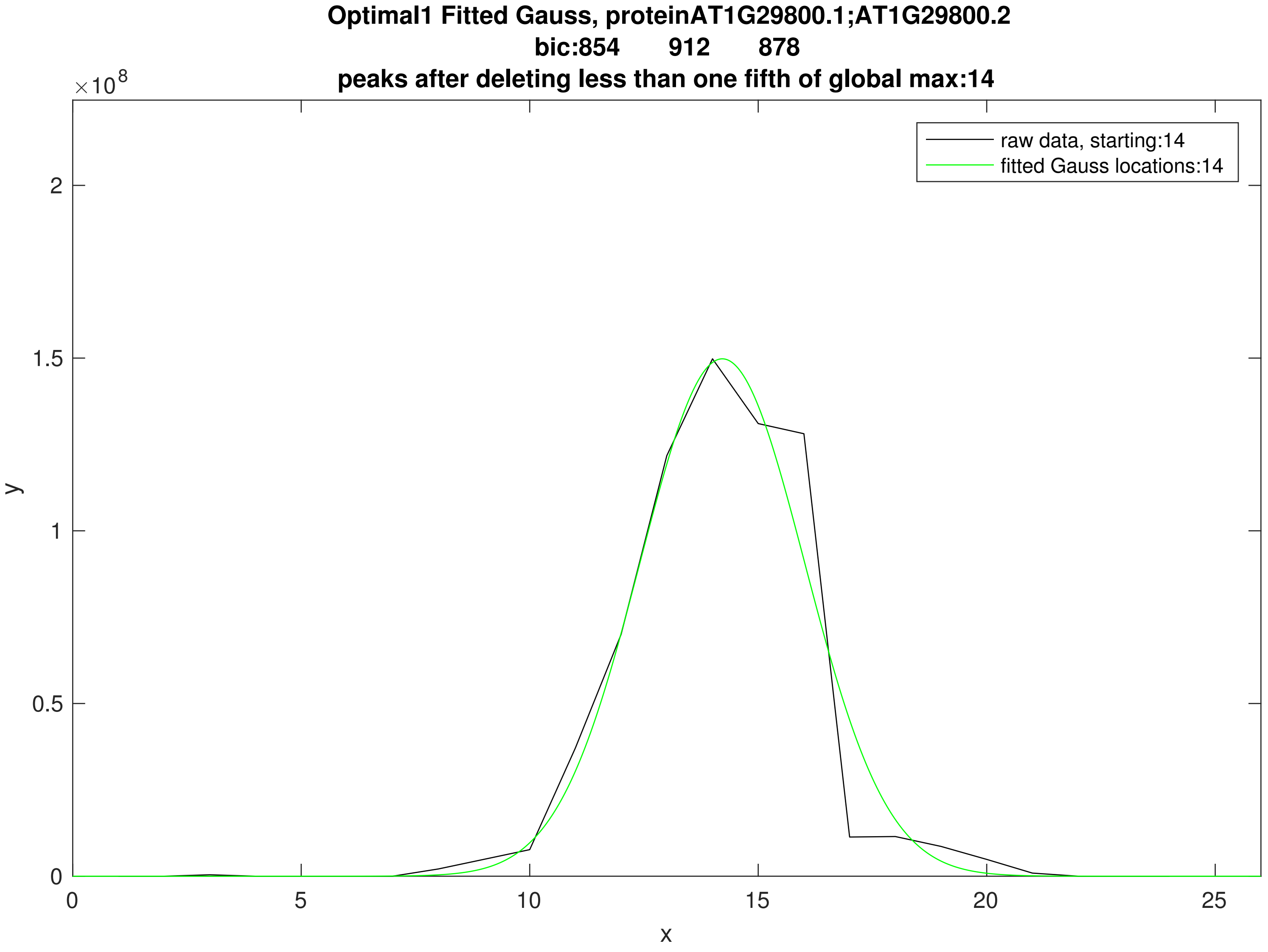

Optimal1 Fitted Gauss, proteinAT1G29800.1;AT1G29800.2
bic:854 912 878
peaks after deleting less than one fifth of global max:14
×10^8
raw data, starting:14
fitted Gauss locations:14
y
x
0
0.5
1
1.5
2
0
5
10
15
20
25

**Optimal2 Fitted Gauss, proteinAT1G29840.1**
**bic:803 801**
**peaks after deleting less than one fifth of global max:18**

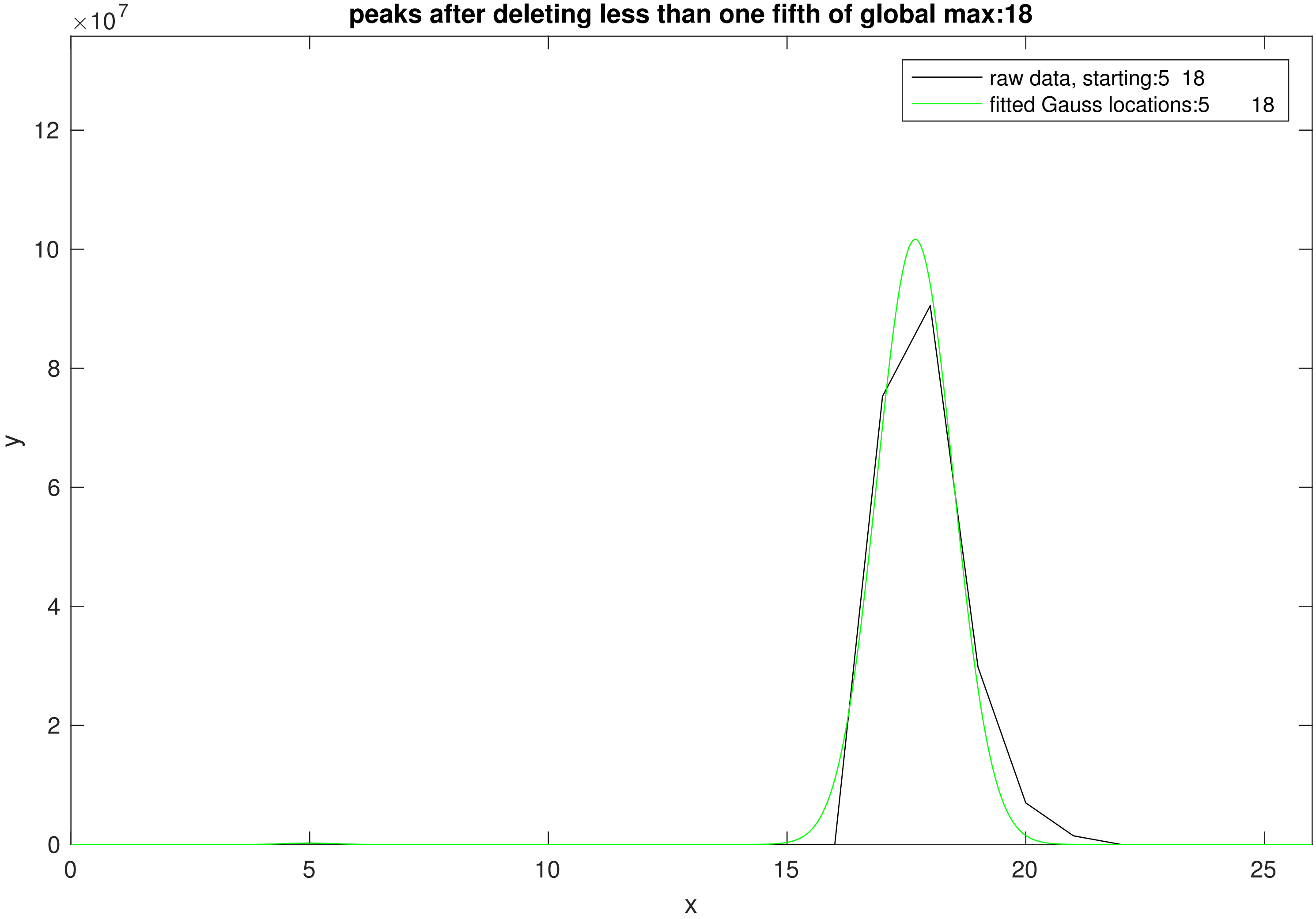

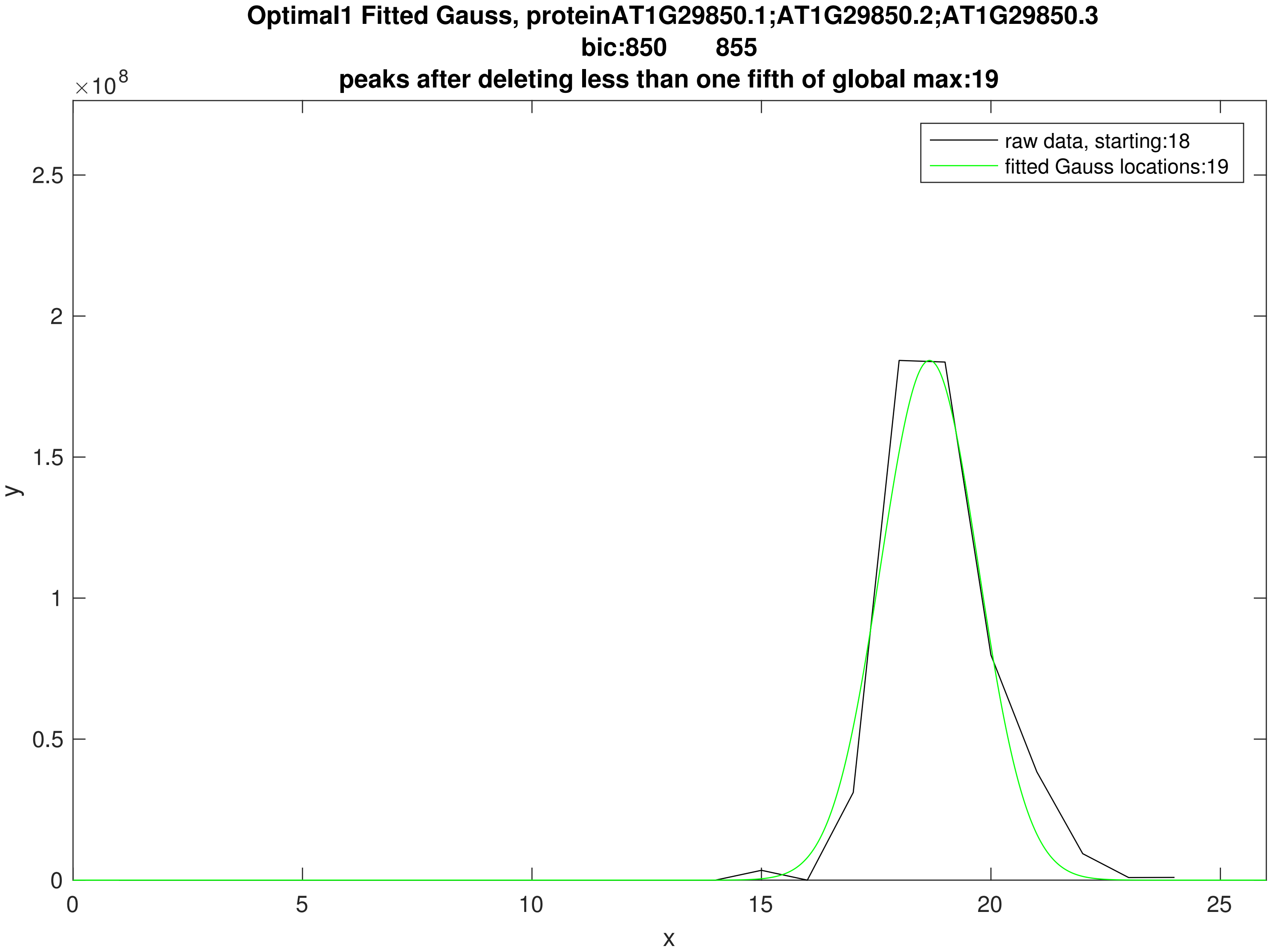

Optimal1 Fitted Gauss, proteinAT1G29850.1;AT1G29850.2;AT1G29850.3
bic:850 855
peaks after deleting less than one fifth of global max:19
×10^8
2.5
2
1.5
1
0.5
0
y
0
5
10
15
20
25
x
raw data, starting:18
fitted Gauss locations:19

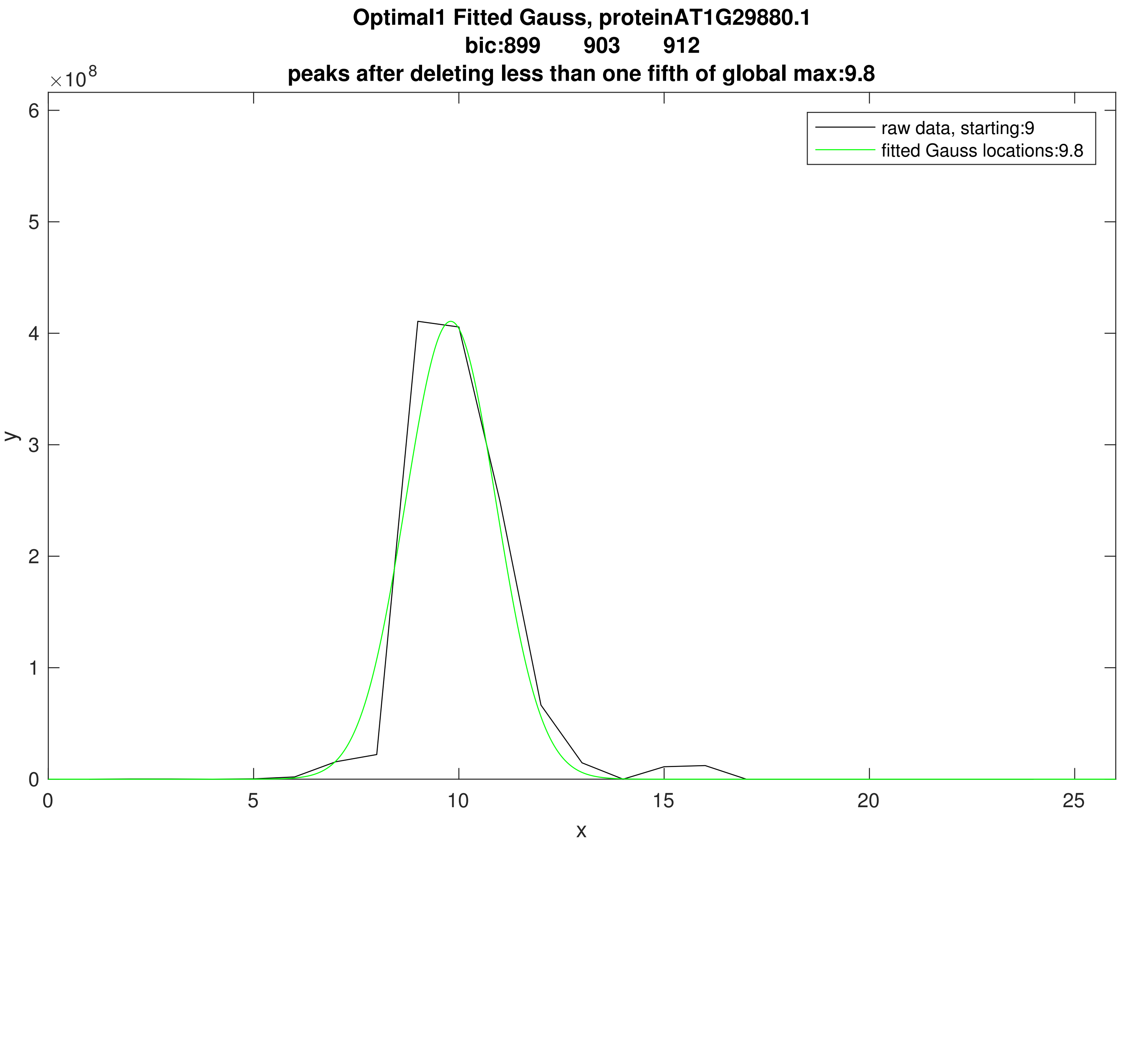
Optimal1 Fitted Gauss, proteinAT1G29880.1
bic:899 903 912
peaks after deleting less than one fifth of global max:9.8
×10^8
y
x
raw data, starting:9
fitted Gauss locations:9.8
0
1
2
3
4
5
6
0
5
10
15
20
25

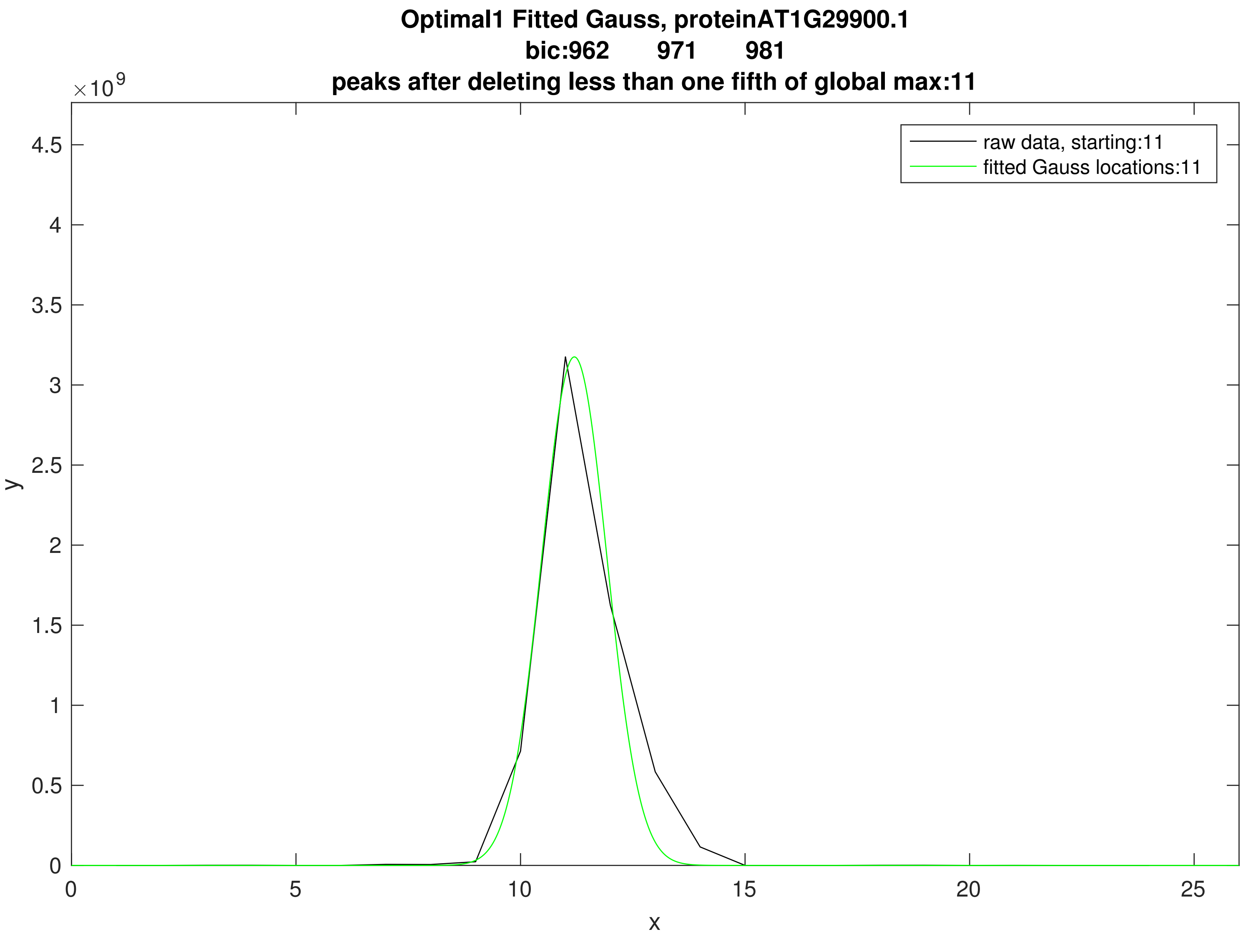
Optimal1 Fitted Gauss, proteinAT1G29900.1
bic:962 971 981
peaks after deleting less than one fifth of global max:11
×10^9
raw data, starting:11
fitted Gauss locations:11
y
x
0
0.5
1
1.5
2
2.5
3
3.5
4
4.5
0
5
10
15
20
25

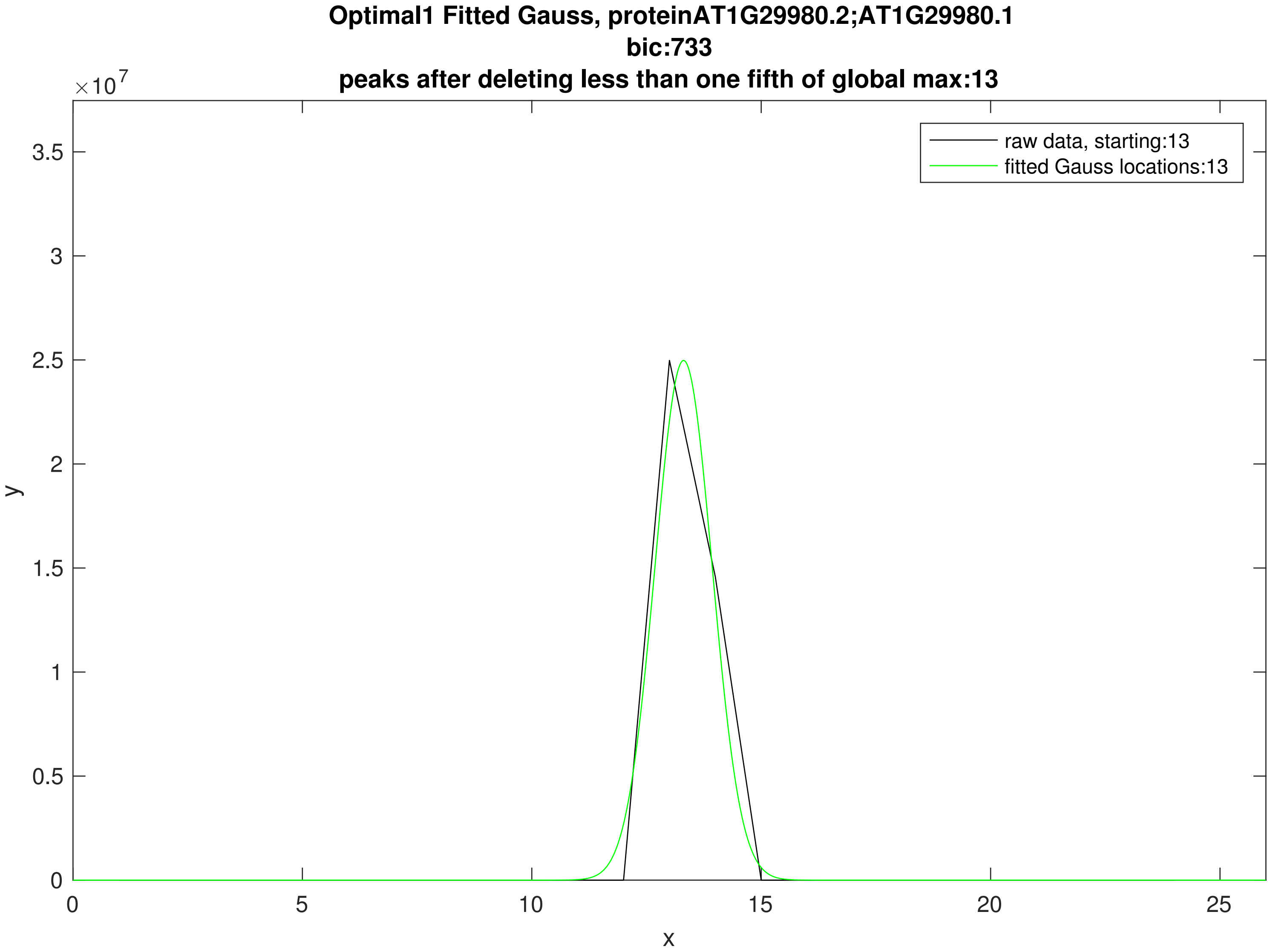

Optimal1 Fitted Gauss, proteinAT1G29980.2;AT1G29980.1
bic:733
peaks after deleting less than one fifth of global max:13
×10^7
3.5
3
2.5
2
1.5
1
0.5
0
y
0
5
10
15
20
25
x
raw data, starting:13
fitted Gauss locations:13

**Optimal2 Fitted Gauss, proteinAT1G29990.1**
**bic:709 701**
**peaks after deleting less than one fifth of global max:12 18**

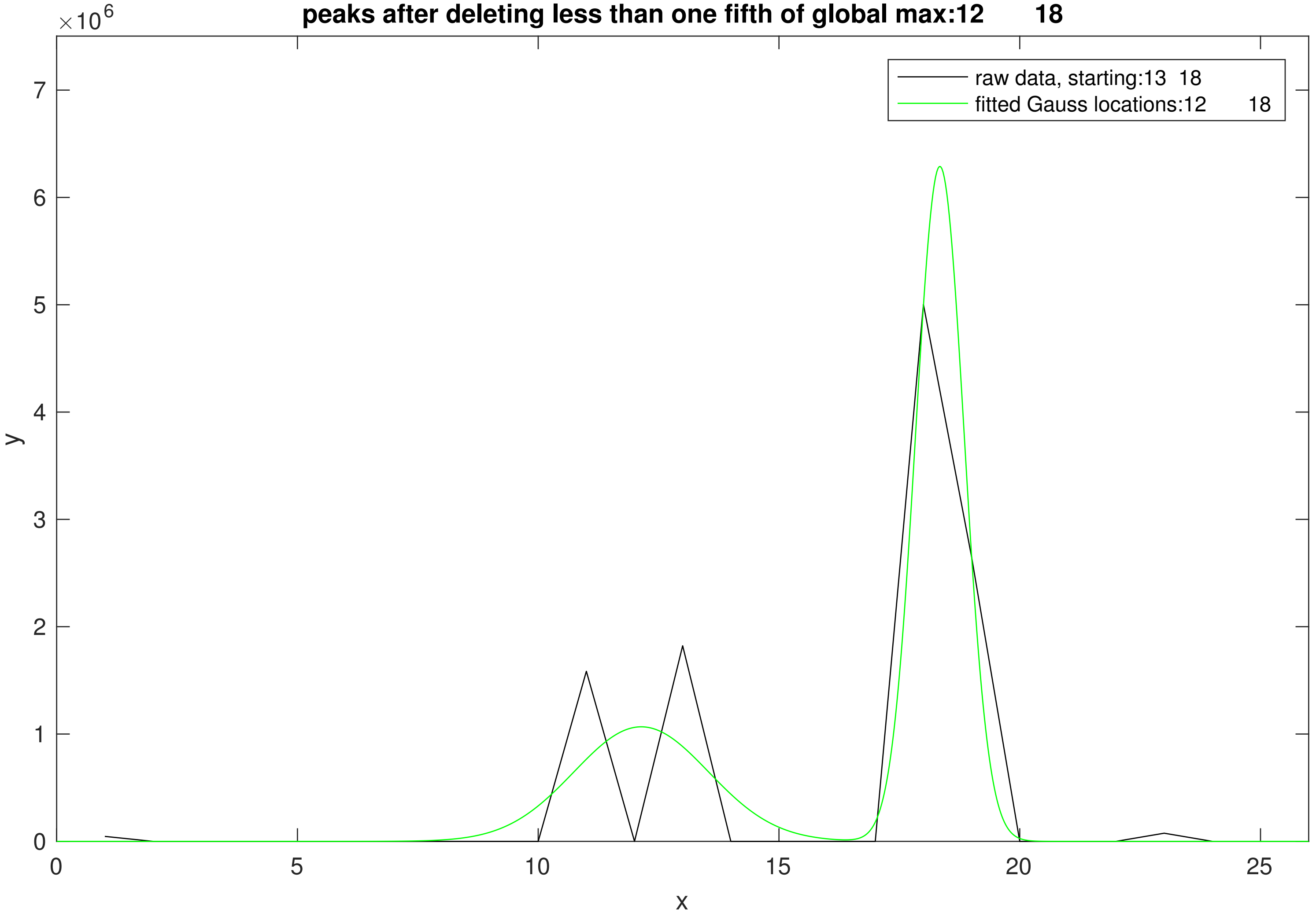

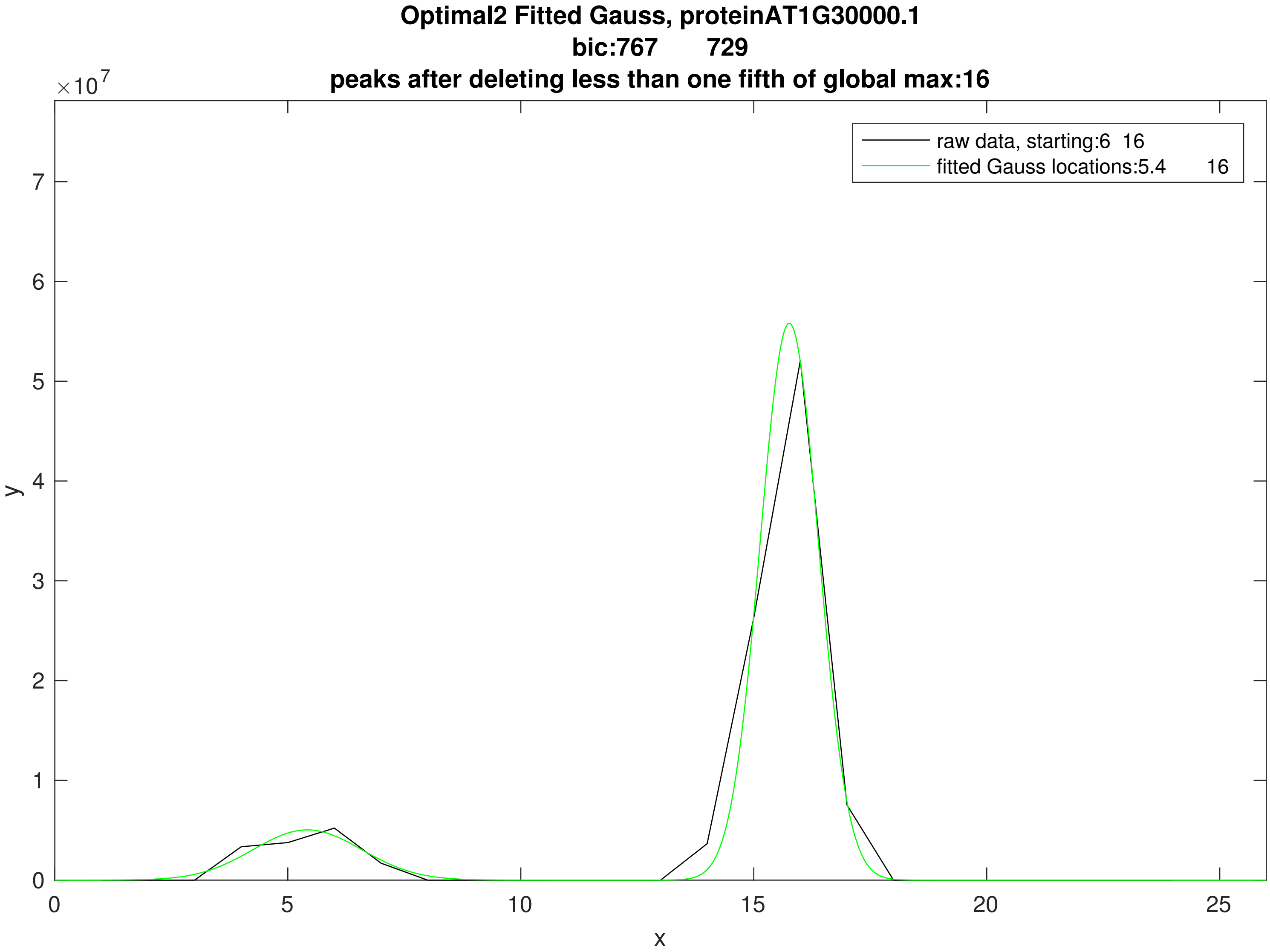
Optimal2 Fitted Gauss, proteinAT1G30000.1
bic:767 729
peaks after deleting less than one fifth of global max:16
×10^7
raw data, starting:6 16
fitted Gauss locations:5.4 16
y
x
0
1
2
3
4
5
6
7
0
5
10
15
20
25

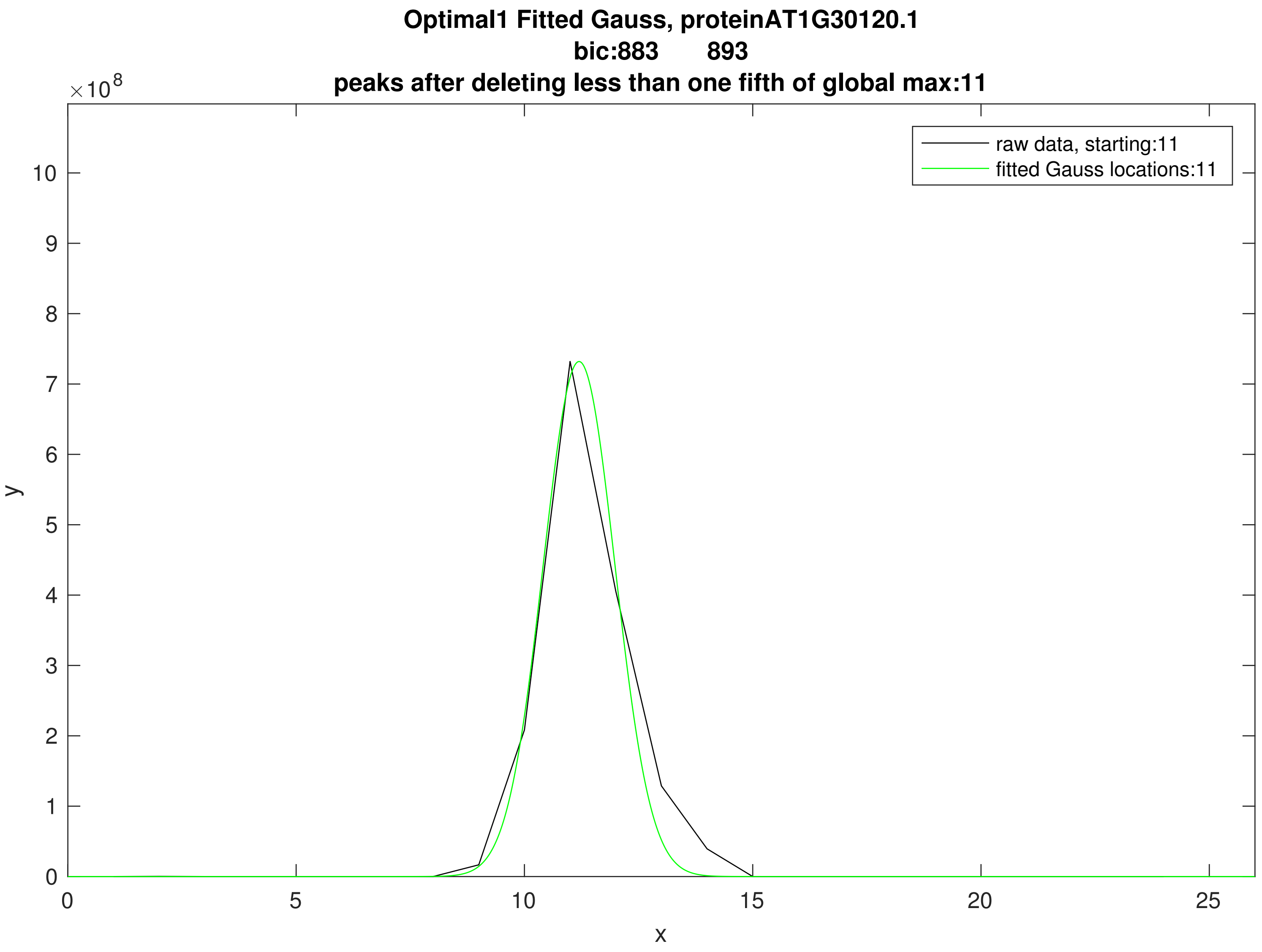
Optimal1 Fitted Gauss, proteinAT1G30120.1
bic:883 893
peaks after deleting less than one fifth of global max:11
×10^8
raw data, starting:11
fitted Gauss locations:11
y
x
0
1
2
3
4
5
6
7
8
9
10
0
5
10
15
20
25

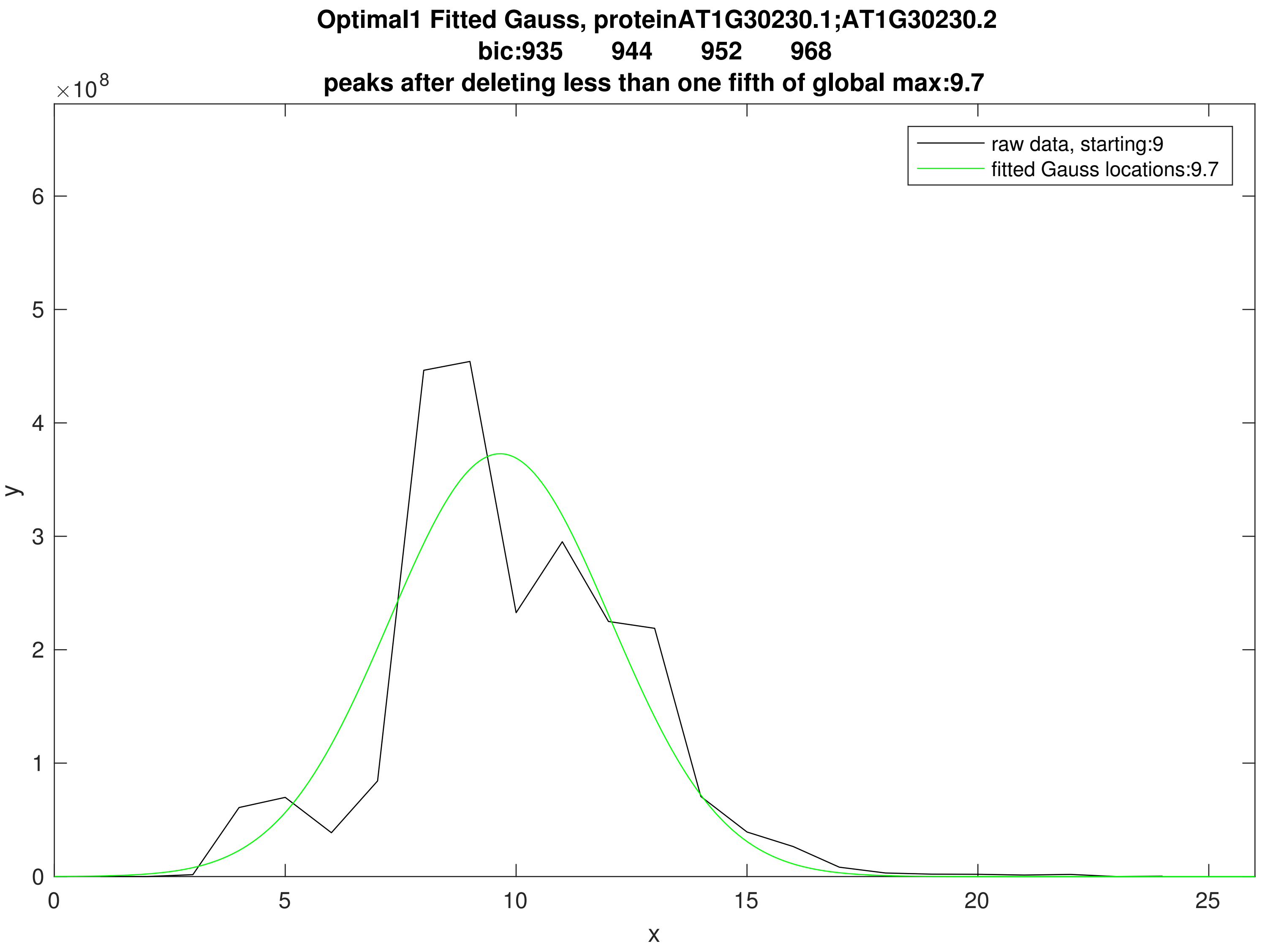

Optimal1 Fitted Gauss, proteinAT1G30230.1;AT1G30230.2
bic:935 944 952 968
peaks after deleting less than one fifth of global max:9.7
×10^8
raw data, starting:9
fitted Gauss locations:9.7
y
x
0
1
2
3
4
5
6
0
5
10
15
20
25

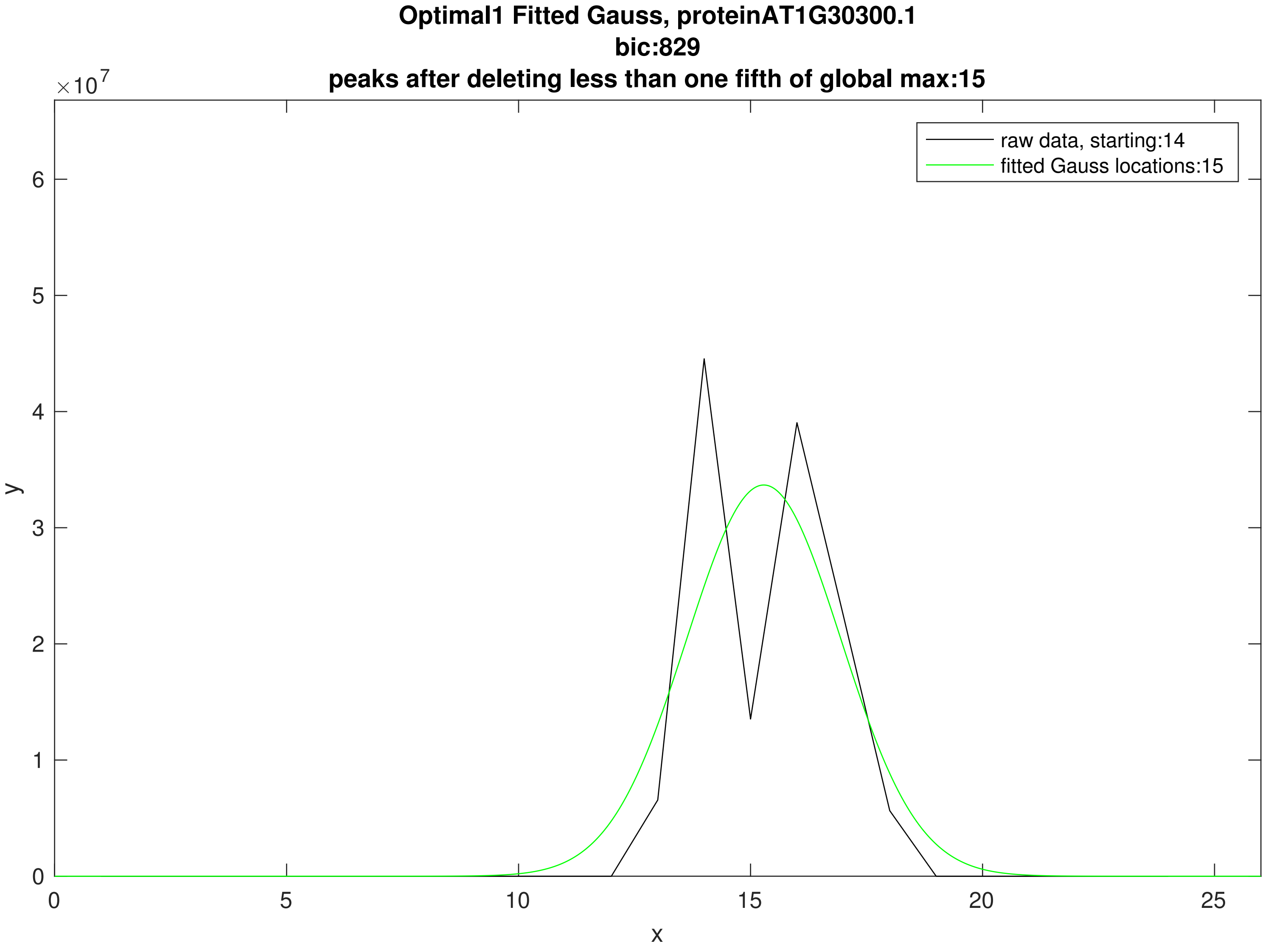
Optimal1 Fitted Gauss, proteinAT1G30300.1
bic:829
peaks after deleting less than one fifth of global max:15
×10^7
raw data, starting:14
fitted Gauss locations:15
y
x
0
1
2
3
4
5
6
0
5
10
15
20
25

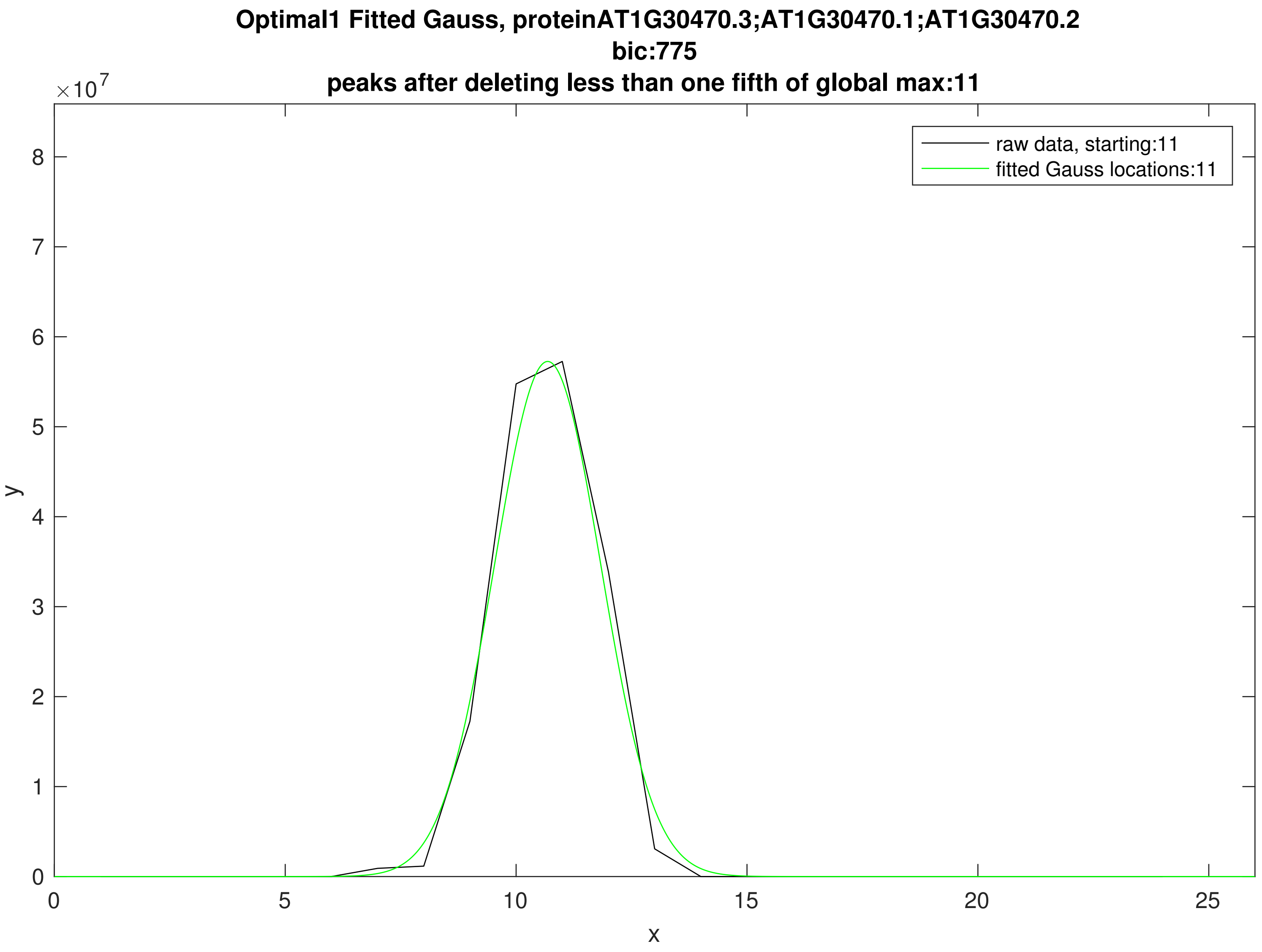

Optimal1 Fitted Gauss, proteinAT1G30470.3;AT1G30470.1;AT1G30470.2
bic:775
peaks after deleting less than one fifth of global max:11
×10^7
raw data, starting:11
fitted Gauss locations:11
y
x
0
1
2
3
4
5
6
7
8
0
5
10
15
20
25

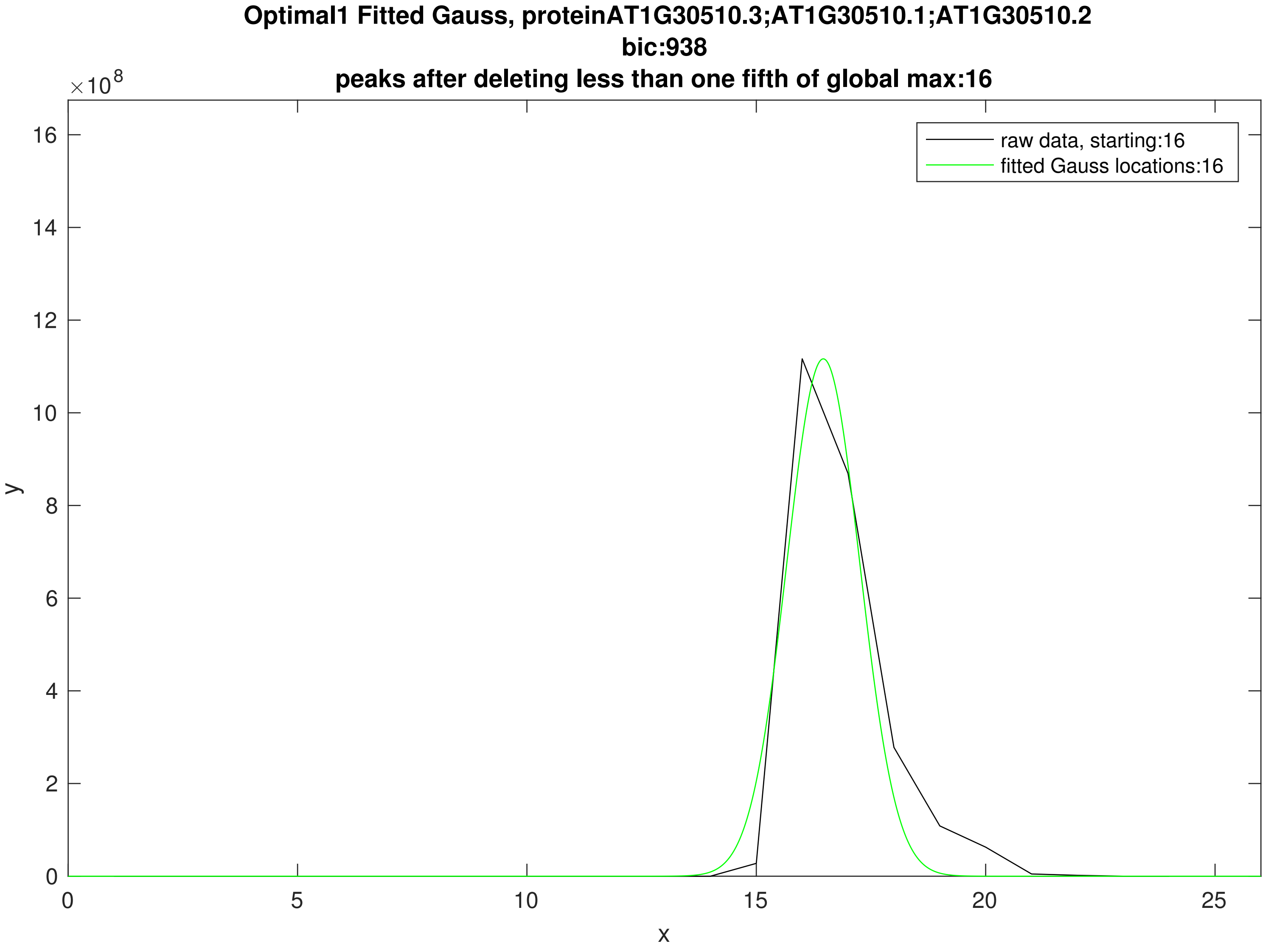

Optimal1 Fitted Gauss, proteinAT1G30510.3;AT1G30510.1;AT1G30510.2
bic:938
peaks after deleting less than one fifth of global max:16
×10^8
raw data, starting:16
fitted Gauss locations:16
y
x
0
2
4
6
8
10
12
14
16
0
5
10
15
20
25

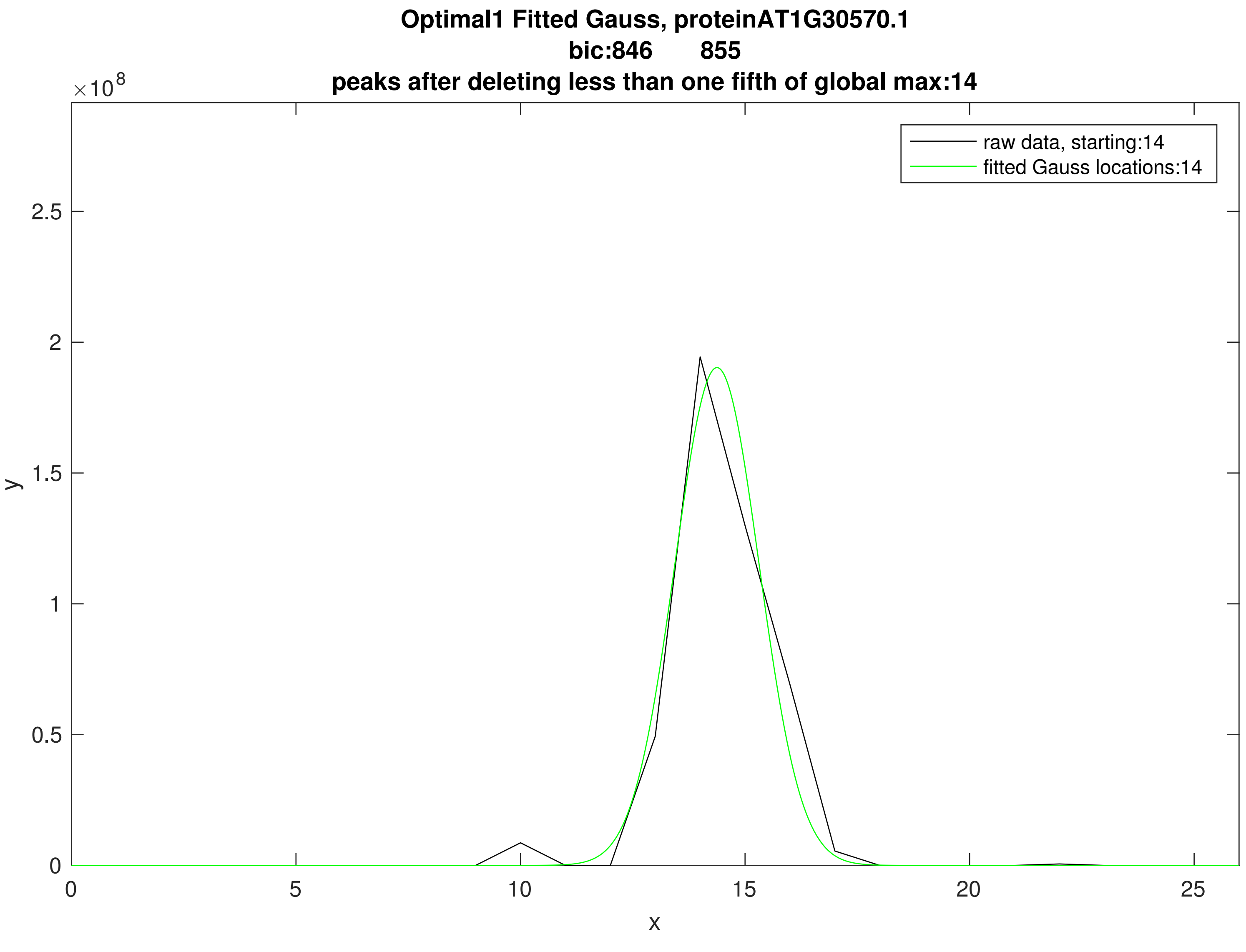
Optimal1 Fitted Gauss, proteinAT1G30570.1
bic:846 855
peaks after deleting less than one fifth of global max:14
×10^8
raw data, starting:14
fitted Gauss locations:14
y
x
0
0.5
1
1.5
2
2.5
0
5
10
15
20
25

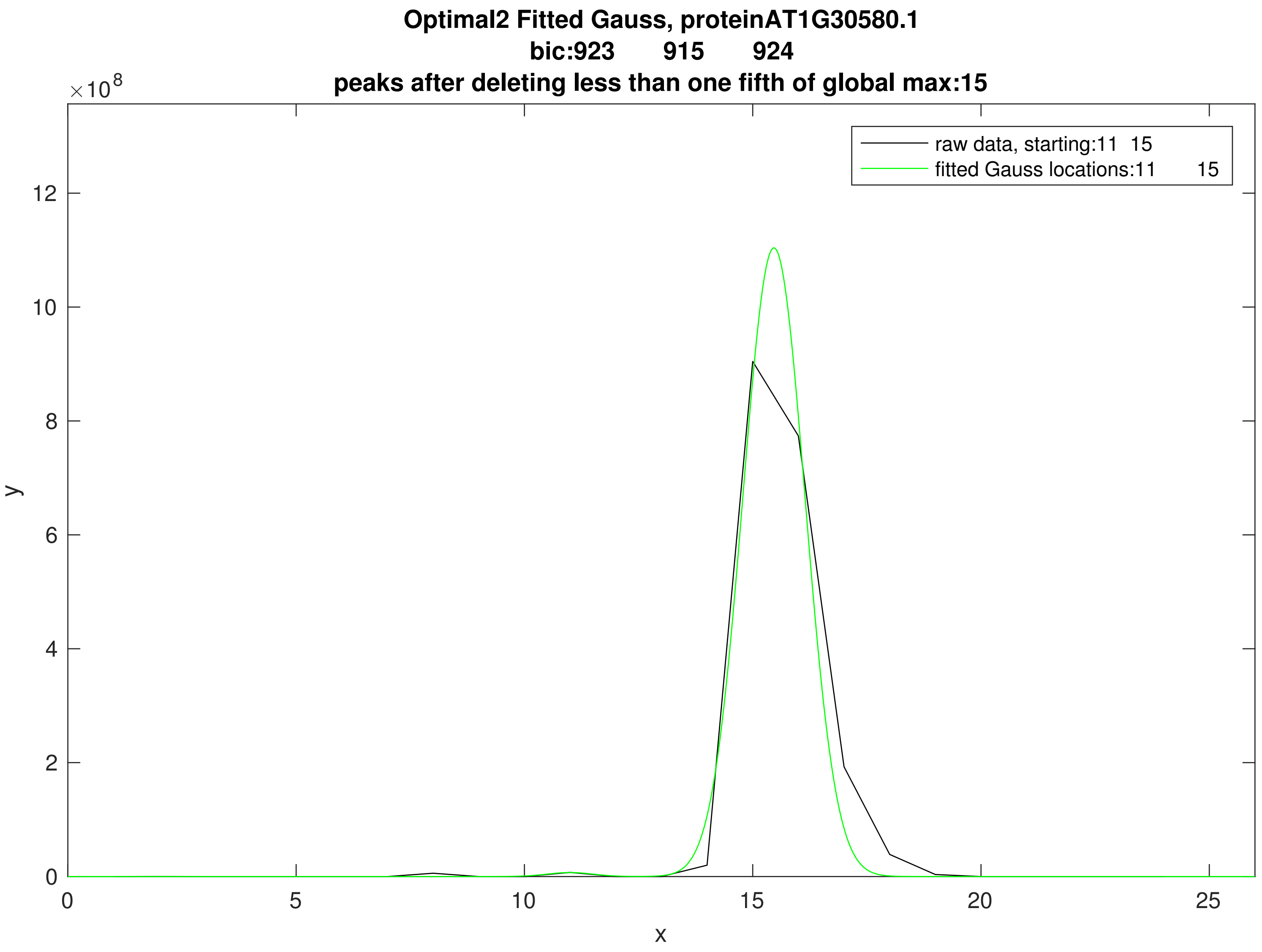

Optimal2 Fitted Gauss, proteinAT1G30580.1
bic:923 915 924
peaks after deleting less than one fifth of global max:15
×10^8
raw data, starting:11 15
fitted Gauss locations:11 15
12
10
8
6
4
2
0
y
0
5
10
15
20
25
x

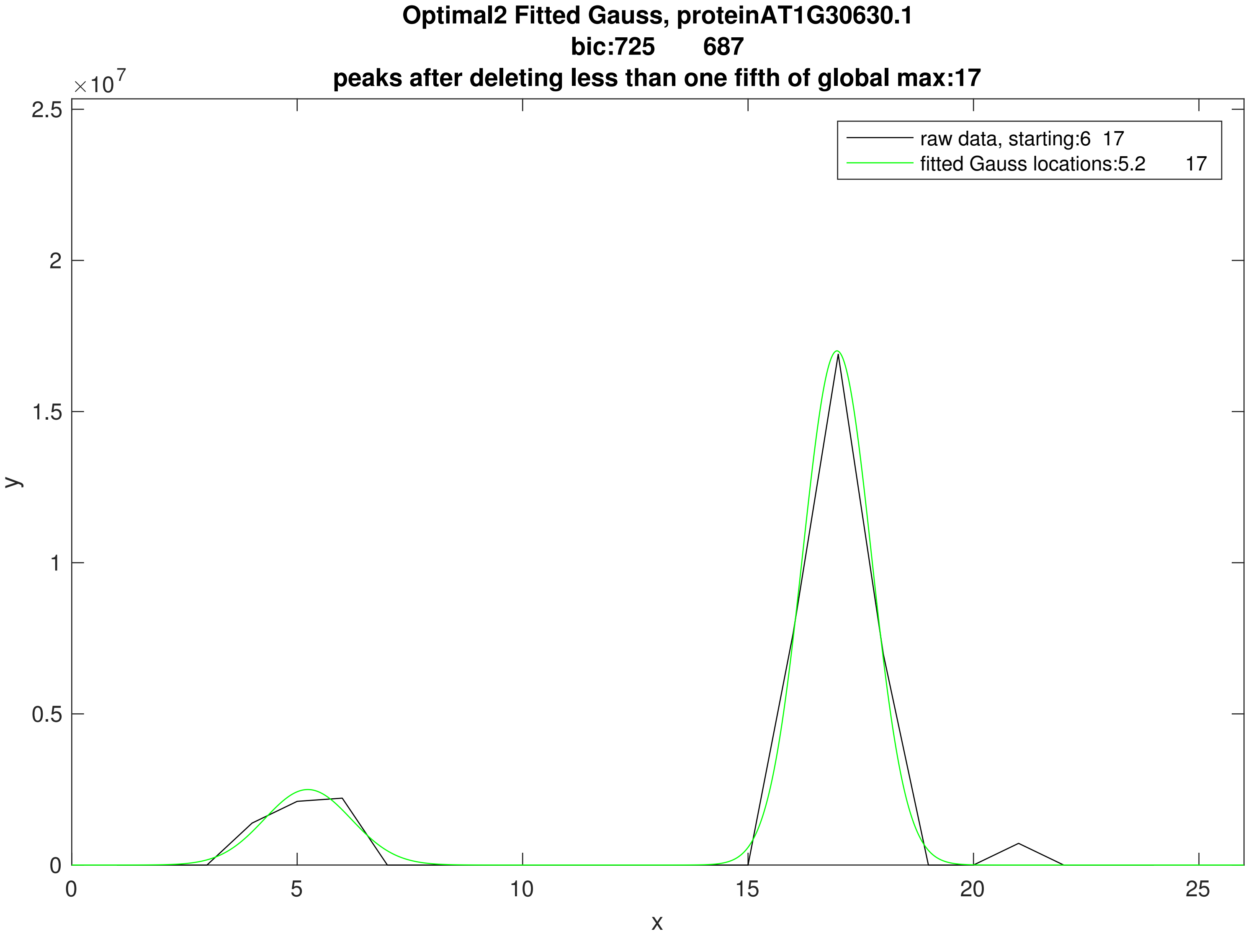
Optimal2 Fitted Gauss, proteinAT1G30630.1
bic:725 687
peaks after deleting less than one fifth of global max:17
×10^7
2.5
2
1.5
1
0.5
0
y
0
5
10
15
20
25
x
raw data, starting:6 17
fitted Gauss locations:5.2 17

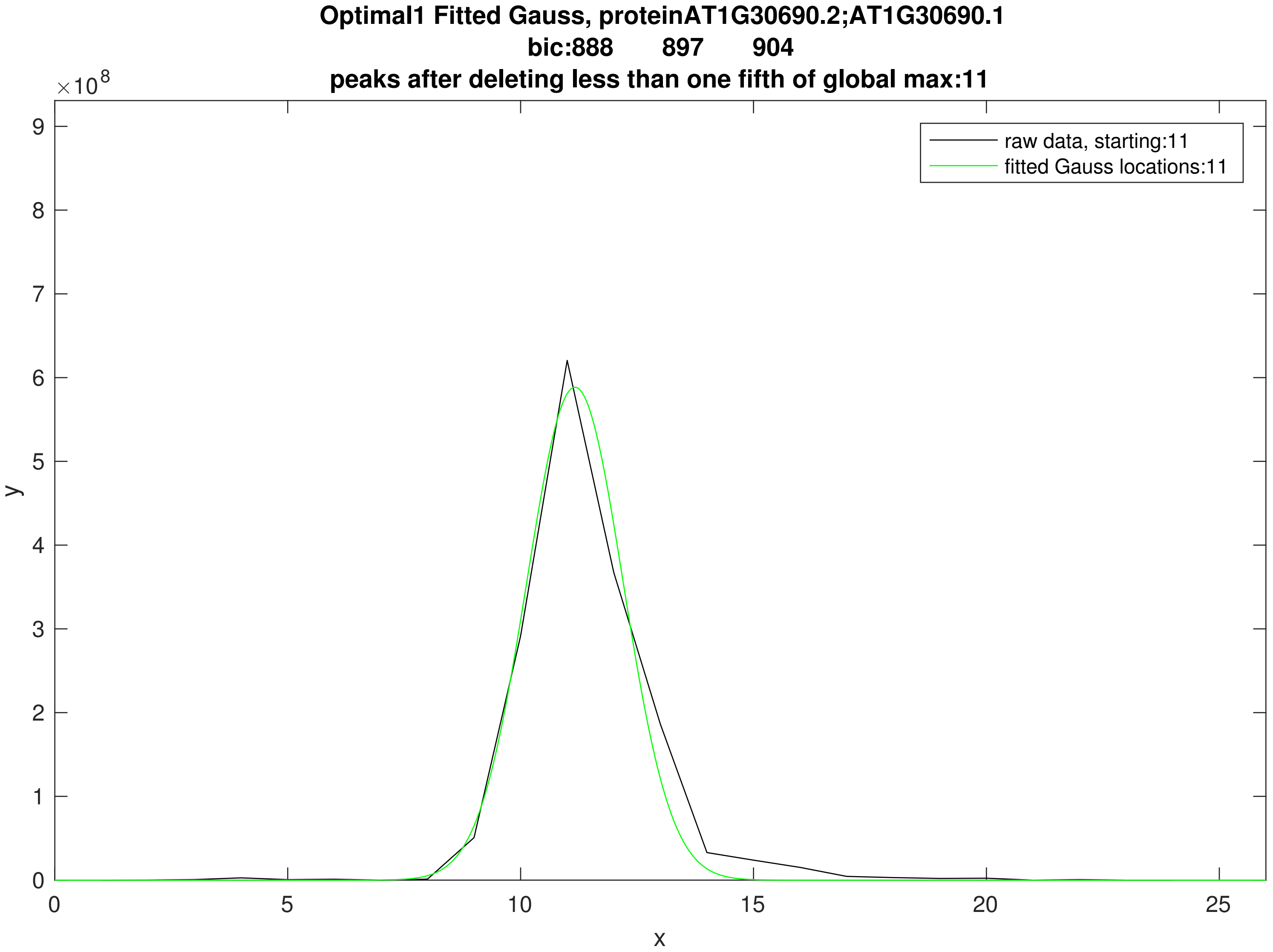
Optimal1 Fitted Gauss, proteinAT1G30690.2;AT1G30690.1
bic:888 897 904
peaks after deleting less than one fifth of global max:11
×10^8
raw data, starting:11
fitted Gauss locations:11
y
x
0
1
2
3
4
5
6
7
8
9
0
5
10
15
20
25

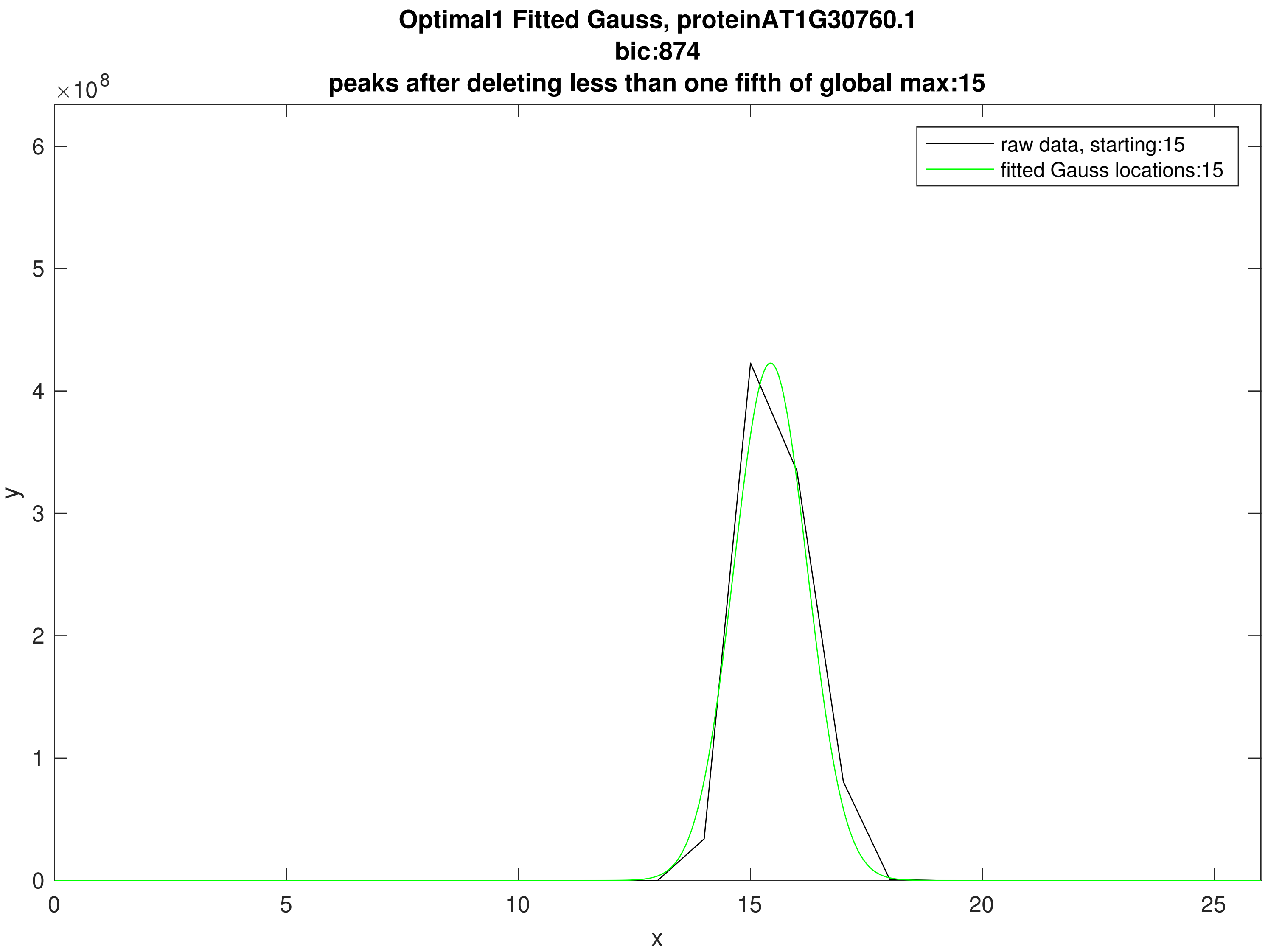

Optimal1 Fitted Gauss, proteinAT1G30760.1
bic:874
peaks after deleting less than one fifth of global max:15
×10^8
raw data, starting:15
fitted Gauss locations:15
y
x
0
1
2
3
4
5
6
0
5
10
15
20
25

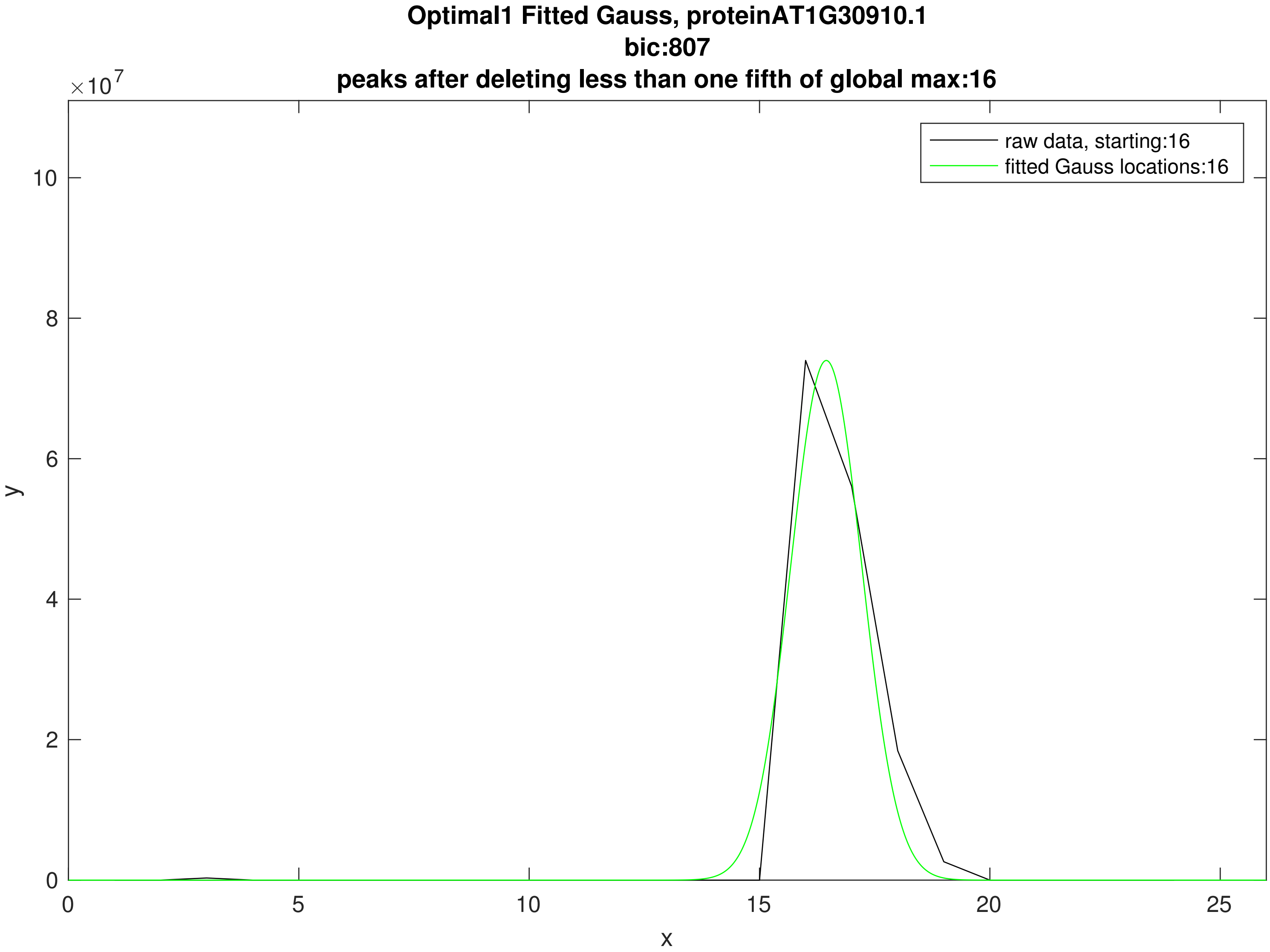

Optimal1 Fitted Gauss, proteinAT1G30910.1
bic:807
peaks after deleting less than one fifth of global max:16
×10^7
raw data, starting:16
fitted Gauss locations:16
y
x
0
2
4
6
8
10
0
5
10
15
20
25

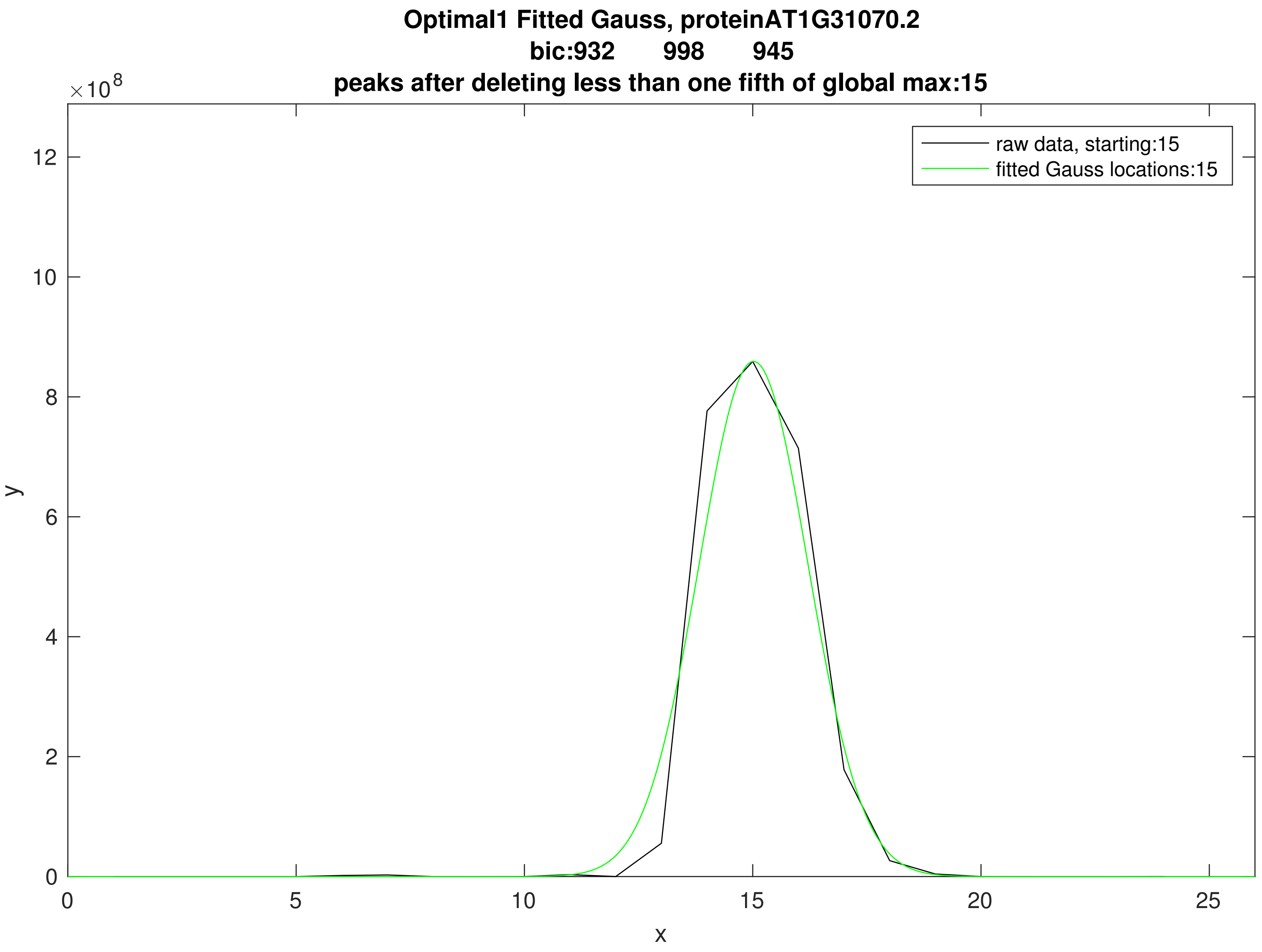

Optimal1 Fitted Gauss, proteinAT1G31070.2
bic:932 998 945
peaks after deleting less than one fifth of global max:15
×10^8
12
10
8
6
4
2
0
y
0
5
10
15
20
25
x
raw data, starting:15
fitted Gauss locations:15

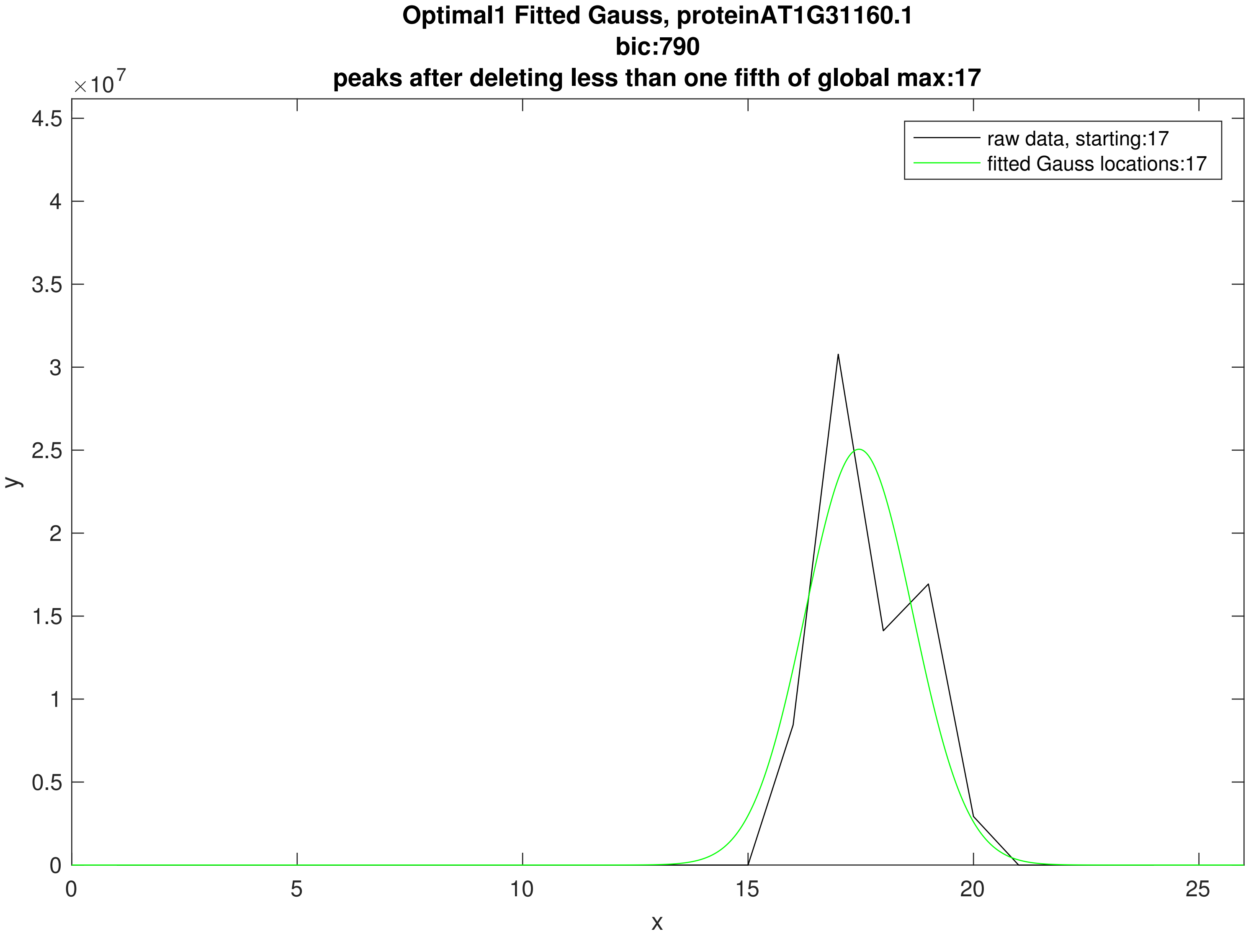

Optimal1 Fitted Gauss, proteinAT1G31160.1
bic:790
peaks after deleting less than one fifth of global max:17
×10^7
4.5
4
3.5
3
2.5
2
1.5
1
0.5
0
y
0
5
10
15
20
25
x
raw data, starting:17
fitted Gauss locations:17

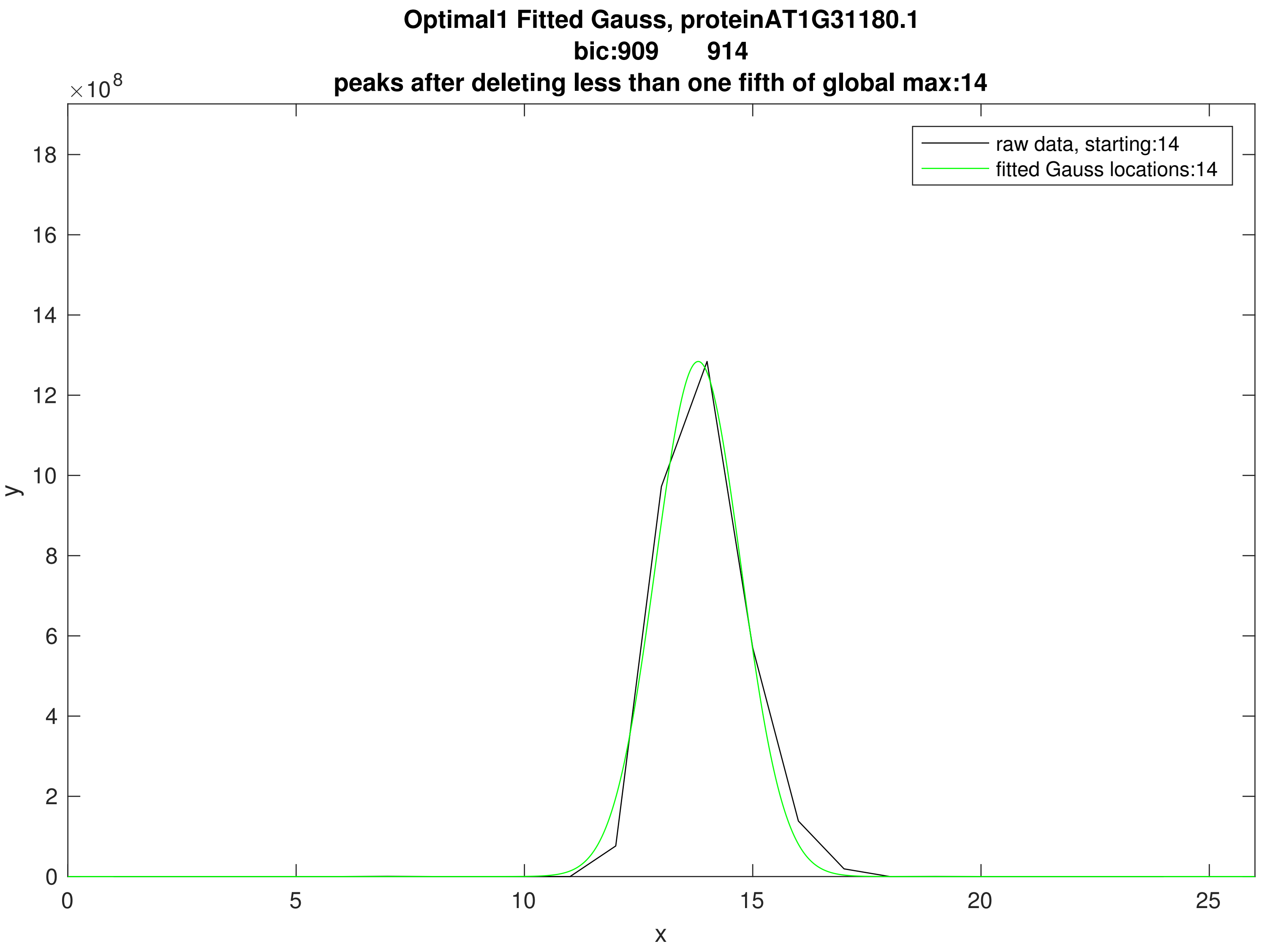

Optimal1 Fitted Gauss, proteinAT1G31180.1
bic:909 914
peaks after deleting less than one fifth of global max:14
×10^8
raw data, starting:14
fitted Gauss locations:14
y
x
0
2
4
6
8
10
12
14
16
18
0
5
10
15
20
25

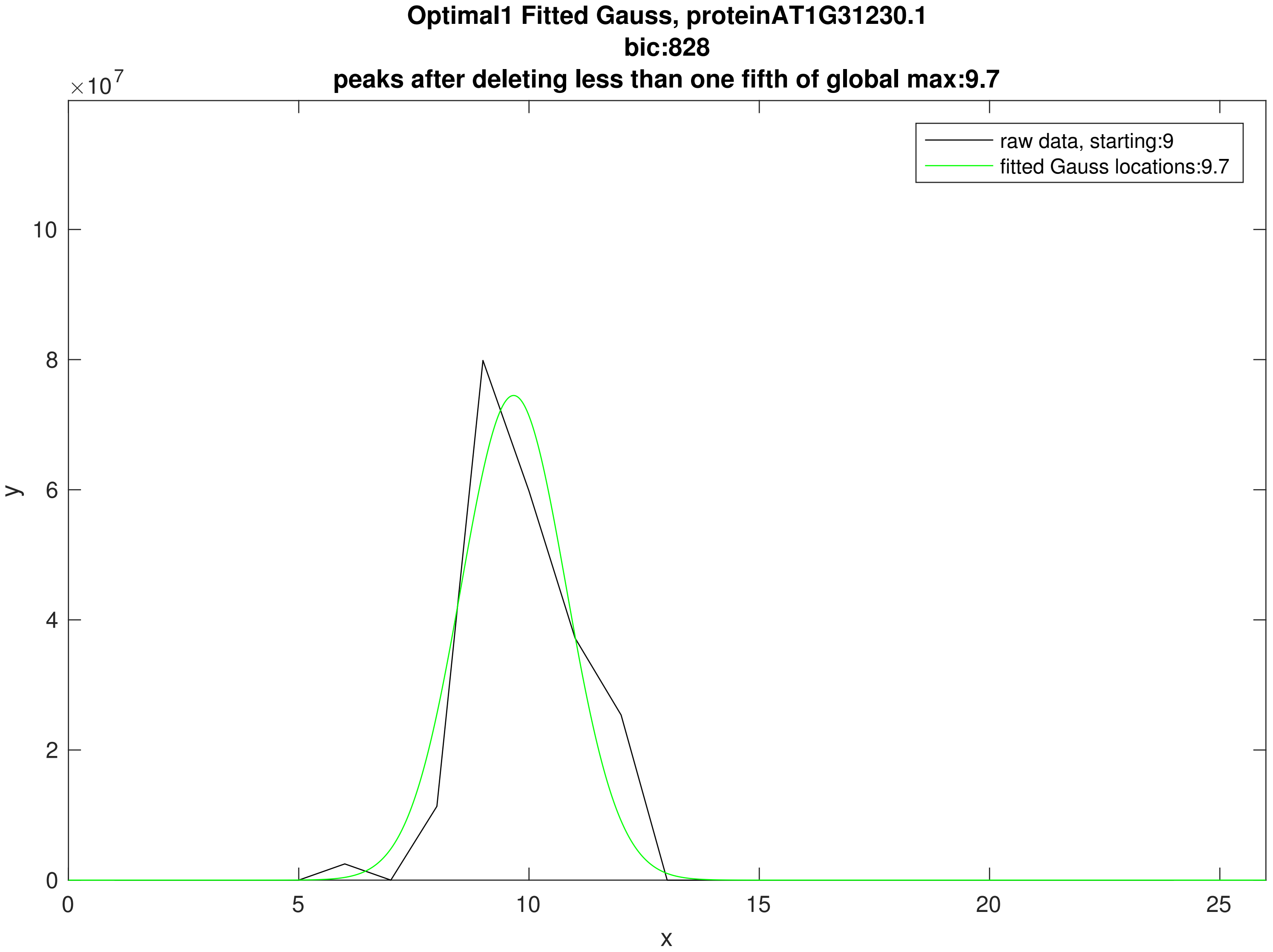
Optimal1 Fitted Gauss, proteinAT1G31230.1
bic:828
peaks after deleting less than one fifth of global max:9.7
×10^7
raw data, starting:9
fitted Gauss locations:9.7
y
x
0
2
4
6
8
10
5
15
20
25

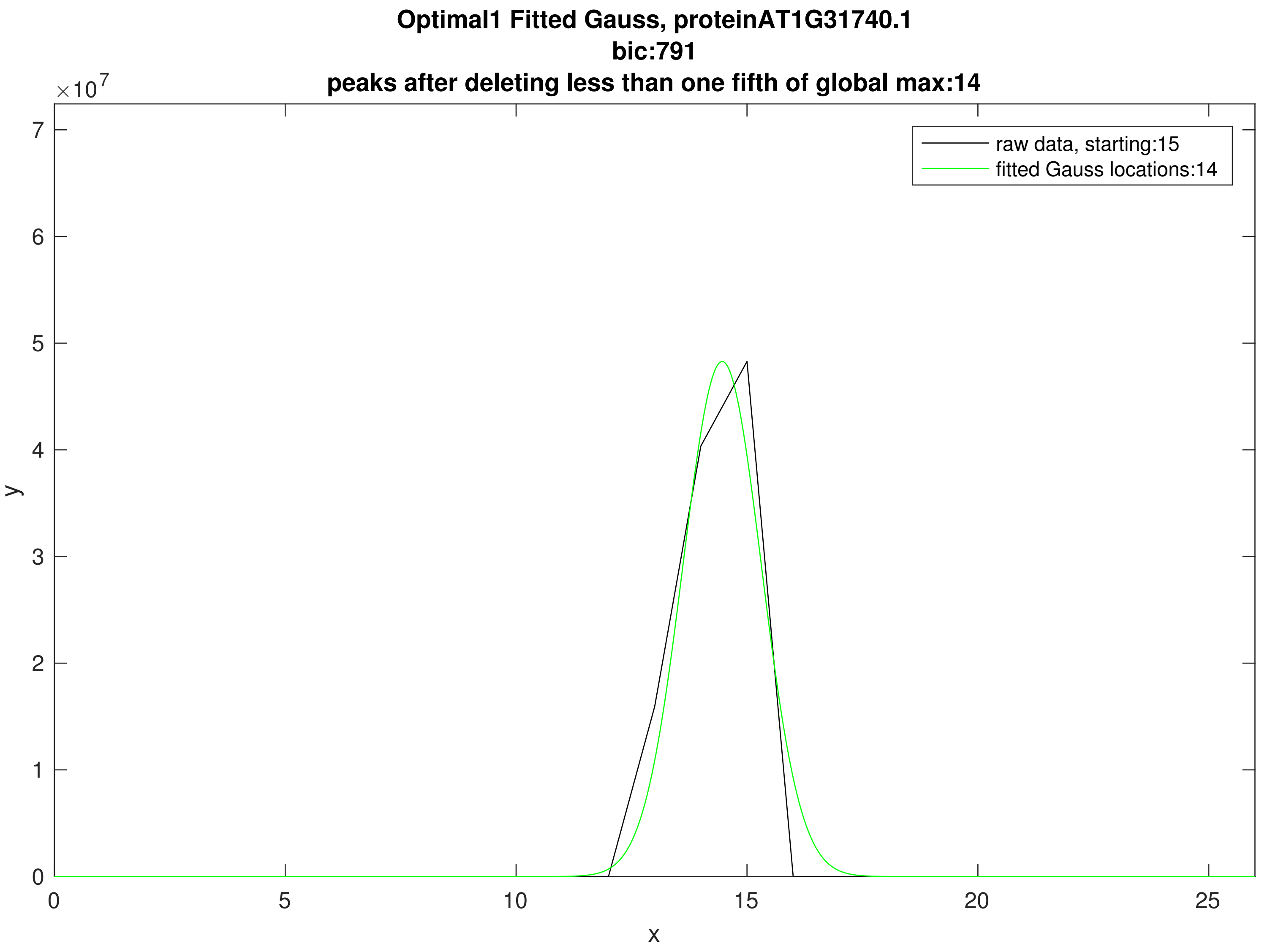
Optimal1 Fitted Gauss, proteinAT1G31740.1
bic:791
peaks after deleting less than one fifth of global max:14
×10^7
raw data, starting:15
fitted Gauss locations:14
y
x
0
1
2
3
4
5
6
7
0
5
10
15
20
25

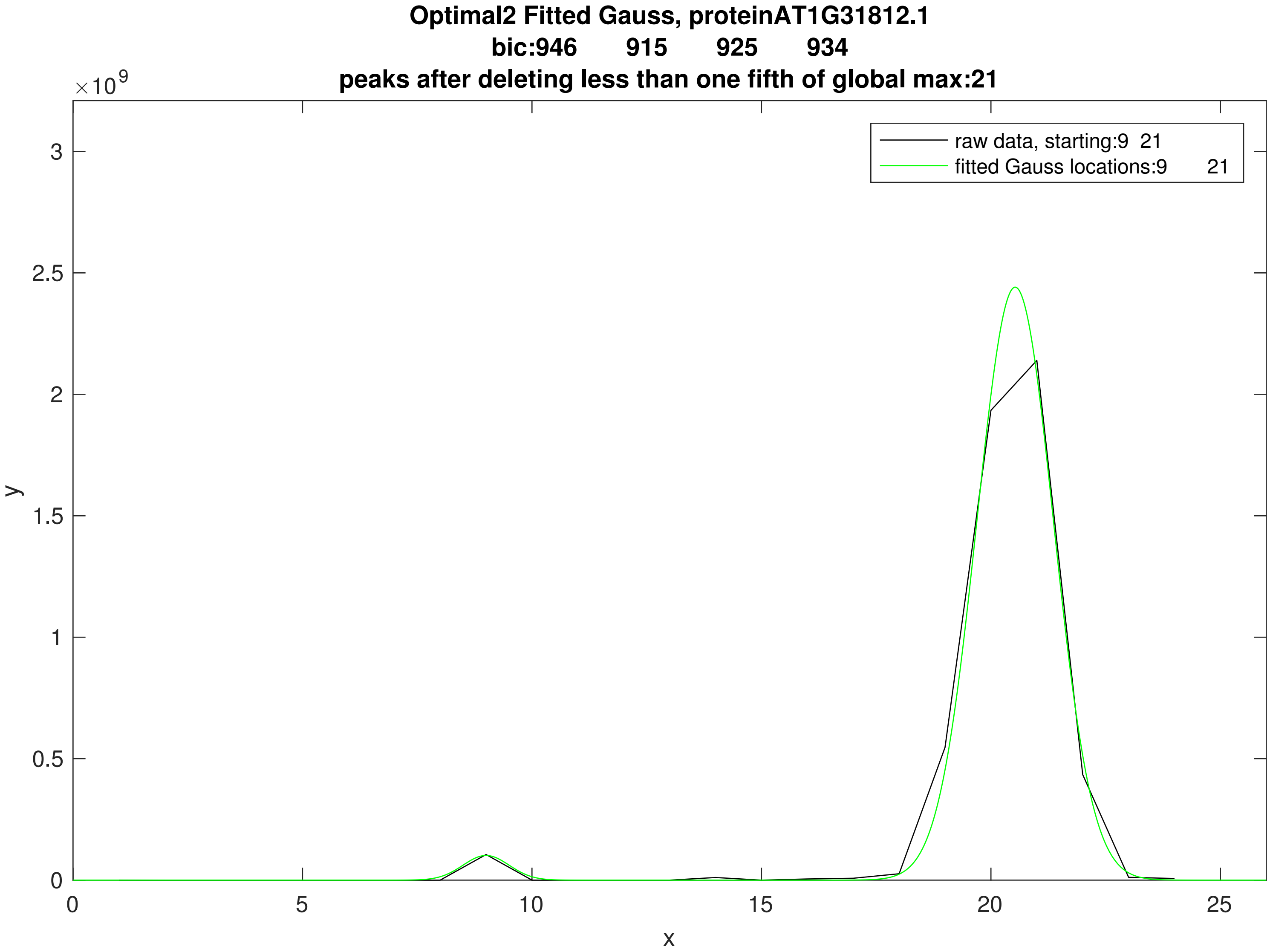
Optimal2 Fitted Gauss, proteinAT1G31812.1
bic:946 915 925 934
peaks after deleting less than one fifth of global max:21
×10^9
3
2.5
2
1.5
1
0.5
0
y
0
5
10
15
20
25
x
raw data, starting:9 21
fitted Gauss locations:9 21

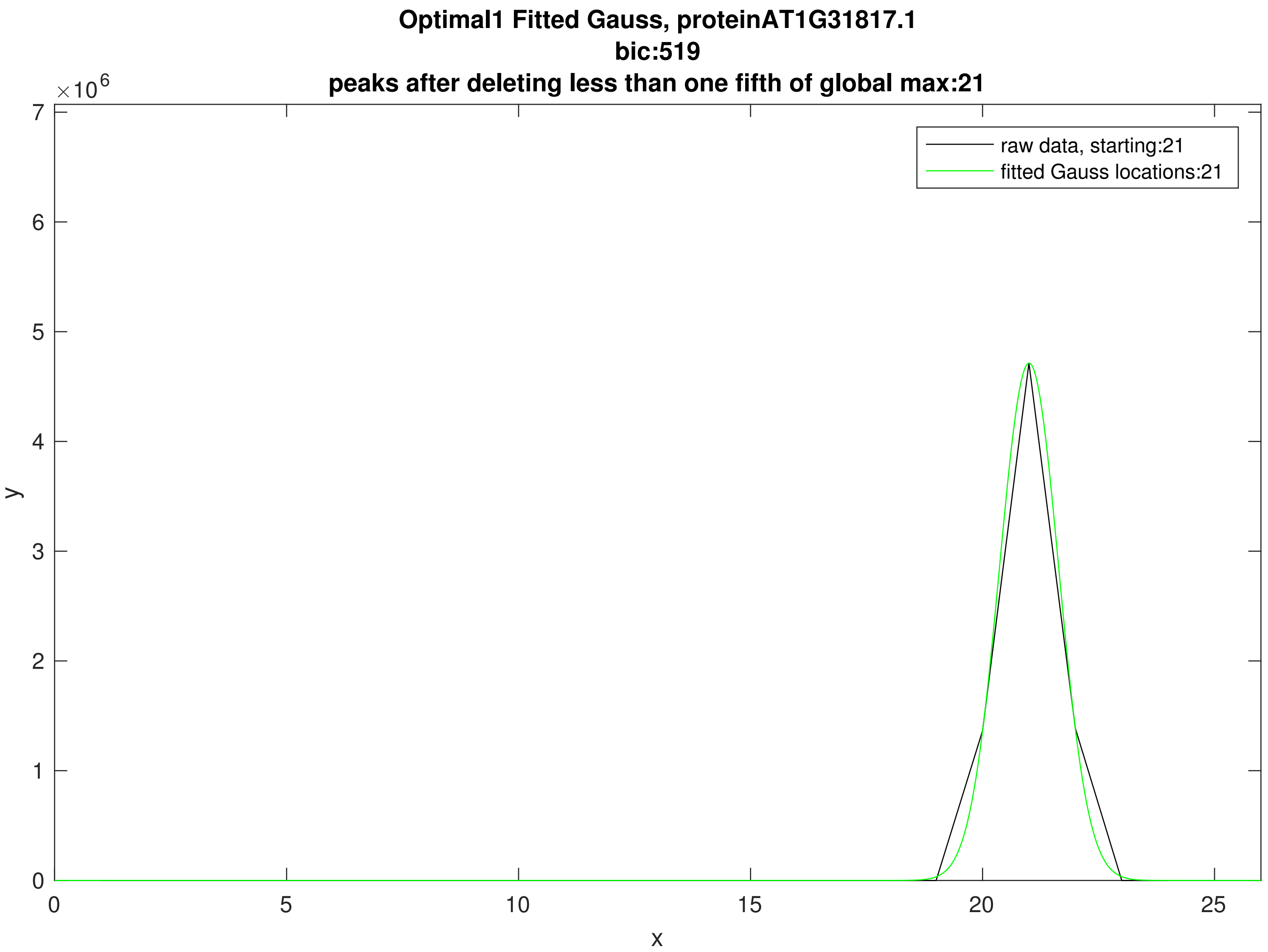

Optimal1 Fitted Gauss, proteinAT1G31817.1
bic:519
peaks after deleting less than one fifth of global max:21
×10^6
raw data, starting:21
fitted Gauss locations:21
y
x

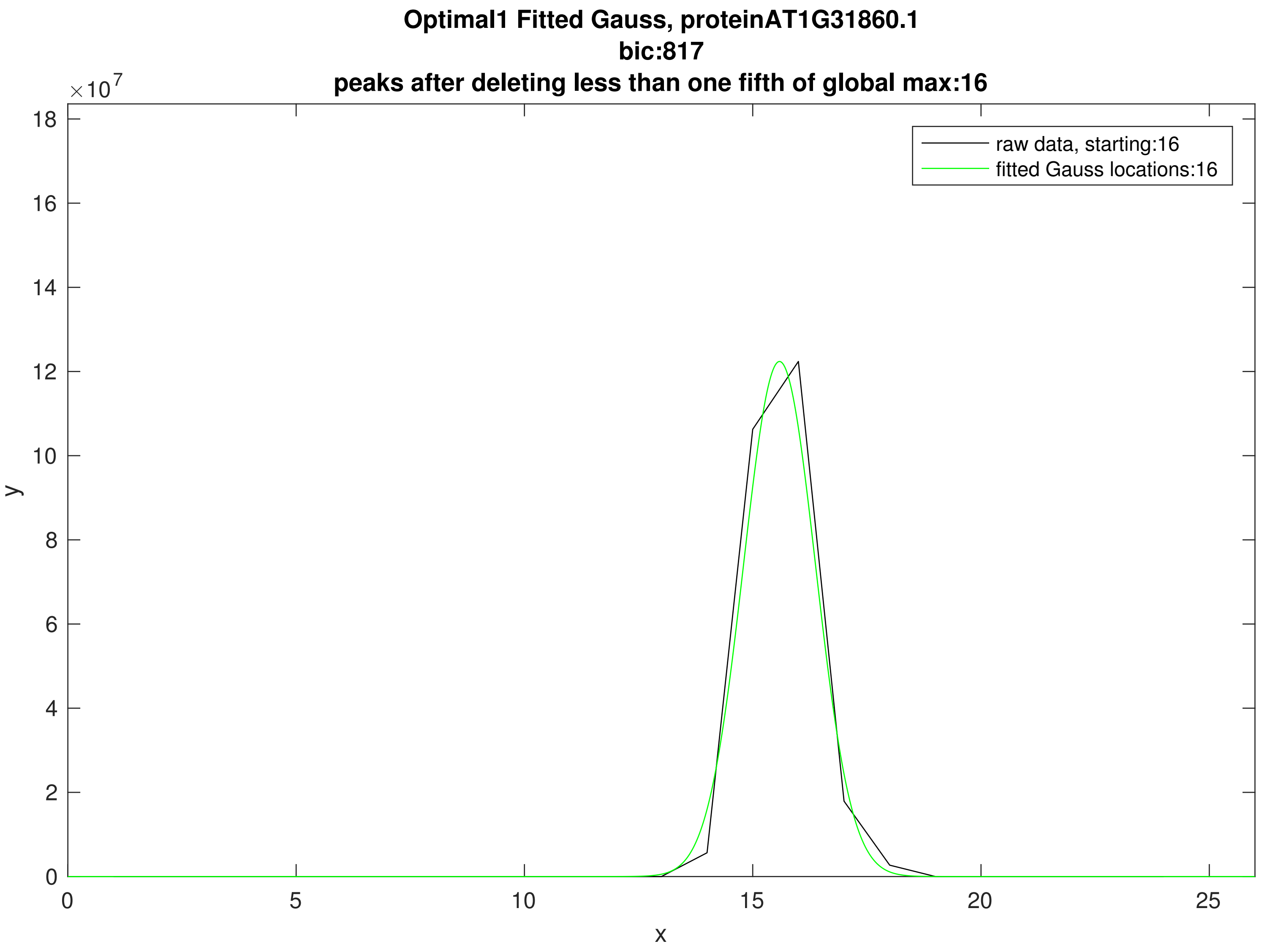

Optimal1 Fitted Gauss, proteinAT1G31860.1
bic:817
peaks after deleting less than one fifth of global max:16
×10^7
y
x
raw data, starting:16
fitted Gauss locations:16

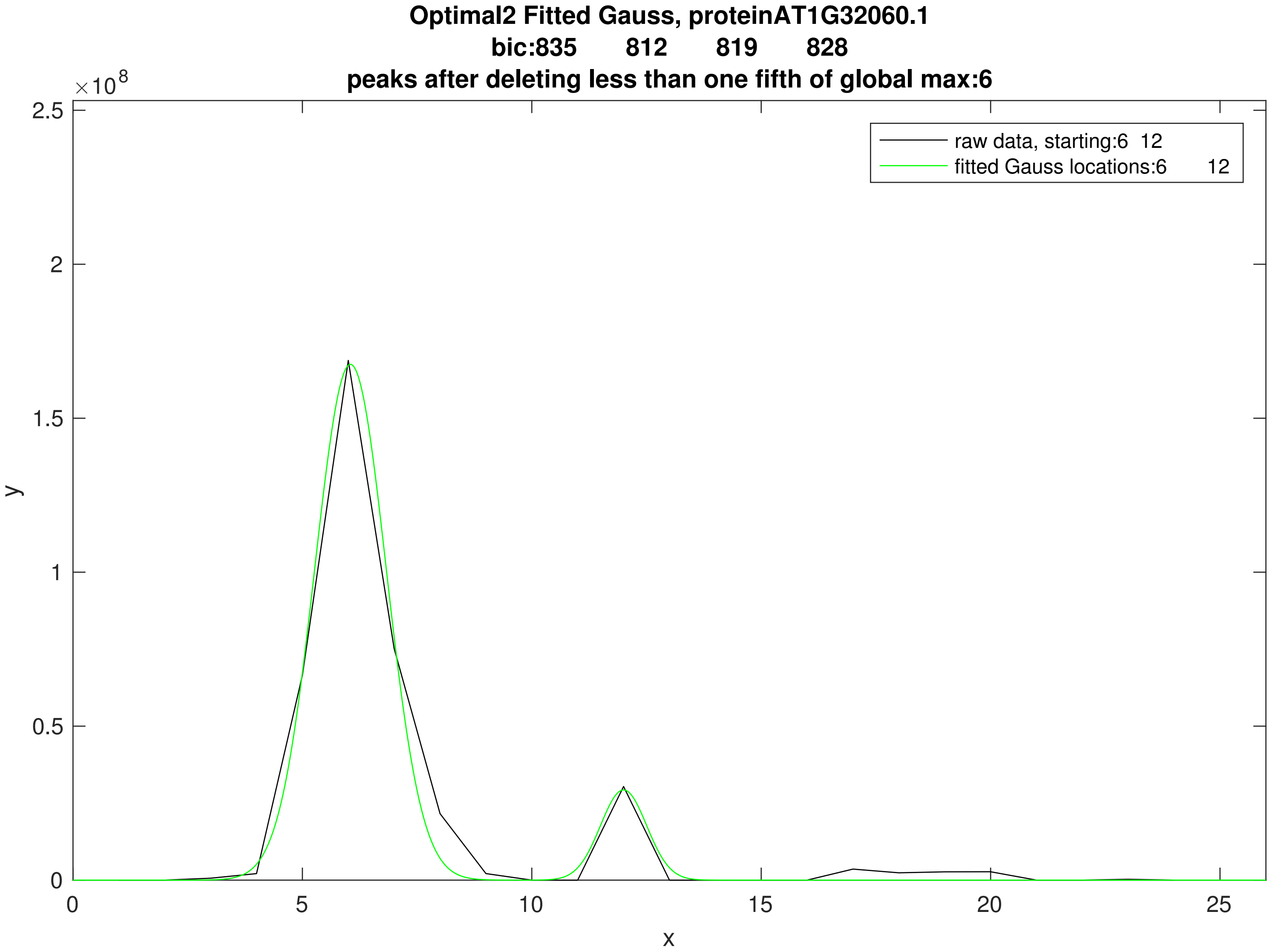
Optimal2 Fitted Gauss, proteinAT1G32060.1
bic:835 812 819 828
peaks after deleting less than one fifth of global max:6
×10^8
2.5
2
1.5
1
0.5
0
y
0
5
10
15
20
25
x
raw data, starting:6 12
fitted Gauss locations:6 12

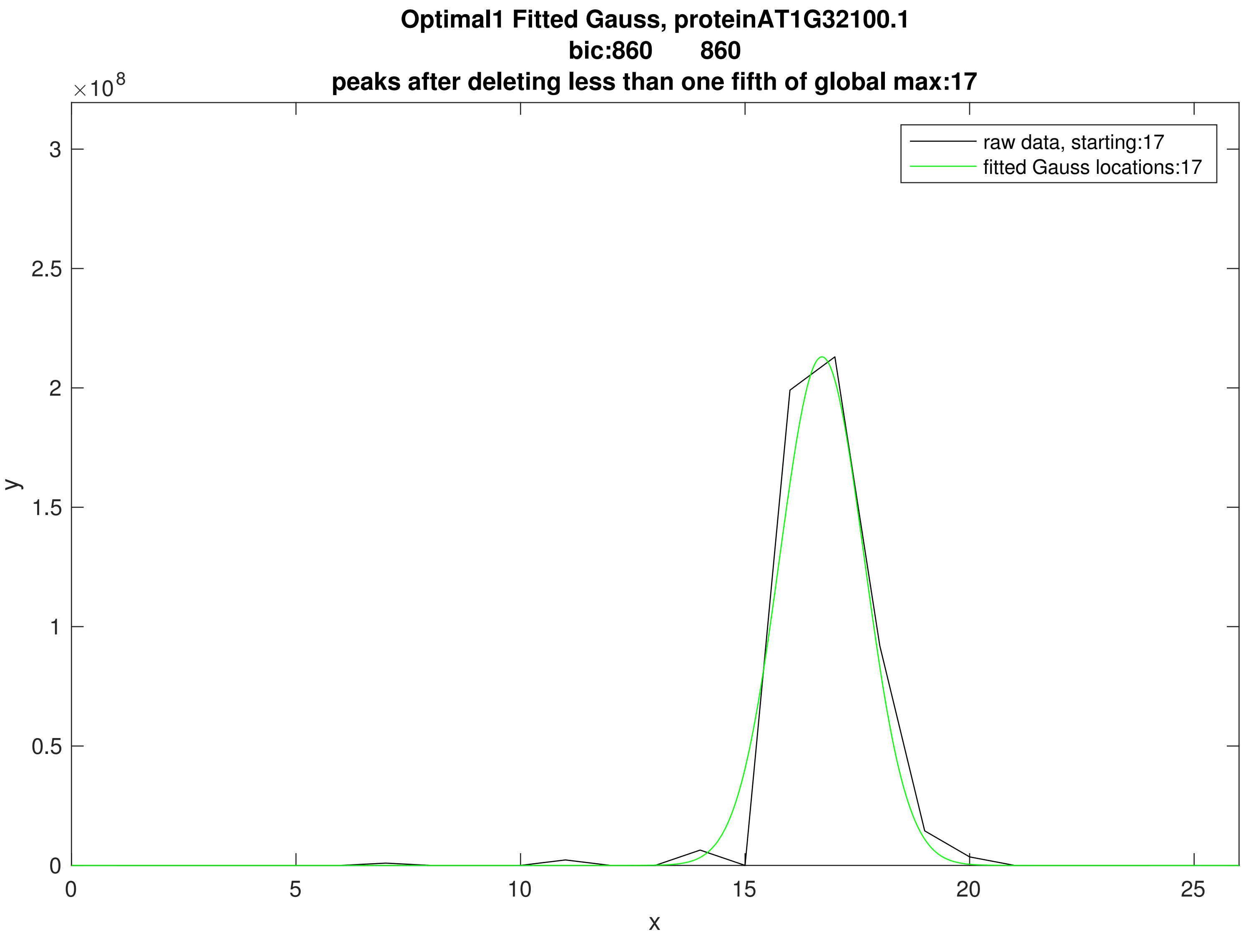
Optimal1 Fitted Gauss, proteinAT1G32100.1
bic:860 860
peaks after deleting less than one fifth of global max:17
×10^8
3
2.5
2
1.5
1
0.5
0
y
0
5
10
15
20
25
x
raw data, starting:17
fitted Gauss locations:17

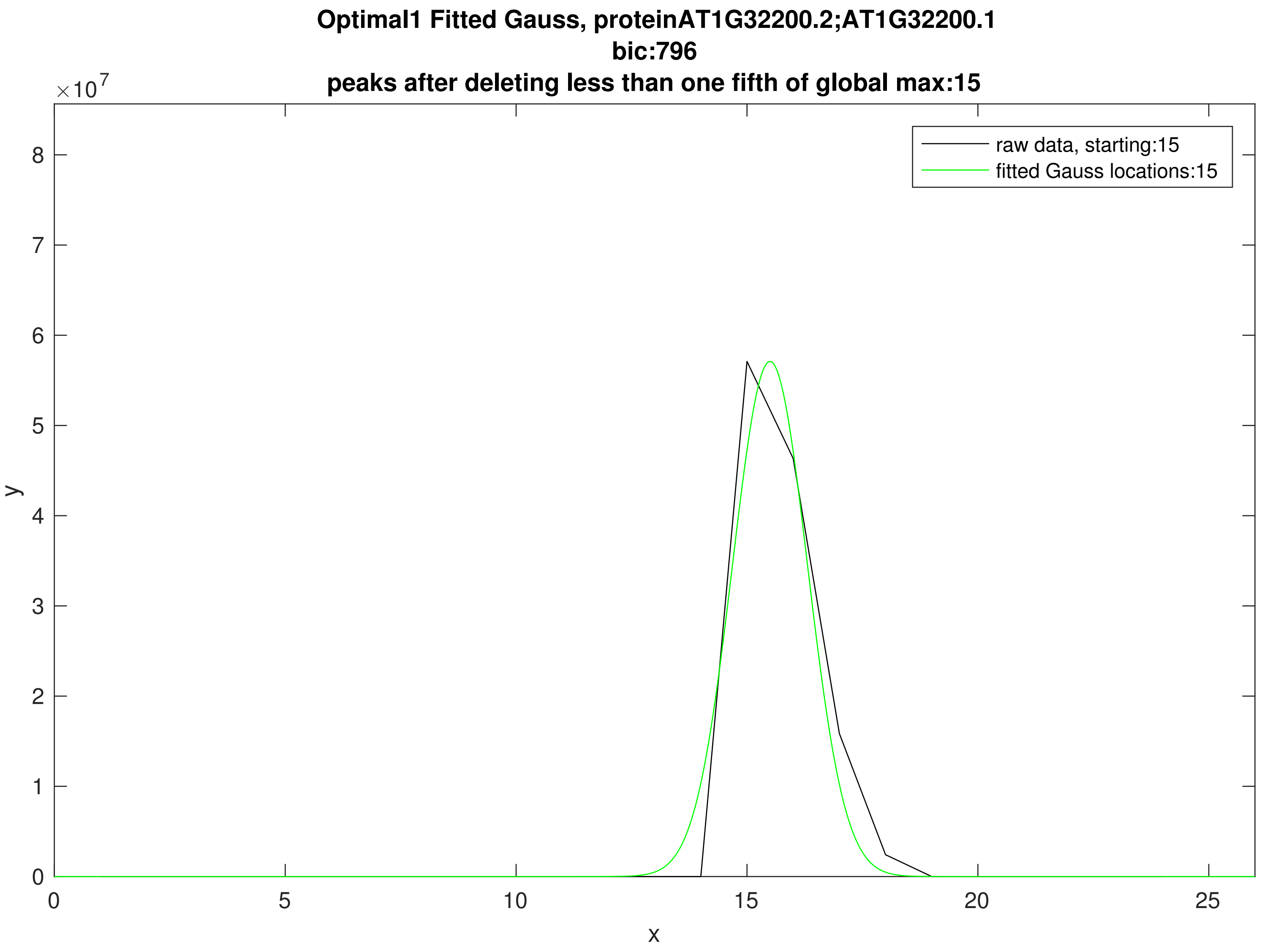
Optimal1 Fitted Gauss, proteinAT1G32200.2;AT1G32200.1
bic:796
peaks after deleting less than one fifth of global max:15
×10^7
8
7
6
5
4
3
2
1
0
y
0
5
10
15
20
25
x
raw data, starting:15
fitted Gauss locations:15

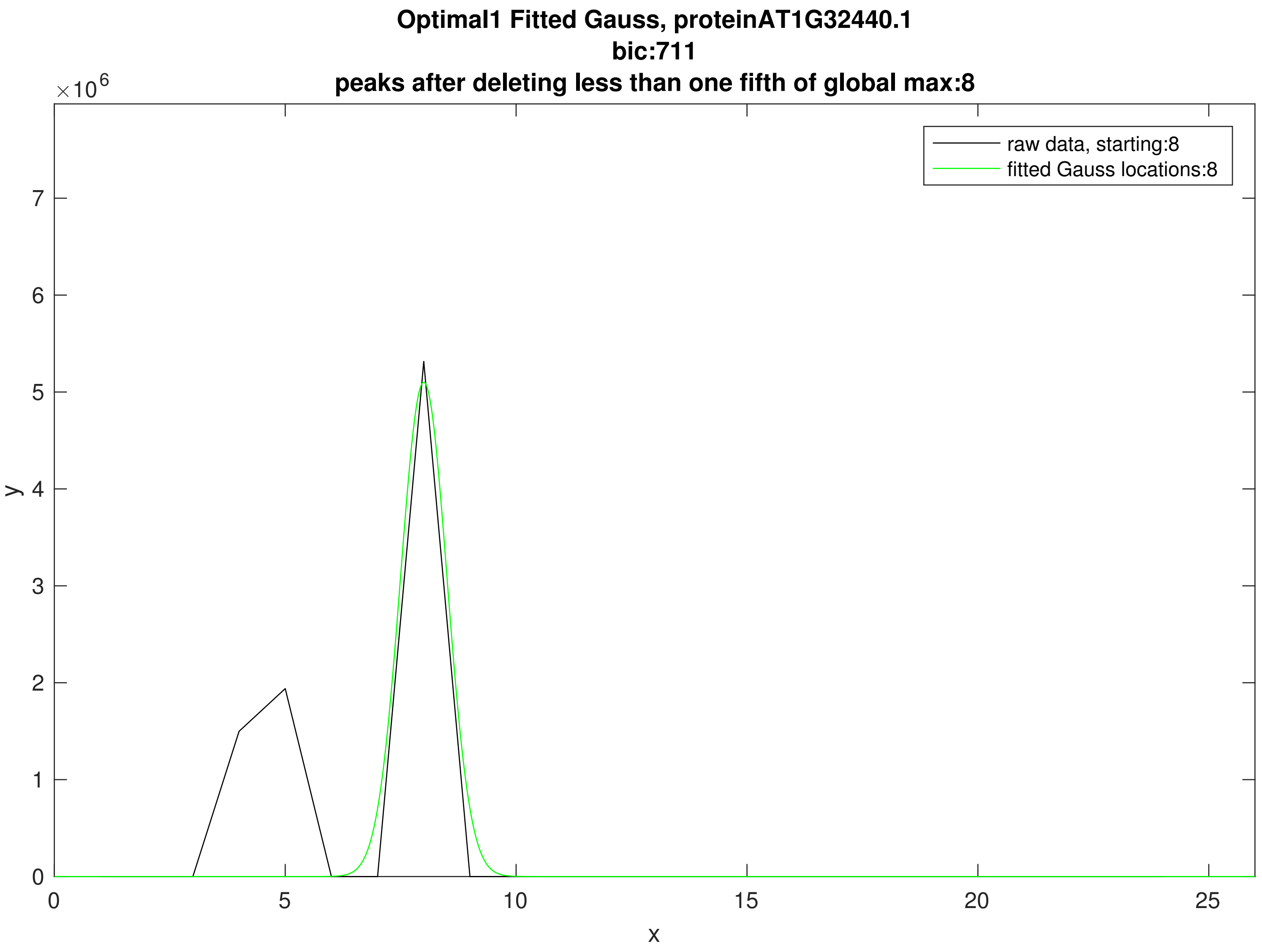
Optimal1 Fitted Gauss, proteinAT1G32440.1
bic:711
peaks after deleting less than one fifth of global max:8
×10^6
raw data, starting:8
fitted Gauss locations:8
y
x
0
1
2
3
4
5
6
7
0
5
10
15
20
25

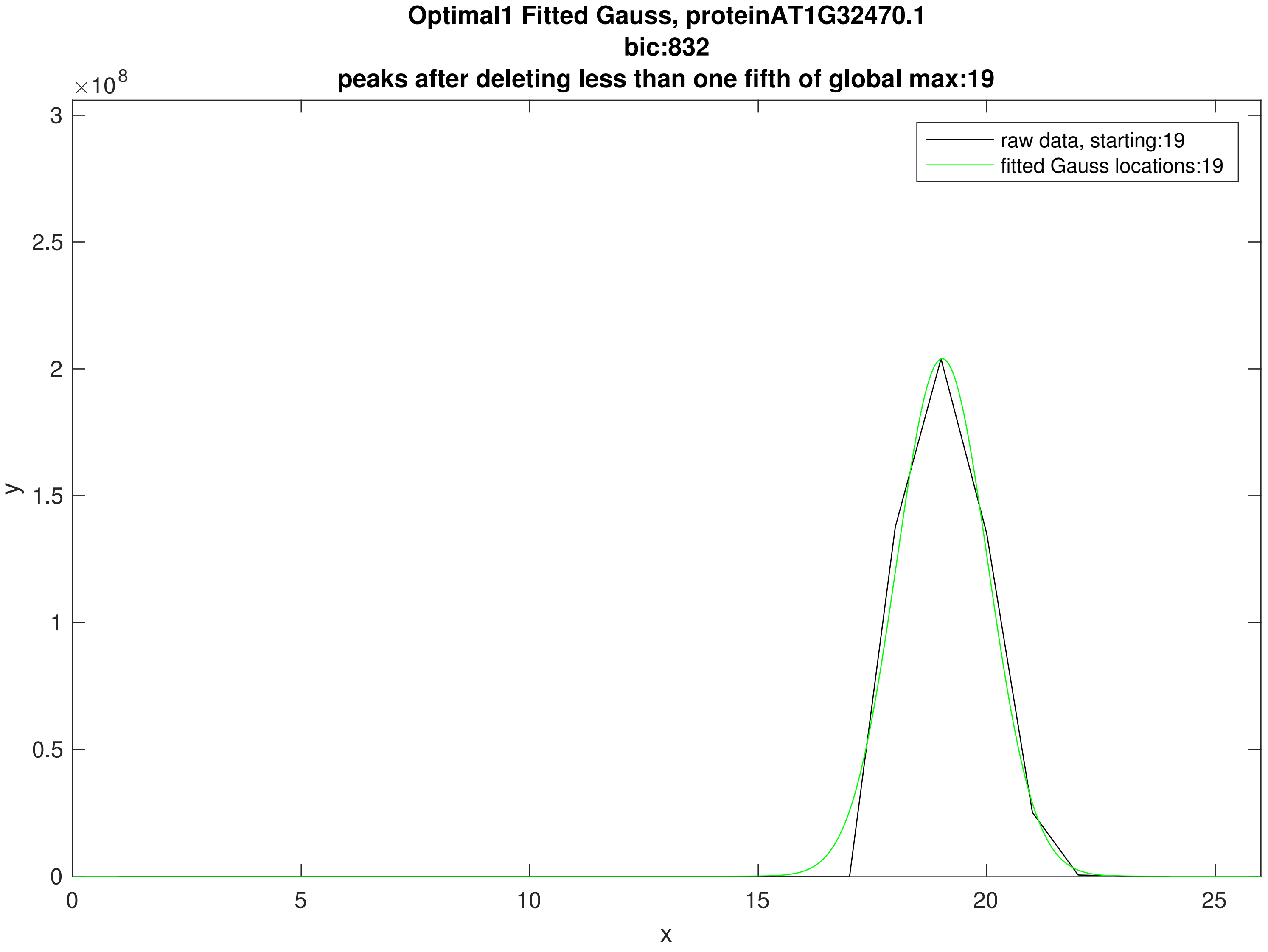

Optimal1 Fitted Gauss, proteinAT1G32470.1
bic:832
peaks after deleting less than one fifth of global max:19
×10^8
raw data, starting:19
fitted Gauss locations:19
y
x

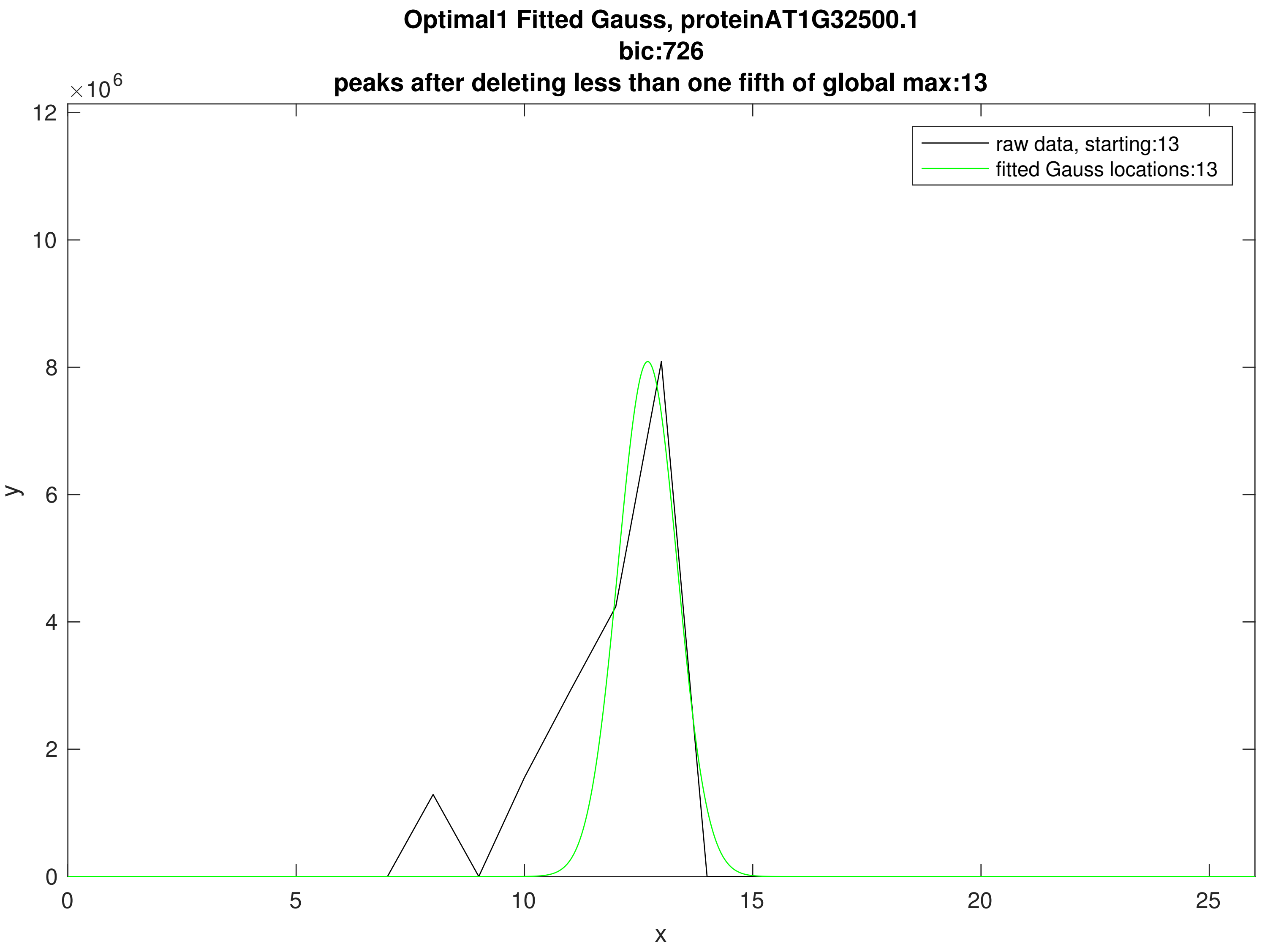
Optimal1 Fitted Gauss, proteinAT1G32500.1
bic:726
peaks after deleting less than one fifth of global max:13
×10^6
12
10
8
6
4
2
0
y
0
5
10
15
20
25
x
raw data, starting:13
fitted Gauss locations:13

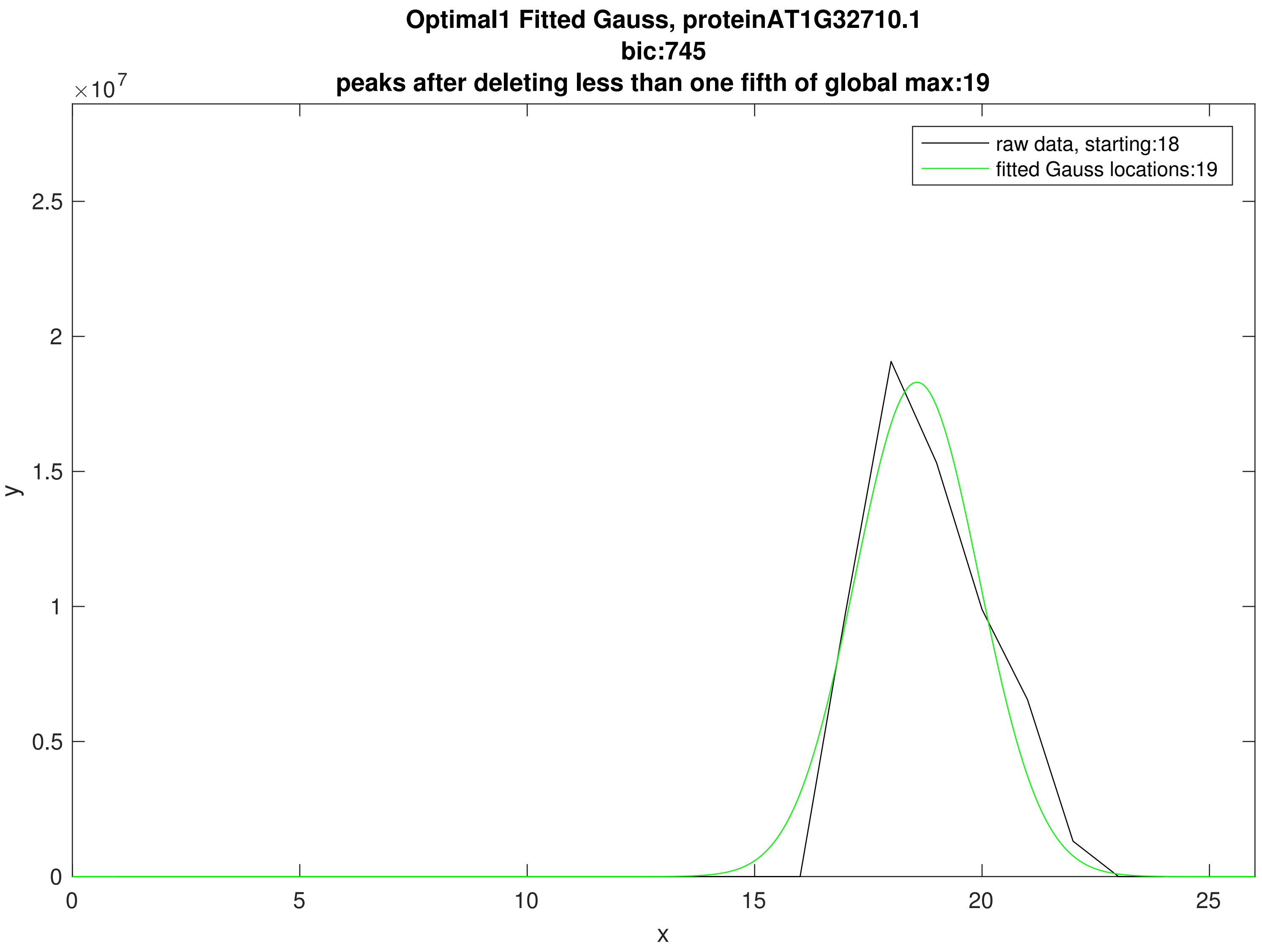

Optimal1 Fitted Gauss, proteinAT1G32710.1
bic:745
peaks after deleting less than one fifth of global max:19
×10^7
raw data, starting:18
fitted Gauss locations:19
y
x
2.5
2
1.5
1
0.5
0
5
10
15
20
25

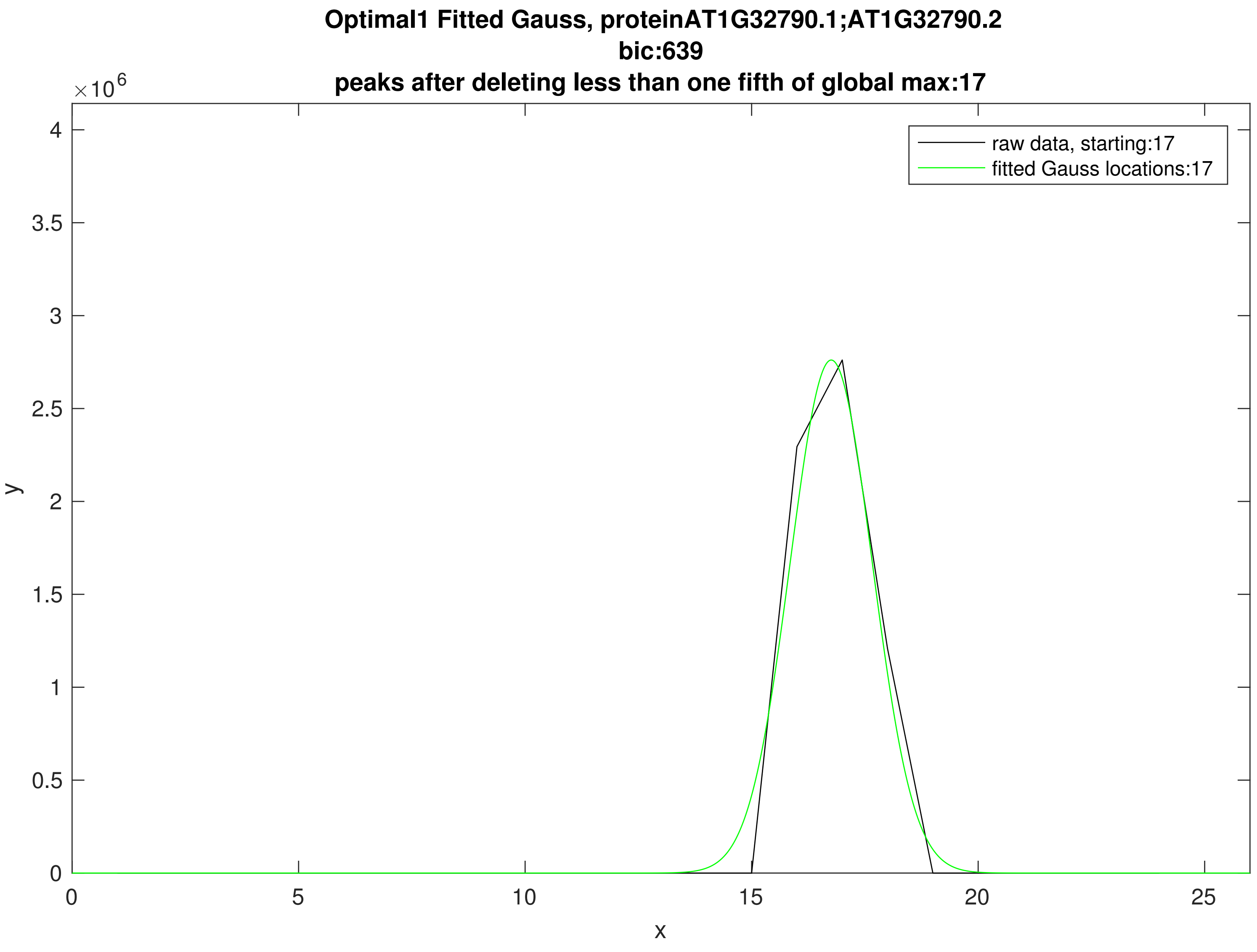
Optimal1 Fitted Gauss, proteinAT1G32790.1;AT1G32790.2
bic:639
peaks after deleting less than one fifth of global max:17
×10^6
raw data, starting:17
fitted Gauss locations:17
y
x
0
0.5
1
1.5
2
2.5
3
3.5
4
5
10
15
20
25

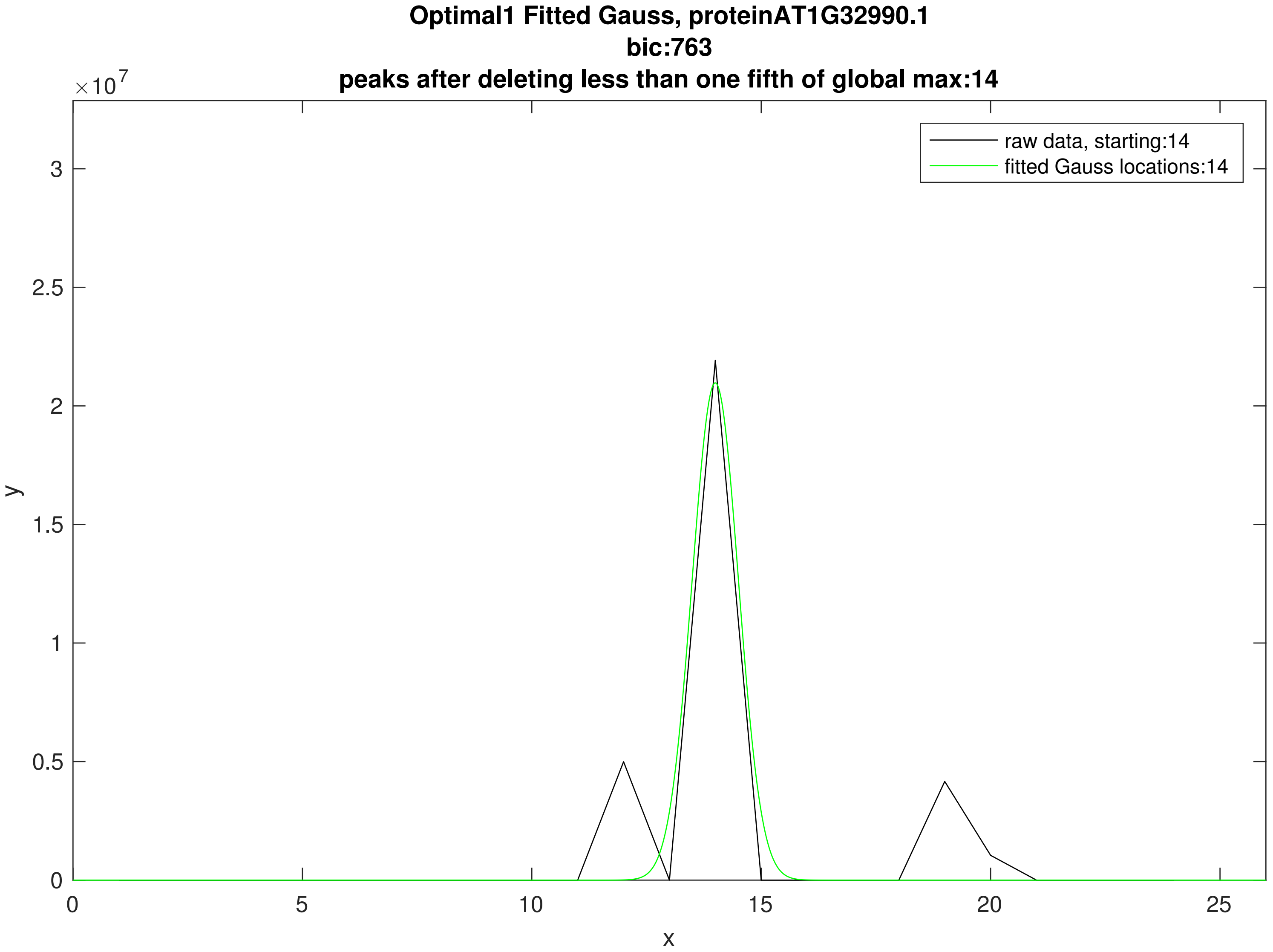

Optimal1 Fitted Gauss, proteinAT1G32990.1
bic:763
peaks after deleting less than one fifth of global max:14
×10^7
raw data, starting:14
fitted Gauss locations:14
3
2.5
2
1.5
1
0.5
0
y
0
5
10
15
20
25
x

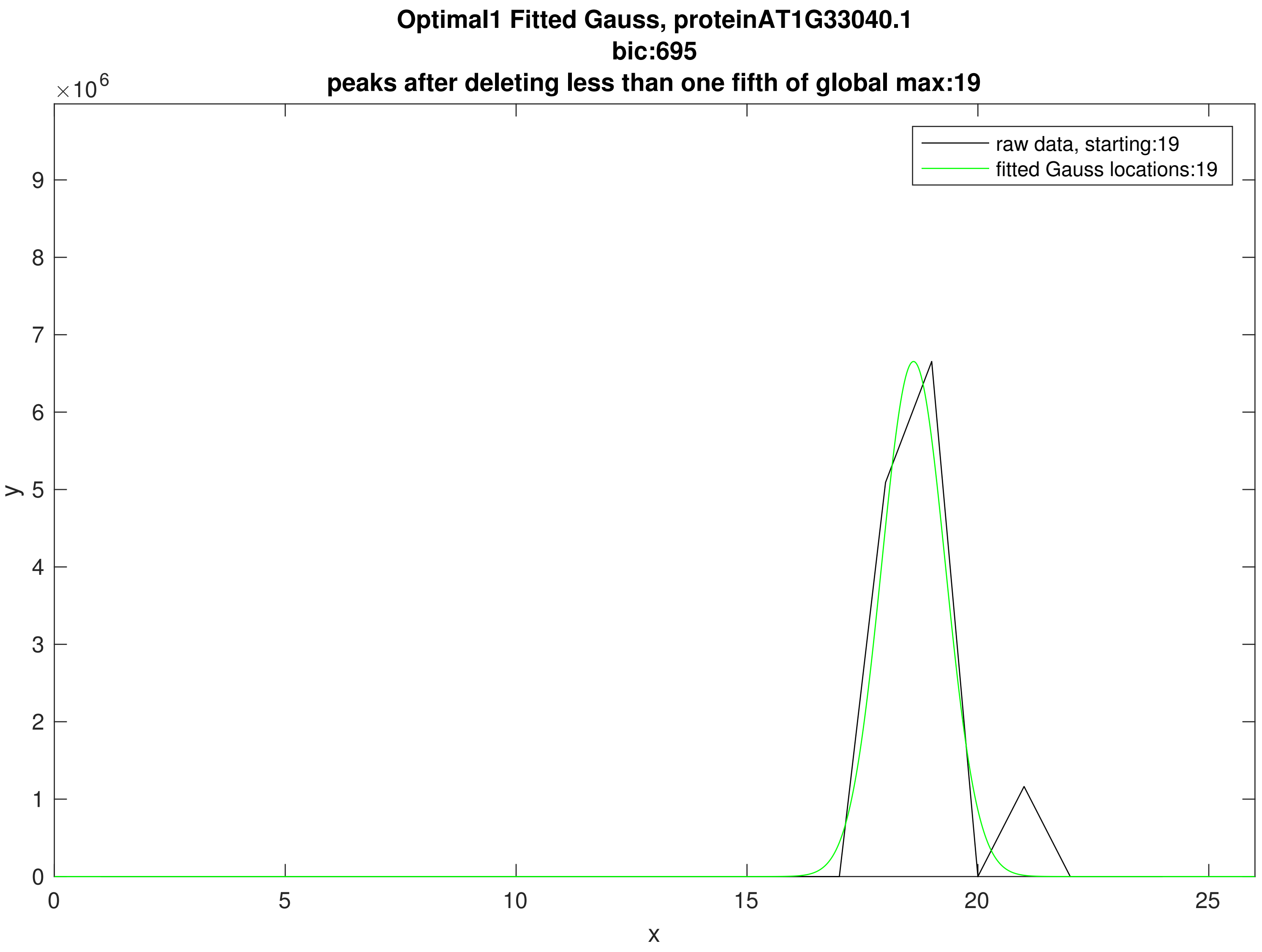
Optimal1 Fitted Gauss, proteinAT1G33040.1
bic:695
peaks after deleting less than one fifth of global max:19
×10^6
y
x
raw data, starting:19
fitted Gauss locations:19
0
1
2
3
4
5
6
7
8
9
0
5
10
15
20
25

Optimal1 Fitted Gauss, proteinAT1G33140.1;AT1G33120.1
bic:807 816 825
peaks after deleting less than one fifth of global max:3.7

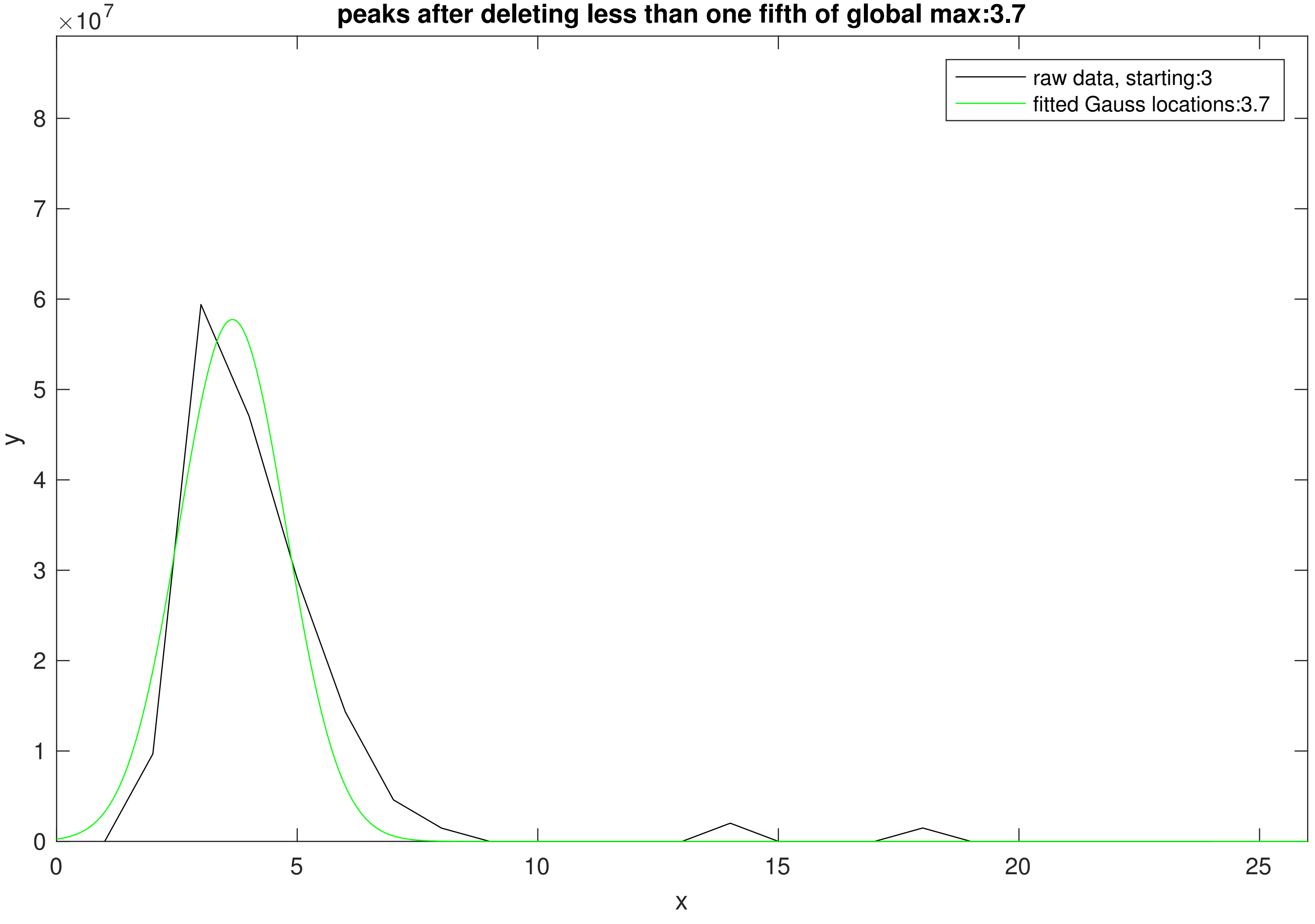

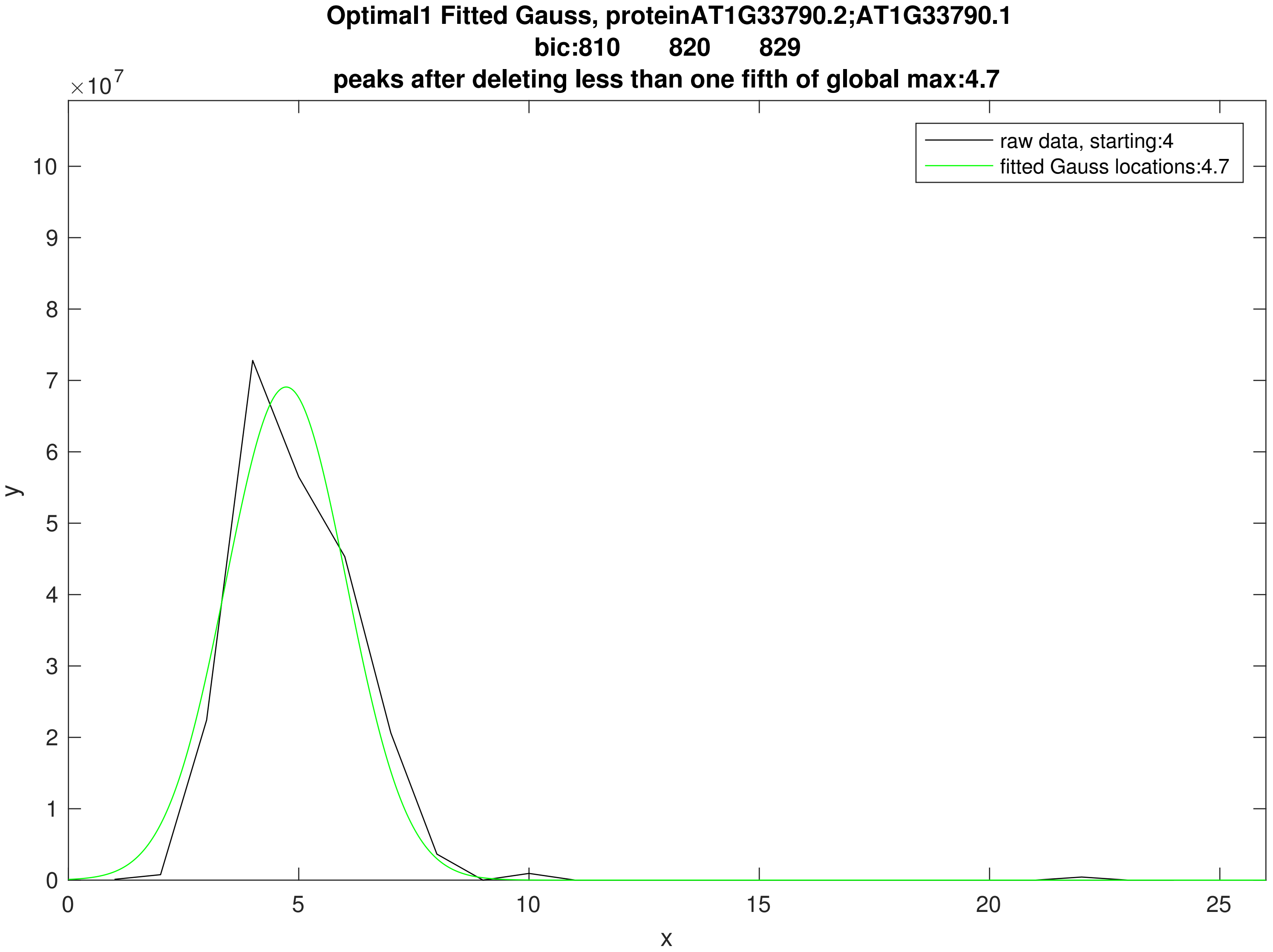
Optimal1 Fitted Gauss, proteinAT1G33790.2;AT1G33790.1
bic:810 820 829
peaks after deleting less than one fifth of global max:4.7
×10^7
raw data, starting:4
fitted Gauss locations:4.7
y
x
0
1
2
3
4
5
6
7
8
9
10
0
5
10
15
20
25

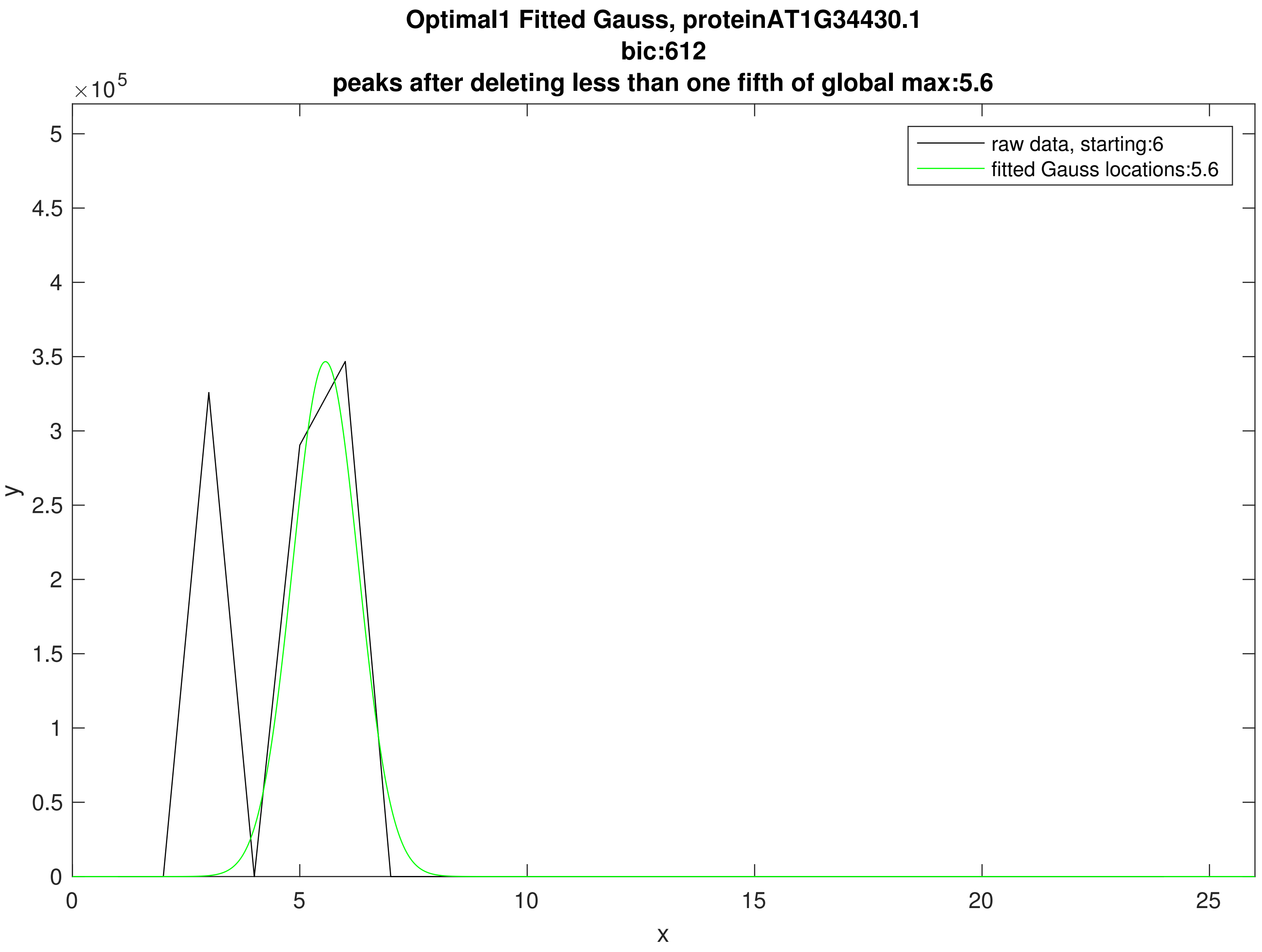
Optimal1 Fitted Gauss, proteinAT1G34430.1
bic:612
peaks after deleting less than one fifth of global max:5.6
×10^5
raw data, starting:6
fitted Gauss locations:5.6
y
x
0
0.5
1
1.5
2
2.5
3
3.5
4
4.5
5
0
5
10
15
20
25

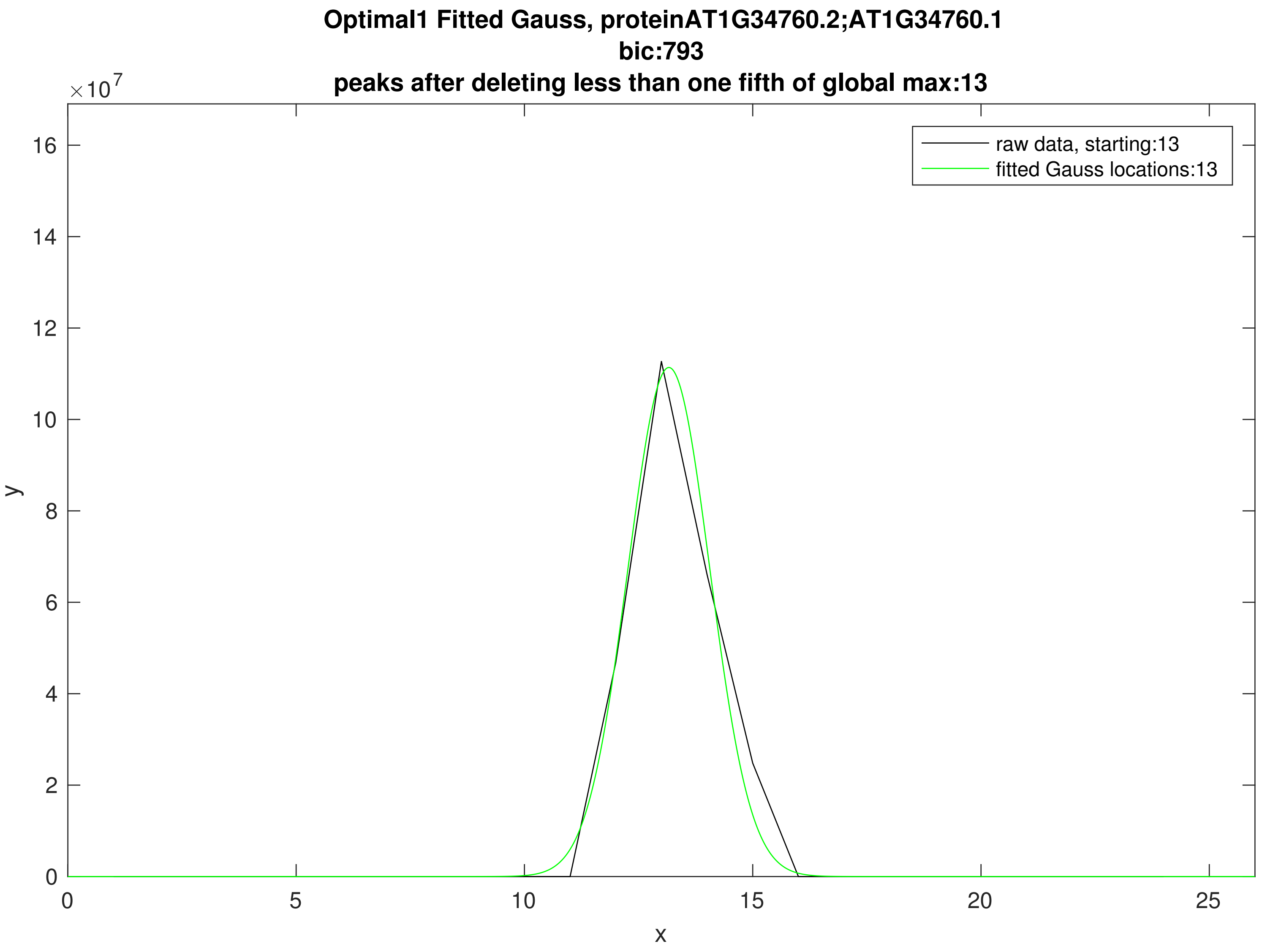

Optimal1 Fitted Gauss, proteinAT1G34760.2;AT1G34760.1
bic:793
peaks after deleting less than one fifth of global max:13
×10^7
16
14
12
10
8
6
4
2
0
y
0
5
10
15
20
25
x
raw data, starting:13
fitted Gauss locations:13

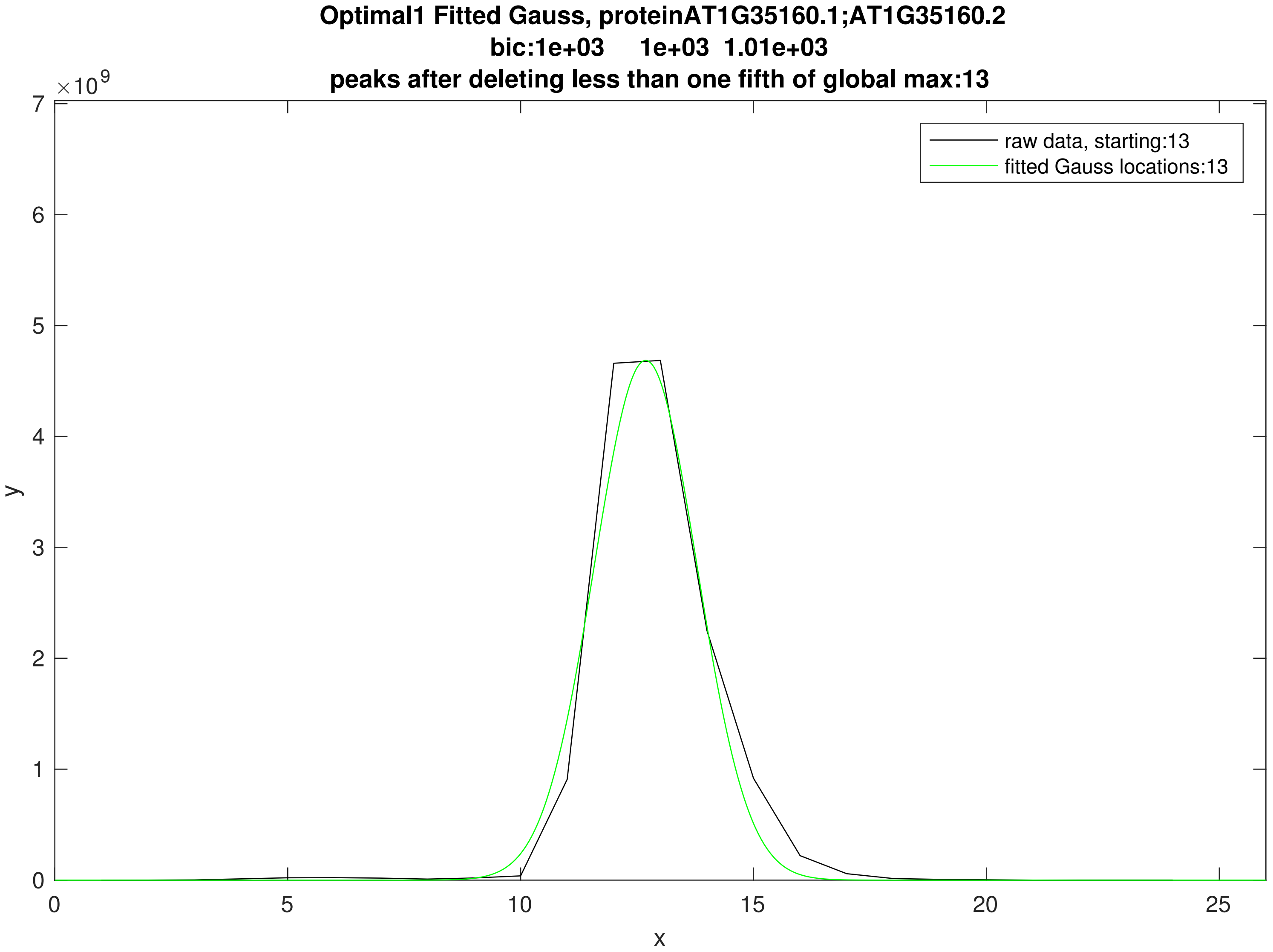
Optimal1 Fitted Gauss, proteinAT1G35160.1;AT1G35160.2
bic:1e+03 1e+03 1.01e+03
peaks after deleting less than one fifth of global max:13
×10^9
raw data, starting:13
fitted Gauss locations:13
y
x
0
1
2
3
4
5
6
7
0
5
10
15
20
25

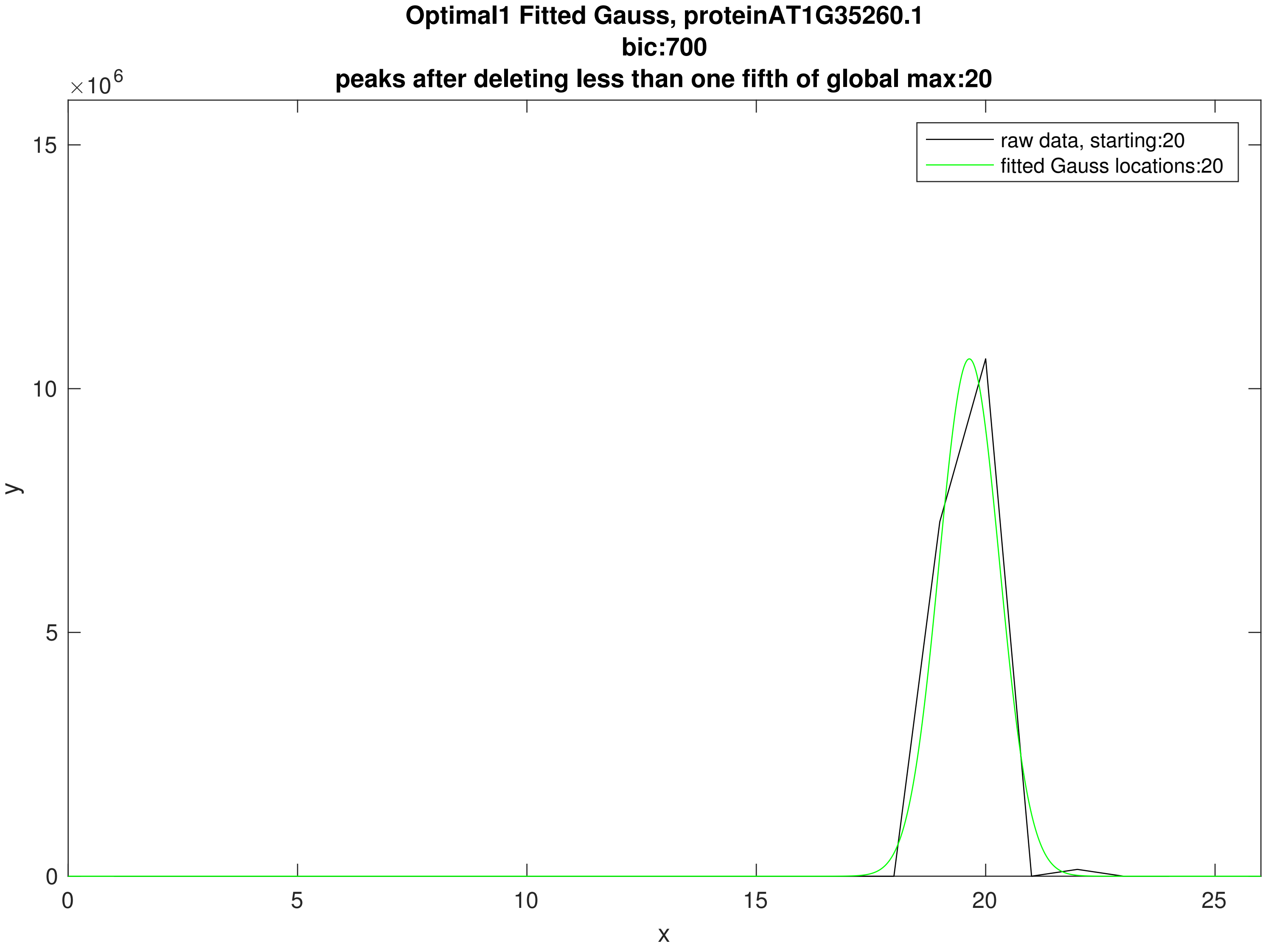
Optimal1 Fitted Gauss, proteinAT1G35260.1
bic:700
peaks after deleting less than one fifth of global max:20
×10^6
raw data, starting:20
fitted Gauss locations:20
y
x
0
5
10
15
20
25
0
5
10
15

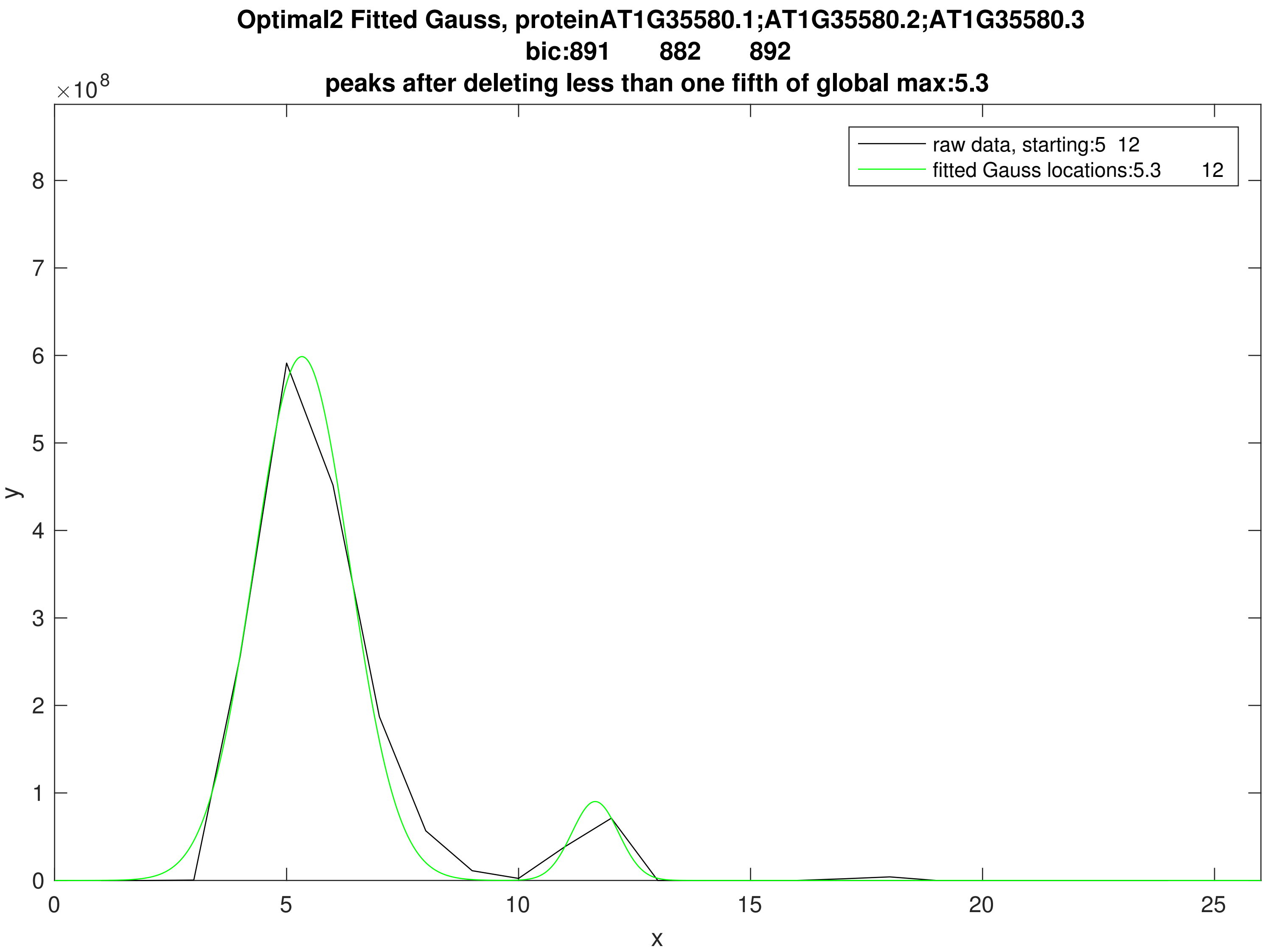
Optimal2 Fitted Gauss, proteinAT1G35580.1;AT1G35580.2;AT1G35580.3
bic:891 882 892
peaks after deleting less than one fifth of global max:5.3
×10^8
raw data, starting:5 12
fitted Gauss locations:5.3 12
y
x
0
1
2
3
4
5
6
7
8
0
5
10
15
20
25

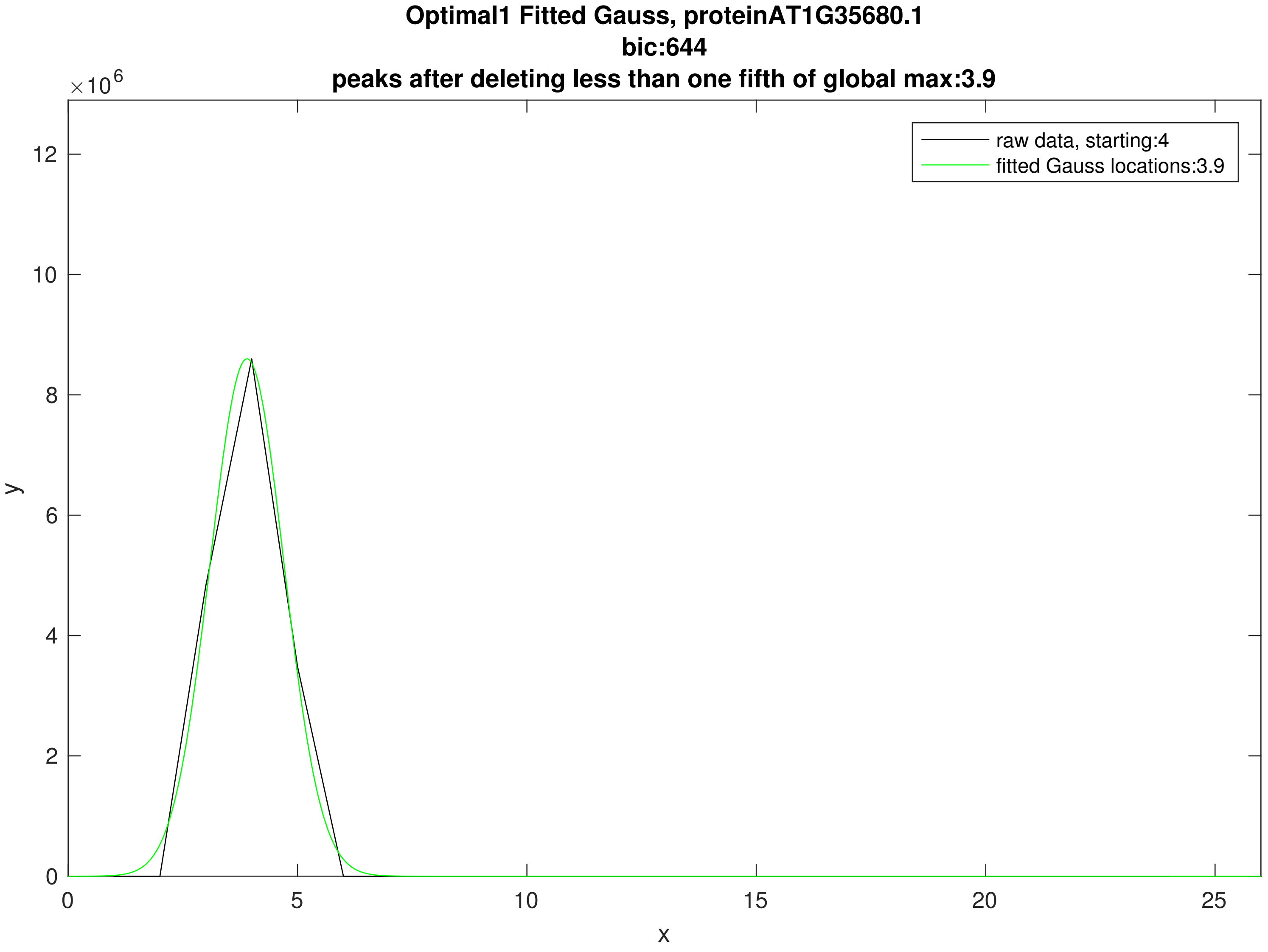

Optimal1 Fitted Gauss, proteinAT1G35680.1
bic:644
peaks after deleting less than one fifth of global max:3.9
×10^6
raw data, starting:4
fitted Gauss locations:3.9
y
x
0
2
4
6
8
10
12
0
5
10
15
20
25

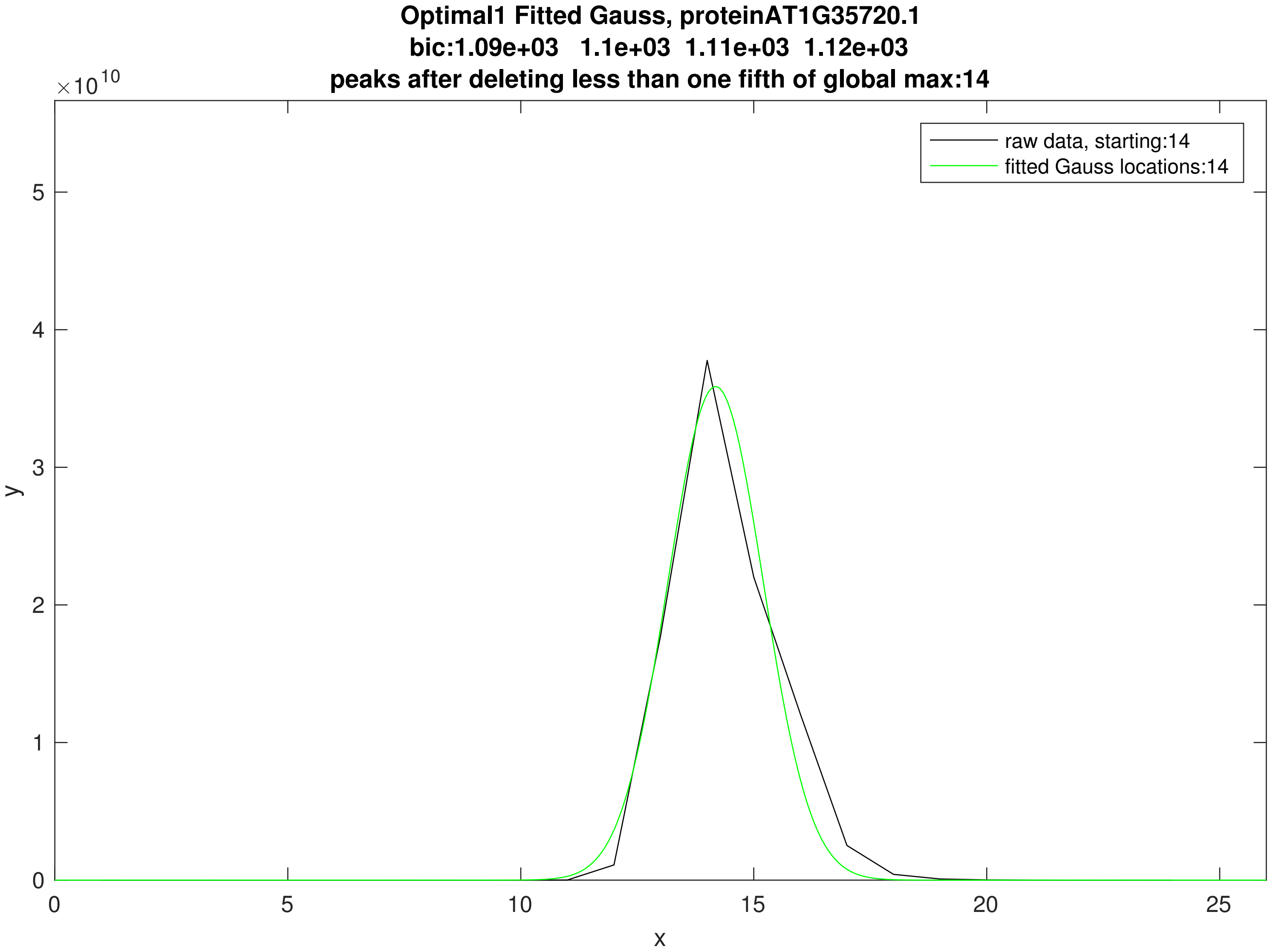

Optimal1 Fitted Gauss, proteinAT1G35720.1
bic:1.09e+03 1.1e+03 1.11e+03 1.12e+03
peaks after deleting less than one fifth of global max:14
×10^10
raw data, starting:14
fitted Gauss locations:14
y
x
0
1
2
3
4
5
0
5
10
15
20
25

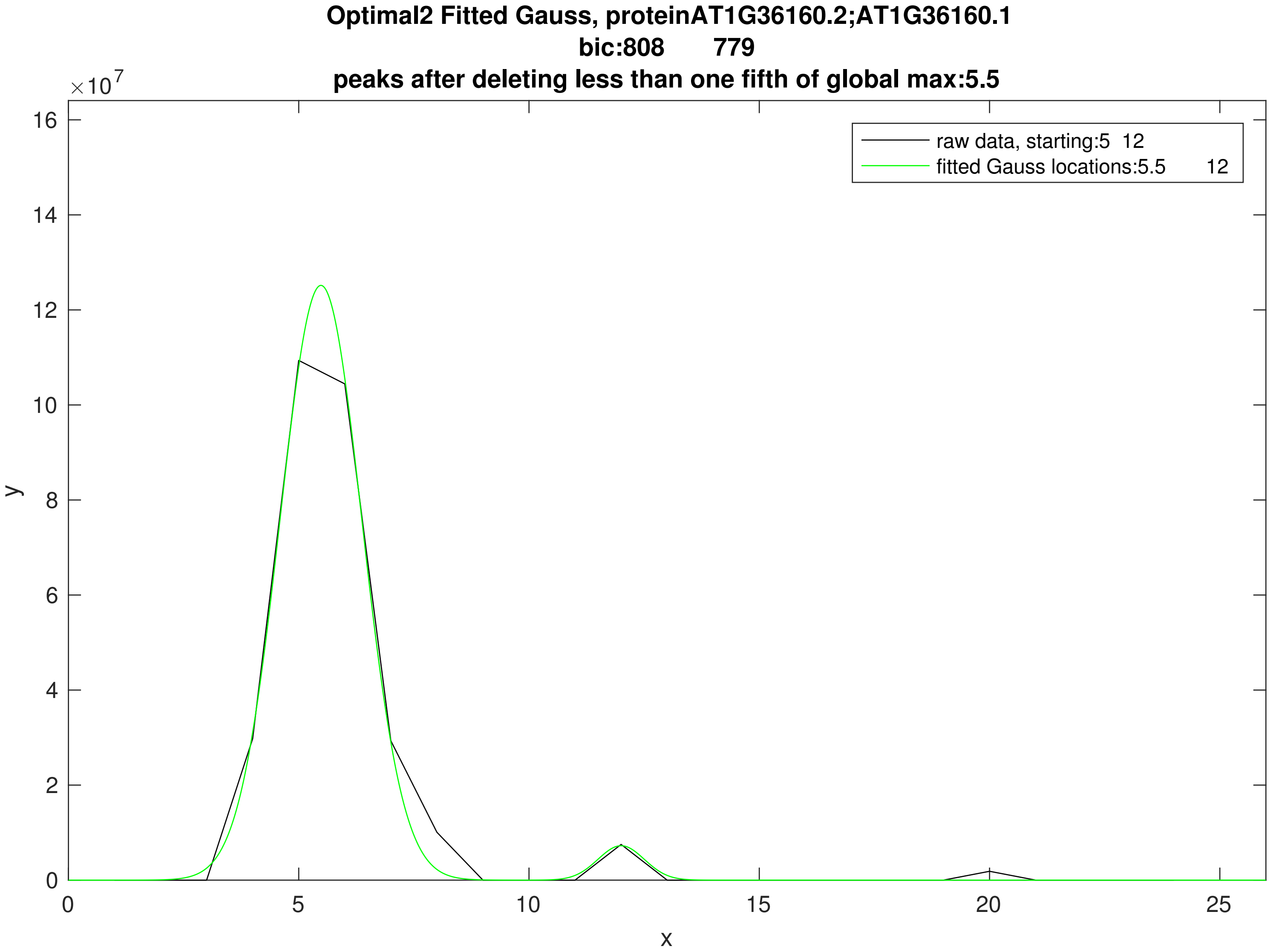
Optimal2 Fitted Gauss, proteinAT1G36160.2;AT1G36160.1
bic:808 779
peaks after deleting less than one fifth of global max:5.5
×10^7
raw data, starting:5 12
fitted Gauss locations:5.5 12
y
x
0
2
4
6
8
10
12
14
16
0
5
10
15
20
25

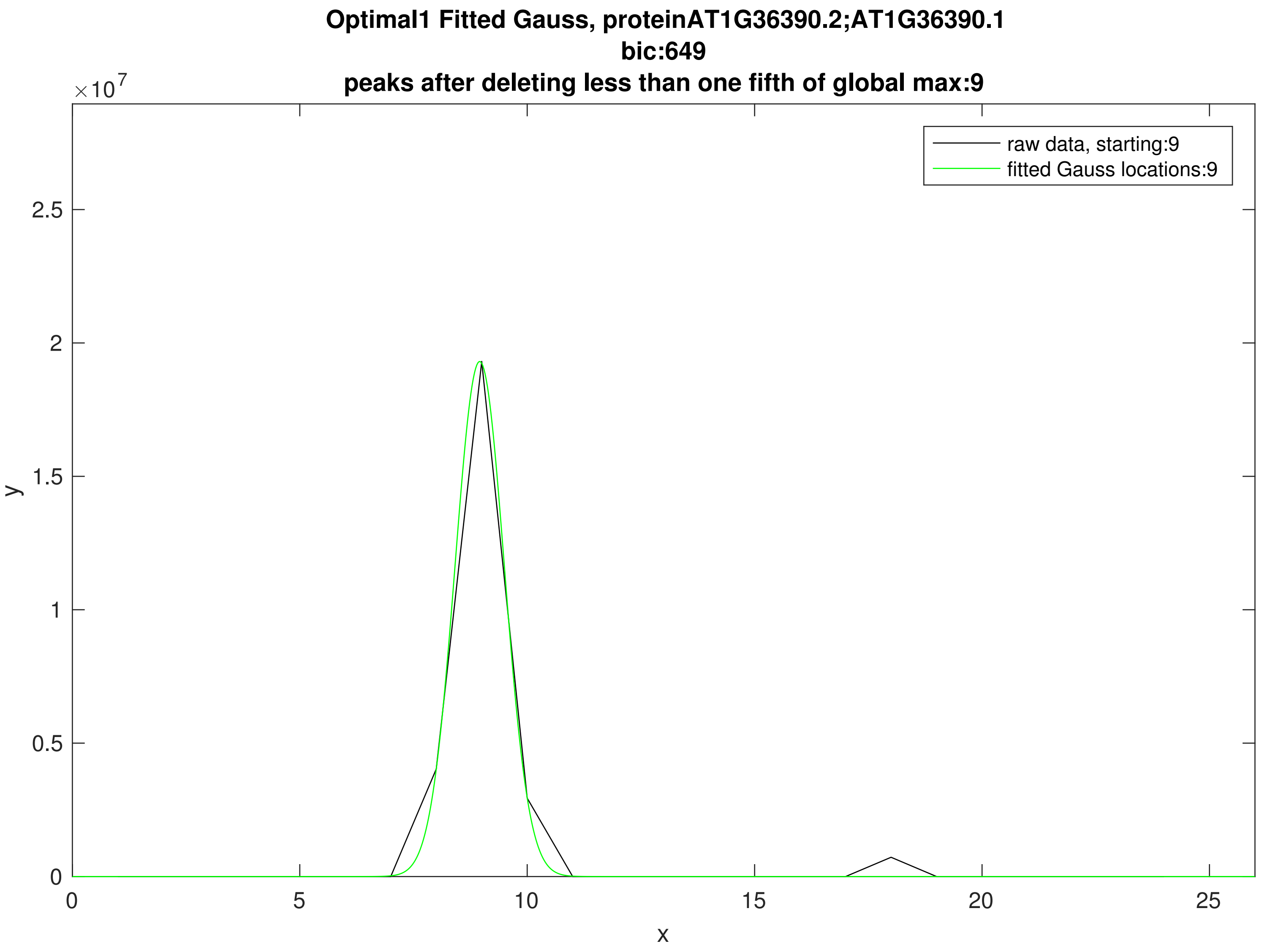

Optimal1 Fitted Gauss, proteinAT1G36390.2;AT1G36390.1
bic:649
peaks after deleting less than one fifth of global max:9
×10^7
raw data, starting:9
fitted Gauss locations:9
2.5
2
1.5
1
0.5
0
y
0
5
10
15
20
25
x

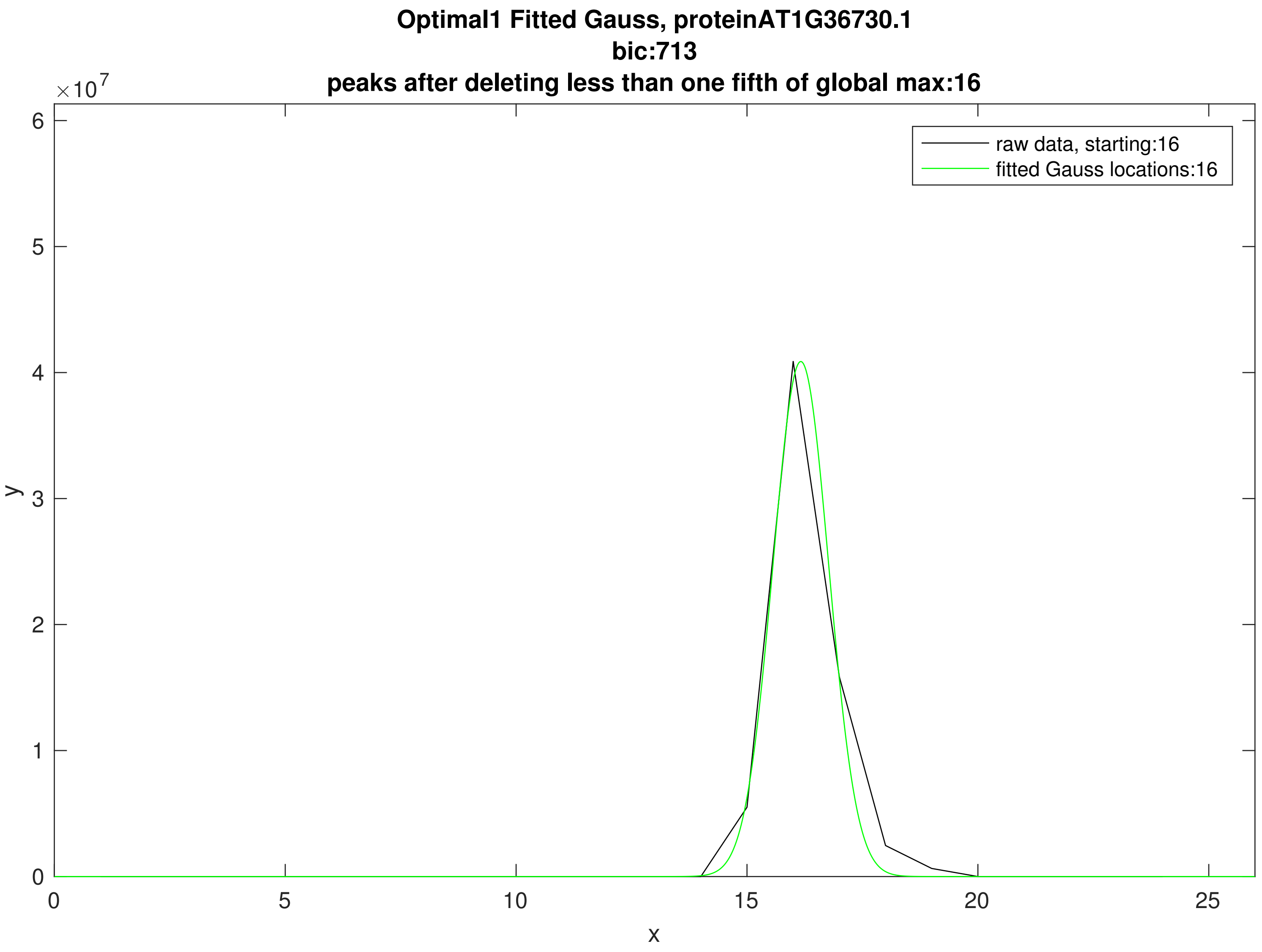

Optimal1 Fitted Gauss, proteinAT1G36730.1
bic:713
peaks after deleting less than one fifth of global max:16
×10^7
raw data, starting:16
fitted Gauss locations:16
y
x
0
1
2
3
4
5
6
0
5
10
15
20
25

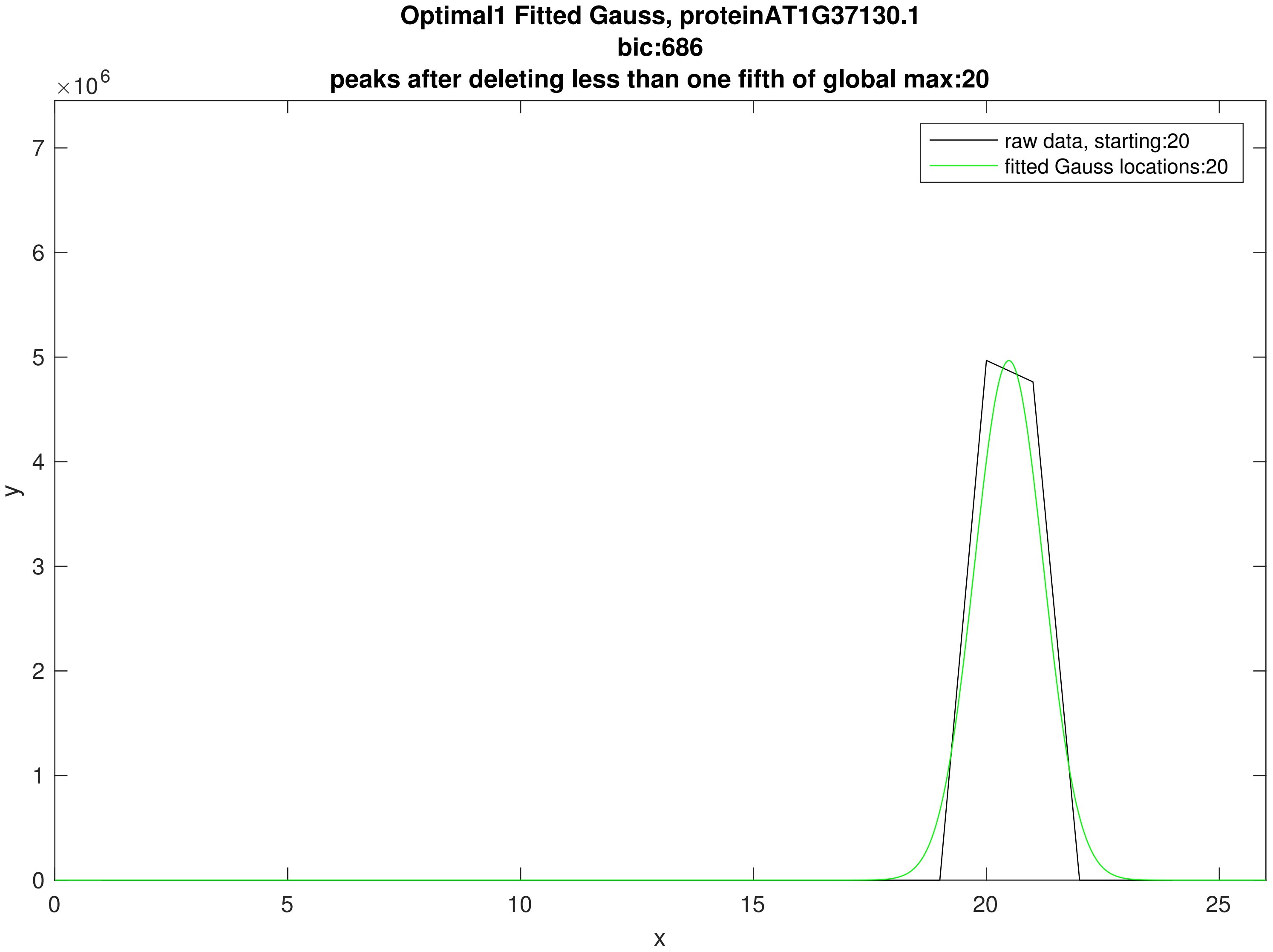

Optimal1 Fitted Gauss, proteinAT1G37130.1
bic:686
peaks after deleting less than one fifth of global max:20
×10^6
raw data, starting:20
fitted Gauss locations:20
y
x
0
1
2
3
4
5
6
7
0
5
10
15
20
25

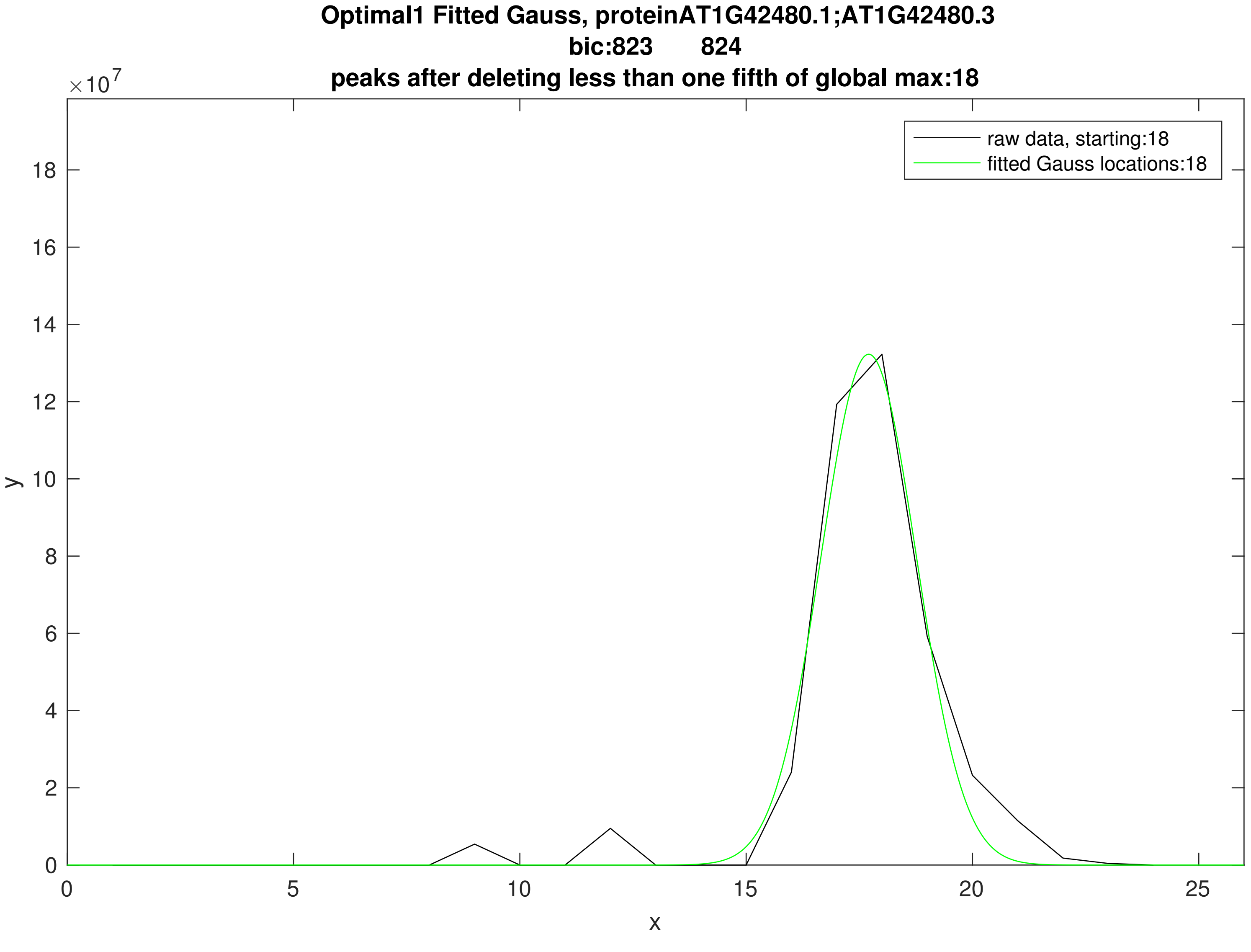
Optimal1 Fitted Gauss, proteinAT1G42480.1;AT1G42480.3
bic:823 824
peaks after deleting less than one fifth of global max:18
×10^7
raw data, starting:18
fitted Gauss locations:18
y
x
0
2
4
6
8
10
12
14
16
18
0
5
10
15
20
25

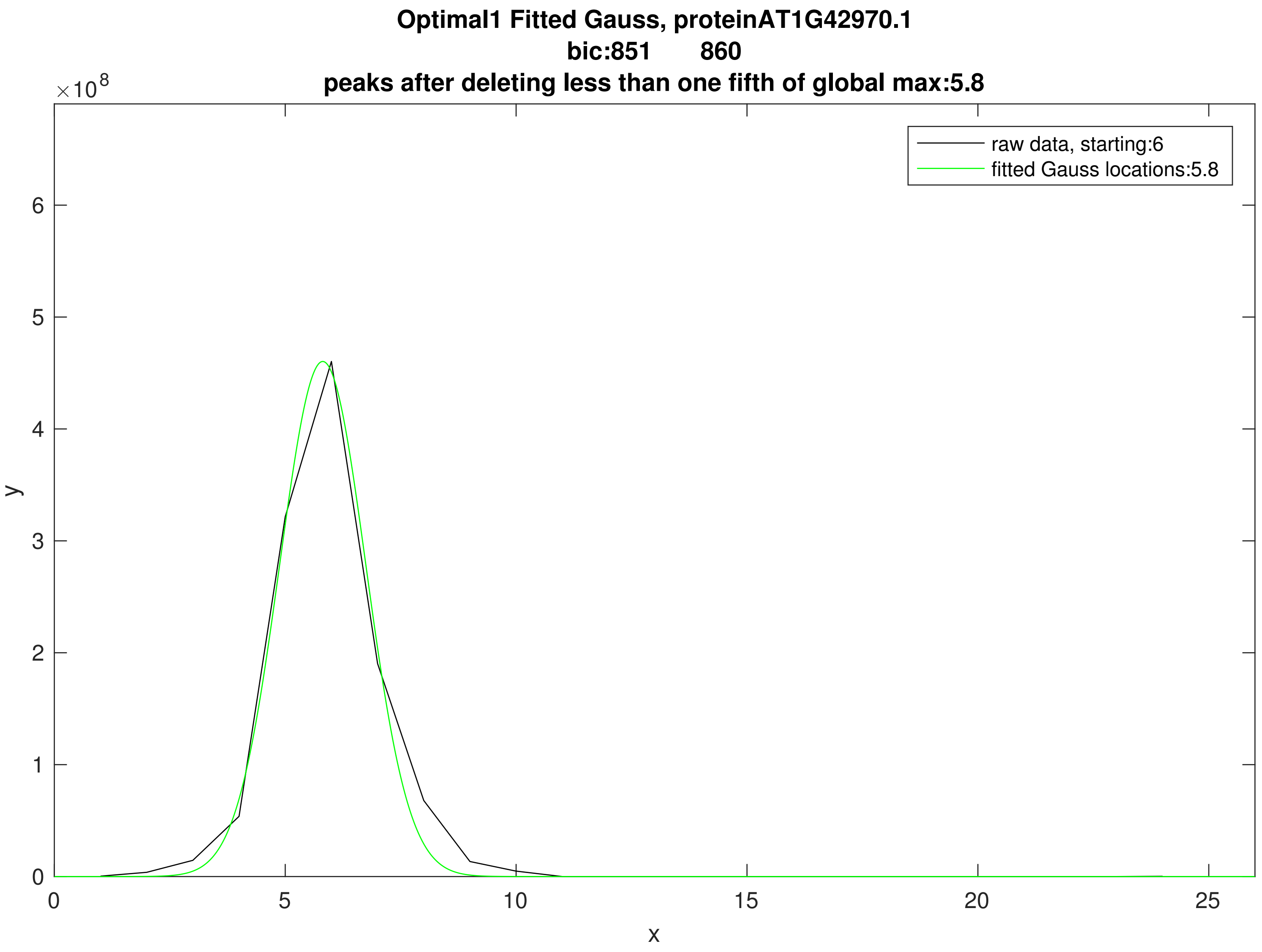

Optimal1 Fitted Gauss, proteinAT1G42970.1
bic:851 860
peaks after deleting less than one fifth of global max:5.8
×10^8
raw data, starting:6
fitted Gauss locations:5.8
y
x
0
1
2
3
4
5
6
0
5
10
15
20
25

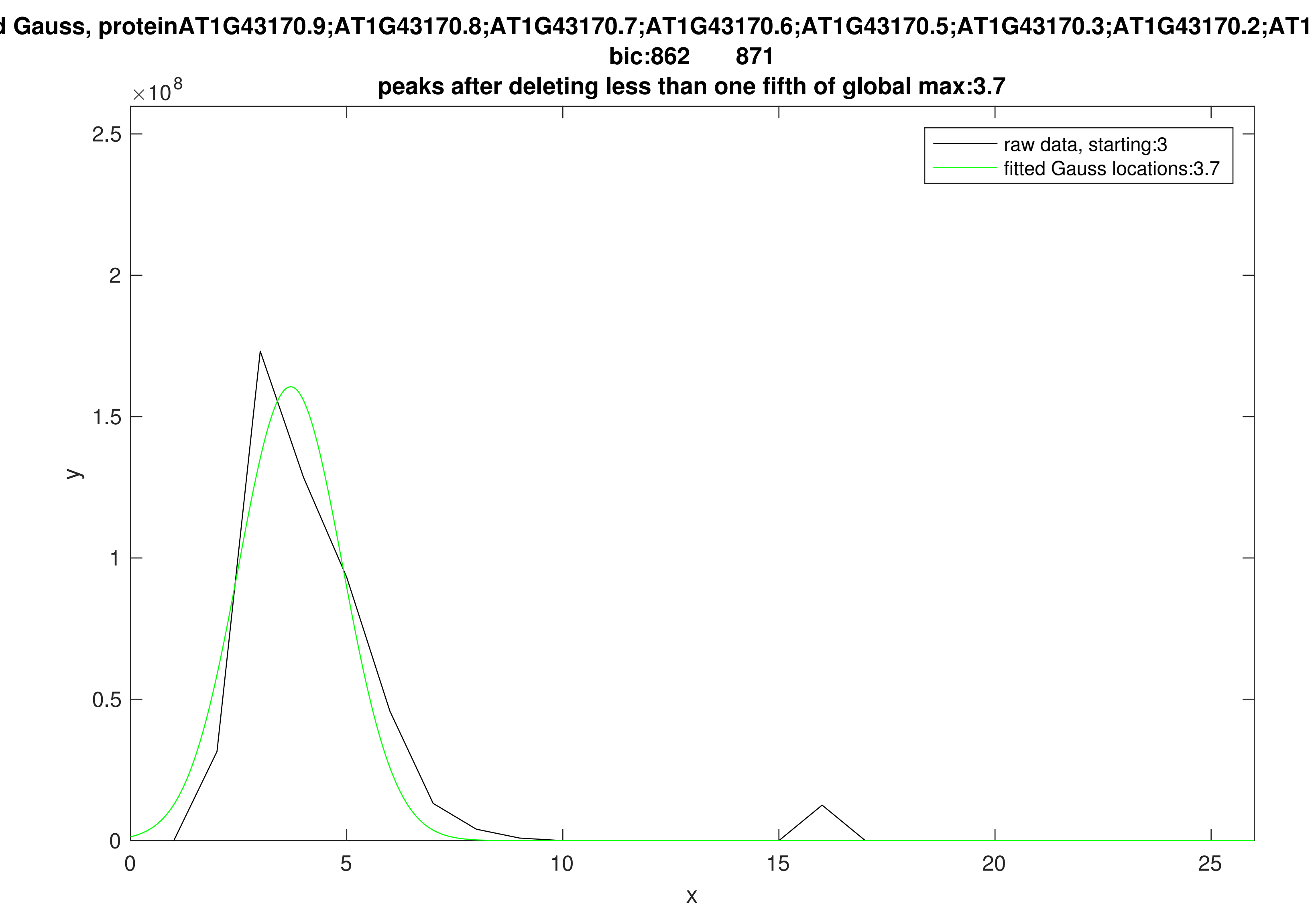

d Gauss, proteinAT1G43170.9;AT1G43170.8;AT1G43170.7;AT1G43170.6;AT1G43170.5;AT1G43170.3;AT1G43170.2;AT1
bic:862 871
peaks after deleting less than one fifth of global max:3.7
×10^8
raw data, starting:3
fitted Gauss locations:3.7
y
x

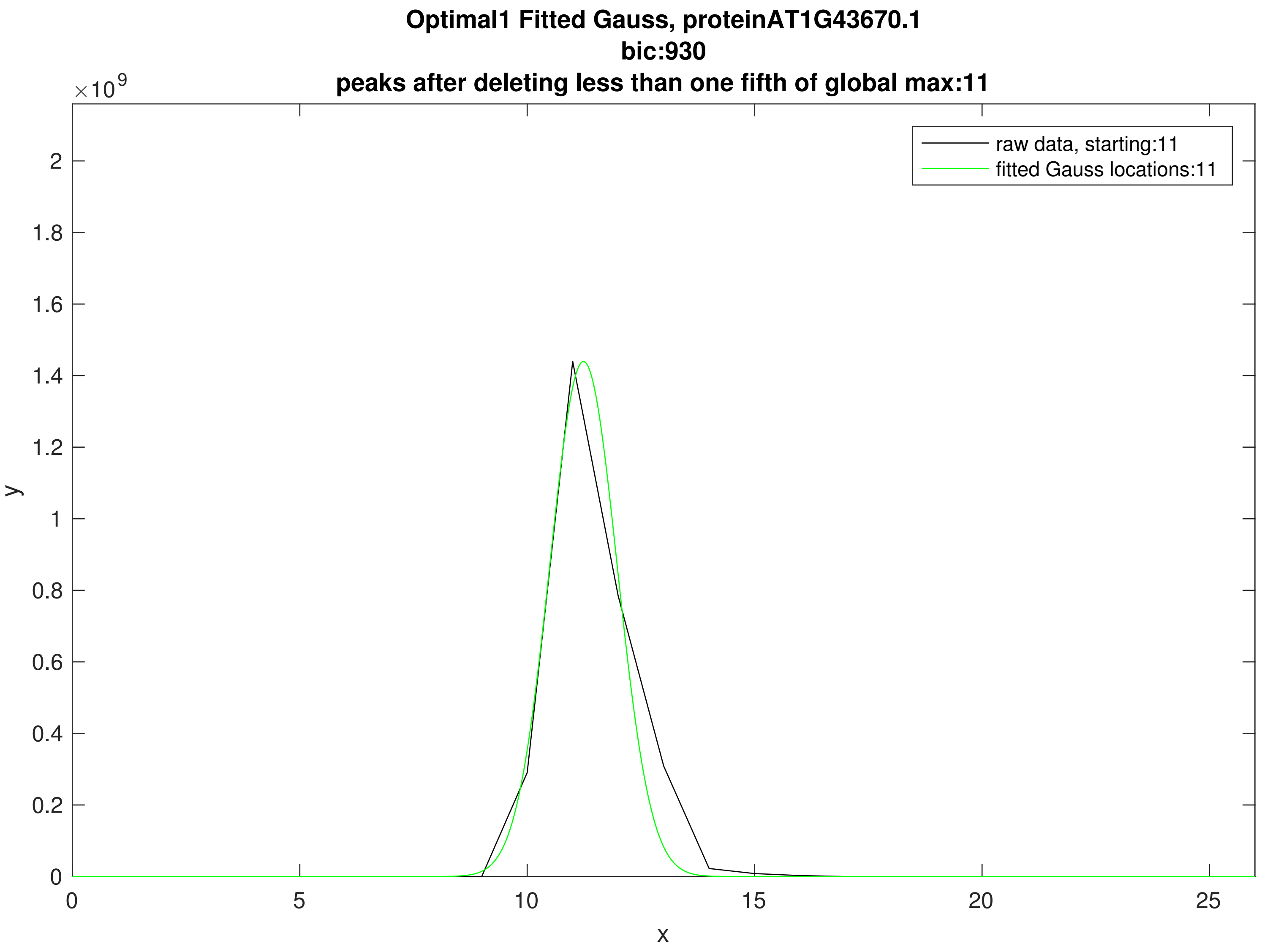
Optimal1 Fitted Gauss, proteinAT1G43670.1
bic:930
peaks after deleting less than one fifth of global max:11
×10^9
raw data, starting:11
fitted Gauss locations:11
y
0
0.2
0.4
0.6
0.8
1
1.2
1.4
1.6
1.8
2
x
0
5
10
15
20
25

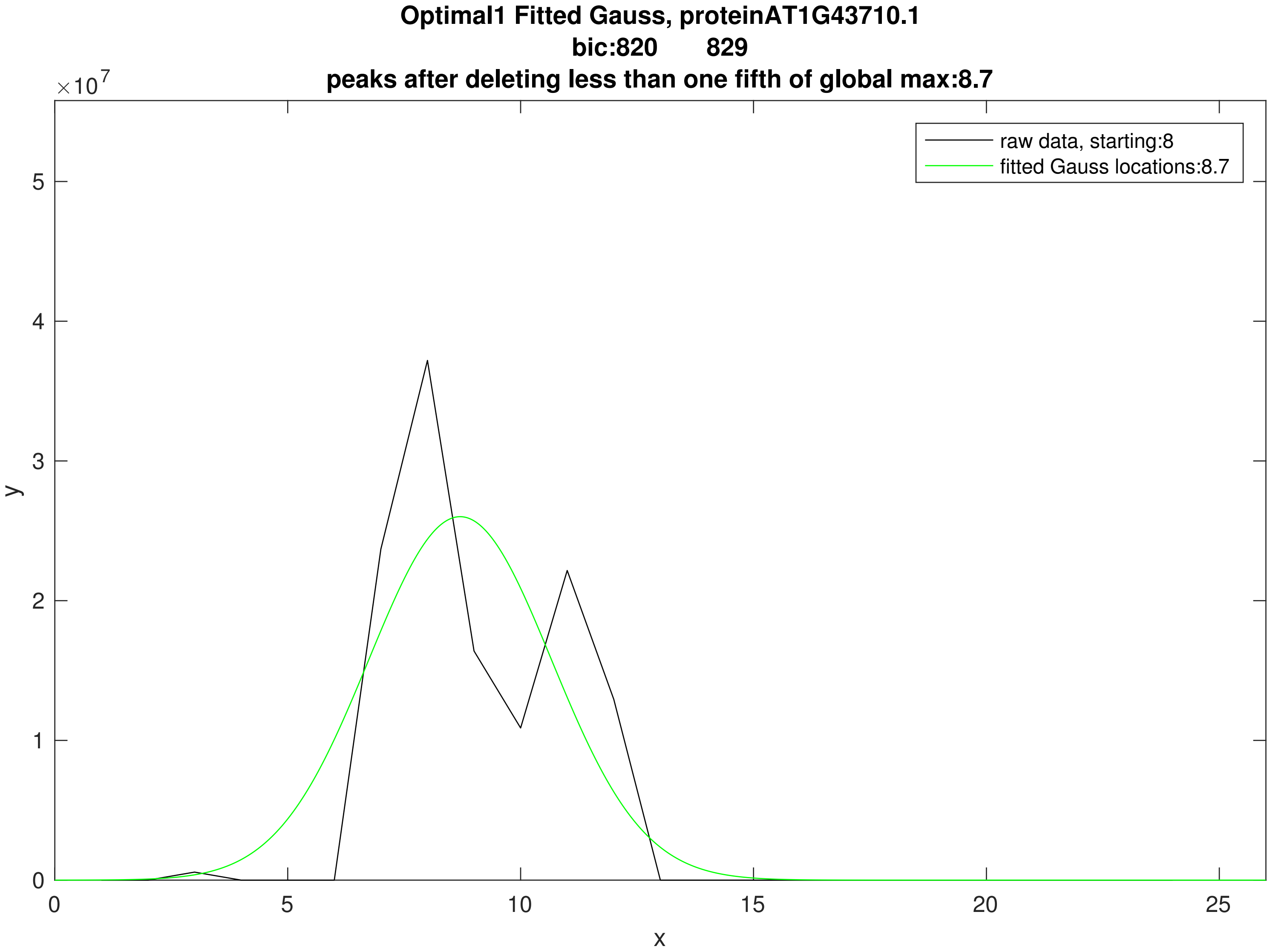
Optimal1 Fitted Gauss, proteinAT1G43710.1
bic:820 829
peaks after deleting less than one fifth of global max:8.7
×10^7
raw data, starting:8
fitted Gauss locations:8.7
y
x
0
1
2
3
4
5
0
5
10
15
20
25

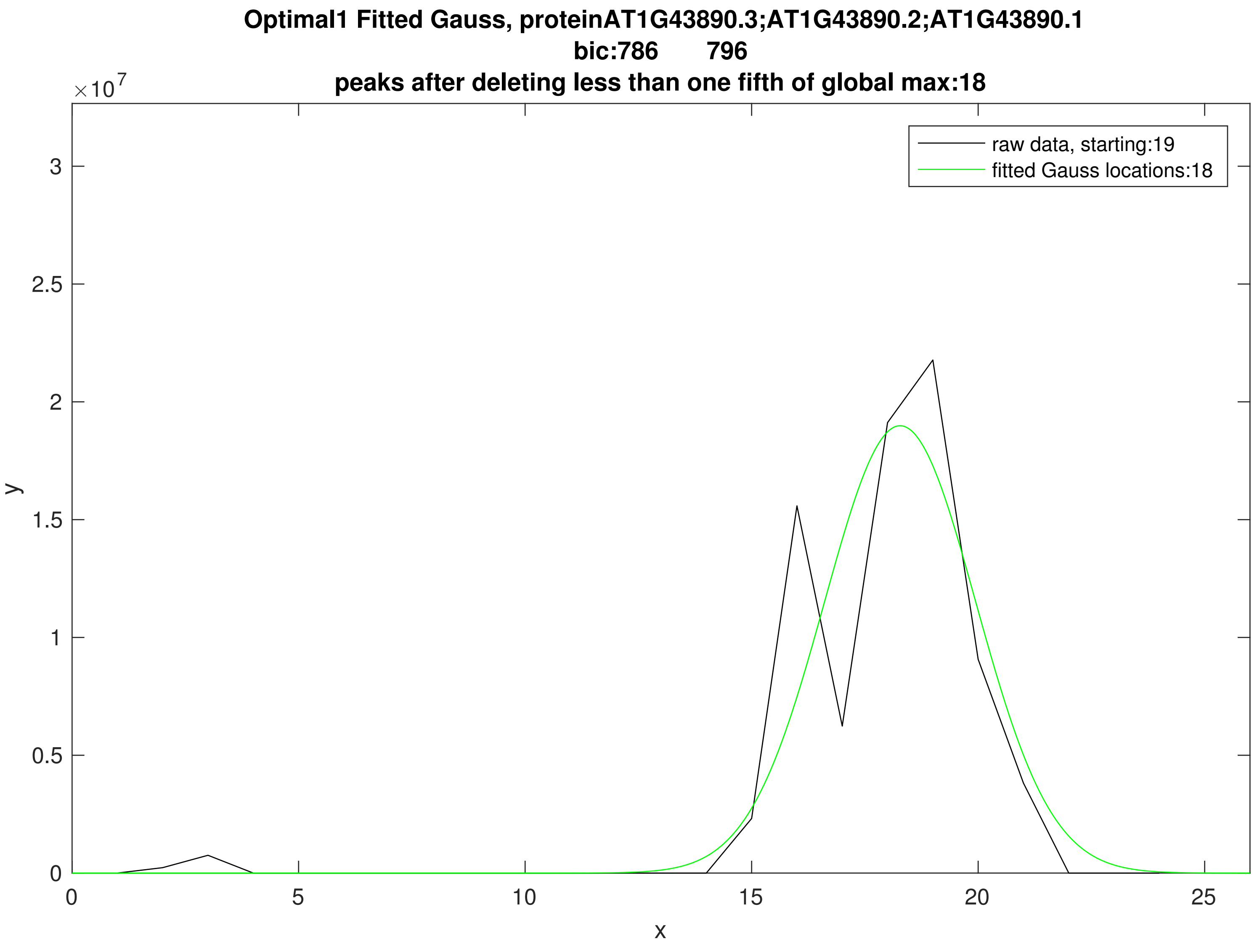

Optimal1 Fitted Gauss, proteinAT1G43890.3;AT1G43890.2;AT1G43890.1
bic:786 796
peaks after deleting less than one fifth of global max:18
×10^7
raw data, starting:19
fitted Gauss locations:18
y
x
0
0.5
1
1.5
2
2.5
3
5
10
15
20
25

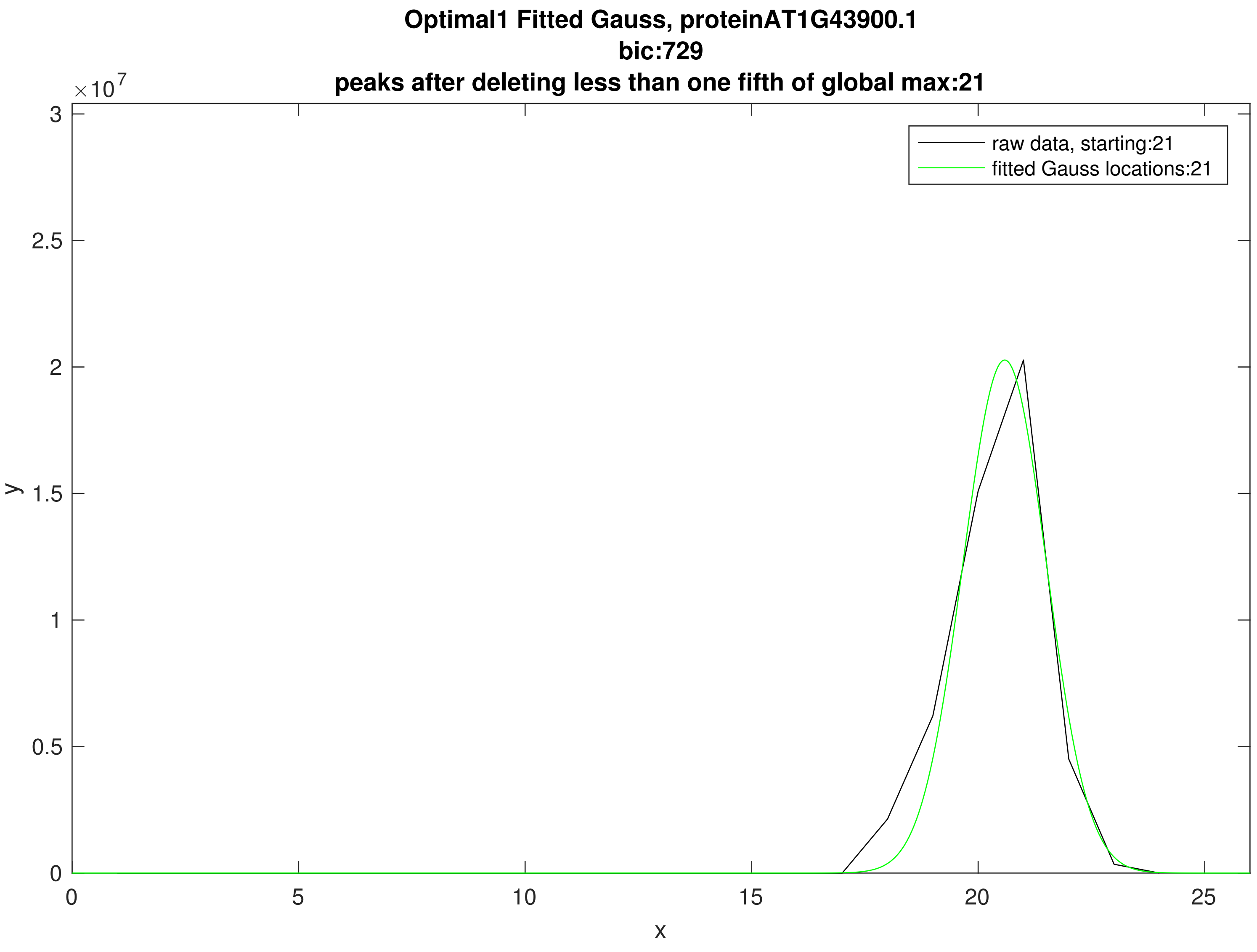
Optimal1 Fitted Gauss, proteinAT1G43900.1
bic:729
peaks after deleting less than one fifth of global max:21
×10^7
raw data, starting:21
fitted Gauss locations:21
3
2.5
2
1.5
1
0.5
0
y
0
5
10
15
20
25
x

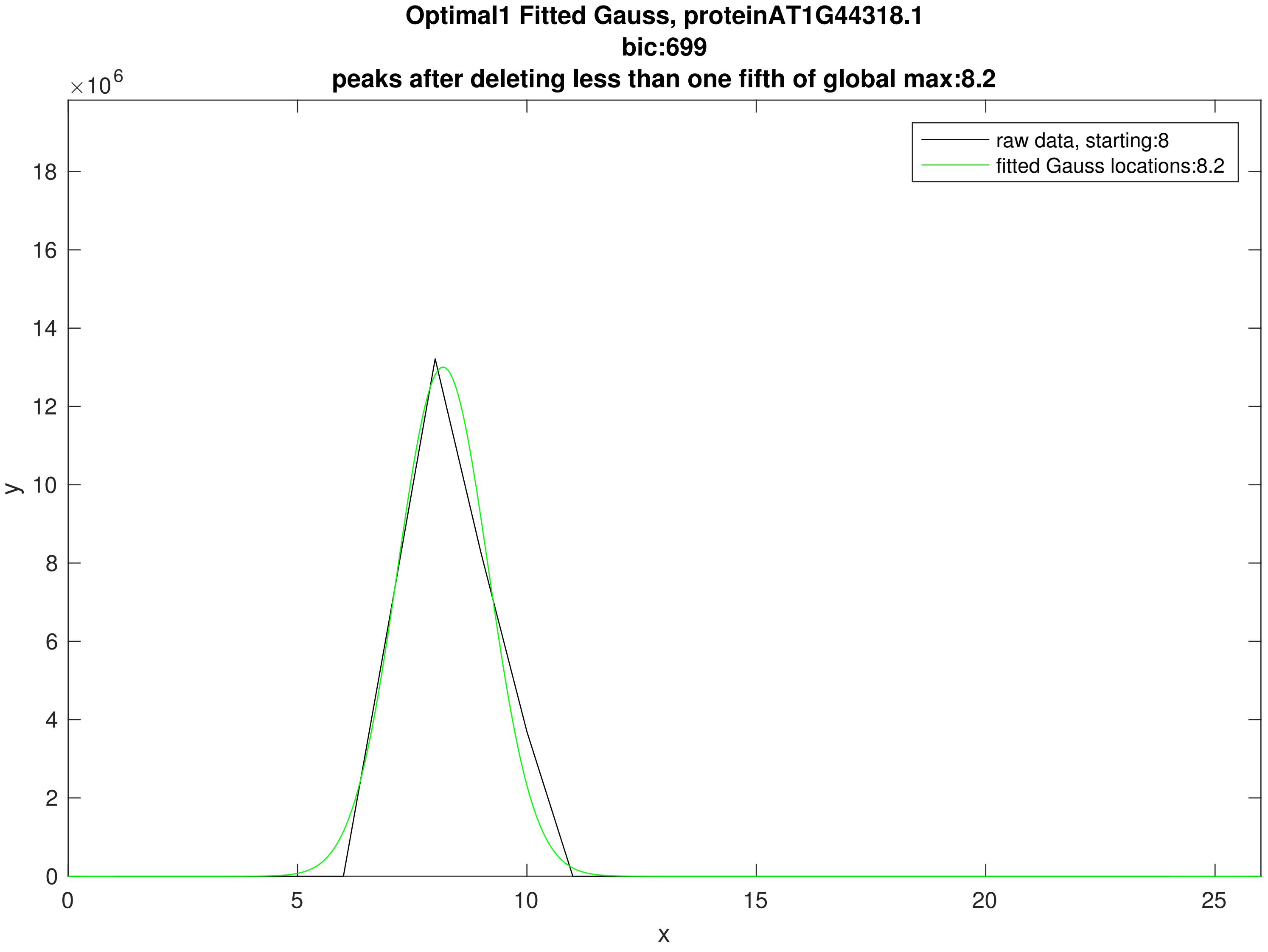

Optimal1 Fitted Gauss, proteinAT1G44318.1
bic:699
peaks after deleting less than one fifth of global max:8.2
×10^6
raw data, starting:8
fitted Gauss locations:8.2
y
x
0
2
4
6
8
10
12
14
16
18
0
5
10
15
20
25

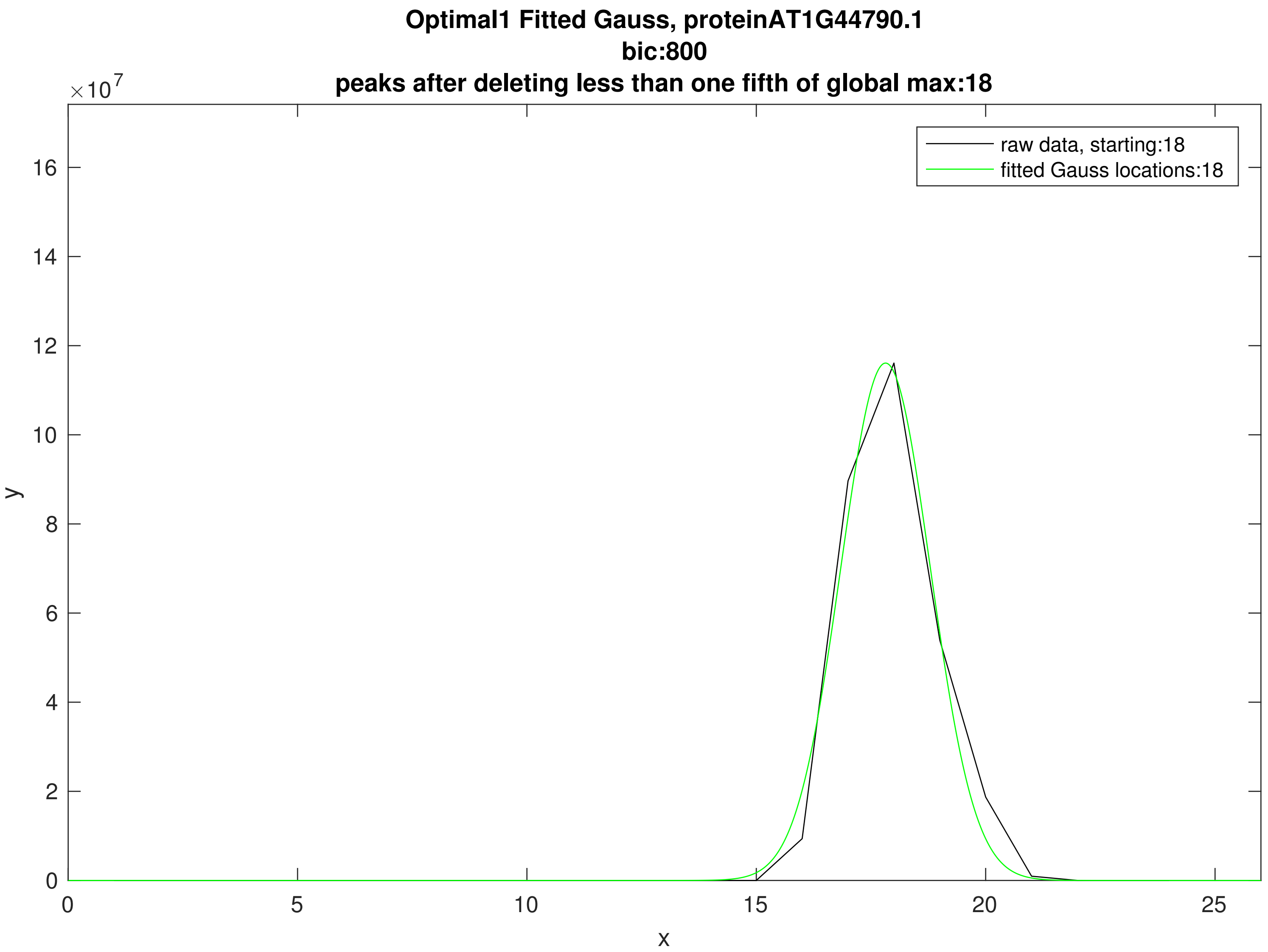
Optimal1 Fitted Gauss, proteinAT1G44790.1
bic:800
peaks after deleting less than one fifth of global max:18
×10^7
raw data, starting:18
fitted Gauss locations:18
y
x
0
2
4
6
8
10
12
14
16
0
5
10
15
20
25

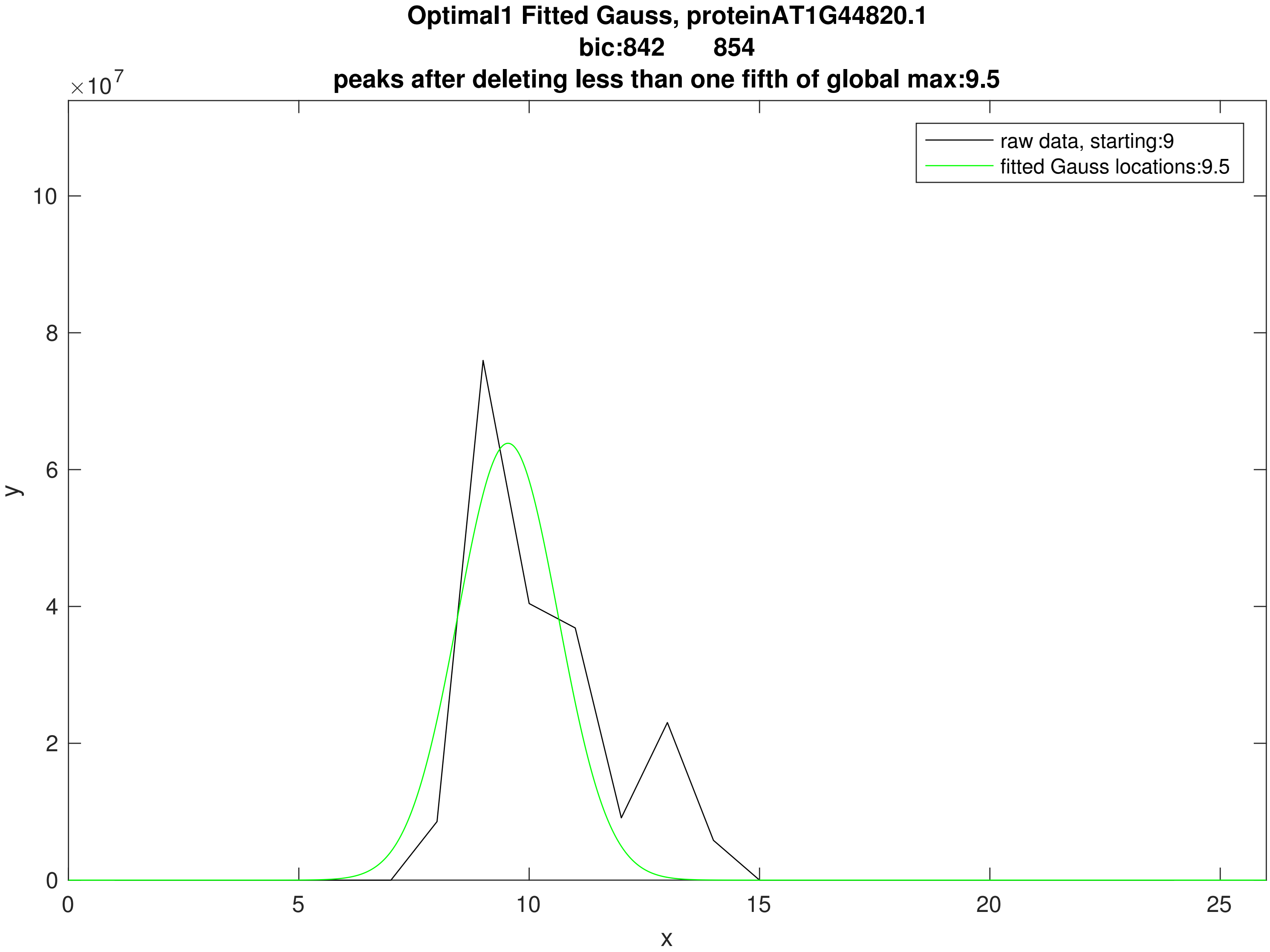

Optimal1 Fitted Gauss, proteinAT1G44820.1
bic:842 854
peaks after deleting less than one fifth of global max:9.5
×10^7
raw data, starting:9
fitted Gauss locations:9.5
0
2
4
6
8
10
y
0
5
10
15
20
25
x

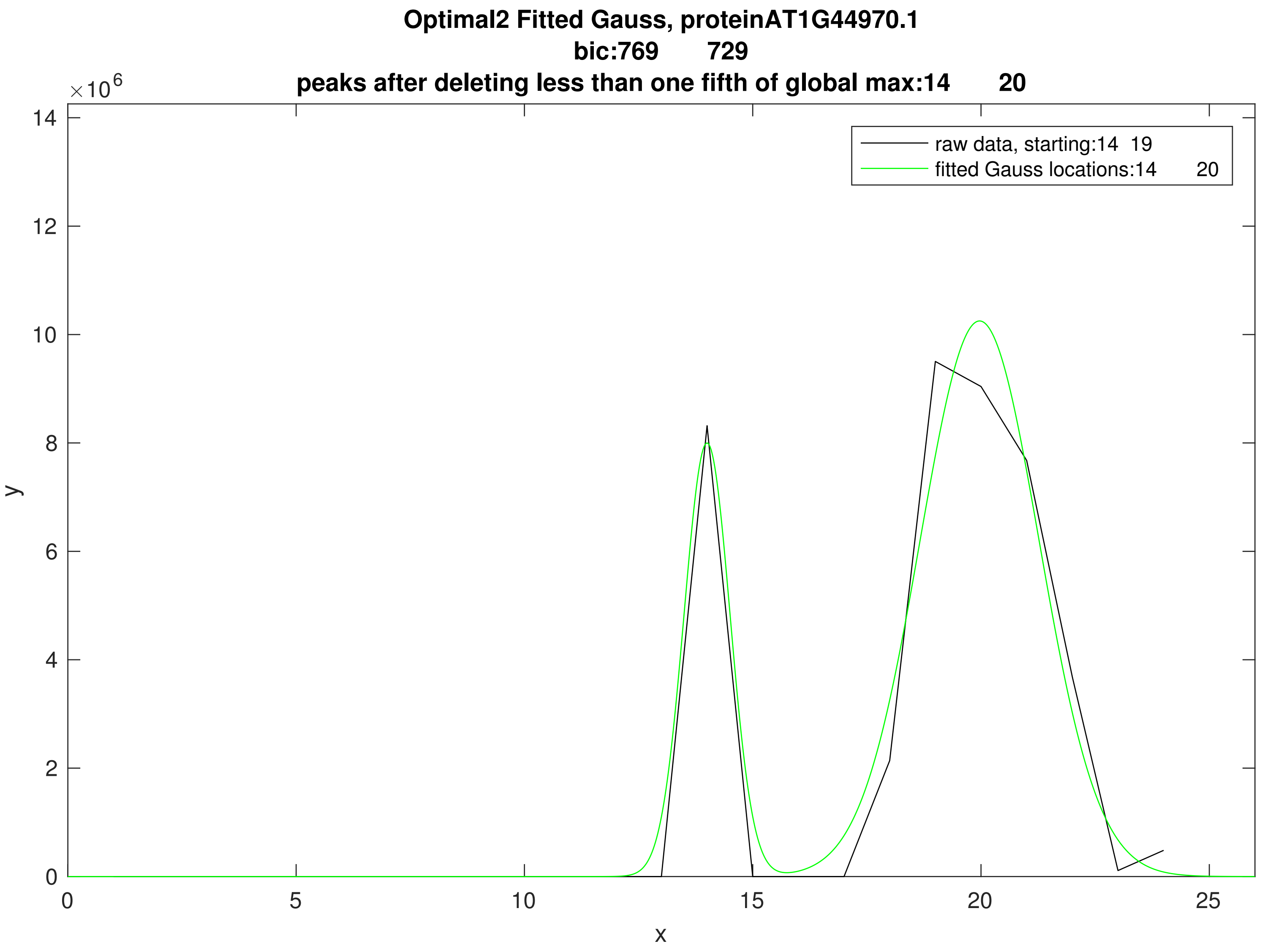
Optimal2 Fitted Gauss, proteinAT1G44970.1
bic:769 729
peaks after deleting less than one fifth of global max:14 20
×10^6
raw data, starting:14 19
fitted Gauss locations:14 20
y
x
0
2
4
6
8
10
12
14
0
5
10
15
20
25

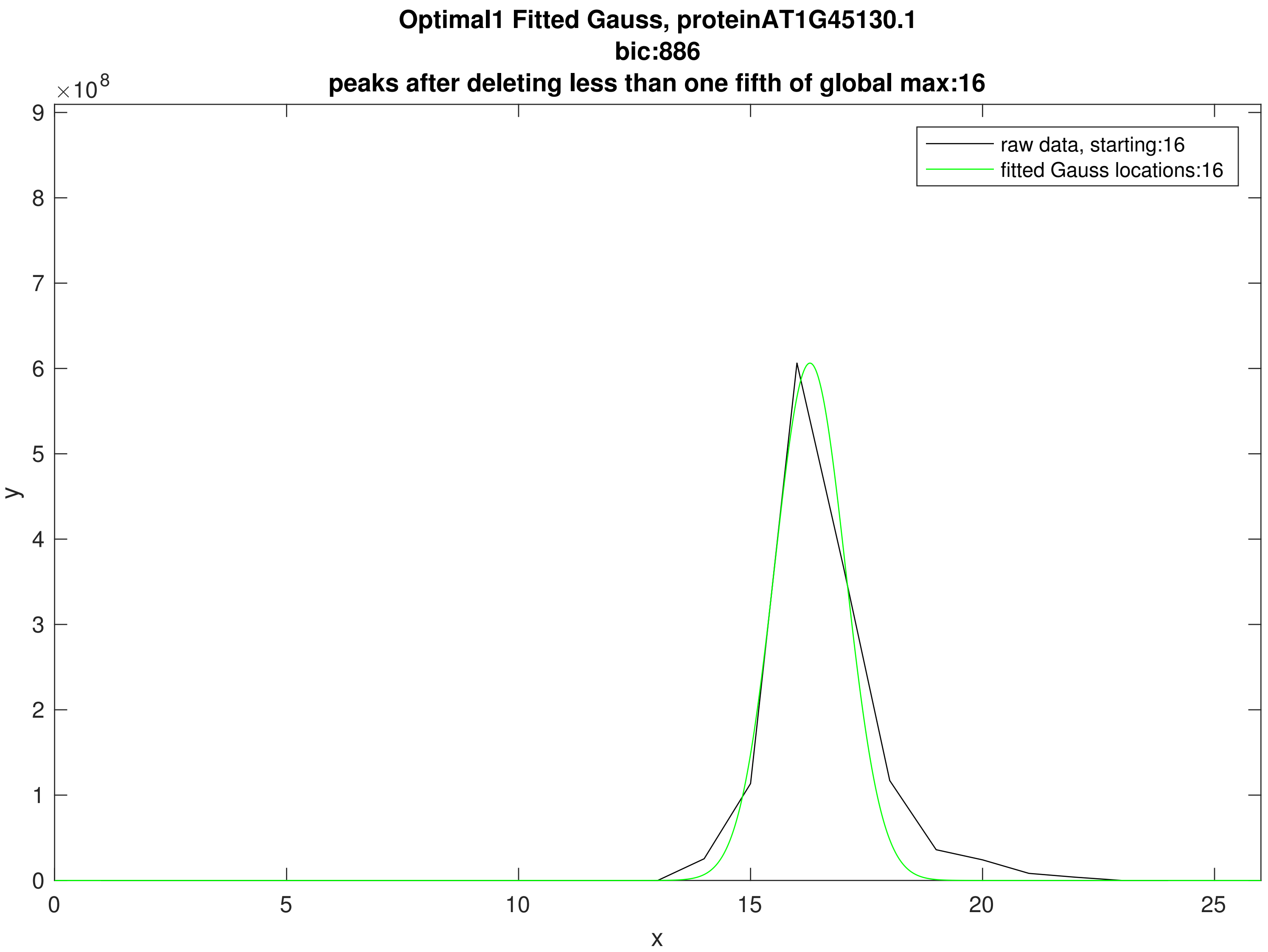

Optimal1 Fitted Gauss, proteinAT1G45130.1
bic:886
peaks after deleting less than one fifth of global max:16
×10^8
raw data, starting:16
fitted Gauss locations:16
y
x
0
1
2
3
4
5
6
7
8
9
0
5
10
15
20
25

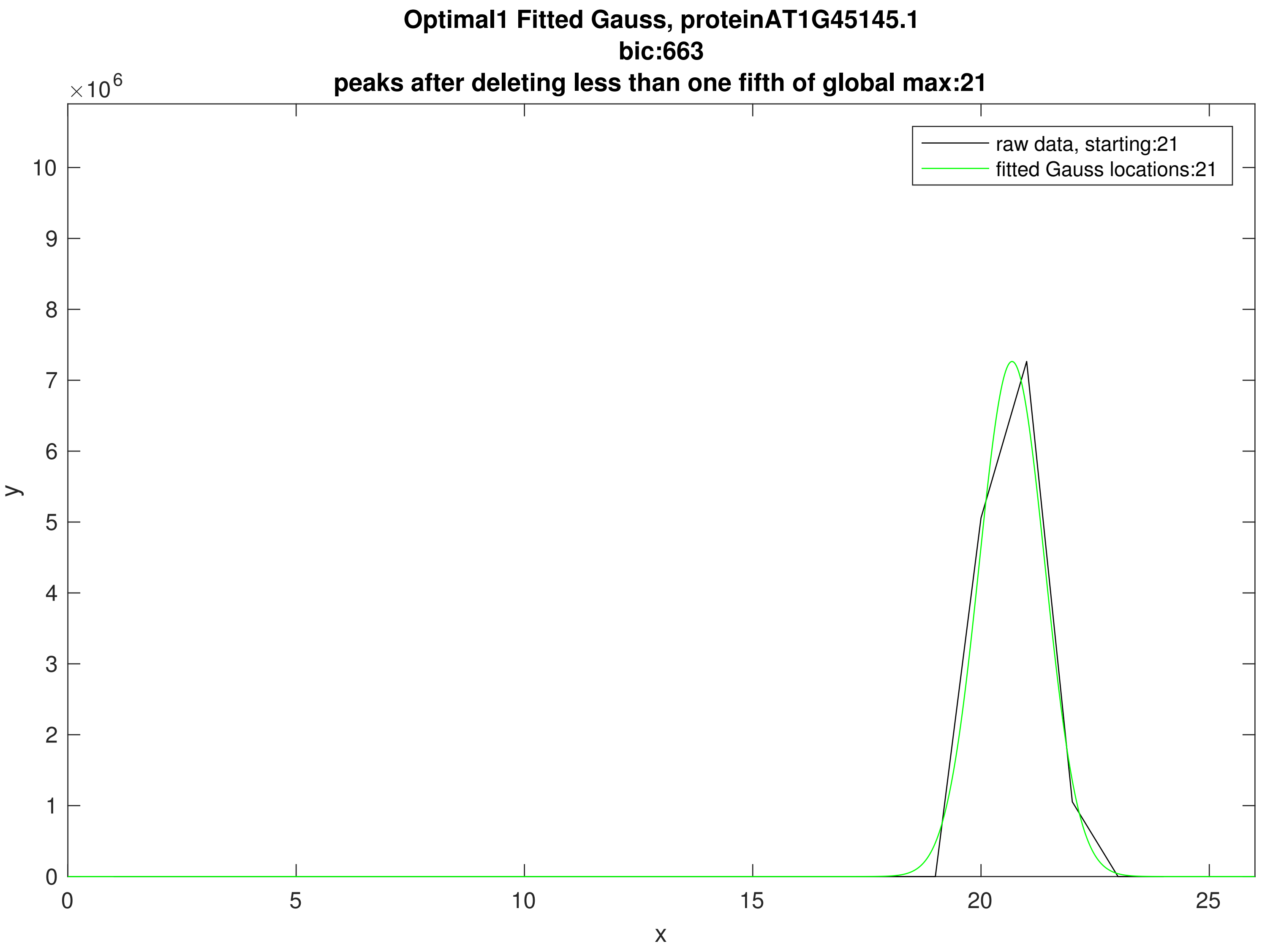

Optimal1 Fitted Gauss, proteinAT1G45145.1
bic:663
peaks after deleting less than one fifth of global max:21
×10^6
raw data, starting:21
fitted Gauss locations:21
10
9
8
7
6
5
4
3
2
1
0
y
0
5
10
15
20
25
x

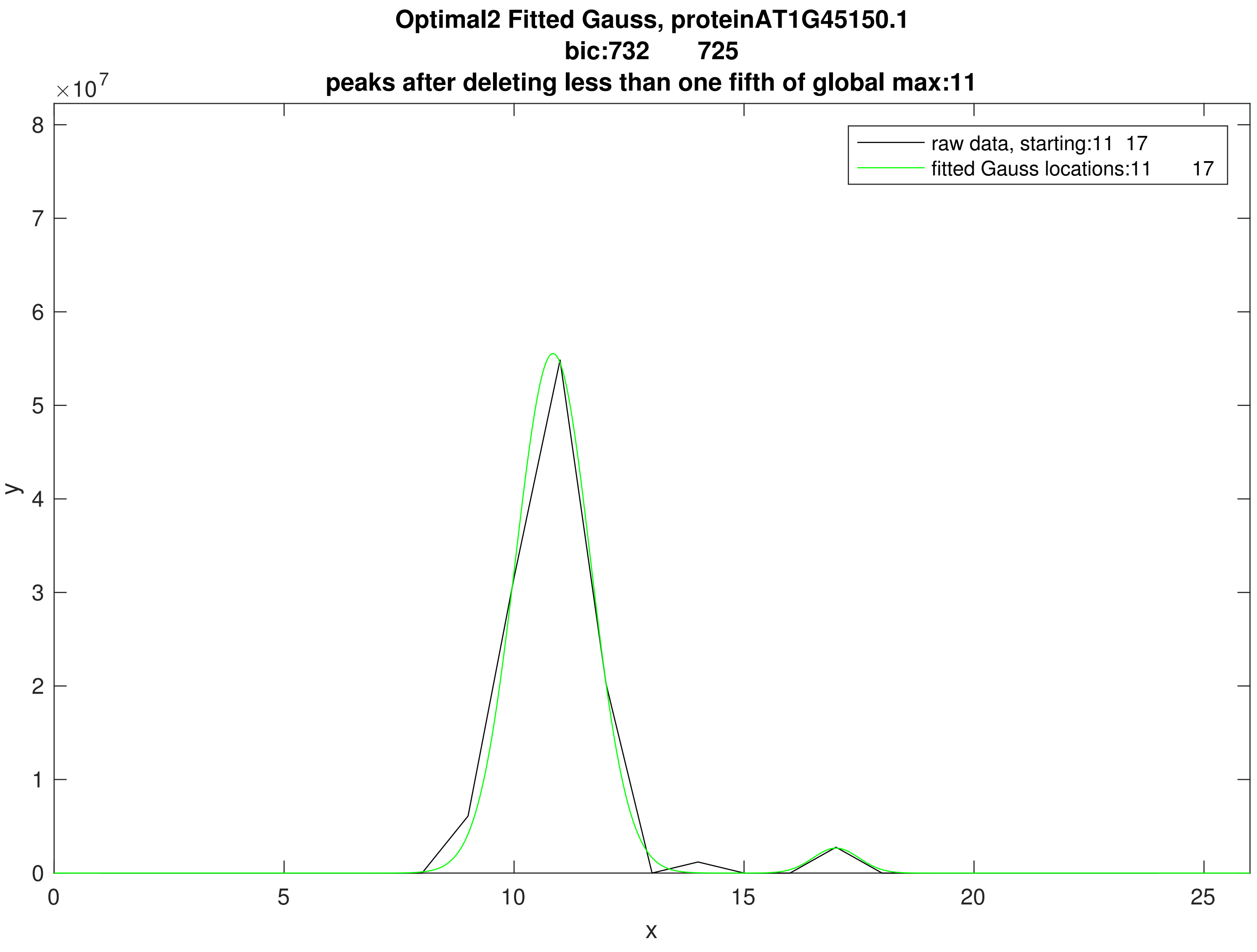
Optimal2 Fitted Gauss, proteinAT1G45150.1
bic:732 725
peaks after deleting less than one fifth of global max:11
×10^7
raw data, starting:11 17
fitted Gauss locations:11 17
y
x
0
1
2
3
4
5
6
7
8
0
5
10
15
20
25

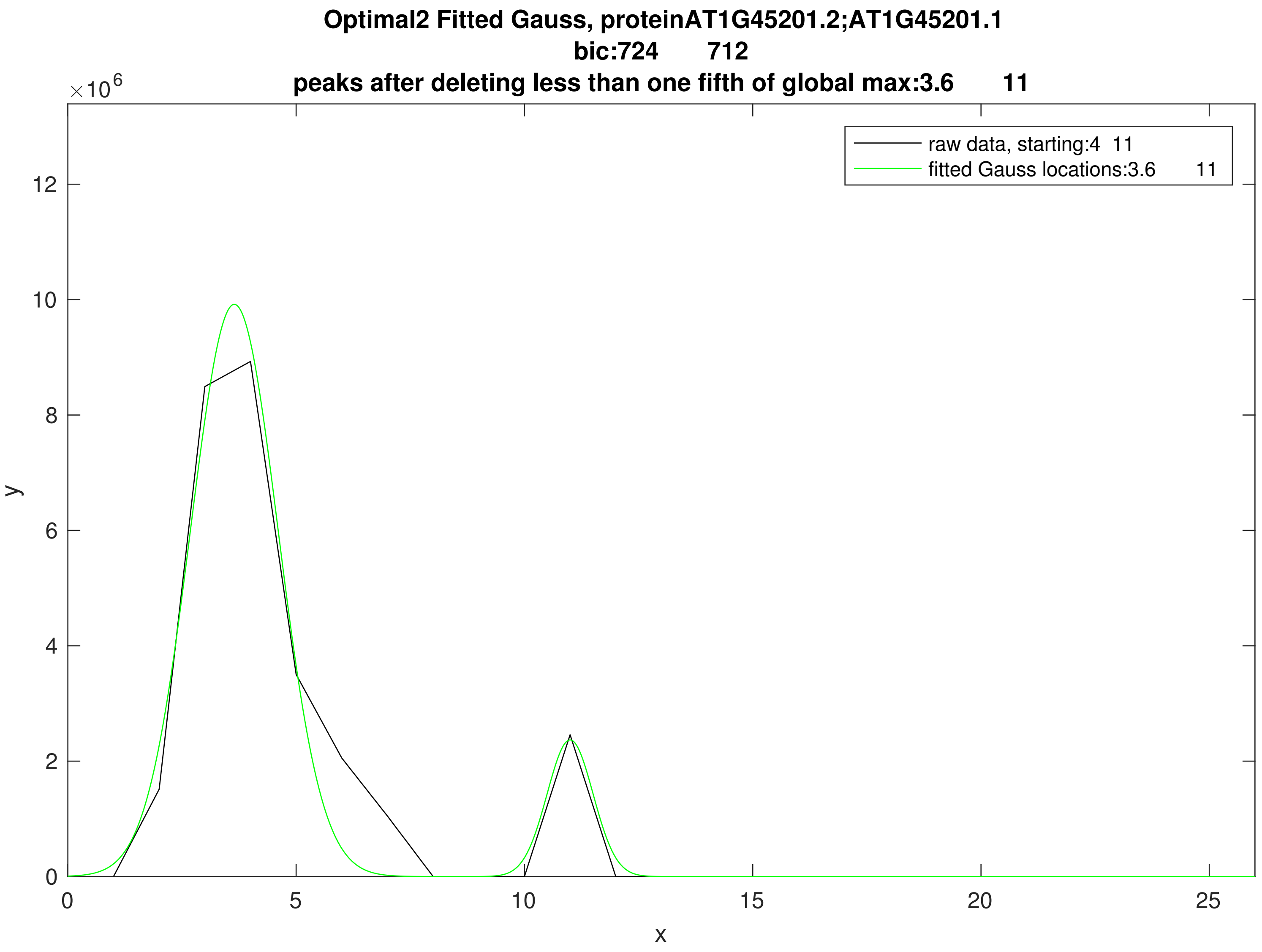
Optimal2 Fitted Gauss, proteinAT1G45201.2;AT1G45201.1
bic:724 712
peaks after deleting less than one fifth of global max:3.6 11
×10^6
raw data, starting:4 11
fitted Gauss locations:3.6 11
y
x
0
2
4
6
8
10
12
5
15
20
25

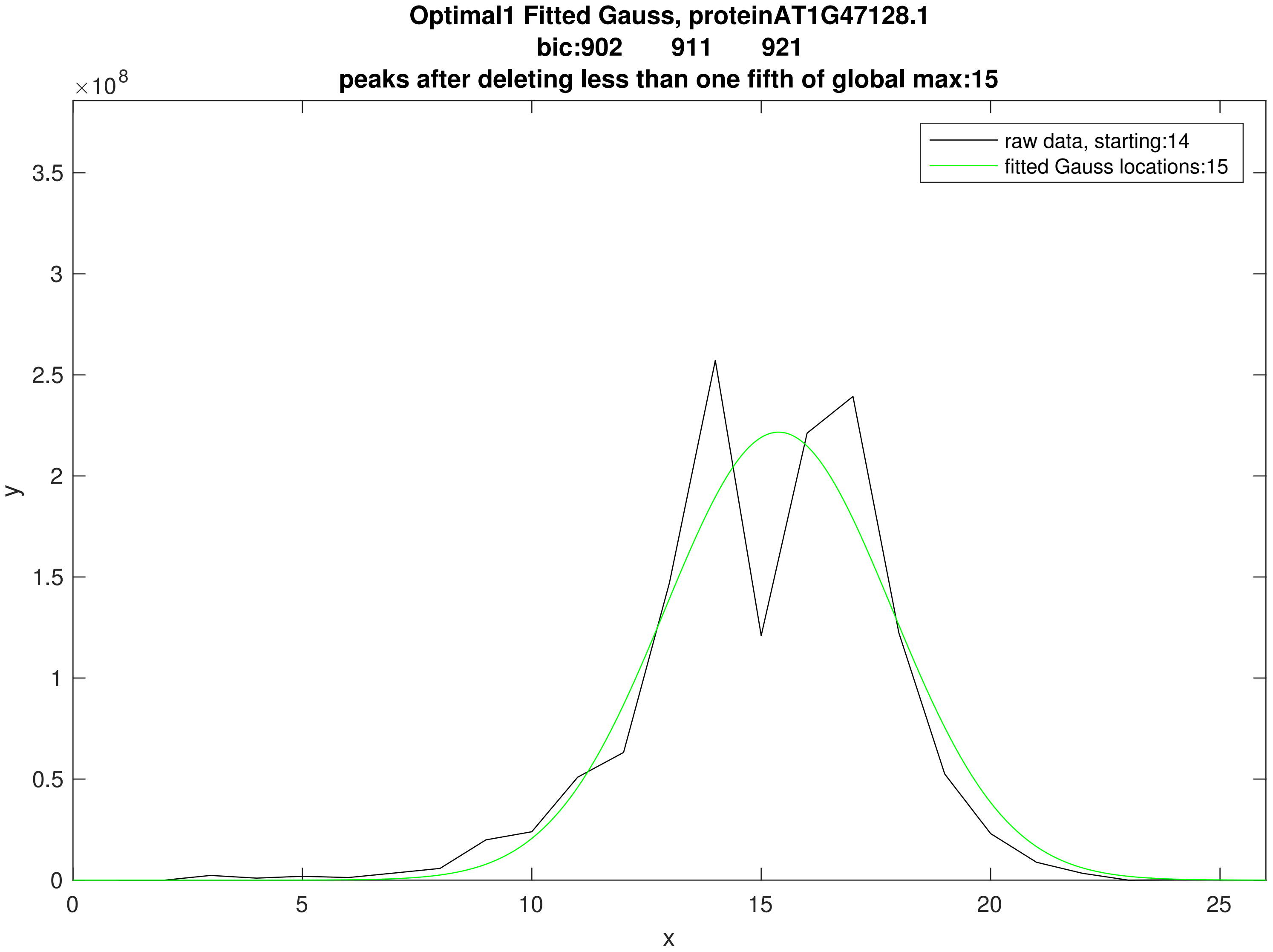

Optimal1 Fitted Gauss, proteinAT1G47128.1
bic:902 911 921
peaks after deleting less than one fifth of global max:15
×10^8
raw data, starting:14
fitted Gauss locations:15
y
x

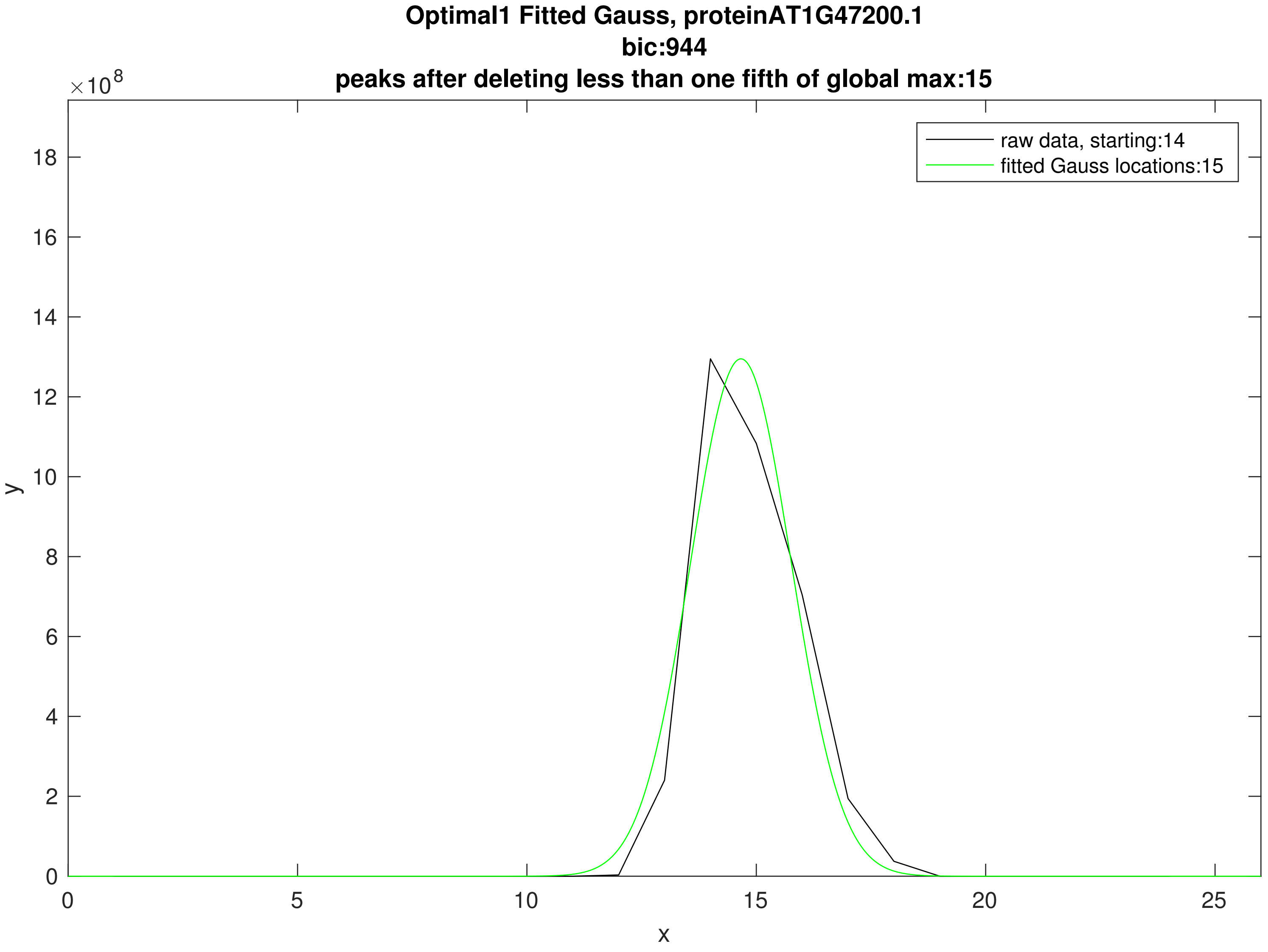

Optimal1 Fitted Gauss, proteinAT1G47200.1
bic:944
peaks after deleting less than one fifth of global max:15
×10^8
raw data, starting:14
fitted Gauss locations:15
18
16
14
12
10
8
6
4
2
0
y
0
5
10
15
20
25
x

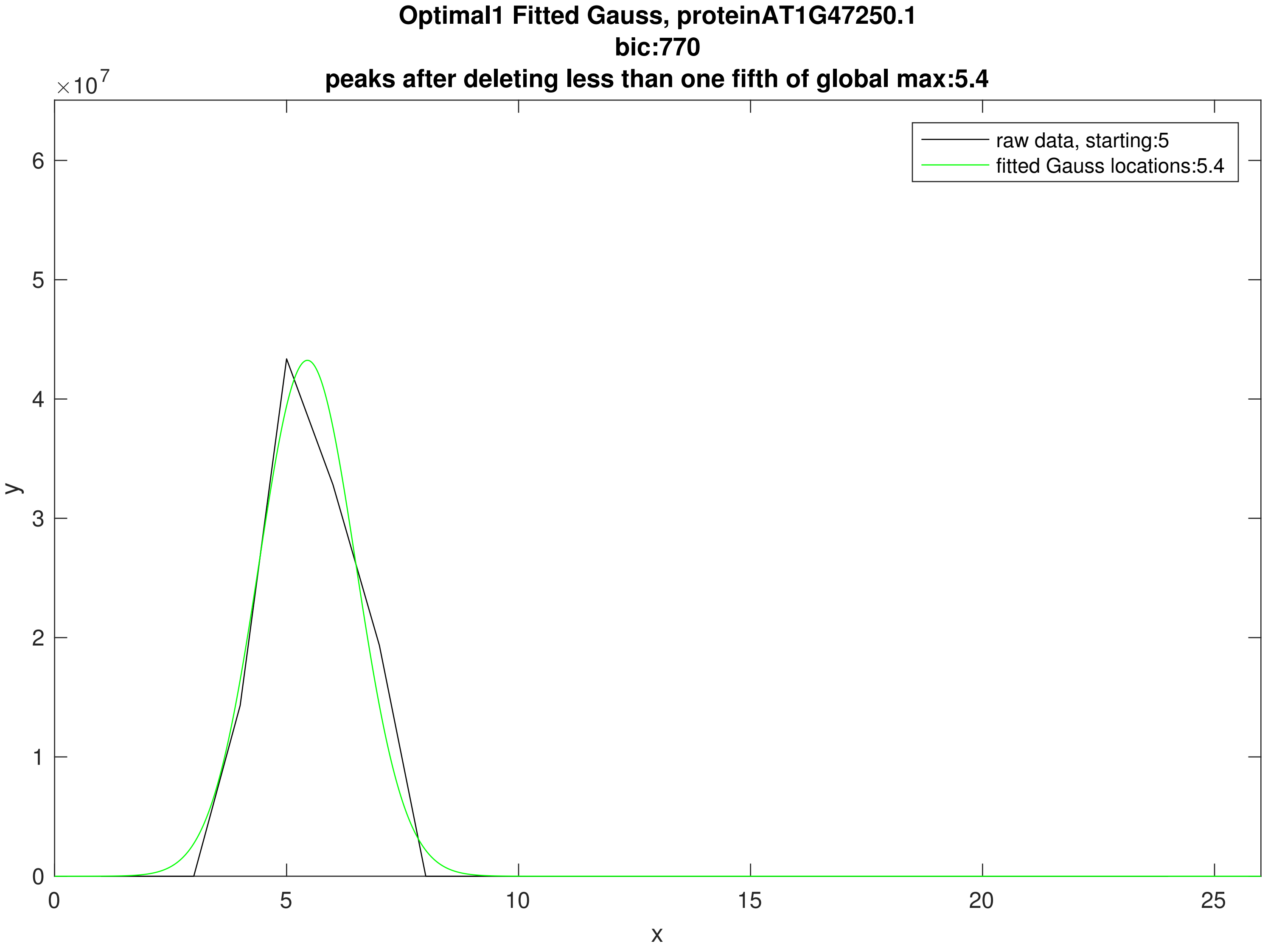

Optimal1 Fitted Gauss, proteinAT1G47250.1
bic:770
peaks after deleting less than one fifth of global max:5.4
×10^7
raw data, starting:5
fitted Gauss locations:5.4
y
x
0
1
2
3
4
5
6
0
5
10
15
20
25

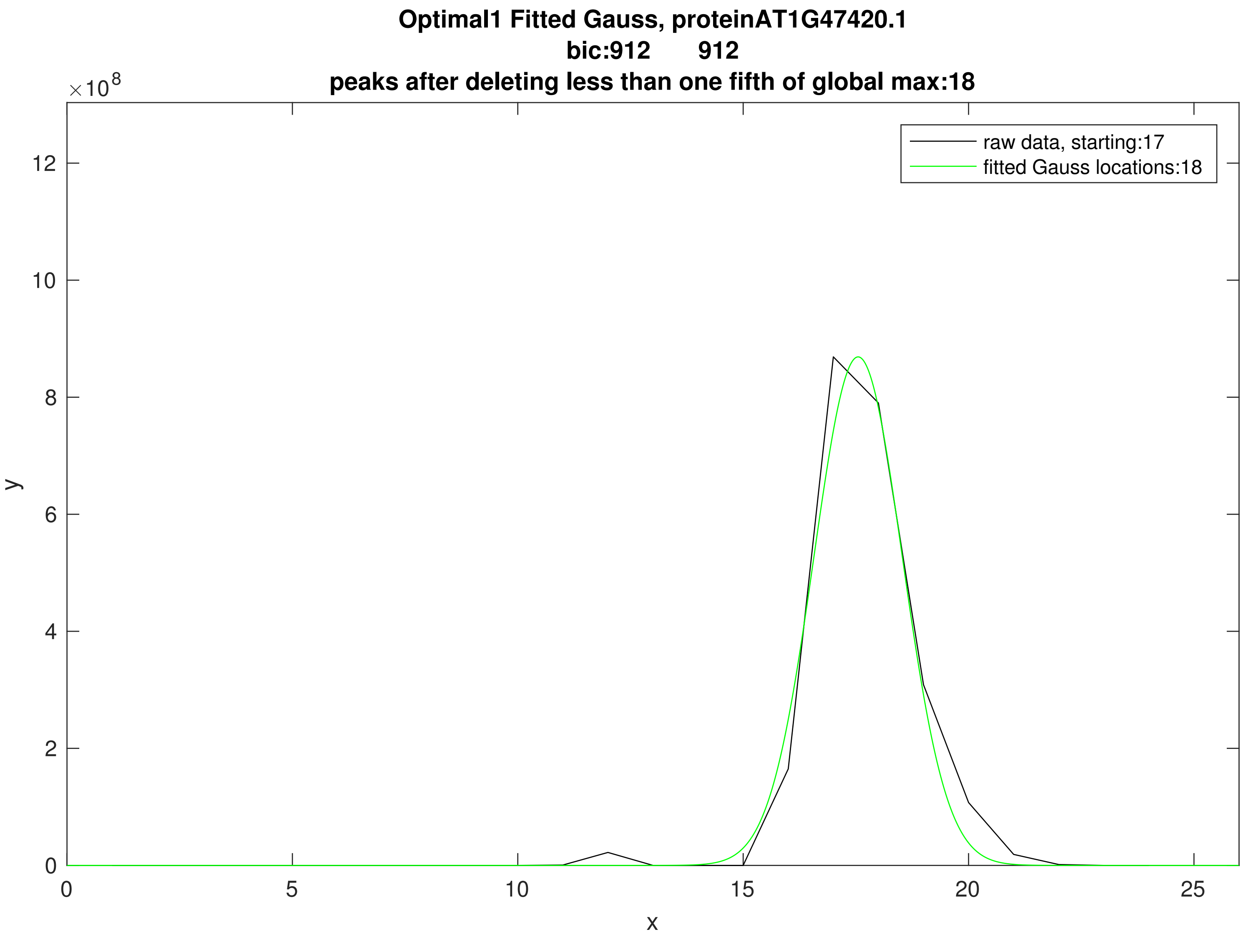
Optimal1 Fitted Gauss, proteinAT1G47420.1
bic:912 912
peaks after deleting less than one fifth of global max:18
×10^8
raw data, starting:17
fitted Gauss locations:18
12
10
8
6
4
2
0
y
0
5
10
15
20
25
x

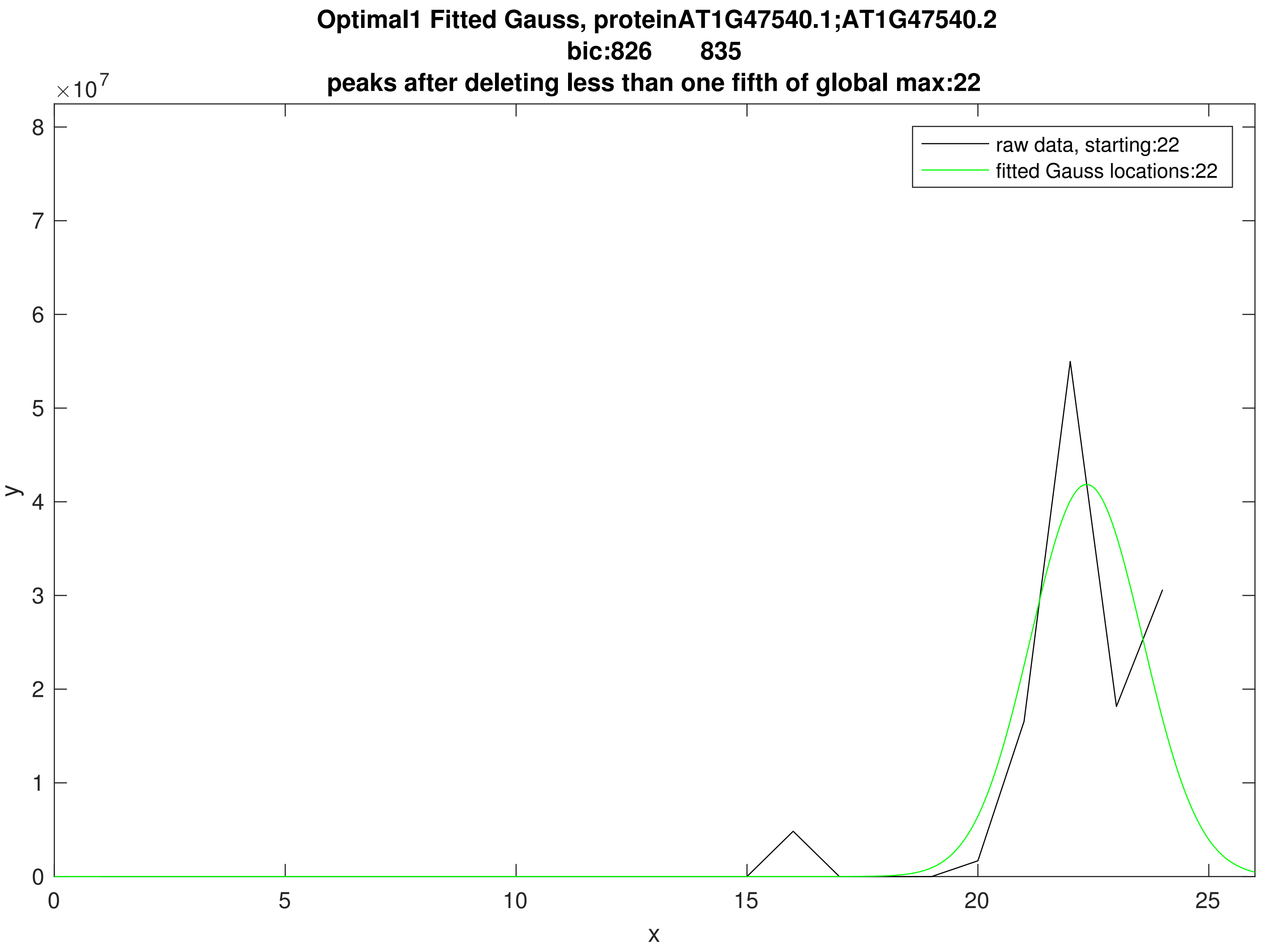

Optimal1 Fitted Gauss, proteinAT1G47540.1;AT1G47540.2
bic:826 835
peaks after deleting less than one fifth of global max:22
×10^7
raw data, starting:22
fitted Gauss locations:22
y
x
0
1
2
3
4
5
6
7
8
0
5
10
15
20
25

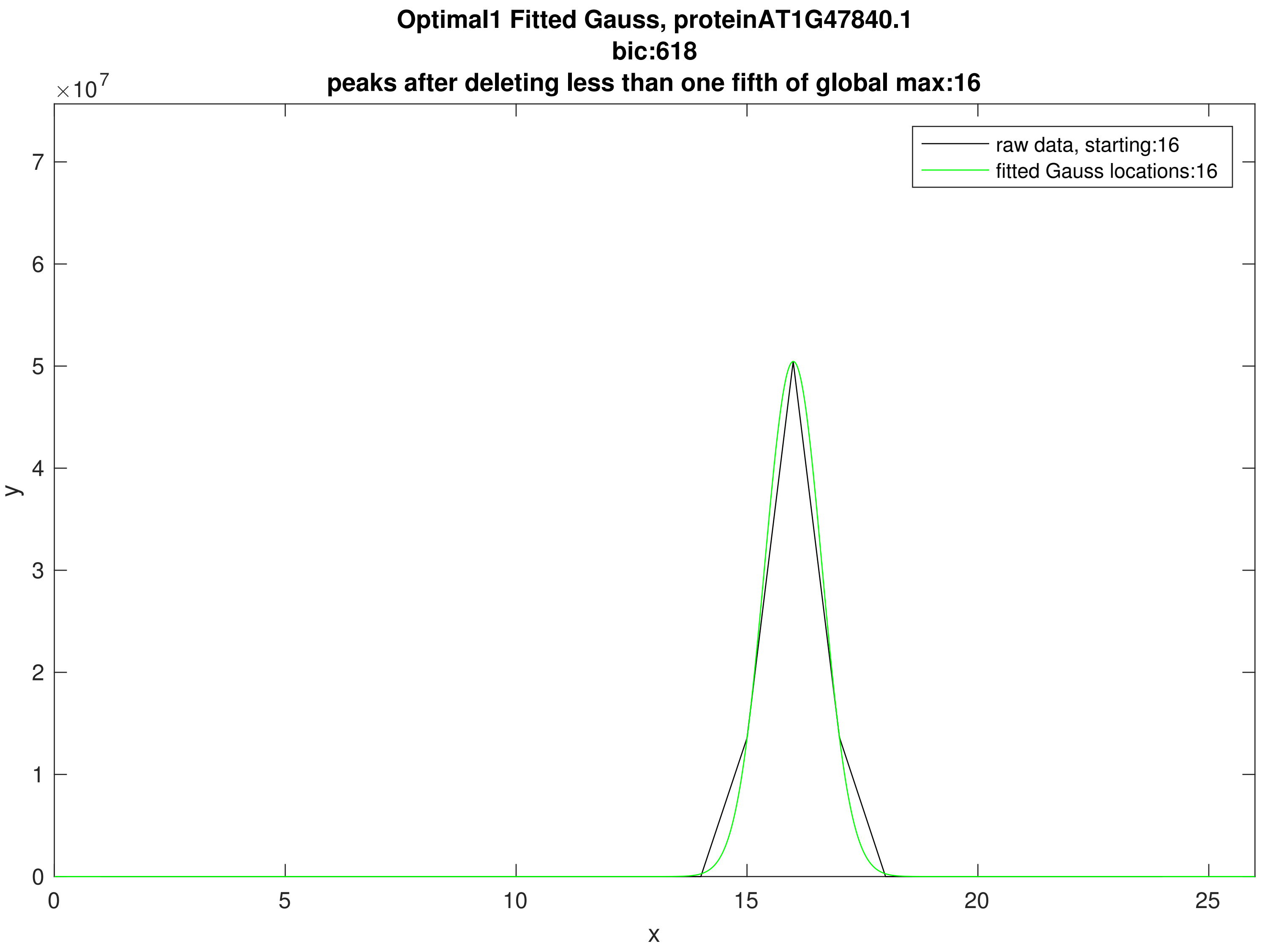
Optimal1 Fitted Gauss, proteinAT1G47840.1
bic:618
peaks after deleting less than one fifth of global max:16
×10^7
raw data, starting:16
fitted Gauss locations:16
y
x
0
1
2
3
4
5
6
7
0
5
10
15
20
25

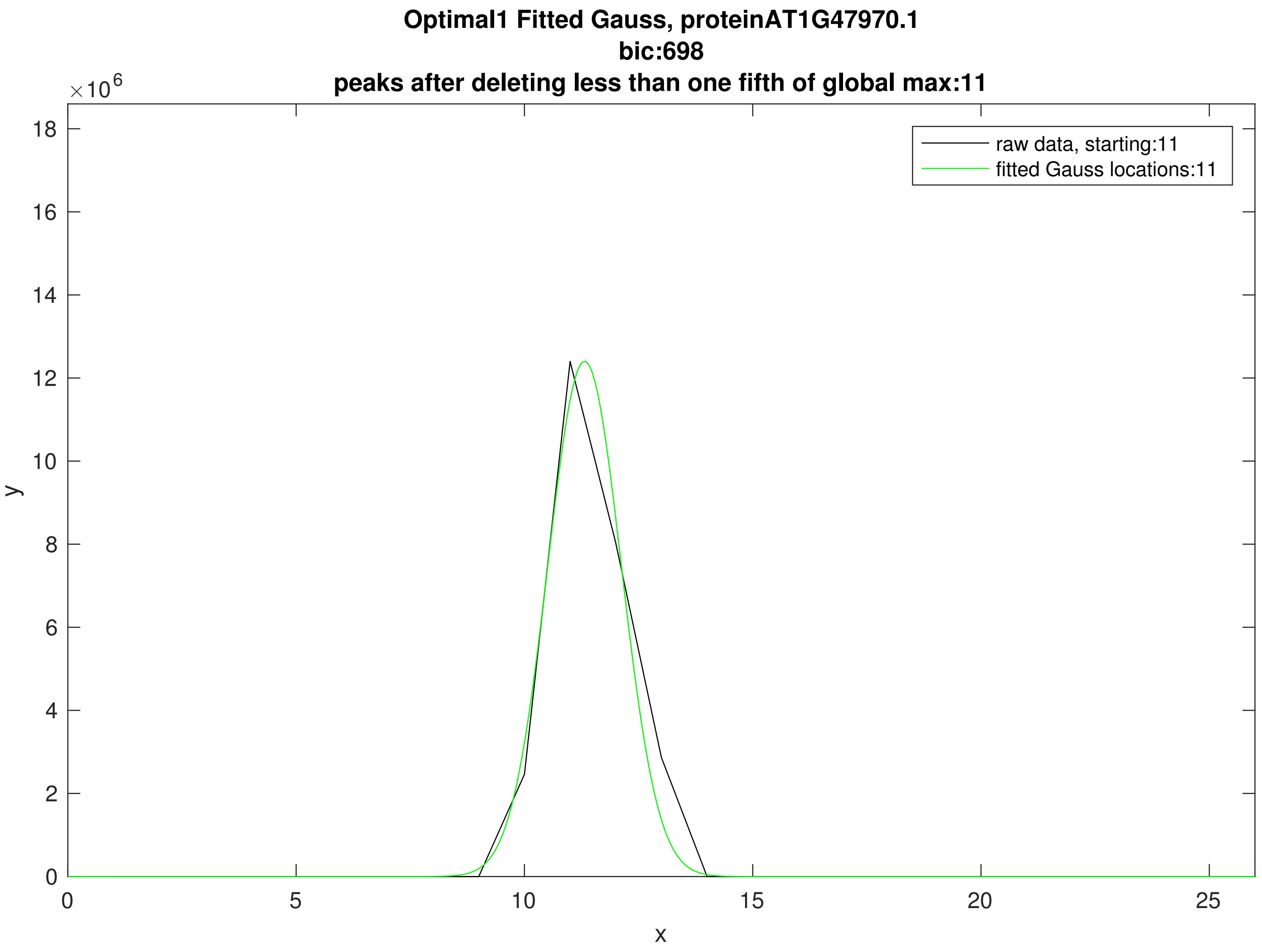

Optimal1 Fitted Gauss, proteinAT1G47970.1
bic:698
peaks after deleting less than one fifth of global max:11
×10^6
18
16
14
12
10
8
6
4
2
0
y
0
5
10
15
20
25
x
raw data, starting:11
fitted Gauss locations:11

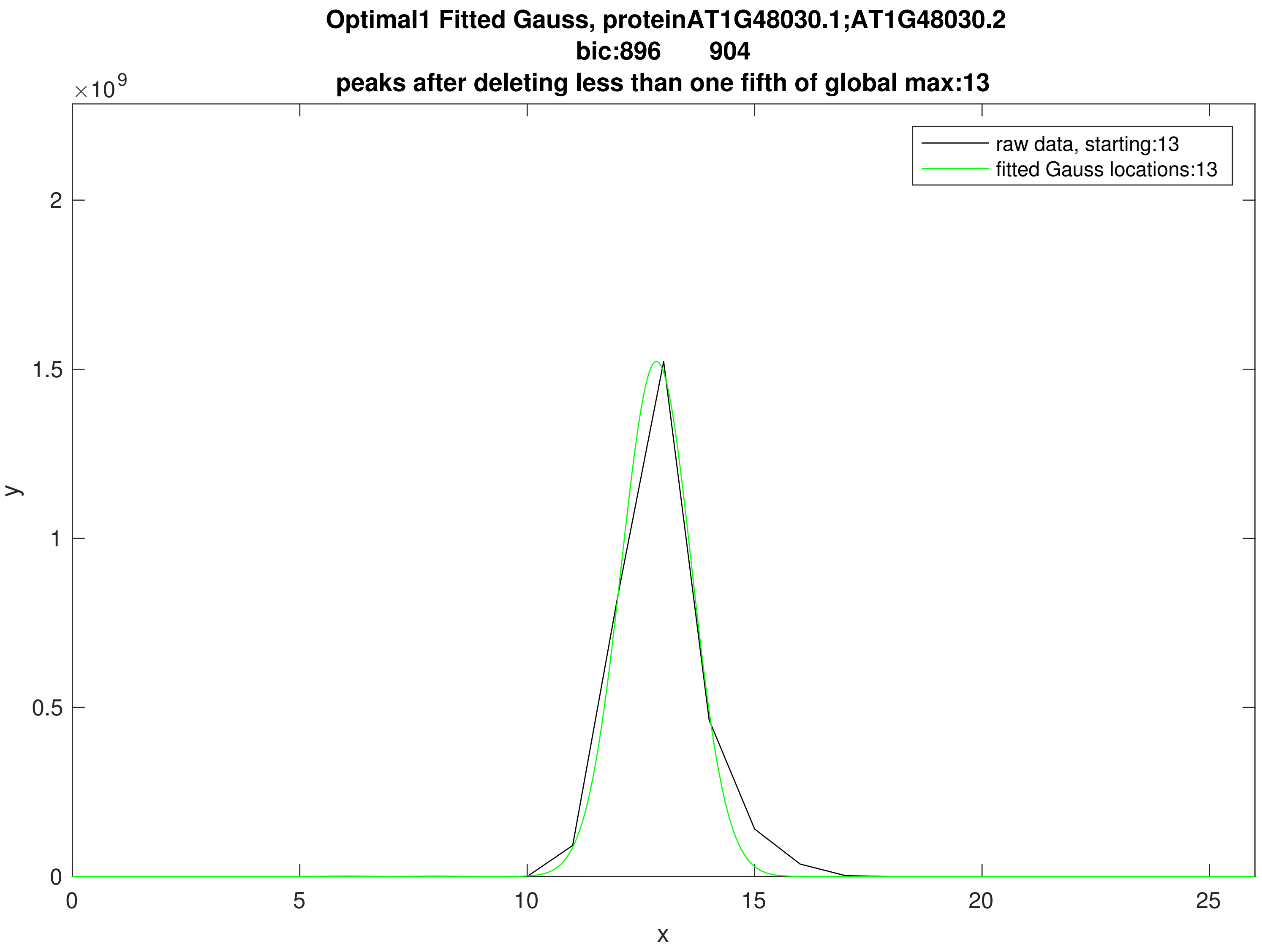

Optimal1 Fitted Gauss, proteinAT1G48030.1;AT1G48030.2
bic:896 904
peaks after deleting less than one fifth of global max:13
×10^9
raw data, starting:13
fitted Gauss locations:13
2
1.5
1
0.5
0
y
0
5
10
15
20
25
x

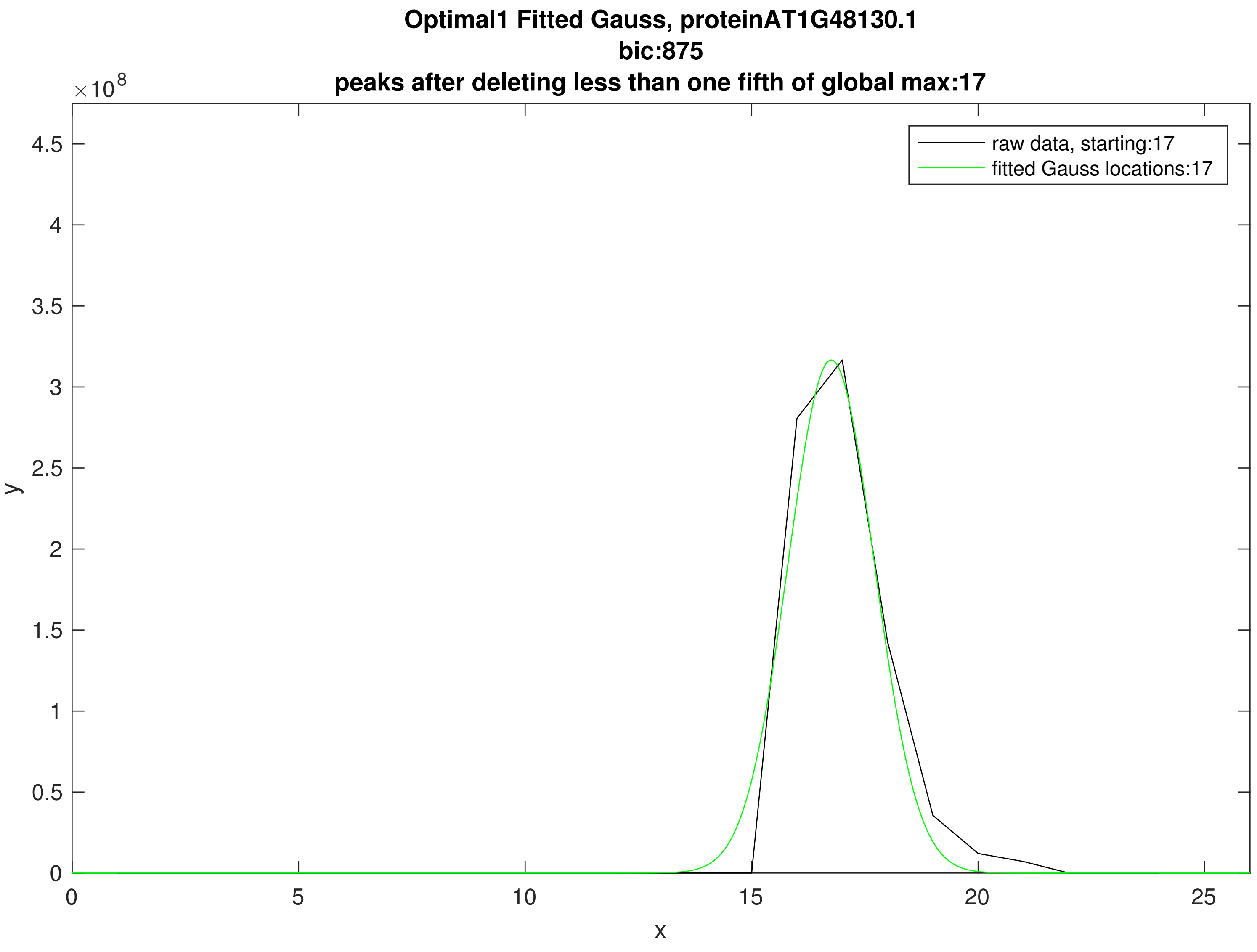
Optimal1 Fitted Gauss, proteinAT1G48130.1
bic:875
peaks after deleting less than one fifth of global max:17
raw data, starting:17
fitted Gauss locations:17
y
x

**Optimal1 Fitted Gauss, proteinAT1G48160.1**
**bic:677**
**peaks after deleting less than one fifth of global max:8.4**

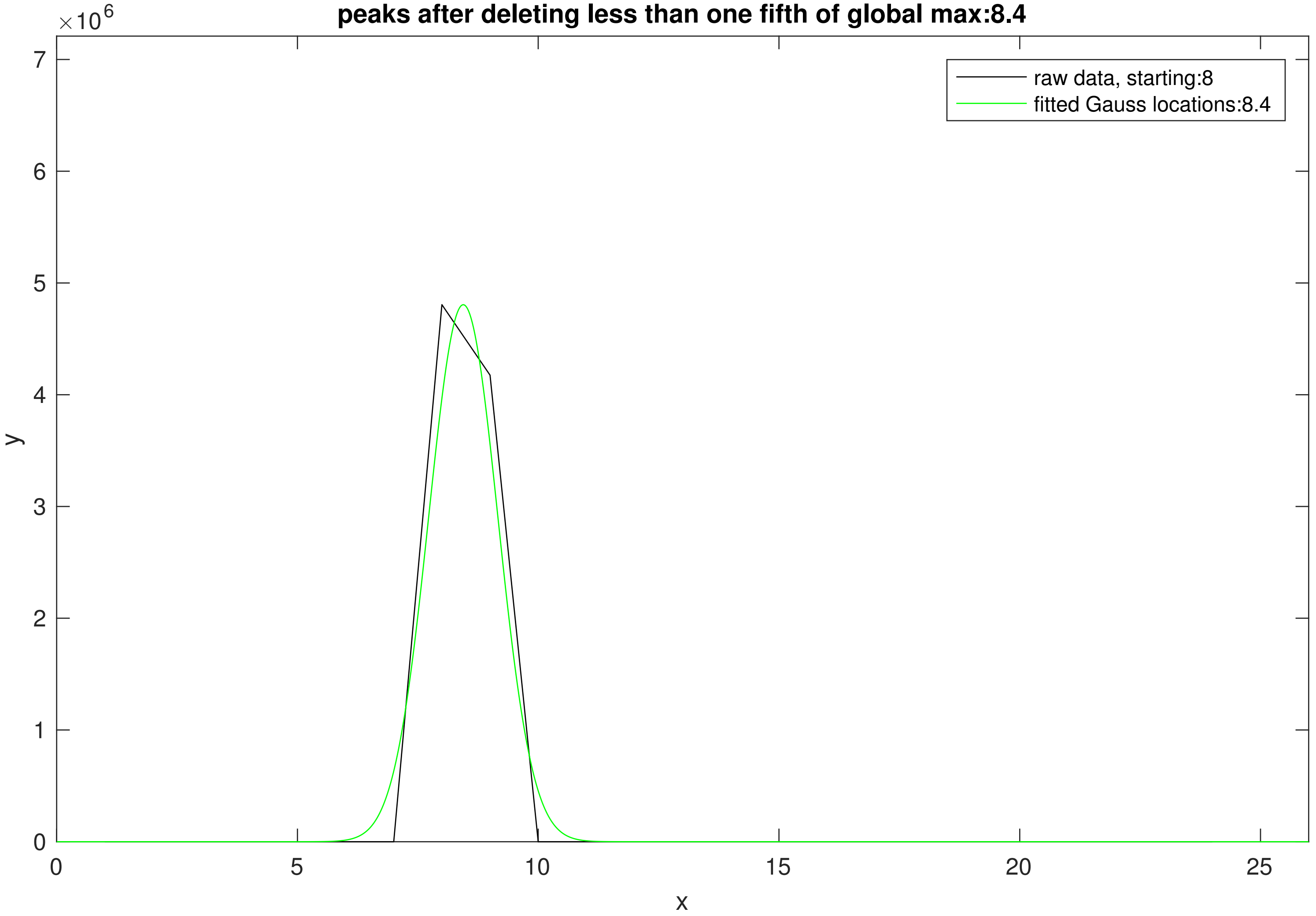

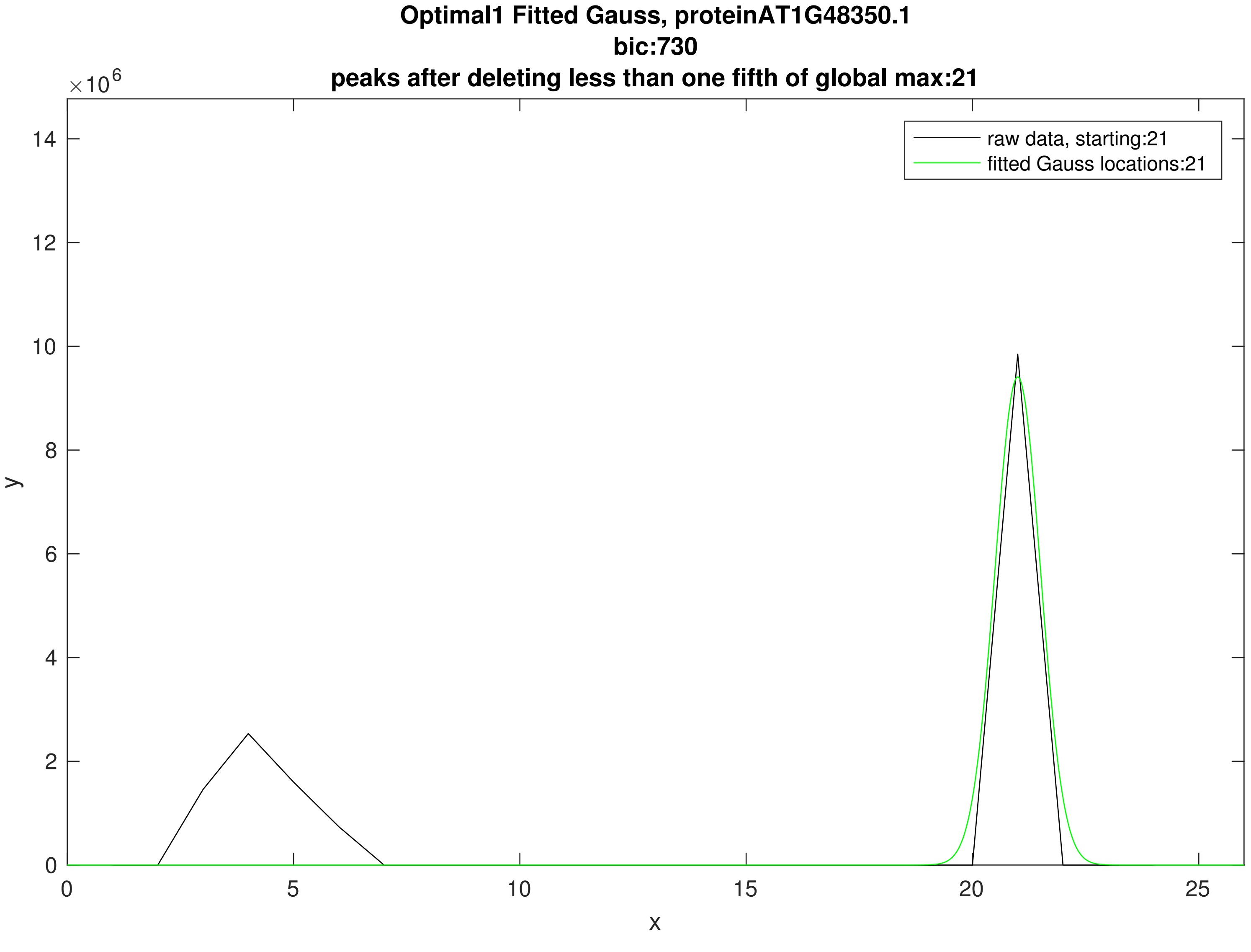

Optimal1 Fitted Gauss, proteinAT1G48350.1
bic:730
peaks after deleting less than one fifth of global max:21
×10^6
raw data, starting:21
fitted Gauss locations:21
y
x
0
2
4
6
8
10
12
14
0
5
10
15
20
25

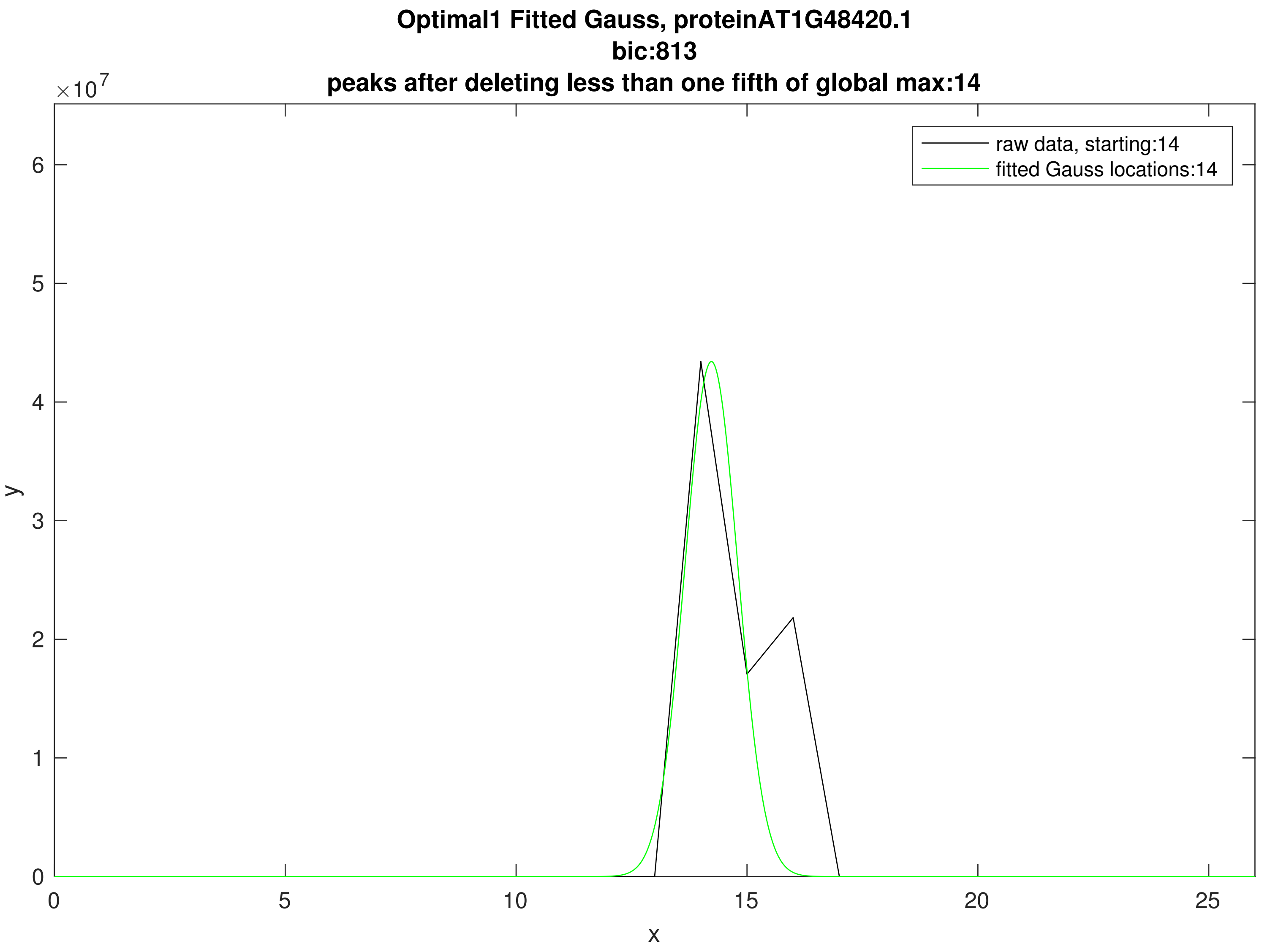

Optimal1 Fitted Gauss, proteinAT1G48420.1
bic:813
peaks after deleting less than one fifth of global max:14
×10^7
raw data, starting:14
fitted Gauss locations:14
y
x
0
1
2
3
4
5
6
0
5
10
15
20
25

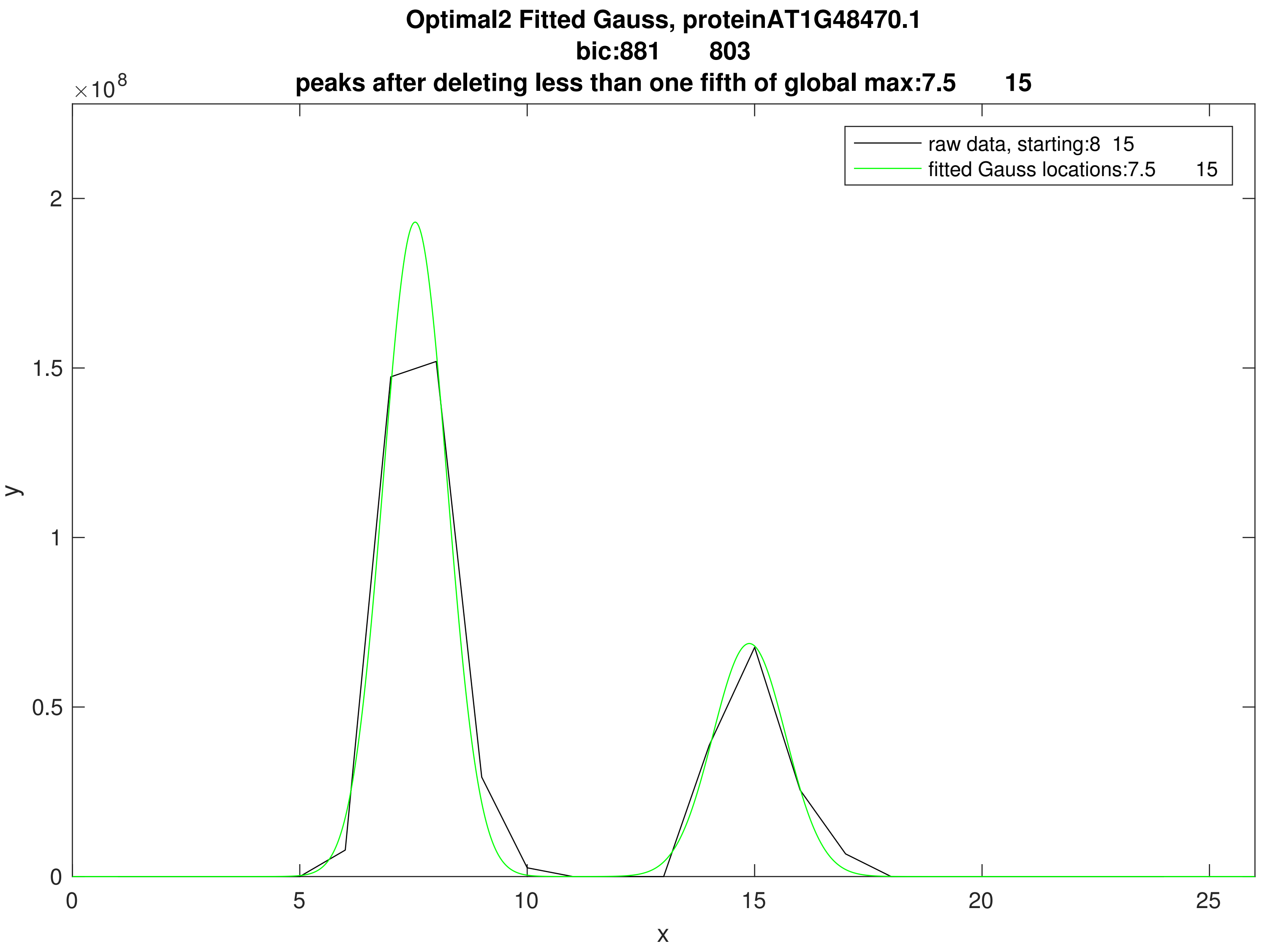

Optimal2 Fitted Gauss, proteinAT1G48470.1
bic:881 803
peaks after deleting less than one fifth of global max:7.5 15
×10^8
raw data, starting:8 15
fitted Gauss locations:7.5 15
2
1.5
1
0.5
0
y
0
5
10
15
20
25
x

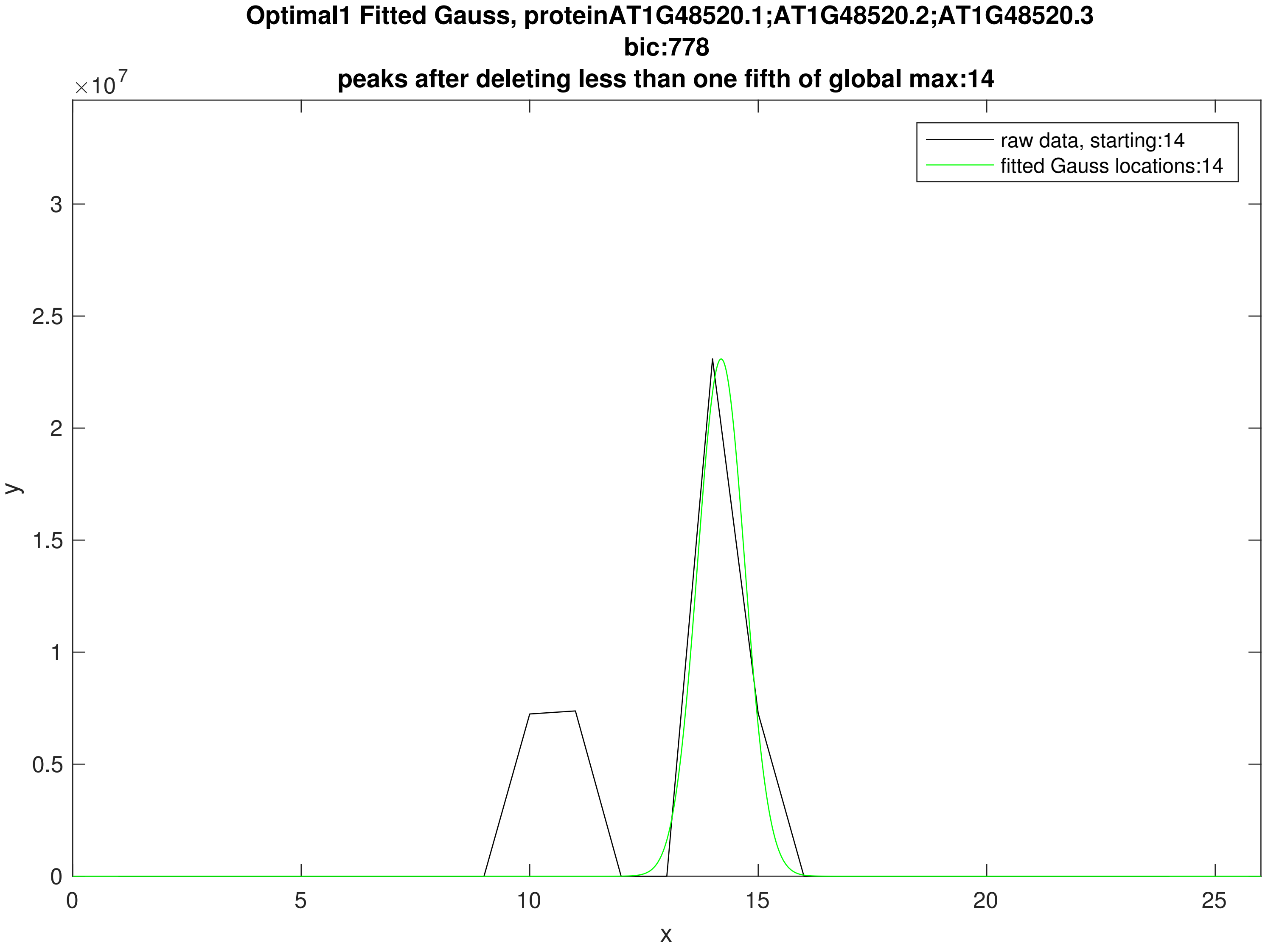
Optimal1 Fitted Gauss, proteinAT1G48520.1;AT1G48520.2;AT1G48520.3
bic:778
peaks after deleting less than one fifth of global max:14
×10^7
raw data, starting:14
fitted Gauss locations:14
y
x
0
0.5
1
1.5
2
2.5
3
0
5
10
15
20
25

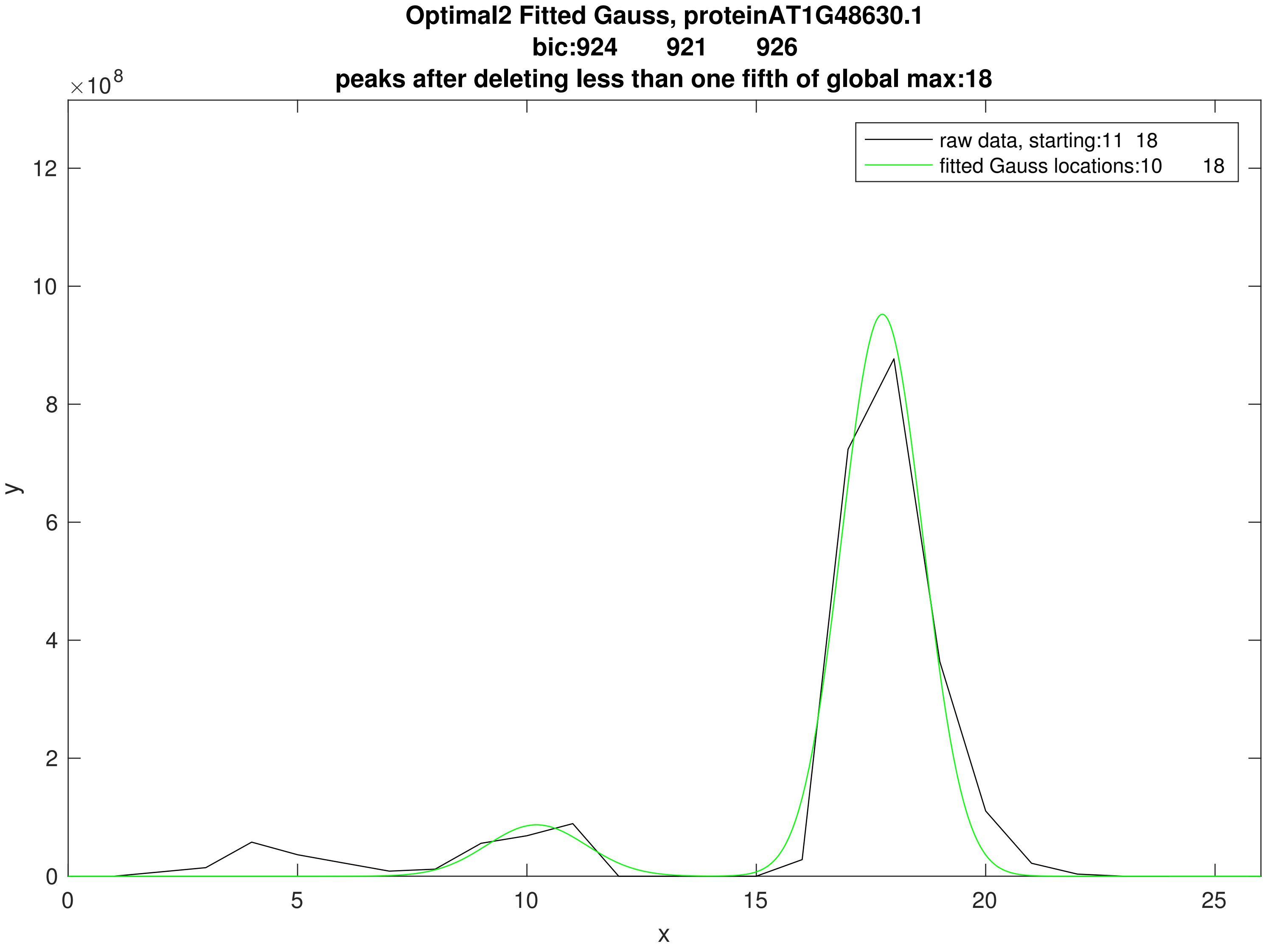
Optimal2 Fitted Gauss, proteinAT1G48630.1
bic:924 921 926
peaks after deleting less than one fifth of global max:18
×10^8
raw data, starting:11 18
fitted Gauss locations:10 18
y
x
0
2
4
6
8
10
12
0
5
10
15
20
25

Optimal1 Fitted Gauss, proteinAT1G48660.1
bic:806
peaks after deleting less than one fifth of global max:14

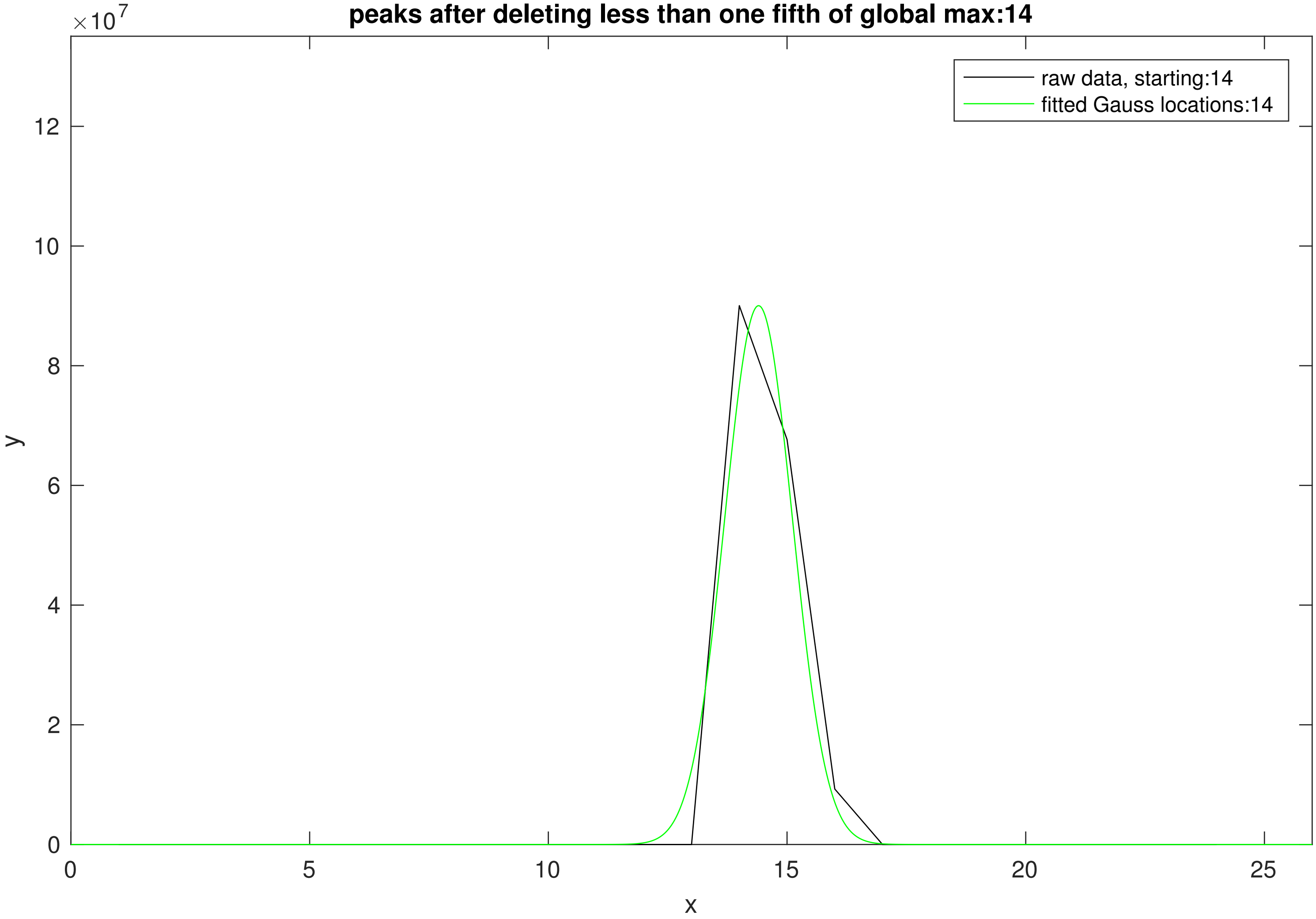

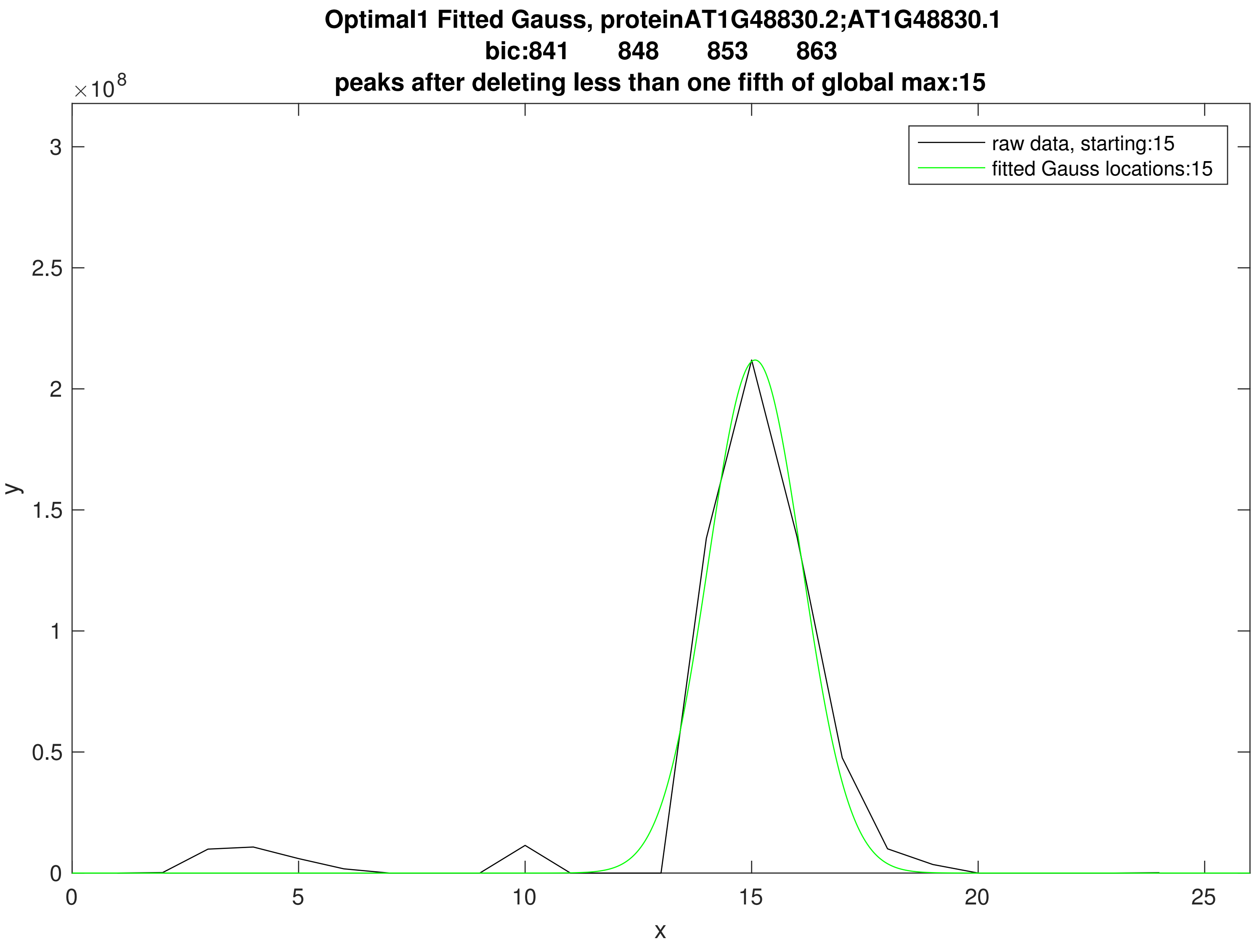
Optimal1 Fitted Gauss, proteinAT1G48830.2;AT1G48830.1
bic:841 848 853 863
peaks after deleting less than one fifth of global max:15
×10^8
raw data, starting:15
fitted Gauss locations:15
y
x
0
0.5
1
1.5
2
2.5
3
0
5
10
15
20
25

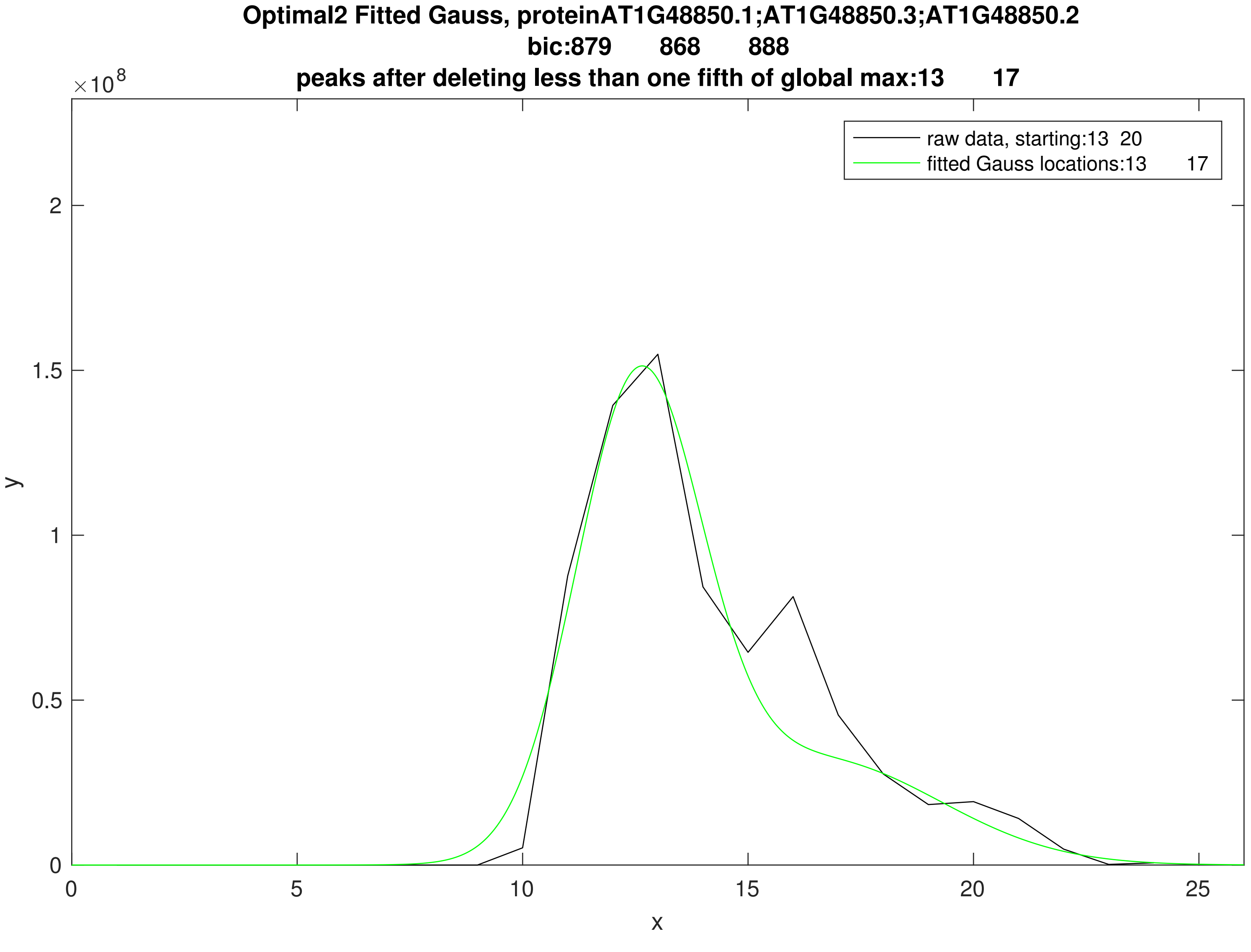
Optimal2 Fitted Gauss, proteinAT1G48850.1;AT1G48850.3;AT1G48850.2
bic:879 868 888
peaks after deleting less than one fifth of global max:13 17
raw data, starting:13 20
fitted Gauss locations:13 17
y
x

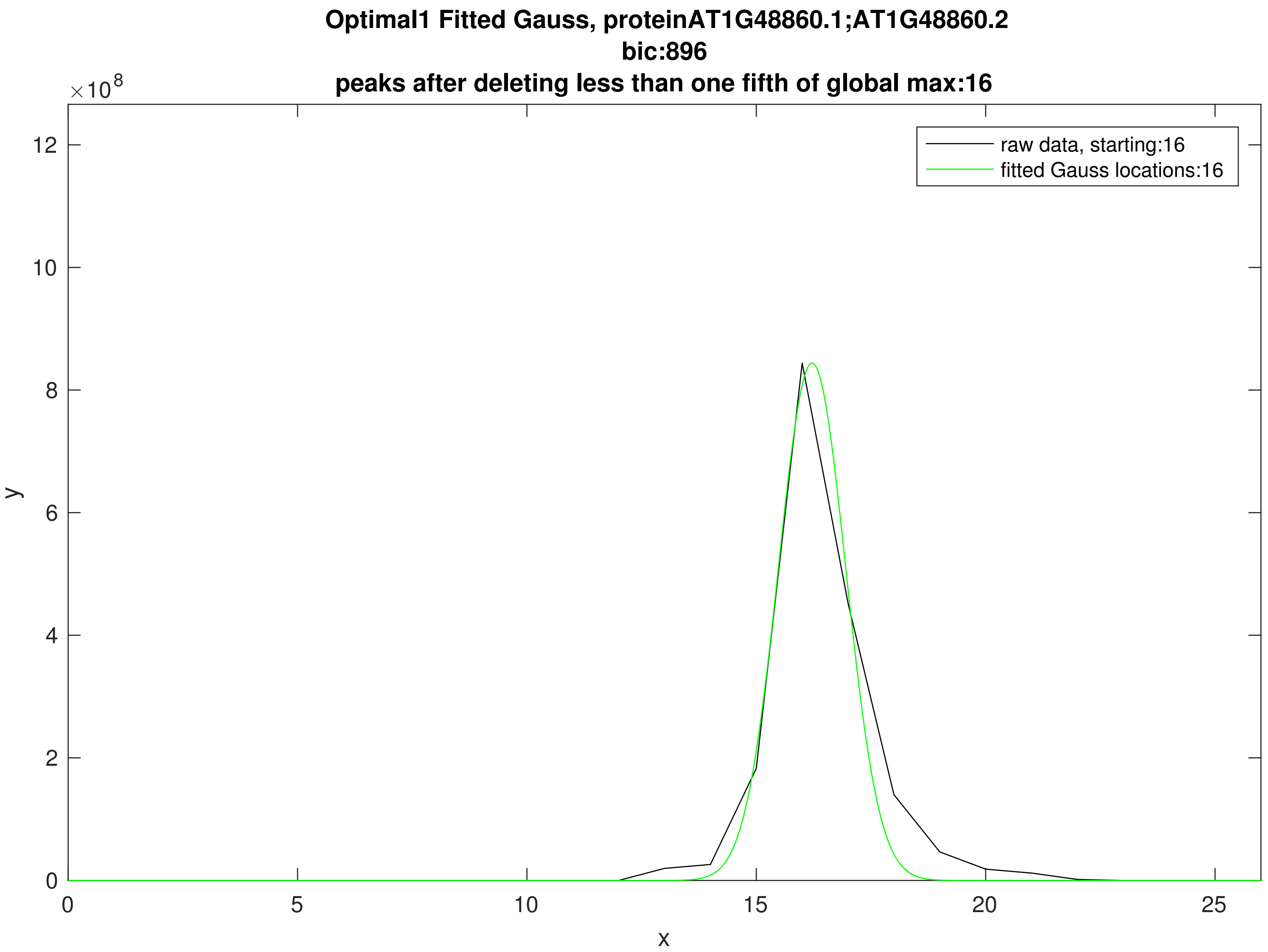
Optimal1 Fitted Gauss, proteinAT1G48860.1;AT1G48860.2
bic:896
peaks after deleting less than one fifth of global max:16
×10^8
raw data, starting:16
fitted Gauss locations:16
y
x
0
2
4
6
8
10
12
0
5
10
15
20
25

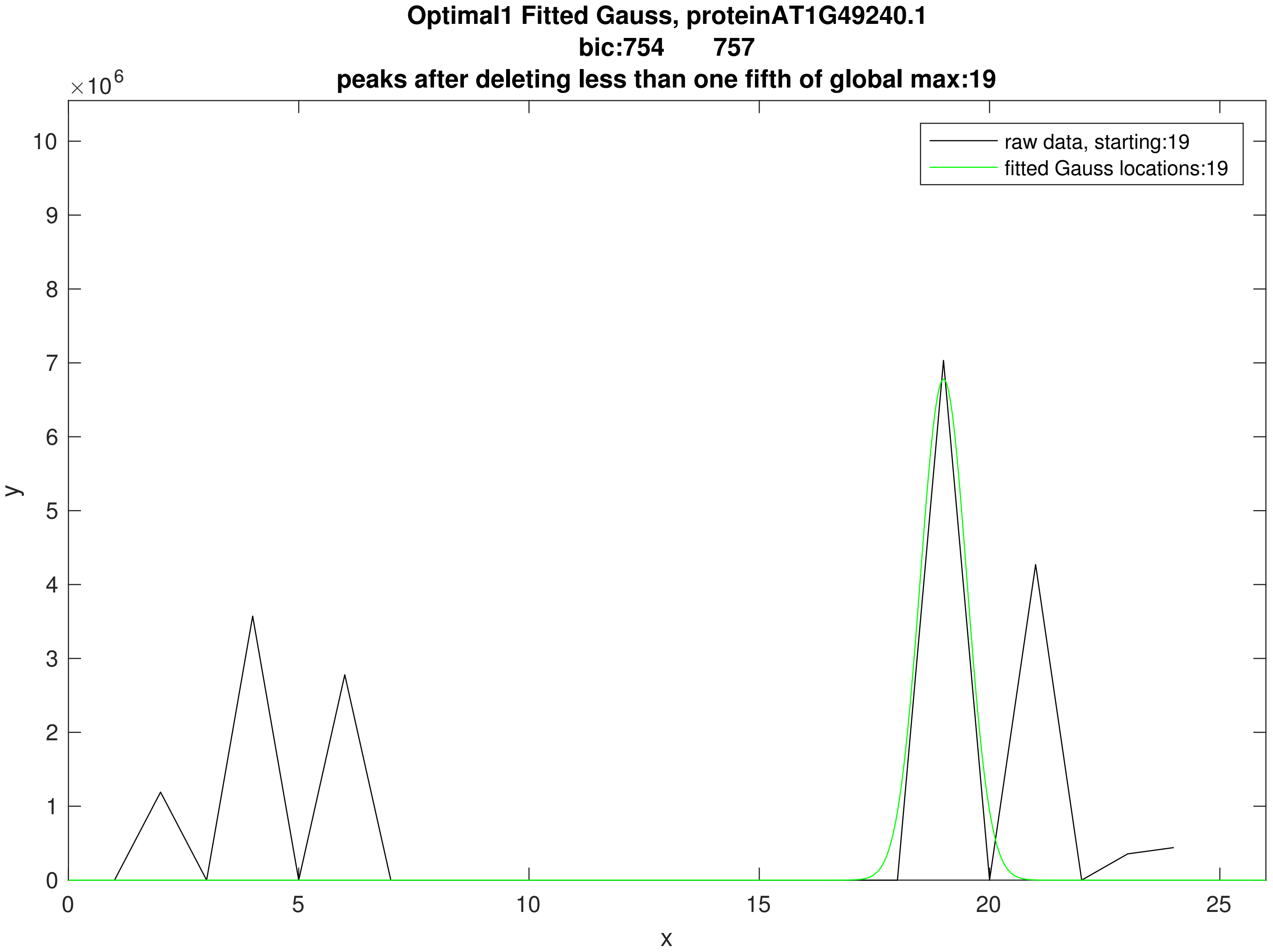
Optimal1 Fitted Gauss, proteinAT1G49240.1
bic:754 757
peaks after deleting less than one fifth of global max:19
×10^6
raw data, starting:19
fitted Gauss locations:19
y
x
0
1
2
3
4
5
6
7
8
9
10
0
5
10
15
20
25

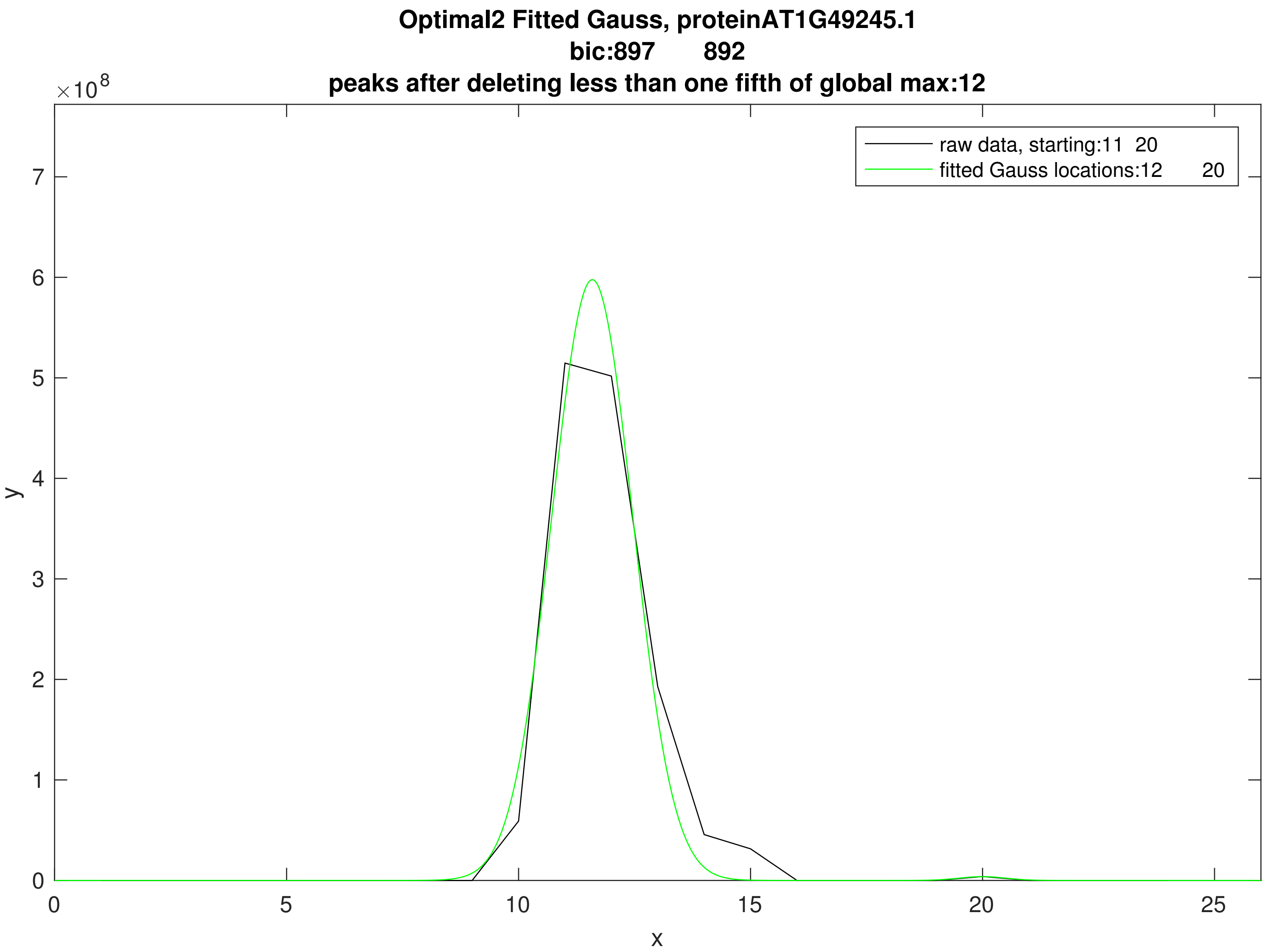

Optimal2 Fitted Gauss, proteinAT1G49245.1
bic:897 892
peaks after deleting less than one fifth of global max:12
×10^8
raw data, starting:11 20
fitted Gauss locations:12 20
y
x
0
1
2
3
4
5
6
7
0
5
10
15
20
25

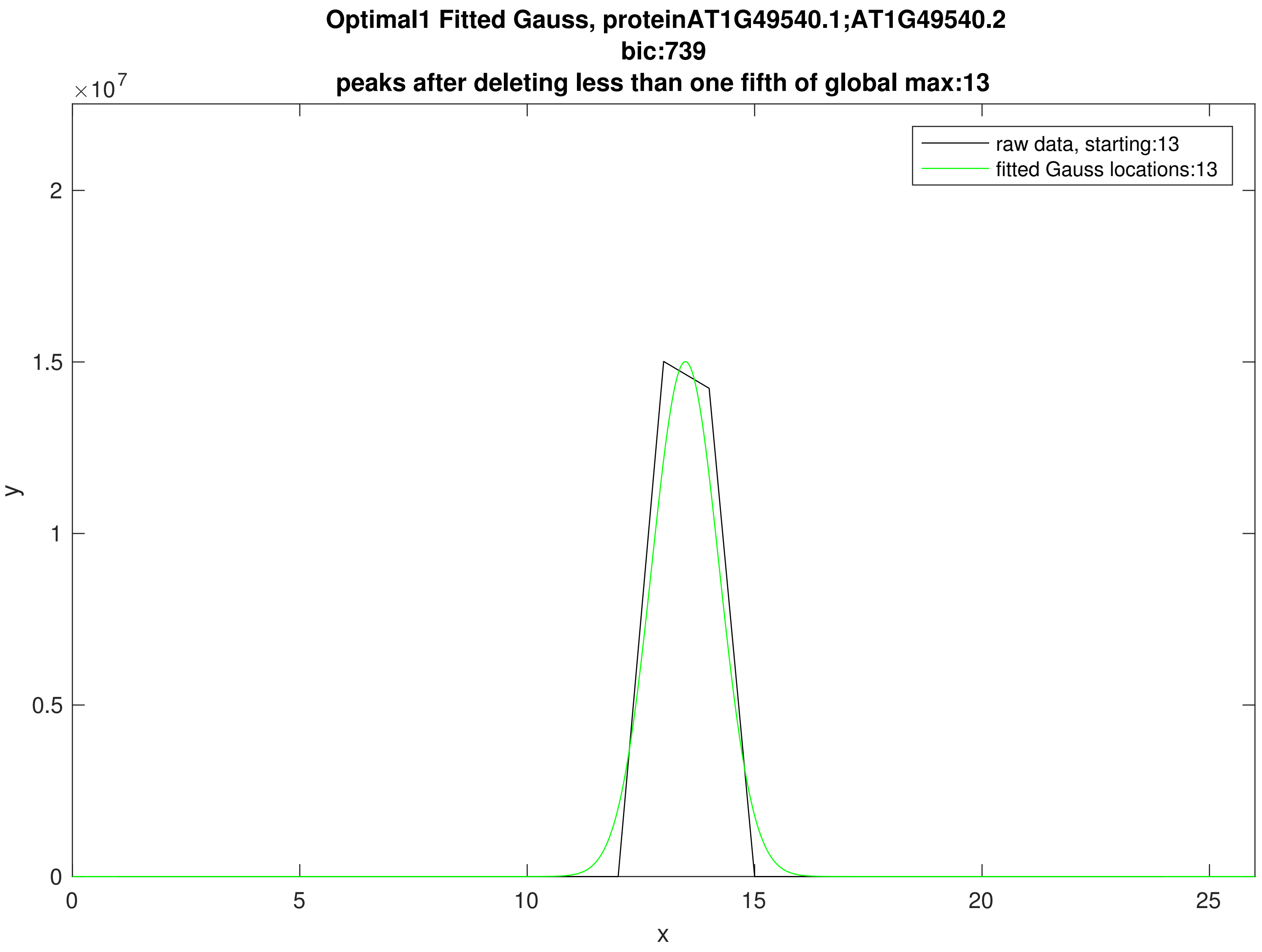
Optimal1 Fitted Gauss, proteinAT1G49540.1;AT1G49540.2
bic:739
peaks after deleting less than one fifth of global max:13
×10^7
raw data, starting:13
fitted Gauss locations:13
y
x
0
0.5
1
1.5
2
5
10
15
20
25

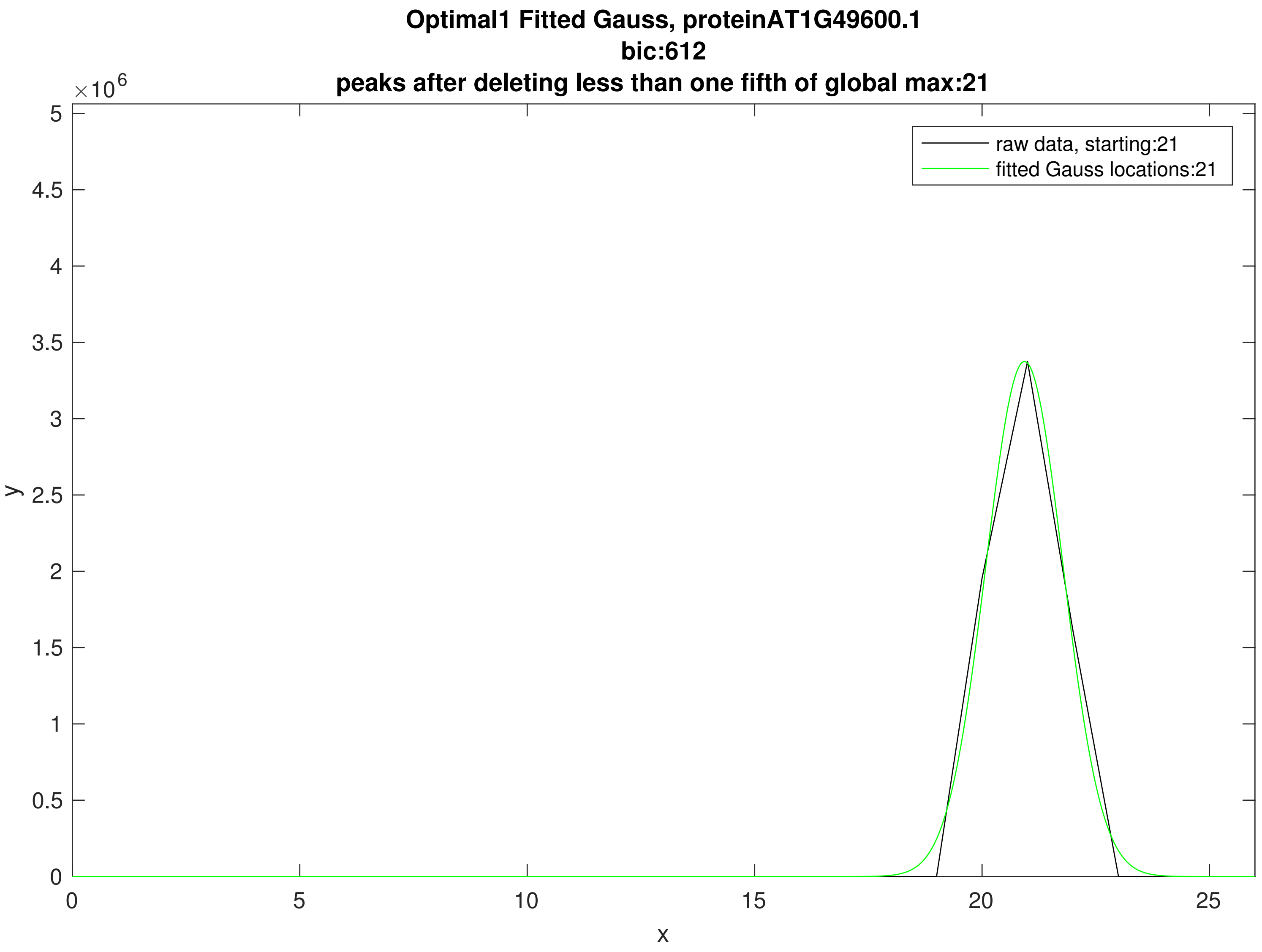
Optimal1 Fitted Gauss, proteinAT1G49600.1
bic:612
peaks after deleting less than one fifth of global max:21
×10^6
5
4.5
4
3.5
3
2.5
2
1.5
1
0.5
0
y
0
5
10
15
20
25
x
raw data, starting:21
fitted Gauss locations:21

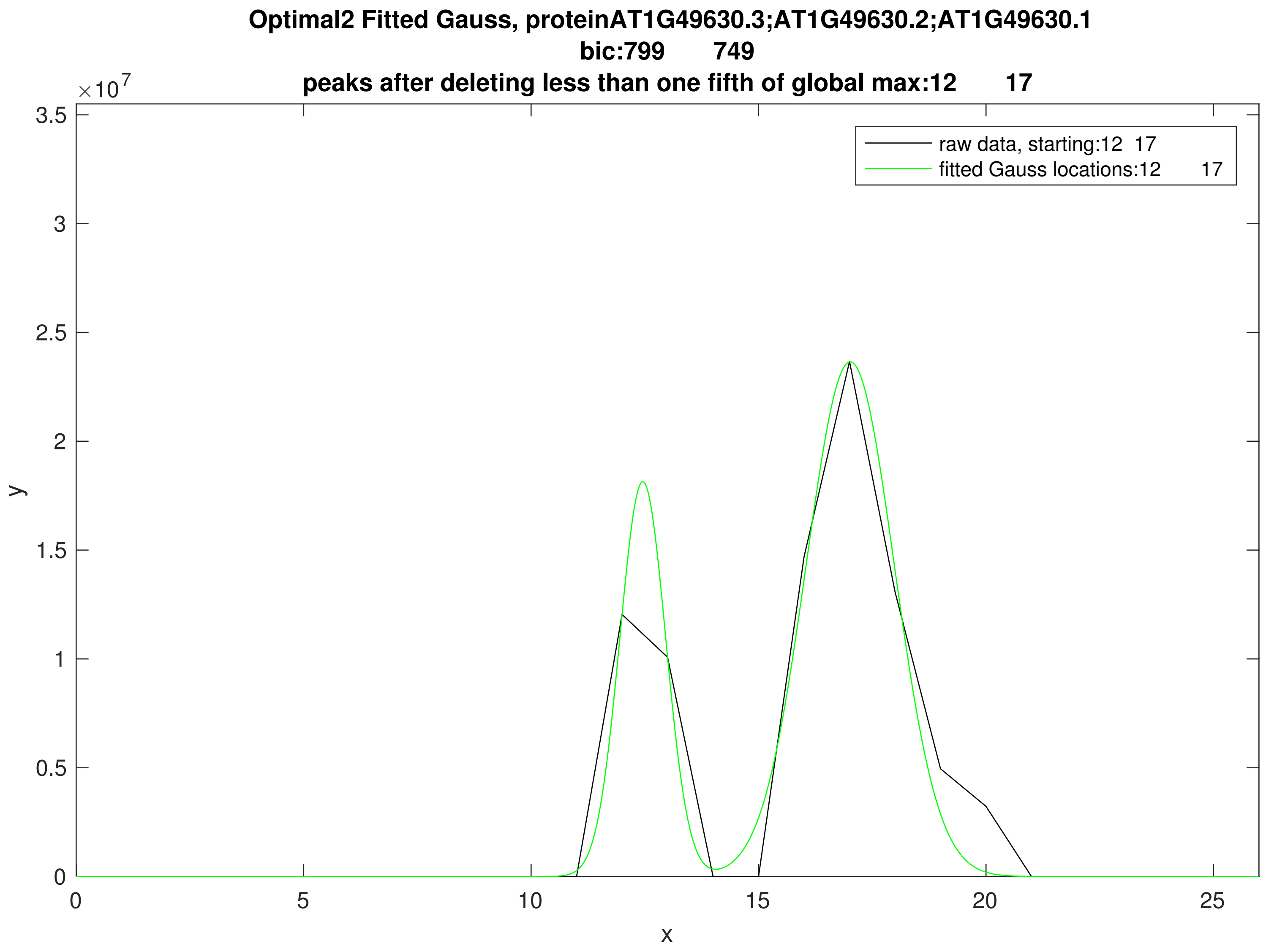
Optimal2 Fitted Gauss, proteinAT1G49630.3;AT1G49630.2;AT1G49630.1
bic:799 749
peaks after deleting less than one fifth of global max:12 17
raw data, starting:12 17
fitted Gauss locations:12 17
×10^7
y
x

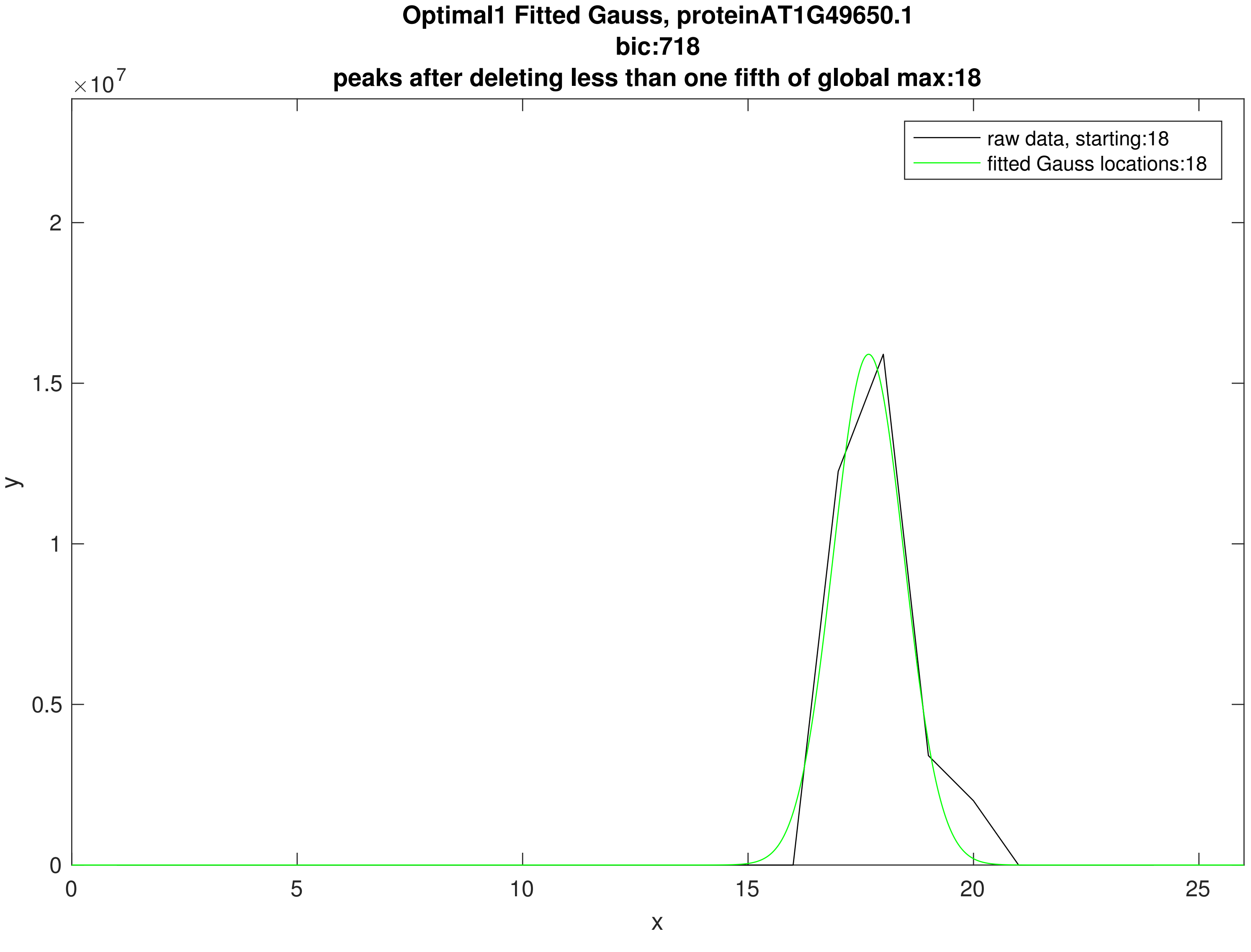
Optimal1 Fitted Gauss, proteinAT1G49650.1
bic:718
peaks after deleting less than one fifth of global max:18
×10^7
raw data, starting:18
fitted Gauss locations:18
2
1.5
1
0.5
0
y
0
5
10
15
20
25
x

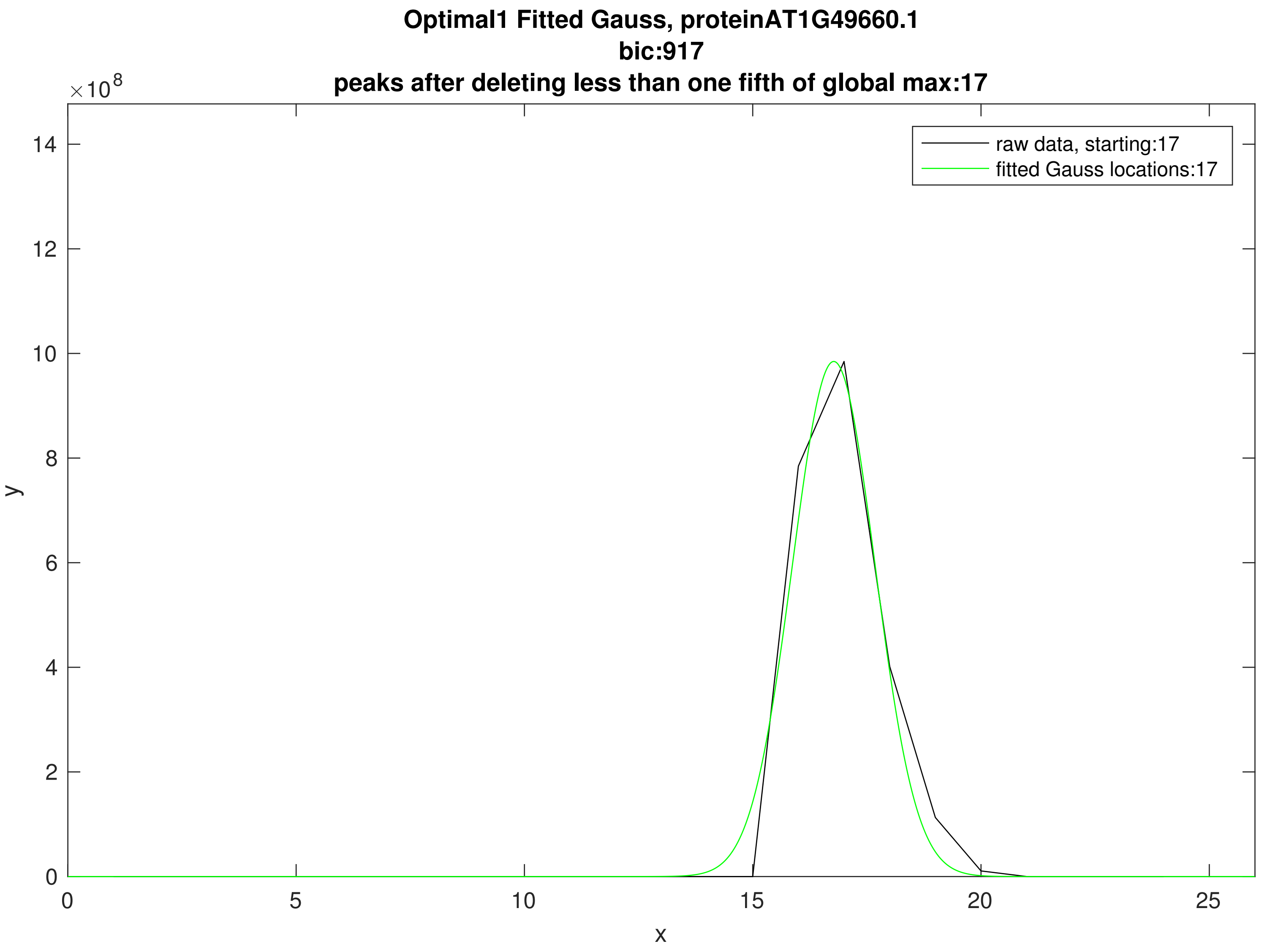
Optimal1 Fitted Gauss, proteinAT1G49660.1
bic:917
peaks after deleting less than one fifth of global max:17
×10^8
raw data, starting:17
fitted Gauss locations:17
y
x
0
2
4
6
8
10
12
14
0
5
10
15
20
25

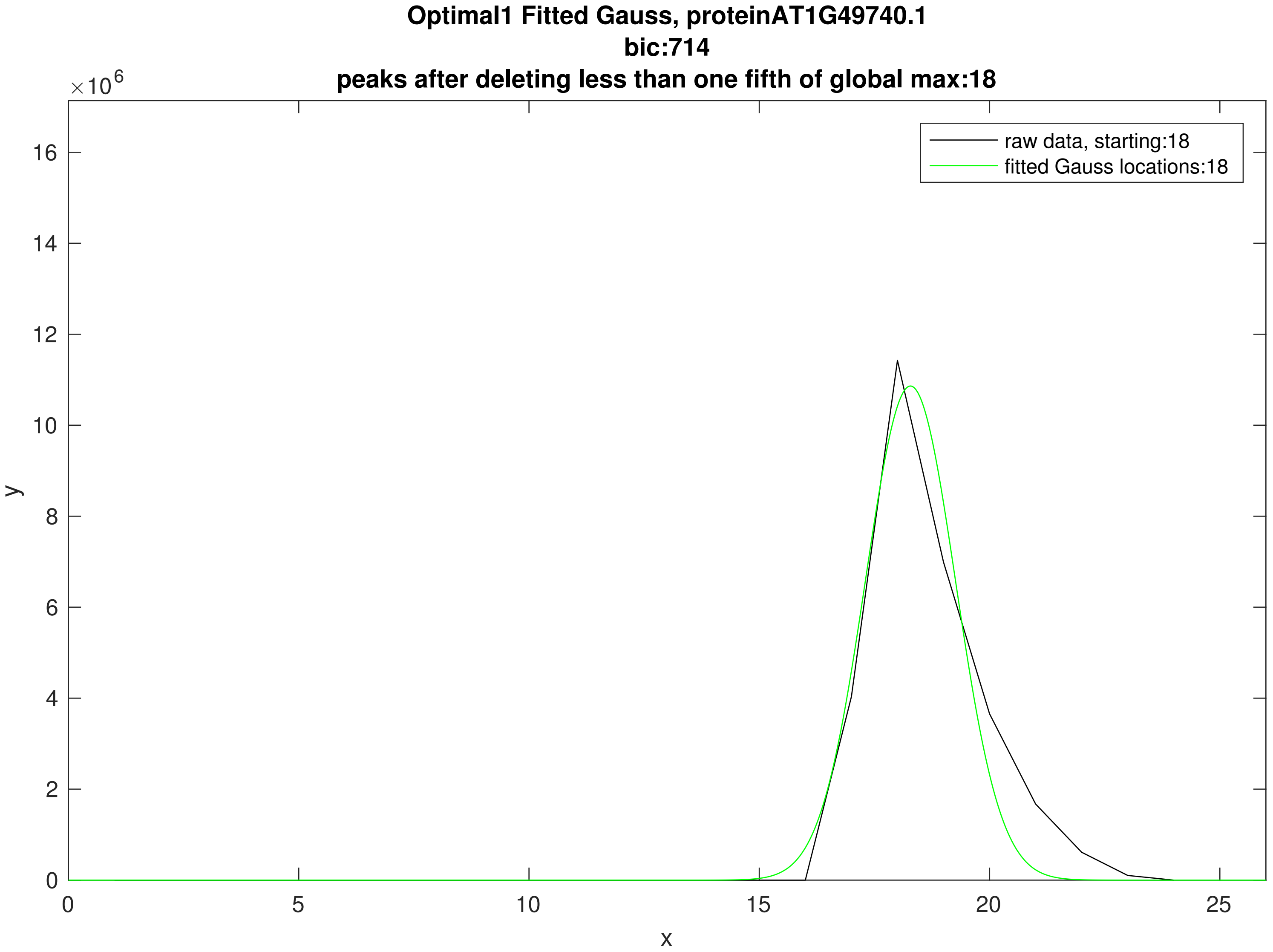
Optimal1 Fitted Gauss, proteinAT1G49740.1
bic:714
peaks after deleting less than one fifth of global max:18
×10^6
raw data, starting:18
fitted Gauss locations:18
y
x
0
2
4
6
8
10
12
14
16
0
5
10
15
20
25

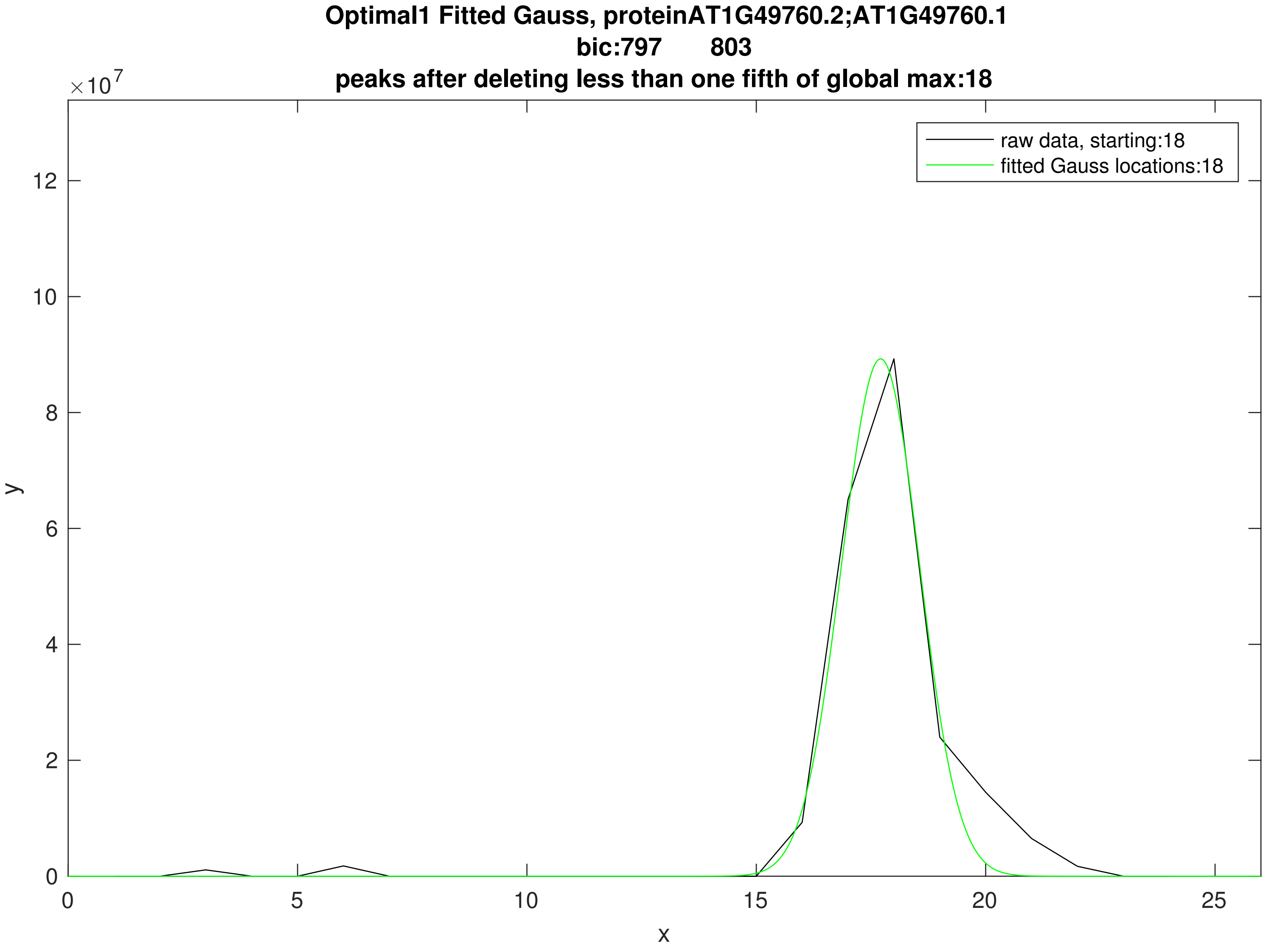
Optimal1 Fitted Gauss, proteinAT1G49760.2;AT1G49760.1
bic:797 803
peaks after deleting less than one fifth of global max:18
×10^7
raw data, starting:18
fitted Gauss locations:18
y
x
0
2
4
6
8
10
12
0
5
10
15
20
25

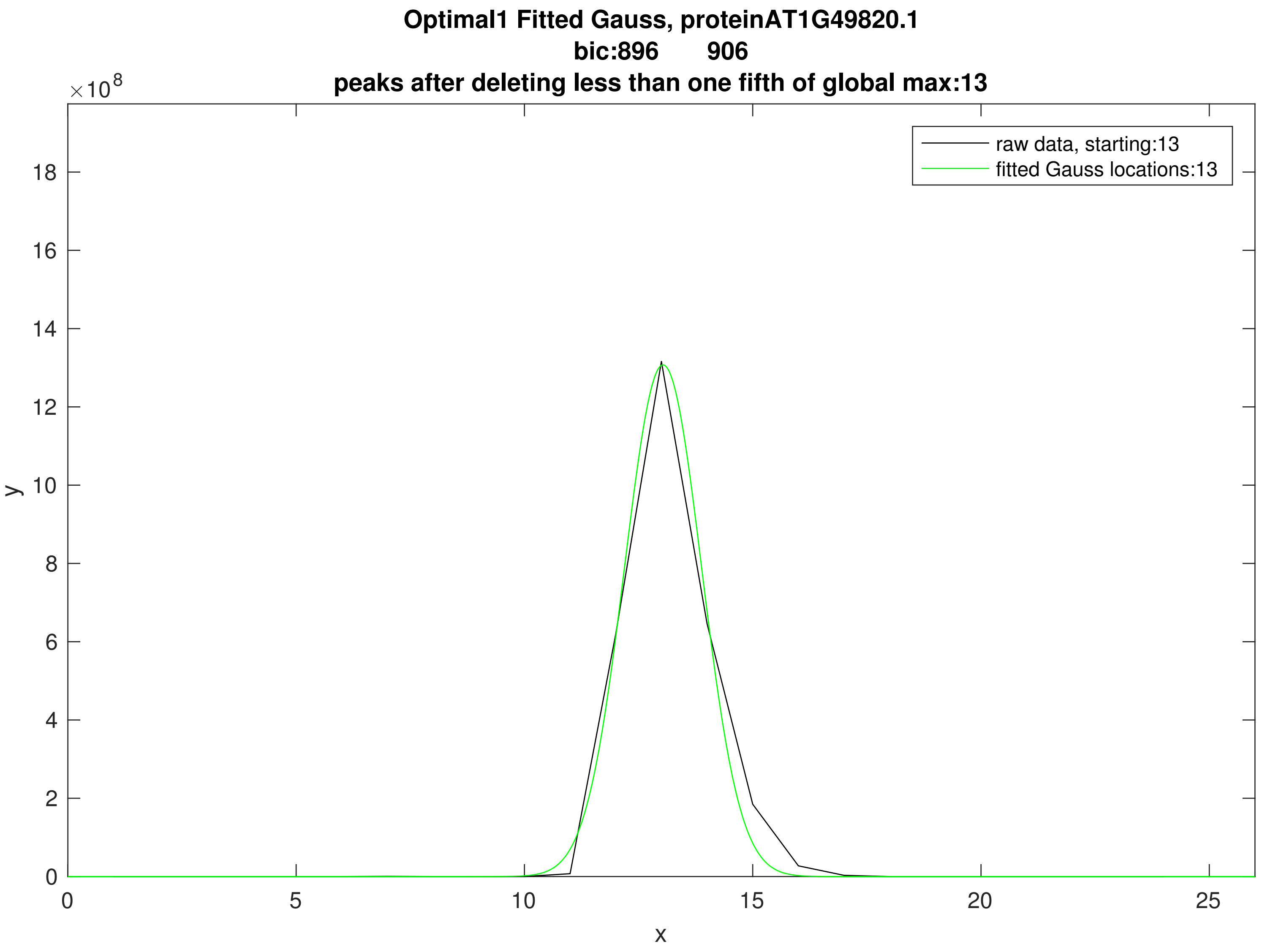
Optimal1 Fitted Gauss, proteinAT1G49820.1
bic:896 906
peaks after deleting less than one fifth of global max:13
×10^8
raw data, starting:13
fitted Gauss locations:13
y
x
0
2
4
6
8
10
12
14
16
18
0
5
10
15
20
25

**Optimal1 Fitted Gauss, proteinAT1G49970.1**
**bic:814 823**
**peaks after deleting less than one fifth of global max:8.8**

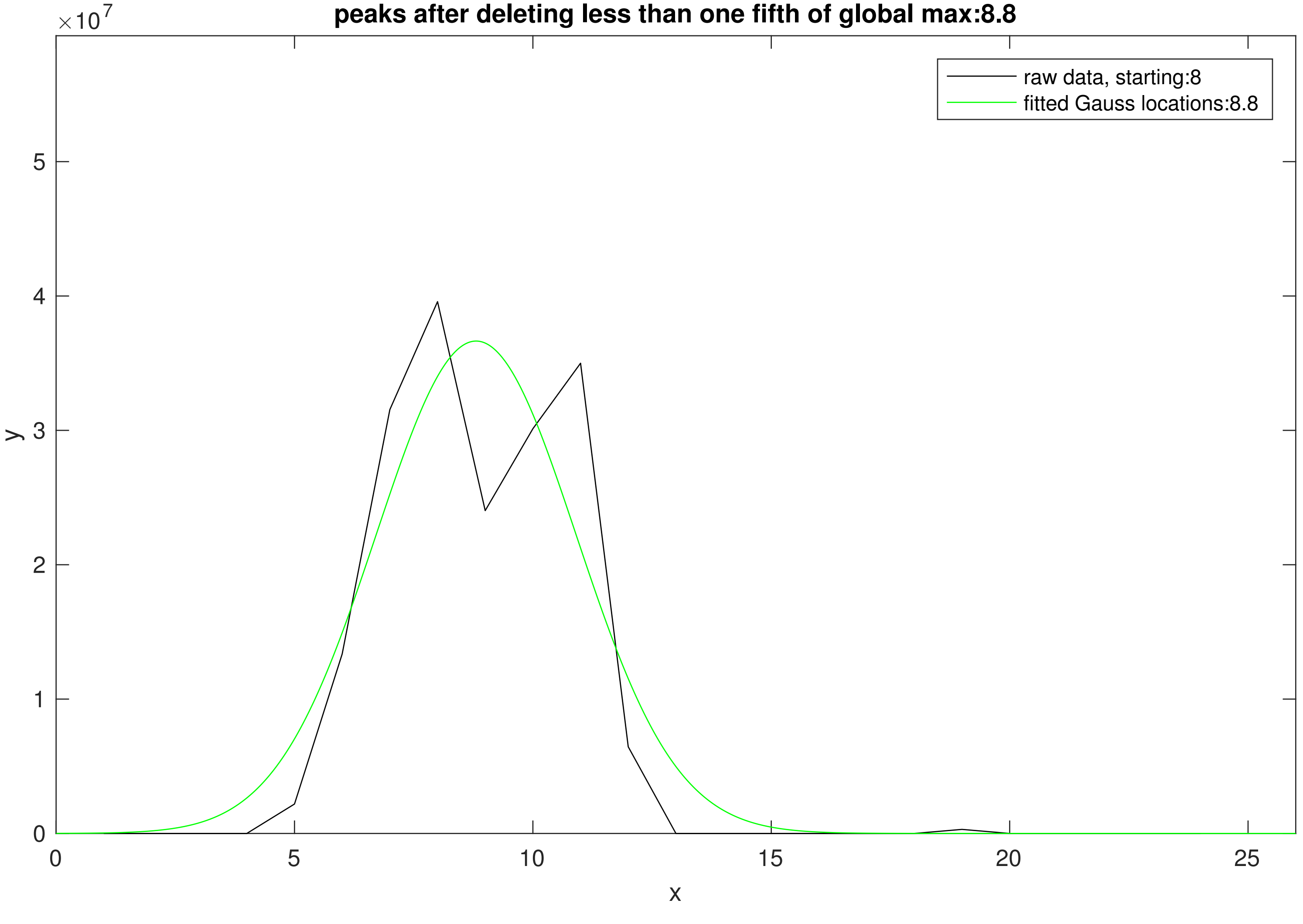

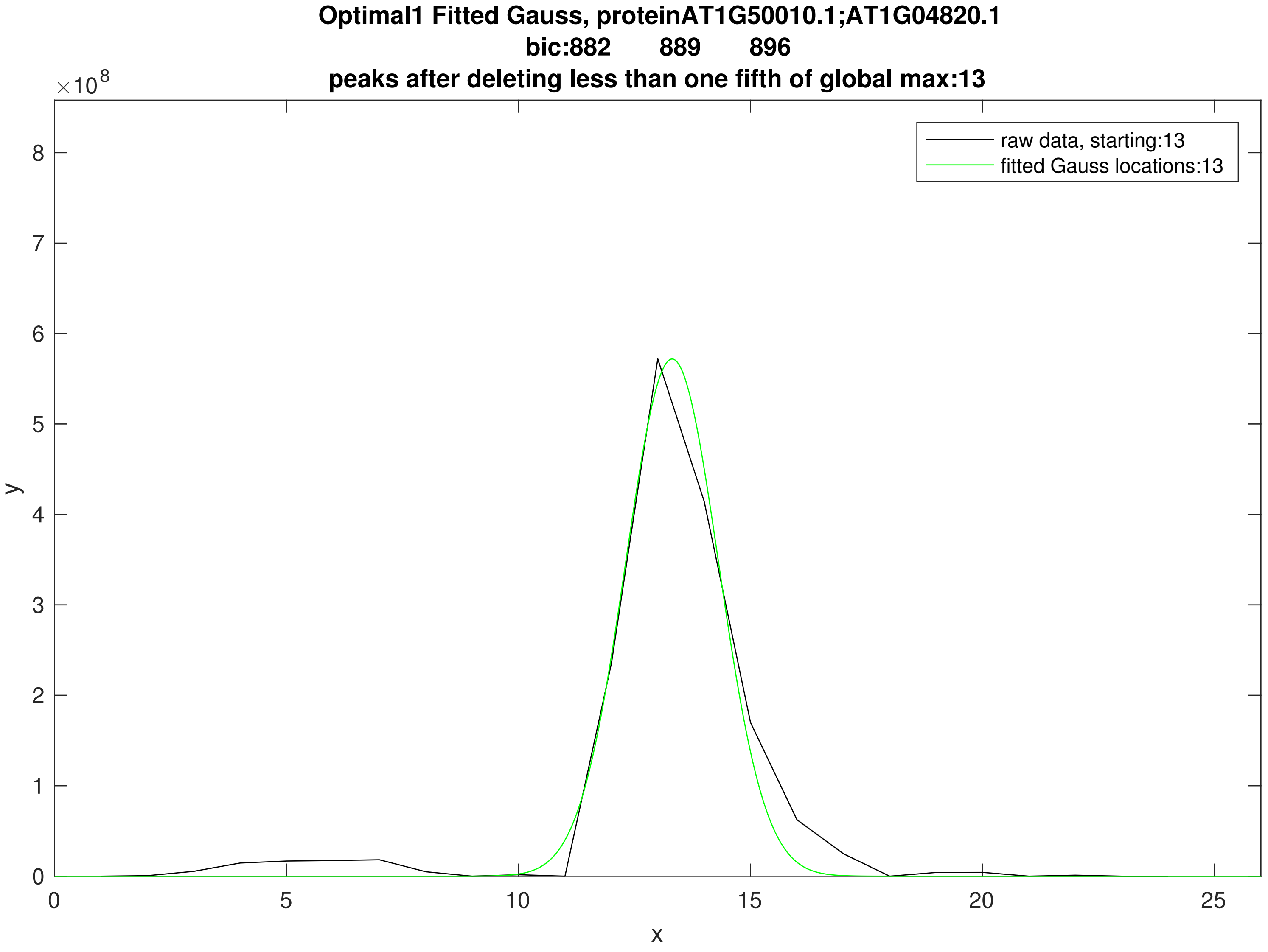
Optimal1 Fitted Gauss, proteinAT1G50010.1;AT1G04820.1
bic:882 889 896
peaks after deleting less than one fifth of global max:13
×10^8
raw data, starting:13
fitted Gauss locations:13
y
x
0
1
2
3
4
5
6
7
8
0
5
10
15
20
25

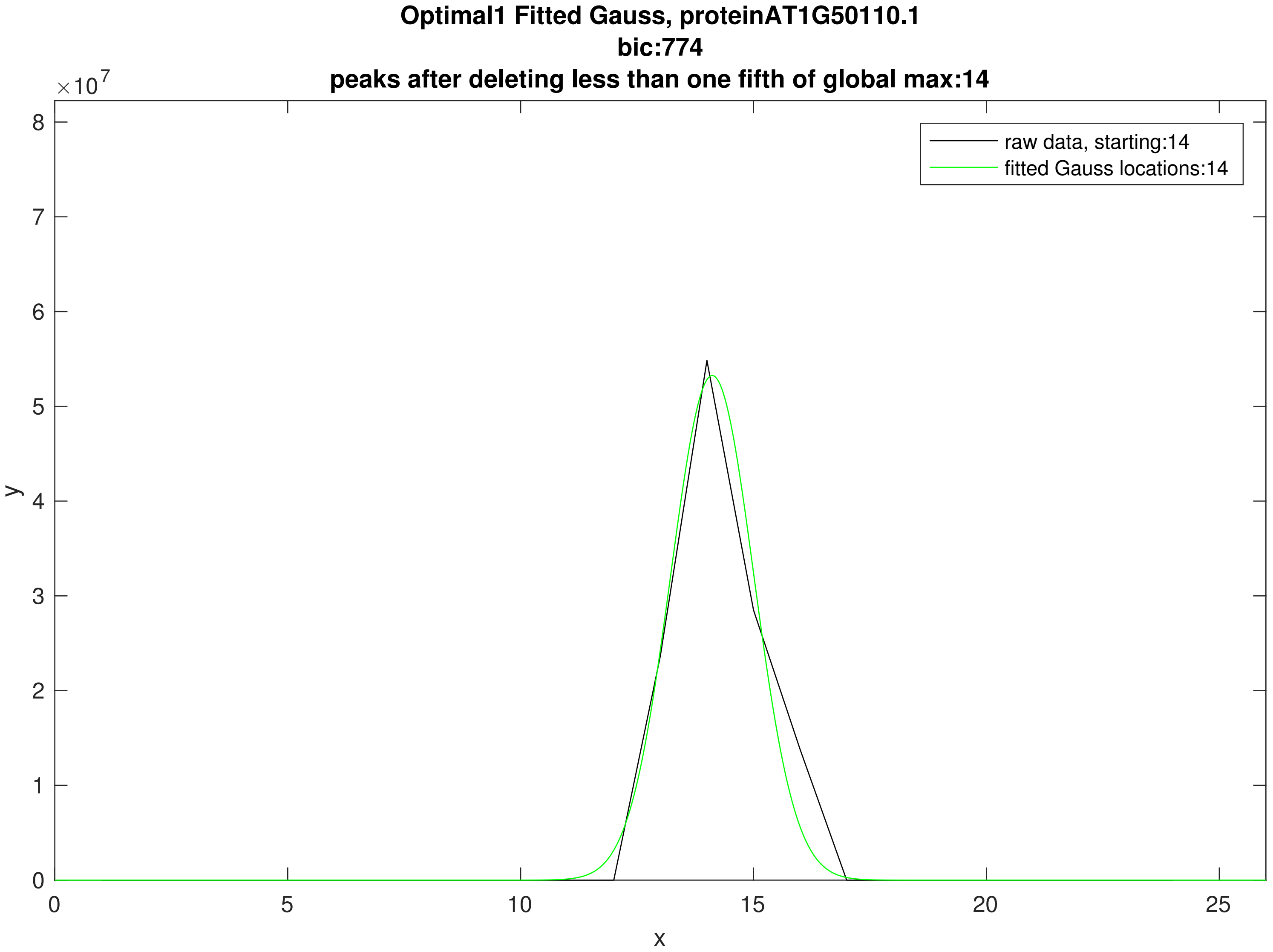

Optimal1 Fitted Gauss, proteinAT1G50110.1
bic:774
peaks after deleting less than one fifth of global max:14
×10^7
raw data, starting:14
fitted Gauss locations:14
y
x
0
1
2
3
4
5
6
7
8
0
5
10
15
20
25

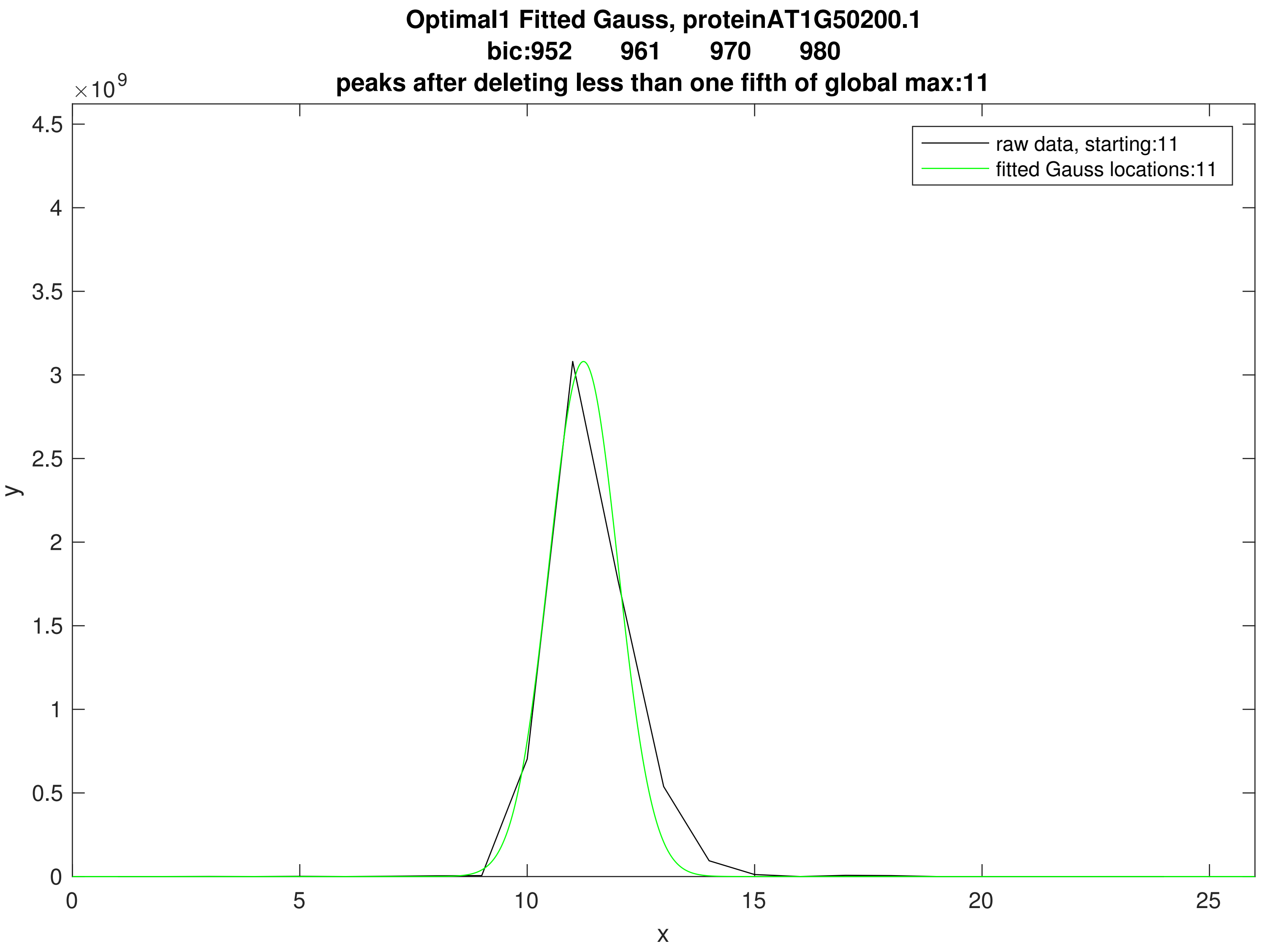

Optimal1 Fitted Gauss, proteinAT1G50200.1
bic:952 961 970 980
peaks after deleting less than one fifth of global max:11
×10^9
4.5
4
3.5
3
2.5
2
1.5
1
0.5
0
y
0
5
10
15
20
25
x
raw data, starting:11
fitted Gauss locations:11

Optimal1 Fitted Gauss, proteinAT1G50200.2
bic:652
peaks after deleting less than one fifth of global max:11

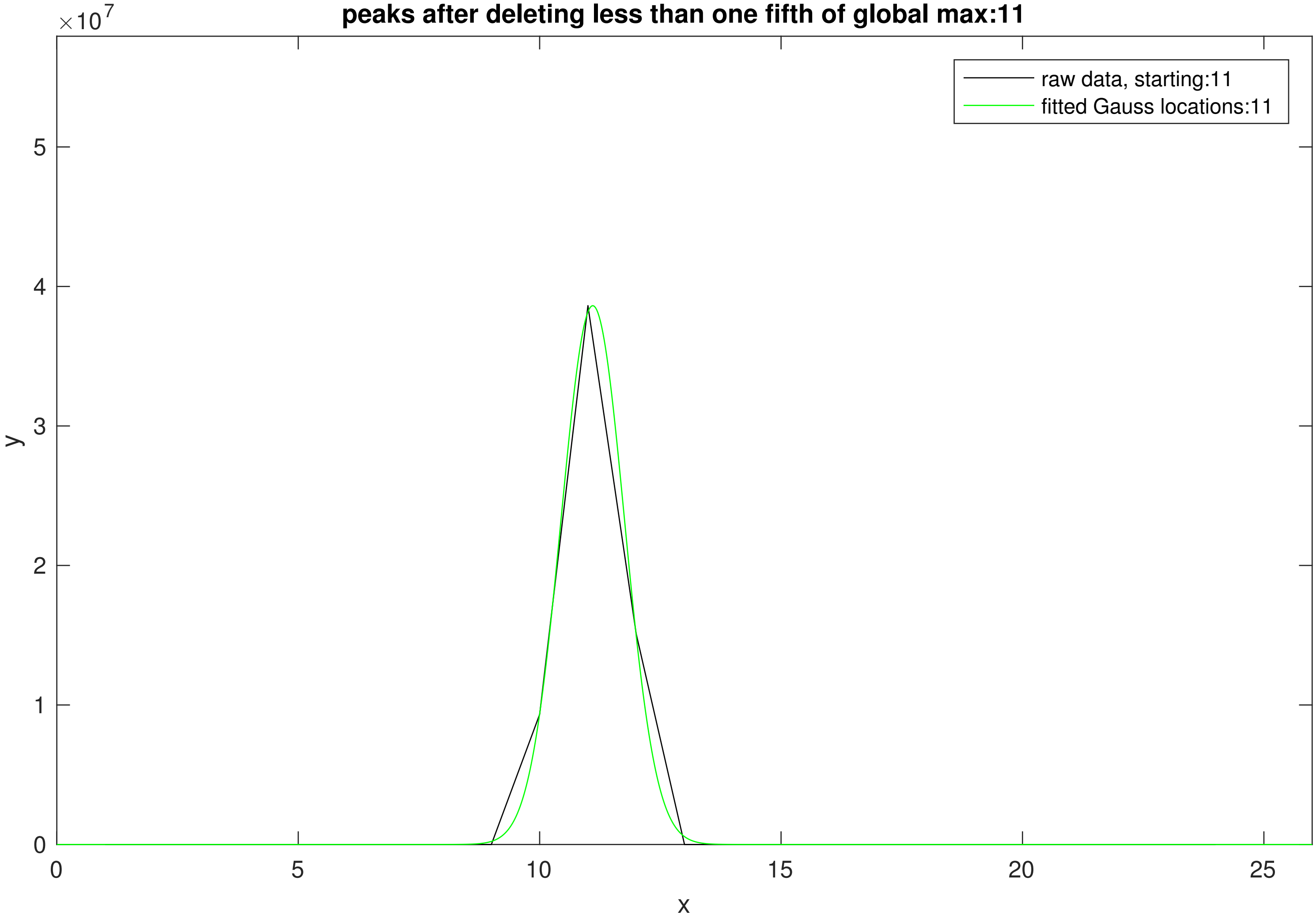

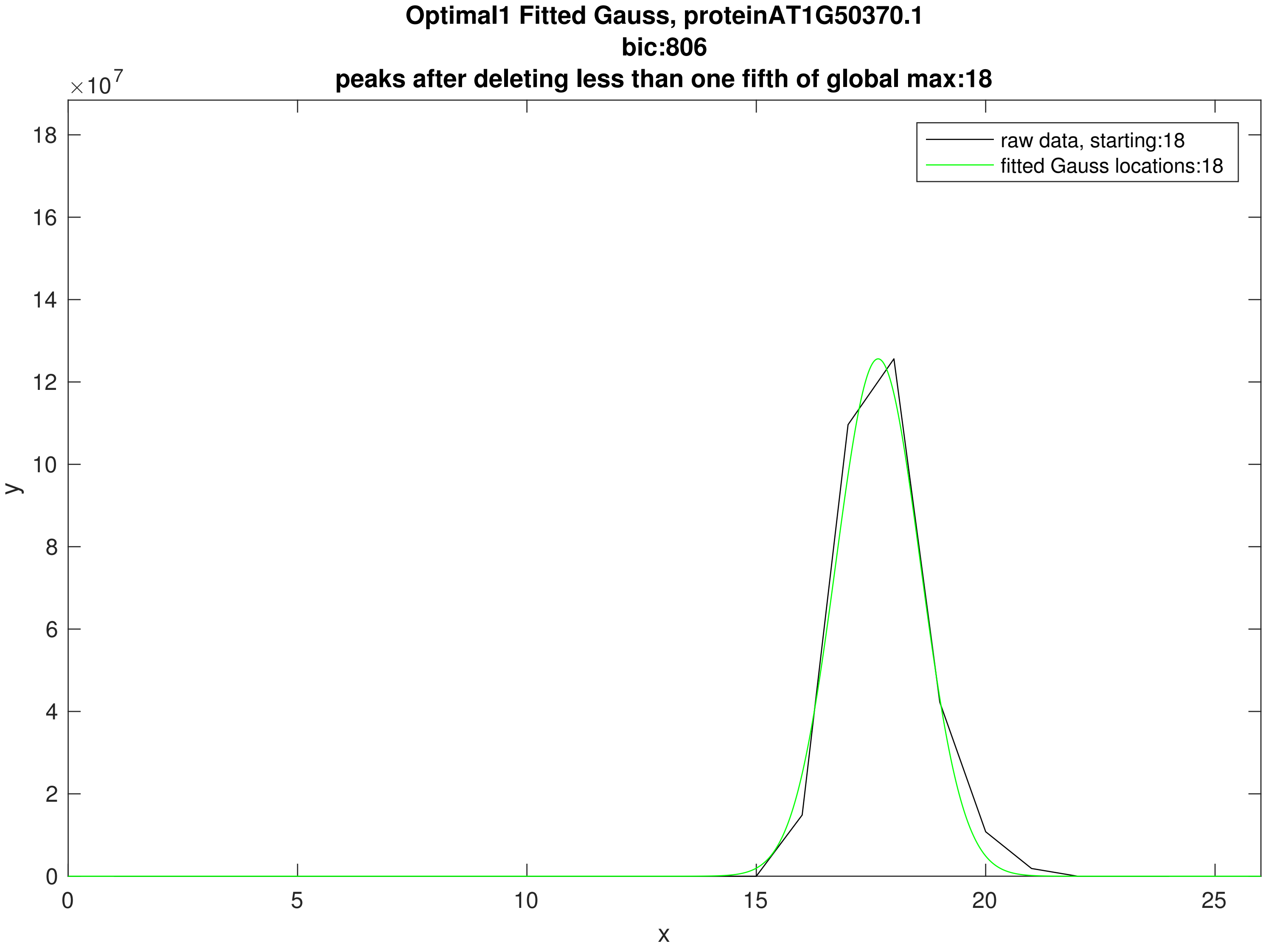
Optimal1 Fitted Gauss, proteinAT1G50370.1
bic:806
peaks after deleting less than one fifth of global max:18
×10^7
18
16
14
12
10
8
6
4
2
0
y
0
5
10
15
20
25
x
raw data, starting:18
fitted Gauss locations:18

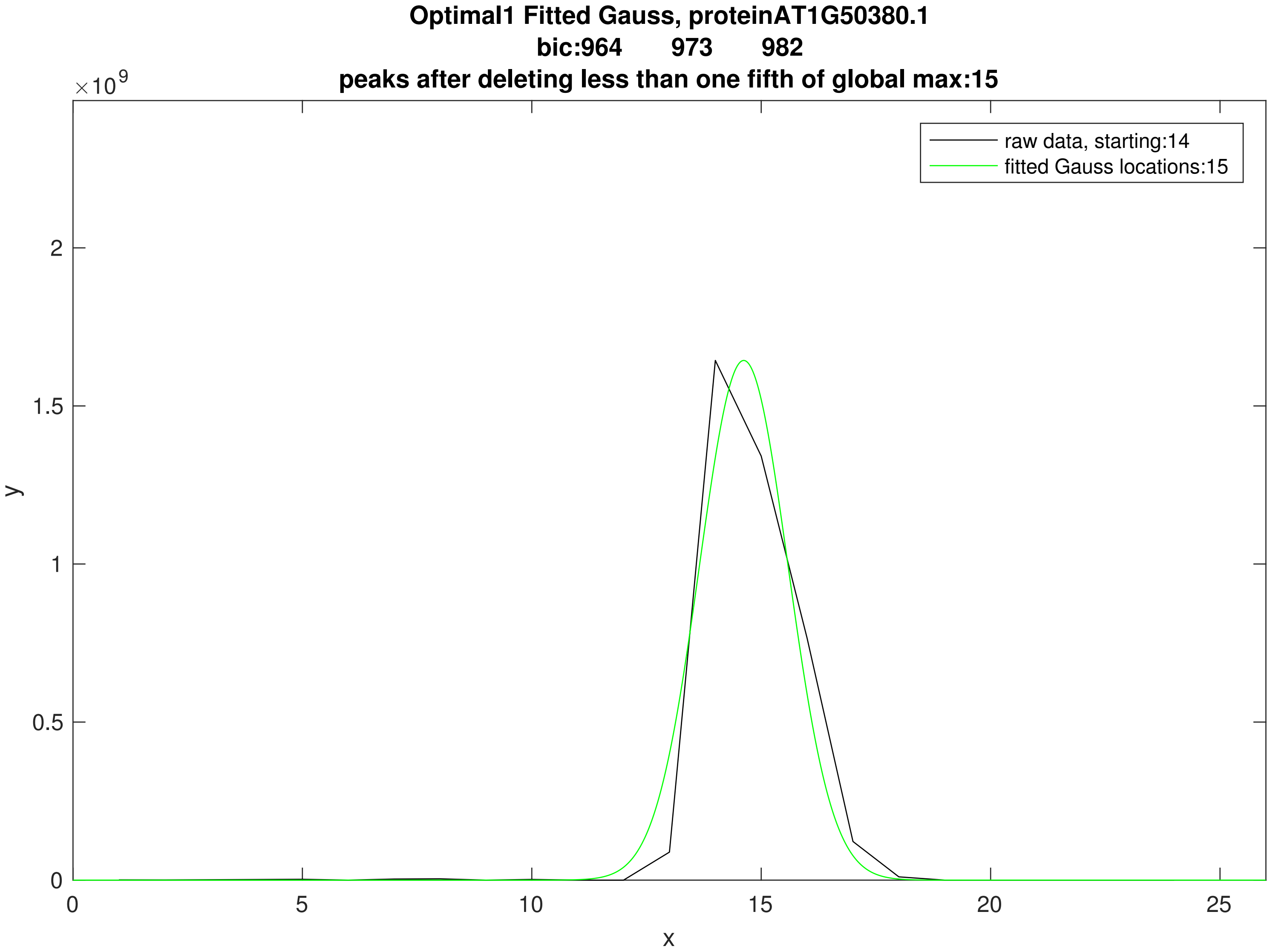

Optimal1 Fitted Gauss, proteinAT1G50380.1
bic:964 973 982
peaks after deleting less than one fifth of global max:15
×10^9
raw data, starting:14
fitted Gauss locations:15
y
x
0
0.5
1
1.5
2
0
5
10
15
20
25

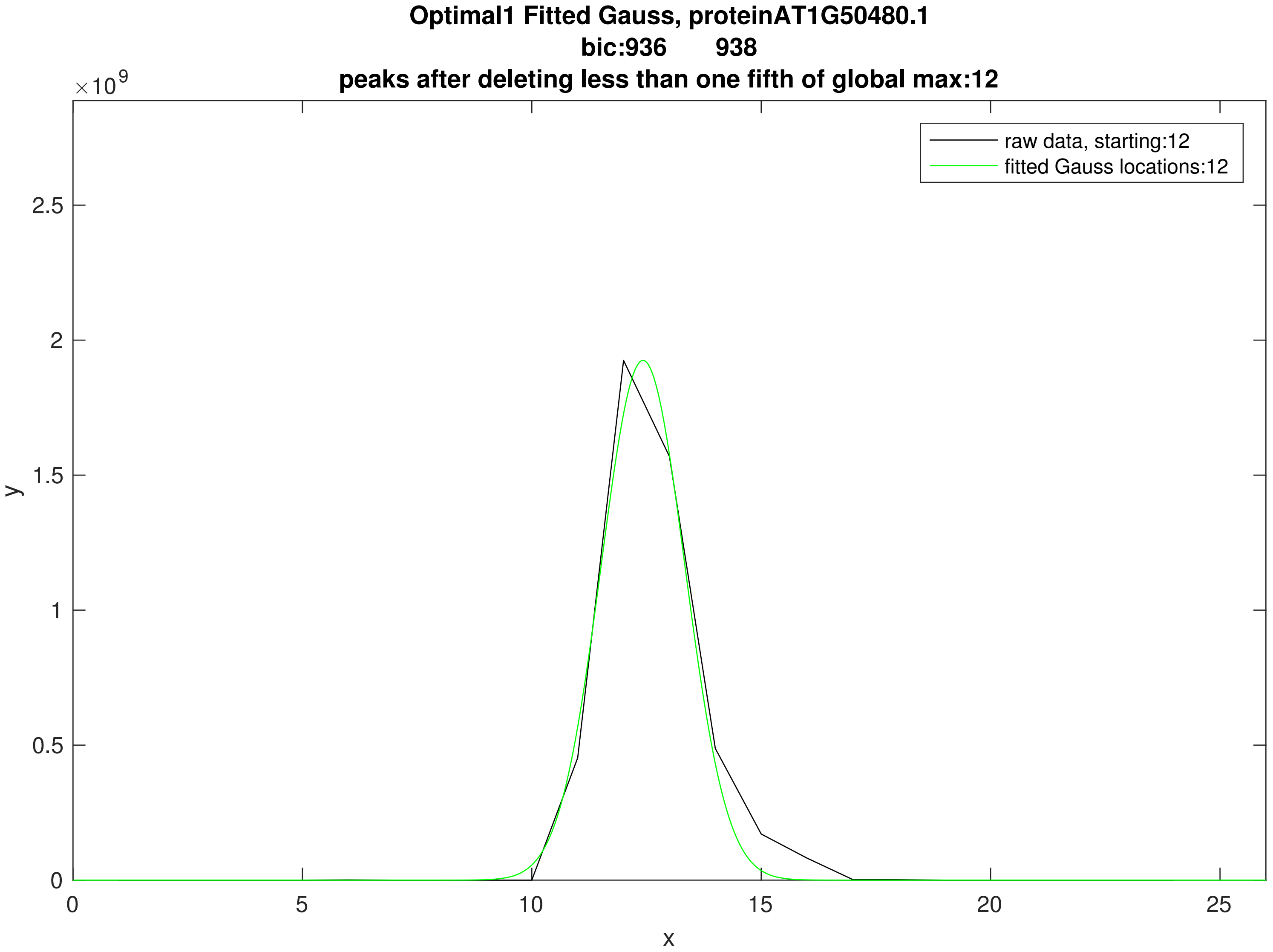
Optimal1 Fitted Gauss, proteinAT1G50480.1
bic:936 938
peaks after deleting less than one fifth of global max:12
×10^9
raw data, starting:12
fitted Gauss locations:12
2.5
2
1.5
1
0.5
0
y
0
5
10
15
20
25
x

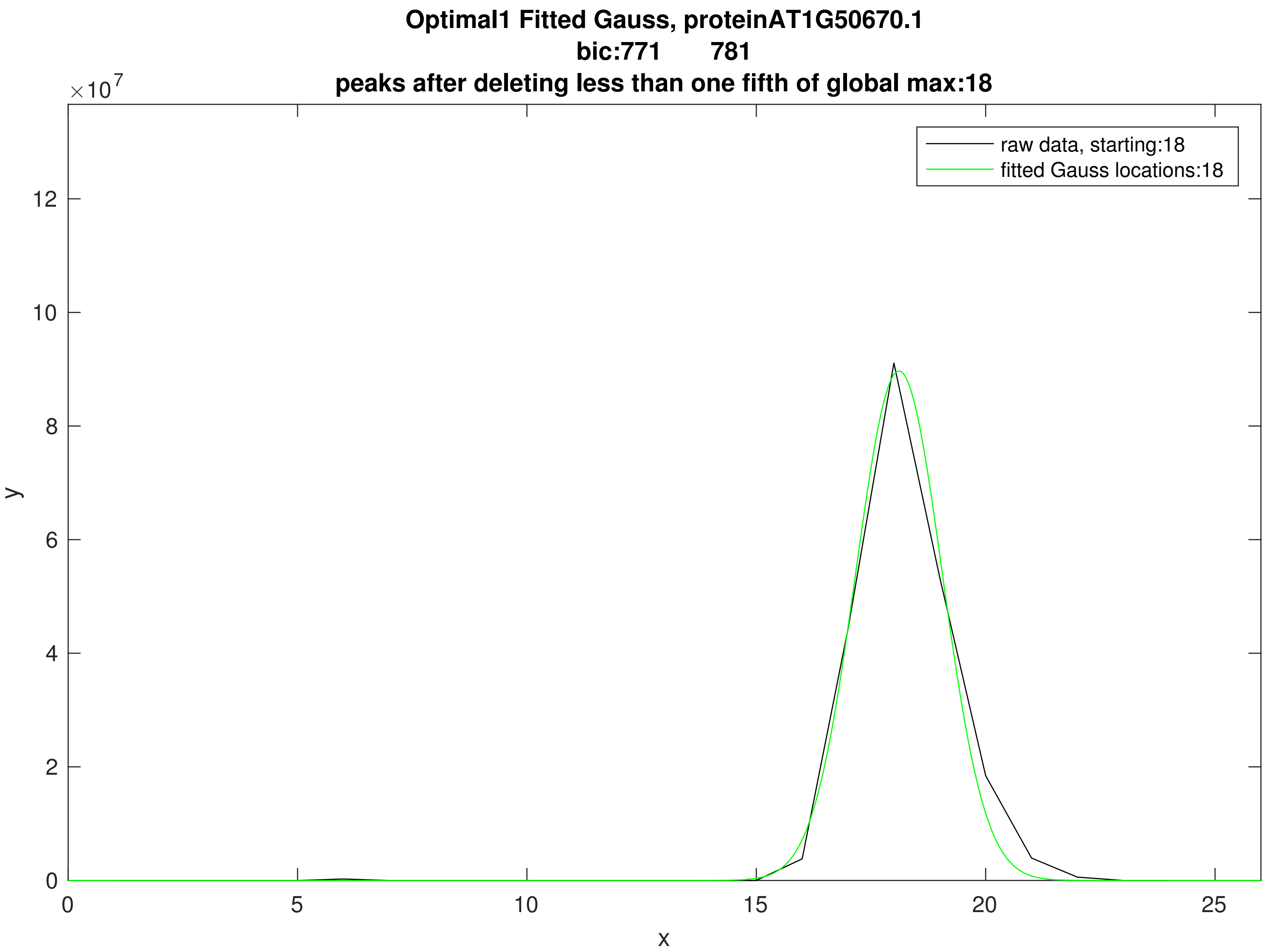
Optimal1 Fitted Gauss, proteinAT1G50670.1
bic:771 781
peaks after deleting less than one fifth of global max:18
×10^7
raw data, starting:18
fitted Gauss locations:18
12
10
8
6
4
2
0
y
0
5
10
15
20
25
x

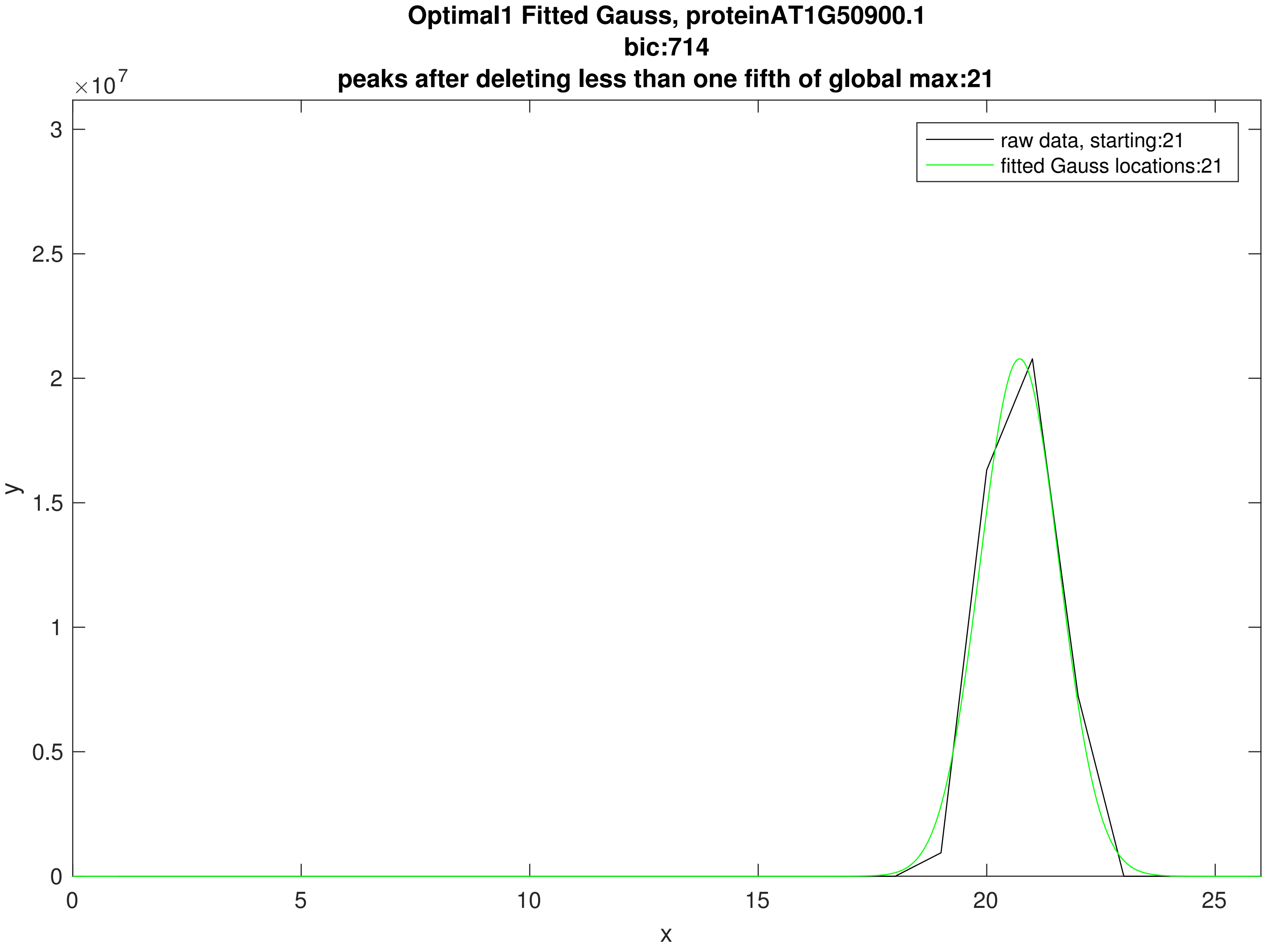
Optimal1 Fitted Gauss, proteinAT1G50900.1
bic:714
peaks after deleting less than one fifth of global max:21
×10^7
3
2.5
2
1.5
1
0.5
0
y
0
5
10
15
20
25
x
raw data, starting:21
fitted Gauss locations:21

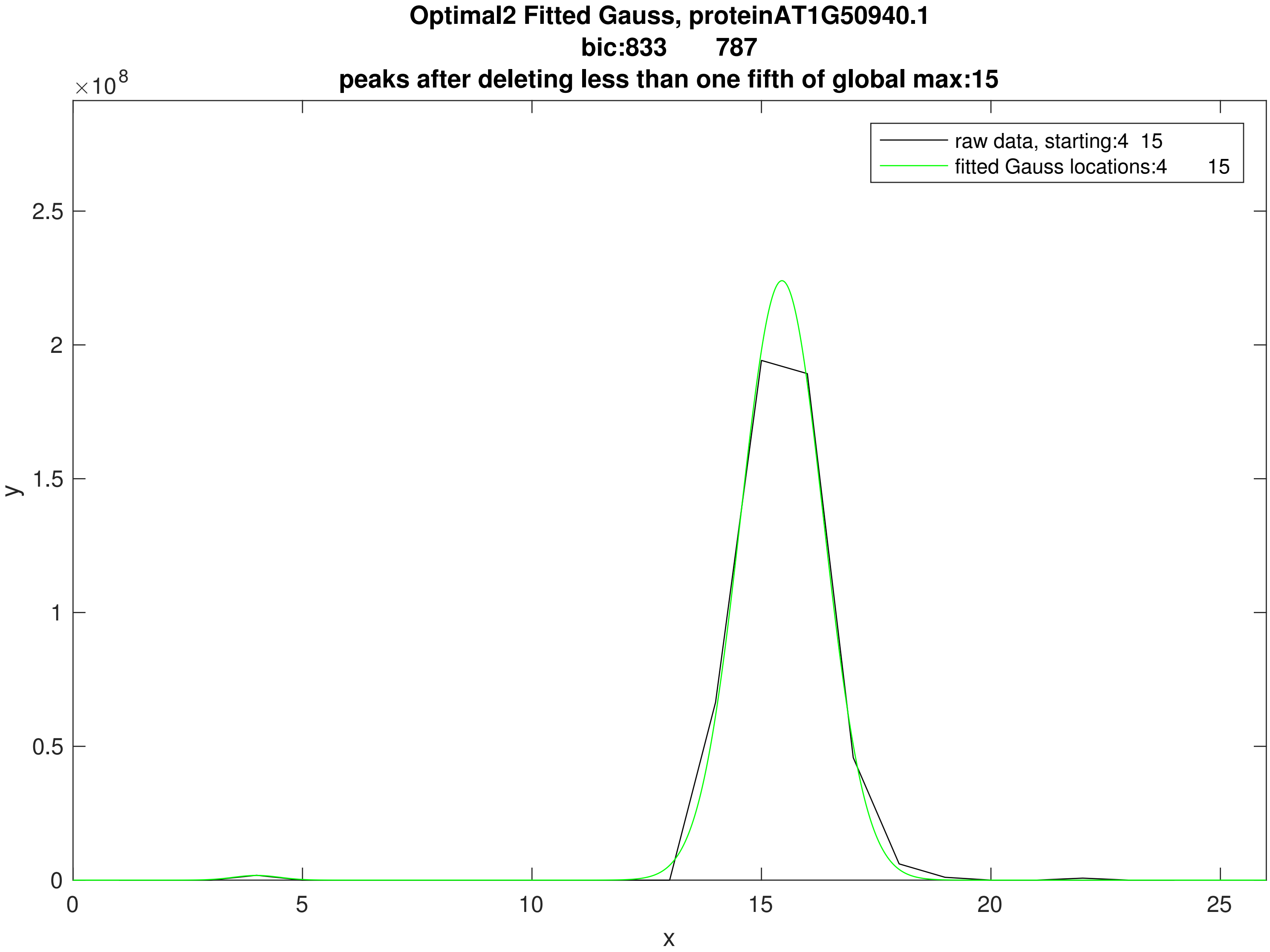
Optimal2 Fitted Gauss, proteinAT1G50940.1
bic:833 787
peaks after deleting less than one fifth of global max:15
×10^8
raw data, starting:4 15
fitted Gauss locations:4 15
2.5
2
1.5
1
0.5
0
y
0
5
10
15
20
25
x

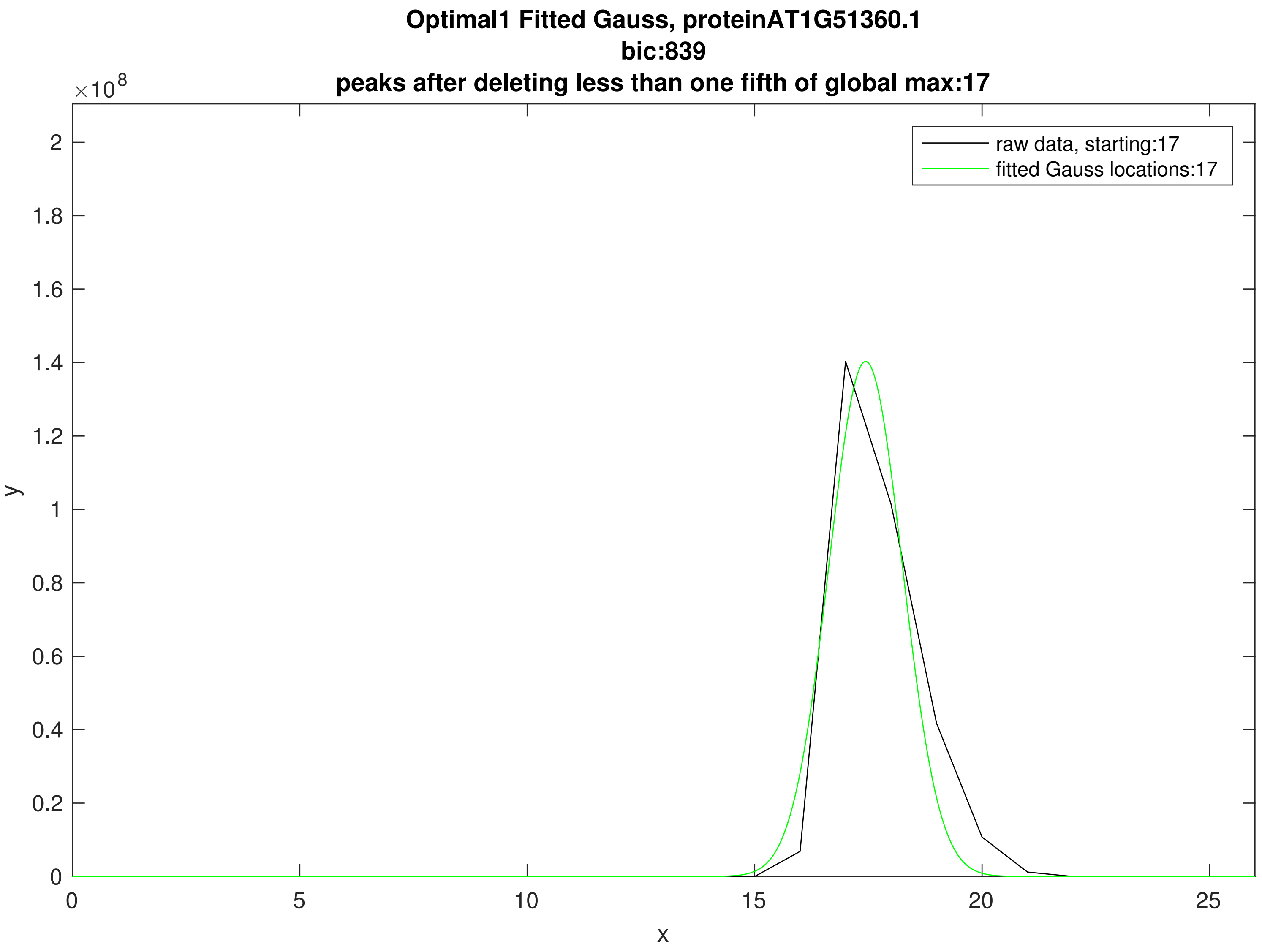
Optimal1 Fitted Gauss, proteinAT1G51360.1
bic:839
peaks after deleting less than one fifth of global max:17
×10^8
raw data, starting:17
fitted Gauss locations:17
2
1.8
1.6
1.4
1.2
1
0.8
0.6
0.4
0.2
0
y
0
5
10
15
20
25
x

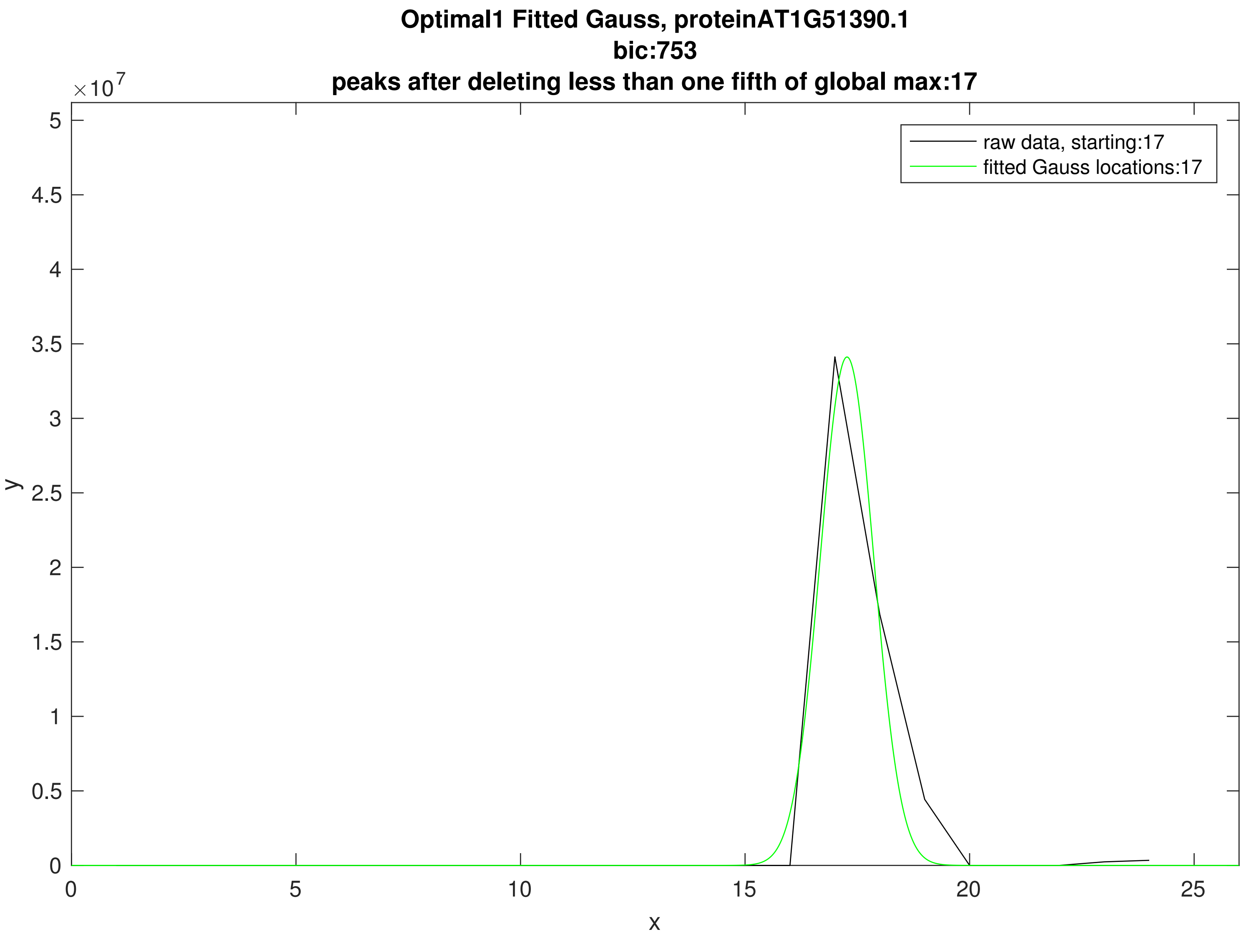
Optimal1 Fitted Gauss, proteinAT1G51390.1
bic:753
peaks after deleting less than one fifth of global max:17
×10^7
y
x
raw data, starting:17
fitted Gauss locations:17
0
0.5
1
1.5
2
2.5
3
3.5
4
4.5
5
0
5
10
15
20
25

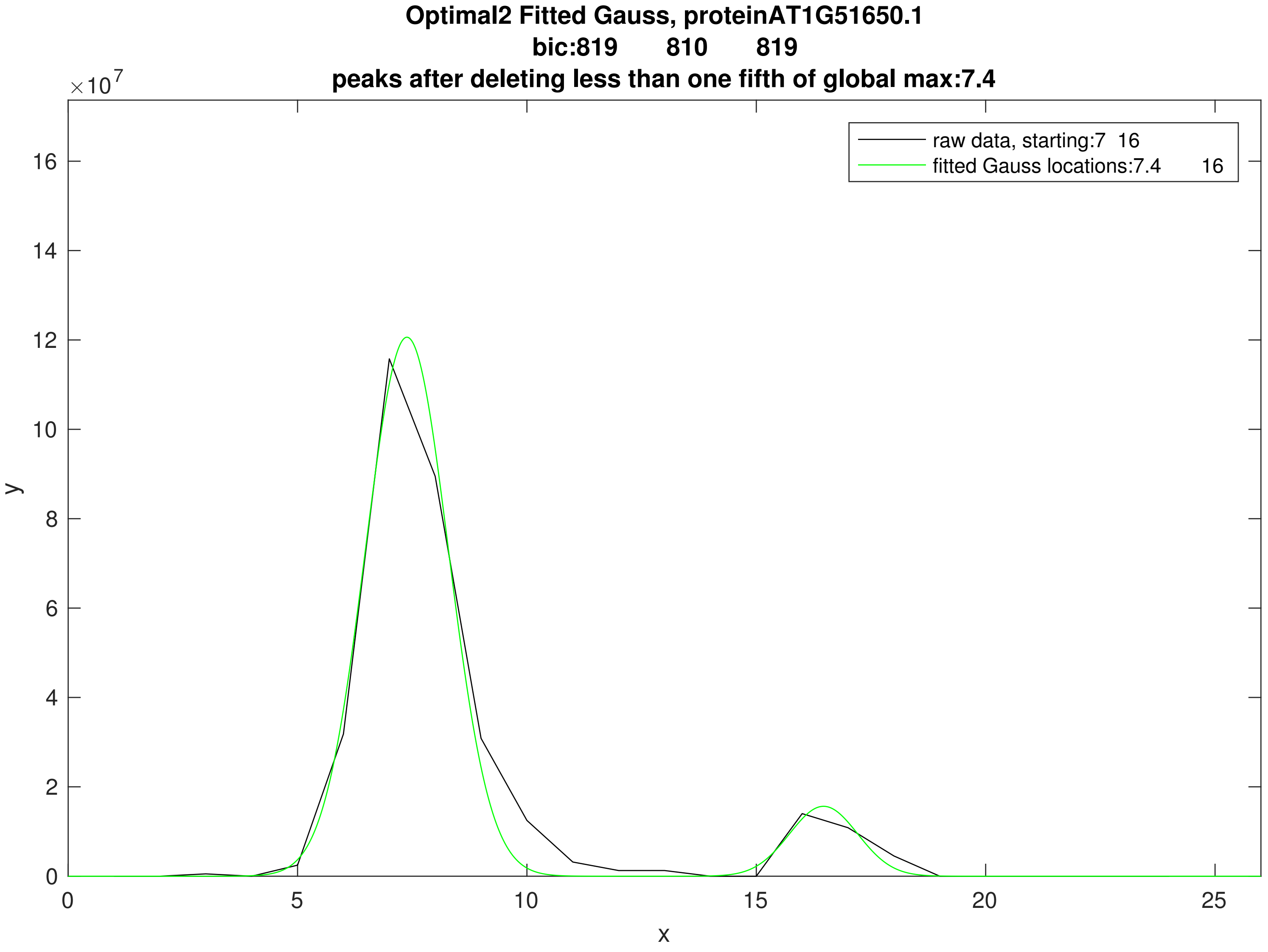
Optimal2 Fitted Gauss, proteinAT1G51650.1
bic:819 810 819
peaks after deleting less than one fifth of global max:7.4
×10^7
raw data, starting:7 16
fitted Gauss locations:7.4 16
y
x
0
2
4
6
8
10
12
14
16
5
15
20
25

#### Optimal1 Fitted Gauss, proteinAT1G51680.1;AT1G51680.3;AT1G51680.2
#### bic:906
#### peaks after deleting less than one fifth of global max:14

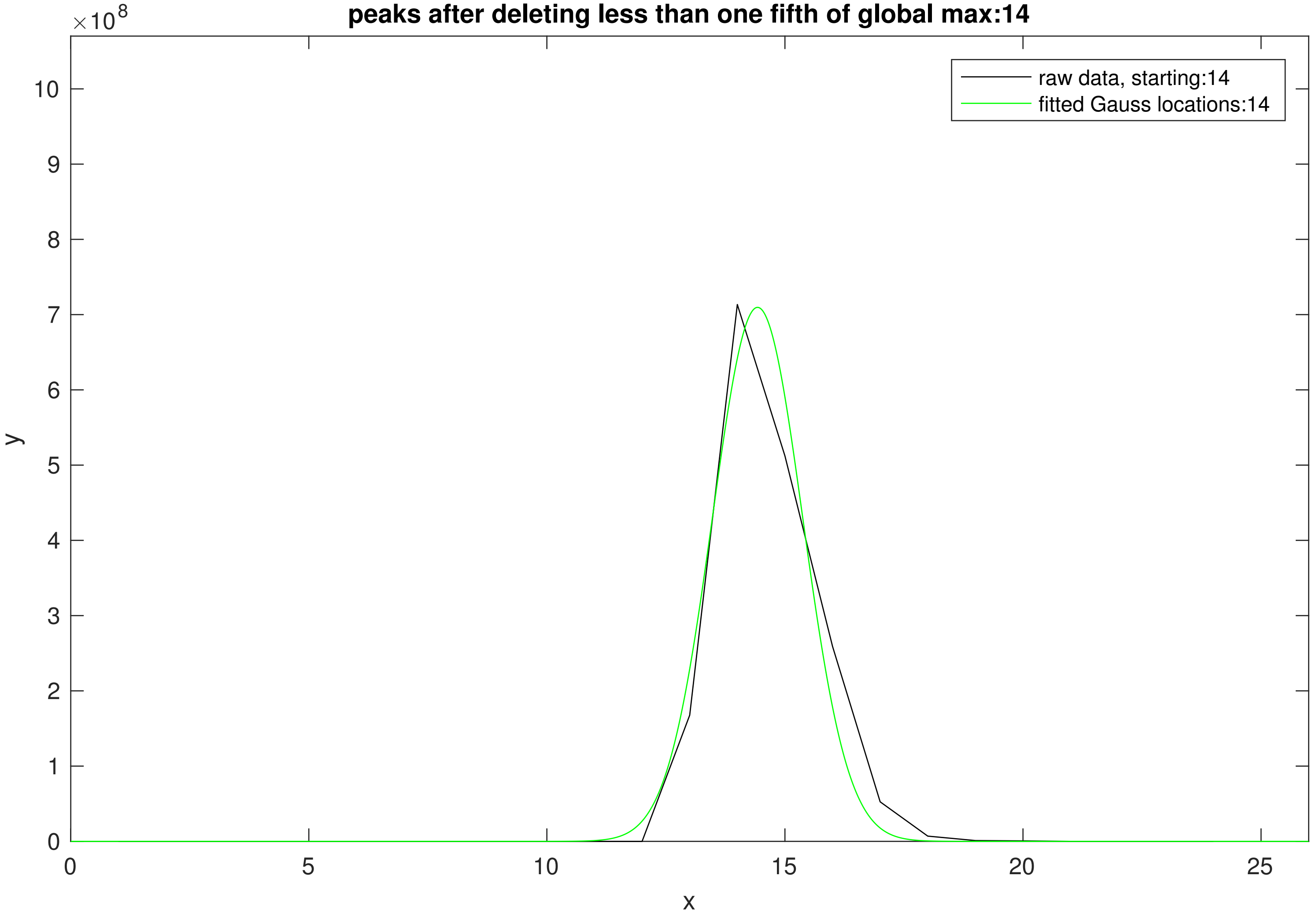

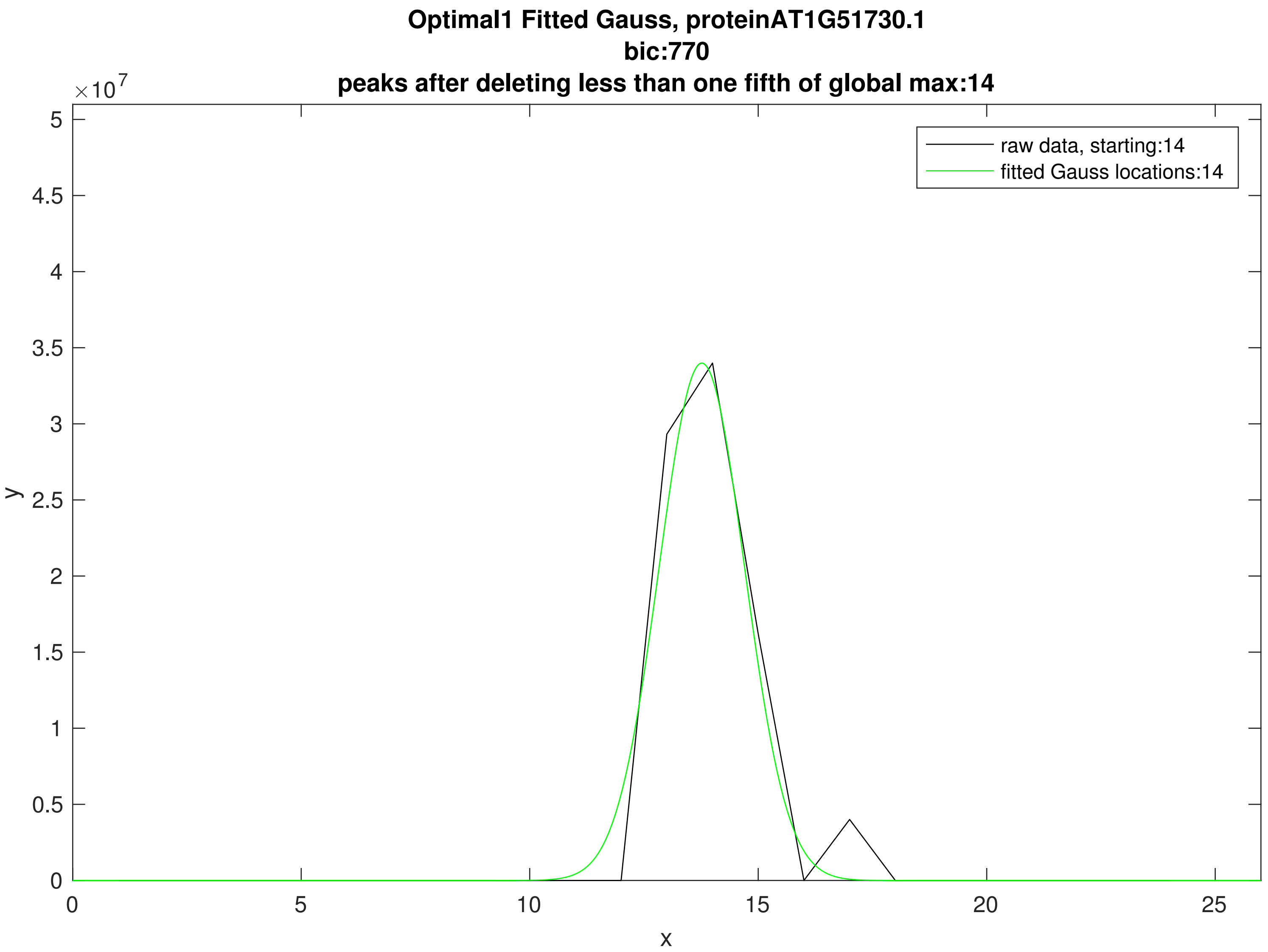
Optimal1 Fitted Gauss, proteinAT1G51730.1
bic:770
peaks after deleting less than one fifth of global max:14
×10^7
raw data, starting:14
fitted Gauss locations:14
y
x
0
0.5
1
1.5
2
2.5
3
3.5
4
4.5
5
0
5
10
15
20
25

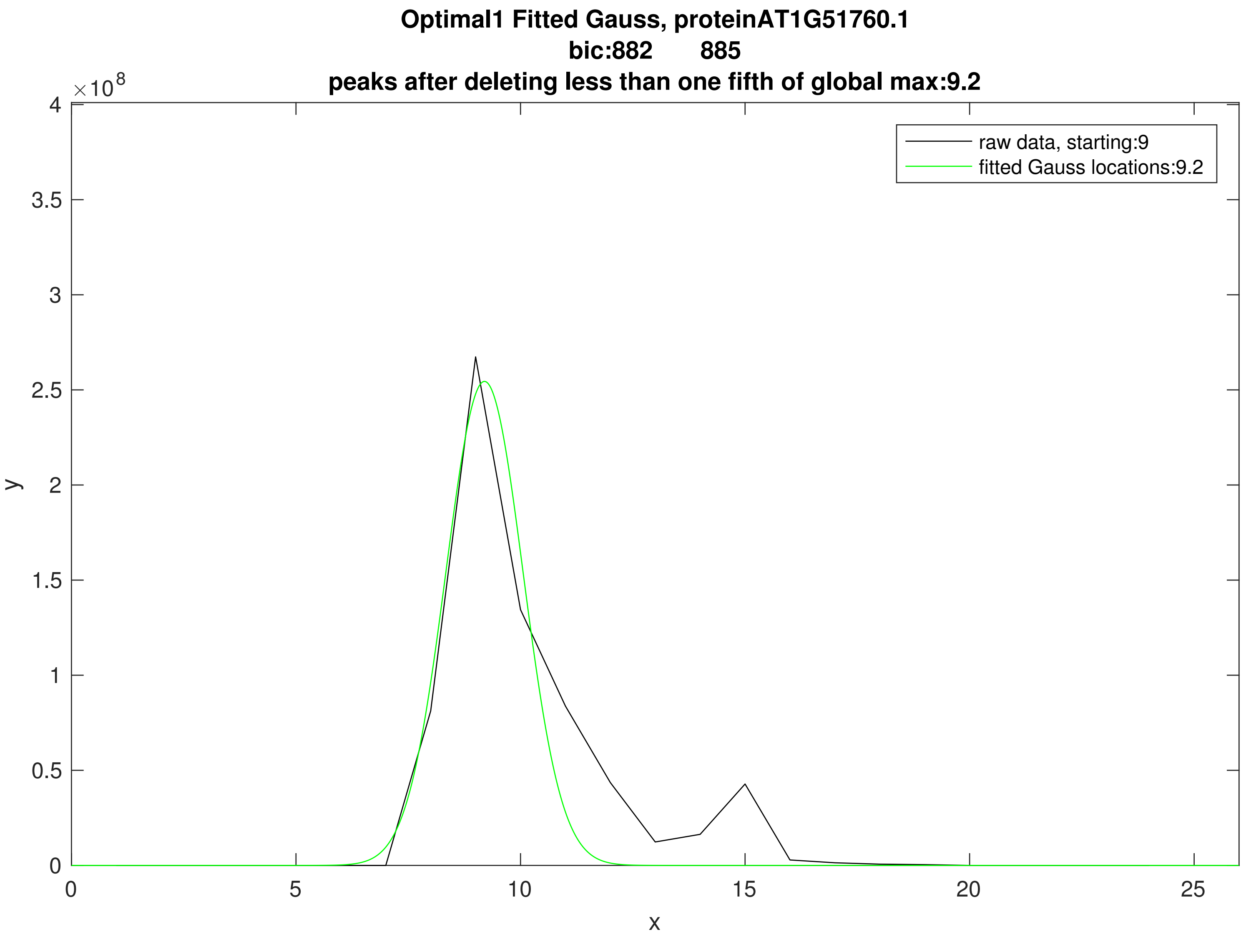

Optimal1 Fitted Gauss, proteinAT1G51760.1
bic:882 885
peaks after deleting less than one fifth of global max:9.2
×10^8
raw data, starting:9
fitted Gauss locations:9.2
y
x
0
0.5
1
1.5
2
2.5
3
3.5
4
0
5
10
15
20
25

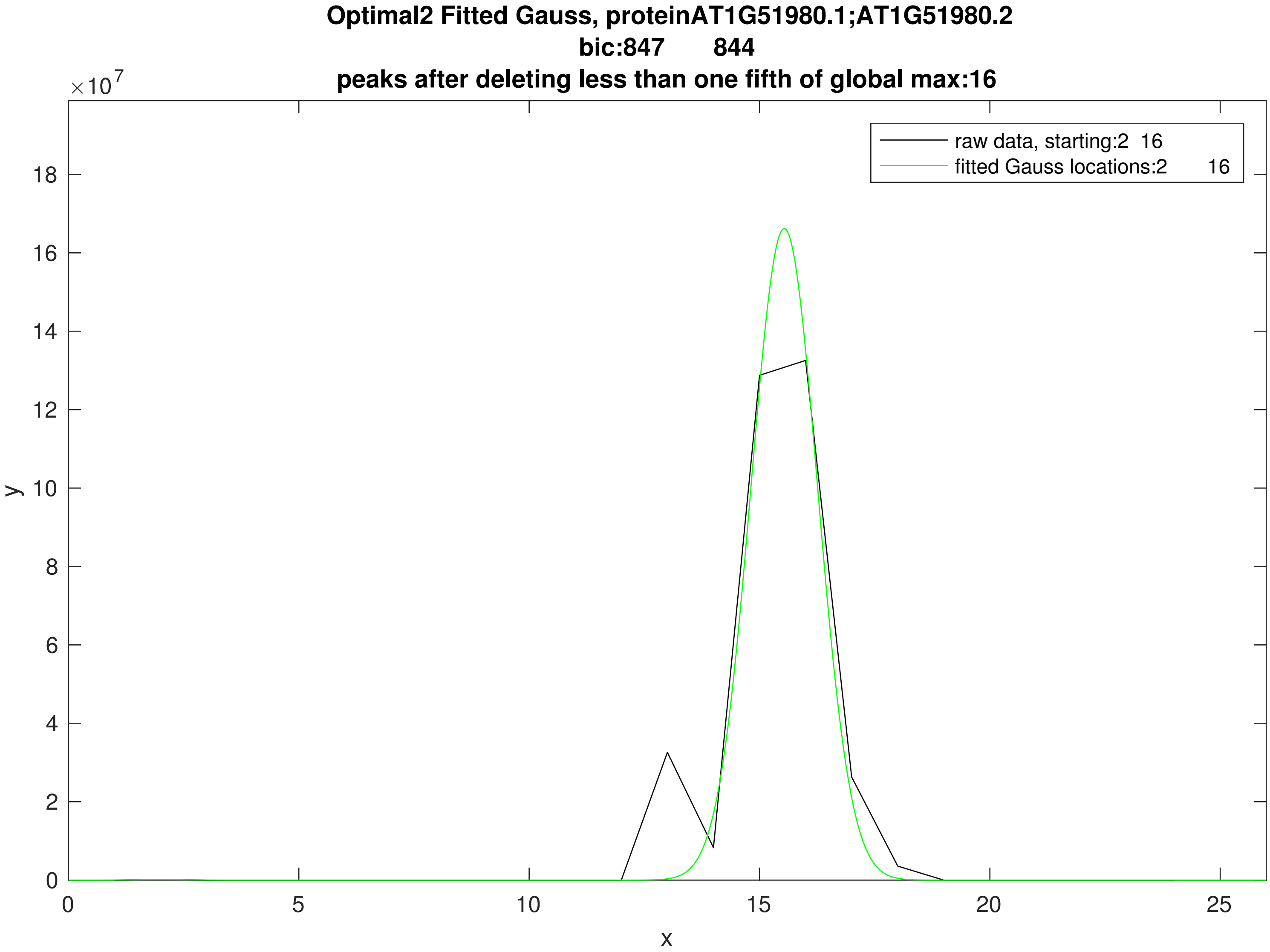

Optimal2 Fitted Gauss, proteinAT1G51980.1;AT1G51980.2
bic:847 844
peaks after deleting less than one fifth of global max:16
×10^7
raw data, starting:2 16
fitted Gauss locations:2 16
y
x
0
2
4
6
8
10
12
14
16
18
0
5
10
15
20
25

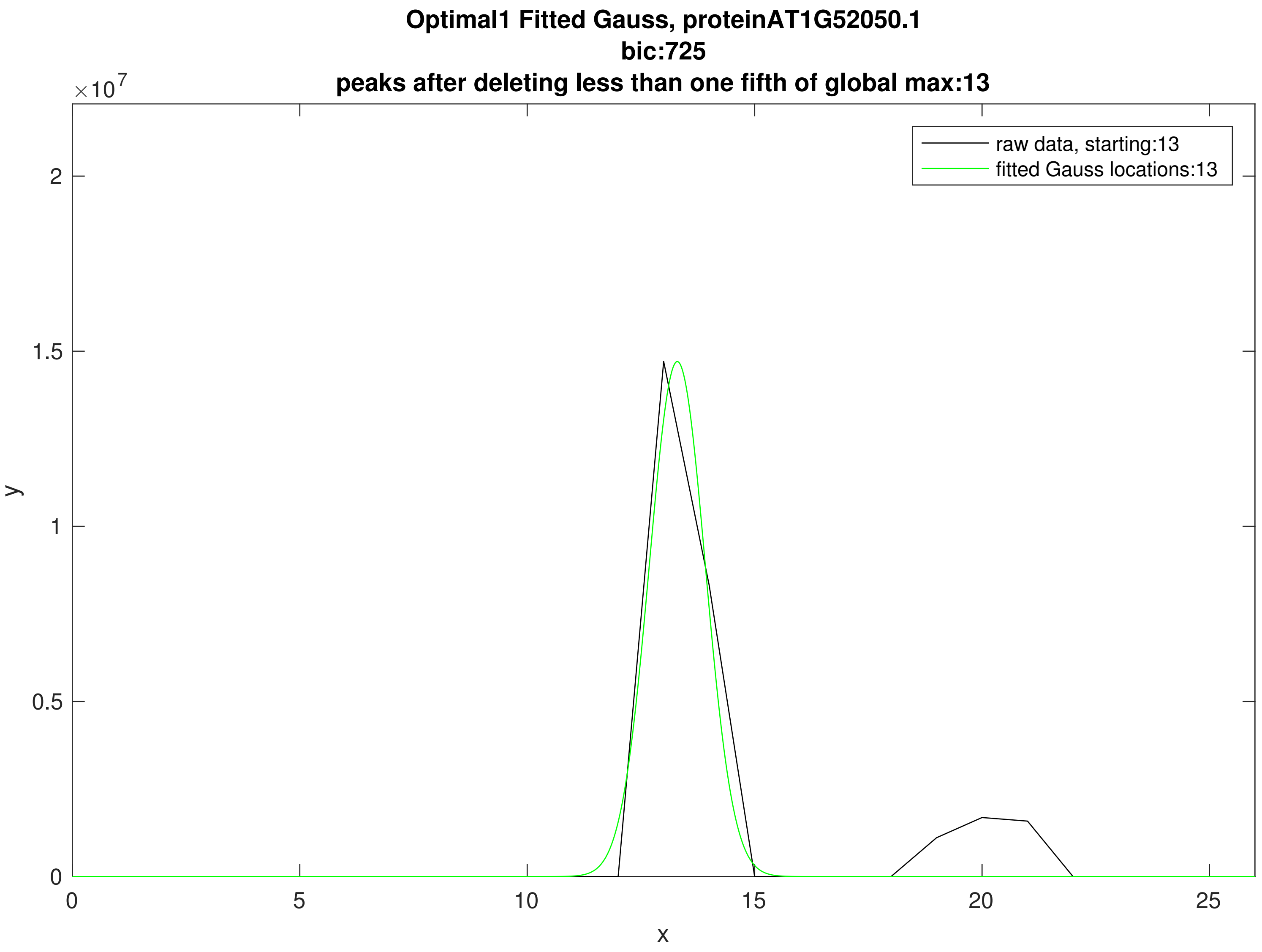
Optimal1 Fitted Gauss, proteinAT1G52050.1
bic:725
peaks after deleting less than one fifth of global max:13
×10^7
raw data, starting:13
fitted Gauss locations:13
y
x
0
0.5
1
1.5
2
5
10
15
20
25

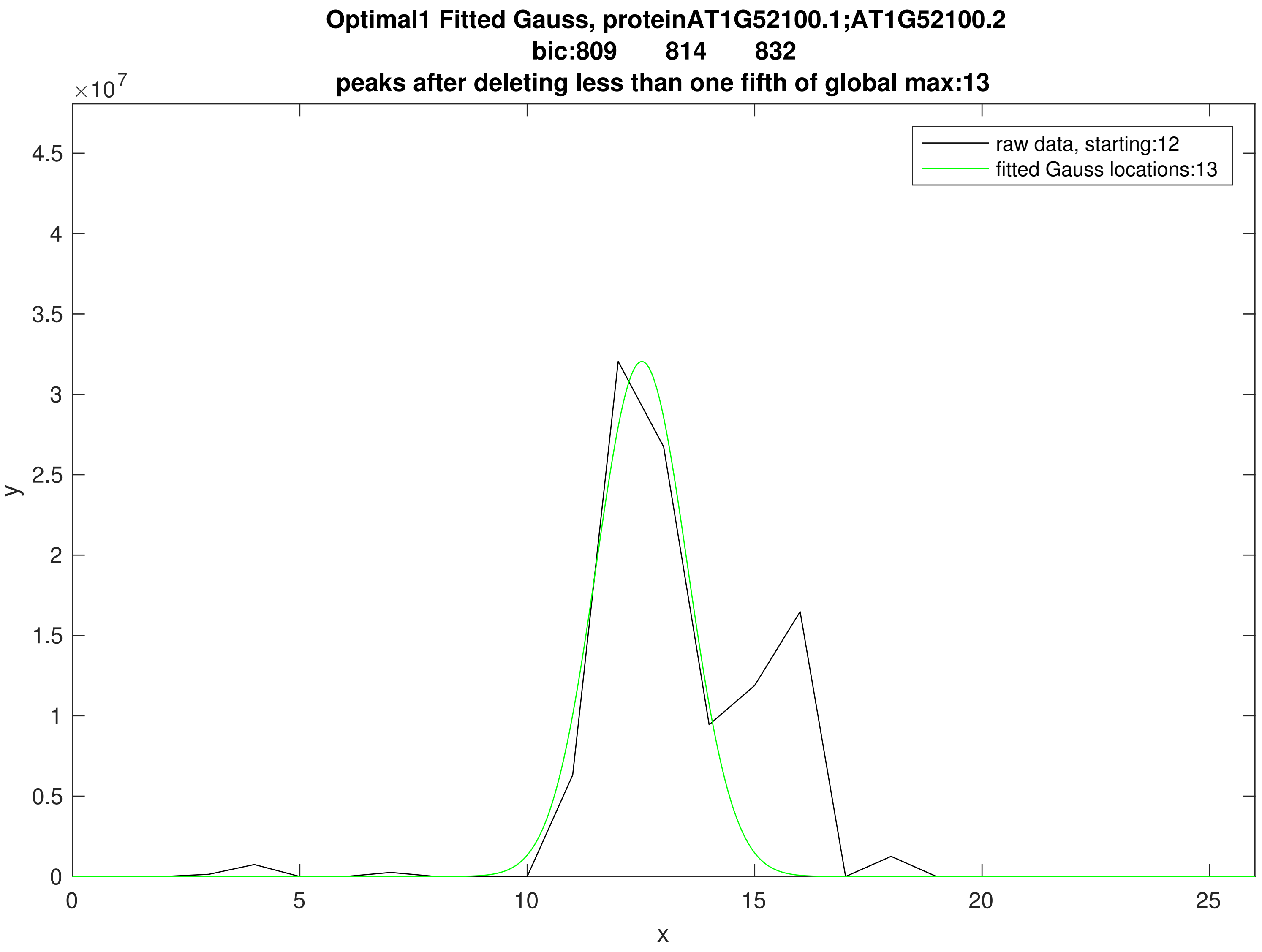

Optimal1 Fitted Gauss, proteinAT1G52100.1;AT1G52100.2
bic:809 814 832
peaks after deleting less than one fifth of global max:13
×10^7
4.5
4
3.5
3
2.5
2
1.5
1
0.5
0
y
0
5
10
15
20
25
x
raw data, starting:12
fitted Gauss locations:13

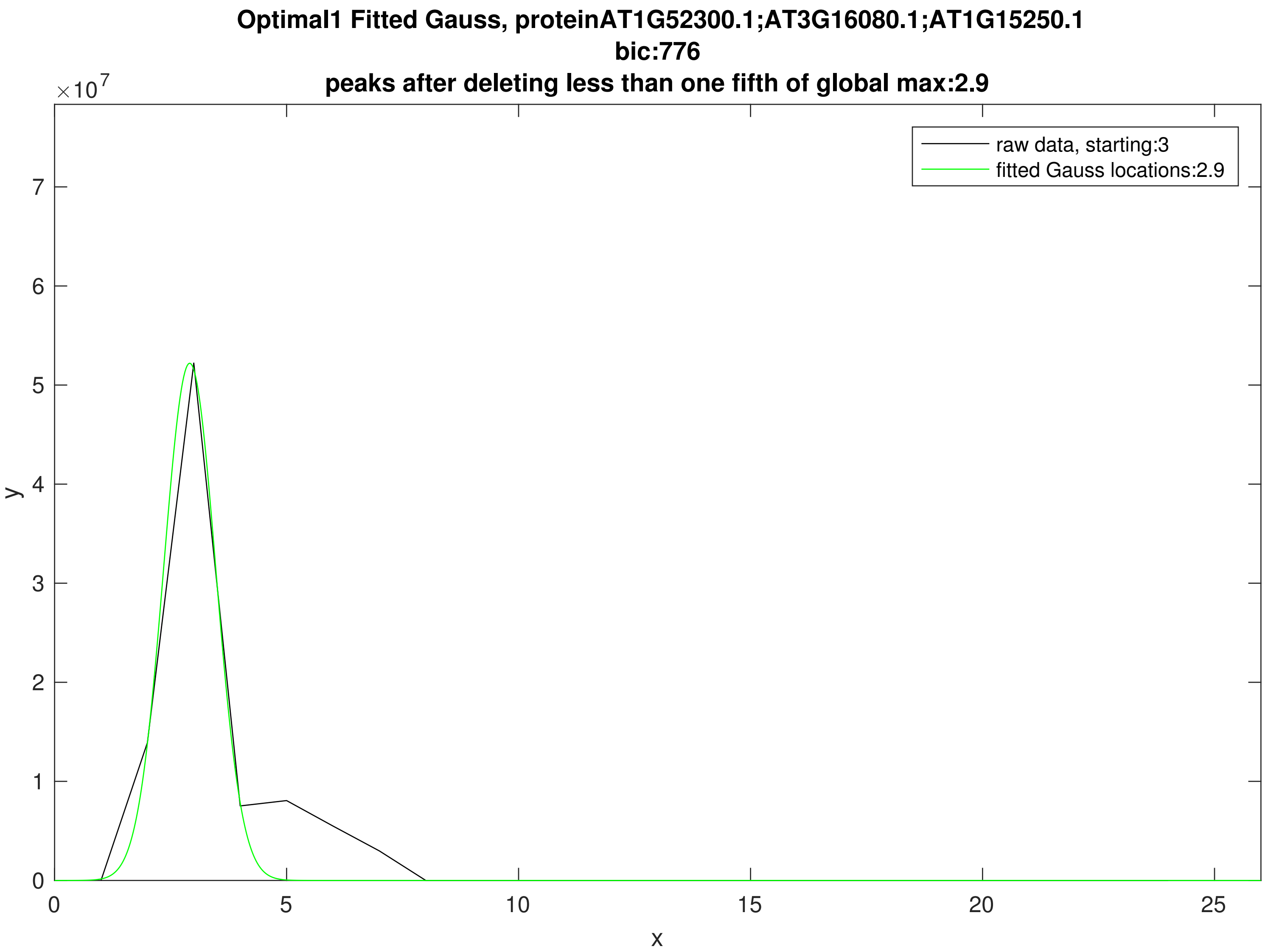

Optimal1 Fitted Gauss, proteinAT1G52300.1;AT3G16080.1;AT1G15250.1
bic:776
peaks after deleting less than one fifth of global max:2.9
×10^7
raw data, starting:3
fitted Gauss locations:2.9
y
x
0
1
2
3
4
5
6
7
0
5
10
15
20
25

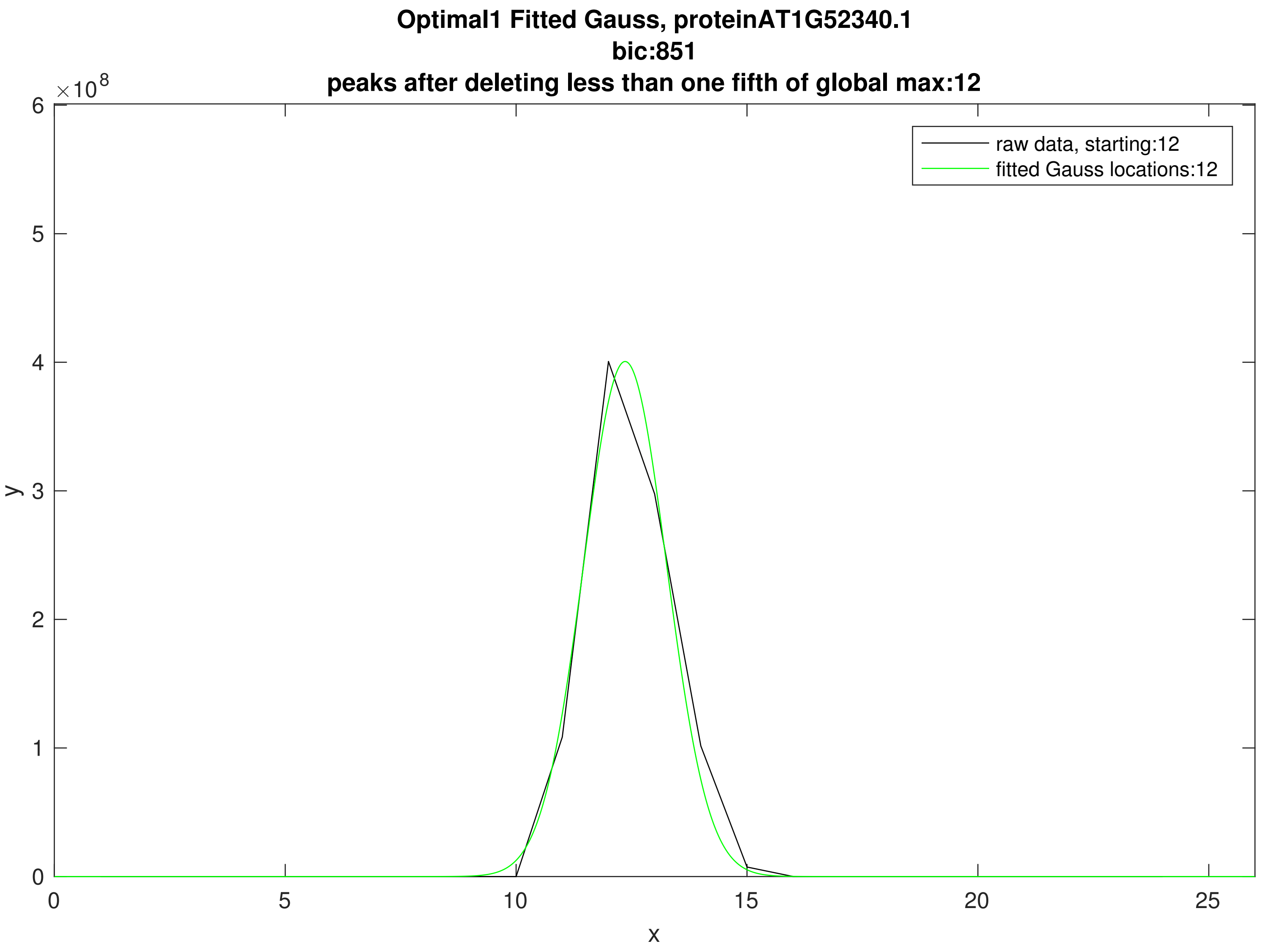

Optimal1 Fitted Gauss, proteinAT1G52340.1
bic:851
peaks after deleting less than one fifth of global max:12
×10^8
raw data, starting:12
fitted Gauss locations:12
y
x
0
1
2
3
4
5
6
0
5
10
15
20
25

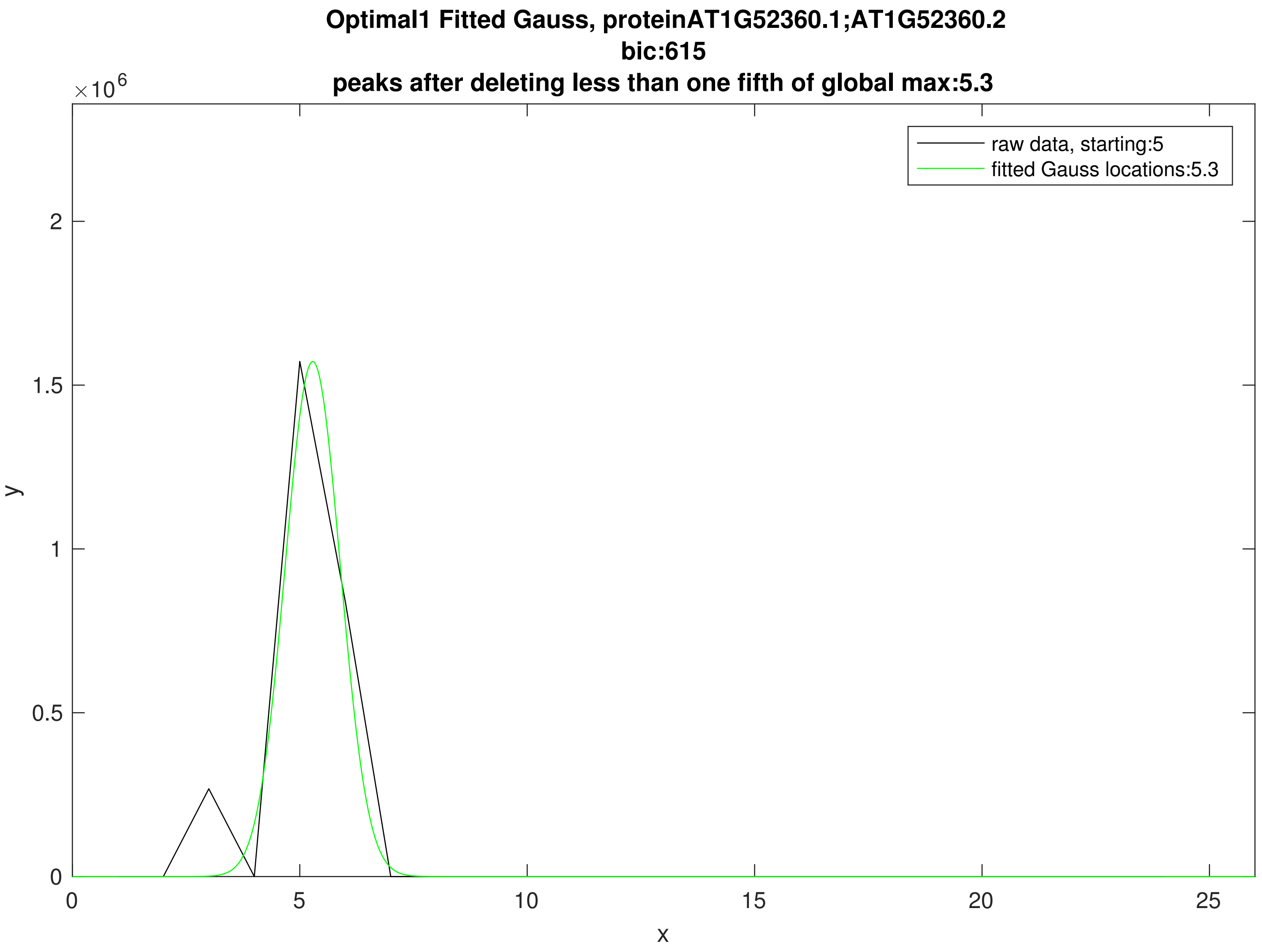

Optimal1 Fitted Gauss, proteinAT1G52360.1;AT1G52360.2
bic:615
peaks after deleting less than one fifth of global max:5.3
×10^6
raw data, starting:5
fitted Gauss locations:5.3
y
x
0
0.5
1
1.5
2
5
10
15
20
25

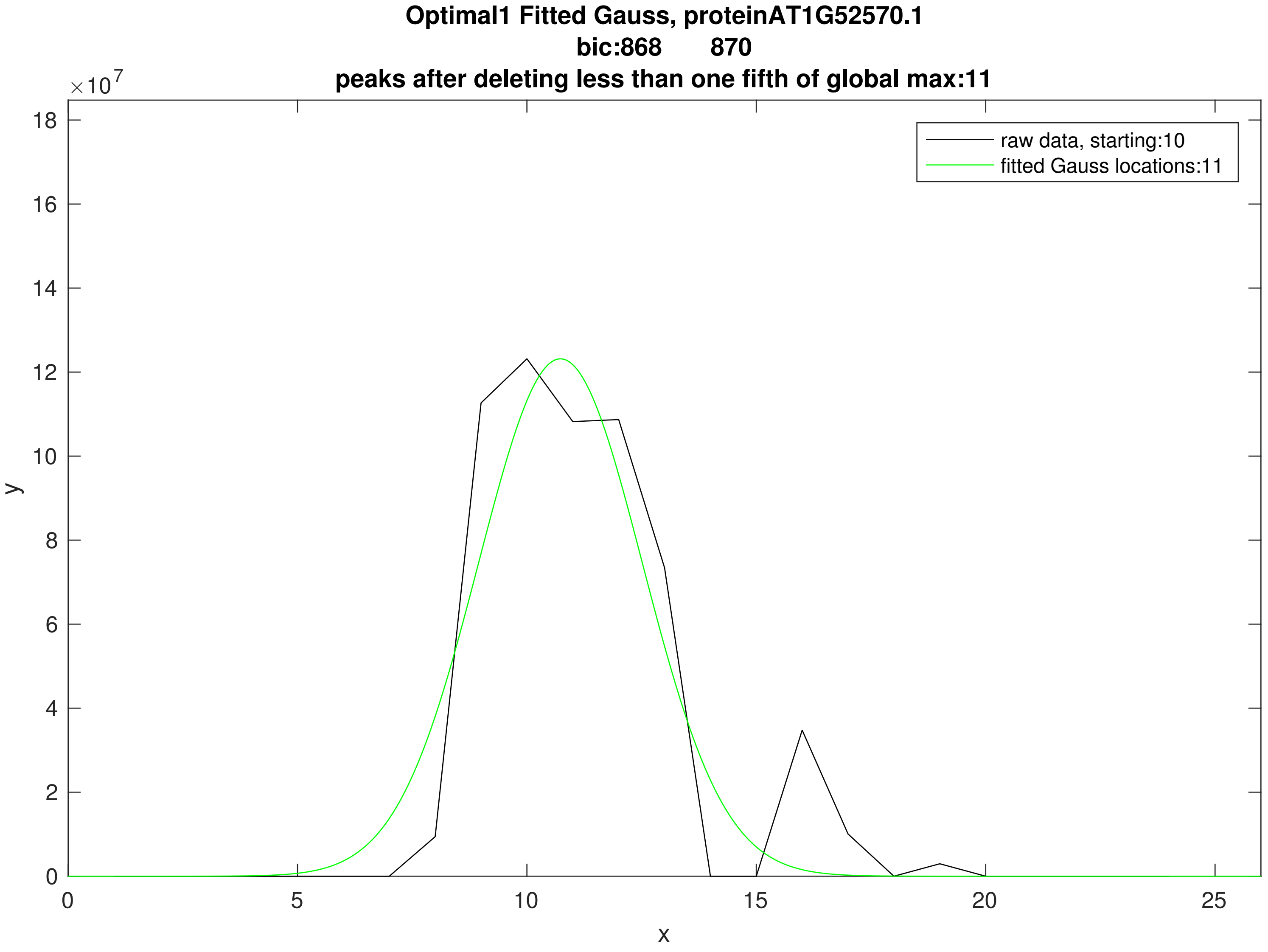
Optimal1 Fitted Gauss, proteinAT1G52570.1
bic:868 870
peaks after deleting less than one fifth of global max:11
×10^7
raw data, starting:10
fitted Gauss locations:11
y
x
0
2
4
6
8
10
12
14
16
18
0
5
10
15
20
25

#### Optimal1 Fitted Gauss, proteinAT1G52670.1
#### bic:669
#### peaks after deleting less than one fifth of global max:17

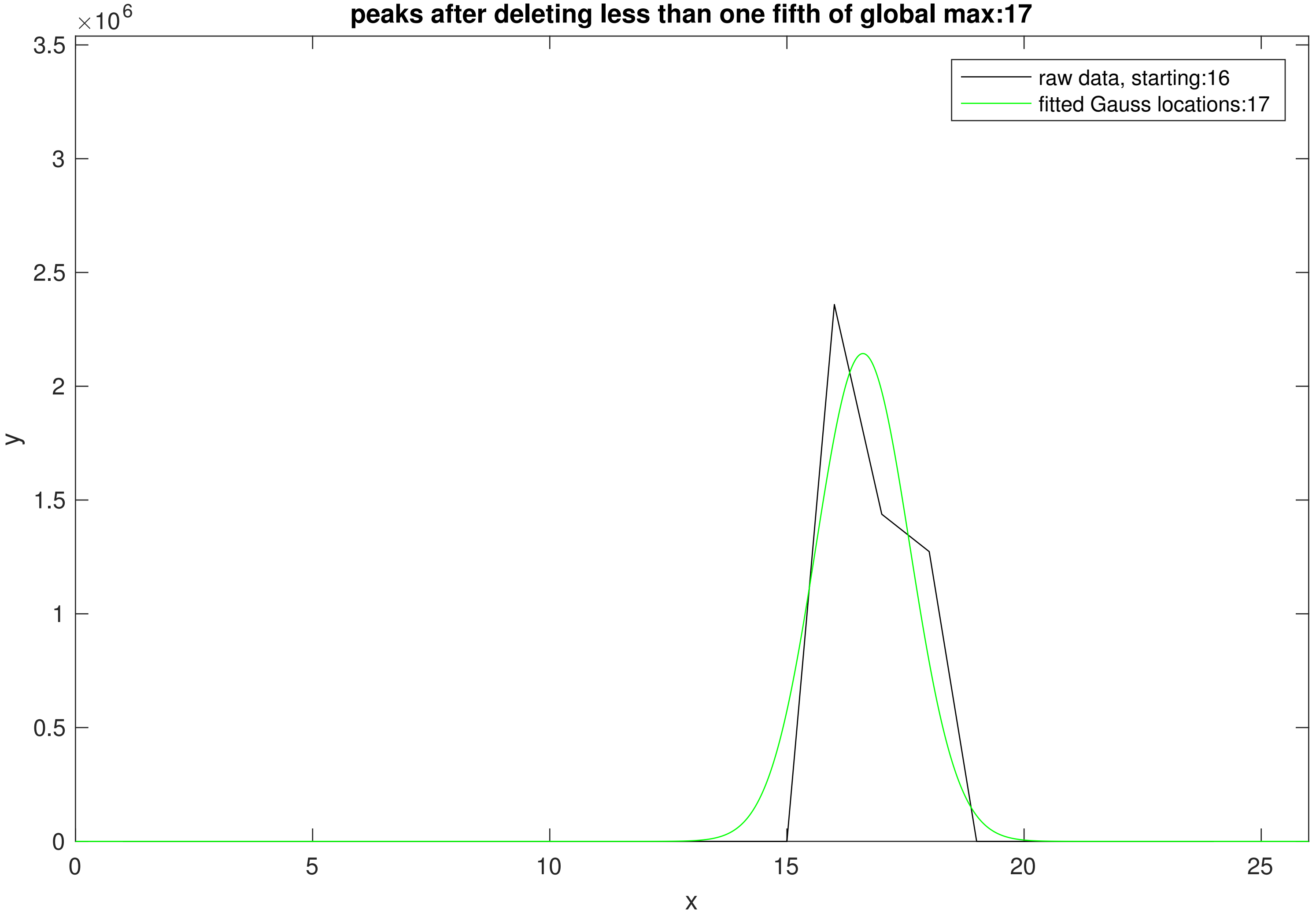

Optimal1 Fitted Gauss, proteinAT1G52690.2;AT1G52690.1
bic:738
peaks after deleting less than one fifth of global max:15

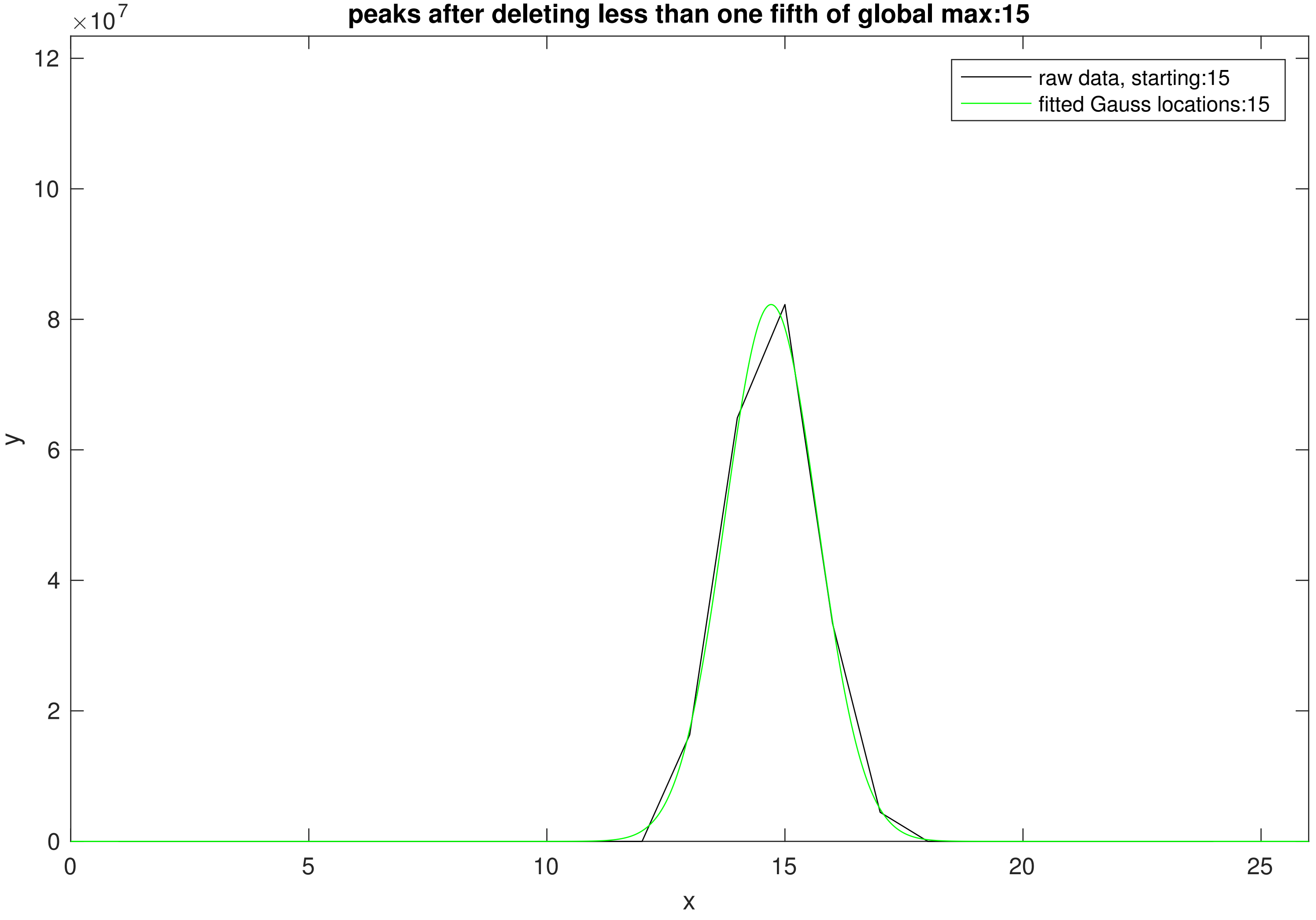

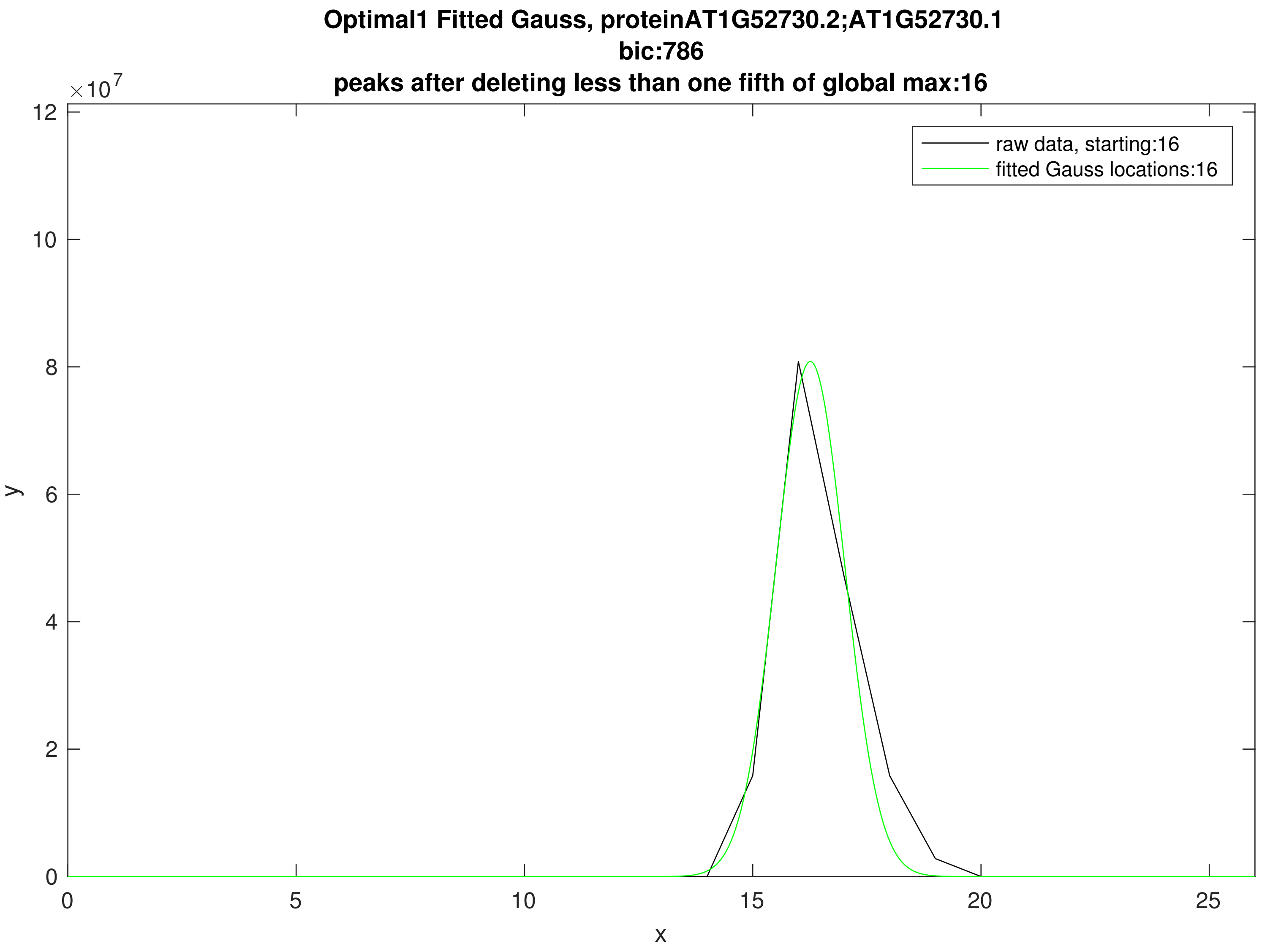

Optimal1 Fitted Gauss, proteinAT1G52730.2;AT1G52730.1
bic:786
peaks after deleting less than one fifth of global max:16
×10^7
raw data, starting:16
fitted Gauss locations:16
y
x
0
2
4
6
8
10
12
0
5
10
15
20
25

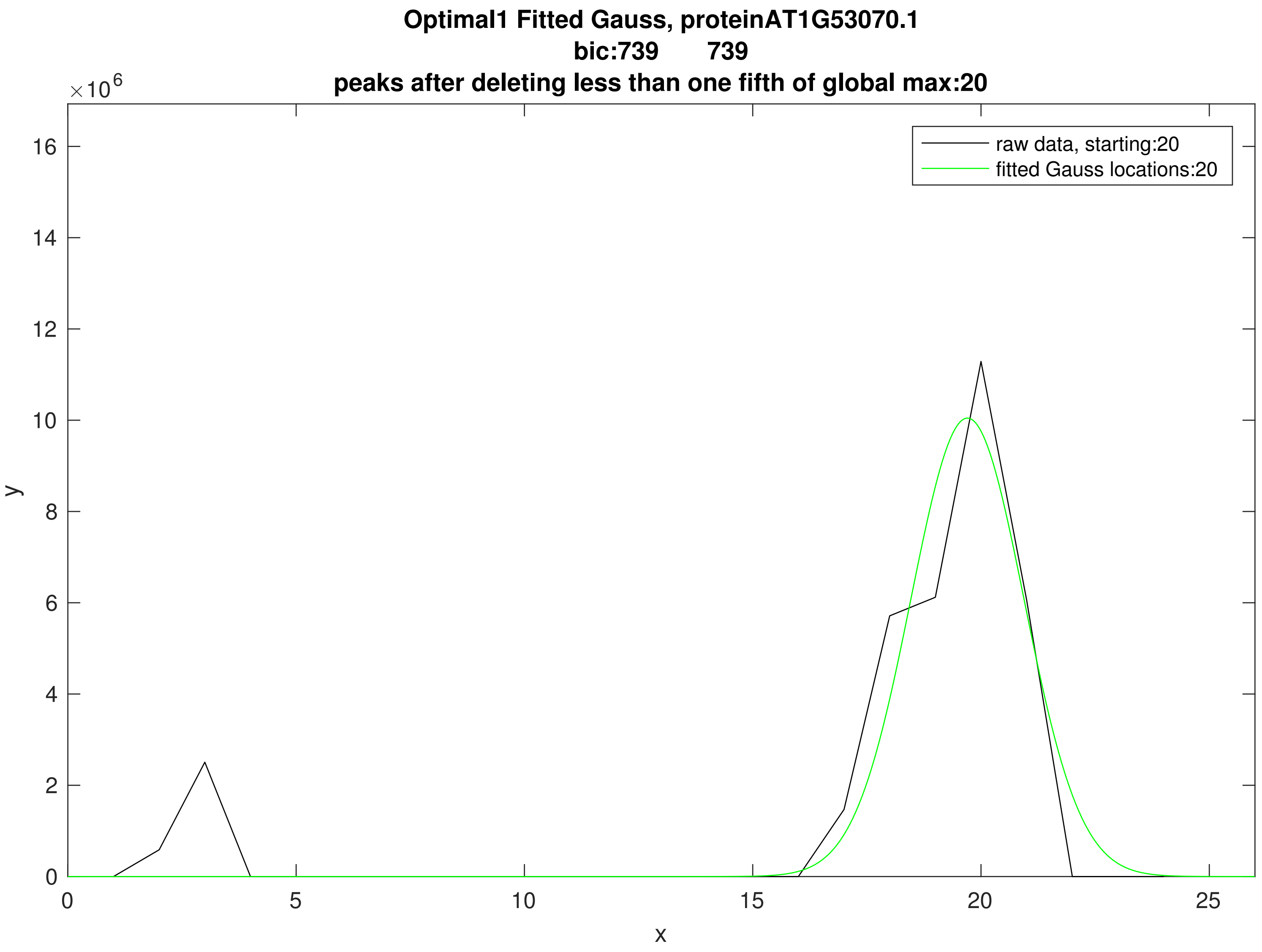
Optimal1 Fitted Gauss, proteinAT1G53070.1
bic:739 739
peaks after deleting less than one fifth of global max:20
×10^6
raw data, starting:20
fitted Gauss locations:20
y
x
0
2
4
6
8
10
12
14
16
0
5
10
15
20
25

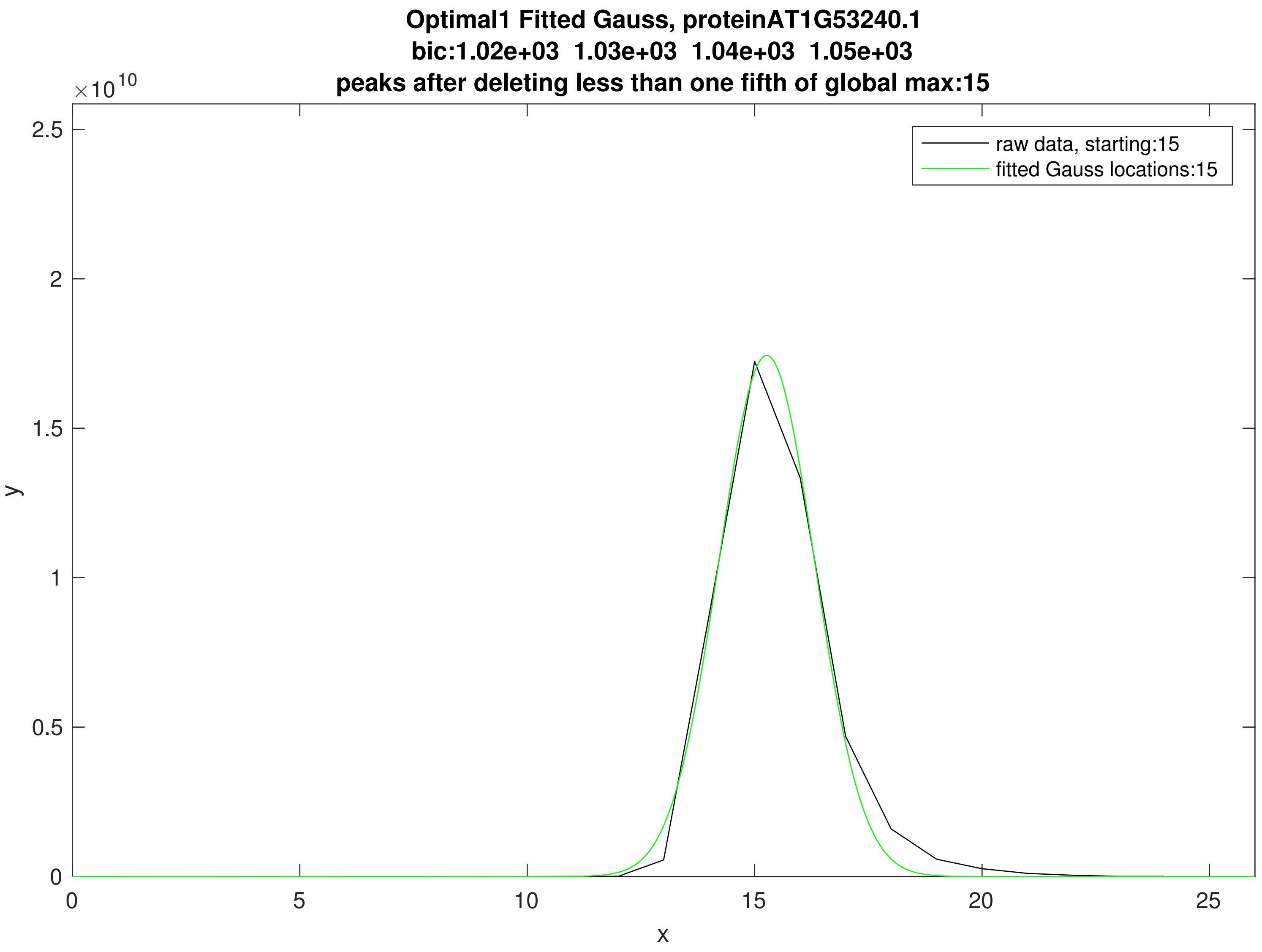
Optimal1 Fitted Gauss, proteinAT1G53240.1
bic:1.02e+03  1.03e+03  1.04e+03  1.05e+03
peaks after deleting less than one fifth of global max:15
×10^10
2.5
2
1.5
1
0.5
0
y
0
5
10
15
20
25
x
raw data, starting:15
fitted Gauss locations:15

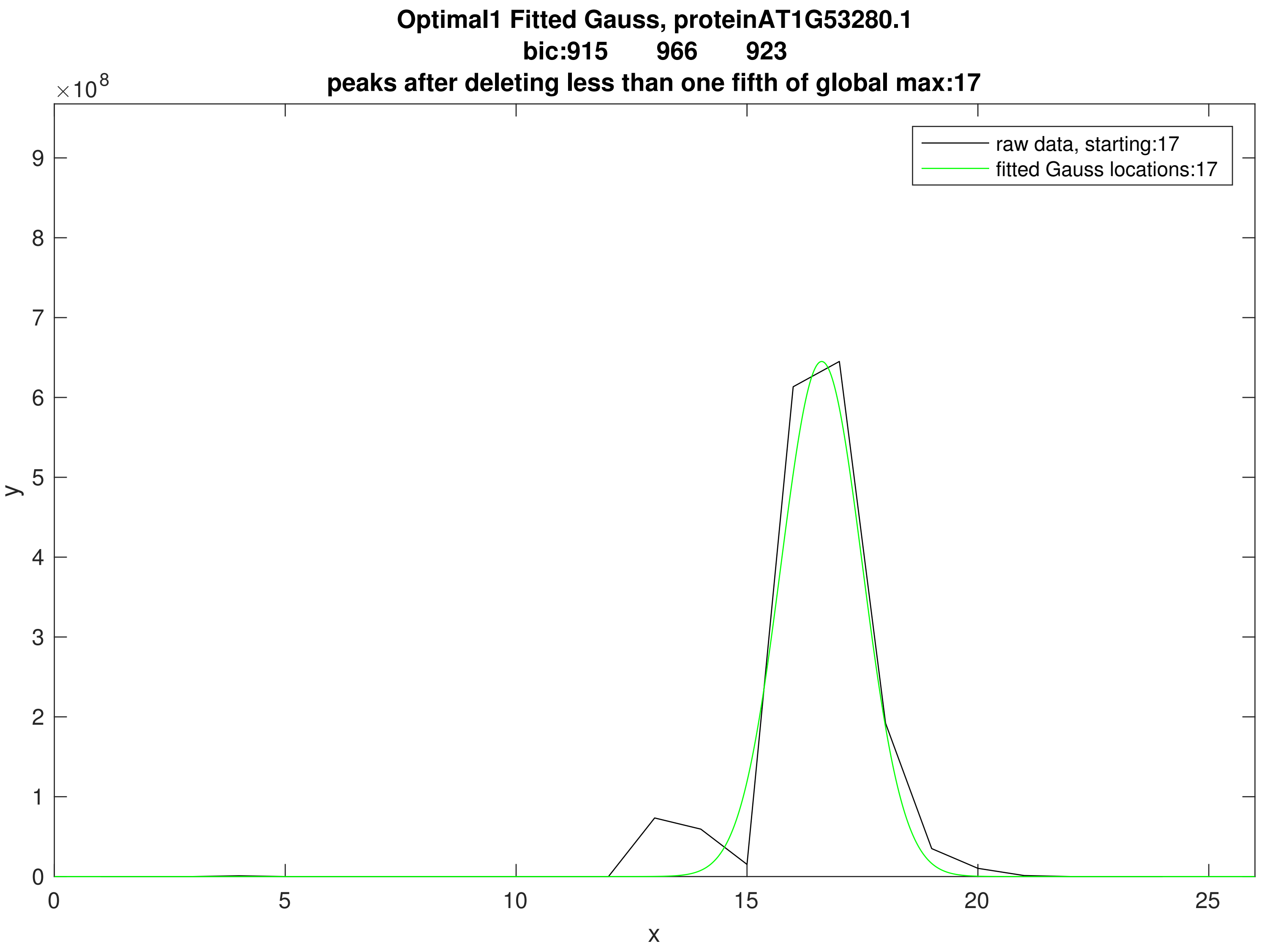

Optimal1 Fitted Gauss, proteinAT1G53280.1
bic:915 966 923
peaks after deleting less than one fifth of global max:17
×10^8
raw data, starting:17
fitted Gauss locations:17
y
x
0
1
2
3
4
5
6
7
8
9
10
15
20
25

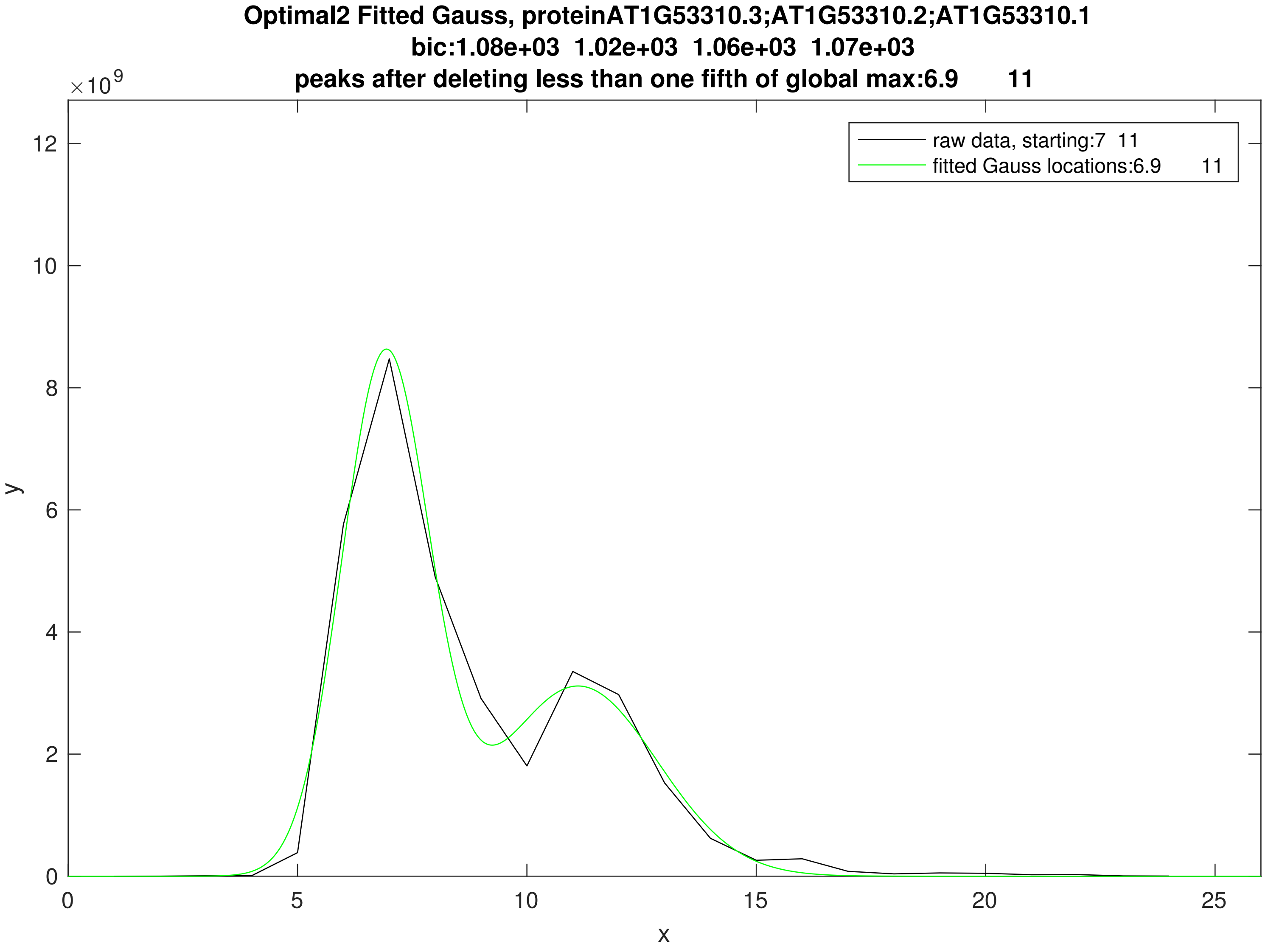

Optimal2 Fitted Gauss, proteinAT1G53310.3;AT1G53310.2;AT1G53310.1
bic:1.08e+03 1.02e+03 1.06e+03 1.07e+03
peaks after deleting less than one fifth of global max:6.9 11
×10^9
raw data, starting:7 11
fitted Gauss locations:6.9 11
y
x
0
2
4
6
8
10
12
0
5
10
15
20
25

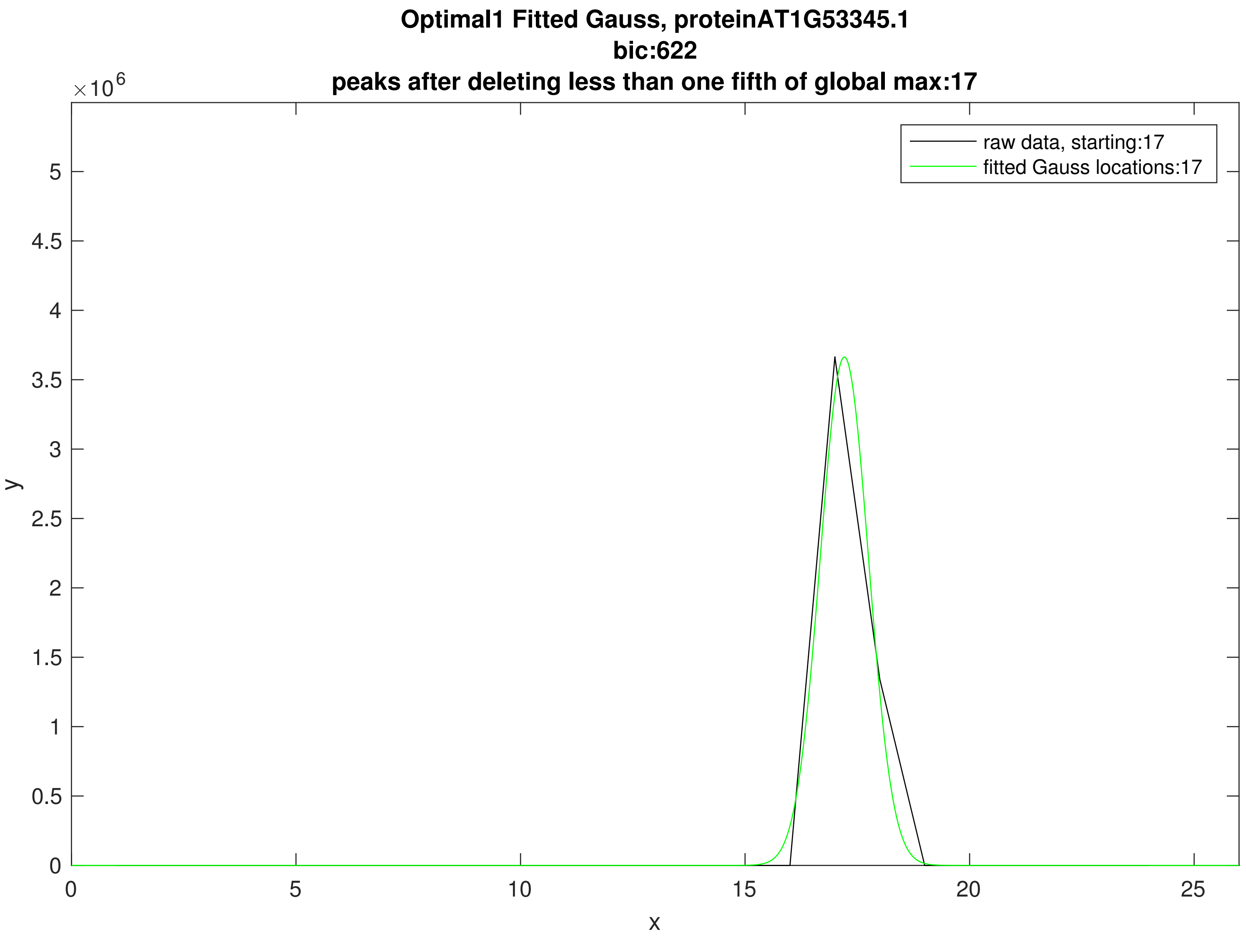
Optimal1 Fitted Gauss, proteinAT1G53345.1
bic:622
peaks after deleting less than one fifth of global max:17
×10^6
raw data, starting:17
fitted Gauss locations:17
y
x
0
0.5
1
1.5
2
2.5
3
3.5
4
4.5
5
5
10
15
20
25

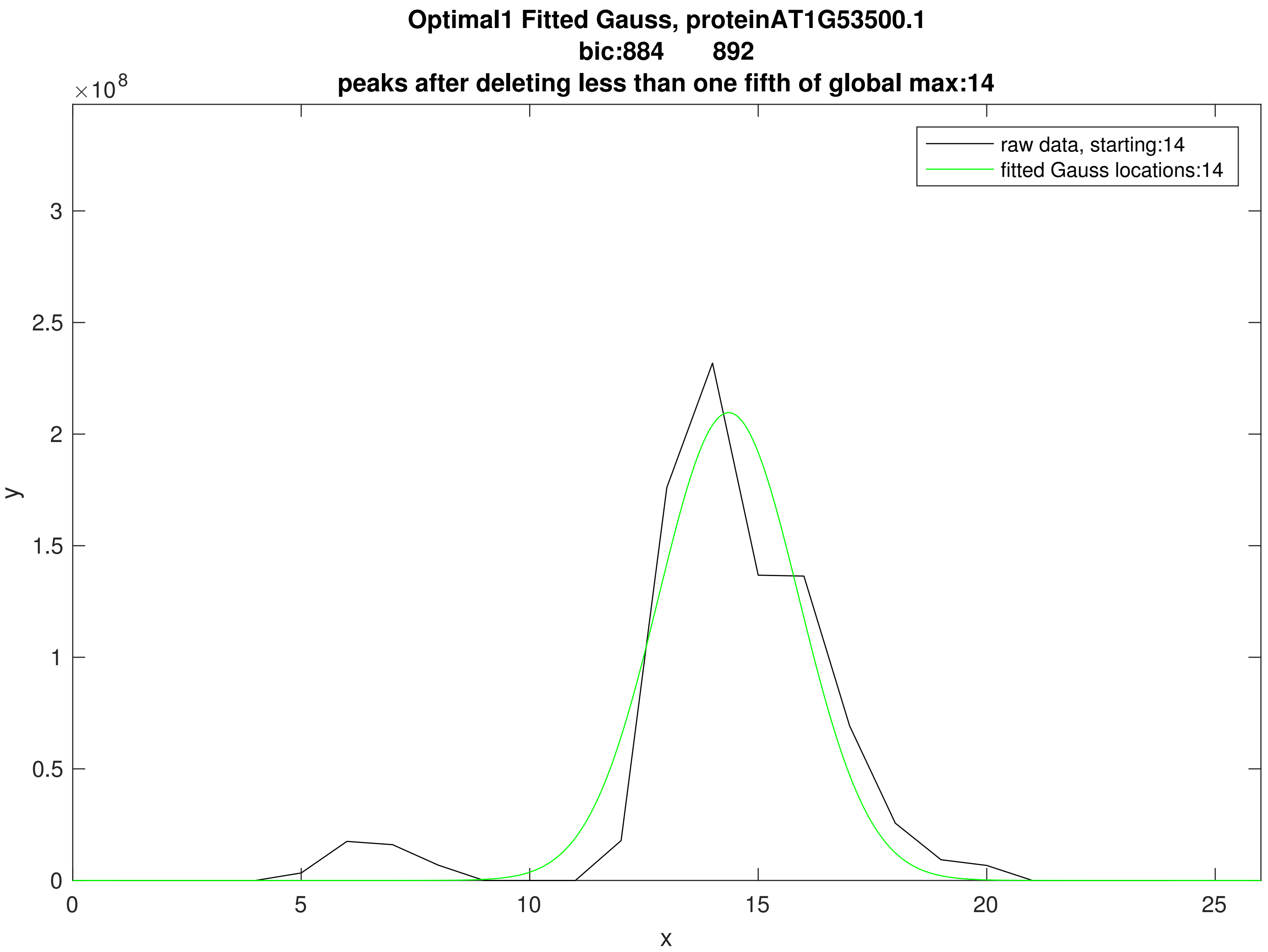
Optimal1 Fitted Gauss, proteinAT1G53500.1
bic:884 892
peaks after deleting less than one fifth of global max:14
×10^8
raw data, starting:14
fitted Gauss locations:14
y
x
0
0.5
1
1.5
2
2.5
3
0
5
10
15
20
25

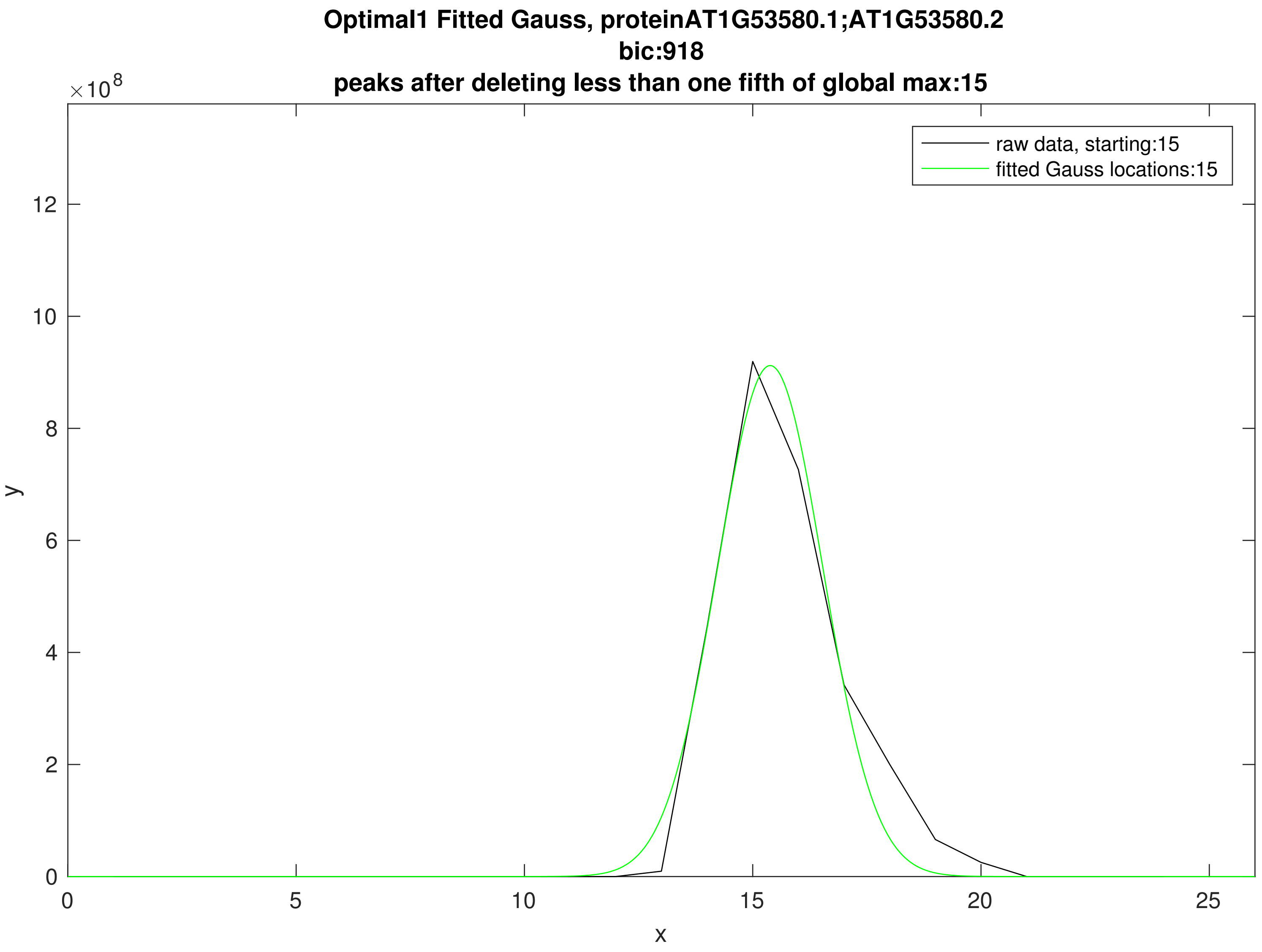
Optimal1 Fitted Gauss, proteinAT1G53580.1;AT1G53580.2
bic:918
peaks after deleting less than one fifth of global max:15
×10^8
raw data, starting:15
fitted Gauss locations:15
y
x
0
2
4
6
8
10
12
0
5
10
15
20
25

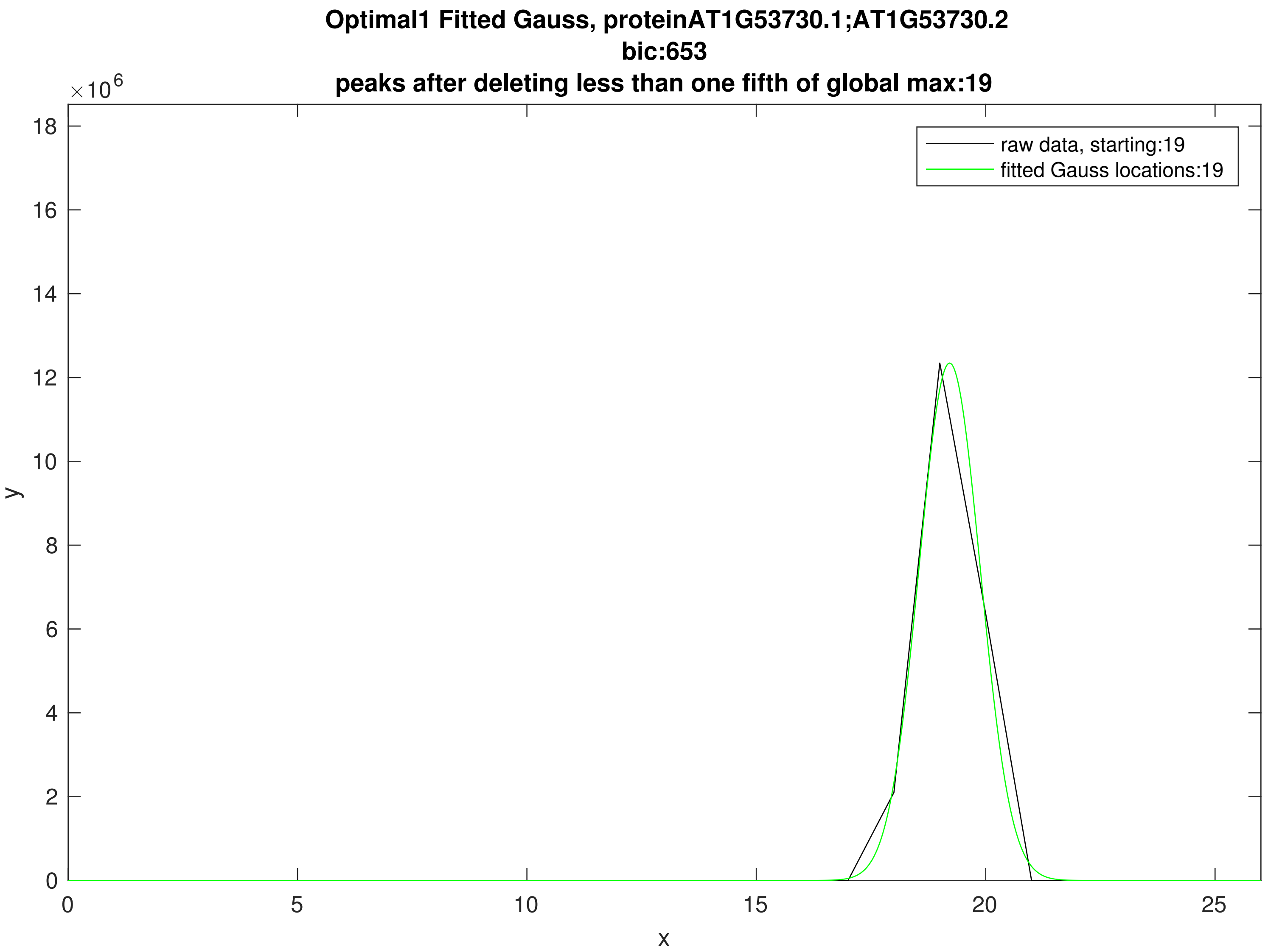

Optimal1 Fitted Gauss, proteinAT1G53730.1;AT1G53730.2
bic:653
peaks after deleting less than one fifth of global max:19
×10^6
raw data, starting:19
fitted Gauss locations:19
y
x

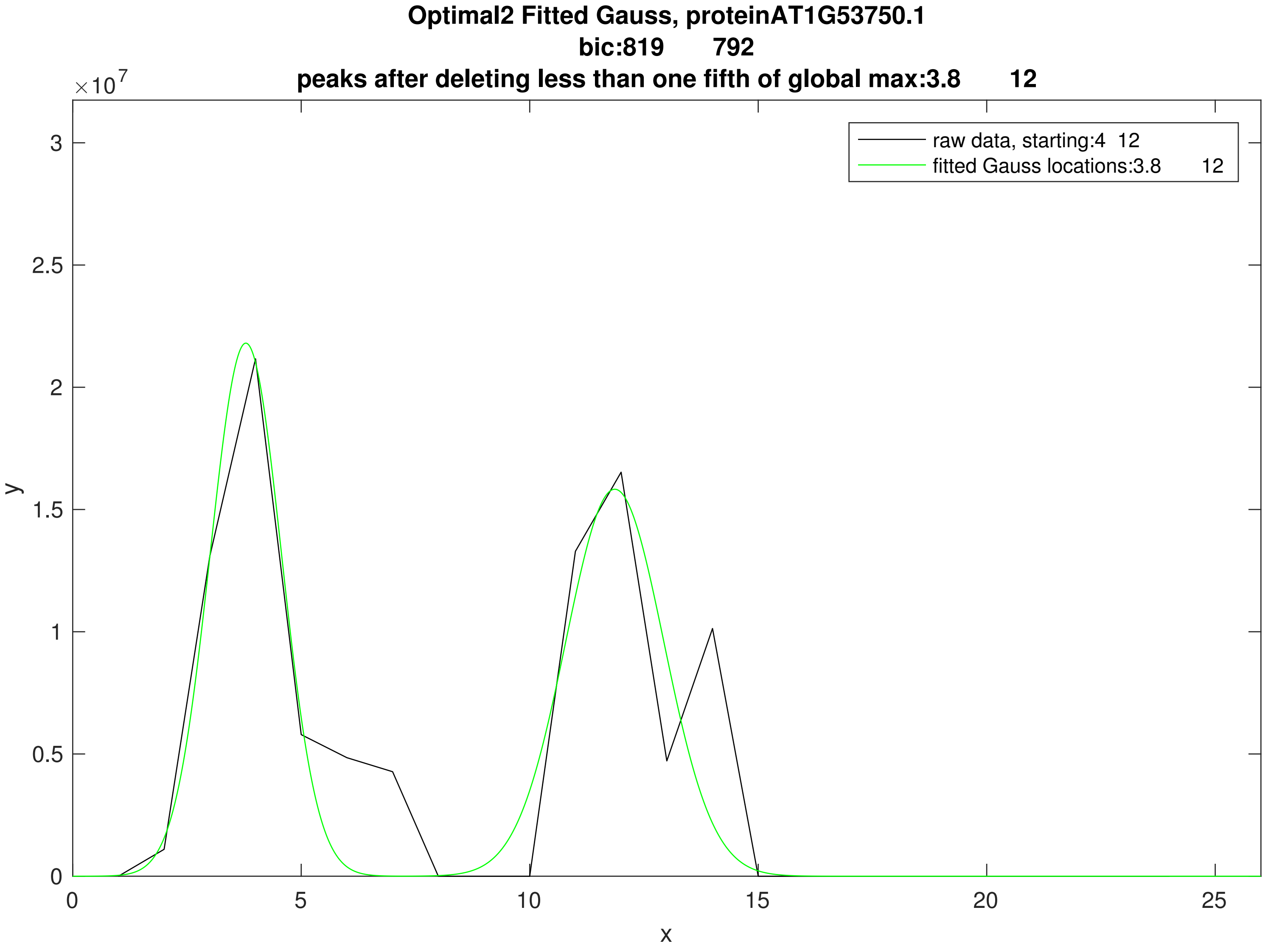

Optimal2 Fitted Gauss, proteinAT1G53750.1
bic:819 792
peaks after deleting less than one fifth of global max:3.8 12
×10^7
raw data, starting:4 12
fitted Gauss locations:3.8 12
y
x
0
0.5
1
1.5
2
2.5
3
5
10
15
20
25

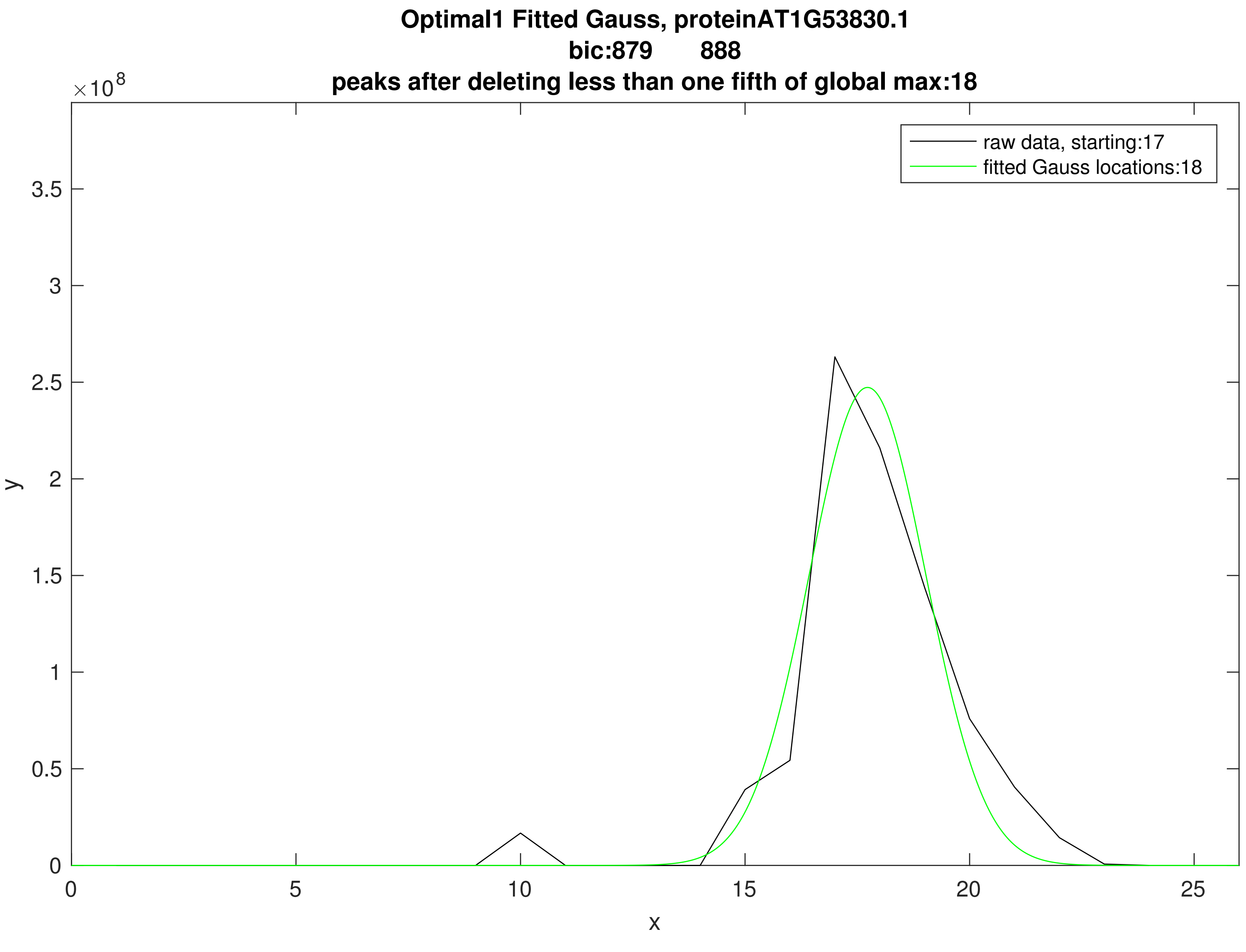

Optimal1 Fitted Gauss, proteinAT1G53830.1
bic:879 888
peaks after deleting less than one fifth of global max:18
×10^8
raw data, starting:17
fitted Gauss locations:18
3.5
3
2.5
2
1.5
1
0.5
0
y
0
5
10
15
20
25
x

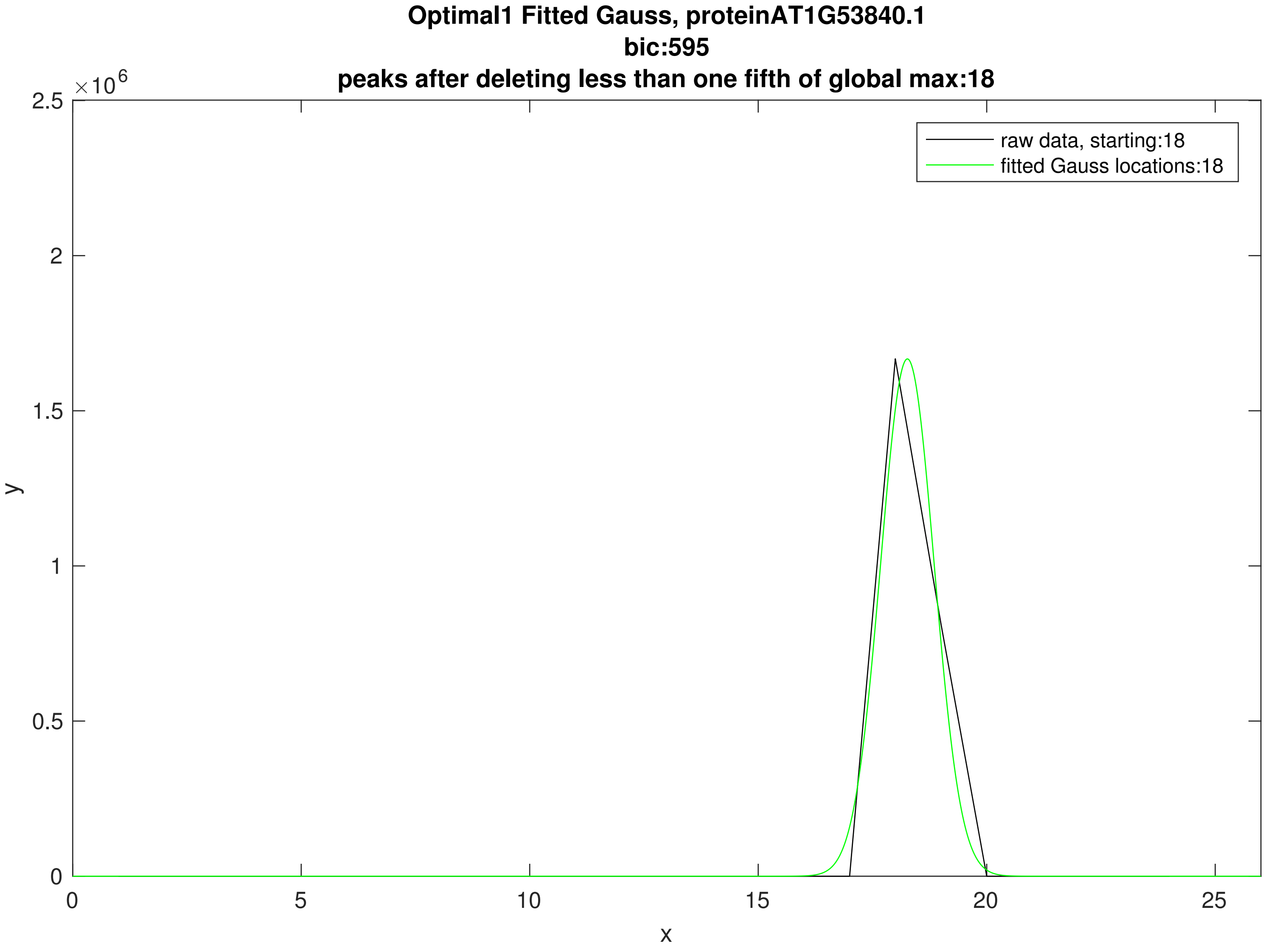

Optimal1 Fitted Gauss, proteinAT1G53840.1
bic:595
peaks after deleting less than one fifth of global max:18
×10^6
2.5
2
1.5
1
0.5
0
y
0
5
10
15
20
25
x
raw data, starting:18
fitted Gauss locations:18

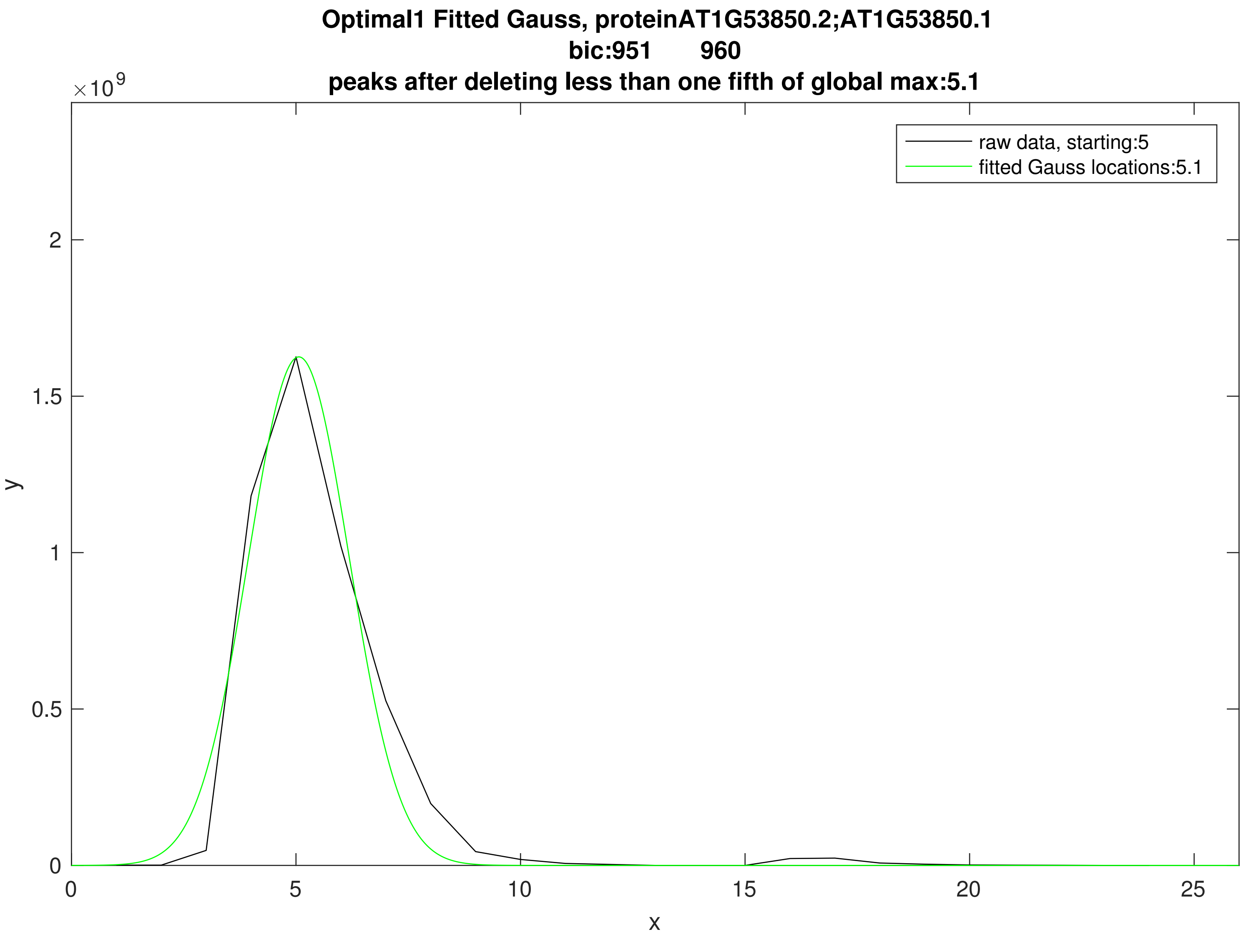
Optimal1 Fitted Gauss, proteinAT1G53850.2;AT1G53850.1
bic:951 960
peaks after deleting less than one fifth of global max:5.1
×10^9
raw data, starting:5
fitted Gauss locations:5.1
y
x
0
0.5
1
1.5
2
0
5
10
15
20
25

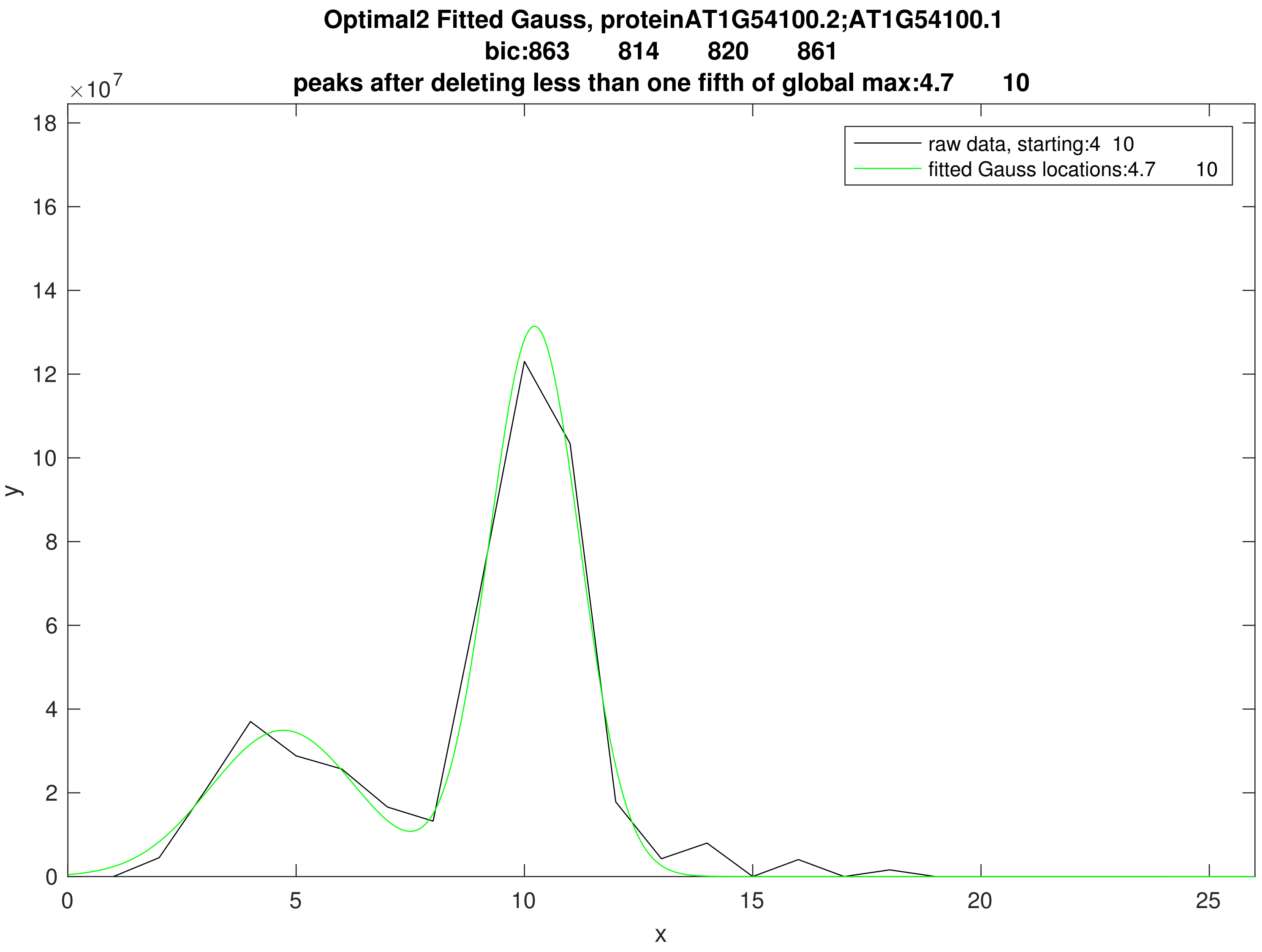
Optimal2 Fitted Gauss, proteinAT1G54100.2;AT1G54100.1
bic:863 814 820 861
peaks after deleting less than one fifth of global max:4.7 10
×10^7
raw data, starting:4 10
fitted Gauss locations:4.7 10
y
x
0
2
4
6
8
10
12
14
16
18
0
5
10
15
20
25

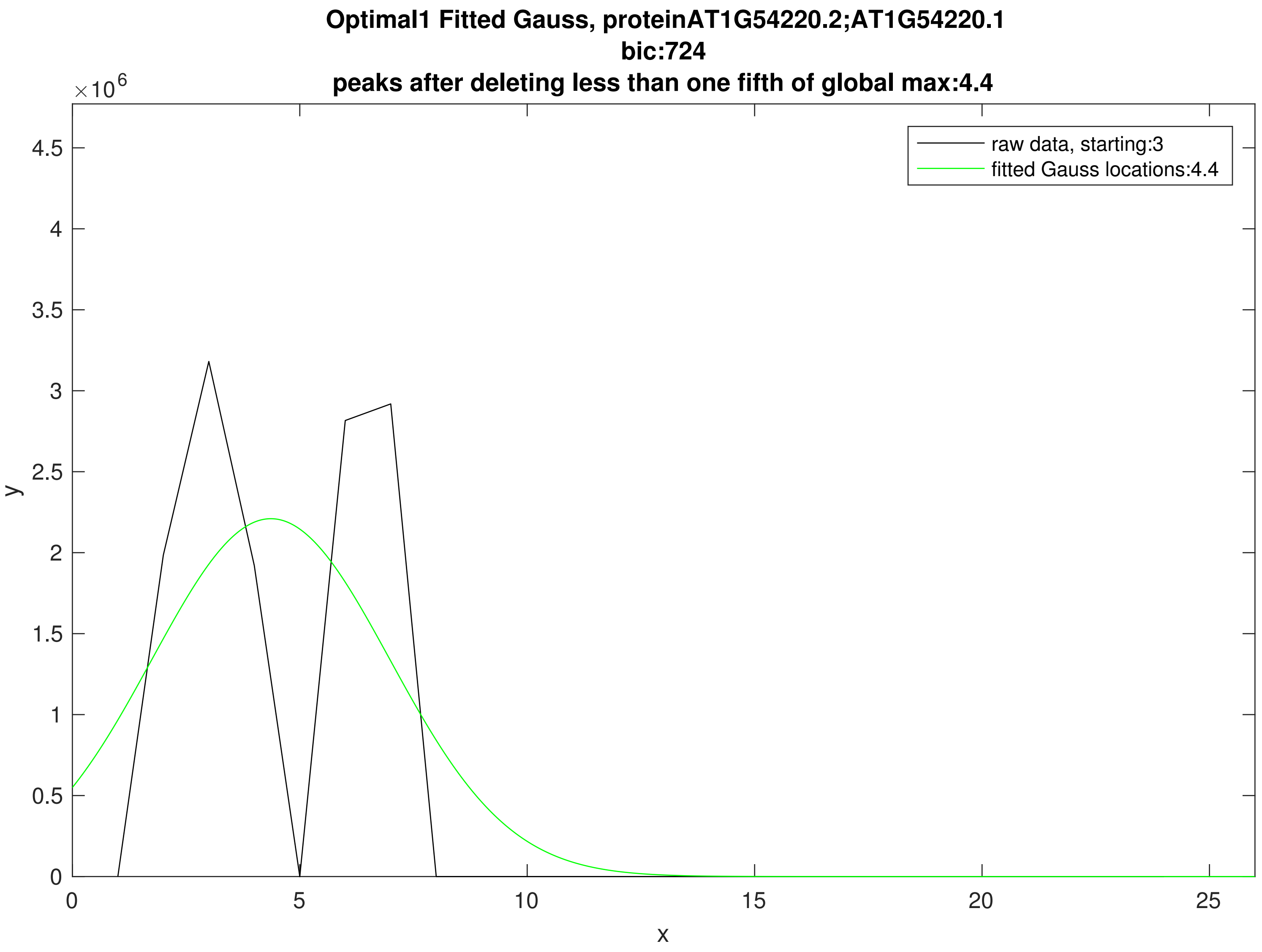
Optimal1 Fitted Gauss, proteinAT1G54220.2;AT1G54220.1
bic:724
peaks after deleting less than one fifth of global max:4.4
×10^6
raw data, starting:3
fitted Gauss locations:4.4
y
x
0
0.5
1
1.5
2
2.5
3
3.5
4
4.5
5
10
15
20
25

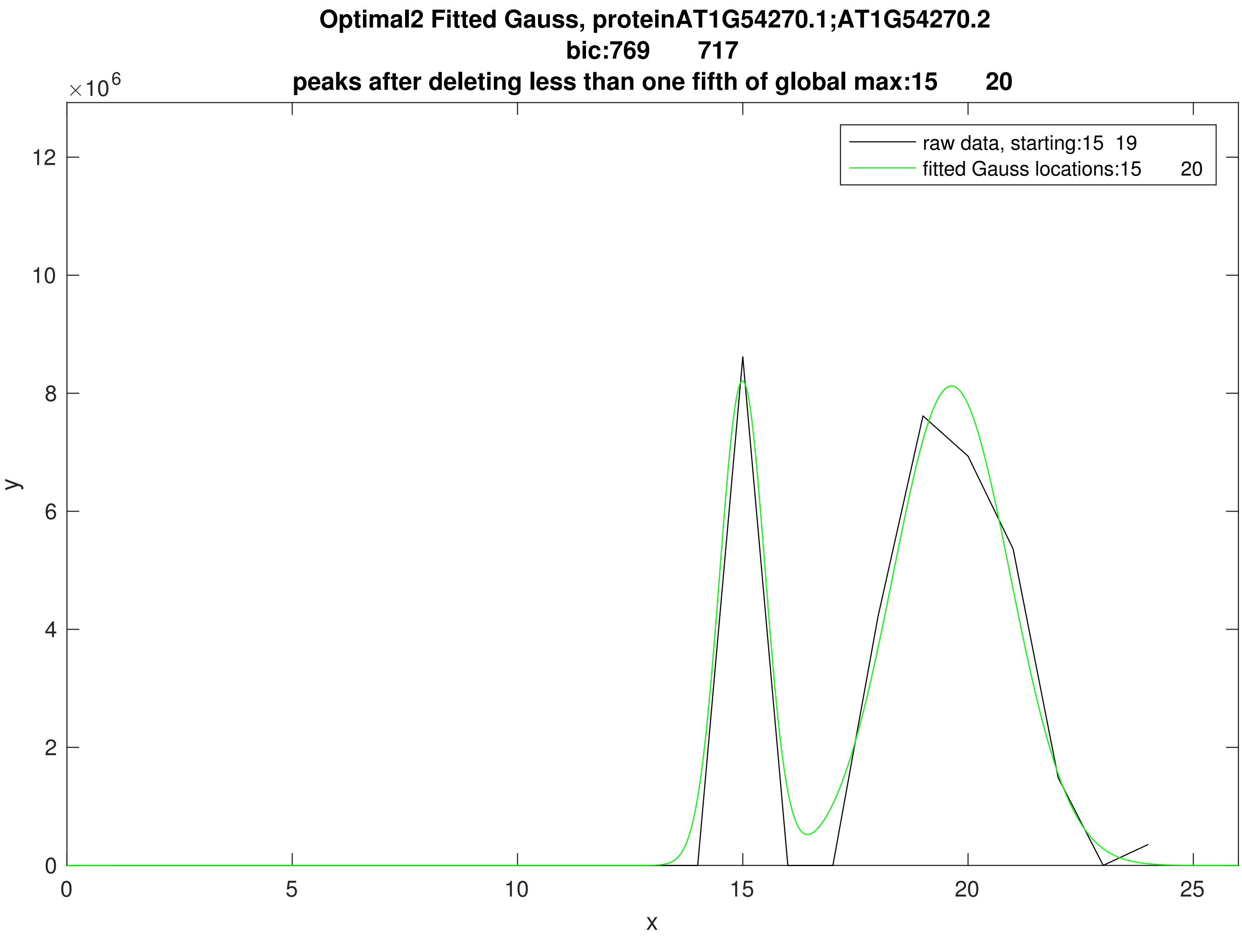

Optimal2 Fitted Gauss, proteinAT1G54270.1;AT1G54270.2
bic:769 717
peaks after deleting less than one fifth of global max:15 20
×10^6
raw data, starting:15 19
fitted Gauss locations:15 20
y
x
0
2
4
6
8
10
12
0
5
10
15
20
25

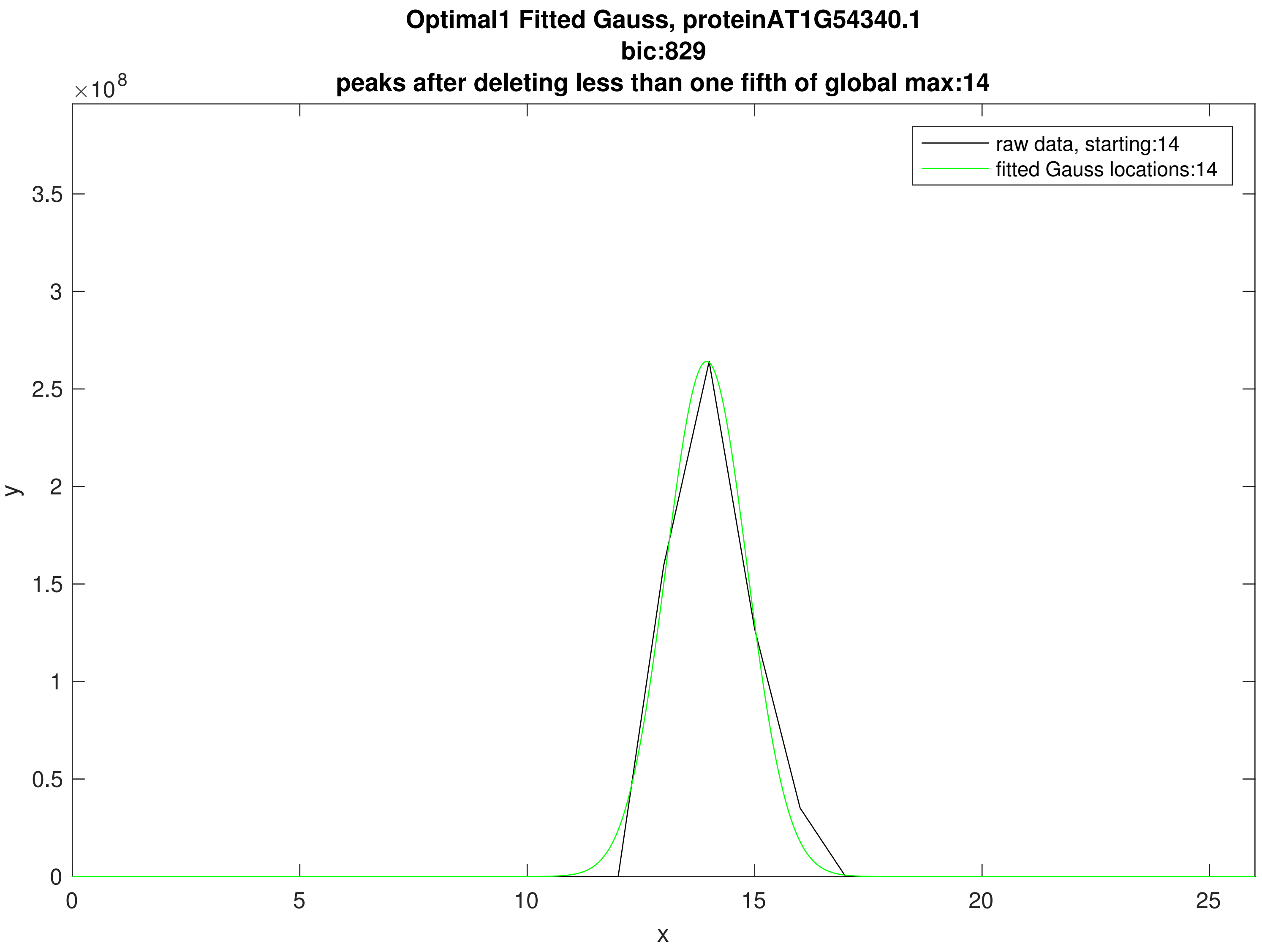

Optimal1 Fitted Gauss, proteinAT1G54340.1
bic:829
peaks after deleting less than one fifth of global max:14
×10^8
raw data, starting:14
fitted Gauss locations:14
3.5
3
2.5
2
1.5
1
0.5
0
y
0
5
10
15
20
25
x

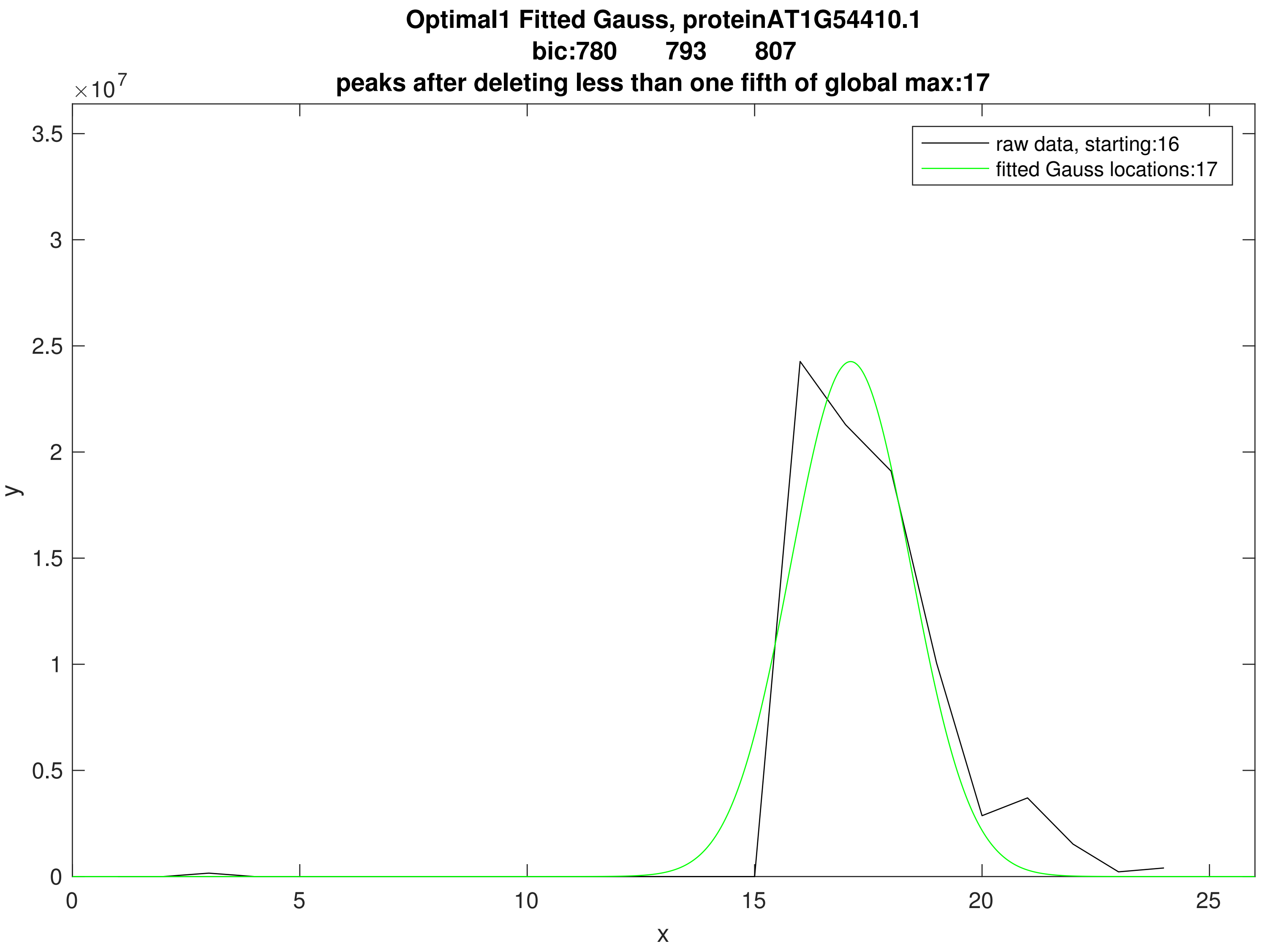

Optimal1 Fitted Gauss, proteinAT1G54410.1
bic:780 793 807
peaks after deleting less than one fifth of global max:17
×10^7
raw data, starting:16
fitted Gauss locations:17
y
x
0
0.5
1
1.5
2
2.5
3
3.5
0
5
10
15
20
25

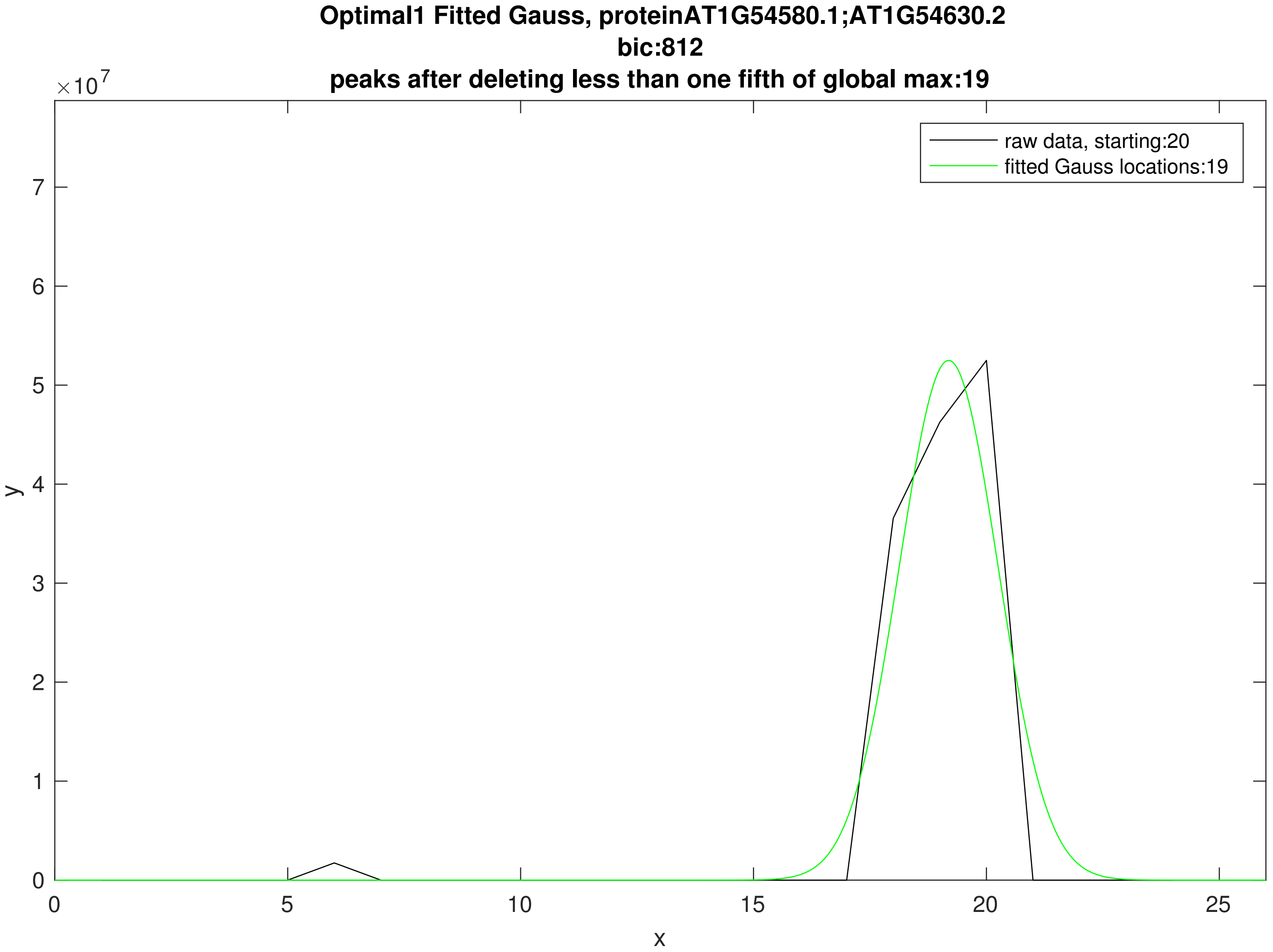
Optimal1 Fitted Gauss, proteinAT1G54580.1;AT1G54630.2
bic:812
peaks after deleting less than one fifth of global max:19
×10^7
raw data, starting:20
fitted Gauss locations:19
y
x
0
1
2
3
4
5
6
7
0
5
10
15
20
25

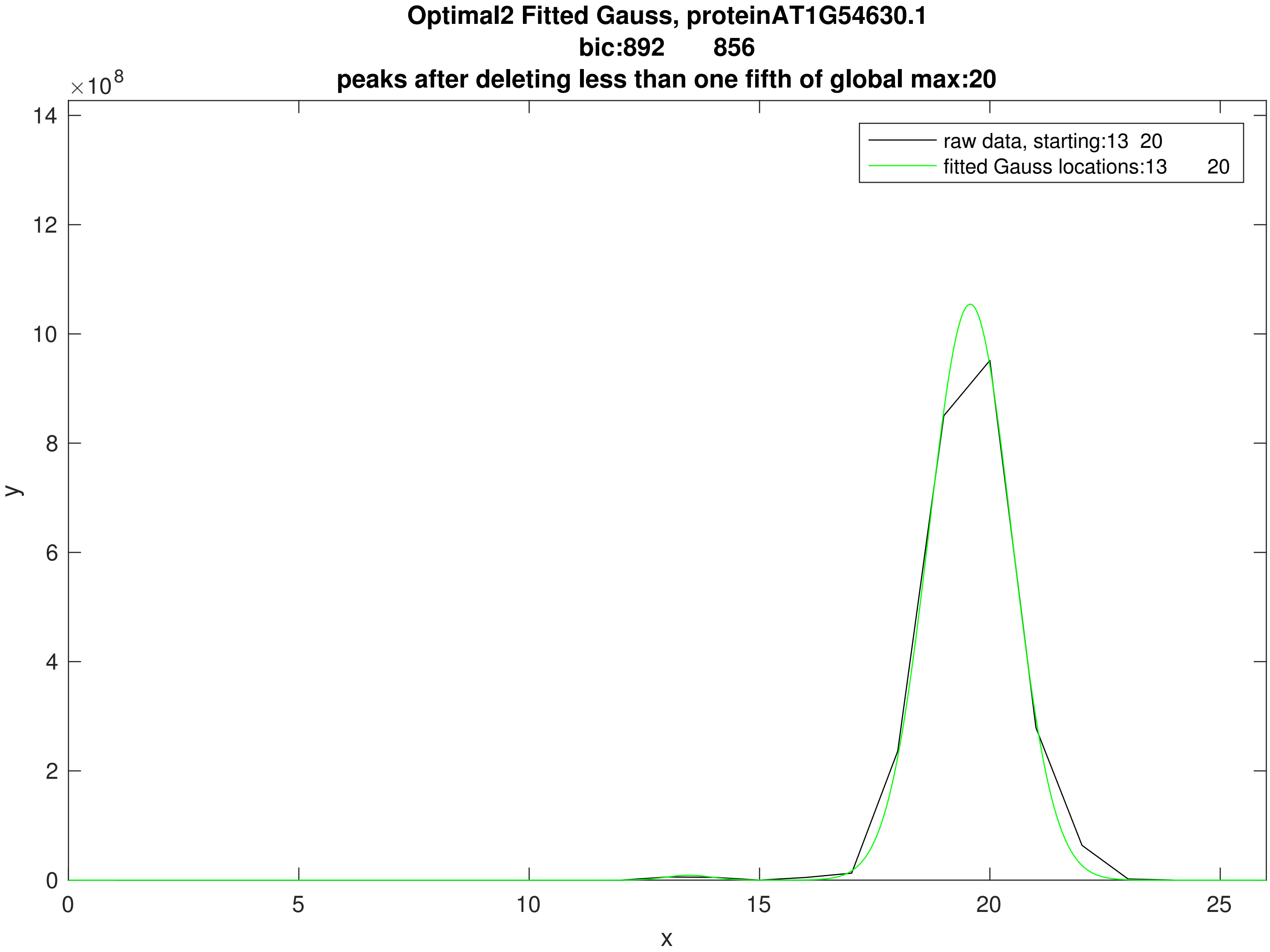
Optimal2 Fitted Gauss, proteinAT1G54630.1
bic:892 856
peaks after deleting less than one fifth of global max:20
×10^8
14
12
10
8
6
4
2
0
y
0
5
10
15
20
25
x
raw data, starting:13 20
fitted Gauss locations:13 20

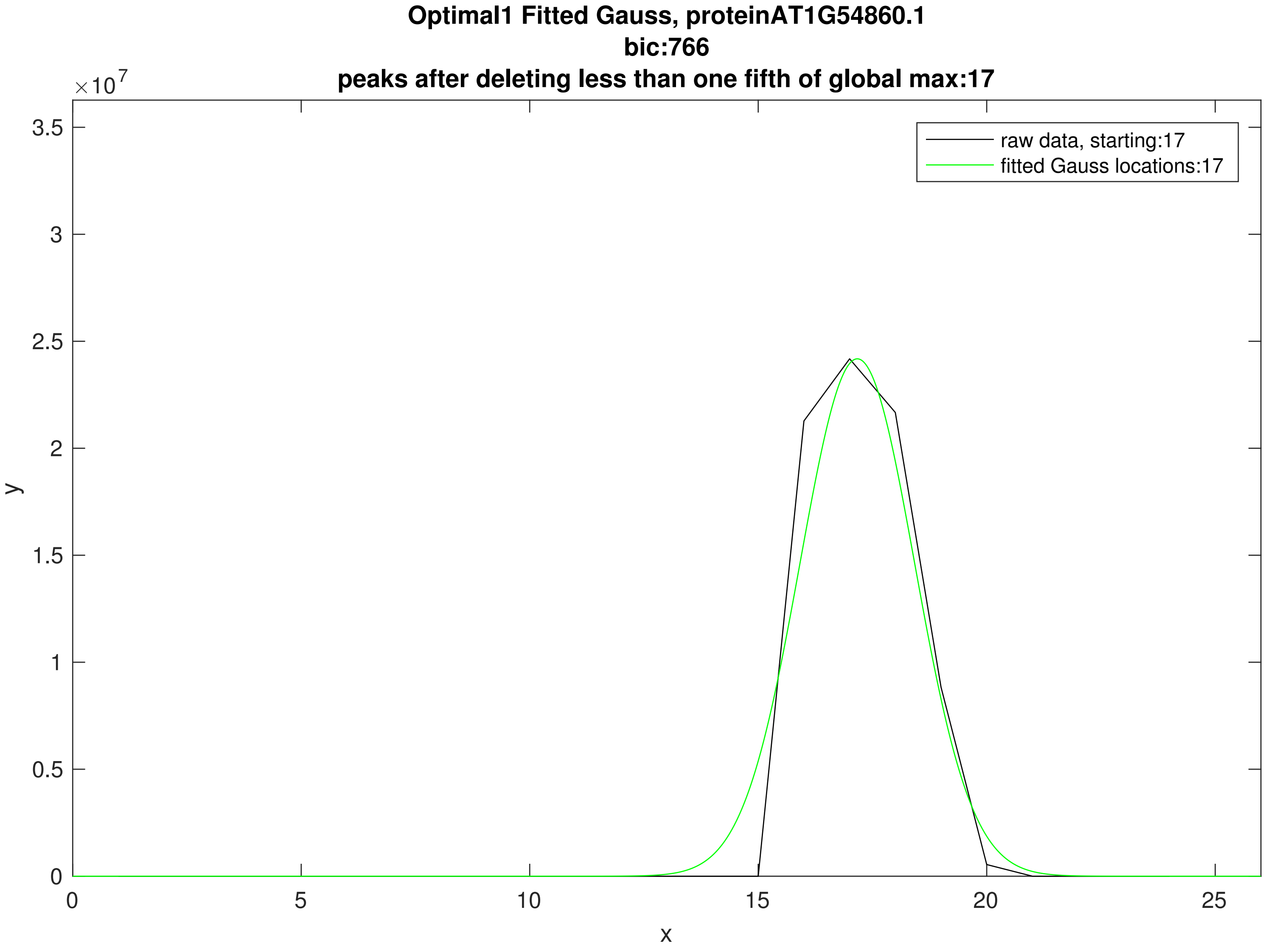
Optimal1 Fitted Gauss, proteinAT1G54860.1
bic:766
peaks after deleting less than one fifth of global max:17
×10^7
raw data, starting:17
fitted Gauss locations:17
y
x
0
0.5
1
1.5
2
2.5
3
3.5
0
5
10
15
20
25

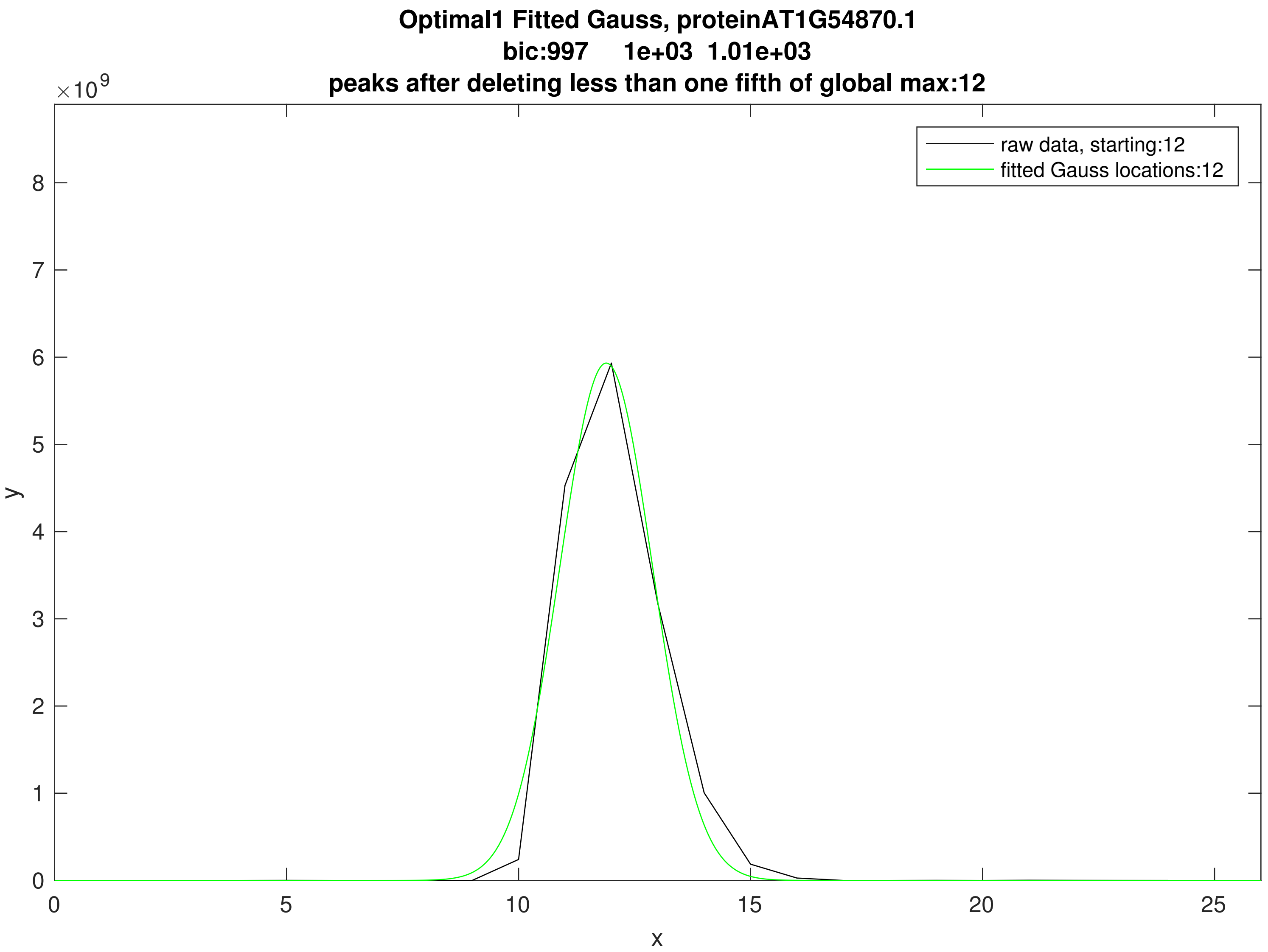
Optimal1 Fitted Gauss, proteinAT1G54870.1
bic:997 1e+03 1.01e+03
peaks after deleting less than one fifth of global max:12
×10^9
raw data, starting:12
fitted Gauss locations:12
y
x
0
1
2
3
4
5
6
7
8
0
5
10
15
20
25

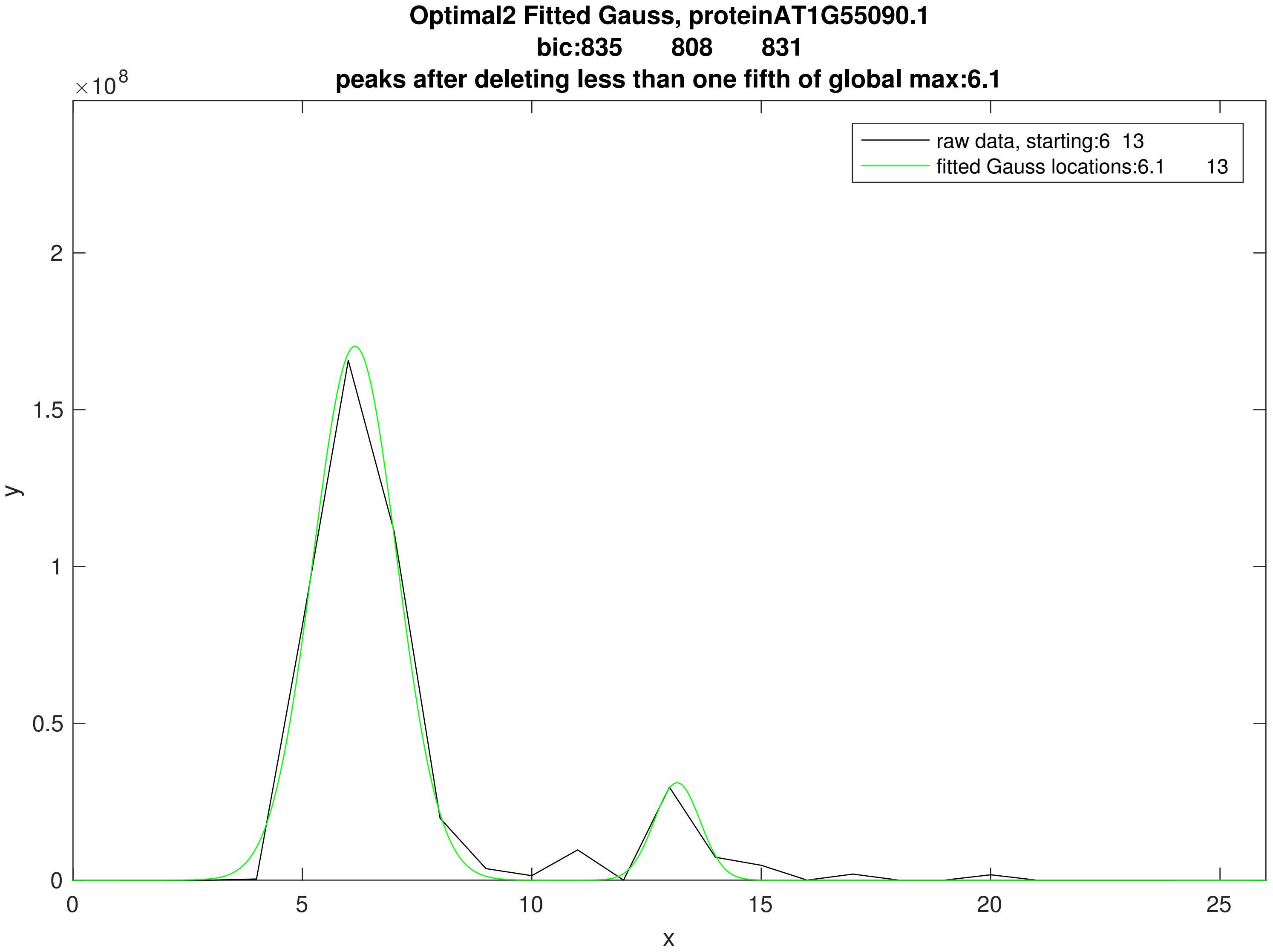

Optimal2 Fitted Gauss, proteinAT1G55090.1
bic:835 808 831
peaks after deleting less than one fifth of global max:6.1
×10^8
raw data, starting:6 13
fitted Gauss locations:6.1 13
2
1.5
1
0.5
0
y
0
5
10
15
20
25
x

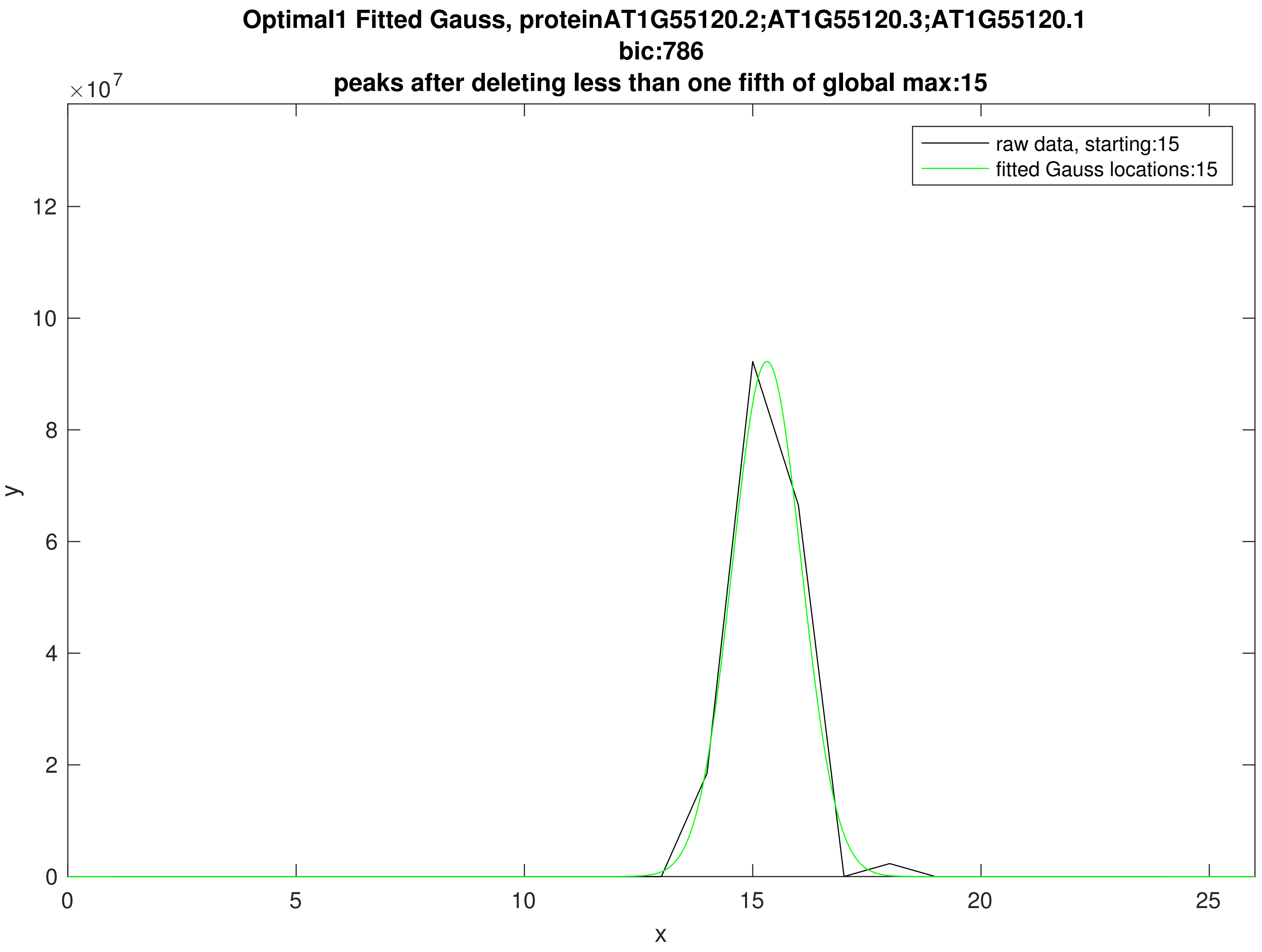

Optimal1 Fitted Gauss, proteinAT1G55120.2;AT1G55120.3;AT1G55120.1
bic:786
peaks after deleting less than one fifth of global max:15
×10^7
raw data, starting:15
fitted Gauss locations:15
y
x
0
2
4
6
8
10
12
0
5
10
15
20
25

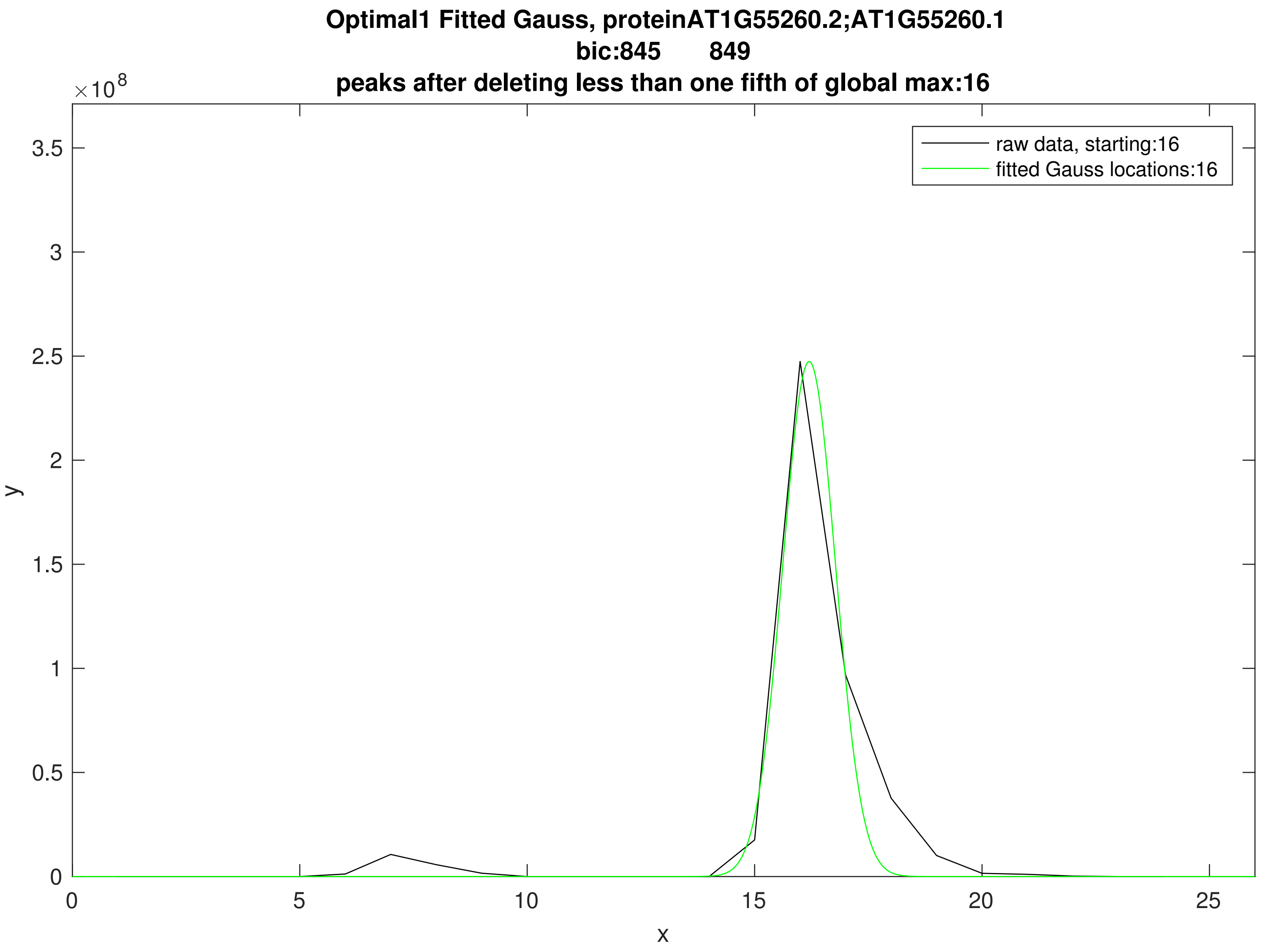
Optimal1 Fitted Gauss, proteinAT1G55260.2;AT1G55260.1
bic:845 849
peaks after deleting less than one fifth of global max:16
×10^8
raw data, starting:16
fitted Gauss locations:16
y
x
0
0.5
1
1.5
2
2.5
3
3.5
0
5
10
15
20
25

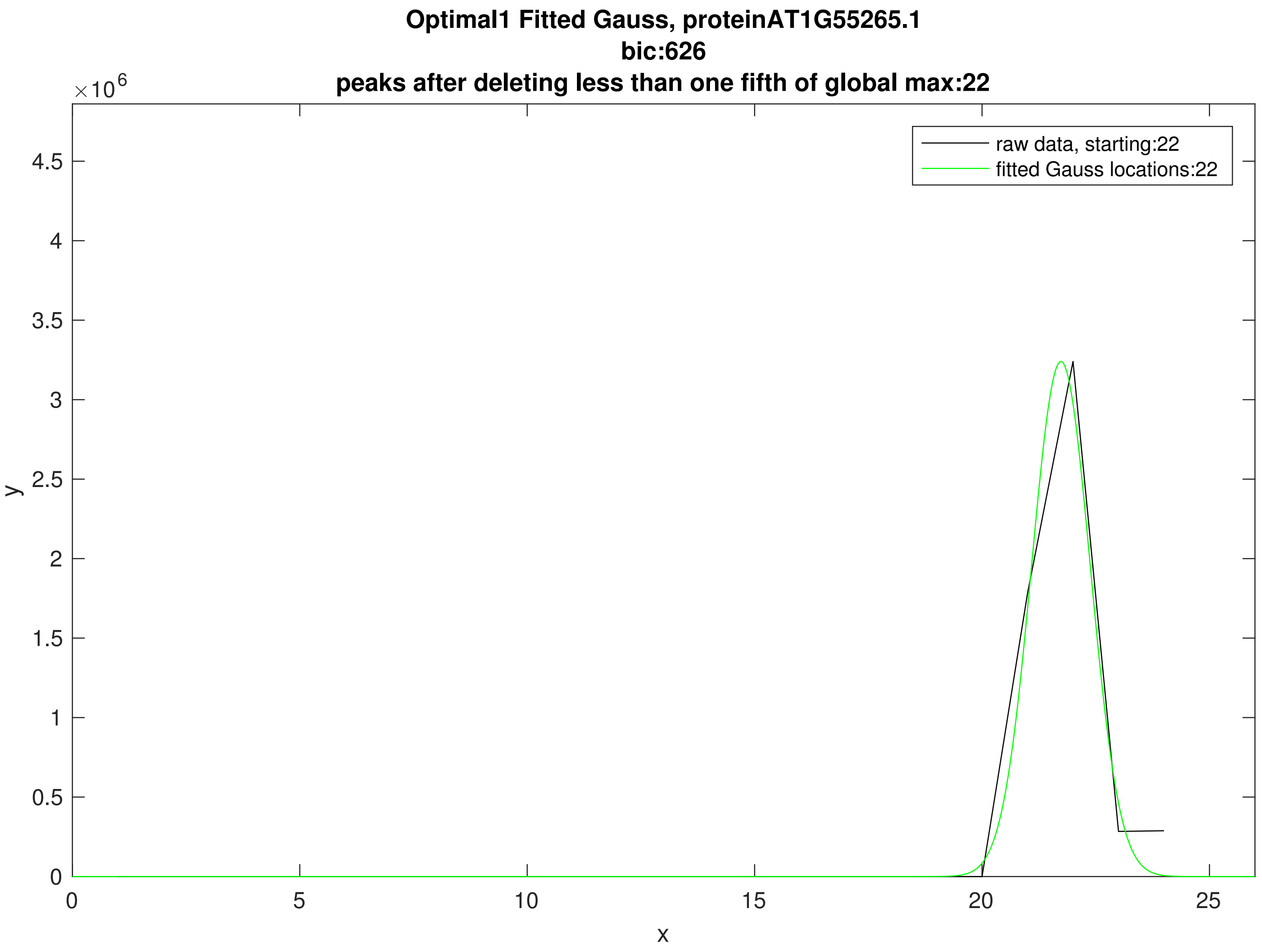
Optimal1 Fitted Gauss, proteinAT1G55265.1
bic:626
peaks after deleting less than one fifth of global max:22
×10^6
raw data, starting:22
fitted Gauss locations:22
y
x
0
0.5
1
1.5
2
2.5
3
3.5
4
4.5
5
10
15
20
25

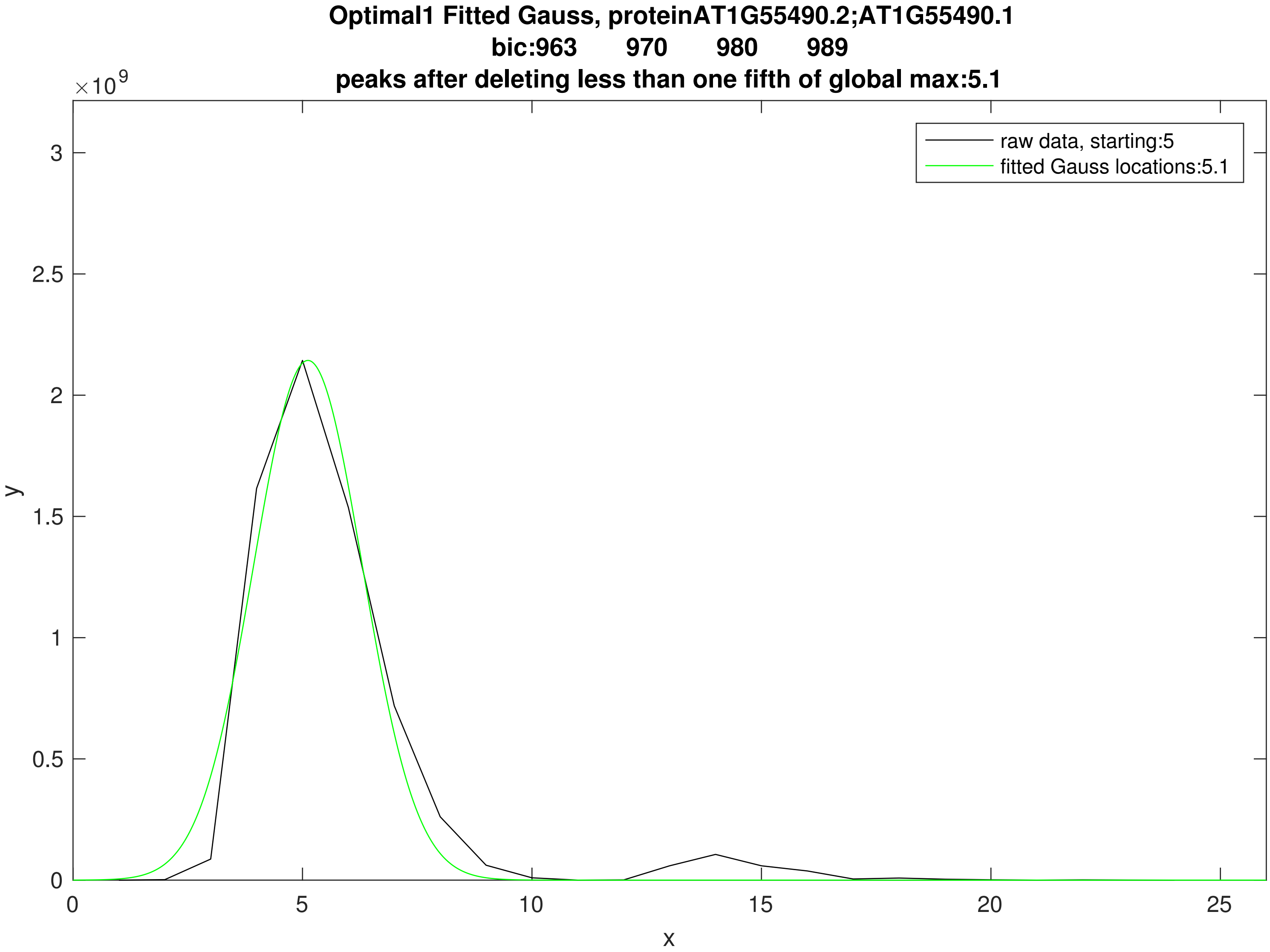
Optimal1 Fitted Gauss, proteinAT1G55490.2;AT1G55490.1
bic:963 970 980 989
peaks after deleting less than one fifth of global max:5.1
×10^9
3
2.5
2
1.5
1
0.5
0
y
0
5
10
15
20
25
x
raw data, starting:5
fitted Gauss locations:5.1

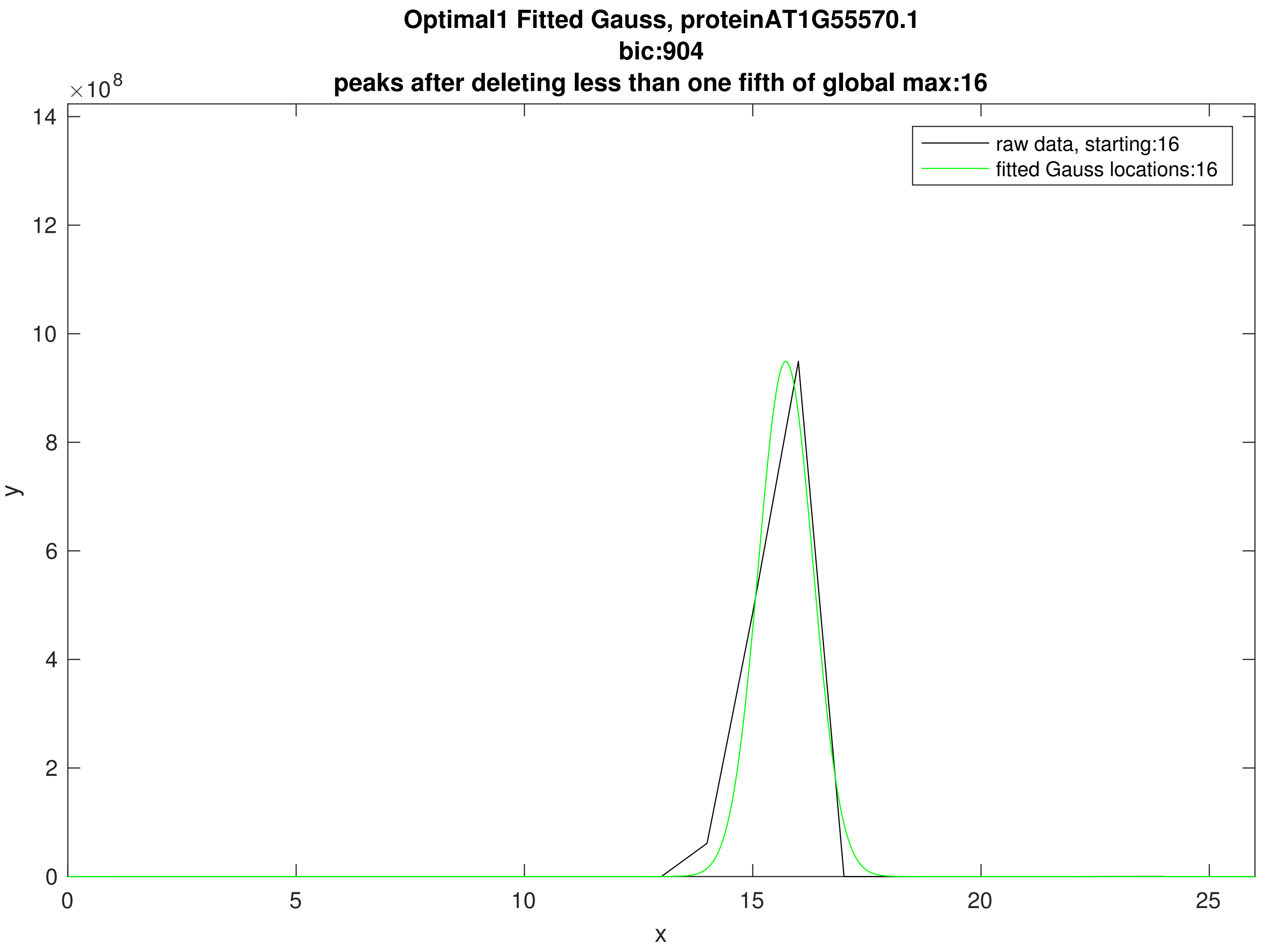

Optimal1 Fitted Gauss, proteinAT1G55570.1
bic:904
peaks after deleting less than one fifth of global max:16
×10^8
14
12
10
8
6
4
2
0
y
0
5
10
15
20
25
x
raw data, starting:16
fitted Gauss locations:16

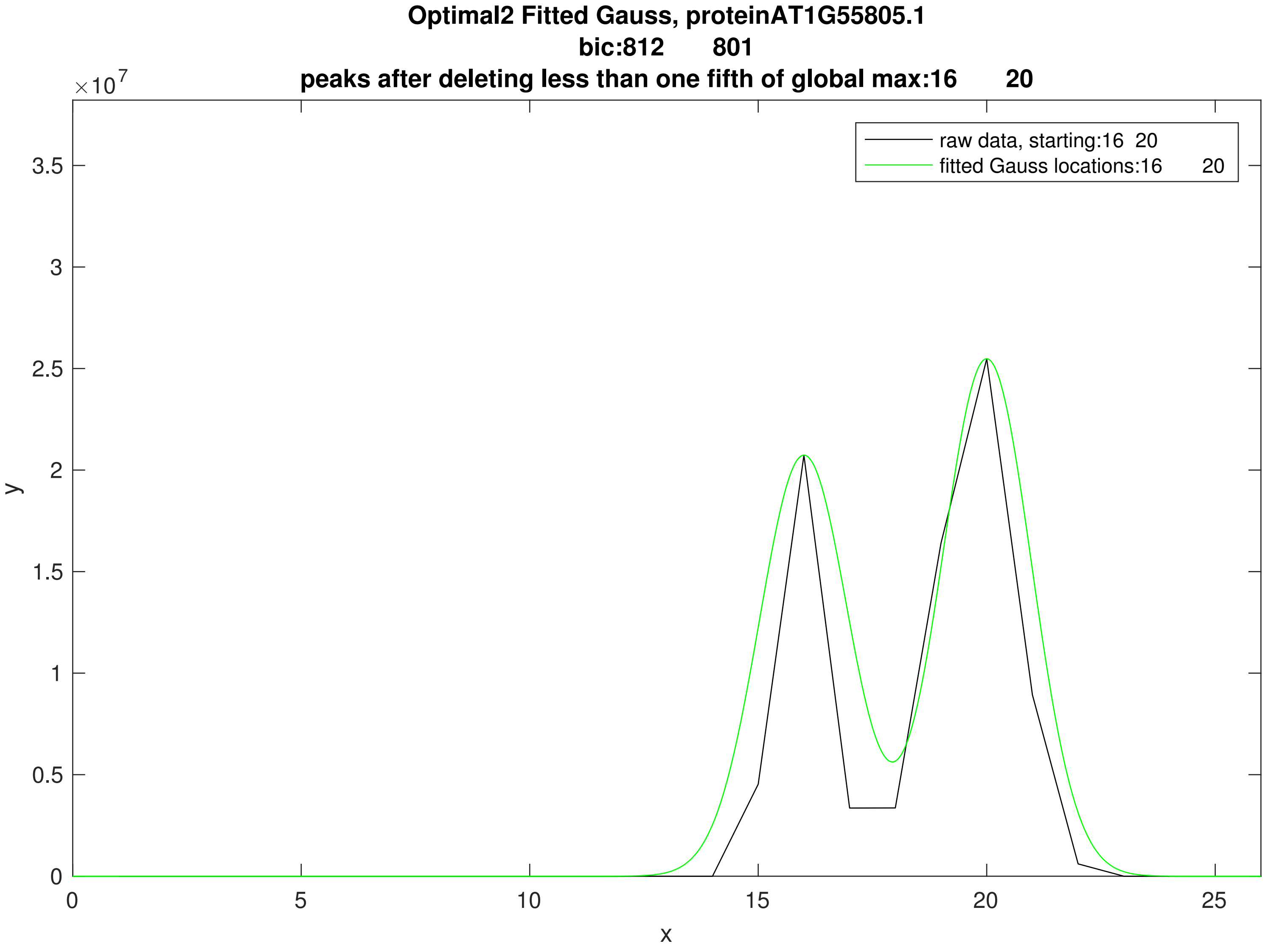
Optimal2 Fitted Gauss, proteinAT1G55805.1
bic:812 801
peaks after deleting less than one fifth of global max:16 20
×10^7
raw data, starting:16 20
fitted Gauss locations:16 20
y
x
0
0.5
1
1.5
2
2.5
3
3.5
5
10
15
20
25

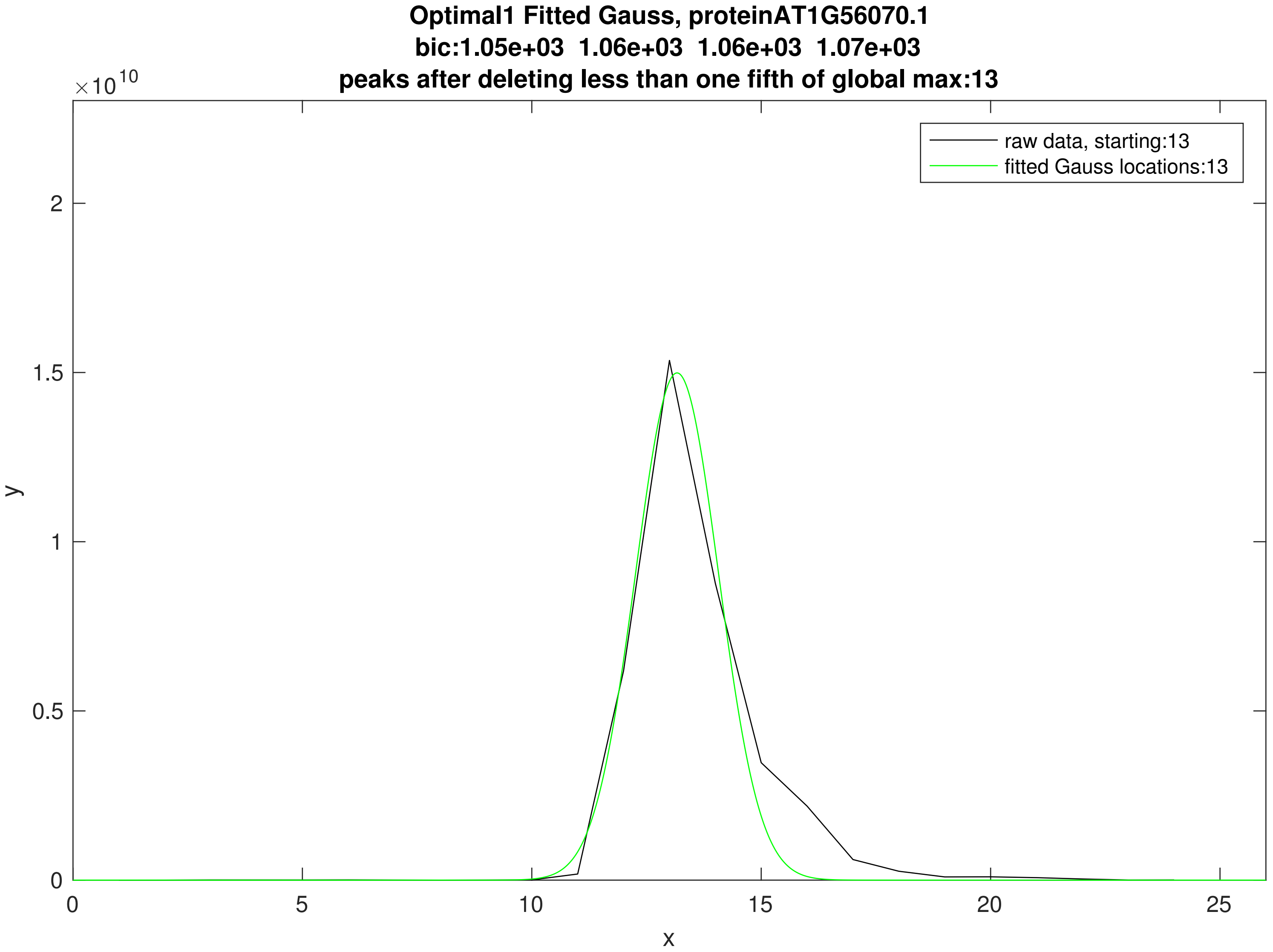
Optimal1 Fitted Gauss, proteinAT1G56070.1
bic:1.05e+03  1.06e+03  1.06e+03  1.07e+03
peaks after deleting less than one fifth of global max:13
×10^10
raw data, starting:13
fitted Gauss locations:13
y
x
0
0.5
1
1.5
2
0
5
10
15
20
25

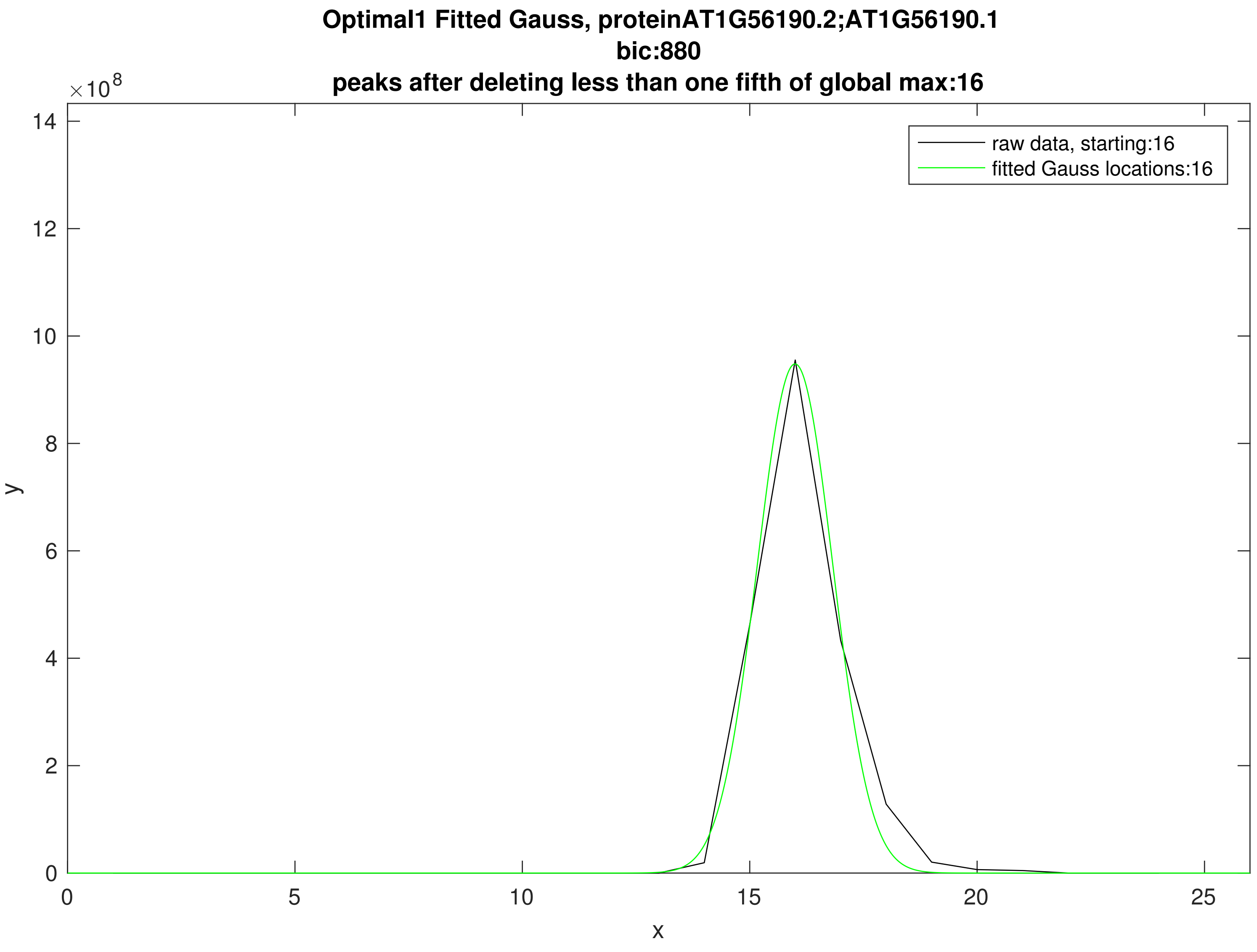
Optimal1 Fitted Gauss, proteinAT1G56190.2;AT1G56190.1
bic:880
peaks after deleting less than one fifth of global max:16
×10^8
raw data, starting:16
fitted Gauss locations:16
y
x
0
2
4
6
8
10
12
14
0
5
10
15
20
25

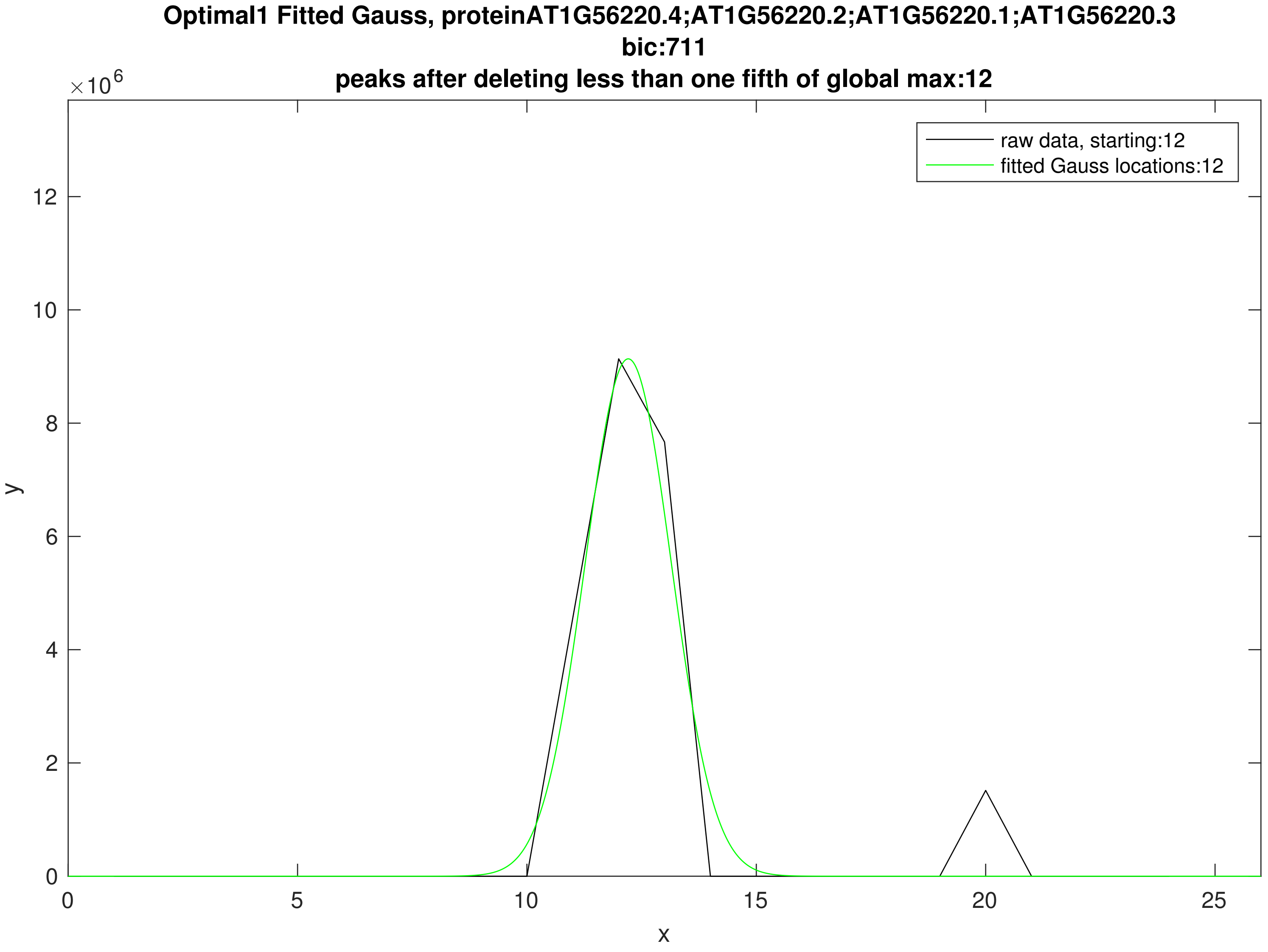
Optimal1 Fitted Gauss, proteinAT1G56220.4;AT1G56220.2;AT1G56220.1;AT1G56220.3
bic:711
peaks after deleting less than one fifth of global max:12
×10^6
raw data, starting:12
fitted Gauss locations:12
y
x
0
2
4
6
8
10
12
0
5
10
15
20
25

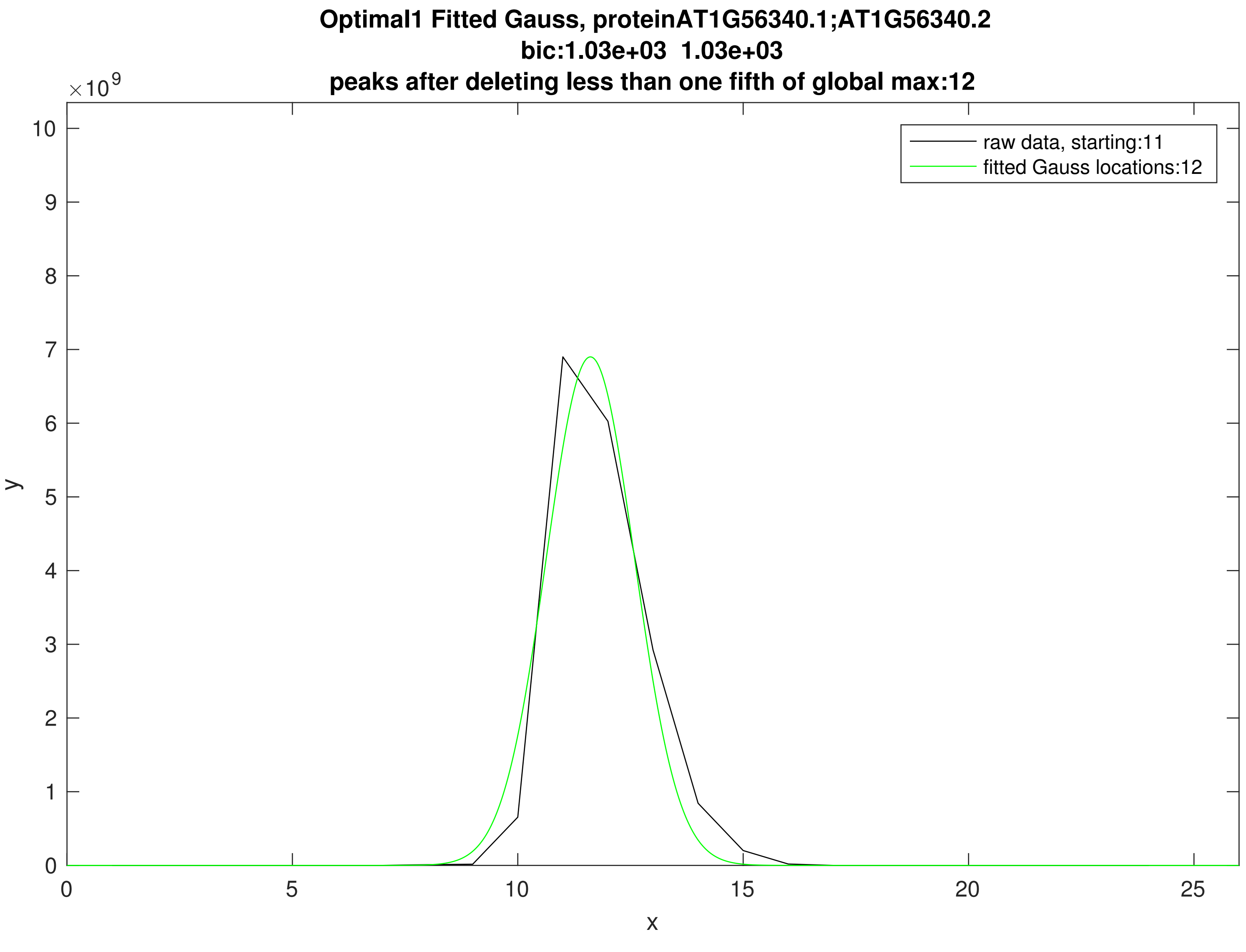
Optimal1 Fitted Gauss, proteinAT1G56340.1;AT1G56340.2
bic:1.03e+03  1.03e+03
peaks after deleting less than one fifth of global max:12
×10^9
raw data, starting:11
fitted Gauss locations:12
y
x
0
1
2
3
4
5
6
7
8
9
10
0
5
10
15
20
25

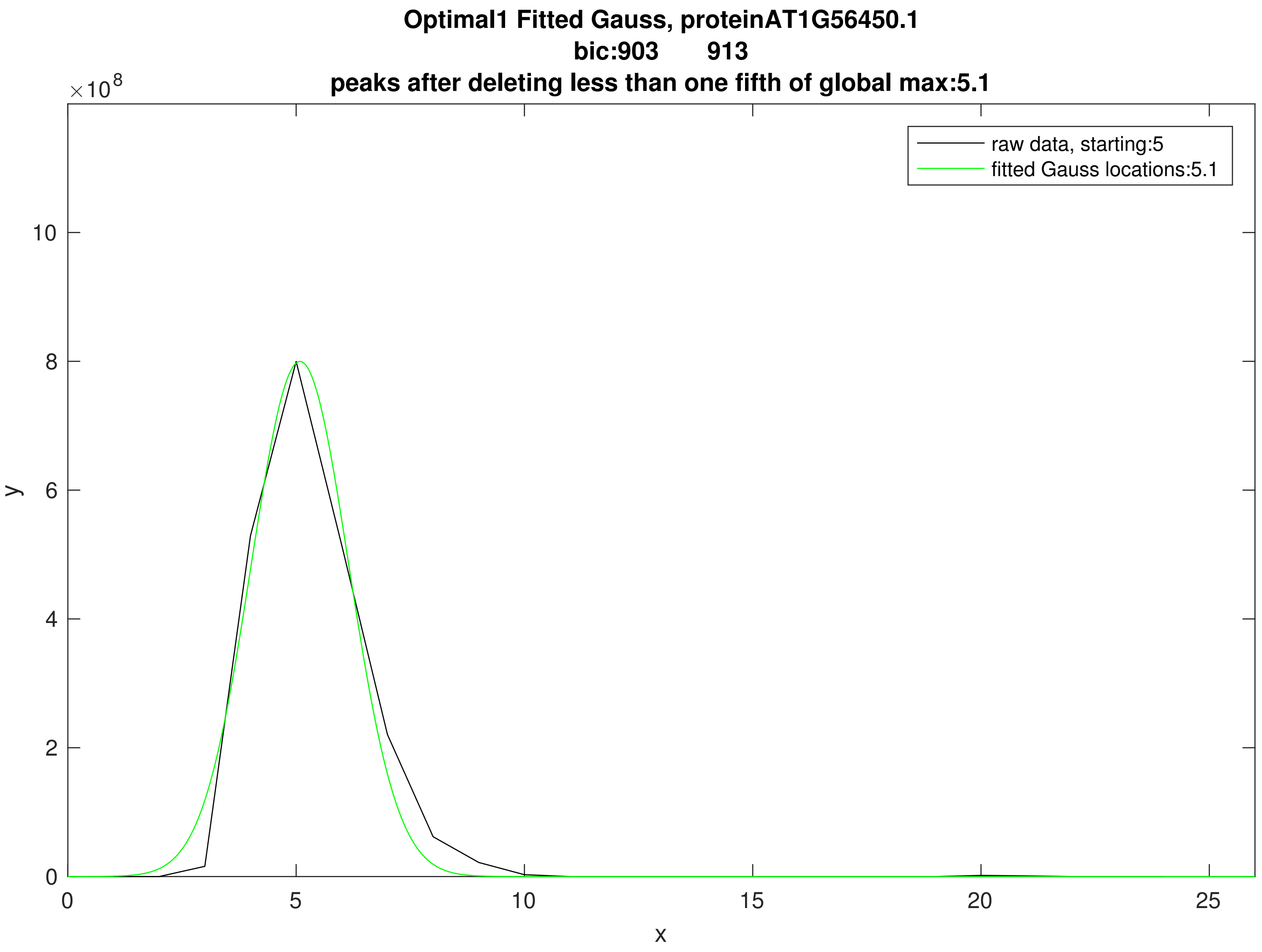
Optimal1 Fitted Gauss, proteinAT1G56450.1
bic:903 913
peaks after deleting less than one fifth of global max:5.1
×10^8
raw data, starting:5
fitted Gauss locations:5.1
y
x
0
2
4
6
8
10
0
5
10
15
20
25

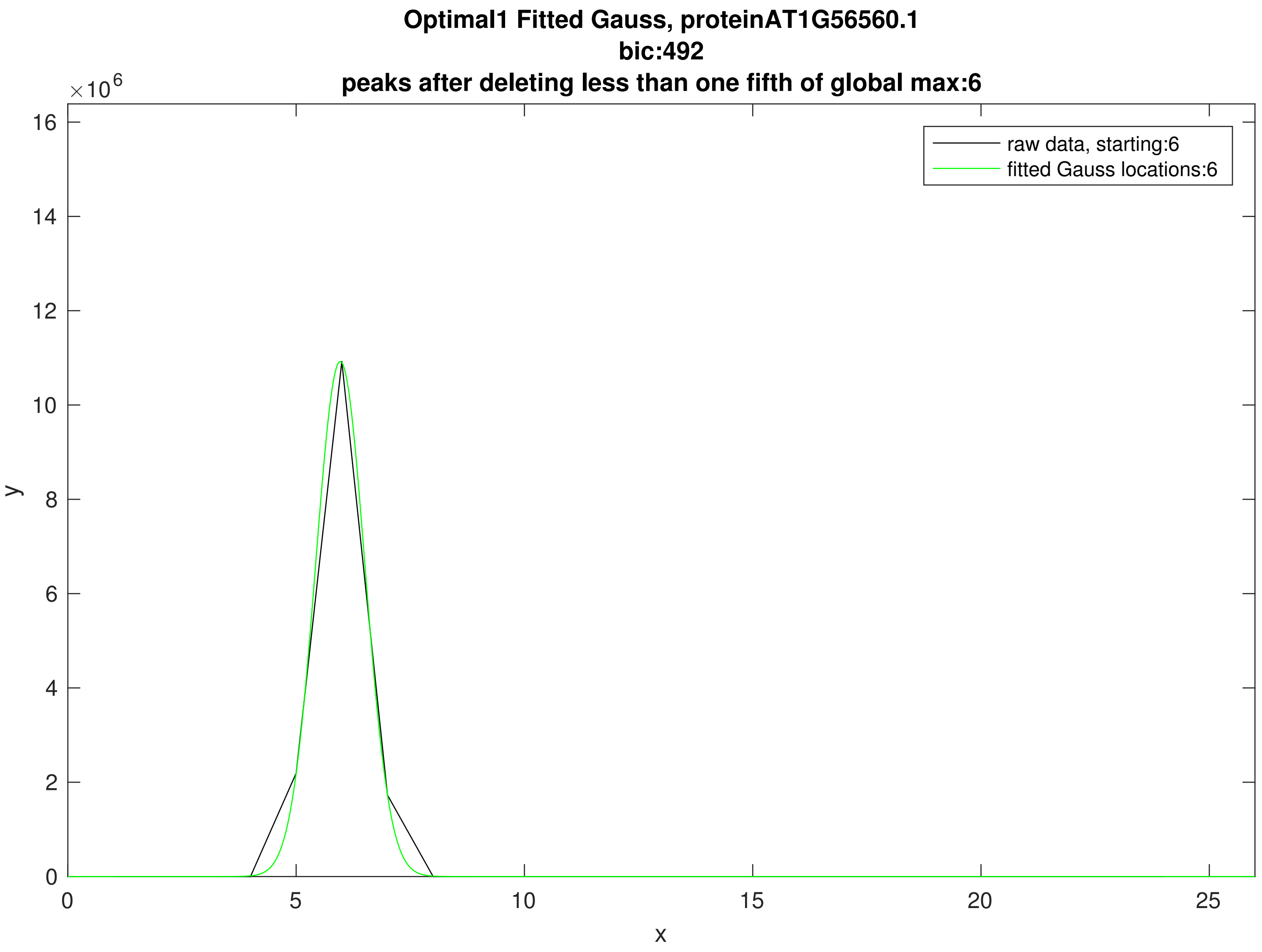
Optimal1 Fitted Gauss, proteinAT1G56560.1
bic:492
peaks after deleting less than one fifth of global max:6
×10^6
16
14
12
10
8
6
4
2
0
y
0
5
10
15
20
25
x
raw data, starting:6
fitted Gauss locations:6

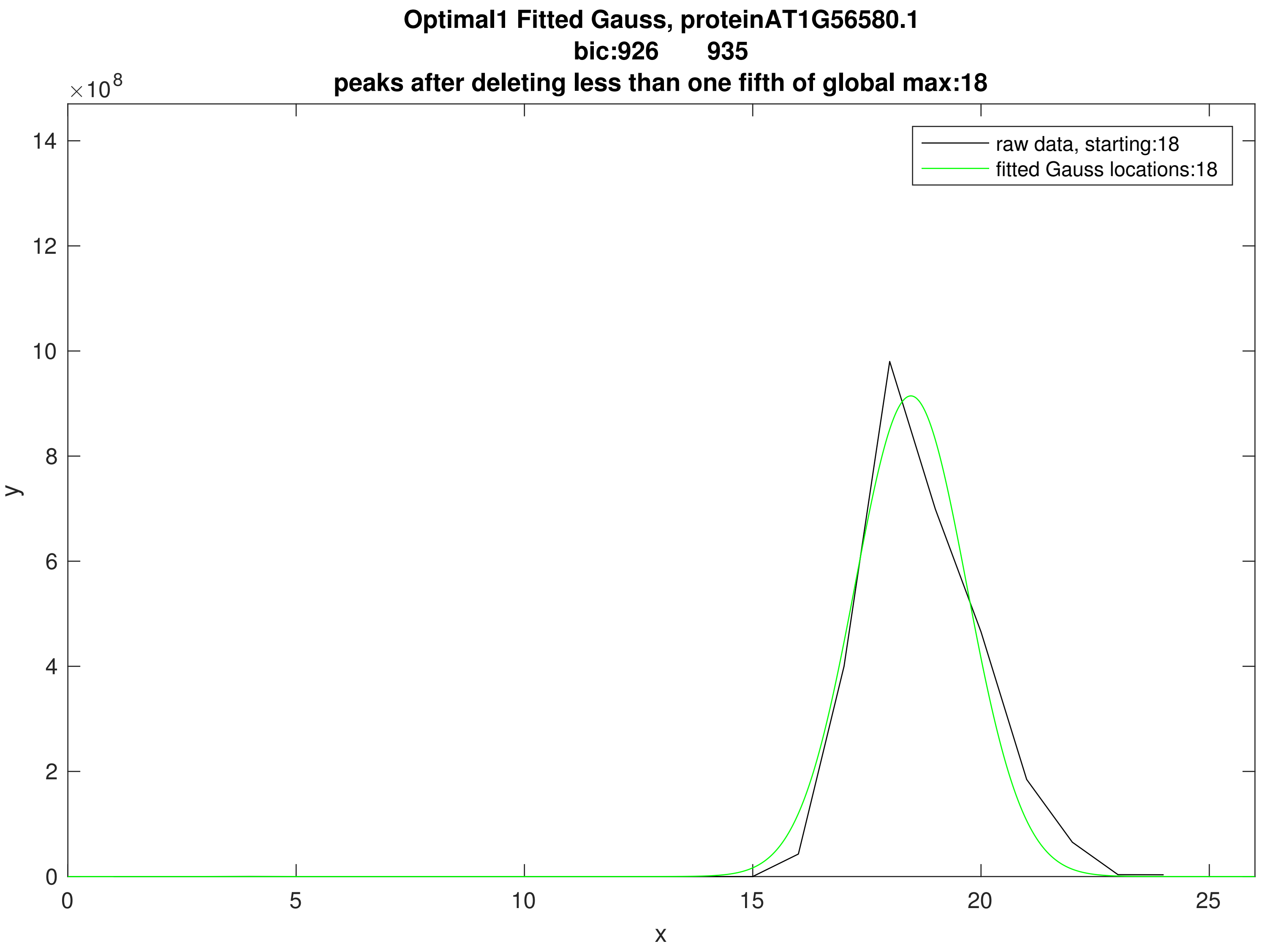

Optimal1 Fitted Gauss, proteinAT1G56580.1
bic:926 935
peaks after deleting less than one fifth of global max:18
×10^8
14
12
10
8
6
4
2
0
y
0
5
10
15
20
25
x
raw data, starting:18
fitted Gauss locations:18

**Optimal1 Fitted Gauss, proteinAT1G56700.1;AT1G56700.2;AT1G56700.3**
**bic:671**
**peaks after deleting less than one fifth of global max:19**

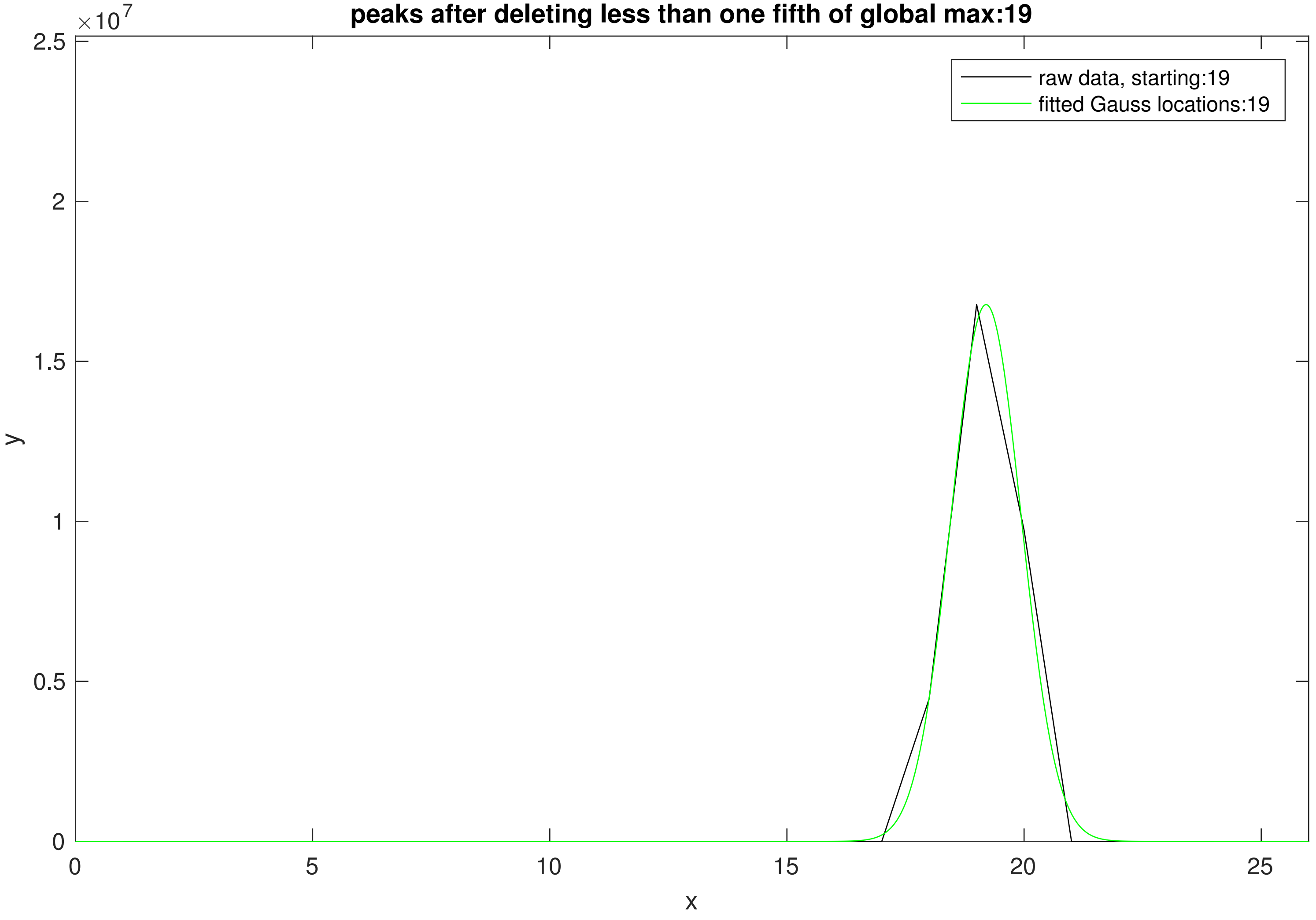

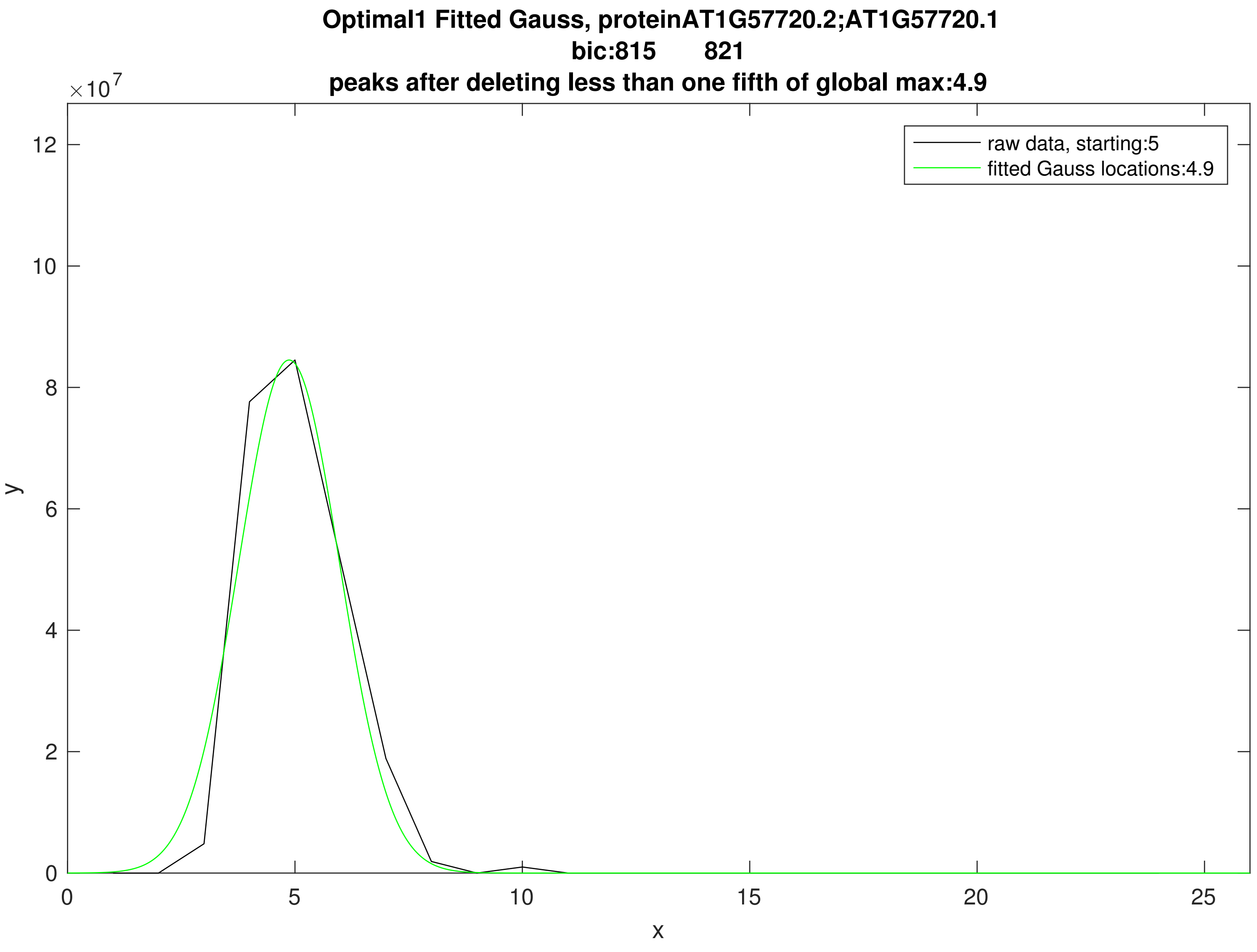
Optimal1 Fitted Gauss, proteinAT1G57720.2;AT1G57720.1
bic:815 821
peaks after deleting less than one fifth of global max:4.9
×10^7
12
10
8
6
4
2
0
y
0
5
10
15
20
25
x
raw data, starting:5
fitted Gauss locations:4.9

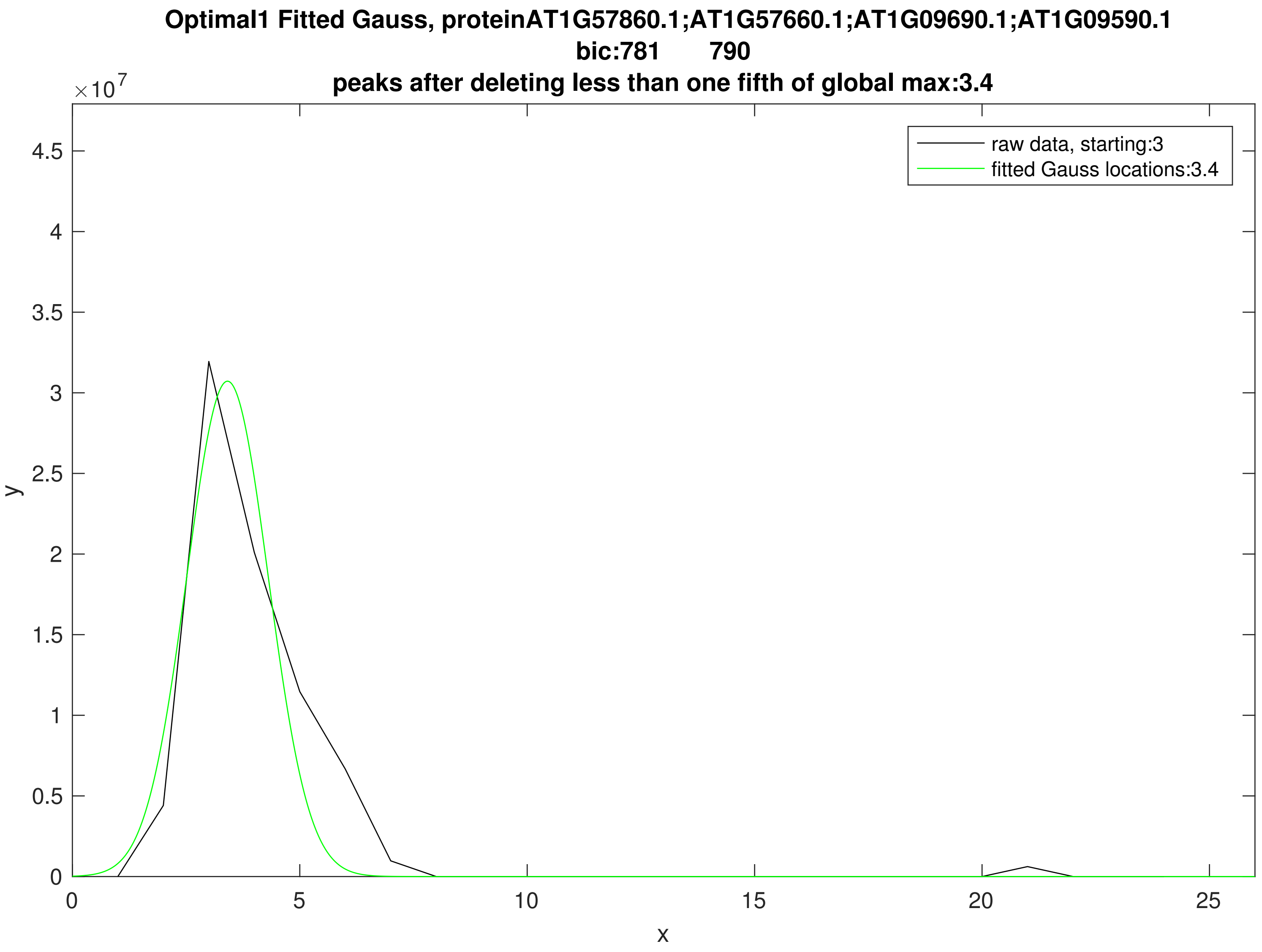
Optimal1 Fitted Gauss, proteinAT1G57860.1;AT1G57660.1;AT1G09690.1;AT1G09590.1
bic:781 790
peaks after deleting less than one fifth of global max:3.4
×10^7
raw data, starting:3
fitted Gauss locations:3.4
y
x
0
0.5
1
1.5
2
2.5
3
3.5
4
4.5
0
5
10
15
20
25

**Optimal1 Fitted Gauss, proteinAT1G58080.1**
**bic:822 826**
**peaks after deleting less than one fifth of global max:11**

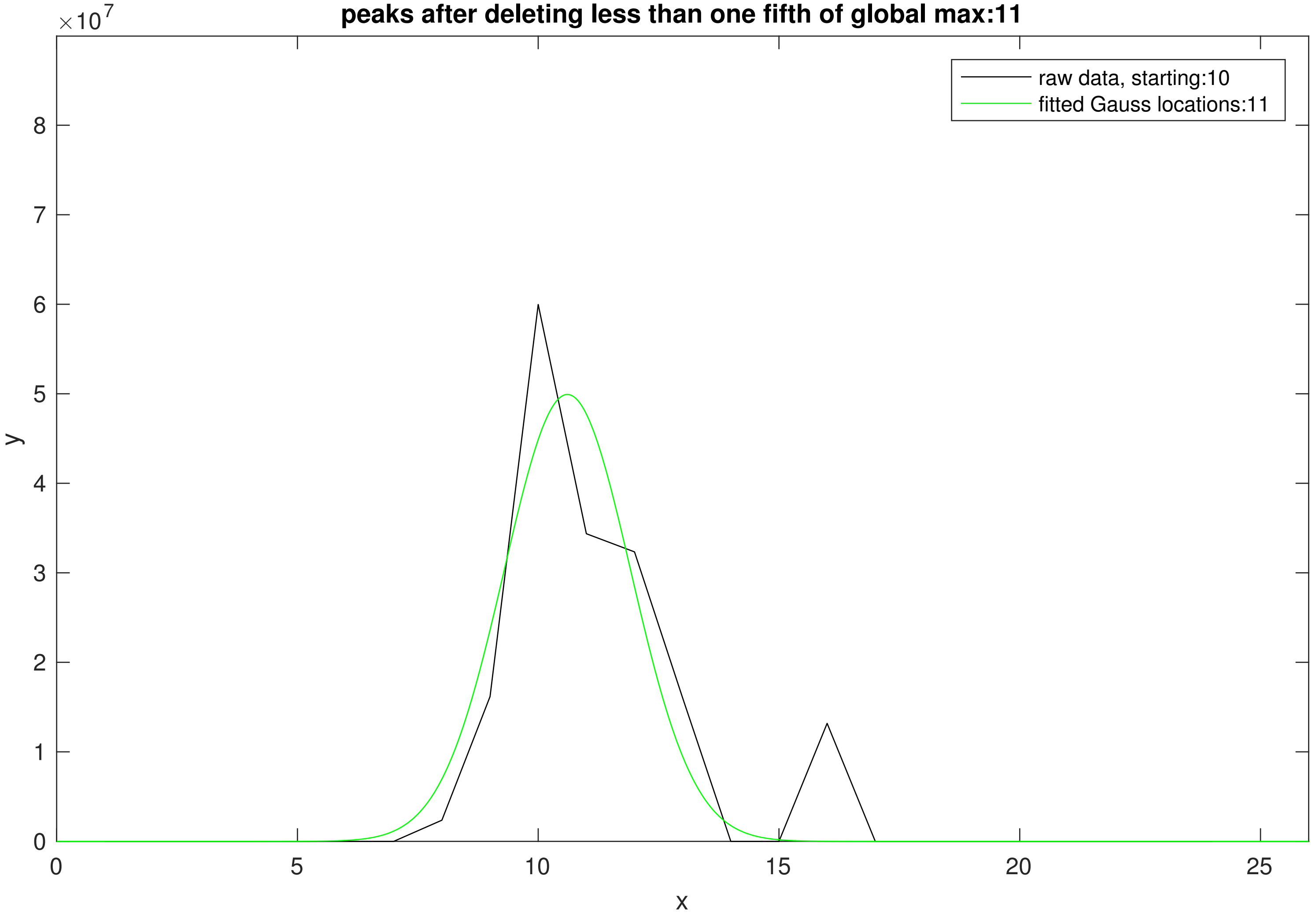

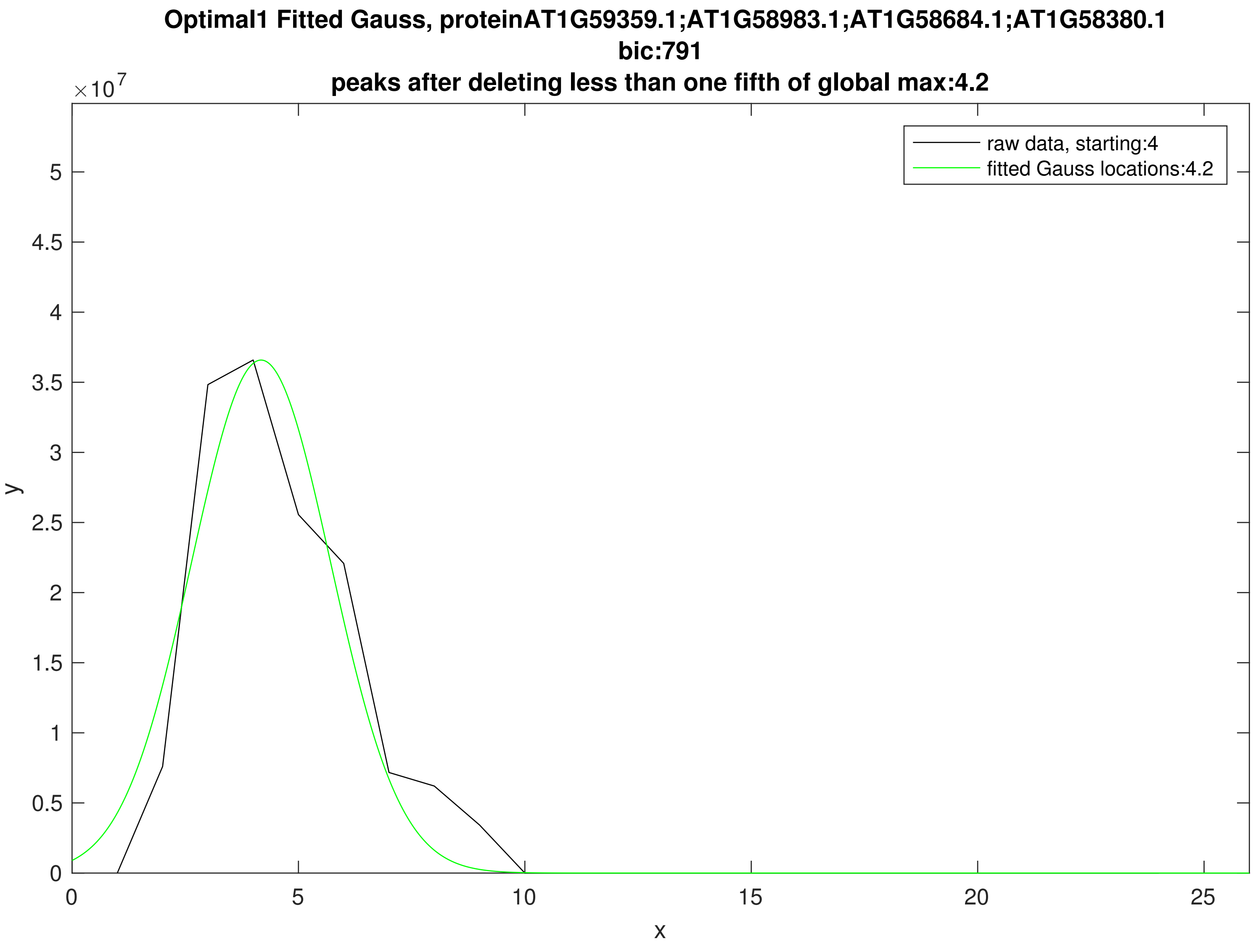

Optimal1 Fitted Gauss, proteinAT1G59359.1;AT1G58983.1;AT1G58684.1;AT1G58380.1
bic:791
peaks after deleting less than one fifth of global max:4.2
×10^7
raw data, starting:4
fitted Gauss locations:4.2
y
x
0
0.5
1
1.5
2
2.5
3
3.5
4
4.5
5
0
5
10
15
20
25

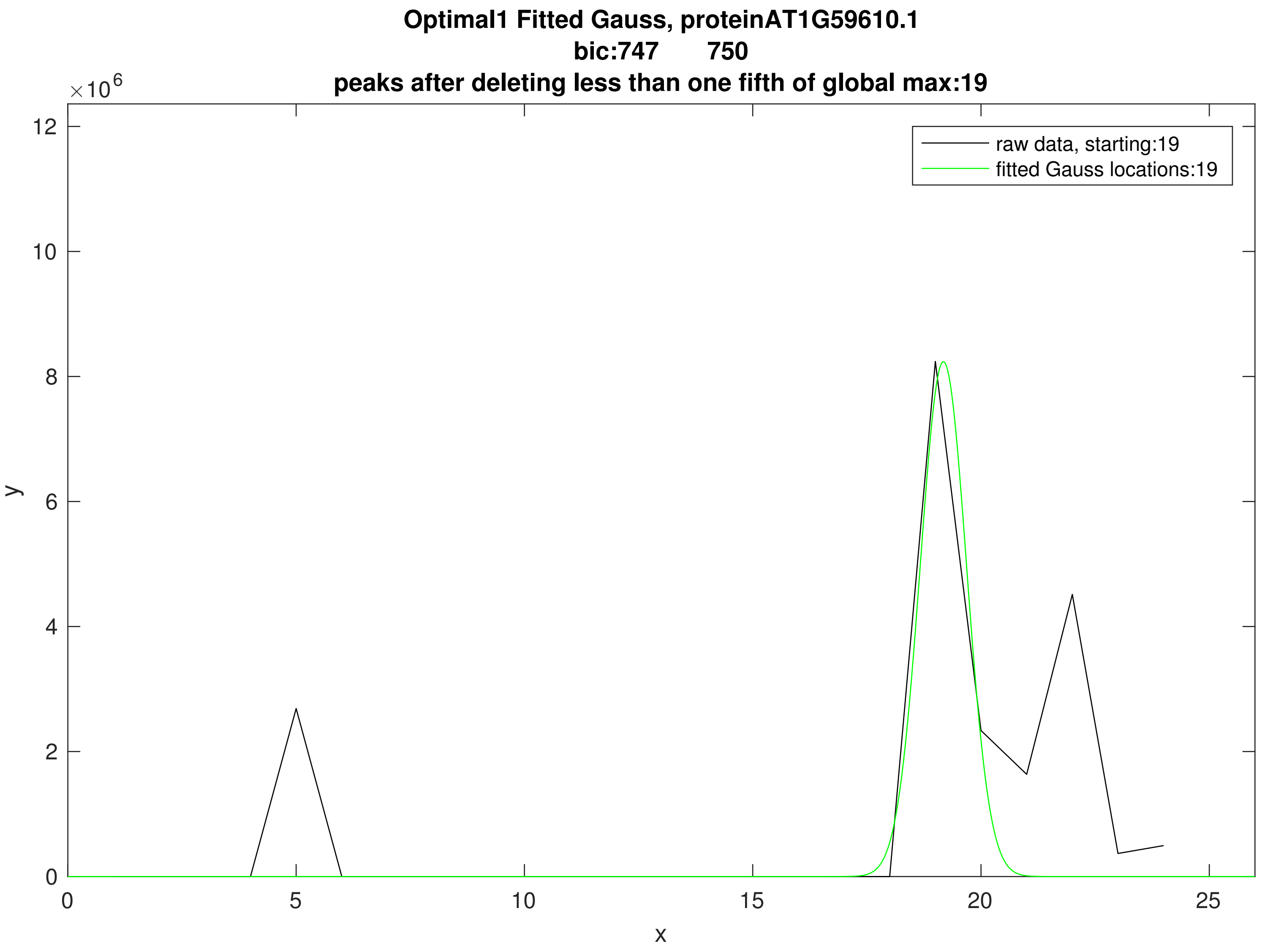
Optimal1 Fitted Gauss, proteinAT1G59610.1
bic:747 750
peaks after deleting less than one fifth of global max:19
×10^6
raw data, starting:19
fitted Gauss locations:19
y
x
0
2
4
6
8
10
12
0
5
10
15
20
25

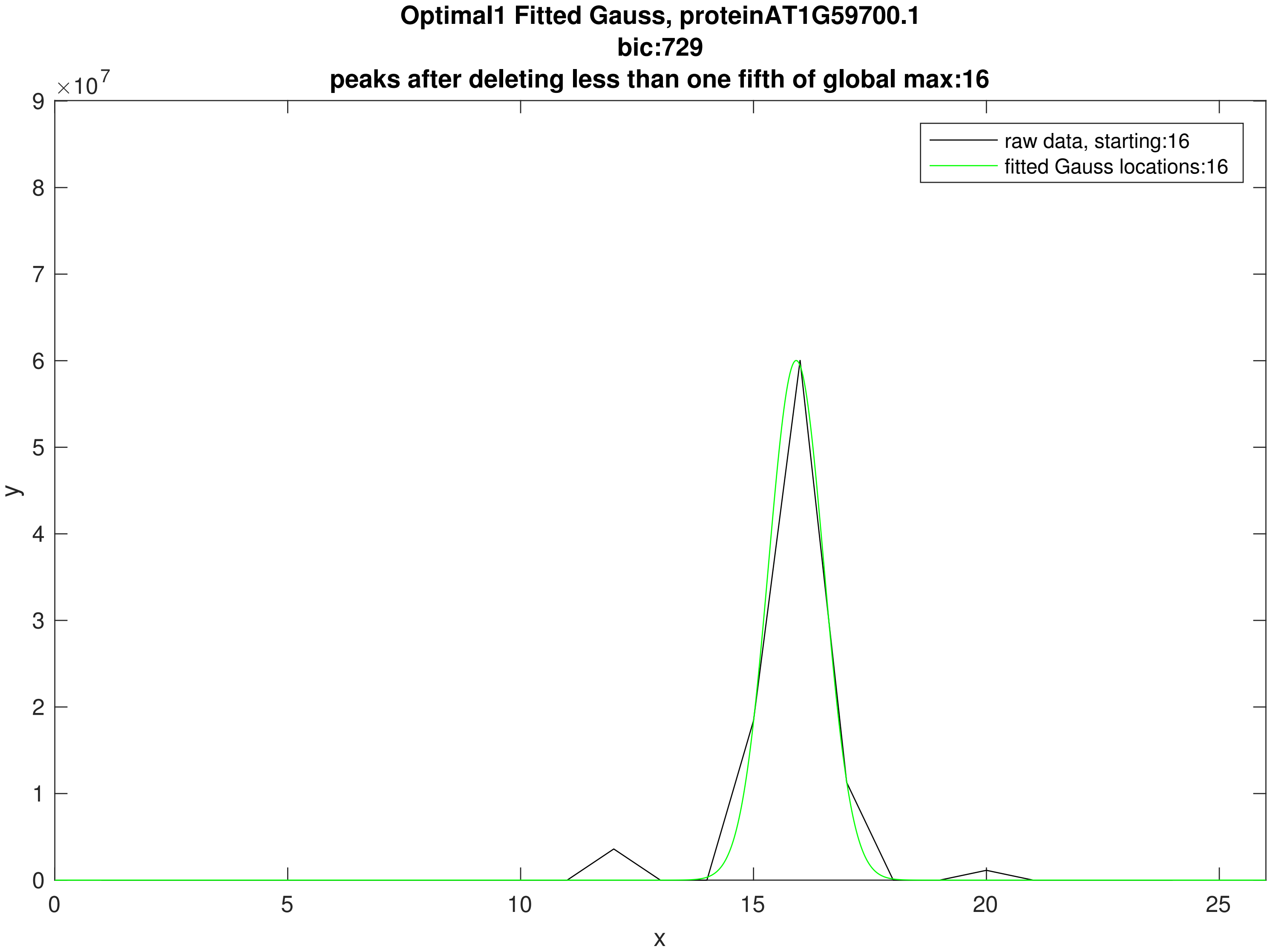
Optimal1 Fitted Gauss, proteinAT1G59700.1
bic:729
peaks after deleting less than one fifth of global max:16
×10^7
9
8
7
6
5
4
3
2
1
0
y
0
5
10
15
20
25
x
raw data, starting:16
fitted Gauss locations:16

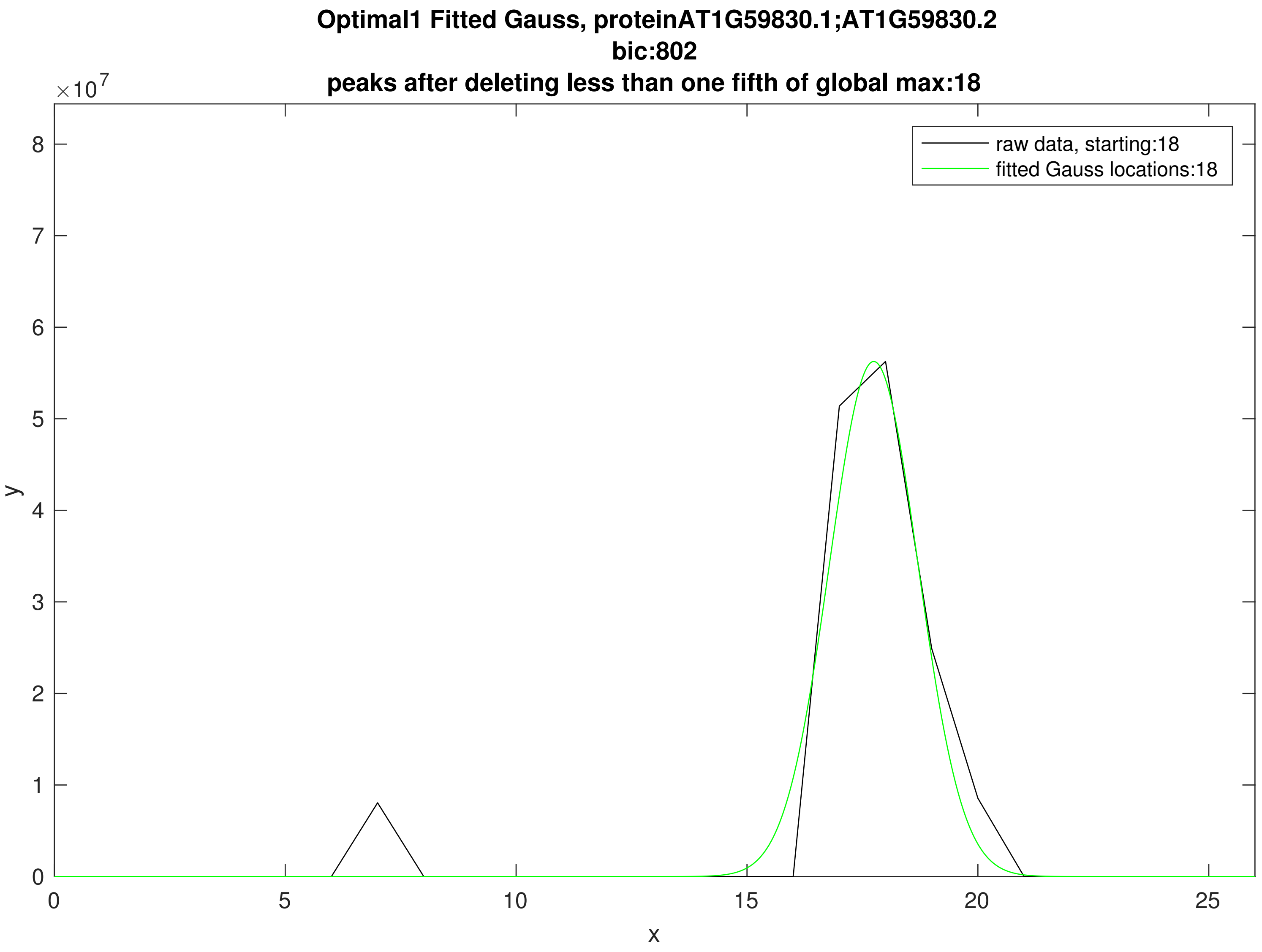

Optimal1 Fitted Gauss, proteinAT1G59830.1;AT1G59830.2
bic:802
peaks after deleting less than one fifth of global max:18
×10^7
raw data, starting:18
fitted Gauss locations:18
y
x
0
1
2
3
4
5
6
7
8
0
5
10
15
20
25

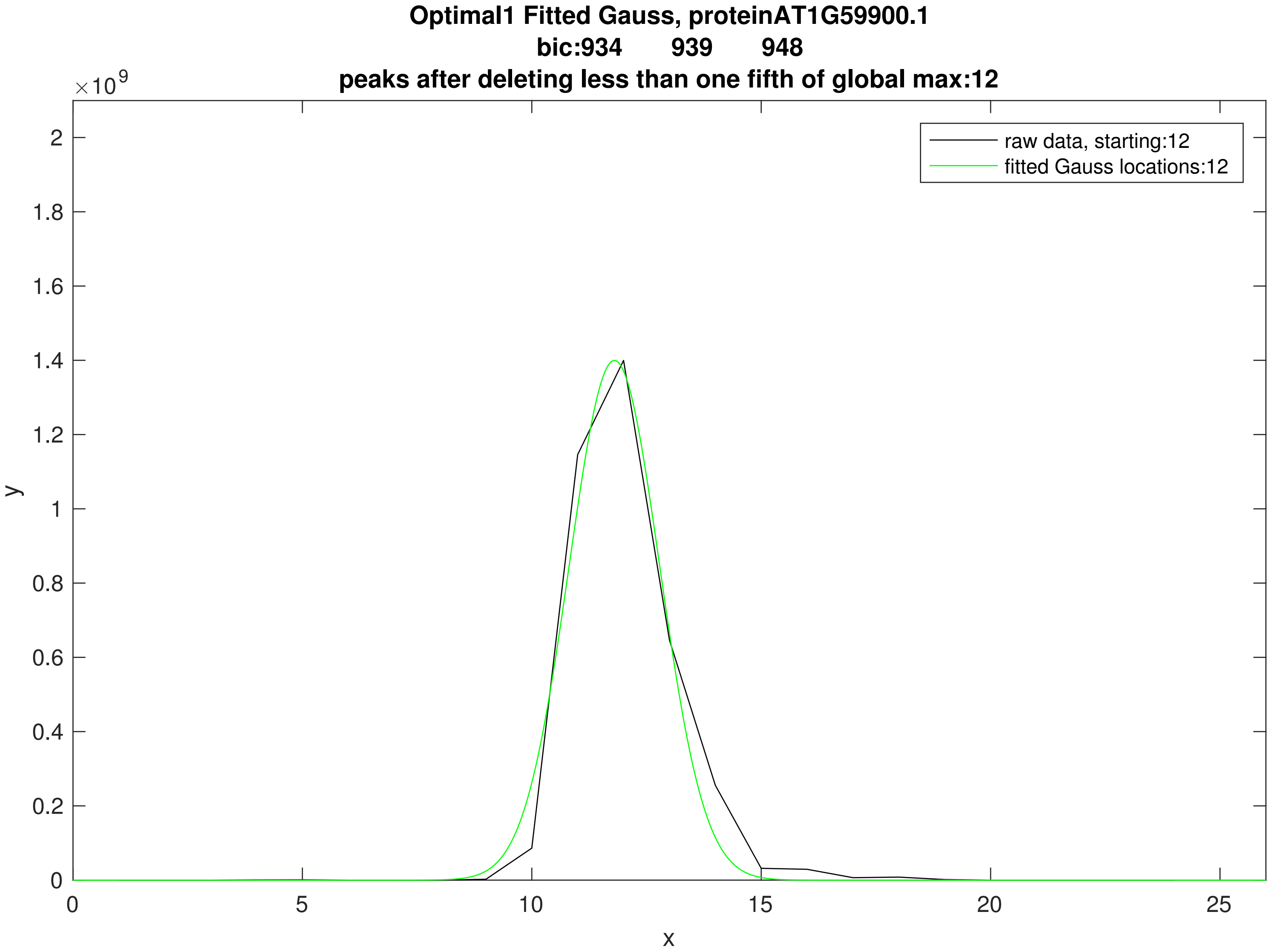
Optimal1 Fitted Gauss, proteinAT1G59900.1
bic:934 939 948
peaks after deleting less than one fifth of global max:12
×10^9
raw data, starting:12
fitted Gauss locations:12
y
x
0
0.2
0.4
0.6
0.8
1
1.2
1.4
1.6
1.8
2
0
5
10
15
20
25

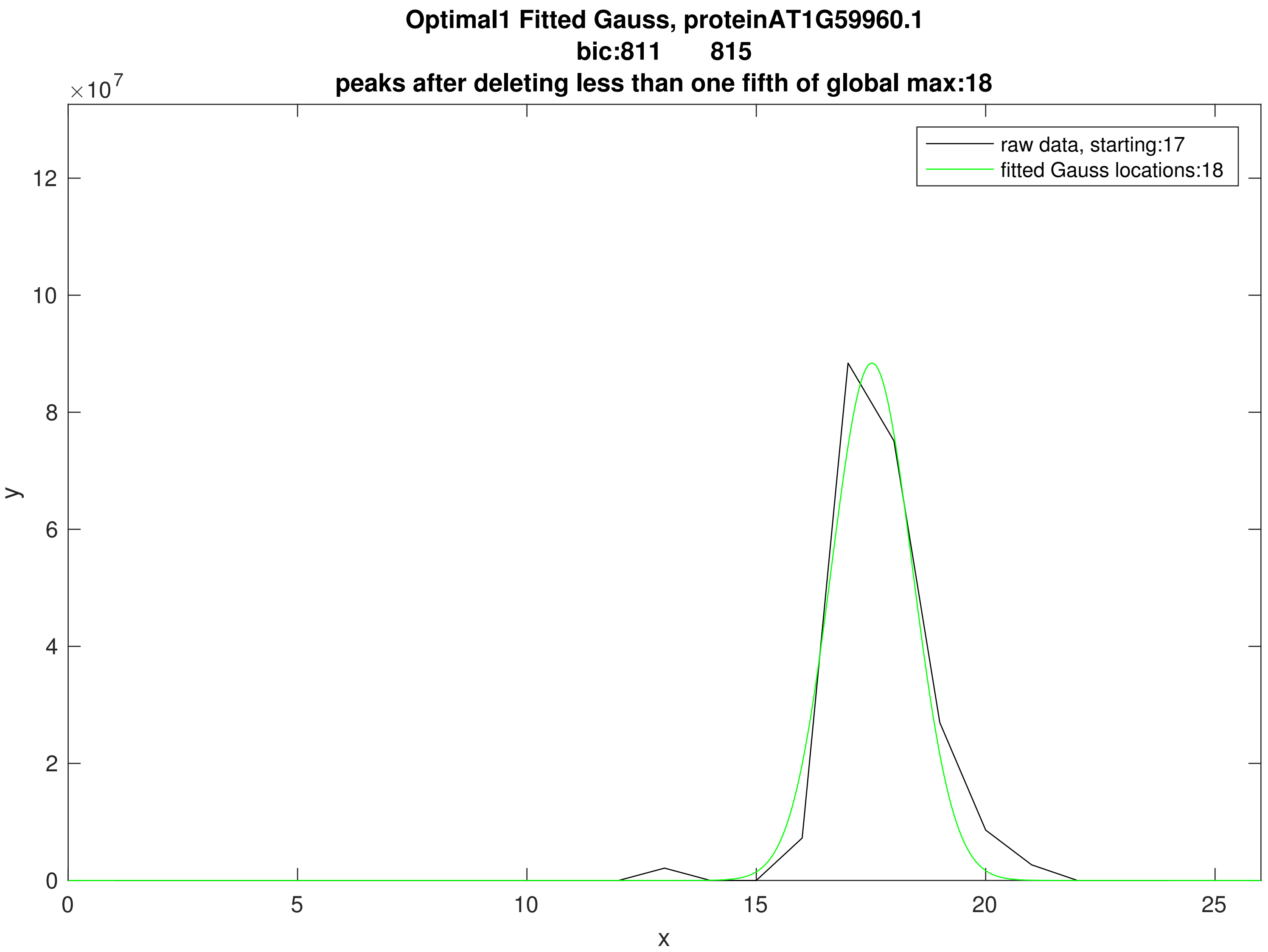
Optimal1 Fitted Gauss, proteinAT1G59960.1
bic:811 815
peaks after deleting less than one fifth of global max:18
×10^7
raw data, starting:17
fitted Gauss locations:18
y
x
0
2
4
6
8
10
12
0
5
10
15
20
25

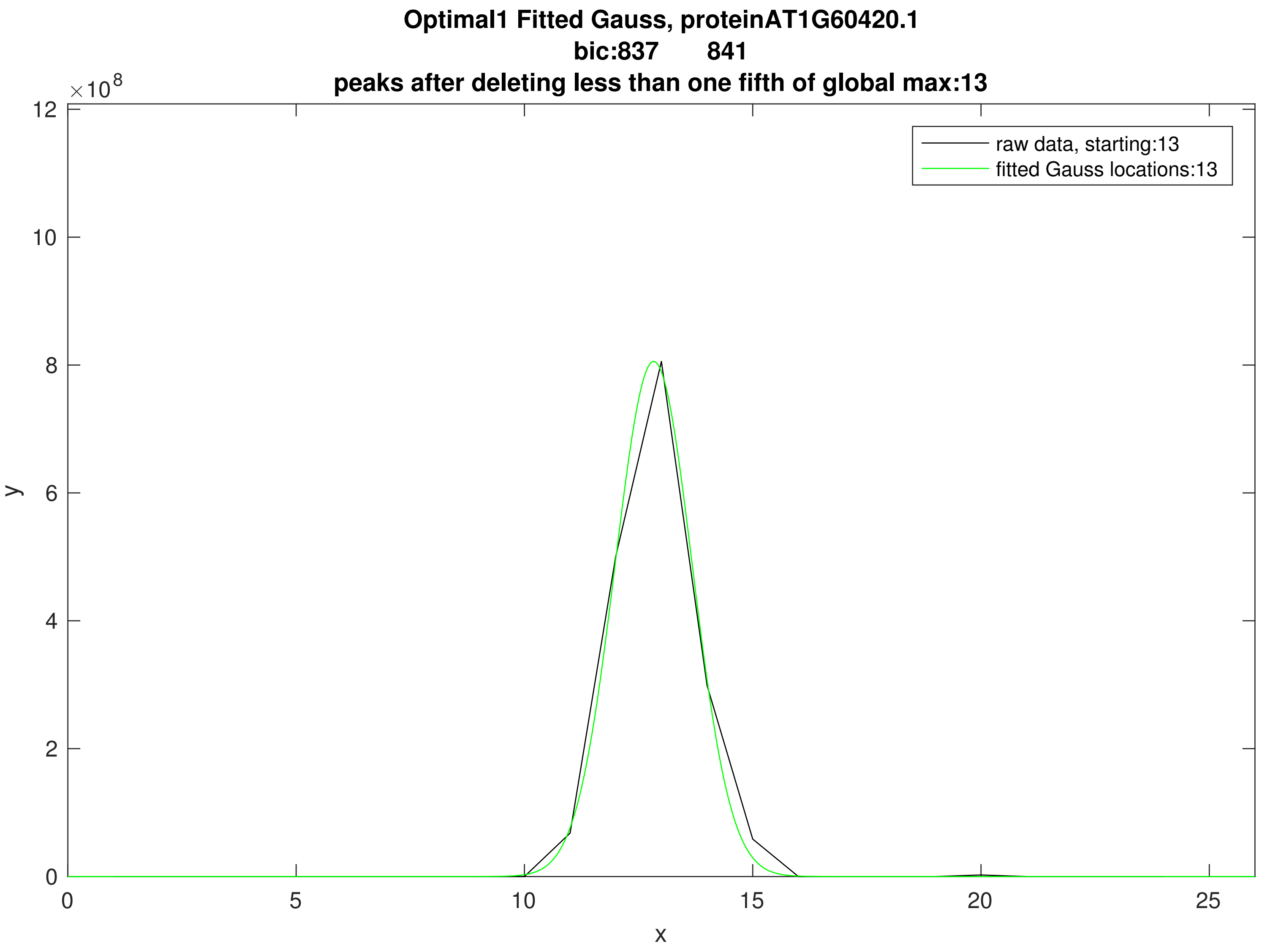
Optimal1 Fitted Gauss, proteinAT1G60420.1
bic:837 841
peaks after deleting less than one fifth of global max:13
×10^8
12
10
8
6
4
2
0
y
0
5
10
15
20
25
x
raw data, starting:13
fitted Gauss locations:13

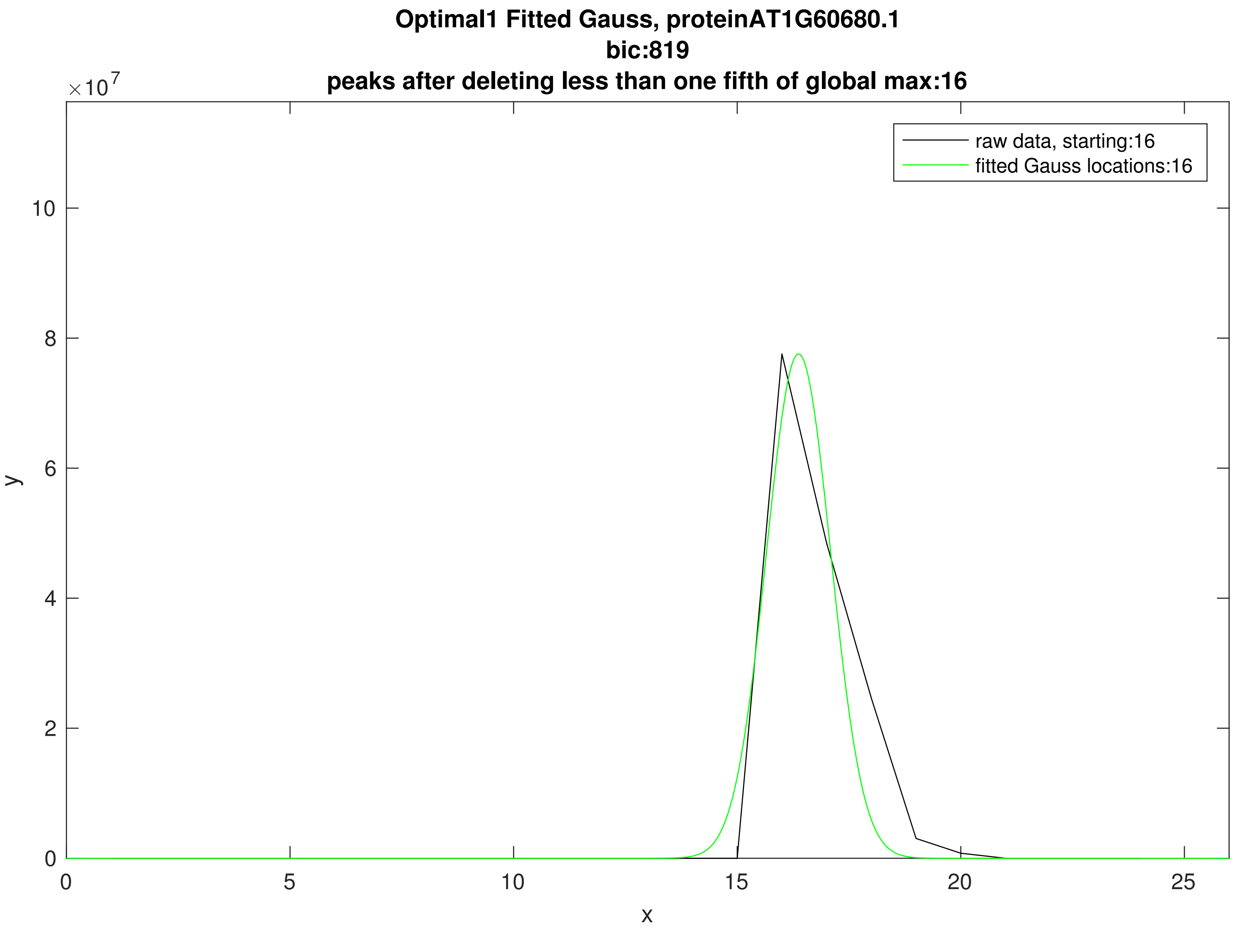

Optimal1 Fitted Gauss, proteinAT1G60680.1
bic:819
peaks after deleting less than one fifth of global max:16
×10^7
raw data, starting:16
fitted Gauss locations:16
10
8
6
4
2
0
y
0
5
10
15
20
25
x

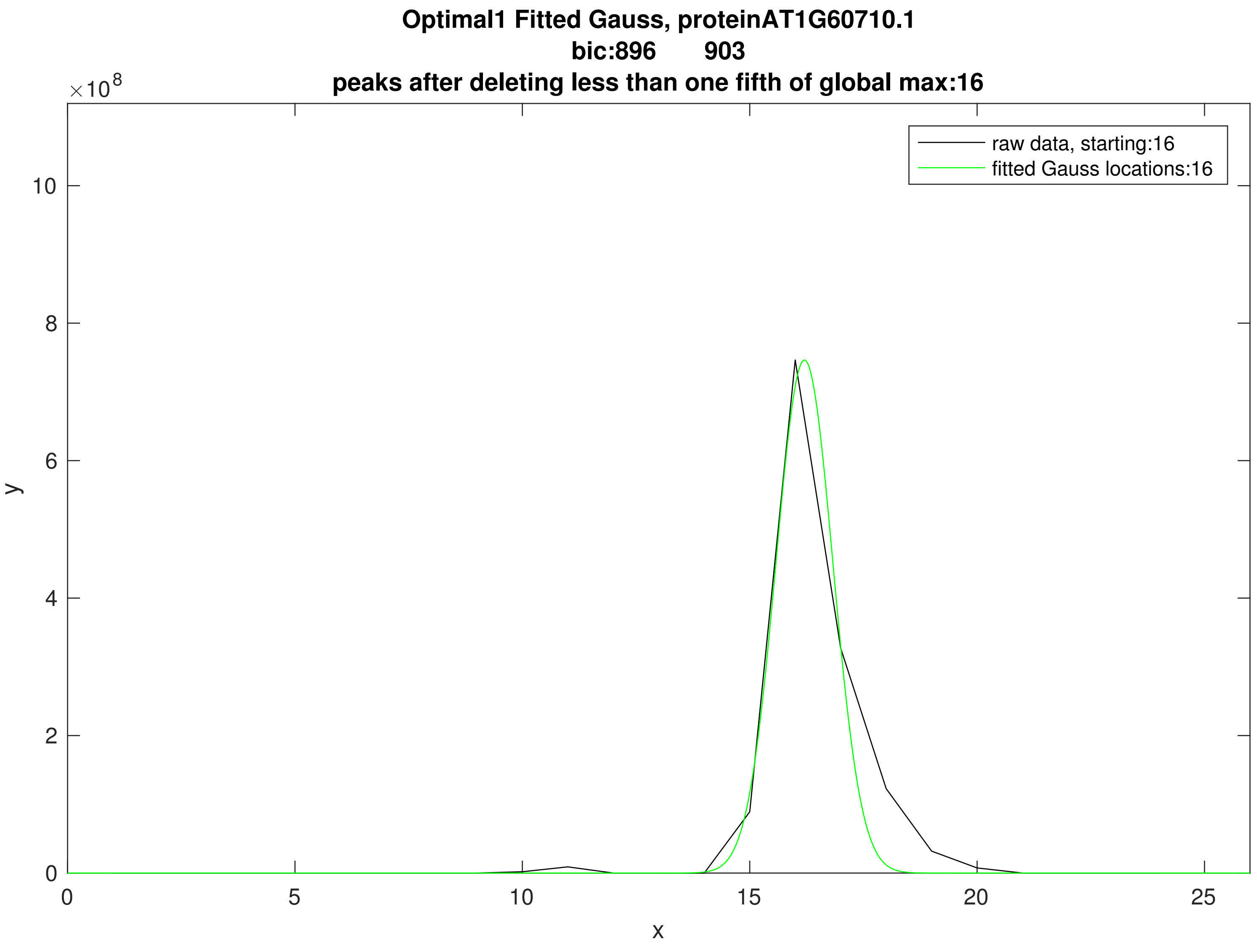
Optimal1 Fitted Gauss, proteinAT1G60710.1
bic:896 903
peaks after deleting less than one fifth of global max:16
×10^8
raw data, starting:16
fitted Gauss locations:16
y
x
0
2
4
6
8
10
0
5
10
15
20
25

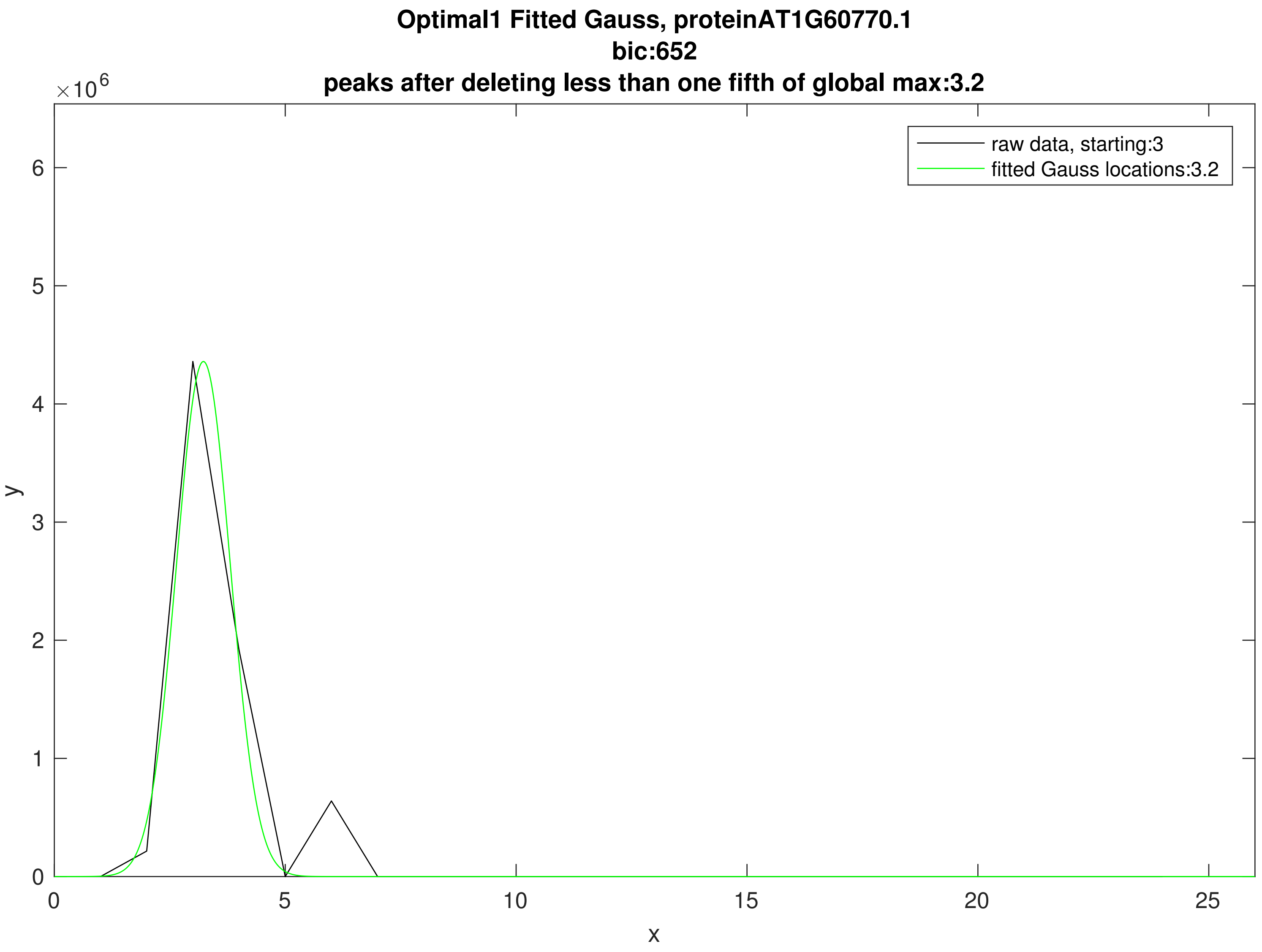
Optimal1 Fitted Gauss, proteinAT1G60770.1
bic:652
peaks after deleting less than one fifth of global max:3.2
×10^6
raw data, starting:3
fitted Gauss locations:3.2
y
x
0
1
2
3
4
5
6
0
5
10
15
20
25

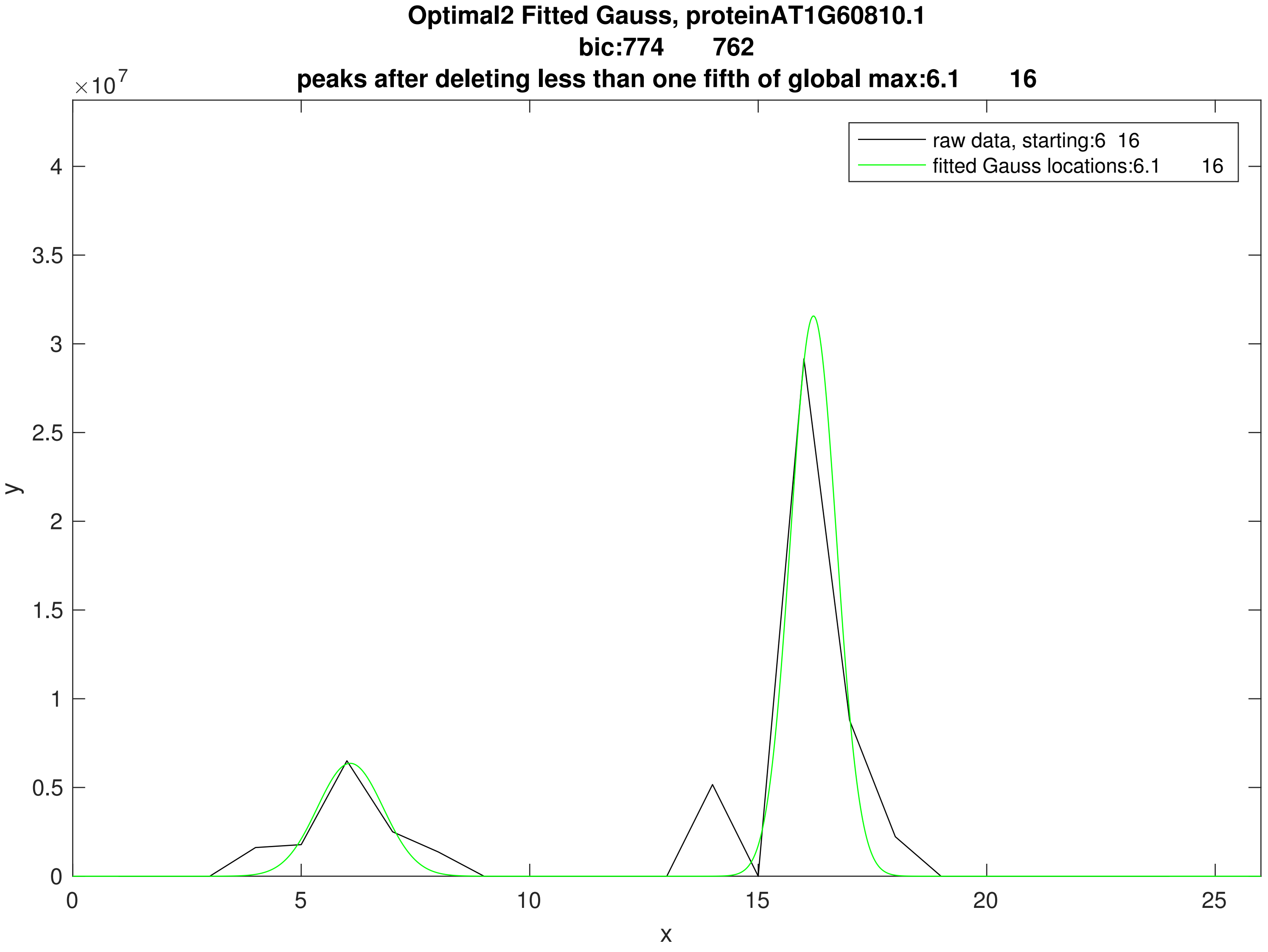
Optimal2 Fitted Gauss, proteinAT1G60810.1
bic:774 762
peaks after deleting less than one fifth of global max:6.1 16
×10^7
raw data, starting:6 16
fitted Gauss locations:6.1 16
y
x
0
0.5
1
1.5
2
2.5
3
3.5
4
0
5
10
15
20
25

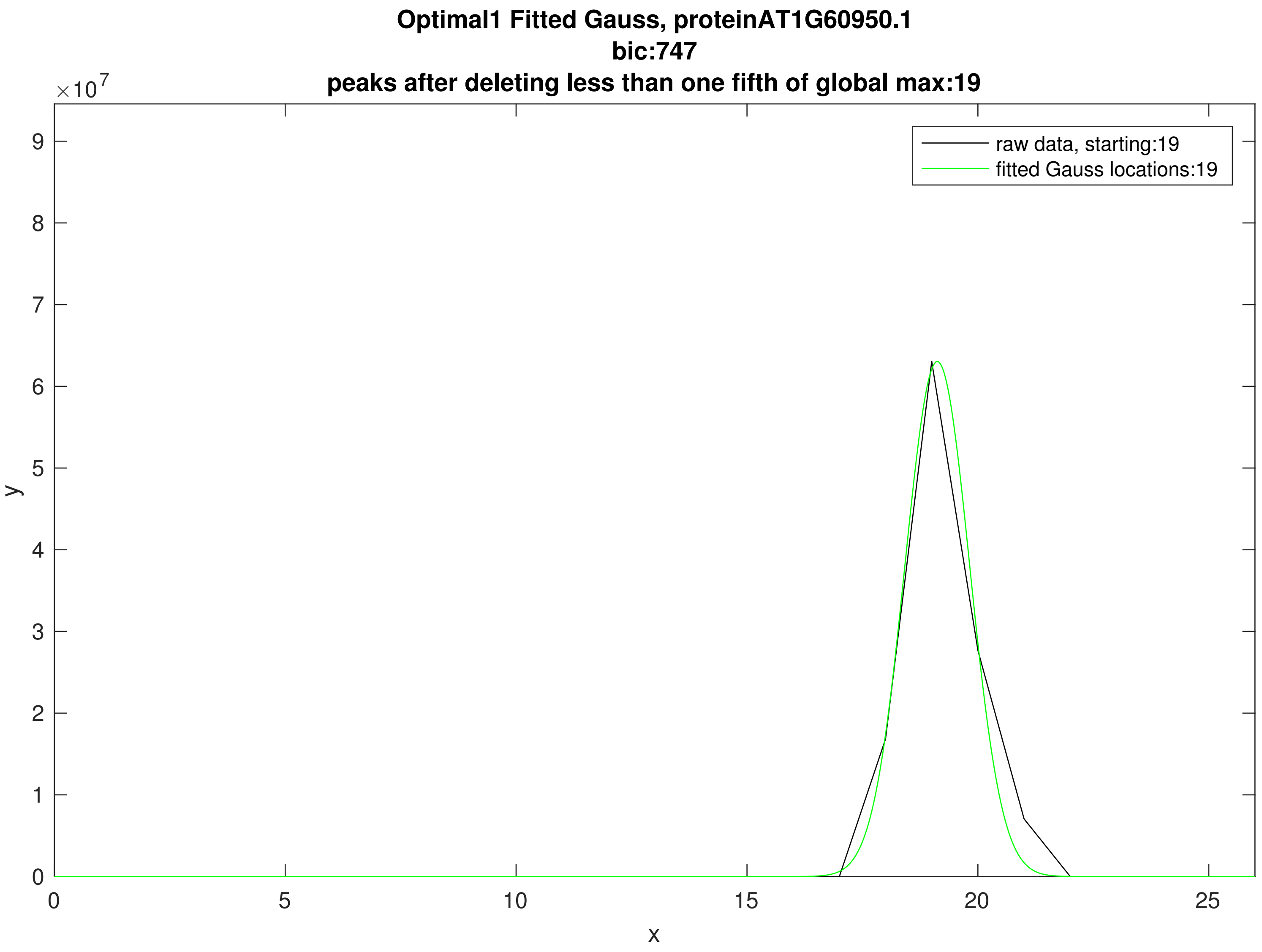
Optimal1 Fitted Gauss, proteinAT1G60950.1
bic:747
peaks after deleting less than one fifth of global max:19
×10^7
raw data, starting:19
fitted Gauss locations:19
y
x
0
1
2
3
4
5
6
7
8
9
0
5
10
15
20
25

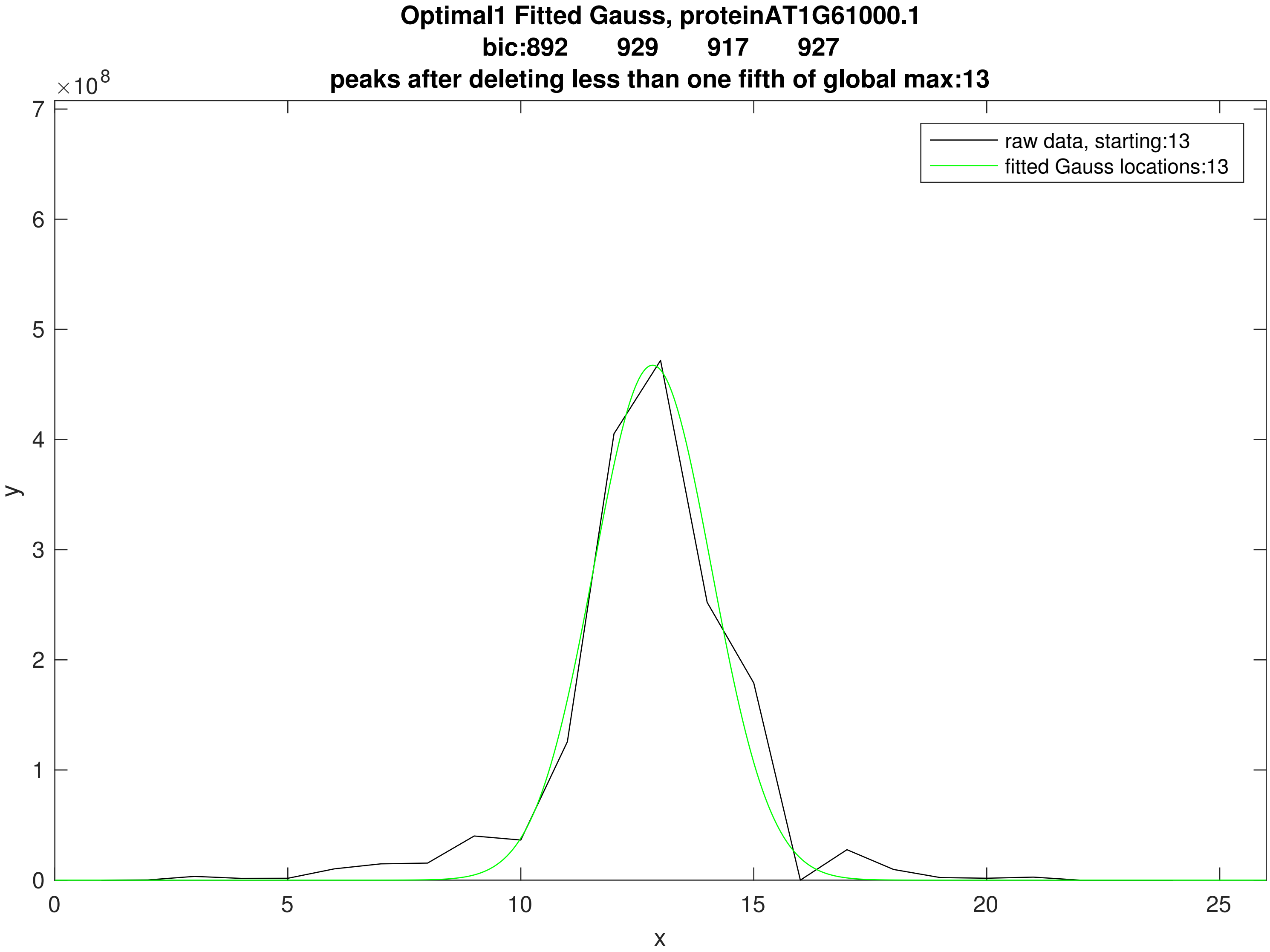
Optimal1 Fitted Gauss, proteinAT1G61000.1
bic:892 929 917 927
peaks after deleting less than one fifth of global max:13
×10^8
raw data, starting:13
fitted Gauss locations:13
y
x

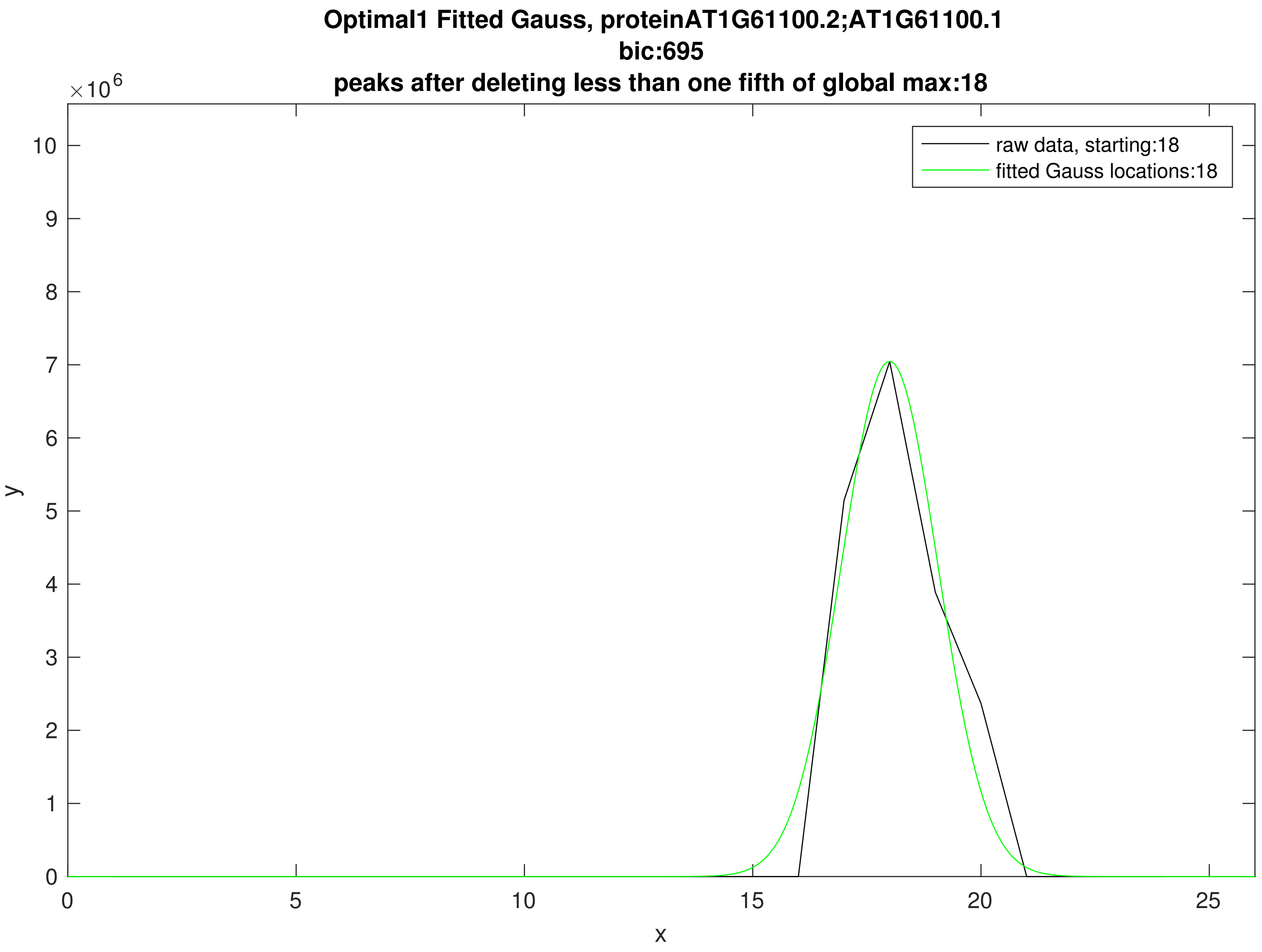
Optimal1 Fitted Gauss, proteinAT1G61100.2;AT1G61100.1
bic:695
peaks after deleting less than one fifth of global max:18
×10^6
raw data, starting:18
fitted Gauss locations:18
y
x
0
1
2
3
4
5
6
7
8
9
10
0
5
10
15
20
25

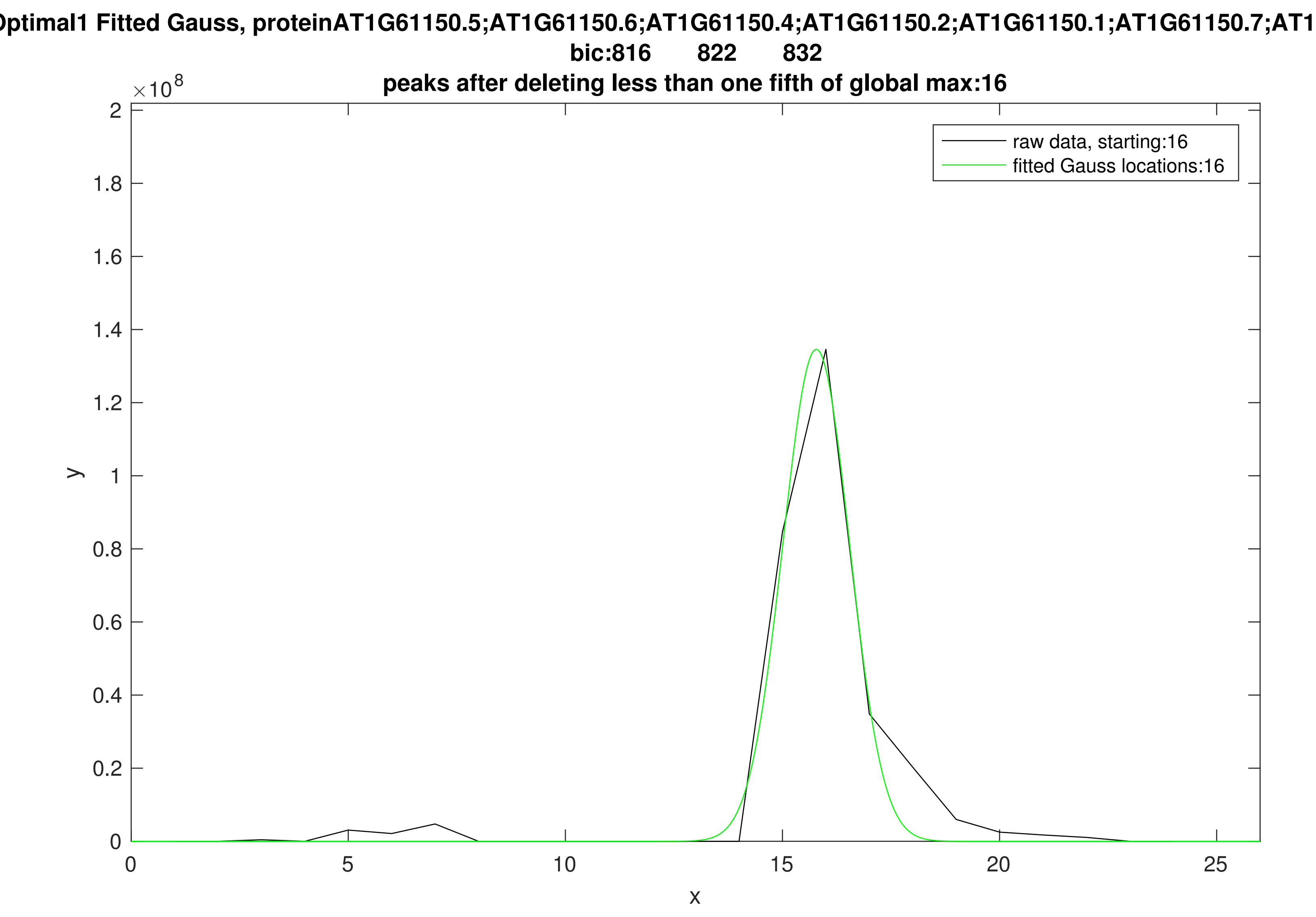
Optimal1 Fitted Gauss, proteinAT1G61150.5;AT1G61150.6;AT1G61150.4;AT1G61150.2;AT1G61150.1;AT1G61150.7;AT1
bic:816 822 832
peaks after deleting less than one fifth of global max:16
×10^8
2
1.8
1.6
1.4
1.2
1
0.8
0.6
0.4
0.2
0
y
0
5
10
15
20
25
x
raw data, starting:16
fitted Gauss locations:16

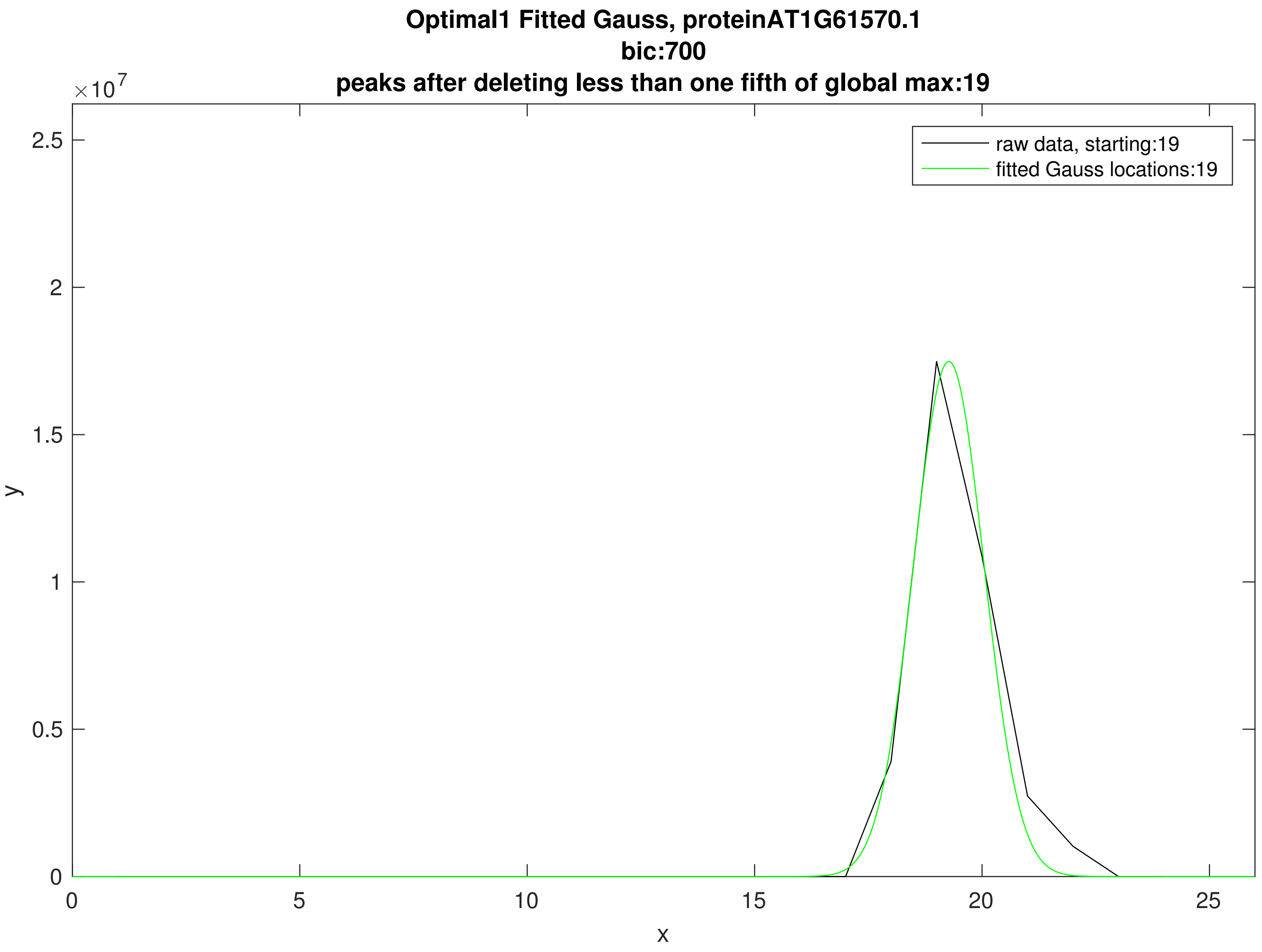
Optimal1 Fitted Gauss, proteinAT1G61570.1
bic:700
peaks after deleting less than one fifth of global max:19
×10^7
2.5
2
1.5
1
0.5
0
y
0
5
10
15
20
25
x
raw data, starting:19
fitted Gauss locations:19

#### Optimal1 Fitted Gauss, proteinAT1G61820.1
#### bic:807
#### peaks after deleting less than one fifth of global max:16

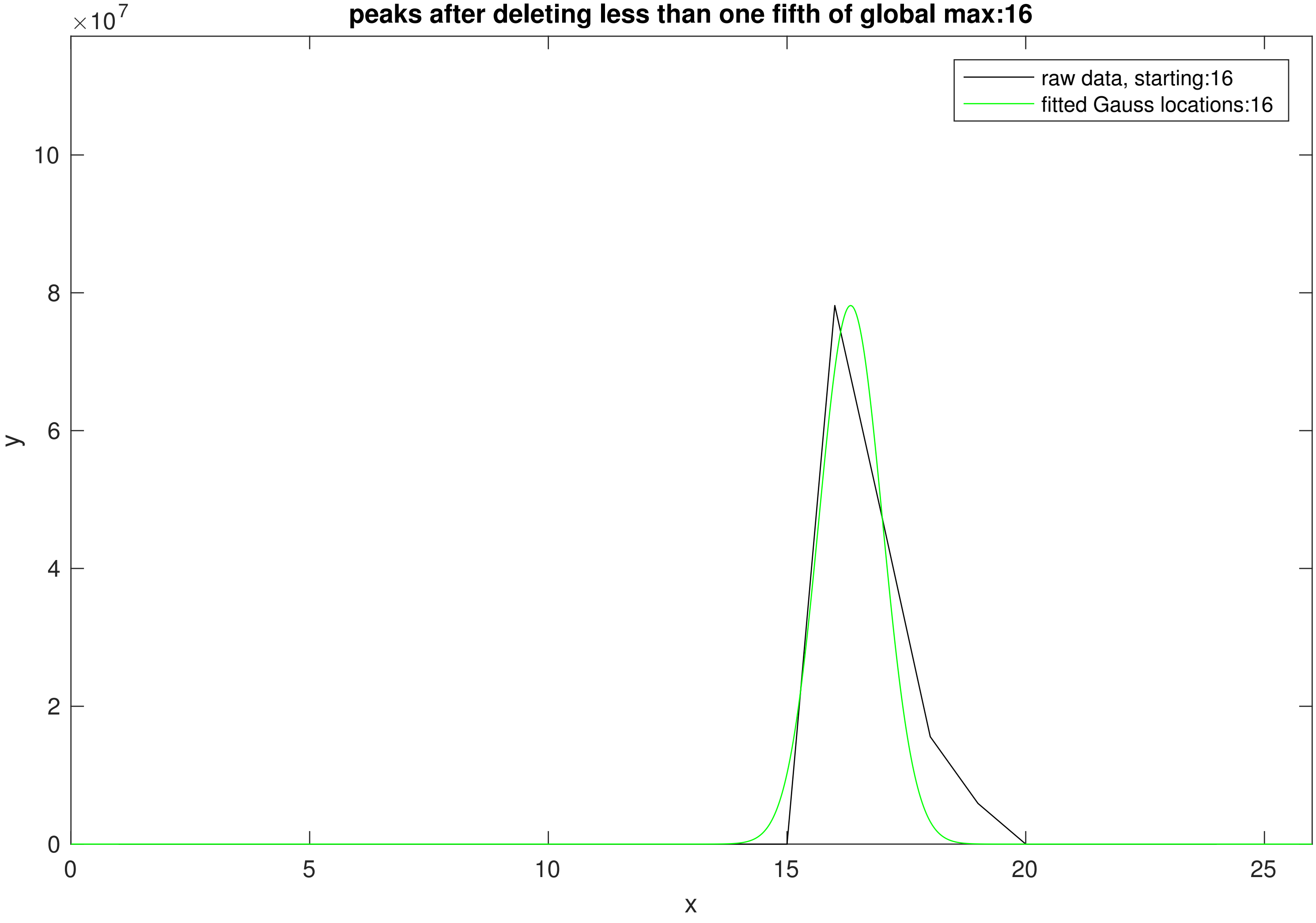

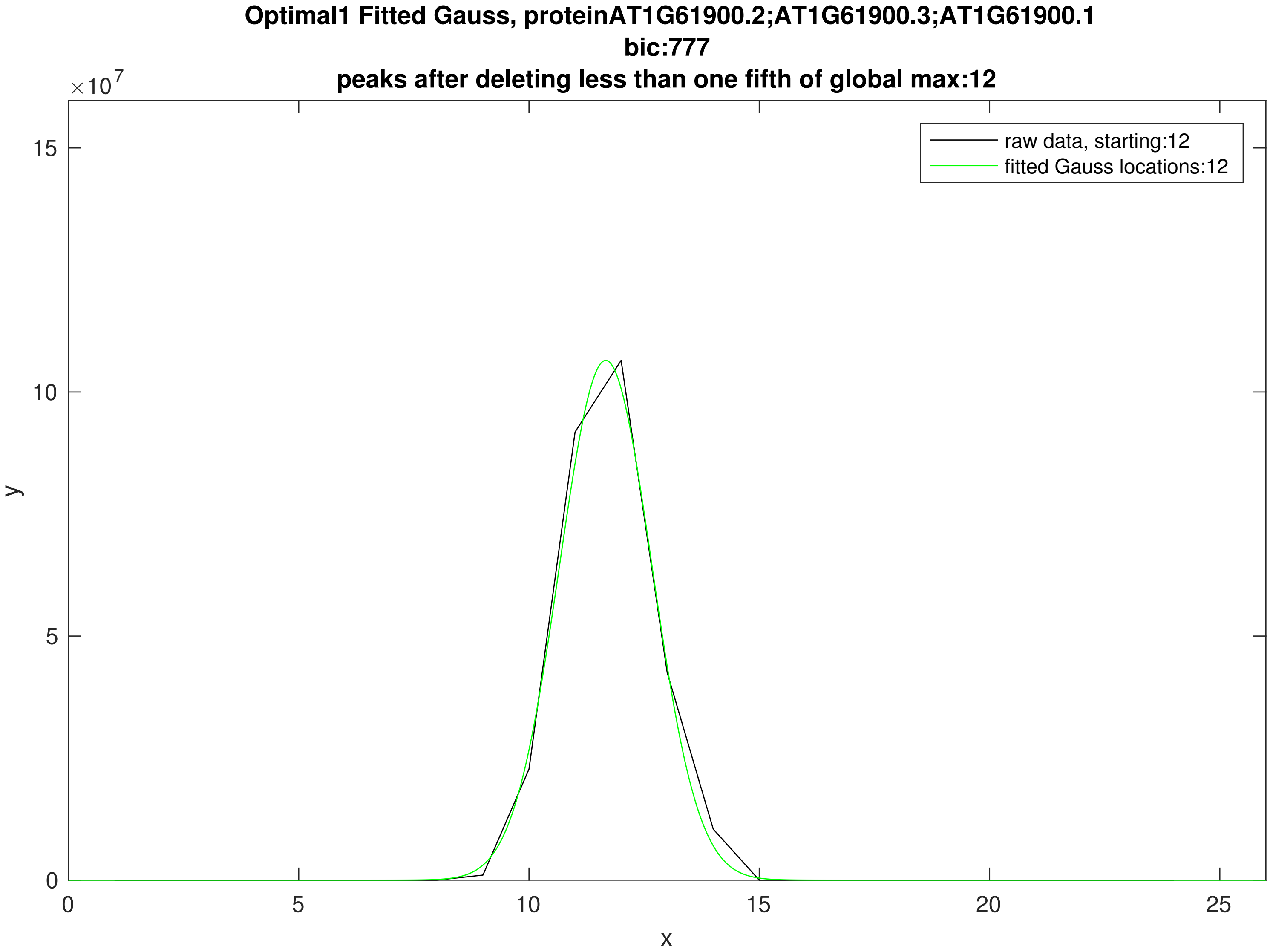

Optimal1 Fitted Gauss, proteinAT1G61900.2;AT1G61900.3;AT1G61900.1
bic:777
peaks after deleting less than one fifth of global max:12
×10^7
15
10
5
0
0
5
10
15
20
25
x
y
raw data, starting:12
fitted Gauss locations:12

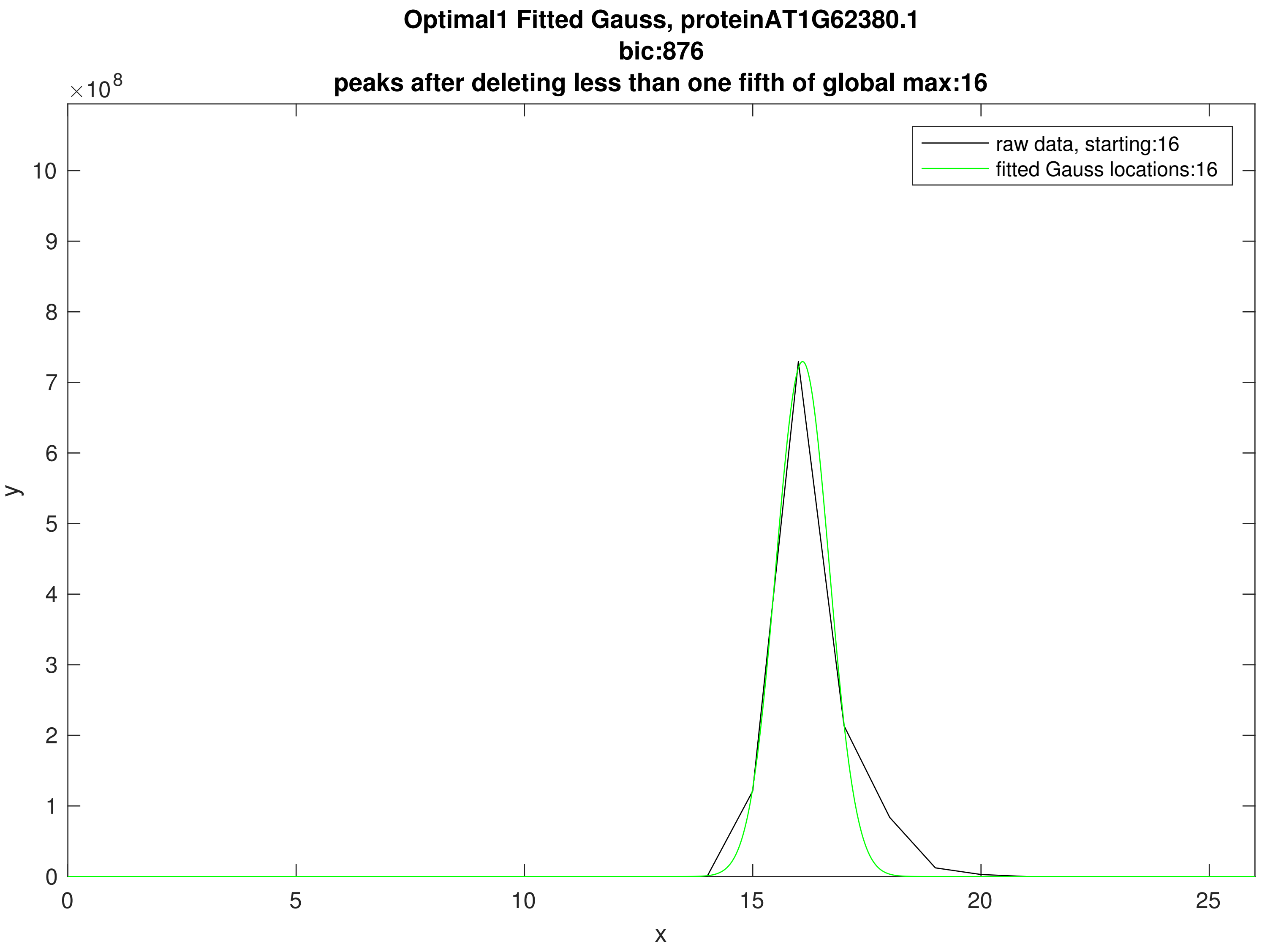
Optimal1 Fitted Gauss, proteinAT1G62380.1
bic:876
peaks after deleting less than one fifth of global max:16
×10^8
raw data, starting:16
fitted Gauss locations:16
y
x
0
1
2
3
4
5
6
7
8
9
10
0
5
10
15
20
25

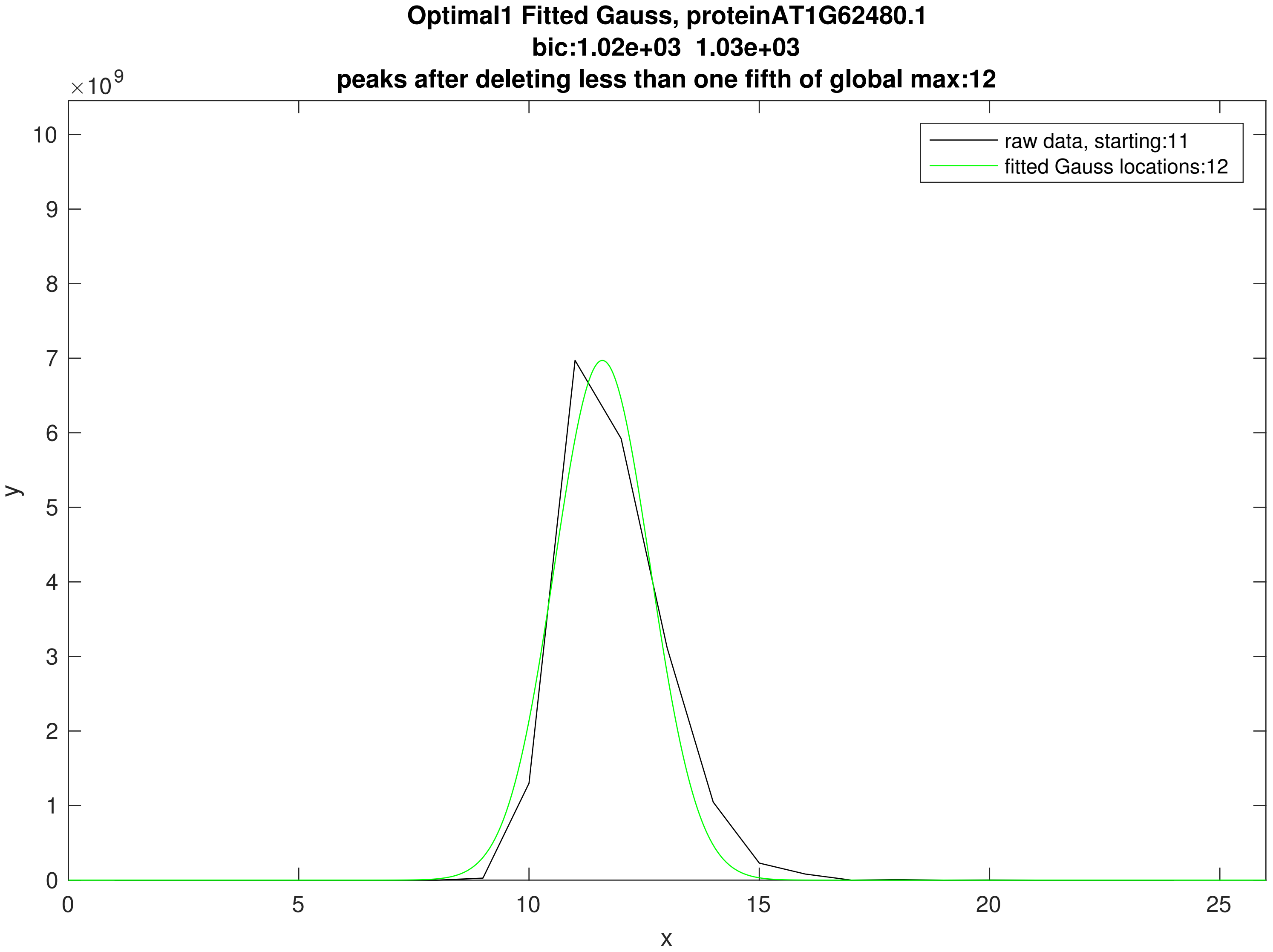
Optimal1 Fitted Gauss, proteinAT1G62480.1
bic:1.02e+03  1.03e+03
peaks after deleting less than one fifth of global max:12
×10^9
10
9
8
7
6
5
4
3
2
1
0
y
0
5
10
15
20
25
x
raw data, starting:11
fitted Gauss locations:12

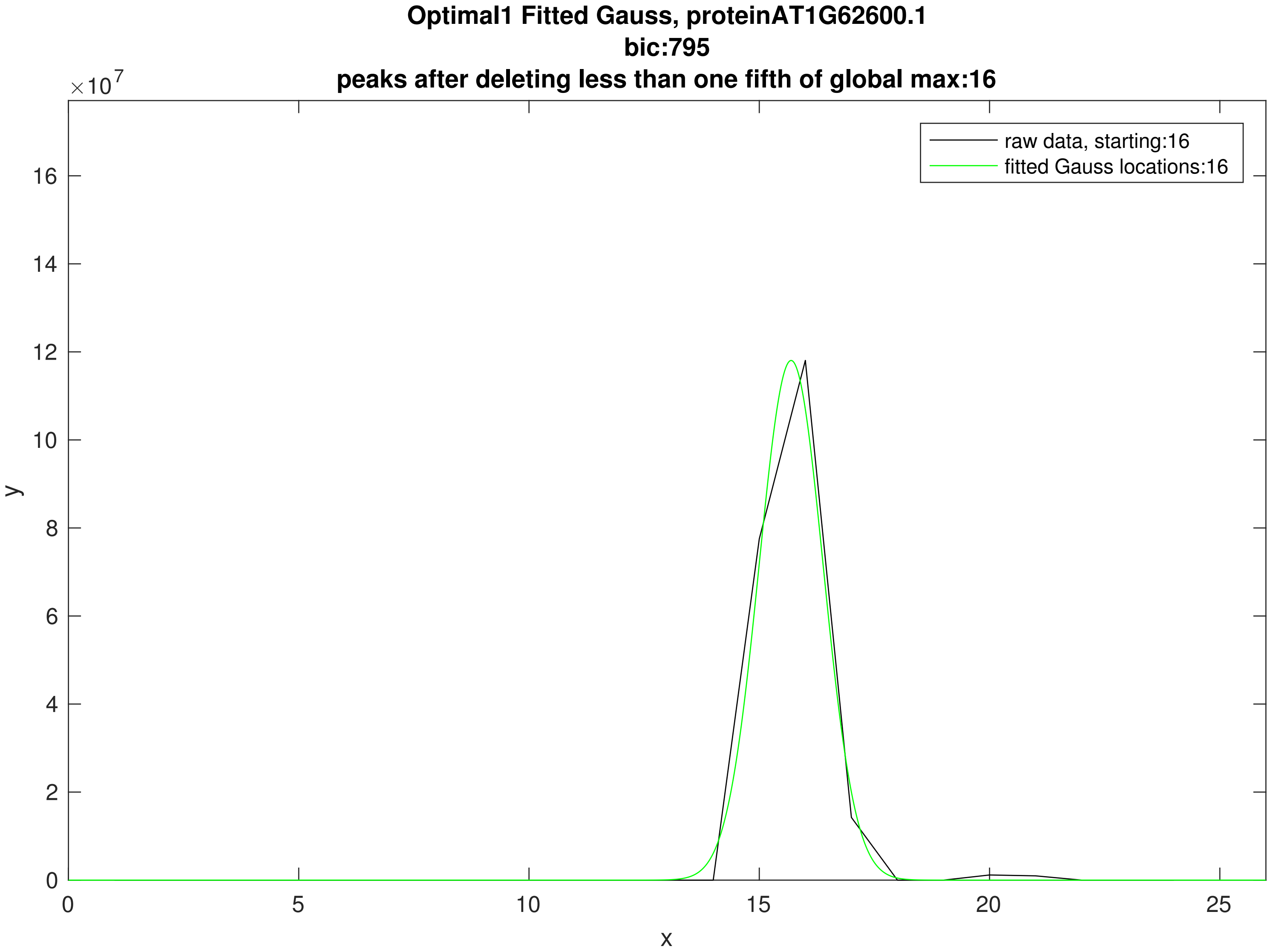
Optimal1 Fitted Gauss, proteinAT1G62600.1
bic:795
peaks after deleting less than one fifth of global max:16
×10^7
raw data, starting:16
fitted Gauss locations:16
y
x
0
2
4
6
8
10
12
14
16
0
5
10
15
20
25

**Optimal1 Fitted Gauss, proteinAT1G62640.2;AT1G62640.1**
**bic:898**
**peaks after deleting less than one fifth of global max:16**

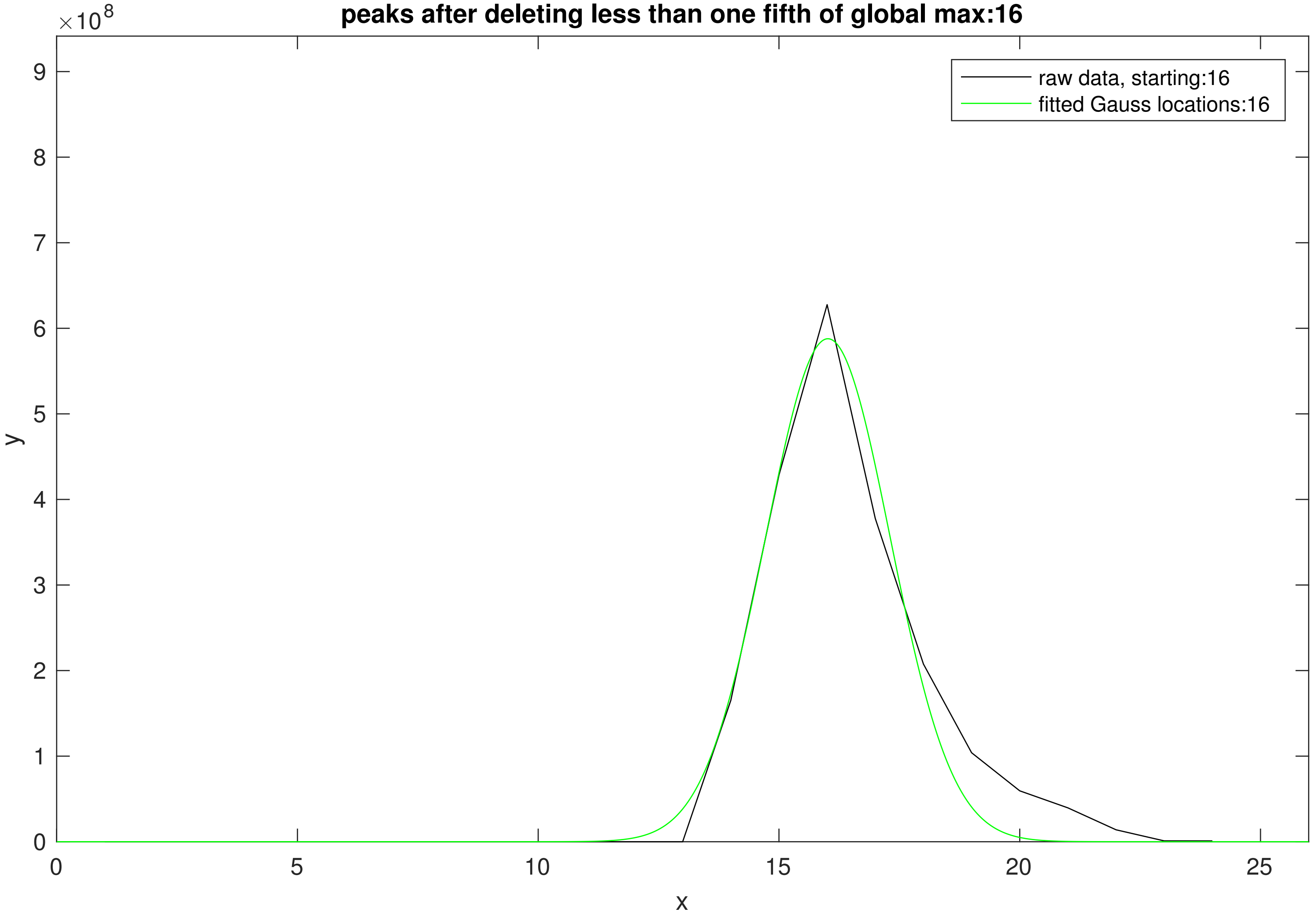

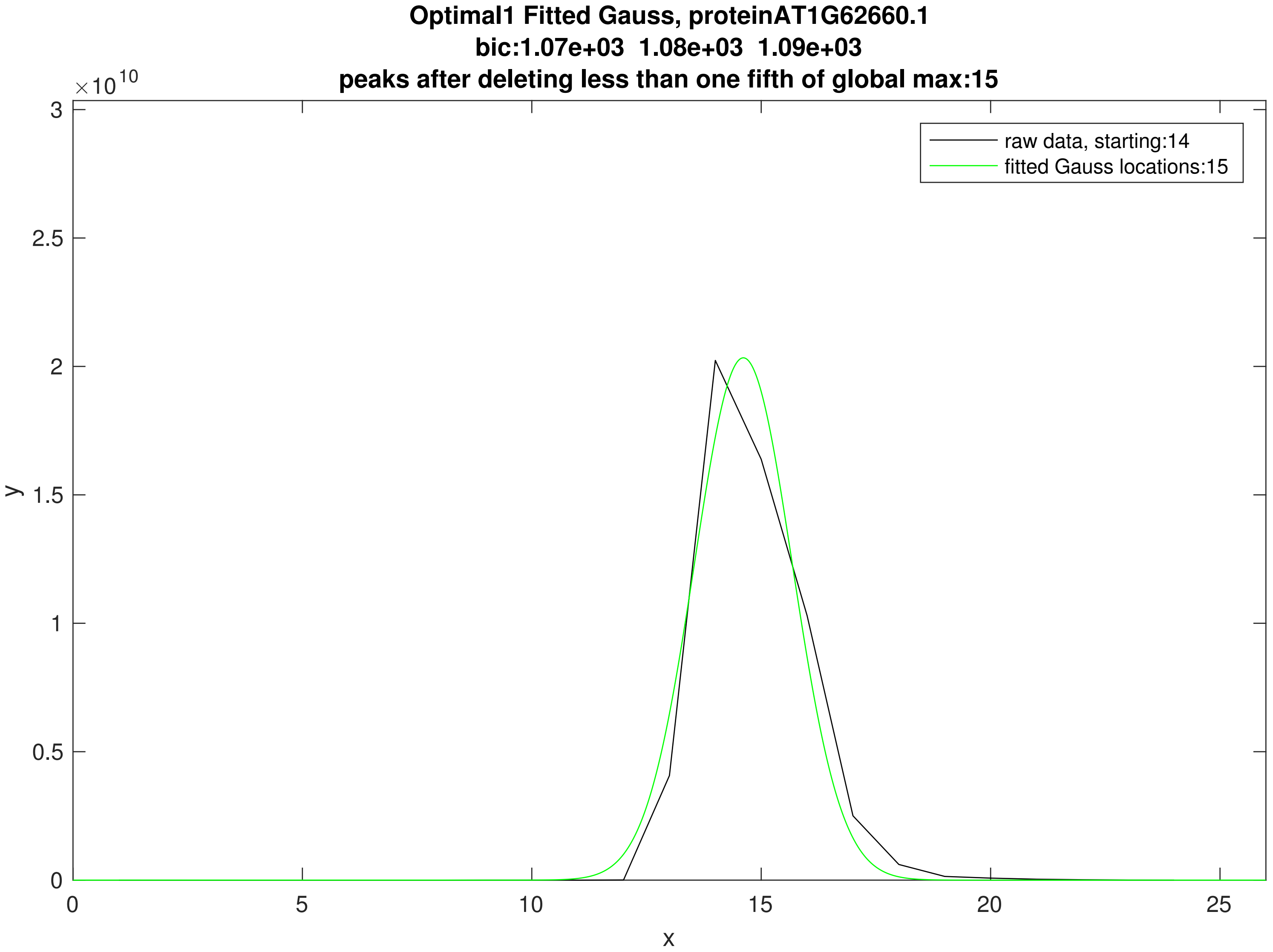

Optimal1 Fitted Gauss, proteinAT1G62660.1
bic:1.07e+03 1.08e+03 1.09e+03
peaks after deleting less than one fifth of global max:15
×10^10
raw data, starting:14
fitted Gauss locations:15
y
x
0
0.5
1
1.5
2
2.5
3
0
5
10
15
20
25

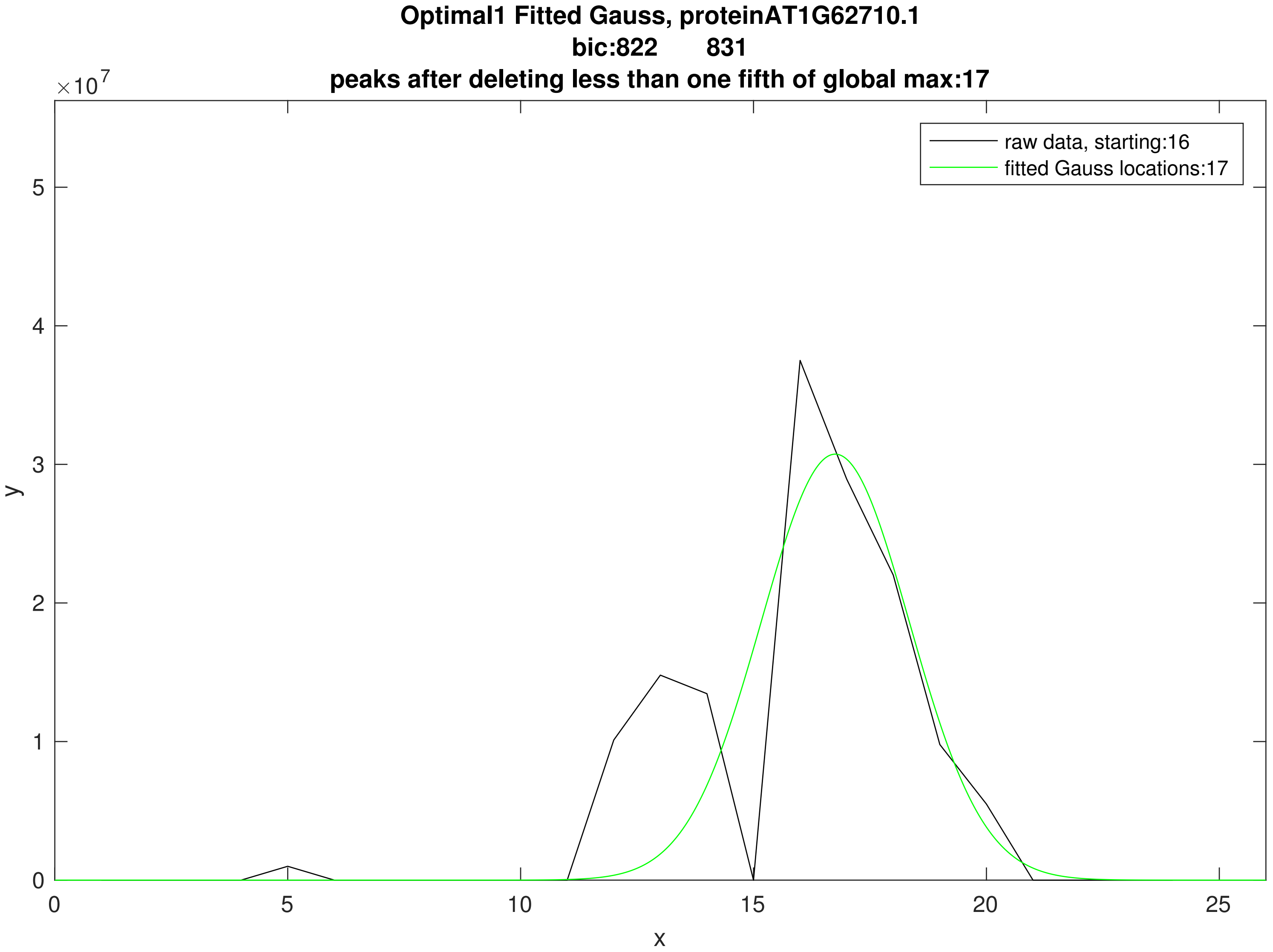

Optimal1 Fitted Gauss, proteinAT1G62710.1
bic:822 831
peaks after deleting less than one fifth of global max:17
×10^7
raw data, starting:16
fitted Gauss locations:17
y
x
0
1
2
3
4
5
0
5
10
15
20
25

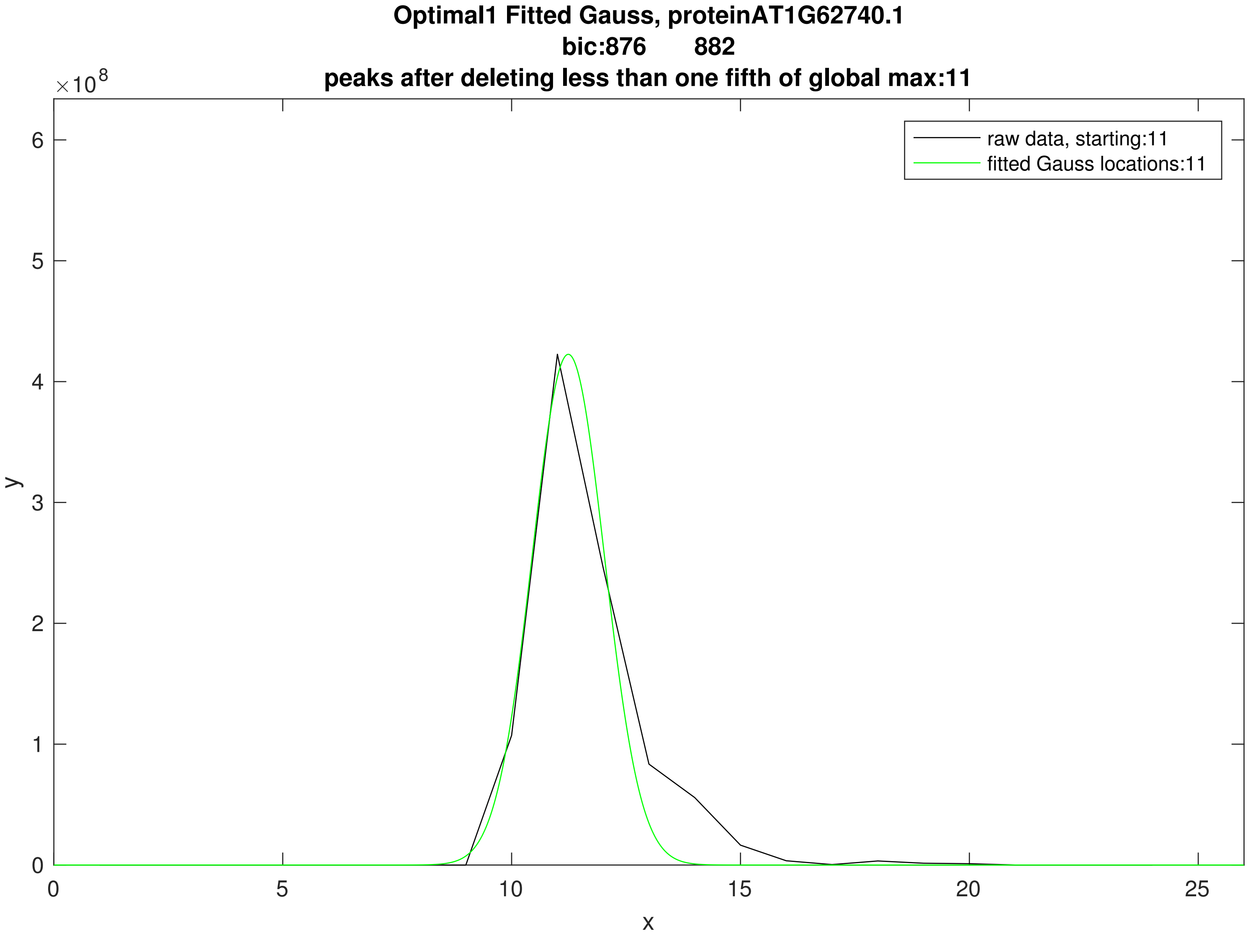
Optimal1 Fitted Gauss, proteinAT1G62740.1
bic:876 882
peaks after deleting less than one fifth of global max:11
×10^8
raw data, starting:11
fitted Gauss locations:11
y
x
0
1
2
3
4
5
6
0
5
10
15
20
25

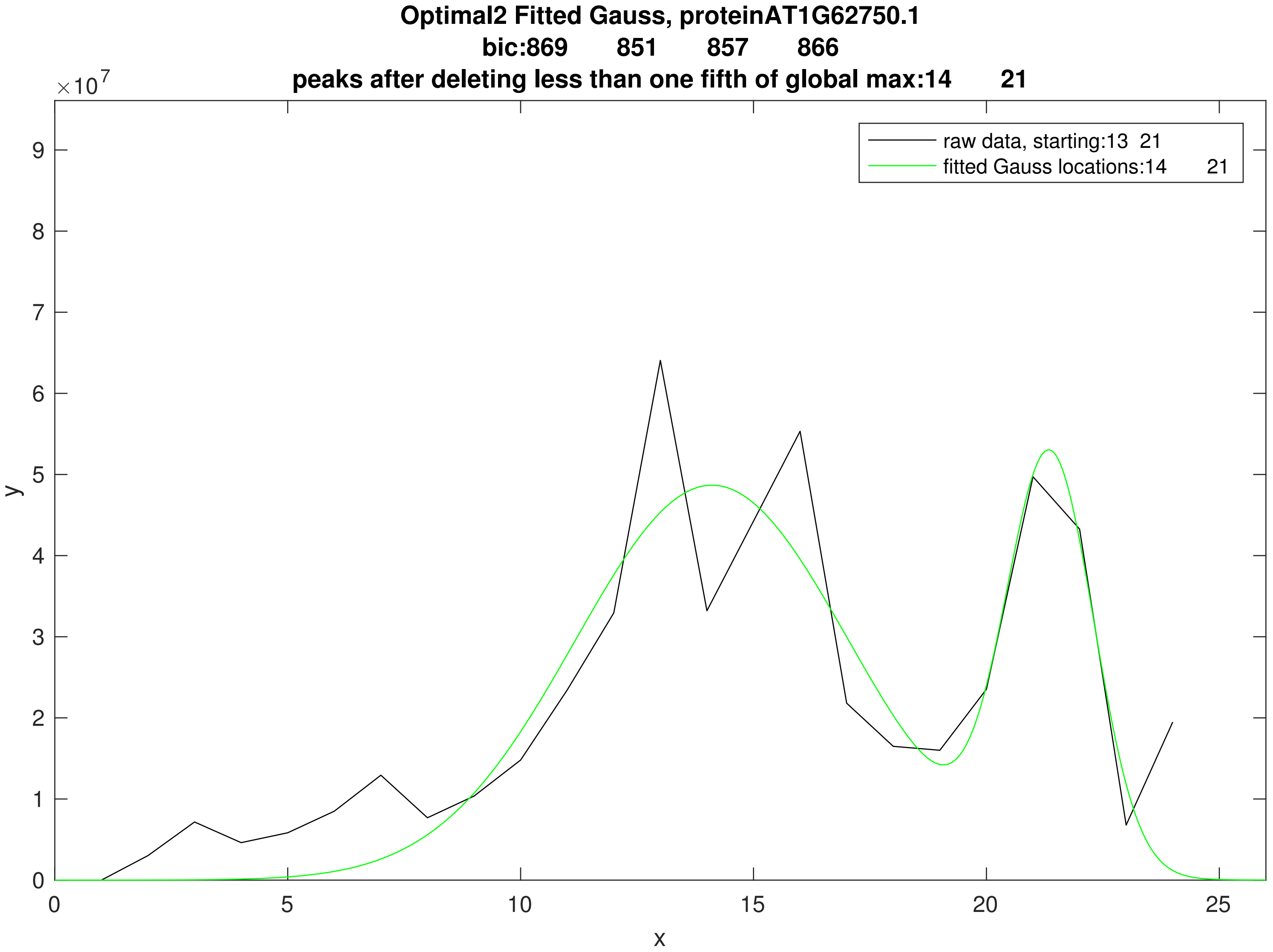

Optimal2 Fitted Gauss, proteinAT1G62750.1
bic:869 851 857 866
peaks after deleting less than one fifth of global max:14 21
×10^7
raw data, starting:13 21
fitted Gauss locations:14 21
y
x

**Optimal1 Fitted Gauss, proteinAT1G62780.1**
**bic:721**
**peaks after deleting less than one fifth of global max:17**

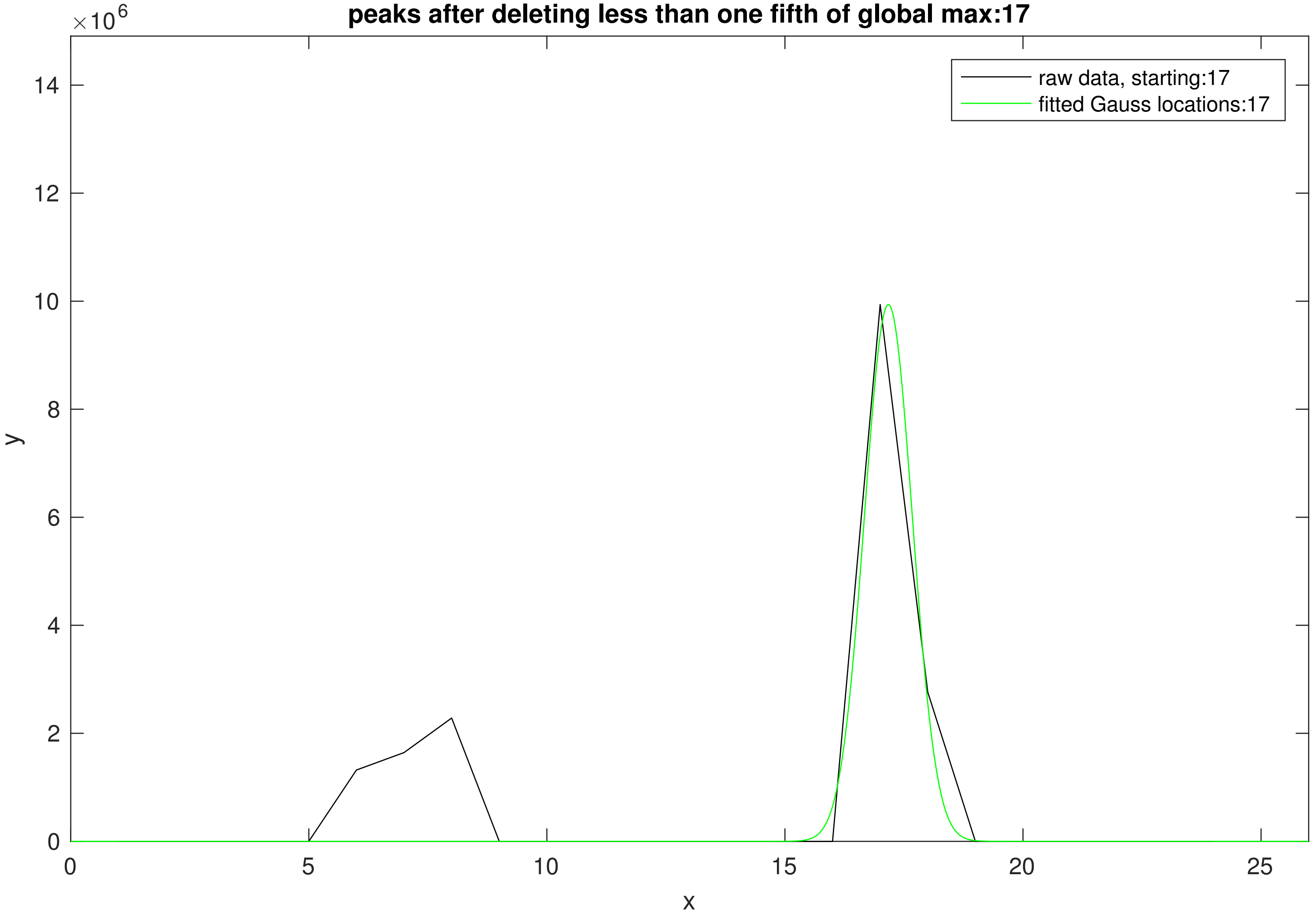

**Optimal1 Fitted Gauss, proteinAT1G62800.2;AT1G62800.1**
**bic:862 871**
**peaks after deleting less than one fifth of global max:14**

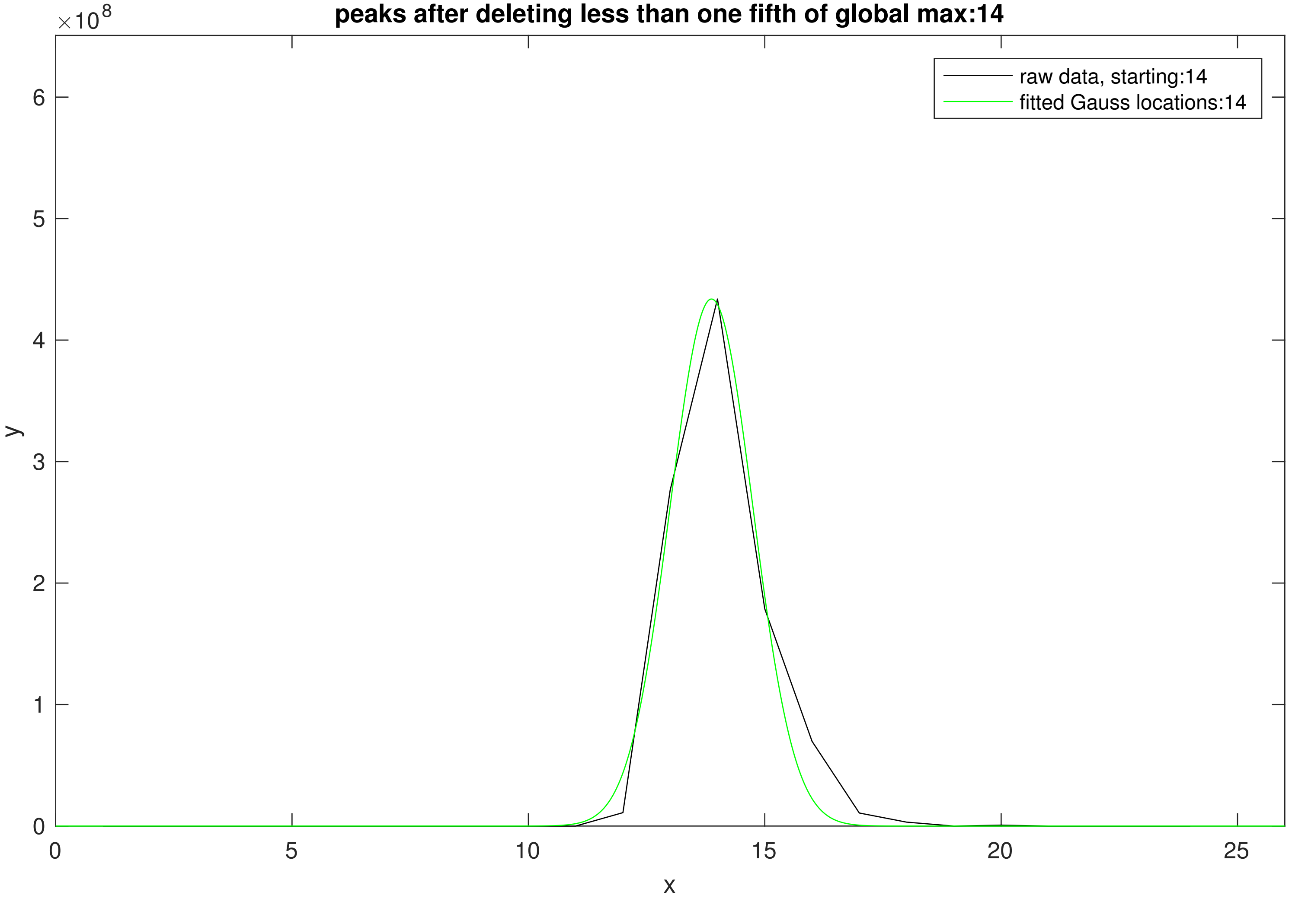

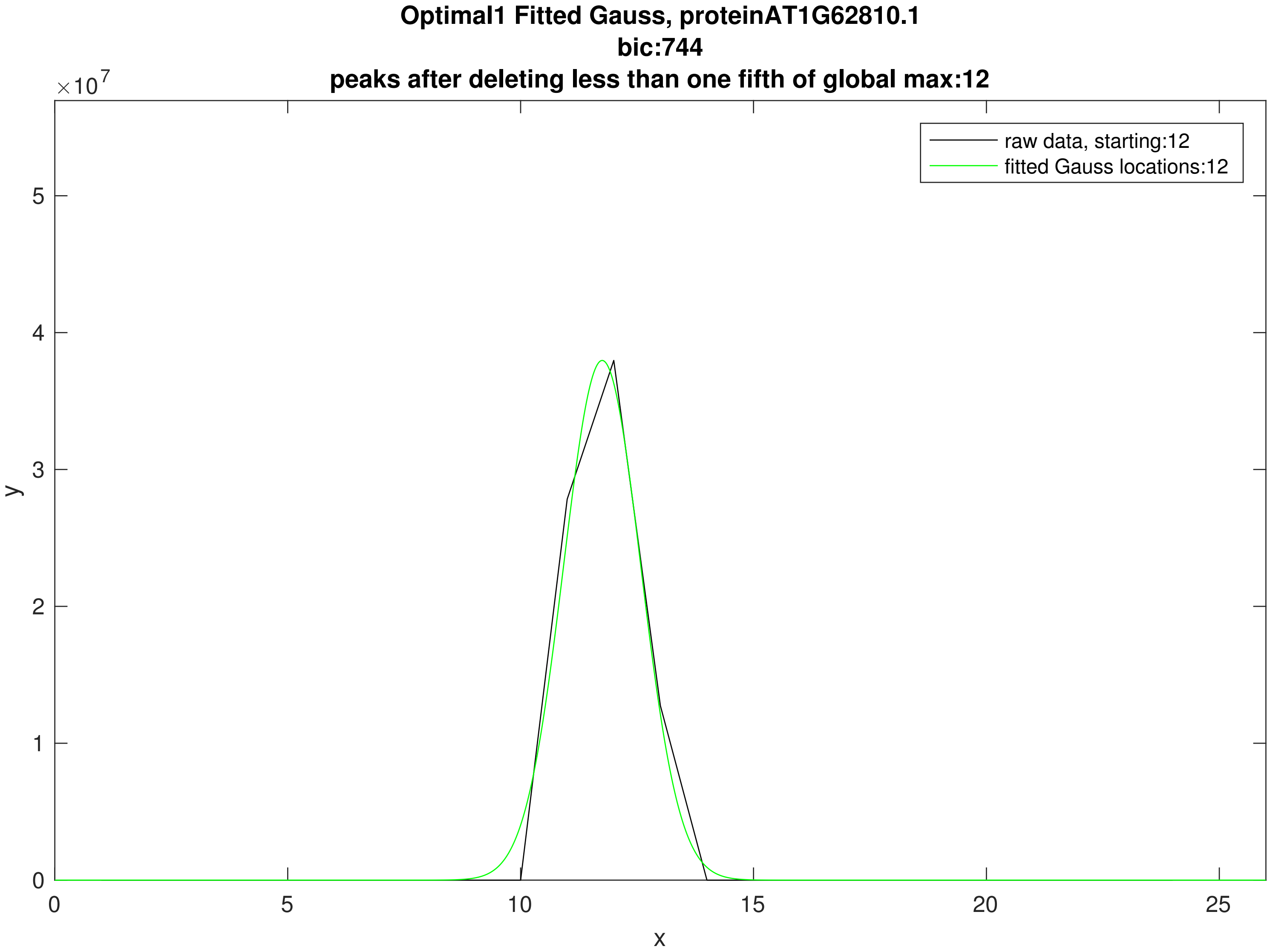
Optimal1 Fitted Gauss, proteinAT1G62810.1
bic:744
peaks after deleting less than one fifth of global max:12
×10^7
raw data, starting:12
fitted Gauss locations:12
y
x
0
1
2
3
4
5
0
5
10
15
20
25

**Optimal2 Fitted Gauss, proteinAT1G62820.1**
**bic:858 841**
**peaks after deleting less than one fifth of global max:19**

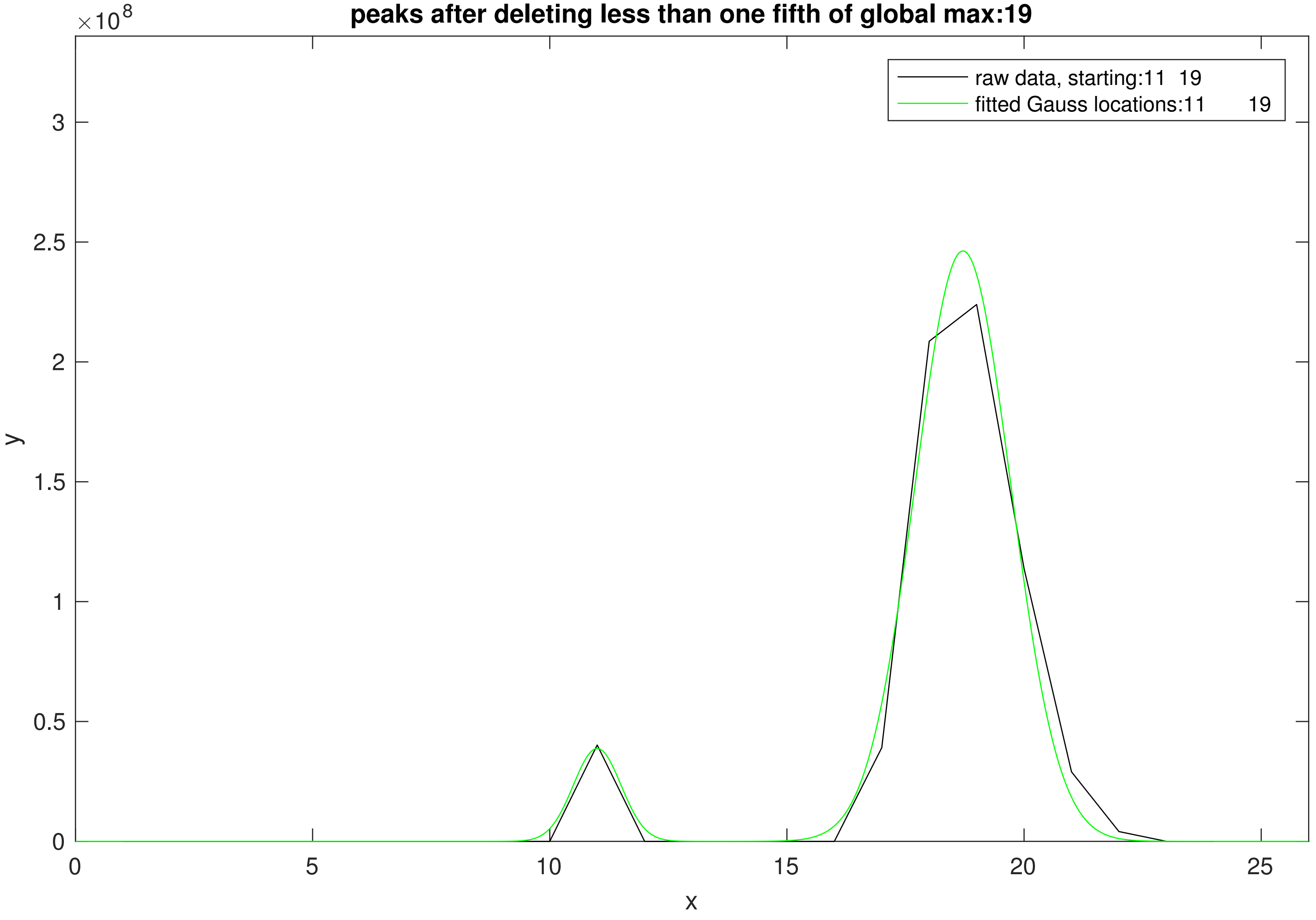

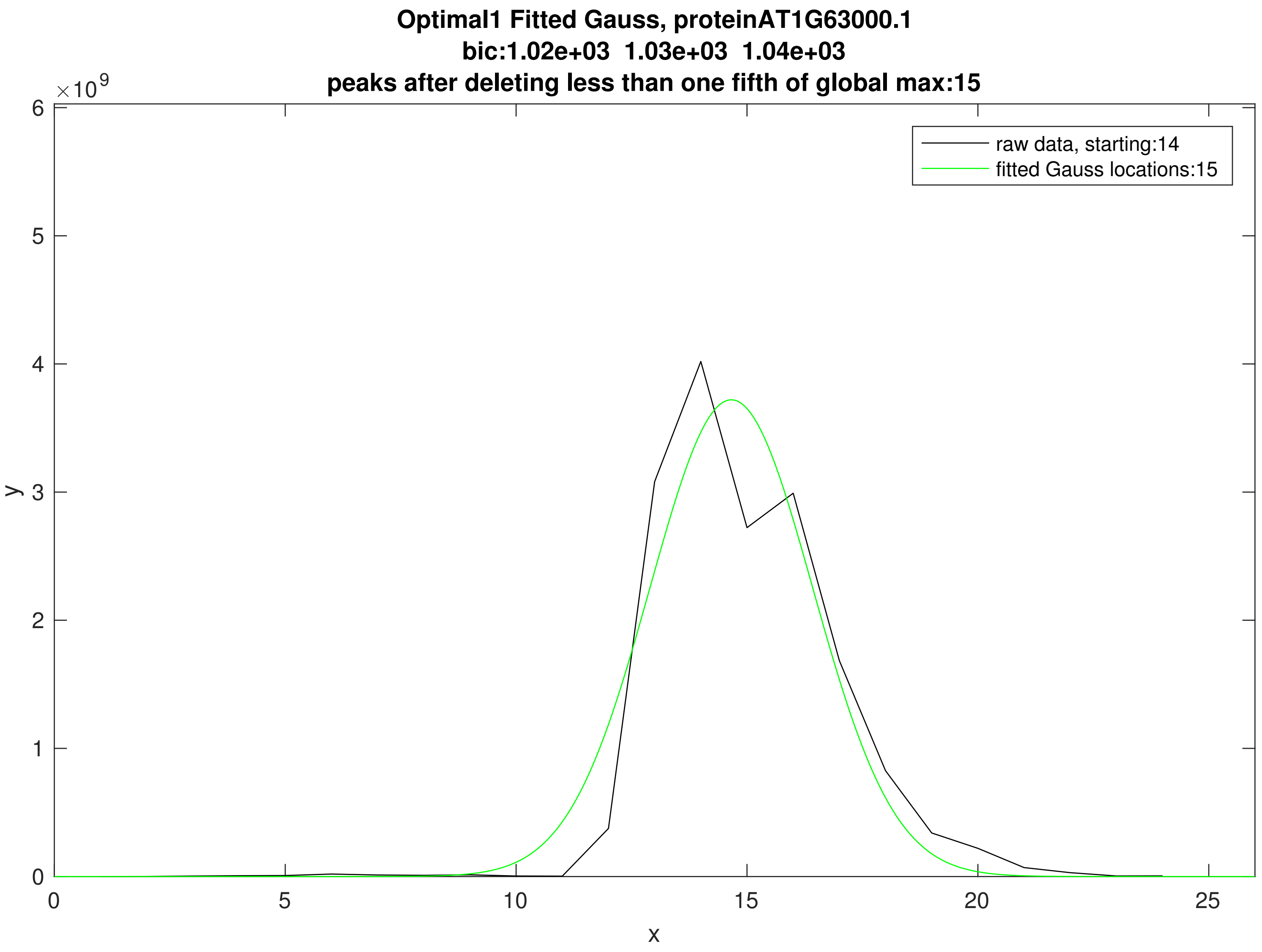
Optimal1 Fitted Gauss, proteinAT1G63000.1
bic:1.02e+03  1.03e+03  1.04e+03
peaks after deleting less than one fifth of global max:15
raw data, starting:14
fitted Gauss locations:15
y
x
×10^9
0
1
2
3
4
5
6
0
5
10
15
20
25

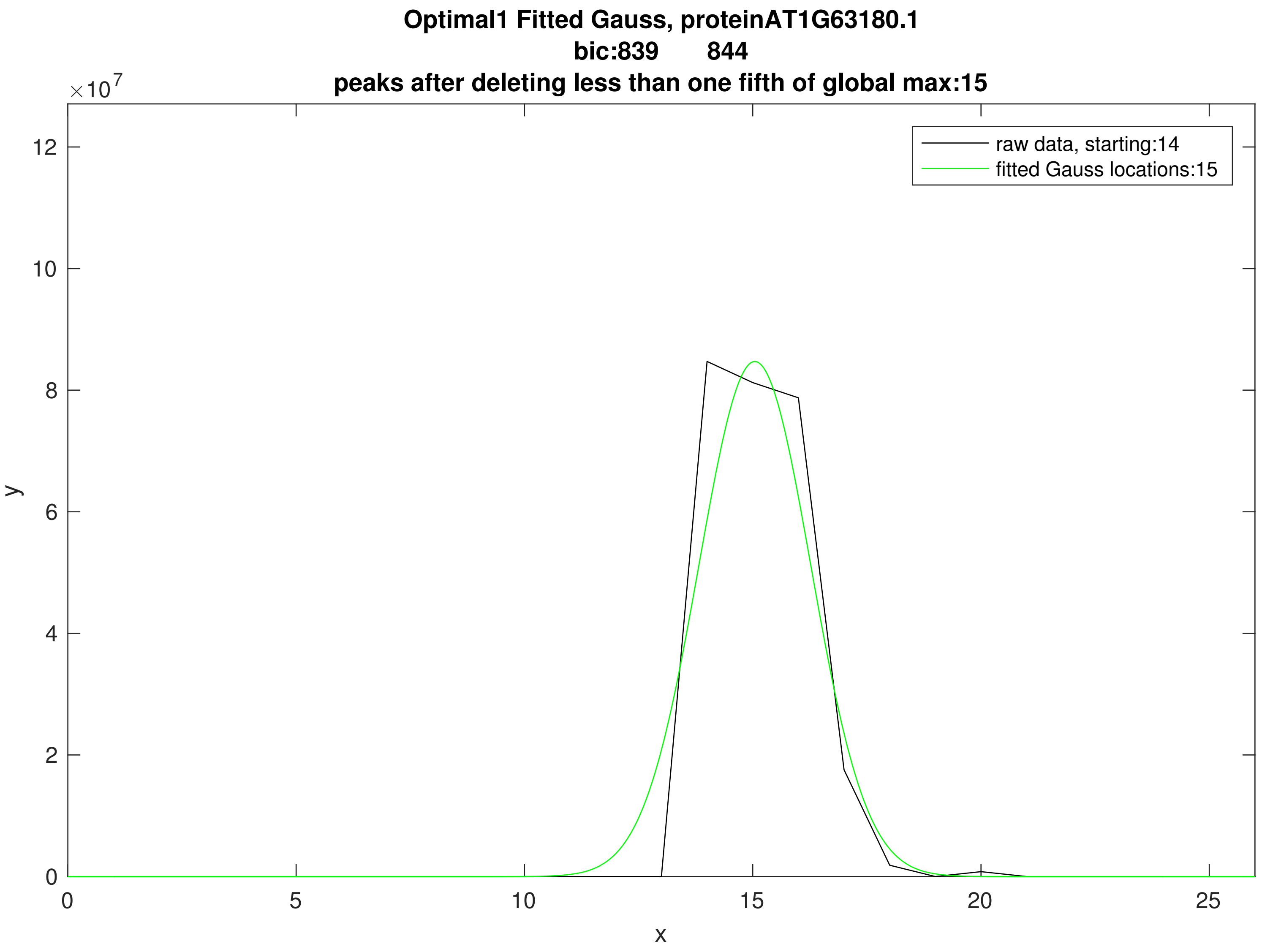
Optimal1 Fitted Gauss, proteinAT1G63180.1
bic:839 844
peaks after deleting less than one fifth of global max:15
×10^7
raw data, starting:14
fitted Gauss locations:15
y
x
0
2
4
6
8
10
12
5
15
20
25

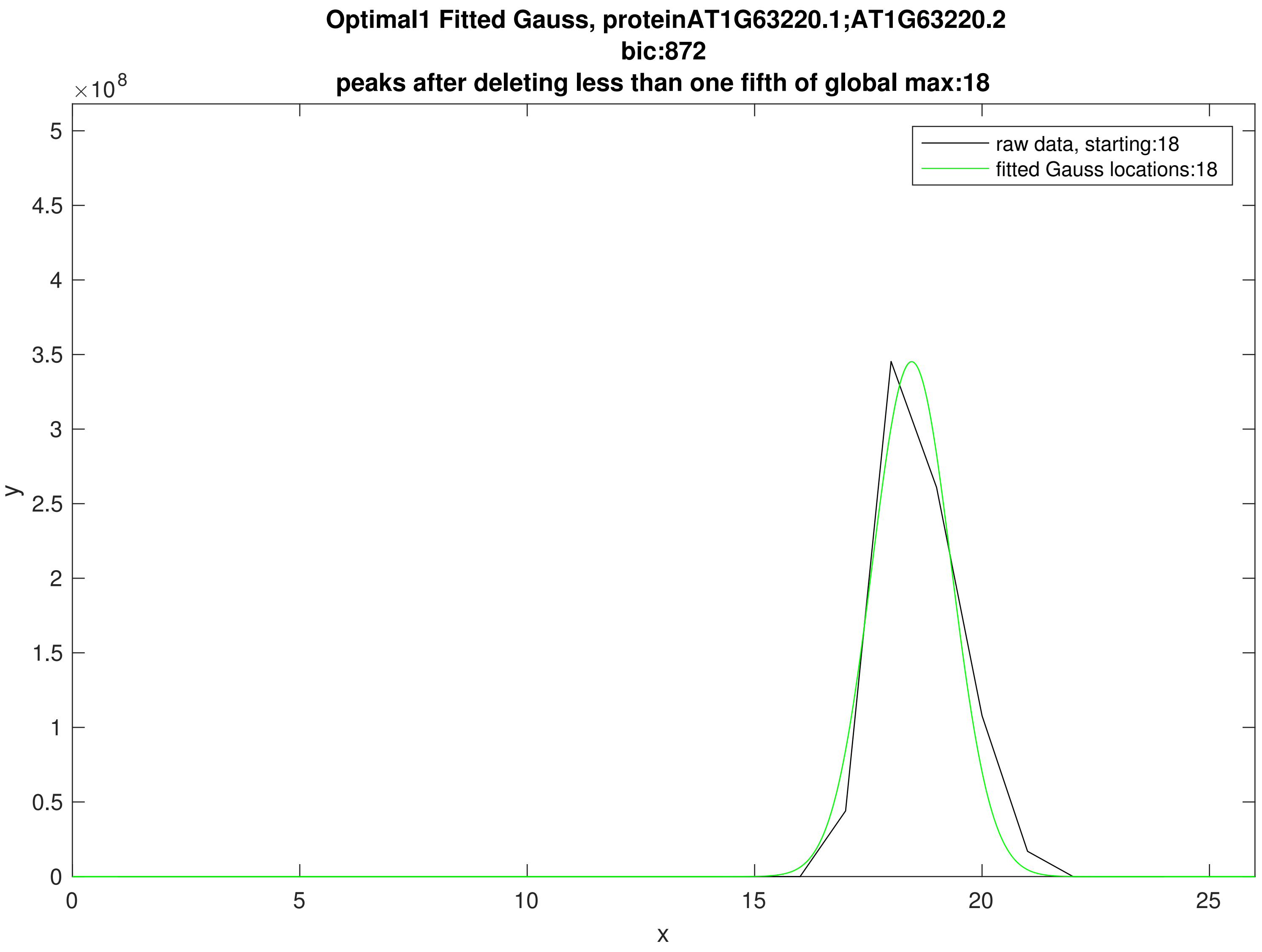

Optimal1 Fitted Gauss, proteinAT1G63220.1;AT1G63220.2
bic:872
peaks after deleting less than one fifth of global max:18
×10^8
raw data, starting:18
fitted Gauss locations:18
y
x
0
0.5
1
1.5
2
2.5
3
3.5
4
4.5
5
0
5
10
15
20
25

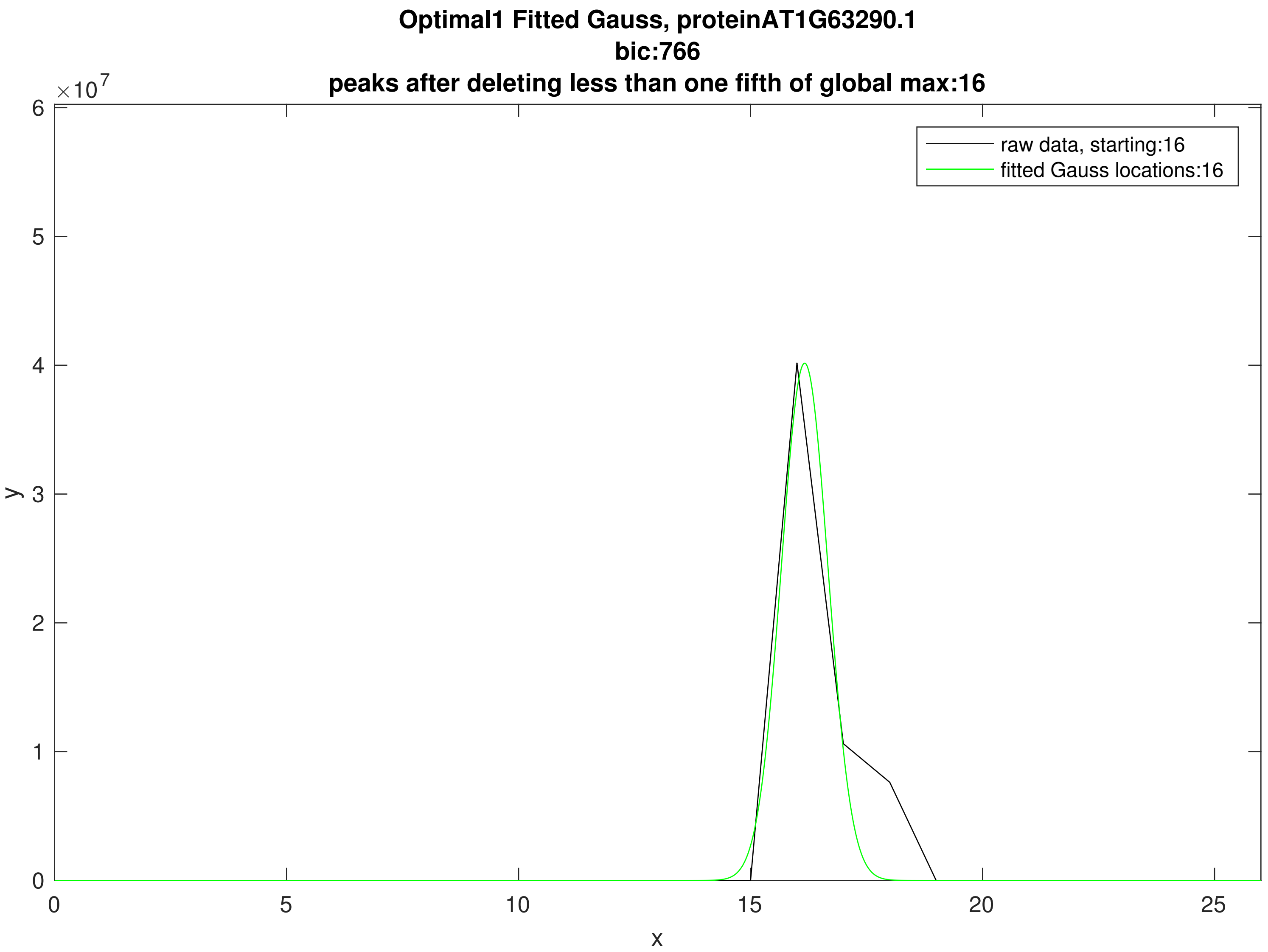

Optimal1 Fitted Gauss, proteinAT1G63290.1
bic:766
peaks after deleting less than one fifth of global max:16
×10^7
raw data, starting:16
fitted Gauss locations:16
y
x
0
1
2
3
4
5
6
0
5
10
15
20
25

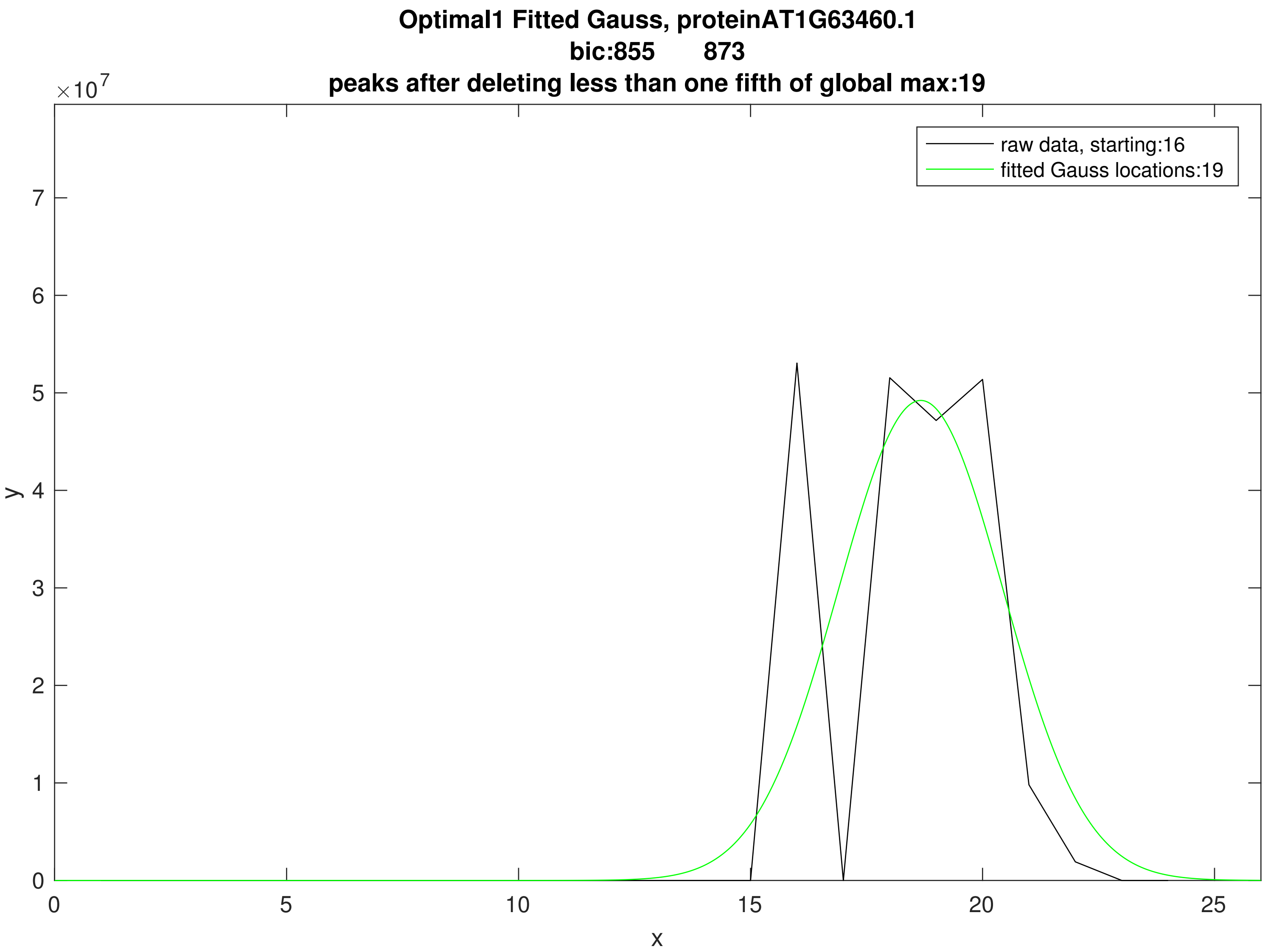
Optimal1 Fitted Gauss, proteinAT1G63460.1
bic:855 873
peaks after deleting less than one fifth of global max:19
×10^7
raw data, starting:16
fitted Gauss locations:19
y
x
0
1
2
3
4
5
6
7
0
5
10
15
20
25

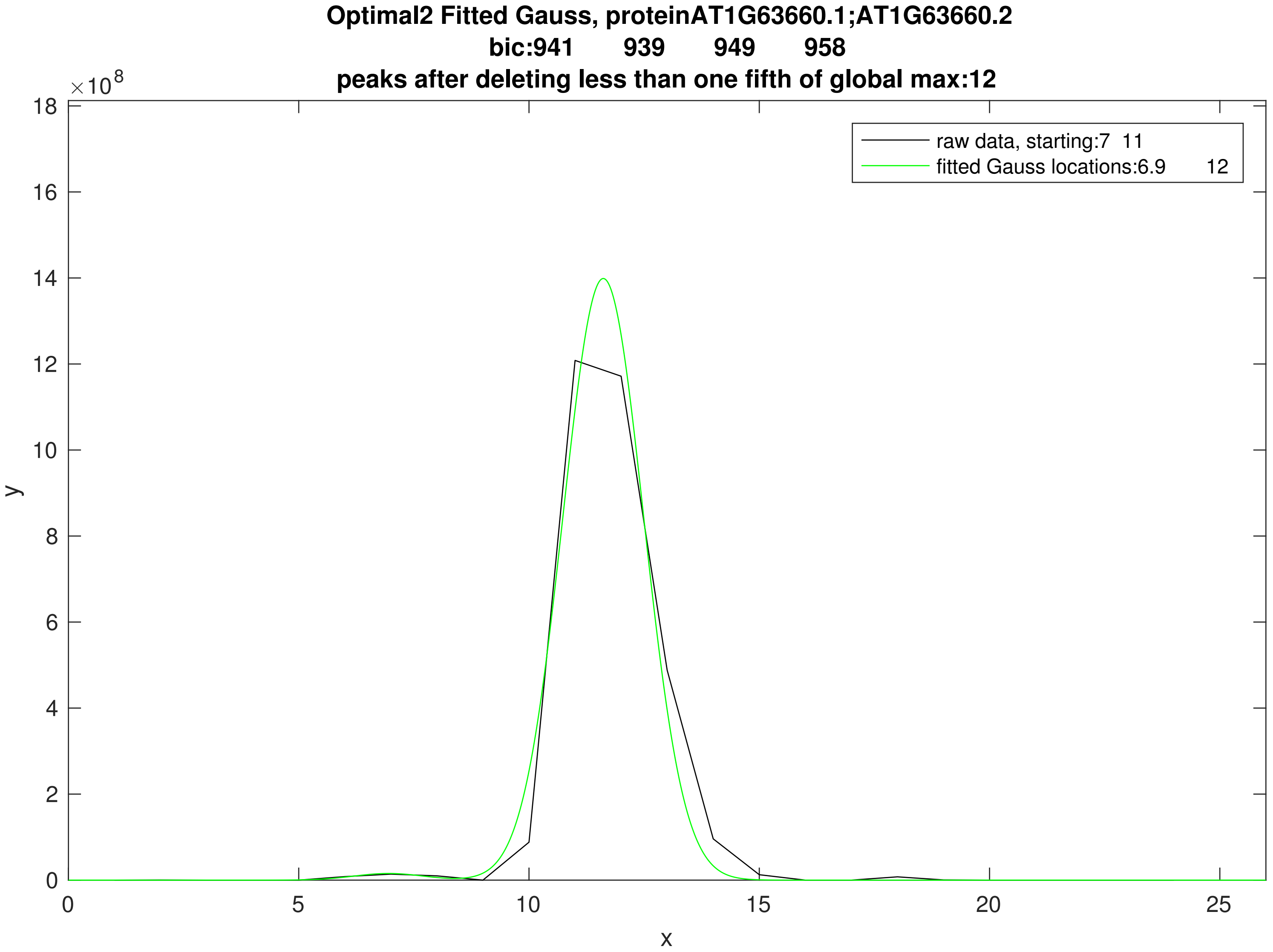

Optimal2 Fitted Gauss, proteinAT1G63660.1;AT1G63660.2
bic:941 939 949 958
peaks after deleting less than one fifth of global max:12
×10^8
18
16
14
12
10
8
6
4
2
0
y
0
5
10
15
20
25
x
raw data, starting:7 11
fitted Gauss locations:6.9 12

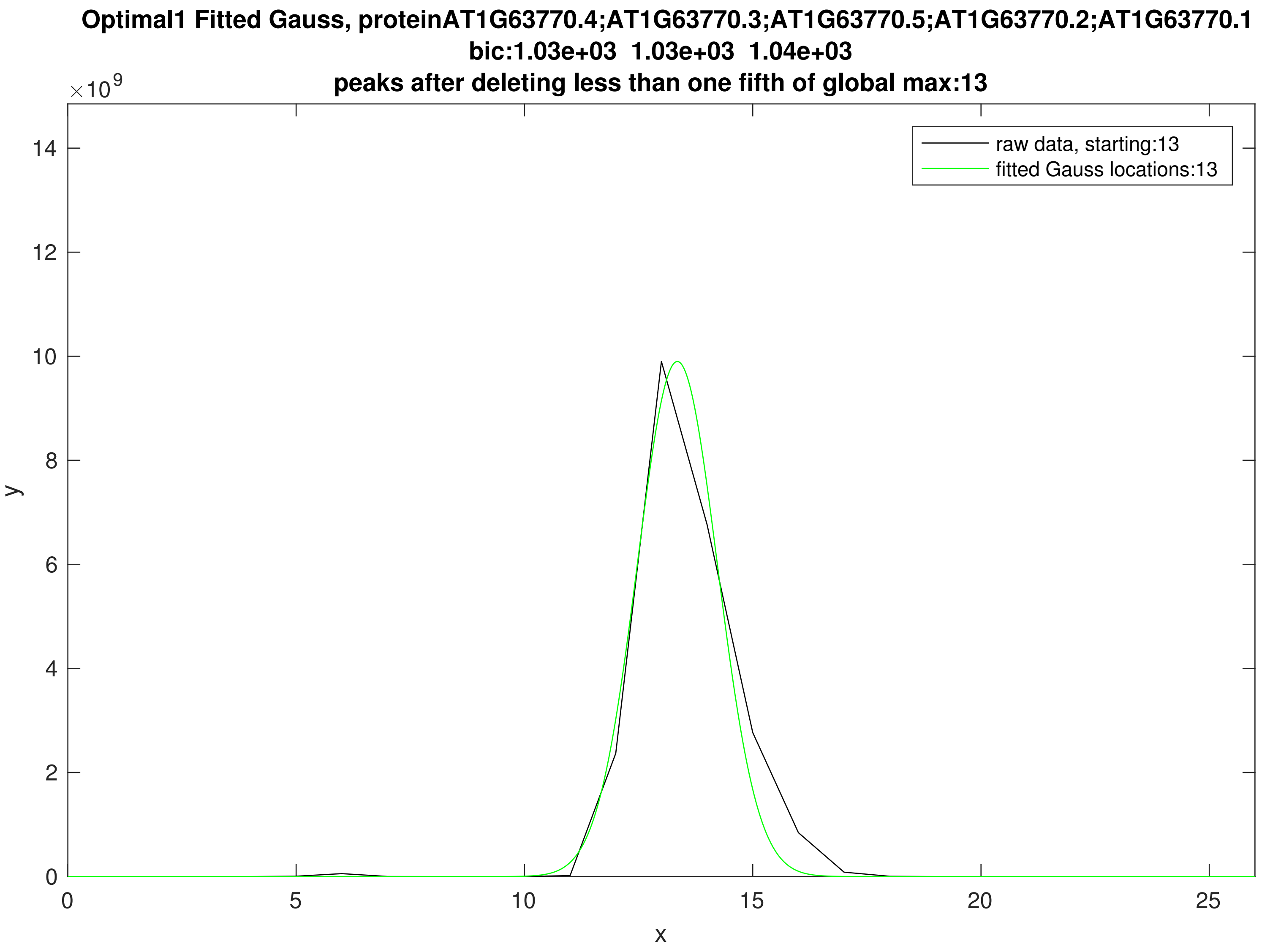
Optimal1 Fitted Gauss, proteinAT1G63770.4;AT1G63770.3;AT1G63770.5;AT1G63770.2;AT1G63770.1
bic:1.03e+03 1.03e+03 1.04e+03
peaks after deleting less than one fifth of global max:13
×10^9
raw data, starting:13
fitted Gauss locations:13
y
x
0
2
4
6
8
10
12
14
0
5
10
15
20
25

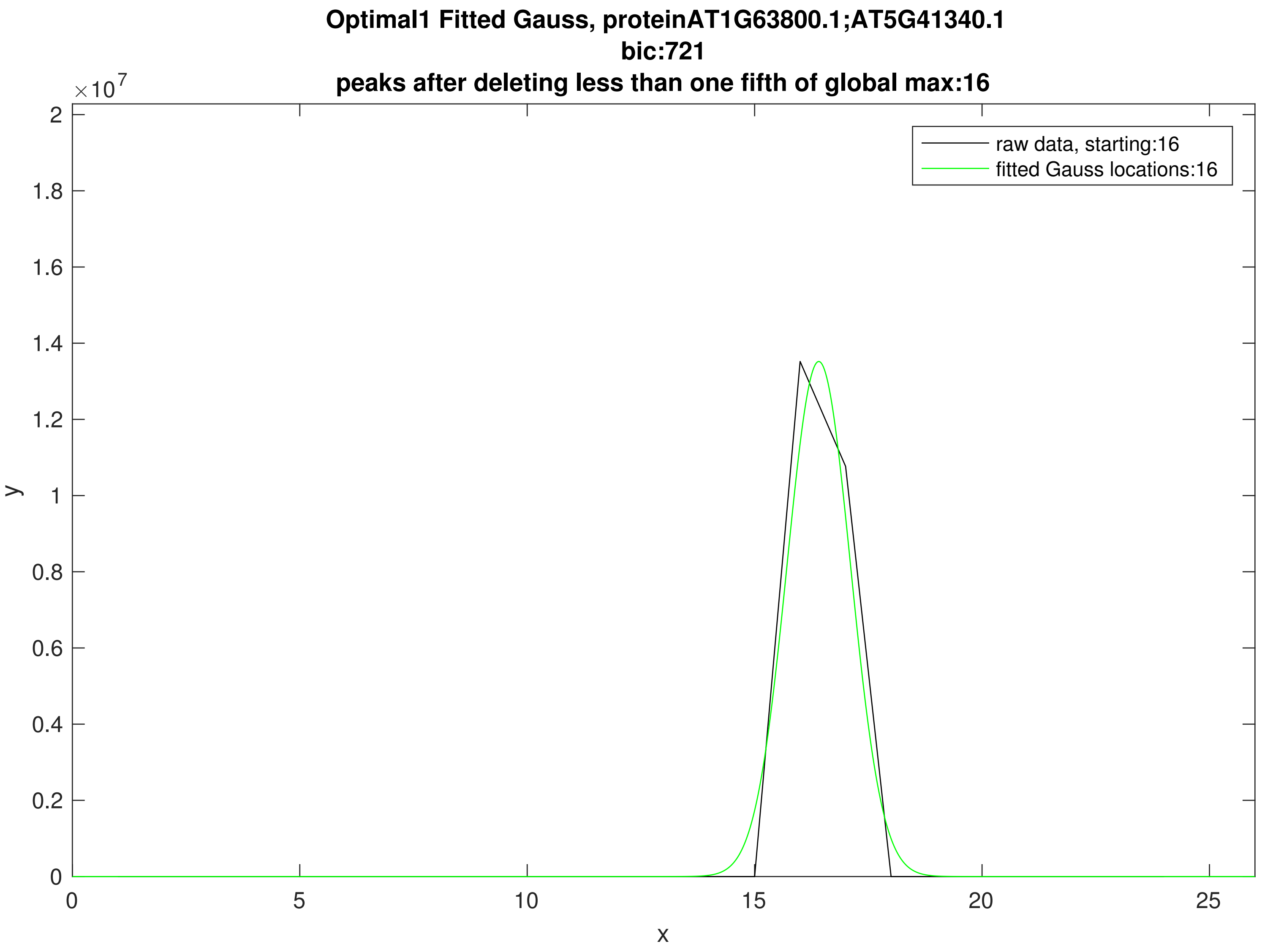

Optimal1 Fitted Gauss, proteinAT1G63800.1;AT5G41340.1
bic:721
peaks after deleting less than one fifth of global max:16
×10^7
2
1.8
1.6
1.4
1.2
1
0.8
0.6
0.4
0.2
0
y
0
5
10
15
20
25
x
raw data, starting:16
fitted Gauss locations:16

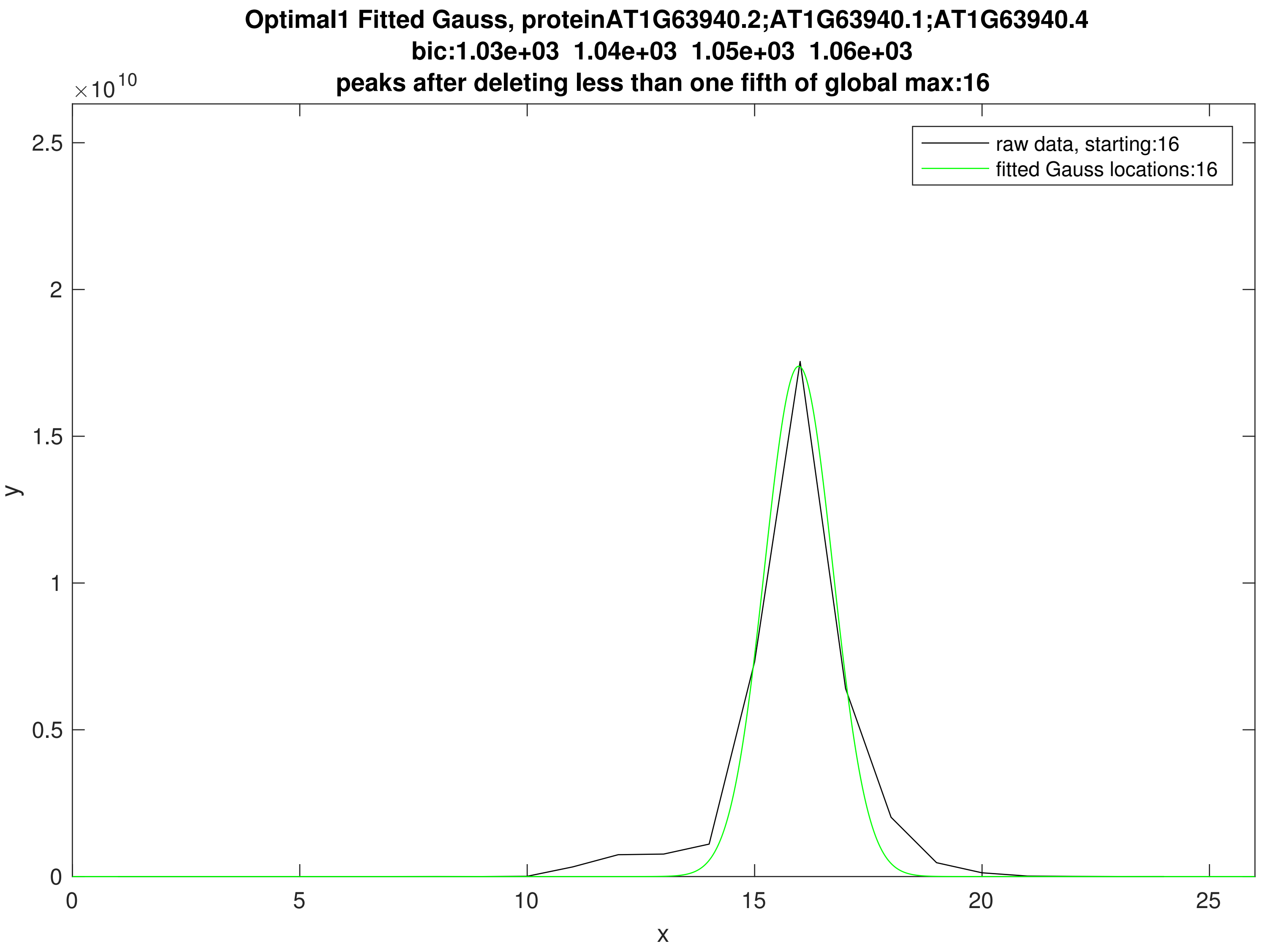
Optimal1 Fitted Gauss, proteinAT1G63940.2;AT1G63940.1;AT1G63940.4
bic:1.03e+03 1.04e+03 1.05e+03 1.06e+03
peaks after deleting less than one fifth of global max:16
×10^10
2.5
2
1.5
1
0.5
0
y
0
5
10
15
20
25
x
raw data, starting:16
fitted Gauss locations:16

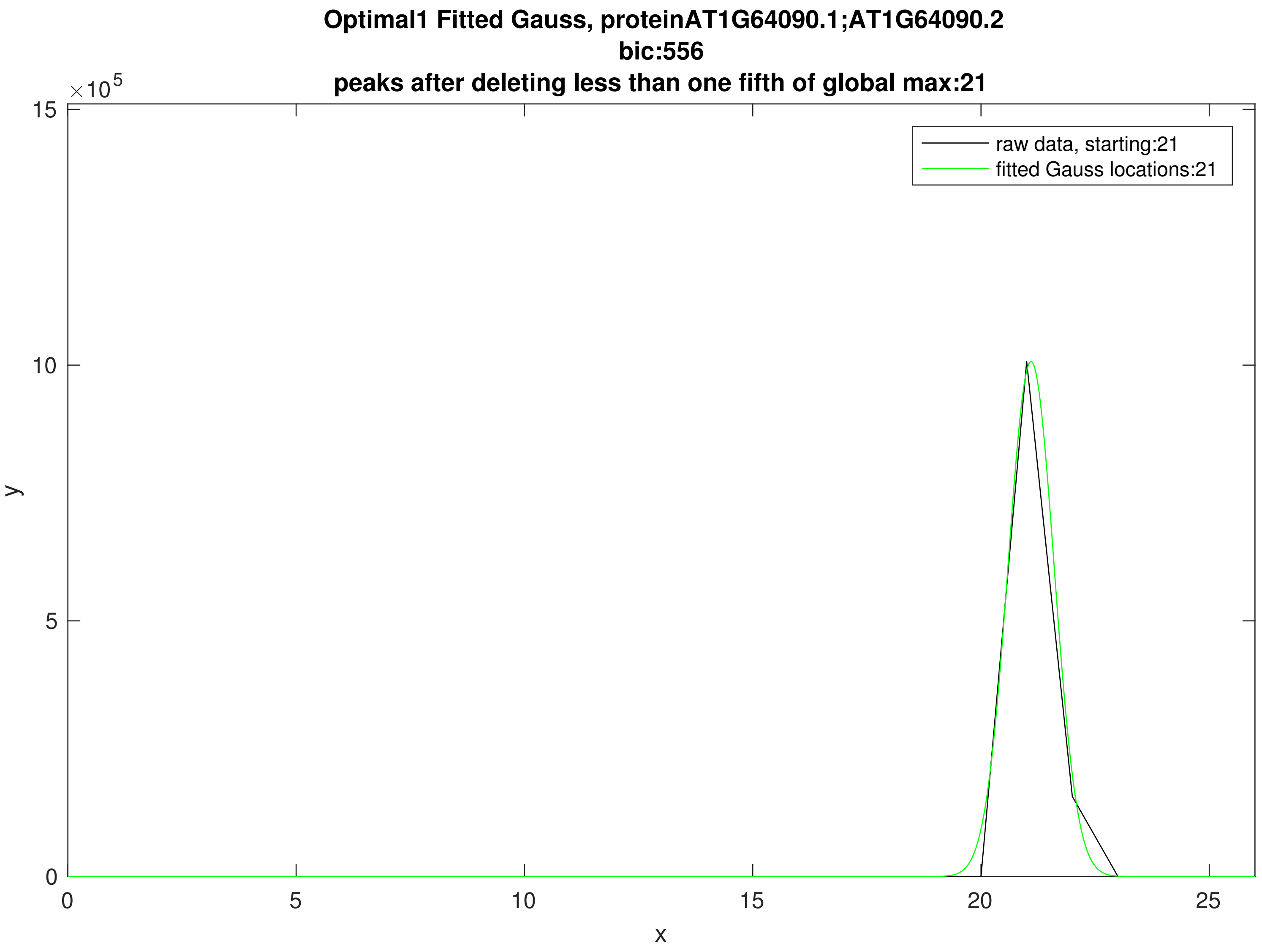
Optimal1 Fitted Gauss, proteinAT1G64090.1;AT1G64090.2
bic:556
peaks after deleting less than one fifth of global max:21
×10^5
15
10
5
0
y
0
5
10
15
20
25
x
raw data, starting:21
fitted Gauss locations:21

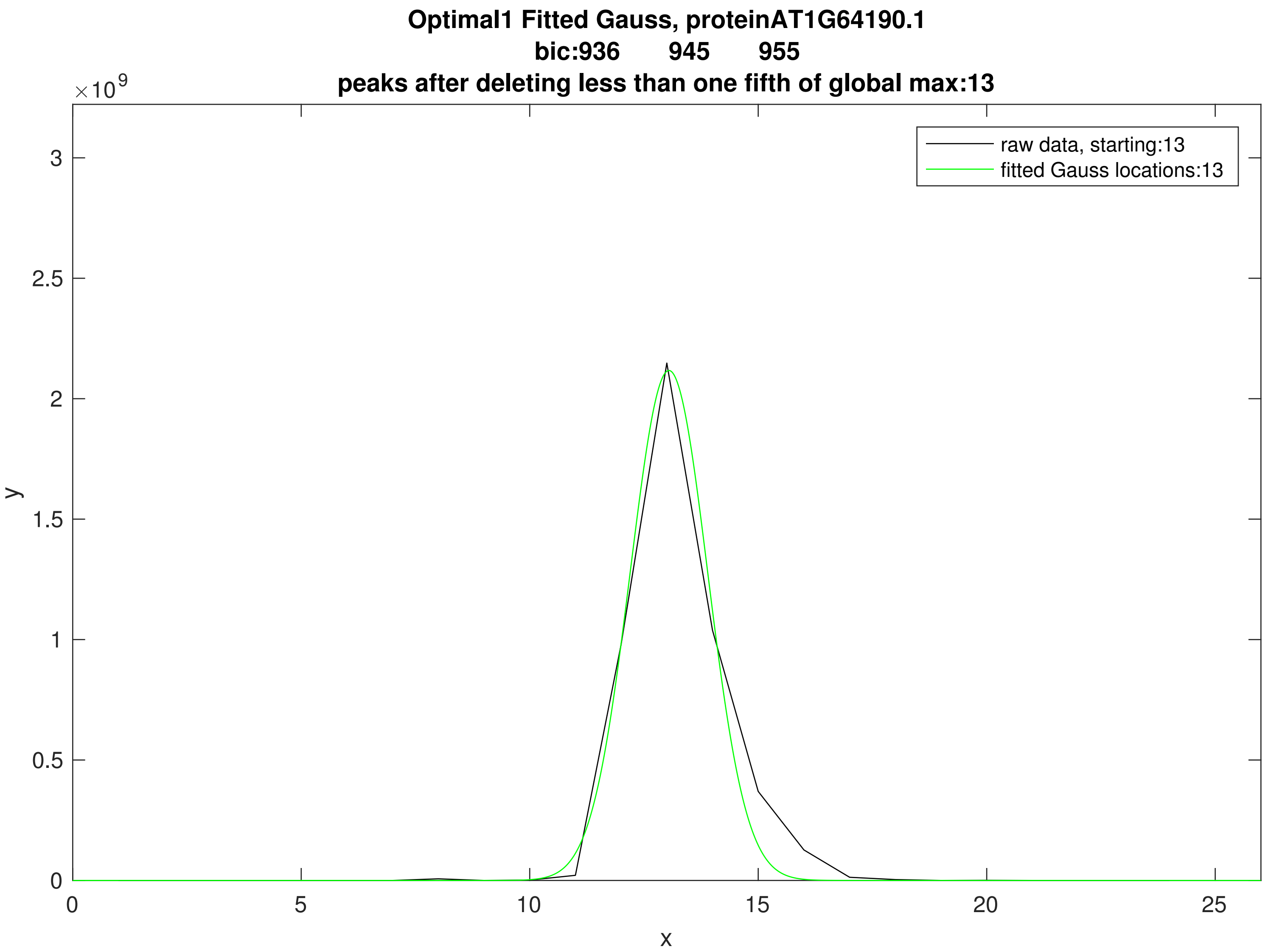
Optimal1 Fitted Gauss, proteinAT1G64190.1
bic:936 945 955
peaks after deleting less than one fifth of global max:13
×10^9
raw data, starting:13
fitted Gauss locations:13
y
x
0
0.5
1
1.5
2
2.5
3
0
5
10
15
20
25

**Optimal1 Fitted Gauss, proteinAT1G64440.1**
**bic:903 912 922**
**peaks after deleting less than one fifth of global max:15**

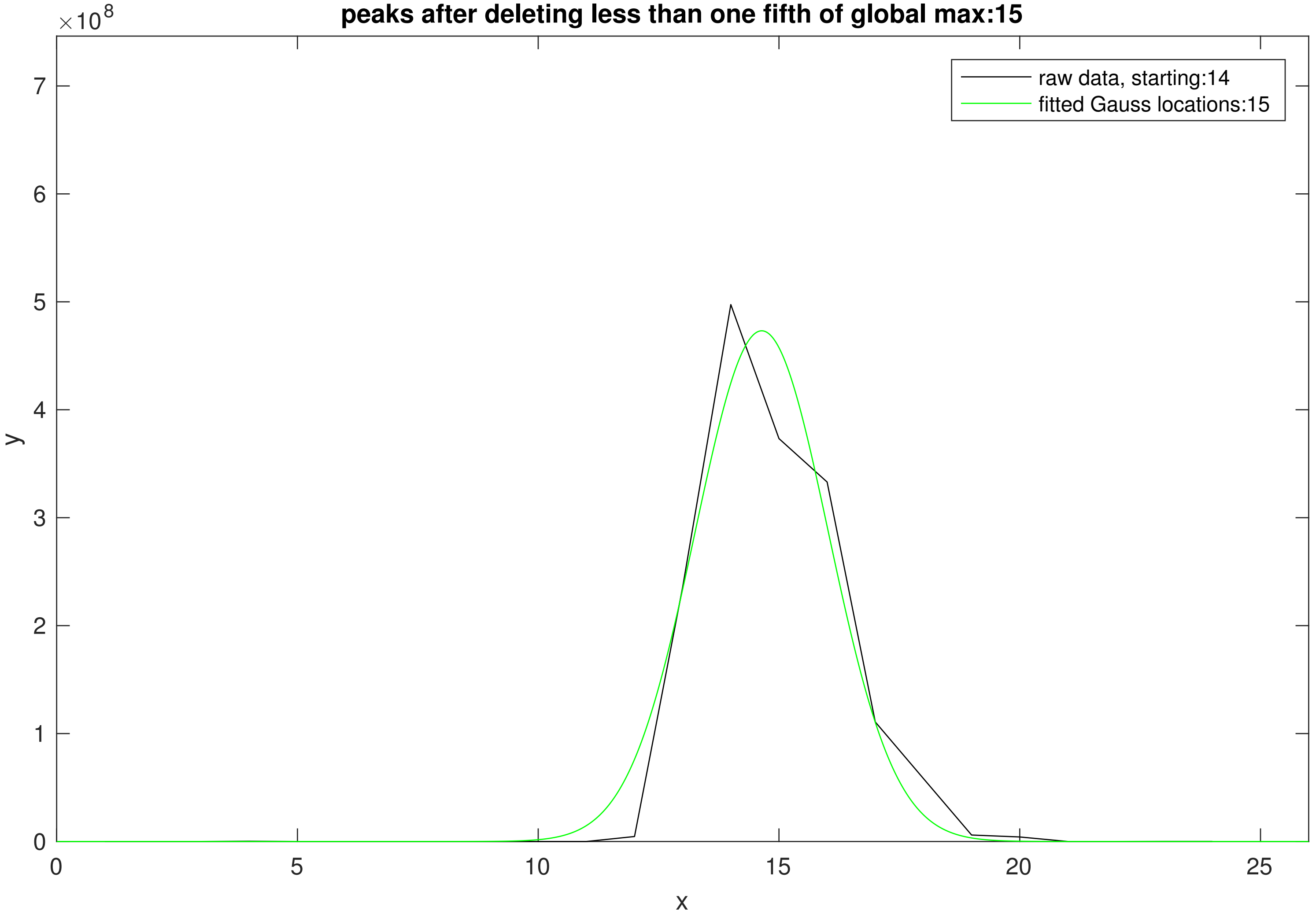

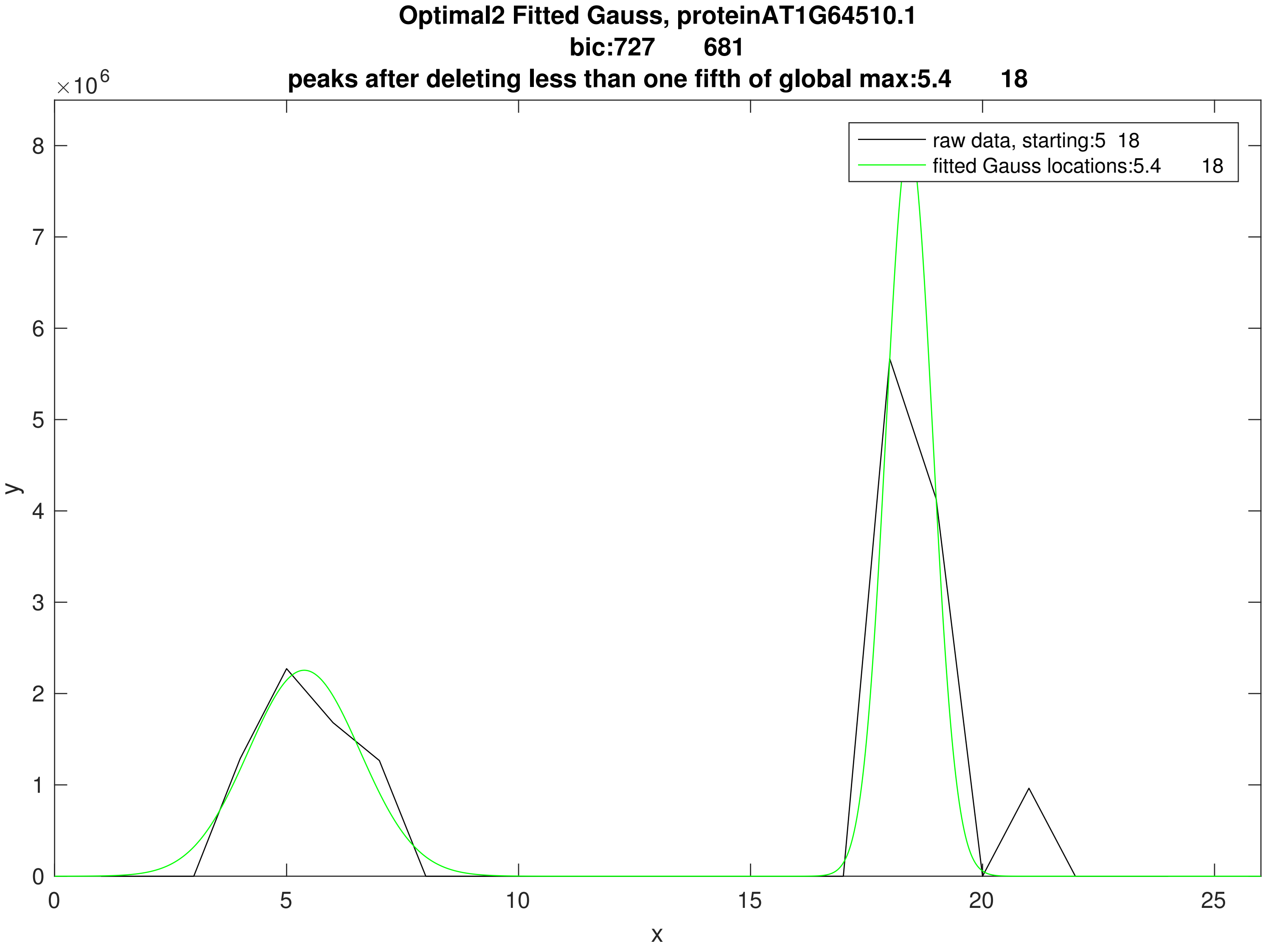
Optimal2 Fitted Gauss, proteinAT1G64510.1
bic:727 681
peaks after deleting less than one fifth of global max:5.4 18
×10^6
raw data, starting:5 18
fitted Gauss locations:5.4 18
y
x

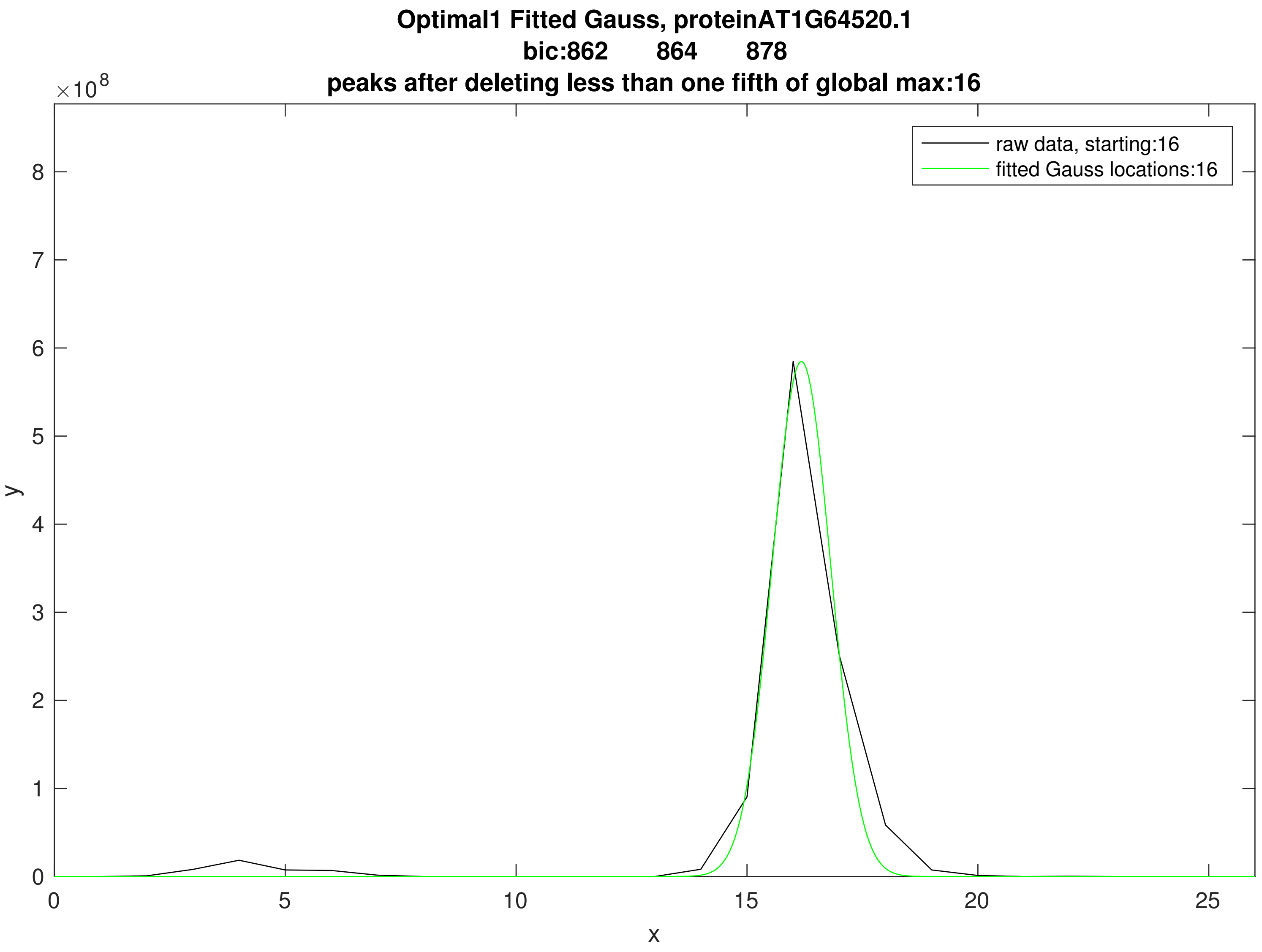
Optimal1 Fitted Gauss, proteinAT1G64520.1
bic:862 864 878
peaks after deleting less than one fifth of global max:16
×10^8
raw data, starting:16
fitted Gauss locations:16
y
x
0
1
2
3
4
5
6
7
8
0
5
10
15
20
25

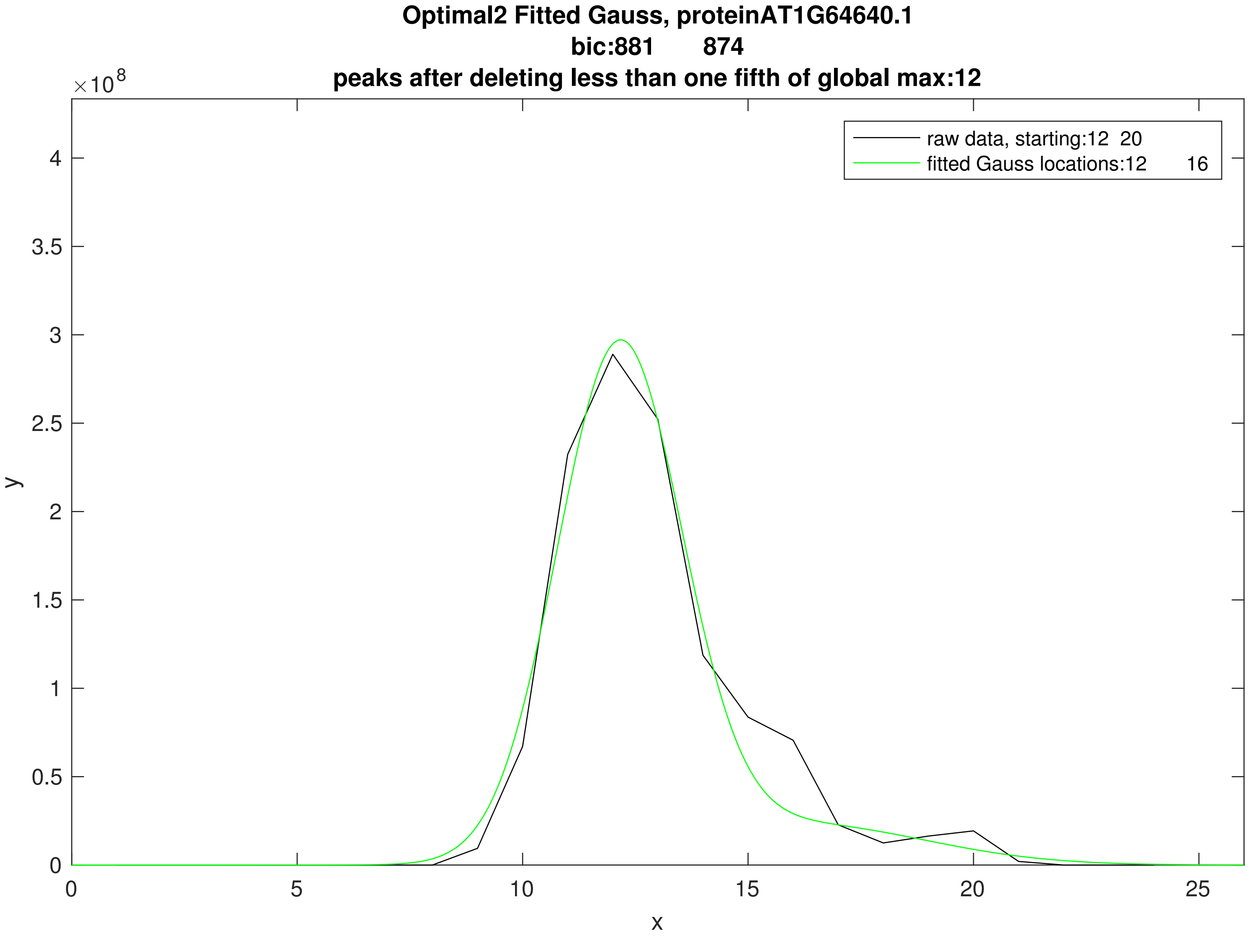

Optimal2 Fitted Gauss, proteinAT1G64640.1
bic:881 874
peaks after deleting less than one fifth of global max:12
×10^8
raw data, starting:12 20
fitted Gauss locations:12 16
y
x
0
0.5
1
1.5
2
2.5
3
3.5
4
5
10
15
20
25

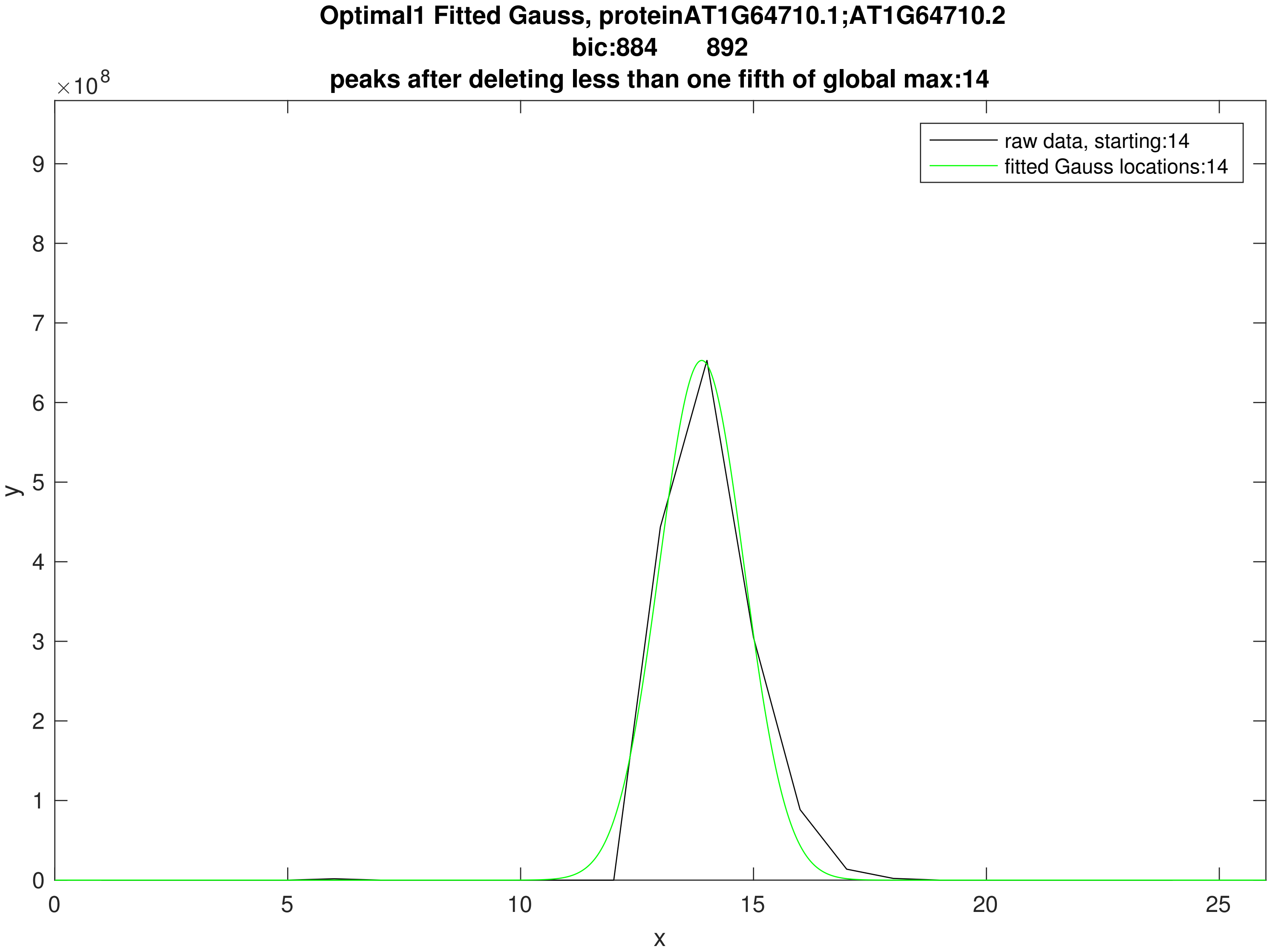
Optimal1 Fitted Gauss, proteinAT1G64710.1;AT1G64710.2
bic:884 892
peaks after deleting less than one fifth of global max:14
×10^8
raw data, starting:14
fitted Gauss locations:14
y
x
0
1
2
3
4
5
6
7
8
9
0
5
10
15
20
25

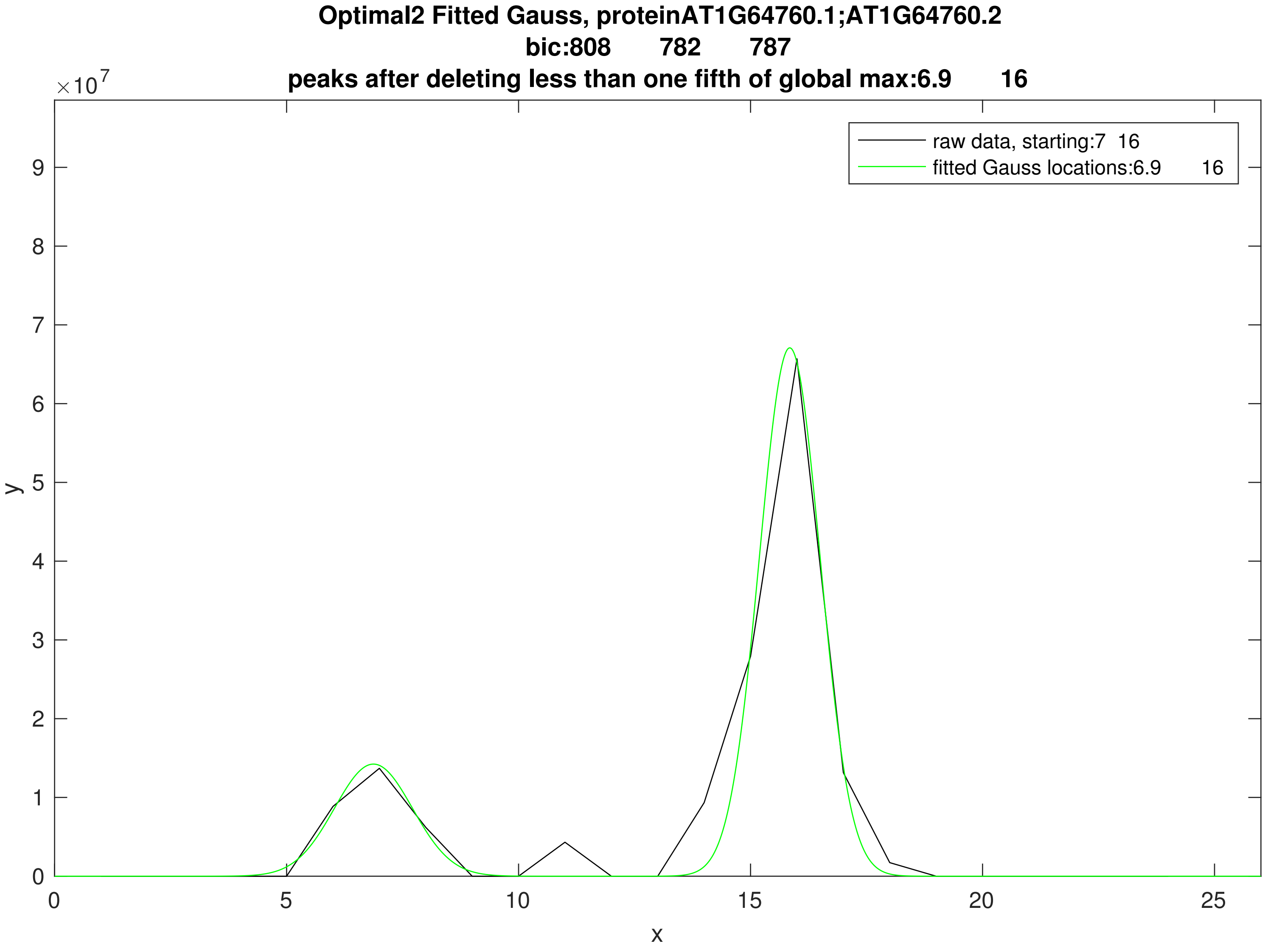

Optimal2 Fitted Gauss, proteinAT1G64760.1;AT1G64760.2
bic:808 782 787
peaks after deleting less than one fifth of global max:6.9 16
×10^7
raw data, starting:7 16
fitted Gauss locations:6.9 16
y
x

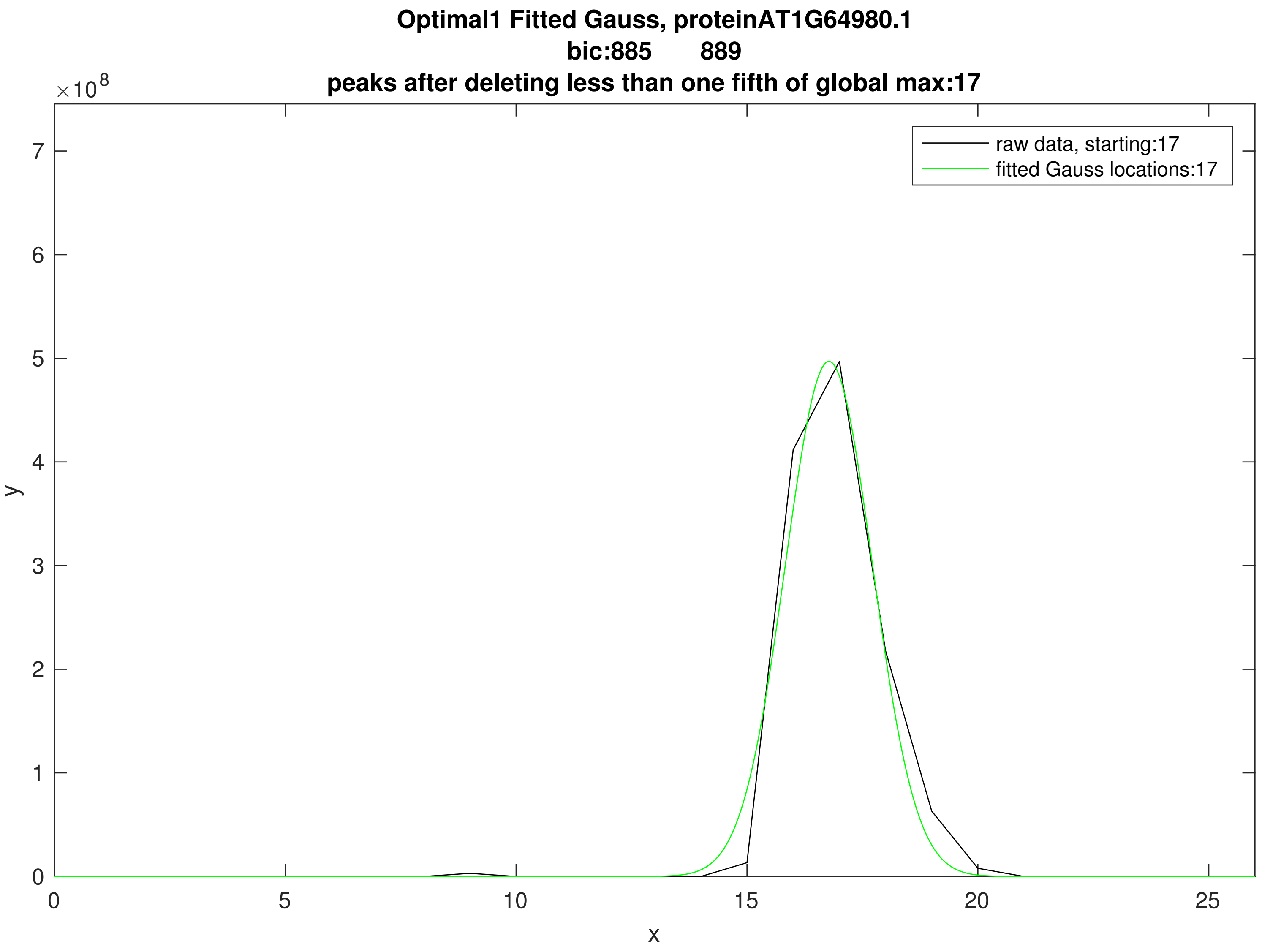

Optimal1 Fitted Gauss, proteinAT1G64980.1
bic:885 889
peaks after deleting less than one fifth of global max:17
×10^8
raw data, starting:17
fitted Gauss locations:17
y
x
0
1
2
3
4
5
6
7
0
5
10
15
20
25

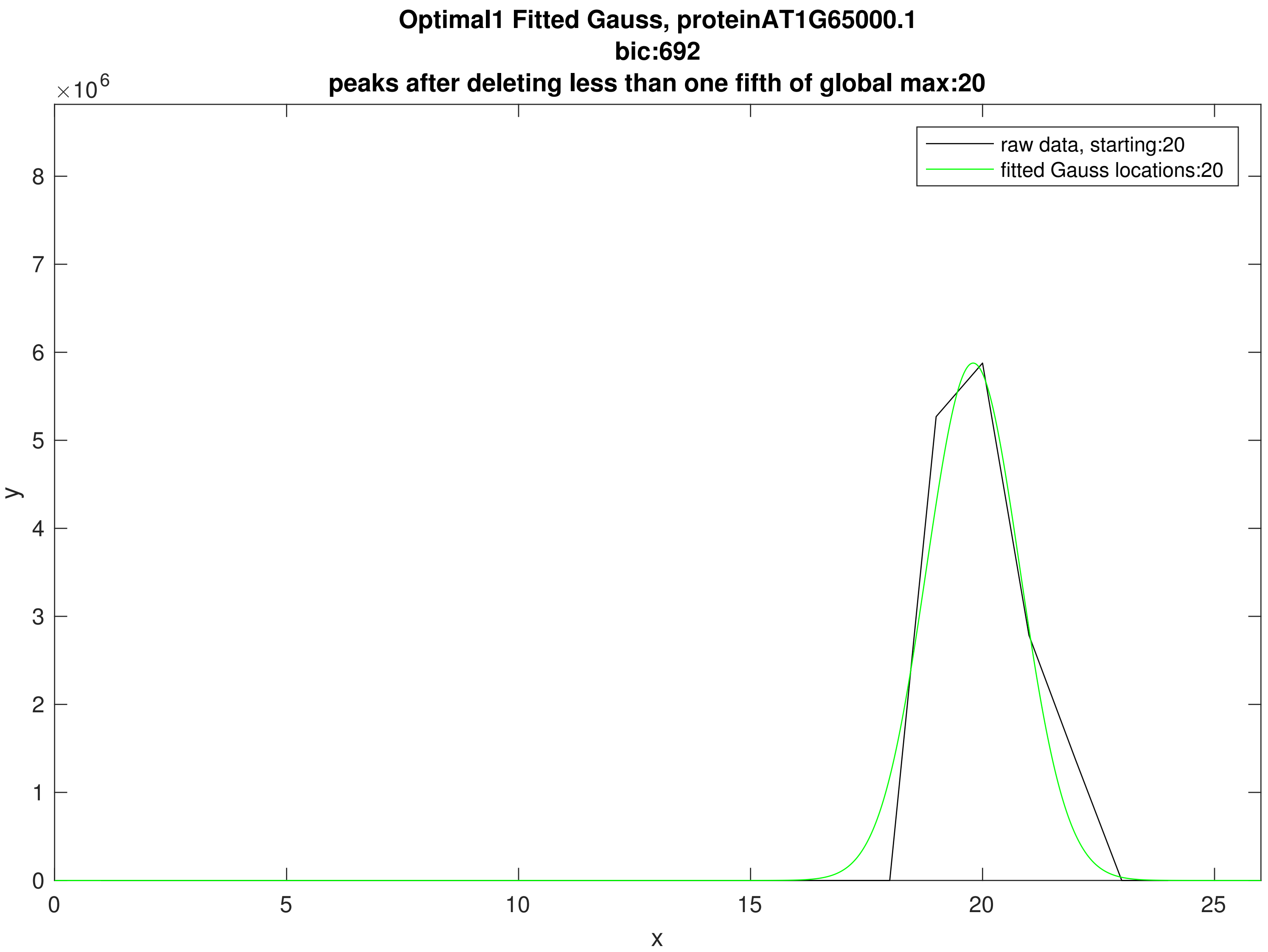

Optimal1 Fitted Gauss, proteinAT1G65000.1
bic:692
peaks after deleting less than one fifth of global max:20
×10^6
raw data, starting:20
fitted Gauss locations:20
y
x
0
1
2
3
4
5
6
7
8
0
5
10
15
20
25

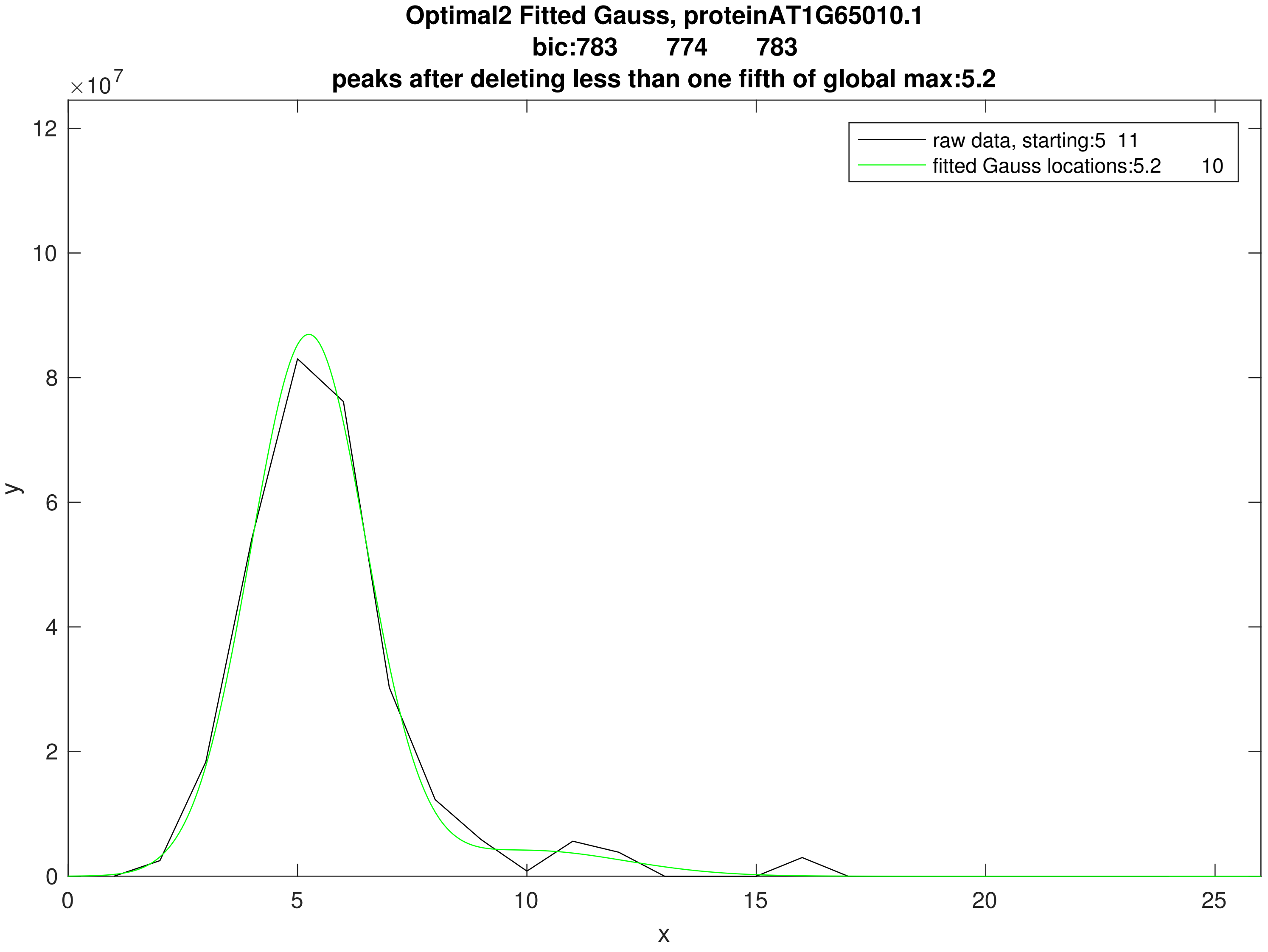

Optimal2 Fitted Gauss, proteinAT1G65010.1
bic:783 774 783
peaks after deleting less than one fifth of global max:5.2
×10^7
raw data, starting:5 11
fitted Gauss locations:5.2 10
y
x
0
2
4
6
8
10
12
0
5
10
15
20
25

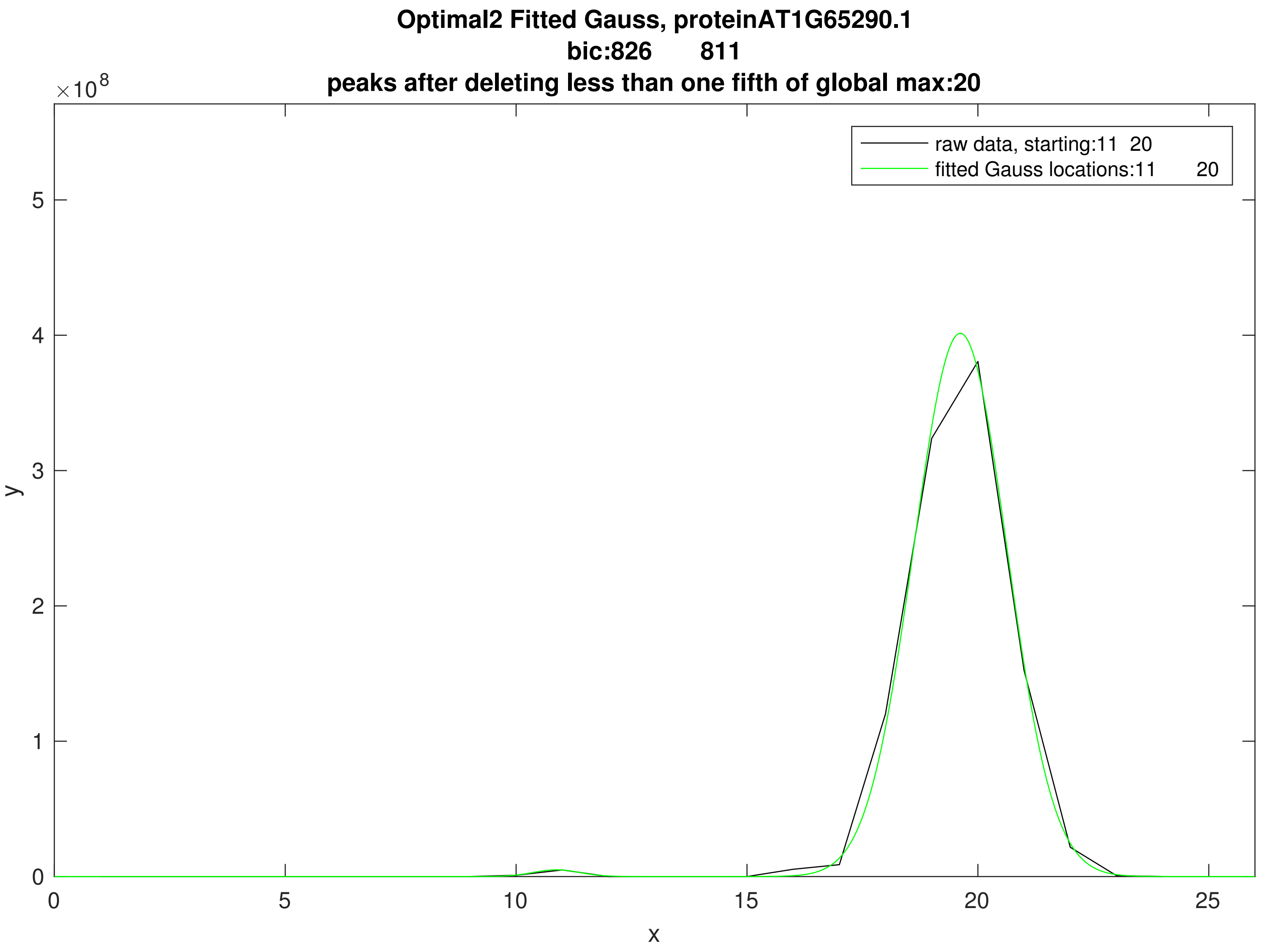
Optimal2 Fitted Gauss, proteinAT1G65290.1
bic:826 811
peaks after deleting less than one fifth of global max:20
×10^8
raw data, starting:11 20
fitted Gauss locations:11 20
y
x
0
1
2
3
4
5
0
5
10
15
20
25

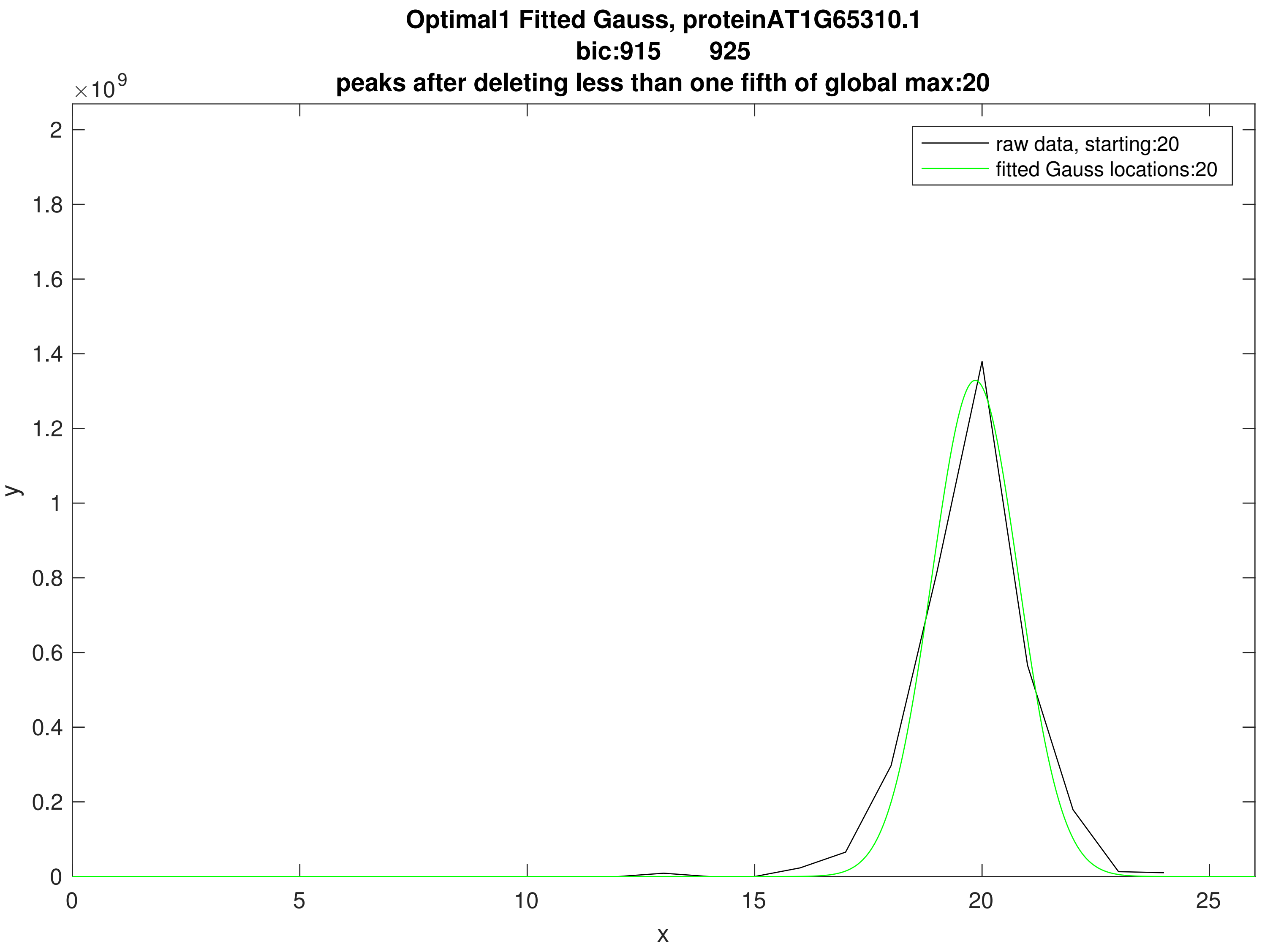

Optimal1 Fitted Gauss, proteinAT1G65310.1
bic:915 925
peaks after deleting less than one fifth of global max:20
×10^9
raw data, starting:20
fitted Gauss locations:20
y
x
0
0.2
0.4
0.6
0.8
1
1.2
1.4
1.6
1.8
2
0
5
10
15
20
25

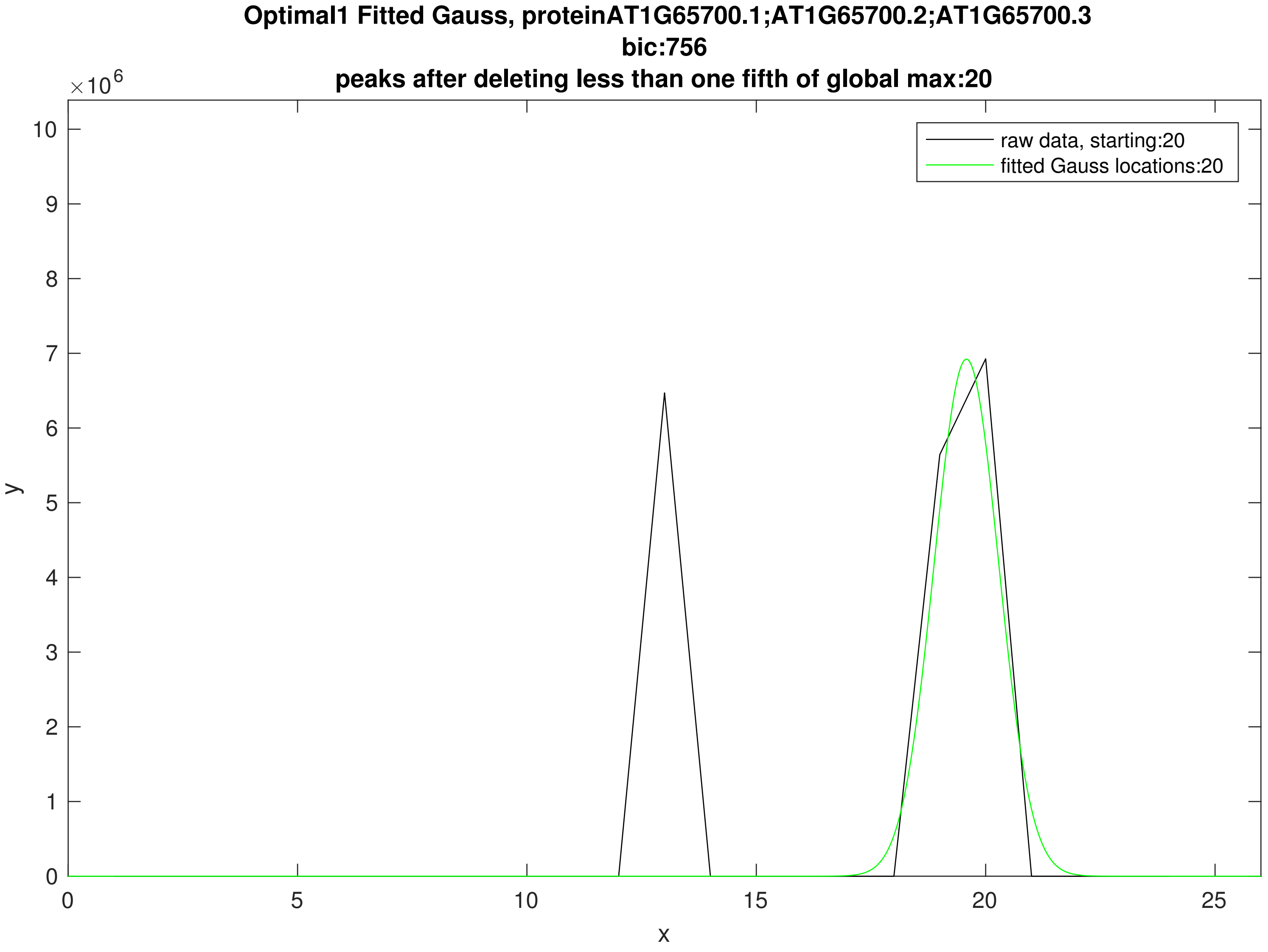

Optimal1 Fitted Gauss, proteinAT1G65700.1;AT1G65700.2;AT1G65700.3
bic:756
peaks after deleting less than one fifth of global max:20
×10^6
raw data, starting:20
fitted Gauss locations:20
y
x
0
1
2
3
4
5
6
7
8
9
10
0
5
10
15
20
25

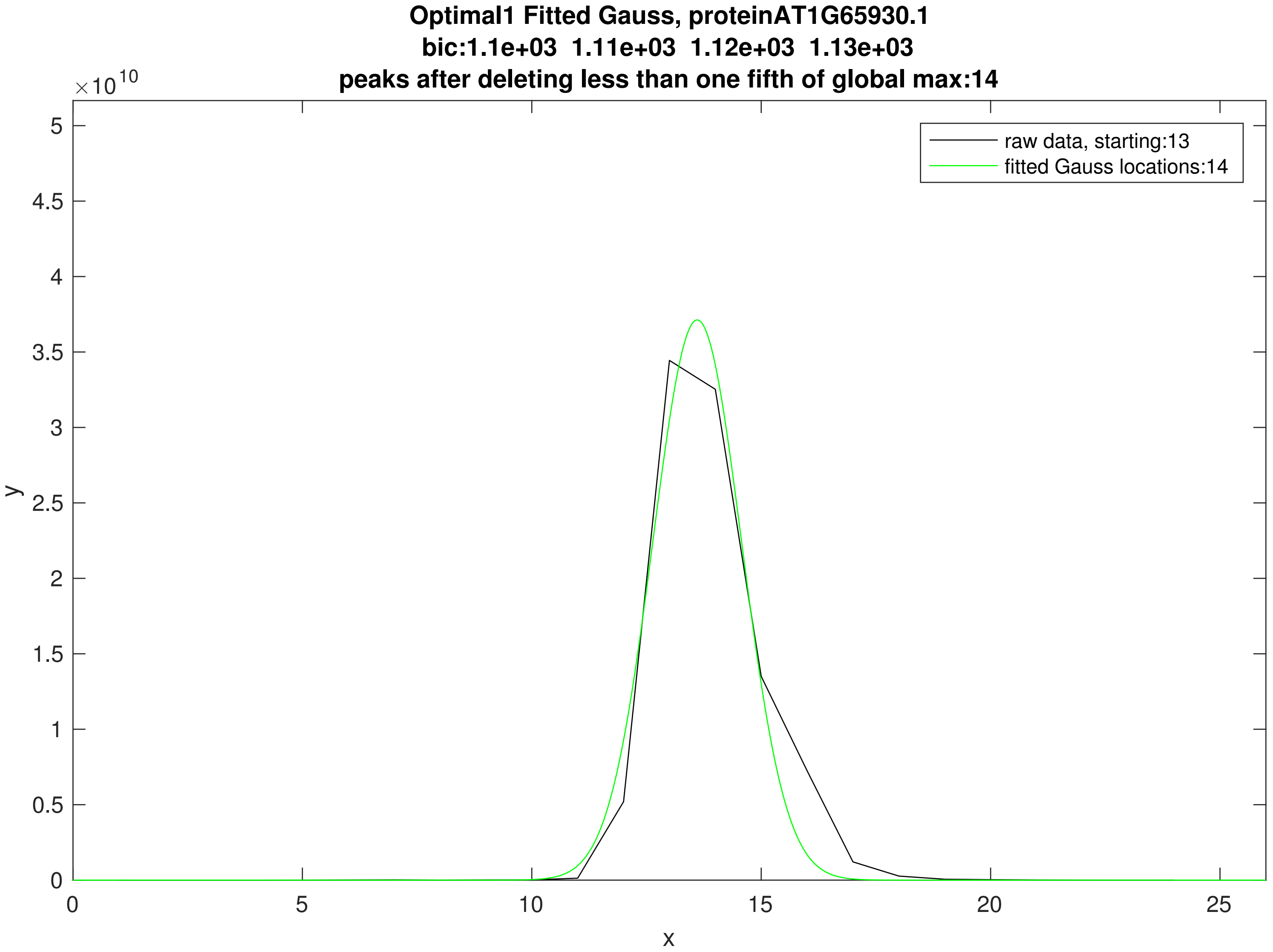
Optimal1 Fitted Gauss, proteinAT1G65930.1
bic:1.1e+03  1.11e+03  1.12e+03  1.13e+03
peaks after deleting less than one fifth of global max:14
×10^10
raw data, starting:13
fitted Gauss locations:14
y
x
0
0.5
1
1.5
2
2.5
3
3.5
4
4.5
5
0
5
10
15
20
25

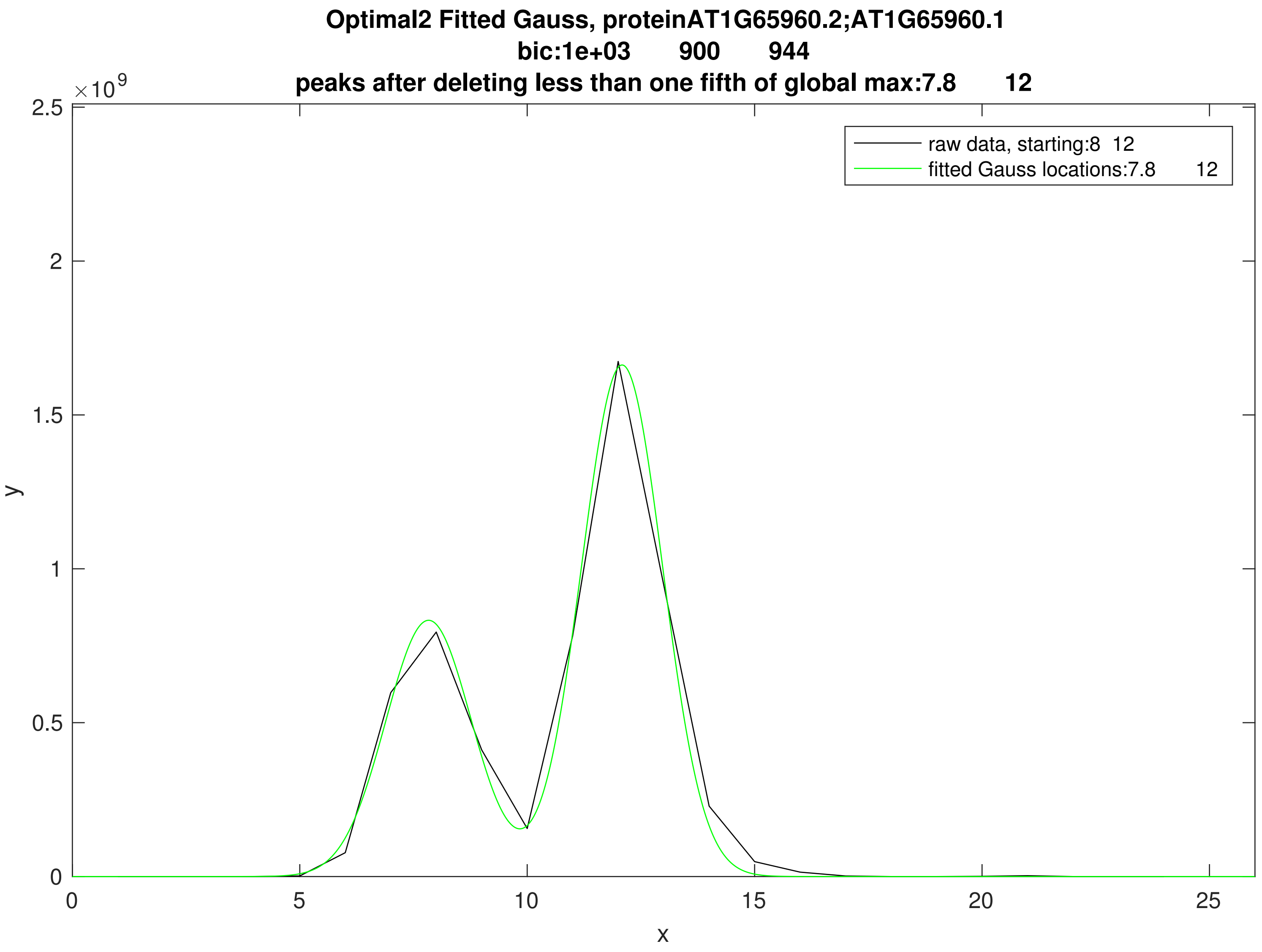
Optimal2 Fitted Gauss, proteinAT1G65960.2;AT1G65960.1
bic:1e+03 900 944
peaks after deleting less than one fifth of global max:7.8 12
×10^9
2.5
2
1.5
1
0.5
0
y
0
5
10
15
20
25
x
raw data, starting:8 12
fitted Gauss locations:7.8 12

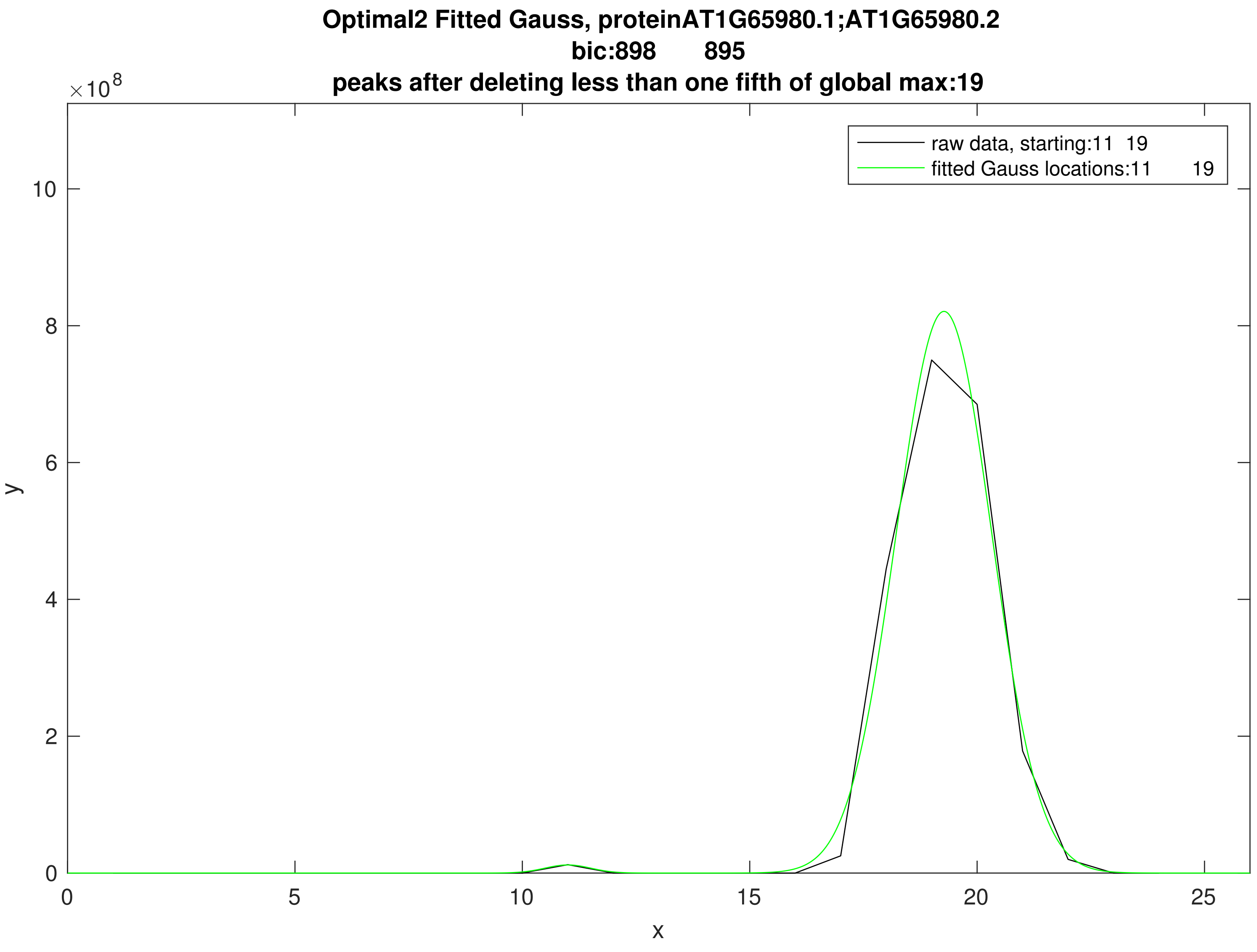
Optimal2 Fitted Gauss, proteinAT1G65980.1;AT1G65980.2
bic:898 895
peaks after deleting less than one fifth of global max:19
×10^8
raw data, starting:11 19
fitted Gauss locations:11 19
y
x
0
2
4
6
8
10
0
5
10
15
20
25

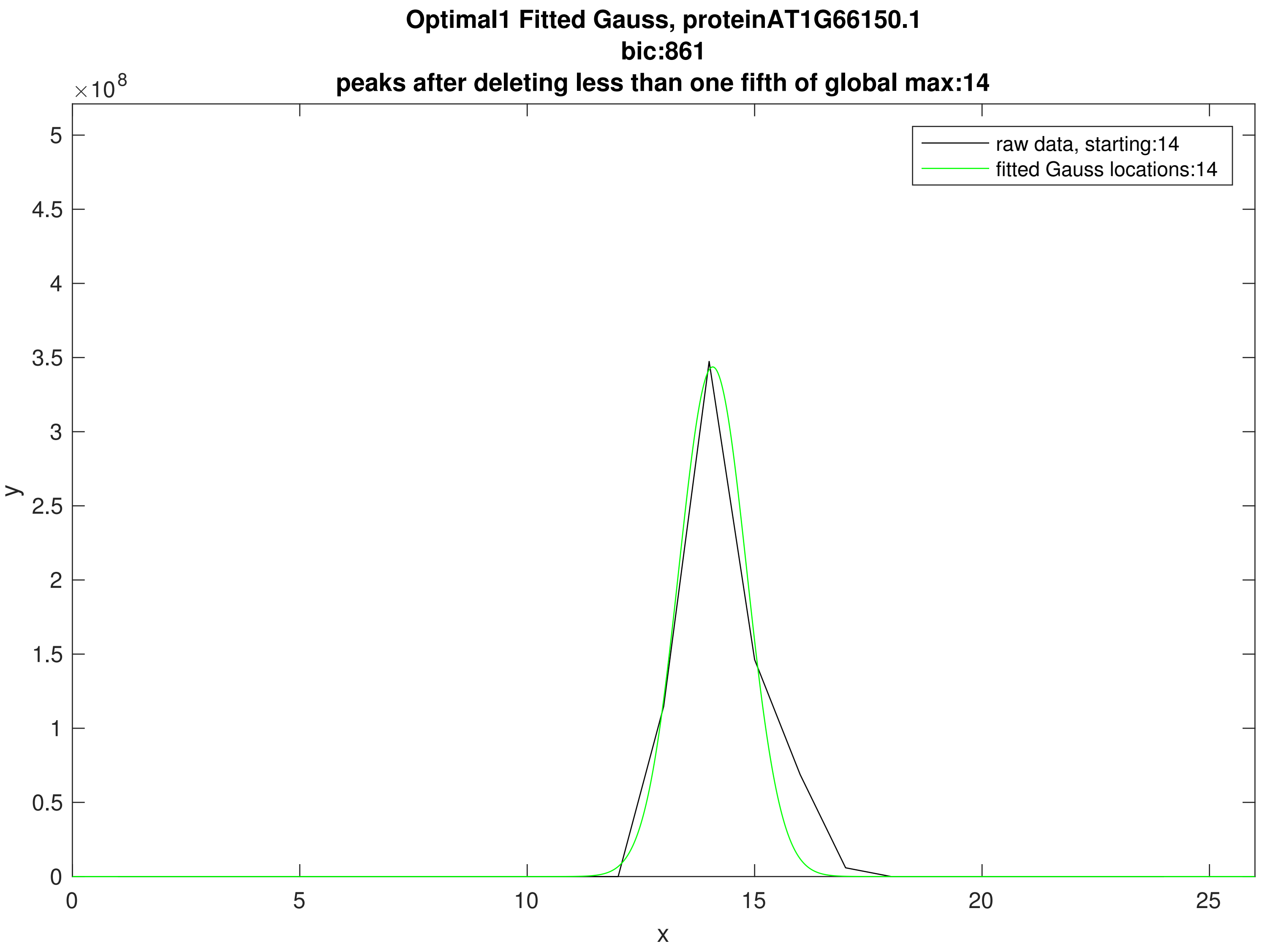
Optimal1 Fitted Gauss, proteinAT1G66150.1
bic:861
peaks after deleting less than one fifth of global max:14
×10^8
y
x
raw data, starting:14
fitted Gauss locations:14
0
0.5
1
1.5
2
2.5
3
3.5
4
4.5
5
0
5
10
15
20
25

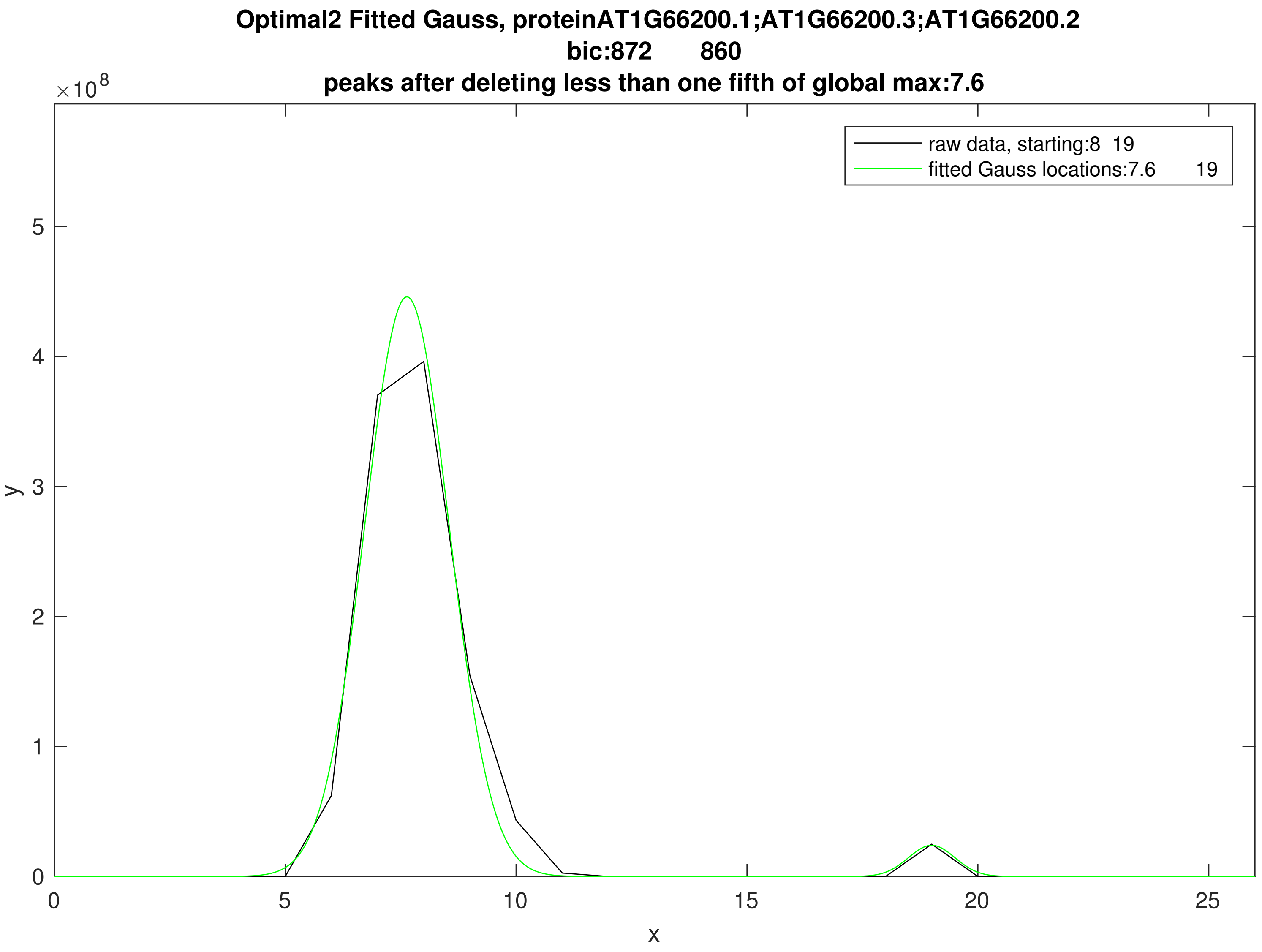
Optimal2 Fitted Gauss, proteinAT1G66200.1;AT1G66200.3;AT1G66200.2
bic:872 860
peaks after deleting less than one fifth of global max:7.6
×10^8
raw data, starting:8 19
fitted Gauss locations:7.6 19
y
x
0
1
2
3
4
5
0
5
10
15
20
25

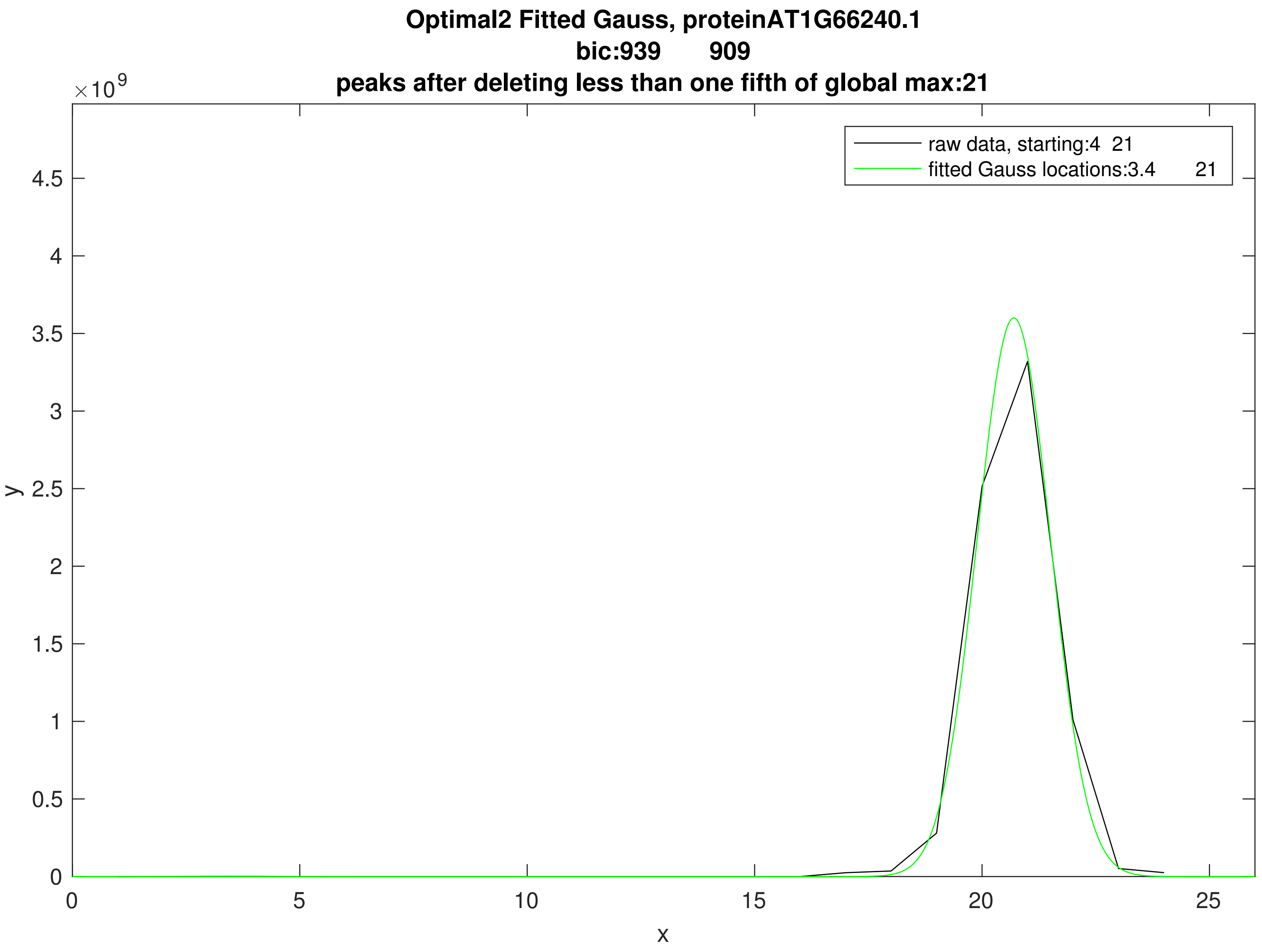

Optimal2 Fitted Gauss, proteinAT1G66240.1
bic:939 909
peaks after deleting less than one fifth of global max:21
×10^9
raw data, starting:4 21
fitted Gauss locations:3.4 21
4.5
4
3.5
3
2.5
2
1.5
1
0.5
0
y
0
5
10
15
20
25
x

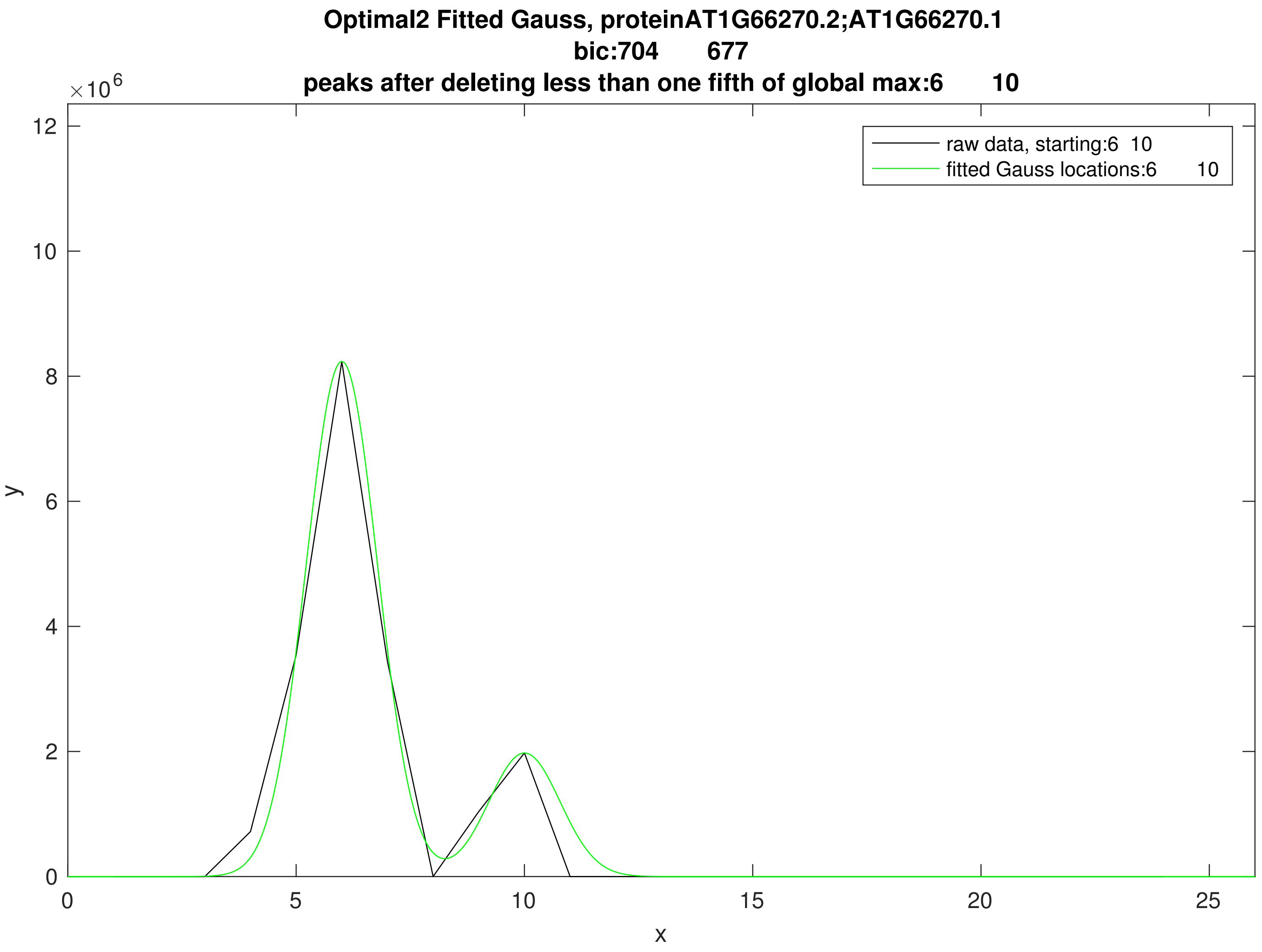

Optimal2 Fitted Gauss, proteinAT1G66270.2;AT1G66270.1
bic:704 677
peaks after deleting less than one fifth of global max:6 10
×10^6
raw data, starting:6 10
fitted Gauss locations:6 10
y
x
0
2
4
6
8
10
12
0
5
10
15
20
25

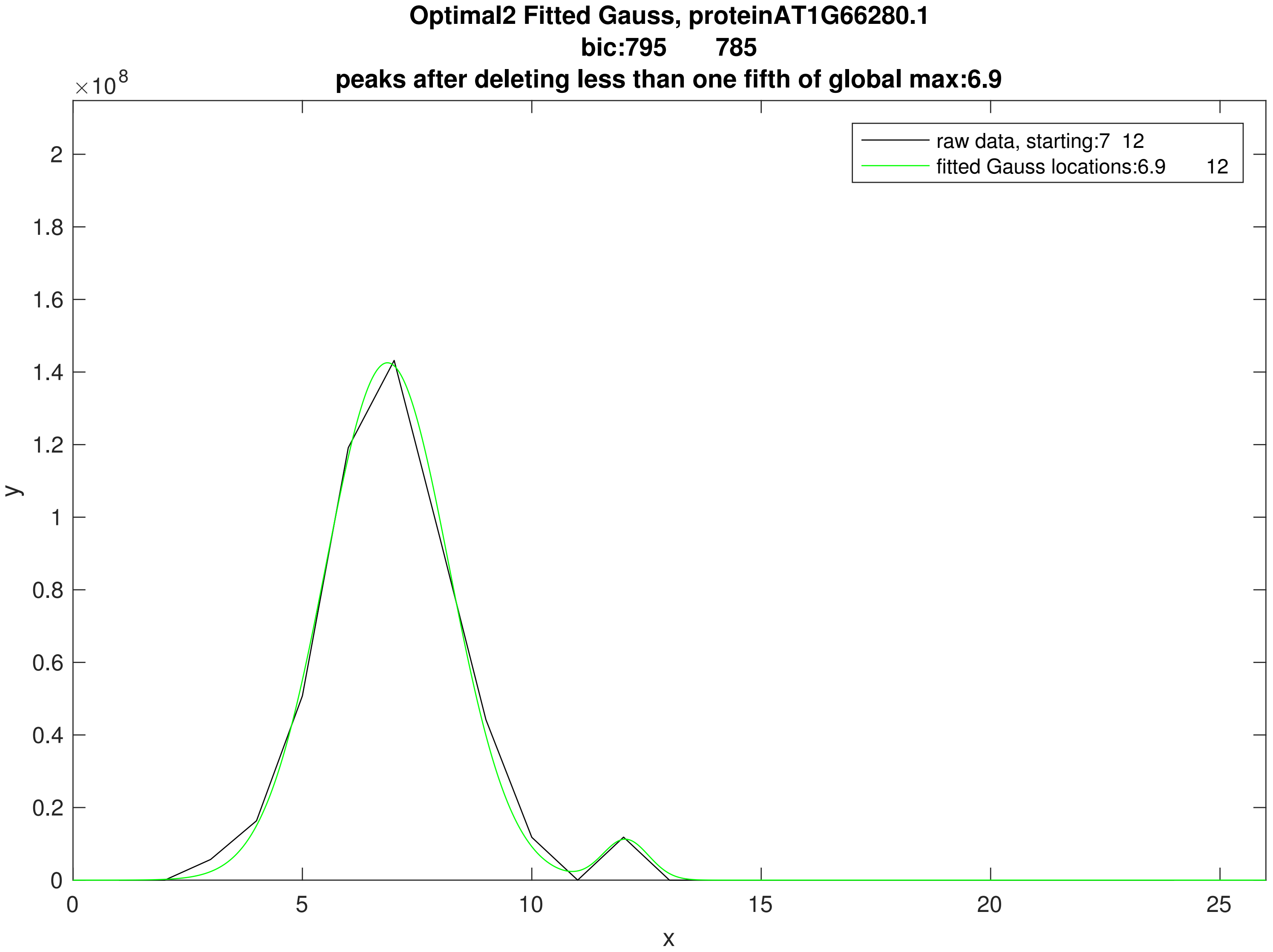

Optimal2 Fitted Gauss, proteinAT1G66280.1
bic:795 785
peaks after deleting less than one fifth of global max:6.9
$\times 10^8$
raw data, starting:7 12
fitted Gauss locations:6.9 12
2
1.8
1.6
1.4
1.2
1
0.8
0.6
0.4
0.2
0
y
0
5
10
15
20
25
x

Optimal1 Fitted Gauss, proteinAT1G66400.1
bic:672
peaks after deleting less than one fifth of global max:19

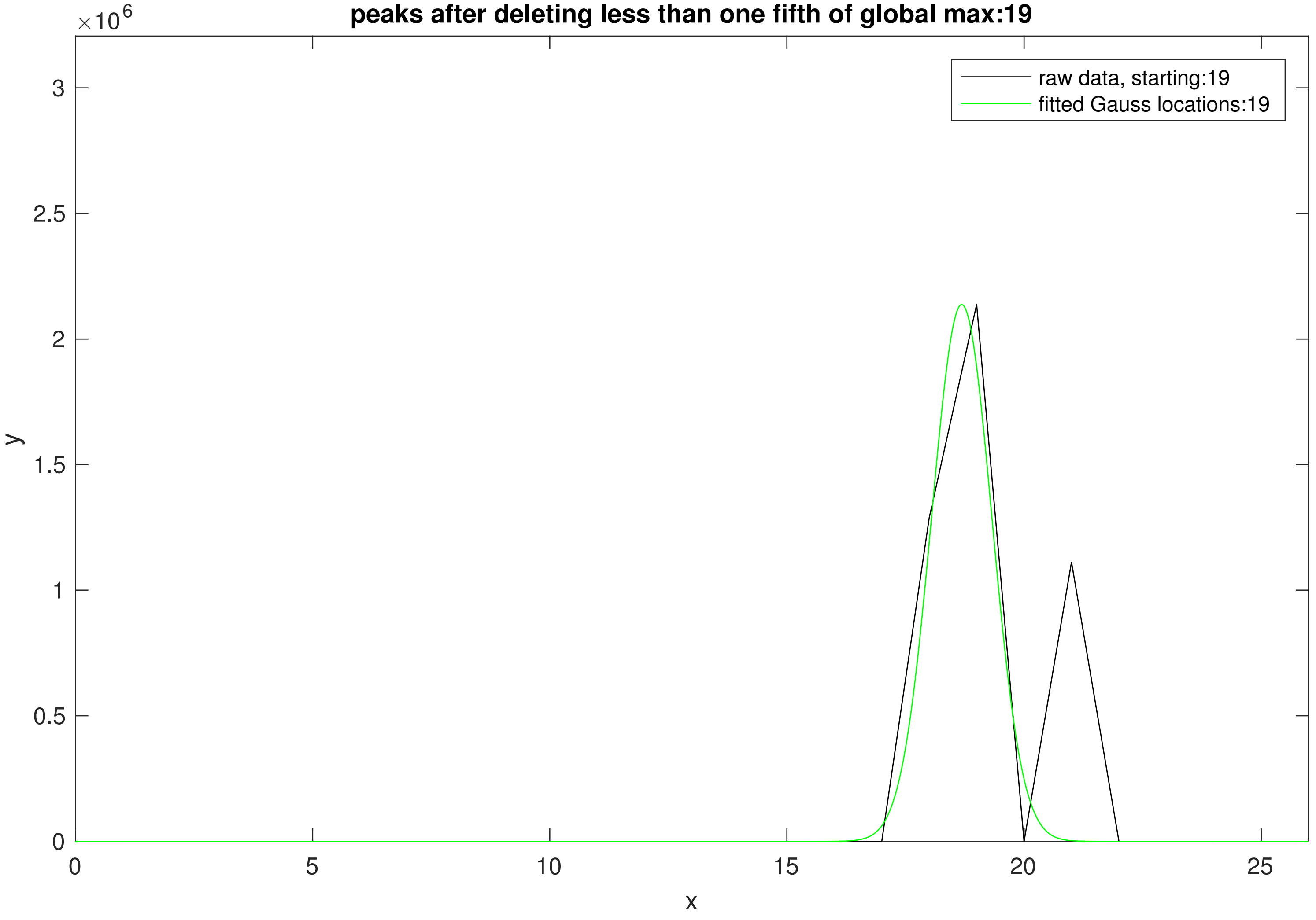

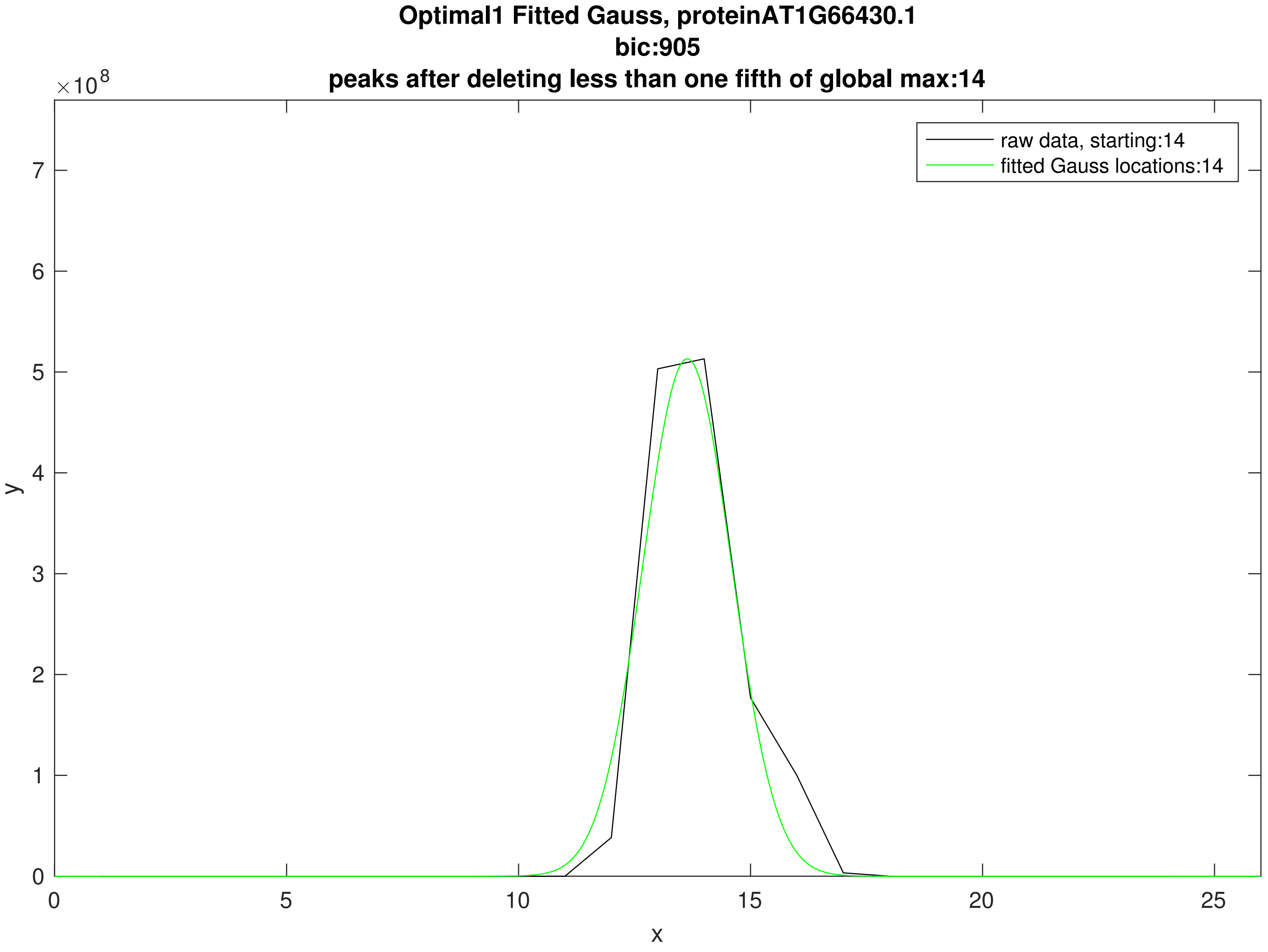

Optimal1 Fitted Gauss, proteinAT1G66430.1
bic:905
peaks after deleting less than one fifth of global max:14
×10^8
raw data, starting:14
fitted Gauss locations:14
y
x
0
1
2
3
4
5
6
7
0
5
10
15
20
25

**Optimal1 Fitted Gauss, proteinAT1G66580.1**
**bic:740**
**peaks after deleting less than one fifth of global max:3.3**

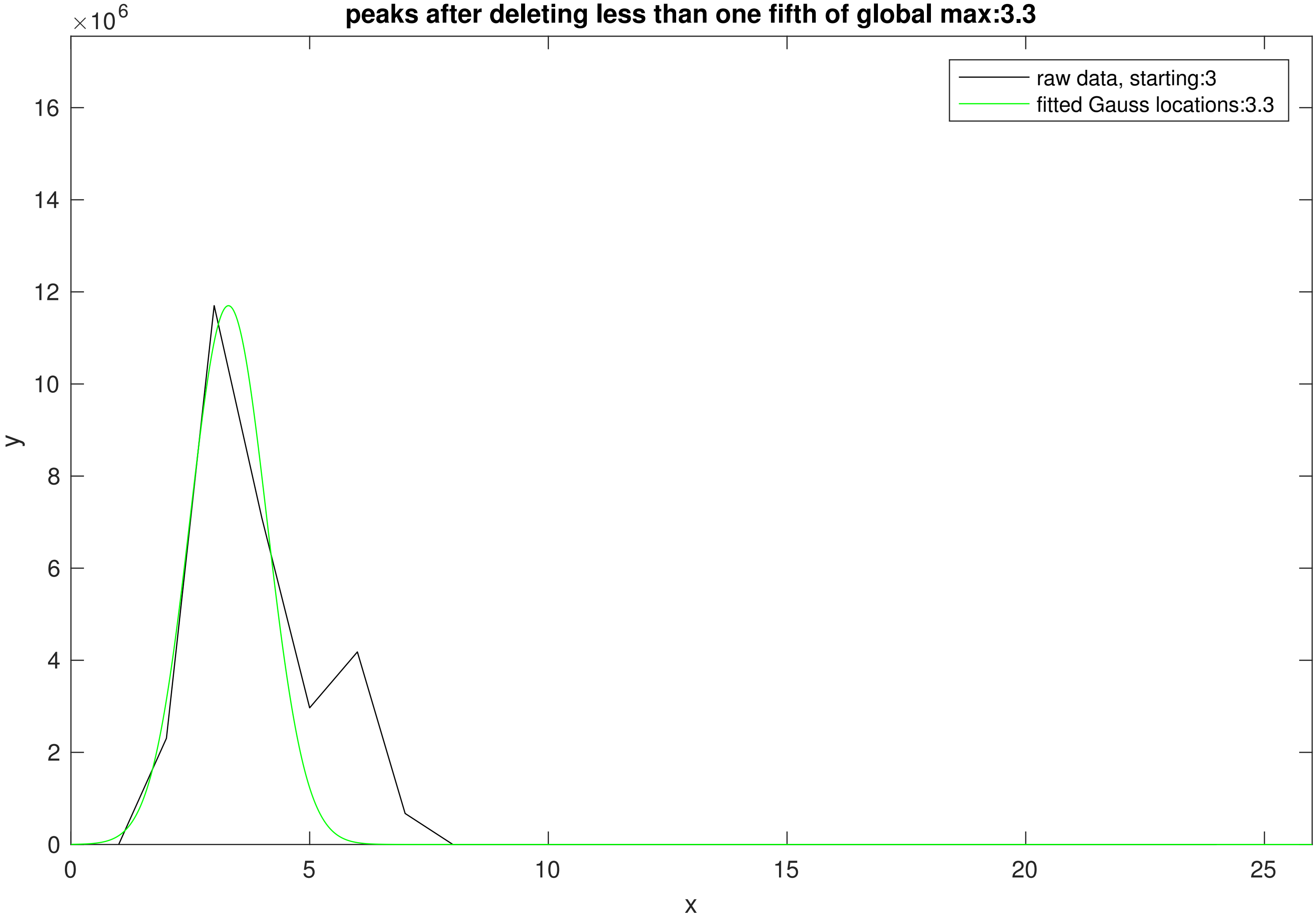

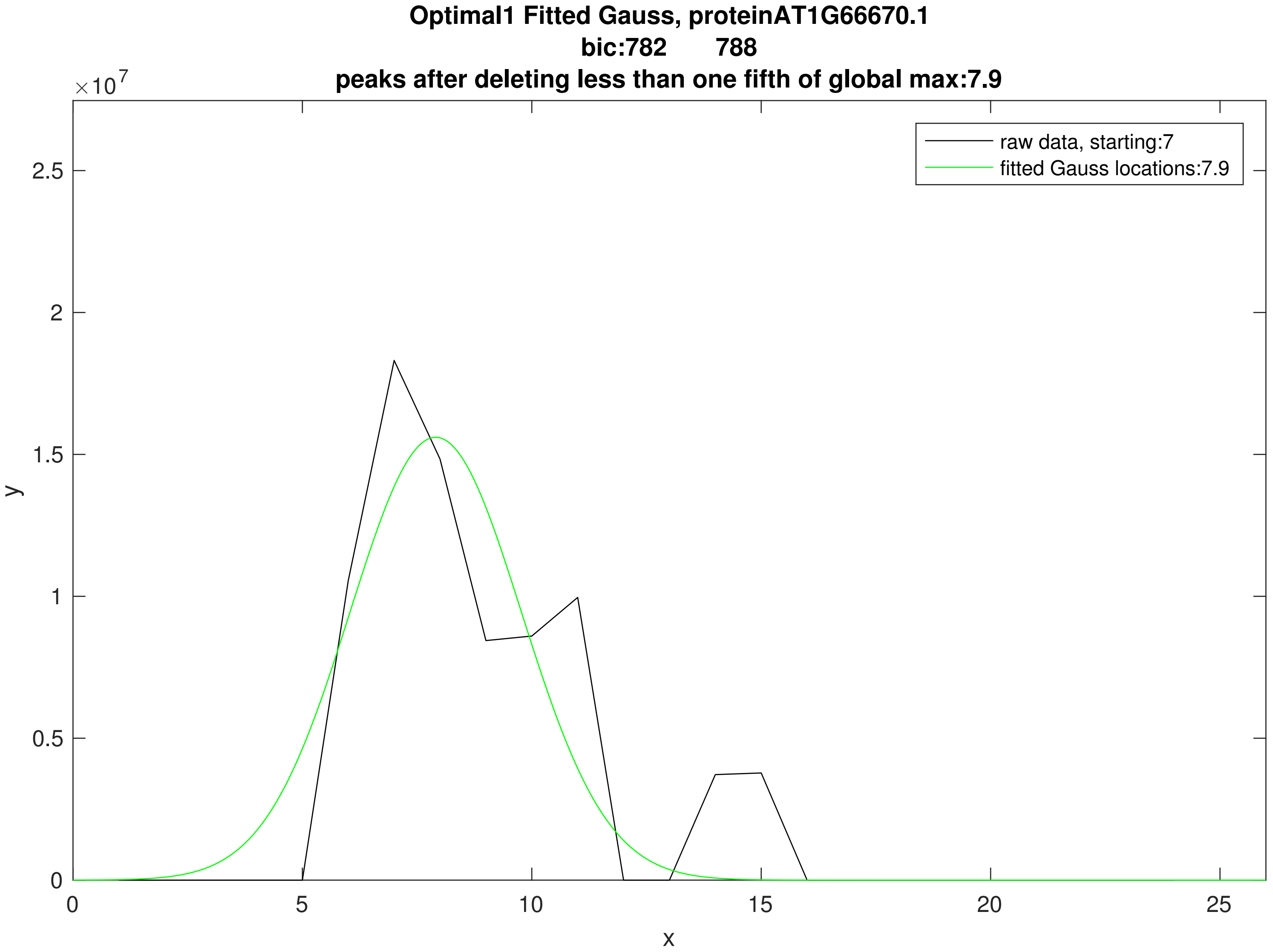
Optimal1 Fitted Gauss, proteinAT1G66670.1
bic:782 788
peaks after deleting less than one fifth of global max:7.9
×10^7
raw data, starting:7
fitted Gauss locations:7.9
y
x
0
0.5
1
1.5
2
2.5
5
10
15
20
25

**Optimal1 Fitted Gauss, proteinAT1G66680.1**
**bic:750**
**peaks after deleting less than one fifth of global max:17**

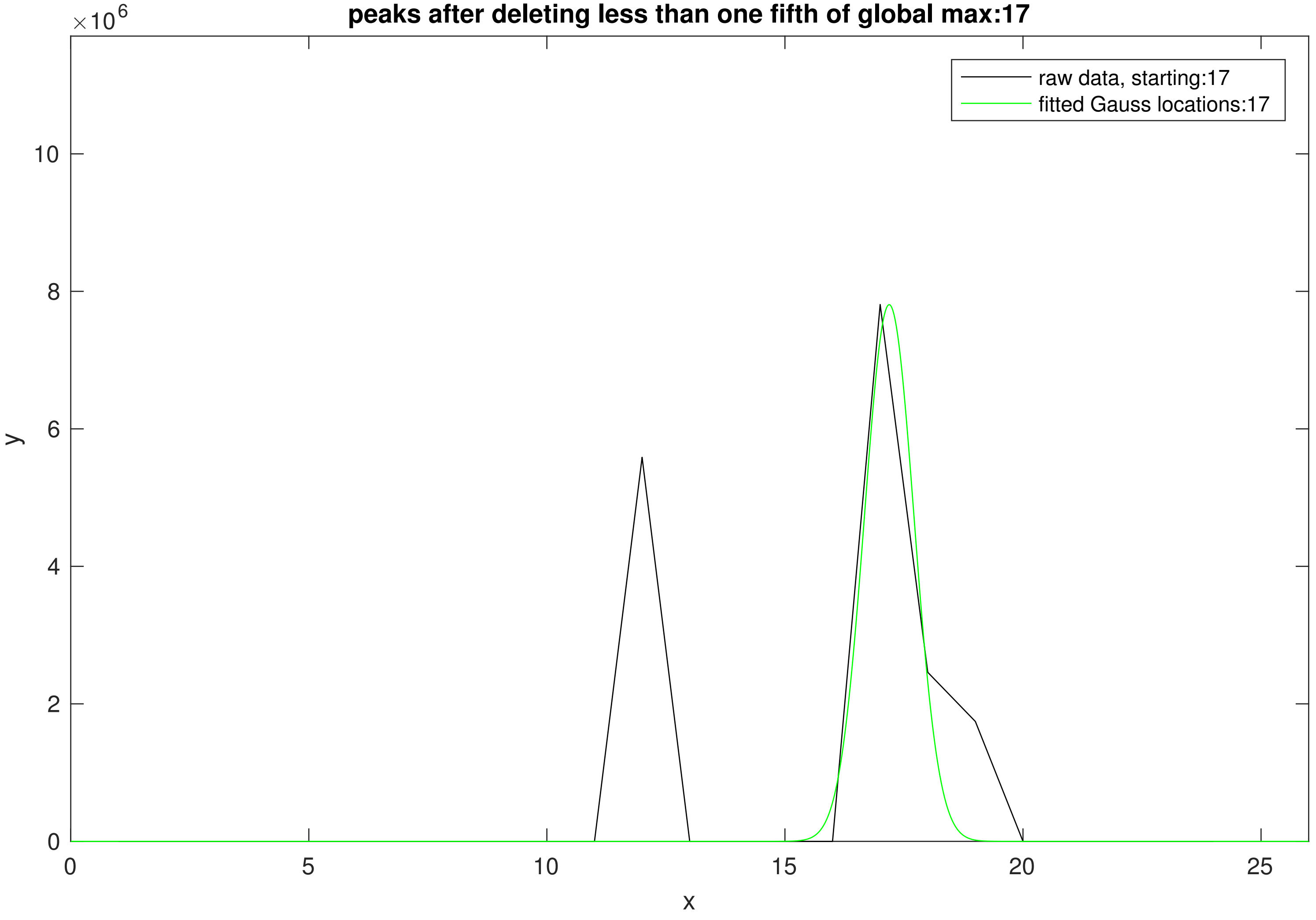

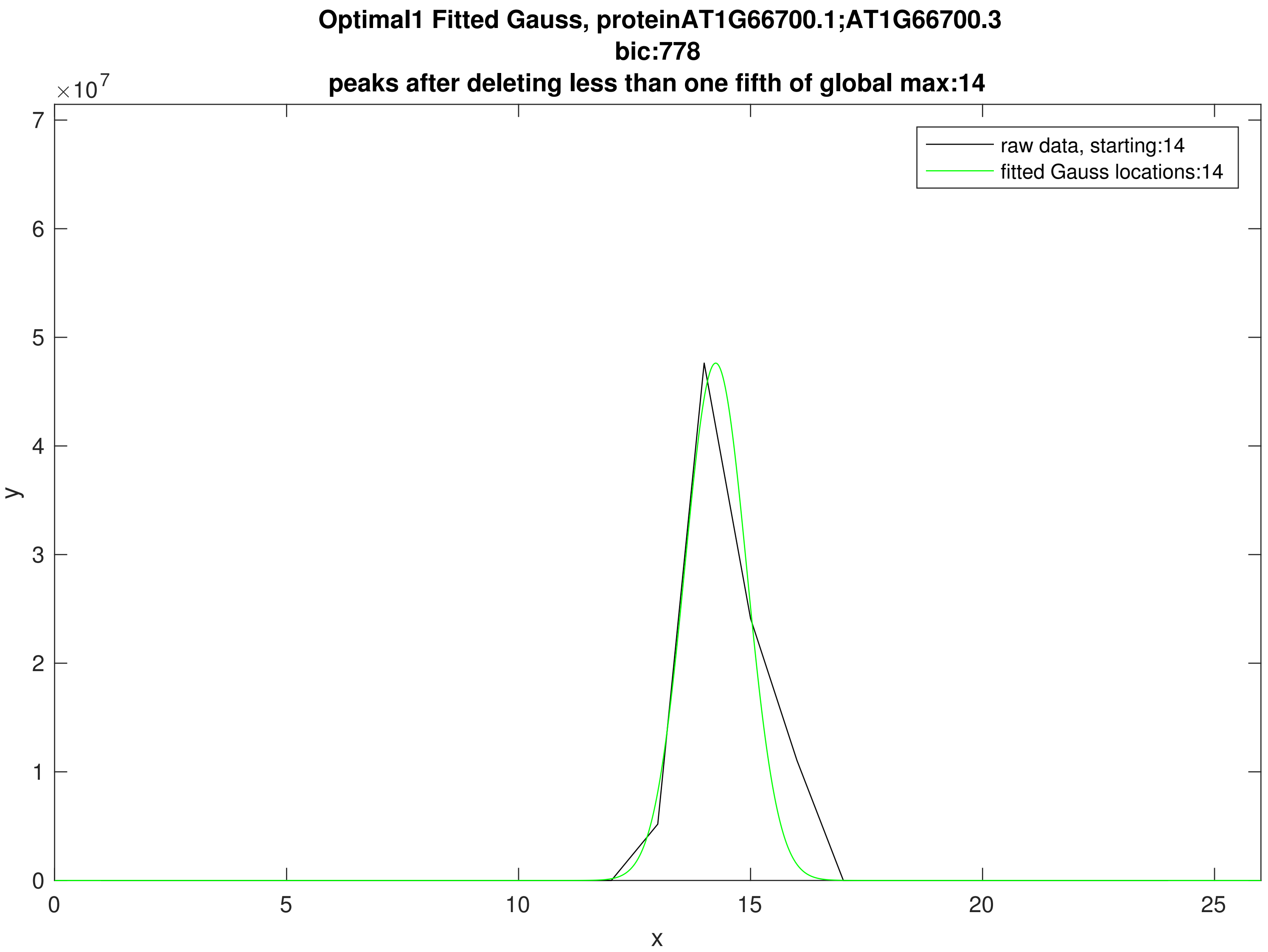
Optimal1 Fitted Gauss, proteinAT1G66700.1;AT1G66700.3
bic:778
peaks after deleting less than one fifth of global max:14
×10^7
raw data, starting:14
fitted Gauss locations:14
y
x
0
1
2
3
4
5
6
7
0
5
10
15
20
25

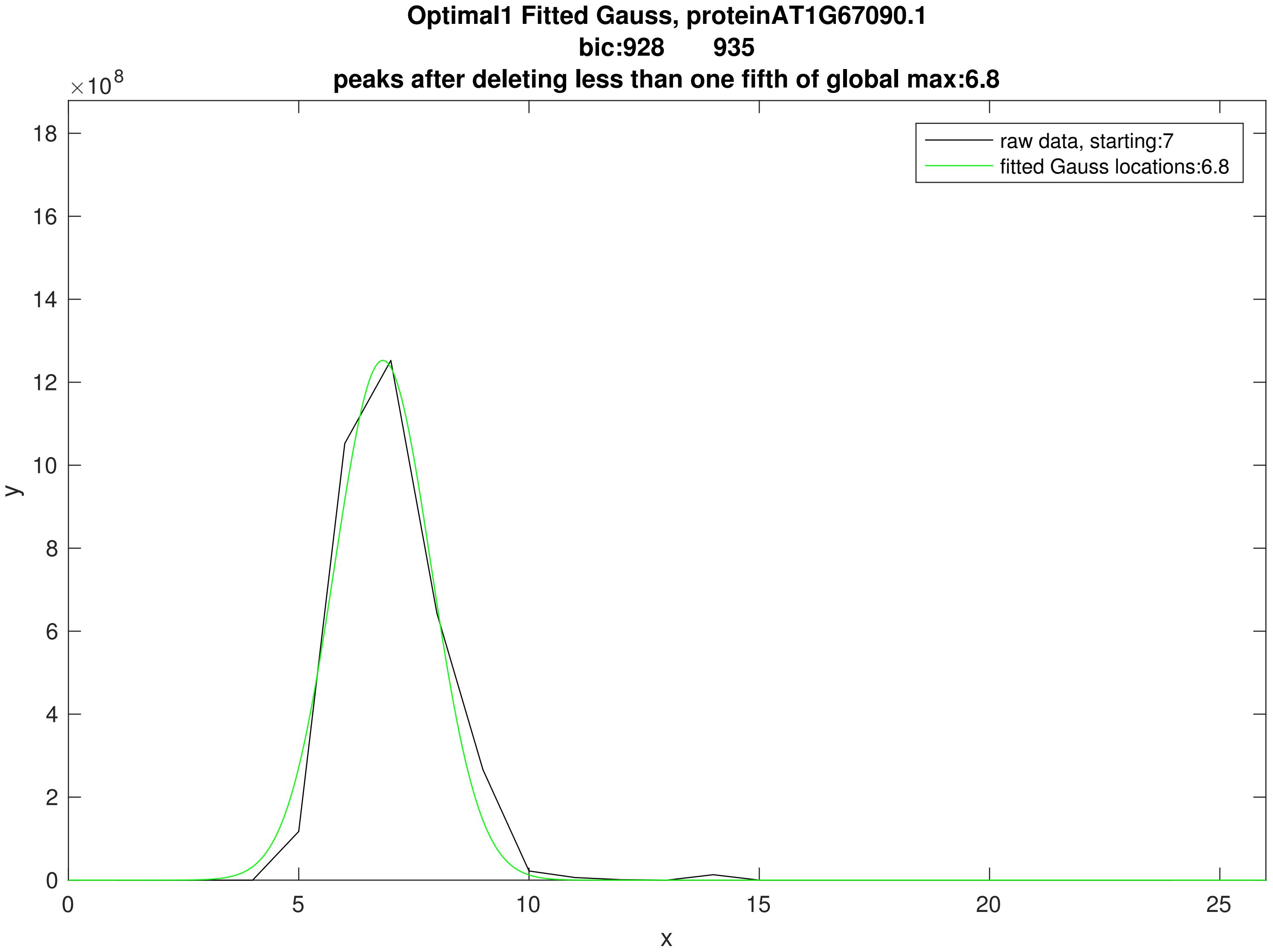
Optimal1 Fitted Gauss, proteinAT1G67090.1
bic:928 935
peaks after deleting less than one fifth of global max:6.8
×10^8
raw data, starting:7
fitted Gauss locations:6.8
y
x
0
2
4
6
8
10
12
14
16
18
0
5
10
15
20
25

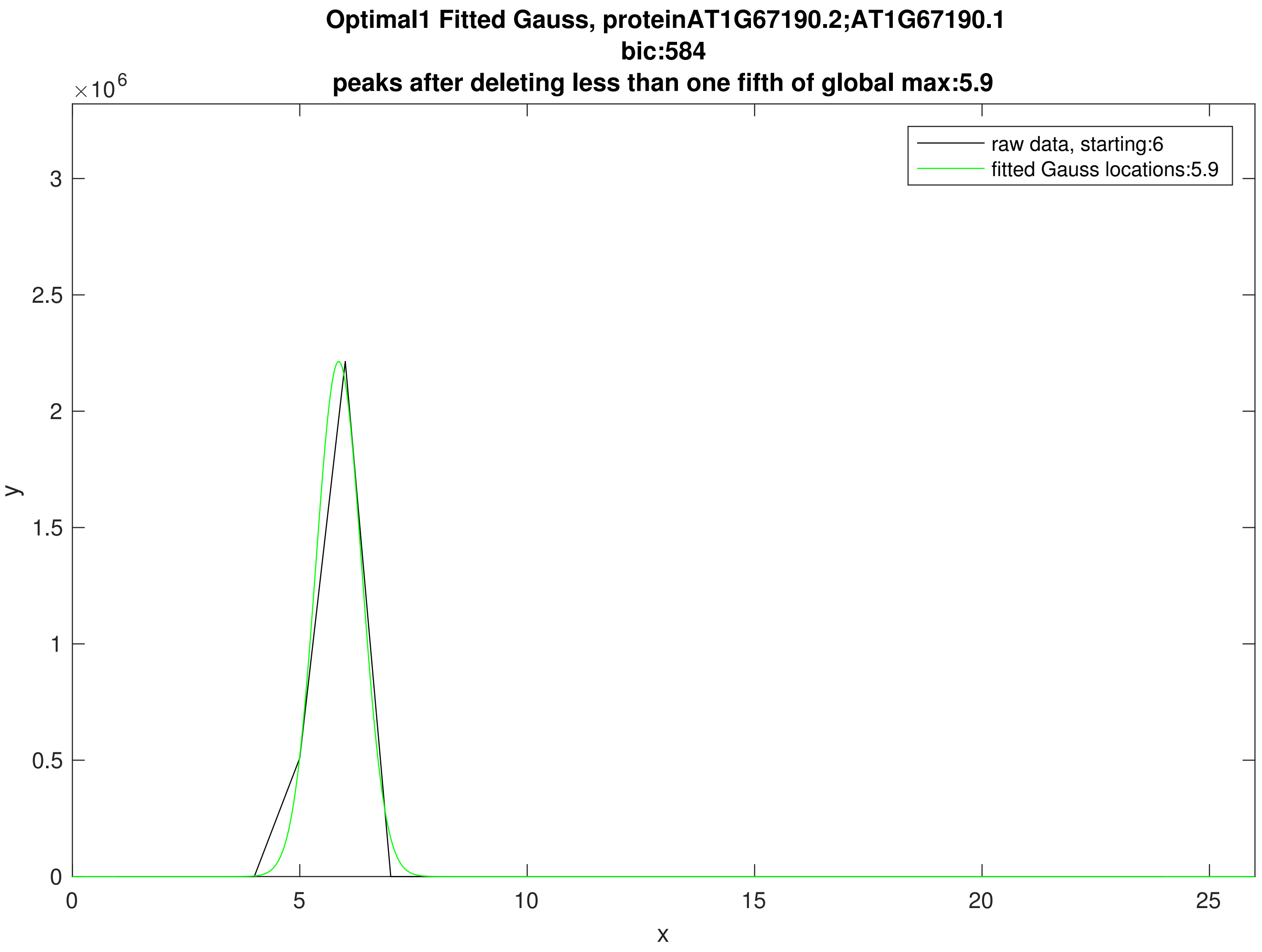
Optimal1 Fitted Gauss, proteinAT1G67190.2;AT1G67190.1
bic:584
peaks after deleting less than one fifth of global max:5.9
×10^6
raw data, starting:6
fitted Gauss locations:5.9
y
x
0
0.5
1
1.5
2
2.5
3
0
5
10
15
20
25

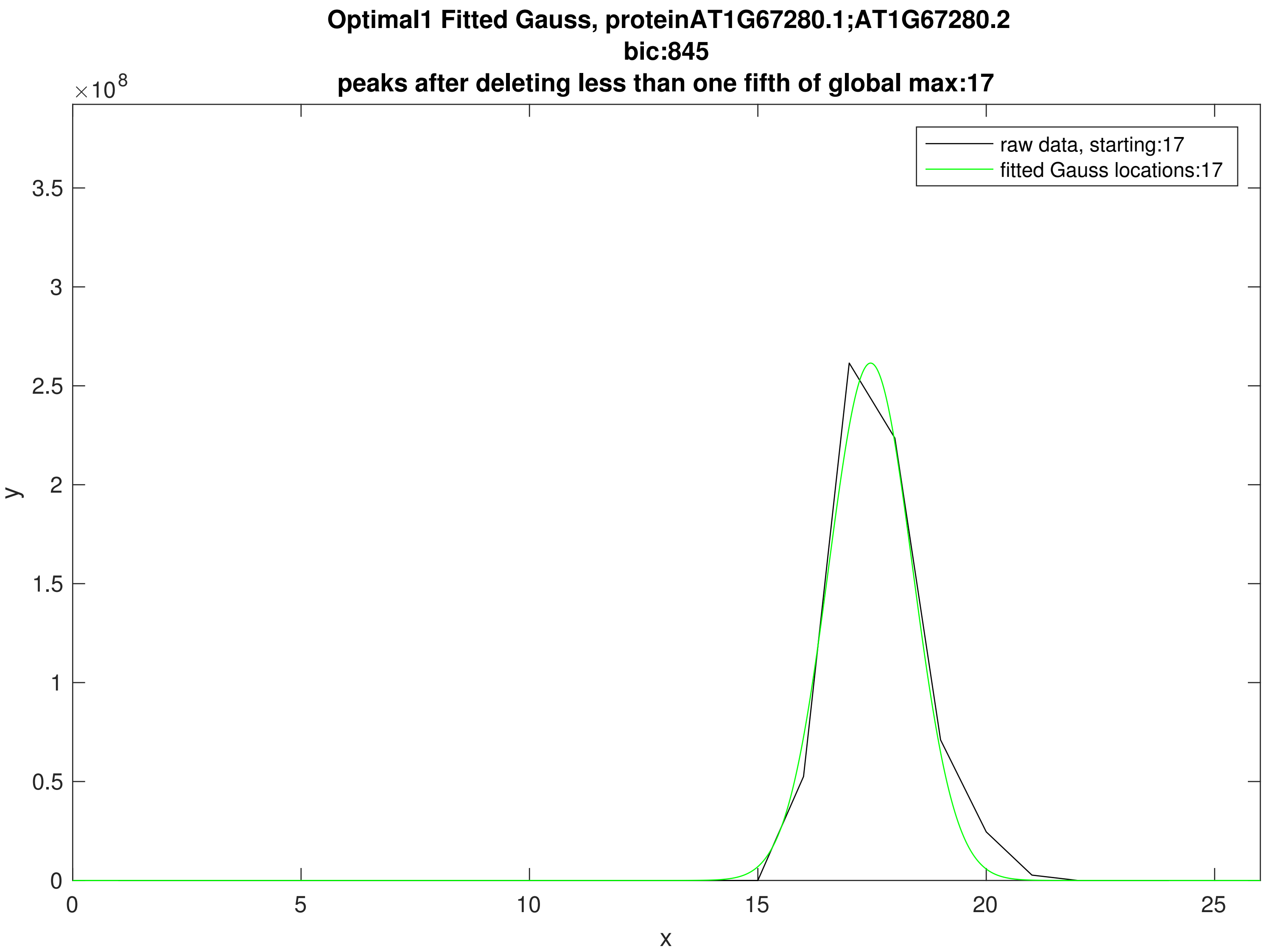
Optimal1 Fitted Gauss, proteinAT1G67280.1;AT1G67280.2
bic:845
peaks after deleting less than one fifth of global max:17
×10^8
3.5
3
2.5
2
1.5
1
0.5
0
y
0
5
10
15
20
25
x
raw data, starting:17
fitted Gauss locations:17

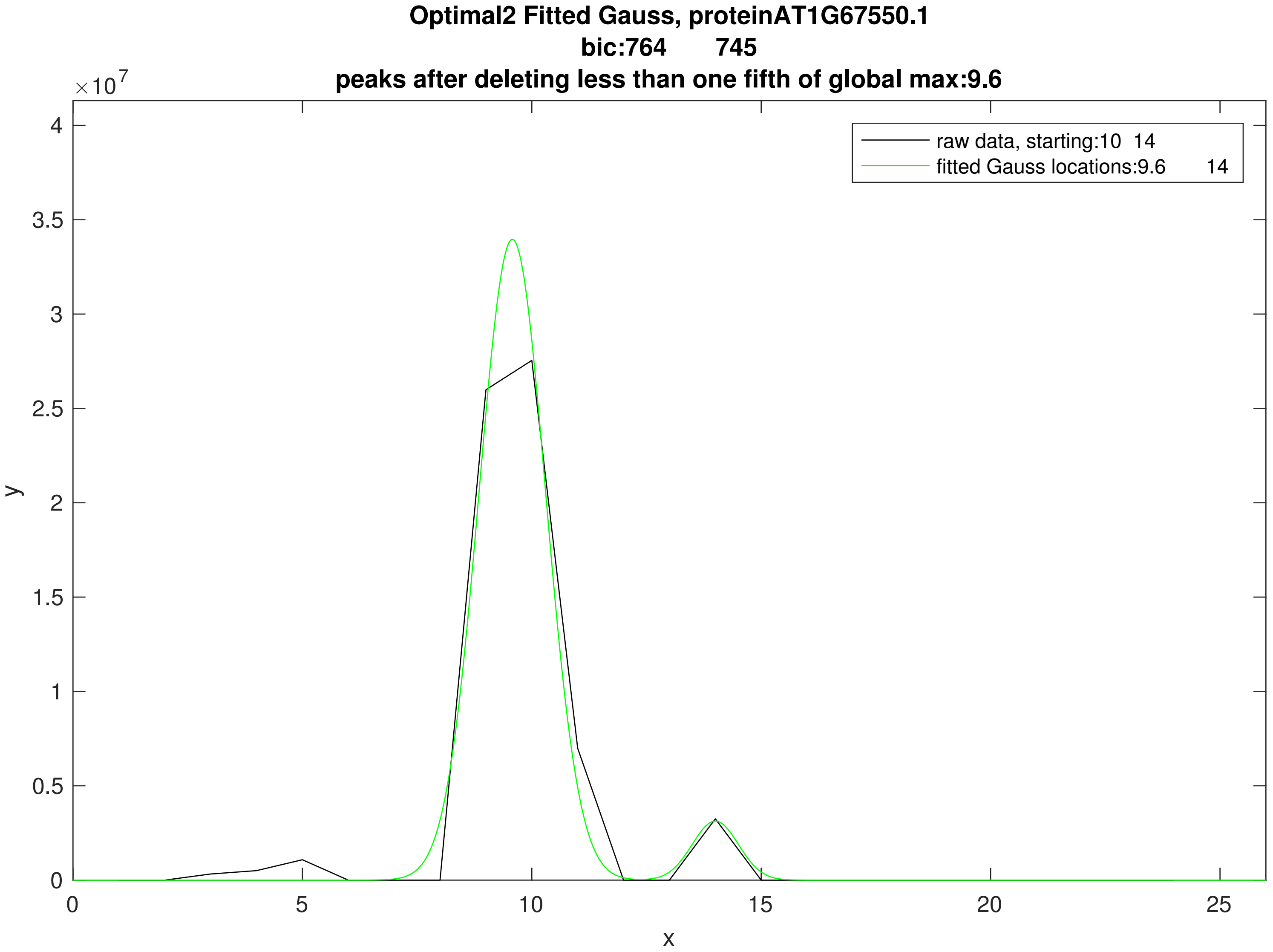
Optimal2 Fitted Gauss, proteinAT1G67550.1
bic:764 745
peaks after deleting less than one fifth of global max:9.6
×10^7
raw data, starting:10 14
fitted Gauss locations:9.6 14
y
x
0
0.5
1
1.5
2
2.5
3
3.5
4
0
5
10
15
20
25

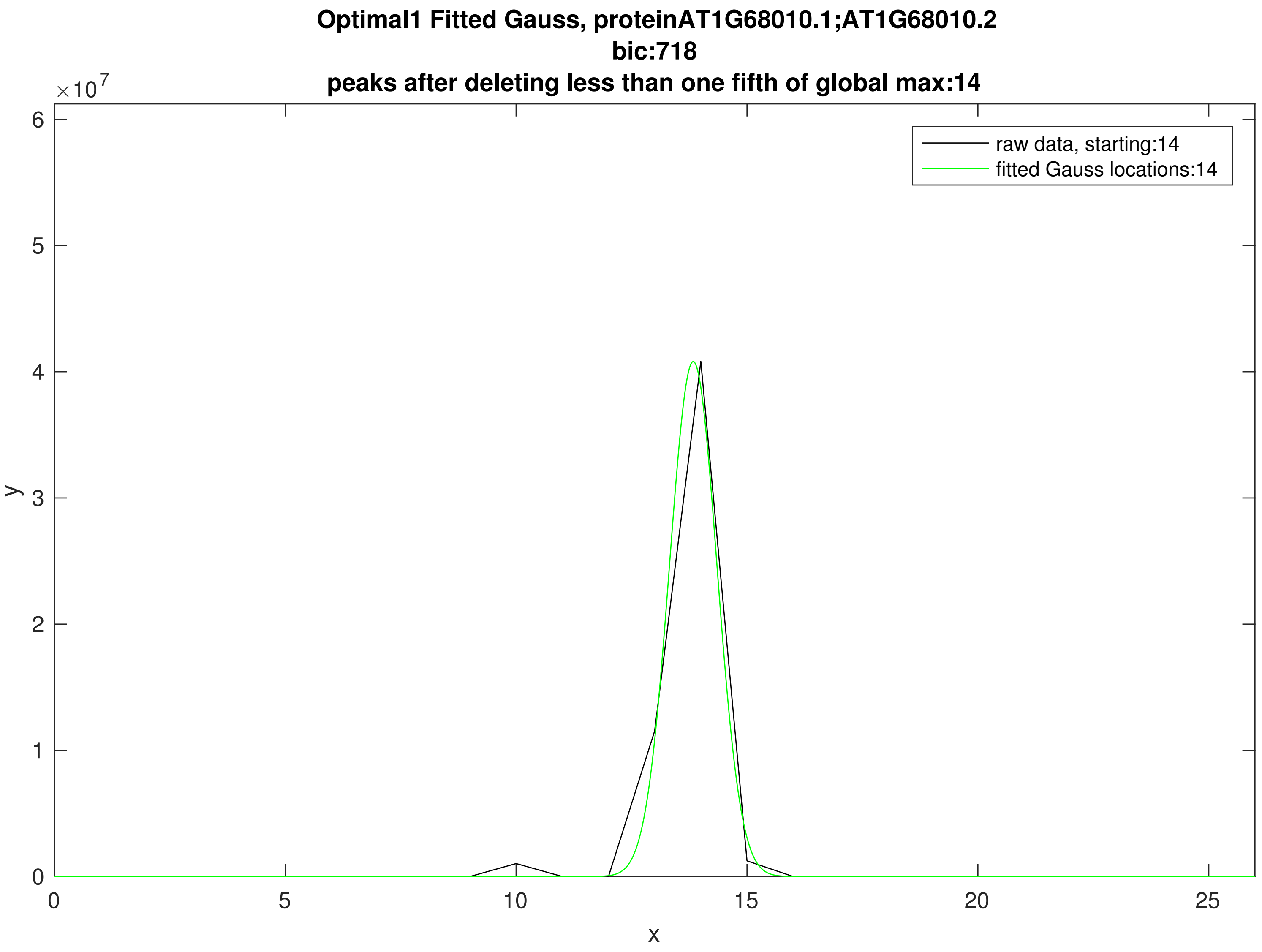

Optimal1 Fitted Gauss, proteinAT1G68010.1;AT1G68010.2
bic:718
peaks after deleting less than one fifth of global max:14
×10^7
6
5
4
3
2
1
0
y
0
5
10
15
20
25
x
raw data, starting:14
fitted Gauss locations:14

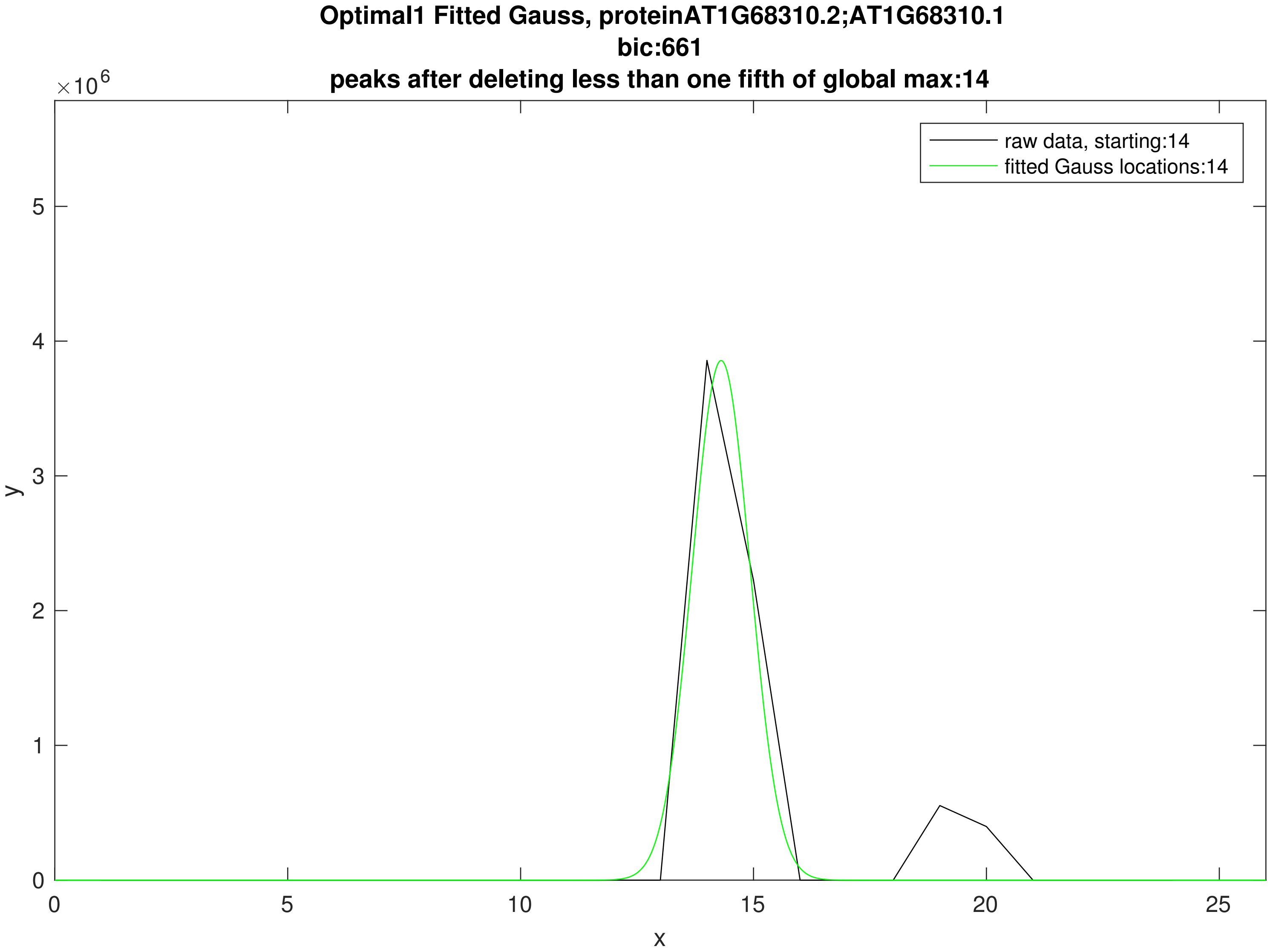
Optimal1 Fitted Gauss, proteinAT1G68310.2;AT1G68310.1
bic:661
peaks after deleting less than one fifth of global max:14
×10^6
raw data, starting:14
fitted Gauss locations:14
y
x
0
1
2
3
4
5
0
5
10
15
20
25

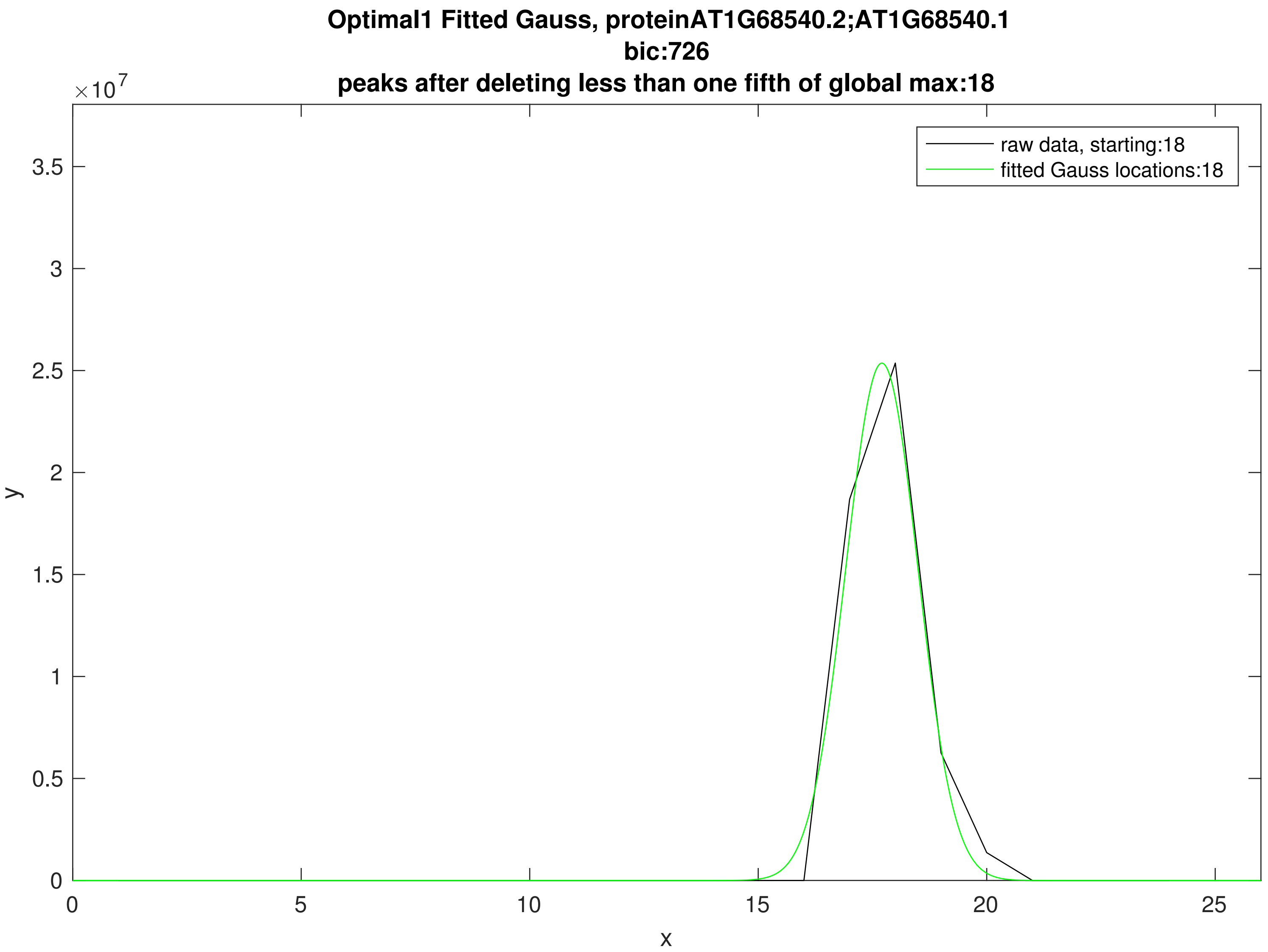
Optimal1 Fitted Gauss, proteinAT1G68540.2;AT1G68540.1
bic:726
peaks after deleting less than one fifth of global max:18
×10^7
3.5
3
2.5
2
1.5
1
0.5
0
y
0
5
10
15
20
25
x
raw data, starting:18
fitted Gauss locations:18

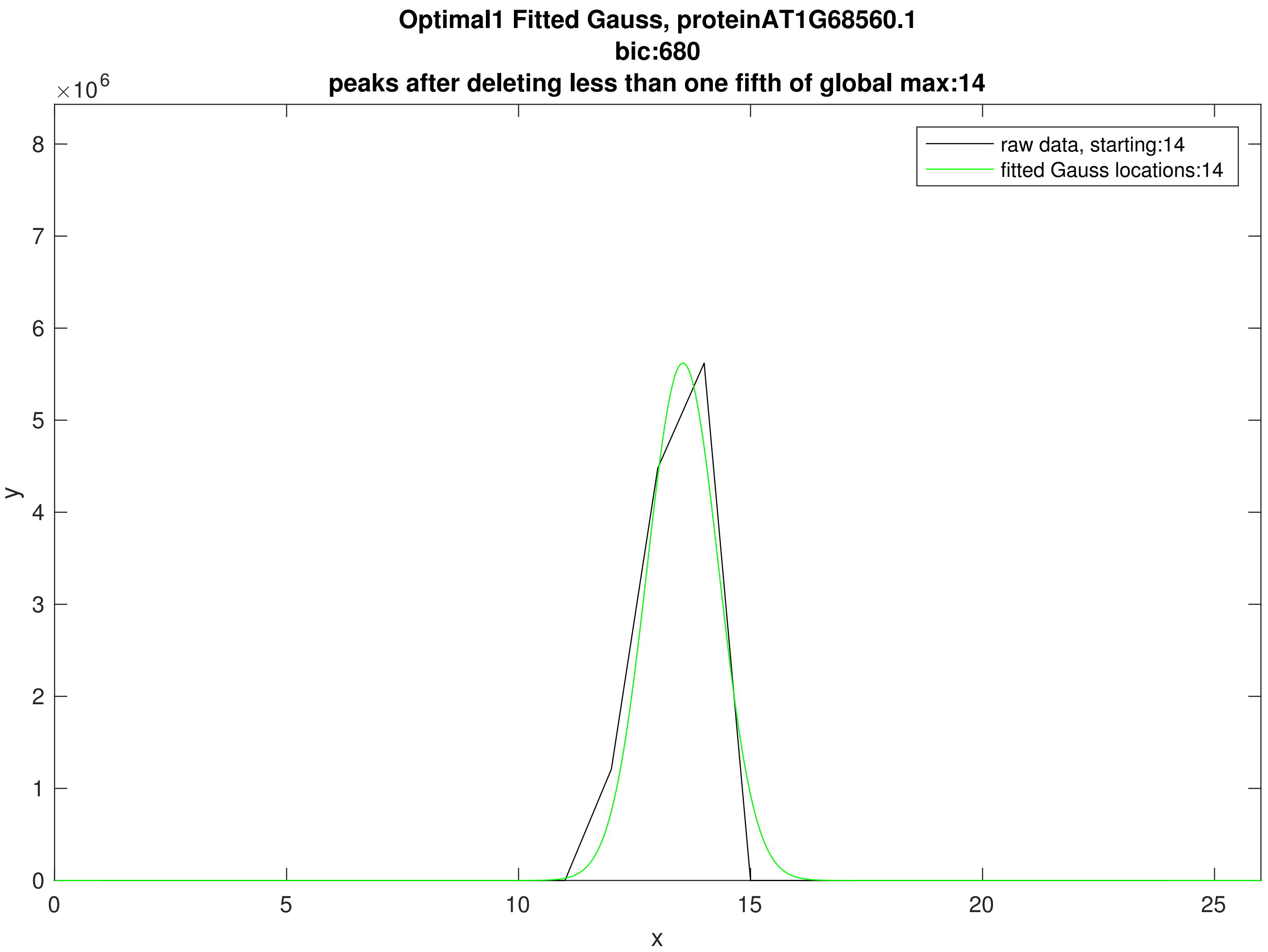
Optimal1 Fitted Gauss, proteinAT1G68560.1
bic:680
peaks after deleting less than one fifth of global max:14
×10^6
raw data, starting:14
fitted Gauss locations:14
y
x
0
1
2
3
4
5
6
7
8
0
5
10
15
20
25

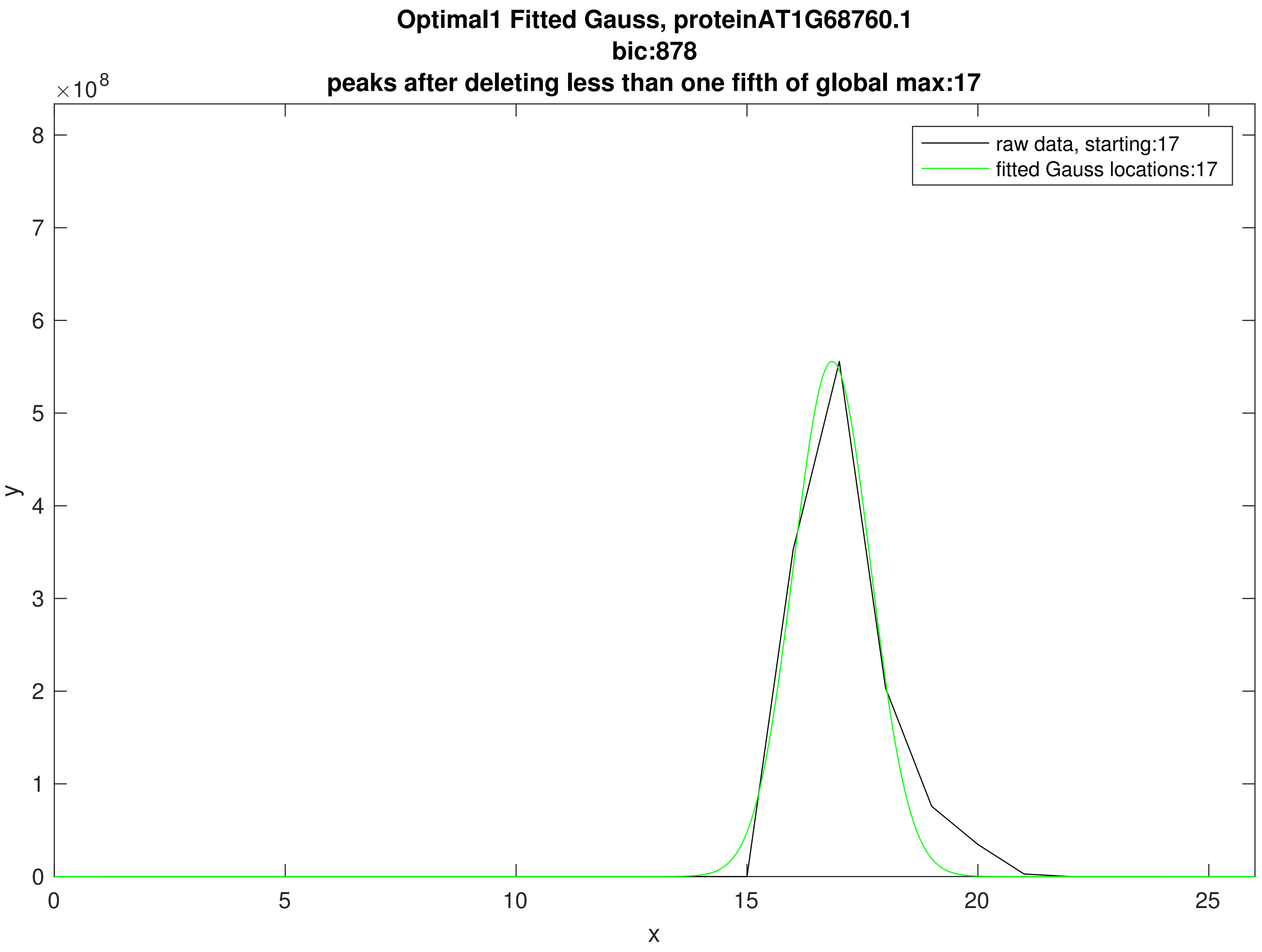

Optimal1 Fitted Gauss, proteinAT1G68760.1
bic:878
peaks after deleting less than one fifth of global max:17
×10^8
raw data, starting:17
fitted Gauss locations:17
y
x
0
1
2
3
4
5
6
7
8
0
5
10
15
20
25

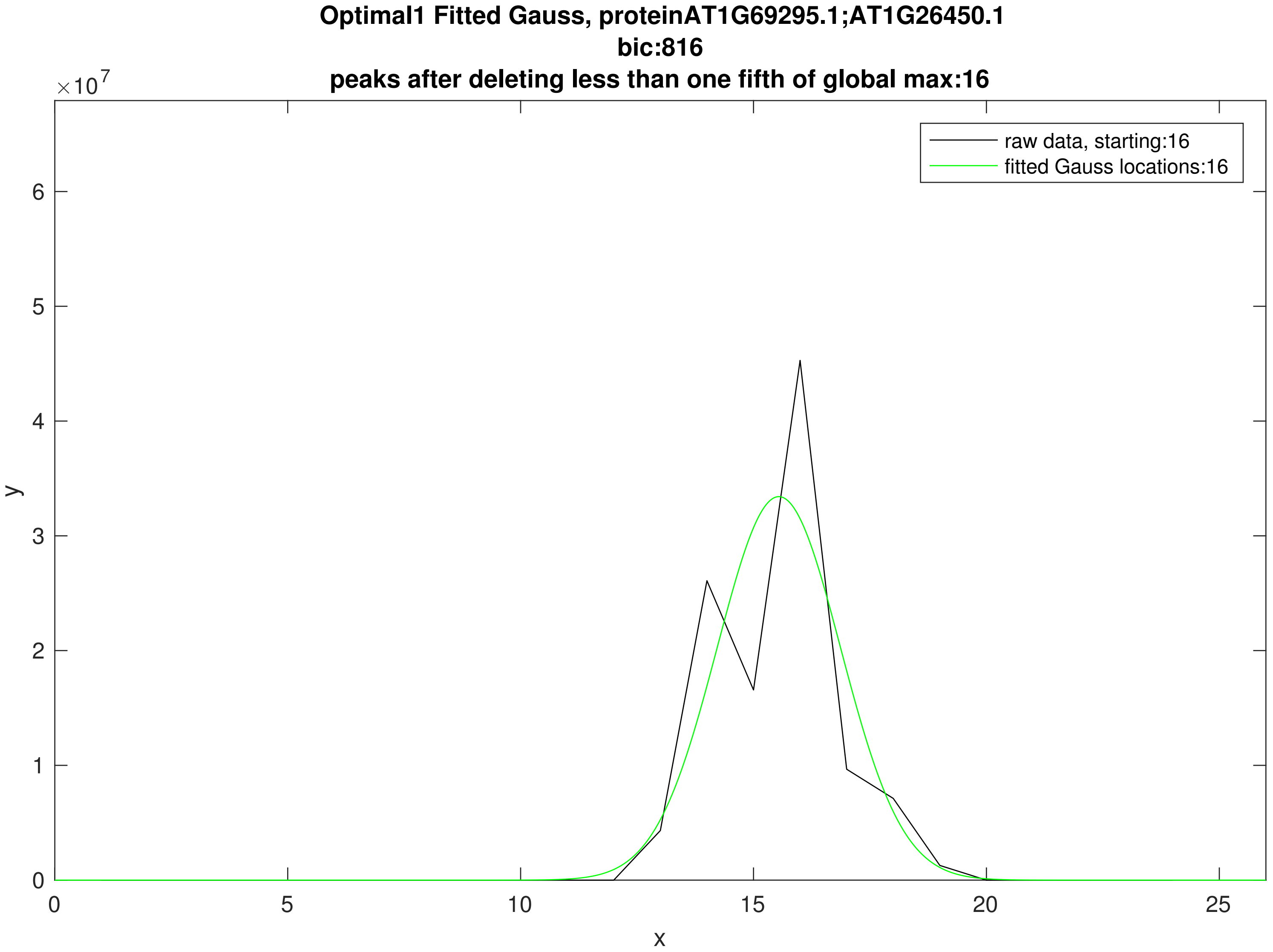
Optimal1 Fitted Gauss, proteinAT1G69295.1;AT1G26450.1
bic:816
peaks after deleting less than one fifth of global max:16
×10^7
raw data, starting:16
fitted Gauss locations:16
y
x
0
1
2
3
4
5
6
0
5
10
15
20
25

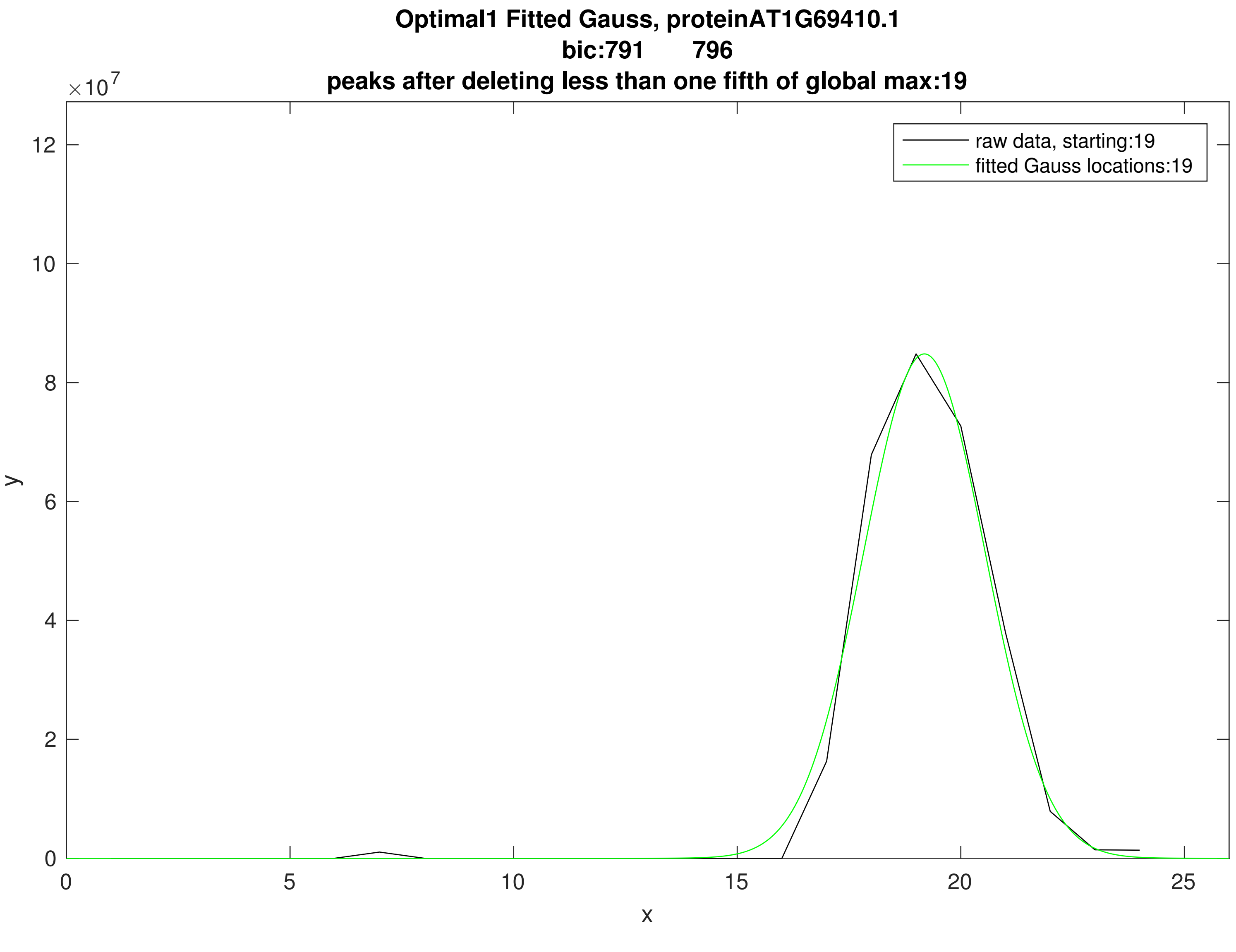

Optimal1 Fitted Gauss, proteinAT1G69410.1
bic:791 796
peaks after deleting less than one fifth of global max:19
×10^7
12
10
8
6
4
2
0
y
0
5
10
15
20
25
x
raw data, starting:19
fitted Gauss locations:19

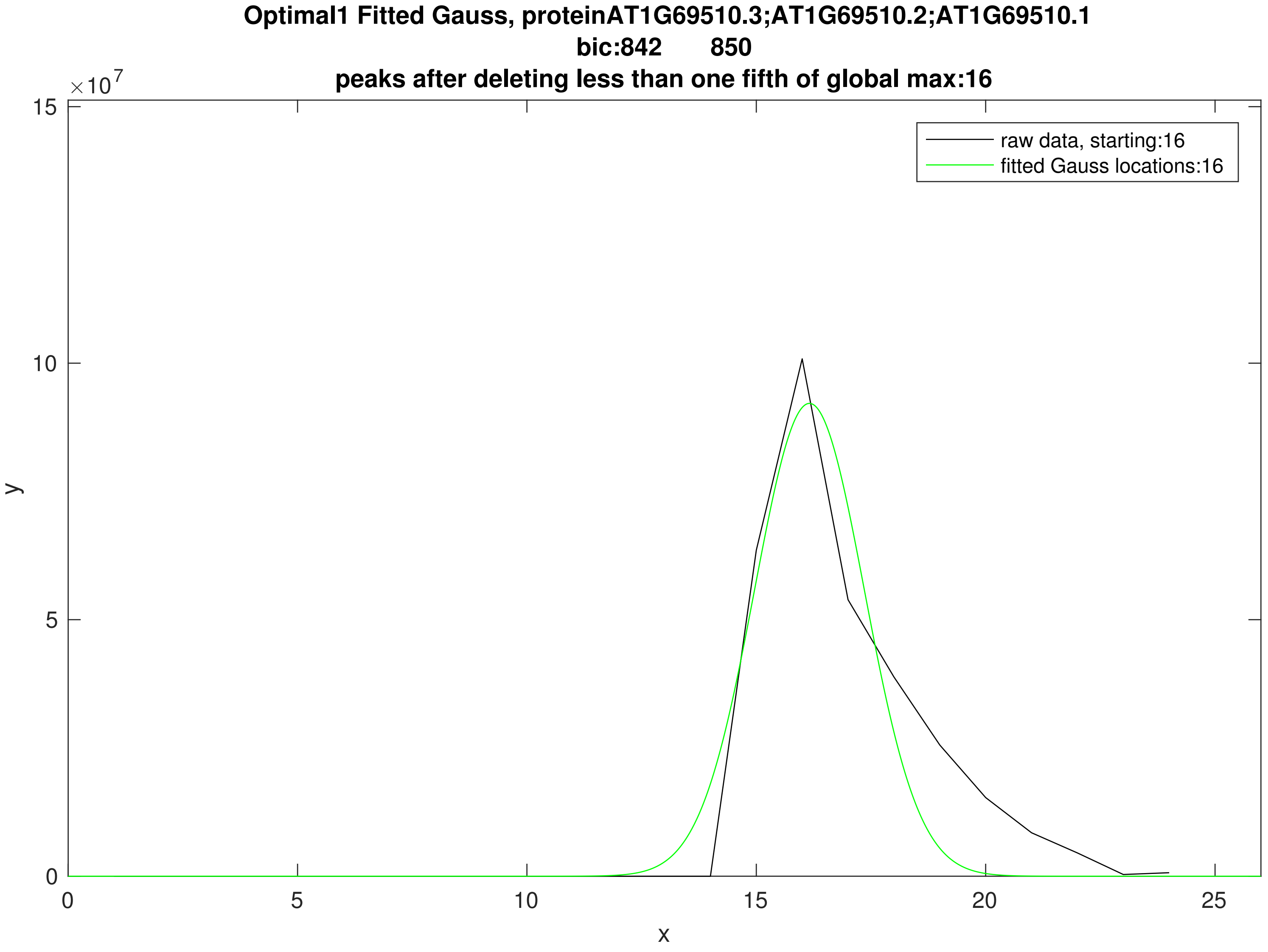
Optimal1 Fitted Gauss, proteinAT1G69510.3;AT1G69510.2;AT1G69510.1
bic:842 850
peaks after deleting less than one fifth of global max:16
×10^7
raw data, starting:16
fitted Gauss locations:16
y
x
0
5
10
15
20
25

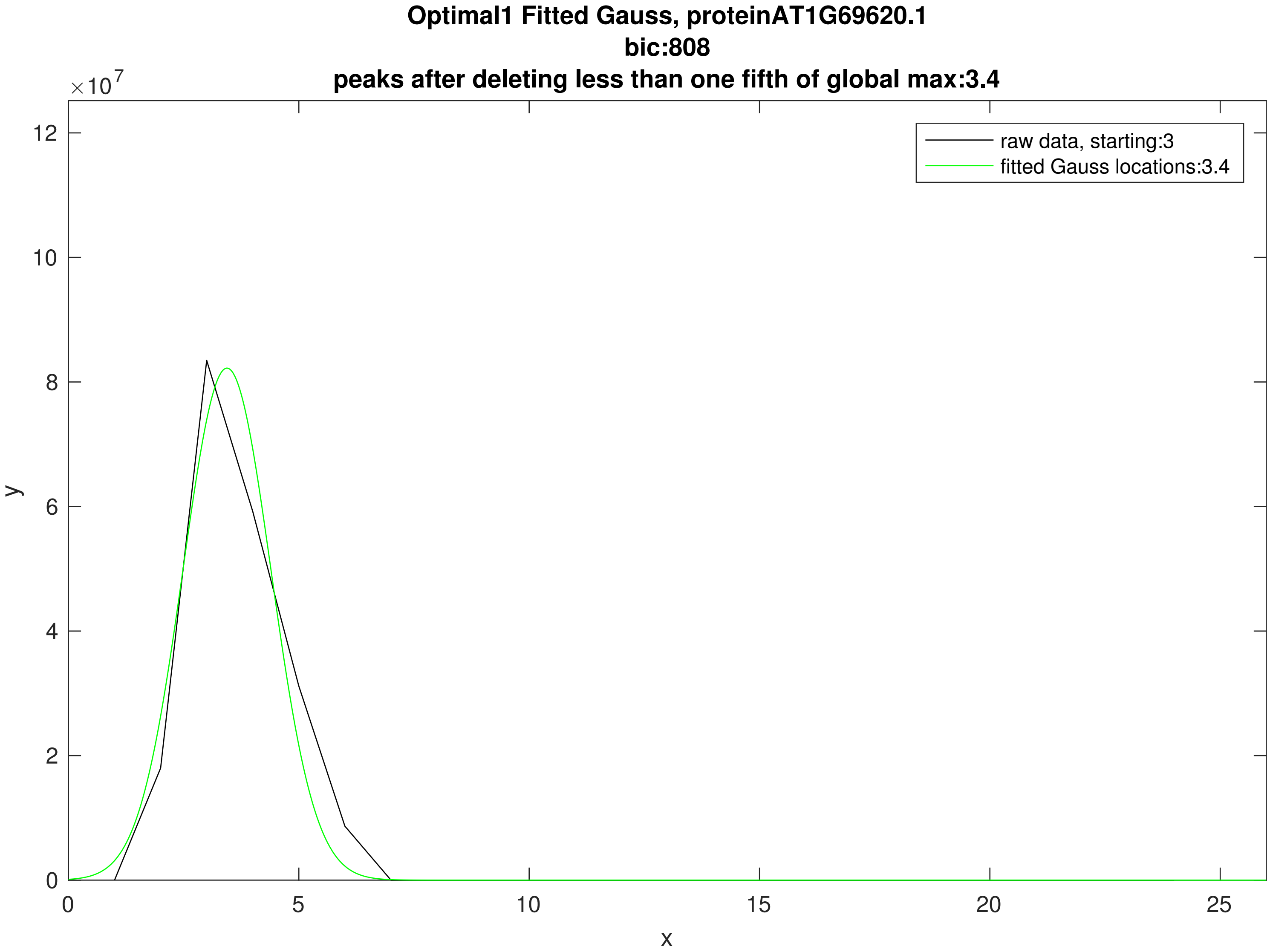
Optimal1 Fitted Gauss, proteinAT1G69620.1
bic:808
peaks after deleting less than one fifth of global max:3.4
×10^7
12
10
8
6
4
2
0
y
0
5
10
15
20
25
x
raw data, starting:3
fitted Gauss locations:3.4

**Optimal2 Fitted Gauss, proteinAT1G69740.2;AT1G69740.1**
**bic:885 879 888**
**peaks after deleting less than one fifth of global max:8.1**

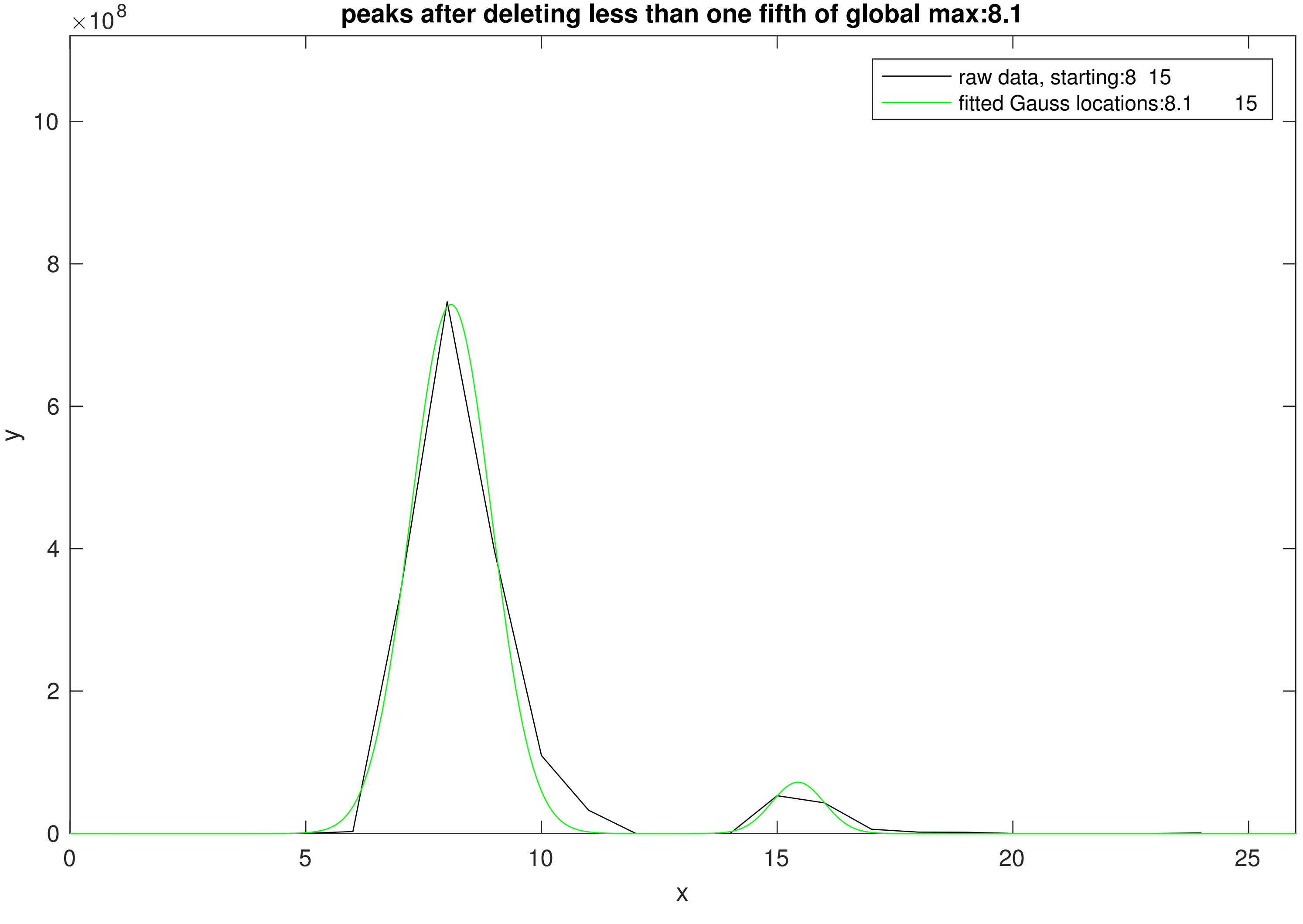

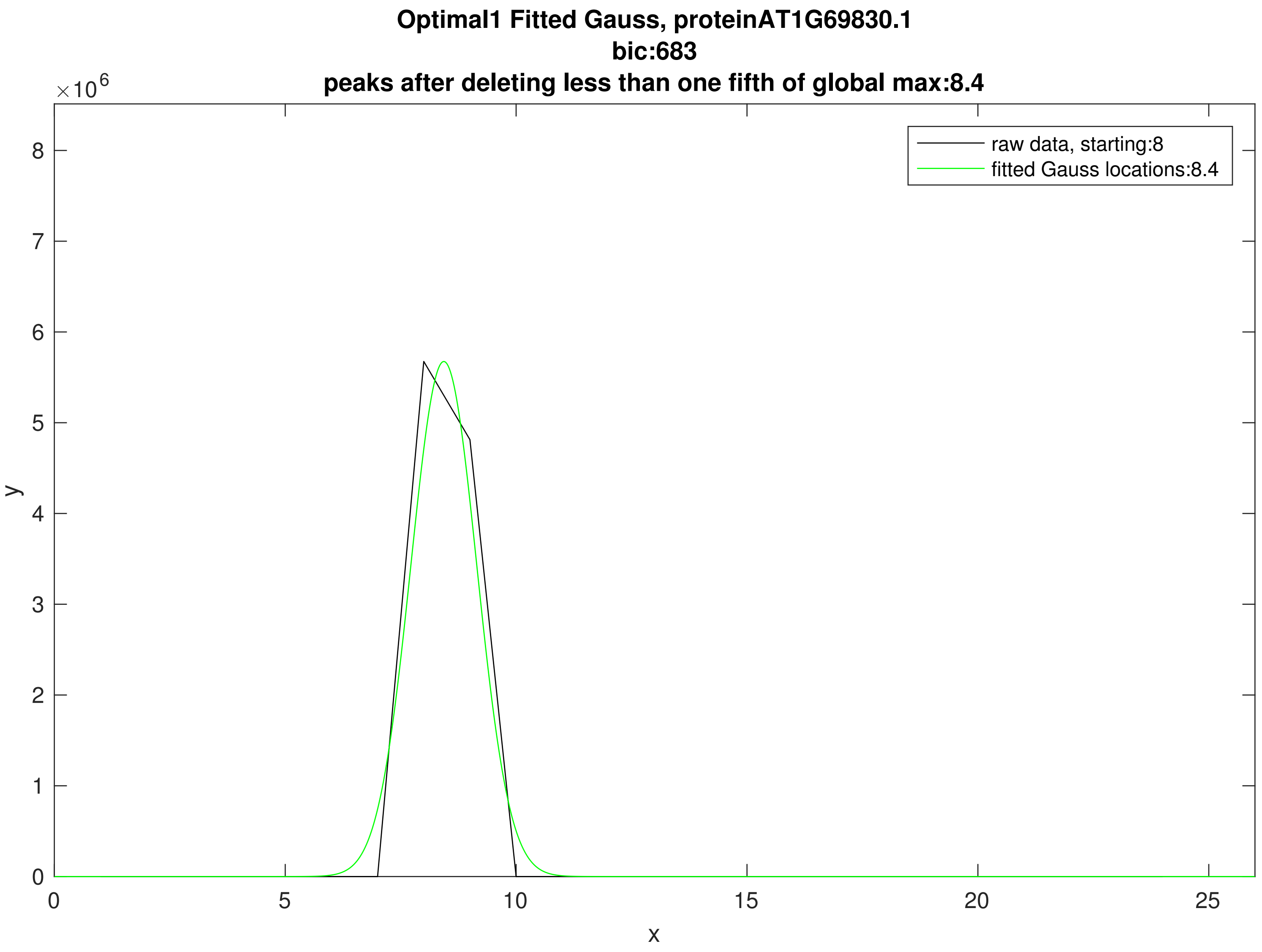

Optimal1 Fitted Gauss, proteinAT1G69830.1
bic:683
peaks after deleting less than one fifth of global max:8.4
×10^6
8
7
6
5
4
3
2
1
0
y
0
5
10
15
20
25
x
raw data, starting:8
fitted Gauss locations:8.4

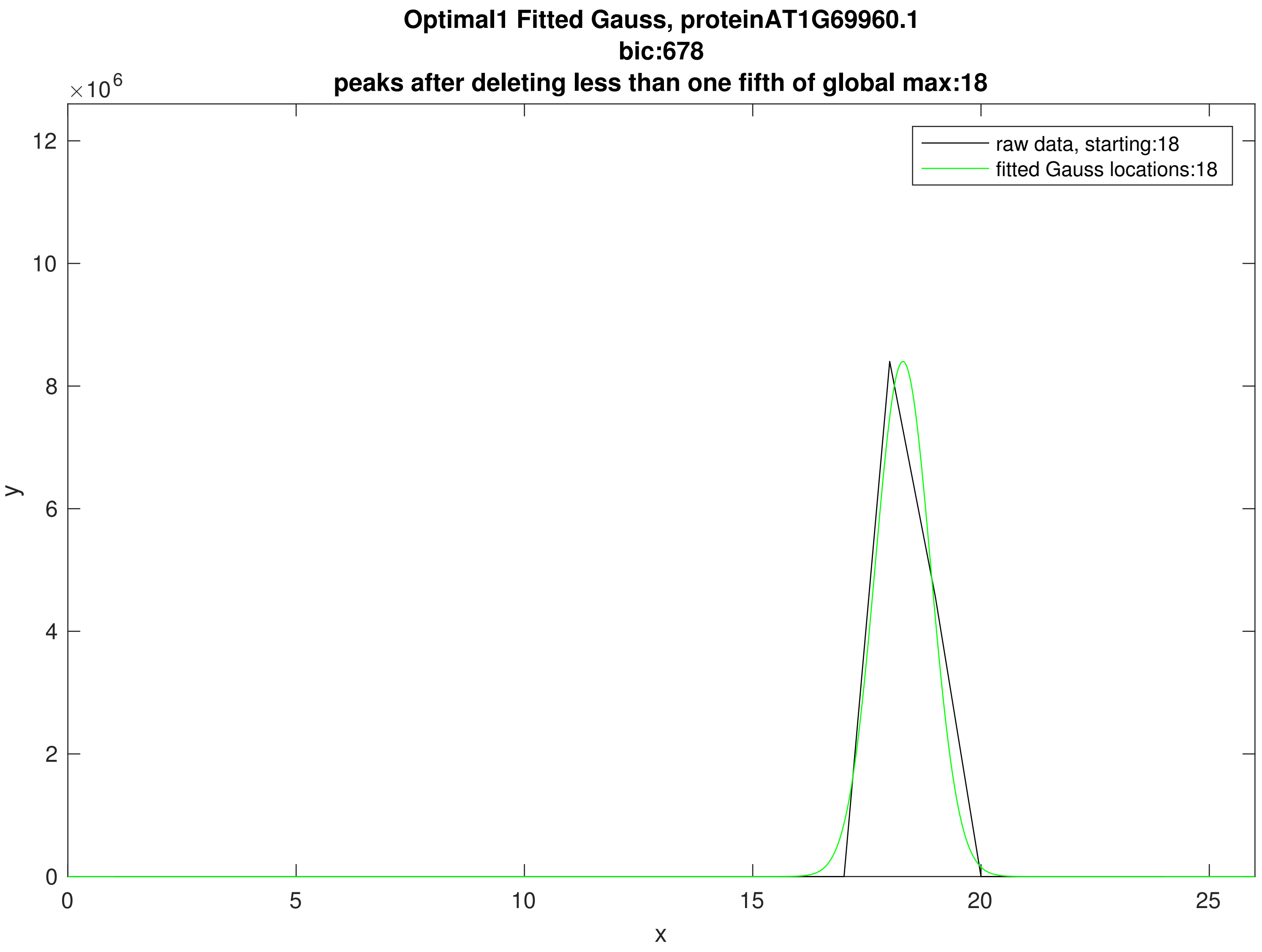

Optimal1 Fitted Gauss, proteinAT1G69960.1
bic:678
peaks after deleting less than one fifth of global max:18
×10^6
raw data, starting:18
fitted Gauss locations:18
y
x
0
2
4
6
8
10
12
0
5
10
15
20
25

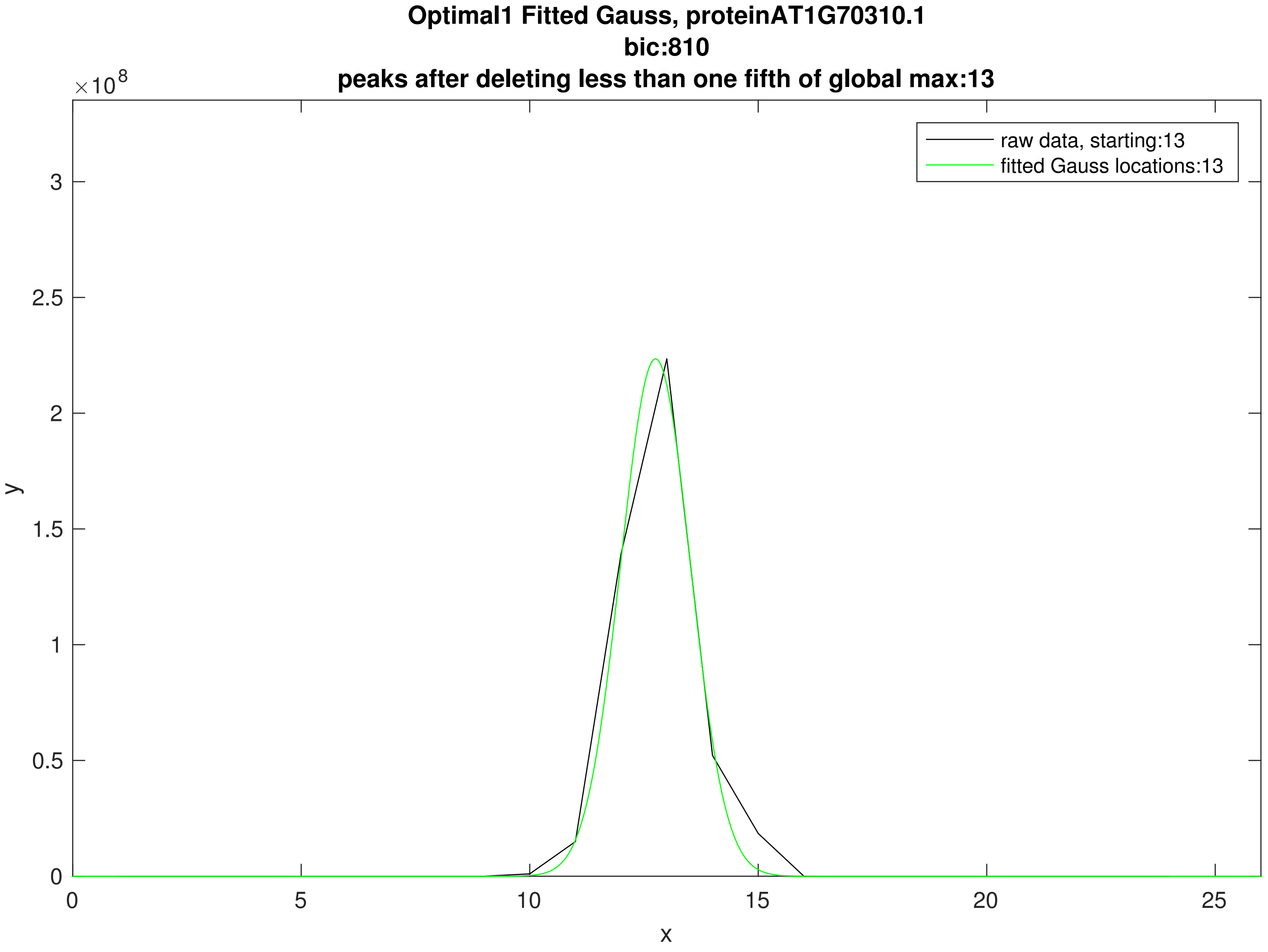

Optimal1 Fitted Gauss, proteinAT1G70310.1
bic:810
peaks after deleting less than one fifth of global max:13
×10^8
raw data, starting:13
fitted Gauss locations:13
3
2.5
2
1.5
1
0.5
0
y
0
5
10
15
20
25
x

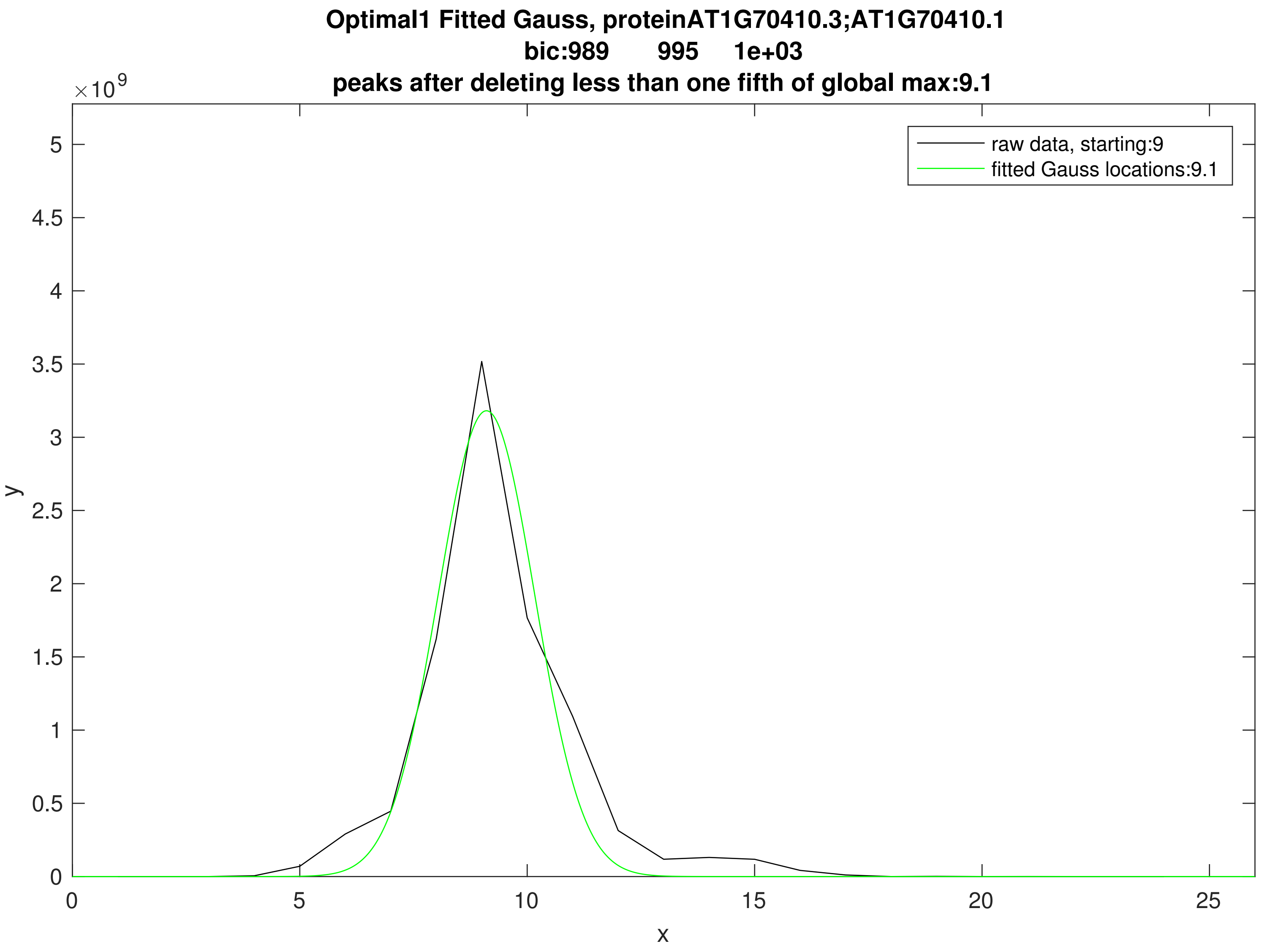

Optimal1 Fitted Gauss, proteinAT1G70410.3;AT1G70410.1
bic:989 995 1e+03
peaks after deleting less than one fifth of global max:9.1
×10^9
raw data, starting:9
fitted Gauss locations:9.1
y
x
0
0.5
1
1.5
2
2.5
3
3.5
4
4.5
5
0
5
10
15
20
25

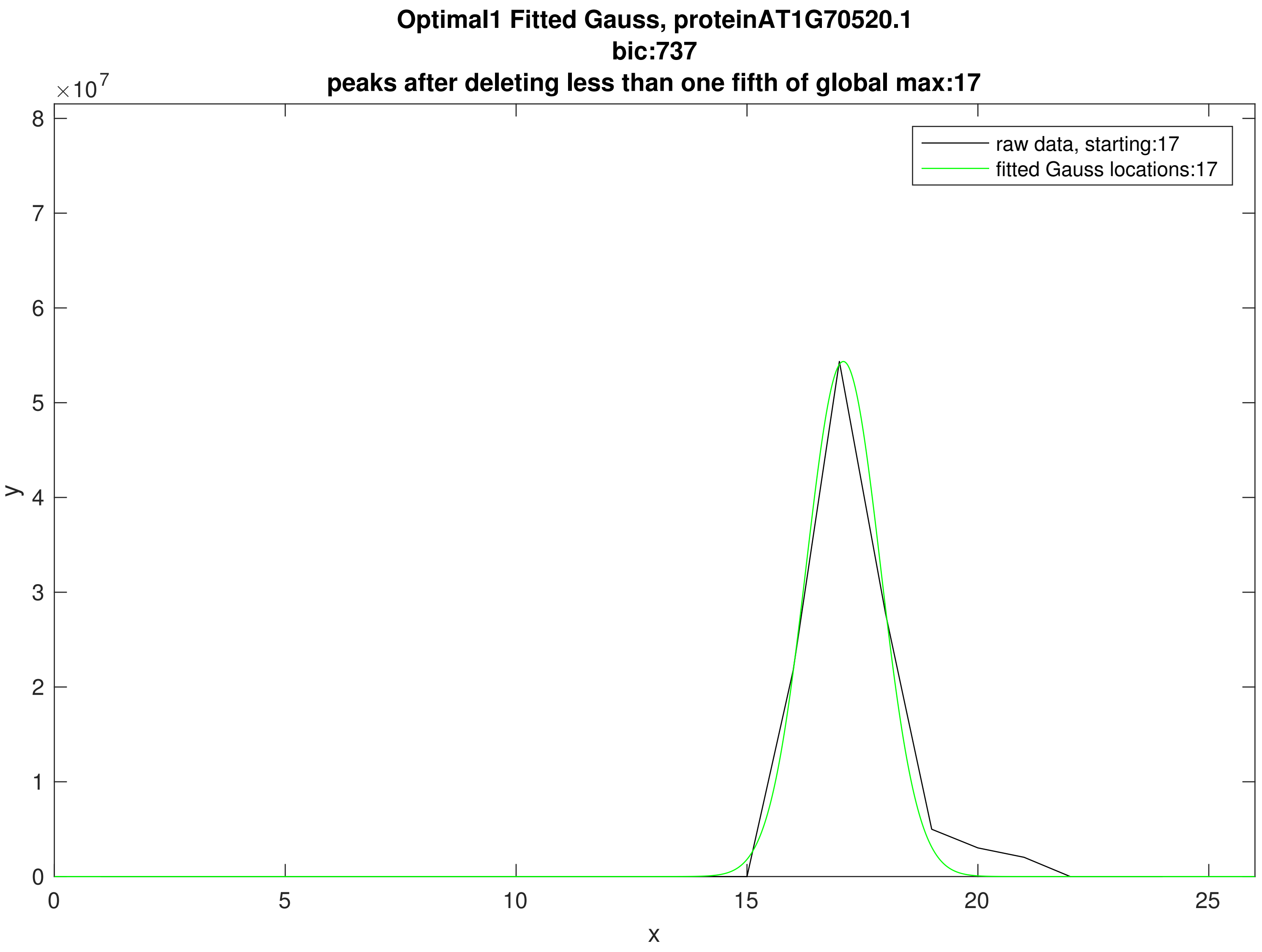

Optimal1 Fitted Gauss, proteinAT1G70520.1
bic:737
peaks after deleting less than one fifth of global max:17
×10^7
8
7
6
5
4
3
2
1
0
y
0
5
10
15
20
25
x
raw data, starting:17
fitted Gauss locations:17

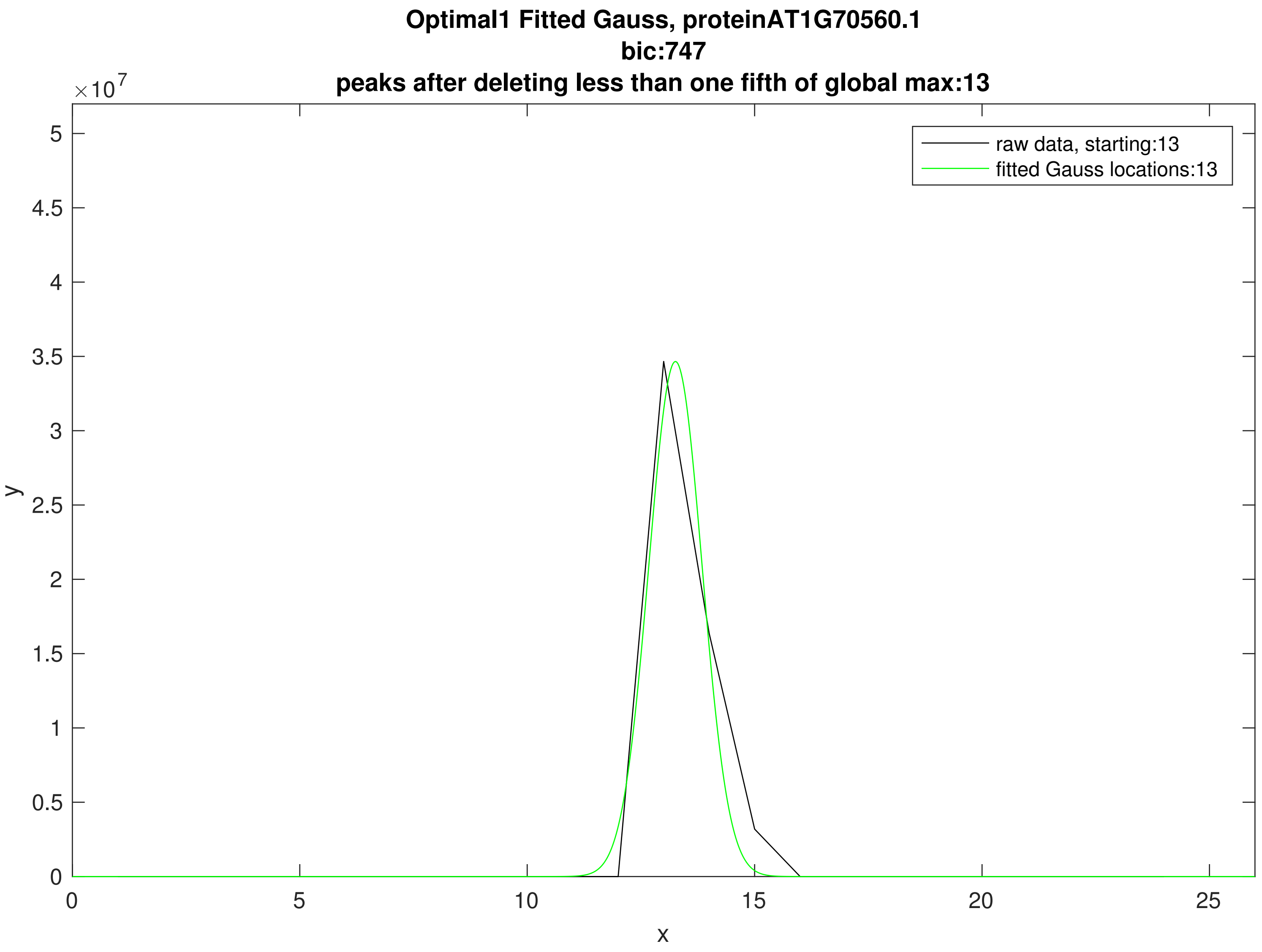
Optimal1 Fitted Gauss, proteinAT1G70560.1
bic:747
peaks after deleting less than one fifth of global max:13
×10^7
y
x
raw data, starting:13
fitted Gauss locations:13
0
0.5
1
1.5
2
2.5
3
3.5
4
4.5
5
0
5
10
15
20
25

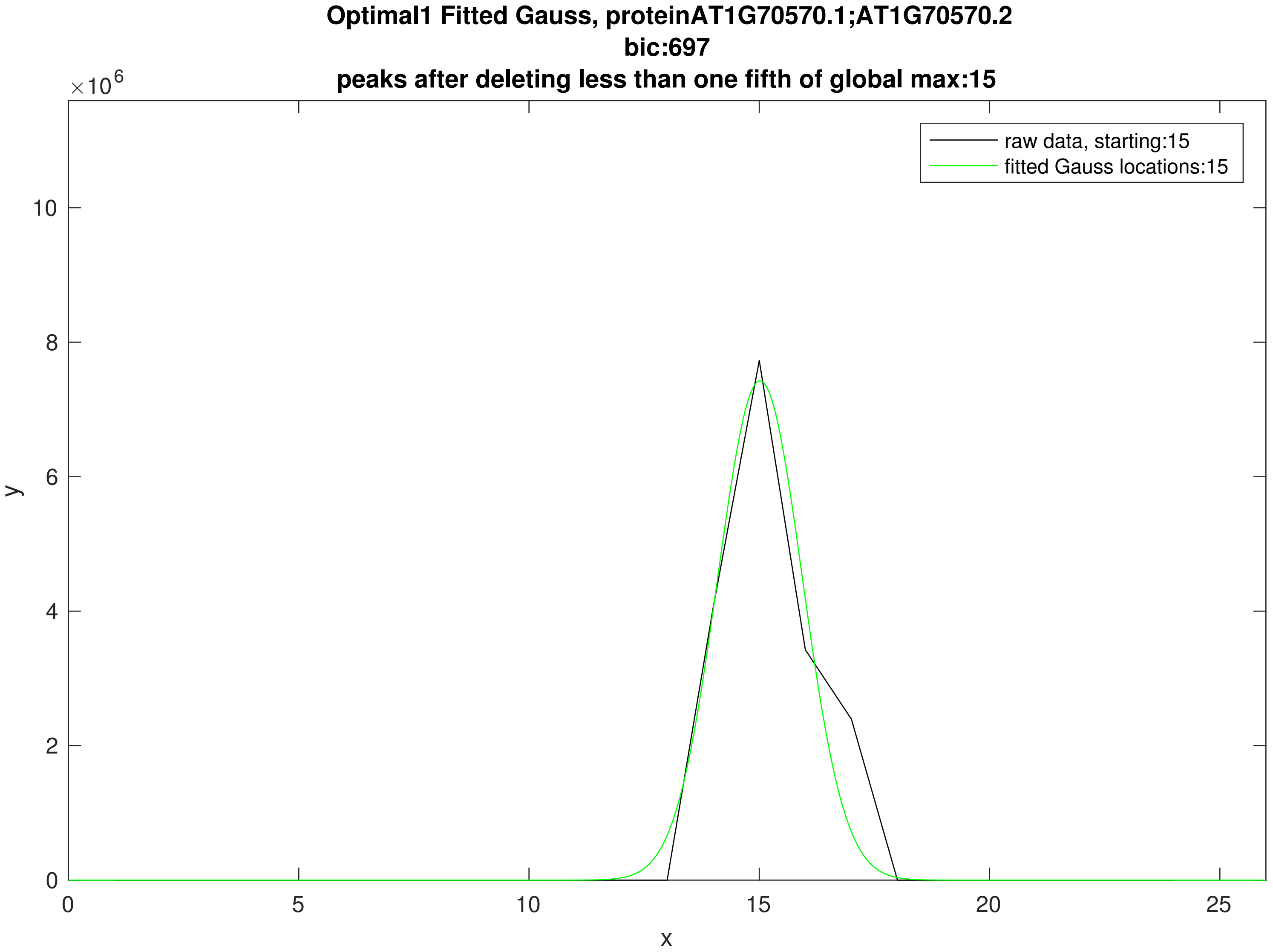
Optimal1 Fitted Gauss, proteinAT1G70570.1;AT1G70570.2
bic:697
peaks after deleting less than one fifth of global max:15
$\times 10^6$
raw data, starting:15
fitted Gauss locations:15
10
8
6
4
2
0
y
0
5
10
15
20
25
x

#### Optimal1 Fitted Gauss, proteinAT1G70580.4;AT1G70580.3;AT1G70580.2;AT1G70580.1
#### bic:875
#### peaks after deleting less than one fifth of global max:14

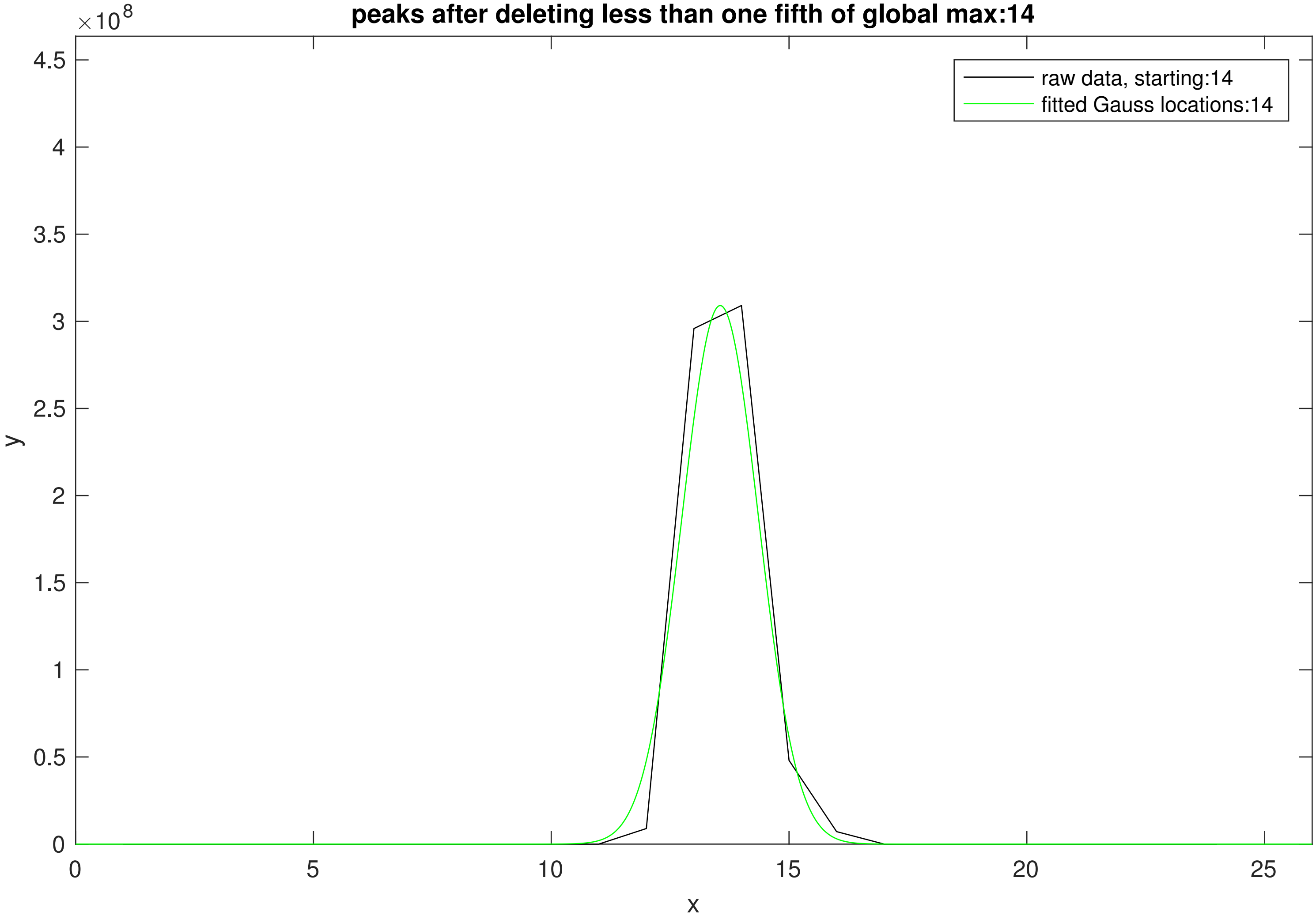

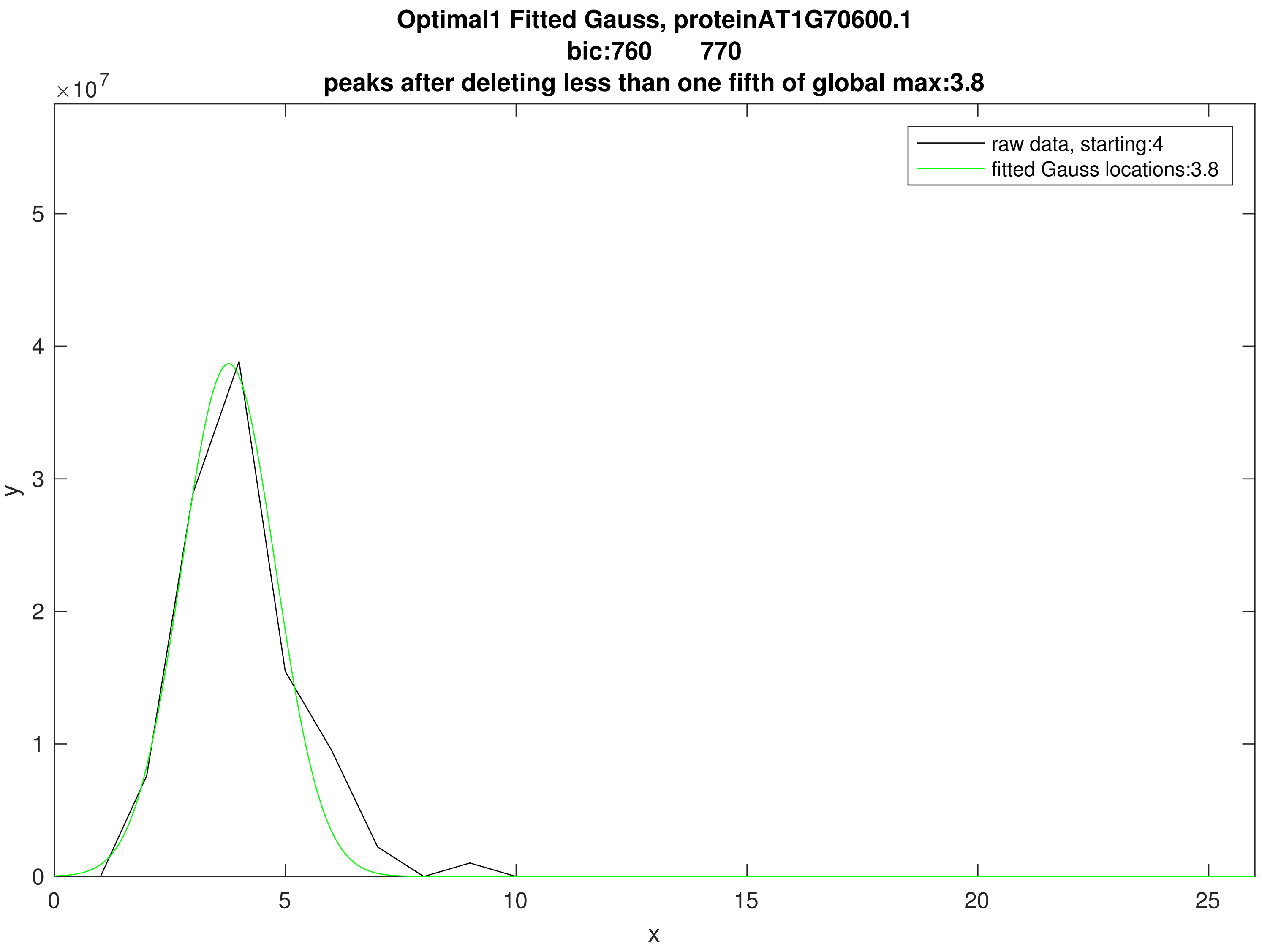
Optimal1 Fitted Gauss, proteinAT1G70600.1
bic:760 770
peaks after deleting less than one fifth of global max:3.8
×10^7
raw data, starting:4
fitted Gauss locations:3.8
y
0
1
2
3
4
5
0
5
10
15
20
25
x

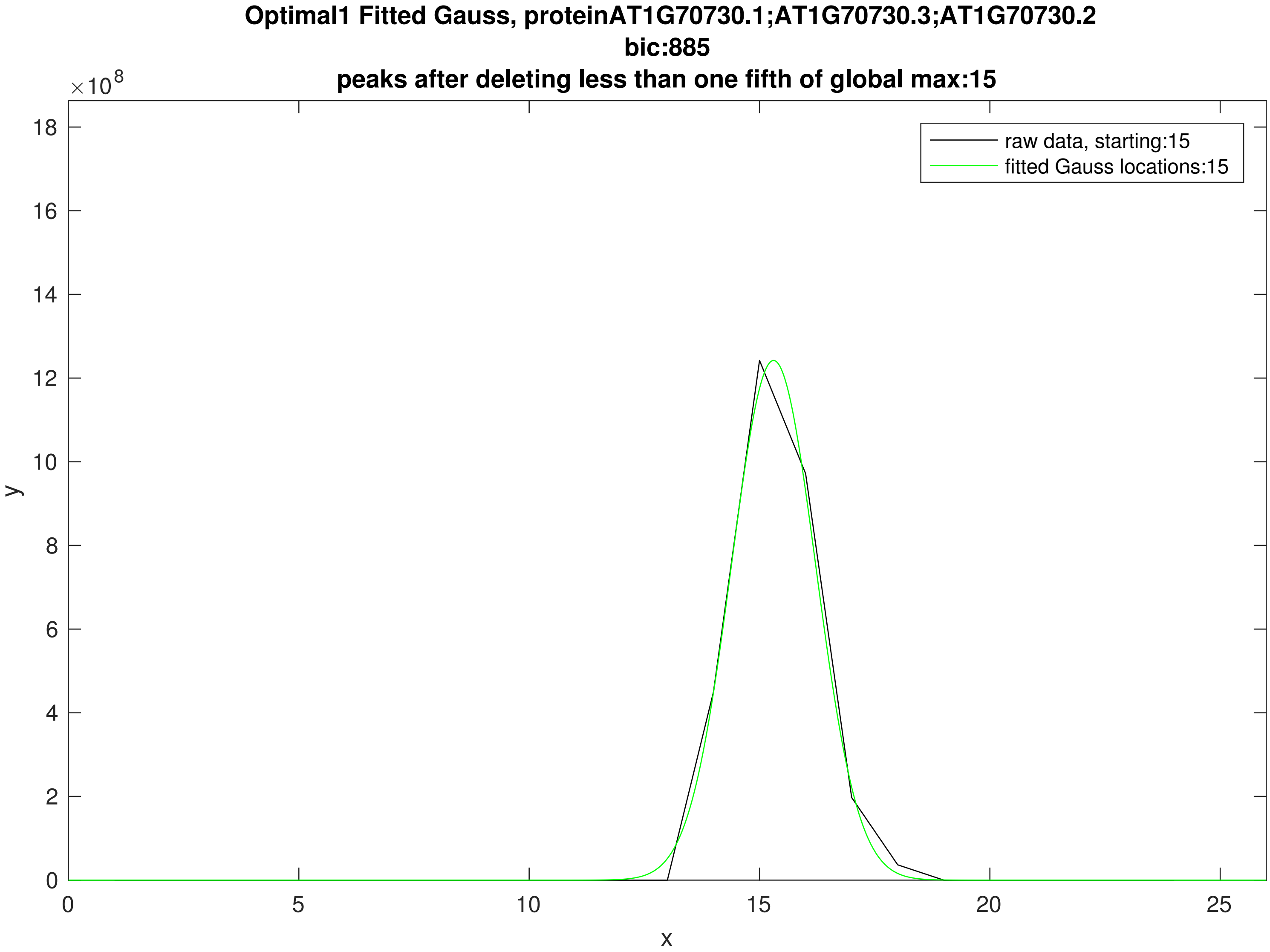

Optimal1 Fitted Gauss, proteinAT1G70730.1;AT1G70730.3;AT1G70730.2
bic:885
peaks after deleting less than one fifth of global max:15
×10^8
raw data, starting:15
fitted Gauss locations:15
y
x
0
2
4
6
8
10
12
14
16
18
0
5
10
15
20
25

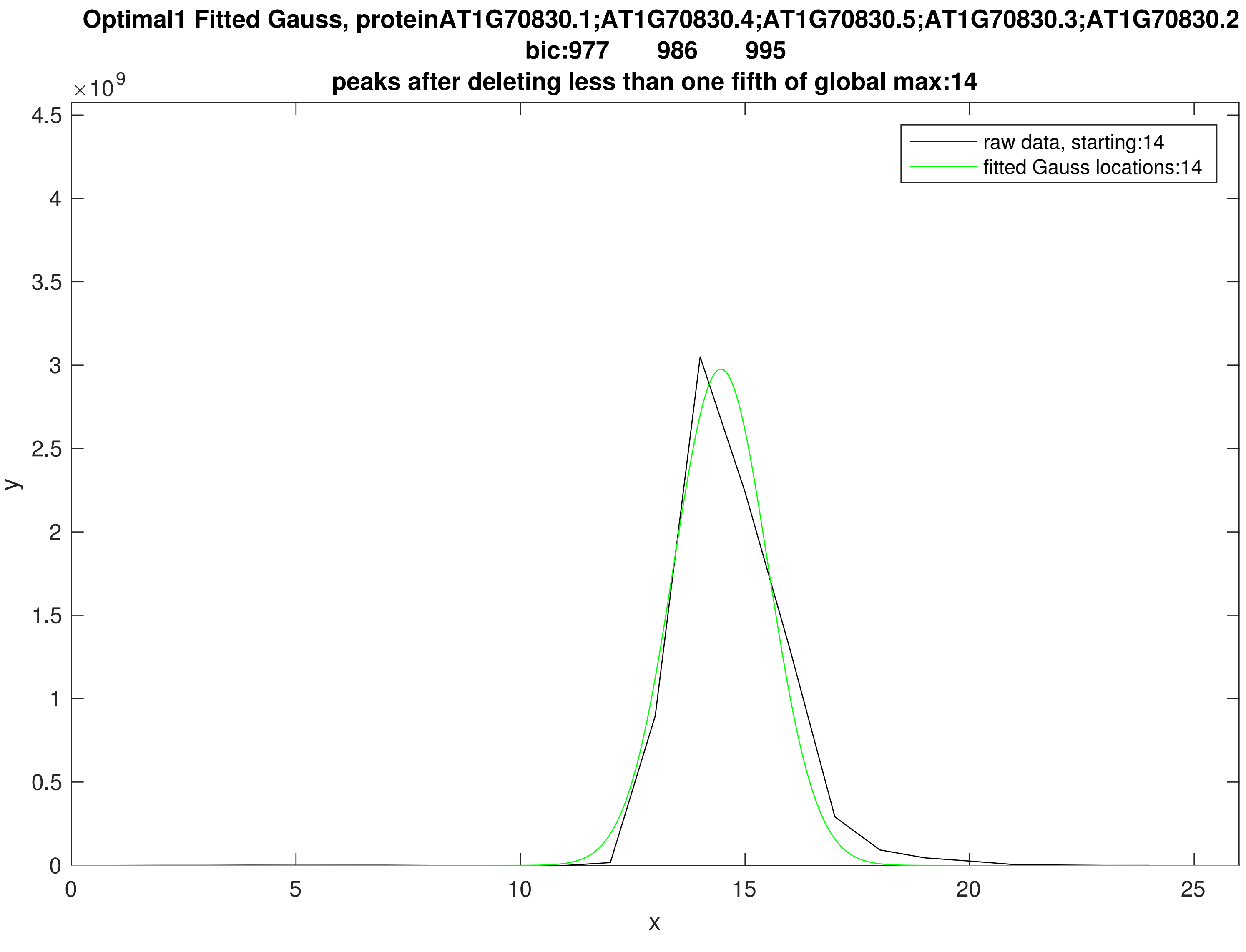
Optimal1 Fitted Gauss, proteinAT1G70830.1;AT1G70830.4;AT1G70830.5;AT1G70830.3;AT1G70830.2
bic:977 986 995
peaks after deleting less than one fifth of global max:14
×10^9
4.5
4
3.5
3
2.5
2
1.5
1
0.5
0
y
0
5
10
15
20
25
x
raw data, starting:14
fitted Gauss locations:14

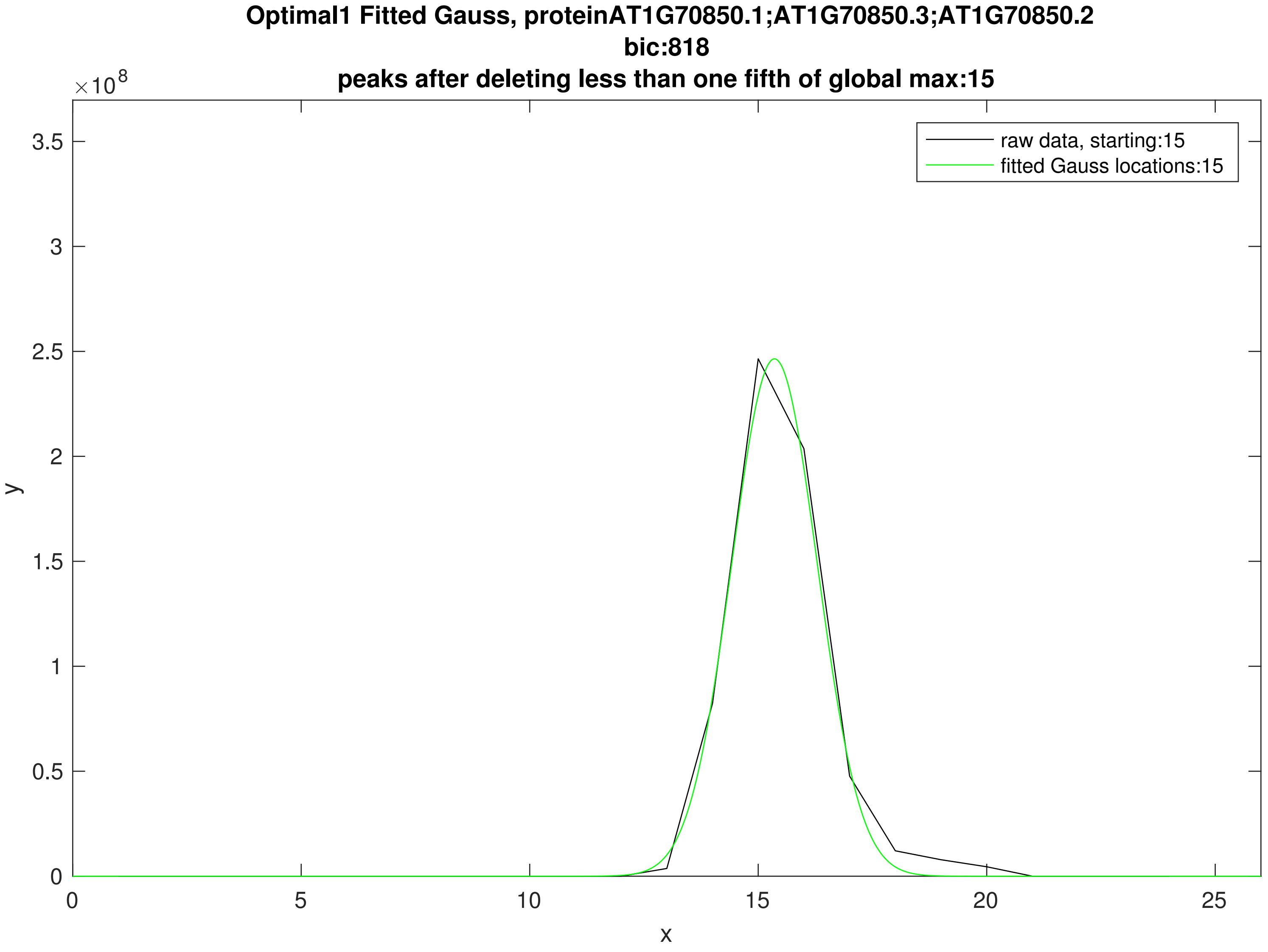
Optimal1 Fitted Gauss, proteinAT1G70850.1;AT1G70850.3;AT1G70850.2
bic:818
peaks after deleting less than one fifth of global max:15
×10^8
3.5
3
2.5
2
1.5
1
0.5
0
y
0
5
10
15
20
25
x
raw data, starting:15
fitted Gauss locations:15

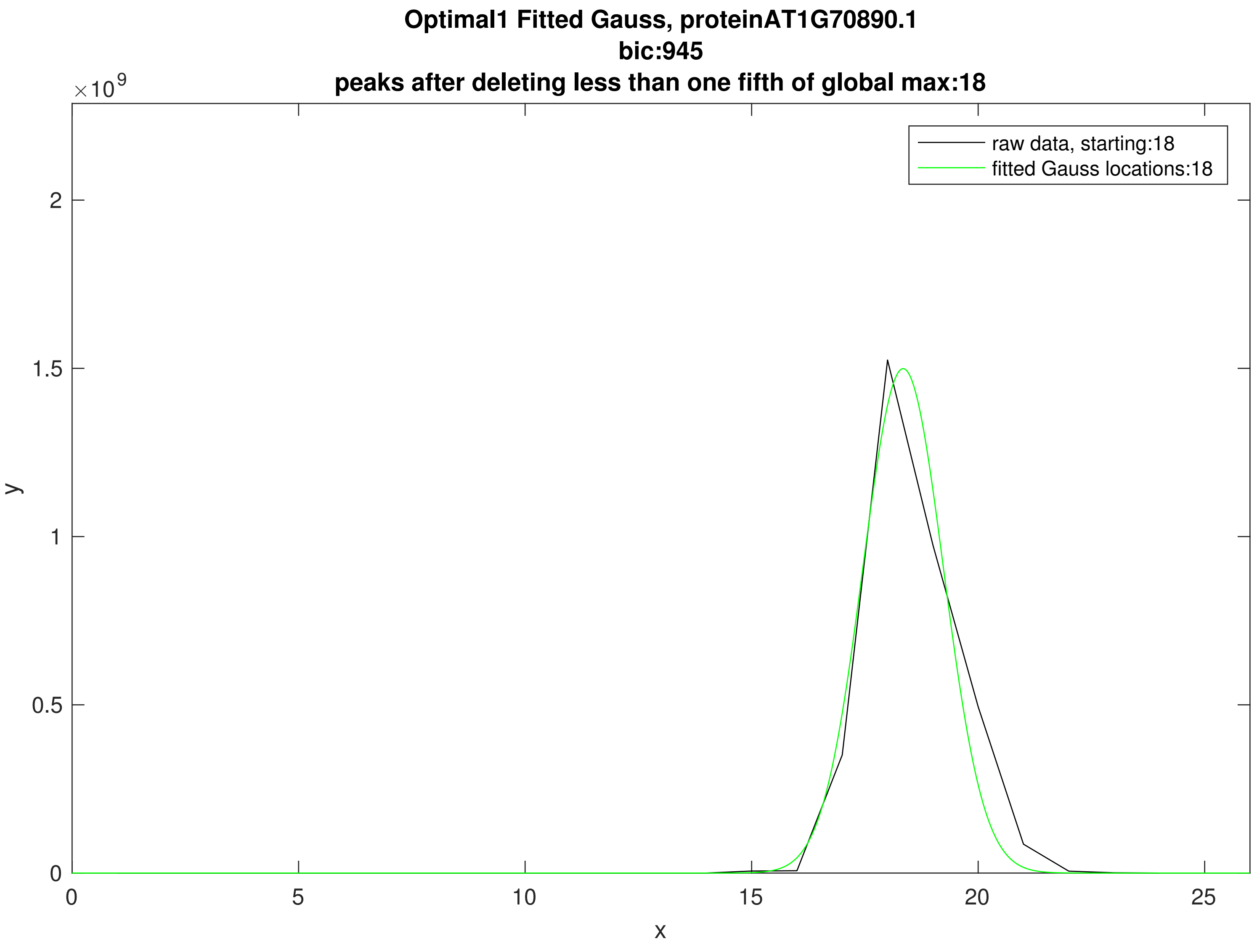

Optimal1 Fitted Gauss, proteinAT1G70890.1
bic:945
peaks after deleting less than one fifth of global max:18
×10^9
raw data, starting:18
fitted Gauss locations:18
y
x
0
0.5
1
1.5
2
0
5
10
15
20
25

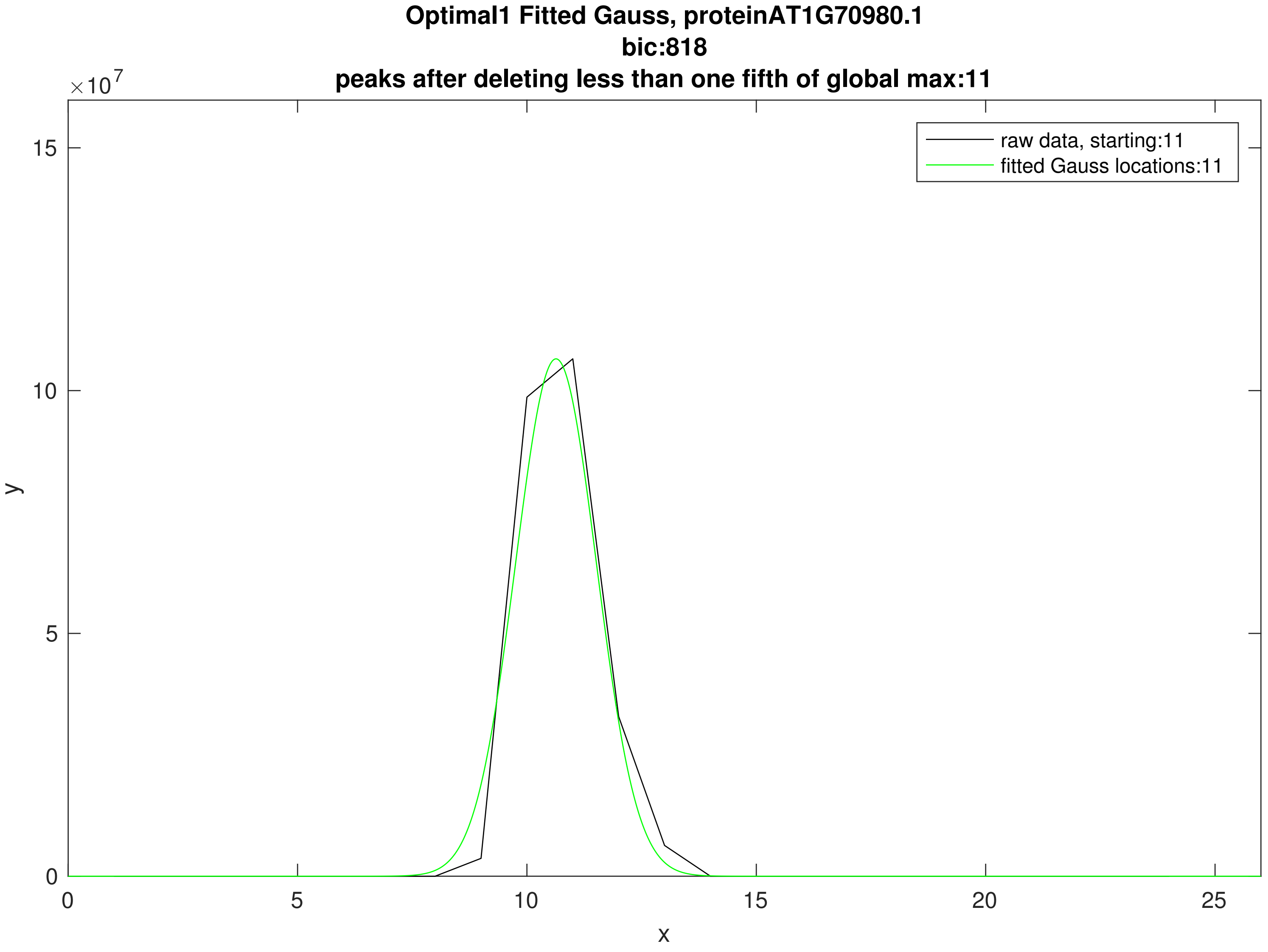
Optimal1 Fitted Gauss, proteinAT1G70980.1
bic:818
peaks after deleting less than one fifth of global max:11
×10^7
raw data, starting:11
fitted Gauss locations:11
y
x
0
5
10
15
20
25

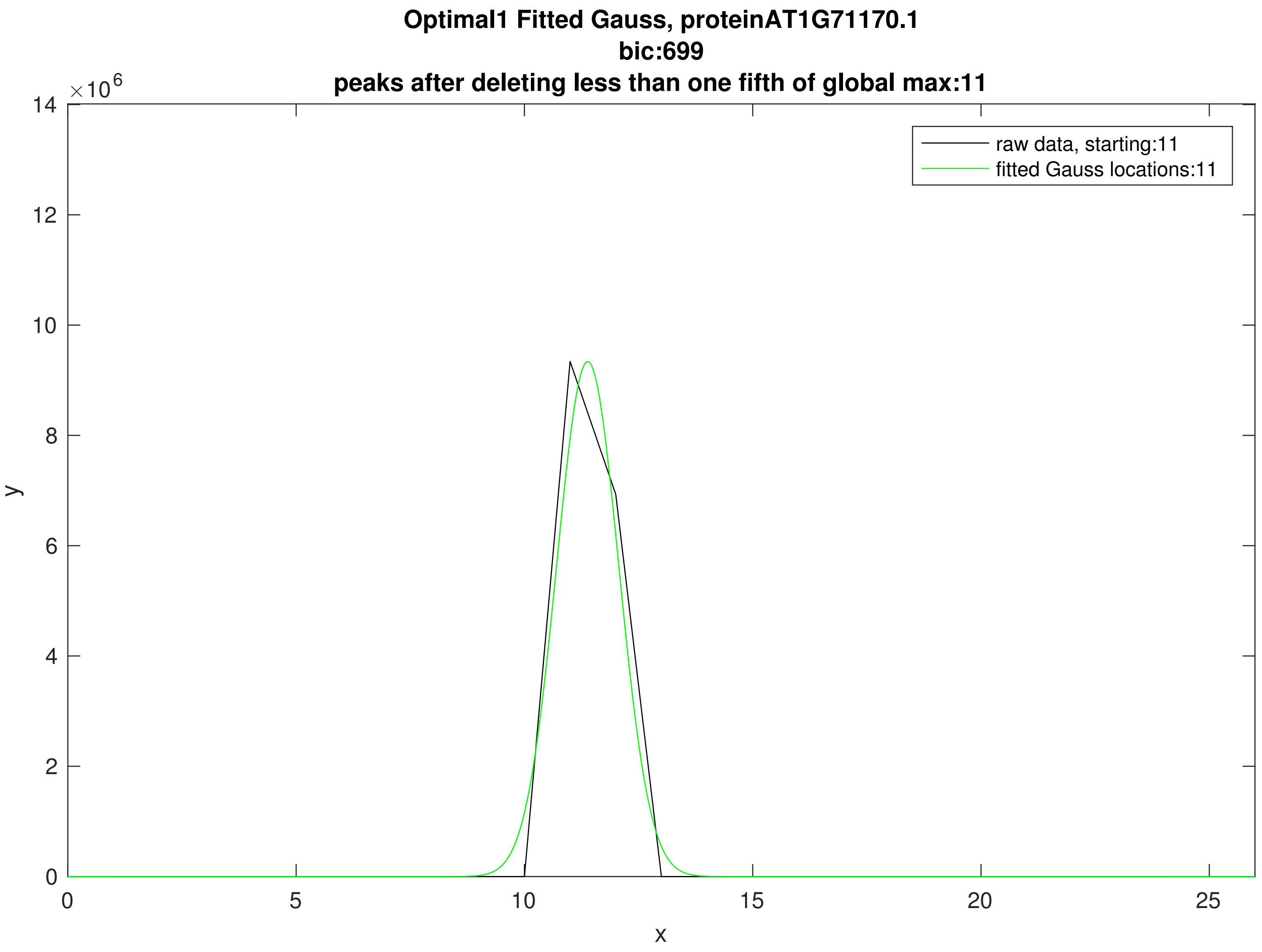
Optimal1 Fitted Gauss, proteinAT1G71170.1
bic:699
peaks after deleting less than one fifth of global max:11
×10^6
14
12
10
8
6
4
2
0
y
0
5
10
15
20
25
x
raw data, starting:11
fitted Gauss locations:11

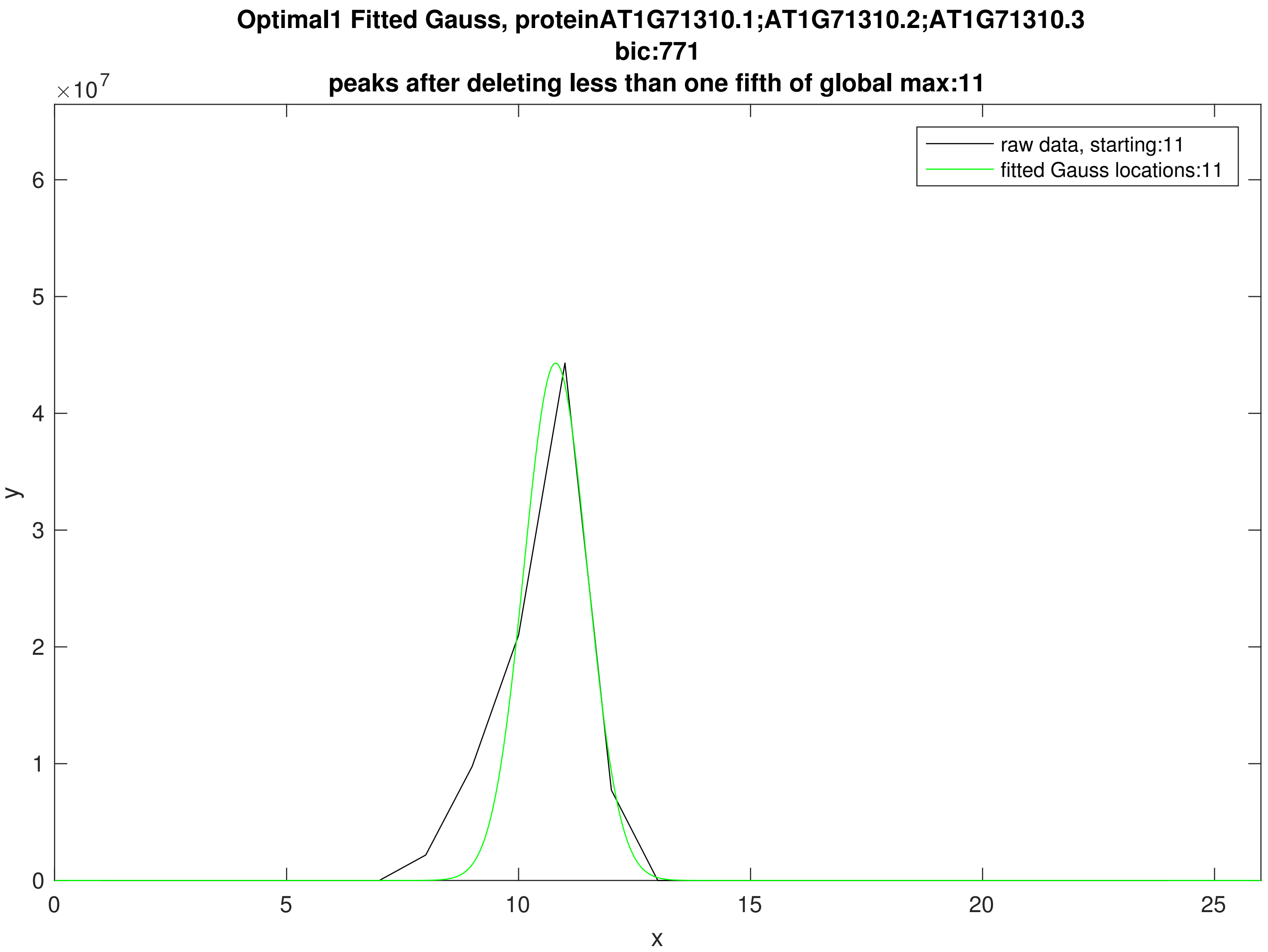
Optimal1 Fitted Gauss, proteinAT1G71310.1;AT1G71310.2;AT1G71310.3
bic:771
peaks after deleting less than one fifth of global max:11
×10^7
raw data, starting:11
fitted Gauss locations:11
y
x
0
1
2
3
4
5
6
0
5
10
15
20
25

Optimal1 Fitted Gauss, proteinAT1G71695.1
bic:669
peaks after deleting less than one fifth of global max:5.7

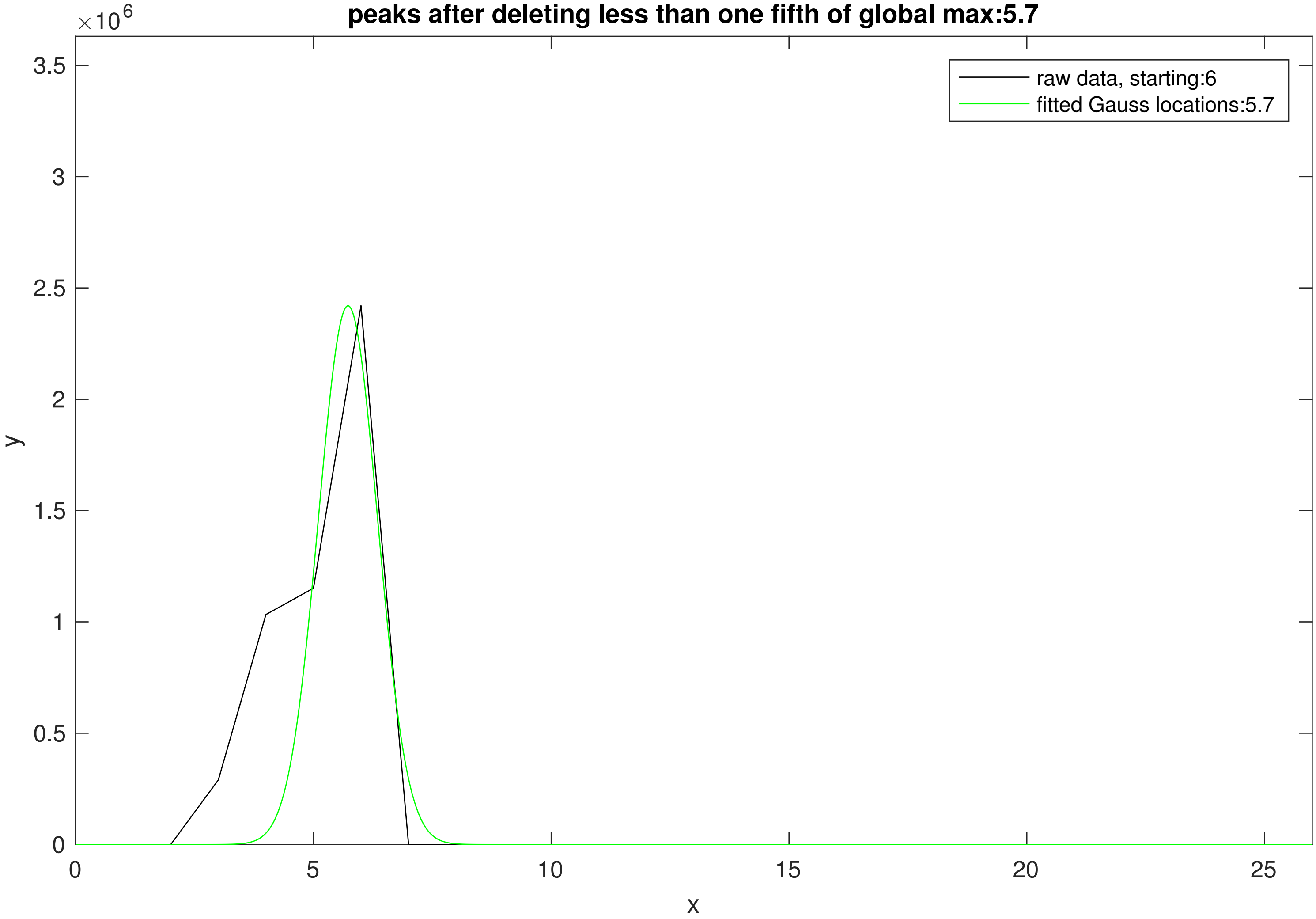

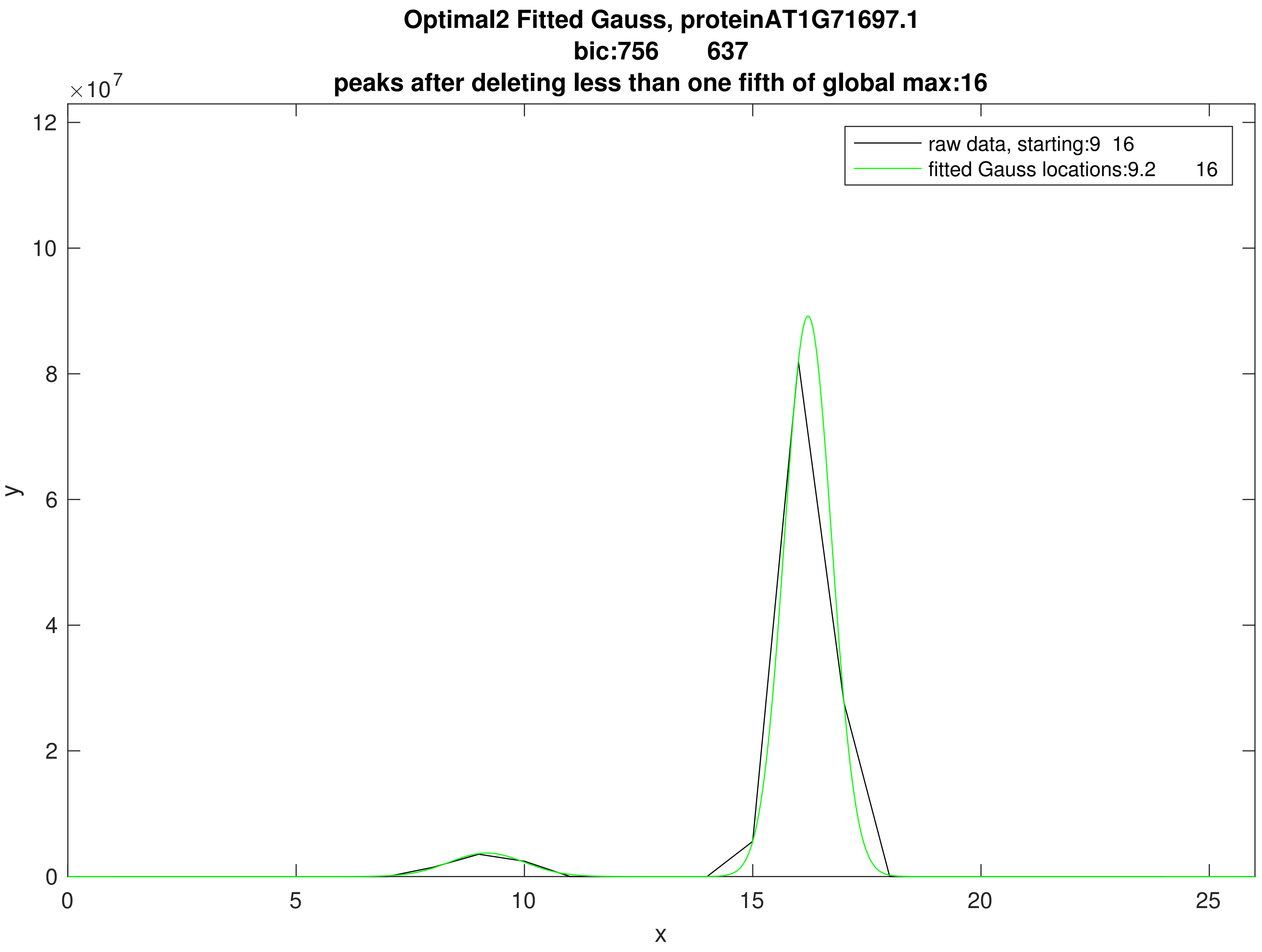

Optimal2 Fitted Gauss, proteinAT1G71697.1
bic:756 637
peaks after deleting less than one fifth of global max:16
×10^7
12
10
8
6
4
2
0
y
0
5
10
15
20
25
x
raw data, starting:9 16
fitted Gauss locations:9.2 16

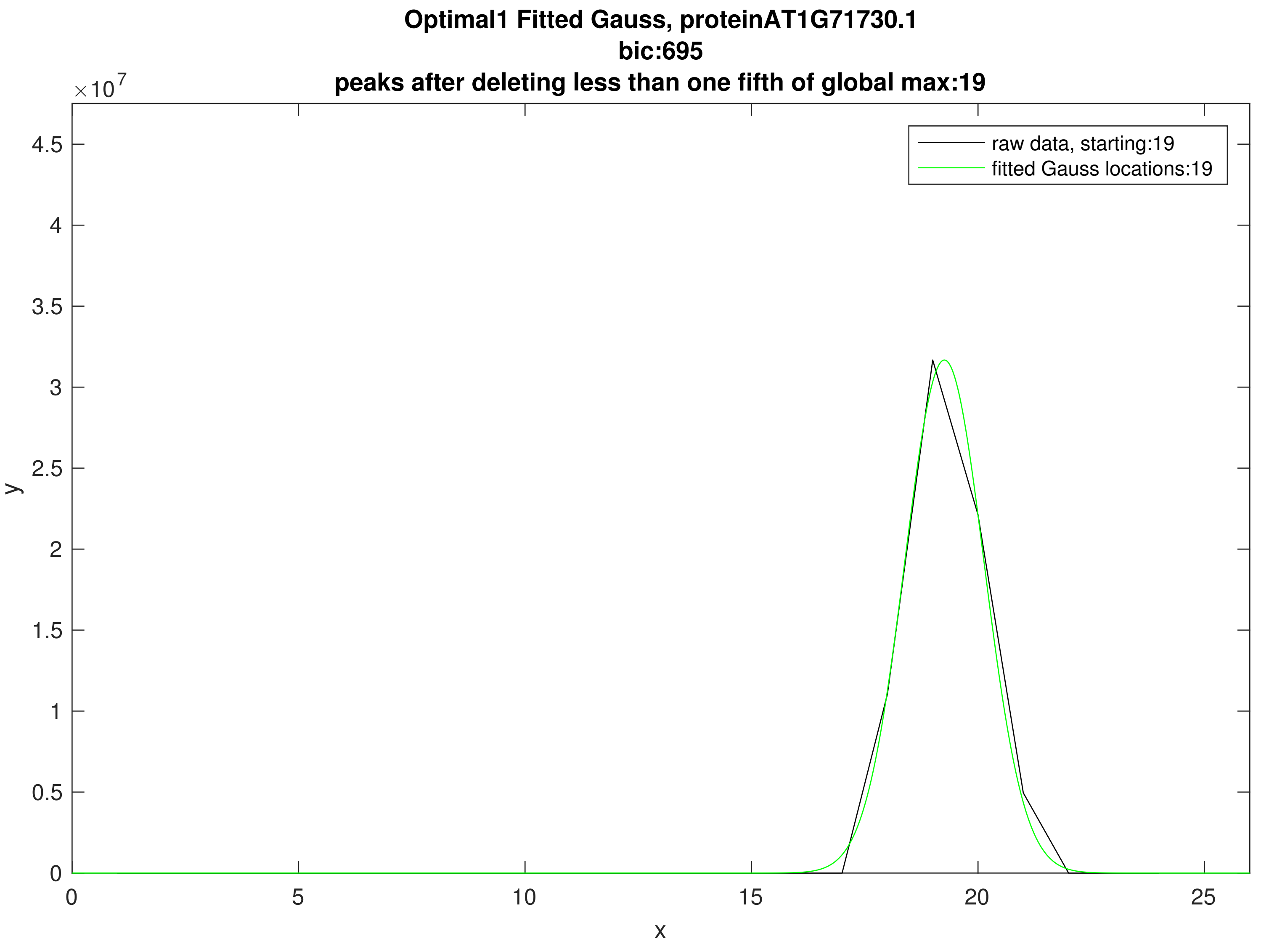
Optimal1 Fitted Gauss, proteinAT1G71730.1
bic:695
peaks after deleting less than one fifth of global max:19
×10^7
raw data, starting:19
fitted Gauss locations:19
y
x
0
0.5
1
1.5
2
2.5
3
3.5
4
4.5
0
5
10
15
20
25

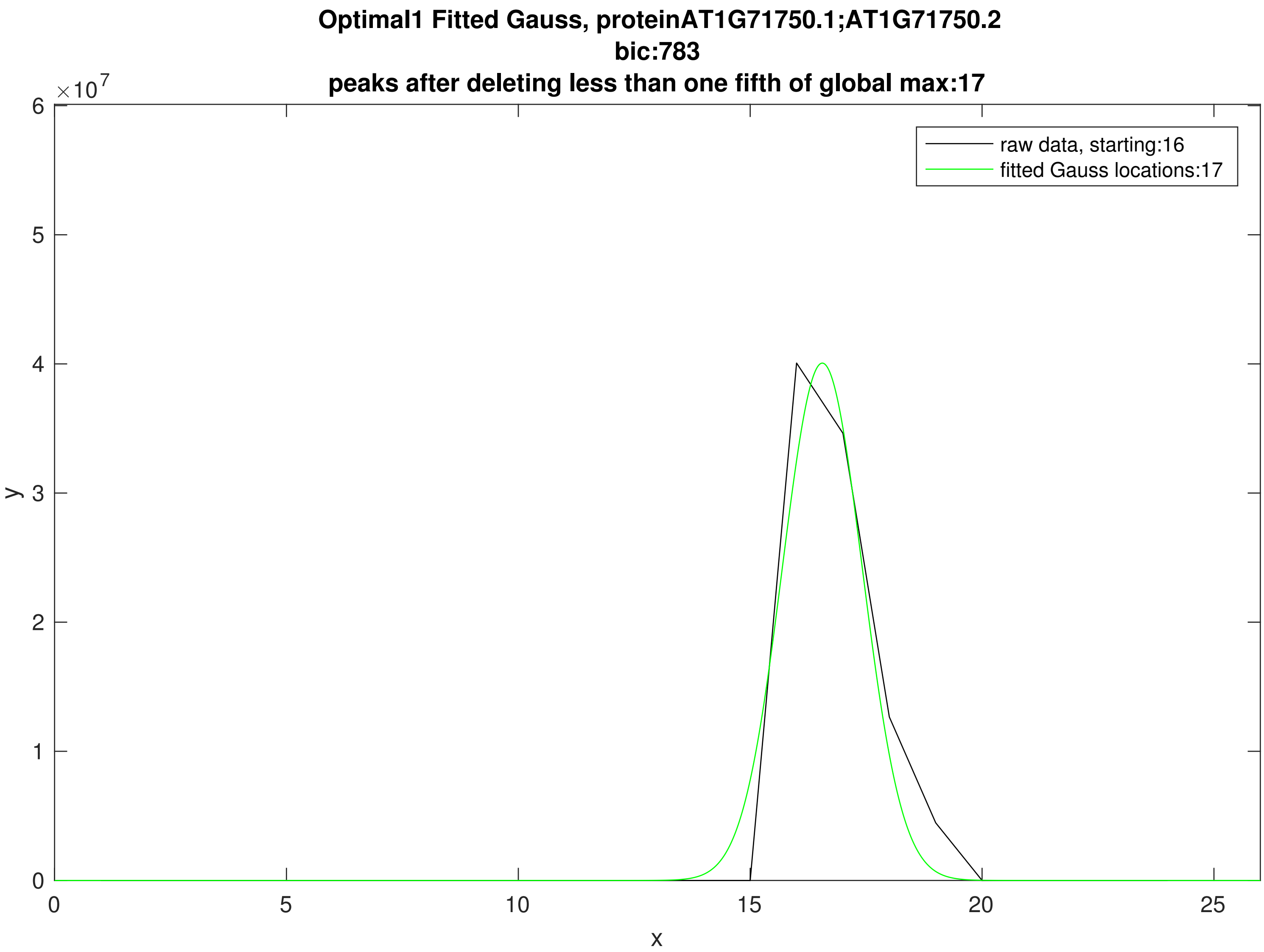

Optimal1 Fitted Gauss, proteinAT1G71750.1;AT1G71750.2
bic:783
peaks after deleting less than one fifth of global max:17
×10^7
raw data, starting:16
fitted Gauss locations:17
y
x
0
1
2
3
4
5
6
0
5
10
15
20
25

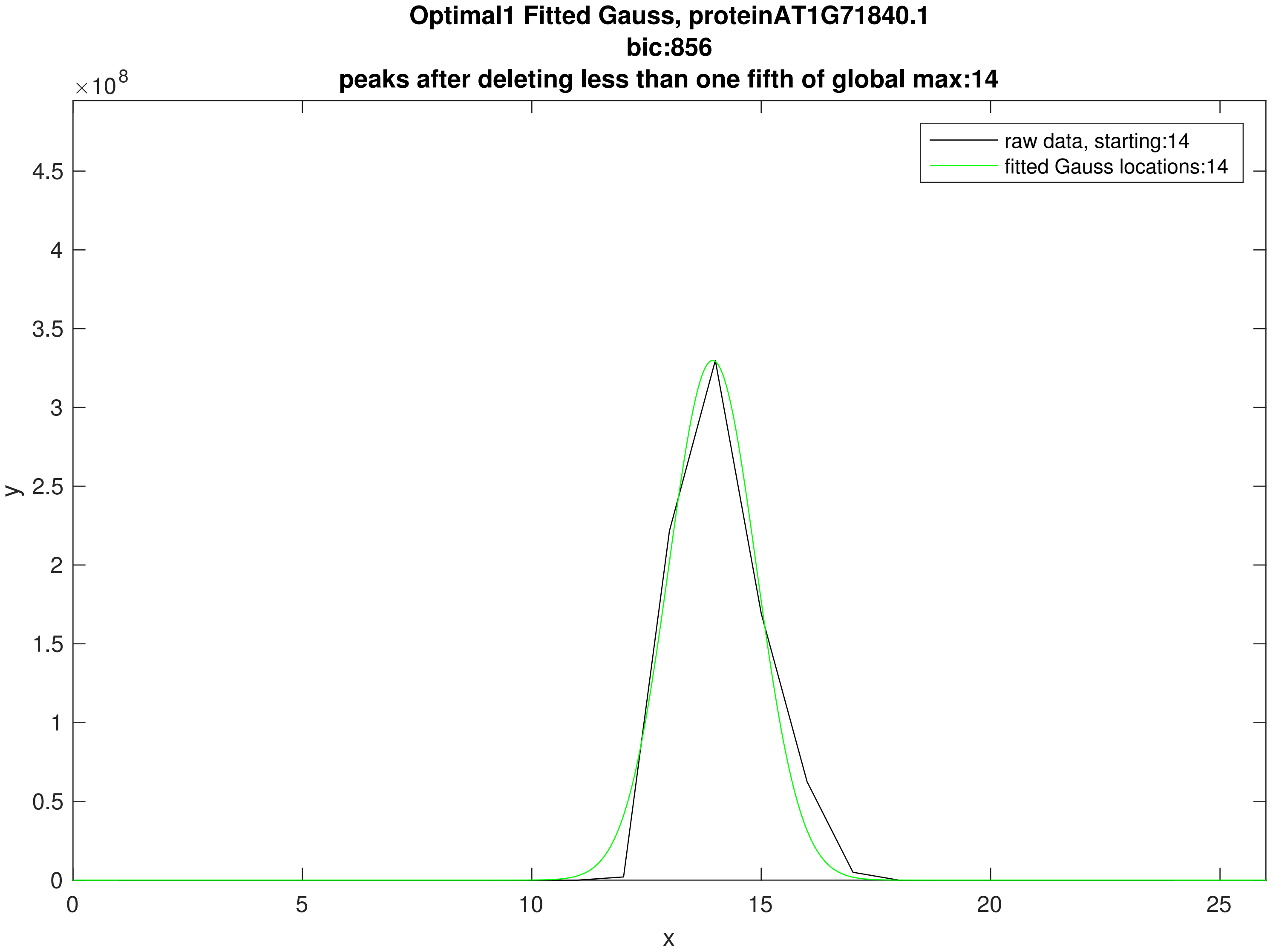
Optimal1 Fitted Gauss, proteinAT1G71840.1
bic:856
peaks after deleting less than one fifth of global max:14
×10^8
raw data, starting:14
fitted Gauss locations:14
y
x
4.5
4
3.5
3
2.5
2
1.5
1
0.5
0
0
5
10
15
20
25

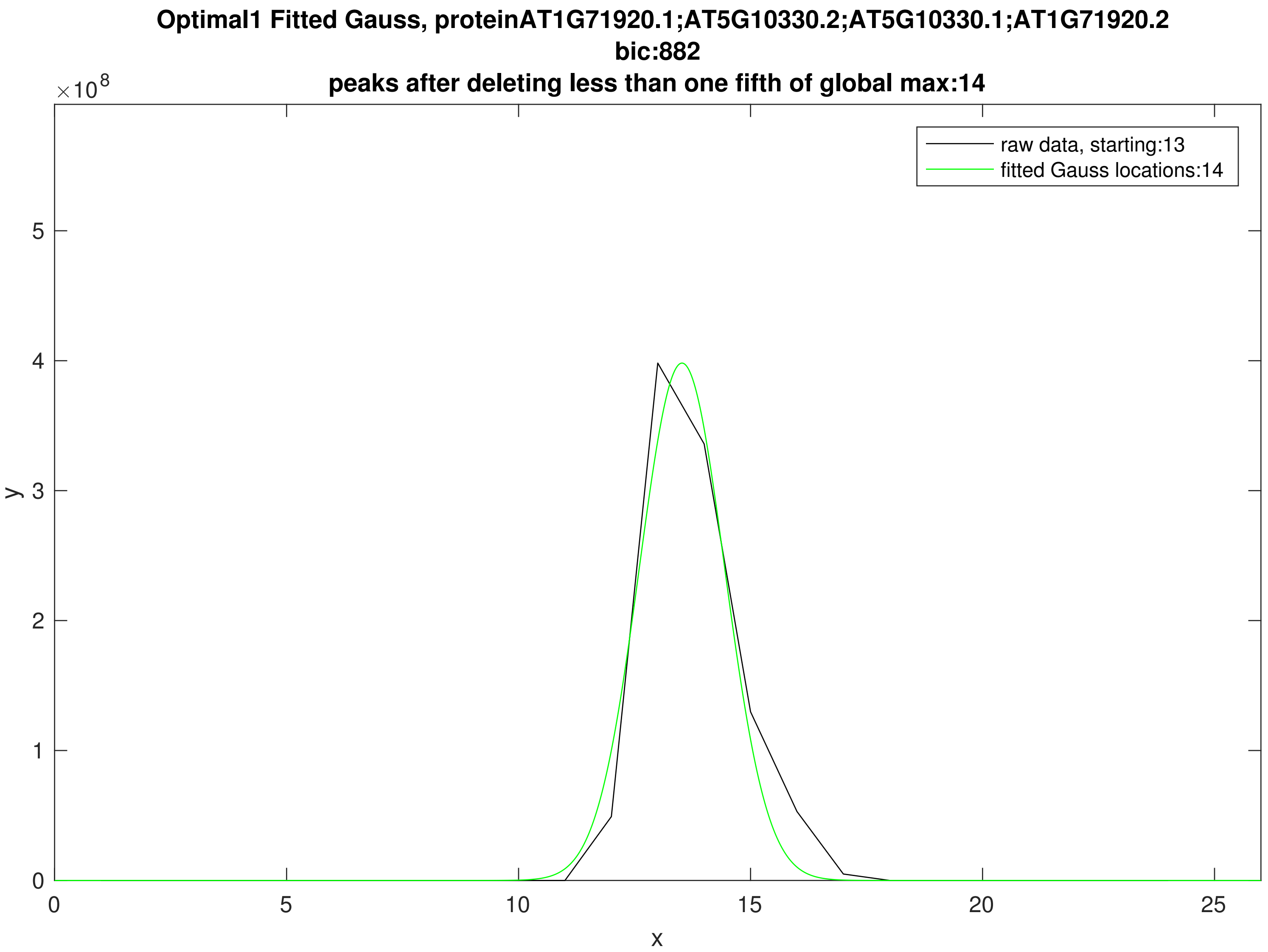

Optimal1 Fitted Gauss, proteinAT1G71920.1;AT5G10330.2;AT5G10330.1;AT1G71920.2
bic:882
peaks after deleting less than one fifth of global max:14
×10^8
raw data, starting:13
fitted Gauss locations:14
0
1
2
3
4
5
y
0
5
10
15
20
25
x

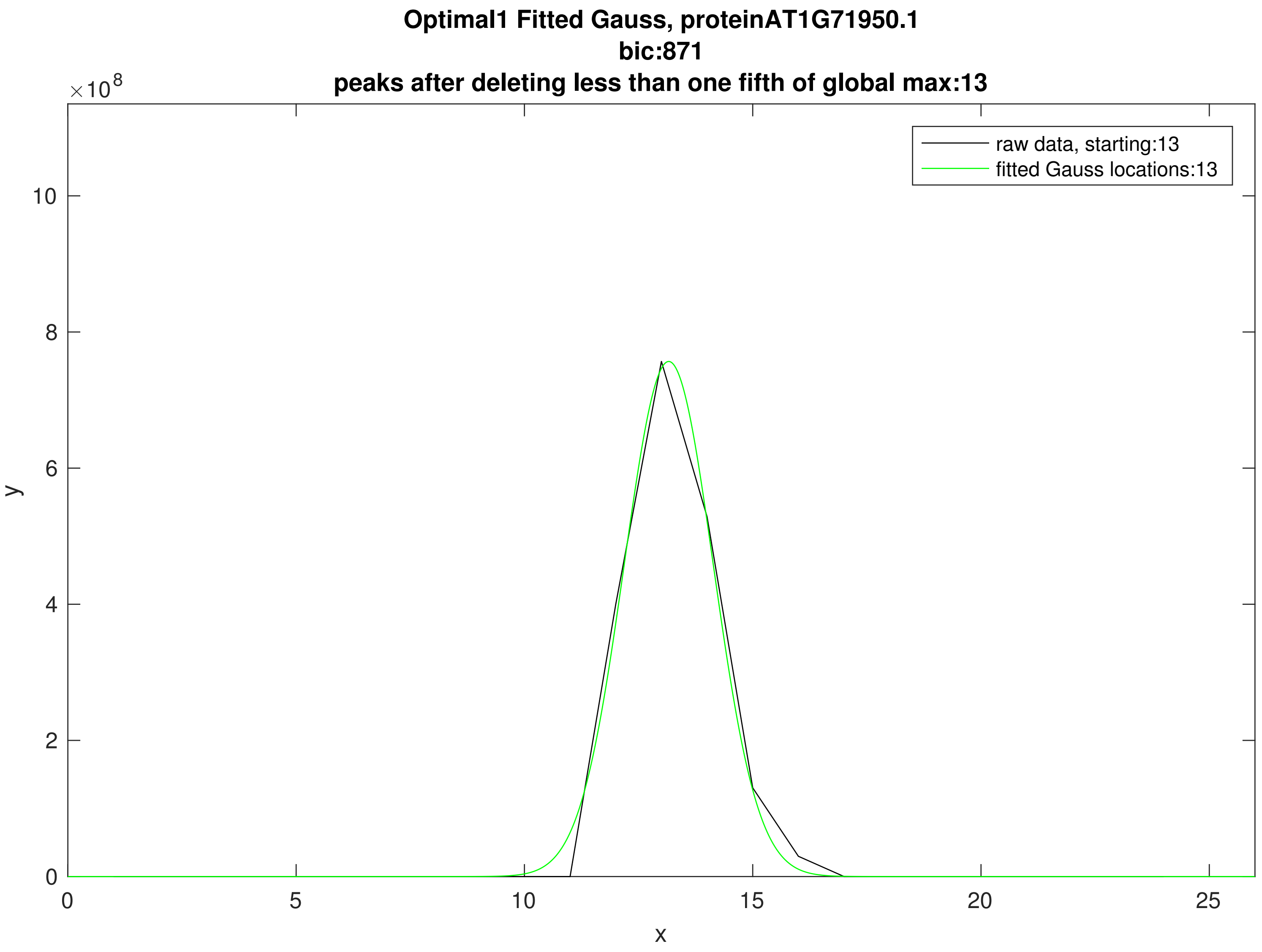

Optimal1 Fitted Gauss, proteinAT1G71950.1
bic:871
peaks after deleting less than one fifth of global max:13
×10^8
raw data, starting:13
fitted Gauss locations:13
y
x
0
2
4
6
8
10
0
5
10
15
20
25

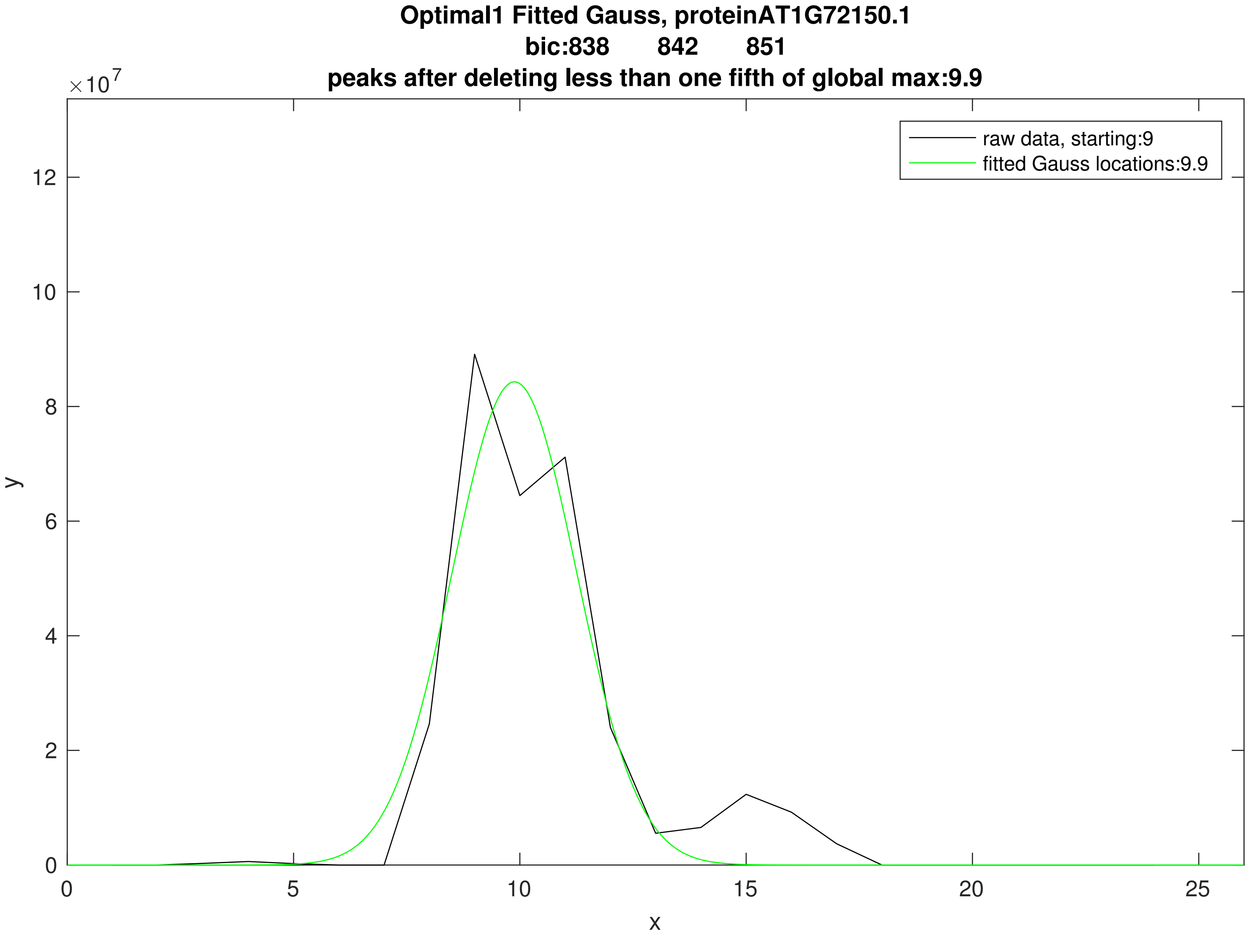

Optimal1 Fitted Gauss, proteinAT1G72150.1
bic:838 842 851
peaks after deleting less than one fifth of global max:9.9
×10^7
raw data, starting:9
fitted Gauss locations:9.9
y
x

Optimal1 Fitted Gauss, proteinAT1G72230.1
bic:813
peaks after deleting less than one fifth of global max:18

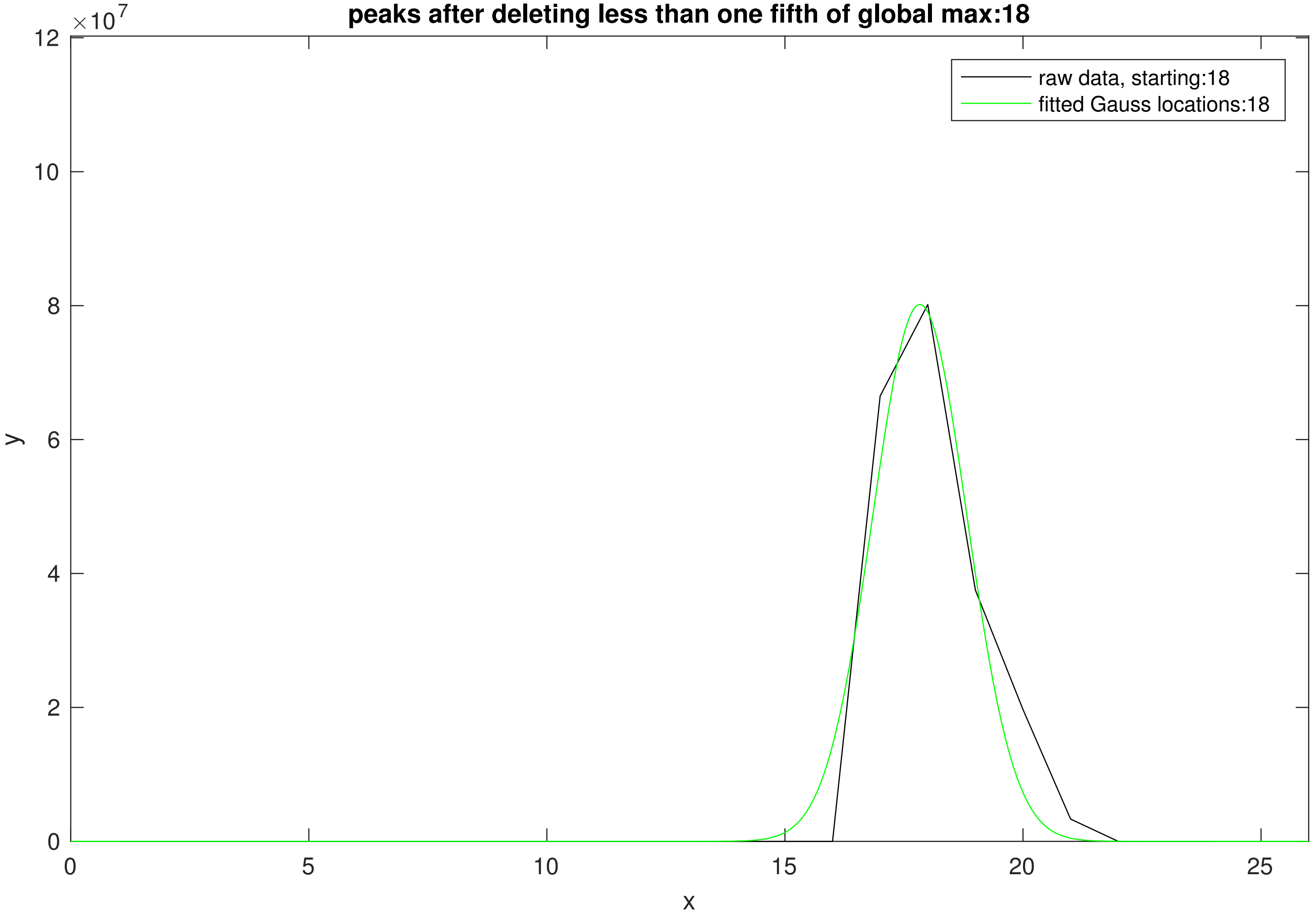

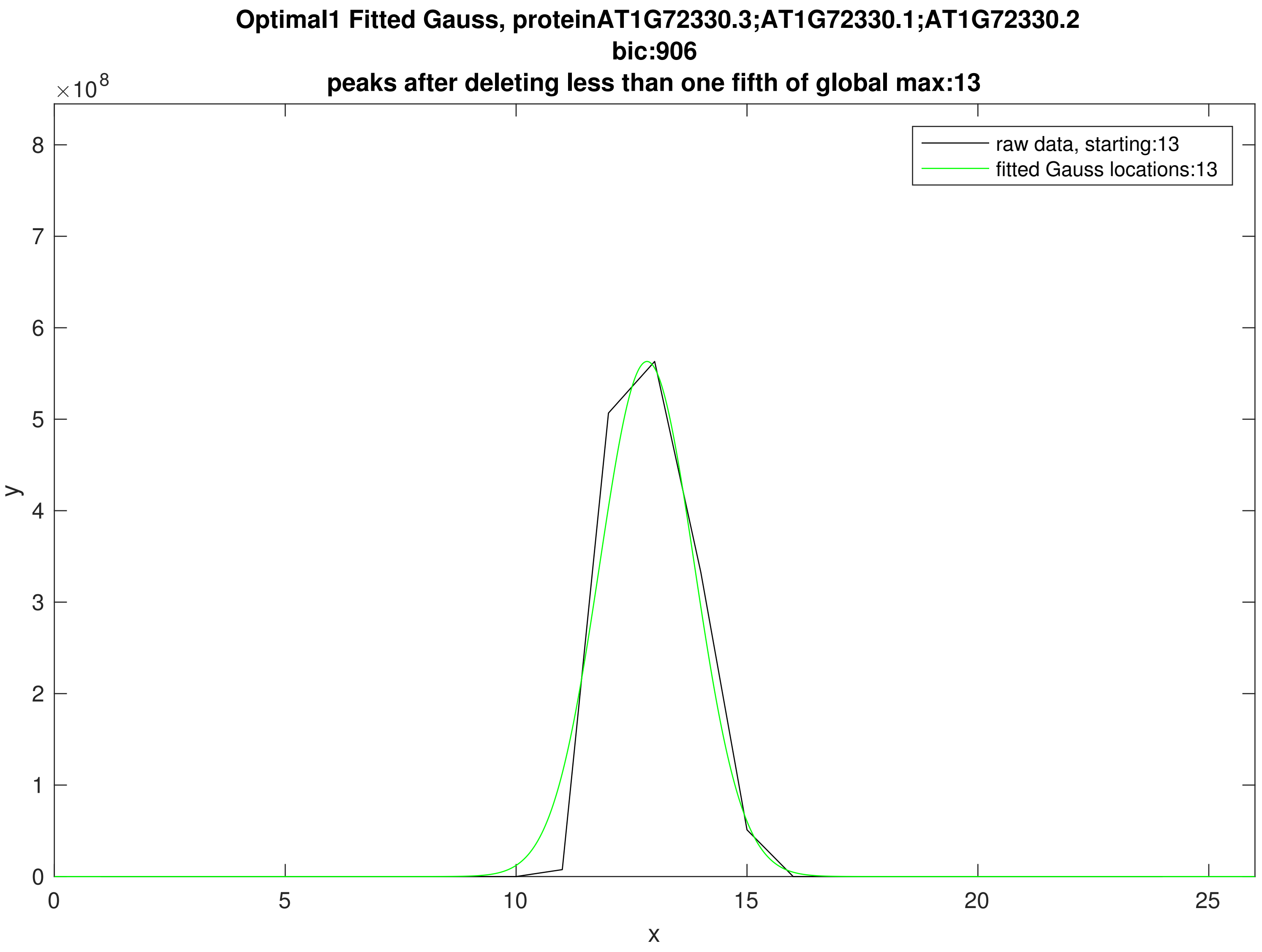

Optimal1 Fitted Gauss, proteinAT1G72330.3;AT1G72330.1;AT1G72330.2
bic:906
peaks after deleting less than one fifth of global max:13
×10^8
raw data, starting:13
fitted Gauss locations:13
y
x
0
1
2
3
4
5
6
7
8
0
5
10
15
20
25

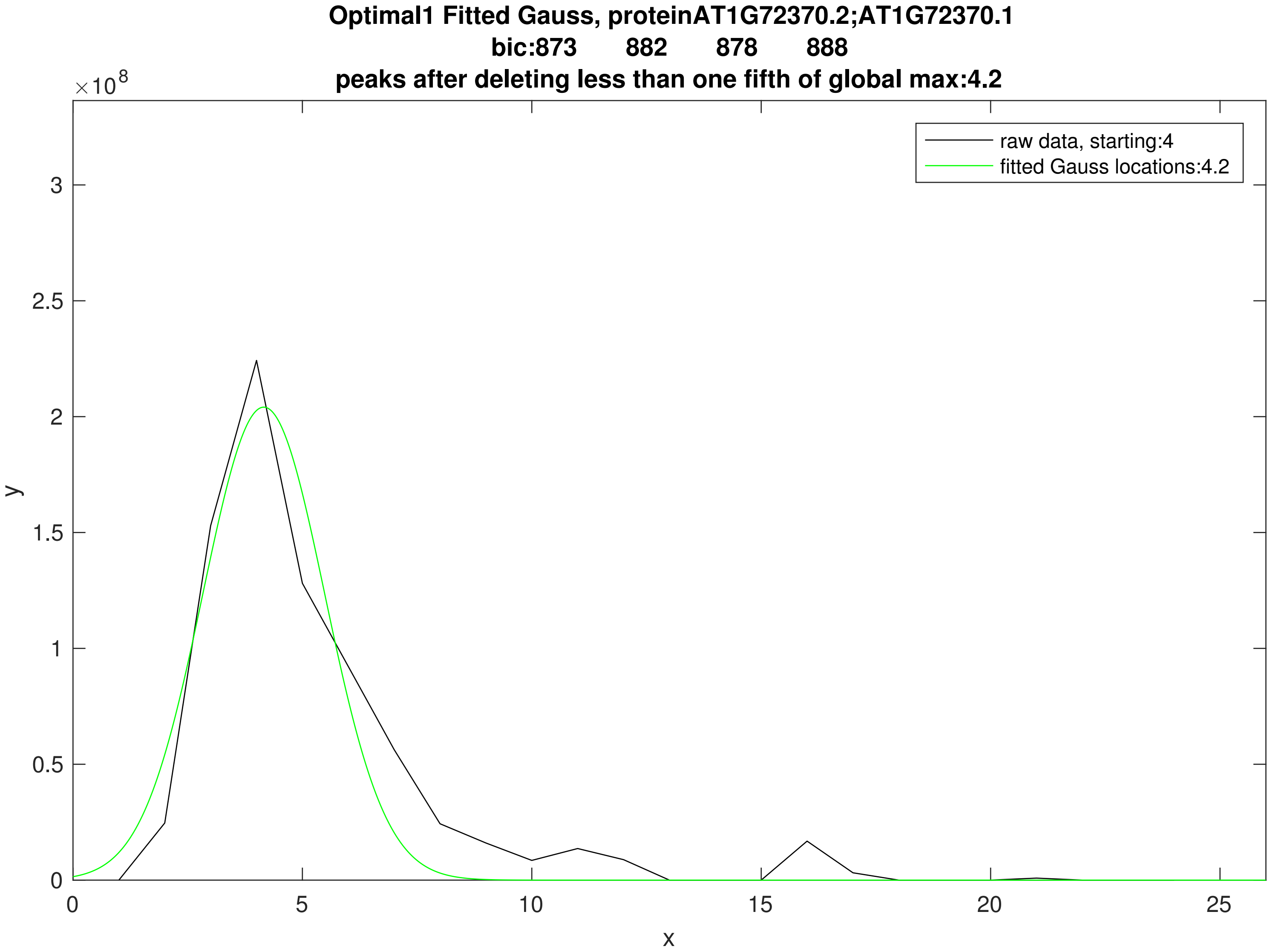

Optimal1 Fitted Gauss, proteinAT1G72370.2;AT1G72370.1
bic:873 882 878 888
peaks after deleting less than one fifth of global max:4.2
×10^8
raw data, starting:4
fitted Gauss locations:4.2
y
x
0
0.5
1
1.5
2
2.5
3
5
10
15
20
25

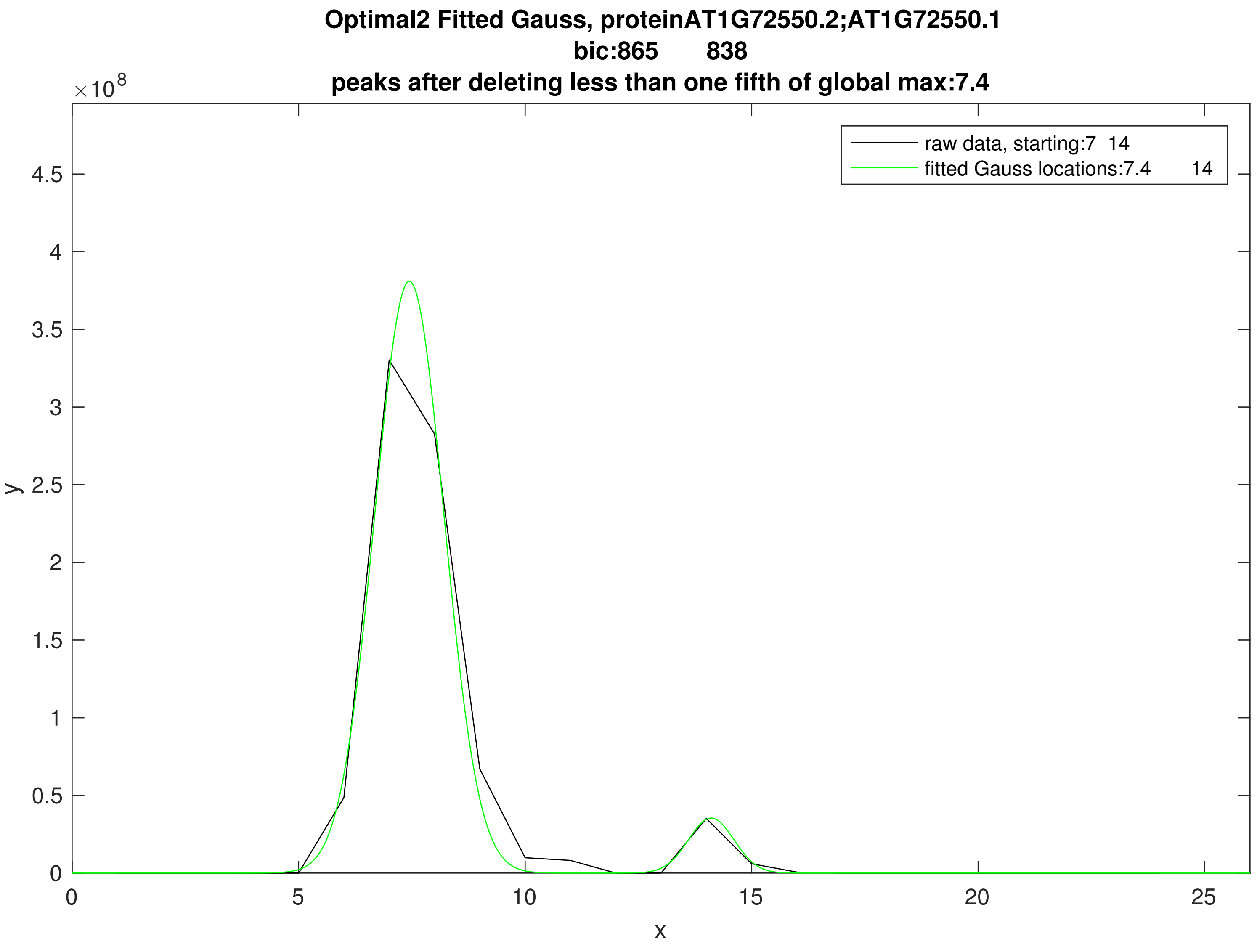

Optimal2 Fitted Gauss, proteinAT1G72550.2;AT1G72550.1
bic:865 838
peaks after deleting less than one fifth of global max:7.4
raw data, starting:7 14
fitted Gauss locations:7.4 14
×10^8
y
x

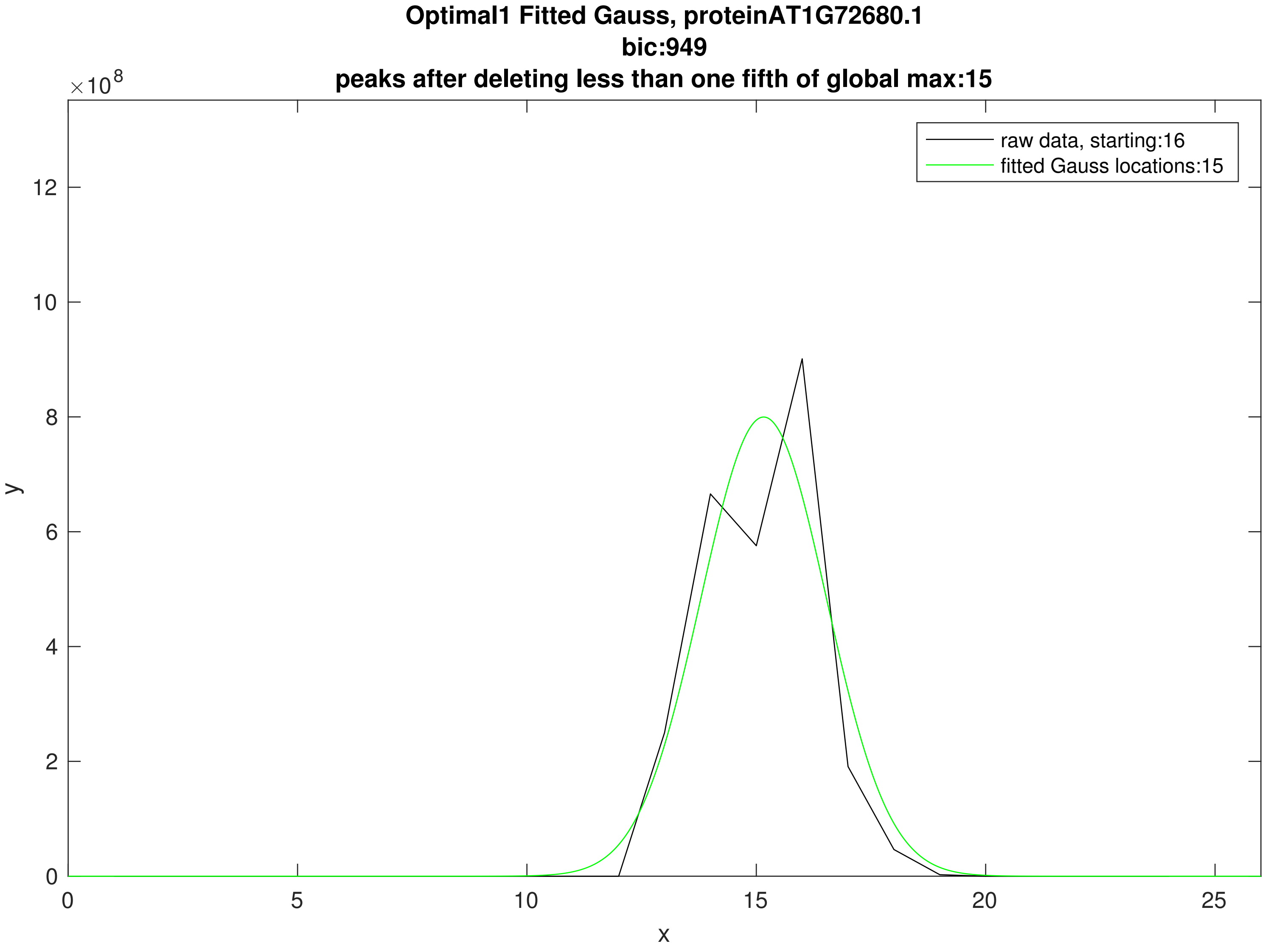

Optimal1 Fitted Gauss, proteinAT1G72680.1
bic:949
peaks after deleting less than one fifth of global max:15
×10^8
raw data, starting:16
fitted Gauss locations:15
y
x
0
2
4
6
8
10
12
0
5
10
15
20
25

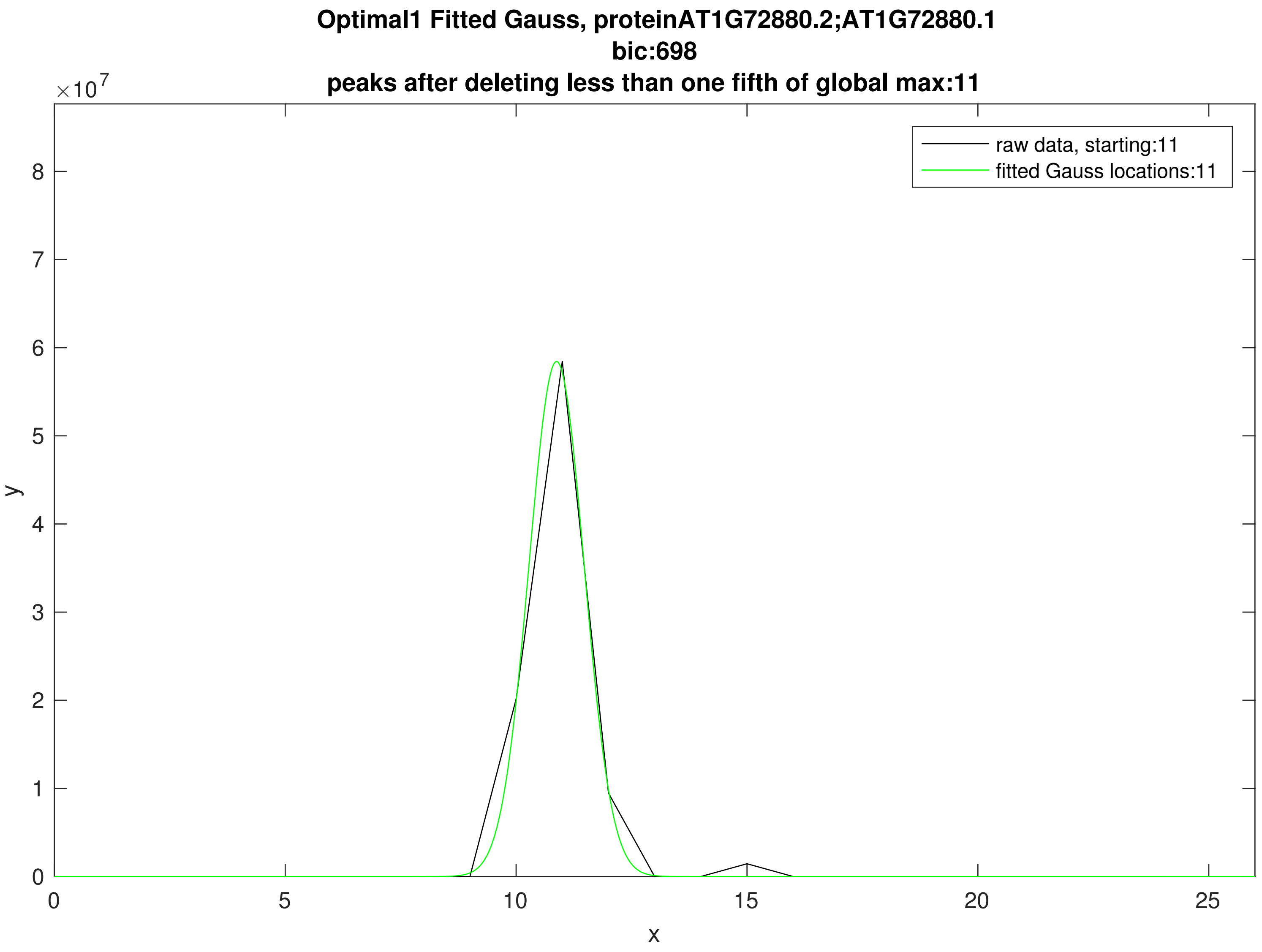
Optimal1 Fitted Gauss, proteinAT1G72880.2;AT1G72880.1
bic:698
peaks after deleting less than one fifth of global max:11
×10^7
raw data, starting:11
fitted Gauss locations:11
y
x
0
1
2
3
4
5
6
7
8
0
5
10
15
20
25

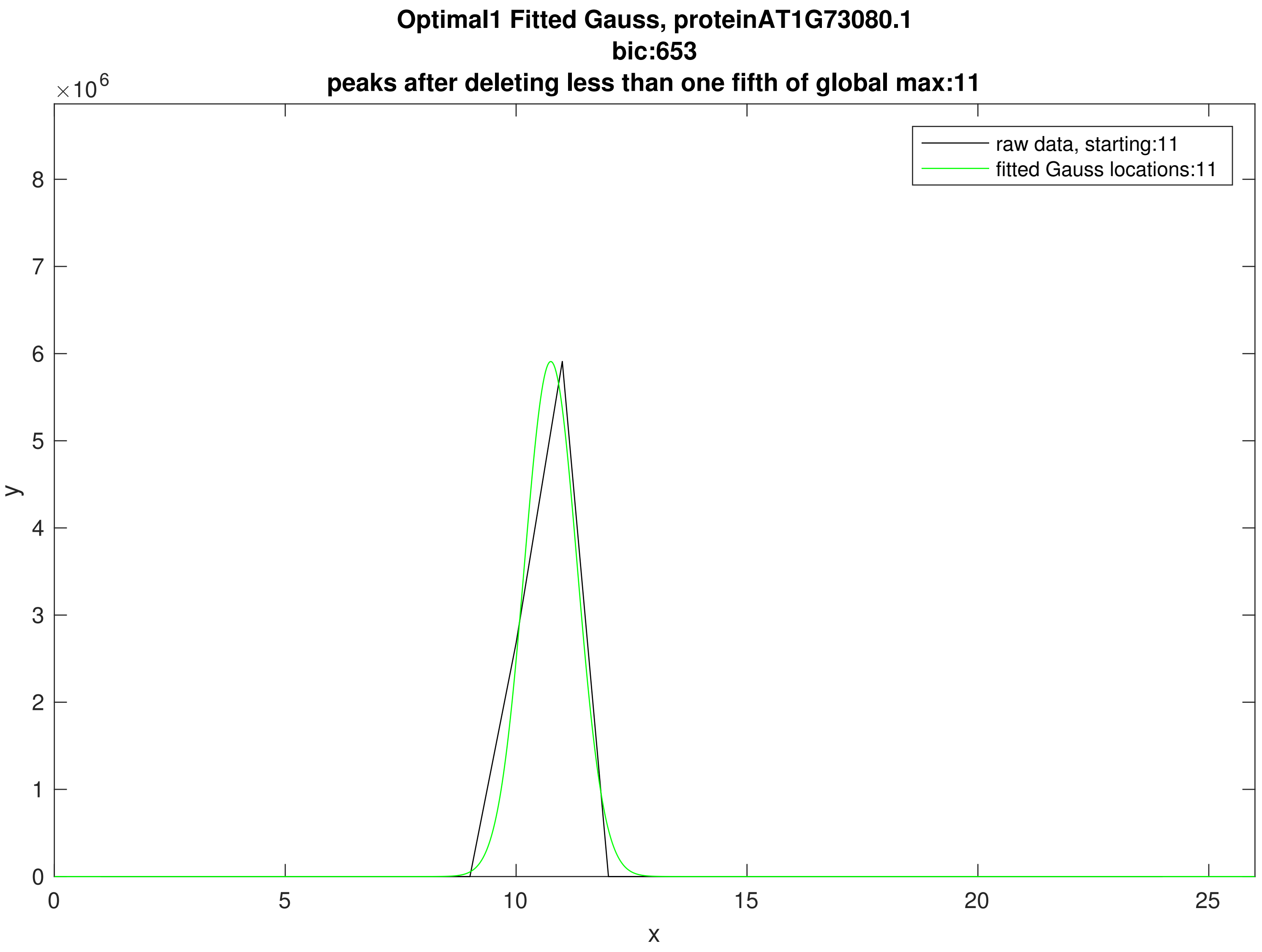
Optimal1 Fitted Gauss, proteinAT1G73080.1
bic:653
peaks after deleting less than one fifth of global max:11
×10^6
raw data, starting:11
fitted Gauss locations:11
y
x
0
1
2
3
4
5
6
7
8
0
5
10
15
20
25

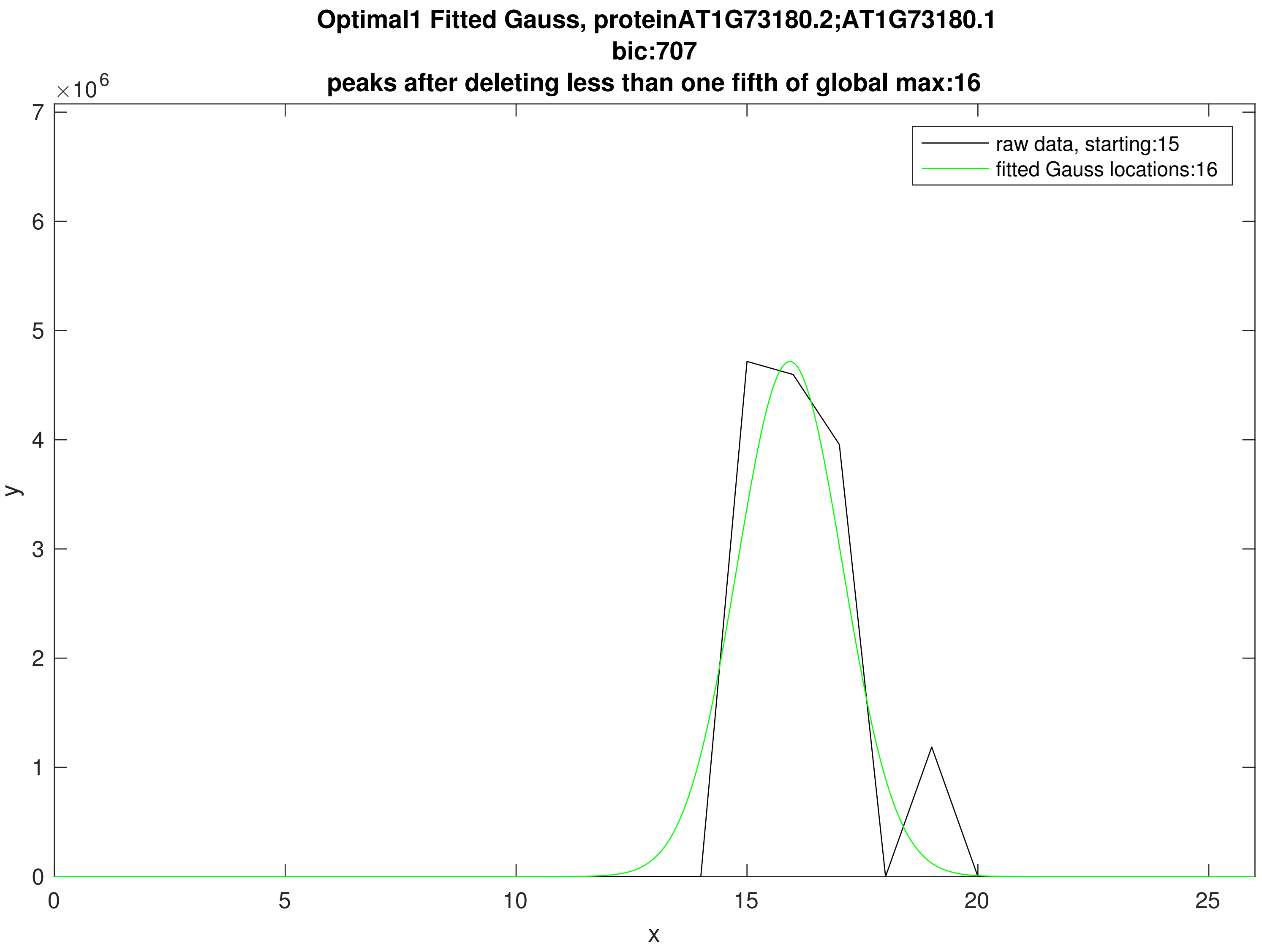

Optimal1 Fitted Gauss, proteinAT1G73180.2;AT1G73180.1
bic:707
peaks after deleting less than one fifth of global max:16
×10^6
raw data, starting:15
fitted Gauss locations:16
y
x
0
1
2
3
4
5
6
7
0
5
10
15
20
25

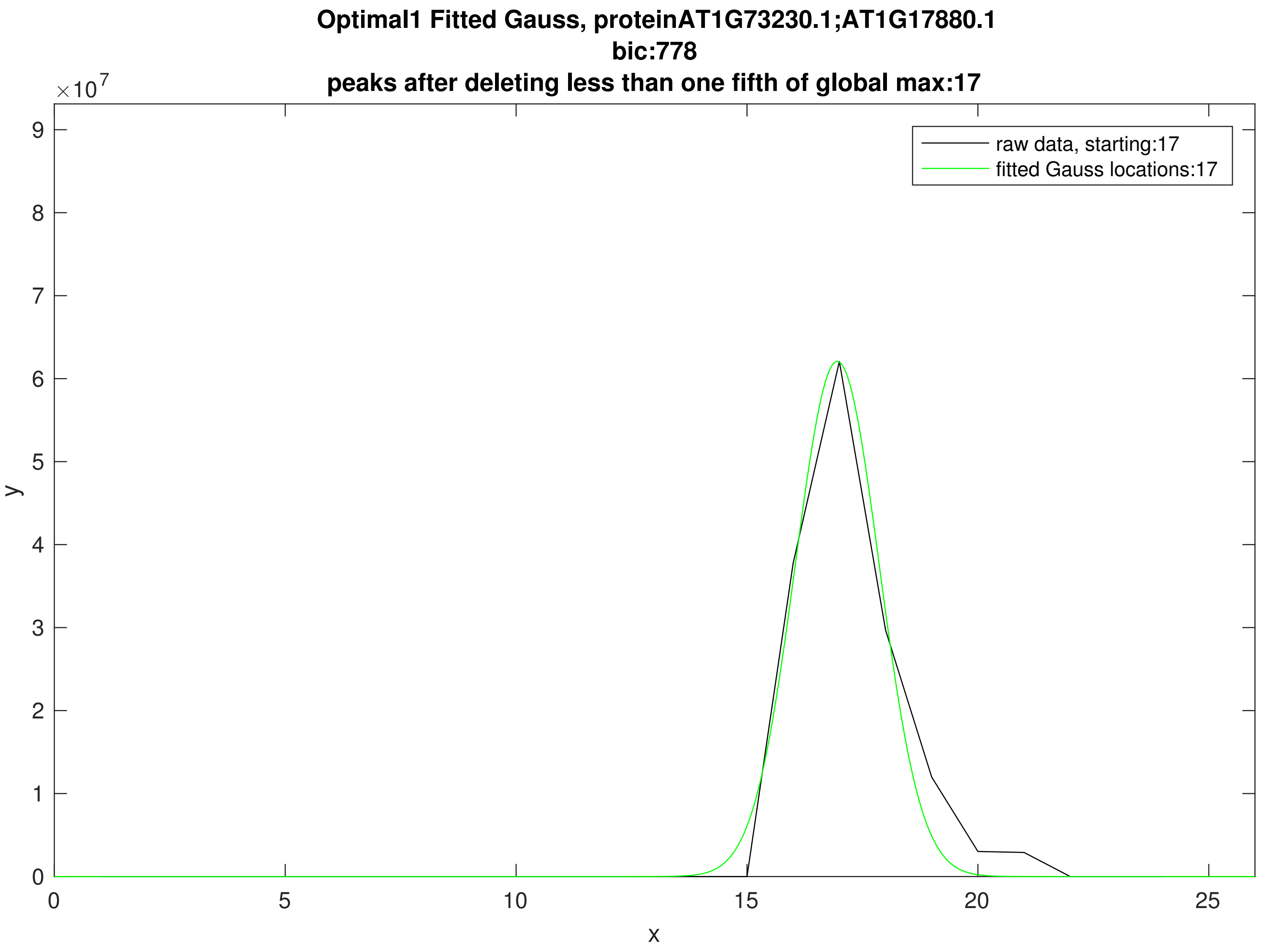
Optimal1 Fitted Gauss, proteinAT1G73230.1;AT1G17880.1
bic:778
peaks after deleting less than one fifth of global max:17
×10^7
raw data, starting:17
fitted Gauss locations:17
y
x
0
1
2
3
4
5
6
7
8
9
0
5
10
15
20
25

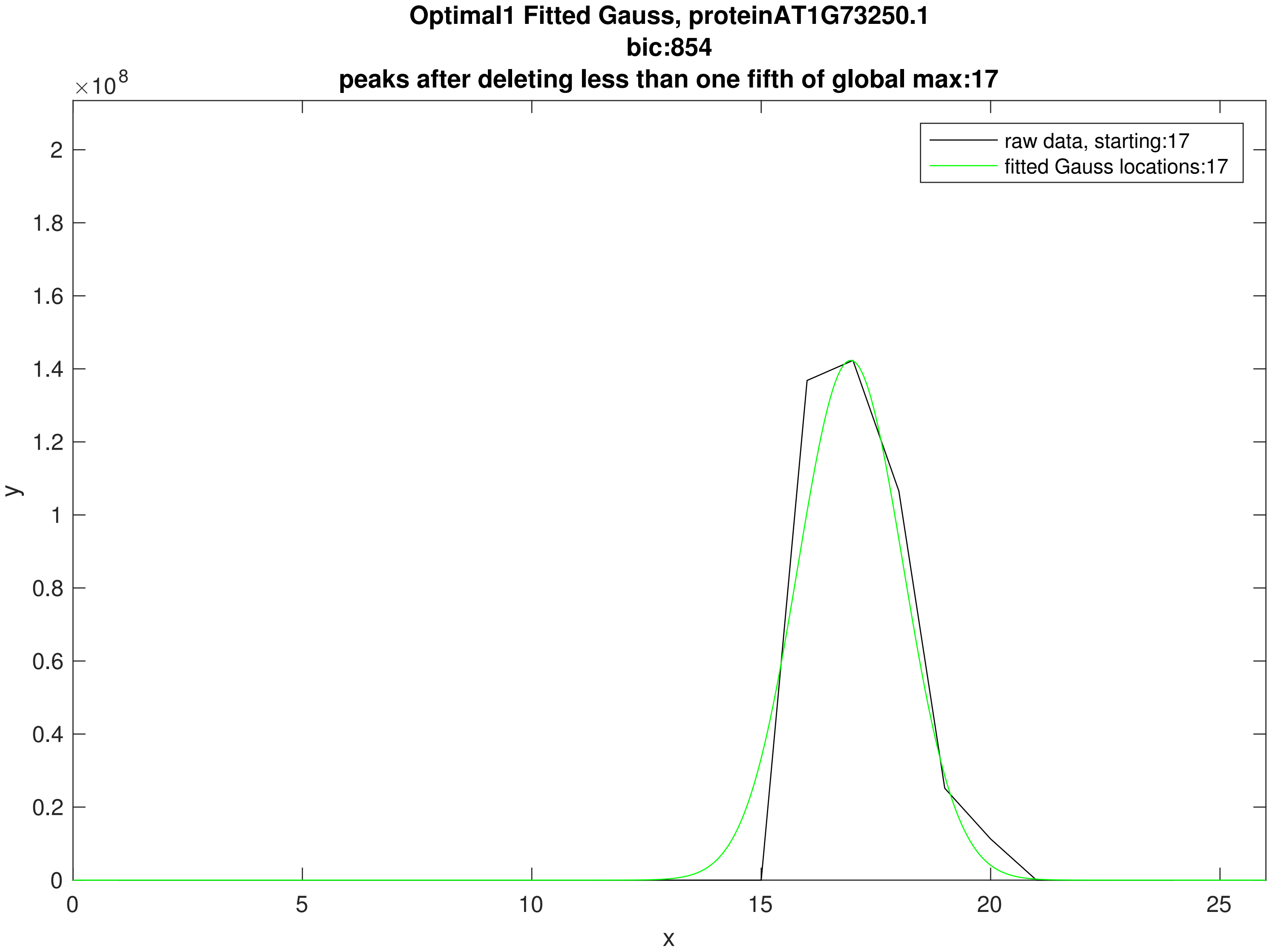

Optimal1 Fitted Gauss, proteinAT1G73250.1
bic:854
peaks after deleting less than one fifth of global max:17
×10^8
raw data, starting:17
fitted Gauss locations:17
y
x
0
0.2
0.4
0.6
0.8
1
1.2
1.4
1.6
1.8
2
0
5
10
15
20
25

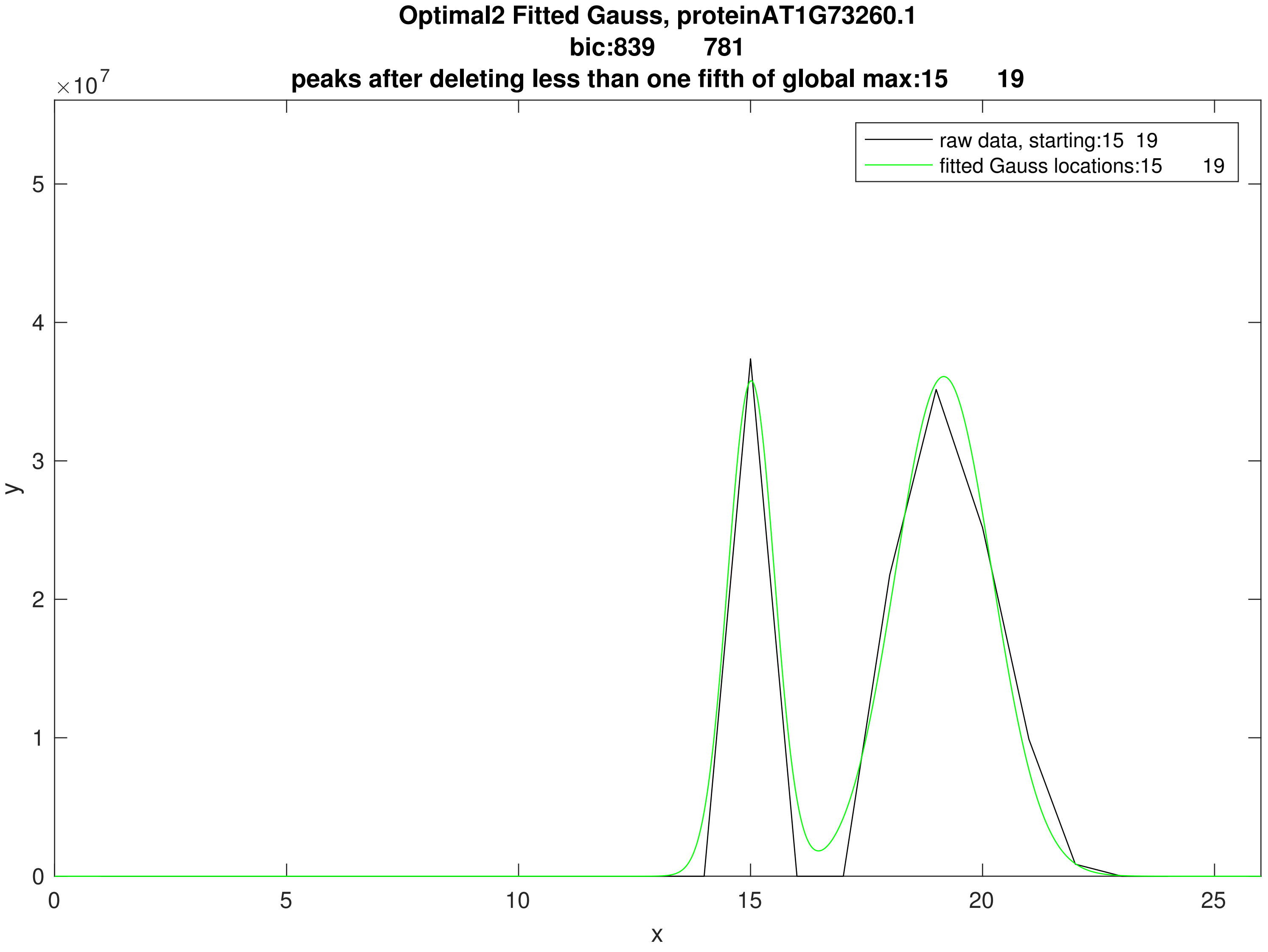
Optimal2 Fitted Gauss, proteinAT1G73260.1
bic:839 781
peaks after deleting less than one fifth of global max:15 19
×10^7
raw data, starting:15 19
fitted Gauss locations:15 19
y
x
0
1
2
3
4
5
0
5
10
15
20
25

#### Optimal1 Fitted Gauss, proteinAT1G73330.1
#### bic:647
#### peaks after deleting less than one fifth of global max:20

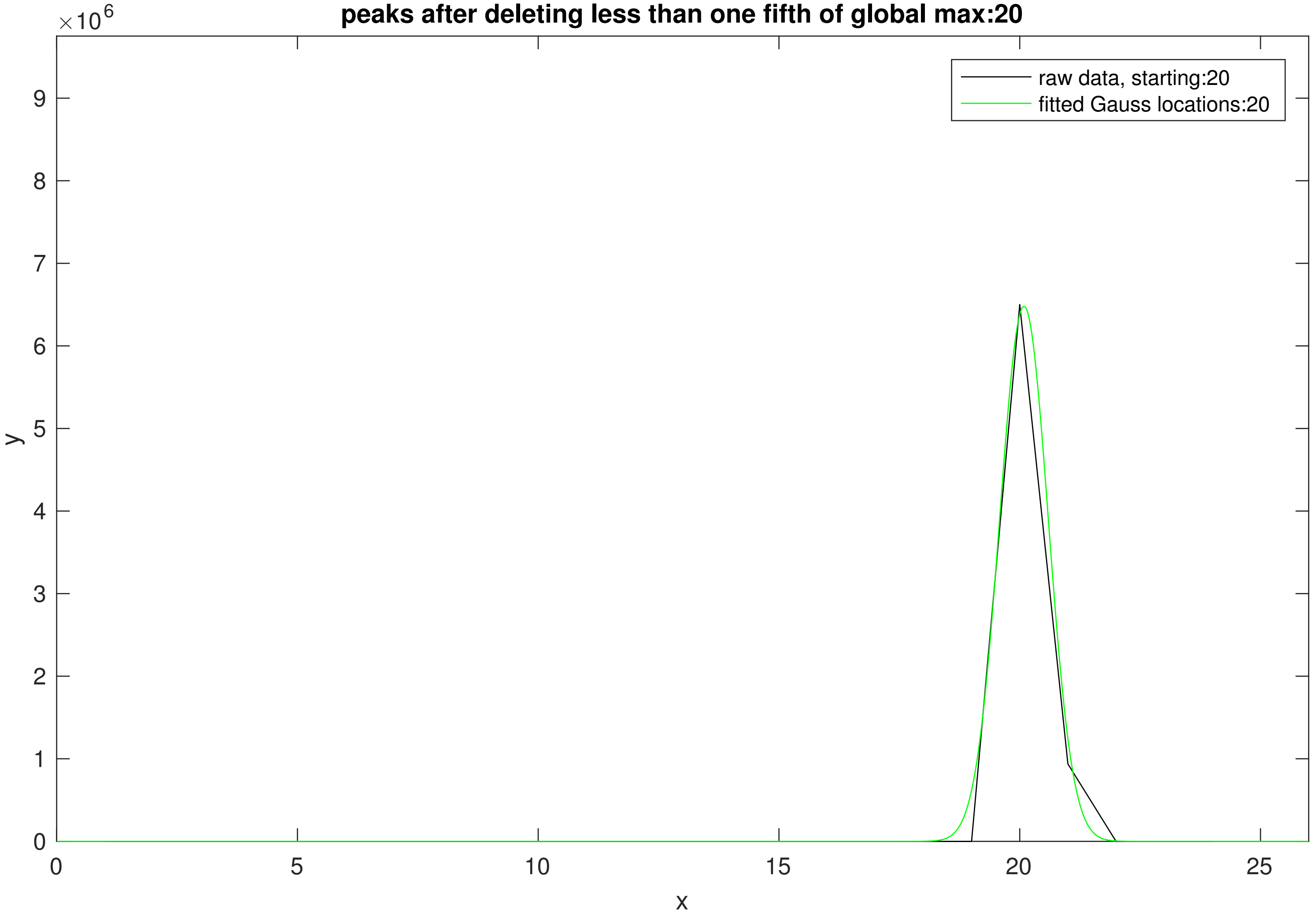

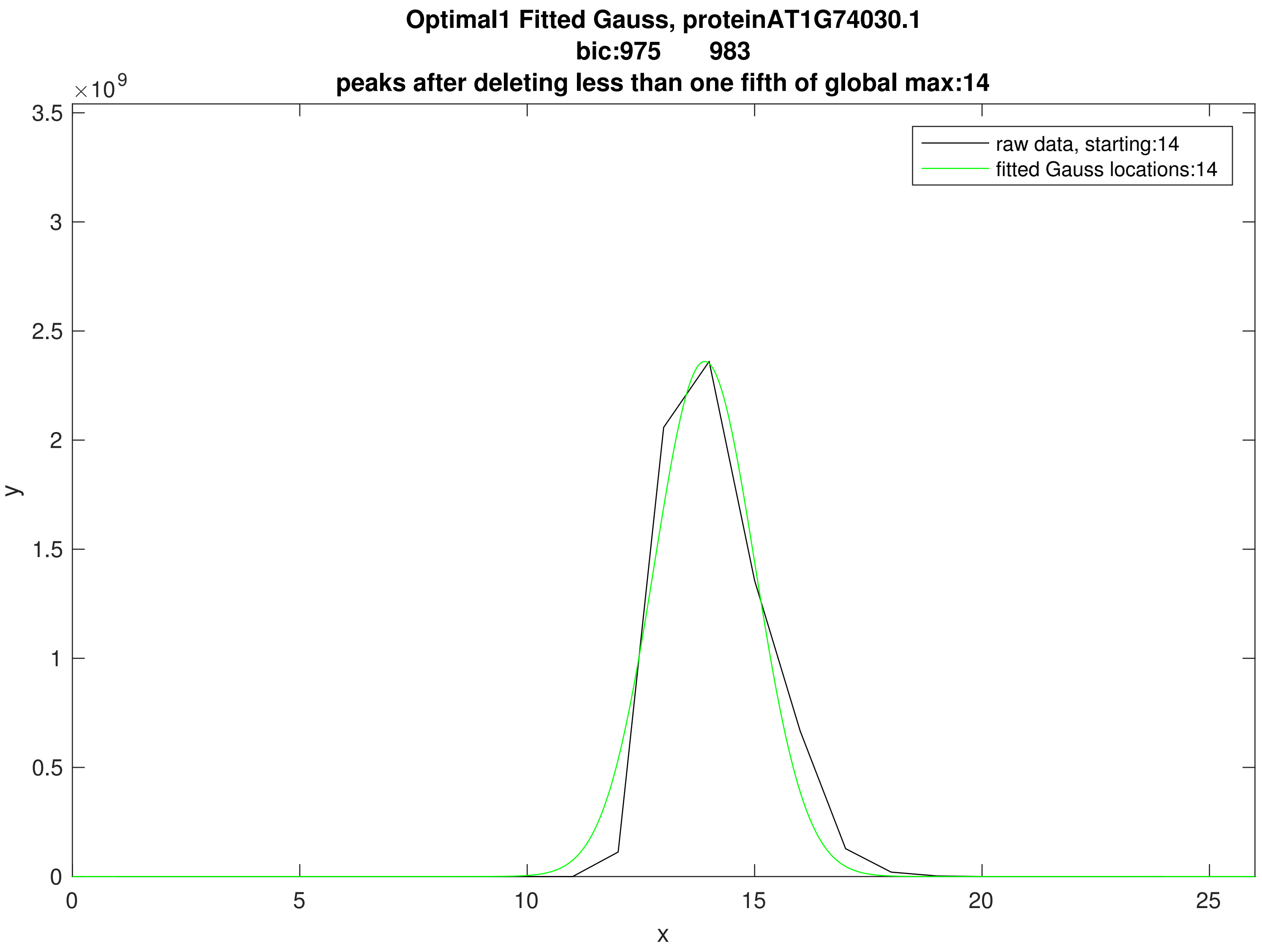
Optimal1 Fitted Gauss, proteinAT1G74030.1
bic:975 983
peaks after deleting less than one fifth of global max:14
×10^9
3.5
3
2.5
2
1.5
1
0.5
0
y
0
5
10
15
20
25
x
raw data, starting:14
fitted Gauss locations:14

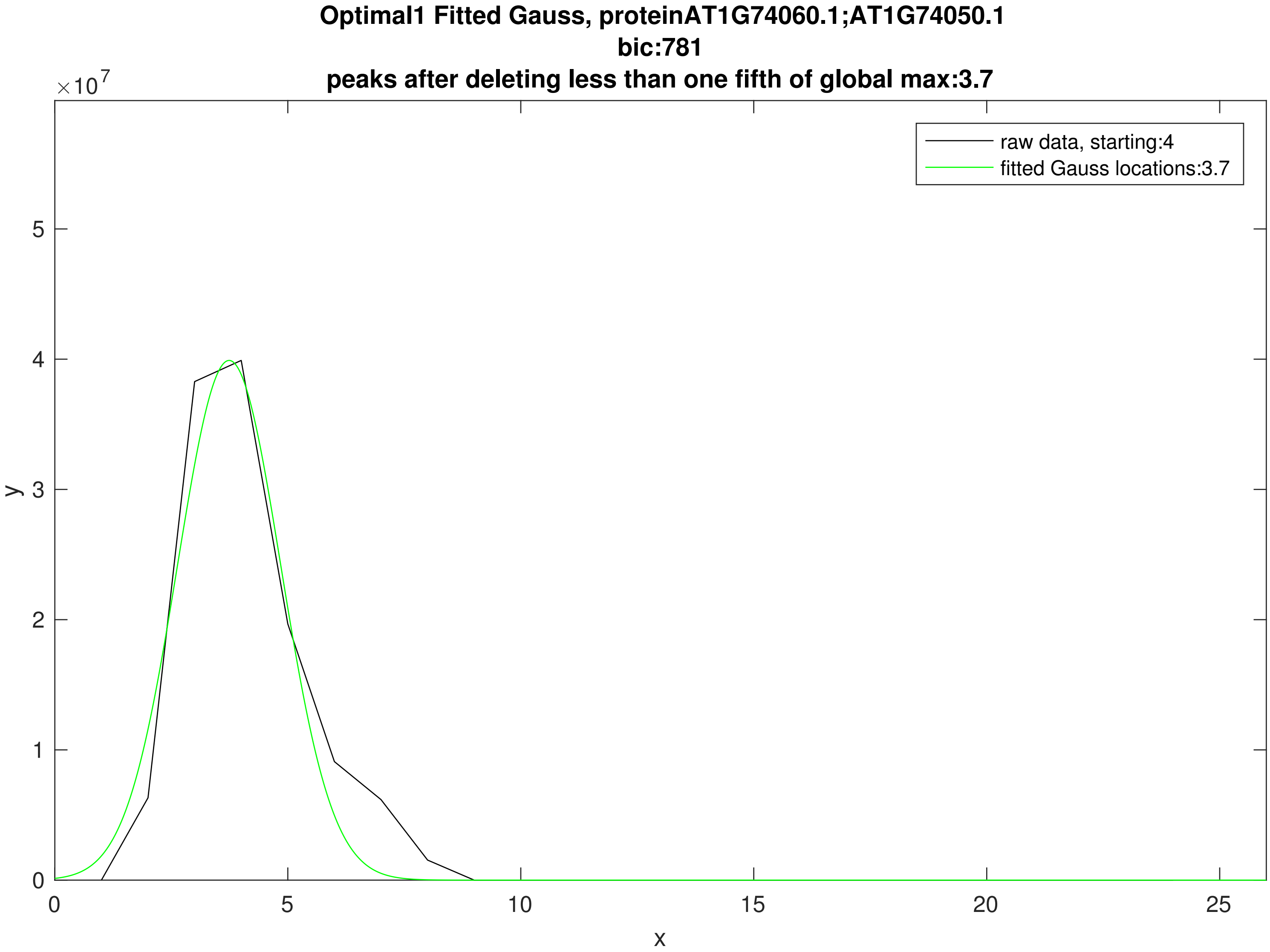
Optimal1 Fitted Gauss, proteinAT1G74060.1;AT1G74050.1
bic:781
peaks after deleting less than one fifth of global max:3.7
×10^7
raw data, starting:4
fitted Gauss locations:3.7
y
0
1
2
3
4
5
0
5
10
15
20
25
x

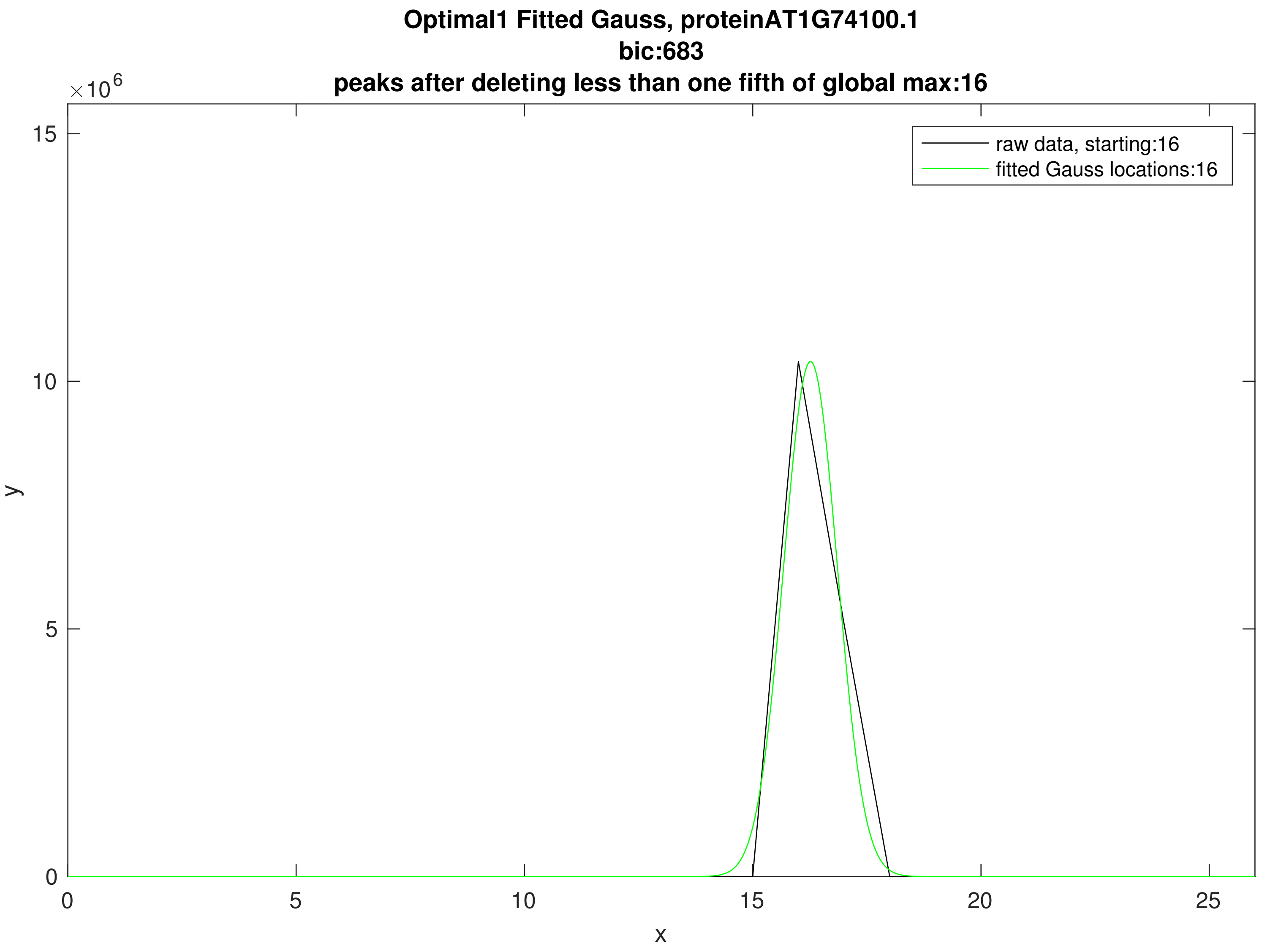

Optimal1 Fitted Gauss, proteinAT1G74100.1
bic:683
peaks after deleting less than one fifth of global max:16
×10^6
15
10
5
0
y
0
5
10
15
20
25
x
raw data, starting:16
fitted Gauss locations:16

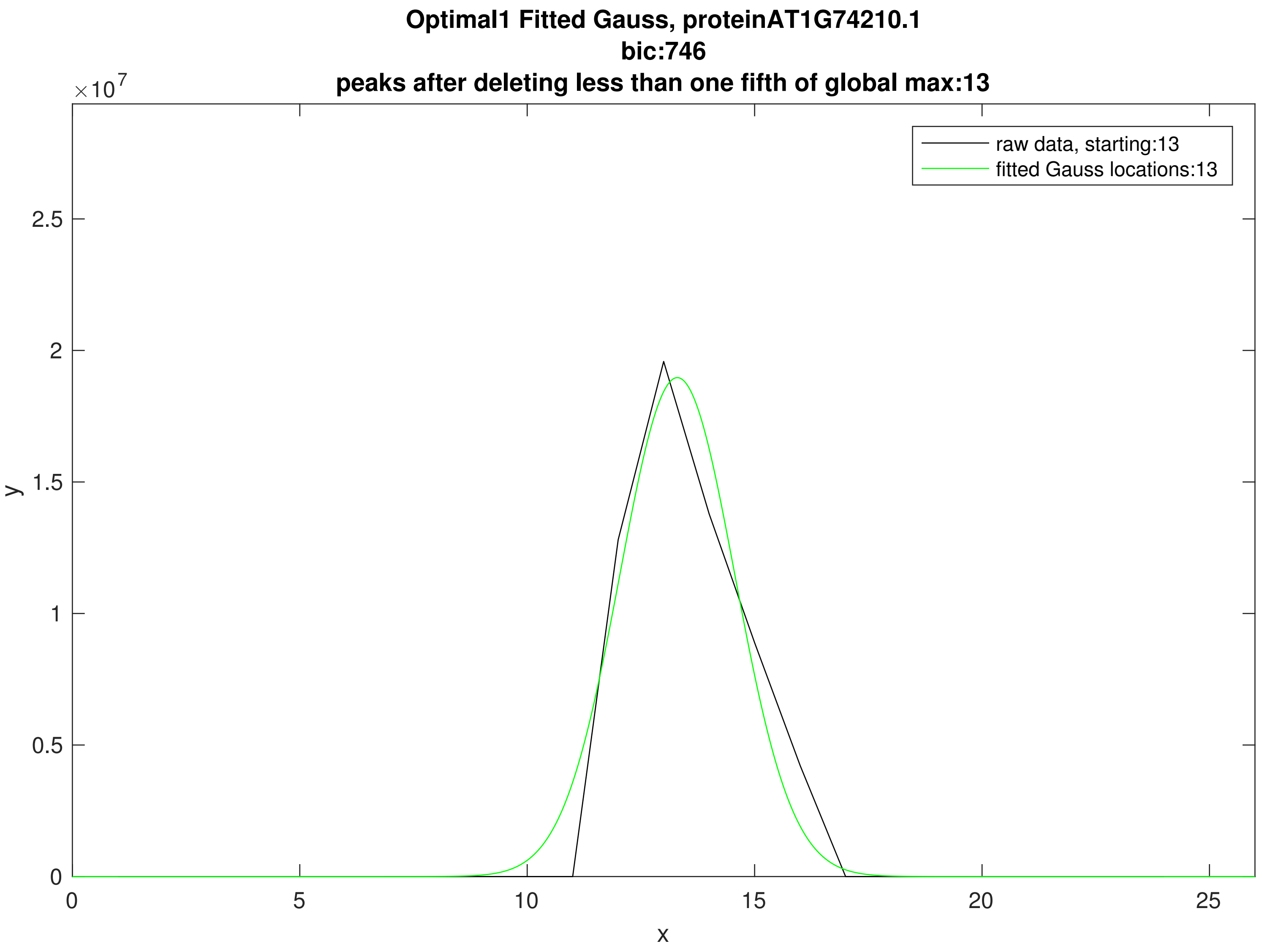

Optimal1 Fitted Gauss, proteinAT1G74210.1
bic:746
peaks after deleting less than one fifth of global max:13
×10^7
raw data, starting:13
fitted Gauss locations:13
2.5
2
1.5
1
0.5
0
y
0
5
10
15
20
25
x

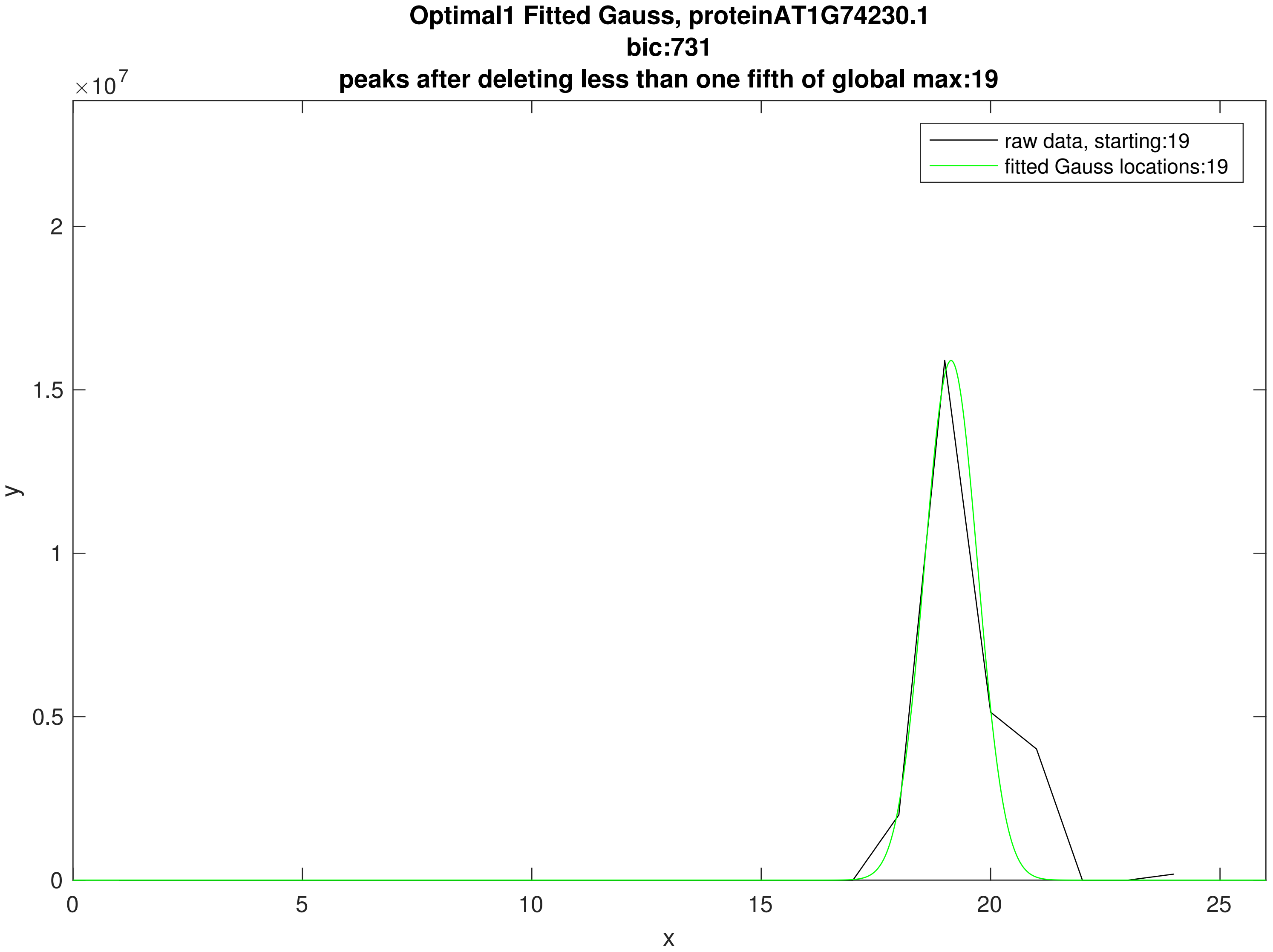
Optimal1 Fitted Gauss, proteinAT1G74230.1
bic:731
peaks after deleting less than one fifth of global max:19
×10^7
raw data, starting:19
fitted Gauss locations:19
y
x
0
0.5
1
1.5
2
0
5
10
15
20
25

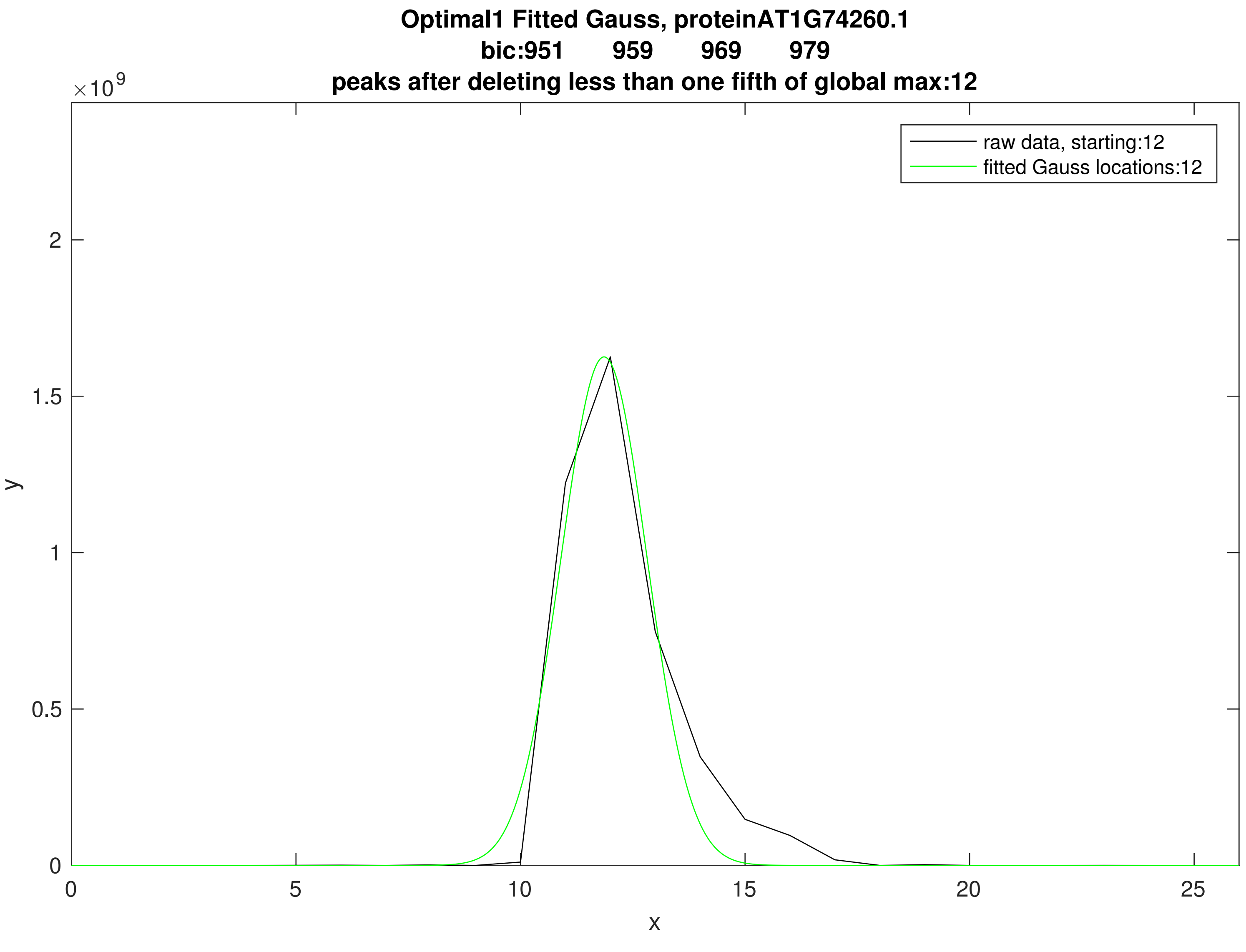

Optimal1 Fitted Gauss, proteinAT1G74260.1
bic:951 959 969 979
peaks after deleting less than one fifth of global max:12
×10^9
raw data, starting:12
fitted Gauss locations:12
y
x
0
0.5
1
1.5
2
0
5
10
15
20
25

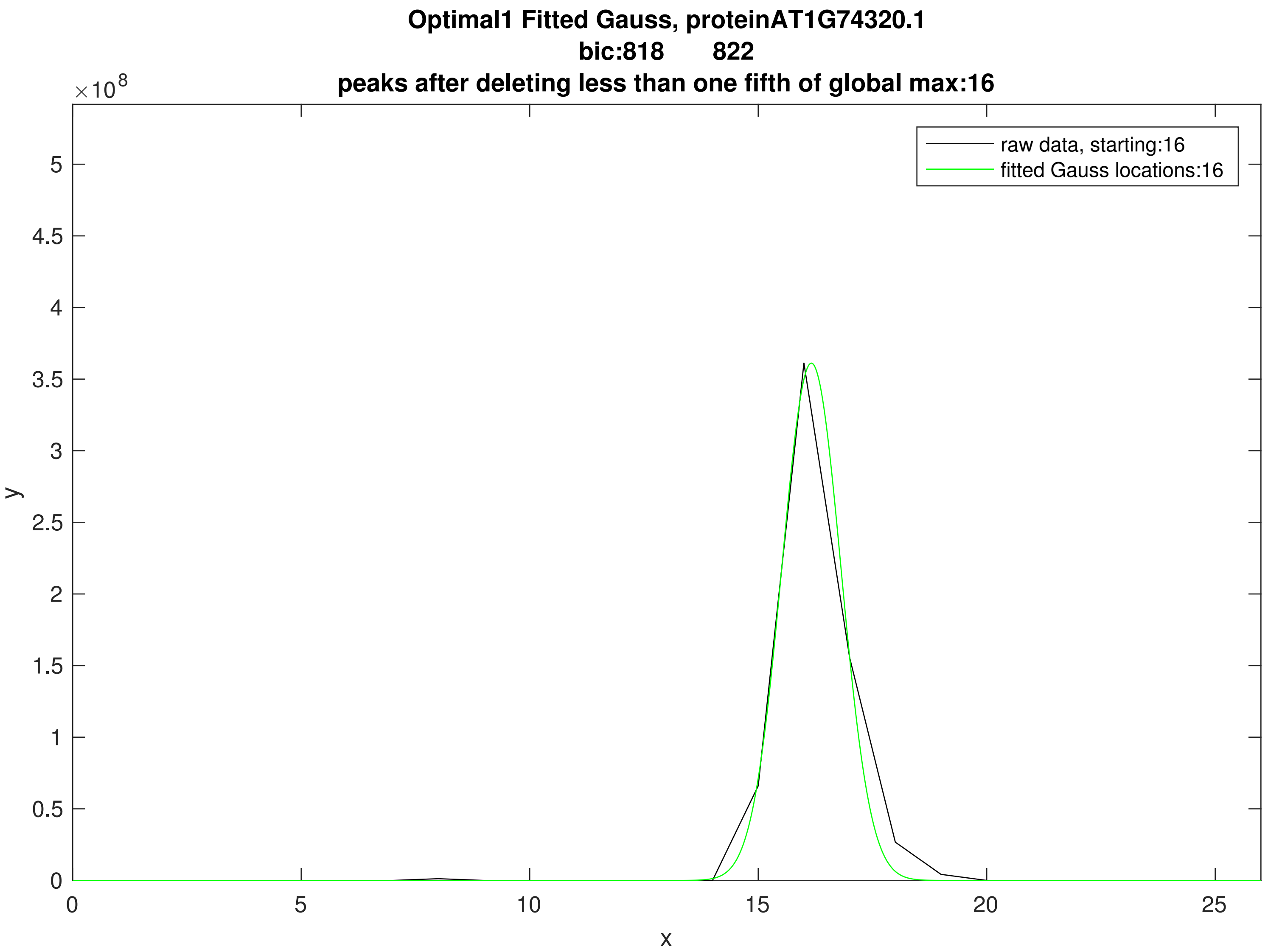

Optimal1 Fitted Gauss, proteinAT1G74320.1
bic:818 822
peaks after deleting less than one fifth of global max:16
×10^8
raw data, starting:16
fitted Gauss locations:16
y
x

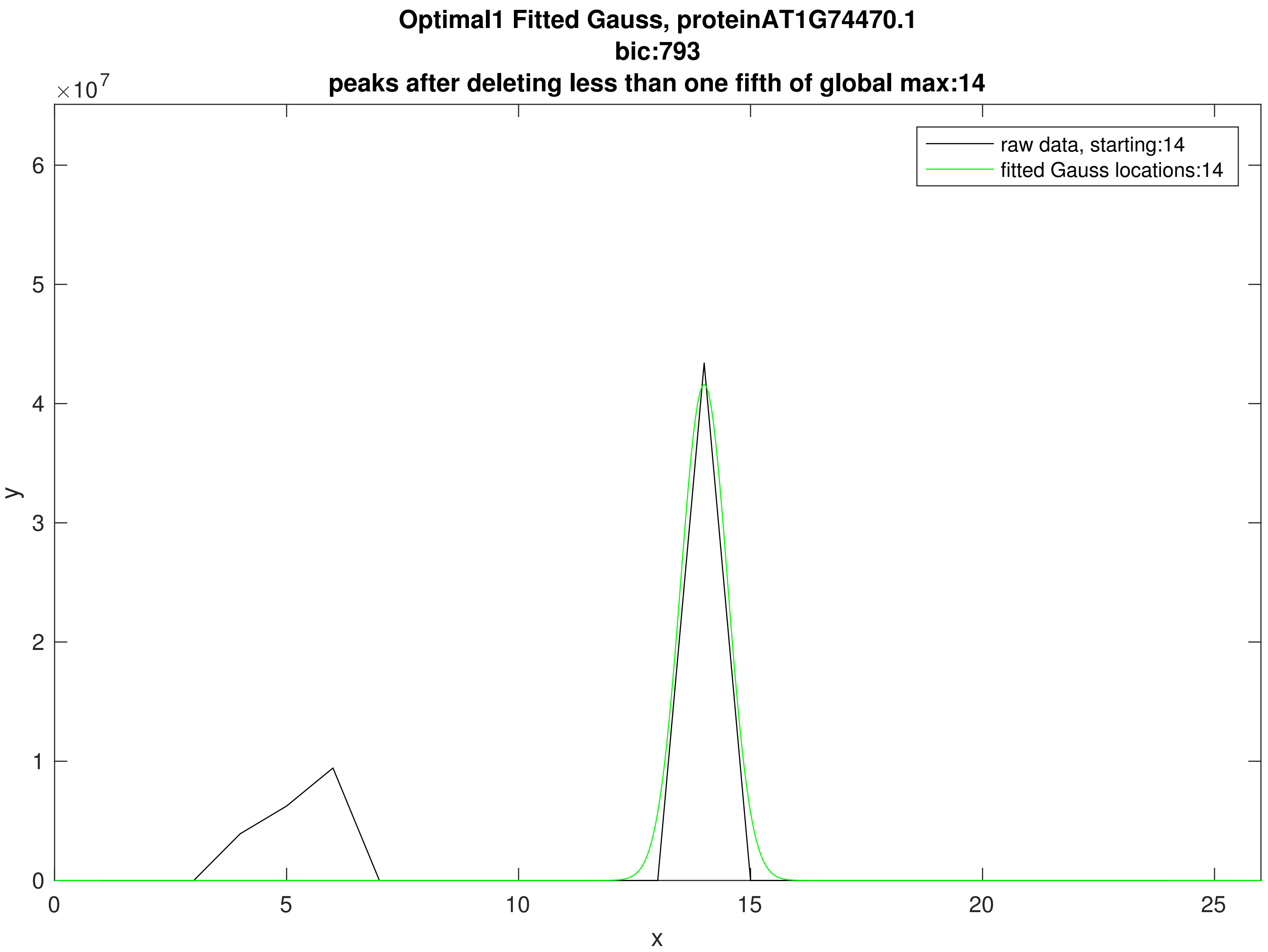
Optimal1 Fitted Gauss, proteinAT1G74470.1
bic:793
peaks after deleting less than one fifth of global max:14
×10^7
raw data, starting:14
fitted Gauss locations:14
y
x
0
1
2
3
4
5
6
0
5
10
15
20
25

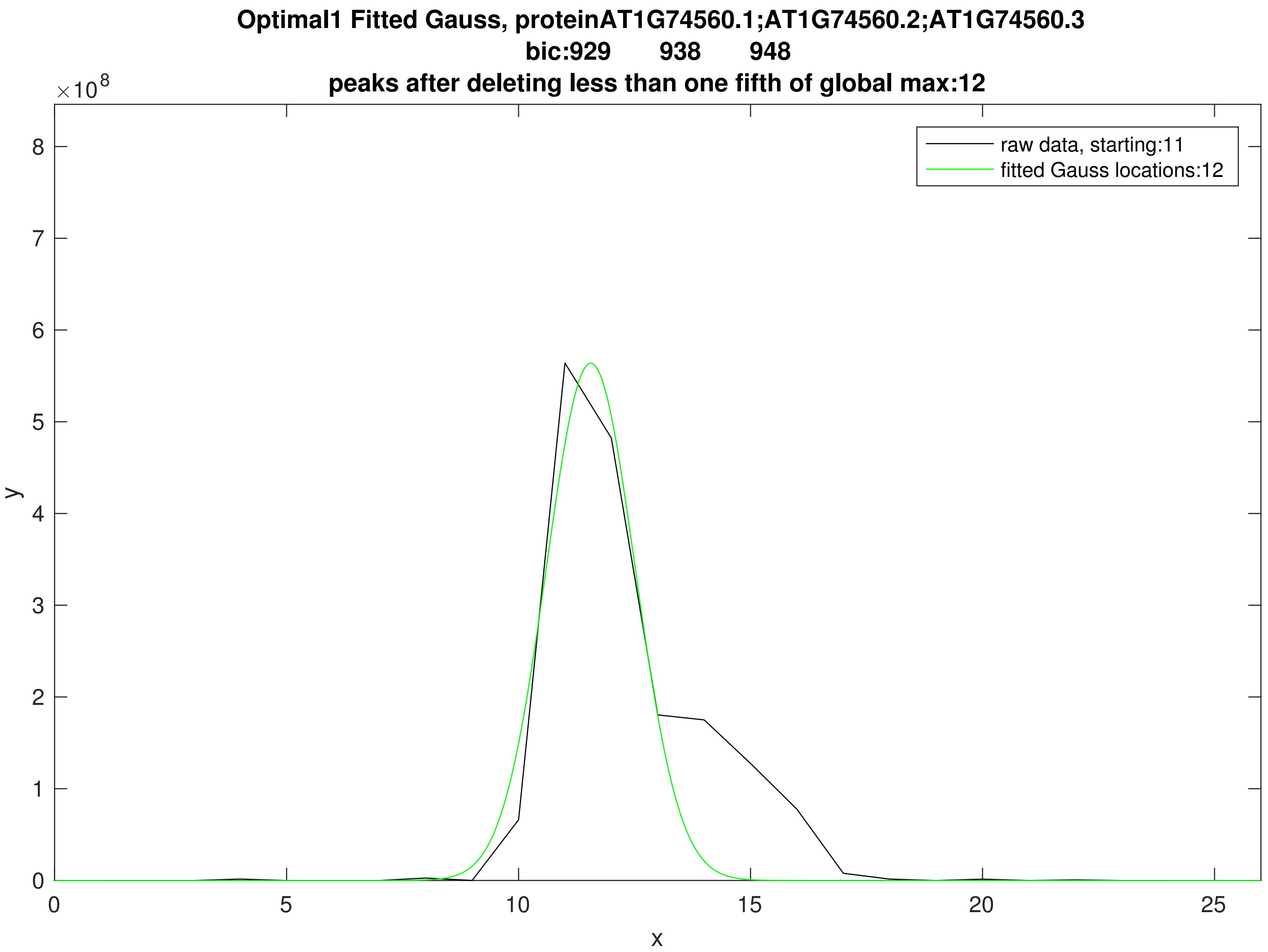
Optimal1 Fitted Gauss, proteinAT1G74560.1;AT1G74560.2;AT1G74560.3
bic:929 938 948
peaks after deleting less than one fifth of global max:12
×10^8
raw data, starting:11
fitted Gauss locations:12
y
x
0
1
2
3
4
5
6
7
8
0
5
10
15
20
25

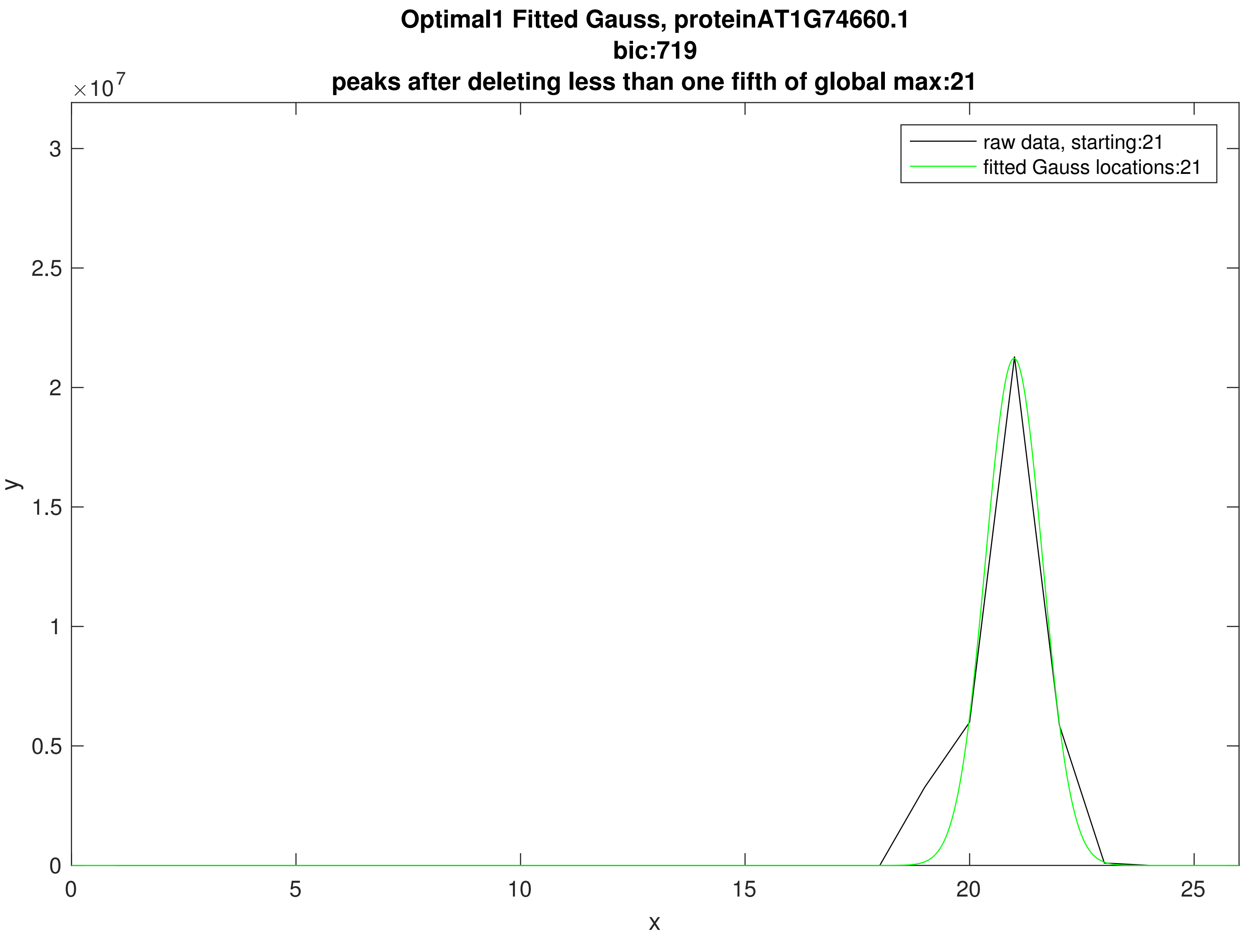
Optimal1 Fitted Gauss, proteinAT1G74660.1
bic:719
peaks after deleting less than one fifth of global max:21
×10^7
raw data, starting:21
fitted Gauss locations:21
y
x
0
0.5
1
1.5
2
2.5
3
0
5
10
15
20
25

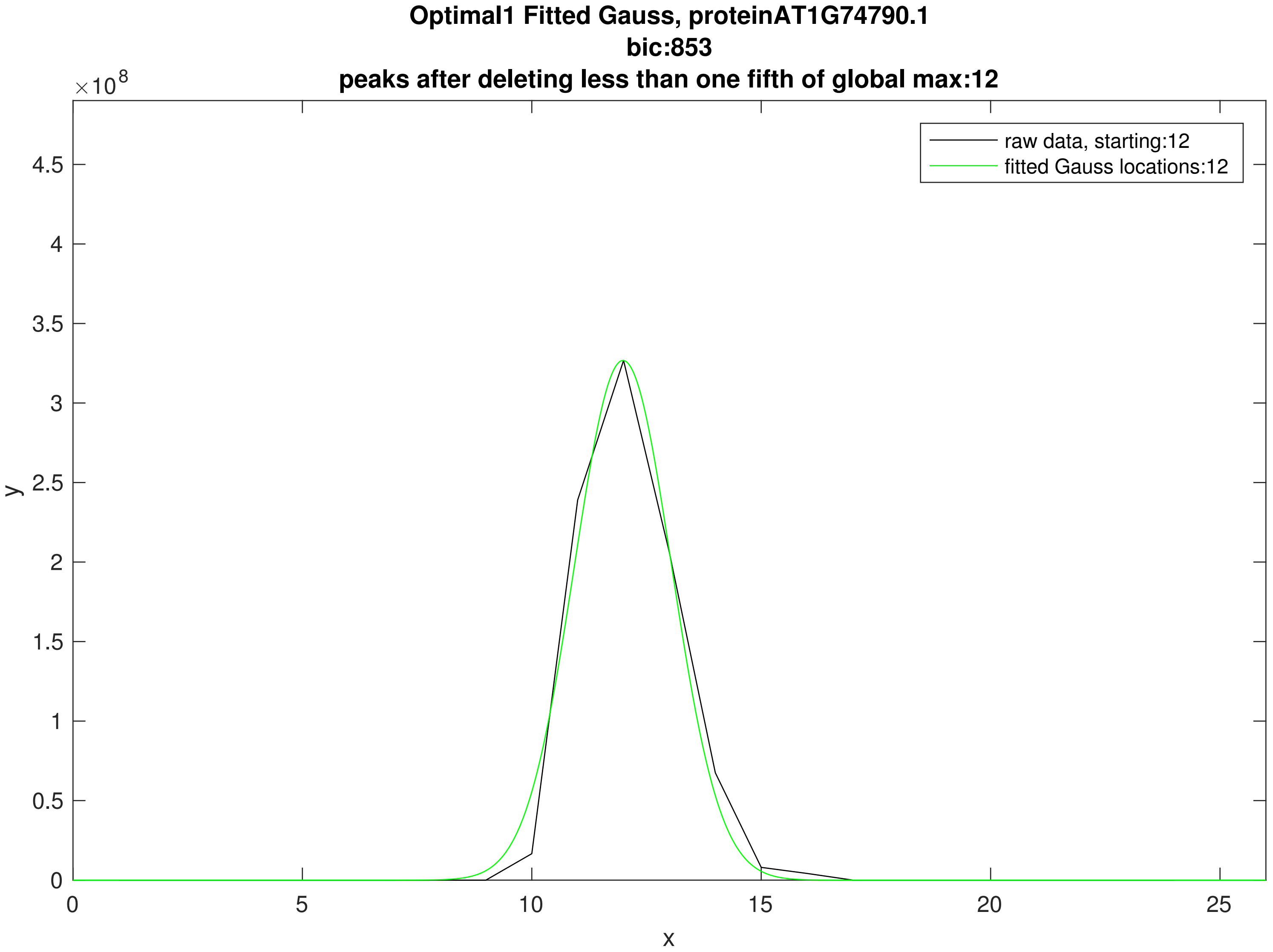
Optimal1 Fitted Gauss, proteinAT1G74790.1
bic:853
peaks after deleting less than one fifth of global max:12
×10^8
4.5
4
3.5
3
2.5
2
1.5
1
0.5
0
y
0
5
10
15
20
25
x
raw data, starting:12
fitted Gauss locations:12

Optimal1 Fitted Gauss, proteinAT1G74910.1;AT1G74910.2;AT1G74910.3
bic:954  1.01e+03
peaks after deleting less than one fifth of global max:14

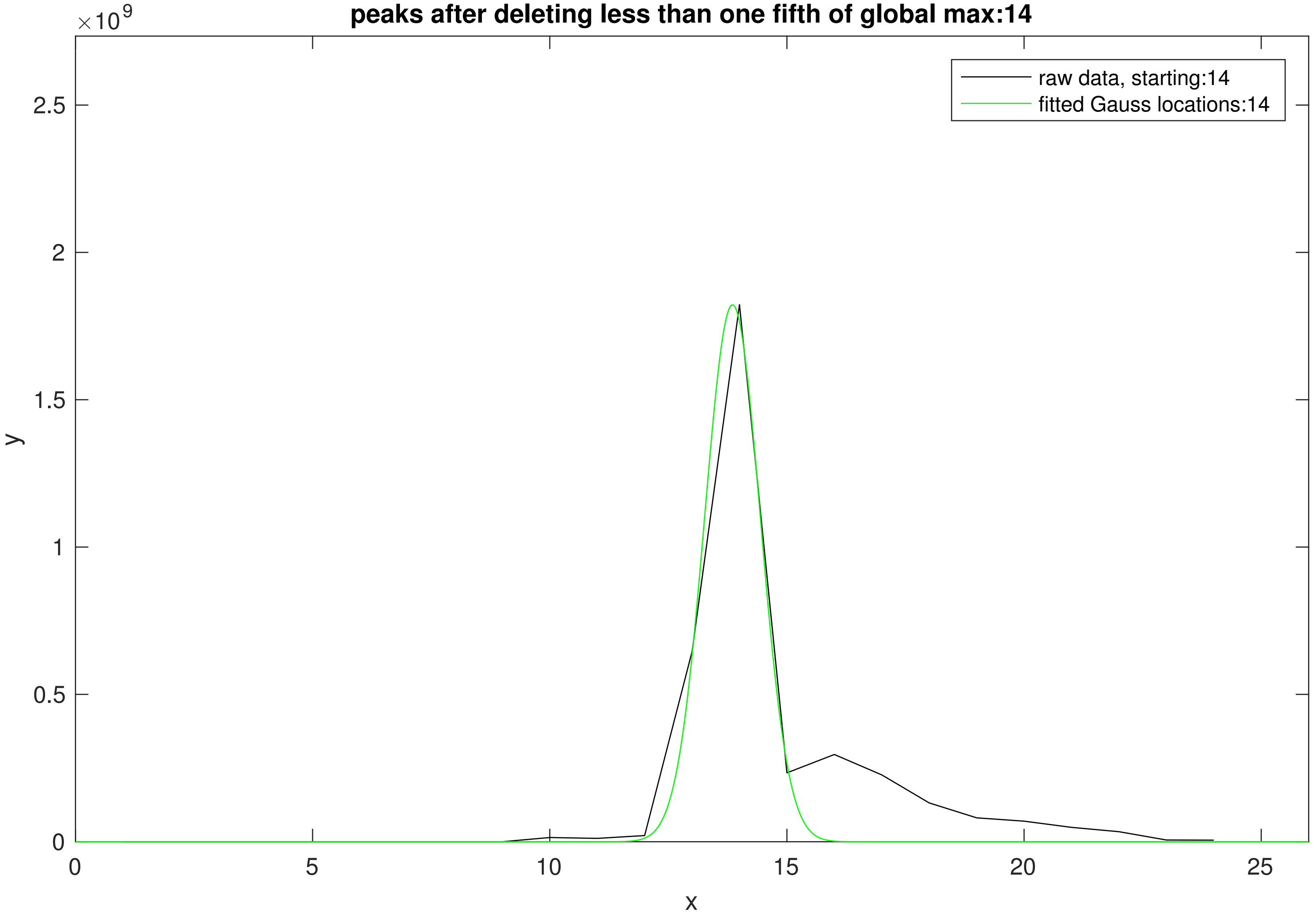

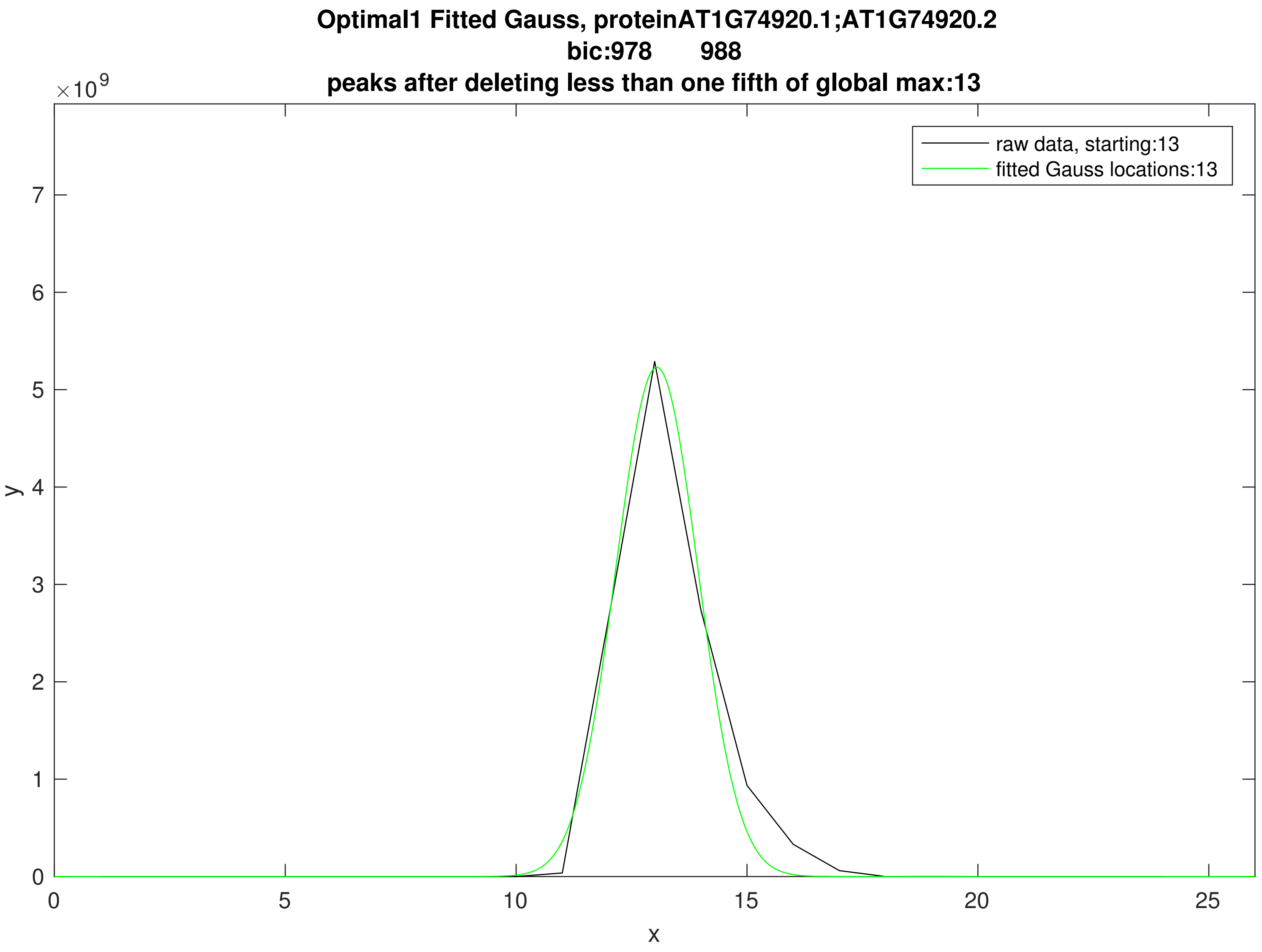

Optimal1 Fitted Gauss, proteinAT1G74920.1;AT1G74920.2
bic:978 988
peaks after deleting less than one fifth of global max:13
×10^9
raw data, starting:13
fitted Gauss locations:13
y
x
0
1
2
3
4
5
6
7
0
5
10
15
20
25

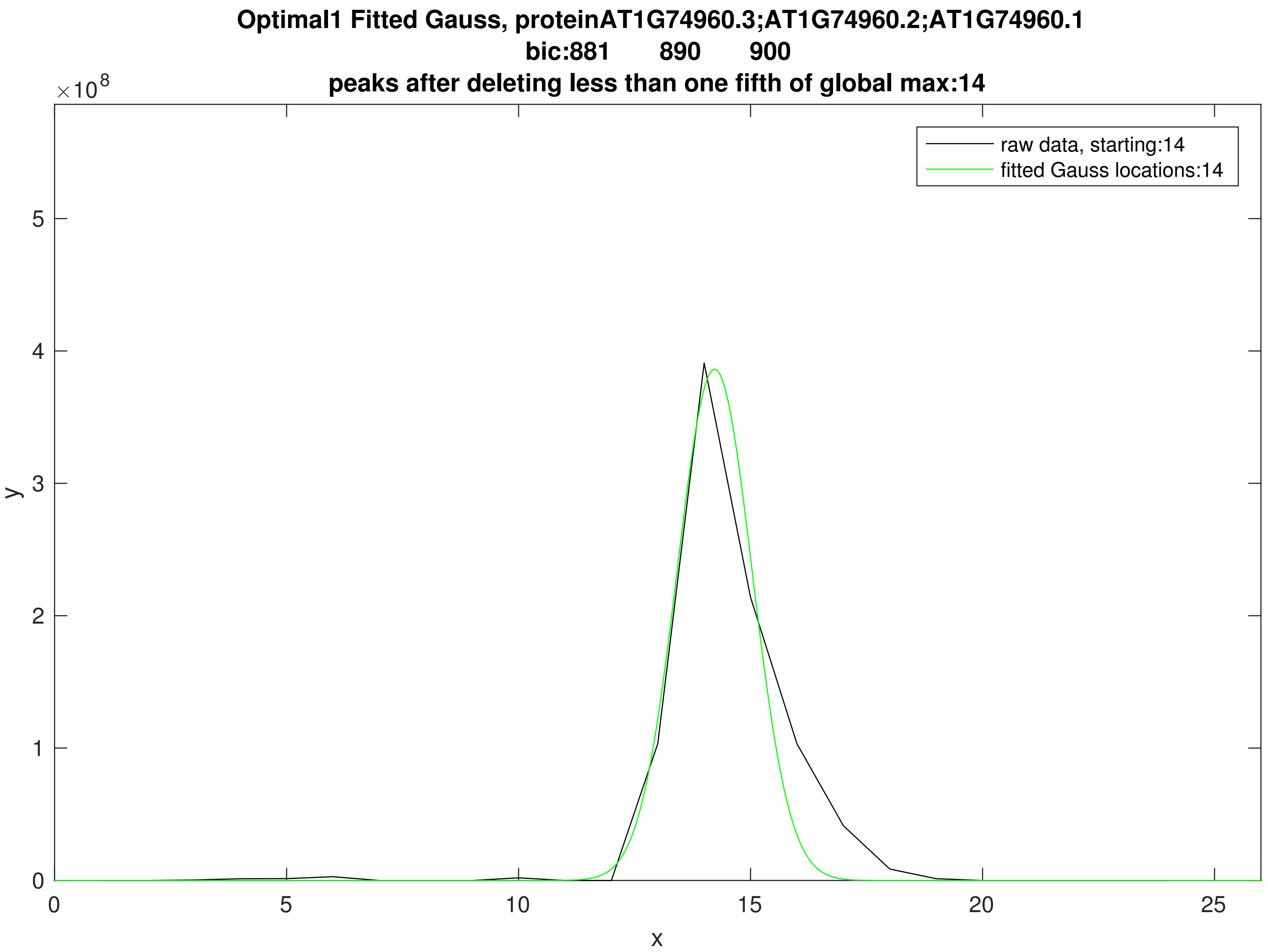
Optimal1 Fitted Gauss, proteinAT1G74960.3;AT1G74960.2;AT1G74960.1
bic:881 890 900
peaks after deleting less than one fifth of global max:14
×10^8
raw data, starting:14
fitted Gauss locations:14
y
x
0
1
2
3
4
5
0
5
10
15
20
25

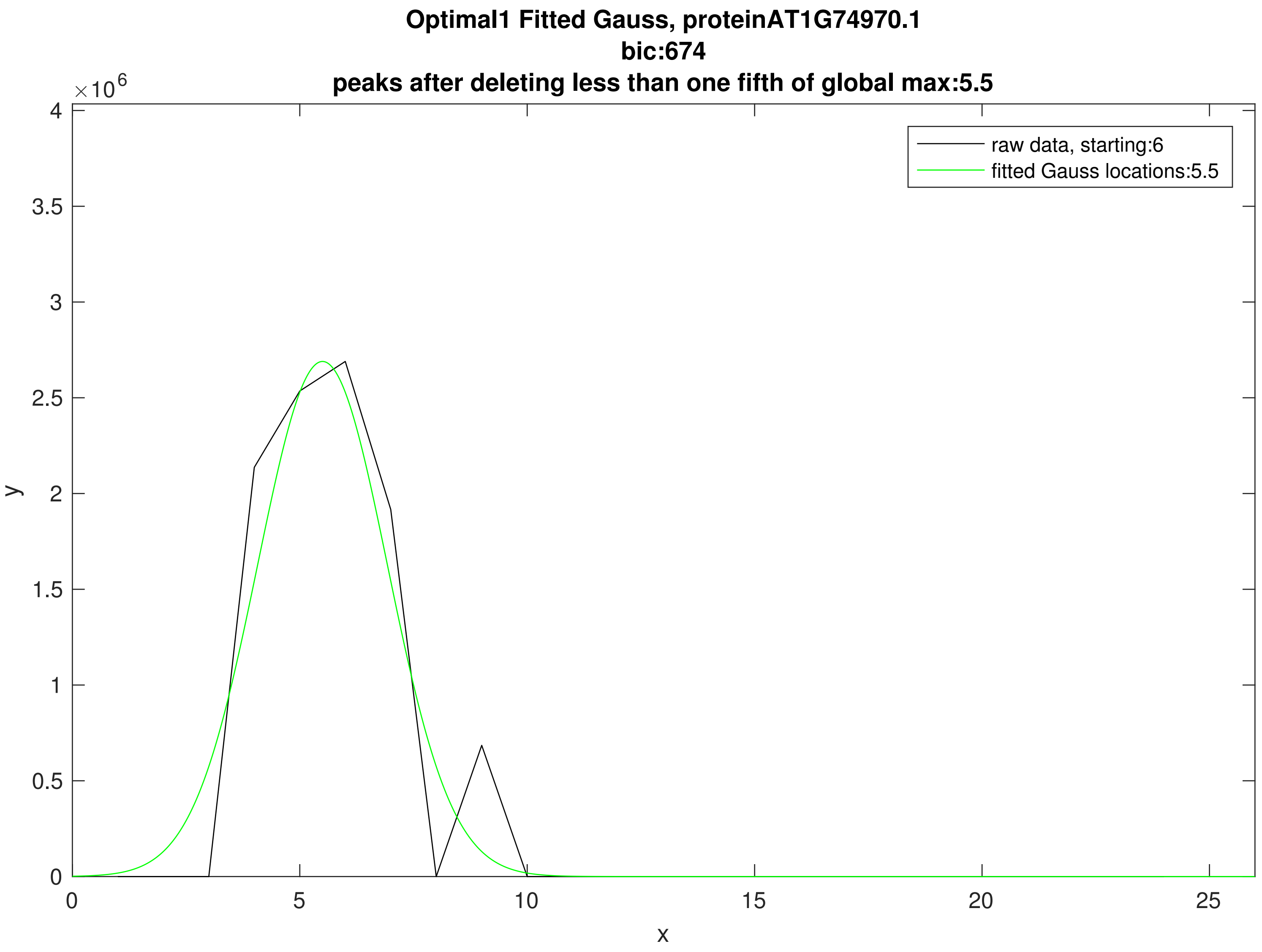
Optimal1 Fitted Gauss, proteinAT1G74970.1
bic:674
peaks after deleting less than one fifth of global max:5.5
×10^6
raw data, starting:6
fitted Gauss locations:5.5
y
x
0
0.5
1
1.5
2
2.5
3
3.5
4
5
10
15
20
25

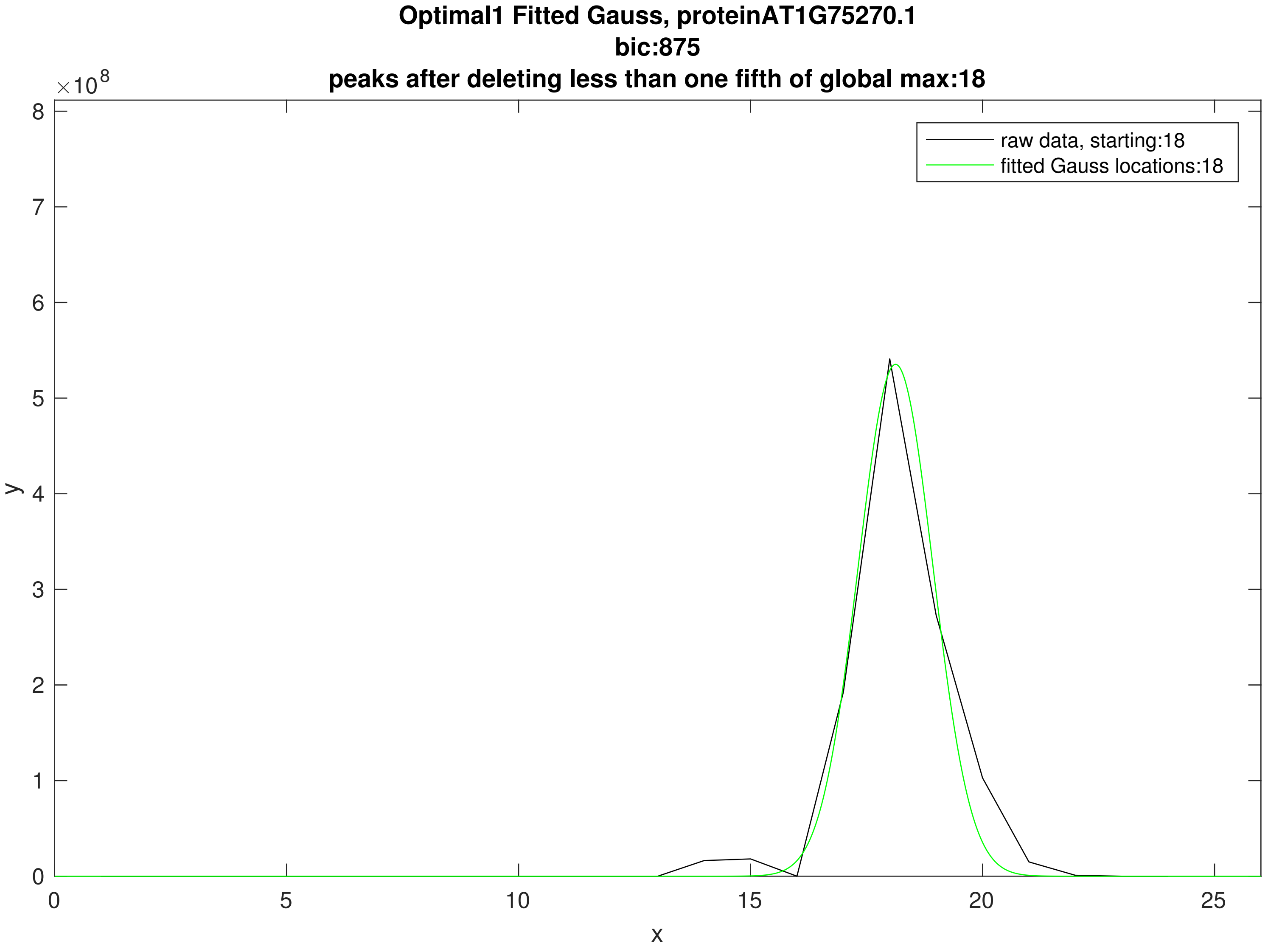

Optimal1 Fitted Gauss, proteinAT1G75270.1
bic:875
peaks after deleting less than one fifth of global max:18
×10^8
y
x
raw data, starting:18
fitted Gauss locations:18

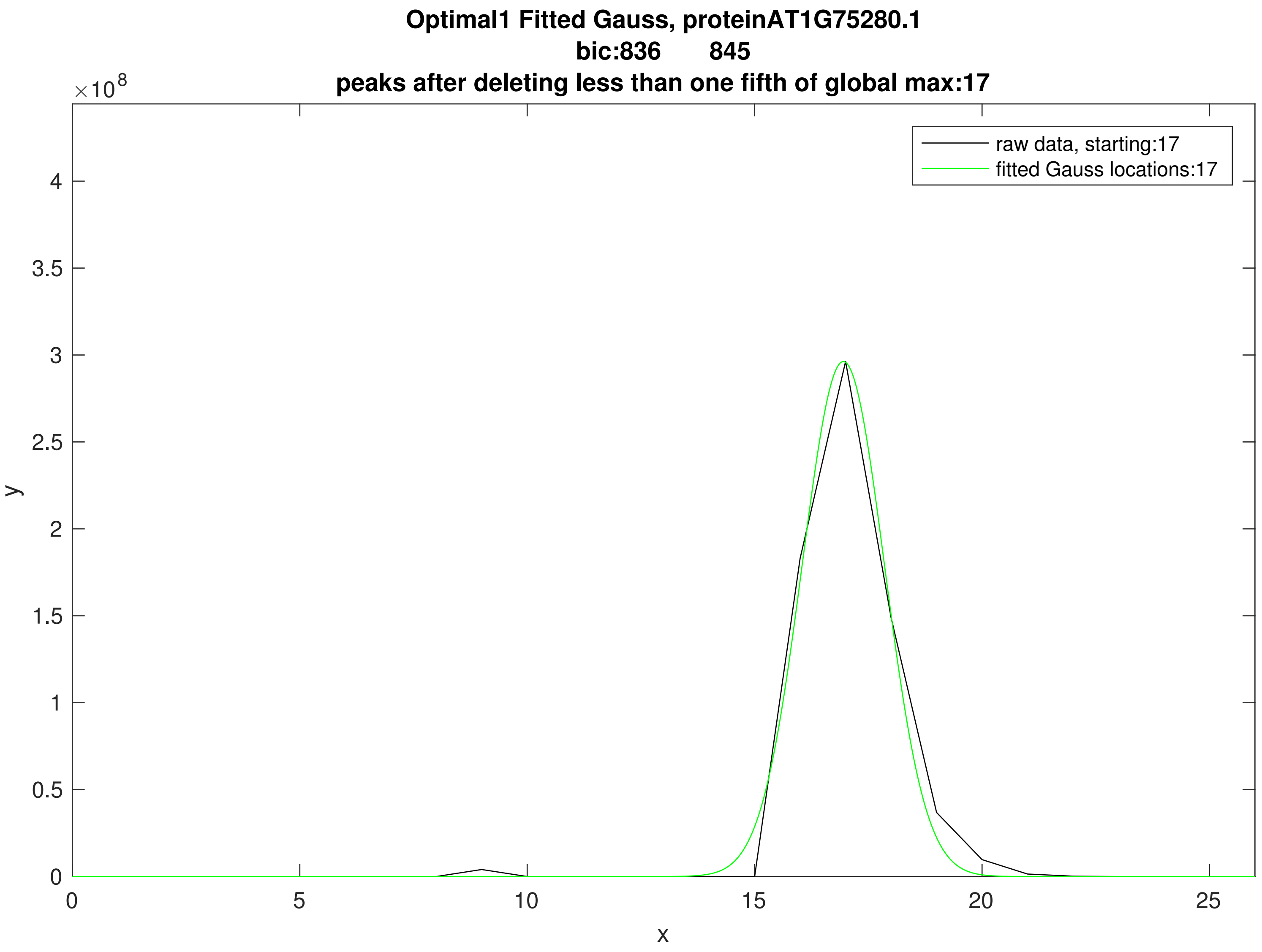
Optimal1 Fitted Gauss, proteinAT1G75280.1
bic:836 845
peaks after deleting less than one fifth of global max:17
×10^8
raw data, starting:17
fitted Gauss locations:17
y
x
0
0.5
1
1.5
2
2.5
3
3.5
4
0
5
10
15
20
25

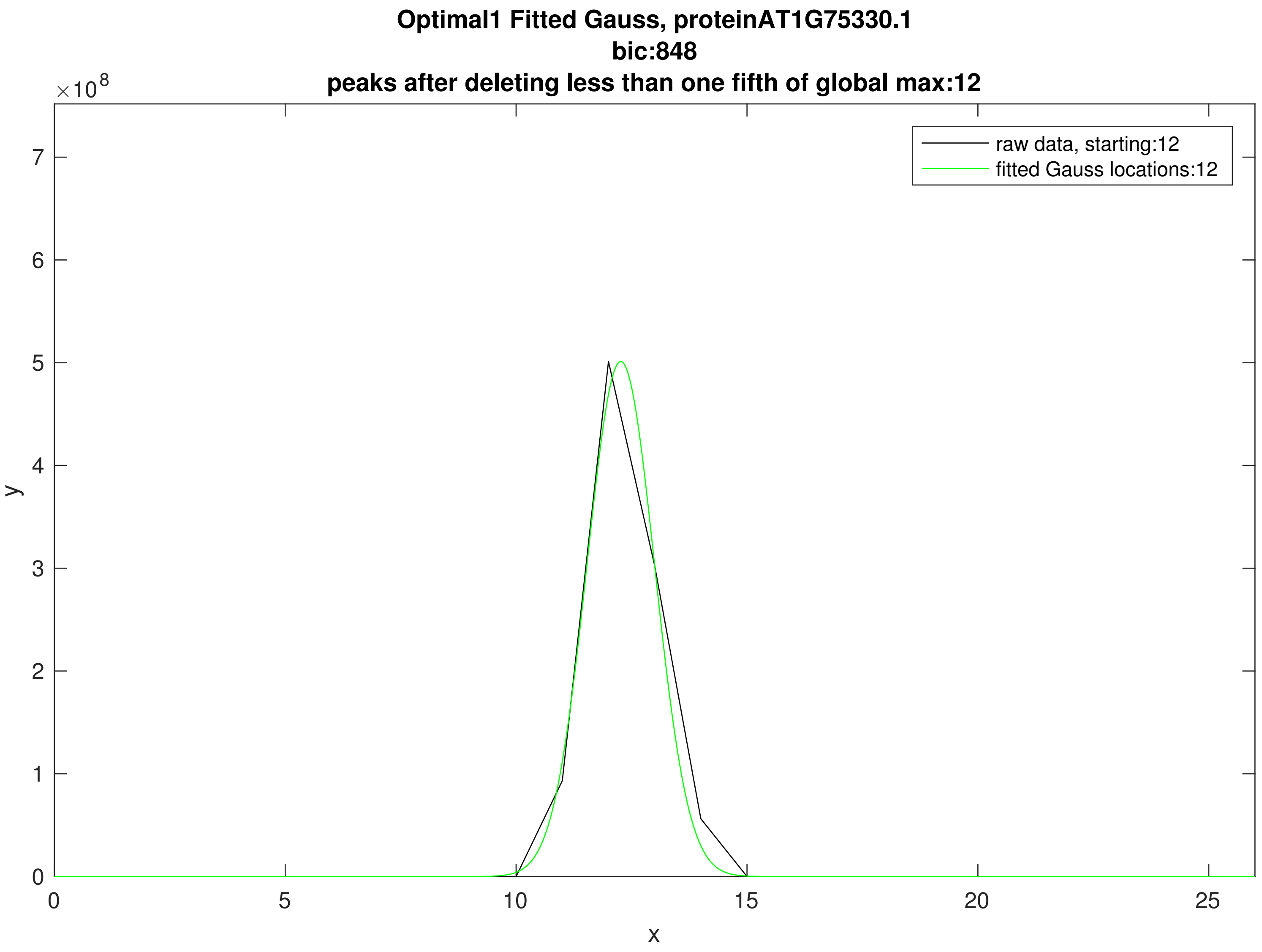
Optimal1 Fitted Gauss, proteinAT1G75330.1
bic:848
peaks after deleting less than one fifth of global max:12
×10^8
raw data, starting:12
fitted Gauss locations:12
y
x
0
1
2
3
4
5
6
7
0
5
10
15
20
25

**Optimal1 Fitted Gauss, proteinAT1G75780.1**
**bic:828 838**
**peaks after deleting less than one fifth of global max:13**

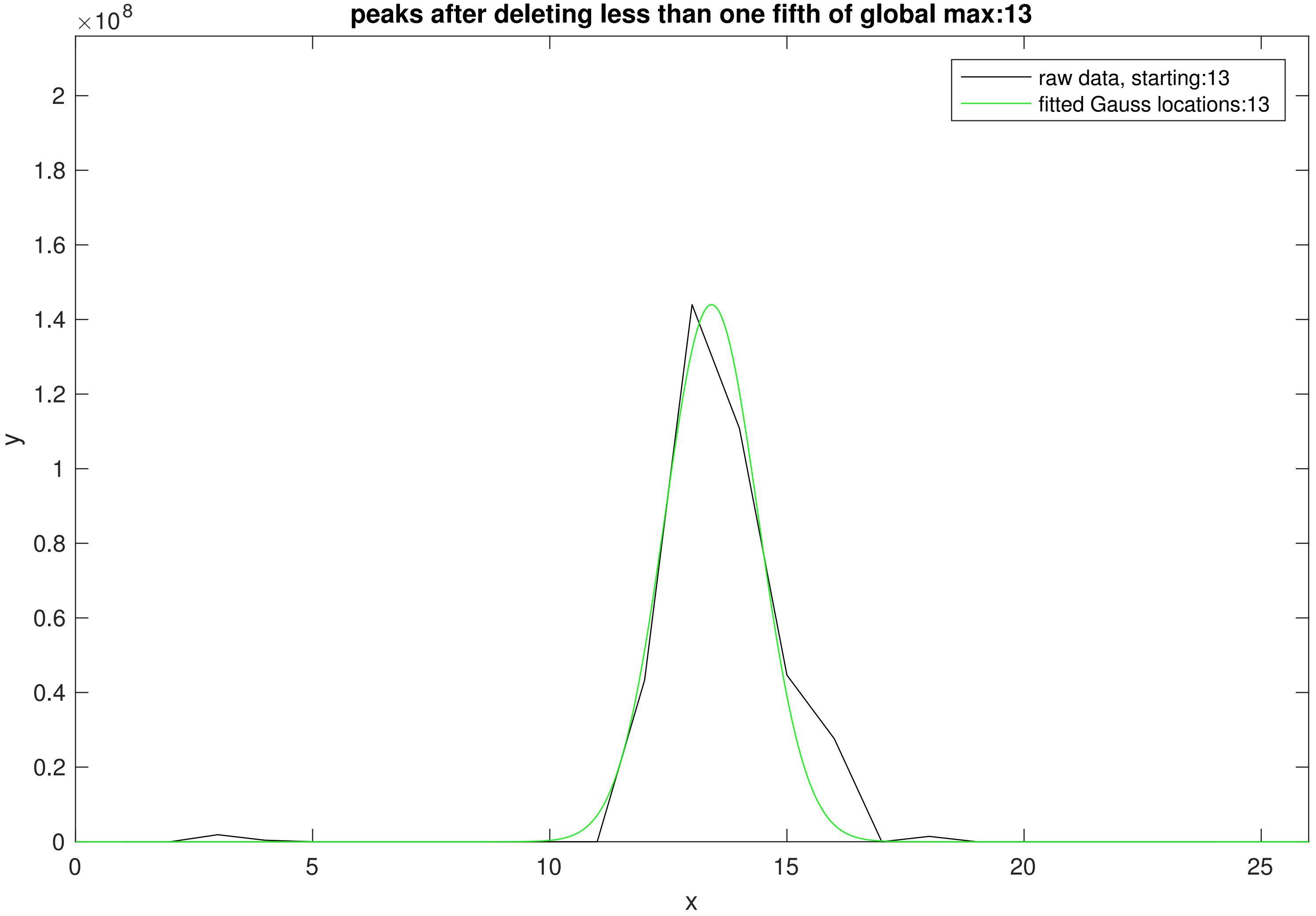

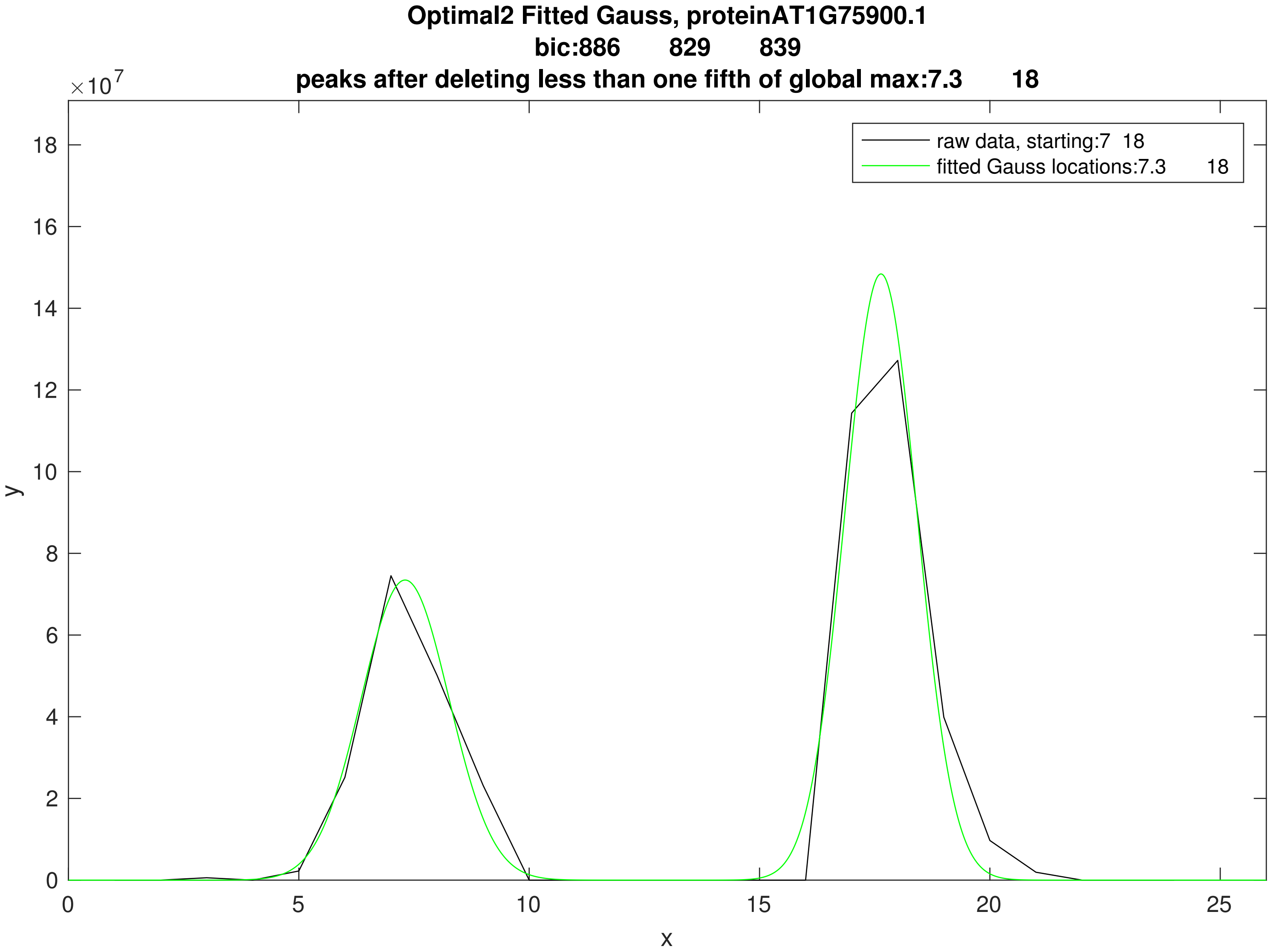
Optimal2 Fitted Gauss, proteinAT1G75900.1
bic:886 829 839
peaks after deleting less than one fifth of global max:7.3 18
×10^7
raw data, starting:7 18
fitted Gauss locations:7.3 18
y
x
0
2
4
6
8
10
12
14
16
18
0
5
10
15
20
25

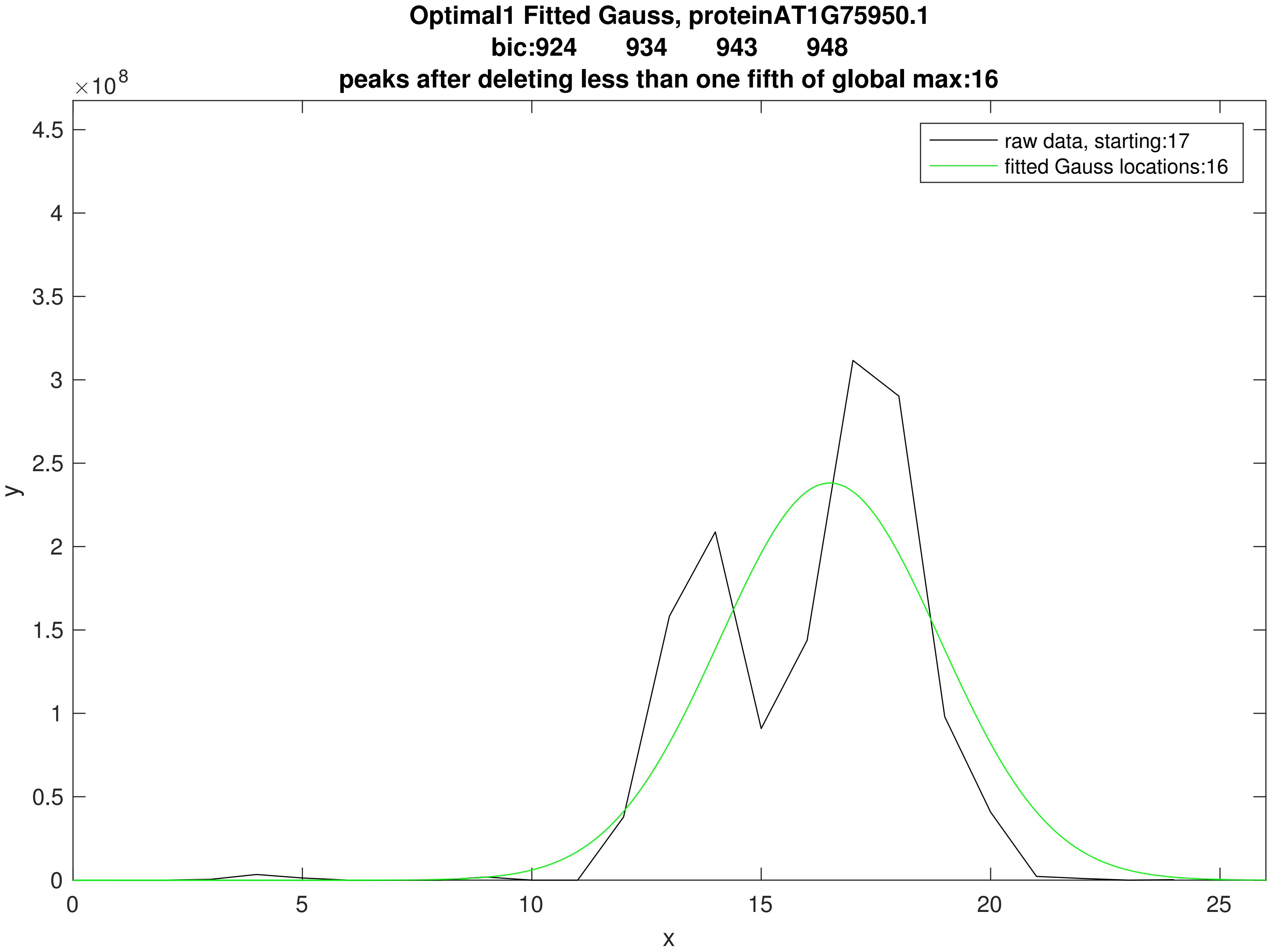
Optimal1 Fitted Gauss, proteinAT1G75950.1
bic:924 934 943 948
peaks after deleting less than one fifth of global max:16
×10^8
raw data, starting:17
fitted Gauss locations:16
y
x
0
0.5
1
1.5
2
2.5
3
3.5
4
4.5
5
10
15
20
25

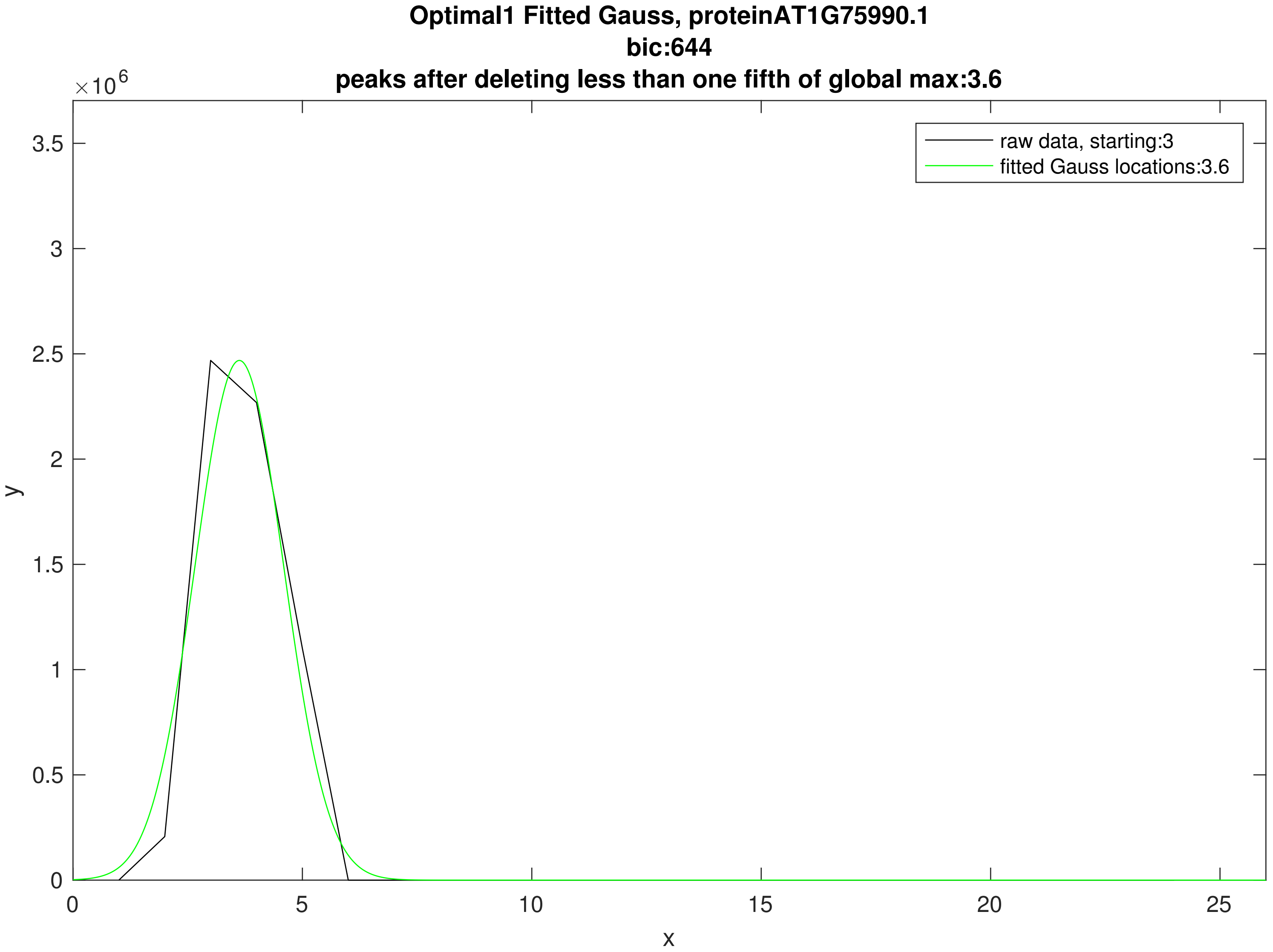
Optimal1 Fitted Gauss, proteinAT1G75990.1
bic:644
peaks after deleting less than one fifth of global max:3.6
×10^6
raw data, starting:3
fitted Gauss locations:3.6
y
x
0
0.5
1
1.5
2
2.5
3
3.5
0
5
10
15
20
25

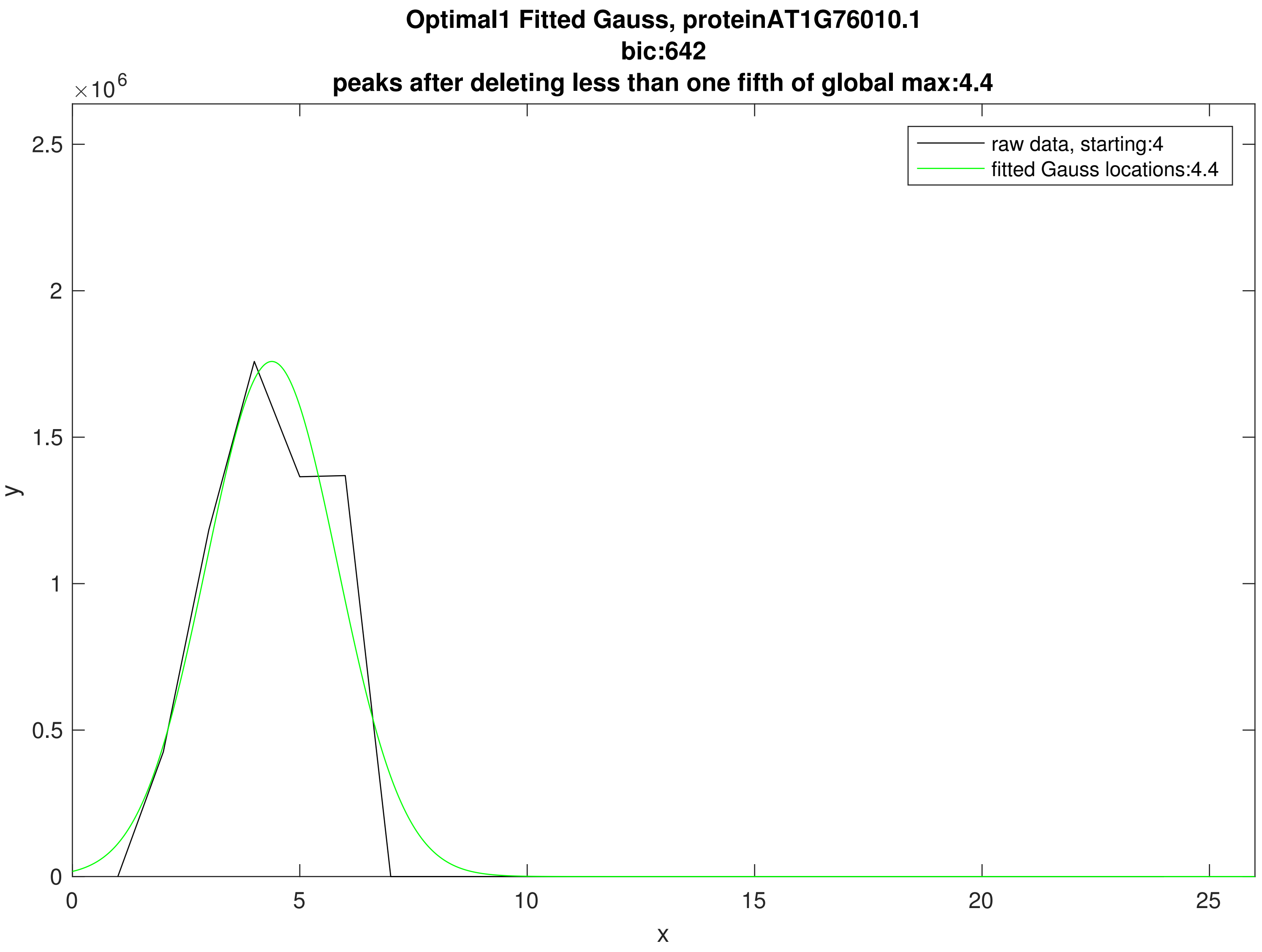

Optimal1 Fitted Gauss, proteinAT1G76010.1
bic:642
peaks after deleting less than one fifth of global max:4.4
×10^6
raw data, starting:4
fitted Gauss locations:4.4
2.5
2
1.5
1
0.5
0
y
0
5
10
15
20
25
x

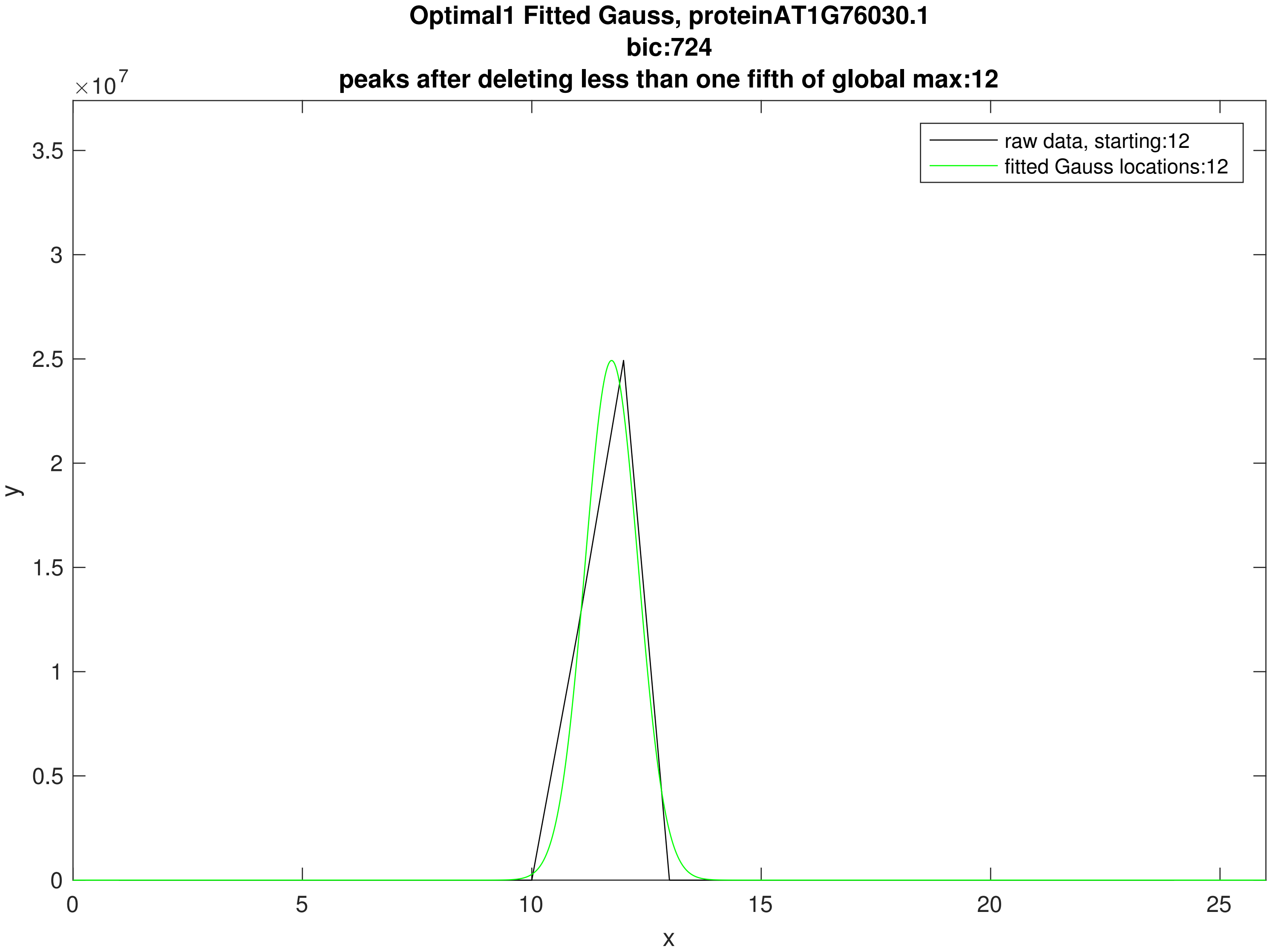
Optimal1 Fitted Gauss, proteinAT1G76030.1
bic:724
peaks after deleting less than one fifth of global max:12
×10^7
3.5
3
2.5
2
1.5
1
0.5
0
y
0
5
10
15
20
25
x
raw data, starting:12
fitted Gauss locations:12

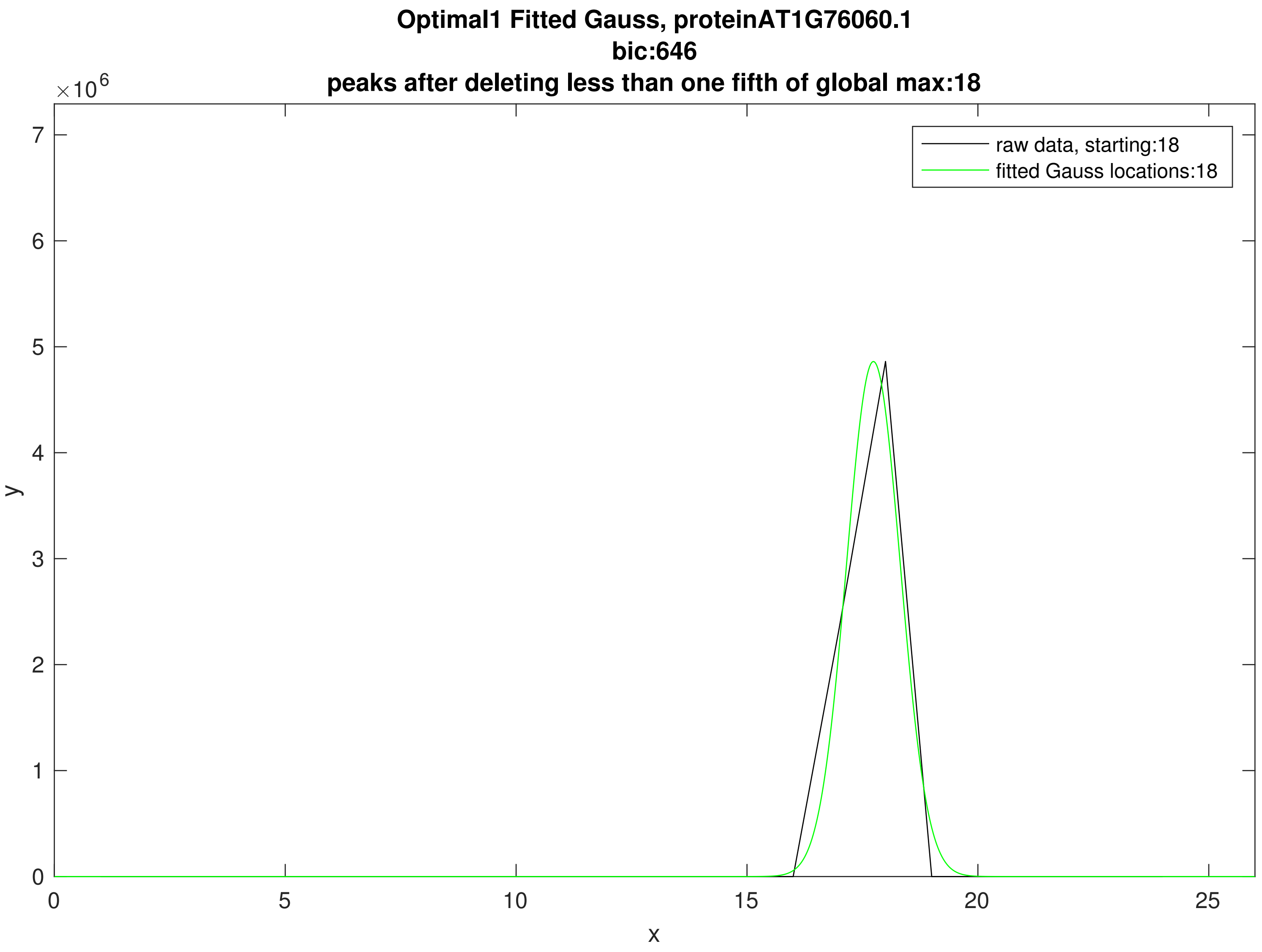
Optimal1 Fitted Gauss, proteinAT1G76060.1
bic:646
peaks after deleting less than one fifth of global max:18
×10^6
raw data, starting:18
fitted Gauss locations:18
y
x
0
1
2
3
4
5
6
7
0
5
10
15
20
25

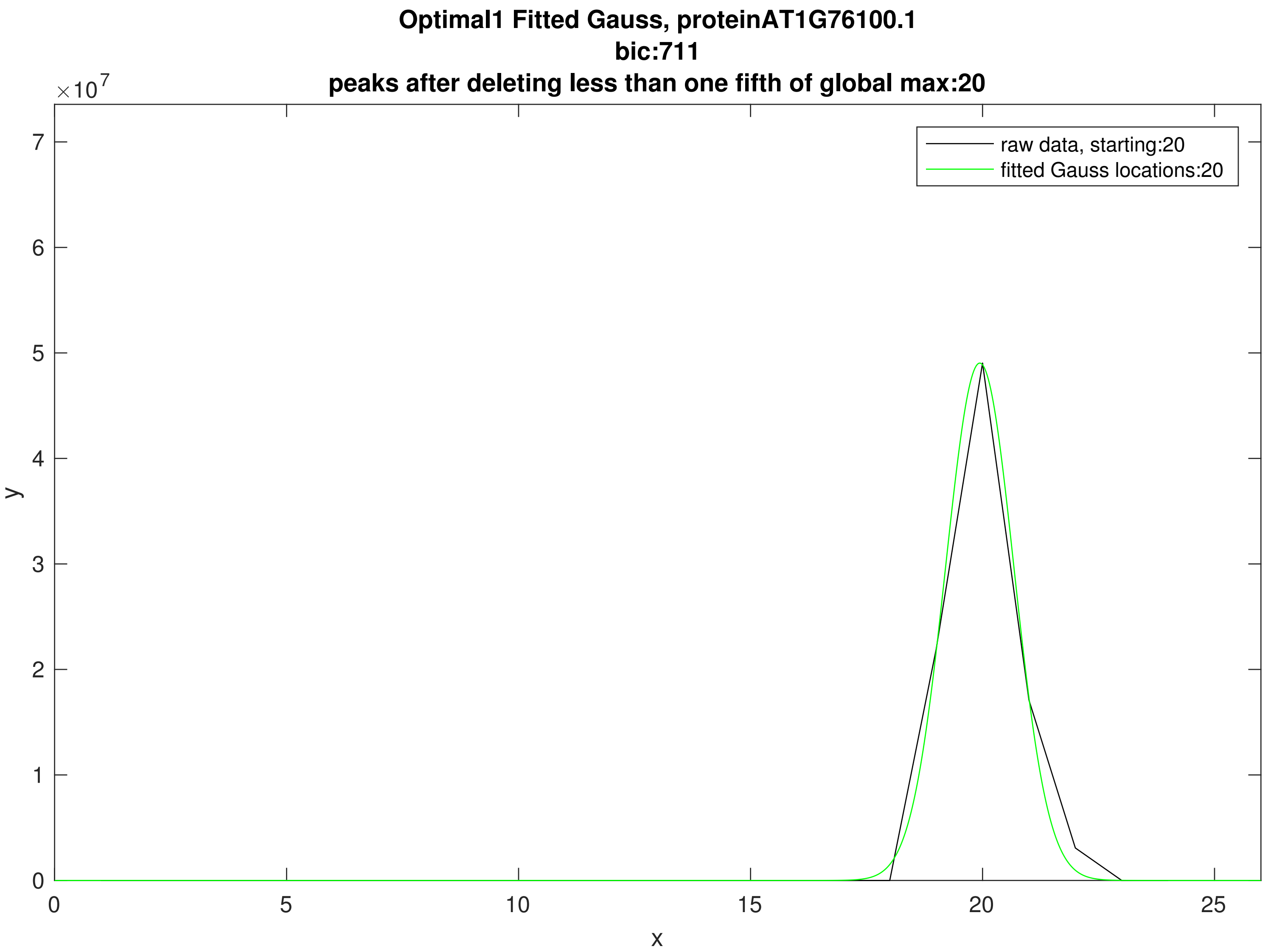
Optimal1 Fitted Gauss, proteinAT1G76100.1
bic:711
peaks after deleting less than one fifth of global max:20
$\times 10^7$
raw data, starting:20
fitted Gauss locations:20
y
x
0
1
2
3
4
5
6
7
0
5
10
15
20
25

##### Optimal1 Fitted Gauss, proteinAT1G76140.1;AT1G76140.2
##### bic:913
##### peaks after deleting less than one fifth of global max:14

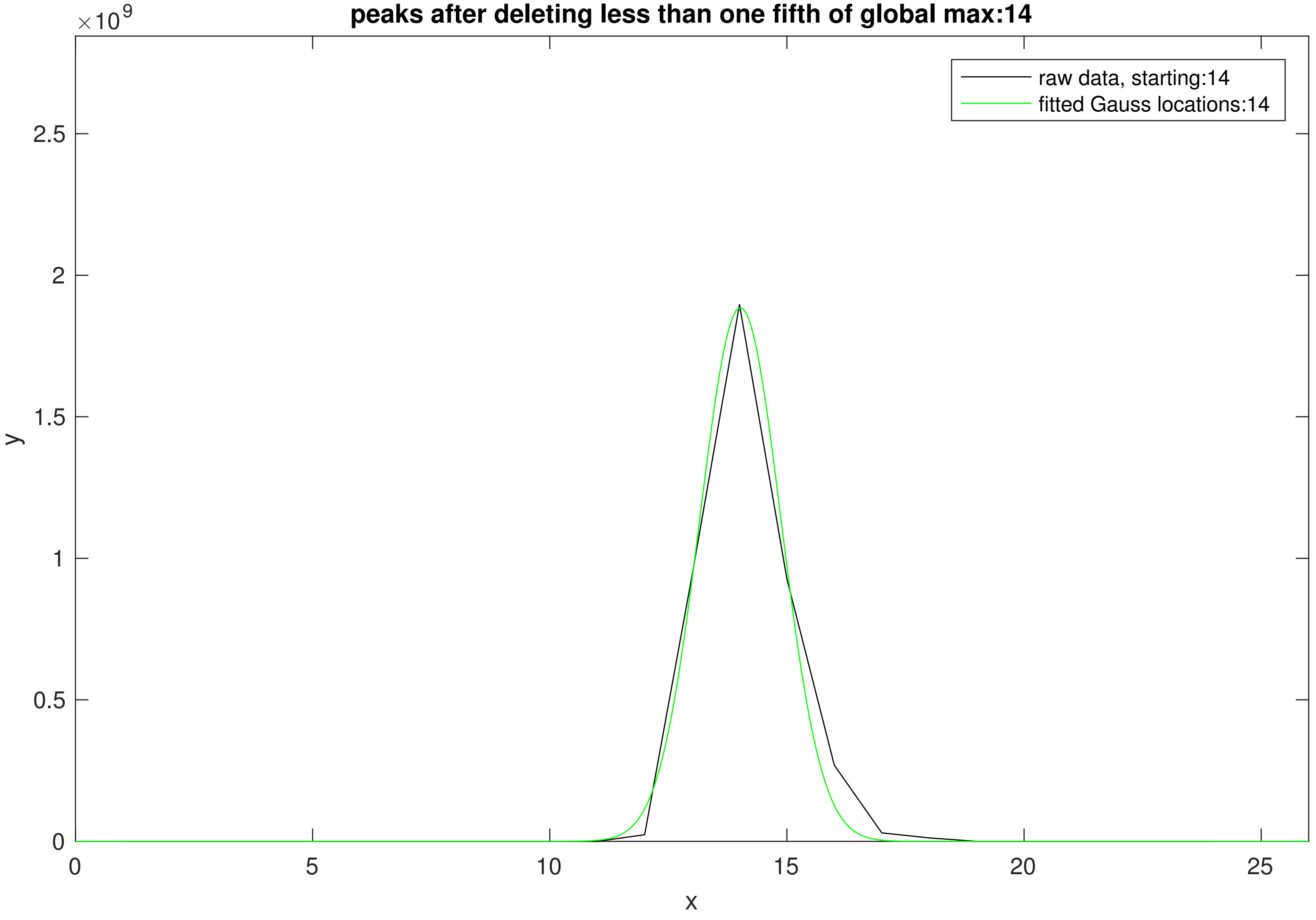

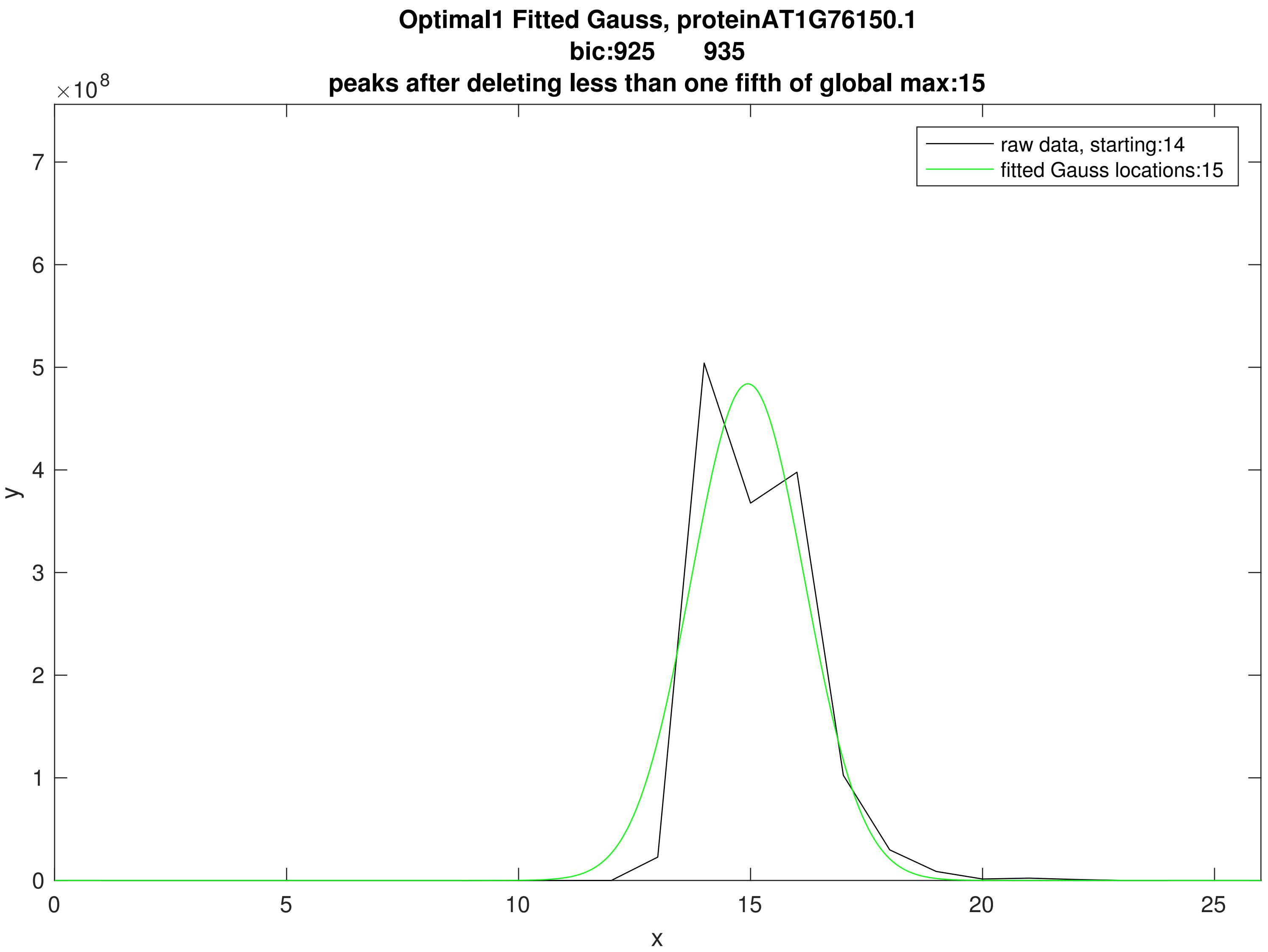
Optimal1 Fitted Gauss, proteinAT1G76150.1
bic:925 935
peaks after deleting less than one fifth of global max:15
×10^8
raw data, starting:14
fitted Gauss locations:15
y
x
0
1
2
3
4
5
6
7
0
5
10
15
20
25

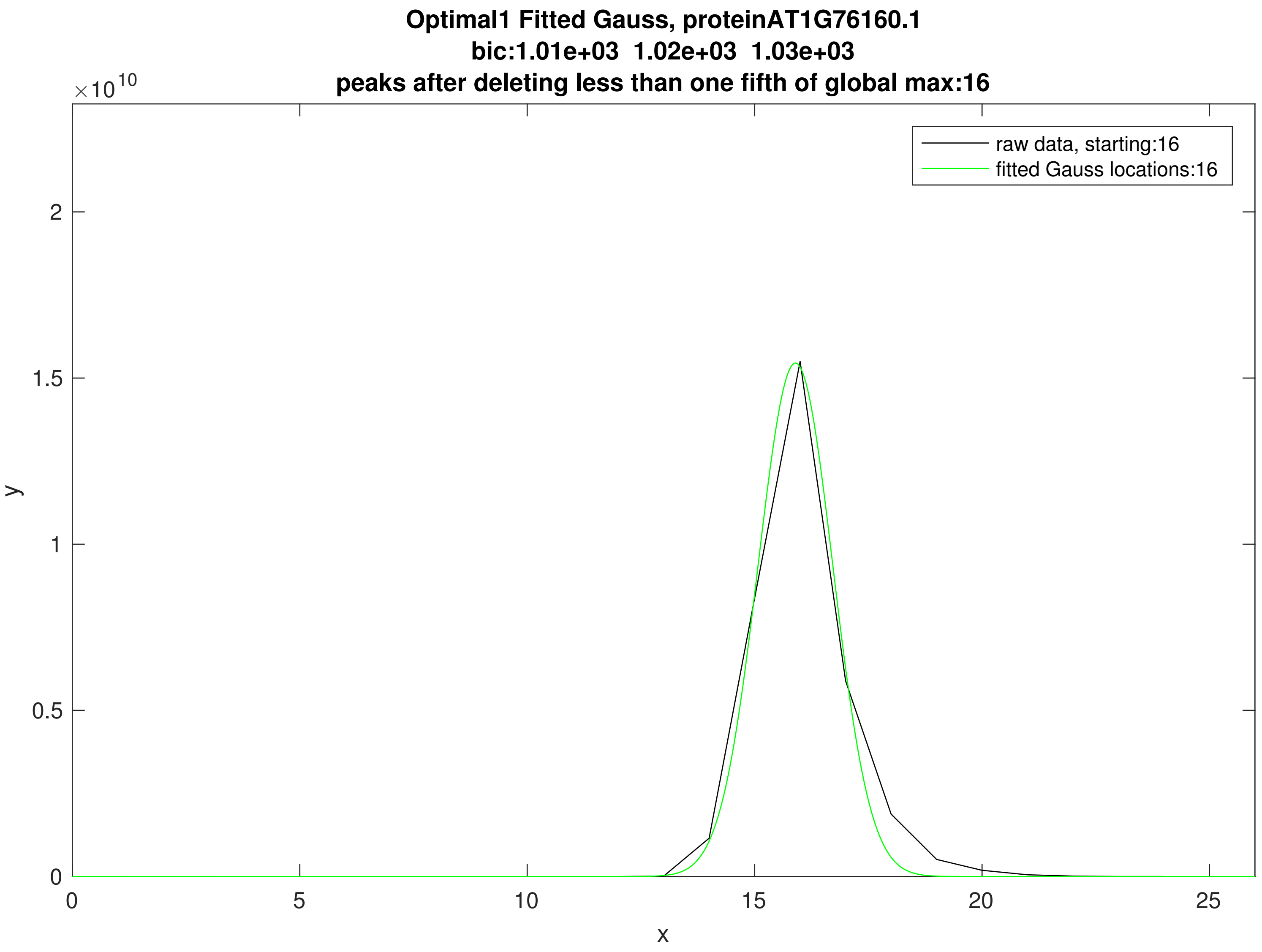

Optimal1 Fitted Gauss, proteinAT1G76160.1
bic:1.01e+03 1.02e+03 1.03e+03
peaks after deleting less than one fifth of global max:16
×10^10
raw data, starting:16
fitted Gauss locations:16
2
1.5
1
0.5
0
y
0
5
10
15
20
25
x

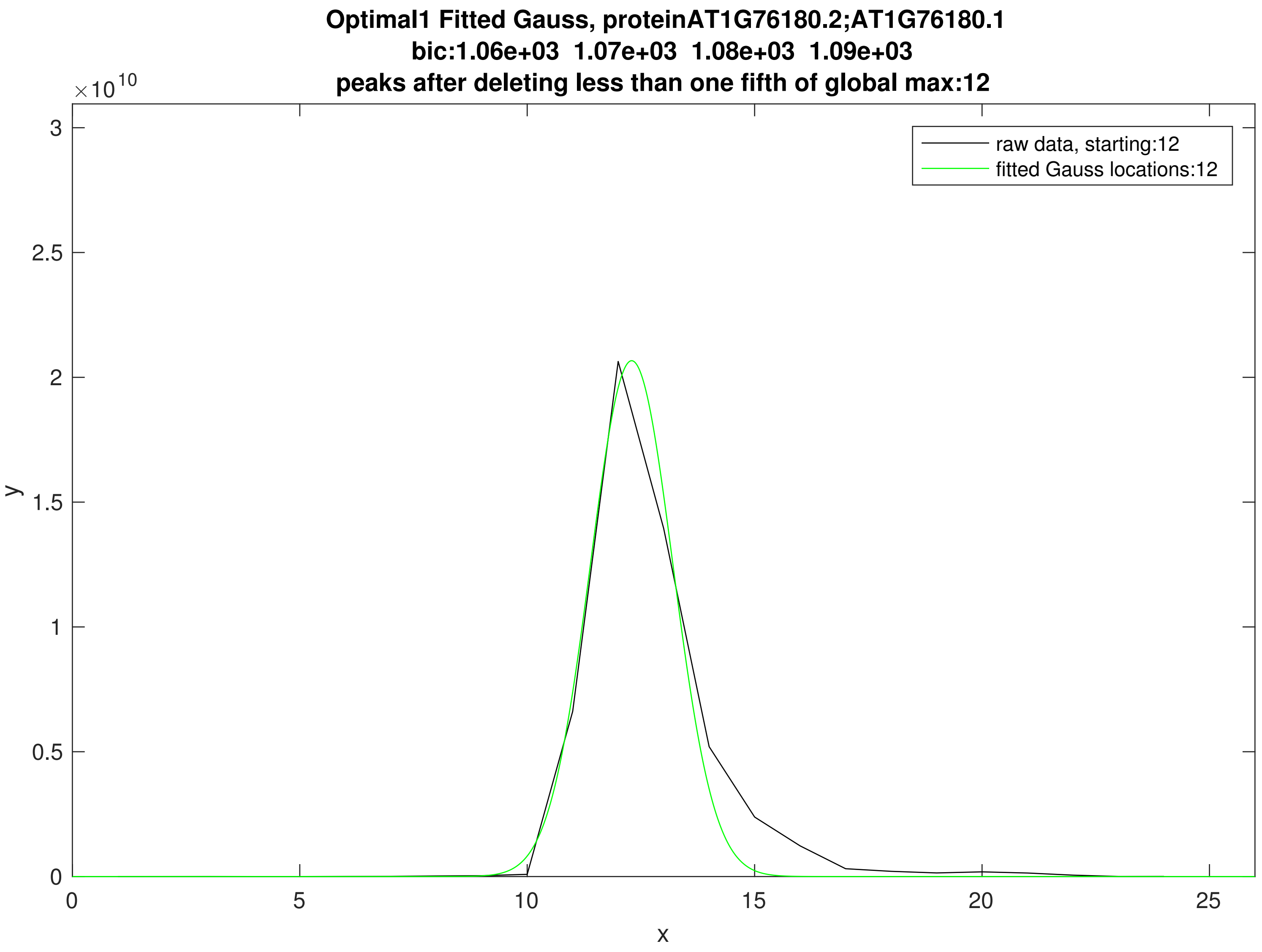
Optimal1 Fitted Gauss, proteinAT1G76180.2;AT1G76180.1
bic:1.06e+03  1.07e+03  1.08e+03  1.09e+03
peaks after deleting less than one fifth of global max:12
×10^10
raw data, starting:12
fitted Gauss locations:12
y
0
0.5
1
1.5
2
2.5
3
x
0
5
10
15
20
25

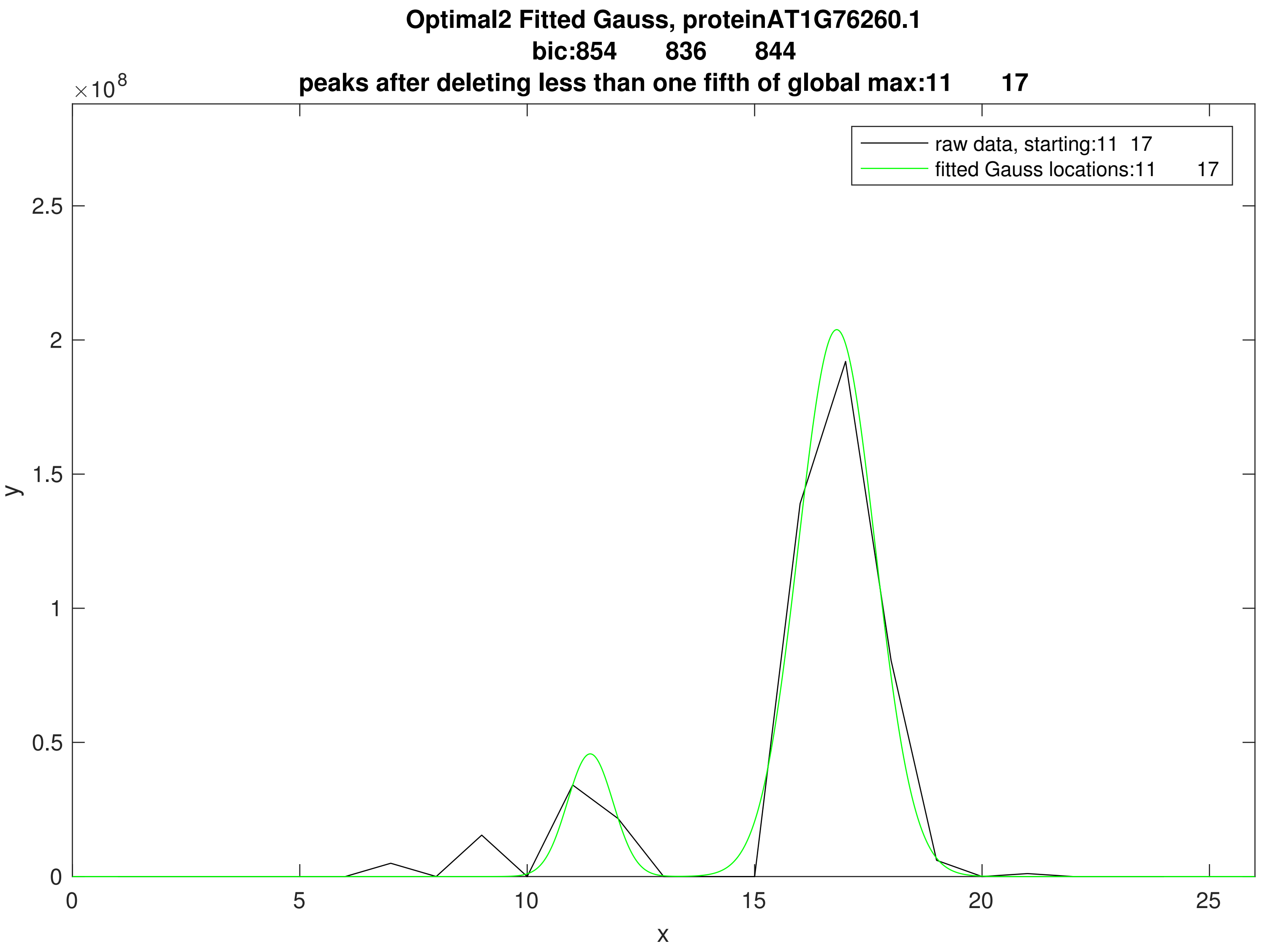

Optimal2 Fitted Gauss, proteinAT1G76260.1
bic:854 836 844
peaks after deleting less than one fifth of global max:11 17
×10^8
raw data, starting:11 17
fitted Gauss locations:11 17
y
x
0
0.5
1
1.5
2
2.5
0
5
10
15
20
25

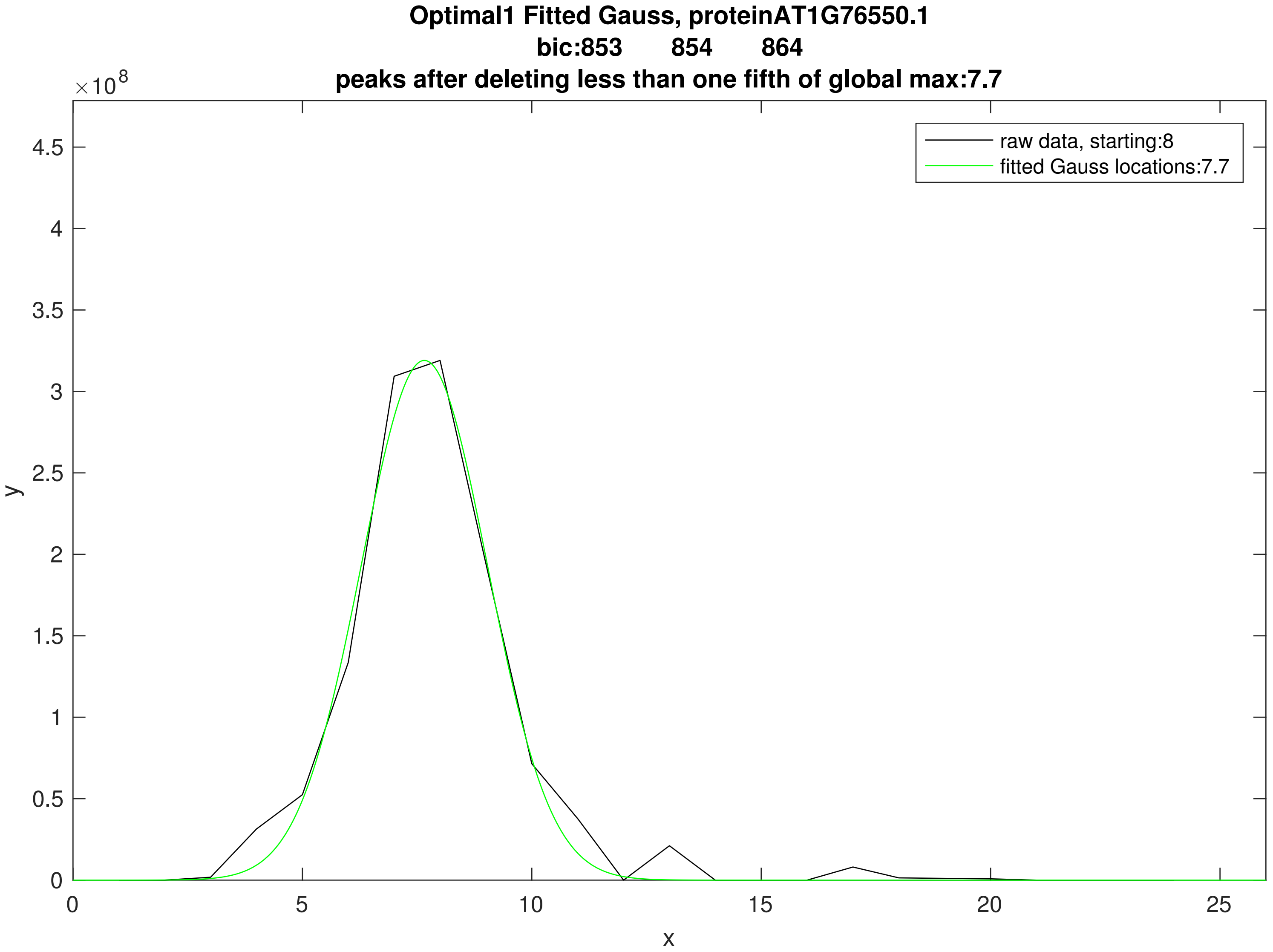

Optimal1 Fitted Gauss, proteinAT1G76550.1
bic:853 854 864
peaks after deleting less than one fifth of global max:7.7
×10^8
4.5
4
3.5
3
2.5
2
1.5
1
0.5
0
y
0
5
10
15
20
25
x
raw data, starting:8
fitted Gauss locations:7.7

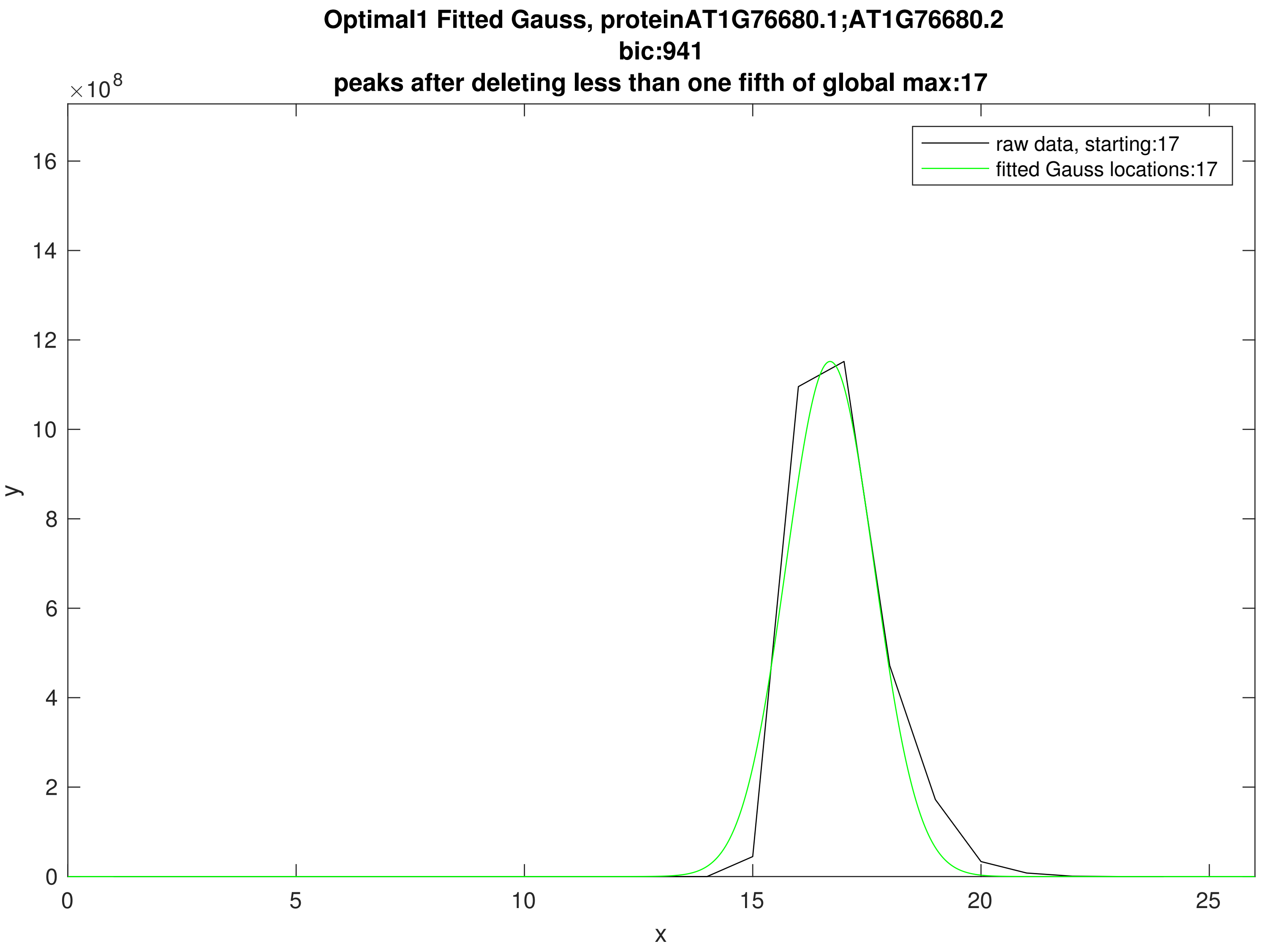
Optimal1 Fitted Gauss, proteinAT1G76680.1;AT1G76680.2
bic:941
peaks after deleting less than one fifth of global max:17
×10^8
16
14
12
10
8
6
4
2
0
y
0
5
10
15
20
25
x
raw data, starting:17
fitted Gauss locations:17

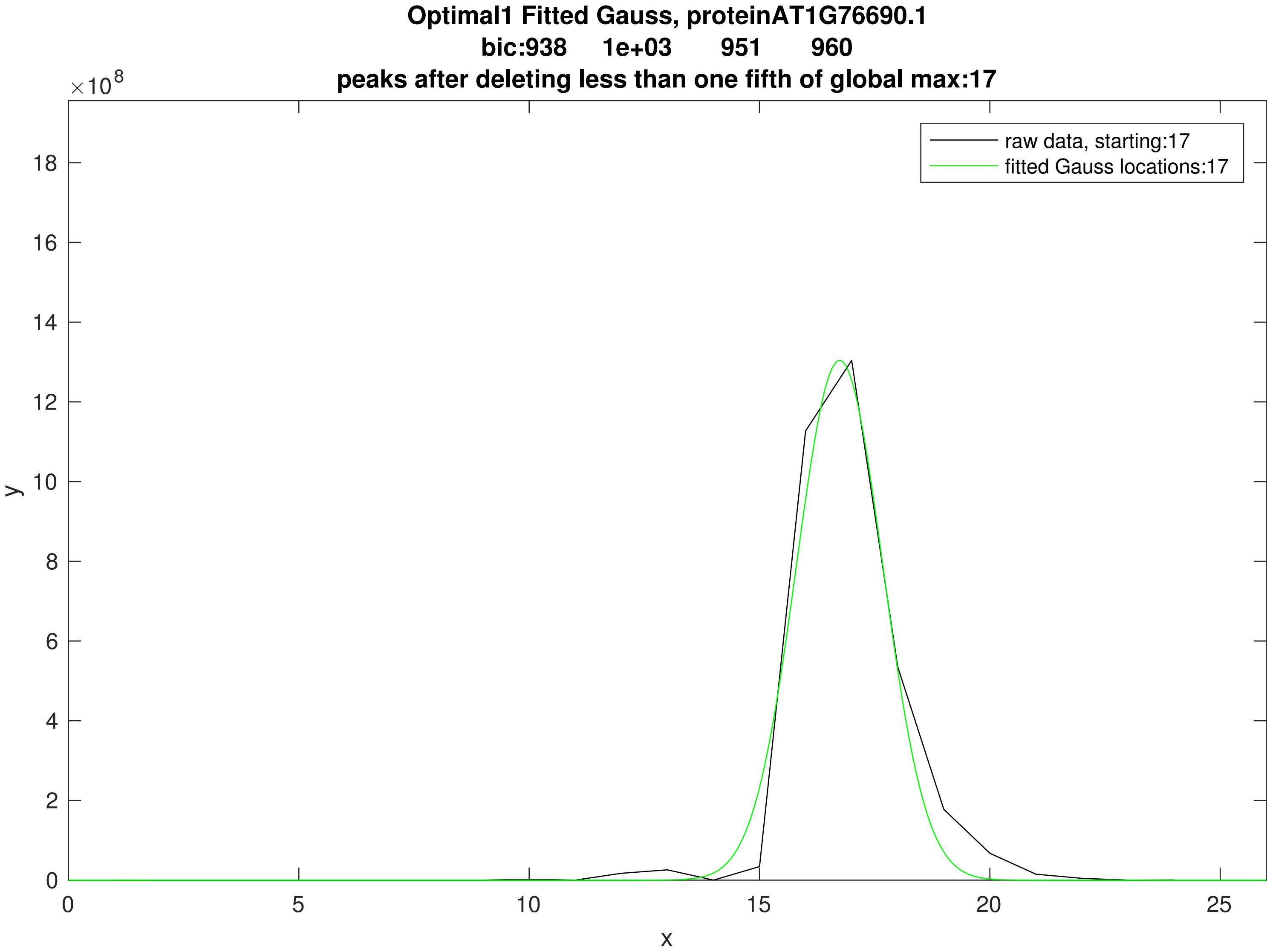

Optimal1 Fitted Gauss, proteinAT1G76690.1
bic:938 1e+03 951 960
peaks after deleting less than one fifth of global max:17
×10^8
raw data, starting:17
fitted Gauss locations:17
y
x
0
2
4
6
8
10
12
14
16
18
0
5
10
15
20
25

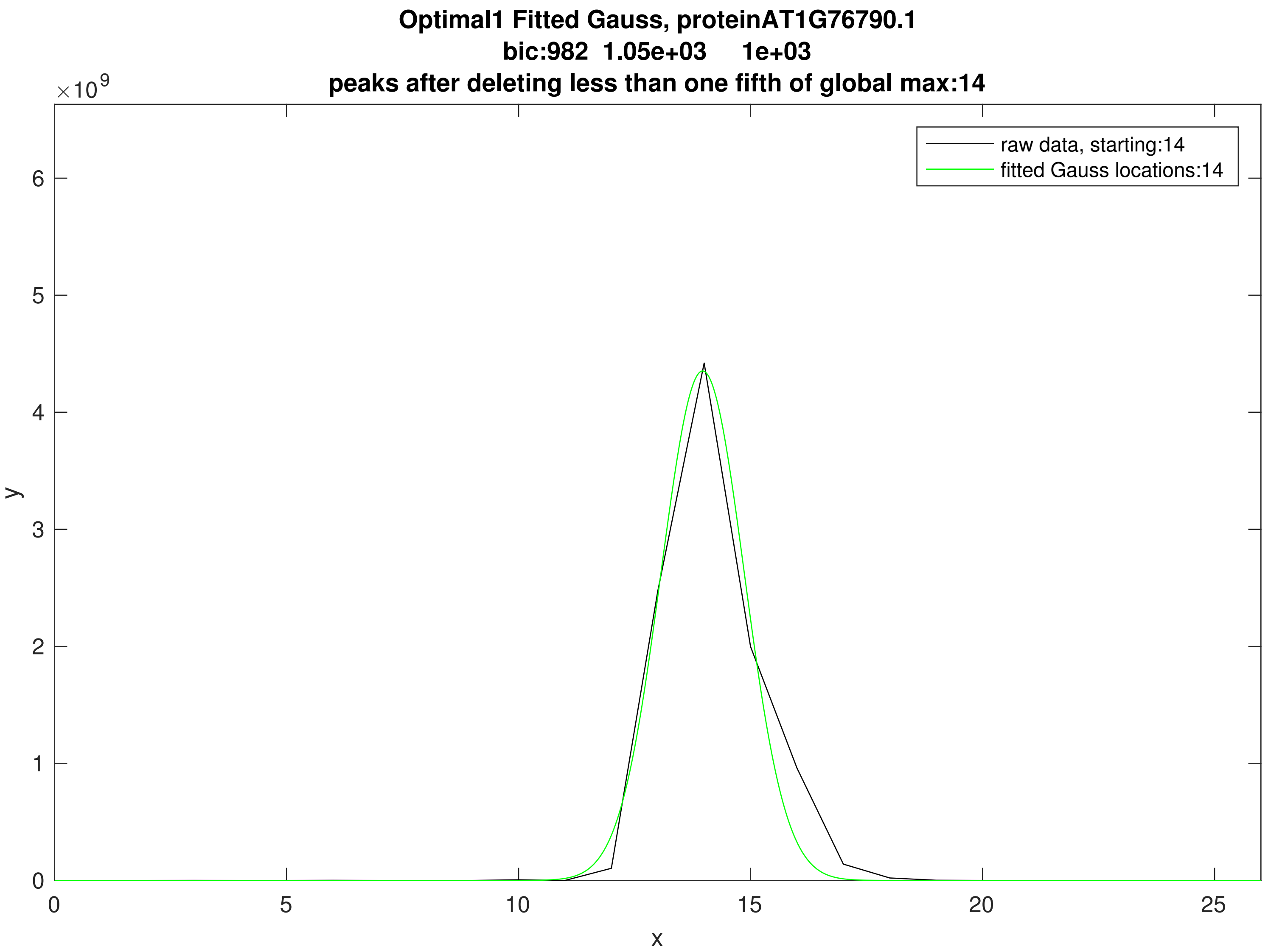
Optimal1 Fitted Gauss, proteinAT1G76790.1
bic:982 1.05e+03 1e+03
peaks after deleting less than one fifth of global max:14
×10^9
raw data, starting:14
fitted Gauss locations:14
y
x
0
1
2
3
4
5
6
0
5
10
15
20
25

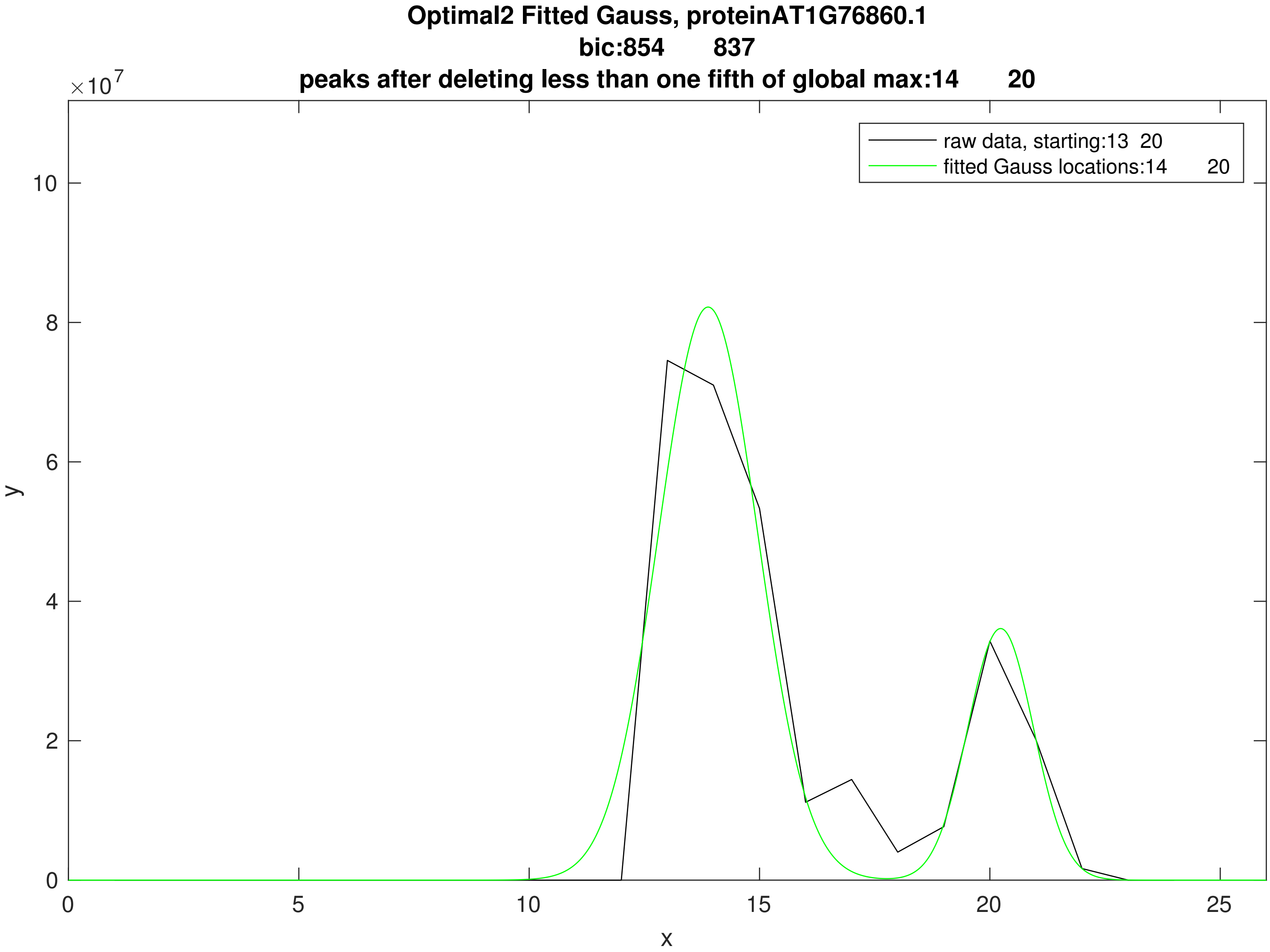
Optimal2 Fitted Gauss, proteinAT1G76860.1
bic:854 837
peaks after deleting less than one fifth of global max:14 20
×10^7
raw data, starting:13 20
fitted Gauss locations:14 20
10
8
6
4
2
0
y
0
5
10
15
20
25
x

**Optimal2 Fitted Gauss, proteinAT1G77060.1**
**bic:787 772**
**peaks after deleting less than one fifth of global max:12 17**

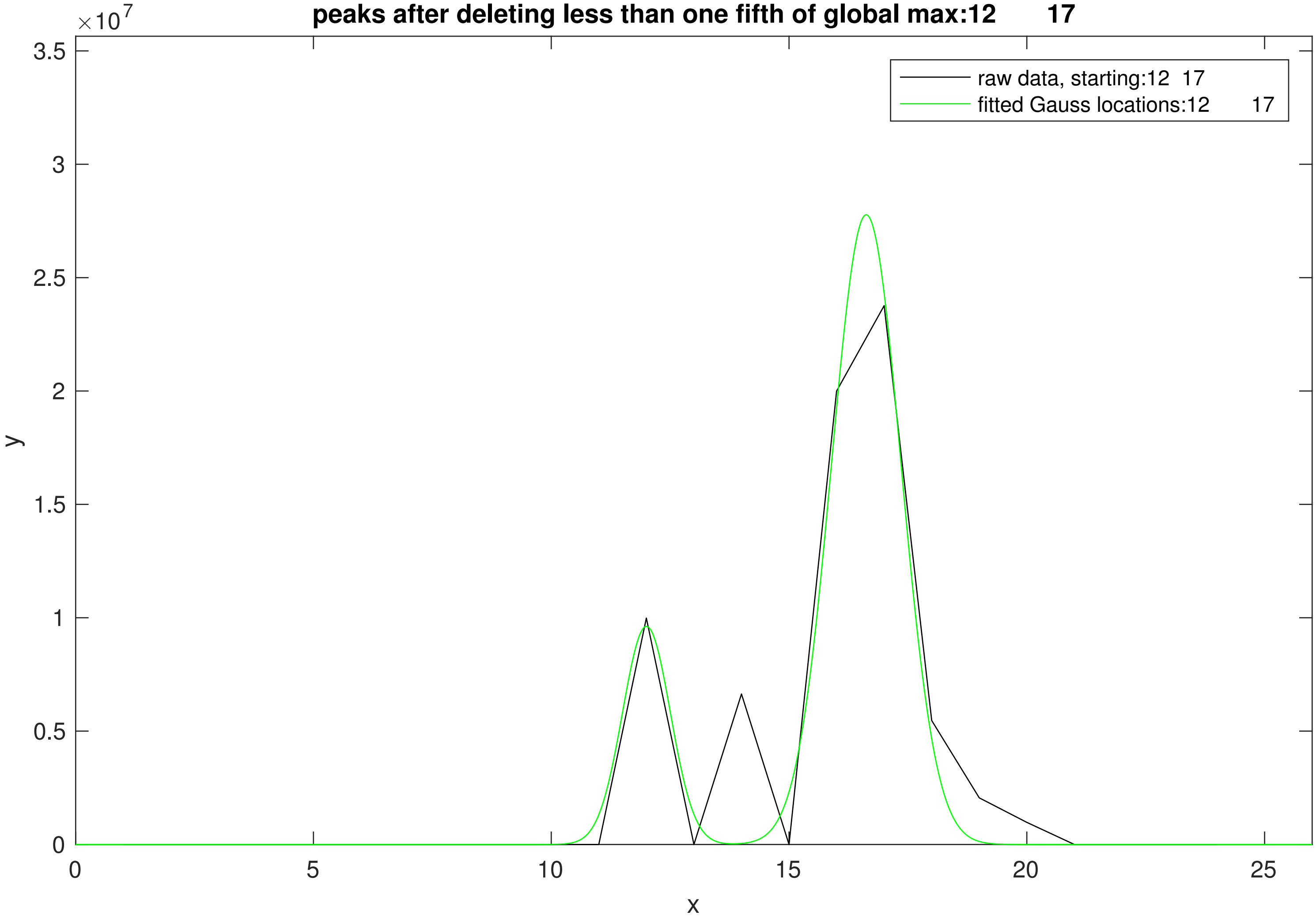

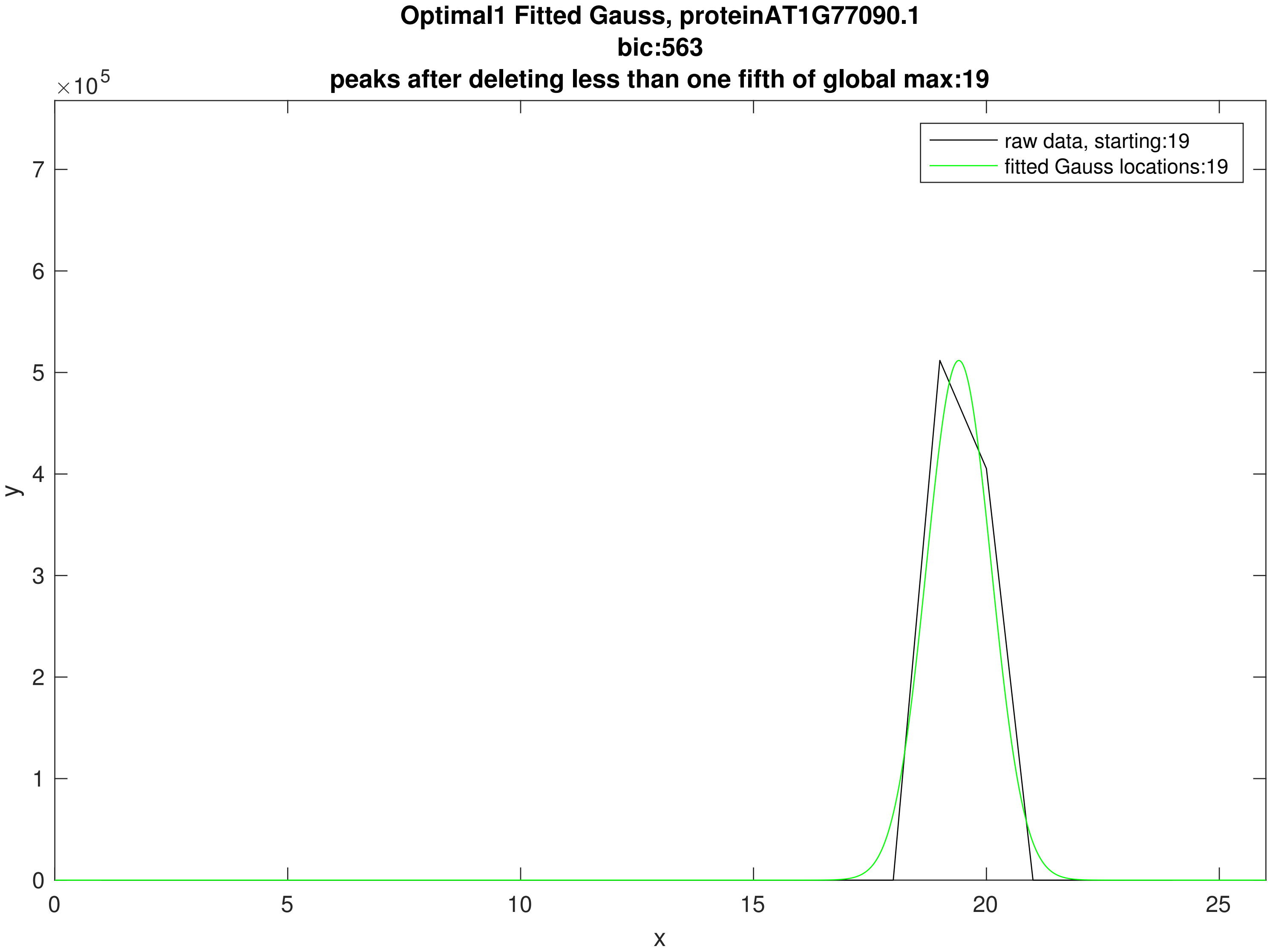
Optimal1 Fitted Gauss, proteinAT1G77090.1
bic:563
peaks after deleting less than one fifth of global max:19
×10^5
raw data, starting:19
fitted Gauss locations:19
0
1
2
3
4
5
6
7
y
0
5
10
15
20
25
x

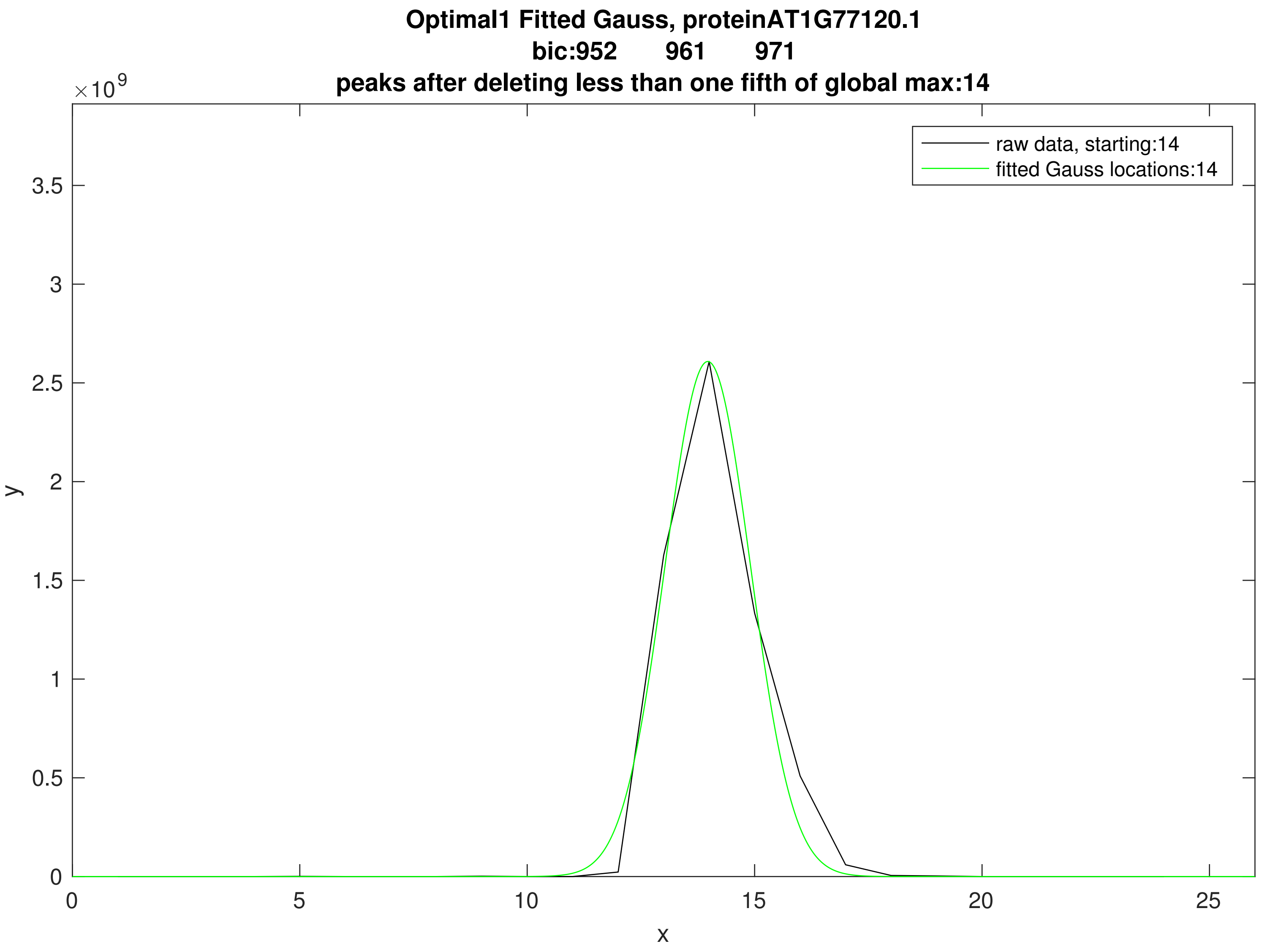
Optimal1 Fitted Gauss, proteinAT1G77120.1
bic:952 961 971
peaks after deleting less than one fifth of global max:14
raw data, starting:14
fitted Gauss locations:14
×10^9
y
x

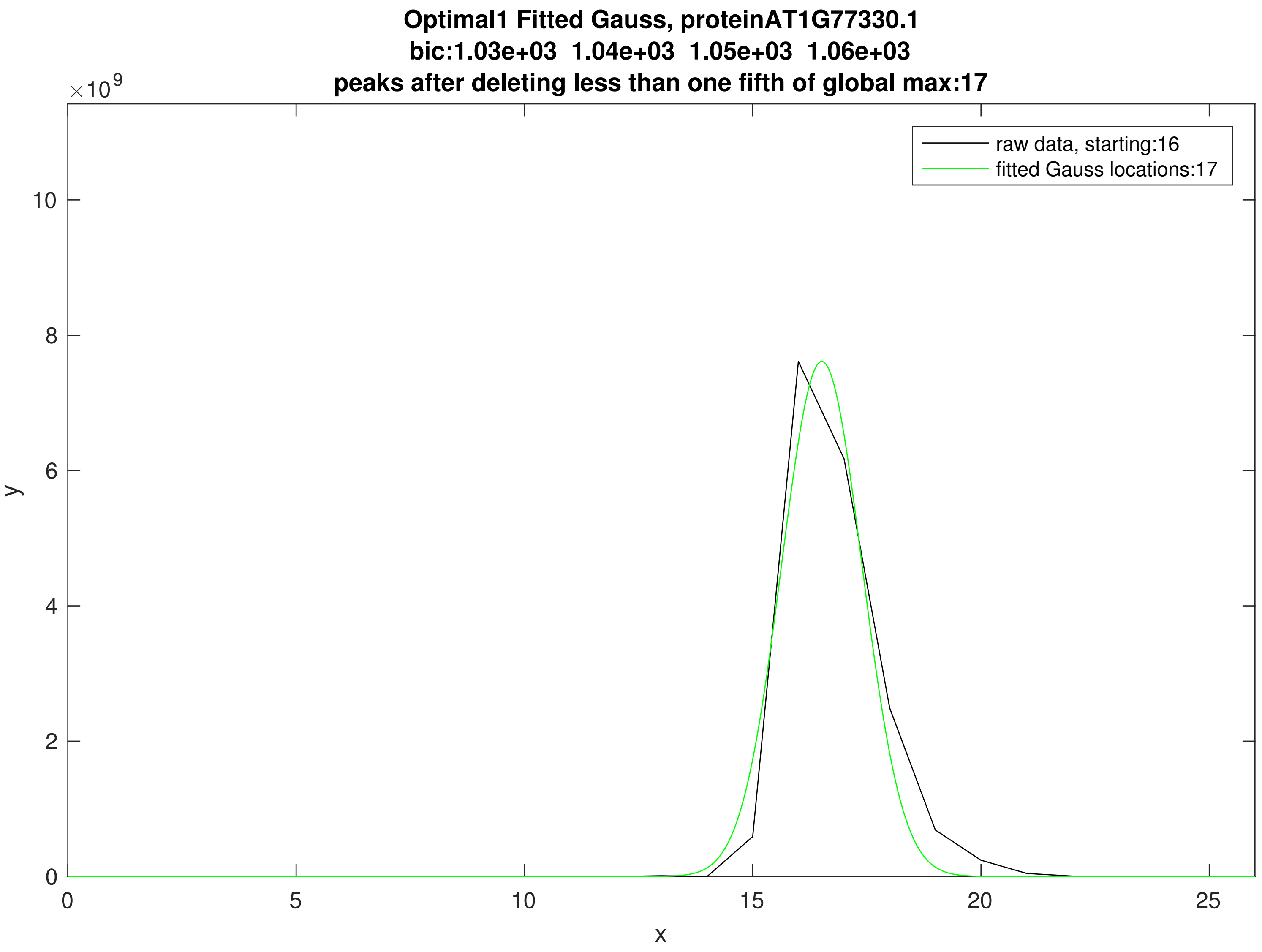

Optimal1 Fitted Gauss, proteinAT1G77330.1
bic:1.03e+03 1.04e+03 1.05e+03 1.06e+03
peaks after deleting less than one fifth of global max:17
×10^9
raw data, starting:16
fitted Gauss locations:17
y
x
0
2
4
6
8
10
0
5
10
15
20
25

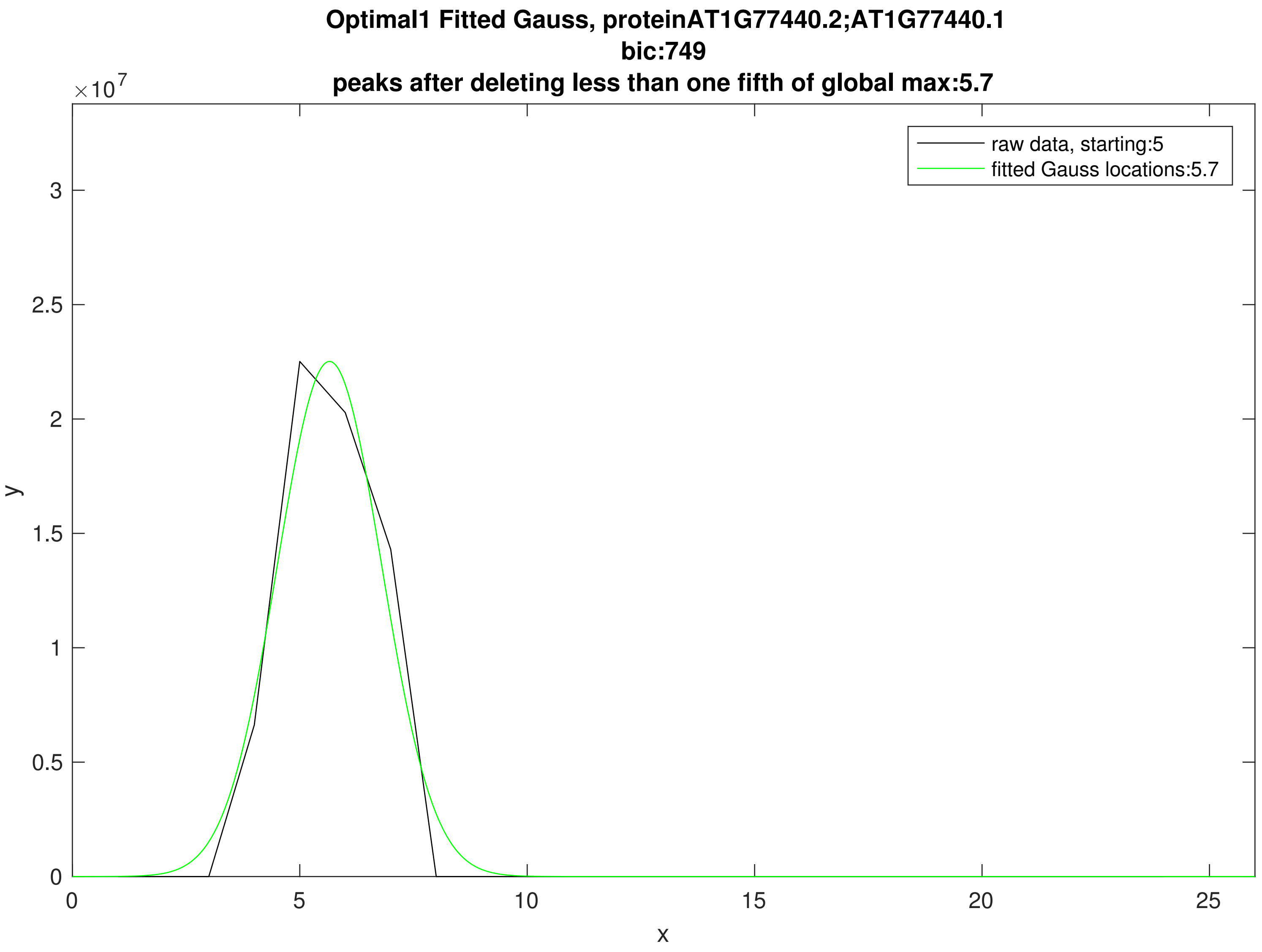

Optimal1 Fitted Gauss, proteinAT1G77440.2;AT1G77440.1
bic:749
peaks after deleting less than one fifth of global max:5.7
×10^7
3
2.5
2
1.5
1
0.5
0
y
0
5
10
15
20
25
x
raw data, starting:5
fitted Gauss locations:5.7

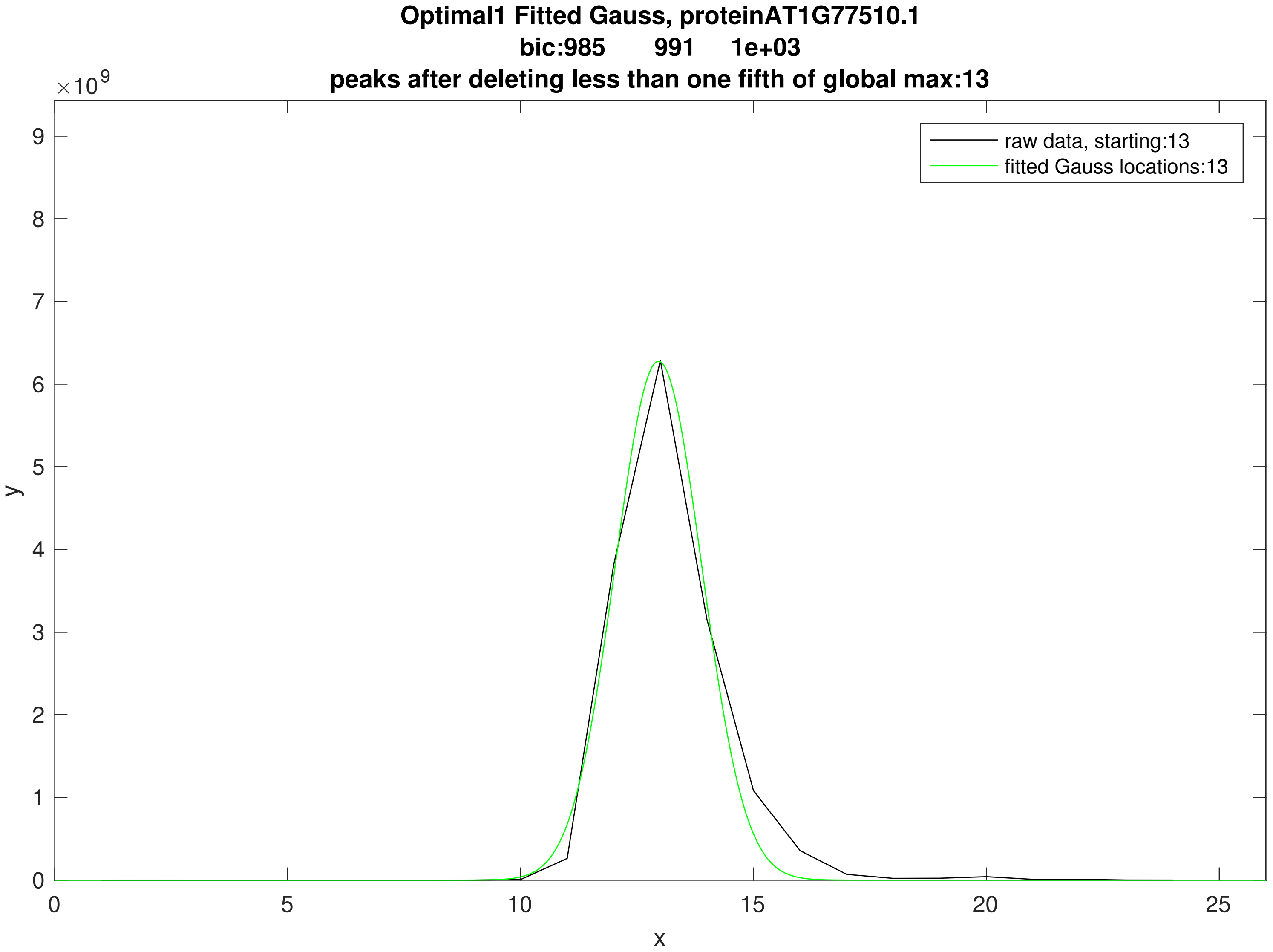

Optimal1 Fitted Gauss, proteinAT1G77510.1
bic:985 991 1e+03
peaks after deleting less than one fifth of global max:13
×10^9
raw data, starting:13
fitted Gauss locations:13
y
x
0
1
2
3
4
5
6
7
8
9
0
5
10
15
20
25

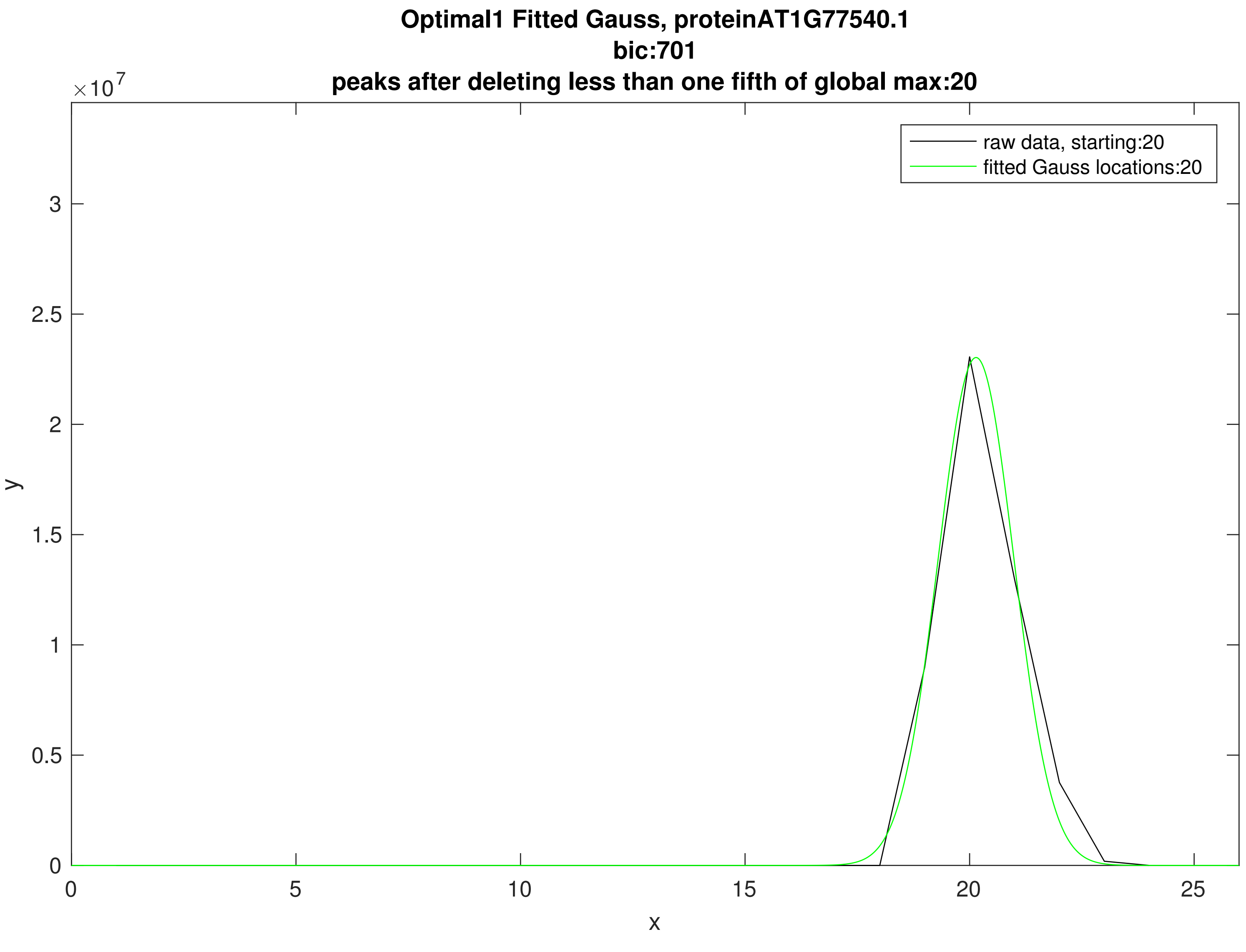

Optimal1 Fitted Gauss, proteinAT1G77540.1
bic:701
peaks after deleting less than one fifth of global max:20
×10^7
raw data, starting:20
fitted Gauss locations:20
y
x
0
0.5
1
1.5
2
2.5
3
0
5
10
15
20
25

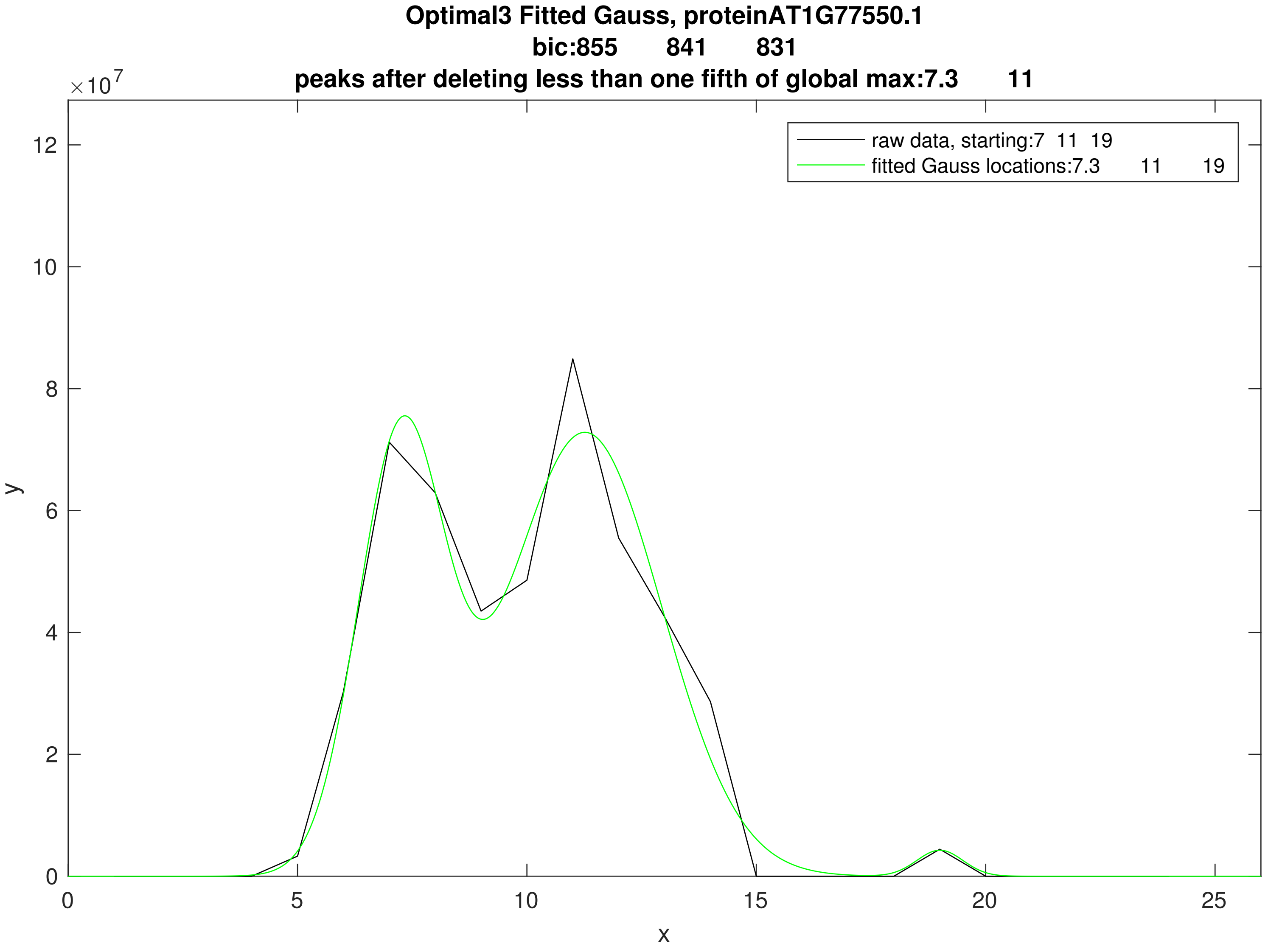
Optimal3 Fitted Gauss, proteinAT1G77550.1
bic:855 841 831
peaks after deleting less than one fifth of global max:7.3 11
×10^7
raw data, starting:7 11 19
fitted Gauss locations:7.3 11 19
y
x
0
2
4
6
8
10
12
0
5
10
15
20
25

**Optimal1 Fitted Gauss, proteinAT1G77670.1**
**bic:849**
**peaks after deleting less than one fifth of global max:14**

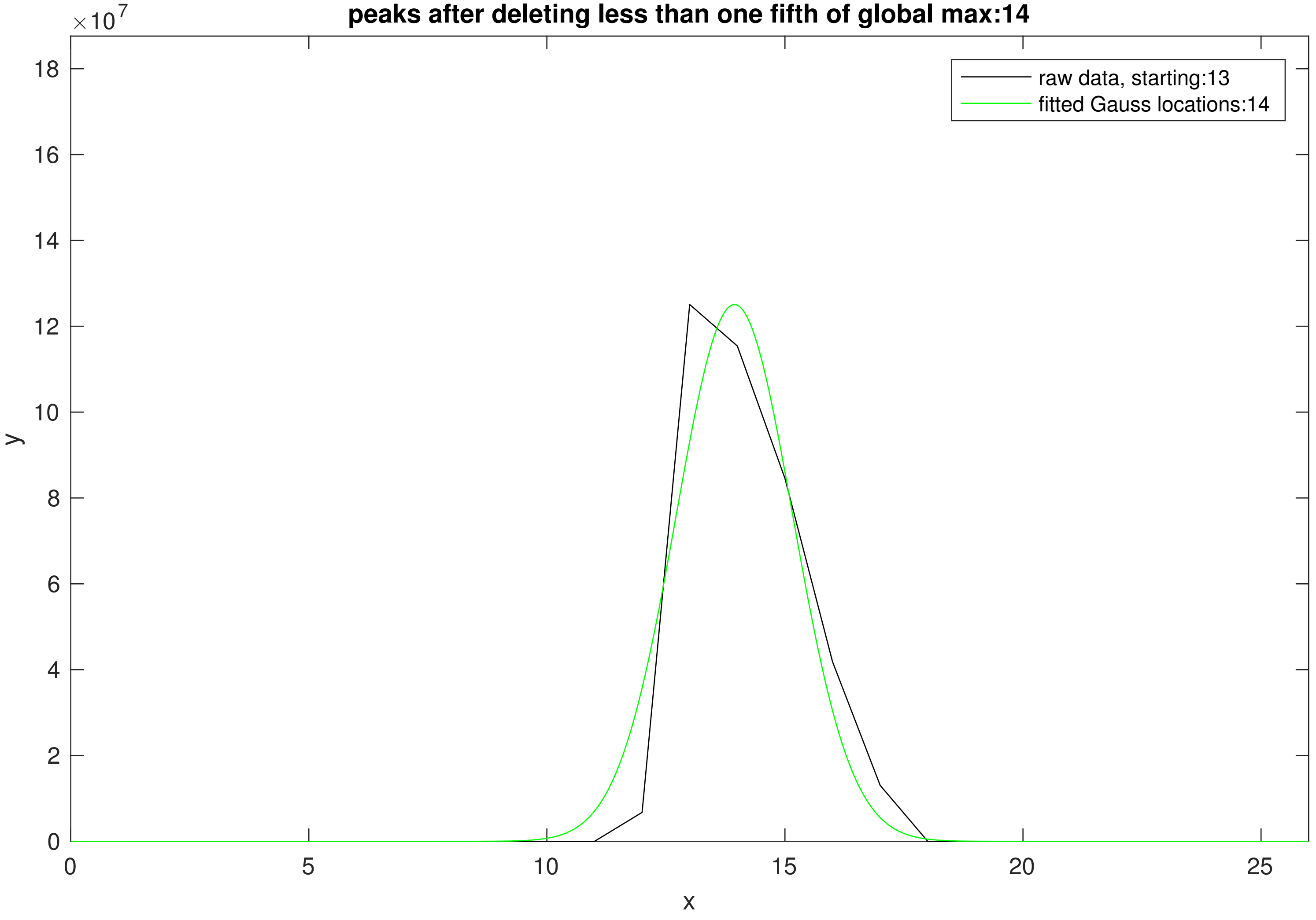

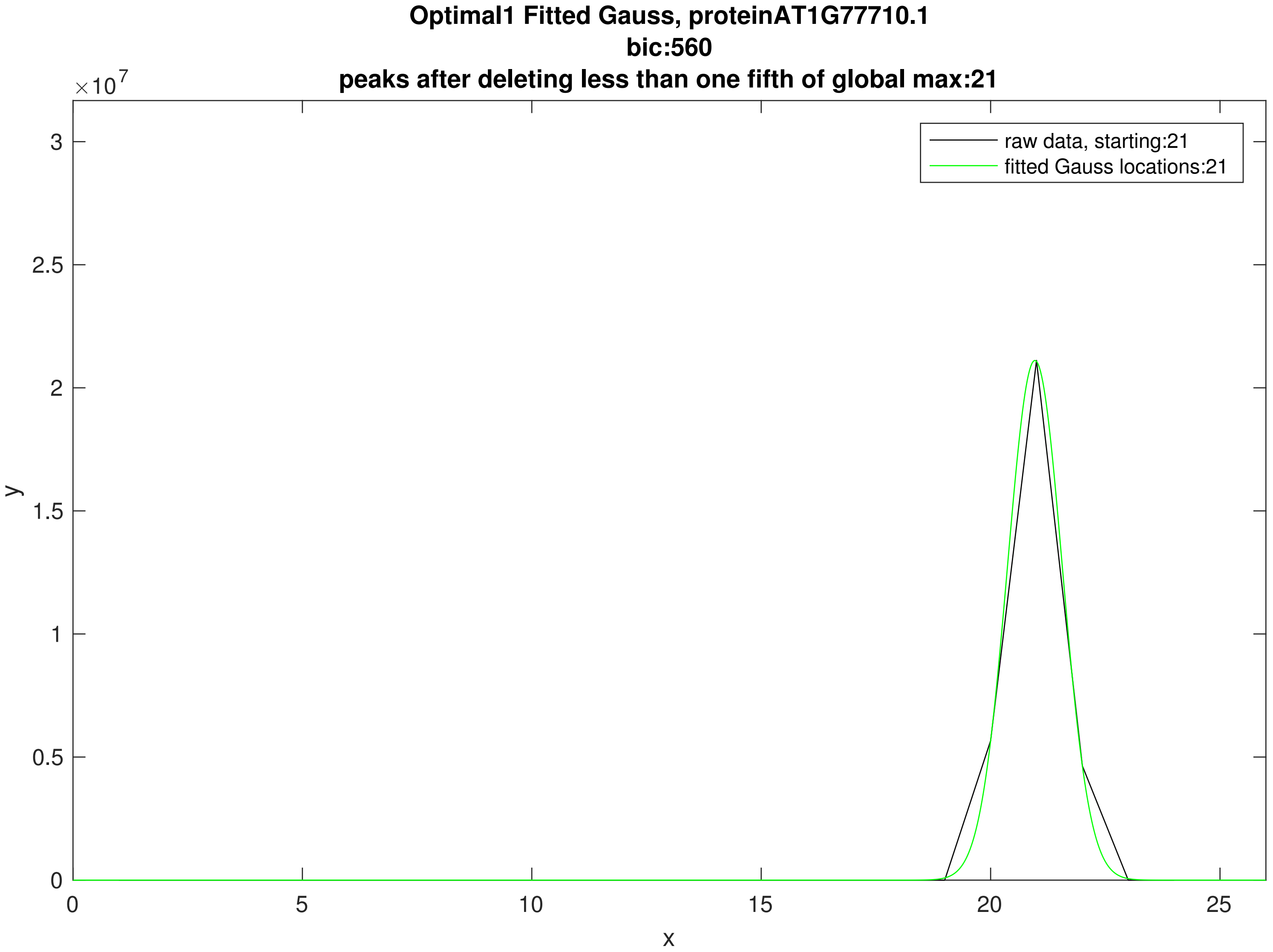

Optimal1 Fitted Gauss, proteinAT1G77710.1
bic:560
peaks after deleting less than one fifth of global max:21
×10^7
3
2.5
2
1.5
1
0.5
0
y
0
5
10
15
20
25
x
raw data, starting:21
fitted Gauss locations:21

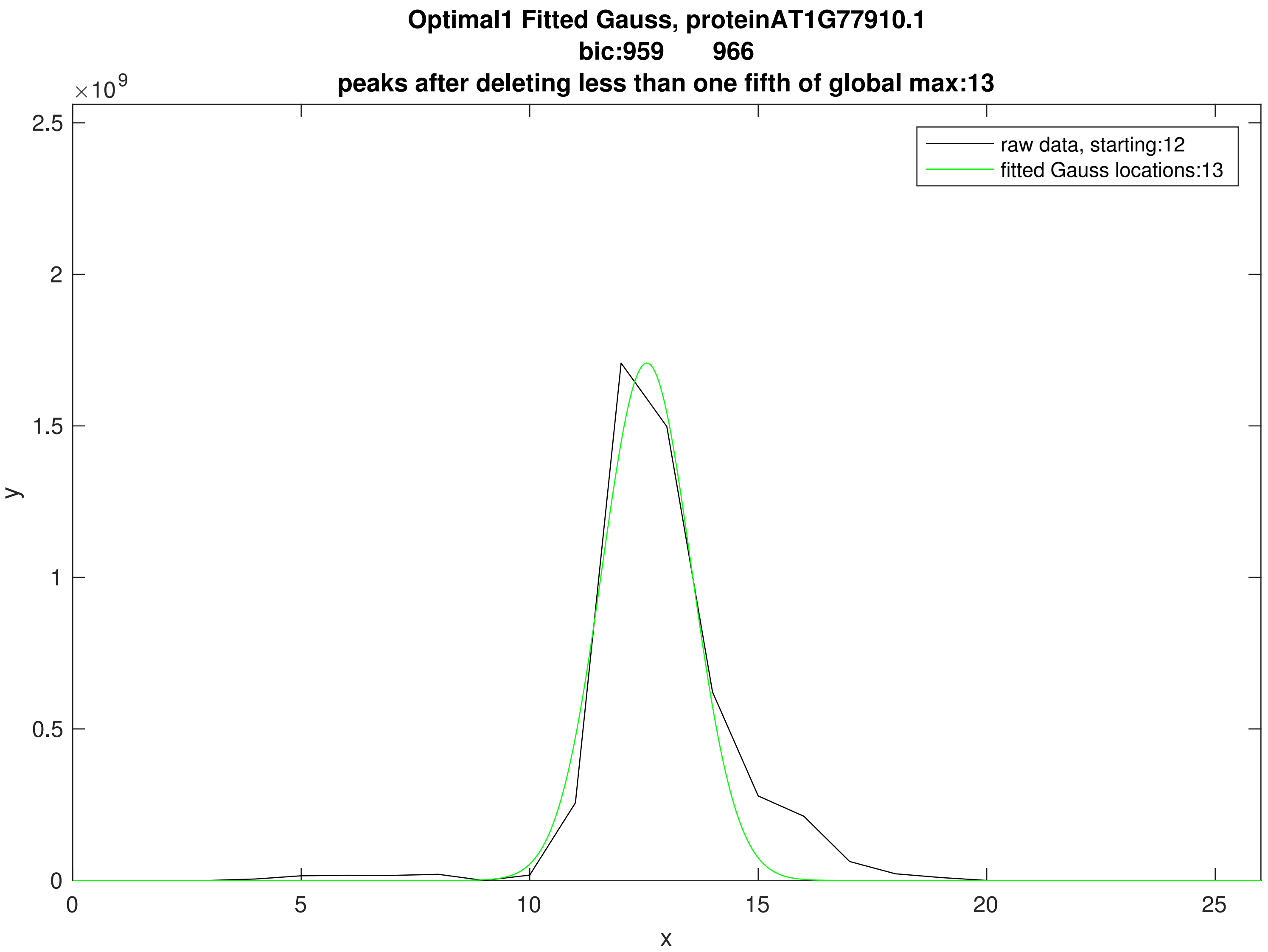
Optimal1 Fitted Gauss, proteinAT1G77910.1
bic:959 966
peaks after deleting less than one fifth of global max:13
×10^9
2.5
2
1.5
1
0.5
0
y
0
5
10
15
20
25
x
raw data, starting:12
fitted Gauss locations:13

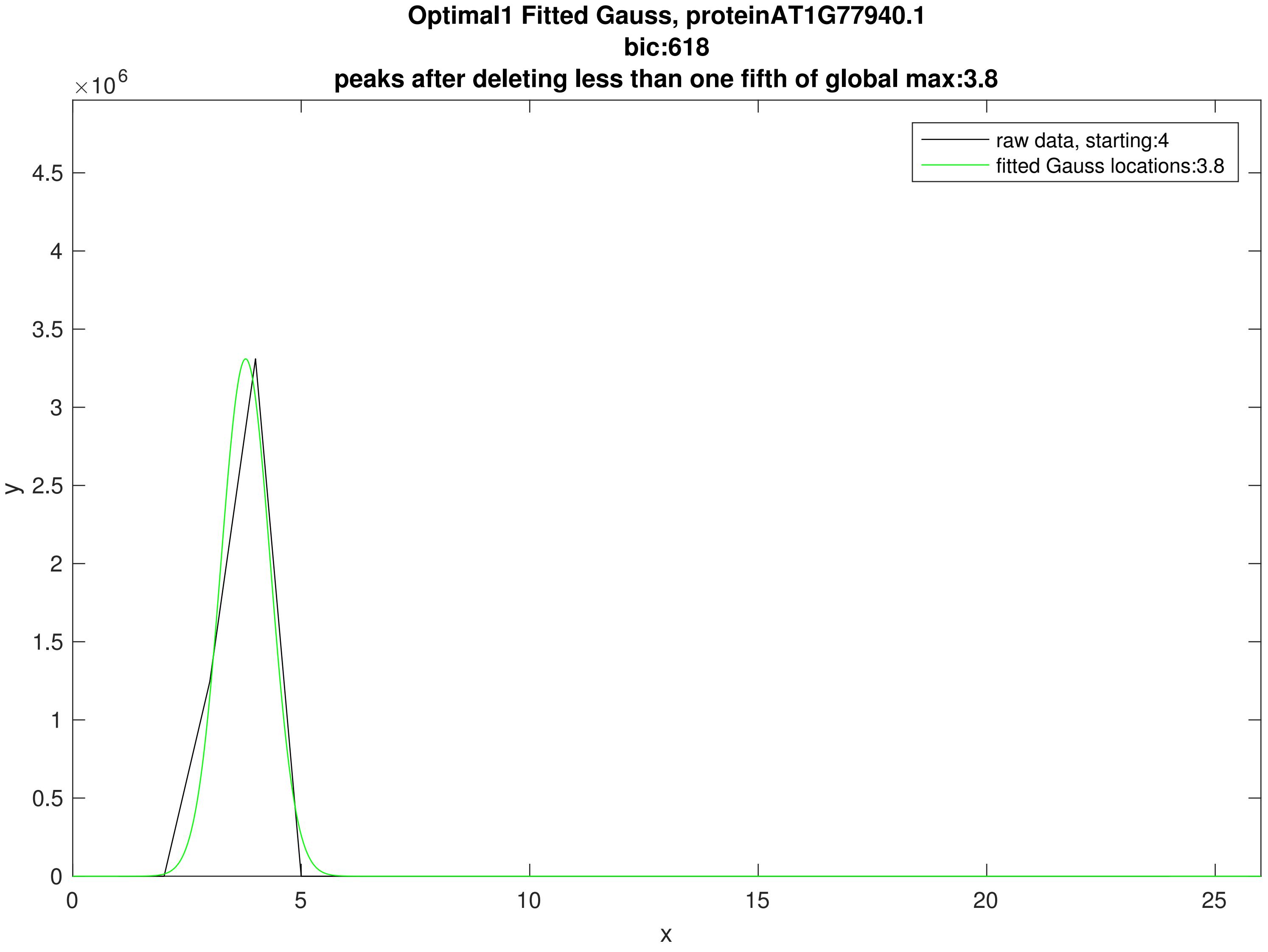
Optimal1 Fitted Gauss, proteinAT1G77940.1
bic:618
peaks after deleting less than one fifth of global max:3.8
×10^6
raw data, starting:4
fitted Gauss locations:3.8
y
x
0
0.5
1
1.5
2
2.5
3
3.5
4
4.5
5
10
15
20
25

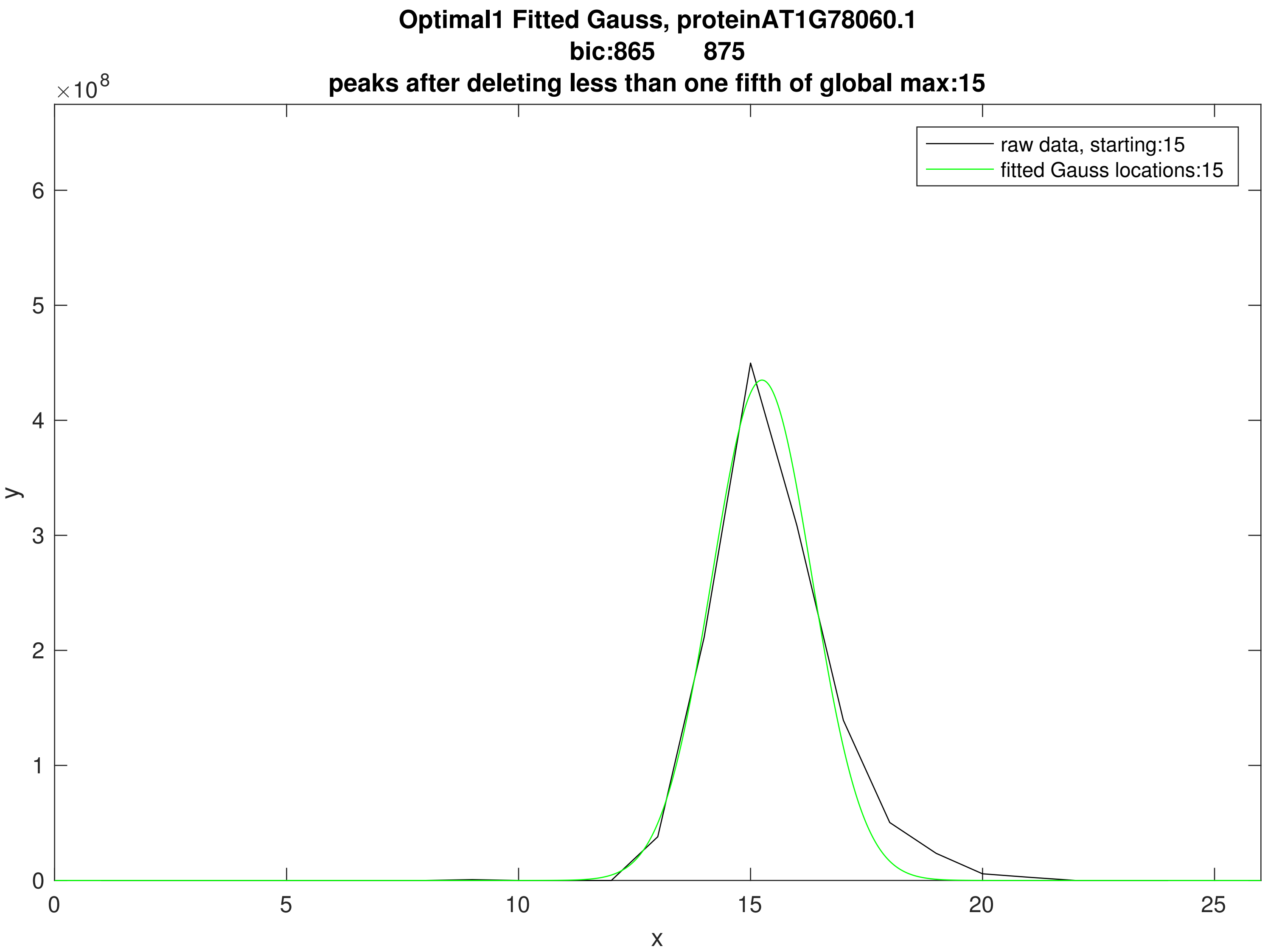
Optimal1 Fitted Gauss, proteinAT1G78060.1
bic:865 875
peaks after deleting less than one fifth of global max:15
×10^8
raw data, starting:15
fitted Gauss locations:15
6
5
4
3
2
1
0
y
0
5
10
15
20
25
x

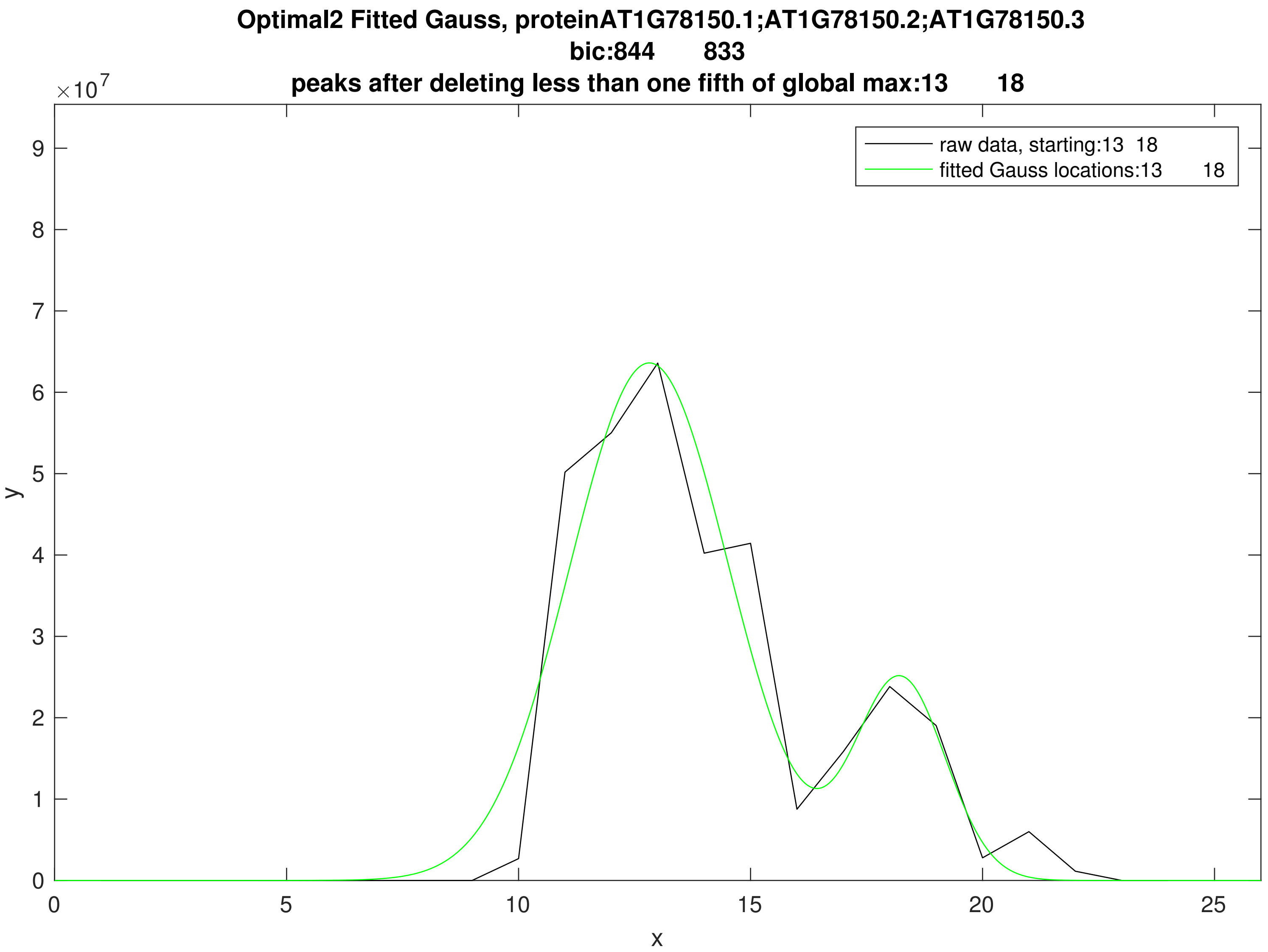

Optimal2 Fitted Gauss, proteinAT1G78150.1;AT1G78150.2;AT1G78150.3
bic:844 833
peaks after deleting less than one fifth of global max:13 18
×10^7
raw data, starting:13 18
fitted Gauss locations:13 18
y
x

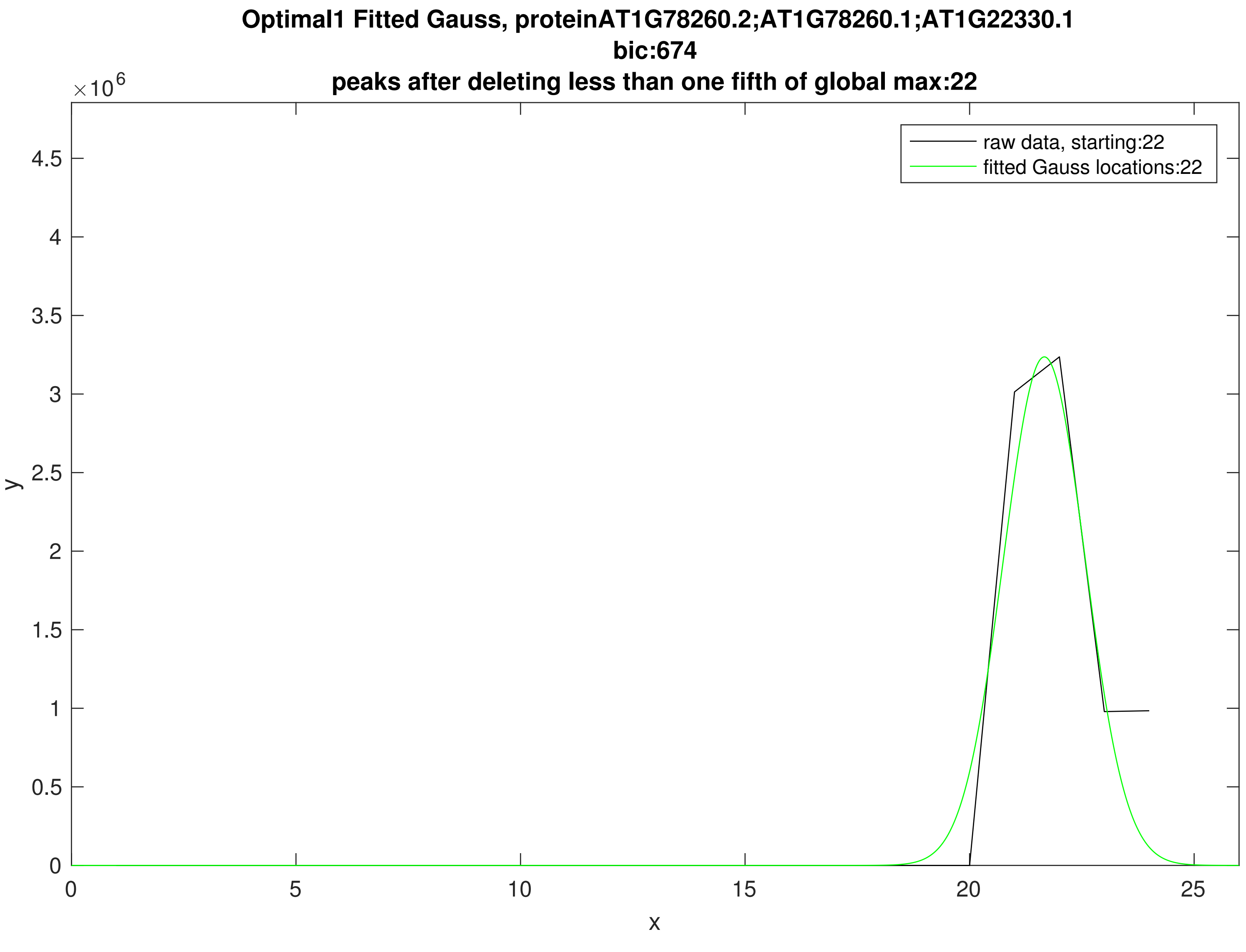
Optimal1 Fitted Gauss, proteinAT1G78260.2;AT1G78260.1;AT1G22330.1
bic:674
peaks after deleting less than one fifth of global max:22
×10^6
raw data, starting:22
fitted Gauss locations:22
y
x
0
0.5
1
1.5
2
2.5
3
3.5
4
4.5
0
5
10
15
20
25

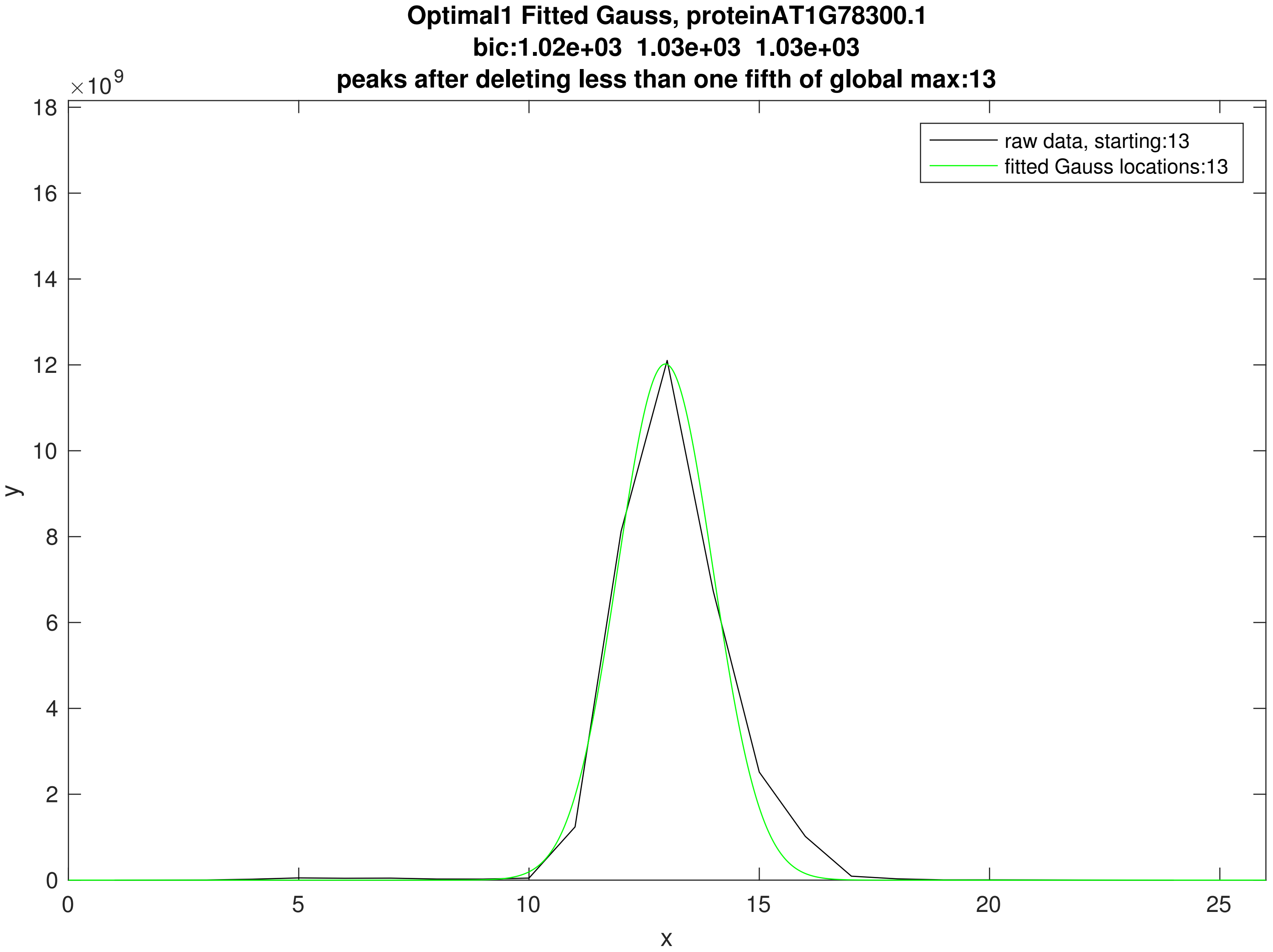
Optimal1 Fitted Gauss, proteinAT1G78300.1
bic:1.02e+03  1.03e+03  1.03e+03
peaks after deleting less than one fifth of global max:13
×10^9
raw data, starting:13
fitted Gauss locations:13
y
x
0
2
4
6
8
10
12
14
16
18
0
5
10
15
20
25

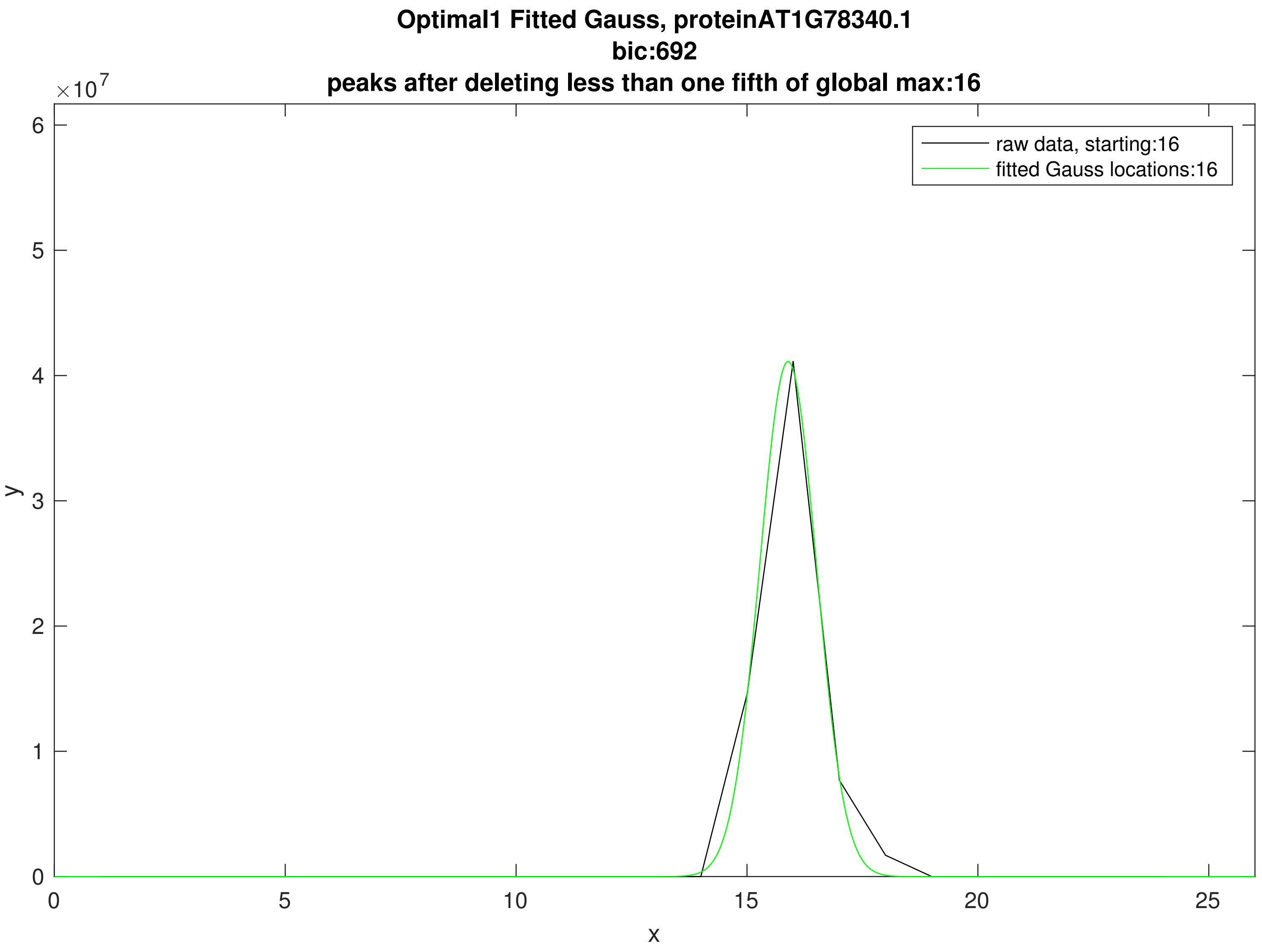

Optimal1 Fitted Gauss, proteinAT1G78340.1
bic:692
peaks after deleting less than one fifth of global max:16
×10^7
raw data, starting:16
fitted Gauss locations:16
y
x
0
1
2
3
4
5
6
0
5
10
15
20
25

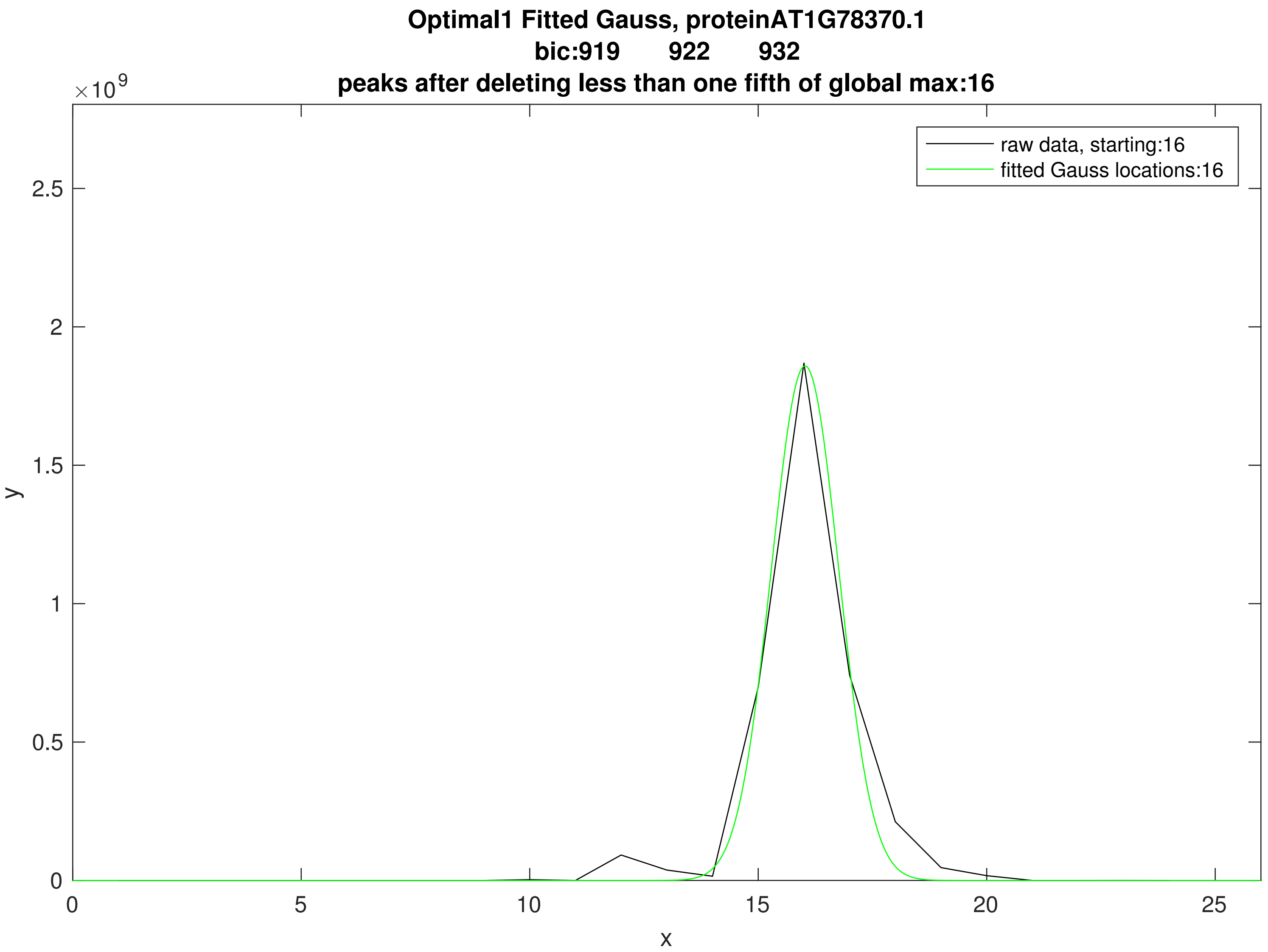
Optimal1 Fitted Gauss, proteinAT1G78370.1
bic:919 922 932
peaks after deleting less than one fifth of global max:16
×10^9
2.5
2
1.5
1
0.5
0
y
0
5
10
15
20
25
x
raw data, starting:16
fitted Gauss locations:16

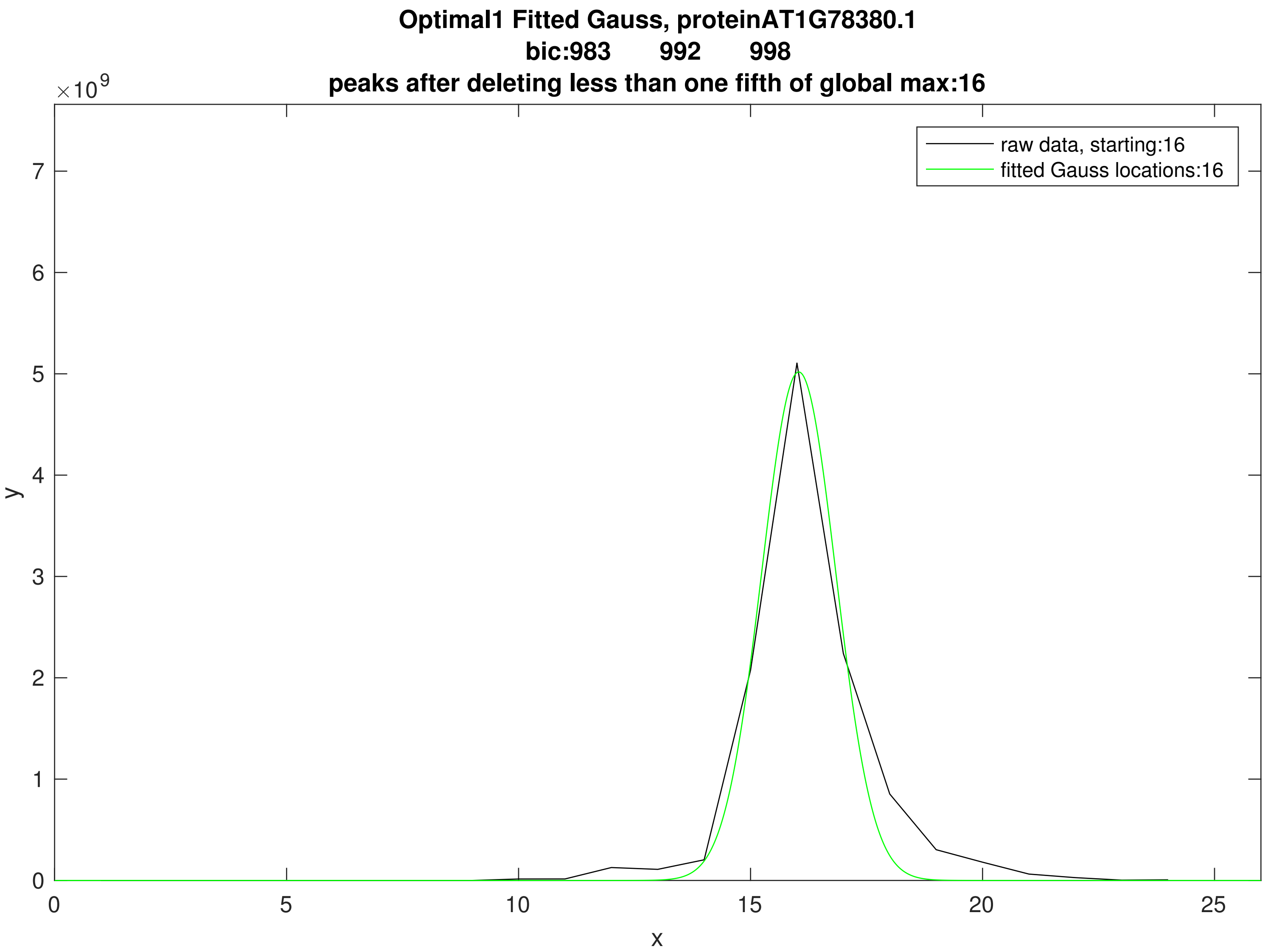
Optimal1 Fitted Gauss, proteinAT1G78380.1
bic:983 992 998
peaks after deleting less than one fifth of global max:16
×10^9
raw data, starting:16
fitted Gauss locations:16
y
x
0
1
2
3
4
5
6
7
0
5
10
15
20
25

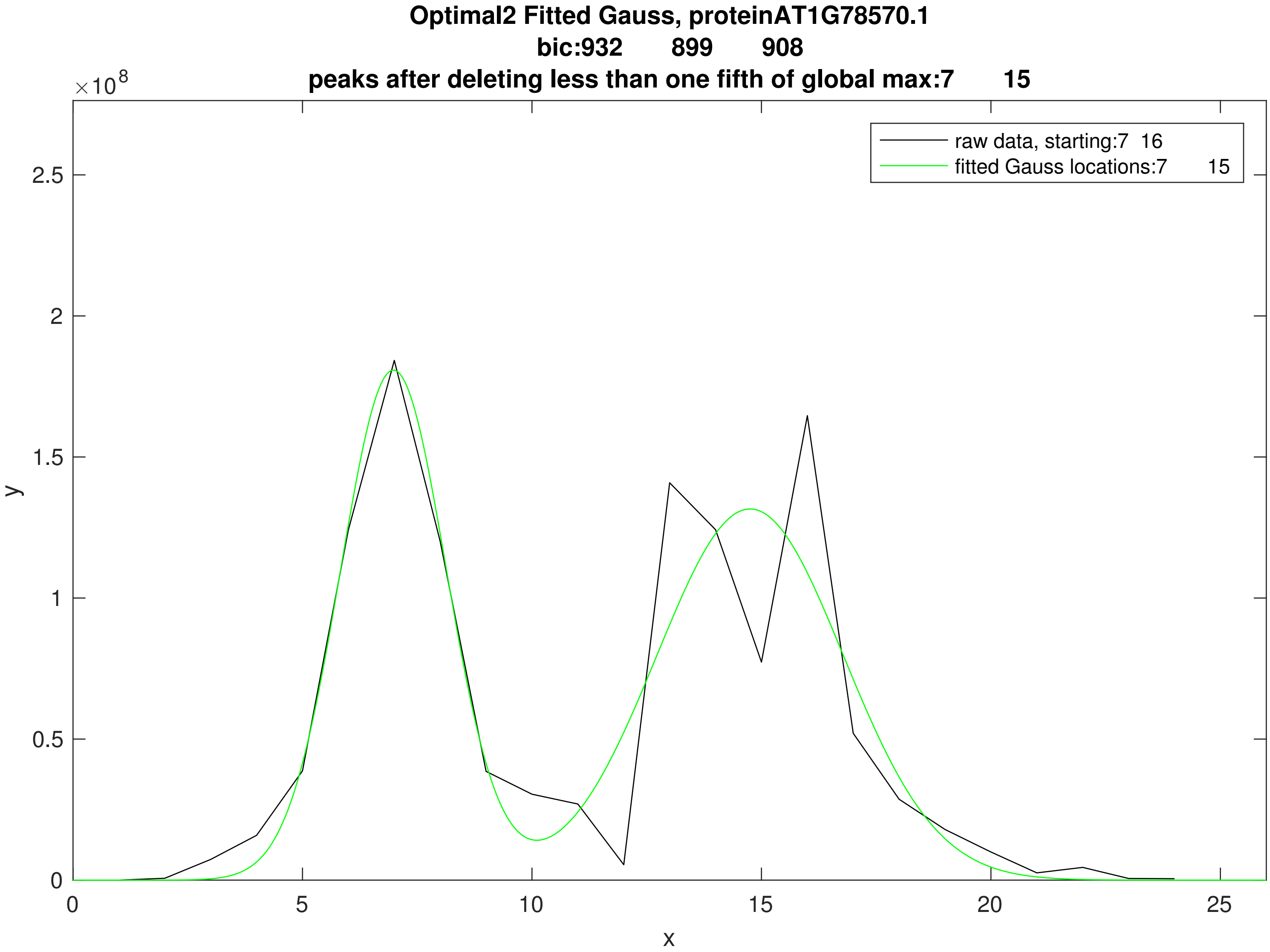
Optimal2 Fitted Gauss, proteinAT1G78570.1
bic:932 899 908
peaks after deleting less than one fifth of global max:7 15
×10^8
raw data, starting:7 16
fitted Gauss locations:7 15
y
x
0
0.5
1
1.5
2
2.5
5
10
15
20
25

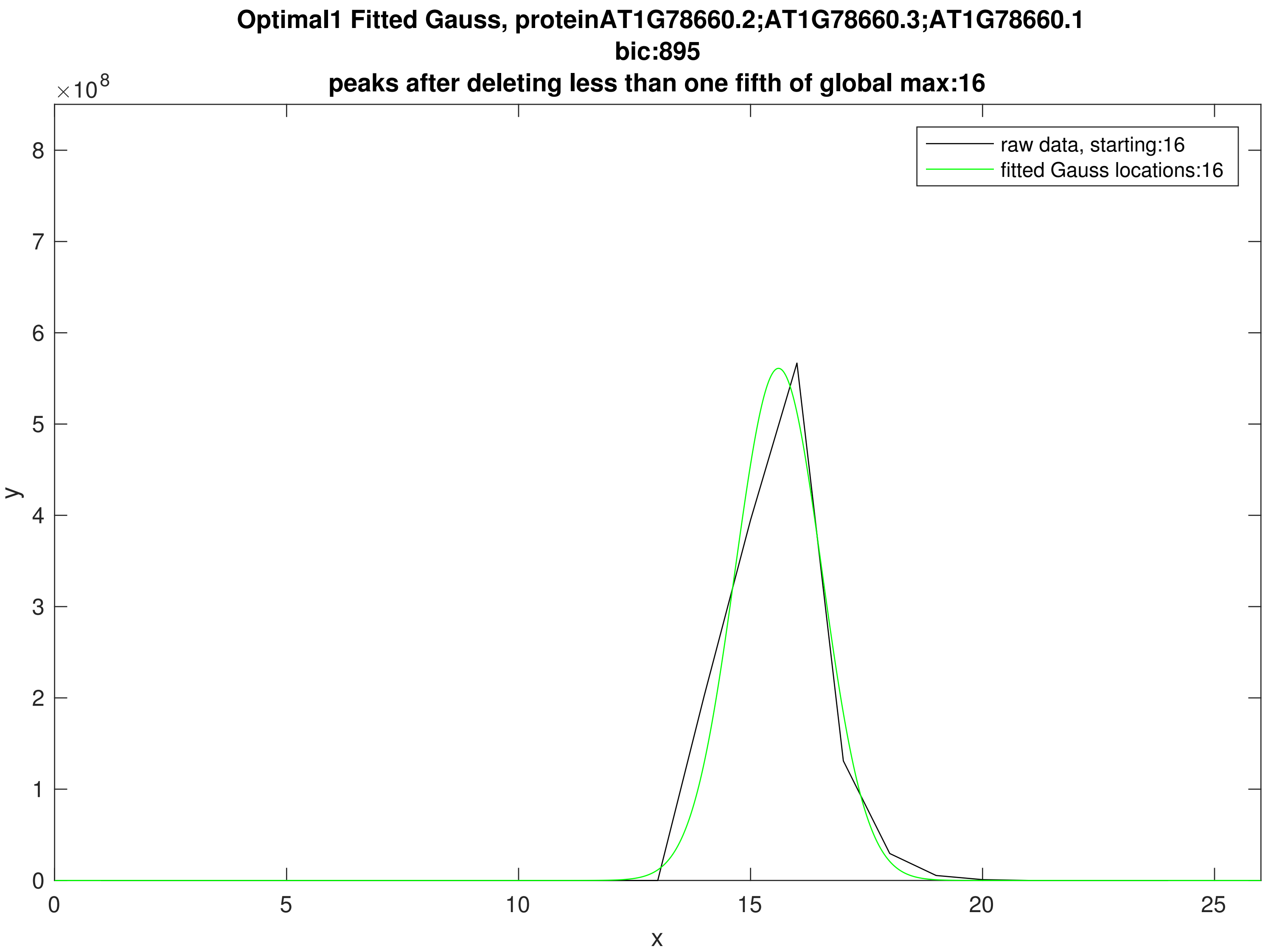
Optimal1 Fitted Gauss, proteinAT1G78660.2;AT1G78660.3;AT1G78660.1
bic:895
peaks after deleting less than one fifth of global max:16
×10^8
raw data, starting:16
fitted Gauss locations:16
y
x
0
1
2
3
4
5
6
7
8
0
5
10
15
20
25

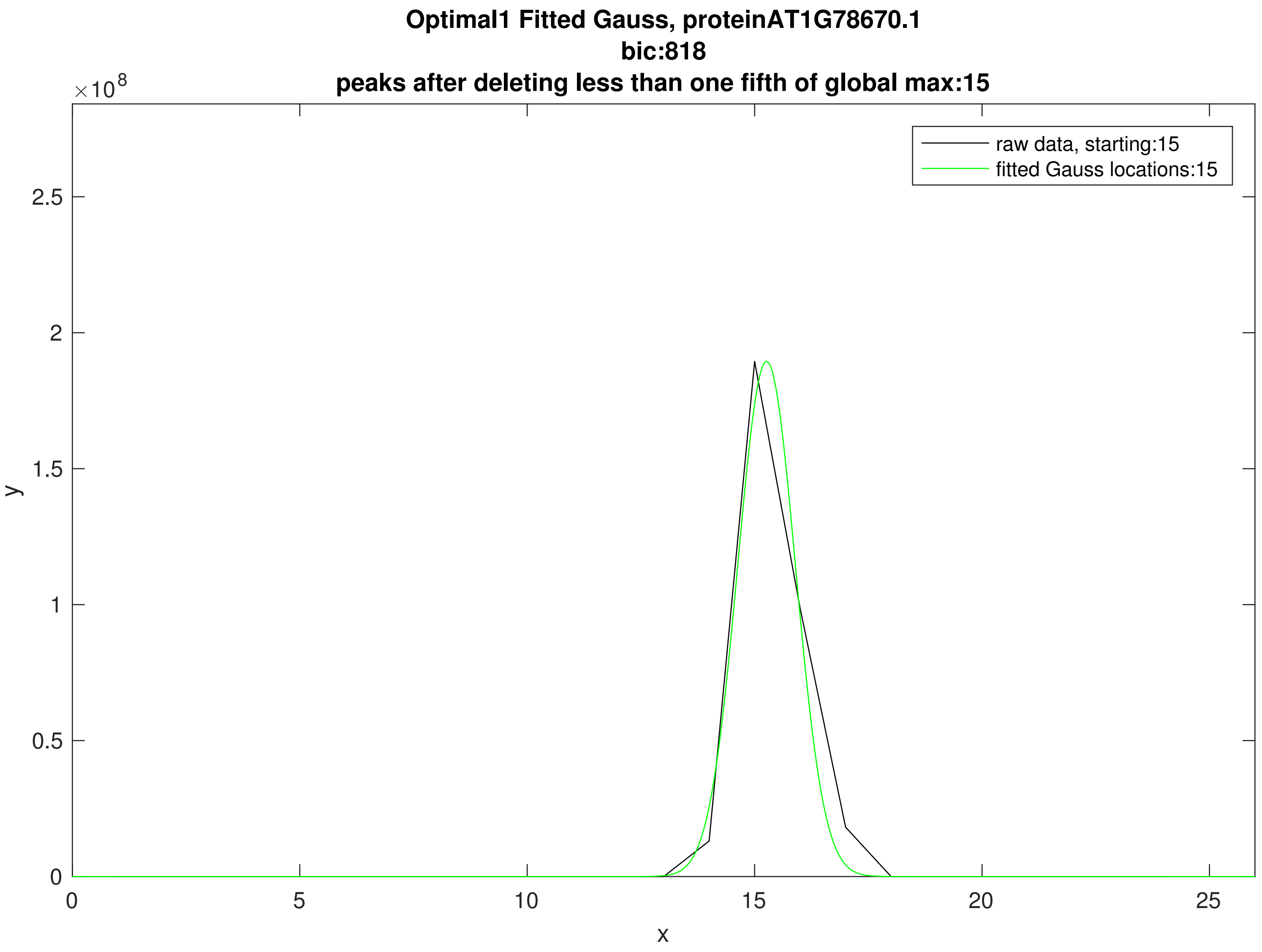
Optimal1 Fitted Gauss, proteinAT1G78670.1
bic:818
peaks after deleting less than one fifth of global max:15
×10^8
raw data, starting:15
fitted Gauss locations:15
y
x
0
0.5
1
1.5
2
2.5
0
5
10
15
20
25

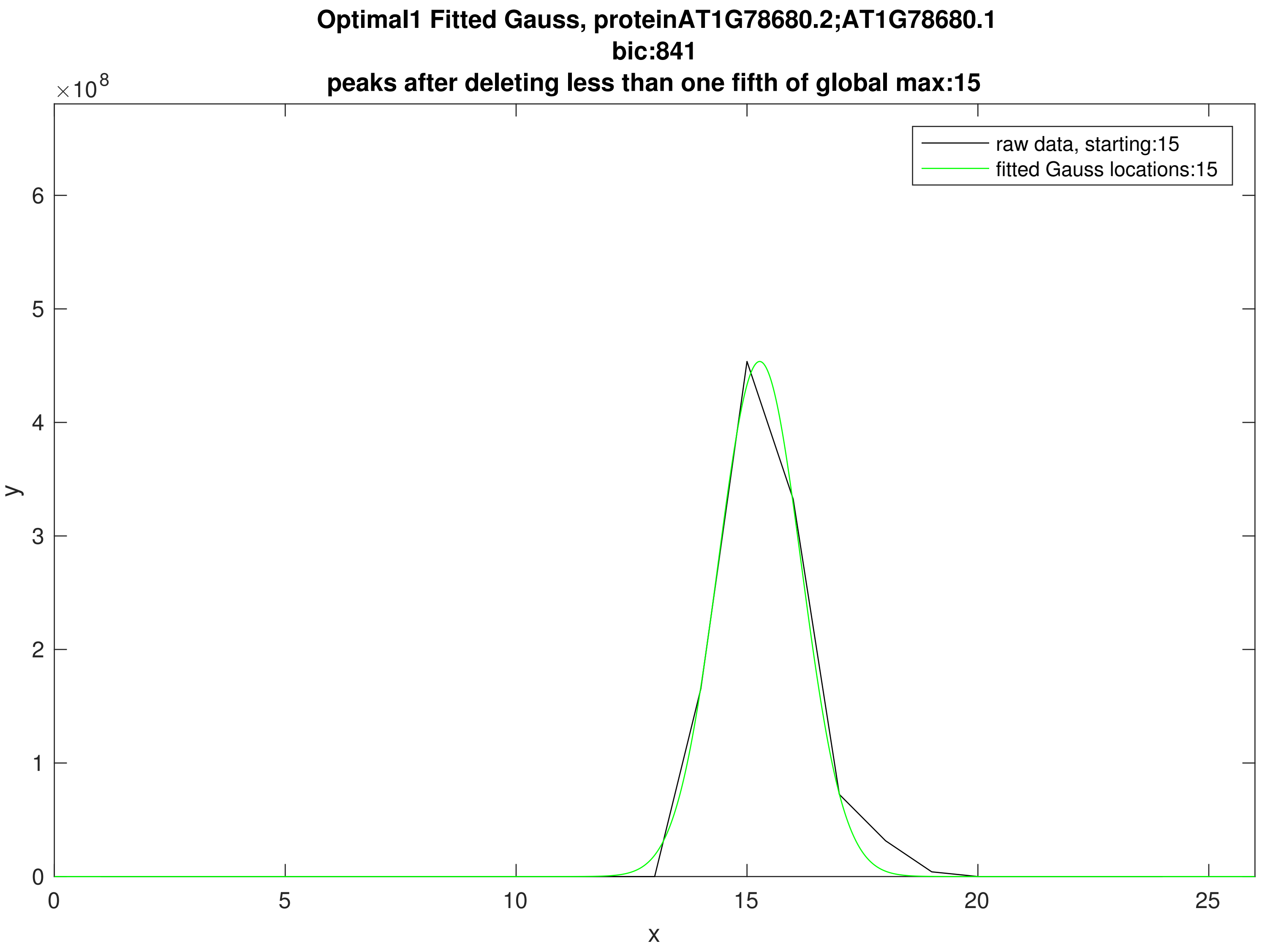
Optimal1 Fitted Gauss, proteinAT1G78680.2;AT1G78680.1
bic:841
peaks after deleting less than one fifth of global max:15
×10^8
raw data, starting:15
fitted Gauss locations:15
y
x
0
1
2
3
4
5
6
0
5
10
15
20
25

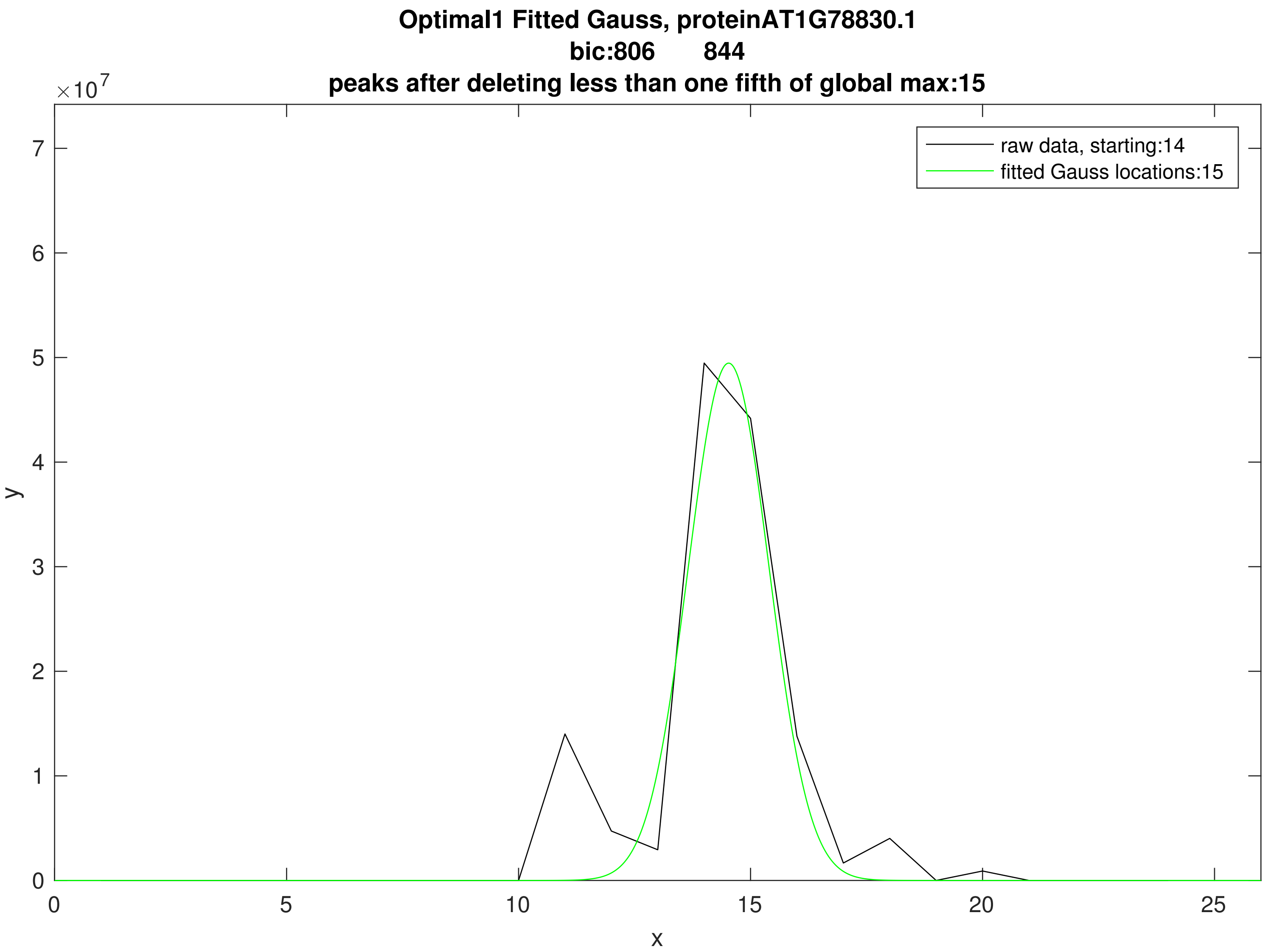
Optimal1 Fitted Gauss, proteinAT1G78830.1
bic:806 844
peaks after deleting less than one fifth of global max:15
×10^7
raw data, starting:14
fitted Gauss locations:15
y
x
0
1
2
3
4
5
6
7
0
5
10
15
20
25

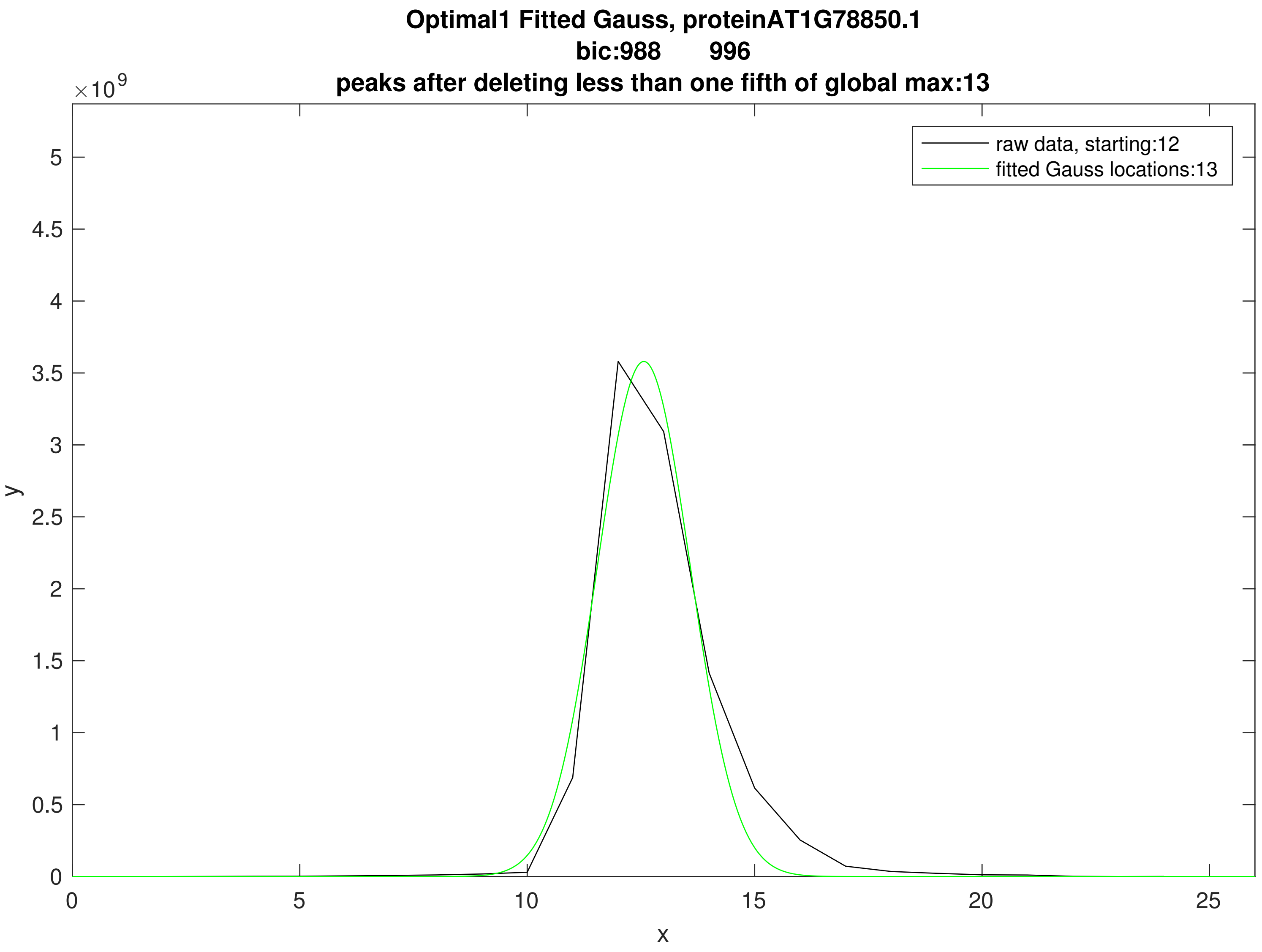

Optimal1 Fitted Gauss, proteinAT1G78850.1
bic:988 996
peaks after deleting less than one fifth of global max:13
×10^9
5
4.5
4
3.5
3
2.5
2
1.5
1
0.5
0
y
0
5
10
15
20
25
x
raw data, starting:12
fitted Gauss locations:13

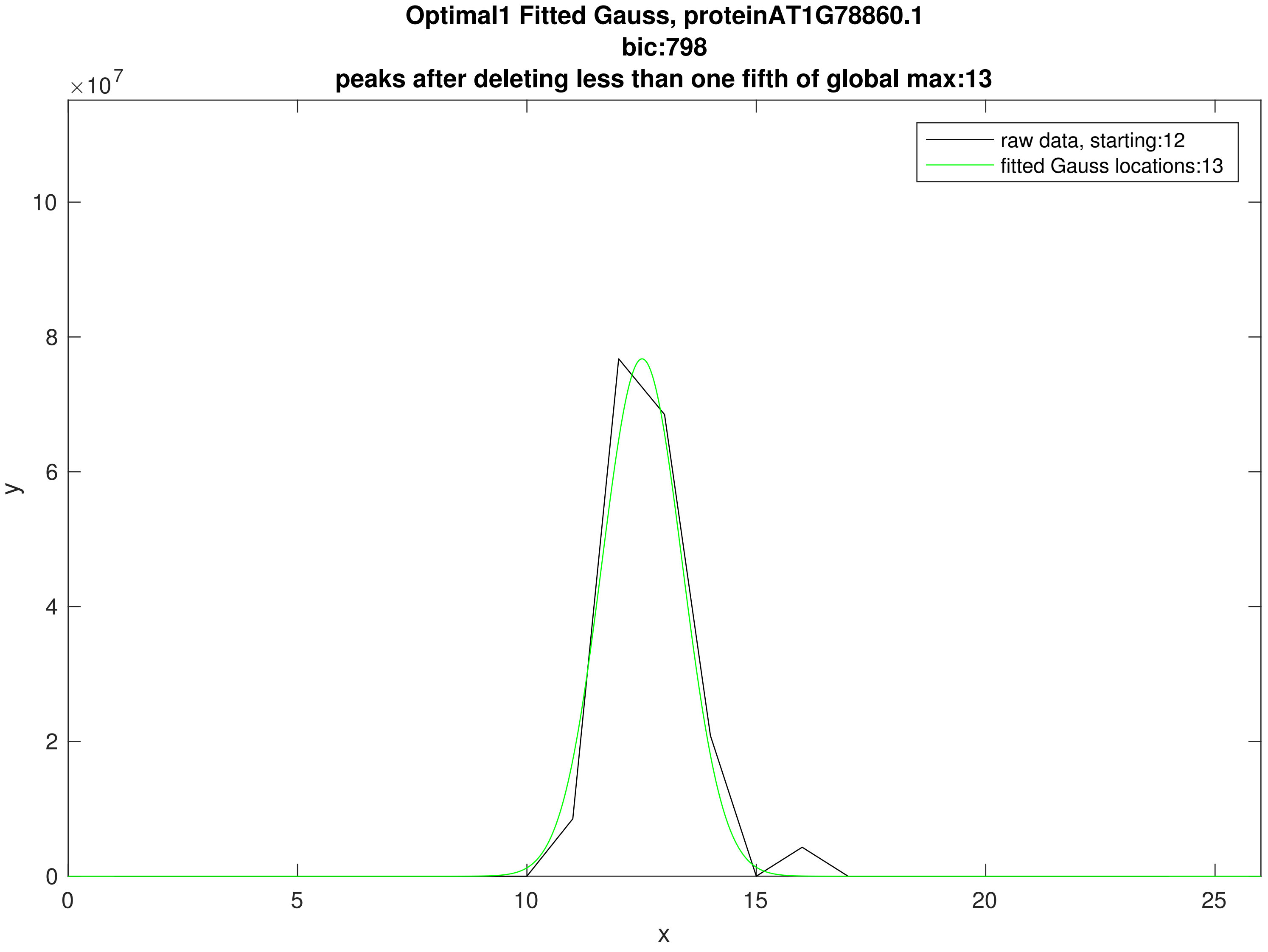

Optimal1 Fitted Gauss, proteinAT1G78860.1
bic:798
peaks after deleting less than one fifth of global max:13
×10^7
raw data, starting:12
fitted Gauss locations:13
y
x
0
2
4
6
8
10
0
5
10
15
20
25

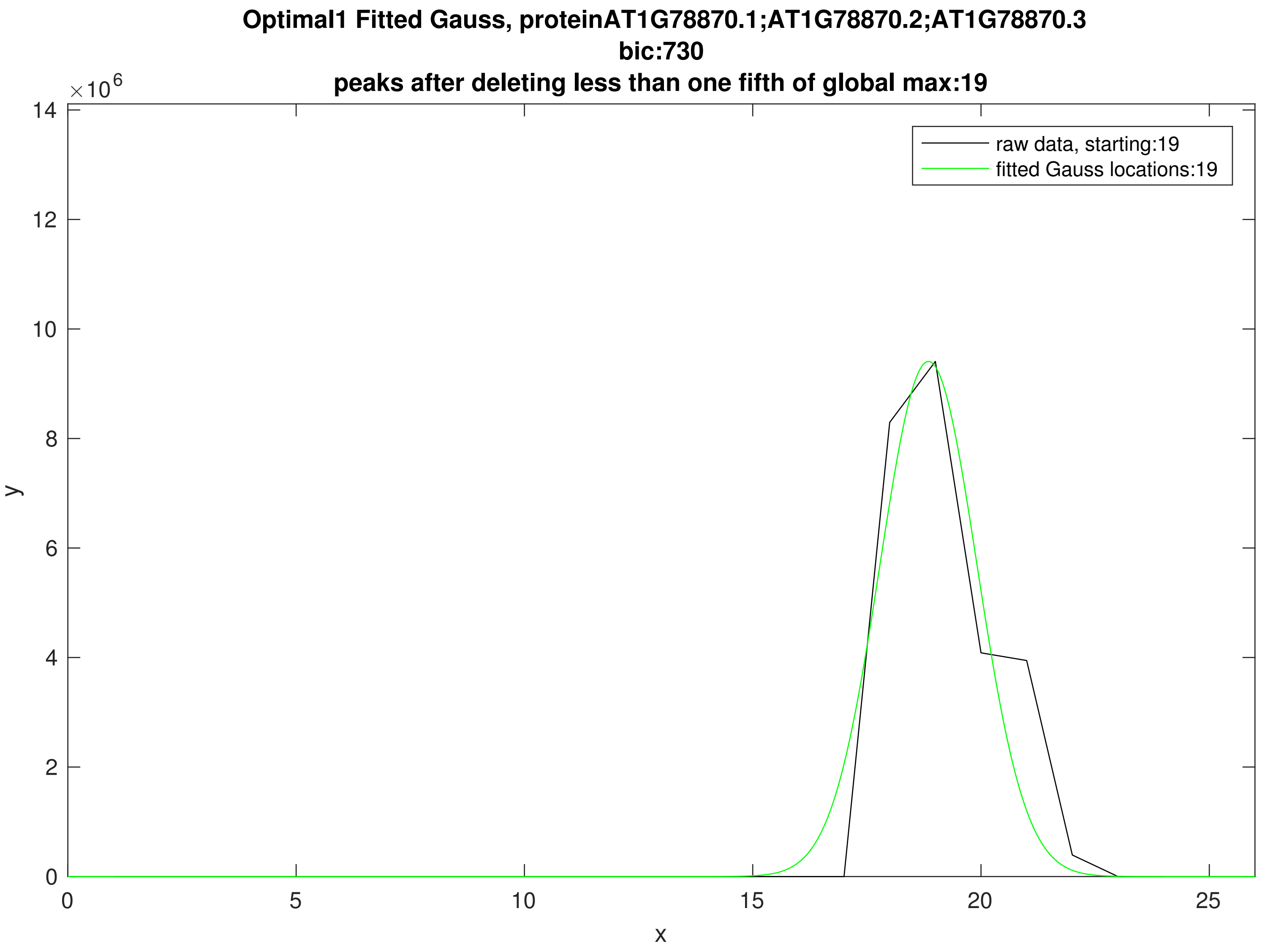
Optimal1 Fitted Gauss, proteinAT1G78870.1;AT1G78870.2;AT1G78870.3
bic:730
peaks after deleting less than one fifth of global max:19
×10^6
14
12
10
8
6
4
2
0
y
0
5
10
15
20
25
x
raw data, starting:19
fitted Gauss locations:19

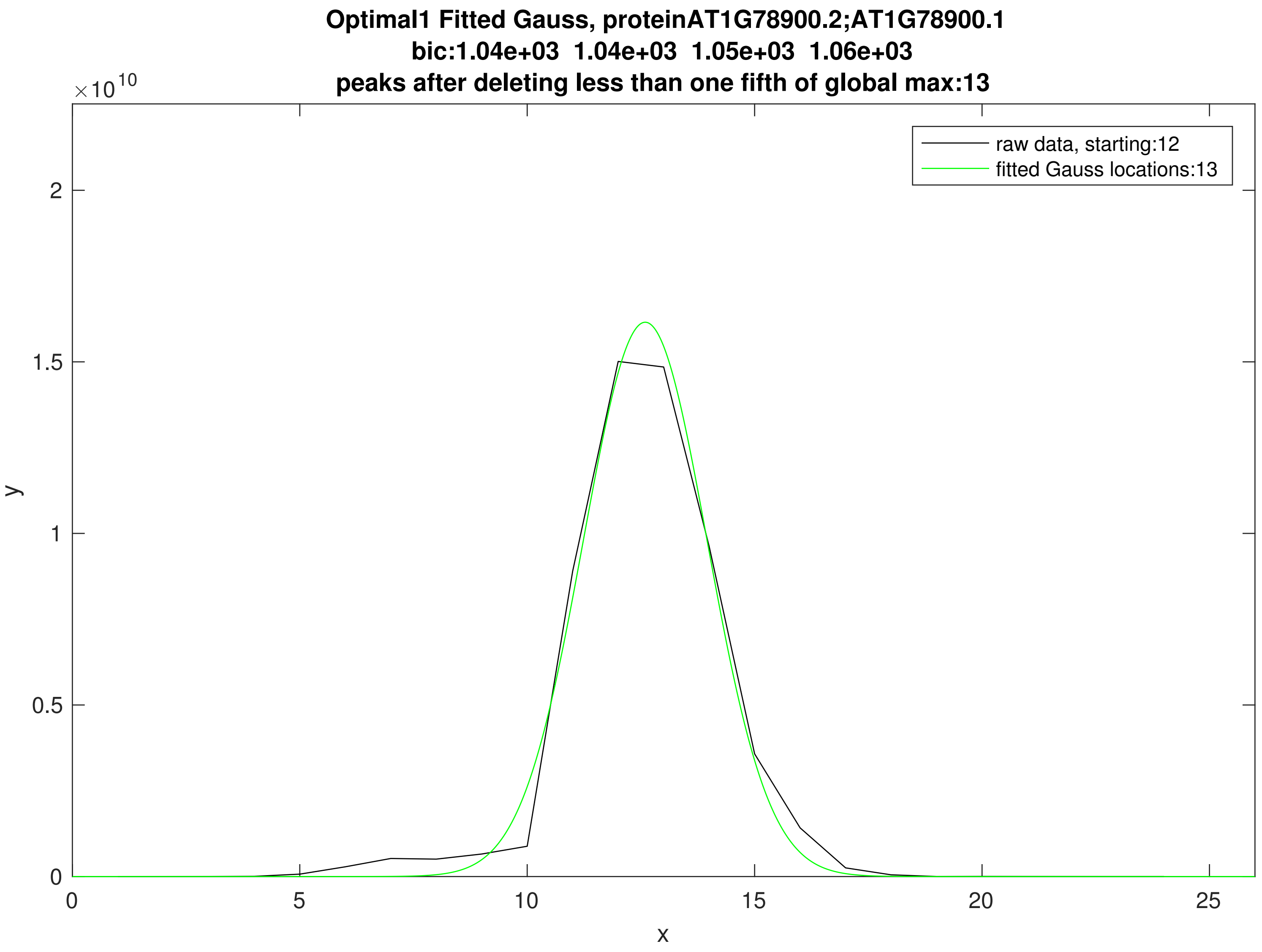
Optimal1 Fitted Gauss, proteinAT1G78900.2;AT1G78900.1
bic:1.04e+03 1.04e+03 1.05e+03 1.06e+03
peaks after deleting less than one fifth of global max:13
×10^10
raw data, starting:12
fitted Gauss locations:13
y
x
0
0.5
1
1.5
2
5
10
15
20
25

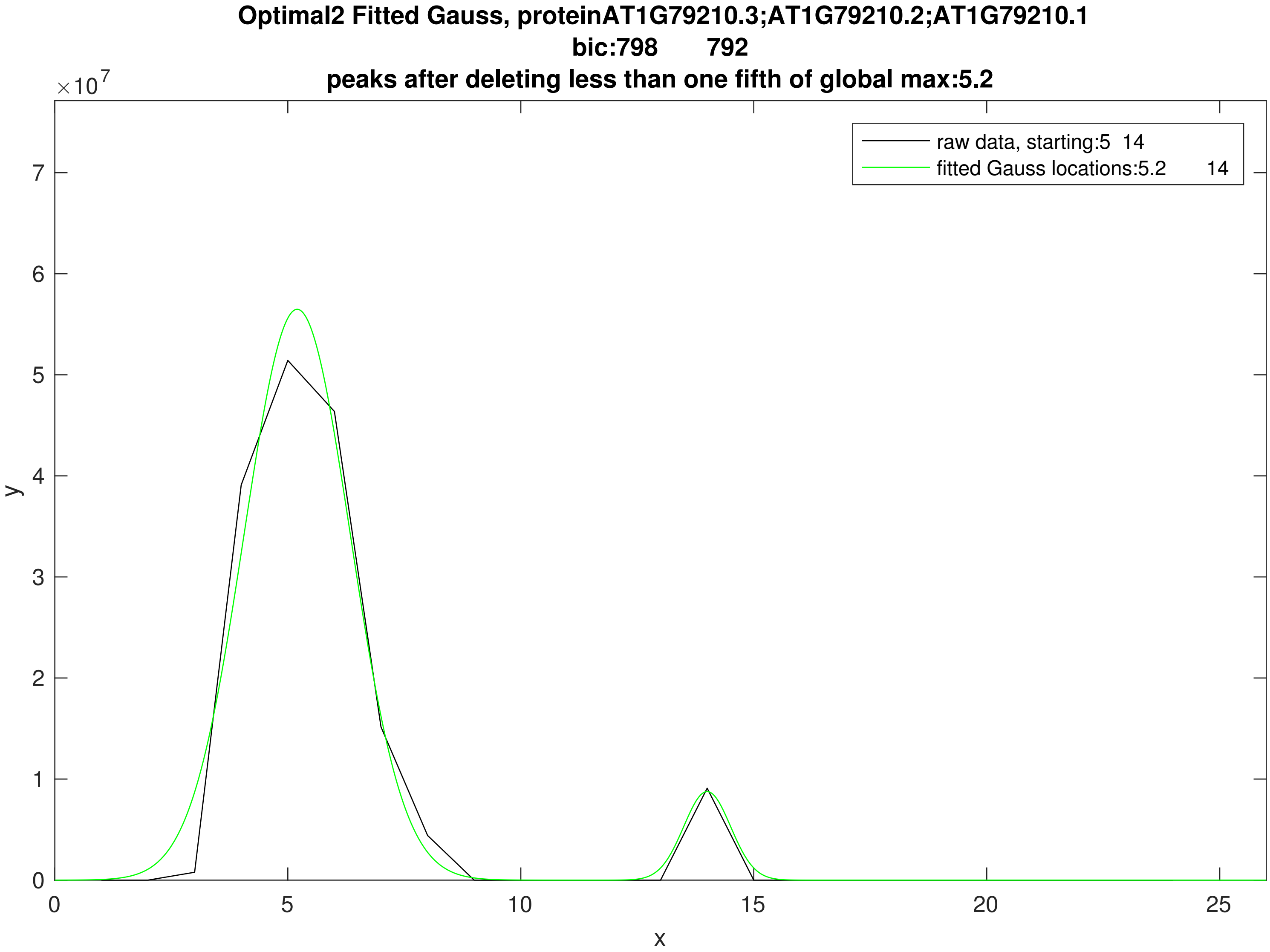

Optimal2 Fitted Gauss, proteinAT1G79210.3;AT1G79210.2;AT1G79210.1
bic:798 792
peaks after deleting less than one fifth of global max:5.2
×10^7
raw data, starting:5 14
fitted Gauss locations:5.2 14
y
x
0
1
2
3
4
5
6
7
0
5
10
15
20
25

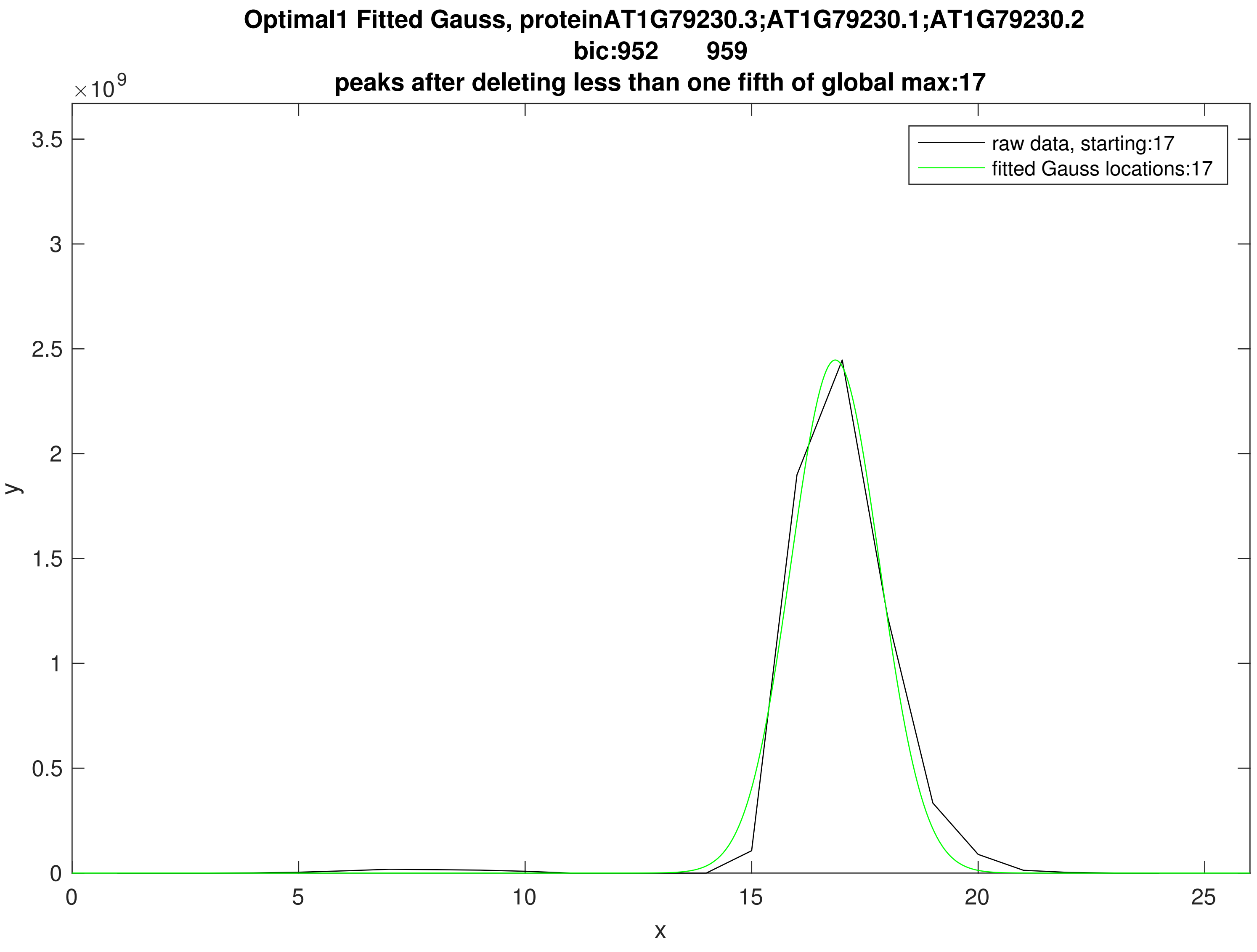
Optimal1 Fitted Gauss, proteinAT1G79230.3;AT1G79230.1;AT1G79230.2
bic:952 959
peaks after deleting less than one fifth of global max:17
×10^9
3.5
3
2.5
2
1.5
1
0.5
0
y
0
5
10
15
20
25
x
raw data, starting:17
fitted Gauss locations:17

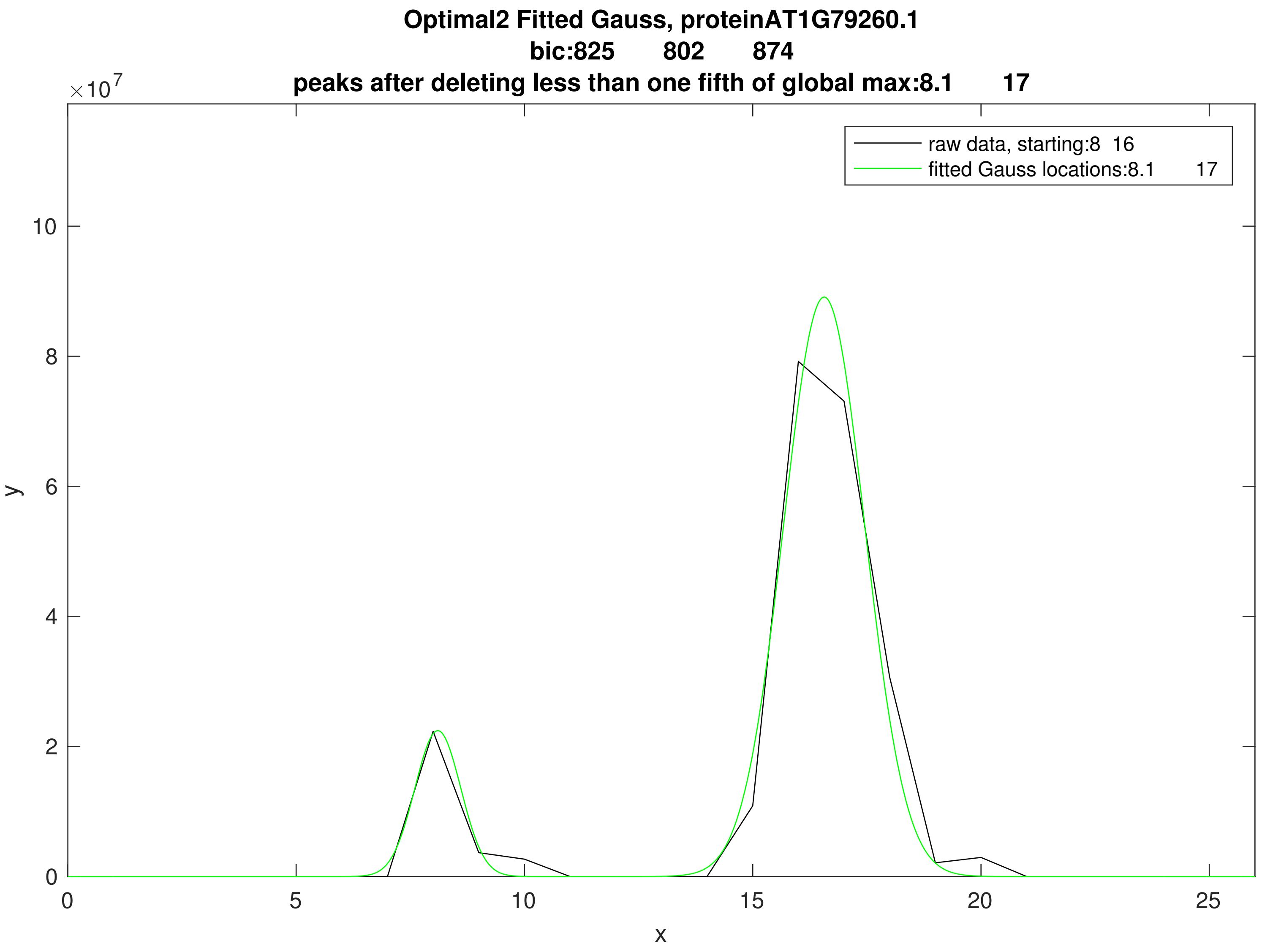
Optimal2 Fitted Gauss, proteinAT1G79260.1
bic:825 802 874
peaks after deleting less than one fifth of global max:8.1 17
×10^7
raw data, starting:8 16
fitted Gauss locations:8.1 17
10
8
6
4
2
0
y
0
5
10
15
20
25
x

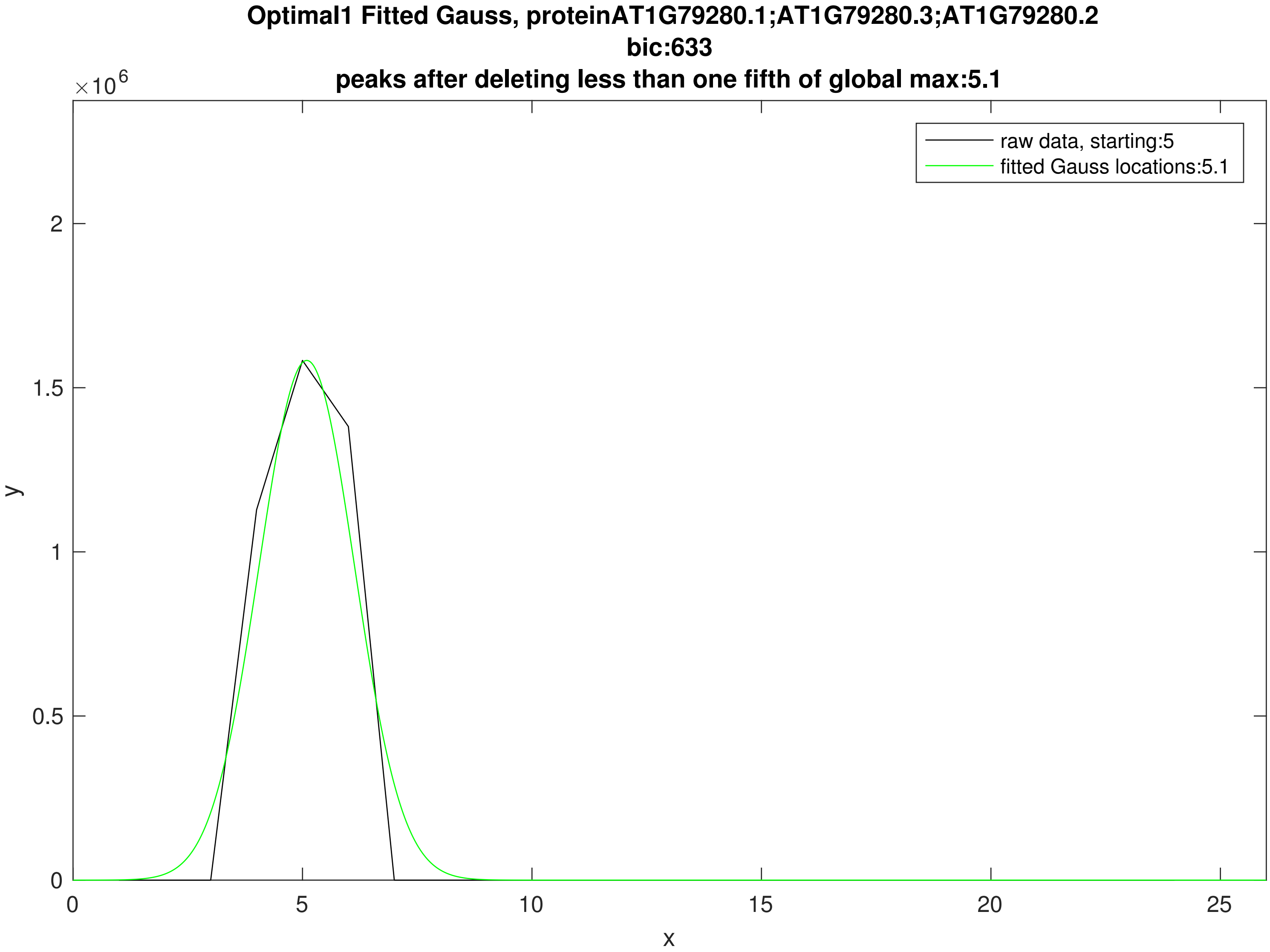
Optimal1 Fitted Gauss, proteinAT1G79280.1;AT1G79280.3;AT1G79280.2
bic:633
peaks after deleting less than one fifth of global max:5.1
×10^6
raw data, starting:5
fitted Gauss locations:5.1
y
x
0
0.5
1
1.5
2
5
10
15
20
25

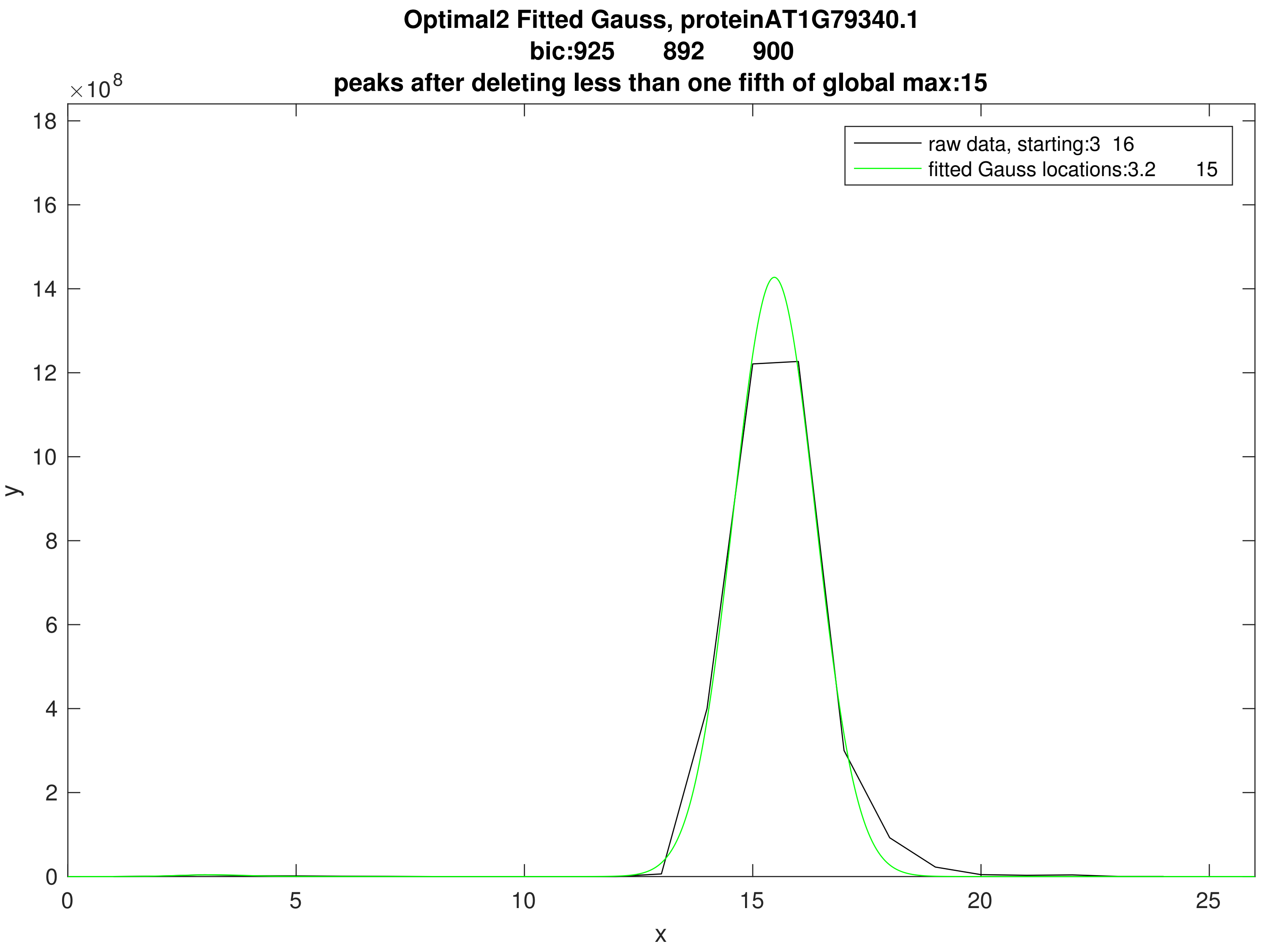
Optimal2 Fitted Gauss, proteinAT1G79340.1
bic:925 892 900
peaks after deleting less than one fifth of global max:15
×10^8
raw data, starting:3 16
fitted Gauss locations:3.2 15
y
x
0
2
4
6
8
10
12
14
16
18
0
5
10
15
20
25

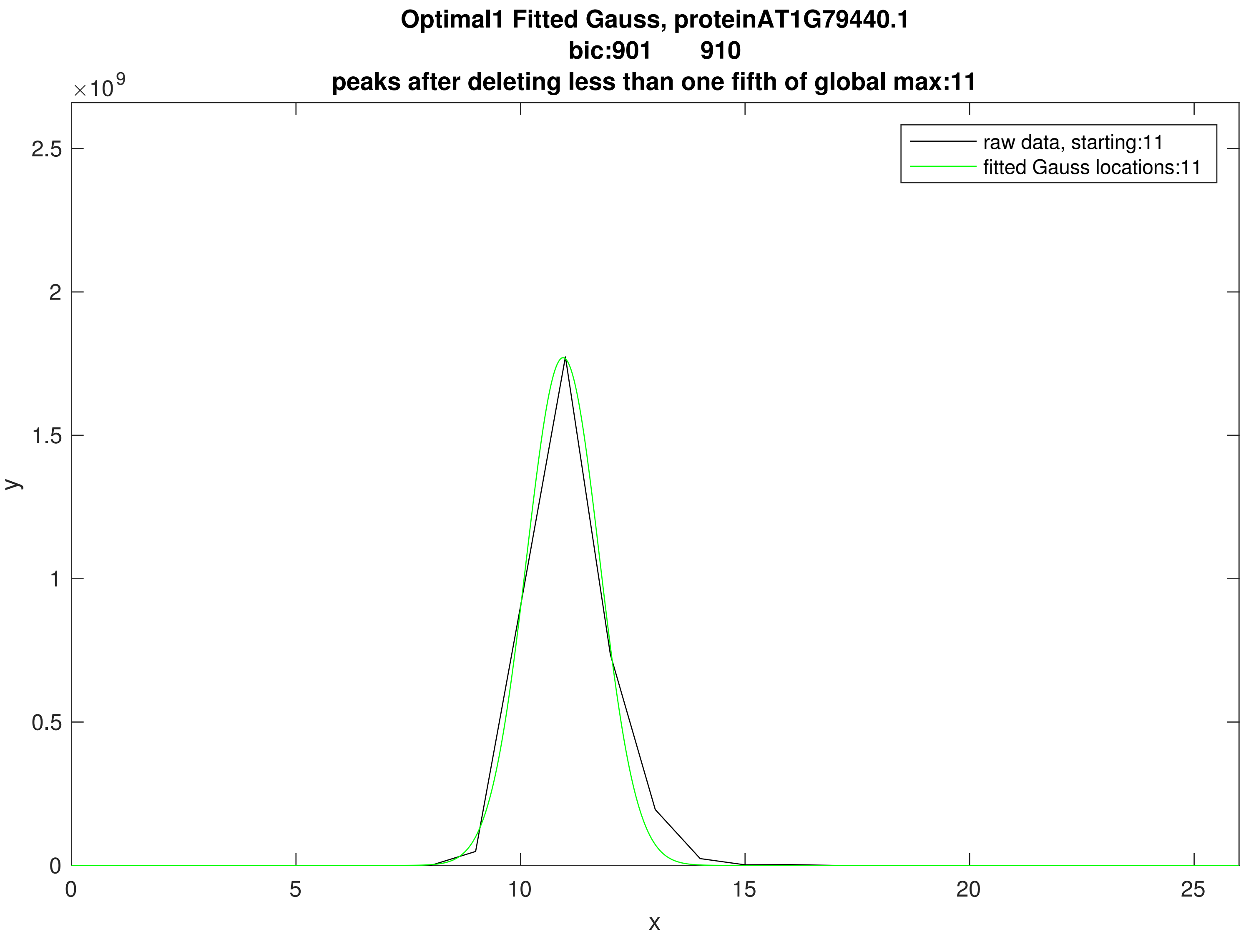
Optimal1 Fitted Gauss, proteinAT1G79440.1
bic:901 910
peaks after deleting less than one fifth of global max:11
×10^9
2.5
2
1.5
1
0.5
0
y
0
5
10
15
20
25
x
raw data, starting:11
fitted Gauss locations:11

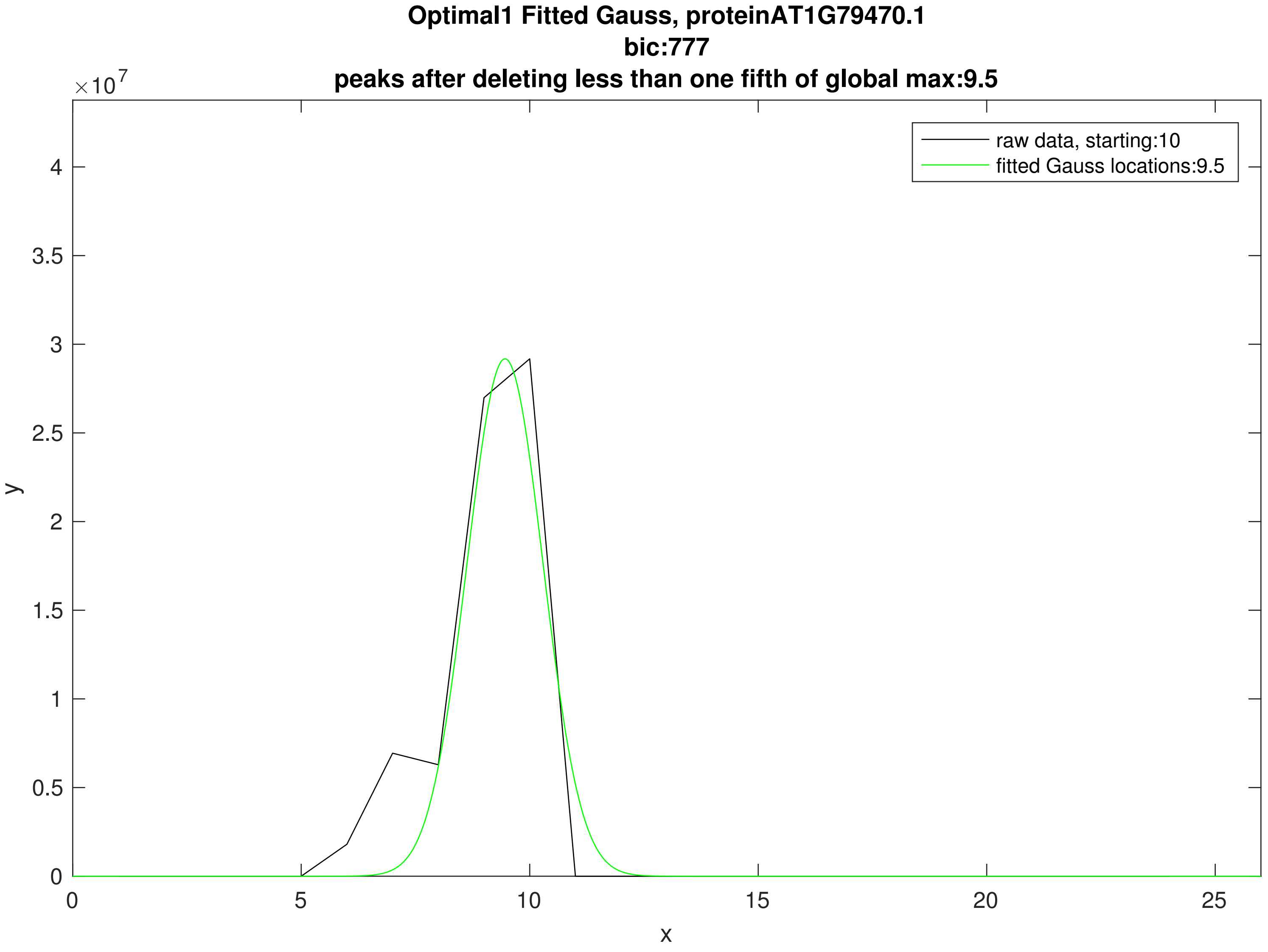
Optimal1 Fitted Gauss, proteinAT1G79470.1
bic:777
peaks after deleting less than one fifth of global max:9.5
×10^7
raw data, starting:10
fitted Gauss locations:9.5
y
x
0
0.5
1
1.5
2
2.5
3
3.5
4
5
10
15
20
25

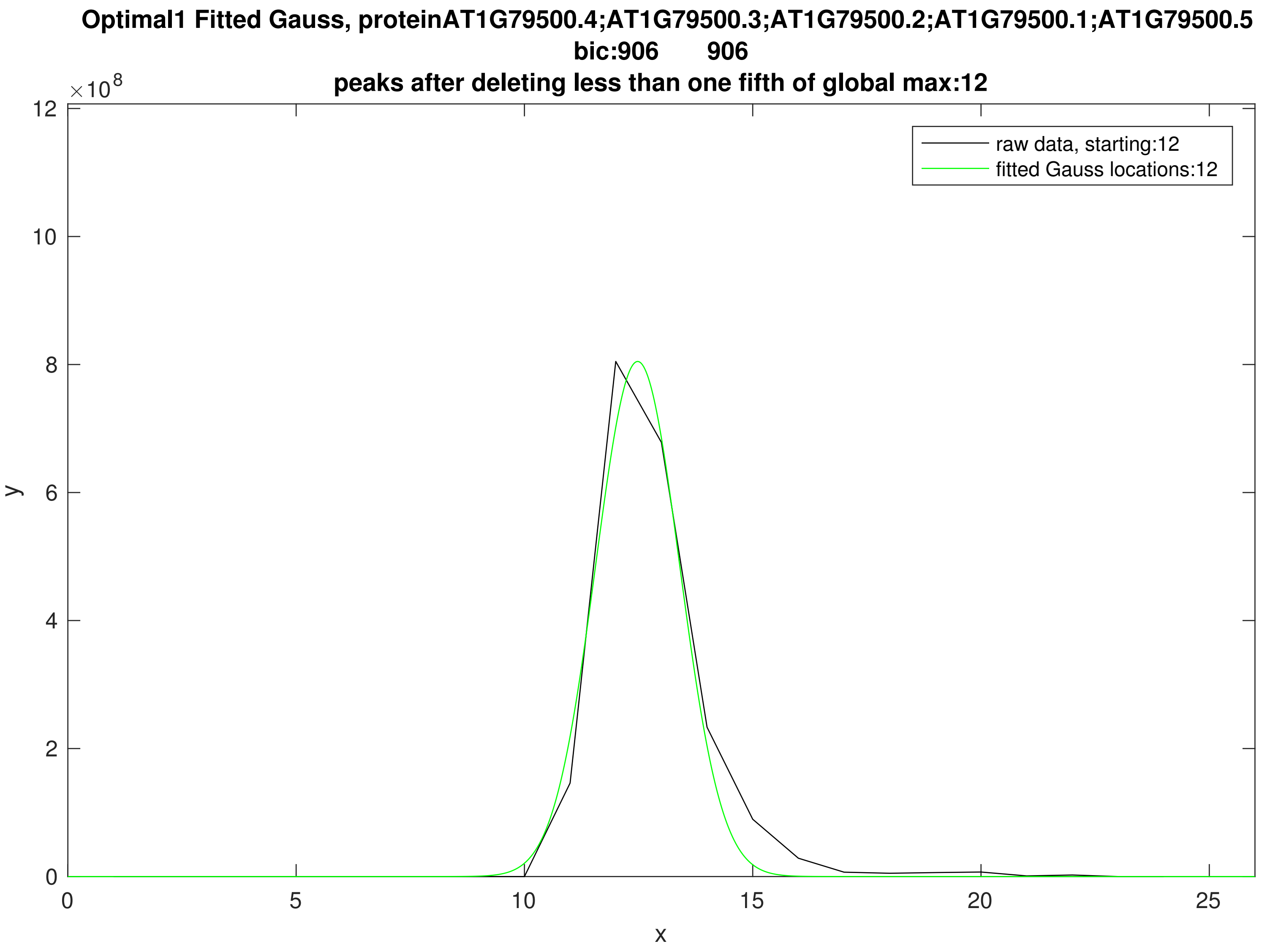
Optimal1 Fitted Gauss, proteinAT1G79500.4;AT1G79500.3;AT1G79500.2;AT1G79500.1;AT1G79500.5
bic:906 906
peaks after deleting less than one fifth of global max:12
raw data, starting:12
fitted Gauss locations:12
×10^8
y
x
0
2
4
6
8
10
12
0
5
10
15
20
25

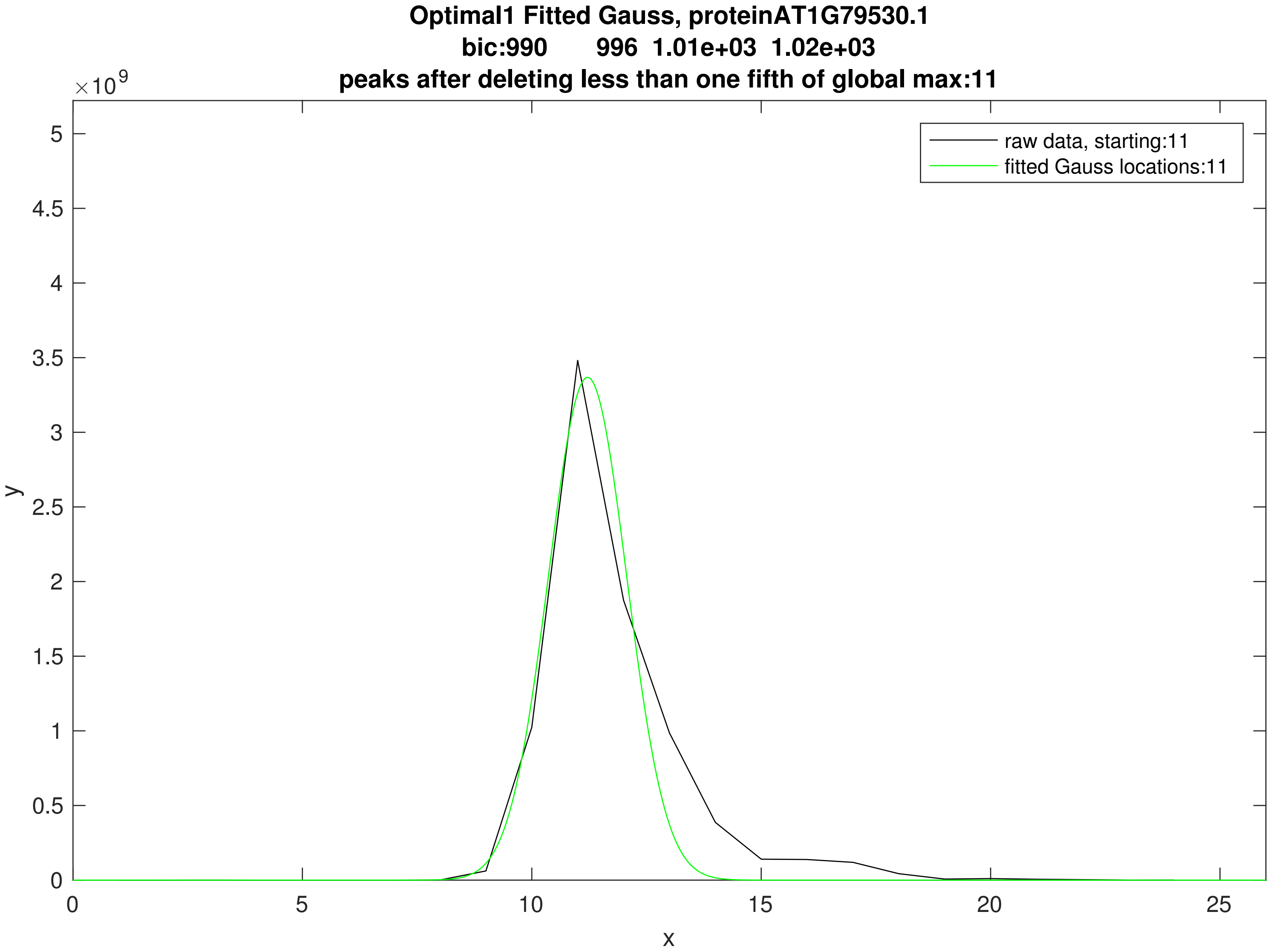

Optimal1 Fitted Gauss, proteinAT1G79530.1
bic:990 996 1.01e+03 1.02e+03
peaks after deleting less than one fifth of global max:11
×10^9
5
4.5
4
3.5
3
2.5
2
1.5
1
0.5
0
y
0
5
10
15
20
25
x
raw data, starting:11
fitted Gauss locations:11

Optimal1 Fitted Gauss, proteinAT1G79550.2;AT1G79550.1
bic:1.08e+03  1.09e+03   1.1e+03  1.11e+03
peaks after deleting less than one fifth of global max:16

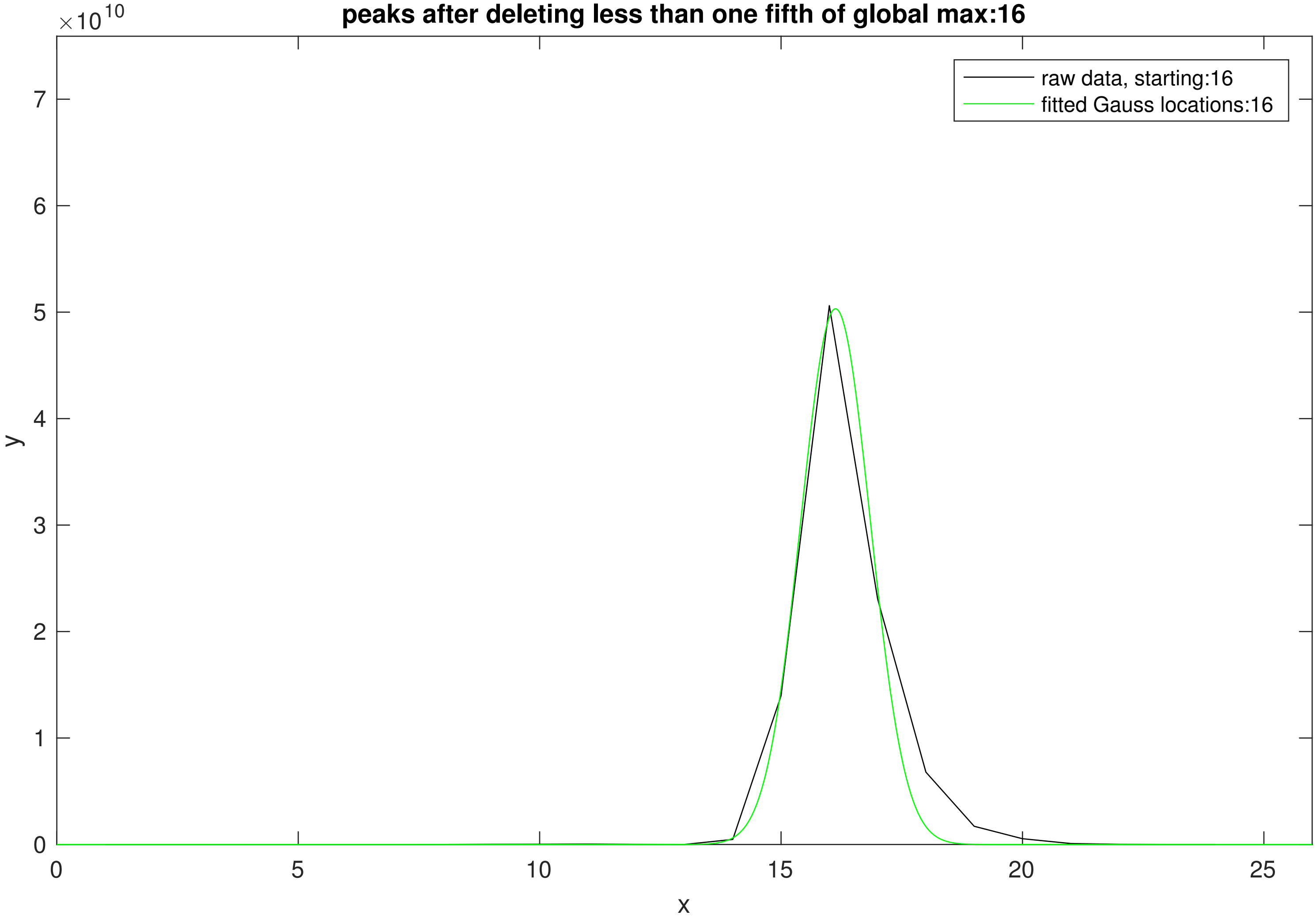

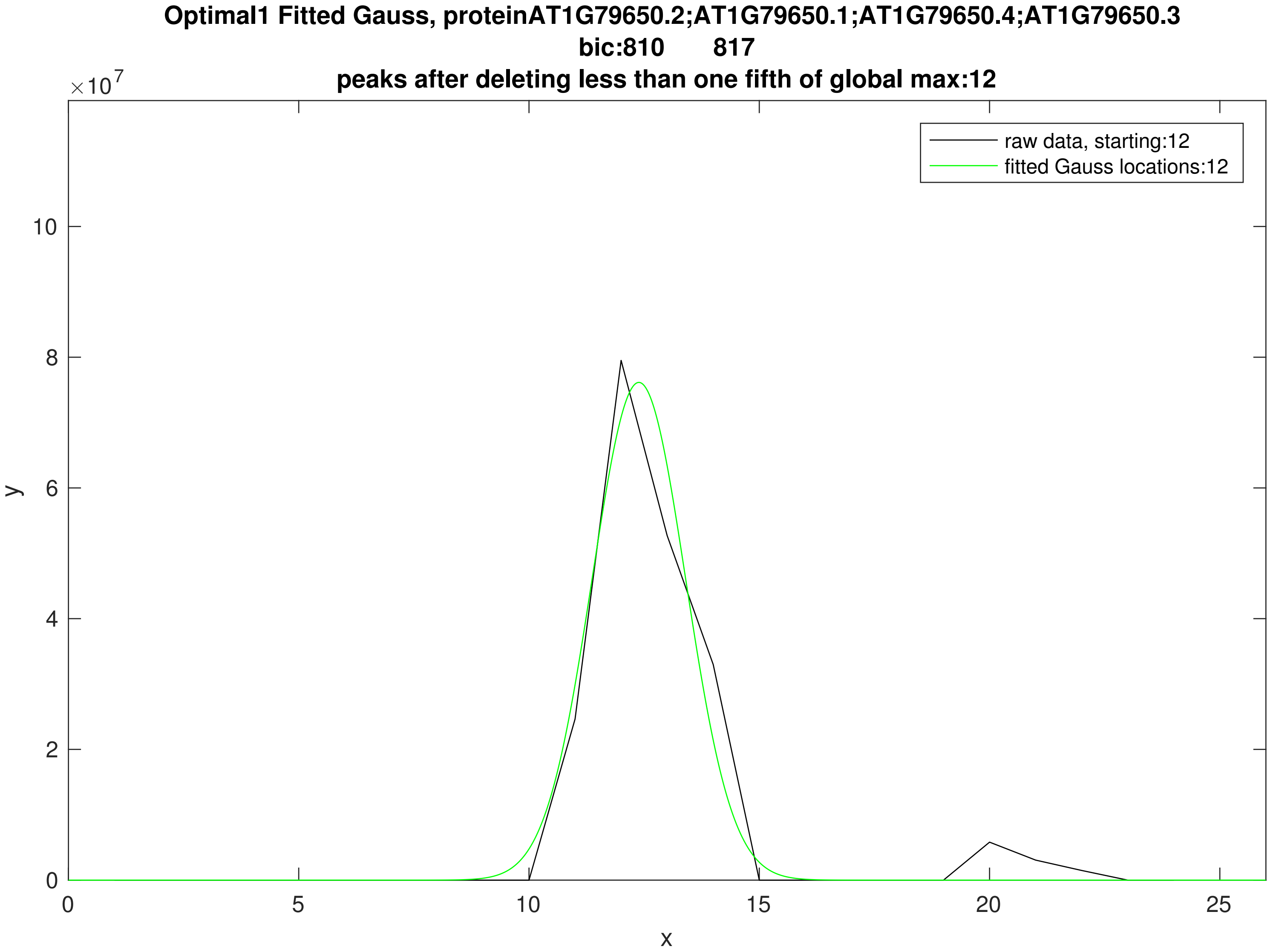
Optimal1 Fitted Gauss, proteinAT1G79650.2;AT1G79650.1;AT1G79650.4;AT1G79650.3
bic:810 817
peaks after deleting less than one fifth of global max:12
×10^7
raw data, starting:12
fitted Gauss locations:12
y
x
0
2
4
6
8
10
0
5
10
15
20
25

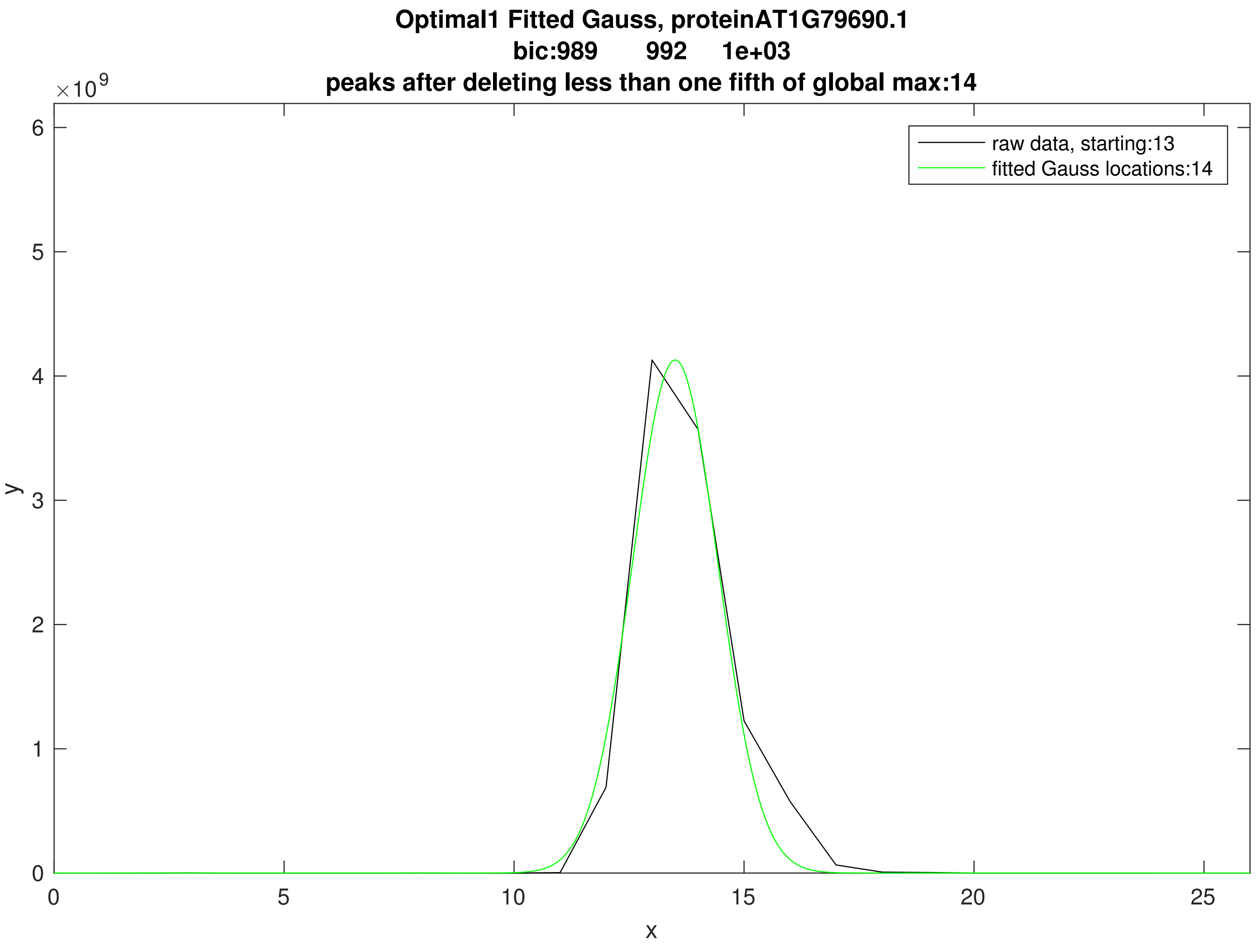

Optimal1 Fitted Gauss, proteinAT1G79690.1
bic:989 992 1e+03
peaks after deleting less than one fifth of global max:14
×10^9
6
5
4
3
2
1
0
y
0
5
10
15
20
25
x
raw data, starting:13
fitted Gauss locations:14

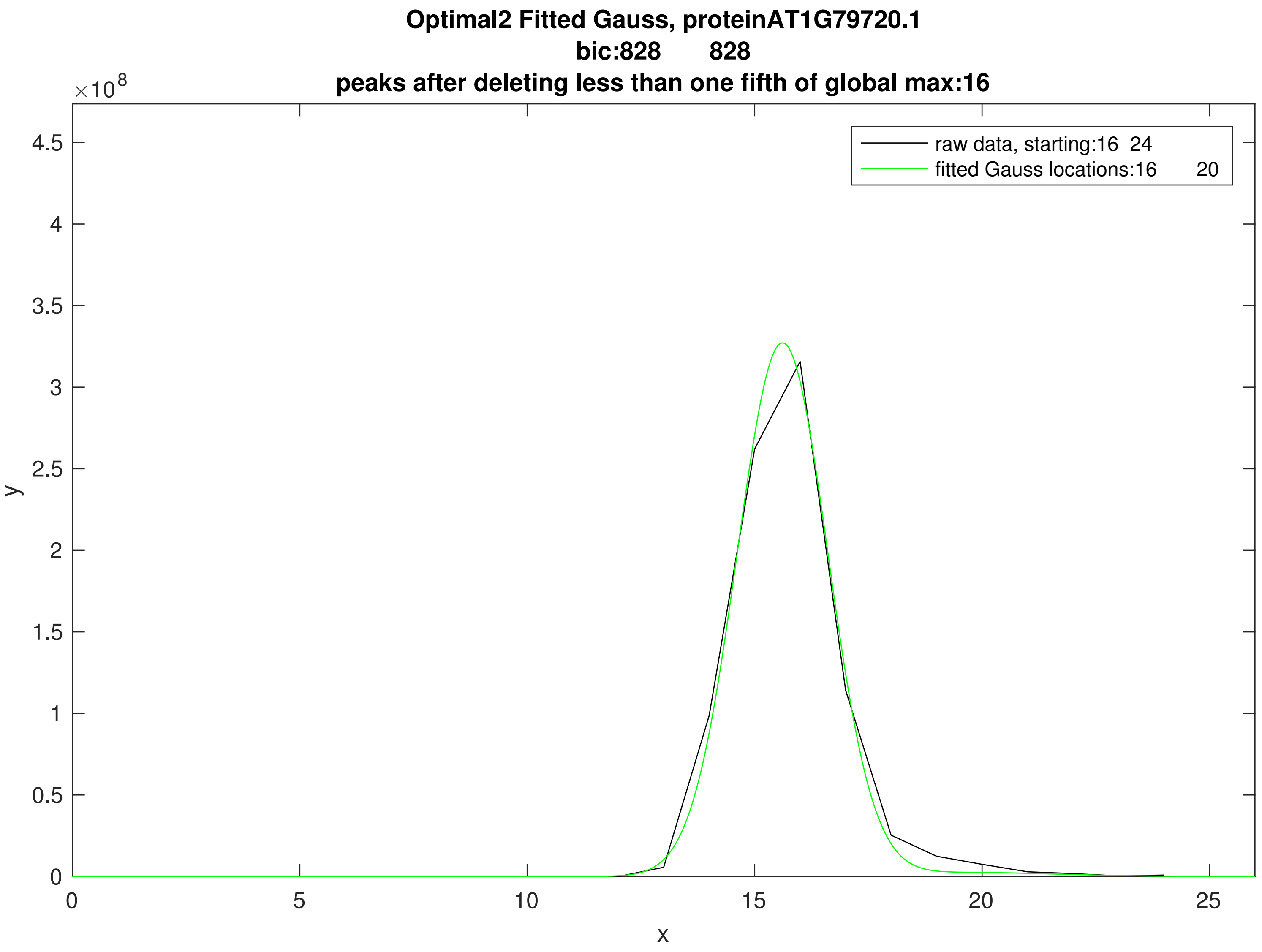

Optimal2 Fitted Gauss, proteinAT1G79720.1
bic:828 828
peaks after deleting less than one fifth of global max:16
×10^8
4.5
4
3.5
3
2.5
2
1.5
1
0.5
0
y
0
5
10
15
20
25
x
raw data, starting:16 24
fitted Gauss locations:16 20

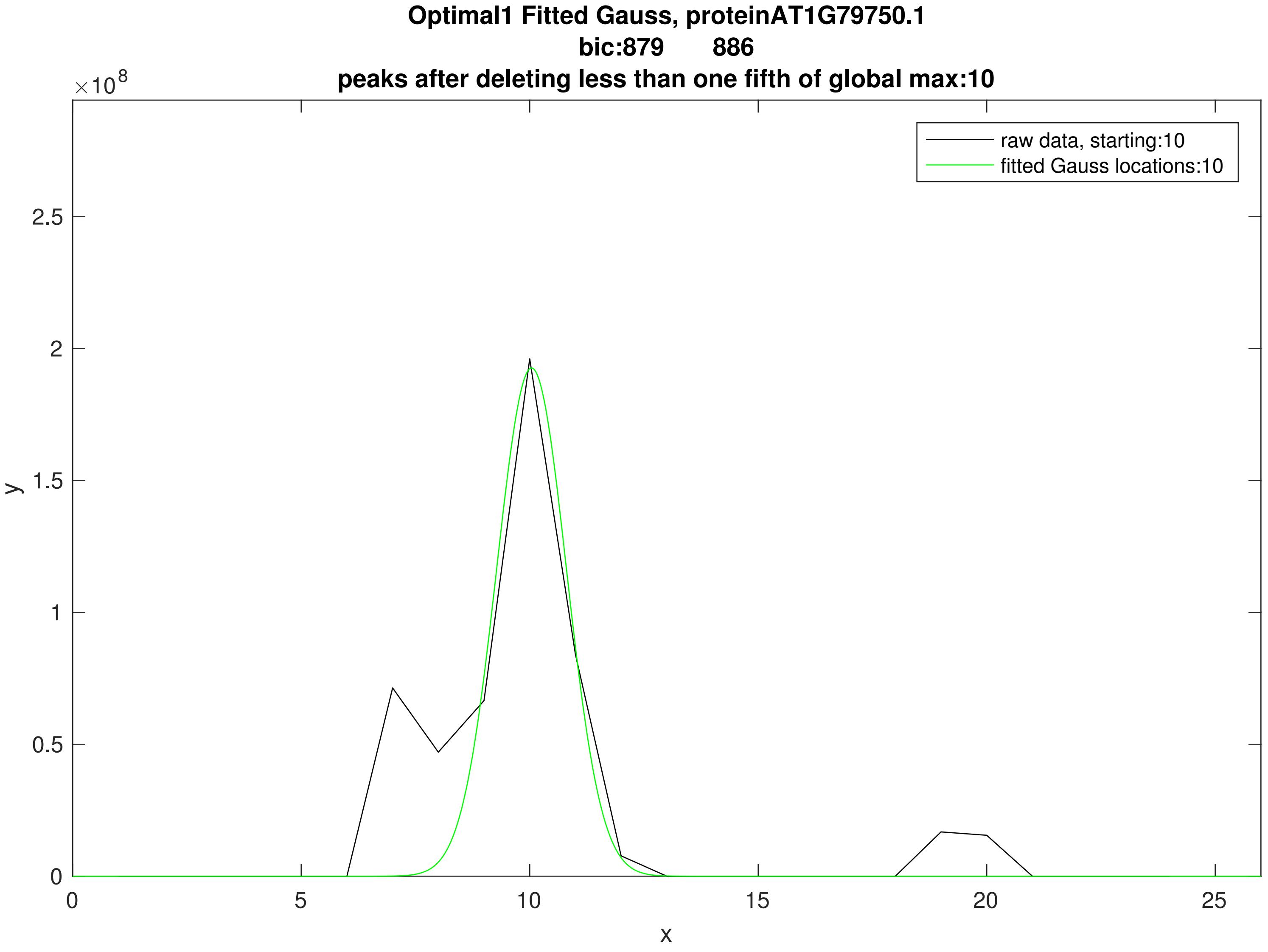

Optimal1 Fitted Gauss, proteinAT1G79750.1
bic:879 886
peaks after deleting less than one fifth of global max:10
×10^8
raw data, starting:10
fitted Gauss locations:10
2.5
2
1.5
1
0.5
0
y
0
5
10
15
20
25
x

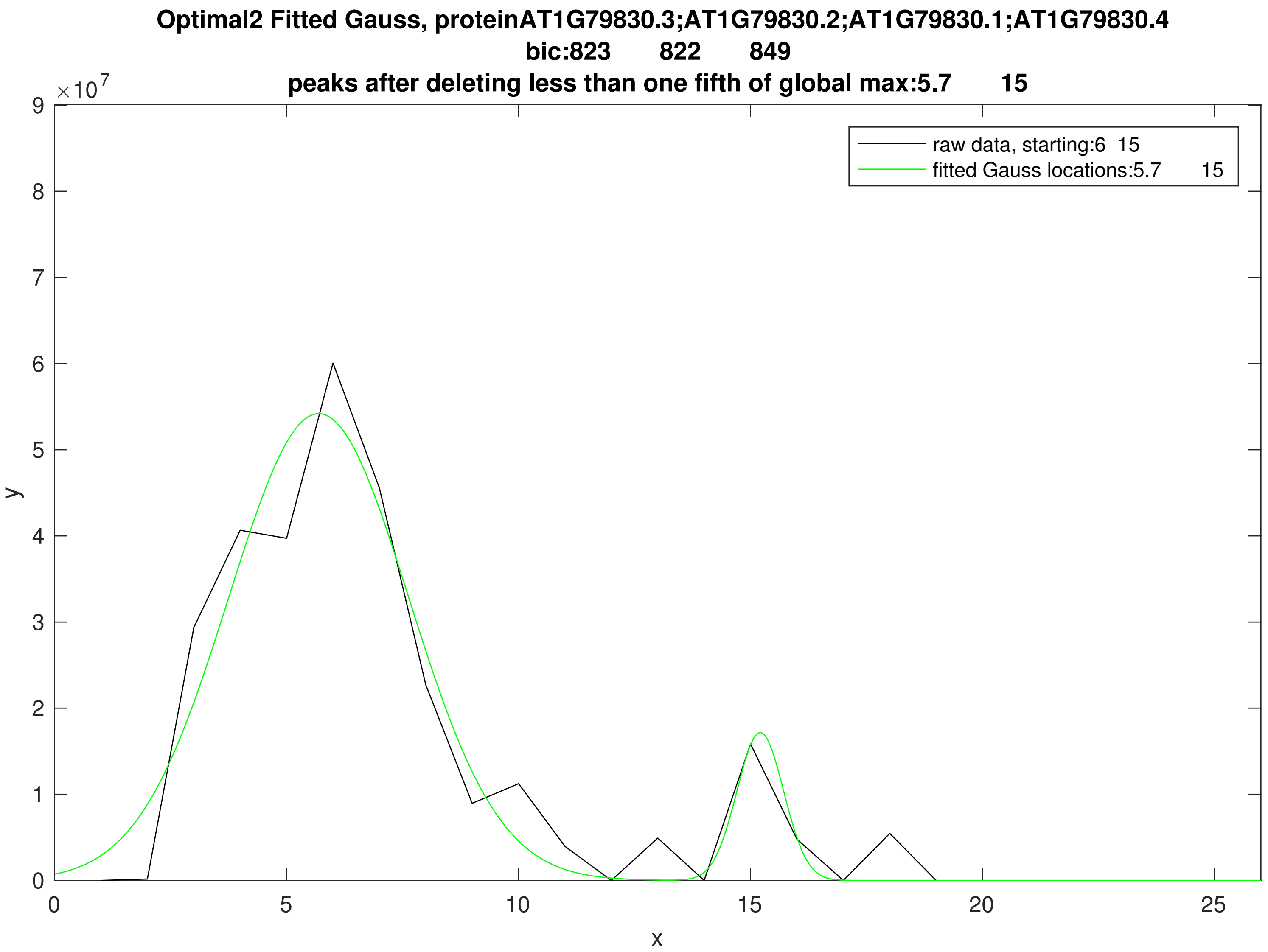

Optimal2 Fitted Gauss, proteinAT1G79830.3;AT1G79830.2;AT1G79830.1;AT1G79830.4
bic:823 822 849
peaks after deleting less than one fifth of global max:5.7 15
×10^7
raw data, starting:6 15
fitted Gauss locations:5.7 15
y
x
0
1
2
3
4
5
6
7
8
9
0
5
10
15
20
25

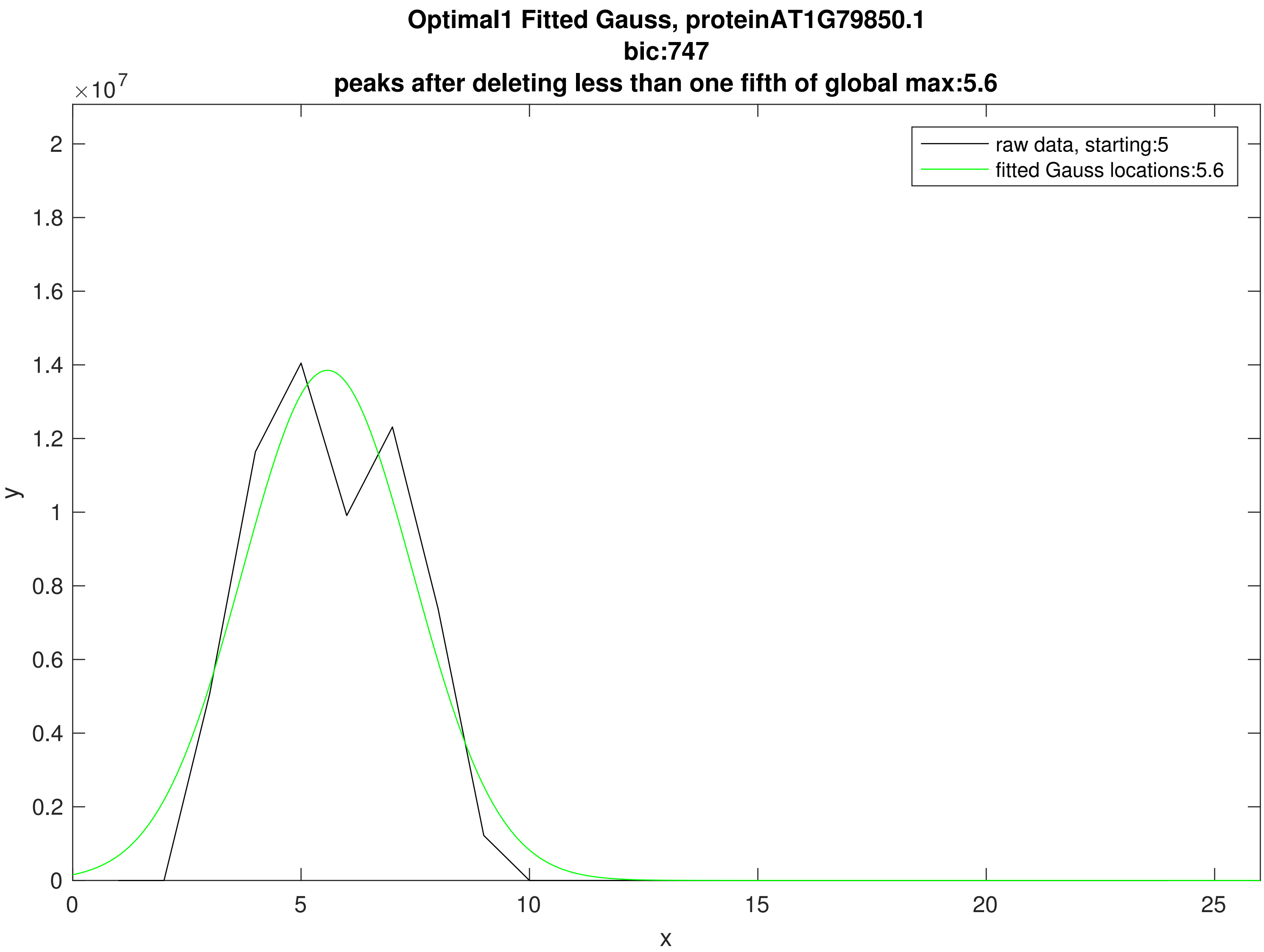
Optimal1 Fitted Gauss, proteinAT1G79850.1
bic:747
peaks after deleting less than one fifth of global max:5.6
×10^7
2
1.8
1.6
1.4
1.2
1
0.8
0.6
0.4
0.2
0
y
0
5
10
15
20
25
x
raw data, starting:5
fitted Gauss locations:5.6

Optimal1 Fitted Gauss, proteinAT1G79870.1;AT1G79870.2
bic:959 968
peaks after deleting less than one fifth of global max:14

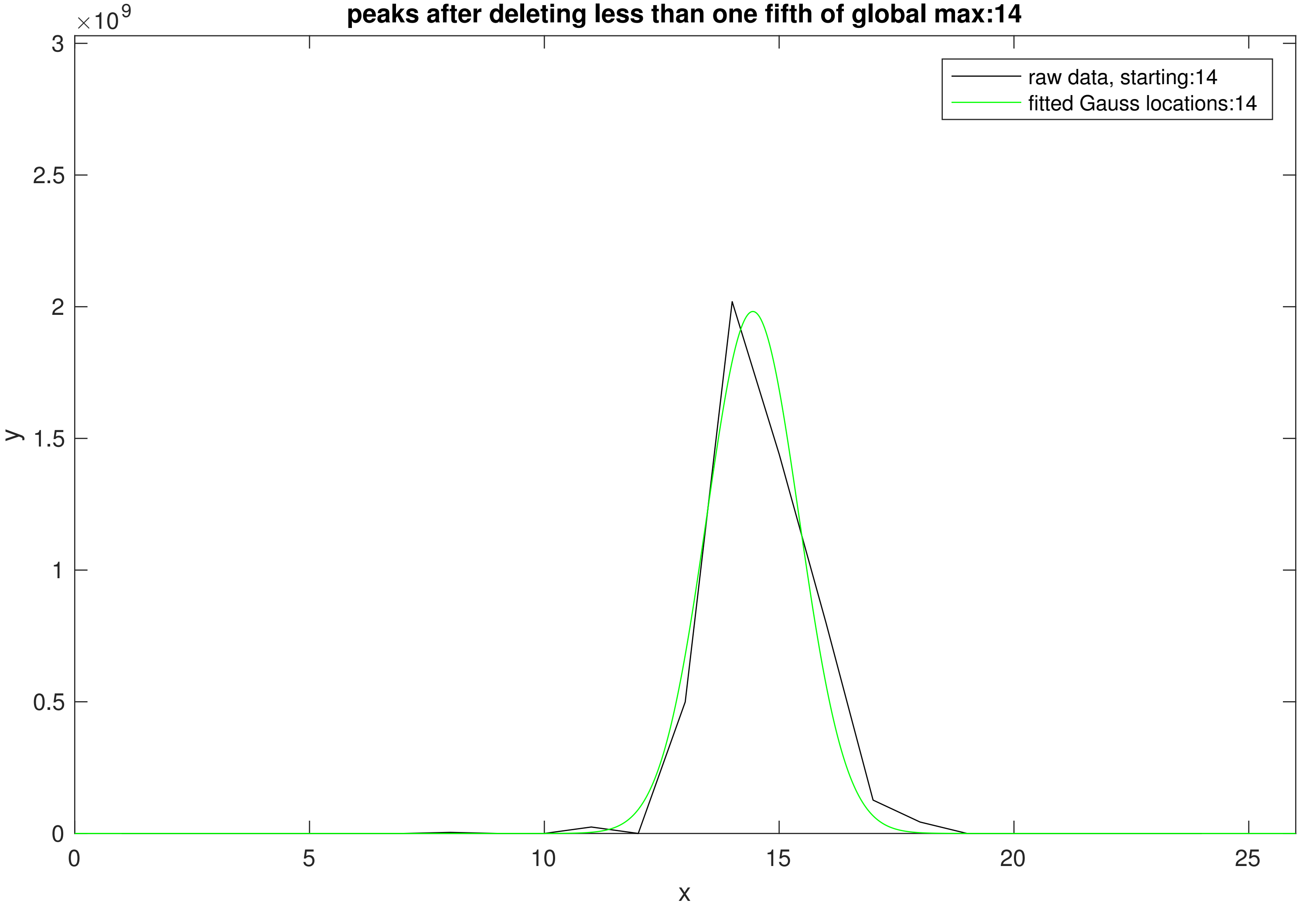

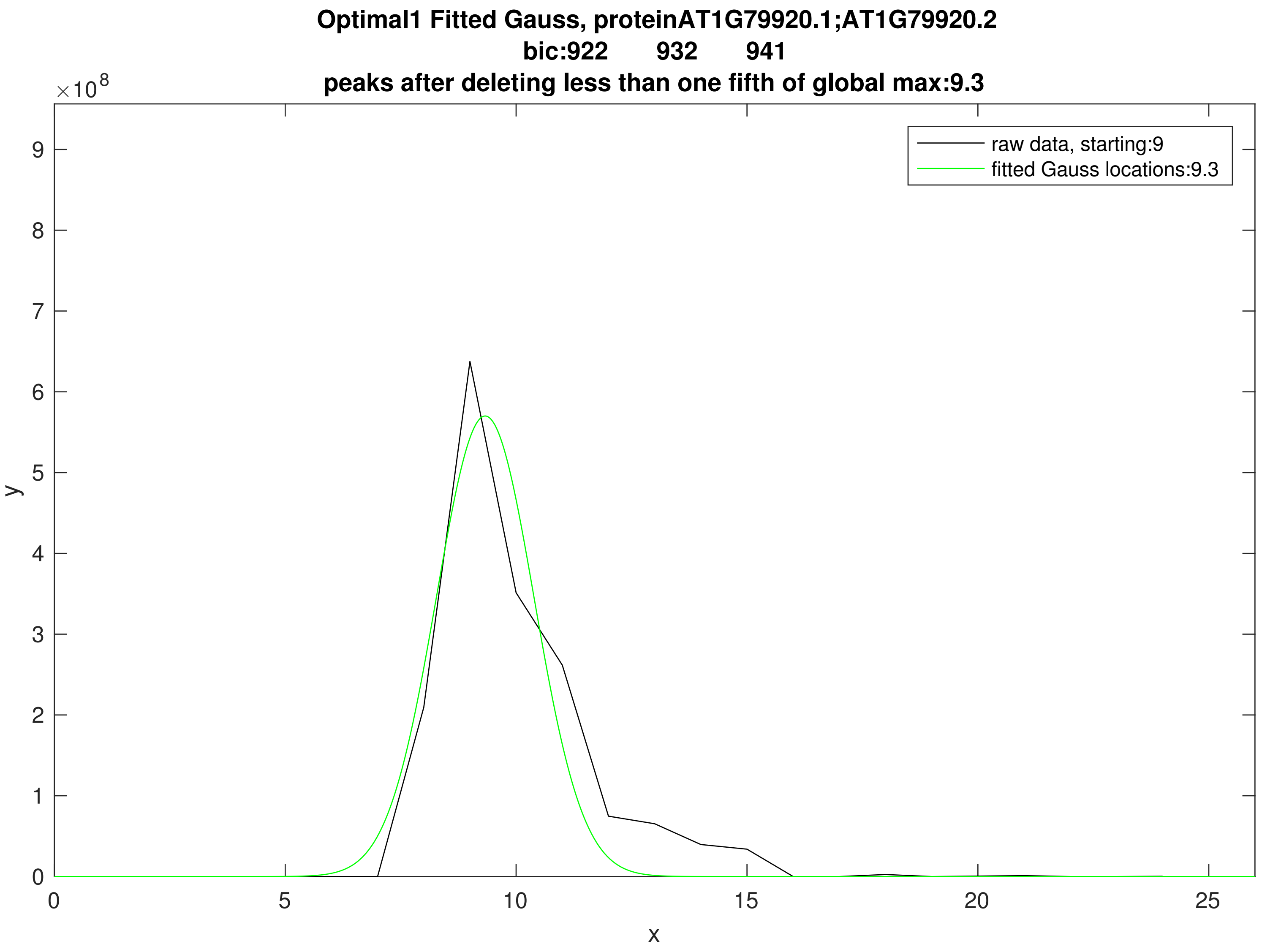

Optimal1 Fitted Gauss, proteinAT1G79920.1;AT1G79920.2
bic:922 932 941
peaks after deleting less than one fifth of global max:9.3
×10^8
raw data, starting:9
fitted Gauss locations:9.3
y
x
0
1
2
3
4
5
6
7
8
9
0
5
10
15
20
25

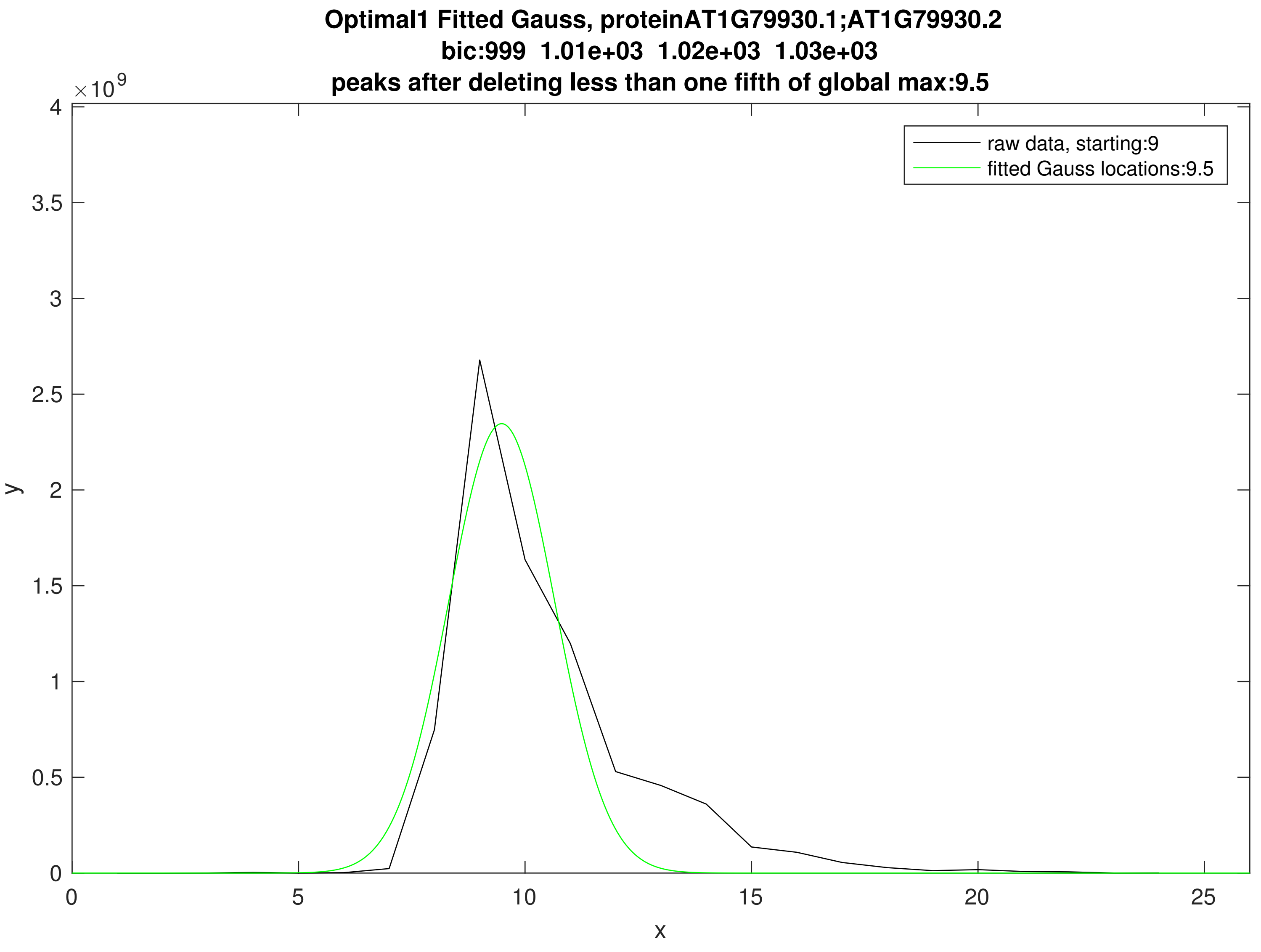

Optimal1 Fitted Gauss, proteinAT1G79930.1;AT1G79930.2
bic:999 1.01e+03 1.02e+03 1.03e+03
peaks after deleting less than one fifth of global max:9.5
×10^9
raw data, starting:9
fitted Gauss locations:9.5
y
x
0
0.5
1
1.5
2
2.5
3
3.5
4
5
10
15
20
25

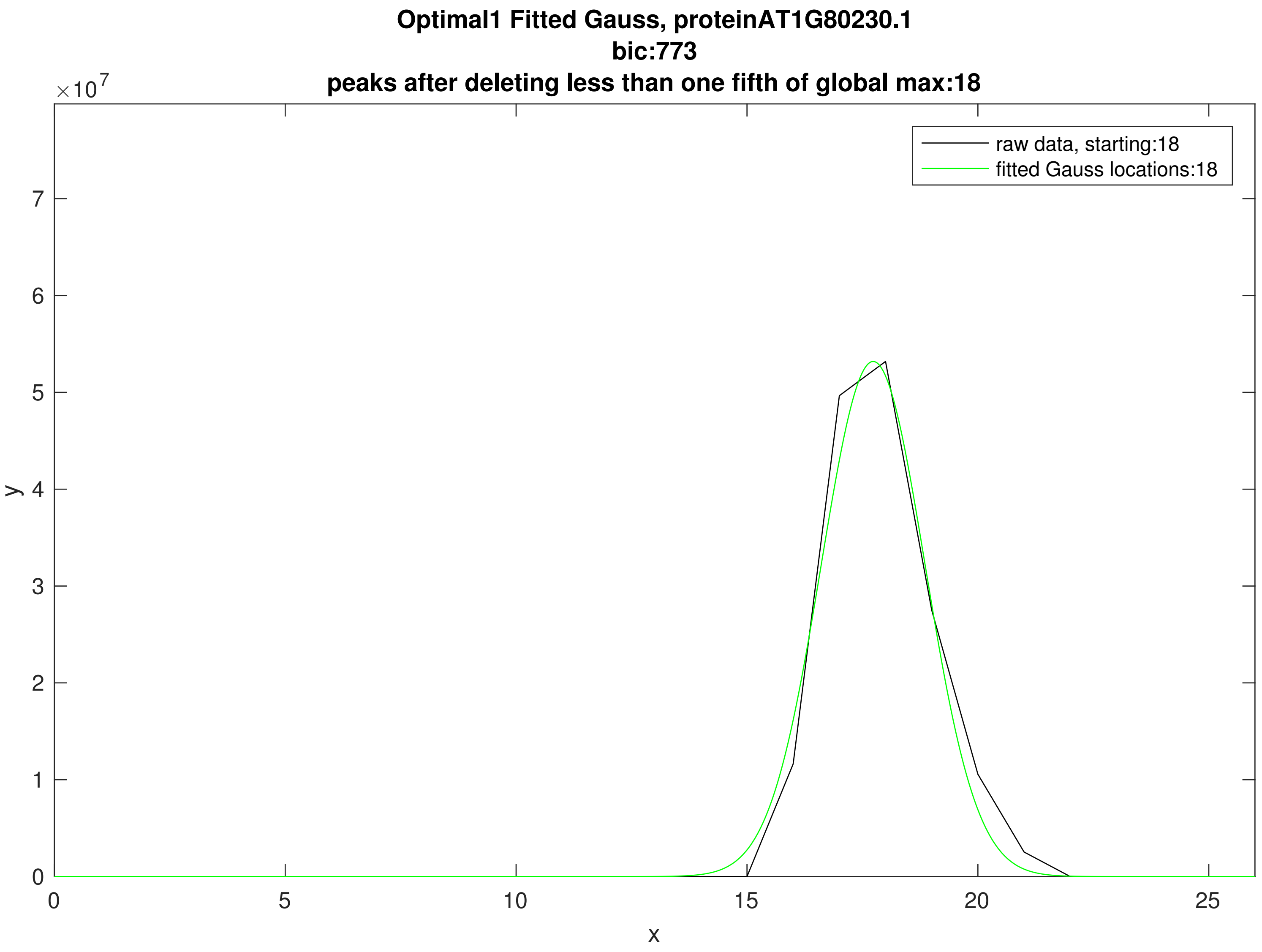

Optimal1 Fitted Gauss, proteinAT1G80230.1
bic:773
peaks after deleting less than one fifth of global max:18
×10^7
raw data, starting:18
fitted Gauss locations:18
y
x
0
1
2
3
4
5
6
7
0
5
10
15
20
25

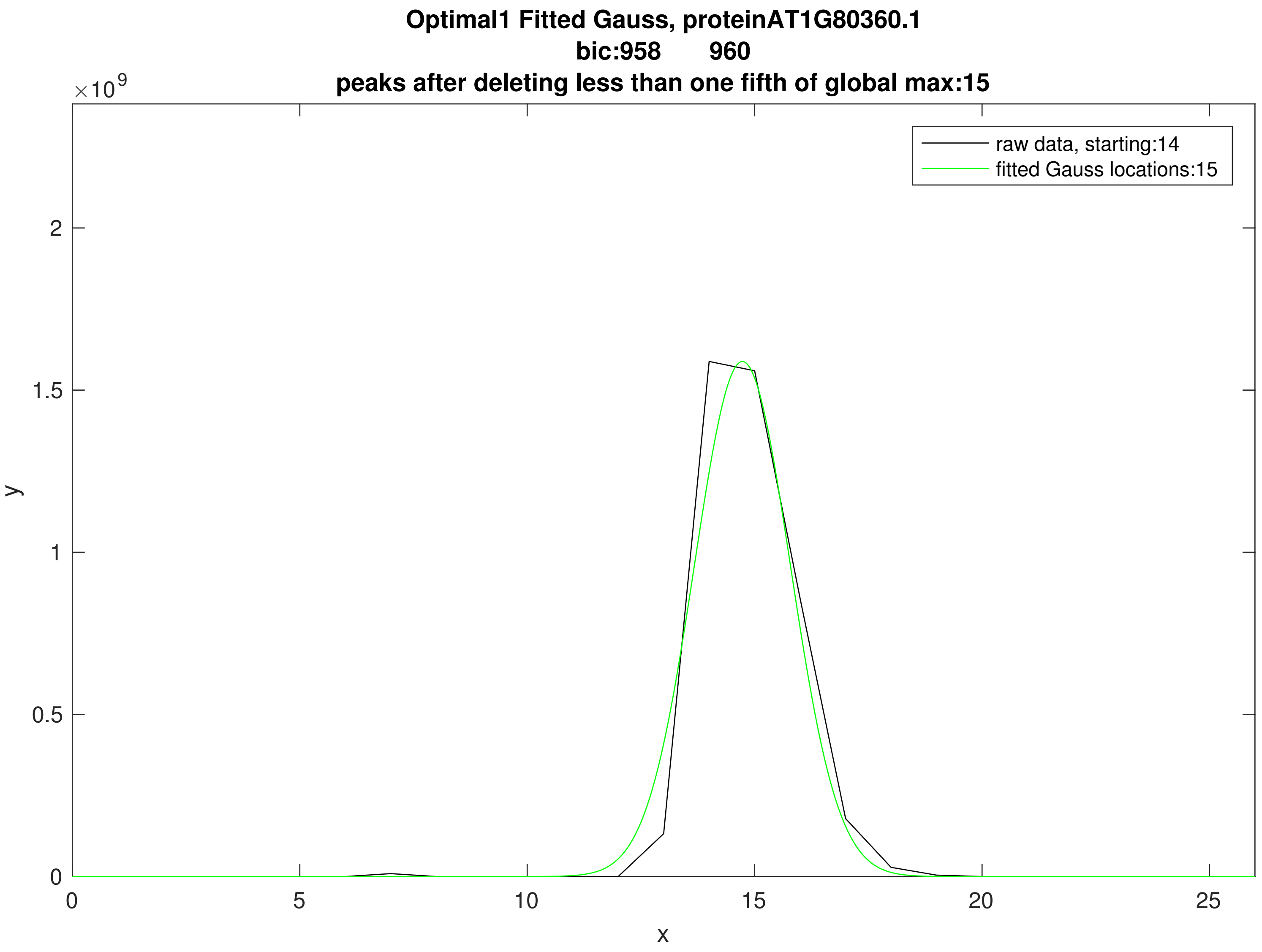

Optimal1 Fitted Gauss, proteinAT1G80360.1
bic:958 960
peaks after deleting less than one fifth of global max:15
×10^9
raw data, starting:14
fitted Gauss locations:15
2
1.5
1
0.5
0
y
0
5
10
15
20
25
x

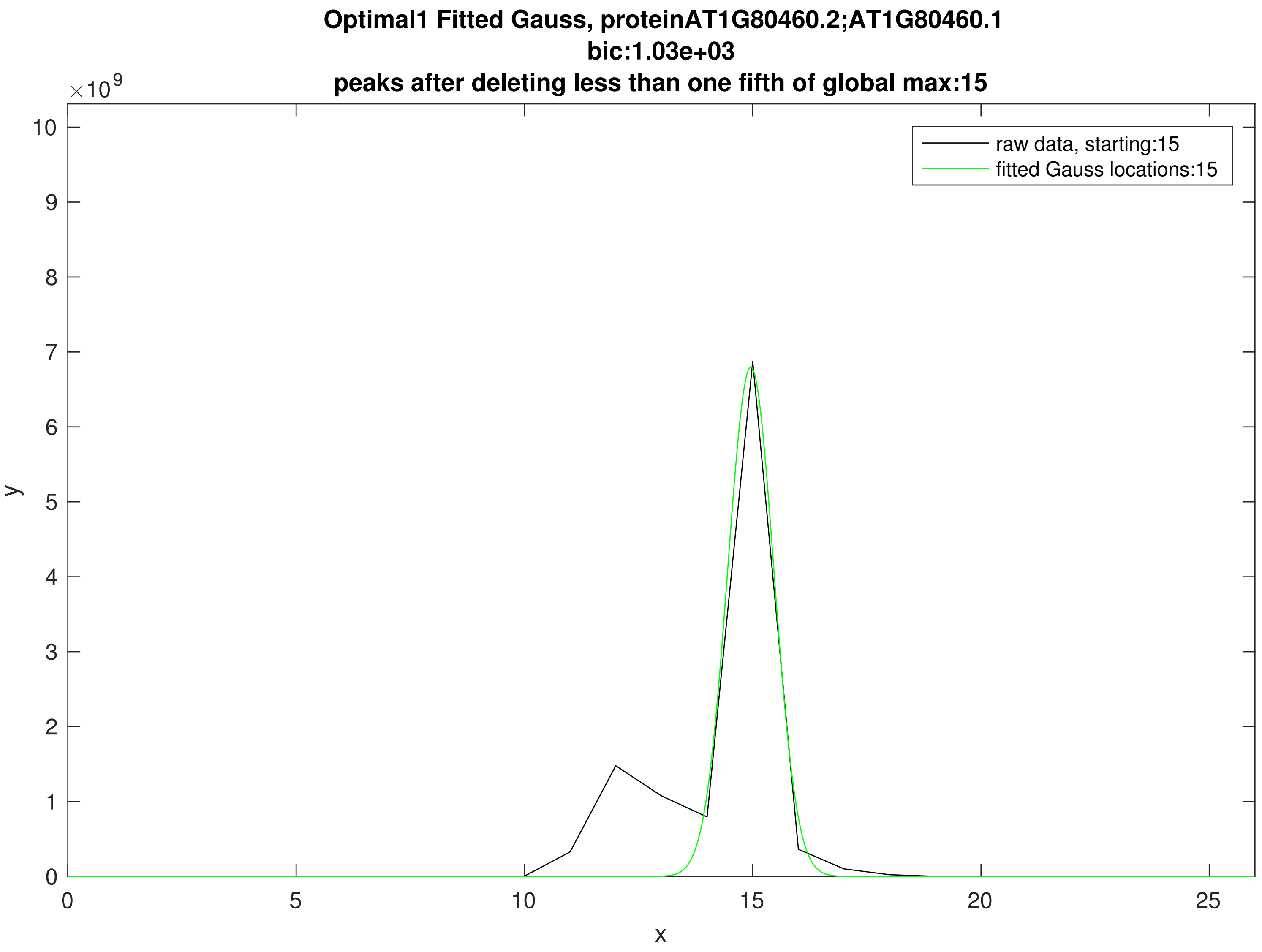
Optimal1 Fitted Gauss, proteinAT1G80460.2;AT1G80460.1
bic:1.03e+03
peaks after deleting less than one fifth of global max:15
×10^9
raw data, starting:15
fitted Gauss locations:15
y
x
0
1
2
3
4
5
6
7
8
9
10
0
5
10
15
20
25

**Optimal1 Fitted Gauss, proteinAT1G80560.1**
**bic:839 845**
**peaks after deleting less than one fifth of global max:14**

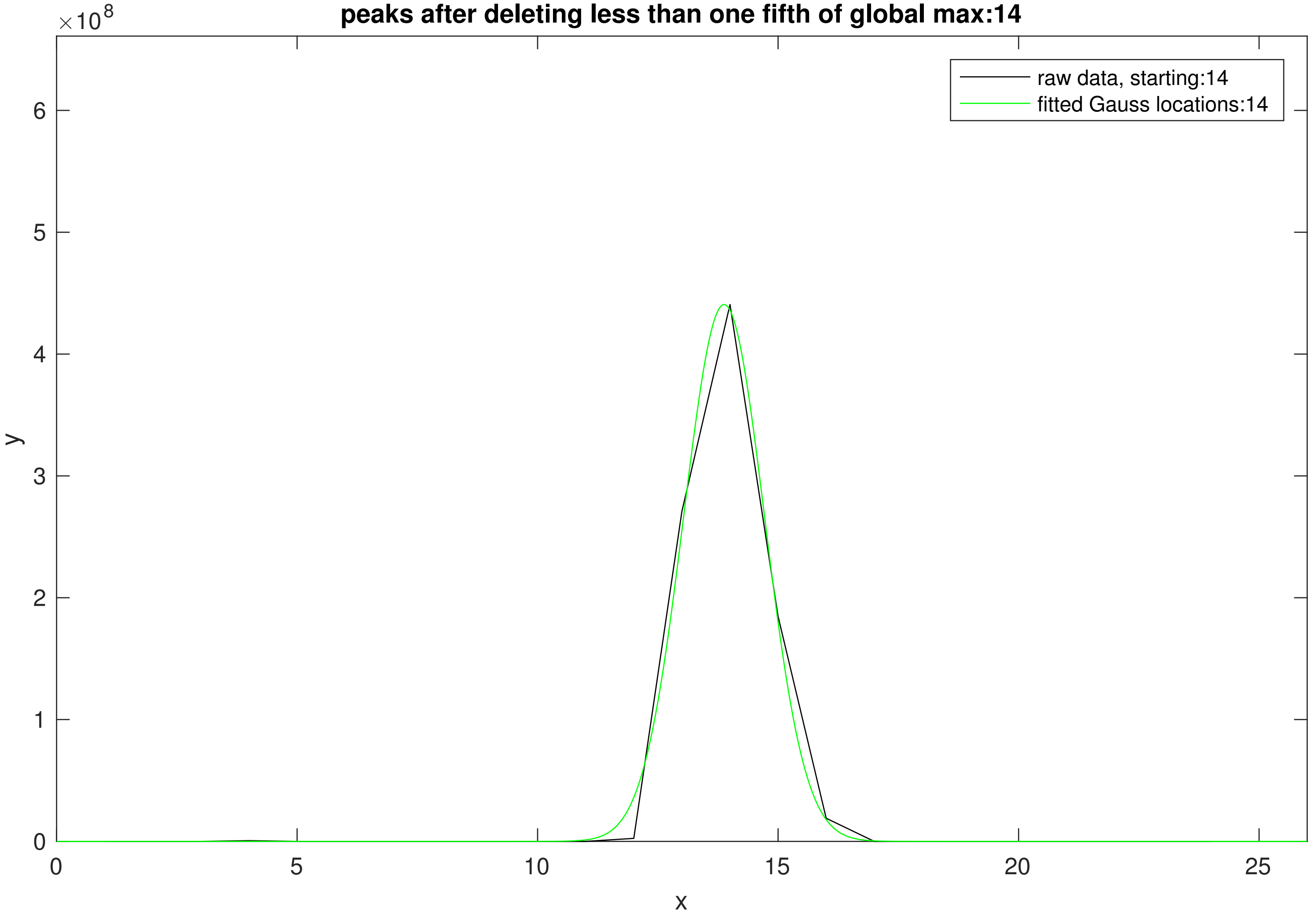

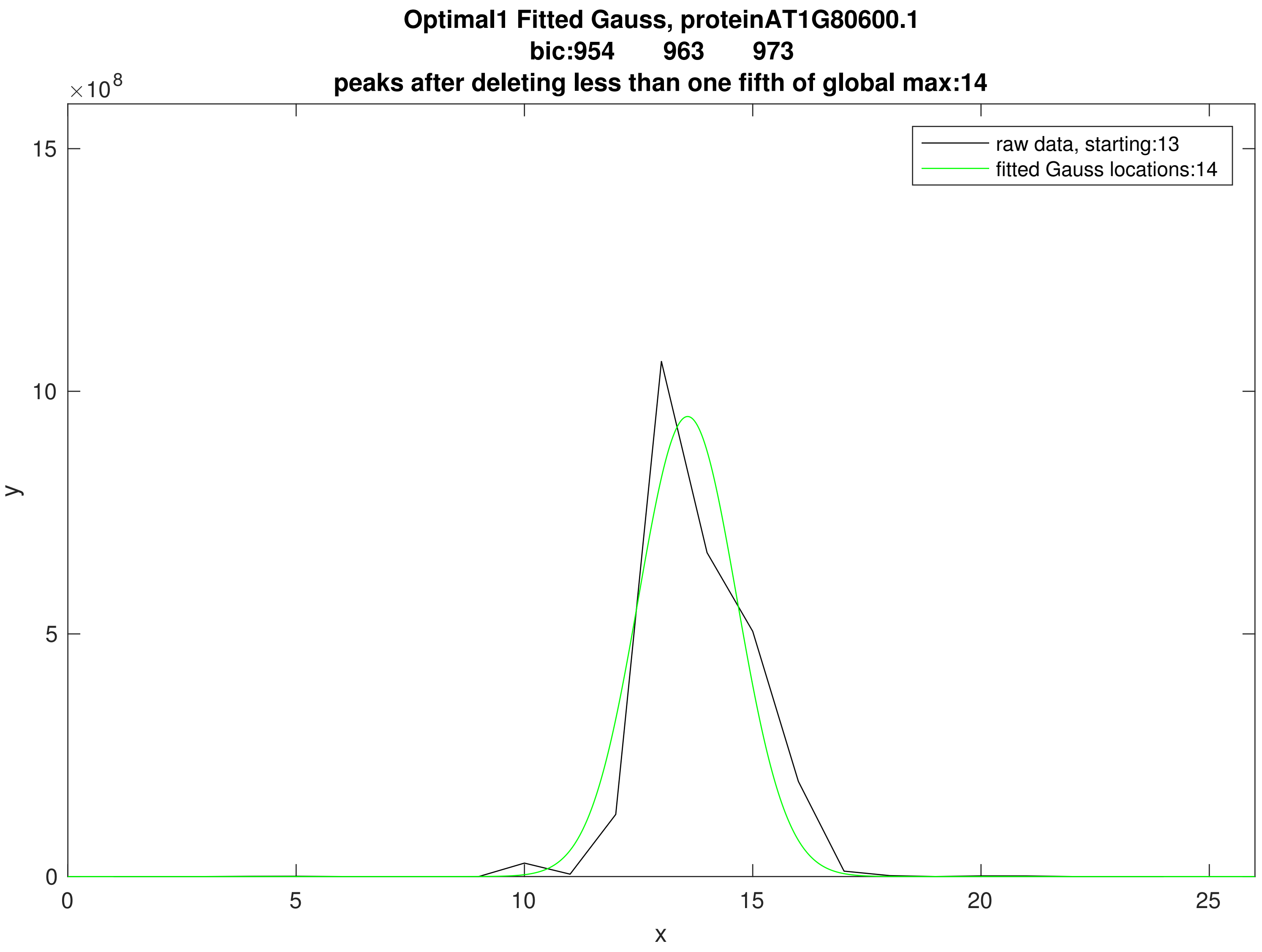
Optimal1 Fitted Gauss, proteinAT1G80600.1
bic:954 963 973
peaks after deleting less than one fifth of global max:14
×10^8
raw data, starting:13
fitted Gauss locations:14
y
x
0
5
10
15
20
25

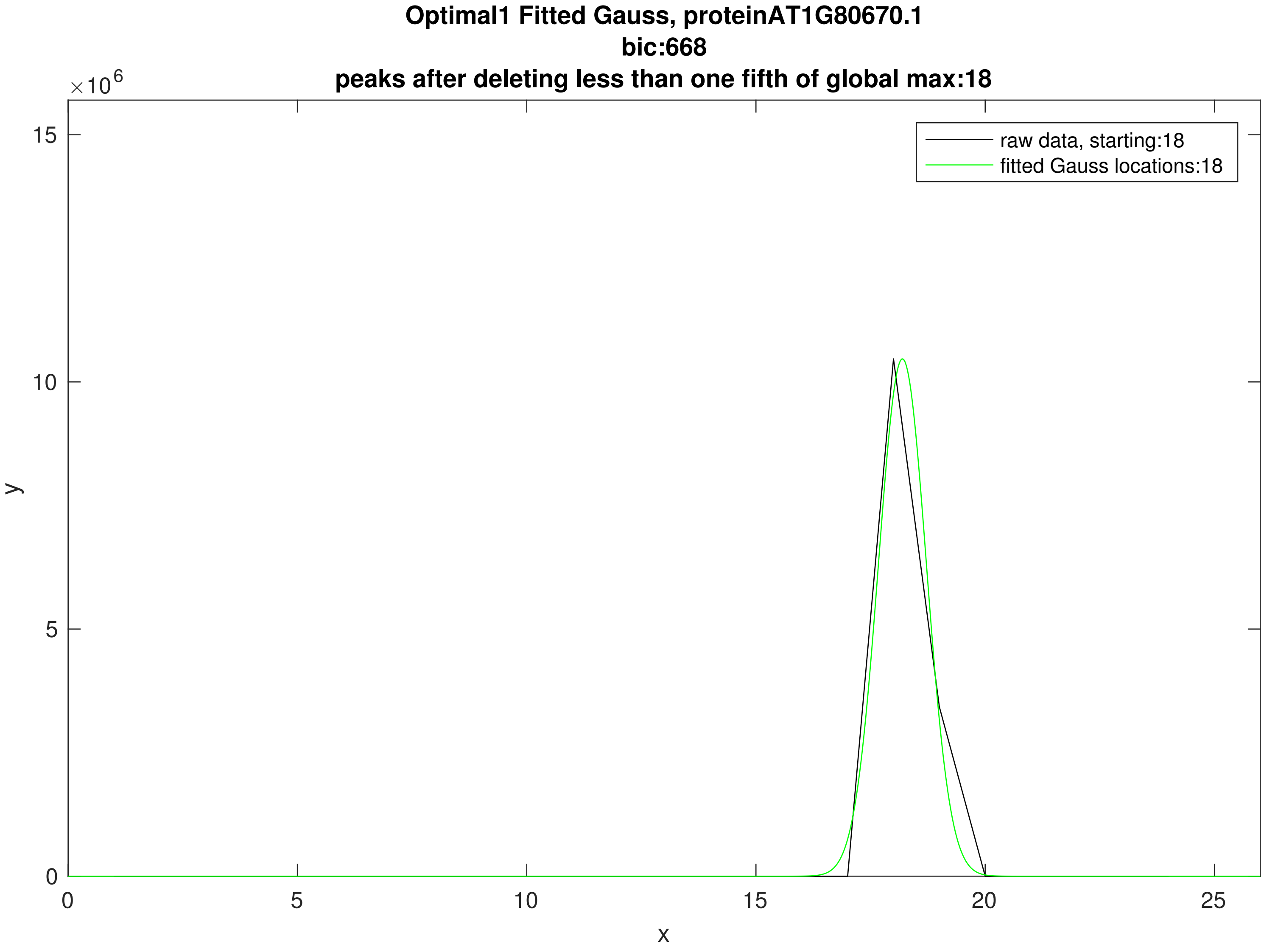

Optimal1 Fitted Gauss, proteinAT1G80670.1
bic:668
peaks after deleting less than one fifth of global max:18
×10^6
15
10
5
0
y
0
5
10
15
20
25
x
raw data, starting:18
fitted Gauss locations:18

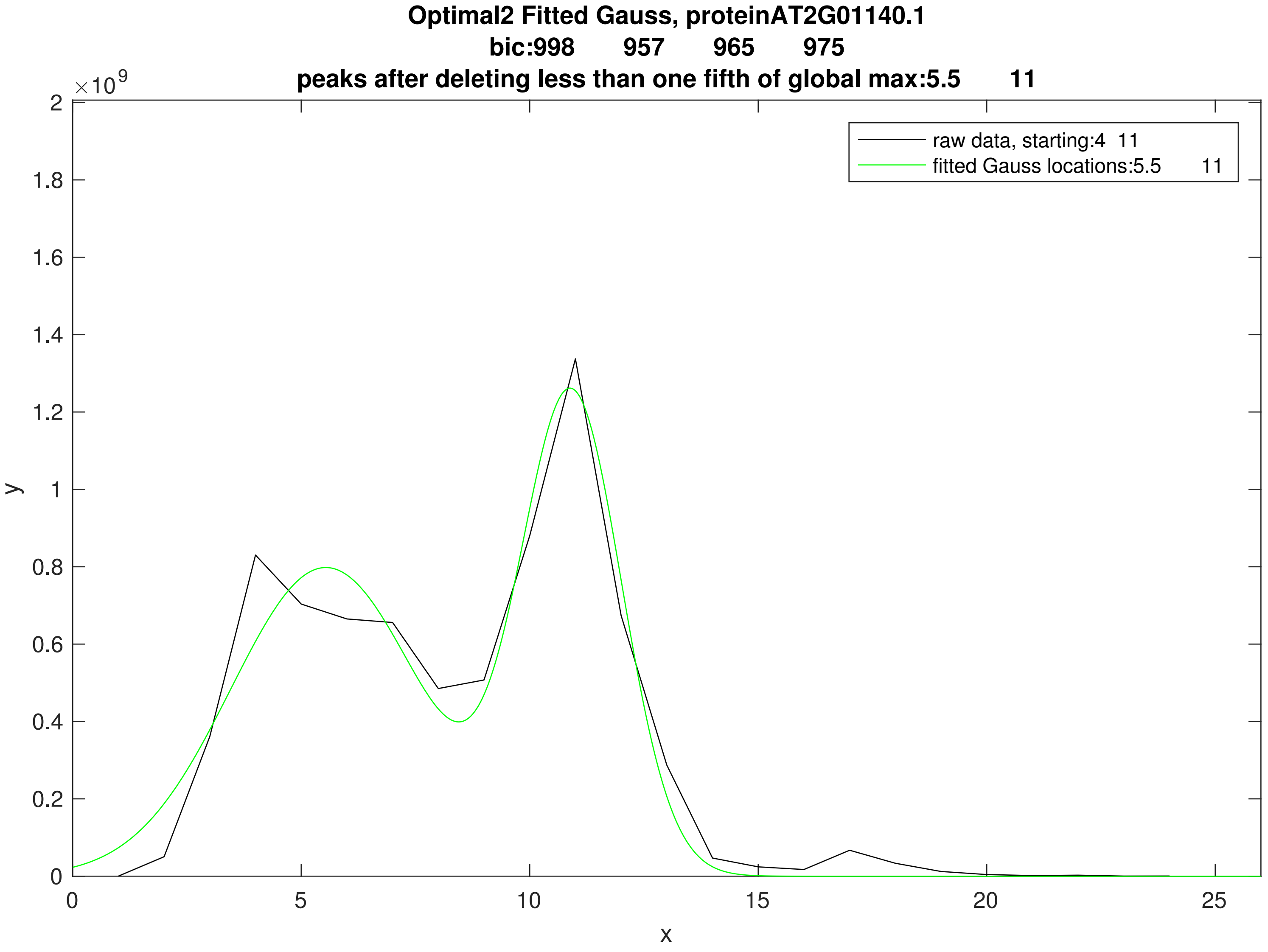

Optimal2 Fitted Gauss, proteinAT2G01140.1
bic:998 957 965 975
peaks after deleting less than one fifth of global max:5.5 11
×10^9
raw data, starting:4 11
fitted Gauss locations:5.5 11
y
x
0
0.2
0.4
0.6
0.8
1
1.2
1.4
1.6
1.8
2
0
5
10
15
20
25

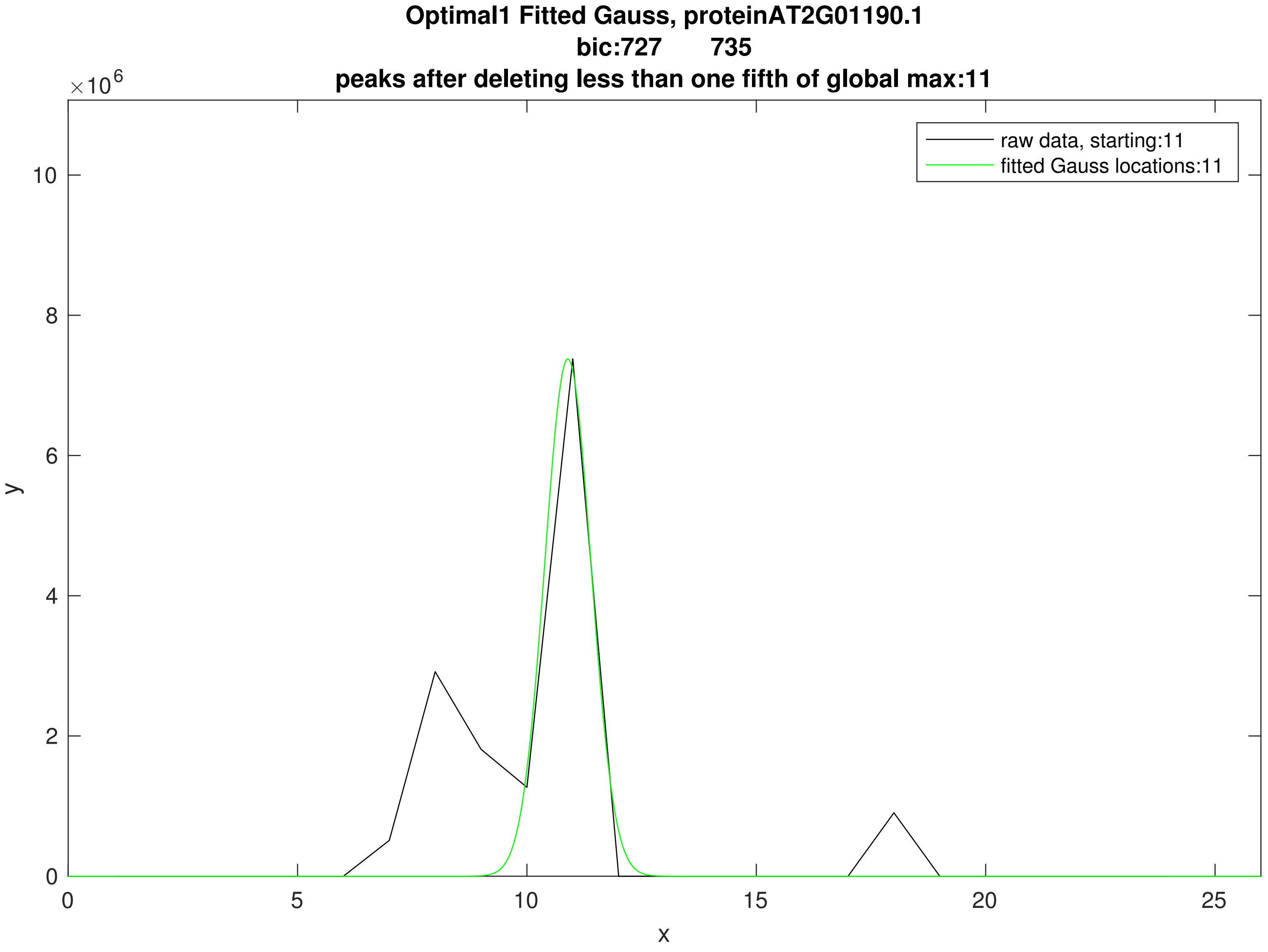

Optimal1 Fitted Gauss, proteinAT2G01190.1
bic:727 735
peaks after deleting less than one fifth of global max:11
×10^6
raw data, starting:11
fitted Gauss locations:11
y
x
0
2
4
6
8
10
0
5
10
15
20
25

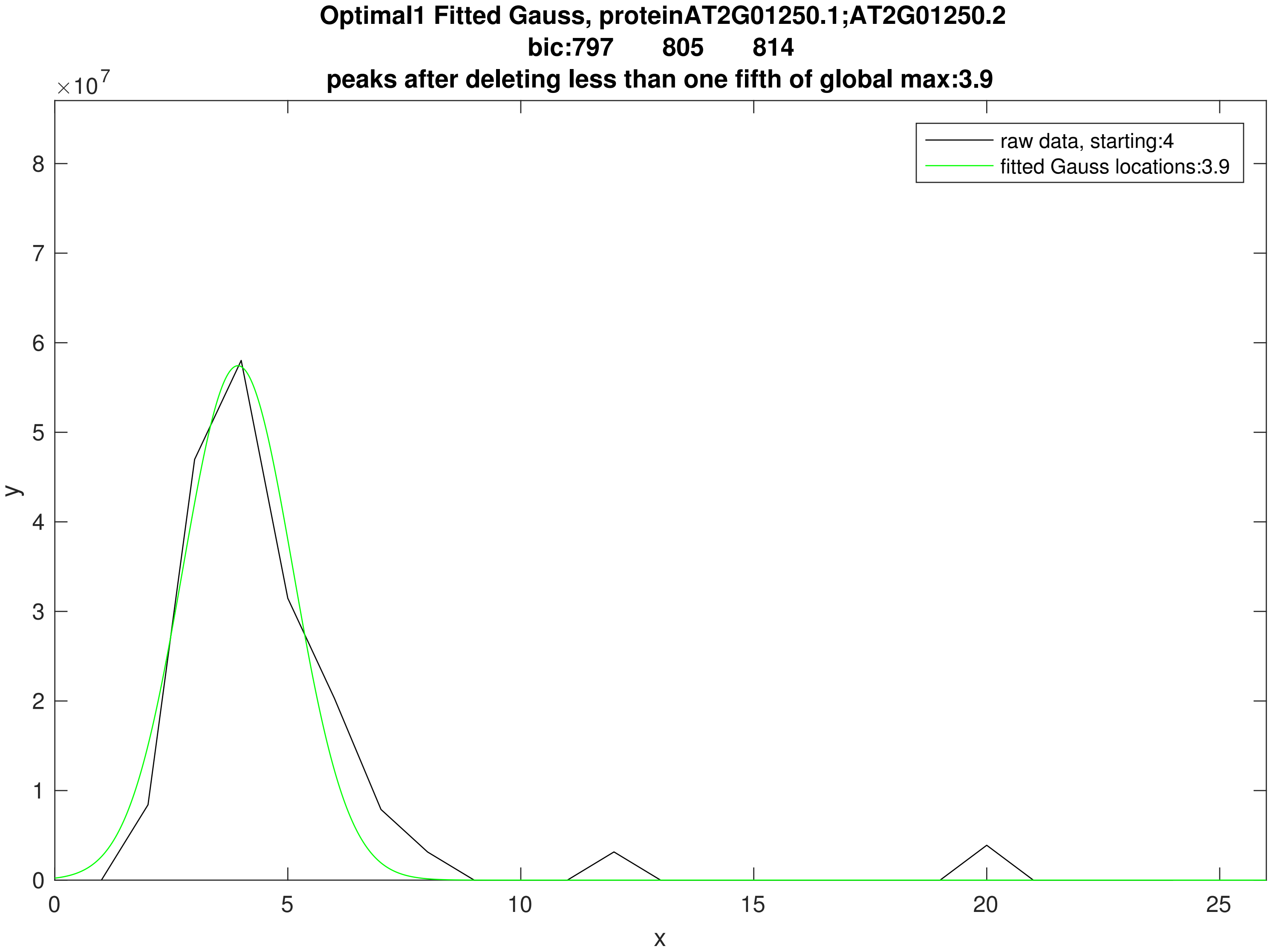
Optimal1 Fitted Gauss, proteinAT2G01250.1;AT2G01250.2
bic:797 805 814
peaks after deleting less than one fifth of global max:3.9
×10^7
raw data, starting:4
fitted Gauss locations:3.9
y
x
0
1
2
3
4
5
6
7
8
0
5
10
15
20
25

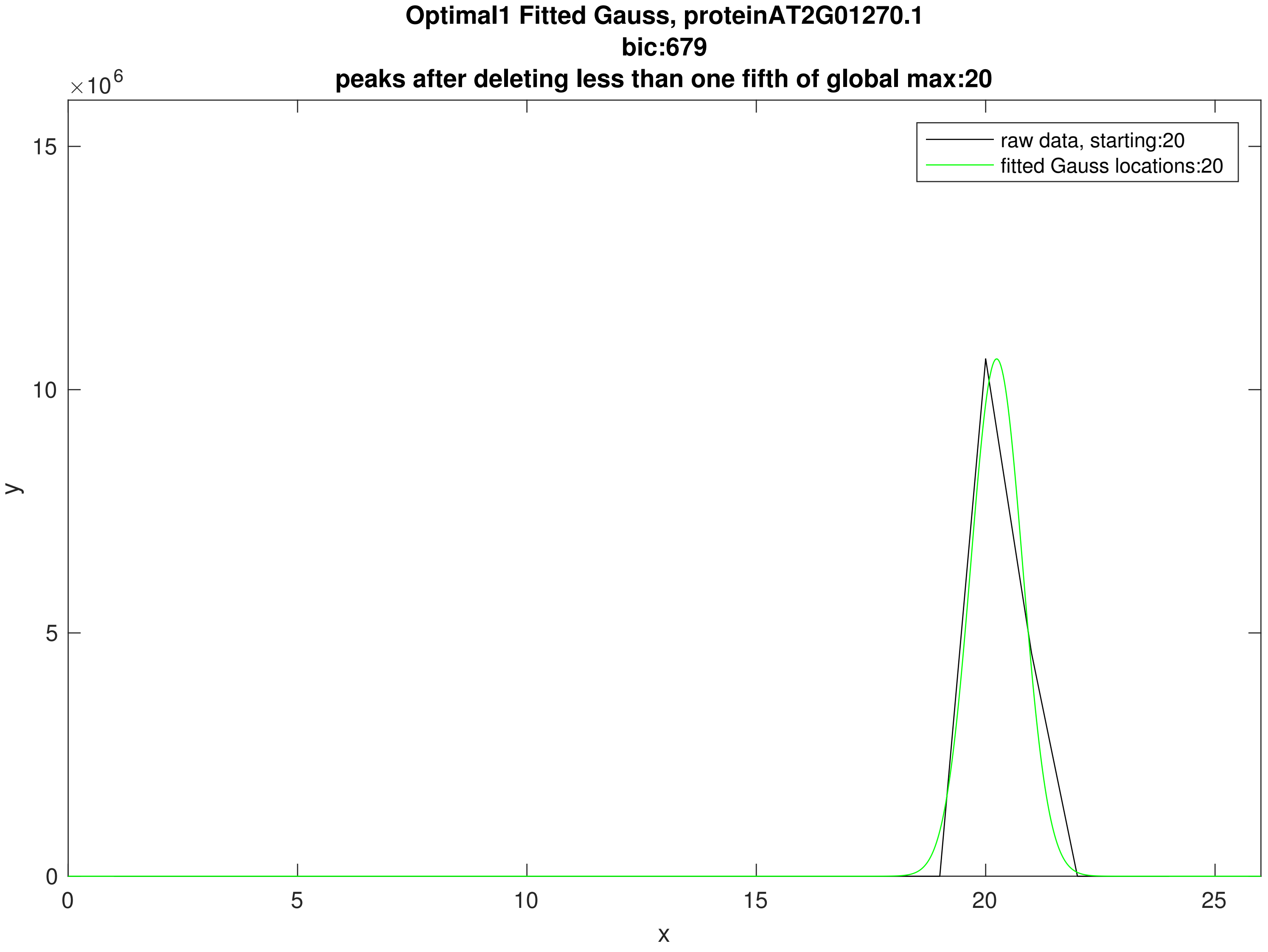

Optimal1 Fitted Gauss, proteinAT2G01270.1
bic:679
peaks after deleting less than one fifth of global max:20
×10^6
15
10
5
0
y
0
5
10
15
20
25
x
raw data, starting:20
fitted Gauss locations:20

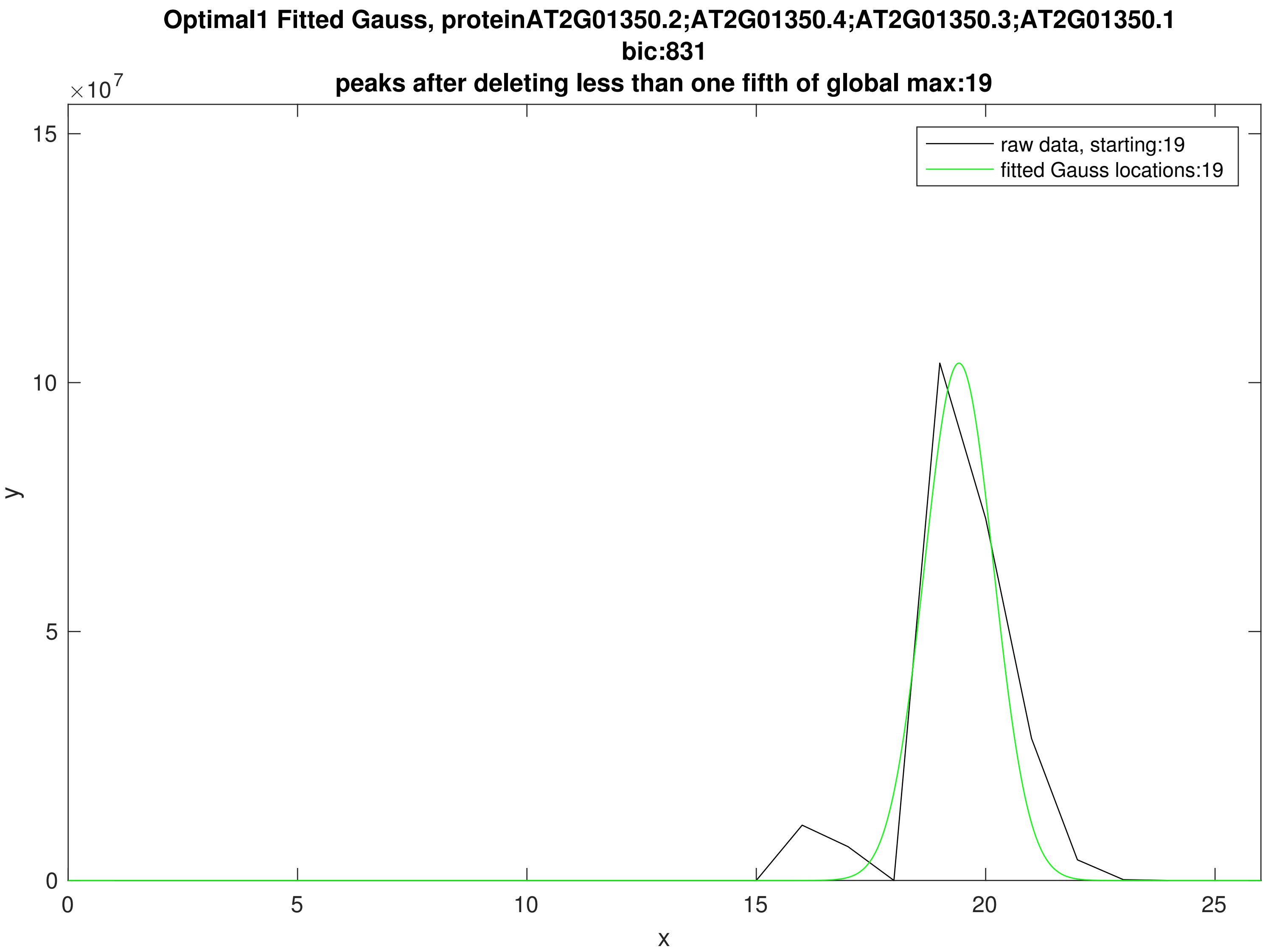

Optimal1 Fitted Gauss, proteinAT2G01350.2;AT2G01350.4;AT2G01350.3;AT2G01350.1
bic:831
peaks after deleting less than one fifth of global max:19
×10^7
15
10
5
0
y
0
5
10
15
20
25
x
raw data, starting:19
fitted Gauss locations:19

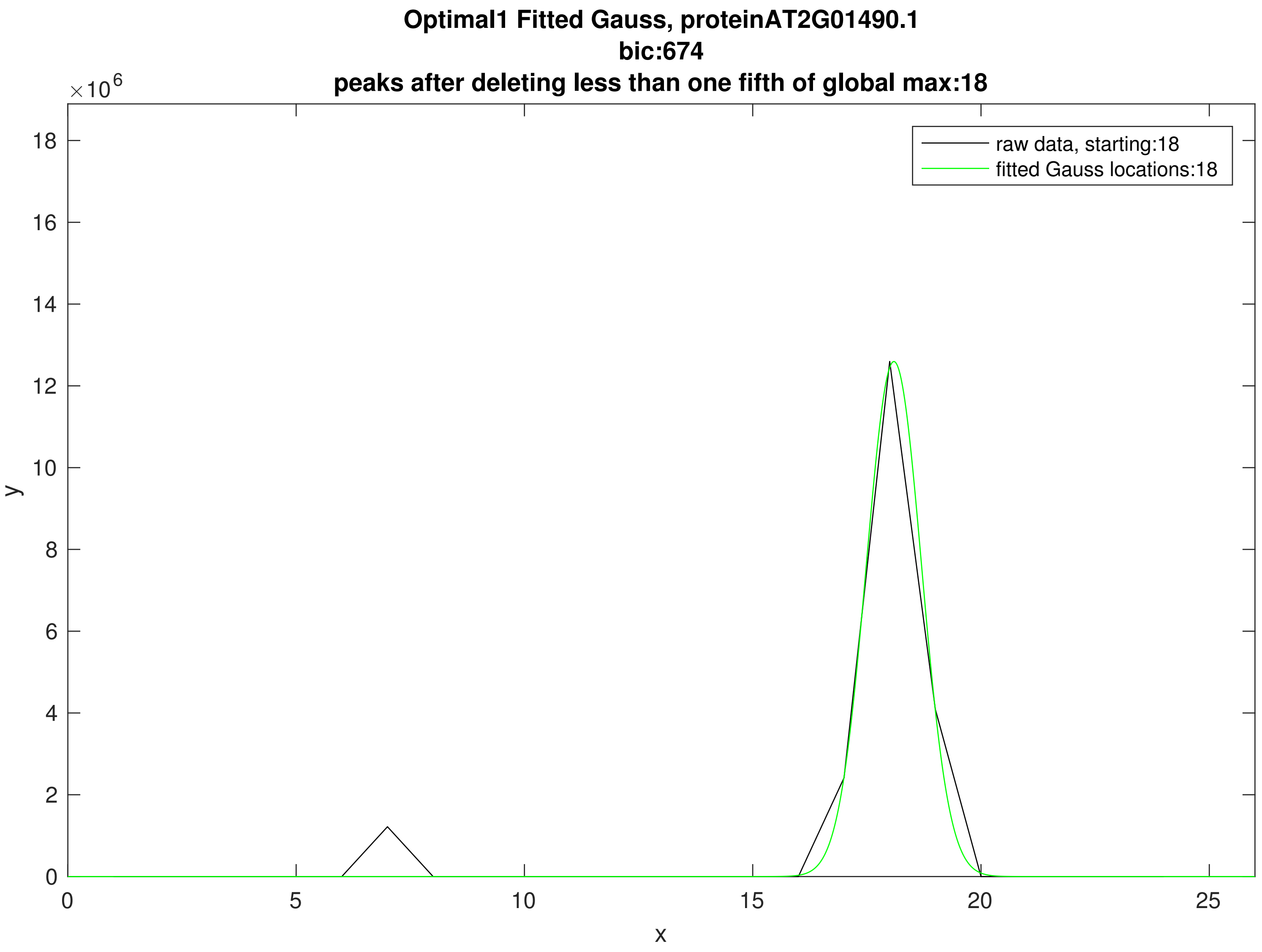
Optimal1 Fitted Gauss, proteinAT2G01490.1
bic:674
peaks after deleting less than one fifth of global max:18
×10^6
raw data, starting:18
fitted Gauss locations:18
y
x
0
2
4
6
8
10
12
14
16
18
0
5
10
15
20
25

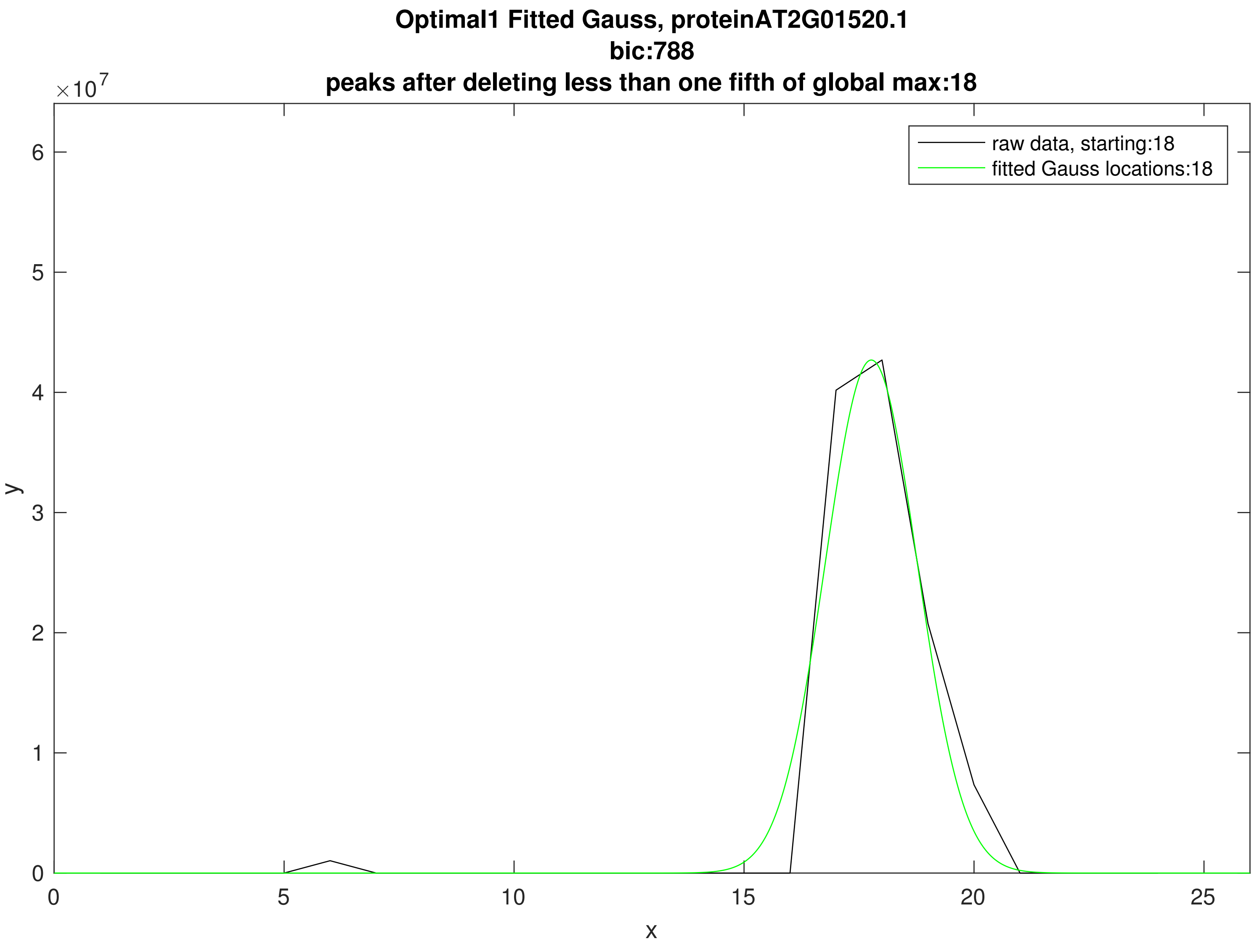

Optimal1 Fitted Gauss, proteinAT2G01520.1
bic:788
peaks after deleting less than one fifth of global max:18
×10^7
raw data, starting:18
fitted Gauss locations:18
y
x
0
1
2
3
4
5
6
0
5
10
15
20
25

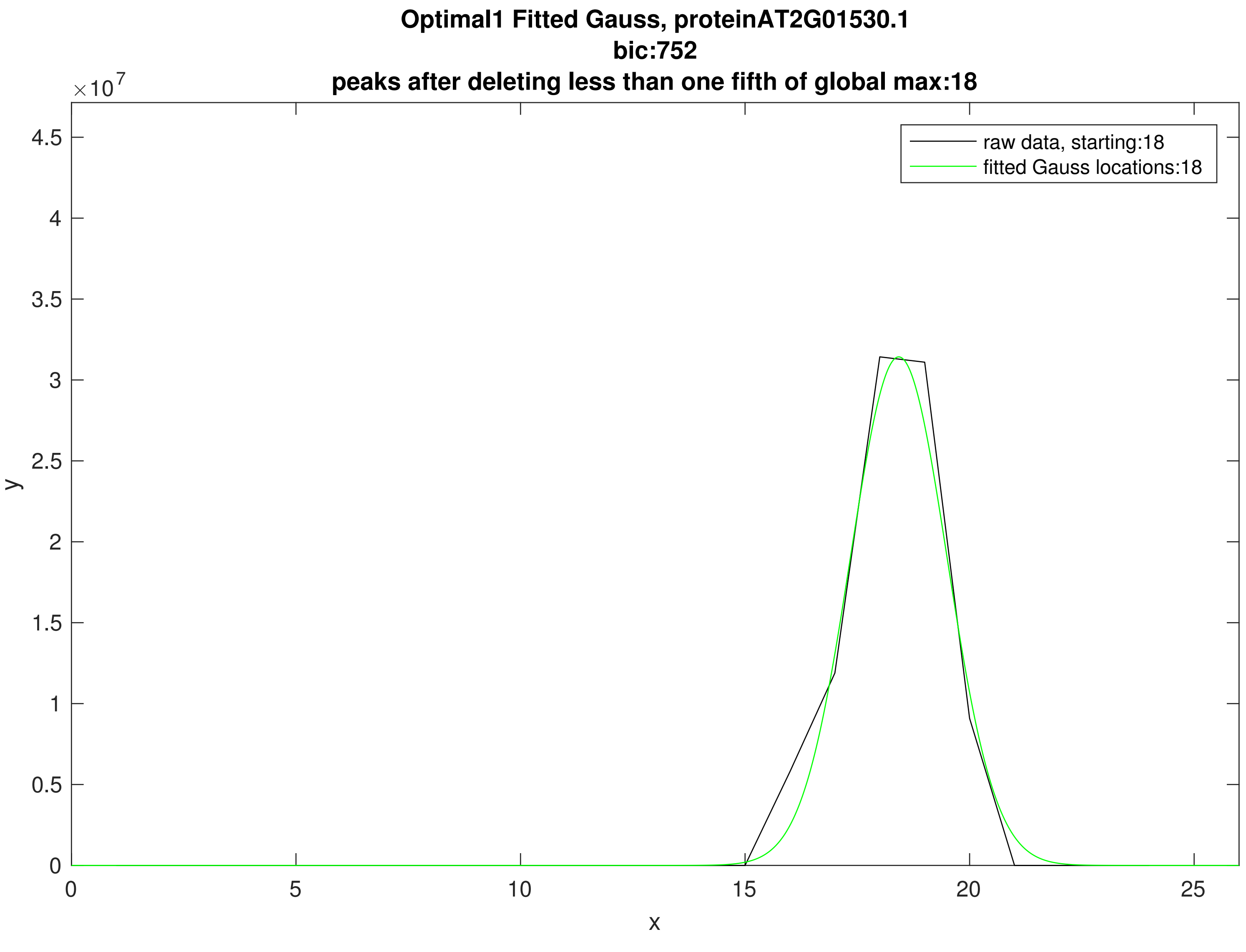
Optimal1 Fitted Gauss, proteinAT2G01530.1
bic:752
peaks after deleting less than one fifth of global max:18
×10^7
4.5
4
3.5
3
2.5
2
1.5
1
0.5
0
y
0
5
10
15
20
25
x
raw data, starting:18
fitted Gauss locations:18

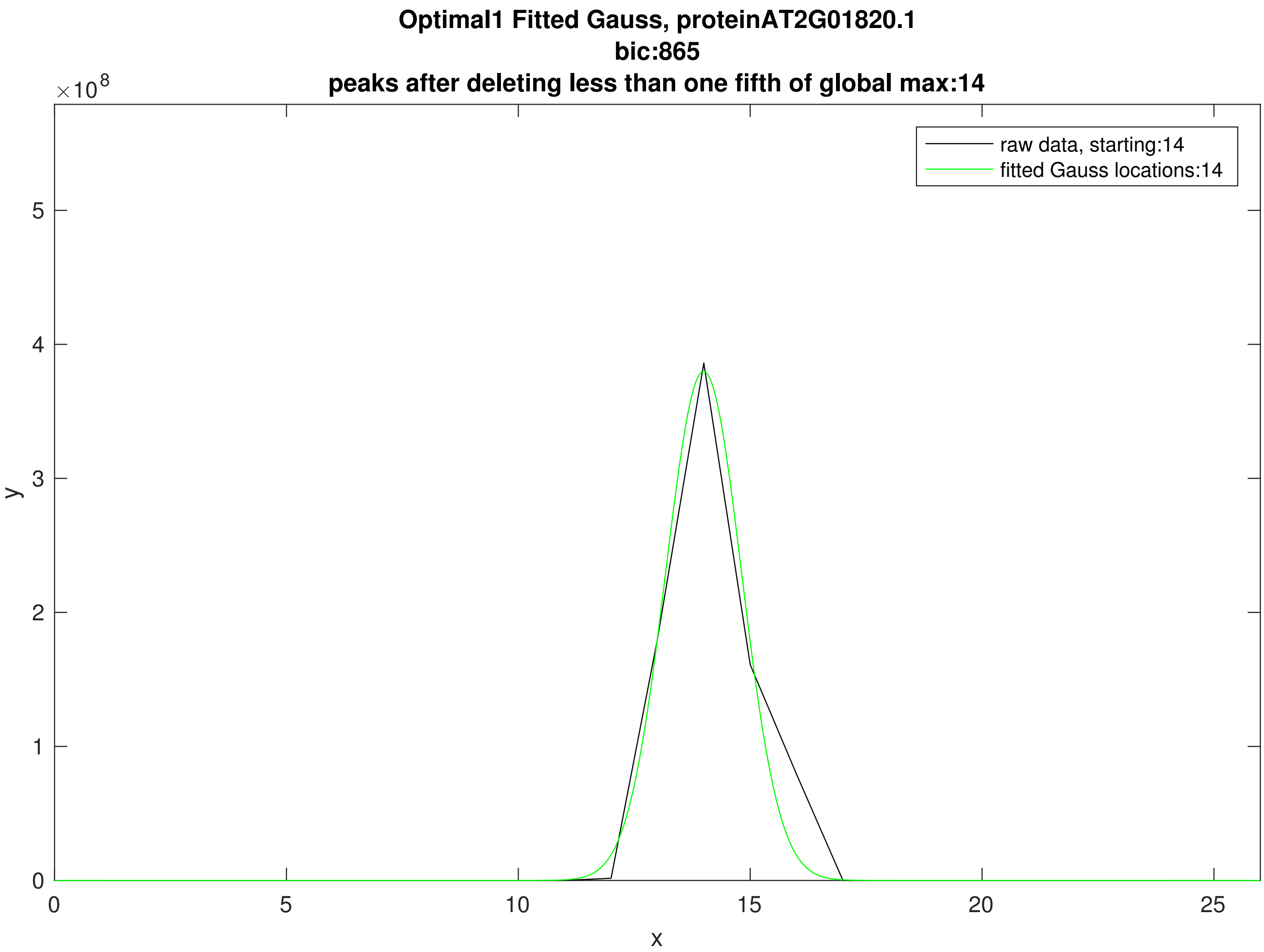
Optimal1 Fitted Gauss, proteinAT2G01820.1
bic:865
peaks after deleting less than one fifth of global max:14
×10^8
raw data, starting:14
fitted Gauss locations:14
y
x
0
1
2
3
4
5
0
5
10
15
20
25

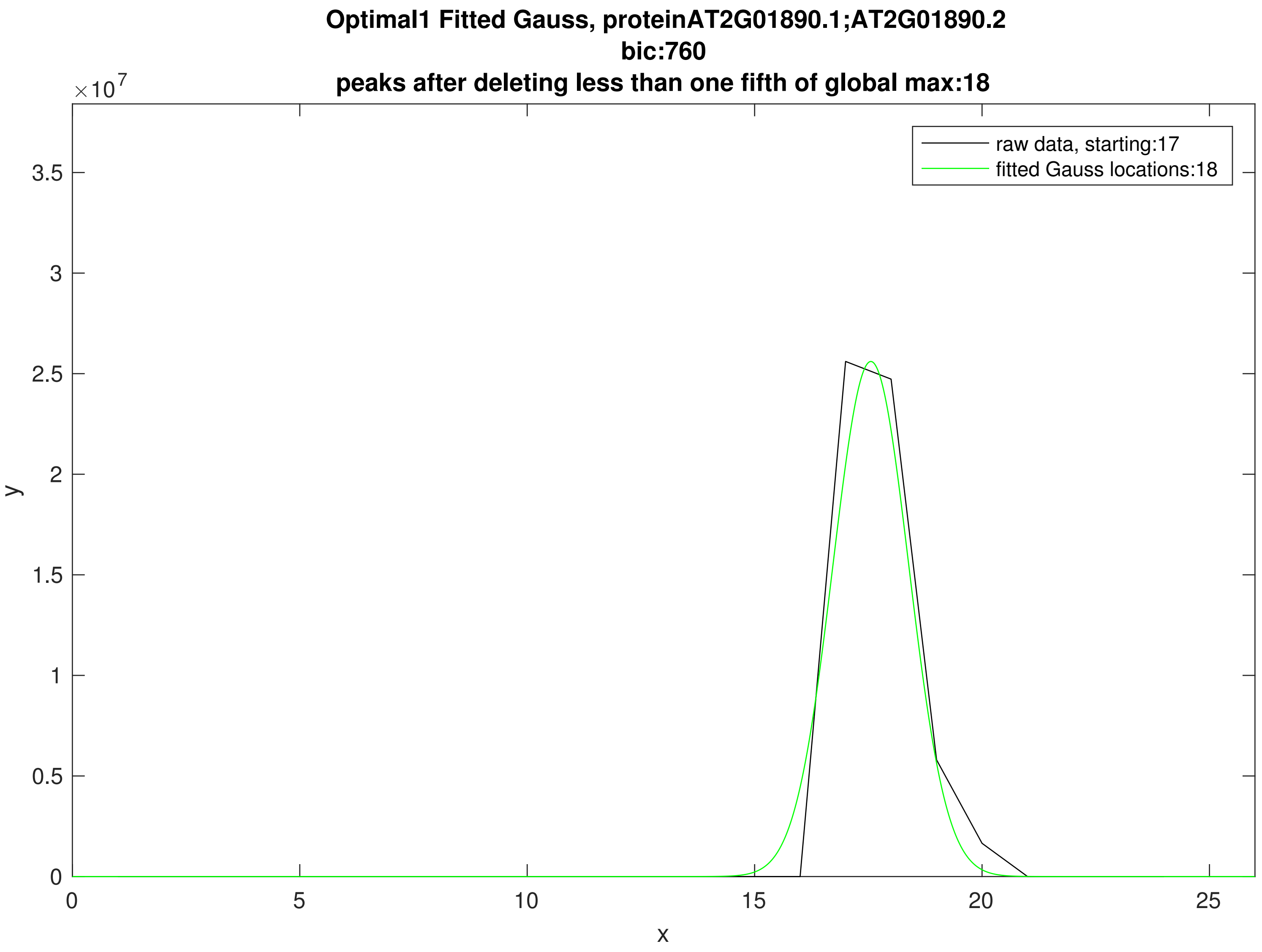

Optimal1 Fitted Gauss, proteinAT2G01890.1;AT2G01890.2
bic:760
peaks after deleting less than one fifth of global max:18
×10^7
raw data, starting:17
fitted Gauss locations:18
3.5
3
2.5
2
1.5
1
0.5
0
y
0
5
10
15
20
25
x

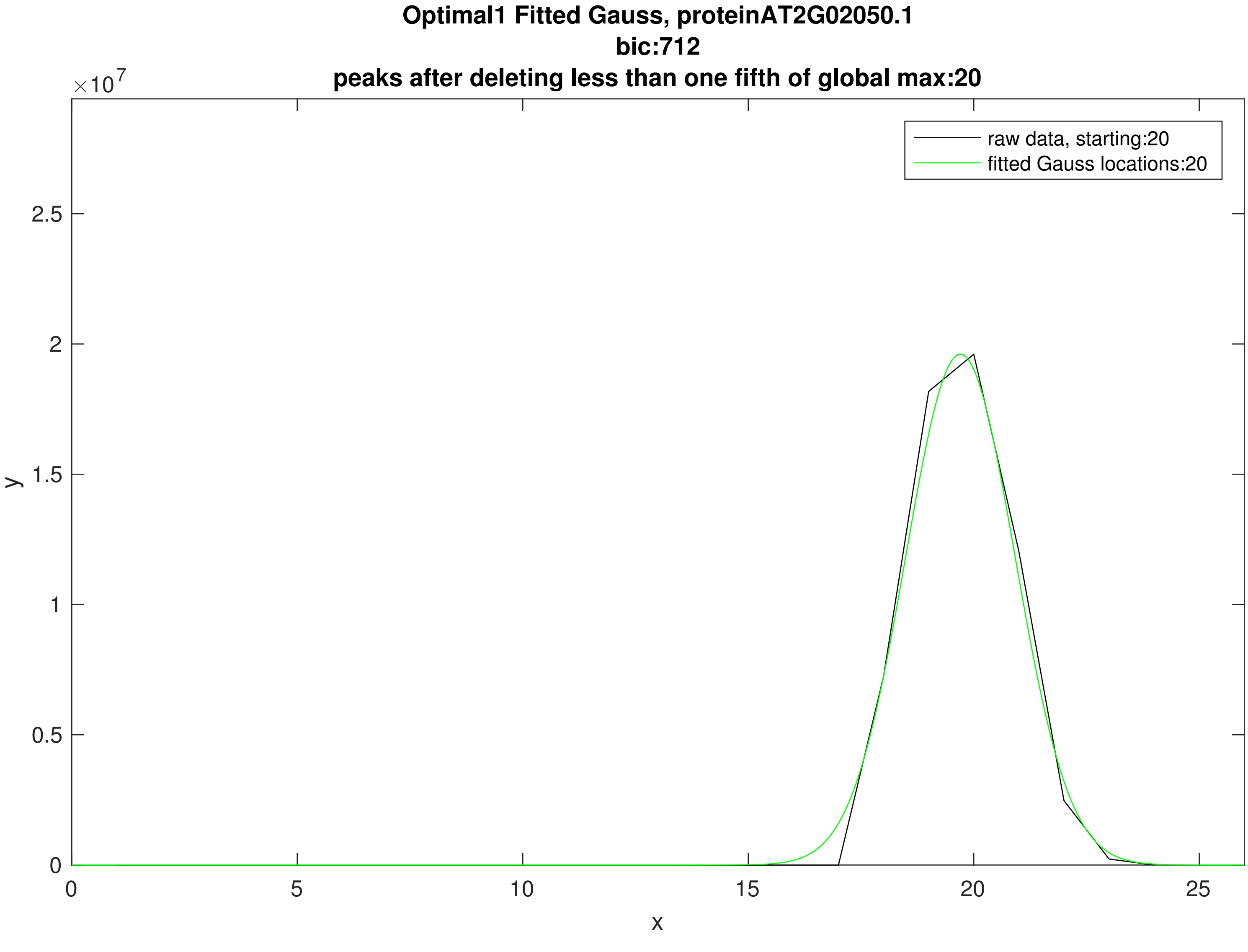
Optimal1 Fitted Gauss, proteinAT2G02050.1
bic:712
peaks after deleting less than one fifth of global max:20
×10^7
raw data, starting:20
fitted Gauss locations:20
y
x
0
0.5
1
1.5
2
2.5
0
5
10
15
20
25

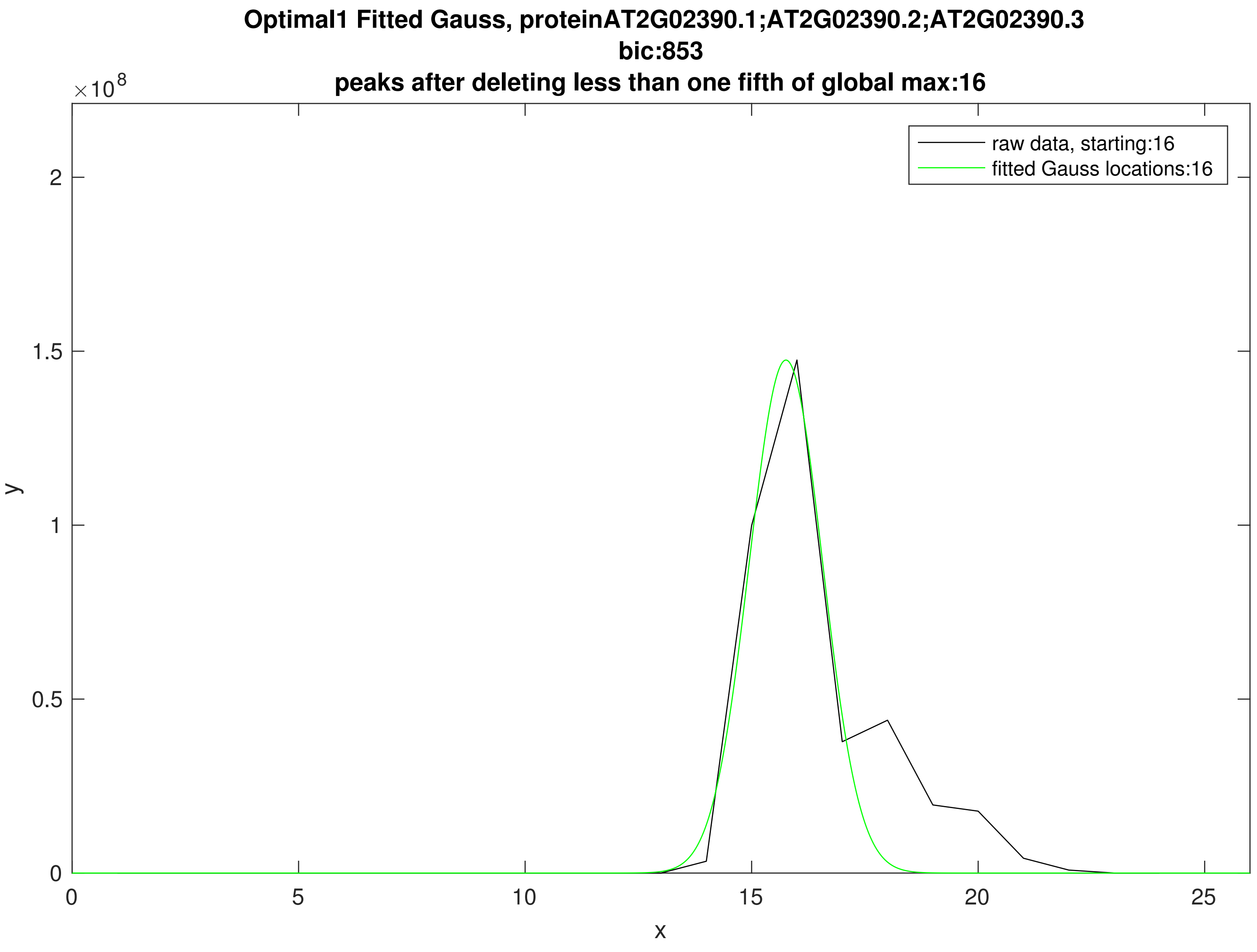

Optimal1 Fitted Gauss, proteinAT2G02390.1;AT2G02390.2;AT2G02390.3
bic:853
peaks after deleting less than one fifth of global max:16
×10^8
raw data, starting:16
fitted Gauss locations:16
y
x
0
0.5
1
1.5
2
0
5
10
15
20
25

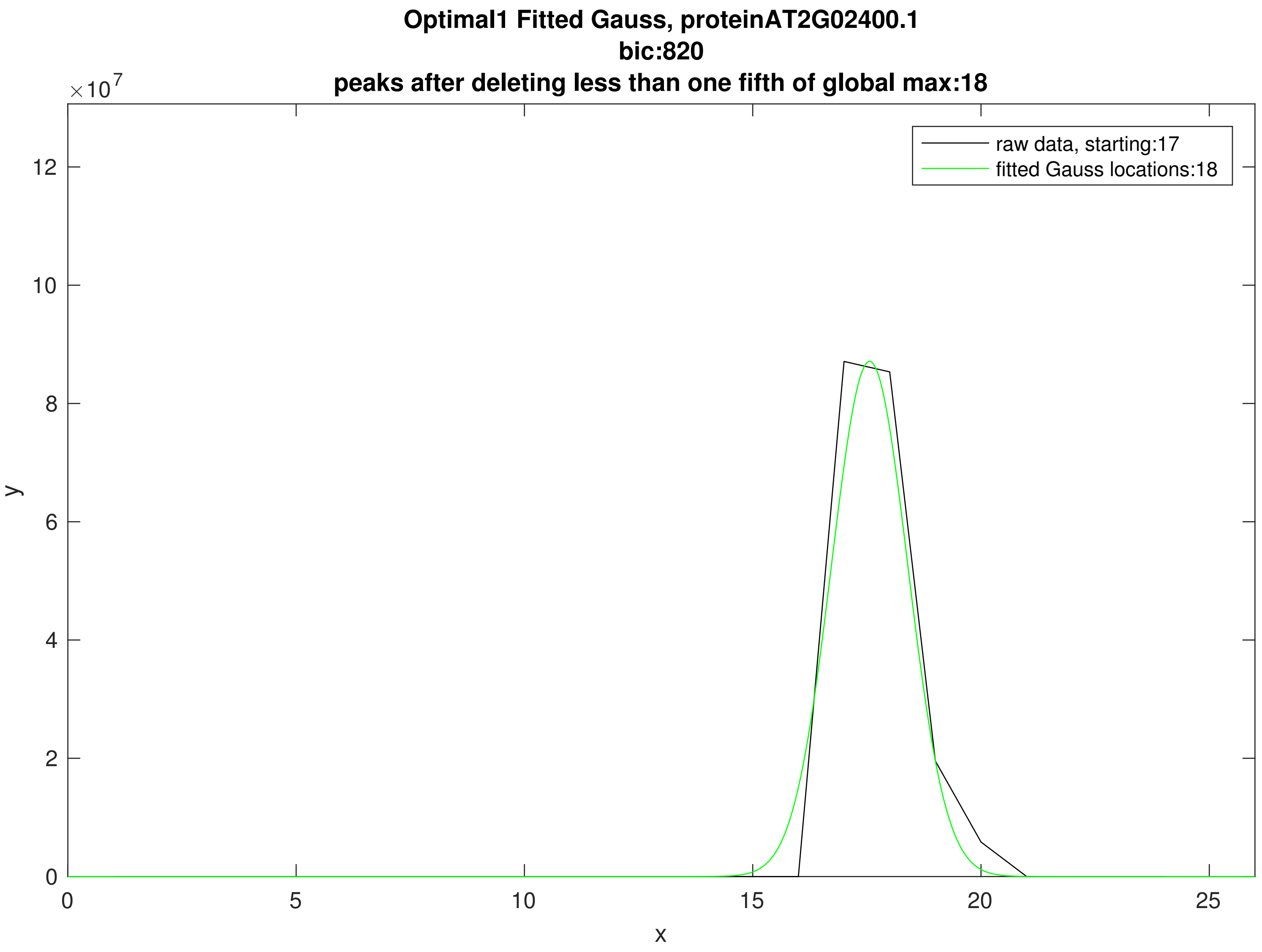
Optimal1 Fitted Gauss, proteinAT2G02400.1
bic:820
peaks after deleting less than one fifth of global max:18
×10^7
raw data, starting:17
fitted Gauss locations:18
y
x
0
2
4
6
8
10
12
0
5
10
15
20
25

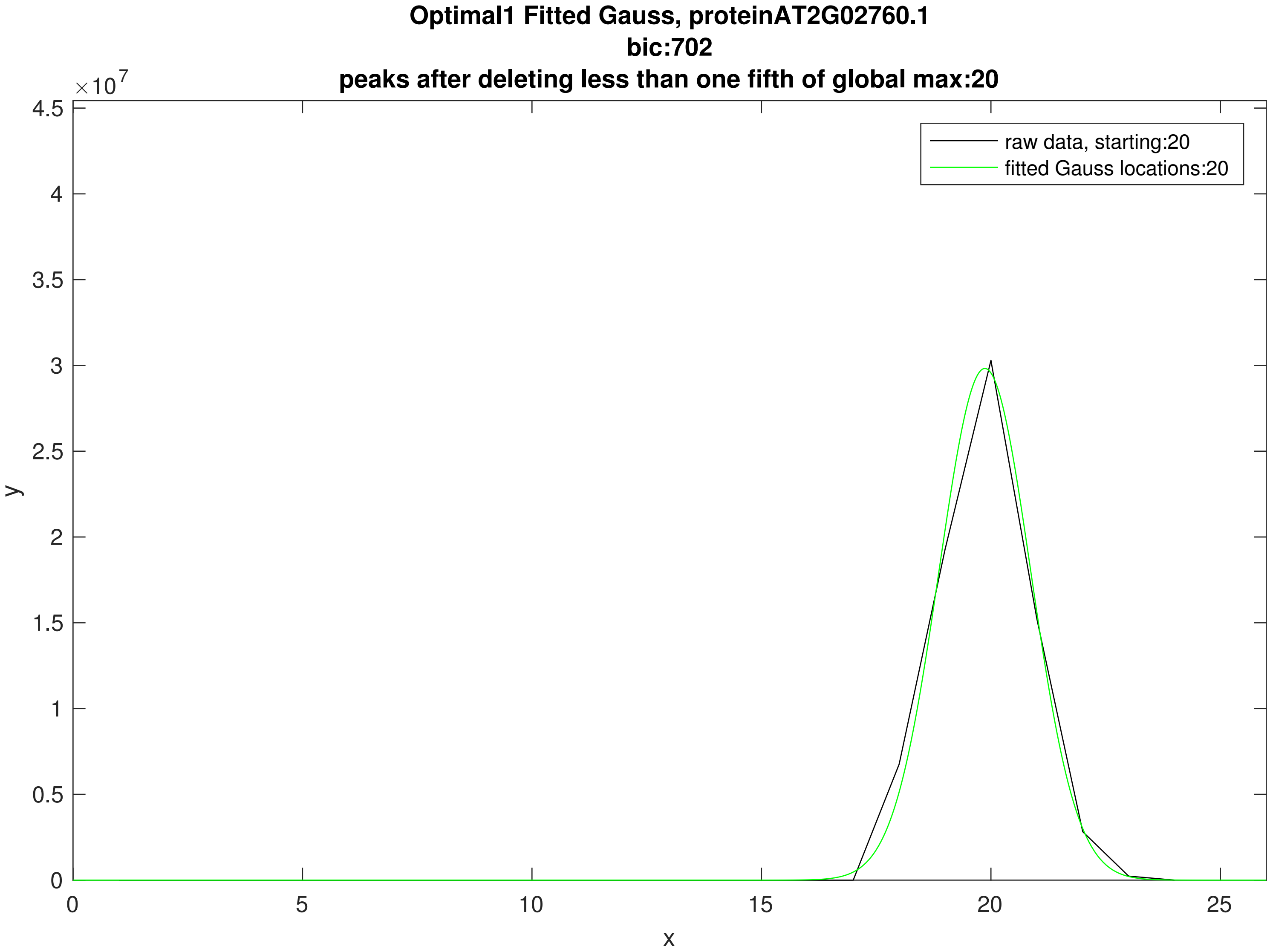

Optimal1 Fitted Gauss, proteinAT2G02760.1
bic:702
peaks after deleting less than one fifth of global max:20
×10^7
4.5
4
3.5
3
2.5
2
1.5
1
0.5
0
y
0
5
10
15
20
25
x
raw data, starting:20
fitted Gauss locations:20

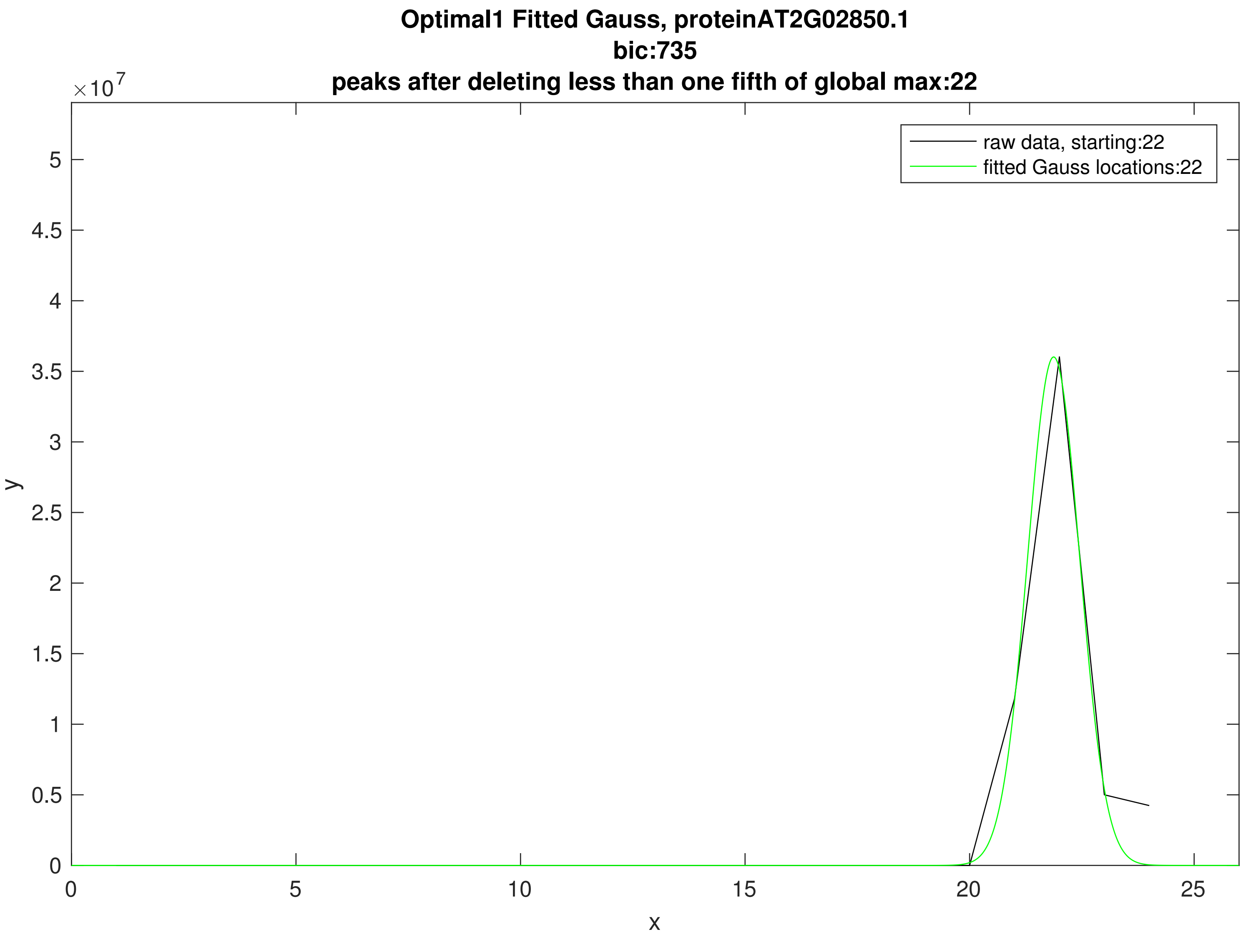

Optimal1 Fitted Gauss, proteinAT2G02850.1
bic:735
peaks after deleting less than one fifth of global max:22
×10^7
raw data, starting:22
fitted Gauss locations:22
y
x
0
0.5
1
1.5
2
2.5
3
3.5
4
4.5
5
0
5
10
15
20
25

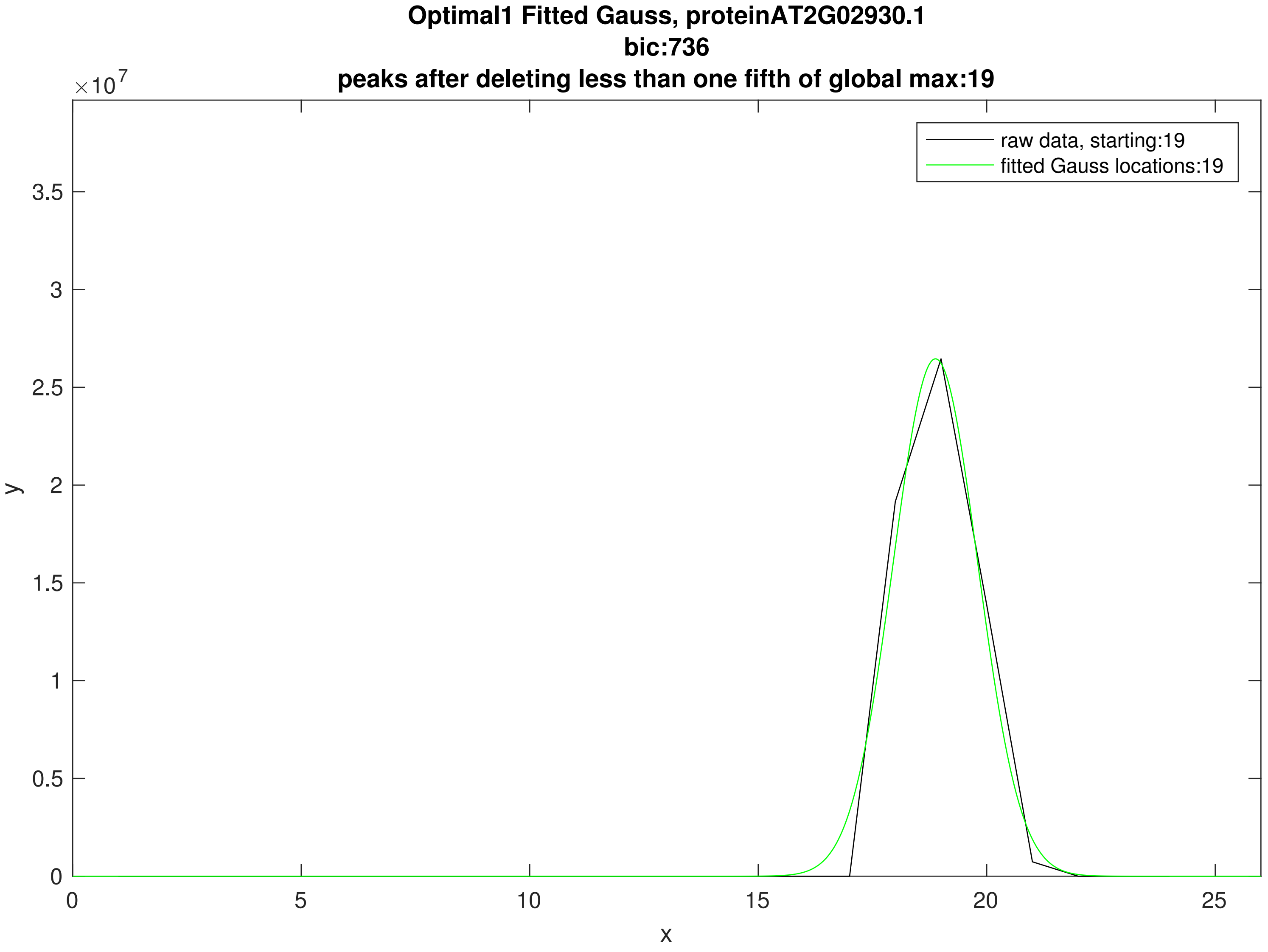
Optimal1 Fitted Gauss, proteinAT2G02930.1
bic:736
peaks after deleting less than one fifth of global max:19
×10^7
3.5
3
2.5
2
1.5
1
0.5
0
y
0
5
10
15
20
25
x
raw data, starting:19
fitted Gauss locations:19

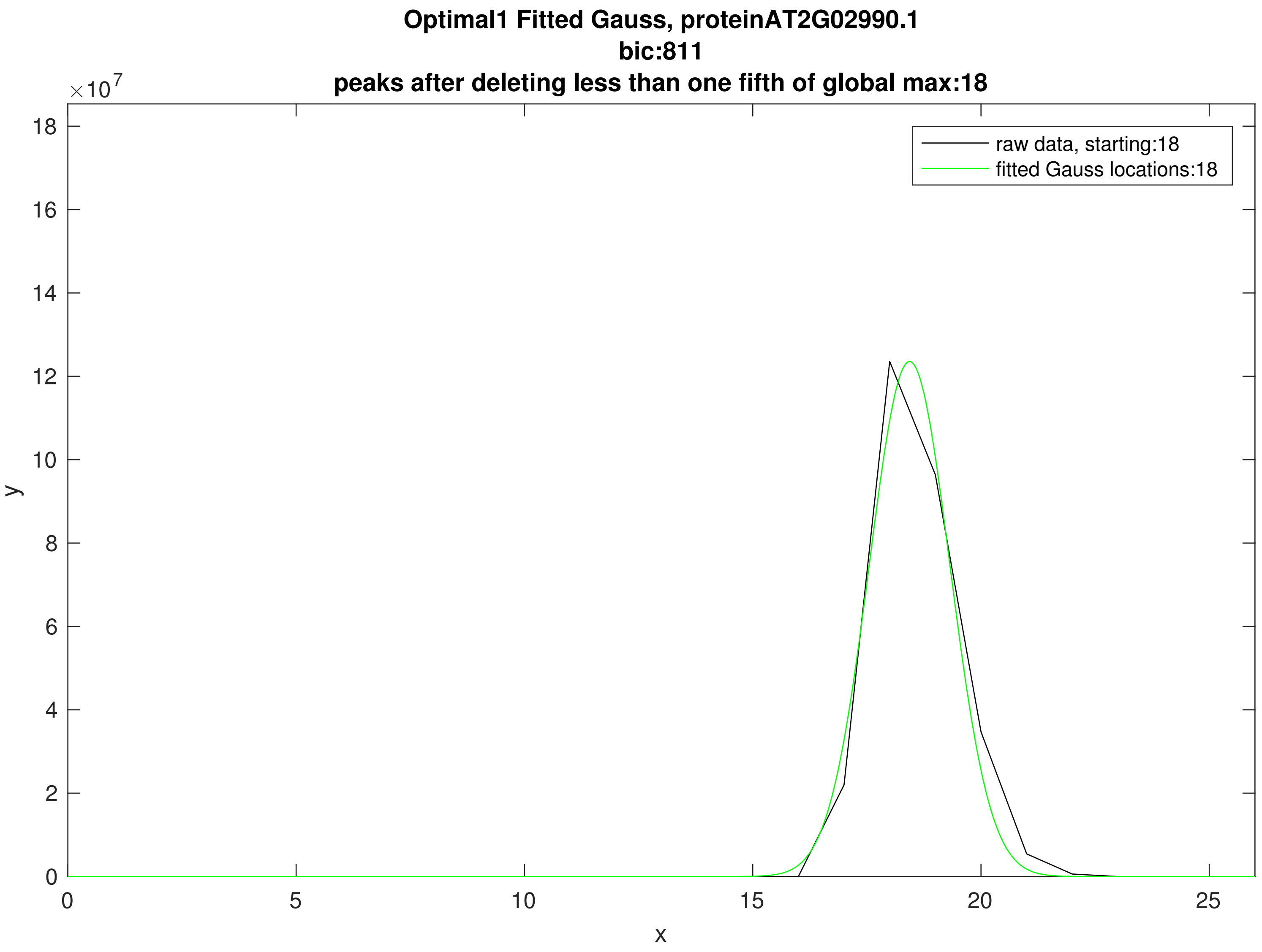

Optimal1 Fitted Gauss, proteinAT2G02990.1
bic:811
peaks after deleting less than one fifth of global max:18
×10^7
18
16
14
12
10
8
6
4
2
0
y
0
5
10
15
20
25
x
raw data, starting:18
fitted Gauss locations:18

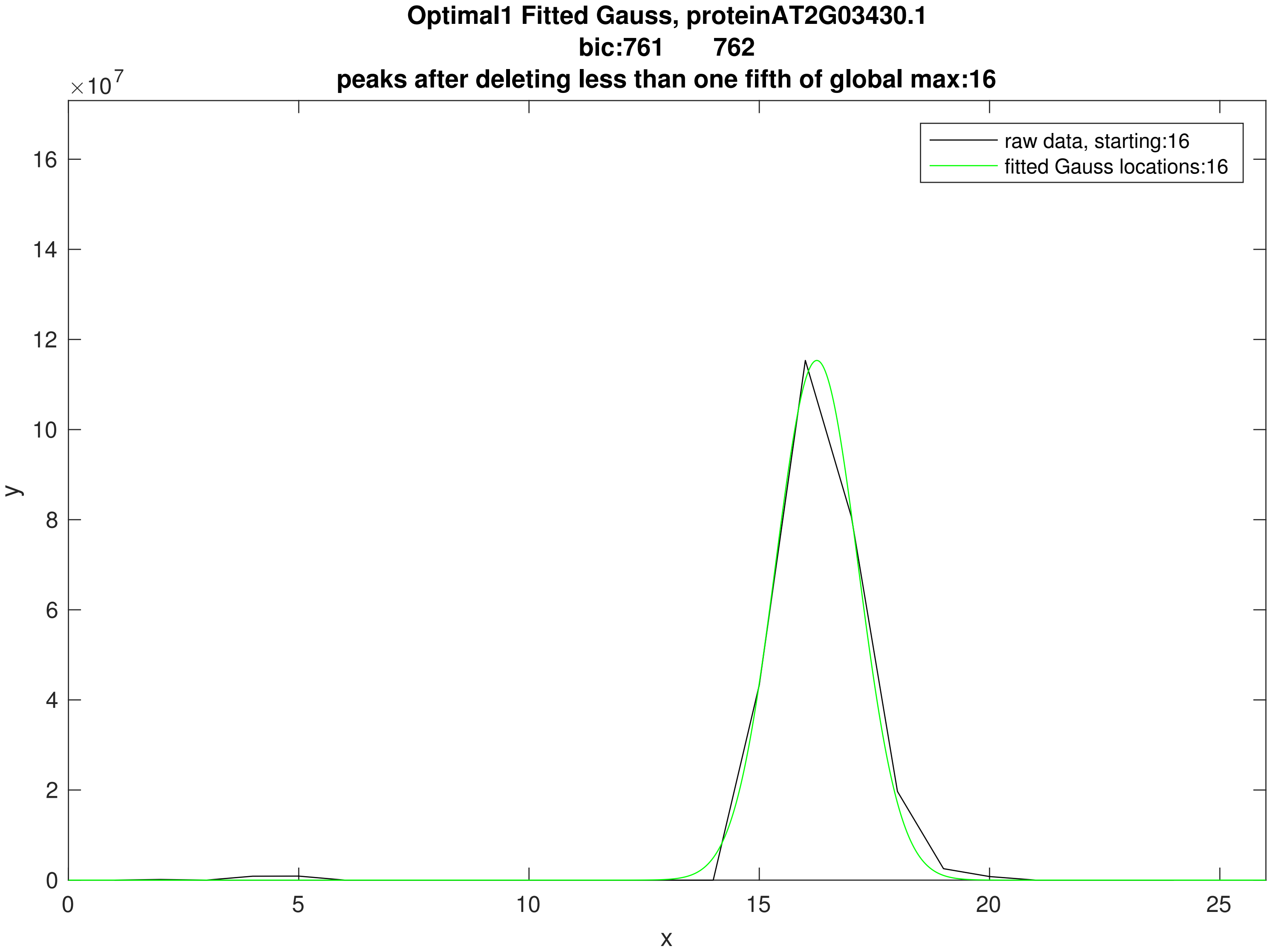

Optimal1 Fitted Gauss, proteinAT2G03430.1
bic:761 762
peaks after deleting less than one fifth of global max:16
×10^7
raw data, starting:16
fitted Gauss locations:16
y
x
0
2
4
6
8
10
12
14
16
5
15
20
25

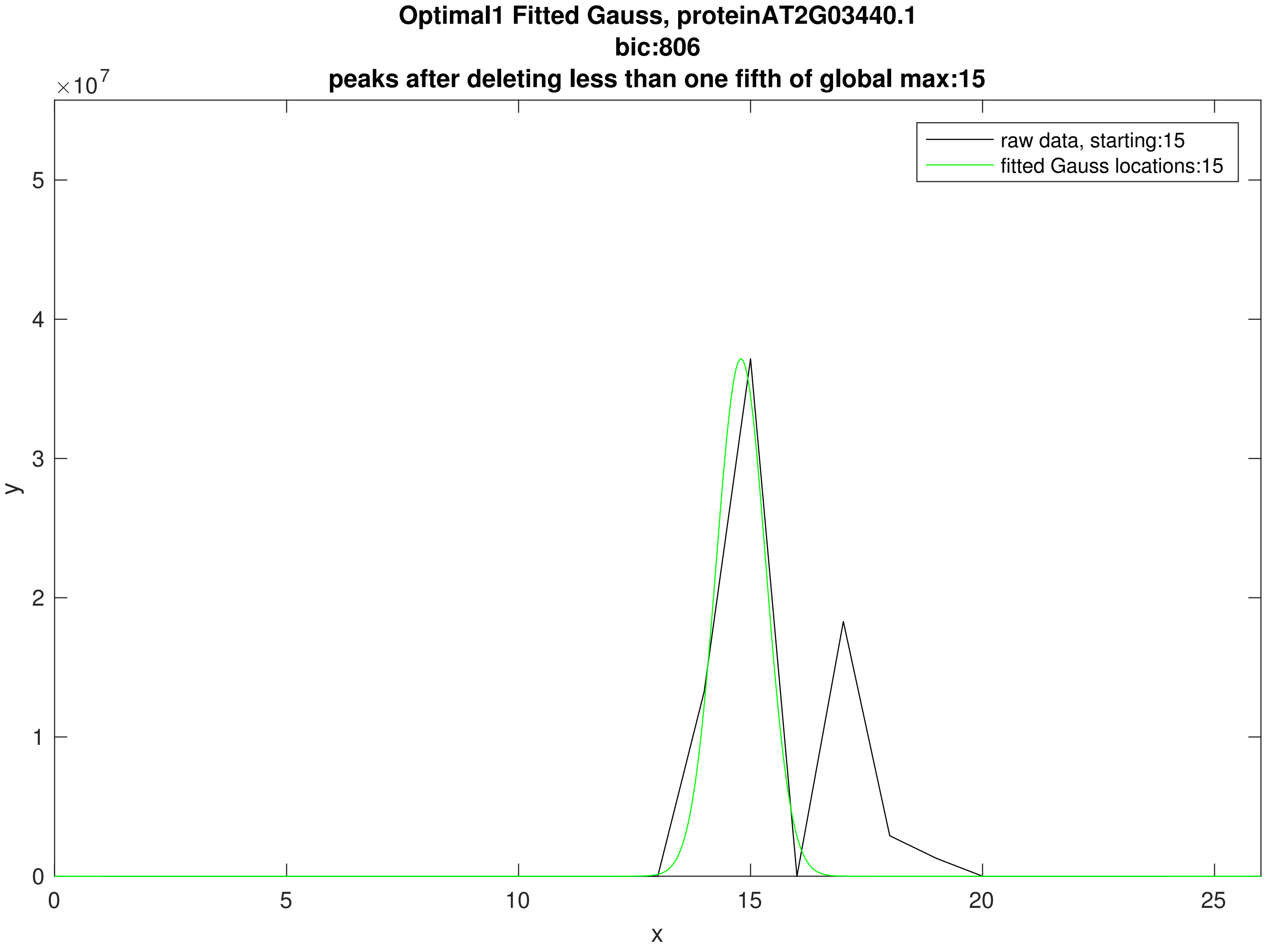
Optimal1 Fitted Gauss, proteinAT2G03440.1
bic:806
peaks after deleting less than one fifth of global max:15
×10^7
raw data, starting:15
fitted Gauss locations:15
y
x
0
1
2
3
4
5
0
5
10
15
20
25

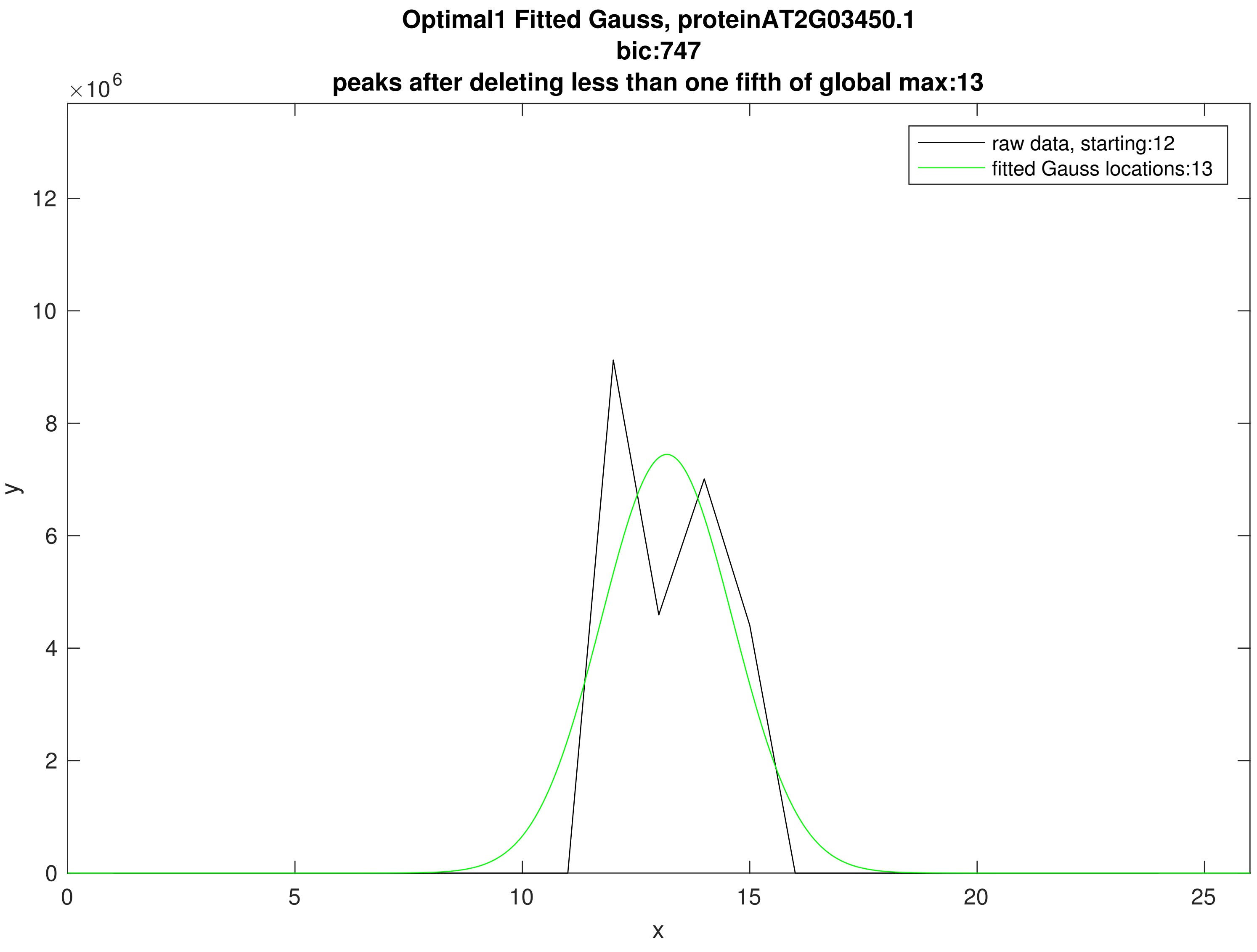
Optimal1 Fitted Gauss, proteinAT2G03450.1
bic:747
peaks after deleting less than one fifth of global max:13
×10^6
raw data, starting:12
fitted Gauss locations:13
y
x
0
2
4
6
8
10
12
0
5
10
15
20
25

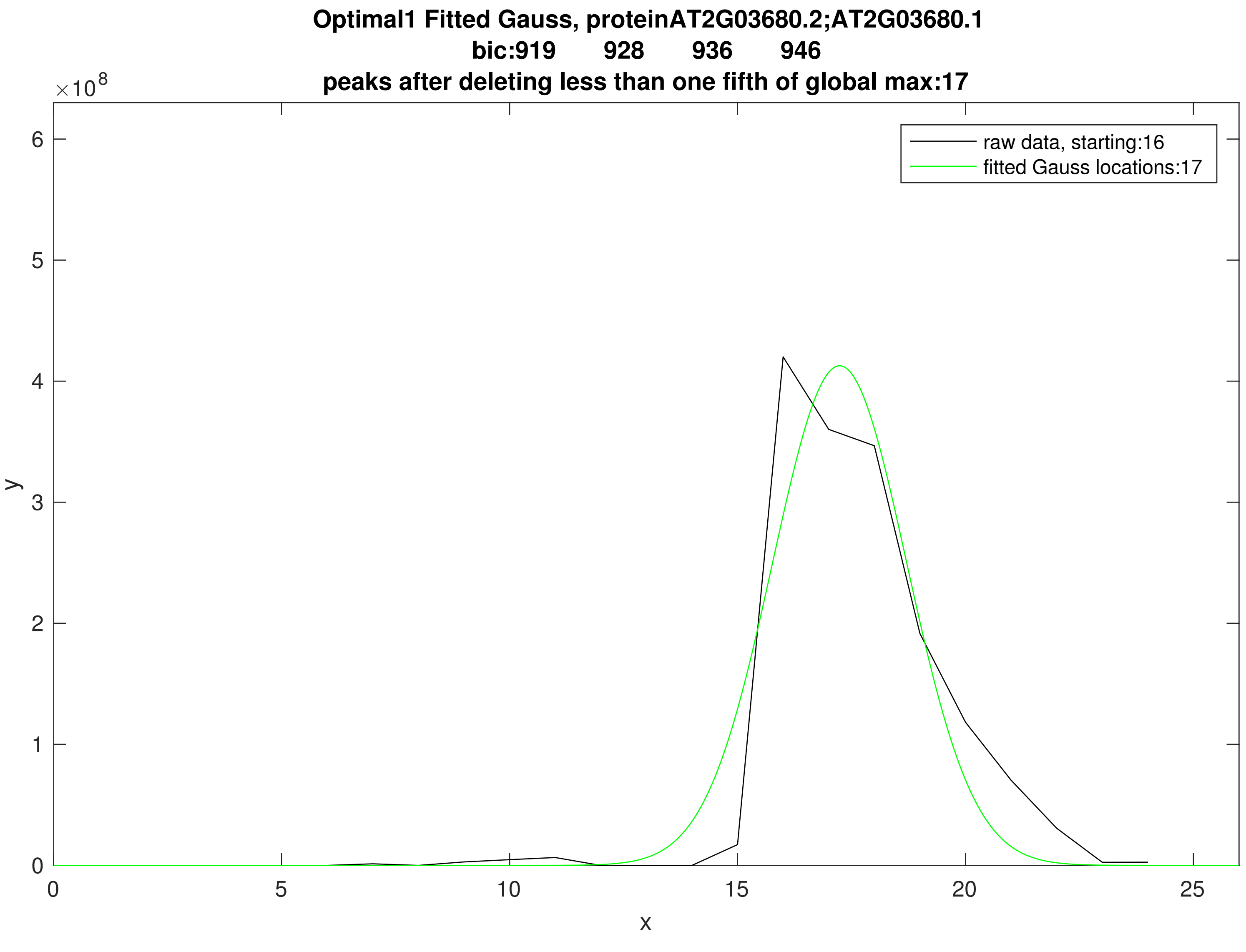

Optimal1 Fitted Gauss, proteinAT2G03680.2;AT2G03680.1
bic:919 928 936 946
peaks after deleting less than one fifth of global max:17
×10^8
raw data, starting:16
fitted Gauss locations:17
y
x
0
1
2
3
4
5
6
0
5
10
15
20
25

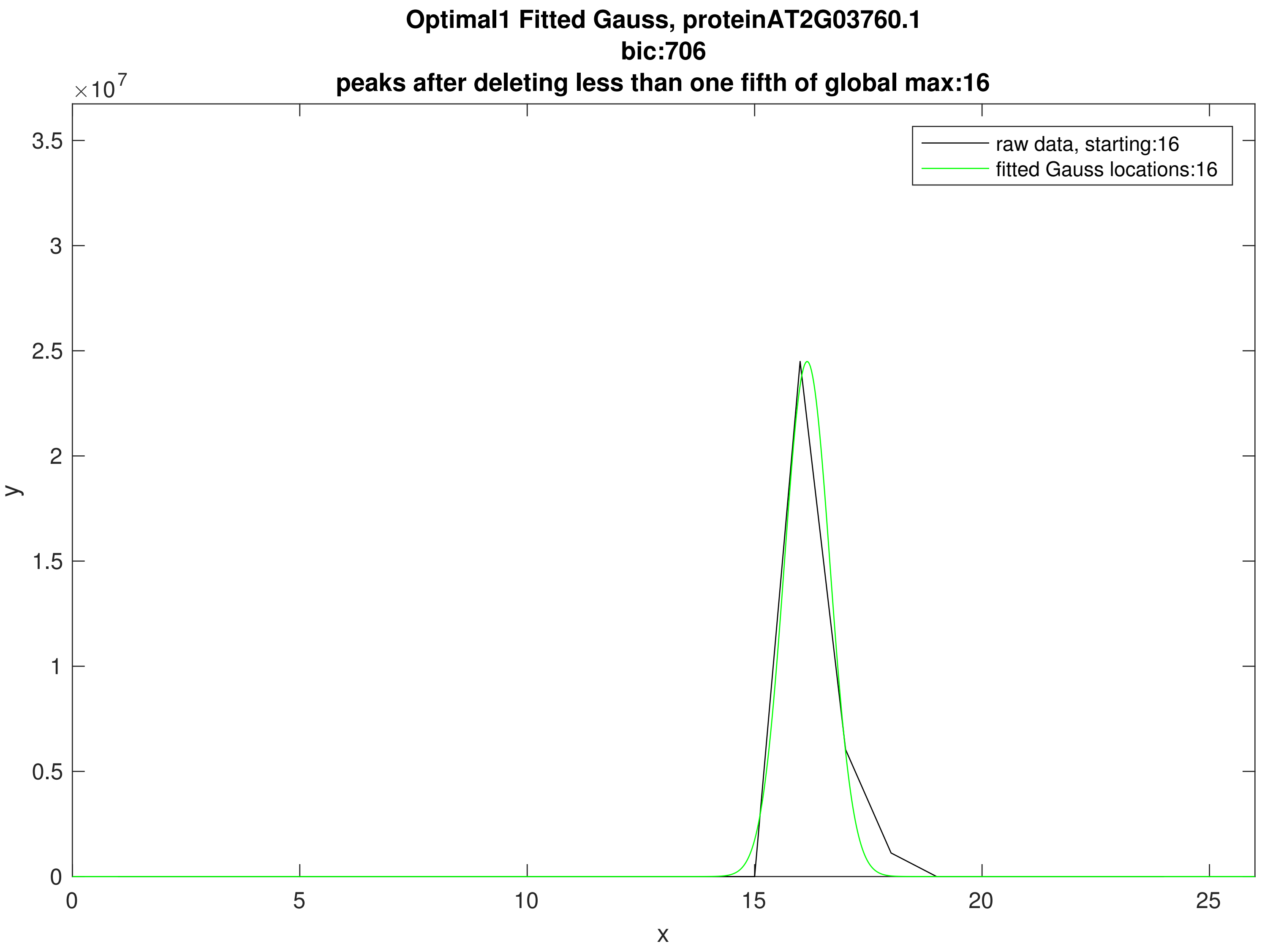

Optimal1 Fitted Gauss, proteinAT2G03760.1
bic:706
peaks after deleting less than one fifth of global max:16
×10^7
raw data, starting:16
fitted Gauss locations:16
y
x
0
0.5
1
1.5
2
2.5
3
3.5
5
10
15
20
25

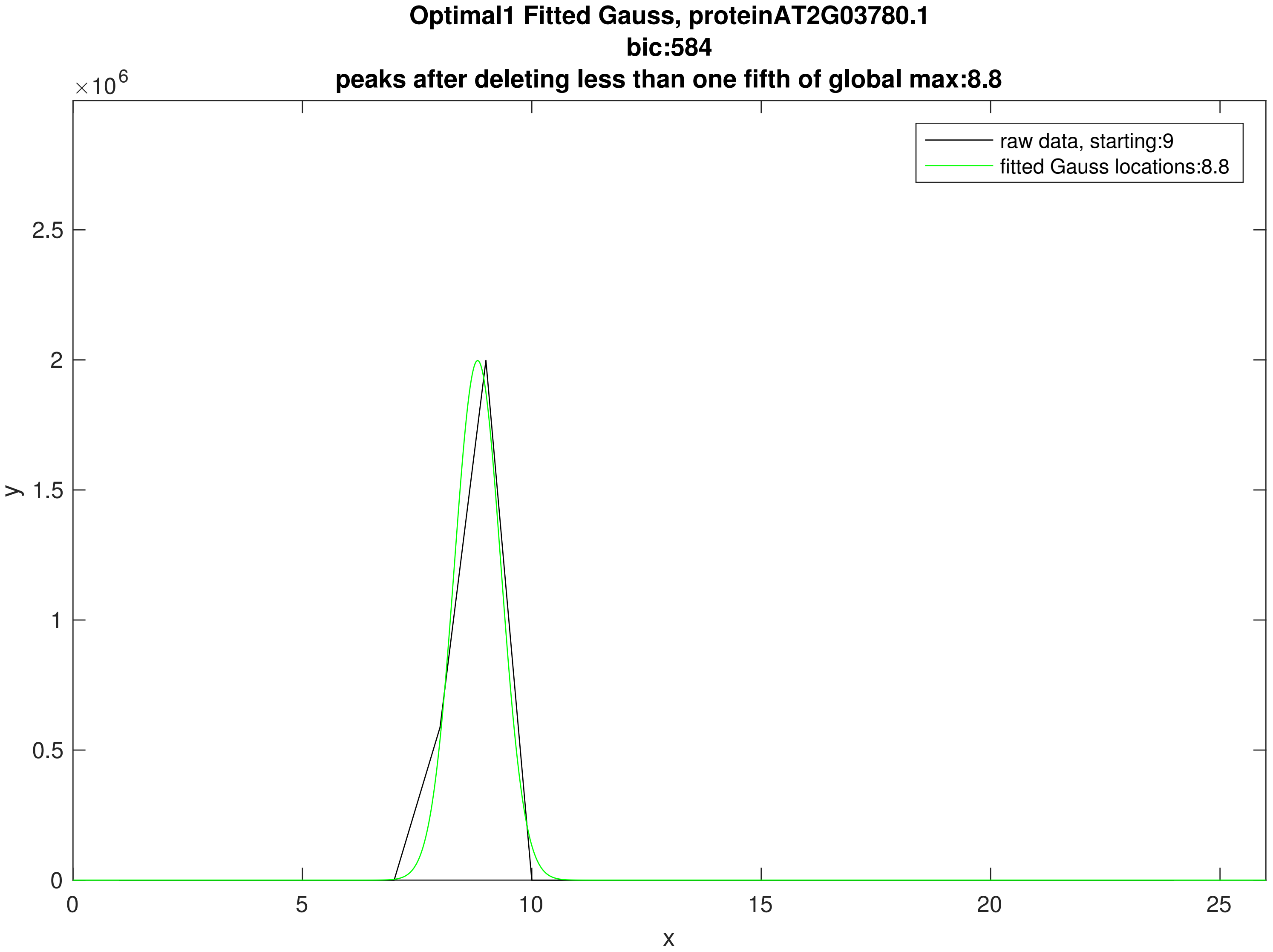

Optimal1 Fitted Gauss, proteinAT2G03780.1
bic:584
peaks after deleting less than one fifth of global max:8.8
×10^6
raw data, starting:9
fitted Gauss locations:8.8
y
x
0
0.5
1
1.5
2
2.5
0
5
10
15
20
25

**Optimal1 Fitted Gauss, proteinAT2G03800.1**
**bic:780**
**peaks after deleting less than one fifth of global max:17**

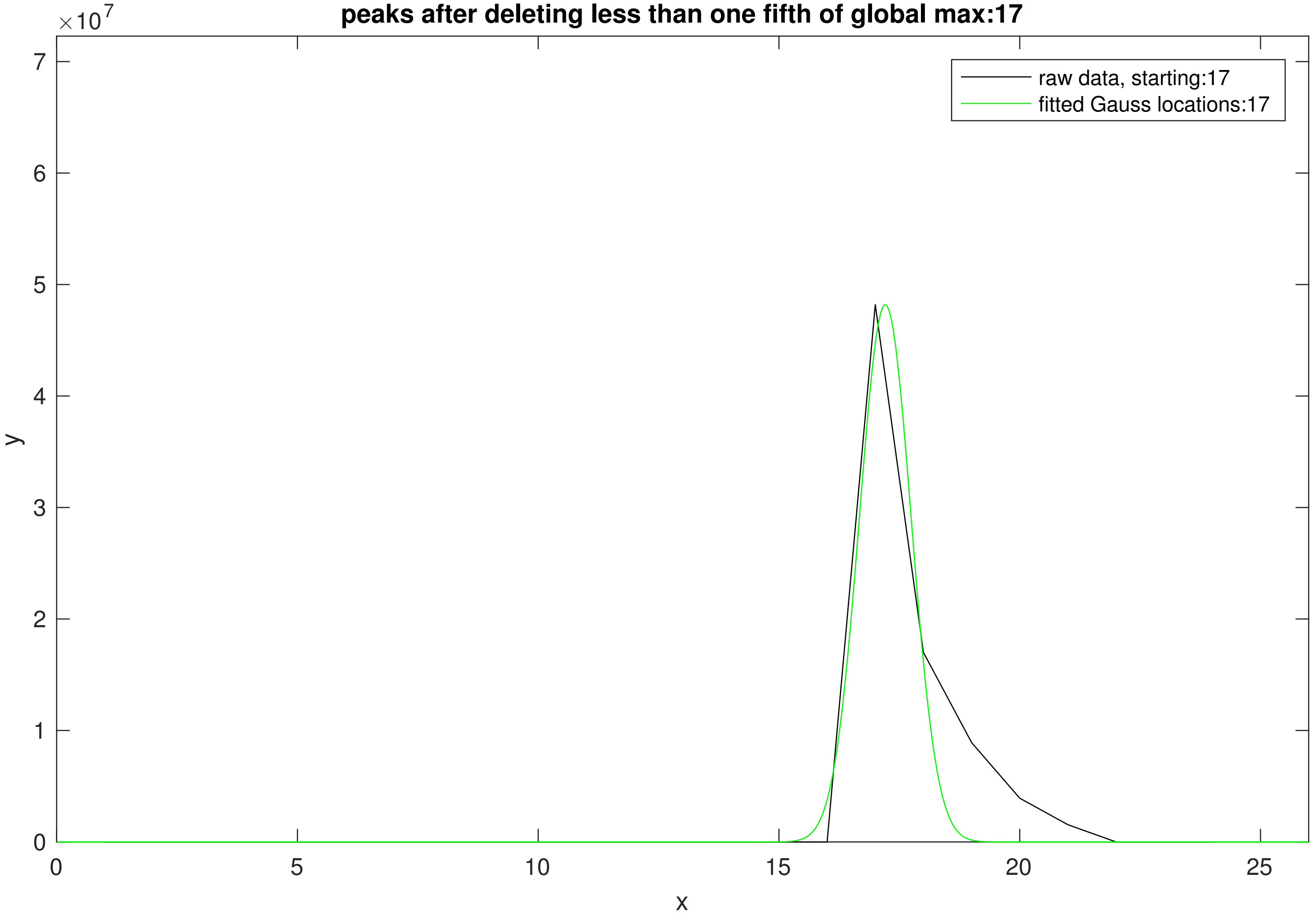

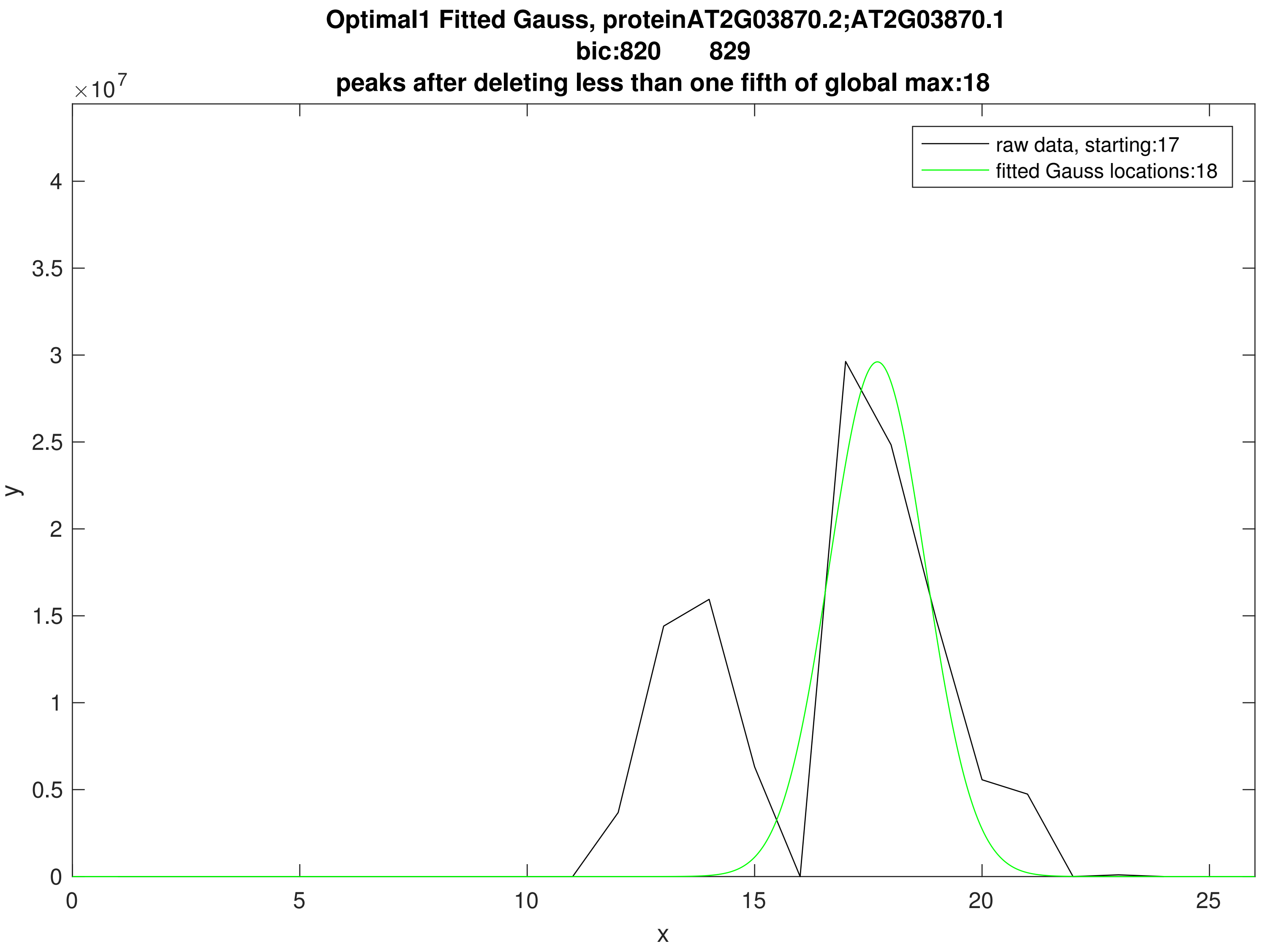
Optimal1 Fitted Gauss, proteinAT2G03870.2;AT2G03870.1
bic:820 829
peaks after deleting less than one fifth of global max:18
×10^7
raw data, starting:17
fitted Gauss locations:18
y
x
0
0.5
1
1.5
2
2.5
3
3.5
4
5
10
15
20
25

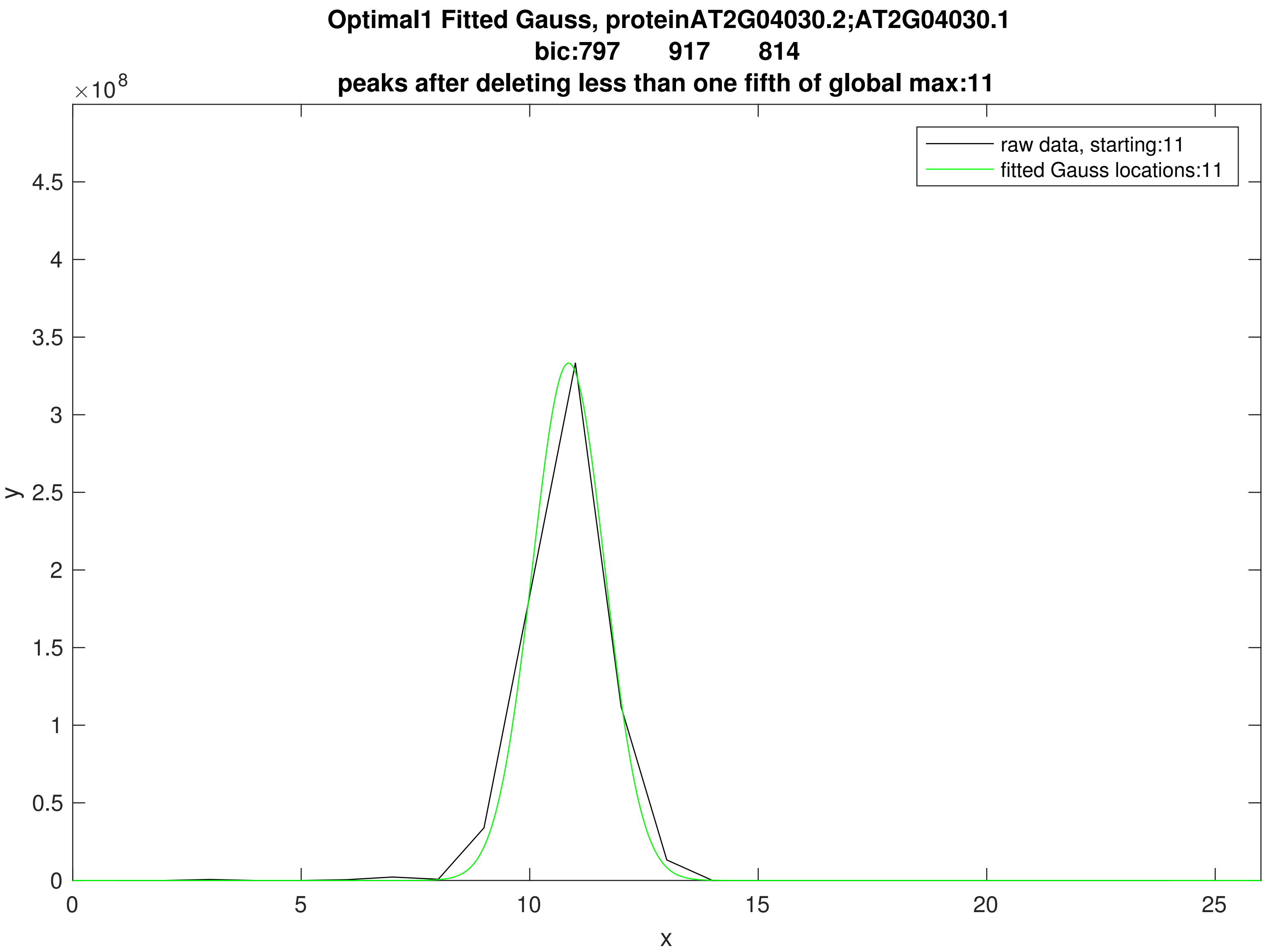
Optimal1 Fitted Gauss, proteinAT2G04030.2;AT2G04030.1
bic:797 917 814
peaks after deleting less than one fifth of global max:11
×10^8
raw data, starting:11
fitted Gauss locations:11
y
x
0
0.5
1
1.5
2
2.5
3
3.5
4
4.5
0
5
10
15
20
25

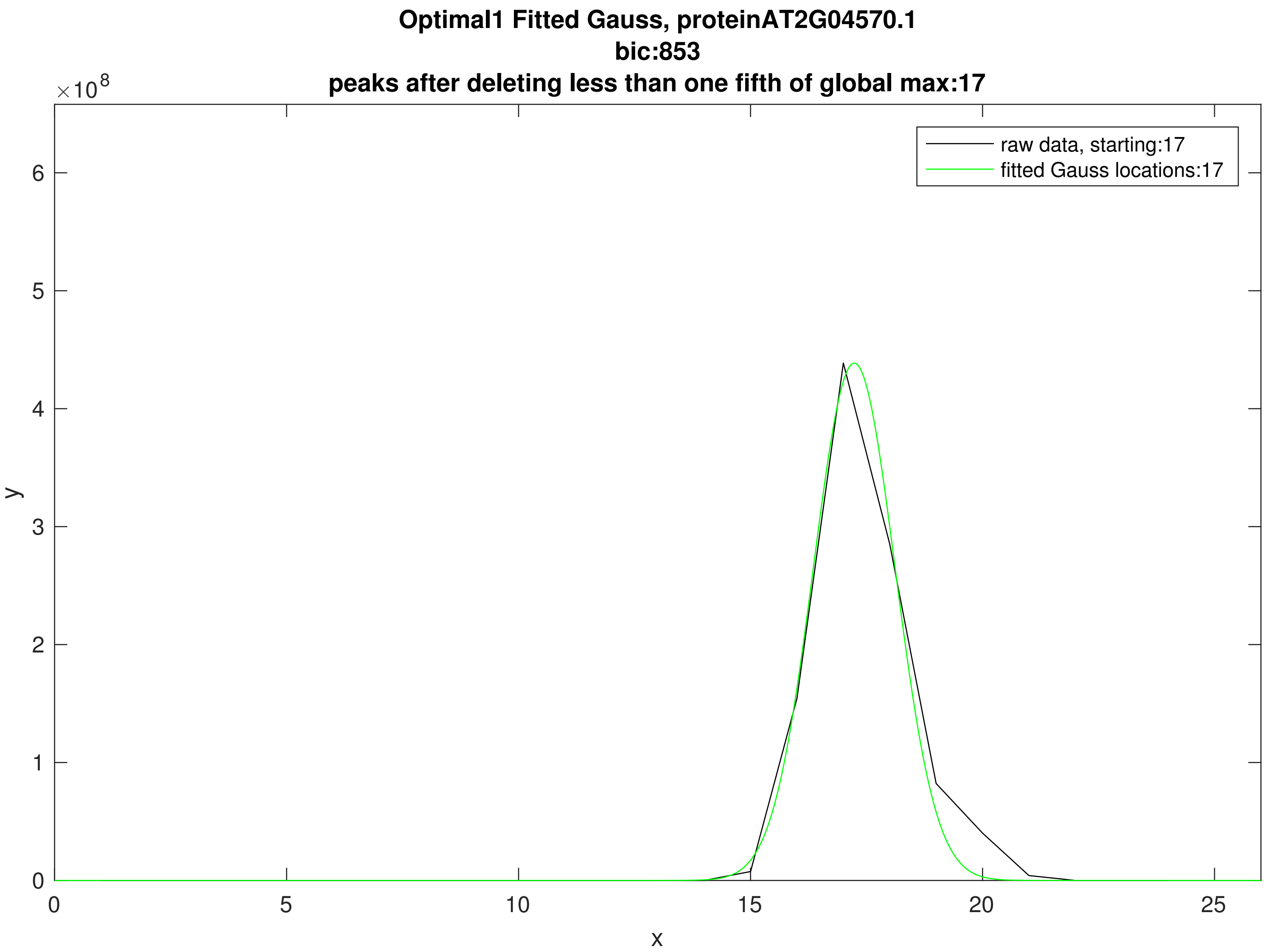
Optimal1 Fitted Gauss, proteinAT2G04570.1
bic:853
peaks after deleting less than one fifth of global max:17
×10^8
6
5
4
3
2
1
0
y
0
5
10
15
20
25
x
raw data, starting:17
fitted Gauss locations:17

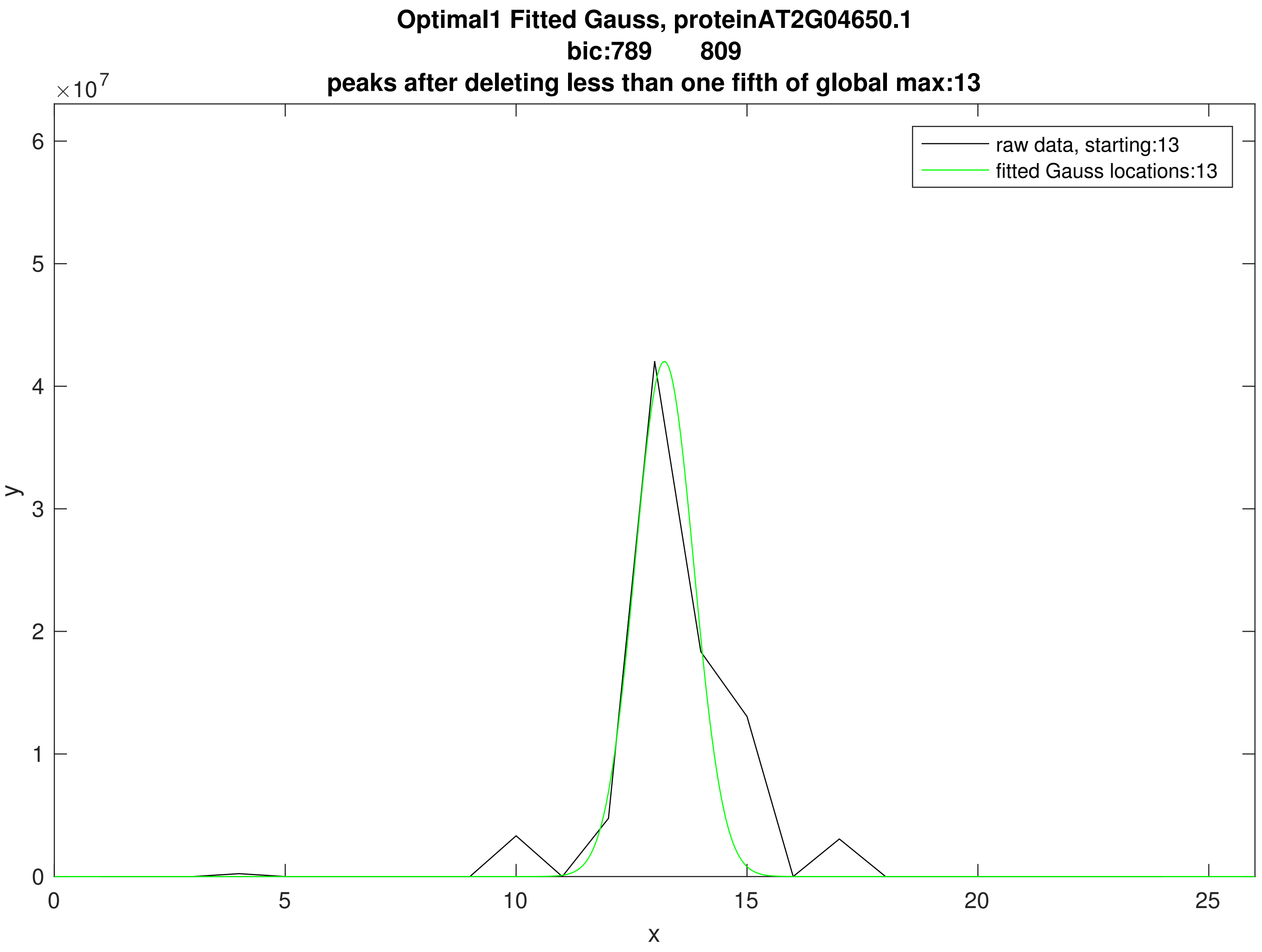
Optimal1 Fitted Gauss, proteinAT2G04650.1
bic:789 809
peaks after deleting less than one fifth of global max:13
×10^7
raw data, starting:13
fitted Gauss locations:13
y
x
0
1
2
3
4
5
6
0
5
10
15
20
25

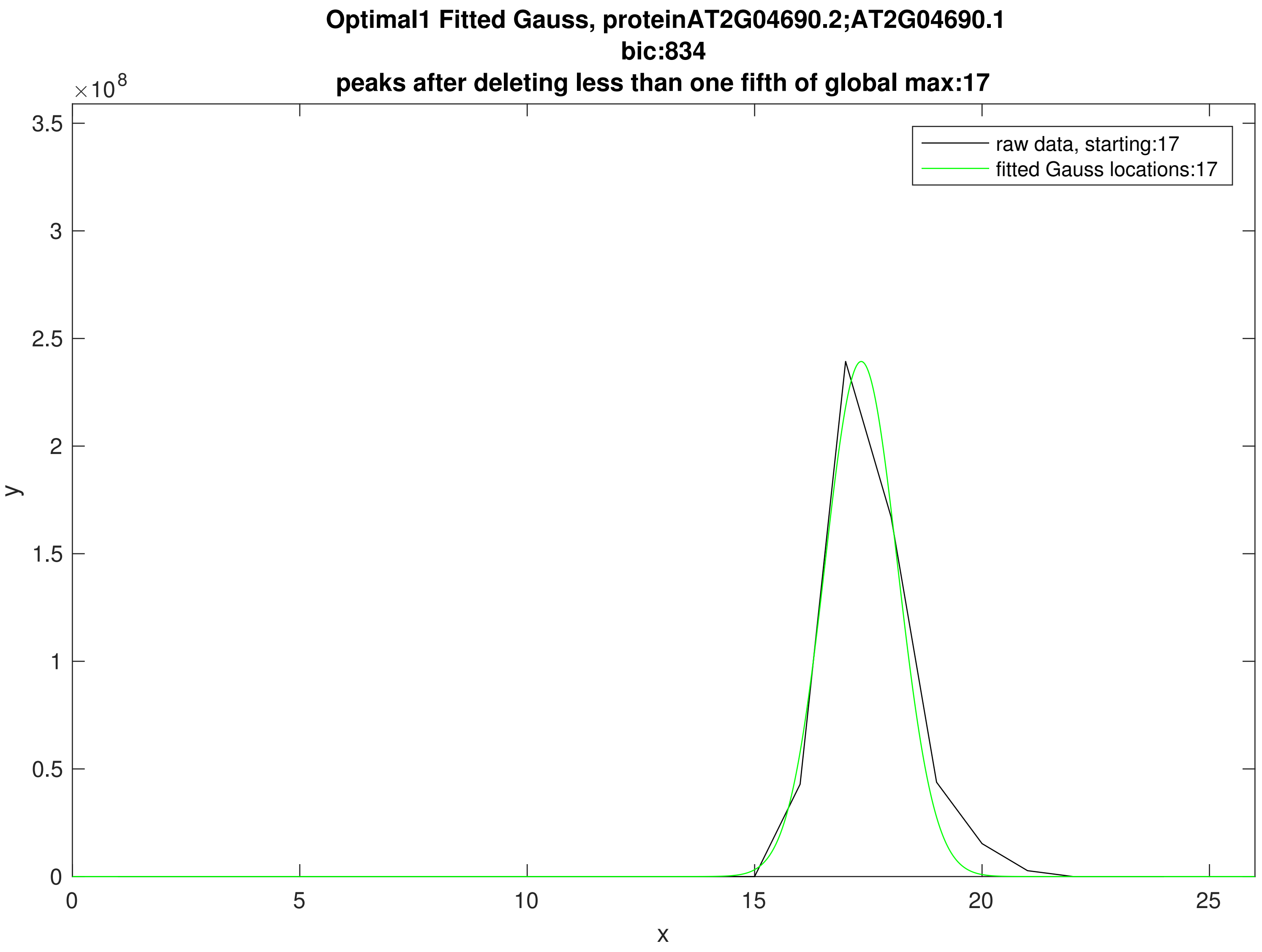
Optimal1 Fitted Gauss, proteinAT2G04690.2;AT2G04690.1
bic:834
peaks after deleting less than one fifth of global max:17
×10^8
3.5
3
2.5
2
1.5
1
0.5
0
y
0
5
10
15
20
25
x
raw data, starting:17
fitted Gauss locations:17

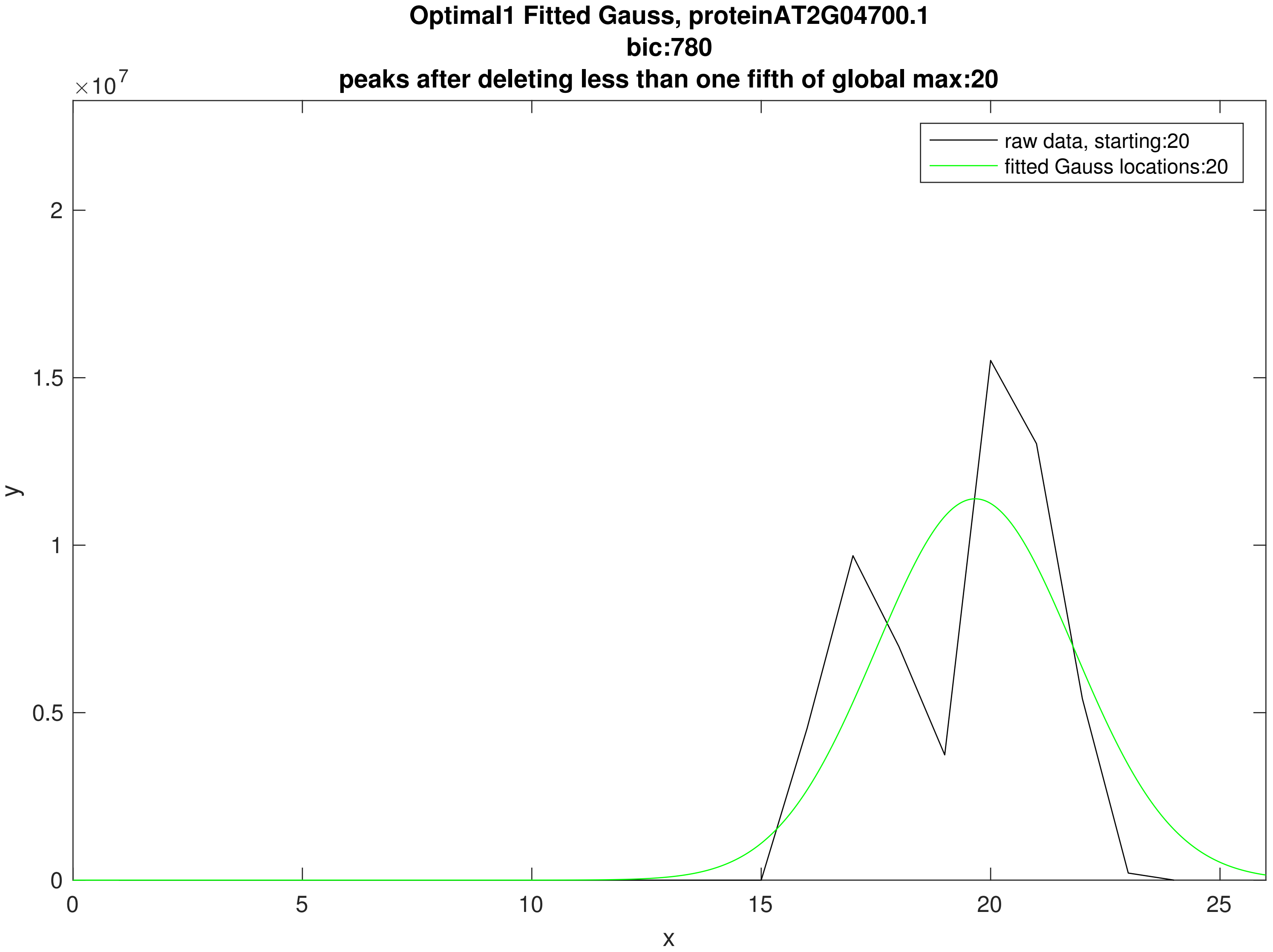
Optimal1 Fitted Gauss, proteinAT2G04700.1
bic:780
peaks after deleting less than one fifth of global max:20
×10^7
raw data, starting:20
fitted Gauss locations:20
y
x
0
0.5
1
1.5
2
0
5
10
15
20
25

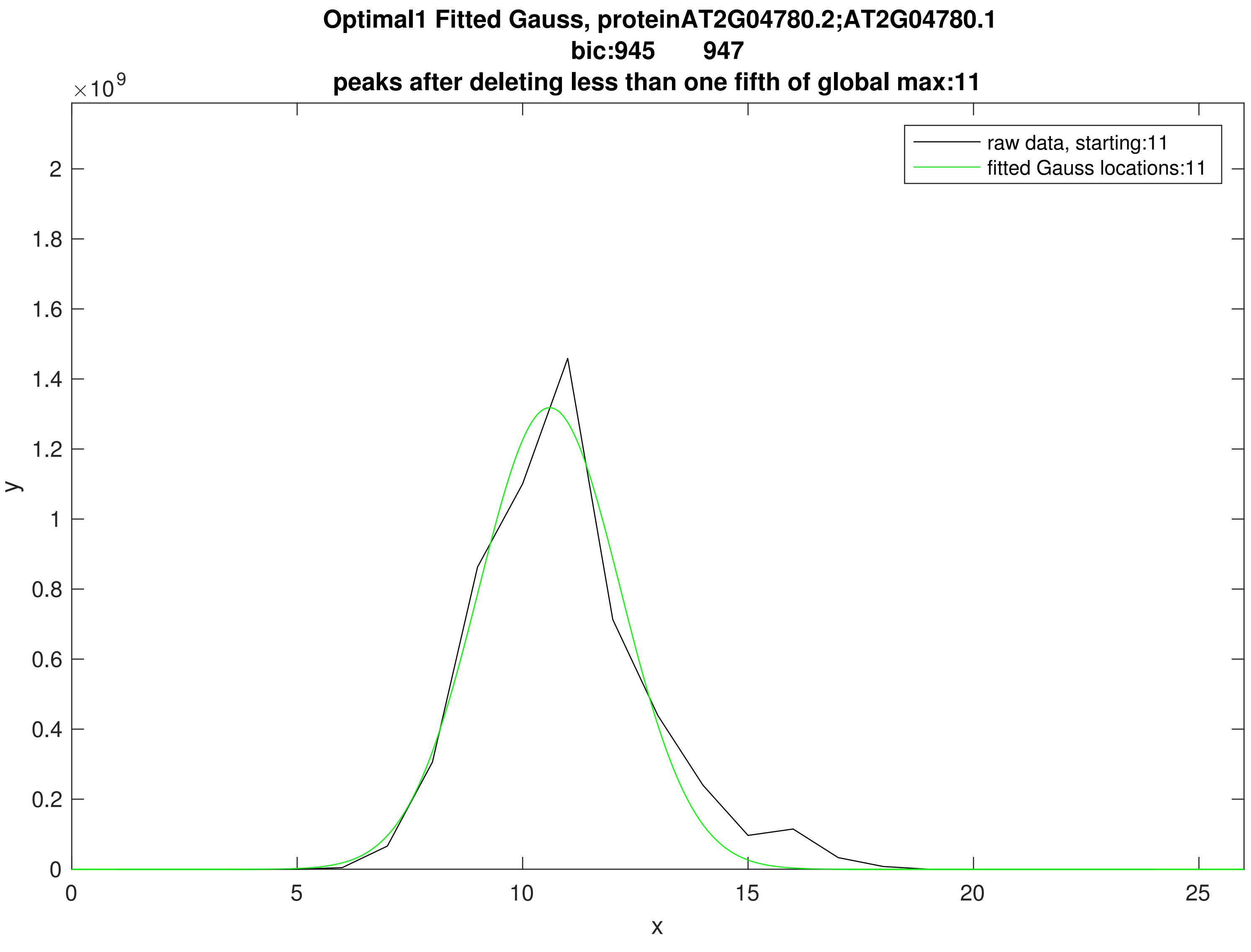

Optimal1 Fitted Gauss, proteinAT2G04780.2;AT2G04780.1
bic:945 947
peaks after deleting less than one fifth of global max:11
×10^9
raw data, starting:11
fitted Gauss locations:11
y
x
0
0.2
0.4
0.6
0.8
1
1.2
1.4
1.6
1.8
2
0
5
10
15
20
25

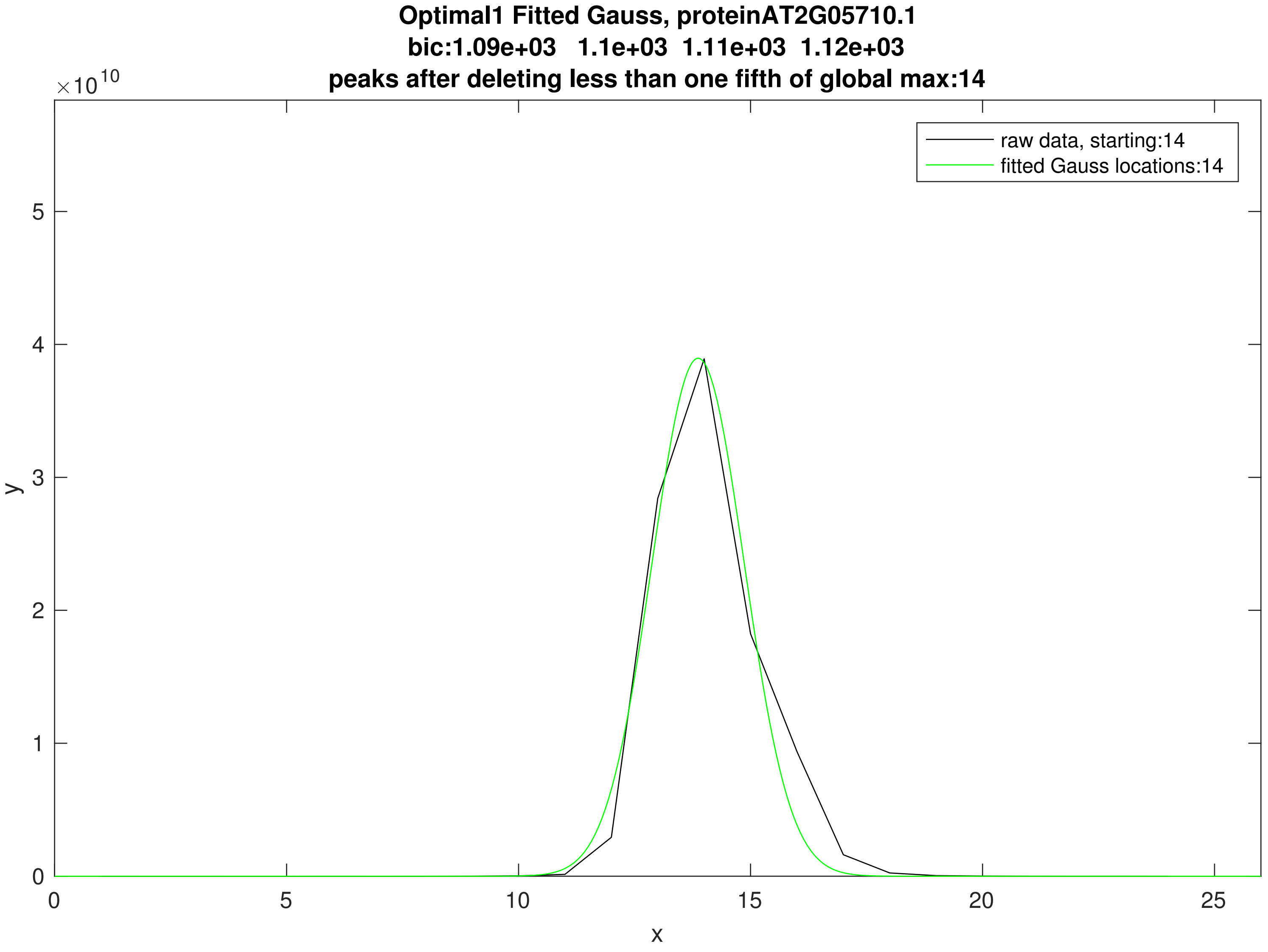
Optimal1 Fitted Gauss, proteinAT2G05710.1
bic:1.09e+03 1.1e+03 1.11e+03 1.12e+03
peaks after deleting less than one fifth of global max:14
×10^10
raw data, starting:14
fitted Gauss locations:14
y
x
0
1
2
3
4
5
5
10
15
20
25

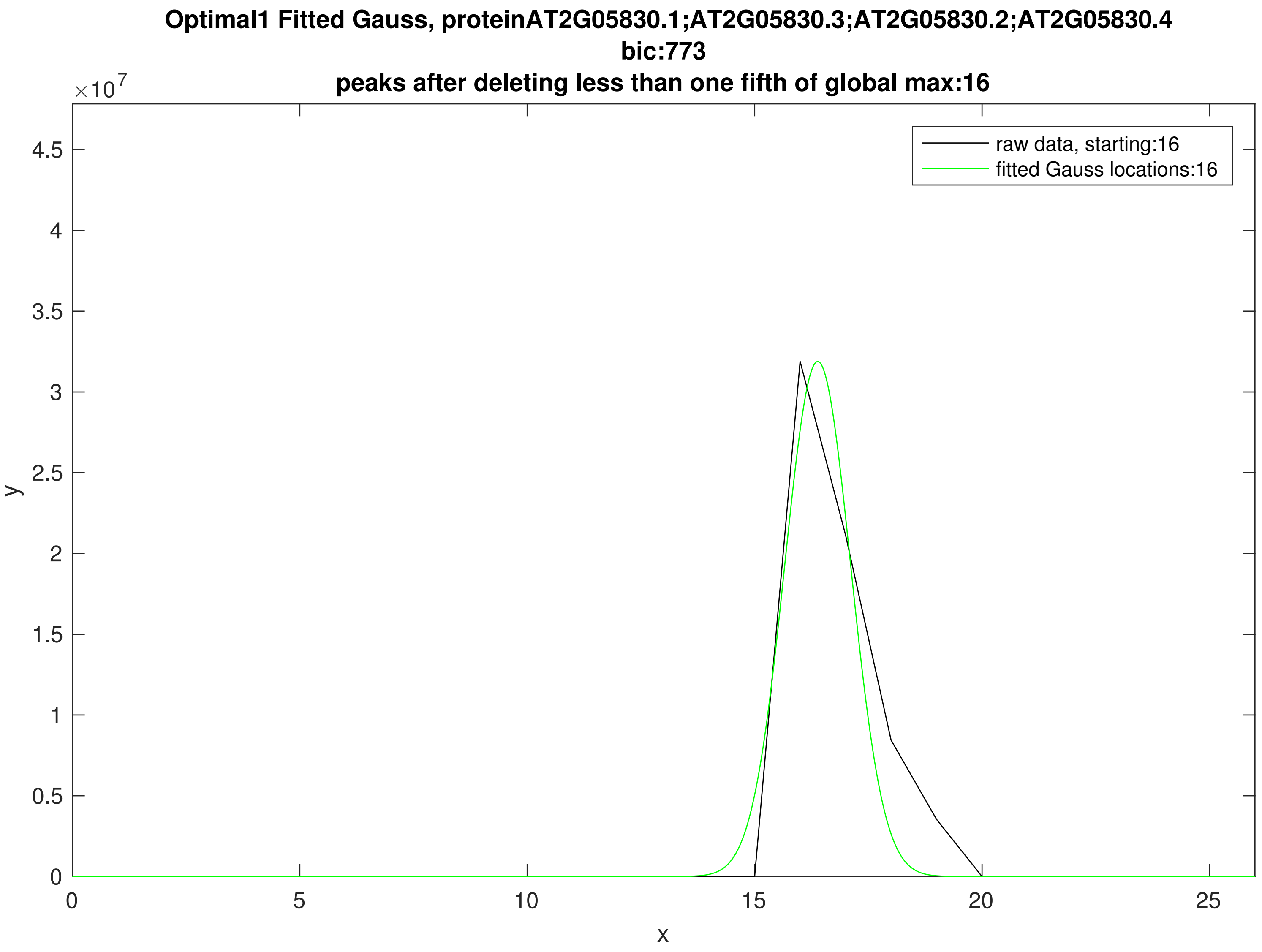
Optimal1 Fitted Gauss, proteinAT2G05830.1;AT2G05830.3;AT2G05830.2;AT2G05830.4
bic:773
peaks after deleting less than one fifth of global max:16
×10^7
raw data, starting:16
fitted Gauss locations:16
y
x
0
0.5
1
1.5
2
2.5
3
3.5
4
4.5
0
5
10
15
20
25

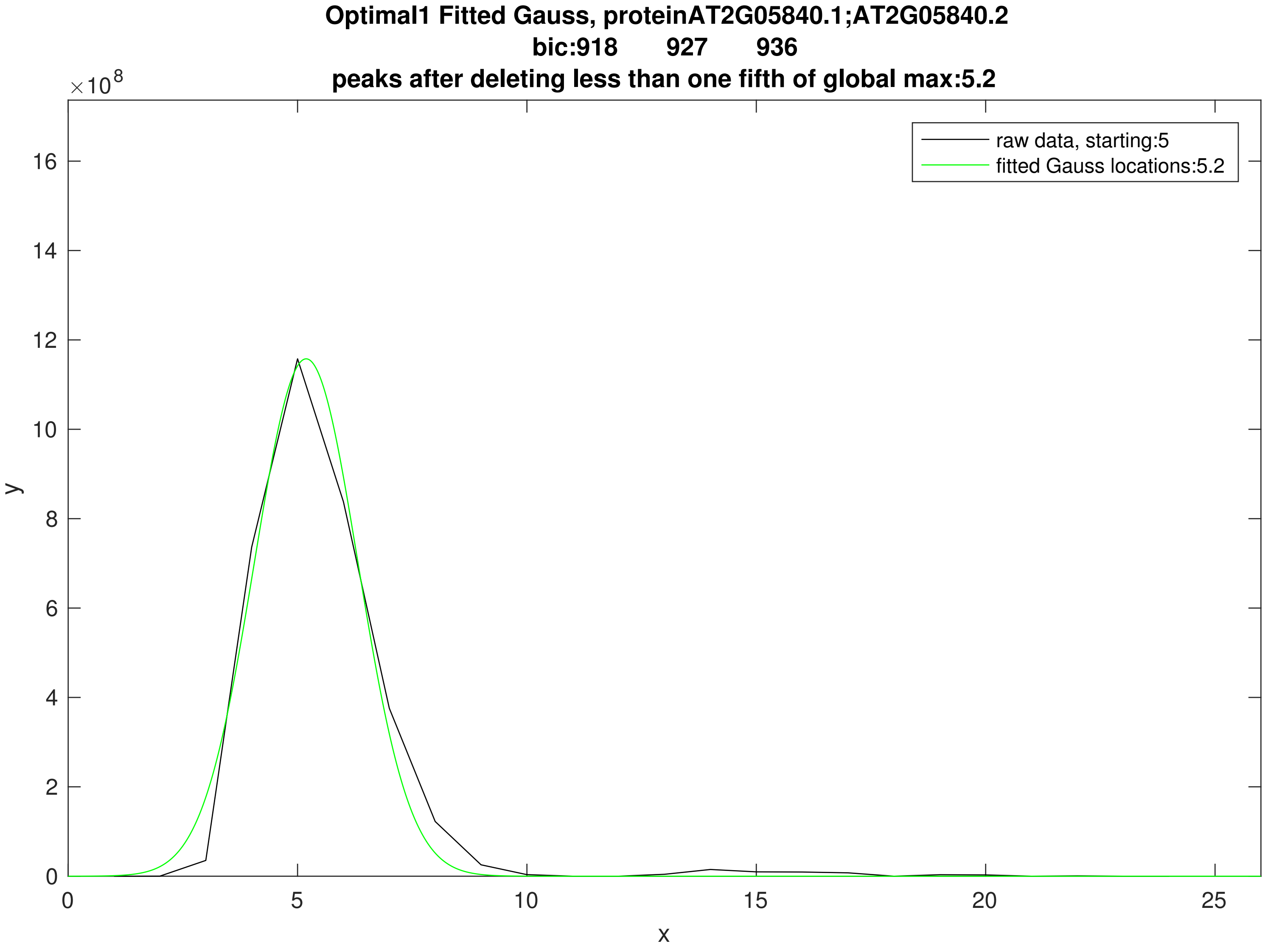
Optimal1 Fitted Gauss, proteinAT2G05840.1;AT2G05840.2
bic:918 927 936
peaks after deleting less than one fifth of global max:5.2
×10^8
raw data, starting:5
fitted Gauss locations:5.2
y
x
0
2
4
6
8
10
12
14
16
0
5
10
15
20
25

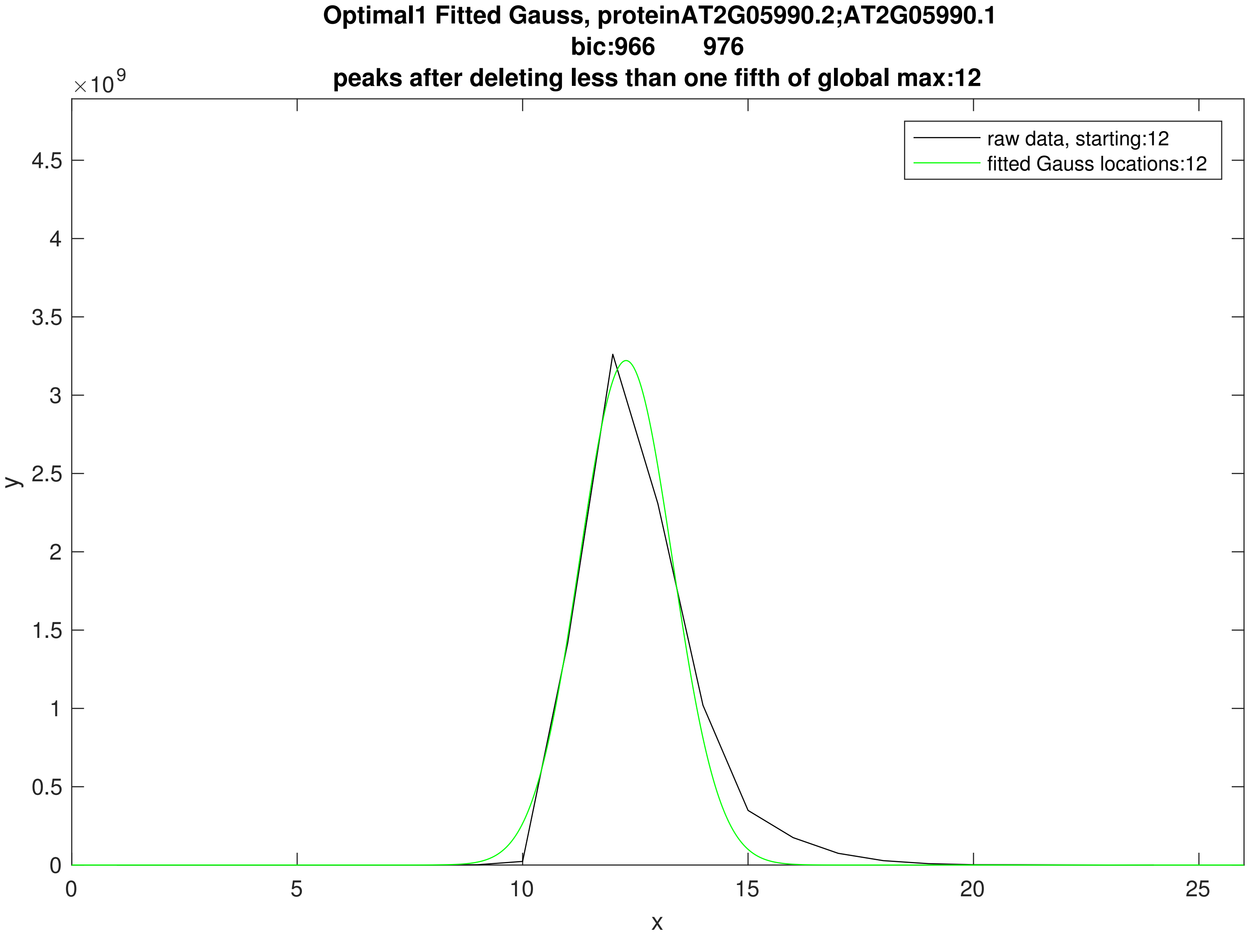

Optimal1 Fitted Gauss, proteinAT2G05990.2;AT2G05990.1
bic:966 976
peaks after deleting less than one fifth of global max:12
×10^9
raw data, starting:12
fitted Gauss locations:12
y
x
0
0.5
1
1.5
2
2.5
3
3.5
4
4.5
5
10
15
20
25

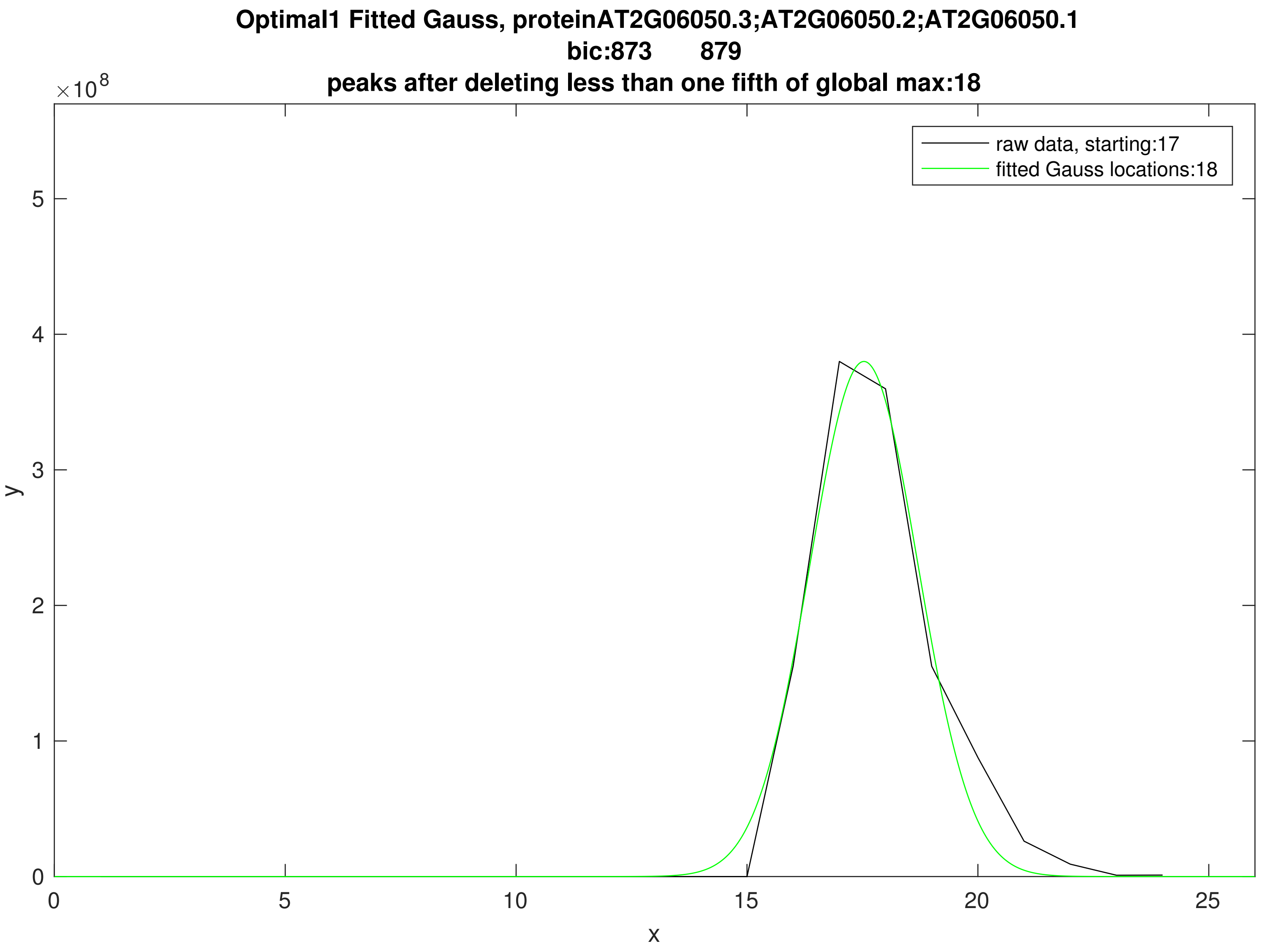
Optimal1 Fitted Gauss, proteinAT2G06050.3;AT2G06050.2;AT2G06050.1
bic:873 879
peaks after deleting less than one fifth of global max:18
×10^8
raw data, starting:17
fitted Gauss locations:18
y
x
0
1
2
3
4
5
0
5
10
15
20
25

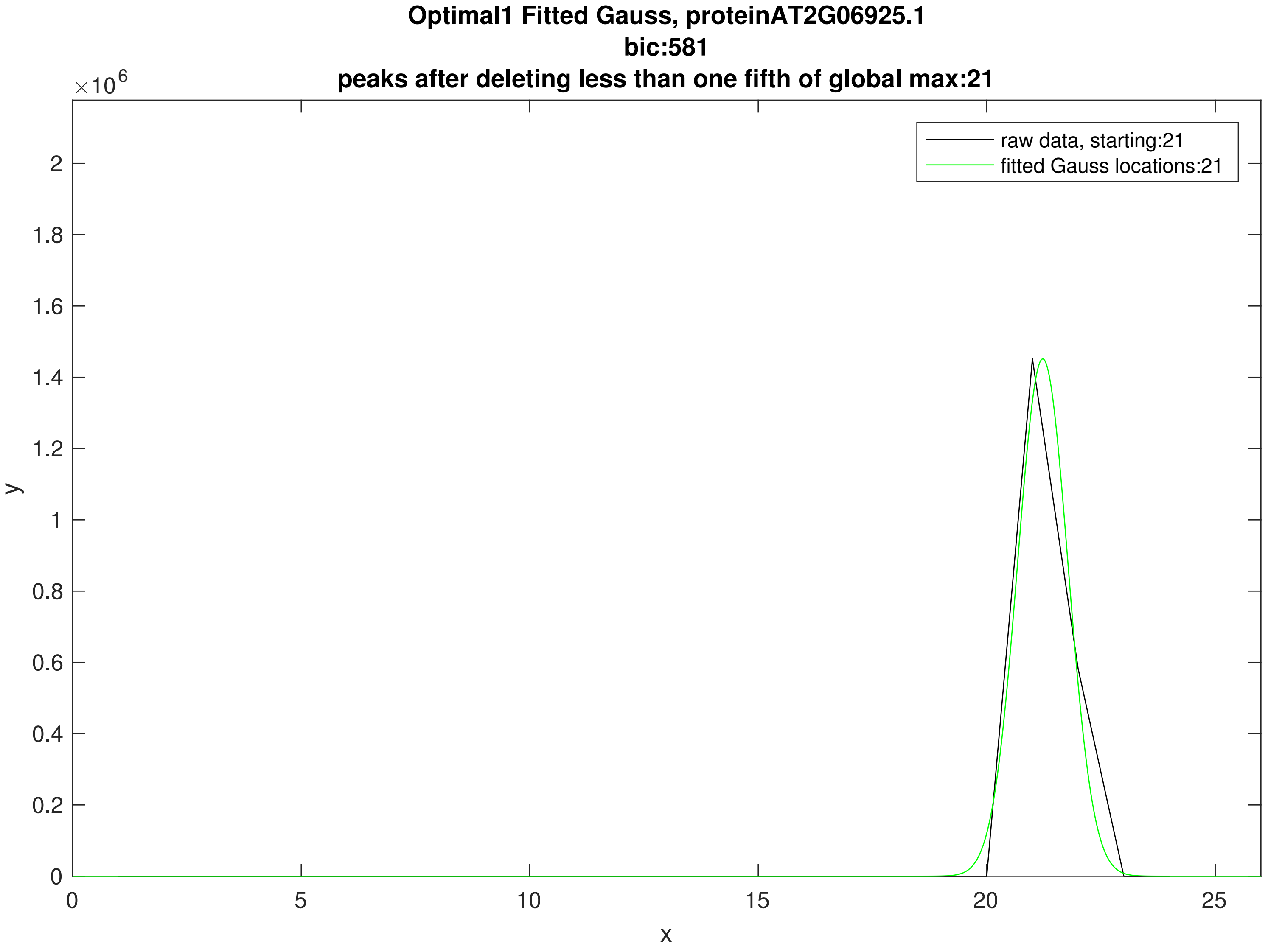
Optimal1 Fitted Gauss, proteinAT2G06925.1
bic:581
peaks after deleting less than one fifth of global max:21
×10^6
2
1.8
1.6
1.4
1.2
1
0.8
0.6
0.4
0.2
0
y
0
5
10
15
20
25
x
raw data, starting:21
fitted Gauss locations:21

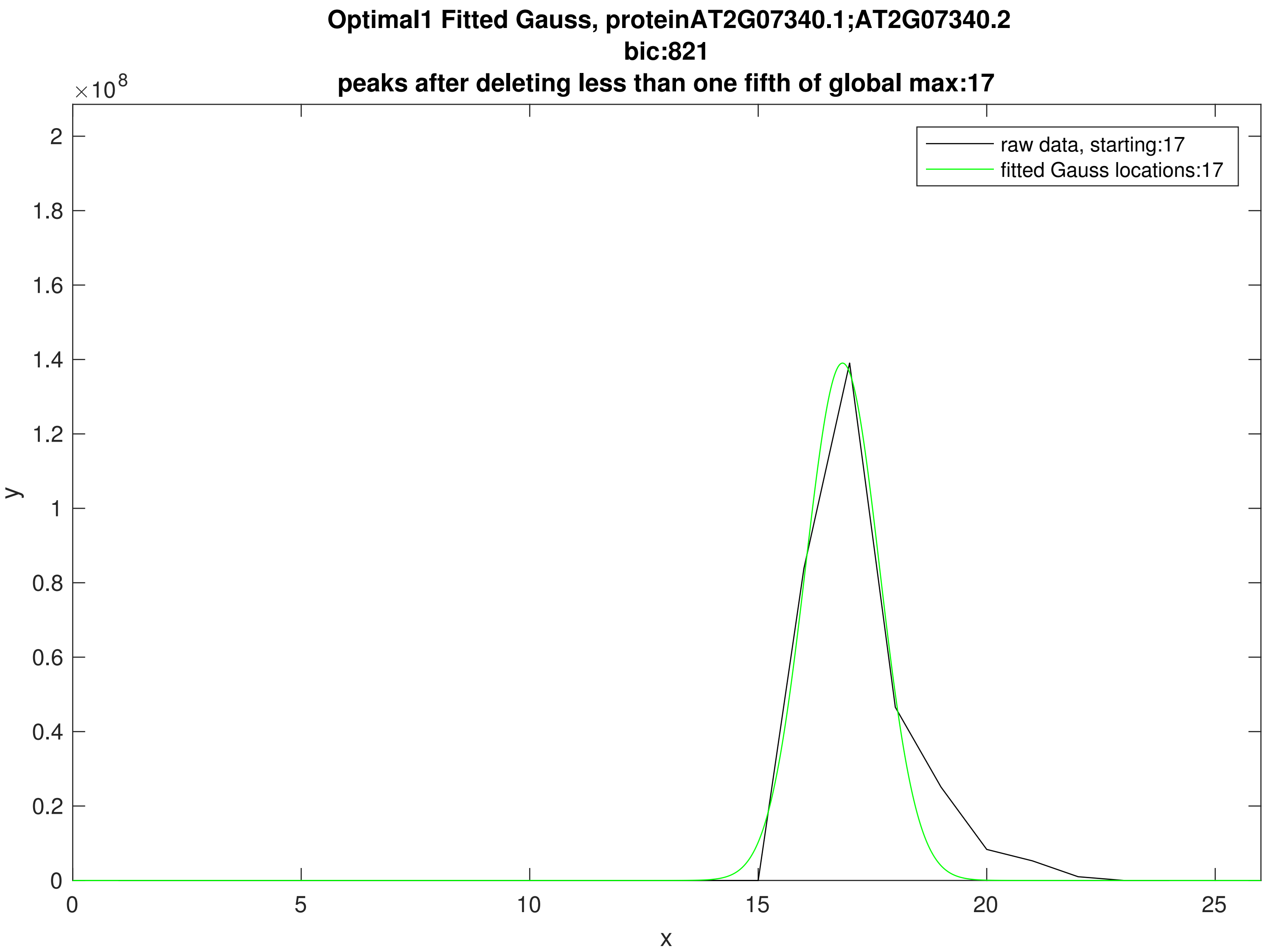
Optimal1 Fitted Gauss, proteinAT2G07340.1;AT2G07340.2
bic:821
peaks after deleting less than one fifth of global max:17
×10^8
raw data, starting:17
fitted Gauss locations:17
y
x
0
0.2
0.4
0.6
0.8
1
1.2
1.4
1.6
1.8
2
0
5
10
15
20
25

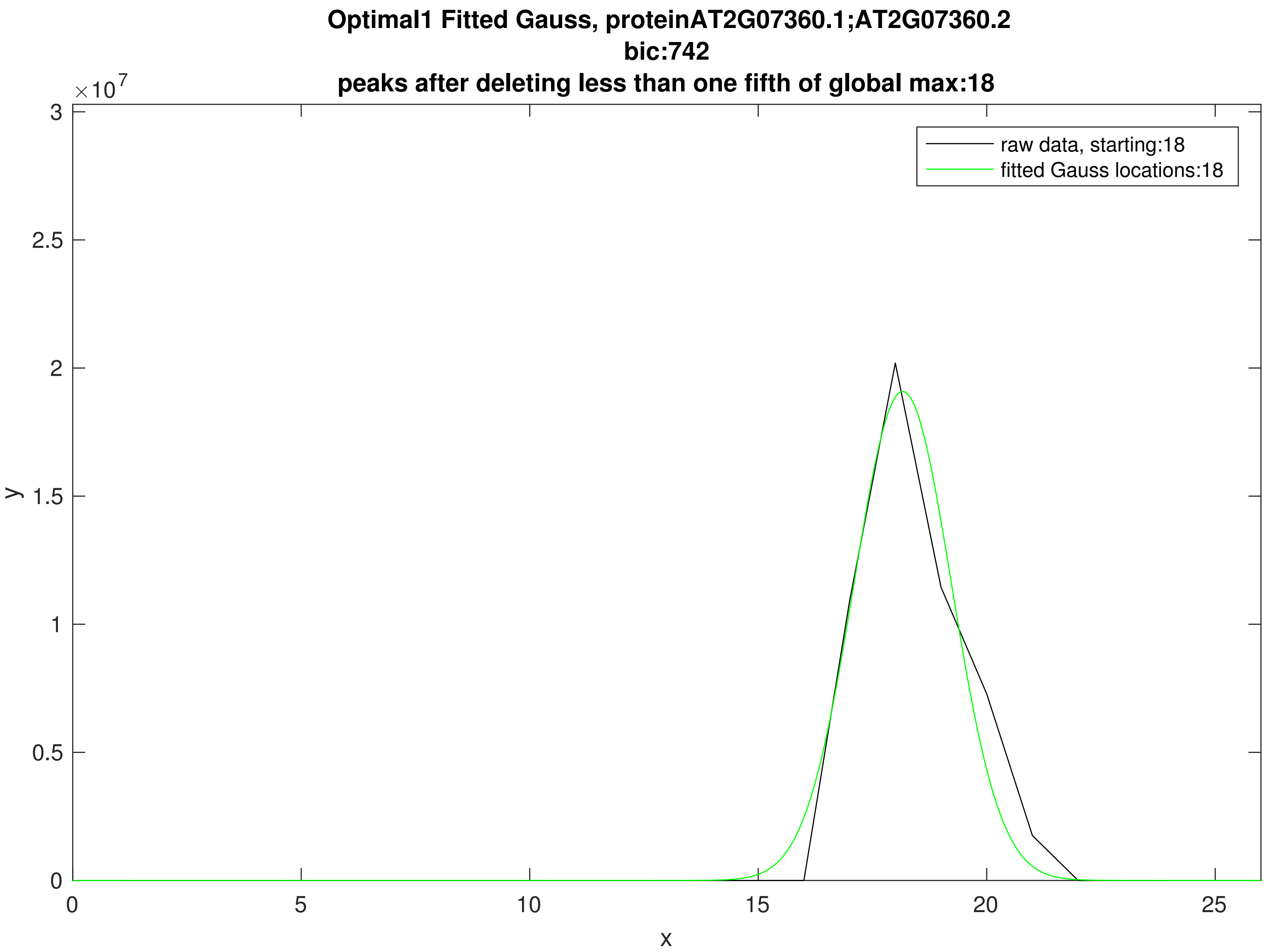
Optimal1 Fitted Gauss, proteinAT2G07360.1;AT2G07360.2
bic:742
peaks after deleting less than one fifth of global max:18
×10^7
3
2.5
2
1.5
1
0.5
0
y
0
5
10
15
20
25
x
raw data, starting:18
fitted Gauss locations:18

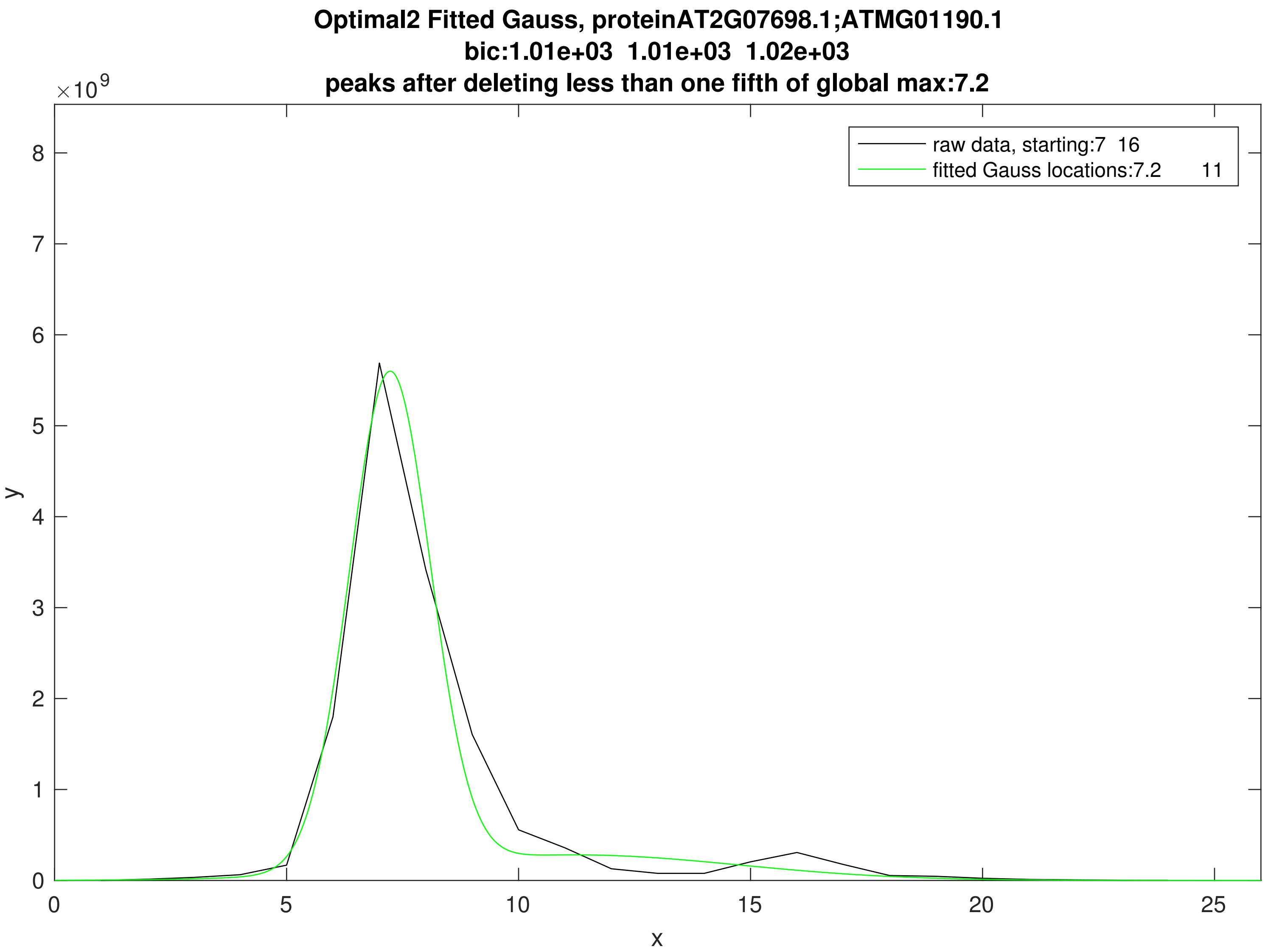
Optimal2 Fitted Gauss, proteinAT2G07698.1;ATMG01190.1
bic:1.01e+03  1.01e+03  1.02e+03
peaks after deleting less than one fifth of global max:7.2
raw data, starting:7  16
fitted Gauss locations:7.2       11
×10^9
y
x
0
1
2
3
4
5
6
7
8
0
5
10
15
20
25

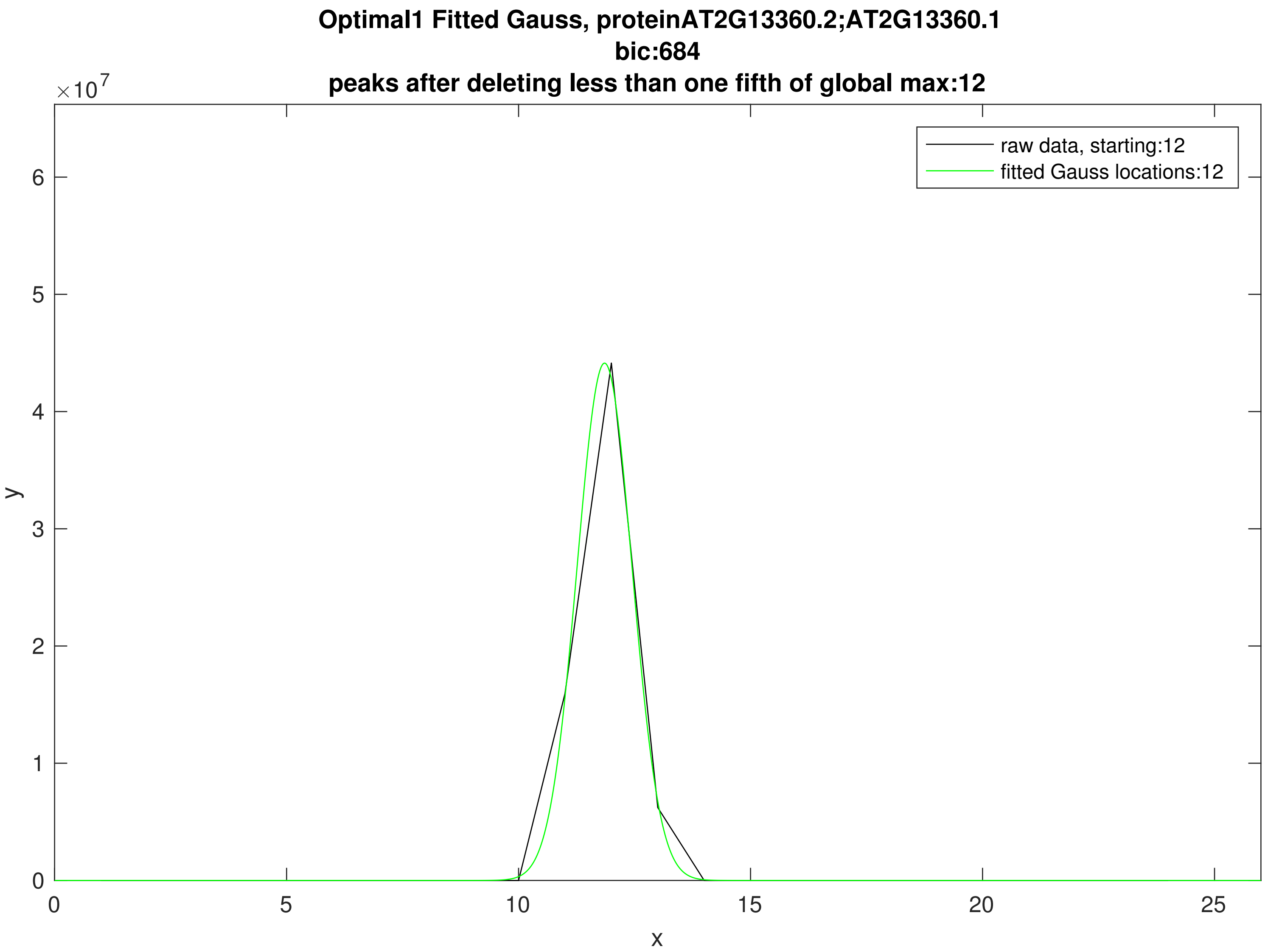
Optimal1 Fitted Gauss, proteinAT2G13360.2;AT2G13360.1
bic:684
peaks after deleting less than one fifth of global max:12
×10^7
raw data, starting:12
fitted Gauss locations:12
y
x
0
1
2
3
4
5
6
0
5
10
15
20
25

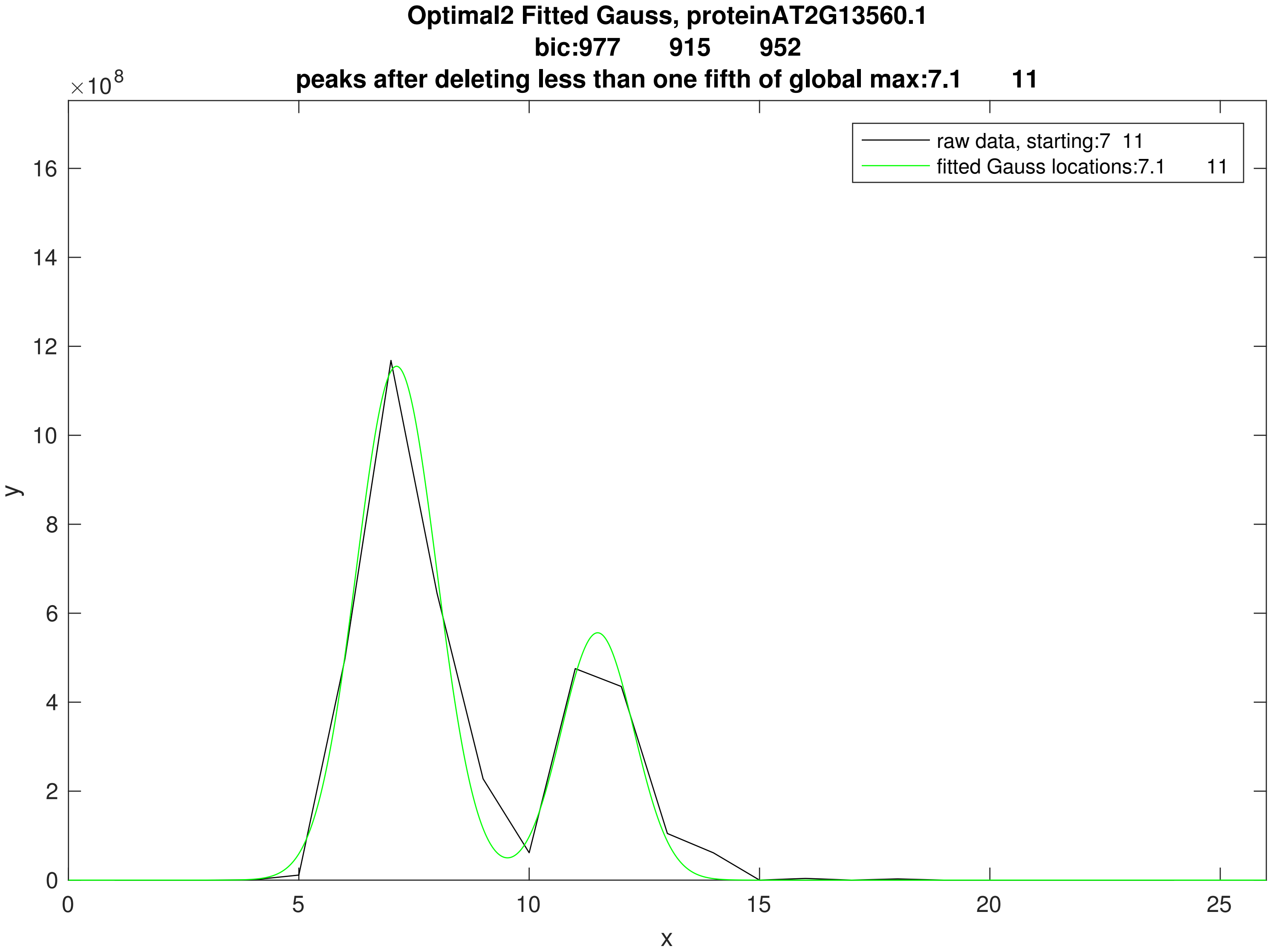
Optimal2 Fitted Gauss, proteinAT2G13560.1
bic:977 915 952
peaks after deleting less than one fifth of global max:7.1 11
×10^8
raw data, starting:7 11
fitted Gauss locations:7.1 11
y
x
0
2
4
6
8
10
12
14
16
0
5
10
15
20
25

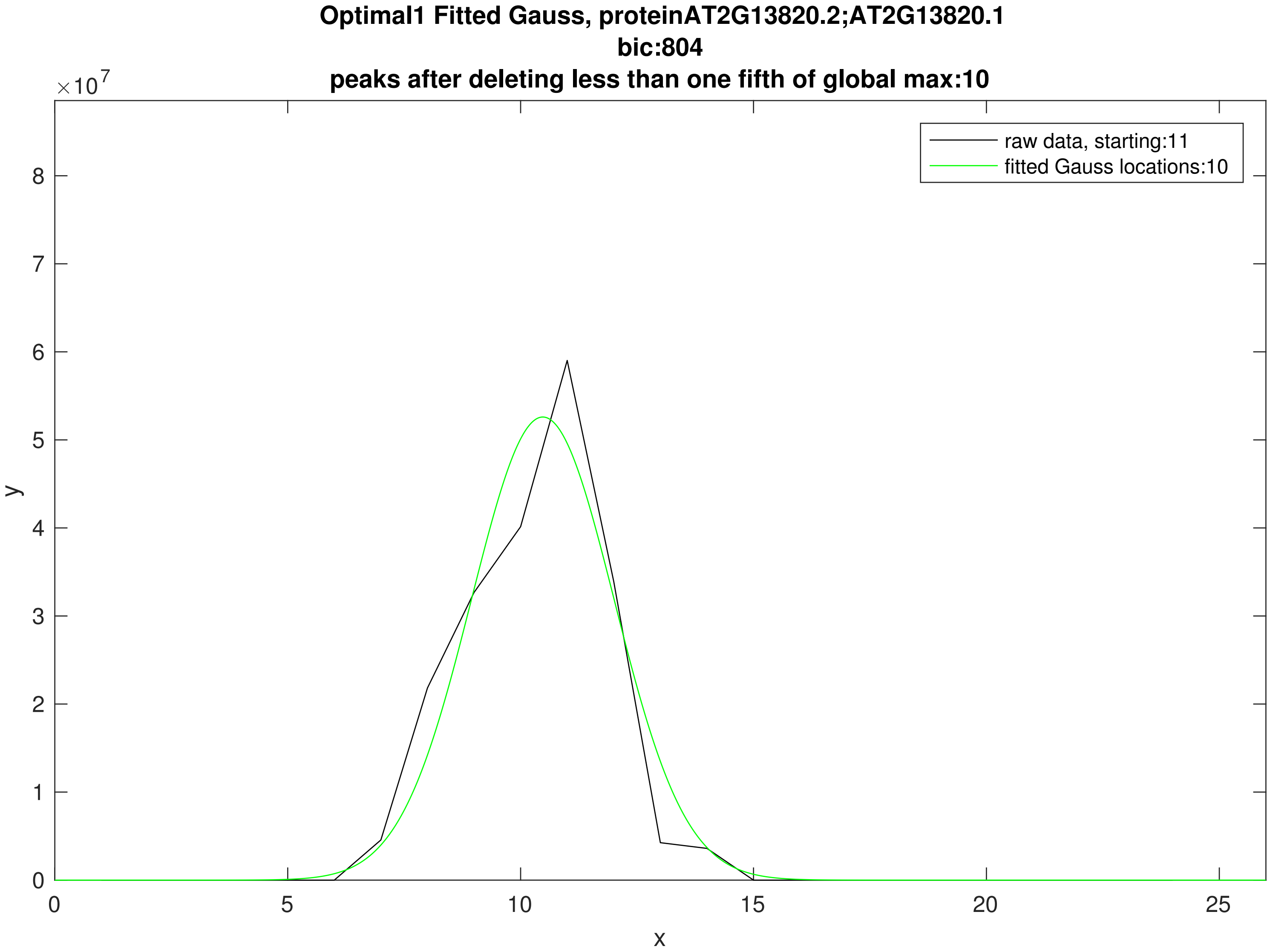
Optimal1 Fitted Gauss, proteinAT2G13820.2;AT2G13820.1
bic:804
peaks after deleting less than one fifth of global max:10
×10^7
8
7
6
5
4
3
2
1
0
y
0
5
10
15
20
25
x
raw data, starting:11
fitted Gauss locations:10

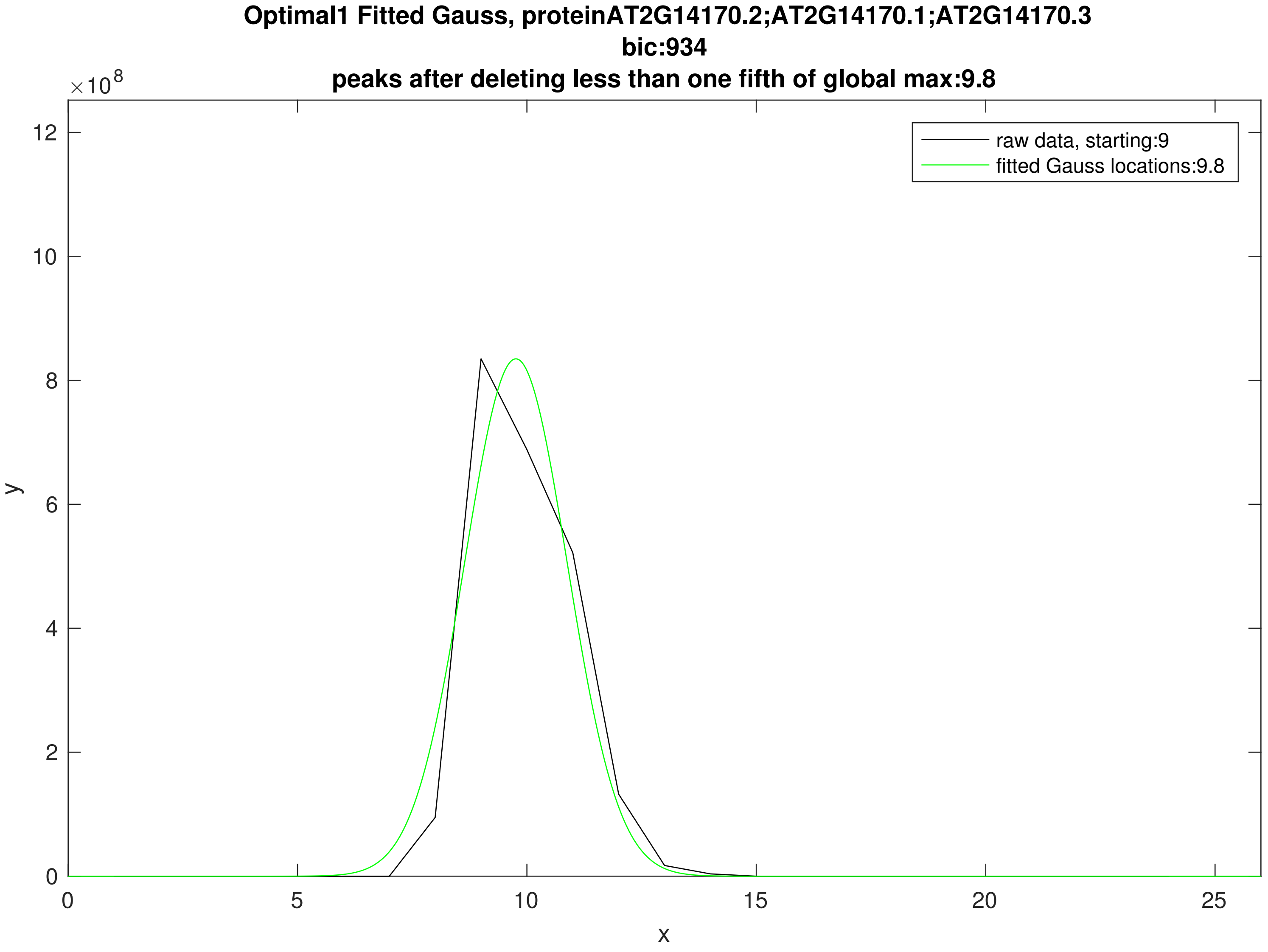
Optimal1 Fitted Gauss, proteinAT2G14170.2;AT2G14170.1;AT2G14170.3
bic:934
peaks after deleting less than one fifth of global max:9.8
×10^8
raw data, starting:9
fitted Gauss locations:9.8
y
x
0
2
4
6
8
10
12
0
5
10
15
20
25

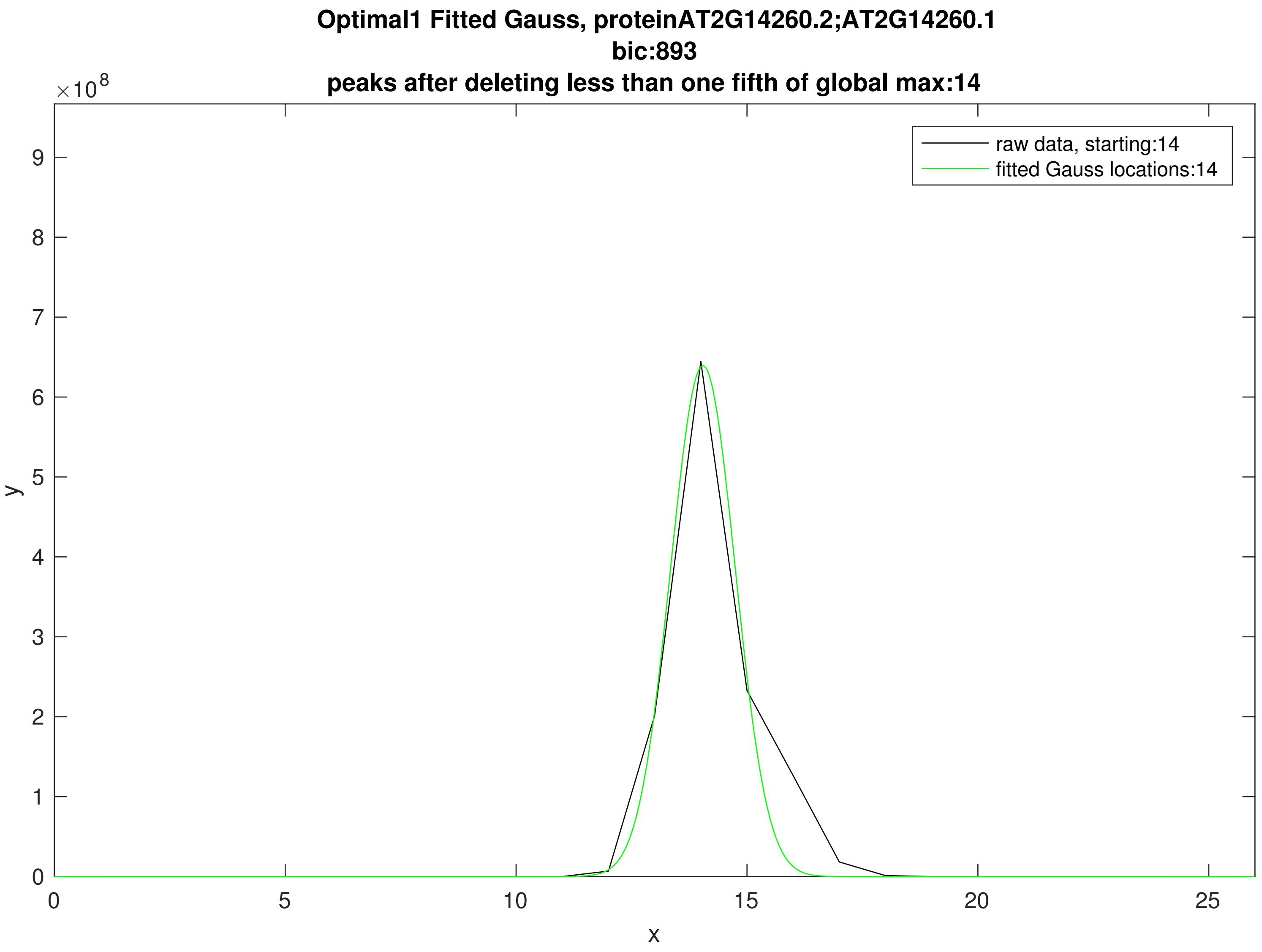

Optimal1 Fitted Gauss, proteinAT2G14260.2;AT2G14260.1
bic:893
peaks after deleting less than one fifth of global max:14
×10^8
raw data, starting:14
fitted Gauss locations:14
y
x
0
1
2
3
4
5
6
7
8
9
0
5
10
15
20
25

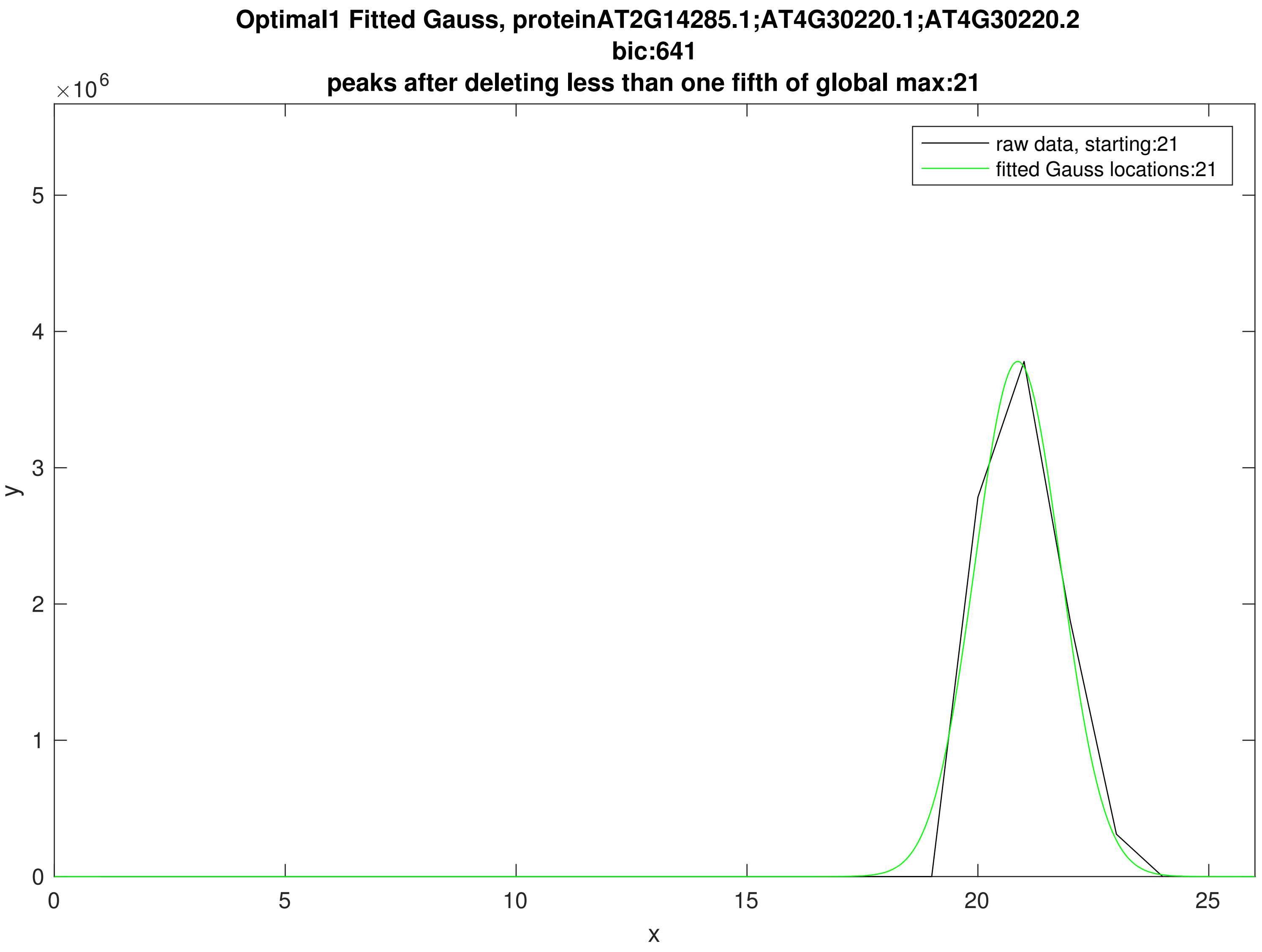
Optimal1 Fitted Gauss, proteinAT2G14285.1;AT4G30220.1;AT4G30220.2
bic:641
peaks after deleting less than one fifth of global max:21
×10^6
5
4
3
2
1
0
y
0
5
10
15
20
25
x
raw data, starting:21
fitted Gauss locations:21

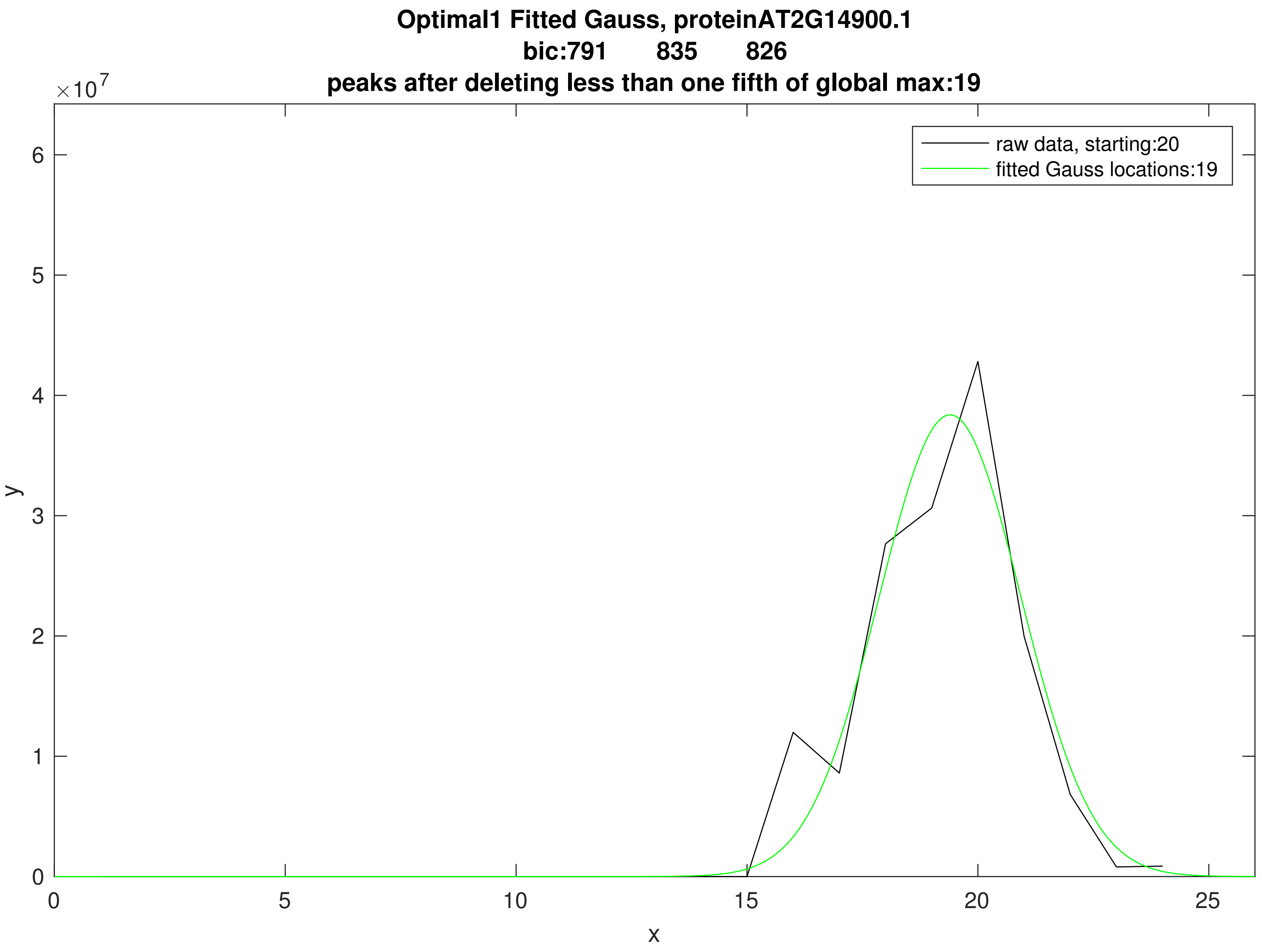

Optimal1 Fitted Gauss, proteinAT2G14900.1
bic:791 835 826
peaks after deleting less than one fifth of global max:19
×10^7
raw data, starting:20
fitted Gauss locations:19
y
x
0
1
2
3
4
5
6
0
5
10
15
20
25

**Optimal1 Fitted Gauss, proteinAT2G15130.2;AT2G15220.1;AT2G15130.1**
**bic:652**
**peaks after deleting less than one fifth of global max:21**

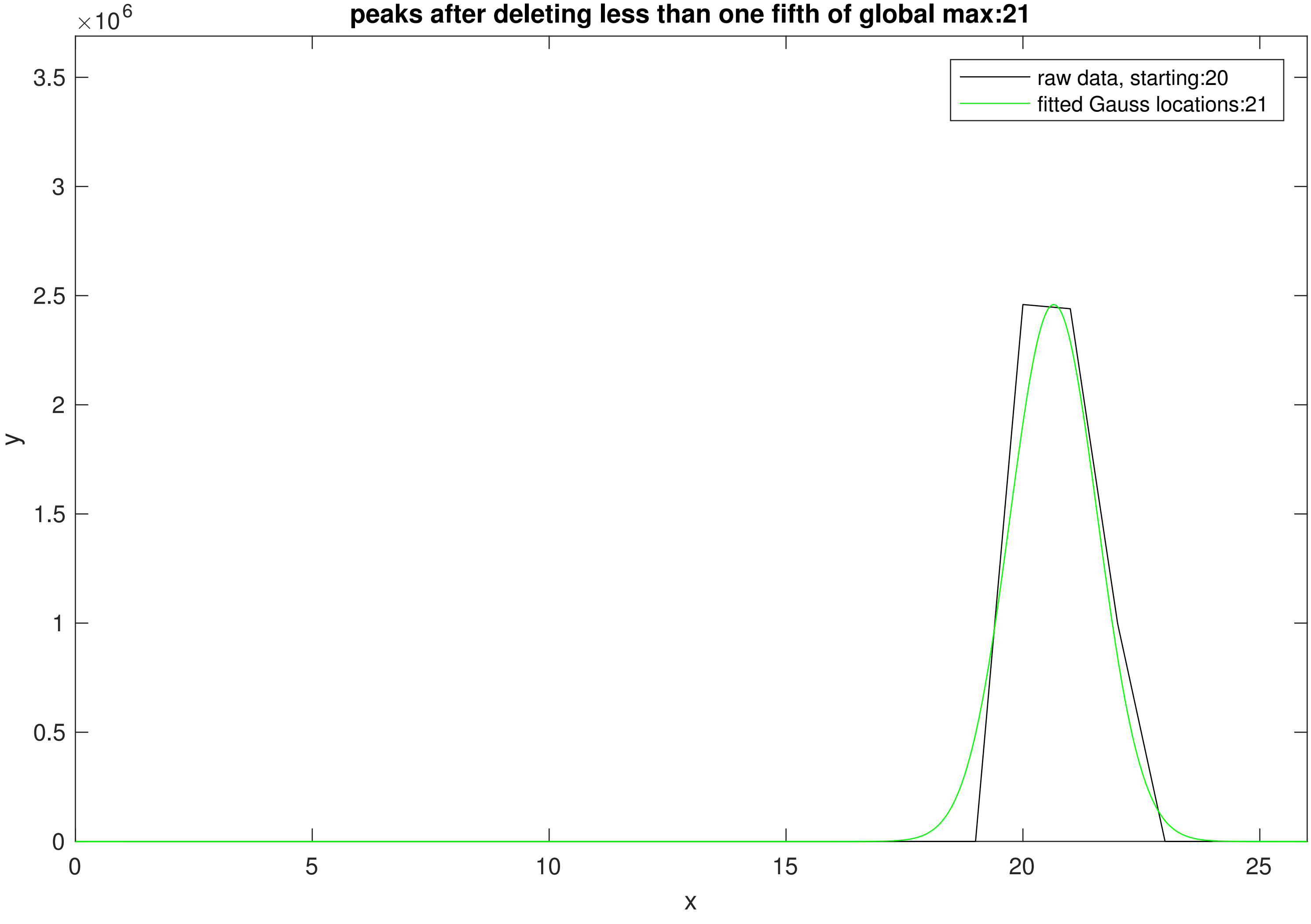

Optimal2 Fitted Gauss, proteinAT2G15430.1
bic:705 637
peaks after deleting less than one fifth of global max:4.4 21

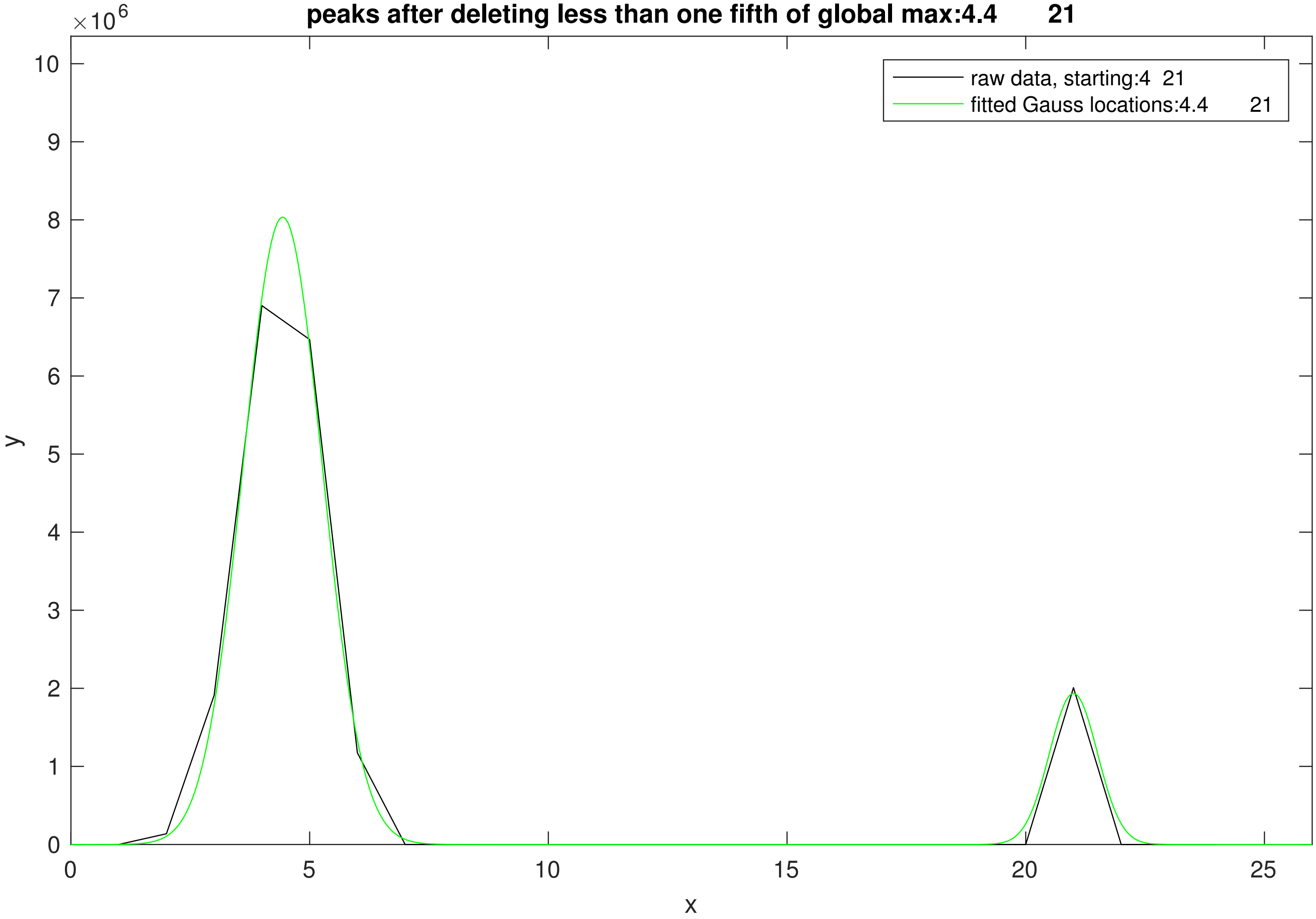

**Optimal1 Fitted Gauss, proteinAT2G15490.1;AT2G15490.3;AT2G15490.2**
**bic:756**
**peaks after deleting less than one fifth of global max:15**

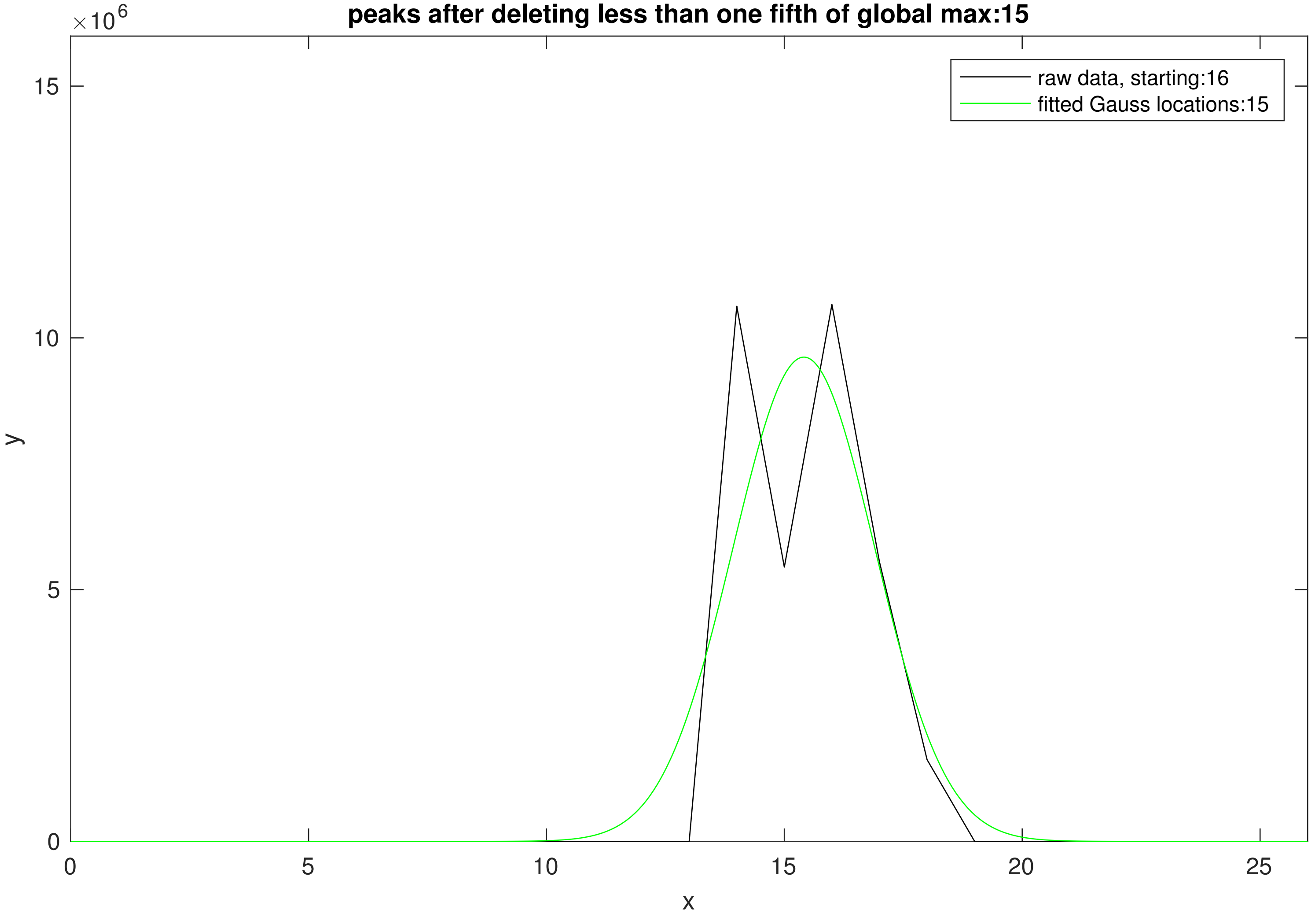

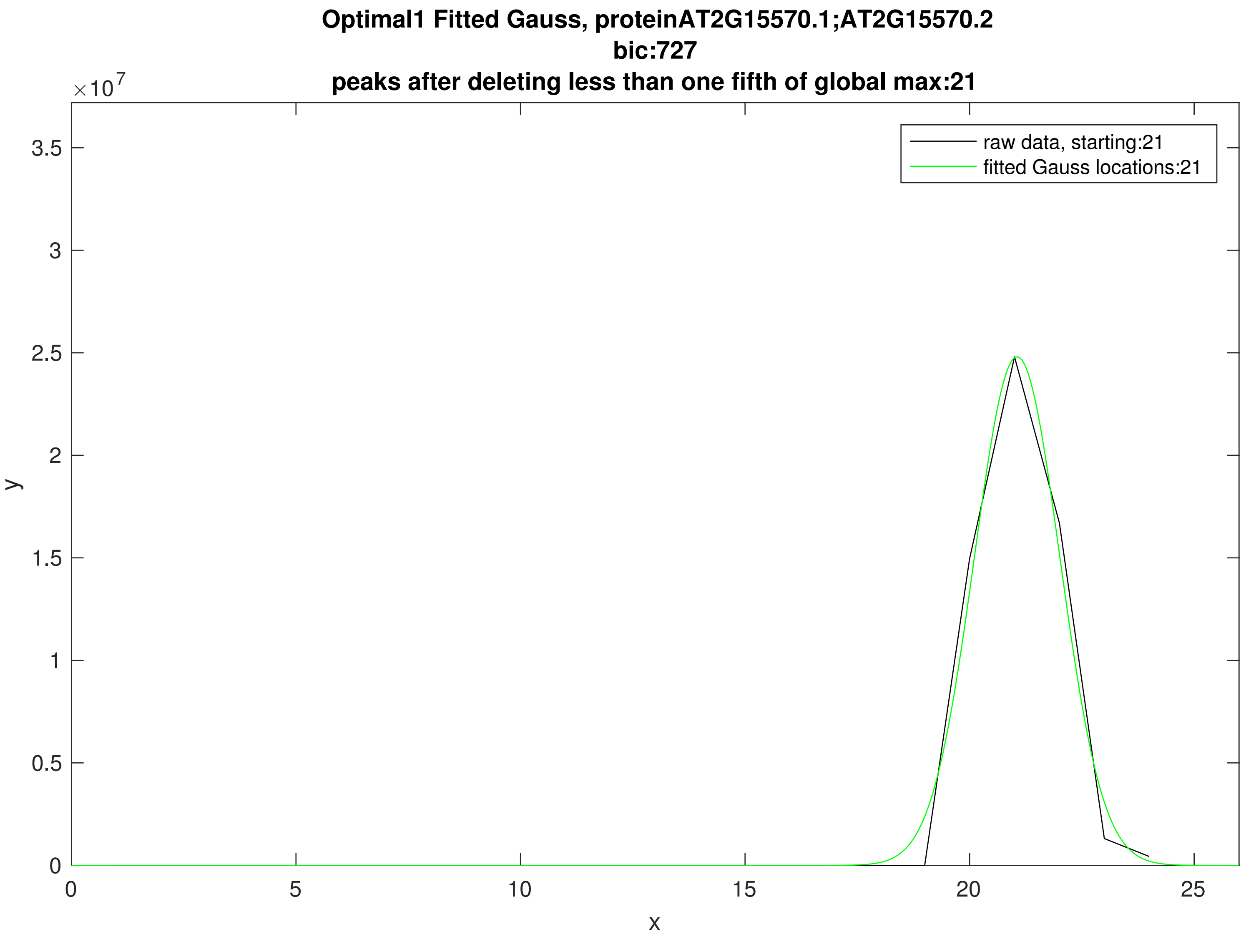

Optimal1 Fitted Gauss, proteinAT2G15570.1;AT2G15570.2
bic:727
peaks after deleting less than one fifth of global max:21
×10^7
3.5
3
2.5
2
1.5
1
0.5
0
y
0
5
10
15
20
25
x
raw data, starting:21
fitted Gauss locations:21

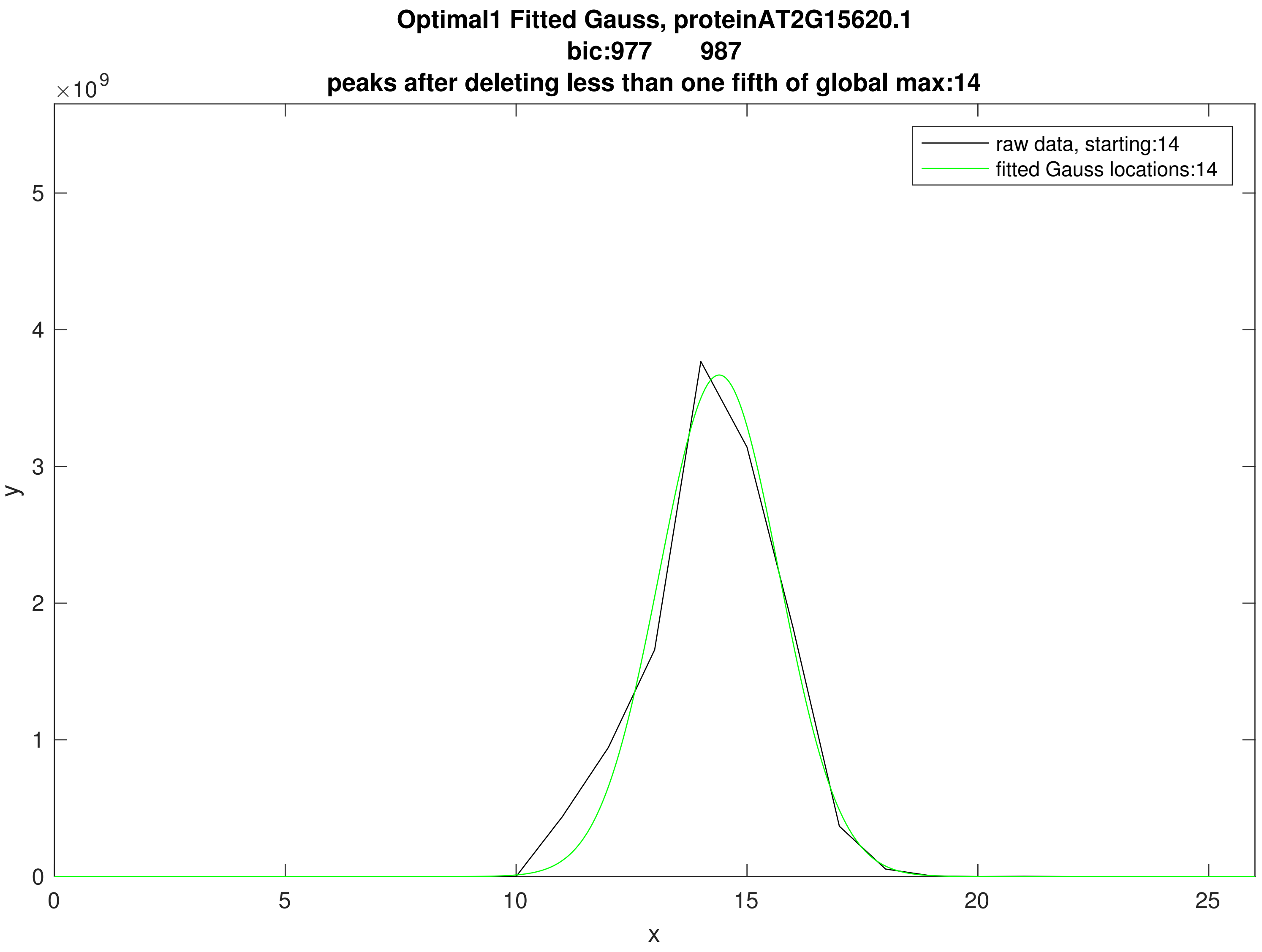
Optimal1 Fitted Gauss, proteinAT2G15620.1
bic:977 987
peaks after deleting less than one fifth of global max:14
×10^9
raw data, starting:14
fitted Gauss locations:14
y
x
0
1
2
3
4
5
0
5
10
15
20
25

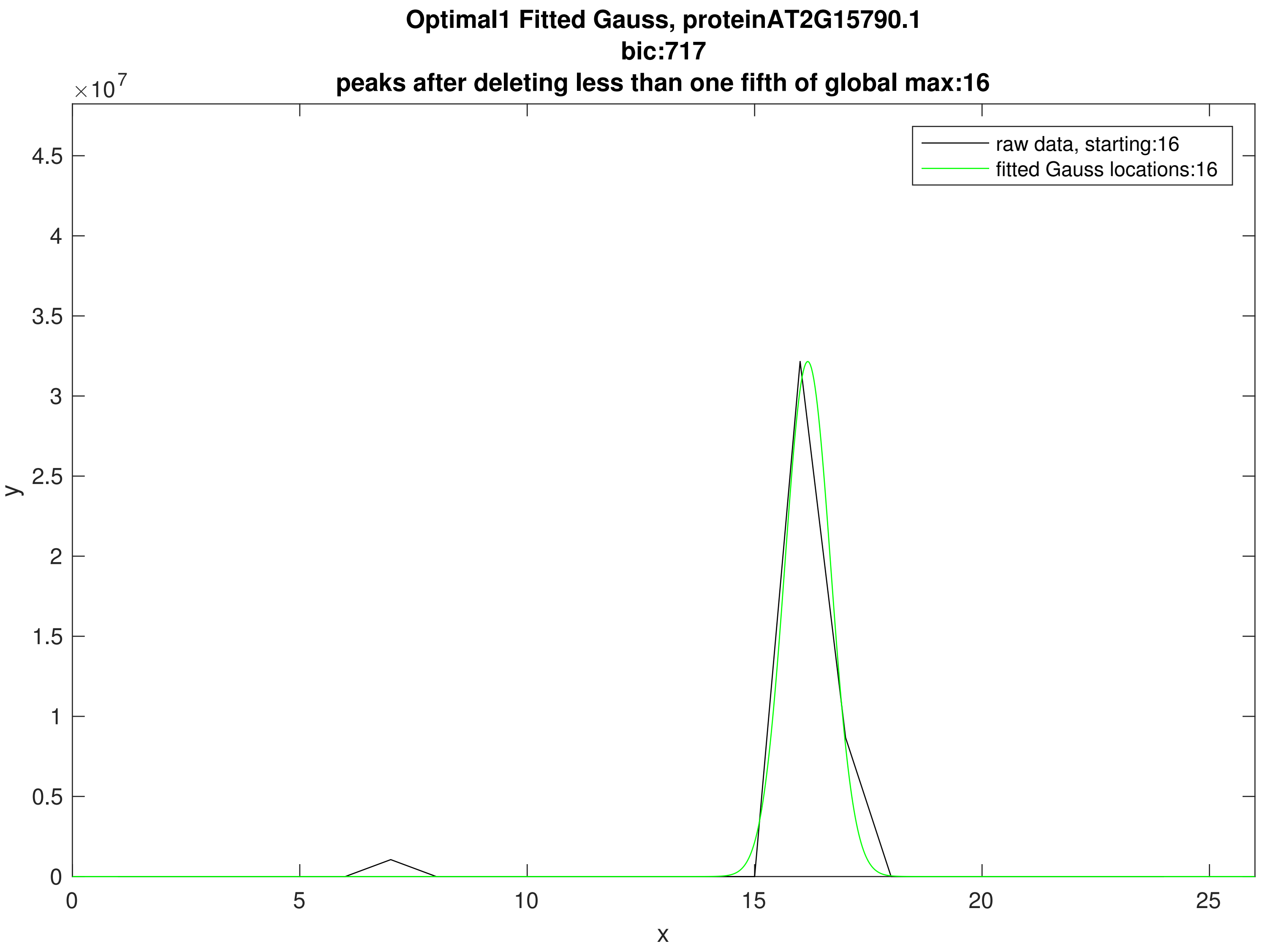

Optimal1 Fitted Gauss, proteinAT2G15790.1
bic:717
peaks after deleting less than one fifth of global max:16
×10^7
4.5
4
3.5
3
2.5
2
1.5
1
0.5
0
y
0
5
10
15
20
25
x
raw data, starting:16
fitted Gauss locations:16

#### Optimal1 Fitted Gauss, proteinAT2G16060.1
#### bic:748
#### peaks after deleting less than one fifth of global max:19

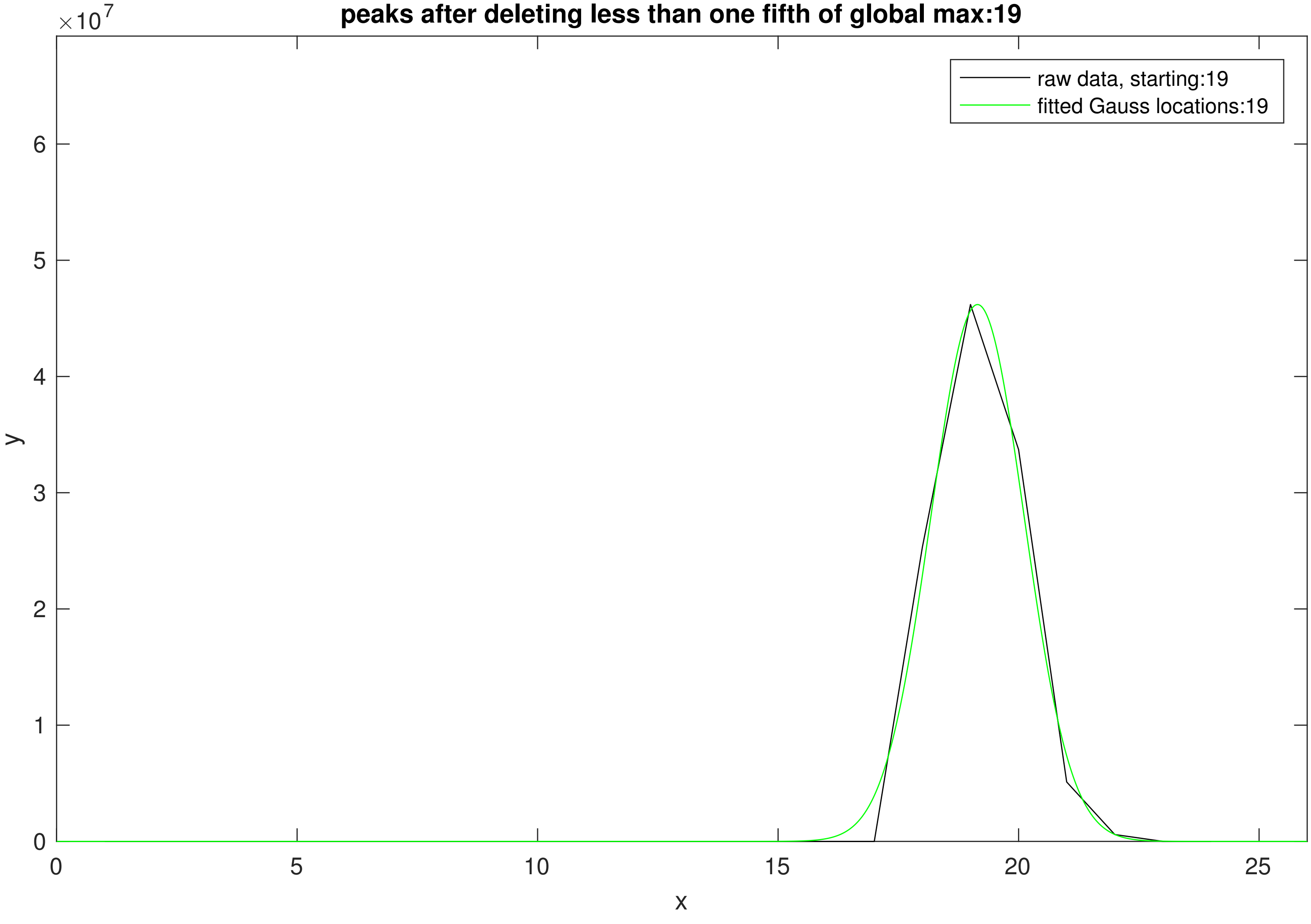

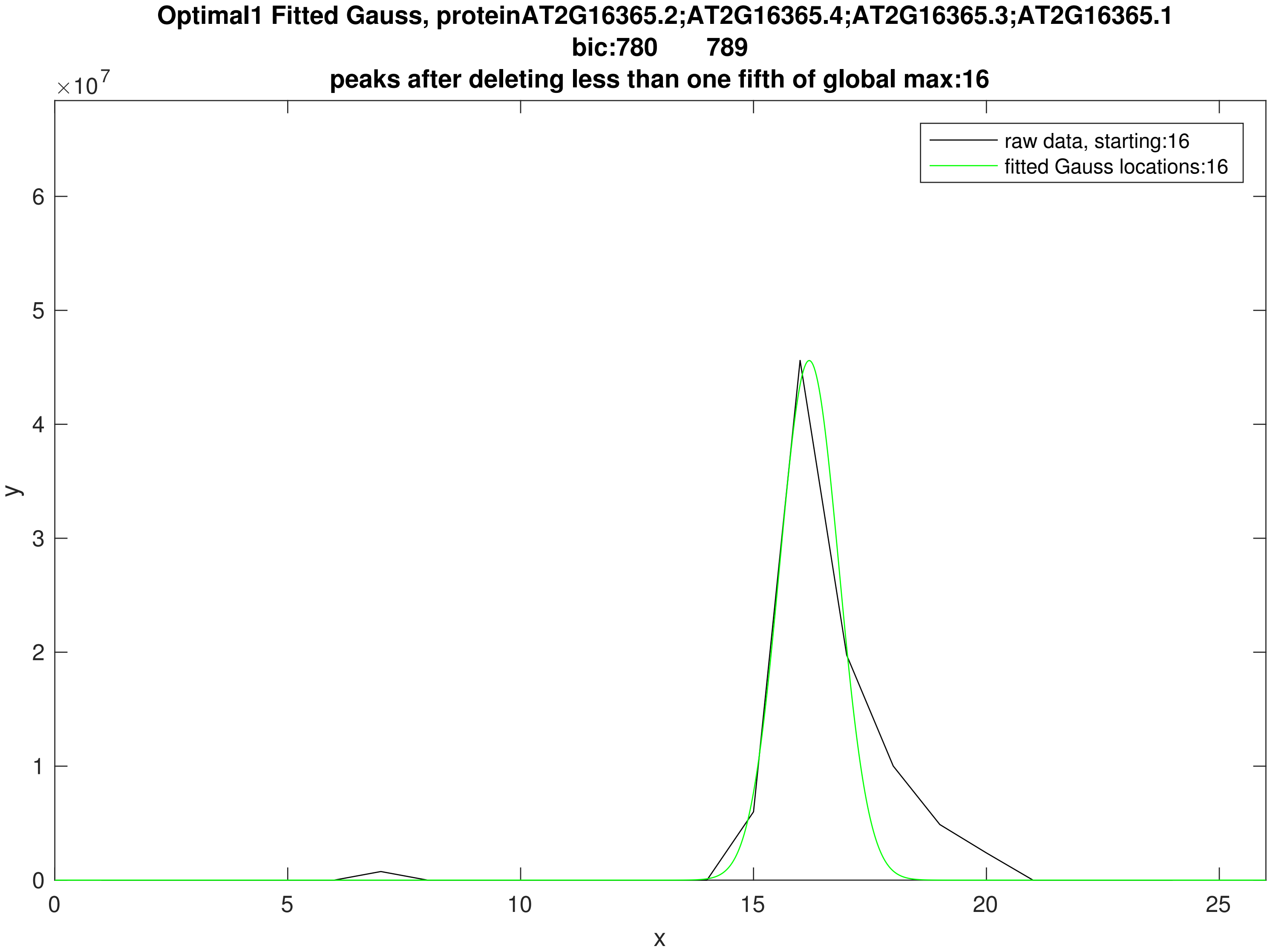
Optimal1 Fitted Gauss, proteinAT2G16365.2;AT2G16365.4;AT2G16365.3;AT2G16365.1
bic:780 789
peaks after deleting less than one fifth of global max:16
×10^7
raw data, starting:16
fitted Gauss locations:16
y
x
0
1
2
3
4
5
6
0
5
10
15
20
25

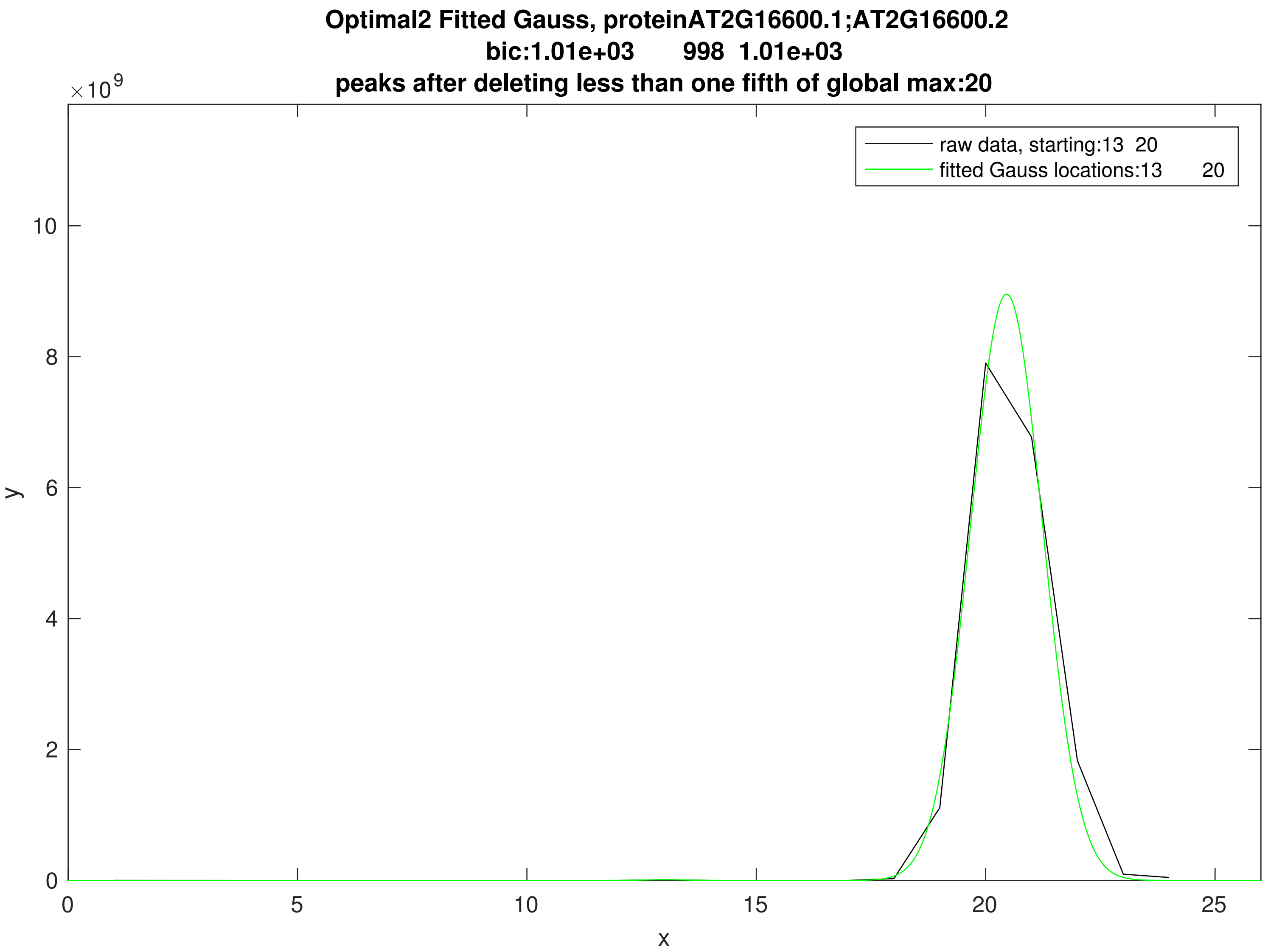
Optimal2 Fitted Gauss, proteinAT2G16600.1;AT2G16600.2
bic:1.01e+03 998 1.01e+03
peaks after deleting less than one fifth of global max:20
×10^9
raw data, starting:13 20
fitted Gauss locations:13 20
y
x
0
2
4
6
8
10
0
5
10
15
20
25

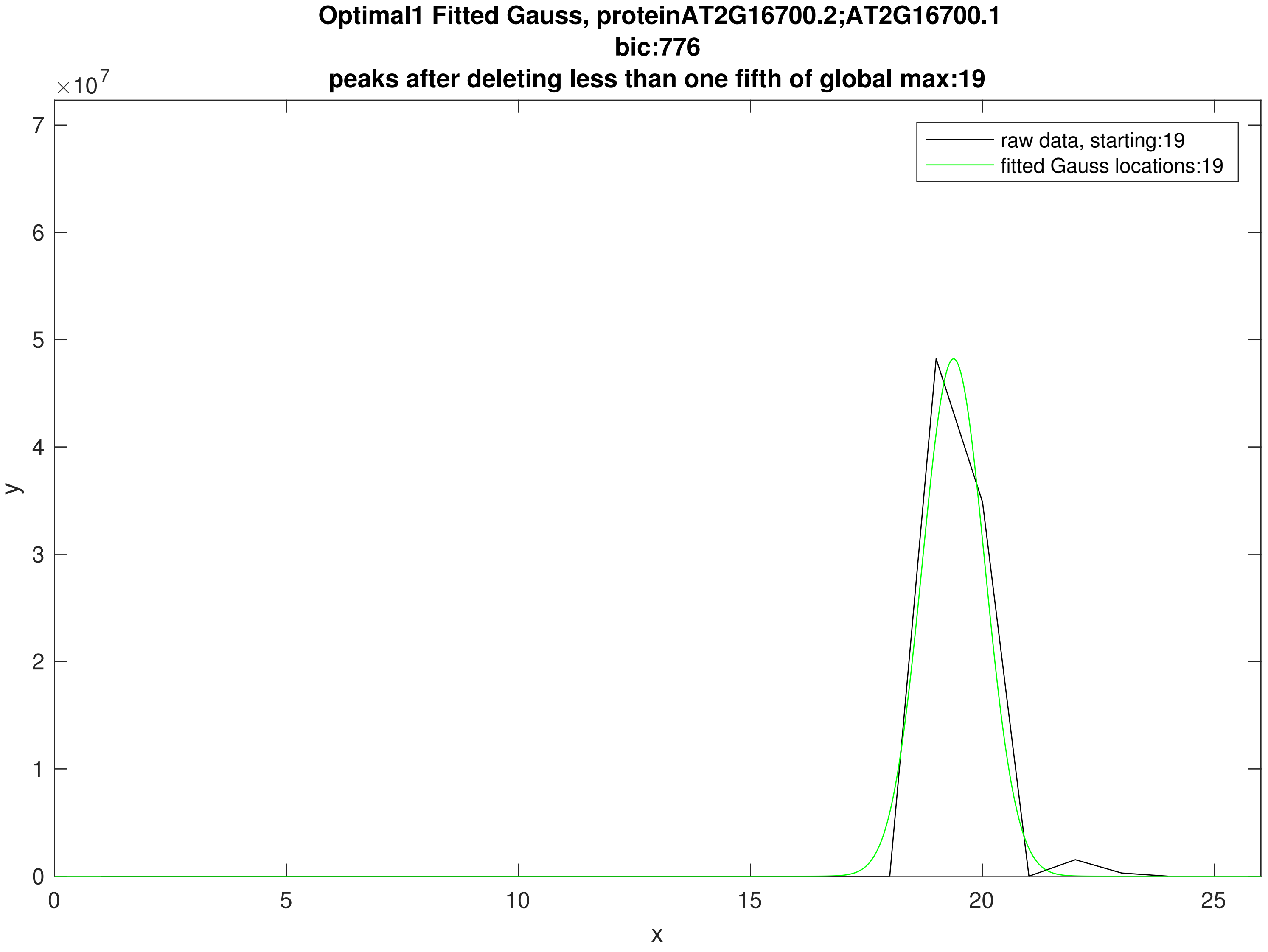
Optimal1 Fitted Gauss, proteinAT2G16700.2;AT2G16700.1
bic:776
peaks after deleting less than one fifth of global max:19
×10^7
raw data, starting:19
fitted Gauss locations:19
y
x
0
1
2
3
4
5
6
7
0
5
10
15
20
25

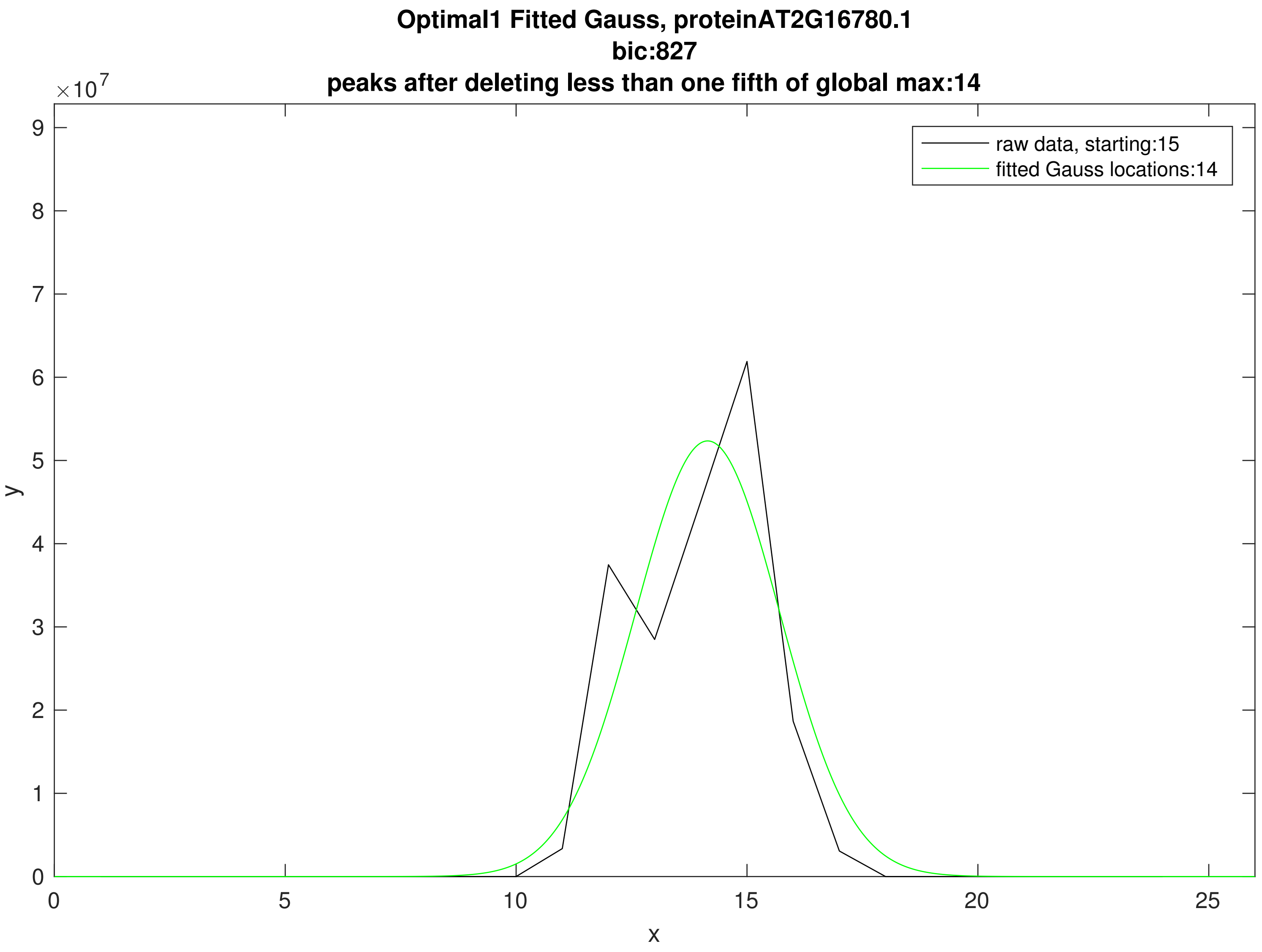

Optimal1 Fitted Gauss, proteinAT2G16780.1
bic:827
peaks after deleting less than one fifth of global max:14
×10^7
raw data, starting:15
fitted Gauss locations:14
y
x
0
1
2
3
4
5
6
7
8
9
0
5
10
15
20
25

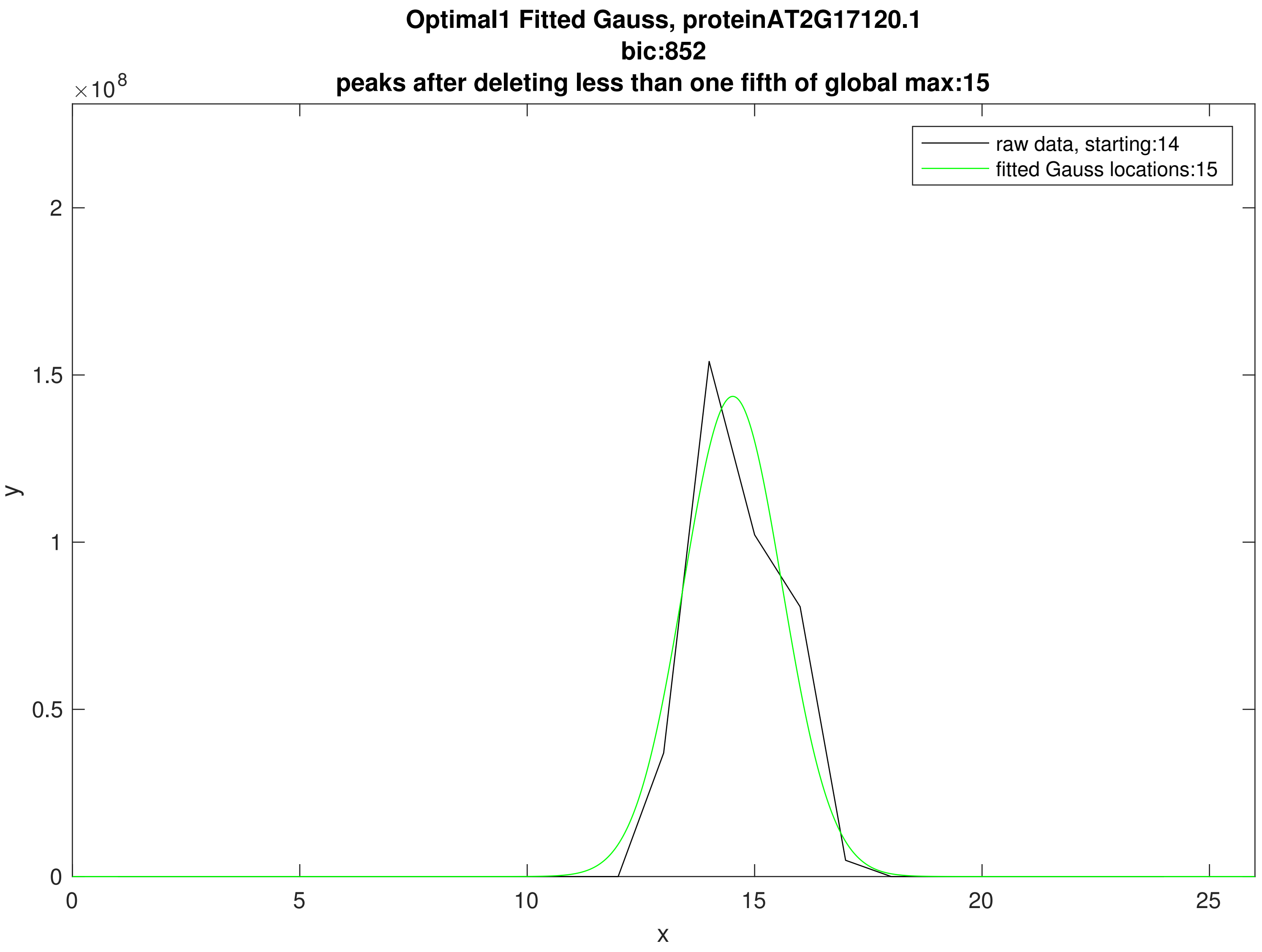
Optimal1 Fitted Gauss, proteinAT2G17120.1
bic:852
peaks after deleting less than one fifth of global max:15
×10^8
raw data, starting:14
fitted Gauss locations:15
y
x
0
0.5
1
1.5
2
0
5
10
15
20
25

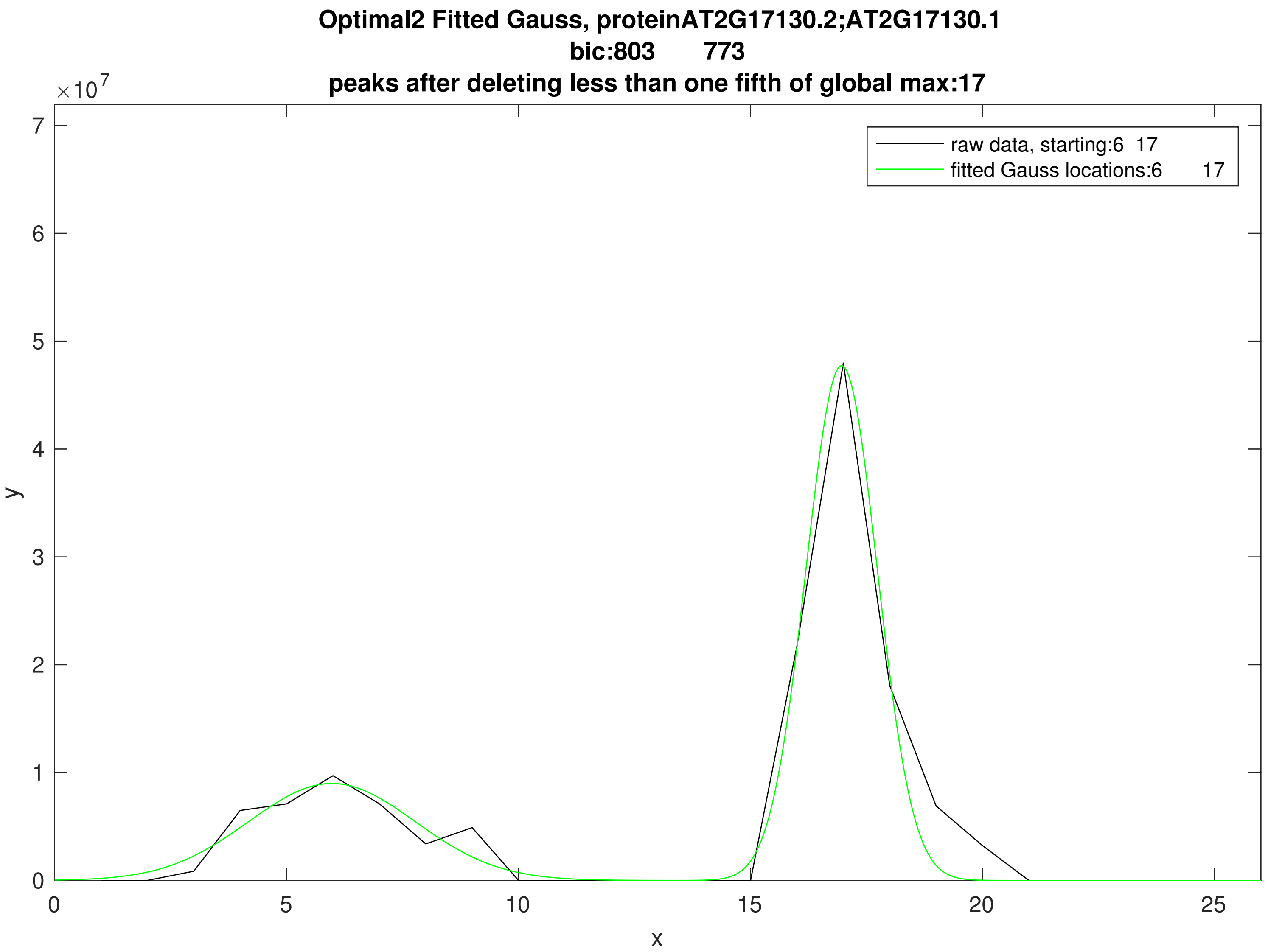
Optimal2 Fitted Gauss, proteinAT2G17130.2;AT2G17130.1
bic:803 773
peaks after deleting less than one fifth of global max:17
×10^7
raw data, starting:6 17
fitted Gauss locations:6 17
y
x
0
1
2
3
4
5
6
7
0
5
10
15
20
25

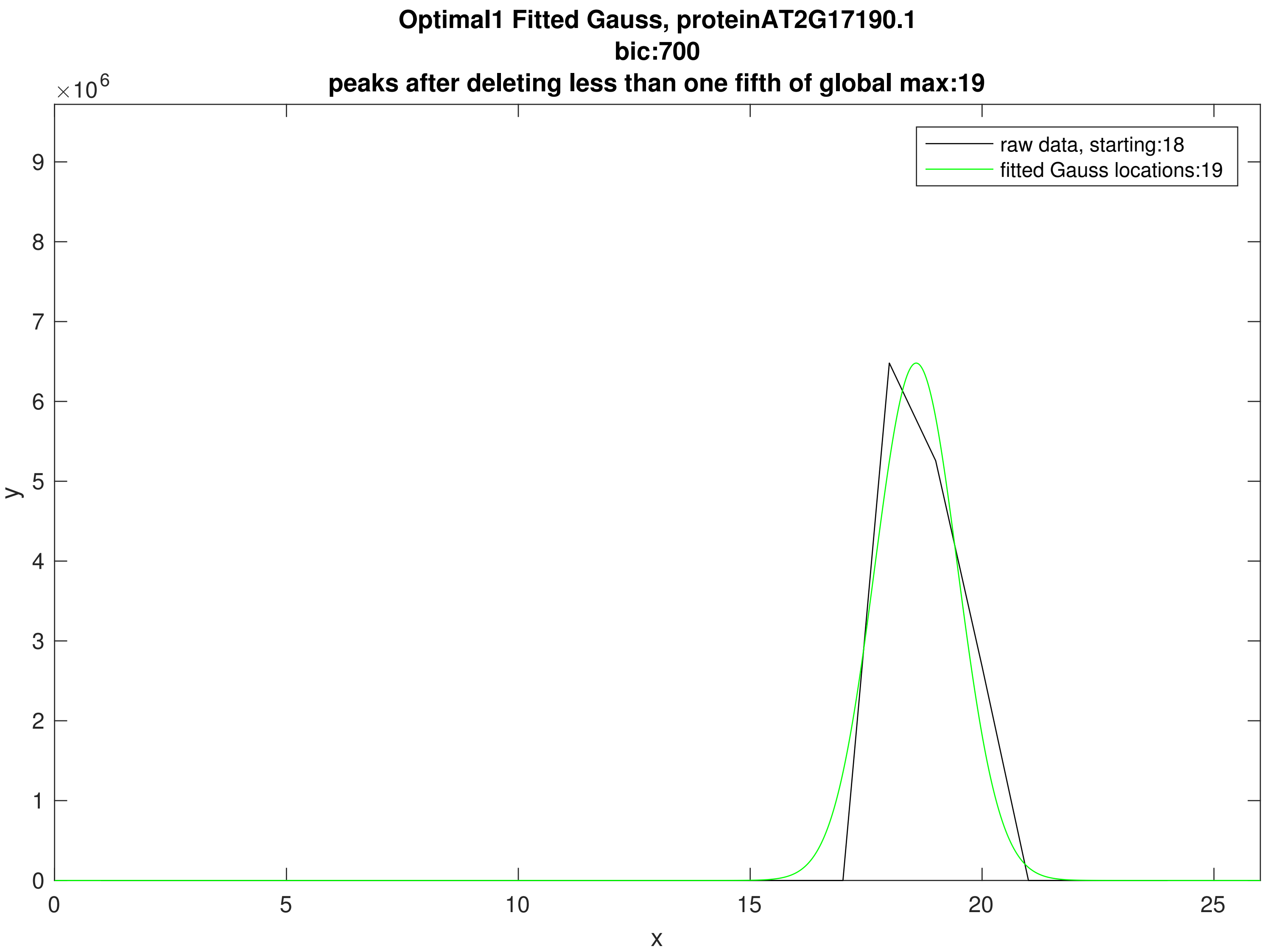

Optimal1 Fitted Gauss, proteinAT2G17190.1
bic:700
peaks after deleting less than one fifth of global max:19
×10^6
raw data, starting:18
fitted Gauss locations:19
y
x
0
1
2
3
4
5
6
7
8
9
0
5
10
15
20
25

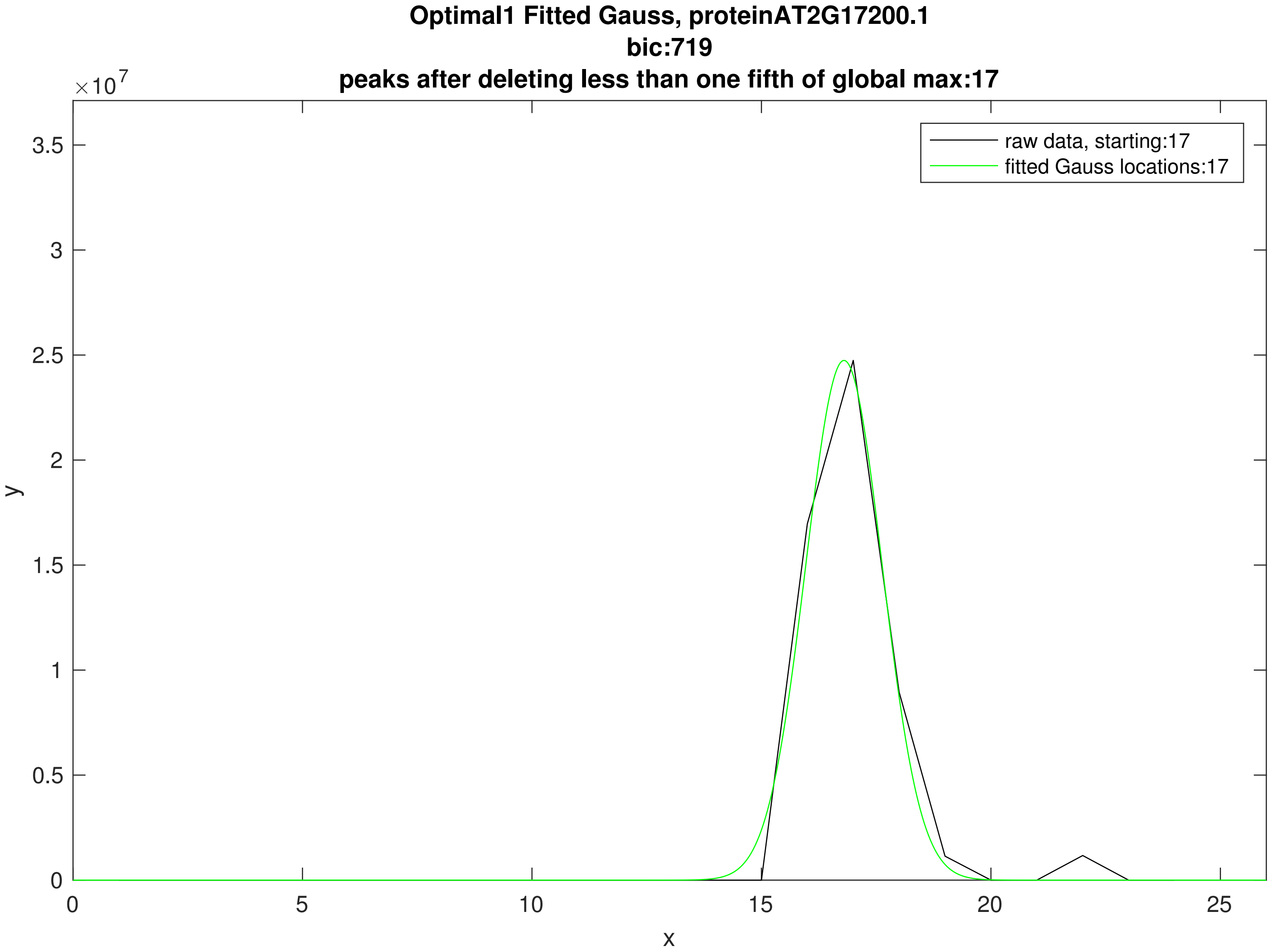
Optimal1 Fitted Gauss, proteinAT2G17200.1
bic:719
peaks after deleting less than one fifth of global max:17
×10^7
3.5
3
2.5
2
1.5
1
0.5
0
y
0
5
10
15
20
25
x
raw data, starting:17
fitted Gauss locations:17

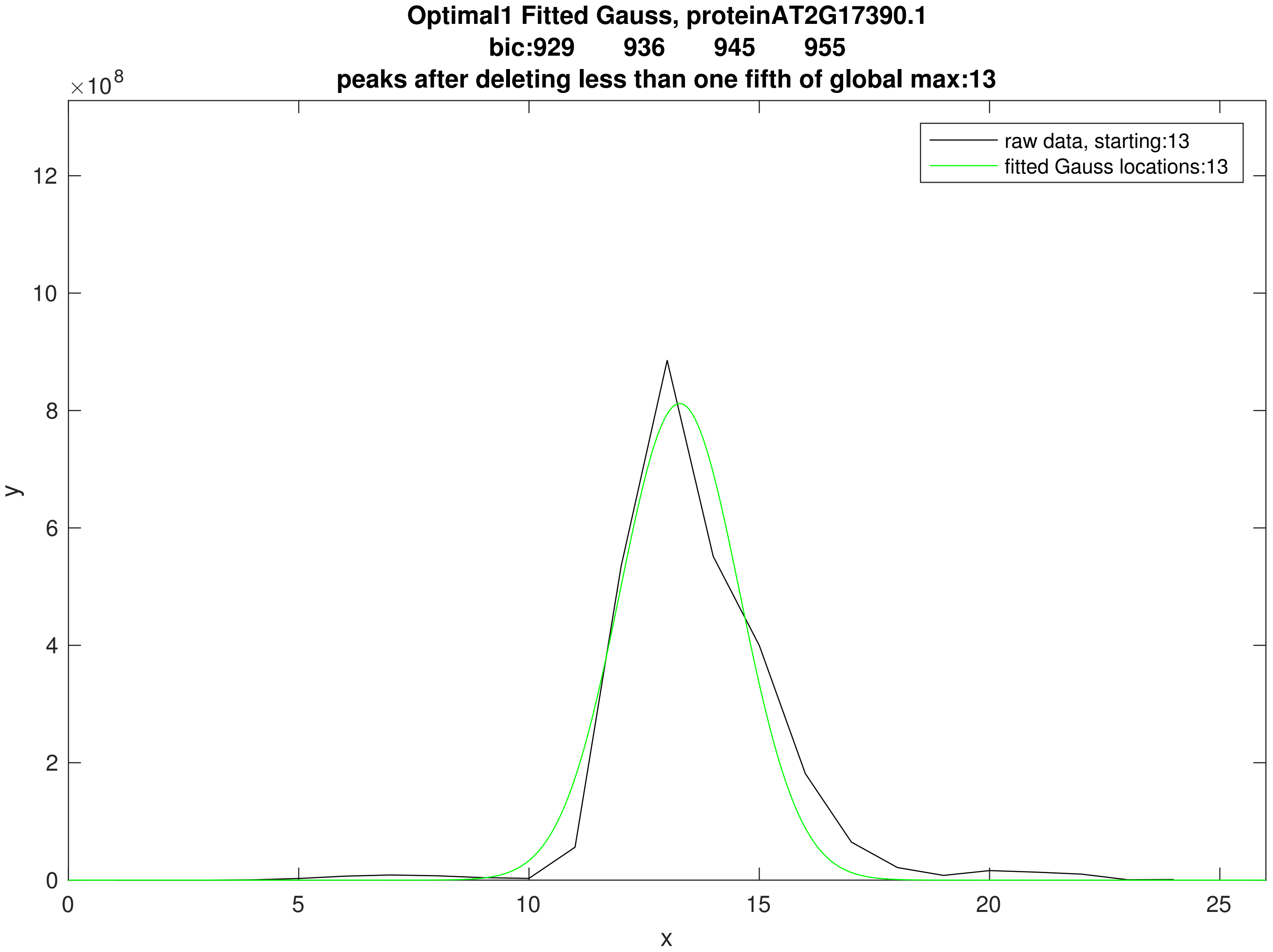

Optimal1 Fitted Gauss, proteinAT2G17390.1
bic:929 936 945 955
peaks after deleting less than one fifth of global max:13
×10^8
raw data, starting:13
fitted Gauss locations:13
y
x
0
2
4
6
8
10
12
0
5
10
15
20
25

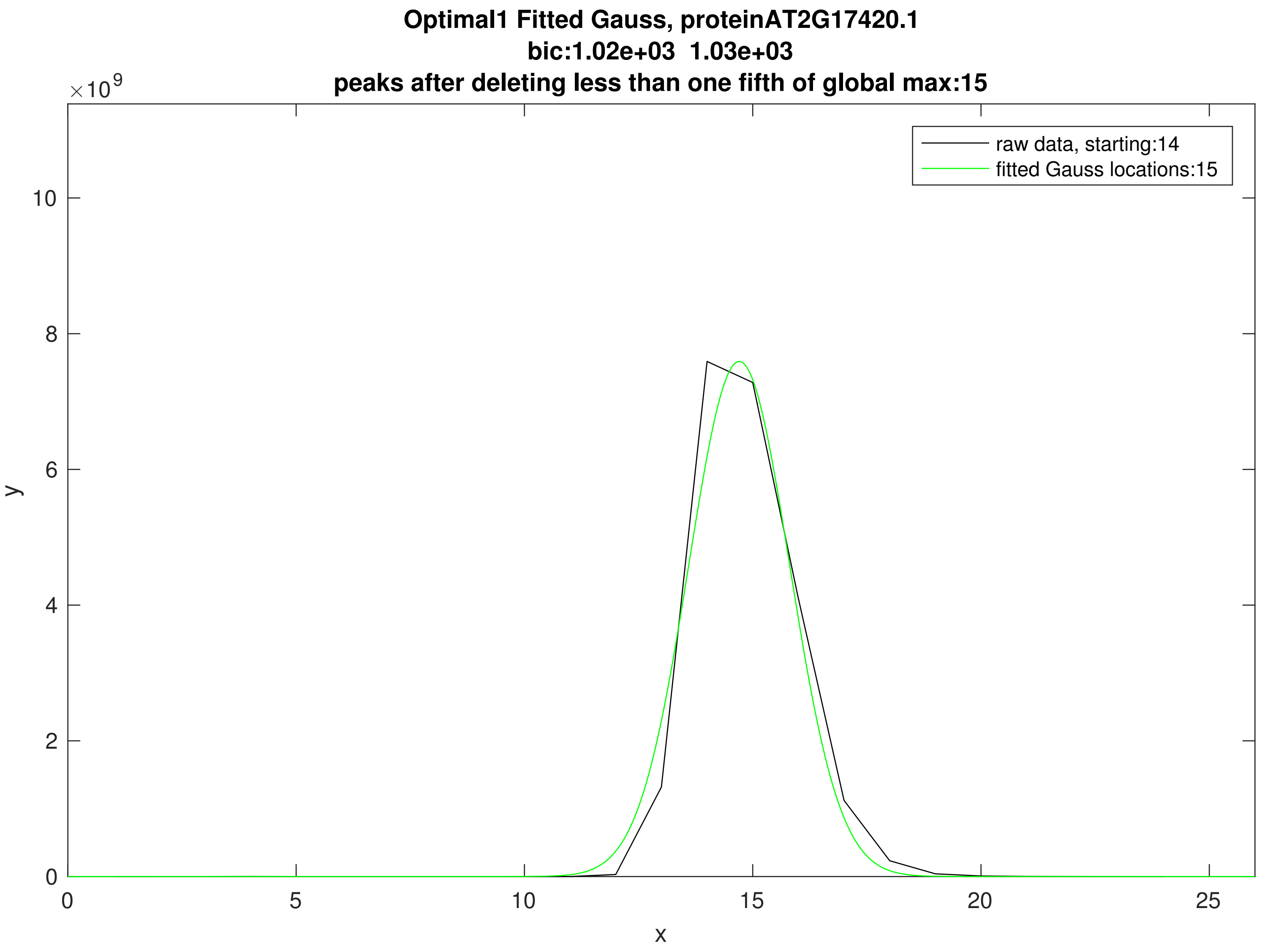

Optimal1 Fitted Gauss, proteinAT2G17420.1
bic:1.02e+03  1.03e+03
peaks after deleting less than one fifth of global max:15
×10^9
raw data, starting:14
fitted Gauss locations:15
y
x
0
2
4
6
8
10
0
5
10
15
20
25

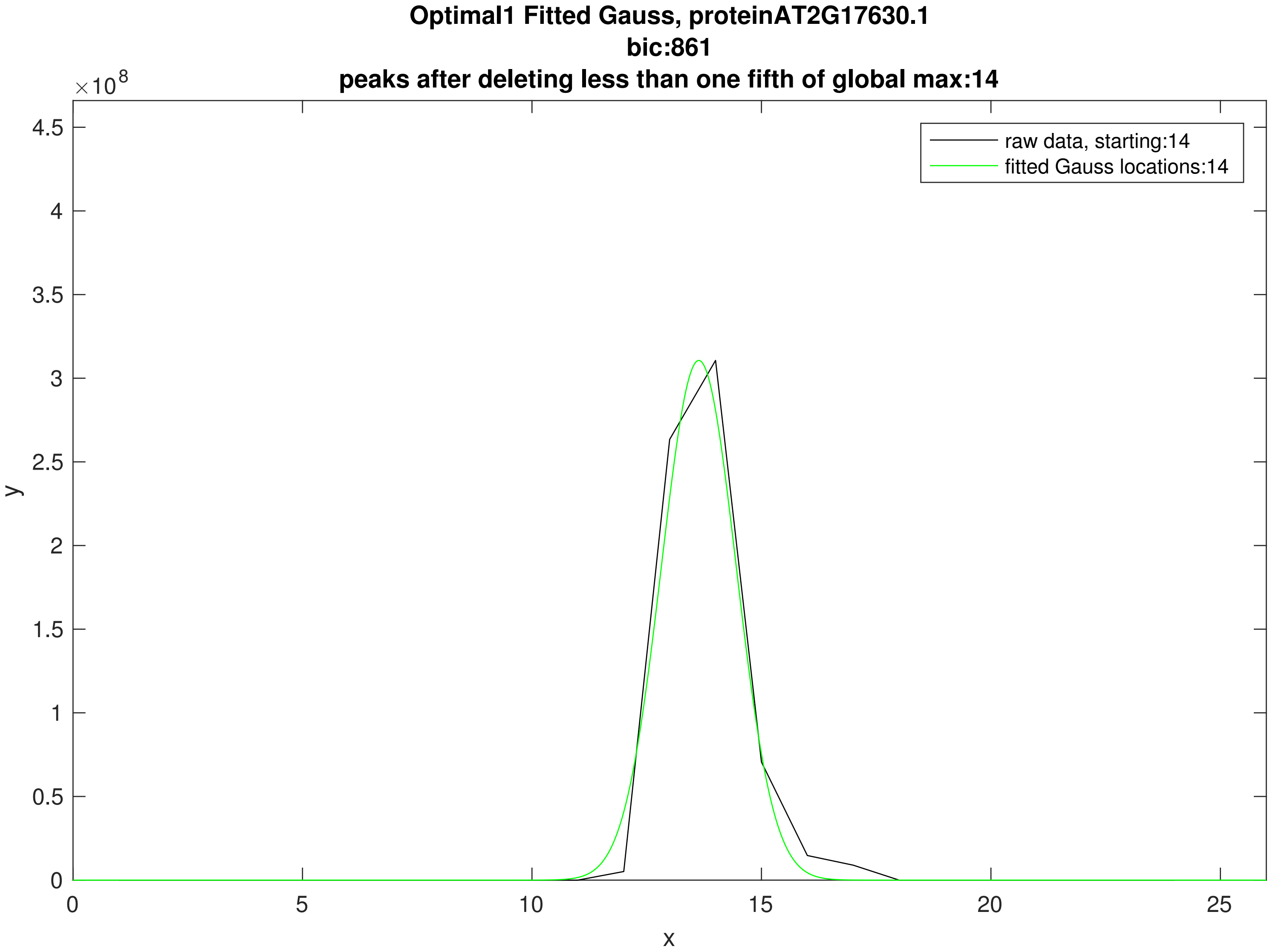

Optimal1 Fitted Gauss, proteinAT2G17630.1
bic:861
peaks after deleting less than one fifth of global max:14
×10^8
4.5
4
3.5
3
2.5
2
1.5
1
0.5
0
y
0
5
10
15
20
25
x
raw data, starting:14
fitted Gauss locations:14

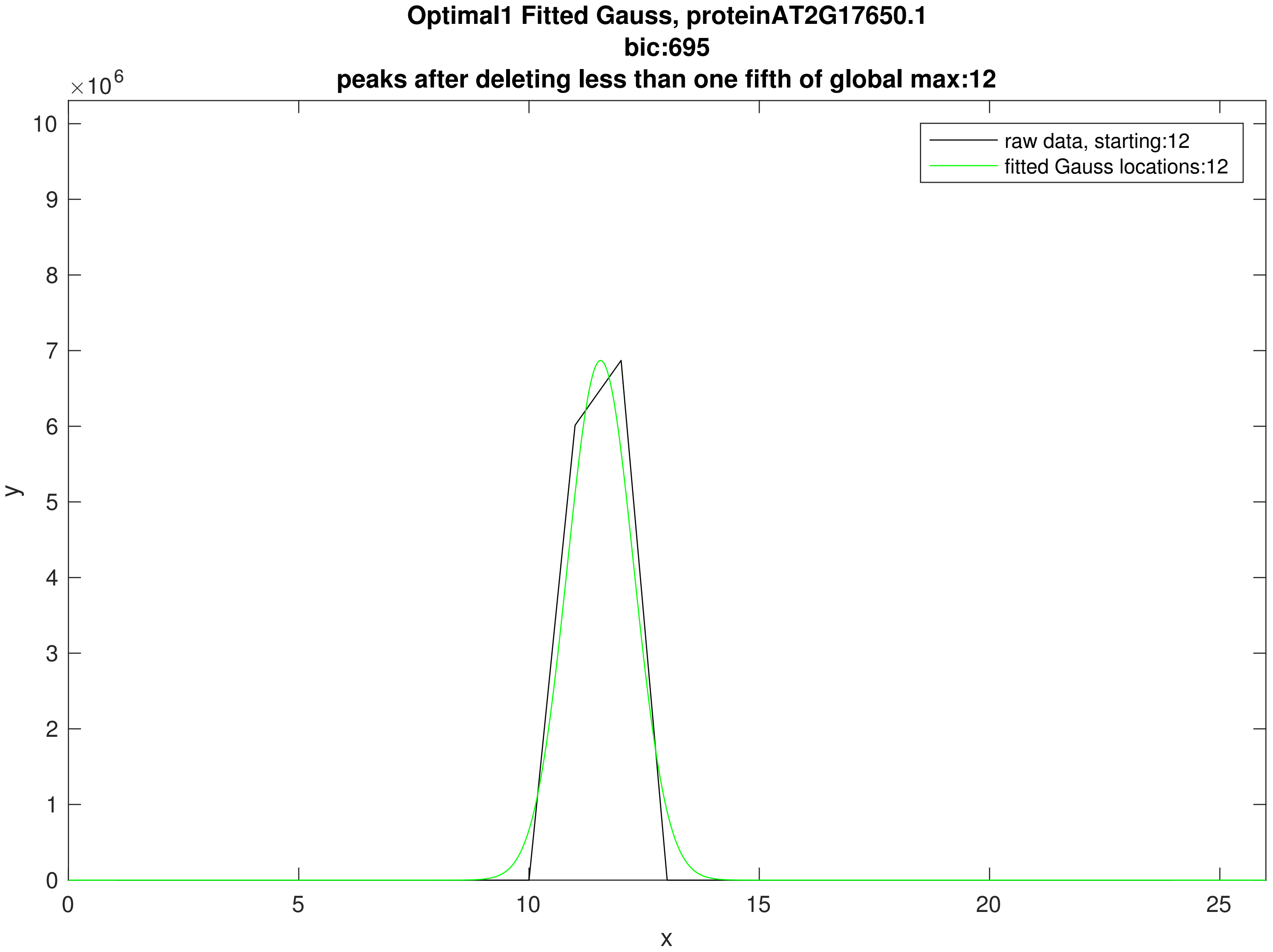

Optimal1 Fitted Gauss, proteinAT2G17650.1
bic:695
peaks after deleting less than one fifth of global max:12
×10^6
raw data, starting:12
fitted Gauss locations:12
y
x
0
1
2
3
4
5
6
7
8
9
10
0
5
10
15
20
25

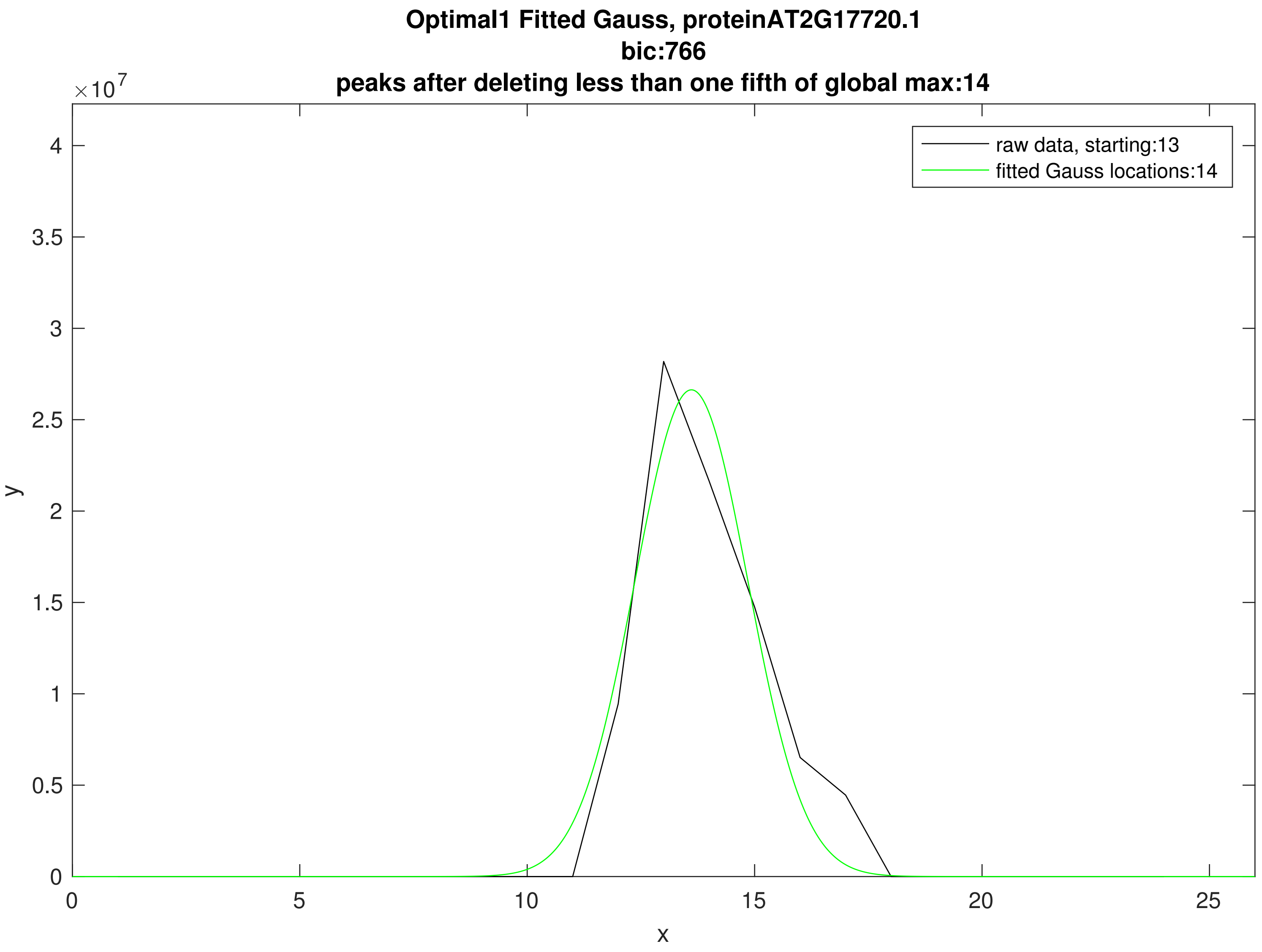
Optimal1 Fitted Gauss, proteinAT2G17720.1
bic:766
peaks after deleting less than one fifth of global max:14
×10^7
raw data, starting:13
fitted Gauss locations:14
y
x
0
0.5
1
1.5
2
2.5
3
3.5
4
0
5
10
15
20
25

**Optimal1 Fitted Gauss, proteinAT2G17760.1**
**bic:698**
**peaks after deleting less than one fifth of global max:14**

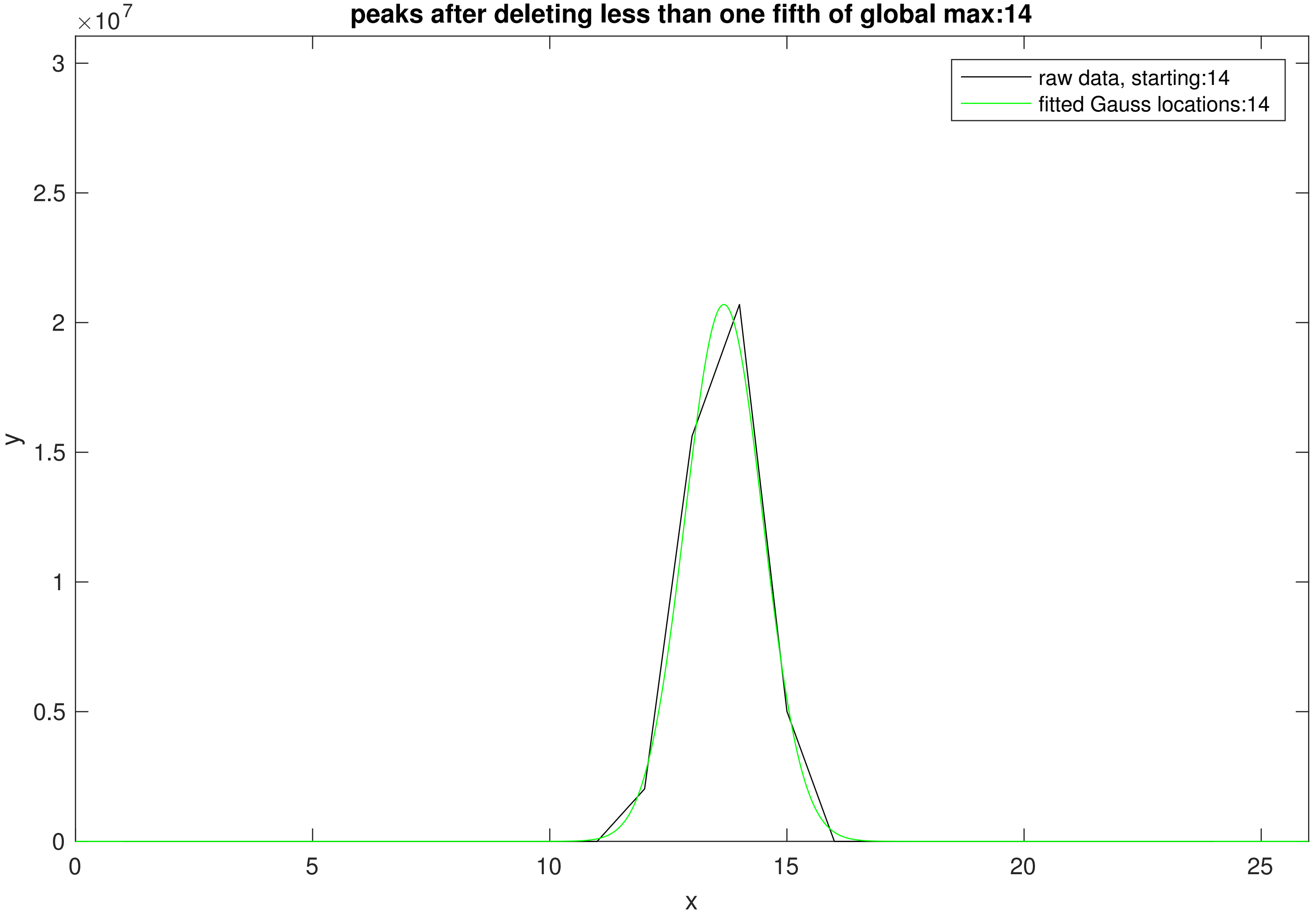

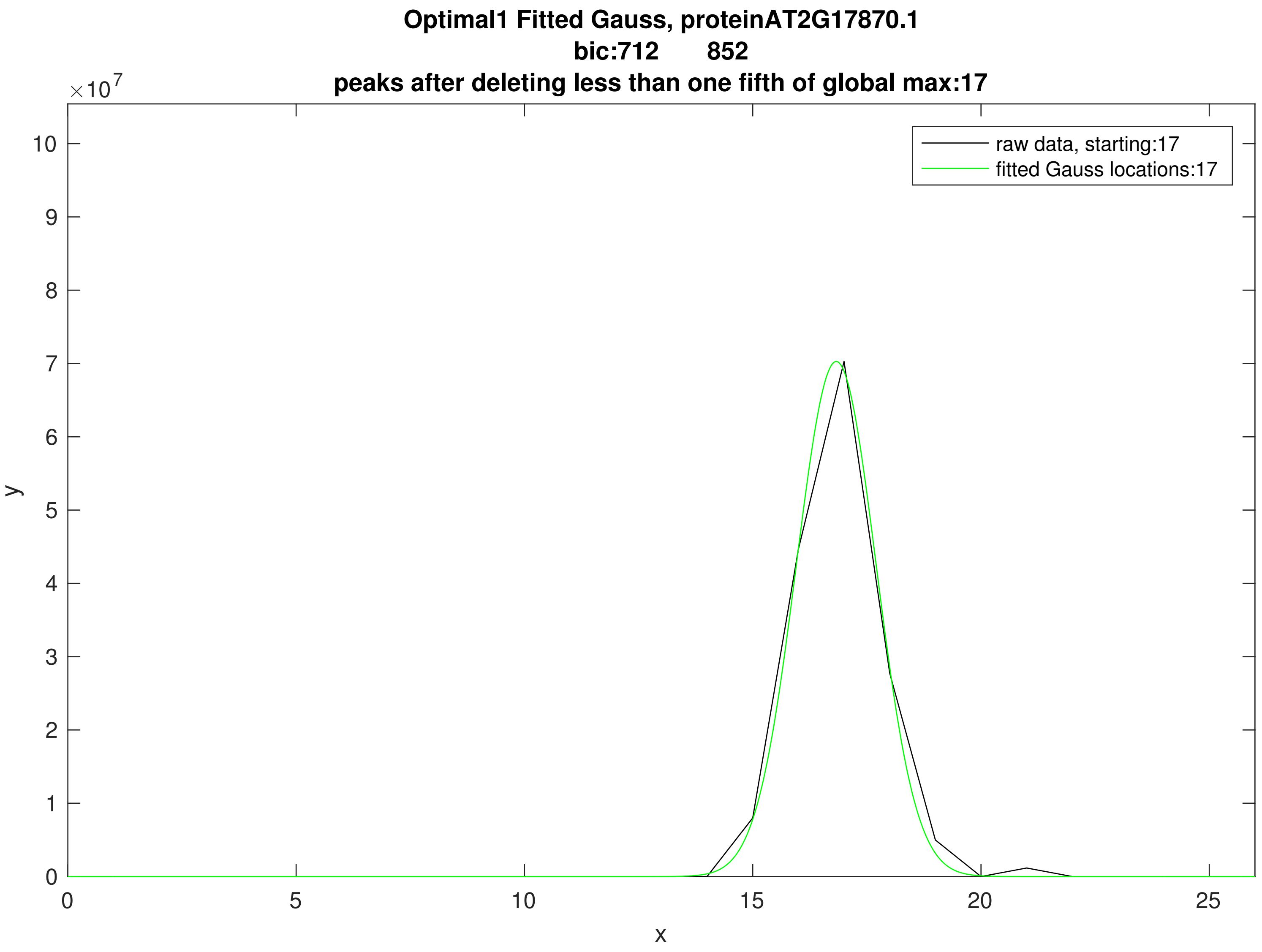

Optimal1 Fitted Gauss, proteinAT2G17870.1
bic:712 852
peaks after deleting less than one fifth of global max:17
×10^7
raw data, starting:17
fitted Gauss locations:17
y
x
0
1
2
3
4
5
6
7
8
9
10
0
5
10
15
20
25

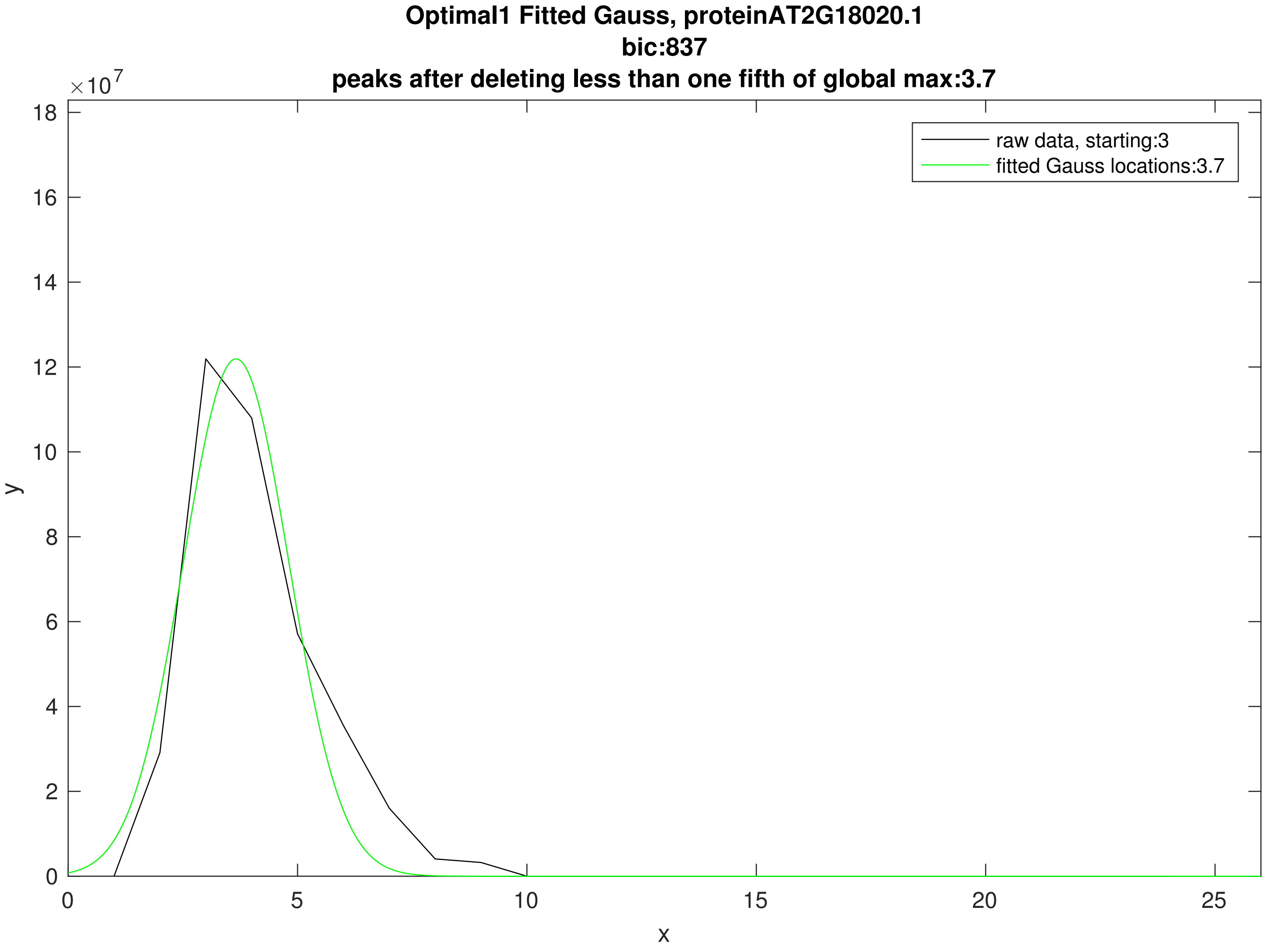
Optimal1 Fitted Gauss, proteinAT2G18020.1
bic:837
peaks after deleting less than one fifth of global max:3.7
raw data, starting:3
fitted Gauss locations:3.7
$\times 10^7$
y
x
0
2
4
6
8
10
12
14
16
18
0
5
10
15
20
25

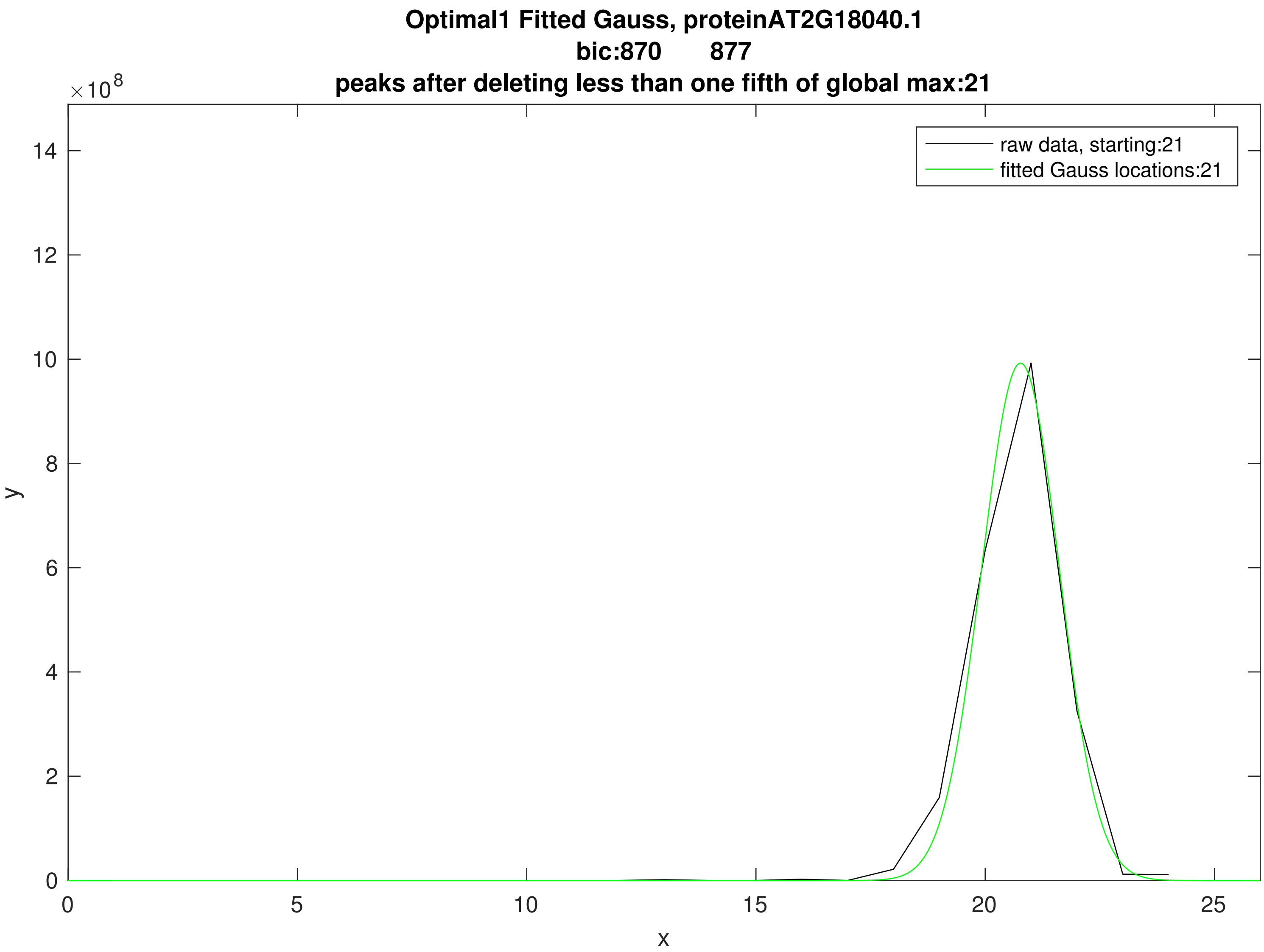

Optimal1 Fitted Gauss, proteinAT2G18040.1
bic:870 877
peaks after deleting less than one fifth of global max:21
×10^8
14
12
10
8
6
4
2
0
y
0
5
10
15
20
25
x
raw data, starting:21
fitted Gauss locations:21

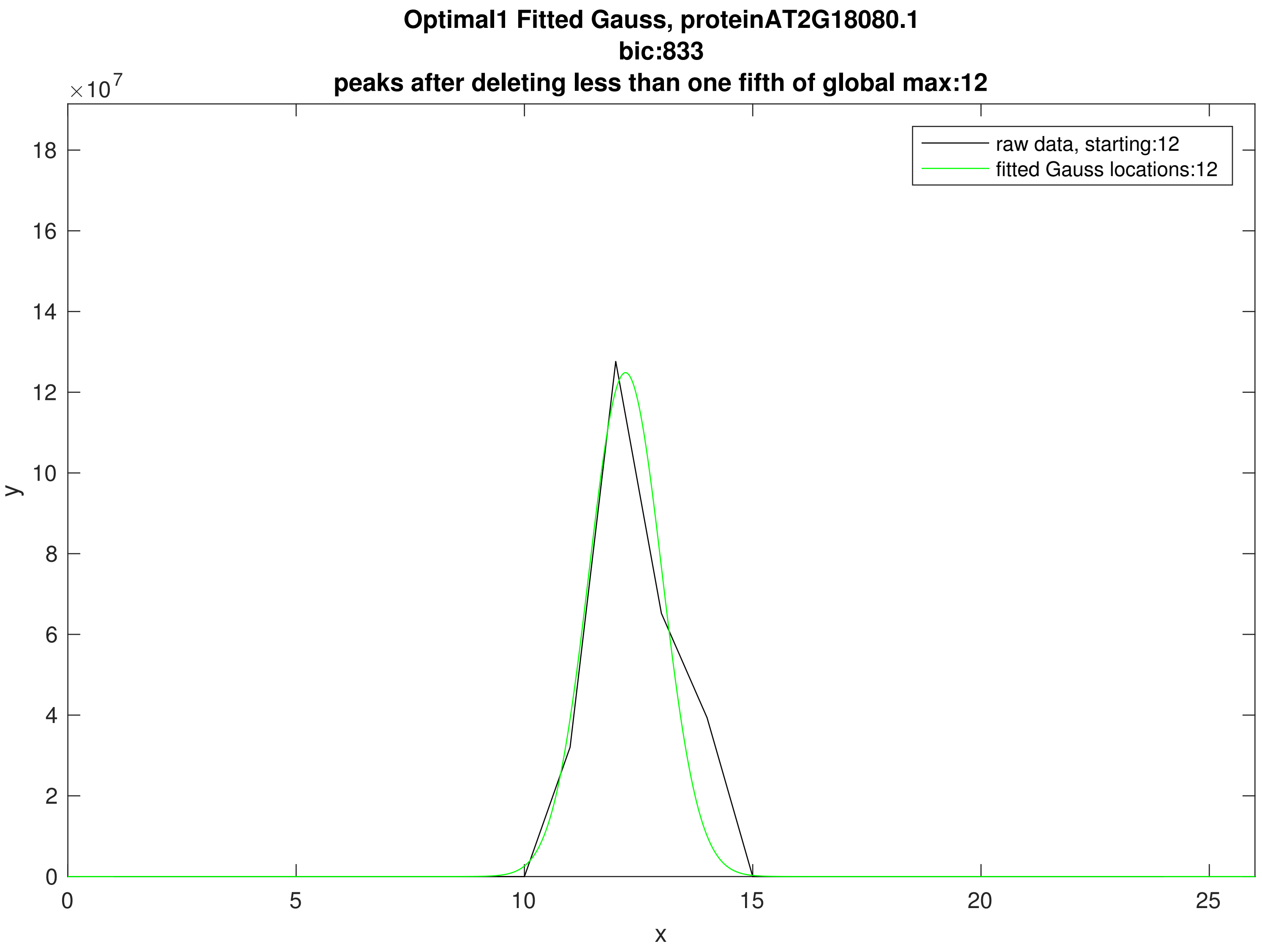
Optimal1 Fitted Gauss, proteinAT2G18080.1
bic:833
peaks after deleting less than one fifth of global max:12
×10^7
raw data, starting:12
fitted Gauss locations:12
y
x
0
2
4
6
8
10
12
14
16
18
0
5
10
15
20
25

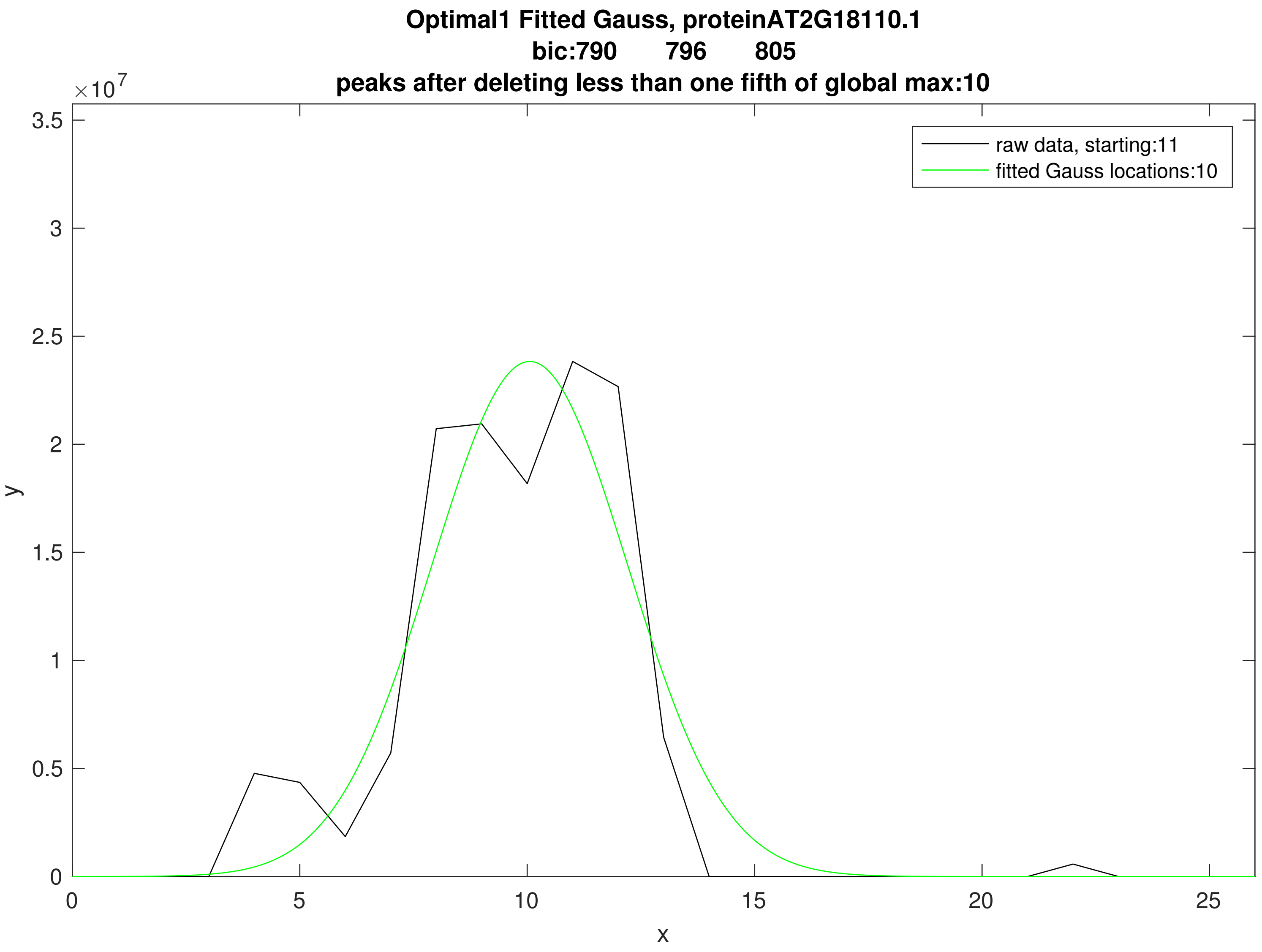

Optimal1 Fitted Gauss, proteinAT2G18110.1
bic:790 796 805
peaks after deleting less than one fifth of global max:10
×10^7
3.5
3
2.5
2
1.5
1
0.5
0
y
0
5
10
15
20
25
x
raw data, starting:11
fitted Gauss locations:10

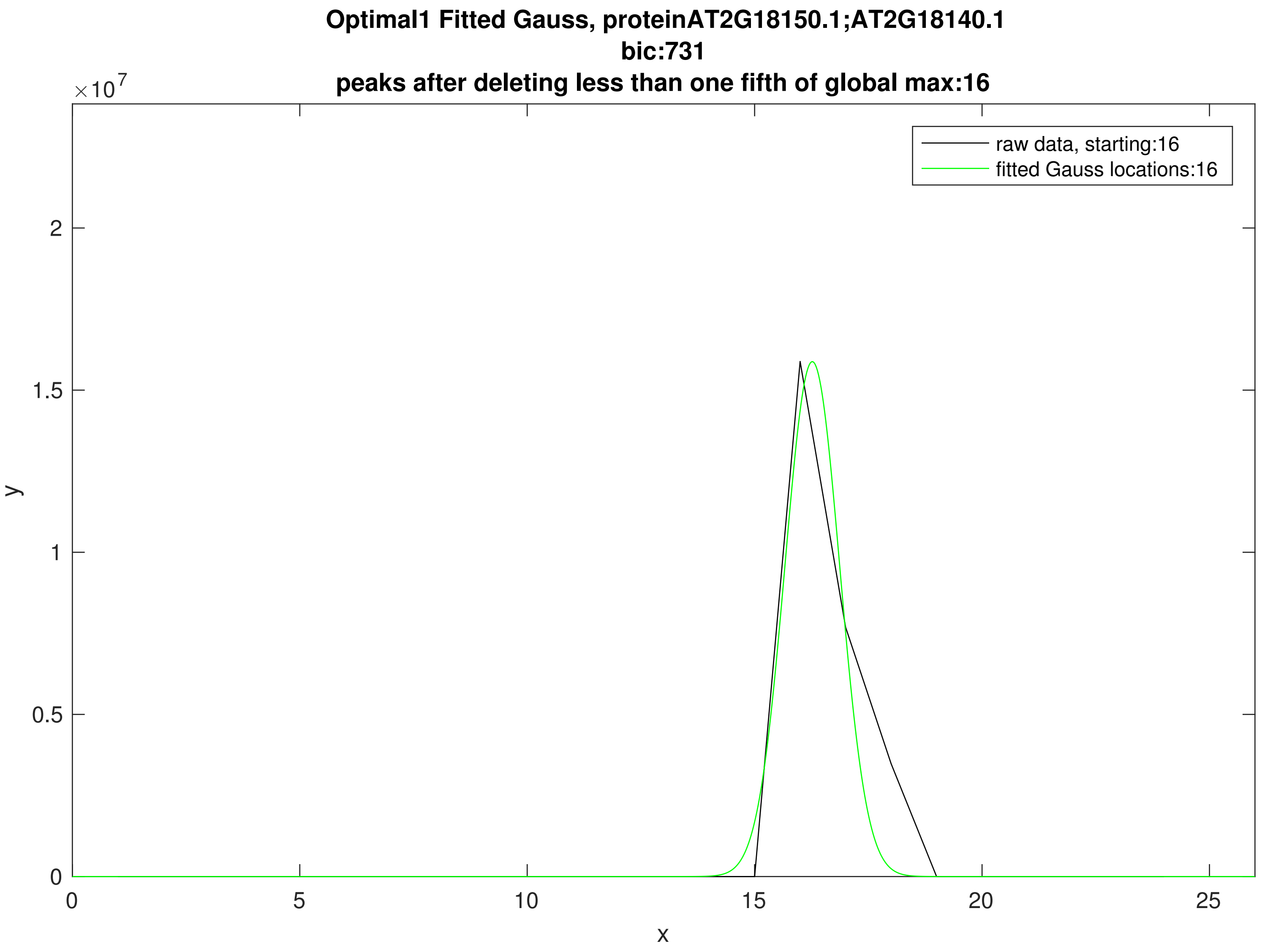
Optimal1 Fitted Gauss, proteinAT2G18150.1;AT2G18140.1
bic:731
peaks after deleting less than one fifth of global max:16
×10^7
raw data, starting:16
fitted Gauss locations:16
2
1.5
1
0.5
0
y
0
5
10
15
20
25
x

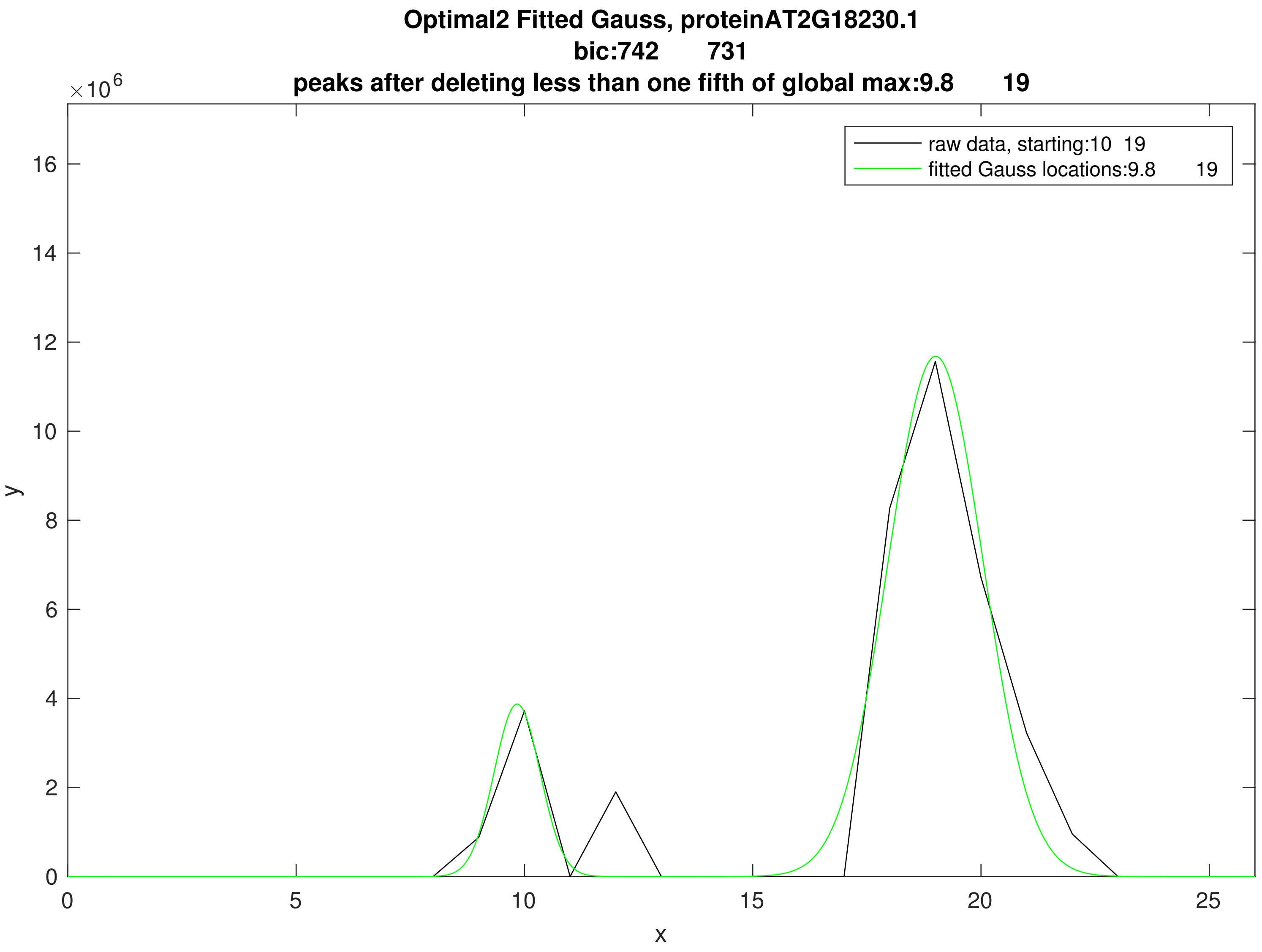
Optimal2 Fitted Gauss, proteinAT2G18230.1
bic:742 731
peaks after deleting less than one fifth of global max:9.8 19
×10^6
raw data, starting:10 19
fitted Gauss locations:9.8 19
y
x
0
2
4
6
8
10
12
14
16
5
15
20
25

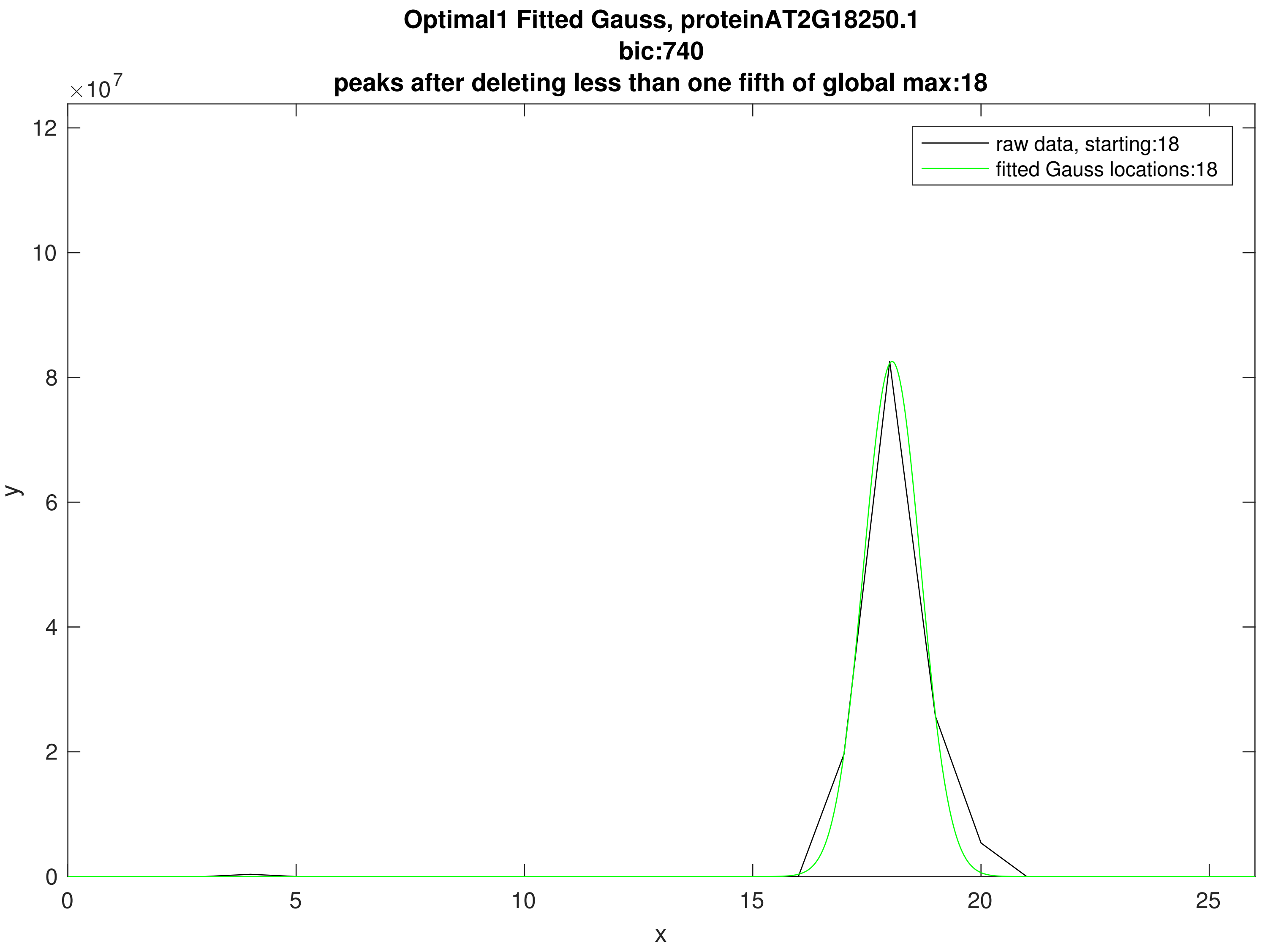

Optimal1 Fitted Gauss, proteinAT2G18250.1
bic:740
peaks after deleting less than one fifth of global max:18
×10^7
12
10
8
6
4
2
0
y
0
5
10
15
20
25
x
raw data, starting:18
fitted Gauss locations:18

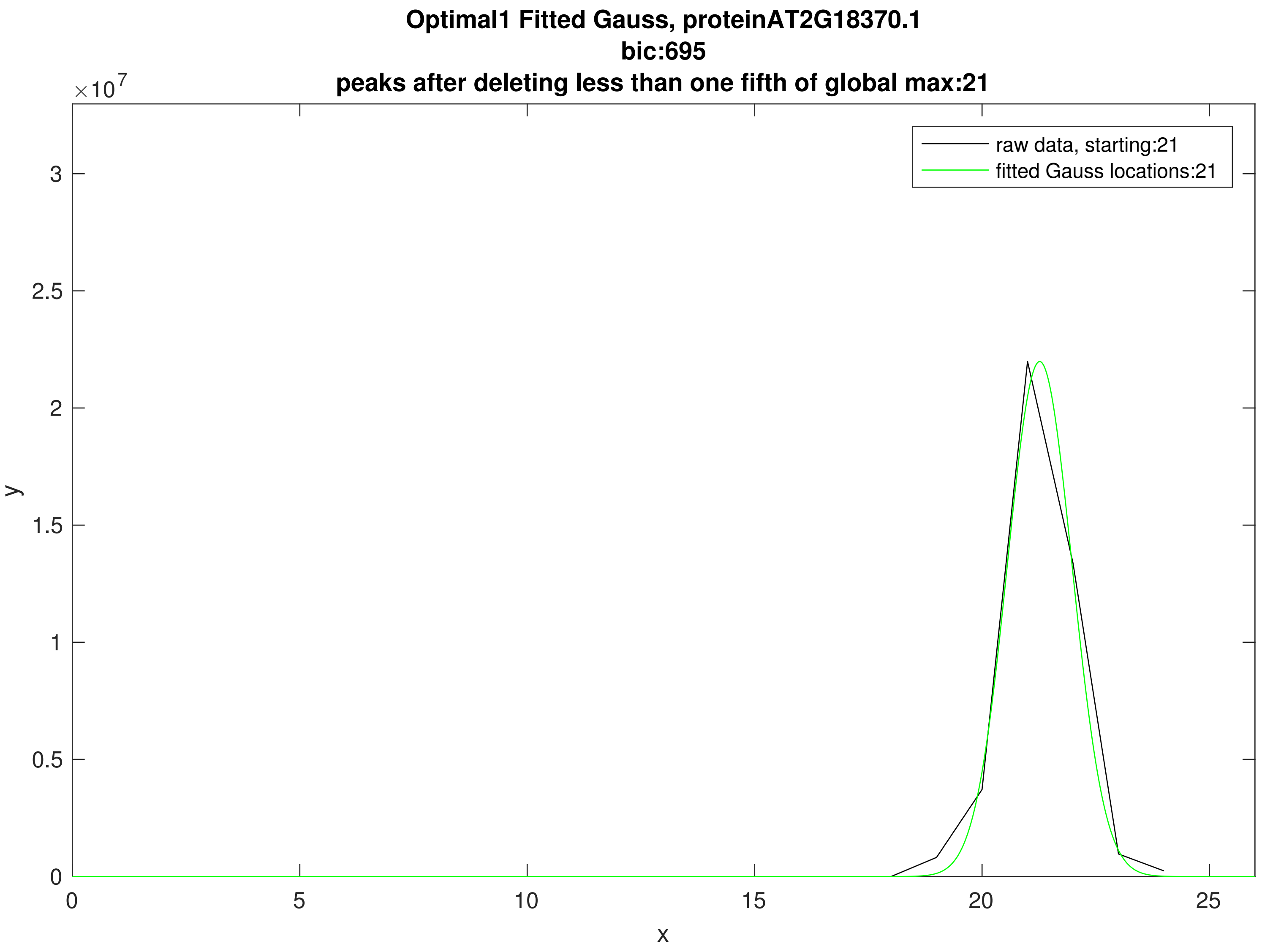

Optimal1 Fitted Gauss, proteinAT2G18370.1
bic:695
peaks after deleting less than one fifth of global max:21
×10^7
3
2.5
2
1.5
1
0.5
0
y
0
5
10
15
20
25
x
raw data, starting:21
fitted Gauss locations:21

**Optimal1 Fitted Gauss, proteinAT2G18990.1;AT3G25580.1**
**bic:647**
**peaks after deleting less than one fifth of global max:17**

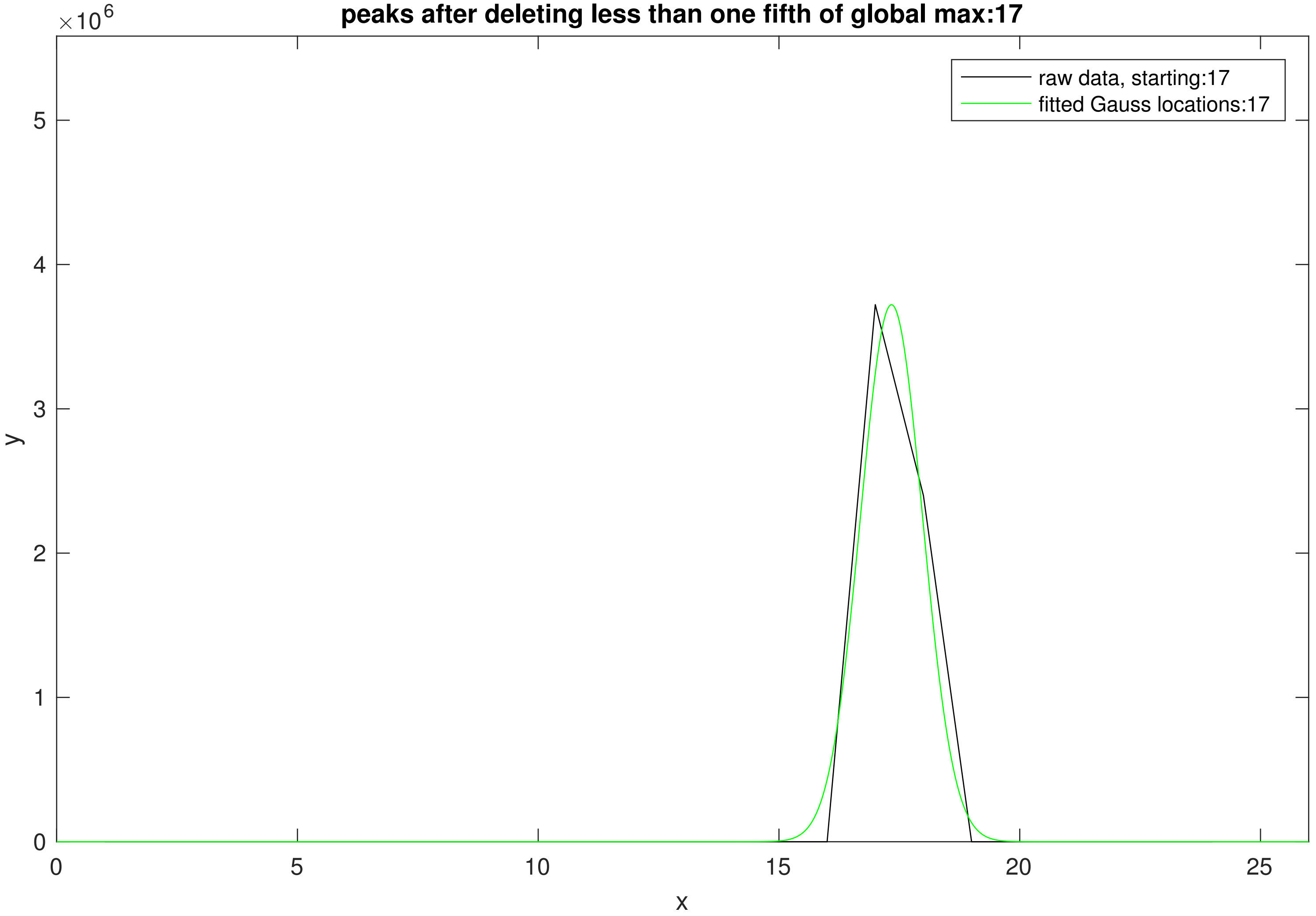

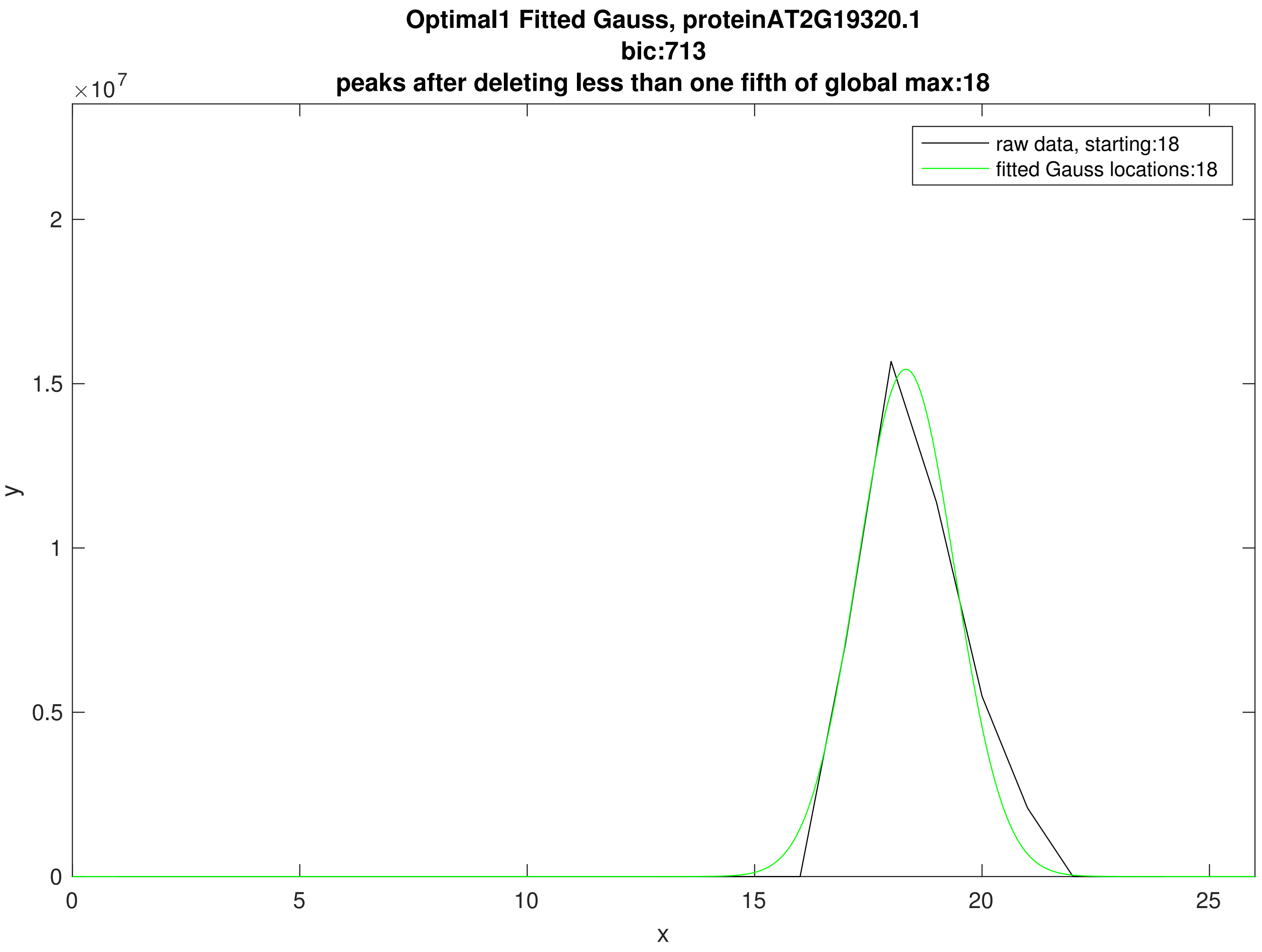

Optimal1 Fitted Gauss, proteinAT2G19320.1
bic:713
peaks after deleting less than one fifth of global max:18
×10^7
raw data, starting:18
fitted Gauss locations:18
y
x
0
0.5
1
1.5
2
0
5
10
15
20
25

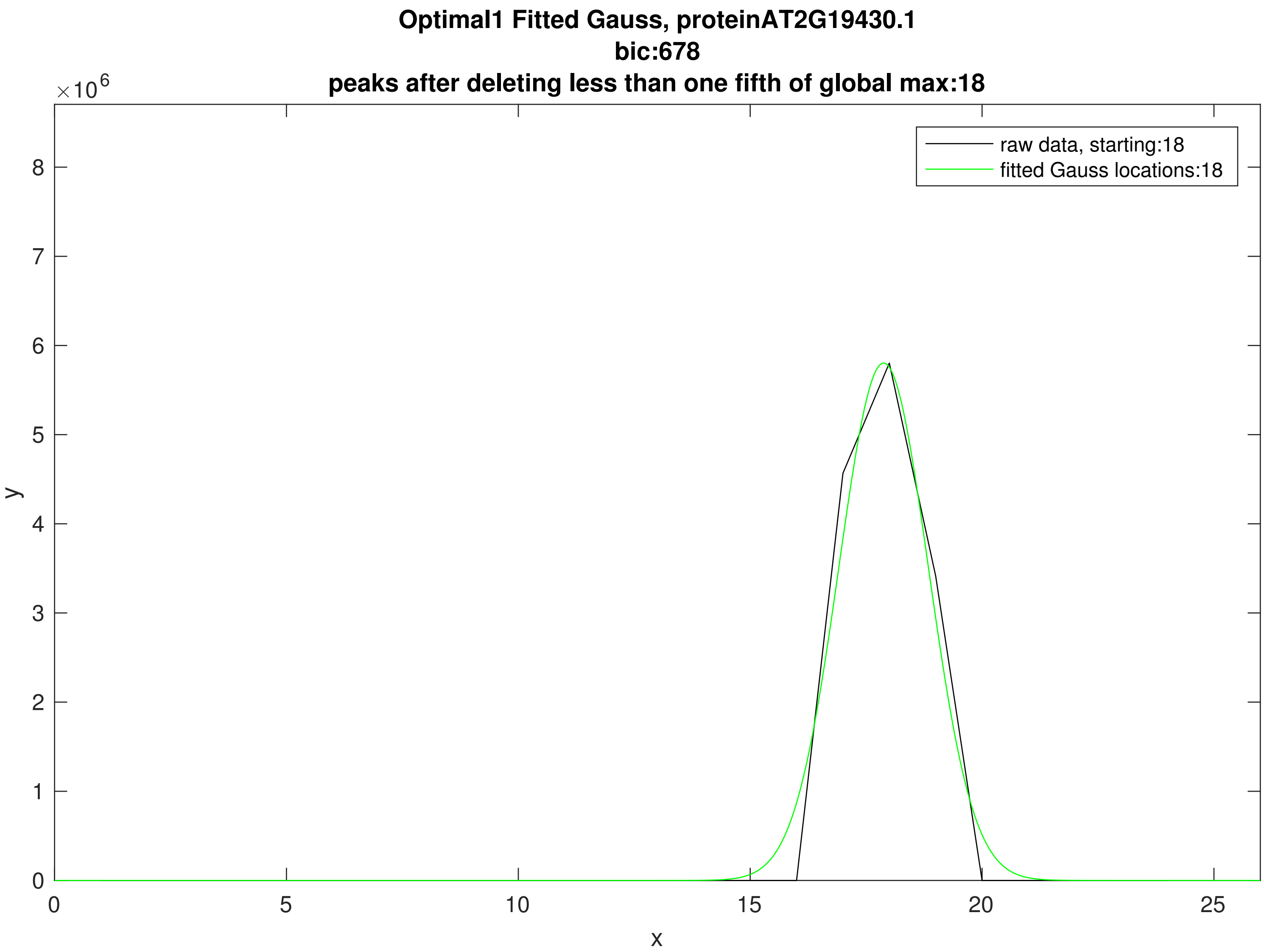
Optimal1 Fitted Gauss, proteinAT2G19430.1
bic:678
peaks after deleting less than one fifth of global max:18
×10^6
raw data, starting:18
fitted Gauss locations:18
y
x
0
1
2
3
4
5
6
7
8
0
5
10
15
20
25

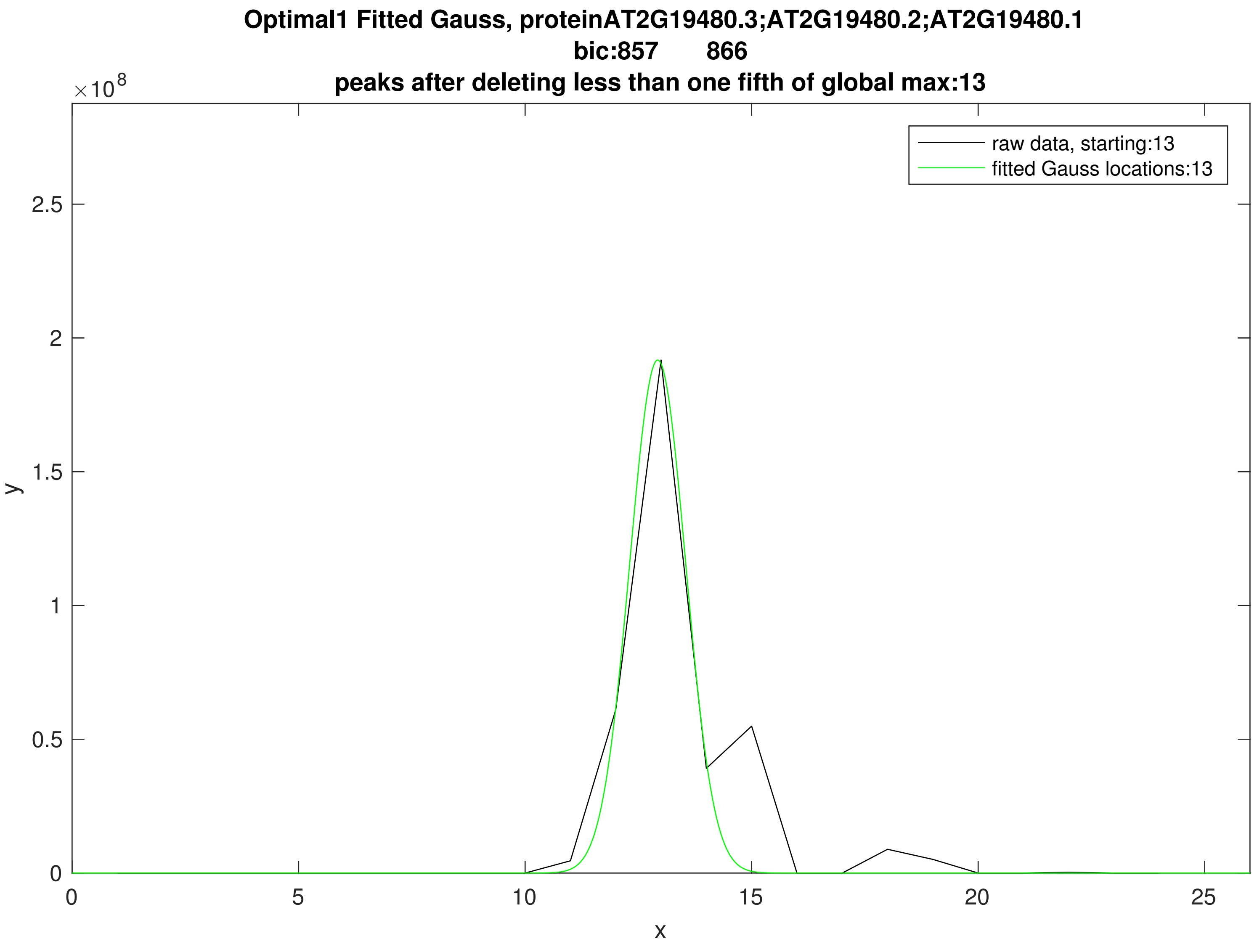

Optimal1 Fitted Gauss, proteinAT2G19480.3;AT2G19480.2;AT2G19480.1
bic:857 866
peaks after deleting less than one fifth of global max:13
×10^8
raw data, starting:13
fitted Gauss locations:13
2.5
2
1.5
1
0.5
0
y
0
5
10
15
20
25
x

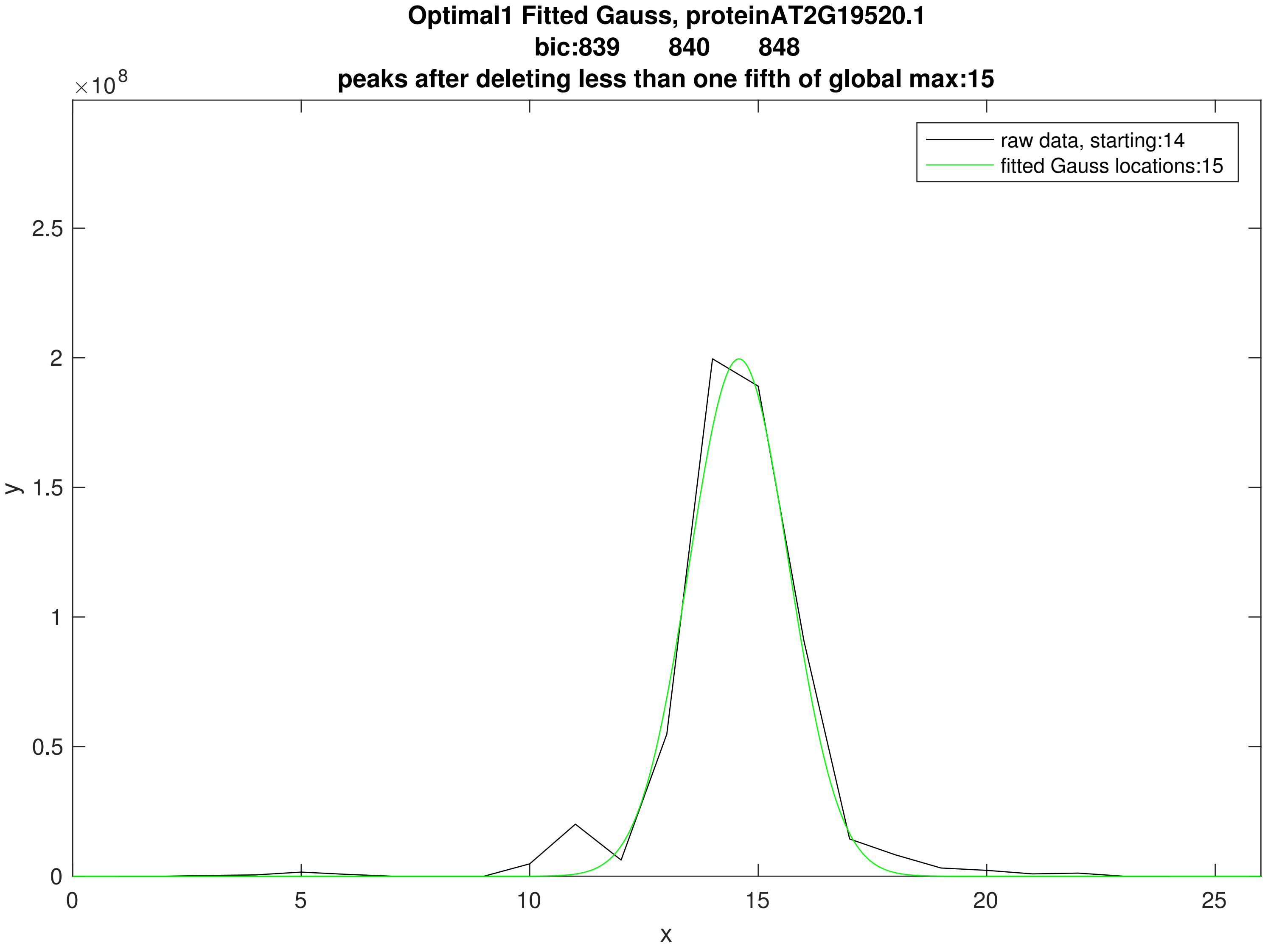
Optimal1 Fitted Gauss, proteinAT2G19520.1
bic:839 840 848
peaks after deleting less than one fifth of global max:15
×10^8
raw data, starting:14
fitted Gauss locations:15
y
x
0
0.5
1
1.5
2
2.5
0
5
10
15
20
25

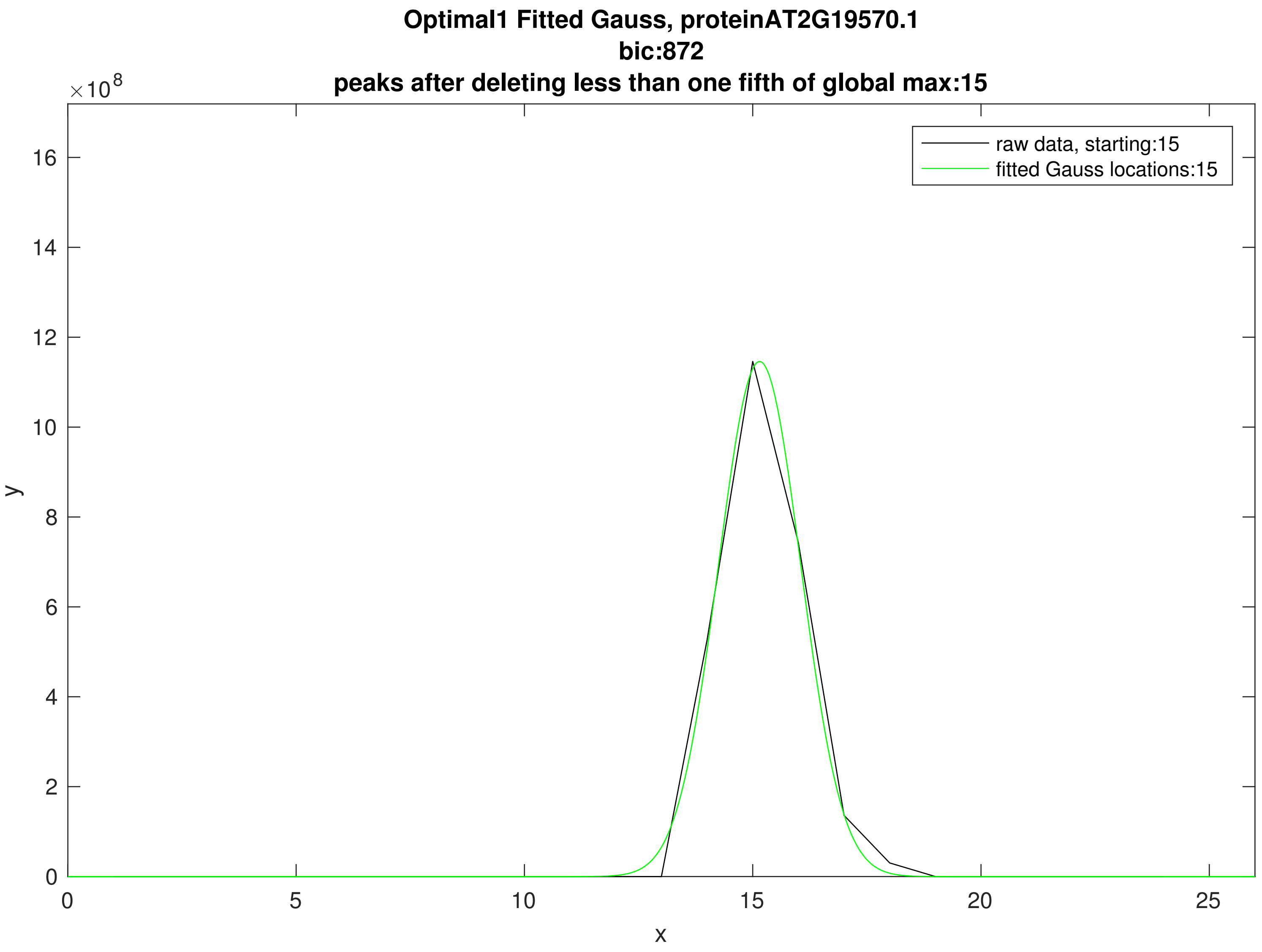

Optimal1 Fitted Gauss, proteinAT2G19570.1
bic:872
peaks after deleting less than one fifth of global max:15
×10^8
raw data, starting:15
fitted Gauss locations:15
y
x
0
2
4
6
8
10
12
14
16
0
5
10
15
20
25

**Optimal1 Fitted Gauss, proteinAT2G19590.1**
**bic:669**
**peaks after deleting less than one fifth of global max:14**

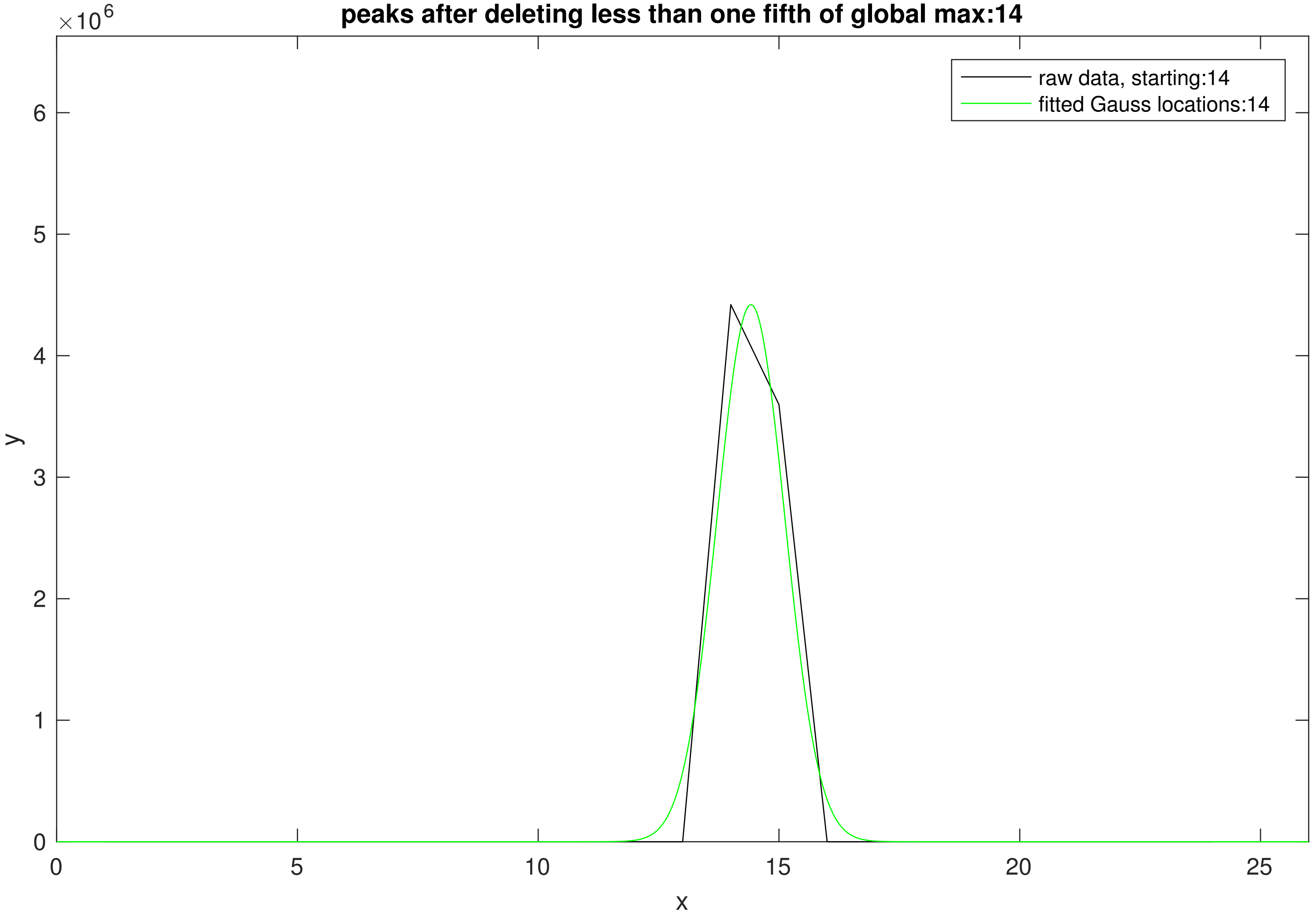

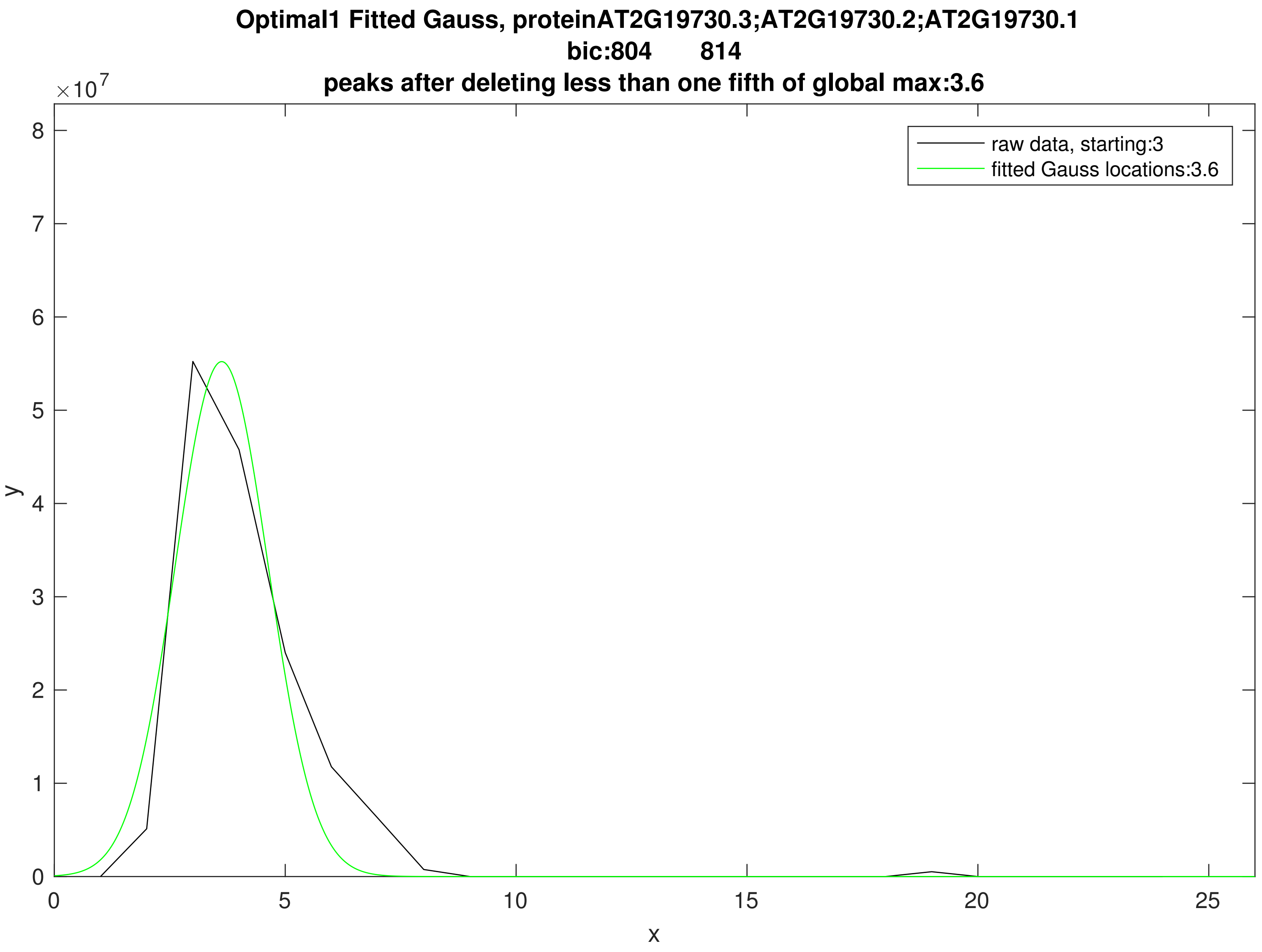
Optimal1 Fitted Gauss, proteinAT2G19730.3;AT2G19730.2;AT2G19730.1
bic:804 814
peaks after deleting less than one fifth of global max:3.6
×10^7
raw data, starting:3
fitted Gauss locations:3.6
y
x
0
1
2
3
4
5
6
7
8
0
5
10
15
20
25

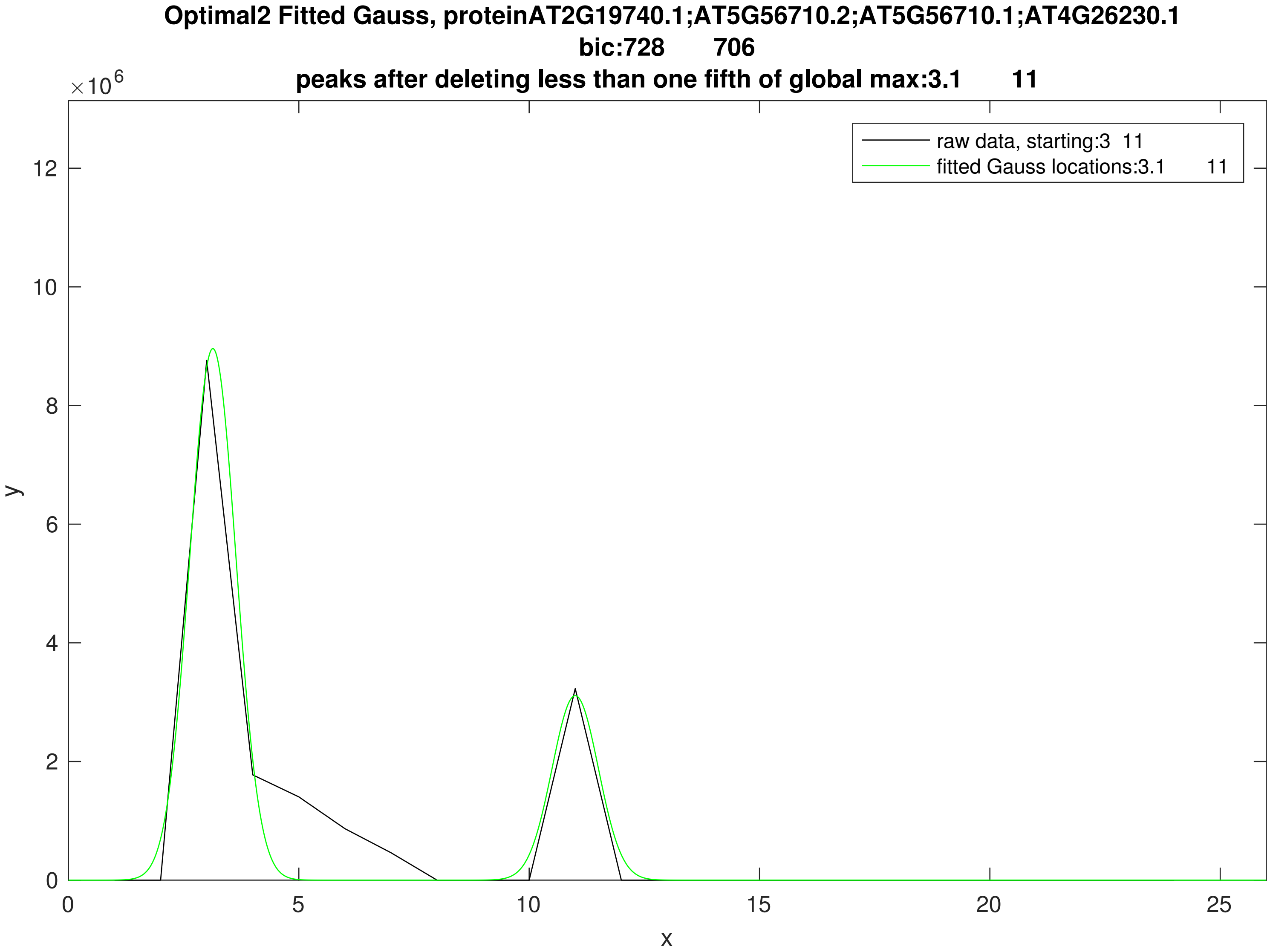

Optimal2 Fitted Gauss, proteinAT2G19740.1;AT5G56710.2;AT5G56710.1;AT4G26230.1
bic:728 706
peaks after deleting less than one fifth of global max:3.1 11
×10^6
raw data, starting:3 11
fitted Gauss locations:3.1 11
y
x
0
2
4
6
8
10
12
5
15
20
25

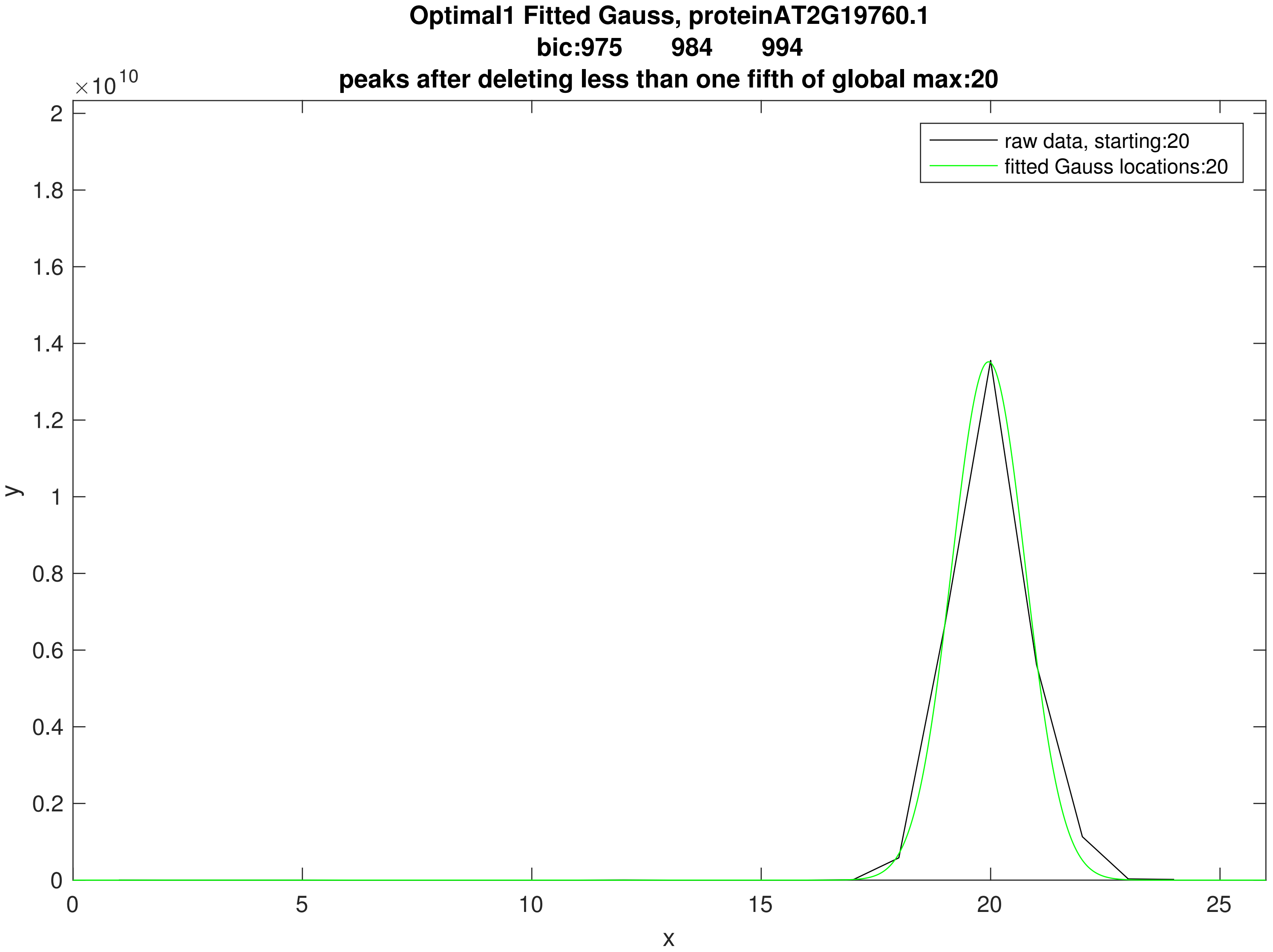
Optimal1 Fitted Gauss, proteinAT2G19760.1
bic:975 984 994
peaks after deleting less than one fifth of global max:20
×10^10
raw data, starting:20
fitted Gauss locations:20
y
x
0
0.2
0.4
0.6
0.8
1
1.2
1.4
1.6
1.8
2
0
5
10
15
20
25

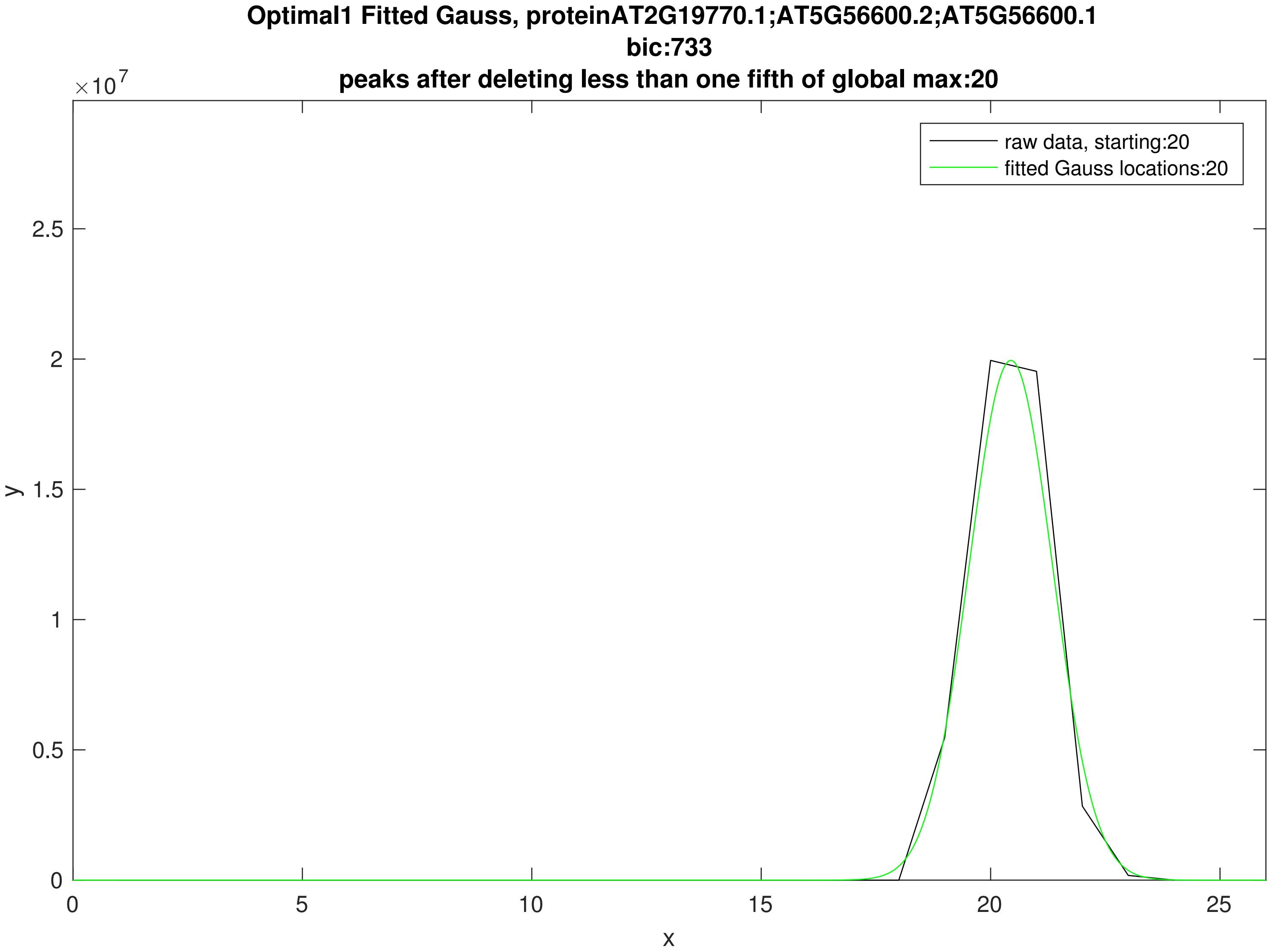
Optimal1 Fitted Gauss, proteinAT2G19770.1;AT5G56600.2;AT5G56600.1
bic:733
peaks after deleting less than one fifth of global max:20
×10^7
raw data, starting:20
fitted Gauss locations:20
y
x
0
0.5
1
1.5
2
2.5
0
5
10
15
20
25

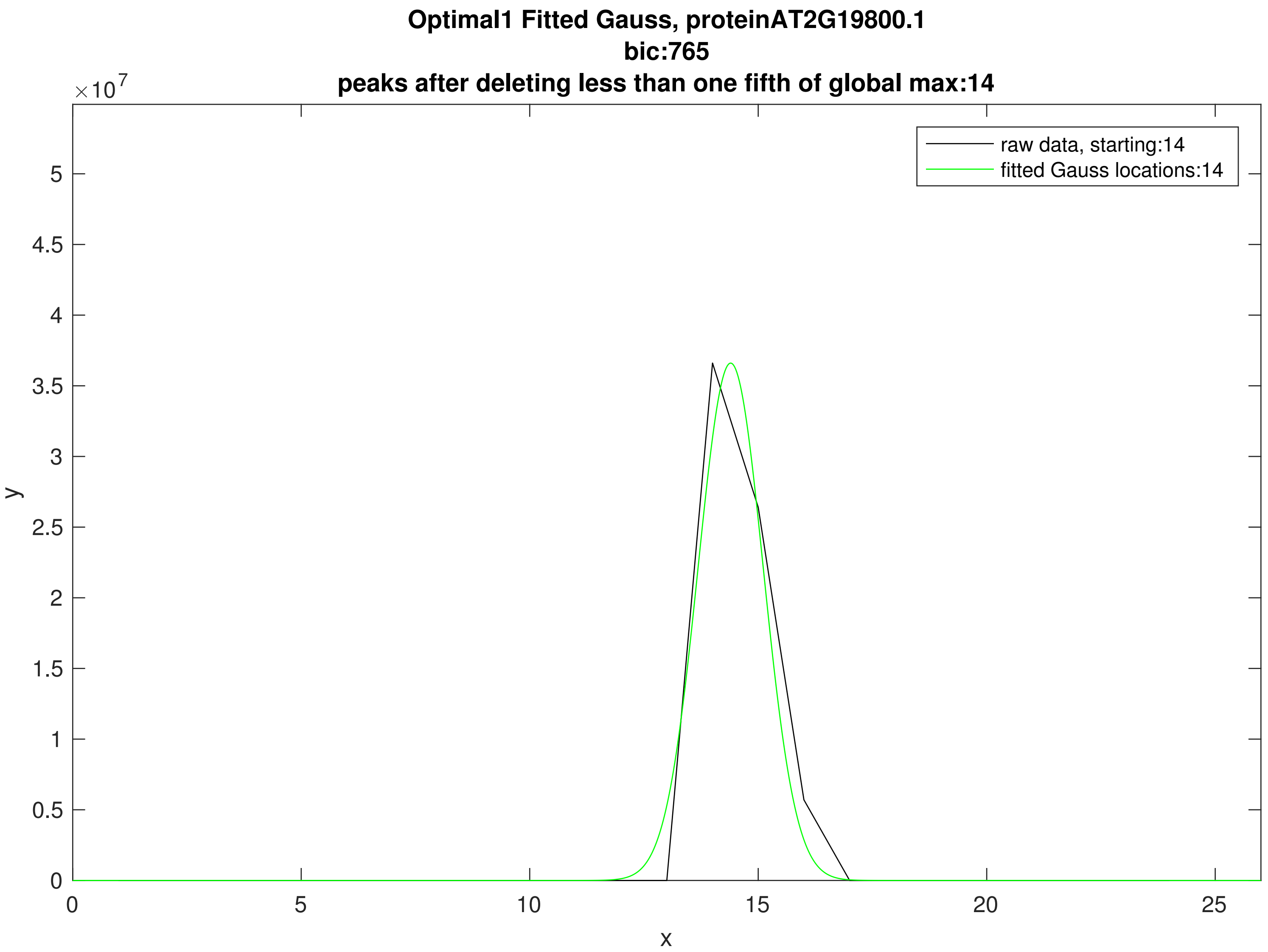
Optimal1 Fitted Gauss, proteinAT2G19800.1
bic:765
peaks after deleting less than one fifth of global max:14
raw data, starting:14
fitted Gauss locations:14
y
x

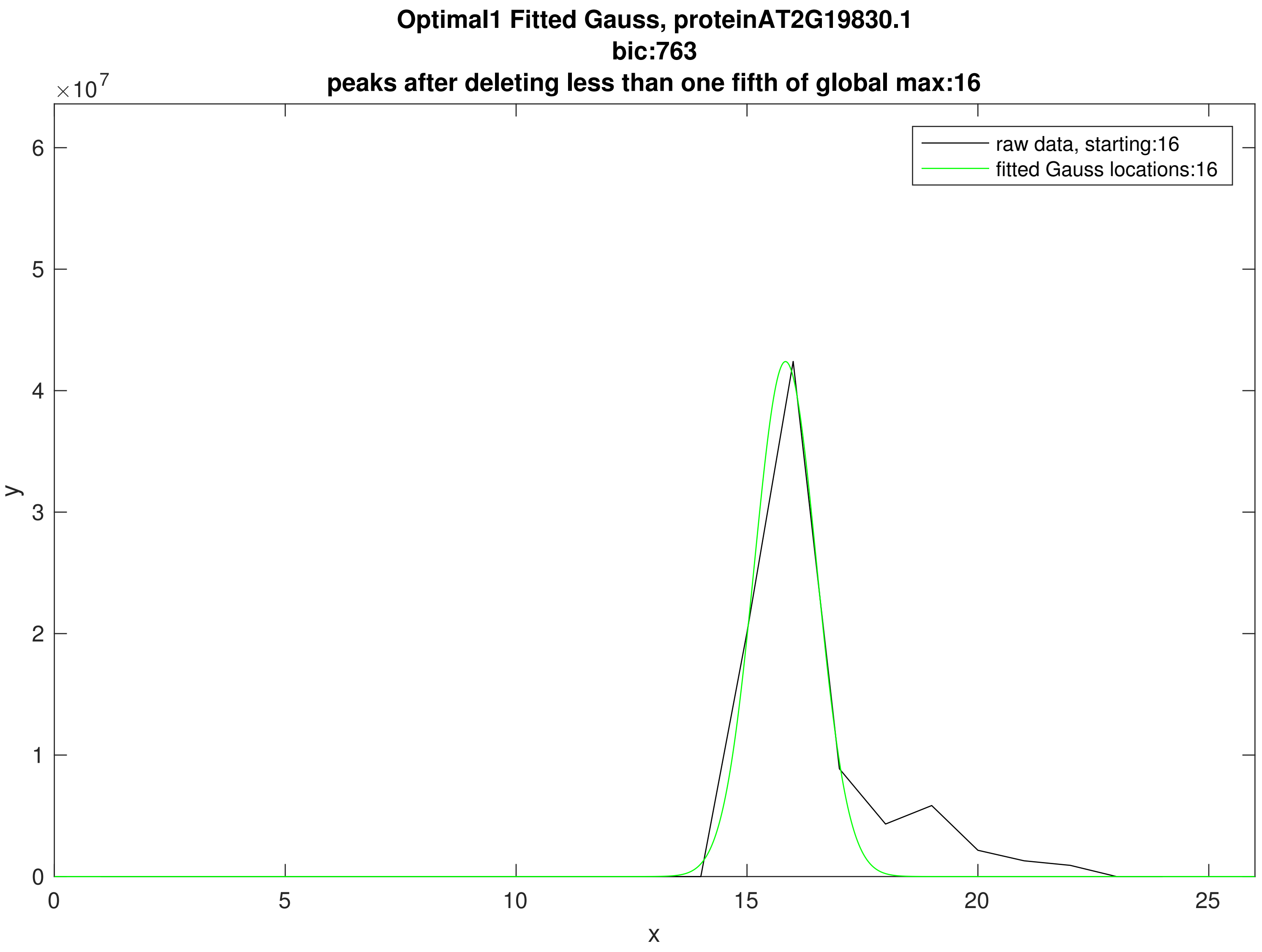
Optimal1 Fitted Gauss, proteinAT2G19830.1
bic:763
peaks after deleting less than one fifth of global max:16
×10^7
raw data, starting:16
fitted Gauss locations:16
y
x
0
1
2
3
4
5
6
0
5
10
15
20
25

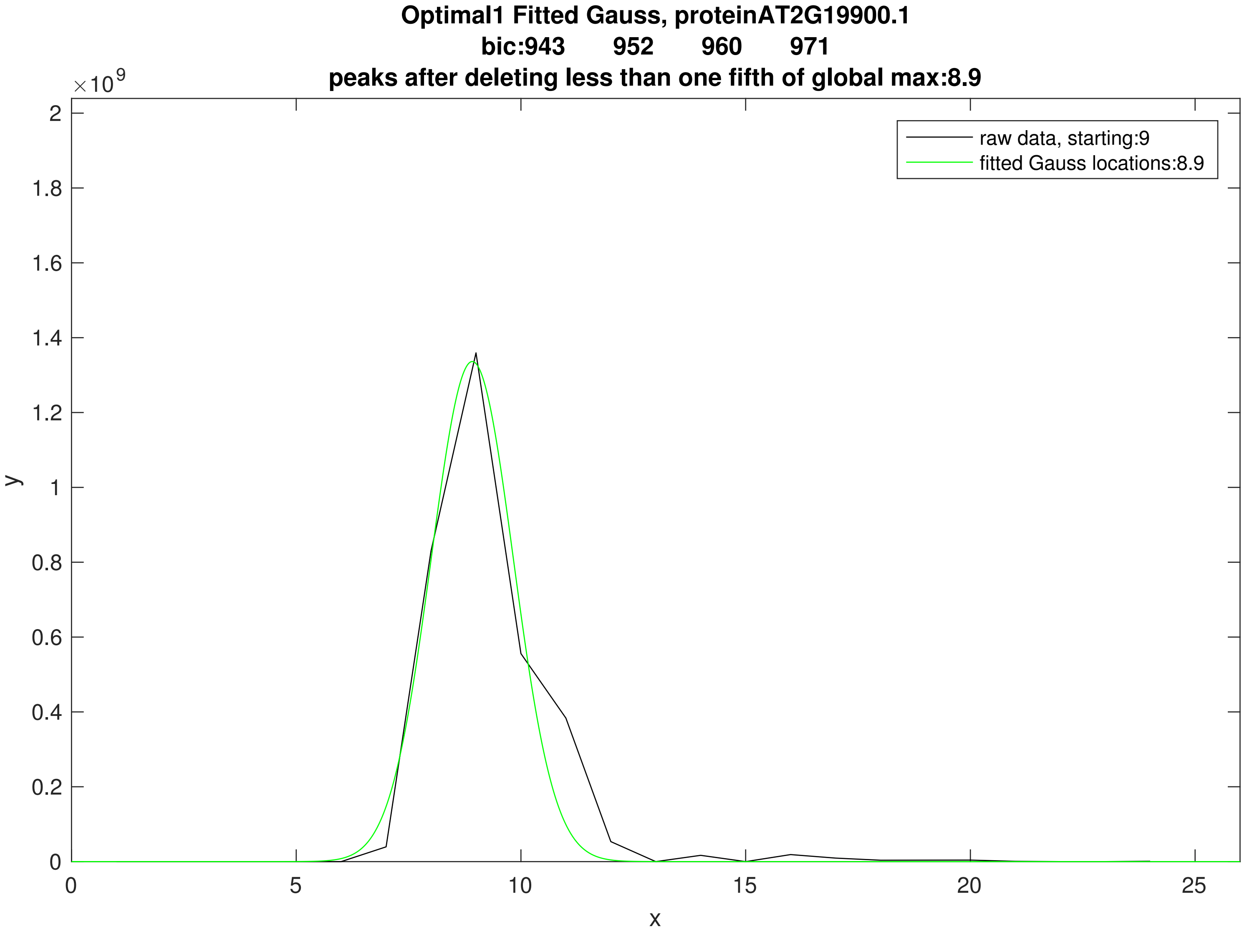
Optimal1 Fitted Gauss, proteinAT2G19900.1
bic:943 952 960 971
peaks after deleting less than one fifth of global max:8.9
×10^9
raw data, starting:9
fitted Gauss locations:8.9
y
x
0
0.2
0.4
0.6
0.8
1
1.2
1.4
1.6
1.8
2
5
10
15
20
25

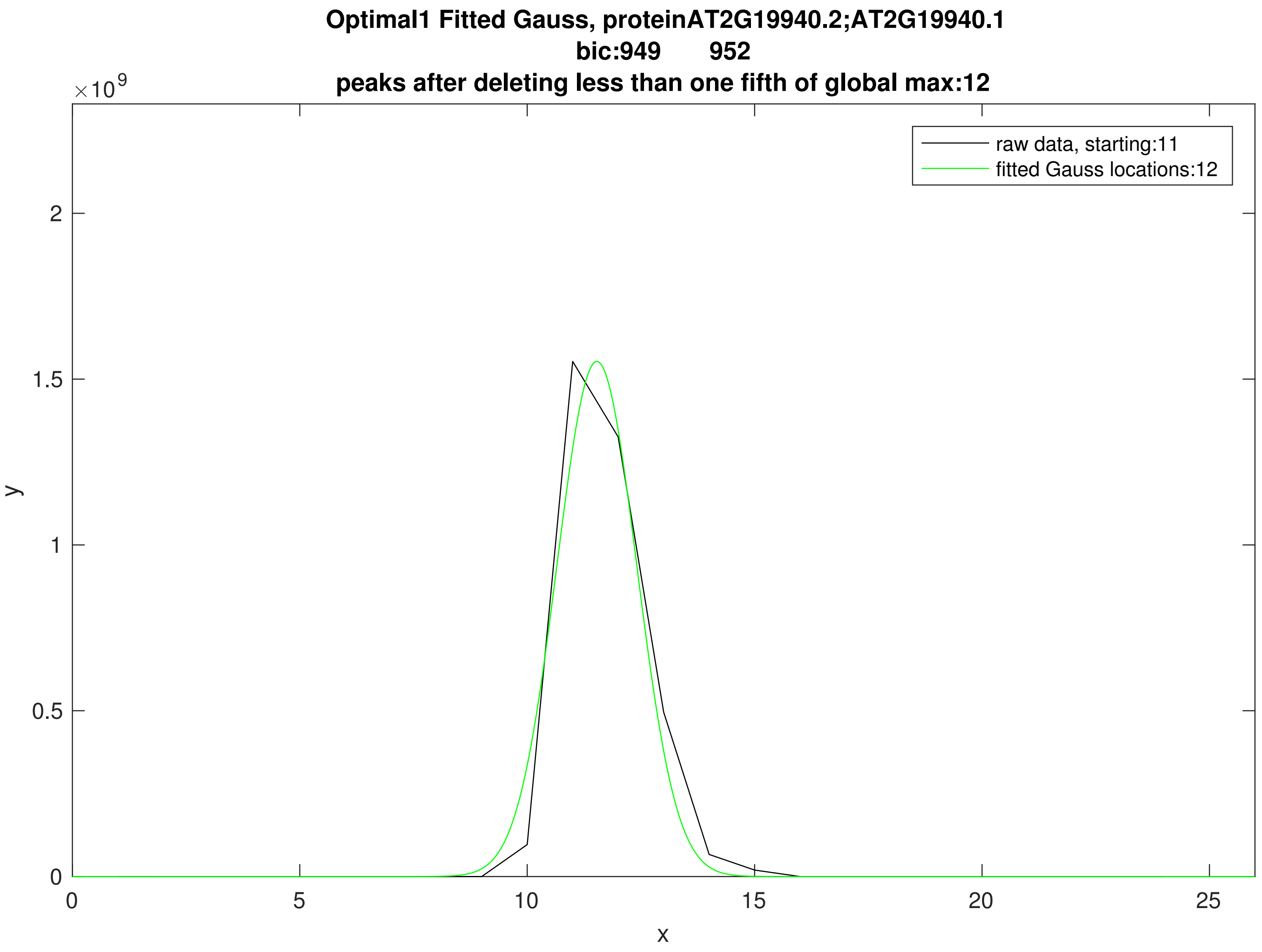
Optimal1 Fitted Gauss, proteinAT2G19940.2;AT2G19940.1
bic:949 952
peaks after deleting less than one fifth of global max:12
×10^9
raw data, starting:11
fitted Gauss locations:12
y
x
0
0.5
1
1.5
2
5
10
15
20
25

**Optimal1 Fitted Gauss, proteinAT2G20260.1**
**bic:738 745 753**
**peaks after deleting less than one fifth of global max:20**

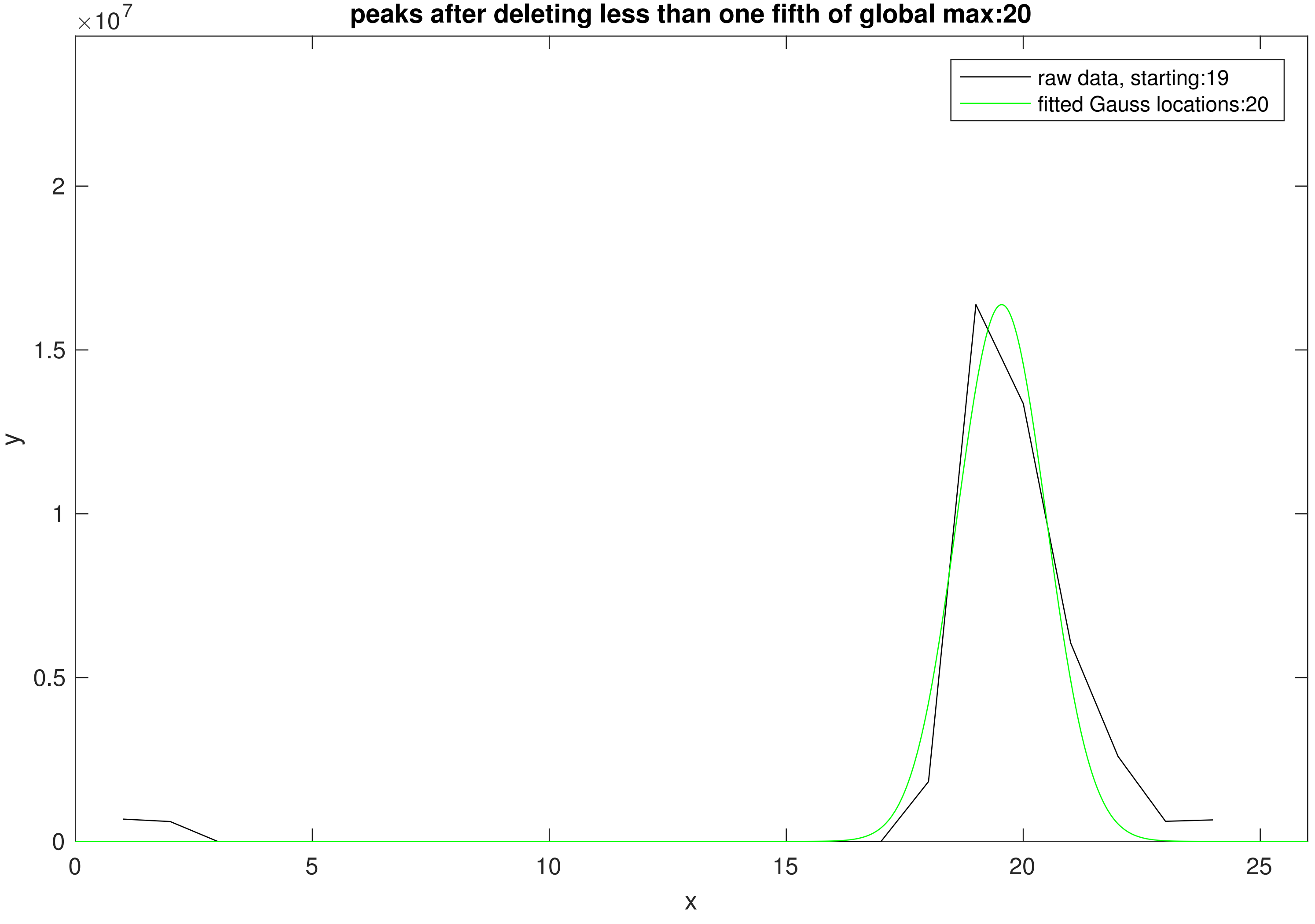

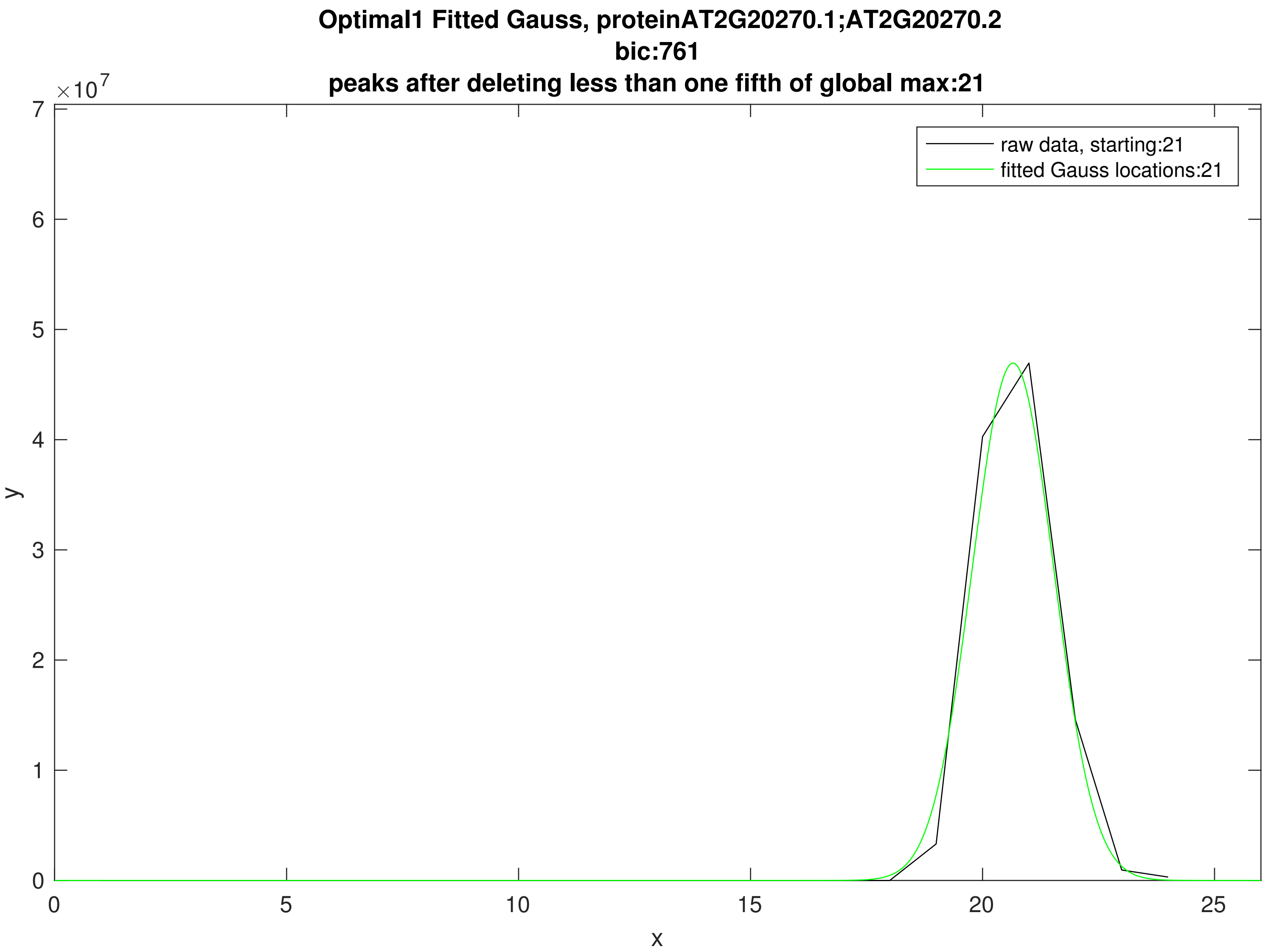

Optimal1 Fitted Gauss, proteinAT2G20270.1;AT2G20270.2
bic:761
peaks after deleting less than one fifth of global max:21
×10^7
raw data, starting:21
fitted Gauss locations:21
y
x
0
1
2
3
4
5
6
7
0
5
10
15
20
25

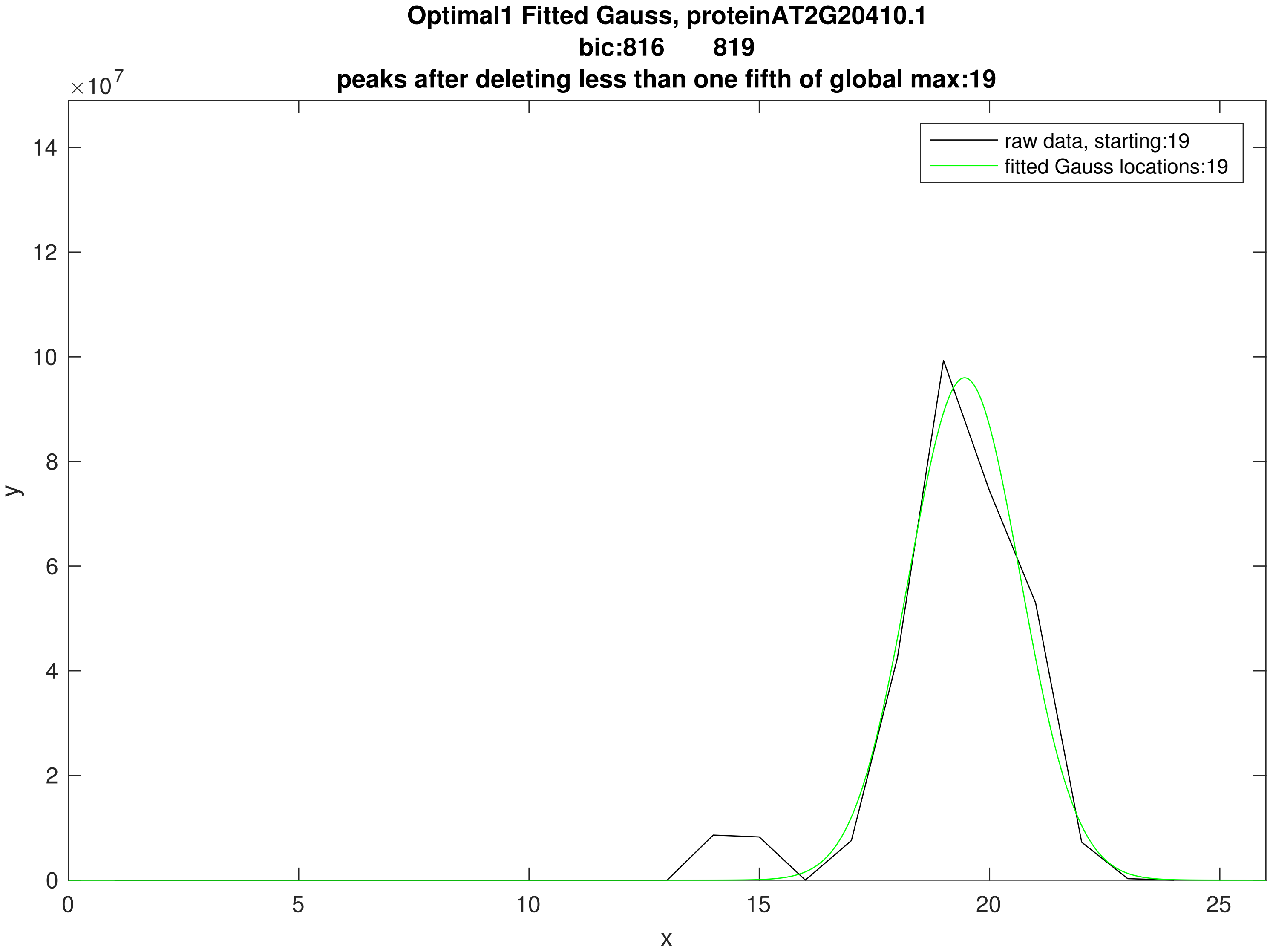

Optimal1 Fitted Gauss, proteinAT2G20410.1
bic:816 819
peaks after deleting less than one fifth of global max:19
×10^7
raw data, starting:19
fitted Gauss locations:19
y
x
0
2
4
6
8
10
12
14
0
5
10
15
20
25

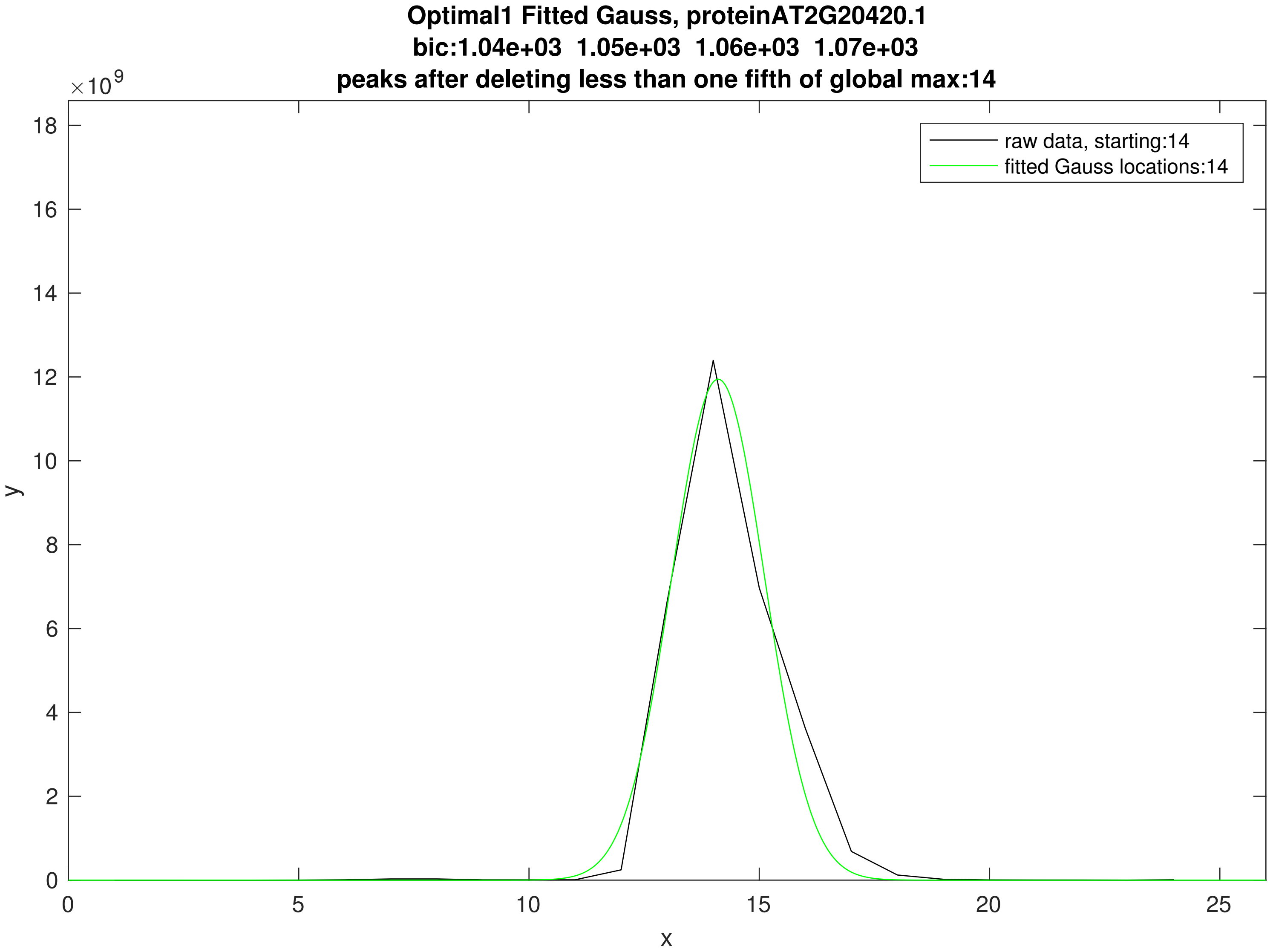

Optimal1 Fitted Gauss, proteinAT2G20420.1
bic:1.04e+03 1.05e+03 1.06e+03 1.07e+03
peaks after deleting less than one fifth of global max:14
×10^9
raw data, starting:14
fitted Gauss locations:14
y
x
0
2
4
6
8
10
12
14
16
18
0
5
10
15
20
25

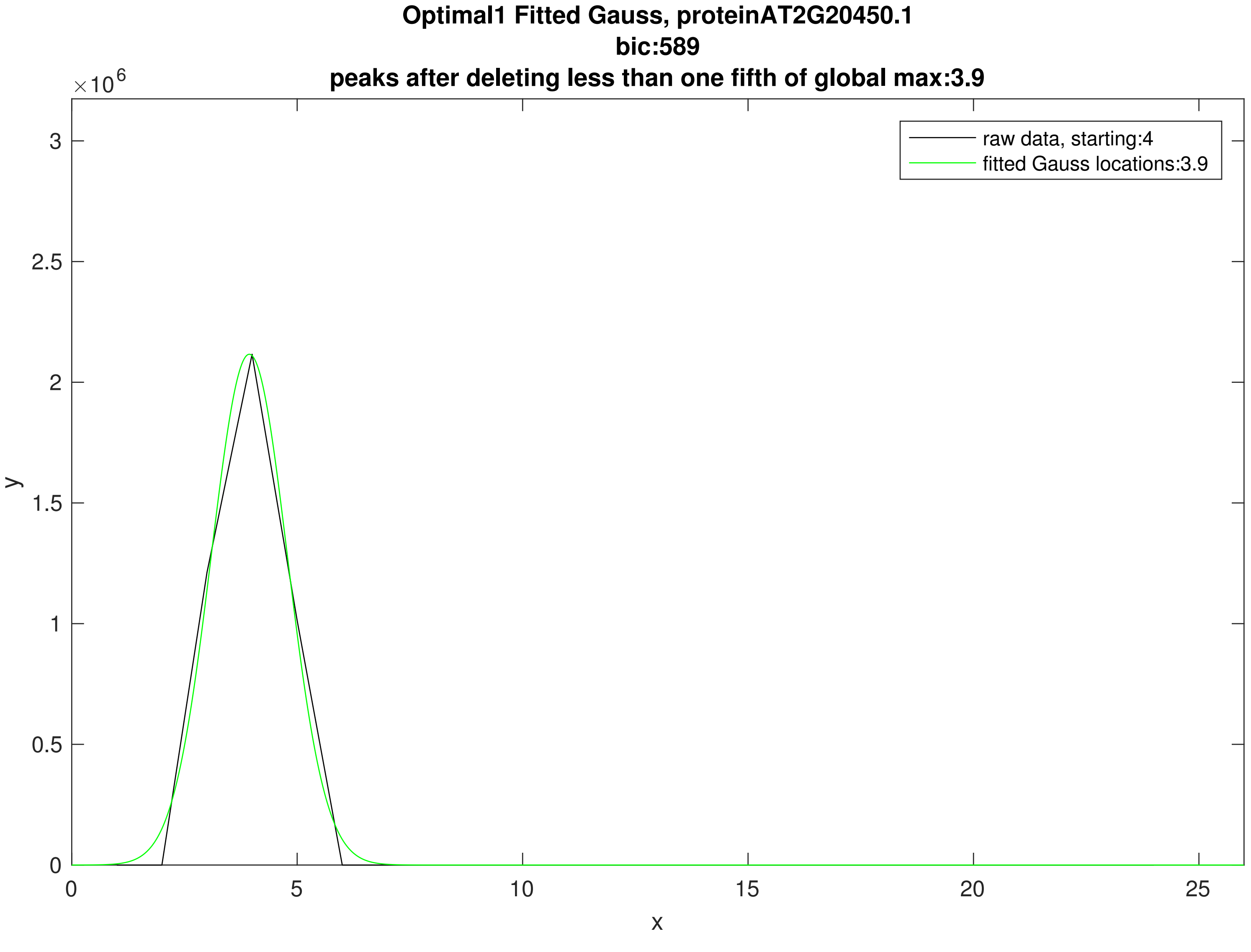

Optimal1 Fitted Gauss, proteinAT2G20450.1
bic:589
peaks after deleting less than one fifth of global max:3.9
×10^6
raw data, starting:4
fitted Gauss locations:3.9
y
x
0
0.5
1
1.5
2
2.5
3
0
5
10
15
20
25

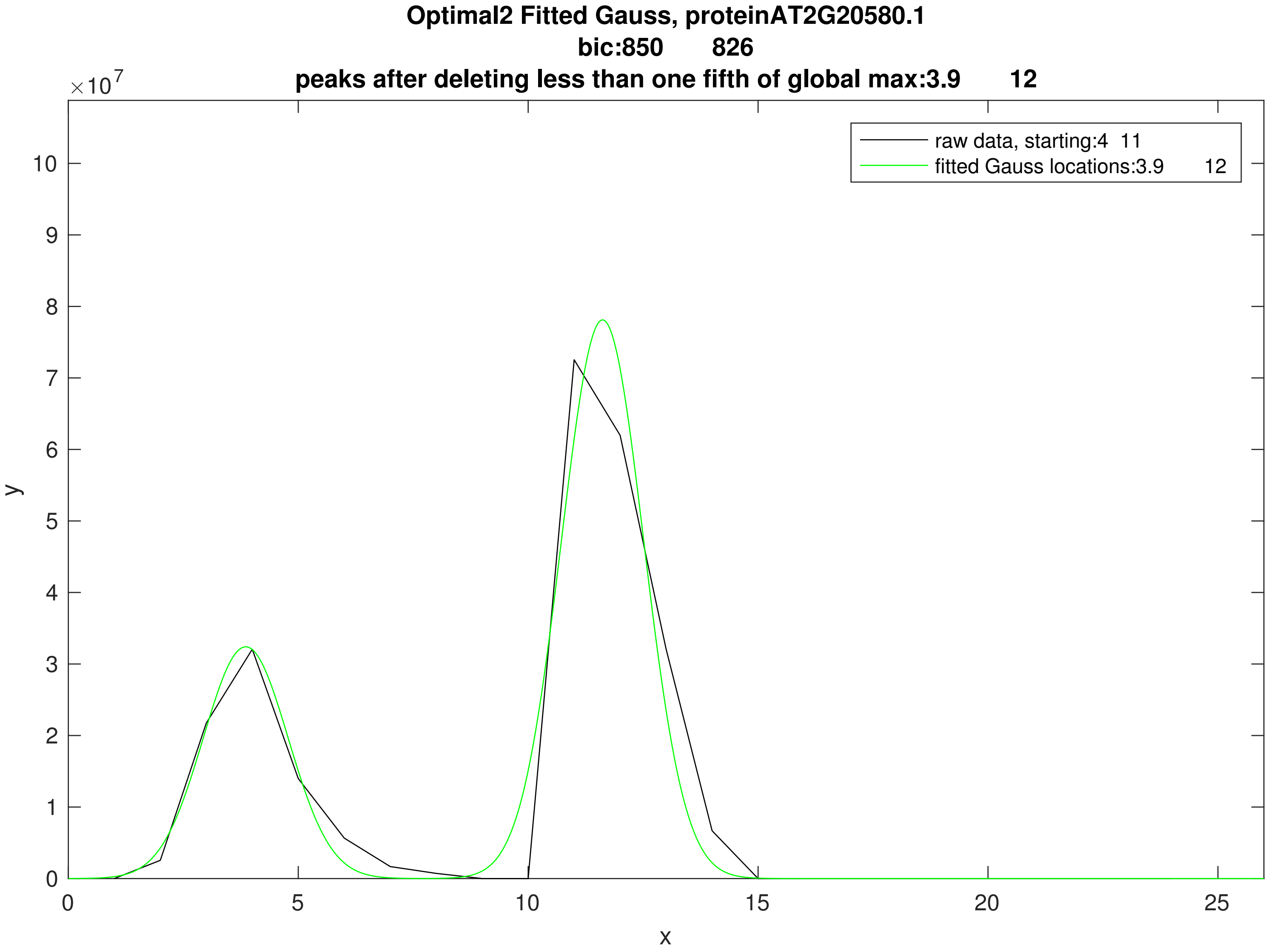
Optimal2 Fitted Gauss, proteinAT2G20580.1
bic:850 826
peaks after deleting less than one fifth of global max:3.9 12
×10^7
raw data, starting:4 11
fitted Gauss locations:3.9 12
y
x
0
1
2
3
4
5
6
7
8
9
10
0
5
10
15
20
25

**Optimal1 Fitted Gauss, proteinAT2G20610.1;AT2G20610.2**
**bic:956 966 975 985**
**peaks after deleting less than one fifth of global max:11**

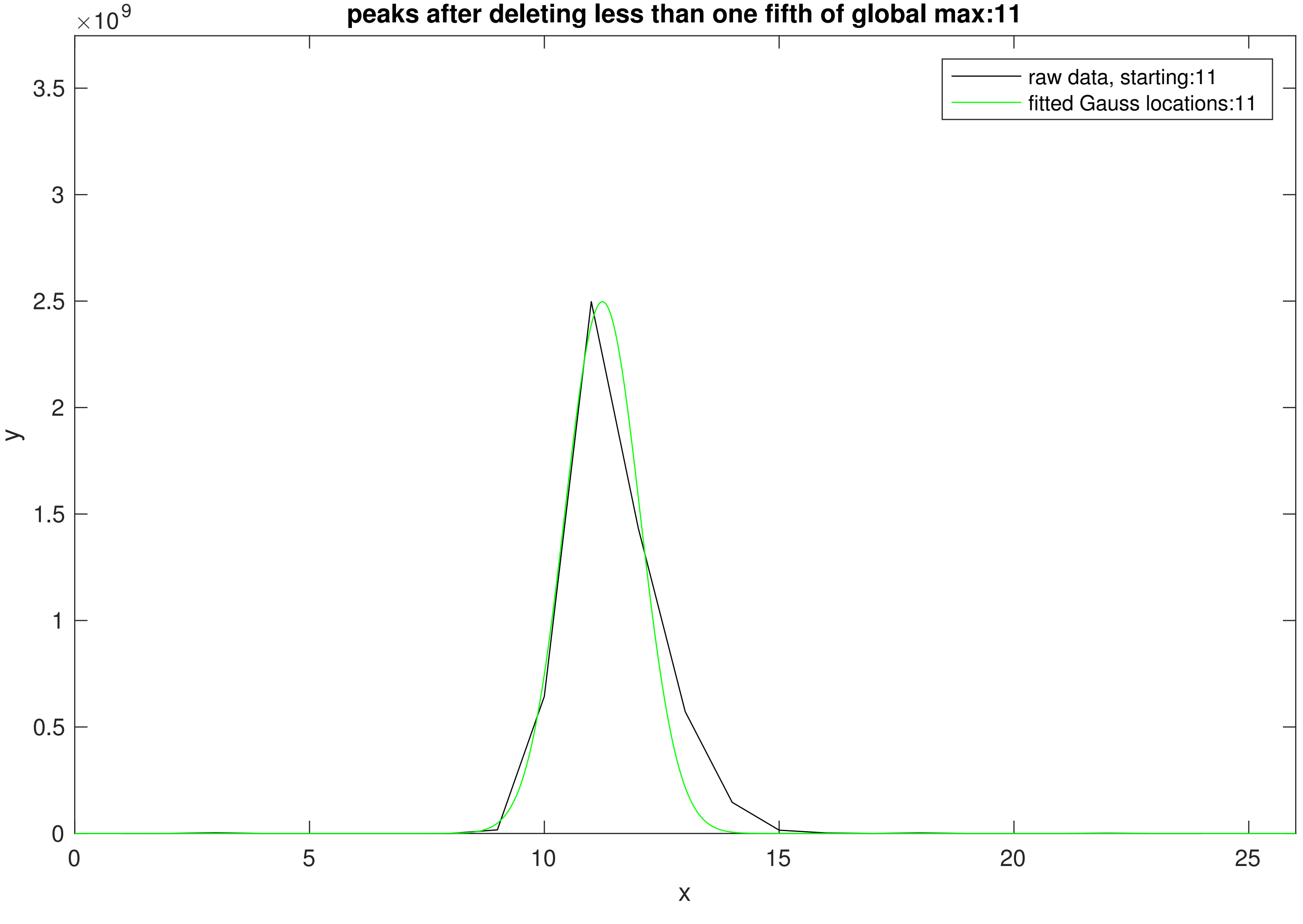

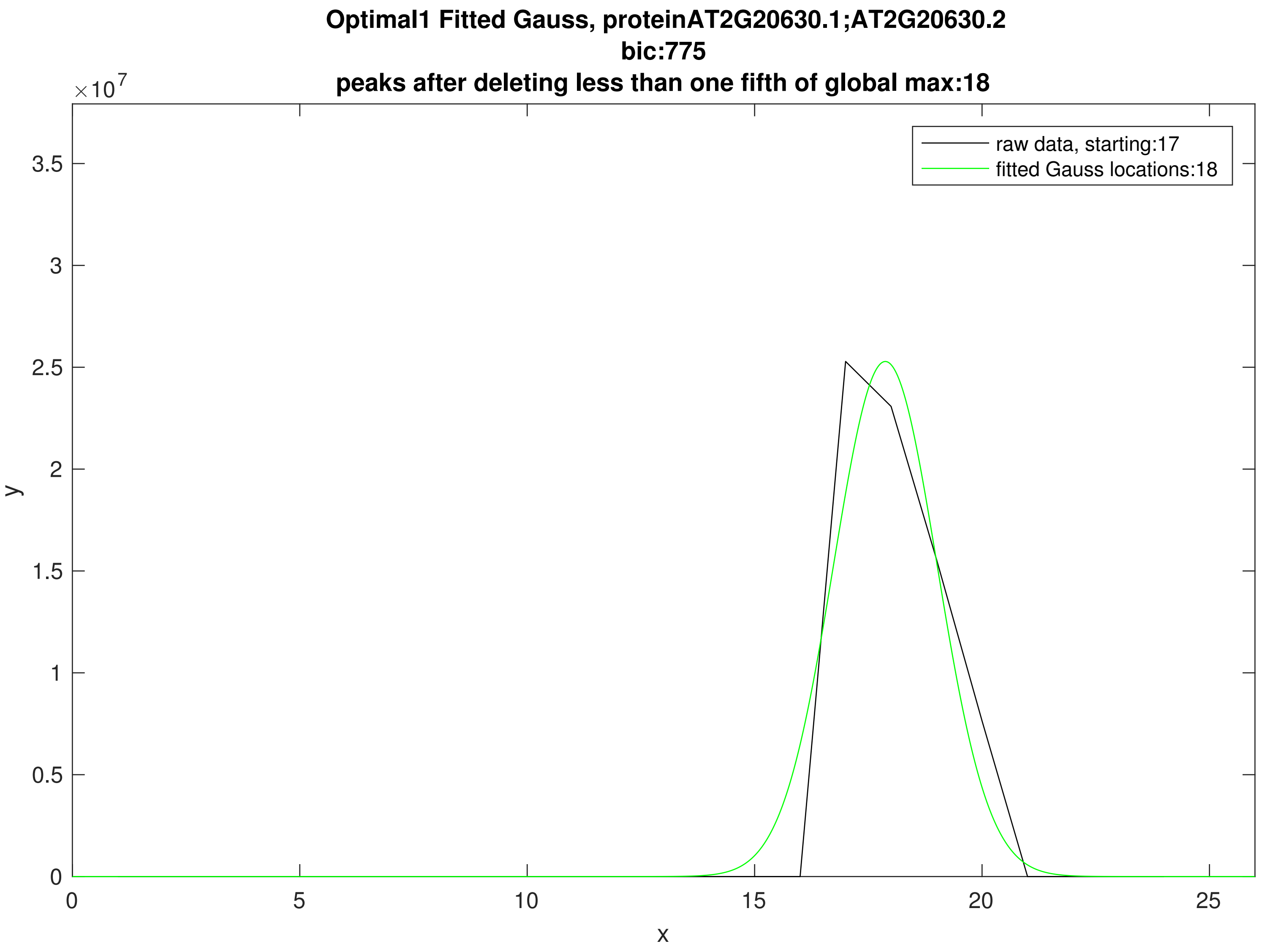
Optimal1 Fitted Gauss, proteinAT2G20630.1;AT2G20630.2
bic:775
peaks after deleting less than one fifth of global max:18
×10^7
raw data, starting:17
fitted Gauss locations:18
3.5
3
2.5
2
1.5
1
0.5
0
y
0
5
10
15
20
25
x

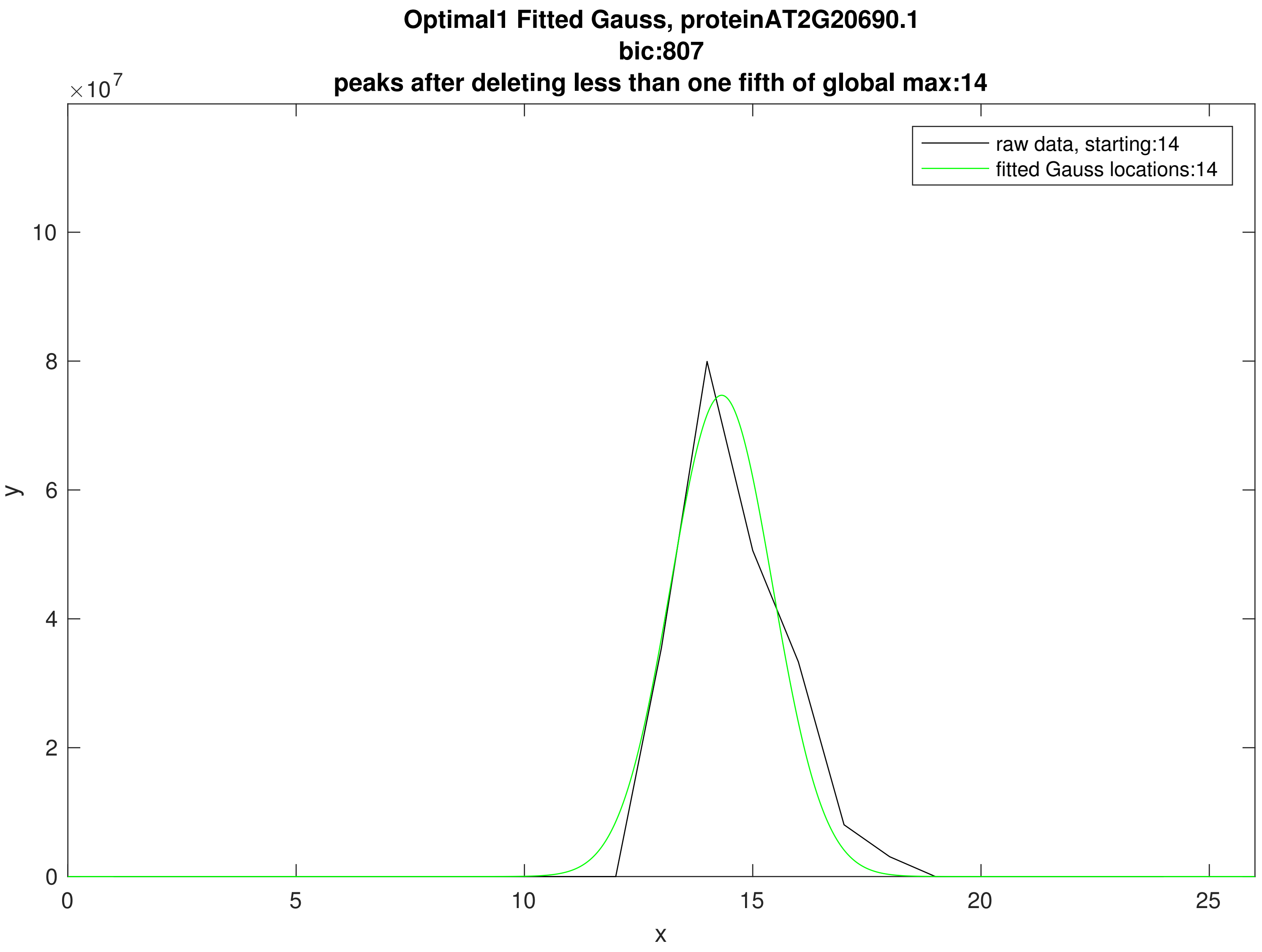

Optimal1 Fitted Gauss, proteinAT2G20690.1
bic:807
peaks after deleting less than one fifth of global max:14
×10^7
raw data, starting:14
fitted Gauss locations:14
y
x
0
2
4
6
8
10
0
5
10
15
20
25

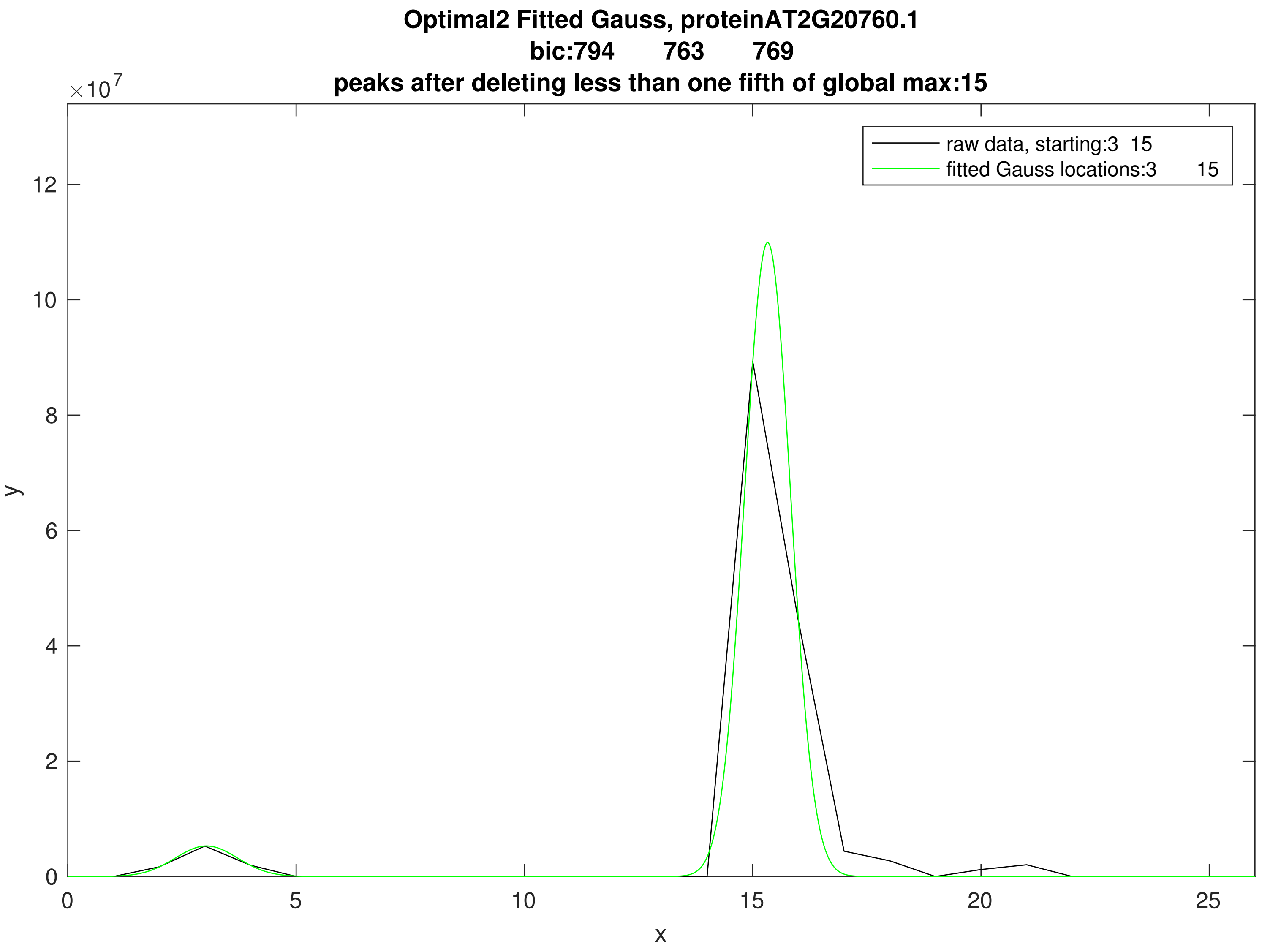

Optimal2 Fitted Gauss, proteinAT2G20760.1
bic:794 763 769
peaks after deleting less than one fifth of global max:15
×10^7
raw data, starting:3 15
fitted Gauss locations:3 15
y
x
0
2
4
6
8
10
12
0
5
10
15
20
25

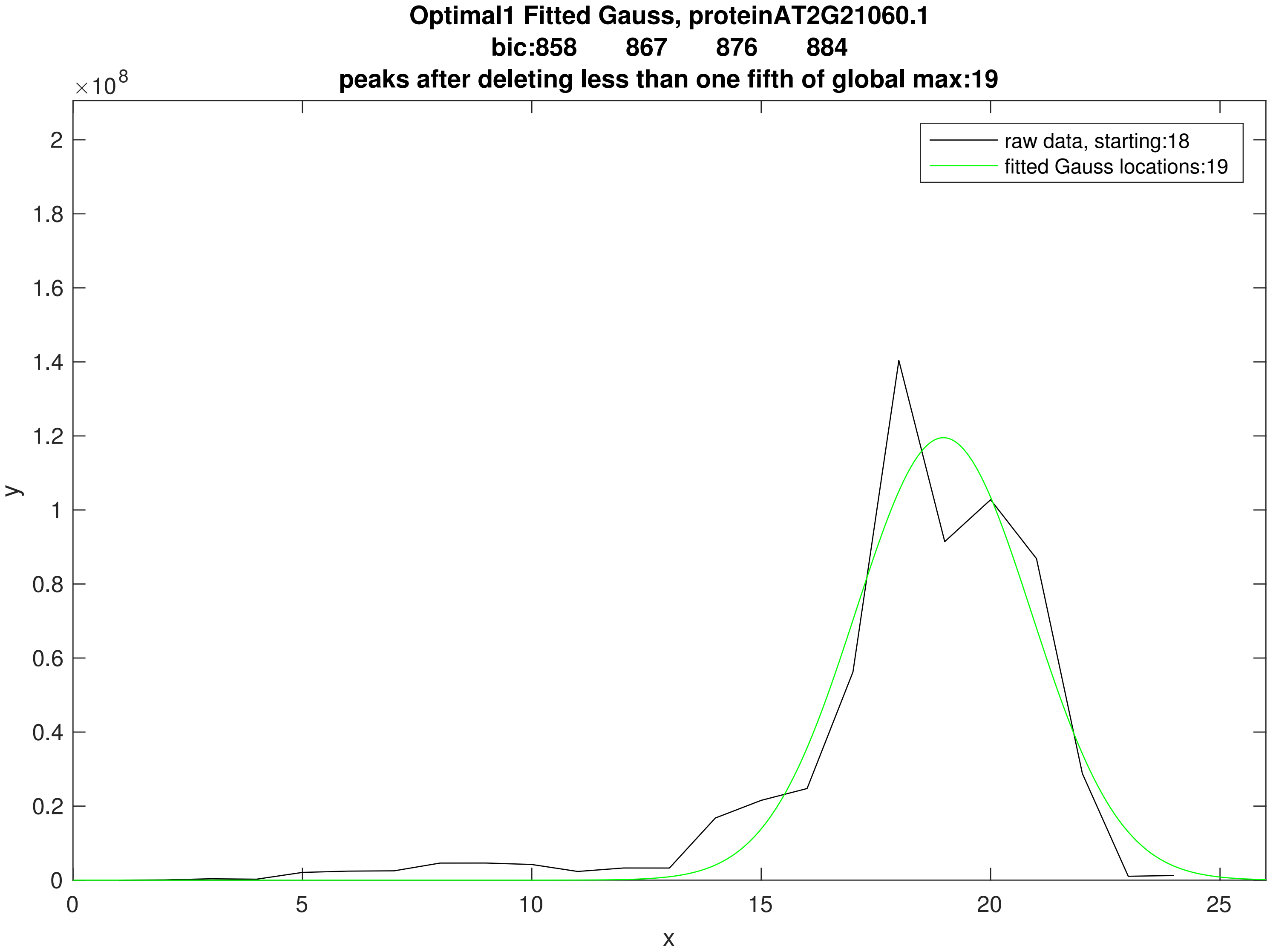
Optimal1 Fitted Gauss, proteinAT2G21060.1
bic:858 867 876 884
peaks after deleting less than one fifth of global max:19
×10^8
raw data, starting:18
fitted Gauss locations:19
y
x
0
0.2
0.4
0.6
0.8
1
1.2
1.4
1.6
1.8
2
0
5
10
15
20
25

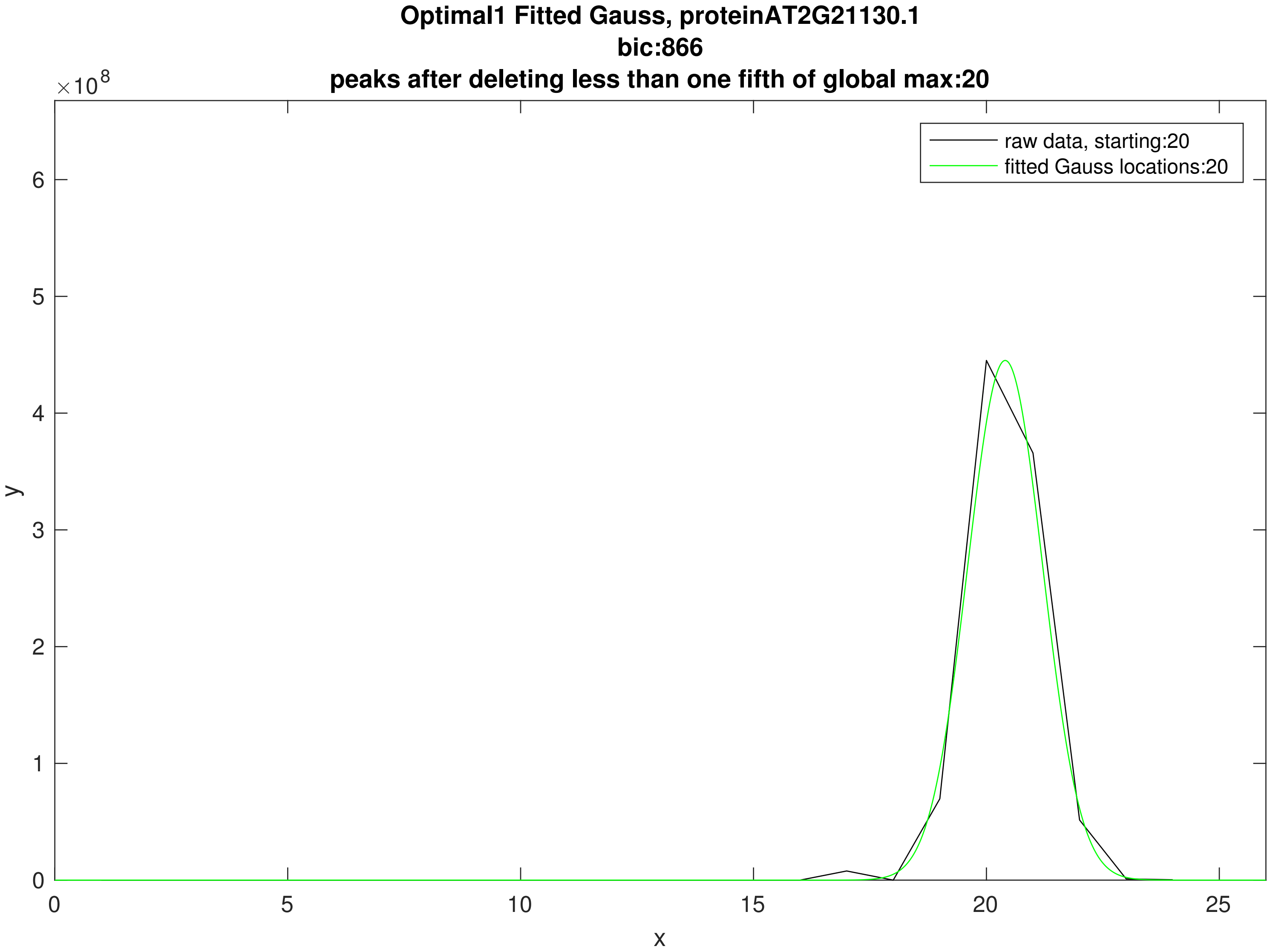

Optimal1 Fitted Gauss, proteinAT2G21130.1
bic:866
peaks after deleting less than one fifth of global max:20
×10^8
raw data, starting:20
fitted Gauss locations:20
6
5
4
3
2
1
0
y
0
5
10
15
20
25
x

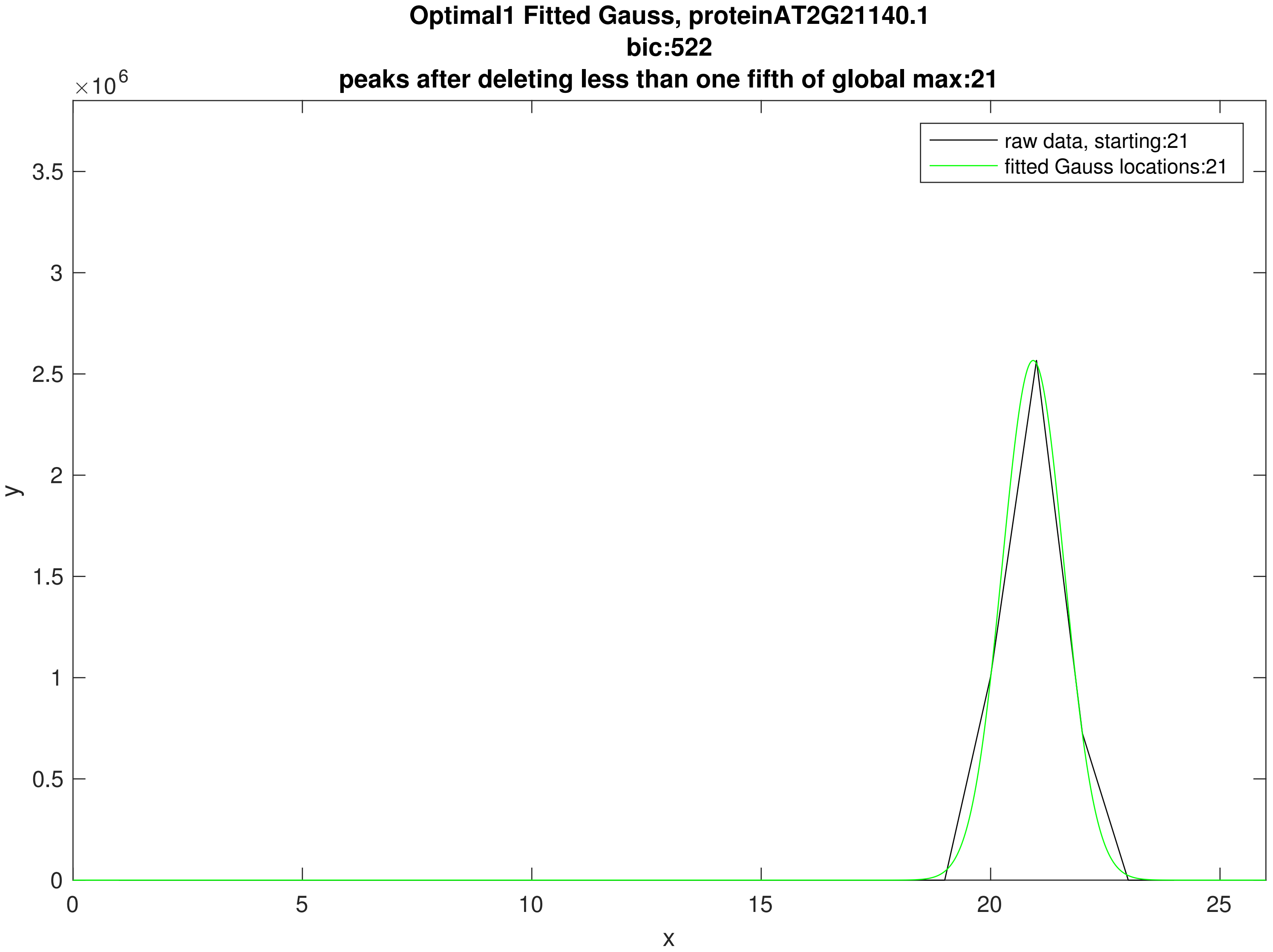
Optimal1 Fitted Gauss, proteinAT2G21140.1
bic:522
peaks after deleting less than one fifth of global max:21
×10^6
raw data, starting:21
fitted Gauss locations:21
y
x
0
0.5
1
1.5
2
2.5
3
3.5
0
5
10
15
20
25

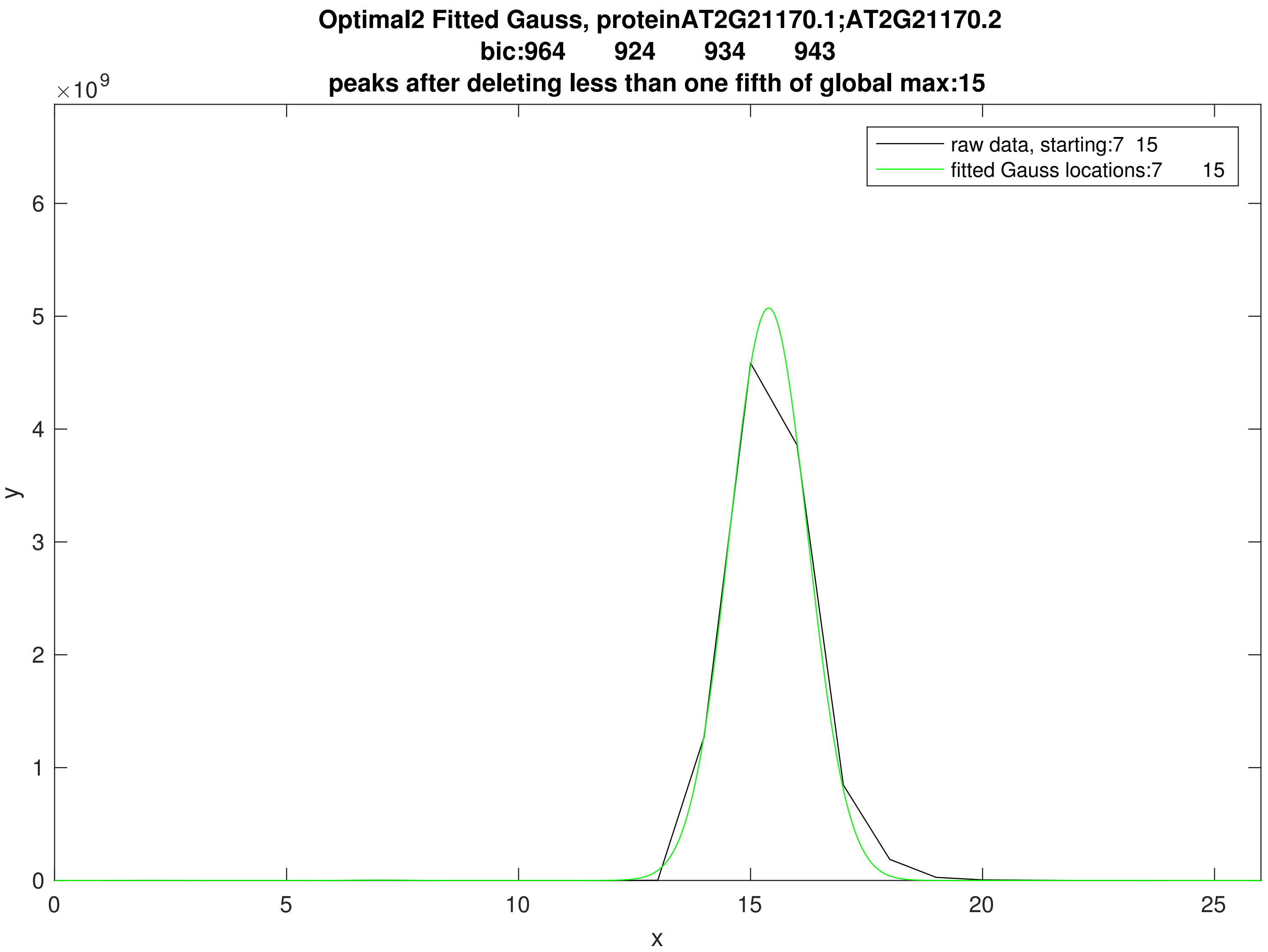

Optimal2 Fitted Gauss, proteinAT2G21170.1;AT2G21170.2
bic:964 924 934 943
peaks after deleting less than one fifth of global max:15
×10^9
raw data, starting:7 15
fitted Gauss locations:7 15
y
x
0
1
2
3
4
5
6
0
5
10
15
20
25

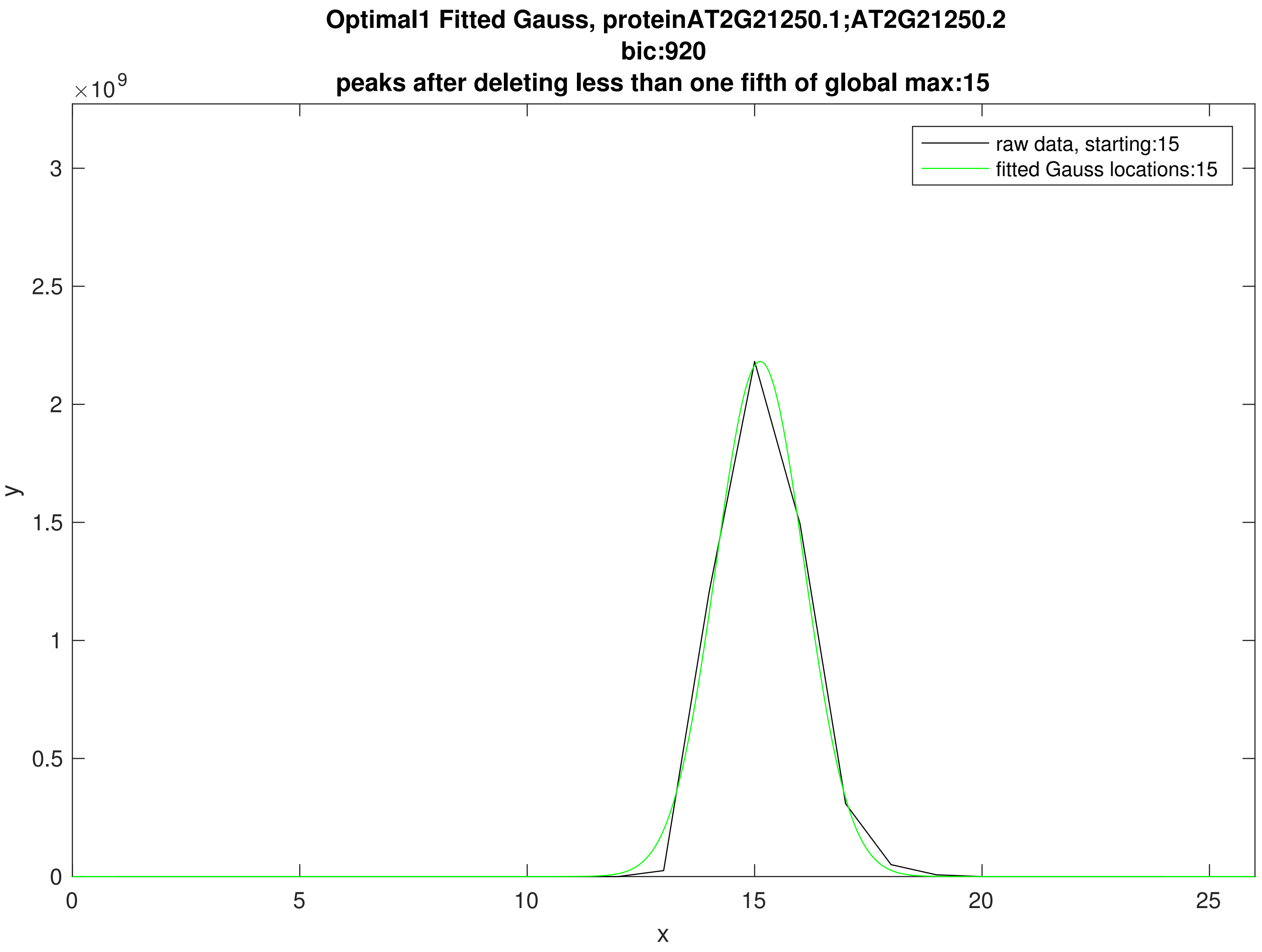
Optimal1 Fitted Gauss, proteinAT2G21250.1;AT2G21250.2
bic:920
peaks after deleting less than one fifth of global max:15
×10^9
3
2.5
2
1.5
1
0.5
0
y
0
5
10
15
20
25
x
raw data, starting:15
fitted Gauss locations:15

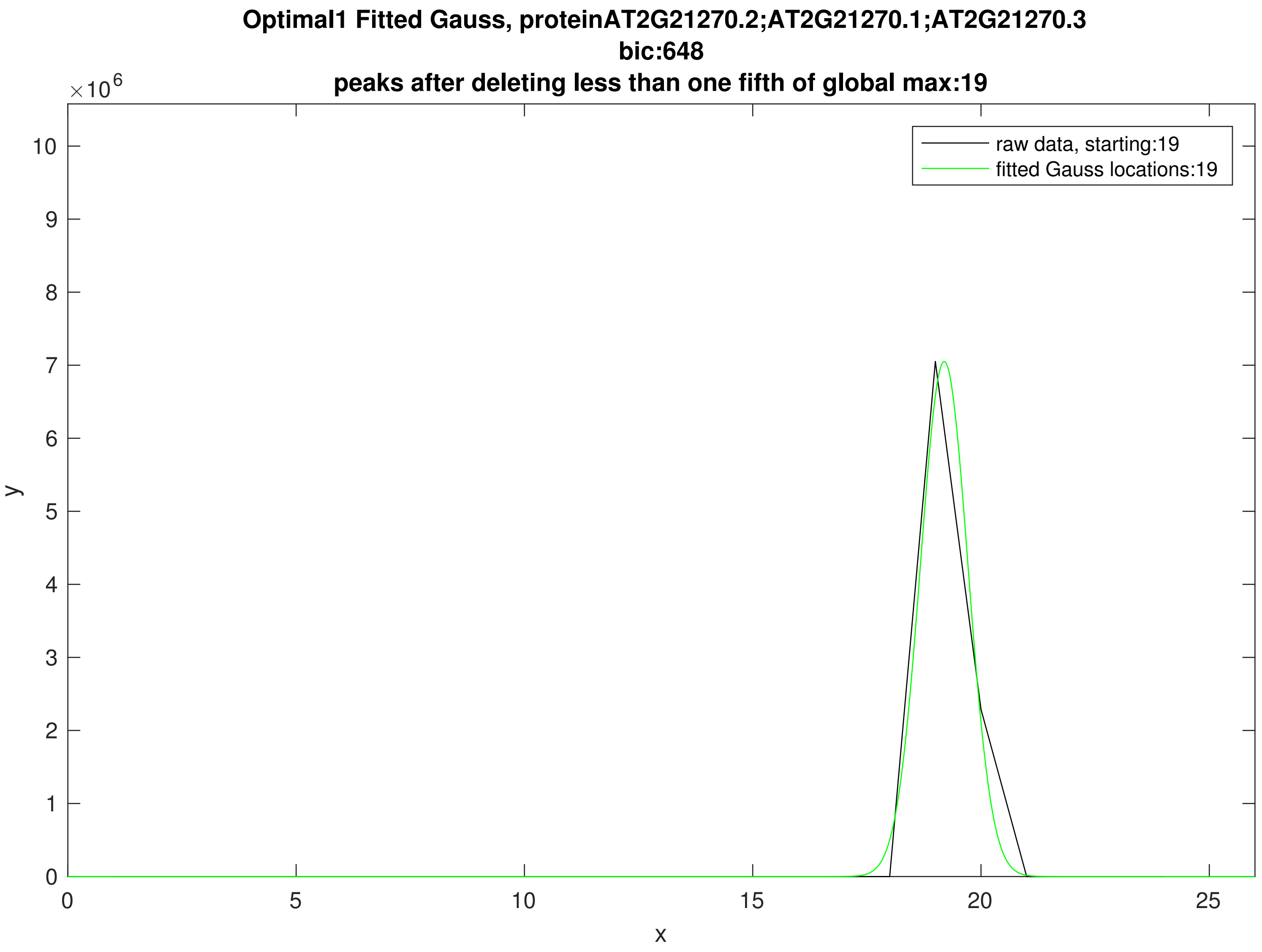

Optimal1 Fitted Gauss, proteinAT2G21270.2;AT2G21270.1;AT2G21270.3
bic:648
peaks after deleting less than one fifth of global max:19
×10^6
raw data, starting:19
fitted Gauss locations:19
y
x
0
1
2
3
4
5
6
7
8
9
10
0
5
10
15
20
25

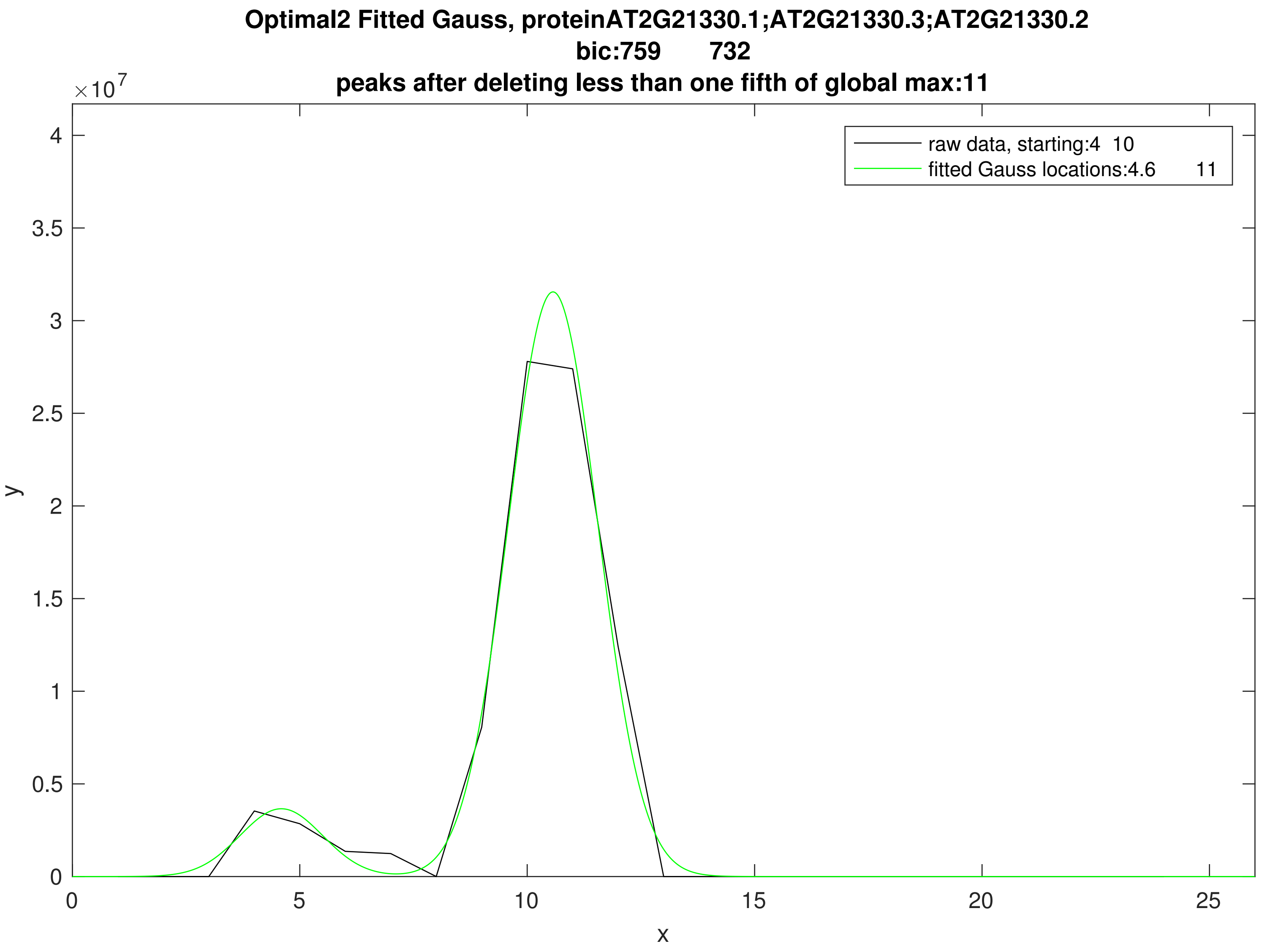

Optimal2 Fitted Gauss, proteinAT2G21330.1;AT2G21330.3;AT2G21330.2
bic:759 732
peaks after deleting less than one fifth of global max:11
×10^7
raw data, starting:4 10
fitted Gauss locations:4.6 11
y
x
0
0.5
1
1.5
2
2.5
3
3.5
4
0
5
10
15
20
25

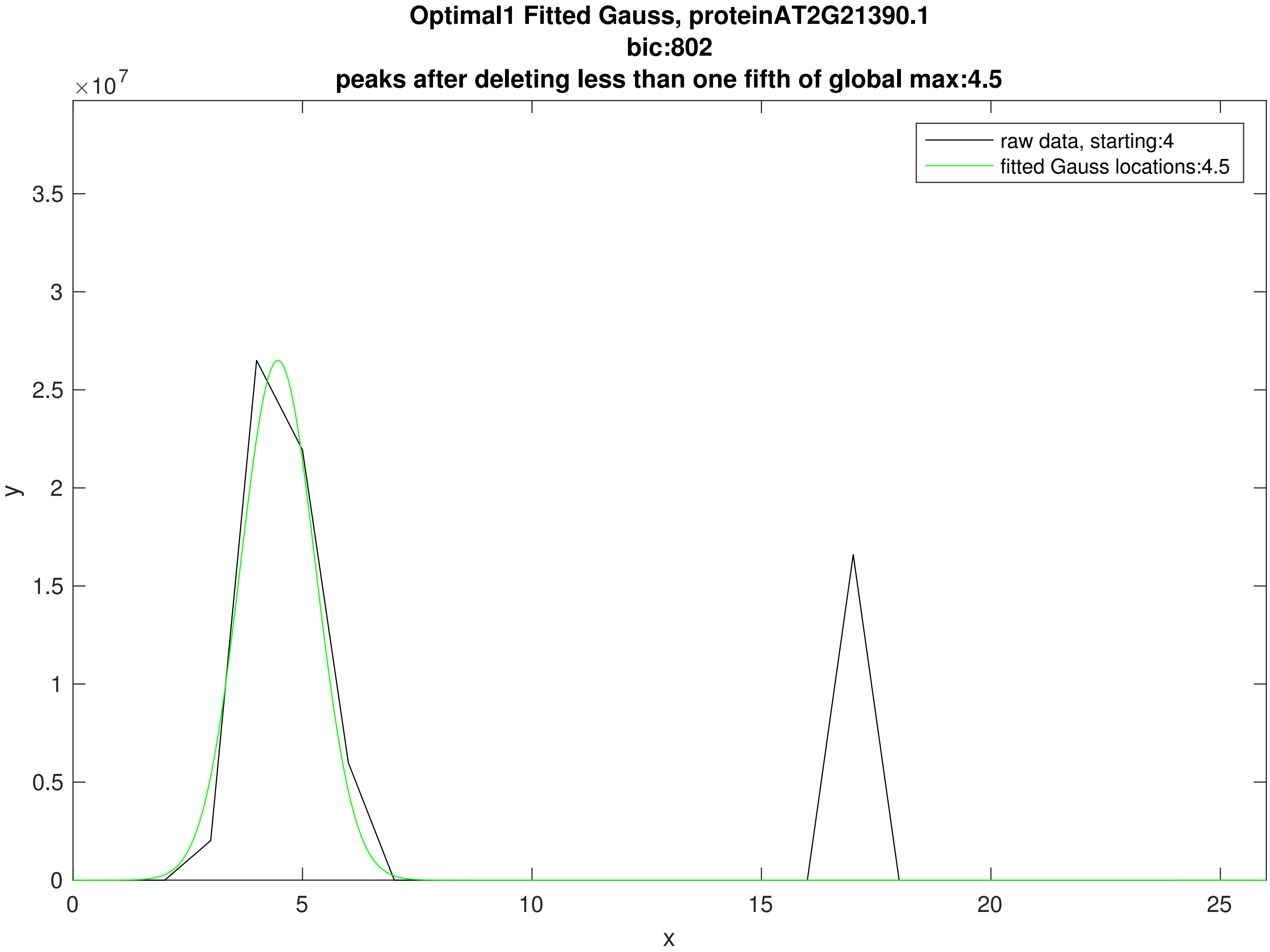

Optimal1 Fitted Gauss, proteinAT2G21390.1
bic:802
peaks after deleting less than one fifth of global max:4.5
×10^7
3.5
3
2.5
2
1.5
1
0.5
0
y
0
5
10
15
20
25
x
raw data, starting:4
fitted Gauss locations:4.5

**Optimal1 Fitted Gauss, proteinAT2G21470.1;AT2G21470.3;AT2G21470.2**
**bic:689**
**peaks after deleting less than one fifth of global max:11**

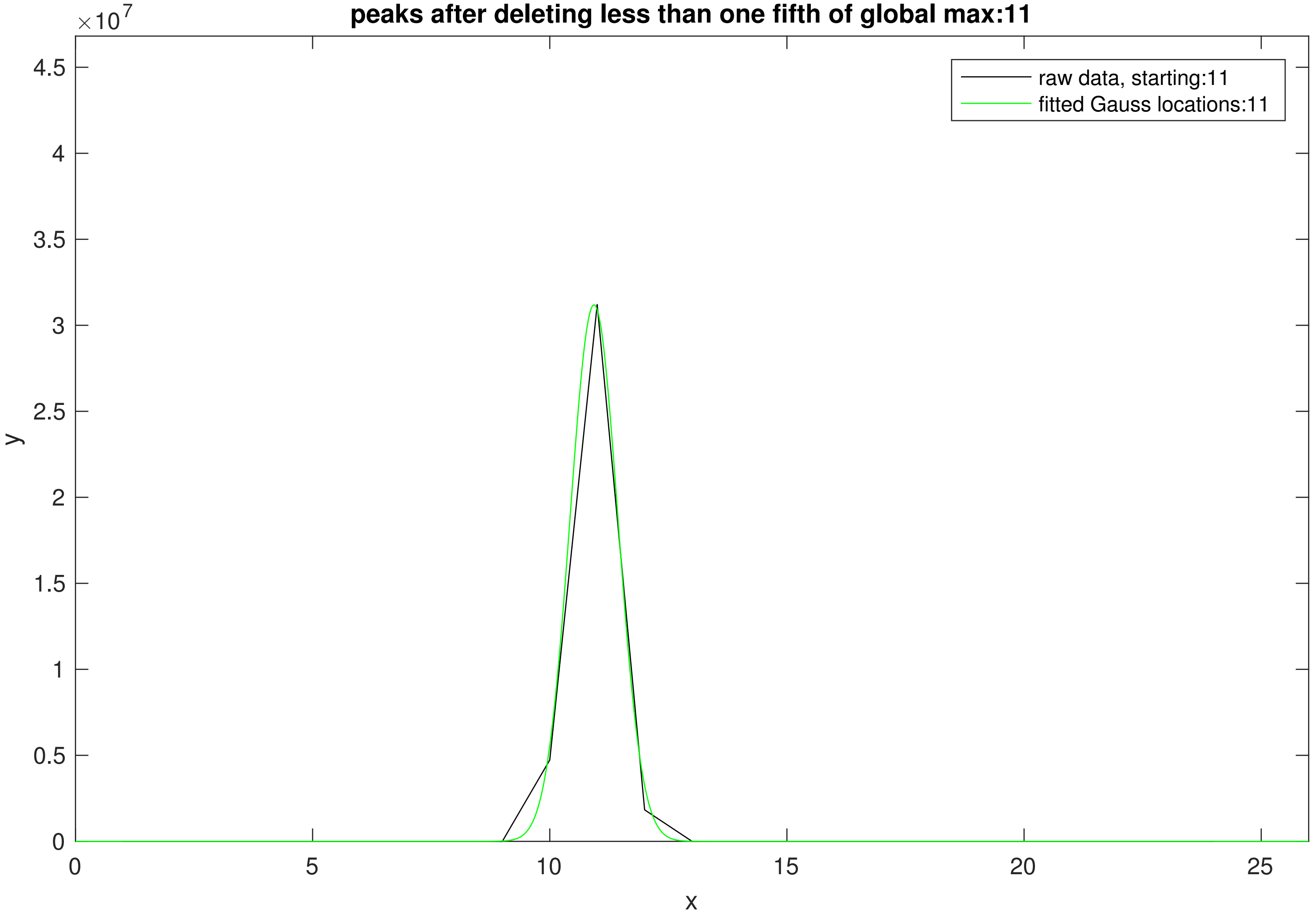

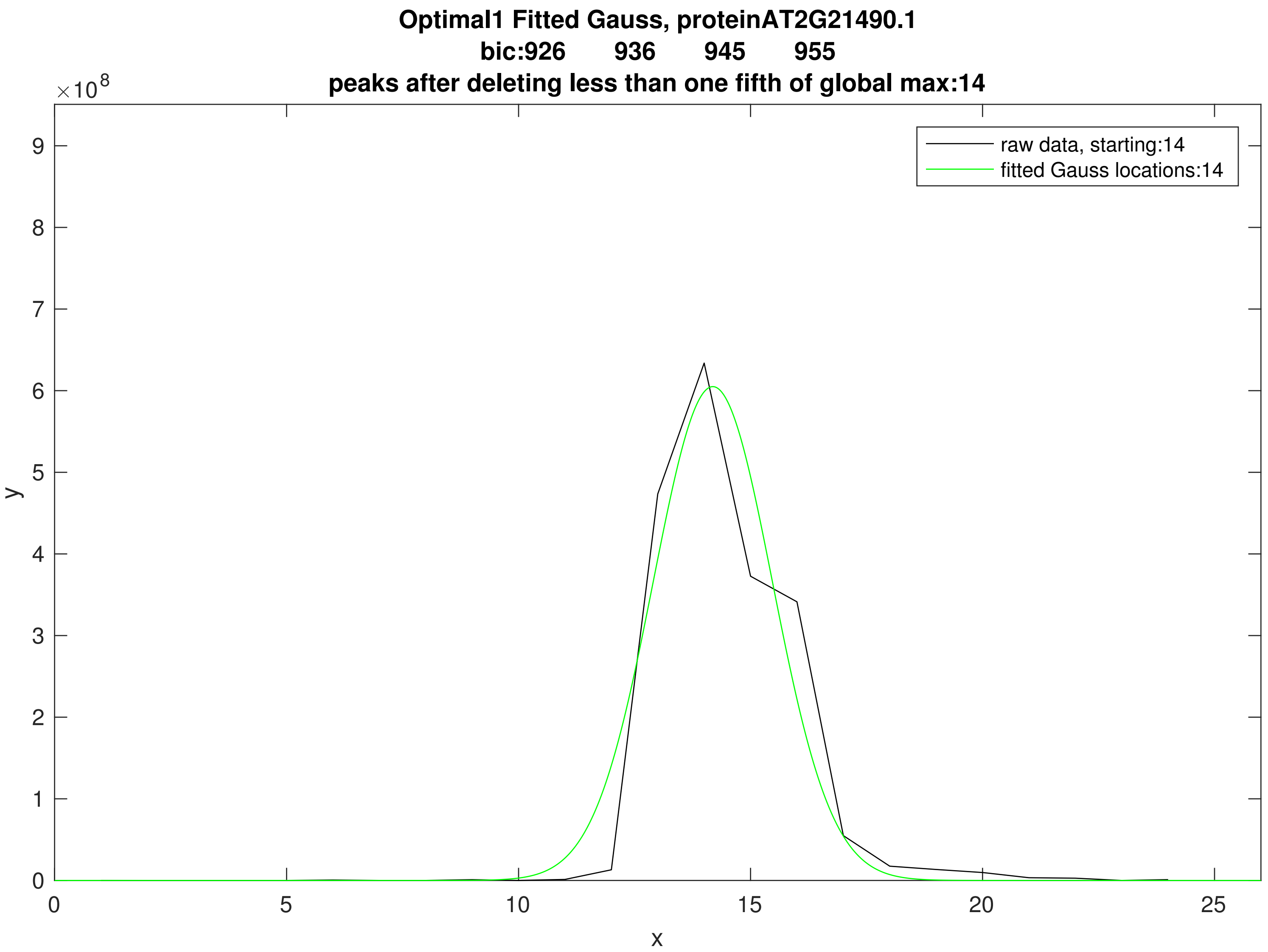
Optimal1 Fitted Gauss, proteinAT2G21490.1
bic:926 936 945 955
peaks after deleting less than one fifth of global max:14
×10^8
raw data, starting:14
fitted Gauss locations:14
0
1
2
3
4
5
6
7
8
9
y
0
5
10
15
20
25
x

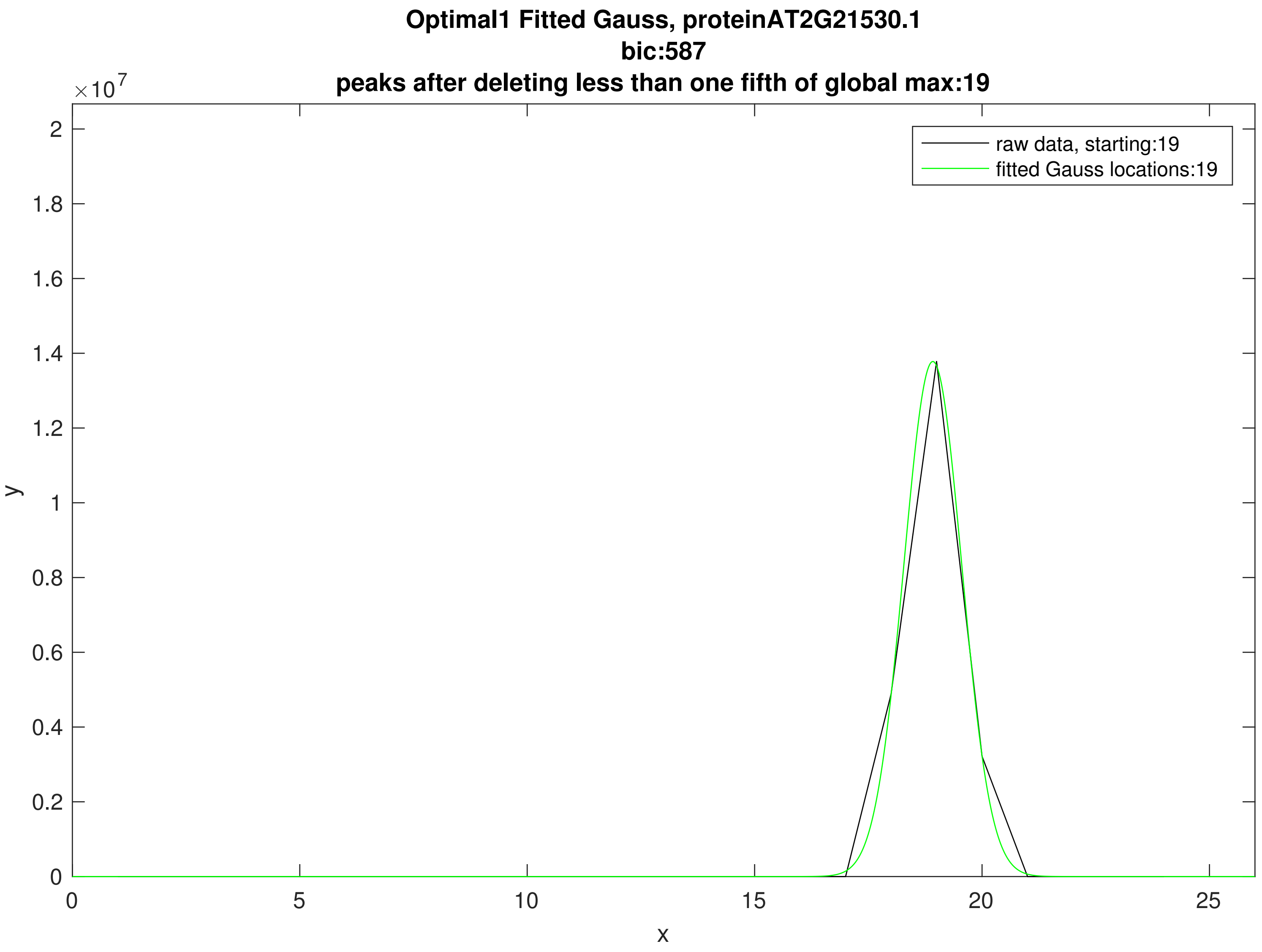

Optimal1 Fitted Gauss, proteinAT2G21530.1
bic:587
peaks after deleting less than one fifth of global max:19
×10^7
2
1.8
1.6
1.4
1.2
1
0.8
0.6
0.4
0.2
0
y
0
5
10
15
20
25
x
raw data, starting:19
fitted Gauss locations:19

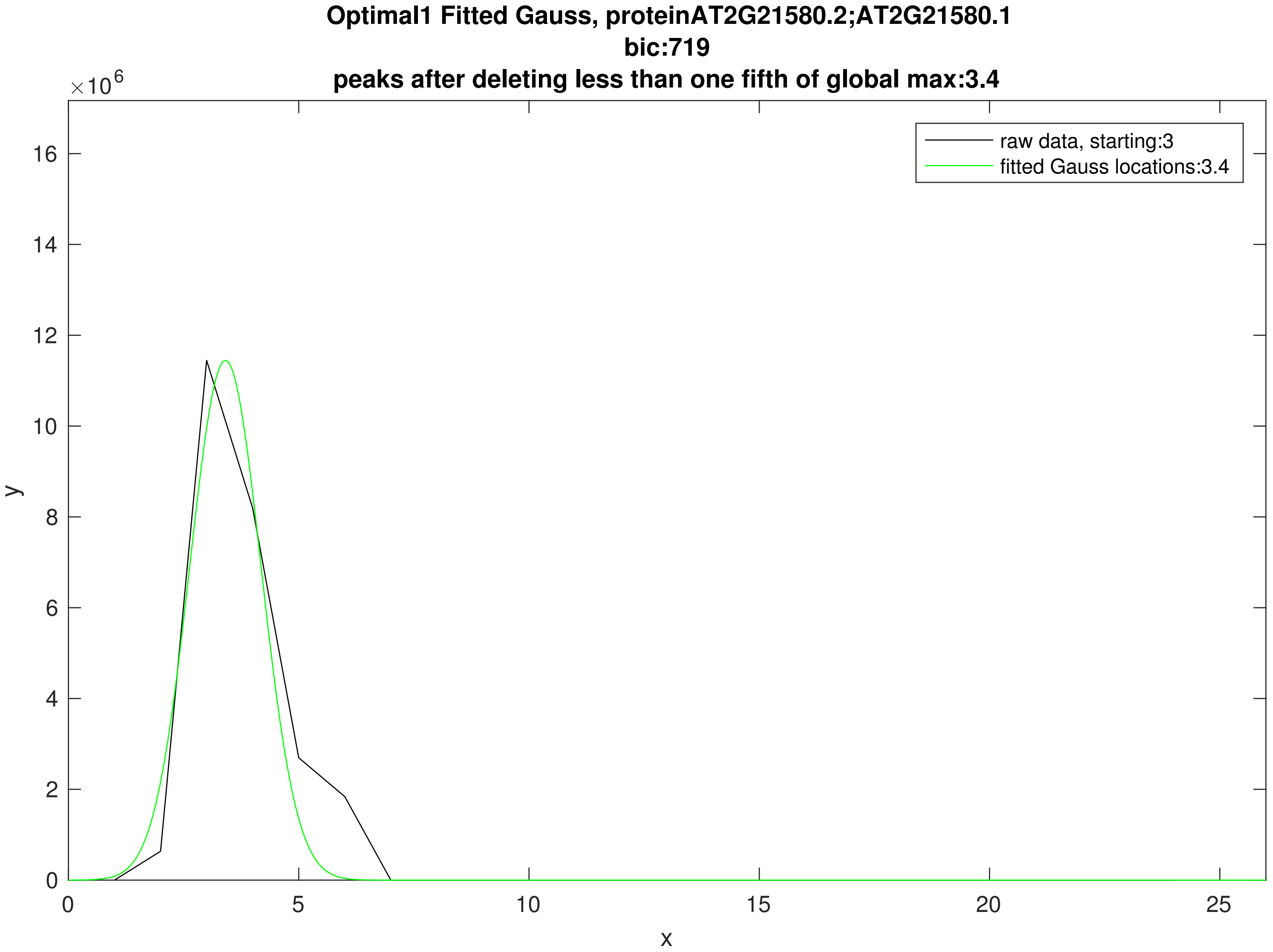
Optimal1 Fitted Gauss, proteinAT2G21580.2;AT2G21580.1
bic:719
peaks after deleting less than one fifth of global max:3.4
×10^6
raw data, starting:3
fitted Gauss locations:3.4
y
x
0
2
4
6
8
10
12
14
16
0
5
10
15
20
25

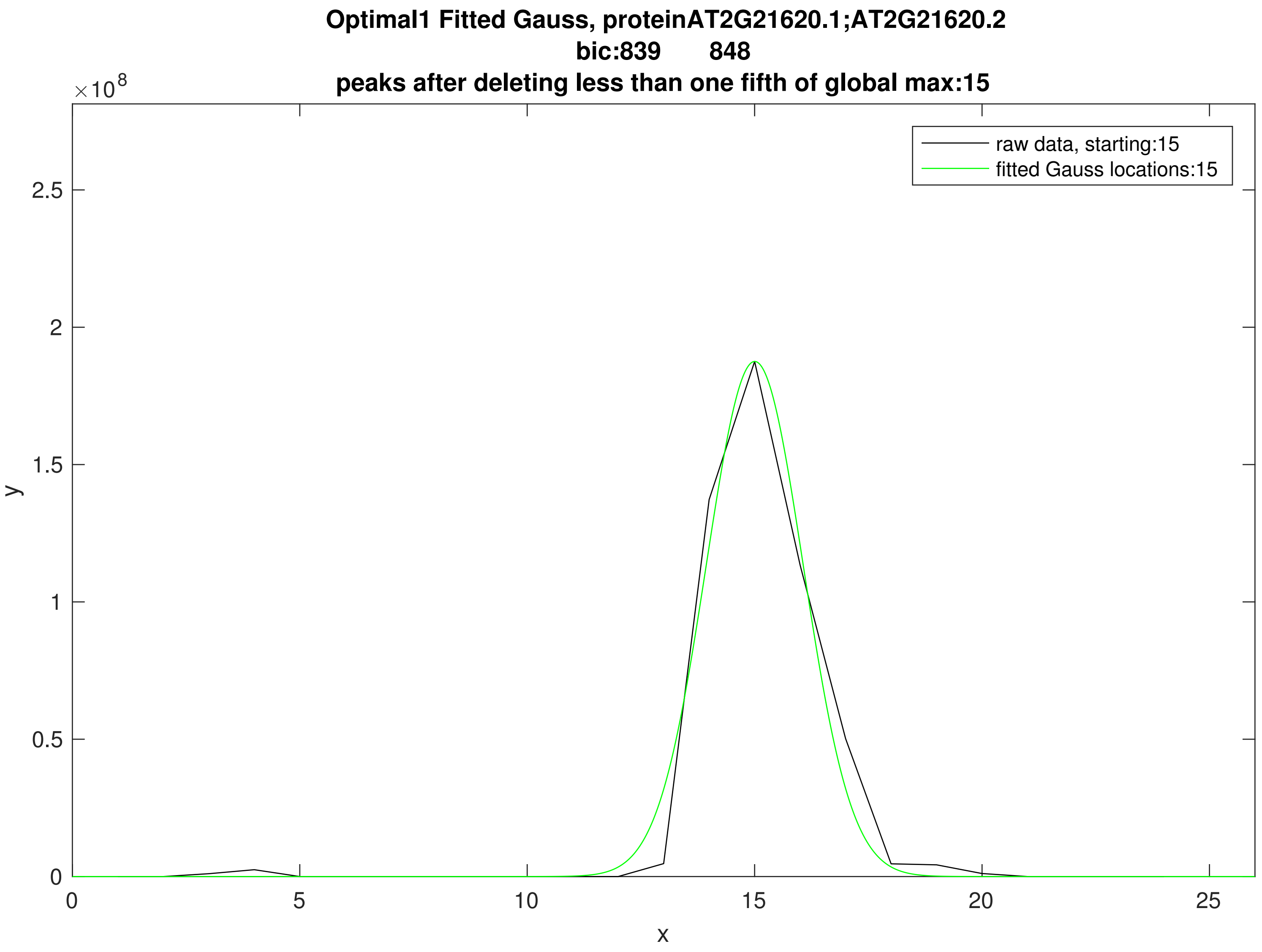
Optimal1 Fitted Gauss, proteinAT2G21620.1;AT2G21620.2
bic:839 848
peaks after deleting less than one fifth of global max:15
×10^8
2.5
2
1.5
1
0.5
0
y
0
5
10
15
20
25
x
raw data, starting:15
fitted Gauss locations:15

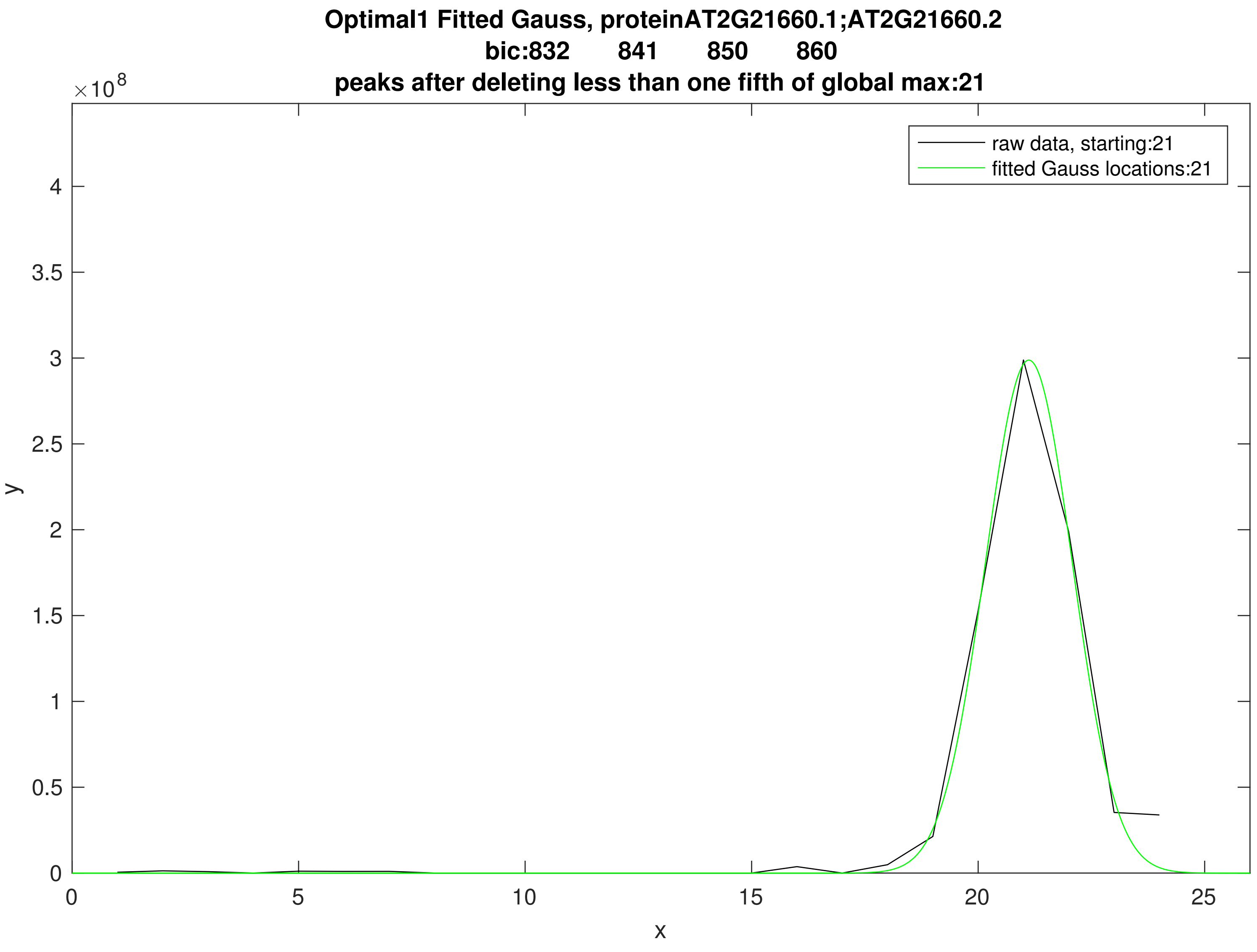

Optimal1 Fitted Gauss, proteinAT2G21660.1;AT2G21660.2
bic:832 841 850 860
peaks after deleting less than one fifth of global max:21
×10^8
raw data, starting:21
fitted Gauss locations:21
4
3.5
3
2.5
2
1.5
1
0.5
0
y
0
5
10
15
20
25
x

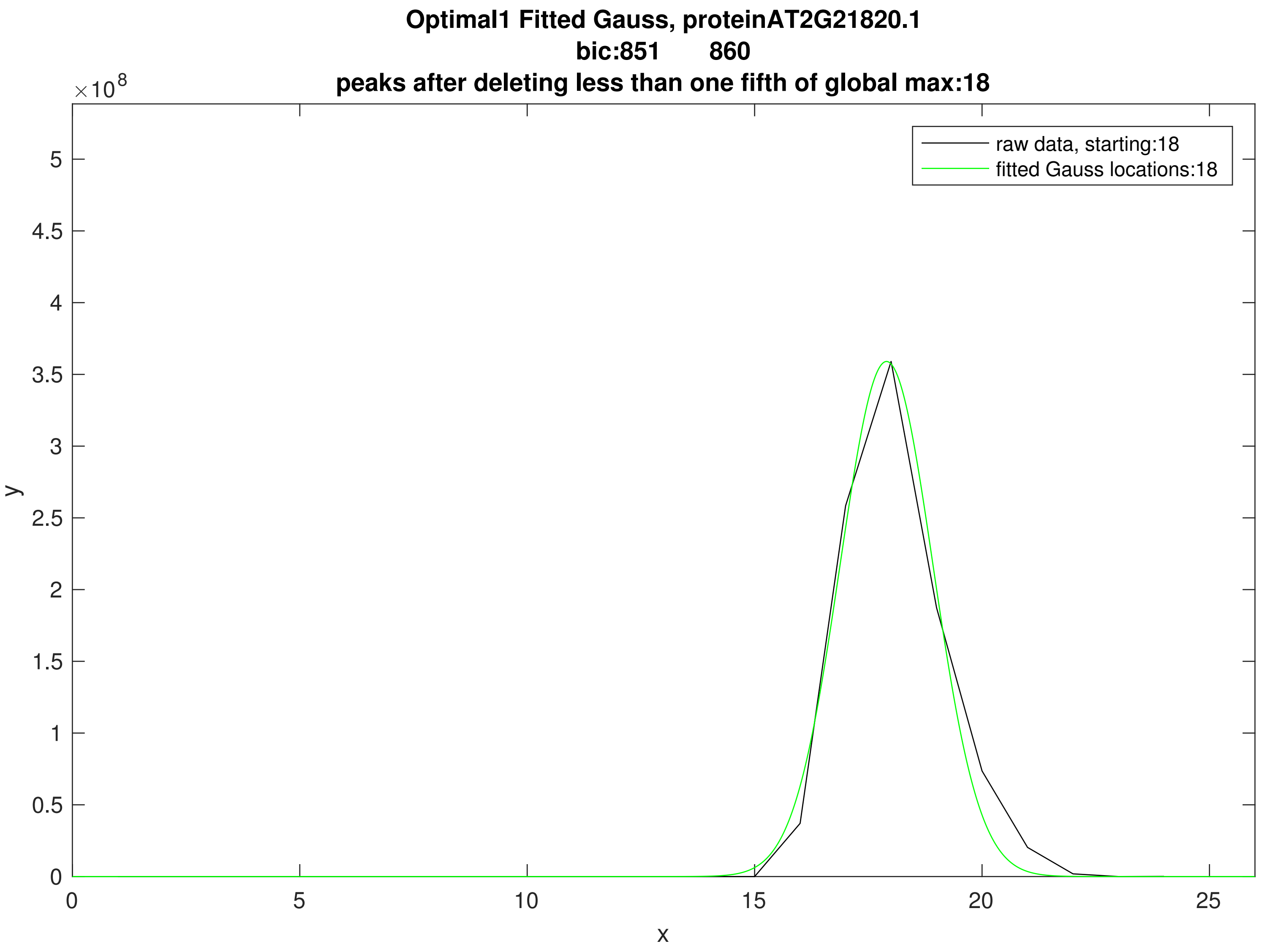
Optimal1 Fitted Gauss, proteinAT2G21820.1
bic:851 860
peaks after deleting less than one fifth of global max:18
×10^8
raw data, starting:18
fitted Gauss locations:18
y
x
0
0.5
1
1.5
2
2.5
3
3.5
4
4.5
5
0
5
10
15
20
25

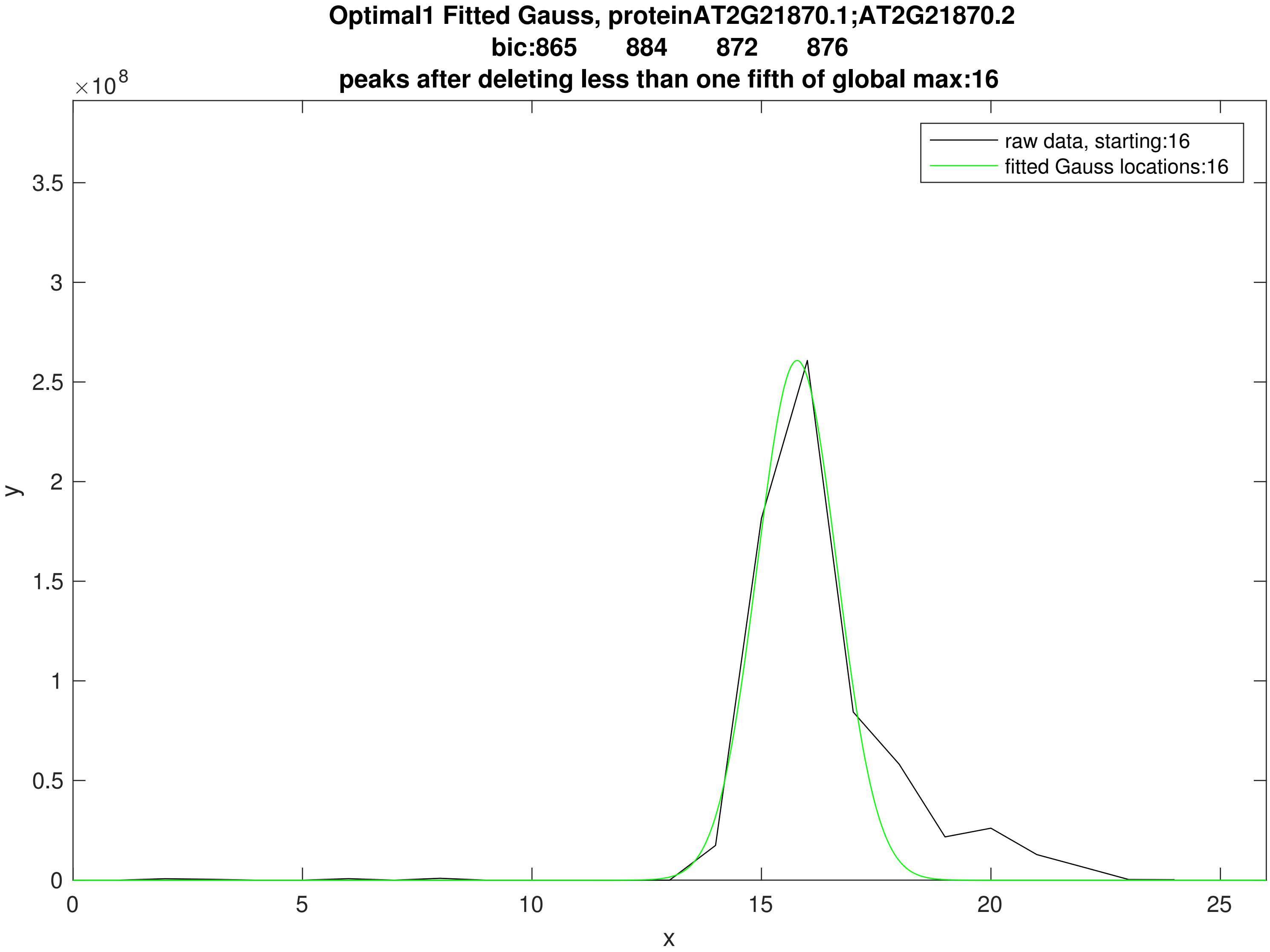

Optimal1 Fitted Gauss, proteinAT2G21870.1;AT2G21870.2
bic:865 884 872 876
peaks after deleting less than one fifth of global max:16
×10^8
raw data, starting:16
fitted Gauss locations:16
y
x
0
0.5
1
1.5
2
2.5
3
3.5
5
10
15
20
25

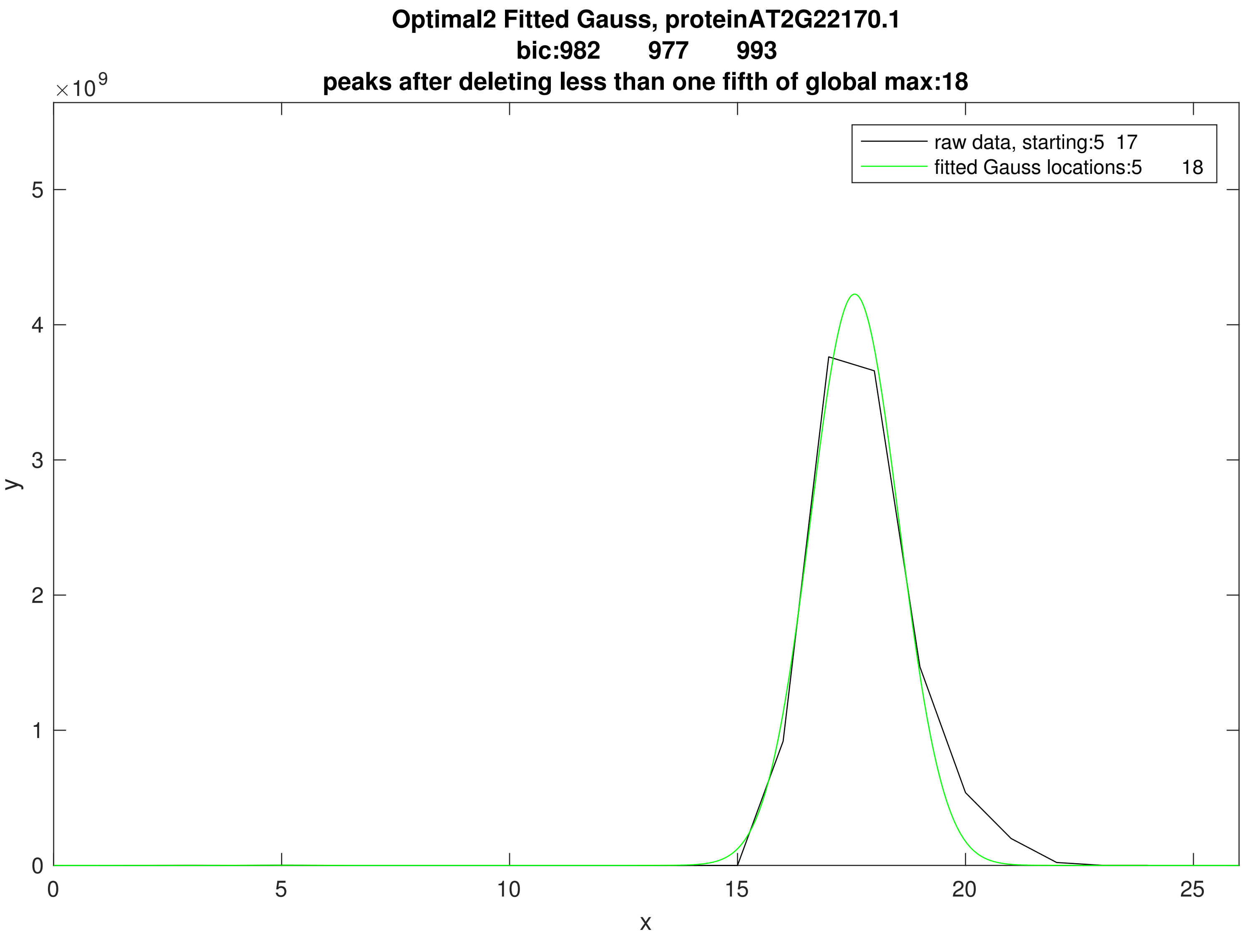

Optimal2 Fitted Gauss, proteinAT2G22170.1
bic:982 977 993
peaks after deleting less than one fifth of global max:18
×10^9
raw data, starting:5 17
fitted Gauss locations:5 18
y
x
0
1
2
3
4
5
0
5
10
15
20
25

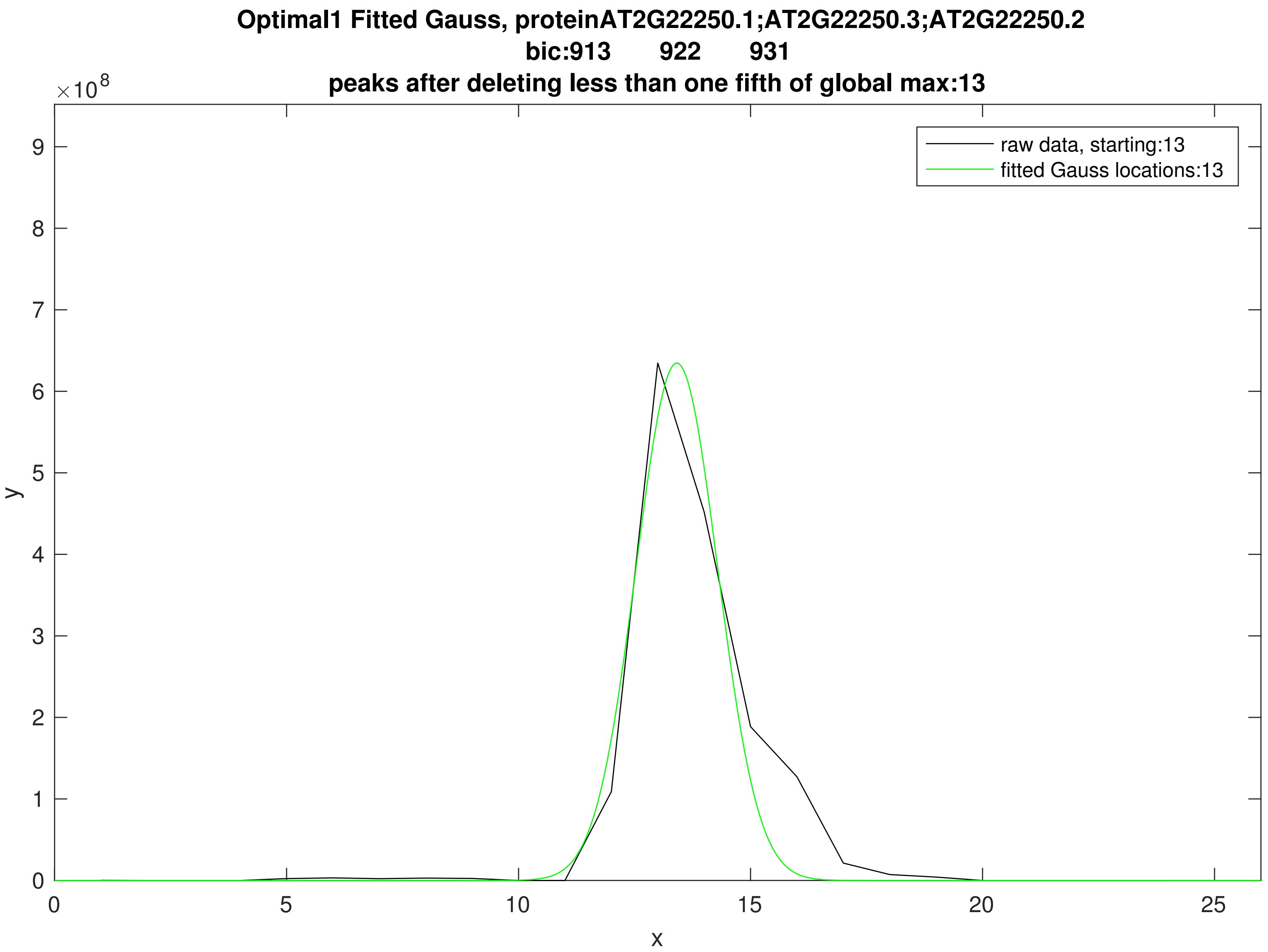

Optimal1 Fitted Gauss, proteinAT2G22250.1;AT2G22250.3;AT2G22250.2
bic:913 922 931
peaks after deleting less than one fifth of global max:13
×10^8
raw data, starting:13
fitted Gauss locations:13
y
x
0
1
2
3
4
5
6
7
8
9
10
15
20
25

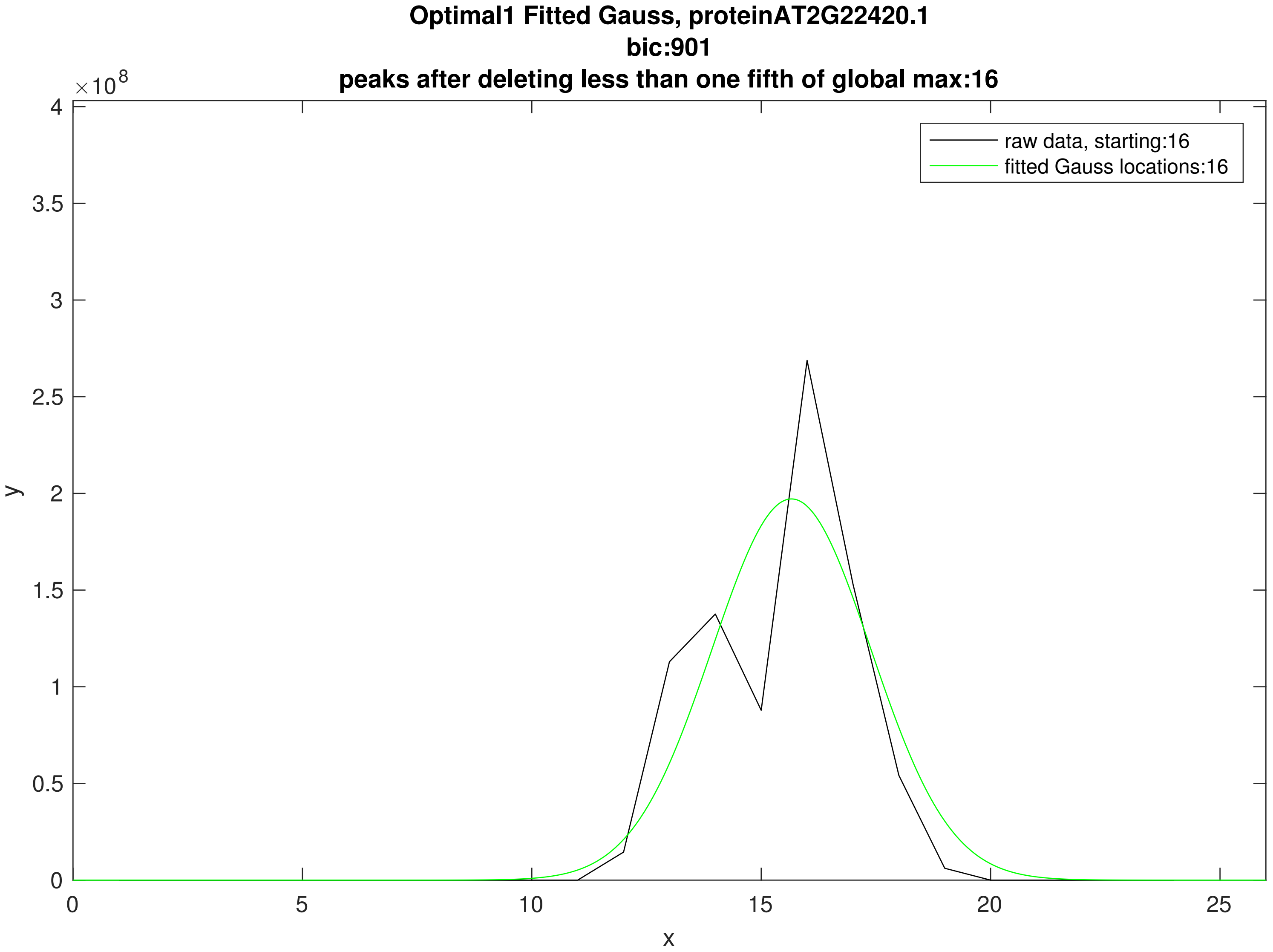
Optimal1 Fitted Gauss, proteinAT2G22420.1
bic:901
peaks after deleting less than one fifth of global max:16
raw data, starting:16
fitted Gauss locations:16
y
x

**Optimal1 Fitted Gauss, proteinAT2G22450.1**
**bic:751**
**peaks after deleting less than one fifth of global max:11**

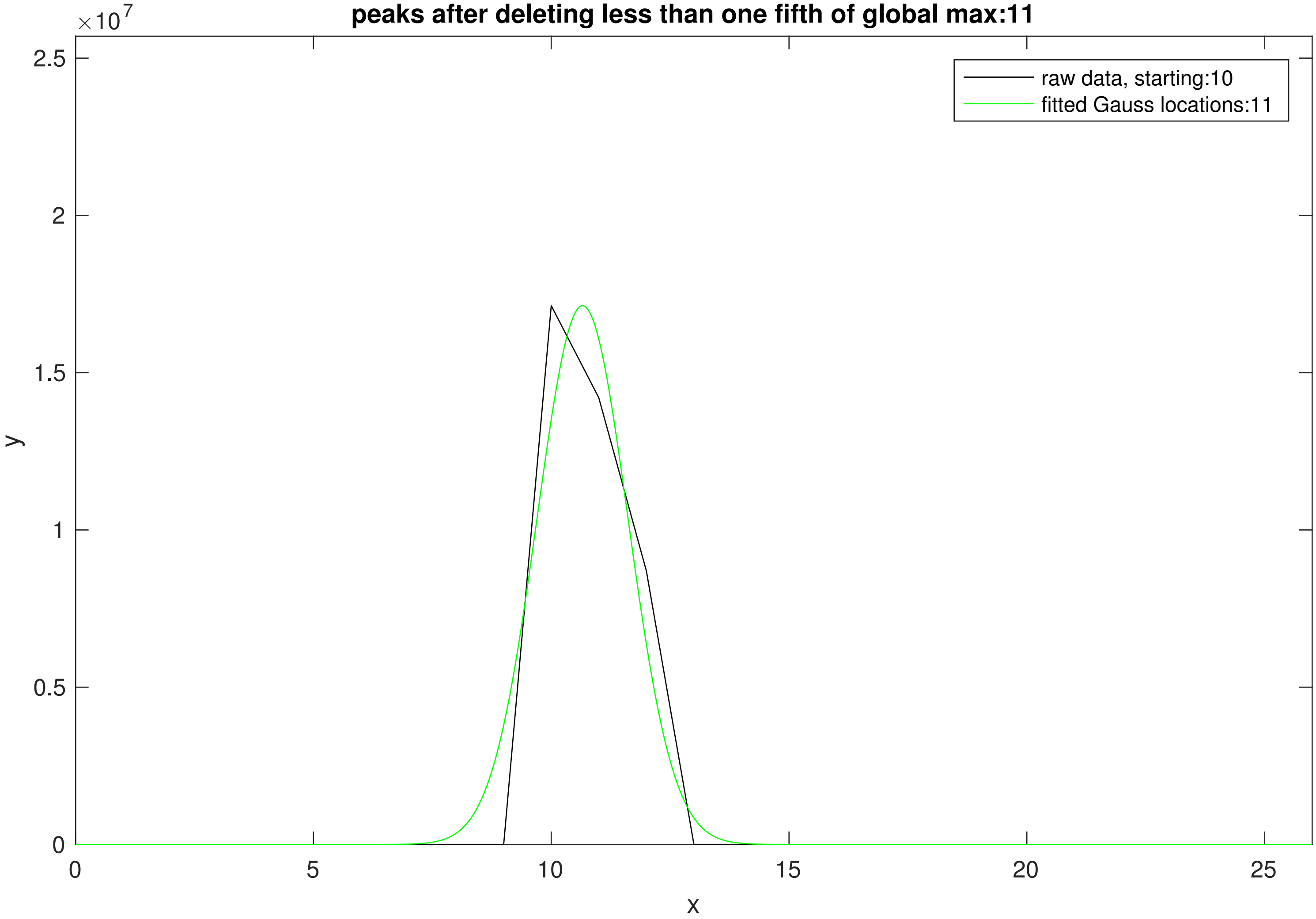

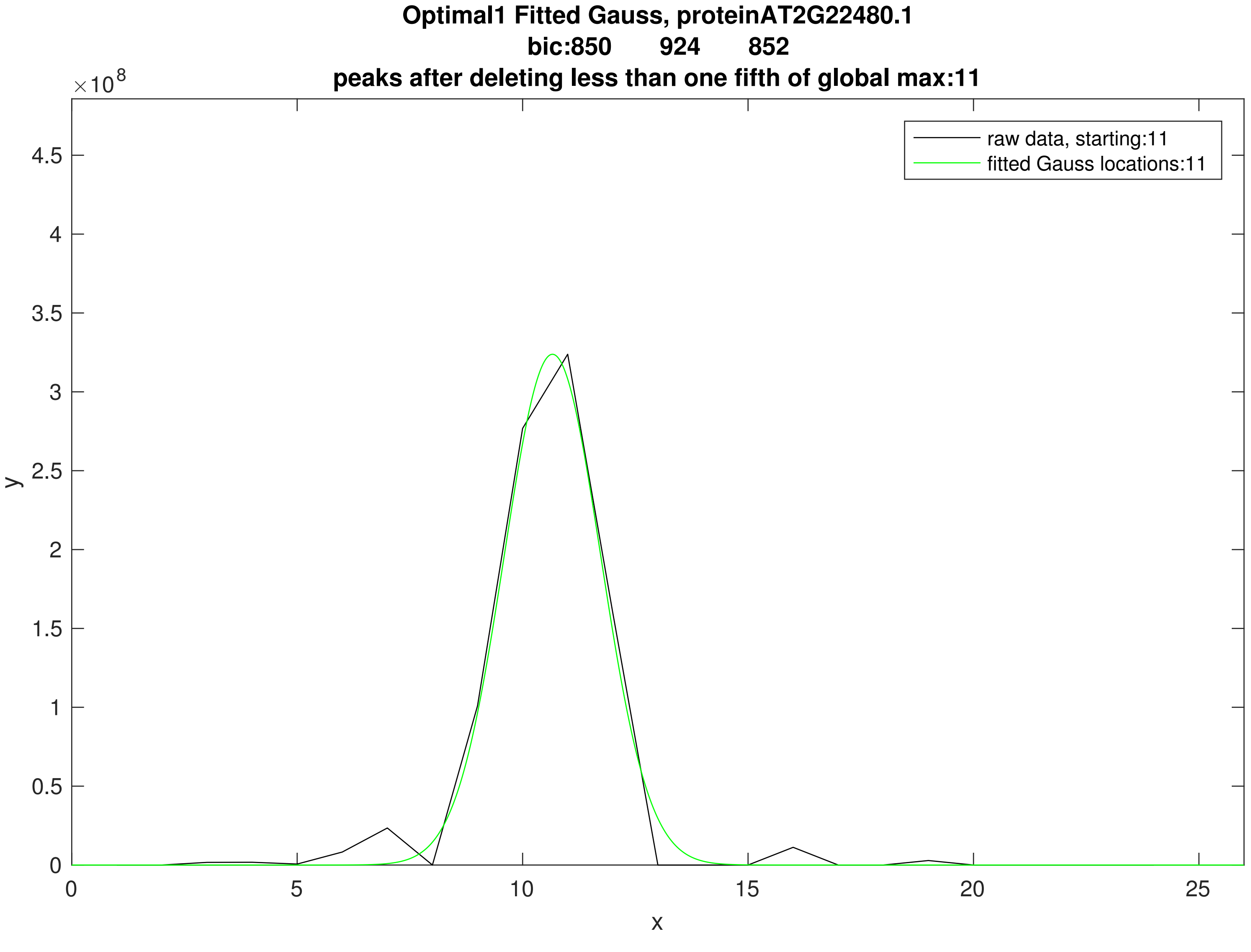
Optimal1 Fitted Gauss, proteinAT2G22480.1
bic:850 924 852
peaks after deleting less than one fifth of global max:11
×10^8
raw data, starting:11
fitted Gauss locations:11
y
x
0
0.5
1
1.5
2
2.5
3
3.5
4
4.5
5
10
15
20
25

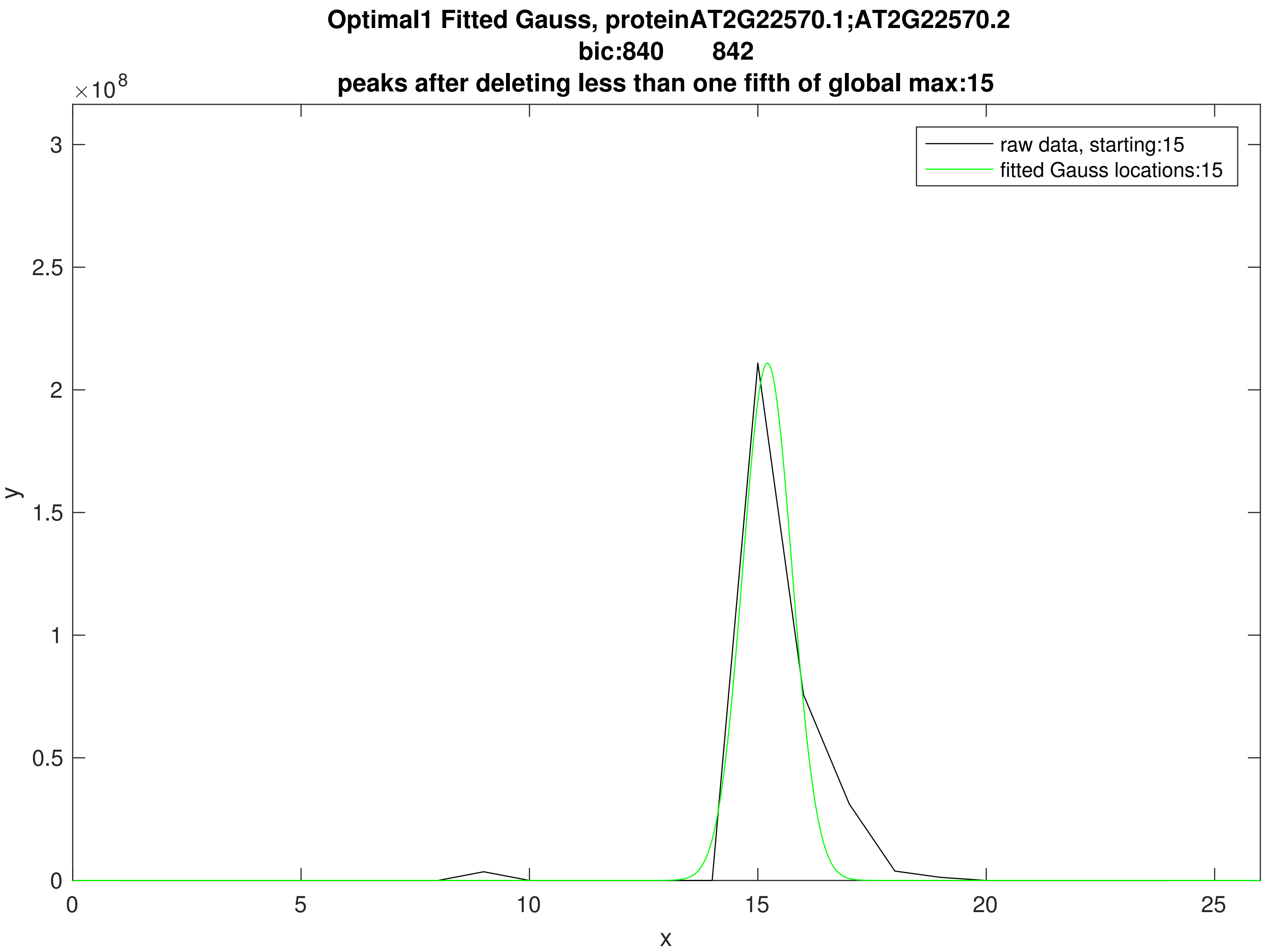

Optimal1 Fitted Gauss, proteinAT2G22570.1;AT2G22570.2
bic:840 842
peaks after deleting less than one fifth of global max:15
×10^8
raw data, starting:15
fitted Gauss locations:15
y
x
0
0.5
1
1.5
2
2.5
3
5
10
15
20
25

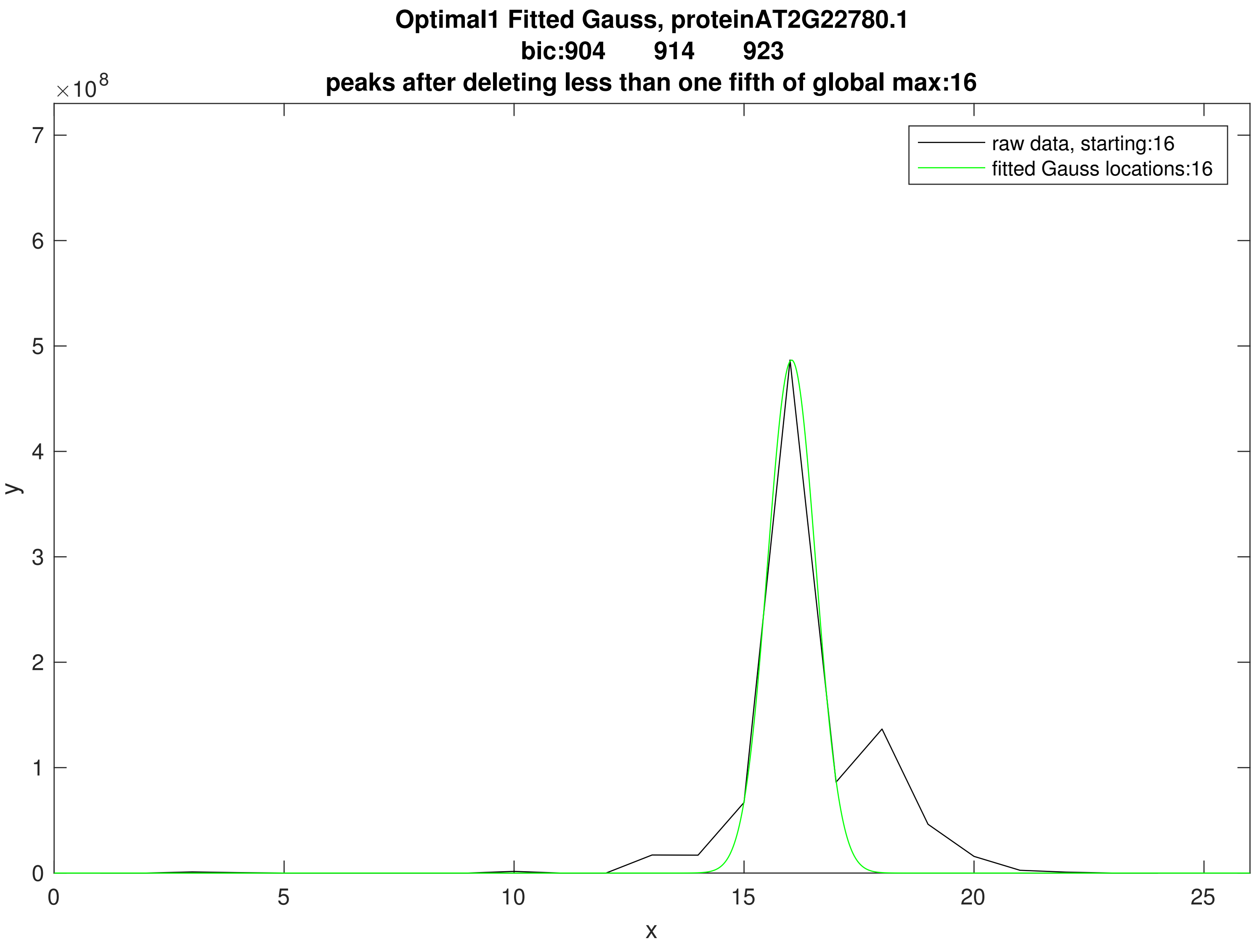

Optimal1 Fitted Gauss, proteinAT2G22780.1
bic:904 914 923
peaks after deleting less than one fifth of global max:16
×10^8
y
x
raw data, starting:16
fitted Gauss locations:16

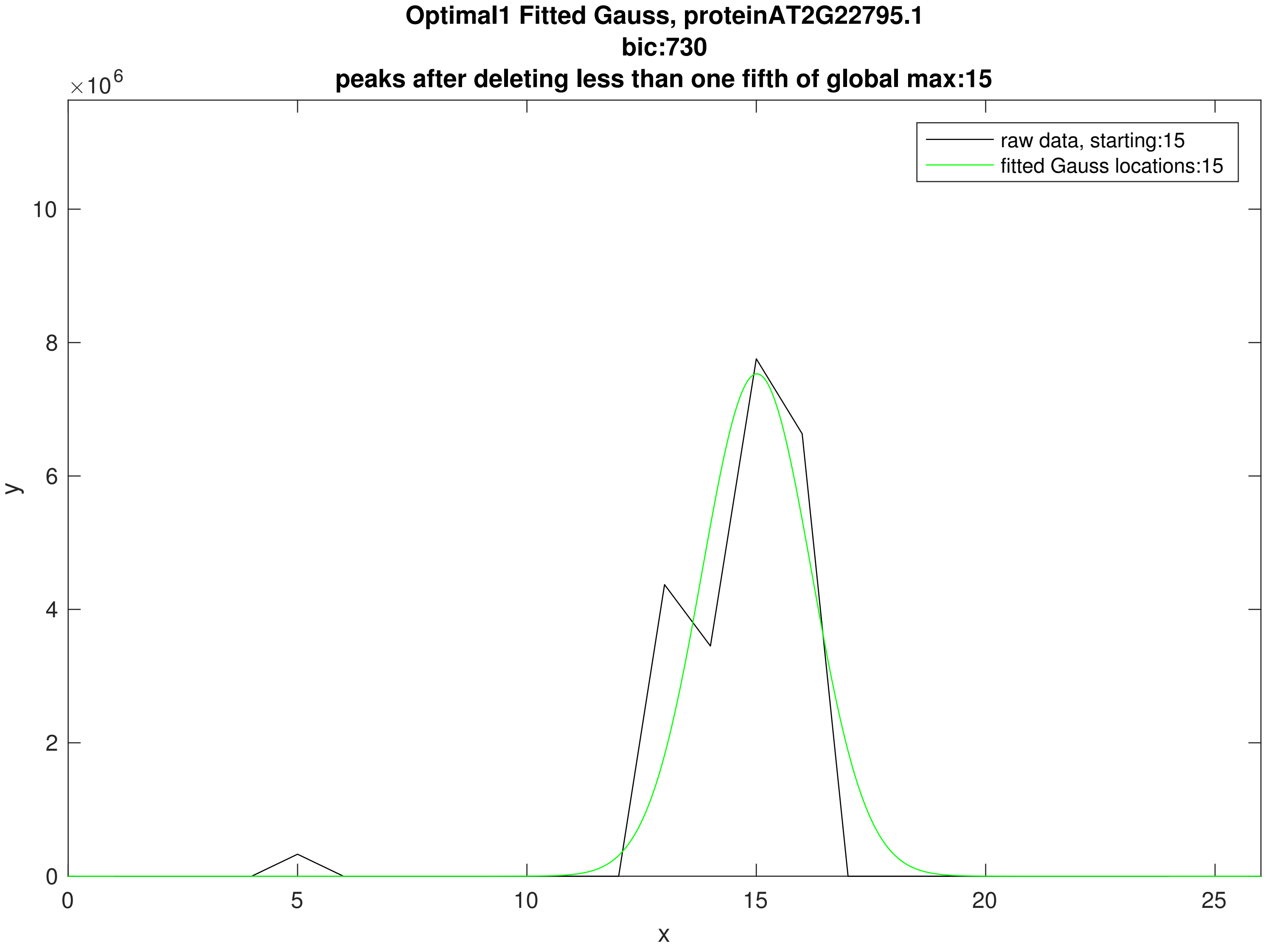
Optimal1 Fitted Gauss, proteinAT2G22795.1
bic:730
peaks after deleting less than one fifth of global max:15
×10^6
raw data, starting:15
fitted Gauss locations:15
y
0
2
4
6
8
10
x
0
5
10
15
20
25

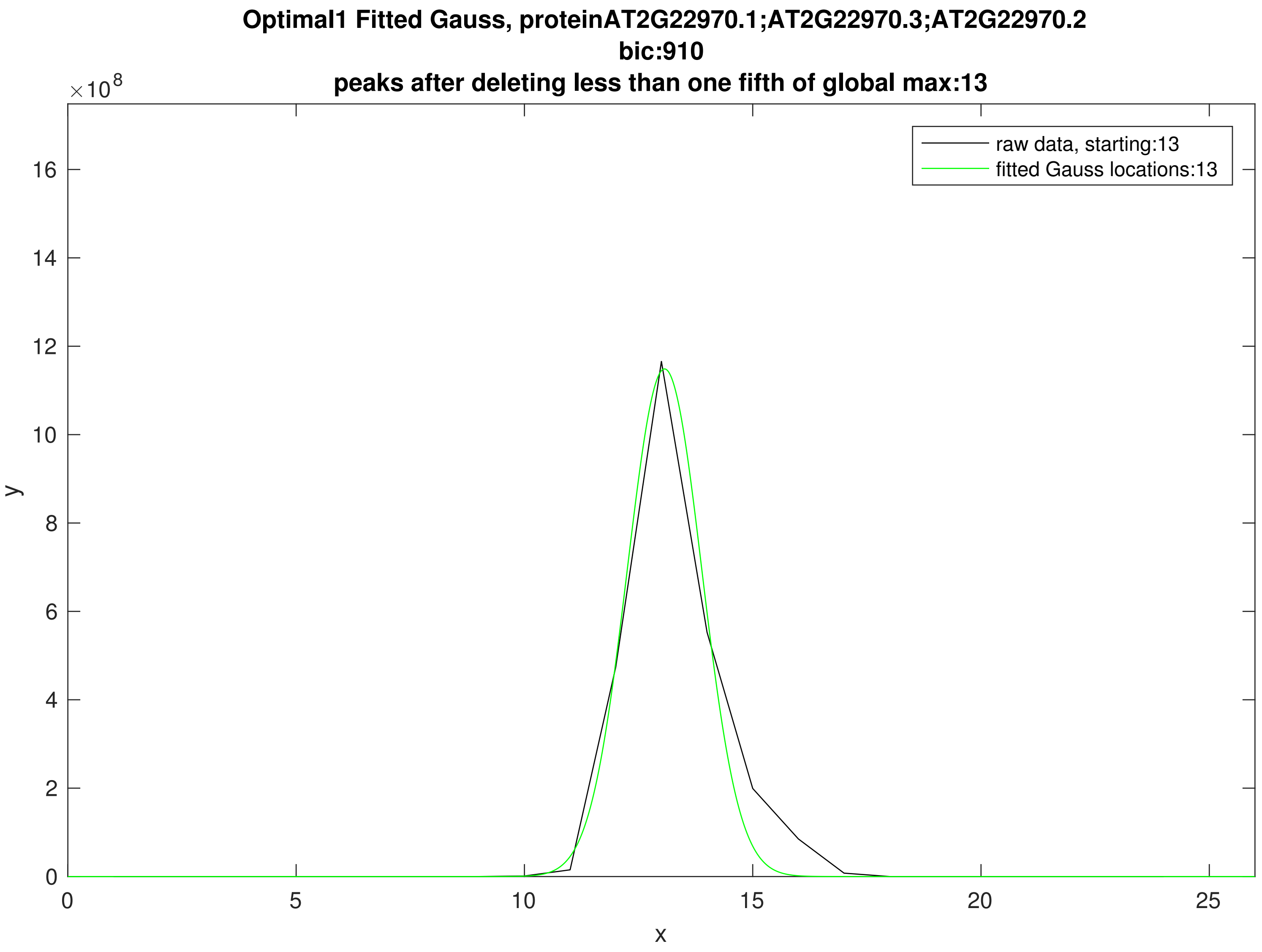
Optimal1 Fitted Gauss, proteinAT2G22970.1;AT2G22970.3;AT2G22970.2
bic:910
peaks after deleting less than one fifth of global max:13
×10^8
raw data, starting:13
fitted Gauss locations:13
16
14
12
10
8
6
4
2
0
y
0
5
10
15
20
25
x

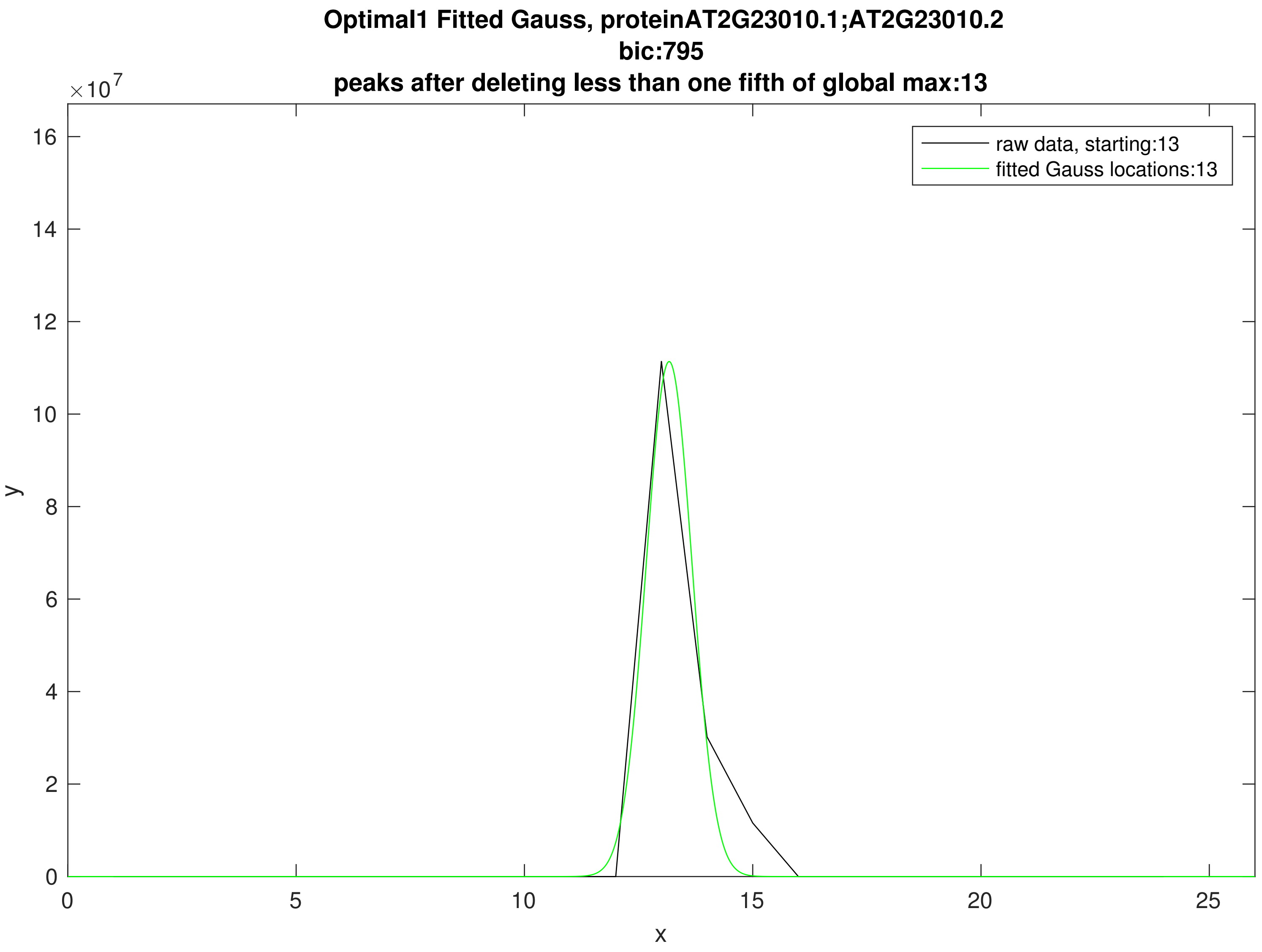

Optimal1 Fitted Gauss, proteinAT2G23010.1;AT2G23010.2
bic:795
peaks after deleting less than one fifth of global max:13
×10^7
16
14
12
10
8
6
4
2
0
y
0
5
10
15
20
25
x
raw data, starting:13
fitted Gauss locations:13

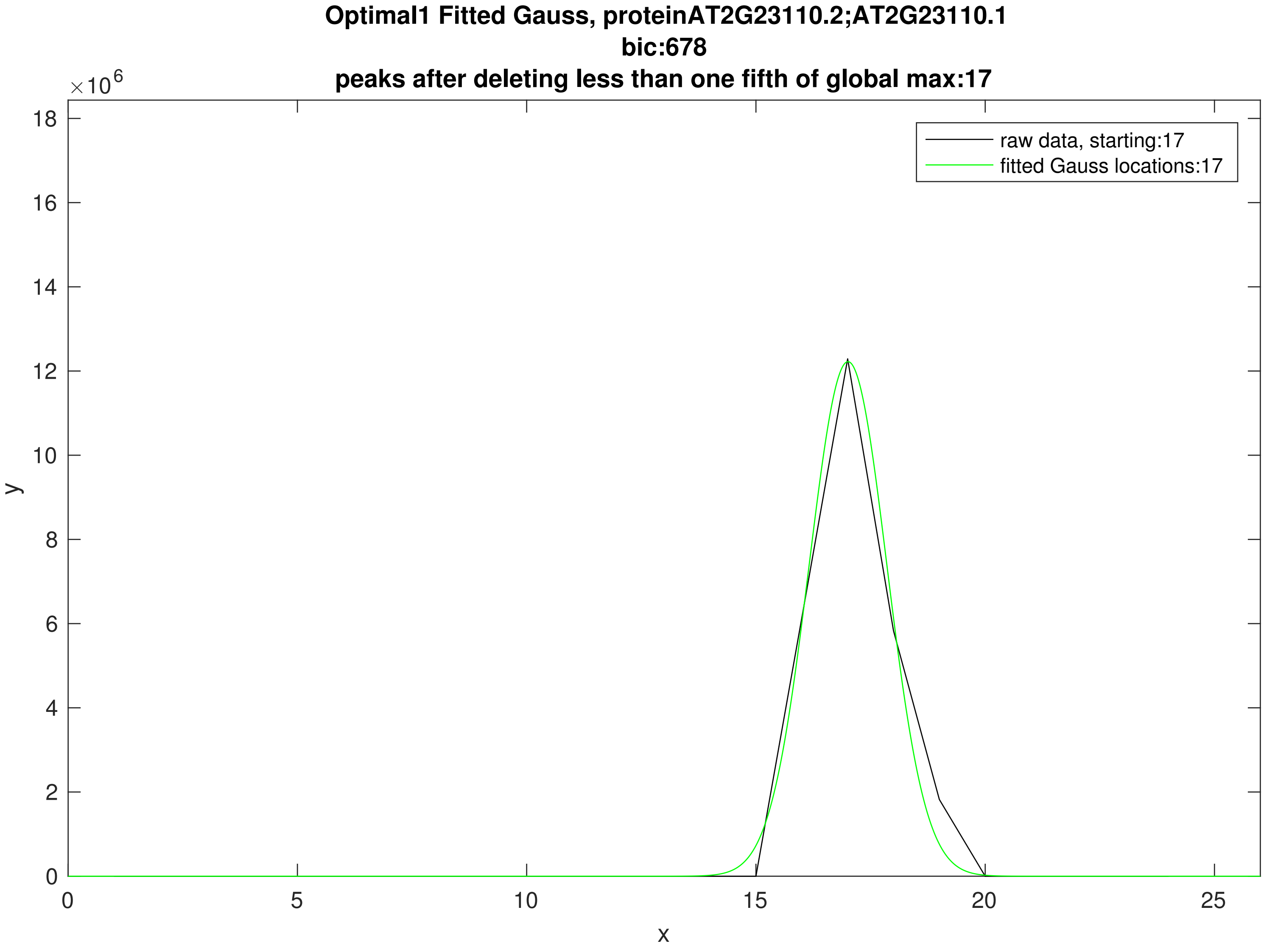
Optimal1 Fitted Gauss, proteinAT2G23110.2;AT2G23110.1
bic:678
peaks after deleting less than one fifth of global max:17
×10^6
18
16
14
12
10
8
6
4
2
0
y
0
5
10
15
20
25
x
raw data, starting:17
fitted Gauss locations:17

**Optimal1 Fitted Gauss, proteinAT2G23120.1**
**bic:886**
**peaks after deleting less than one fifth of global max:17**

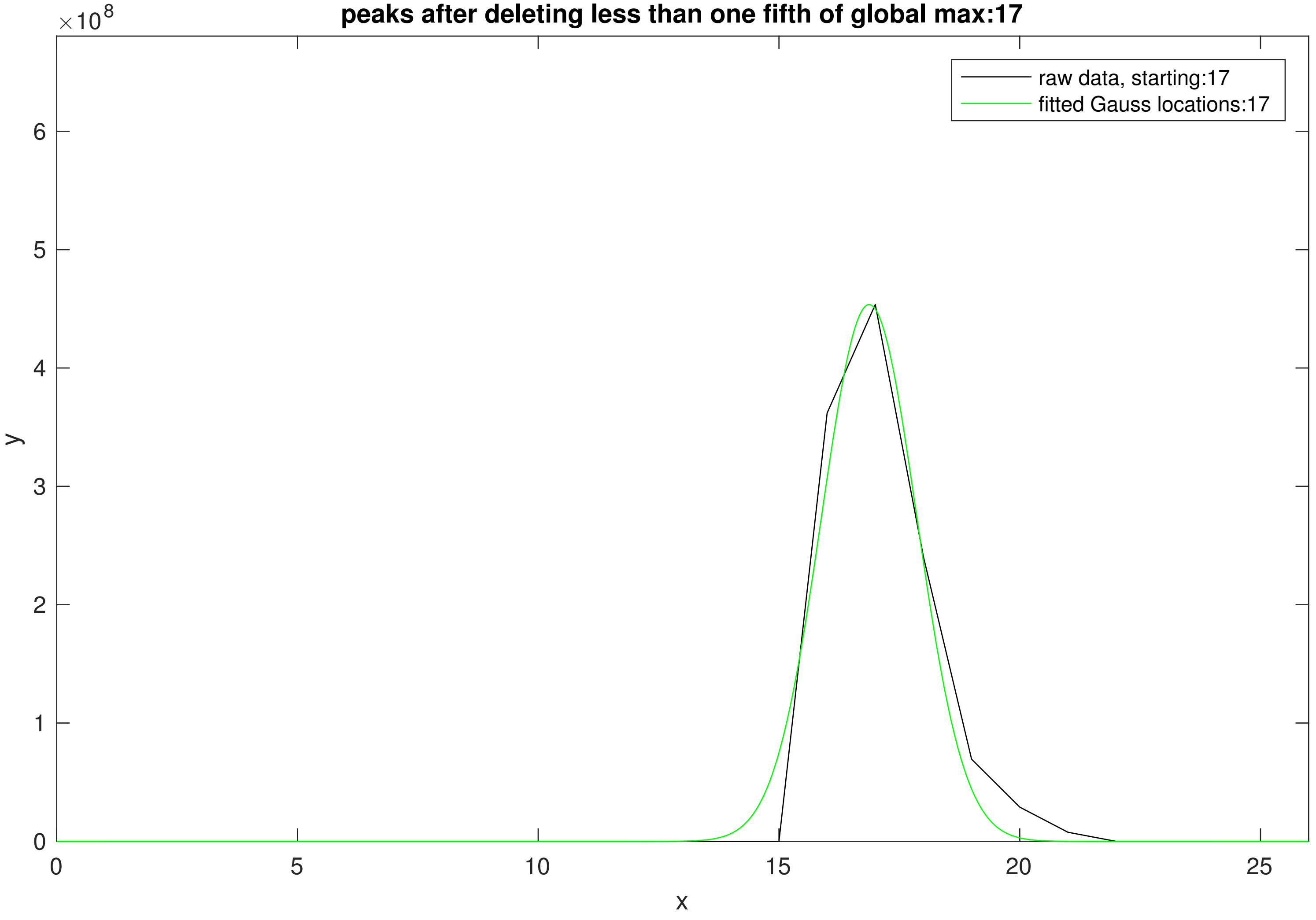

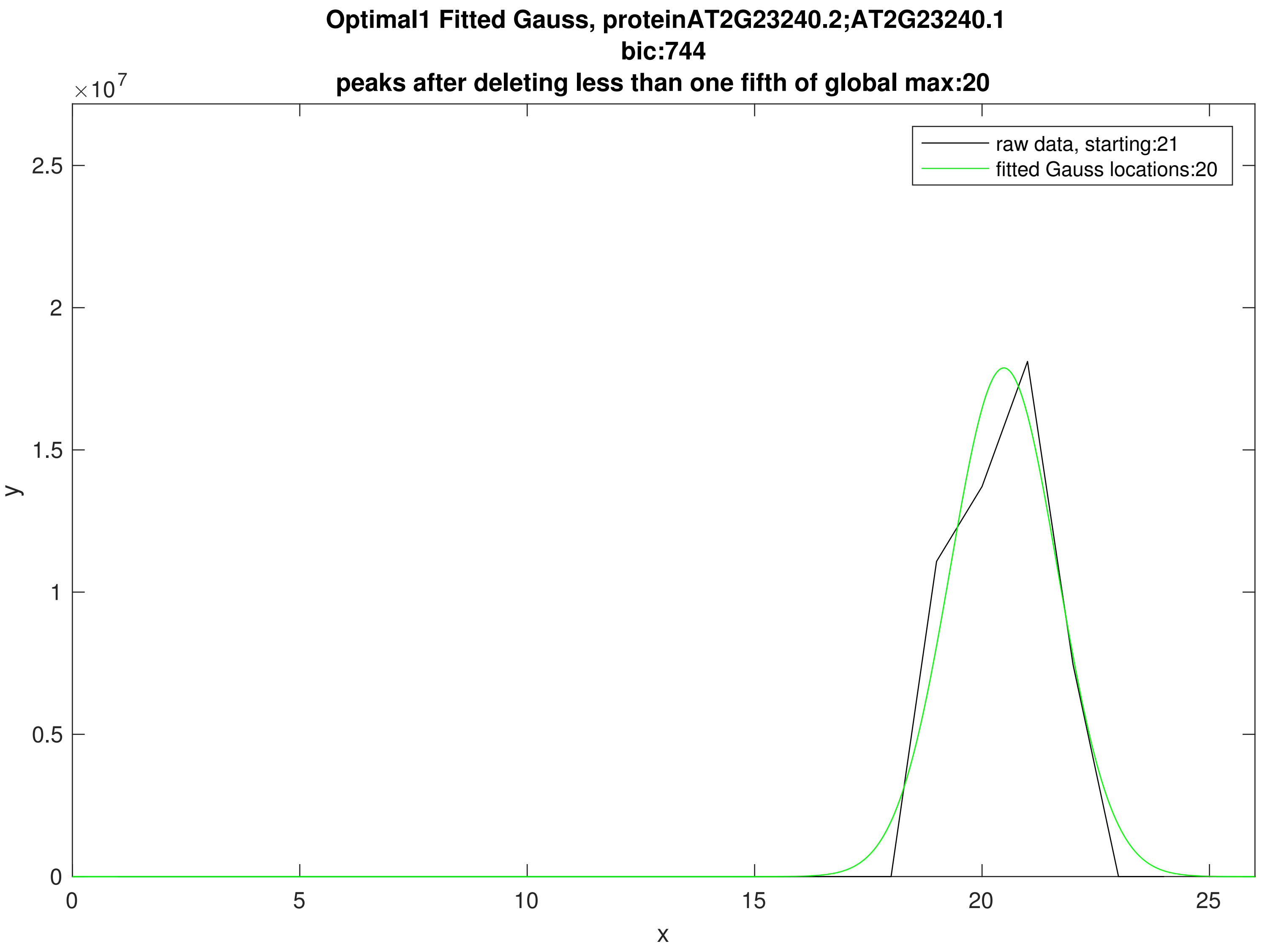
Optimal1 Fitted Gauss, proteinAT2G23240.2;AT2G23240.1
bic:744
peaks after deleting less than one fifth of global max:20
×10^7
2.5
2
1.5
1
0.5
0
y
0
5
10
15
20
25
x
raw data, starting:21
fitted Gauss locations:20

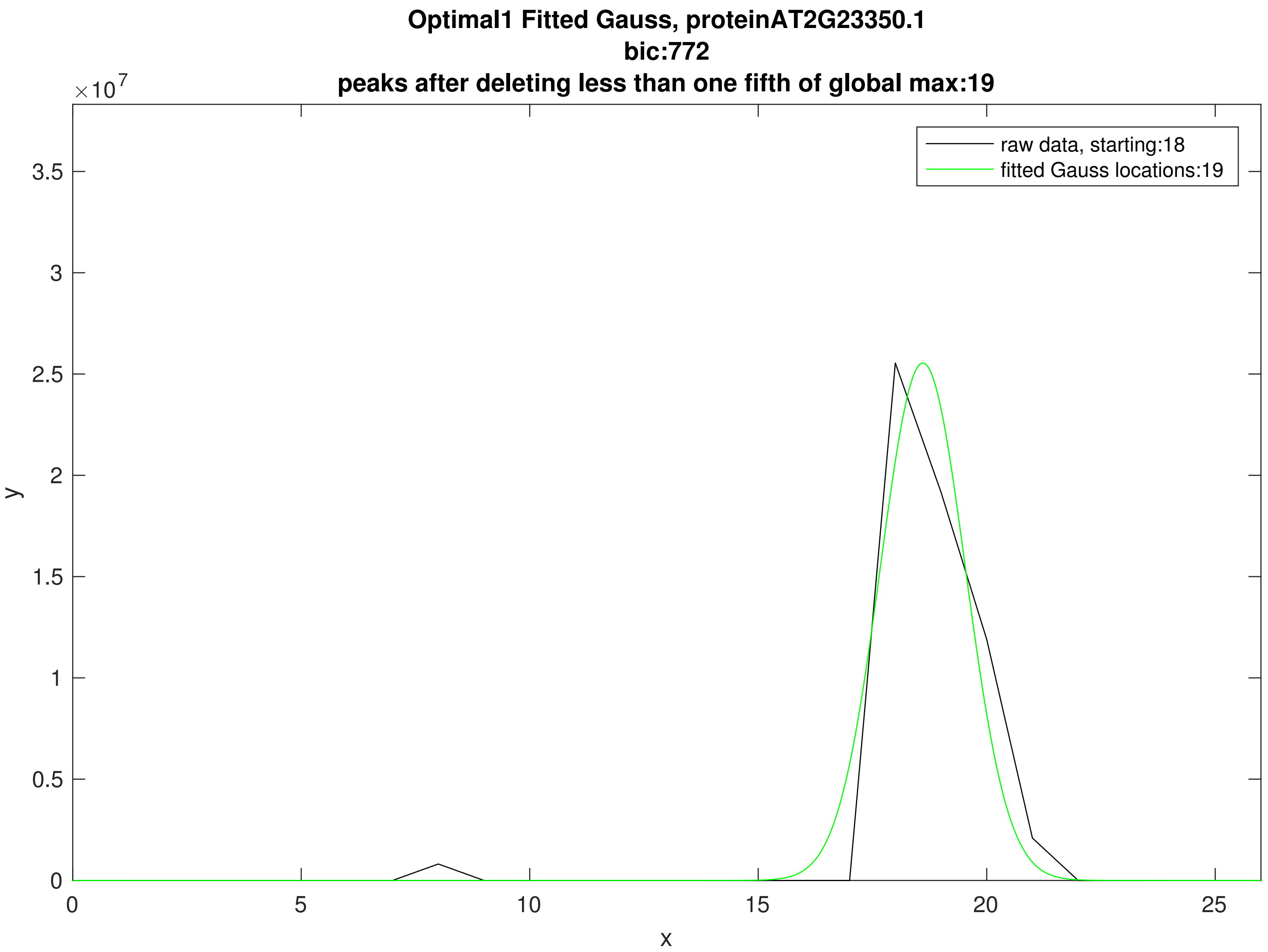
Optimal1 Fitted Gauss, proteinAT2G23350.1
bic:772
peaks after deleting less than one fifth of global max:19
×10^7
raw data, starting:18
fitted Gauss locations:19
y
x
3.5
3
2.5
2
1.5
1
0.5
0
0
5
10
15
20
25

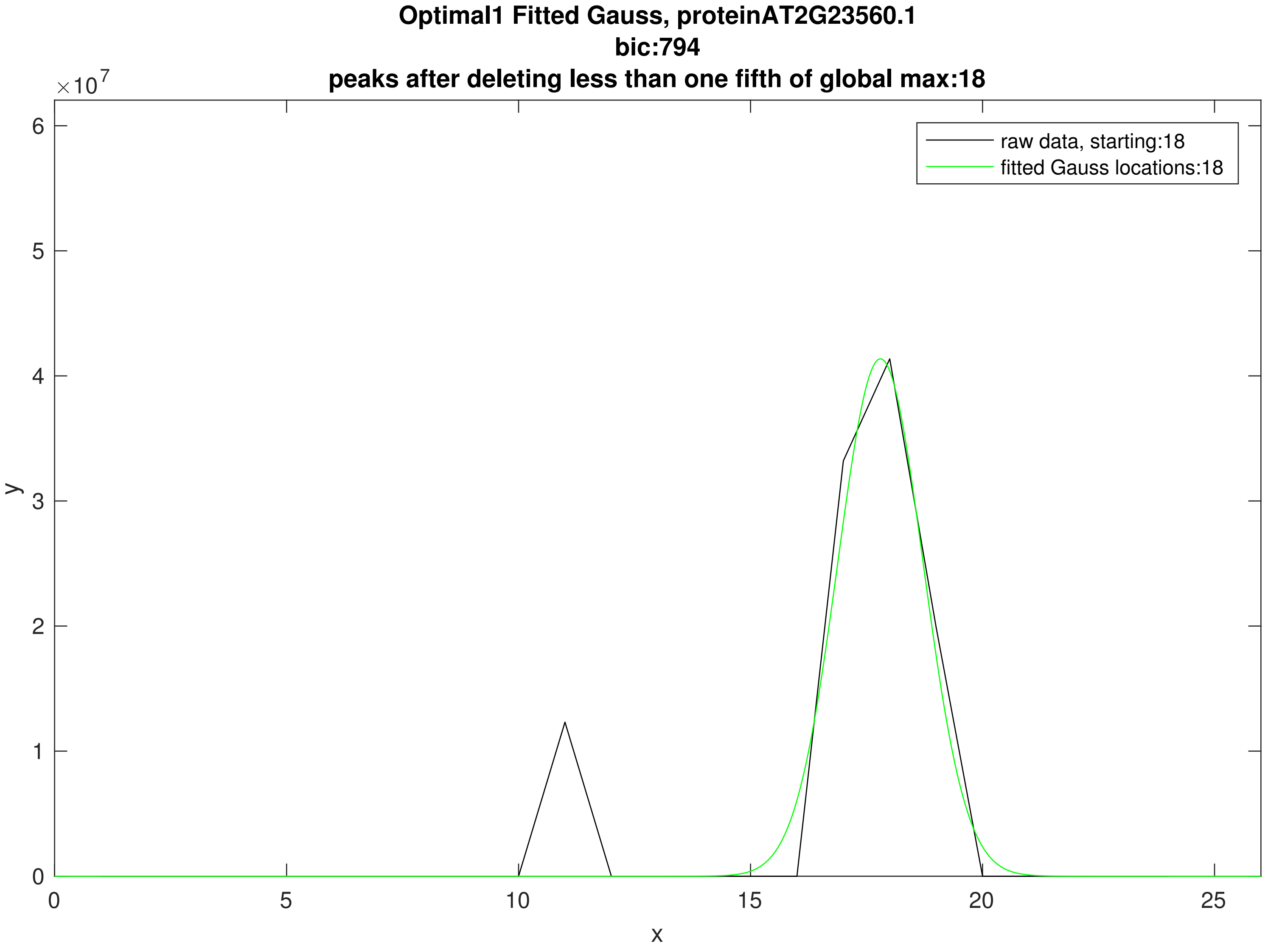
Optimal1 Fitted Gauss, proteinAT2G23560.1
bic:794
peaks after deleting less than one fifth of global max:18
×10^7
raw data, starting:18
fitted Gauss locations:18
y
x
0
1
2
3
4
5
6
0
5
10
15
20
25

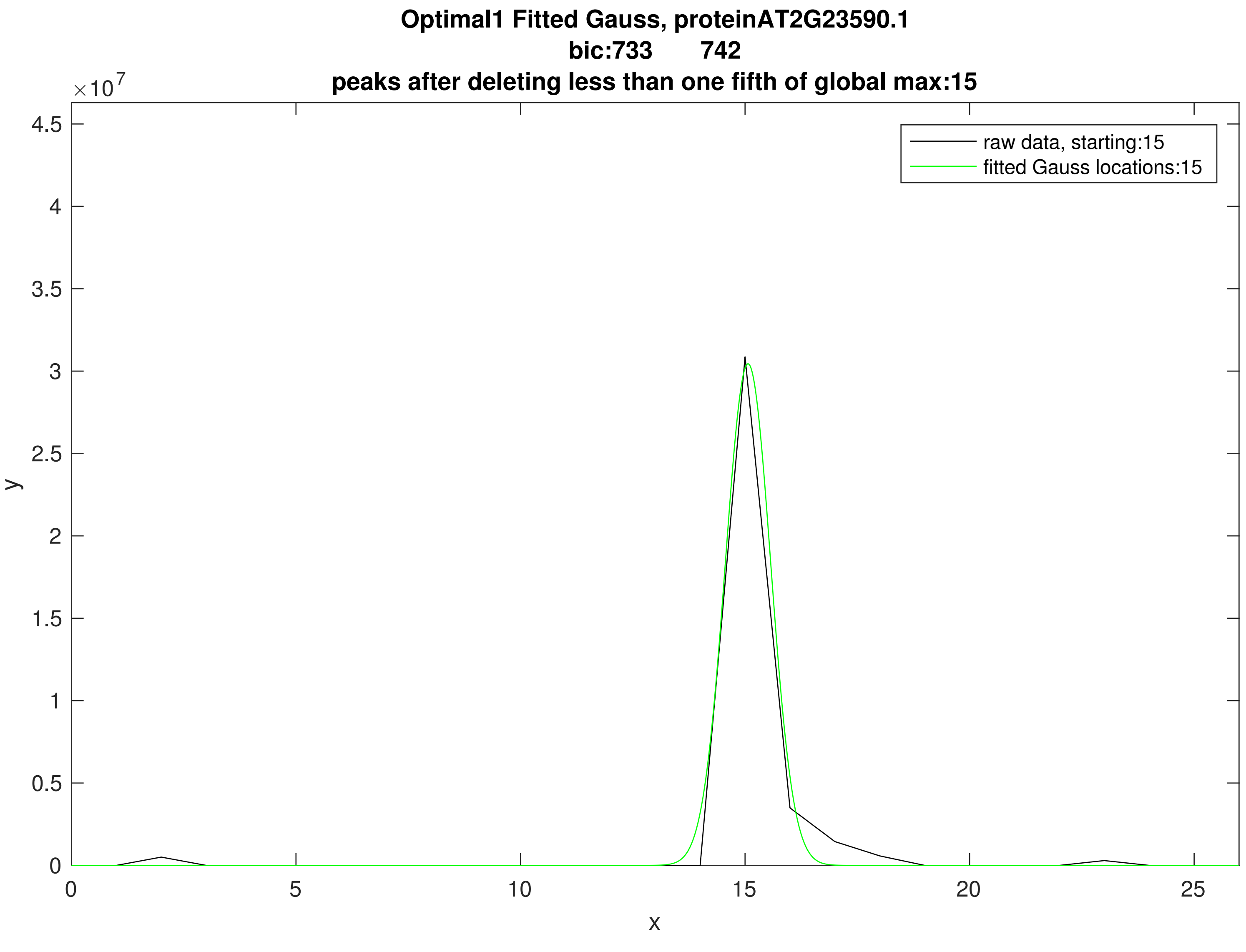
Optimal1 Fitted Gauss, proteinAT2G23590.1
bic:733 742
peaks after deleting less than one fifth of global max:15
×10^7
raw data, starting:15
fitted Gauss locations:15
y
x
0
0.5
1
1.5
2
2.5
3
3.5
4
4.5
0
5
10
15
20
25

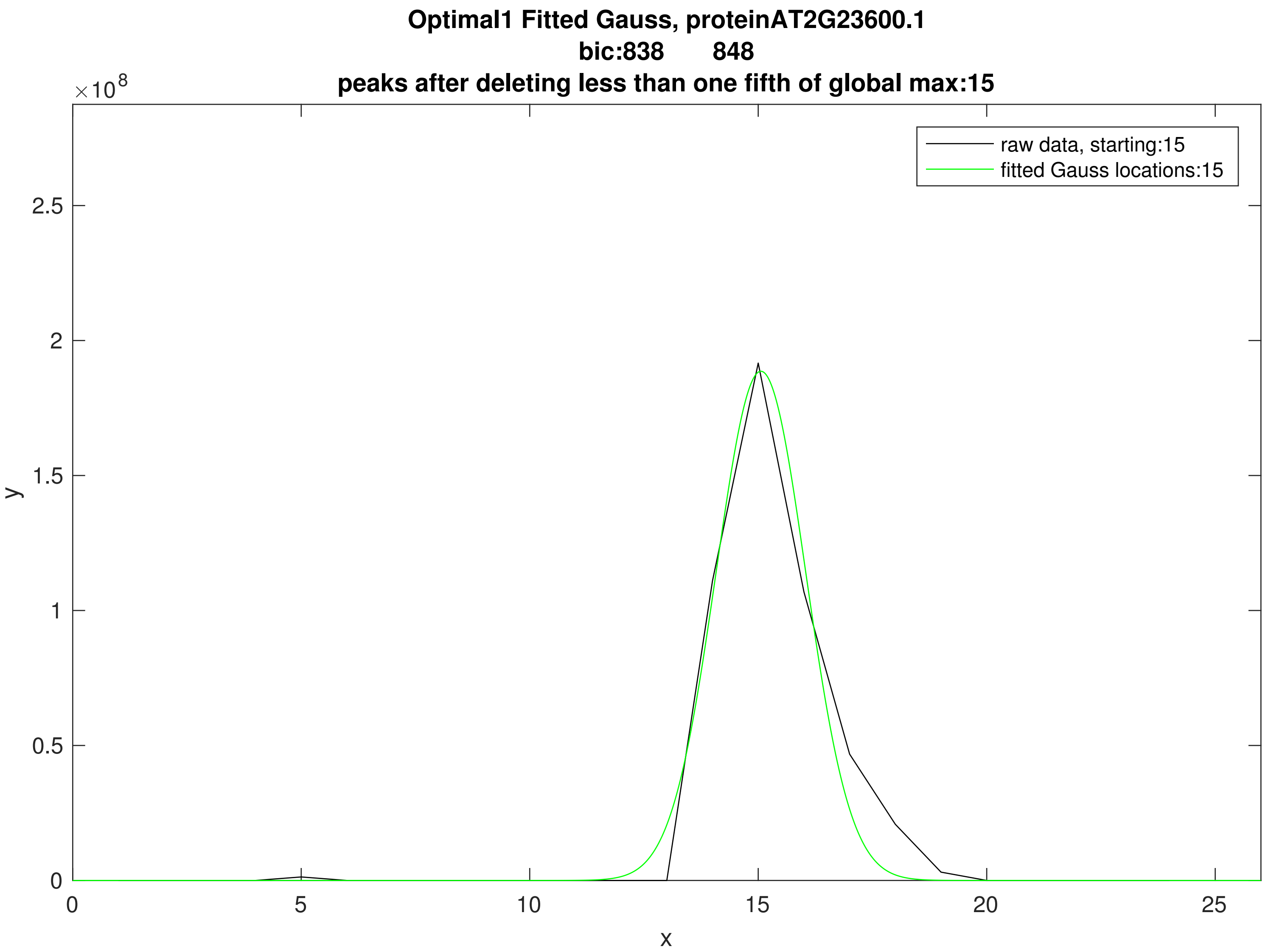
Optimal1 Fitted Gauss, proteinAT2G23600.1
bic:838 848
peaks after deleting less than one fifth of global max:15
raw data, starting:15
fitted Gauss locations:15
×10^8
y
x
0
0.5
1
1.5
2
2.5
0
5
10
15
20
25

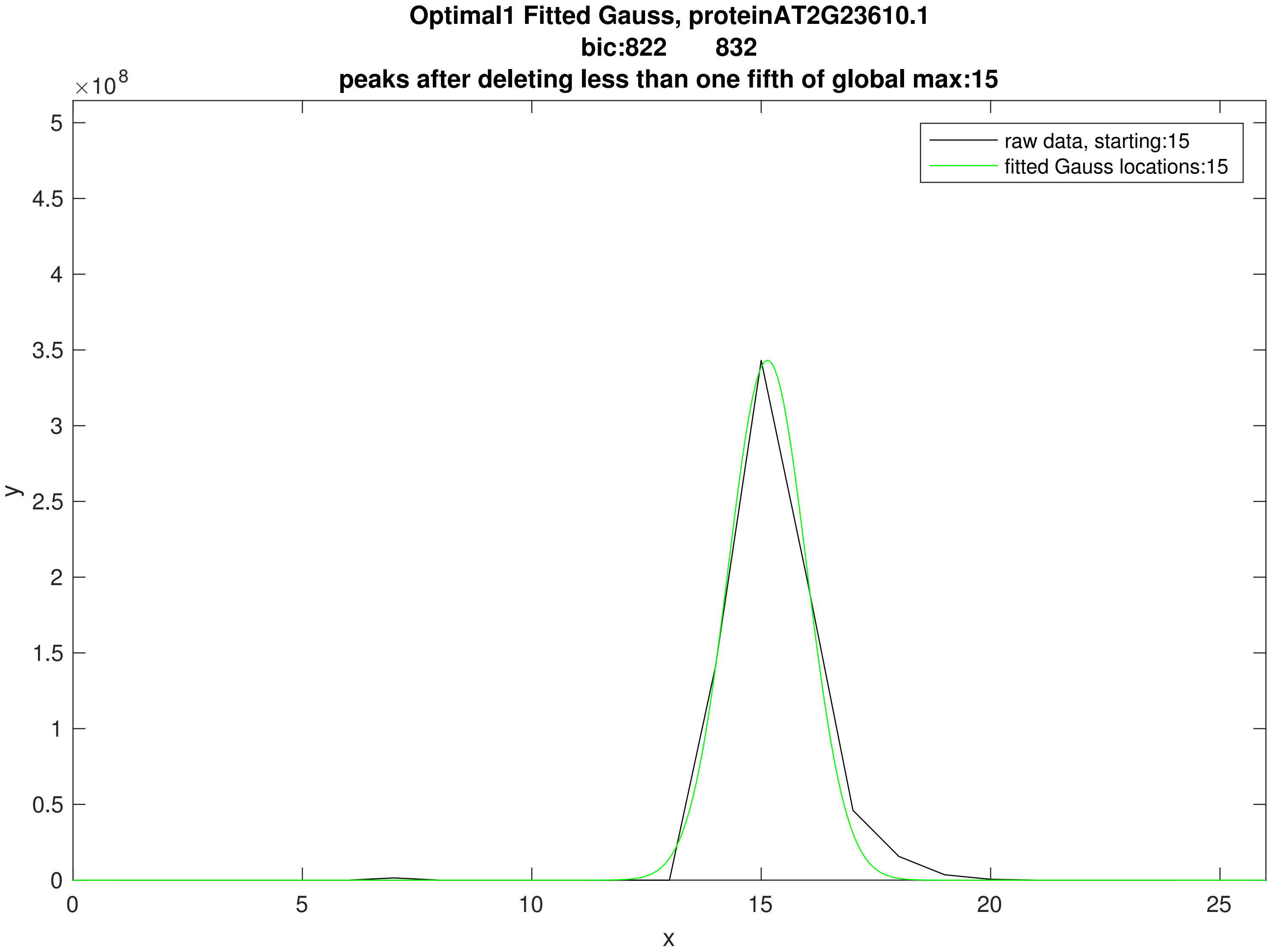
Optimal1 Fitted Gauss, proteinAT2G23610.1
bic:822 832
peaks after deleting less than one fifth of global max:15
×10^8
raw data, starting:15
fitted Gauss locations:15
y
x
0
0.5
1
1.5
2
2.5
3
3.5
4
4.5
5
0
5
10
15
20
25

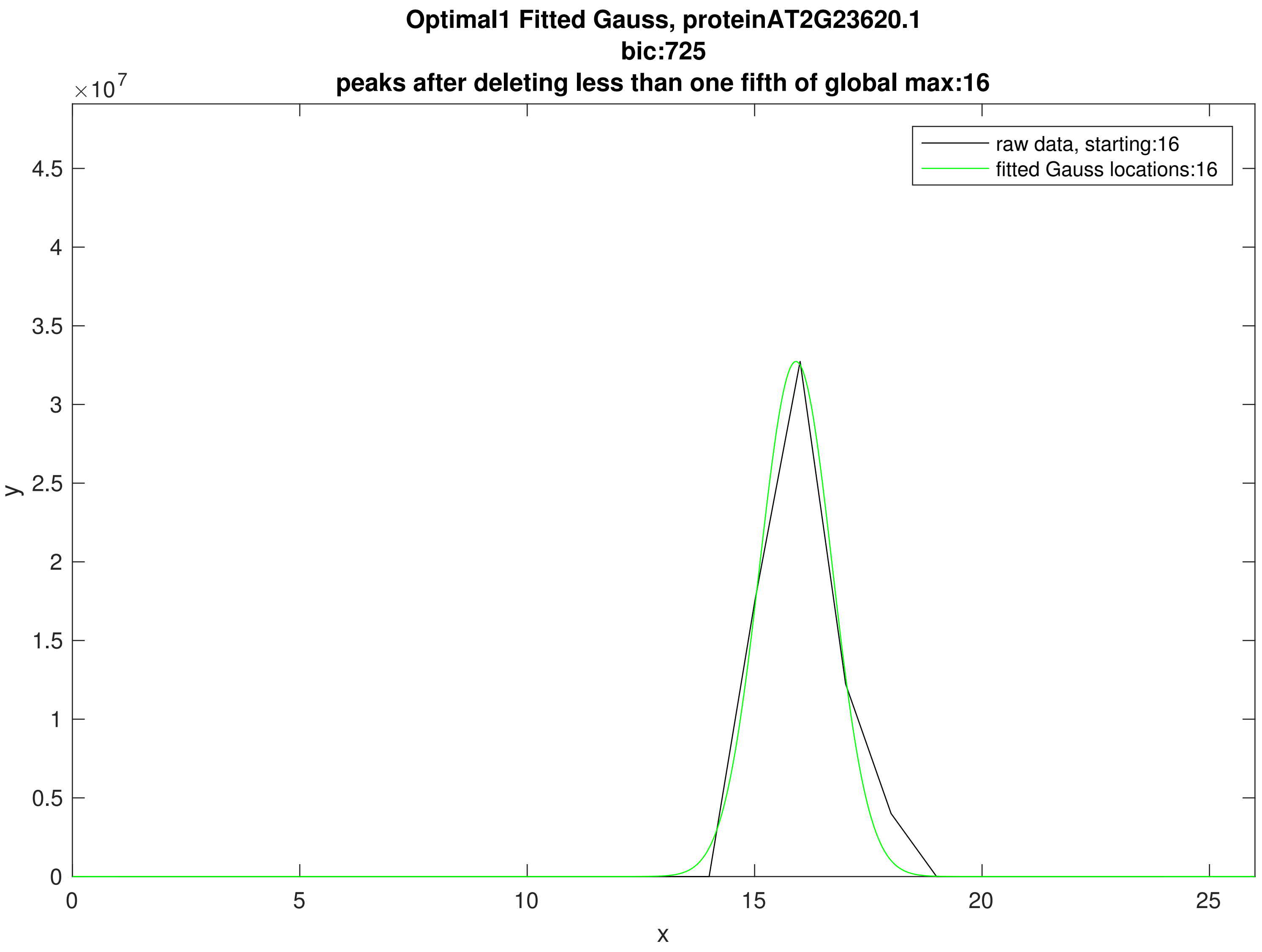
Optimal1 Fitted Gauss, proteinAT2G23620.1
bic:725
peaks after deleting less than one fifth of global max:16
×10^7
raw data, starting:16
fitted Gauss locations:16
y
x
4.5
4
3.5
3
2.5
2
1.5
1
0.5
0
0
5
10
15
20
25

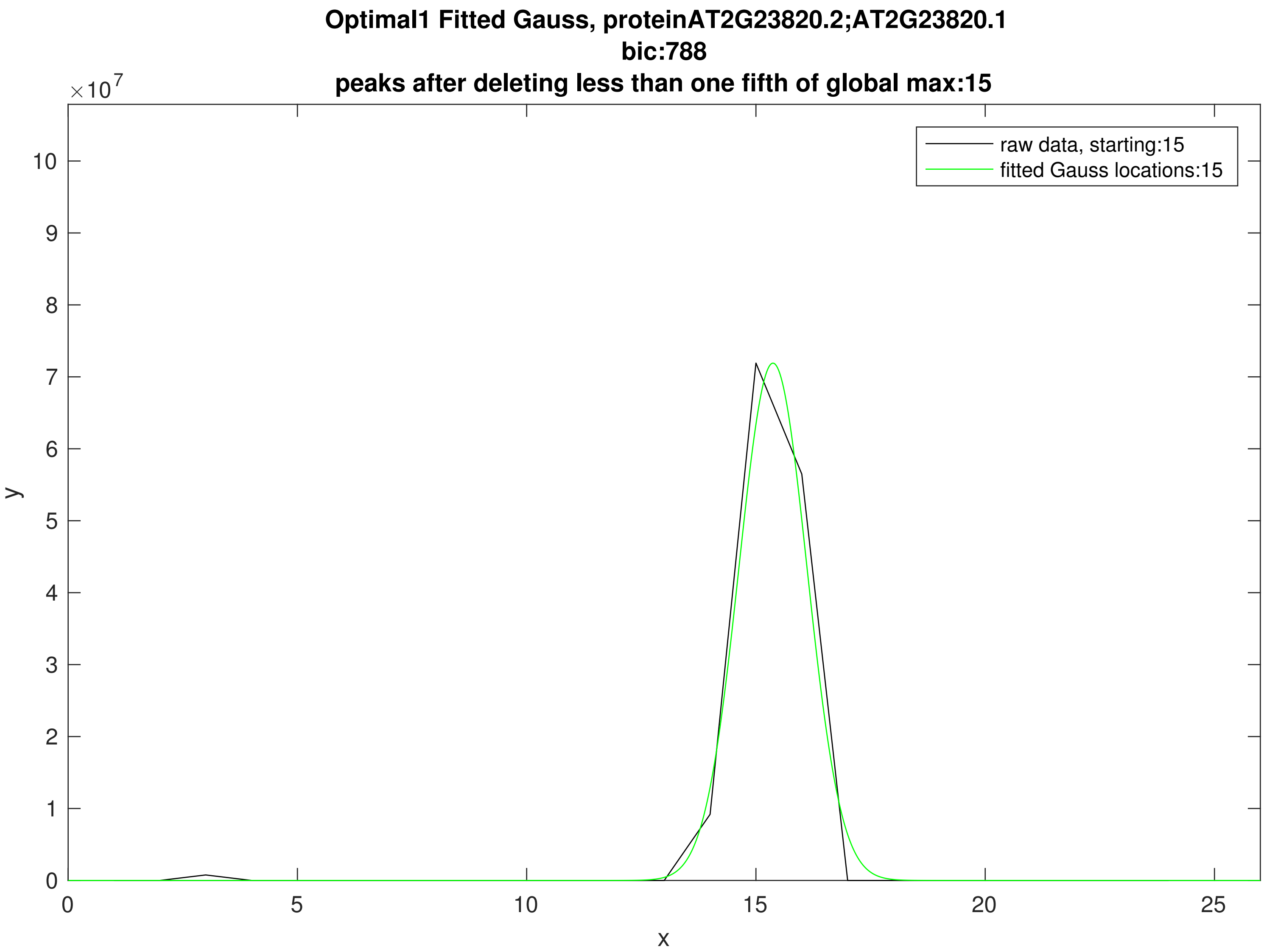

Optimal1 Fitted Gauss, proteinAT2G23820.2;AT2G23820.1
bic:788
peaks after deleting less than one fifth of global max:15
×10^7
raw data, starting:15
fitted Gauss locations:15
y
x
0
1
2
3
4
5
6
7
8
9
10
0
5
10
15
20
25

Optimal1 Fitted Gauss, proteinAT2G24020.1;AT2G24020.2
bic:783
peaks after deleting less than one fifth of global max:17

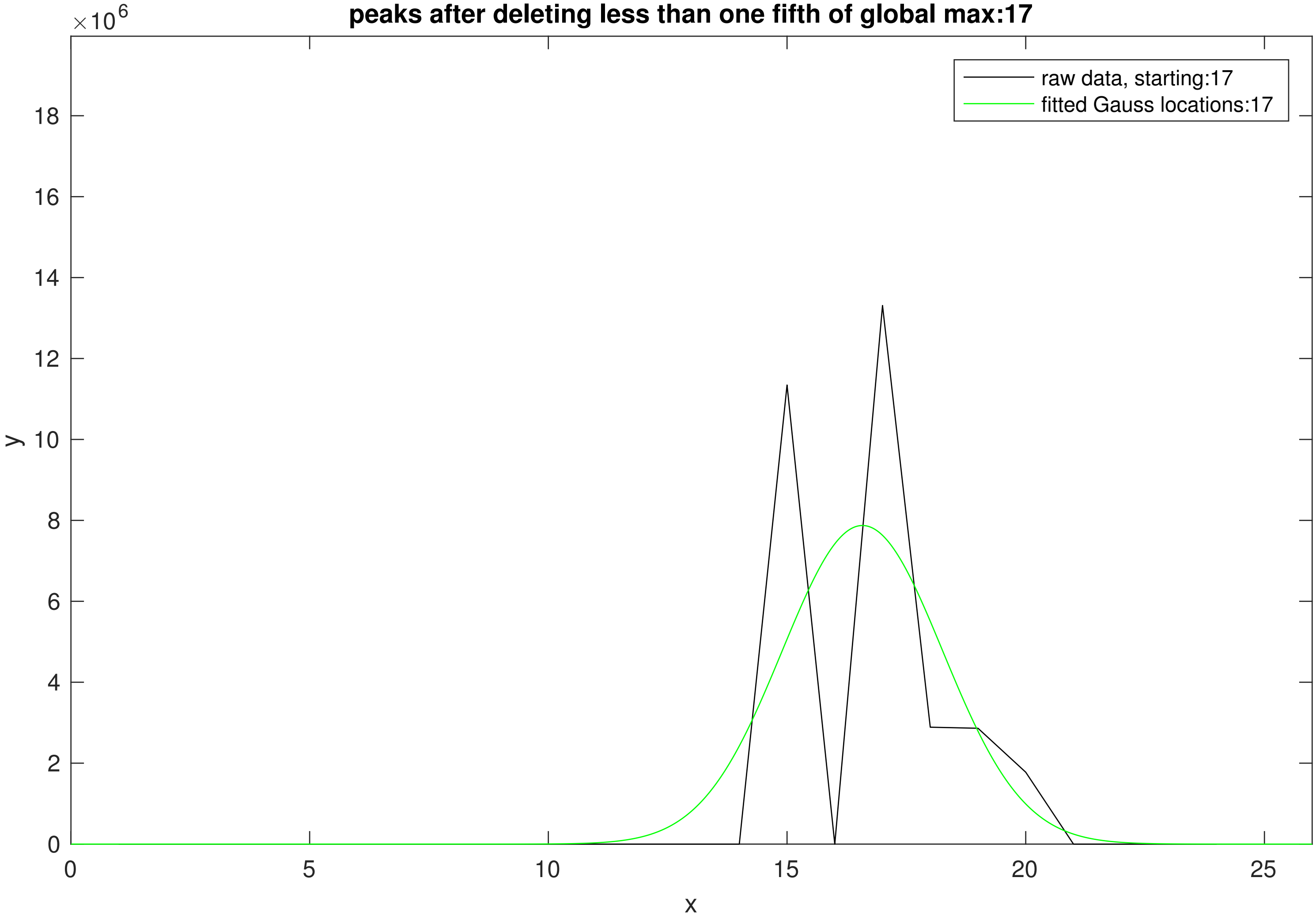

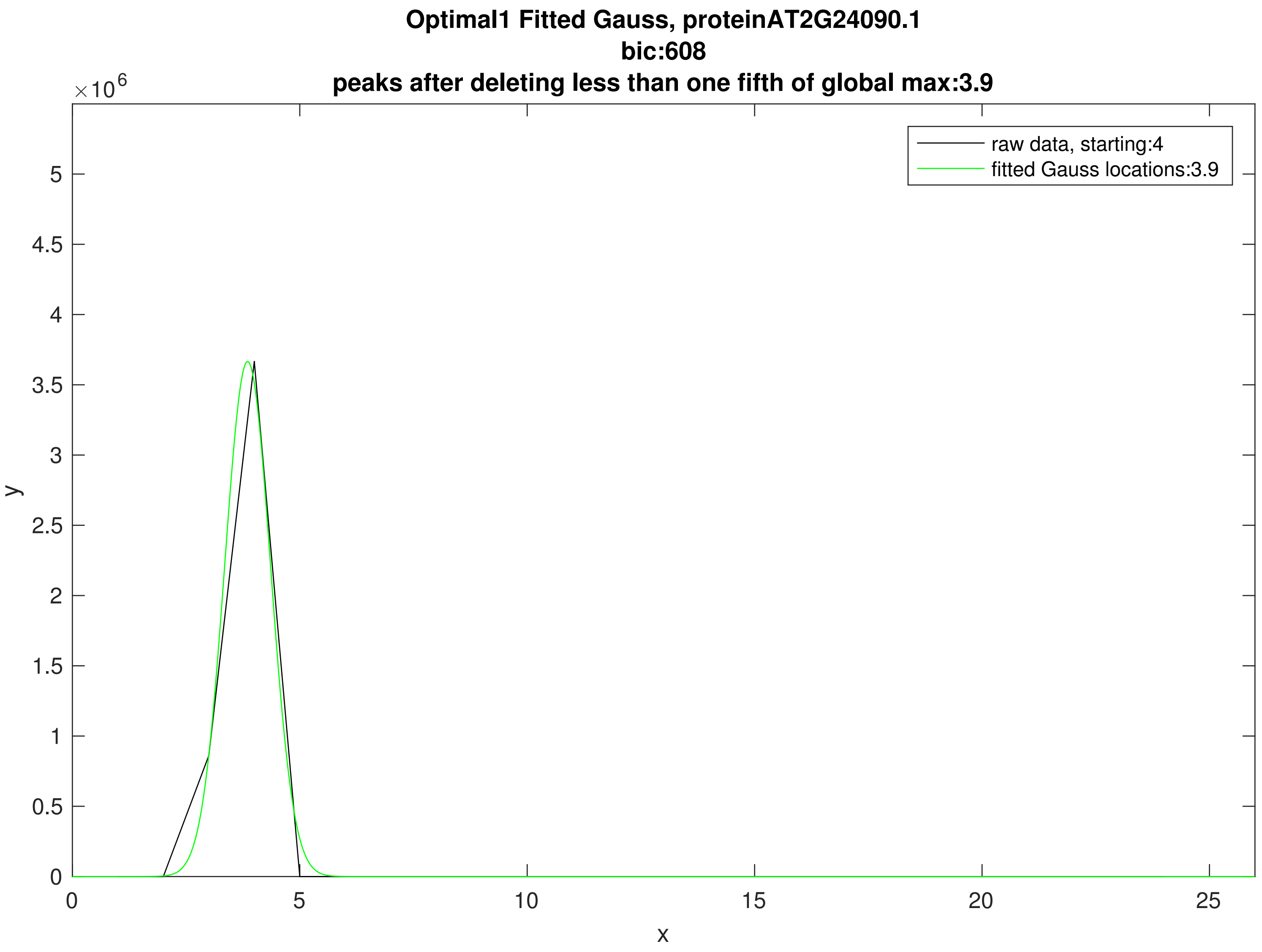

Optimal1 Fitted Gauss, proteinAT2G24090.1
bic:608
peaks after deleting less than one fifth of global max:3.9
×10^6
5
4.5
4
3.5
3
2.5
2
1.5
1
0.5
0
y
0
5
10
15
20
25
x
raw data, starting:4
fitted Gauss locations:3.9

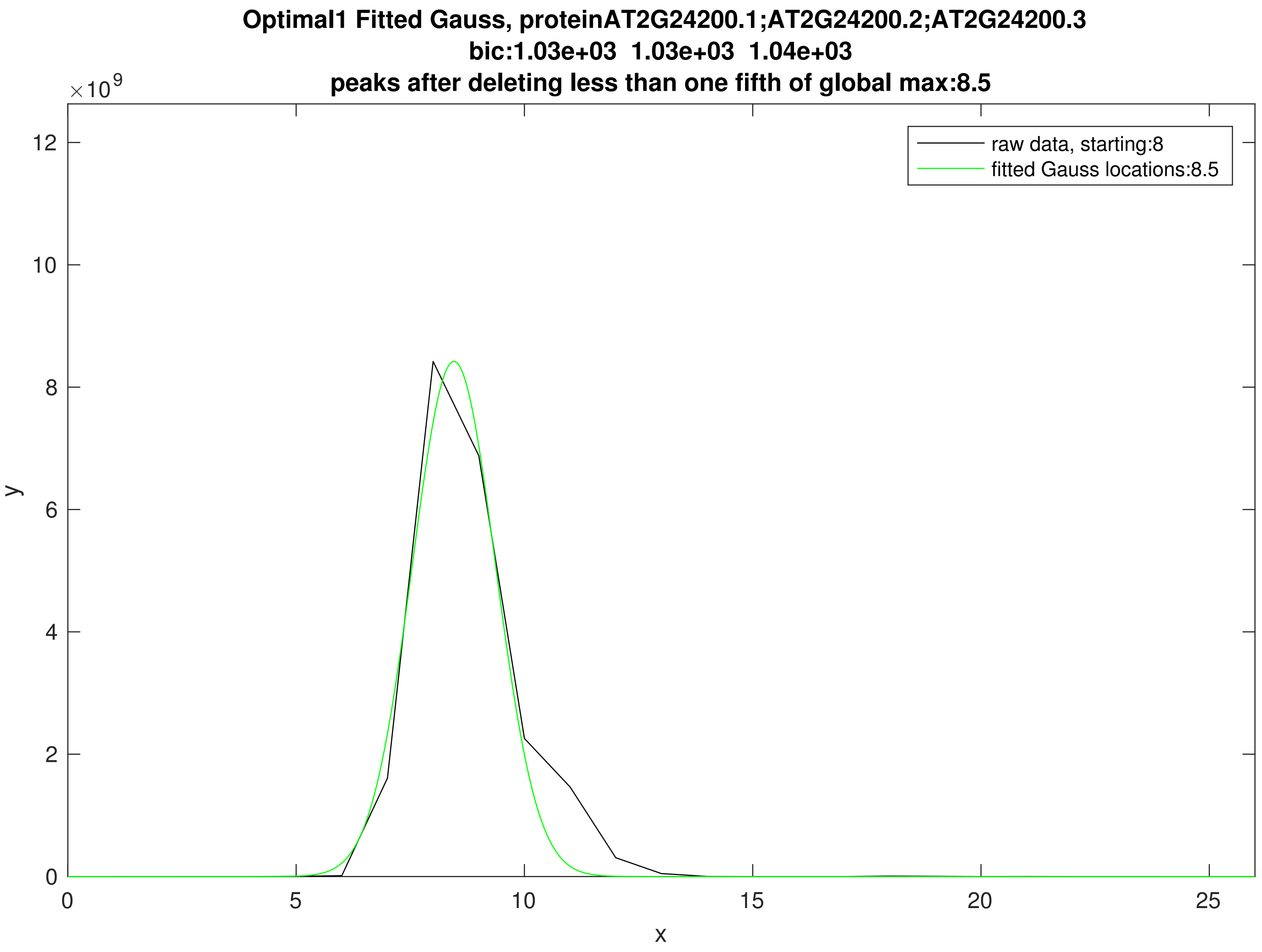
Optimal1 Fitted Gauss, proteinAT2G24200.1;AT2G24200.2;AT2G24200.3
bic:1.03e+03  1.03e+03  1.04e+03
peaks after deleting less than one fifth of global max:8.5
×10^9
raw data, starting:8
fitted Gauss locations:8.5
y
x
0
2
4
6
8
10
12
0
5
10
15
20
25

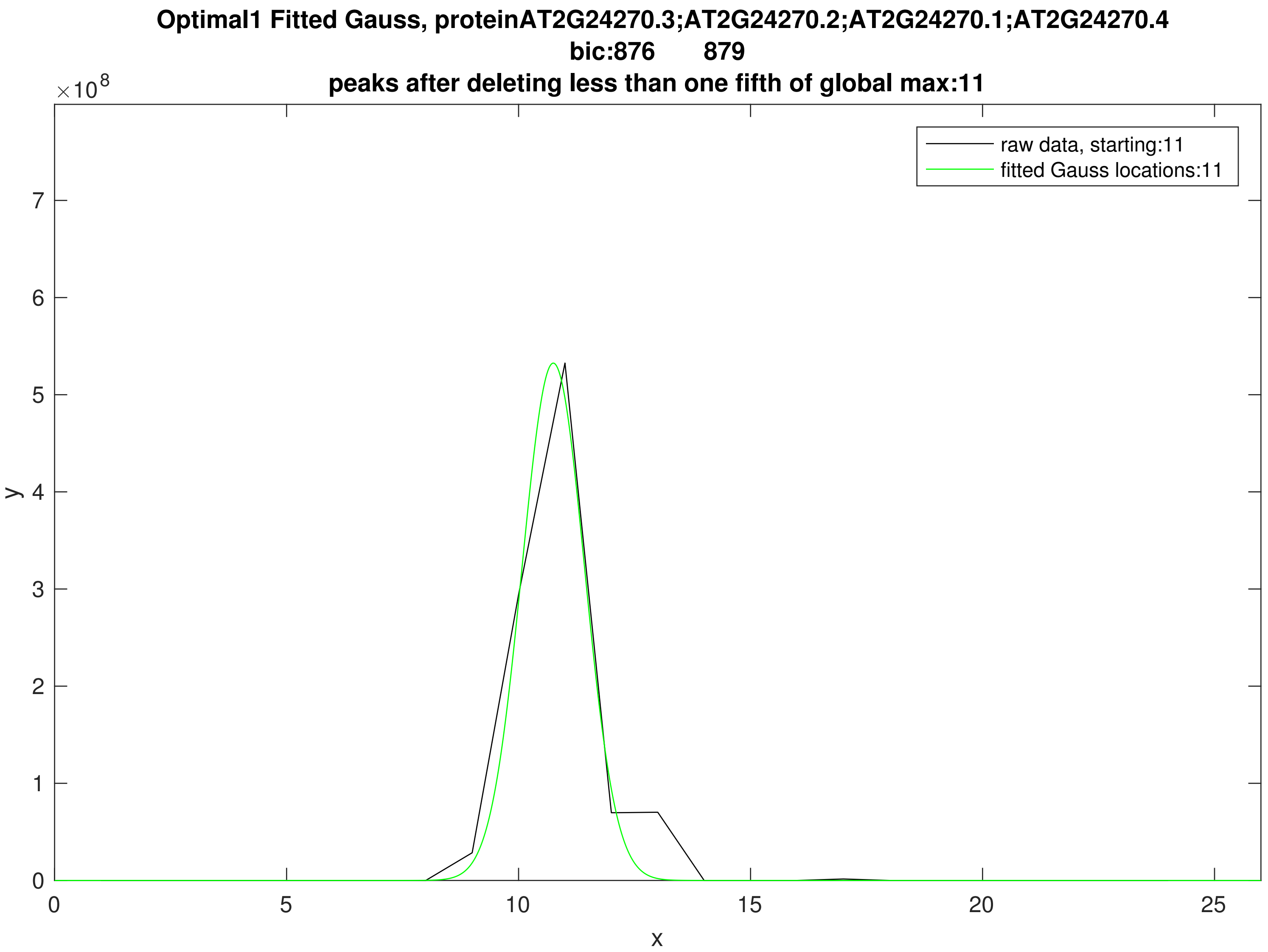
Optimal1 Fitted Gauss, proteinAT2G24270.3;AT2G24270.2;AT2G24270.1;AT2G24270.4
bic:876 879
peaks after deleting less than one fifth of global max:11
×10^8
raw data, starting:11
fitted Gauss locations:11
y
x
0
1
2
3
4
5
6
7
0
5
10
15
20
25

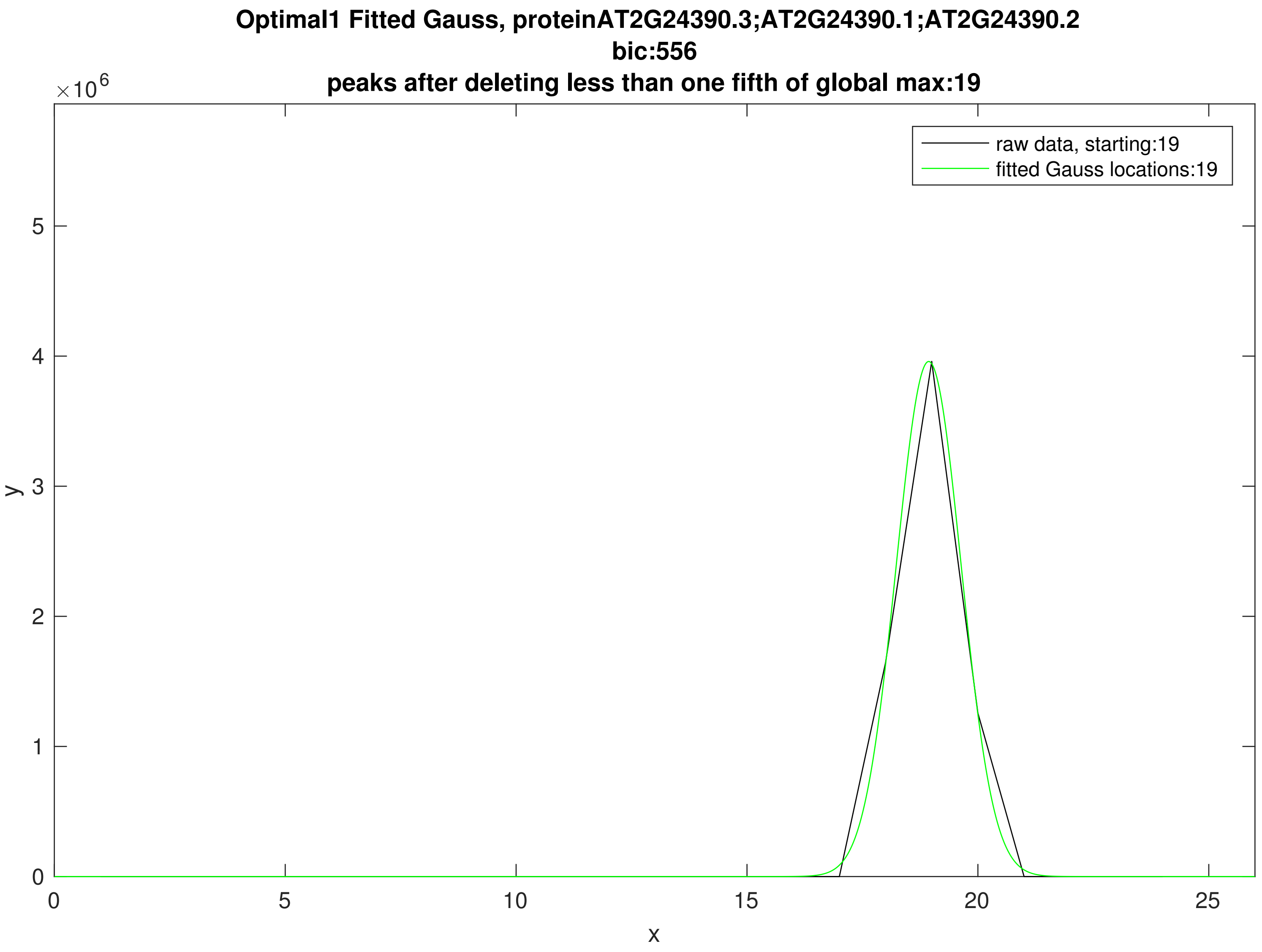
Optimal1 Fitted Gauss, proteinAT2G24390.3;AT2G24390.1;AT2G24390.2
bic:556
peaks after deleting less than one fifth of global max:19
×10^6
raw data, starting:19
fitted Gauss locations:19
0
1
2
3
4
5
y
0
5
10
15
20
25
x

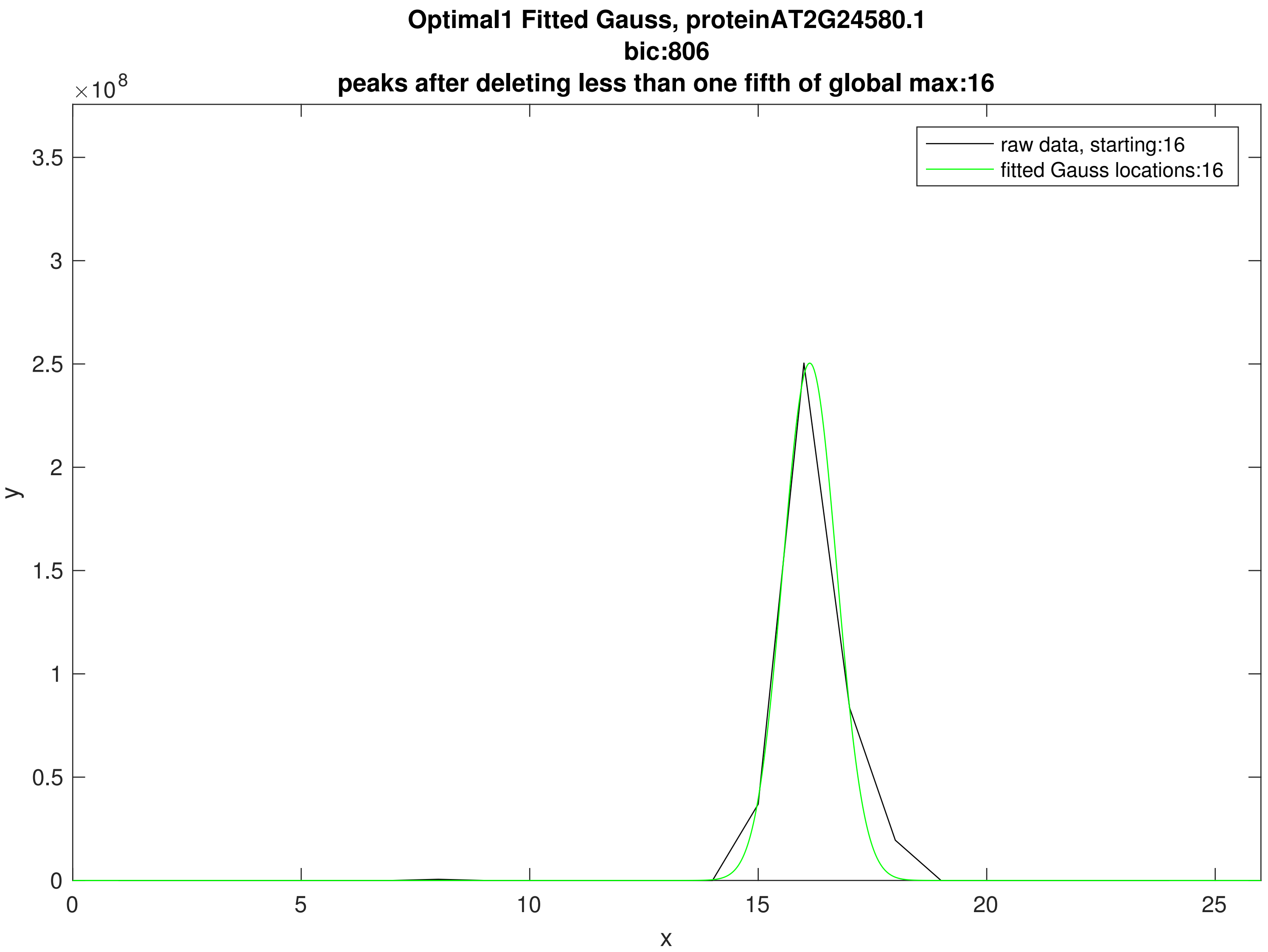

Optimal1 Fitted Gauss, proteinAT2G24580.1
bic:806
peaks after deleting less than one fifth of global max:16
×10^8
raw data, starting:16
fitted Gauss locations:16
y
x
0
0.5
1
1.5
2
2.5
3
3.5
0
5
10
15
20
25

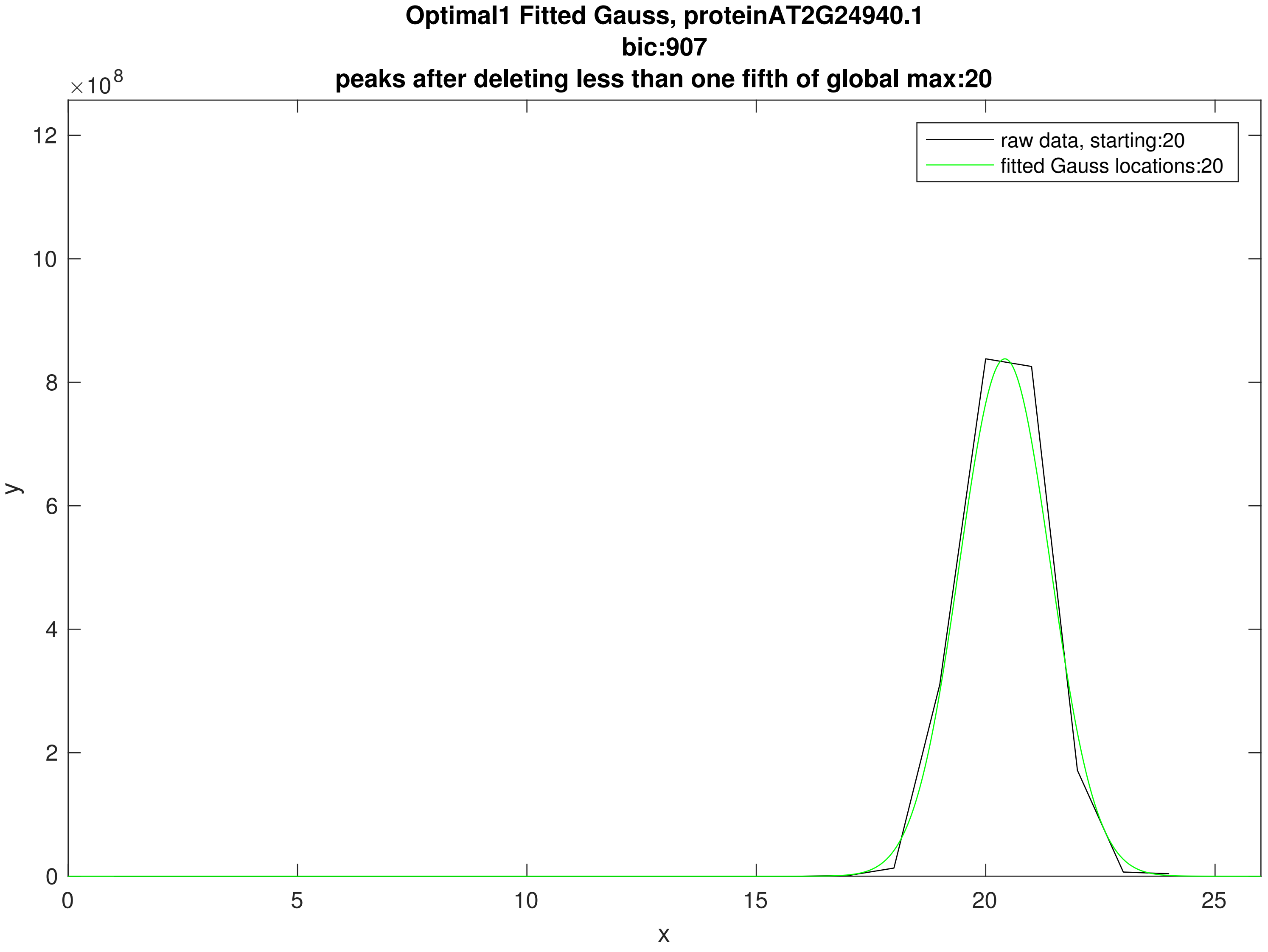
Optimal1 Fitted Gauss, proteinAT2G24940.1
bic:907
peaks after deleting less than one fifth of global max:20
×10^8
12
10
8
6
4
2
0
y
0
5
10
15
20
25
x
raw data, starting:20
fitted Gauss locations:20

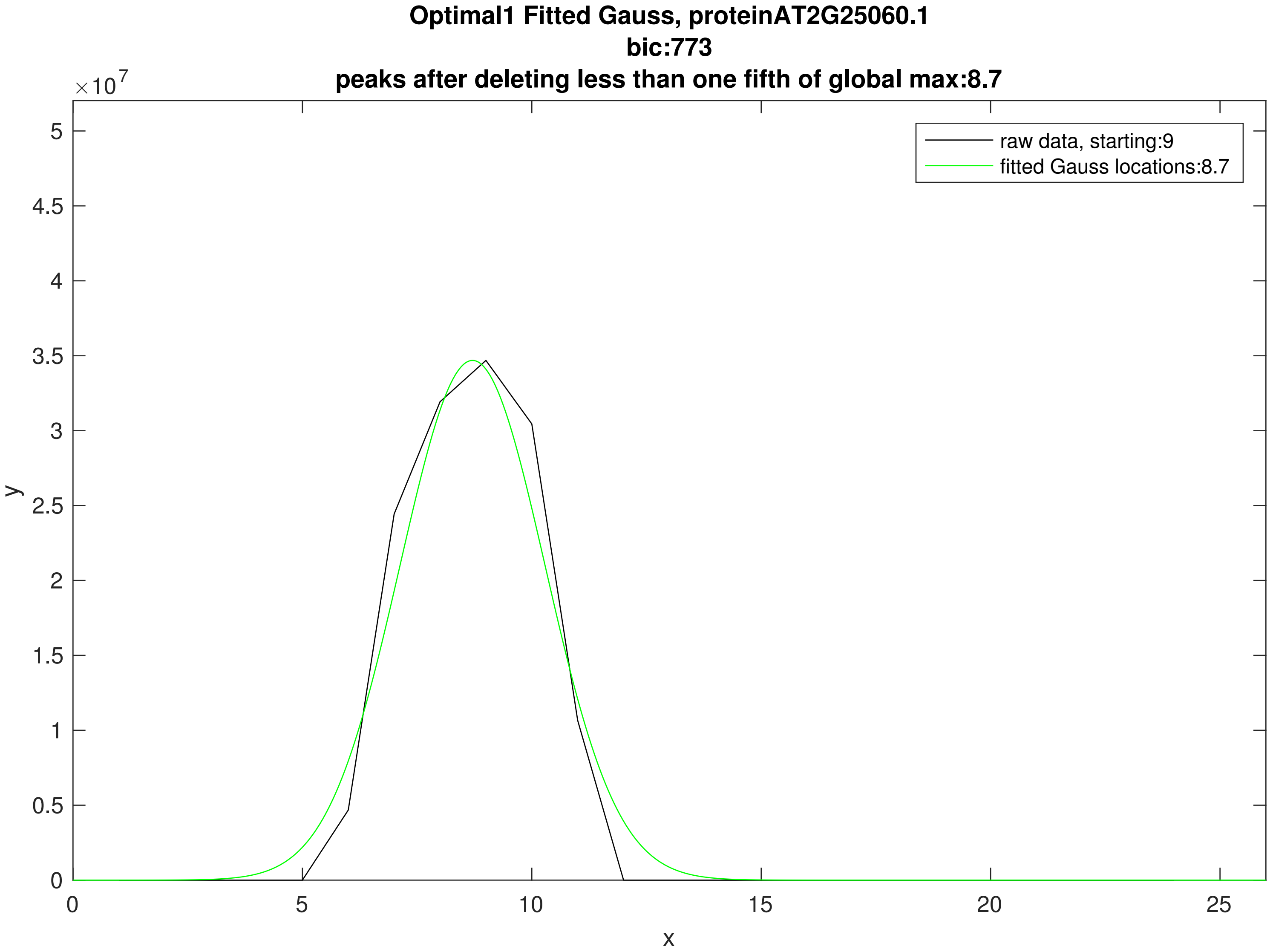
Optimal1 Fitted Gauss, proteinAT2G25060.1
bic:773
peaks after deleting less than one fifth of global max:8.7
×10^7
raw data, starting:9
fitted Gauss locations:8.7
y
x
0
0.5
1
1.5
2
2.5
3
3.5
4
4.5
5
0
5
10
15
20
25

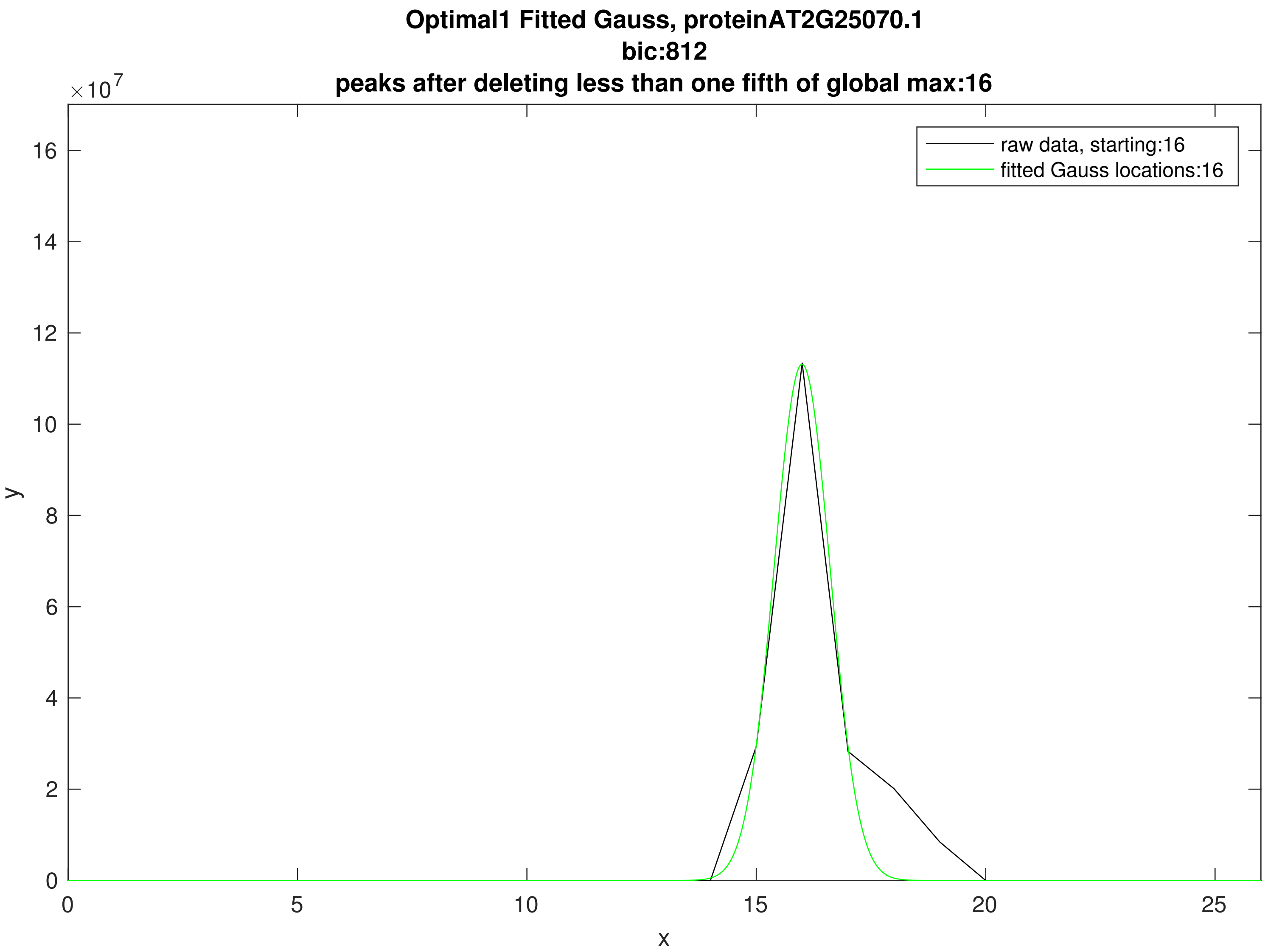
Optimal1 Fitted Gauss, proteinAT2G25070.1
bic:812
peaks after deleting less than one fifth of global max:16
×10^7
16
14
12
10
8
6
4
2
0
y
0
5
10
15
20
25
x
raw data, starting:16
fitted Gauss locations:16

Optimal1 Fitted Gauss, proteinAT2G25110.1
bic:699
peaks after deleting less than one fifth of global max:20

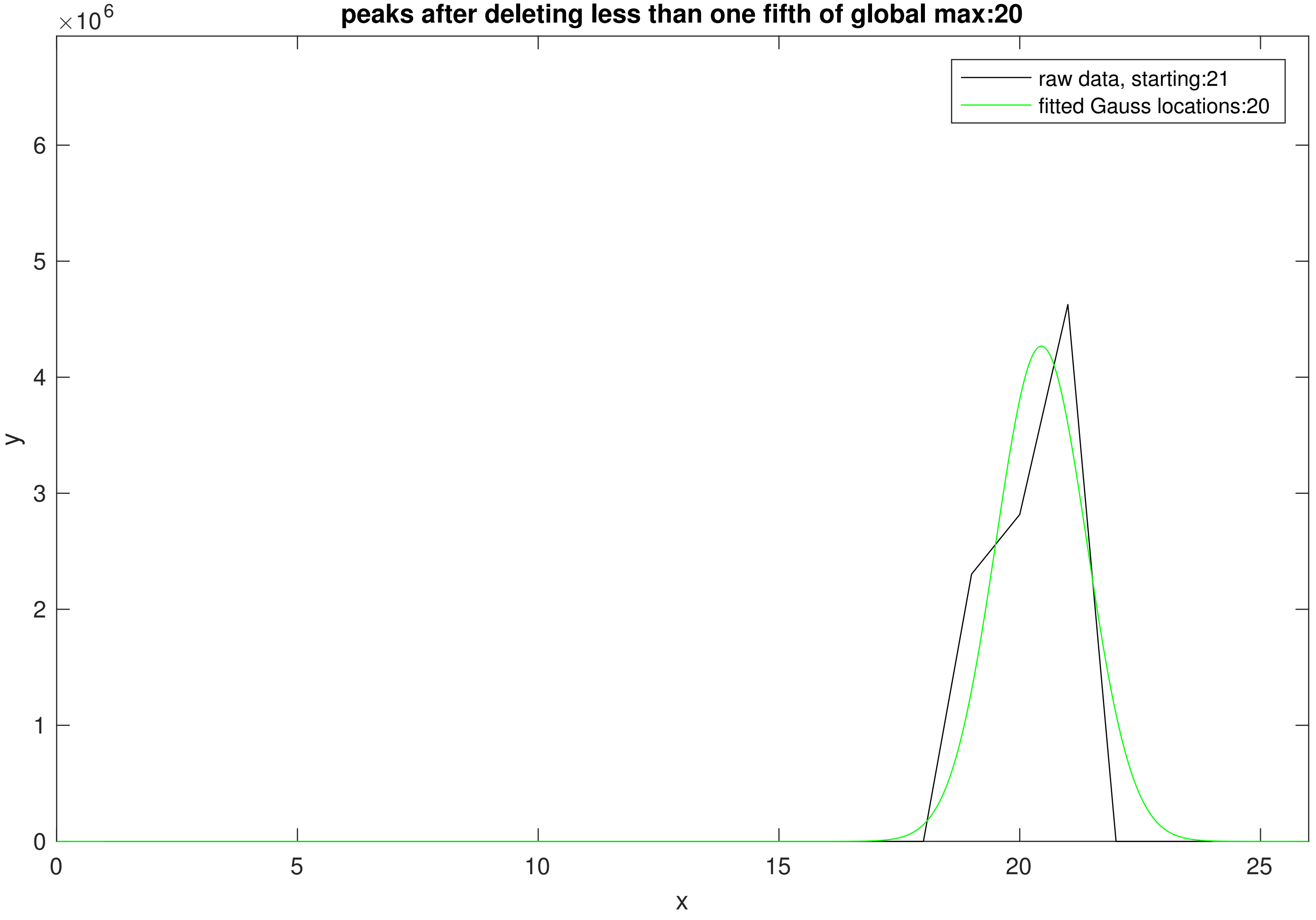

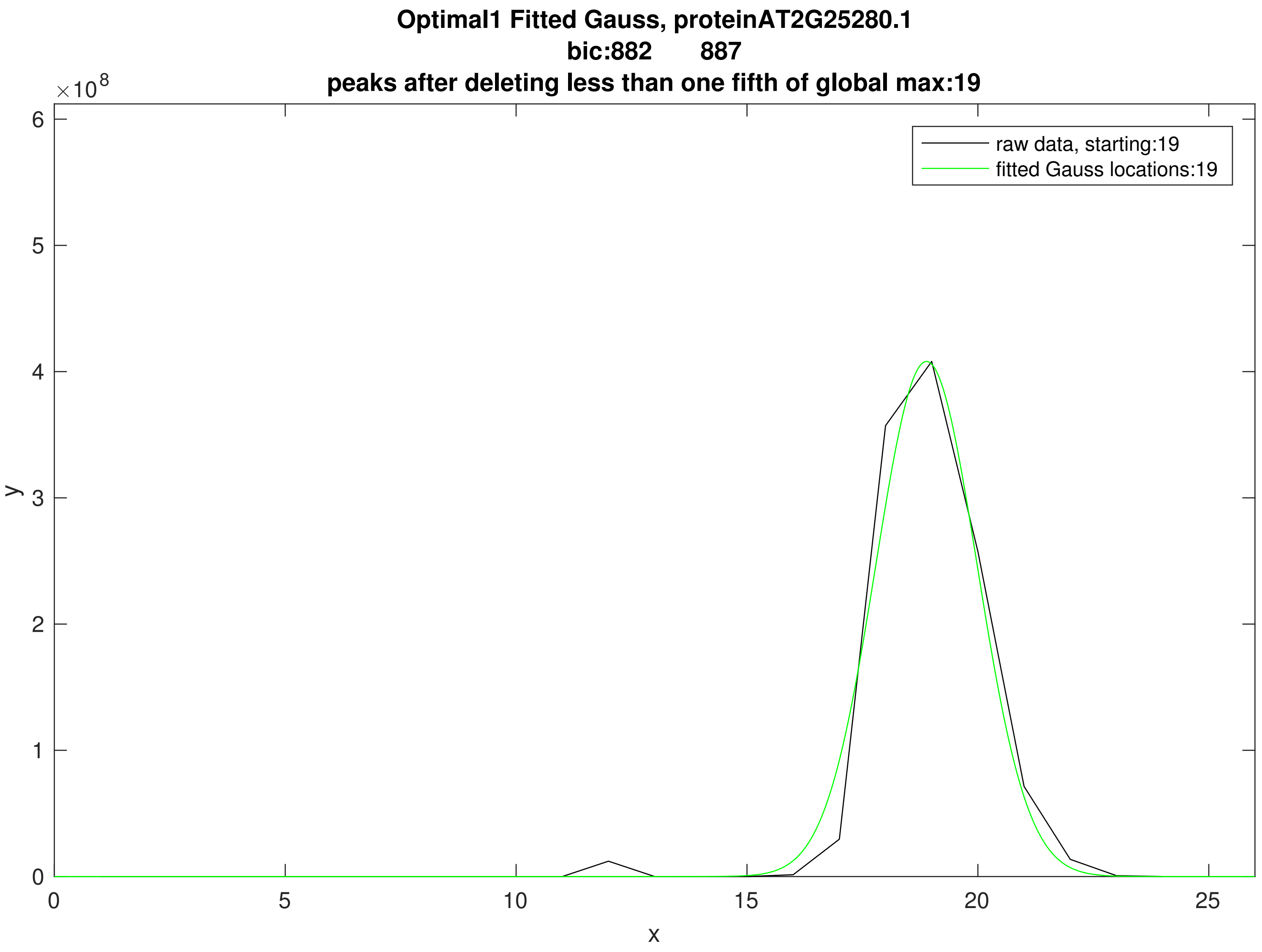

Optimal1 Fitted Gauss, proteinAT2G25280.1
bic:882 887
peaks after deleting less than one fifth of global max:19
×10^8
6
5
4
3
2
1
0
y
0
5
10
15
20
25
x
raw data, starting:19
fitted Gauss locations:19

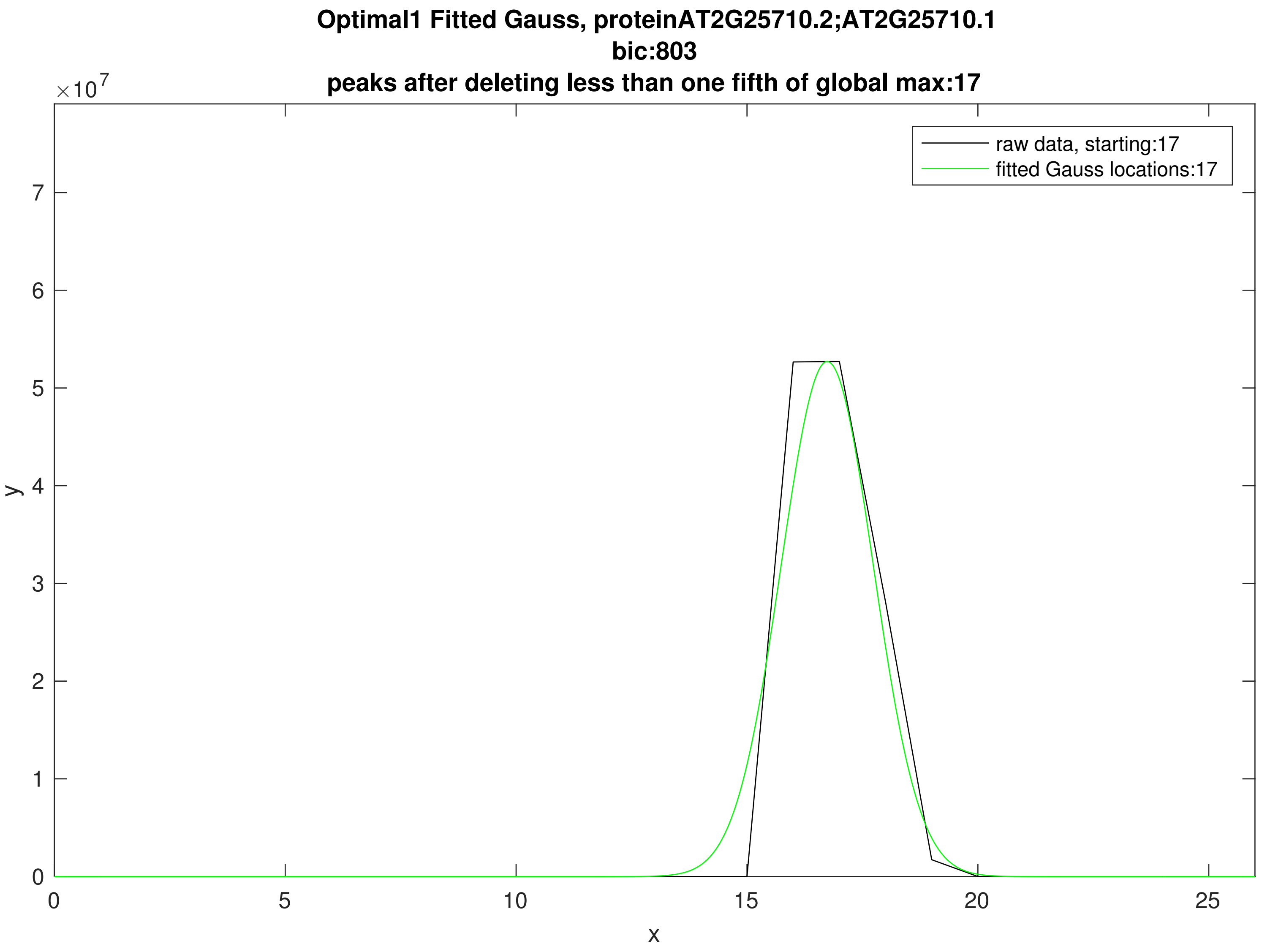

Optimal1 Fitted Gauss, proteinAT2G25710.2;AT2G25710.1
bic:803
peaks after deleting less than one fifth of global max:17
×10^7
raw data, starting:17
fitted Gauss locations:17
y
x
0
1
2
3
4
5
6
7
0
5
10
15
20
25

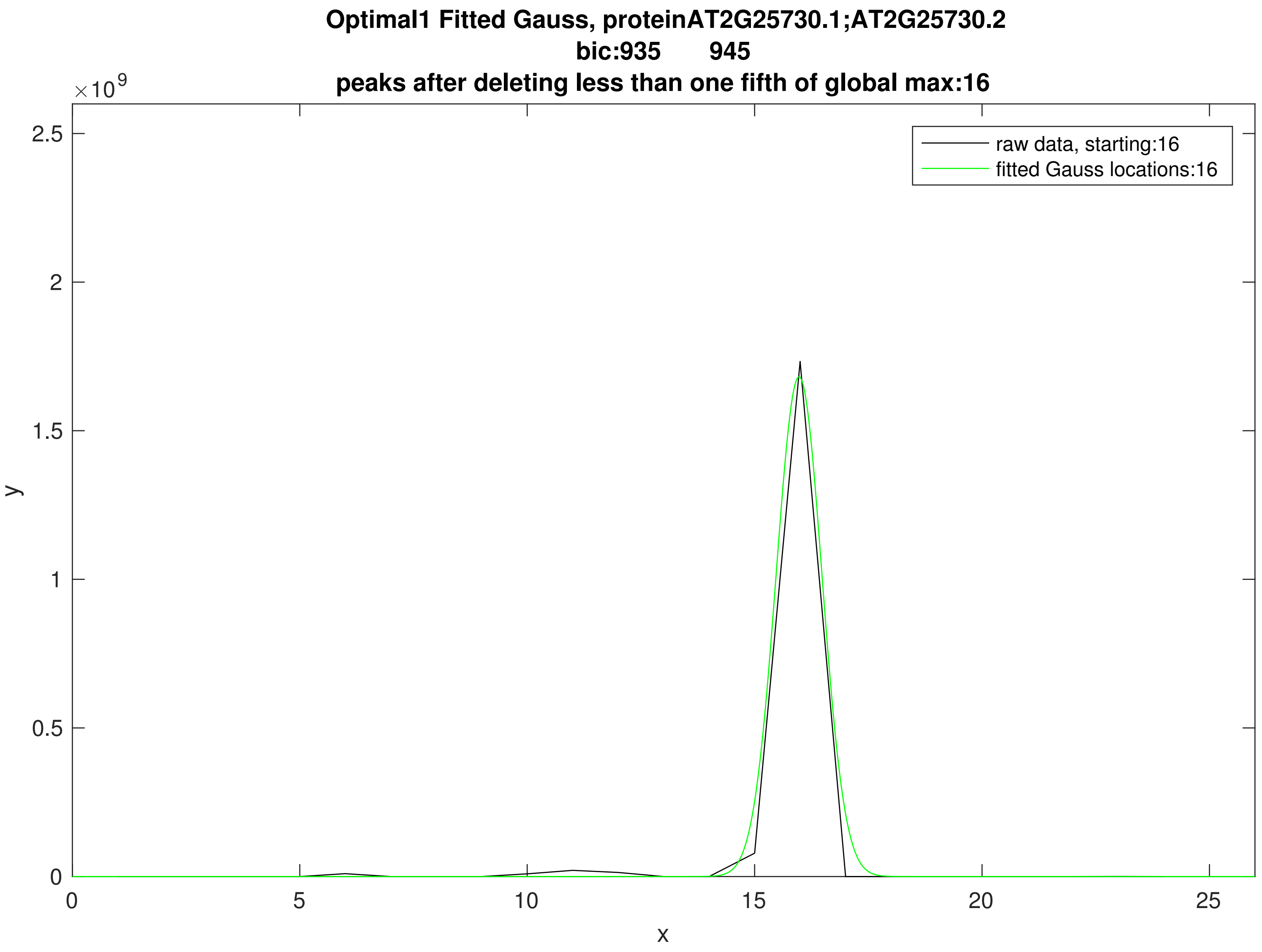
Optimal1 Fitted Gauss, proteinAT2G25730.1;AT2G25730.2
bic:935 945
peaks after deleting less than one fifth of global max:16
×10^9
2.5
2
1.5
1
0.5
0
y
0
5
10
15
20
25
x
raw data, starting:16
fitted Gauss locations:16

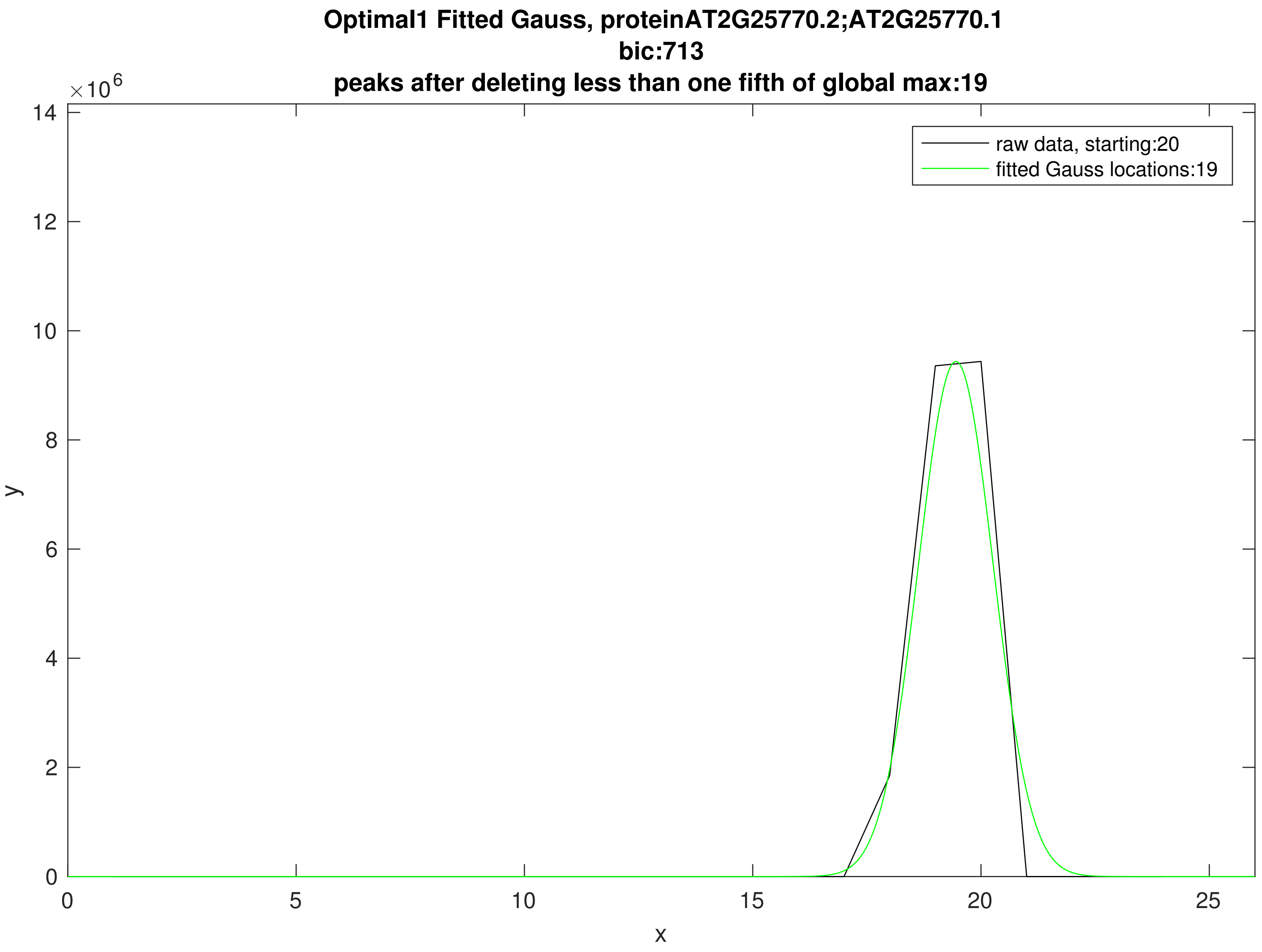
Optimal1 Fitted Gauss, proteinAT2G25770.2;AT2G25770.1
bic:713
peaks after deleting less than one fifth of global max:19
×10^6
raw data, starting:20
fitted Gauss locations:19
y
x
0
2
4
6
8
10
12
14
5
10
15
20
25

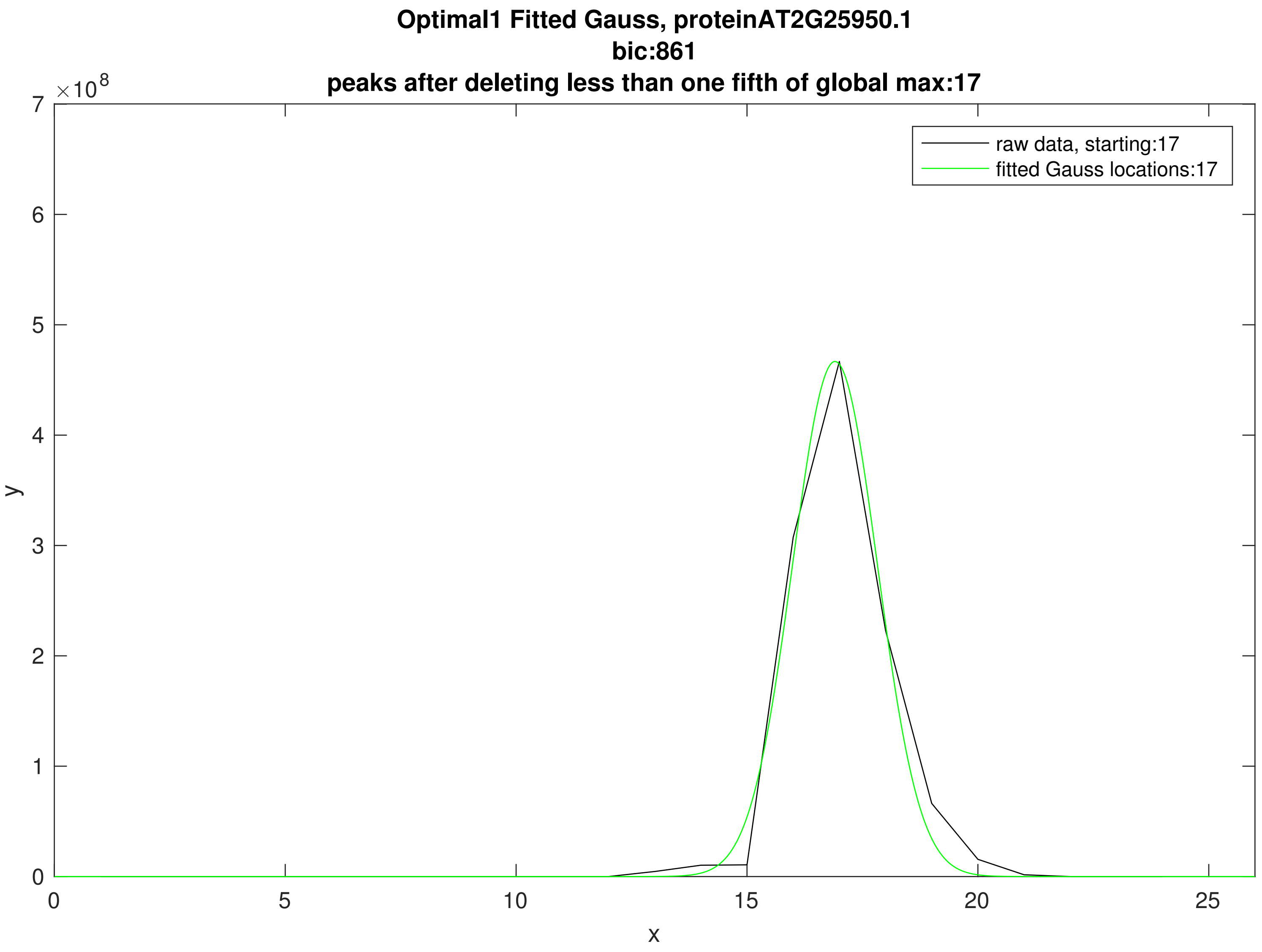
Optimal1 Fitted Gauss, proteinAT2G25950.1
bic:861
peaks after deleting less than one fifth of global max:17
×10^8
raw data, starting:17
fitted Gauss locations:17
y
x
0
1
2
3
4
5
6
7
0
5
10
15
20
25

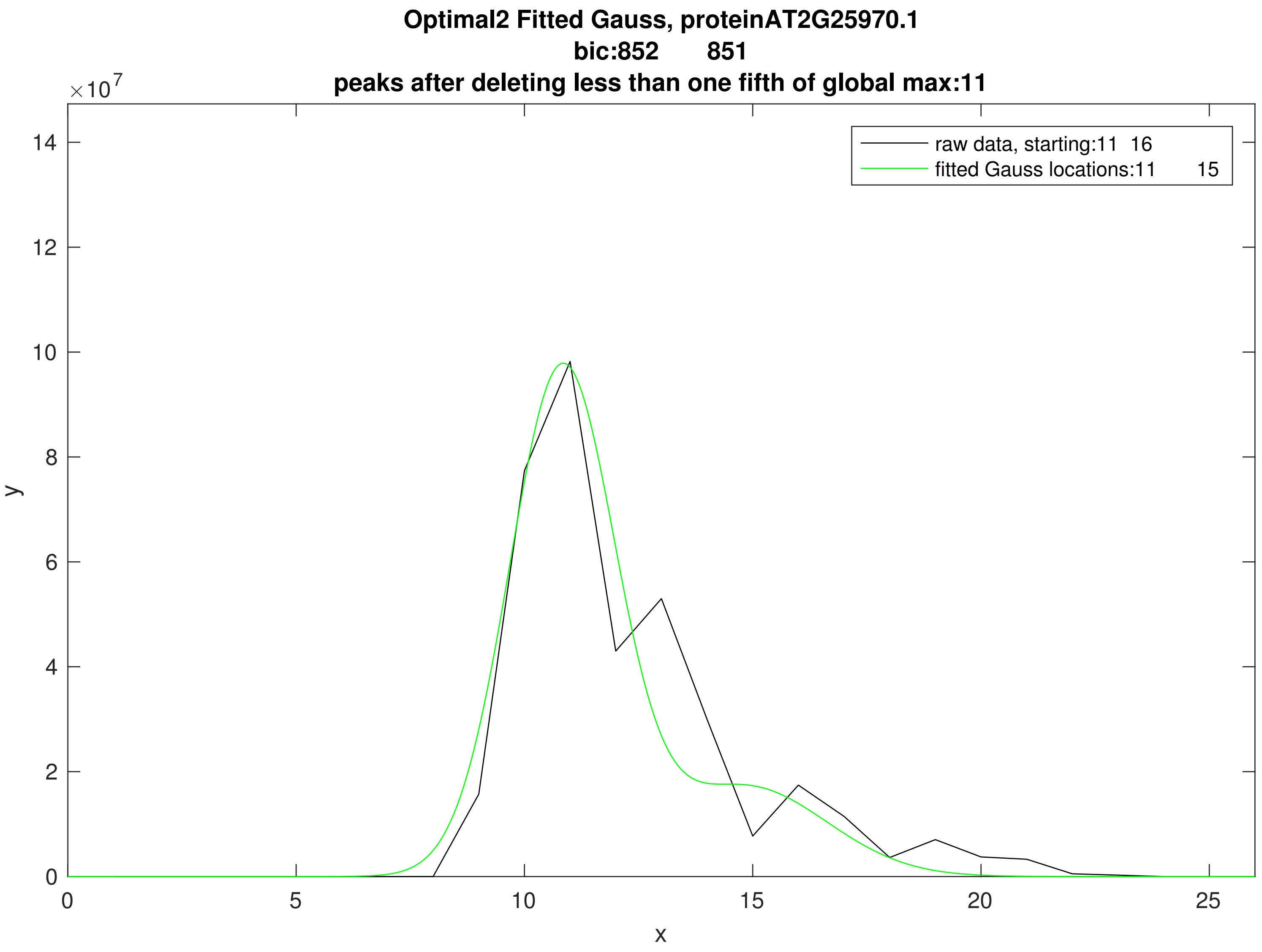
Optimal2 Fitted Gauss, proteinAT2G25970.1
bic:852 851
peaks after deleting less than one fifth of global max:11
×10^7
raw data, starting:11 16
fitted Gauss locations:11 15
y
x
0
2
4
6
8
10
12
14
0
5
10
15
20
25

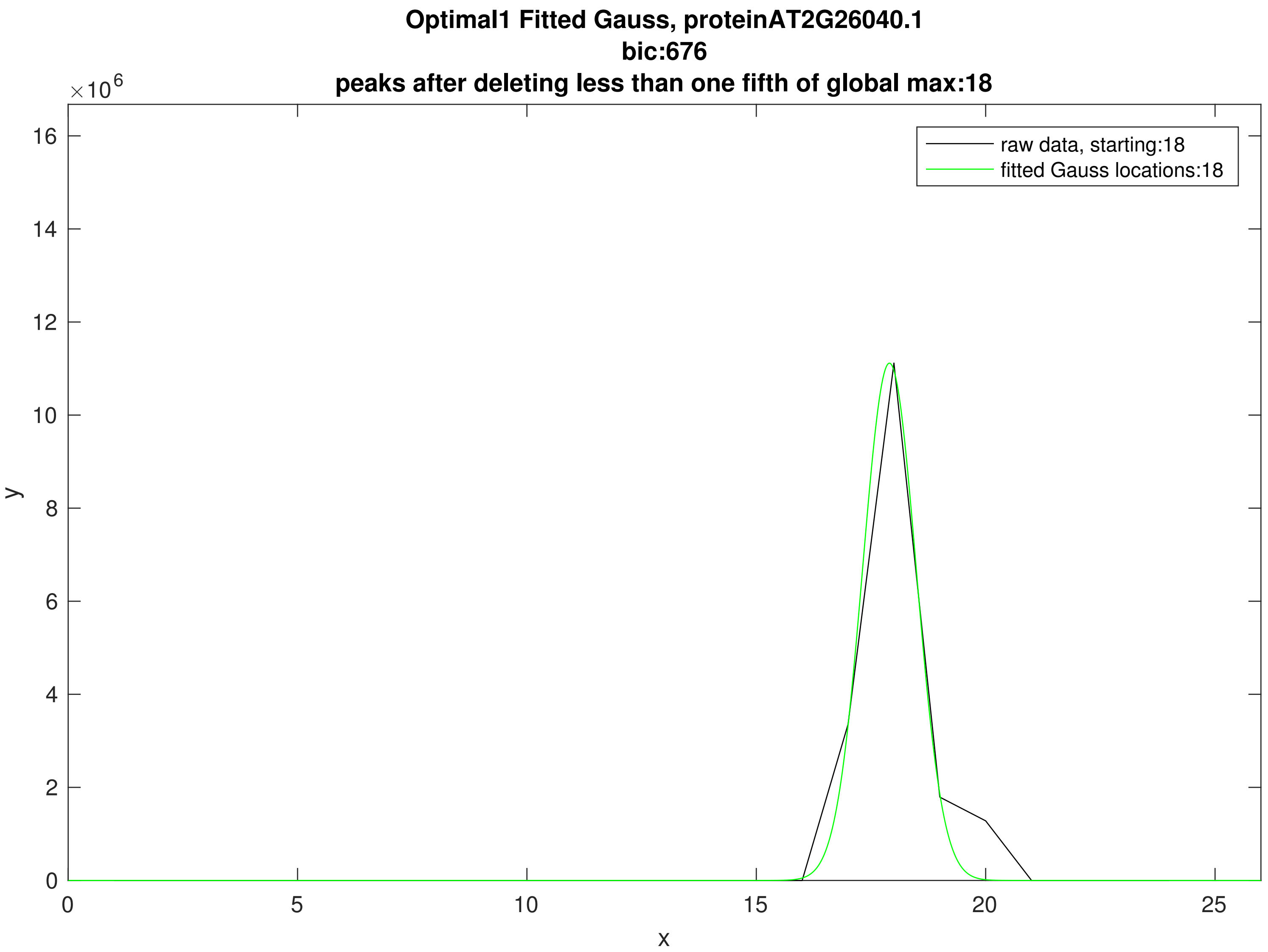

Optimal1 Fitted Gauss, proteinAT2G26040.1
bic:676
peaks after deleting less than one fifth of global max:18
×10^6
16
14
12
10
8
6
4
2
0
y
0
5
10
15
20
25
x
raw data, starting:18
fitted Gauss locations:18

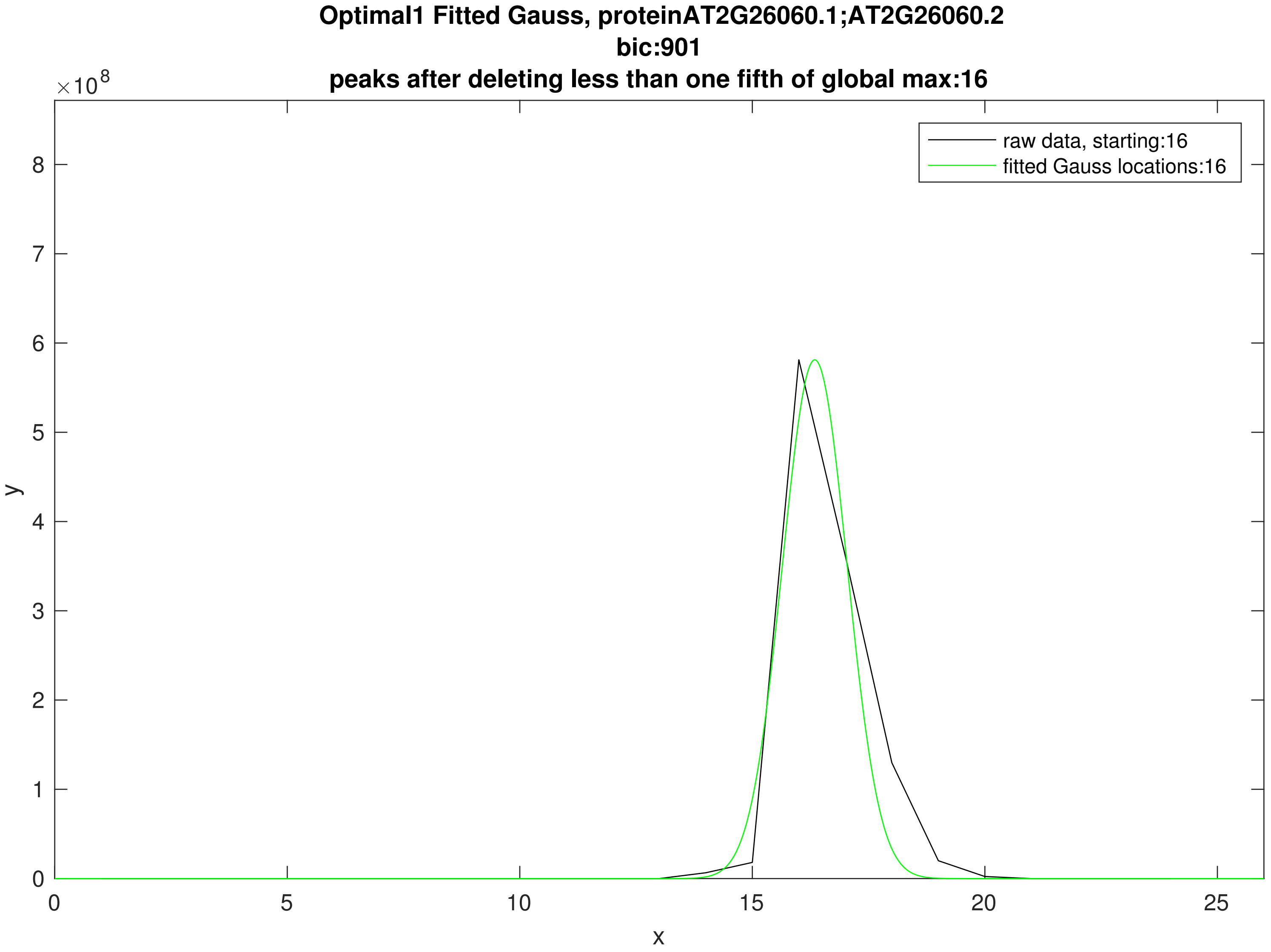
Optimal1 Fitted Gauss, proteinAT2G26060.1;AT2G26060.2
bic:901
peaks after deleting less than one fifth of global max:16
×10^8
raw data, starting:16
fitted Gauss locations:16
y
x
0
1
2
3
4
5
6
7
8
0
5
10
15
20
25

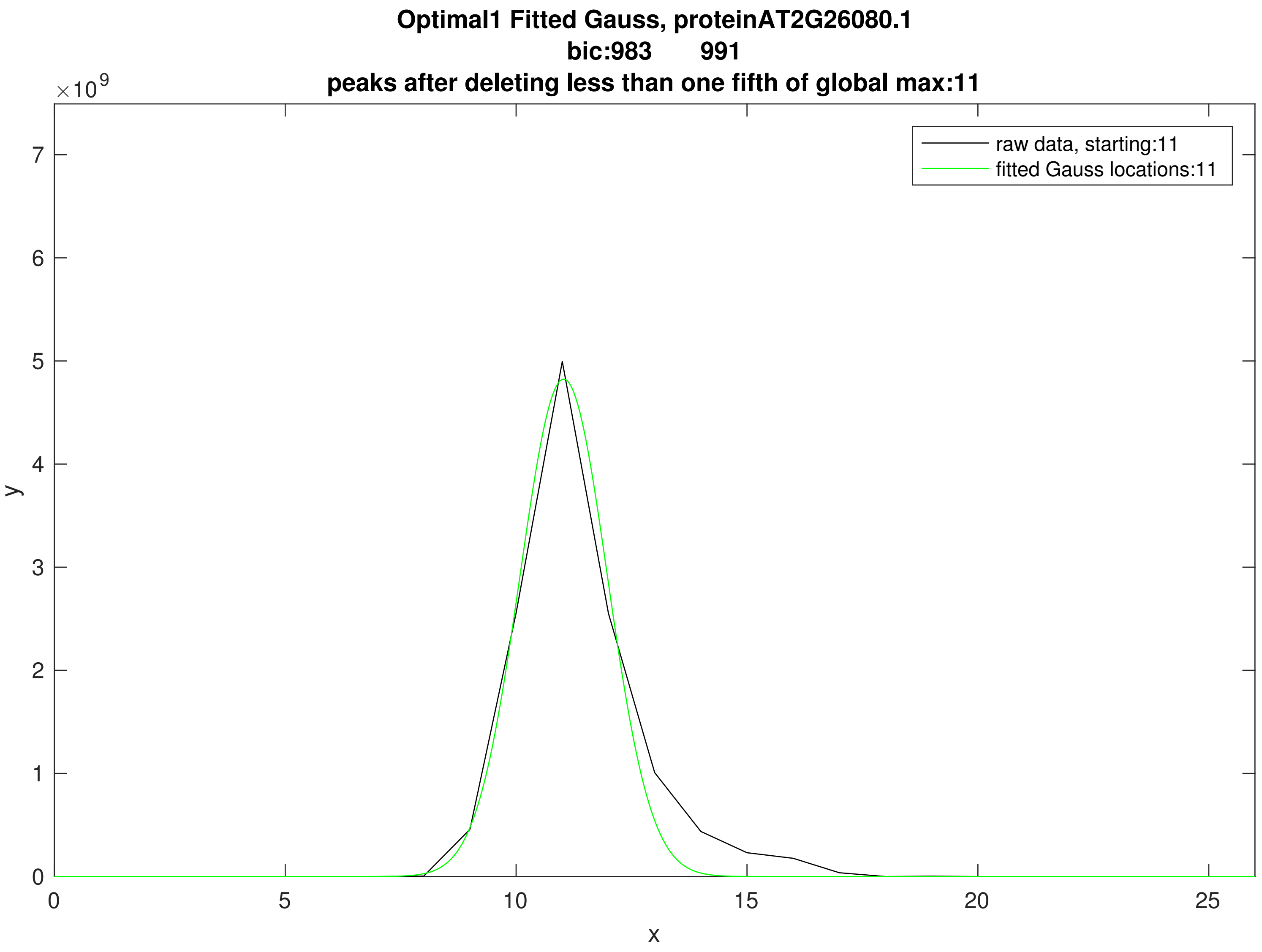
Optimal1 Fitted Gauss, proteinAT2G26080.1
bic:983 991
peaks after deleting less than one fifth of global max:11
×10^9
raw data, starting:11
fitted Gauss locations:11
y
x
0
1
2
3
4
5
6
7
0
5
10
15
20
25

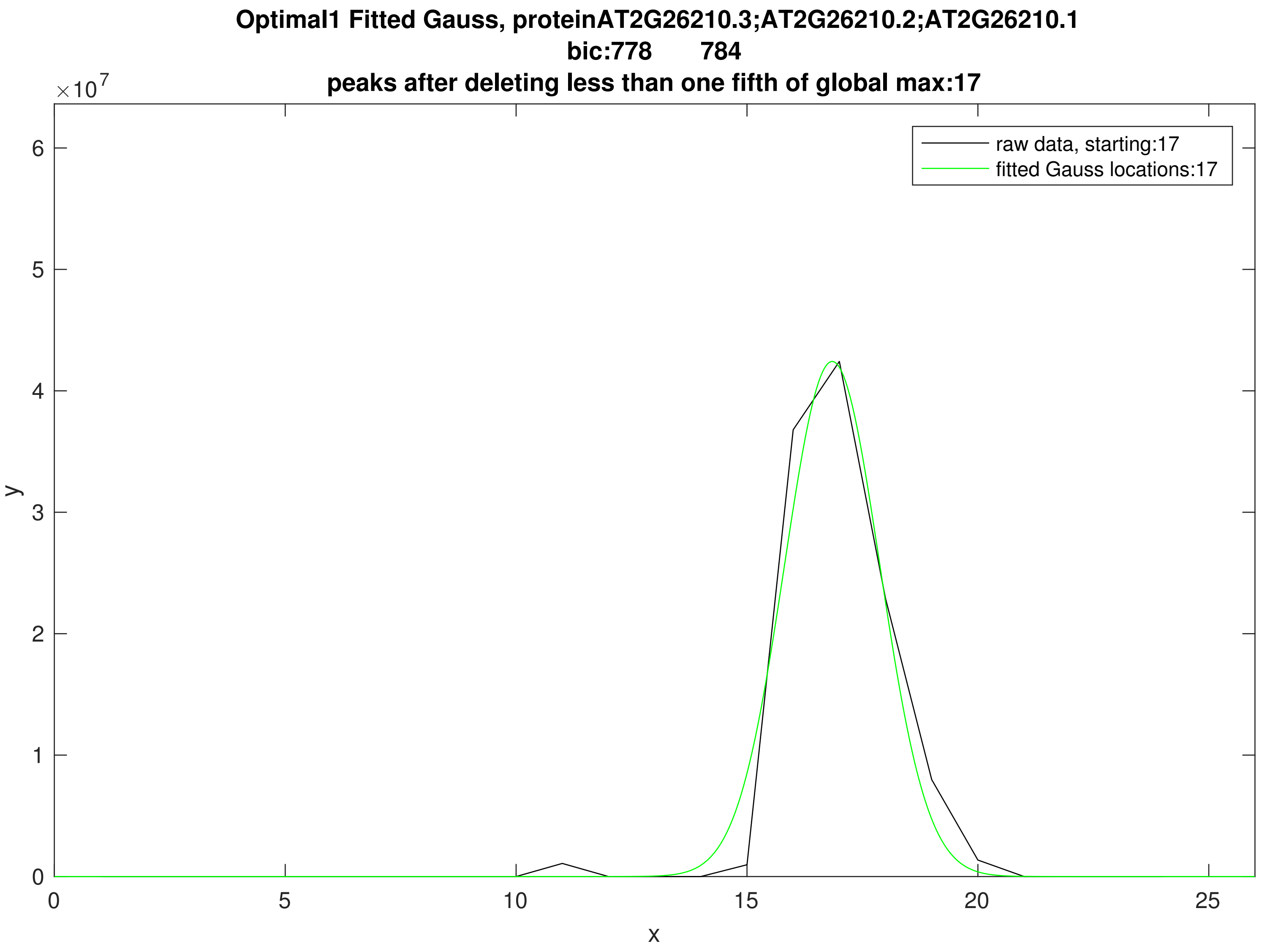
Optimal1 Fitted Gauss, proteinAT2G26210.3;AT2G26210.2;AT2G26210.1
bic:778 784
peaks after deleting less than one fifth of global max:17
×10^7
raw data, starting:17
fitted Gauss locations:17
y
x
0
1
2
3
4
5
6
0
5
10
15
20
25

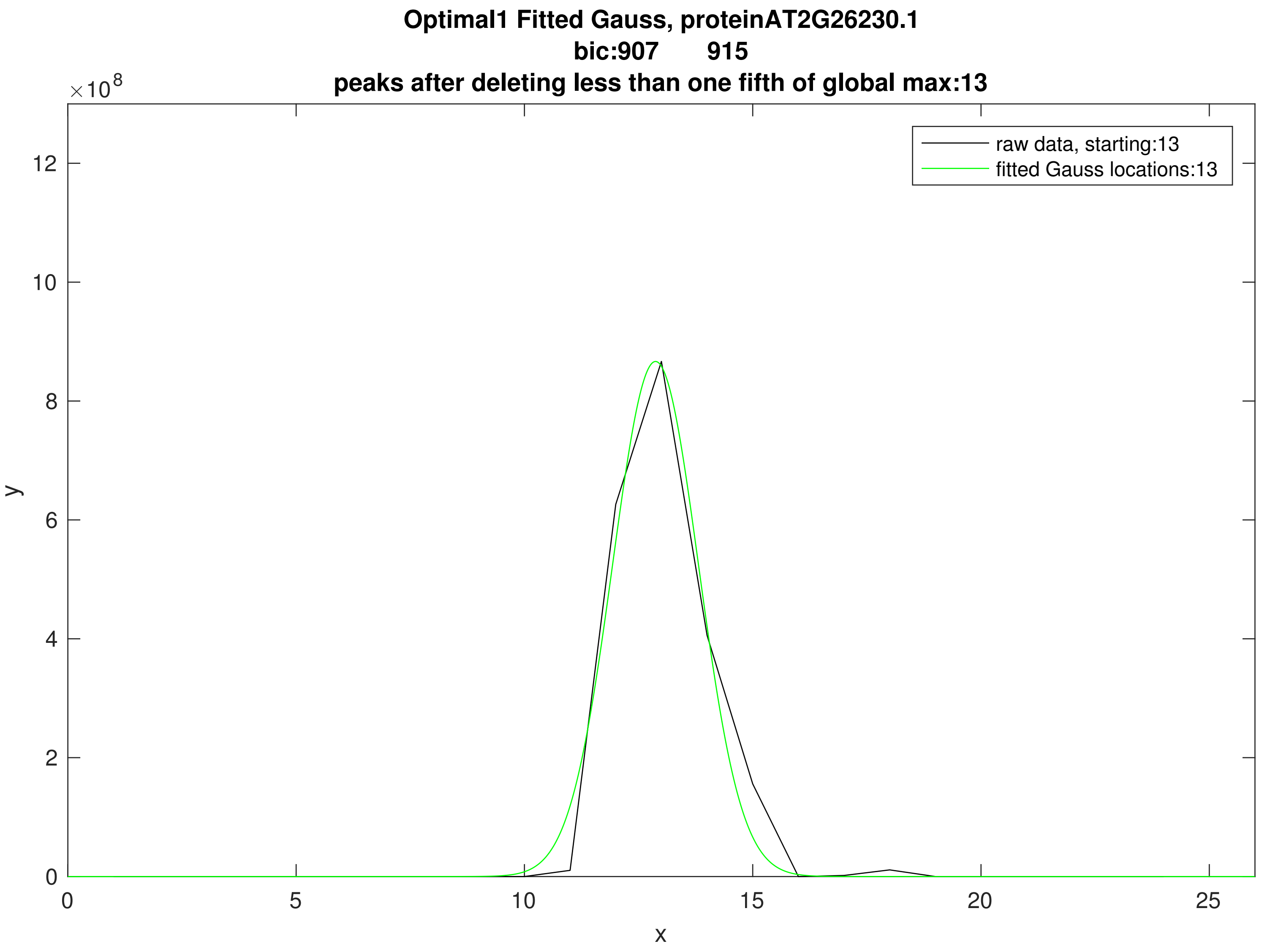
Optimal1 Fitted Gauss, proteinAT2G26230.1
bic:907 915
peaks after deleting less than one fifth of global max:13
×10^8
12
10
8
6
4
2
0
y
0
5
10
15
20
25
x
raw data, starting:13
fitted Gauss locations:13

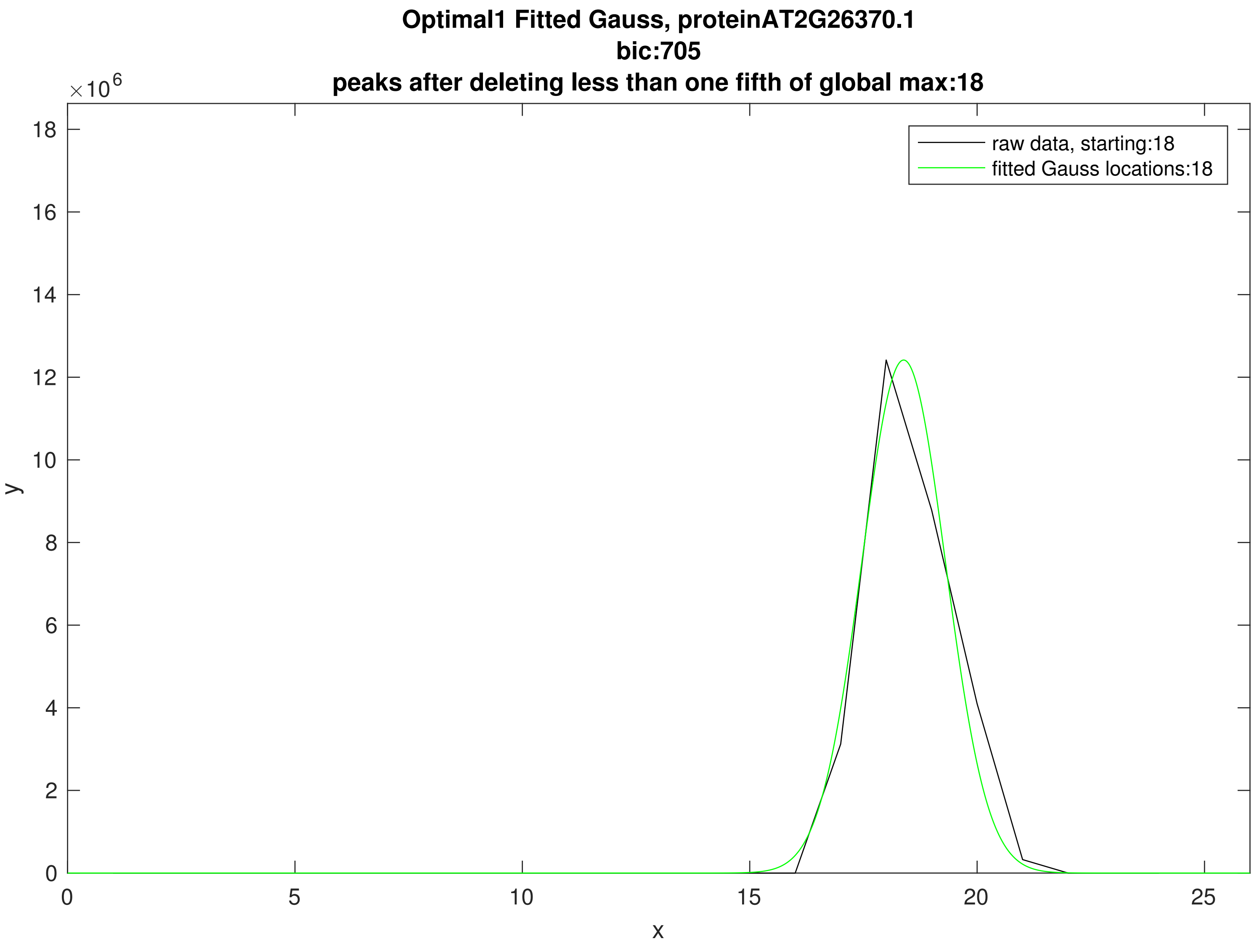

Optimal1 Fitted Gauss, proteinAT2G26370.1
bic:705
peaks after deleting less than one fifth of global max:18
×10^6
raw data, starting:18
fitted Gauss locations:18
y
x
0
2
4
6
8
10
12
14
16
18
0
5
10
15
20
25

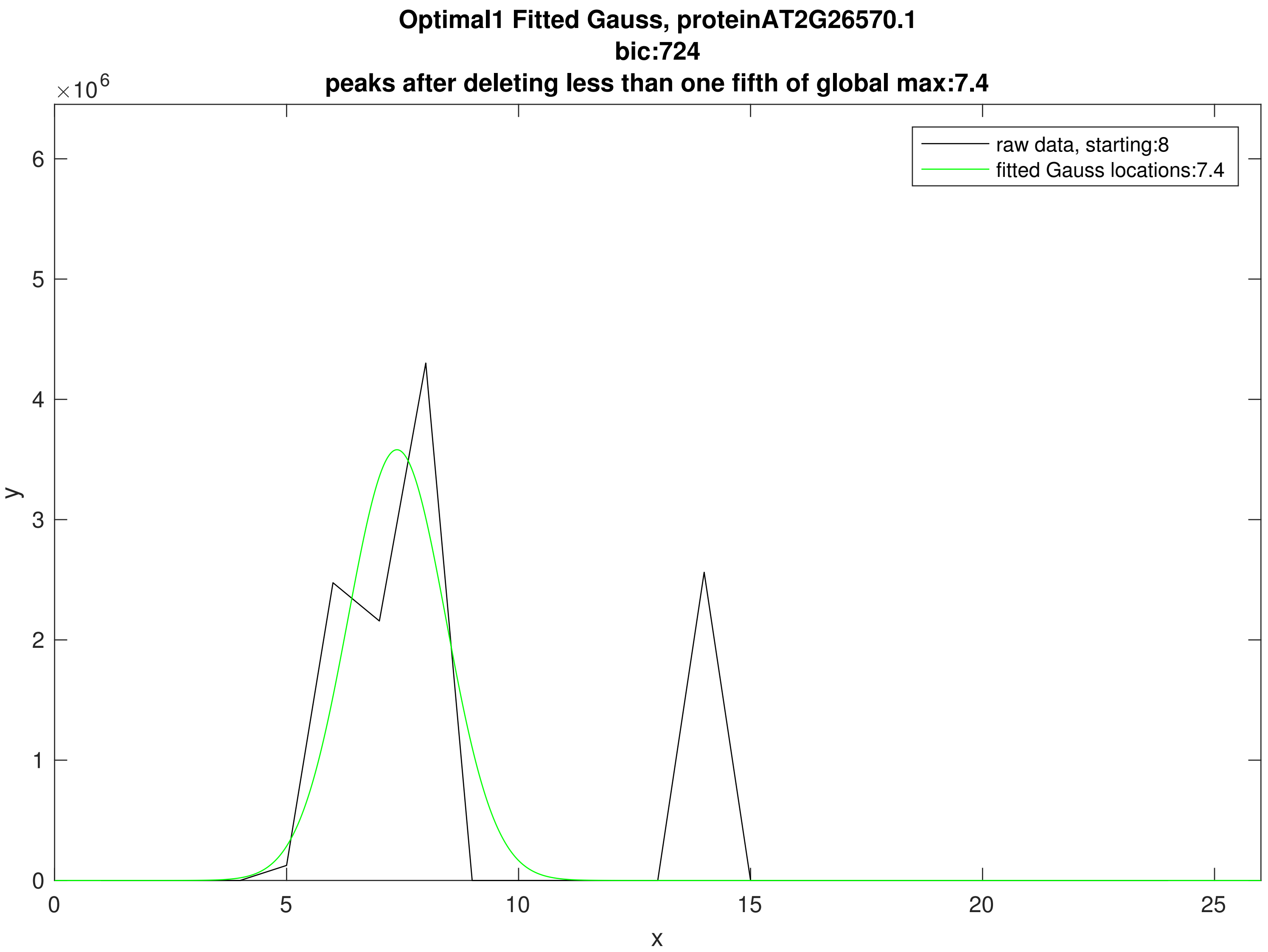

Optimal1 Fitted Gauss, proteinAT2G26570.1
bic:724
peaks after deleting less than one fifth of global max:7.4
×10^6
raw data, starting:8
fitted Gauss locations:7.4
y
x
0
1
2
3
4
5
6
0
5
10
15
20
25

**Optimal1 Fitted Gauss, proteinAT2G26590.3;AT2G26590.2;AT2G26590.1**
**bic:805       857**
**peaks after deleting less than one fifth of global max:14**

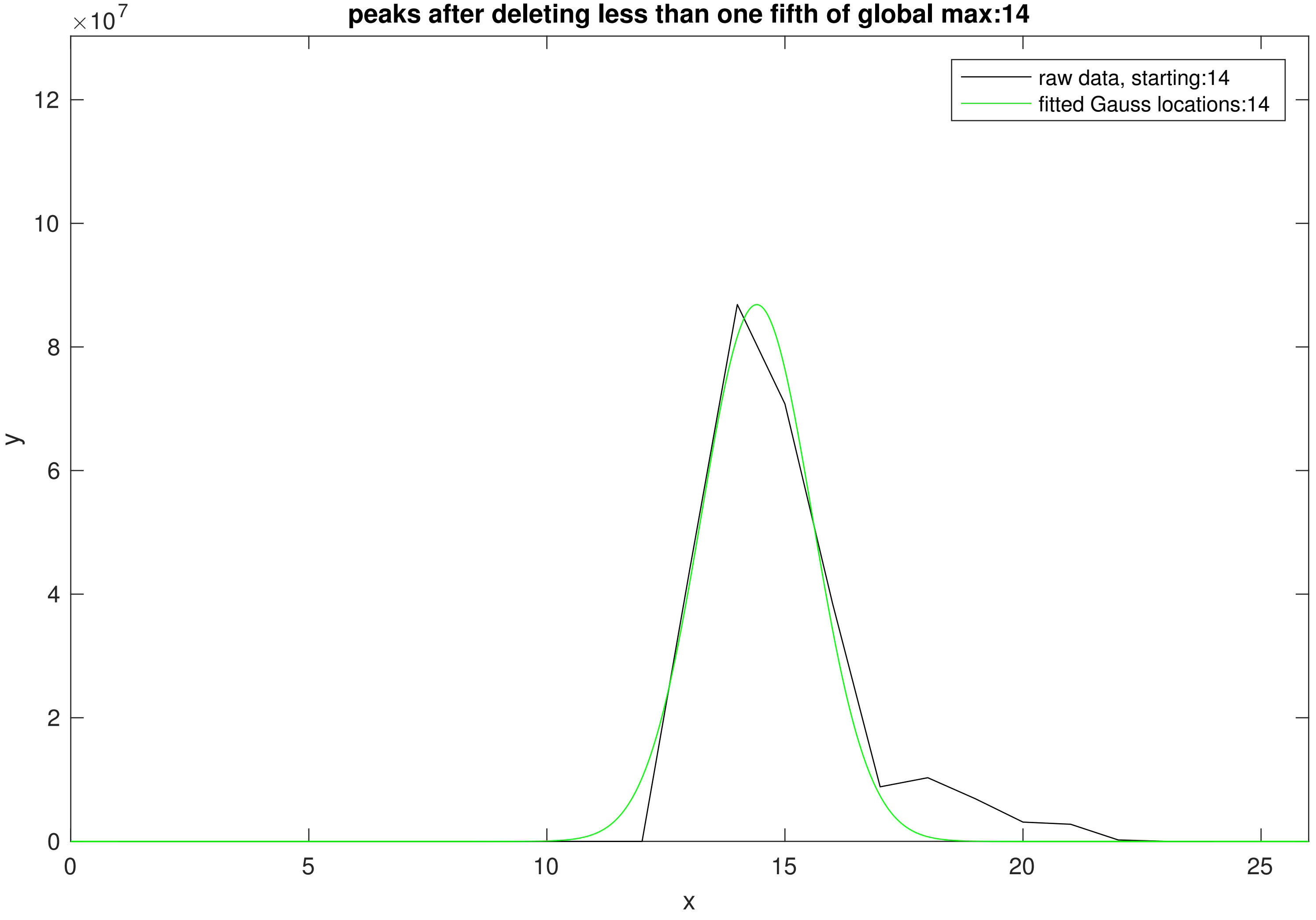

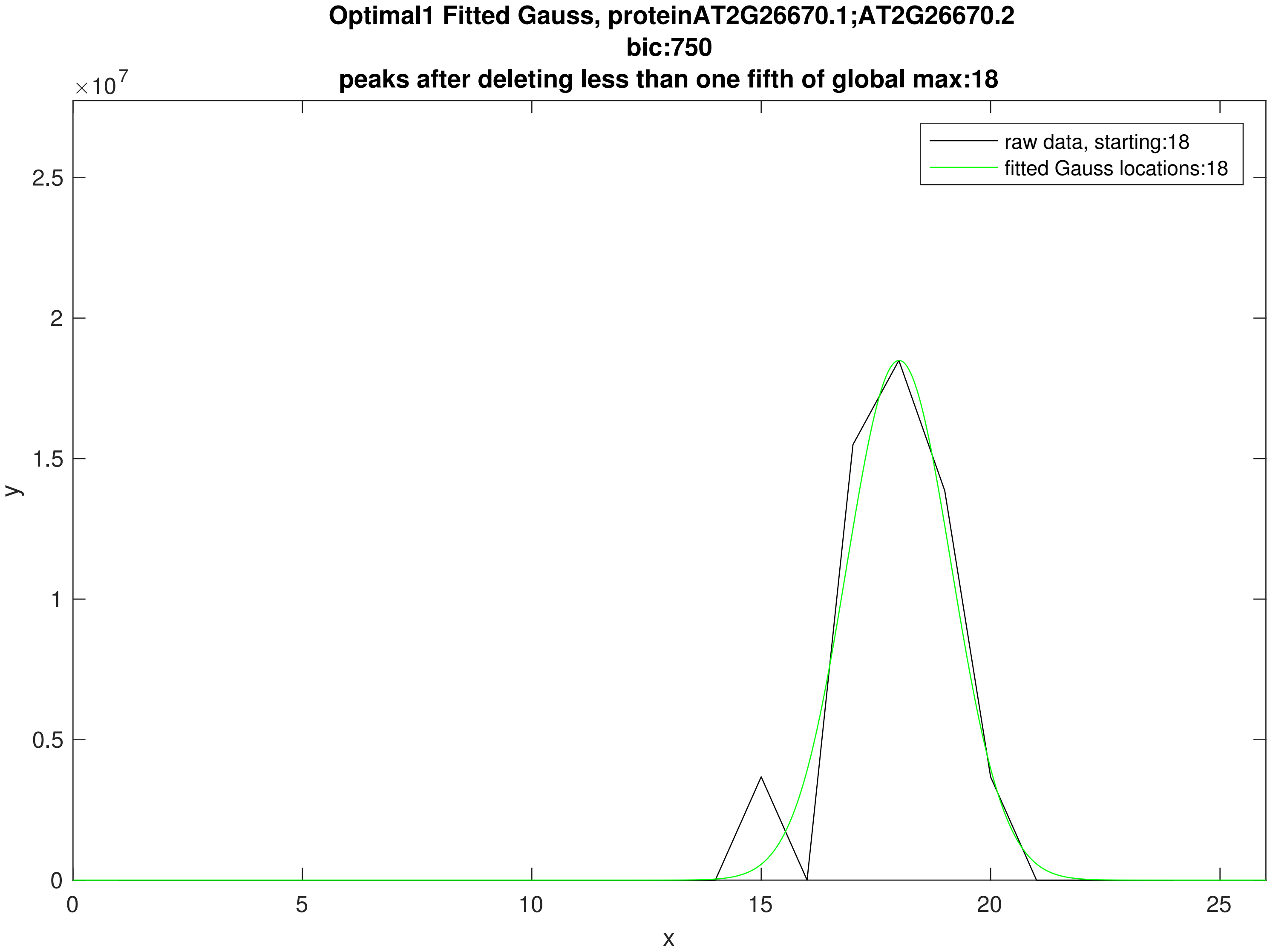
Optimal1 Fitted Gauss, proteinAT2G26670.1;AT2G26670.2
bic:750
peaks after deleting less than one fifth of global max:18
×10^7
raw data, starting:18
fitted Gauss locations:18
y
x
0
0.5
1
1.5
2
2.5
0
5
10
15
20
25

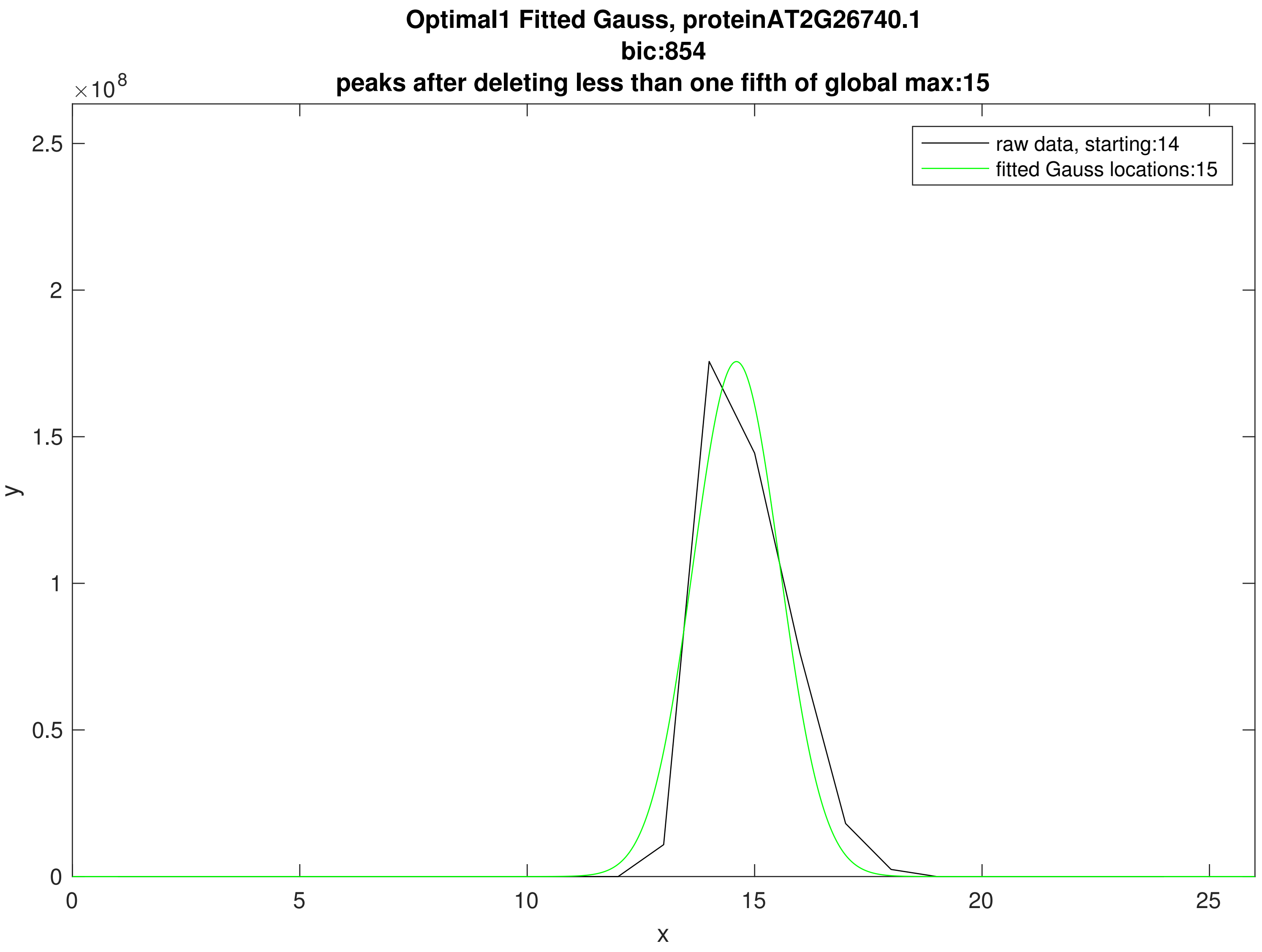
Optimal1 Fitted Gauss, proteinAT2G26740.1
bic:854
peaks after deleting less than one fifth of global max:15
×10^8
raw data, starting:14
fitted Gauss locations:15
y
x
0
0.5
1
1.5
2
2.5
0
5
10
15
20
25

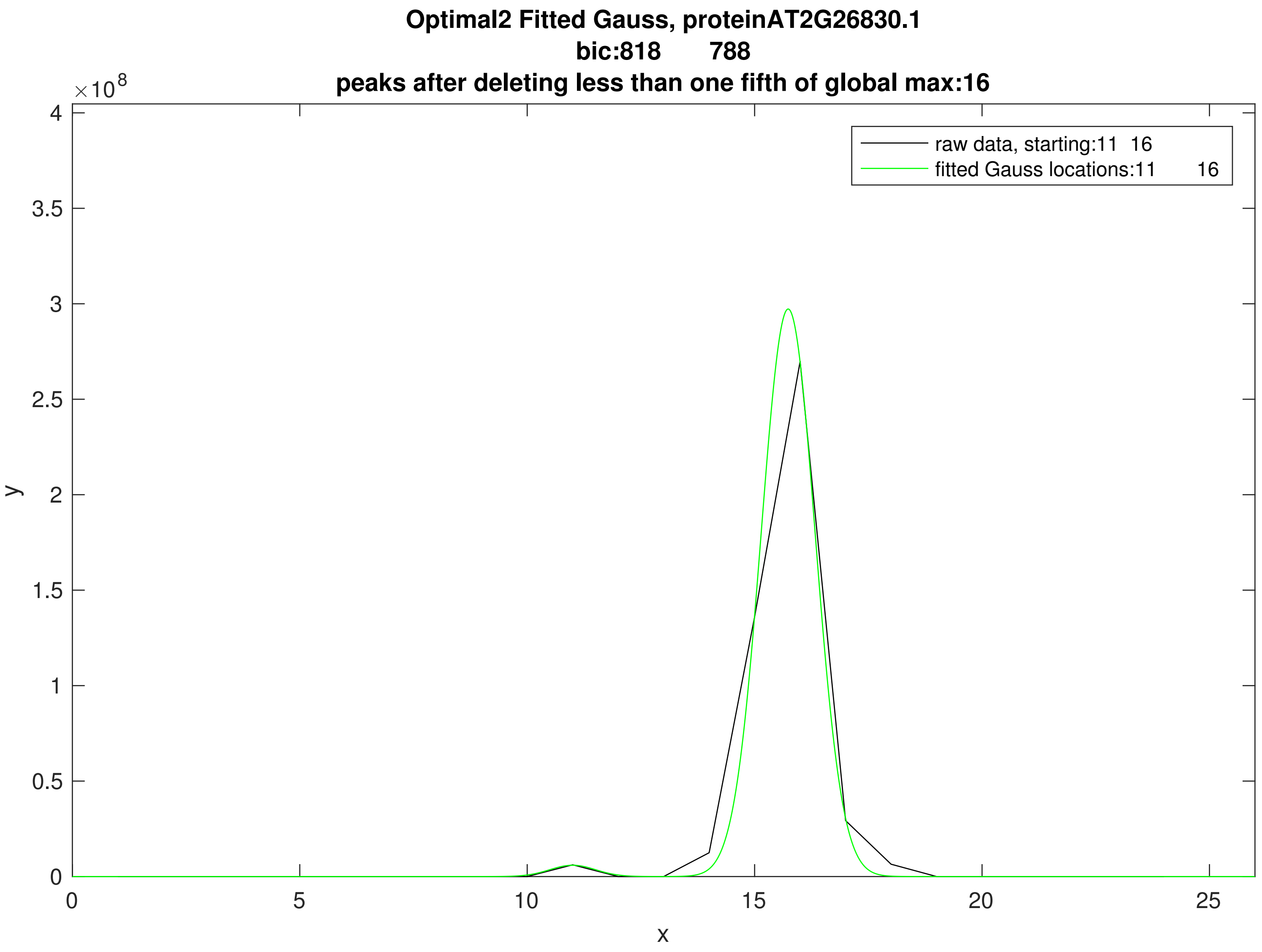
Optimal2 Fitted Gauss, proteinAT2G26830.1
bic:818 788
peaks after deleting less than one fifth of global max:16
×10^8
4
3.5
3
2.5
2
1.5
1
0.5
0
y
0
5
10
15
20
25
x
raw data, starting:11 16
fitted Gauss locations:11 16

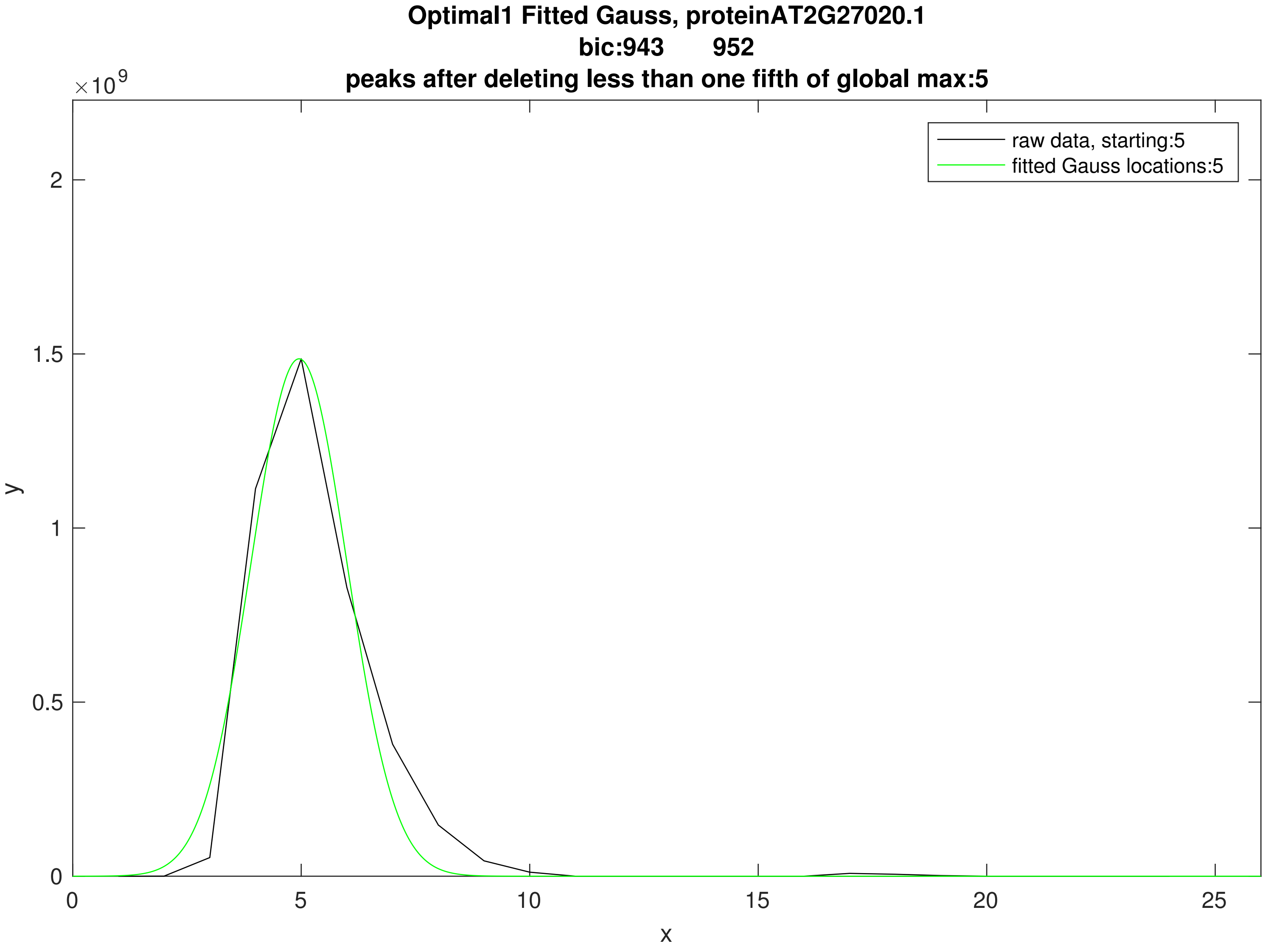
Optimal1 Fitted Gauss, proteinAT2G27020.1
bic:943 952
peaks after deleting less than one fifth of global max:5
×10^9
raw data, starting:5
fitted Gauss locations:5
y
x
0
0.5
1
1.5
2
5
10
15
20
25

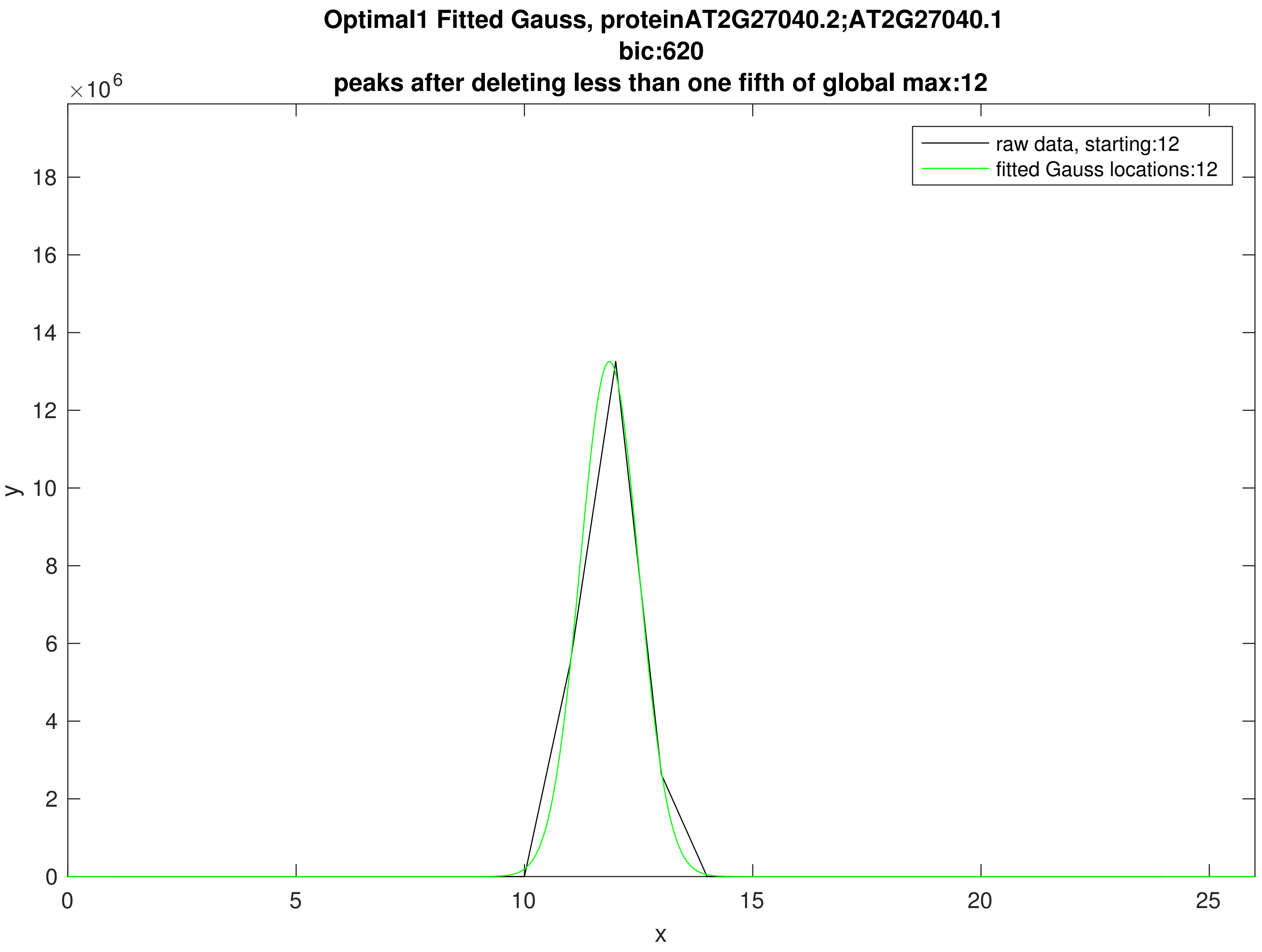

Optimal1 Fitted Gauss, proteinAT2G27040.2;AT2G27040.1
bic:620
peaks after deleting less than one fifth of global max:12
×10^6
raw data, starting:12
fitted Gauss locations:12
y
x
0
2
4
6
8
10
12
14
16
18
0
5
10
15
20
25

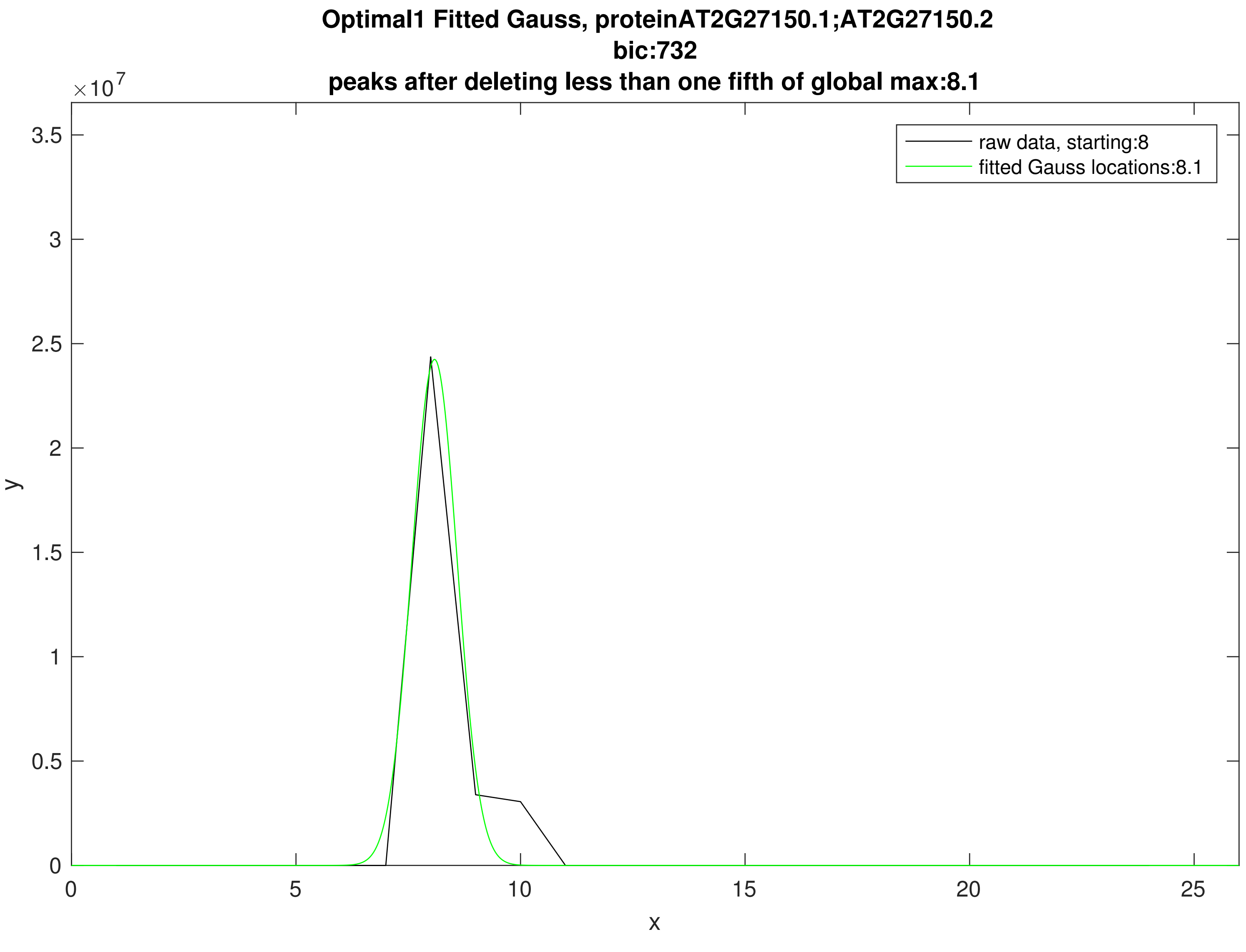
Optimal1 Fitted Gauss, proteinAT2G27150.1;AT2G27150.2
bic:732
peaks after deleting less than one fifth of global max:8.1
×10^7
3.5
3
2.5
2
1.5
1
0.5
0
y
0
5
10
15
20
25
x
raw data, starting:8
fitted Gauss locations:8.1

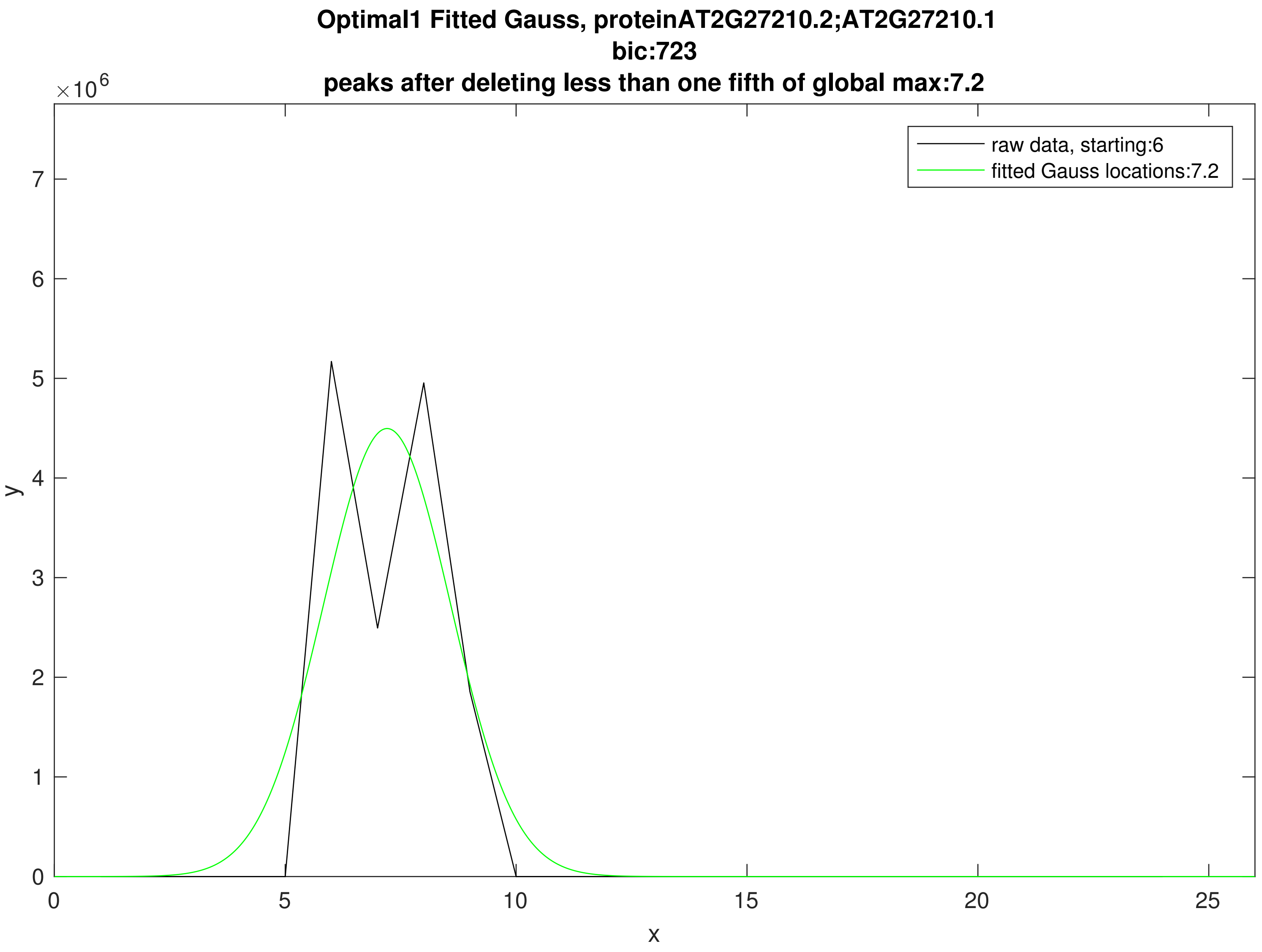
Optimal1 Fitted Gauss, proteinAT2G27210.2;AT2G27210.1
bic:723
peaks after deleting less than one fifth of global max:7.2
×10^6
raw data, starting:6
fitted Gauss locations:7.2
y
x
0
1
2
3
4
5
6
7
0
5
10
15
20
25

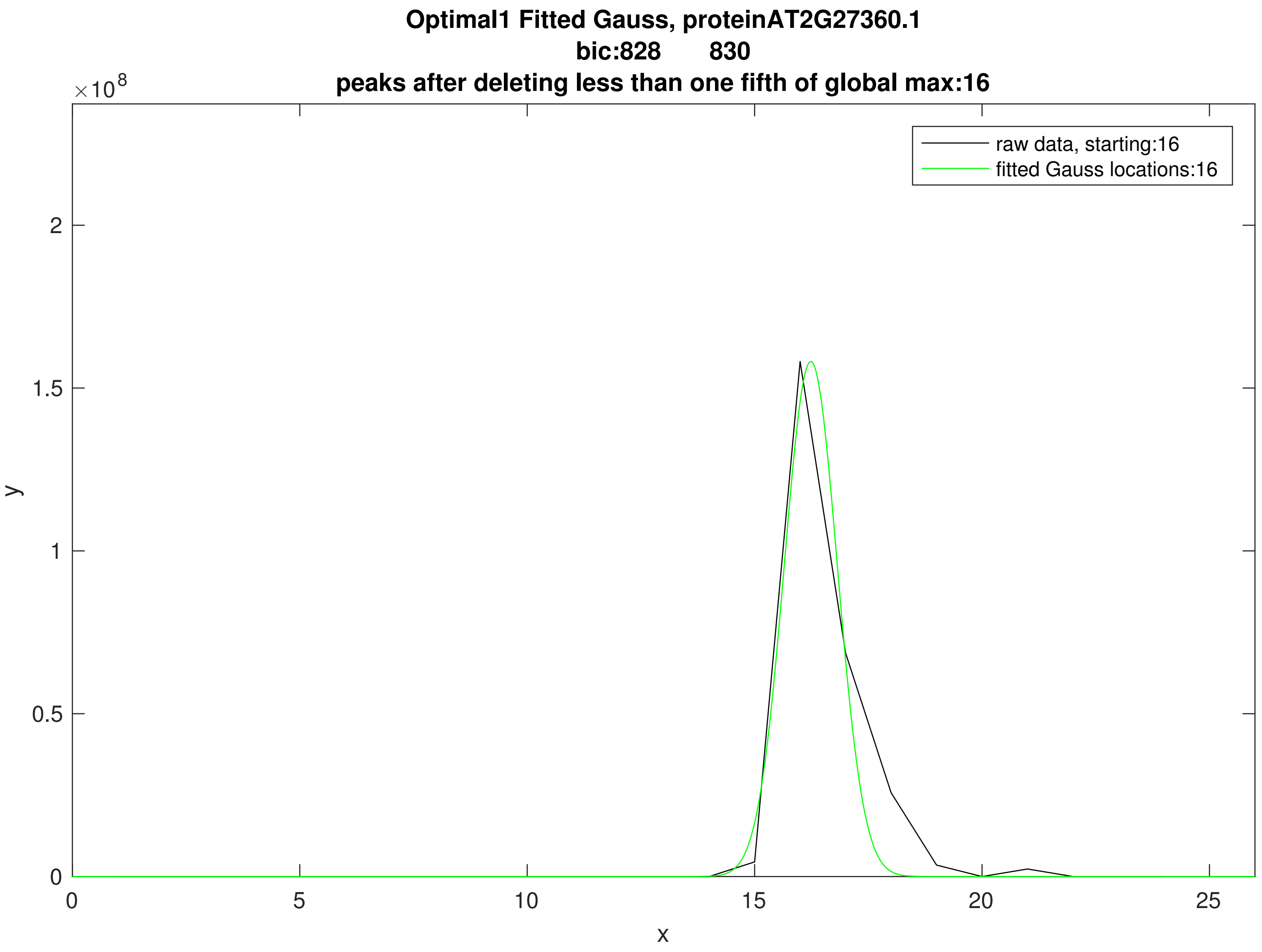

Optimal1 Fitted Gauss, proteinAT2G27360.1
bic:828 830
peaks after deleting less than one fifth of global max:16
×10^8
raw data, starting:16
fitted Gauss locations:16
y
x
0
0.5
1
1.5
2
0
5
10
15
20
25

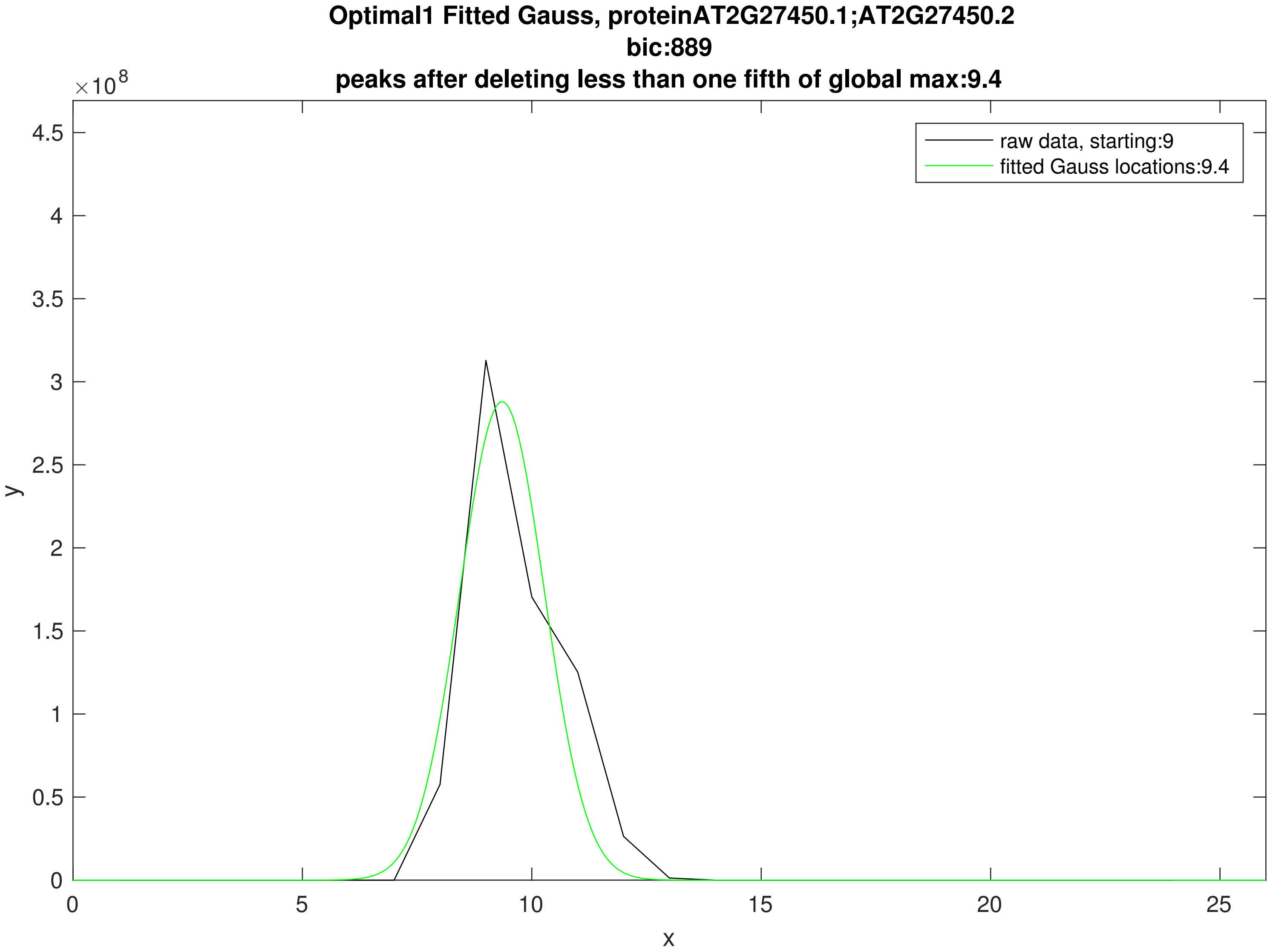

Optimal1 Fitted Gauss, proteinAT2G27450.1;AT2G27450.2
bic:889
peaks after deleting less than one fifth of global max:9.4
×10^8
4.5
4
3.5
3
2.5
2
1.5
1
0.5
0
y
0
5
10
15
20
25
x
raw data, starting:9
fitted Gauss locations:9.4

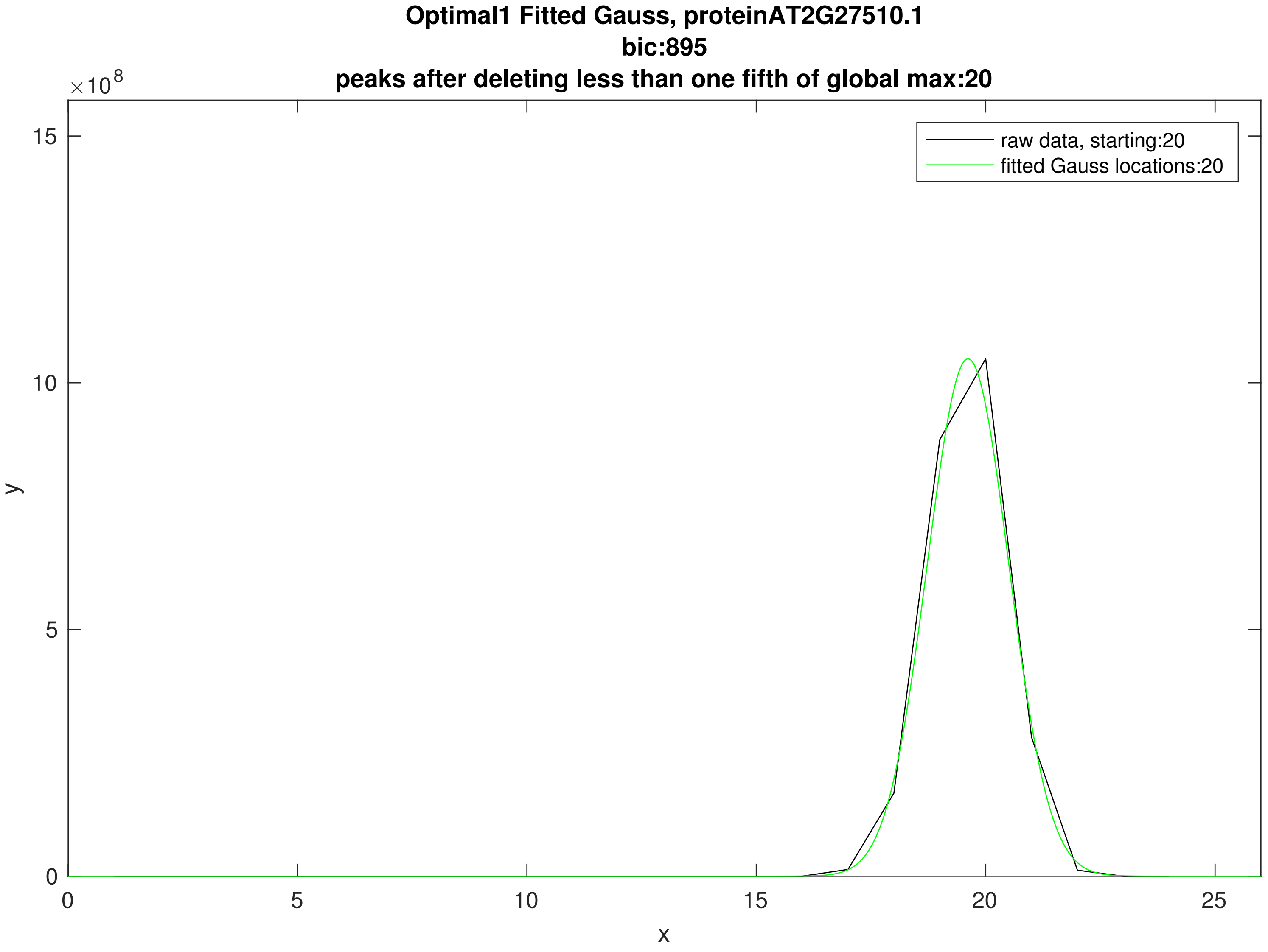

Optimal1 Fitted Gauss, proteinAT2G27510.1
bic:895
peaks after deleting less than one fifth of global max:20
×10^8
15
10
5
0
y
0
5
10
15
20
25
x
raw data, starting:20
fitted Gauss locations:20

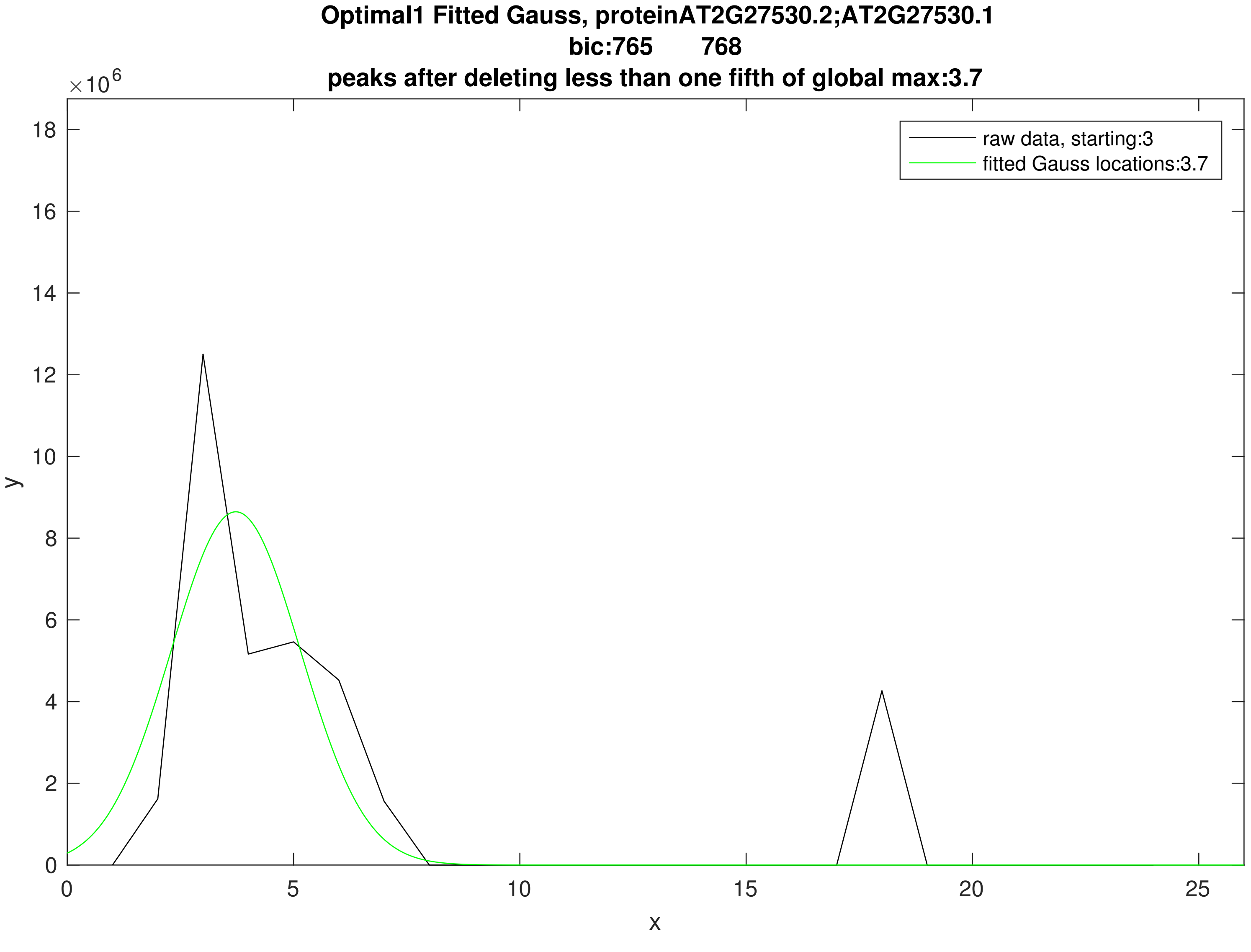

Optimal1 Fitted Gauss, proteinAT2G27530.2;AT2G27530.1
bic:765 768
peaks after deleting less than one fifth of global max:3.7
×10^6
raw data, starting:3
fitted Gauss locations:3.7
y
x
0
2
4
6
8
10
12
14
16
18
5
15
20
25

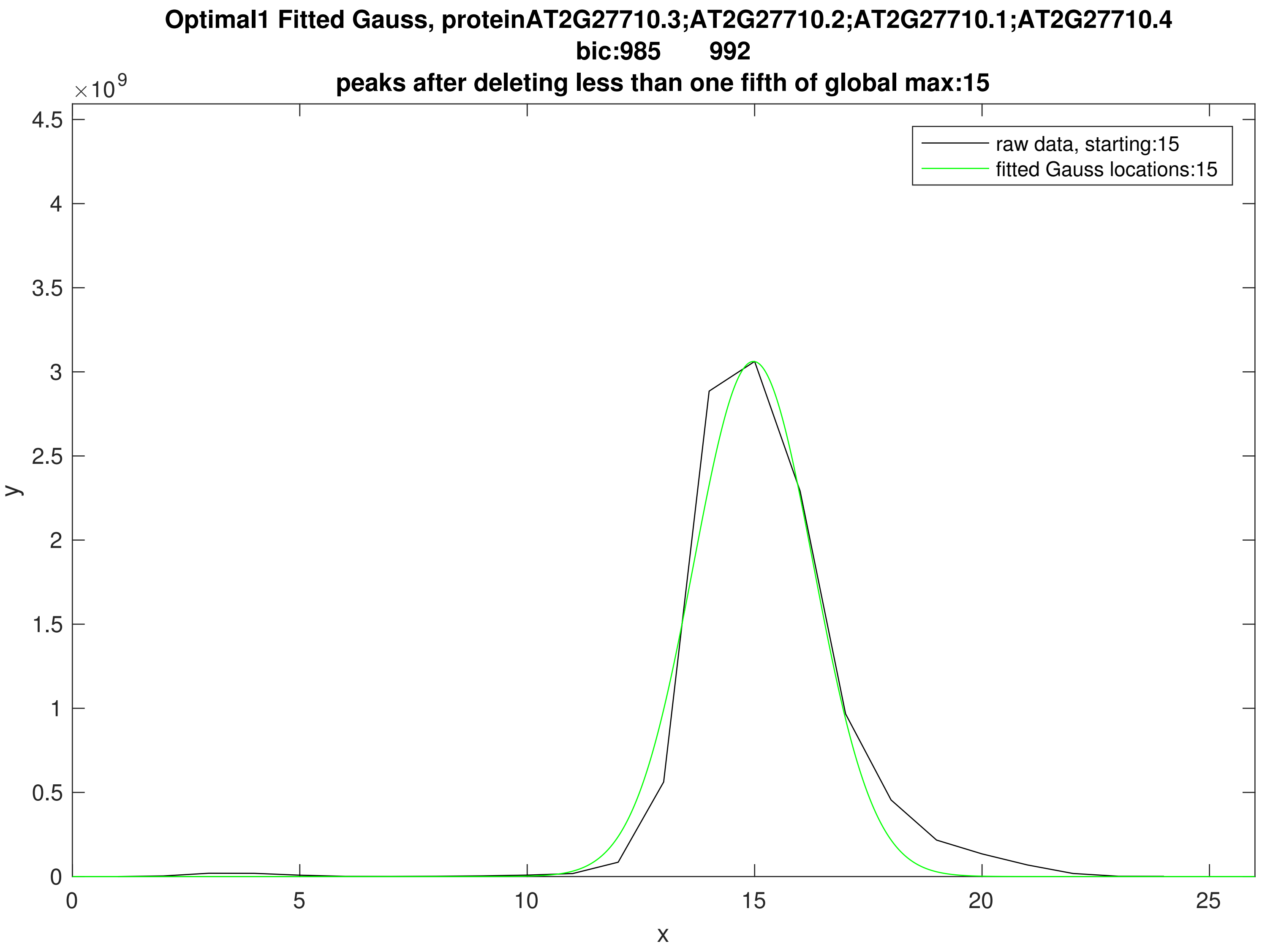
Optimal1 Fitted Gauss, proteinAT2G27710.3;AT2G27710.2;AT2G27710.1;AT2G27710.4
bic:985 992
peaks after deleting less than one fifth of global max:15
×10^9
4.5
4
3.5
3
2.5
2
1.5
1
0.5
0
y
0
5
10
15
20
25
x
raw data, starting:15
fitted Gauss locations:15

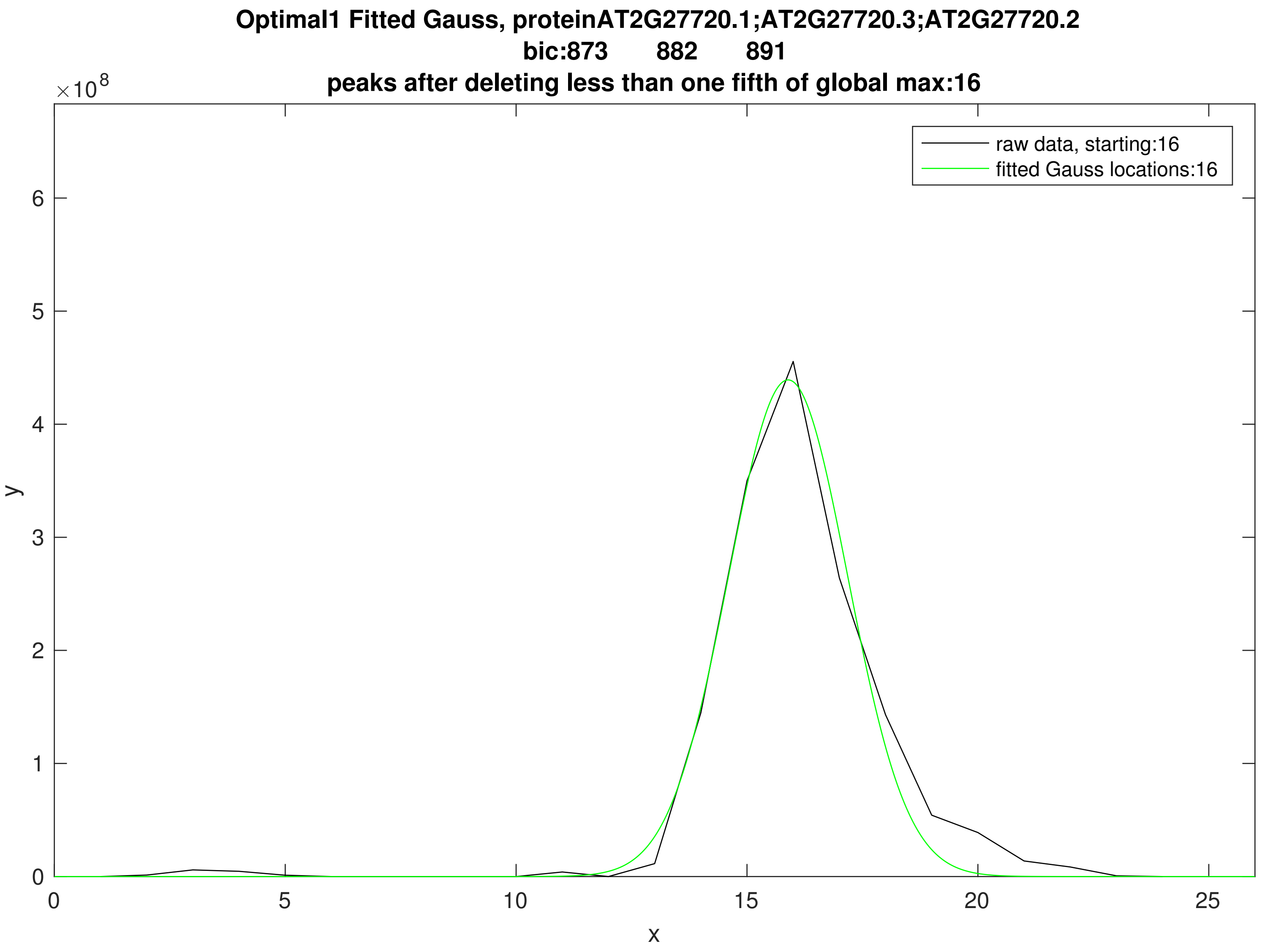

Optimal1 Fitted Gauss, proteinAT2G27720.1;AT2G27720.3;AT2G27720.2
bic:873 882 891
peaks after deleting less than one fifth of global max:16
×10^8
raw data, starting:16
fitted Gauss locations:16
y
x
0
1
2
3
4
5
6
0
5
10
15
20
25

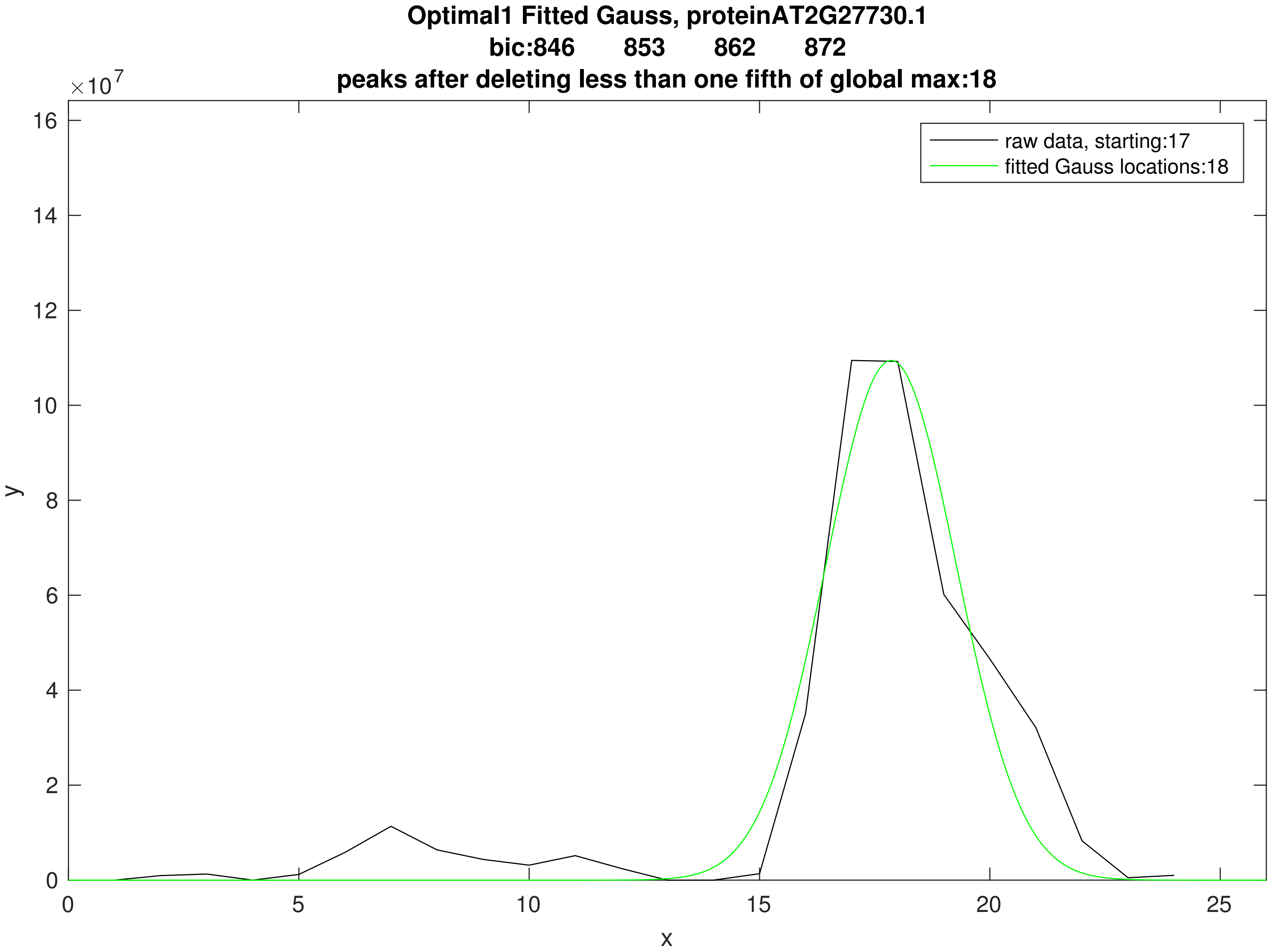
Optimal1 Fitted Gauss, proteinAT2G27730.1
bic:846 853 862 872
peaks after deleting less than one fifth of global max:18
×10^7
16
14
12
10
8
6
4
2
0
y
0
5
10
15
20
25
x
raw data, starting:17
fitted Gauss locations:18

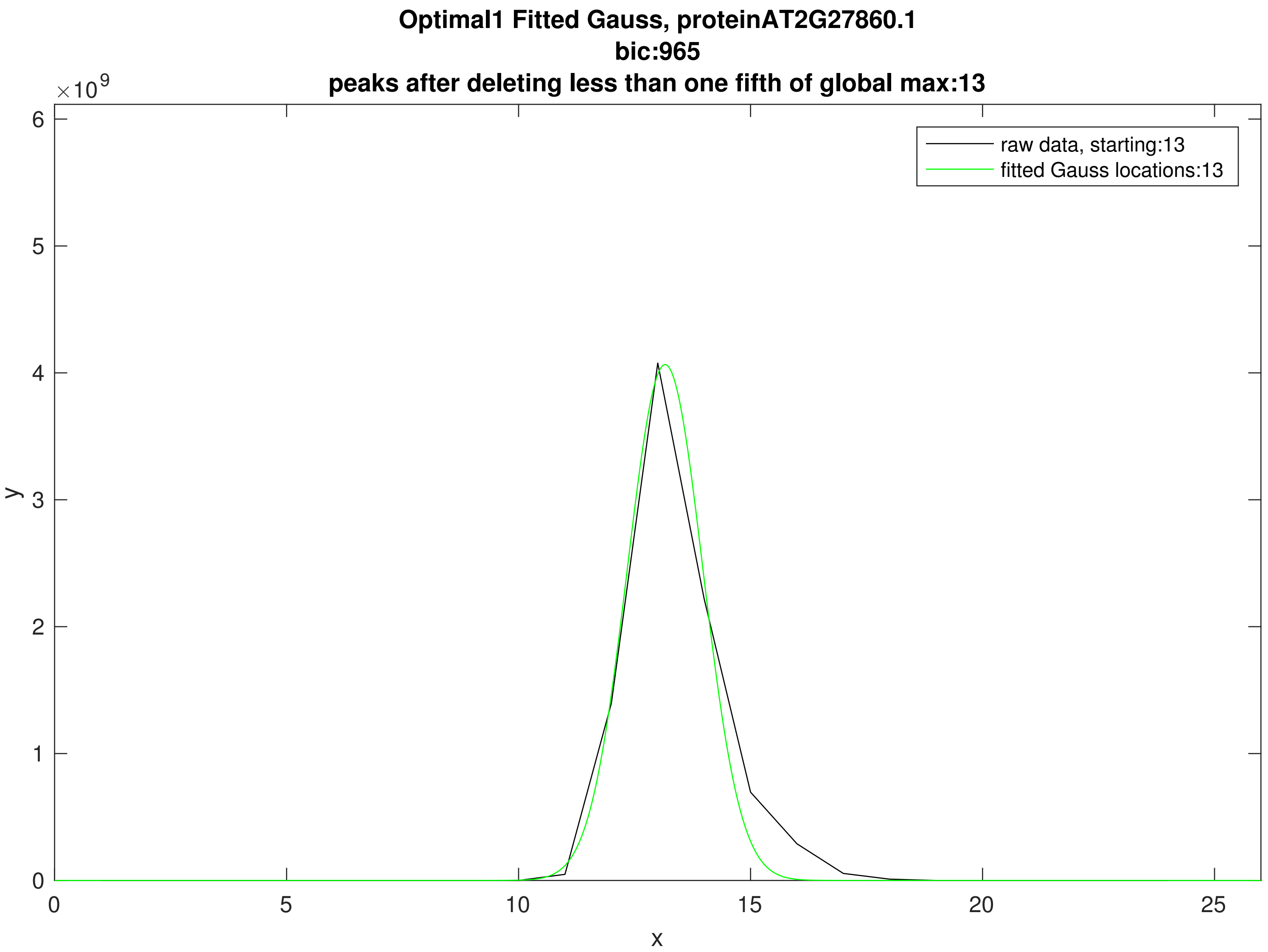
Optimal1 Fitted Gauss, proteinAT2G27860.1
bic:965
peaks after deleting less than one fifth of global max:13
×10^9
6
5
4
3
2
1
0
y
0
5
10
15
20
25
x
raw data, starting:13
fitted Gauss locations:13

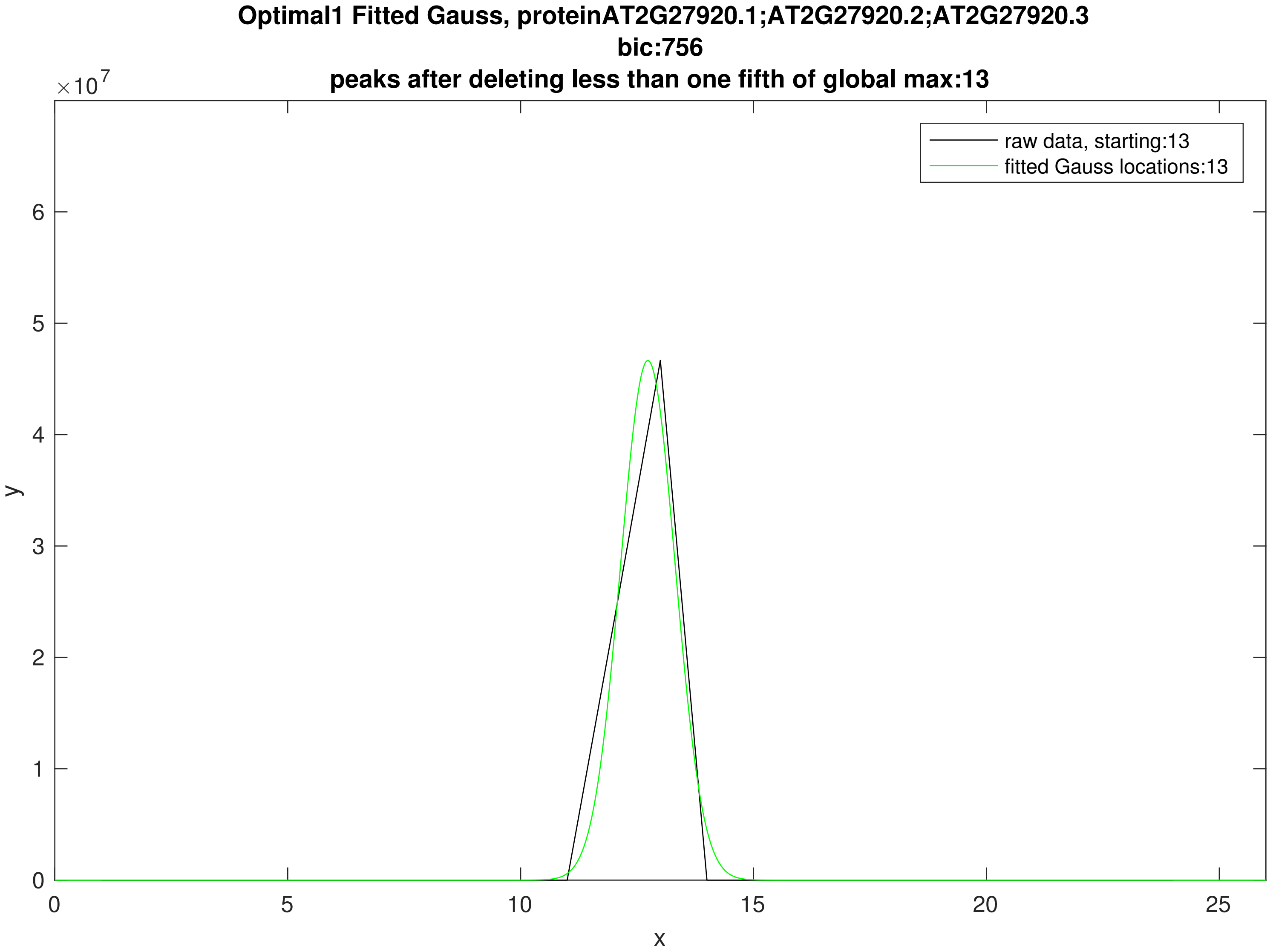

Optimal1 Fitted Gauss, proteinAT2G27920.1;AT2G27920.2;AT2G27920.3
bic:756
peaks after deleting less than one fifth of global max:13
×10^7
raw data, starting:13
fitted Gauss locations:13
y
x
0
1
2
3
4
5
6
0
5
10
15
20
25

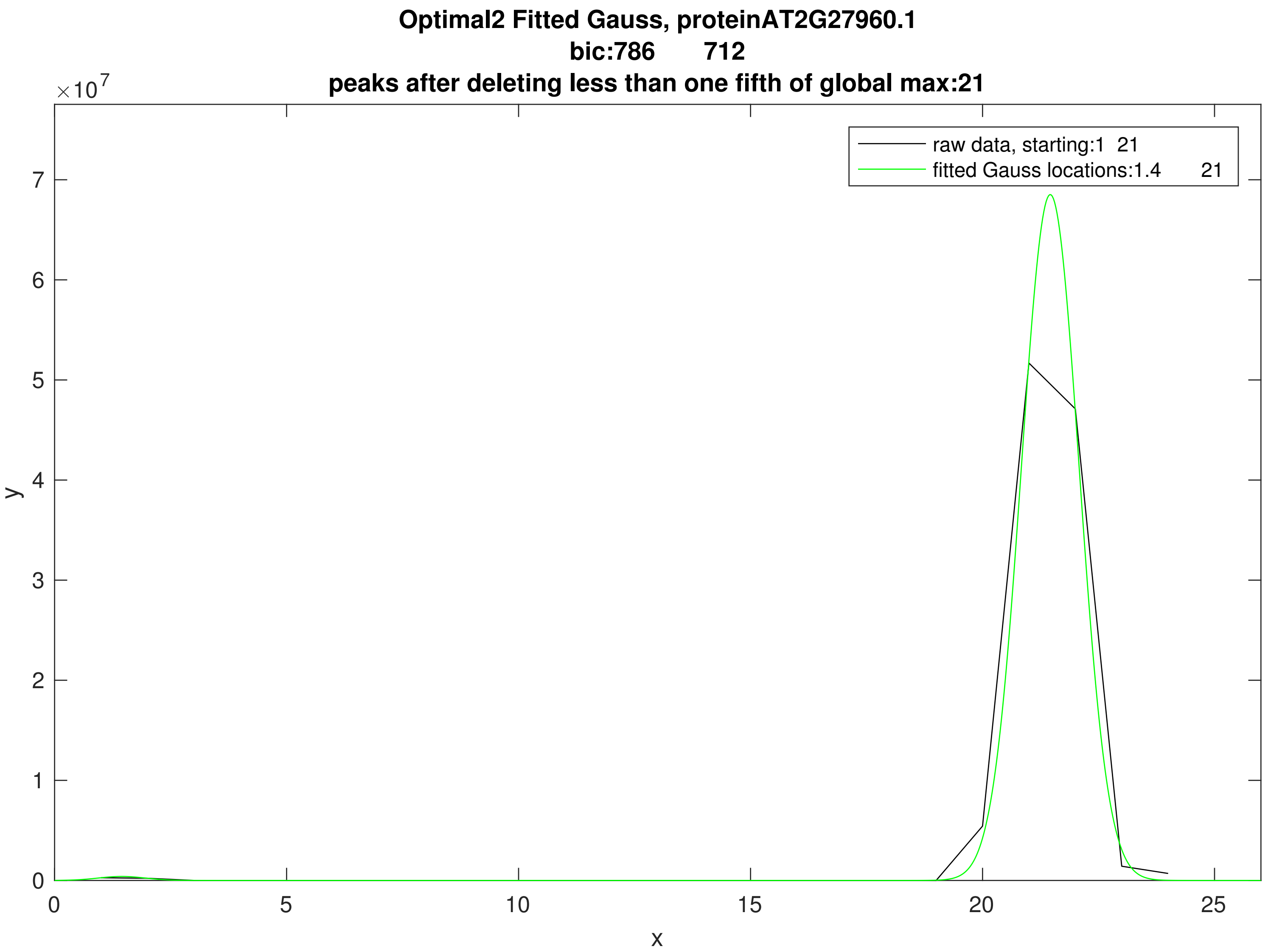
Optimal2 Fitted Gauss, proteinAT2G27960.1
bic:786 712
peaks after deleting less than one fifth of global max:21
×10^7
raw data, starting:1 21
fitted Gauss locations:1.4 21
y
x
0
1
2
3
4
5
6
7
0
5
10
15
20
25

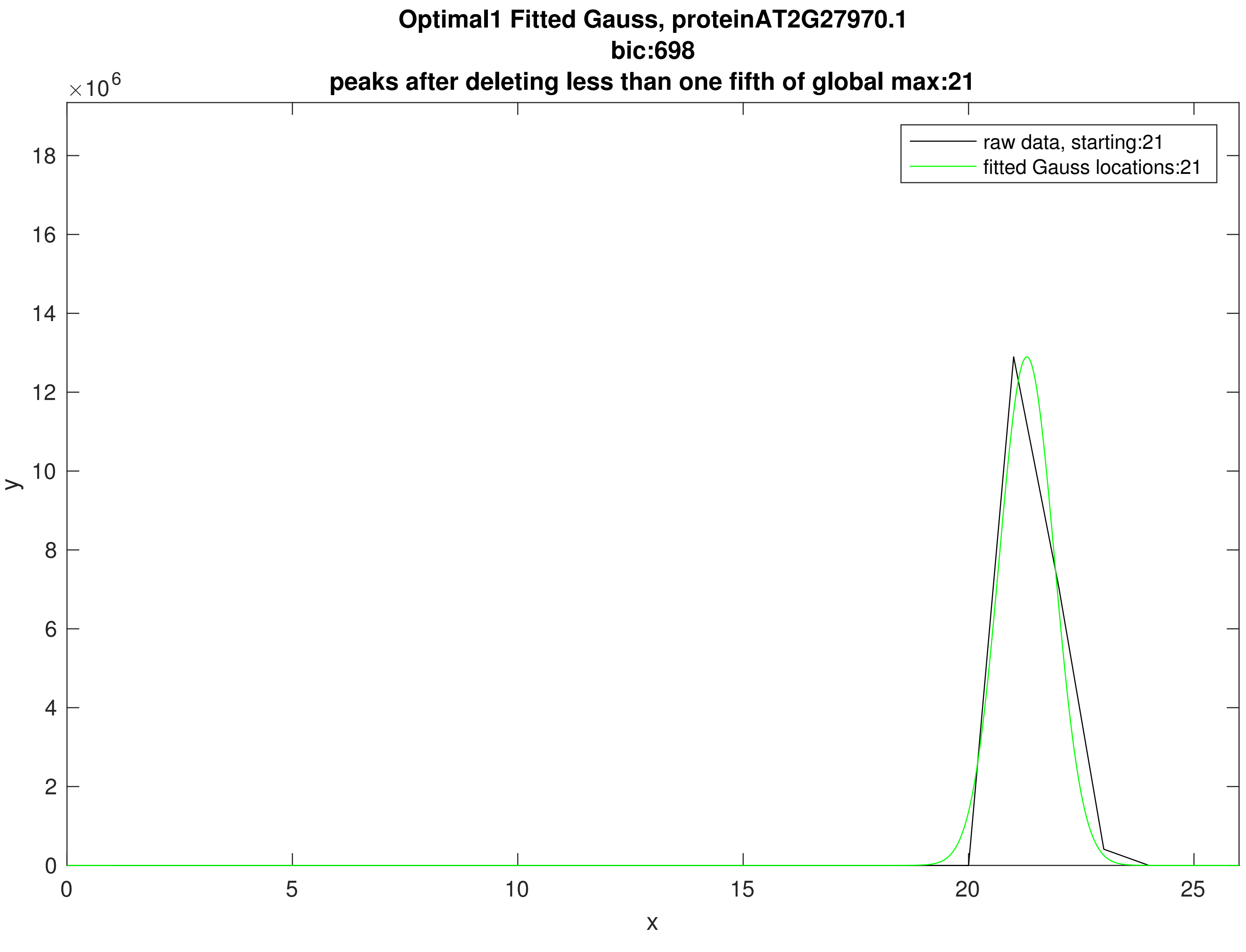

Optimal1 Fitted Gauss, proteinAT2G27970.1
bic:698
peaks after deleting less than one fifth of global max:21
×10^6
18
16
14
12
10
8
6
4
2
0
y
0
5
10
15
20
25
x
raw data, starting:21
fitted Gauss locations:21

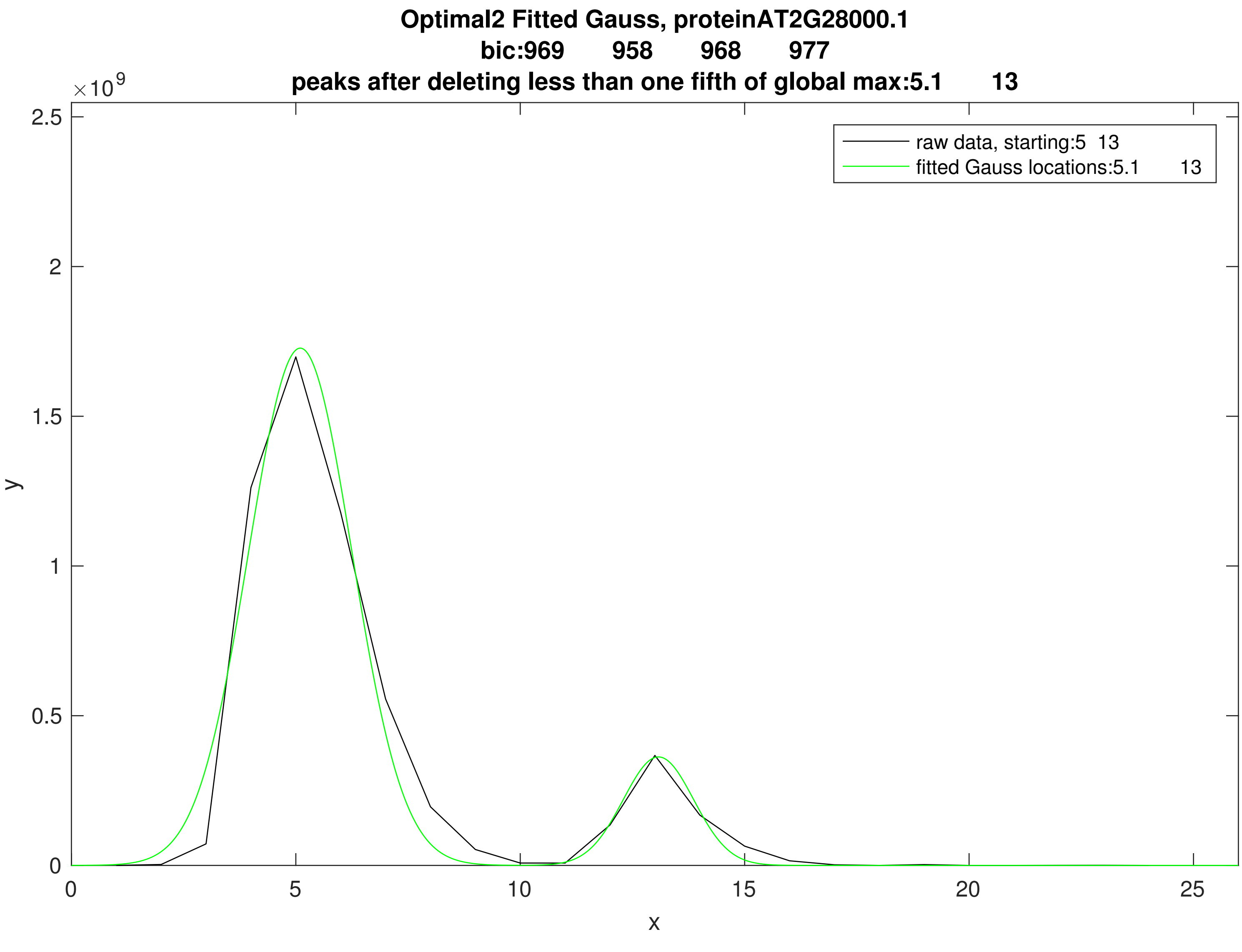

Optimal2 Fitted Gauss, proteinAT2G28000.1
bic:969 958 968 977
peaks after deleting less than one fifth of global max:5.1 13
×10^9
2.5
2
1.5
1
0.5
0
y
0
5
10
15
20
25
x
raw data, starting:5 13
fitted Gauss locations:5.1 13

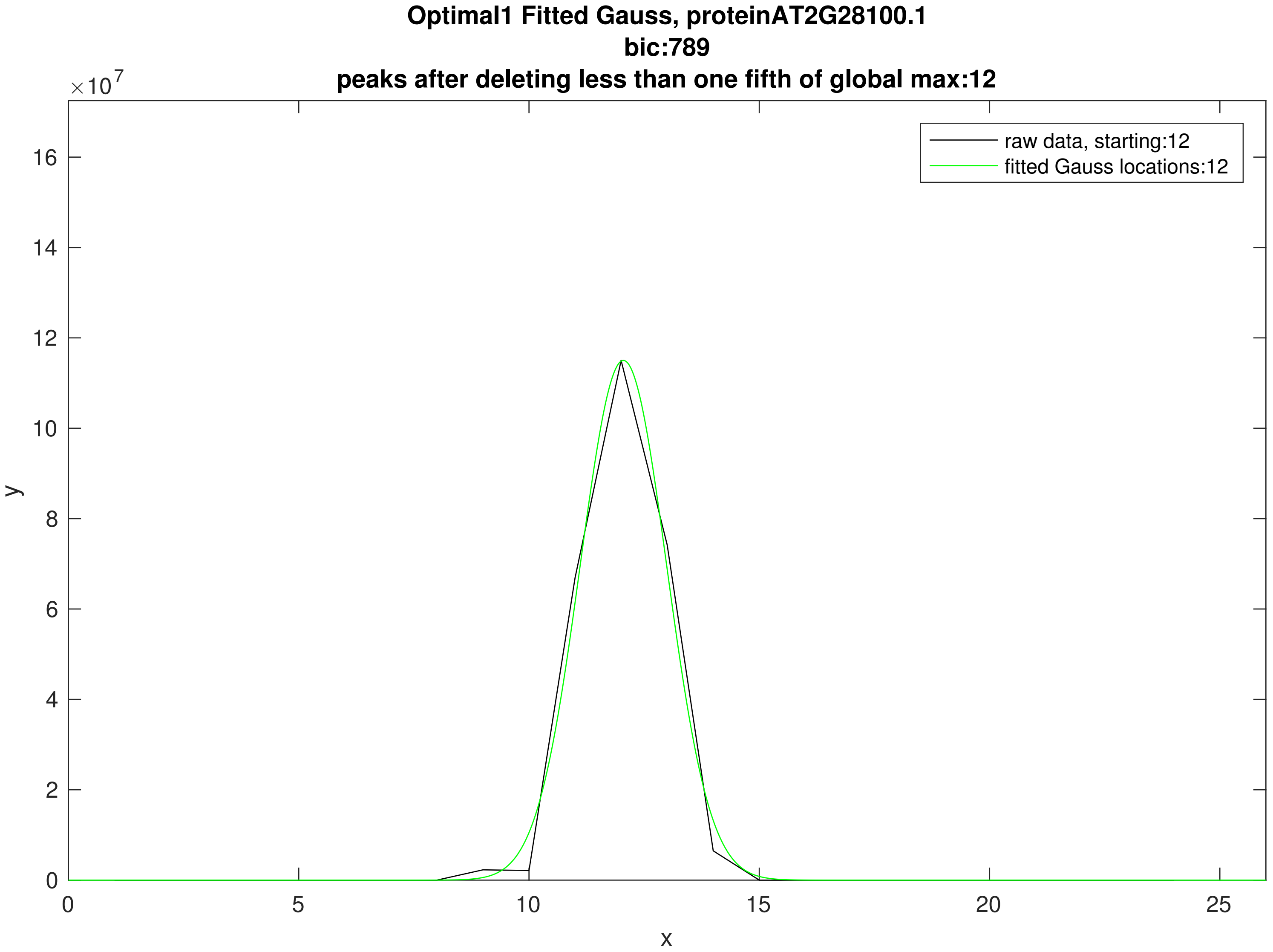
Optimal1 Fitted Gauss, proteinAT2G28100.1
bic:789
peaks after deleting less than one fifth of global max:12
×10^7
raw data, starting:12
fitted Gauss locations:12
y
x
0
2
4
6
8
10
12
14
16
0
5
10
15
20
25

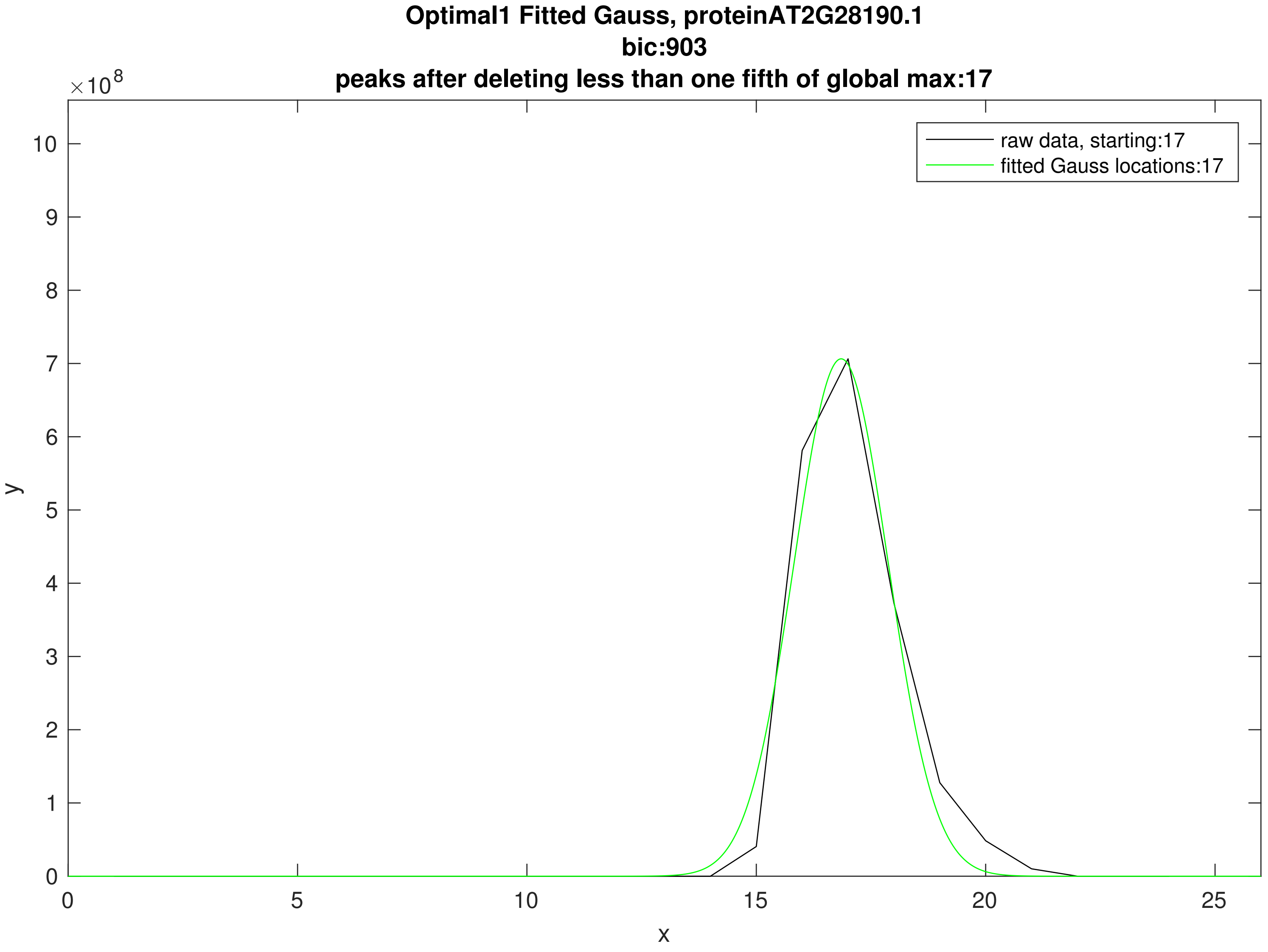

Optimal1 Fitted Gauss, proteinAT2G28190.1
bic:903
peaks after deleting less than one fifth of global max:17
×10^8
raw data, starting:17
fitted Gauss locations:17
y
x
0
1
2
3
4
5
6
7
8
9
10
0
5
10
15
20
25

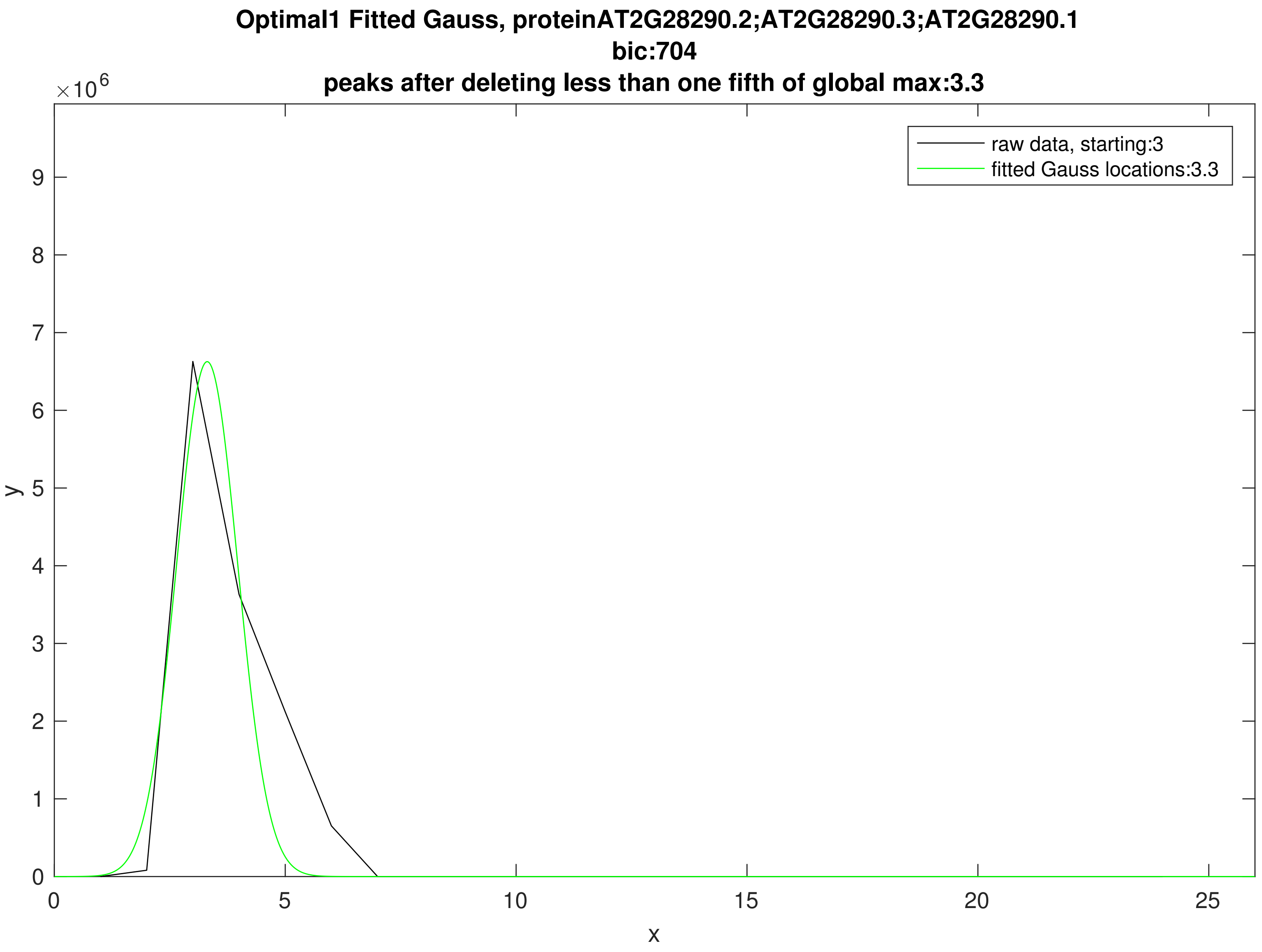
Optimal1 Fitted Gauss, proteinAT2G28290.2;AT2G28290.3;AT2G28290.1
bic:704
peaks after deleting less than one fifth of global max:3.3
×10^6
raw data, starting:3
fitted Gauss locations:3.3
y
x
0
1
2
3
4
5
6
7
8
9
0
5
10
15
20
25

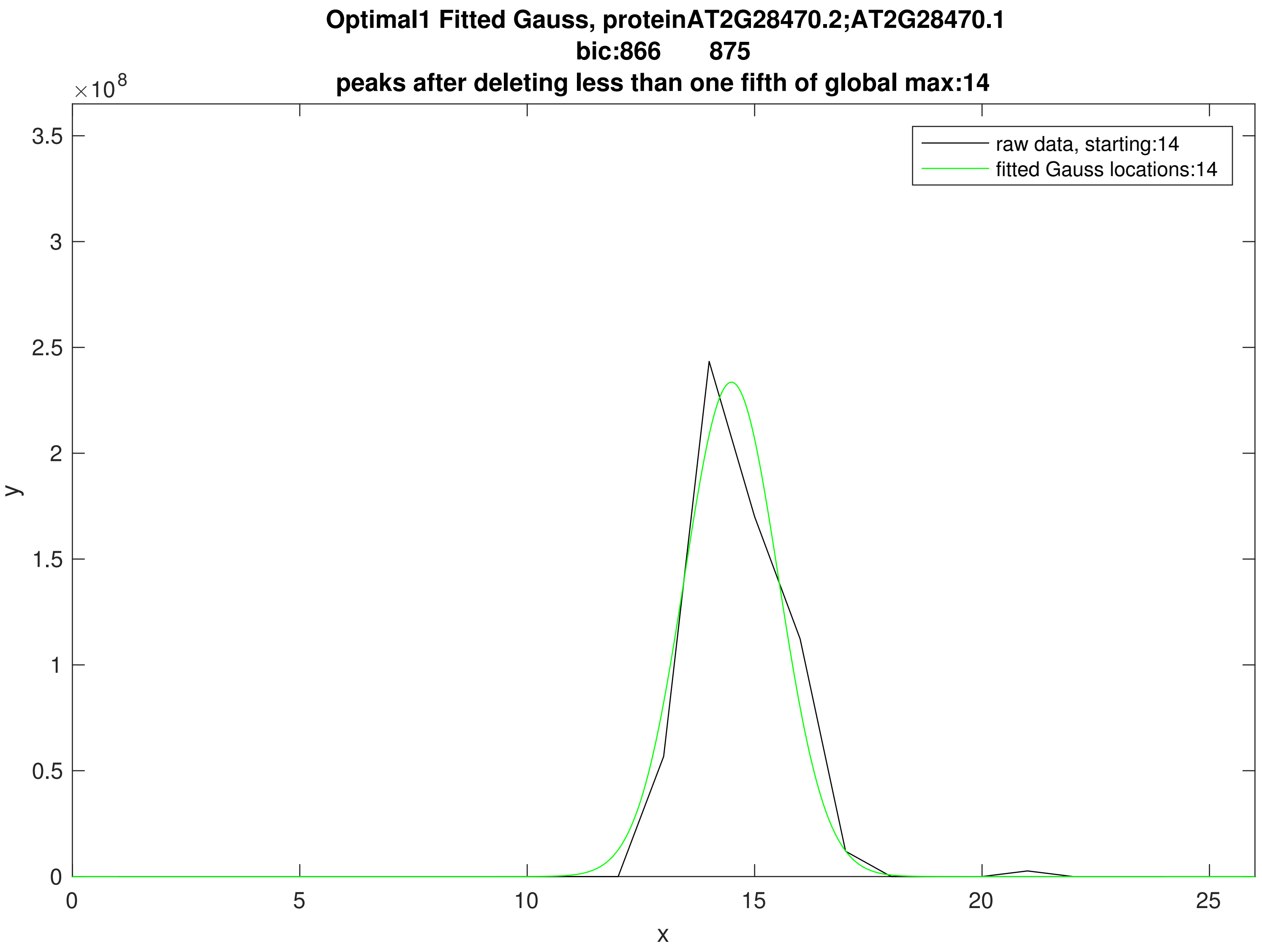

Optimal1 Fitted Gauss, proteinAT2G28470.2;AT2G28470.1
bic:866 875
peaks after deleting less than one fifth of global max:14
×10^8
3.5
3
2.5
2
1.5
1
0.5
0
y
0
5
10
15
20
25
x
raw data, starting:14
fitted Gauss locations:14

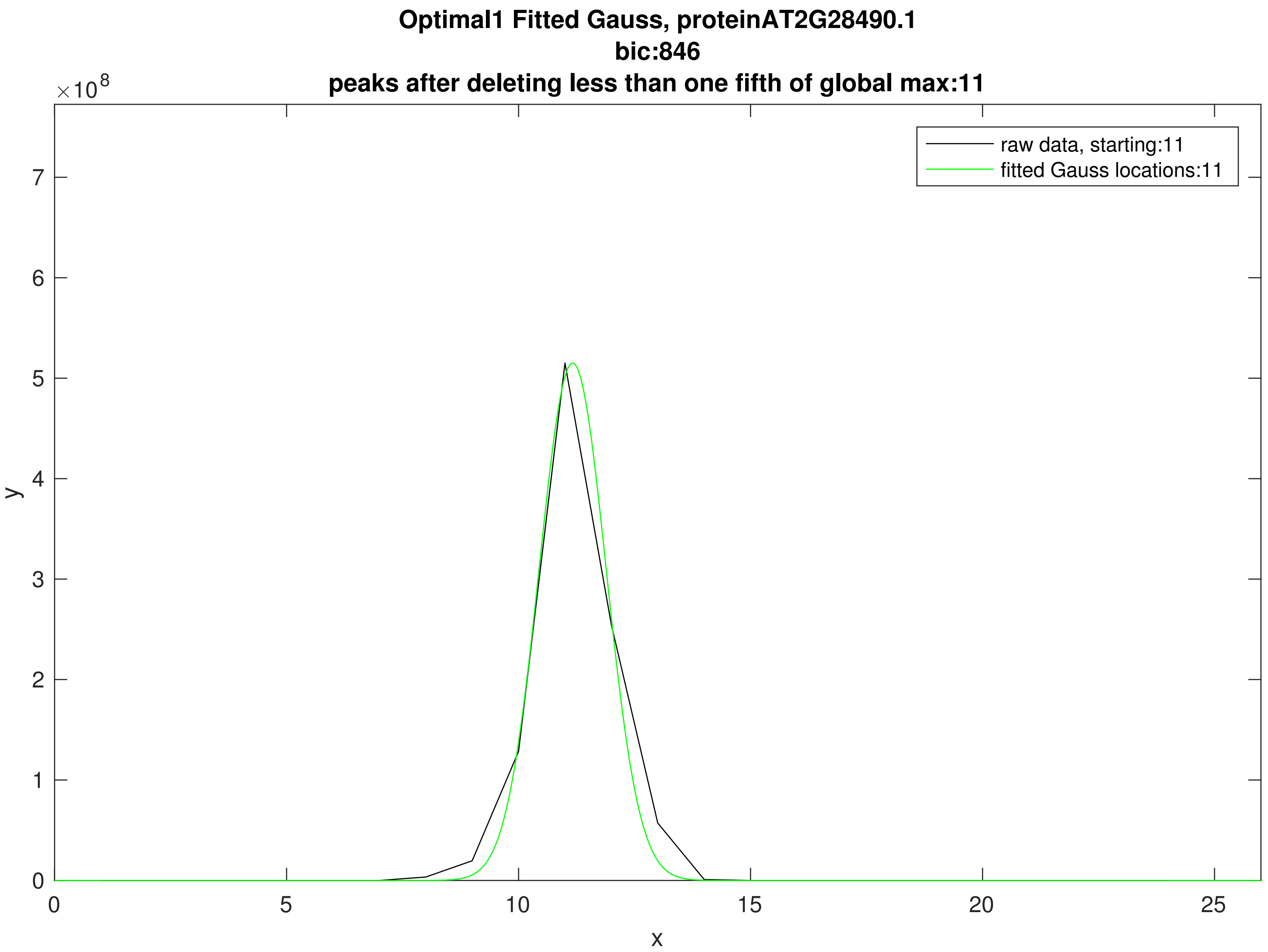
Optimal1 Fitted Gauss, proteinAT2G28490.1
bic:846
peaks after deleting less than one fifth of global max:11
raw data, starting:11
fitted Gauss locations:11
y
x

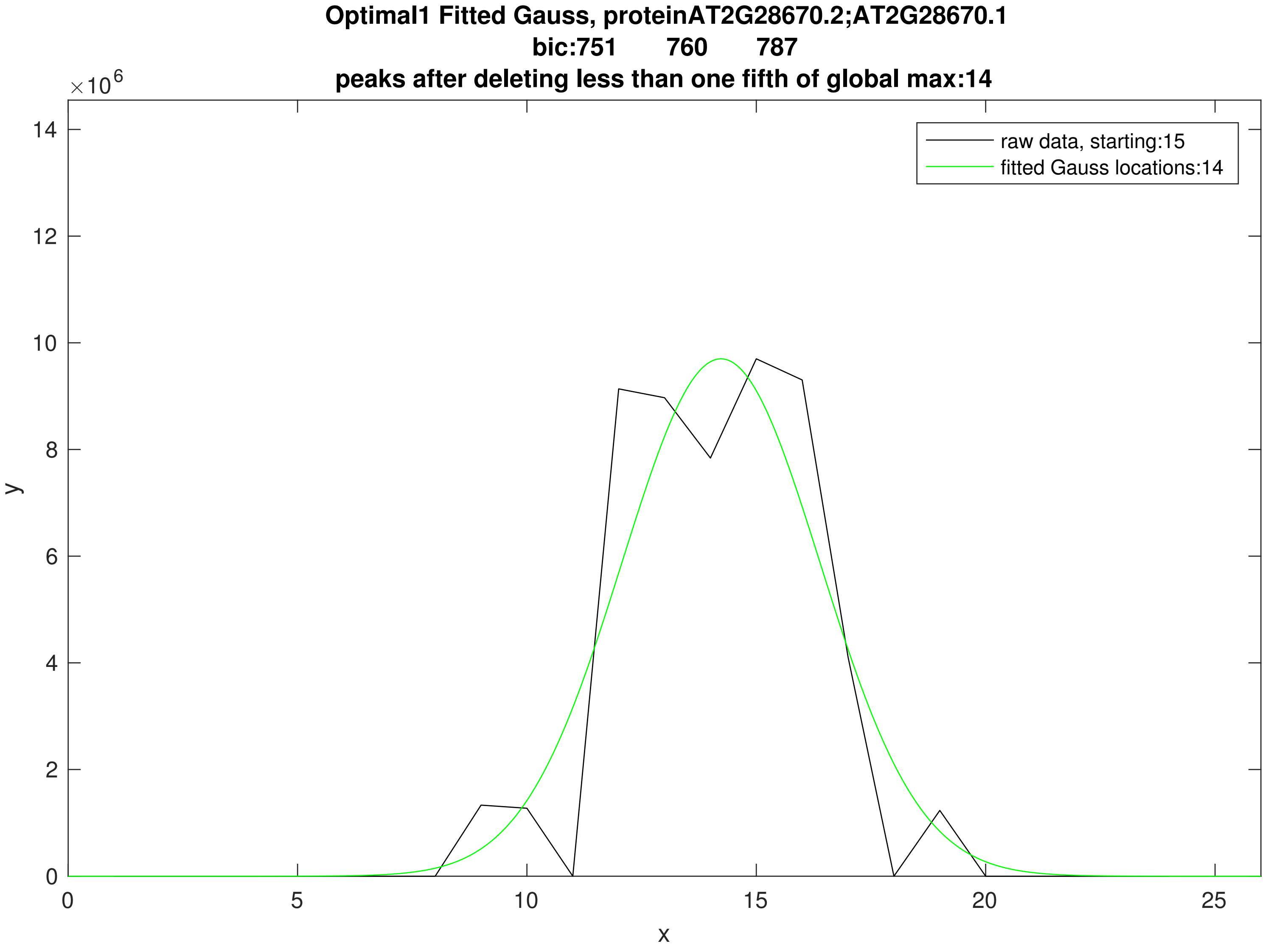

Optimal1 Fitted Gauss, proteinAT2G28670.2;AT2G28670.1
bic:751 760 787
peaks after deleting less than one fifth of global max:14
×10^6
raw data, starting:15
fitted Gauss locations:14
y
x
0
2
4
6
8
10
12
14
5
15
20
25

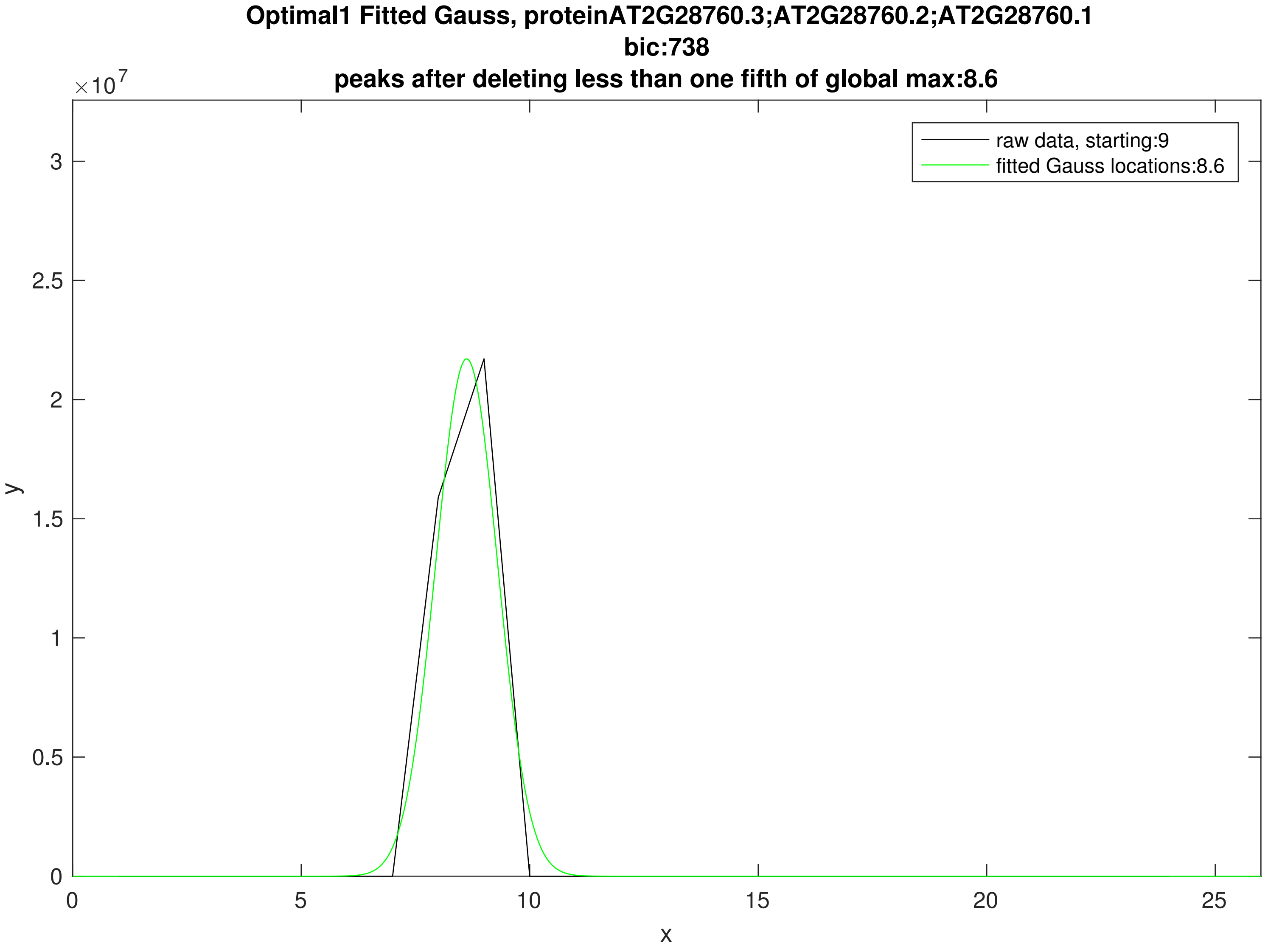
Optimal1 Fitted Gauss, proteinAT2G28760.3;AT2G28760.2;AT2G28760.1
bic:738
peaks after deleting less than one fifth of global max:8.6
×10^7
raw data, starting:9
fitted Gauss locations:8.6
y
x
0
0.5
1
1.5
2
2.5
3
0
5
10
15
20
25

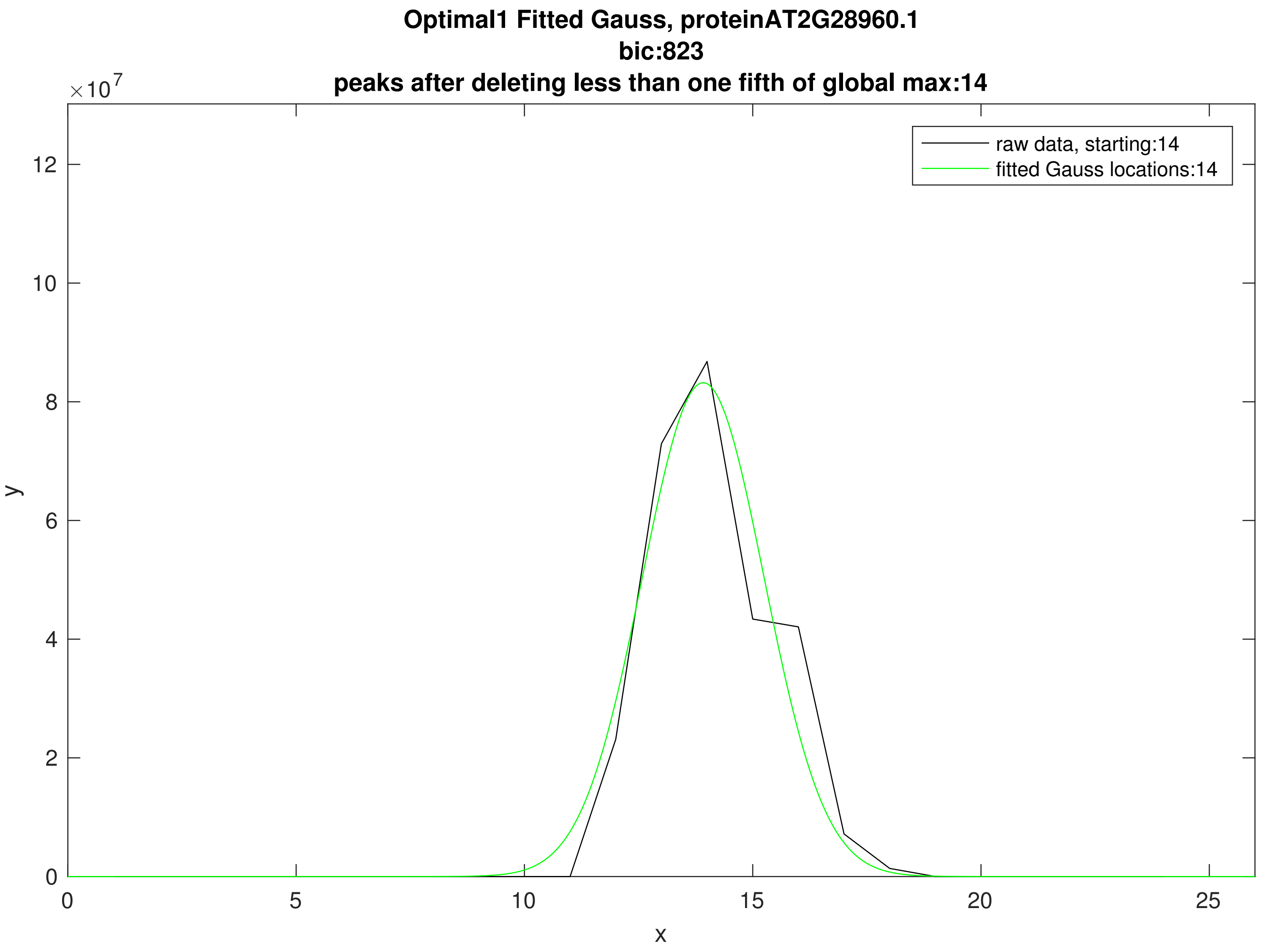
Optimal1 Fitted Gauss, proteinAT2G28960.1
bic:823
peaks after deleting less than one fifth of global max:14
×10^7
raw data, starting:14
fitted Gauss locations:14
12
10
8
6
4
2
0
y
0
5
10
15
20
25
x

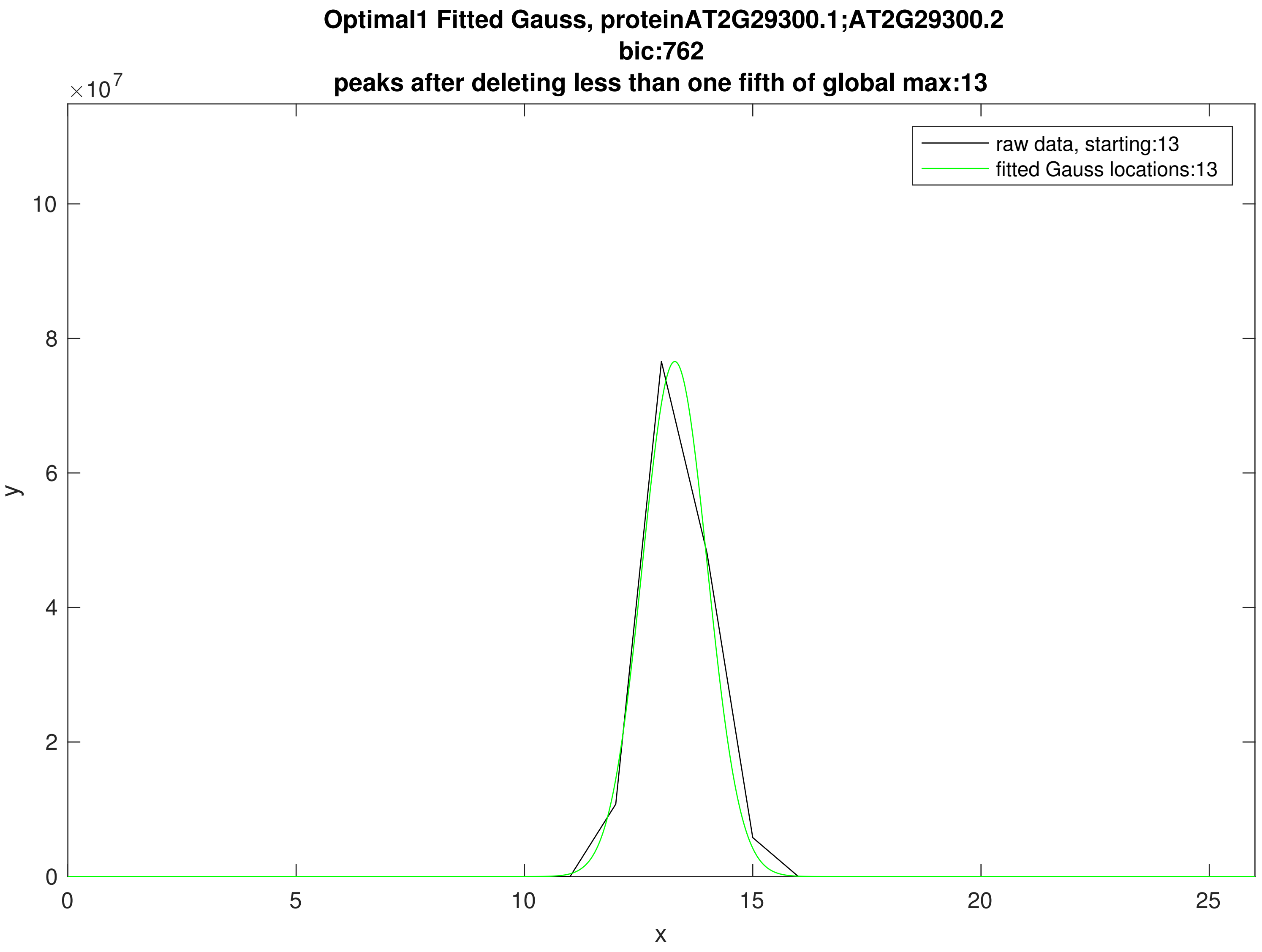
Optimal1 Fitted Gauss, proteinAT2G29300.1;AT2G29300.2
bic:762
peaks after deleting less than one fifth of global max:13
×10^7
raw data, starting:13
fitted Gauss locations:13
y
x
0
2
4
6
8
10
0
5
10
15
20
25

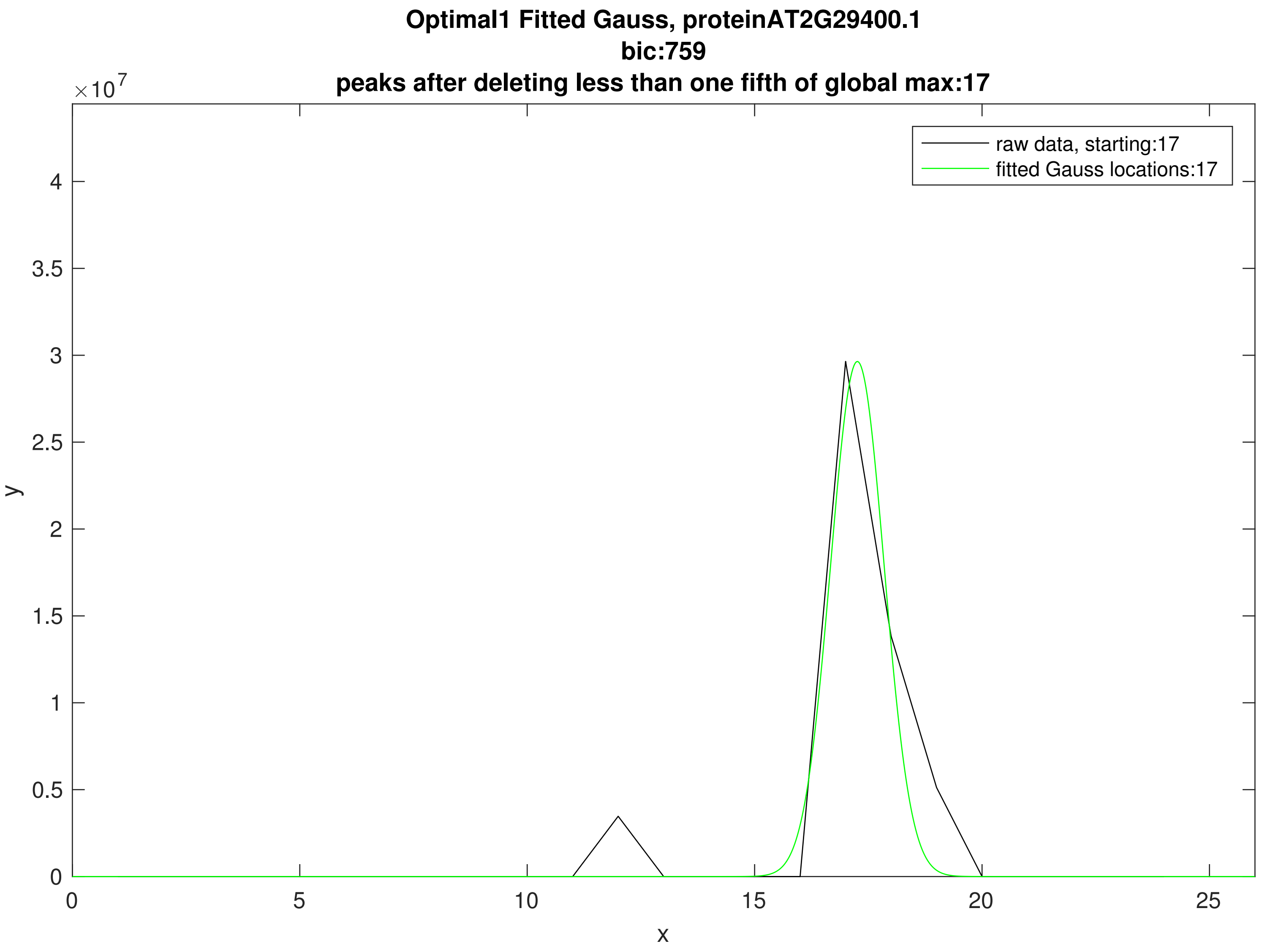
Optimal1 Fitted Gauss, proteinAT2G29400.1
bic:759
peaks after deleting less than one fifth of global max:17
raw data, starting:17
fitted Gauss locations:17
×10^7
y
x

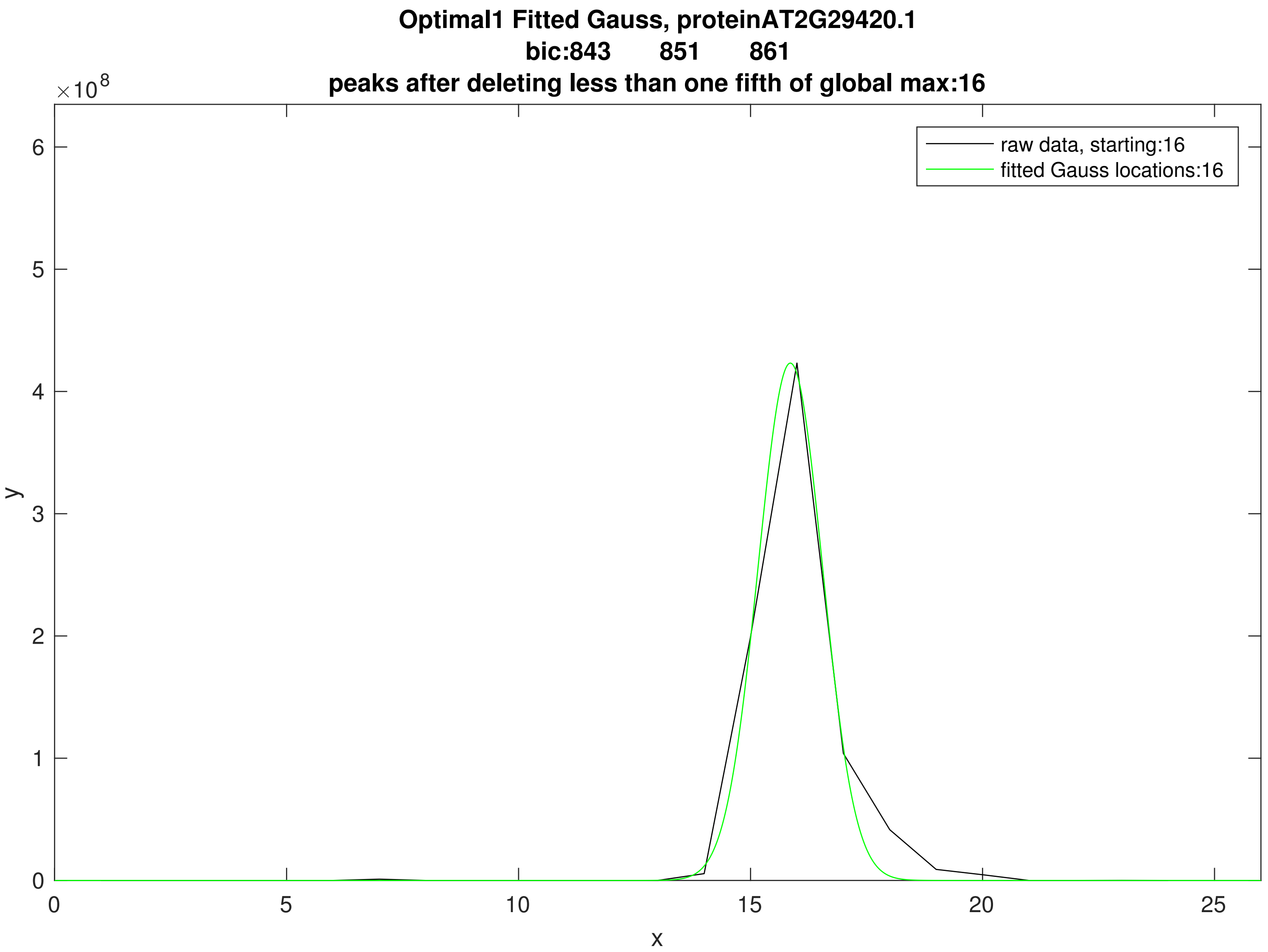
Optimal1 Fitted Gauss, proteinAT2G29420.1
bic:843 851 861
peaks after deleting less than one fifth of global max:16
×10^8
raw data, starting:16
fitted Gauss locations:16
y
x
0
1
2
3
4
5
6
0
5
10
15
20
25

Optimal1 Fitted Gauss, proteinAT2G29440.1
bic:750
peaks after deleting less than one fifth of global max:16

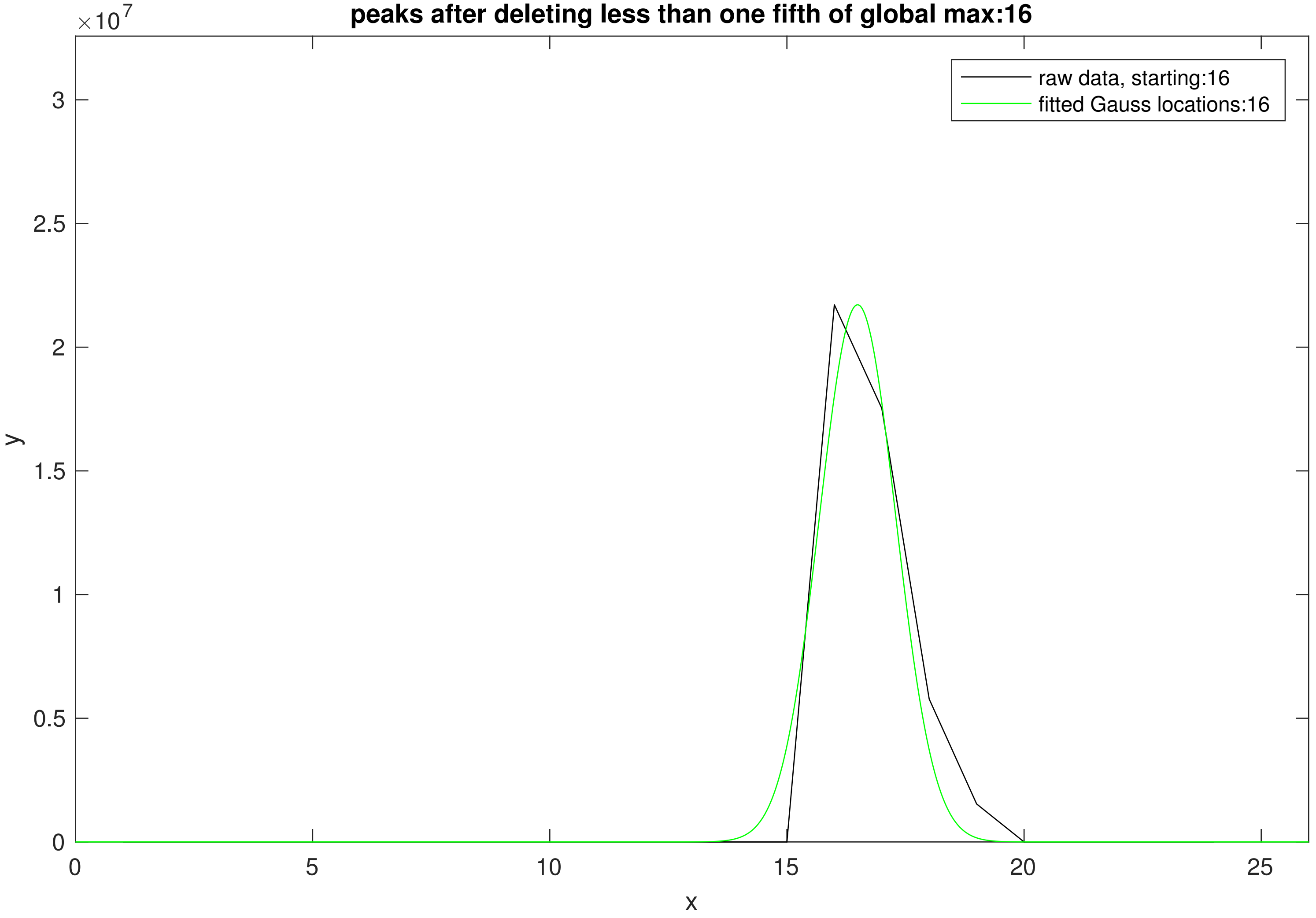

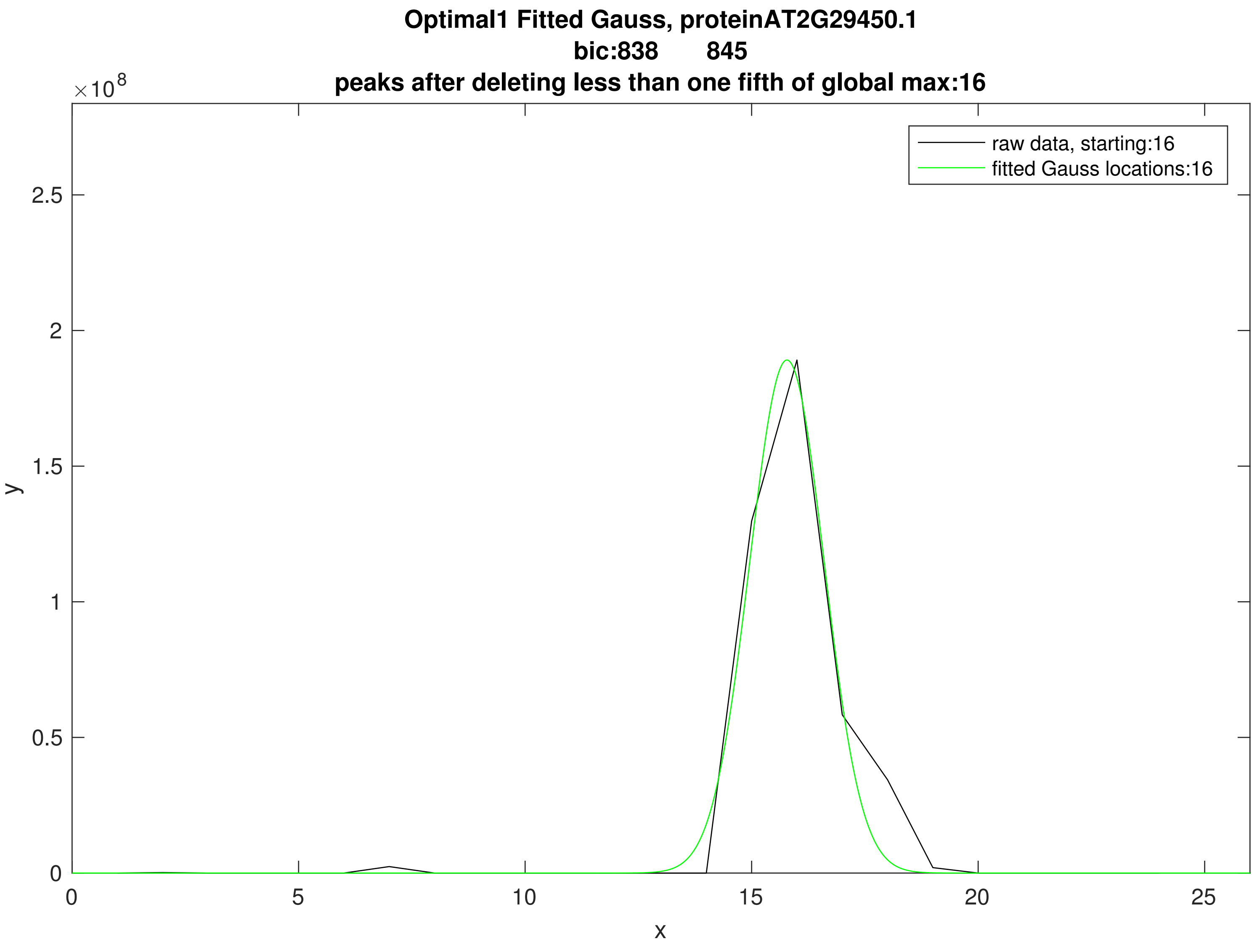

Optimal1 Fitted Gauss, proteinAT2G29450.1
bic:838 845
peaks after deleting less than one fifth of global max:16
×10^8
raw data, starting:16
fitted Gauss locations:16
2.5
2
1.5
1
0.5
0
y
0
5
10
15
20
25
x

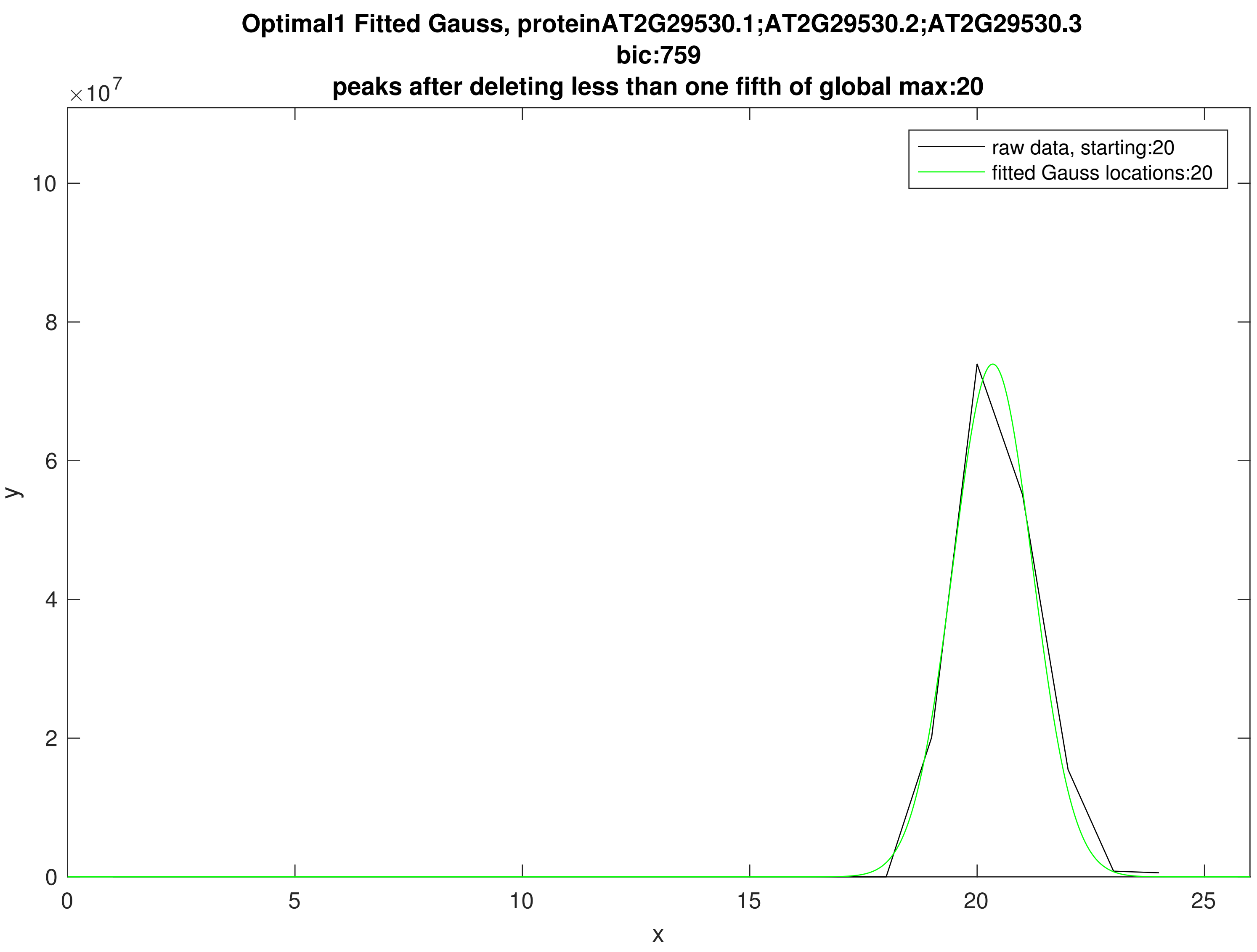
Optimal1 Fitted Gauss, proteinAT2G29530.1;AT2G29530.2;AT2G29530.3
bic:759
peaks after deleting less than one fifth of global max:20
×10^7
raw data, starting:20
fitted Gauss locations:20
y
x
0
2
4
6
8
10
0
5
10
15
20
25

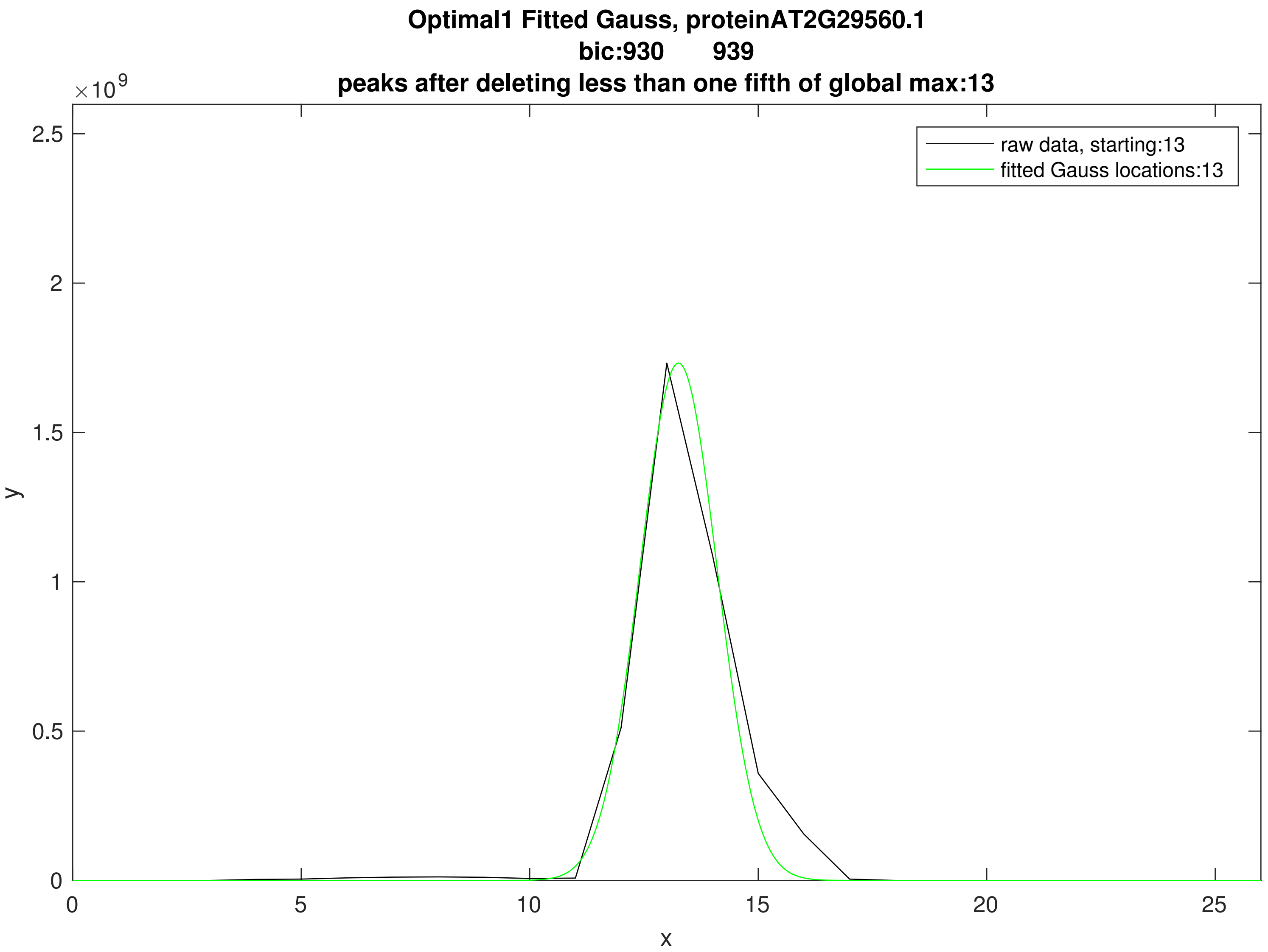
Optimal1 Fitted Gauss, proteinAT2G29560.1
bic:930 939
peaks after deleting less than one fifth of global max:13
×10^9
2.5
2
1.5
1
0.5
0
y
0
5
10
15
20
25
x
raw data, starting:13
fitted Gauss locations:13

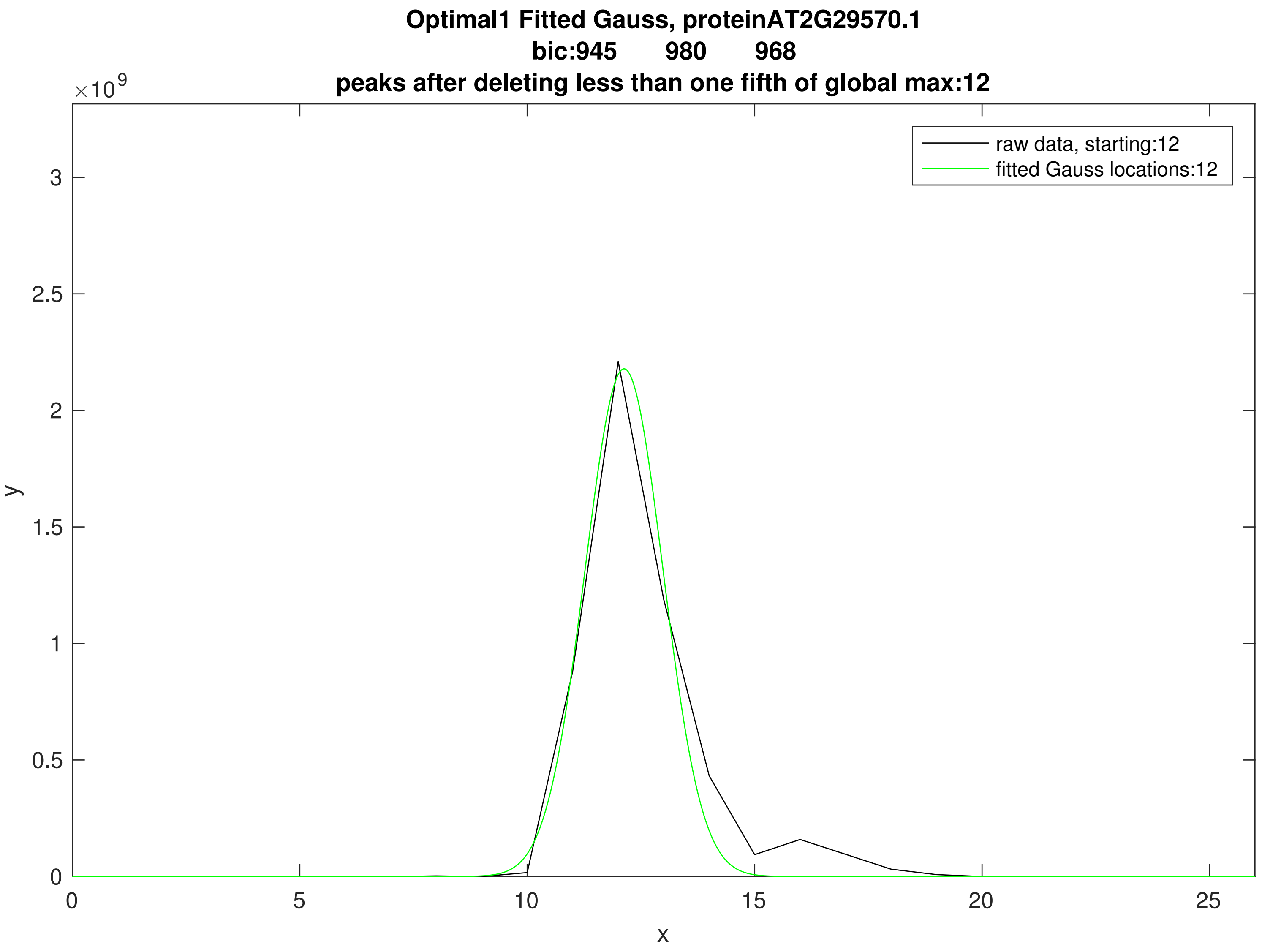
Optimal1 Fitted Gauss, proteinAT2G29570.1
bic:945 980 968
peaks after deleting less than one fifth of global max:12
×10^9
raw data, starting:12
fitted Gauss locations:12
y
x
0
0.5
1
1.5
2
2.5
3
5
10
15
20
25

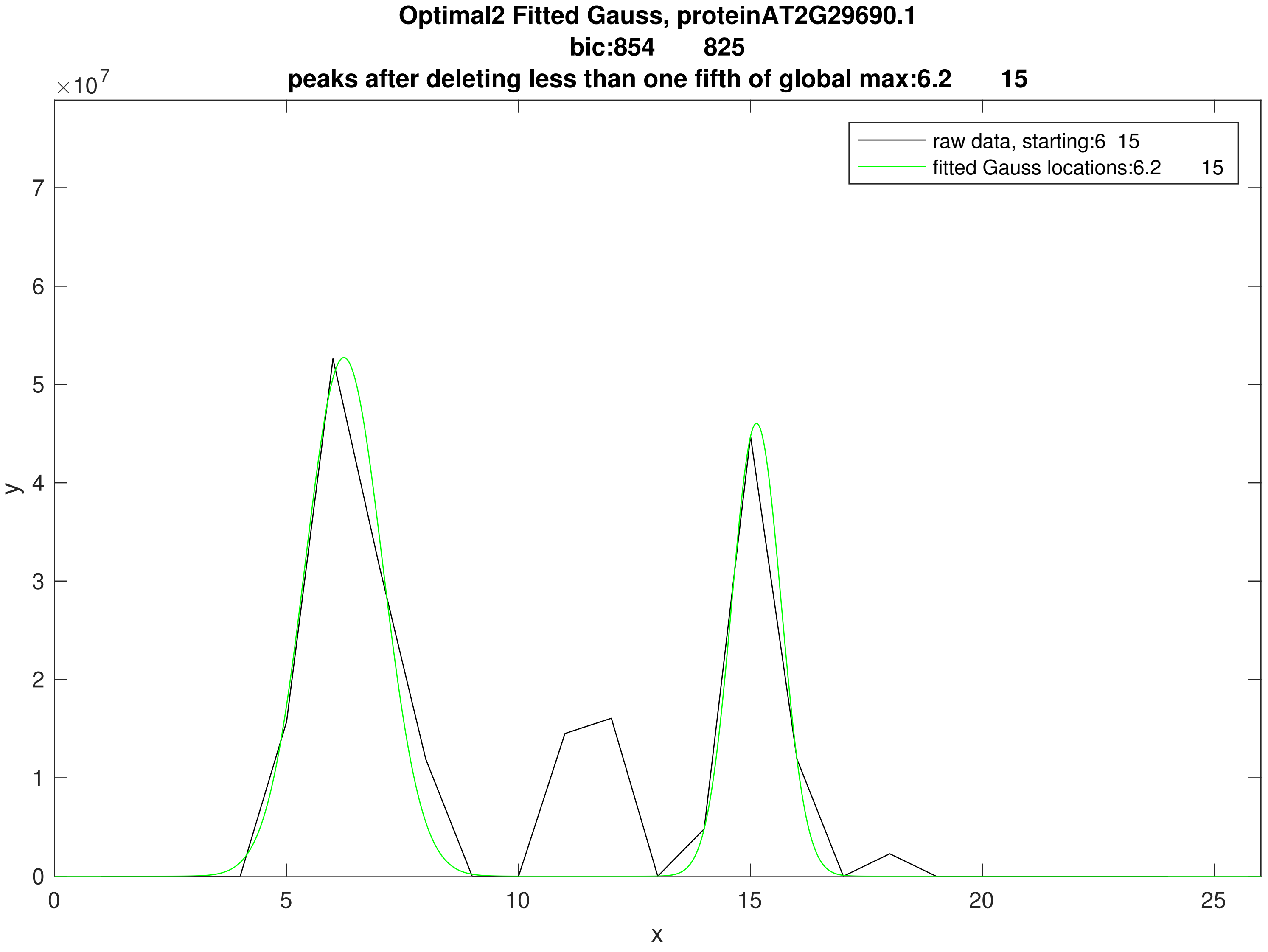
Optimal2 Fitted Gauss, proteinAT2G29690.1
bic:854 825
peaks after deleting less than one fifth of global max:6.2 15
×10^7
raw data, starting:6 15
fitted Gauss locations:6.2 15
y
x
0
1
2
3
4
5
6
7
0
5
10
15
20
25

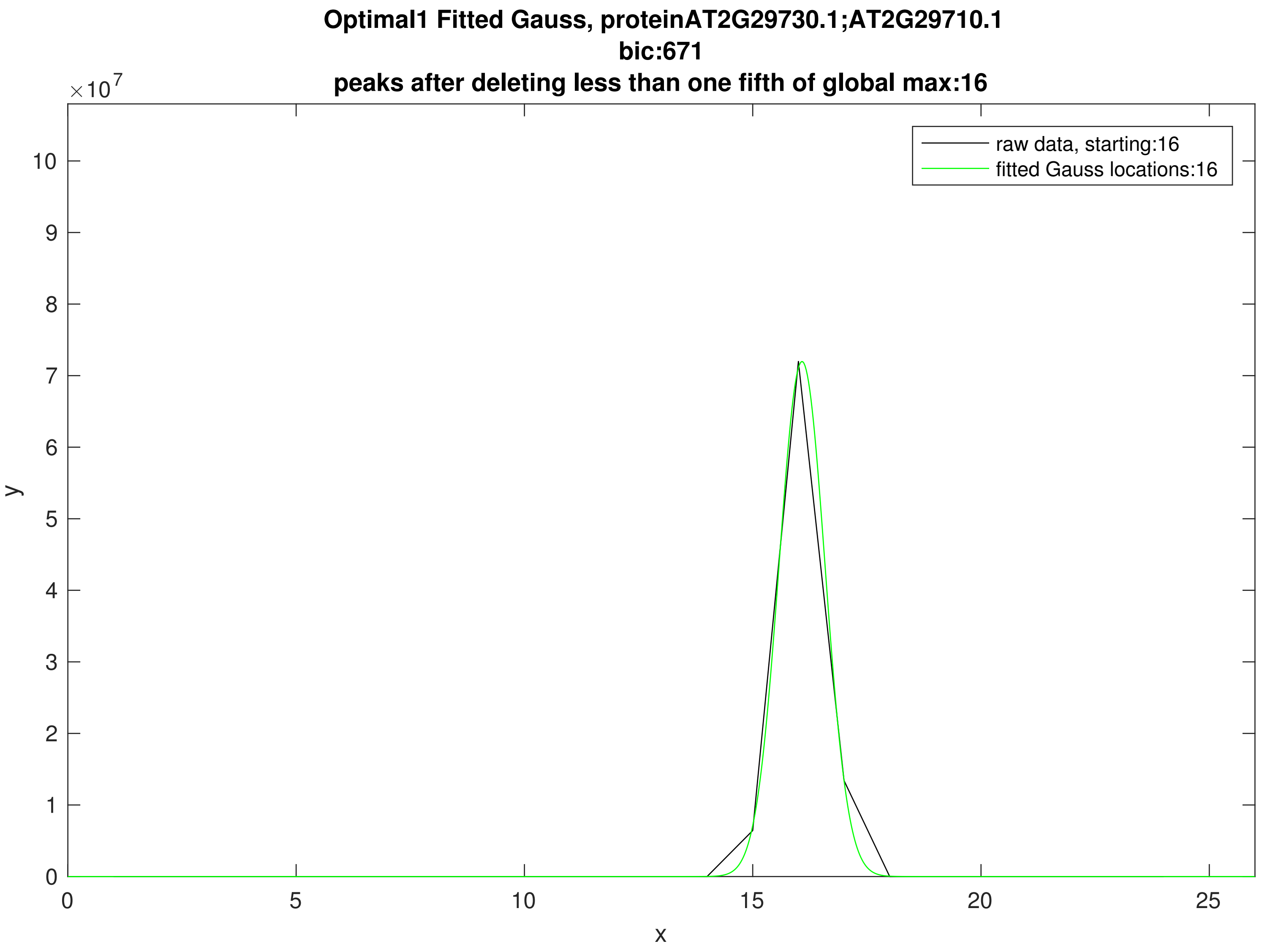

Optimal1 Fitted Gauss, proteinAT2G29730.1;AT2G29710.1
bic:671
peaks after deleting less than one fifth of global max:16
×10^7
raw data, starting:16
fitted Gauss locations:16
y
x
0
1
2
3
4
5
6
7
8
9
10
0
5
10
15
20
25

#### Optimal1 Fitted Gauss, proteinAT2G29750.1
#### bic:870
#### peaks after deleting less than one fifth of global max:16

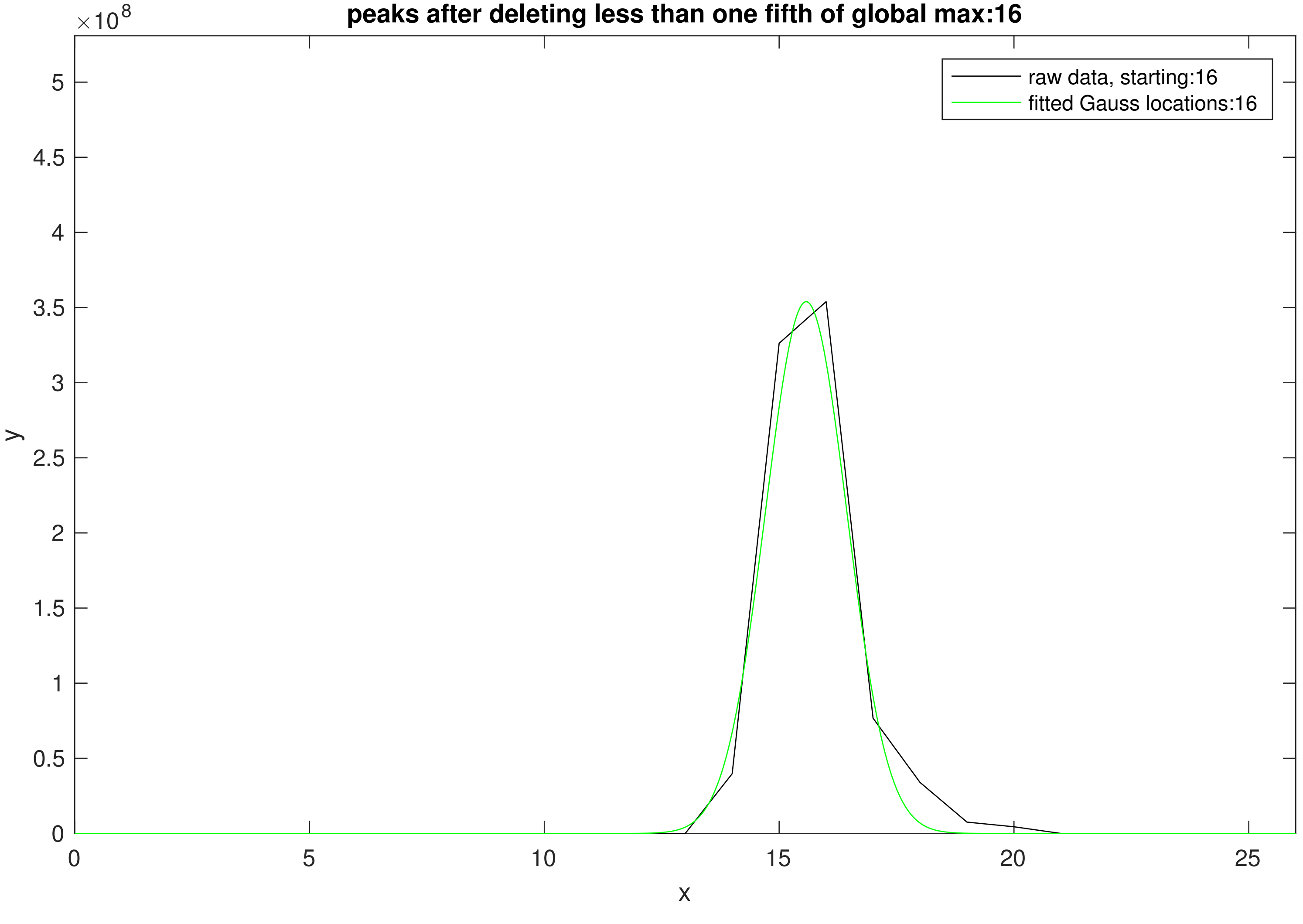

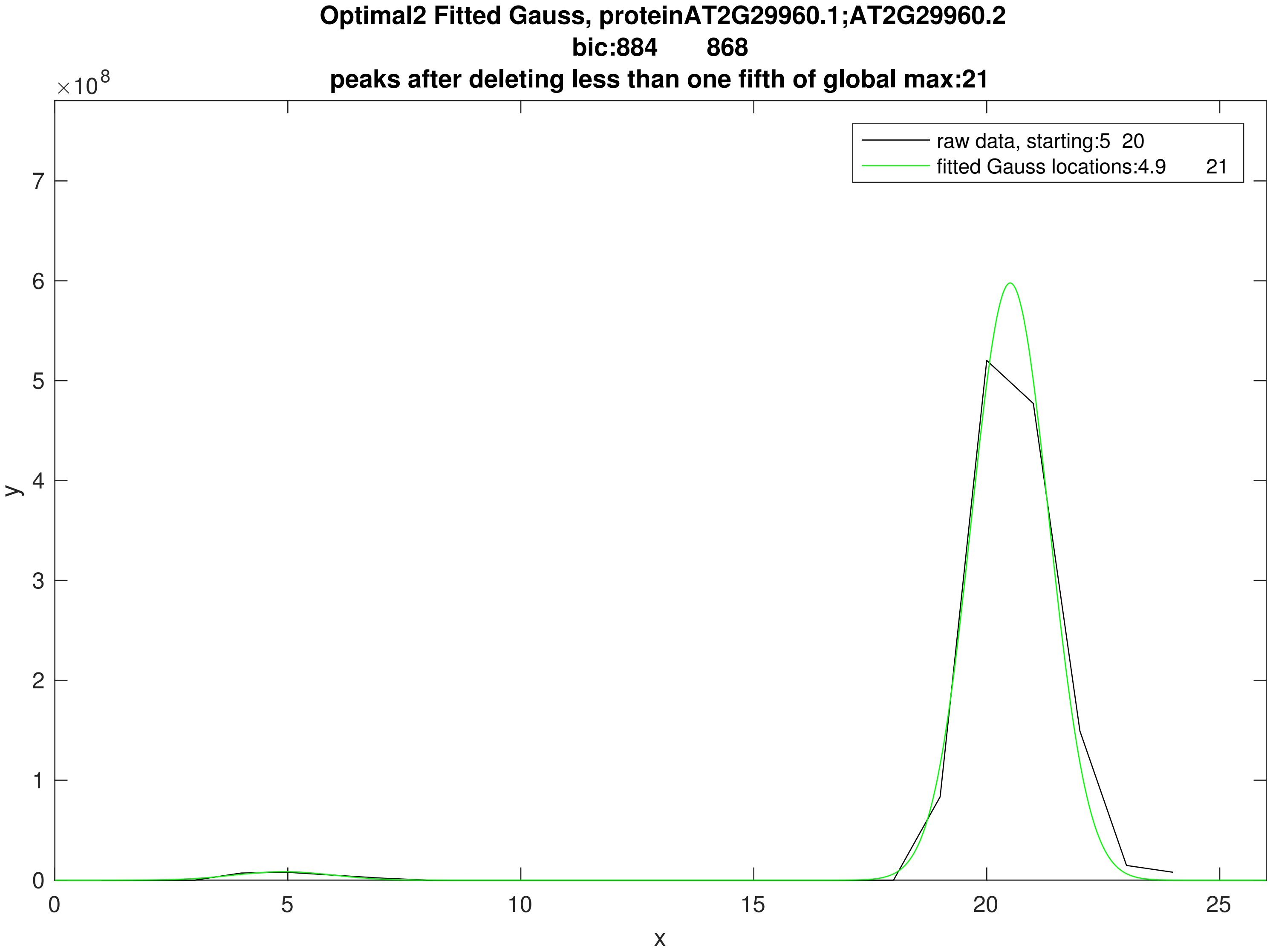

Optimal2 Fitted Gauss, proteinAT2G29960.1;AT2G29960.2
bic:884 868
peaks after deleting less than one fifth of global max:21
×10^8
raw data, starting:5 20
fitted Gauss locations:4.9 21
y
x
0
1
2
3
4
5
6
7
0
5
10
15
20
25

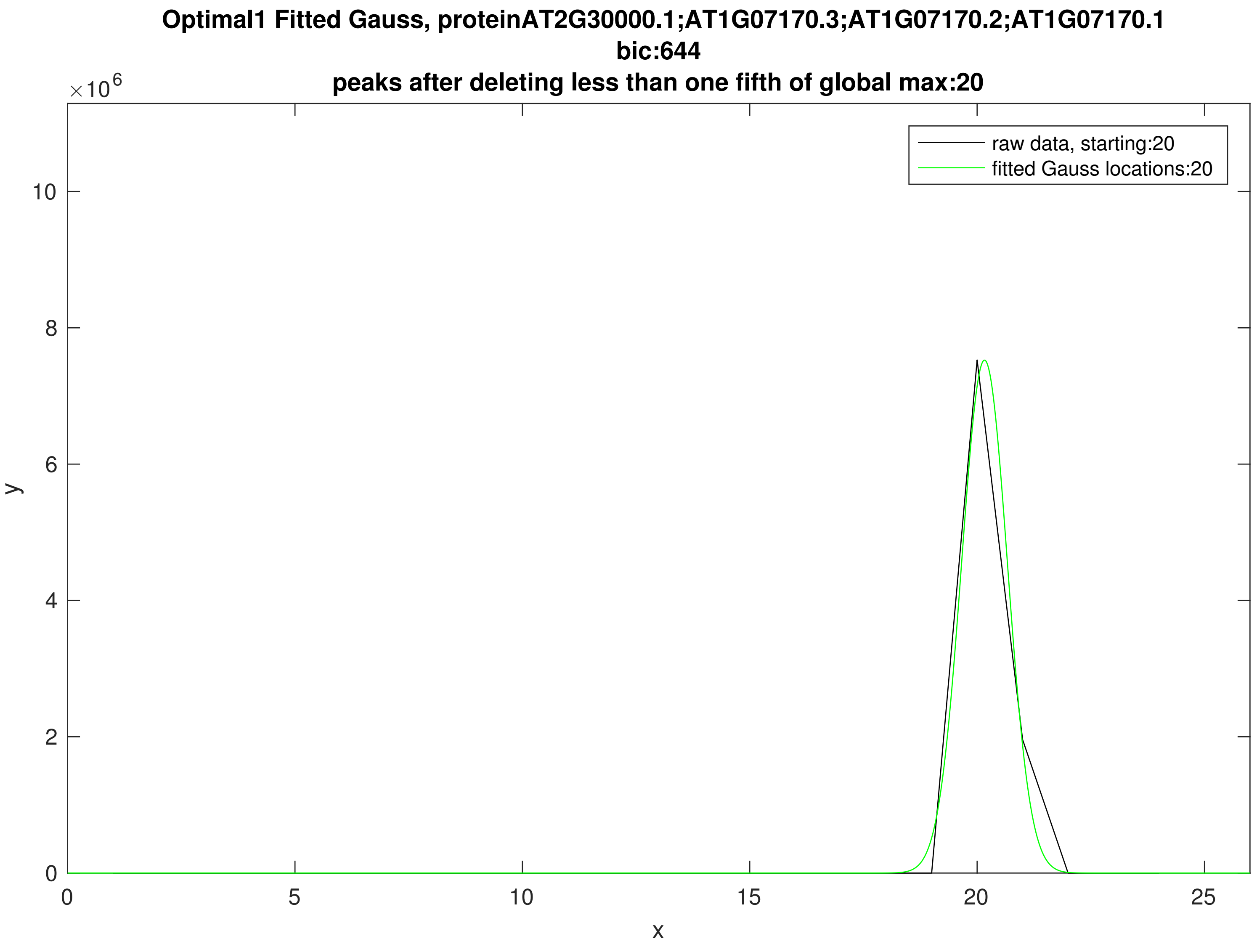
Optimal1 Fitted Gauss, proteinAT2G30000.1;AT1G07170.3;AT1G07170.2;AT1G07170.1
bic:644
peaks after deleting less than one fifth of global max:20
$\times 10^6$
raw data, starting:20
fitted Gauss locations:20
y
x
0
2
4
6
8
10
0
5
10
15
20
25

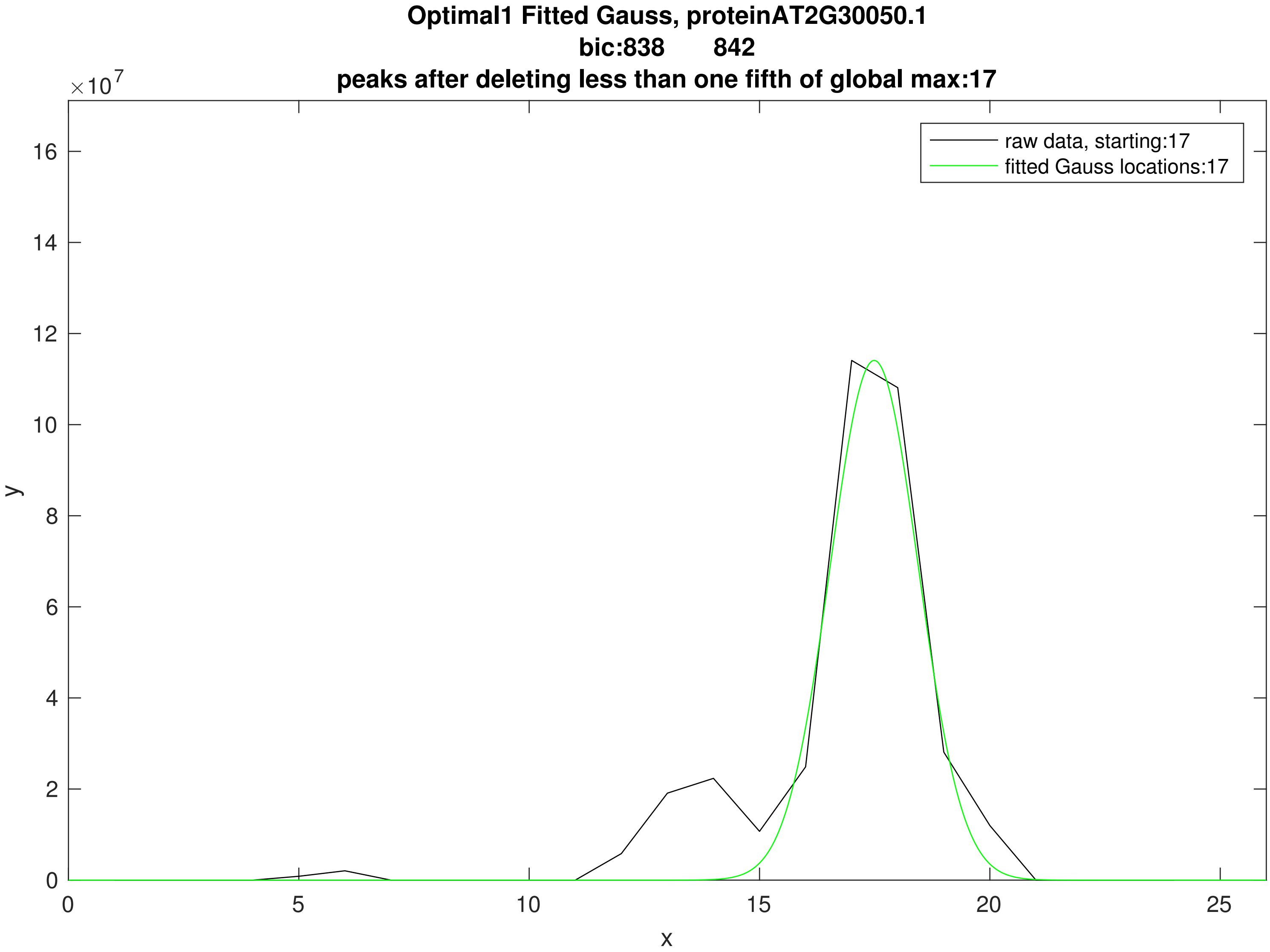
Optimal1 Fitted Gauss, proteinAT2G30050.1
bic:838 842
peaks after deleting less than one fifth of global max:17
×10^7
raw data, starting:17
fitted Gauss locations:17
y
x
0
2
4
6
8
10
12
14
16
0
5
10
15
20
25

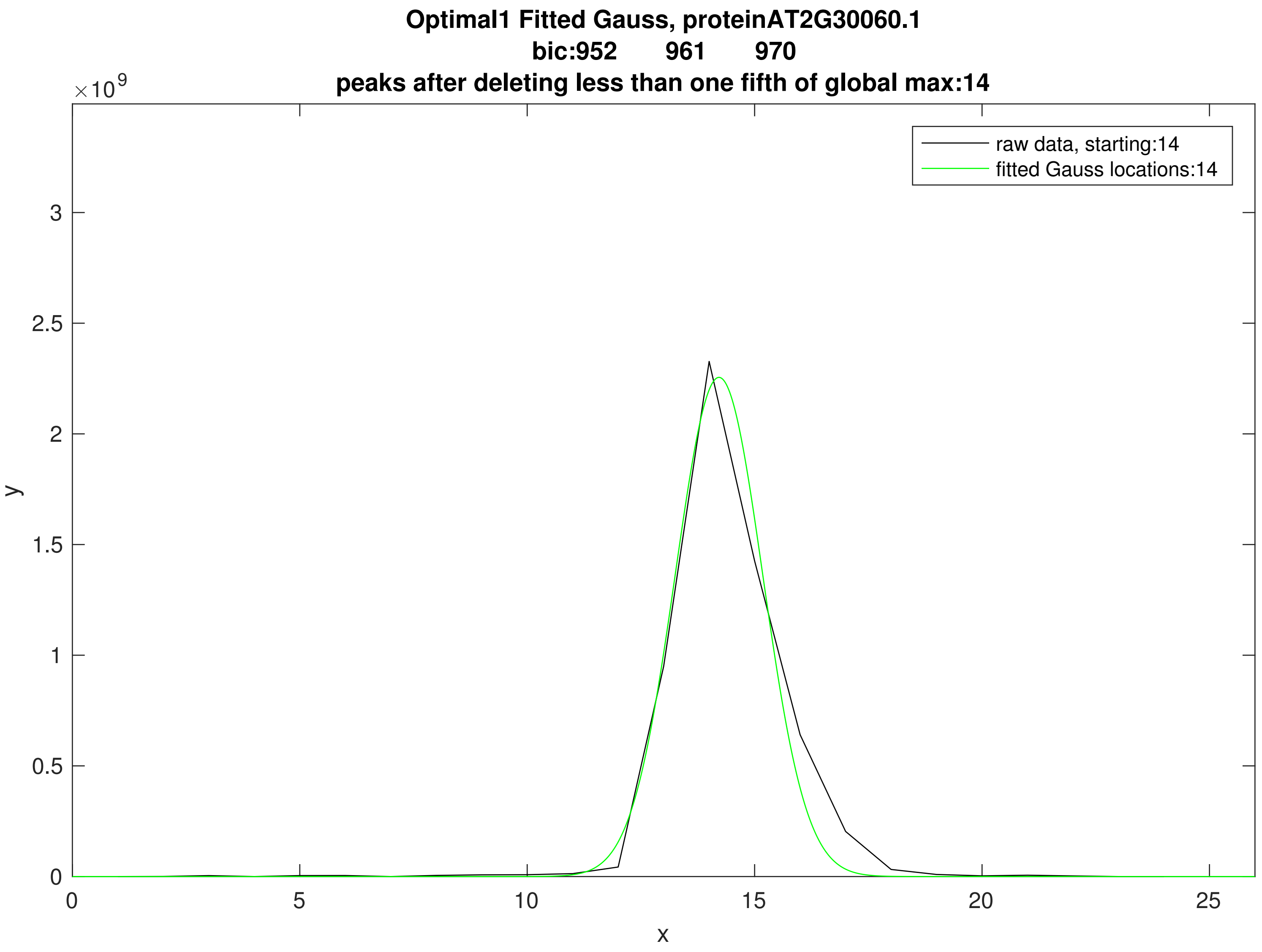

Optimal1 Fitted Gauss, proteinAT2G30060.1
bic:952 961 970
peaks after deleting less than one fifth of global max:14
×10^9
raw data, starting:14
fitted Gauss locations:14
y
x
0
0.5
1
1.5
2
2.5
3
5
10
15
20
25

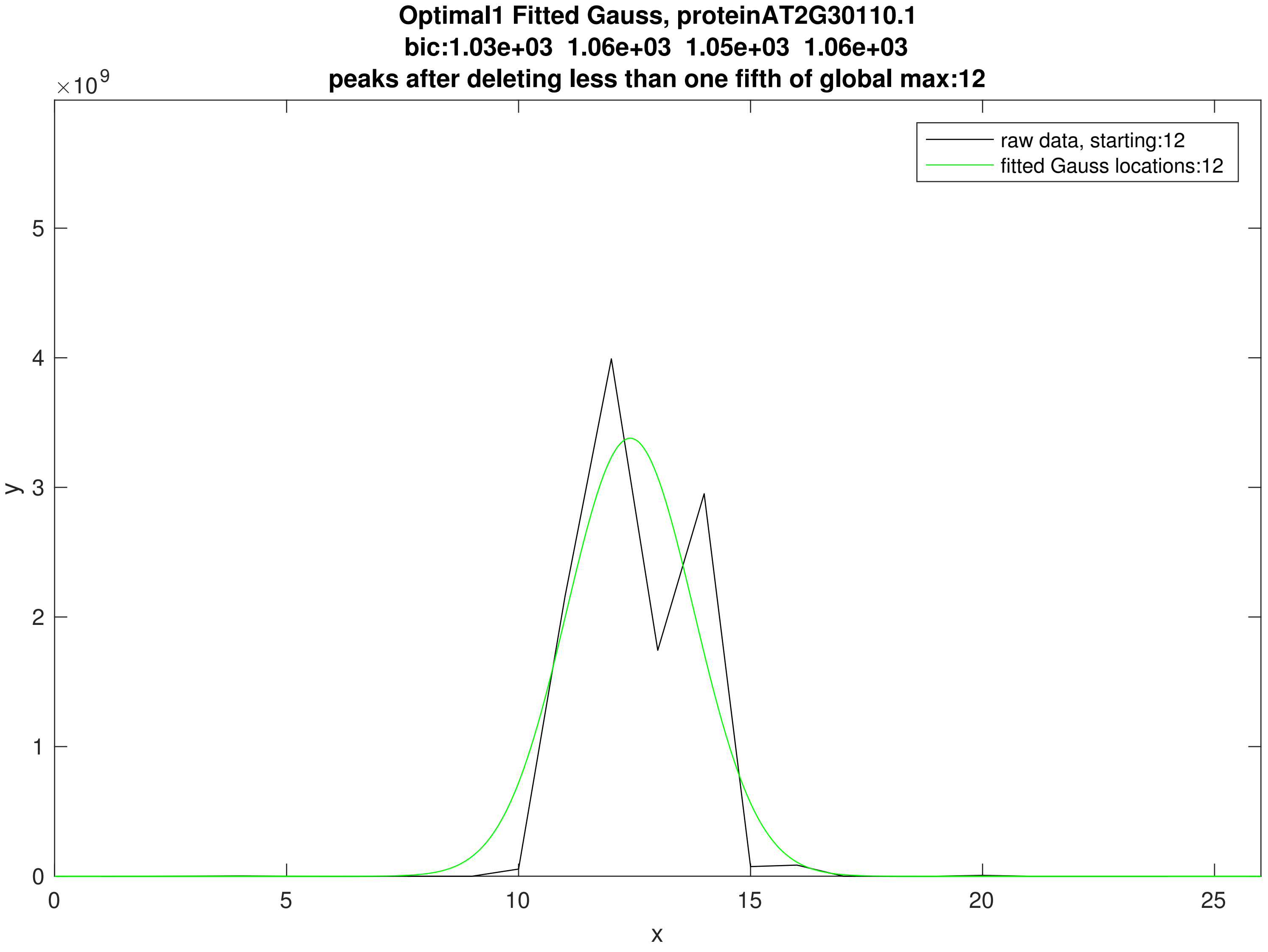

Optimal1 Fitted Gauss, proteinAT2G30110.1
bic:1.03e+03  1.06e+03  1.05e+03  1.06e+03
peaks after deleting less than one fifth of global max:12
×10^9
raw data, starting:12
fitted Gauss locations:12
y
x
0
1
2
3
4
5
10
15
20
25

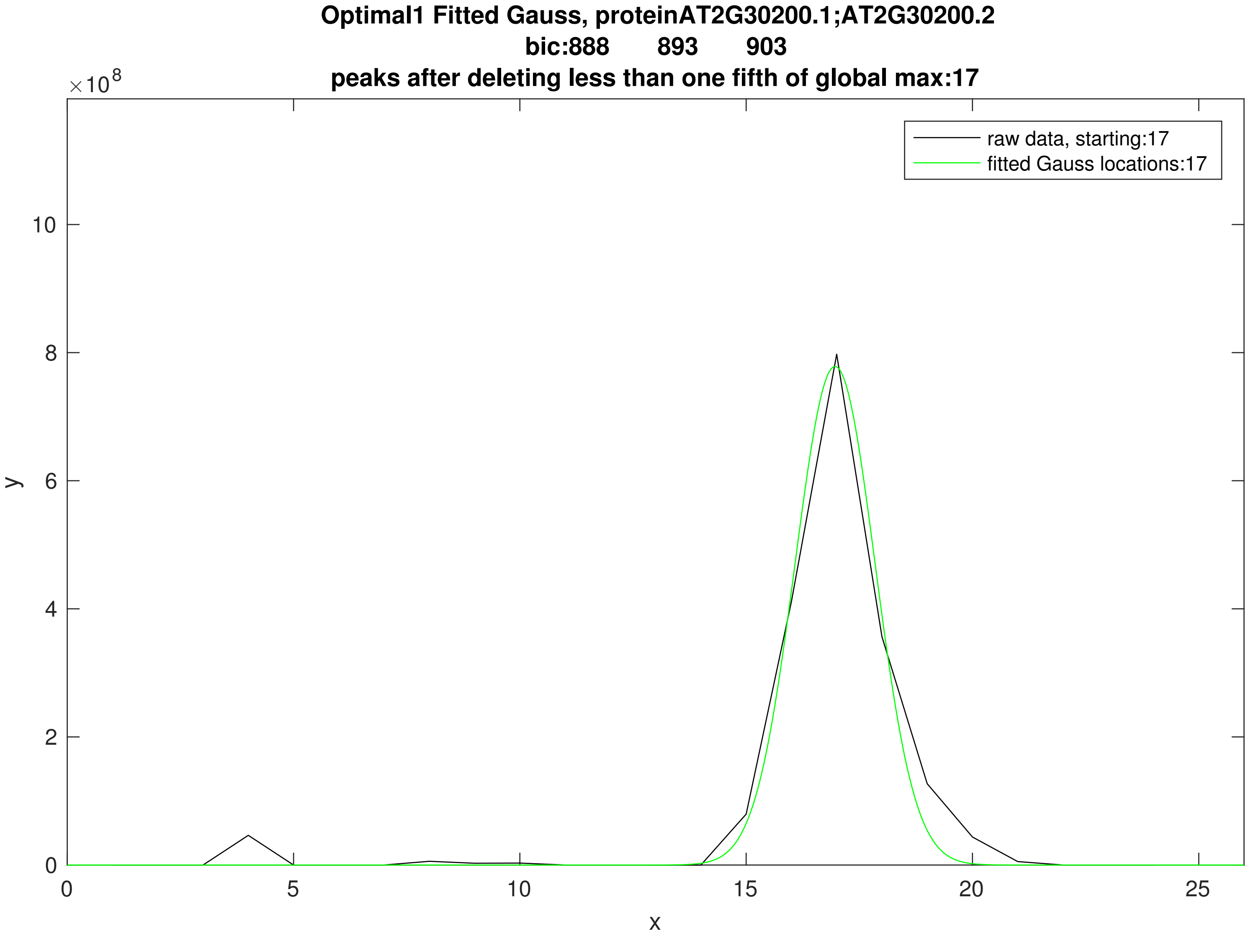

Optimal1 Fitted Gauss, proteinAT2G30200.1;AT2G30200.2
bic:888 893 903
peaks after deleting less than one fifth of global max:17
$\times 10^8$
raw data, starting:17
fitted Gauss locations:17
y
x
0
2
4
6
8
10
0
5
10
15
20
25

**Optimal1 Fitted Gauss, proteinAT2G30410.2;AT2G30410.1**
**bic:937 947**
**peaks after deleting less than one fifth of global max:18**

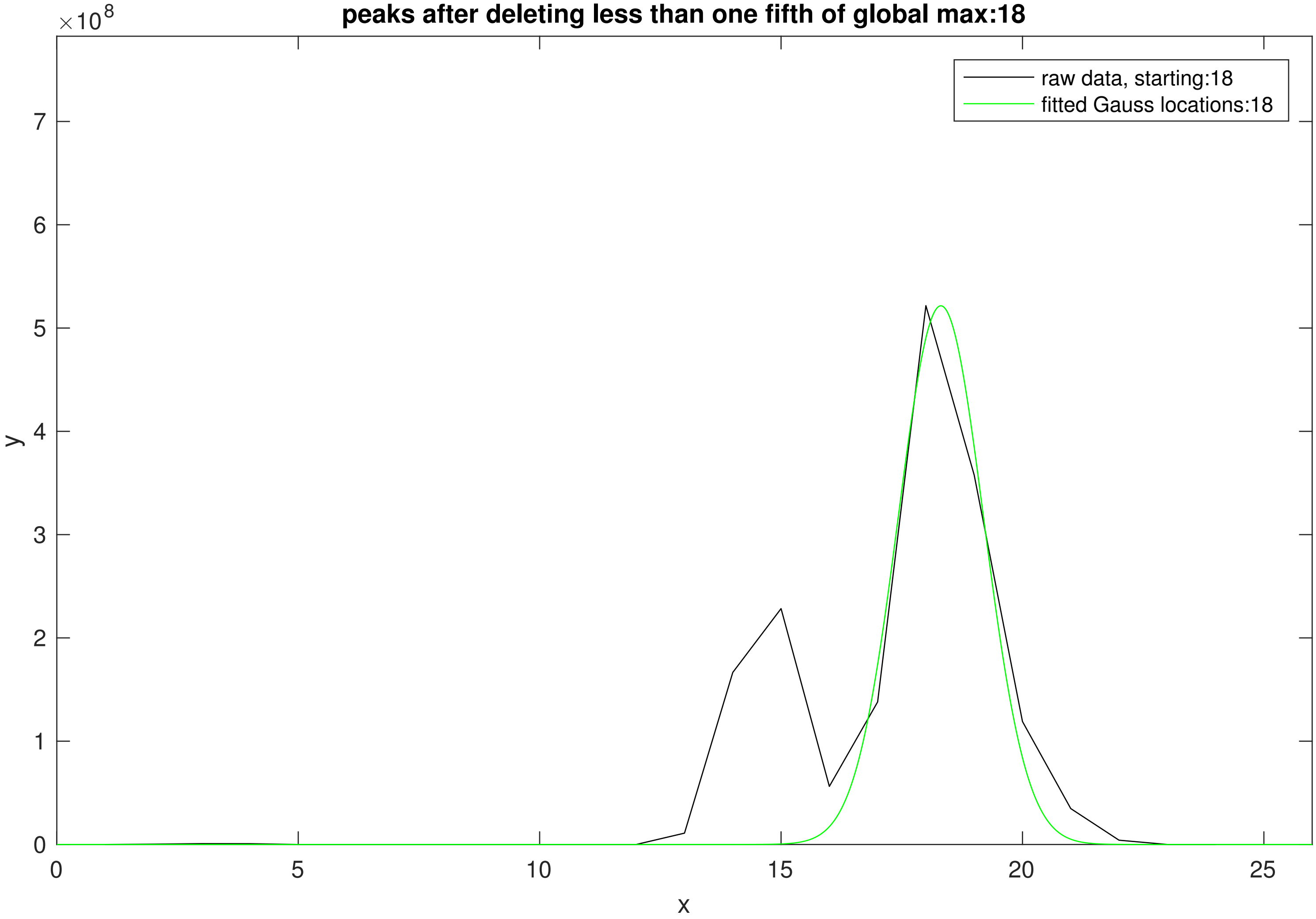

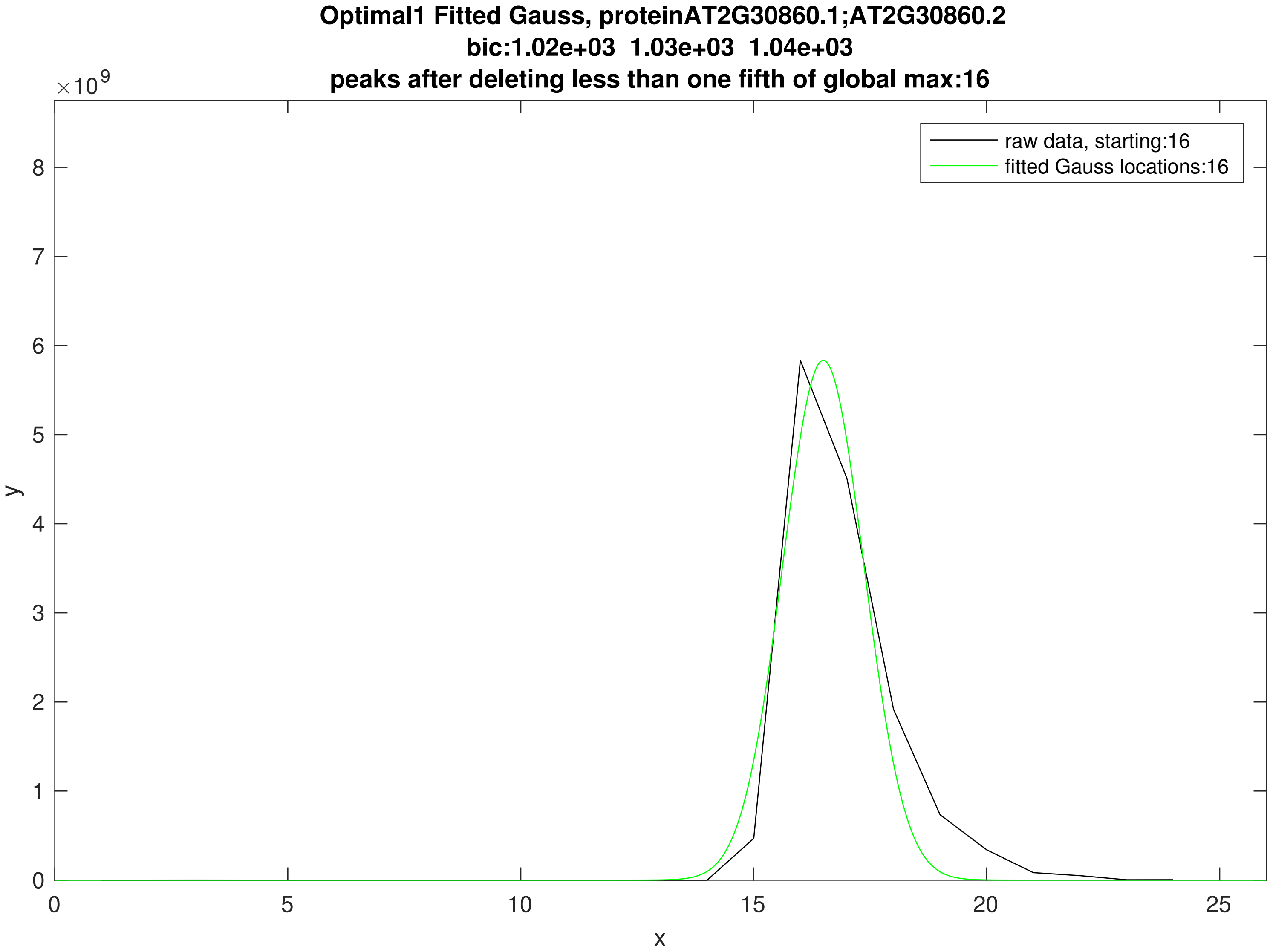

Optimal1 Fitted Gauss, proteinAT2G30860.1;AT2G30860.2
bic:1.02e+03 1.03e+03 1.04e+03
peaks after deleting less than one fifth of global max:16
×10^9
raw data, starting:16
fitted Gauss locations:16
y
x
0
1
2
3
4
5
6
7
8
0
5
10
15
20
25

**Optimal1 Fitted Gauss, proteinAT2G30870.1**
**bic:1.03e+03  1.04e+03  1.05e+03**
**peaks after deleting less than one fifth of global max:16**

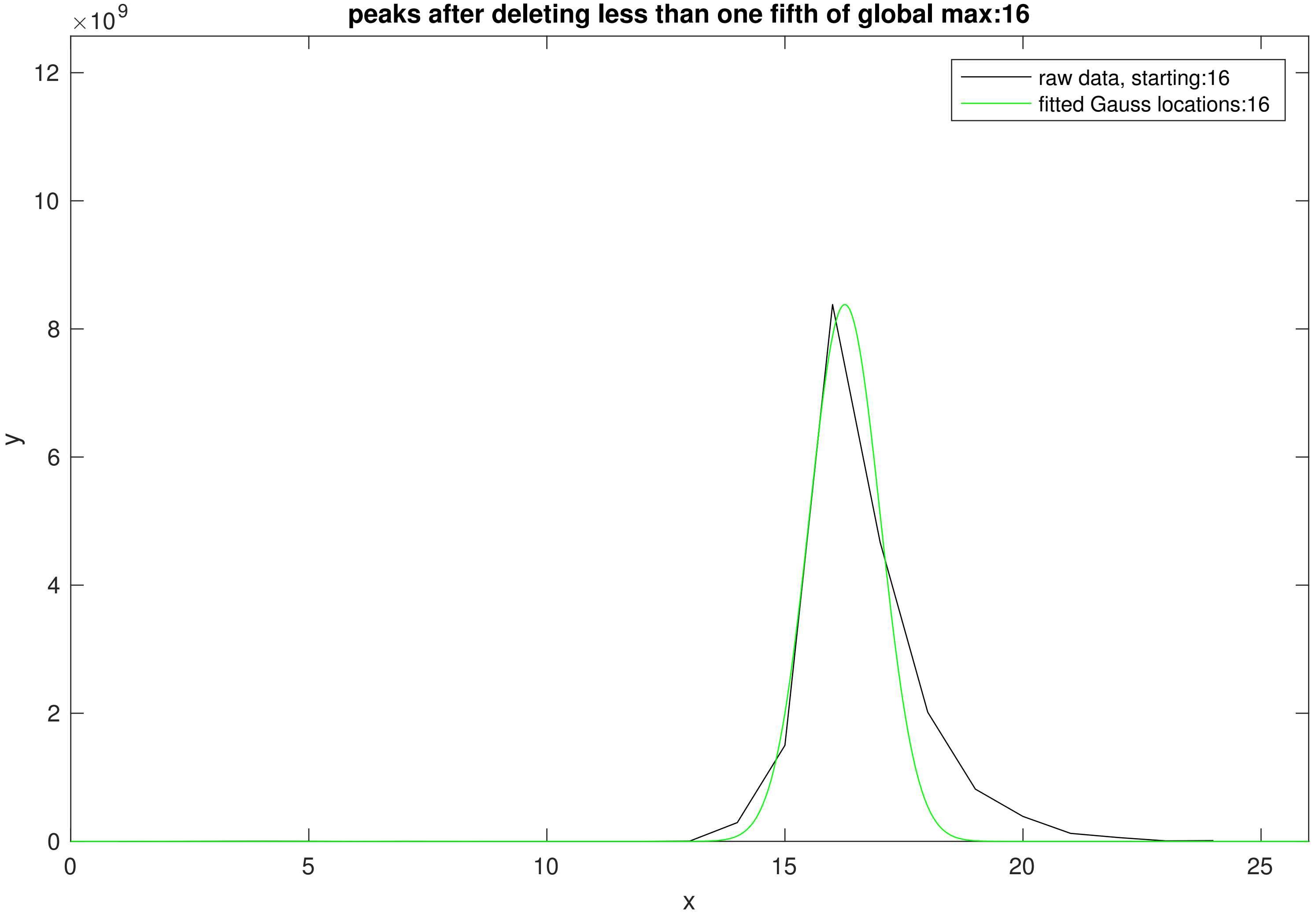

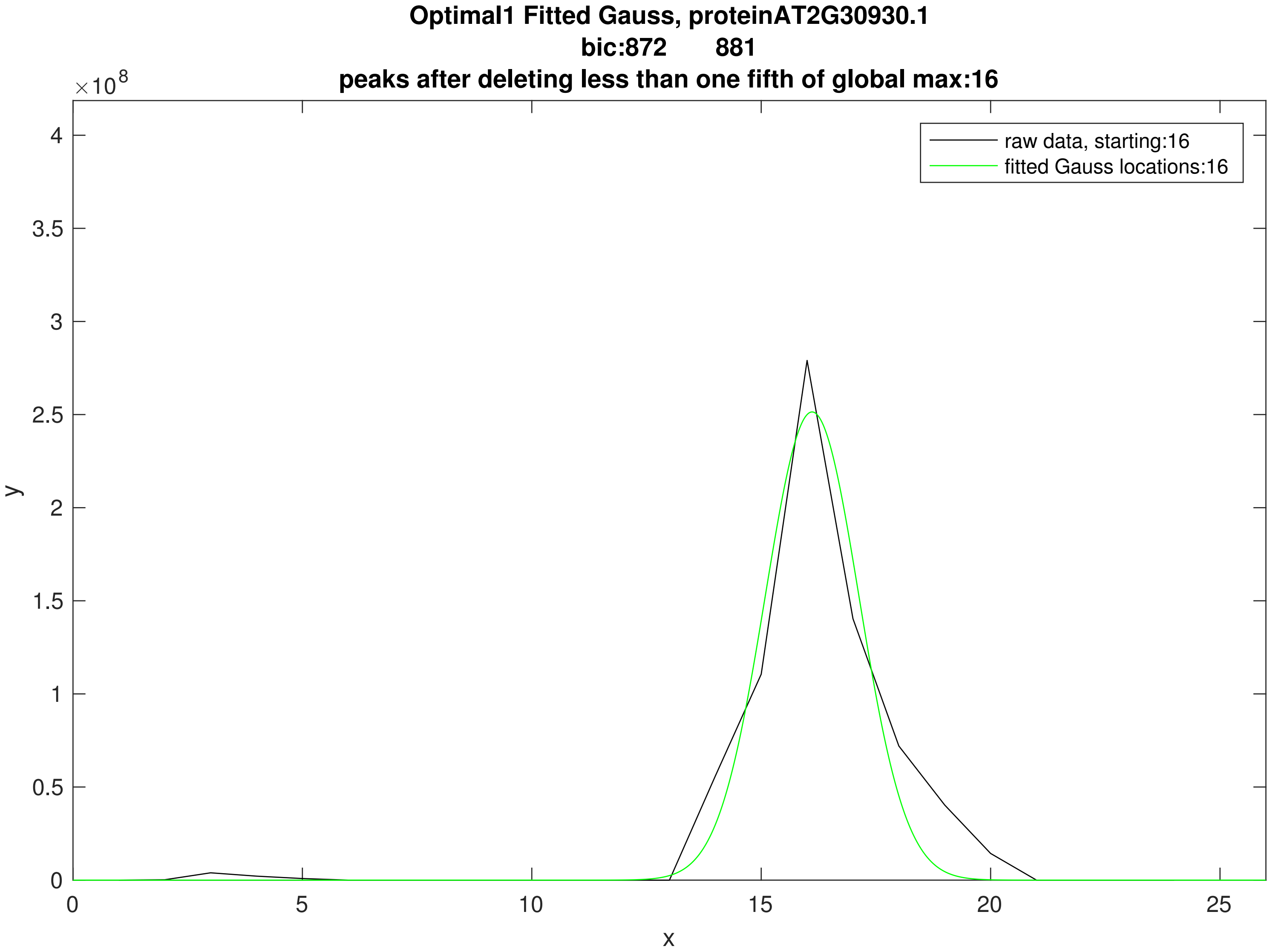
Optimal1 Fitted Gauss, proteinAT2G30930.1
bic:872 881
peaks after deleting less than one fifth of global max:16
×10^8
raw data, starting:16
fitted Gauss locations:16
y
x
0
0.5
1
1.5
2
2.5
3
3.5
4
5
10
15
20
25

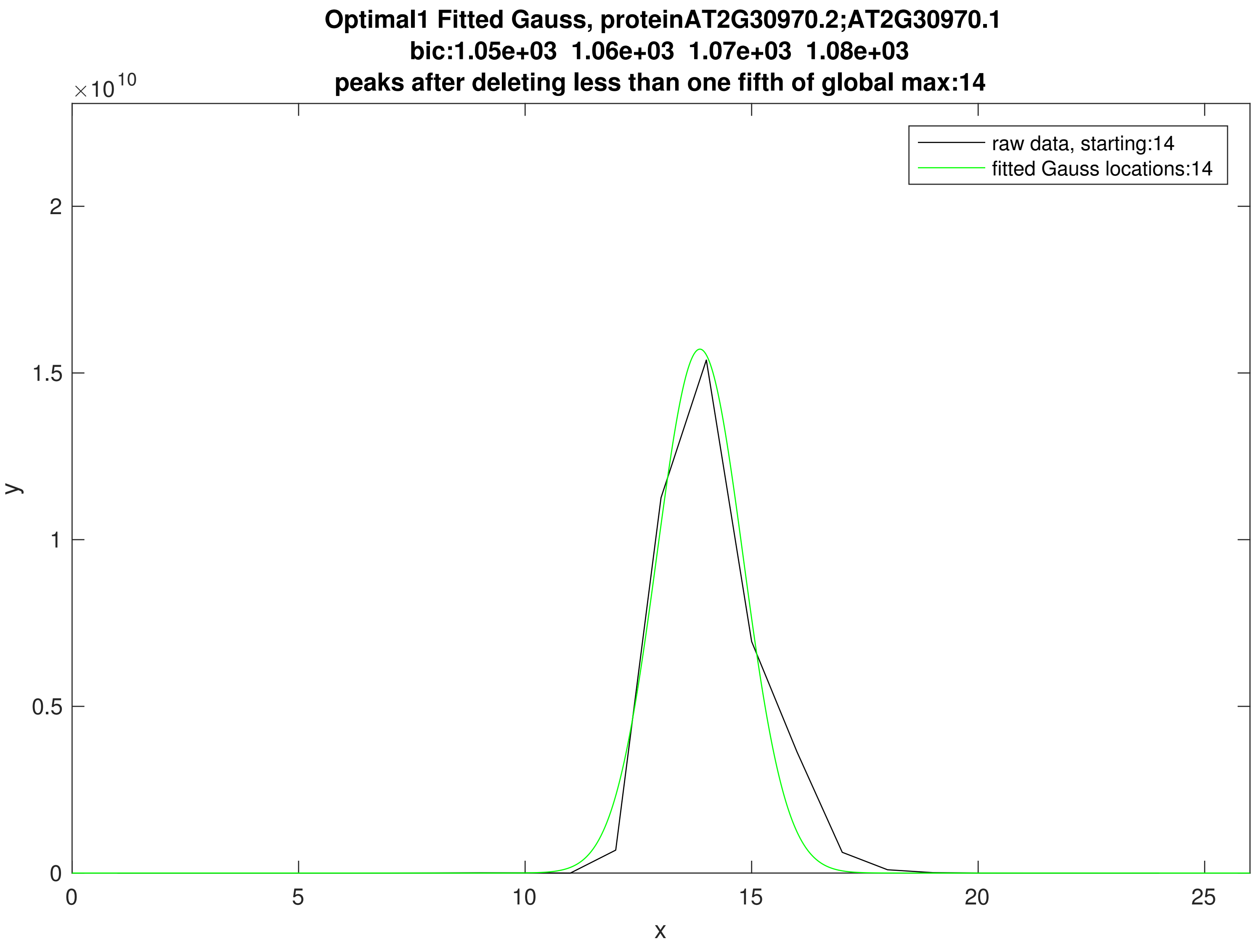

Optimal1 Fitted Gauss, proteinAT2G30970.2;AT2G30970.1
bic:1.05e+03 1.06e+03 1.07e+03 1.08e+03
peaks after deleting less than one fifth of global max:14
×10^10
raw data, starting:14
fitted Gauss locations:14
y
x
0
0.5
1
1.5
2
5
10
15
20
25

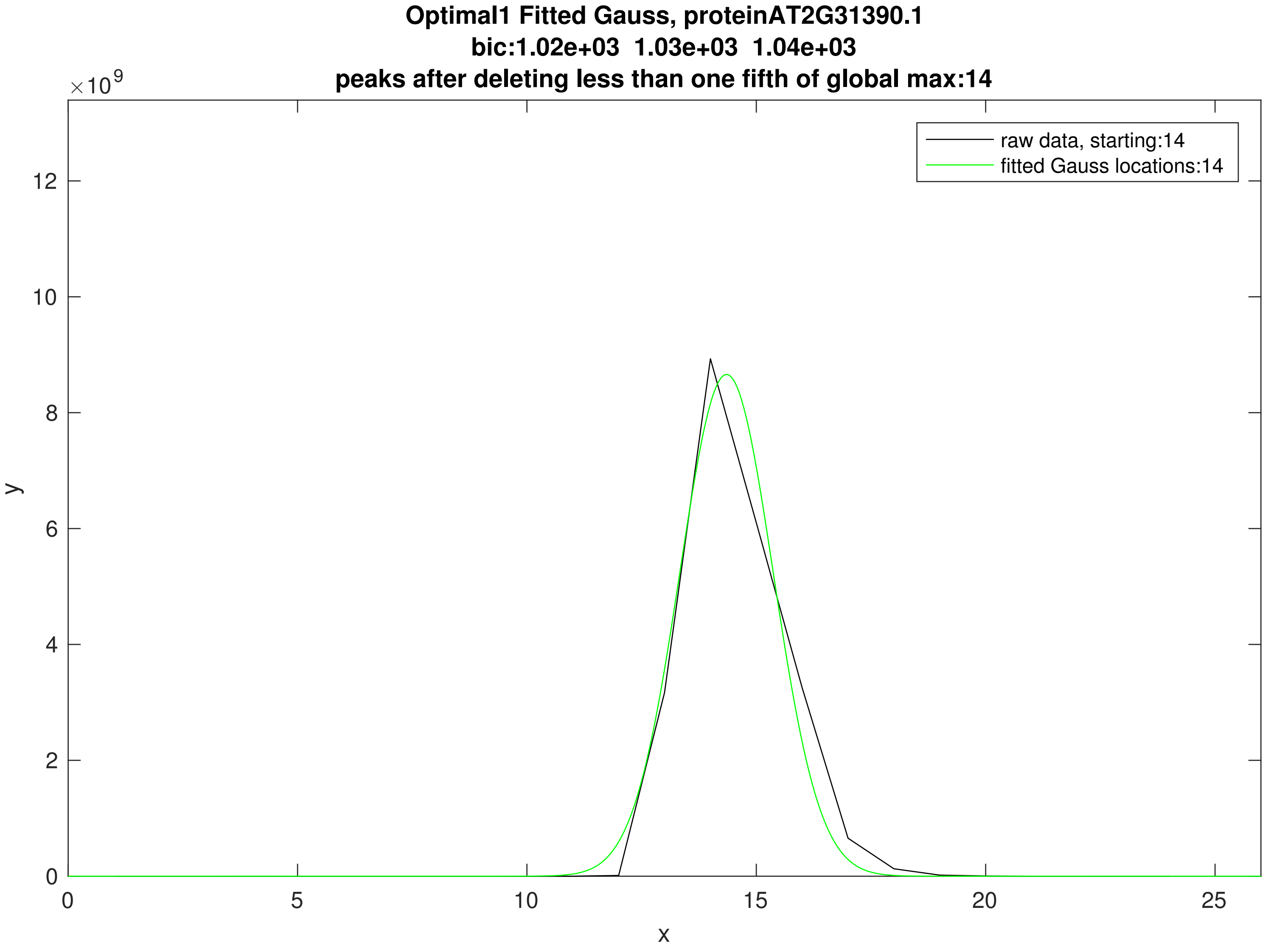
Optimal1 Fitted Gauss, proteinAT2G31390.1
bic:1.02e+03 1.03e+03 1.04e+03
peaks after deleting less than one fifth of global max:14
×10^9
raw data, starting:14
fitted Gauss locations:14
y
x
0
2
4
6
8
10
12
0
5
10
15
20
25

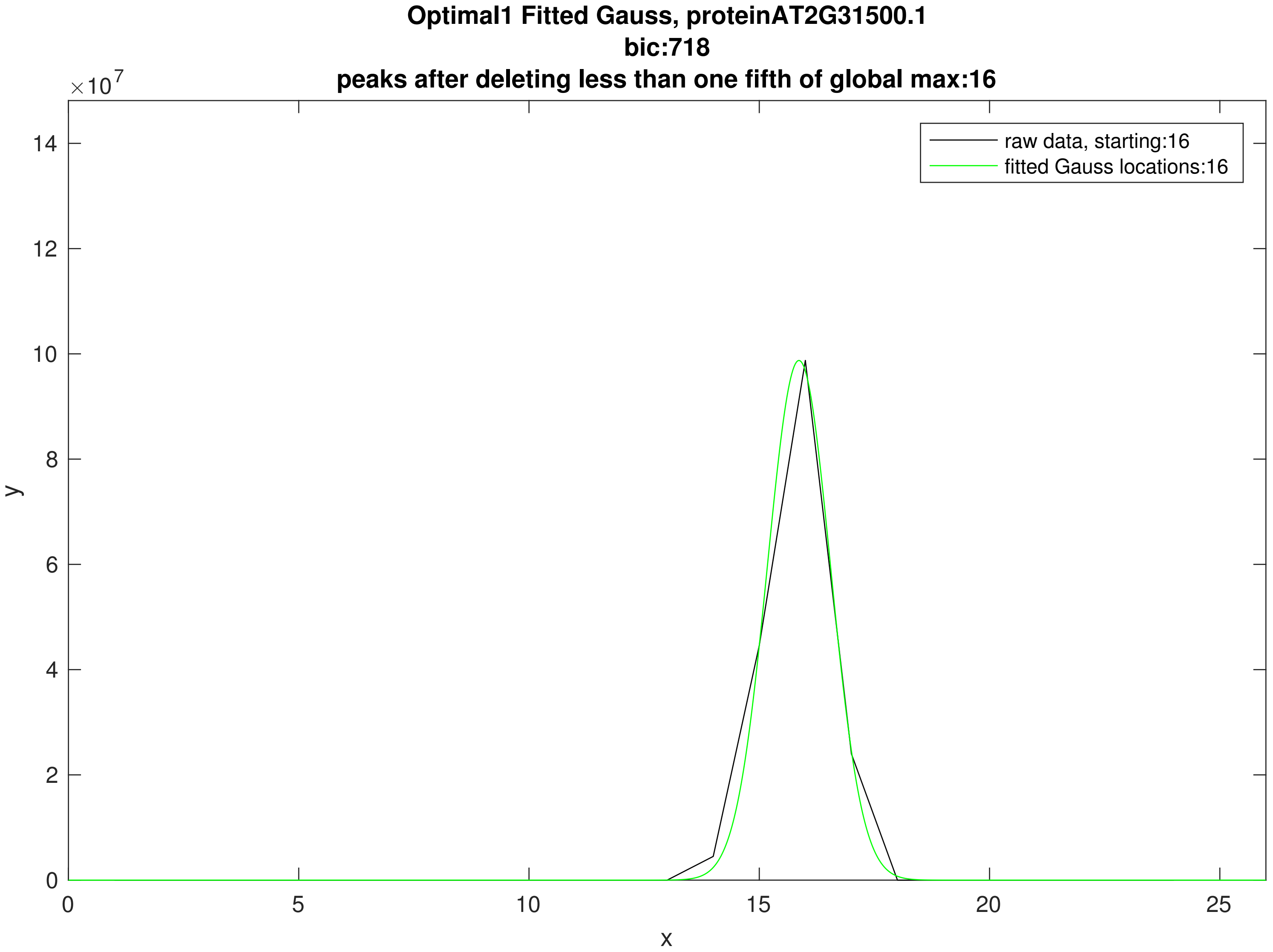
Optimal1 Fitted Gauss, proteinAT2G31500.1
bic:718
peaks after deleting less than one fifth of global max:16
×10^7
14
12
10
8
6
4
2
0
y
0
5
10
15
20
25
x
raw data, starting:16
fitted Gauss locations:16

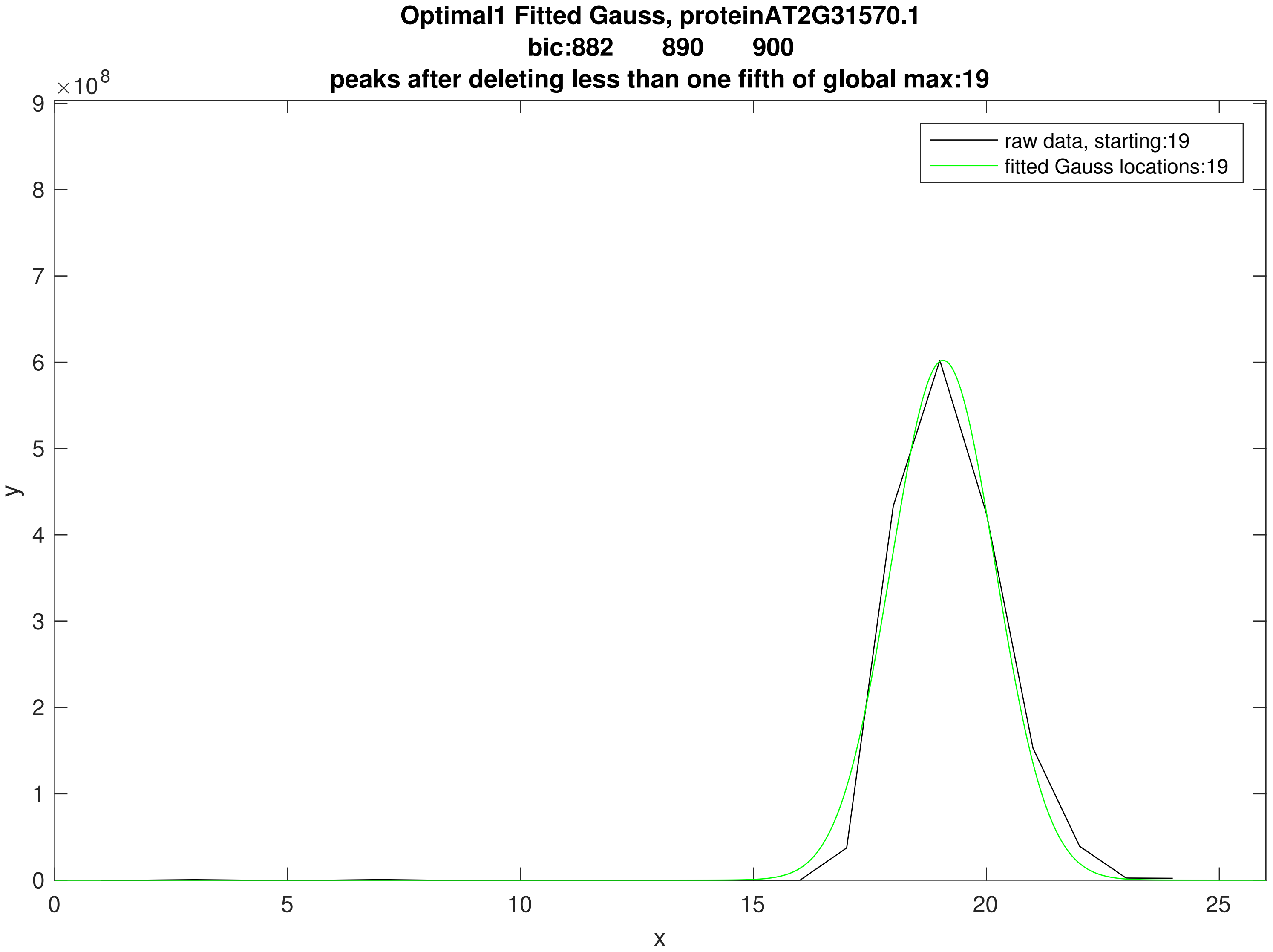
Optimal1 Fitted Gauss, proteinAT2G31570.1
bic:882 890 900
peaks after deleting less than one fifth of global max:19
×10^8
raw data, starting:19
fitted Gauss locations:19
y
x
0
1
2
3
4
5
6
7
8
9
0
5
10
15
20
25

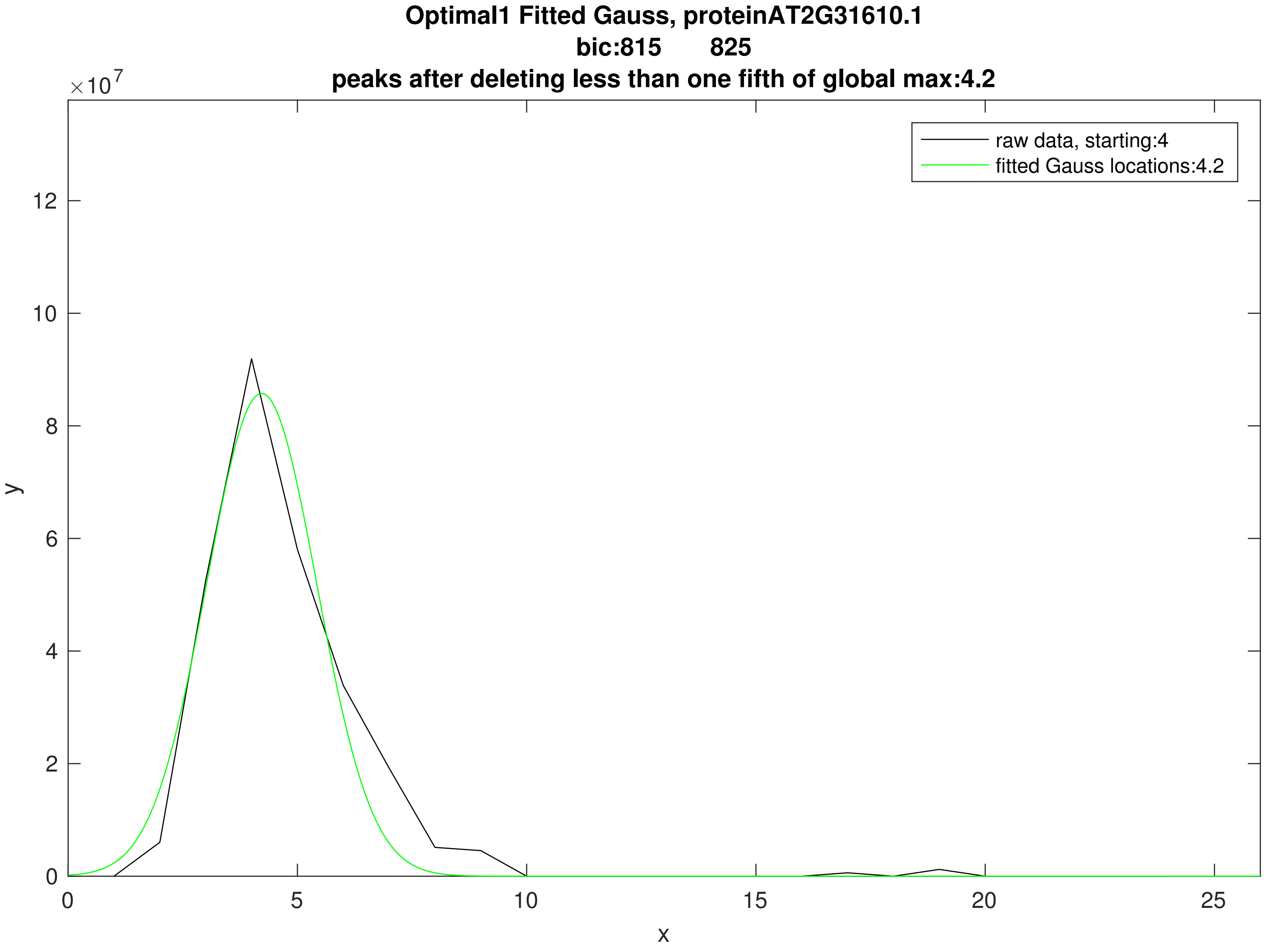
Optimal1 Fitted Gauss, proteinAT2G31610.1
bic:815 825
peaks after deleting less than one fifth of global max:4.2
×10^7
raw data, starting:4
fitted Gauss locations:4.2
12
10
8
6
4
2
0
y
0
5
10
15
20
25
x

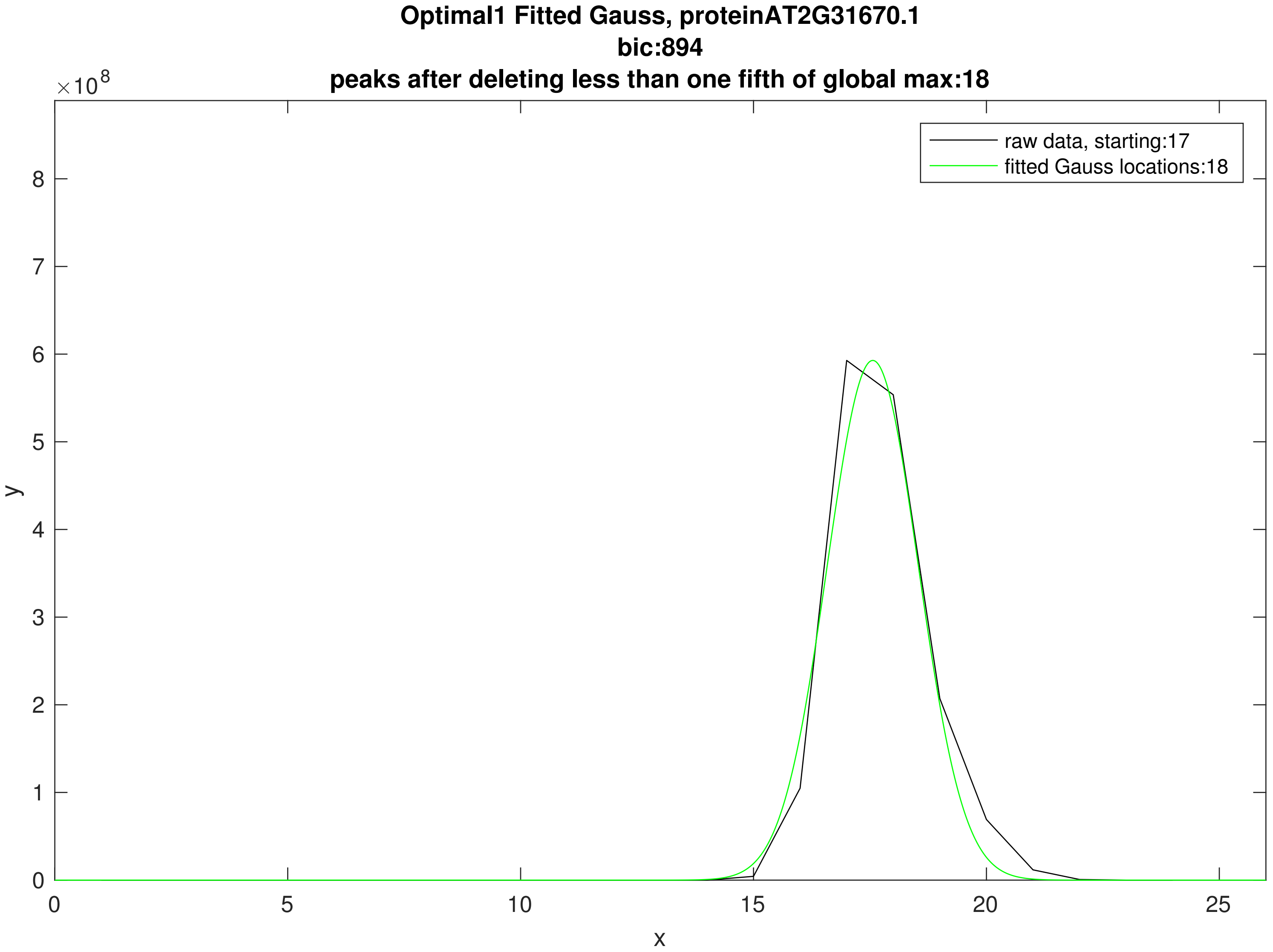
Optimal1 Fitted Gauss, proteinAT2G31670.1
bic:894
peaks after deleting less than one fifth of global max:18
×10^8
raw data, starting:17
fitted Gauss locations:18
y
x
0
1
2
3
4
5
6
7
8
0
5
10
15
20
25

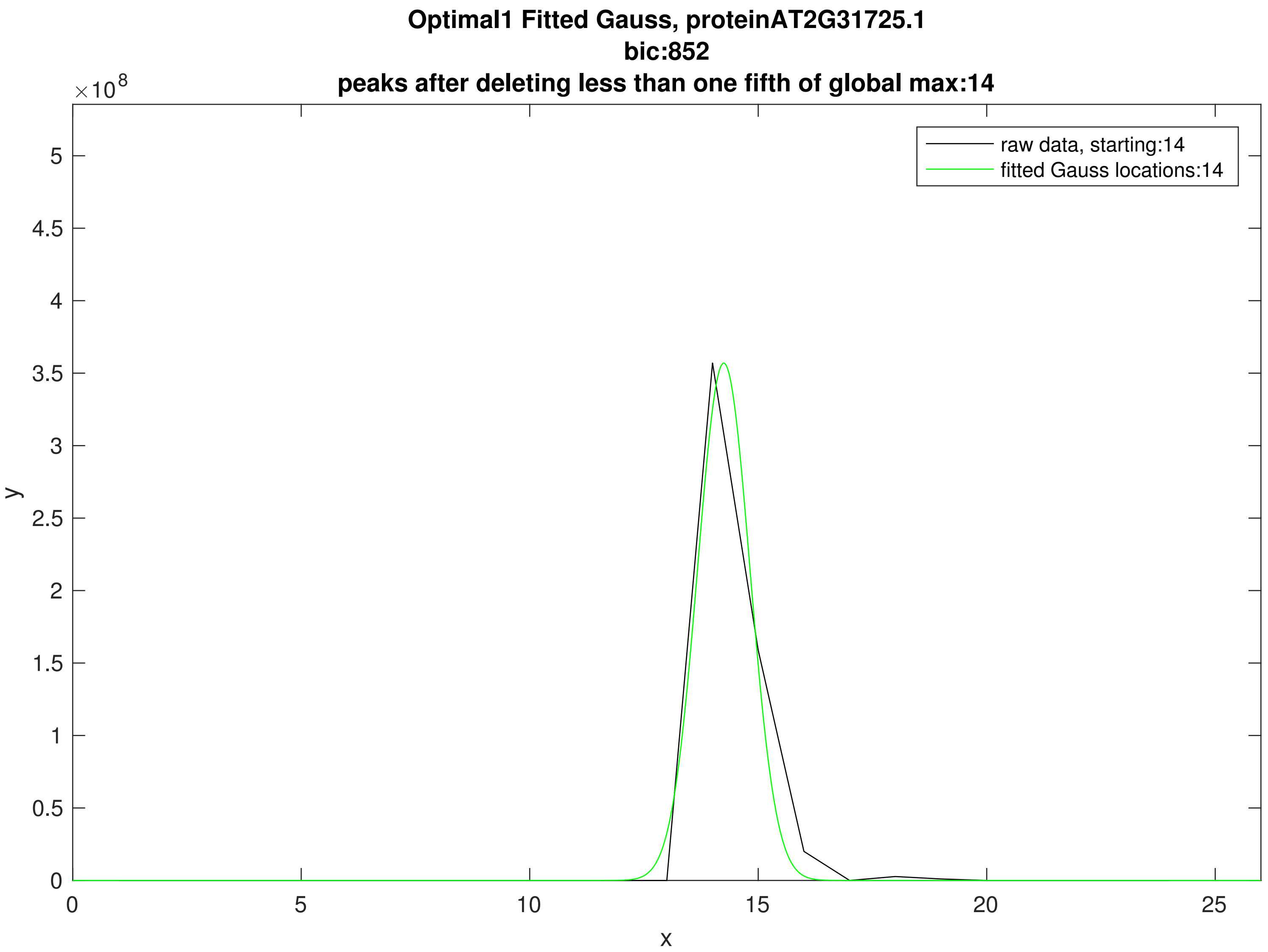
Optimal1 Fitted Gauss, proteinAT2G31725.1
bic:852
peaks after deleting less than one fifth of global max:14
×10^8
raw data, starting:14
fitted Gauss locations:14
y
x
0
0.5
1
1.5
2
2.5
3
3.5
4
4.5
5
0
5
10
15
20
25

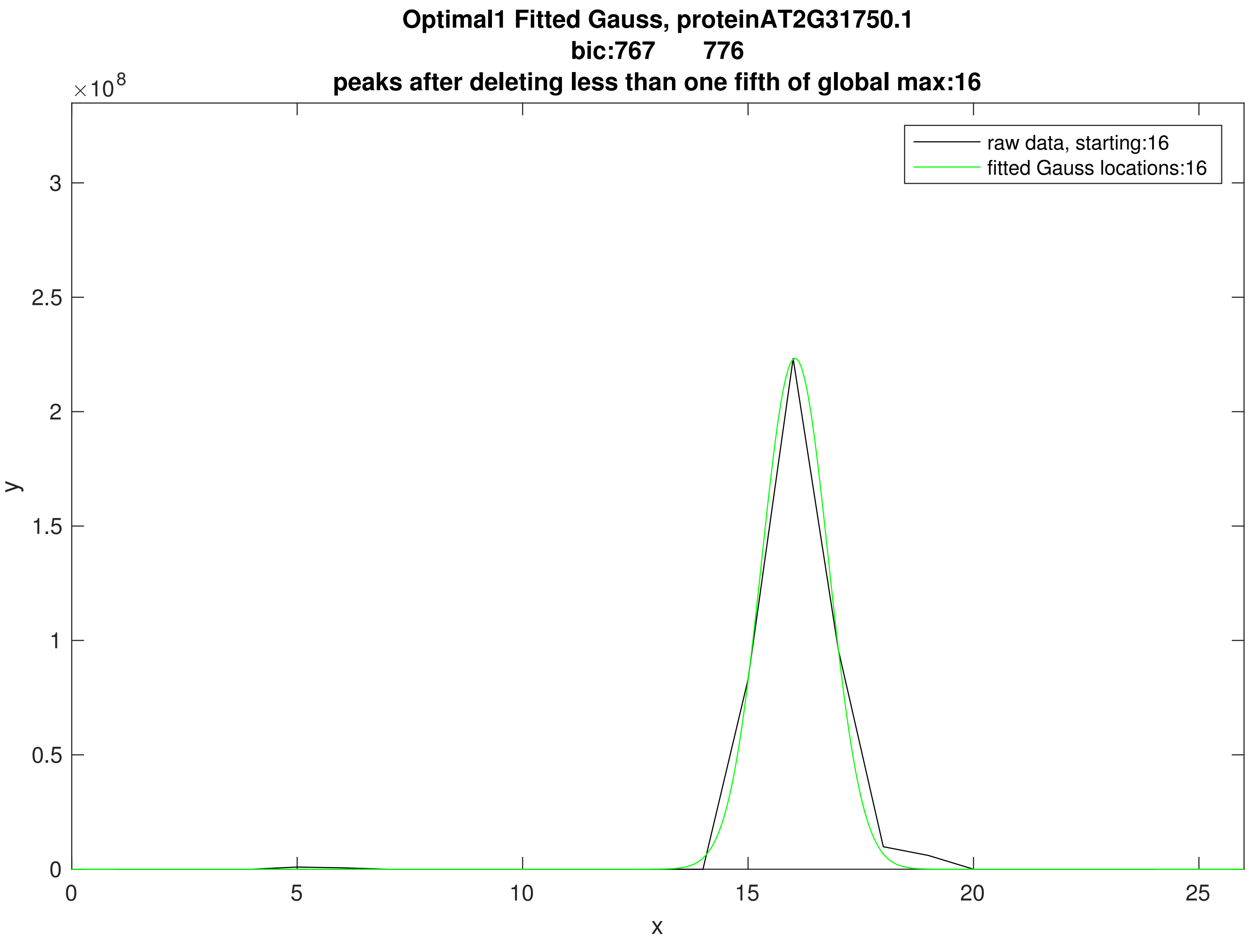
Optimal1 Fitted Gauss, proteinAT2G31750.1
bic:767 776
peaks after deleting less than one fifth of global max:16
×10^8
raw data, starting:16
fitted Gauss locations:16
3
2.5
2
1.5
1
0.5
0
y
0
5
10
15
20
25
x

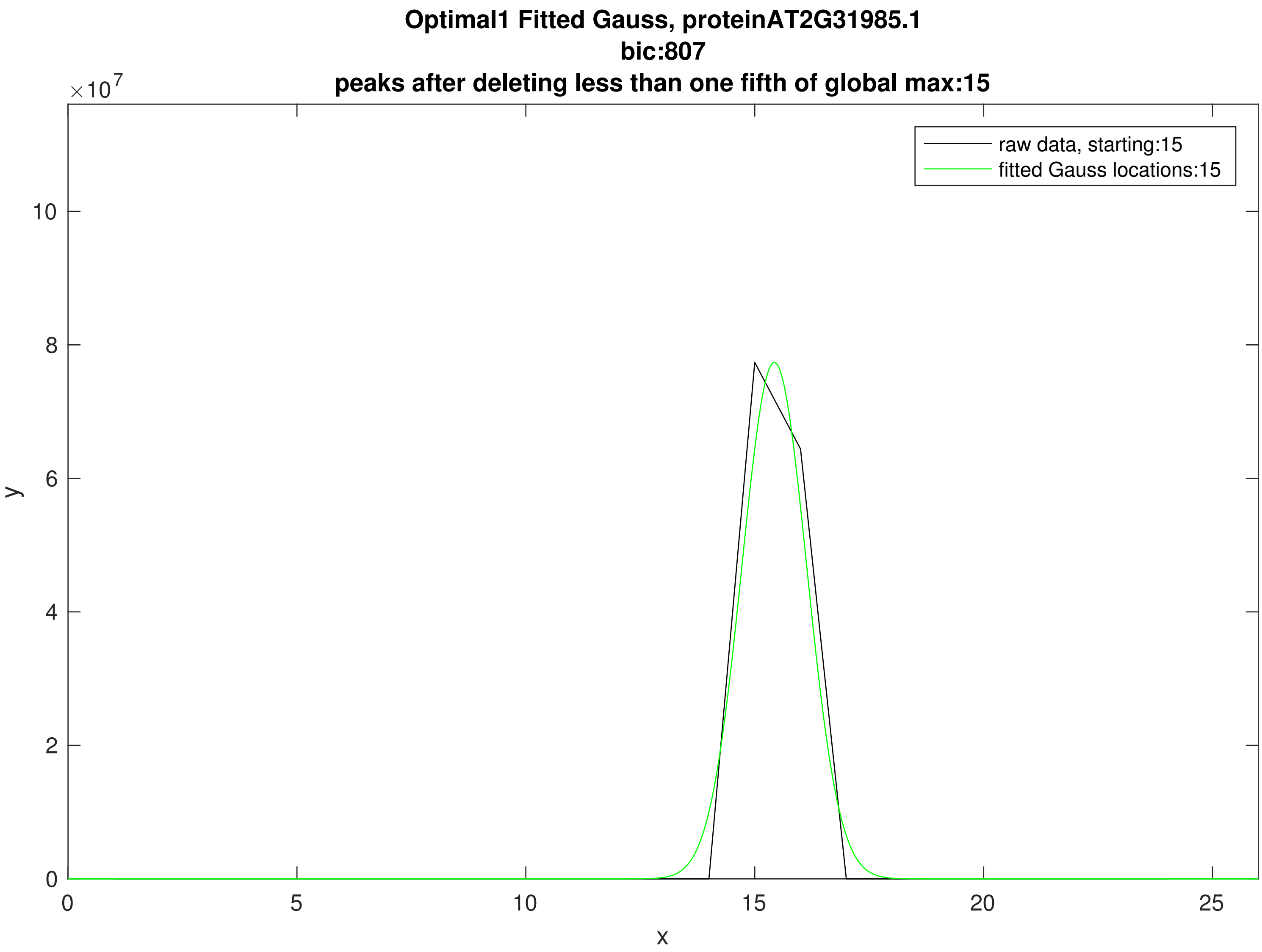
Optimal1 Fitted Gauss, proteinAT2G31985.1
bic:807
peaks after deleting less than one fifth of global max:15
×10^7
raw data, starting:15
fitted Gauss locations:15
10
8
6
4
2
0
y
0
5
10
15
20
25
x

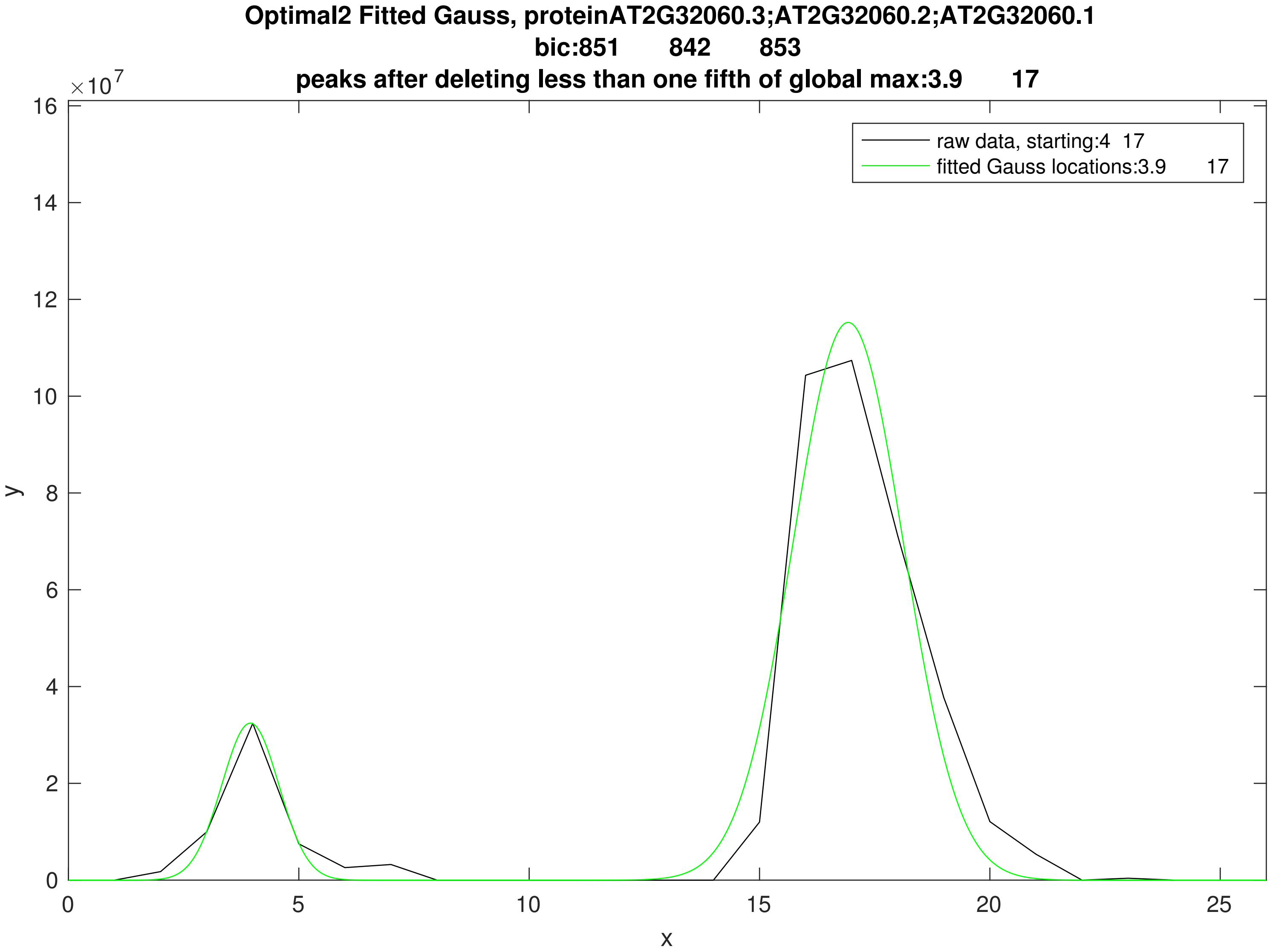

Optimal2 Fitted Gauss, proteinAT2G32060.3;AT2G32060.2;AT2G32060.1
bic:851 842 853
peaks after deleting less than one fifth of global max:3.9 17
×10^7
raw data, starting:4 17
fitted Gauss locations:3.9 17
y
x
0
2
4
6
8
10
12
14
16
5
15
20
25

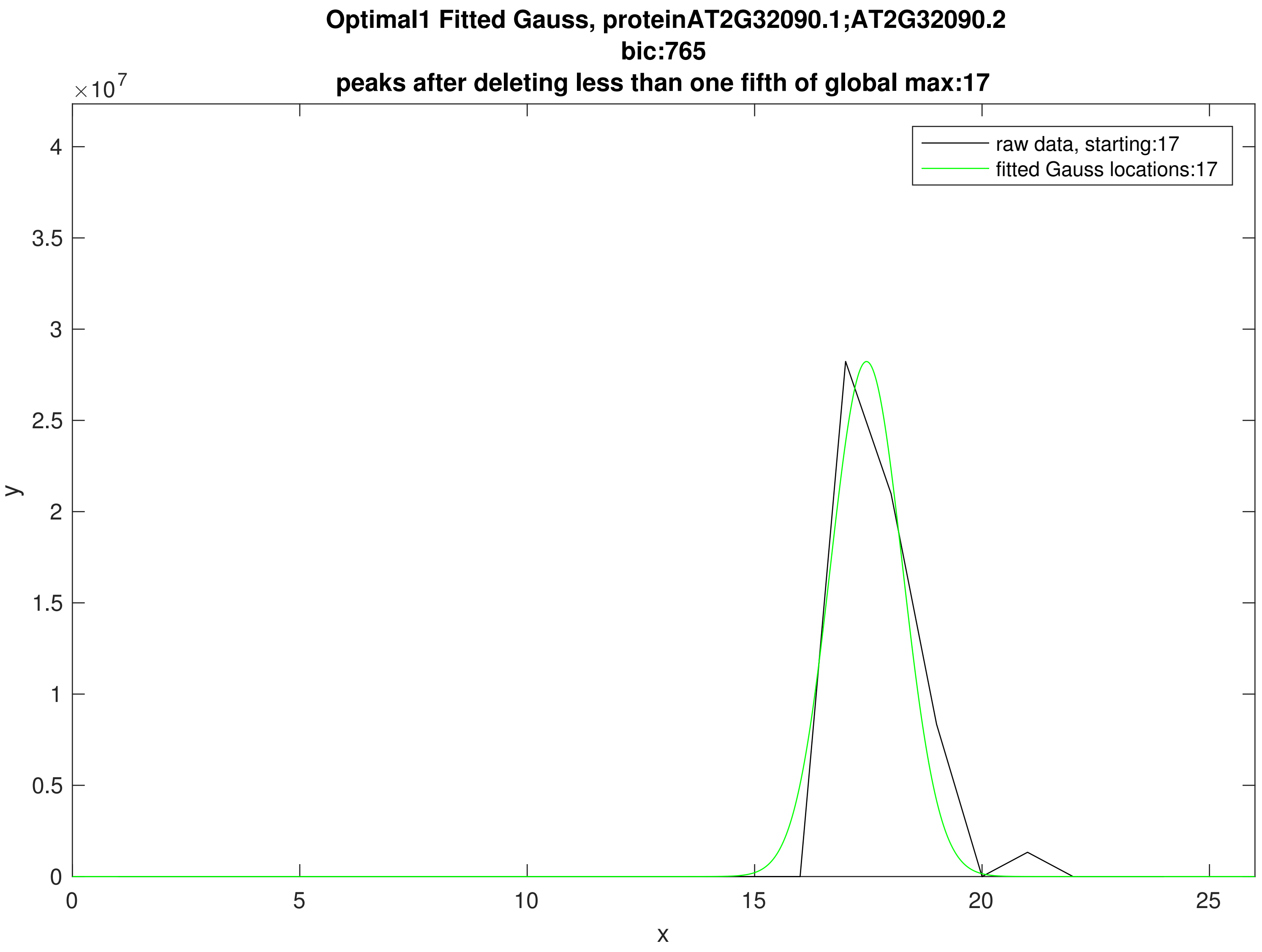
Optimal1 Fitted Gauss, proteinAT2G32090.1;AT2G32090.2
bic:765
peaks after deleting less than one fifth of global max:17
×10^7
raw data, starting:17
fitted Gauss locations:17
4
3.5
3
2.5
2
1.5
1
0.5
0
y
0
5
10
15
20
25
x

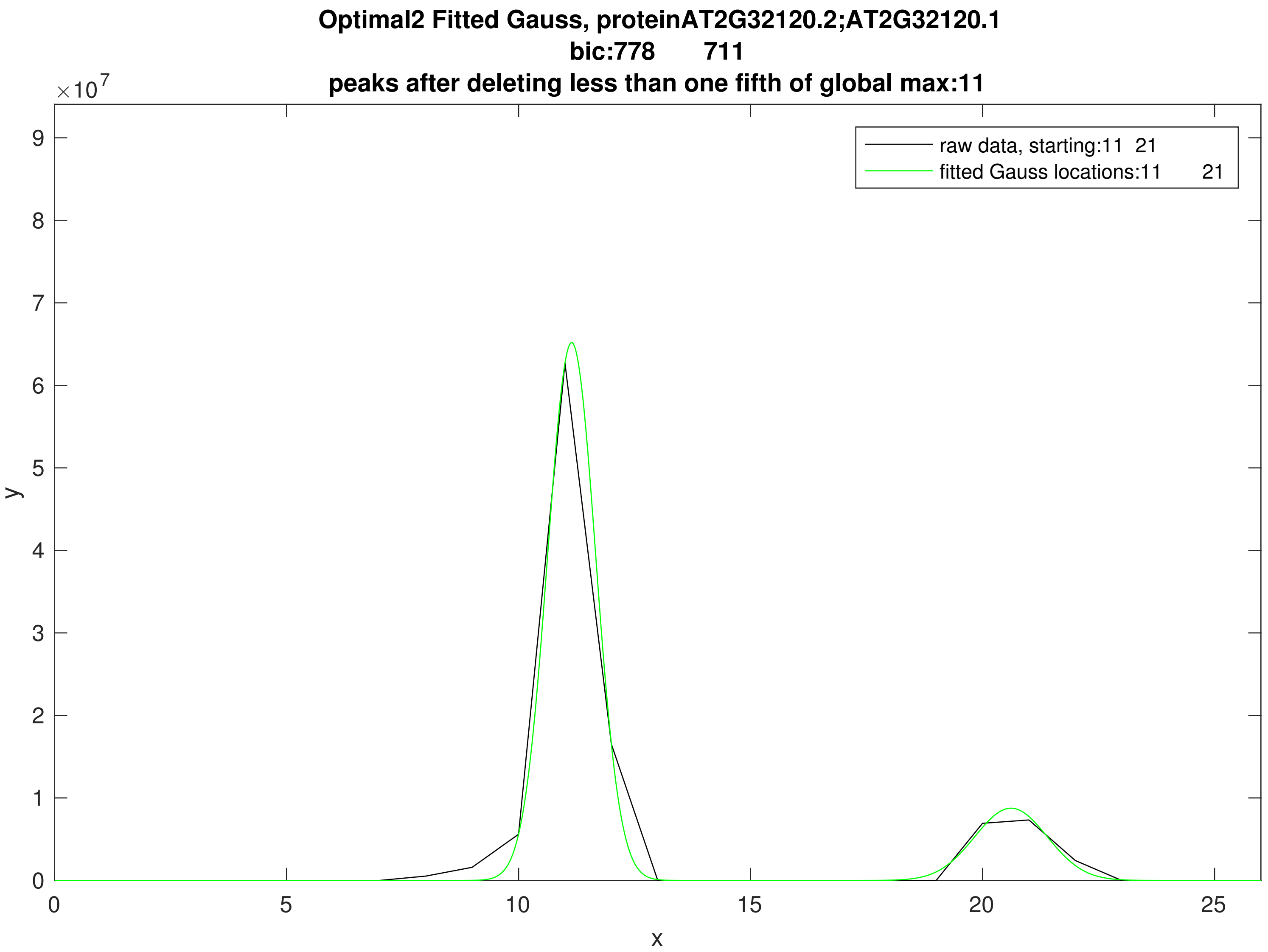
Optimal2 Fitted Gauss, proteinAT2G32120.2;AT2G32120.1
bic:778 711
peaks after deleting less than one fifth of global max:11
×10^7
raw data, starting:11 21
fitted Gauss locations:11 21
y
x
0
1
2
3
4
5
6
7
8
9
0
5
10
15
20
25

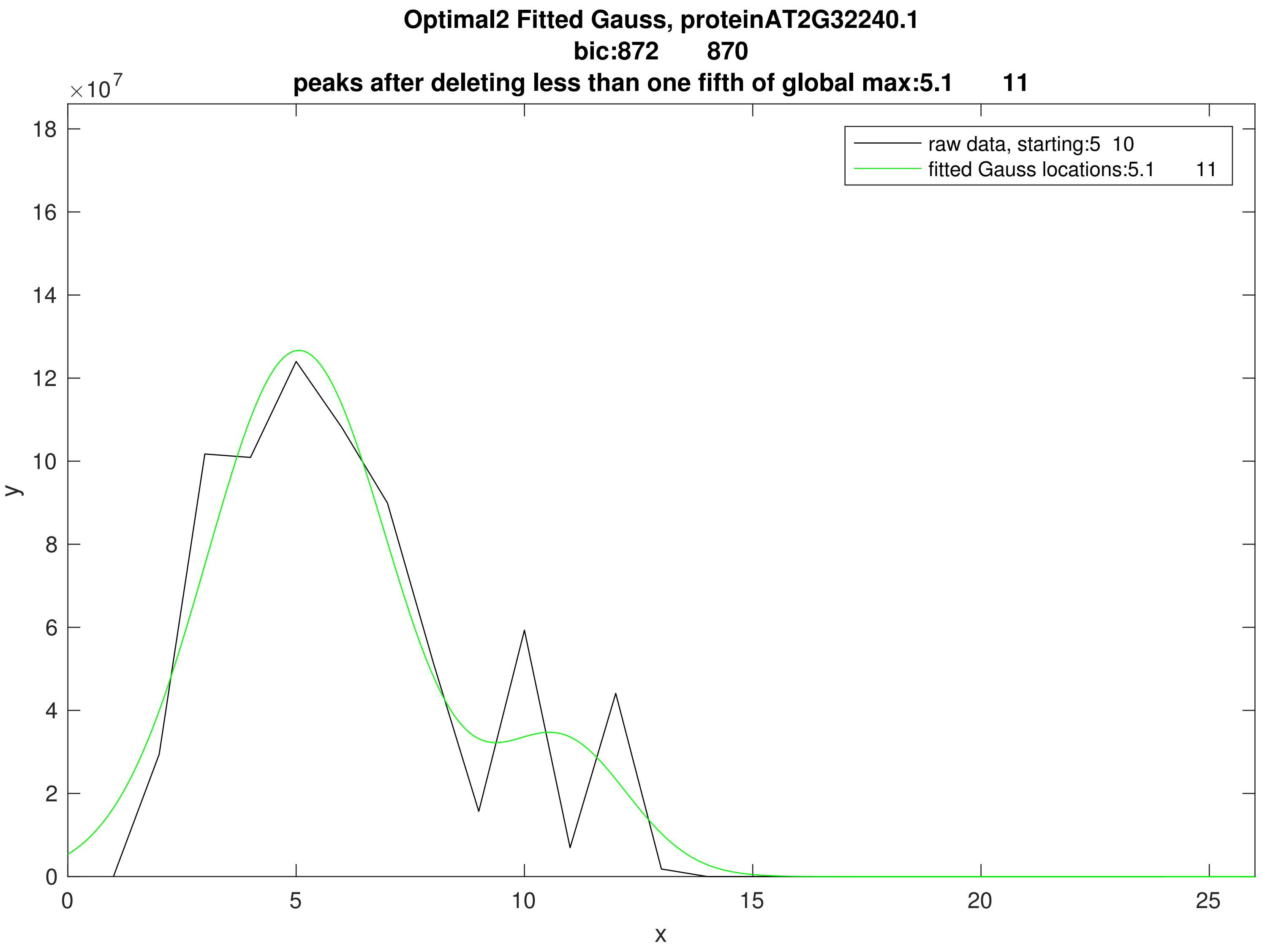

Optimal2 Fitted Gauss, proteinAT2G32240.1
bic:872 870
peaks after deleting less than one fifth of global max:5.1 11
×10^7
raw data, starting:5 10
fitted Gauss locations:5.1 11
y
x
0
2
4
6
8
10
12
14
16
18
0
5
10
15
20
25

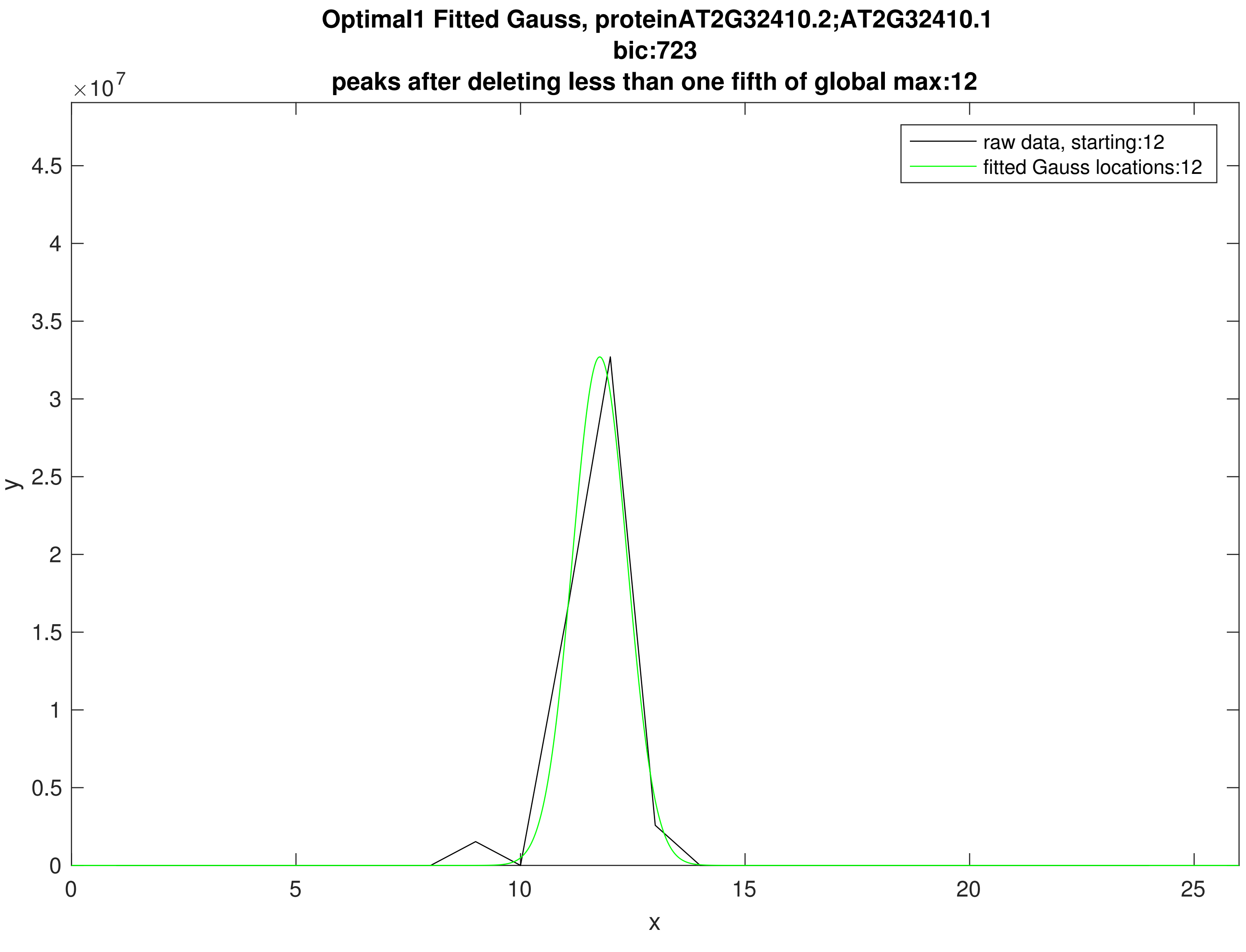
Optimal1 Fitted Gauss, proteinAT2G32410.2;AT2G32410.1
bic:723
peaks after deleting less than one fifth of global max:12
×10^7
4.5
4
3.5
3
2.5
2
1.5
1
0.5
0
y
0
5
10
15
20
25
x
raw data, starting:12
fitted Gauss locations:12

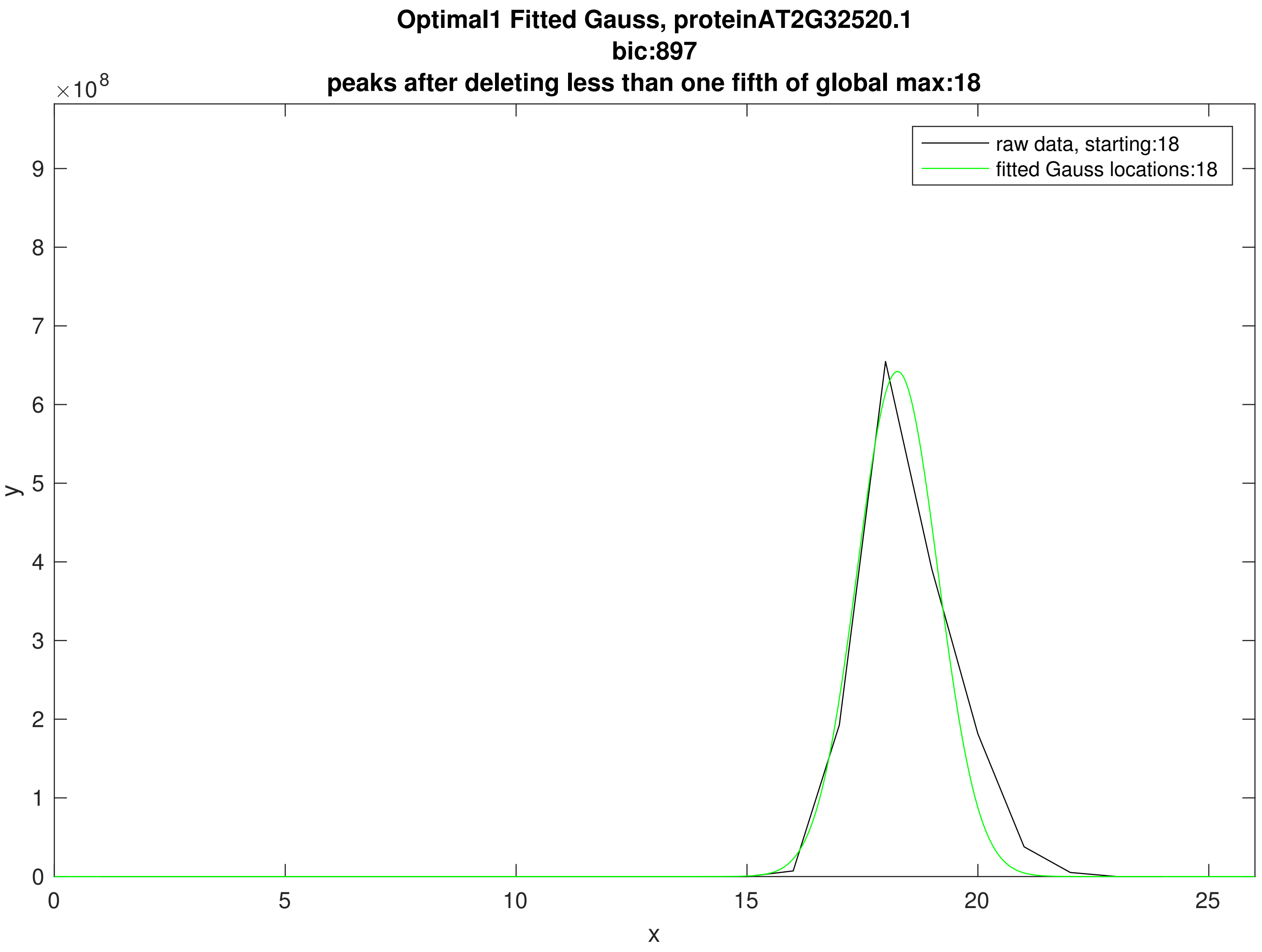
Optimal1 Fitted Gauss, proteinAT2G32520.1
bic:897
peaks after deleting less than one fifth of global max:18
×10^8
raw data, starting:18
fitted Gauss locations:18
y
x
0
1
2
3
4
5
6
7
8
9
0
5
10
15
20
25

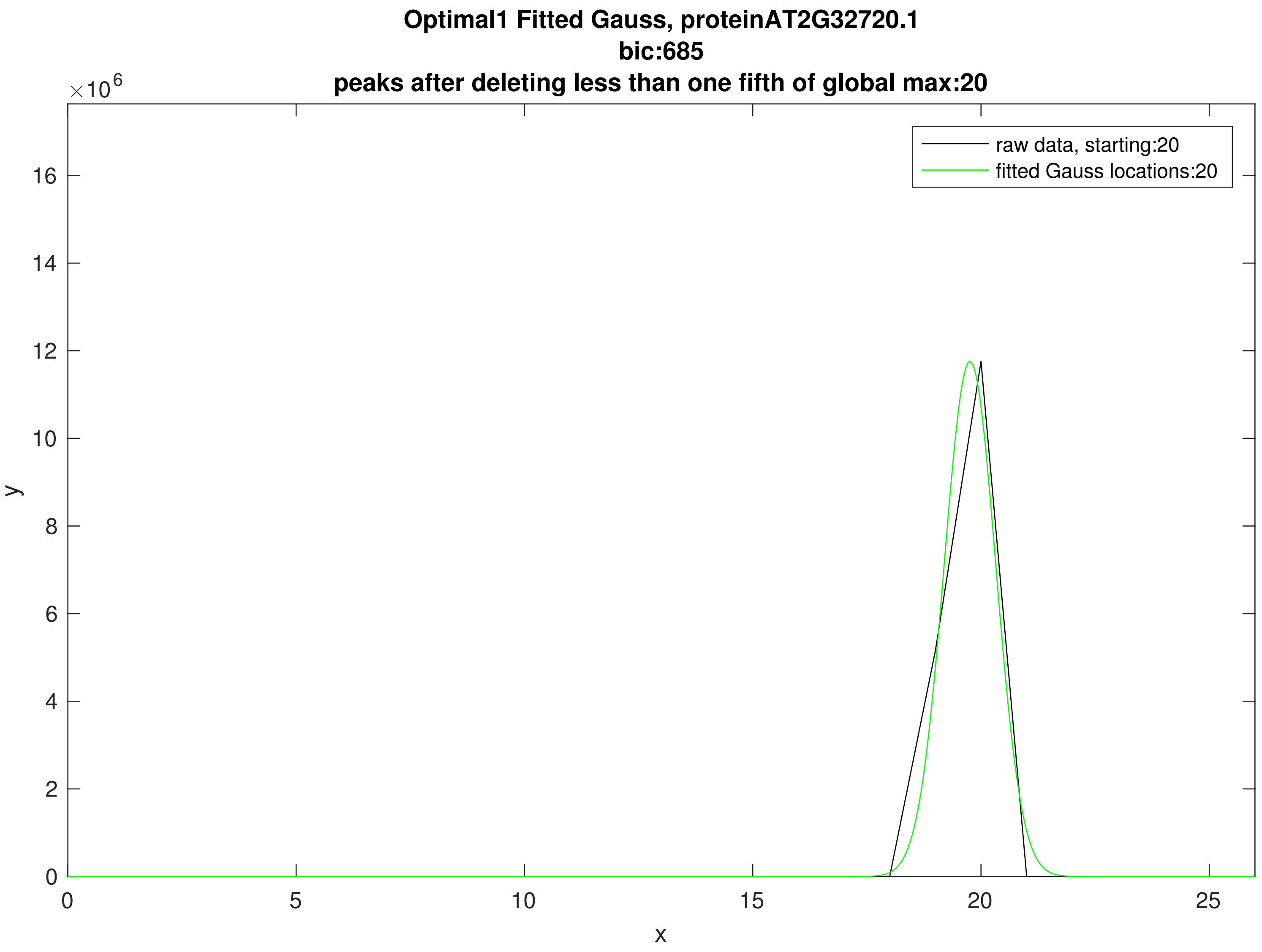
Optimal1 Fitted Gauss, proteinAT2G32720.1
bic:685
peaks after deleting less than one fifth of global max:20
×10^6
raw data, starting:20
fitted Gauss locations:20
y
x
0
2
4
6
8
10
12
14
16
5
15
20
25

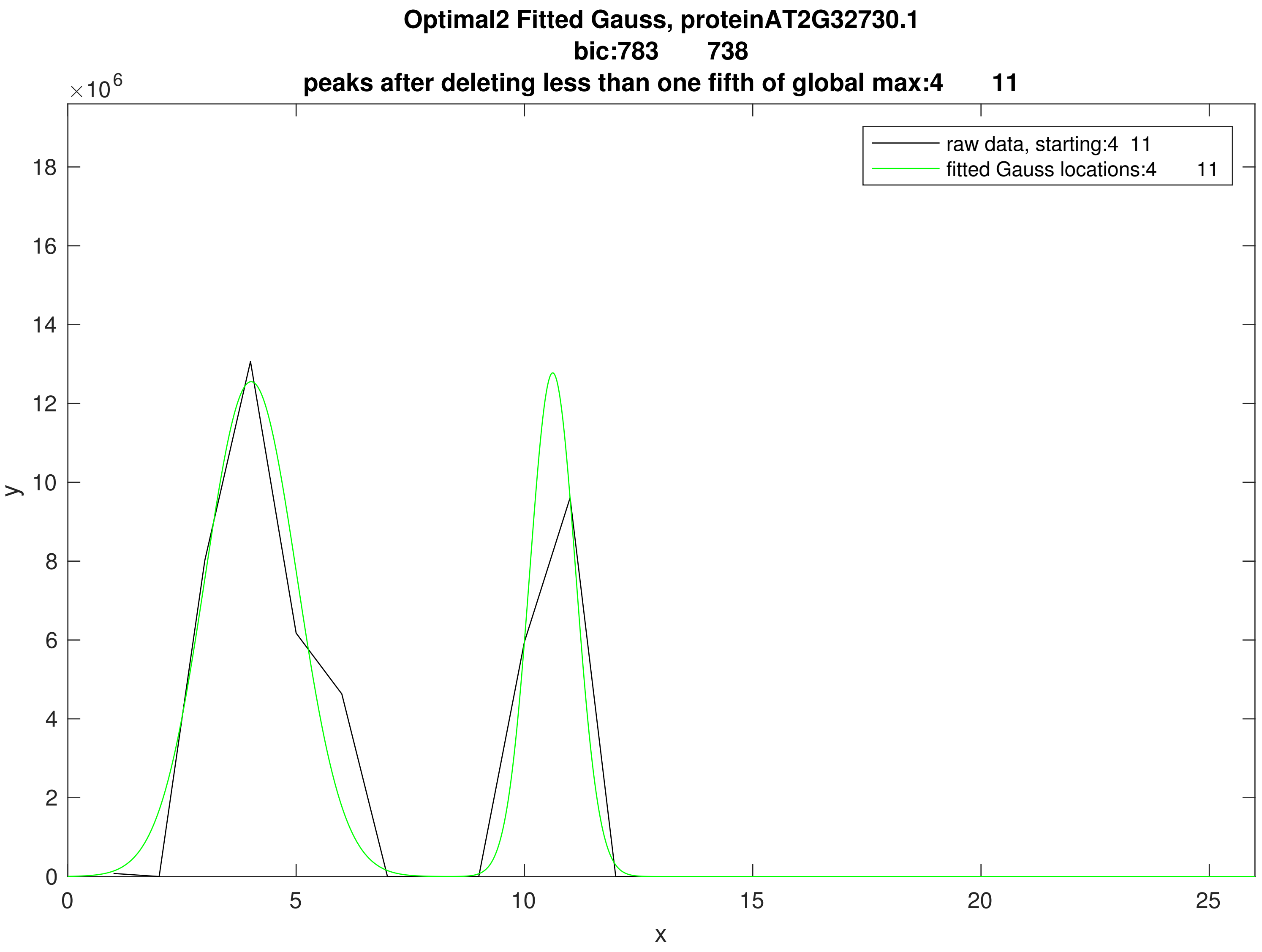
Optimal2 Fitted Gauss, proteinAT2G32730.1
bic:783 738
peaks after deleting less than one fifth of global max:4 11
raw data, starting:4 11
fitted Gauss locations:4 11
×10^6
y
x
0
2
4
6
8
10
12
14
16
18
0
5
10
15
20
25

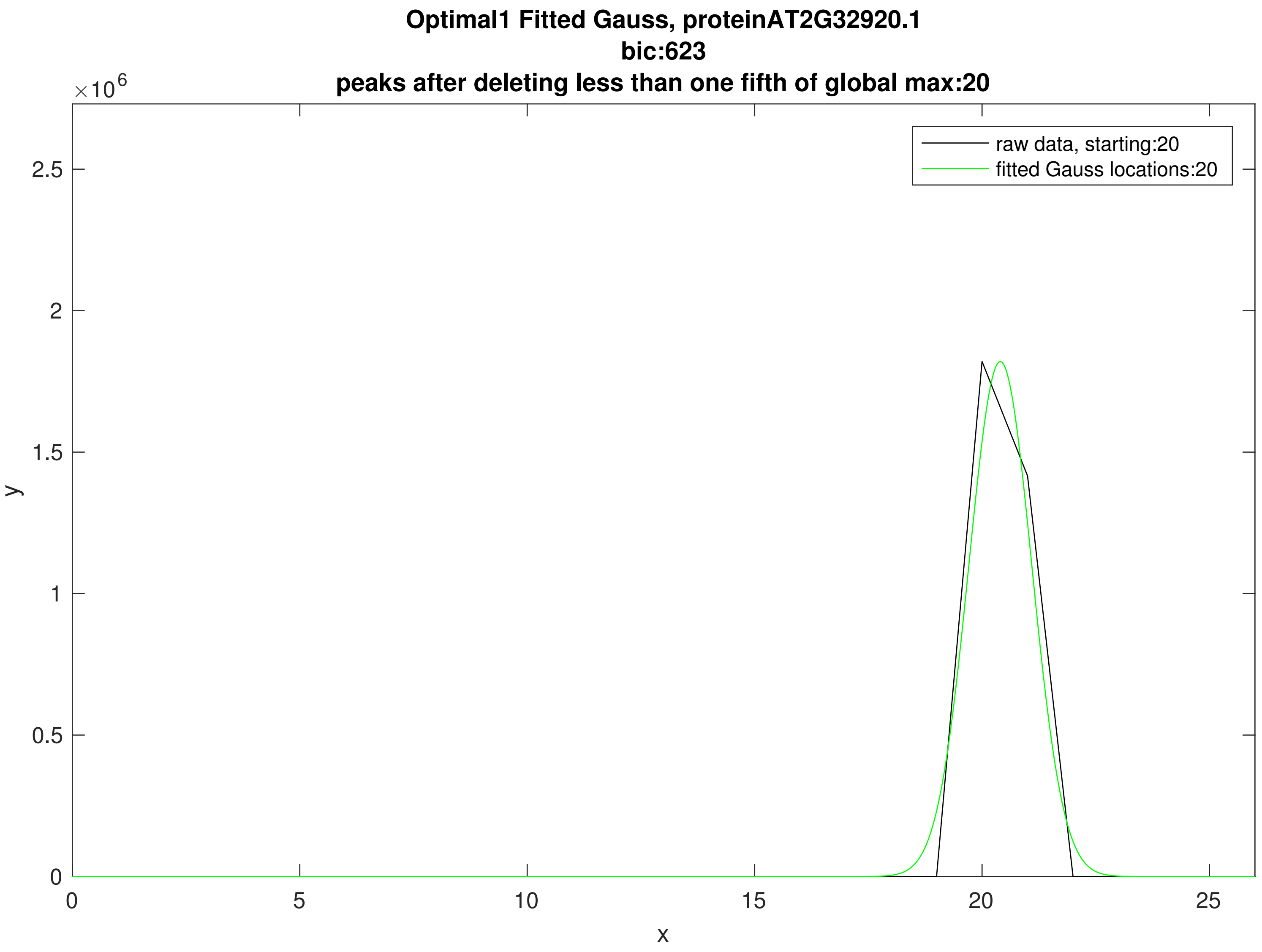

Optimal1 Fitted Gauss, proteinAT2G32920.1
bic:623
peaks after deleting less than one fifth of global max:20
×10^6
2.5
2
1.5
1
0.5
0
y
0
5
10
15
20
25
x
raw data, starting:20
fitted Gauss locations:20

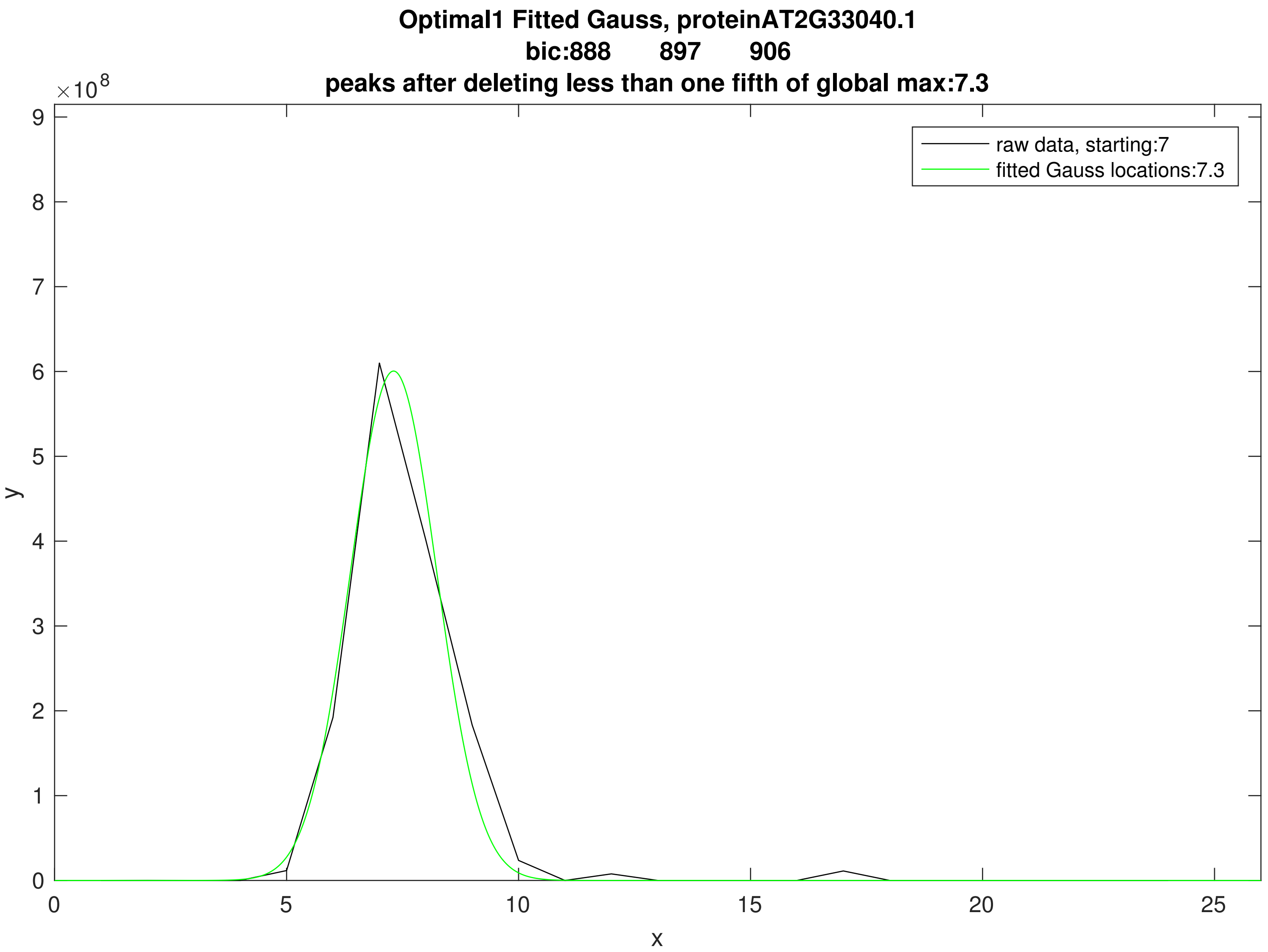

Optimal1 Fitted Gauss, proteinAT2G33040.1
bic:888 897 906
peaks after deleting less than one fifth of global max:7.3
×10^8
raw data, starting:7
fitted Gauss locations:7.3
y
x
0
1
2
3
4
5
6
7
8
9
0
5
10
15
20
25

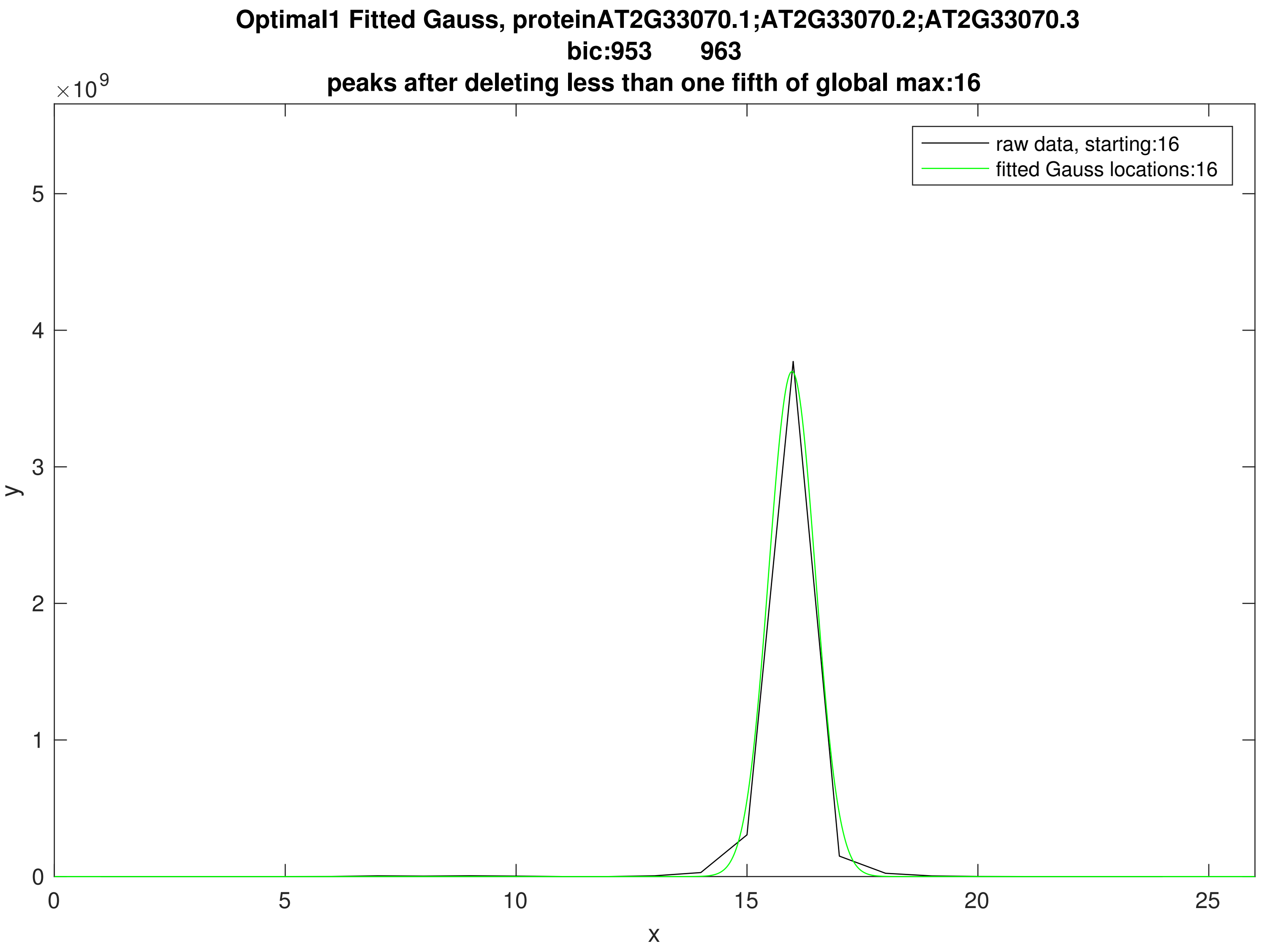

Optimal1 Fitted Gauss, proteinAT2G33070.1;AT2G33070.2;AT2G33070.3
bic:953 963
peaks after deleting less than one fifth of global max:16
×10^9
raw data, starting:16
fitted Gauss locations:16
y
x
0
1
2
3
4
5
0
5
10
15
20
25

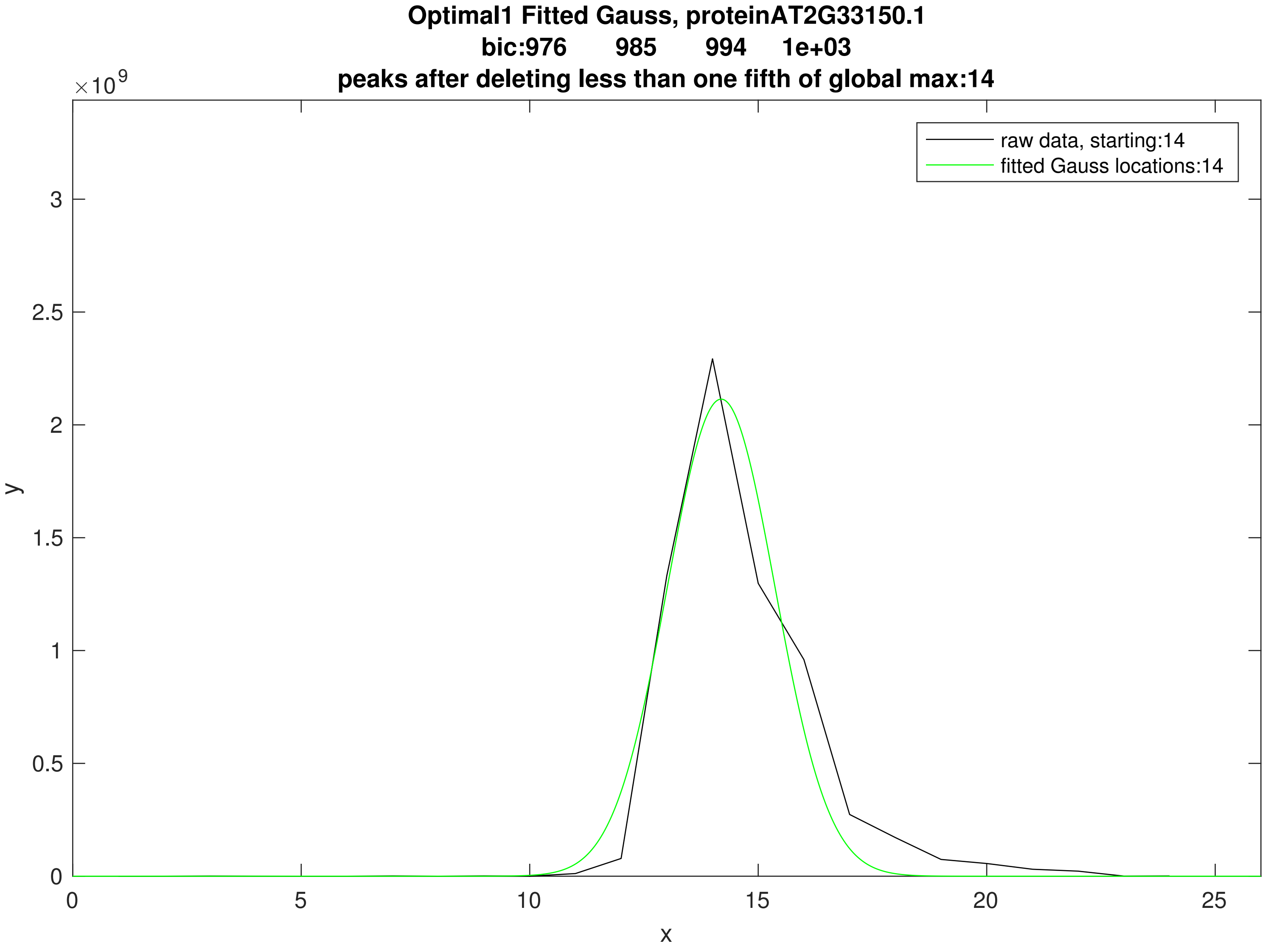

Optimal1 Fitted Gauss, proteinAT2G33150.1
bic:976 985 994 1e+03
peaks after deleting less than one fifth of global max:14
×10^9
3
2.5
2
1.5
1
0.5
0
y
0
5
10
15
20
25
x
raw data, starting:14
fitted Gauss locations:14

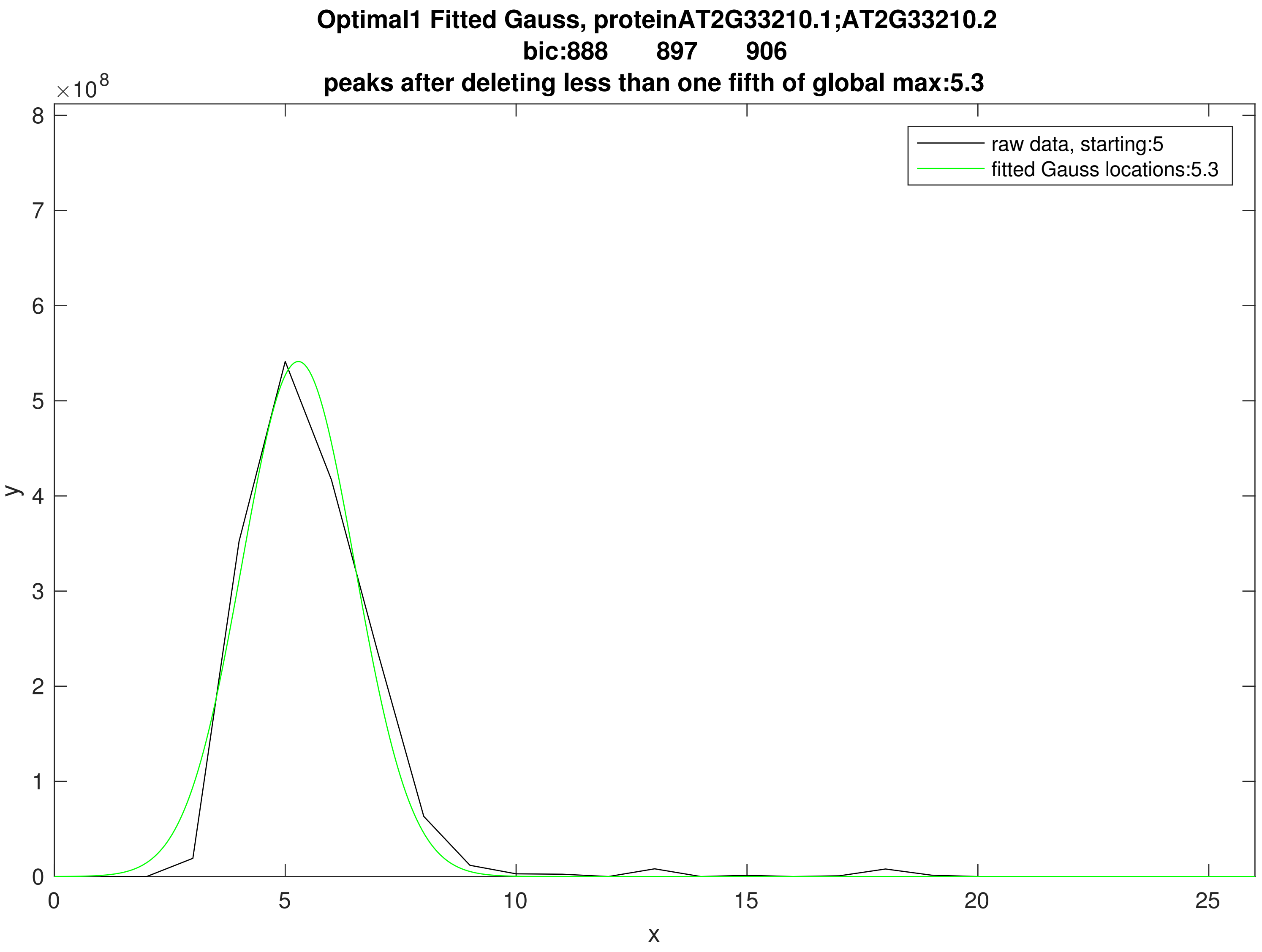

Optimal1 Fitted Gauss, proteinAT2G33210.1;AT2G33210.2
bic:888 897 906
peaks after deleting less than one fifth of global max:5.3
×10^8
raw data, starting:5
fitted Gauss locations:5.3
y
x
0
1
2
3
4
5
6
7
8
10
15
20
25

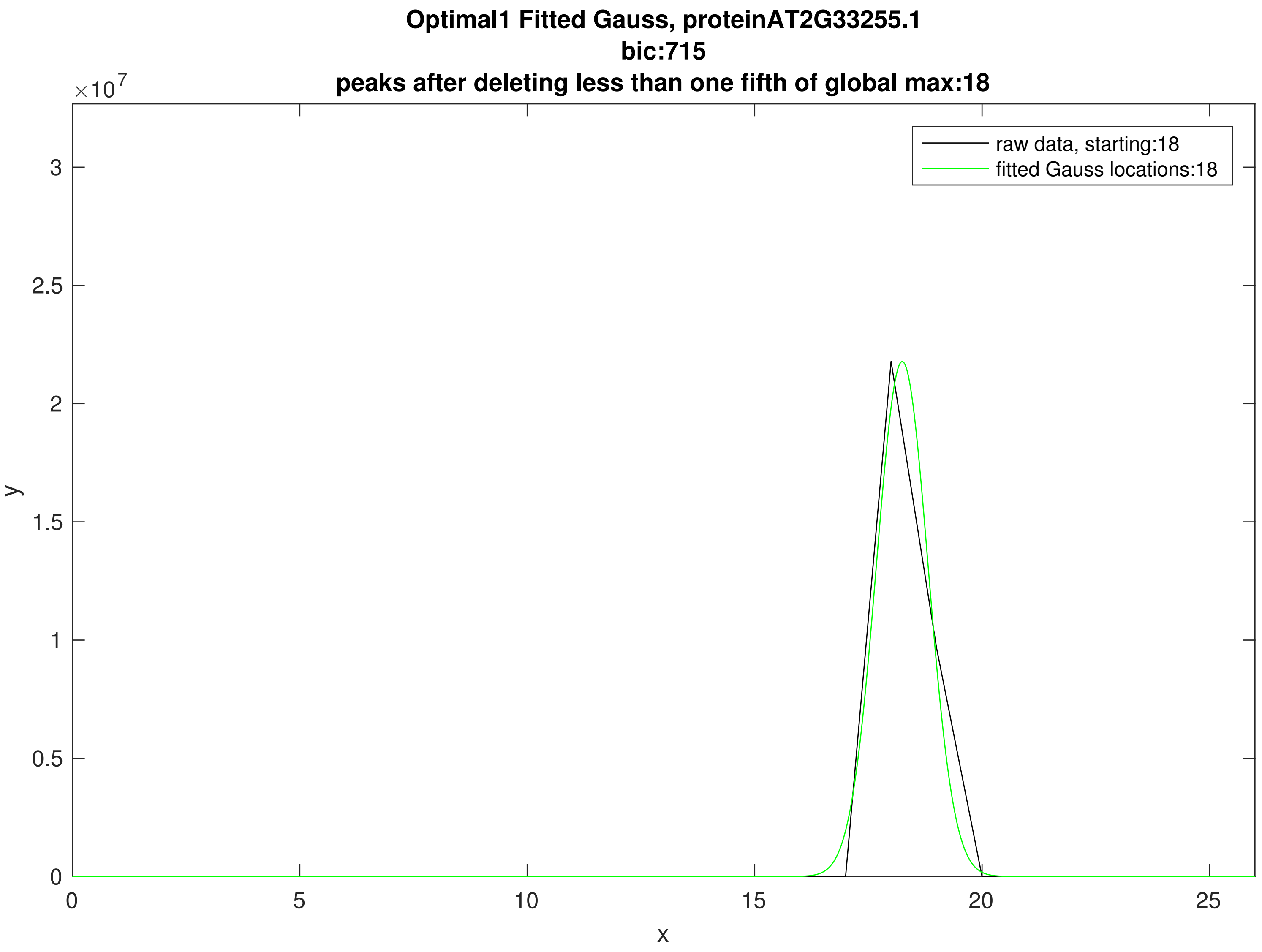
Optimal1 Fitted Gauss, proteinAT2G33255.1
bic:715
peaks after deleting less than one fifth of global max:18
×10^7
raw data, starting:18
fitted Gauss locations:18
y
x
0
0.5
1
1.5
2
2.5
3
0
5
10
15
20
25

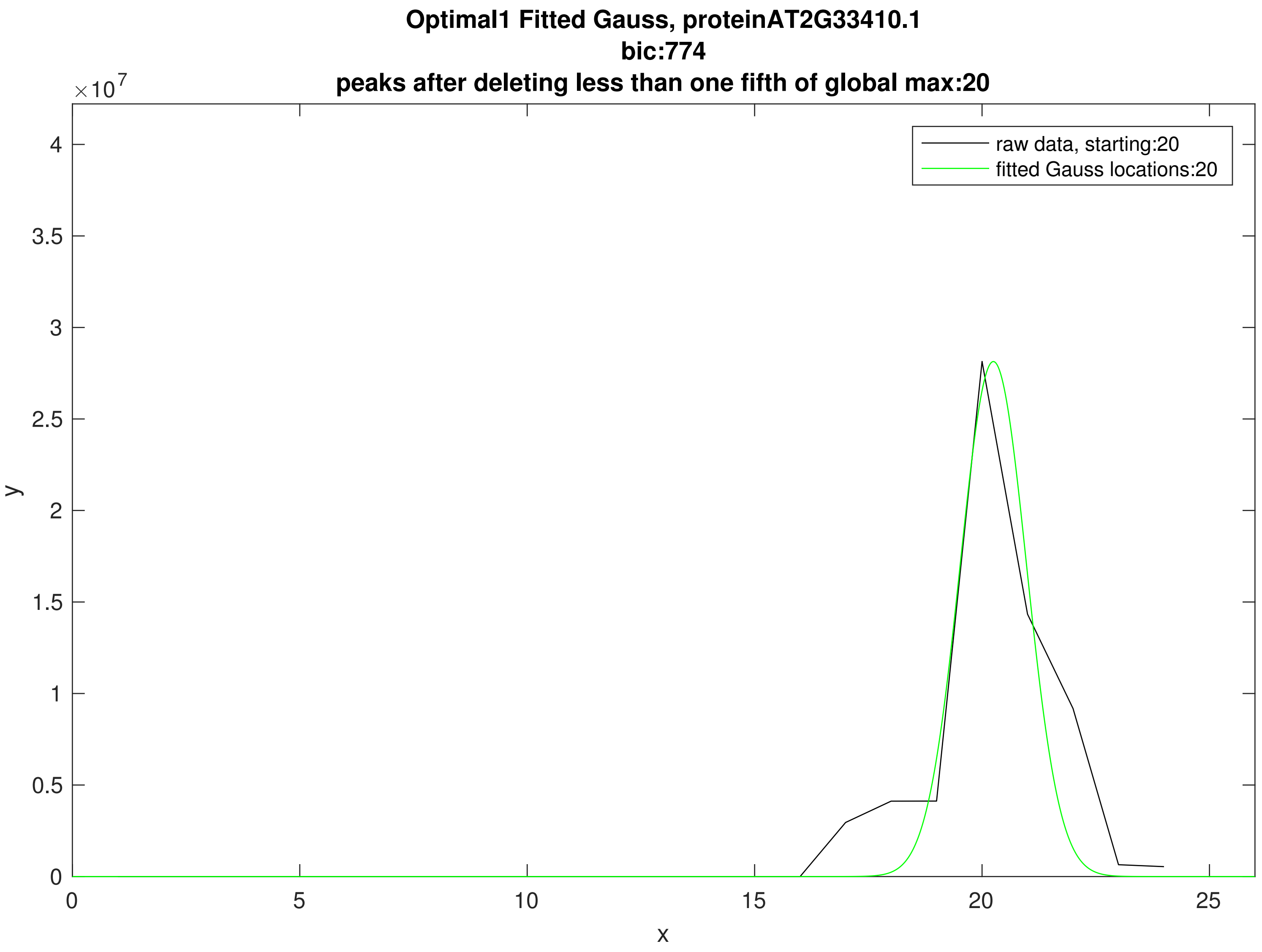

Optimal1 Fitted Gauss, proteinAT2G33410.1
bic:774
peaks after deleting less than one fifth of global max:20
×10^7
raw data, starting:20
fitted Gauss locations:20
y
x
0
0.5
1
1.5
2
2.5
3
3.5
4
0
5
10
15
20
25

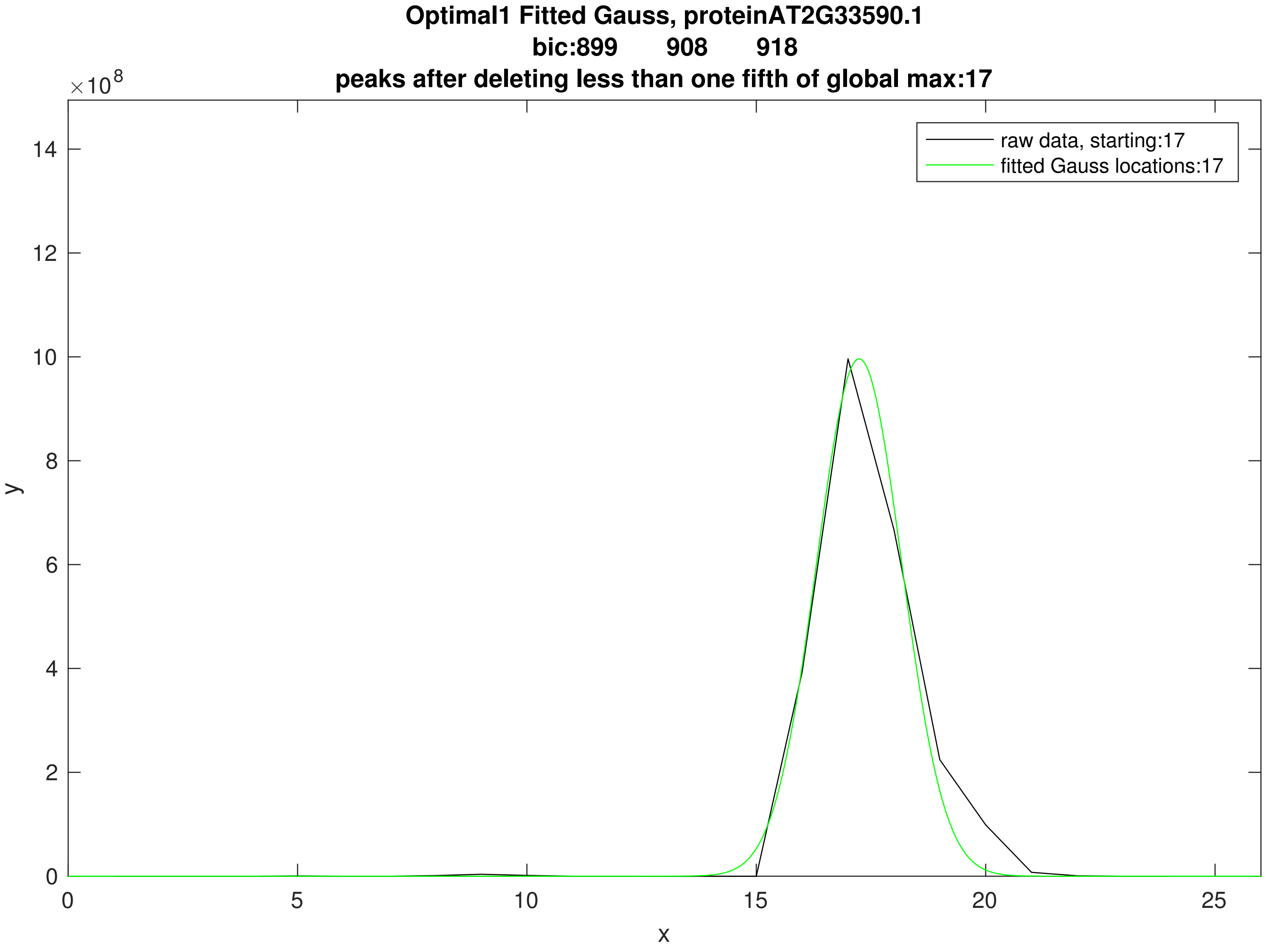

Optimal1 Fitted Gauss, proteinAT2G33590.1
bic:899 908 918
peaks after deleting less than one fifth of global max:17
×10^8
raw data, starting:17
fitted Gauss locations:17
y
x
0
2
4
6
8
10
12
14
5
15
20
25

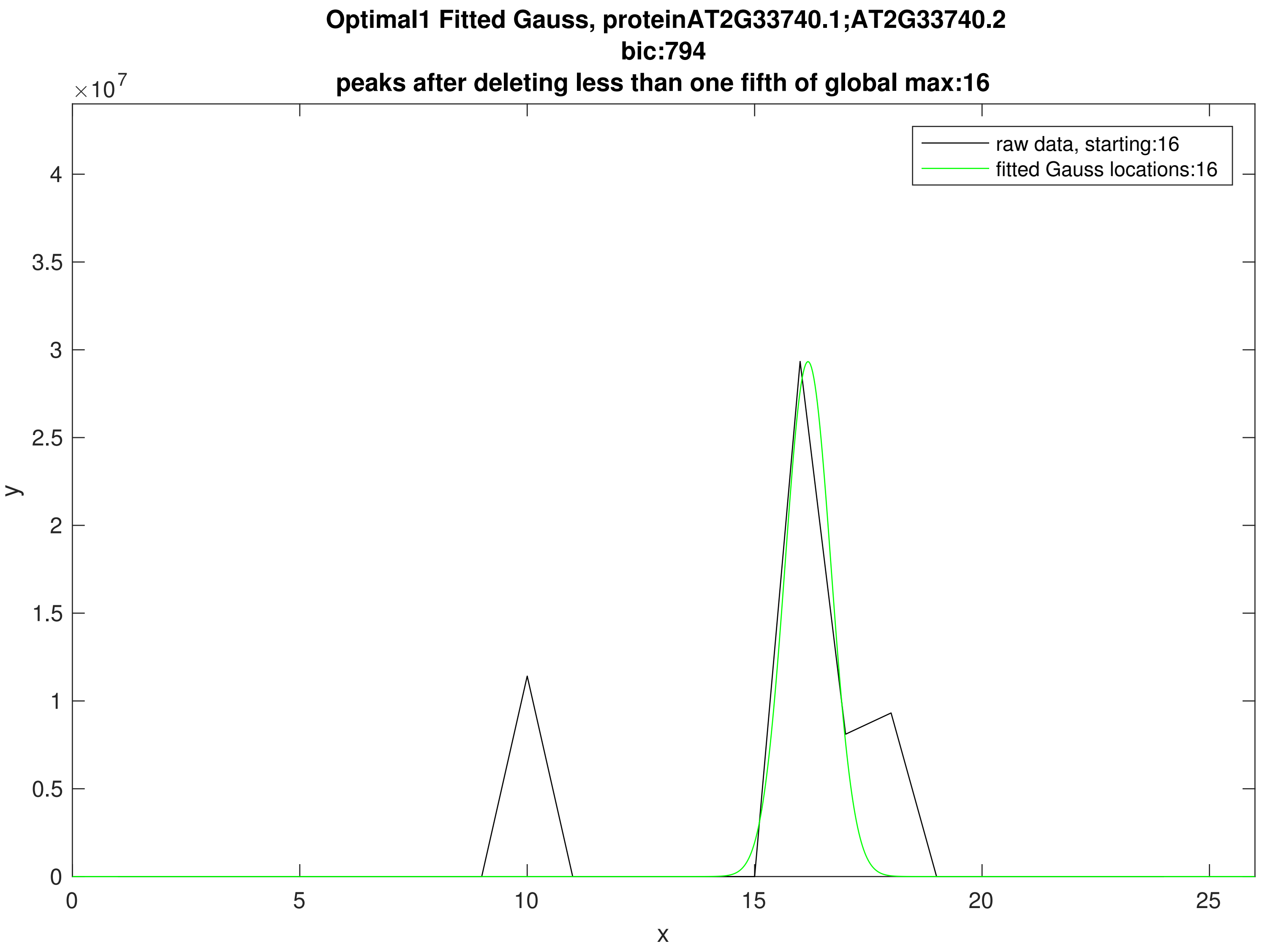
Optimal1 Fitted Gauss, proteinAT2G33740.1;AT2G33740.2
bic:794
peaks after deleting less than one fifth of global max:16
×10^7
raw data, starting:16
fitted Gauss locations:16
y
x
0
0.5
1
1.5
2
2.5
3
3.5
4
5
10
15
20
25

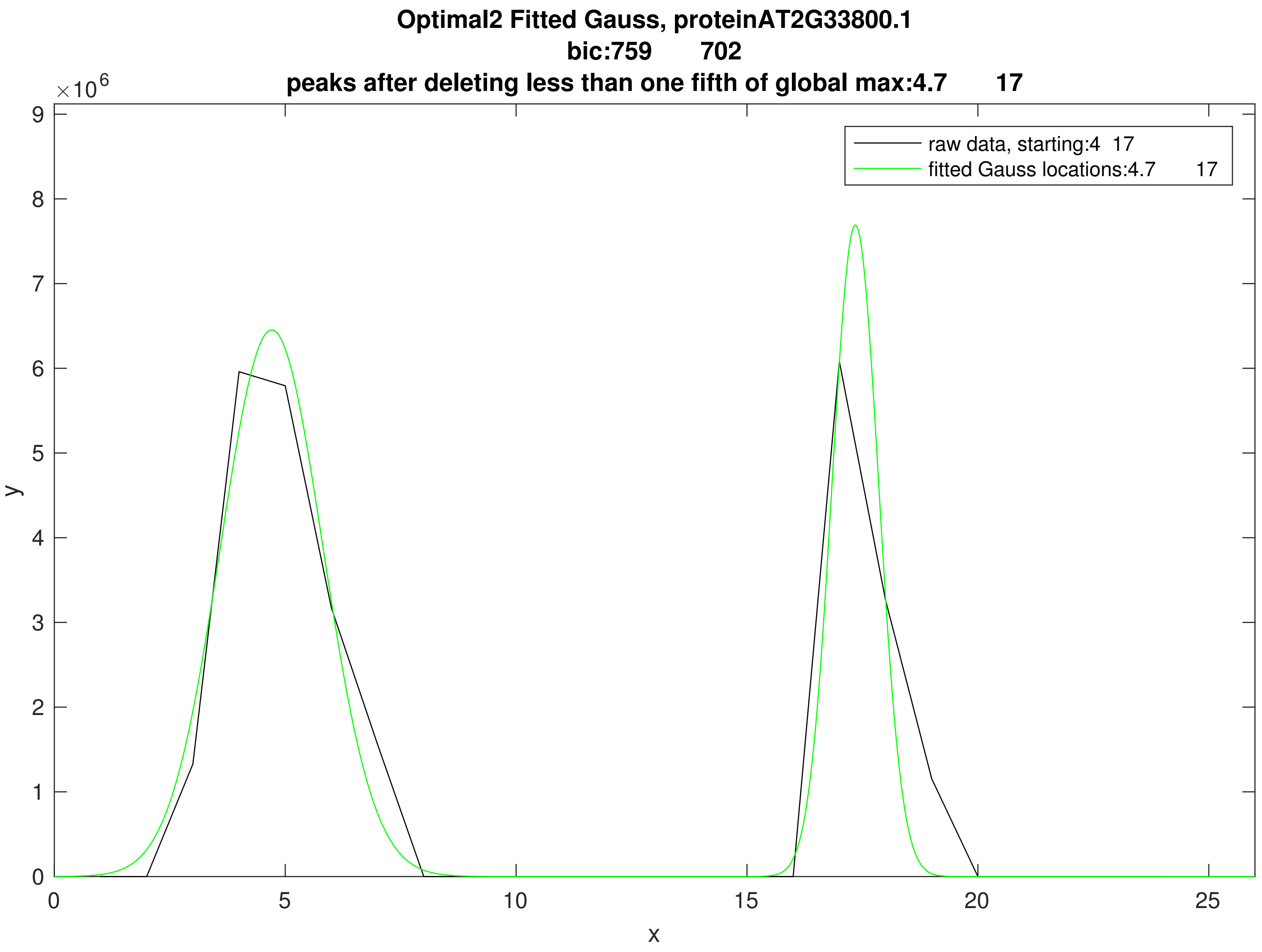
Optimal2 Fitted Gauss, proteinAT2G33800.1
bic:759 702
peaks after deleting less than one fifth of global max:4.7 17
×10^6
raw data, starting:4 17
fitted Gauss locations:4.7 17
y
x
0
1
2
3
4
5
6
7
8
9
0
5
10
15
20
25

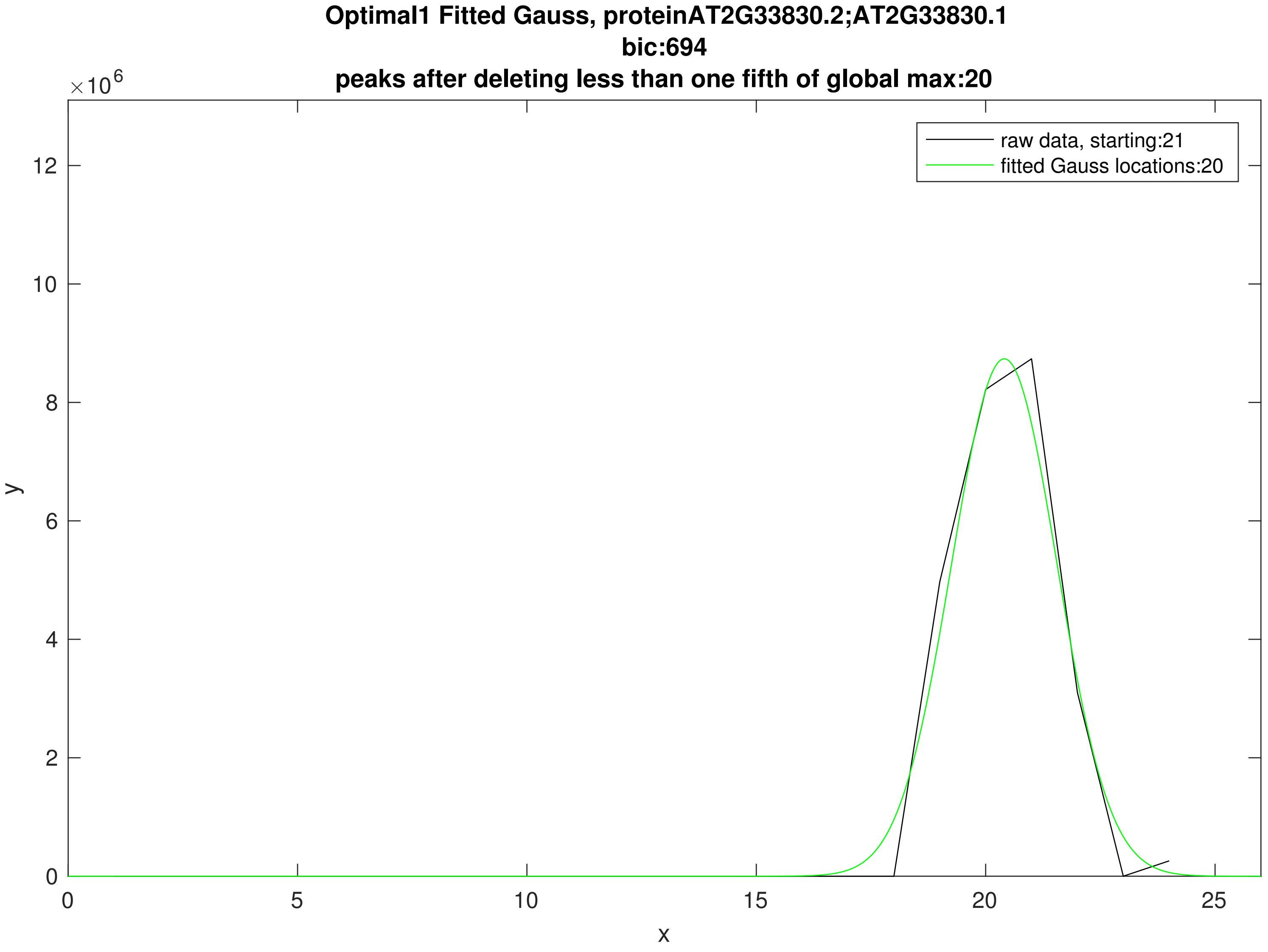

Optimal1 Fitted Gauss, proteinAT2G33830.2;AT2G33830.1
bic:694
peaks after deleting less than one fifth of global max:20
×10^6
12
10
8
6
4
2
0
y
0
5
10
15
20
25
x
raw data, starting:21
fitted Gauss locations:20

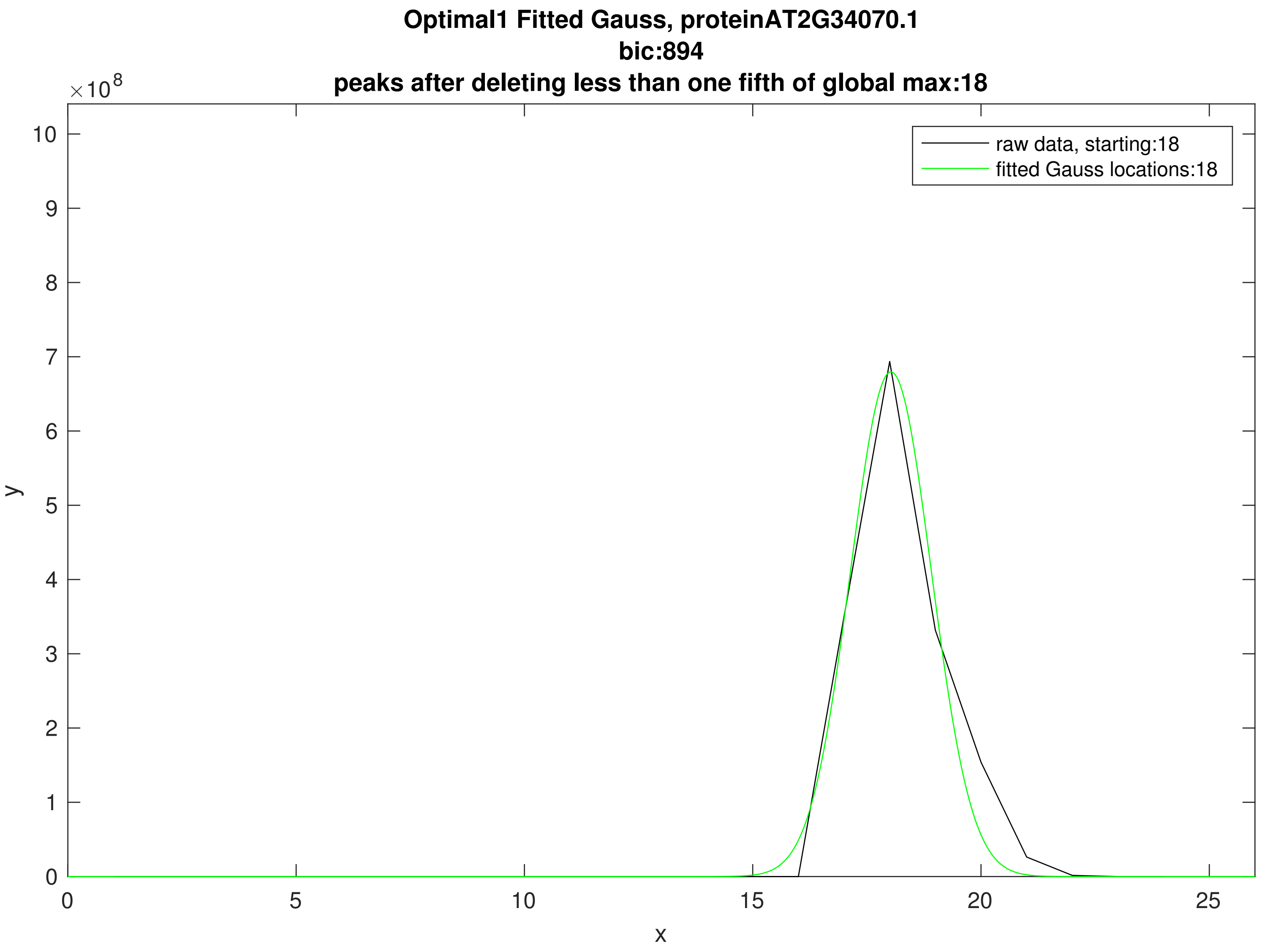

Optimal1 Fitted Gauss, proteinAT2G34070.1
bic:894
peaks after deleting less than one fifth of global max:18
×10^8
raw data, starting:18
fitted Gauss locations:18
y
x
0
1
2
3
4
5
6
7
8
9
10
0
5
10
15
20
25

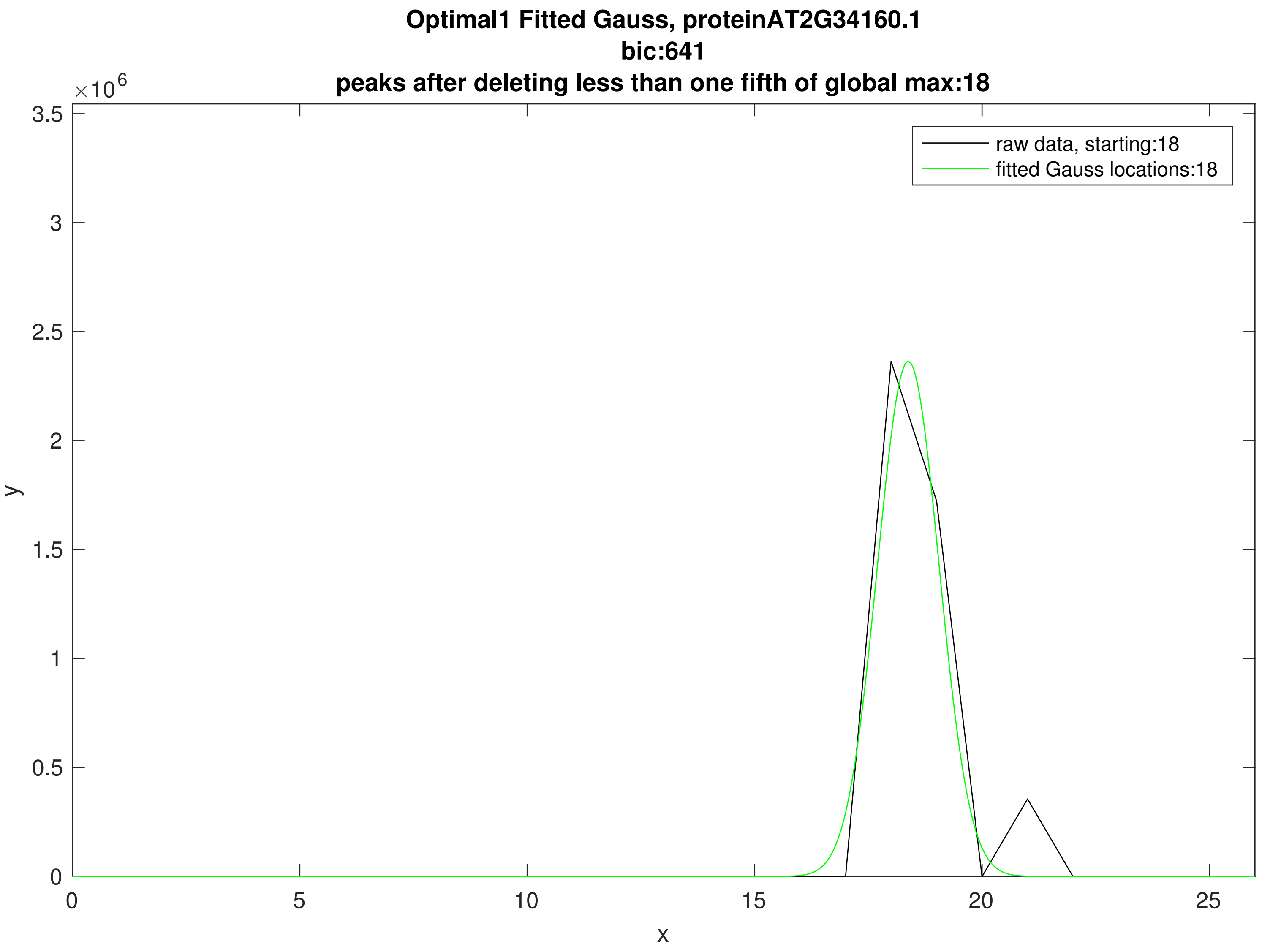
Optimal1 Fitted Gauss, proteinAT2G34160.1
bic:641
peaks after deleting less than one fifth of global max:18
×10^6
3.5
3
2.5
2
1.5
1
0.5
0
y
0
5
10
15
20
25
x
raw data, starting:18
fitted Gauss locations:18

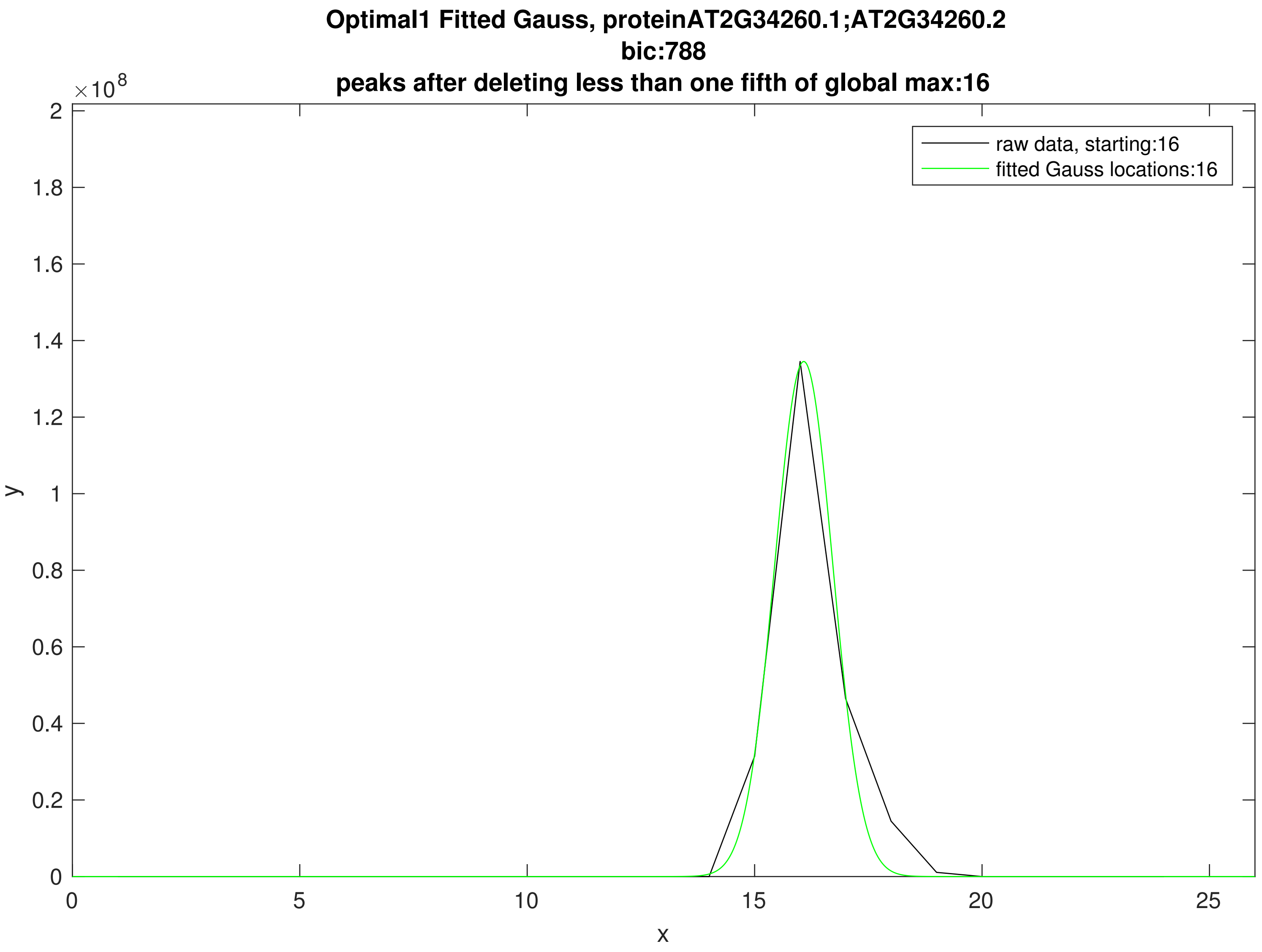
Optimal1 Fitted Gauss, proteinAT2G34260.1;AT2G34260.2
bic:788
peaks after deleting less than one fifth of global max:16
raw data, starting:16
fitted Gauss locations:16
×10^8
y
x

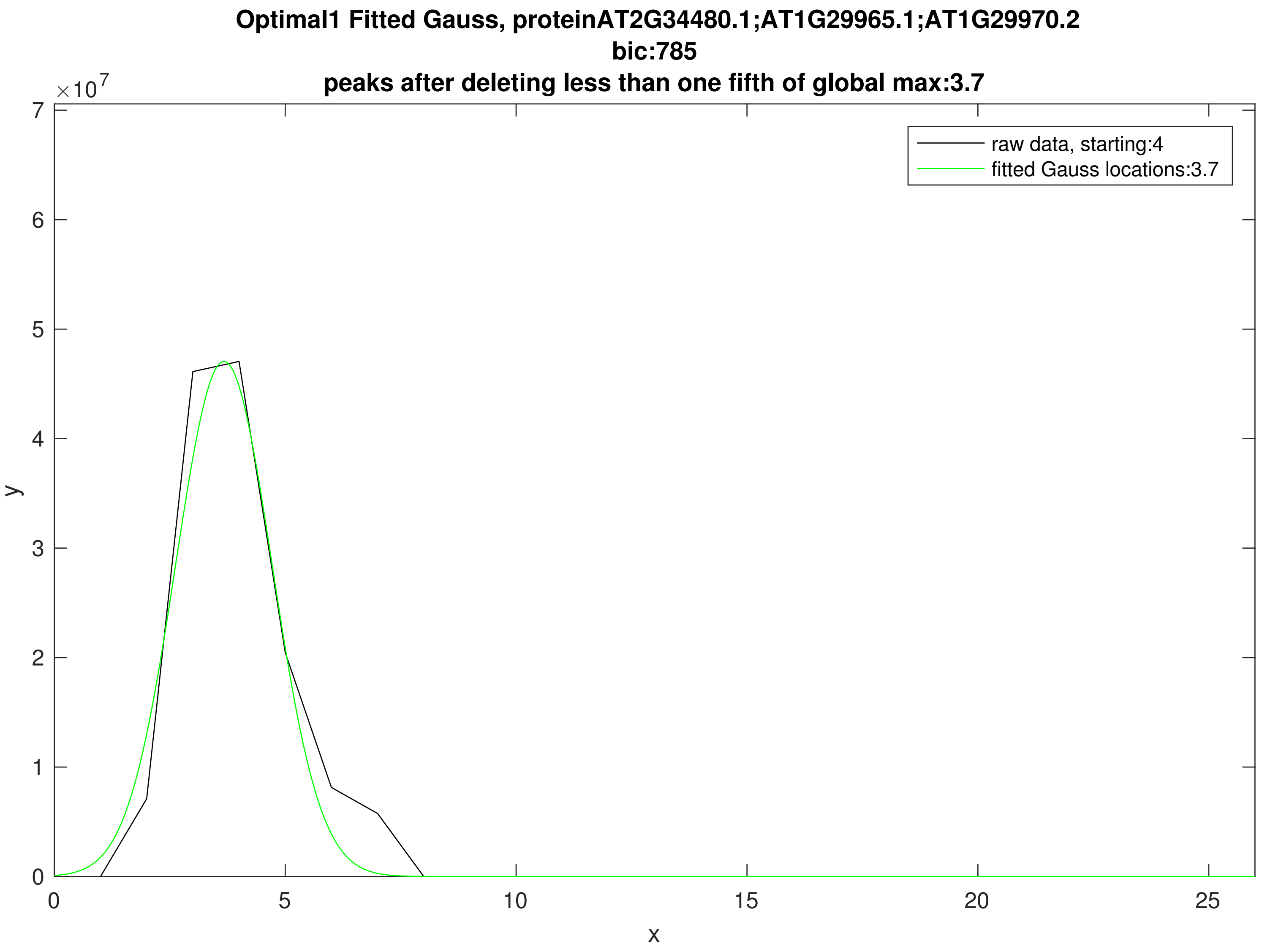

Optimal1 Fitted Gauss, proteinAT2G34480.1;AT1G29965.1;AT1G29970.2
bic:785
peaks after deleting less than one fifth of global max:3.7
×10^7
raw data, starting:4
fitted Gauss locations:3.7
y
x
0
1
2
3
4
5
6
7
0
5
10
15
20
25

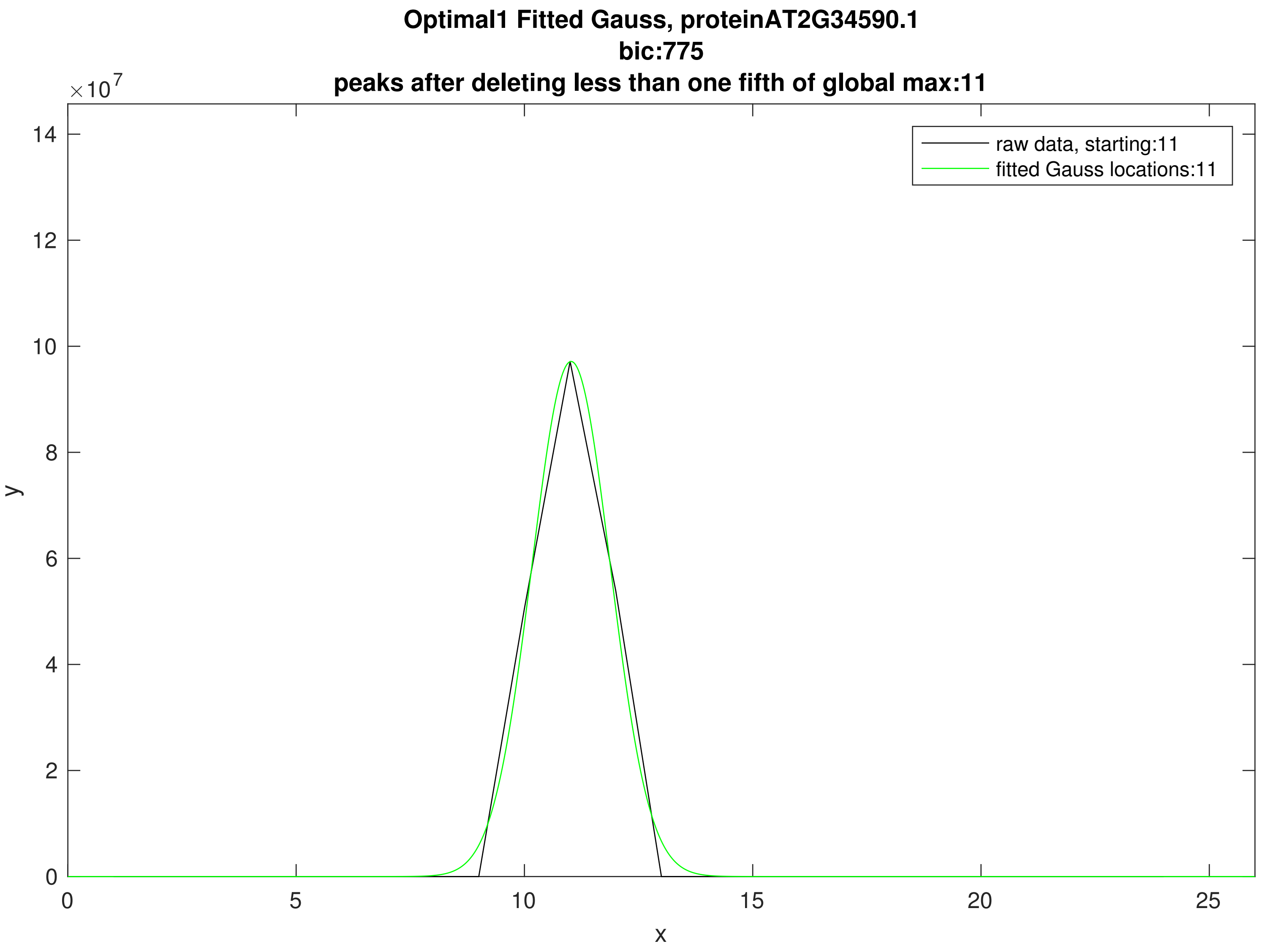
Optimal1 Fitted Gauss, proteinAT2G34590.1
bic:775
peaks after deleting less than one fifth of global max:11
×10^7
14
12
10
8
6
4
2
0
y
0
5
10
15
20
25
x
raw data, starting:11
fitted Gauss locations:11

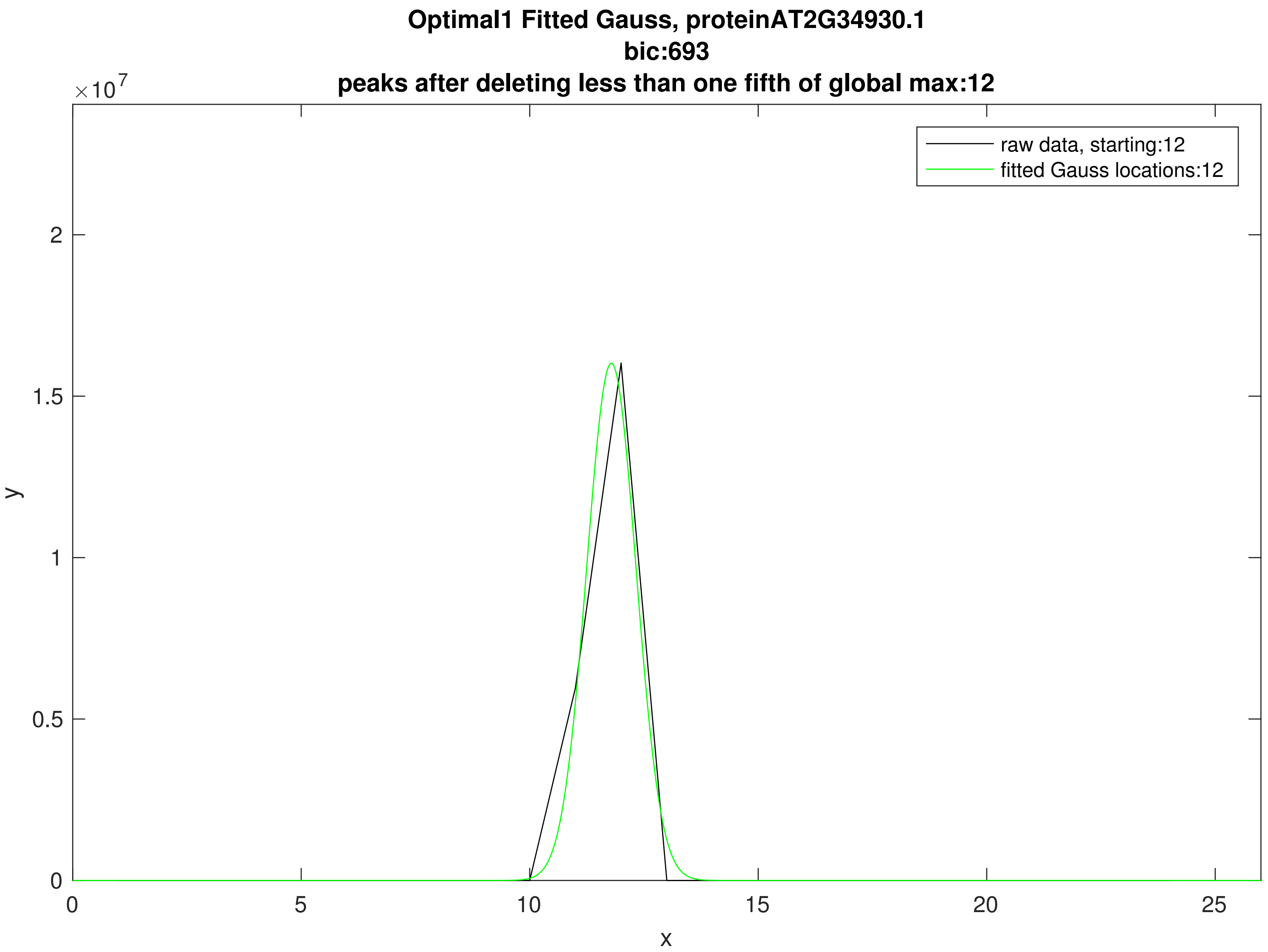
Optimal1 Fitted Gauss, proteinAT2G34930.1
bic:693
peaks after deleting less than one fifth of global max:12
×10^7
raw data, starting:12
fitted Gauss locations:12
2
1.5
1
0.5
0
y
0
5
10
15
20
25
x

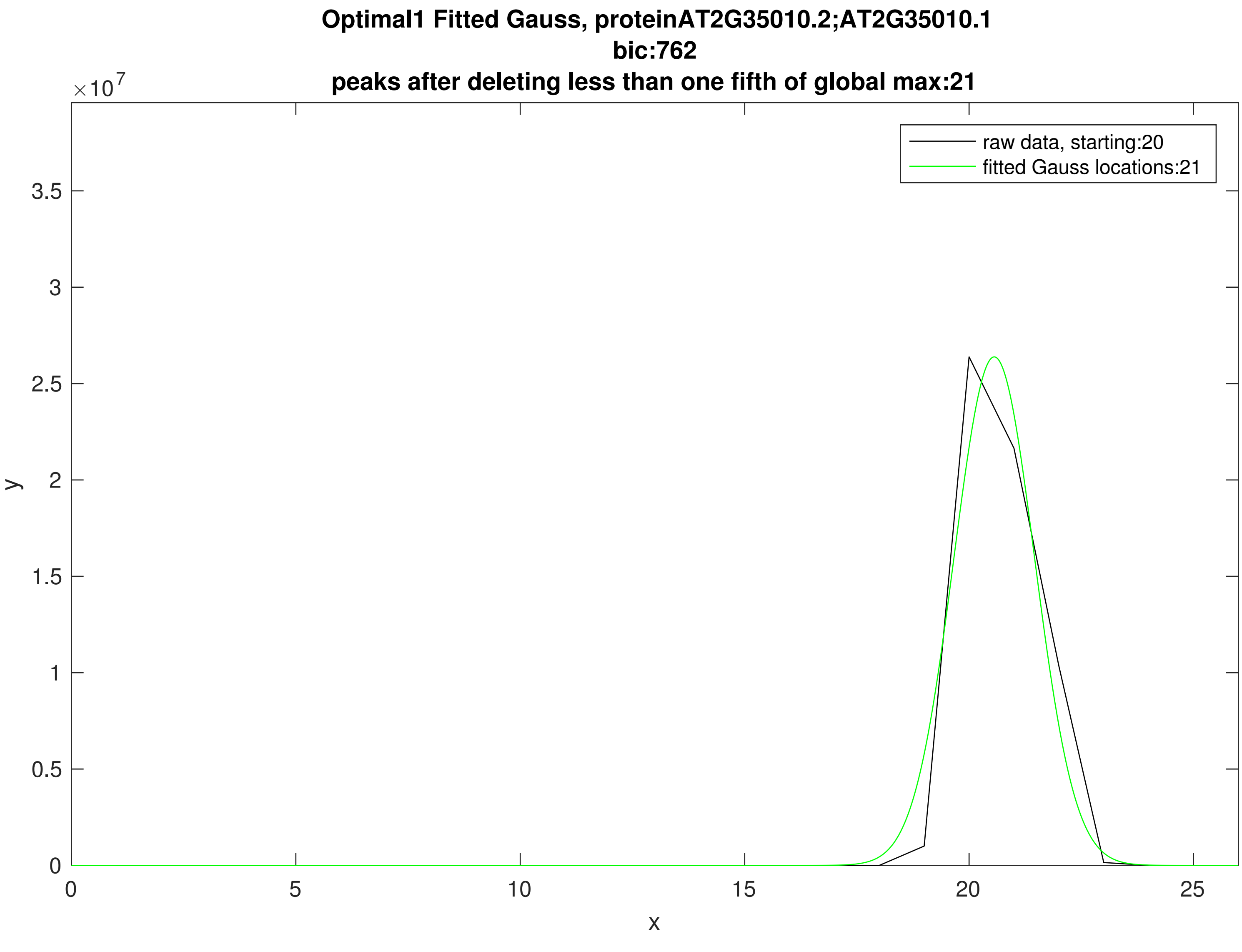

Optimal1 Fitted Gauss, proteinAT2G35010.2;AT2G35010.1
bic:762
peaks after deleting less than one fifth of global max:21
×10^7
raw data, starting:20
fitted Gauss locations:21
3.5
3
2.5
2
1.5
1
0.5
0
y
0
5
10
15
20
25
x

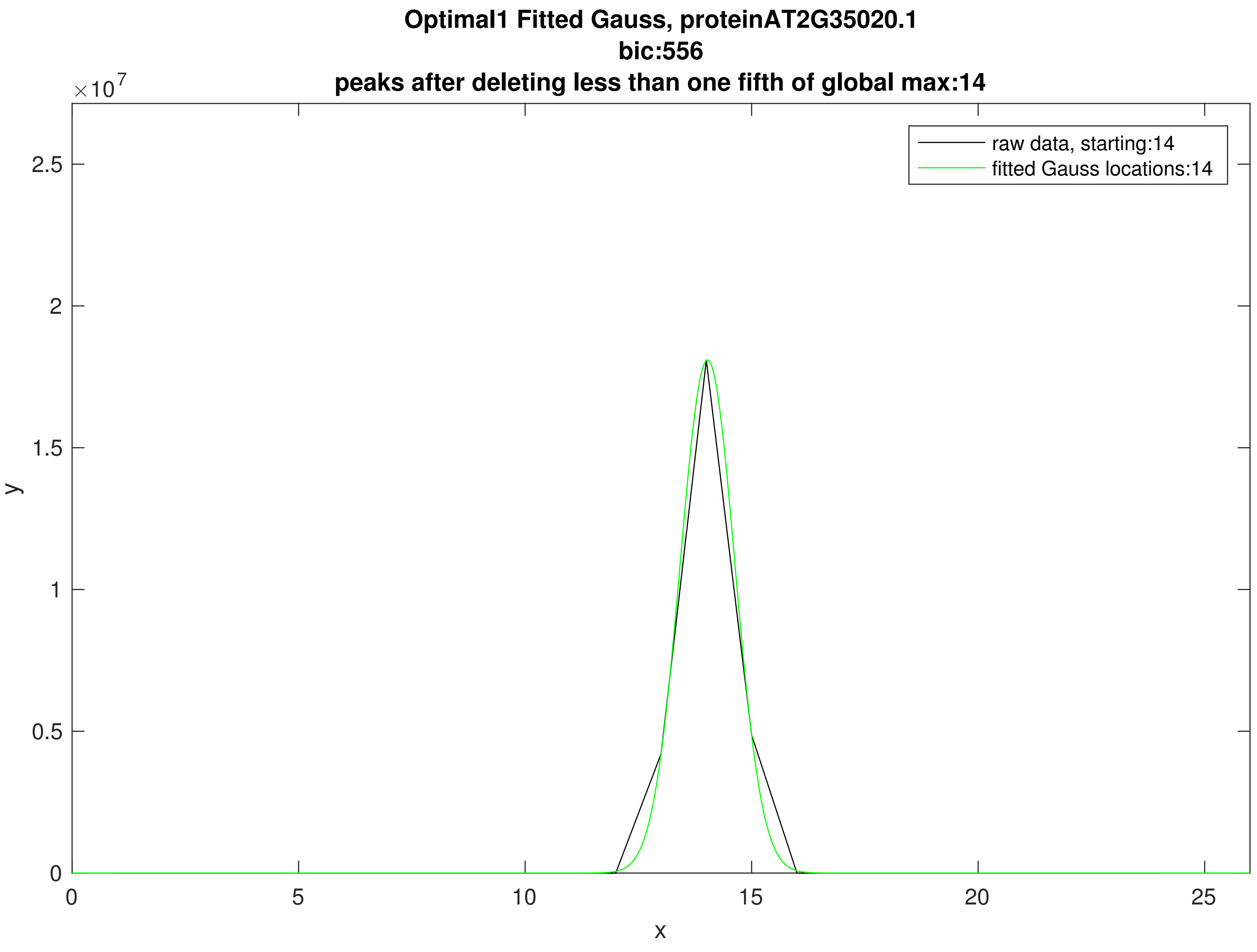

Optimal1 Fitted Gauss, proteinAT2G35020.1
bic:556
peaks after deleting less than one fifth of global max:14
×10^7
raw data, starting:14
fitted Gauss locations:14
y
x
0
0.5
1
1.5
2
2.5
0
5
10
15
20
25

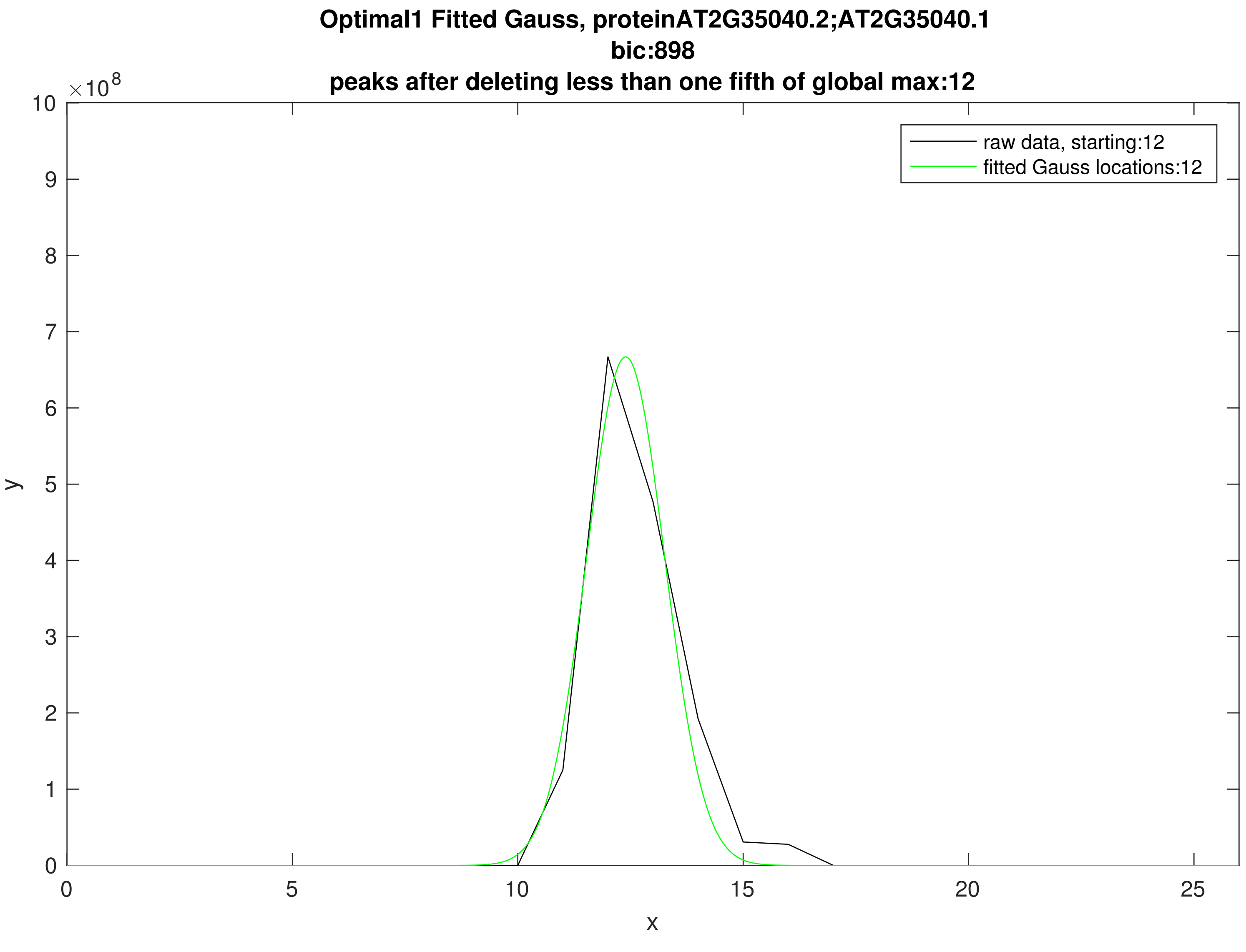

Optimal1 Fitted Gauss, proteinAT2G35040.2;AT2G35040.1
bic:898
peaks after deleting less than one fifth of global max:12
×10^8
raw data, starting:12
fitted Gauss locations:12
y
x
0
1
2
3
4
5
6
7
8
9
10
0
5
10
15
20
25

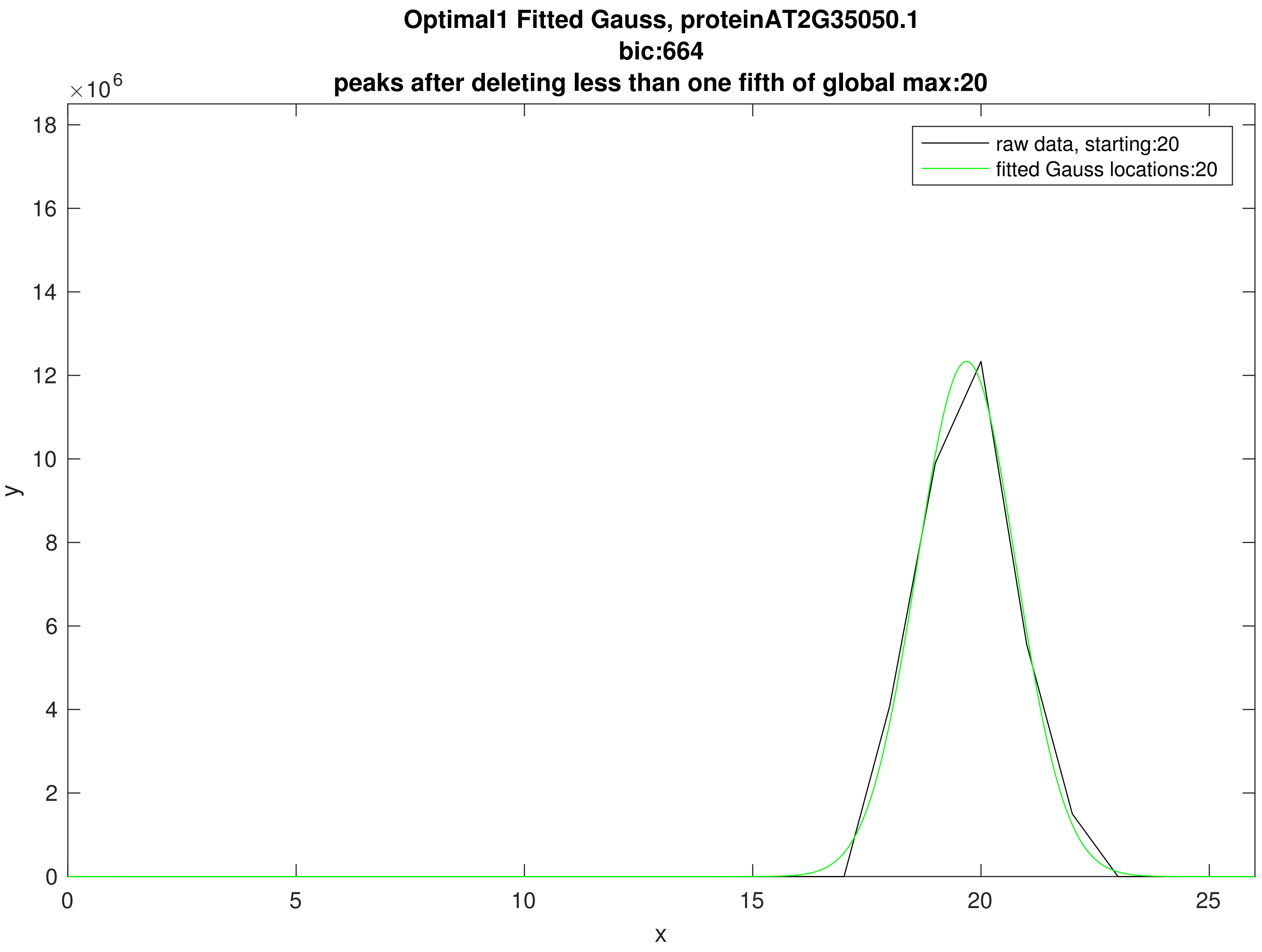

Optimal1 Fitted Gauss, proteinAT2G35050.1
bic:664
peaks after deleting less than one fifth of global max:20
×10^6
raw data, starting:20
fitted Gauss locations:20
18
16
14
12
10
8
6
4
2
0
y
0
5
10
15
20
25
x

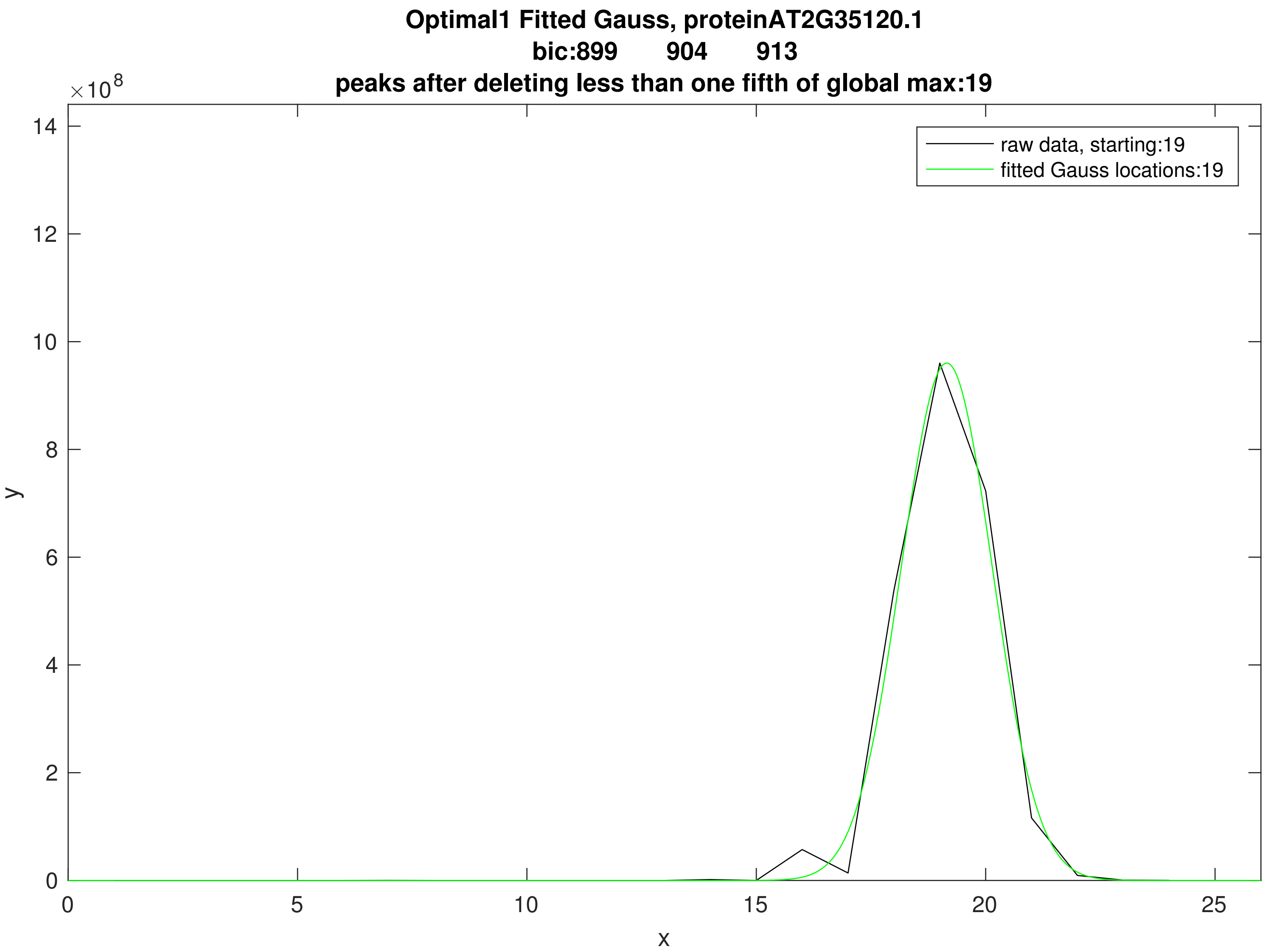

Optimal1 Fitted Gauss, proteinAT2G35120.1
bic:899 904 913
peaks after deleting less than one fifth of global max:19
×10^8
14
12
10
8
6
4
2
0
y
0
5
10
15
20
25
x
raw data, starting:19
fitted Gauss locations:19

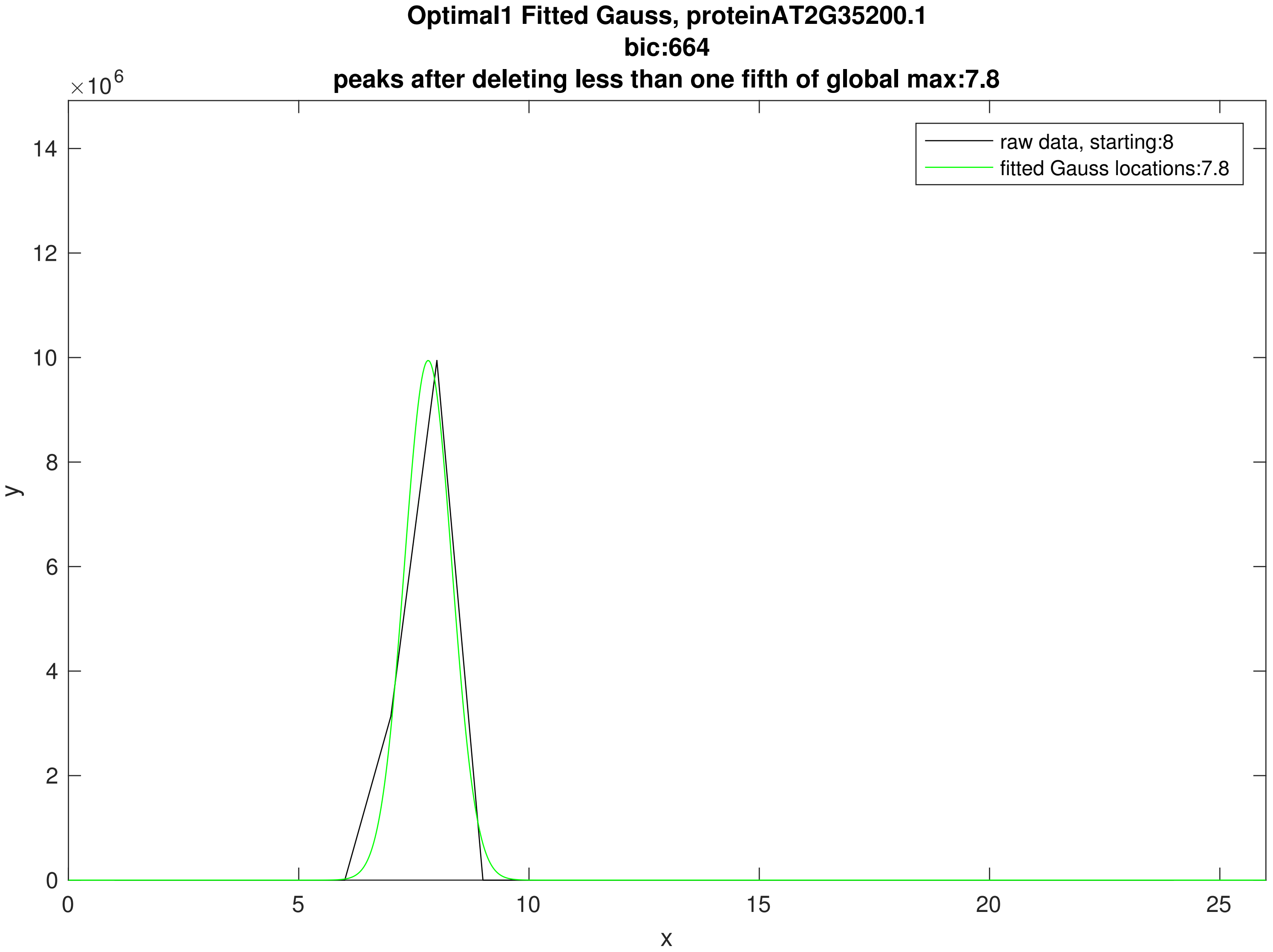
Optimal1 Fitted Gauss, proteinAT2G35200.1
bic:664
peaks after deleting less than one fifth of global max:7.8
×10^6
raw data, starting:8
fitted Gauss locations:7.8
0
2
4
6
8
10
12
14
y
0
5
10
15
20
25
x

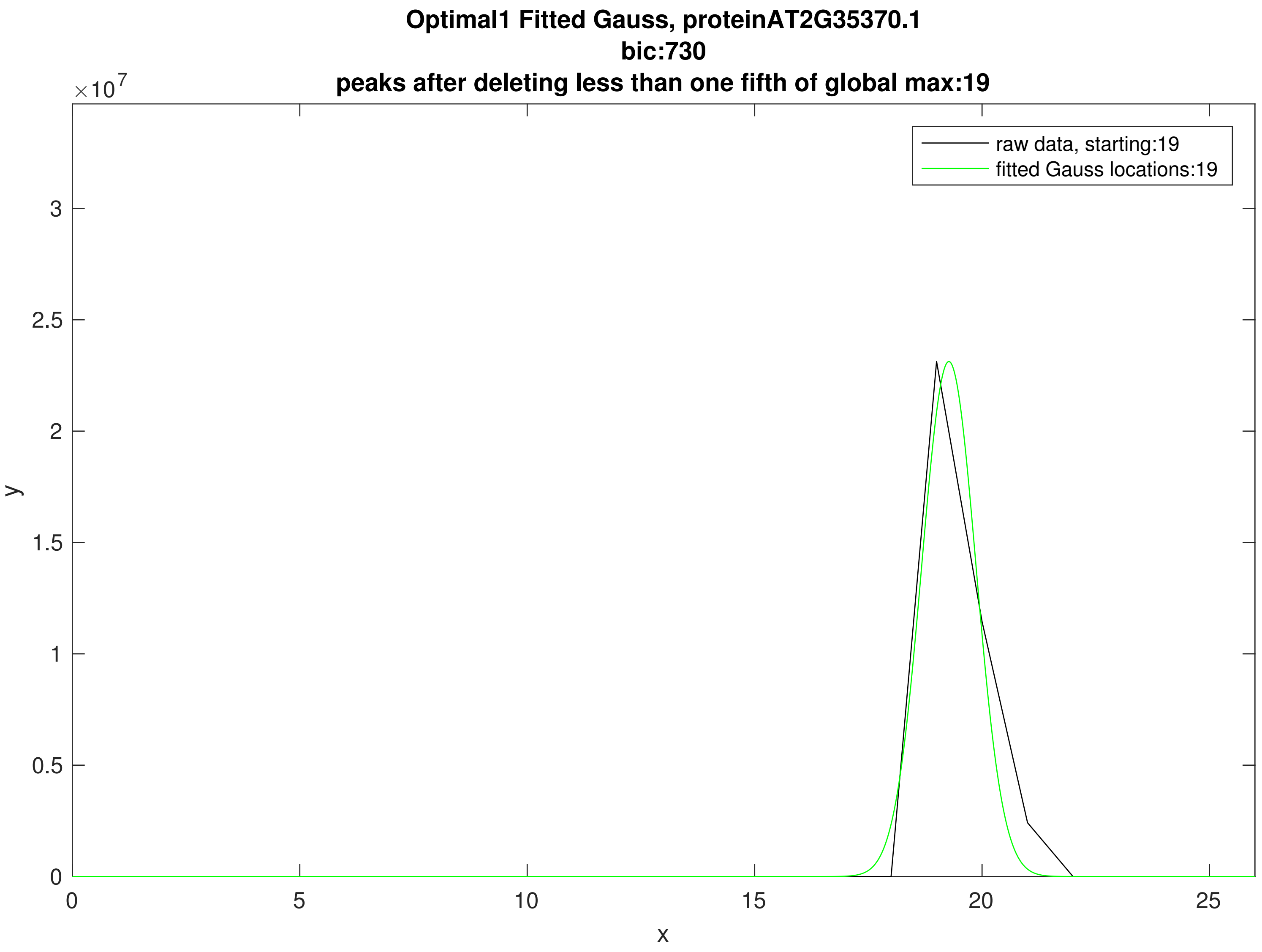
Optimal1 Fitted Gauss, proteinAT2G35370.1
bic:730
peaks after deleting less than one fifth of global max:19
×10^7
3
2.5
2
1.5
1
0.5
0
y
0
5
10
15
20
25
x
raw data, starting:19
fitted Gauss locations:19

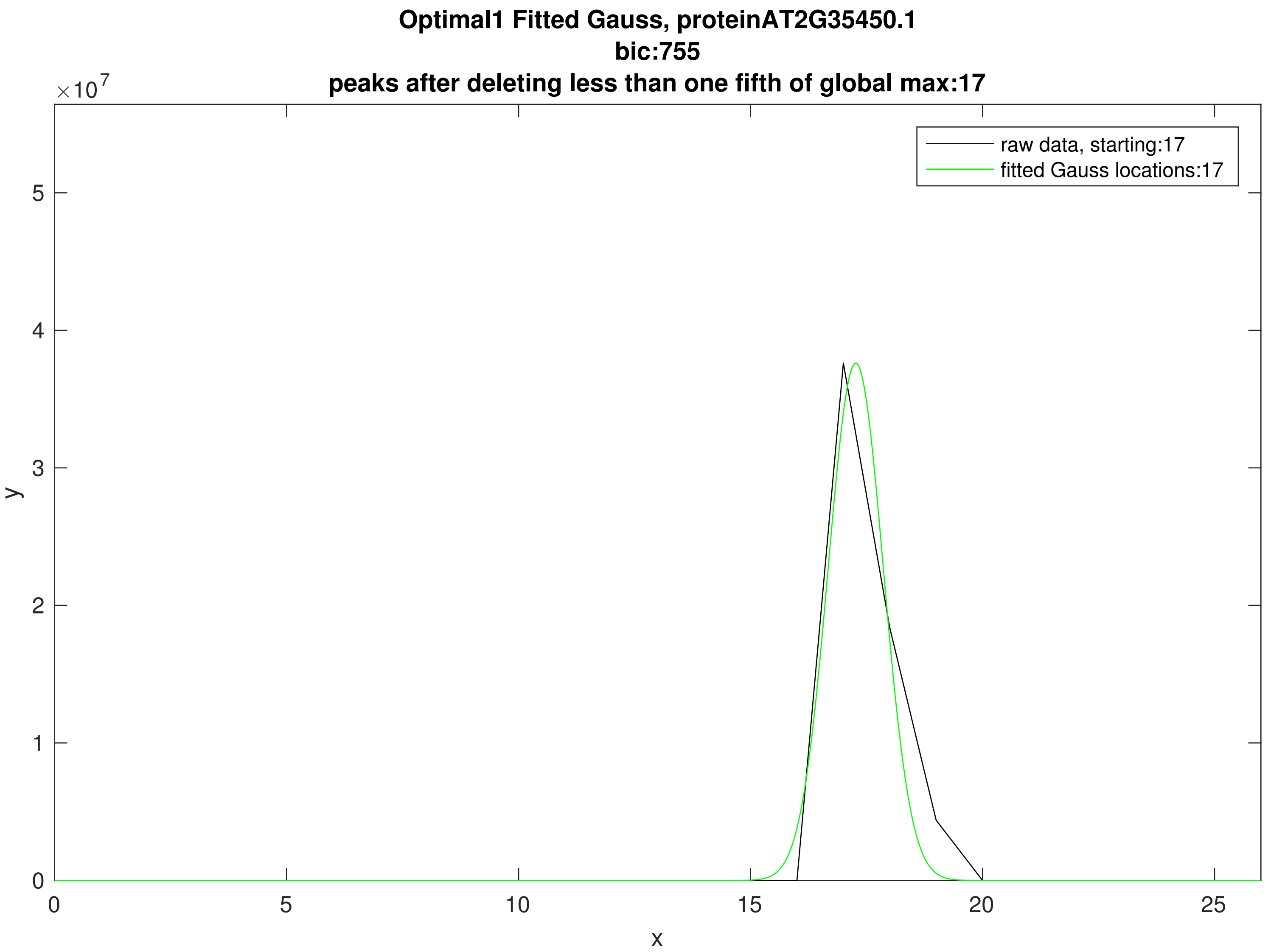
Optimal1 Fitted Gauss, proteinAT2G35450.1
bic:755
peaks after deleting less than one fifth of global max:17
×10^7
raw data, starting:17
fitted Gauss locations:17
y
x
0
1
2
3
4
5
0
5
10
15
20
25

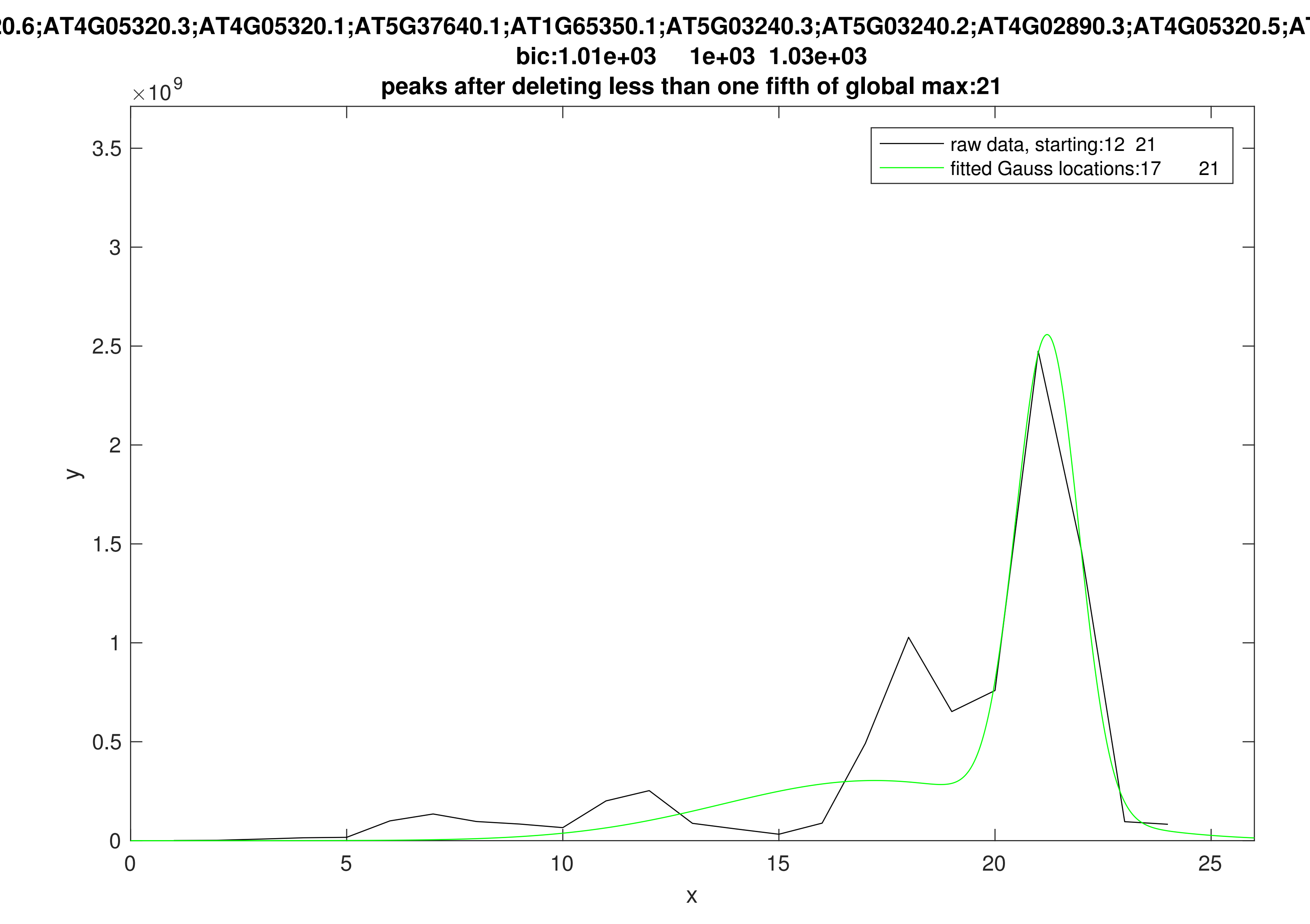
20.6;AT4G05320.3;AT4G05320.1;AT5G37640.1;AT1G65350.1;AT5G03240.3;AT5G03240.2;AT4G02890.3;AT4G05320.5;AT
bic:1.01e+03 1e+03 1.03e+03
peaks after deleting less than one fifth of global max:21
×10^9
raw data, starting:12 21
fitted Gauss locations:17 21
y
x
3.5
3
2.5
2
1.5
1
0.5
0
5
10
15
20
25

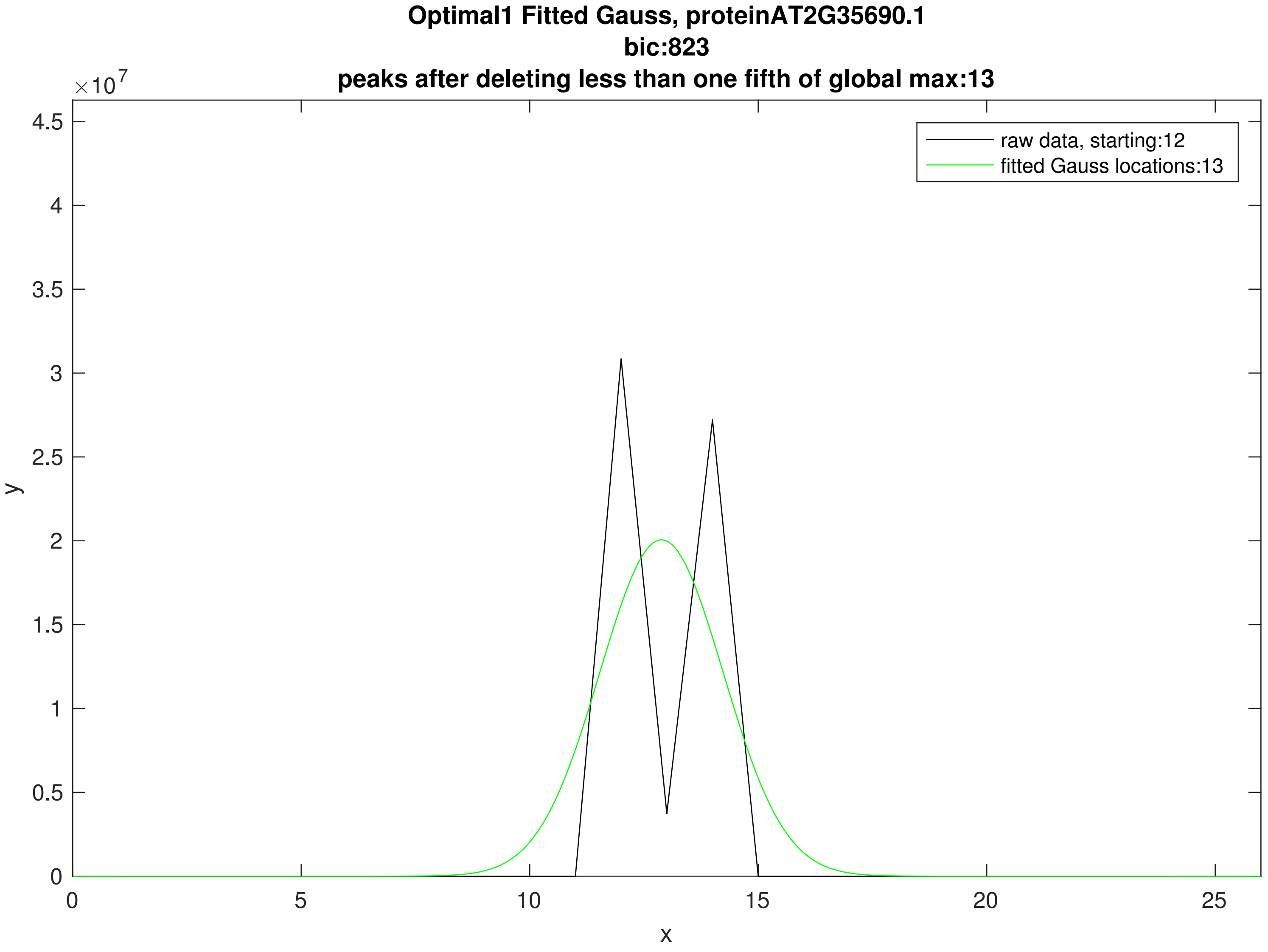
Optimal1 Fitted Gauss, proteinAT2G35690.1
bic:823
peaks after deleting less than one fifth of global max:13
raw data, starting:12
fitted Gauss locations:13
×10^7
y
x
0
0.5
1
1.5
2
2.5
3
3.5
4
4.5
0
5
10
15
20
25

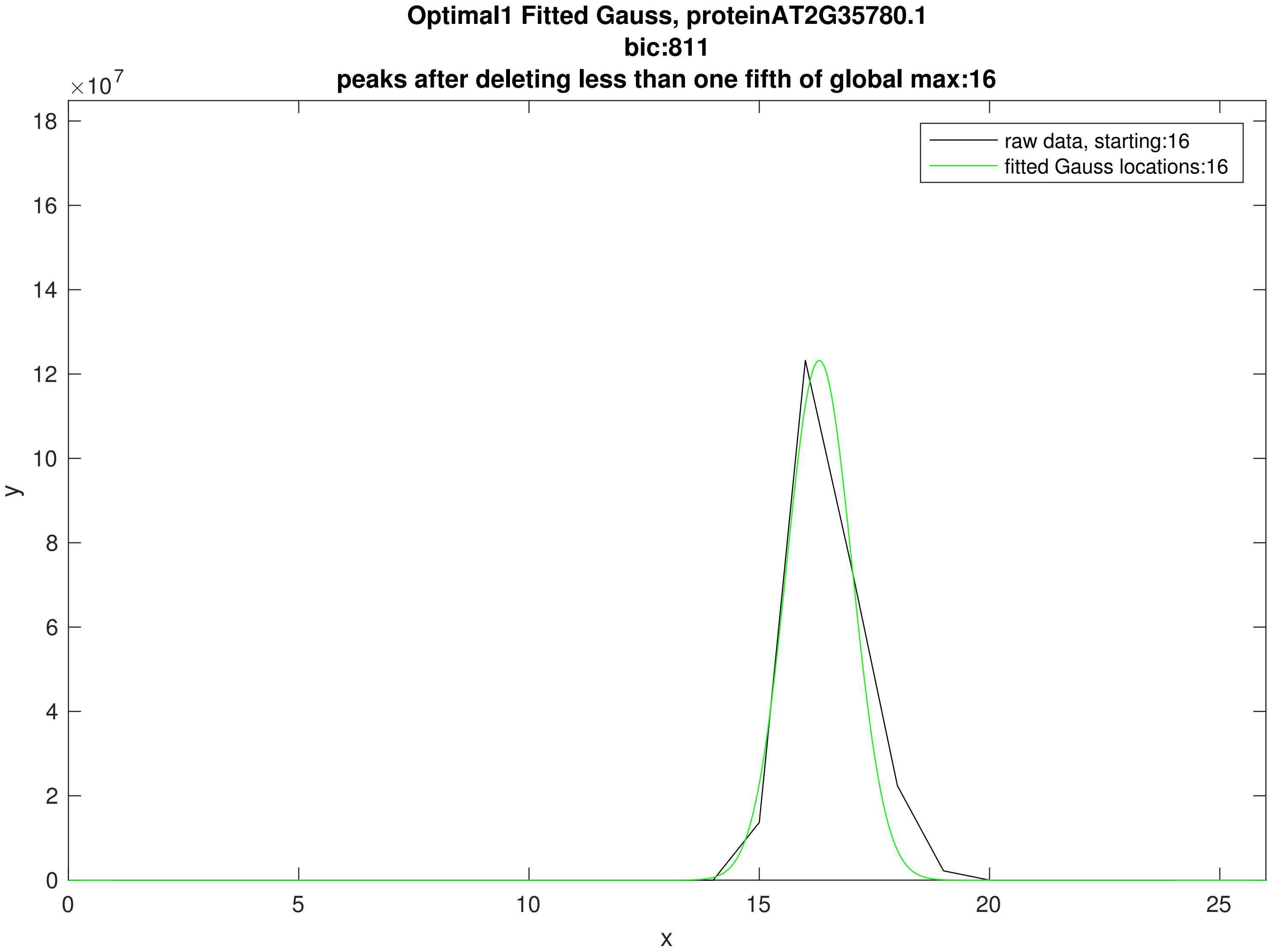
Optimal1 Fitted Gauss, proteinAT2G35780.1
bic:811
peaks after deleting less than one fifth of global max:16
×10^7
y
x
raw data, starting:16
fitted Gauss locations:16
0
2
4
6
8
10
12
14
16
18
0
5
10
15
20
25

#### Optimal1 Fitted Gauss, proteinAT2G35810.1;AT2G35830.1;AT2G35830.2
#### bic:893 902
#### peaks after deleting less than one fifth of global max:16

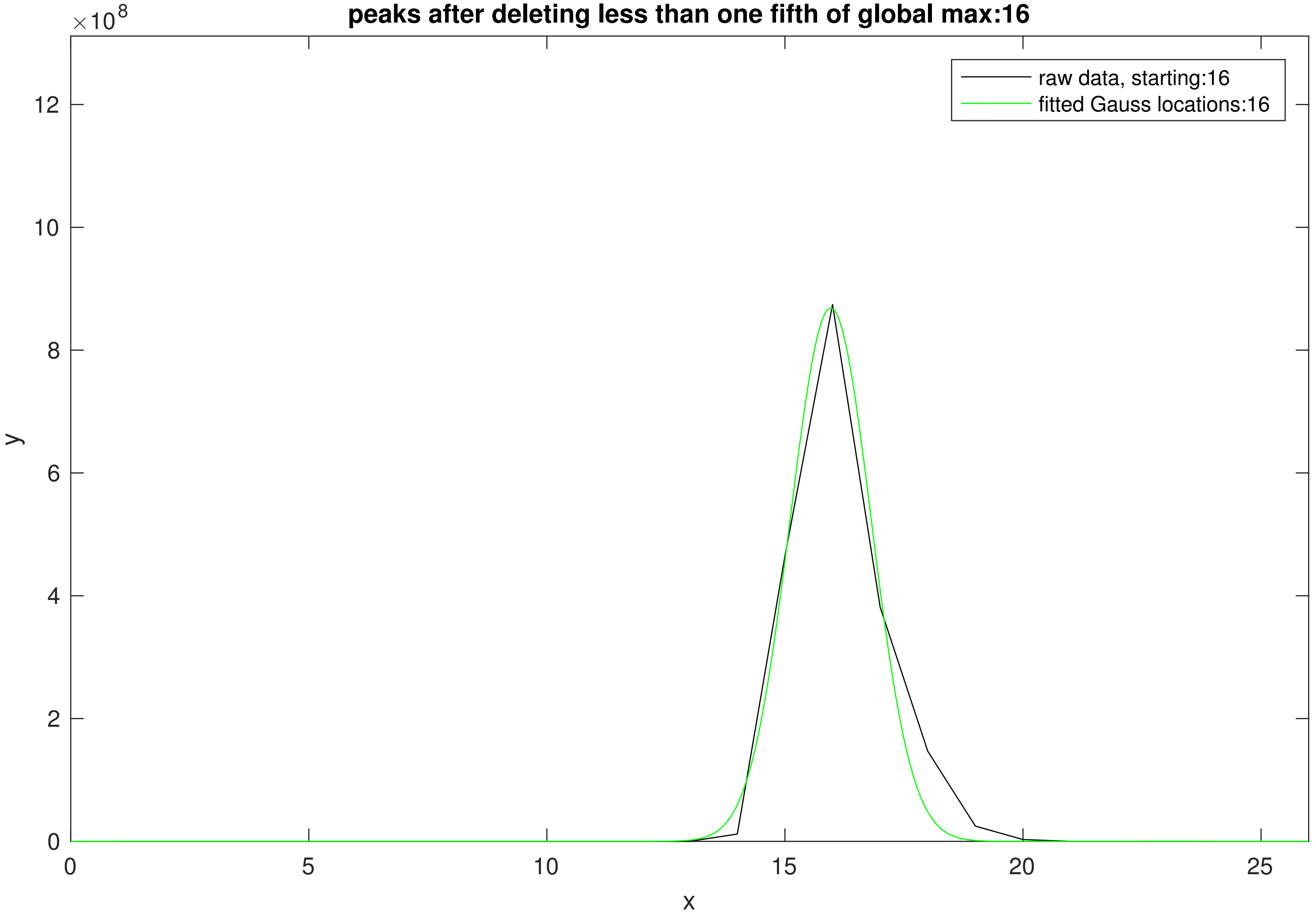

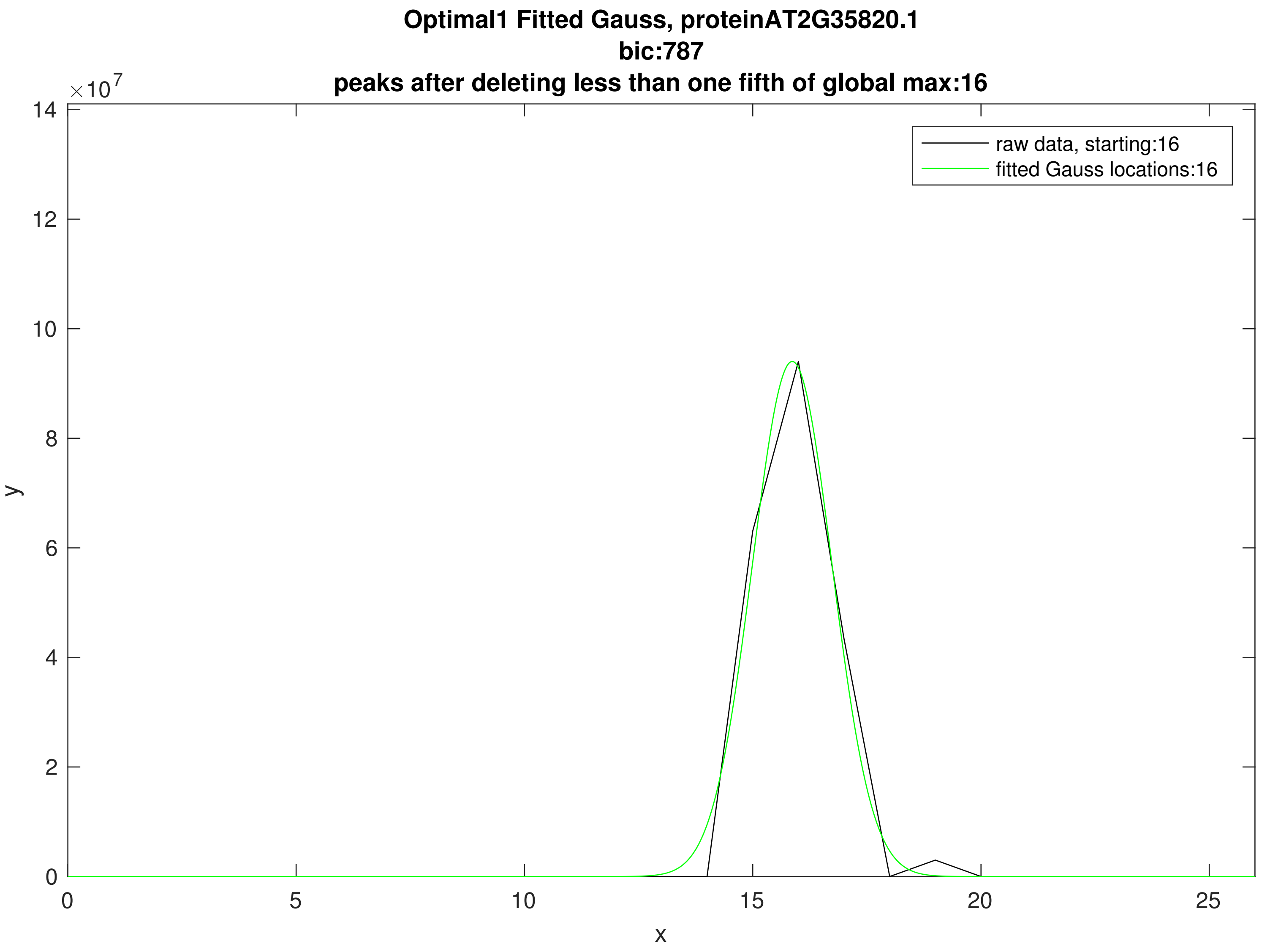
Optimal1 Fitted Gauss, proteinAT2G35820.1
bic:787
peaks after deleting less than one fifth of global max:16
×10^7
14
12
10
8
6
4
2
0
y
0
5
10
15
20
25
x
raw data, starting:16
fitted Gauss locations:16

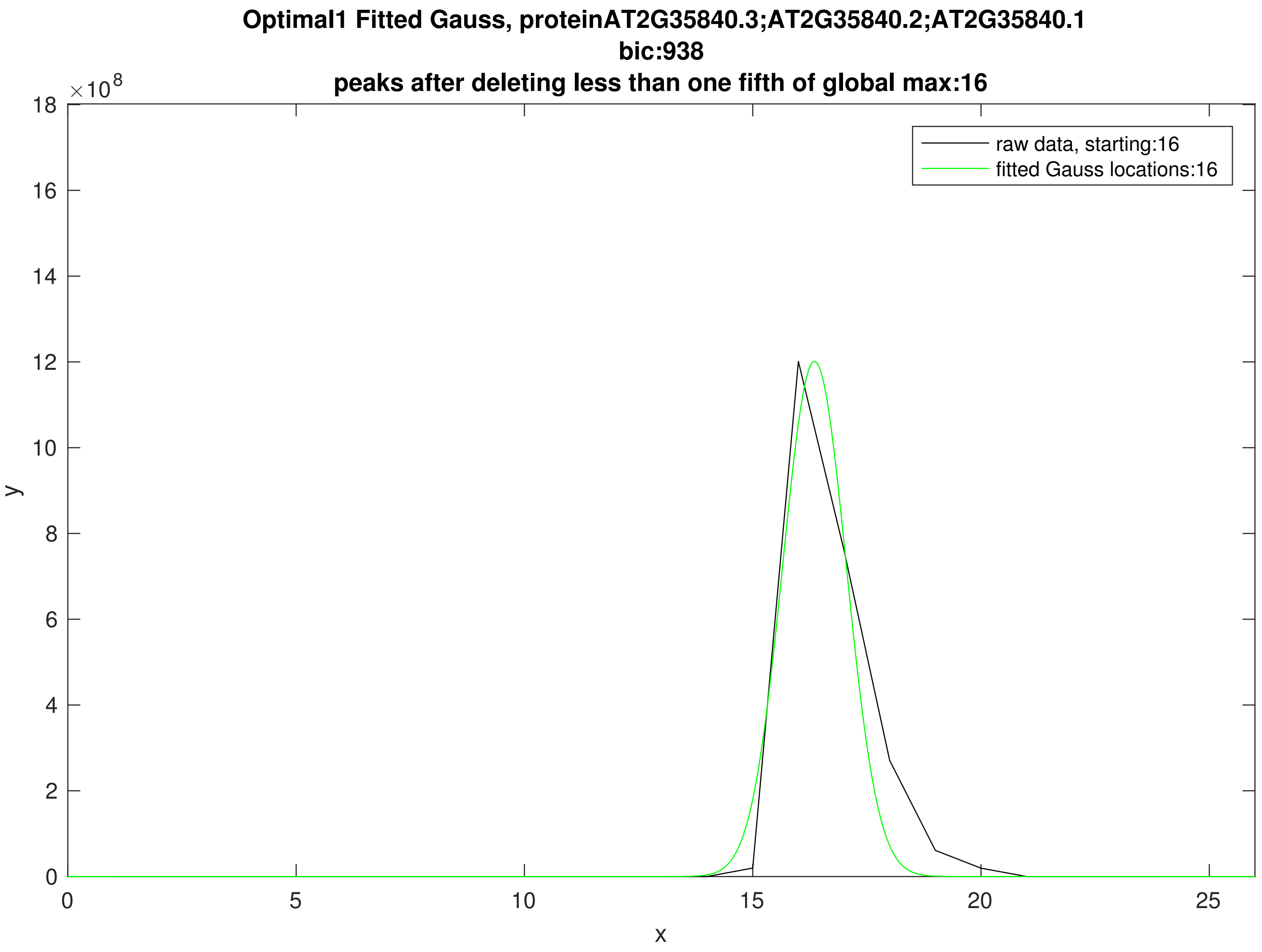
Optimal1 Fitted Gauss, proteinAT2G35840.3;AT2G35840.2;AT2G35840.1
bic:938
peaks after deleting less than one fifth of global max:16
×10^8
raw data, starting:16
fitted Gauss locations:16
y
x

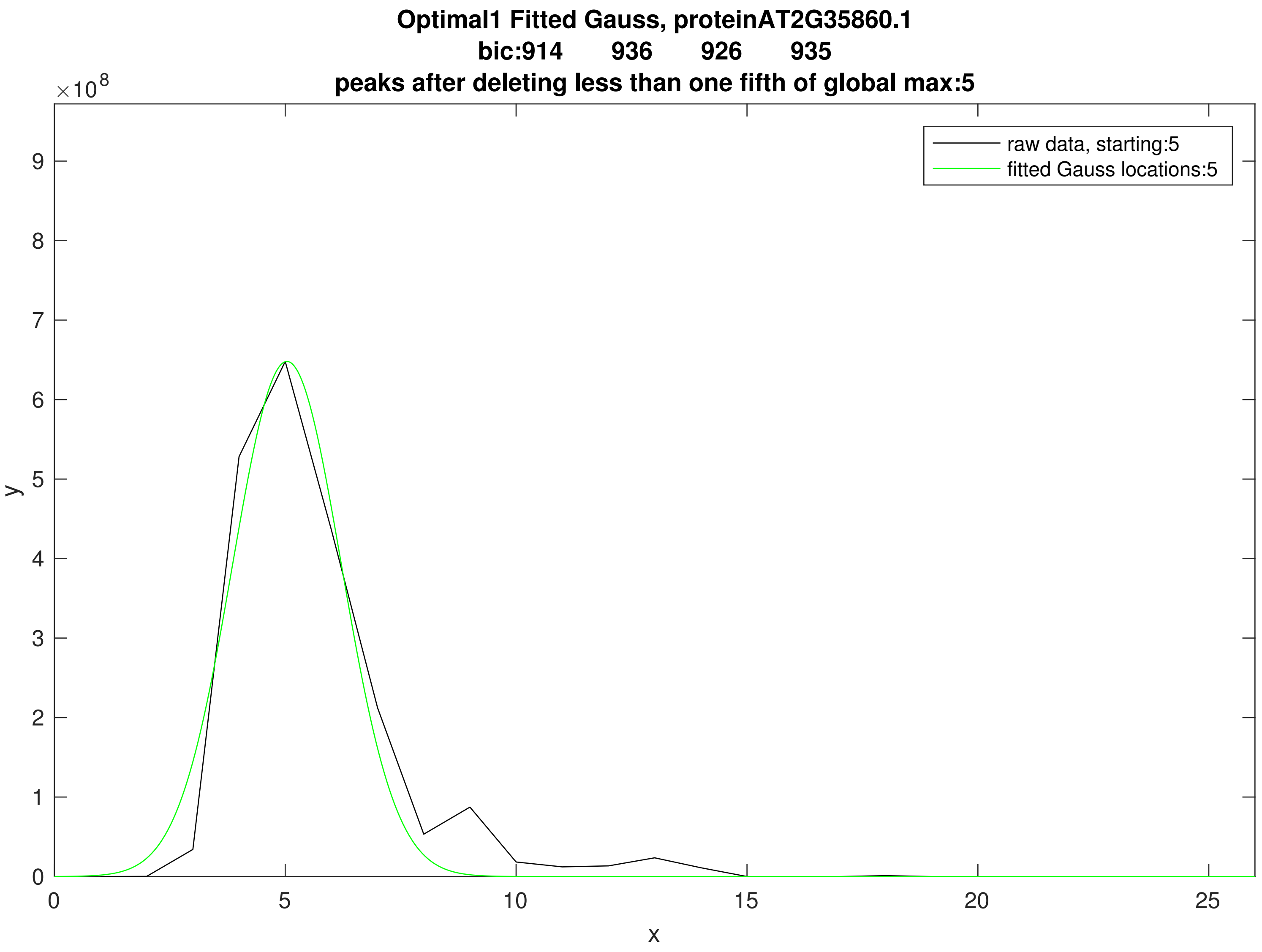
Optimal1 Fitted Gauss, proteinAT2G35860.1
bic:914 936 926 935
peaks after deleting less than one fifth of global max:5
×10^8
raw data, starting:5
fitted Gauss locations:5
y
x
0
1
2
3
4
5
6
7
8
9
0
5
10
15
20
25

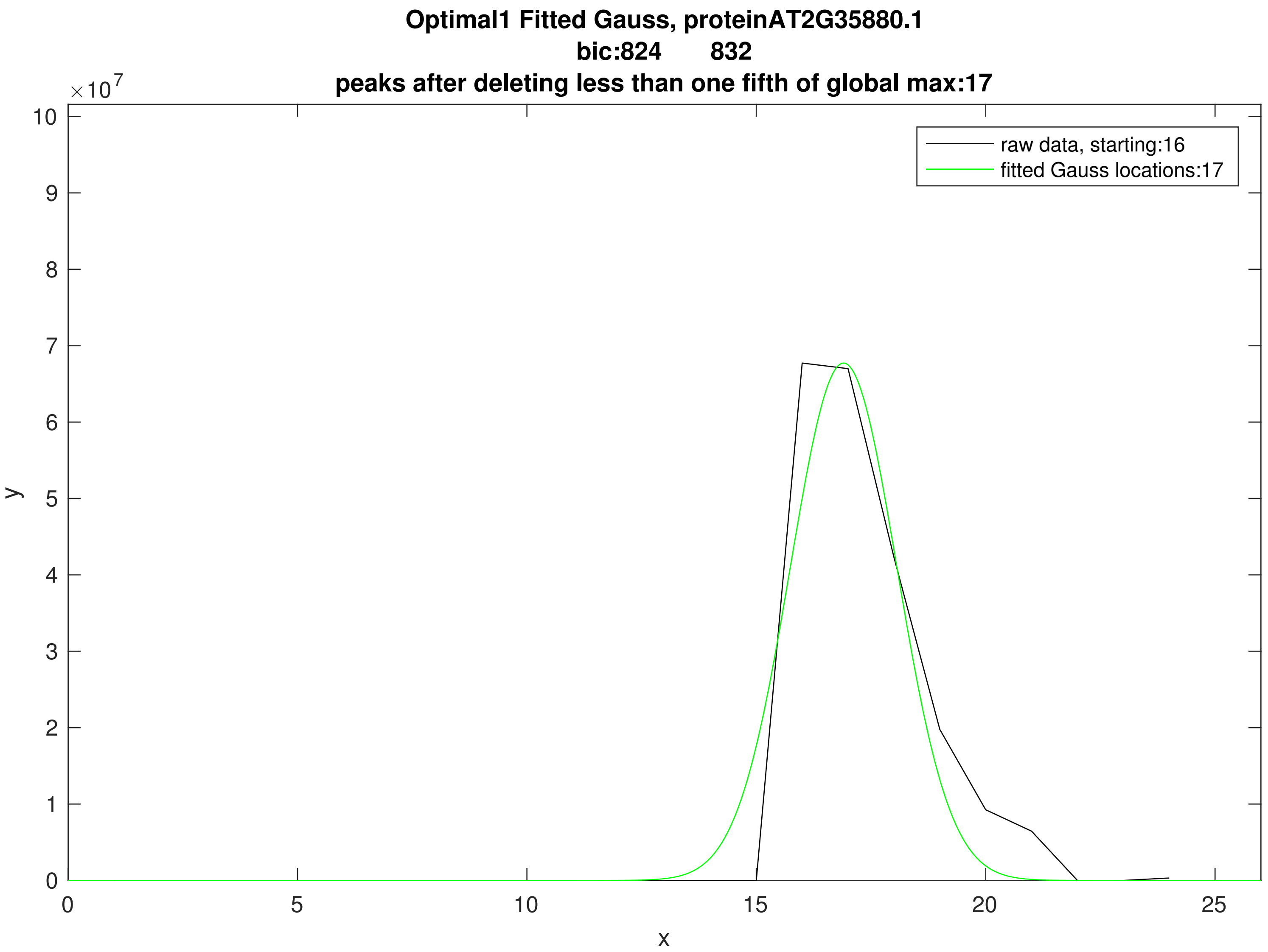

Optimal1 Fitted Gauss, proteinAT2G35880.1
bic:824 832
peaks after deleting less than one fifth of global max:17
×10^7
10
9
8
7
6
5
4
3
2
1
0
y
0
5
10
15
20
25
x
raw data, starting:16
fitted Gauss locations:17

**Optimal1 Fitted Gauss, proteinAT2G36130.1**
**bic:836**
**peaks after deleting less than one fifth of global max:20**

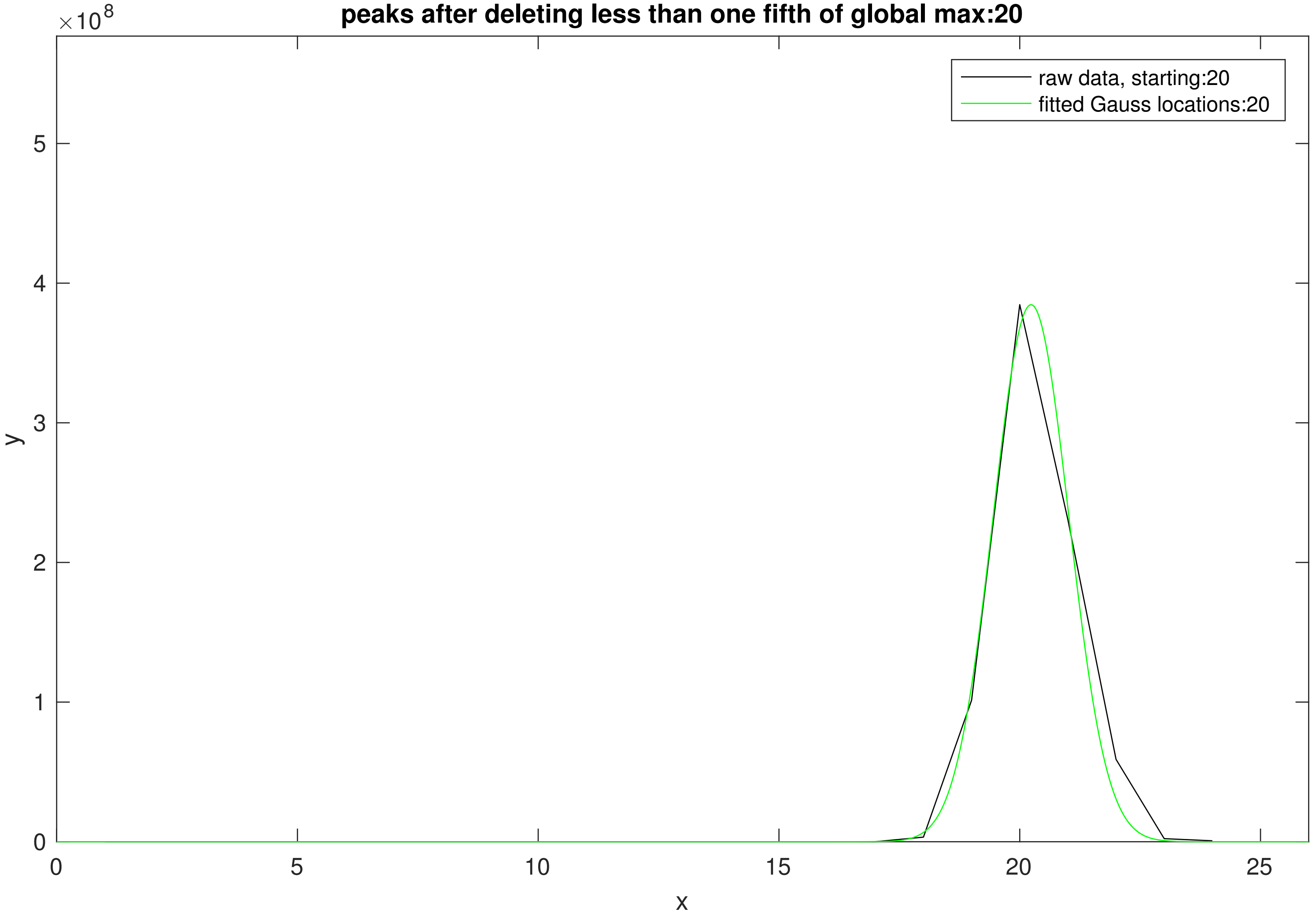

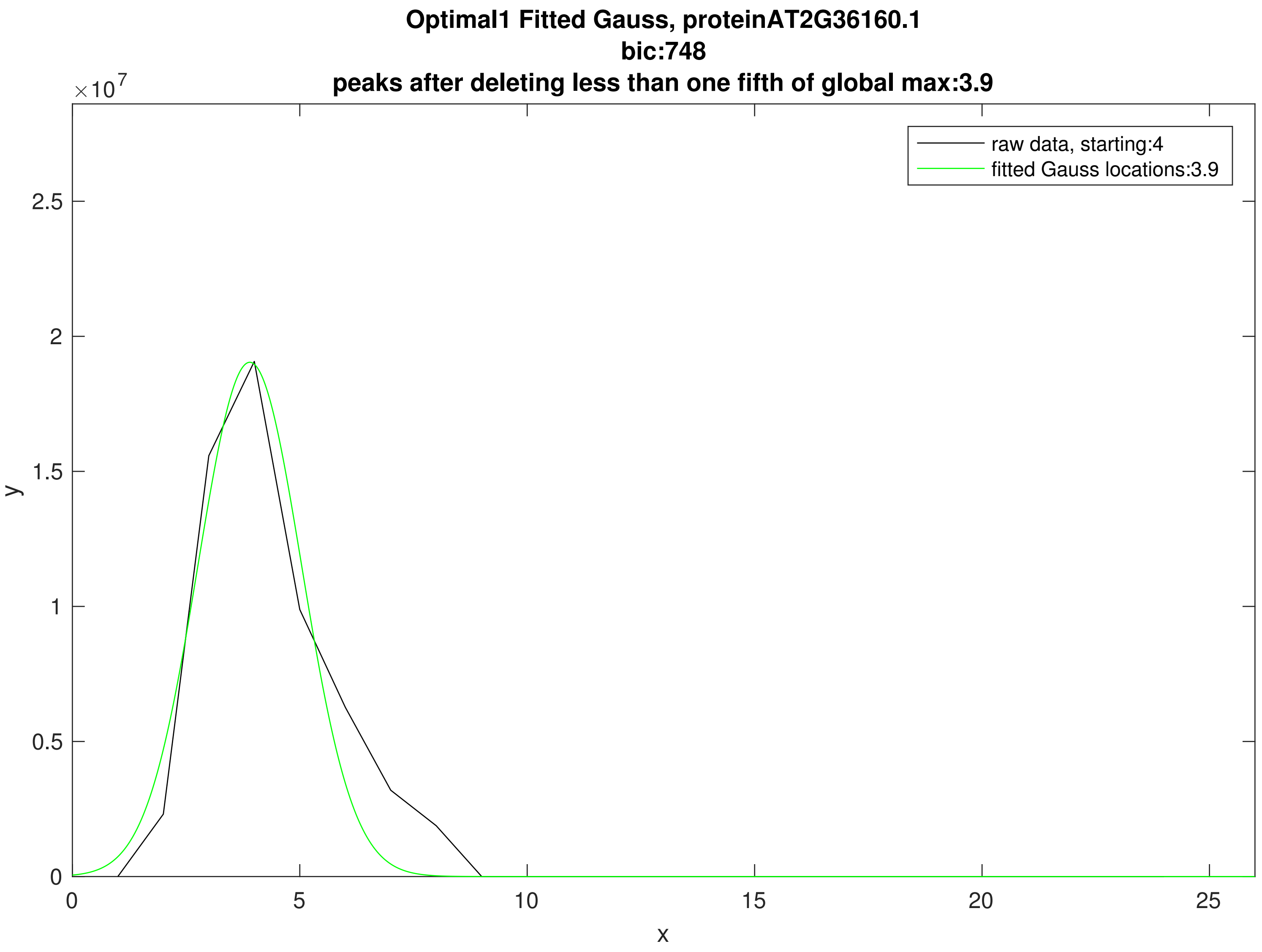
Optimal1 Fitted Gauss, proteinAT2G36160.1
bic:748
peaks after deleting less than one fifth of global max:3.9
×10^7
raw data, starting:4
fitted Gauss locations:3.9
2.5
2
1.5
1
0.5
0
y
0
5
10
15
20
25
x

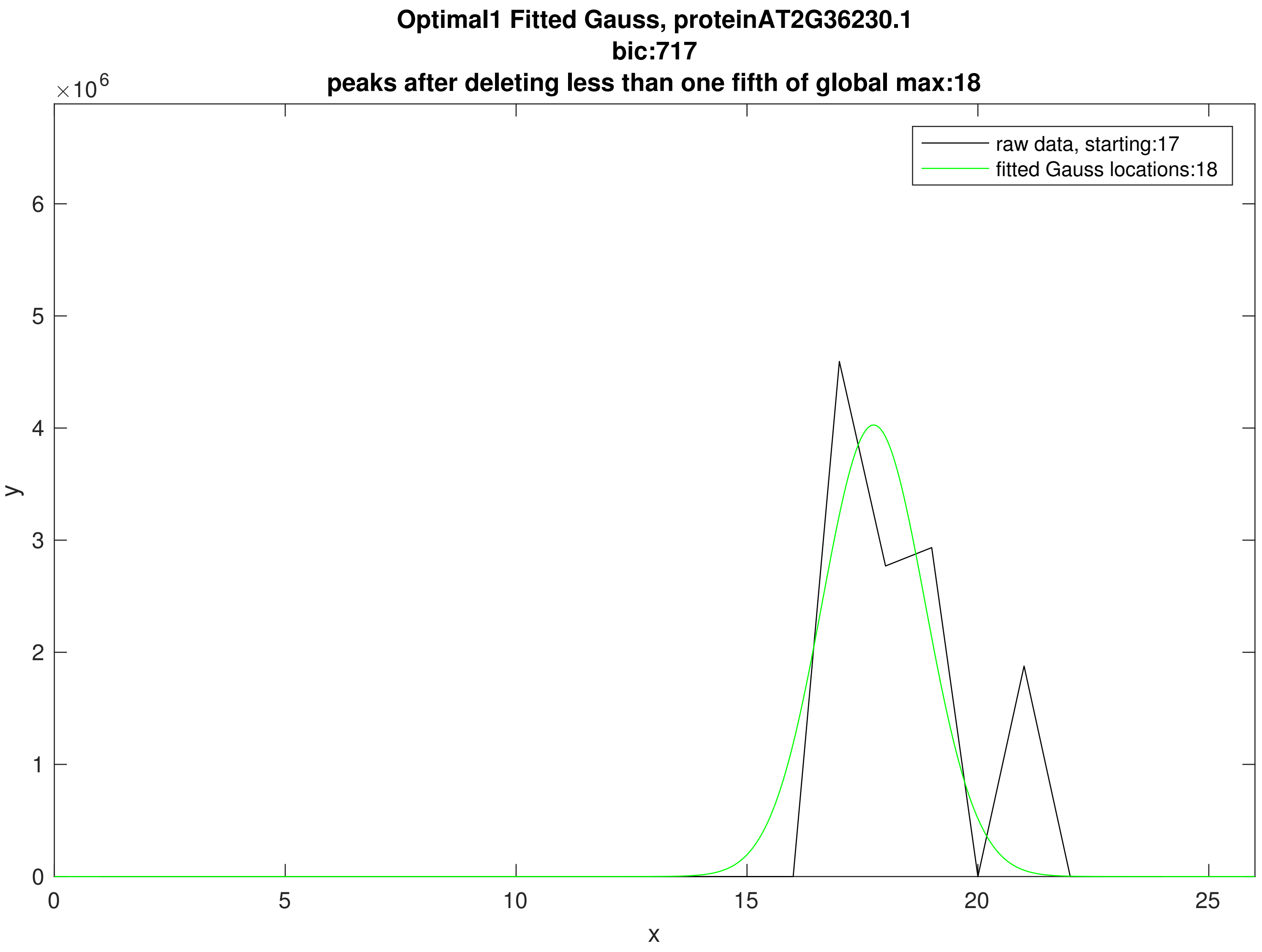
Optimal1 Fitted Gauss, proteinAT2G36230.1
bic:717
peaks after deleting less than one fifth of global max:18
×10^6
raw data, starting:17
fitted Gauss locations:18
y
x
0
1
2
3
4
5
6
0
5
10
15
20
25

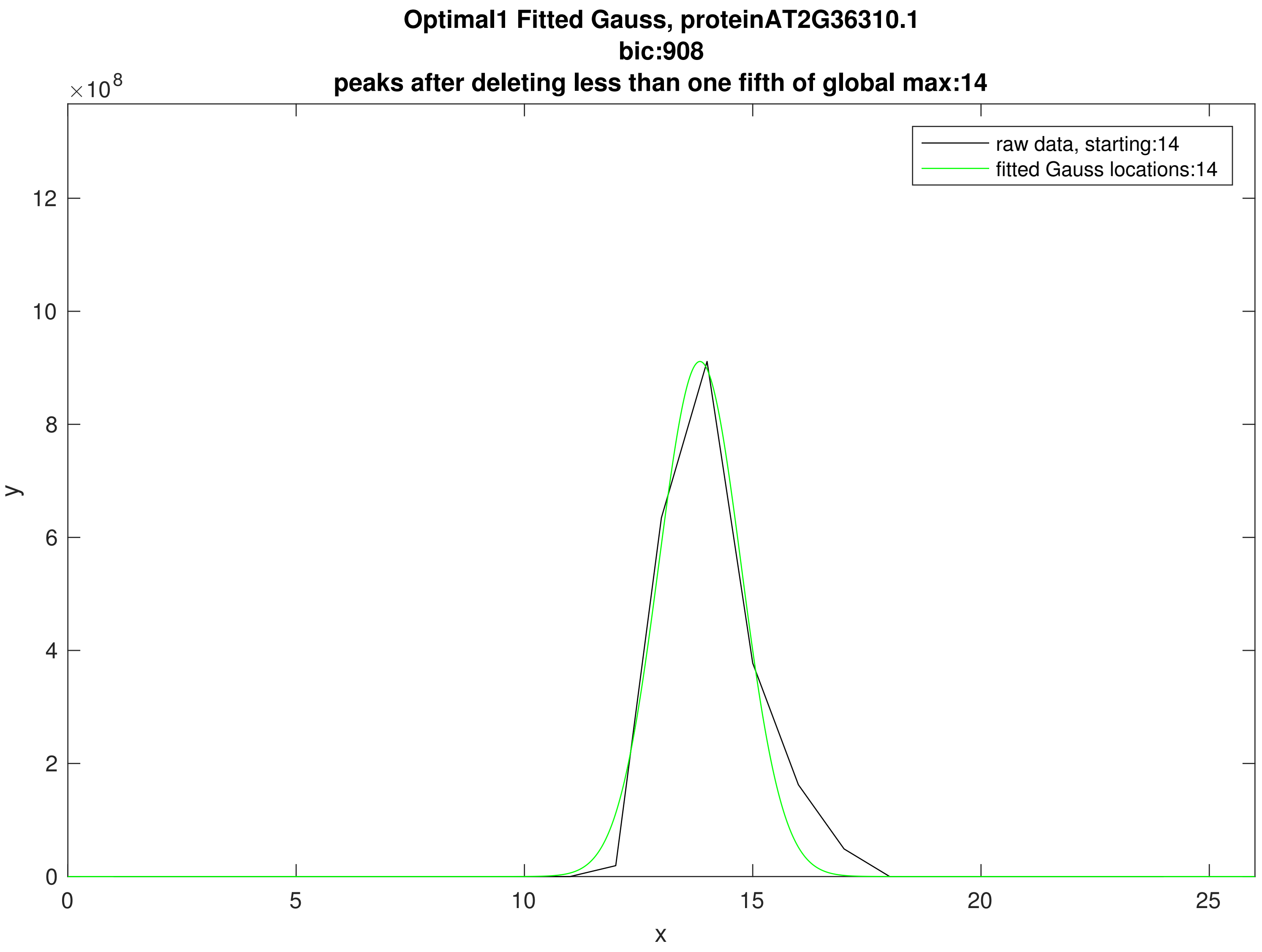
Optimal1 Fitted Gauss, proteinAT2G36310.1
bic:908
peaks after deleting less than one fifth of global max:14
×10^8
raw data, starting:14
fitted Gauss locations:14
y
x
0
2
4
6
8
10
12
0
5
10
15
20
25

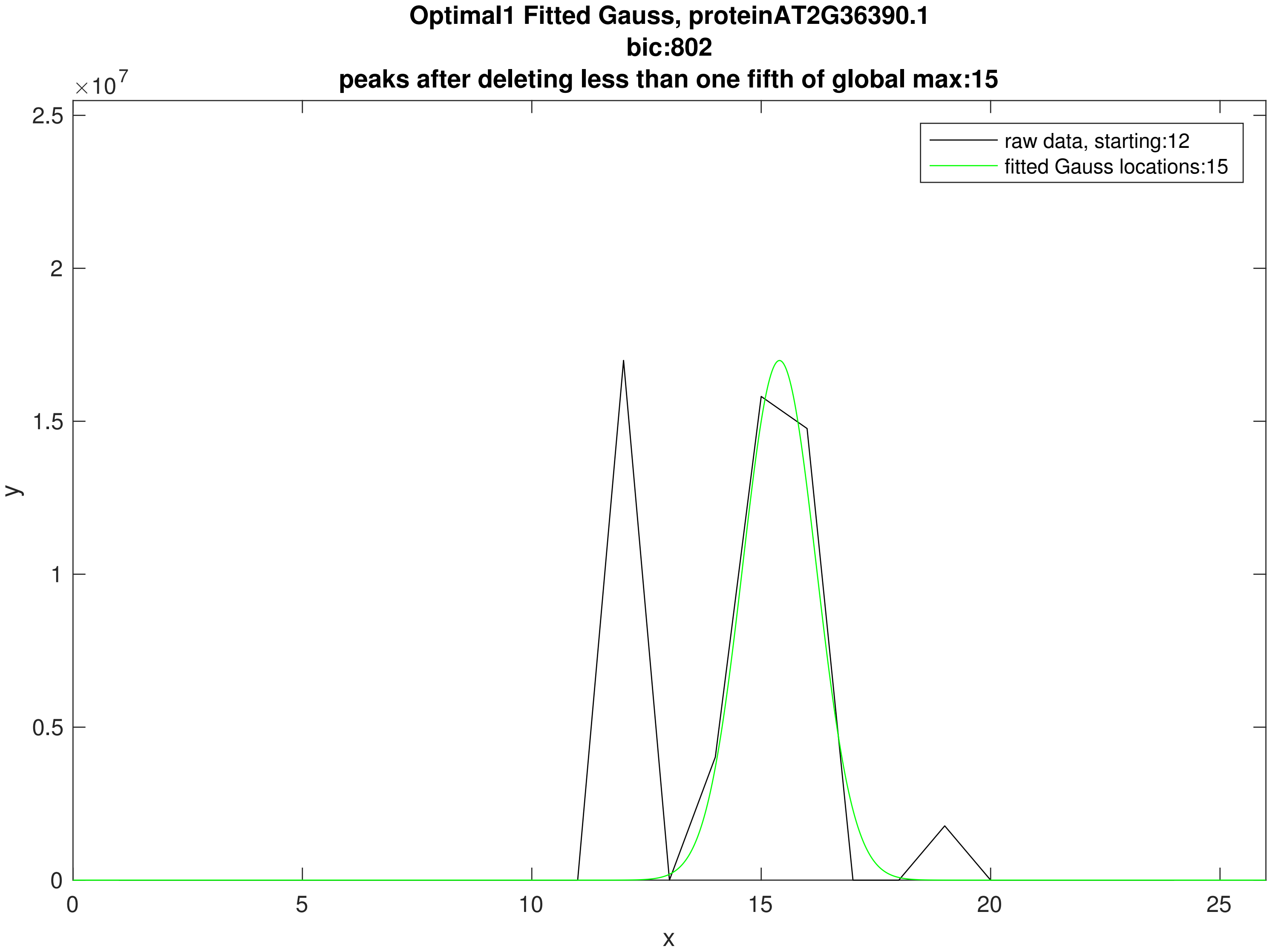
Optimal1 Fitted Gauss, proteinAT2G36390.1
bic:802
peaks after deleting less than one fifth of global max:15
raw data, starting:12
fitted Gauss locations:15
$\times 10^7$
y
x
0
0.5
1
1.5
2
2.5
5
10
15
20
25

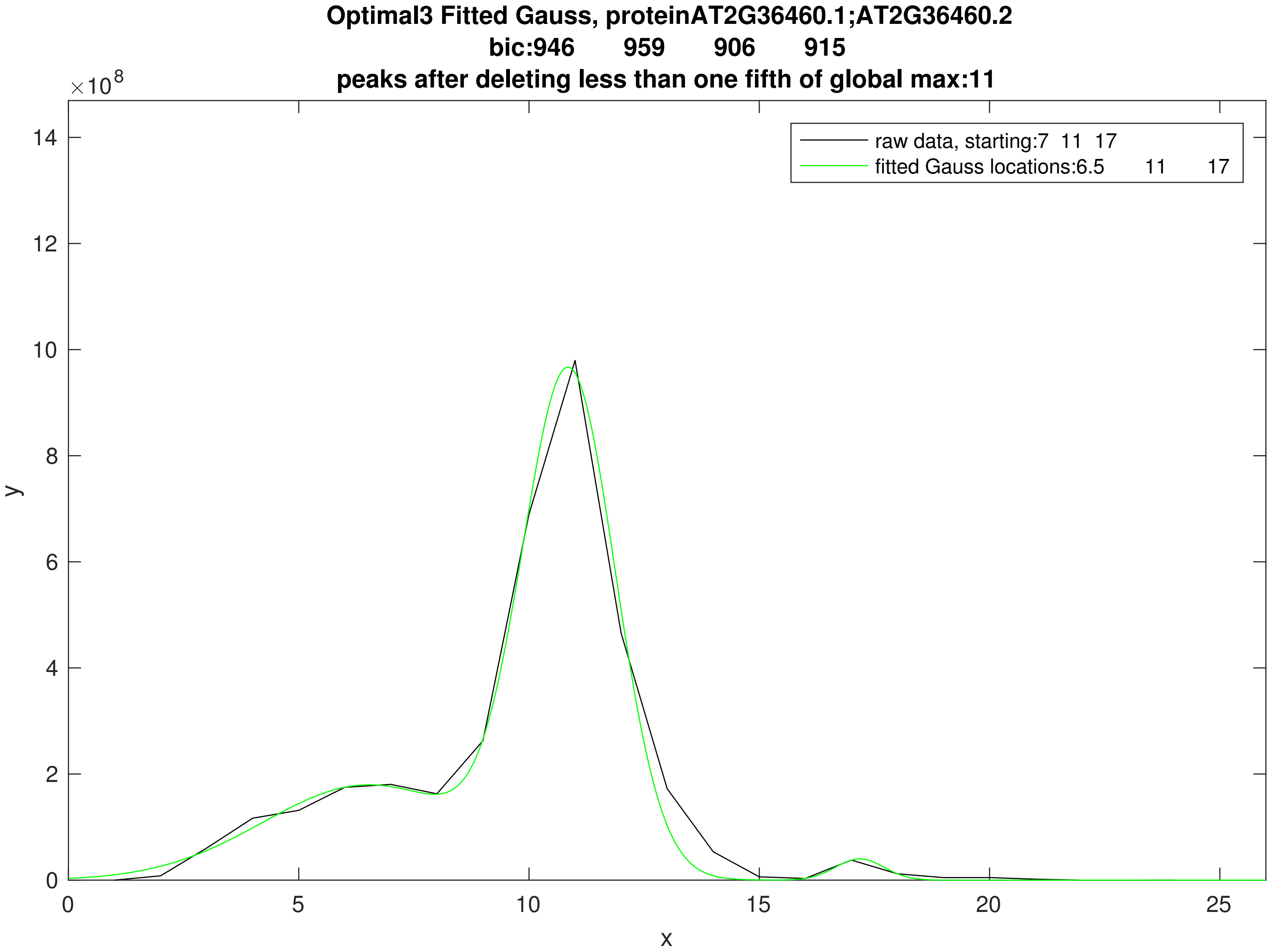
Optimal3 Fitted Gauss, proteinAT2G36460.1;AT2G36460.2
bic:946 959 906 915
peaks after deleting less than one fifth of global max:11
raw data, starting:7 11 17
fitted Gauss locations:6.5 11 17
×10^8
y
x

**Optimal1 Fitted Gauss, proteinAT2G36530.1**
**bic:1.11e+03  1.12e+03  1.13e+03**
**peaks after deleting less than one fifth of global max:14**

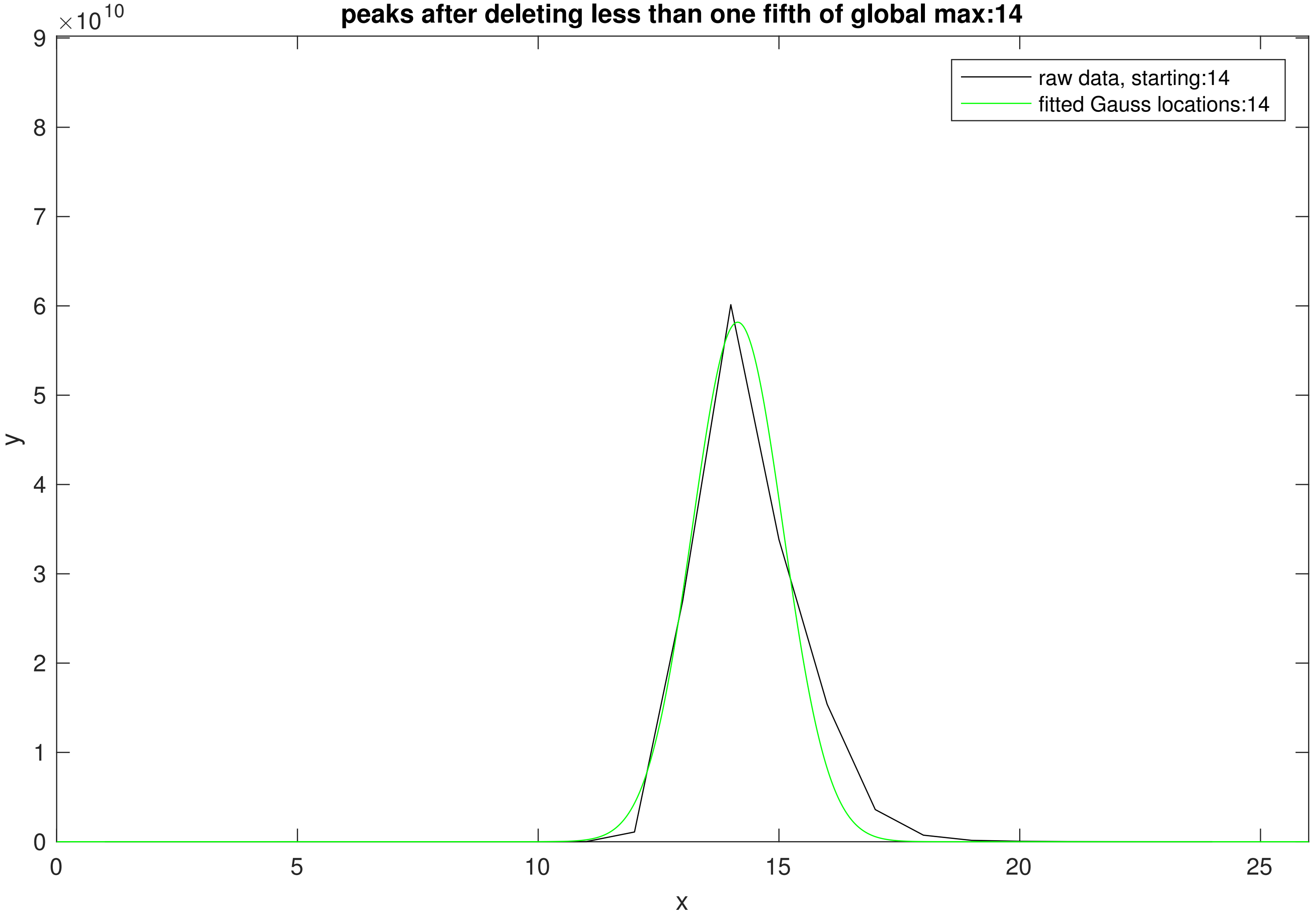

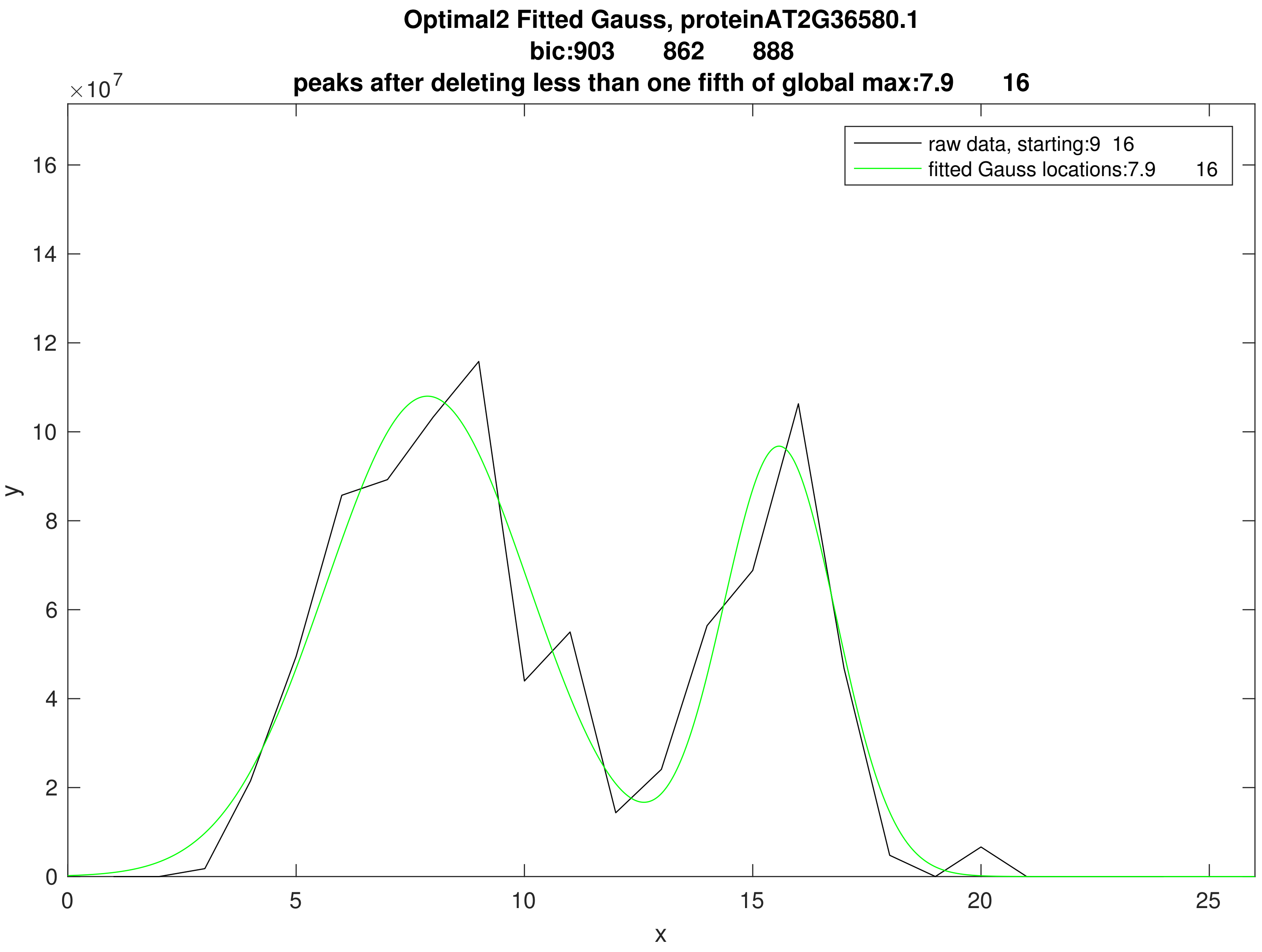

Optimal2 Fitted Gauss, proteinAT2G36580.1
bic:903 862 888
peaks after deleting less than one fifth of global max:7.9 16
×10^7
raw data, starting:9 16
fitted Gauss locations:7.9 16
y
x
0
2
4
6
8
10
12
14
16
0
5
10
15
20
25

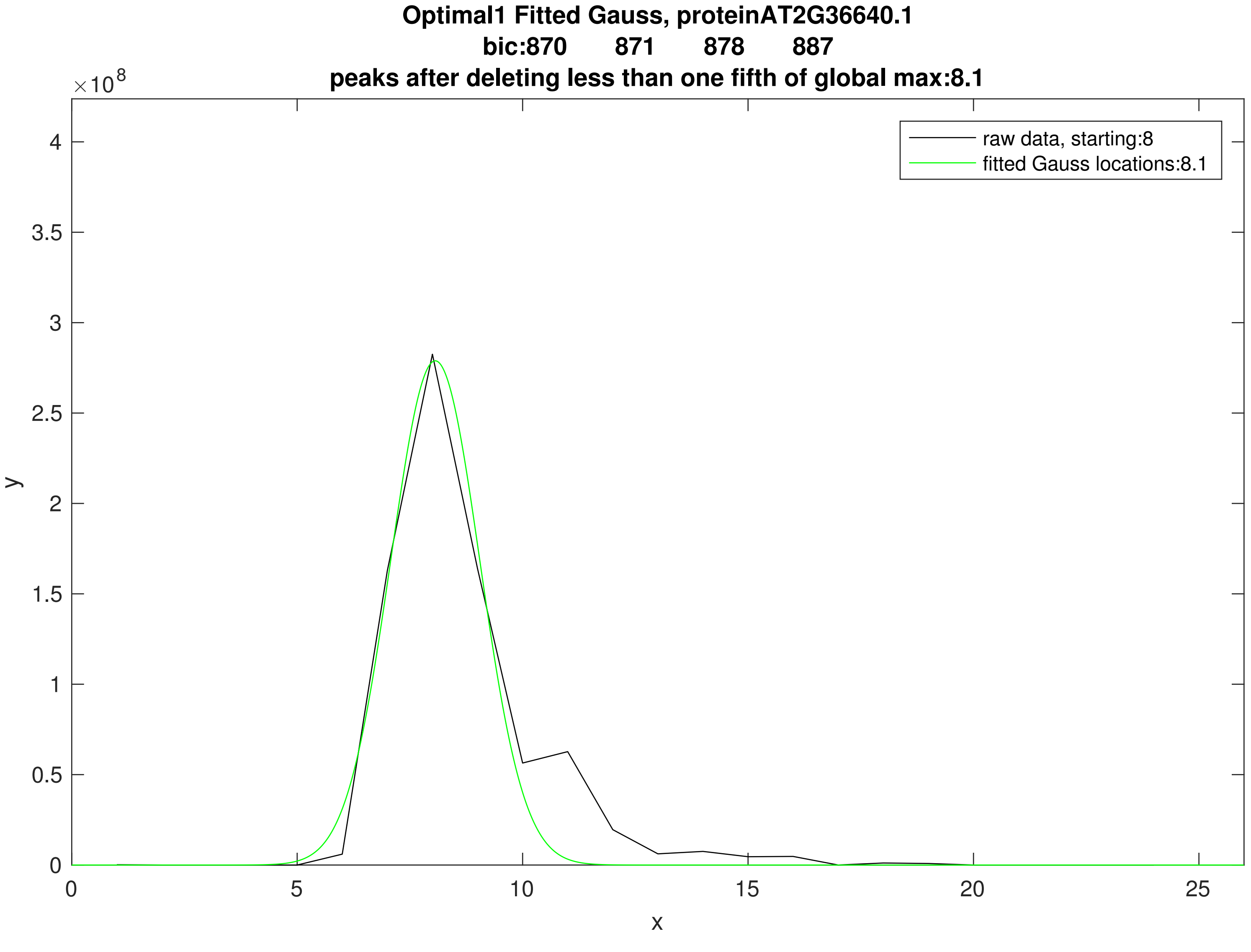

Optimal1 Fitted Gauss, proteinAT2G36640.1
bic:870 871 878 887
peaks after deleting less than one fifth of global max:8.1
×10^8
4
3.5
3
2.5
2
1.5
1
0.5
0
y
0
5
10
15
20
25
x
raw data, starting:8
fitted Gauss locations:8.1

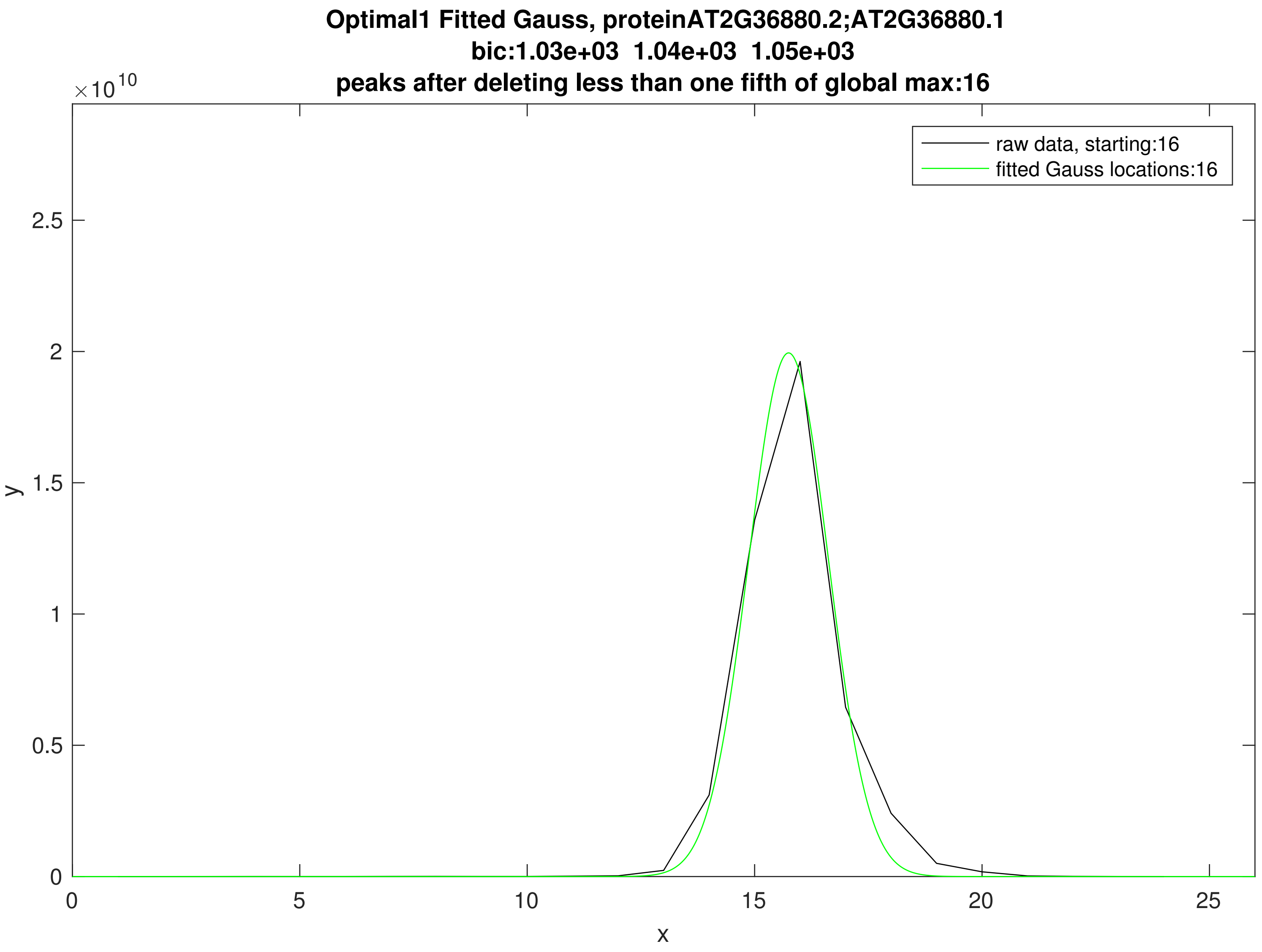
Optimal1 Fitted Gauss, proteinAT2G36880.2;AT2G36880.1
bic:1.03e+03 1.04e+03 1.05e+03
peaks after deleting less than one fifth of global max:16
×10^10
raw data, starting:16
fitted Gauss locations:16
y
x
0
0.5
1
1.5
2
2.5
0
5
10
15
20
25

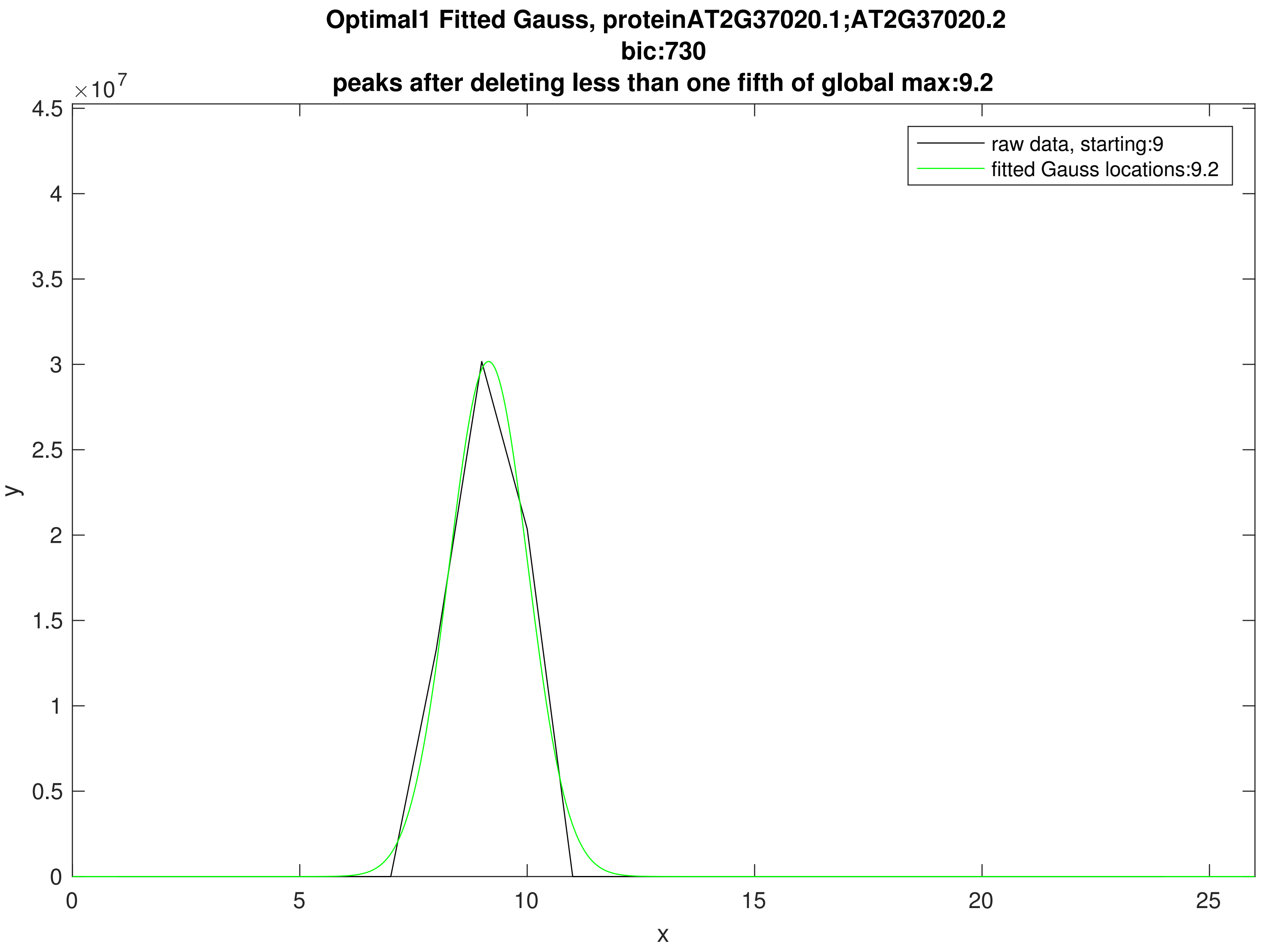
Optimal1 Fitted Gauss, proteinAT2G37020.1;AT2G37020.2
bic:730
peaks after deleting less than one fifth of global max:9.2
×10^7
4.5
4
3.5
3
2.5
2
1.5
1
0.5
0
y
0
5
10
15
20
25
x
raw data, starting:9
fitted Gauss locations:9.2

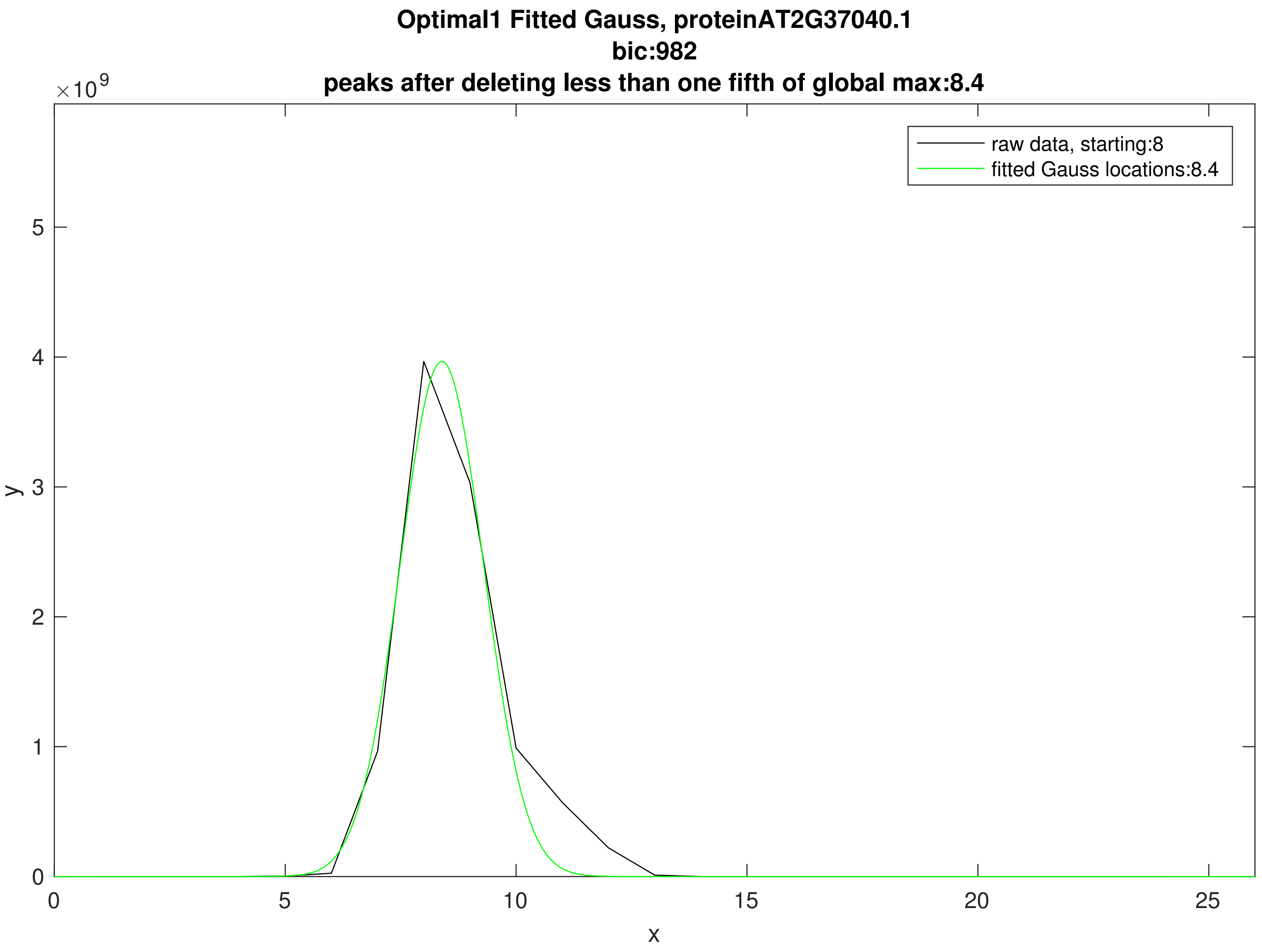
Optimal1 Fitted Gauss, proteinAT2G37040.1
bic:982
peaks after deleting less than one fifth of global max:8.4
×10^9
raw data, starting:8
fitted Gauss locations:8.4
0
1
2
3
4
5
y
0
5
10
15
20
25
x

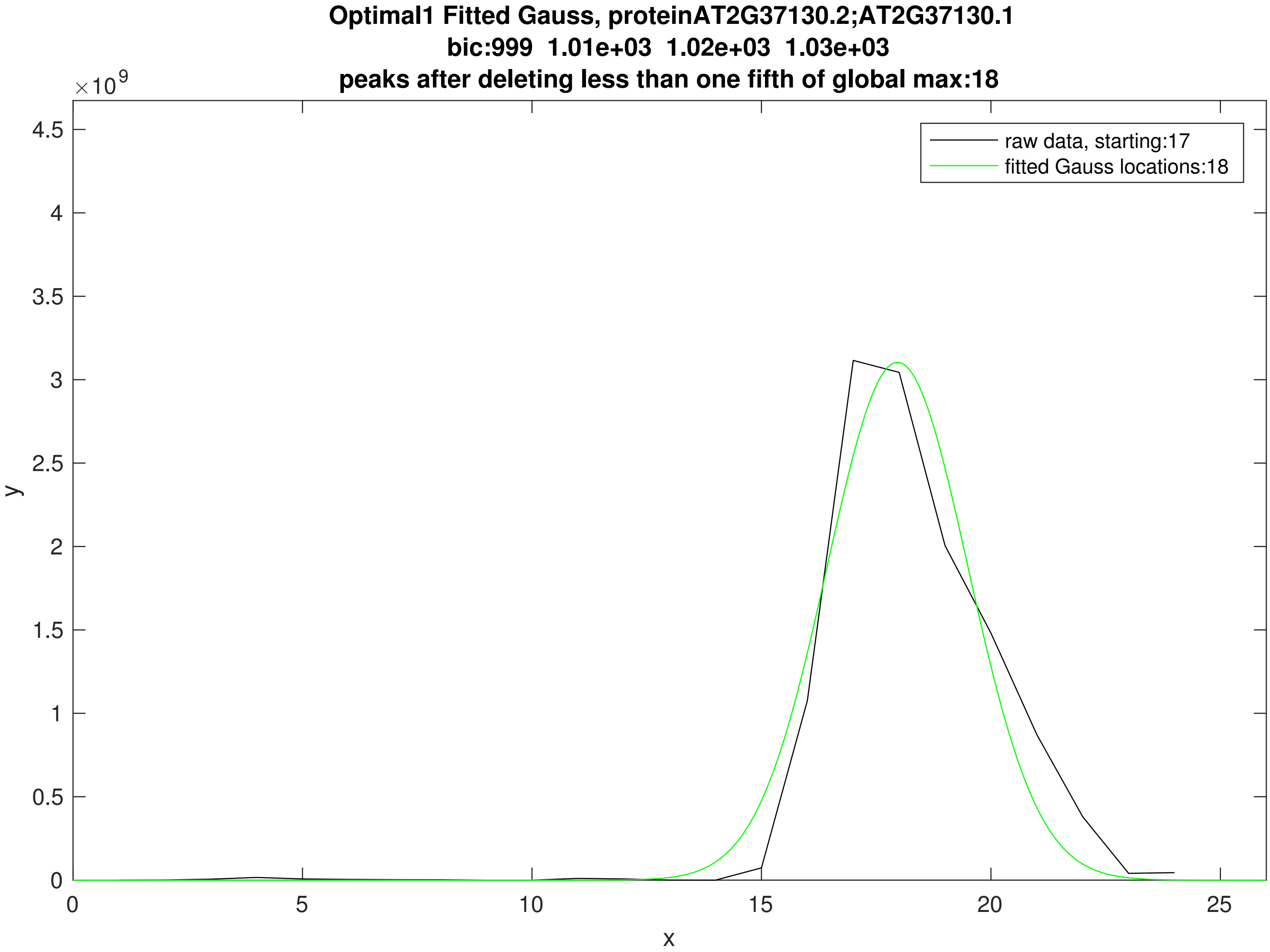
Optimal1 Fitted Gauss, proteinAT2G37130.2;AT2G37130.1
bic:999 1.01e+03 1.02e+03 1.03e+03
peaks after deleting less than one fifth of global max:18
×10^9
4.5
4
3.5
3
2.5
2
1.5
1
0.5
0
y
0
5
10
15
20
25
x
raw data, starting:17
fitted Gauss locations:18

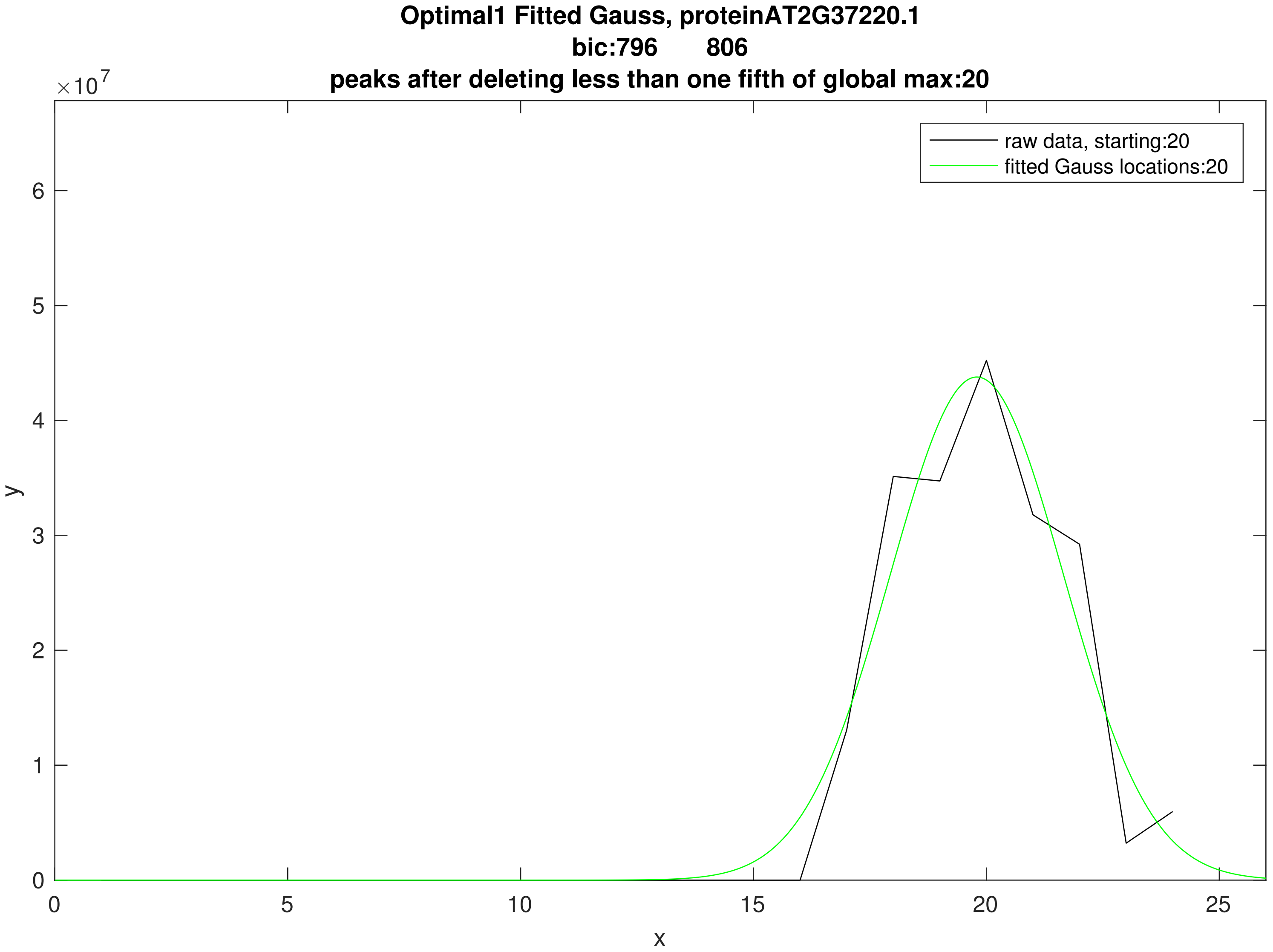

Optimal1 Fitted Gauss, proteinAT2G37220.1
bic:796 806
peaks after deleting less than one fifth of global max:20
×10^7
raw data, starting:20
fitted Gauss locations:20
6
5
4
3
2
1
0
y
0
5
10
15
20
25
x

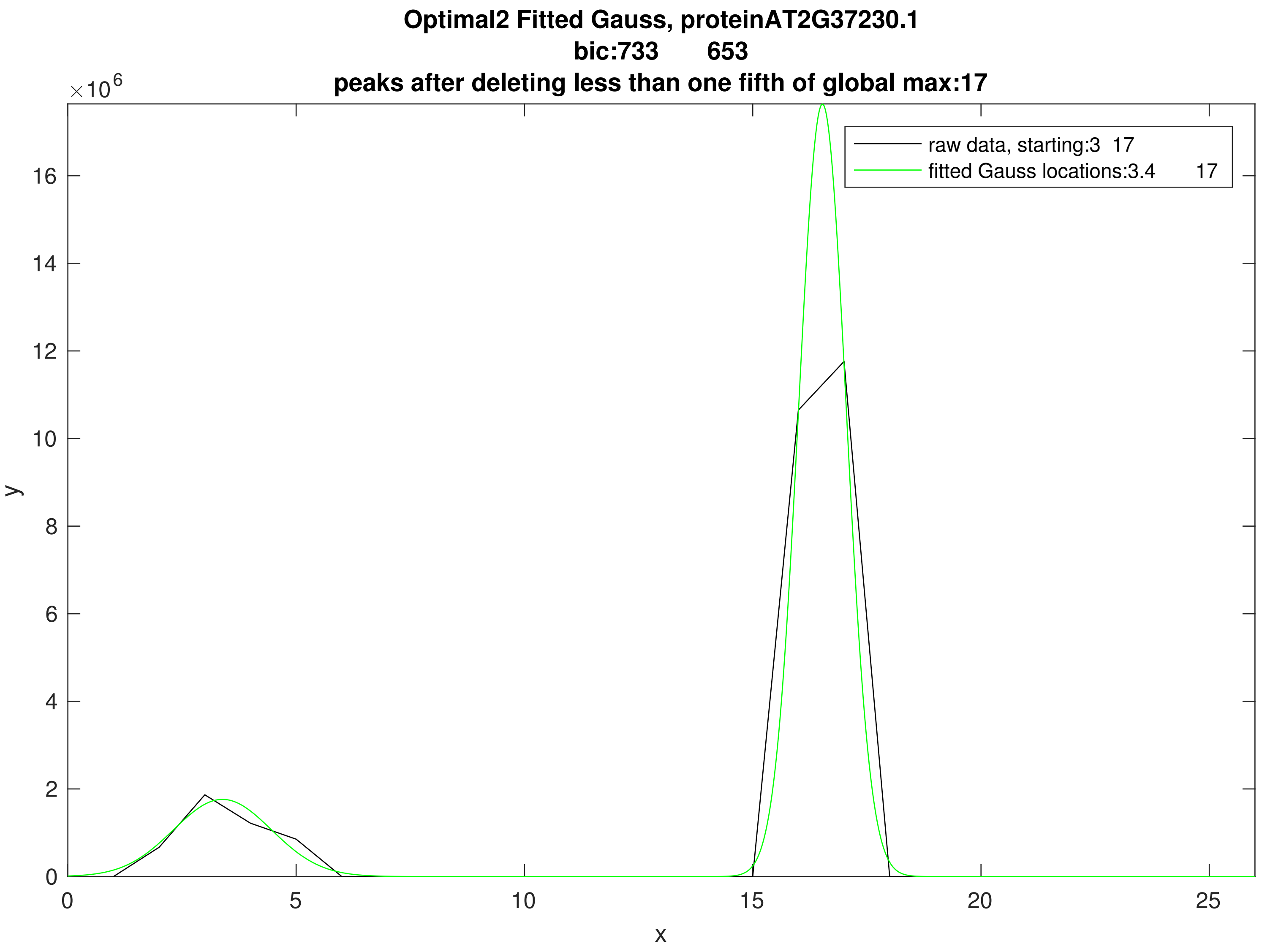
Optimal2 Fitted Gauss, proteinAT2G37230.1
bic:733 653
peaks after deleting less than one fifth of global max:17
×10^6
raw data, starting:3 17
fitted Gauss locations:3.4 17
y
x
0
2
4
6
8
10
12
14
16
0
5
10
15
20
25

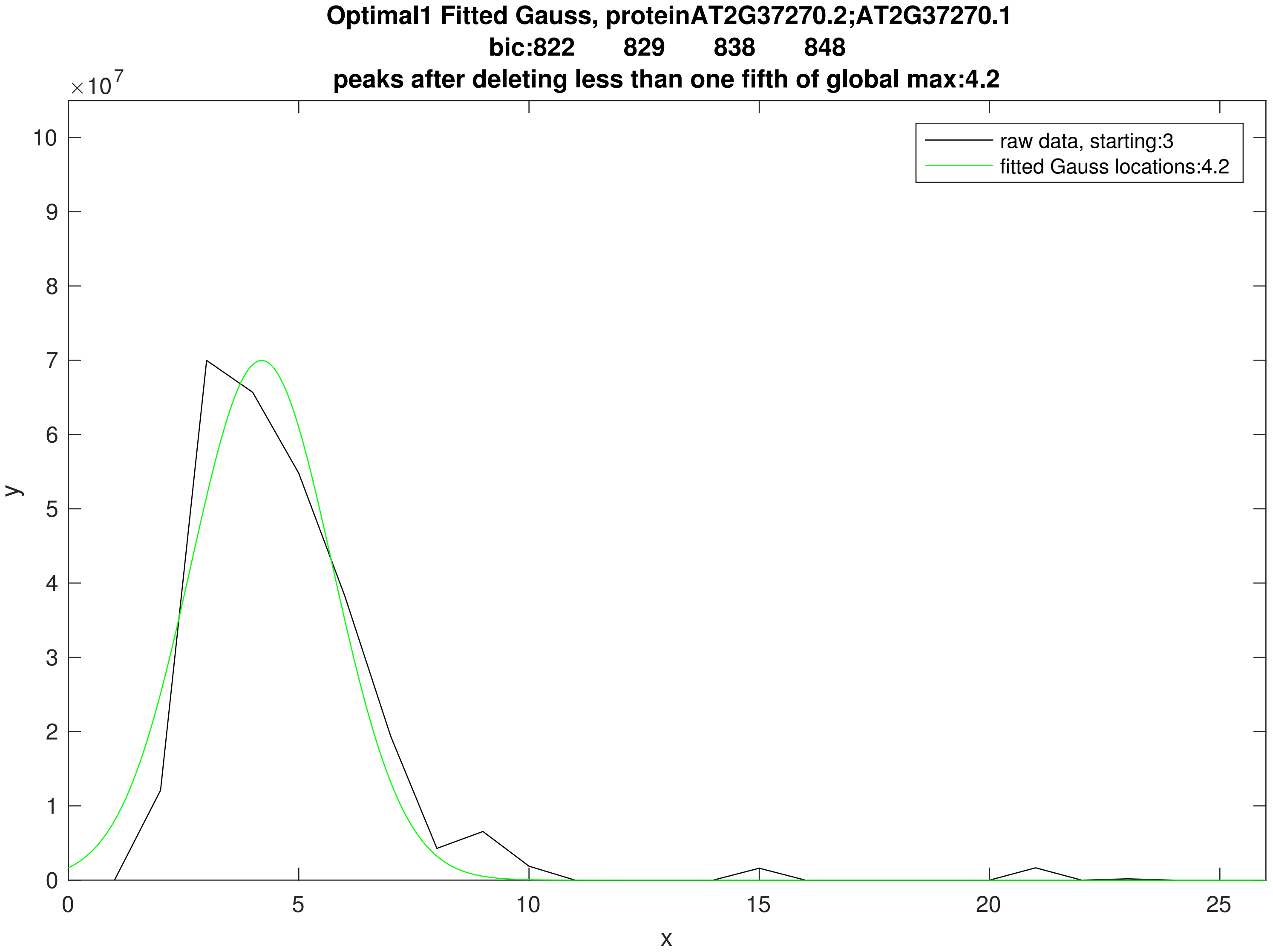

Optimal1 Fitted Gauss, proteinAT2G37270.2;AT2G37270.1
bic:822 829 838 848
peaks after deleting less than one fifth of global max:4.2
×10^7
raw data, starting:3
fitted Gauss locations:4.2
y
x
0
1
2
3
4
5
6
7
8
9
10
15
20
25

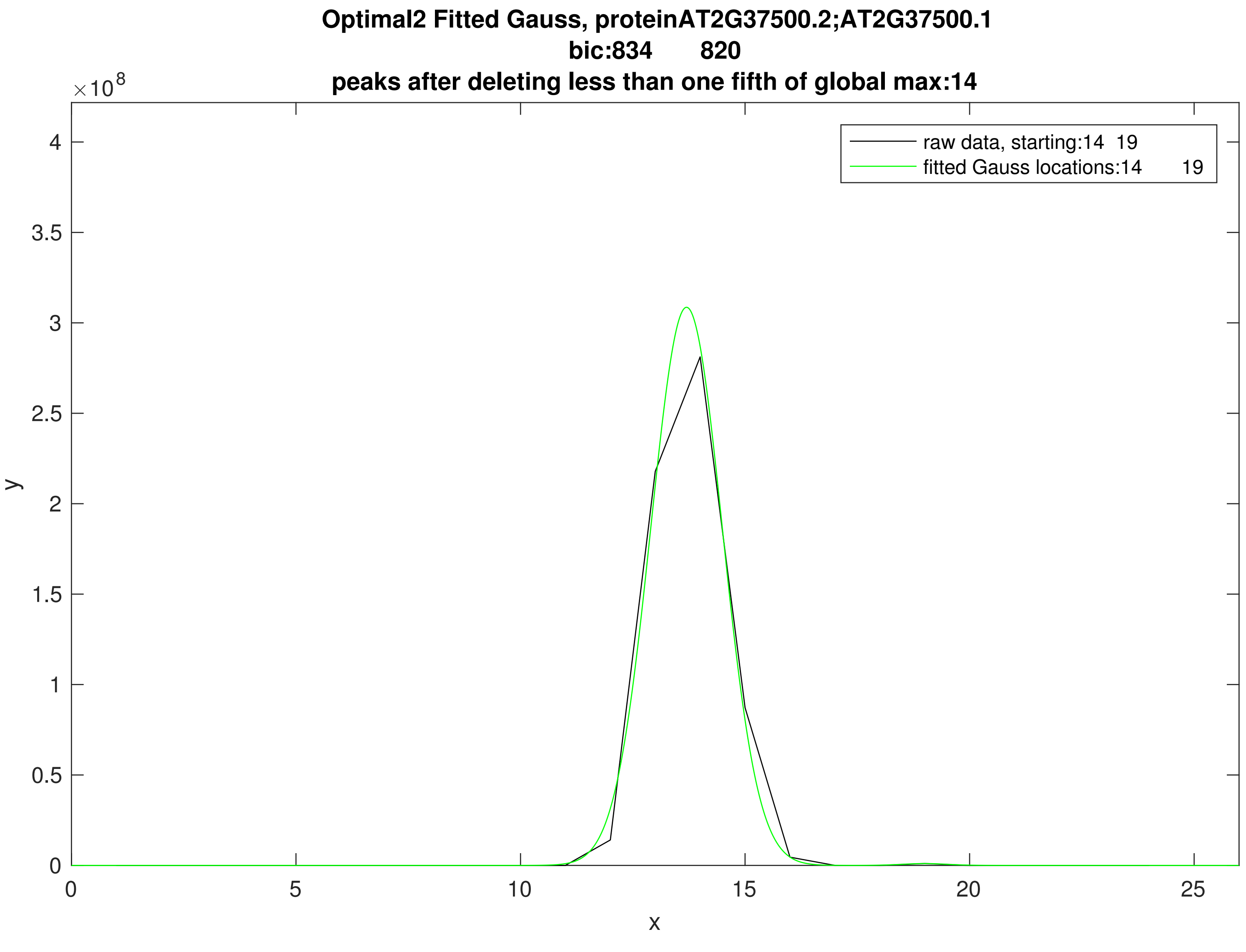

Optimal2 Fitted Gauss, proteinAT2G37500.2;AT2G37500.1
bic:834 820
peaks after deleting less than one fifth of global max:14
×10^8
raw data, starting:14 19
fitted Gauss locations:14 19
4
3.5
3
2.5
2
1.5
1
0.5
0
y
0
5
10
15
20
25
x

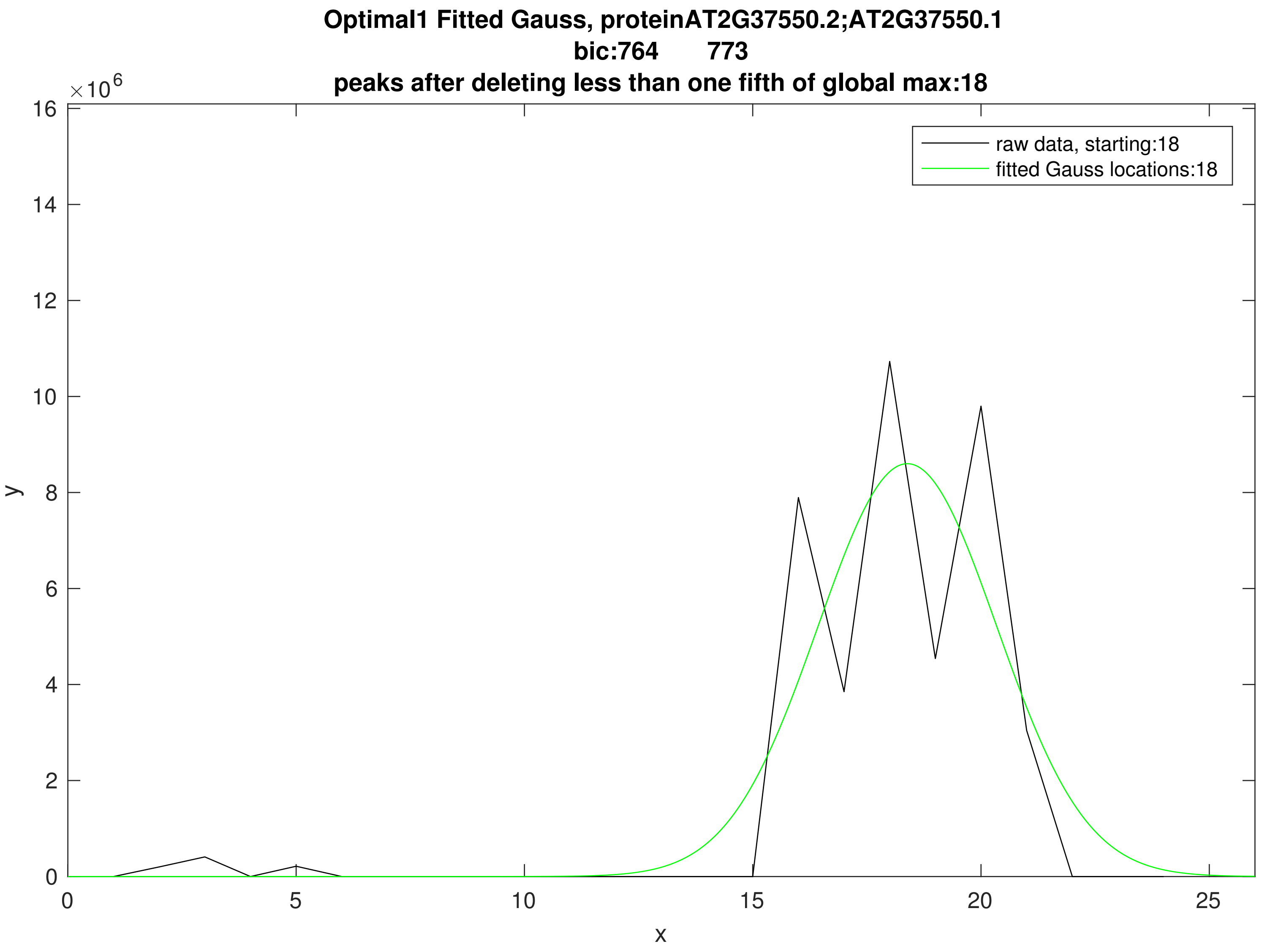
Optimal1 Fitted Gauss, proteinAT2G37550.2;AT2G37550.1
bic:764 773
peaks after deleting less than one fifth of global max:18
×10^6
16
14
12
10
8
6
4
2
0
y
0
5
10
15
20
25
x
raw data, starting:18
fitted Gauss locations:18

Optimal1 Fitted Gauss, proteinAT2G37660.1
bic:891 897
peaks after deleting less than one fifth of global max:18

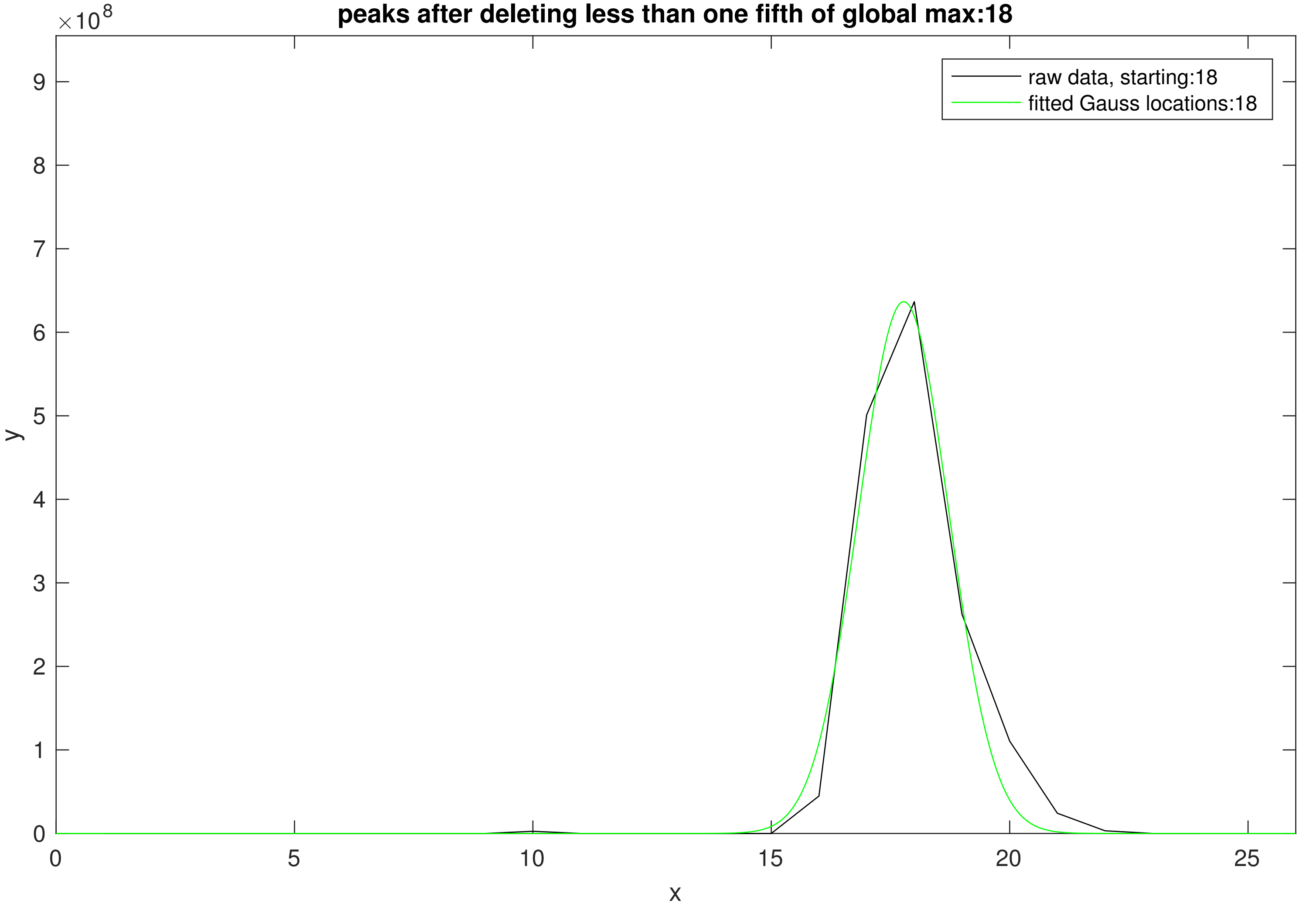

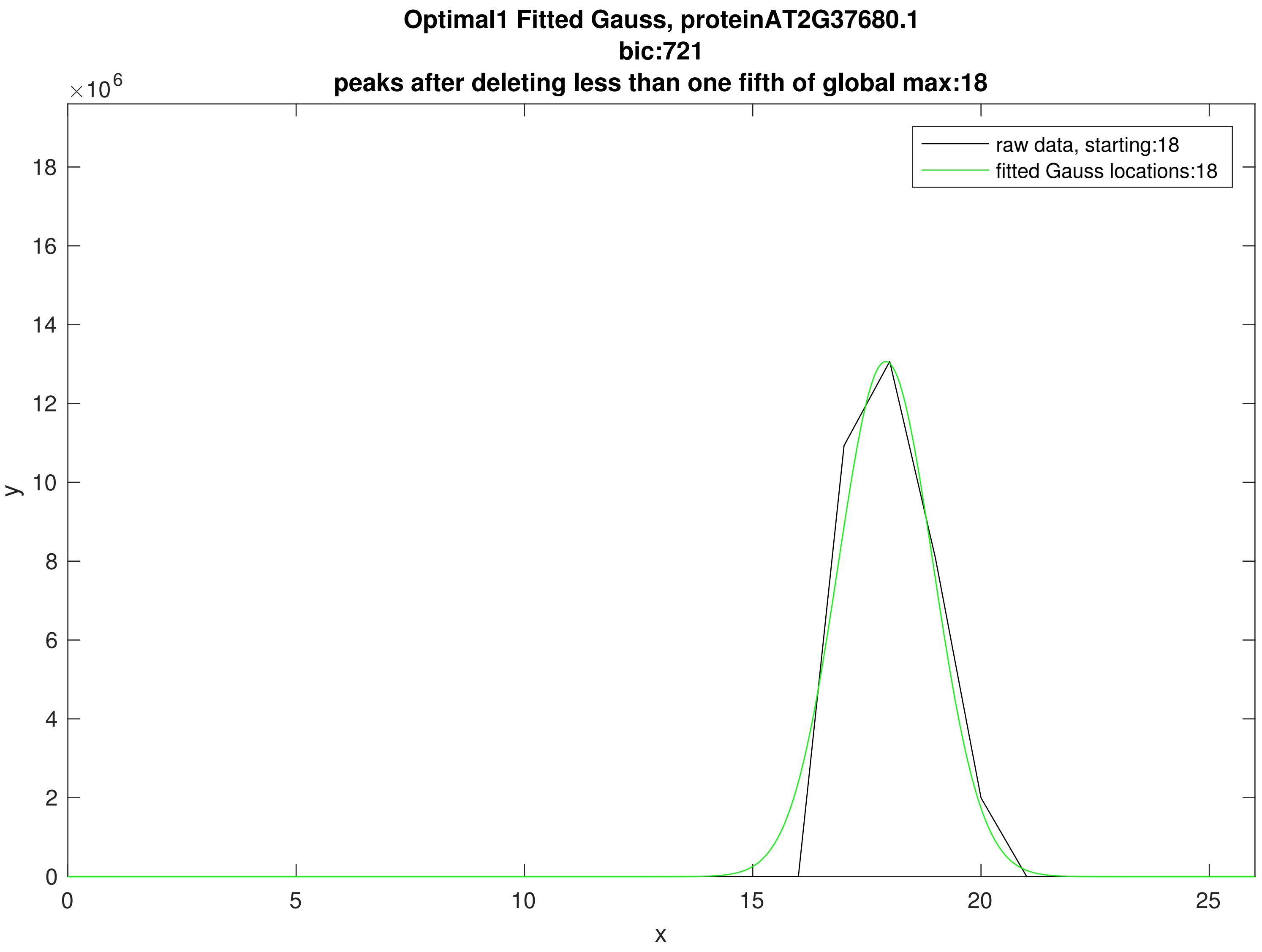

Optimal1 Fitted Gauss, proteinAT2G37680.1
bic:721
peaks after deleting less than one fifth of global max:18
×10^6
raw data, starting:18
fitted Gauss locations:18
y
x
0
2
4
6
8
10
12
14
16
18
0
5
10
15
20
25

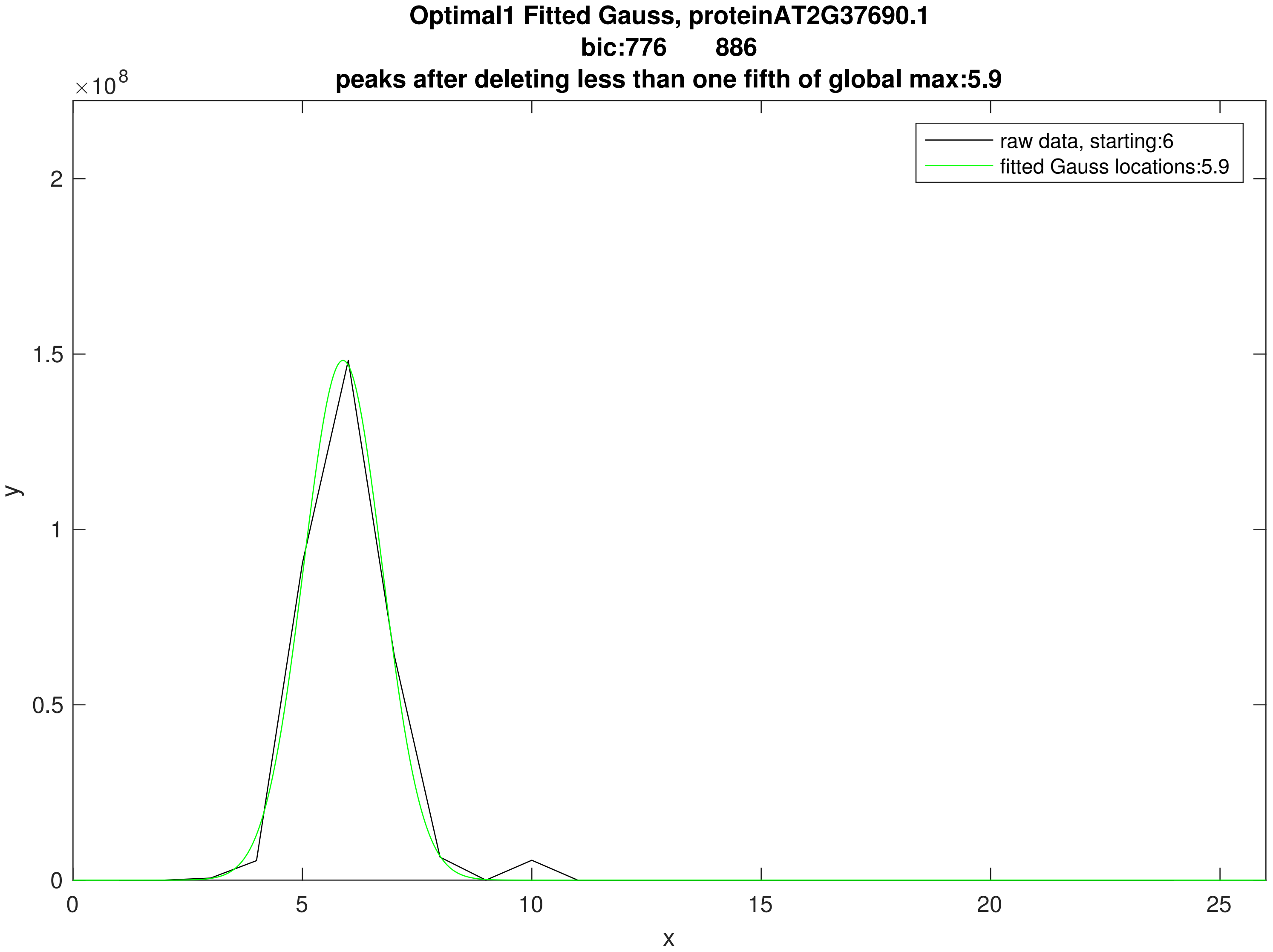

Optimal1 Fitted Gauss, proteinAT2G37690.1
bic:776 886
peaks after deleting less than one fifth of global max:5.9
×10^8
raw data, starting:6
fitted Gauss locations:5.9
y
x
0
0.5
1
1.5
2
5
10
15
20
25

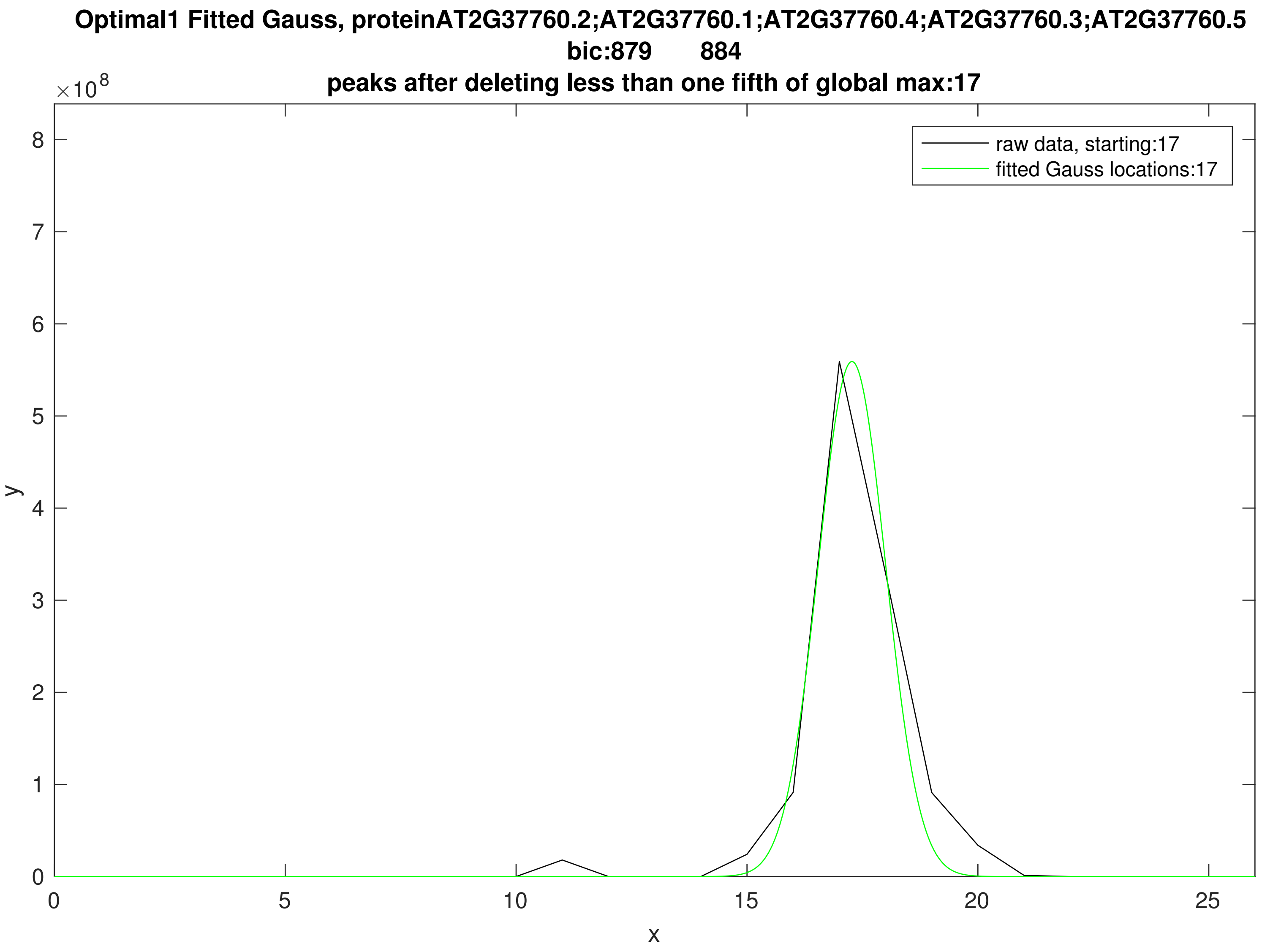
Optimal1 Fitted Gauss, proteinAT2G37760.2;AT2G37760.1;AT2G37760.4;AT2G37760.3;AT2G37760.5
bic:879 884
peaks after deleting less than one fifth of global max:17
×10^8
raw data, starting:17
fitted Gauss locations:17
y
x
0
1
2
3
4
5
6
7
8
0
5
10
15
20
25

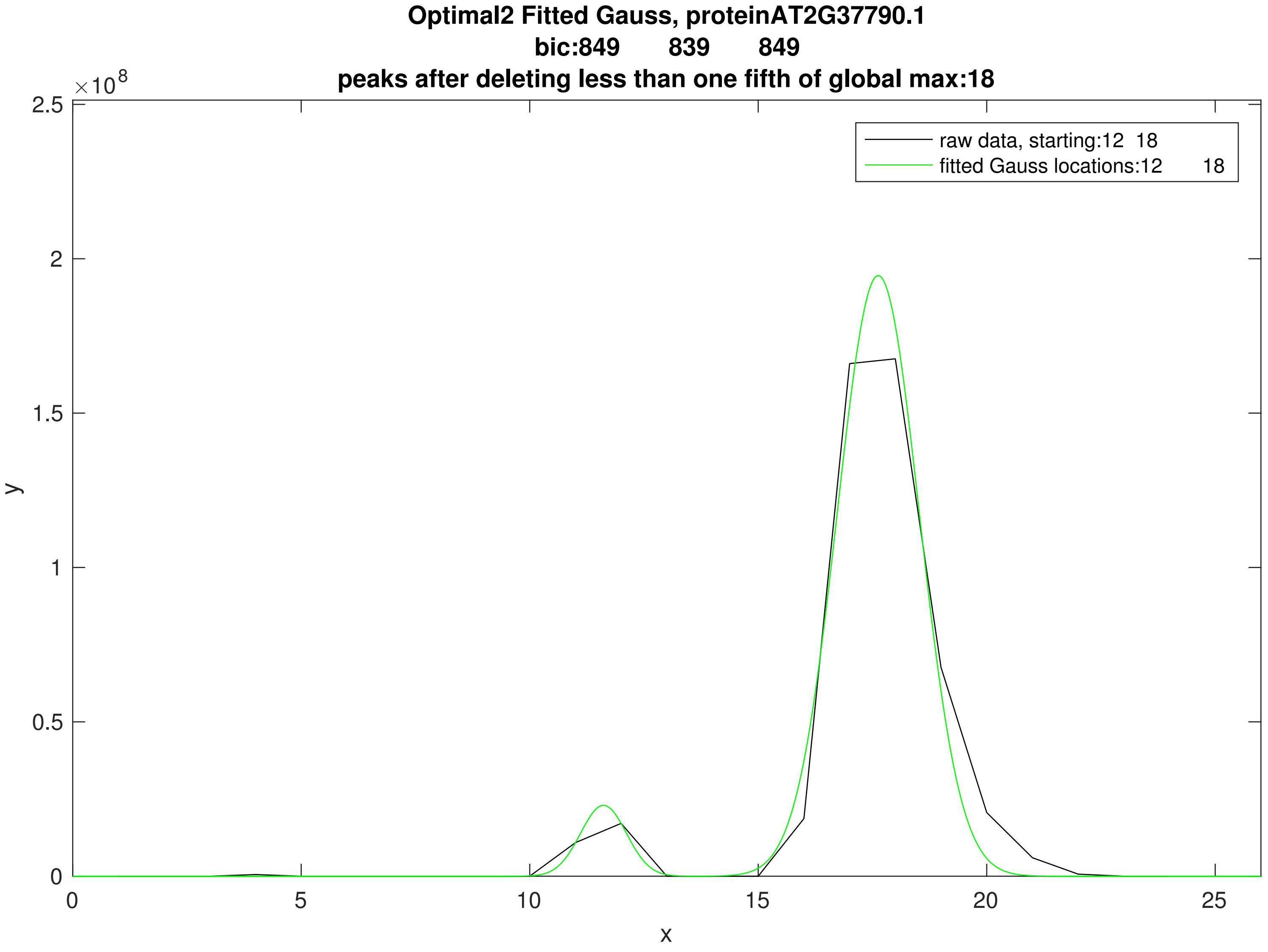
Optimal2 Fitted Gauss, proteinAT2G37790.1
bic:849 839 849
peaks after deleting less than one fifth of global max:18
×10^8
2.5
2
1.5
1
0.5
0
y
0
5
10
15
20
25
x
raw data, starting:12 18
fitted Gauss locations:12 18

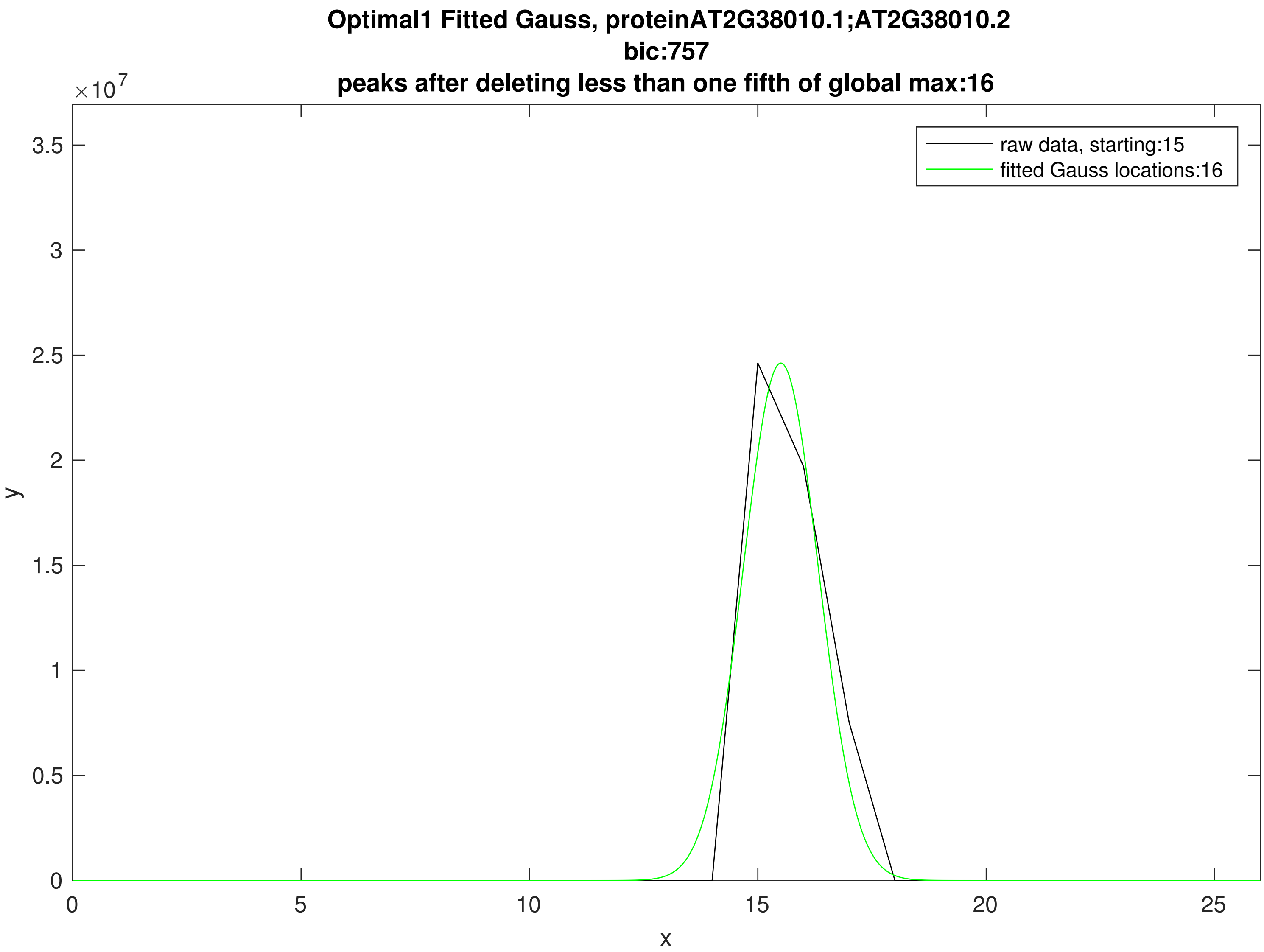

Optimal1 Fitted Gauss, proteinAT2G38010.1;AT2G38010.2
bic:757
peaks after deleting less than one fifth of global max:16
×10^7
3.5
3
2.5
2
1.5
1
0.5
0
y
0
5
10
15
20
25
x
raw data, starting:15
fitted Gauss locations:16

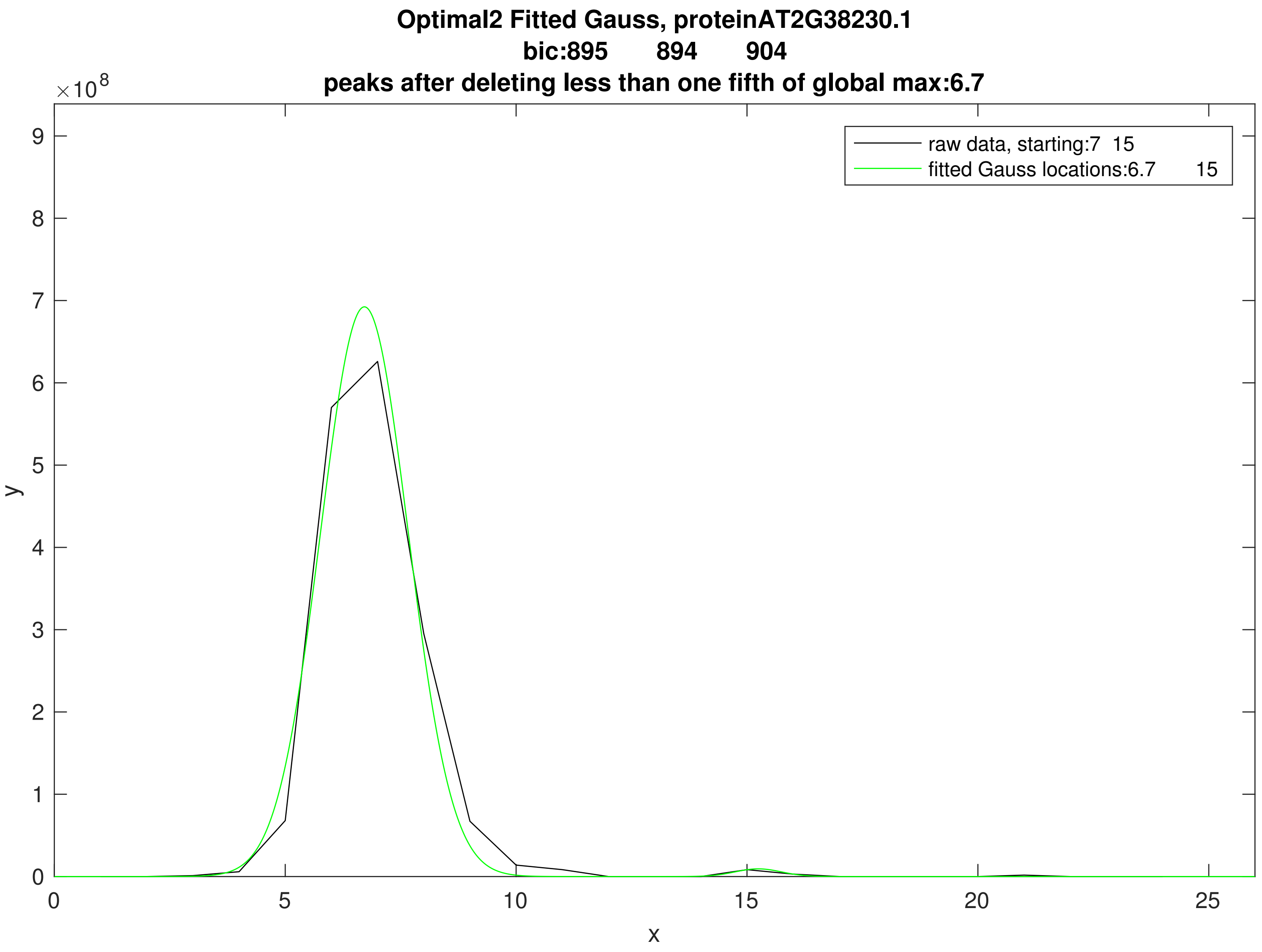
Optimal2 Fitted Gauss, proteinAT2G38230.1
bic:895 894 904
peaks after deleting less than one fifth of global max:6.7
×10^8
raw data, starting:7 15
fitted Gauss locations:6.7 15
y
x
0
1
2
3
4
5
6
7
8
9
0
5
10
15
20
25

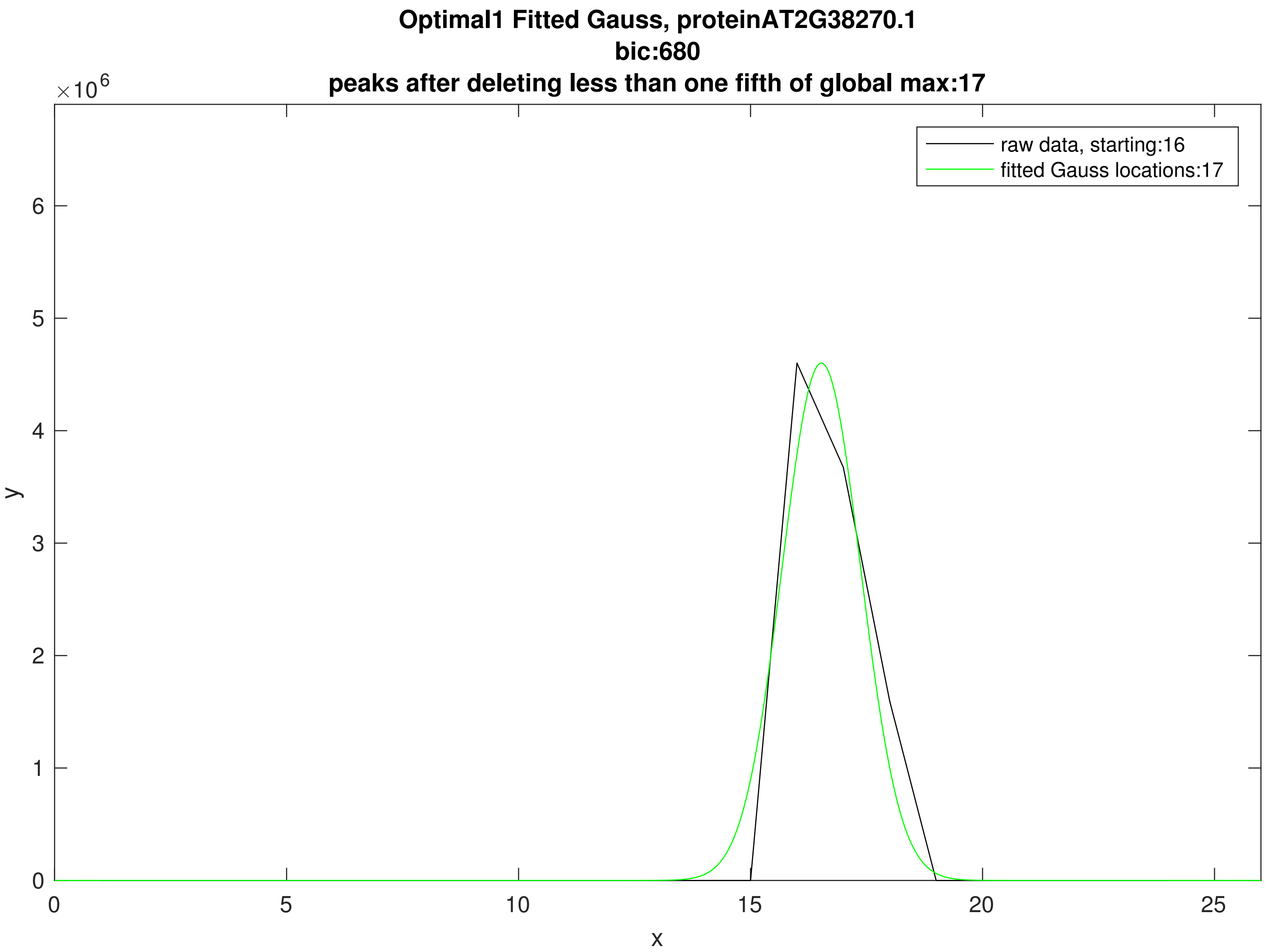

Optimal1 Fitted Gauss, proteinAT2G38270.1
bic:680
peaks after deleting less than one fifth of global max:17
×10^6
raw data, starting:16
fitted Gauss locations:17
y
x
0
1
2
3
4
5
6
0
5
10
15
20
25

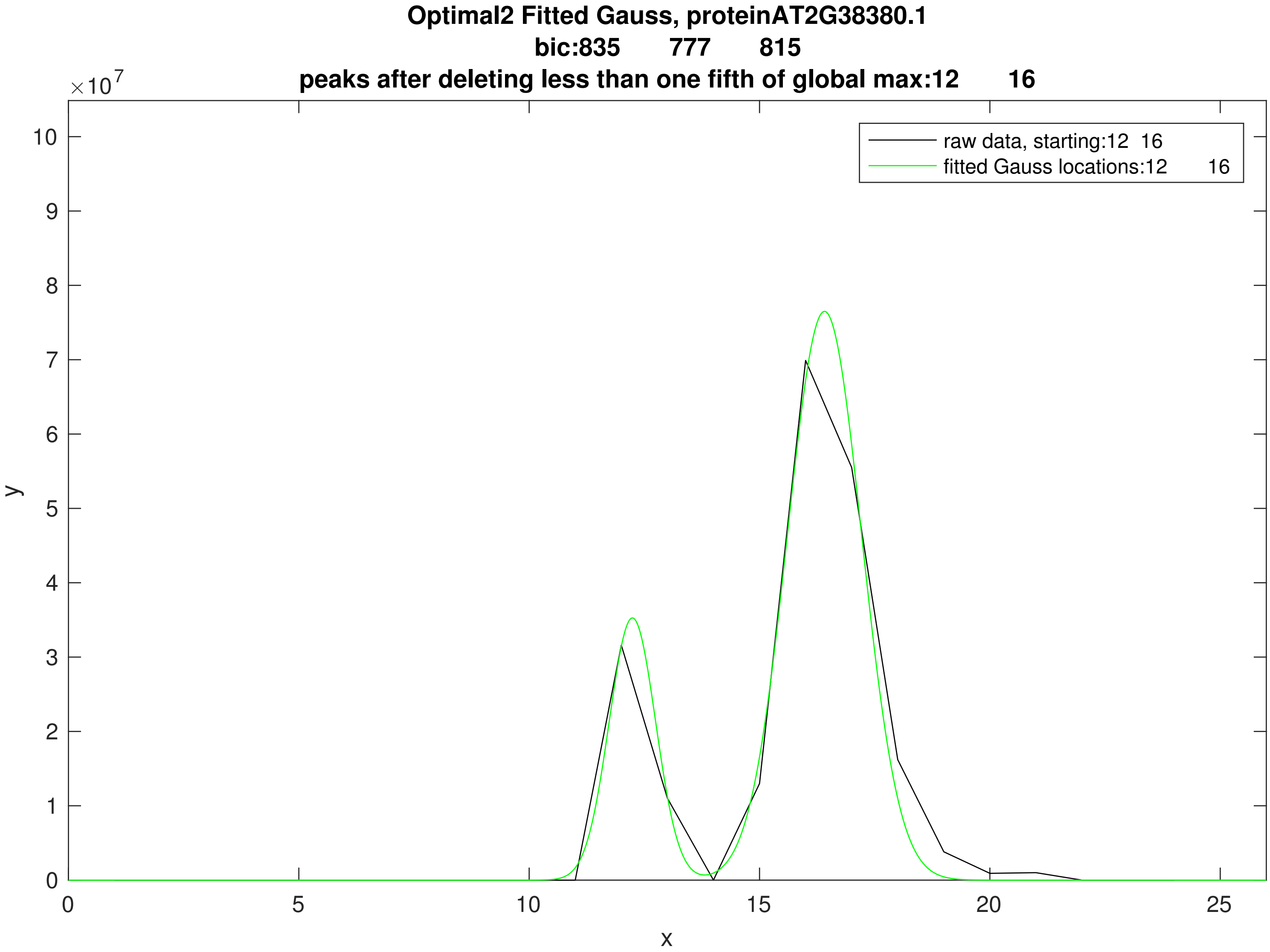

Optimal2 Fitted Gauss, proteinAT2G38380.1
bic:835 777 815
peaks after deleting less than one fifth of global max:12 16
×10^7
raw data, starting:12 16
fitted Gauss locations:12 16
y
x
0
1
2
3
4
5
6
7
8
9
10
0
5
10
15
20
25

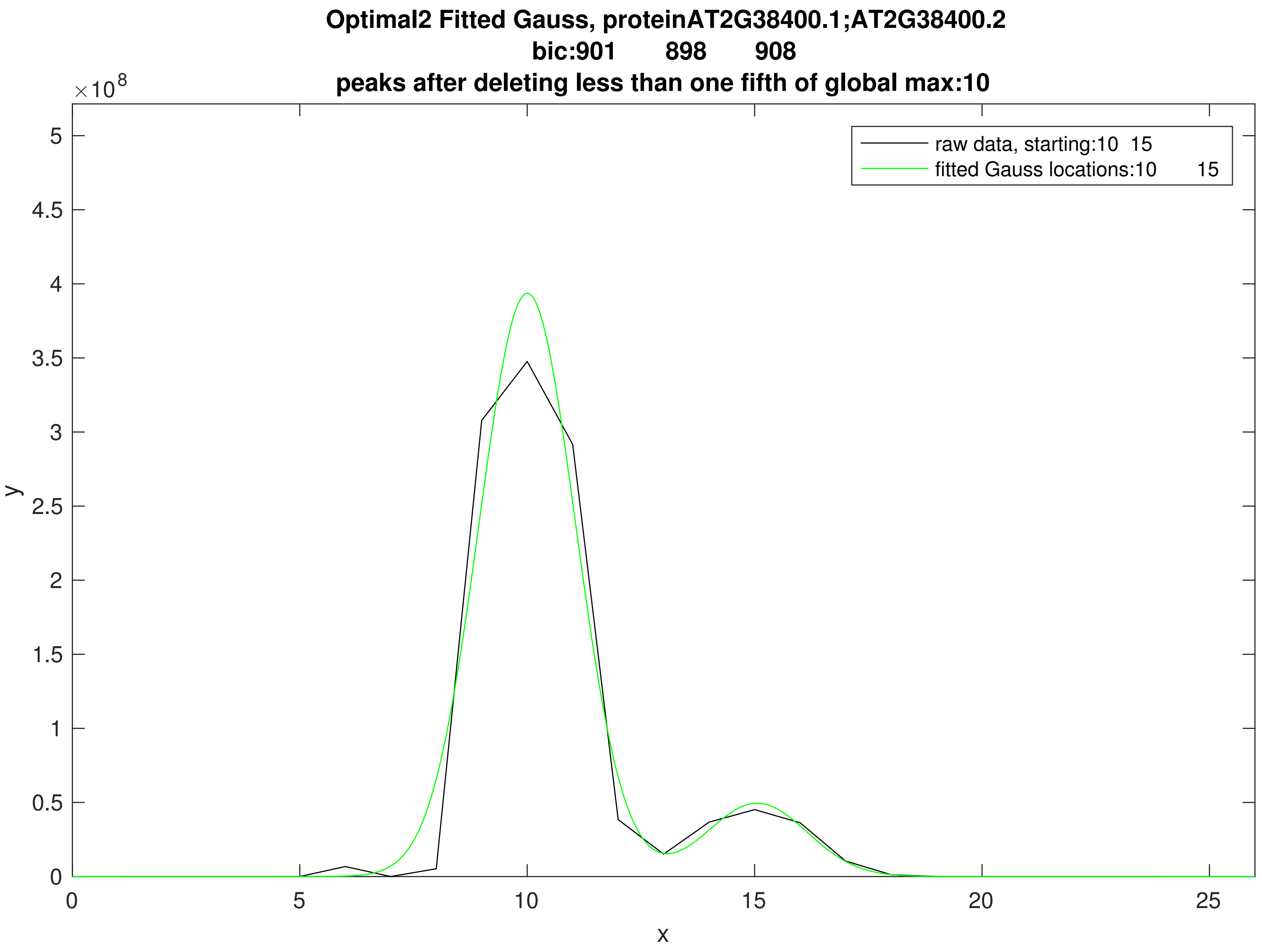
Optimal2 Fitted Gauss, proteinAT2G38400.1;AT2G38400.2
bic:901 898 908
peaks after deleting less than one fifth of global max:10
×10^8
raw data, starting:10 15
fitted Gauss locations:10 15
y
x
0
0.5
1
1.5
2
2.5
3
3.5
4
4.5
5
0
5
10
15
20
25

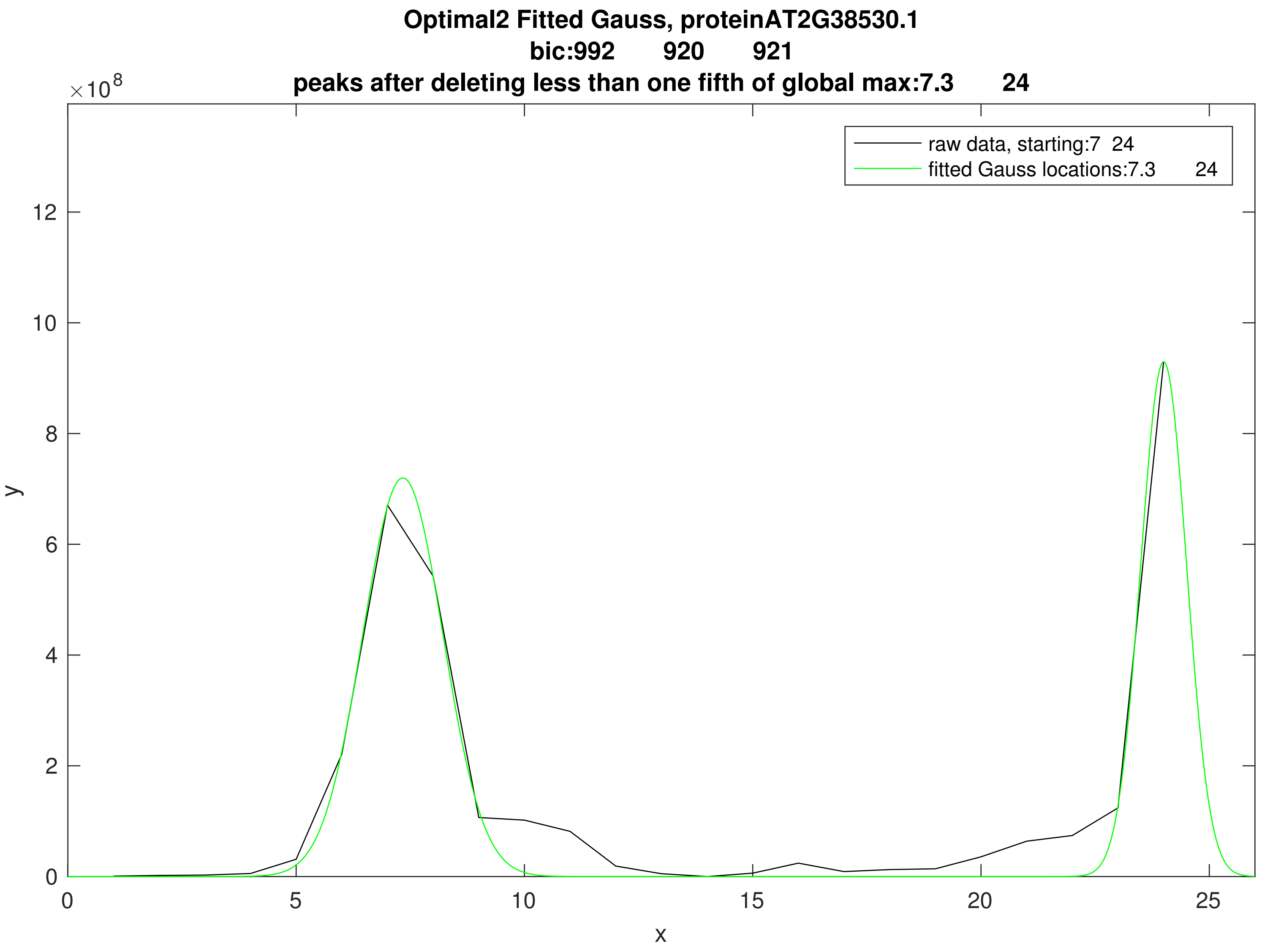
Optimal2 Fitted Gauss, proteinAT2G38530.1
bic:992 920 921
peaks after deleting less than one fifth of global max:7.3 24
×10^8
raw data, starting:7 24
fitted Gauss locations:7.3 24
y
x
0
2
4
6
8
10
12
0
5
10
15
20
25

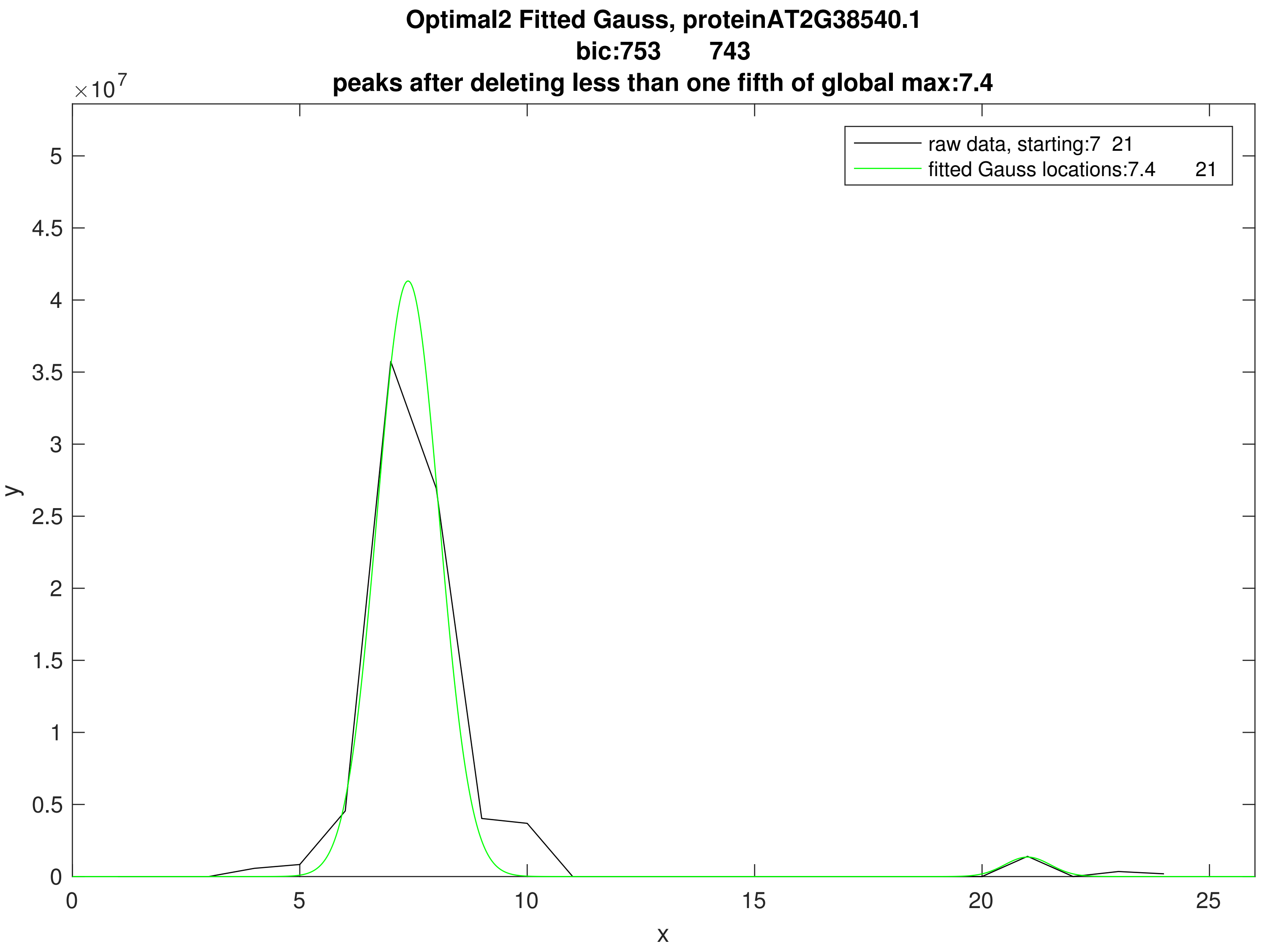
Optimal2 Fitted Gauss, proteinAT2G38540.1
bic:753 743
peaks after deleting less than one fifth of global max:7.4
×10^7
5
4.5
4
3.5
3
2.5
2
1.5
1
0.5
0
y
0
5
10
15
20
25
x
raw data, starting:7 21
fitted Gauss locations:7.4 21

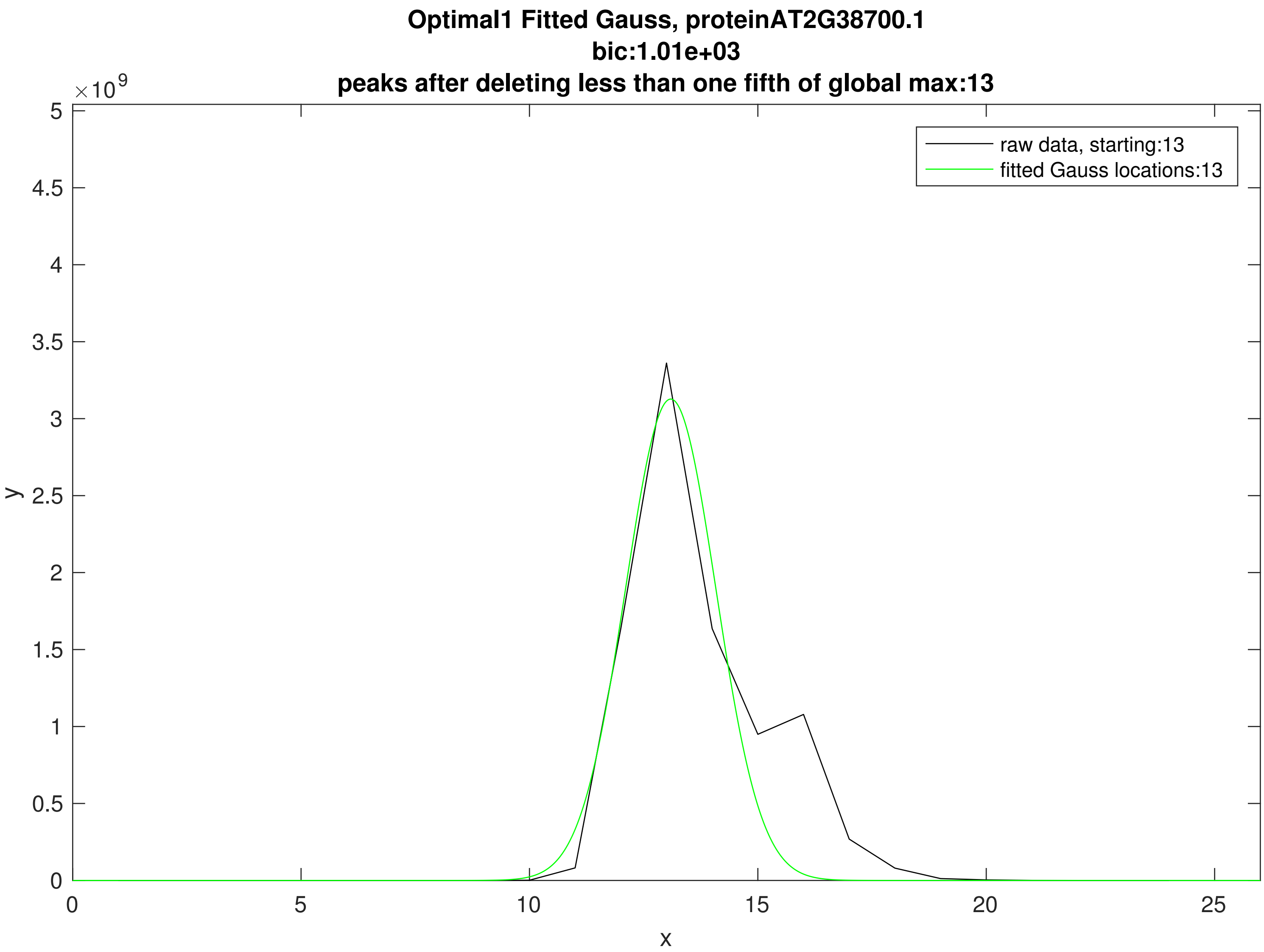

Optimal1 Fitted Gauss, proteinAT2G38700.1
bic:1.01e+03
peaks after deleting less than one fifth of global max:13
×10^9
raw data, starting:13
fitted Gauss locations:13
y
x

Optimal1 Fitted Gauss, proteinAT2G38710.2;AT2G38710.1
bic:751
peaks after deleting less than one fifth of global max:18

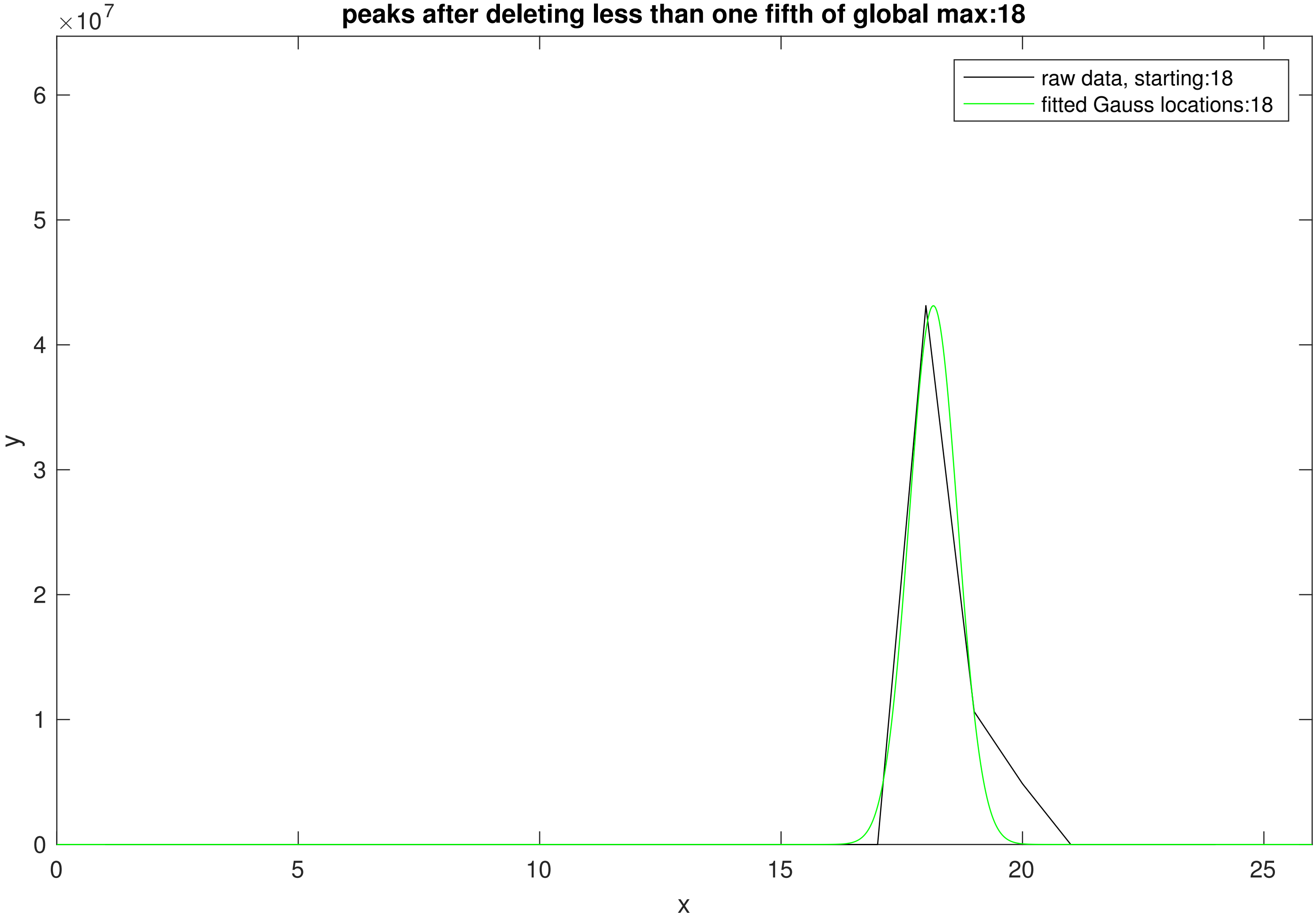

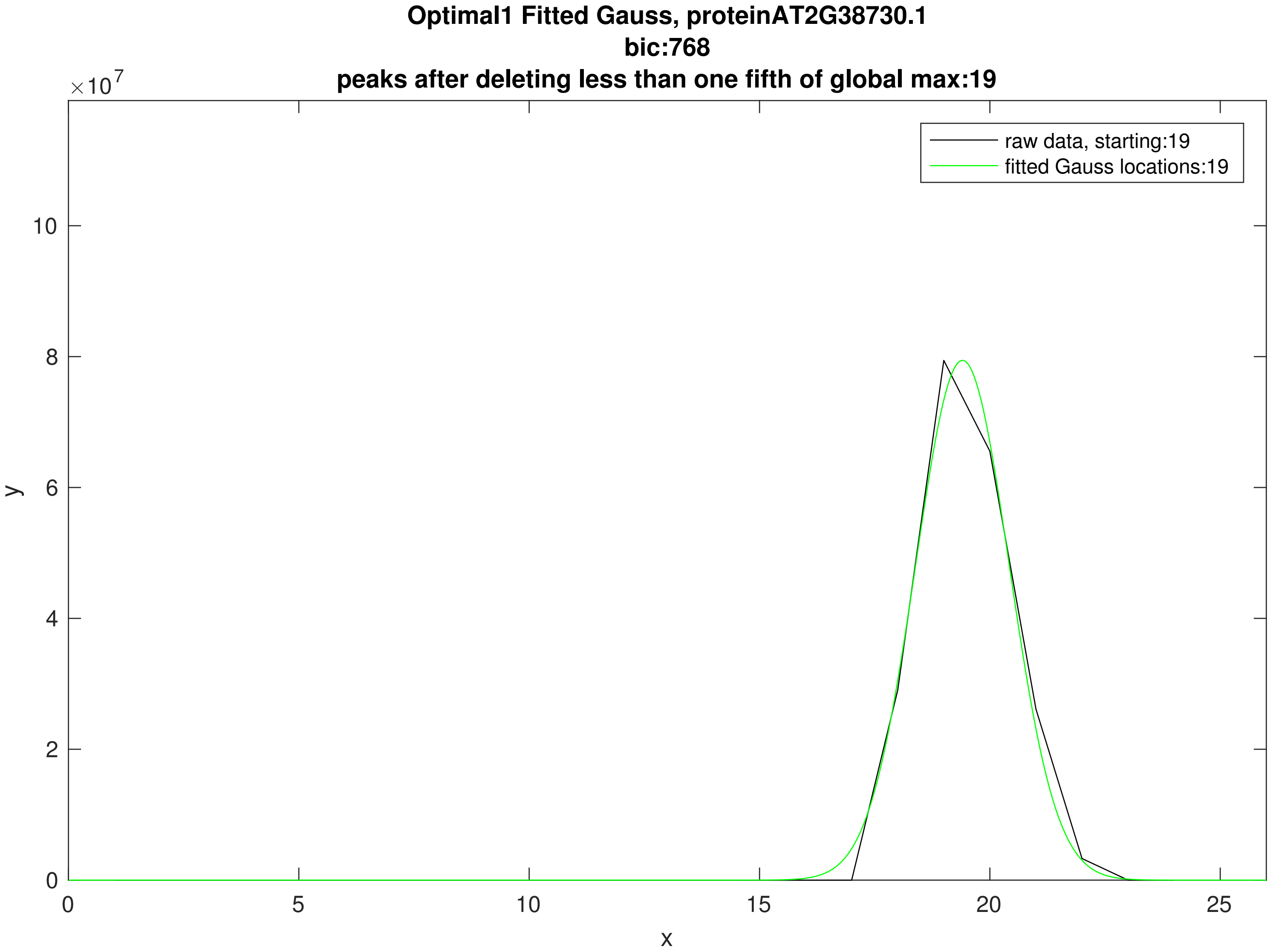
Optimal1 Fitted Gauss, proteinAT2G38730.1
bic:768
peaks after deleting less than one fifth of global max:19
×10^7
raw data, starting:19
fitted Gauss locations:19
y
x
0
2
4
6
8
10
5
15
20
25

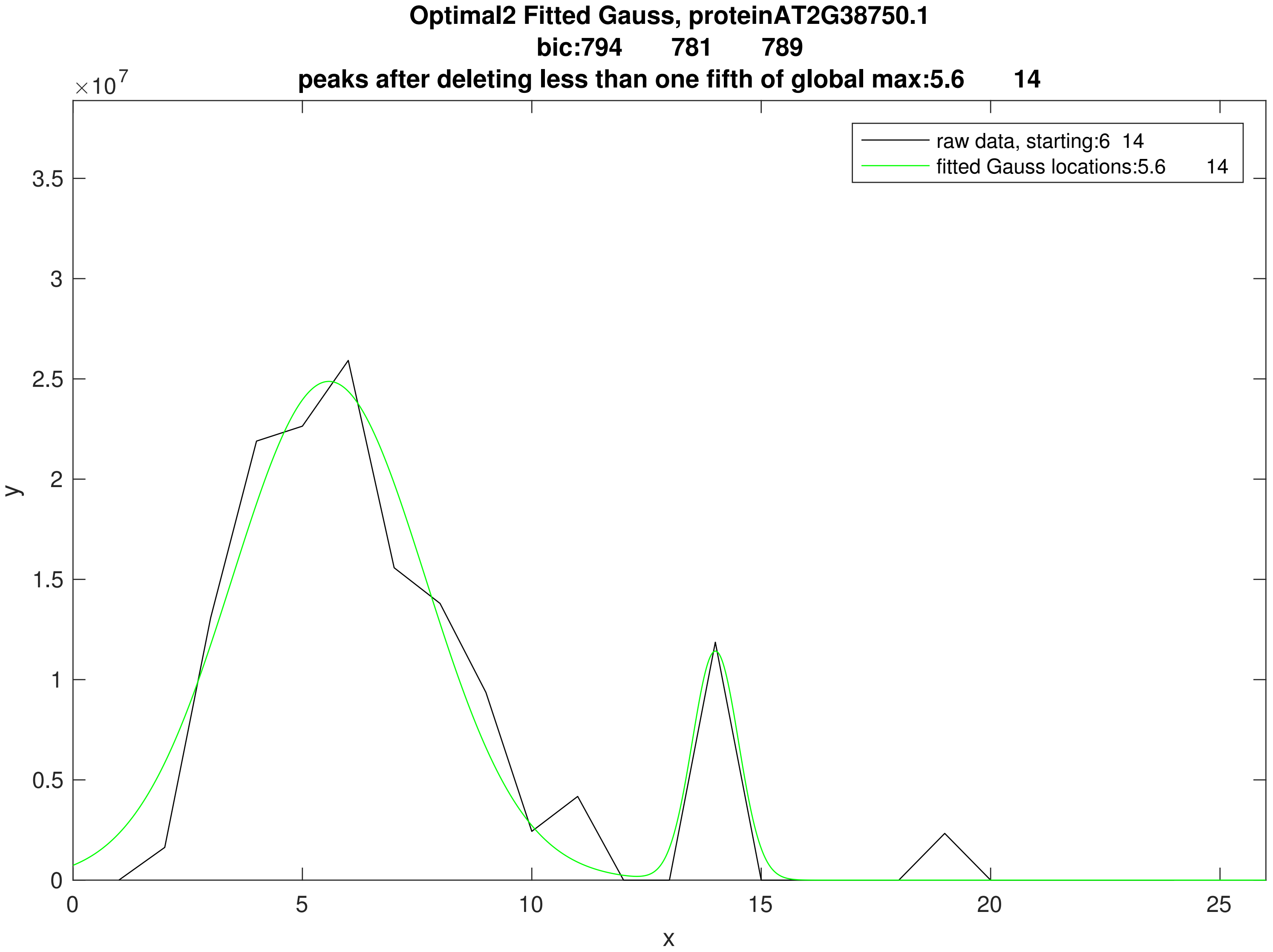
Optimal2 Fitted Gauss, proteinAT2G38750.1
bic:794 781 789
peaks after deleting less than one fifth of global max:5.6 14
×10^7
raw data, starting:6 14
fitted Gauss locations:5.6 14
3.5
3
2.5
2
1.5
1
0.5
0
y
0
5
10
15
20
25
x

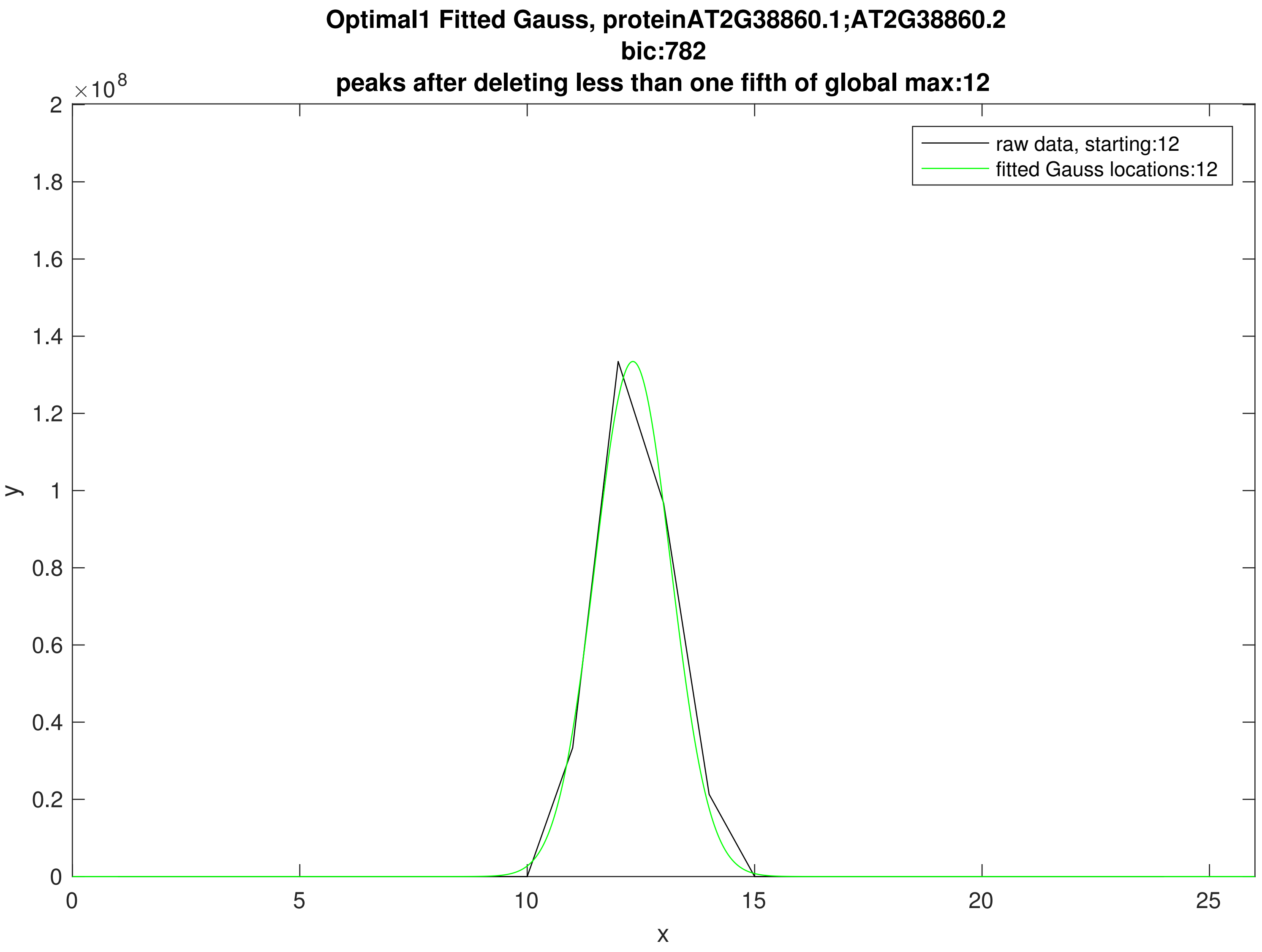

Optimal1 Fitted Gauss, proteinAT2G38860.1;AT2G38860.2
bic:782
peaks after deleting less than one fifth of global max:12
×10^8
2
1.8
1.6
1.4
1.2
1
0.8
0.6
0.4
0.2
0
y
0
5
10
15
20
25
x
raw data, starting:12
fitted Gauss locations:12

Optimal1 Fitted Gauss, proteinAT2G38870.1
bic:682
peaks after deleting less than one fifth of global max:22

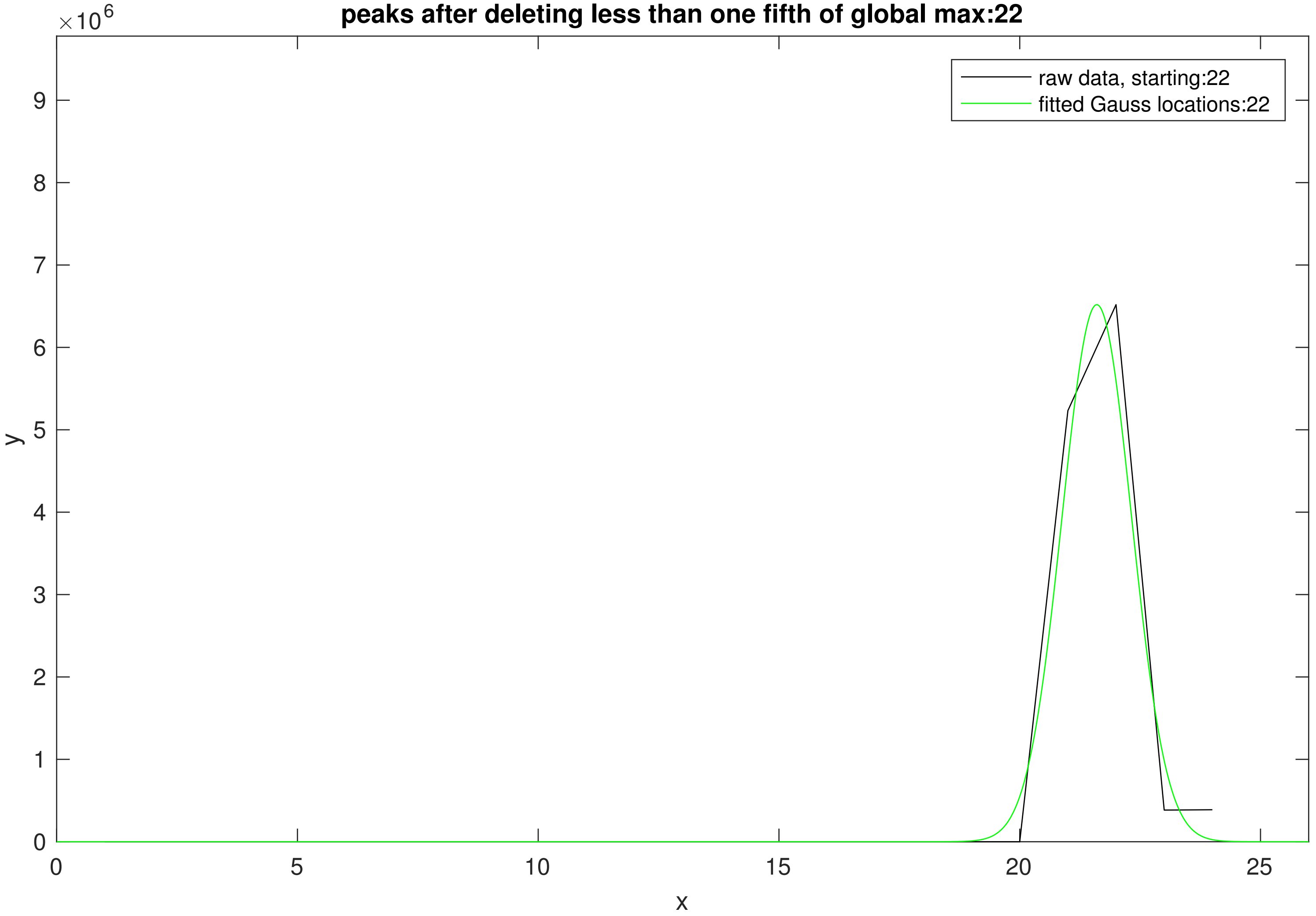

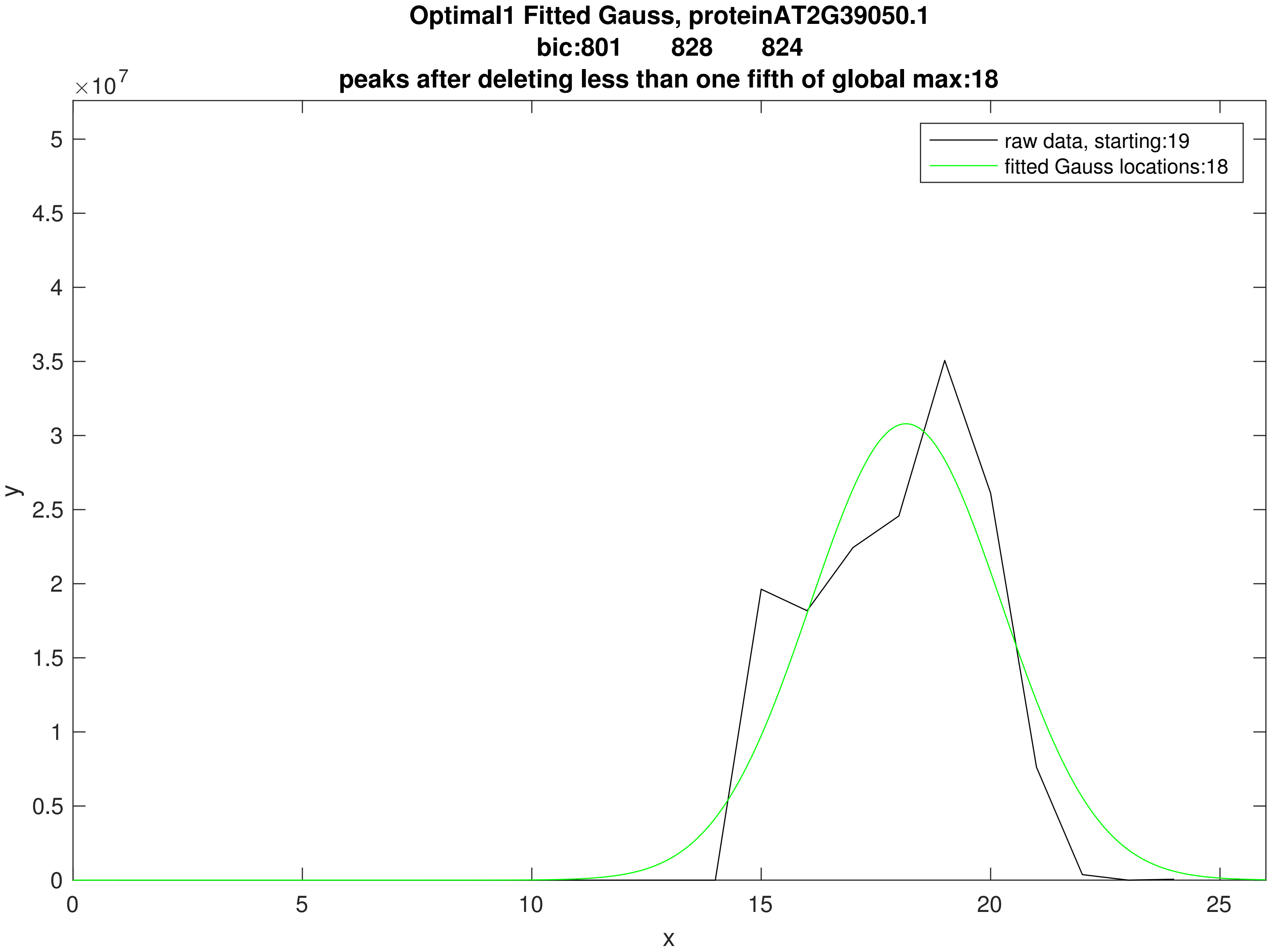

Optimal1 Fitted Gauss, proteinAT2G39050.1
bic:801 828 824
peaks after deleting less than one fifth of global max:18
×10^7
raw data, starting:19
fitted Gauss locations:18
y
x
0
0.5
1
1.5
2
2.5
3
3.5
4
4.5
5
0
5
10
15
20
25

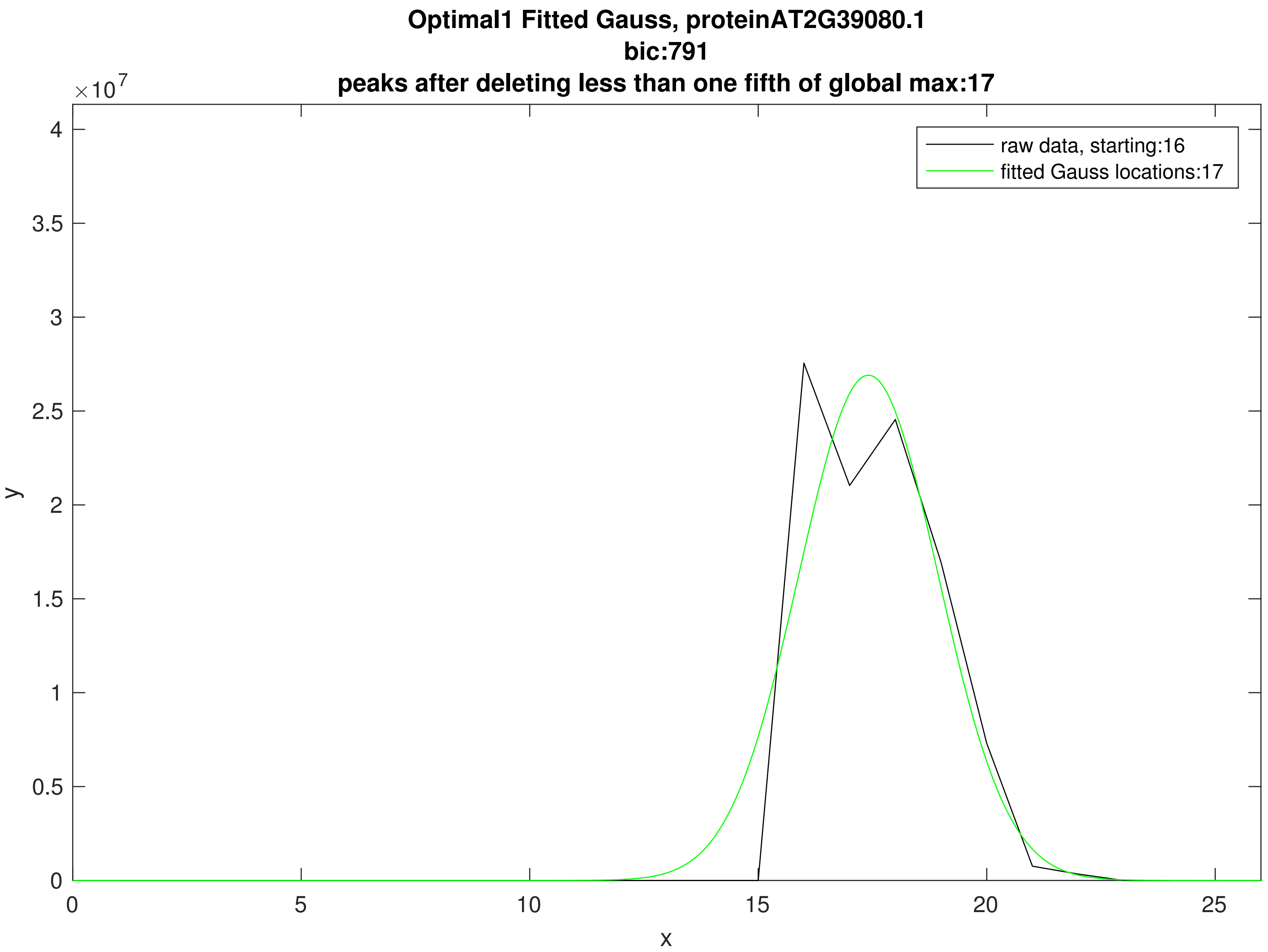
Optimal1 Fitted Gauss, proteinAT2G39080.1
bic:791
peaks after deleting less than one fifth of global max:17
×10^7
raw data, starting:16
fitted Gauss locations:17
y
x
0
0.5
1
1.5
2
2.5
3
3.5
4
0
5
10
15
20
25

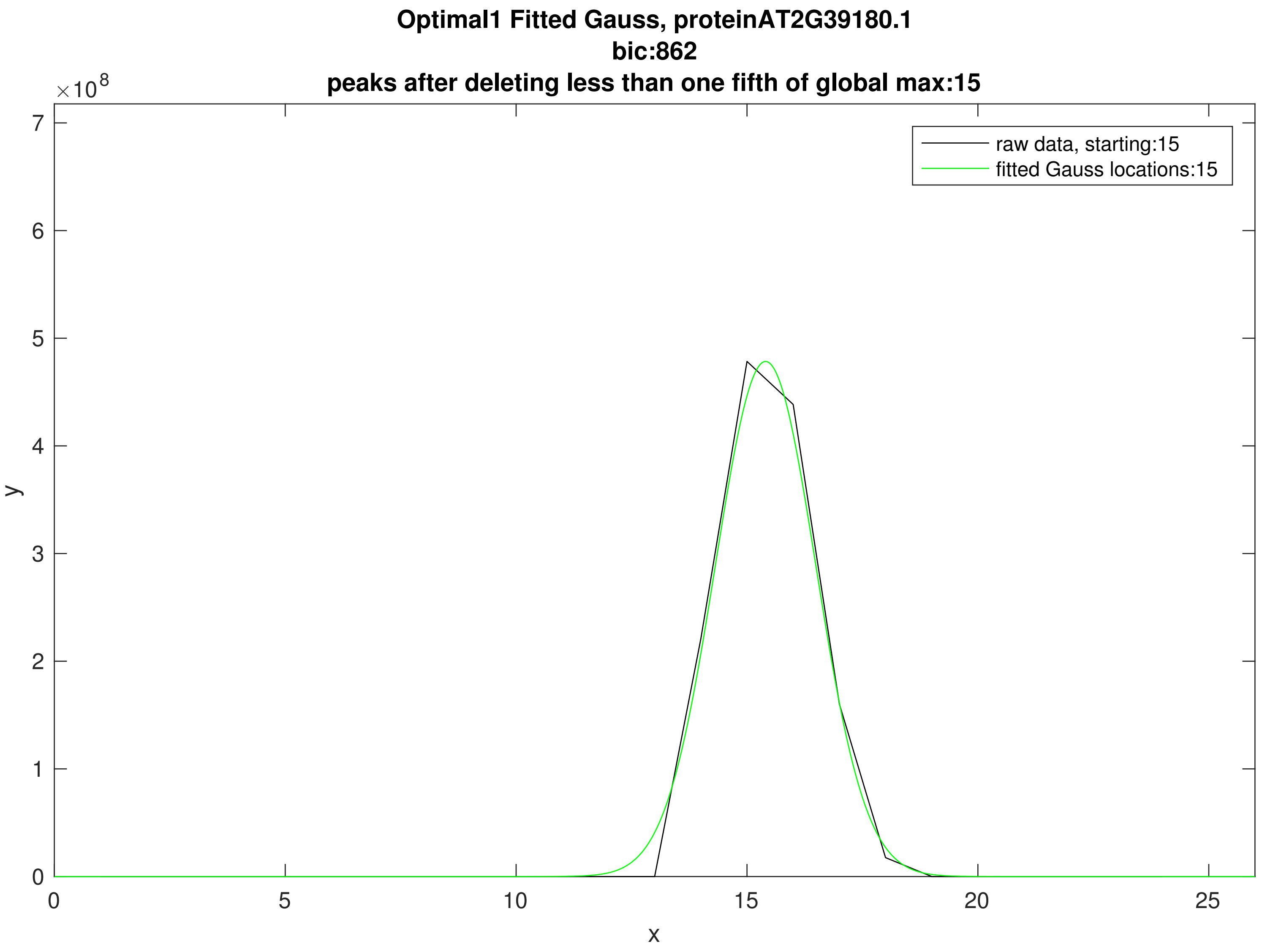

Optimal1 Fitted Gauss, proteinAT2G39180.1
bic:862
peaks after deleting less than one fifth of global max:15
×10^8
raw data, starting:15
fitted Gauss locations:15
y
x
0
1
2
3
4
5
6
7
0
5
10
15
20
25

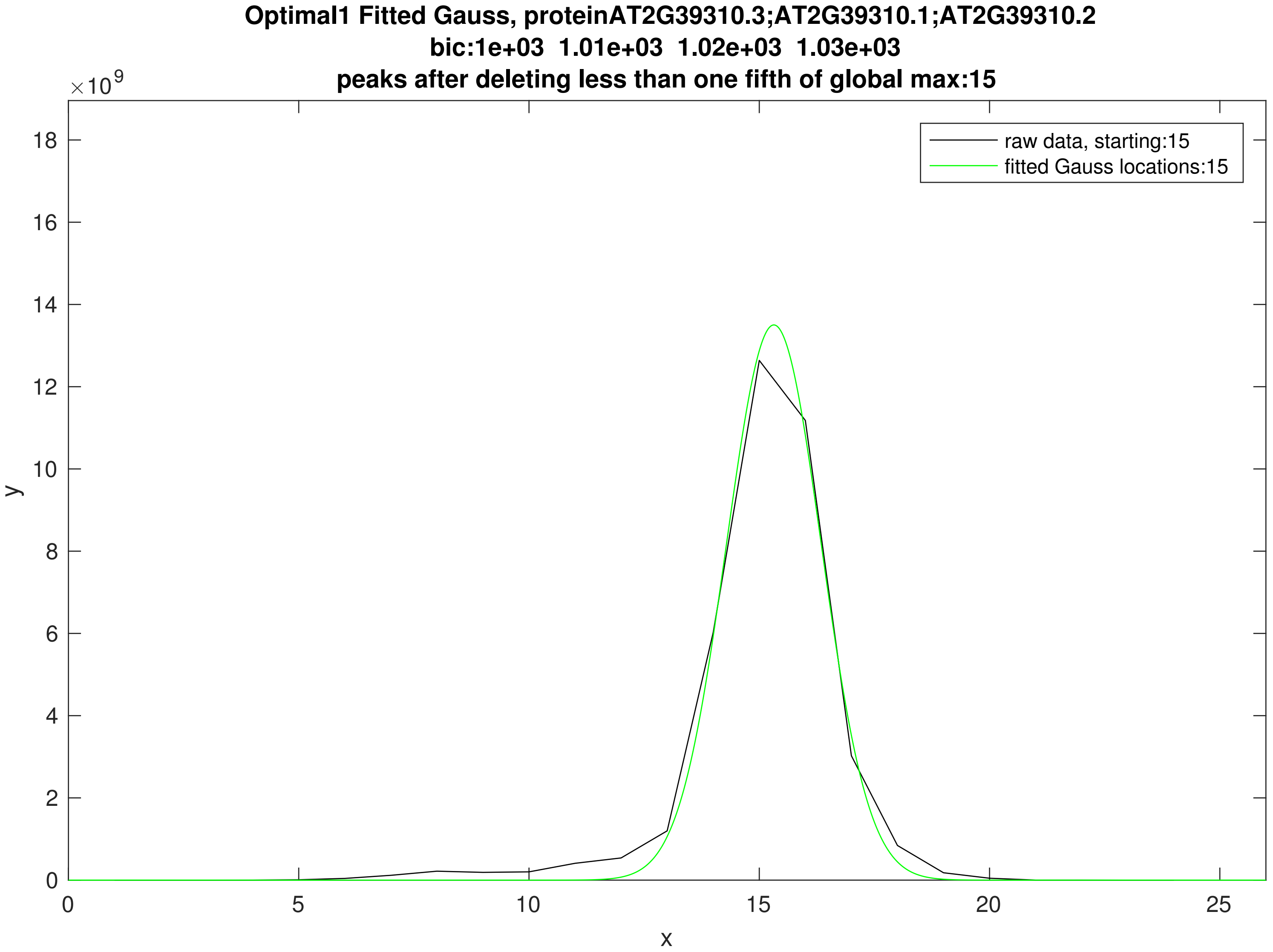
Optimal1 Fitted Gauss, proteinAT2G39310.3;AT2G39310.1;AT2G39310.2
bic:1e+03 1.01e+03 1.02e+03 1.03e+03
peaks after deleting less than one fifth of global max:15
×10^9
raw data, starting:15
fitted Gauss locations:15
y
x
0
2
4
6
8
10
12
14
16
18
0
5
10
15
20
25

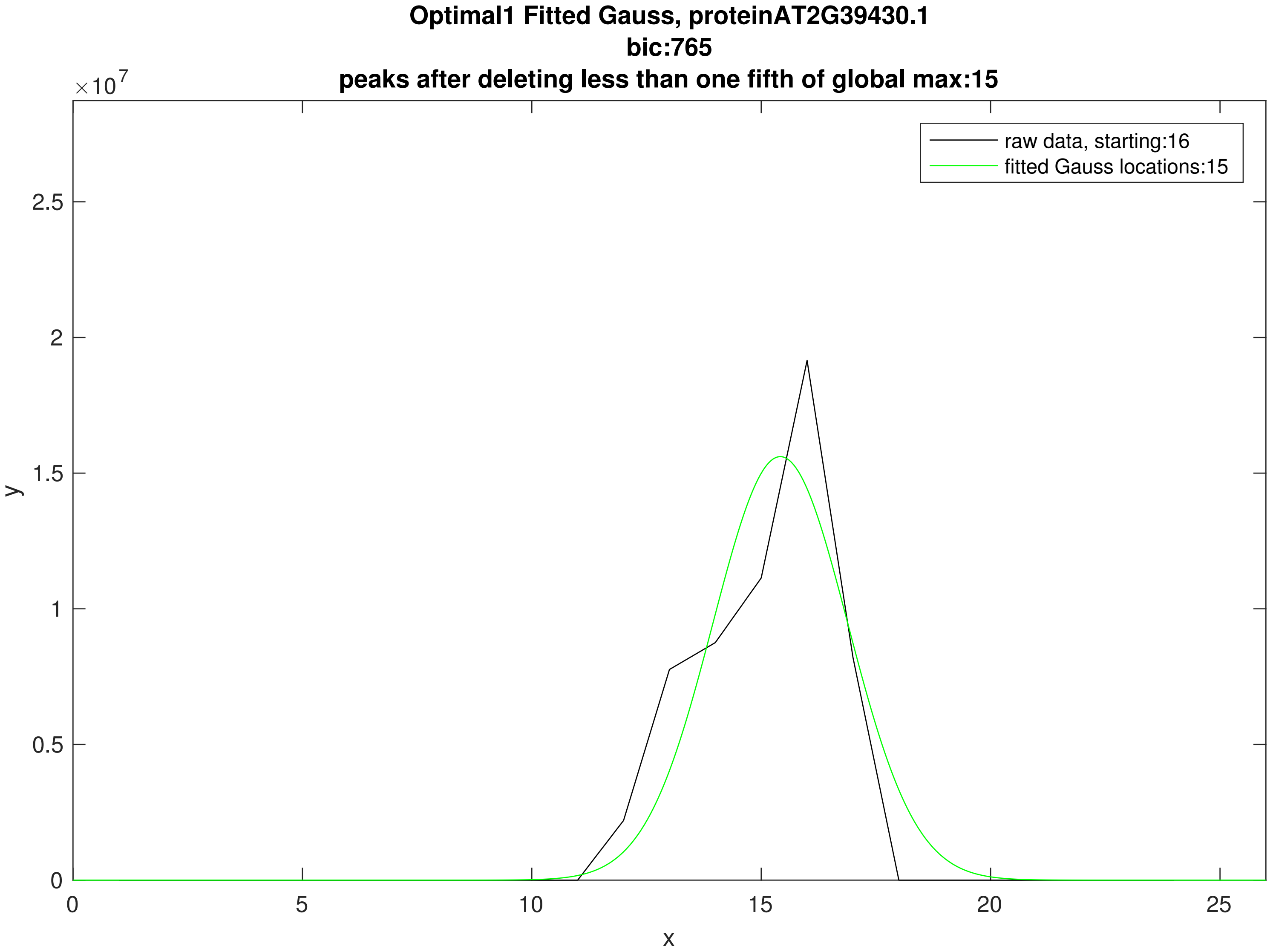
Optimal1 Fitted Gauss, proteinAT2G39430.1
bic:765
peaks after deleting less than one fifth of global max:15
×10^7
raw data, starting:16
fitted Gauss locations:15
2.5
2
1.5
1
0.5
0
y
0
5
10
15
20
25
x

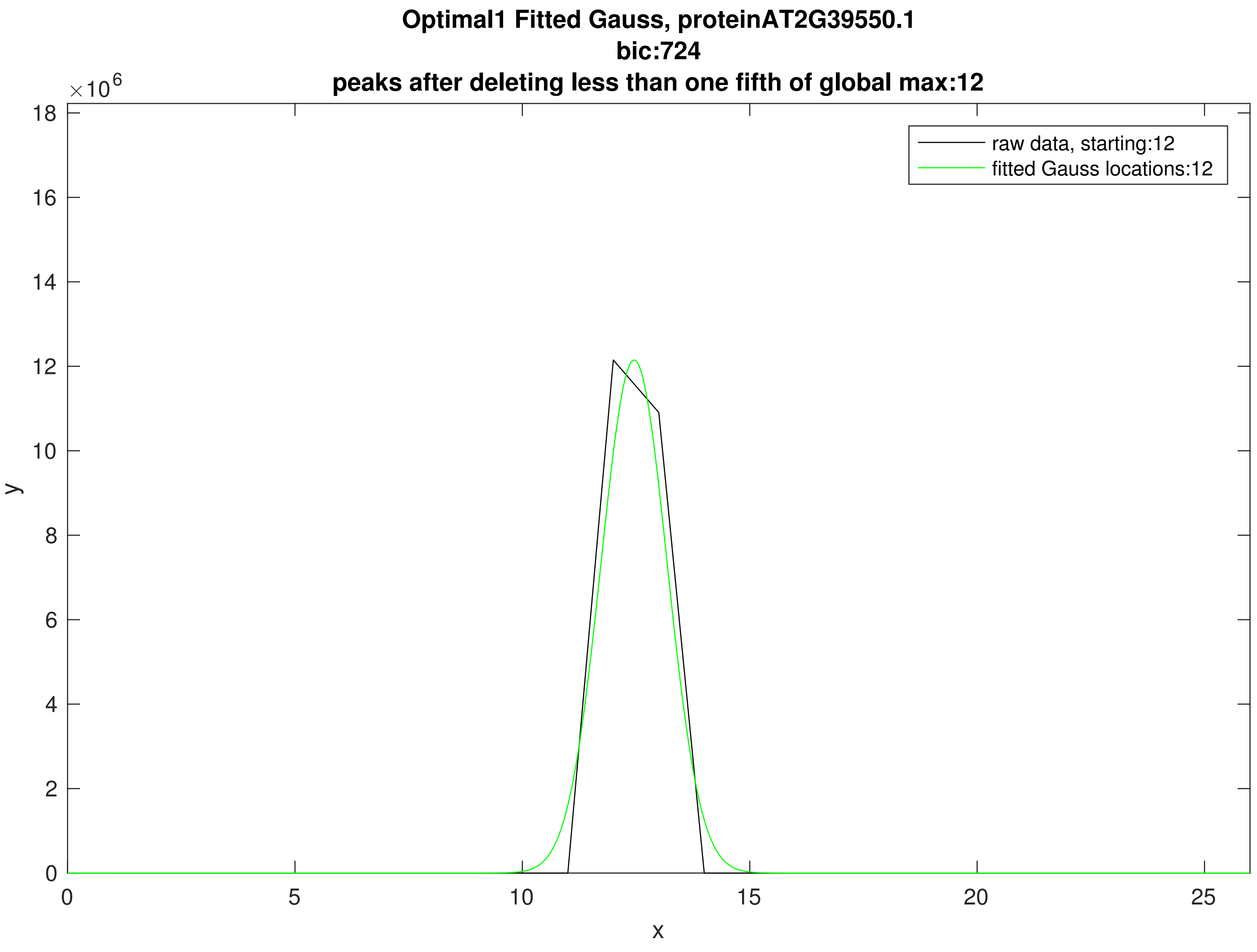
Optimal1 Fitted Gauss, proteinAT2G39550.1
bic:724
peaks after deleting less than one fifth of global max:12
×10^6
raw data, starting:12
fitted Gauss locations:12
y
x
0
2
4
6
8
10
12
14
16
18
0
5
10
15
20
25

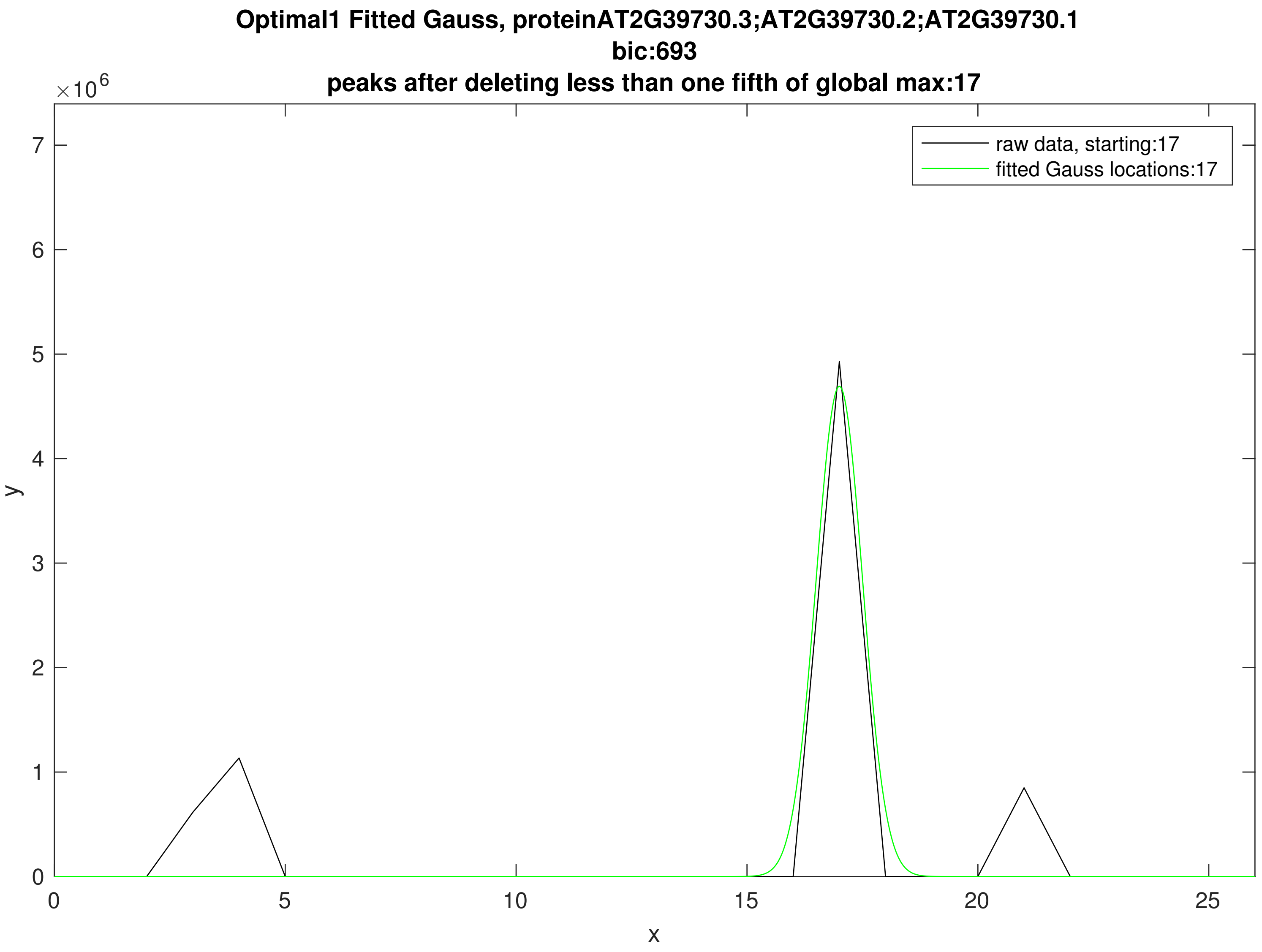
Optimal1 Fitted Gauss, proteinAT2G39730.3;AT2G39730.2;AT2G39730.1
bic:693
peaks after deleting less than one fifth of global max:17
×10^6
raw data, starting:17
fitted Gauss locations:17
y
x
0
1
2
3
4
5
6
7
0
5
10
15
20
25

**Optimal1 Fitted Gauss, proteinAT2G39770.1;AT2G39770.2**
**bic:874**
**peaks after deleting less than one fifth of global max:16**

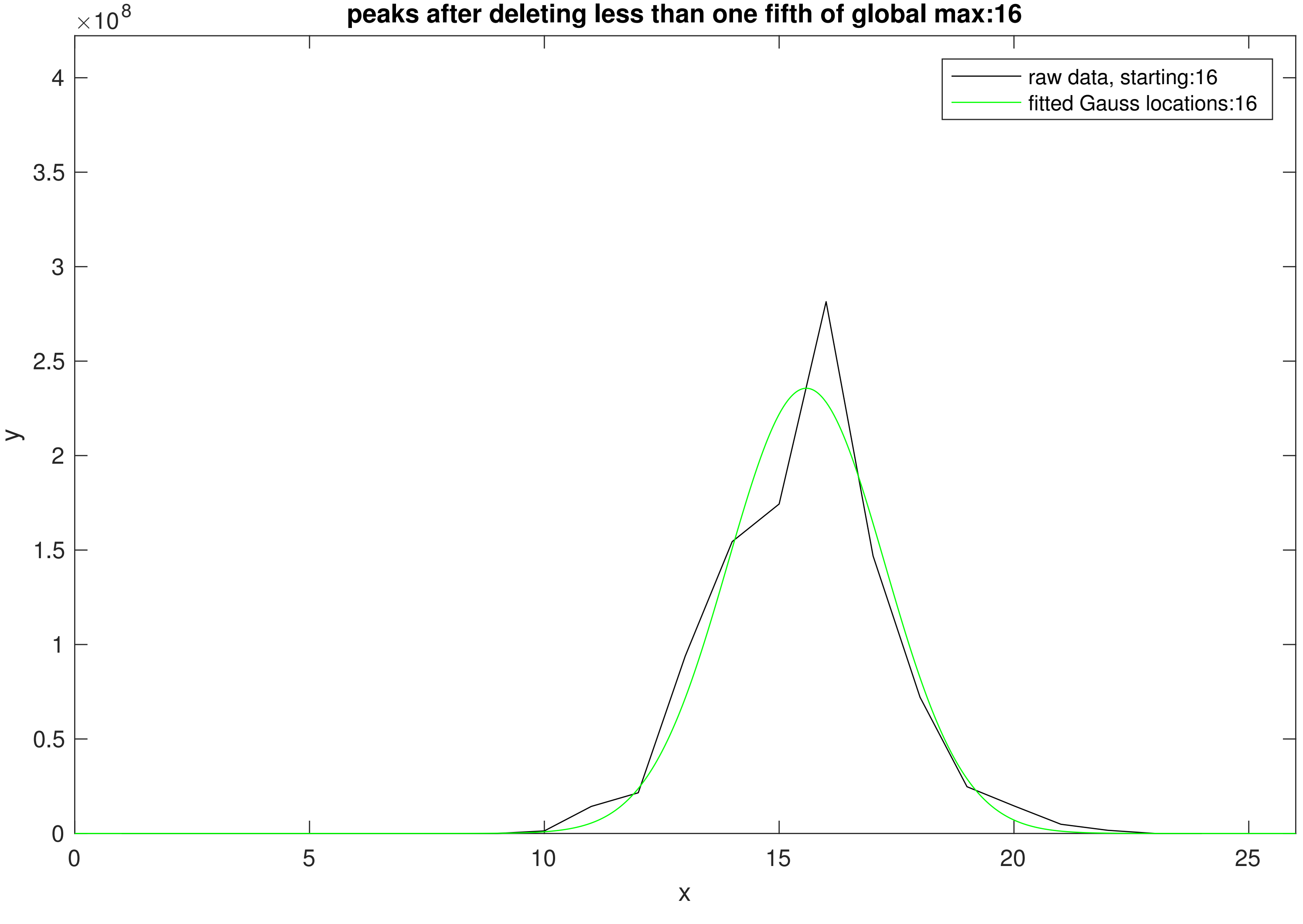

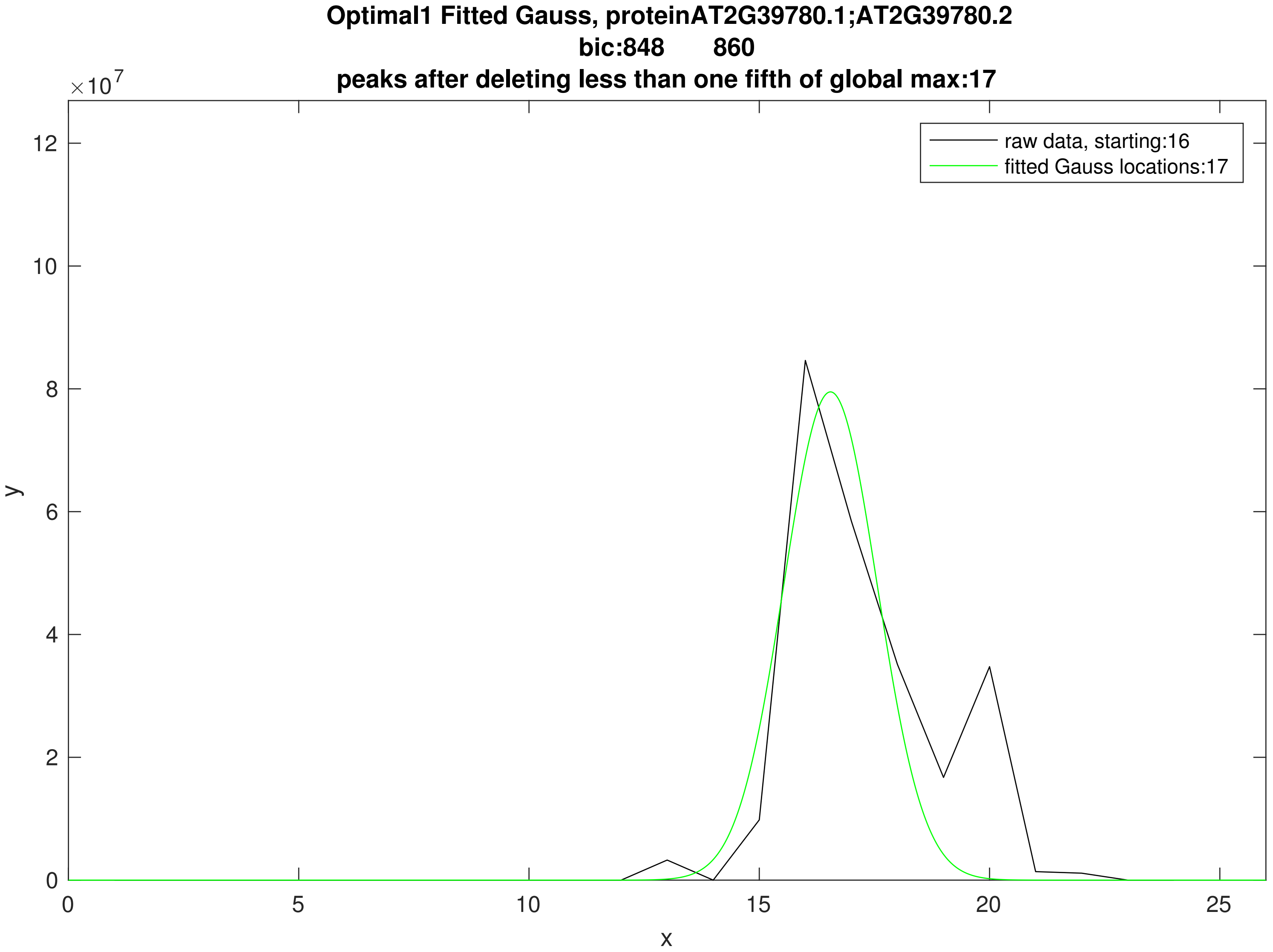

Optimal1 Fitted Gauss, proteinAT2G39780.1;AT2G39780.2
bic:848 860
peaks after deleting less than one fifth of global max:17
×10^7
raw data, starting:16
fitted Gauss locations:17
y
x
0
2
4
6
8
10
12
0
5
10
15
20
25

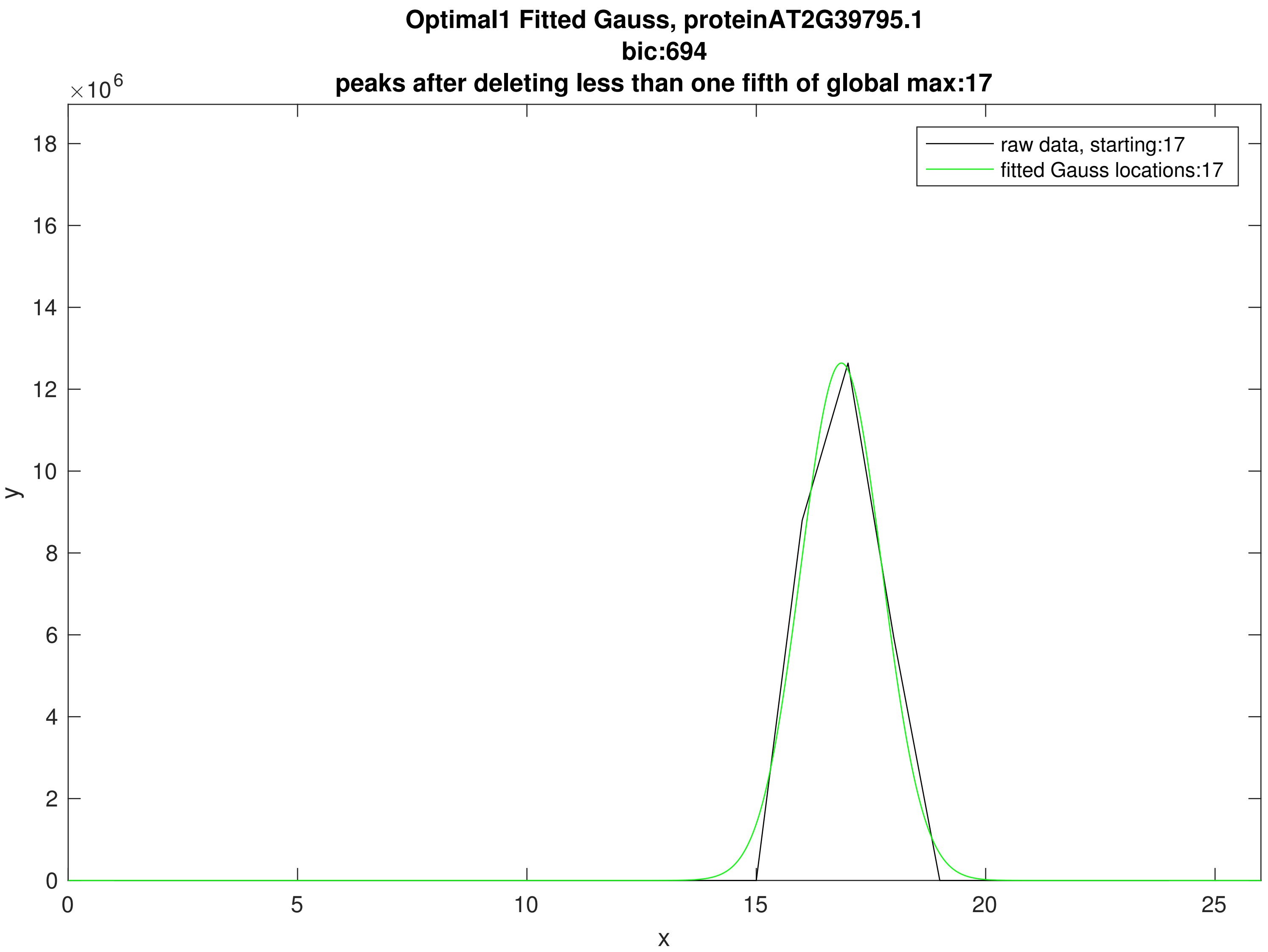

Optimal1 Fitted Gauss, proteinAT2G39795.1
bic:694
peaks after deleting less than one fifth of global max:17
×10^6
raw data, starting:17
fitted Gauss locations:17
y
x
0
2
4
6
8
10
12
14
16
18
0
5
10
15
20
25

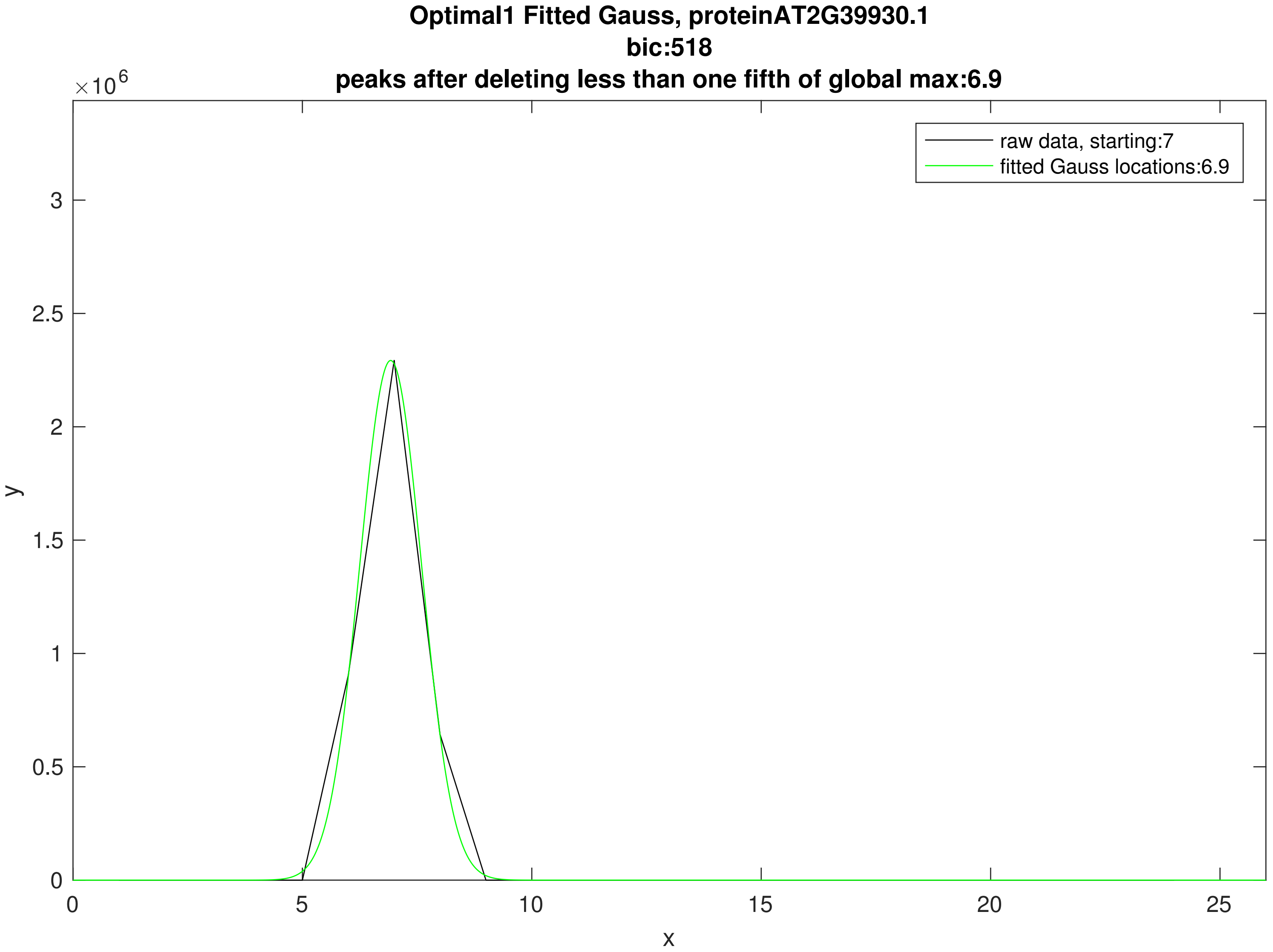

Optimal1 Fitted Gauss, proteinAT2G39930.1
bic:518
peaks after deleting less than one fifth of global max:6.9
×10^6
raw data, starting:7
fitted Gauss locations:6.9
y
x
0
0.5
1
1.5
2
2.5
3
0
5
10
15
20
25

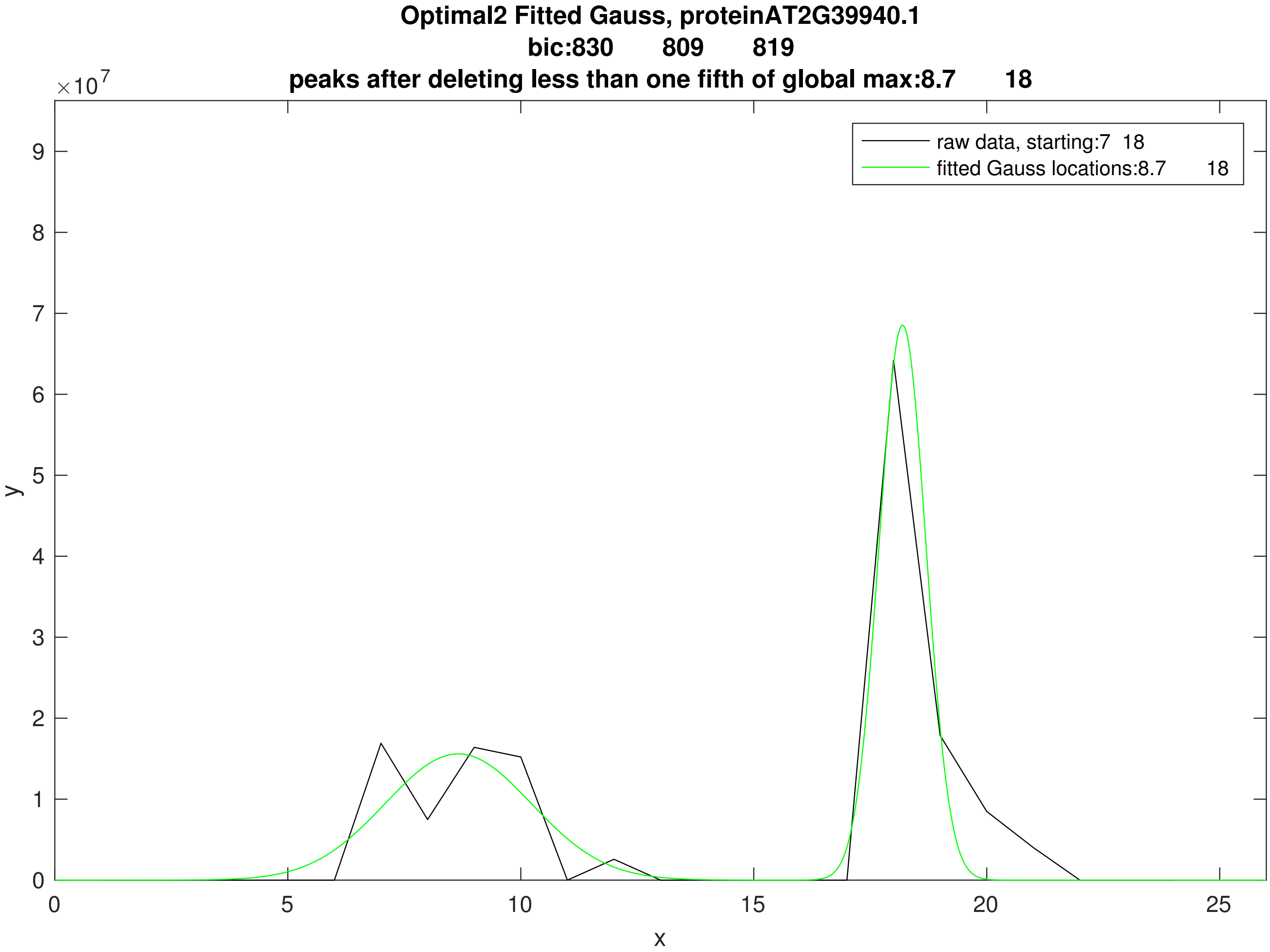

Optimal2 Fitted Gauss, proteinAT2G39940.1
bic:830 809 819
peaks after deleting less than one fifth of global max:8.7 18
×10^7
raw data, starting:7 18
fitted Gauss locations:8.7 18
y
x
0
1
2
3
4
5
6
7
8
9
0
5
10
15
20
25

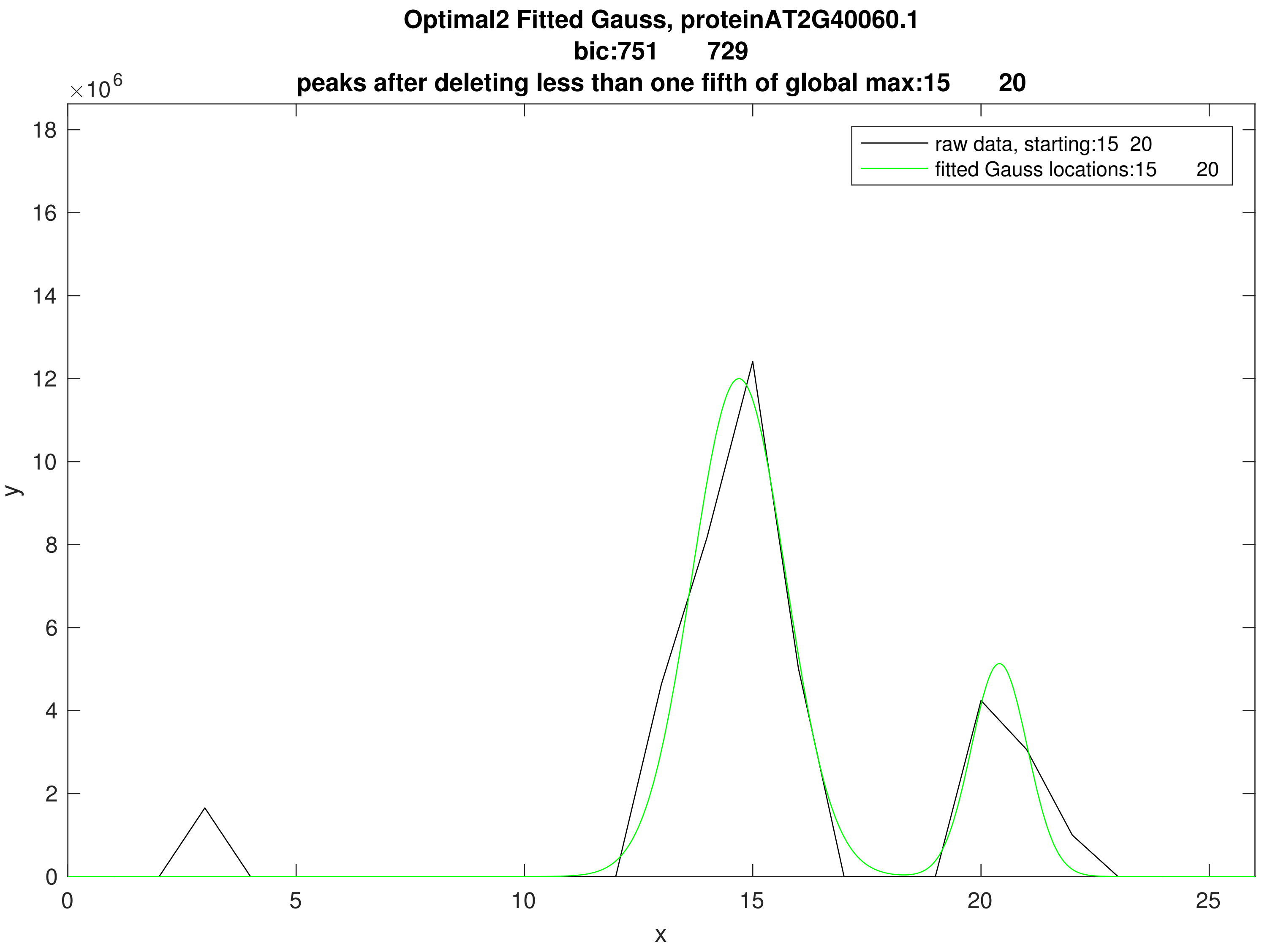

Optimal2 Fitted Gauss, proteinAT2G40060.1
bic:751 729
peaks after deleting less than one fifth of global max:15 20
×10^6
raw data, starting:15 20
fitted Gauss locations:15 20
y
x
0
2
4
6
8
10
12
14
16
18
0
5
10
15
20
25

#### Optimal1 Fitted Gauss, proteinAT2G40170.1
#### bic:721
#### peaks after deleting less than one fifth of global max:18

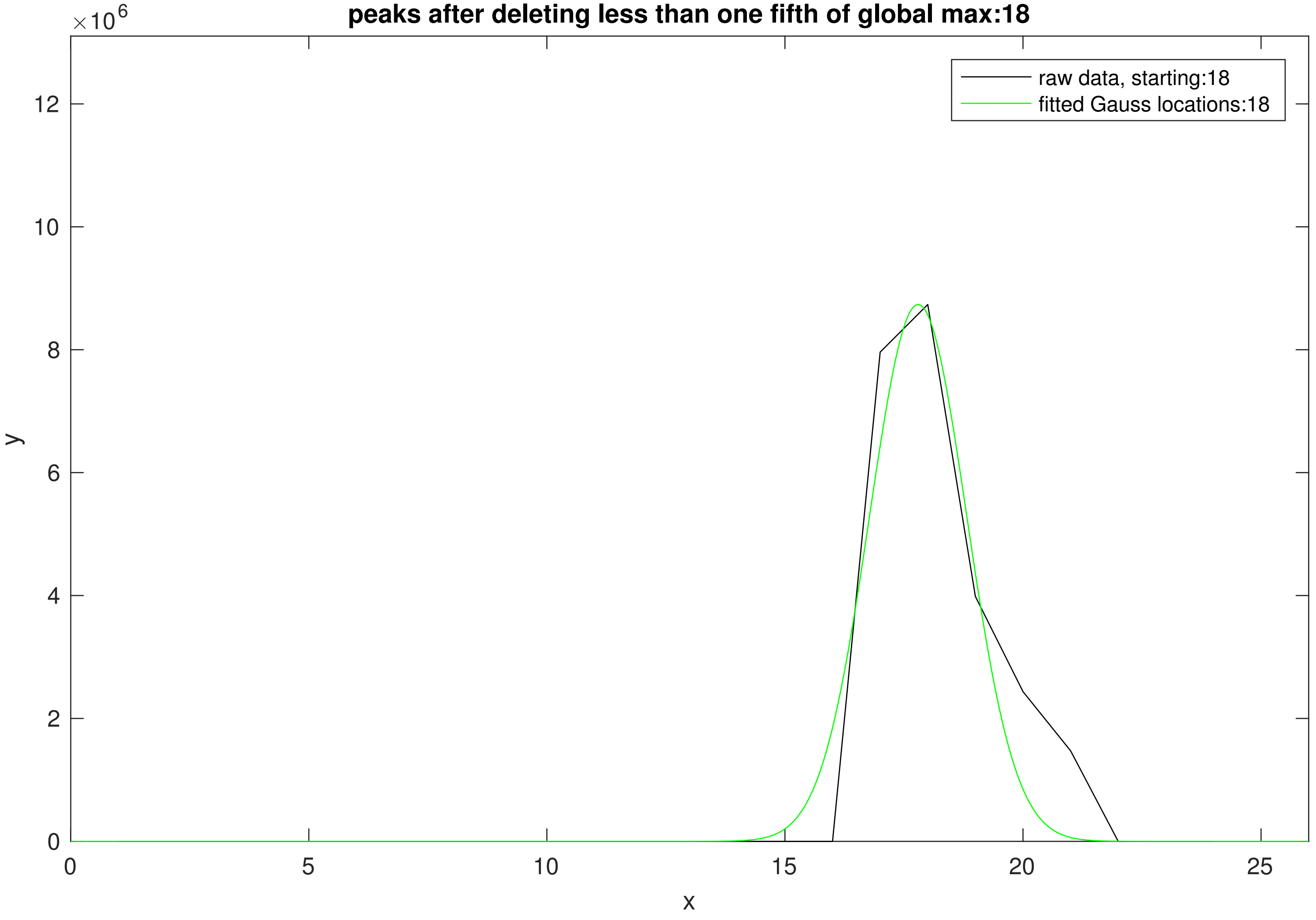

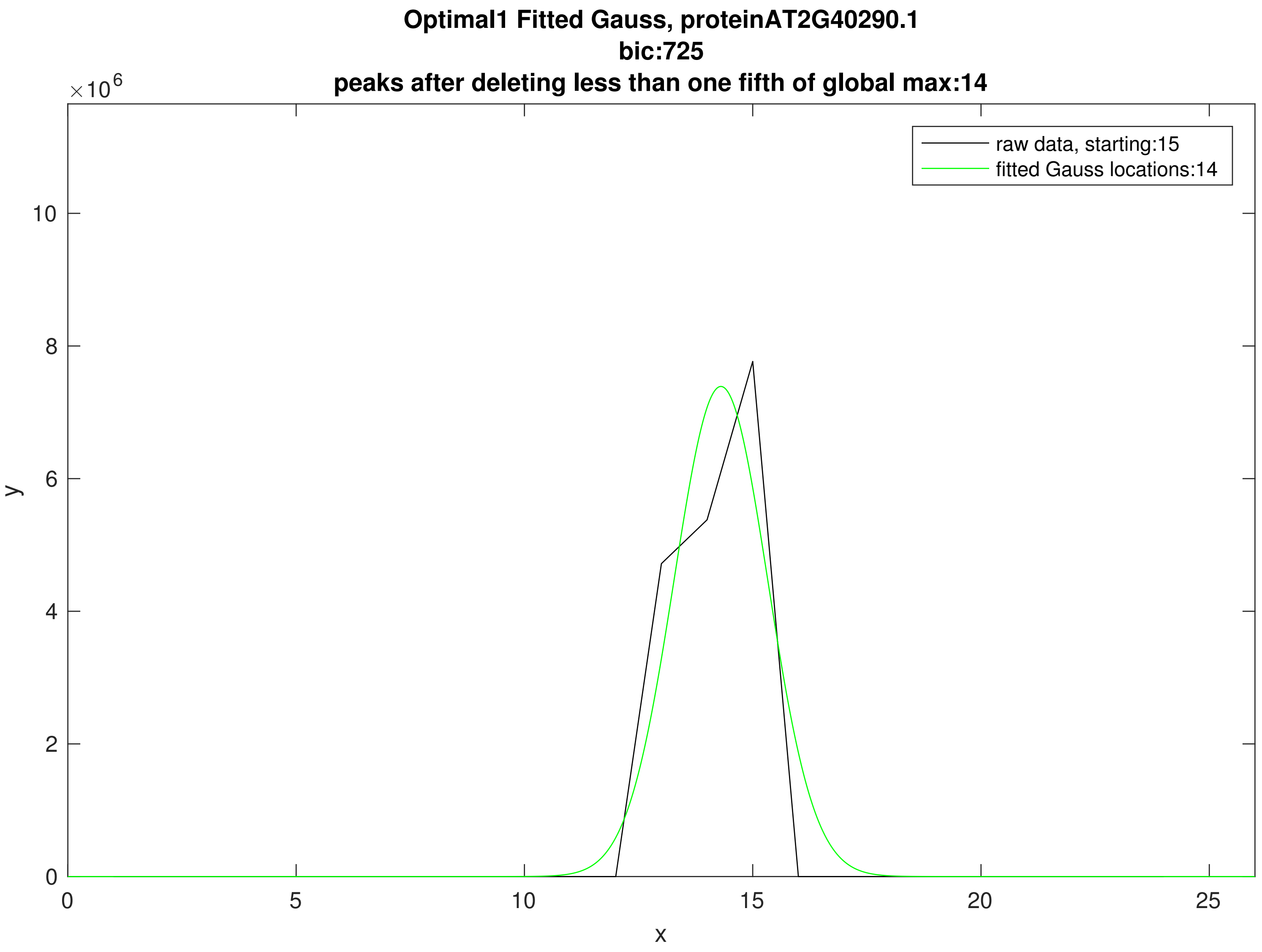
Optimal1 Fitted Gauss, proteinAT2G40290.1
bic:725
peaks after deleting less than one fifth of global max:14
×10^6
raw data, starting:15
fitted Gauss locations:14
10
8
6
4
2
0
y
0
5
10
15
20
25
x

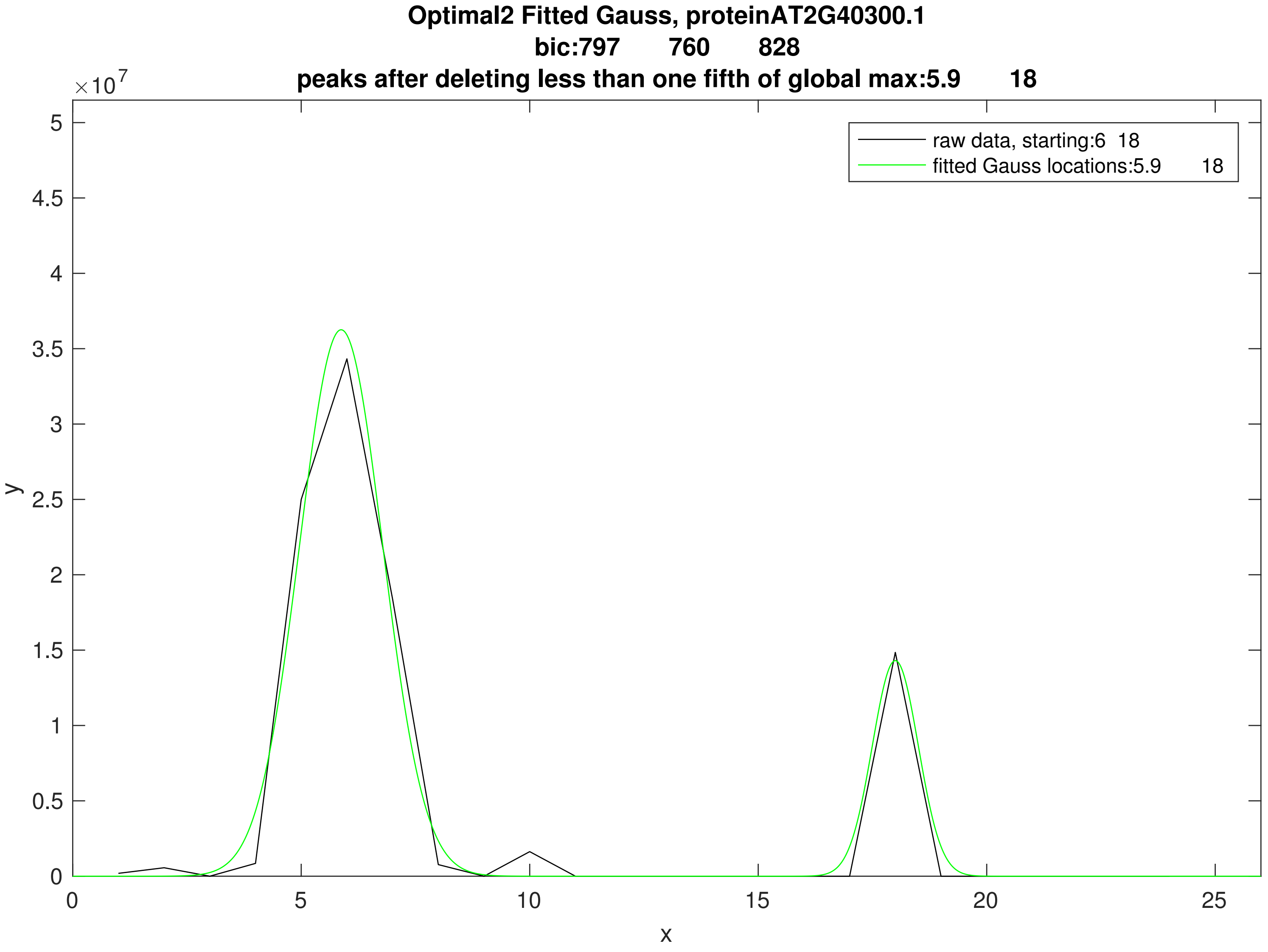
Optimal2 Fitted Gauss, proteinAT2G40300.1
bic:797 760 828
peaks after deleting less than one fifth of global max:5.9 18
raw data, starting:6 18
fitted Gauss locations:5.9 18
×10^7
y
x

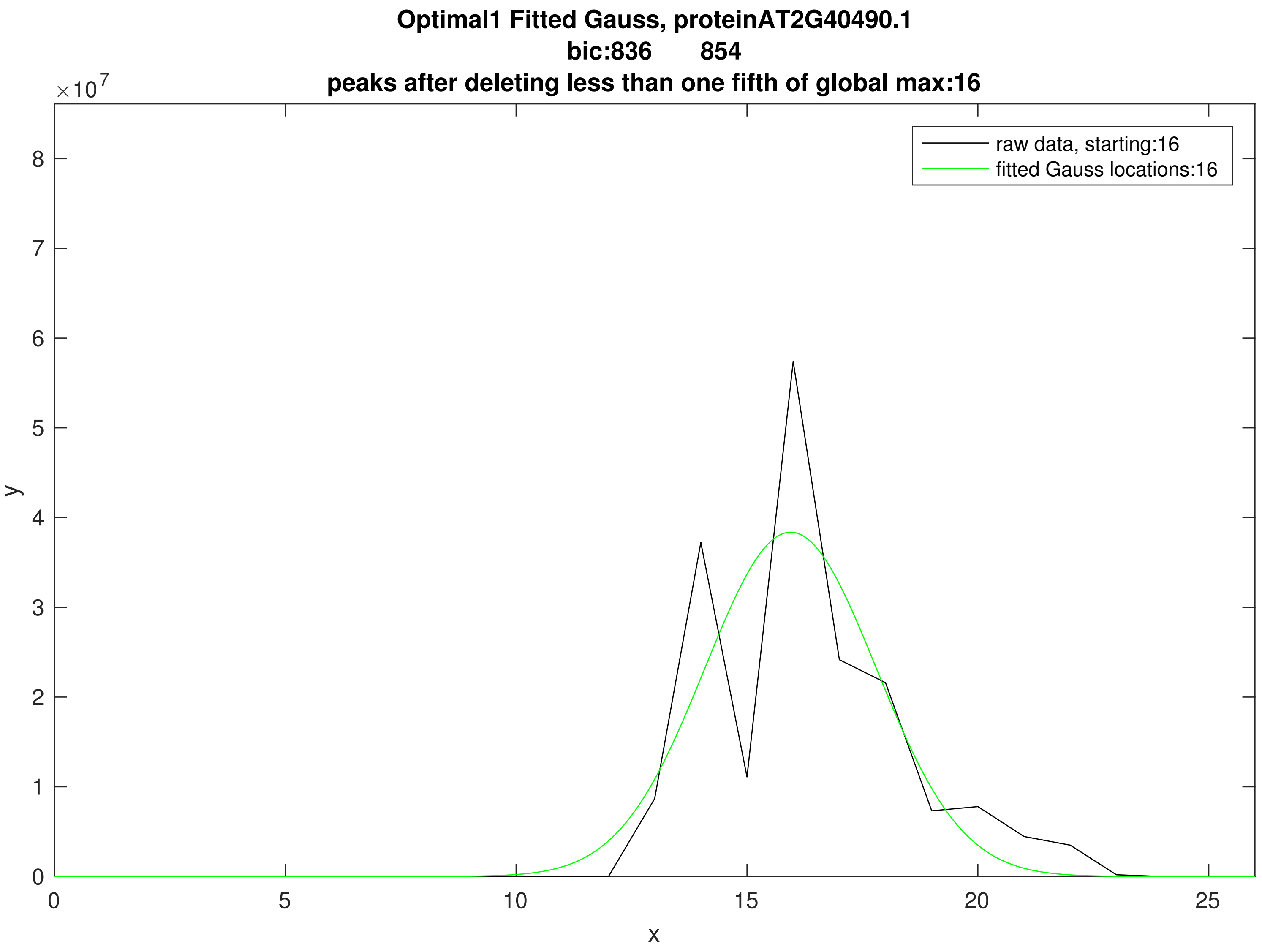

Optimal1 Fitted Gauss, proteinAT2G40490.1
bic:836 854
peaks after deleting less than one fifth of global max:16
×10^7
raw data, starting:16
fitted Gauss locations:16
y
x
0
1
2
3
4
5
6
7
8
0
5
10
15
20
25

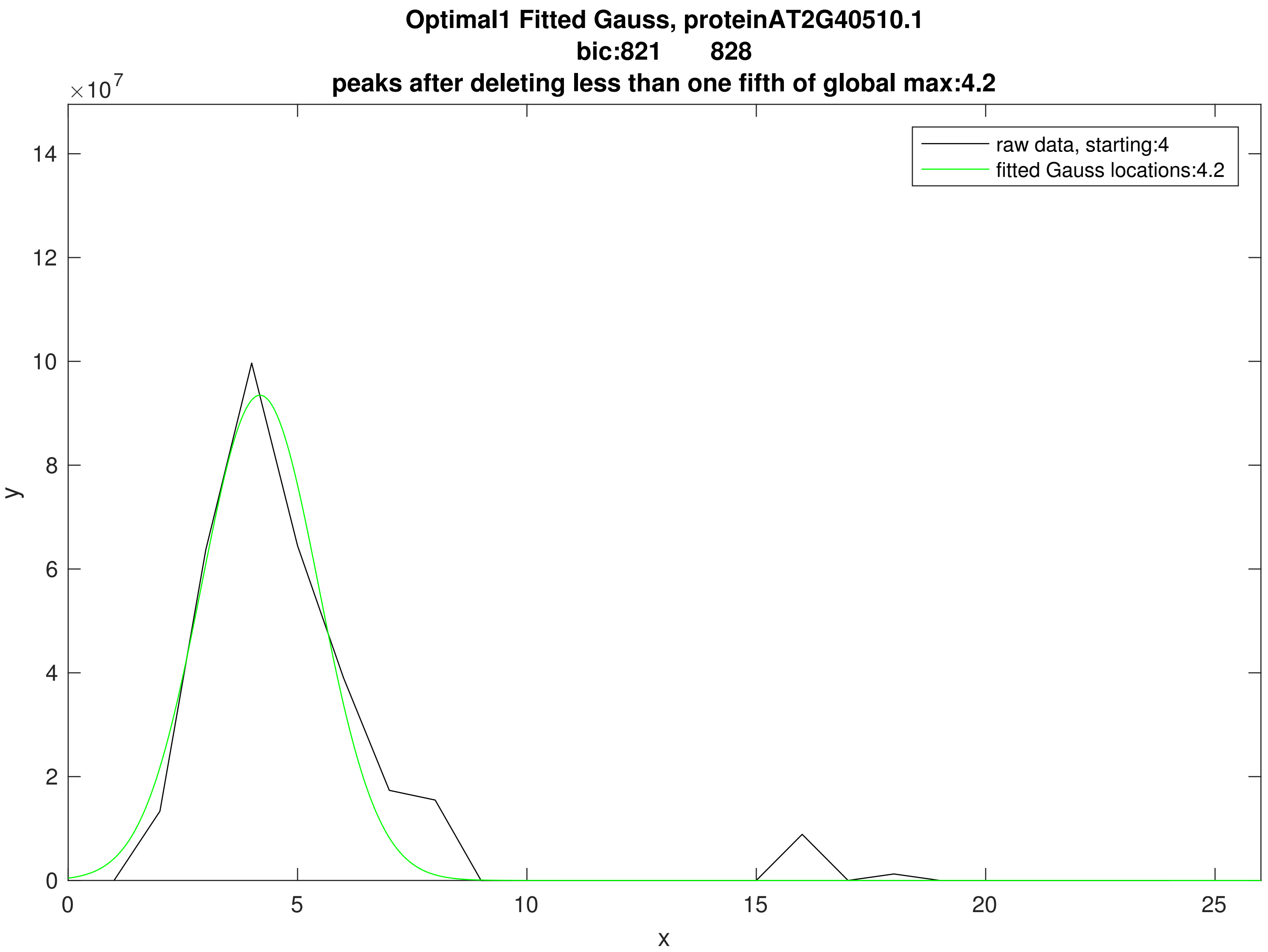

Optimal1 Fitted Gauss, proteinAT2G40510.1
bic:821 828
peaks after deleting less than one fifth of global max:4.2
×10^7
raw data, starting:4
fitted Gauss locations:4.2
y
x
0
2
4
6
8
10
12
14
0
5
10
15
20
25

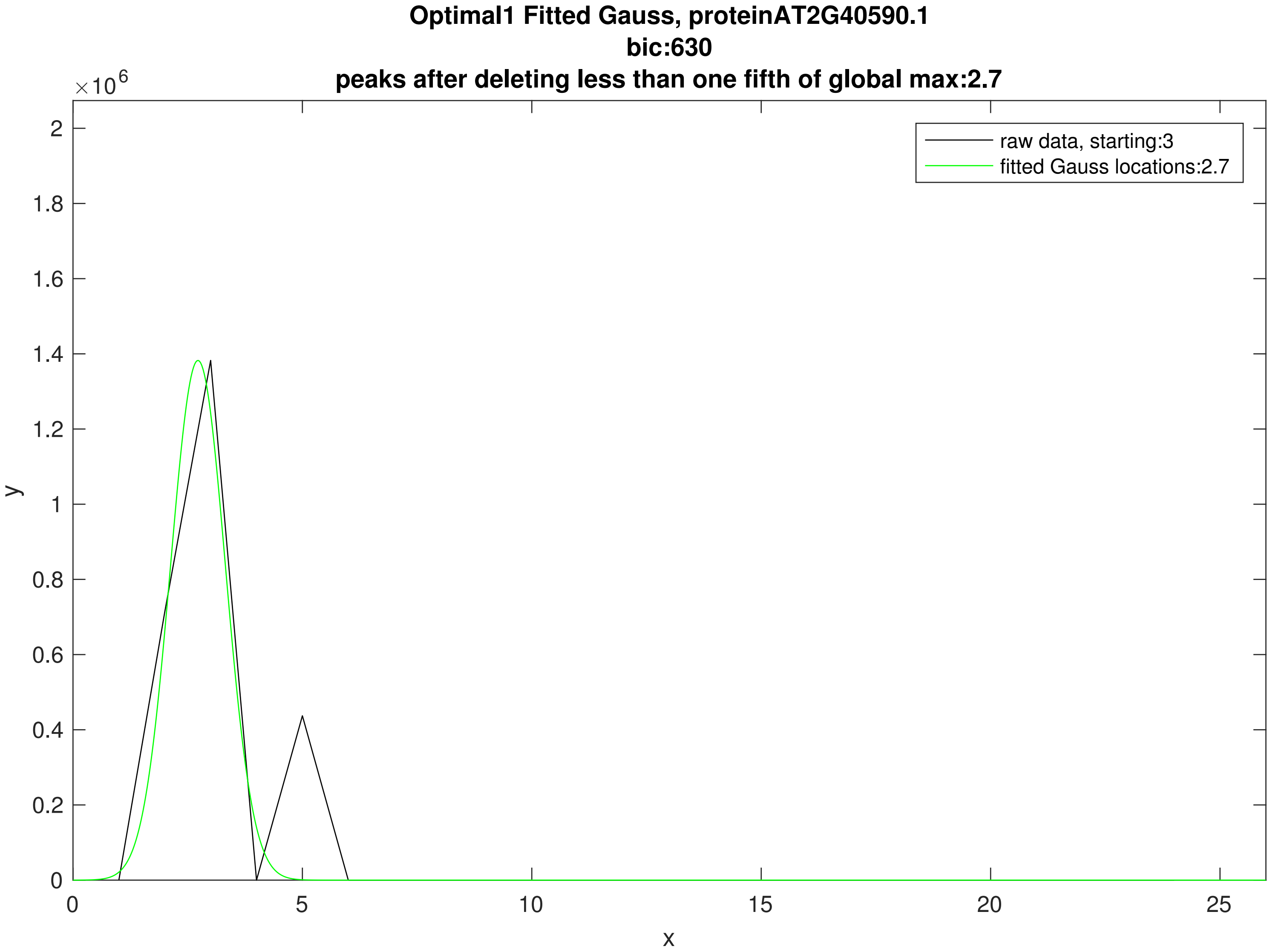

Optimal1 Fitted Gauss, proteinAT2G40590.1
bic:630
peaks after deleting less than one fifth of global max:2.7
×10^6
raw data, starting:3
fitted Gauss locations:2.7
y
x
0
0.2
0.4
0.6
0.8
1
1.2
1.4
1.6
1.8
2
0
5
10
15
20
25

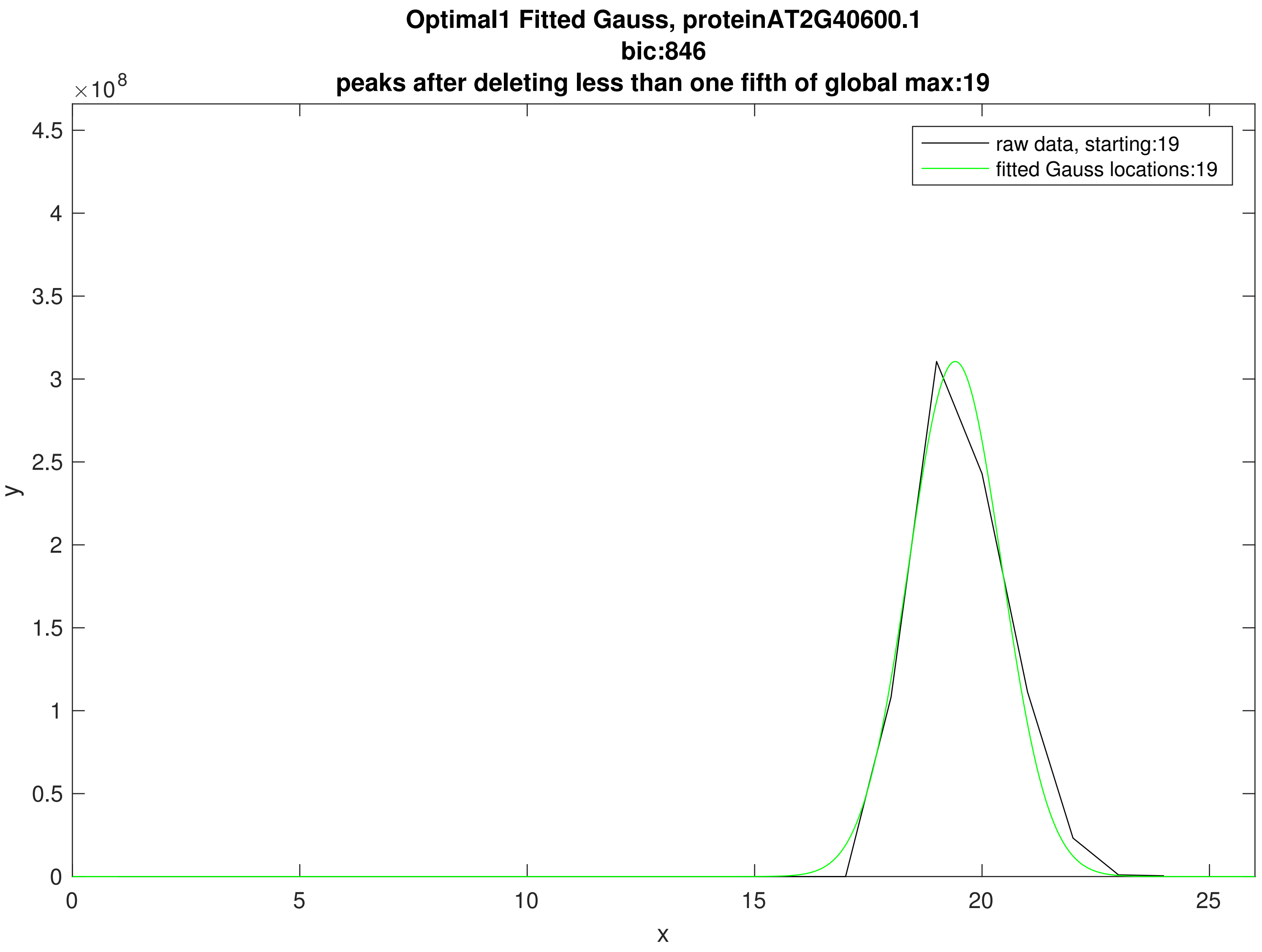

Optimal1 Fitted Gauss, proteinAT2G40600.1
bic:846
peaks after deleting less than one fifth of global max:19
×10^8
4.5
4
3.5
3
2.5
2
1.5
1
0.5
0
y
0
5
10
15
20
25
x
raw data, starting:19
fitted Gauss locations:19

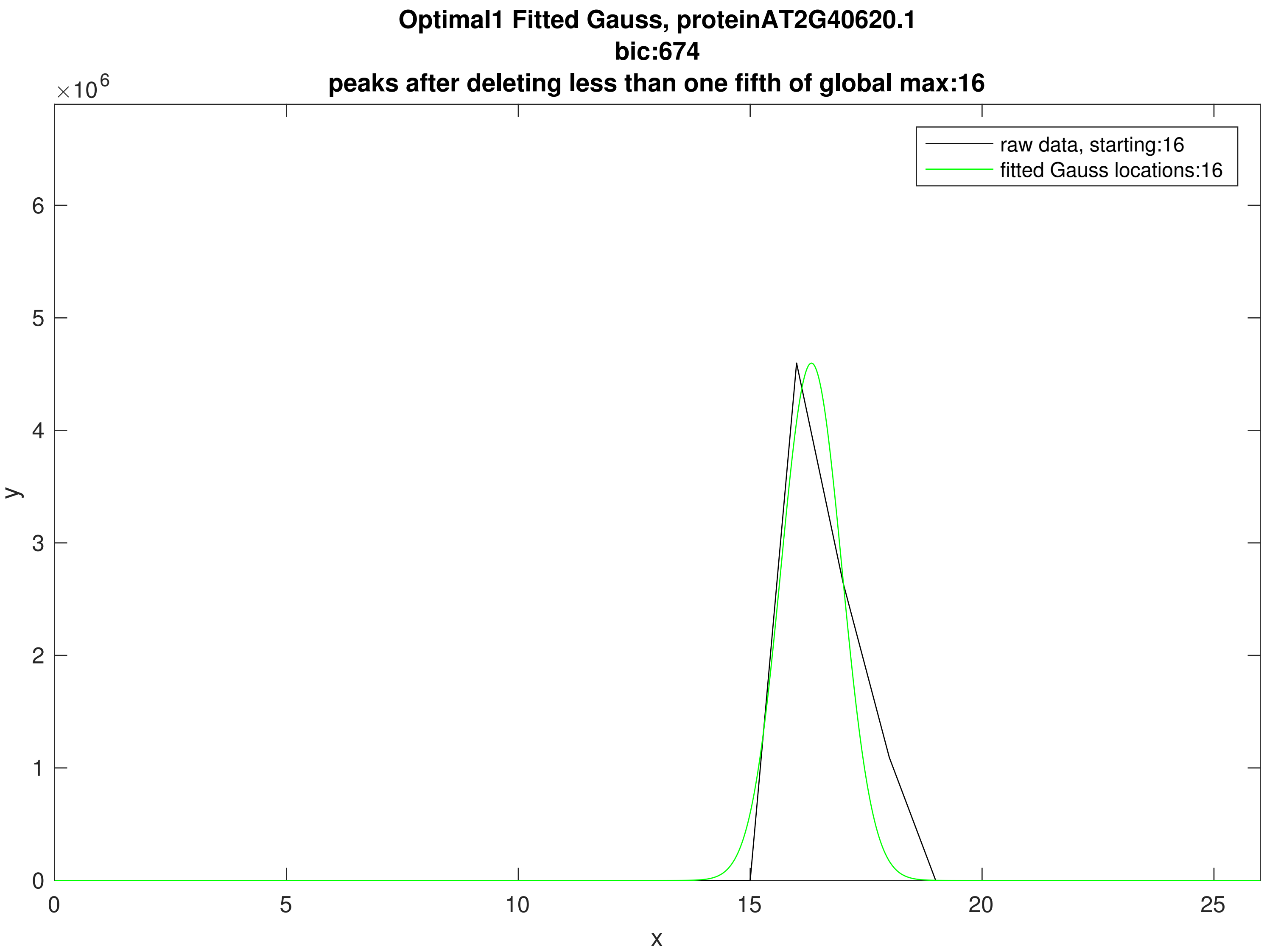

Optimal1 Fitted Gauss, proteinAT2G40620.1
bic:674
peaks after deleting less than one fifth of global max:16
×10^6
raw data, starting:16
fitted Gauss locations:16
y
x
0
1
2
3
4
5
6
0
5
10
15
20
25

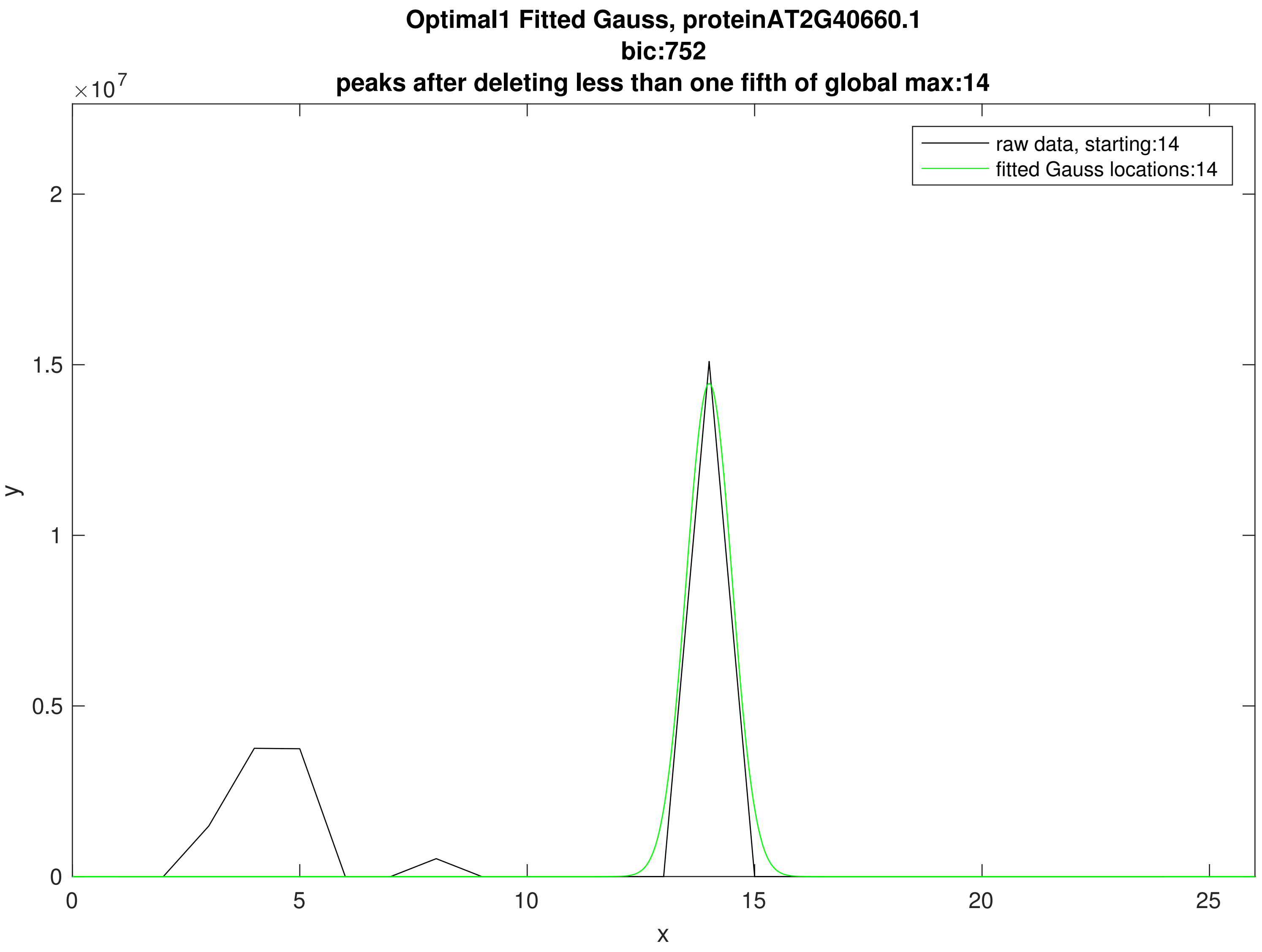

Optimal1 Fitted Gauss, proteinAT2G40660.1
bic:752
peaks after deleting less than one fifth of global max:14
×10^7
raw data, starting:14
fitted Gauss locations:14
y
x
0
0.5
1
1.5
2
5
10
15
20
25

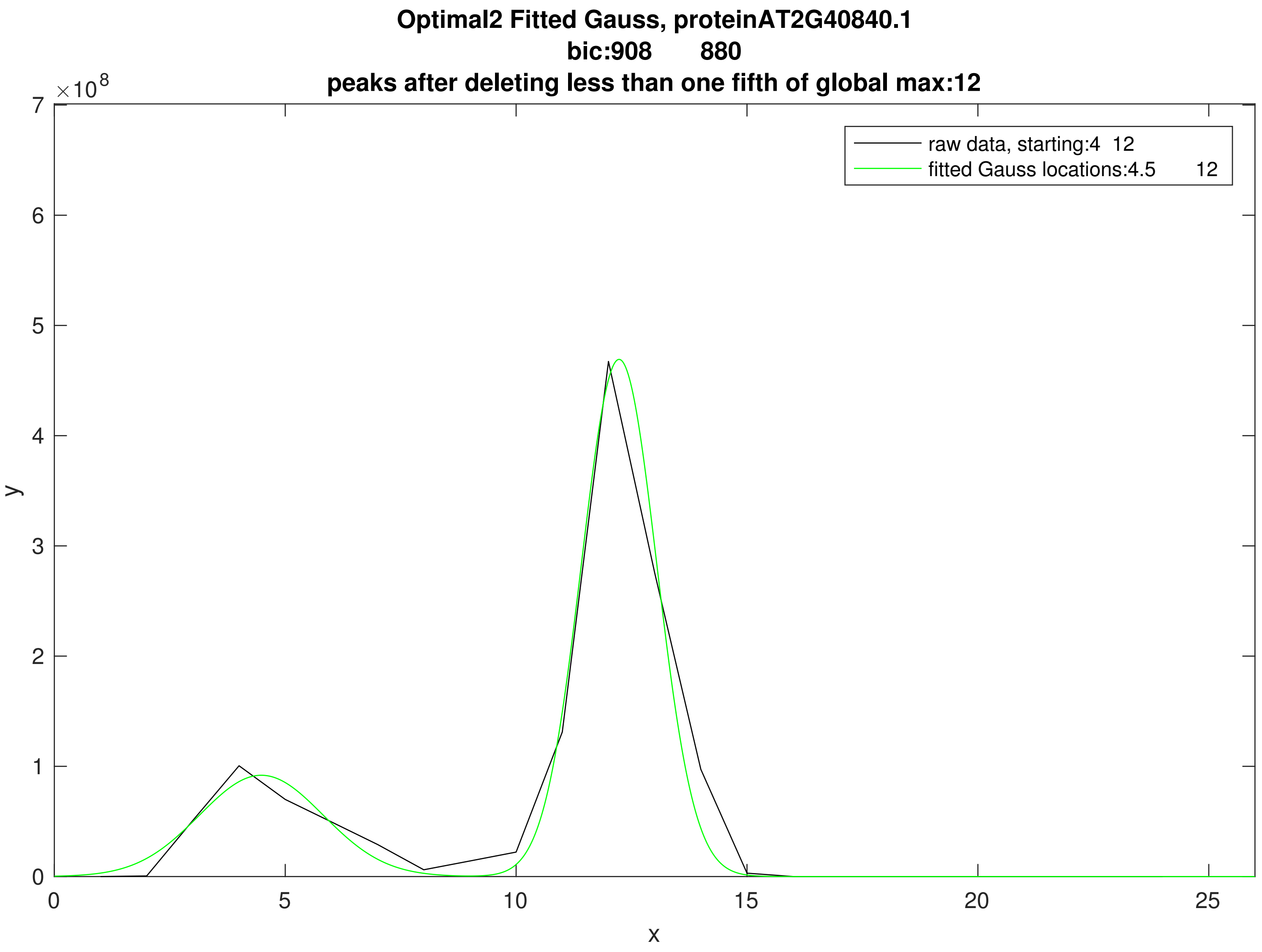
Optimal2 Fitted Gauss, proteinAT2G40840.1
bic:908 880
peaks after deleting less than one fifth of global max:12
×10^8
raw data, starting:4 12
fitted Gauss locations:4.5 12
y
x
0
1
2
3
4
5
6
7
0
5
10
15
20
25

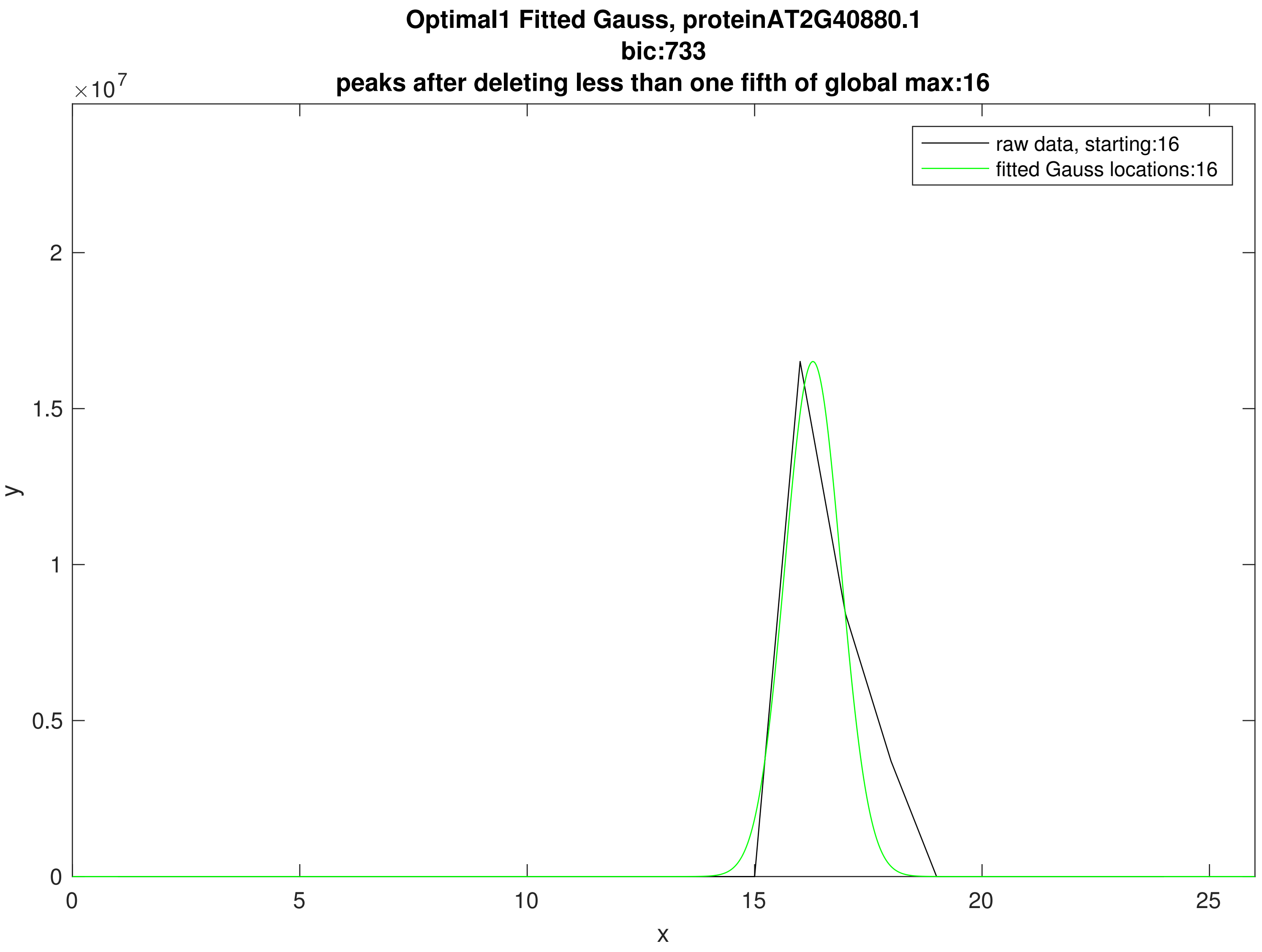

Optimal1 Fitted Gauss, proteinAT2G40880.1
bic:733
peaks after deleting less than one fifth of global max:16
×10^7
raw data, starting:16
fitted Gauss locations:16
2
1.5
1
0.5
0
y
0
5
10
15
20
25
x

**Optimal1 Fitted Gauss, proteinAT2G41100.4;AT2G41100.1**
**bic:762**
**peaks after deleting less than one fifth of global max:18**

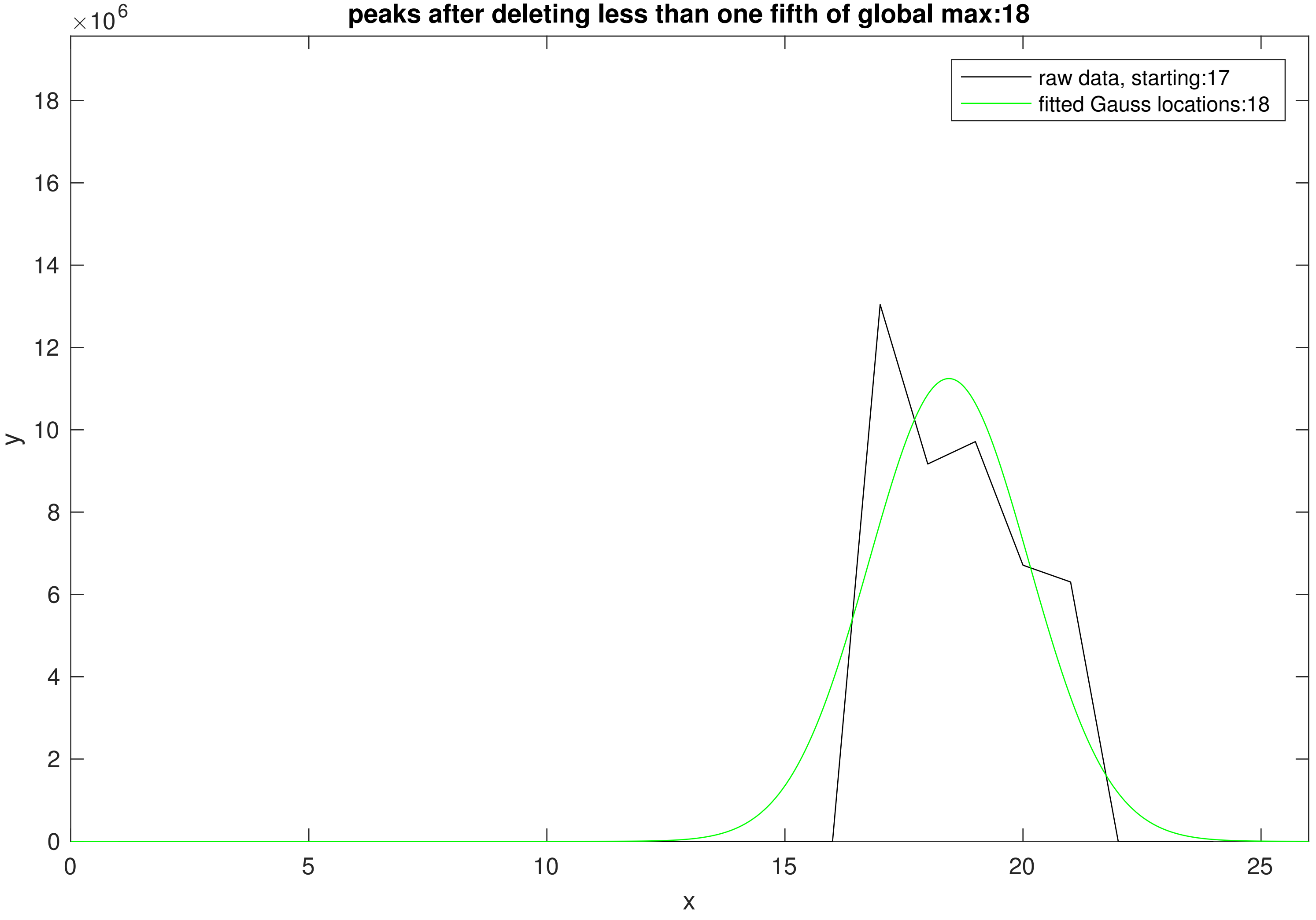

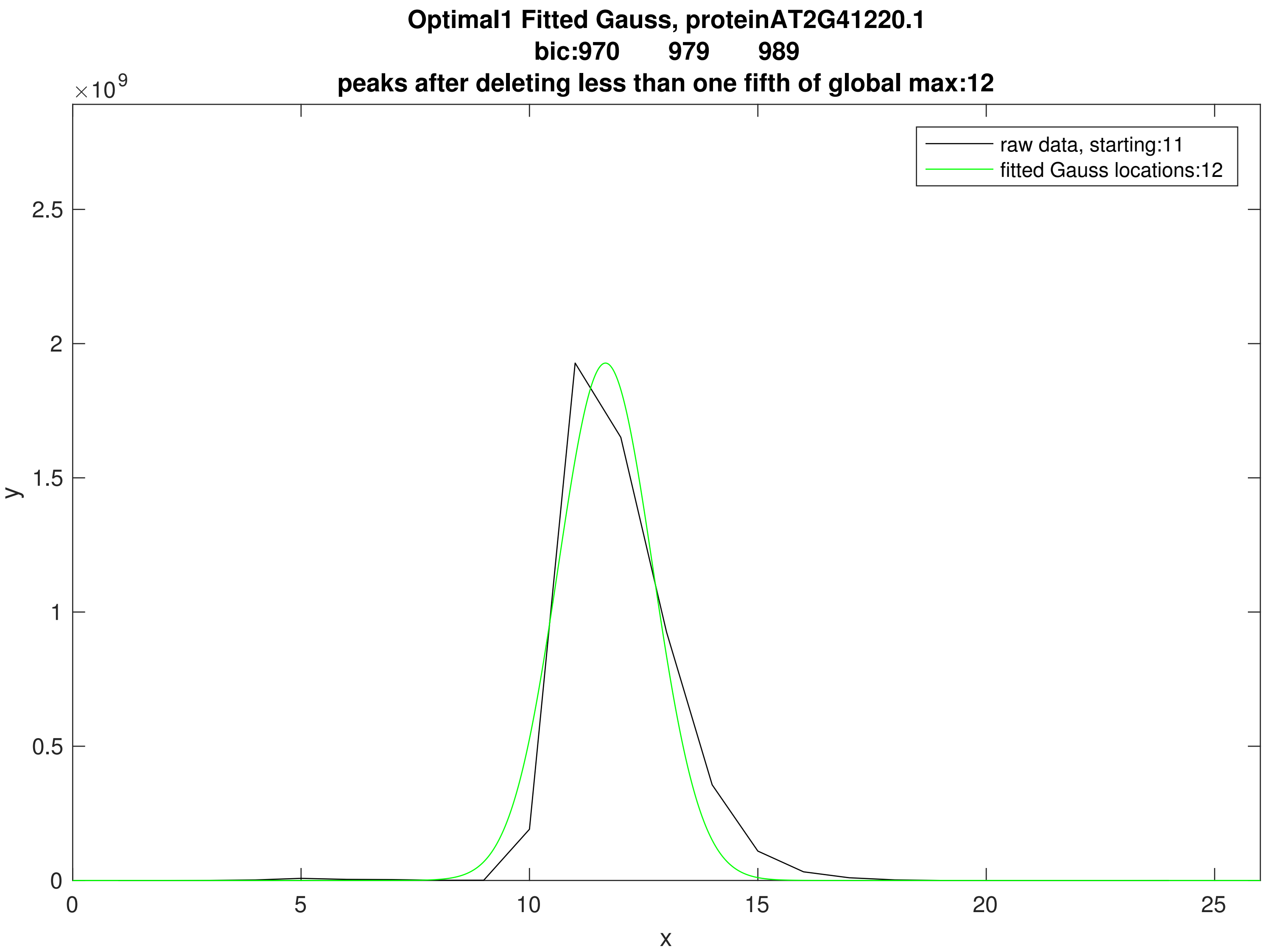

Optimal1 Fitted Gauss, proteinAT2G41220.1
bic:970 979 989
peaks after deleting less than one fifth of global max:12
×10^9
raw data, starting:11
fitted Gauss locations:12
2.5
2
1.5
1
0.5
0
y
0
5
10
15
20
25
x

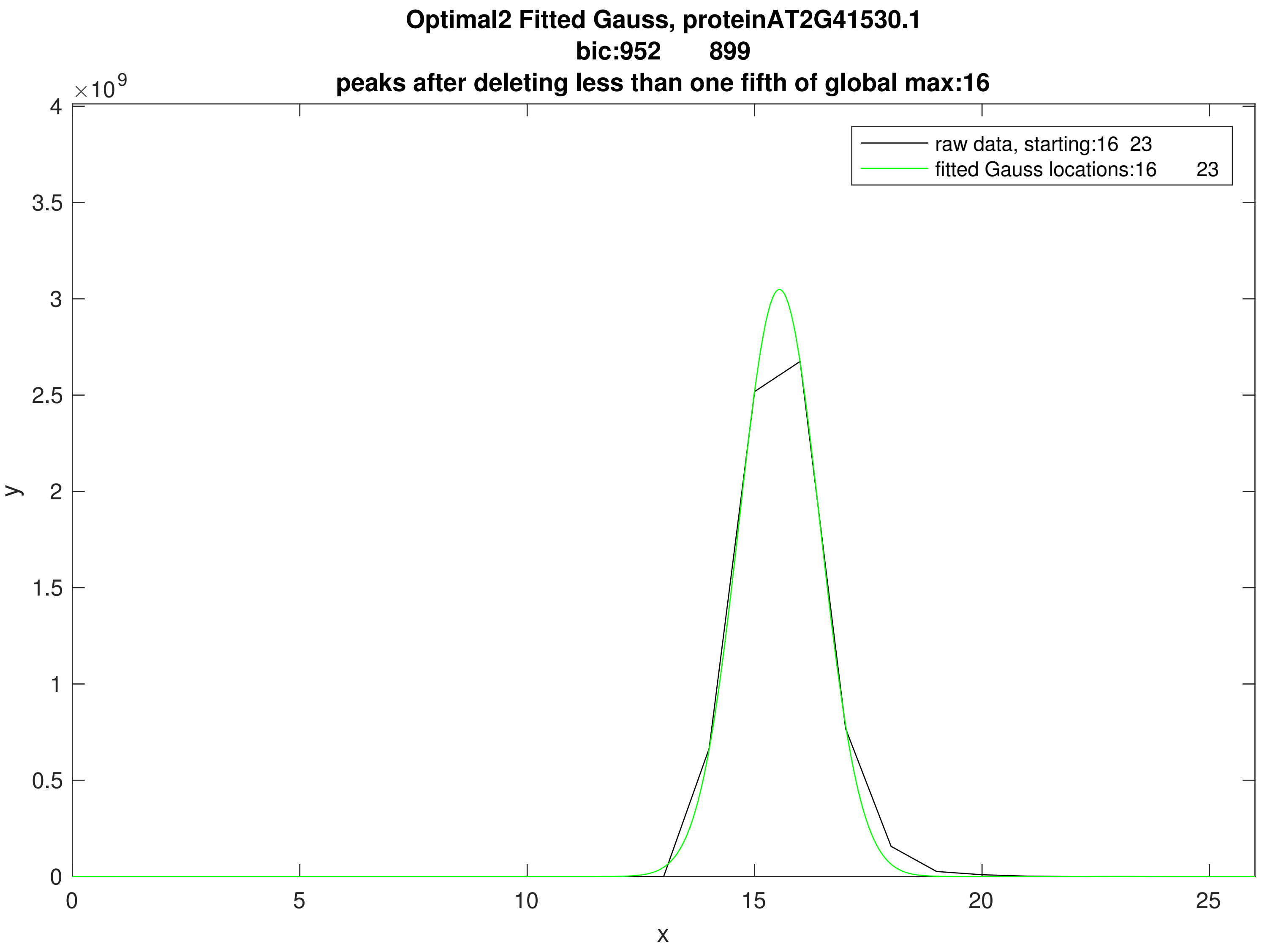

Optimal2 Fitted Gauss, proteinAT2G41530.1
bic:952 899
peaks after deleting less than one fifth of global max:16
×10^9
4
3.5
3
2.5
2
1.5
1
0.5
0
y
0
5
10
15
20
25
x
raw data, starting:16 23
fitted Gauss locations:16 23

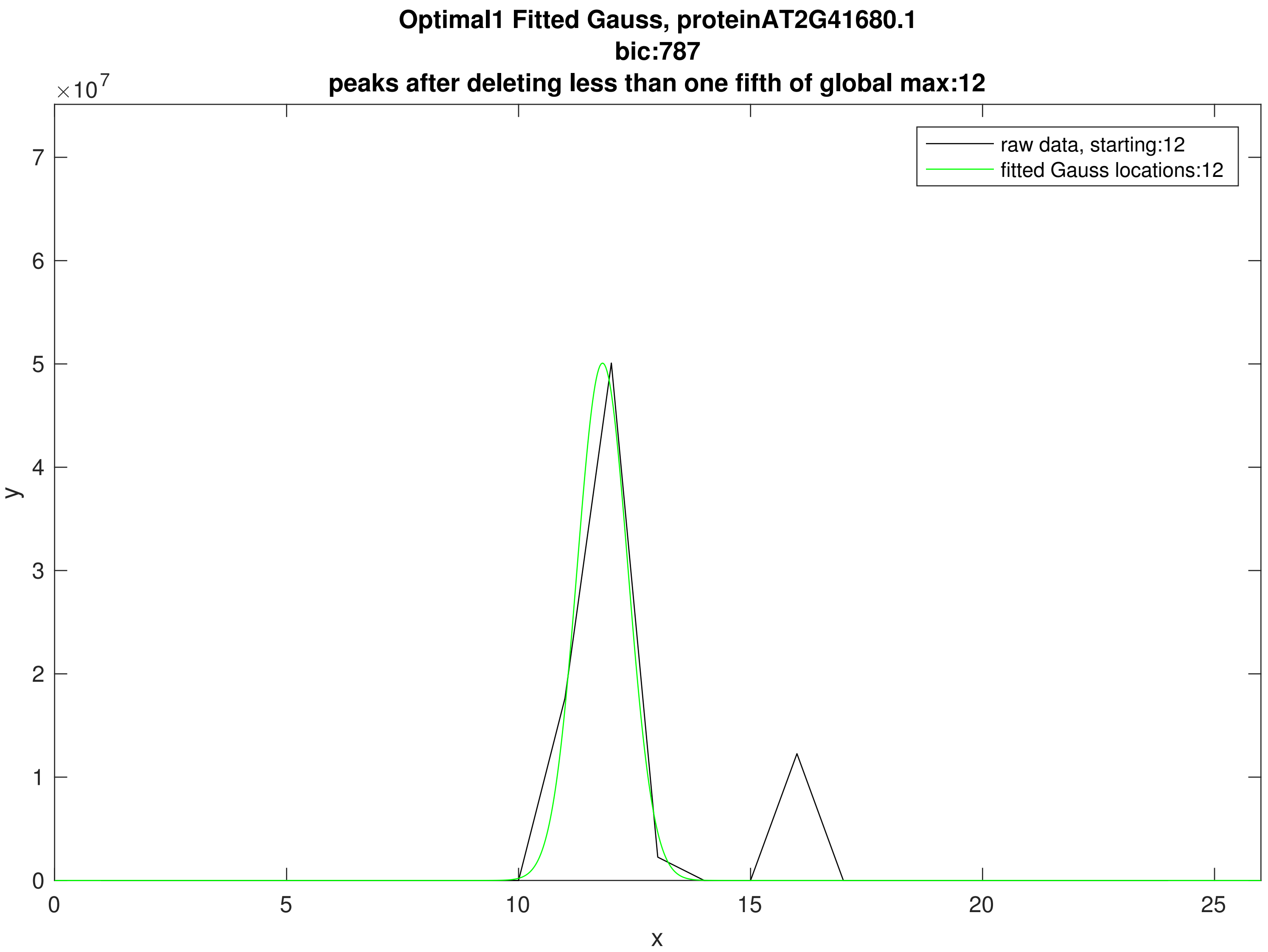
Optimal1 Fitted Gauss, proteinAT2G41680.1
bic:787
peaks after deleting less than one fifth of global max:12
×10^7
raw data, starting:12
fitted Gauss locations:12
y
x
0
1
2
3
4
5
6
7
0
5
10
15
20
25

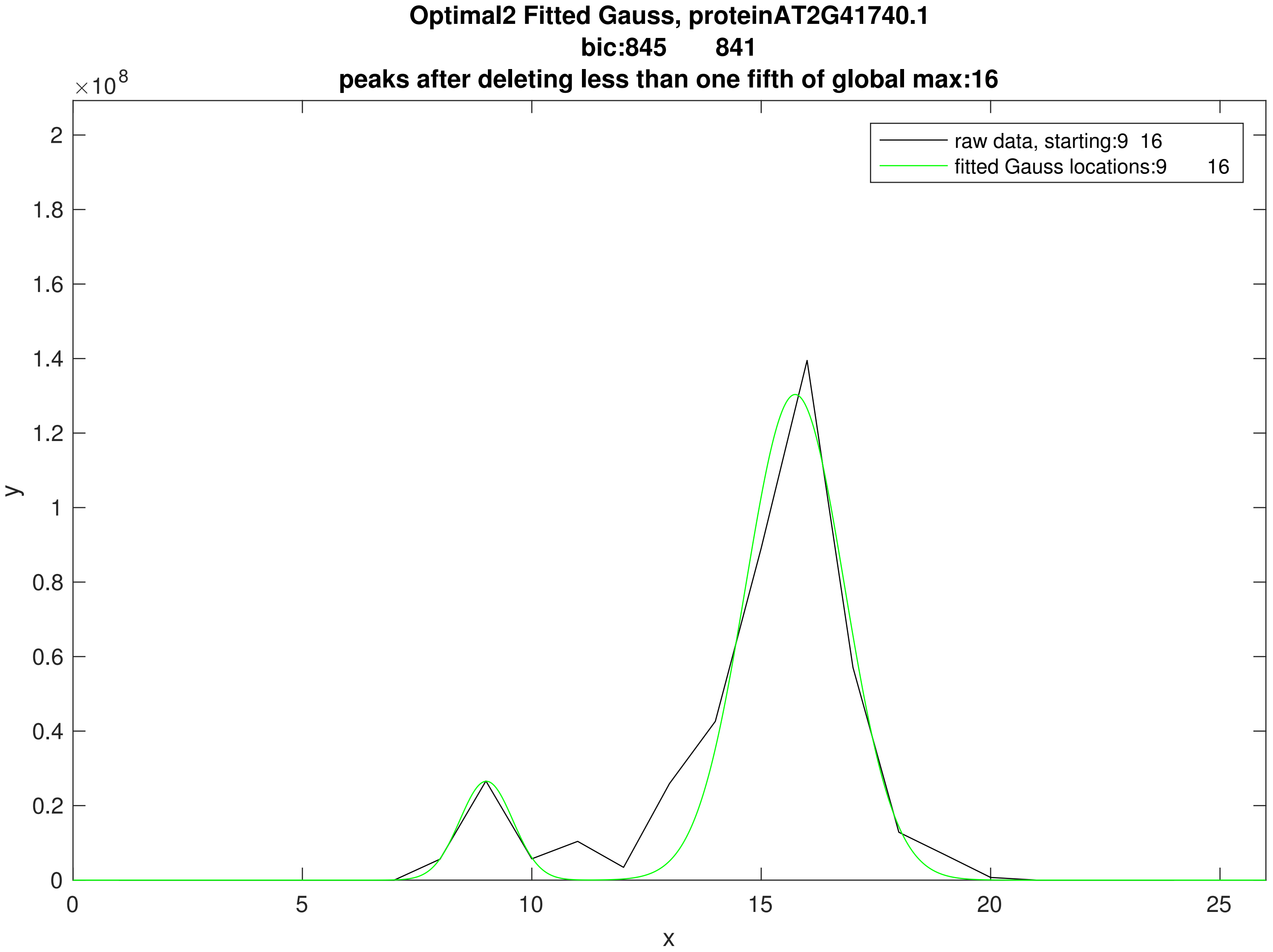
Optimal2 Fitted Gauss, proteinAT2G41740.1
bic:845 841
peaks after deleting less than one fifth of global max:16
×10^8
raw data, starting:9 16
fitted Gauss locations:9 16
y
x
0
0.2
0.4
0.6
0.8
1
1.2
1.4
1.6
1.8
2
0
5
10
15
20
25

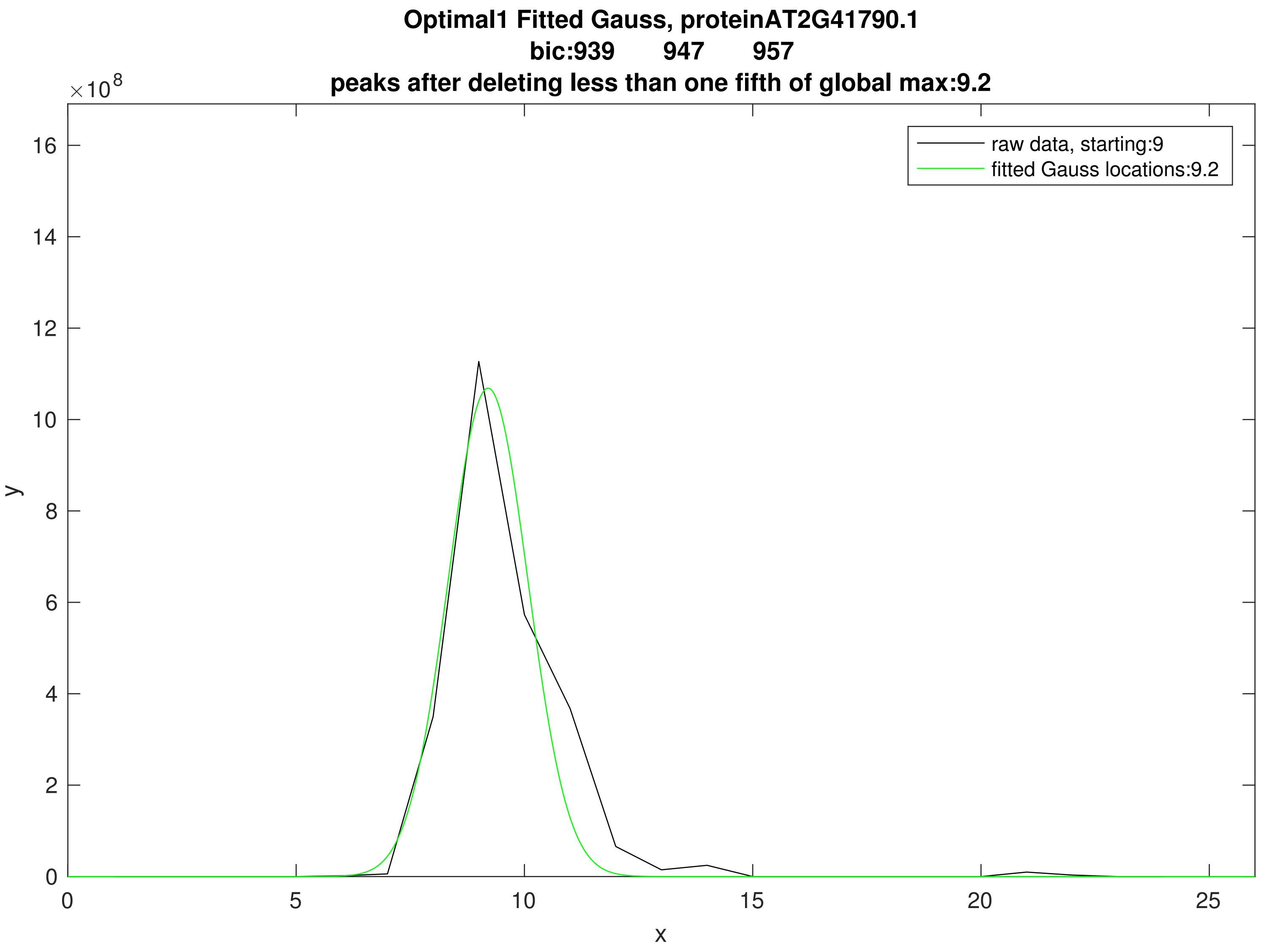

Optimal1 Fitted Gauss, proteinAT2G41790.1
bic:939 947 957
peaks after deleting less than one fifth of global max:9.2
×10^8
16
14
12
10
8
6
4
2
0
y
0
5
10
15
20
25
x
raw data, starting:9
fitted Gauss locations:9.2

**Optimal1 Fitted Gauss, proteinAT2G41840.1**
**bic:857 866 875 885**
**peaks after deleting less than one fifth of global max:4**

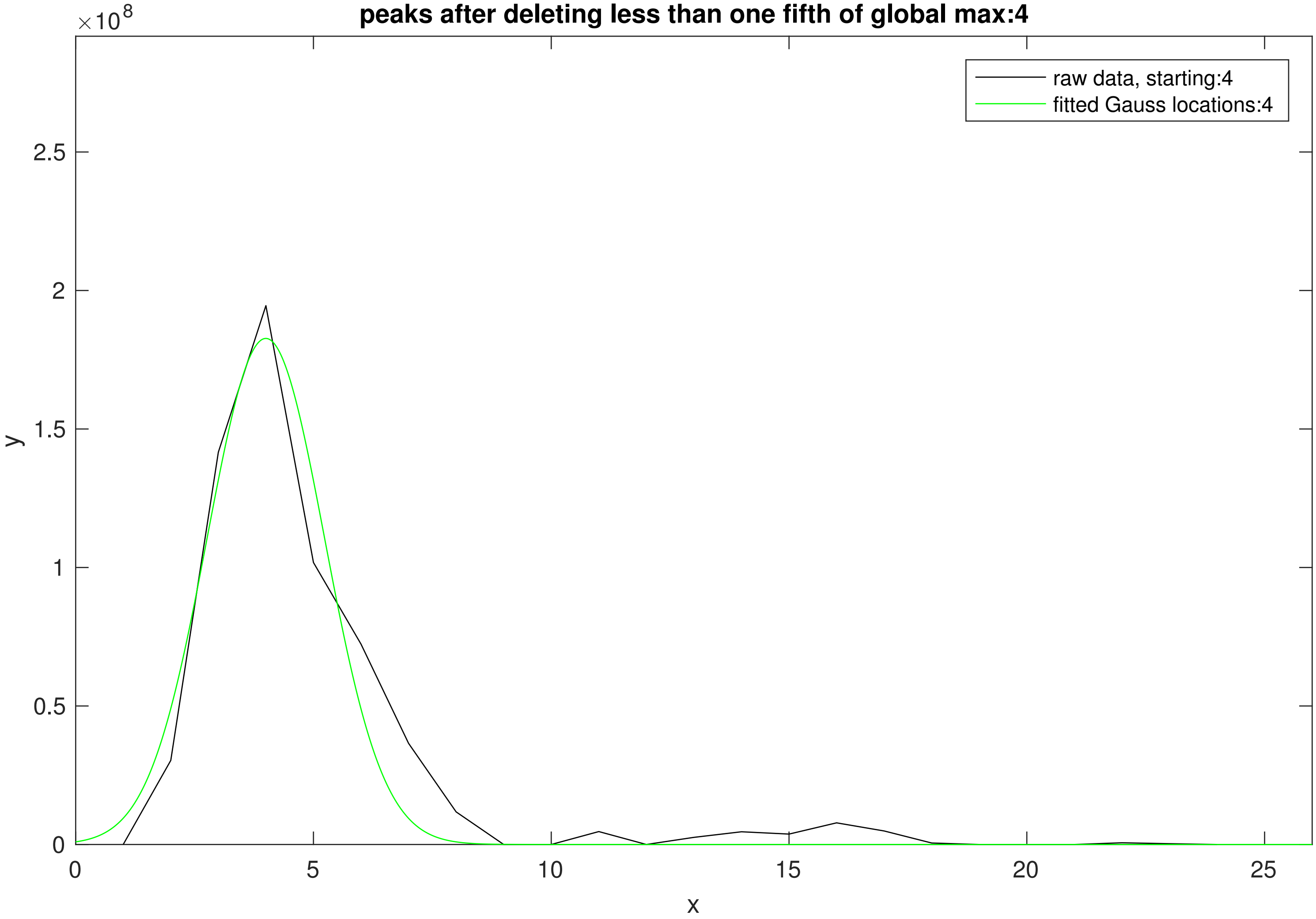

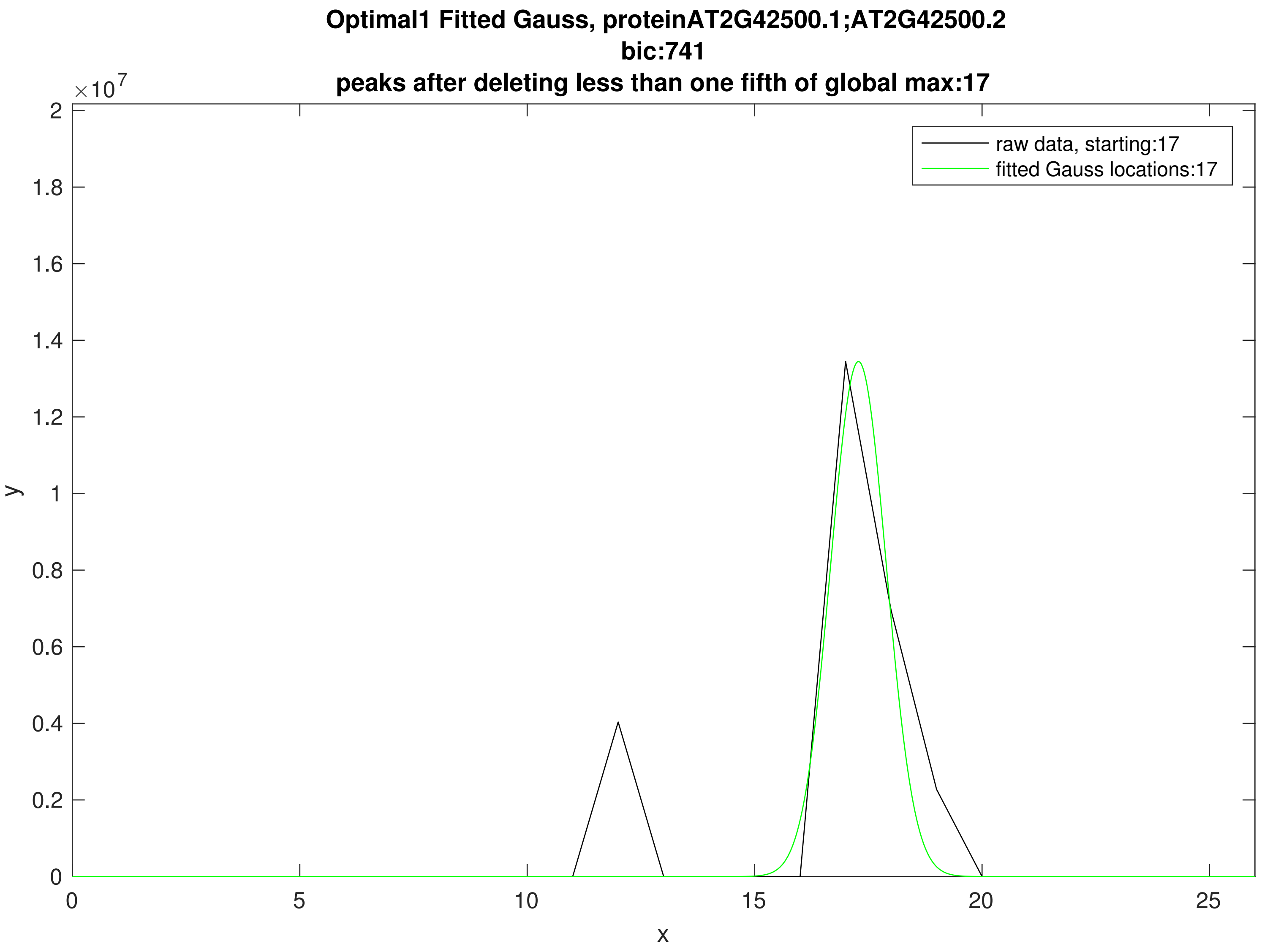
Optimal1 Fitted Gauss, proteinAT2G42500.1;AT2G42500.2
bic:741
peaks after deleting less than one fifth of global max:17
×10^7
2
1.8
1.6
1.4
1.2
1
0.8
0.6
0.4
0.2
0
y
0
5
10
15
20
25
x
raw data, starting:17
fitted Gauss locations:17

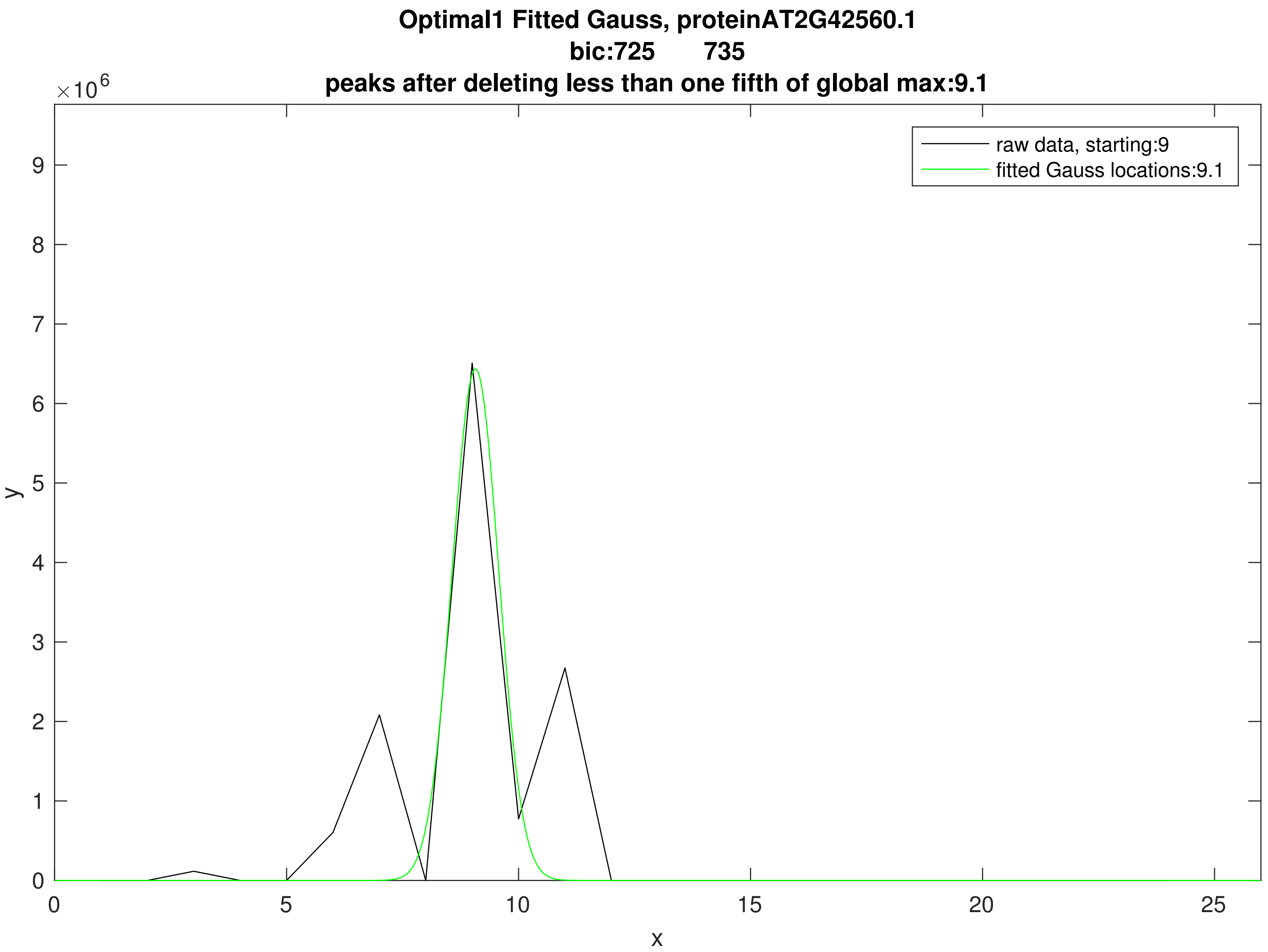

Optimal1 Fitted Gauss, proteinAT2G42560.1
bic:725 735
peaks after deleting less than one fifth of global max:9.1
×10^6
raw data, starting:9
fitted Gauss locations:9.1
y
x
0
1
2
3
4
5
6
7
8
9
10
15
20
25

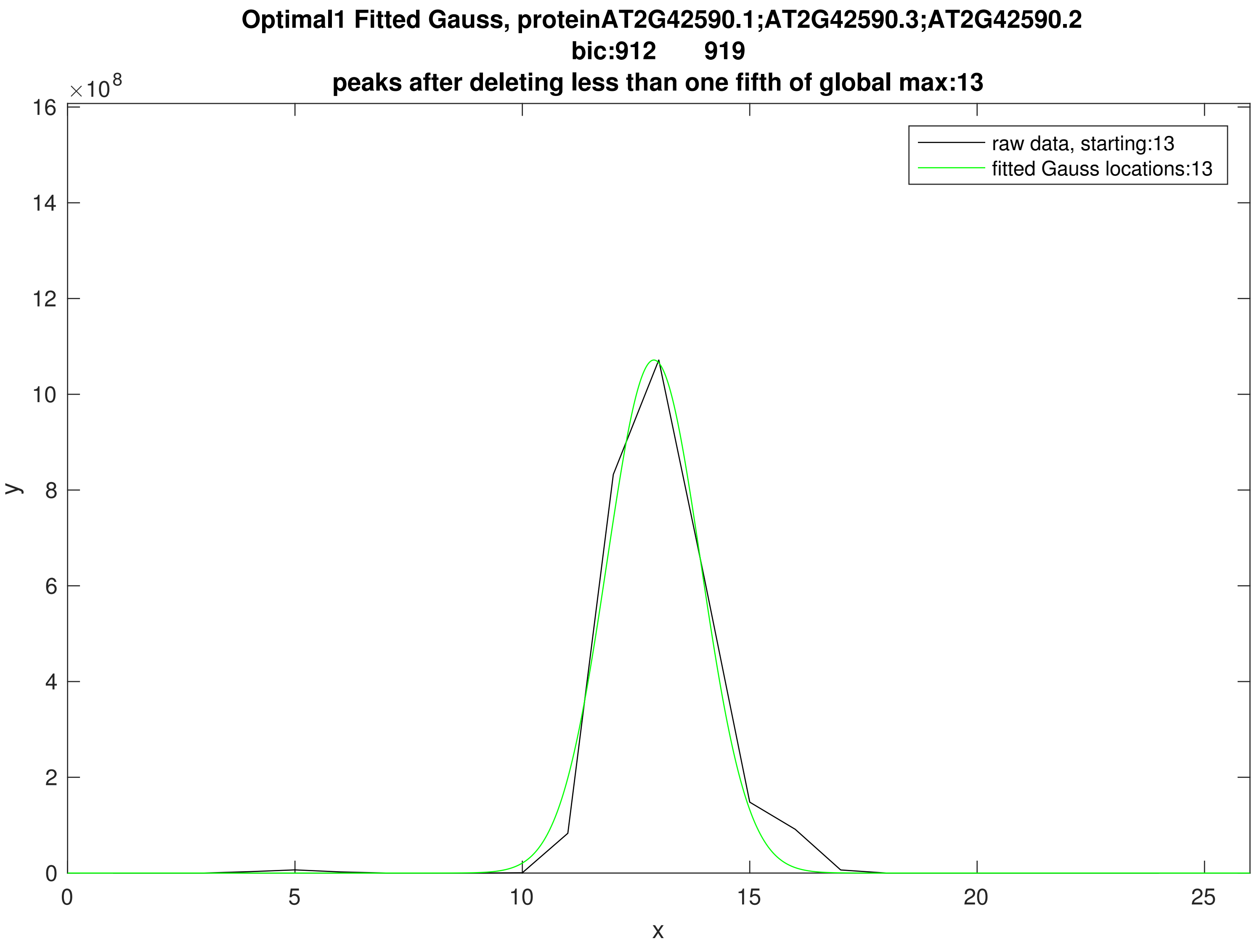
Optimal1 Fitted Gauss, proteinAT2G42590.1;AT2G42590.3;AT2G42590.2
bic:912 919
peaks after deleting less than one fifth of global max:13
×10^8
raw data, starting:13
fitted Gauss locations:13
y
x

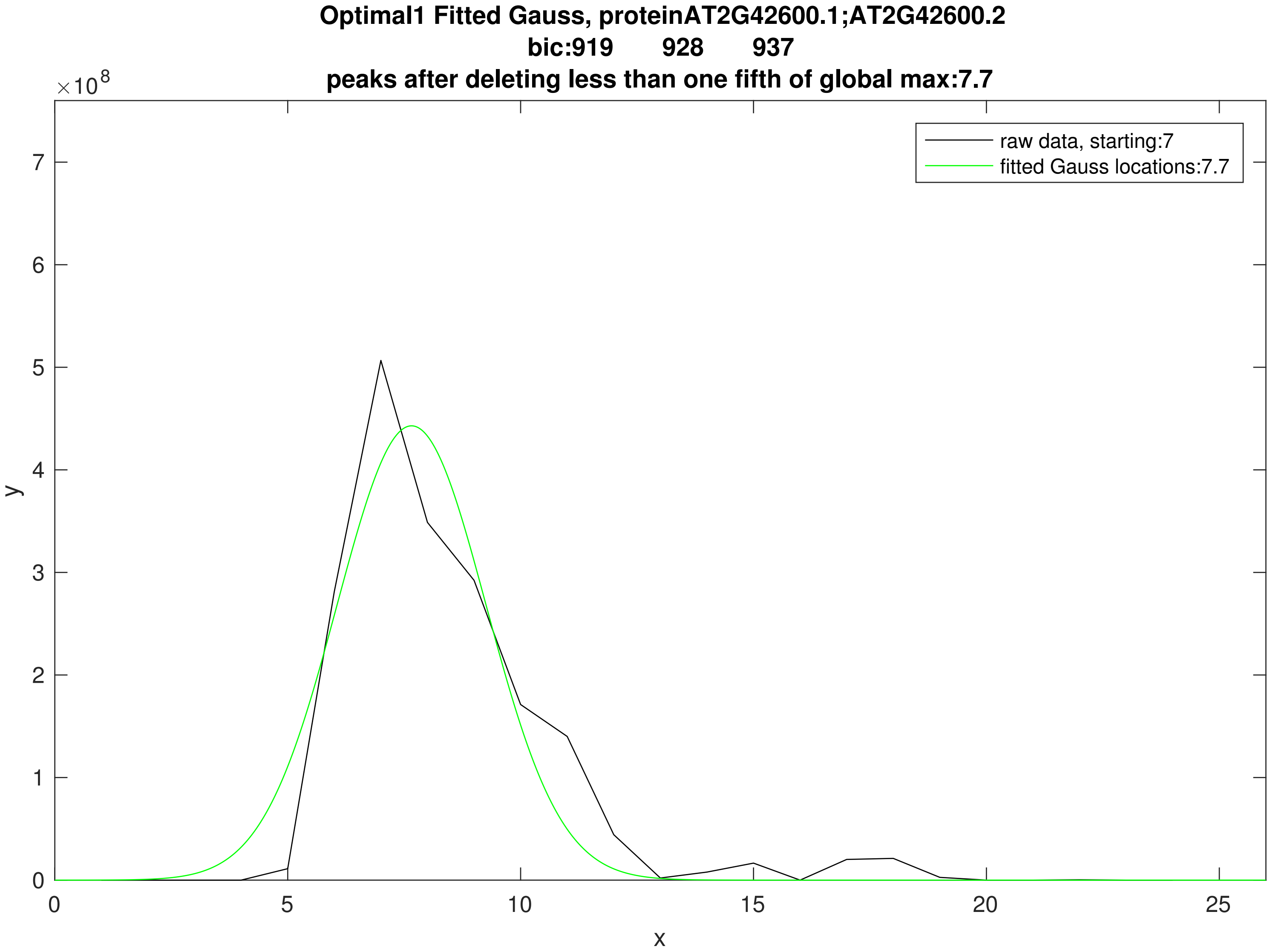
Optimal1 Fitted Gauss, proteinAT2G42600.1;AT2G42600.2
bic:919 928 937
peaks after deleting less than one fifth of global max:7.7
×10^8
raw data, starting:7
fitted Gauss locations:7.7
y
x
0
1
2
3
4
5
6
7
0
5
10
15
20
25

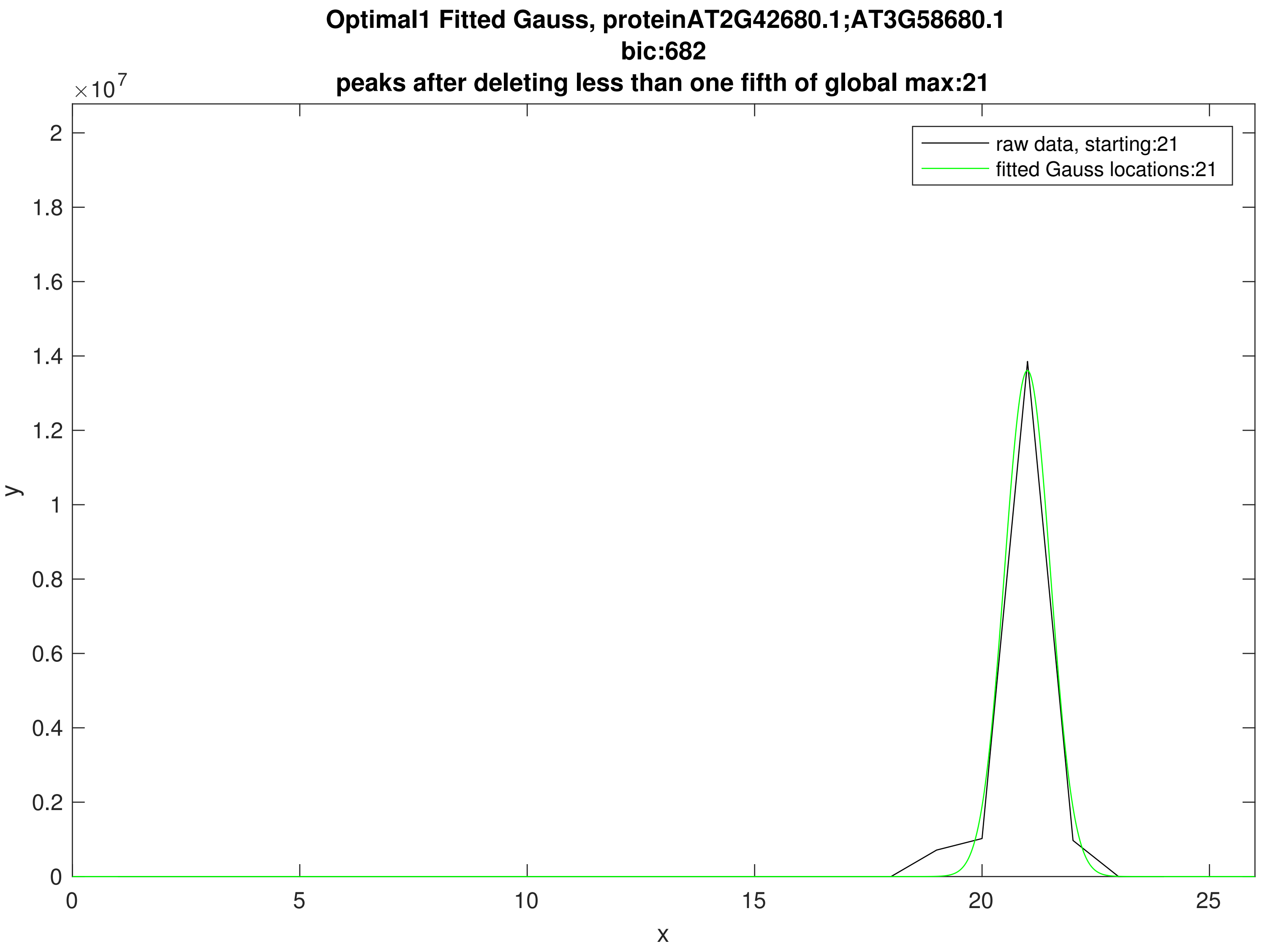
Optimal1 Fitted Gauss, proteinAT2G42680.1;AT3G58680.1
bic:682
peaks after deleting less than one fifth of global max:21
×10^7
raw data, starting:21
fitted Gauss locations:21
y
x
0
0.2
0.4
0.6
0.8
1
1.2
1.4
1.6
1.8
2
0
5
10
15
20
25

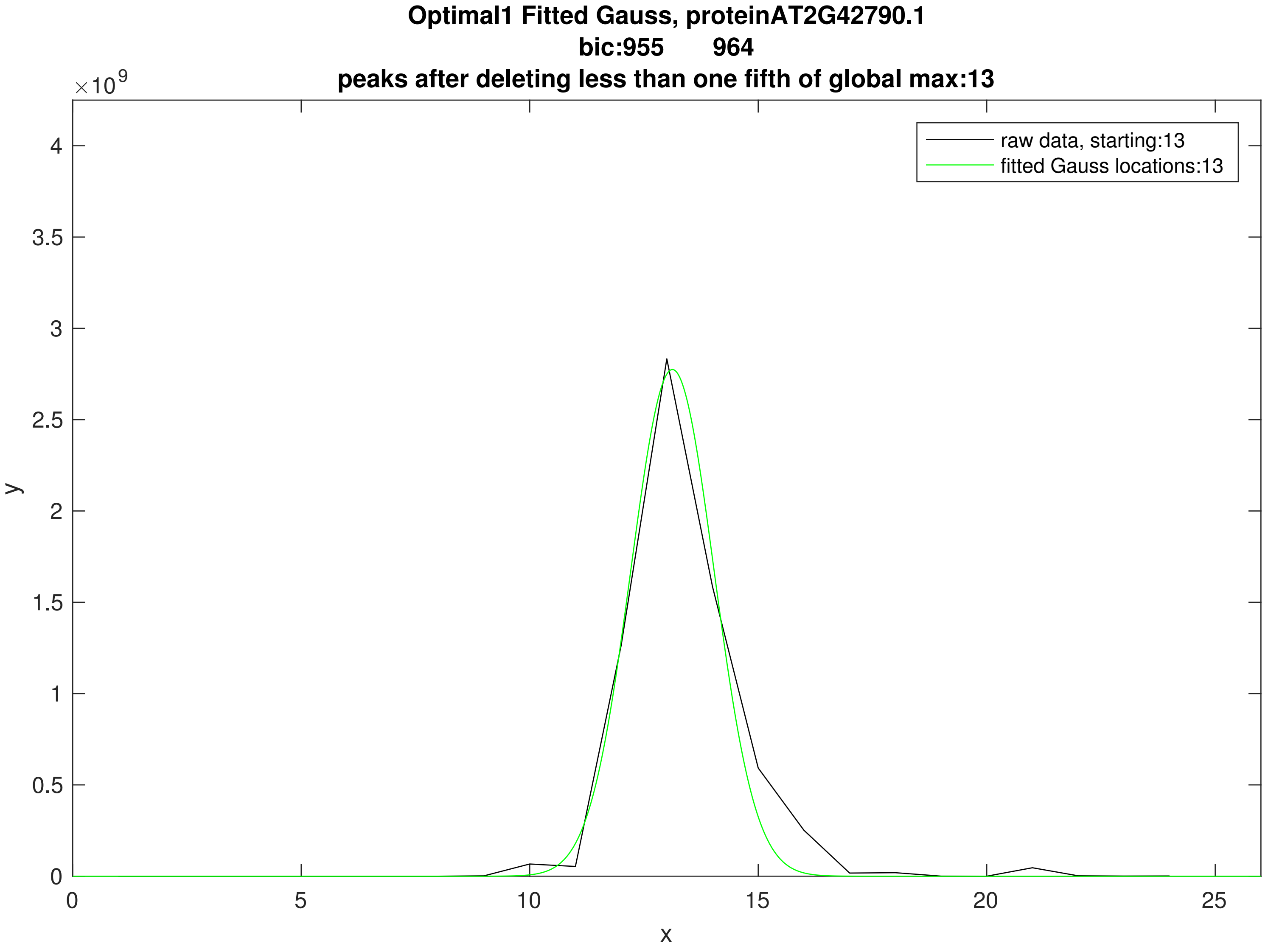
Optimal1 Fitted Gauss, proteinAT2G42790.1
bic:955 964
peaks after deleting less than one fifth of global max:13
×10^9
raw data, starting:13
fitted Gauss locations:13
y
x
0
0.5
1
1.5
2
2.5
3
3.5
4
0
5
10
15
20
25

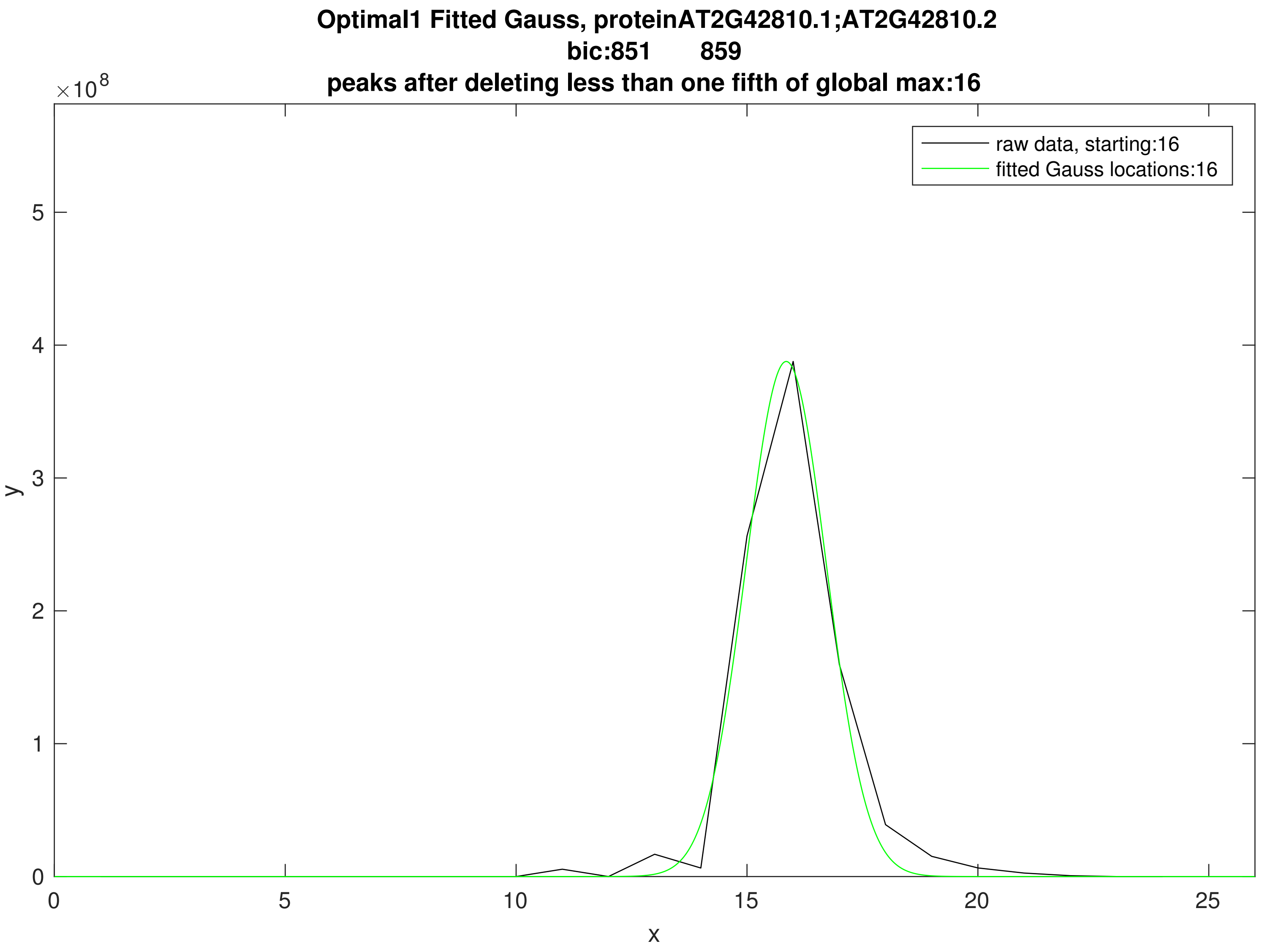
Optimal1 Fitted Gauss, proteinAT2G42810.1;AT2G42810.2
bic:851 859
peaks after deleting less than one fifth of global max:16
×10^8
raw data, starting:16
fitted Gauss locations:16
y
x
0
1
2
3
4
5
0
5
10
15
20
25

#### Optimal2 Fitted Gauss, proteinAT2G42910.1
#### bic:887 820
#### peaks after deleting less than one fifth of global max:6.5 13

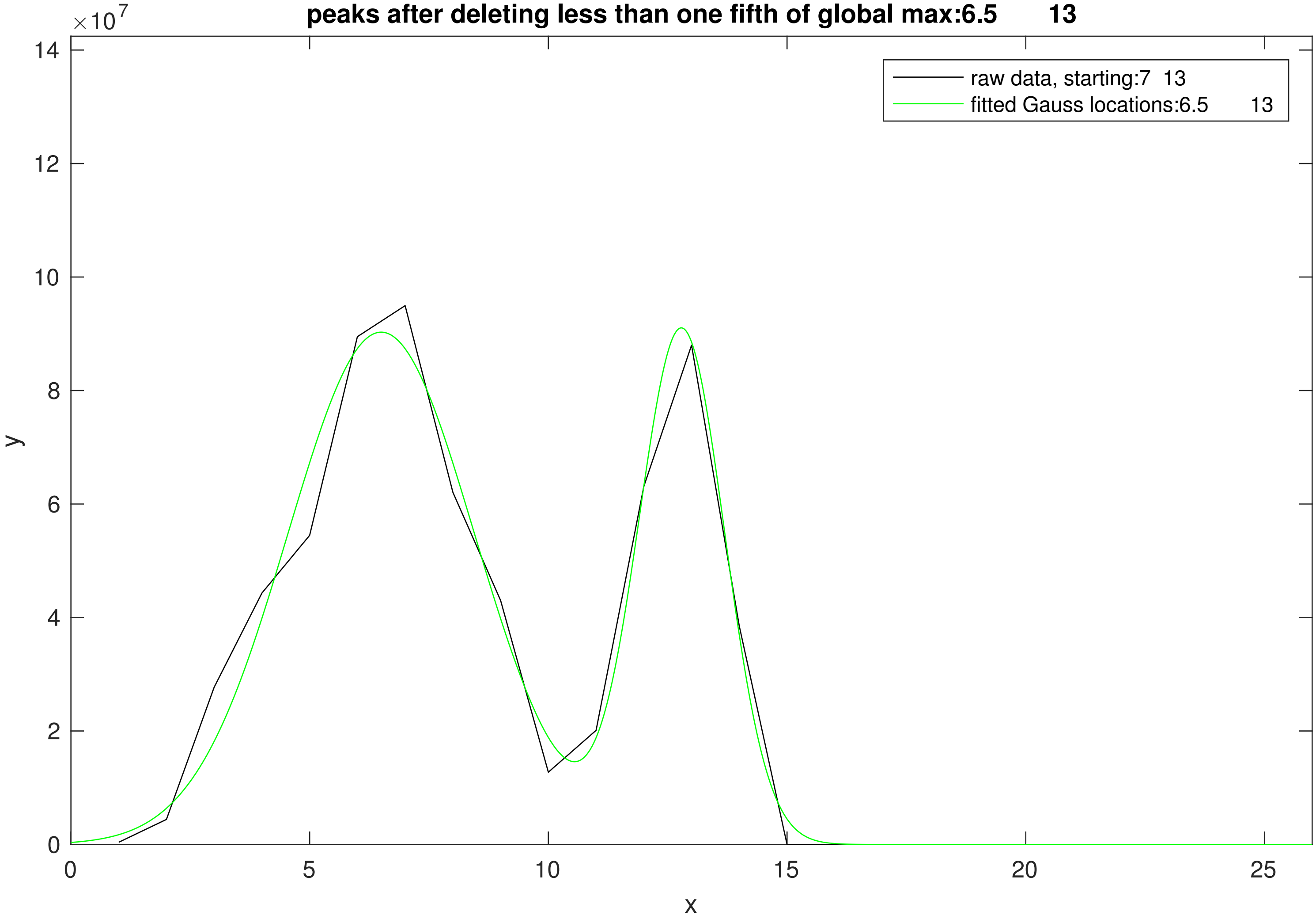

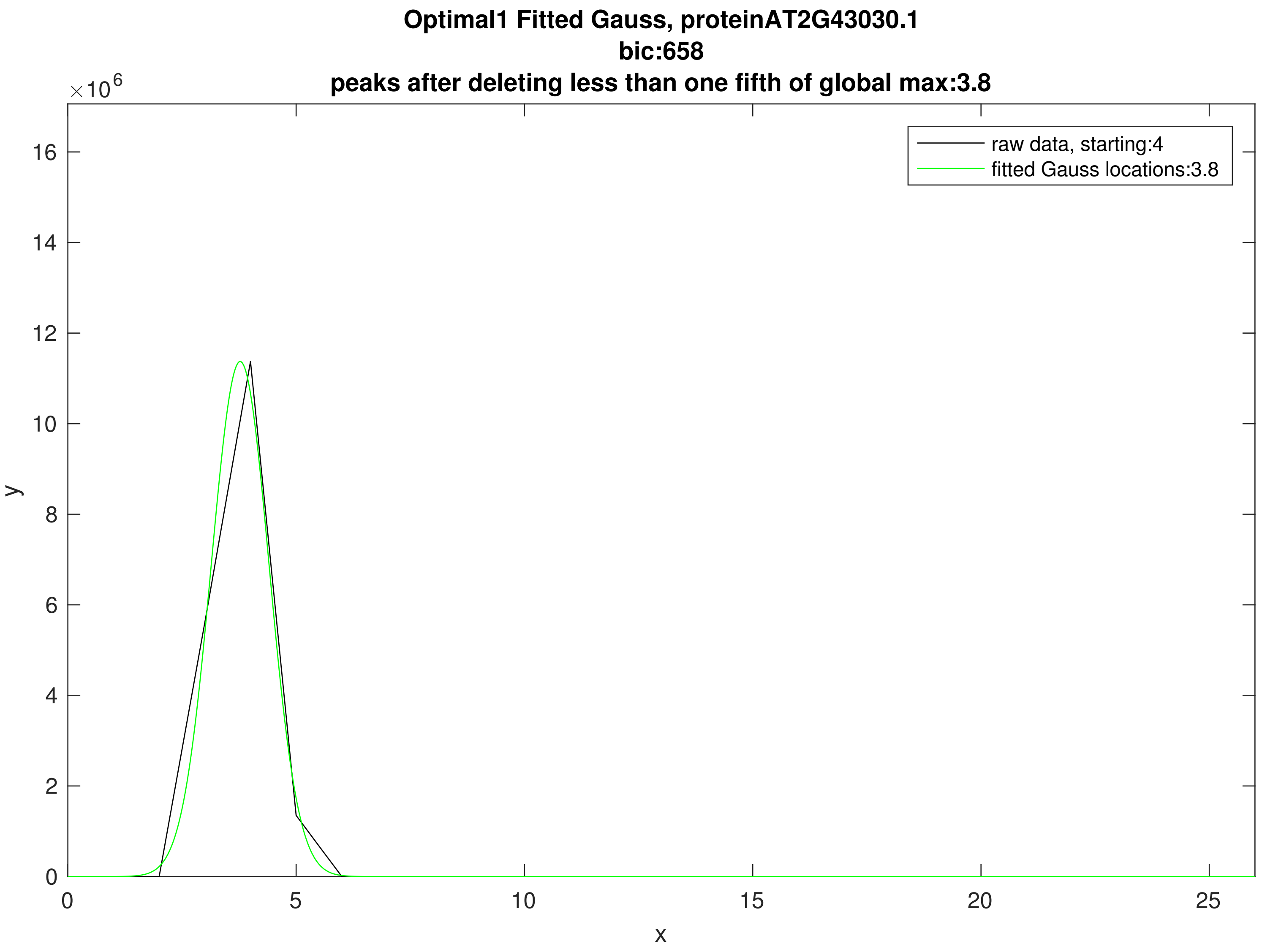
Optimal1 Fitted Gauss, proteinAT2G43030.1
bic:658
peaks after deleting less than one fifth of global max:3.8
×10^6
16
14
12
10
8
6
4
2
0
y
0
5
10
15
20
25
x
raw data, starting:4
fitted Gauss locations:3.8

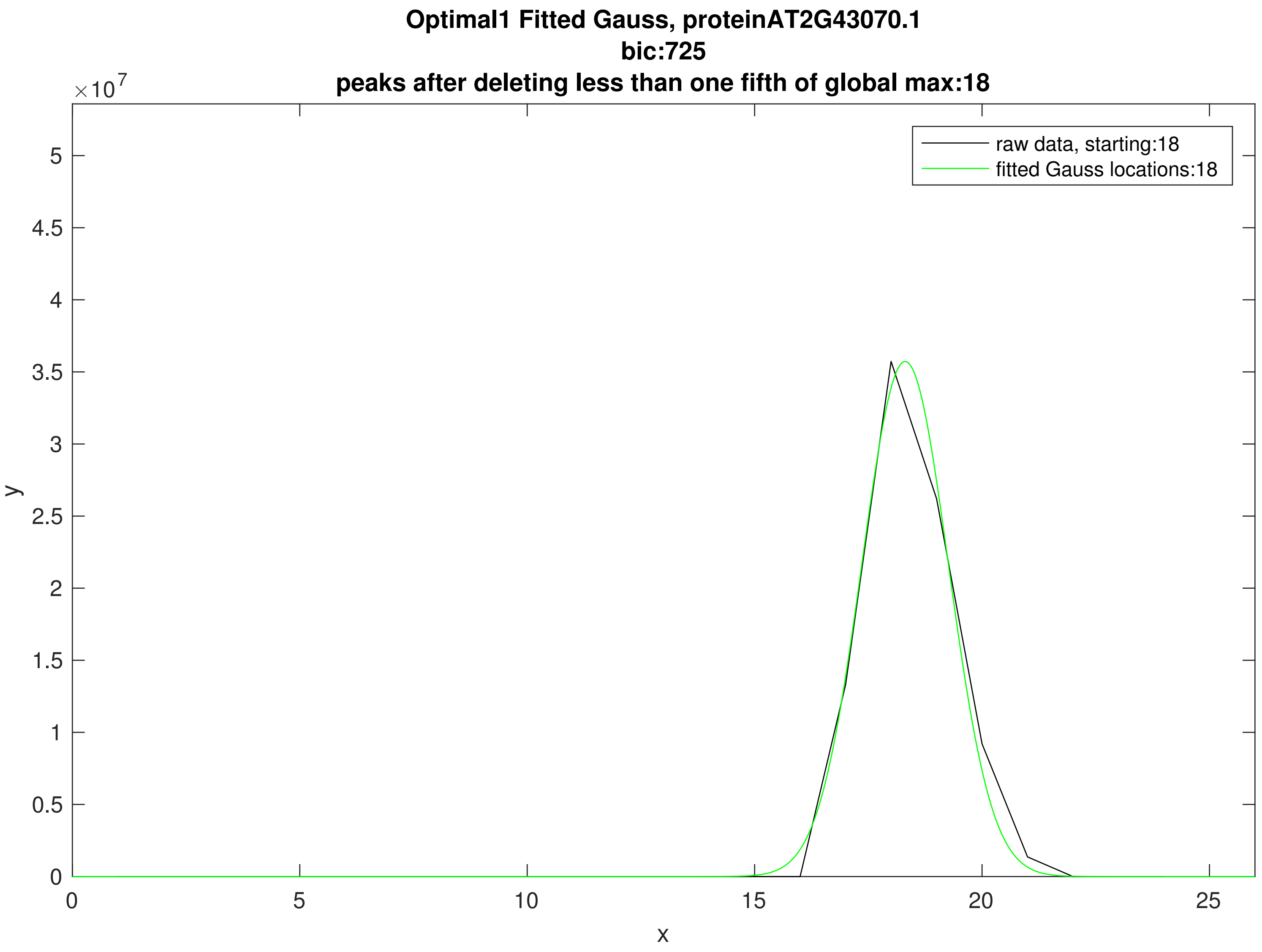

Optimal1 Fitted Gauss, proteinAT2G43070.1
bic:725
peaks after deleting less than one fifth of global max:18
×10^7
5
4.5
4
3.5
3
2.5
2
1.5
1
0.5
0
y
0
5
10
15
20
25
x
raw data, starting:18
fitted Gauss locations:18

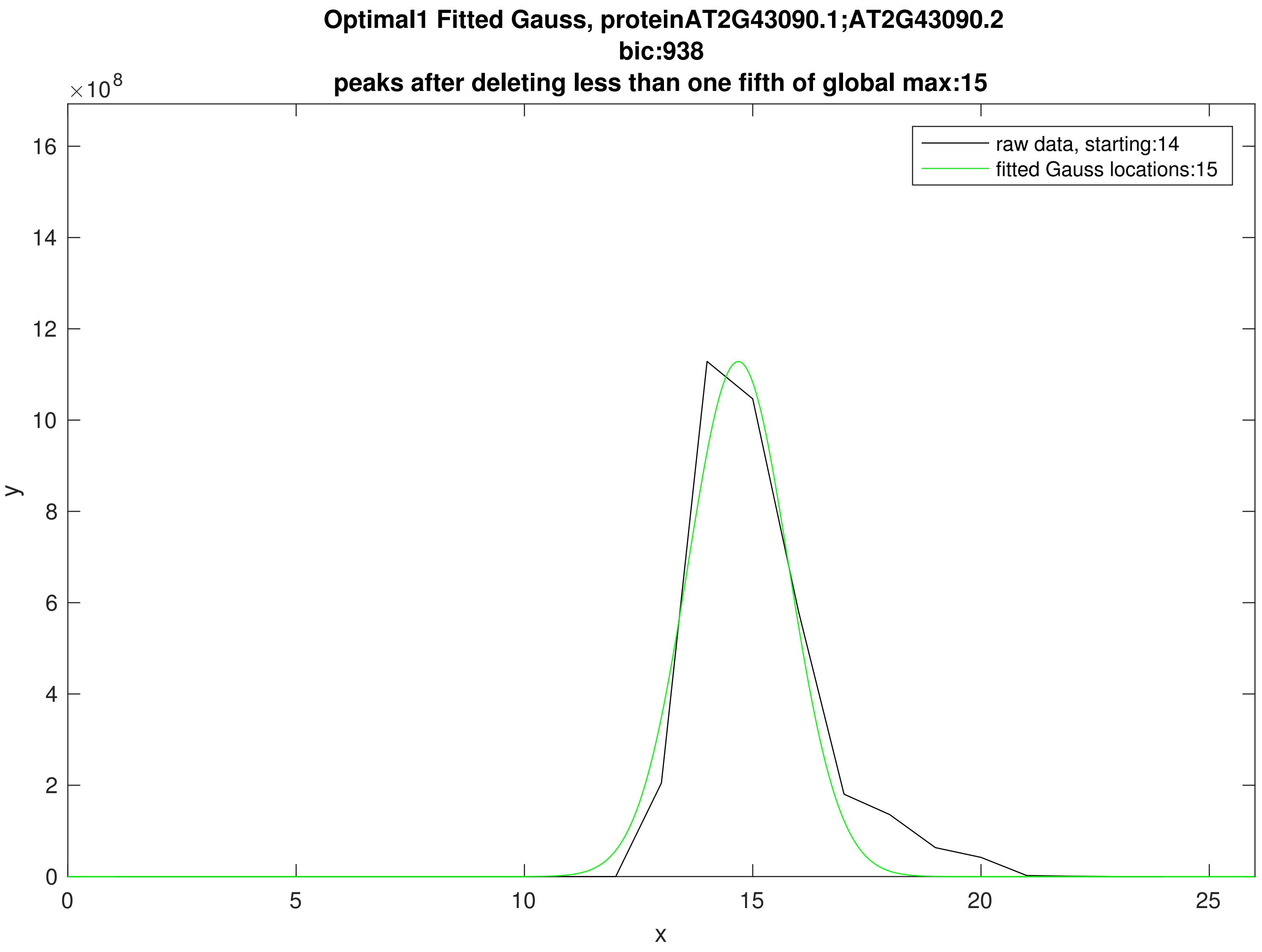
Optimal1 Fitted Gauss, proteinAT2G43090.1;AT2G43090.2
bic:938
peaks after deleting less than one fifth of global max:15
×10^8
raw data, starting:14
fitted Gauss locations:15
16
14
12
10
8
6
4
2
0
y
0
5
10
15
20
25
x

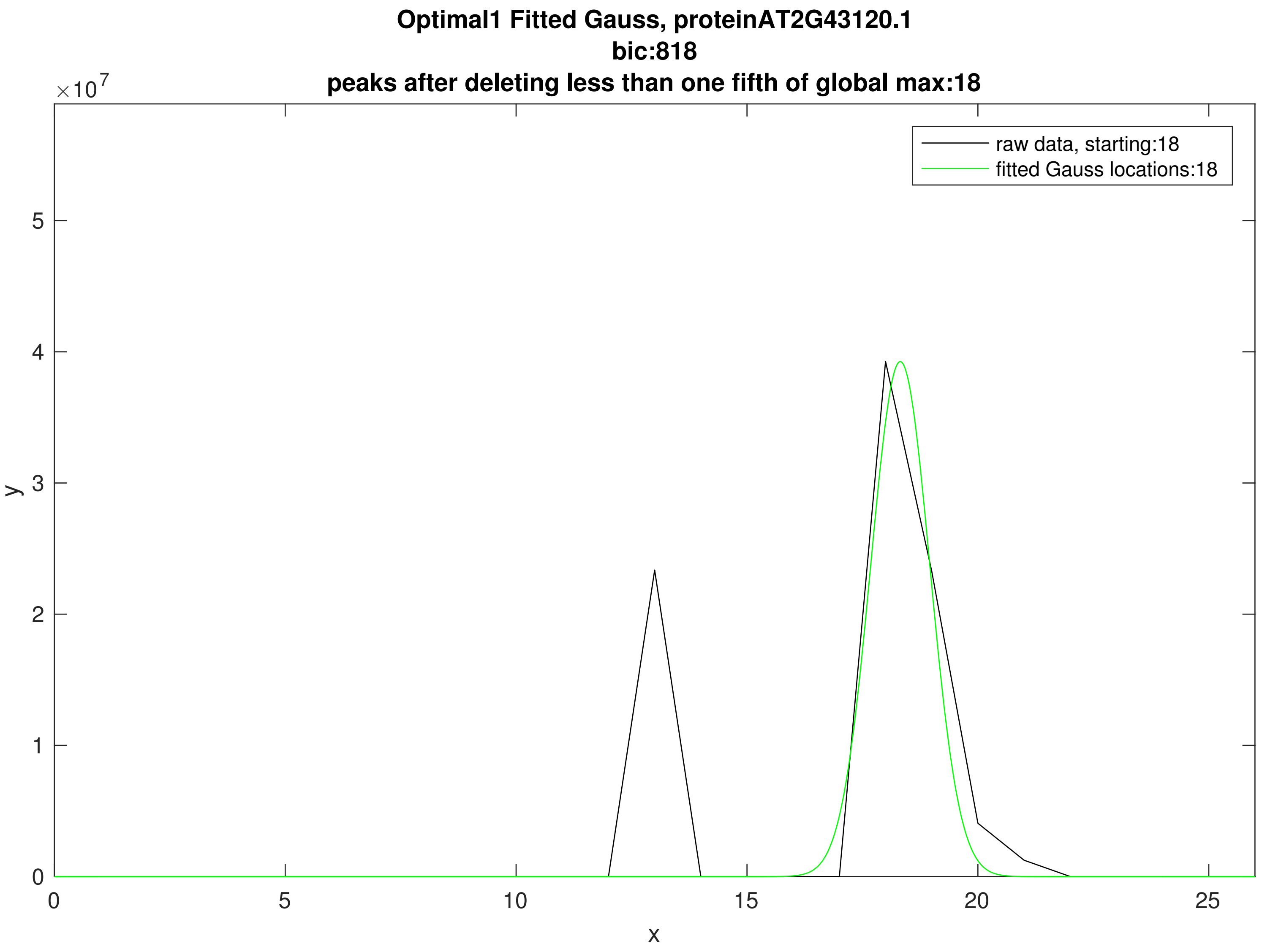
Optimal1 Fitted Gauss, proteinAT2G43120.1
bic:818
peaks after deleting less than one fifth of global max:18
×10^7
raw data, starting:18
fitted Gauss locations:18
y
x
0
1
2
3
4
5
0
5
10
15
20
25

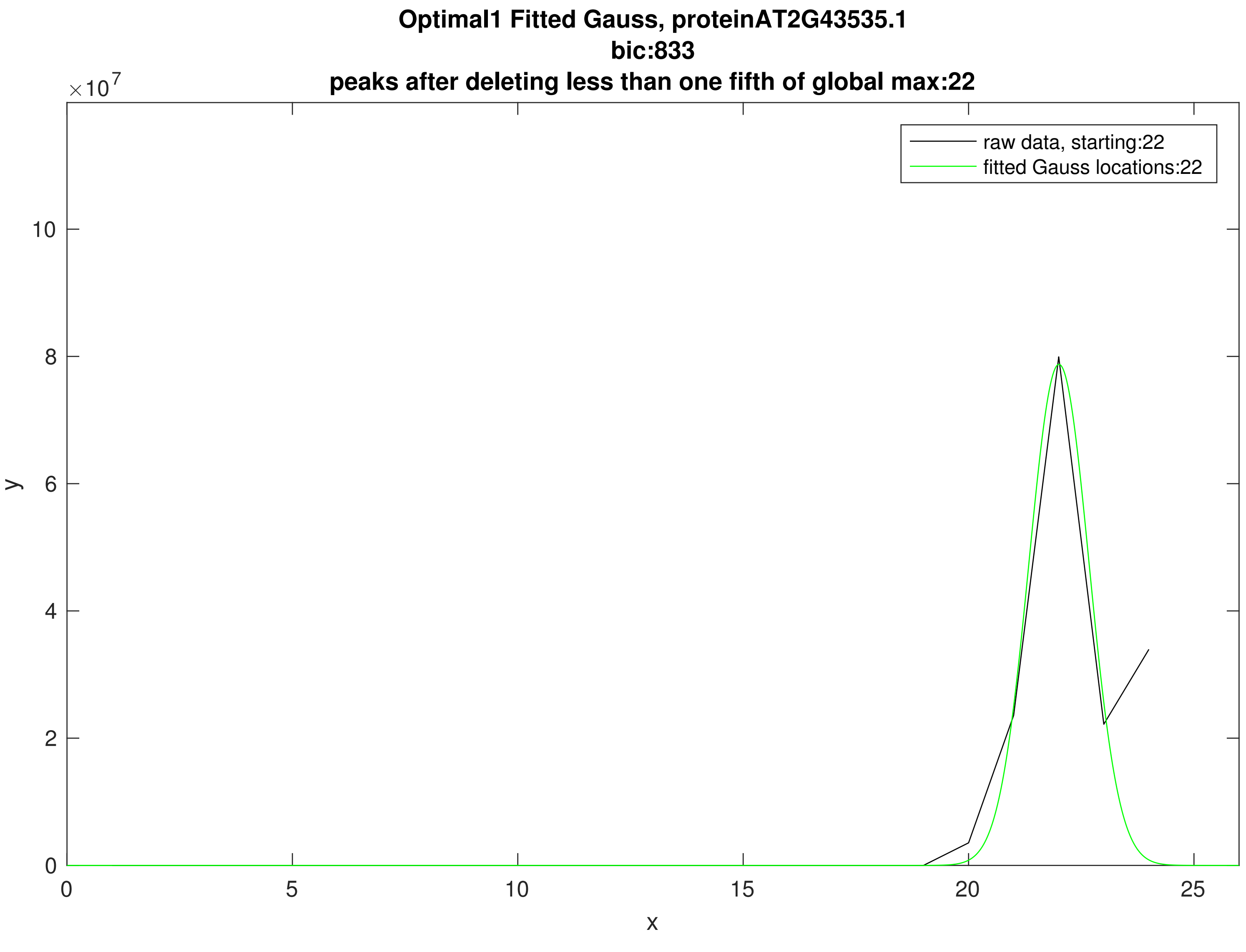
Optimal1 Fitted Gauss, proteinAT2G43535.1
bic:833
peaks after deleting less than one fifth of global max:22
×10^7
raw data, starting:22
fitted Gauss locations:22
y
x
0
2
4
6
8
10
0
5
10
15
20
25

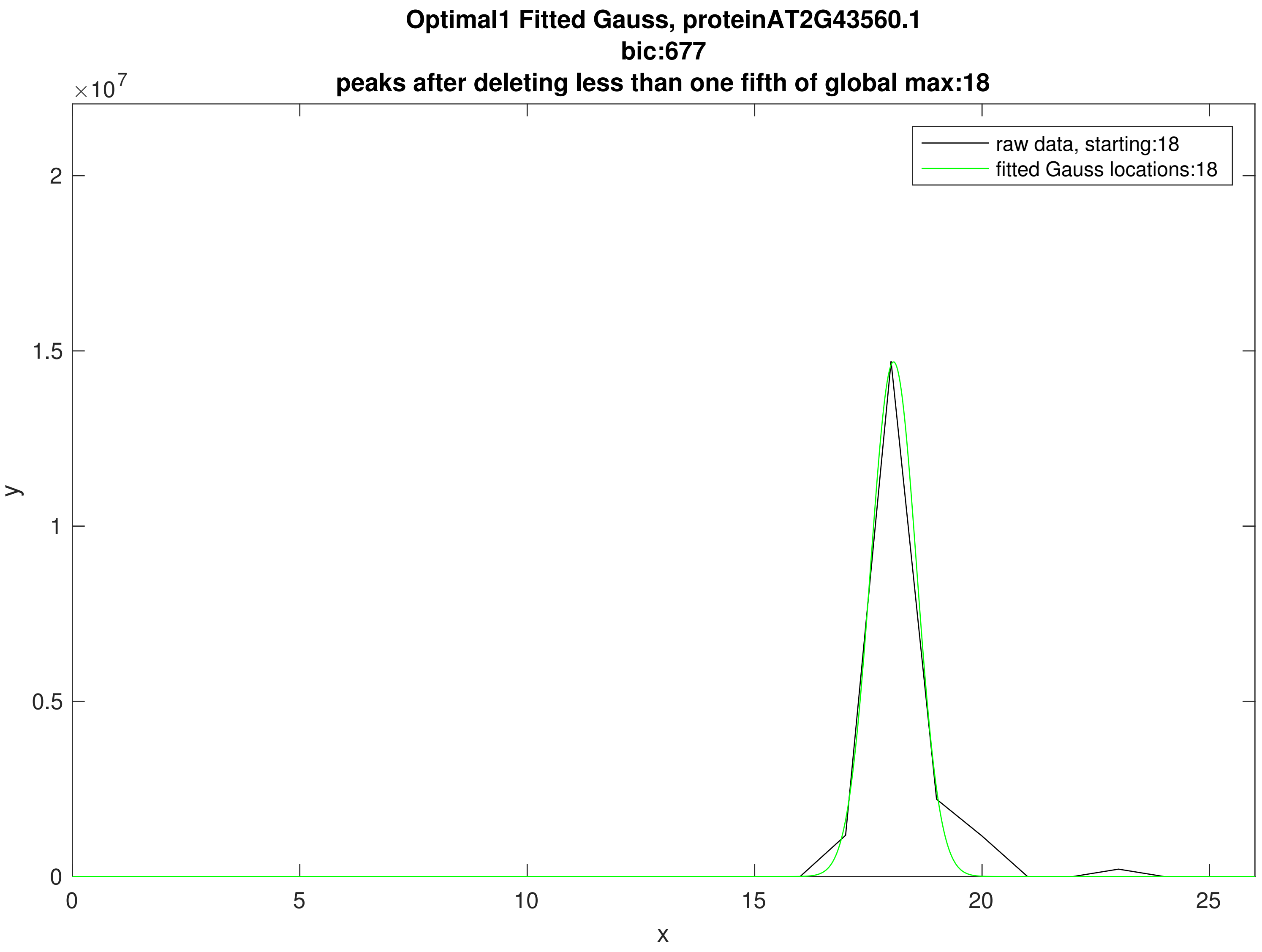
Optimal1 Fitted Gauss, proteinAT2G43560.1
bic:677
peaks after deleting less than one fifth of global max:18
×10^7
2
1.5
1
0.5
0
y
0
5
10
15
20
25
x
raw data, starting:18
fitted Gauss locations:18

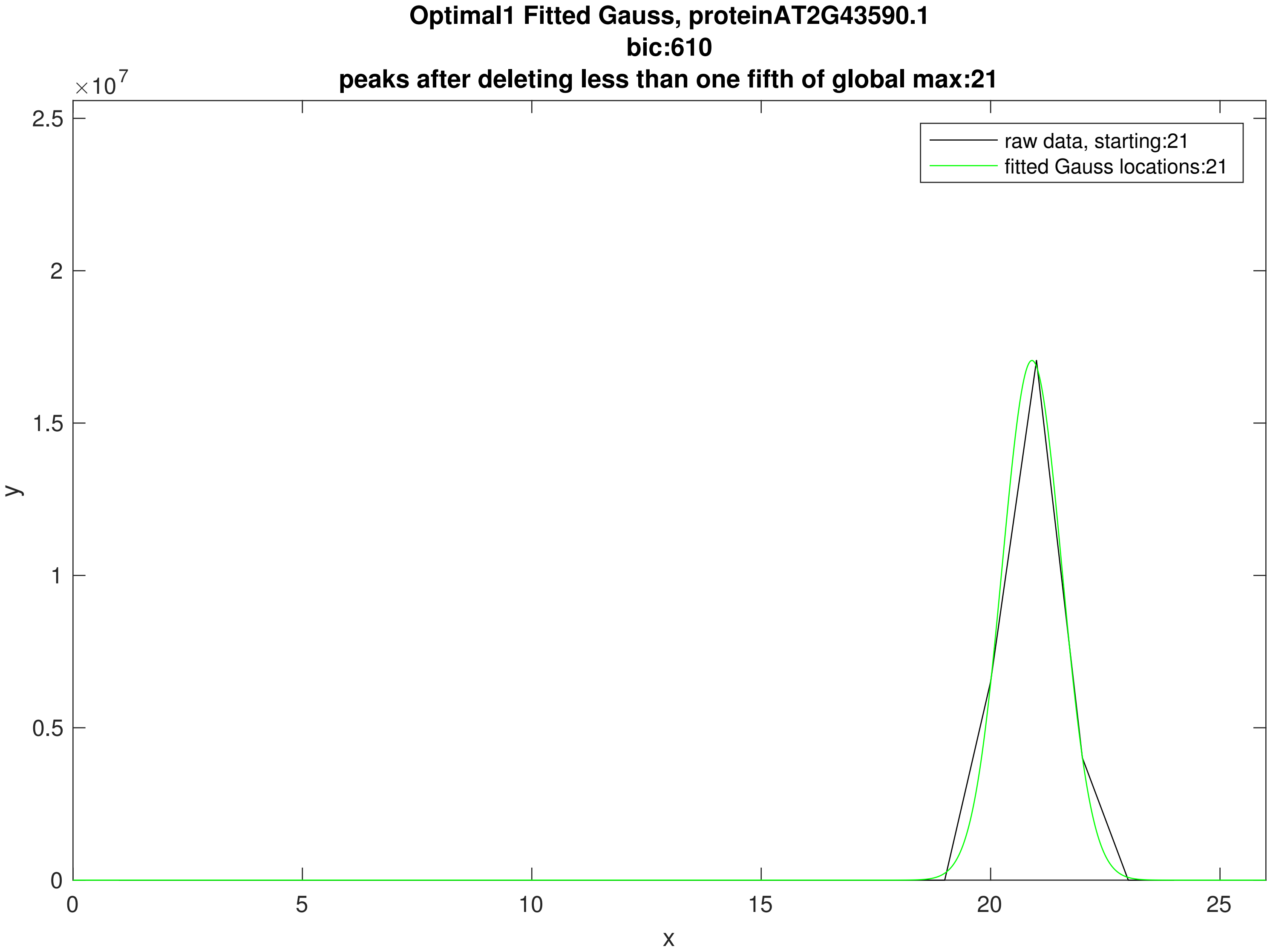
Optimal1 Fitted Gauss, proteinAT2G43590.1
bic:610
peaks after deleting less than one fifth of global max:21
×10^7
2.5
2
1.5
1
0.5
0
y
0
5
10
15
20
25
x
raw data, starting:21
fitted Gauss locations:21

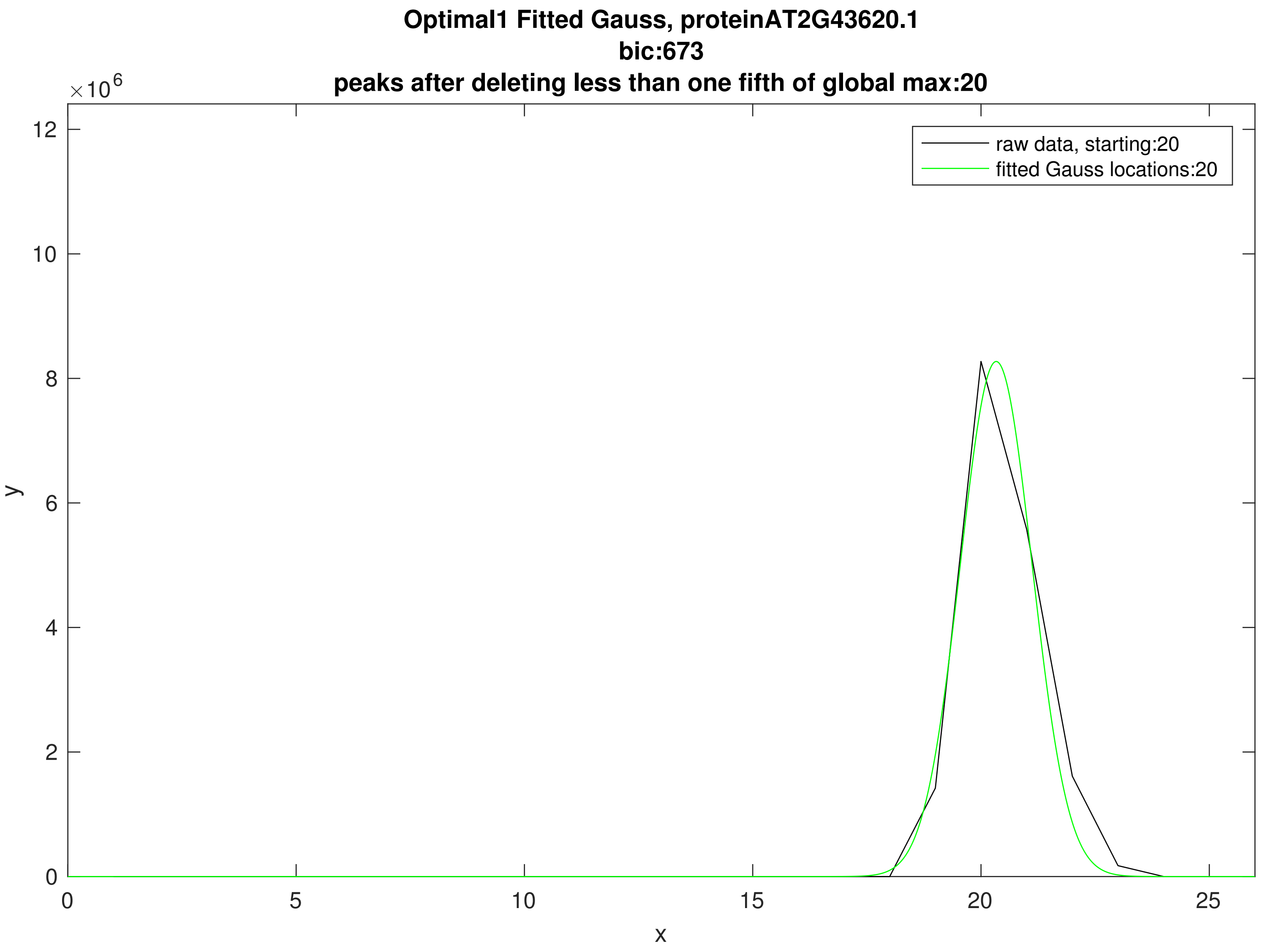
Optimal1 Fitted Gauss, proteinAT2G43620.1
bic:673
peaks after deleting less than one fifth of global max:20
×10^6
raw data, starting:20
fitted Gauss locations:20
y
x
0
2
4
6
8
10
12
0
5
10
15
20
25

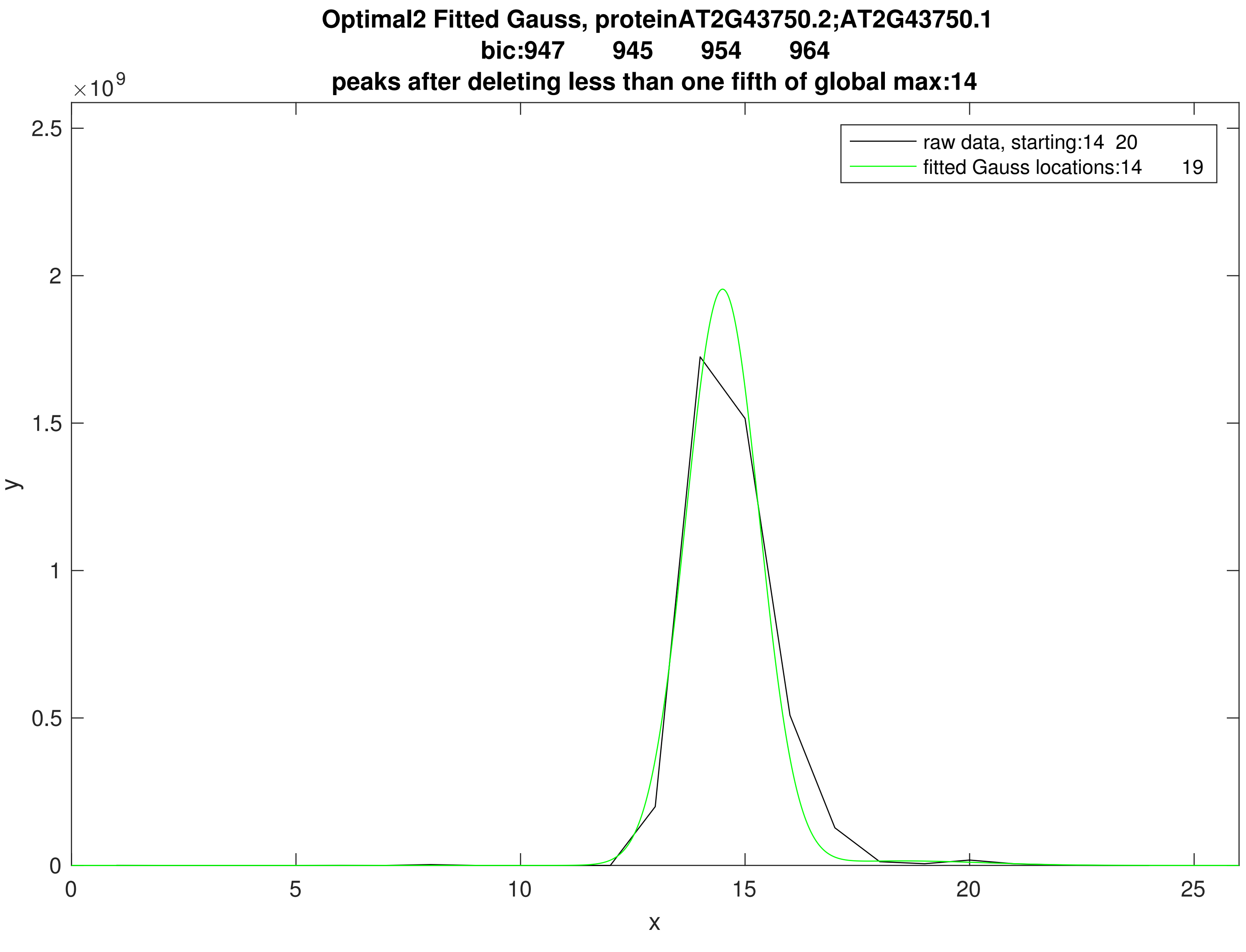
Optimal2 Fitted Gauss, proteinAT2G43750.2;AT2G43750.1
bic:947 945 954 964
peaks after deleting less than one fifth of global max:14
×10^9
raw data, starting:14 20
fitted Gauss locations:14 19
y
x
0
0.5
1
1.5
2
2.5
5
10
15
20
25

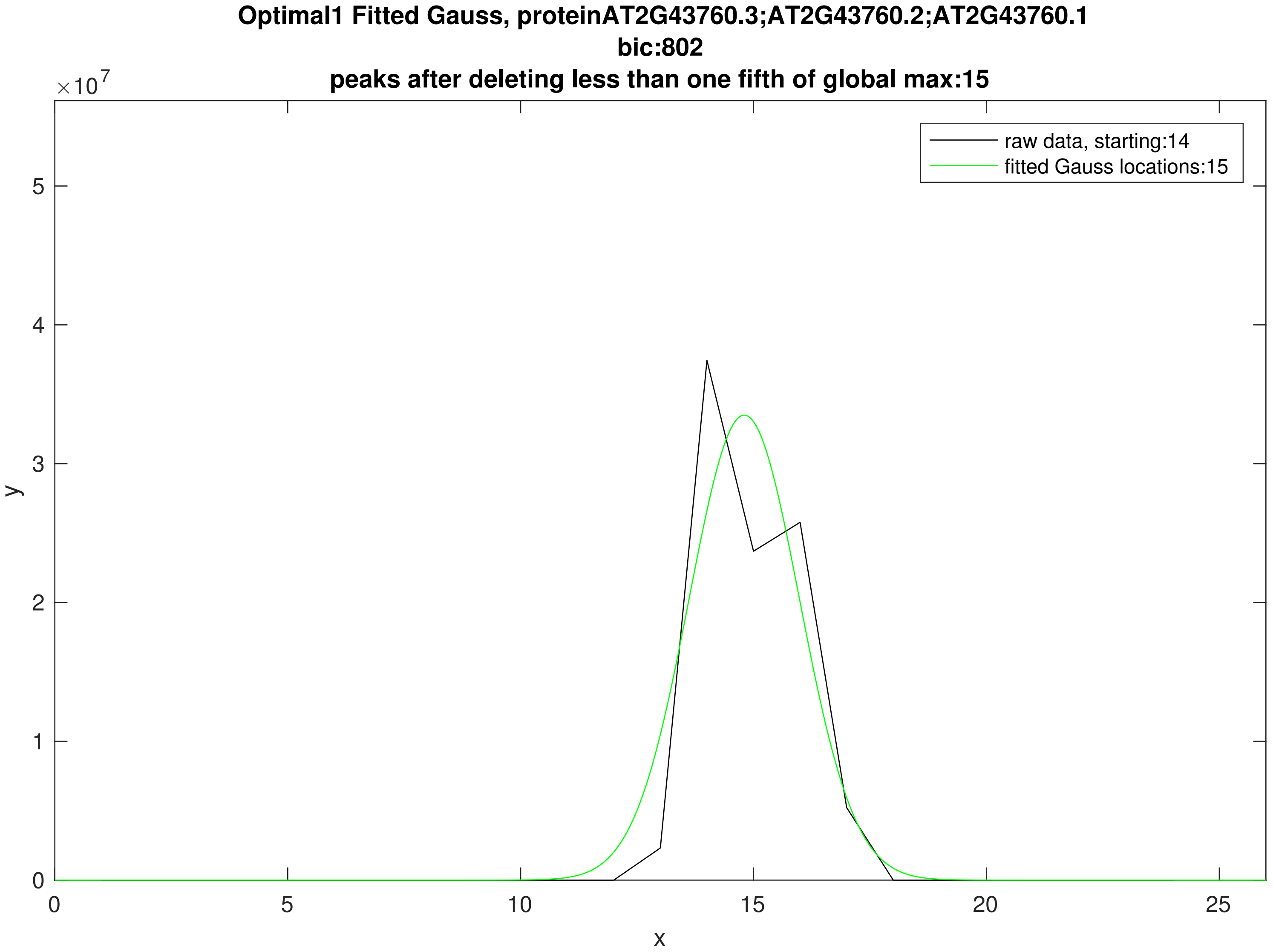
Optimal1 Fitted Gauss, proteinAT2G43760.3;AT2G43760.2;AT2G43760.1
bic:802
peaks after deleting less than one fifth of global max:15
×10^7
raw data, starting:14
fitted Gauss locations:15
y
x
0
1
2
3
4
5
0
5
10
15
20
25

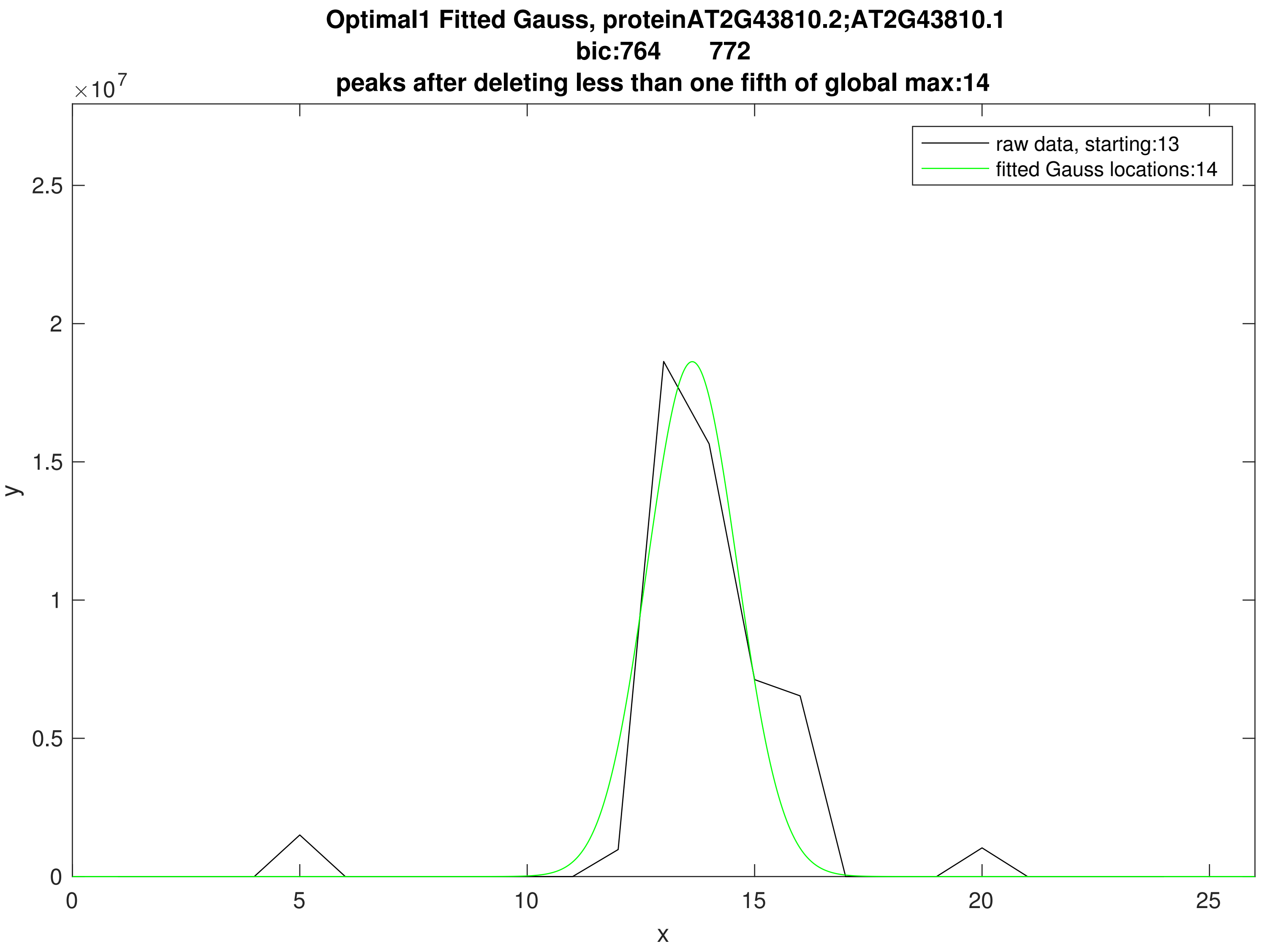

Optimal1 Fitted Gauss, proteinAT2G43810.2;AT2G43810.1
bic:764 772
peaks after deleting less than one fifth of global max:14
×10^7
2.5
2
1.5
1
0.5
0
y
0
5
10
15
20
25
x
raw data, starting:13
fitted Gauss locations:14

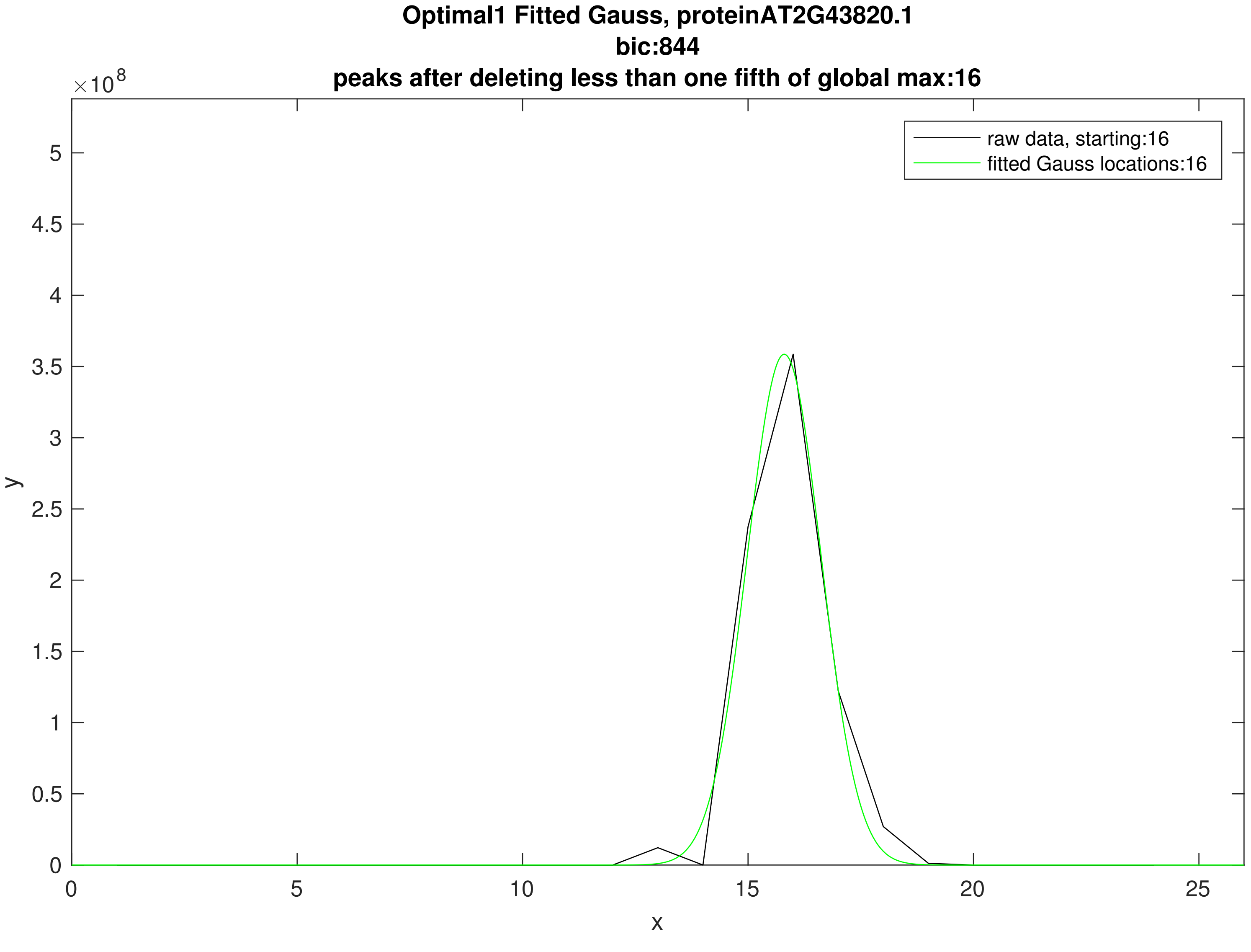
Optimal1 Fitted Gauss, proteinAT2G43820.1
bic:844
peaks after deleting less than one fifth of global max:16
×10^8
y
x
raw data, starting:16
fitted Gauss locations:16
0
0.5
1
1.5
2
2.5
3
3.5
4
4.5
5
0
5
10
15
20
25

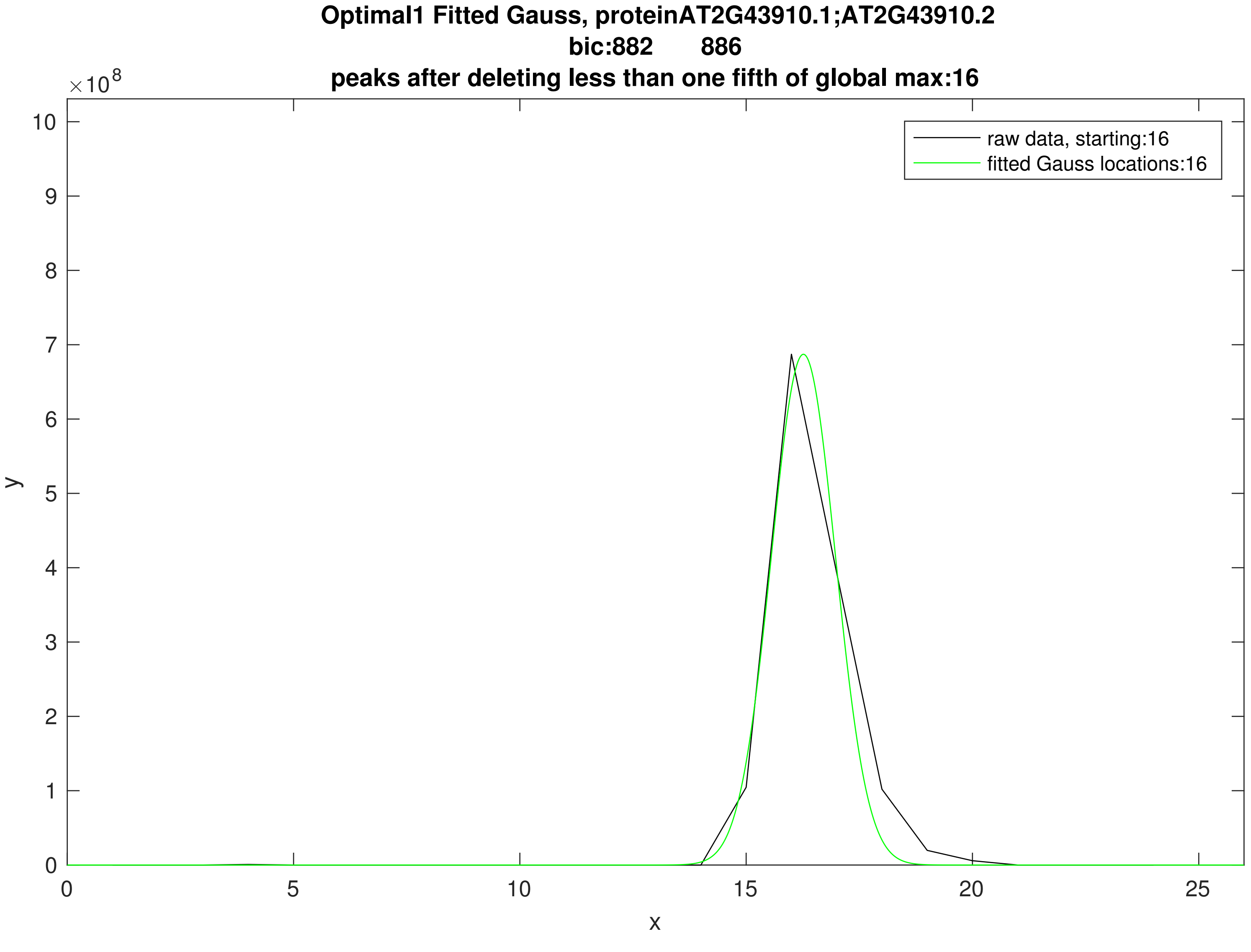
Optimal1 Fitted Gauss, proteinAT2G43910.1;AT2G43910.2
bic:882 886
peaks after deleting less than one fifth of global max:16
×10^8
raw data, starting:16
fitted Gauss locations:16
y
x
0
1
2
3
4
5
6
7
8
9
10
0
5
10
15
20
25

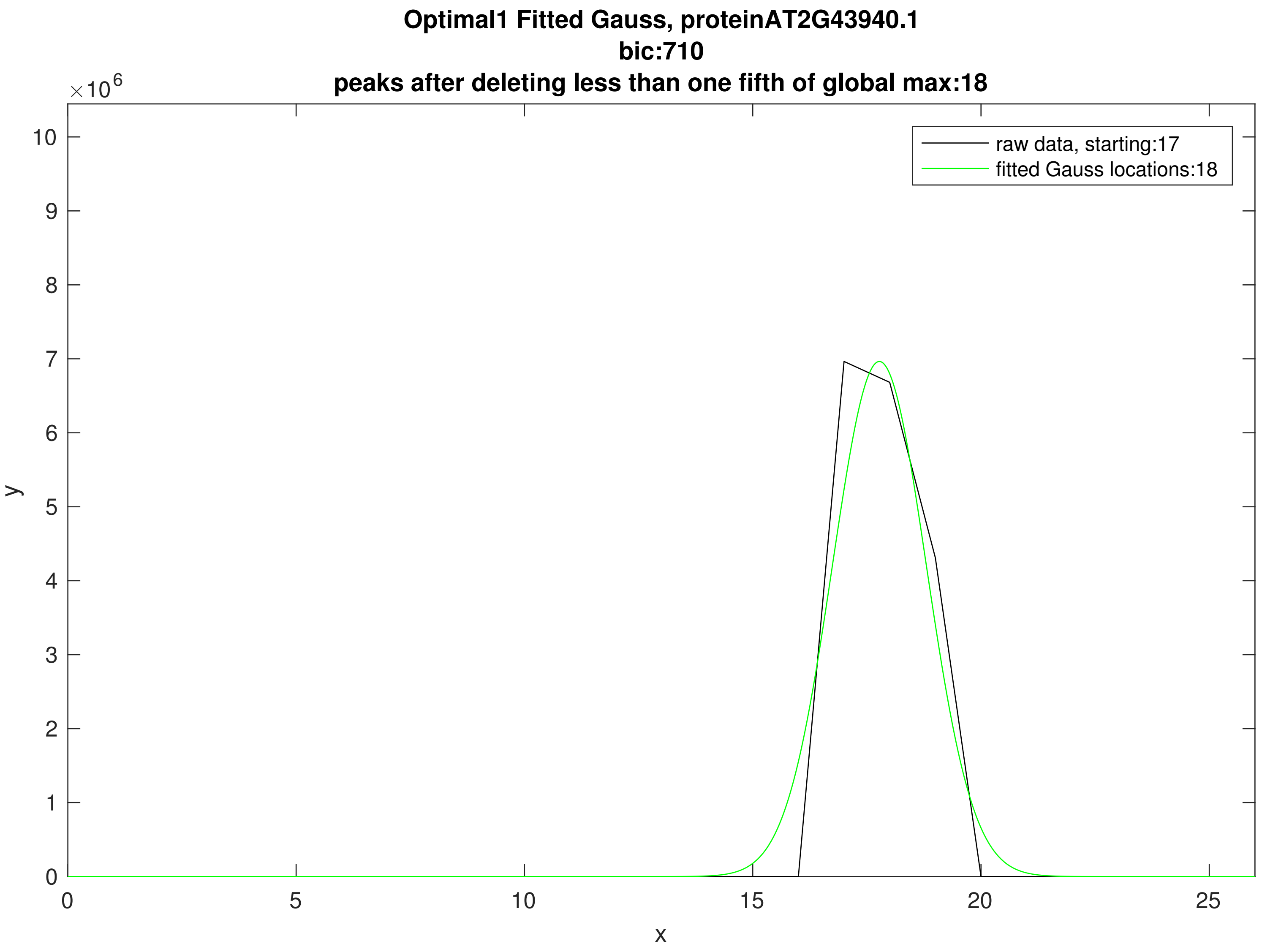
Optimal1 Fitted Gauss, proteinAT2G43940.1
bic:710
peaks after deleting less than one fifth of global max:18
×10^6
raw data, starting:17
fitted Gauss locations:18
y
x
0
1
2
3
4
5
6
7
8
9
10
0
5
10
15
20
25

**Optimal1 Fitted Gauss, proteinAT2G43980.1**
**bic:670**
**peaks after deleting less than one fifth of global max:18**

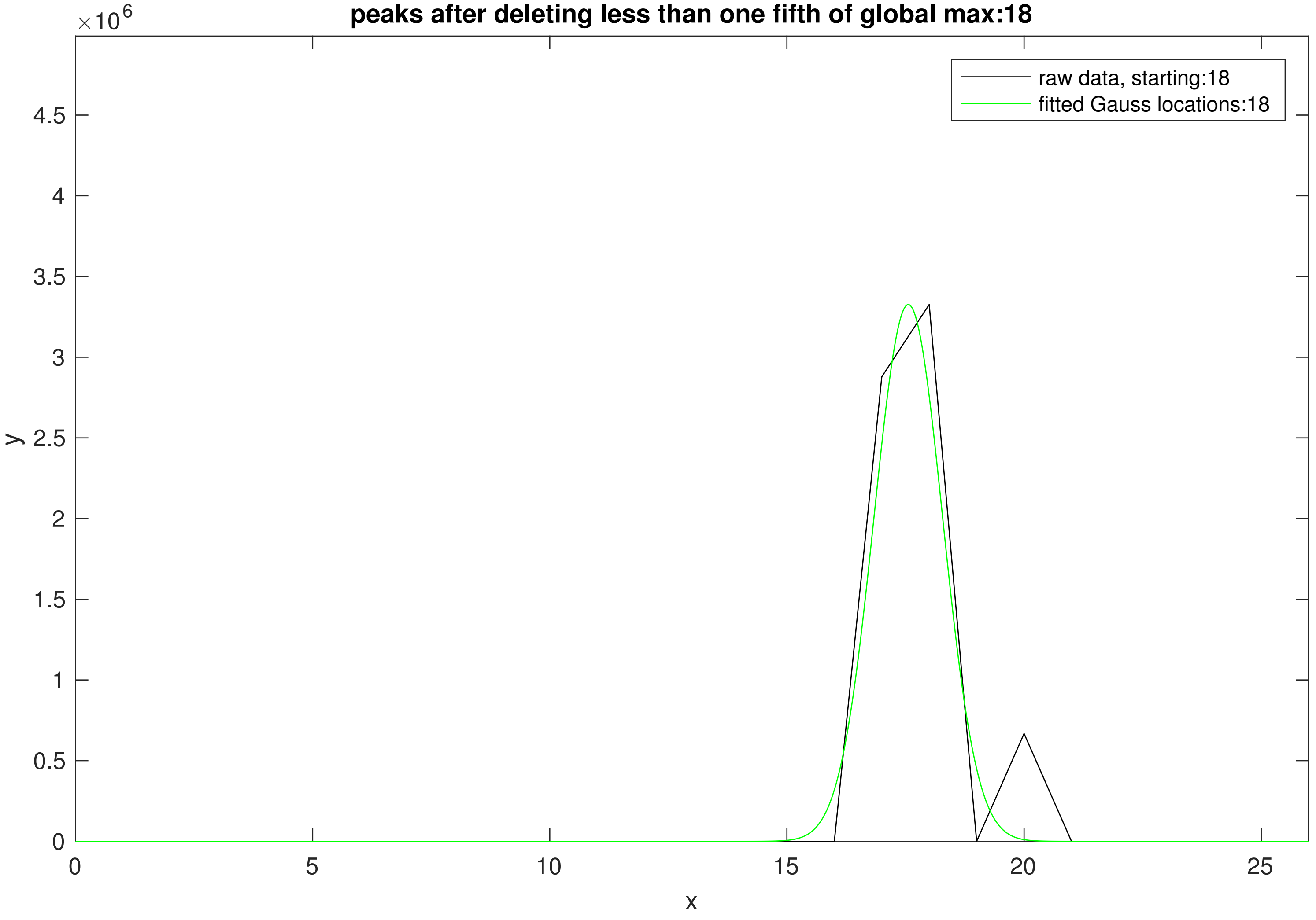

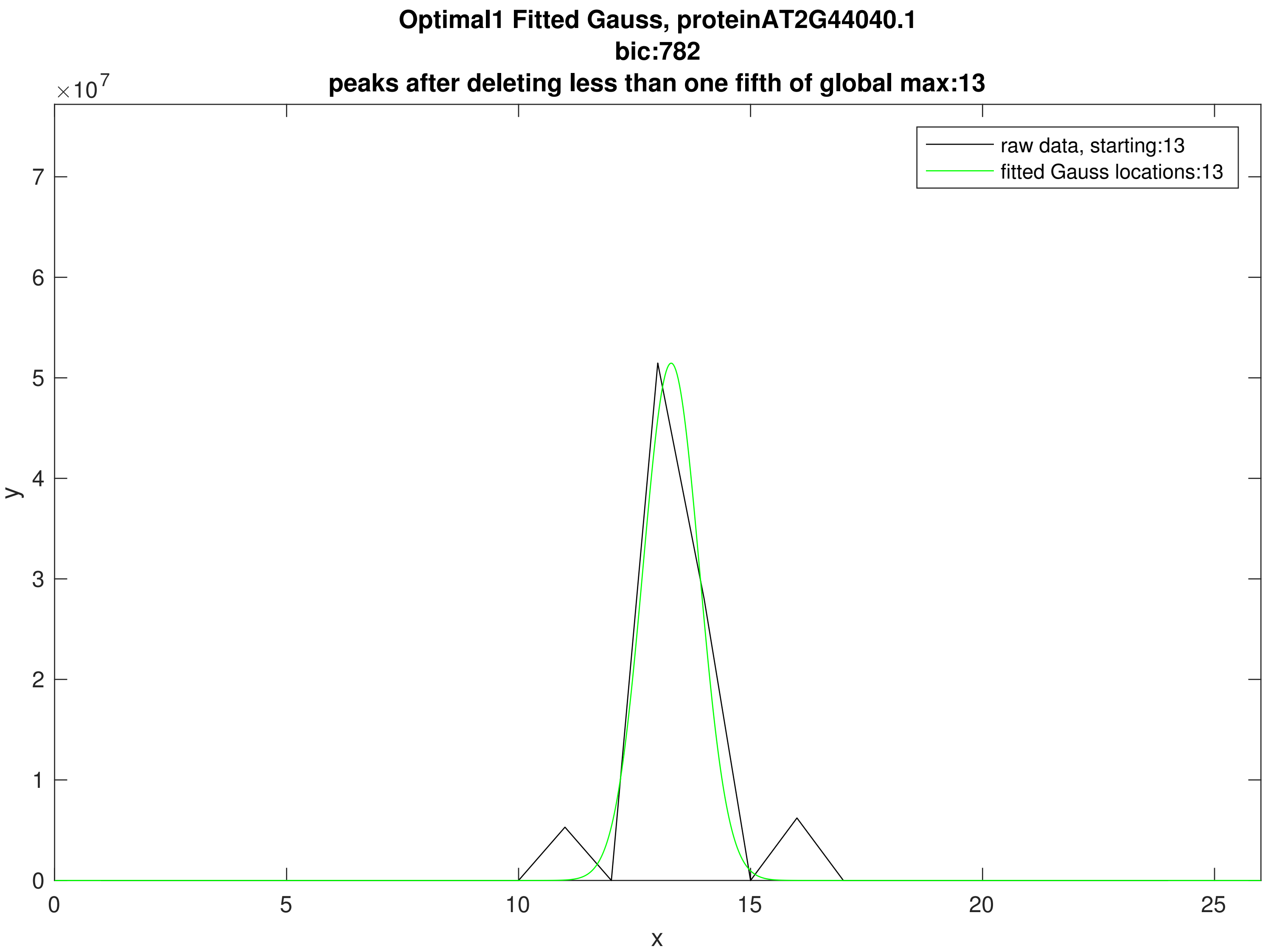
Optimal1 Fitted Gauss, proteinAT2G44040.1
bic:782
peaks after deleting less than one fifth of global max:13
×10^7
7
6
5
4
3
2
1
0
y
0
5
10
15
20
25
x
raw data, starting:13
fitted Gauss locations:13

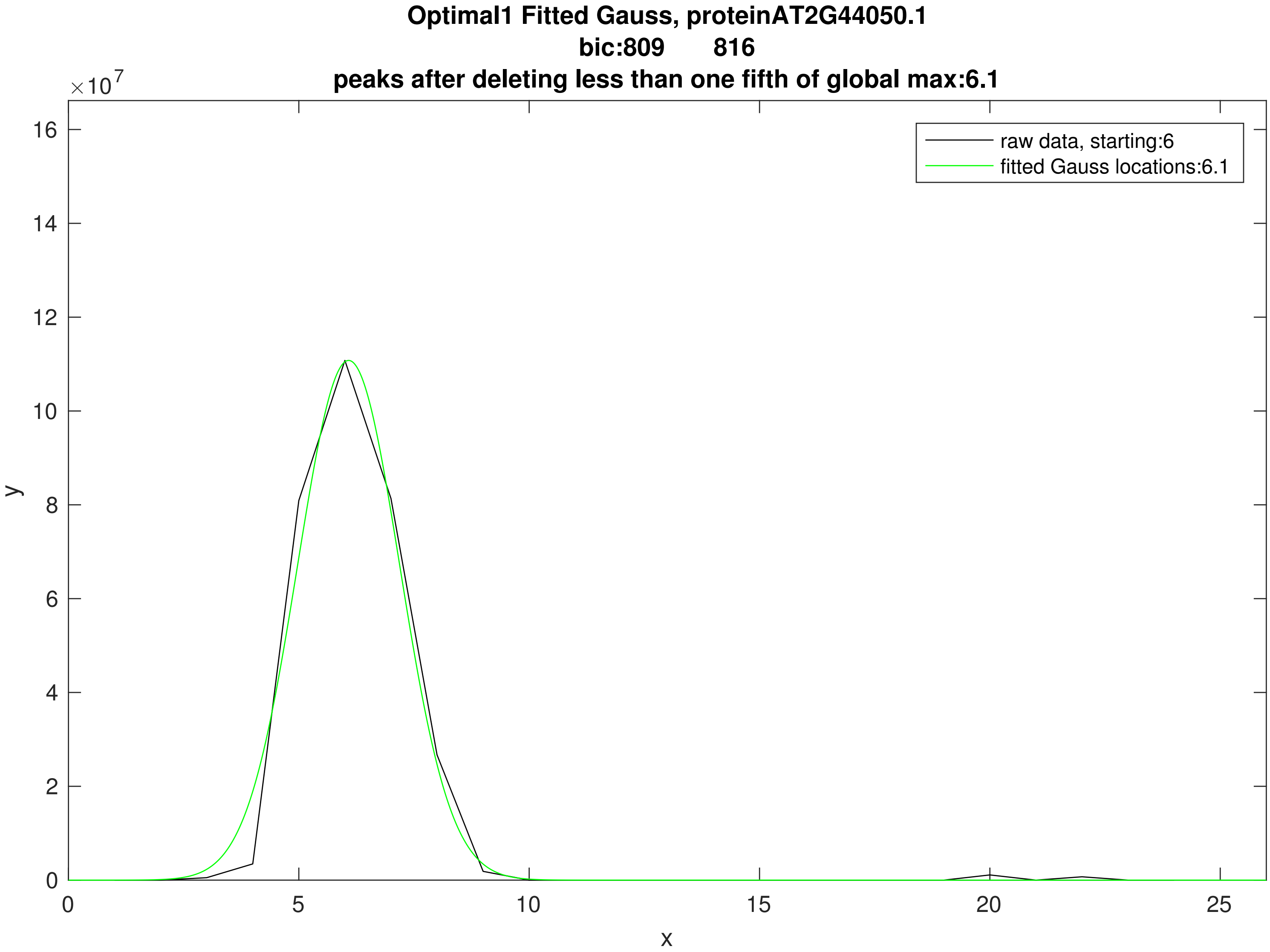

Optimal1 Fitted Gauss, proteinAT2G44050.1
bic:809 816
peaks after deleting less than one fifth of global max:6.1
×10^7
16
14
12
10
8
6
4
2
0
y
0
5
10
15
20
25
x
raw data, starting:6
fitted Gauss locations:6.1

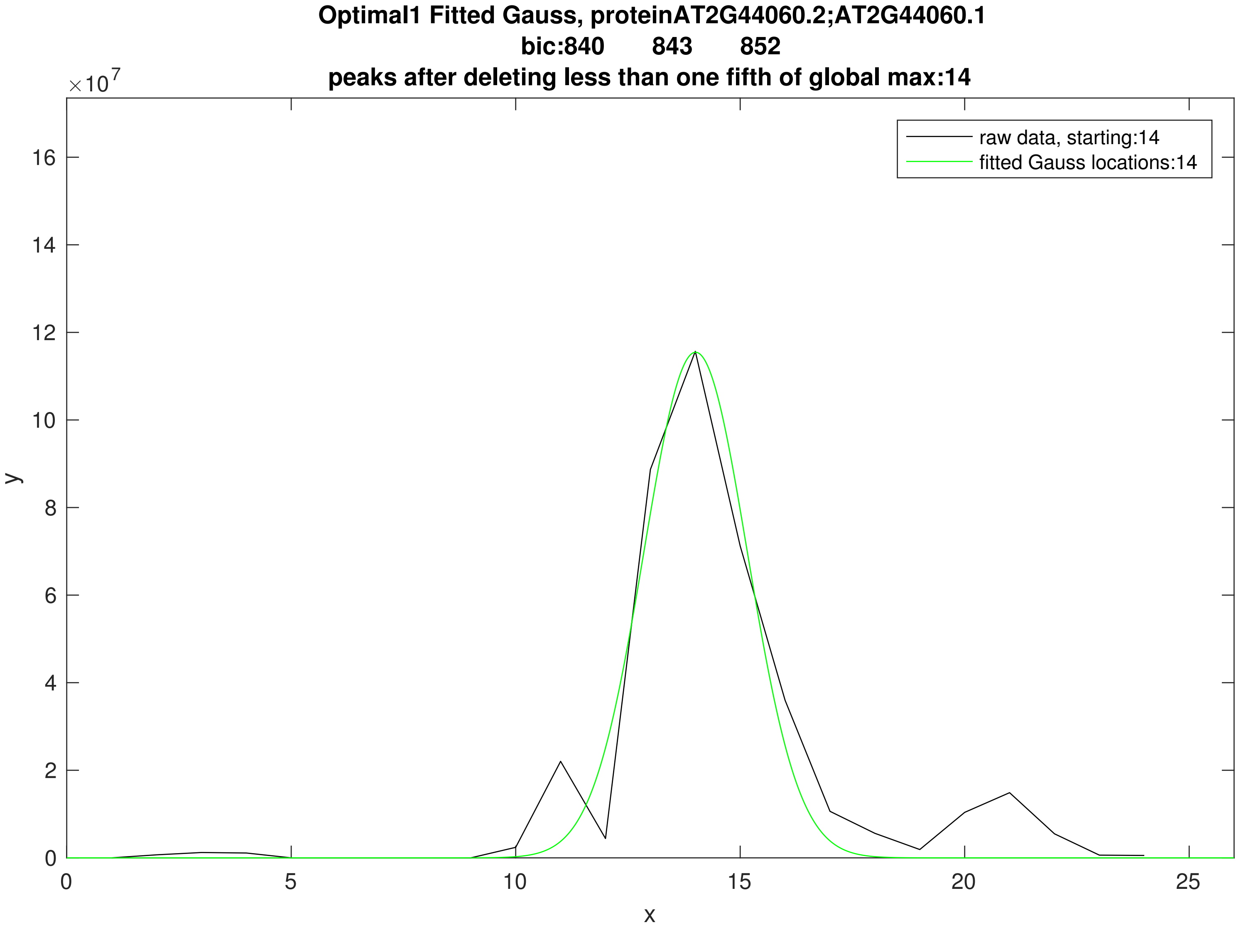
Optimal1 Fitted Gauss, proteinAT2G44060.2;AT2G44060.1
bic:840 843 852
peaks after deleting less than one fifth of global max:14
×10^7
raw data, starting:14
fitted Gauss locations:14
y
x
0
2
4
6
8
10
12
14
16
0
5
10
15
20
25

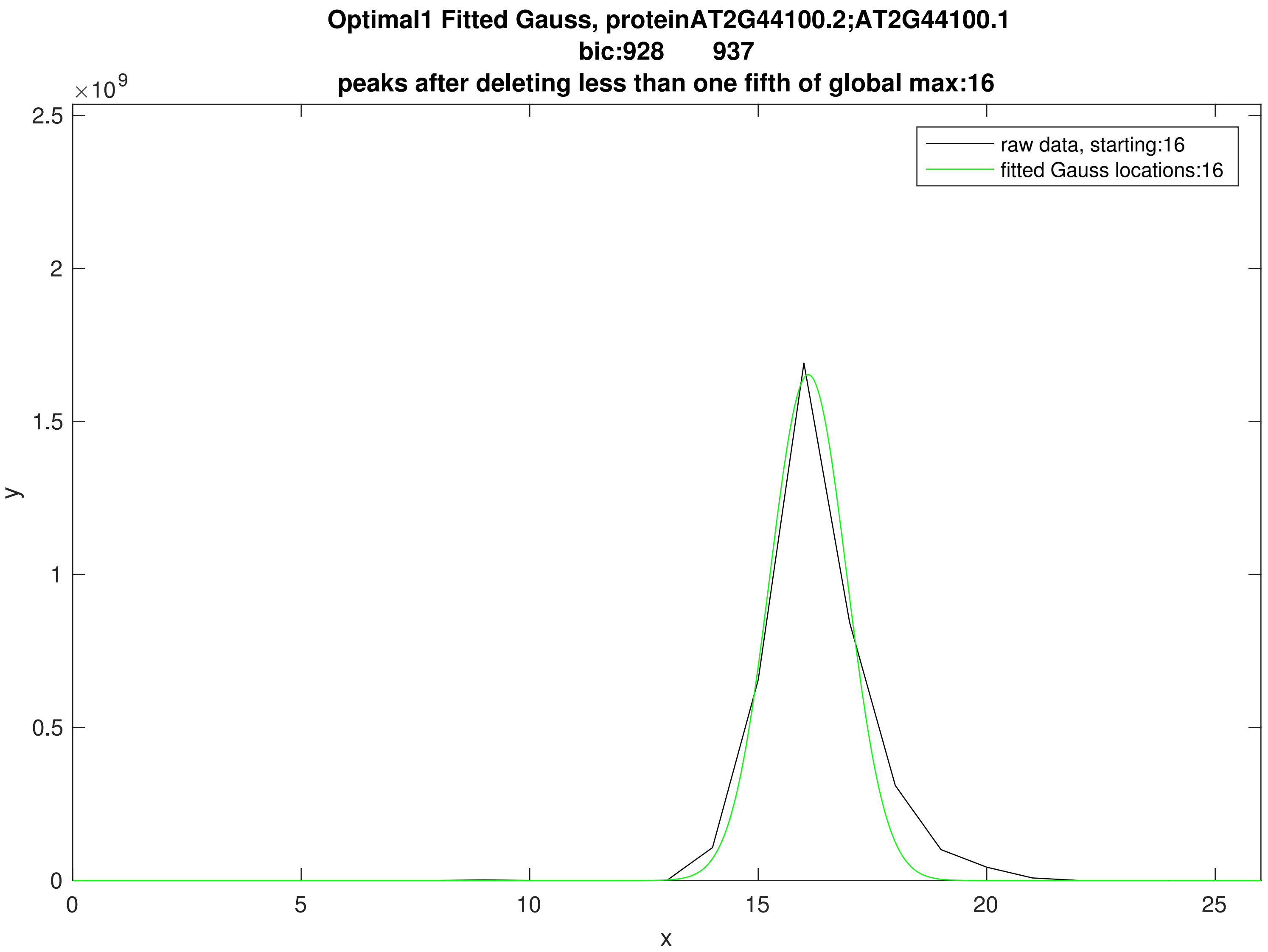
Optimal1 Fitted Gauss, proteinAT2G44100.2;AT2G44100.1
bic:928 937
peaks after deleting less than one fifth of global max:16
×10^9
2.5
2
1.5
1
0.5
0
y
0
5
10
15
20
25
x
raw data, starting:16
fitted Gauss locations:16

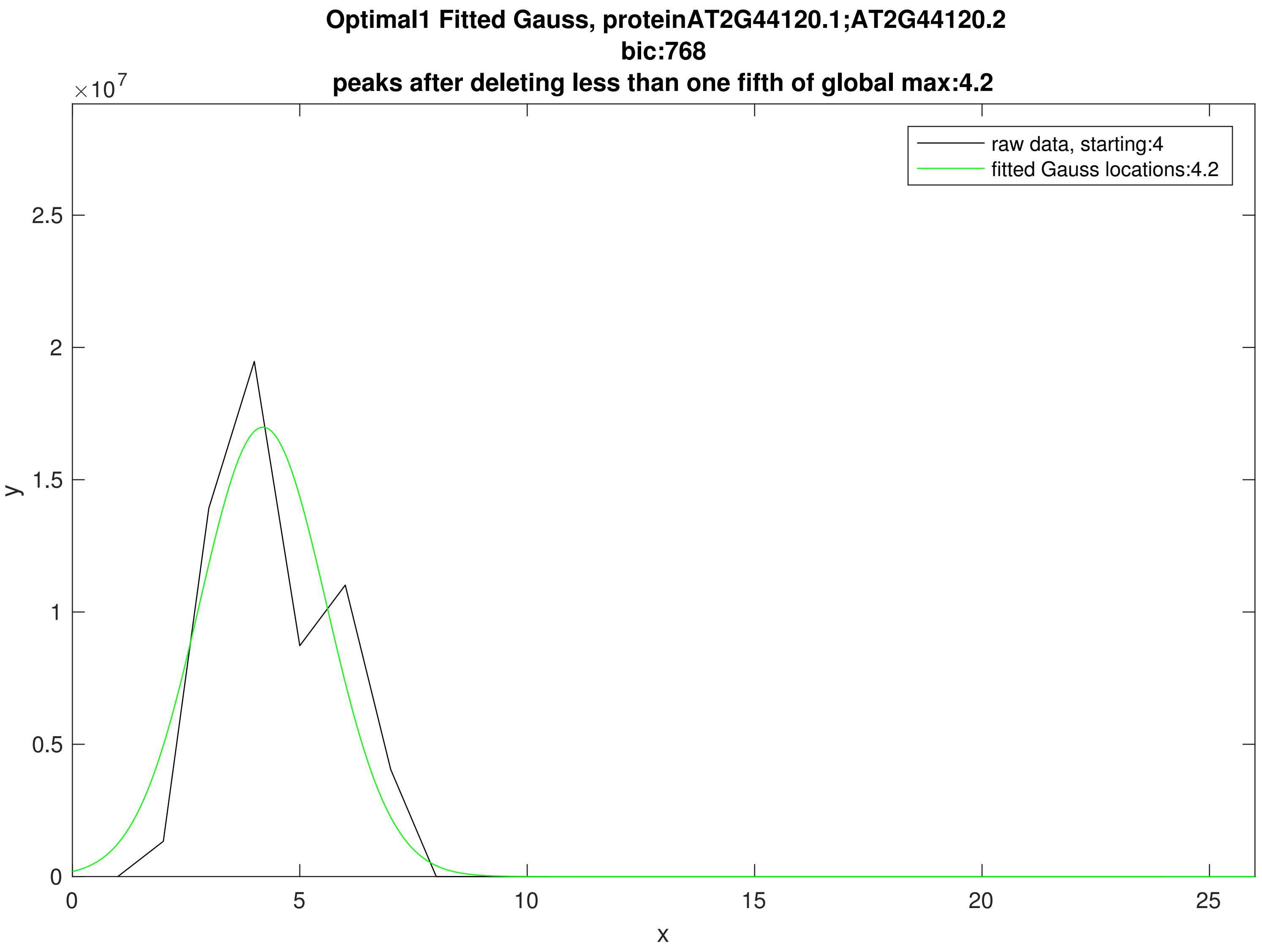

Optimal1 Fitted Gauss, proteinAT2G44120.1;AT2G44120.2
bic:768
peaks after deleting less than one fifth of global max:4.2
×10^7
raw data, starting:4
fitted Gauss locations:4.2
y
x
0
0.5
1
1.5
2
2.5
0
5
10
15
20
25

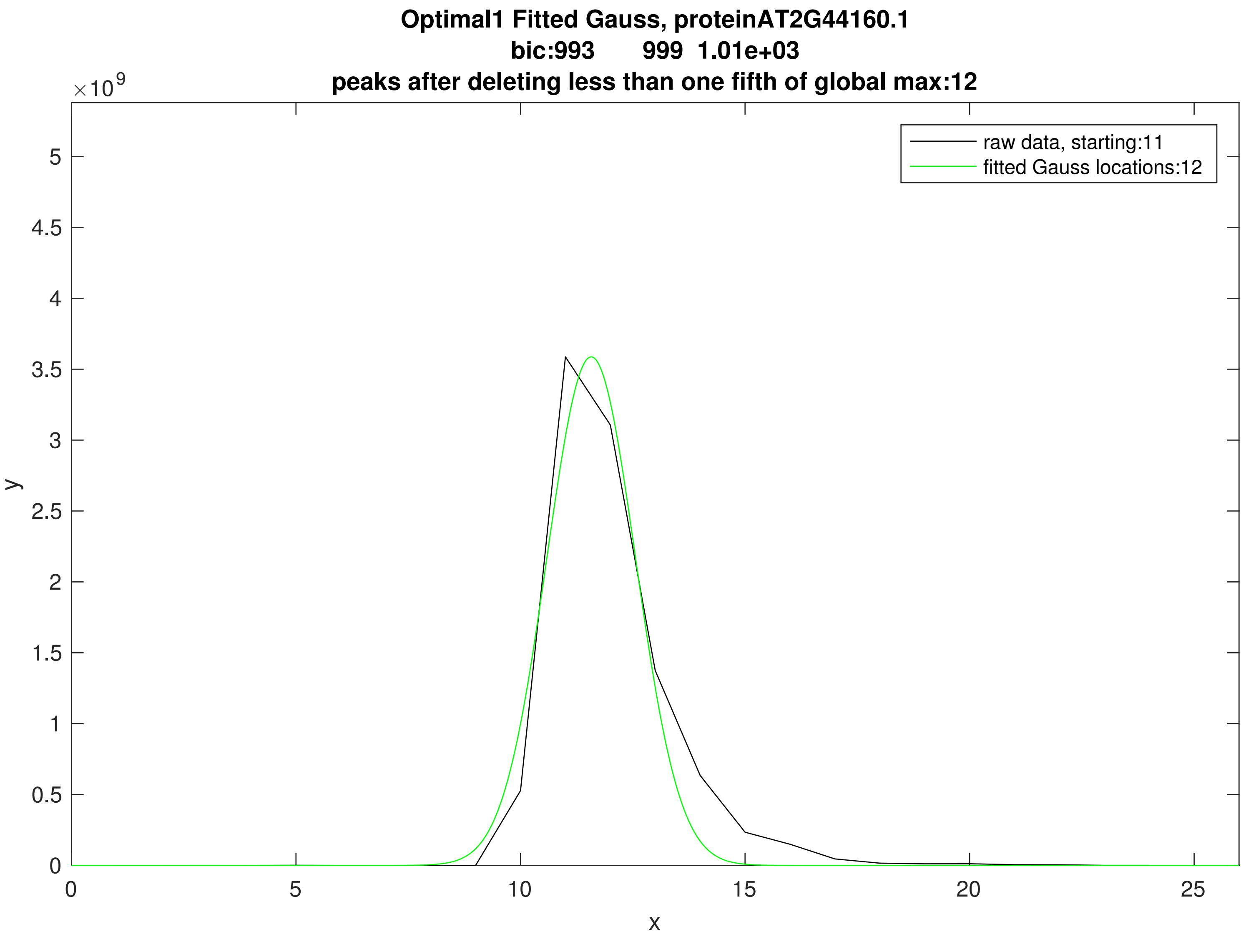
Optimal1 Fitted Gauss, proteinAT2G44160.1
bic:993 999 1.01e+03
peaks after deleting less than one fifth of global max:12
×10^9
raw data, starting:11
fitted Gauss locations:12
y
x
0
0.5
1
1.5
2
2.5
3
3.5
4
4.5
5
0
5
10
15
20
25

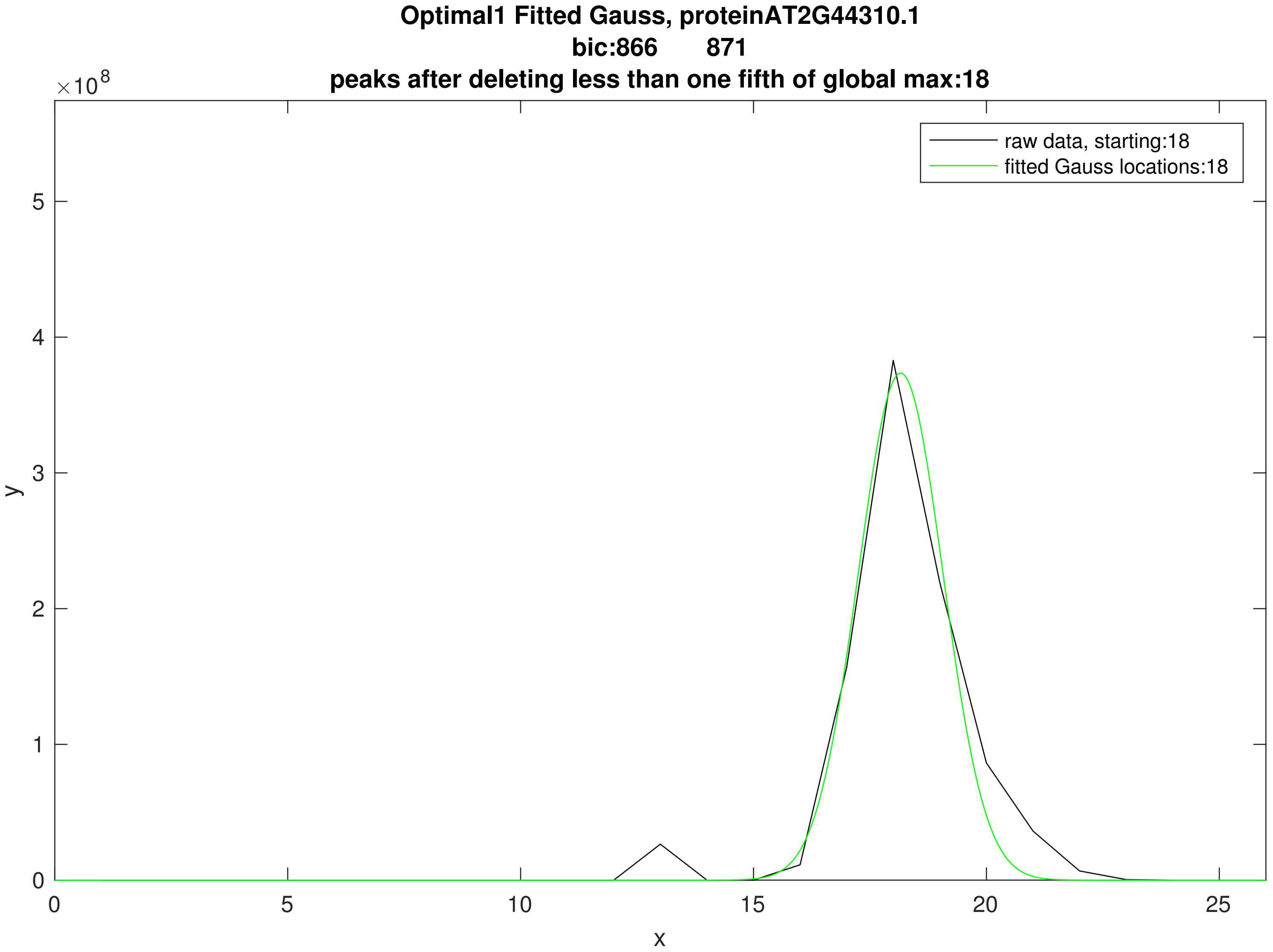
Optimal1 Fitted Gauss, proteinAT2G44310.1
bic:866 871
peaks after deleting less than one fifth of global max:18
raw data, starting:18
fitted Gauss locations:18
y
x

**Optimal1 Fitted Gauss, proteinAT2G44350.2;AT2G44350.1**
**bic:1e+03  1.01e+03**
**peaks after deleting less than one fifth of global max:13**

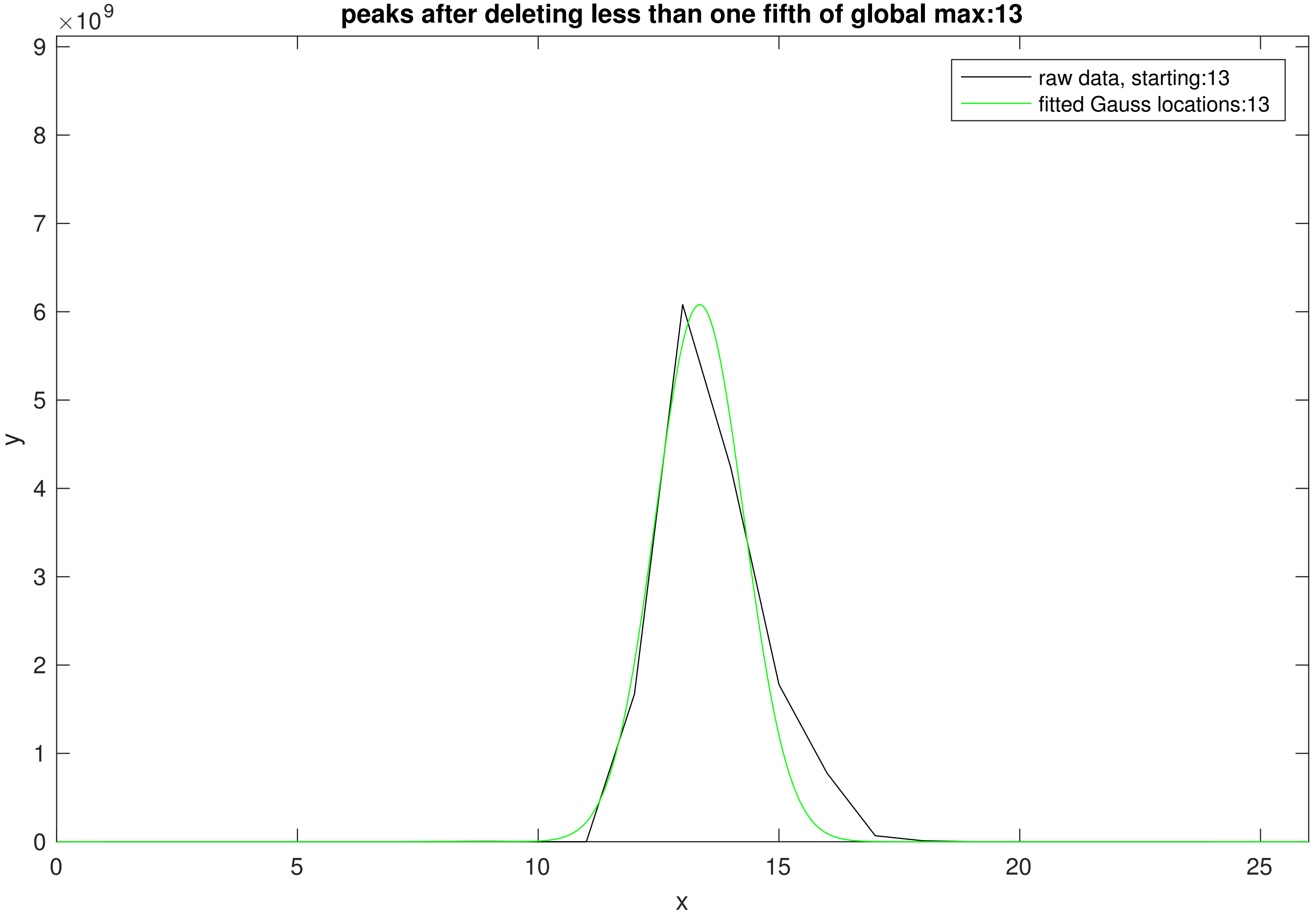

**Optimal1 Fitted Gauss, proteinAT2G44360.1**
**bic:754**
**peaks after deleting less than one fifth of global max:15**

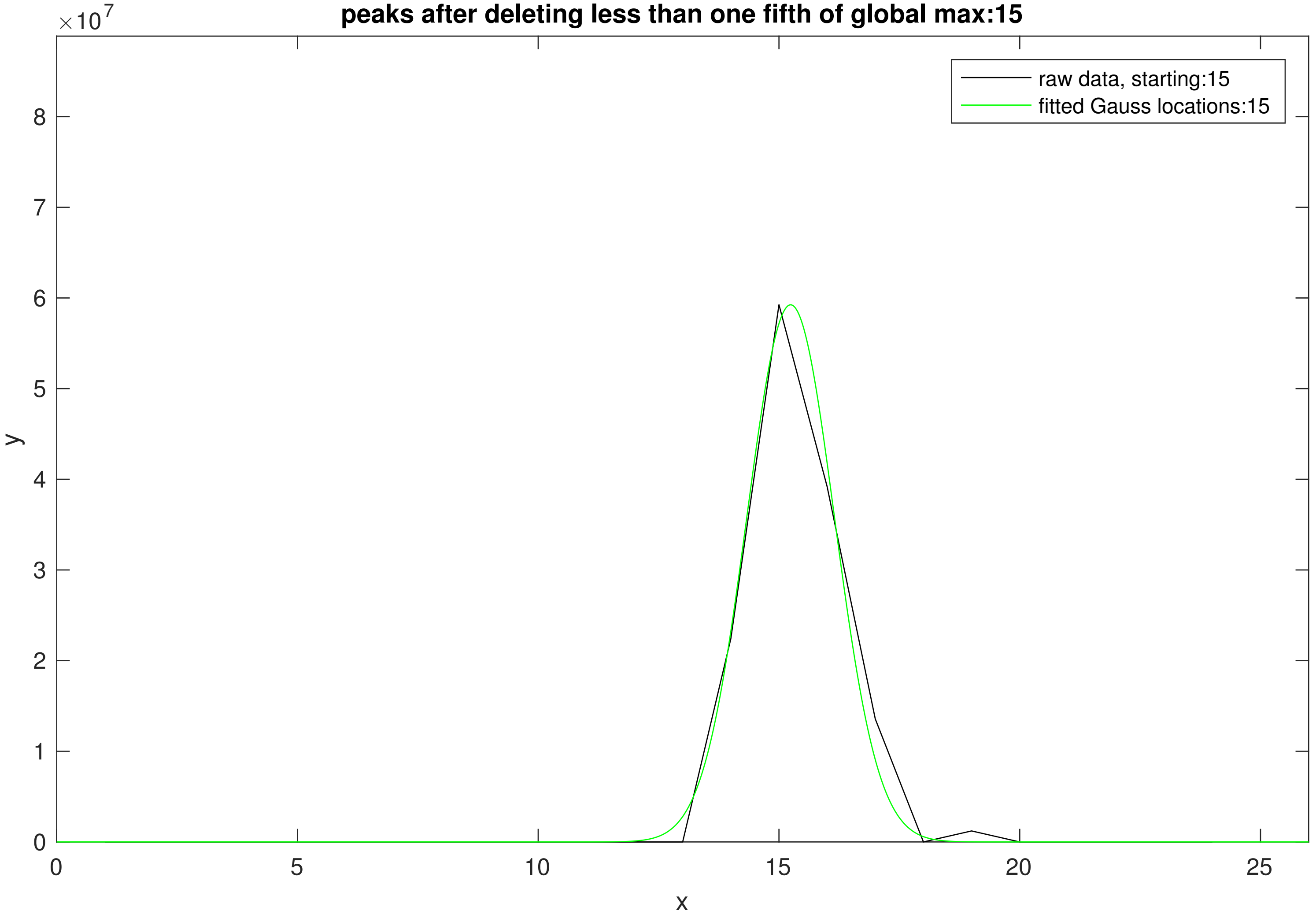

**Optimal1 Fitted Gauss, proteinAT2G44490.1**
**bic:872**
**peaks after deleting less than one fifth of global max:15**

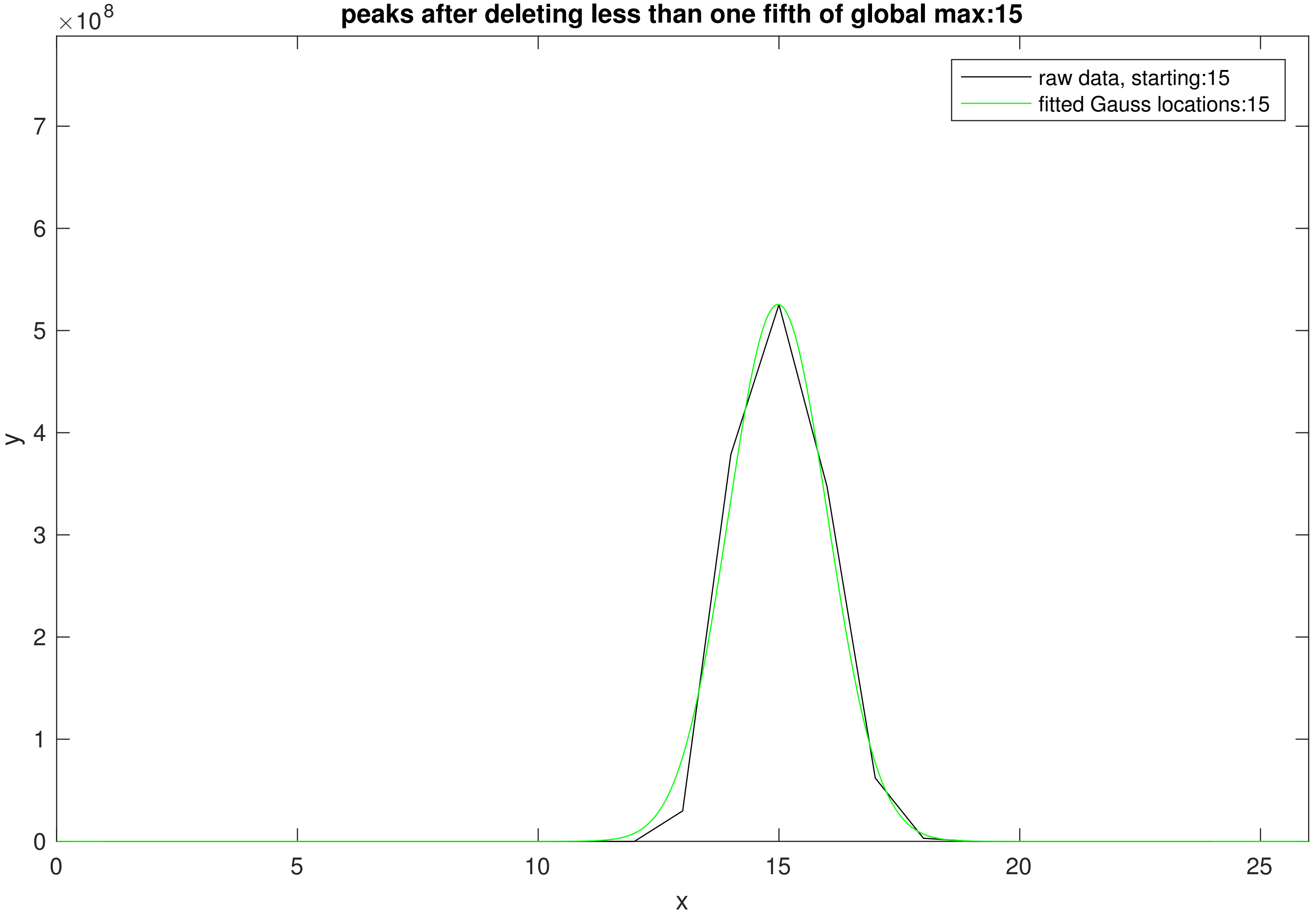

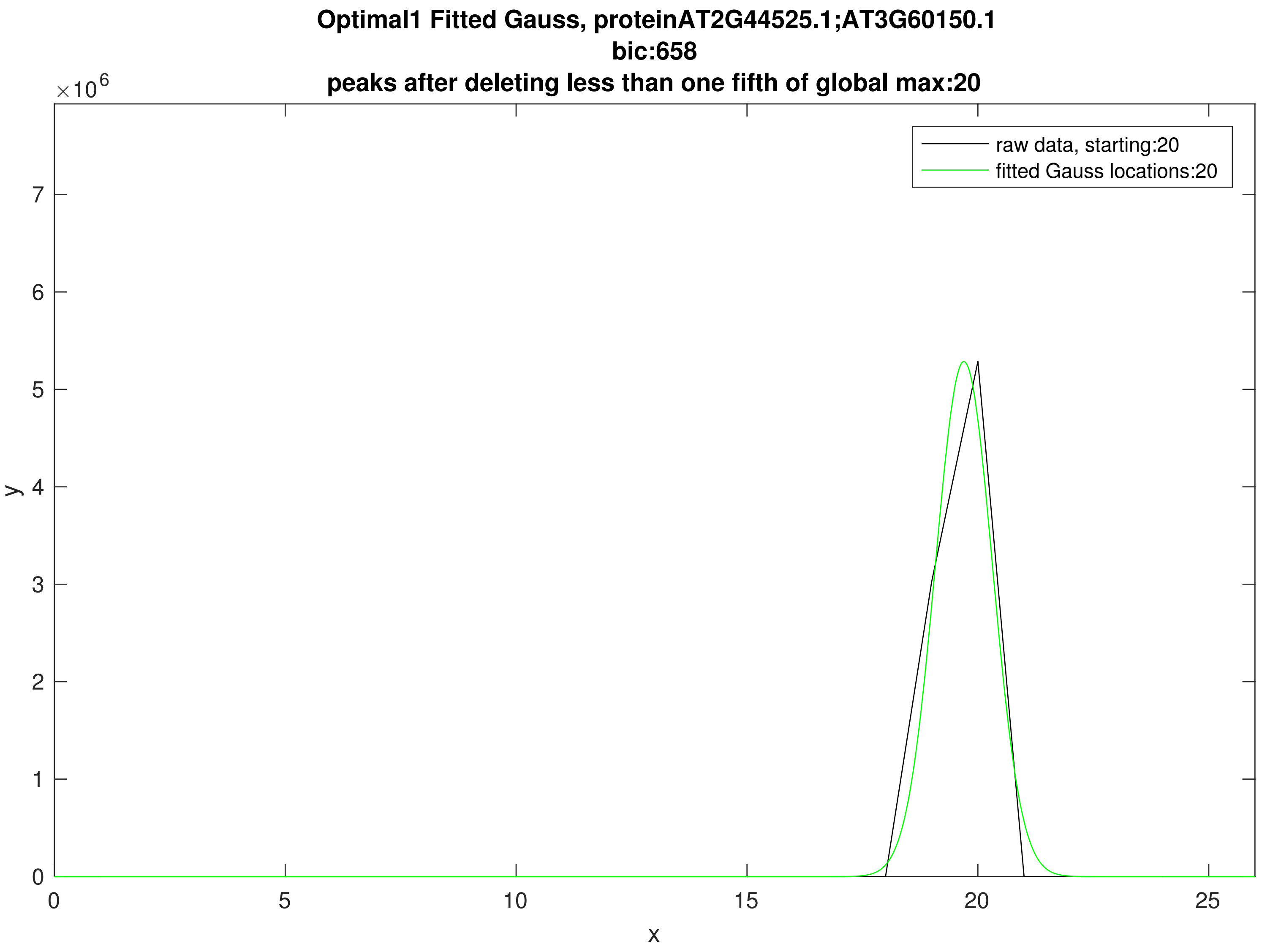

Optimal1 Fitted Gauss, proteinAT2G44525.1;AT3G60150.1
bic:658
peaks after deleting less than one fifth of global max:20
×10^6
raw data, starting:20
fitted Gauss locations:20
7
6
5
4
3
2
1
0
y
0
5
10
15
20
25
x

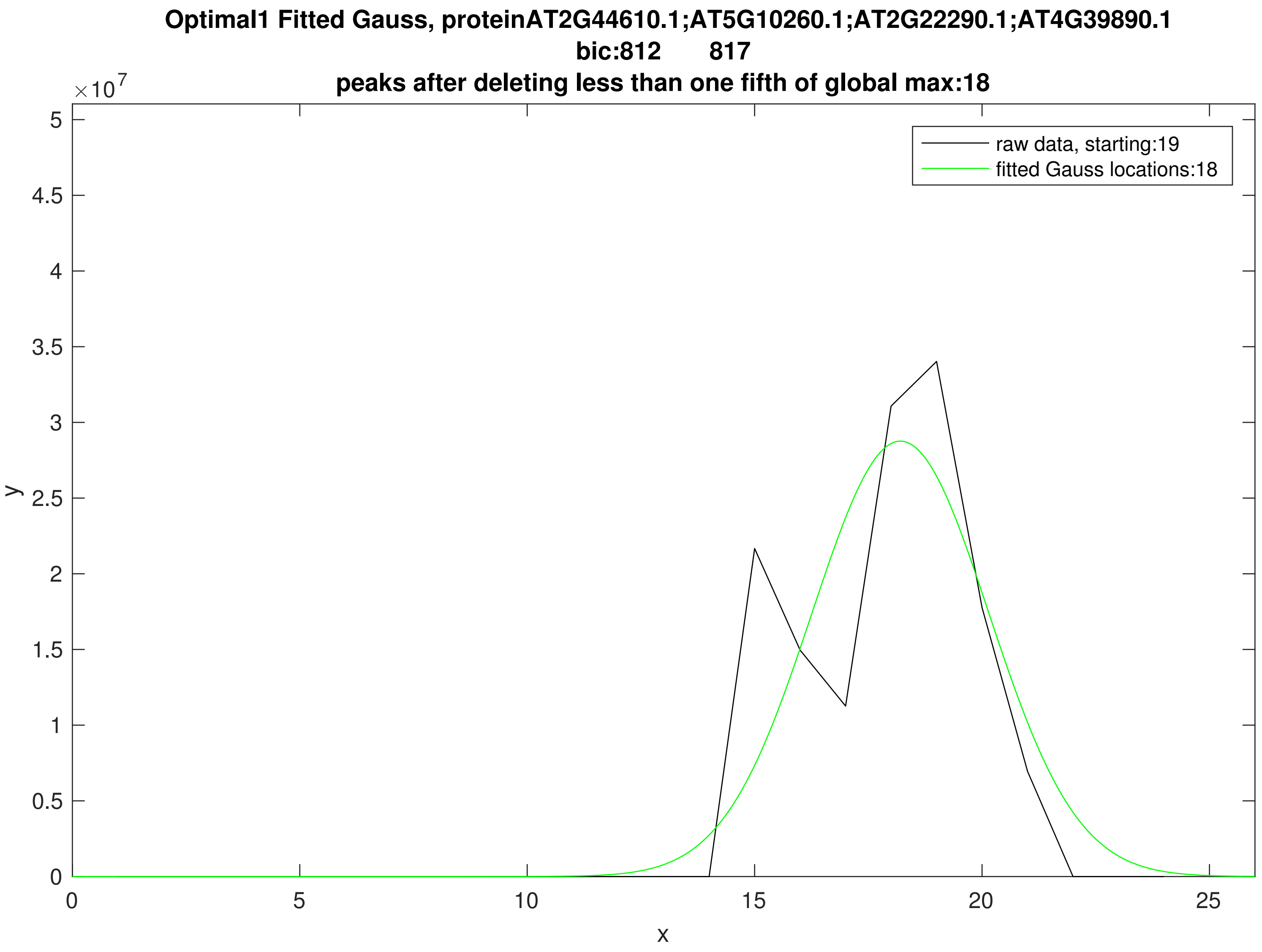
Optimal1 Fitted Gauss, proteinAT2G44610.1;AT5G10260.1;AT2G22290.1;AT4G39890.1
bic:812 817
peaks after deleting less than one fifth of global max:18
×10^7
raw data, starting:19
fitted Gauss locations:18
y
x
0
0.5
1
1.5
2
2.5
3
3.5
4
4.5
5
0
5
10
15
20
25

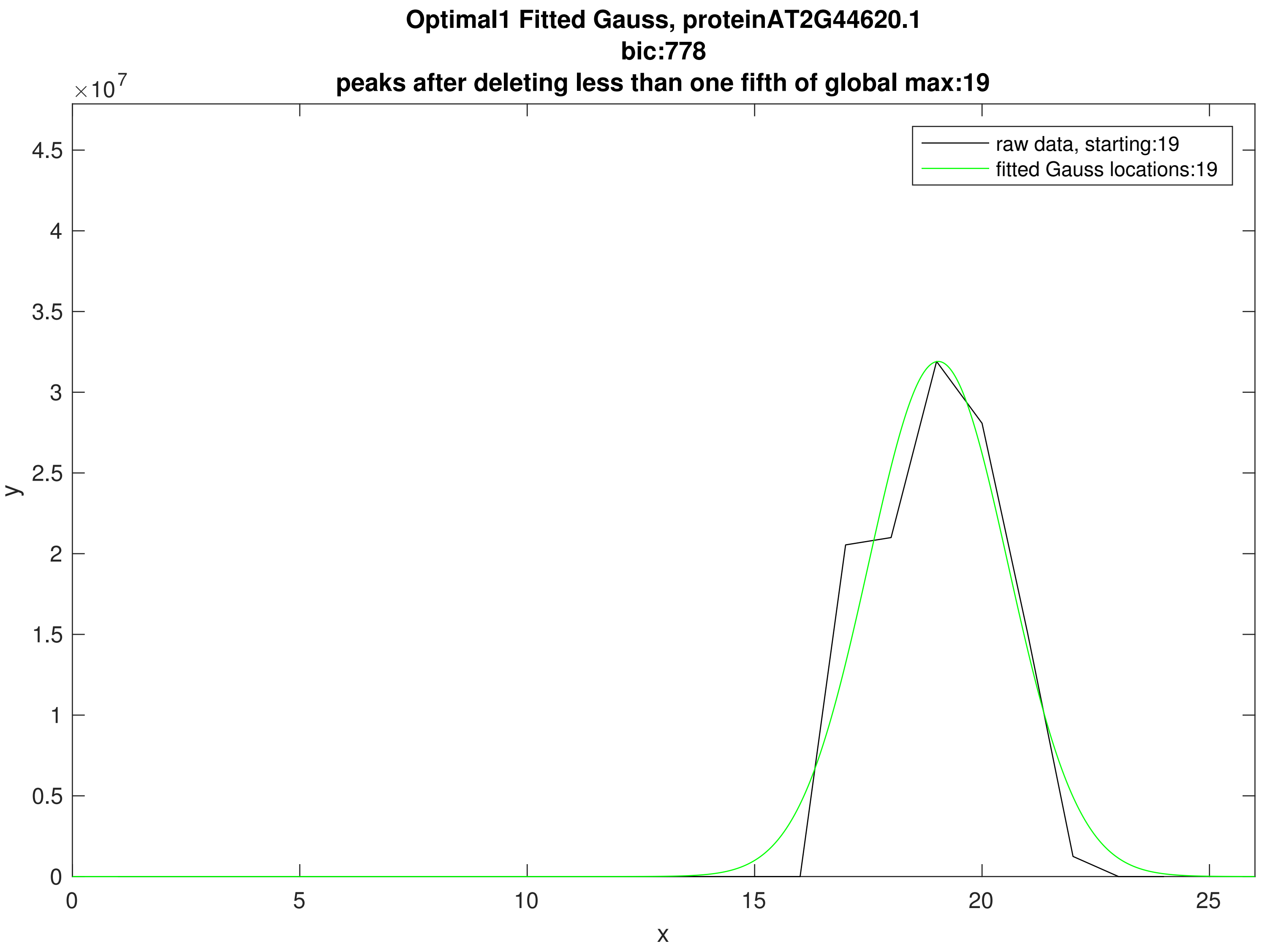
Optimal1 Fitted Gauss, proteinAT2G44620.1
bic:778
peaks after deleting less than one fifth of global max:19
×10^7
raw data, starting:19
fitted Gauss locations:19
4.5
4
3.5
3
2.5
2
1.5
1
0.5
0
y
0
5
10
15
20
25
x

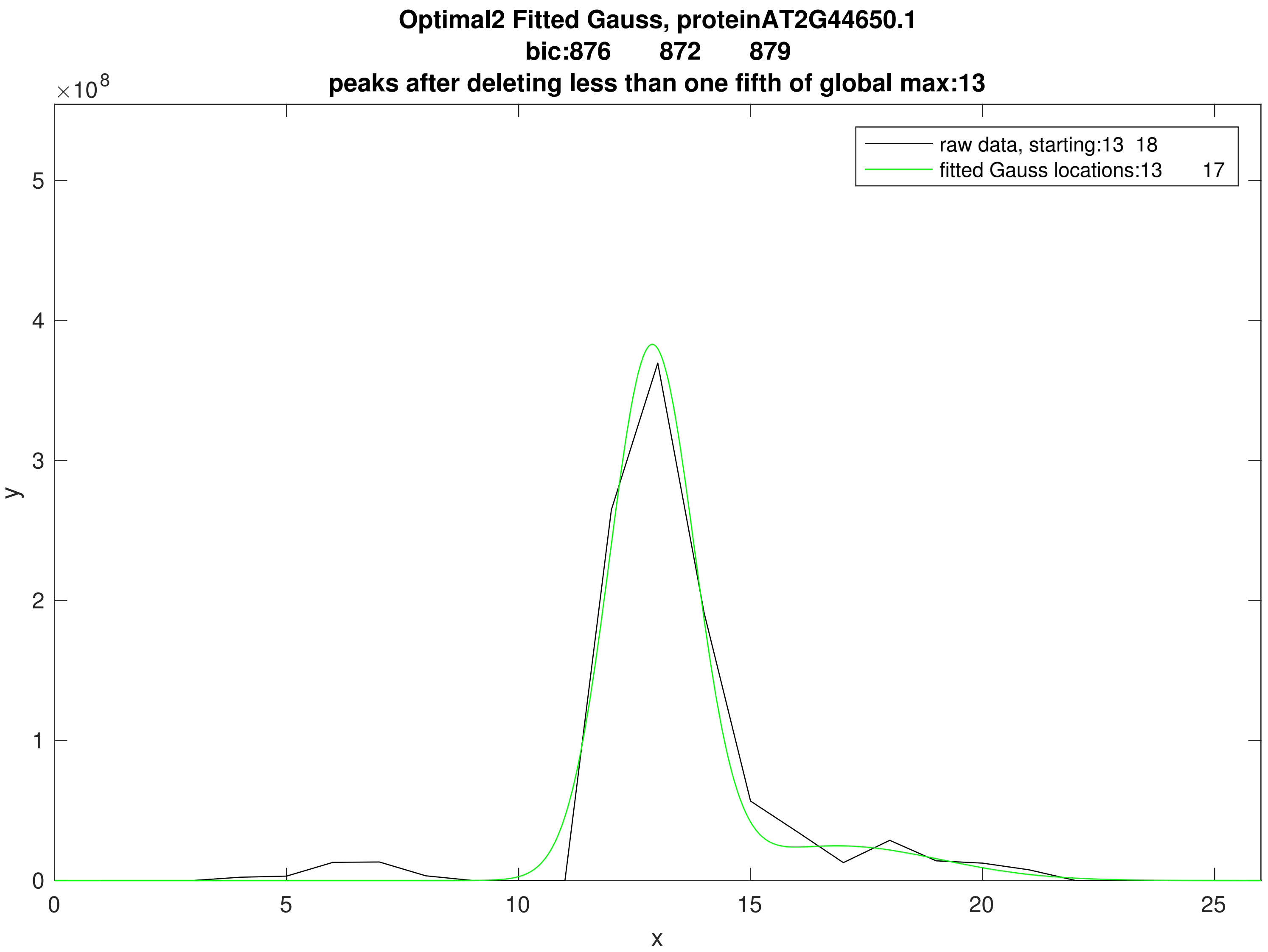

Optimal2 Fitted Gauss, proteinAT2G44650.1
bic:876 872 879
peaks after deleting less than one fifth of global max:13
×10^8
raw data, starting:13 18
fitted Gauss locations:13 17
y
x
0
1
2
3
4
5
0
5
10
15
20
25

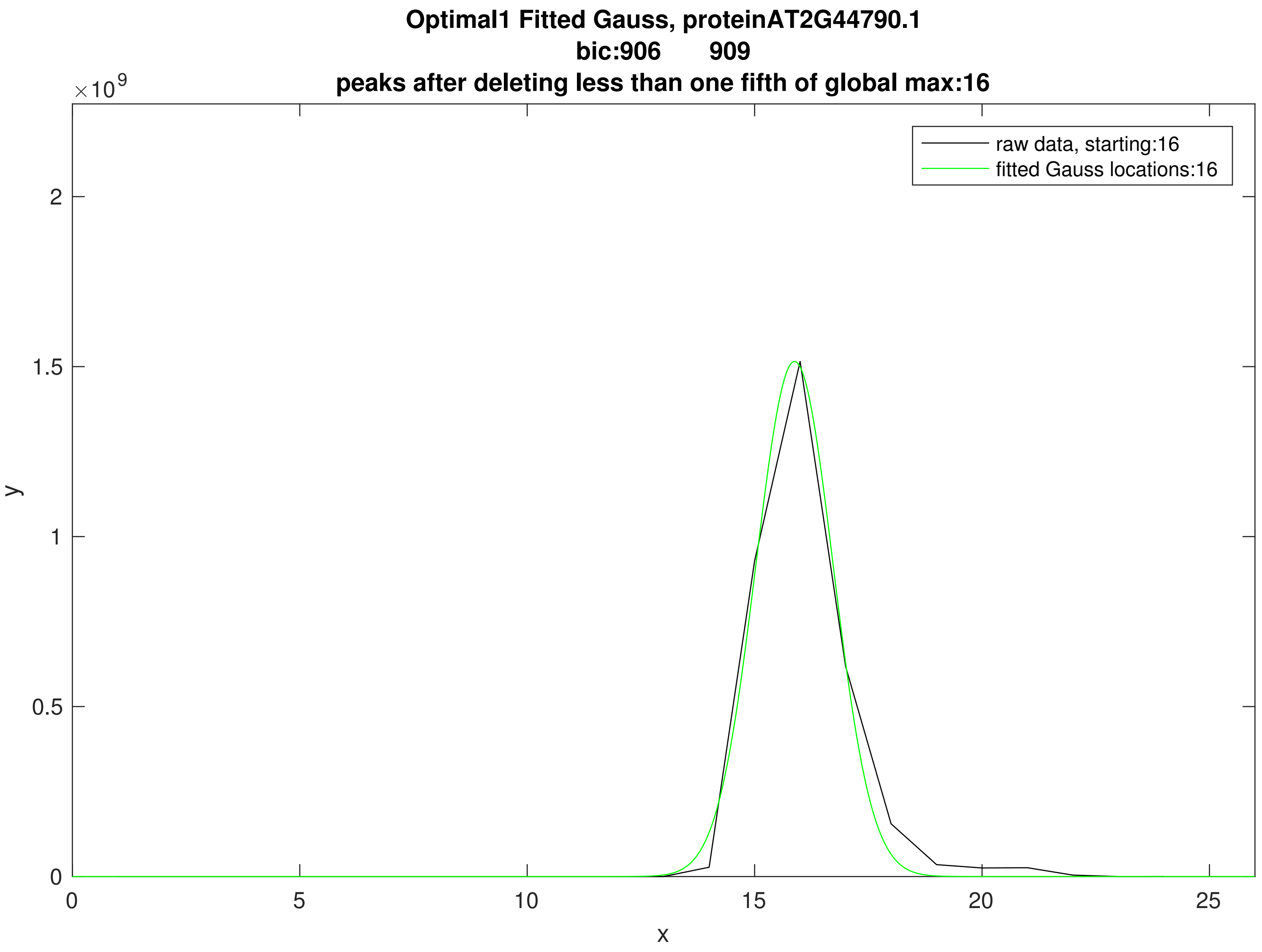

Optimal1 Fitted Gauss, proteinAT2G44790.1
bic:906 909
peaks after deleting less than one fifth of global max:16
×10^9
raw data, starting:16
fitted Gauss locations:16
y
x
0
0.5
1
1.5
2
5
10
15
20
25

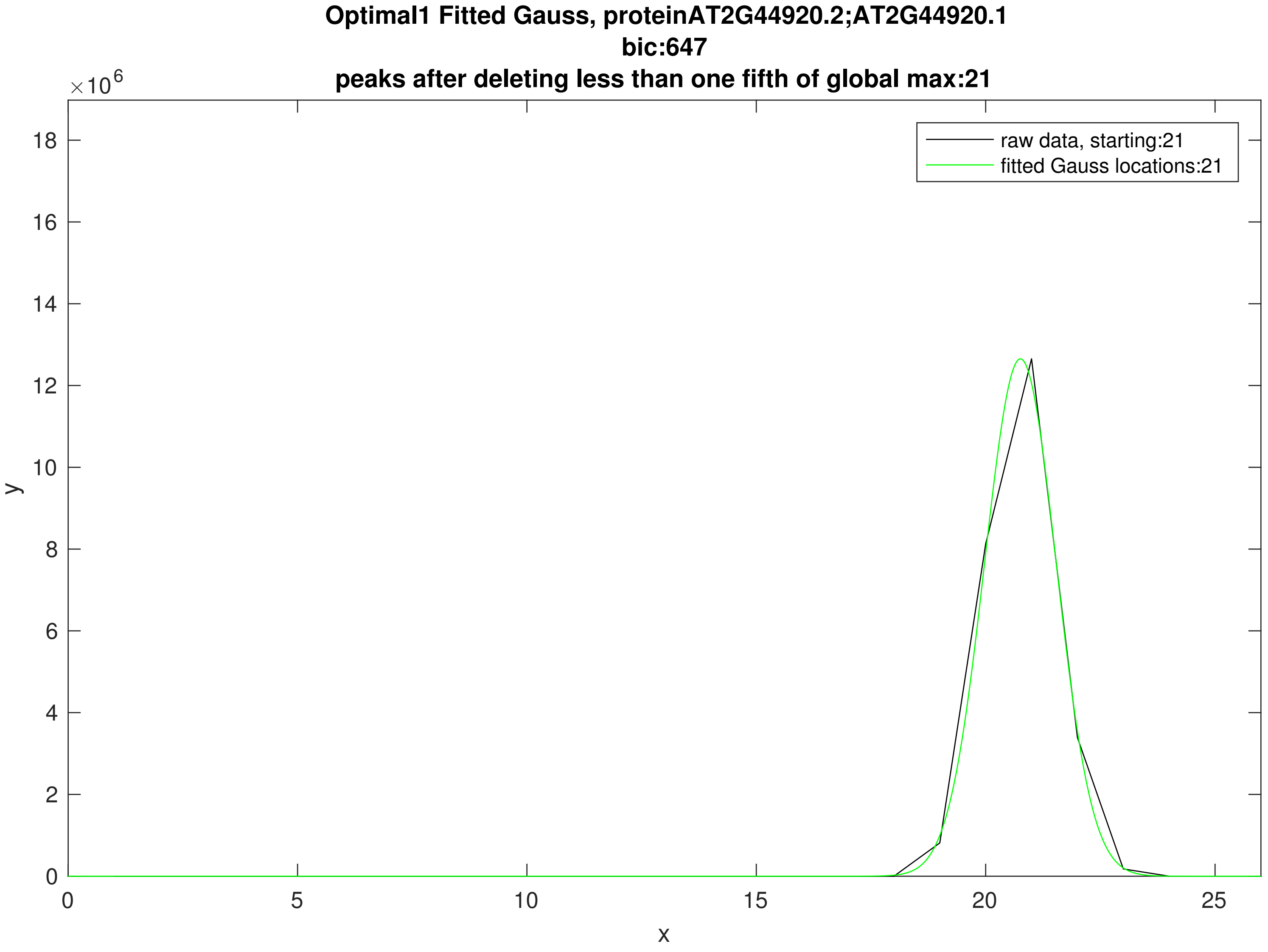

Optimal1 Fitted Gauss, proteinAT2G44920.2;AT2G44920.1
bic:647
peaks after deleting less than one fifth of global max:21
×10^6
raw data, starting:21
fitted Gauss locations:21
y
x
0
2
4
6
8
10
12
14
16
18
0
5
10
15
20
25

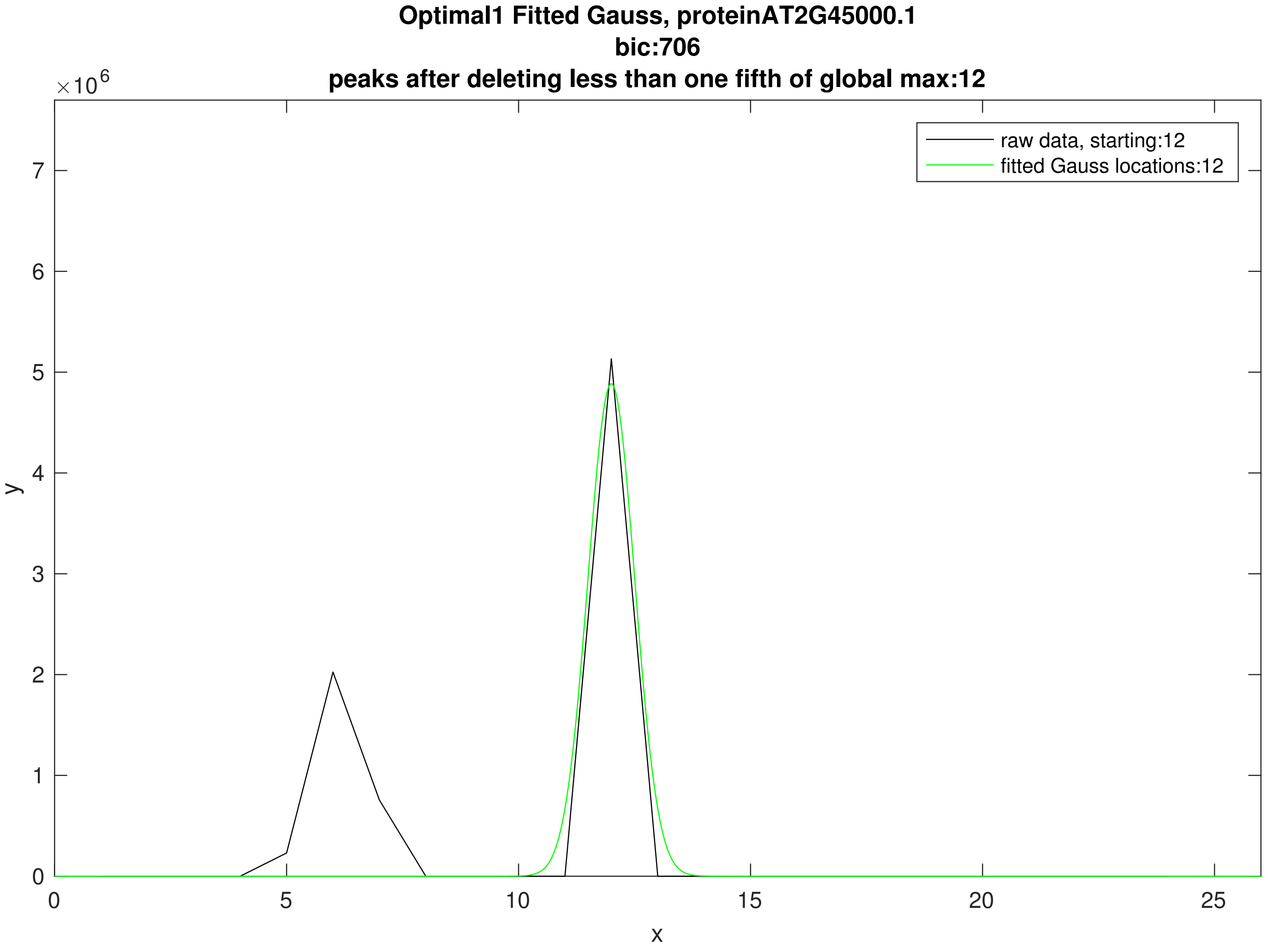
Optimal1 Fitted Gauss, proteinAT2G45000.1
bic:706
peaks after deleting less than one fifth of global max:12
×10^6
raw data, starting:12
fitted Gauss locations:12
y
x
0
1
2
3
4
5
6
7
0
5
10
15
20
25

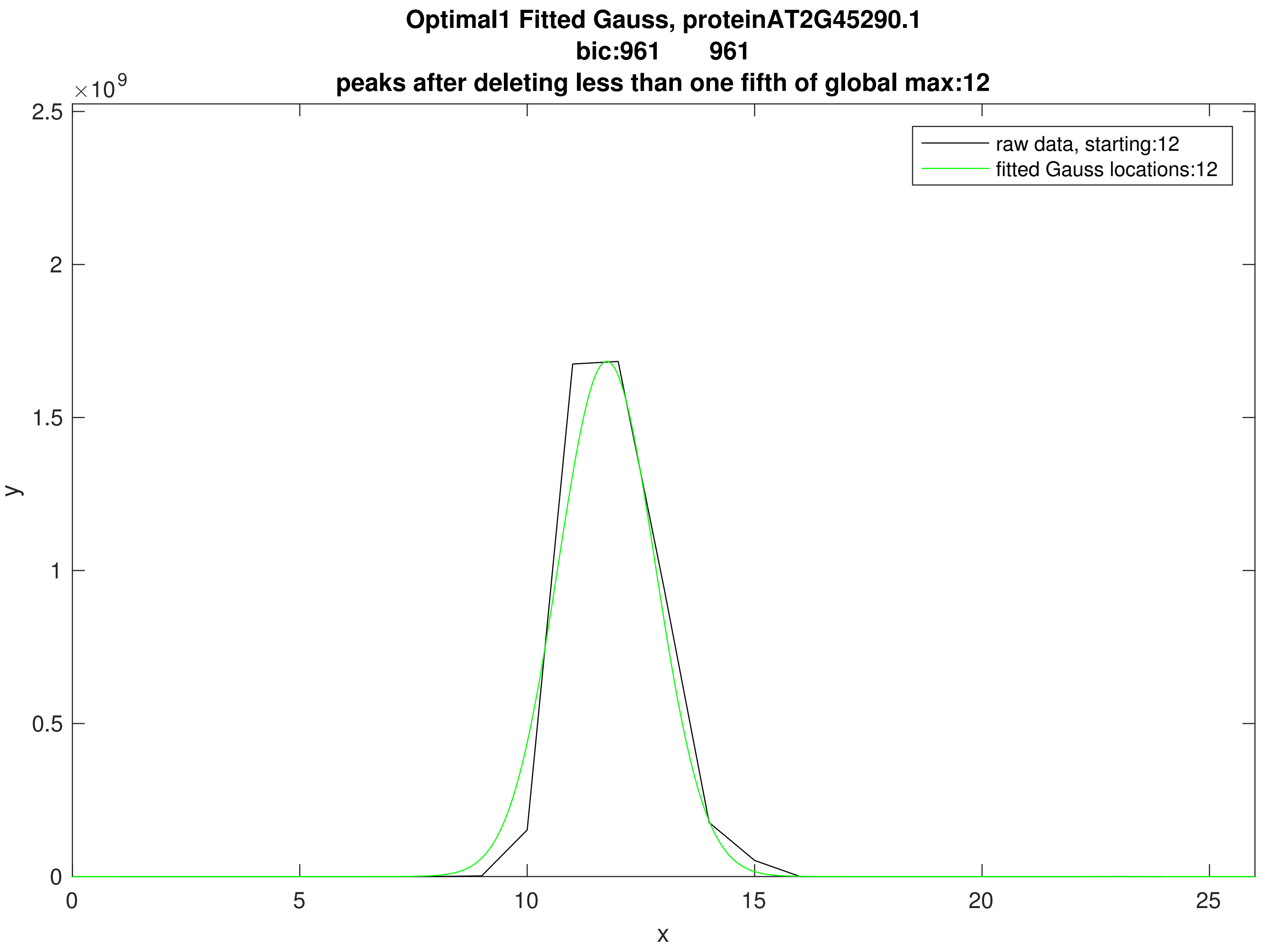
Optimal1 Fitted Gauss, proteinAT2G45290.1
bic:961 961
peaks after deleting less than one fifth of global max:12
×10^9
2.5
2
1.5
1
0.5
0
y
0
5
10
15
20
25
x
raw data, starting:12
fitted Gauss locations:12

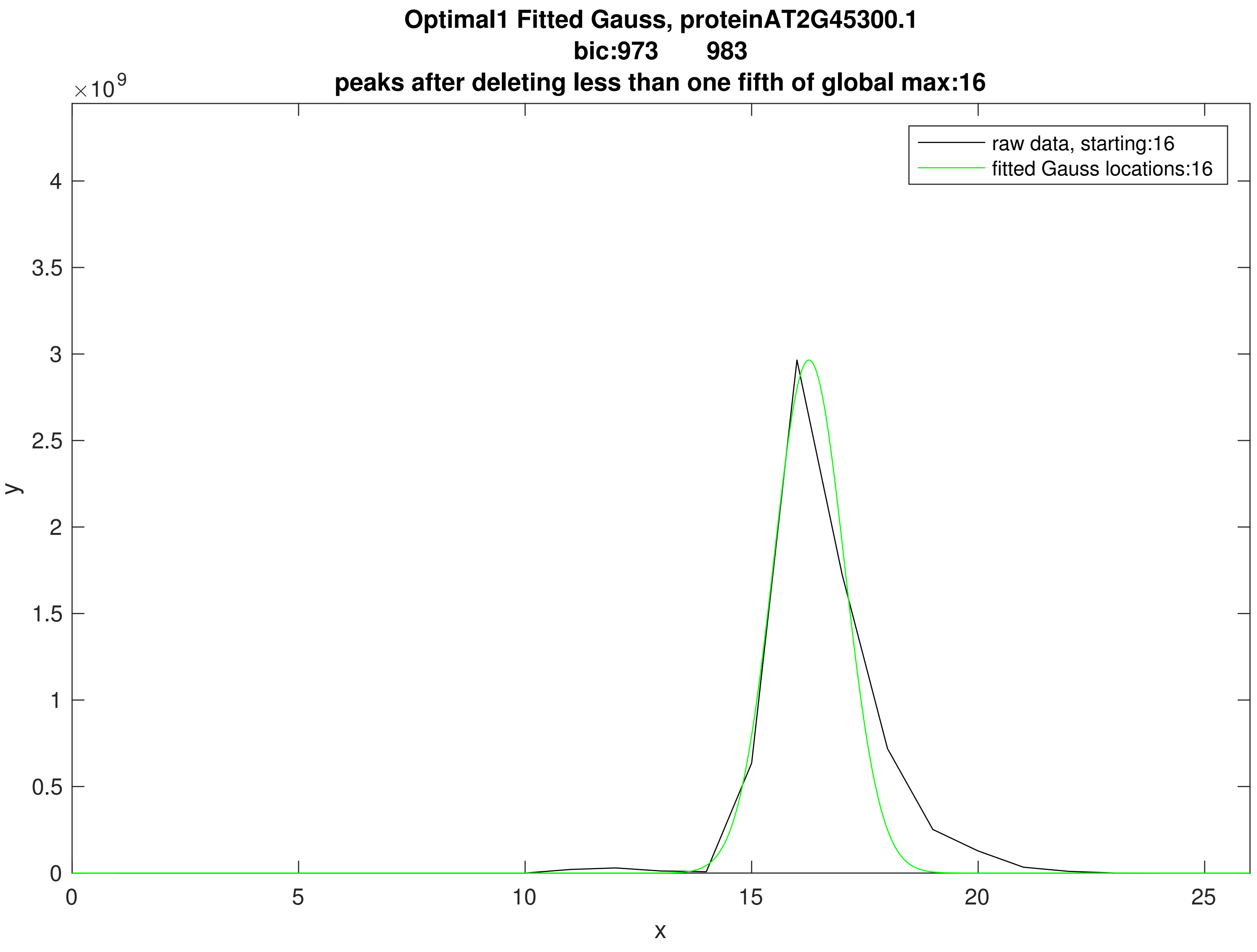

Optimal1 Fitted Gauss, proteinAT2G45300.1
bic:973 983
peaks after deleting less than one fifth of global max:16
×10^9
raw data, starting:16
fitted Gauss locations:16
4
3.5
3
2.5
2
1.5
1
0.5
0
y
0
5
10
15
20
25
x

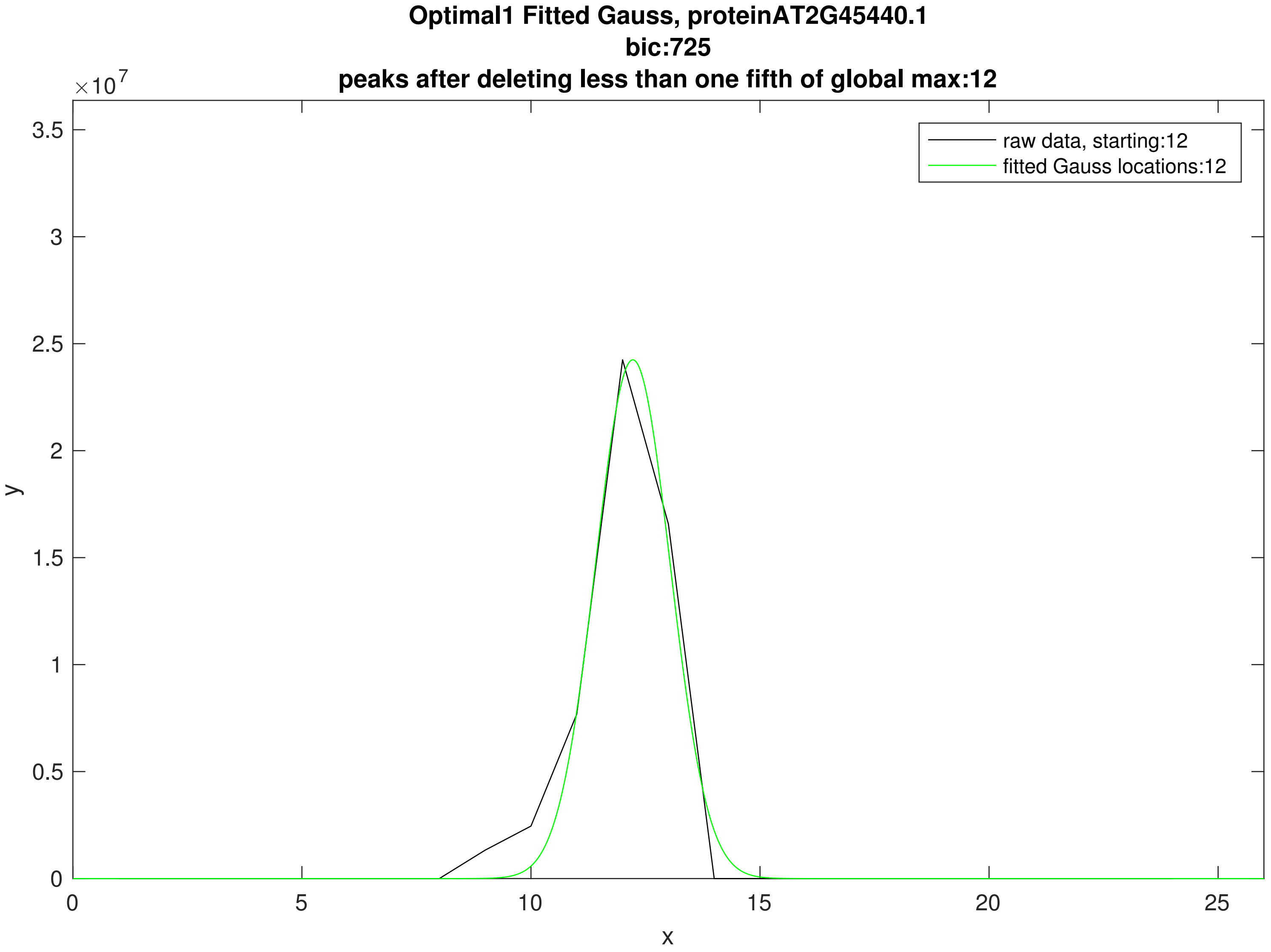
Optimal1 Fitted Gauss, proteinAT2G45440.1
bic:725
peaks after deleting less than one fifth of global max:12
×10^7
raw data, starting:12
fitted Gauss locations:12
y
x
0
0.5
1
1.5
2
2.5
3
3.5
5
10
15
20
25

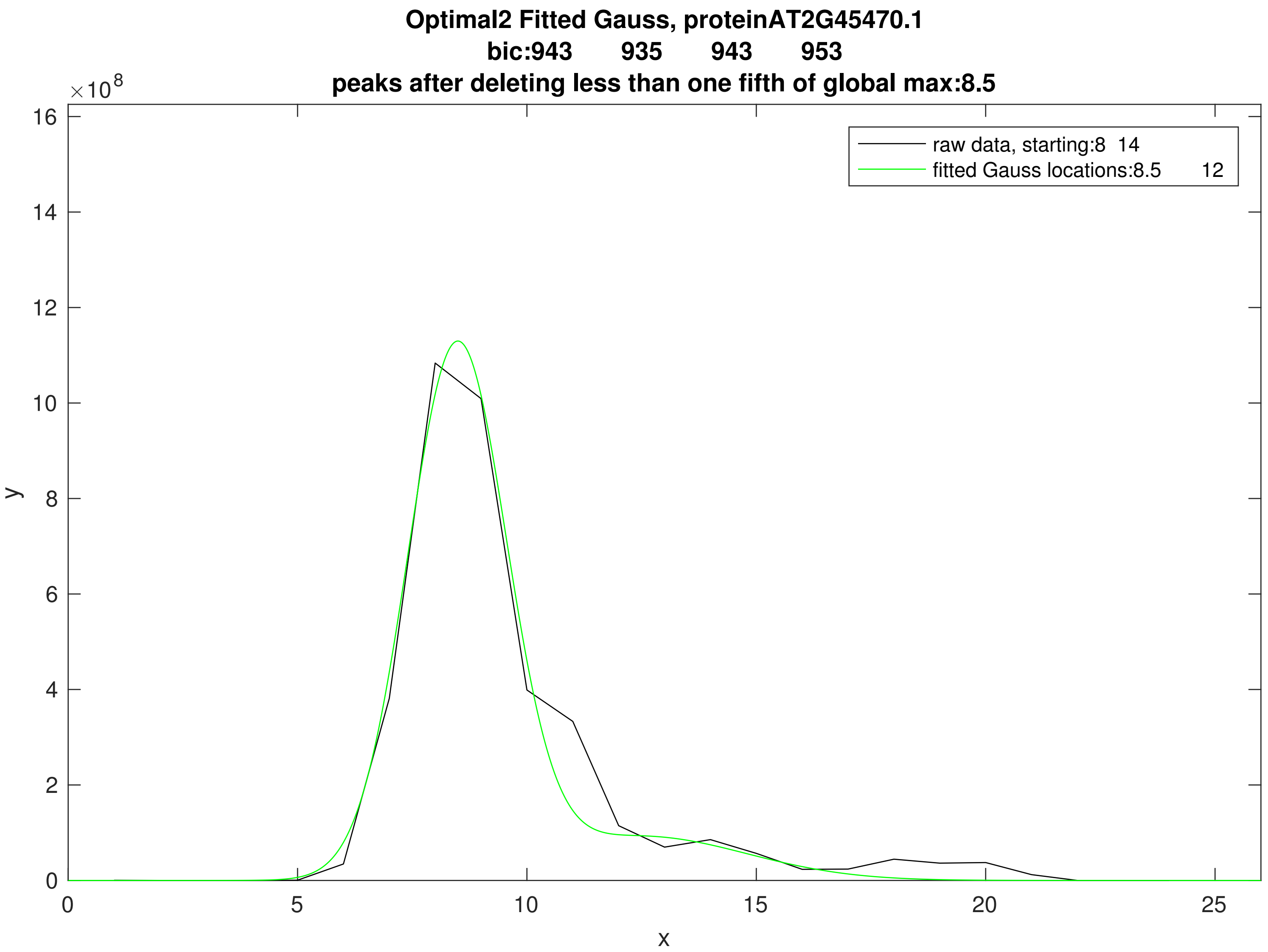
Optimal2 Fitted Gauss, proteinAT2G45470.1
bic:943 935 943 953
peaks after deleting less than one fifth of global max:8.5
×10^8
raw data, starting:8 14
fitted Gauss locations:8.5 12
y
x
0
2
4
6
8
10
12
14
16
0
5
10
15
20
25

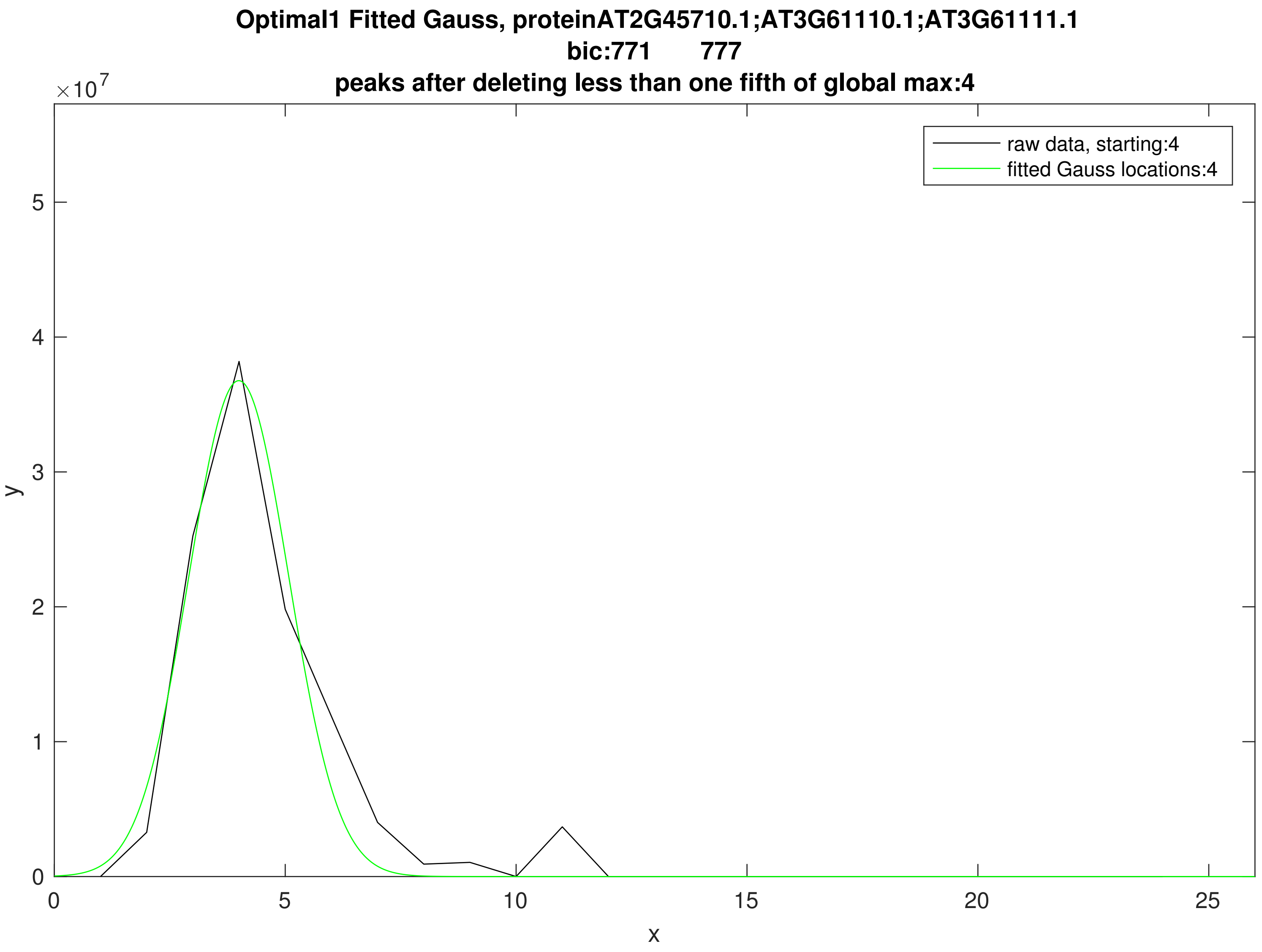

Optimal1 Fitted Gauss, proteinAT2G45710.1;AT3G61110.1;AT3G61111.1
bic:771 777
peaks after deleting less than one fifth of global max:4
×10^7
raw data, starting:4
fitted Gauss locations:4
y
x
0
1
2
3
4
5
0
5
10
15
20
25

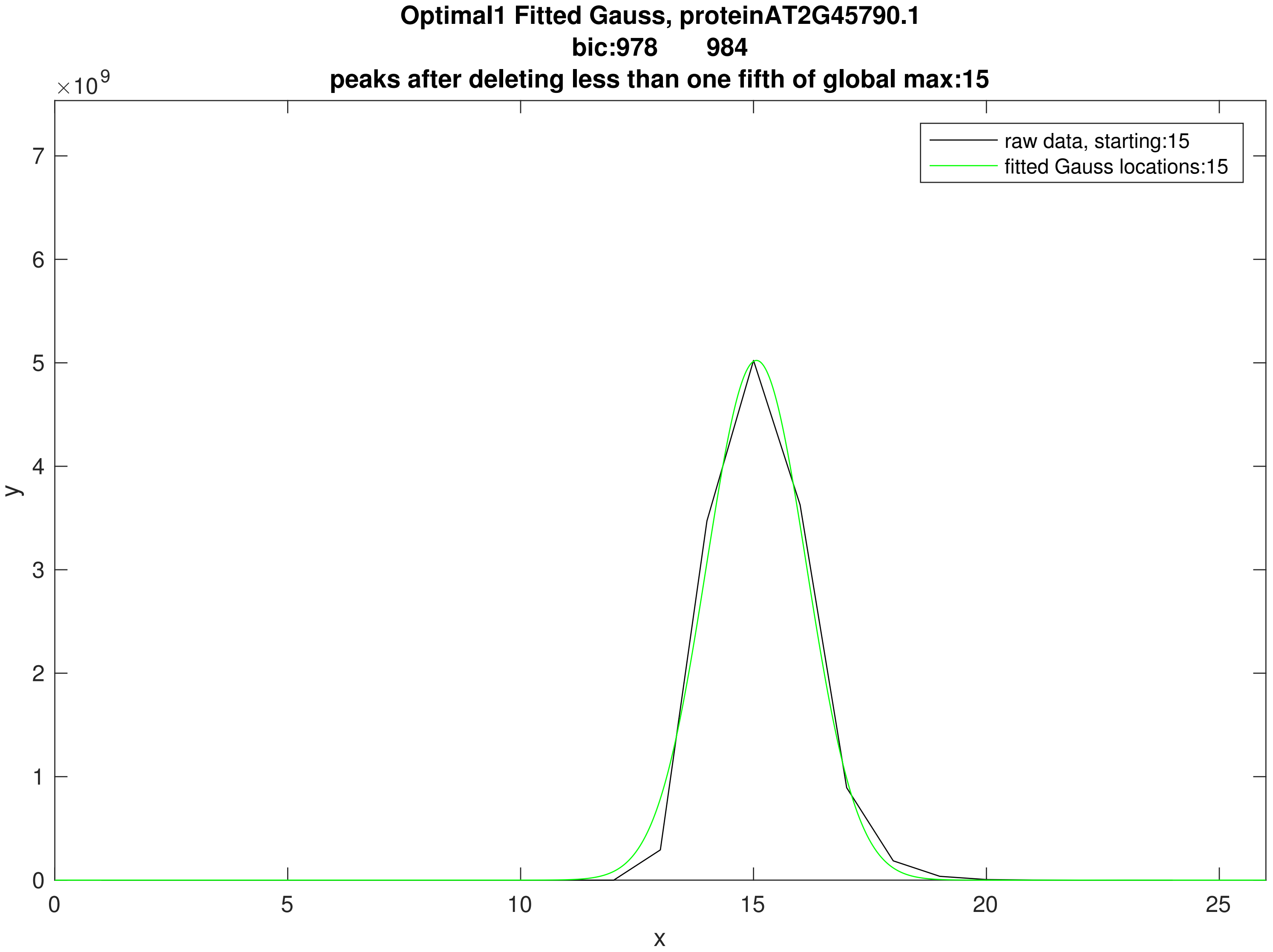
Optimal1 Fitted Gauss, proteinAT2G45790.1
bic:978 984
peaks after deleting less than one fifth of global max:15
×10^9
raw data, starting:15
fitted Gauss locations:15
y
x
0
1
2
3
4
5
6
7
0
5
10
15
20
25

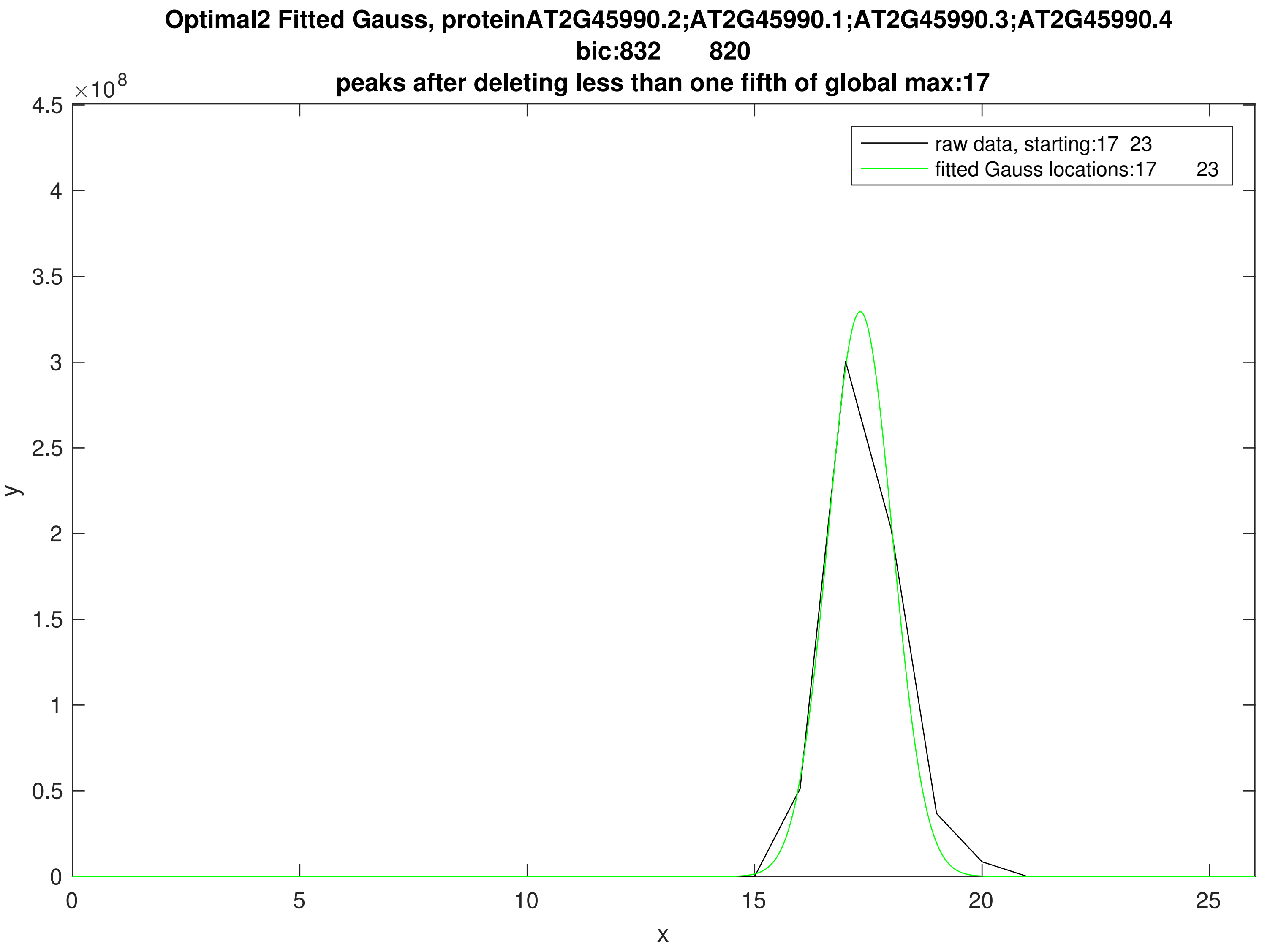

Optimal2 Fitted Gauss, proteinAT2G45990.2;AT2G45990.1;AT2G45990.3;AT2G45990.4
bic:832 820
peaks after deleting less than one fifth of global max:17
×10^8
4.5
4
3.5
3
2.5
2
1.5
1
0.5
0
y
0
5
10
15
20
25
x
raw data, starting:17 23
fitted Gauss locations:17 23

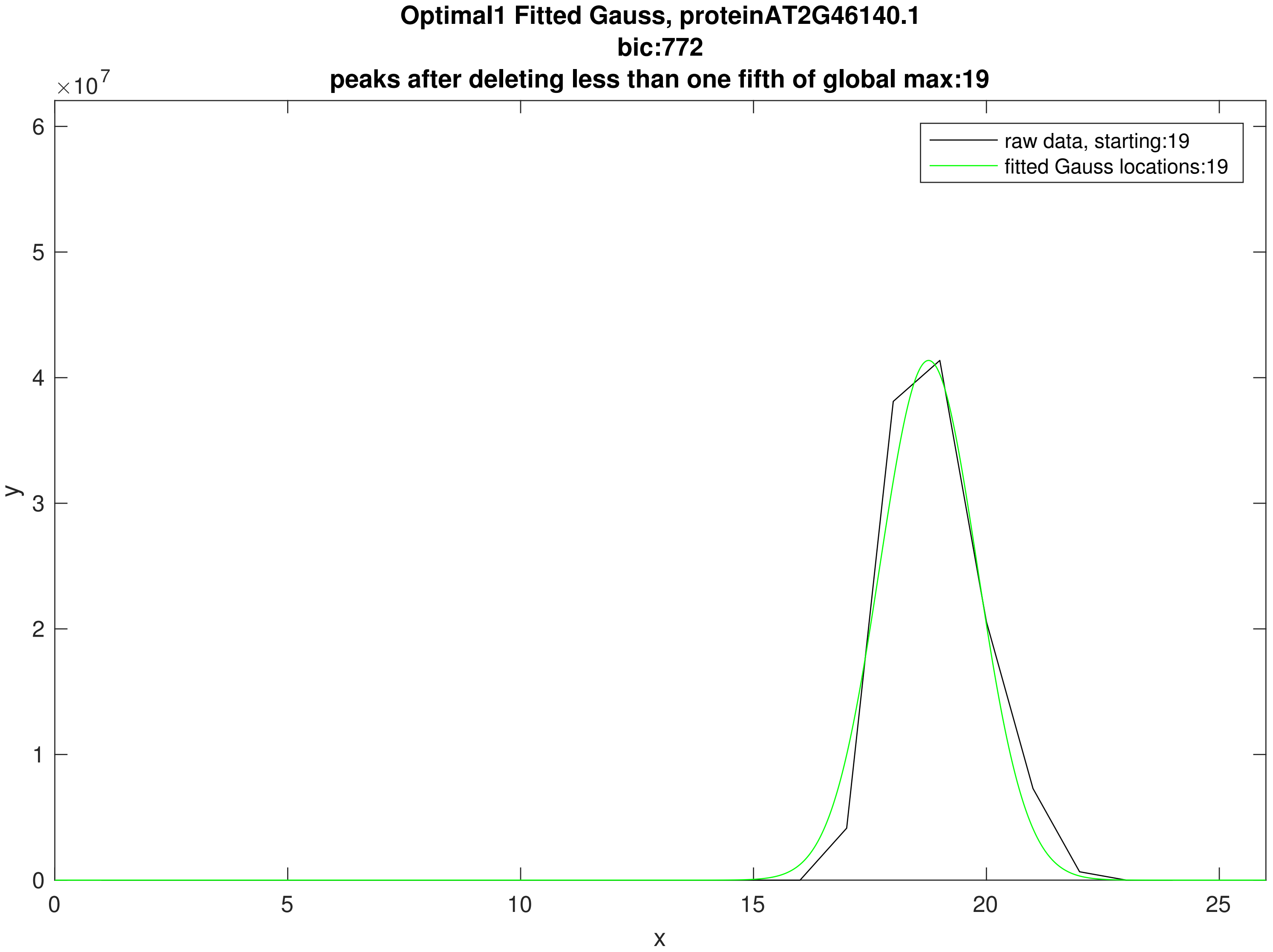
Optimal1 Fitted Gauss, proteinAT2G46140.1
bic:772
peaks after deleting less than one fifth of global max:19
×10^7
raw data, starting:19
fitted Gauss locations:19
y
x
0
1
2
3
4
5
6
0
5
10
15
20
25

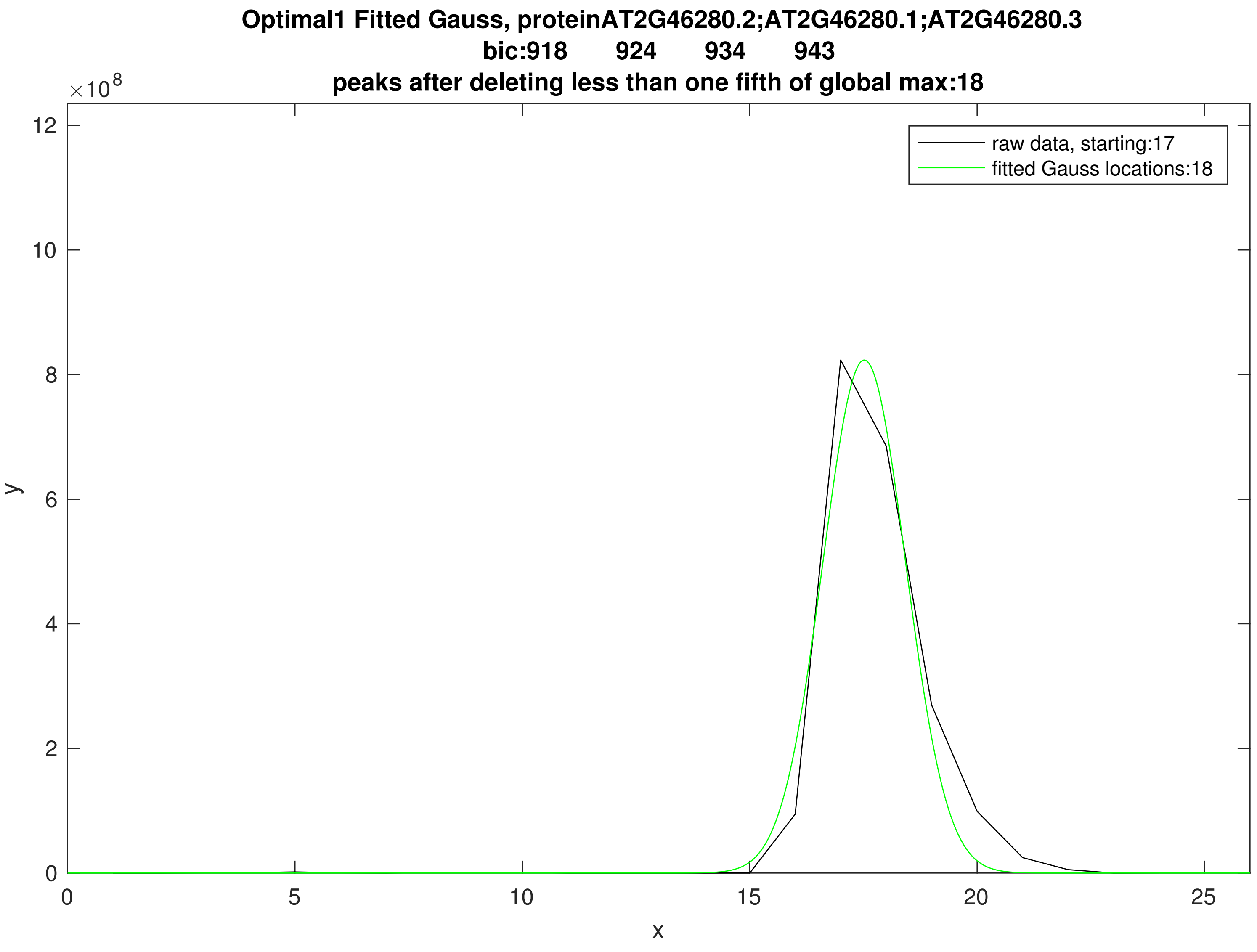

Optimal1 Fitted Gauss, proteinAT2G46280.2;AT2G46280.1;AT2G46280.3
bic:918 924 934 943
peaks after deleting less than one fifth of global max:18
×10^8
raw data, starting:17
fitted Gauss locations:18
y
x
0
2
4
6
8
10
12
5
15
20
25

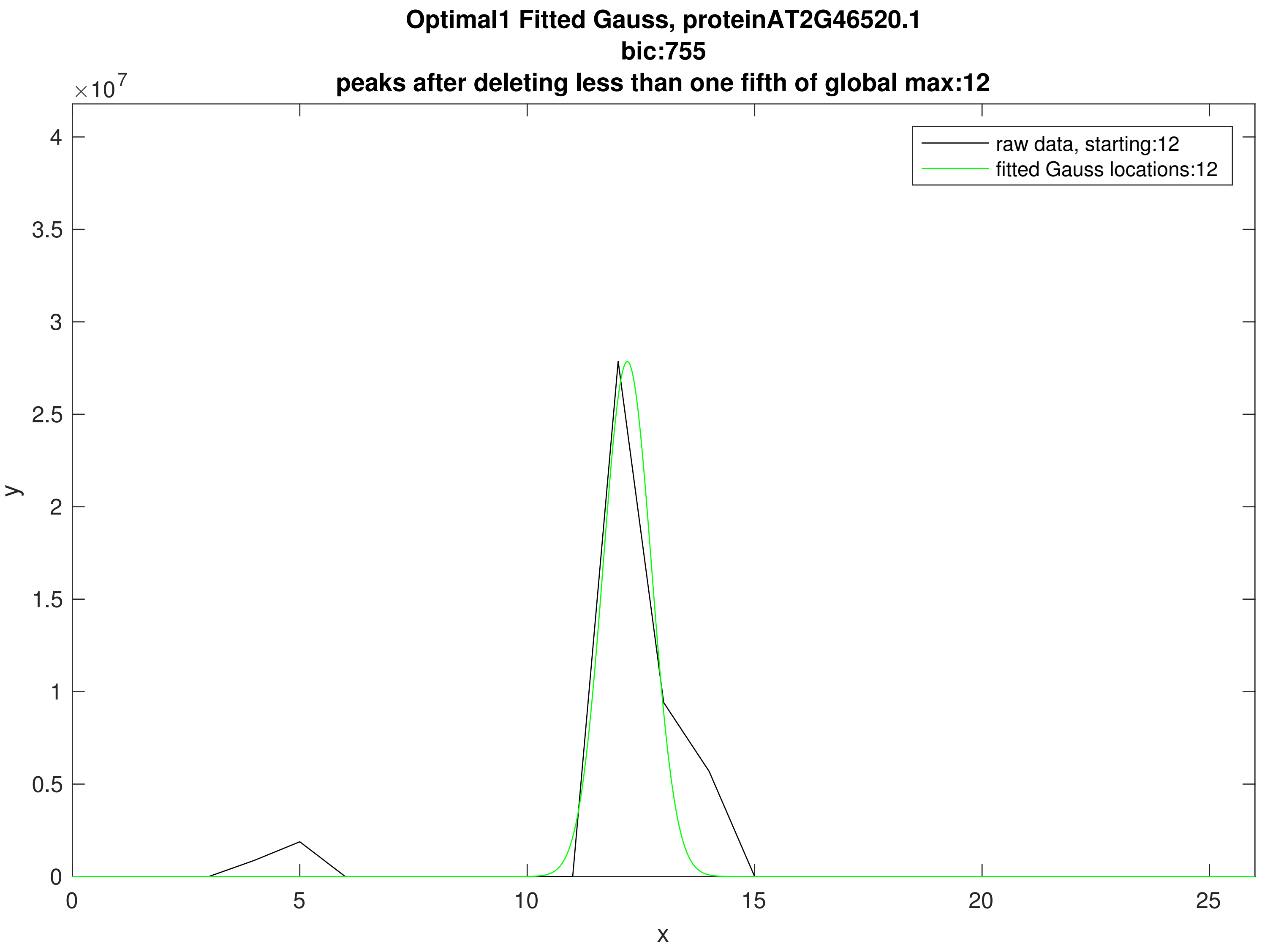
Optimal1 Fitted Gauss, proteinAT2G46520.1
bic:755
peaks after deleting less than one fifth of global max:12
×10^7
4
3.5
3
2.5
2
1.5
1
0.5
0
y
0
5
10
15
20
25
x
raw data, starting:12
fitted Gauss locations:12

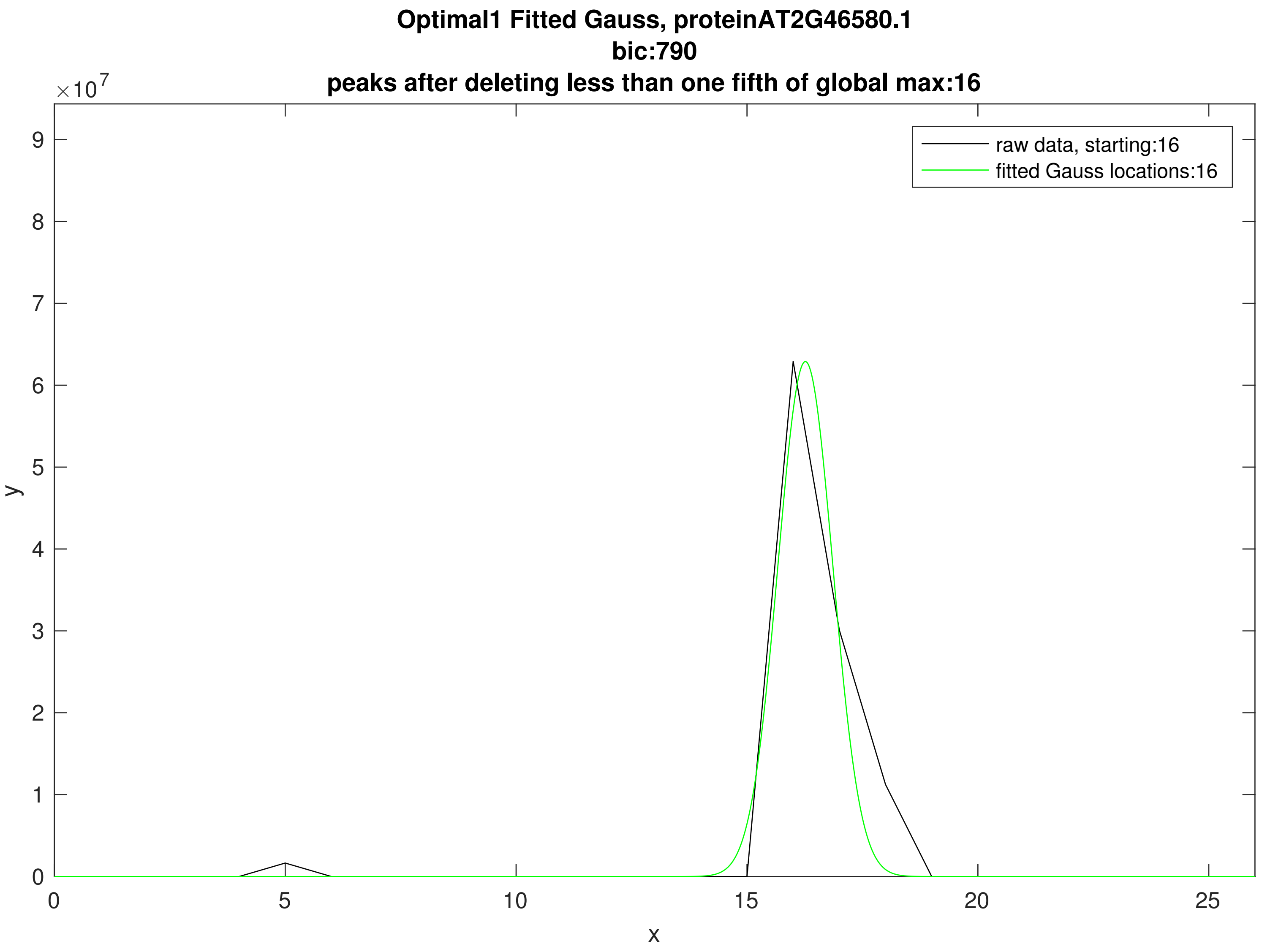
Optimal1 Fitted Gauss, proteinAT2G46580.1
bic:790
peaks after deleting less than one fifth of global max:16
×10^7
raw data, starting:16
fitted Gauss locations:16
y
x
0
1
2
3
4
5
6
7
8
9
0
5
10
15
20
25

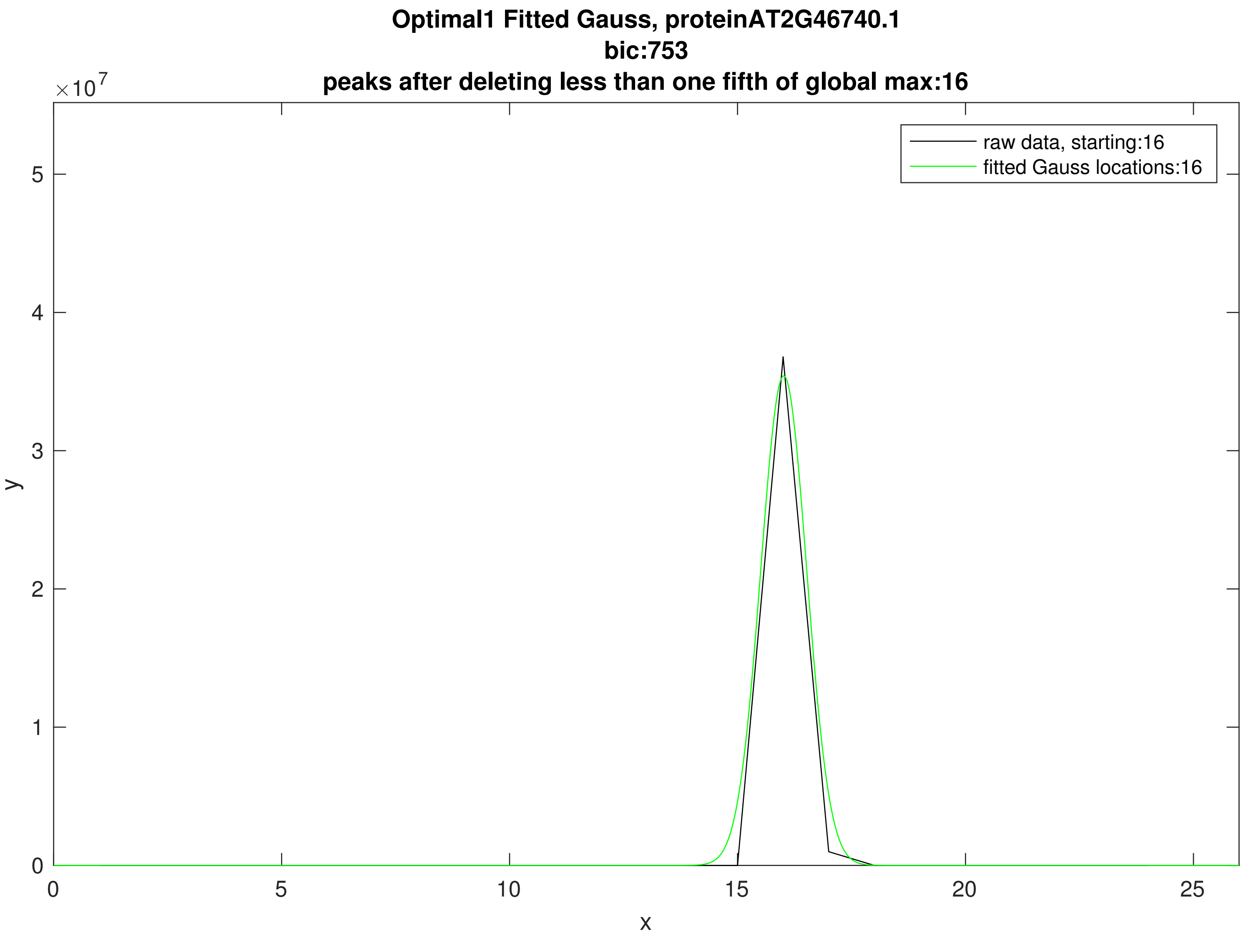
Optimal1 Fitted Gauss, proteinAT2G46740.1
bic:753
peaks after deleting less than one fifth of global max:16
×10^7
y
x
raw data, starting:16
fitted Gauss locations:16
0
1
2
3
4
5
0
5
10
15
20
25

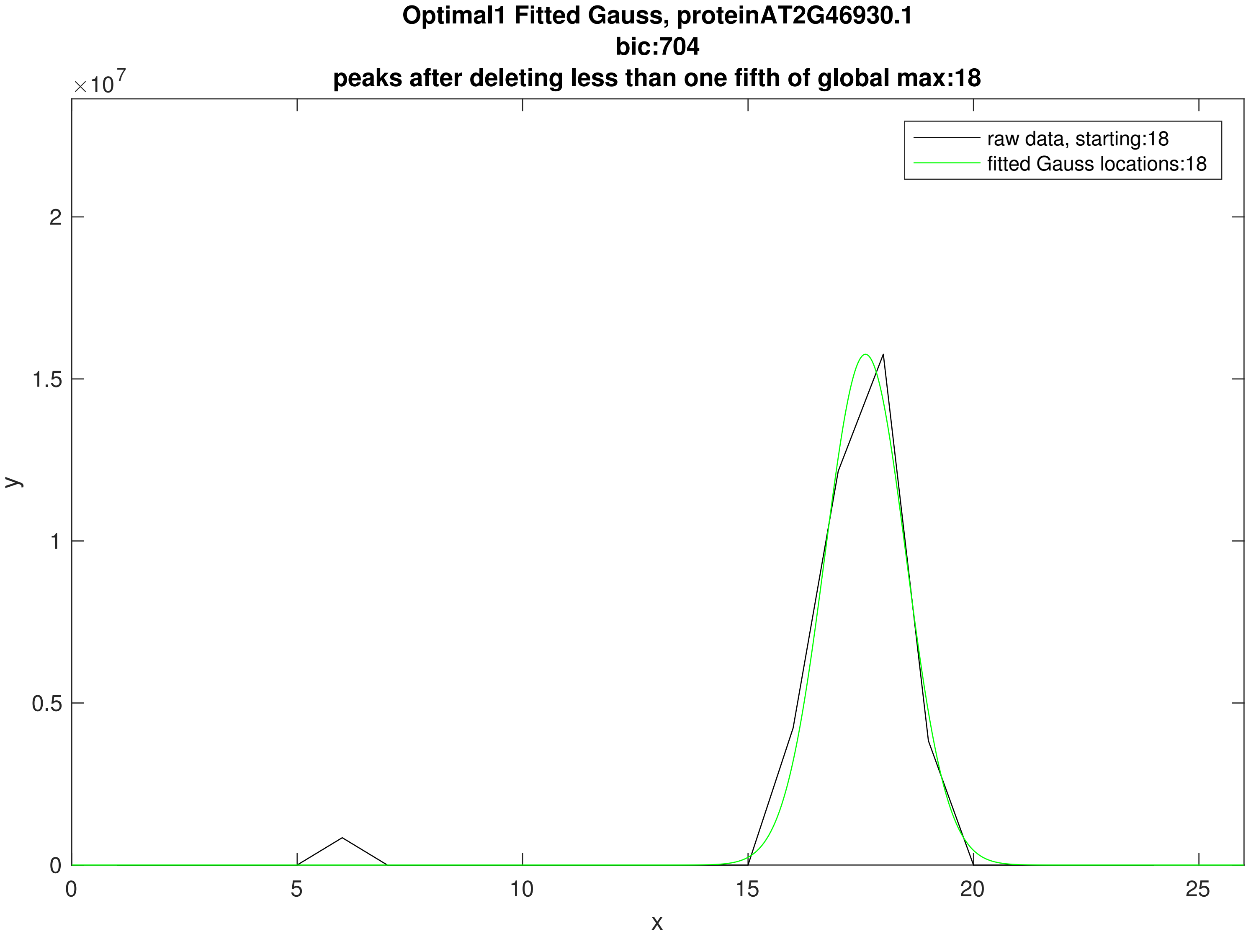

Optimal1 Fitted Gauss, proteinAT2G46930.1
bic:704
peaks after deleting less than one fifth of global max:18
×10^7
raw data, starting:18
fitted Gauss locations:18
y
x
0
0.5
1
1.5
2
0
5
10
15
20
25

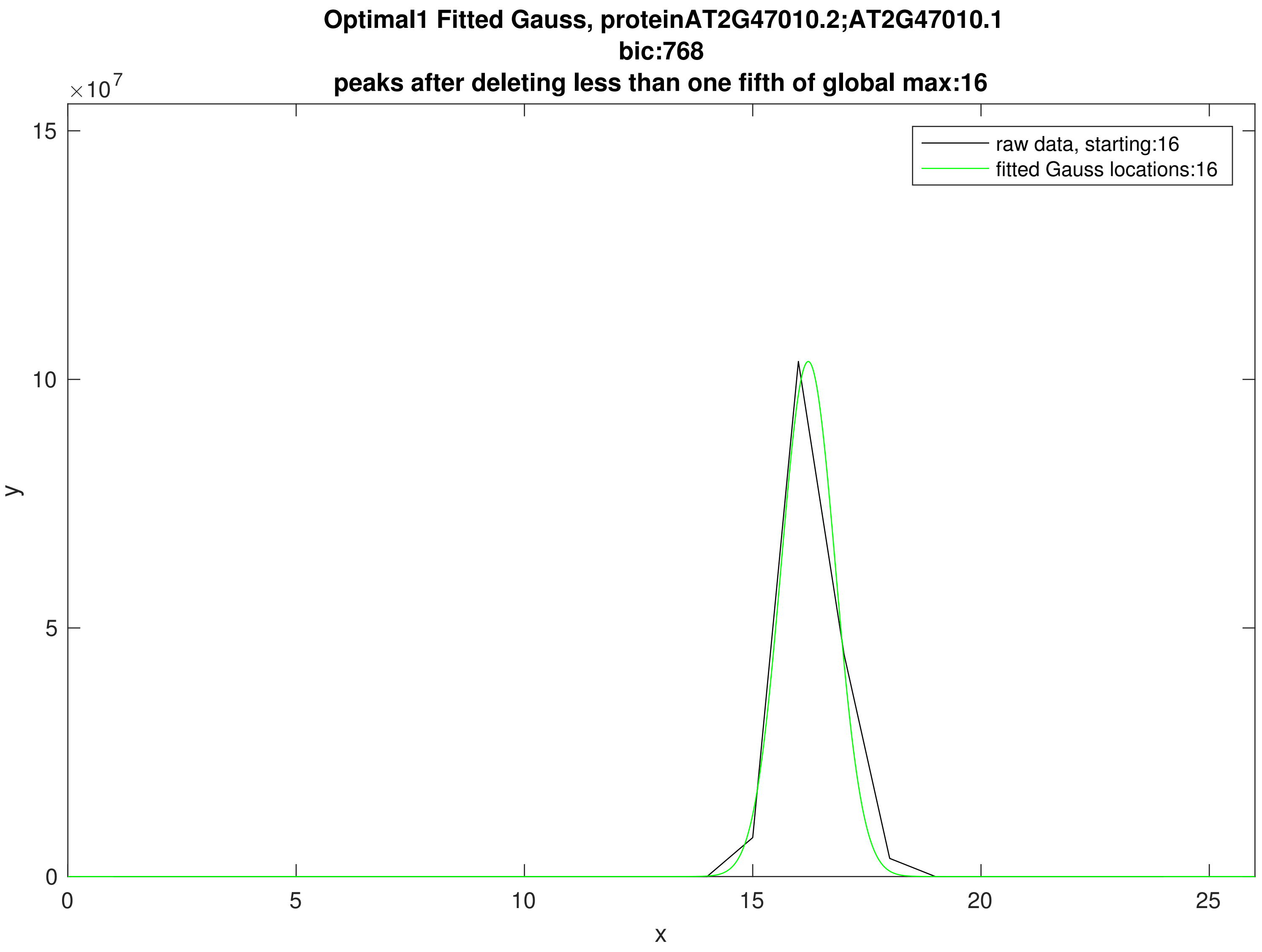
Optimal1 Fitted Gauss, proteinAT2G47010.2;AT2G47010.1
bic:768
peaks after deleting less than one fifth of global max:16
×10^7
15
10
5
0
y
0
5
10
15
20
25
x
raw data, starting:16
fitted Gauss locations:16

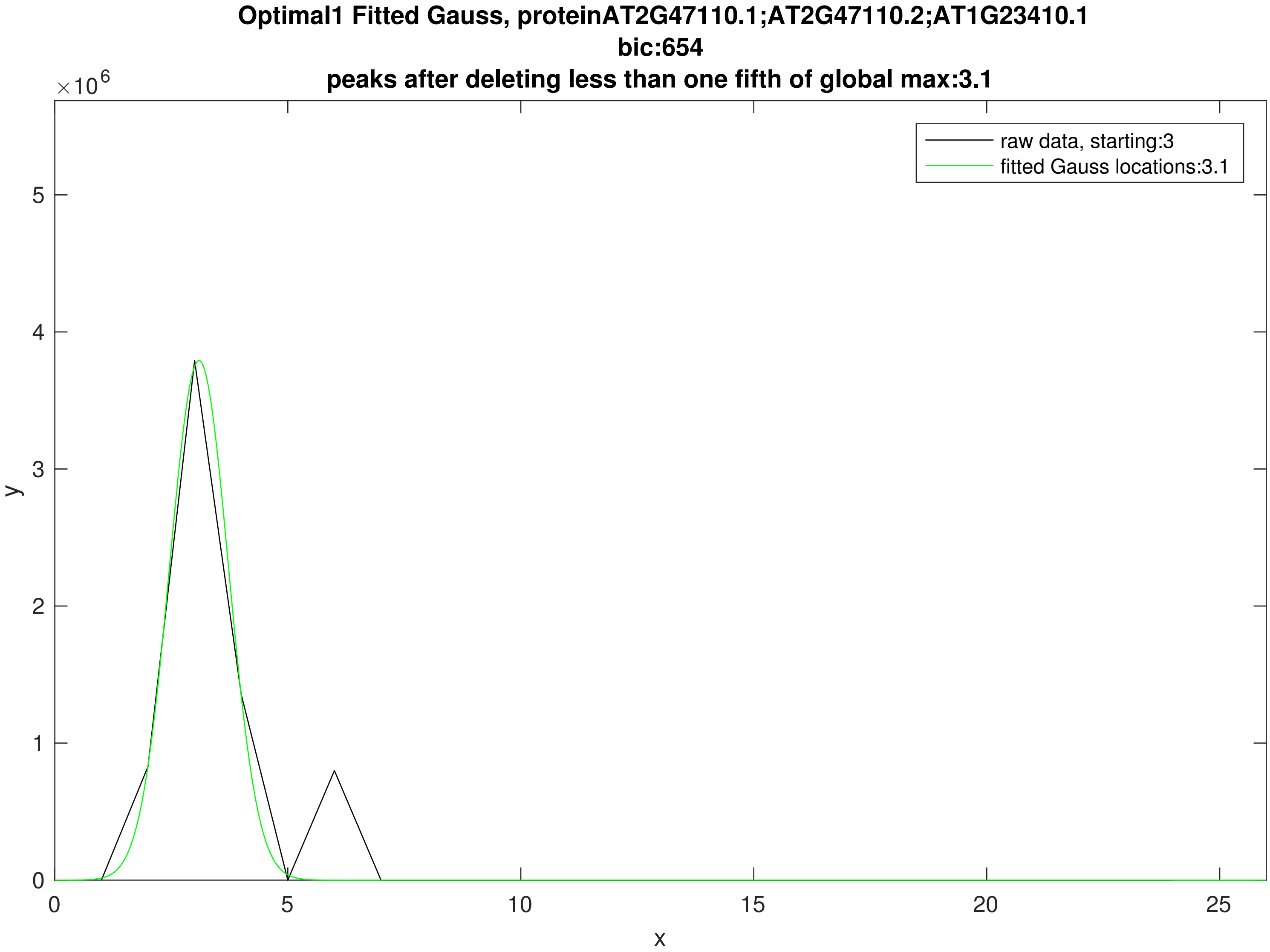
Optimal1 Fitted Gauss, proteinAT2G47110.1;AT2G47110.2;AT1G23410.1
bic:654
peaks after deleting less than one fifth of global max:3.1
×10^6
raw data, starting:3
fitted Gauss locations:3.1
y
x
0
1
2
3
4
5
0
5
10
15
20
25

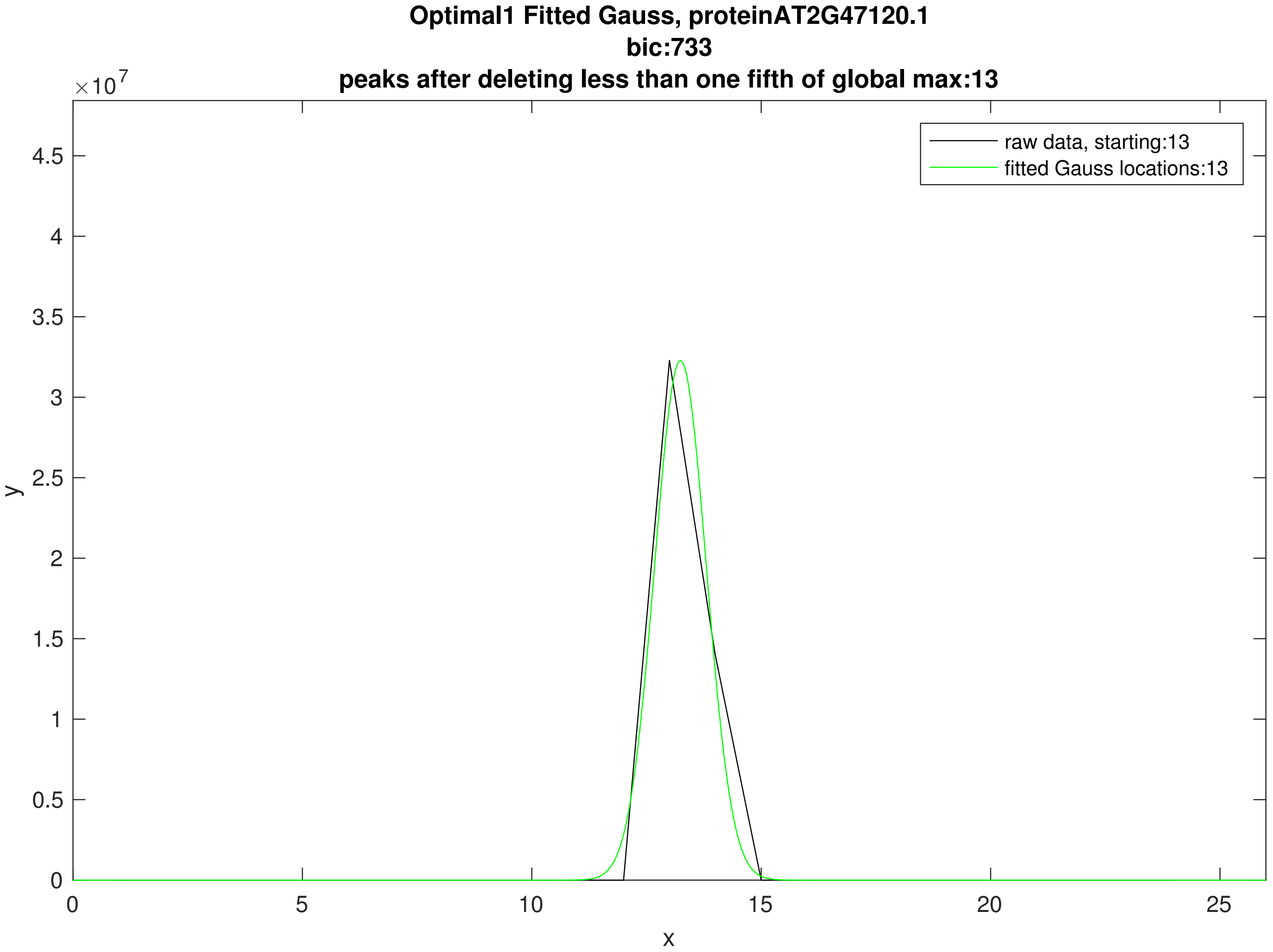

Optimal1 Fitted Gauss, proteinAT2G47120.1
bic:733
peaks after deleting less than one fifth of global max:13
×10^7
4.5
4
3.5
3
2.5
2
1.5
1
0.5
0
y
0
5
10
15
20
25
x
raw data, starting:13
fitted Gauss locations:13

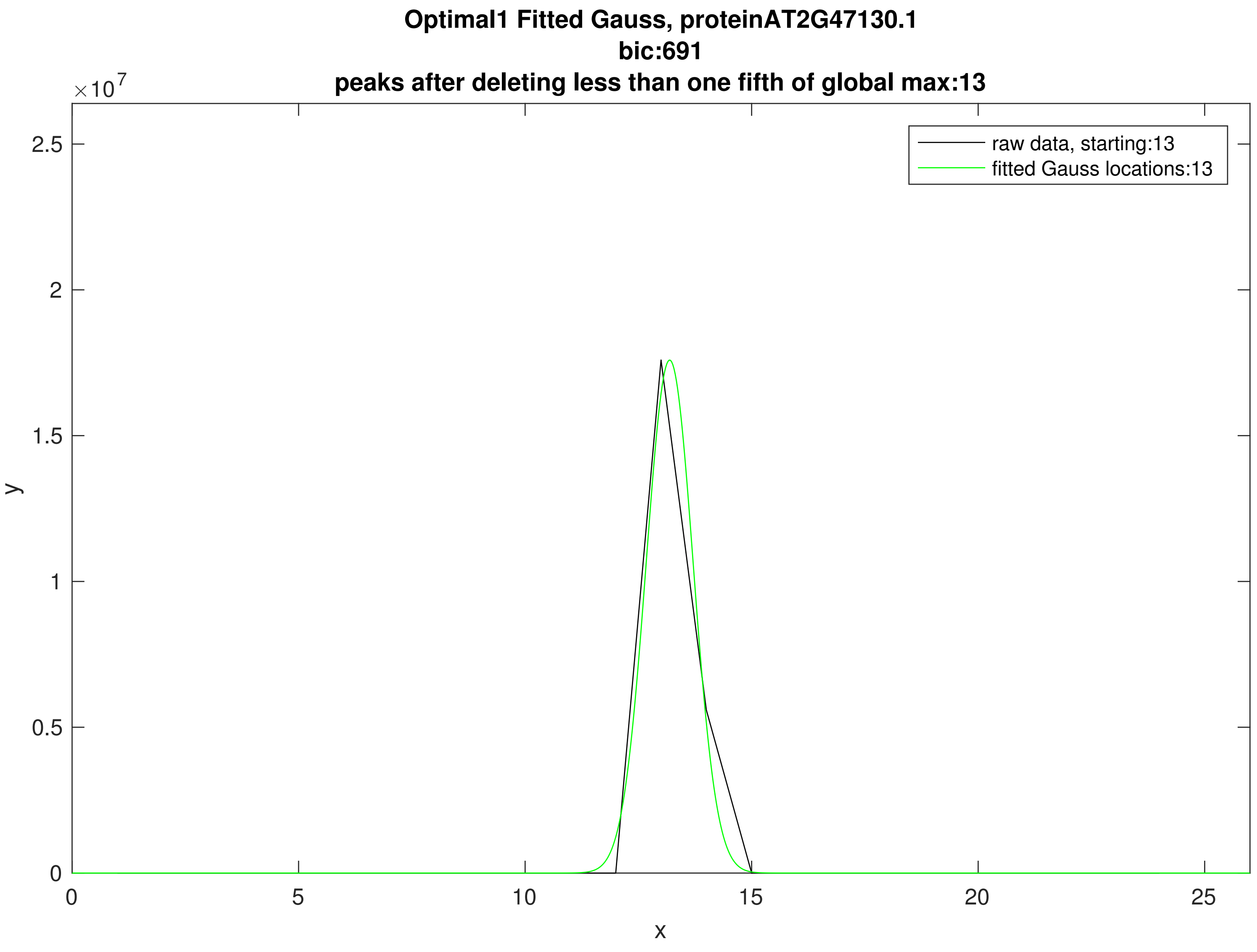
Optimal1 Fitted Gauss, proteinAT2G47130.1
bic:691
peaks after deleting less than one fifth of global max:13
×10^7
2.5
2
1.5
1
0.5
0
y
0
5
10
15
20
25
x
raw data, starting:13
fitted Gauss locations:13

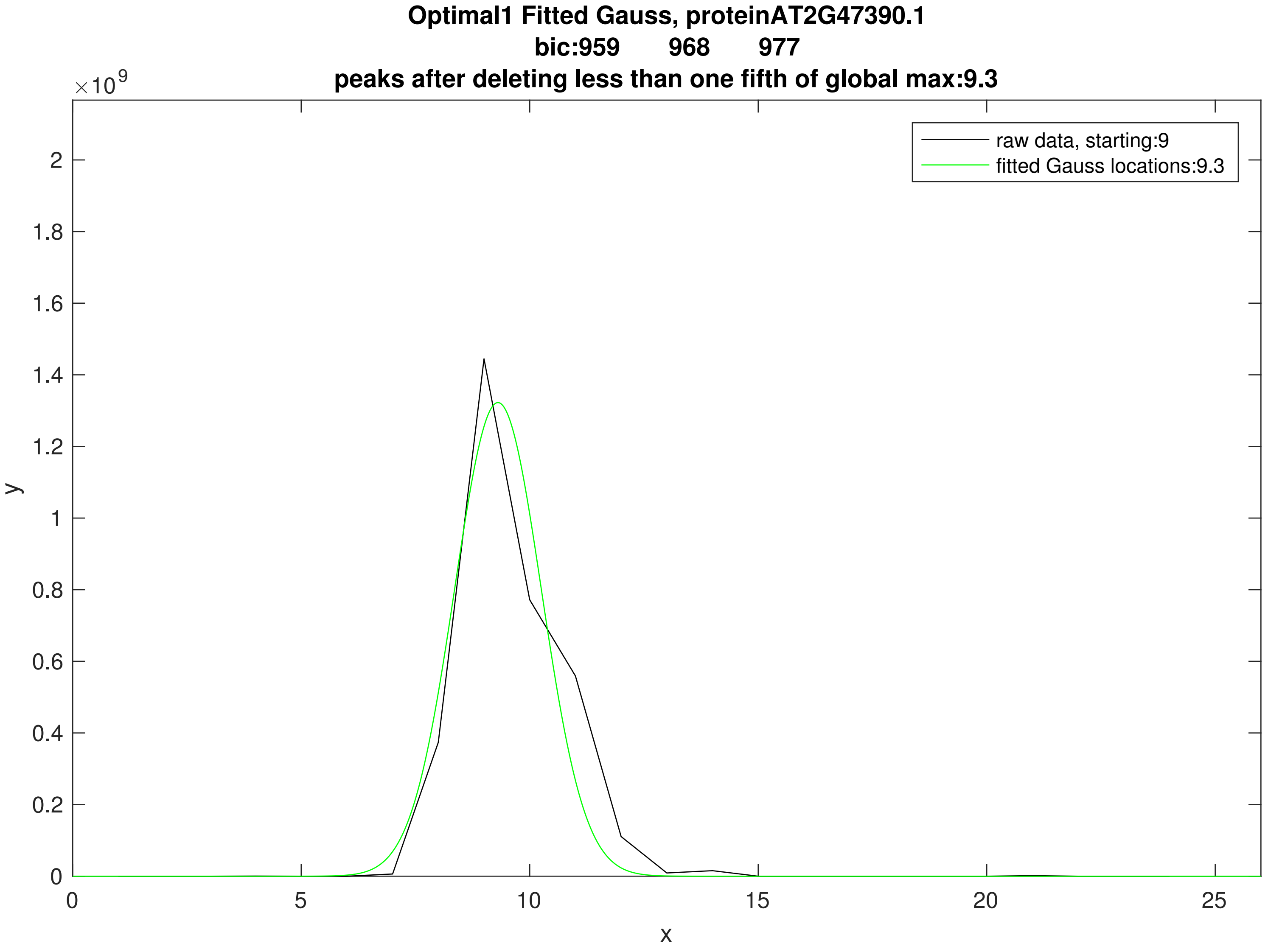
Optimal1 Fitted Gauss, proteinAT2G47390.1
bic:959 968 977
peaks after deleting less than one fifth of global max:9.3
×10^9
raw data, starting:9
fitted Gauss locations:9.3
y
x
0
0.2
0.4
0.6
0.8
1
1.2
1.4
1.6
1.8
2
0
5
10
15
20
25

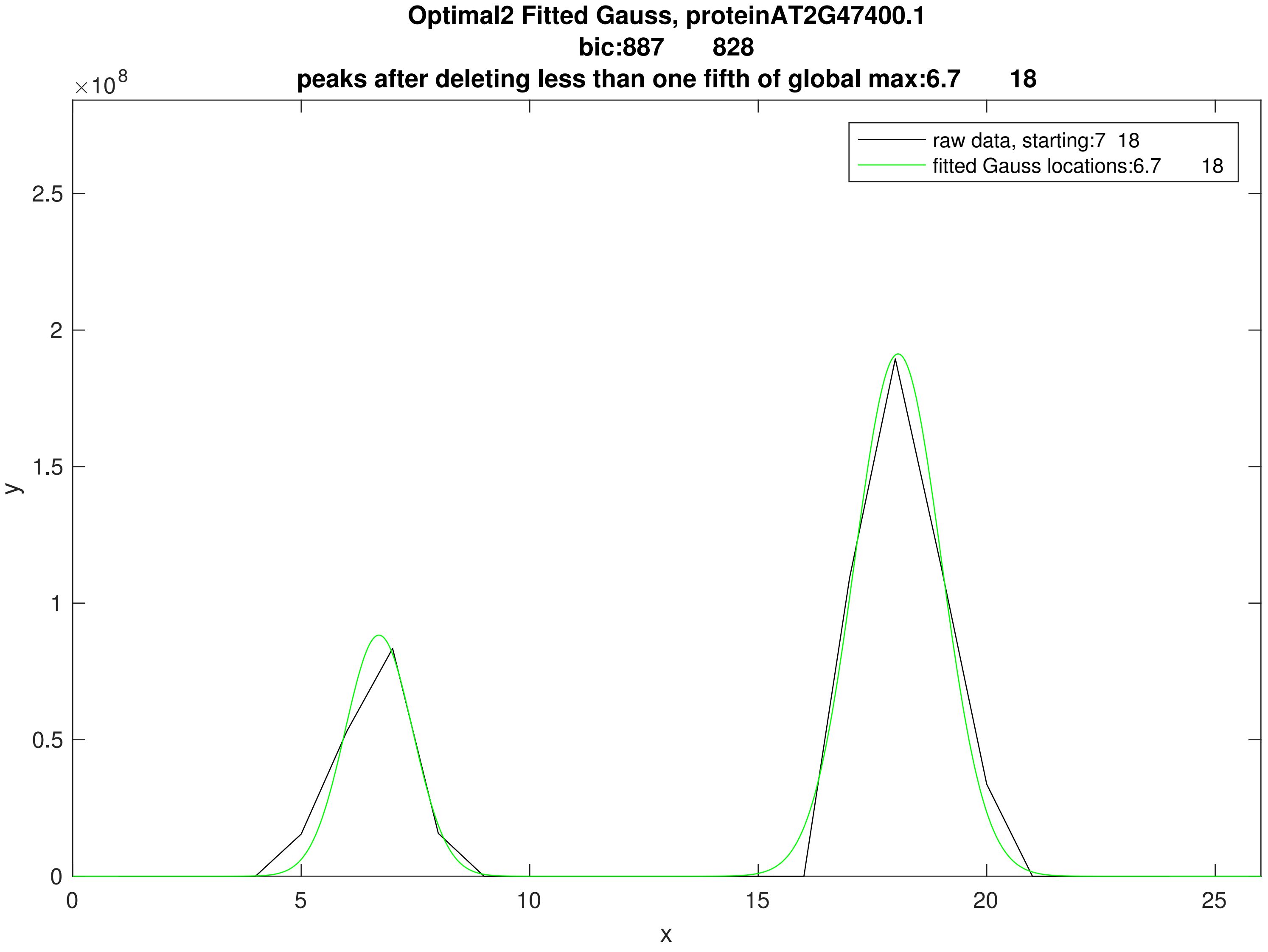
Optimal2 Fitted Gauss, proteinAT2G47400.1
bic:887 828
peaks after deleting less than one fifth of global max:6.7 18
×10^8
raw data, starting:7 18
fitted Gauss locations:6.7 18
y
x
0
0.5
1
1.5
2
2.5
0
5
10
15
20
25

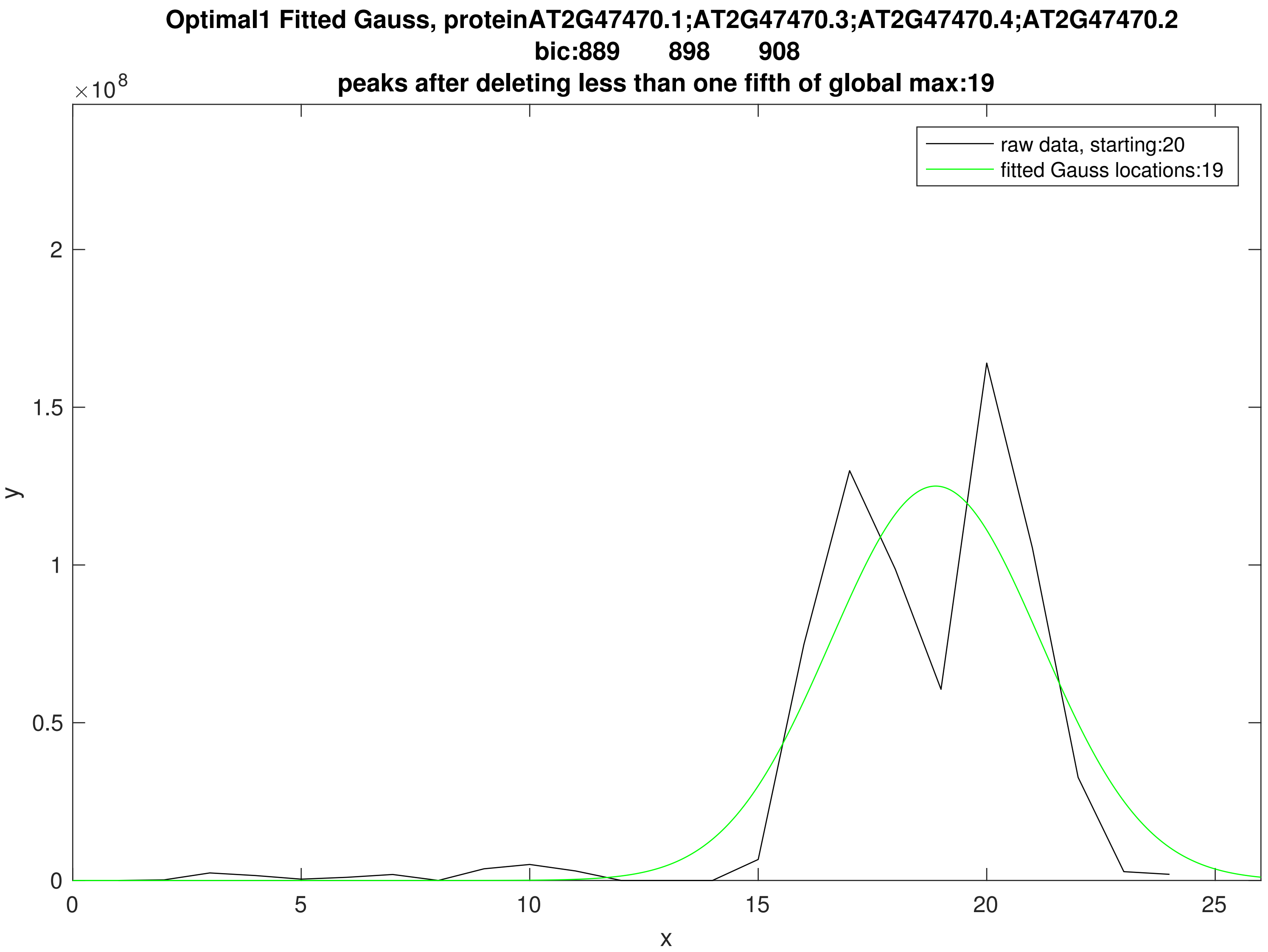
Optimal1 Fitted Gauss, proteinAT2G47470.1;AT2G47470.3;AT2G47470.4;AT2G47470.2
bic:889 898 908
peaks after deleting less than one fifth of global max:19
×10^8
raw data, starting:20
fitted Gauss locations:19
y
x
0
0.5
1
1.5
2
0
5
10
15
20
25

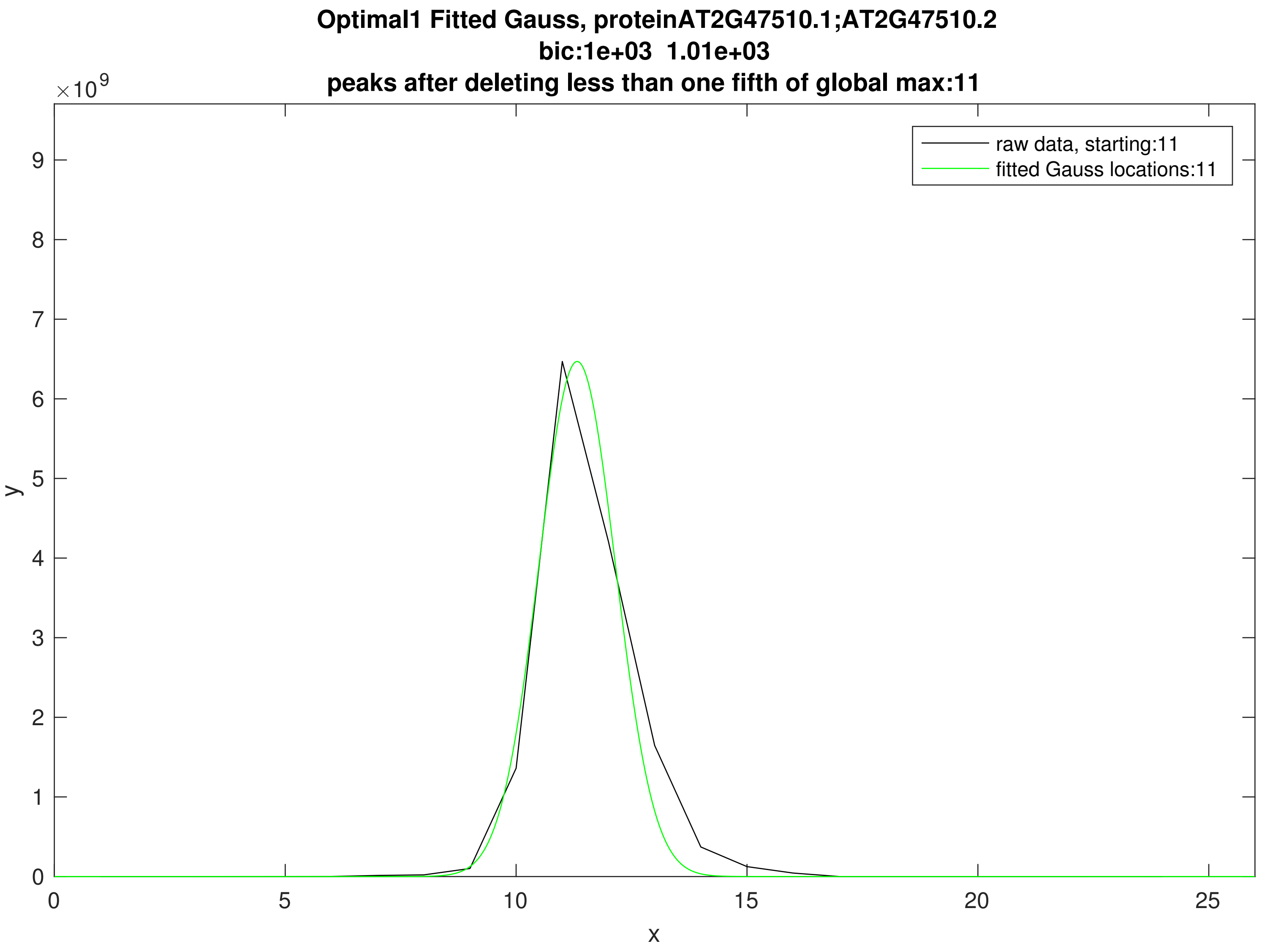
Optimal1 Fitted Gauss, proteinAT2G47510.1;AT2G47510.2
bic:1e+03  1.01e+03
peaks after deleting less than one fifth of global max:11
×10^9
raw data, starting:11
fitted Gauss locations:11
y
x
0
1
2
3
4
5
6
7
8
9
0
5
10
15
20
25

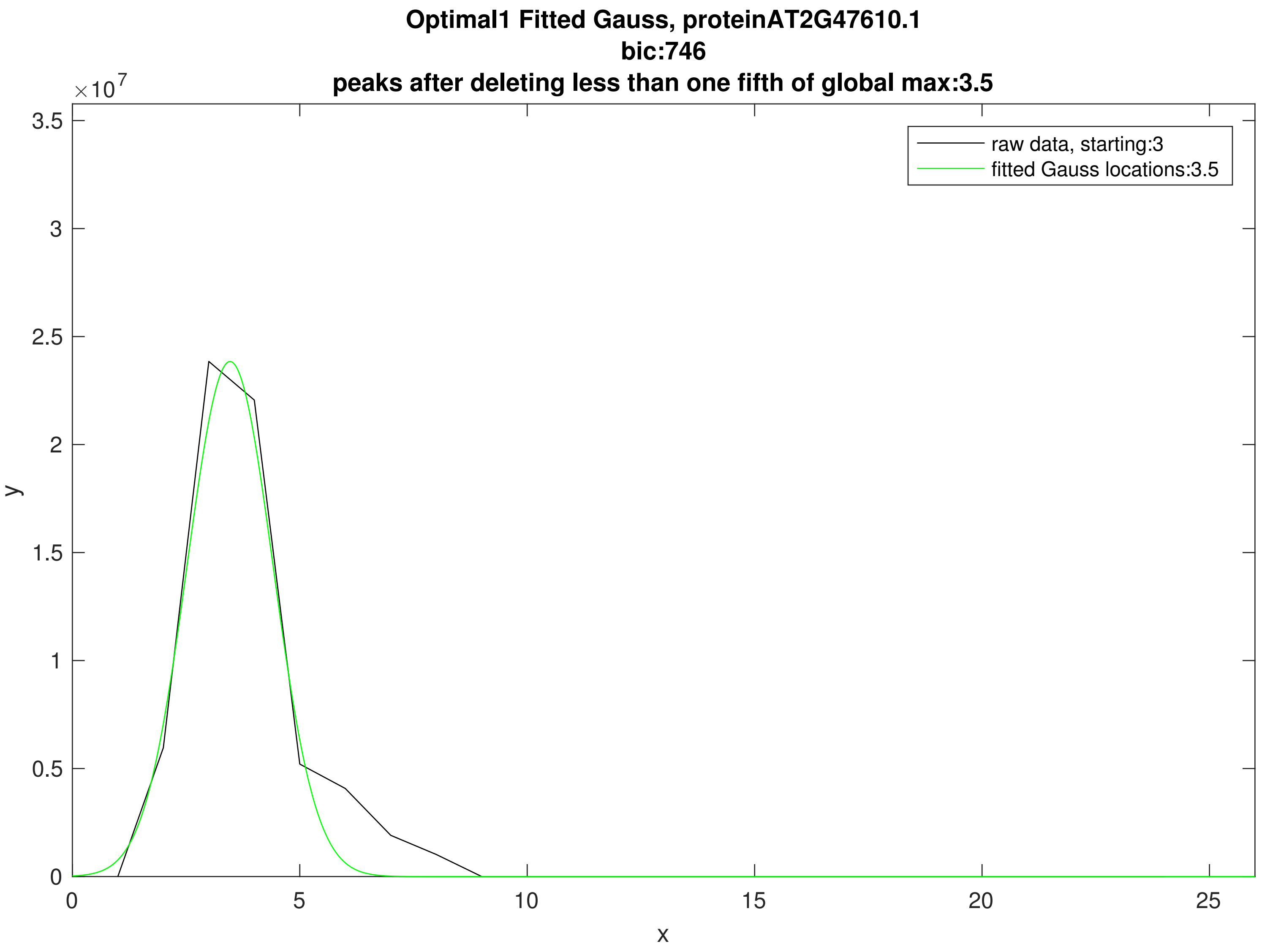

Optimal1 Fitted Gauss, proteinAT2G47610.1
bic:746
peaks after deleting less than one fifth of global max:3.5
×10^7
raw data, starting:3
fitted Gauss locations:3.5
y
x
0
0.5
1
1.5
2
2.5
3
3.5
0
5
10
15
20
25

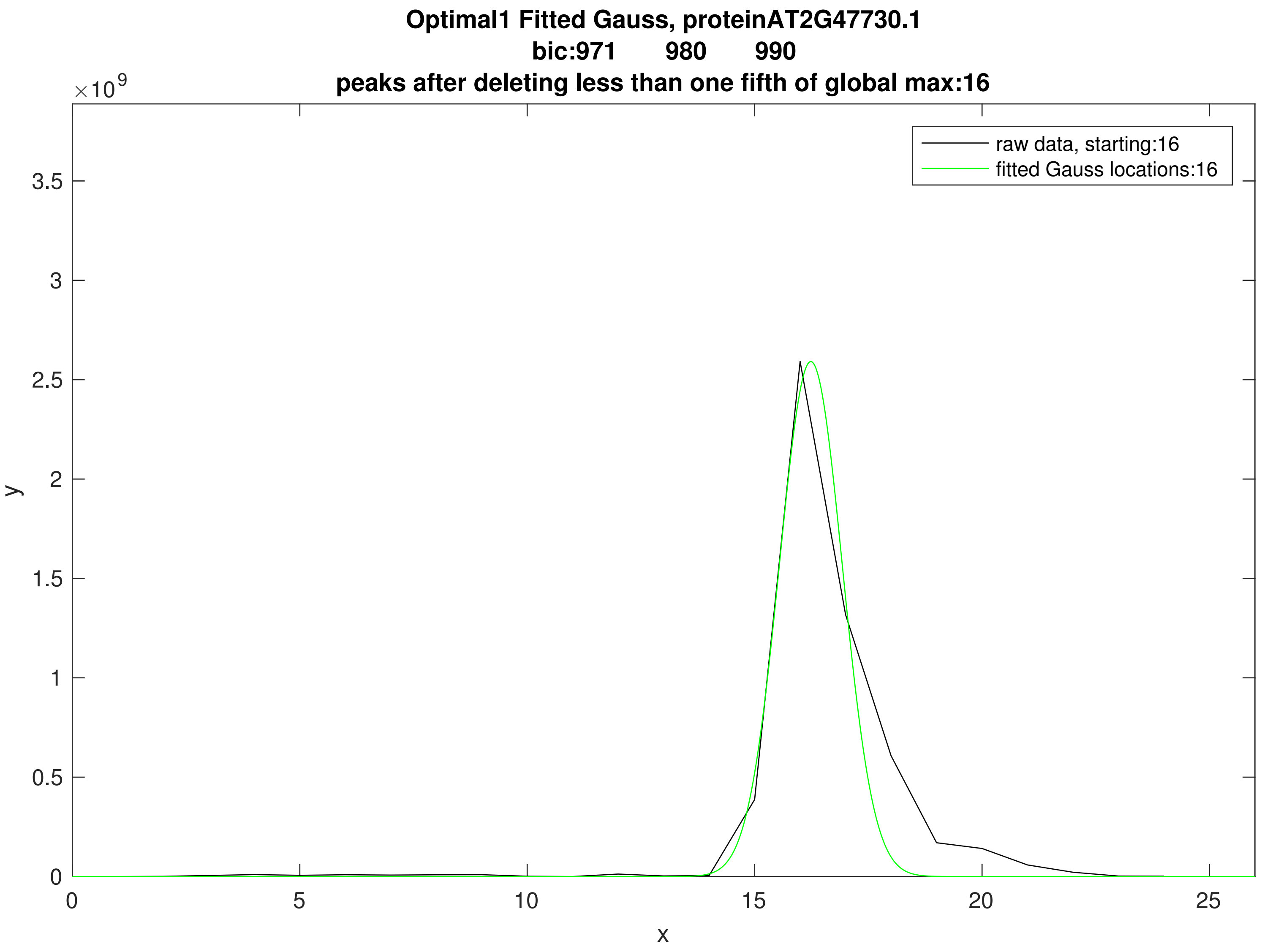
Optimal1 Fitted Gauss, proteinAT2G47730.1
bic:971 980 990
peaks after deleting less than one fifth of global max:16
×10^9
3.5
3
2.5
2
1.5
1
0.5
0
y
0
5
10
15
20
25
x
raw data, starting:16
fitted Gauss locations:16

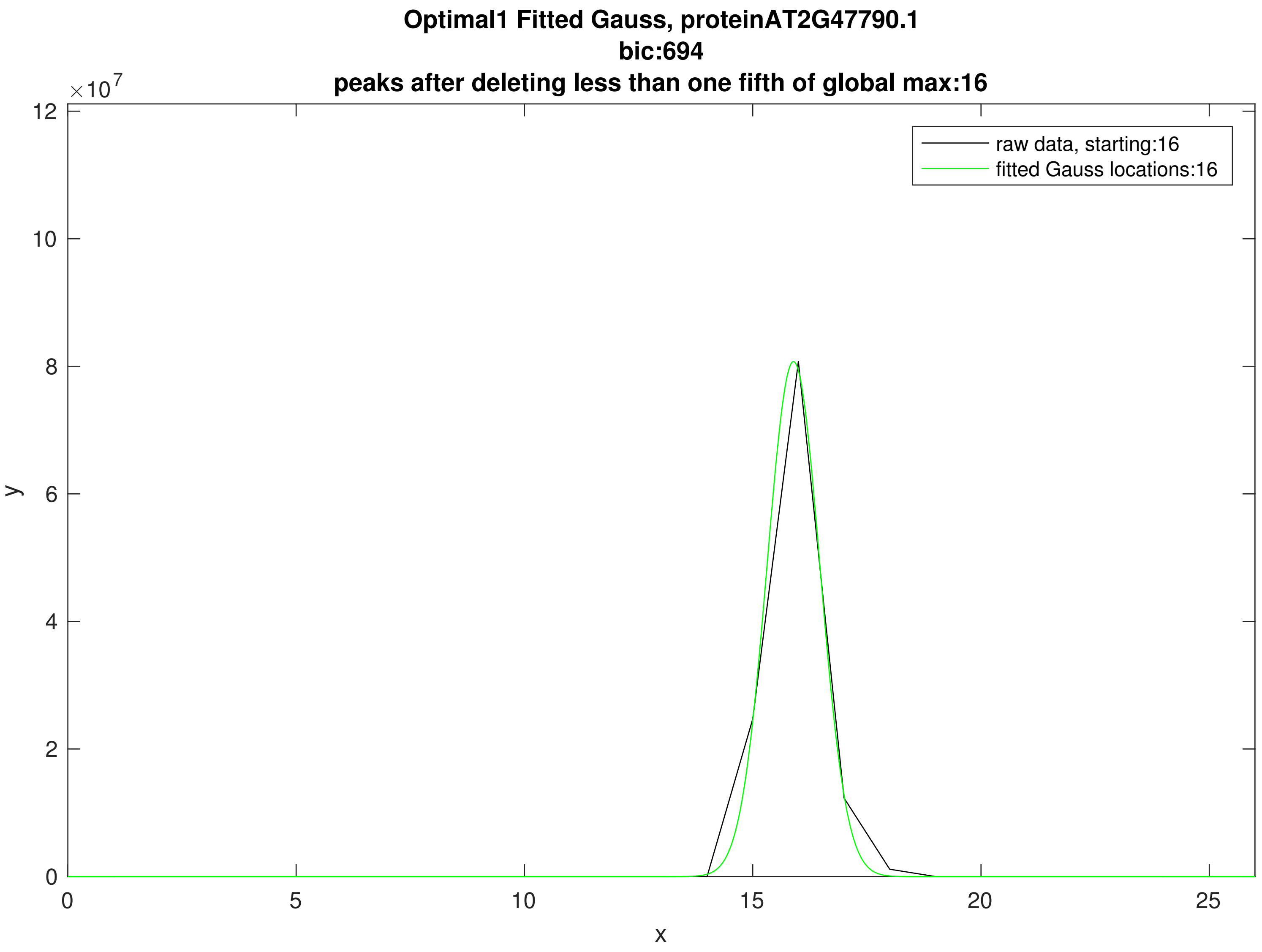

Optimal1 Fitted Gauss, proteinAT2G47790.1
bic:694
peaks after deleting less than one fifth of global max:16
×10^7
raw data, starting:16
fitted Gauss locations:16
y
x

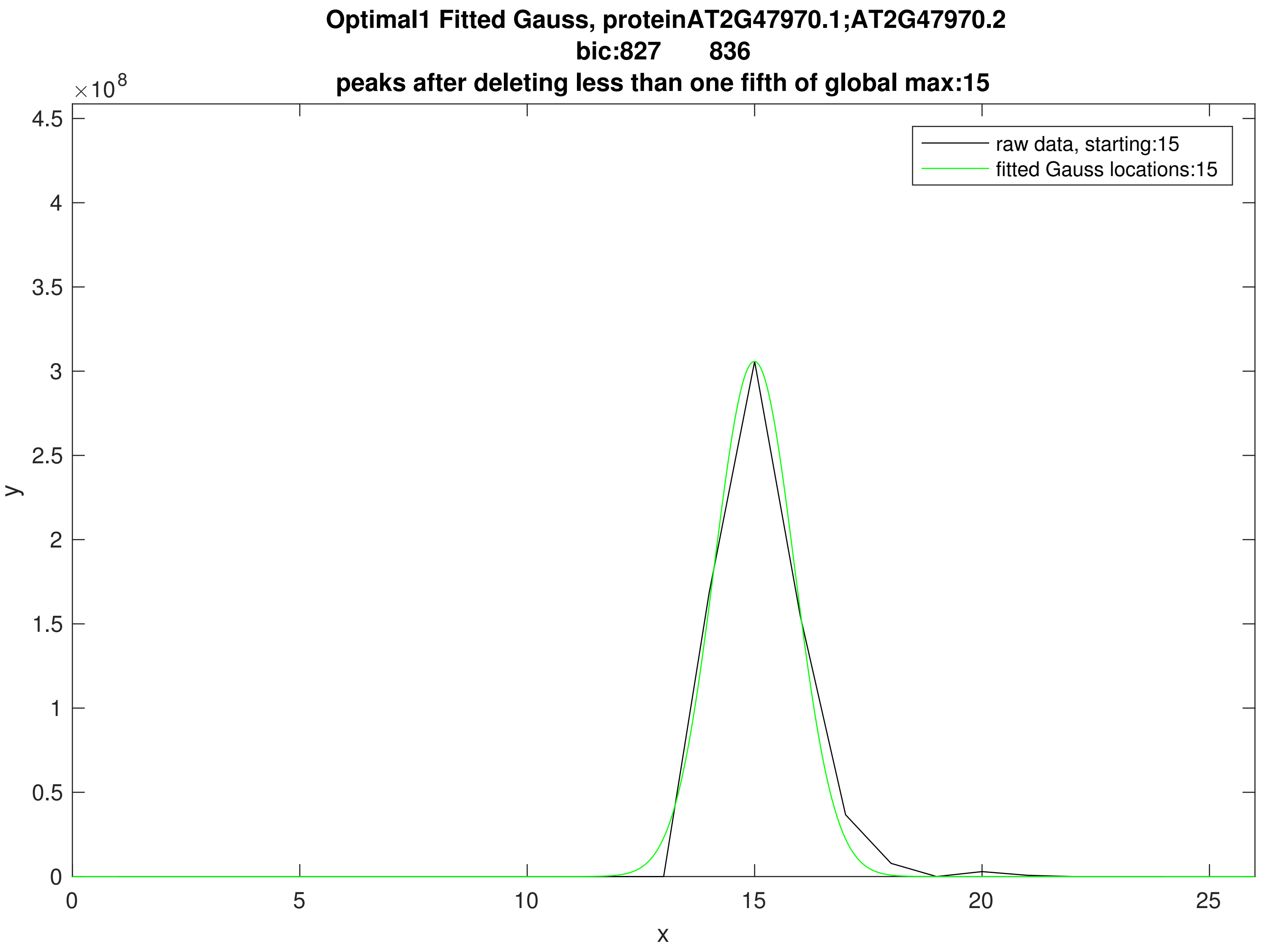
Optimal1 Fitted Gauss, proteinAT2G47970.1;AT2G47970.2
bic:827 836
peaks after deleting less than one fifth of global max:15
×10^8
4.5
4
3.5
3
2.5
2
1.5
1
0.5
0
y
0
5
10
15
20
25
x
raw data, starting:15
fitted Gauss locations:15

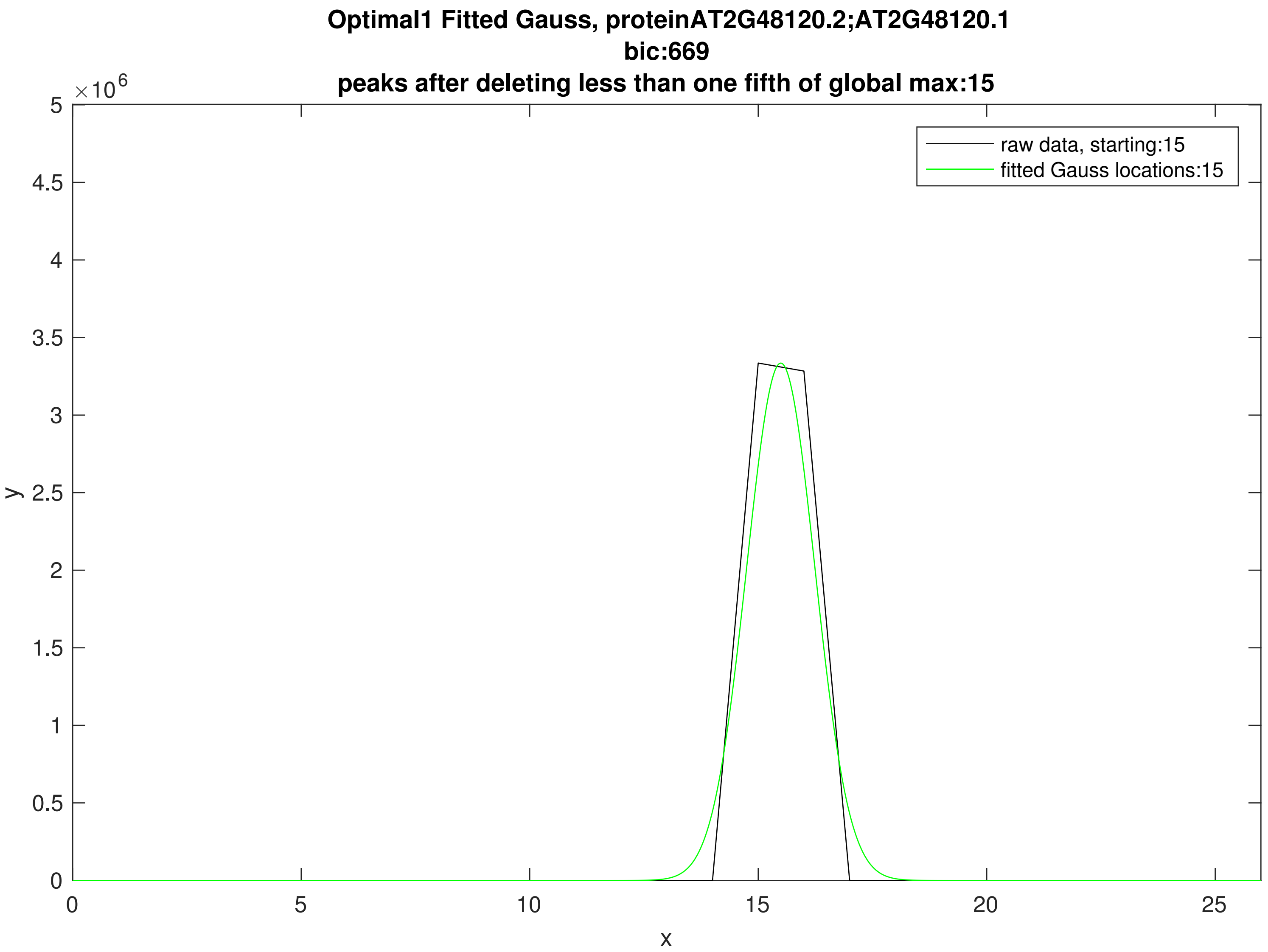
Optimal1 Fitted Gauss, proteinAT2G48120.2;AT2G48120.1
bic:669
peaks after deleting less than one fifth of global max:15
×10^6
raw data, starting:15
fitted Gauss locations:15
y
x
0
0.5
1
1.5
2
2.5
3
3.5
4
4.5
5
0
5
10
15
20
25

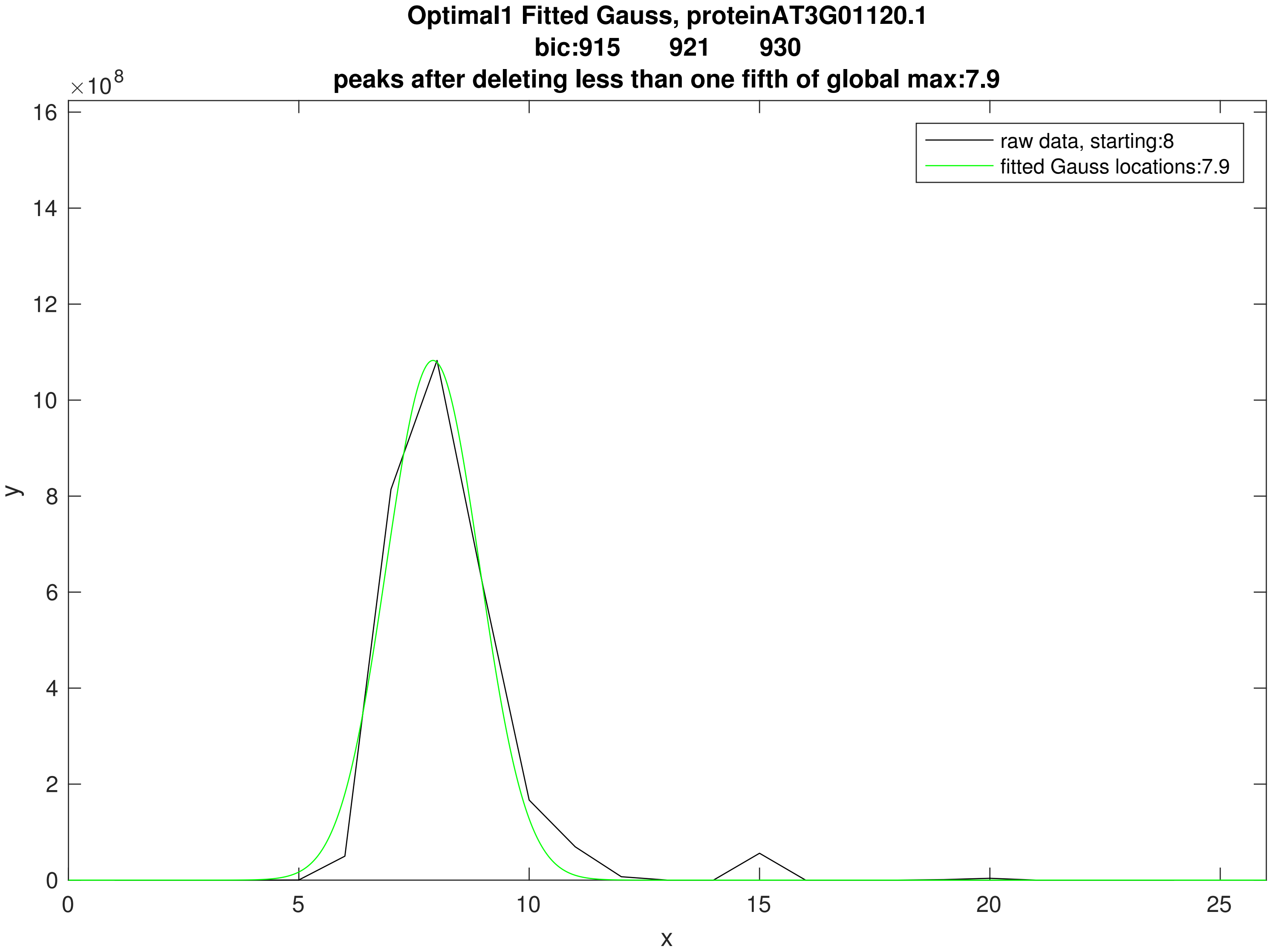
Optimal1 Fitted Gauss, proteinAT3G01120.1
bic:915 921 930
peaks after deleting less than one fifth of global max:7.9
×10^8
16
14
12
10
8
6
4
2
0
y
0
5
10
15
20
25
x
raw data, starting:8
fitted Gauss locations:7.9

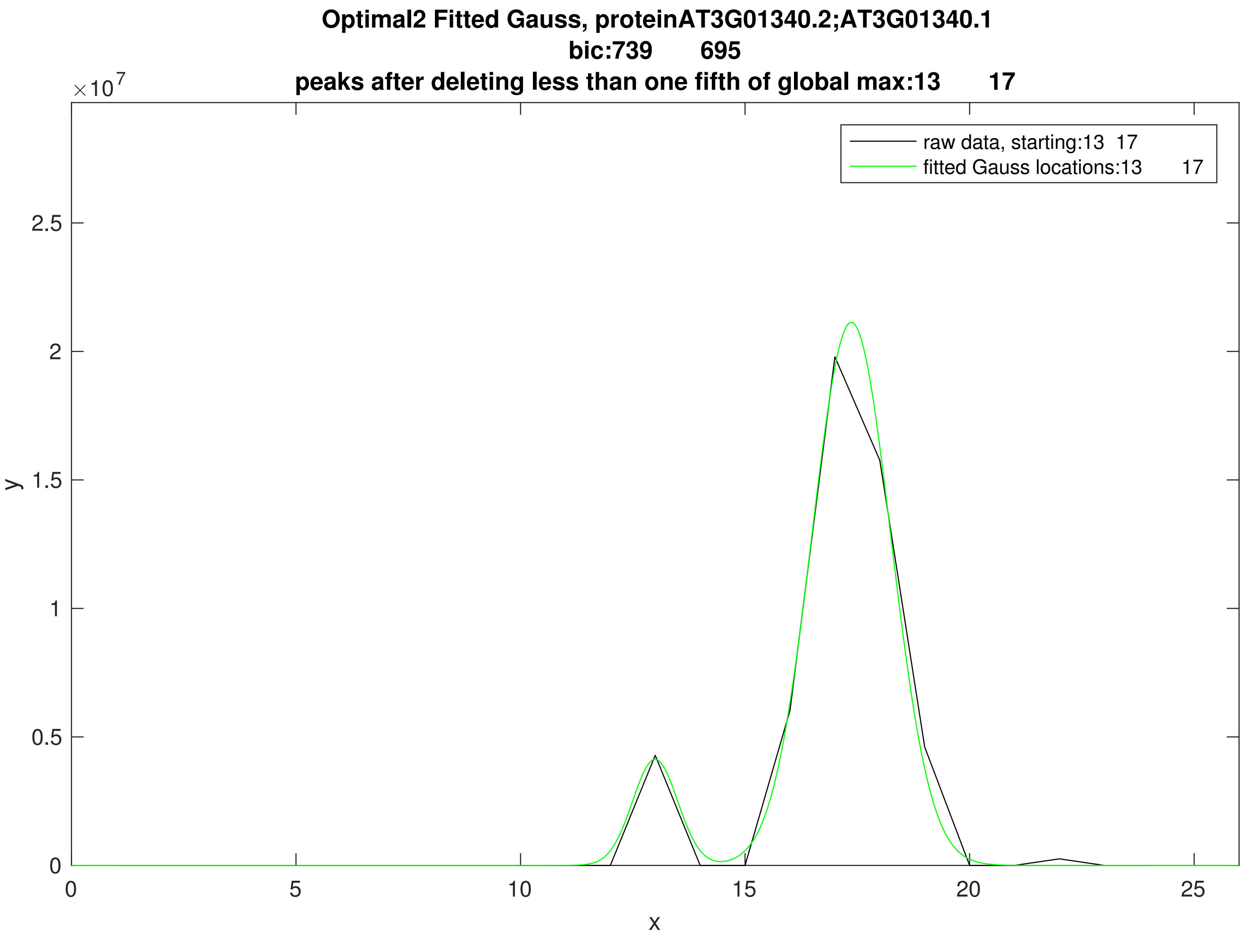
Optimal2 Fitted Gauss, proteinAT3G01340.2;AT3G01340.1
bic:739 695
peaks after deleting less than one fifth of global max:13 17
×10^7
raw data, starting:13 17
fitted Gauss locations:13 17
2.5
2
y 1.5
1
0.5
0
0
5
10
15
20
25
x

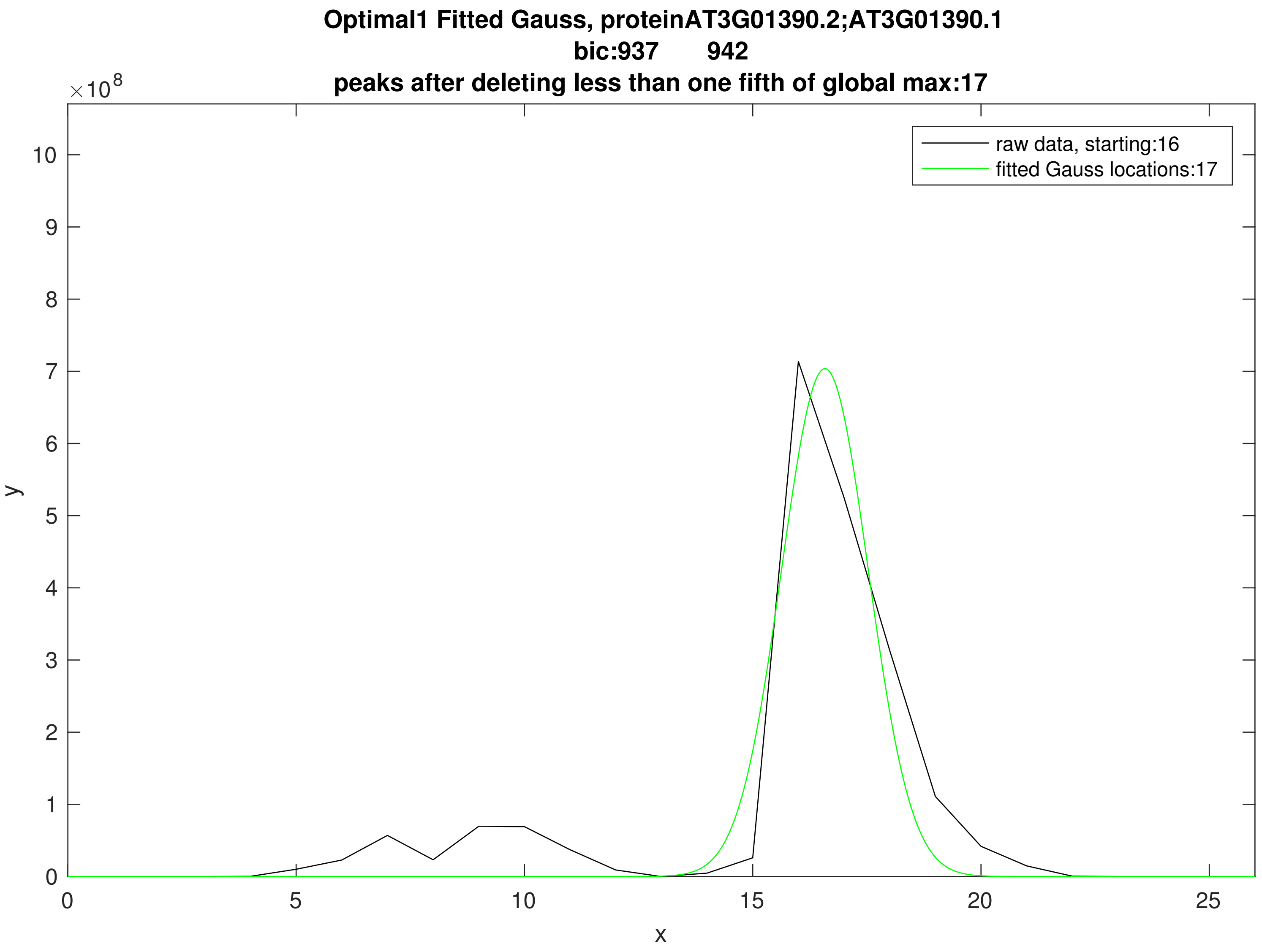
Optimal1 Fitted Gauss, proteinAT3G01390.2;AT3G01390.1
bic:937 942
peaks after deleting less than one fifth of global max:17
×10^8
raw data, starting:16
fitted Gauss locations:17
y
x
0
1
2
3
4
5
6
7
8
9
10
0
5
10
15
20
25

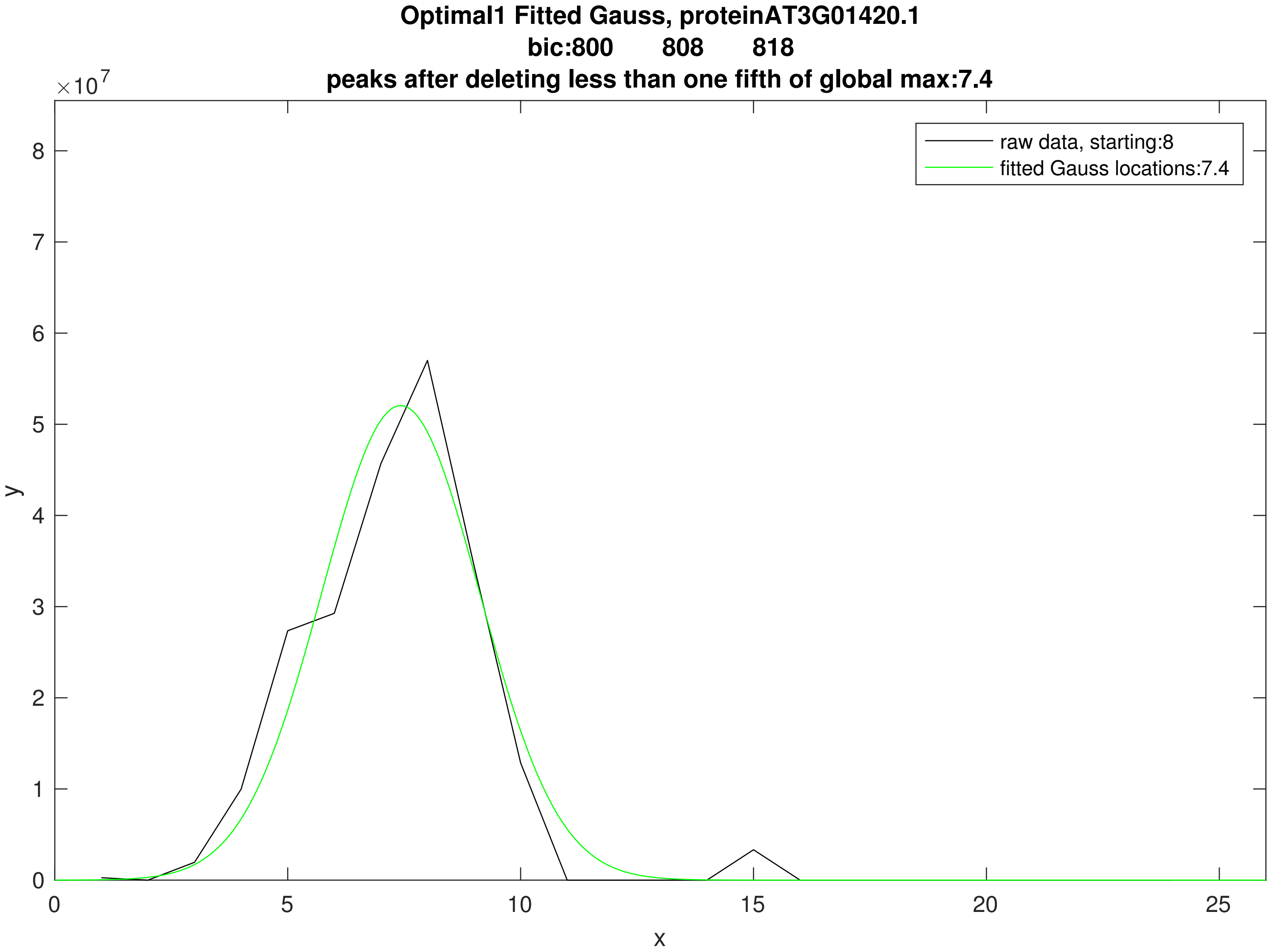
Optimal1 Fitted Gauss, proteinAT3G01420.1
bic:800 808 818
peaks after deleting less than one fifth of global max:7.4
×10^7
raw data, starting:8
fitted Gauss locations:7.4
y
x
0
1
2
3
4
5
6
7
8
0
5
10
15
20
25

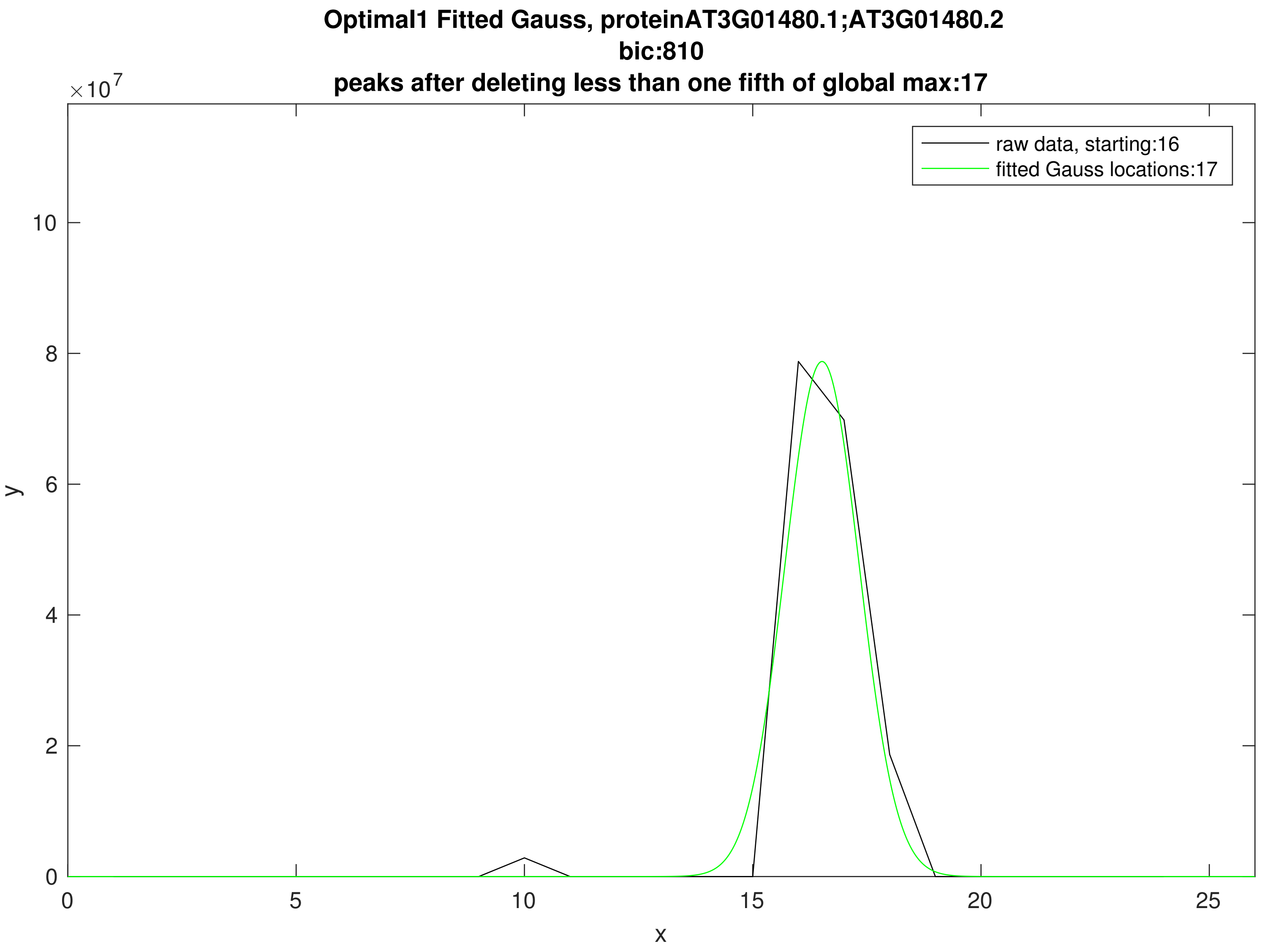
Optimal1 Fitted Gauss, proteinAT3G01480.1;AT3G01480.2
bic:810
peaks after deleting less than one fifth of global max:17
×10^7
raw data, starting:16
fitted Gauss locations:17
y
x
0
2
4
6
8
10
0
5
10
15
20
25

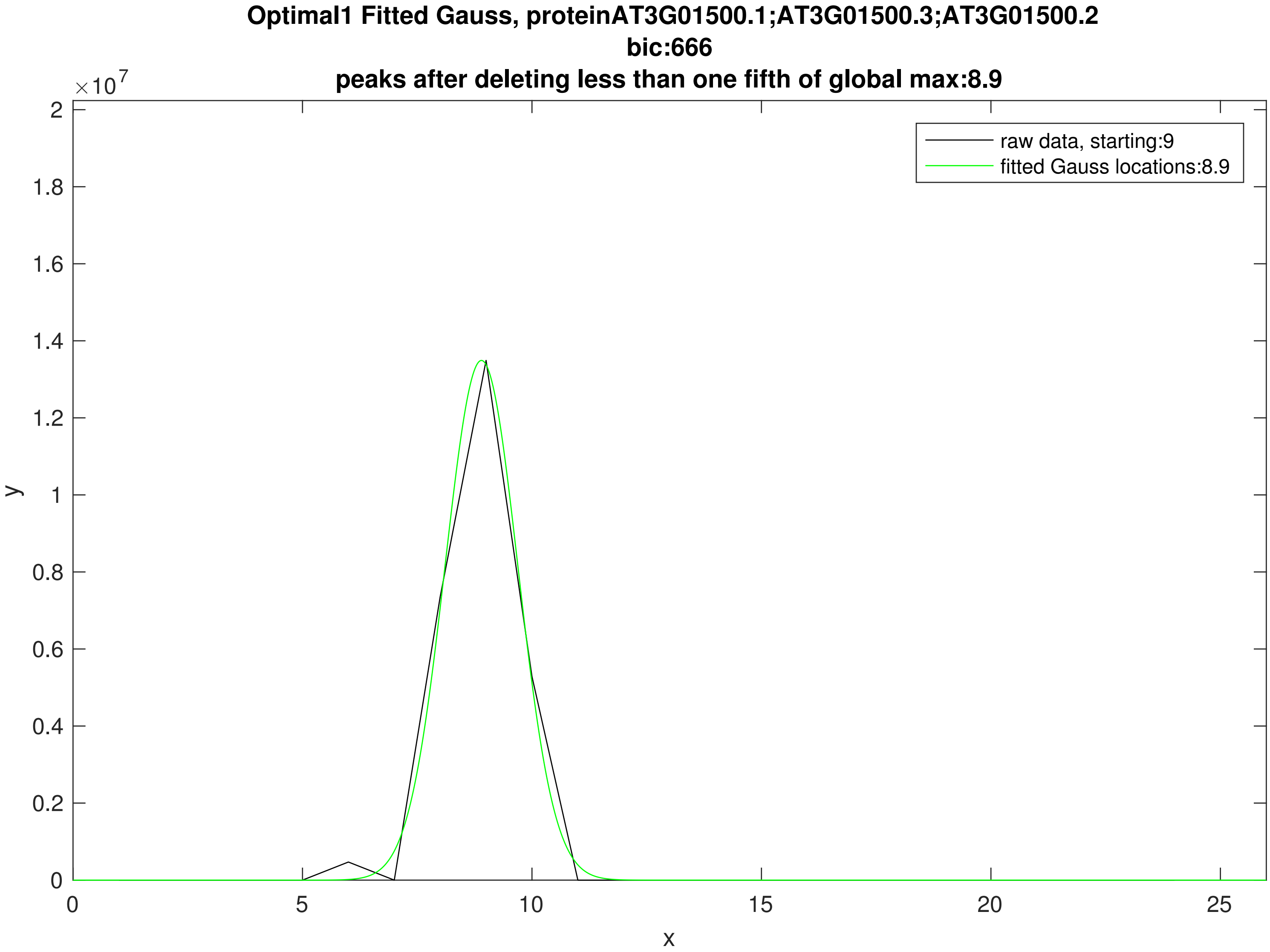
Optimal1 Fitted Gauss, proteinAT3G01500.1;AT3G01500.3;AT3G01500.2
bic:666
peaks after deleting less than one fifth of global max:8.9
$\times 10^7$
2
1.8
1.6
1.4
1.2
1
0.8
0.6
0.4
0.2
0
y
0
5
10
15
20
25
x
raw data, starting:9
fitted Gauss locations:8.9

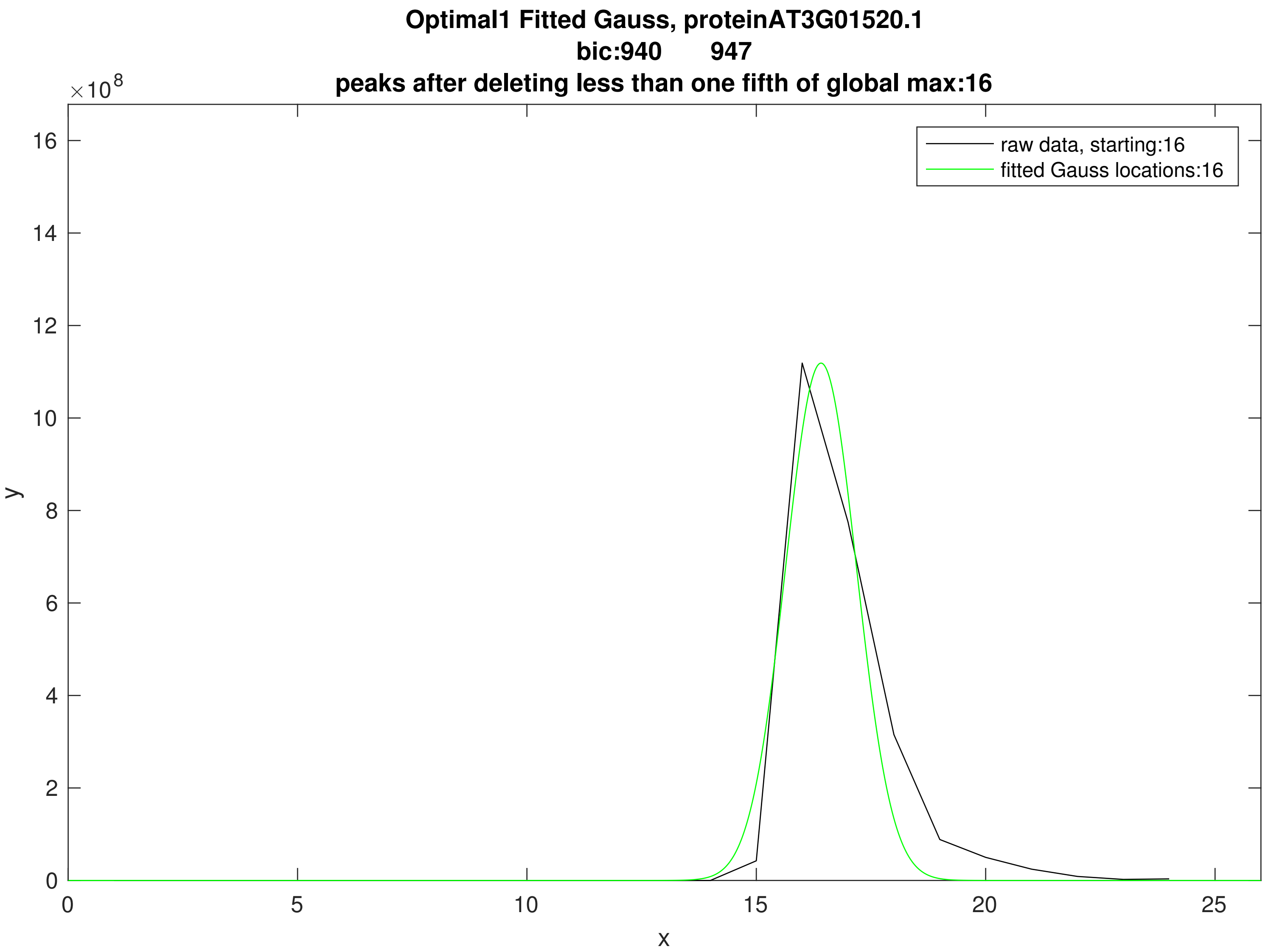
Optimal1 Fitted Gauss, proteinAT3G01520.1
bic:940 947
peaks after deleting less than one fifth of global max:16
raw data, starting:16
fitted Gauss locations:16
×10^8
y
x
0
2
4
6
8
10
12
14
16
0
5
10
15
20
25

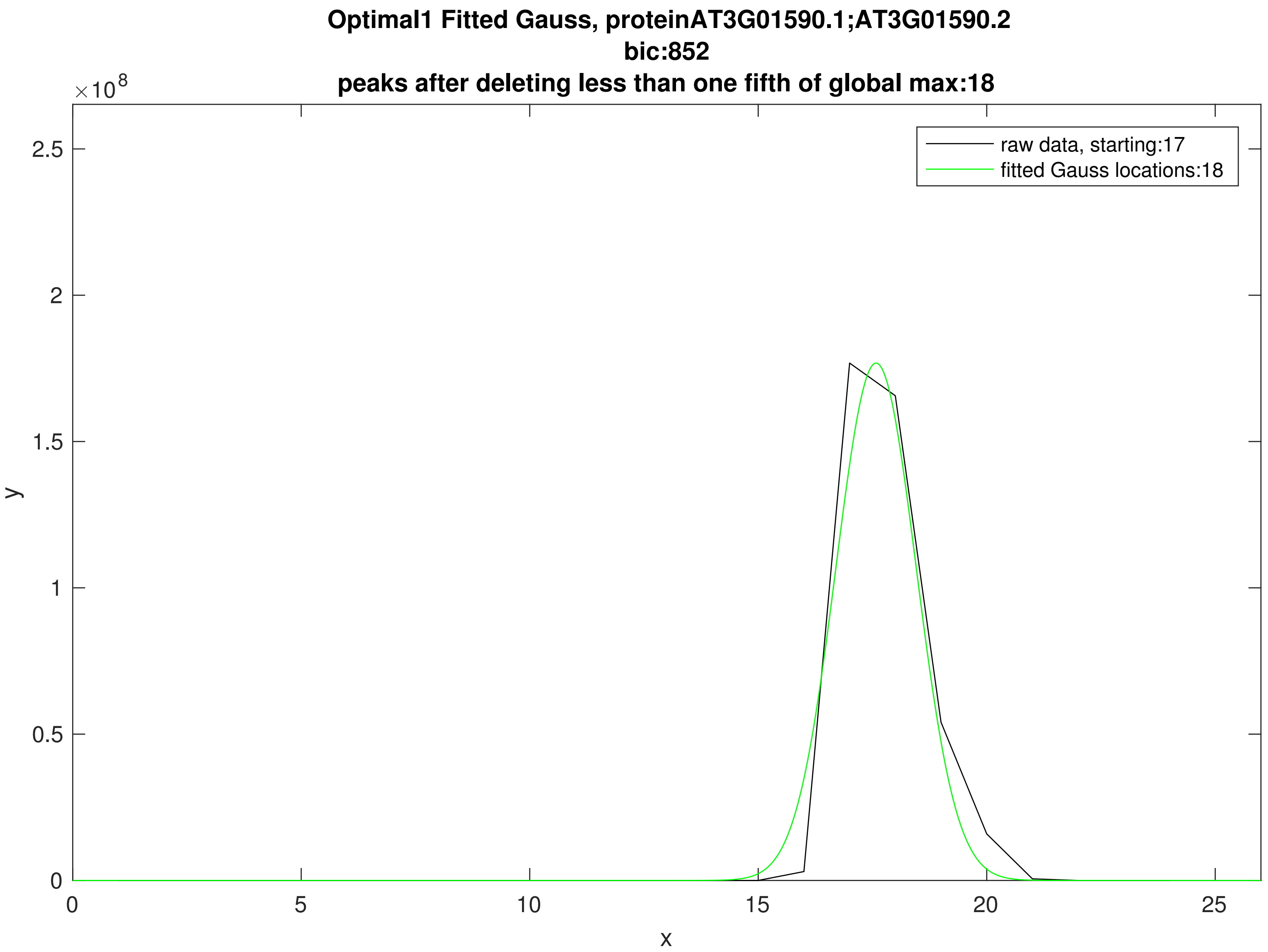

Optimal1 Fitted Gauss, proteinAT3G01590.1;AT3G01590.2
bic:852
peaks after deleting less than one fifth of global max:18
$\times 10^8$
2.5
2
1.5
1
0.5
0
y
0
5
10
15
20
25
x
raw data, starting:17
fitted Gauss locations:18

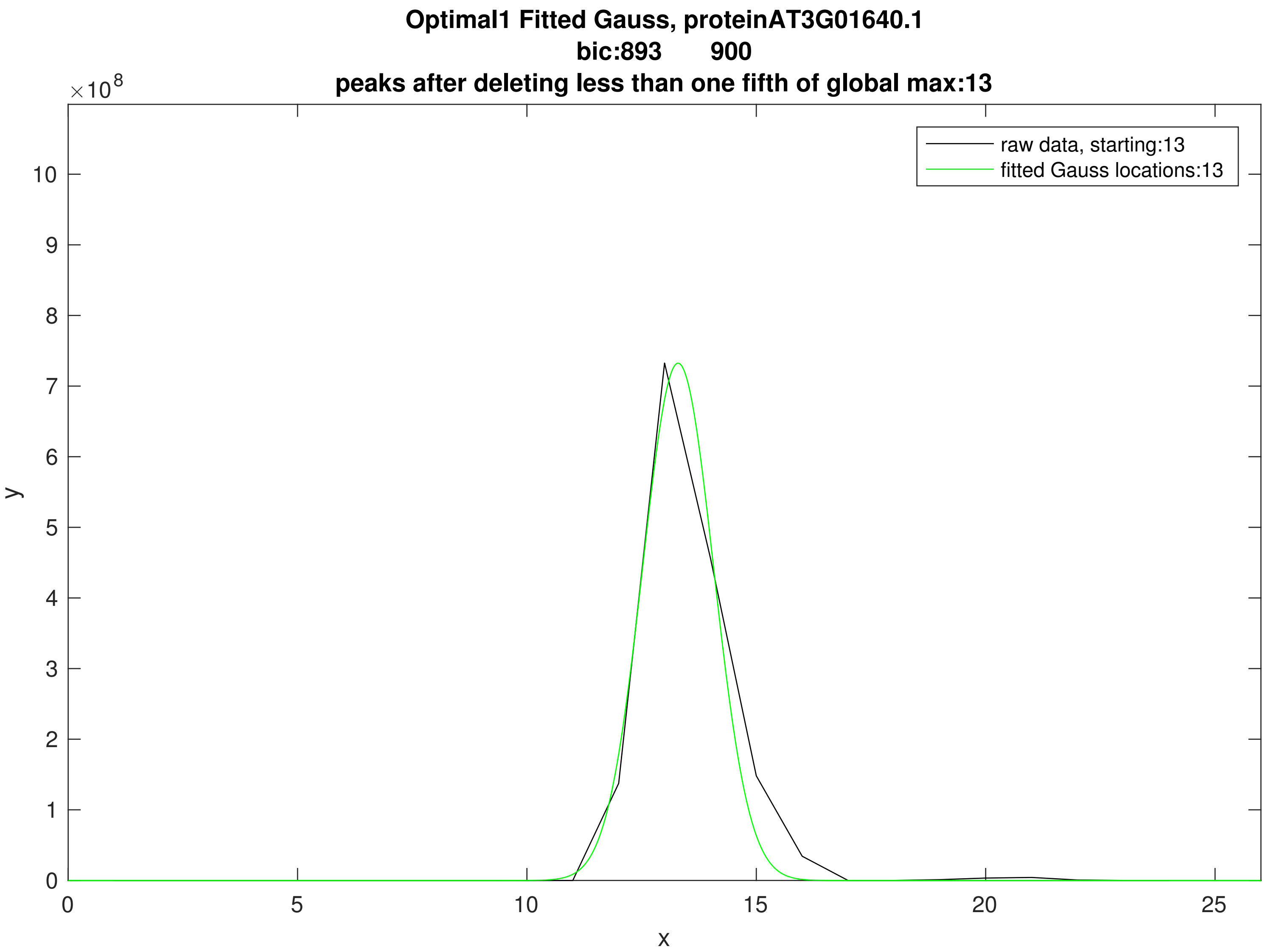
Optimal1 Fitted Gauss, proteinAT3G01640.1
bic:893 900
peaks after deleting less than one fifth of global max:13
$\times 10^8$
raw data, starting:13
fitted Gauss locations:13
y
x
0
1
2
3
4
5
6
7
8
9
10
0
5
10
15
20
25

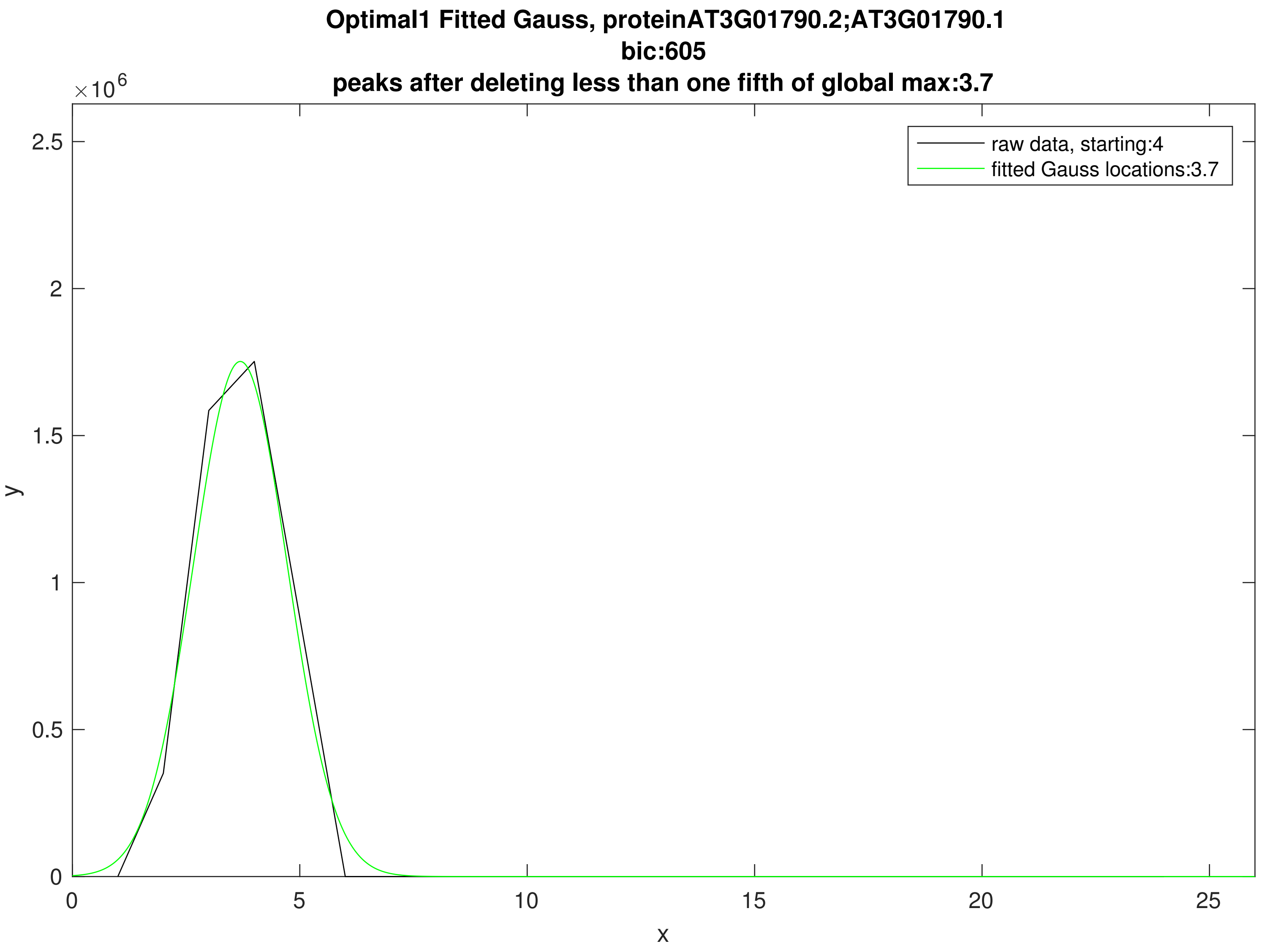
Optimal1 Fitted Gauss, proteinAT3G01790.2;AT3G01790.1
bic:605
peaks after deleting less than one fifth of global max:3.7
×10^6
2.5
2
1.5
1
0.5
0
y
0
5
10
15
20
25
x
raw data, starting:4
fitted Gauss locations:3.7

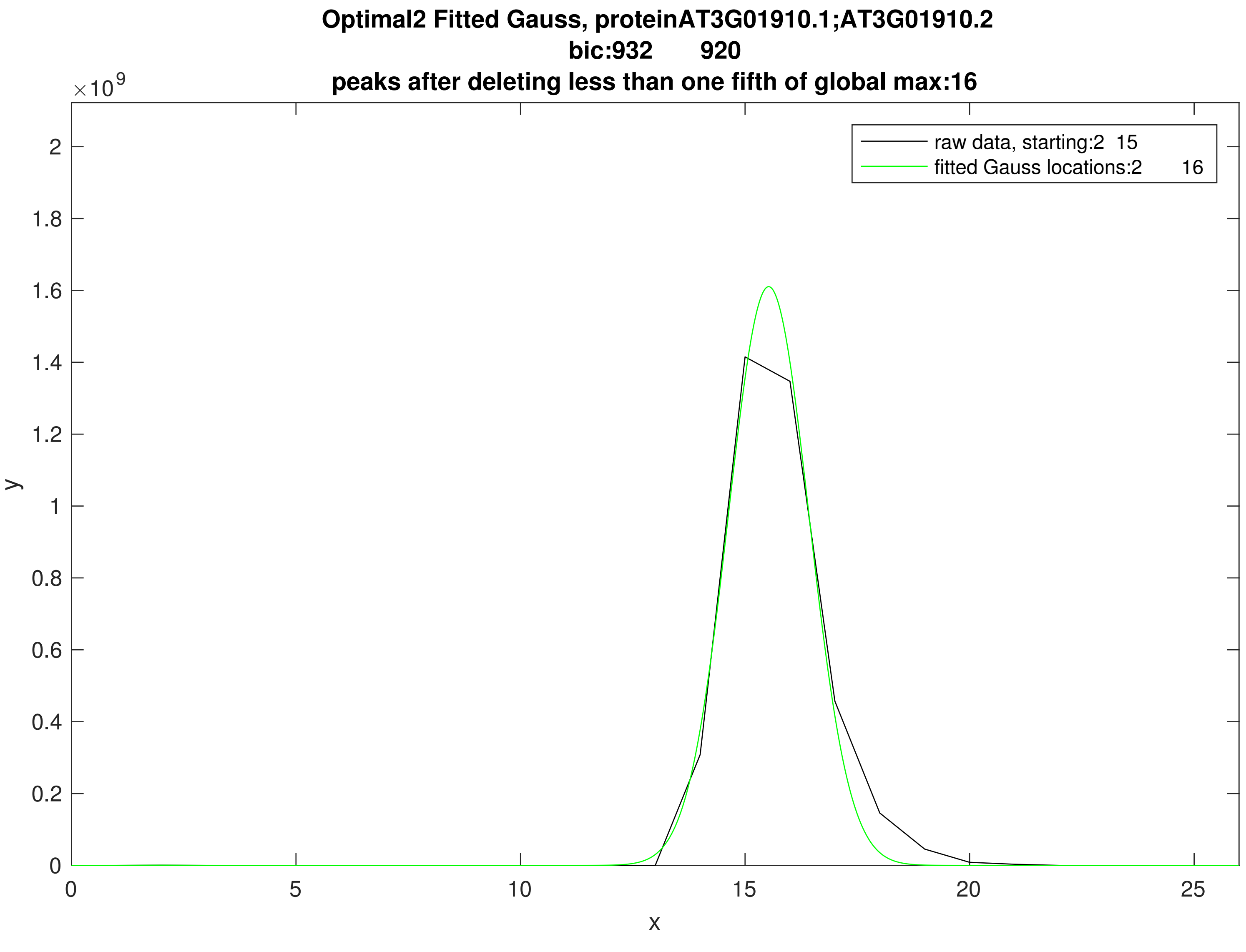
Optimal2 Fitted Gauss, proteinAT3G01910.1;AT3G01910.2
bic:932 920
peaks after deleting less than one fifth of global max:16
×10^9
2
1.8
1.6
1.4
1.2
1
0.8
0.6
0.4
0.2
0
y
raw data, starting:2 15
fitted Gauss locations:2 16
0
5
10
15
20
25
x

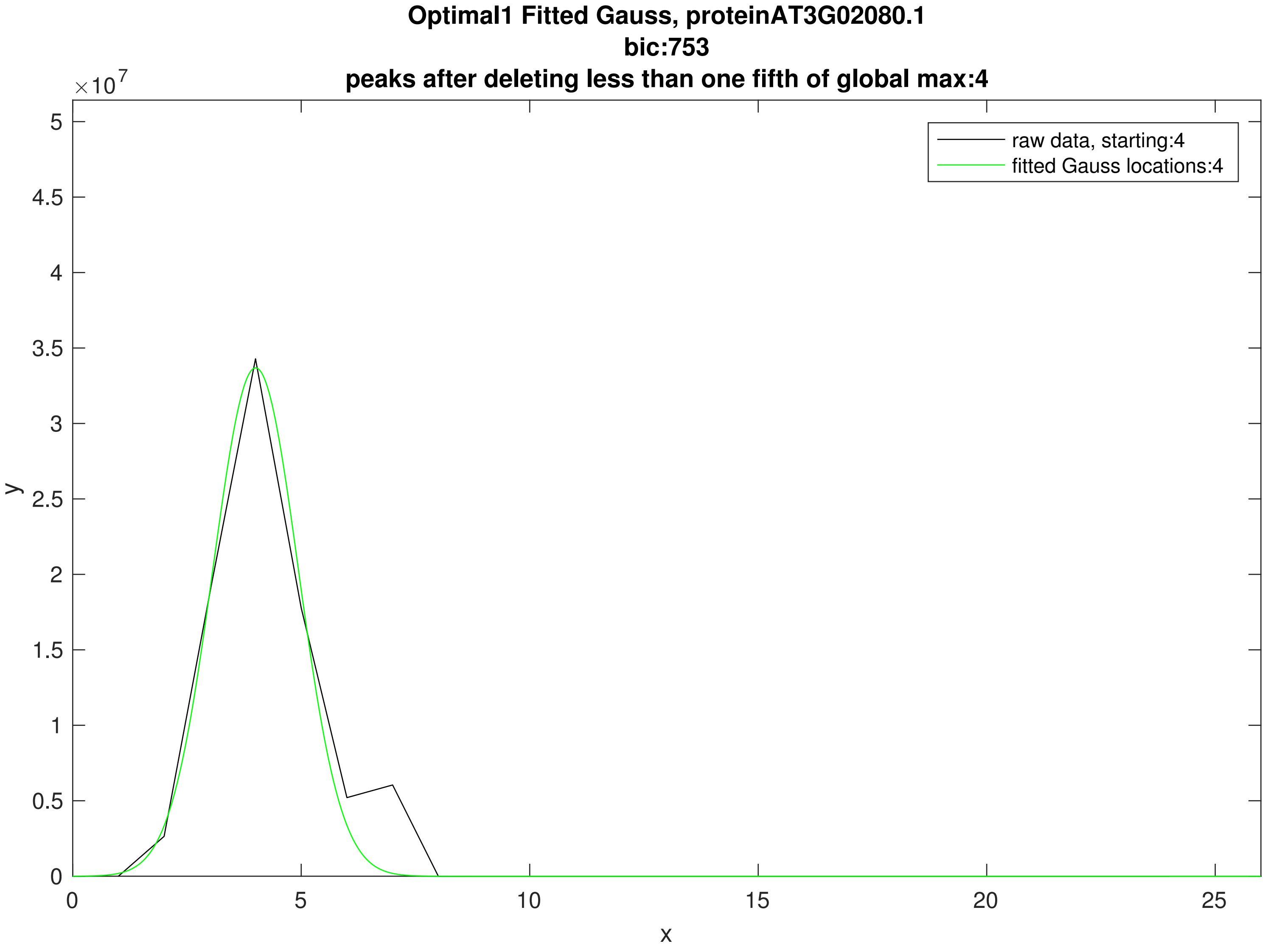
Optimal1 Fitted Gauss, proteinAT3G02080.1
bic:753
peaks after deleting less than one fifth of global max:4
×10^7
5
4.5
4
3.5
3
2.5
2
1.5
1
0.5
0
y
0
5
10
15
20
25
x
raw data, starting:4
fitted Gauss locations:4

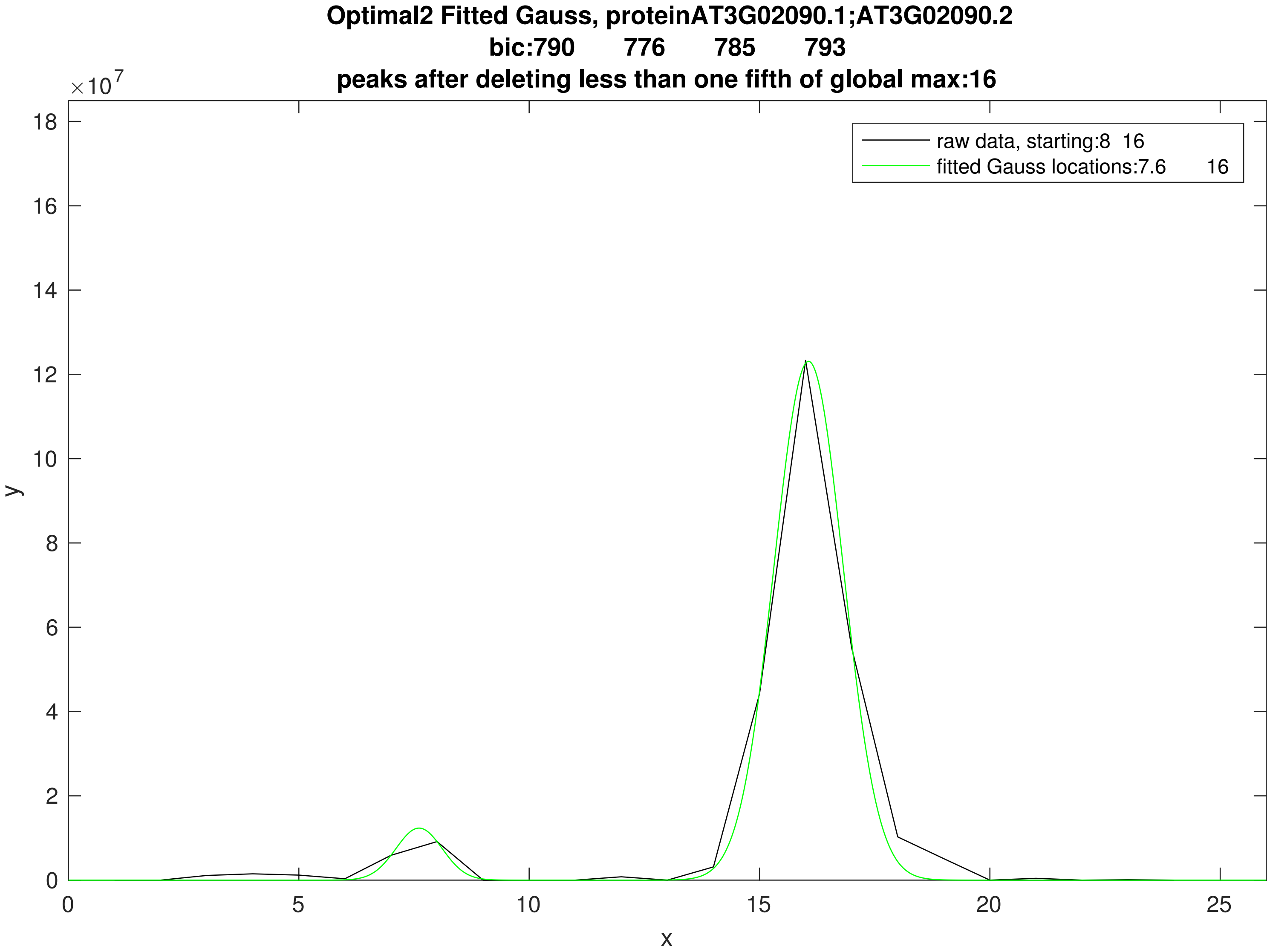
Optimal2 Fitted Gauss, proteinAT3G02090.1;AT3G02090.2
bic:790 776 785 793
peaks after deleting less than one fifth of global max:16
×10^7
raw data, starting:8 16
fitted Gauss locations:7.6 16
y
x
0
2
4
6
8
10
12
14
16
18
0
5
10
15
20
25

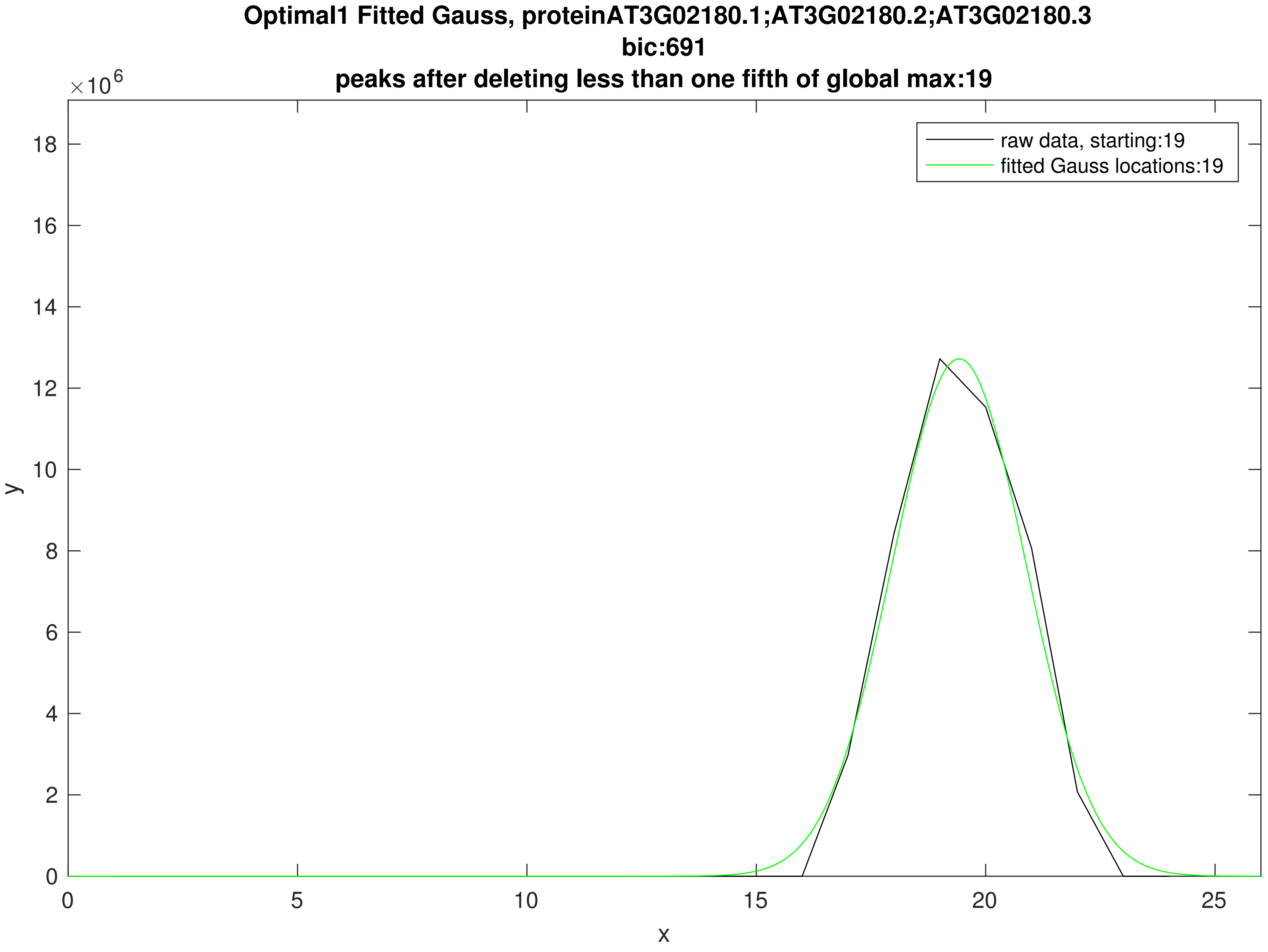
Optimal1 Fitted Gauss, proteinAT3G02180.1;AT3G02180.2;AT3G02180.3
bic:691
peaks after deleting less than one fifth of global max:19
raw data, starting:19
fitted Gauss locations:19
×10^6
y
x
0
2
4
6
8
10
12
14
16
18
0
5
10
15
20
25

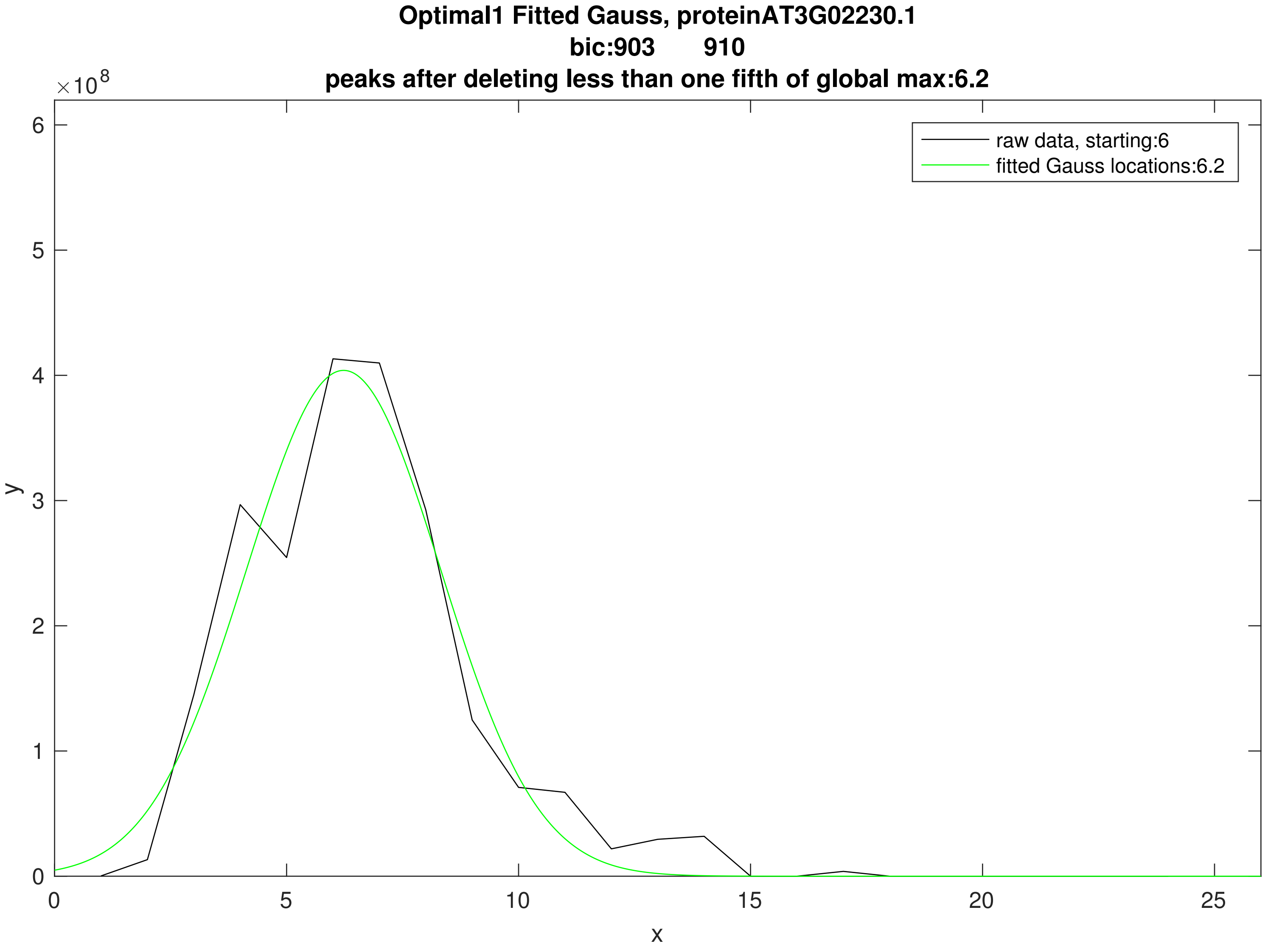
Optimal1 Fitted Gauss, proteinAT3G02230.1
bic:903 910
peaks after deleting less than one fifth of global max:6.2
×10^8
raw data, starting:6
fitted Gauss locations:6.2
y
x
0
1
2
3
4
5
6
0
5
10
15
20
25

Optimal1 Fitted Gauss, proteinAT3G02360.2;AT3G02360.1
bic:1.02e+03  1.03e+03  1.04e+03  1.05e+03
peaks after deleting less than one fifth of global max:13

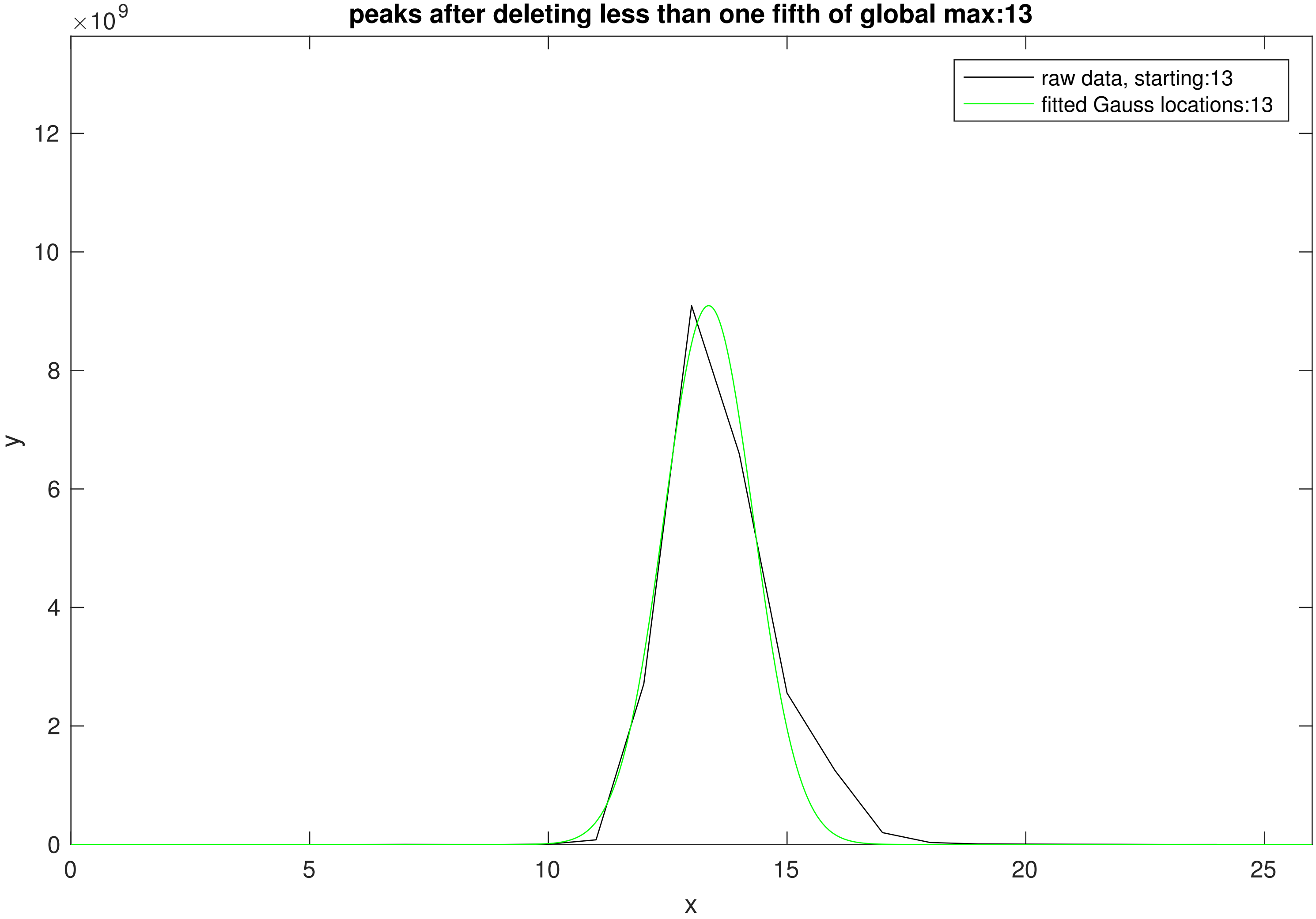

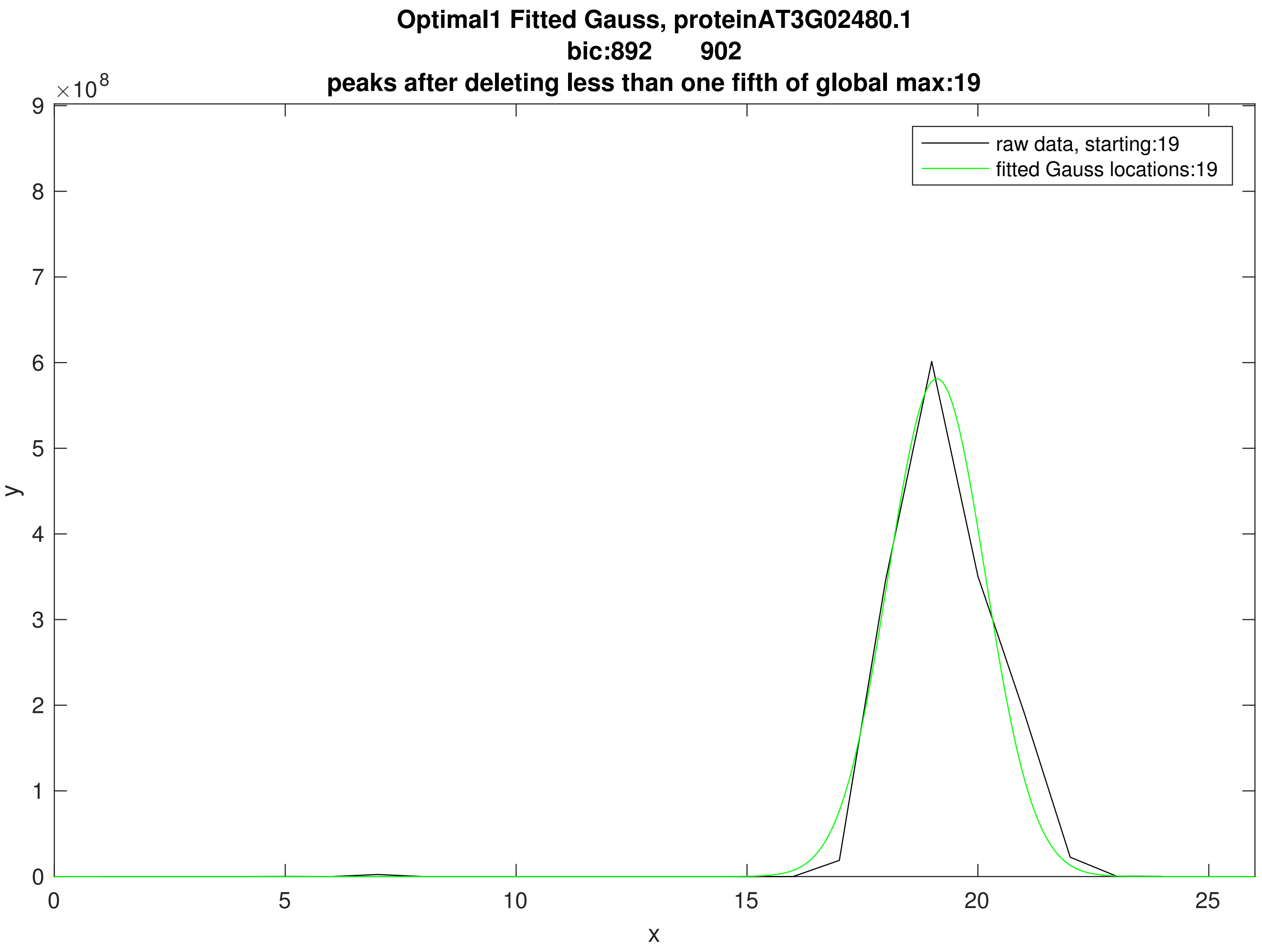
Optimal1 Fitted Gauss, proteinAT3G02480.1
bic:892 902
peaks after deleting less than one fifth of global max:19
×10^8
9
8
7
6
5
4
3
2
1
0
y
0
5
10
15
20
25
x
raw data, starting:19
fitted Gauss locations:19

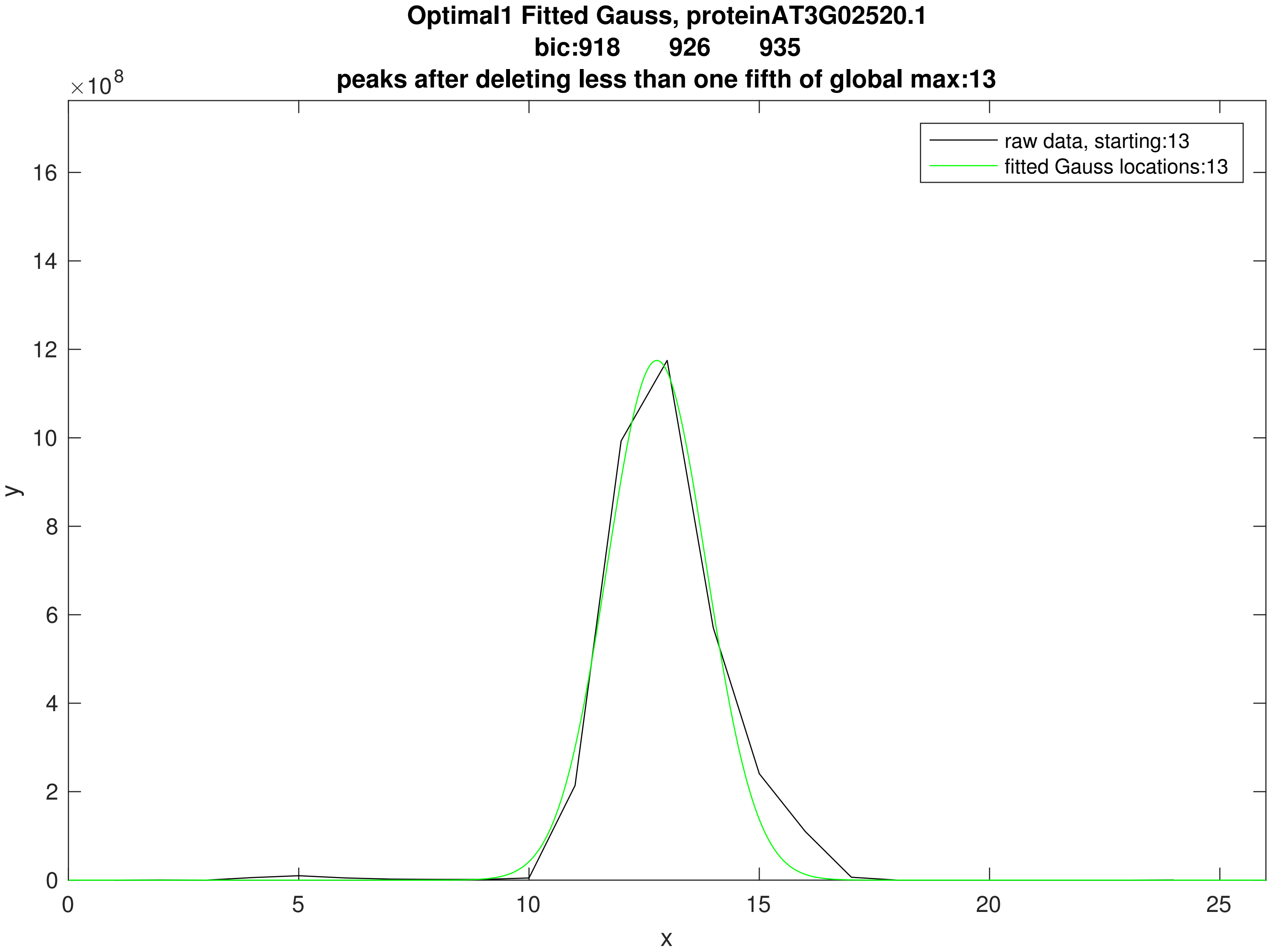
Optimal1 Fitted Gauss, proteinAT3G02520.1
bic:918 926 935
peaks after deleting less than one fifth of global max:13
×10^8
raw data, starting:13
fitted Gauss locations:13
y
x
0
2
4
6
8
10
12
14
16
0
5
10
15
20
25

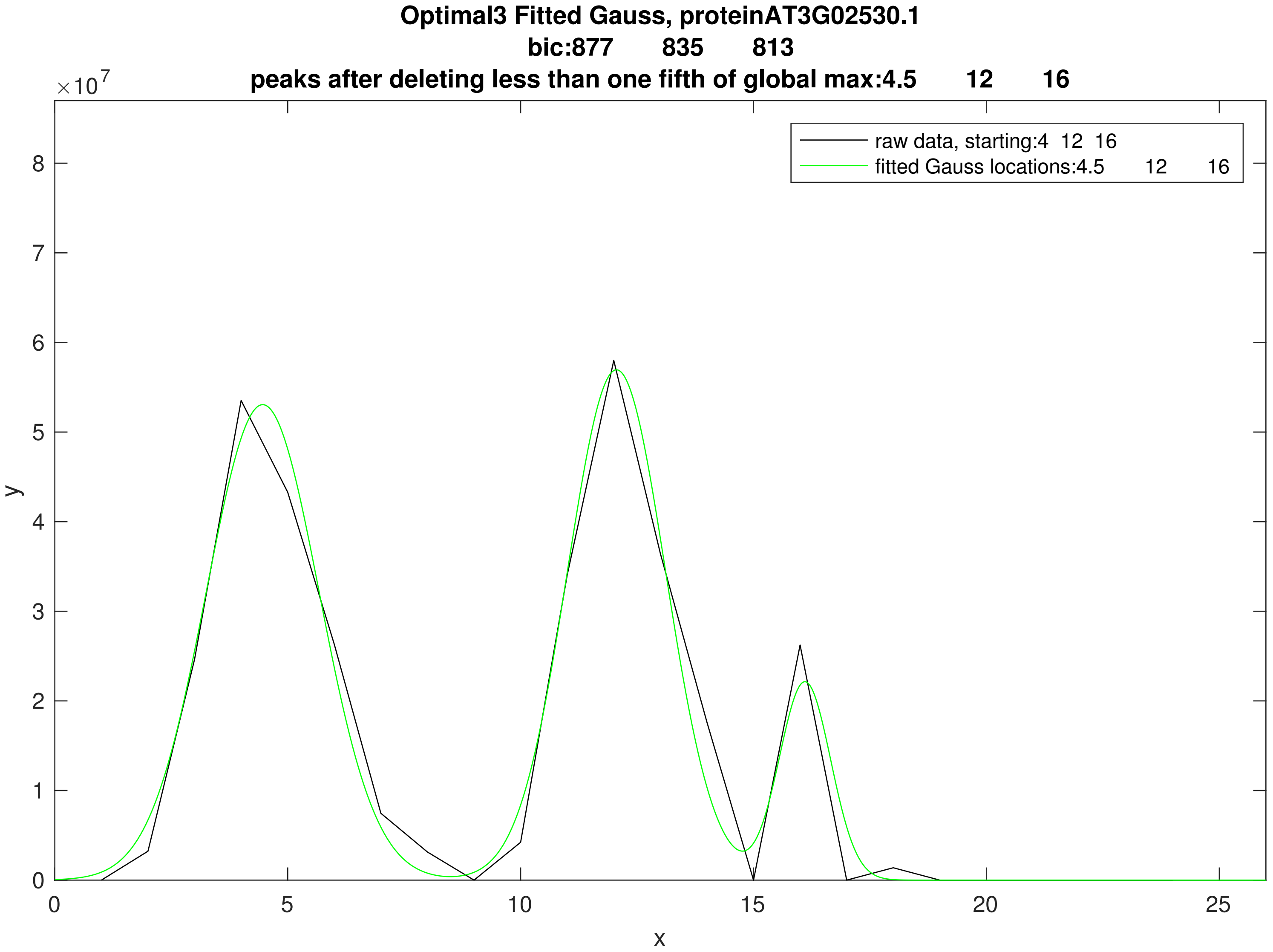

Optimal3 Fitted Gauss, proteinAT3G02530.1
bic:877 835 813
peaks after deleting less than one fifth of global max:4.5 12 16
×10^7
raw data, starting:4 12 16
fitted Gauss locations:4.5 12 16
0
1
2
3
4
5
6
7
8
y
0
5
10
15
20
25
x

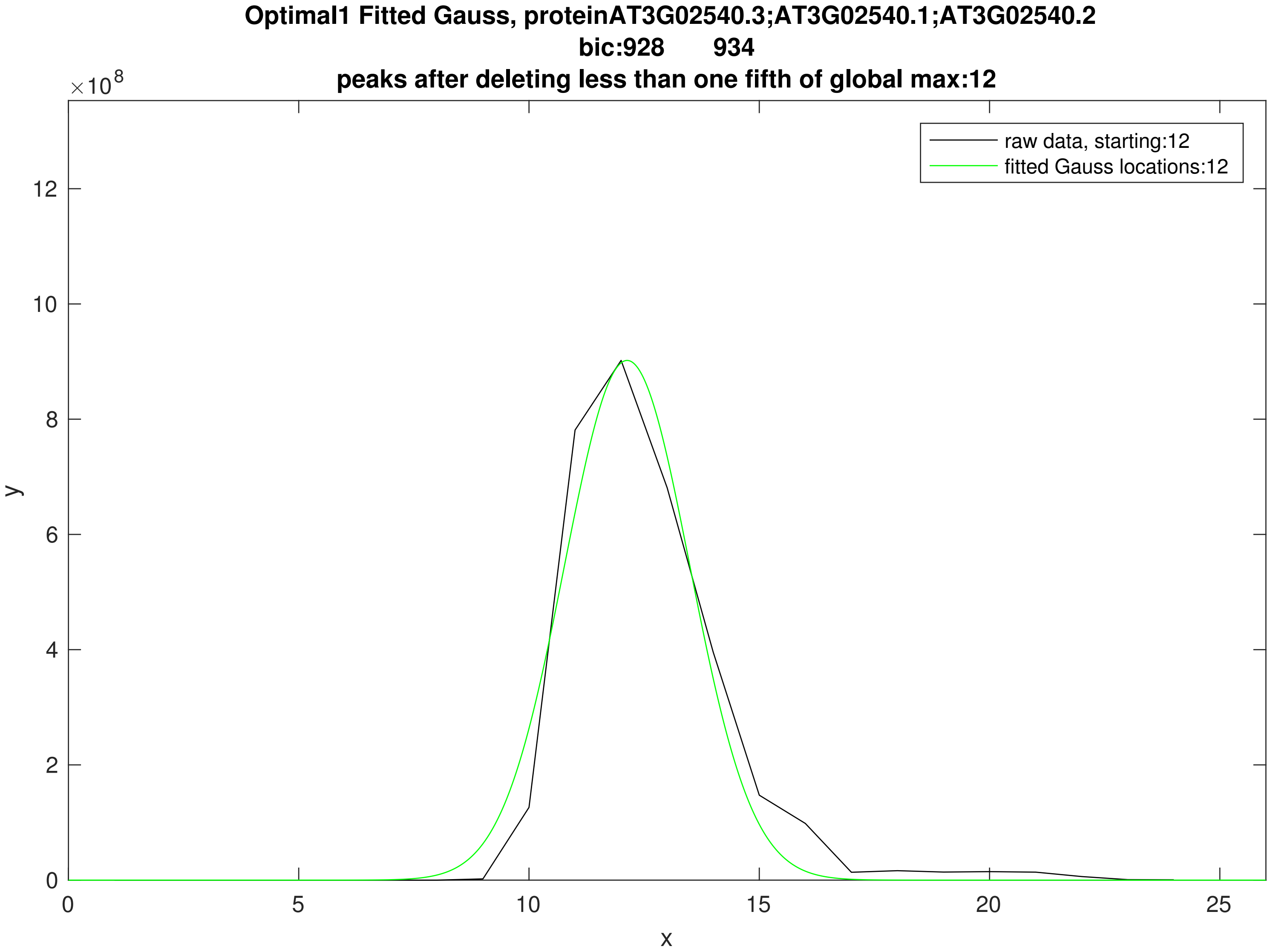
Optimal1 Fitted Gauss, proteinAT3G02540.3;AT3G02540.1;AT3G02540.2
bic:928 934
peaks after deleting less than one fifth of global max:12
×10^8
raw data, starting:12
fitted Gauss locations:12
y
x
0
2
4
6
8
10
12
0
5
10
15
20
25

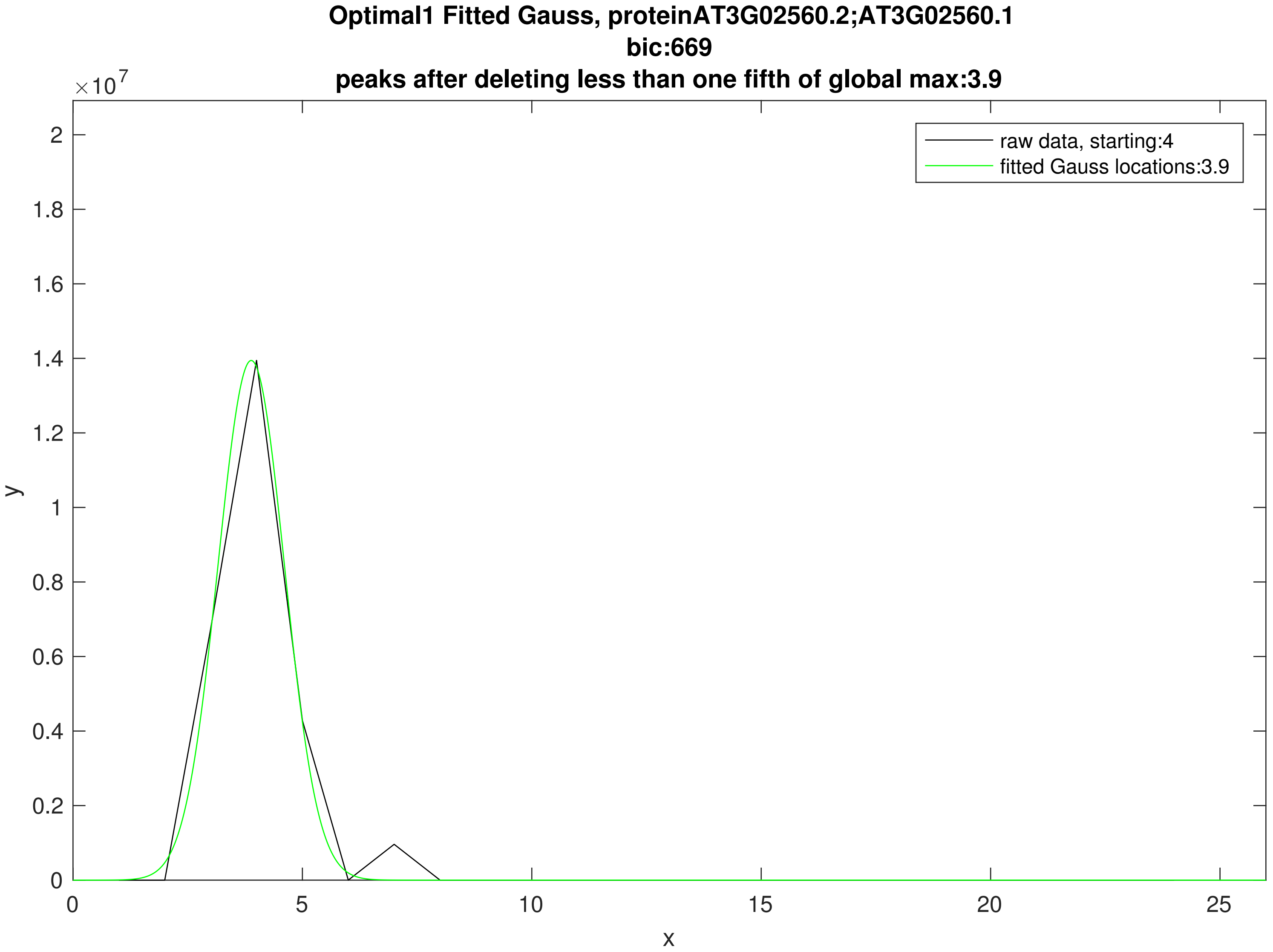

Optimal1 Fitted Gauss, proteinAT3G02560.2;AT3G02560.1
bic:669
peaks after deleting less than one fifth of global max:3.9
×10^7
raw data, starting:4
fitted Gauss locations:3.9
y
x
0
0.2
0.4
0.6
0.8
1
1.2
1.4
1.6
1.8
2
5
10
15
20
25

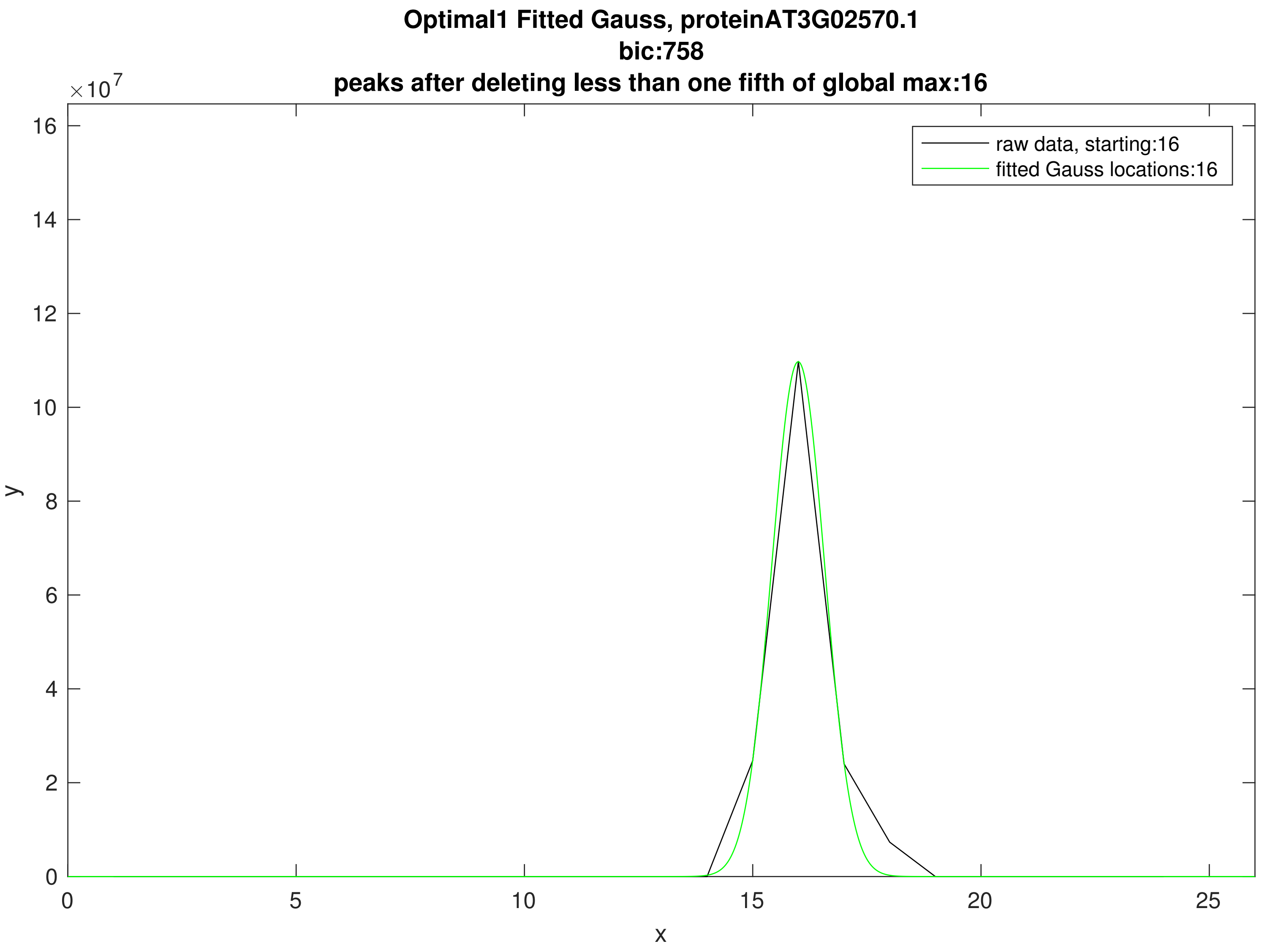
Optimal1 Fitted Gauss, proteinAT3G02570.1
bic:758
peaks after deleting less than one fifth of global max:16
×10^7
raw data, starting:16
fitted Gauss locations:16
y
x
0
2
4
6
8
10
12
14
16
0
5
10
15
20
25

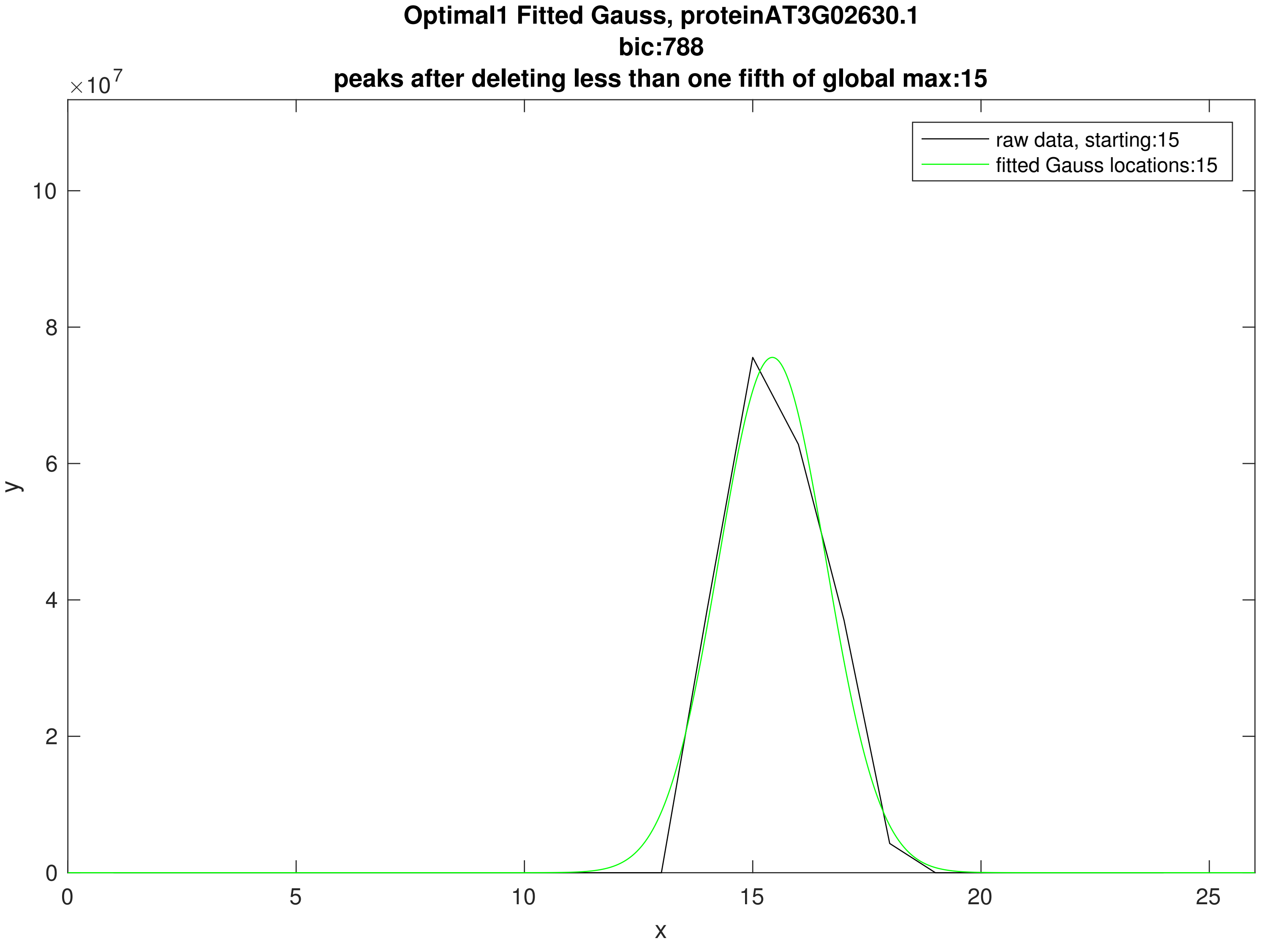
Optimal1 Fitted Gauss, proteinAT3G02630.1
bic:788
peaks after deleting less than one fifth of global max:15
×10^7
raw data, starting:15
fitted Gauss locations:15
y
x
0
2
4
6
8
10
0
5
10
15
20
25

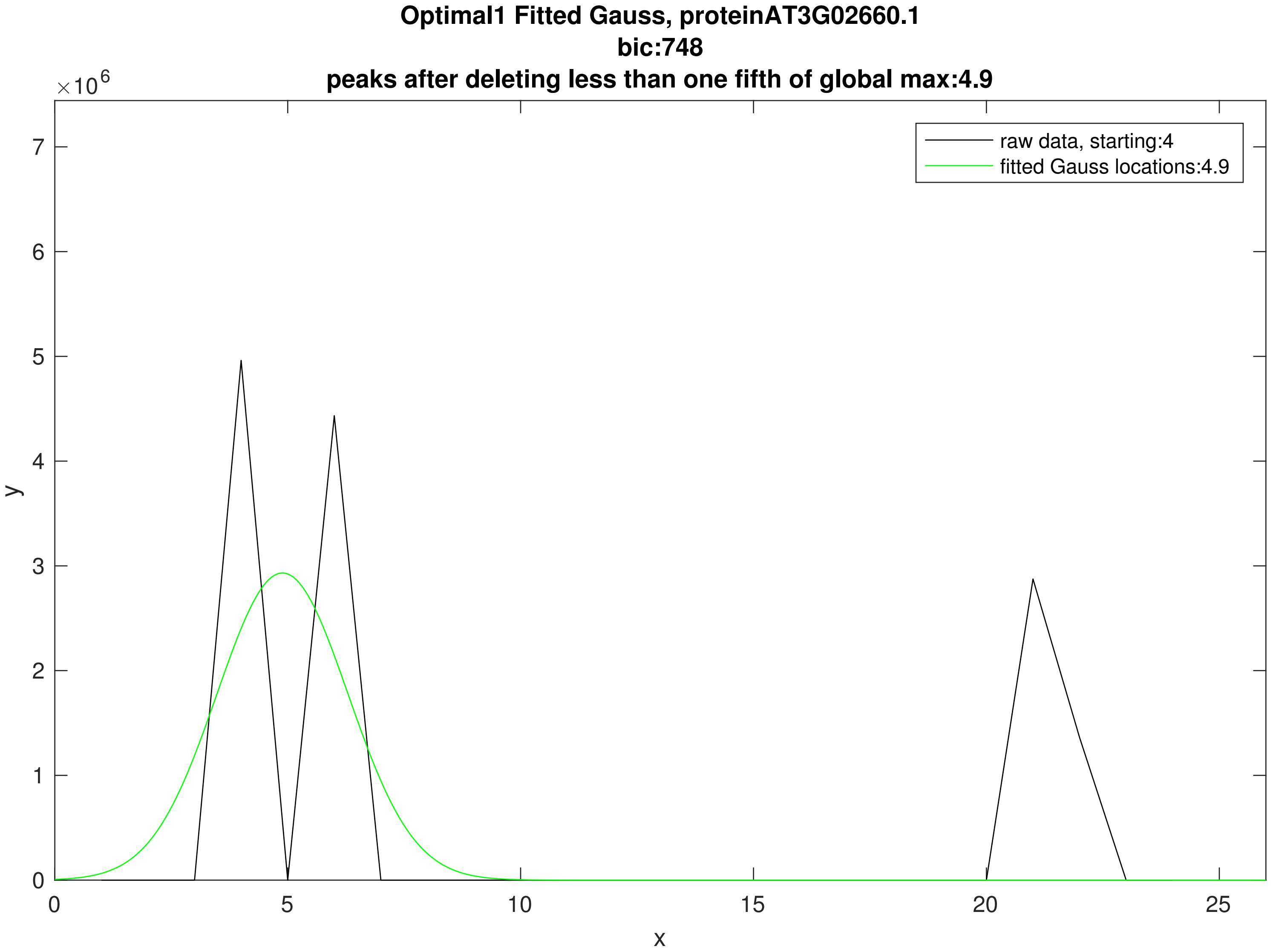
Optimal1 Fitted Gauss, proteinAT3G02660.1
bic:748
peaks after deleting less than one fifth of global max:4.9
×10^6
raw data, starting:4
fitted Gauss locations:4.9
y
x
0
1
2
3
4
5
6
7
0
5
10
15
20
25

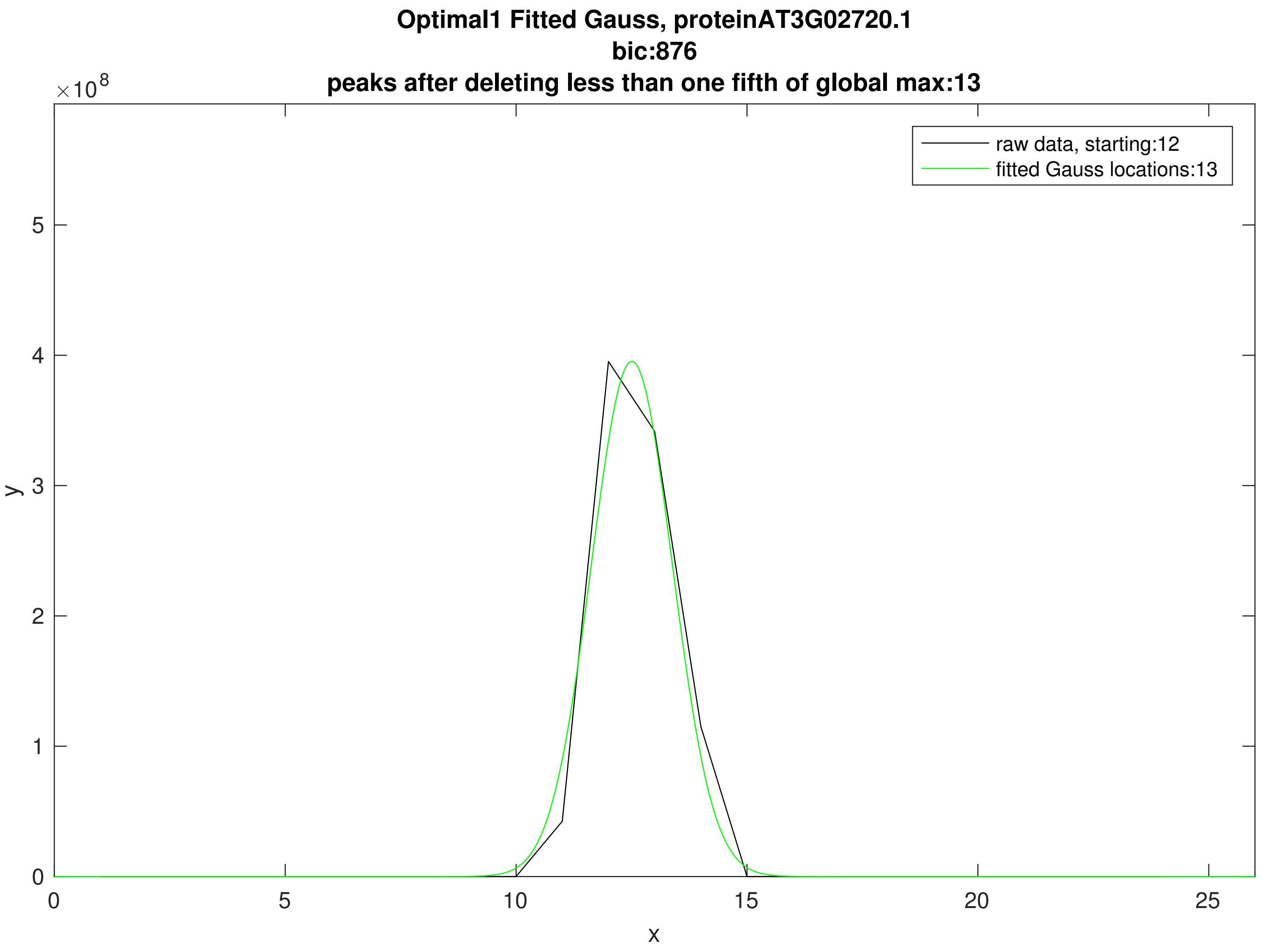
Optimal1 Fitted Gauss, proteinAT3G02720.1
bic:876
peaks after deleting less than one fifth of global max:13
×10^8
raw data, starting:12
fitted Gauss locations:13
y
x
0
1
2
3
4
5
0
5
10
15
20
25

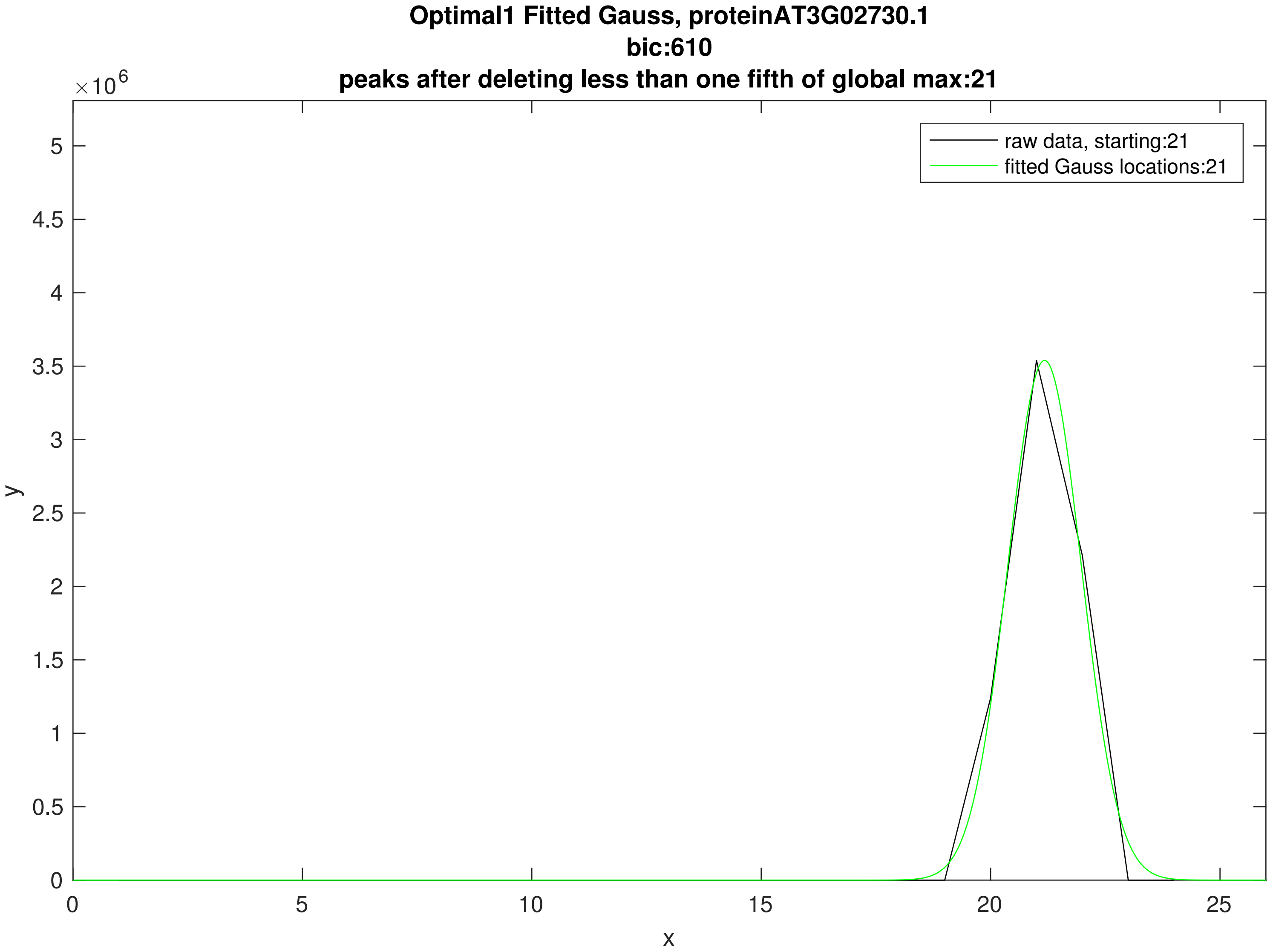
Optimal1 Fitted Gauss, proteinAT3G02730.1
bic:610
peaks after deleting less than one fifth of global max:21
raw data, starting:21
fitted Gauss locations:21
×10^6
y
x
0
0.5
1
1.5
2
2.5
3
3.5
4
4.5
5
5
10
15
20
25

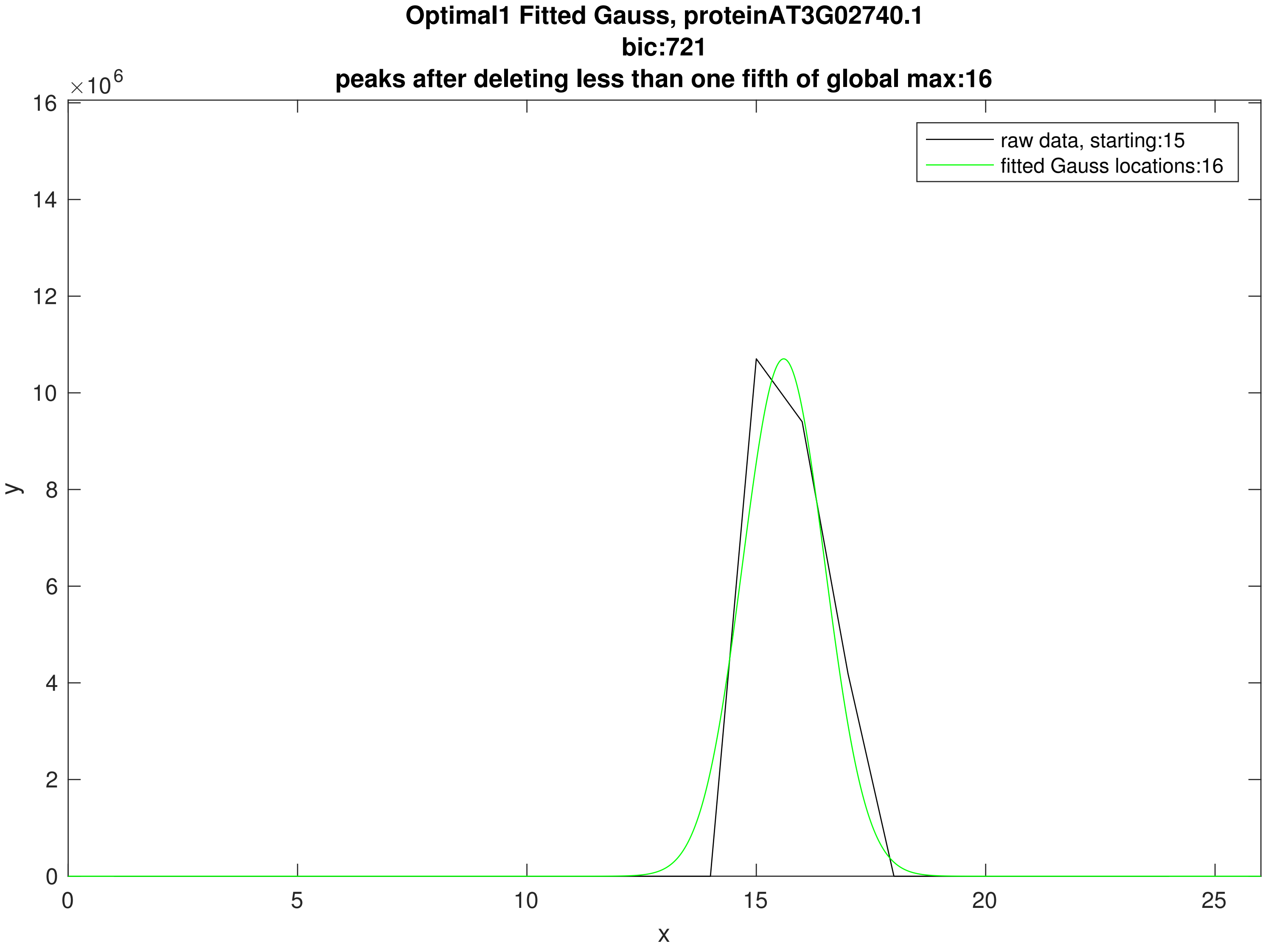
Optimal1 Fitted Gauss, proteinAT3G02740.1
bic:721
peaks after deleting less than one fifth of global max:16
×10^6
raw data, starting:15
fitted Gauss locations:16
y
x
0
2
4
6
8
10
12
14
16
0
5
10
15
20
25

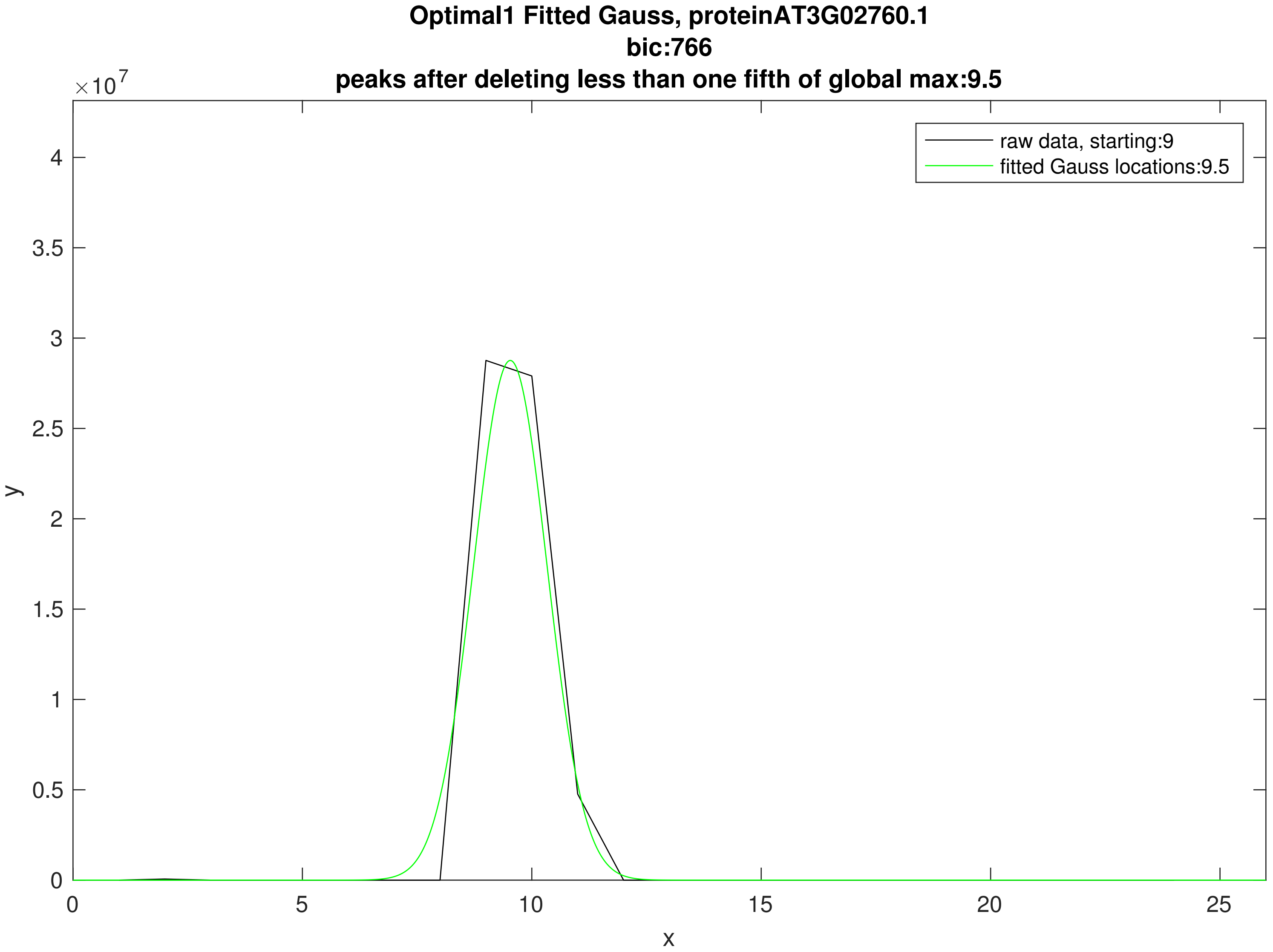

Optimal1 Fitted Gauss, proteinAT3G02760.1
bic:766
peaks after deleting less than one fifth of global max:9.5
×10^7
raw data, starting:9
fitted Gauss locations:9.5
y
x
0
0.5
1
1.5
2
2.5
3
3.5
4
0
5
10
15
20
25

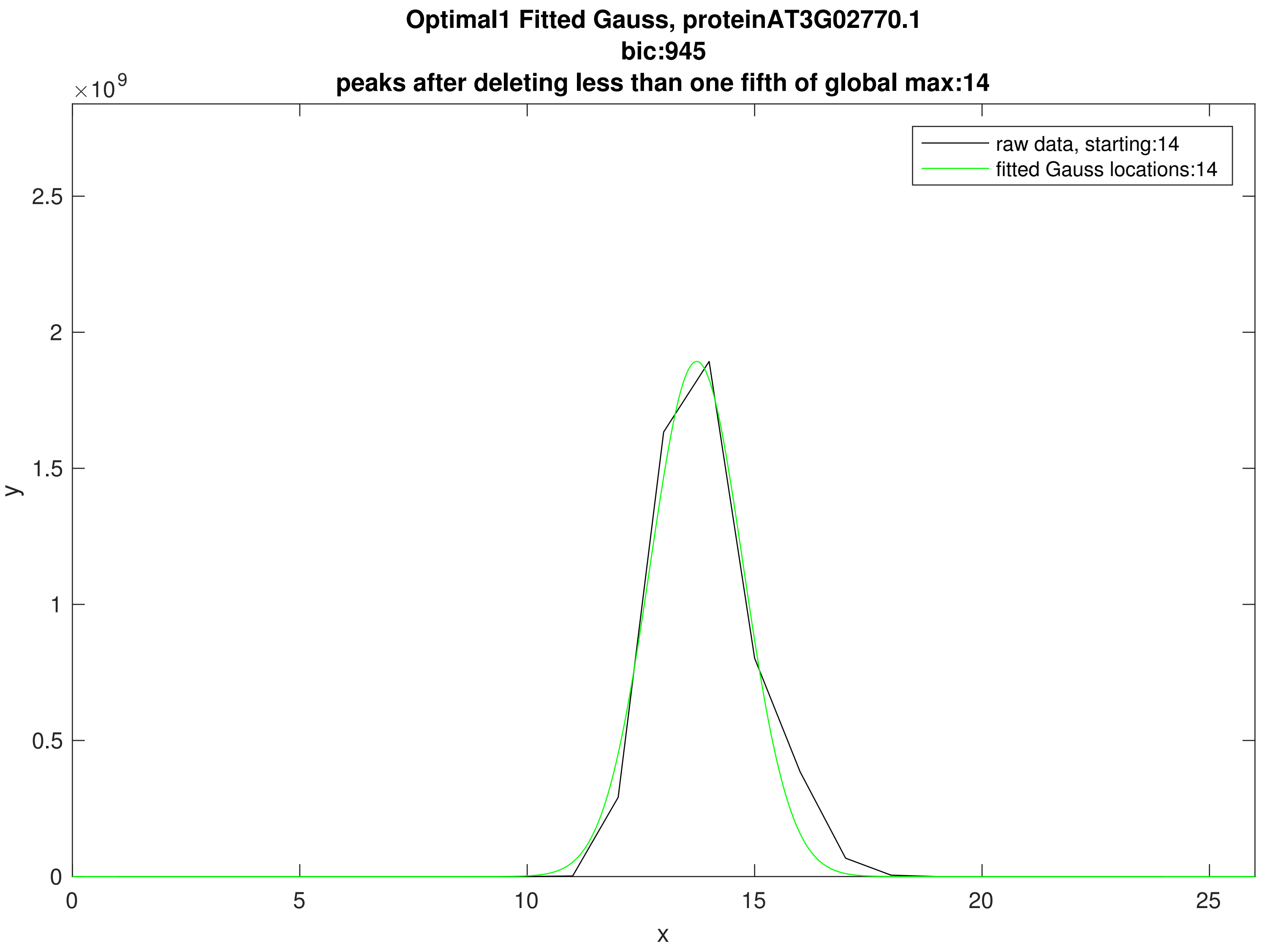
Optimal1 Fitted Gauss, proteinAT3G02770.1
bic:945
peaks after deleting less than one fifth of global max:14
×10^9
2.5
2
1.5
1
0.5
0
y
0
5
10
15
20
25
x
raw data, starting:14
fitted Gauss locations:14

Optimal1 Fitted Gauss, proteinAT3G02780.1
bic:934 939 949
peaks after deleting less than one fifth of global max:17

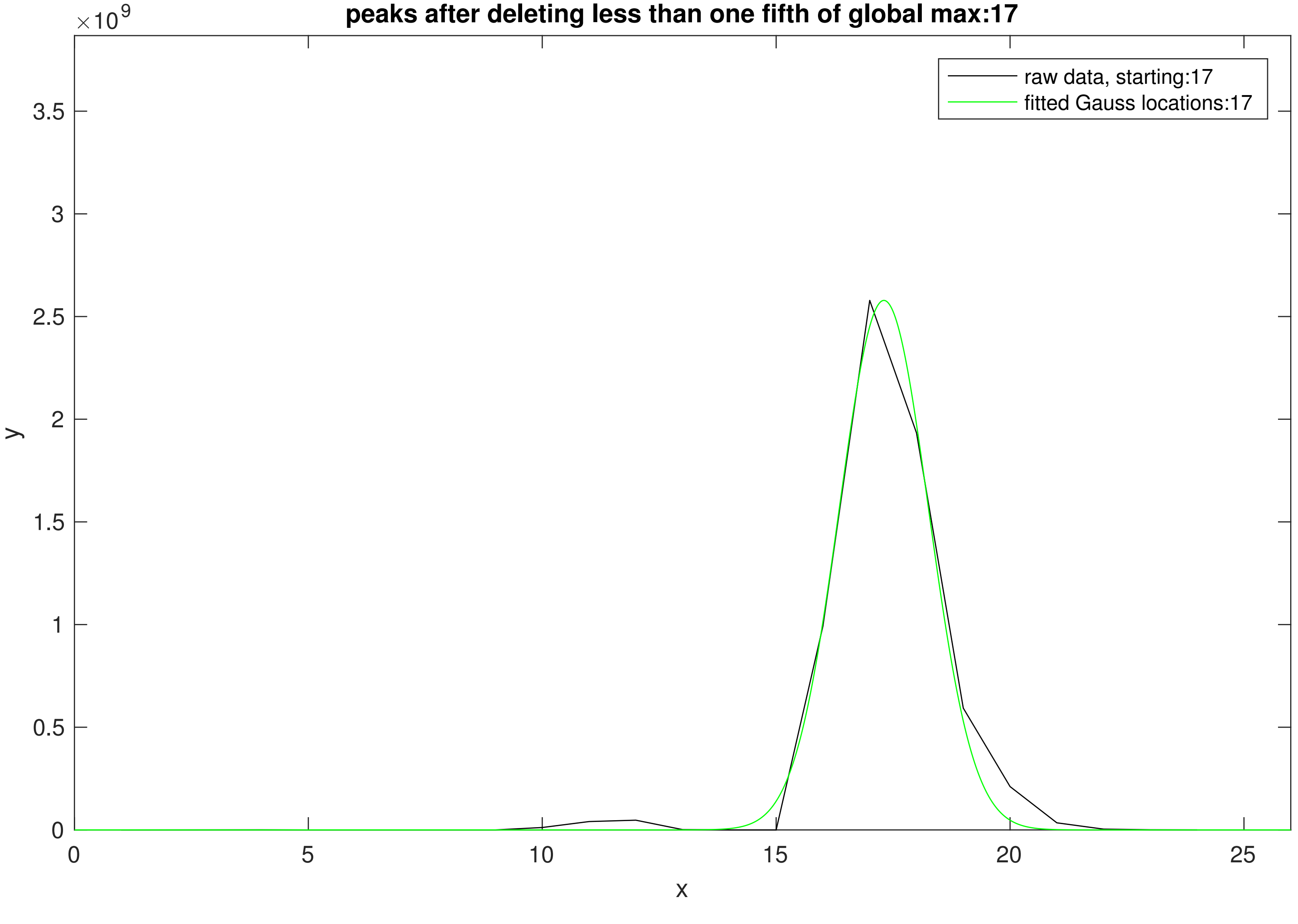

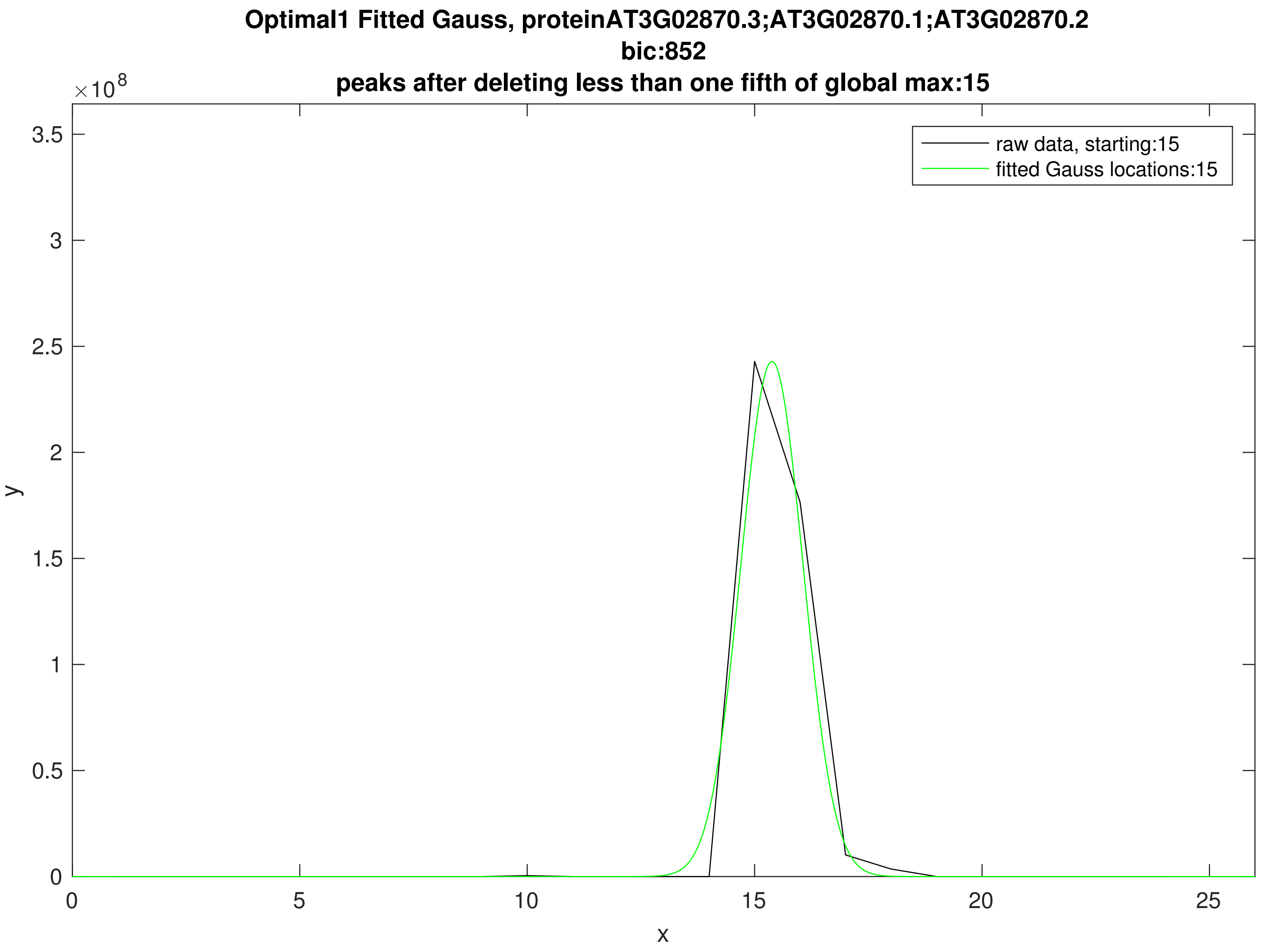

Optimal1 Fitted Gauss, proteinAT3G02870.3;AT3G02870.1;AT3G02870.2
bic:852
peaks after deleting less than one fifth of global max:15
×10^8
3.5
3
2.5
2
1.5
1
0.5
0
y
0
5
10
15
20
25
x
raw data, starting:15
fitted Gauss locations:15

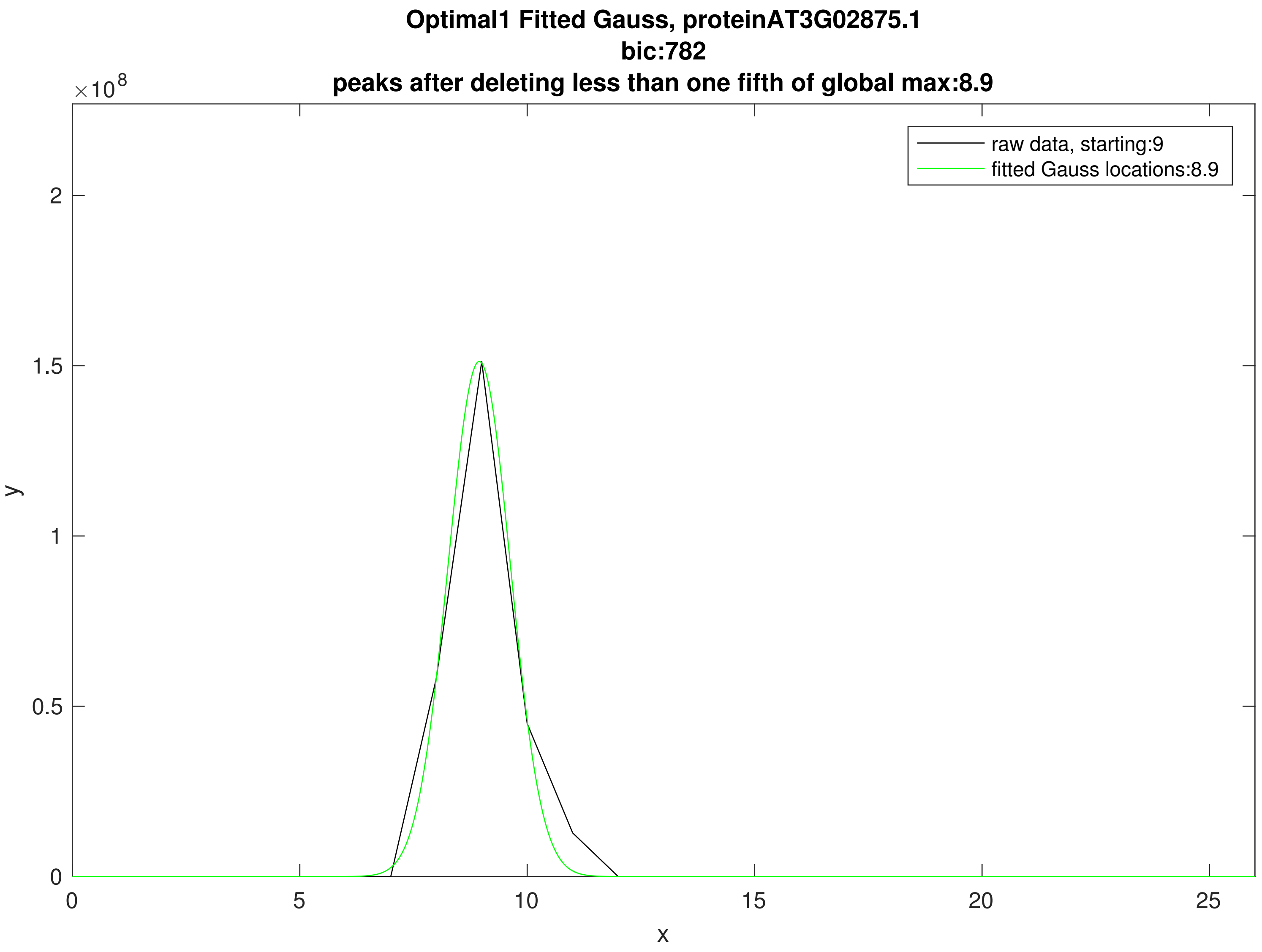

Optimal1 Fitted Gauss, proteinAT3G02875.1
bic:782
peaks after deleting less than one fifth of global max:8.9
×10^8
raw data, starting:9
fitted Gauss locations:8.9
2
1.5
1
0.5
0
y
0
5
10
15
20
25
x

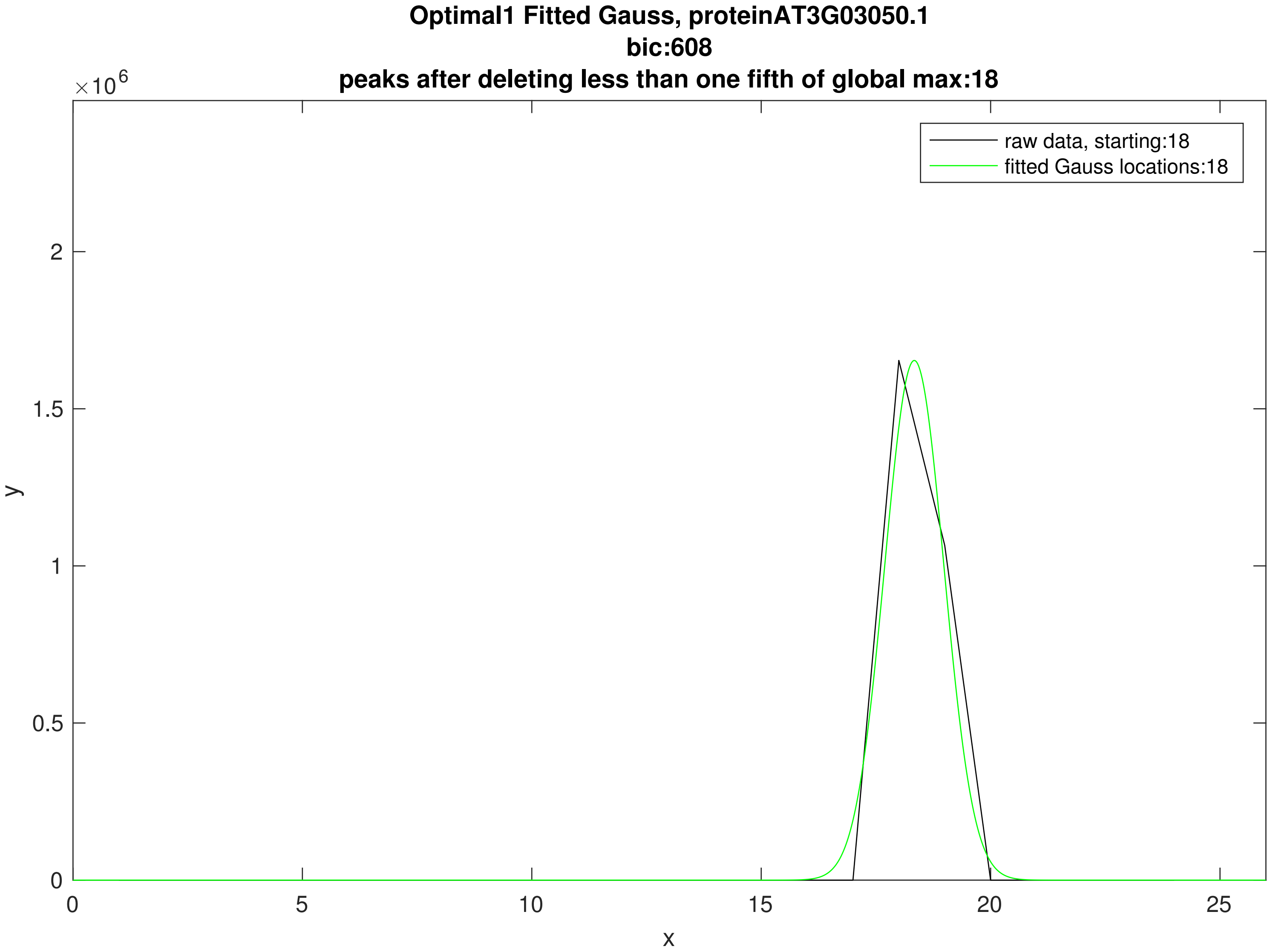
Optimal1 Fitted Gauss, proteinAT3G03050.1
bic:608
peaks after deleting less than one fifth of global max:18
×10^6
raw data, starting:18
fitted Gauss locations:18
y
x
0
0.5
1
1.5
2
0
5
10
15
20
25

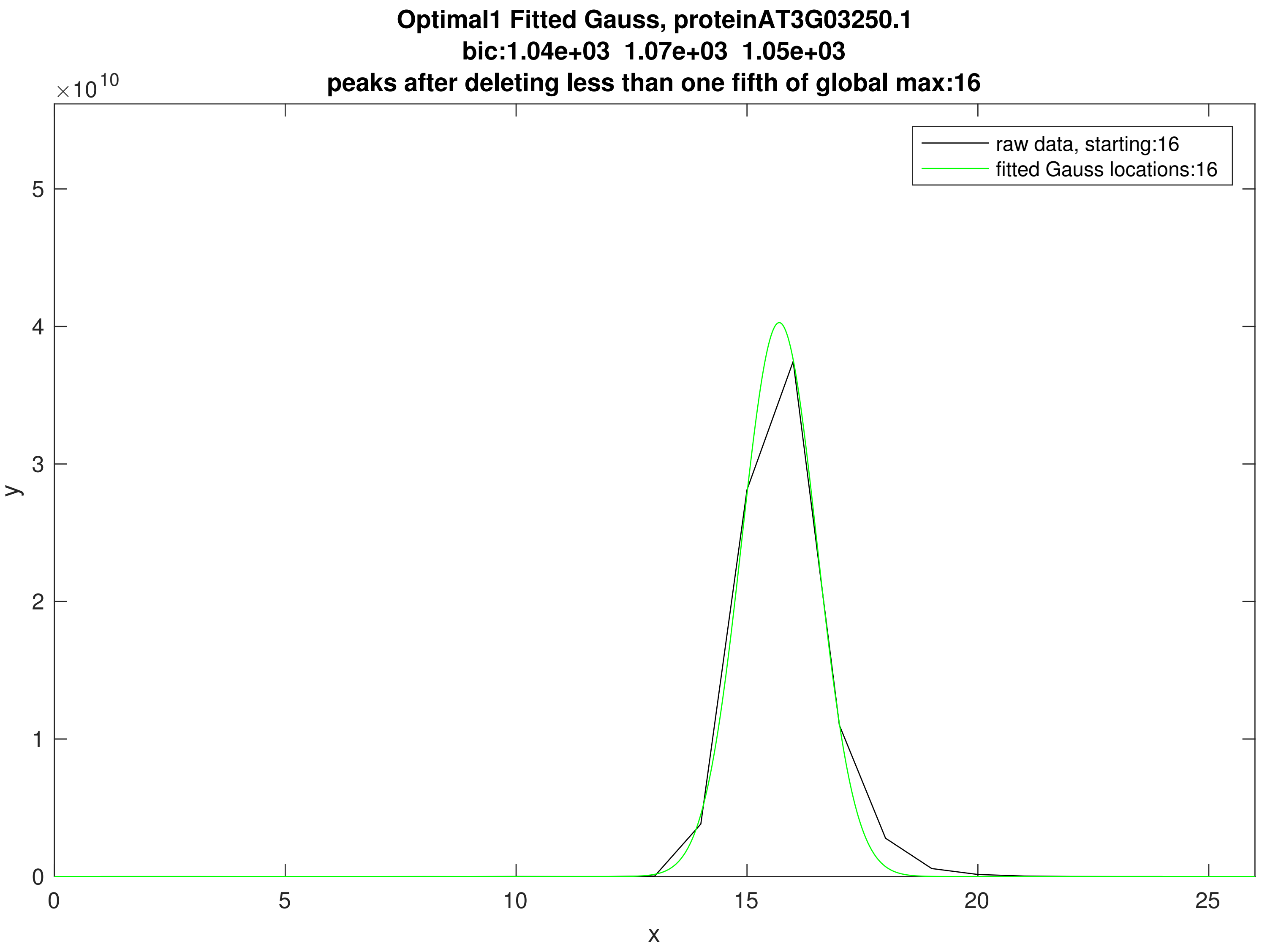

Optimal1 Fitted Gauss, proteinAT3G03250.1
bic:1.04e+03 1.07e+03 1.05e+03
peaks after deleting less than one fifth of global max:16
×10^10
raw data, starting:16
fitted Gauss locations:16
0
1
2
3
4
5
y
0
5
10
15
20
25
x

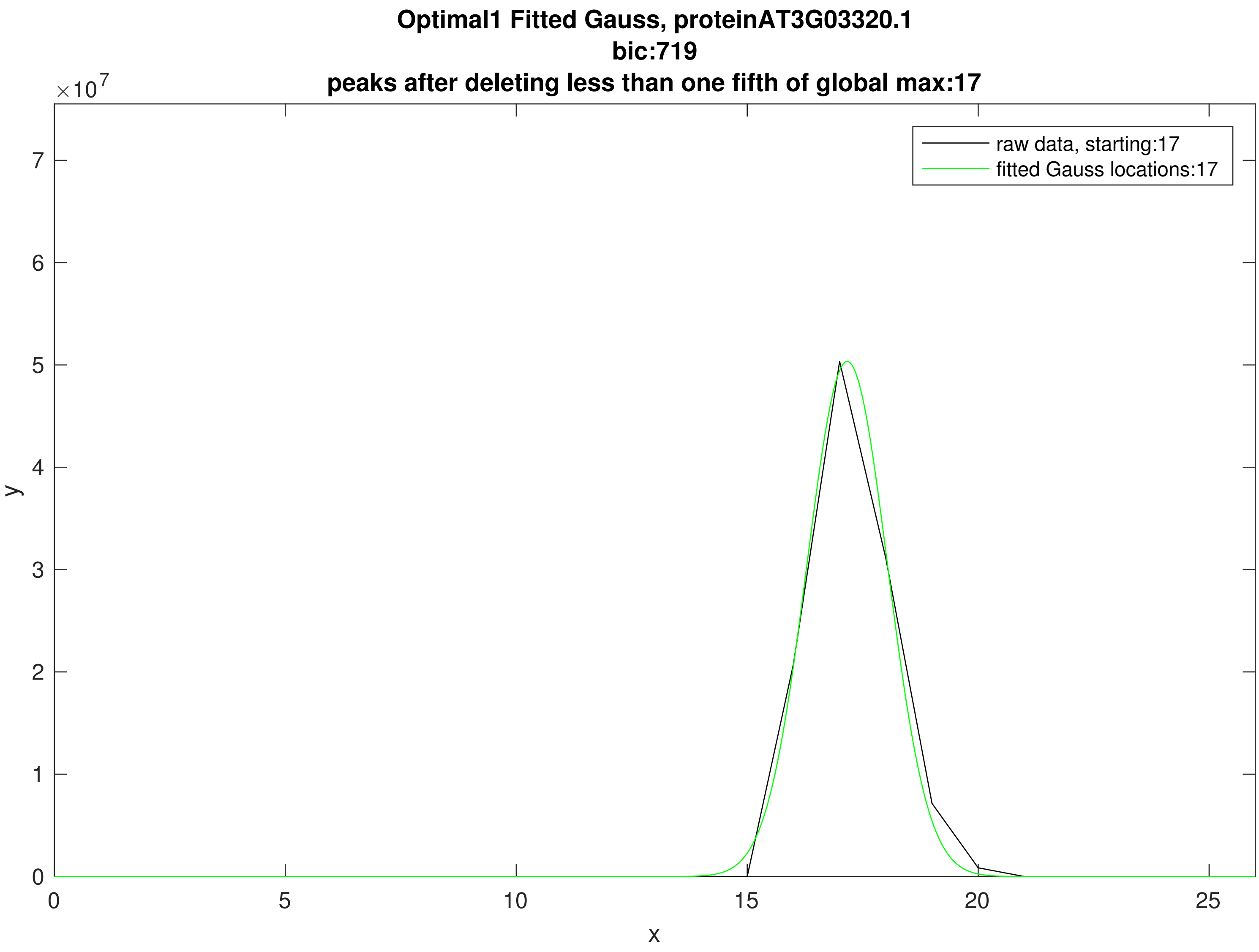

Optimal1 Fitted Gauss, proteinAT3G03320.1
bic:719
peaks after deleting less than one fifth of global max:17
×10^7
raw data, starting:17
fitted Gauss locations:17
y
x
0
1
2
3
4
5
6
7
0
5
10
15
20
25

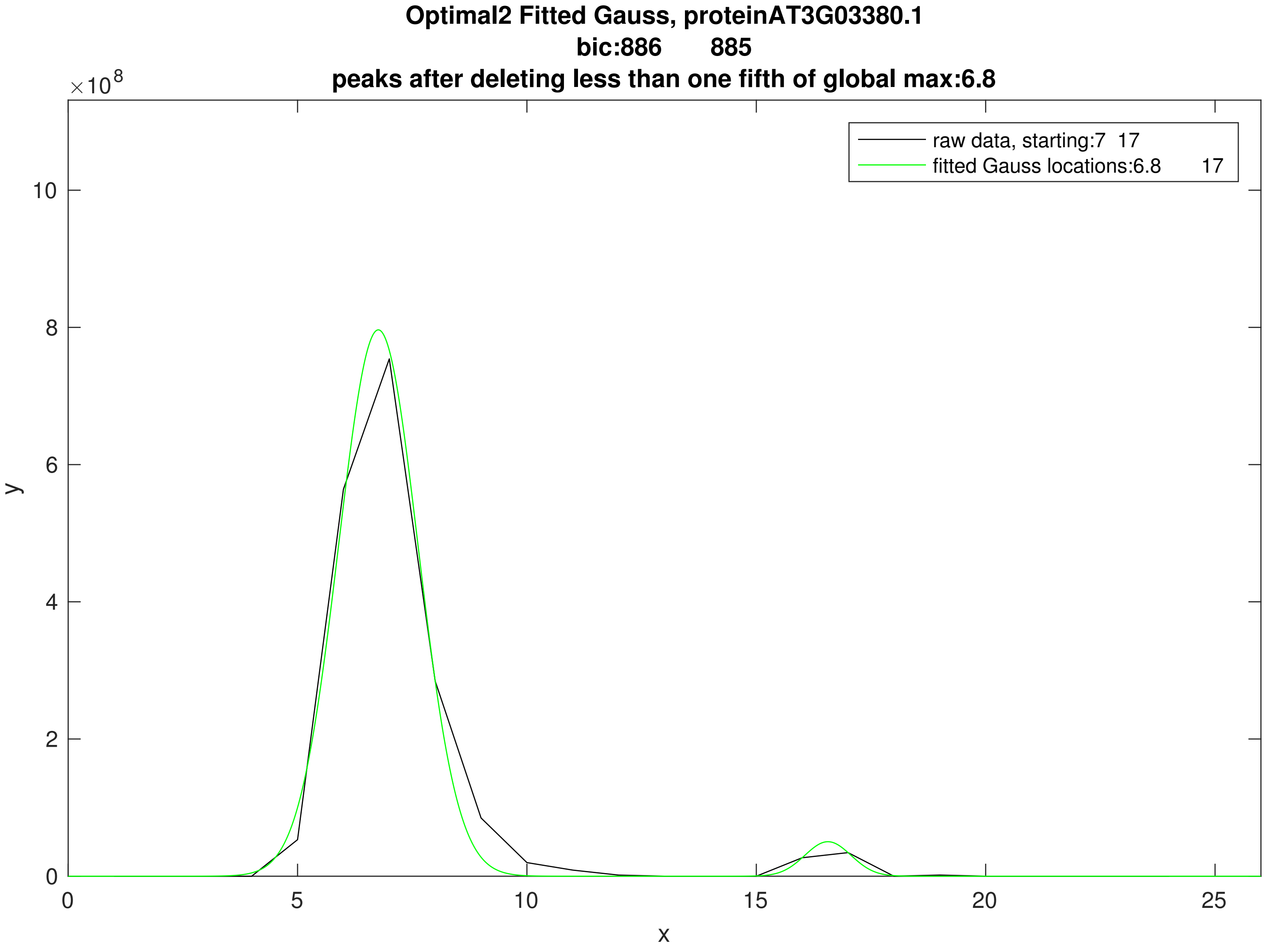

Optimal2 Fitted Gauss, proteinAT3G03380.1
bic:886 885
peaks after deleting less than one fifth of global max:6.8
×10^8
raw data, starting:7 17
fitted Gauss locations:6.8 17
y
x
0
2
4
6
8
10
0
5
10
15
20
25

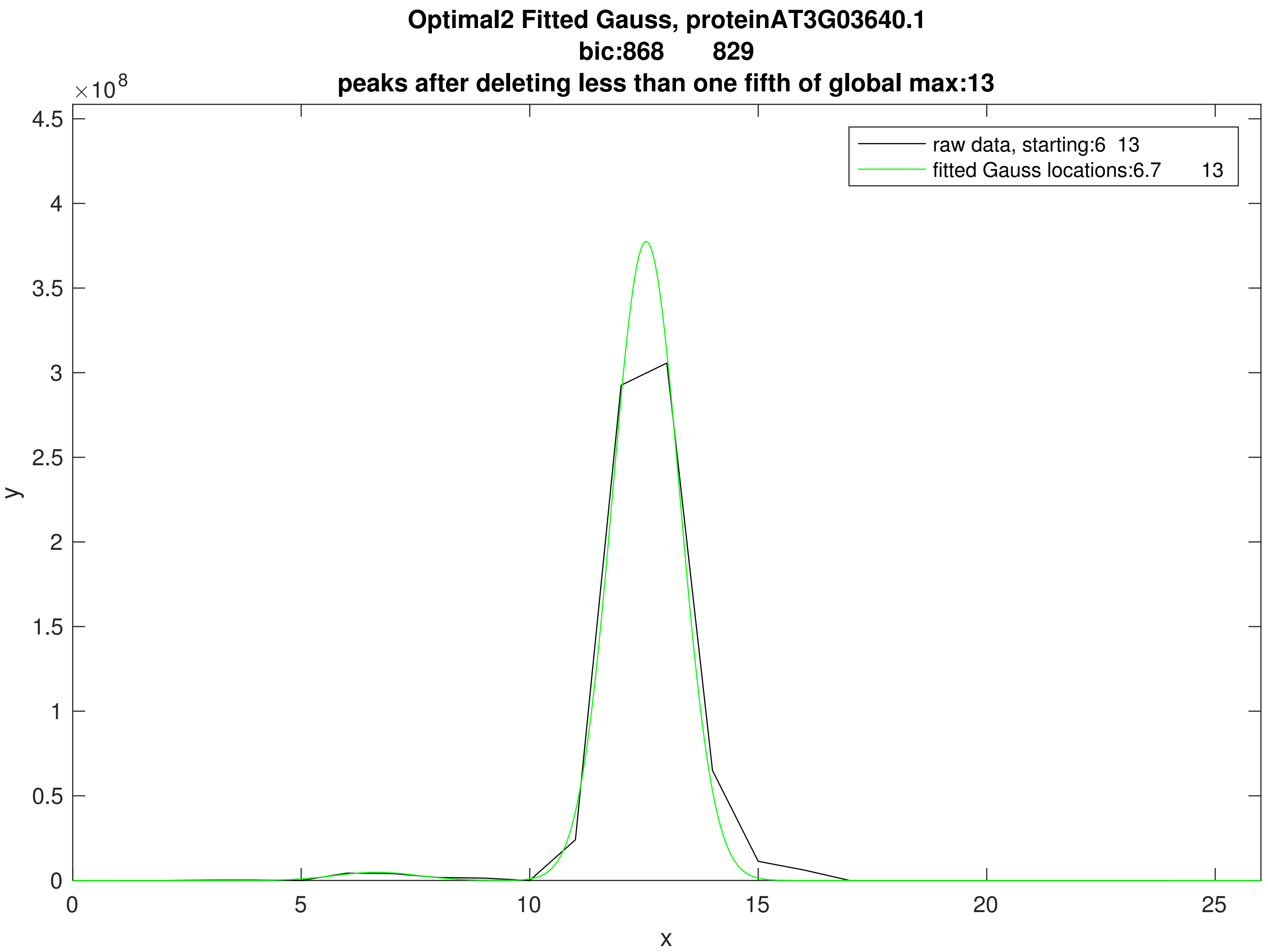
Optimal2 Fitted Gauss, proteinAT3G03640.1
bic:868 829
peaks after deleting less than one fifth of global max:13
×10^8
4.5
4
3.5
3
2.5
2
1.5
1
0.5
0
y
0
5
10
15
20
25
x
raw data, starting:6 13
fitted Gauss locations:6.7 13

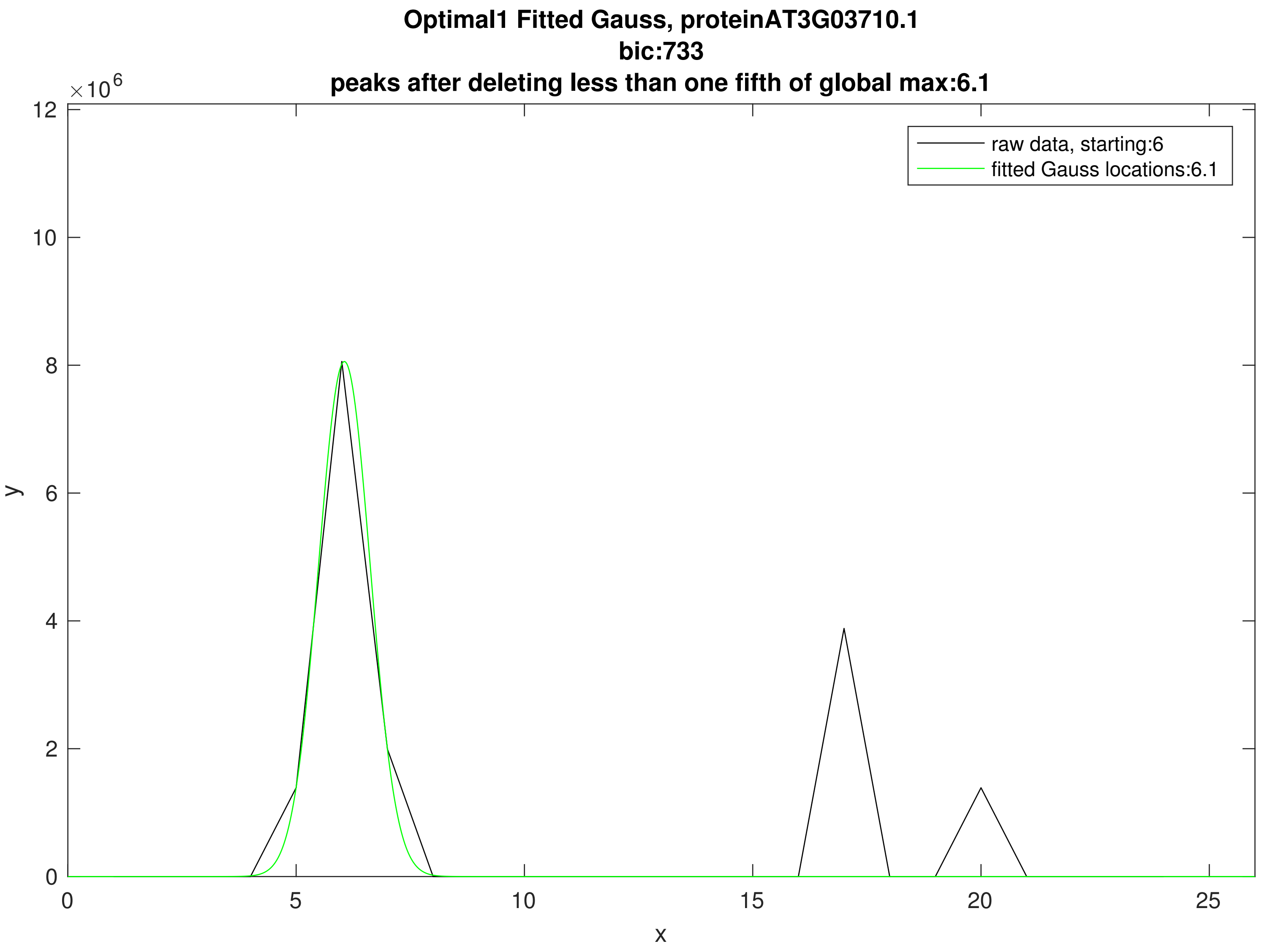

Optimal1 Fitted Gauss, proteinAT3G03710.1
bic:733
peaks after deleting less than one fifth of global max:6.1
×10^6
raw data, starting:6
fitted Gauss locations:6.1
y
x
0
2
4
6
8
10
12
5
15
20
25

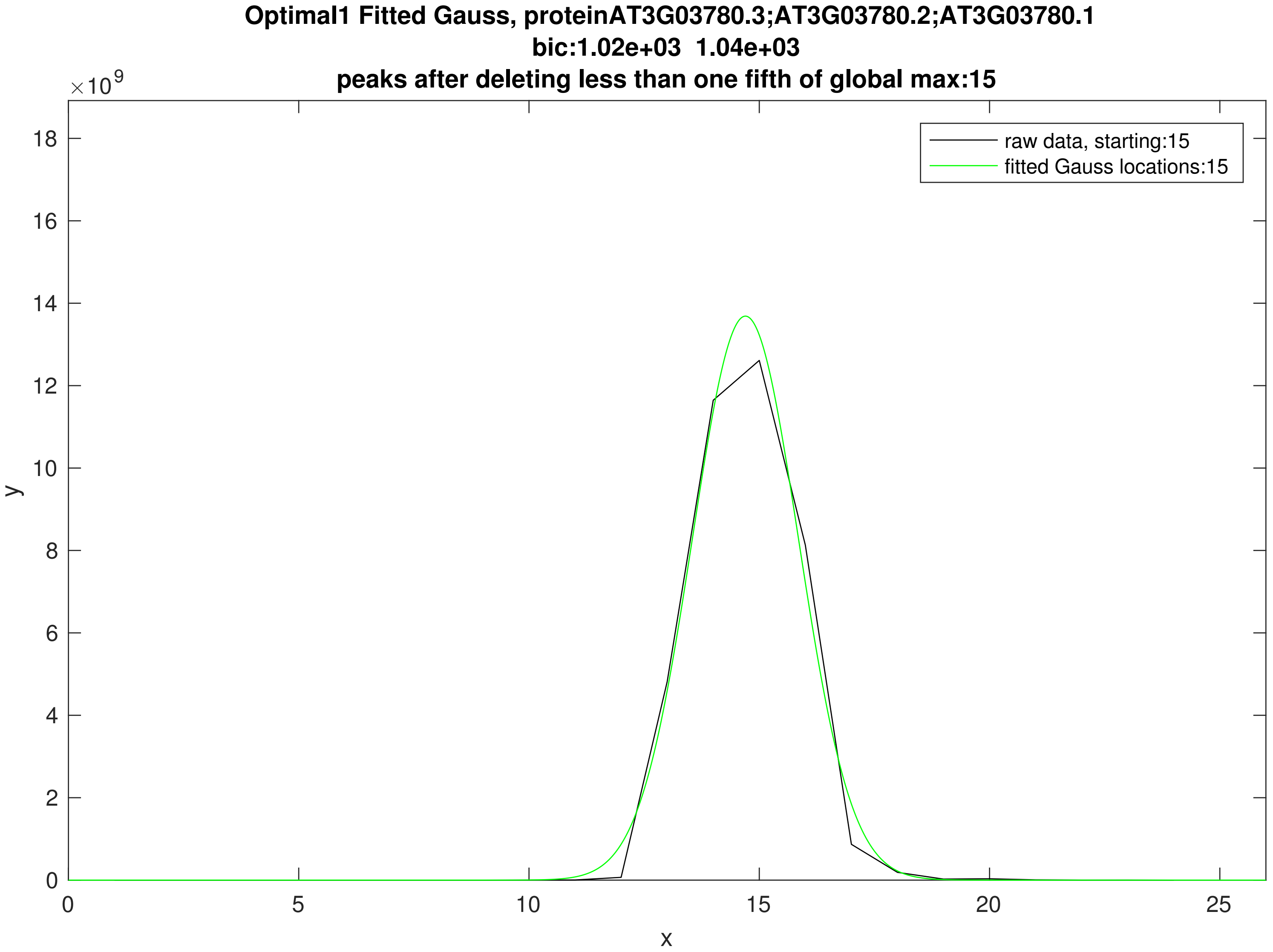

Optimal1 Fitted Gauss, proteinAT3G03780.3;AT3G03780.2;AT3G03780.1
bic:1.02e+03 1.04e+03
peaks after deleting less than one fifth of global max:15
×10^9
raw data, starting:15
fitted Gauss locations:15
y
x
0
2
4
6
8
10
12
14
16
18
0
5
10
15
20
25

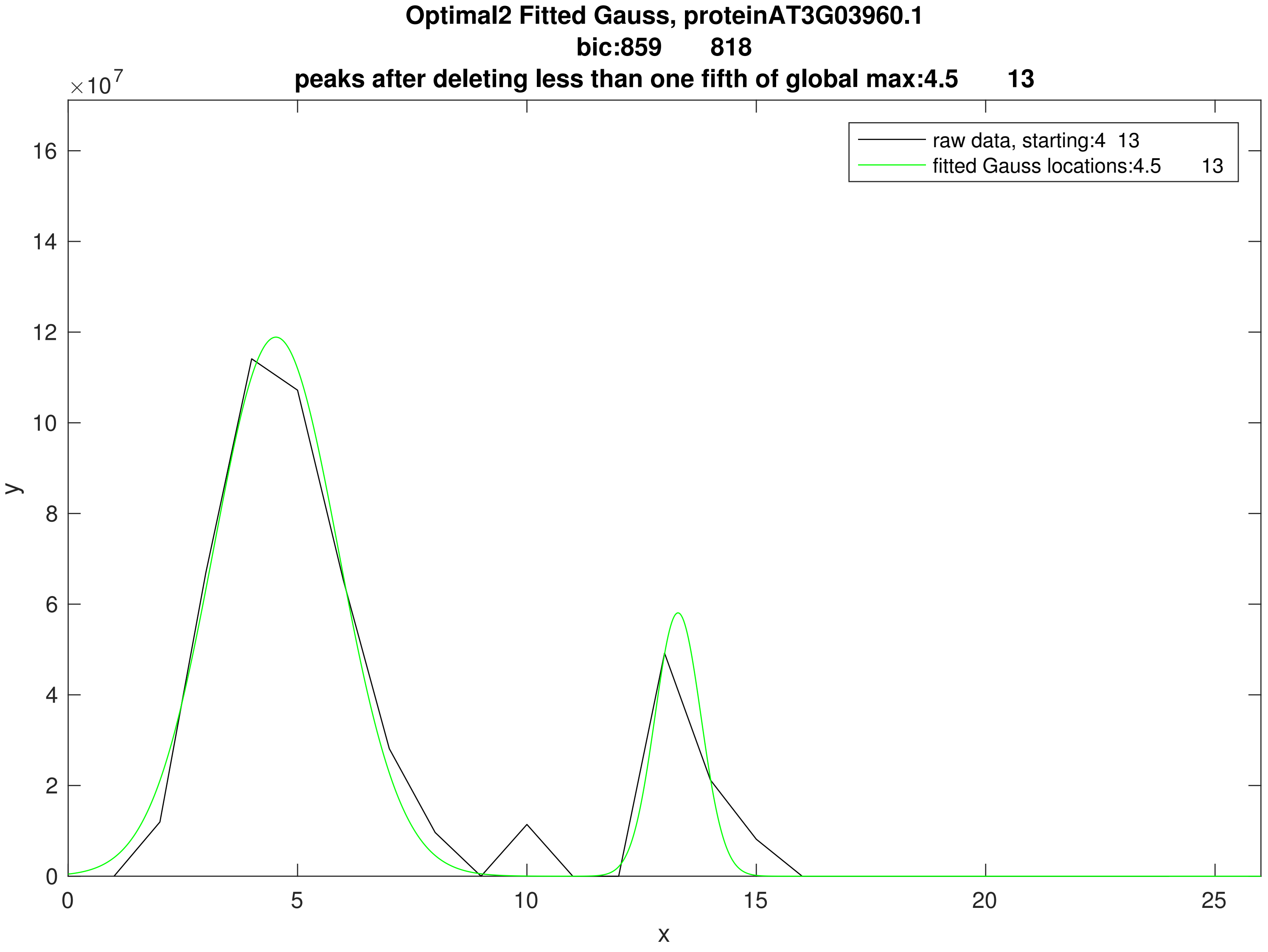
Optimal2 Fitted Gauss, proteinAT3G03960.1
bic:859 818
peaks after deleting less than one fifth of global max:4.5 13
×10^7
raw data, starting:4 13
fitted Gauss locations:4.5 13
y
x
0
2
4
6
8
10
12
14
16
0
5
10
15
20
25

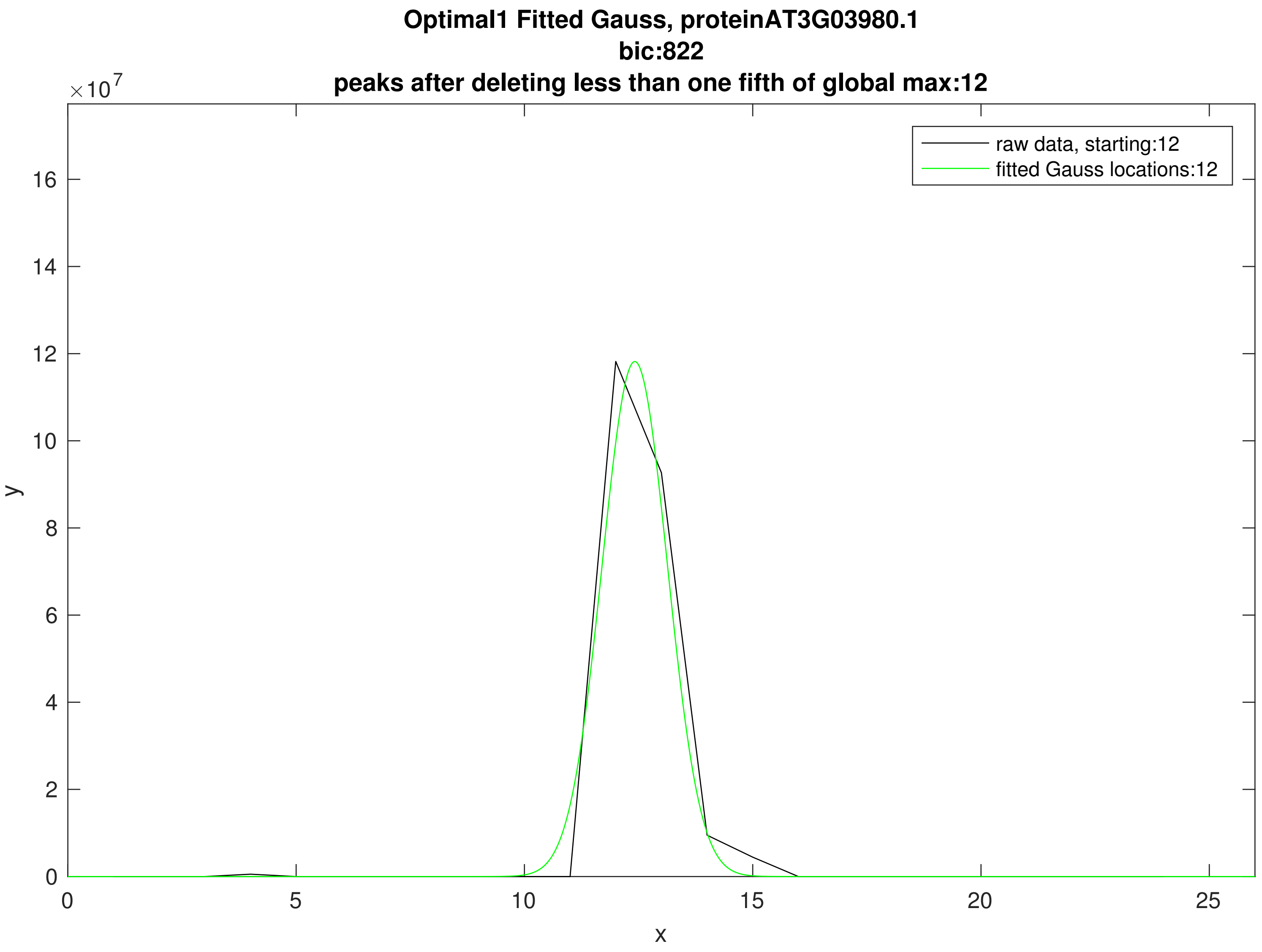
Optimal1 Fitted Gauss, proteinAT3G03980.1
bic:822
peaks after deleting less than one fifth of global max:12
×10^7
raw data, starting:12
fitted Gauss locations:12
16
14
12
10
8
6
4
2
0
y
0
5
10
15
20
25
x

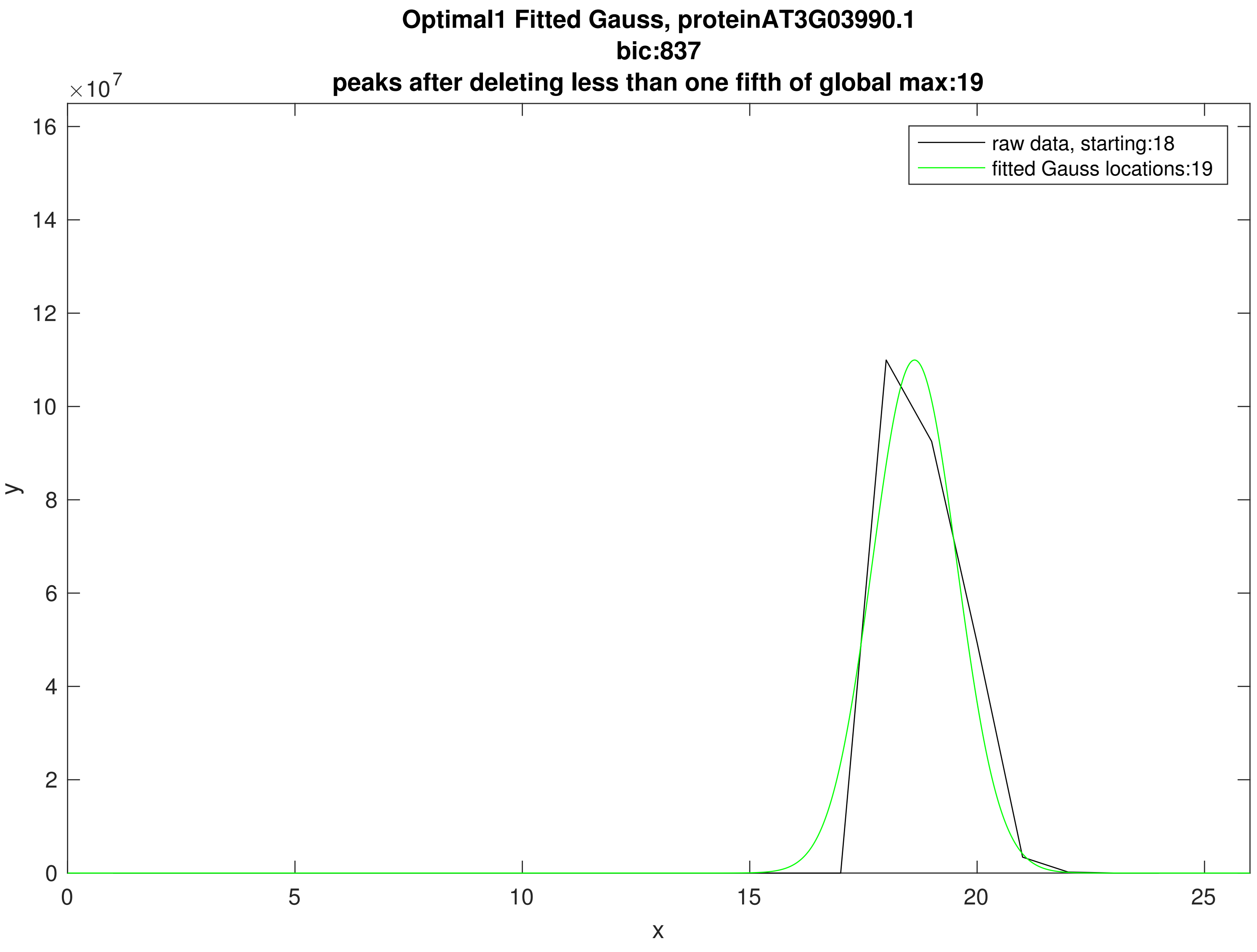

Optimal1 Fitted Gauss, proteinAT3G03990.1
bic:837
peaks after deleting less than one fifth of global max:19
×10^7
16
14
12
10
8
6
4
2
0
y
0
5
10
15
20
25
x
raw data, starting:18
fitted Gauss locations:19

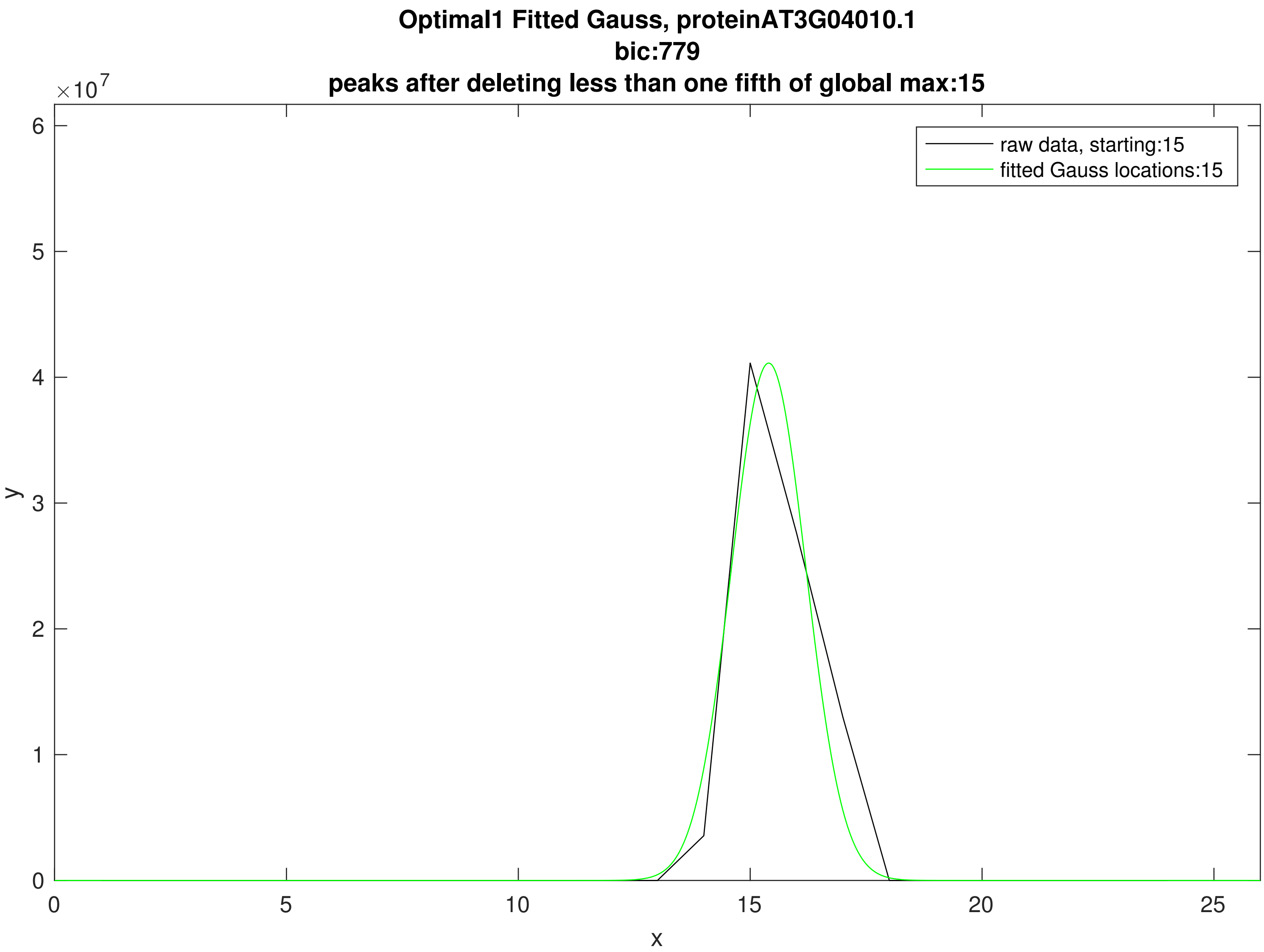

Optimal1 Fitted Gauss, proteinAT3G04010.1
bic:779
peaks after deleting less than one fifth of global max:15
×10^7
6
5
4
3
2
1
0
y
0
5
10
15
20
25
x
raw data, starting:15
fitted Gauss locations:15

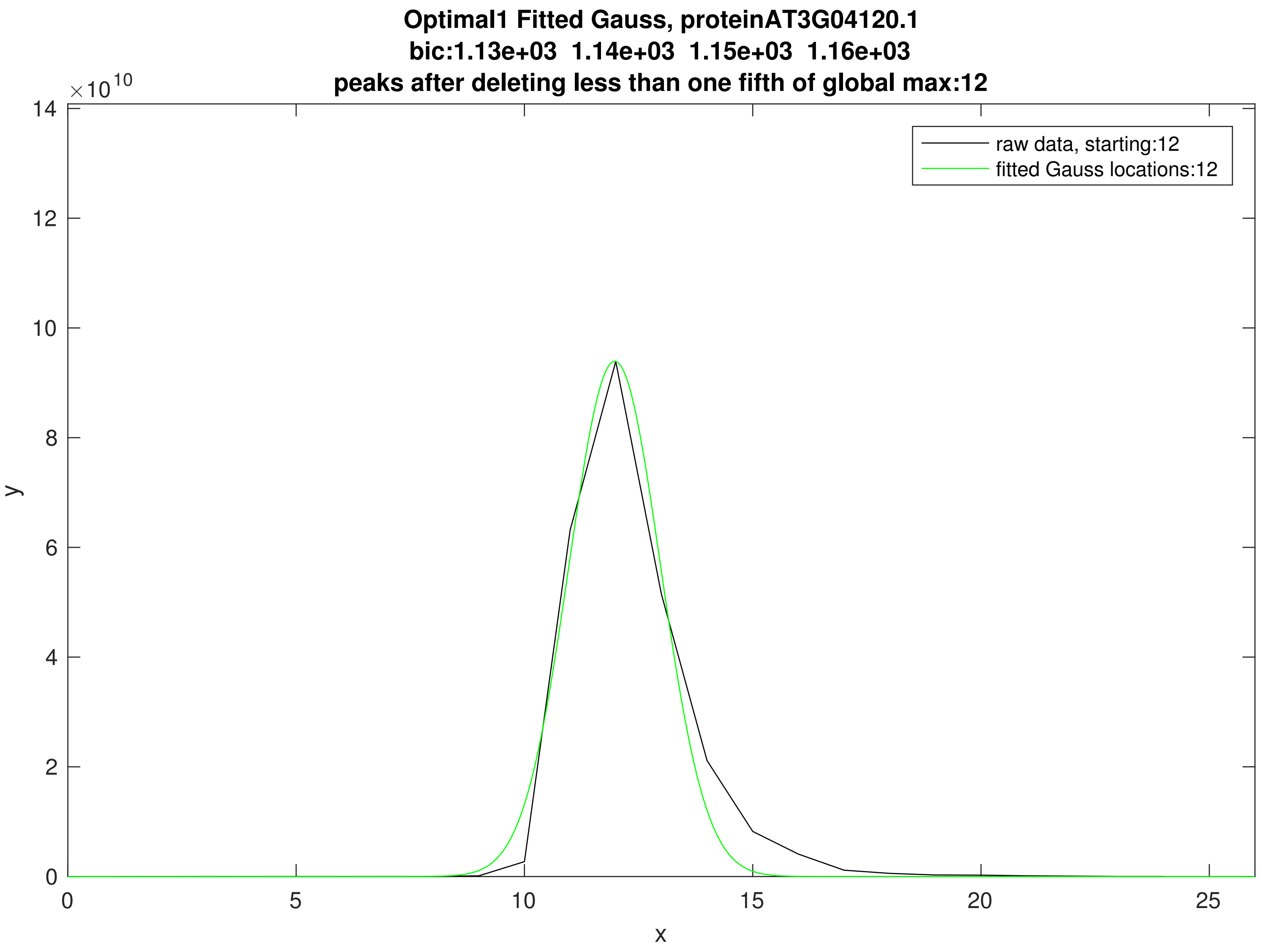
Optimal1 Fitted Gauss, proteinAT3G04120.1
bic:1.13e+03 1.14e+03 1.15e+03 1.16e+03
peaks after deleting less than one fifth of global max:12
×10^10
14
12
10
8
6
4
2
0
y
0
5
10
15
20
25
x
raw data, starting:12
fitted Gauss locations:12

**Optimal1 Fitted Gauss, proteinAT3G04290.1**
**bic:718**
**peaks after deleting less than one fifth of global max:19**

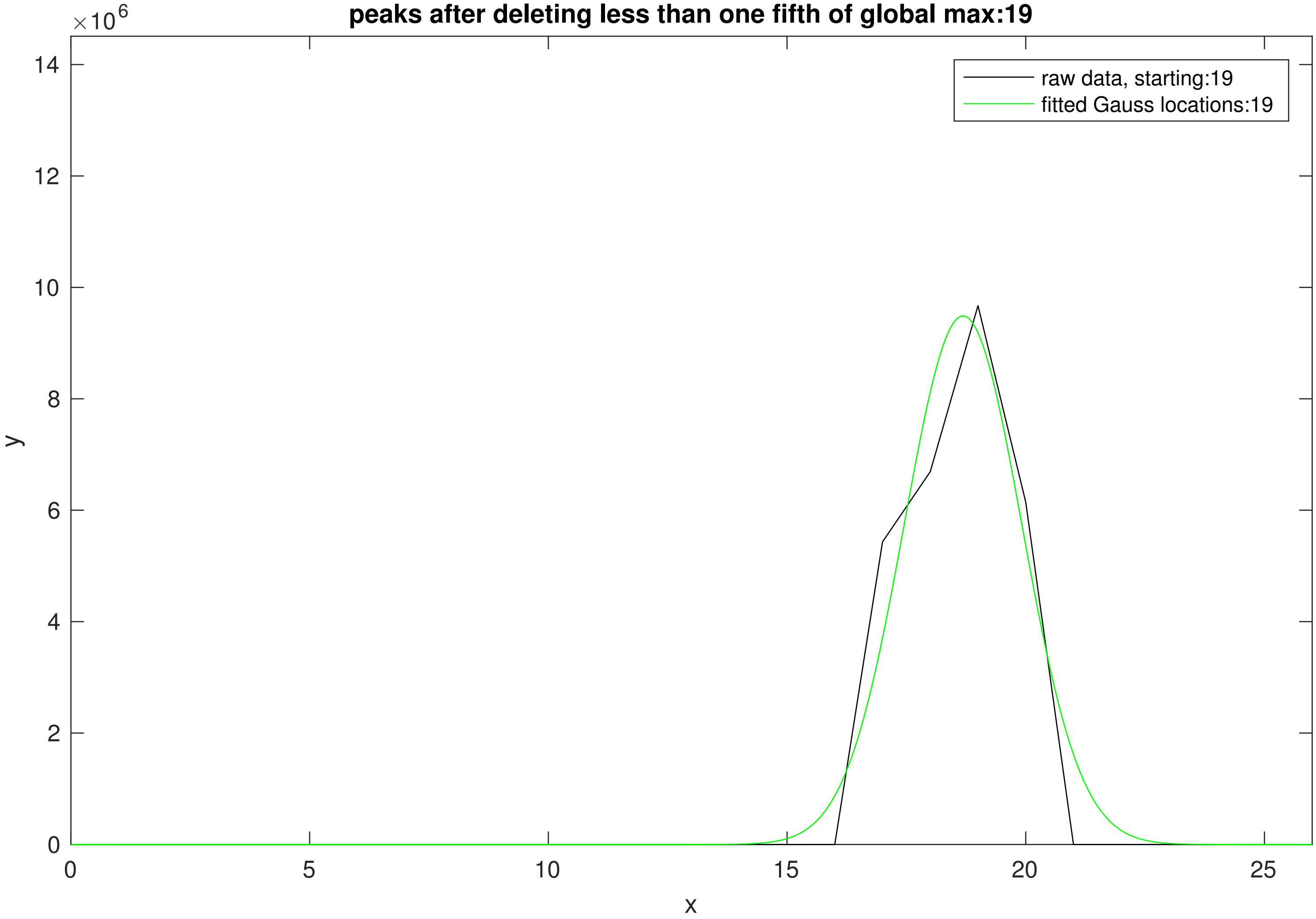

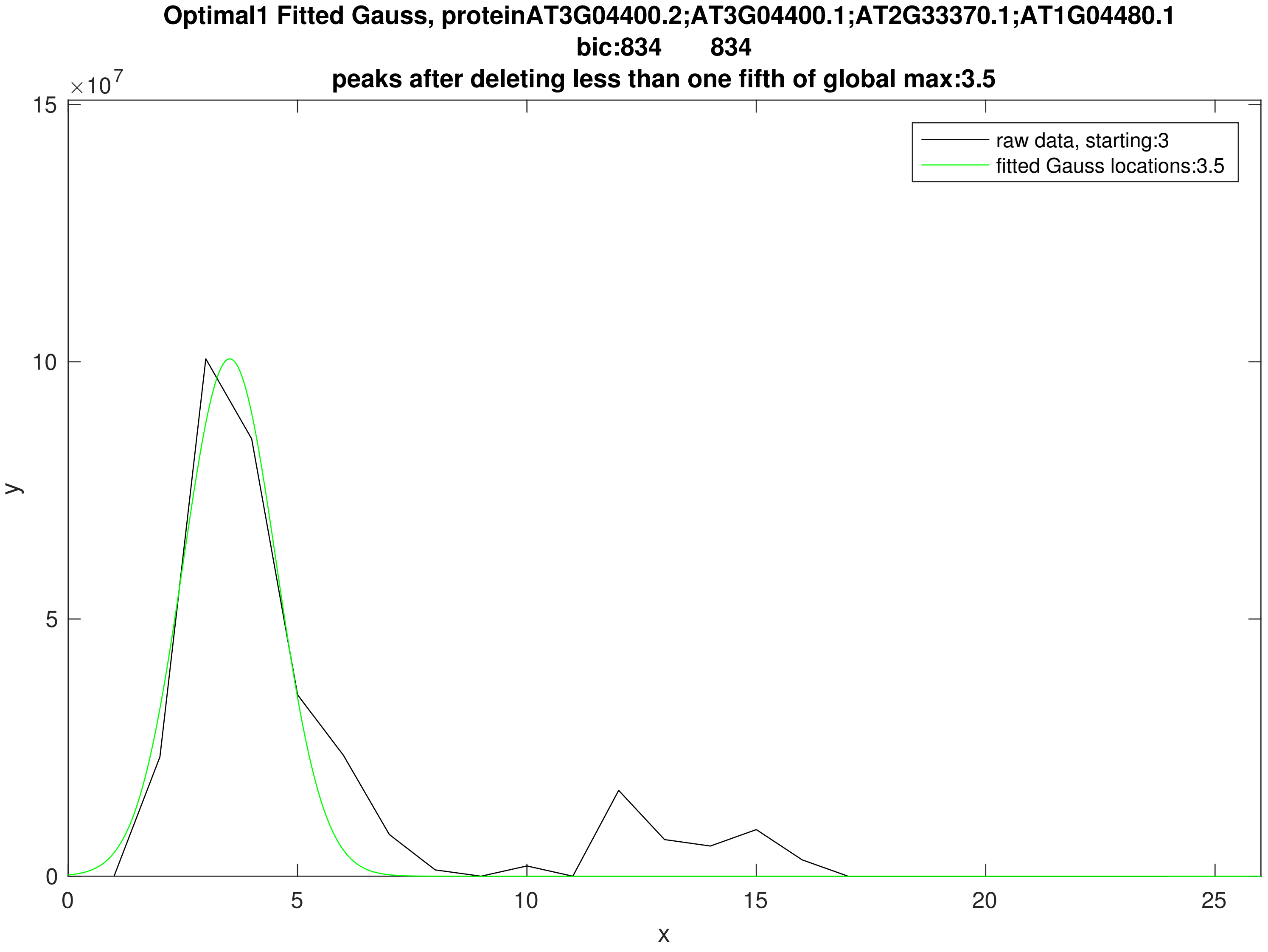
Optimal1 Fitted Gauss, proteinAT3G04400.2;AT3G04400.1;AT2G33370.1;AT1G04480.1
bic:834 834
peaks after deleting less than one fifth of global max:3.5
×10^7
15
10
5
0
y
0
5
10
15
20
25
x
raw data, starting:3
fitted Gauss locations:3.5

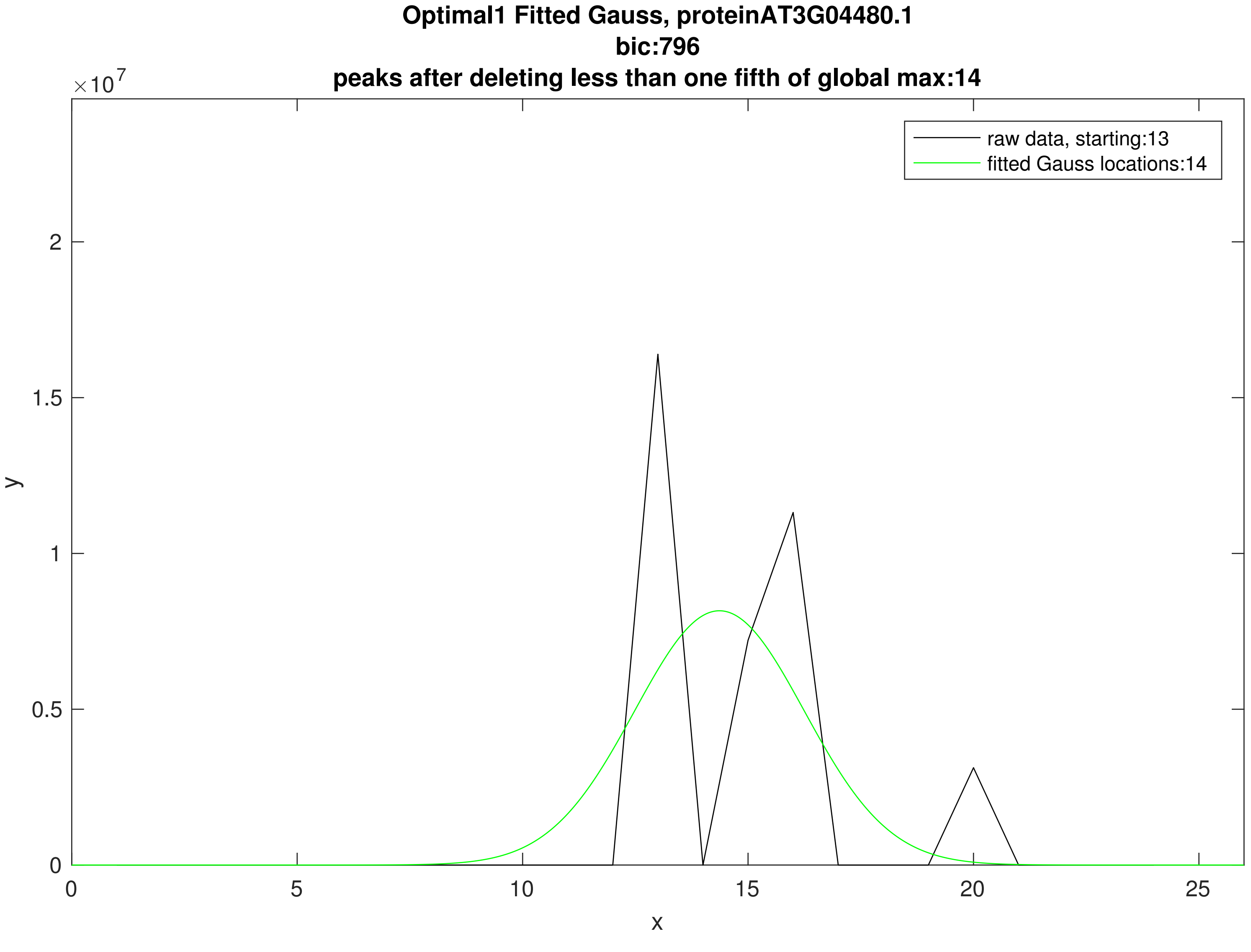

Optimal1 Fitted Gauss, proteinAT3G04480.1
bic:796
peaks after deleting less than one fifth of global max:14
×10^7
raw data, starting:13
fitted Gauss locations:14
2
1.5
1
0.5
0
y
0
5
10
15
20
25
x

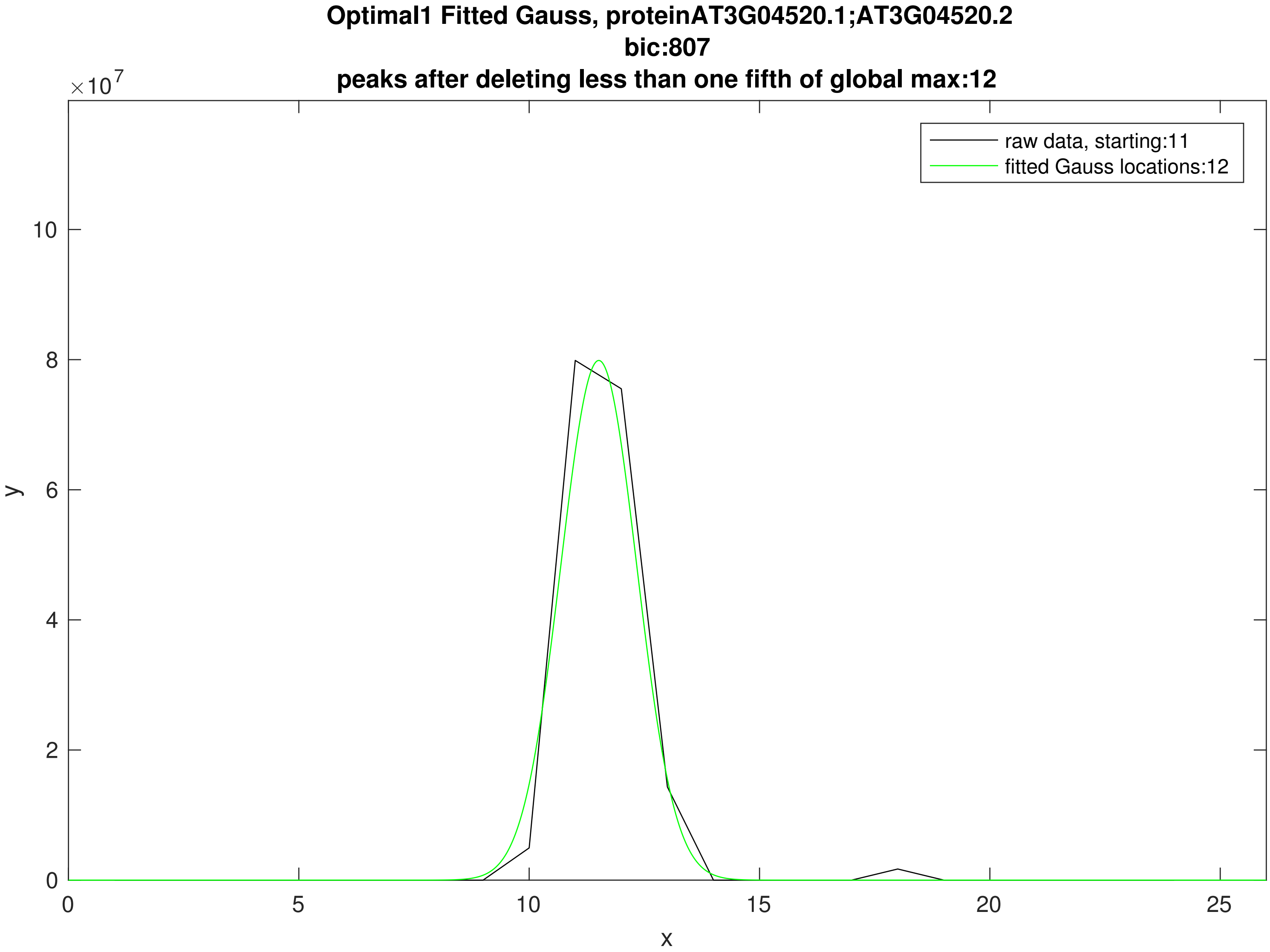

Optimal1 Fitted Gauss, proteinAT3G04520.1;AT3G04520.2
bic:807
peaks after deleting less than one fifth of global max:12
×10^7
raw data, starting:11
fitted Gauss locations:12
y
x
0
2
4
6
8
10
0
5
10
15
20
25

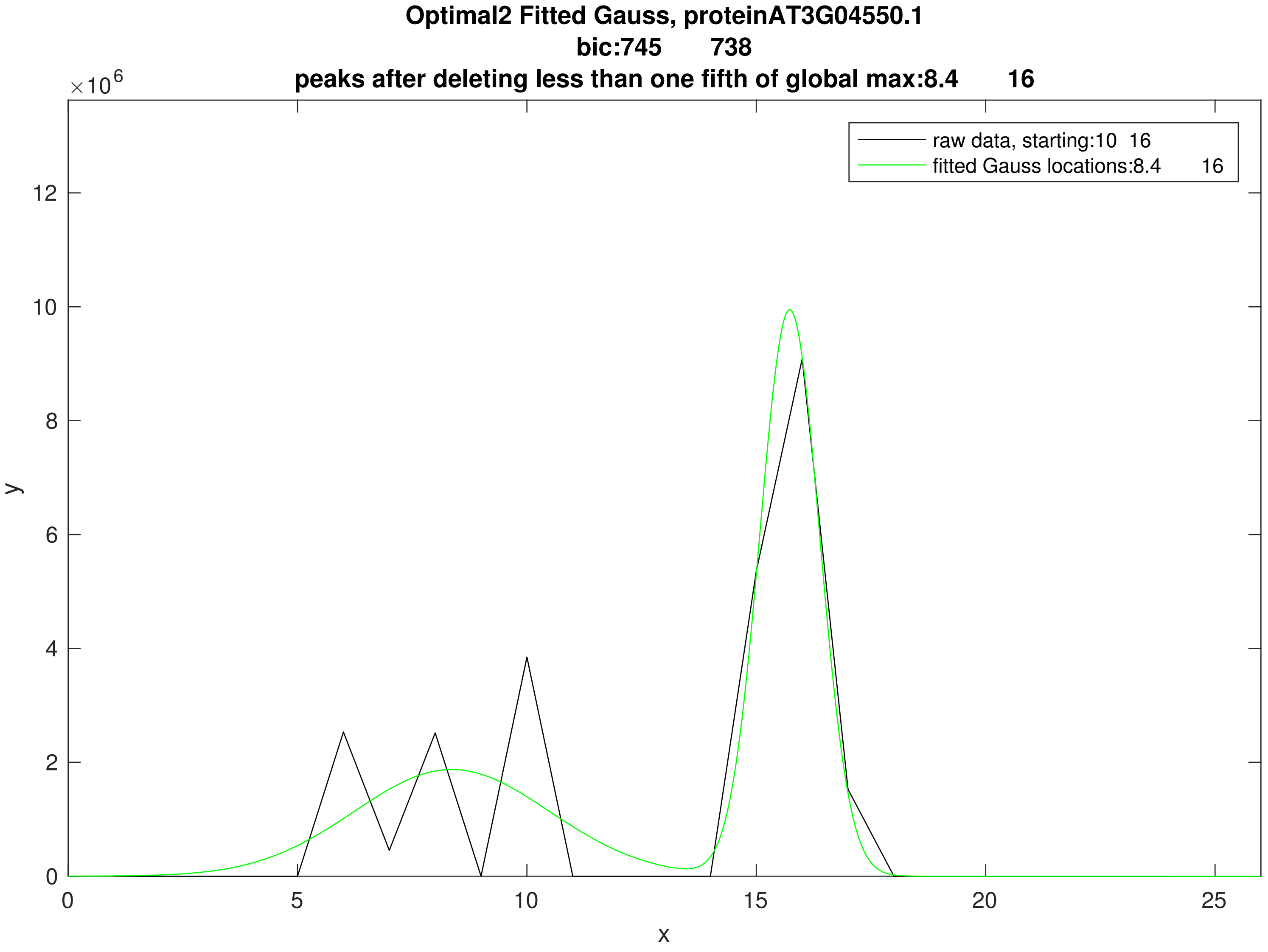
Optimal2 Fitted Gauss, proteinAT3G04550.1
bic:745 738
peaks after deleting less than one fifth of global max:8.4 16
raw data, starting:10 16
fitted Gauss locations:8.4 16
×10^6
y
x
0
2
4
6
8
10
12
5
15
20
25

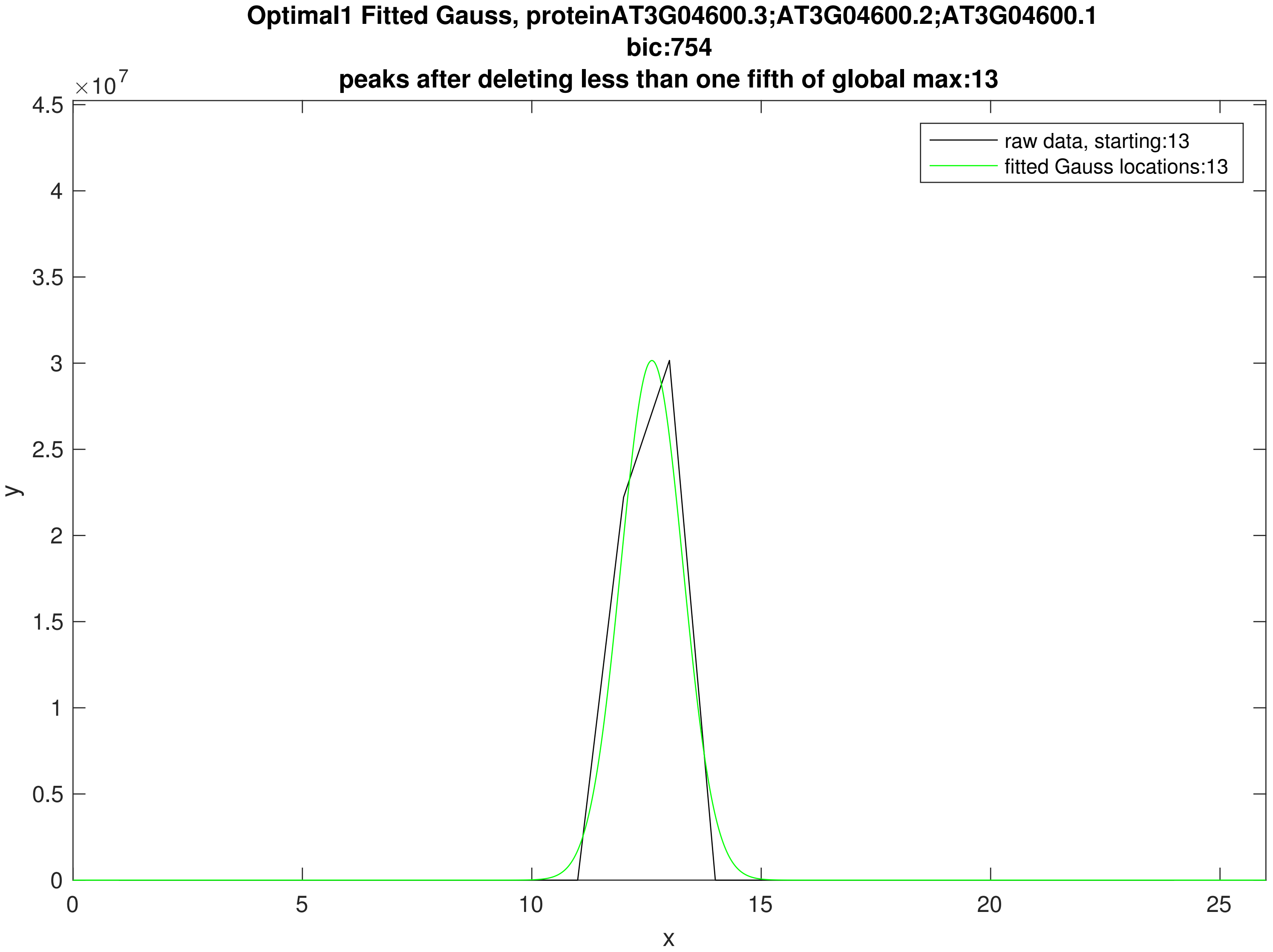
Optimal1 Fitted Gauss, proteinAT3G04600.3;AT3G04600.2;AT3G04600.1
bic:754
peaks after deleting less than one fifth of global max:13
raw data, starting:13
fitted Gauss locations:13
×10^7
4.5
4
3.5
3
2.5
2
1.5
1
0.5
0
y
0
5
10
15
20
25
x

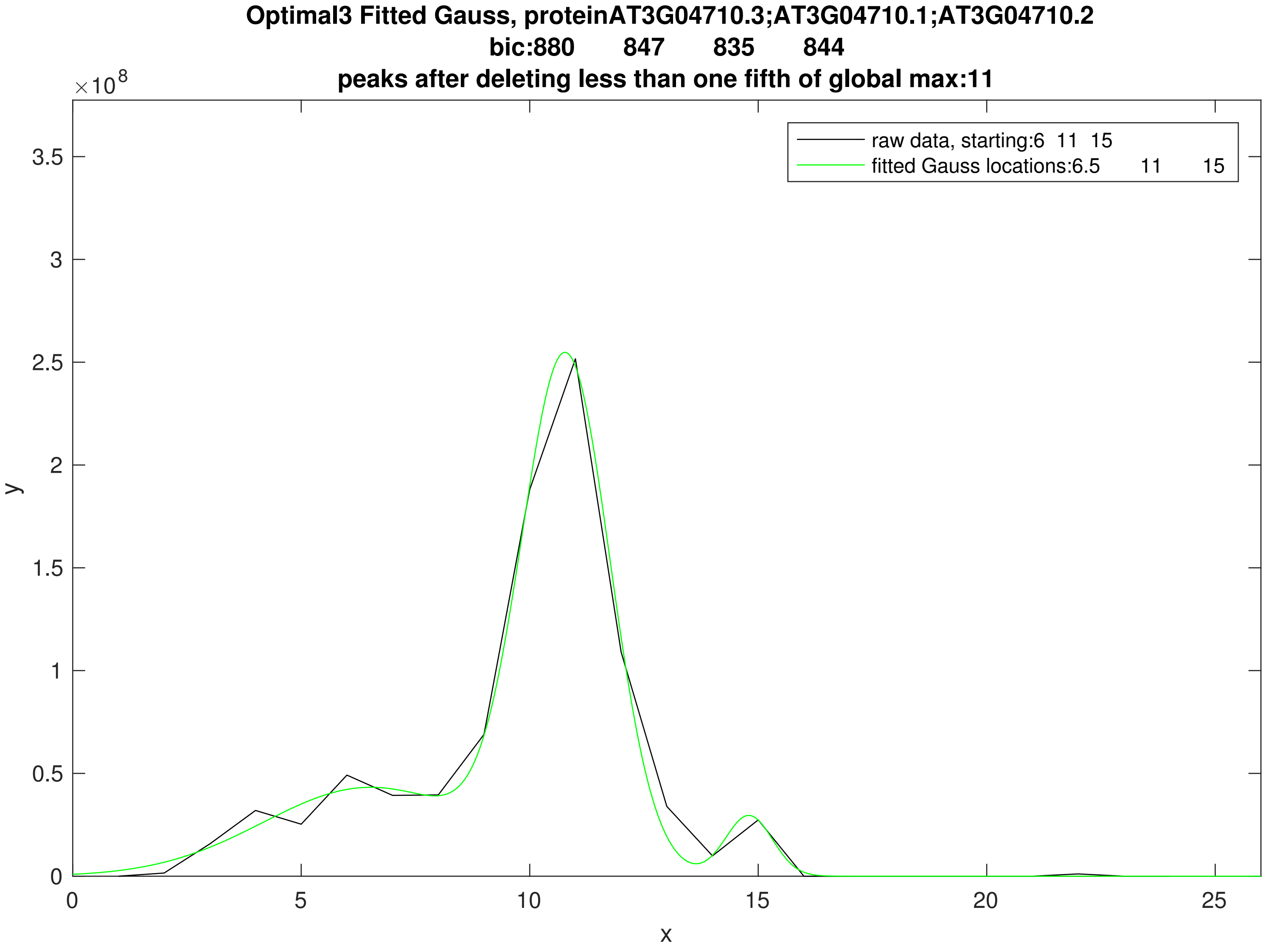

Optimal3 Fitted Gauss, proteinAT3G04710.3;AT3G04710.1;AT3G04710.2
bic:880 847 835 844
peaks after deleting less than one fifth of global max:11
×10^8
3.5
3
2.5
2
1.5
1
0.5
0
y
0
5
10
15
20
25
x
raw data, starting:6 11 15
fitted Gauss locations:6.5 11 15

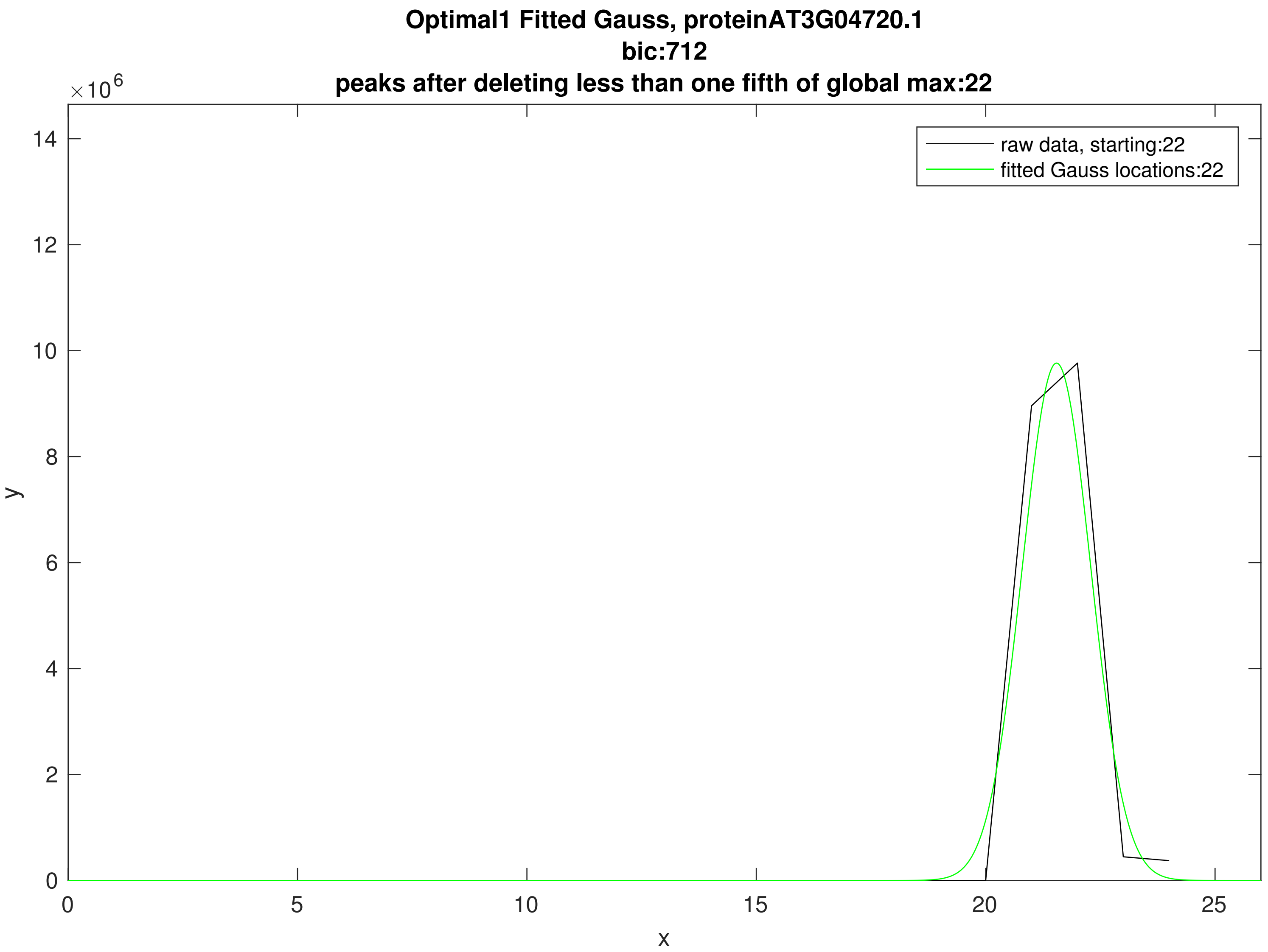
Optimal1 Fitted Gauss, proteinAT3G04720.1
bic:712
peaks after deleting less than one fifth of global max:22
×10^6
raw data, starting:22
fitted Gauss locations:22
y
x
0
2
4
6
8
10
12
14
5
15
20
25

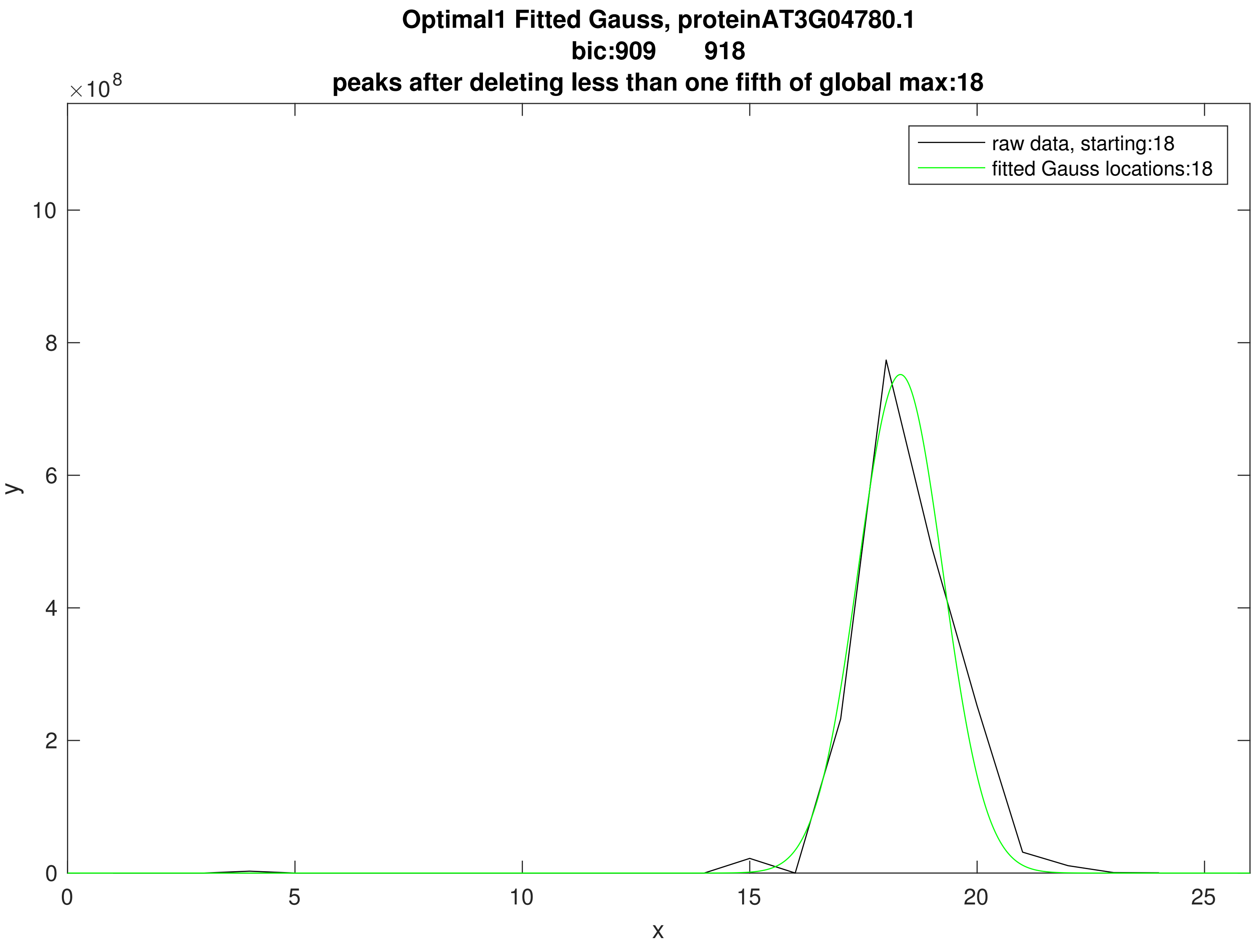

Optimal1 Fitted Gauss, proteinAT3G04780.1
bic:909 918
peaks after deleting less than one fifth of global max:18
×10^8
raw data, starting:18
fitted Gauss locations:18
10
8
6
4
2
0
y
0
5
10
15
20
25
x

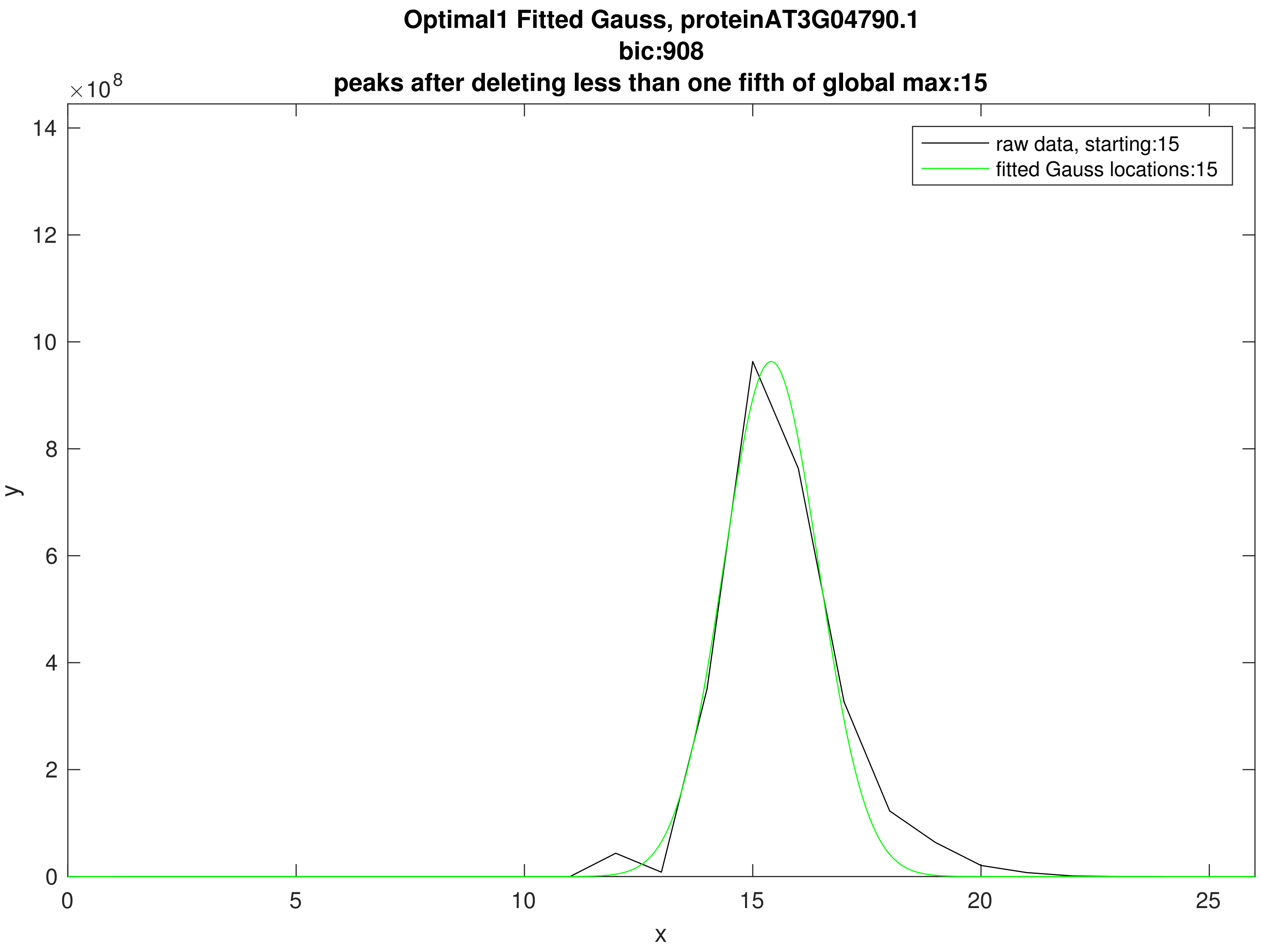

Optimal1 Fitted Gauss, proteinAT3G04790.1
bic:908
peaks after deleting less than one fifth of global max:15
×10^8
14
12
10
8
6
4
2
0
y
0
5
10
15
20
25
x
raw data, starting:15
fitted Gauss locations:15

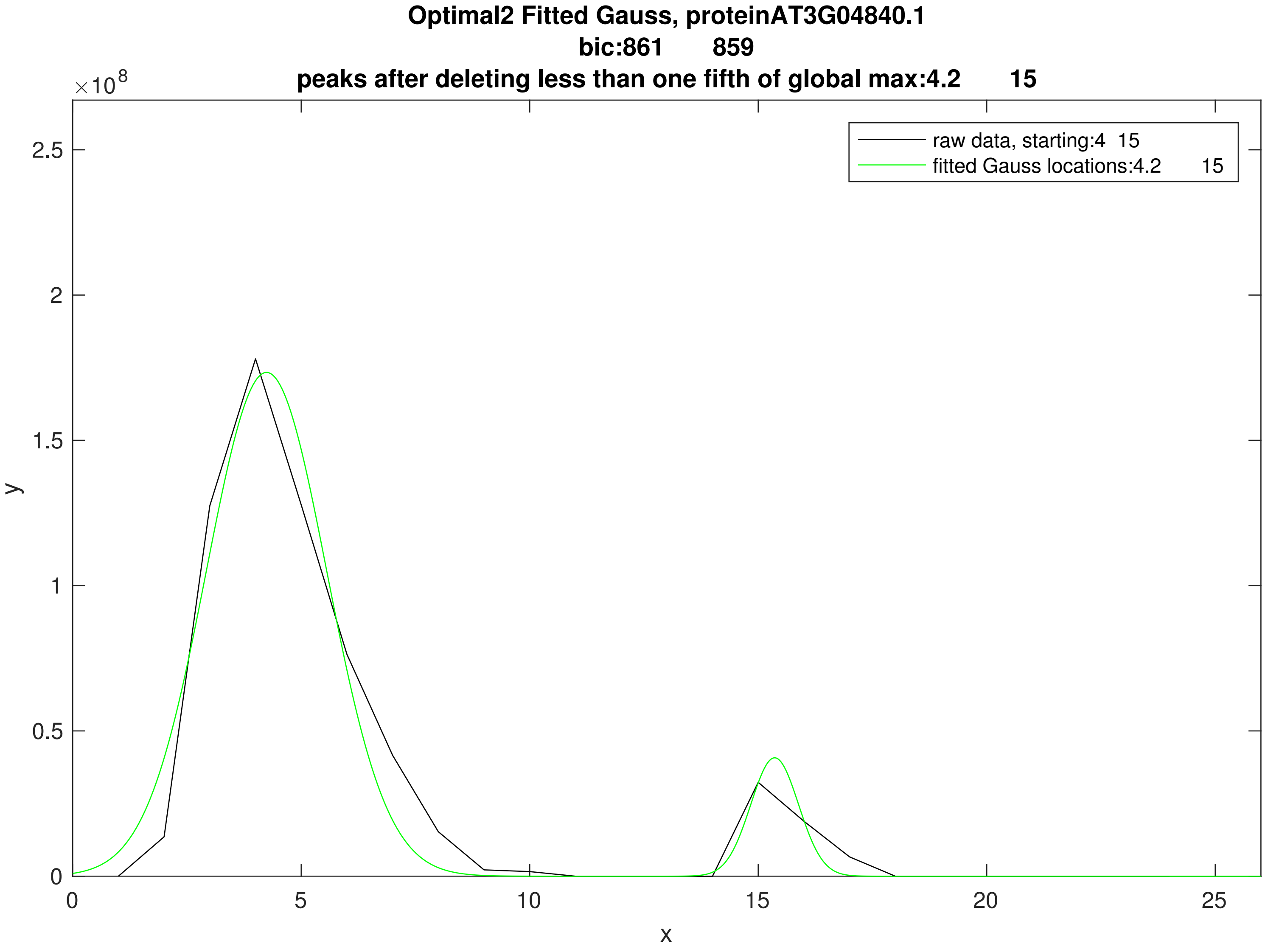
Optimal2 Fitted Gauss, proteinAT3G04840.1
bic:861 859
peaks after deleting less than one fifth of global max:4.2 15
×10^8
raw data, starting:4 15
fitted Gauss locations:4.2 15
y
x
0
0.5
1
1.5
2
2.5
0
5
10
15
20
25

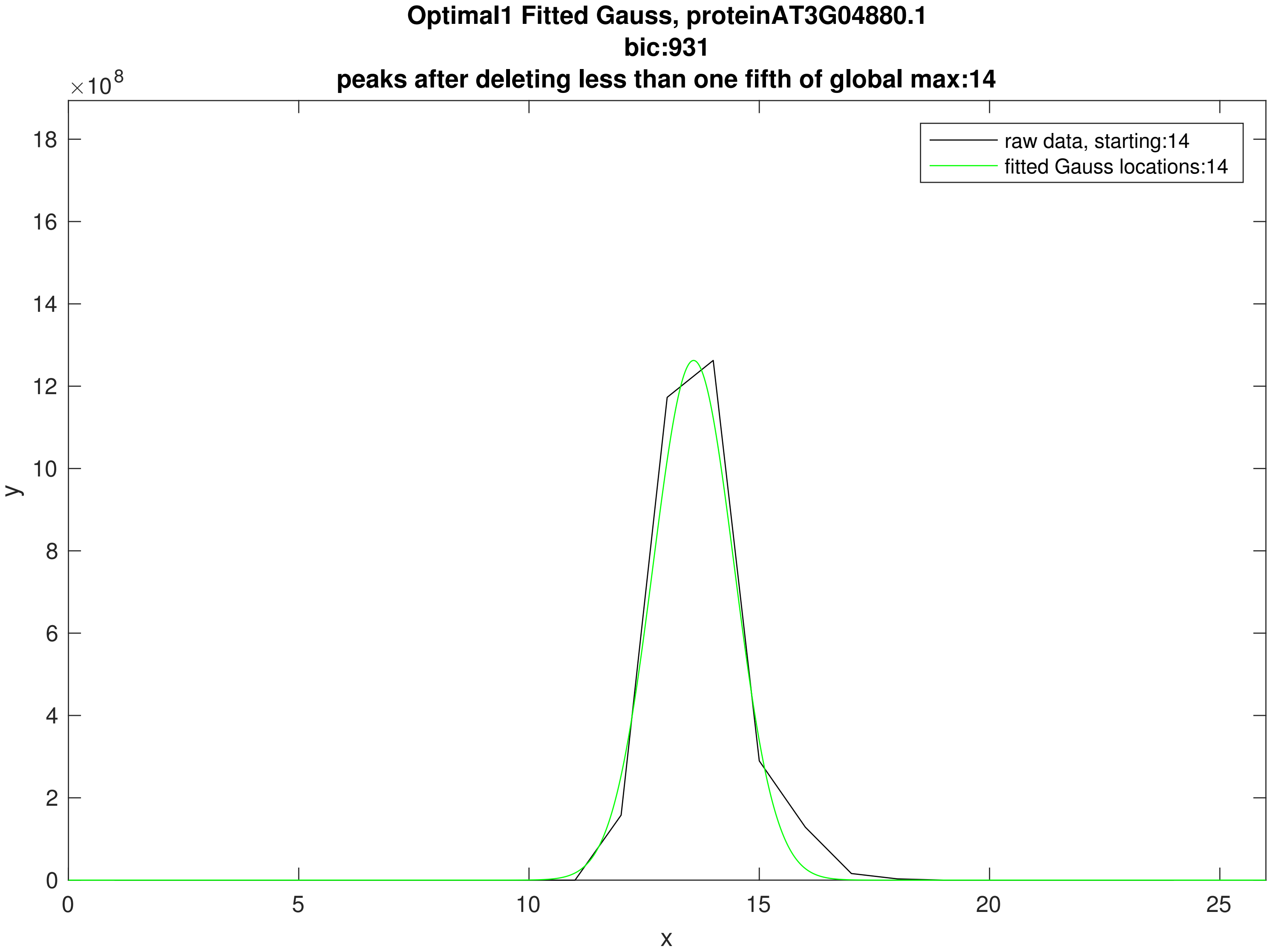

Optimal1 Fitted Gauss, proteinAT3G04880.1
bic:931
peaks after deleting less than one fifth of global max:14
×10^8
raw data, starting:14
fitted Gauss locations:14
y
x
0
2
4
6
8
10
12
14
16
18
0
5
10
15
20
25

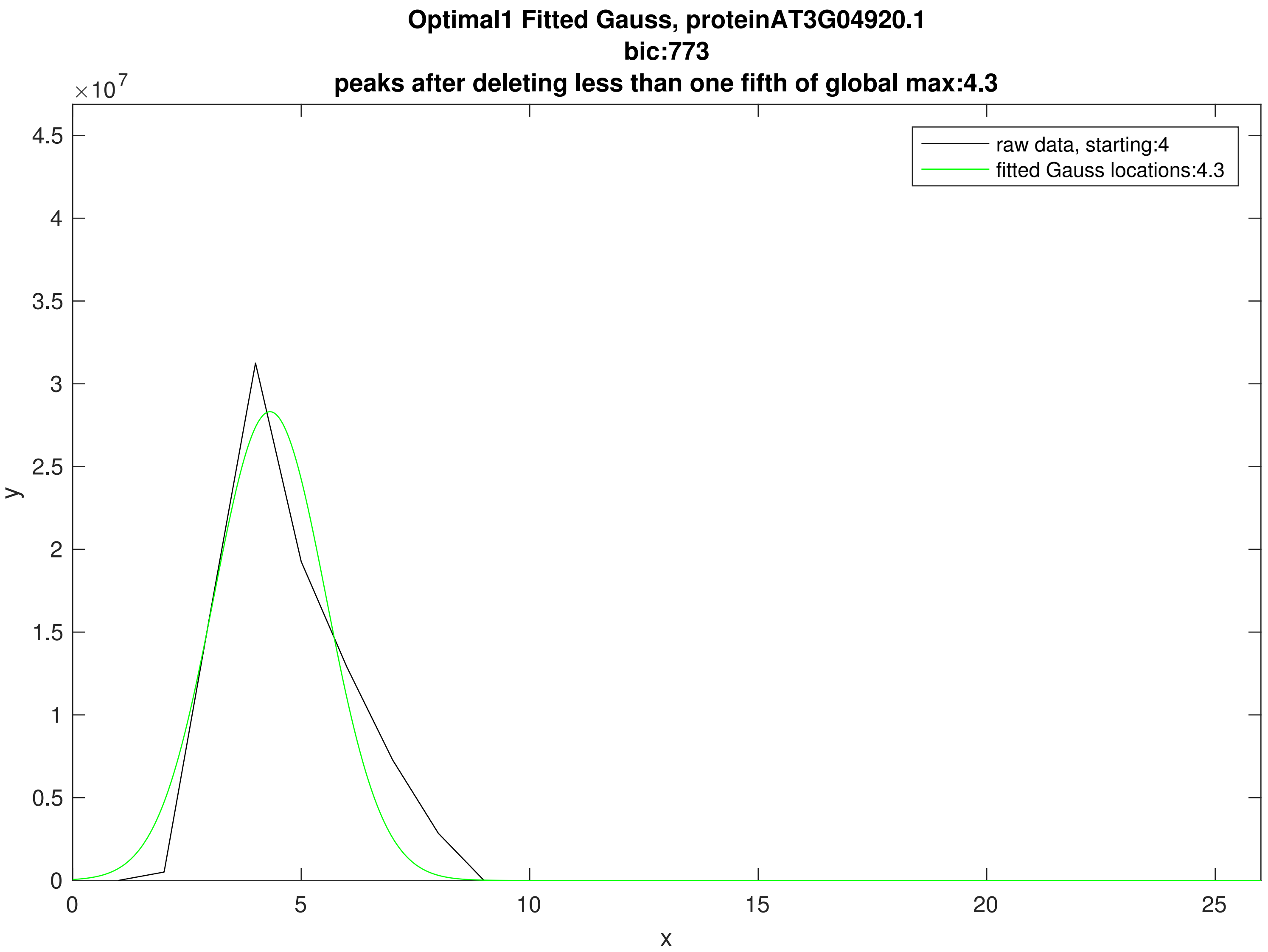
Optimal1 Fitted Gauss, proteinAT3G04920.1
bic:773
peaks after deleting less than one fifth of global max:4.3
×10^7
raw data, starting:4
fitted Gauss locations:4.3
y
x
0
0.5
1
1.5
2
2.5
3
3.5
4
4.5
5
10
15
20
25

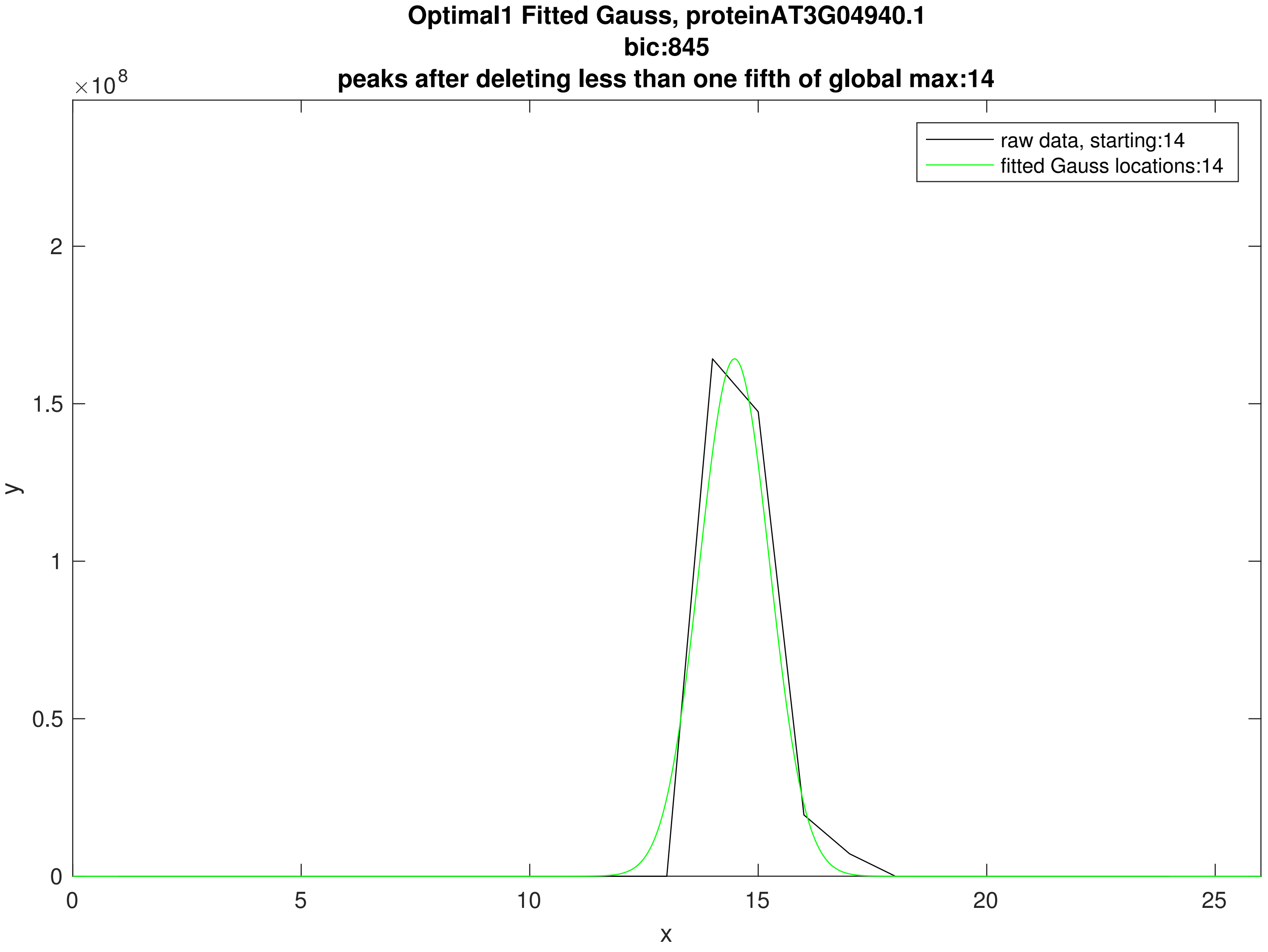
Optimal1 Fitted Gauss, proteinAT3G04940.1
bic:845
peaks after deleting less than one fifth of global max:14
×10^8
raw data, starting:14
fitted Gauss locations:14
2
1.5
1
0.5
0
y
0
5
10
15
20
25
x

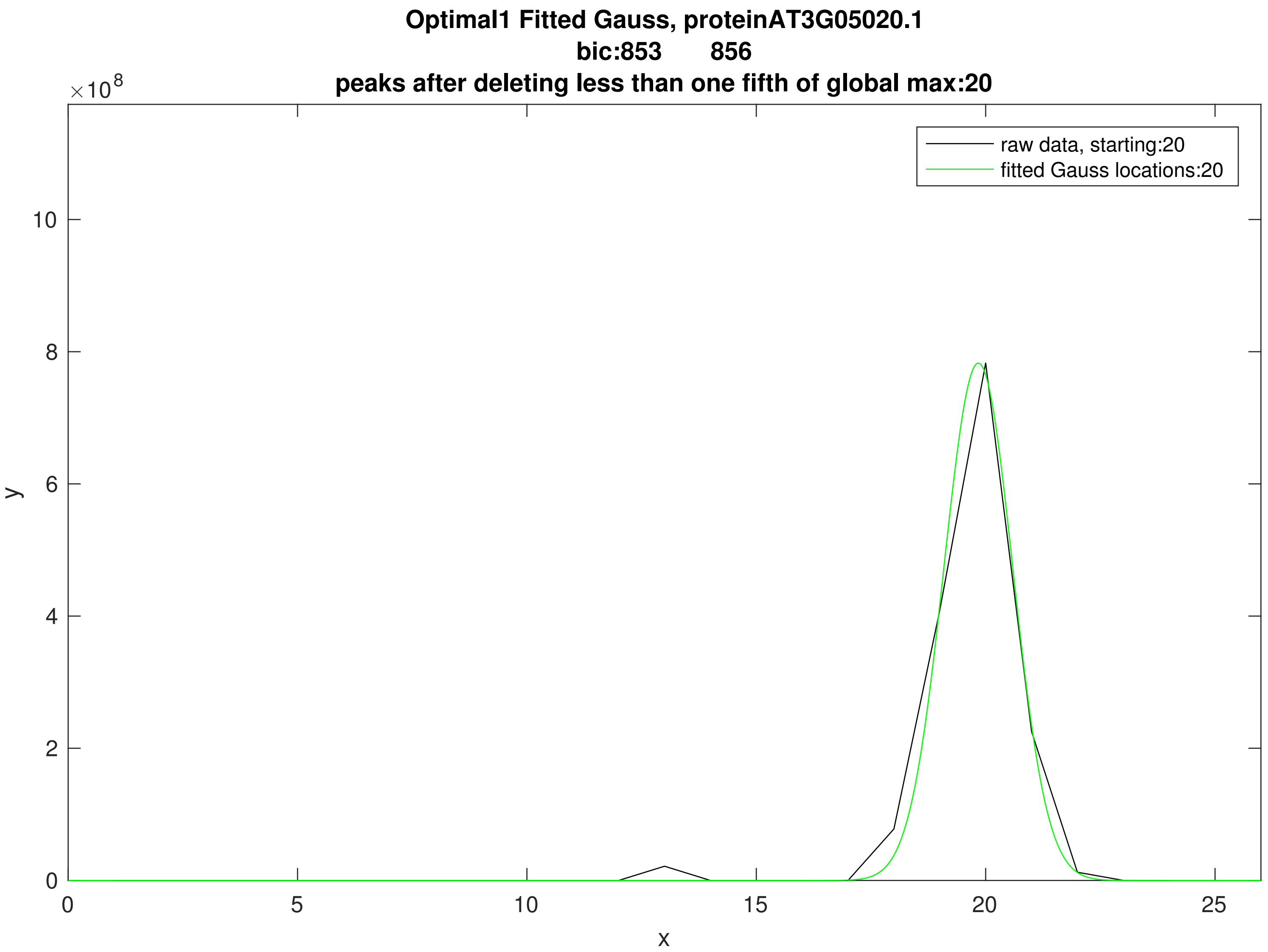
Optimal1 Fitted Gauss, proteinAT3G05020.1
bic:853 856
peaks after deleting less than one fifth of global max:20
×10^8
raw data, starting:20
fitted Gauss locations:20
y
x
0
2
4
6
8
10
0
5
10
15
20
25

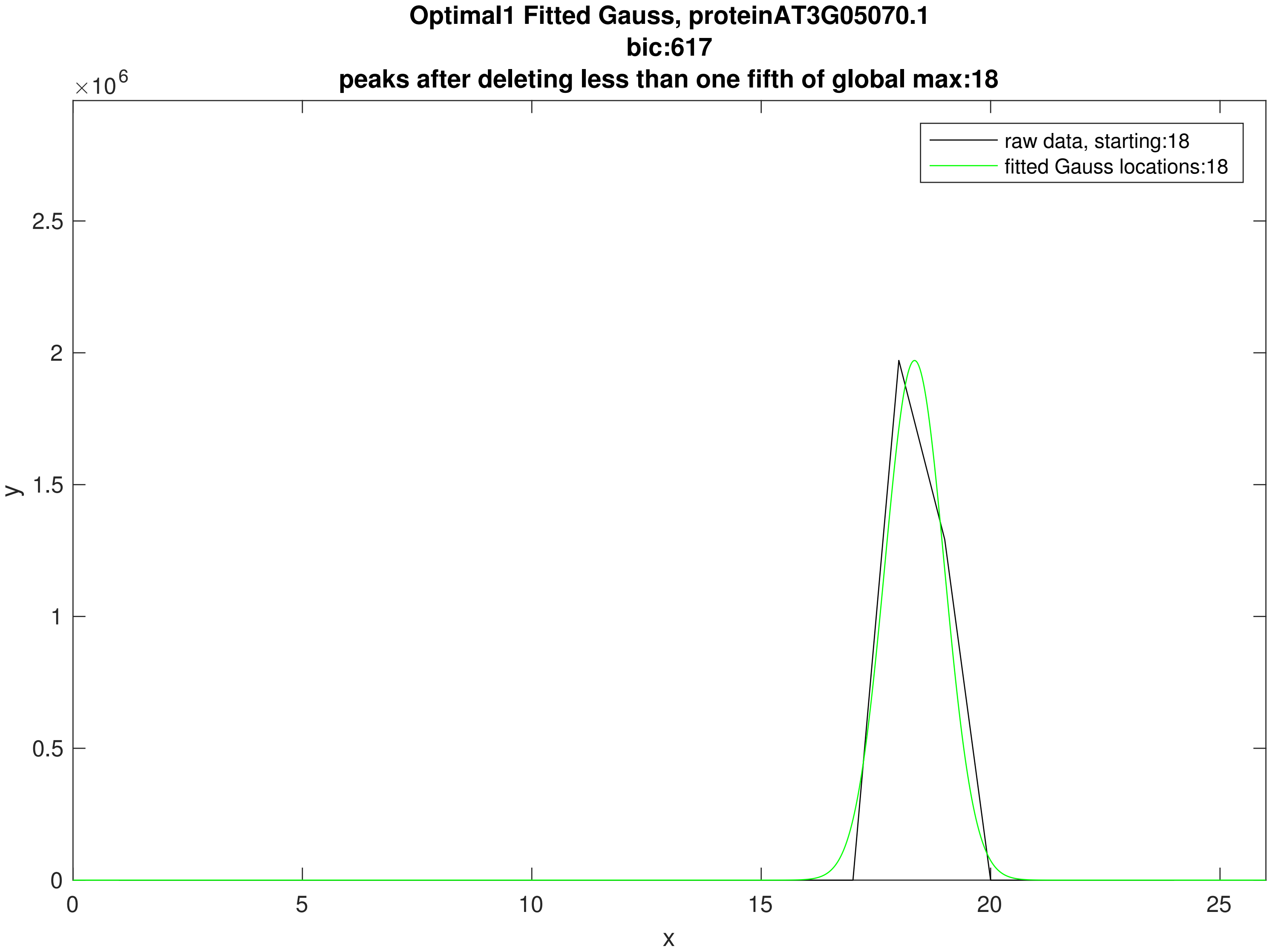
Optimal1 Fitted Gauss, proteinAT3G05070.1
bic:617
peaks after deleting less than one fifth of global max:18
×10^6
raw data, starting:18
fitted Gauss locations:18
y
x
0
0.5
1
1.5
2
2.5
0
5
10
15
20
25

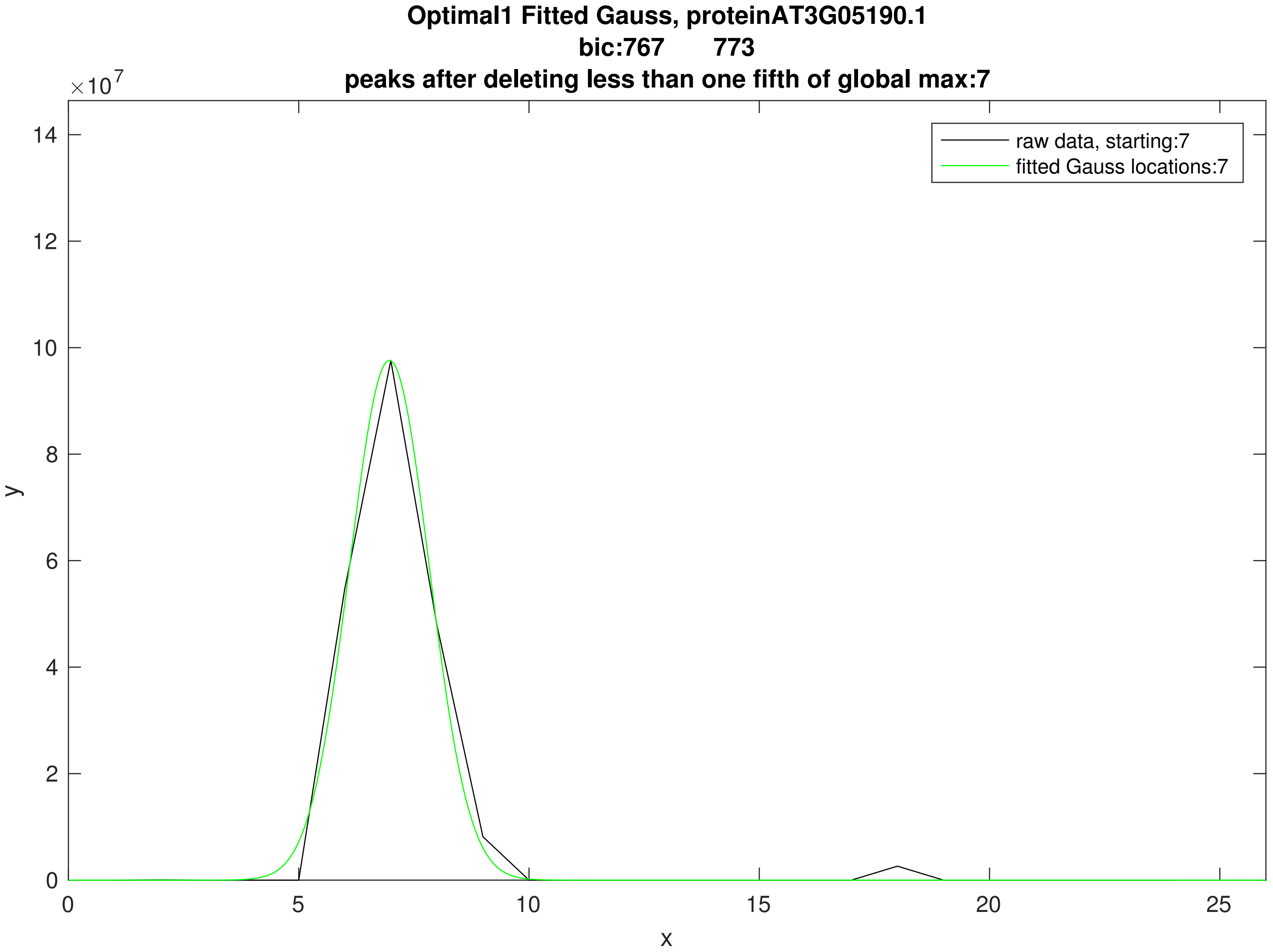
Optimal1 Fitted Gauss, proteinAT3G05190.1
bic:767 773
peaks after deleting less than one fifth of global max:7
×10^7
14
12
10
8
6
4
2
0
y
0
5
10
15
20
25
x
raw data, starting:7
fitted Gauss locations:7

Optimal1 Fitted Gauss, proteinAT3G05260.1
bic:914 915
peaks after deleting less than one fifth of global max:12

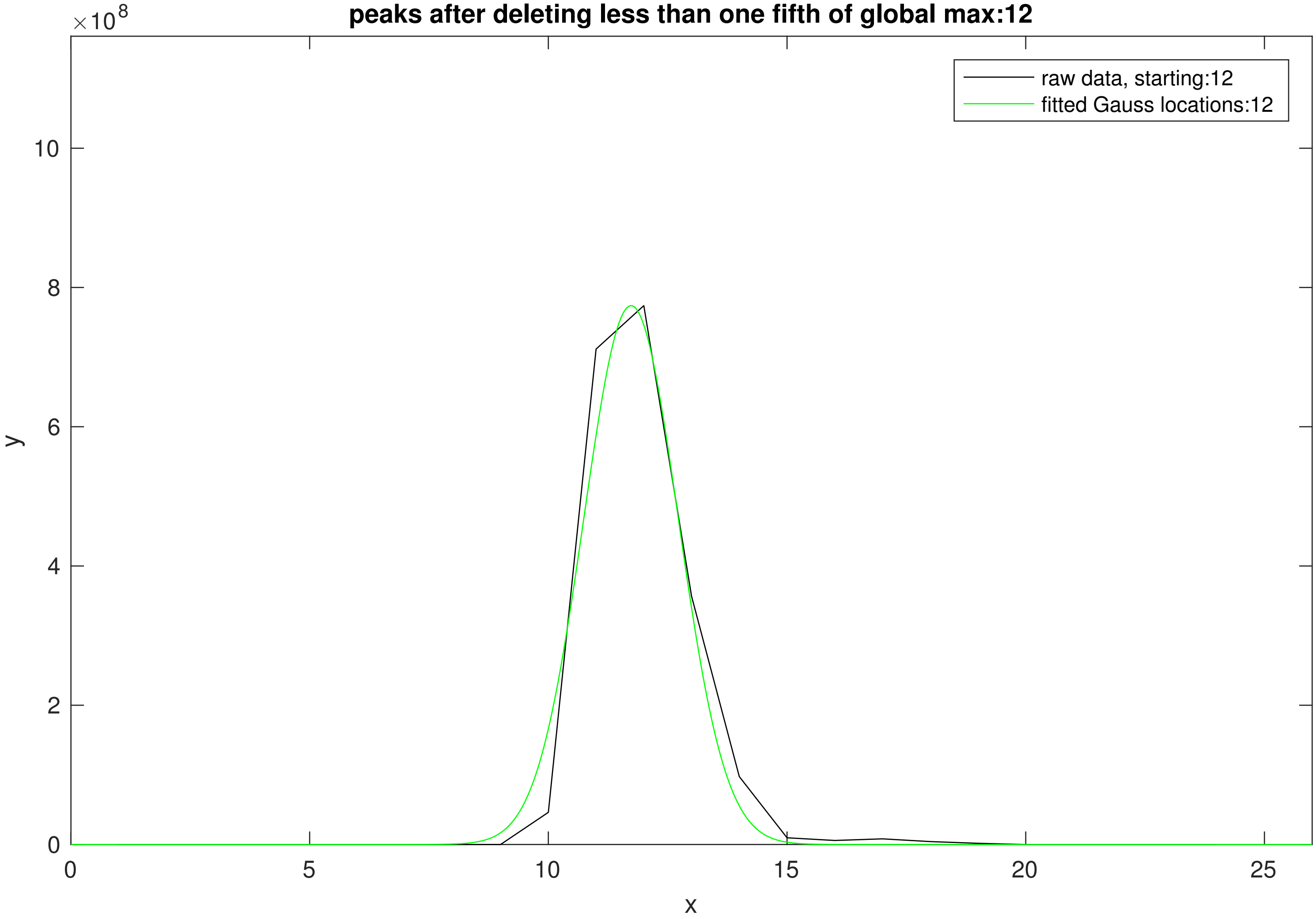

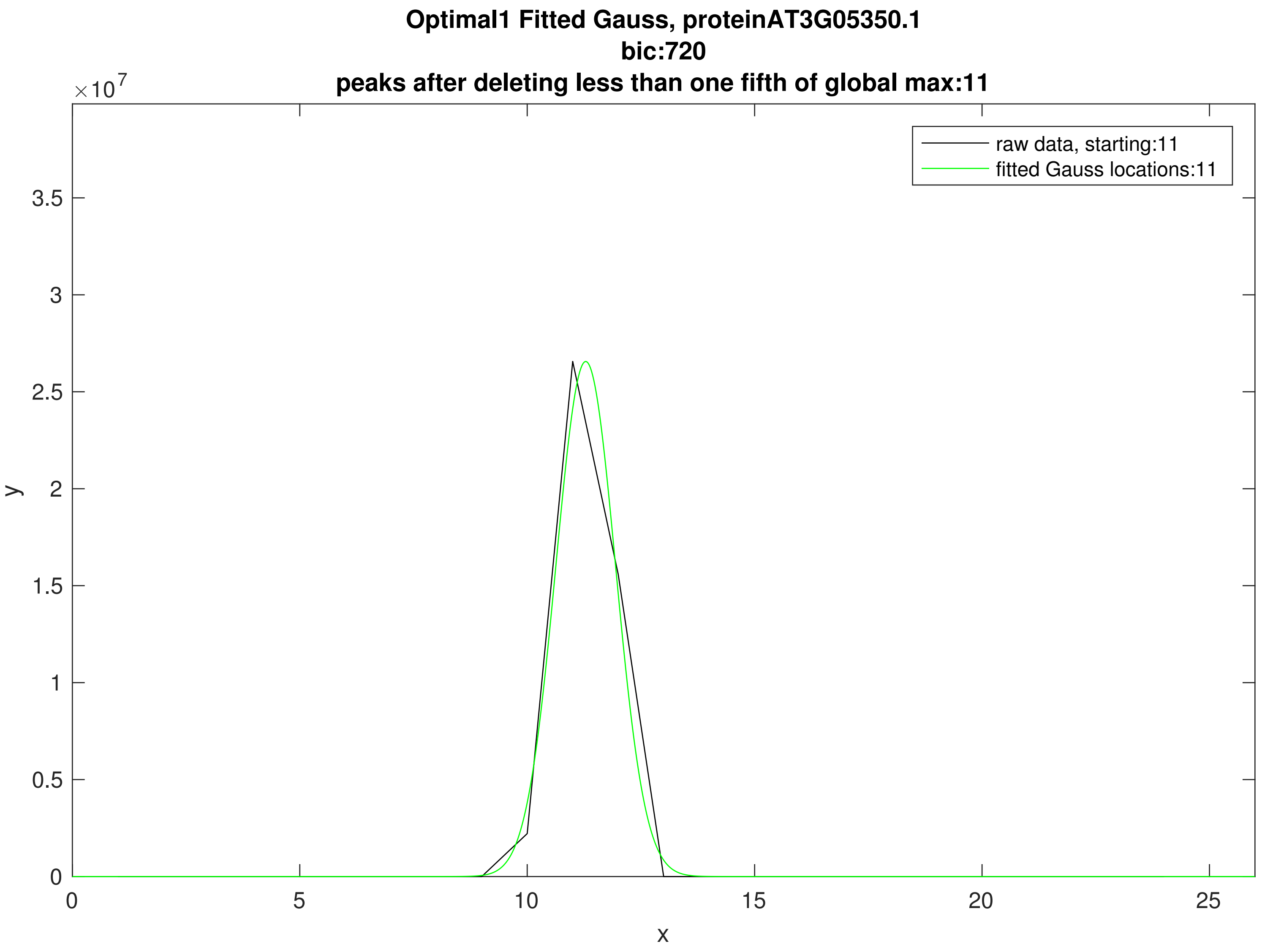
Optimal1 Fitted Gauss, proteinAT3G05350.1
bic:720
peaks after deleting less than one fifth of global max:11
×10^7
raw data, starting:11
fitted Gauss locations:11
y
x
0
0.5
1
1.5
2
2.5
3
3.5
5
10
15
20
25

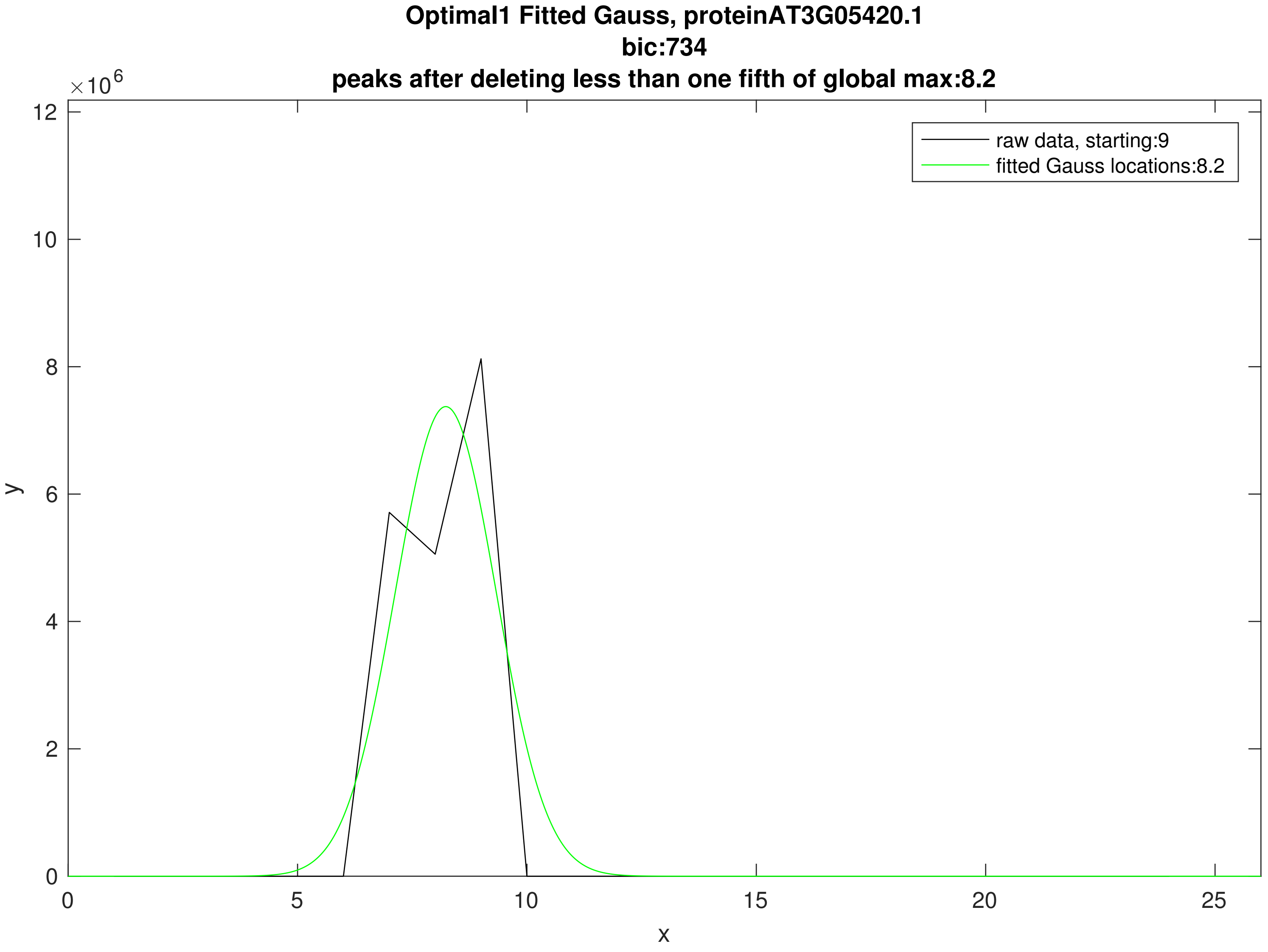

Optimal1 Fitted Gauss, proteinAT3G05420.1
bic:734
peaks after deleting less than one fifth of global max:8.2
×10^6
raw data, starting:9
fitted Gauss locations:8.2
y
x

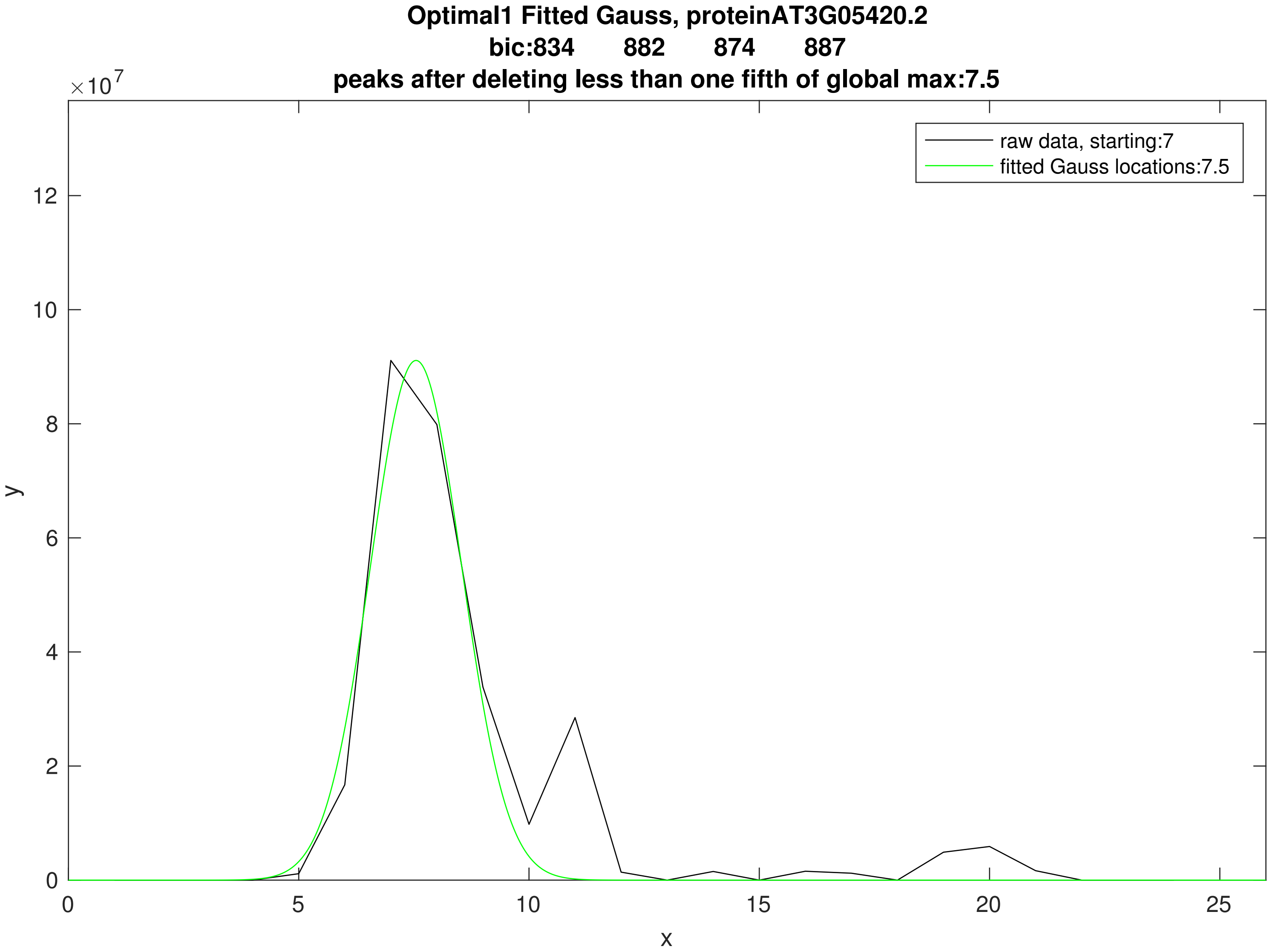
Optimal1 Fitted Gauss, proteinAT3G05420.2
bic:834 882 874 887
peaks after deleting less than one fifth of global max:7.5
×10^7
raw data, starting:7
fitted Gauss locations:7.5
y
x
0
2
4
6
8
10
12
0
5
10
15
20
25

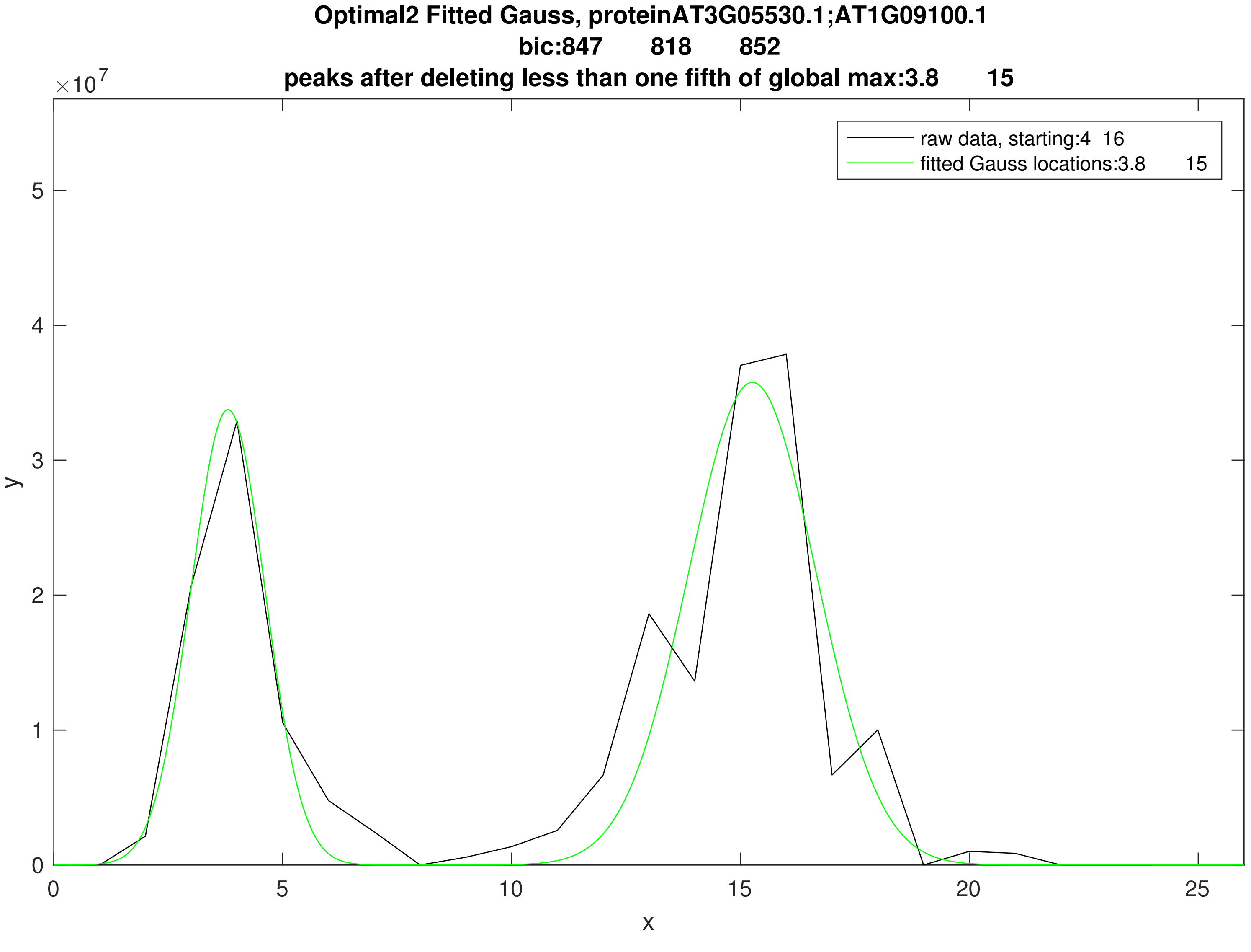
Optimal2 Fitted Gauss, proteinAT3G05530.1;AT1G09100.1
bic:847 818 852
peaks after deleting less than one fifth of global max:3.8 15
×10^7
raw data, starting:4 16
fitted Gauss locations:3.8 15
y
x
0
1
2
3
4
5
0
5
10
15
20
25

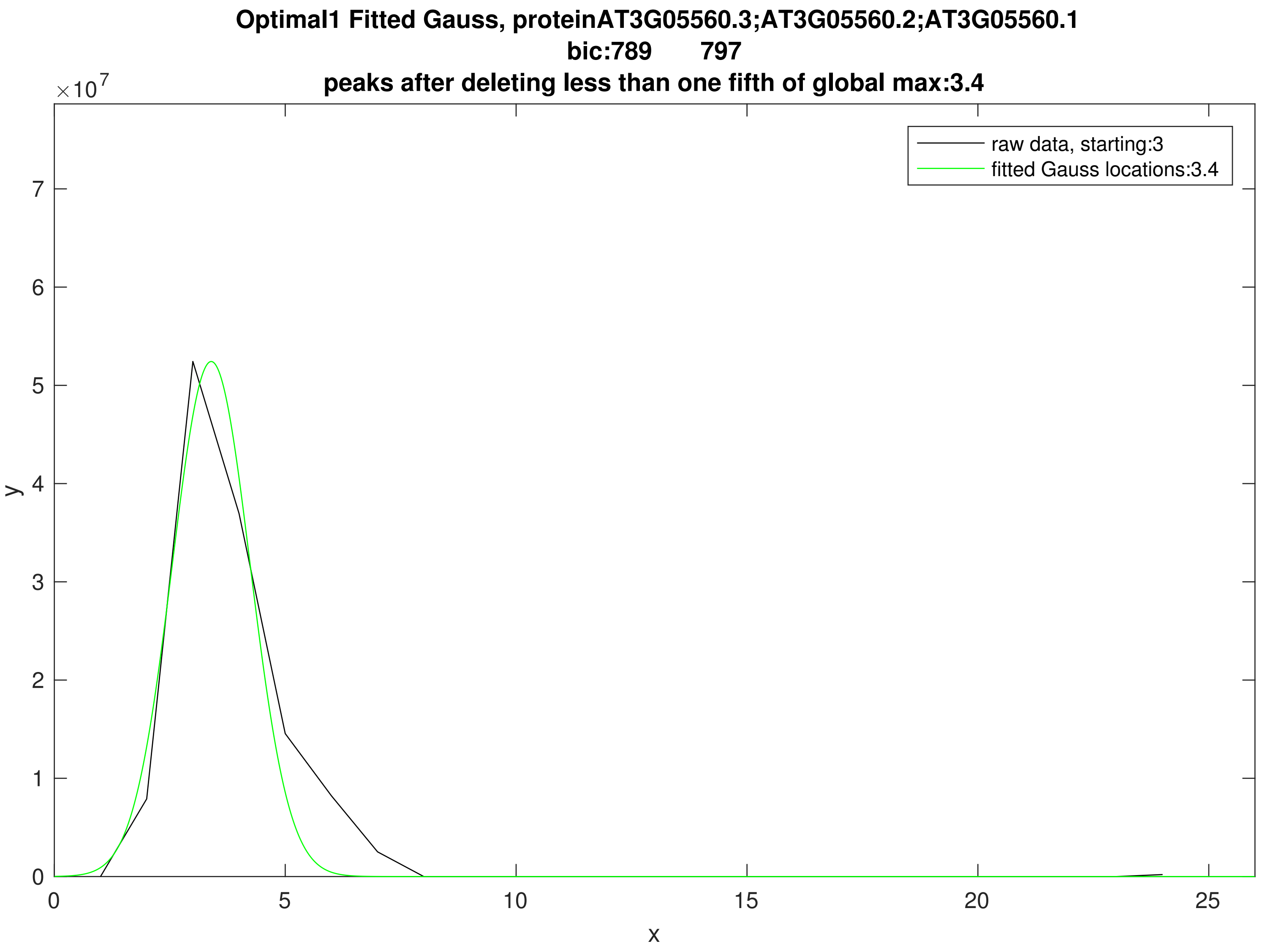
Optimal1 Fitted Gauss, proteinAT3G05560.3;AT3G05560.2;AT3G05560.1
bic:789 797
peaks after deleting less than one fifth of global max:3.4
×10^7
raw data, starting:3
fitted Gauss locations:3.4
y
x
0
1
2
3
4
5
6
7
0
5
10
15
20
25

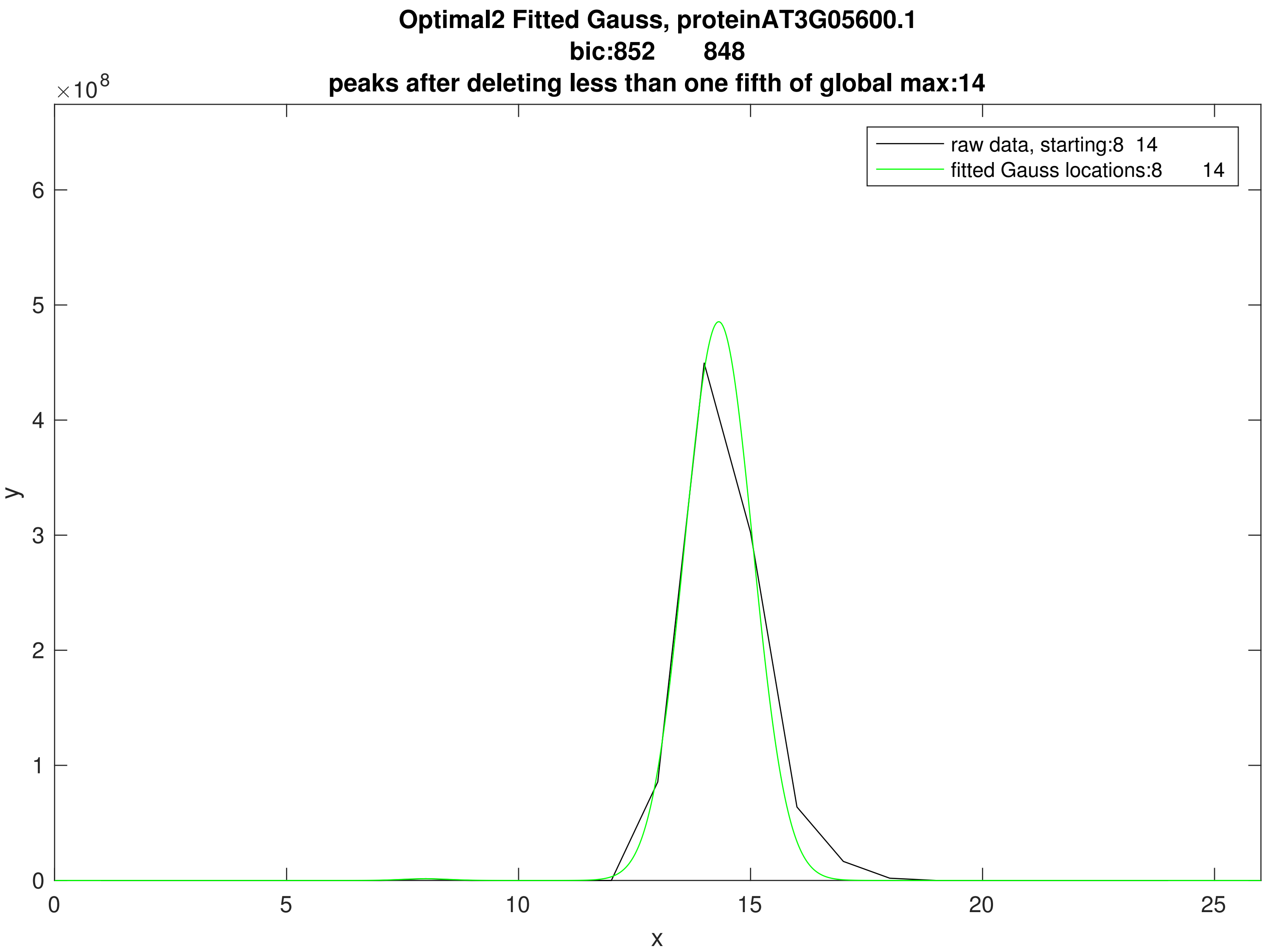
Optimal2 Fitted Gauss, proteinAT3G05600.1
bic:852 848
peaks after deleting less than one fifth of global max:14
raw data, starting:8 14
fitted Gauss locations:8 14
×10^8
y
x

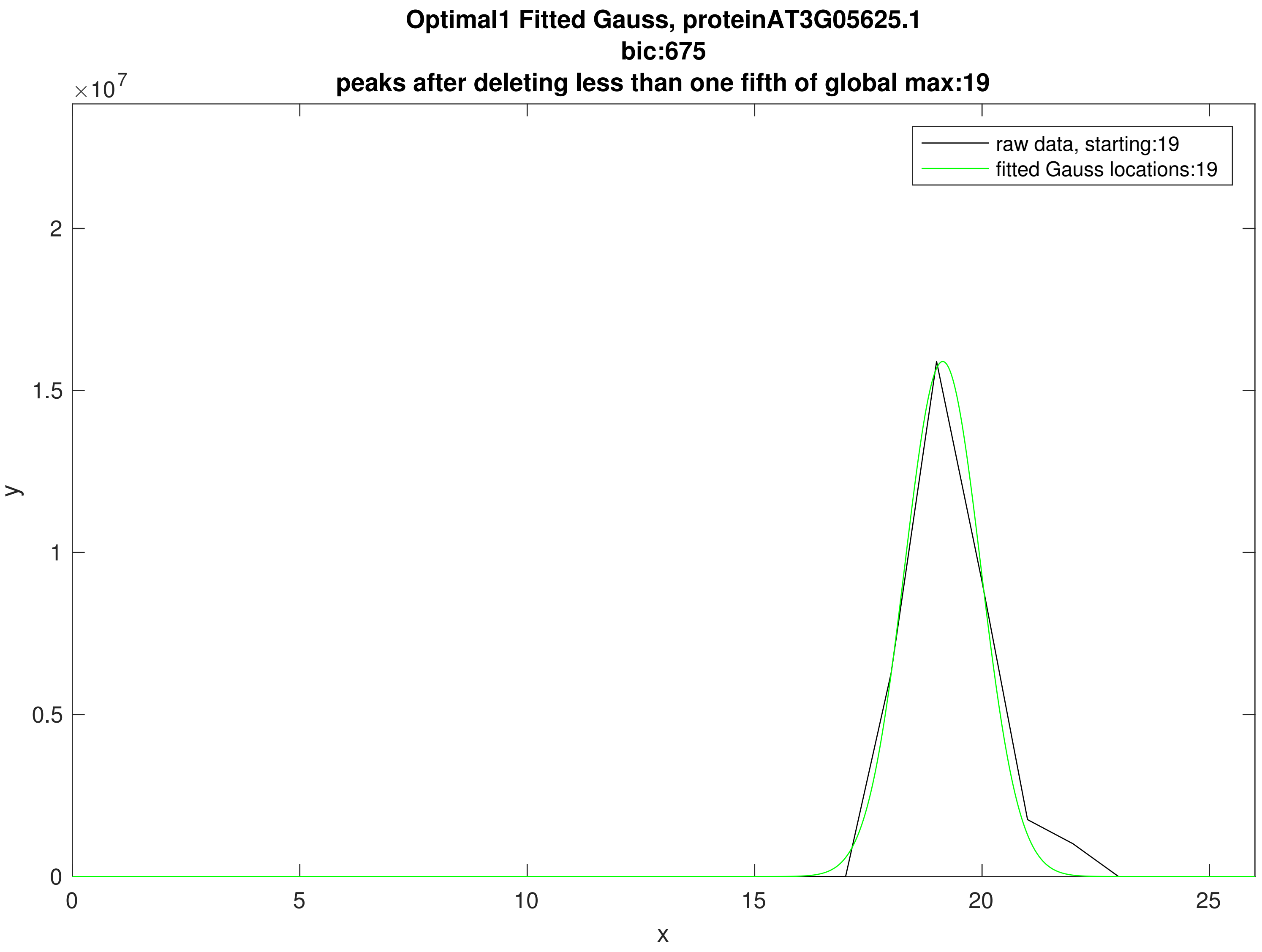

Optimal1 Fitted Gauss, proteinAT3G05625.1
bic:675
peaks after deleting less than one fifth of global max:19
×10^7
raw data, starting:19
fitted Gauss locations:19
2
1.5
1
0.5
0
y
0
5
10
15
20
25
x

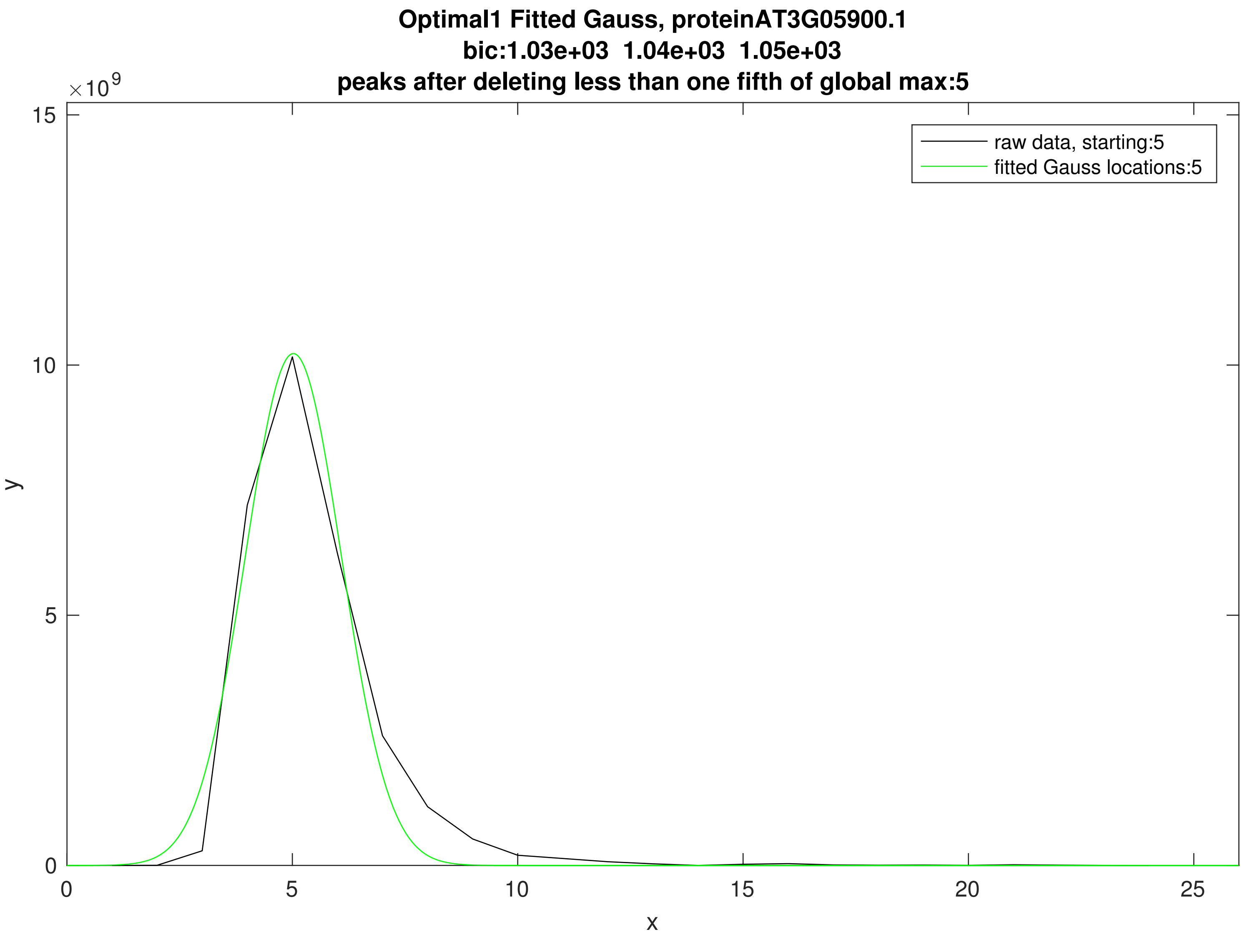
Optimal1 Fitted Gauss, proteinAT3G05900.1
bic:1.03e+03  1.04e+03  1.05e+03
peaks after deleting less than one fifth of global max:5
×10^9
15
10
5
0
y
0
5
10
15
20
25
x
raw data, starting:5
fitted Gauss locations:5

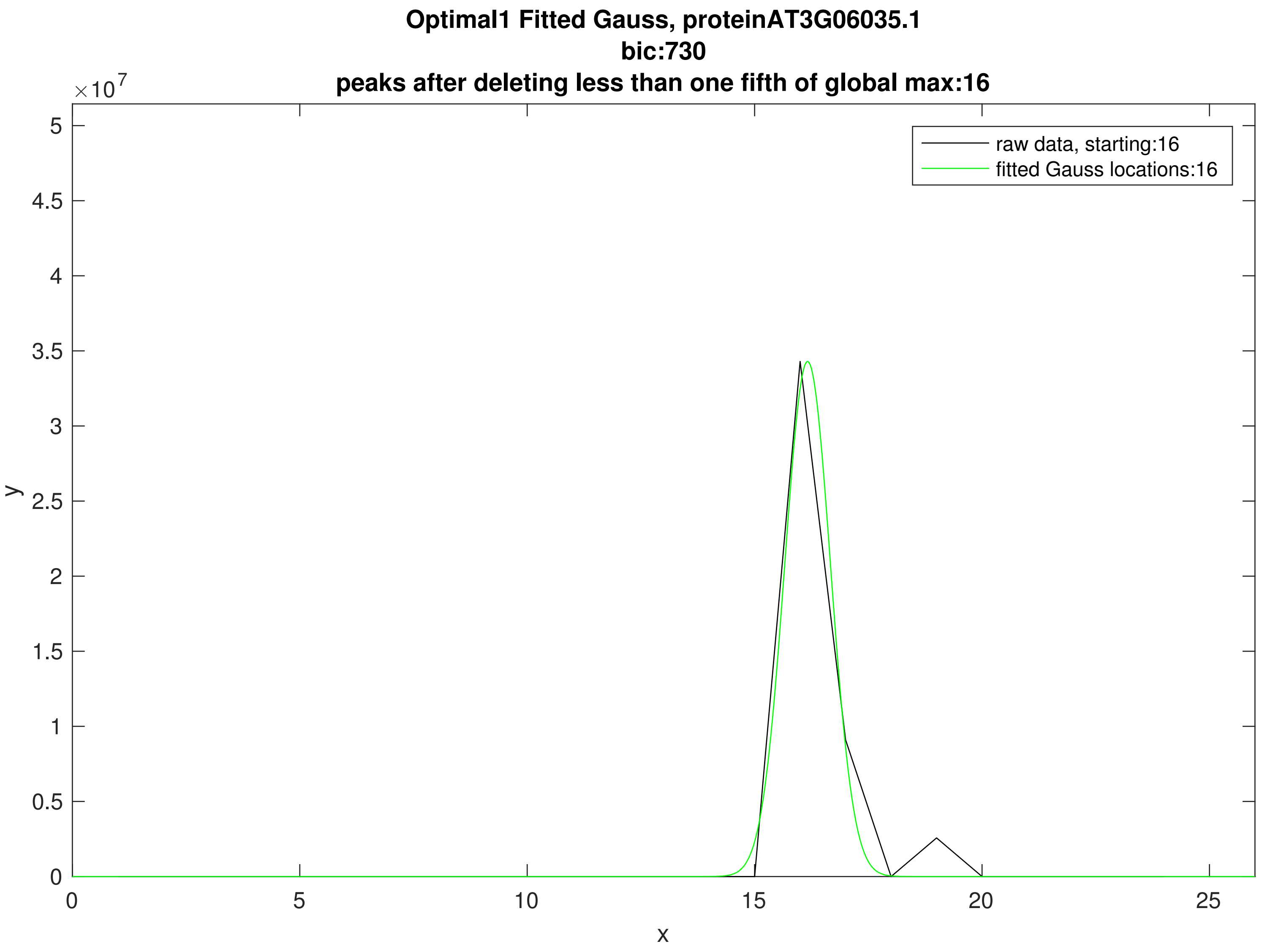

Optimal1 Fitted Gauss, proteinAT3G06035.1
bic:730
peaks after deleting less than one fifth of global max:16
×10^7
y
x
raw data, starting:16
fitted Gauss locations:16
0
0.5
1
1.5
2
2.5
3
3.5
4
4.5
5
0
5
10
15
20
25

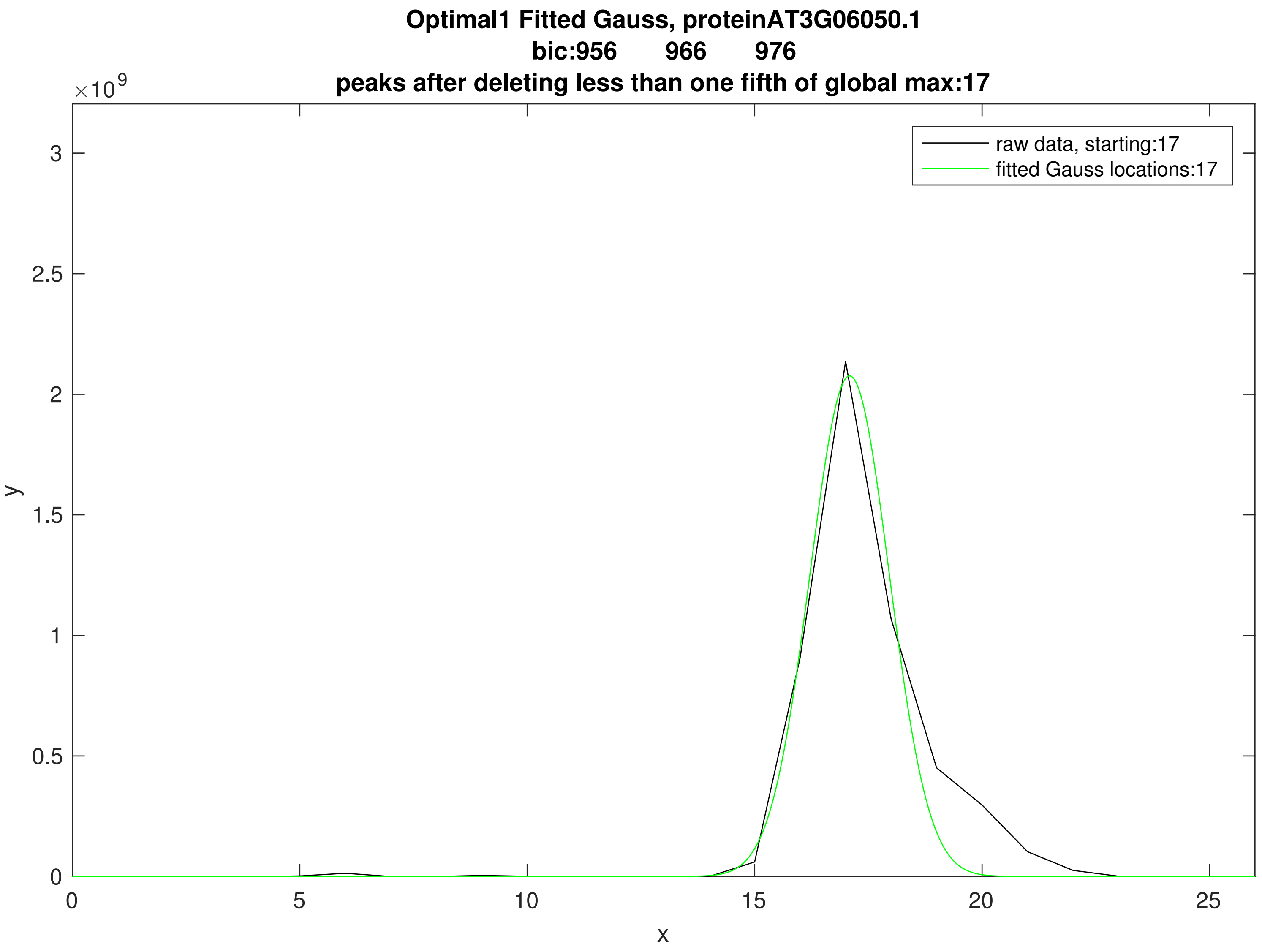

Optimal1 Fitted Gauss, proteinAT3G06050.1
bic:956 966 976
peaks after deleting less than one fifth of global max:17
×10^9
raw data, starting:17
fitted Gauss locations:17
y
x
0
0.5
1
1.5
2
2.5
3
0
5
10
15
20
25

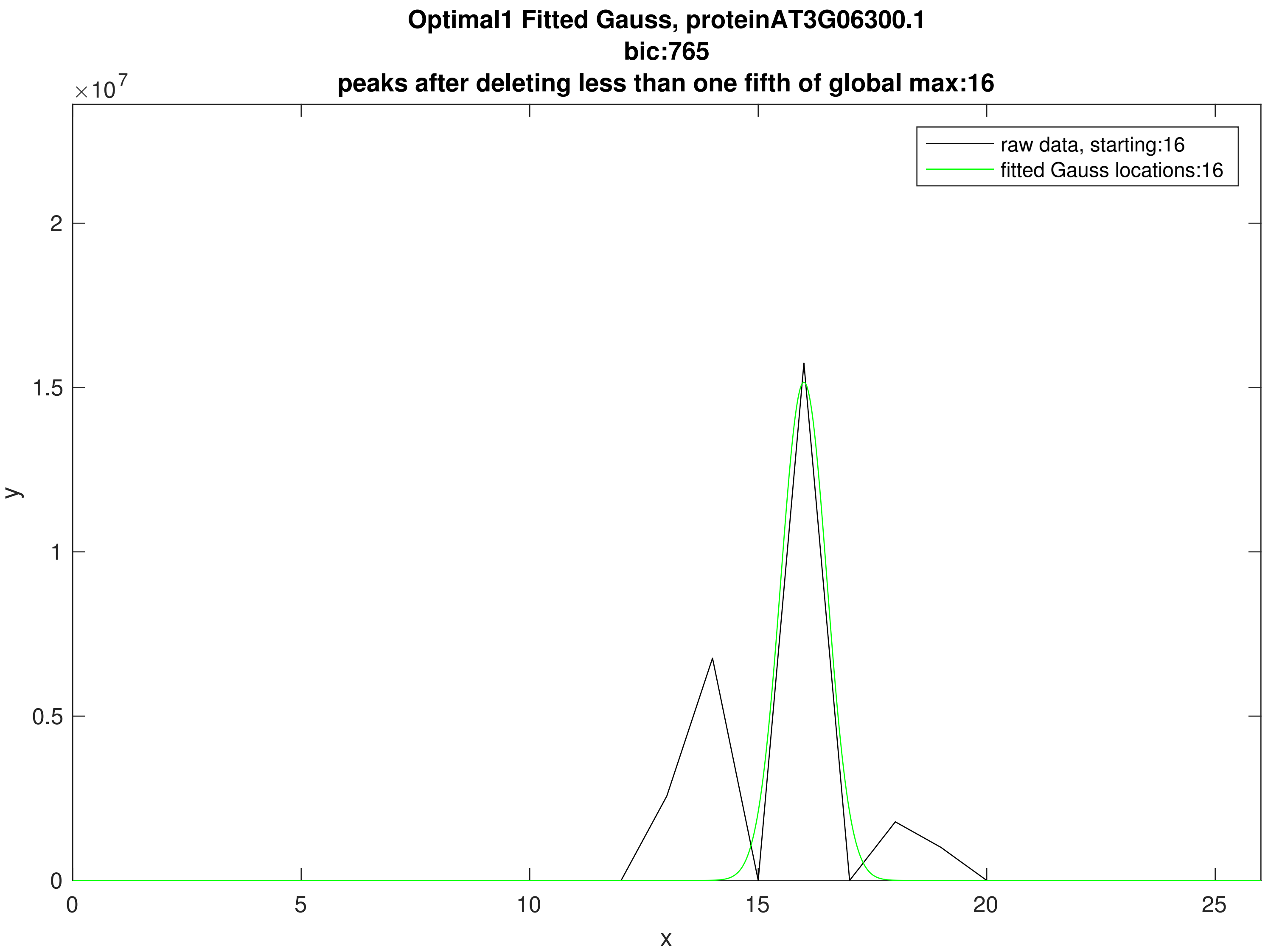

Optimal1 Fitted Gauss, proteinAT3G06300.1
bic:765
peaks after deleting less than one fifth of global max:16
×10^7
raw data, starting:16
fitted Gauss locations:16
y
x
0
0.5
1
1.5
2
0
5
10
15
20
25

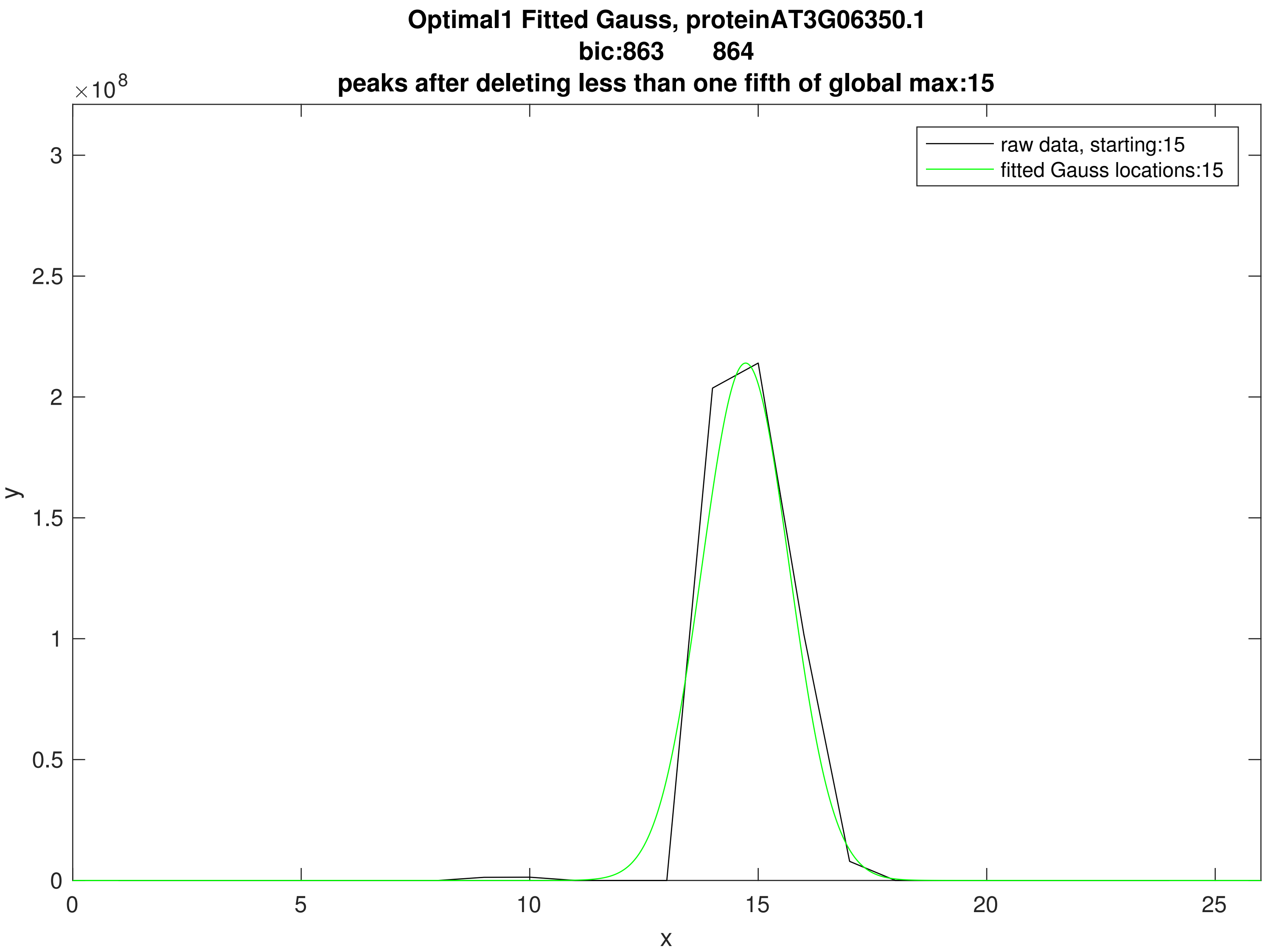
Optimal1 Fitted Gauss, proteinAT3G06350.1
bic:863 864
peaks after deleting less than one fifth of global max:15
×10^8
3
2.5
2
1.5
1
0.5
0
y
0
5
10
15
20
25
x
raw data, starting:15
fitted Gauss locations:15

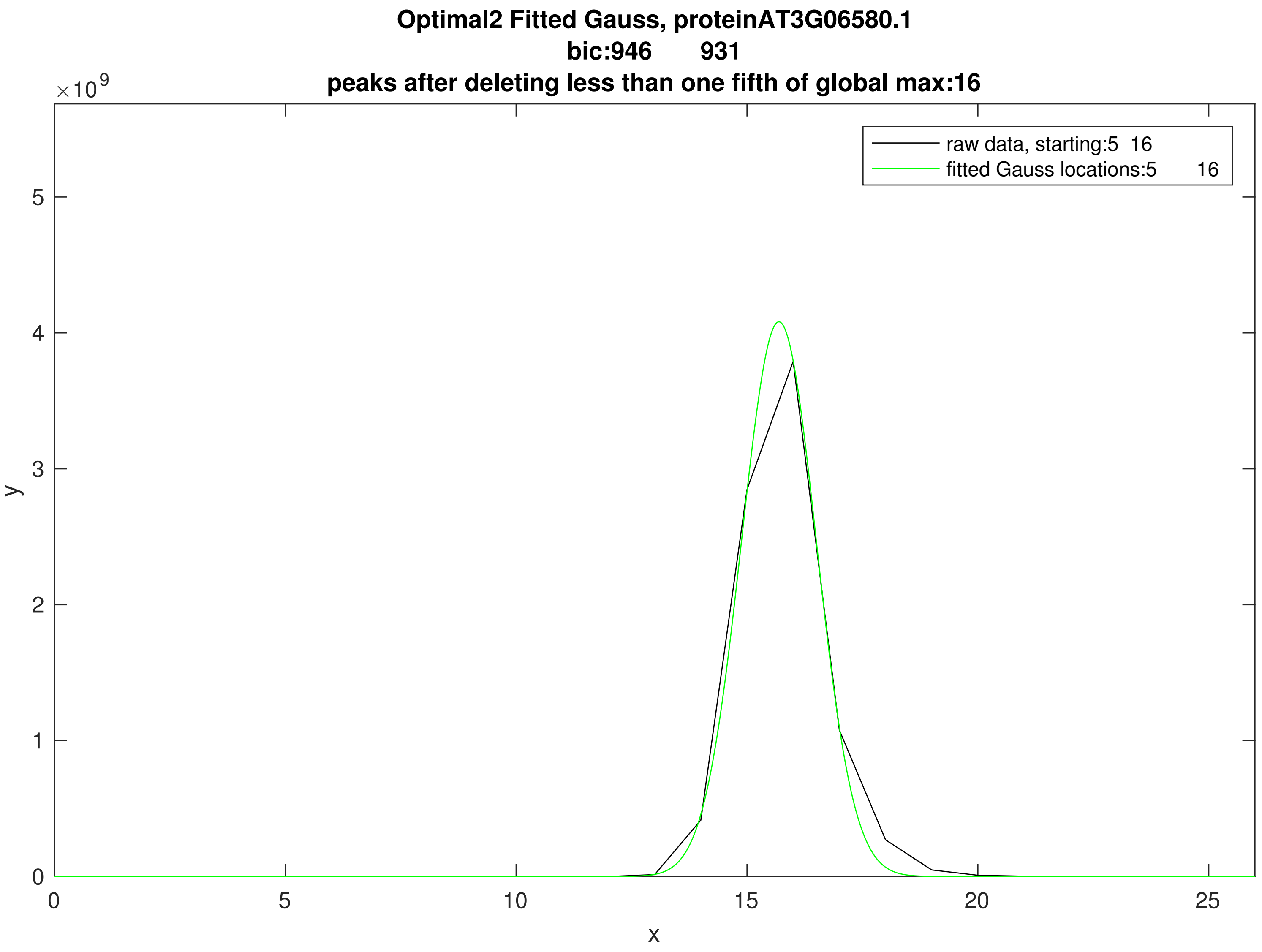

Optimal2 Fitted Gauss, proteinAT3G06580.1
bic:946 931
peaks after deleting less than one fifth of global max:16
×10^9
raw data, starting:5 16
fitted Gauss locations:5 16
y
x
0
1
2
3
4
5
0
5
10
15
20
25

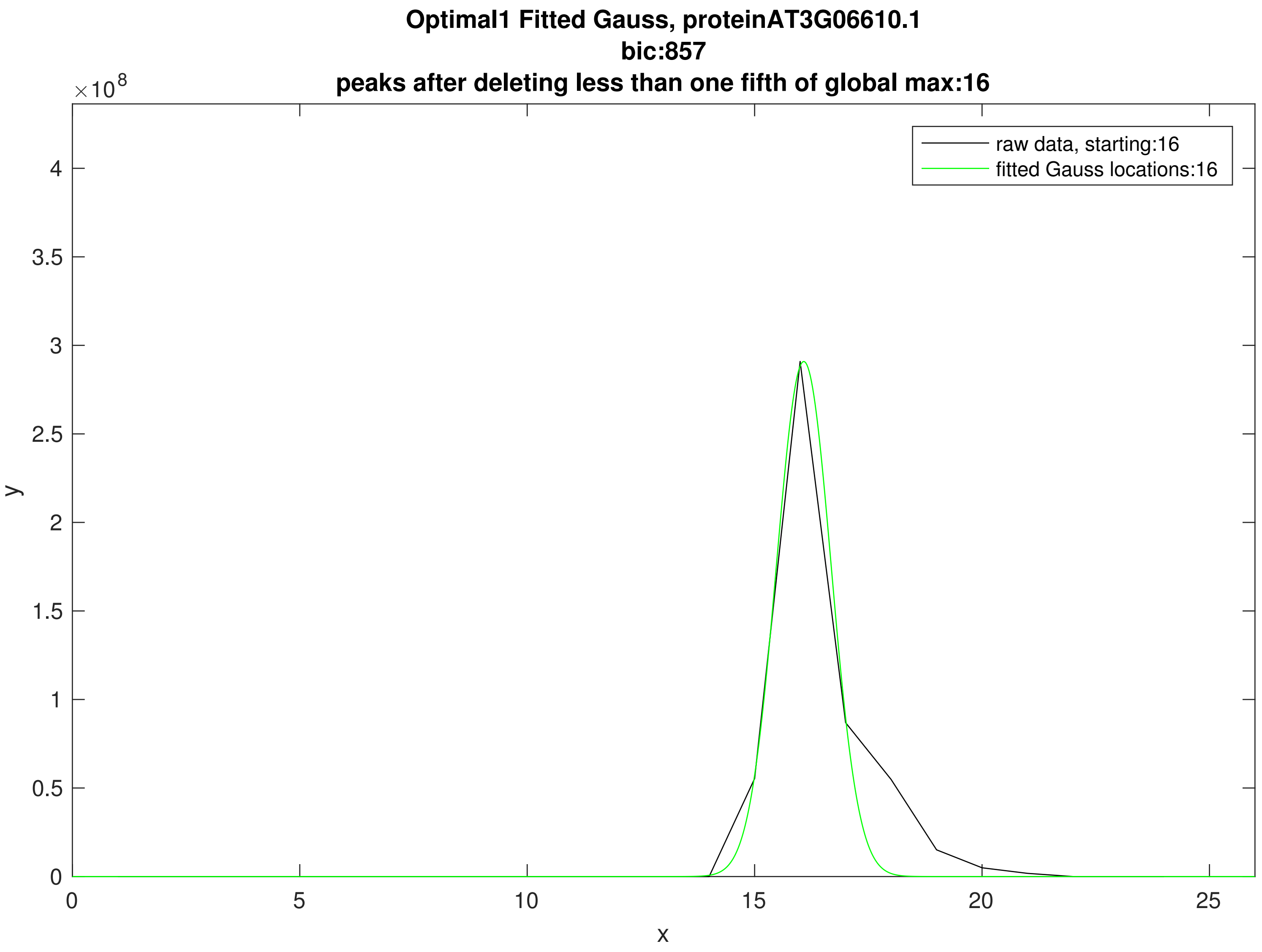
Optimal1 Fitted Gauss, proteinAT3G06610.1
bic:857
peaks after deleting less than one fifth of global max:16
×10^8
4
3.5
3
2.5
2
1.5
1
0.5
0
y
0
5
10
15
20
25
x
raw data, starting:16
fitted Gauss locations:16

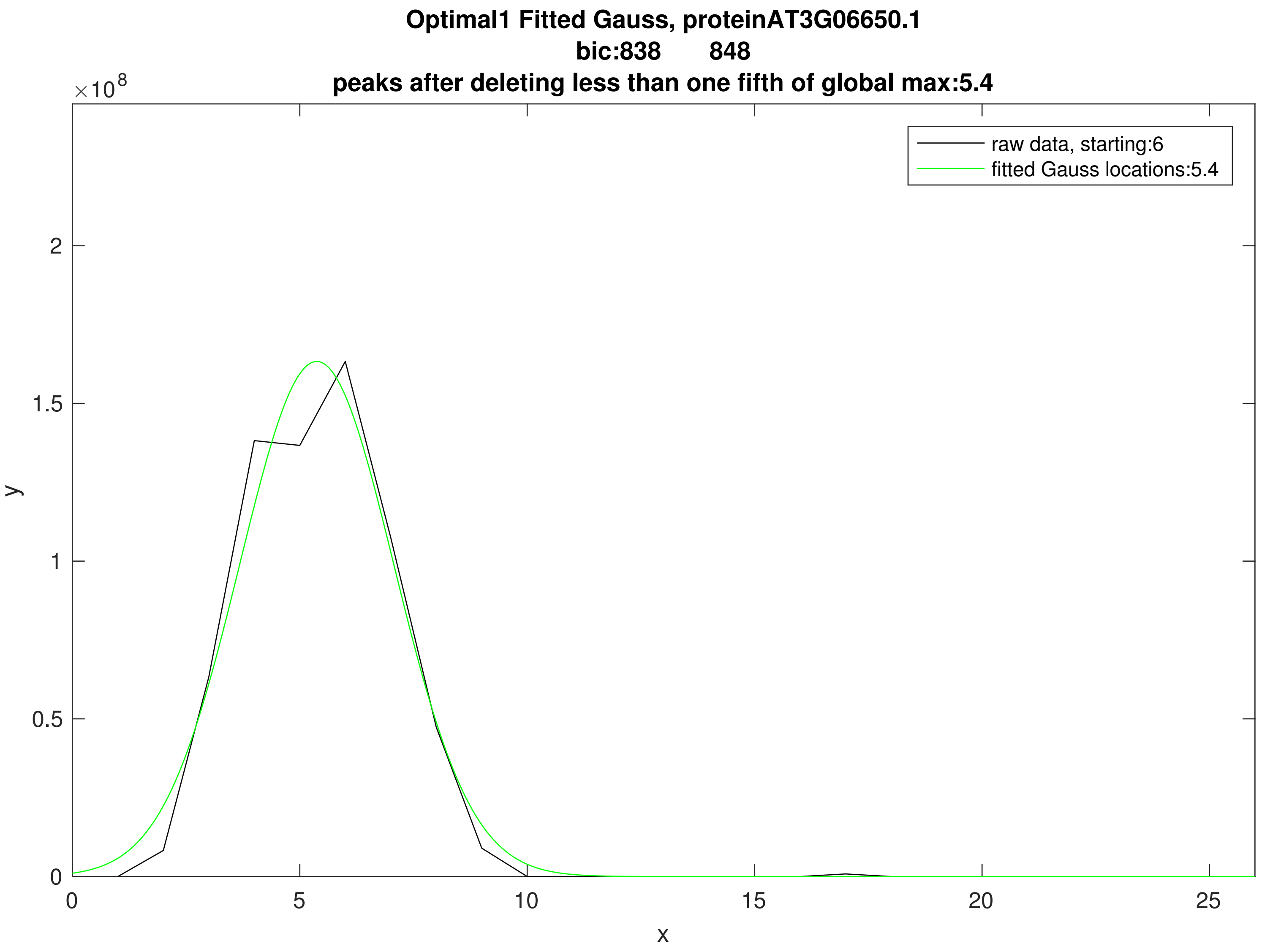
Optimal1 Fitted Gauss, proteinAT3G06650.1
bic:838 848
peaks after deleting less than one fifth of global max:5.4
raw data, starting:6
fitted Gauss locations:5.4
y
x
×10^8
0
0.5
1
1.5
2
0
5
10
15
20
25

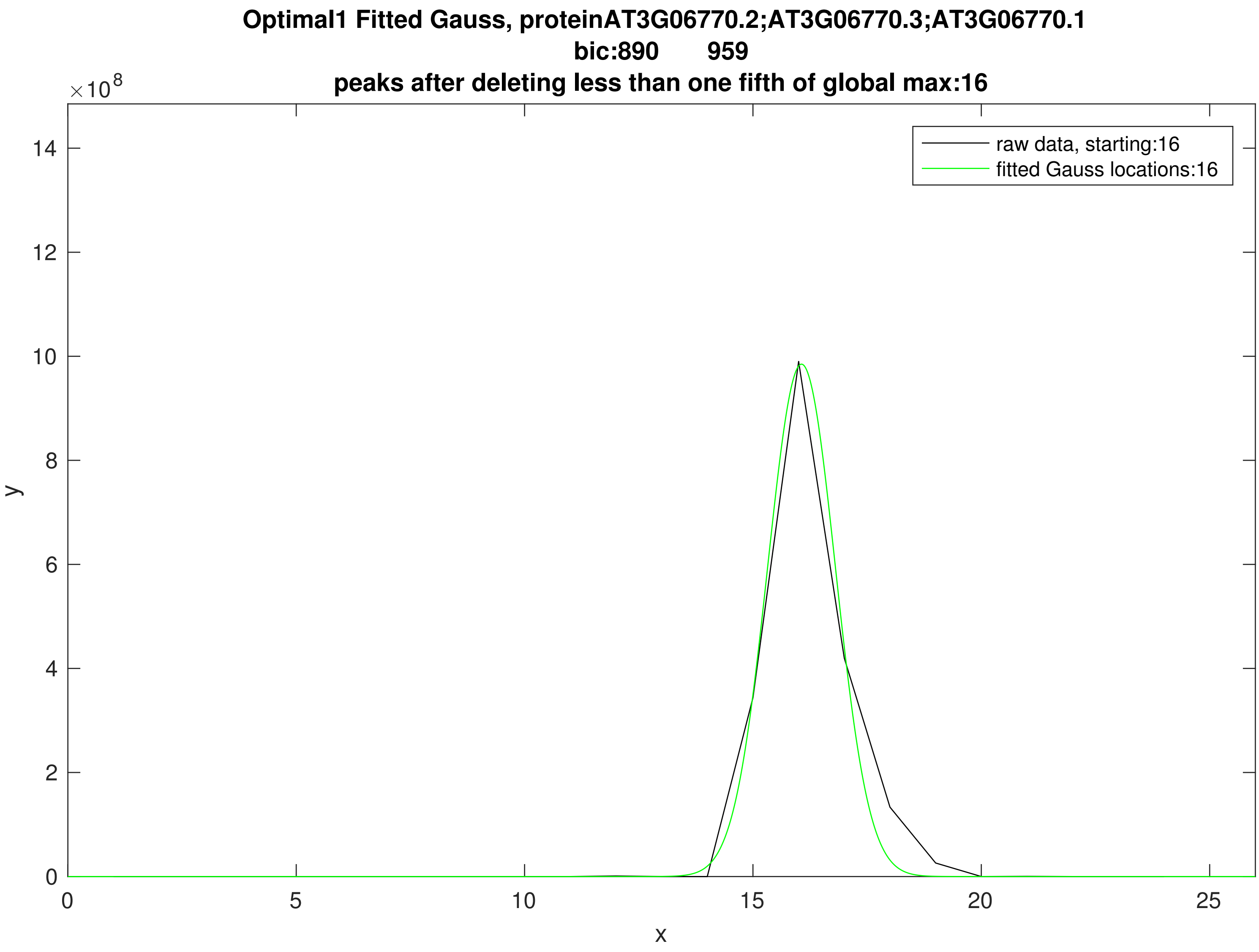

Optimal1 Fitted Gauss, proteinAT3G06770.2;AT3G06770.3;AT3G06770.1
bic:890 959
peaks after deleting less than one fifth of global max:16
×10^8
raw data, starting:16
fitted Gauss locations:16
y
x
0
2
4
6
8
10
12
14
0
5
10
15
20
25

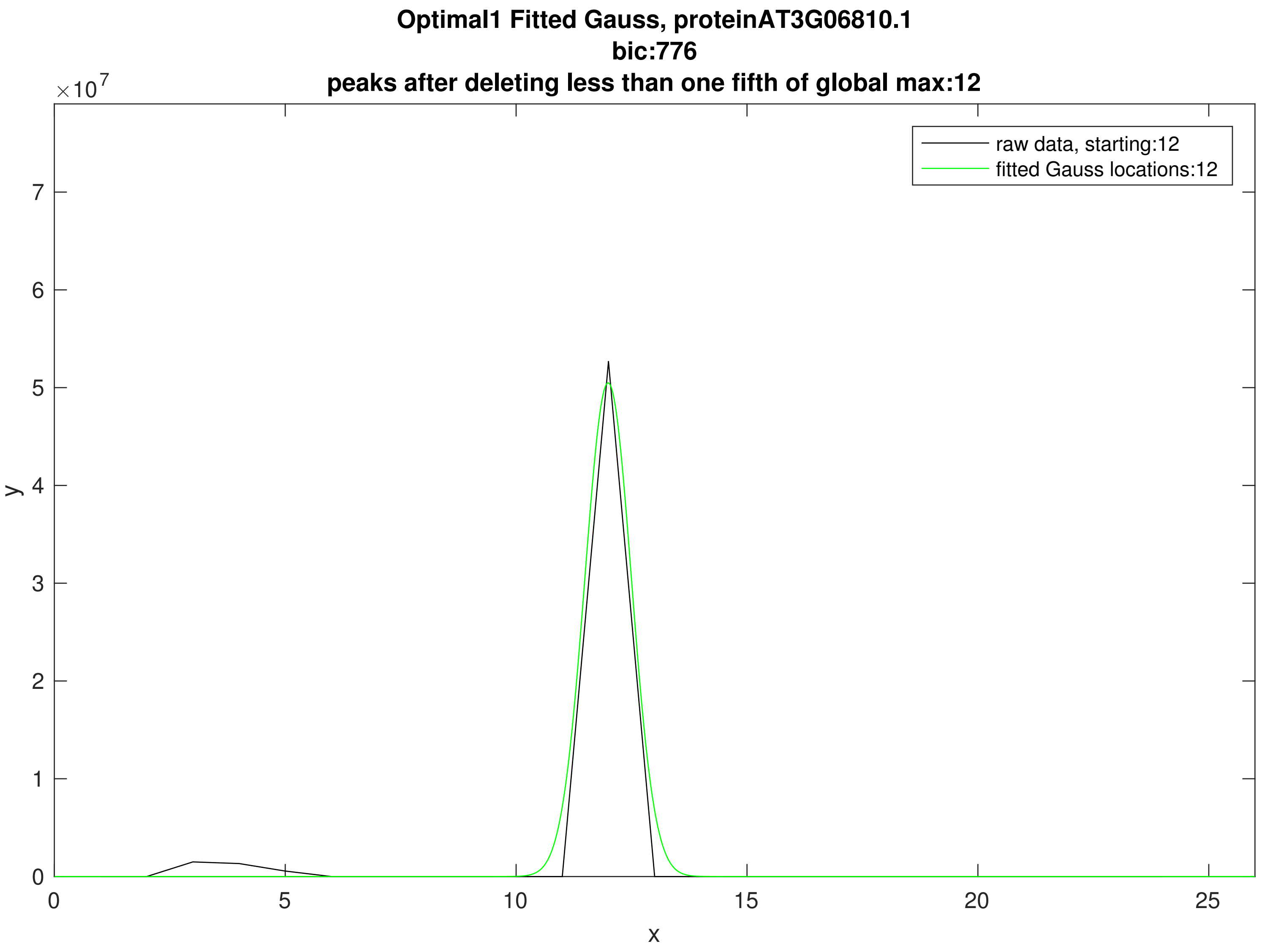
Optimal1 Fitted Gauss, proteinAT3G06810.1
bic:776
peaks after deleting less than one fifth of global max:12
×10^7
raw data, starting:12
fitted Gauss locations:12
y
x
0
1
2
3
4
5
6
7
0
5
10
15
20
25

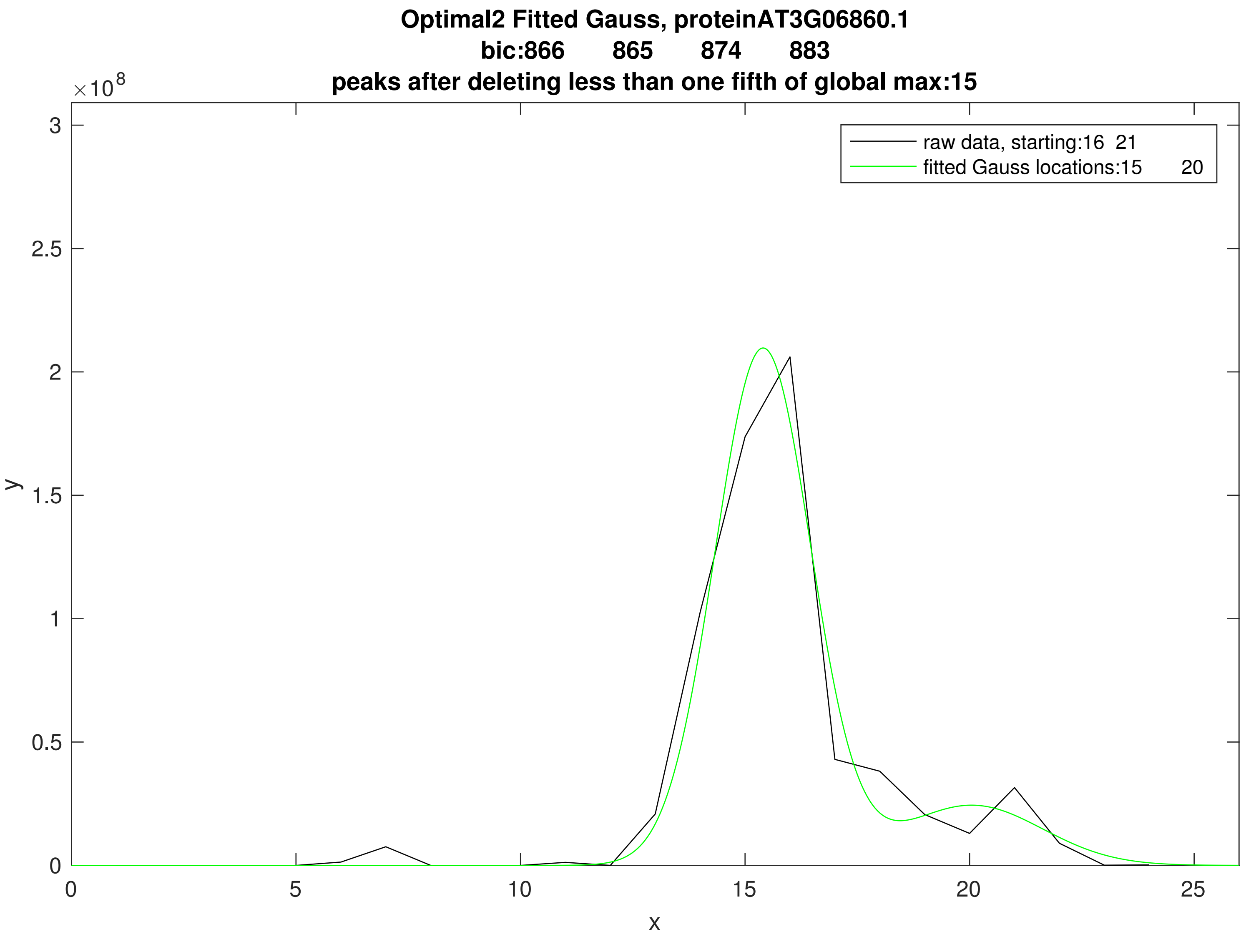
Optimal2 Fitted Gauss, proteinAT3G06860.1
bic:866 865 874 883
peaks after deleting less than one fifth of global max:15
×10^8
raw data, starting:16 21
fitted Gauss locations:15 20
y
x
0
0.5
1
1.5
2
2.5
3
0
5
10
15
20
25

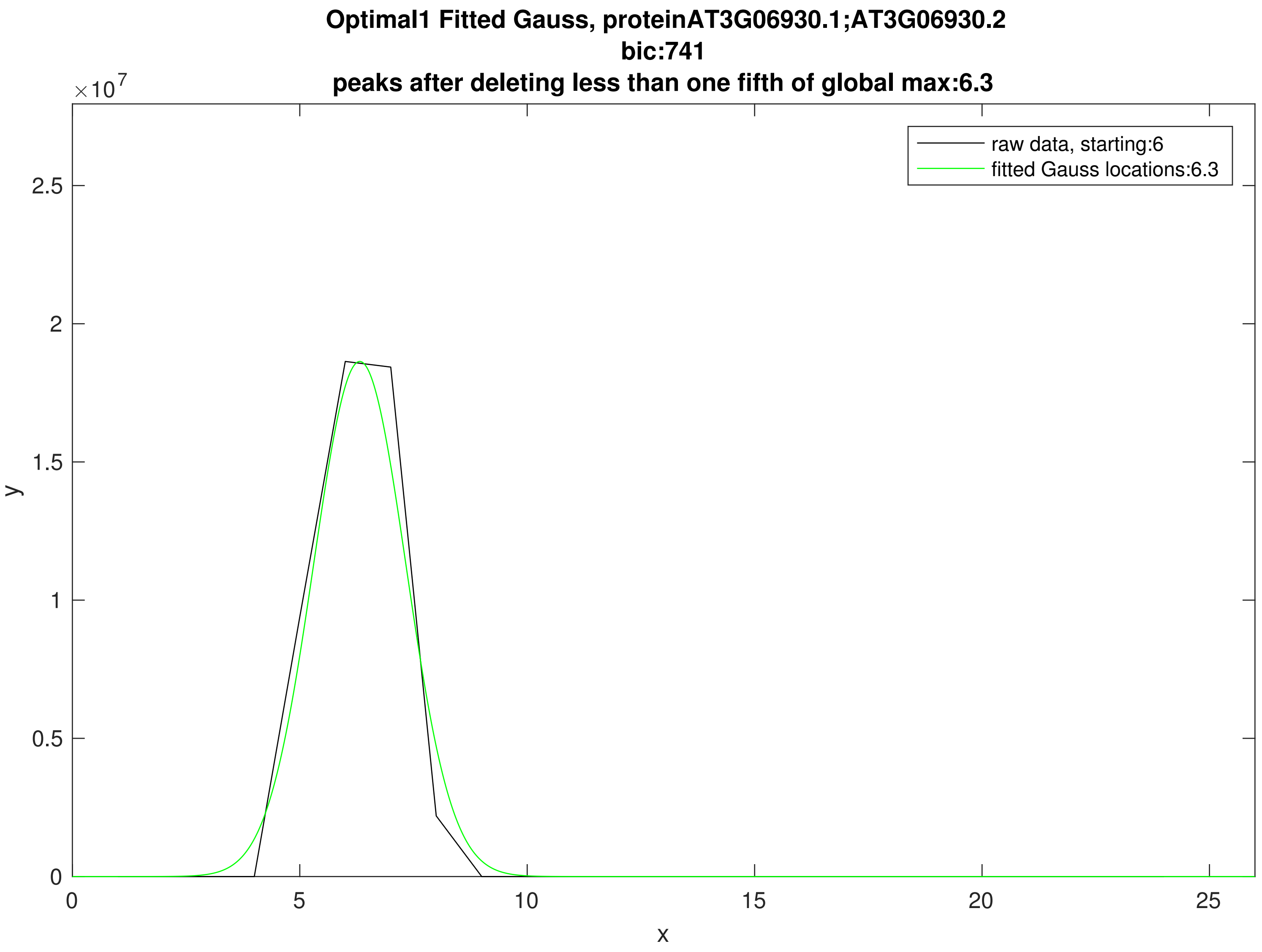
Optimal1 Fitted Gauss, proteinAT3G06930.1;AT3G06930.2
bic:741
peaks after deleting less than one fifth of global max:6.3
×10^7
raw data, starting:6
fitted Gauss locations:6.3
2.5
2
1.5
1
0.5
0
y
0
5
10
15
20
25
x

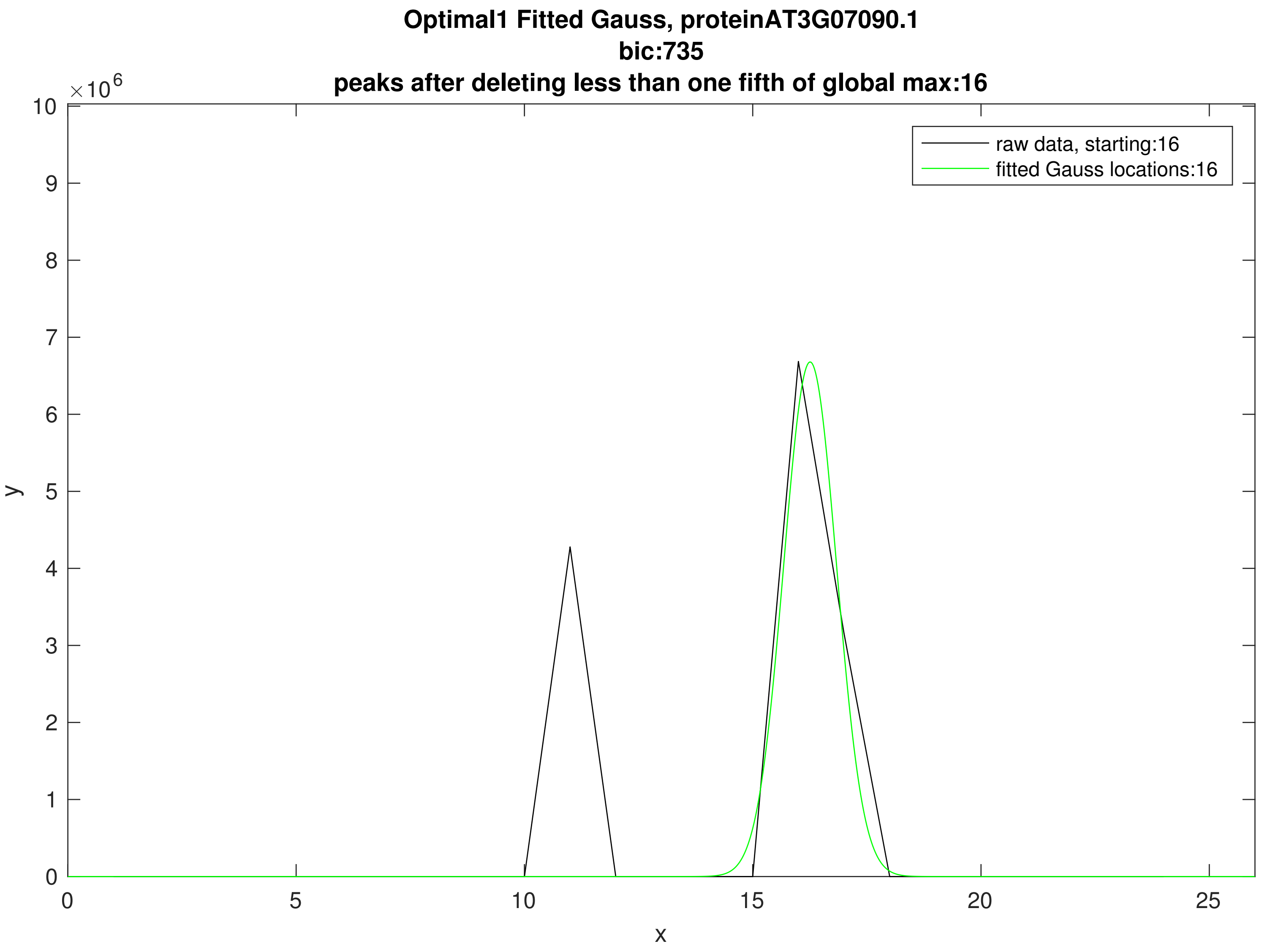
Optimal1 Fitted Gauss, proteinAT3G07090.1
bic:735
peaks after deleting less than one fifth of global max:16
×10^6
10
9
8
7
6
5
4
3
2
1
0
y
0
5
10
15
20
25
x
raw data, starting:16
fitted Gauss locations:16

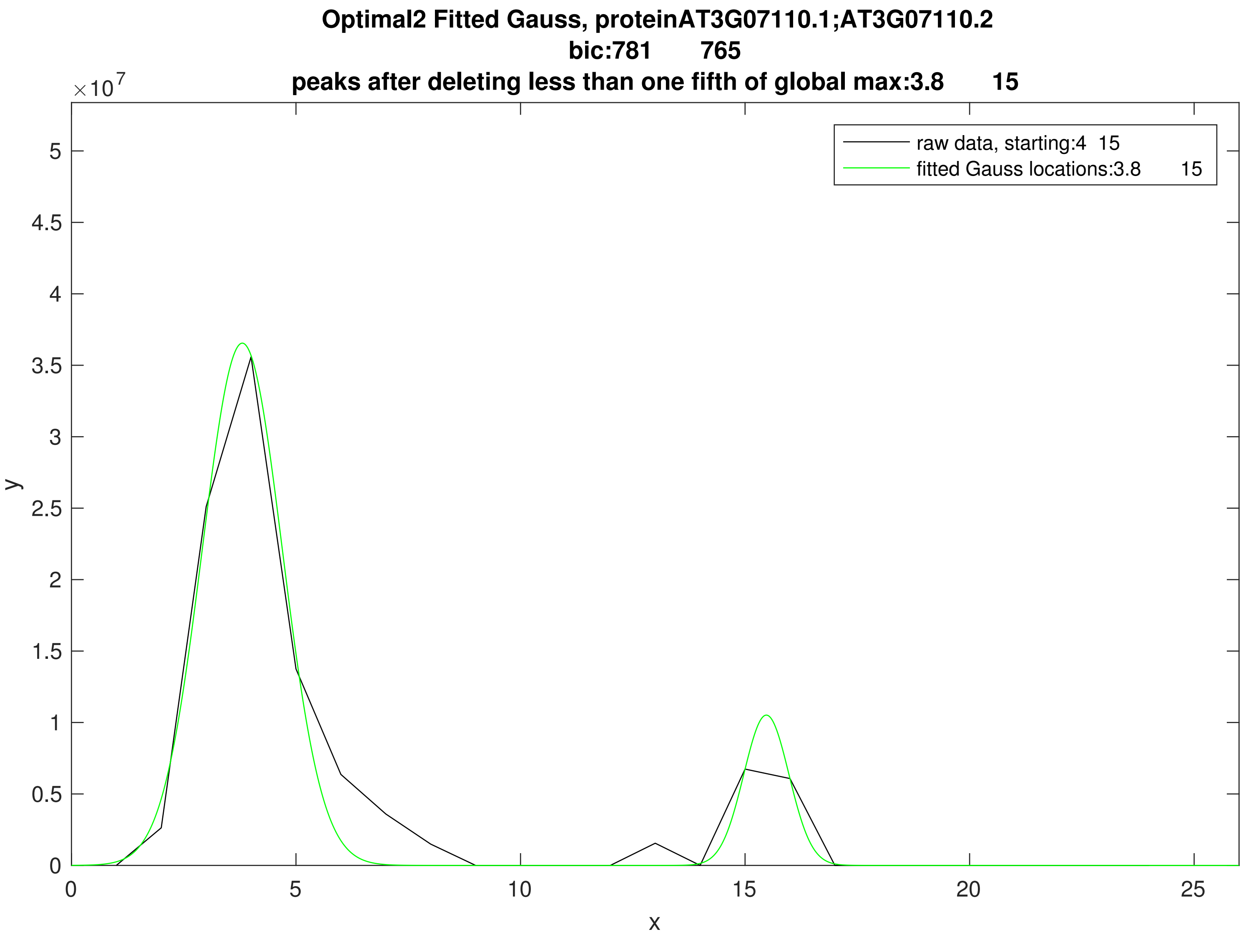

Optimal2 Fitted Gauss, proteinAT3G07110.1;AT3G07110.2
bic:781 765
peaks after deleting less than one fifth of global max:3.8 15
×10^7
raw data, starting:4 15
fitted Gauss locations:3.8 15
y
x
0
0.5
1
1.5
2
2.5
3
3.5
4
4.5
5
0
5
10
15
20
25

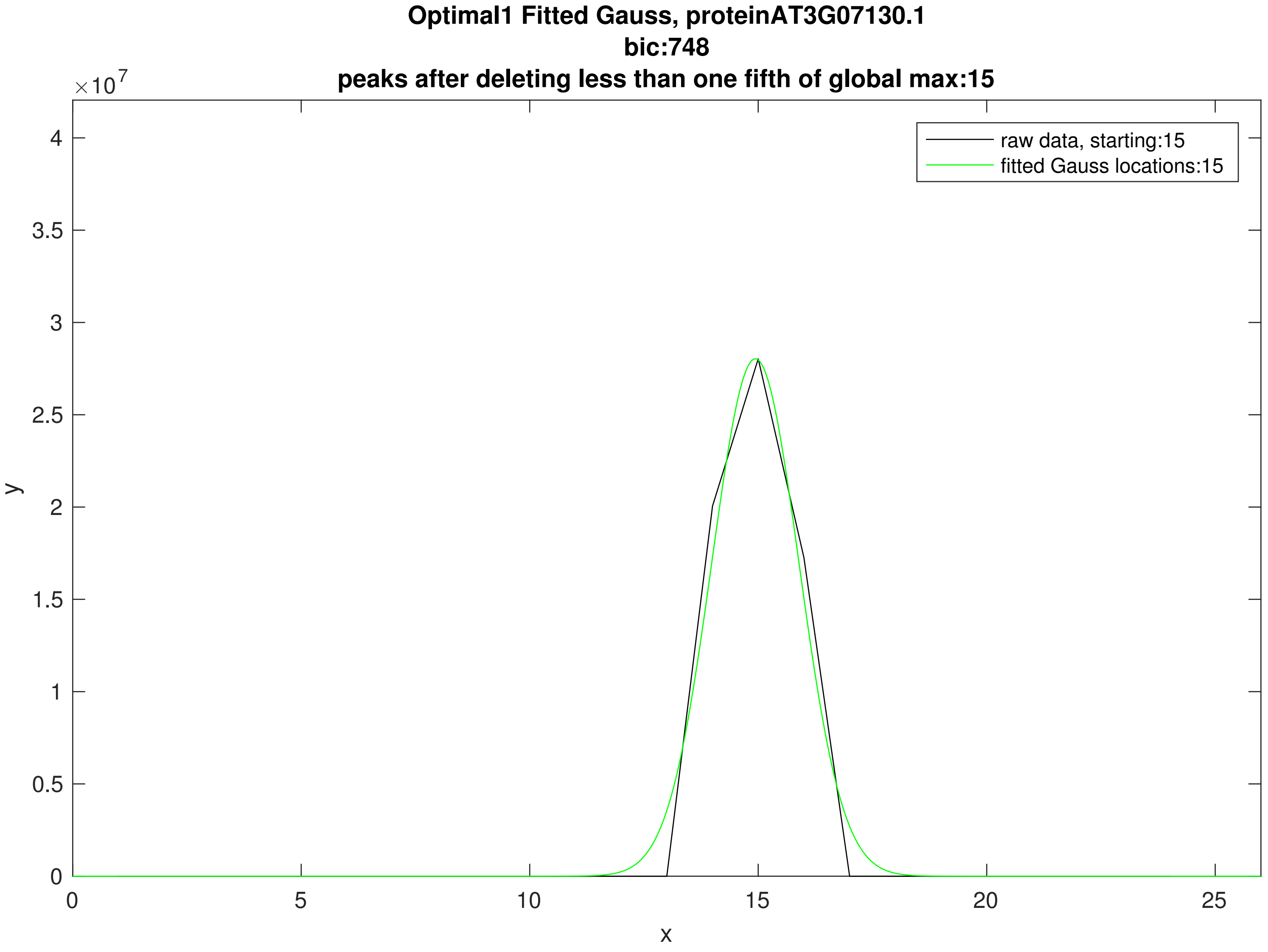
Optimal1 Fitted Gauss, proteinAT3G07130.1
bic:748
peaks after deleting less than one fifth of global max:15
×10^7
raw data, starting:15
fitted Gauss locations:15
y
x
0
0.5
1
1.5
2
2.5
3
3.5
4
0
5
10
15
20
25

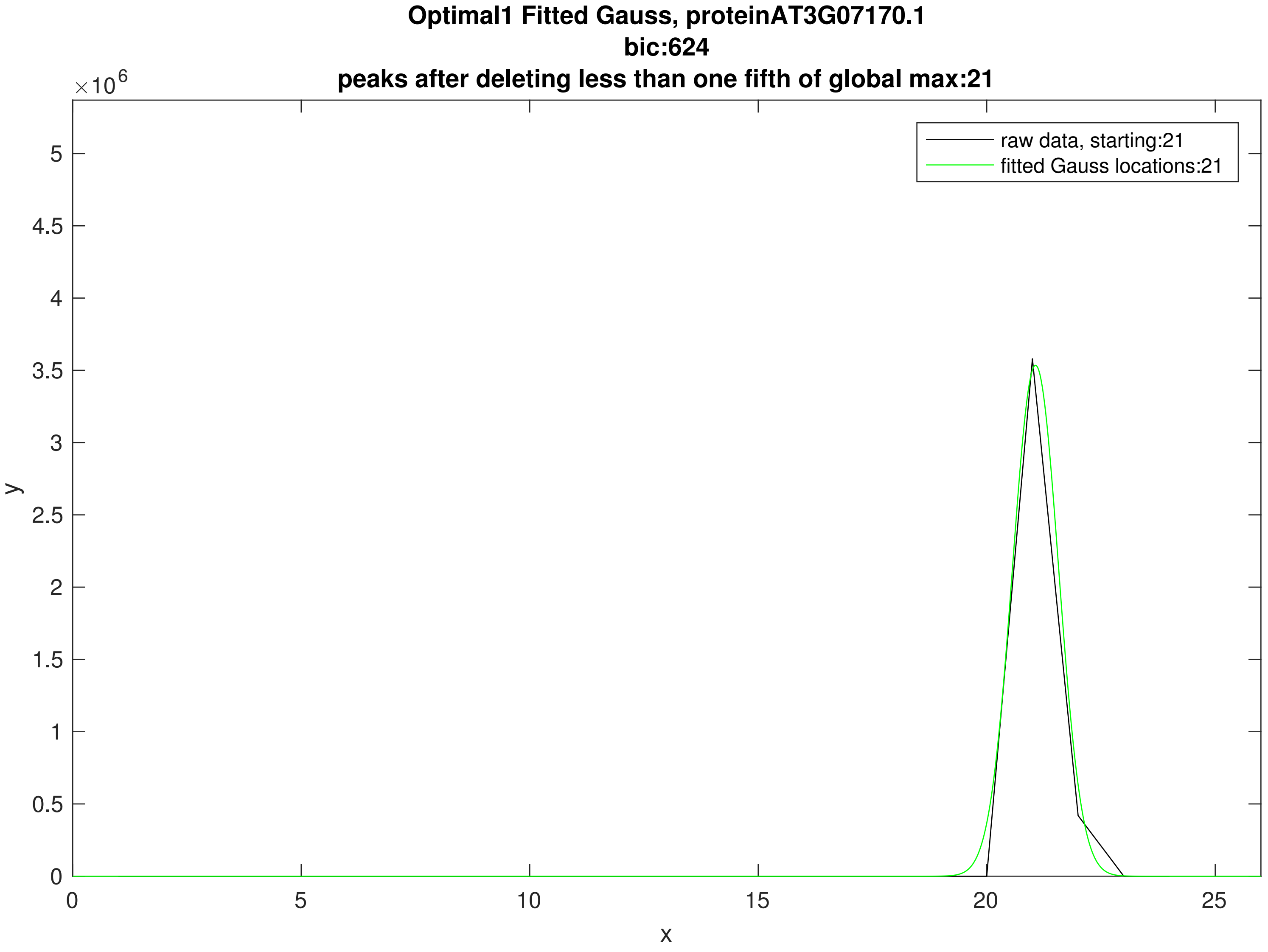
Optimal1 Fitted Gauss, proteinAT3G07170.1
bic:624
peaks after deleting less than one fifth of global max:21
×10^6
raw data, starting:21
fitted Gauss locations:21
y
x
0
0.5
1
1.5
2
2.5
3
3.5
4
4.5
5
0
5
10
15
20
25

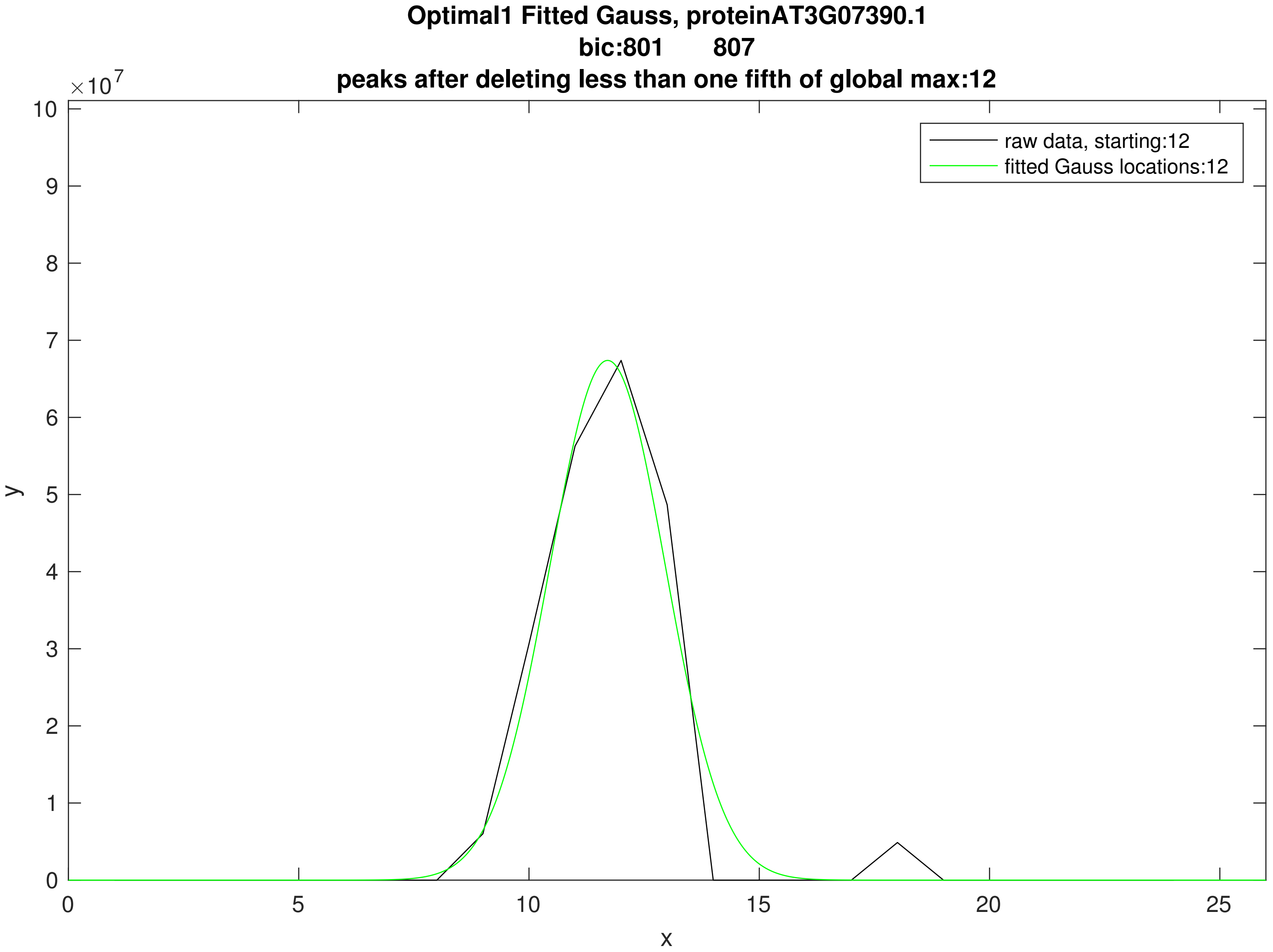
Optimal1 Fitted Gauss, proteinAT3G07390.1
bic:801 807
peaks after deleting less than one fifth of global max:12
raw data, starting:12
fitted Gauss locations:12
y
x
×10^7
0
1
2
3
4
5
6
7
8
9
10
0
5
10
15
20
25

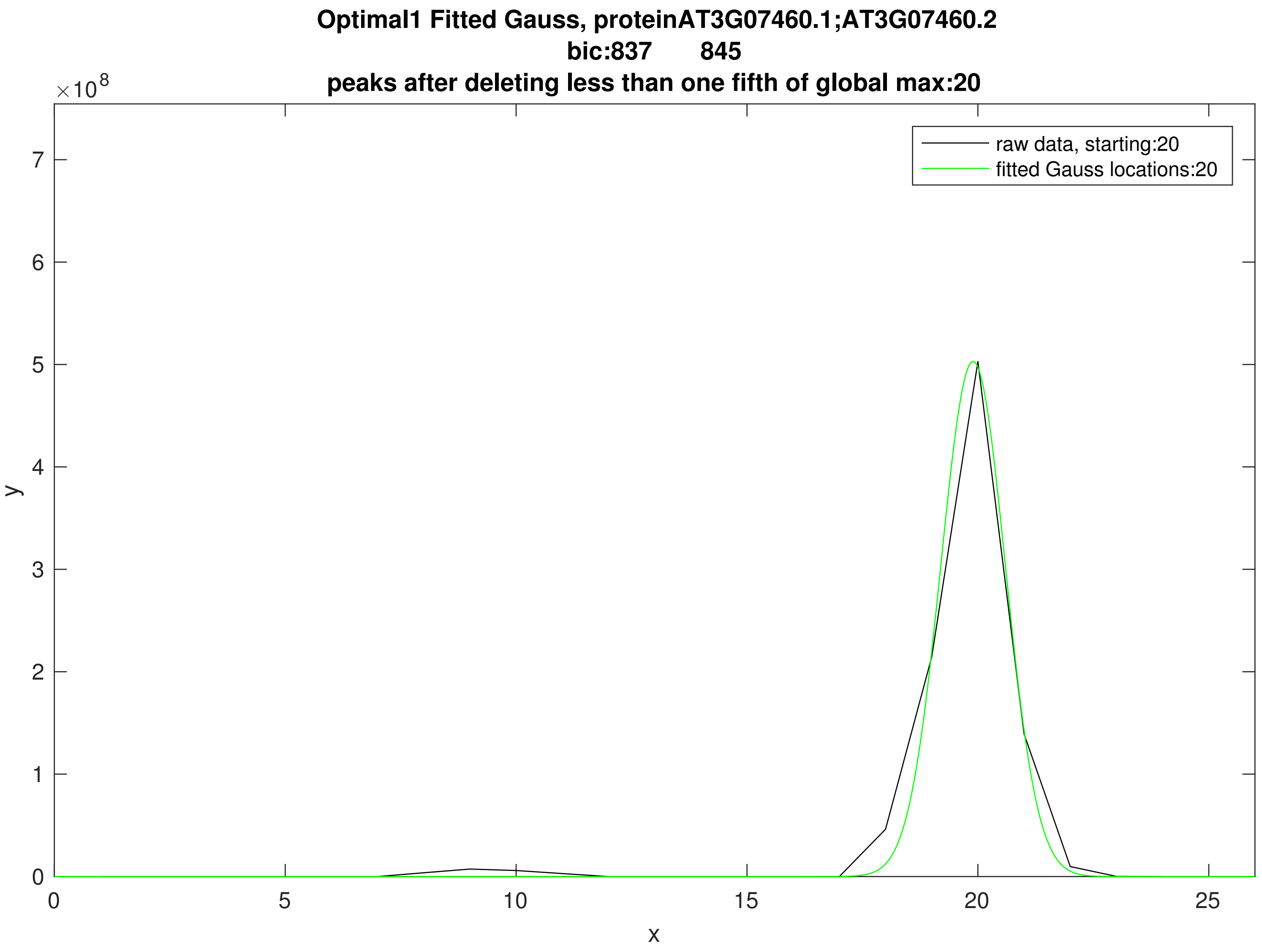

Optimal1 Fitted Gauss, proteinAT3G07460.1;AT3G07460.2
bic:837 845
peaks after deleting less than one fifth of global max:20
×10^8
raw data, starting:20
fitted Gauss locations:20
y
x
0
1
2
3
4
5
6
7
0
5
10
15
20
25

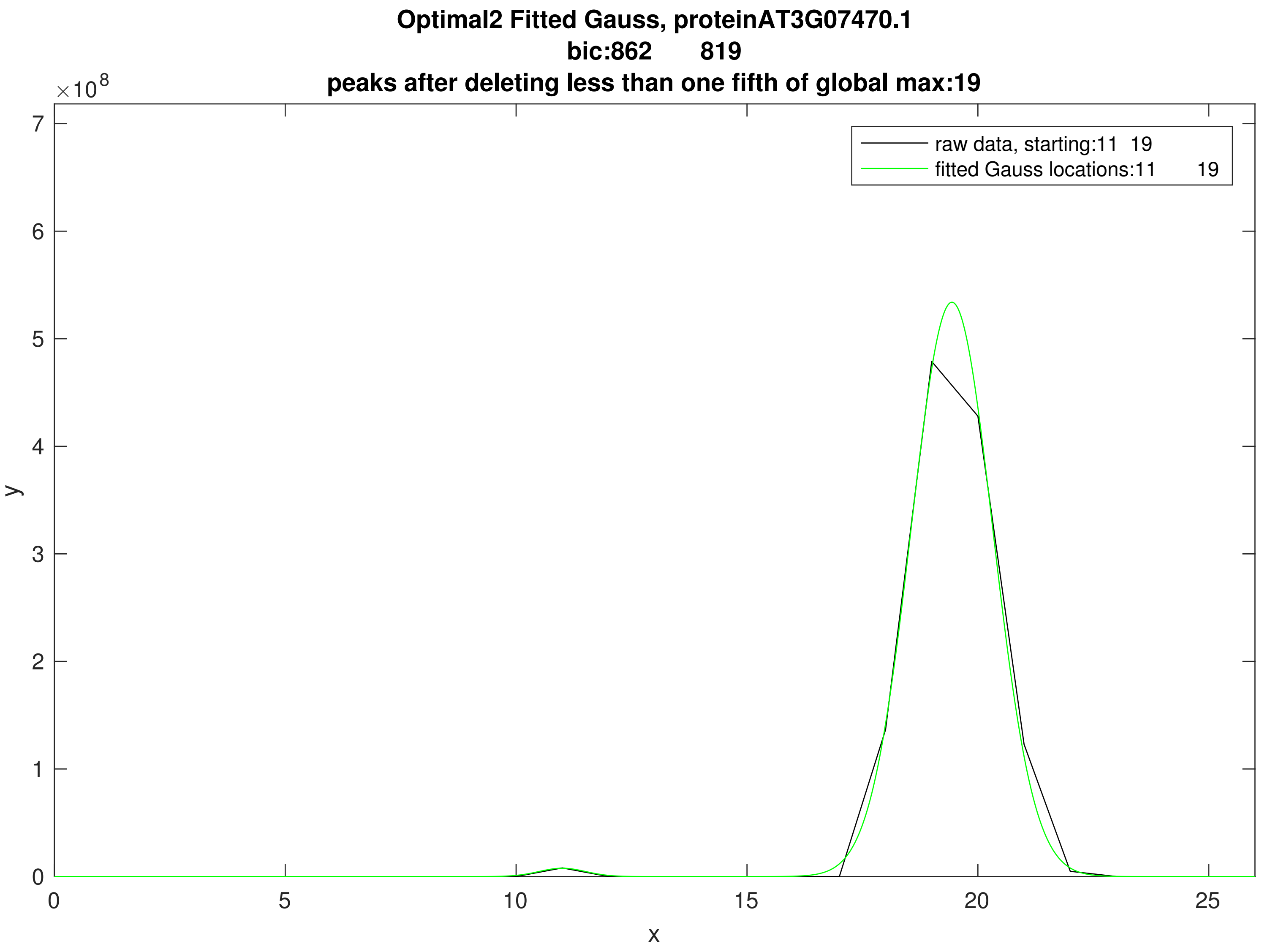

Optimal2 Fitted Gauss, proteinAT3G07470.1
bic:862 819
peaks after deleting less than one fifth of global max:19
×10^8
raw data, starting:11 19
fitted Gauss locations:11 19
y
x
0
1
2
3
4
5
6
7
0
5
10
15
20
25

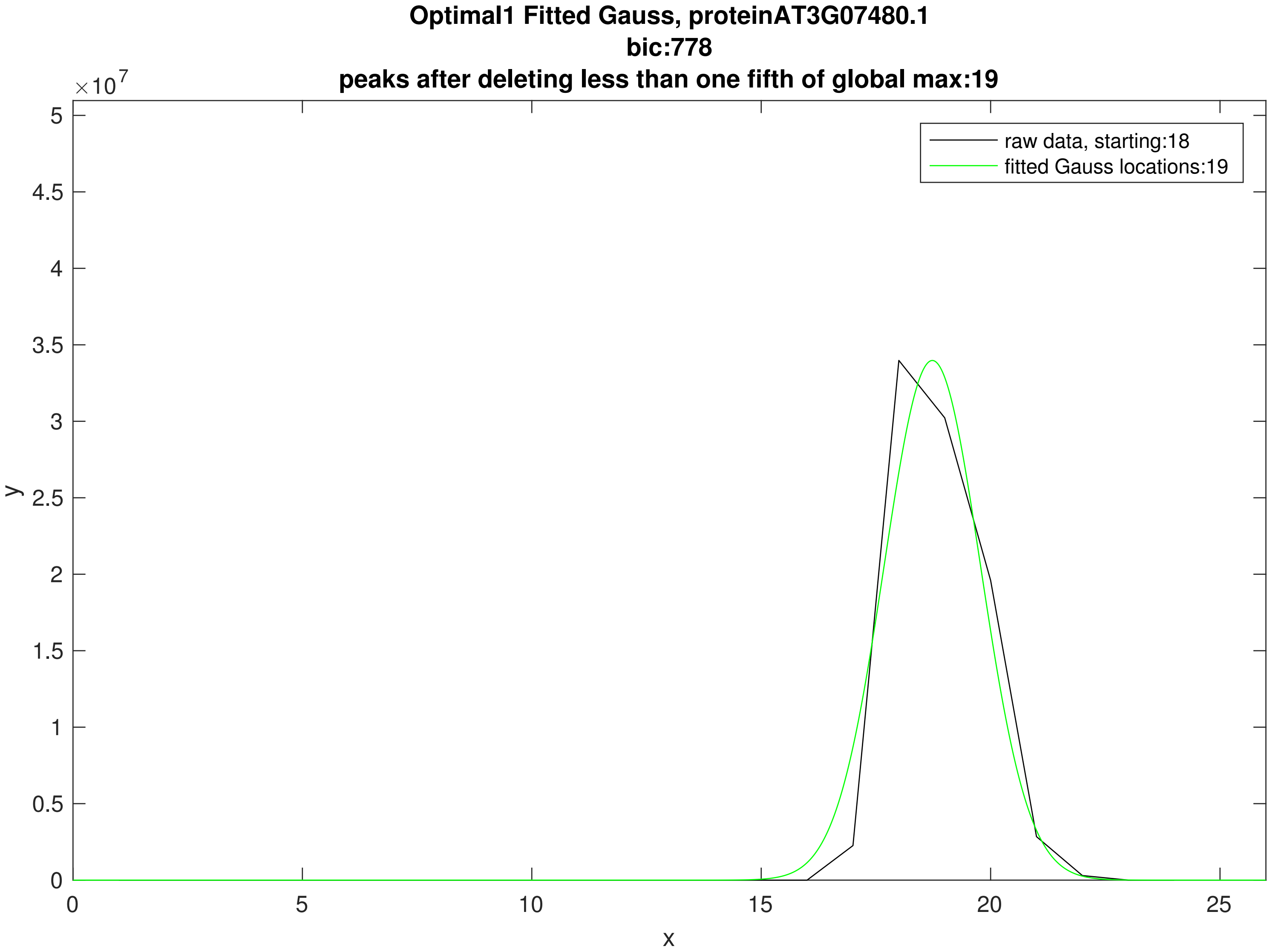

Optimal1 Fitted Gauss, proteinAT3G07480.1
bic:778
peaks after deleting less than one fifth of global max:19
×10^7
raw data, starting:18
fitted Gauss locations:19
y
x
0
0.5
1
1.5
2
2.5
3
3.5
4
4.5
5
0
5
10
15
20
25

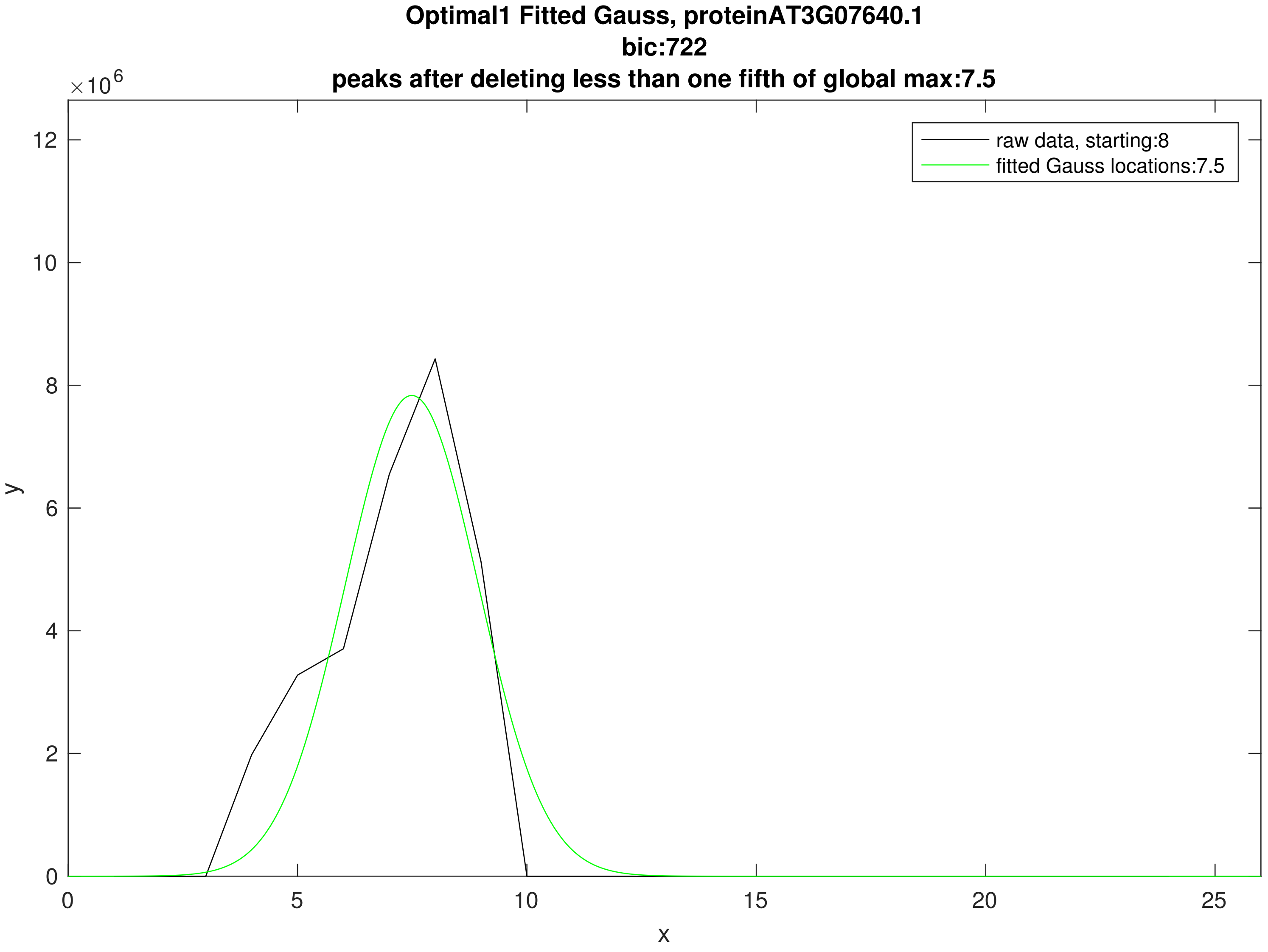
Optimal1 Fitted Gauss, proteinAT3G07640.1
bic:722
peaks after deleting less than one fifth of global max:7.5
×10^6
12
10
8
6
4
2
0
y
0
5
10
15
20
25
x
raw data, starting:8
fitted Gauss locations:7.5

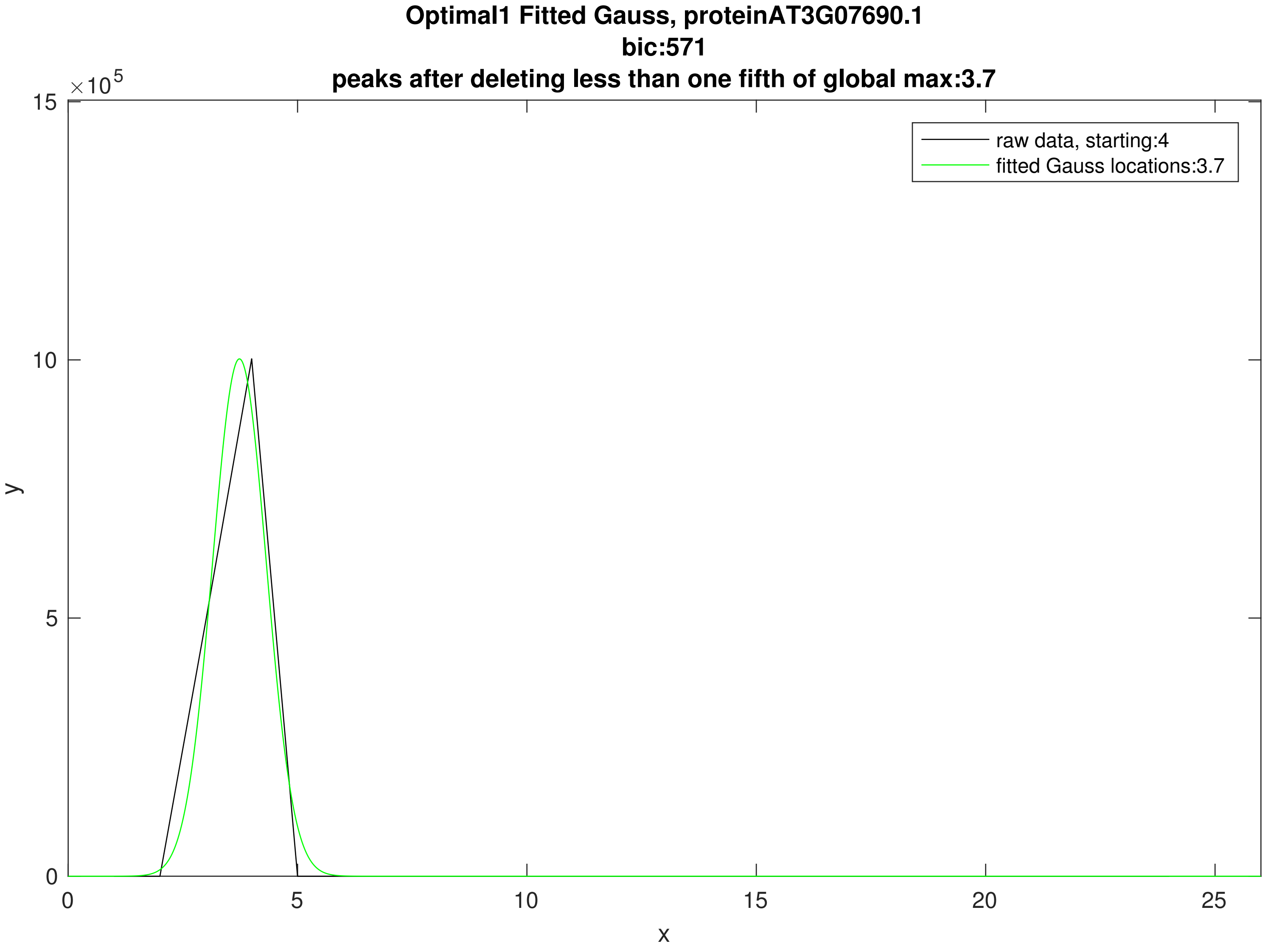

Optimal1 Fitted Gauss, proteinAT3G07690.1
bic:571
peaks after deleting less than one fifth of global max:3.7
×10^5
15
10
5
0
y
0
5
10
15
20
25
x
raw data, starting:4
fitted Gauss locations:3.7

**Optimal1 Fitted Gauss, proteinAT3G07720.1**
**bic:867**
**peaks after deleting less than one fifth of global max:17**

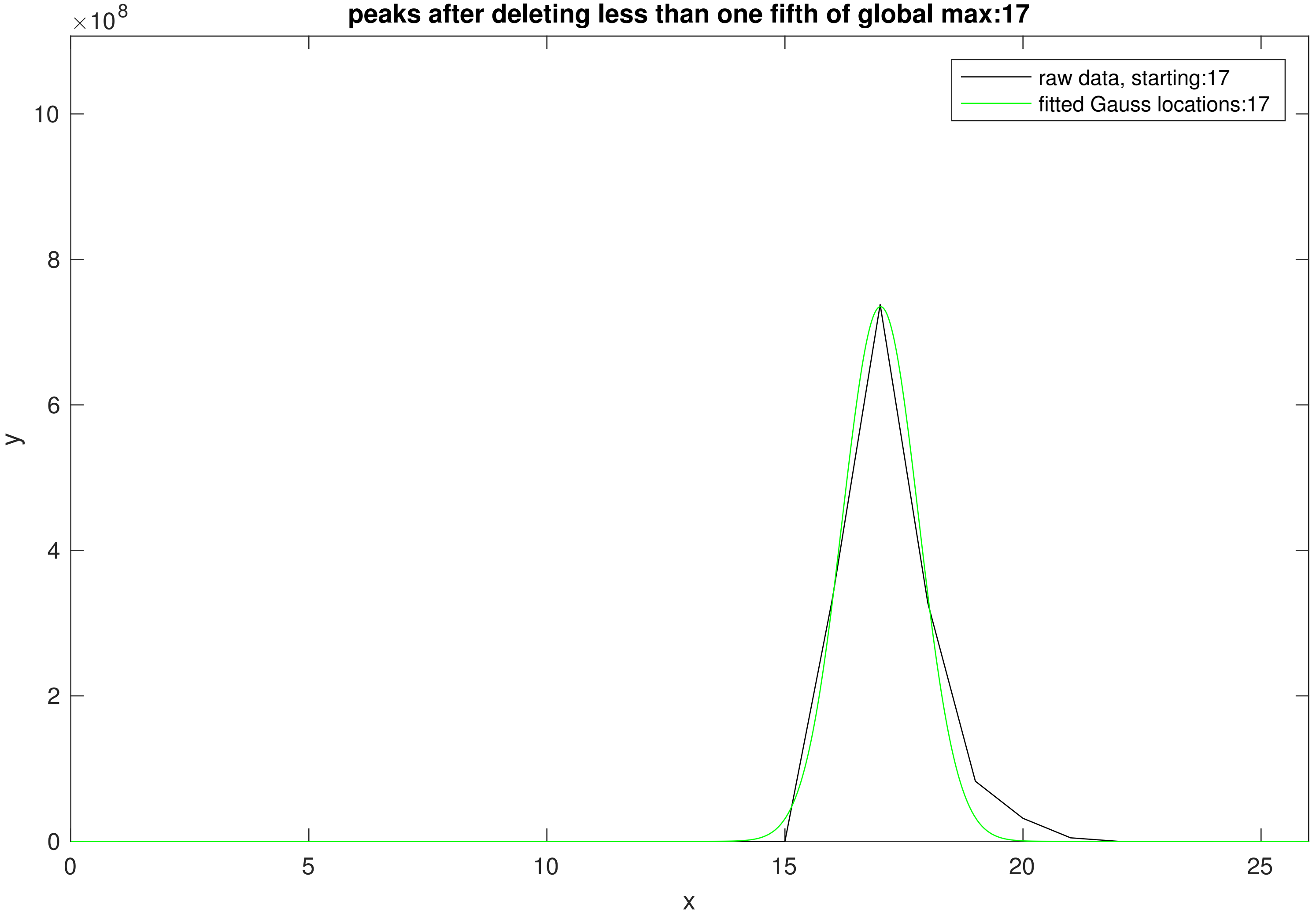

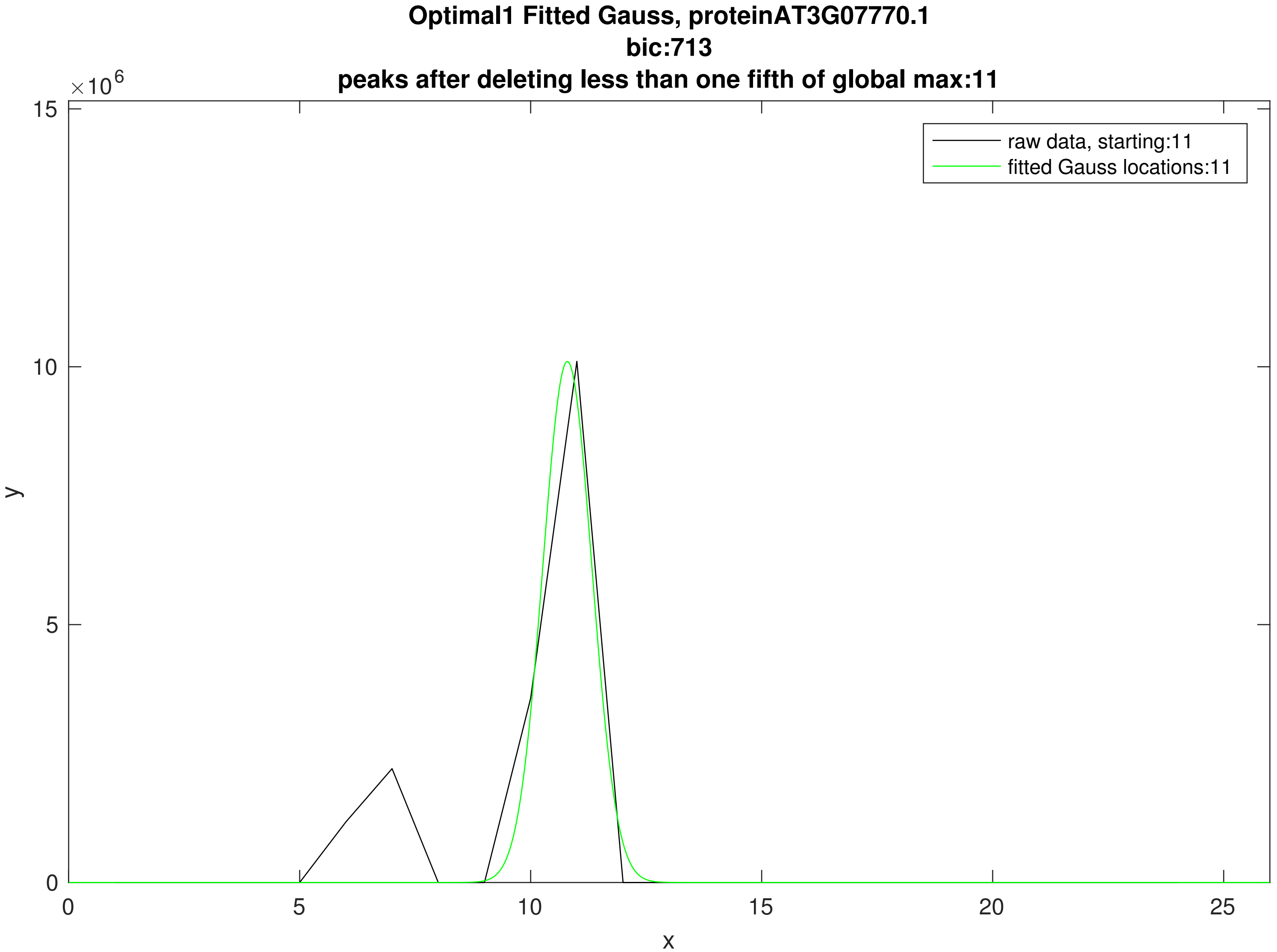

Optimal1 Fitted Gauss, proteinAT3G07770.1
bic:713
peaks after deleting less than one fifth of global max:11
×10^6
15
10
5
0
y
0
5
10
15
20
25
x
raw data, starting:11
fitted Gauss locations:11

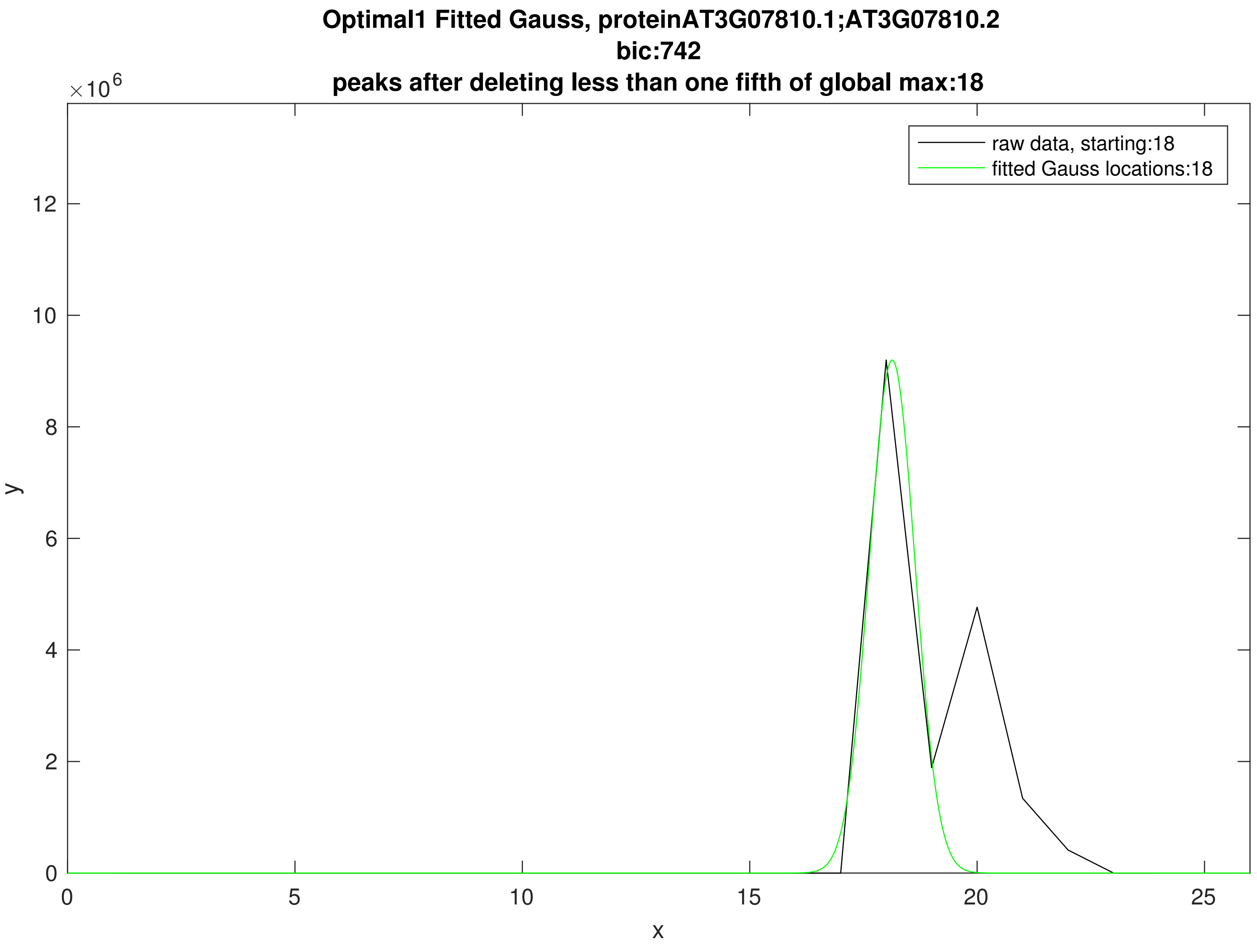

Optimal1 Fitted Gauss, proteinAT3G07810.1;AT3G07810.2
bic:742
peaks after deleting less than one fifth of global max:18
×10^6
raw data, starting:18
fitted Gauss locations:18
y
x
0
2
4
6
8
10
12
0
5
10
15
20
25

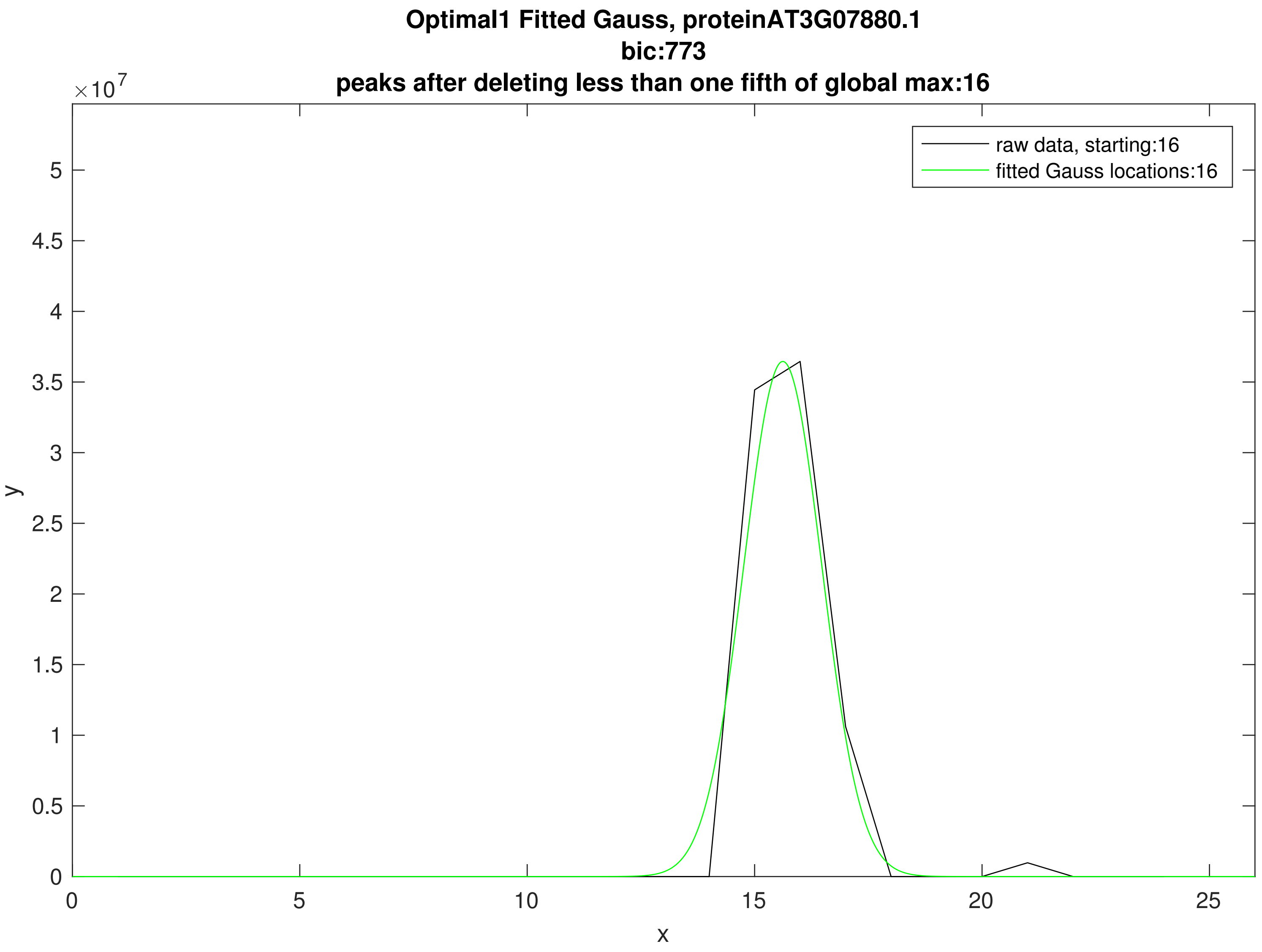

Optimal1 Fitted Gauss, proteinAT3G07880.1
bic:773
peaks after deleting less than one fifth of global max:16
×10^7
raw data, starting:16
fitted Gauss locations:16
y
x
0
0.5
1
1.5
2
2.5
3
3.5
4
4.5
5
0
5
10
15
20
25

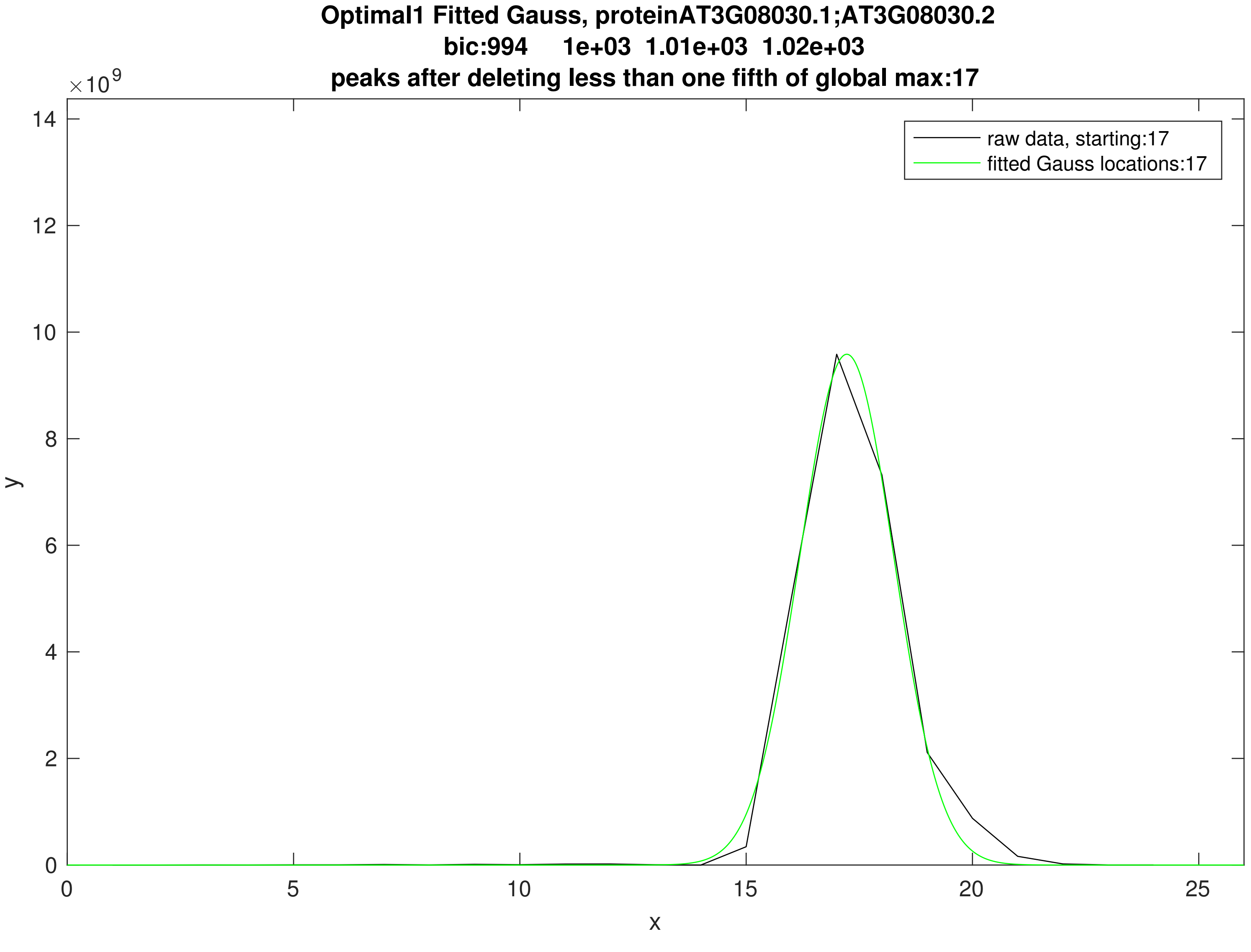
Optimal1 Fitted Gauss, proteinAT3G08030.1;AT3G08030.2
bic:994 1e+03 1.01e+03 1.02e+03
peaks after deleting less than one fifth of global max:17
×10^9
raw data, starting:17
fitted Gauss locations:17
y
x
0
2
4
6
8
10
12
14
5
15
20
25

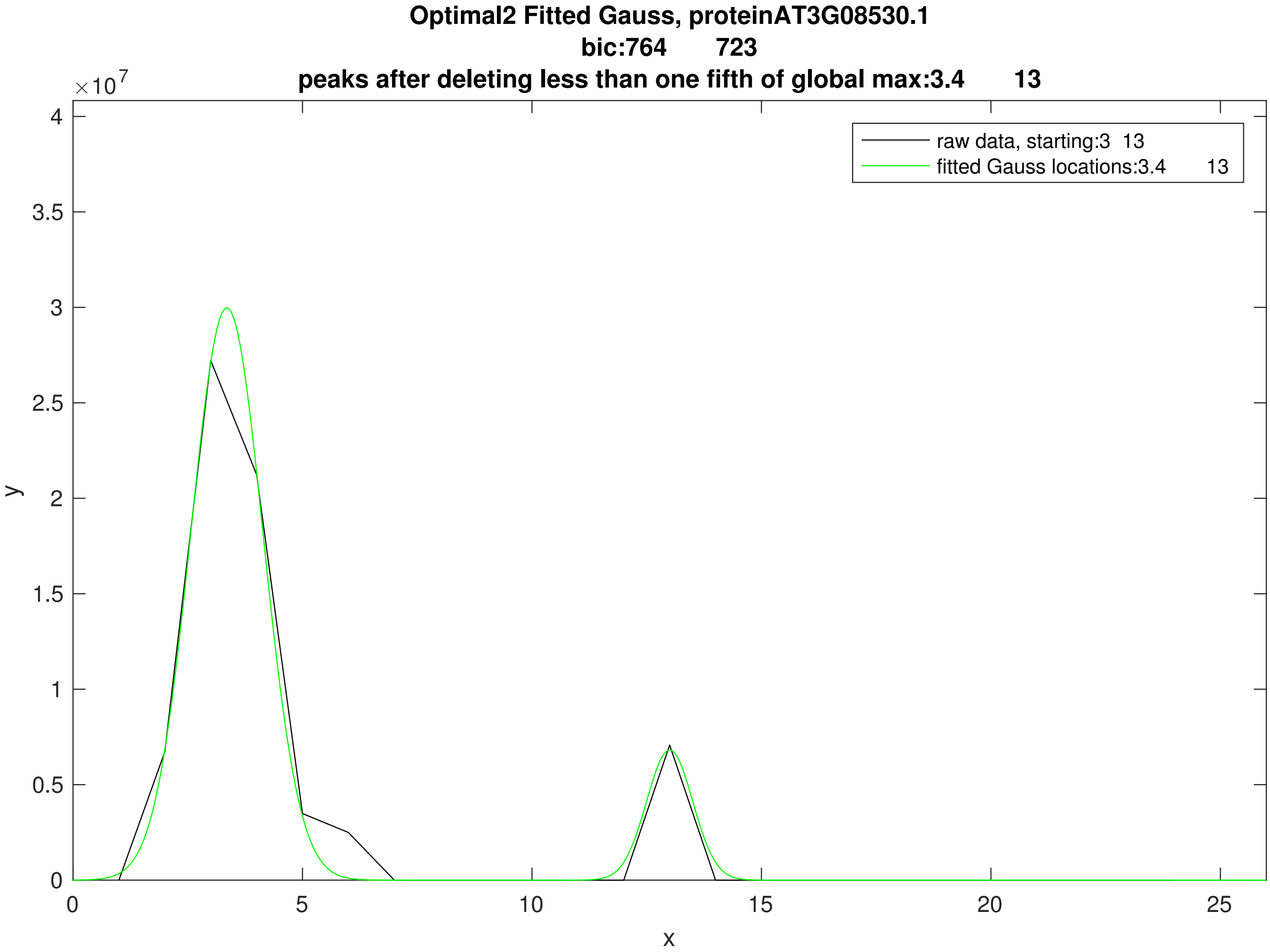
Optimal2 Fitted Gauss, proteinAT3G08530.1
bic:764 723
peaks after deleting less than one fifth of global max:3.4 13
×10^7
raw data, starting:3 13
fitted Gauss locations:3.4 13
y
x
0
0.5
1
1.5
2
2.5
3
3.5
4
5
10
15
20
25

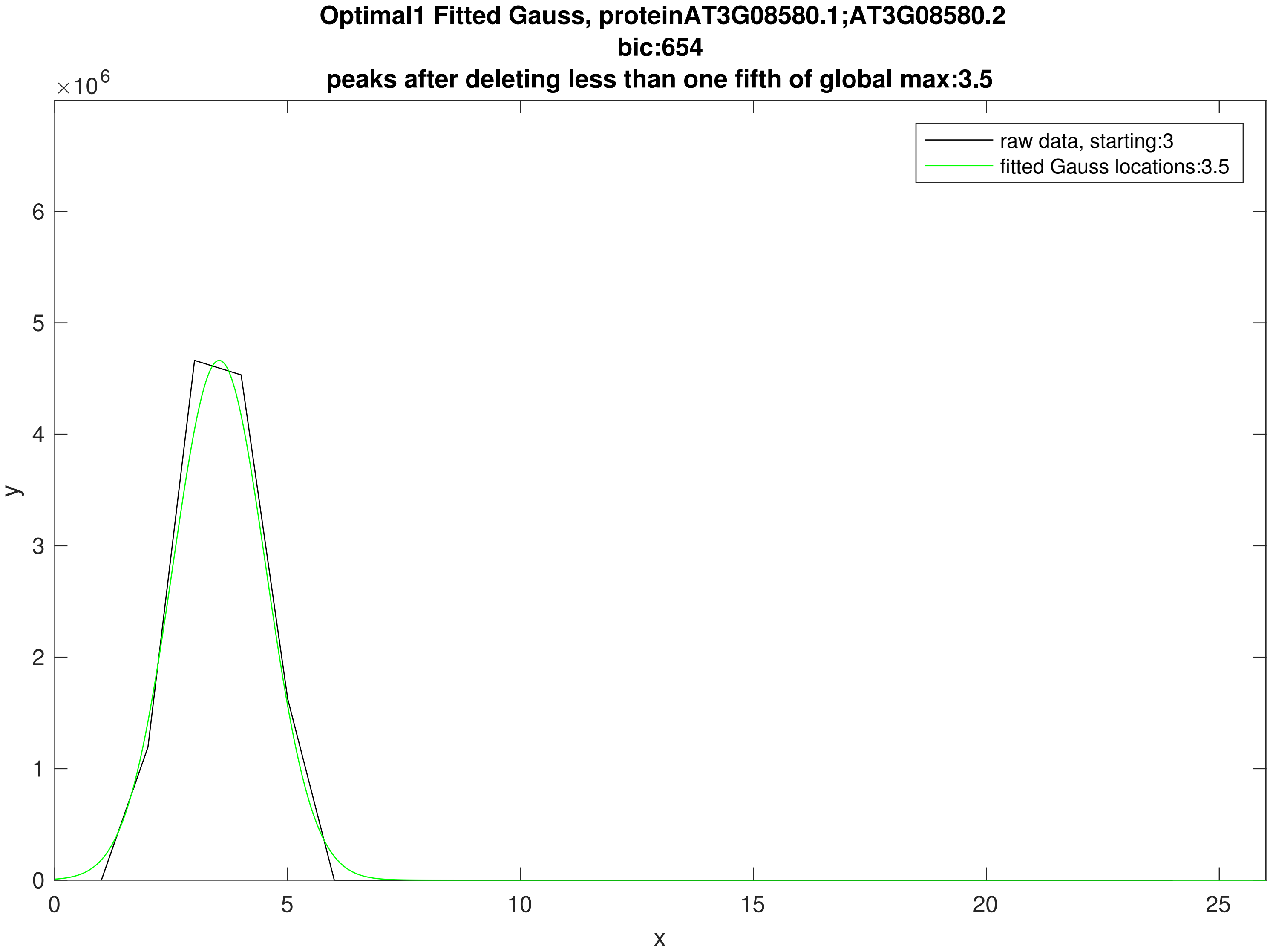
Optimal1 Fitted Gauss, proteinAT3G08580.1;AT3G08580.2
bic:654
peaks after deleting less than one fifth of global max:3.5
×10^6
raw data, starting:3
fitted Gauss locations:3.5
y
x
0
1
2
3
4
5
6
0
5
10
15
20
25

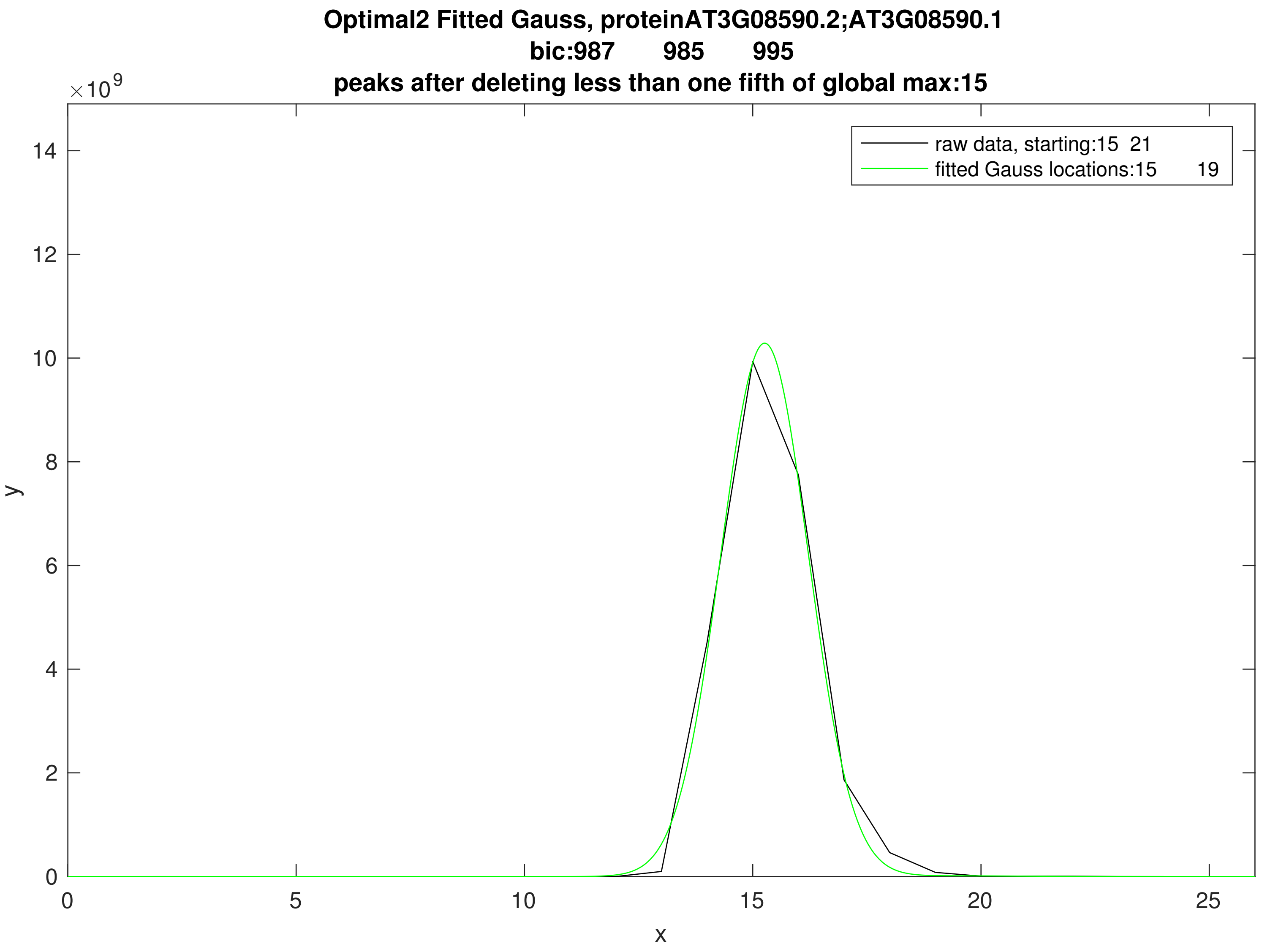
Optimal2 Fitted Gauss, proteinAT3G08590.2;AT3G08590.1
bic:987 985 995
peaks after deleting less than one fifth of global max:15
×10^9
raw data, starting:15 21
fitted Gauss locations:15 19
y
x
0
2
4
6
8
10
12
14
0
5
10
15
20
25

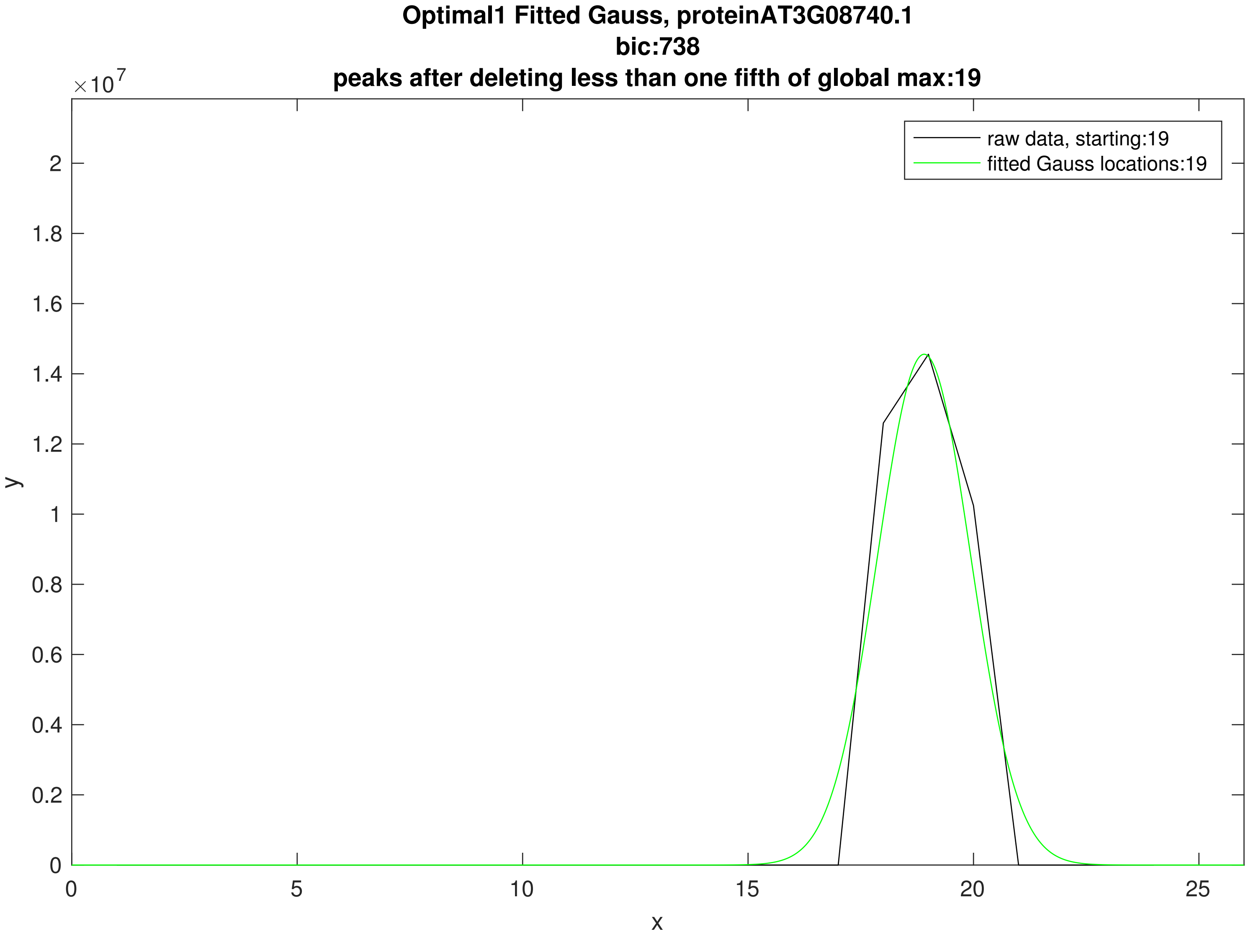
Optimal1 Fitted Gauss, proteinAT3G08740.1
bic:738
peaks after deleting less than one fifth of global max:19
×10^7
raw data, starting:19
fitted Gauss locations:19
y
x
0
0.2
0.4
0.6
0.8
1
1.2
1.4
1.6
1.8
2
5
10
15
20
25

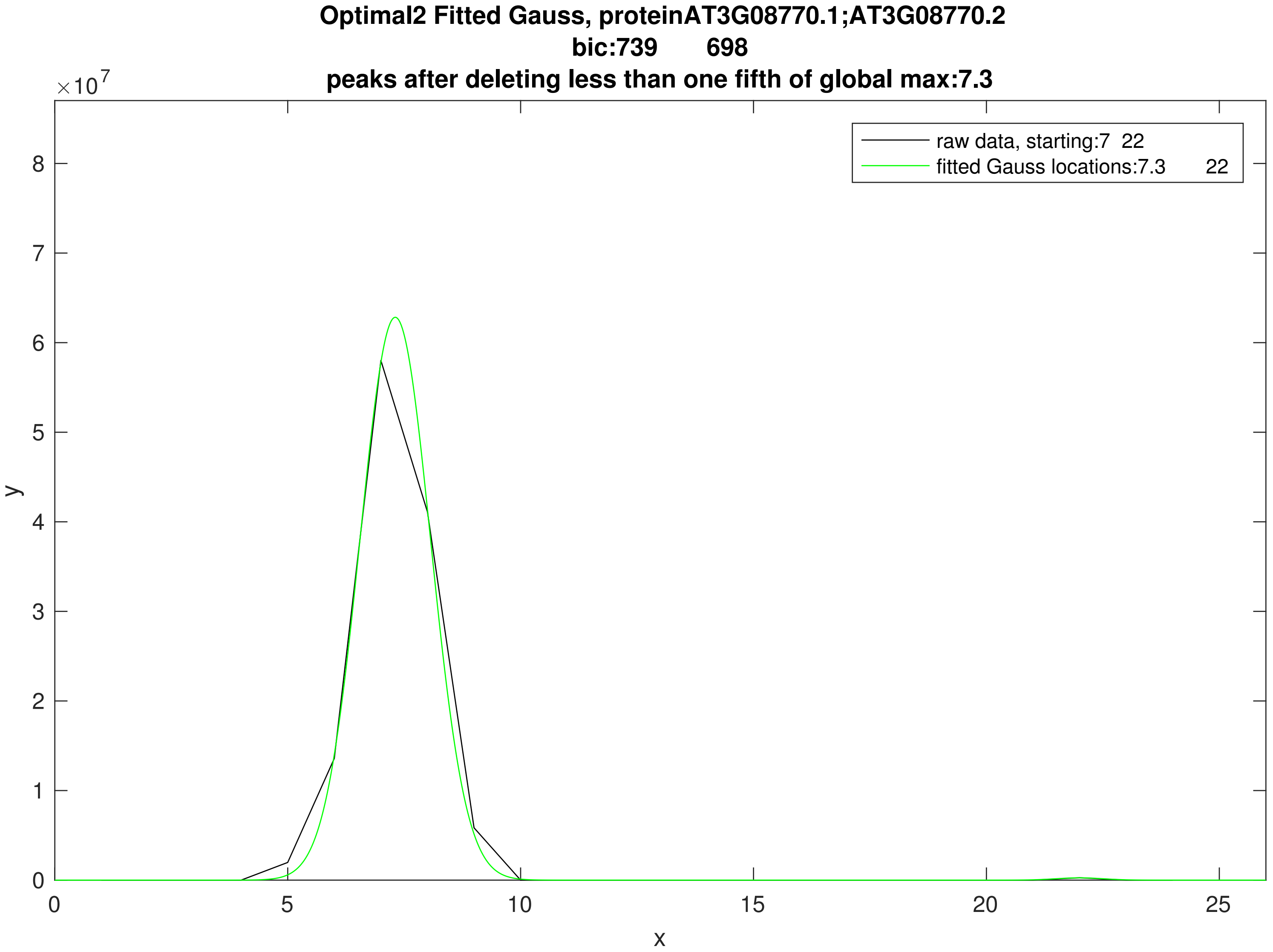
Optimal2 Fitted Gauss, proteinAT3G08770.1;AT3G08770.2
bic:739 698
peaks after deleting less than one fifth of global max:7.3
×10^7
raw data, starting:7 22
fitted Gauss locations:7.3 22
y
x
0
1
2
3
4
5
6
7
8
0
5
10
15
20
25

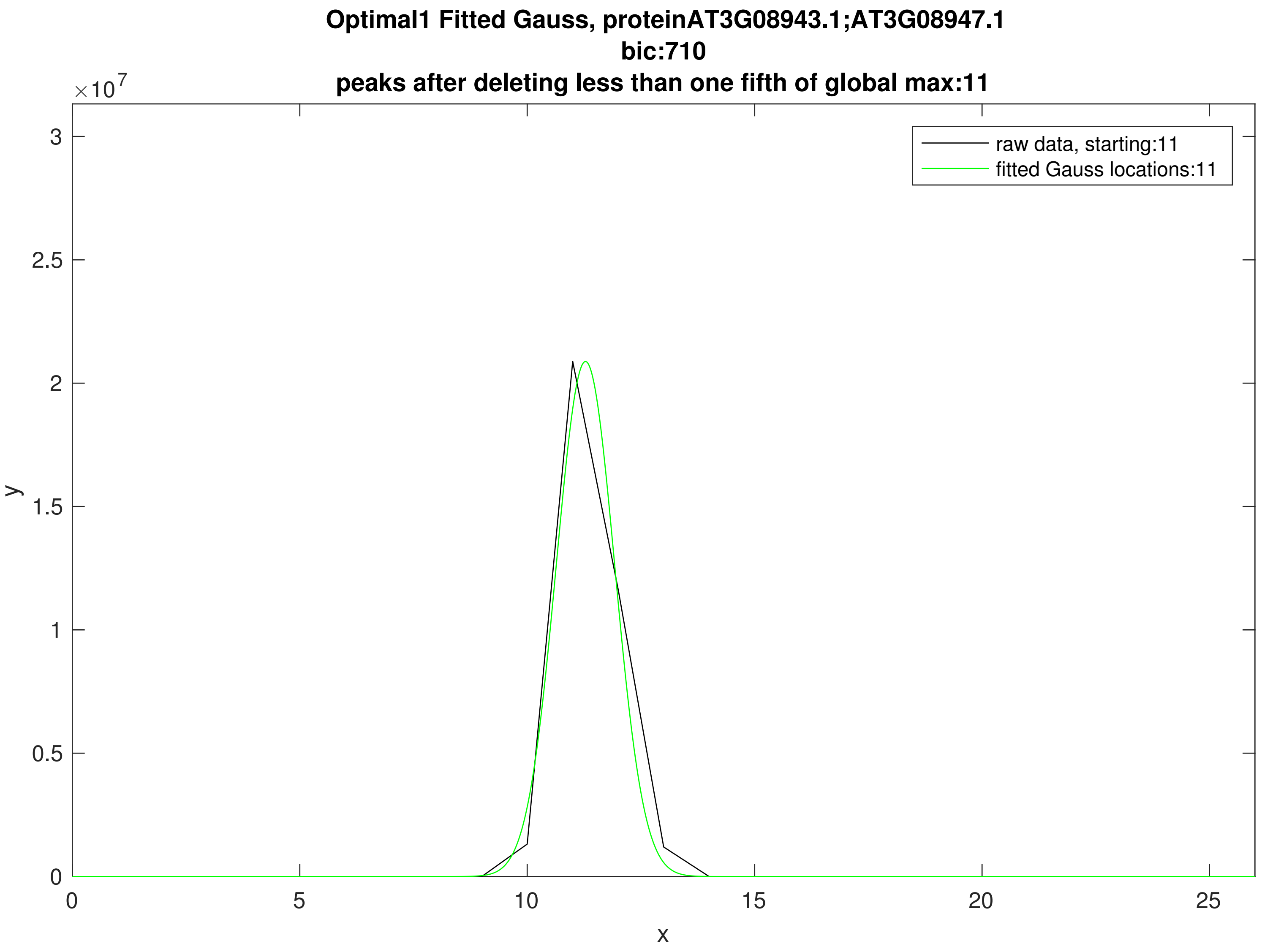

Optimal1 Fitted Gauss, proteinAT3G08943.1;AT3G08947.1
bic:710
peaks after deleting less than one fifth of global max:11
×10^7
3
2.5
2
1.5
1
0.5
0
y
0
5
10
15
20
25
x
raw data, starting:11
fitted Gauss locations:11

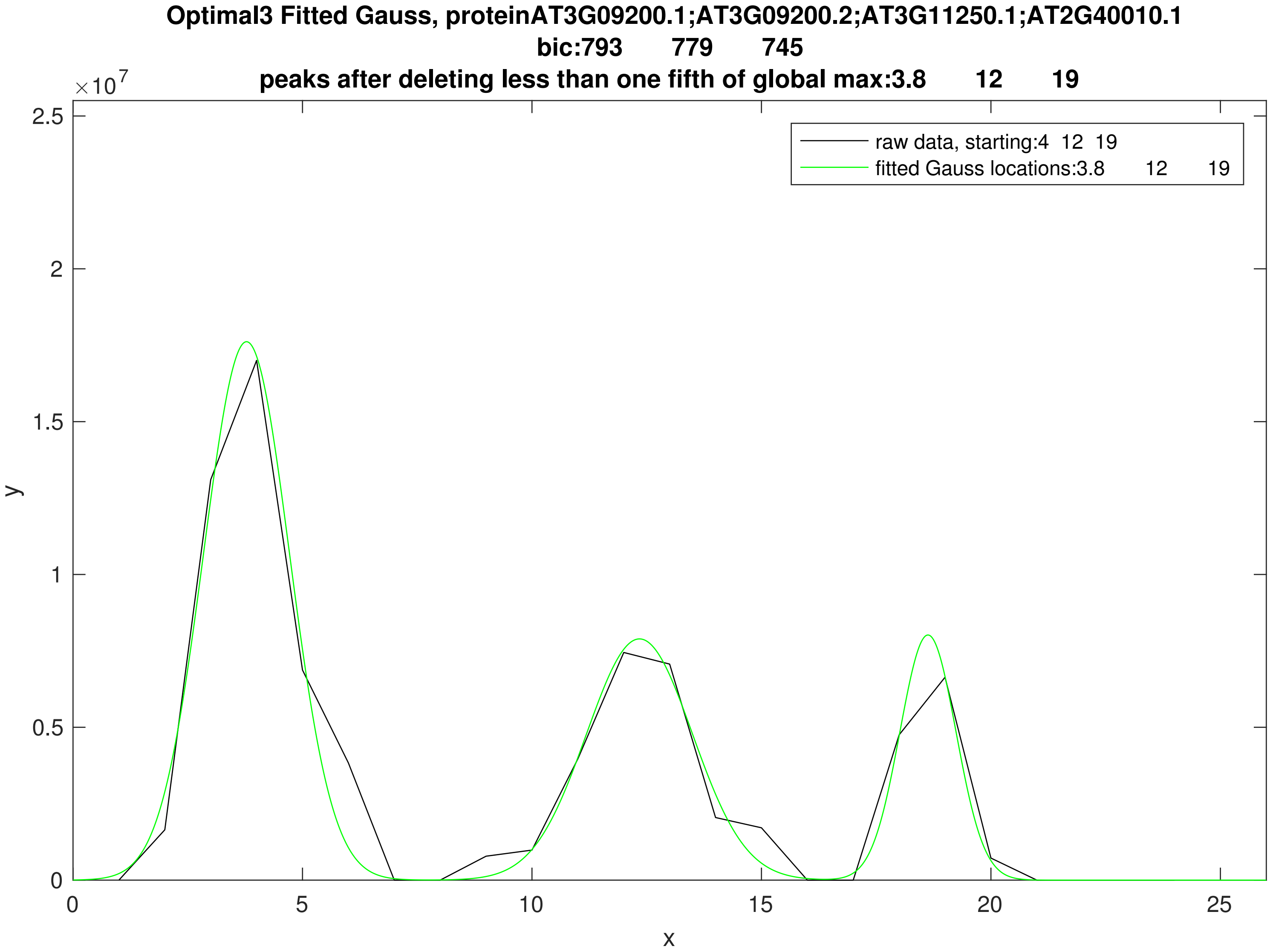

Optimal3 Fitted Gauss, proteinAT3G09200.1;AT3G09200.2;AT3G11250.1;AT2G40010.1
bic:793 779 745
peaks after deleting less than one fifth of global max:3.8 12 19
×10^7
2.5
2
1.5
1
0.5
0
y
0
5
10
15
20
25
x
raw data, starting:4 12 19
fitted Gauss locations:3.8 12 19

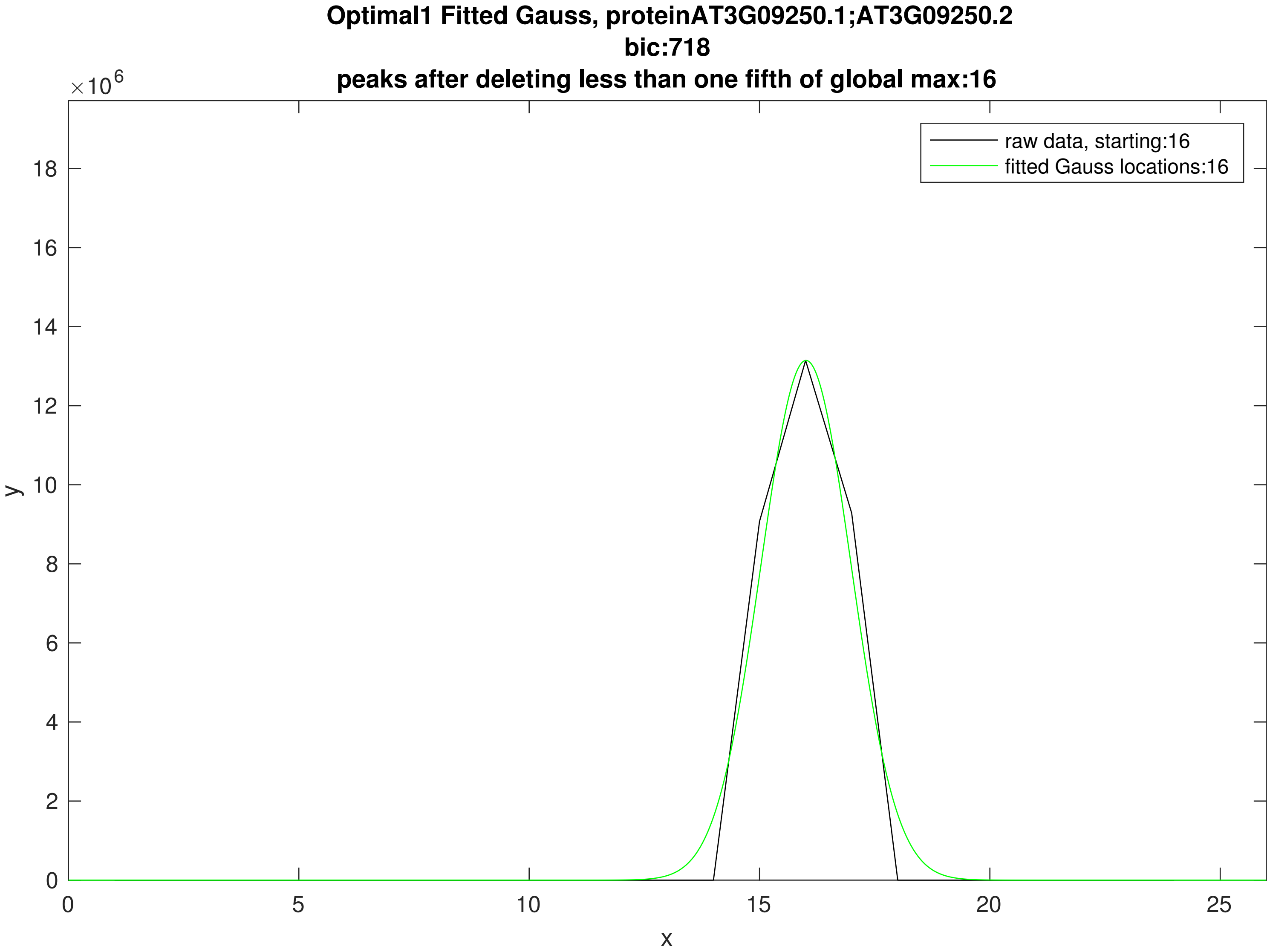

Optimal1 Fitted Gauss, proteinAT3G09250.1;AT3G09250.2
bic:718
peaks after deleting less than one fifth of global max:16
×10^6
raw data, starting:16
fitted Gauss locations:16
18
16
14
12
10
8
6
4
2
0
y
0
5
10
15
20
25
x

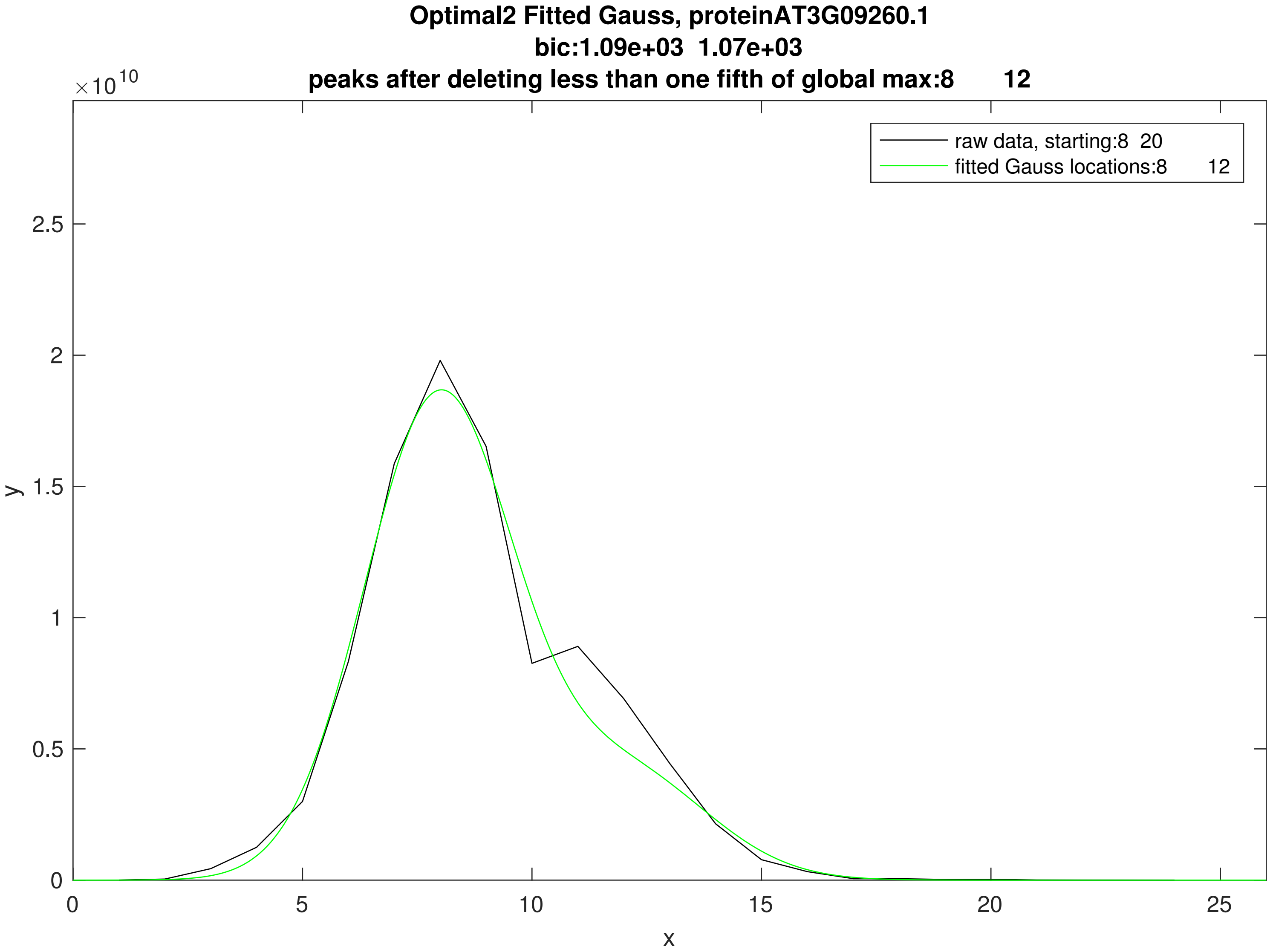

Optimal2 Fitted Gauss, proteinAT3G09260.1
bic:1.09e+03 1.07e+03
peaks after deleting less than one fifth of global max:8 12
×10^10
raw data, starting:8 20
fitted Gauss locations:8 12
2.5
2
1.5
1
0.5
0
y
0
5
10
15
20
25
x

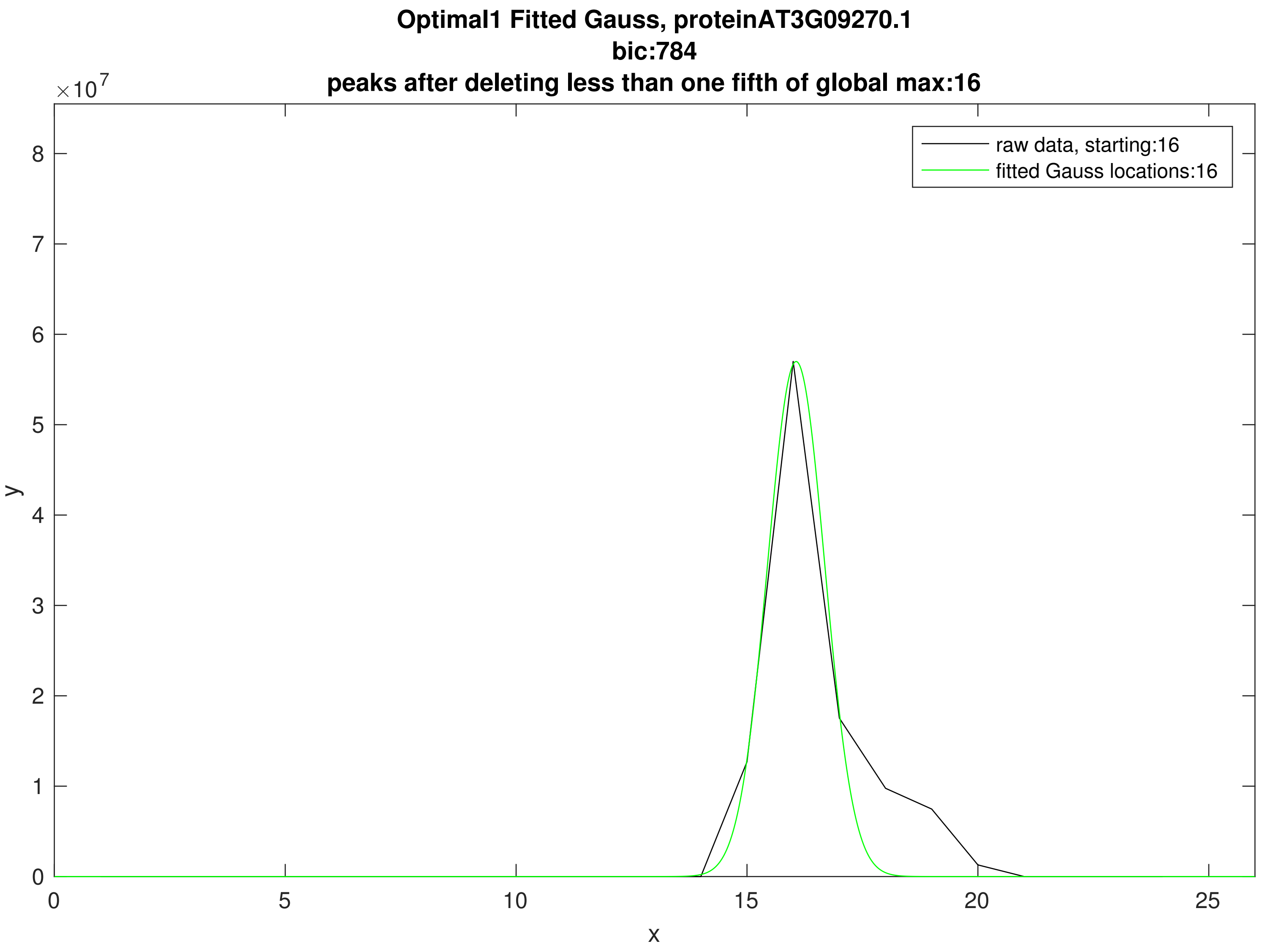
Optimal1 Fitted Gauss, proteinAT3G09270.1
bic:784
peaks after deleting less than one fifth of global max:16
×10^7
raw data, starting:16
fitted Gauss locations:16
y
x
0
1
2
3
4
5
6
7
8
0
5
10
15
20
25

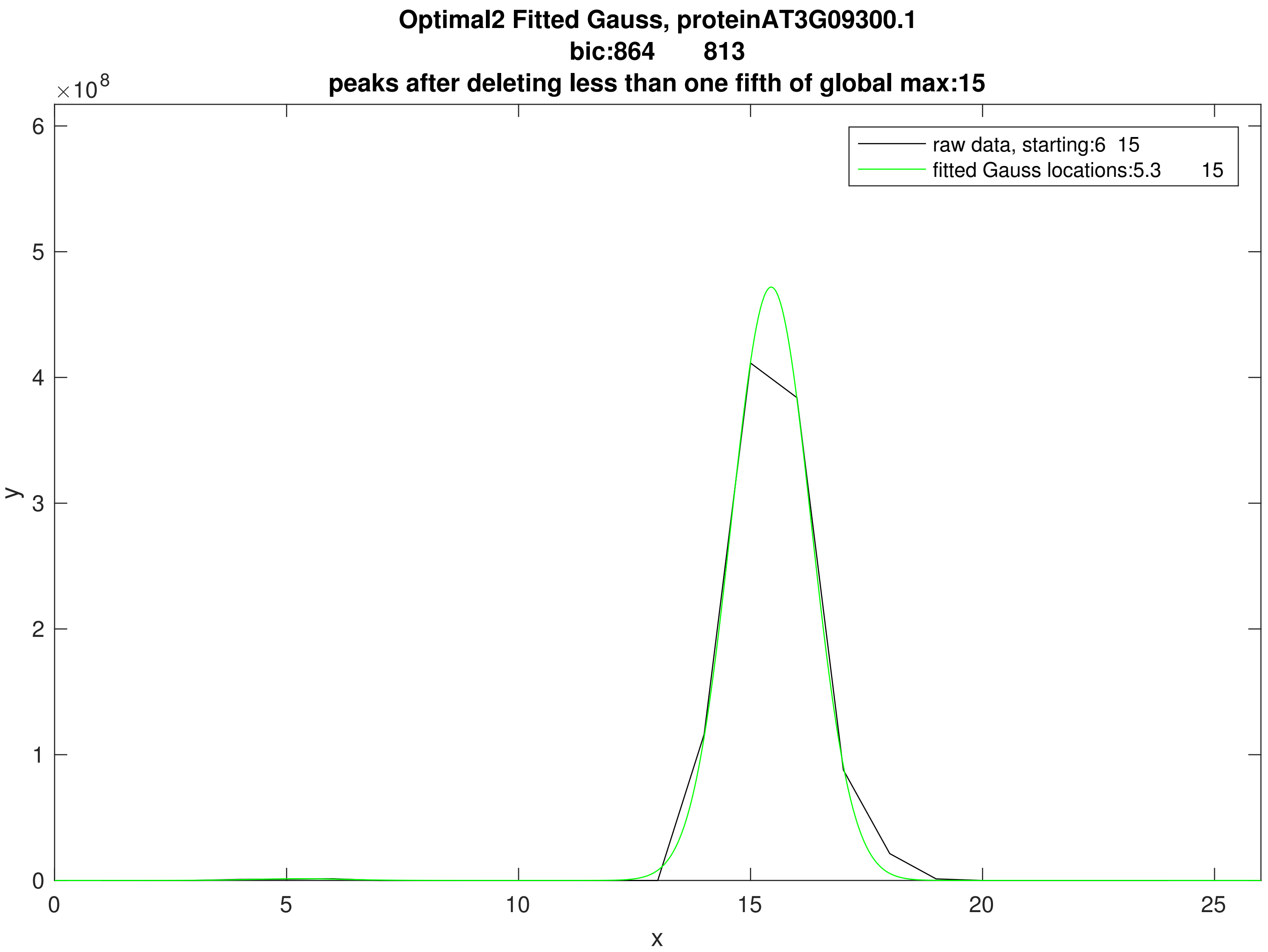
Optimal2 Fitted Gauss, proteinAT3G09300.1
bic:864 813
peaks after deleting less than one fifth of global max:15
×10^8
raw data, starting:6 15
fitted Gauss locations:5.3 15
y
x
0
1
2
3
4
5
6
0
5
10
15
20
25

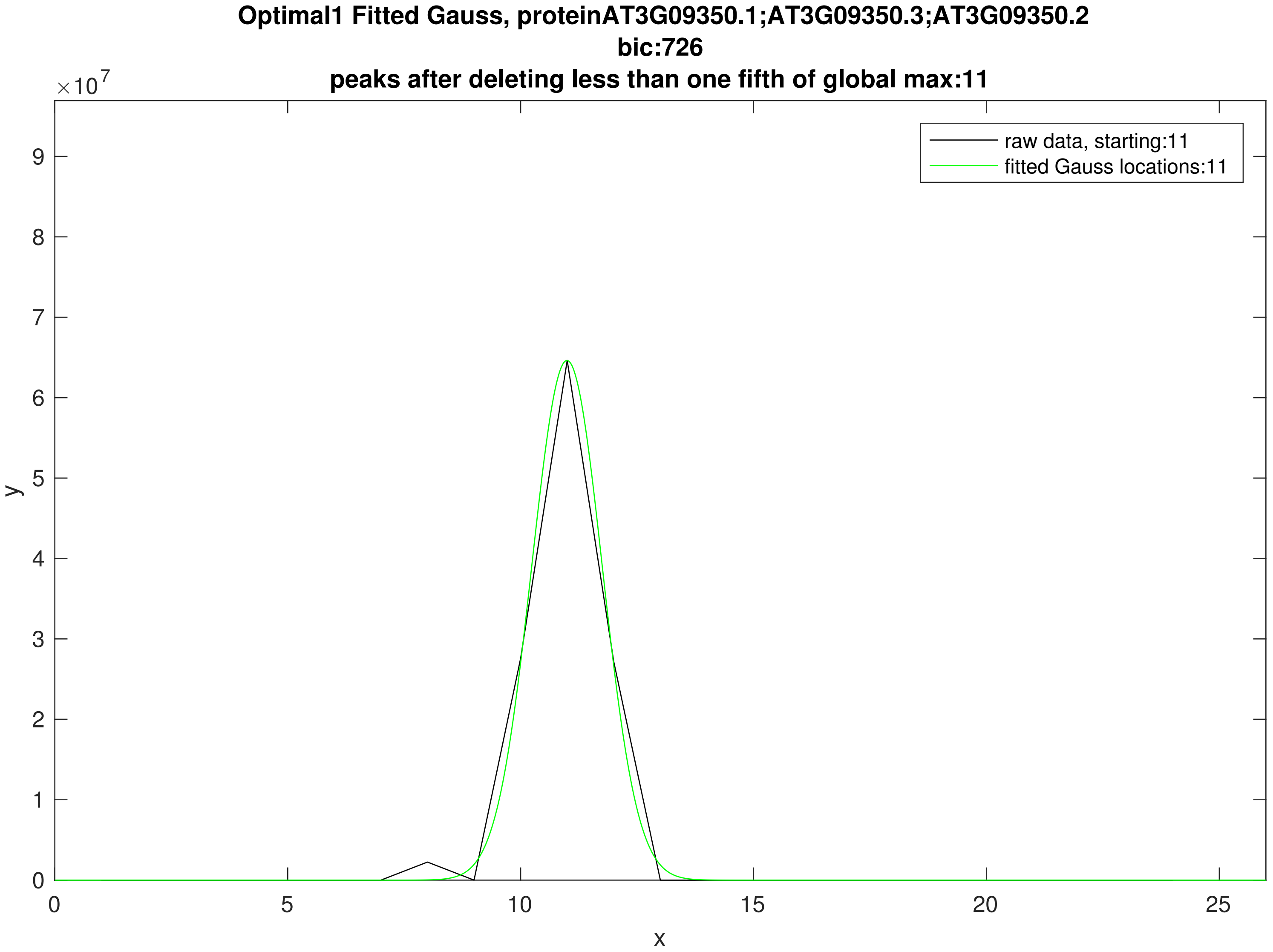
Optimal1 Fitted Gauss, proteinAT3G09350.1;AT3G09350.3;AT3G09350.2
bic:726
peaks after deleting less than one fifth of global max:11
×10^7
raw data, starting:11
fitted Gauss locations:11
y
x
0
1
2
3
4
5
6
7
8
9
0
5
10
15
20
25

Optimal1 Fitted Gauss, proteinAT3G09410.1;AT3G09410.3
bic:812
peaks after deleting less than one fifth of global max:16

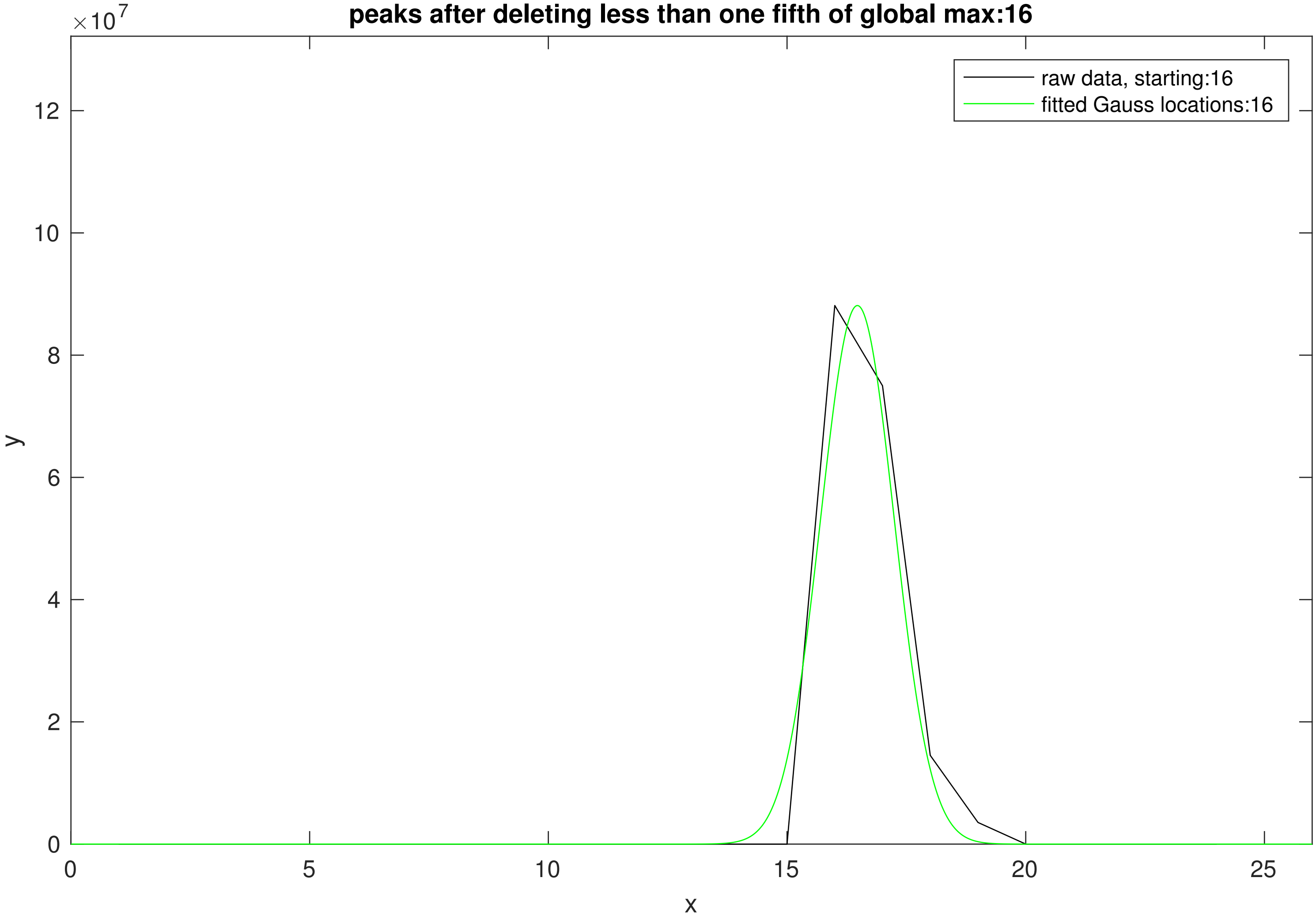

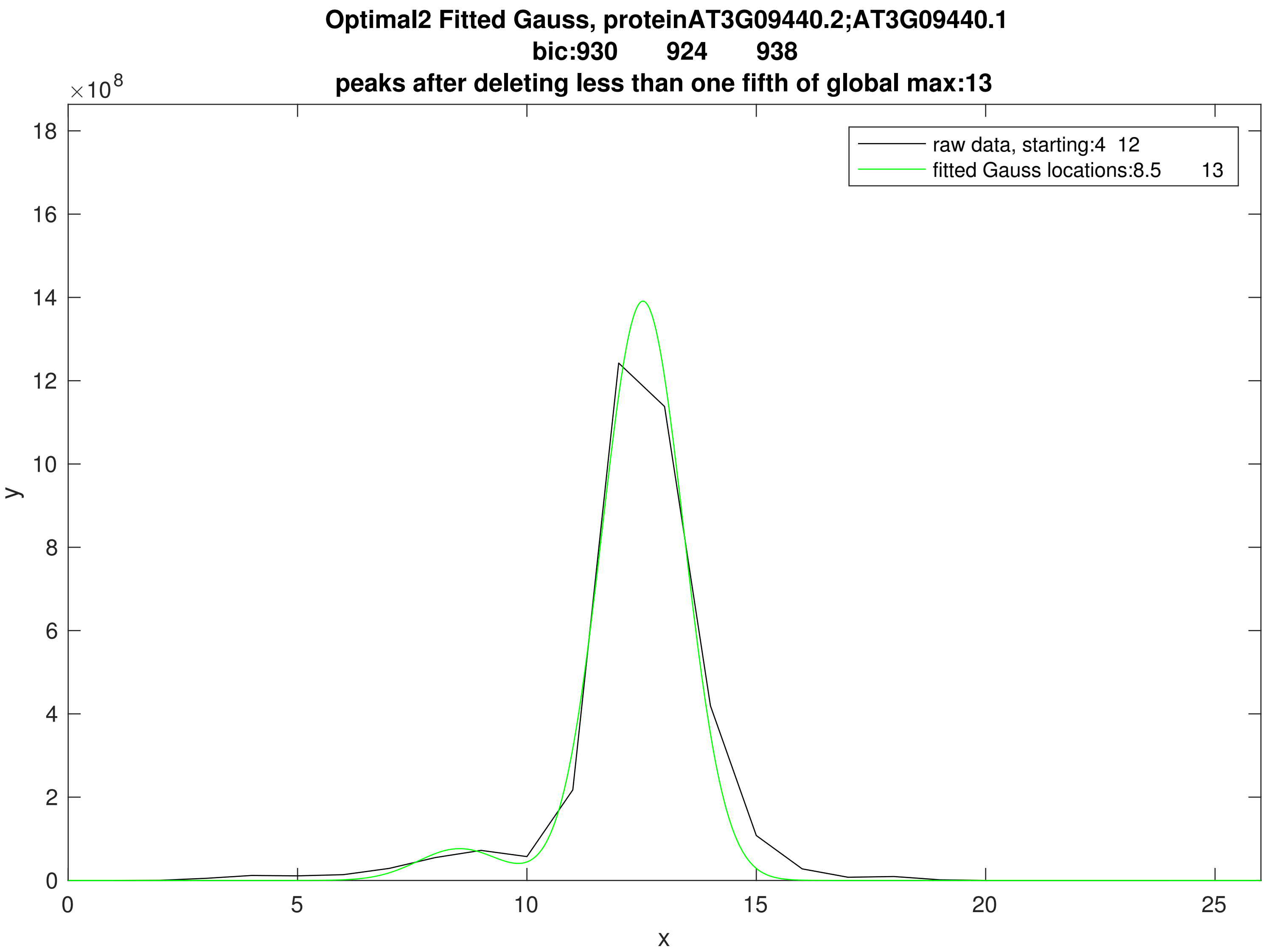

Optimal2 Fitted Gauss, proteinAT3G09440.2;AT3G09440.1
bic:930 924 938
peaks after deleting less than one fifth of global max:13
×10^8
raw data, starting:4 12
fitted Gauss locations:8.5 13
y
x
0
2
4
6
8
10
12
14
16
18
0
5
10
15
20
25

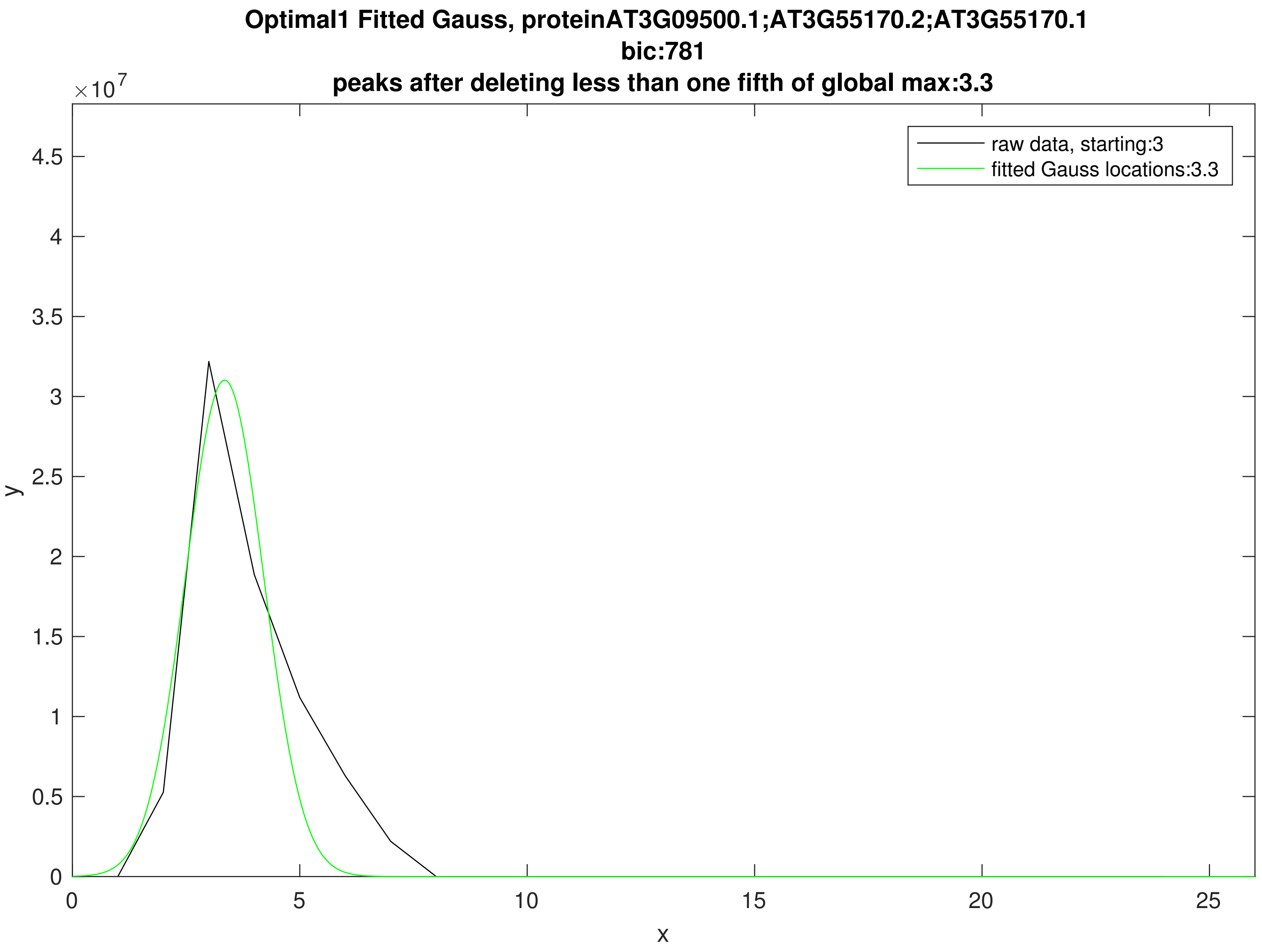
Optimal1 Fitted Gauss, proteinAT3G09500.1;AT3G55170.2;AT3G55170.1
bic:781
peaks after deleting less than one fifth of global max:3.3
×10^7
raw data, starting:3
fitted Gauss locations:3.3
y
x
0
0.5
1
1.5
2
2.5
3
3.5
4
4.5
5
10
15
20
25

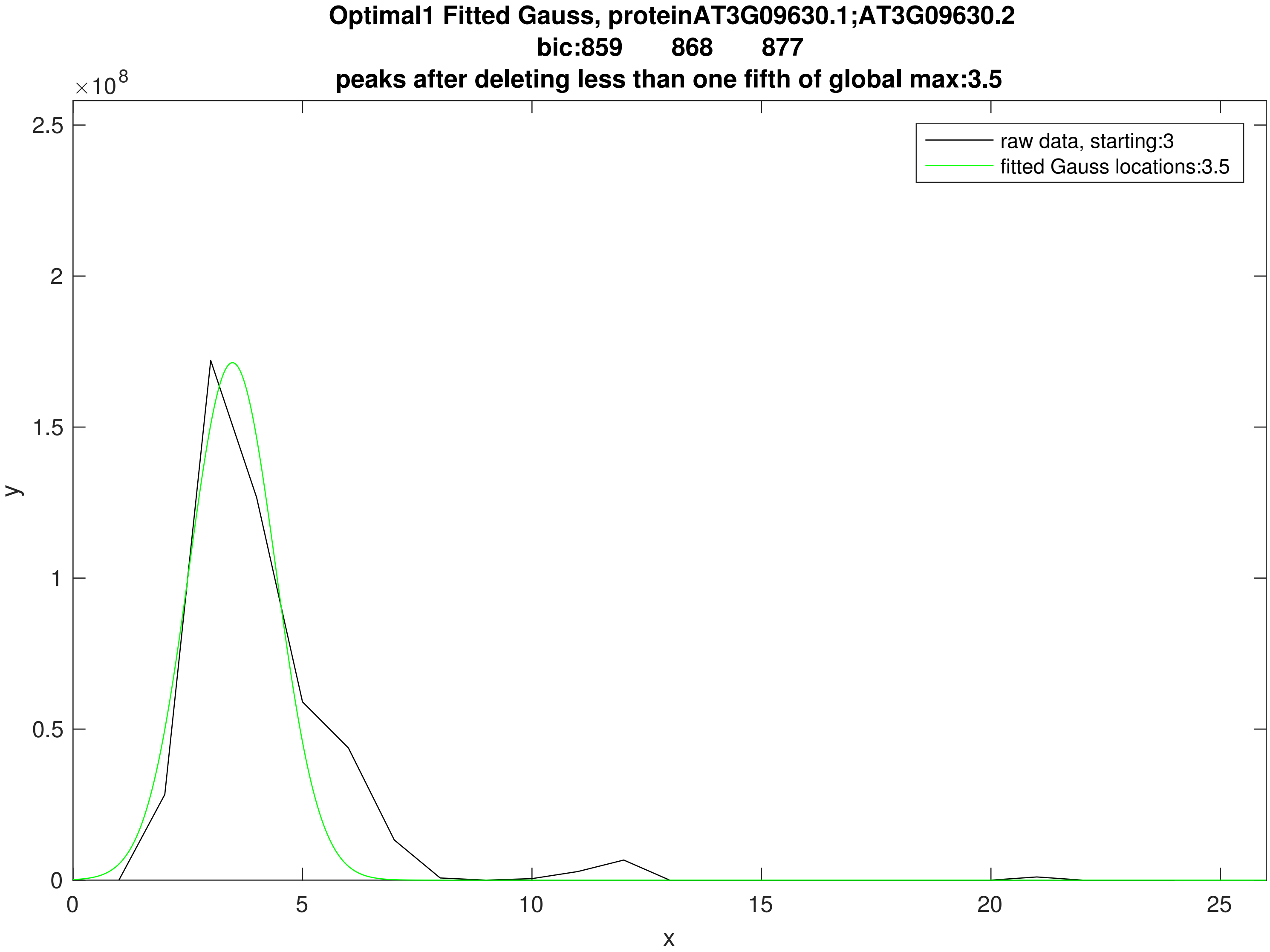
Optimal1 Fitted Gauss, proteinAT3G09630.1;AT3G09630.2
bic:859 868 877
peaks after deleting less than one fifth of global max:3.5
×10^8
2.5
2
1.5
1
0.5
0
y
0
5
10
15
20
25
x
raw data, starting:3
fitted Gauss locations:3.5

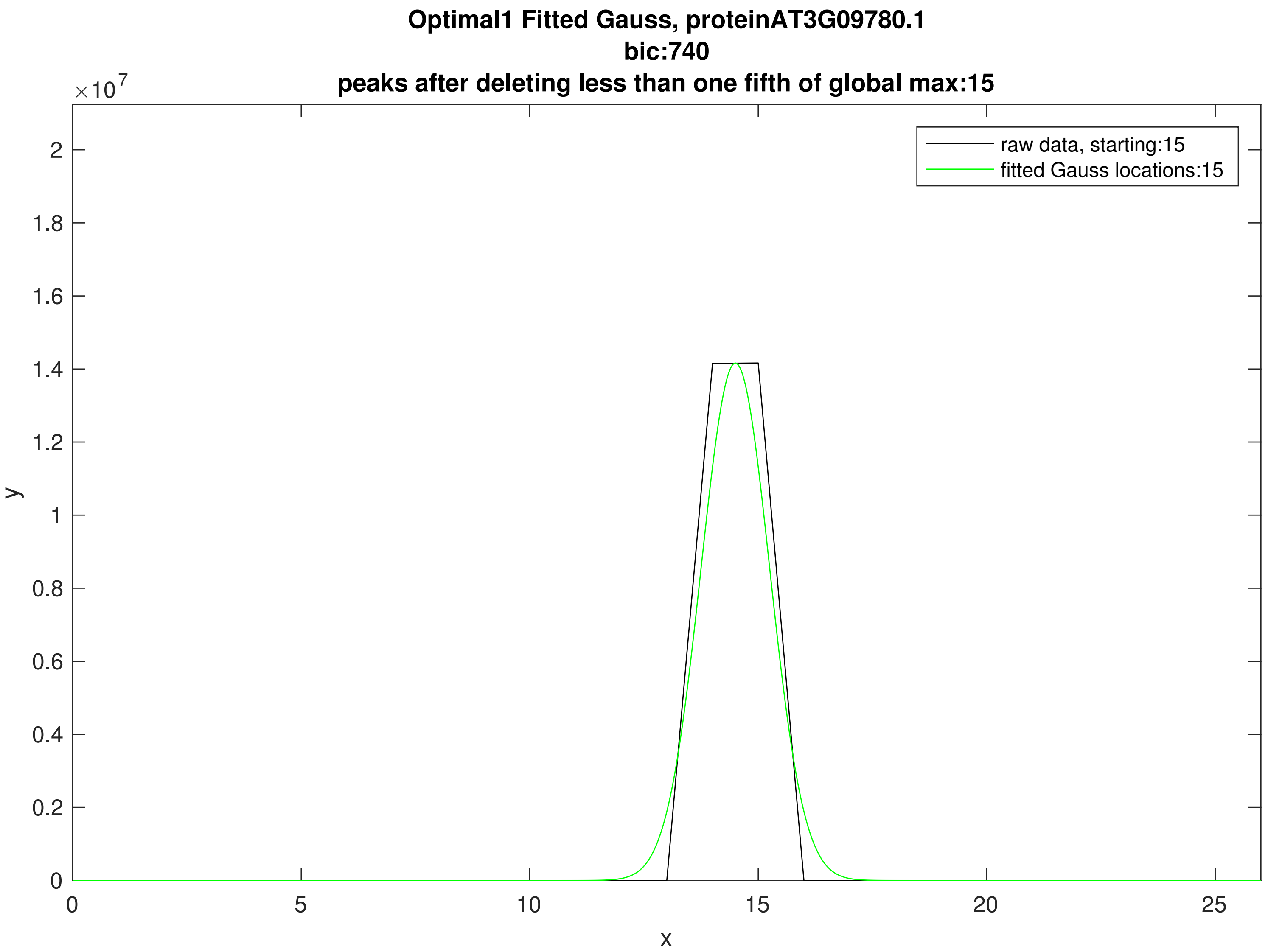
Optimal1 Fitted Gauss, proteinAT3G09780.1
bic:740
peaks after deleting less than one fifth of global max:15
×10^7
raw data, starting:15
fitted Gauss locations:15
y
x
0
0.2
0.4
0.6
0.8
1
1.2
1.4
1.6
1.8
2
0
5
10
15
20
25

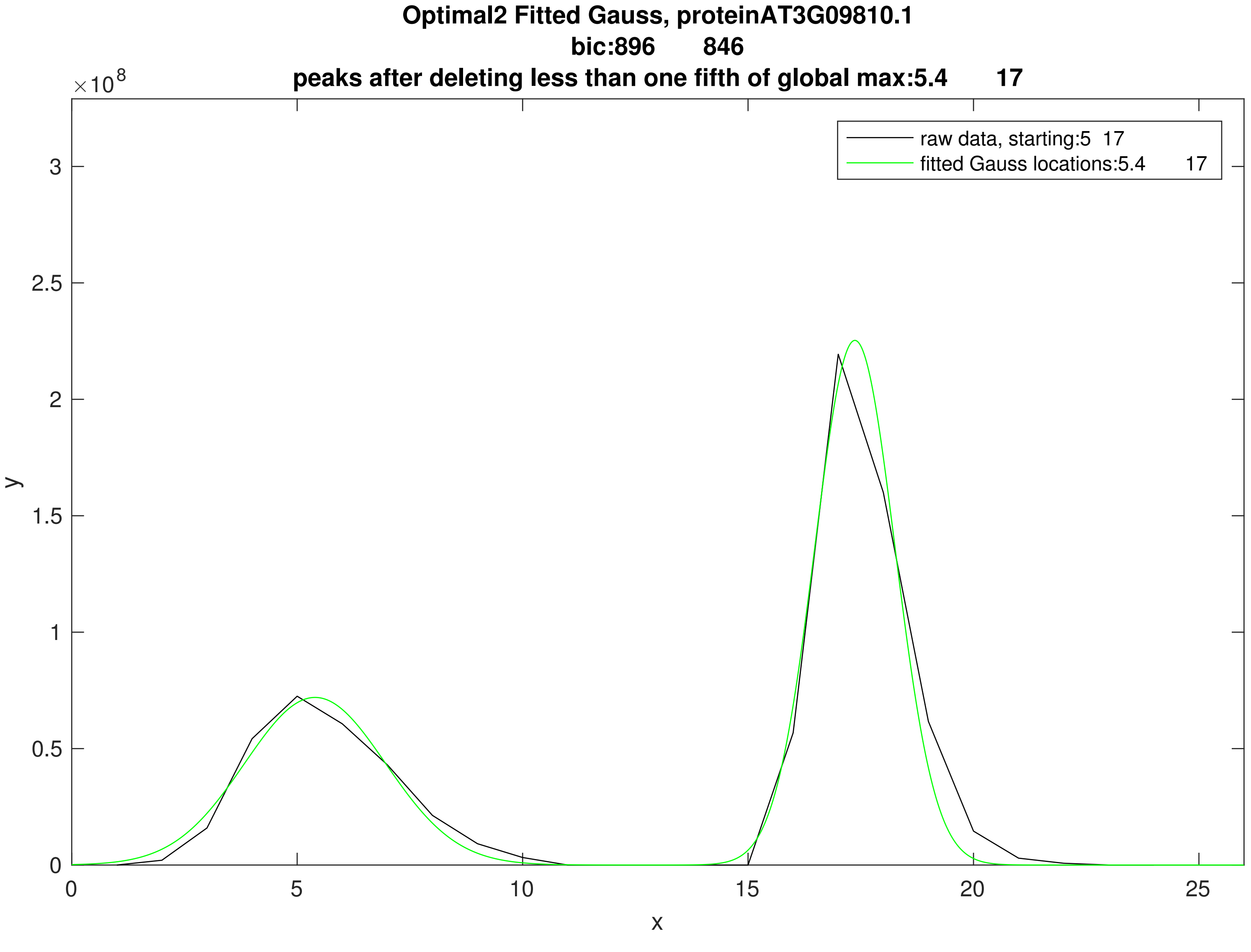
Optimal2 Fitted Gauss, proteinAT3G09810.1
bic:896 846
peaks after deleting less than one fifth of global max:5.4 17
×10^8
raw data, starting:5 17
fitted Gauss locations:5.4 17
y
x
0
0.5
1
1.5
2
2.5
3
0
5
10
15
20
25

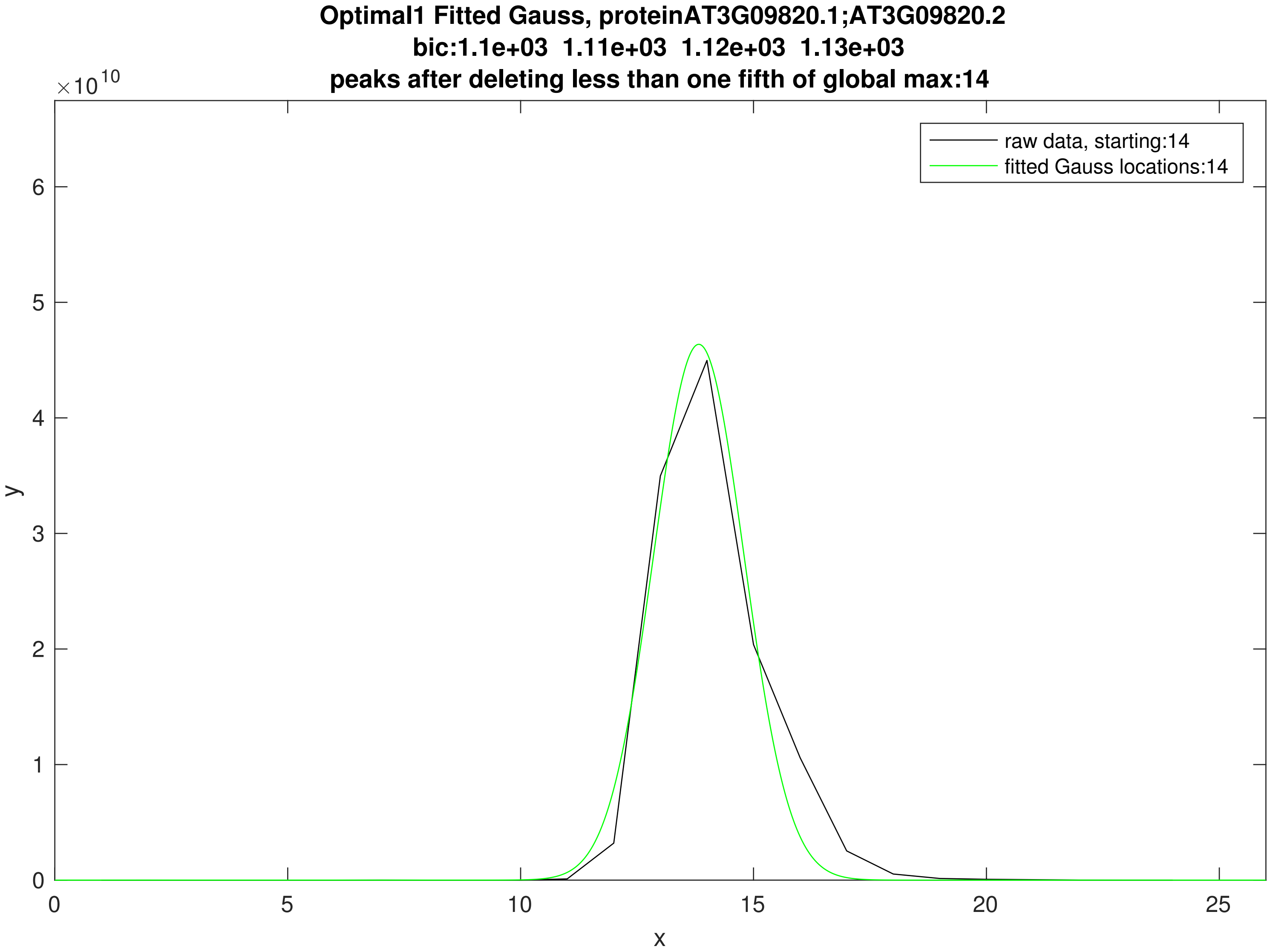
Optimal1 Fitted Gauss, proteinAT3G09820.1;AT3G09820.2
bic:1.1e+03 1.11e+03 1.12e+03 1.13e+03
peaks after deleting less than one fifth of global max:14
×10^10
raw data, starting:14
fitted Gauss locations:14
y
x
0
1
2
3
4
5
6
0
5
10
15
20
25

#### Optimal2 Fitted Gauss, proteinAT3G09840.1
#### bic:928 903 913 922
#### peaks after deleting less than one fifth of global max:5.4

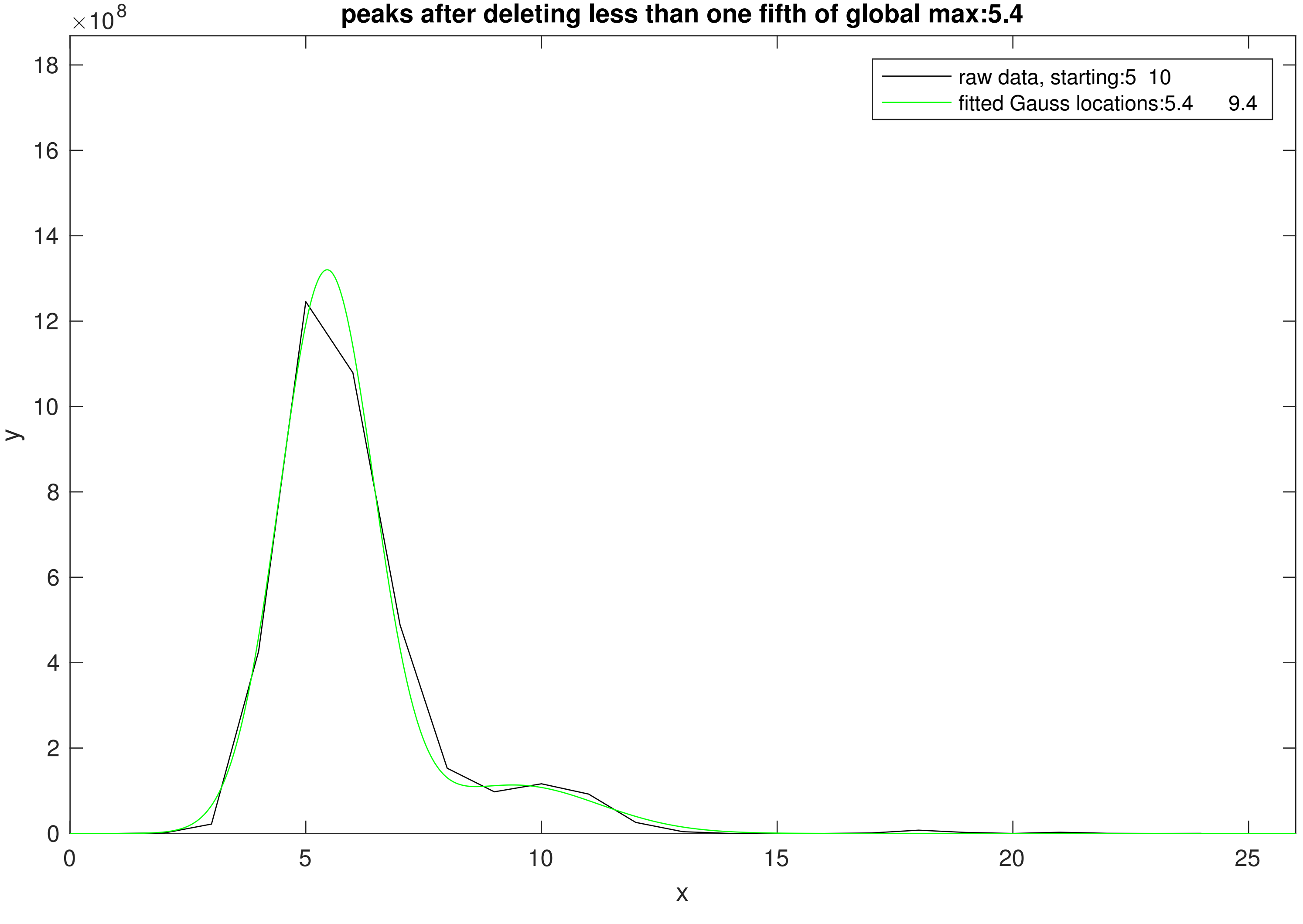

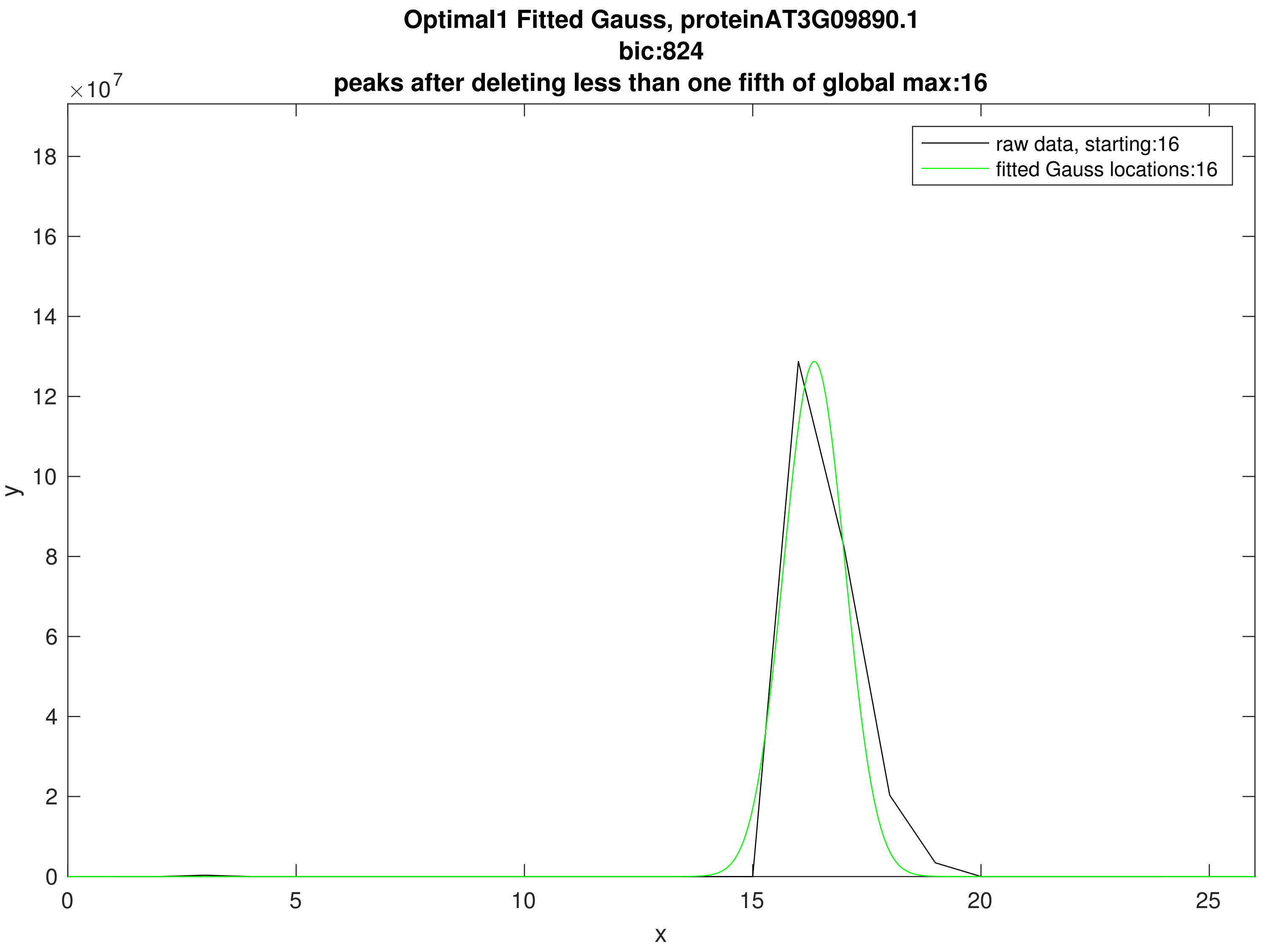

Optimal1 Fitted Gauss, proteinAT3G09890.1
bic:824
peaks after deleting less than one fifth of global max:16
×10^7
18
16
14
12
10
8
6
4
2
0
y
0
5
10
15
20
25
x
raw data, starting:16
fitted Gauss locations:16

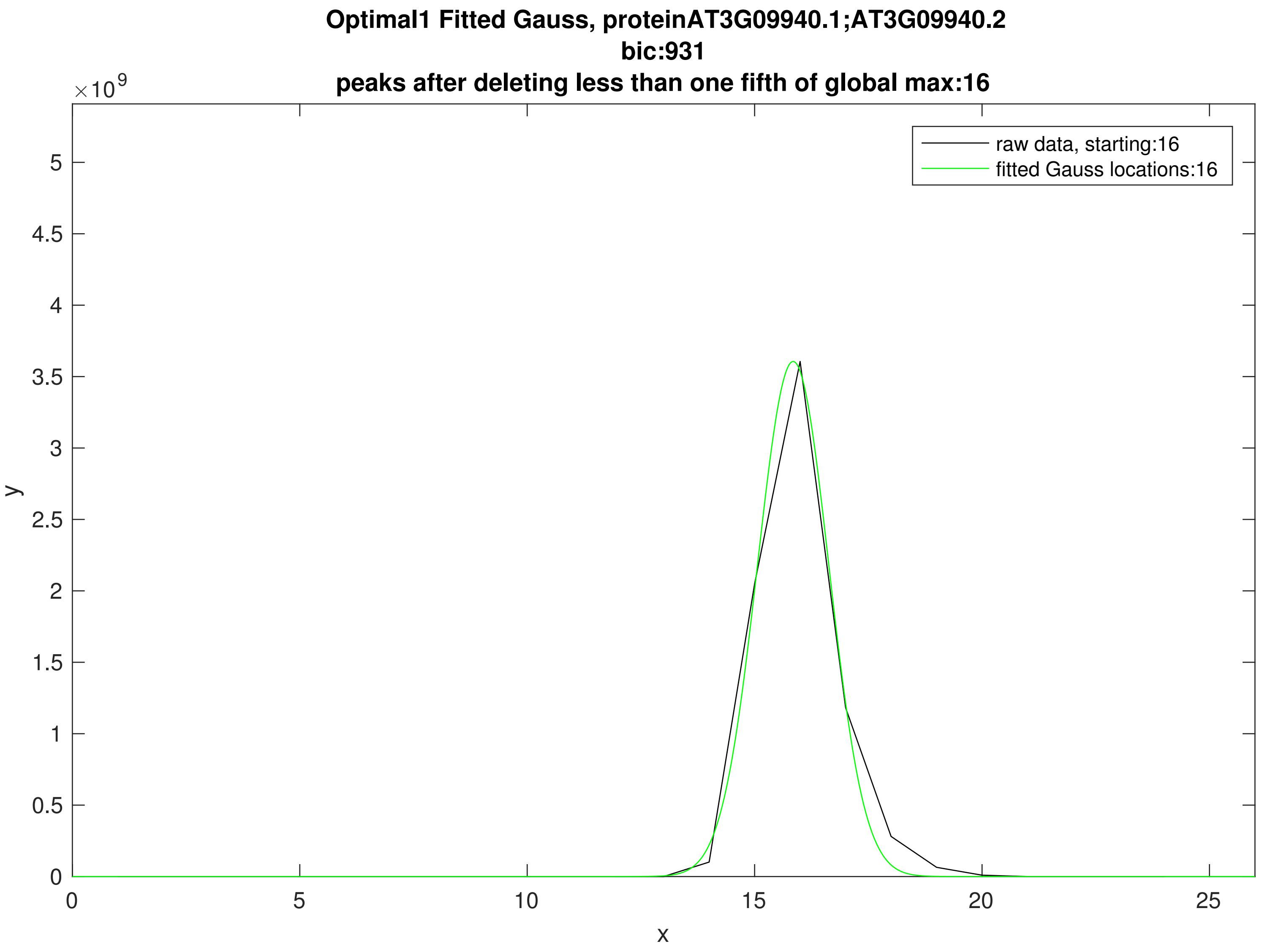
Optimal1 Fitted Gauss, proteinAT3G09940.1;AT3G09940.2
bic:931
peaks after deleting less than one fifth of global max:16
×10^9
raw data, starting:16
fitted Gauss locations:16
y
x
0
0.5
1
1.5
2
2.5
3
3.5
4
4.5
5
0
5
10
15
20
25

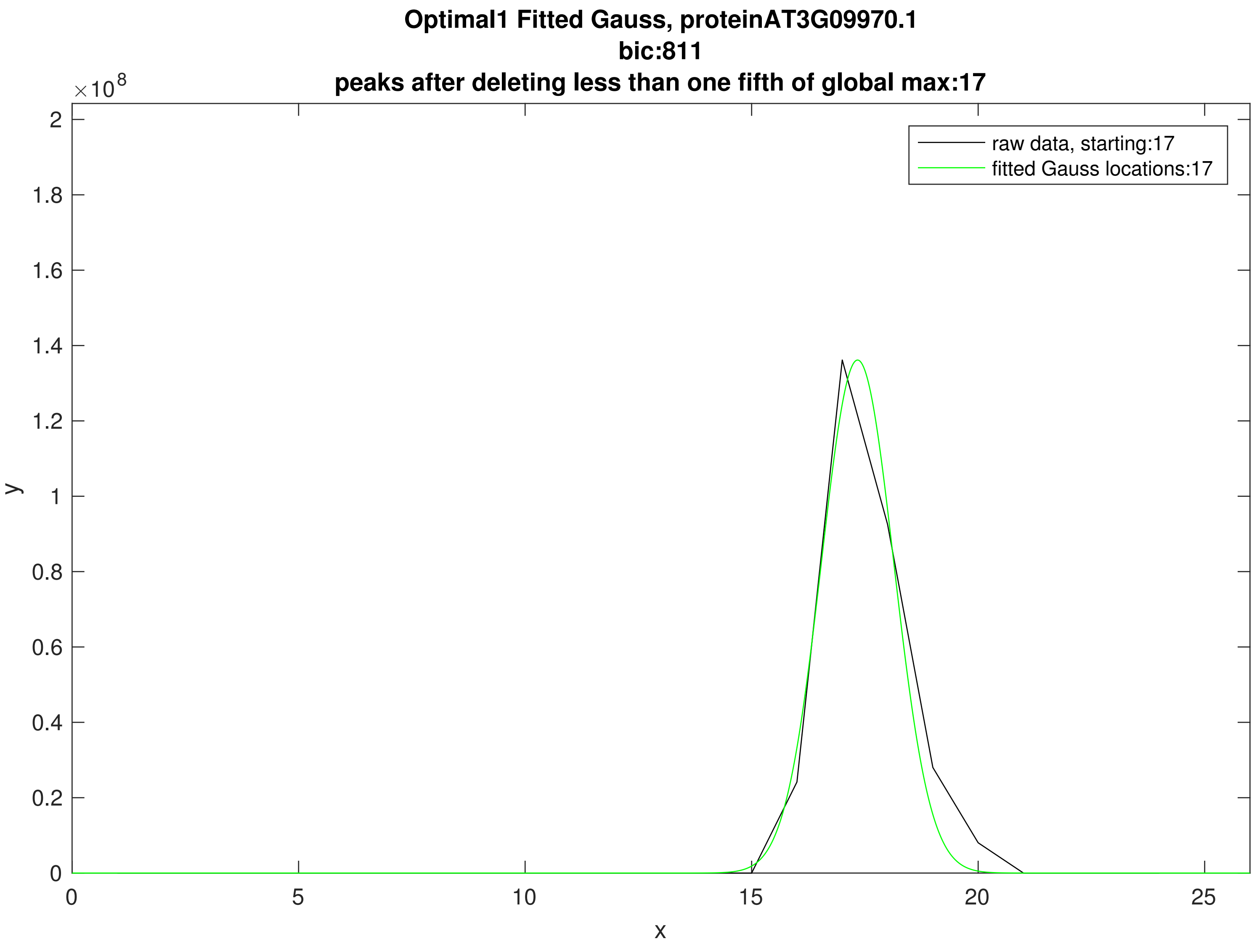

Optimal1 Fitted Gauss, proteinAT3G09970.1
bic:811
peaks after deleting less than one fifth of global max:17
×10^8
raw data, starting:17
fitted Gauss locations:17
y
x
0
0.2
0.4
0.6
0.8
1
1.2
1.4
1.6
1.8
2
0
5
10
15
20
25

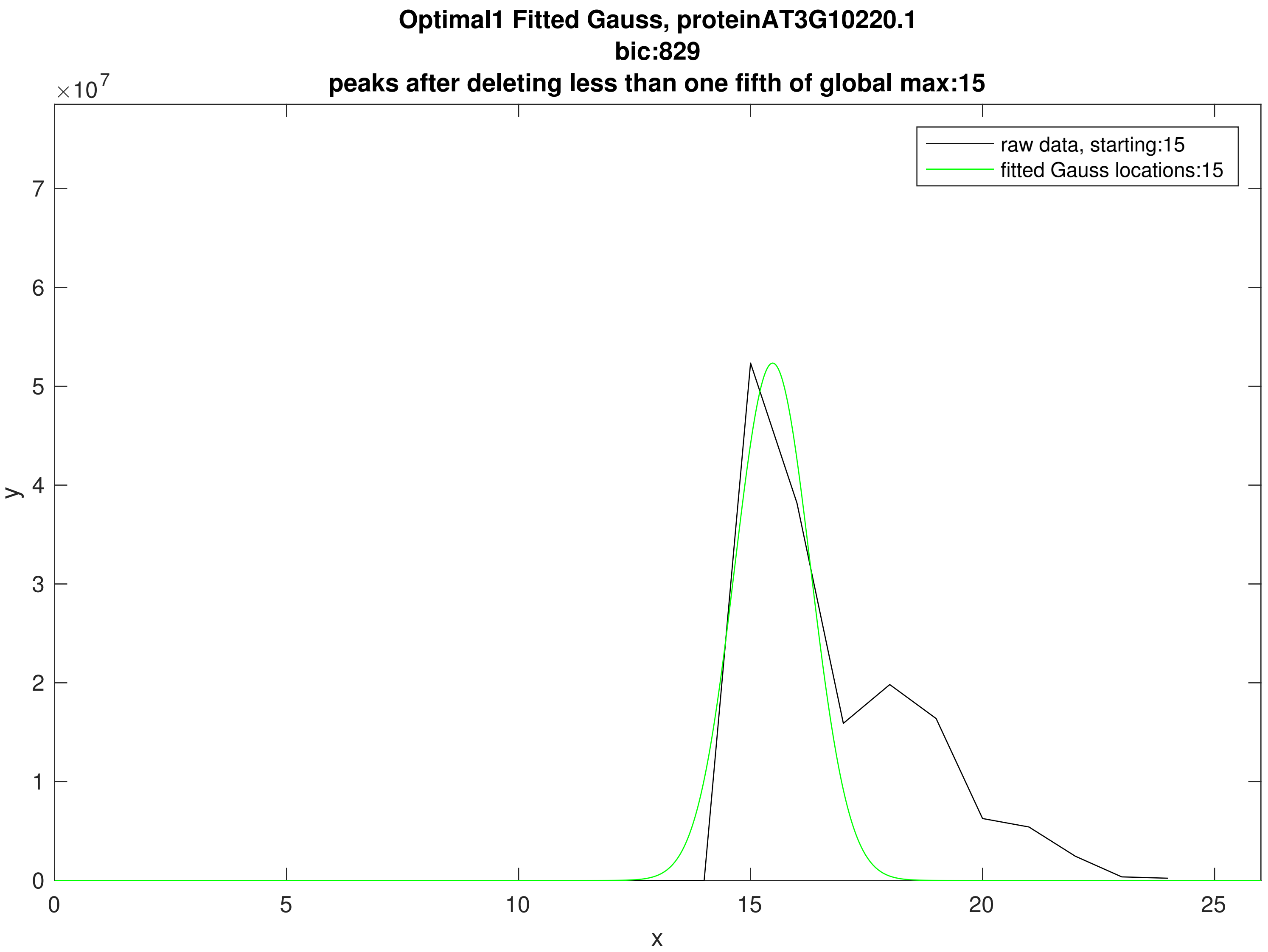
Optimal1 Fitted Gauss, proteinAT3G10220.1
bic:829
peaks after deleting less than one fifth of global max:15
×10^7
raw data, starting:15
fitted Gauss locations:15
y
x
0
1
2
3
4
5
6
7
0
5
10
15
20
25

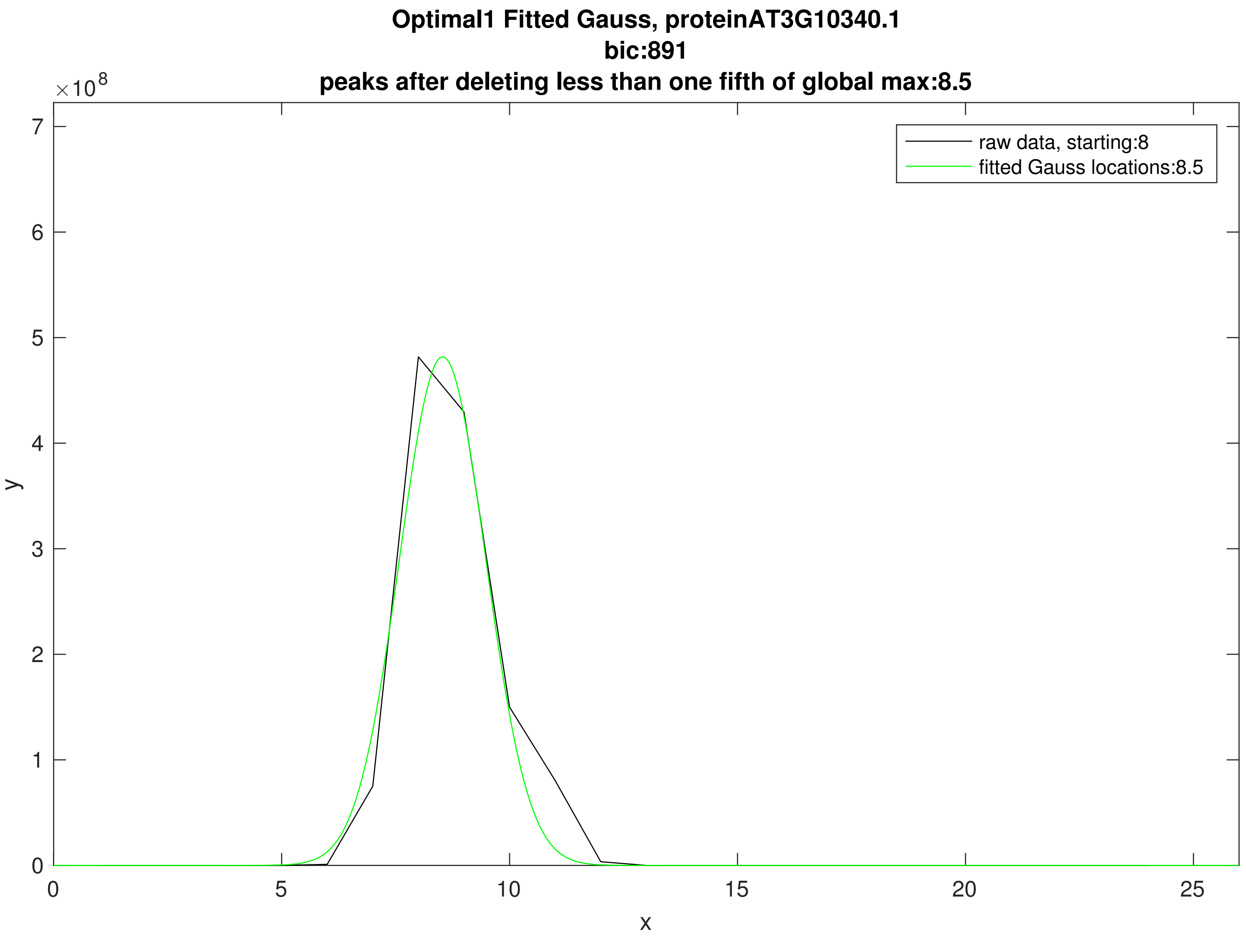

Optimal1 Fitted Gauss, proteinAT3G10340.1
bic:891
peaks after deleting less than one fifth of global max:8.5
×10^8
raw data, starting:8
fitted Gauss locations:8.5
y
x
0
1
2
3
4
5
6
7
0
5
10
15
20
25

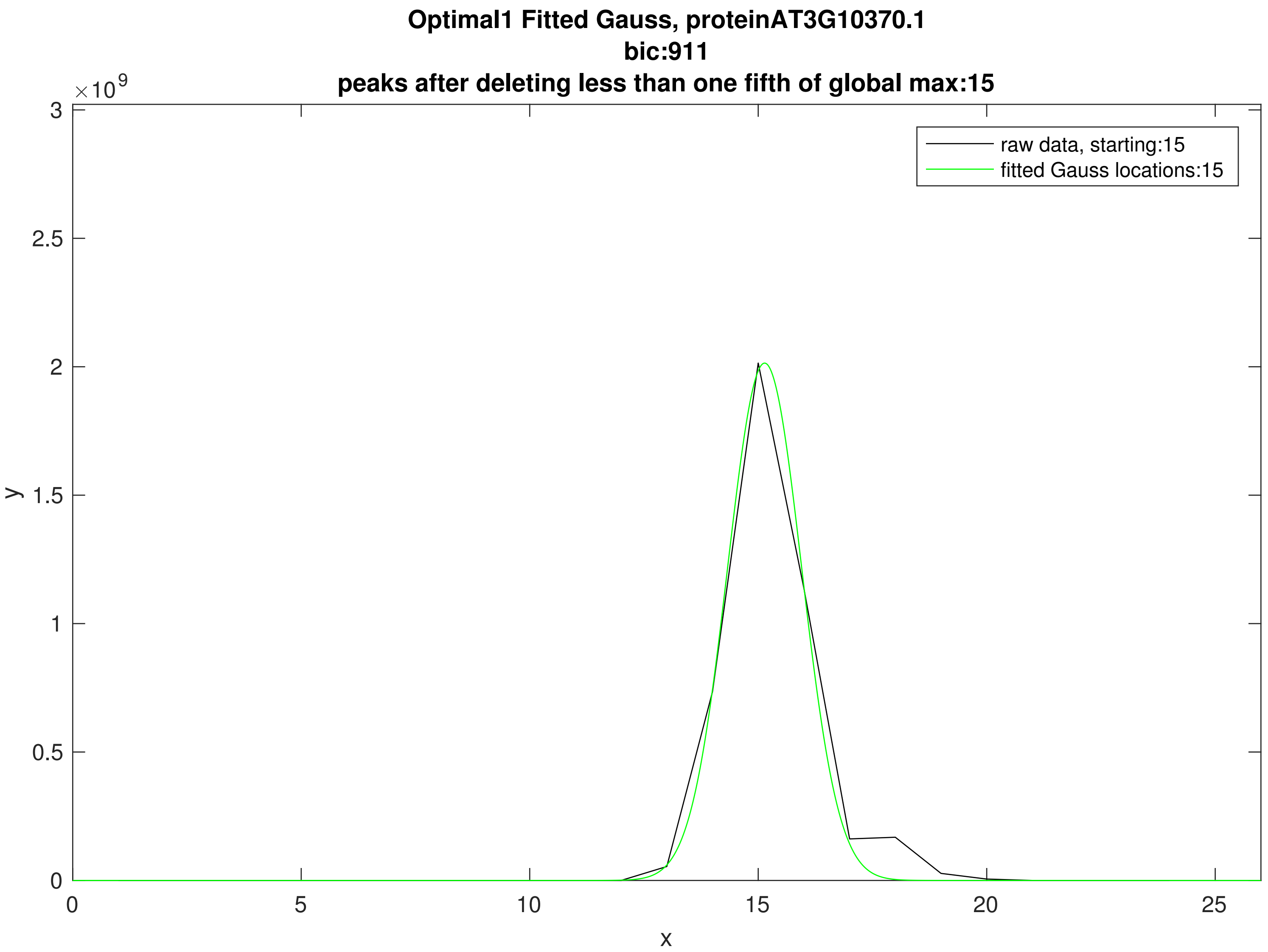
Optimal1 Fitted Gauss, proteinAT3G10370.1
bic:911
peaks after deleting less than one fifth of global max:15
×10^9
raw data, starting:15
fitted Gauss locations:15
y
x
0
0.5
1
1.5
2
2.5
3
0
5
10
15
20
25

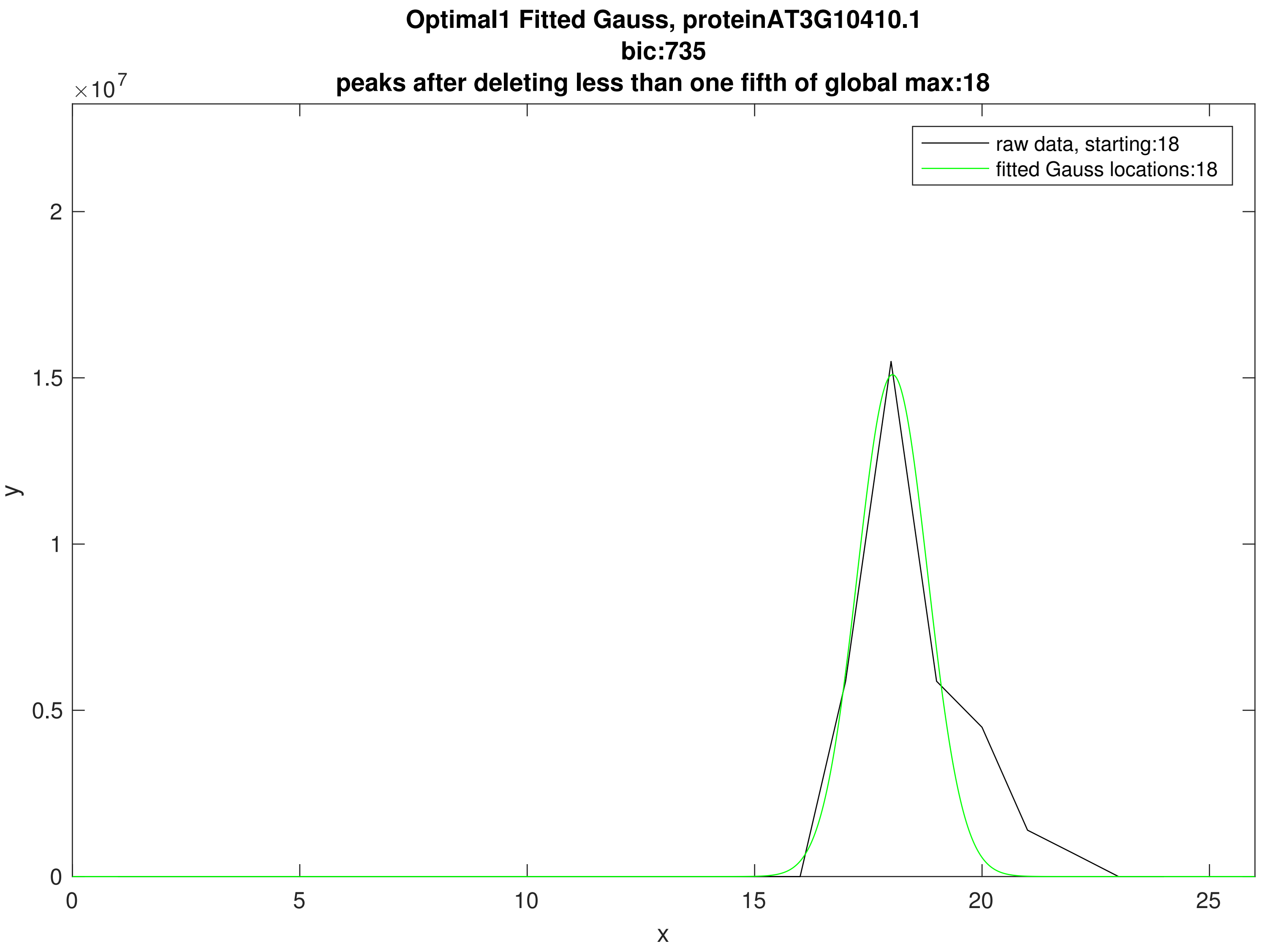
Optimal1 Fitted Gauss, proteinAT3G10410.1
bic:735
peaks after deleting less than one fifth of global max:18
×10^7
raw data, starting:18
fitted Gauss locations:18
y
x
0
0.5
1
1.5
2
0
5
10
15
20
25

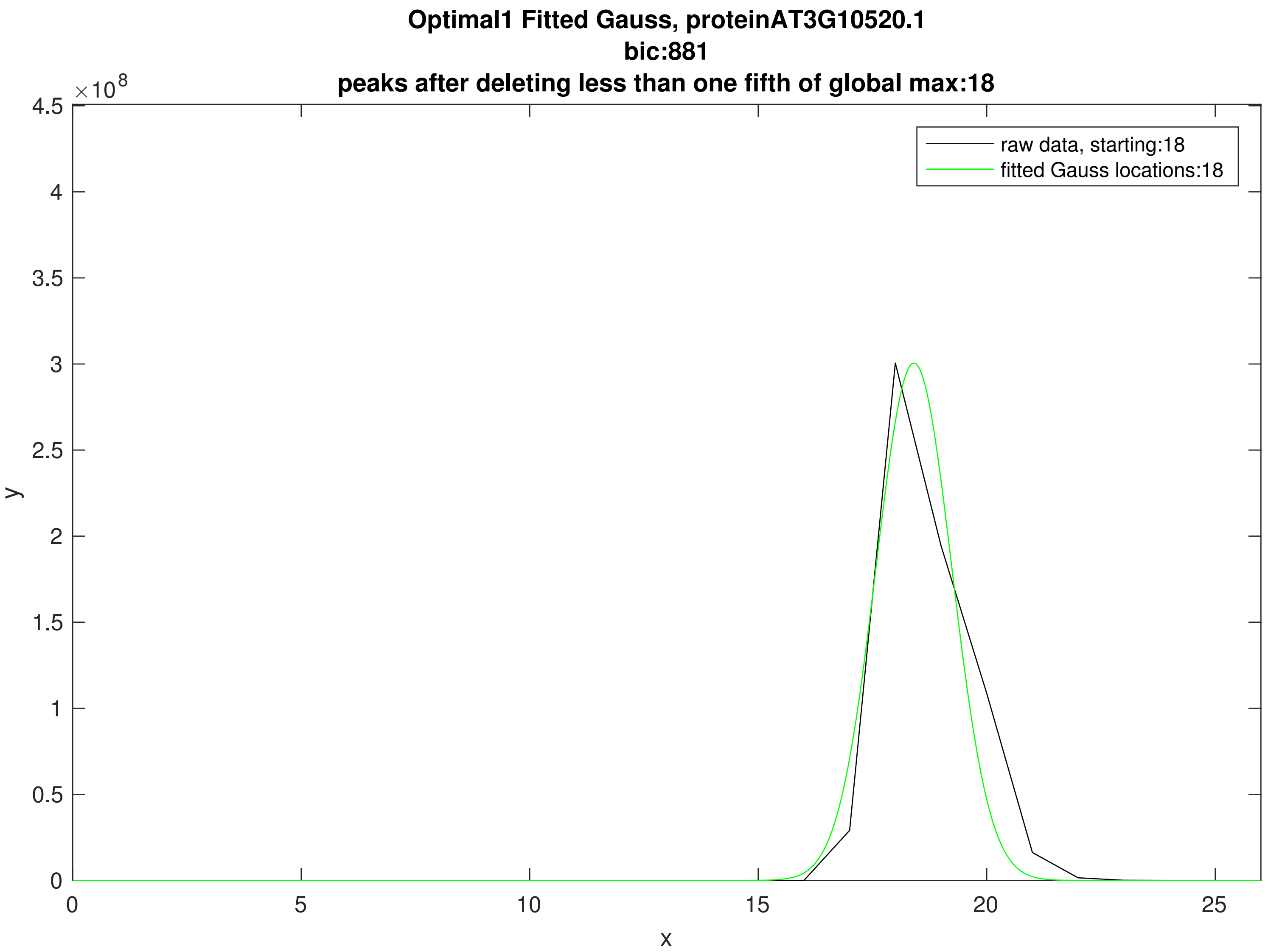

Optimal1 Fitted Gauss, proteinAT3G10520.1
bic:881
peaks after deleting less than one fifth of global max:18
×10^8
4.5
4
3.5
3
2.5
2
1.5
1
0.5
0
y
0
5
10
15
20
25
x
raw data, starting:18
fitted Gauss locations:18

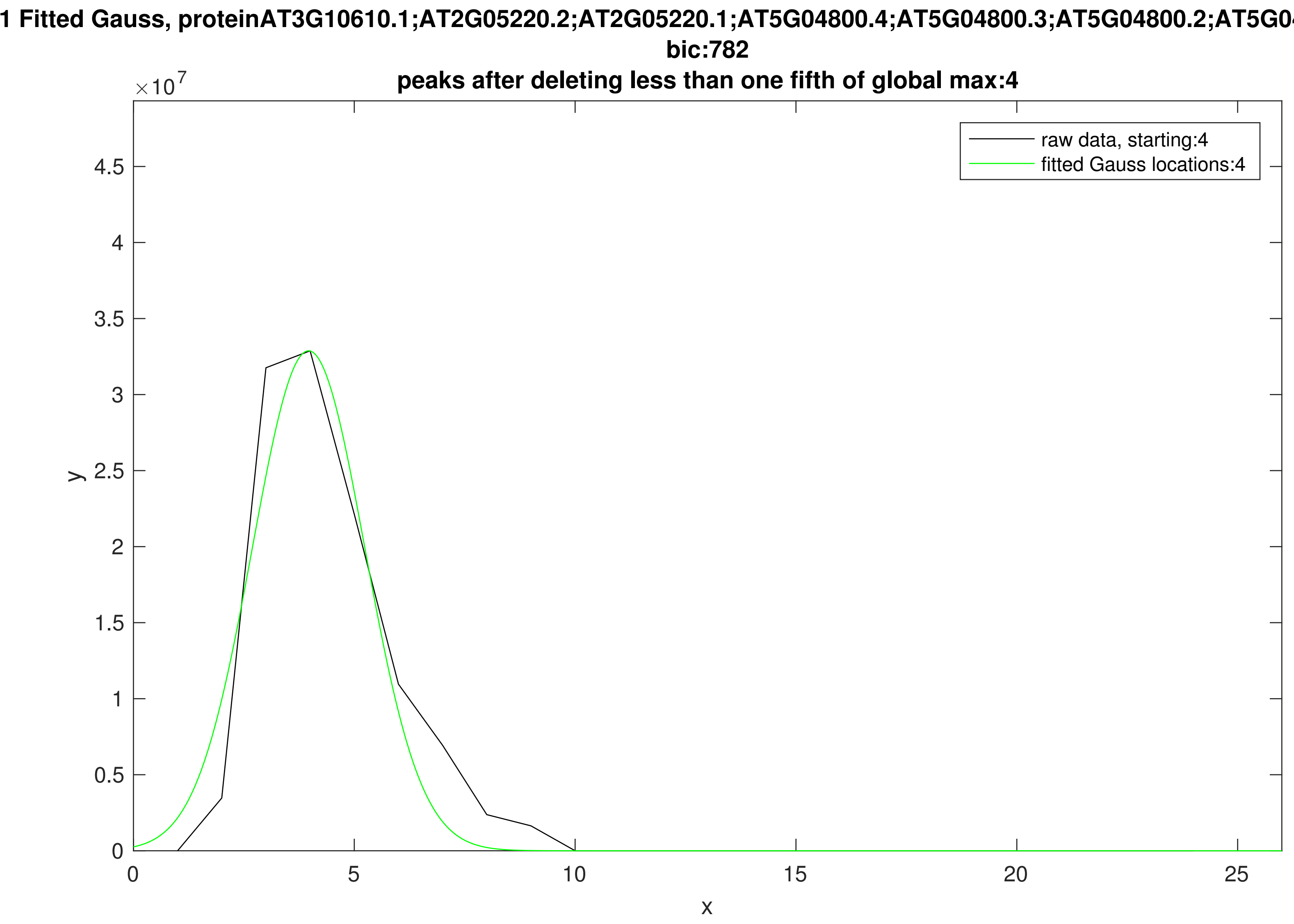
1 Fitted Gauss, proteinAT3G10610.1;AT2G05220.2;AT2G05220.1;AT5G04800.4;AT5G04800.3;AT5G04800.2;AT5G0480
bic:782
peaks after deleting less than one fifth of global max:4
×10^7
raw data, starting:4
fitted Gauss locations:4
y
x
0
0.5
1
1.5
2
2.5
3
3.5
4
4.5
5
10
15
20
25

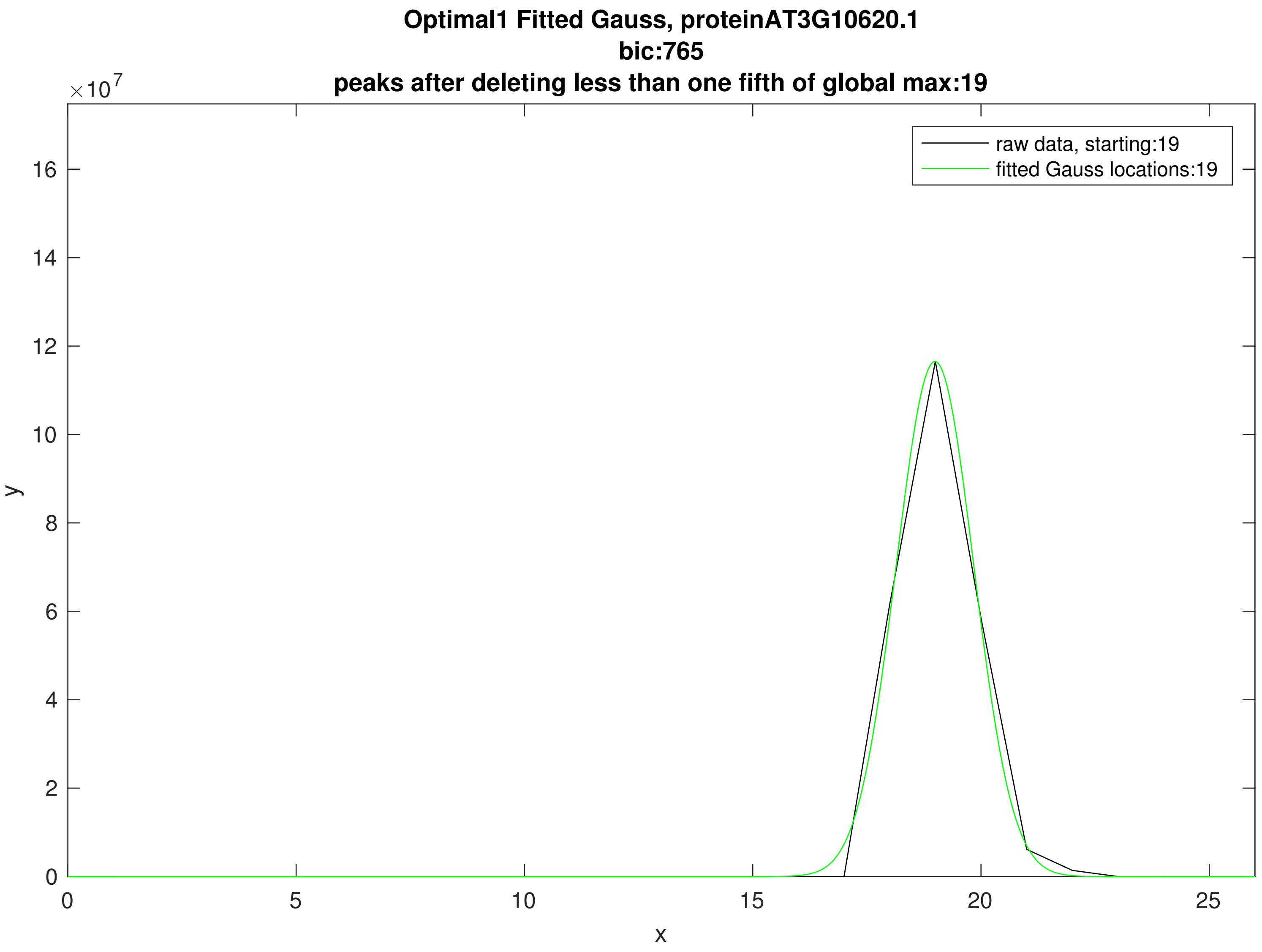
Optimal1 Fitted Gauss, proteinAT3G10620.1
bic:765
peaks after deleting less than one fifth of global max:19
×10^7
raw data, starting:19
fitted Gauss locations:19
y
x
0
2
4
6
8
10
12
14
16
0
5
10
15
20
25

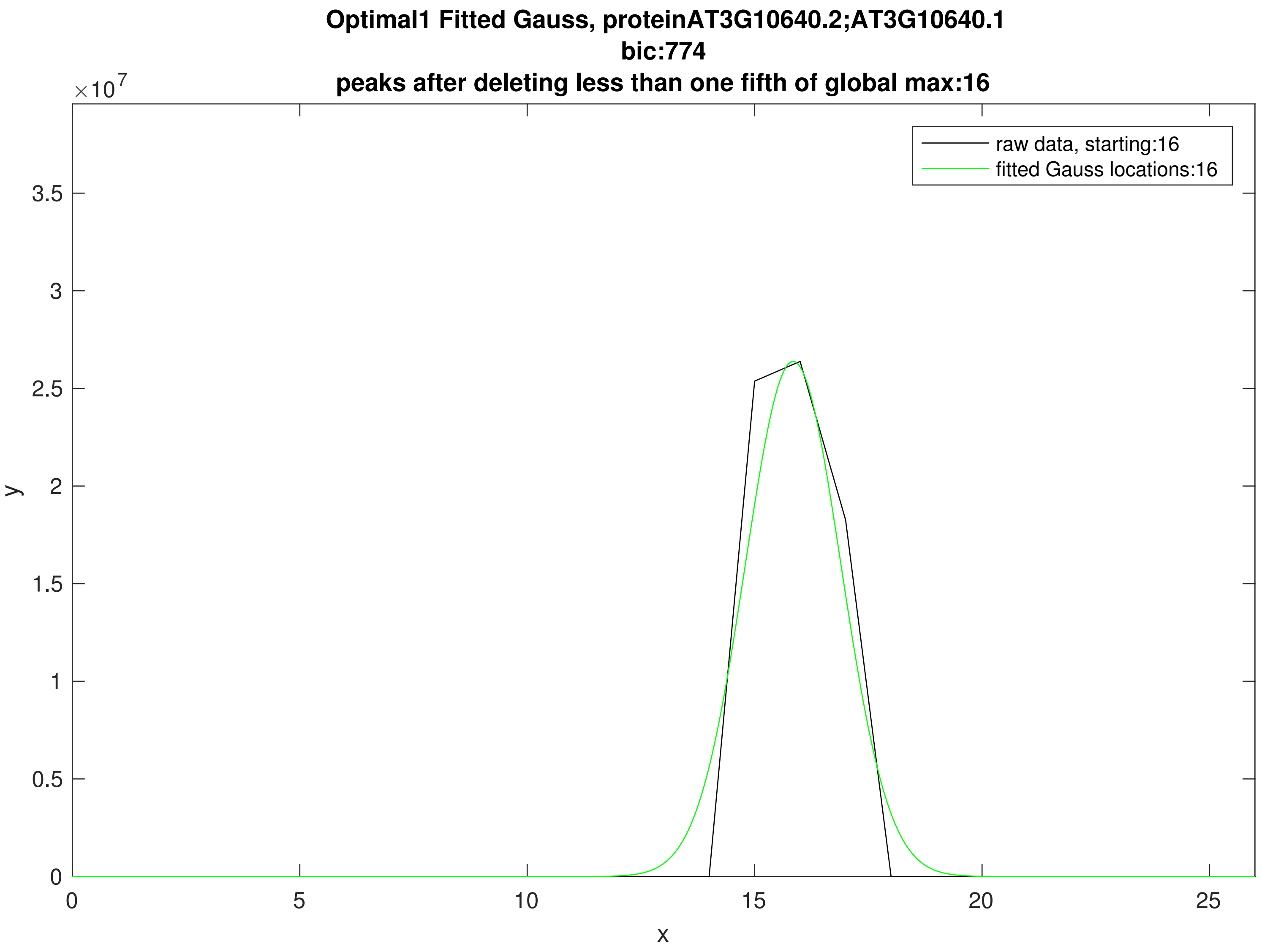

Optimal1 Fitted Gauss, proteinAT3G10640.2;AT3G10640.1
bic:774
peaks after deleting less than one fifth of global max:16
×10^7
raw data, starting:16
fitted Gauss locations:16
y
x
0
0.5
1
1.5
2
2.5
3
3.5
0
5
10
15
20
25

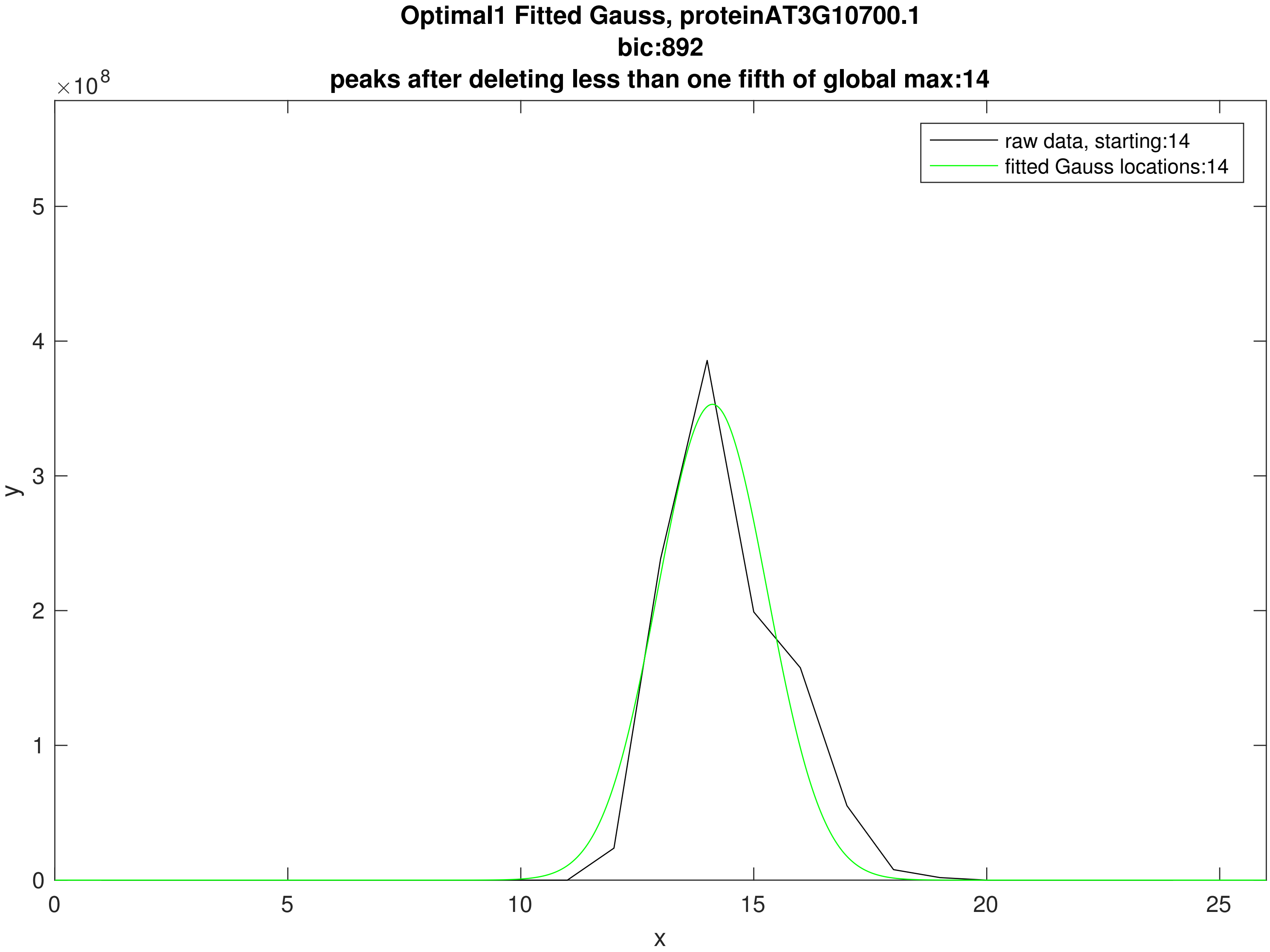
Optimal1 Fitted Gauss, proteinAT3G10700.1
bic:892
peaks after deleting less than one fifth of global max:14
×10^8
raw data, starting:14
fitted Gauss locations:14
y
x
0
1
2
3
4
5
0
5
10
15
20
25

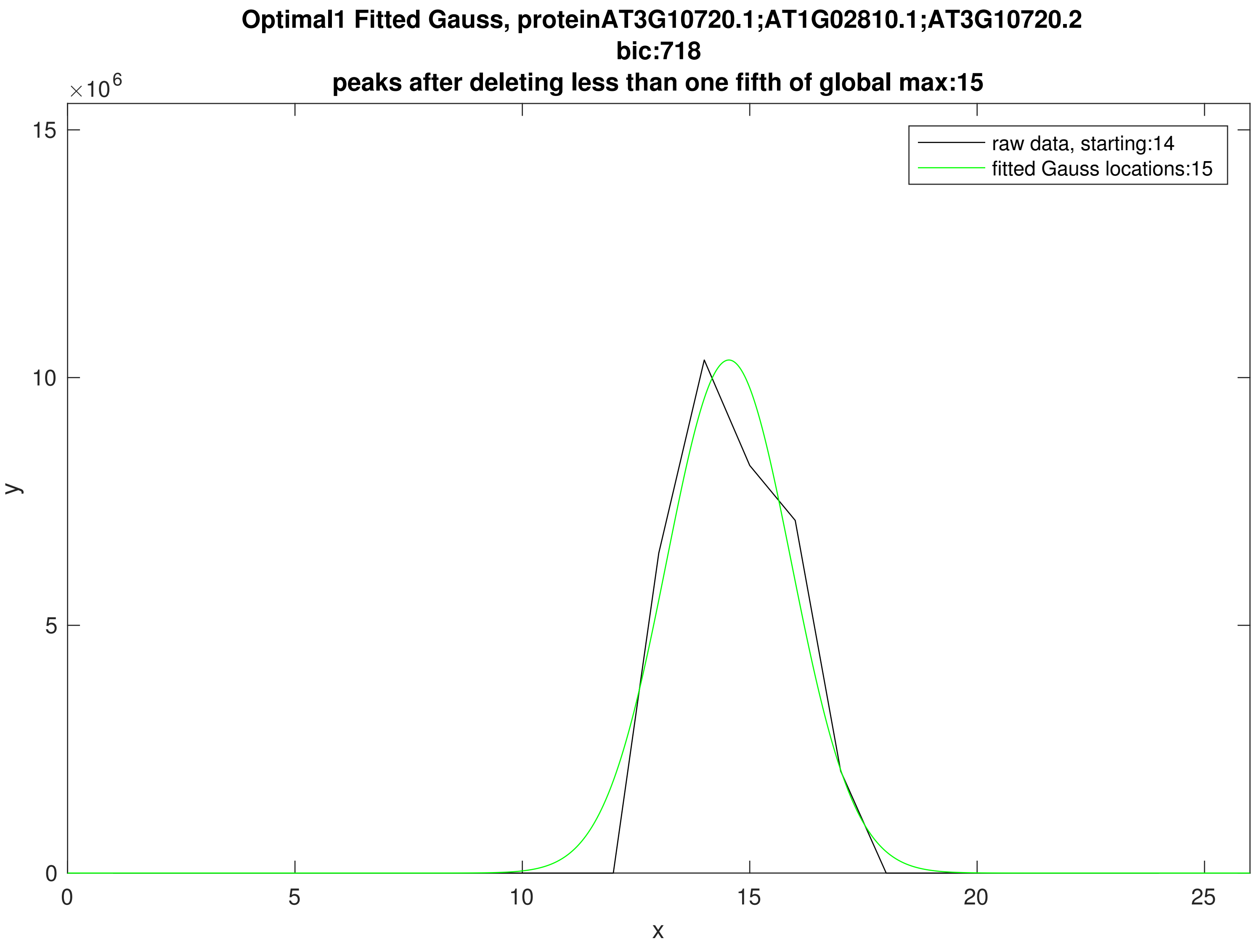
Optimal1 Fitted Gauss, proteinAT3G10720.1;AT1G02810.1;AT3G10720.2
bic:718
peaks after deleting less than one fifth of global max:15
×10^6
15
10
5
0
y
0
5
10
15
20
25
x
raw data, starting:14
fitted Gauss locations:15

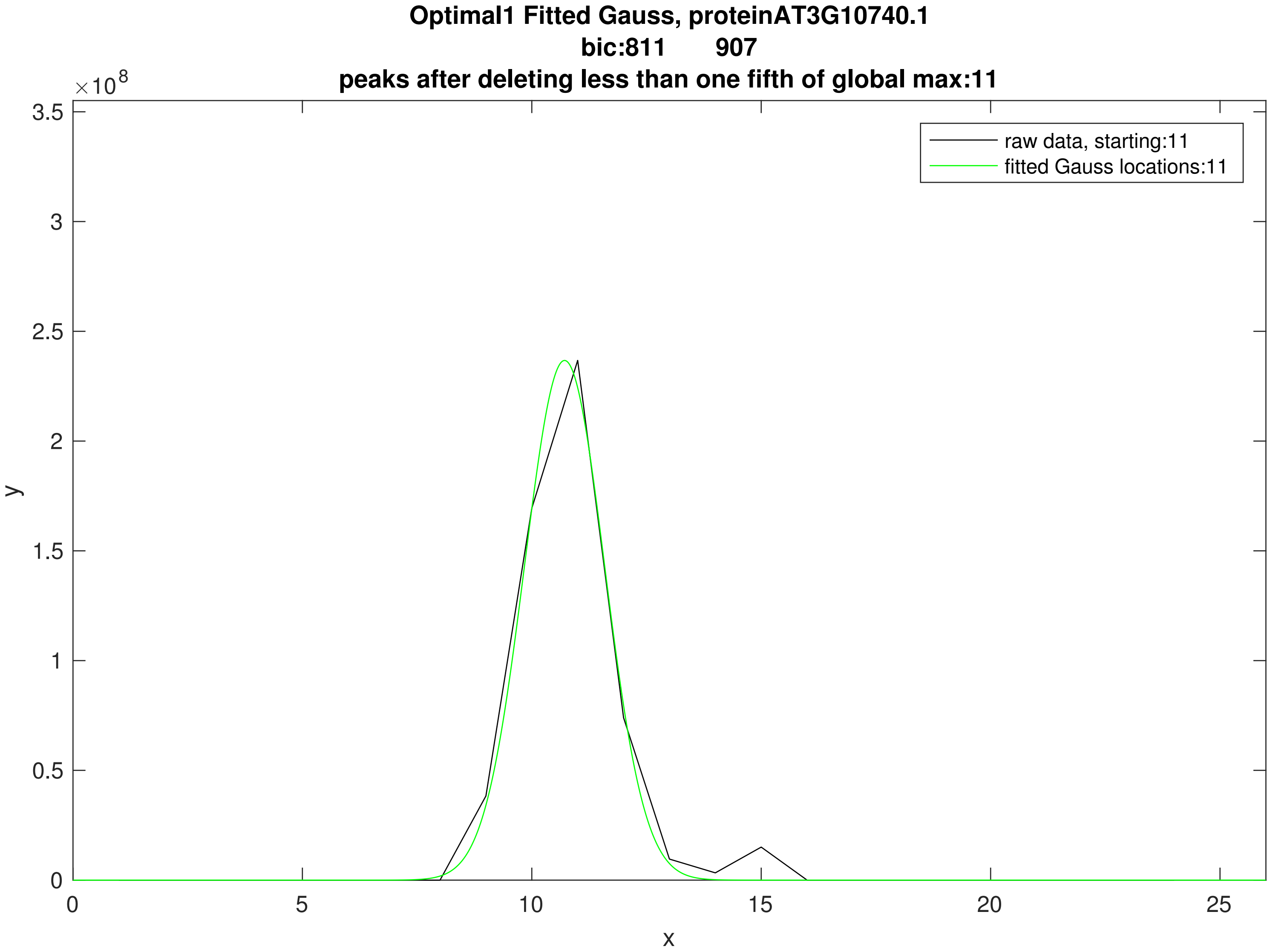
Optimal1 Fitted Gauss, proteinAT3G10740.1
bic:811 907
peaks after deleting less than one fifth of global max:11
×10^8
3.5
3
2.5
2
1.5
1
0.5
0
y
0
5
10
15
20
25
x
raw data, starting:11
fitted Gauss locations:11

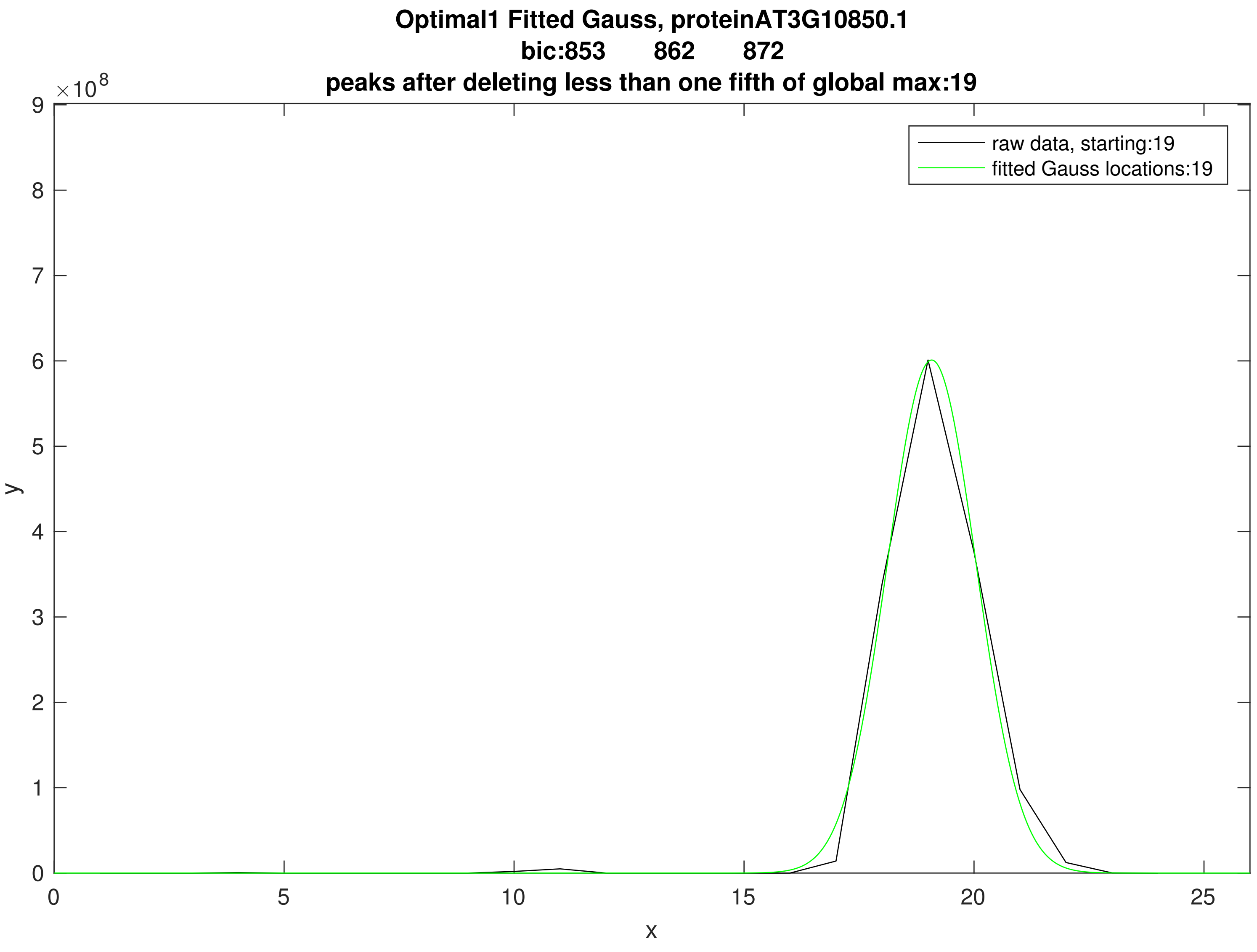
Optimal1 Fitted Gauss, proteinAT3G10850.1
bic:853 862 872
peaks after deleting less than one fifth of global max:19
×10^8
raw data, starting:19
fitted Gauss locations:19
y
x
0
1
2
3
4
5
6
7
8
9
0
5
10
15
20
25

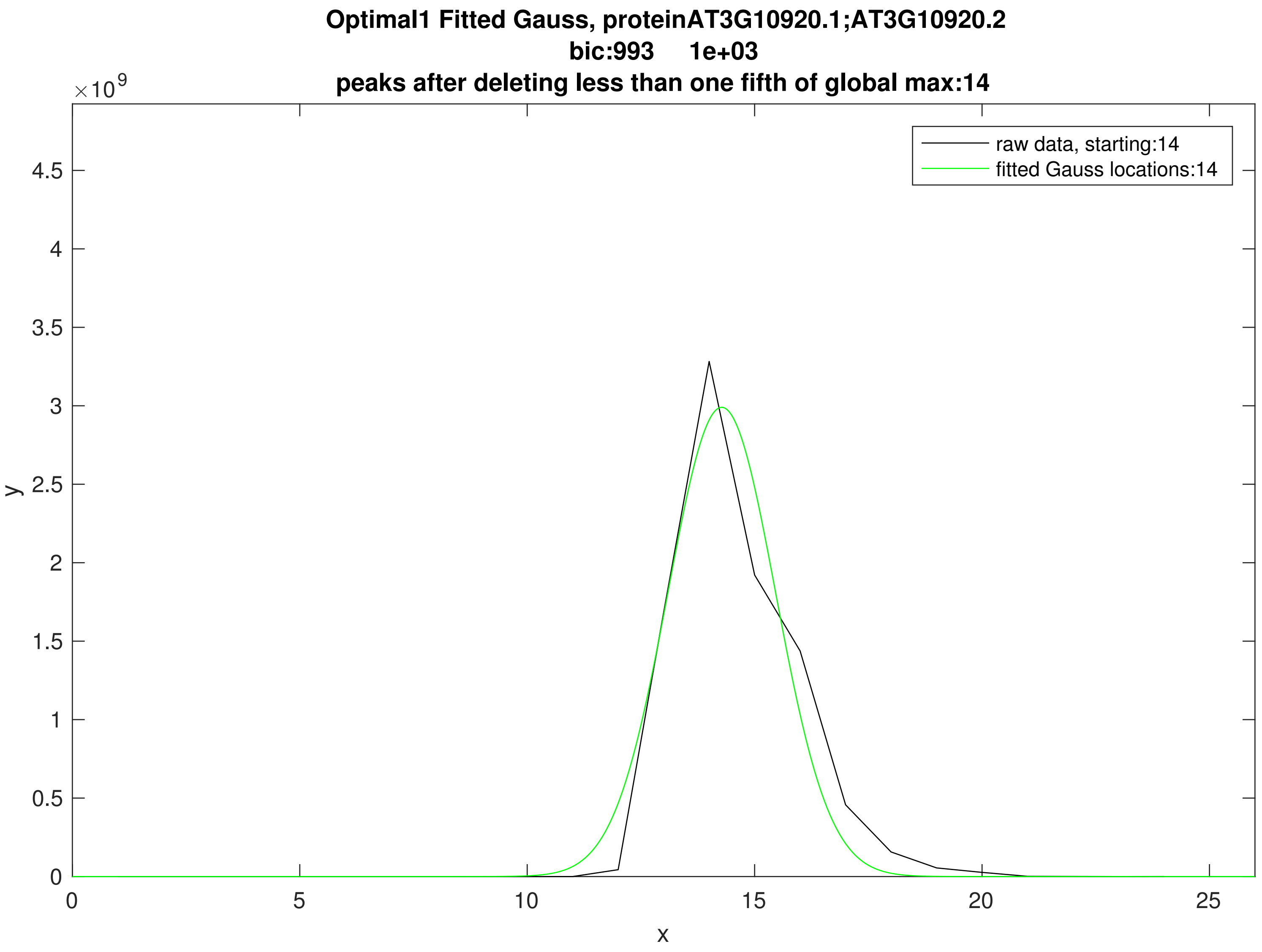

Optimal1 Fitted Gauss, proteinAT3G10920.1;AT3G10920.2
bic:993 1e+03
peaks after deleting less than one fifth of global max:14
×10^9
4.5
4
3.5
3
2.5
2
1.5
1
0.5
0
y
0
5
10
15
20
25
x
raw data, starting:14
fitted Gauss locations:14

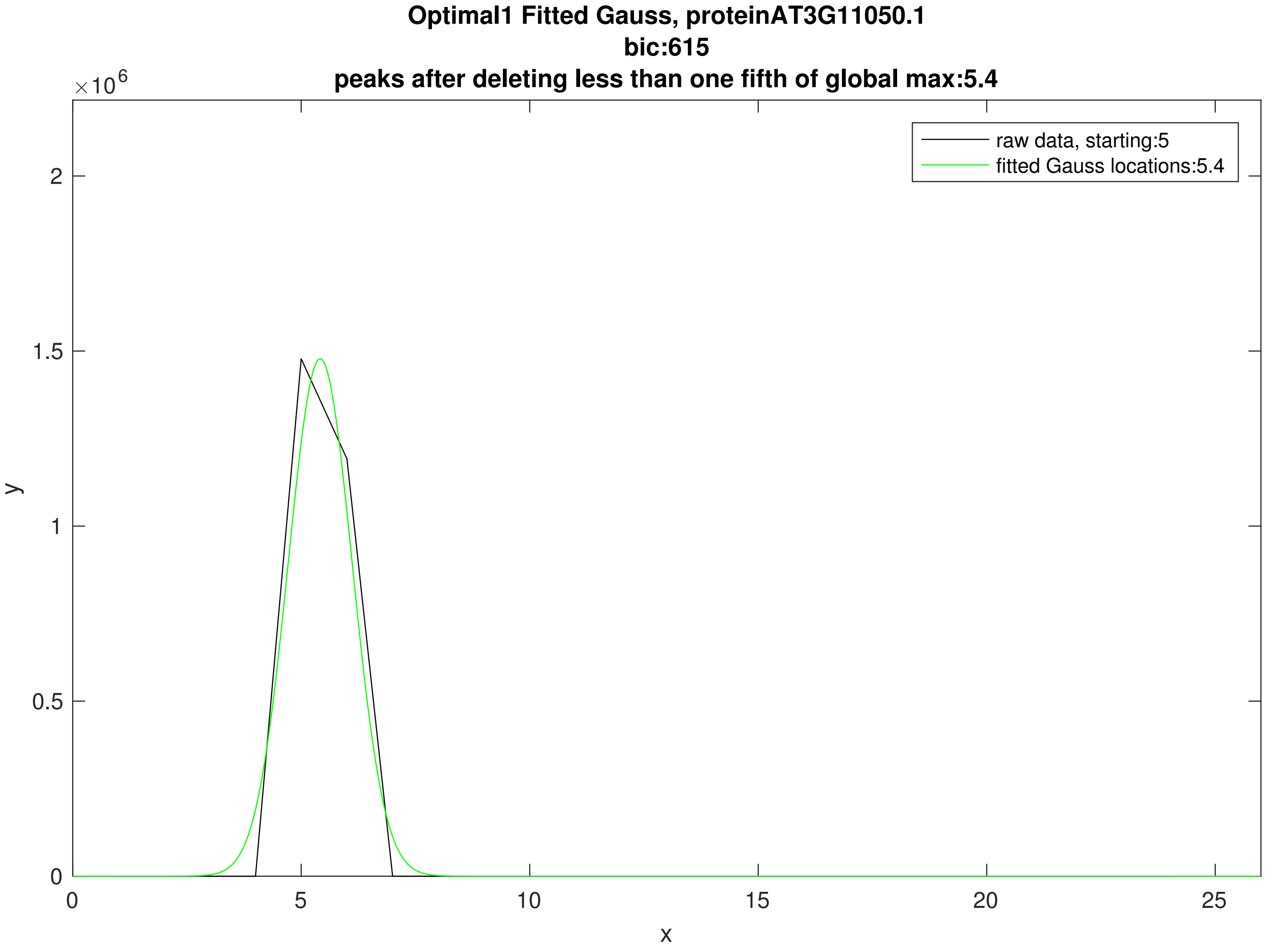

Optimal1 Fitted Gauss, proteinAT3G11050.1
bic:615
peaks after deleting less than one fifth of global max:5.4
×10^6
2
1.5
1
0.5
0
y
0
5
10
15
20
25
x
raw data, starting:5
fitted Gauss locations:5.4

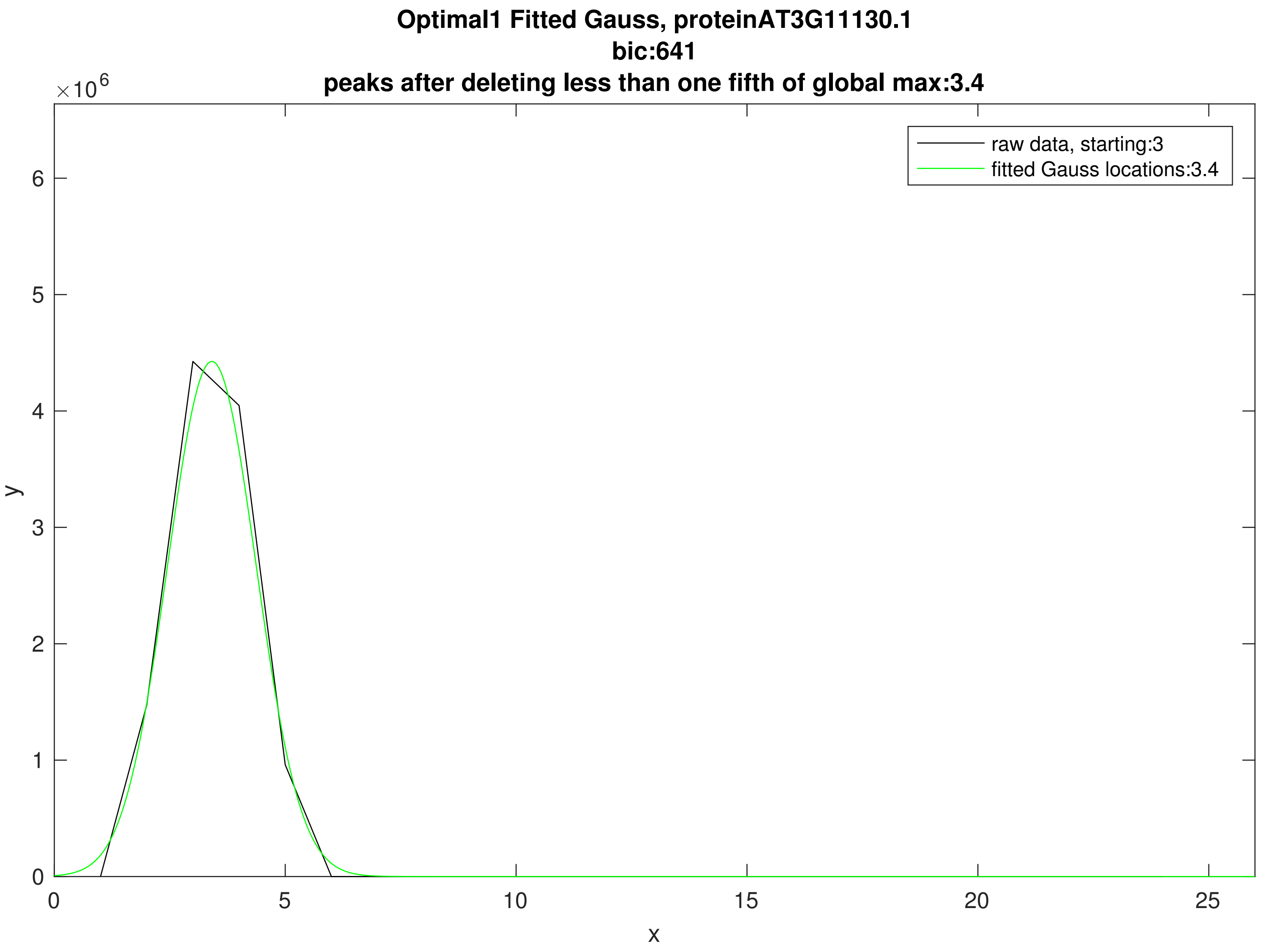
Optimal1 Fitted Gauss, proteinAT3G11130.1
bic:641
peaks after deleting less than one fifth of global max:3.4
×10^6
raw data, starting:3
fitted Gauss locations:3.4
y
x
0
1
2
3
4
5
6
0
5
10
15
20
25

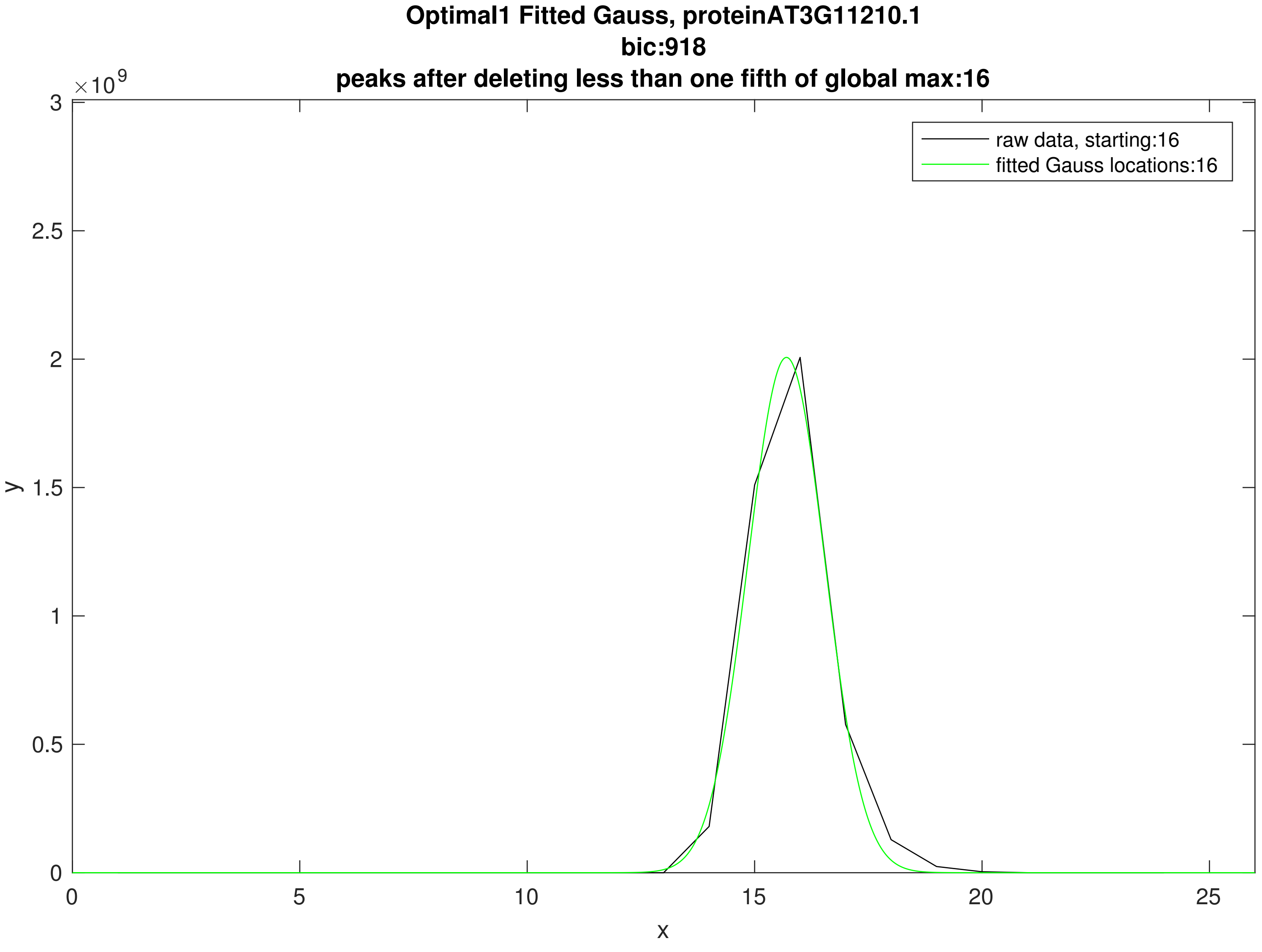

Optimal1 Fitted Gauss, proteinAT3G11210.1
bic:918
peaks after deleting less than one fifth of global max:16
×10^9
3
2.5
2
1.5
1
0.5
0
y
0
5
10
15
20
25
x
raw data, starting:16
fitted Gauss locations:16

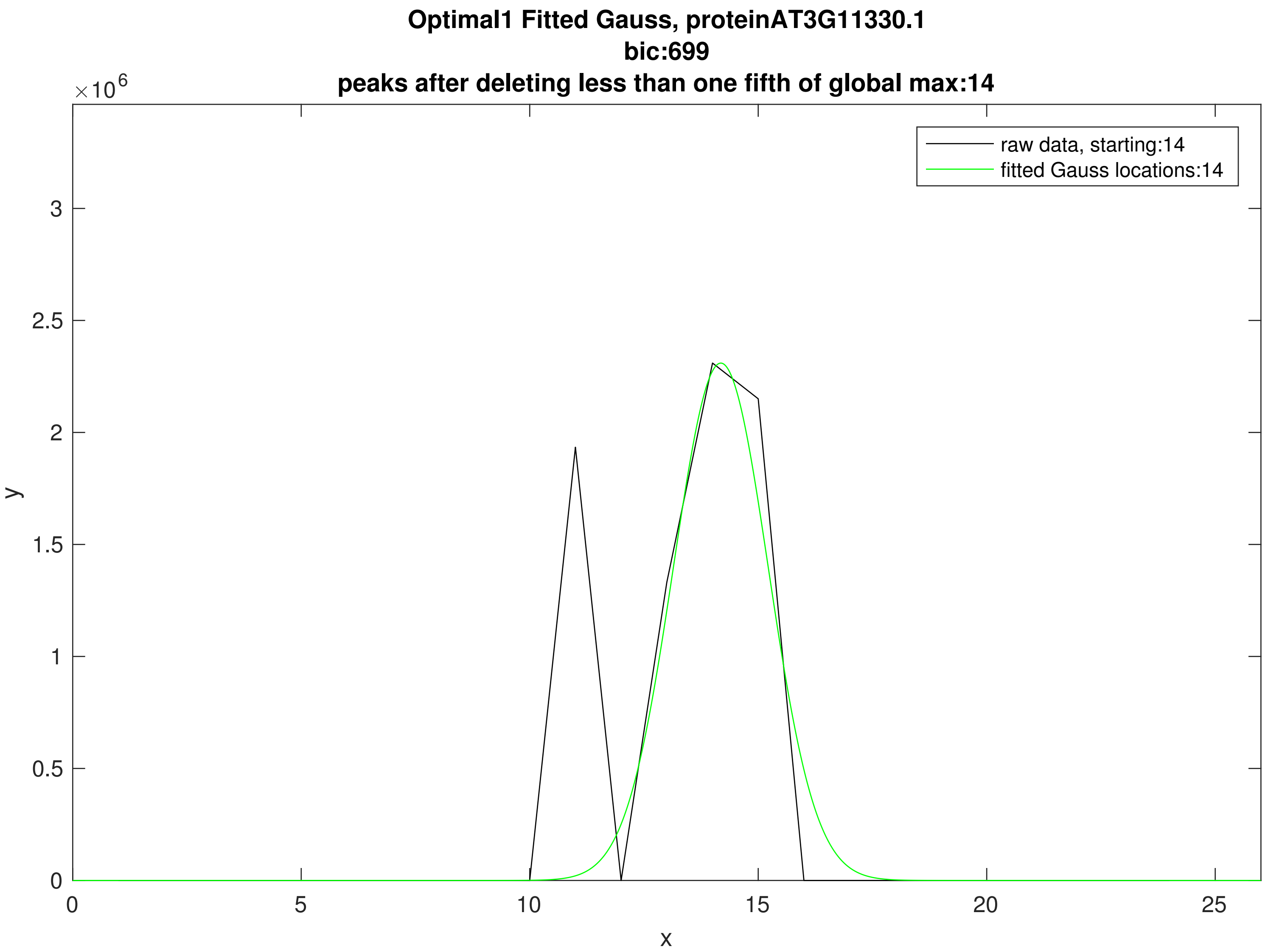
Optimal1 Fitted Gauss, proteinAT3G11330.1
bic:699
peaks after deleting less than one fifth of global max:14
×10^6
raw data, starting:14
fitted Gauss locations:14
y
x
0
0.5
1
1.5
2
2.5
3
0
5
10
15
20
25

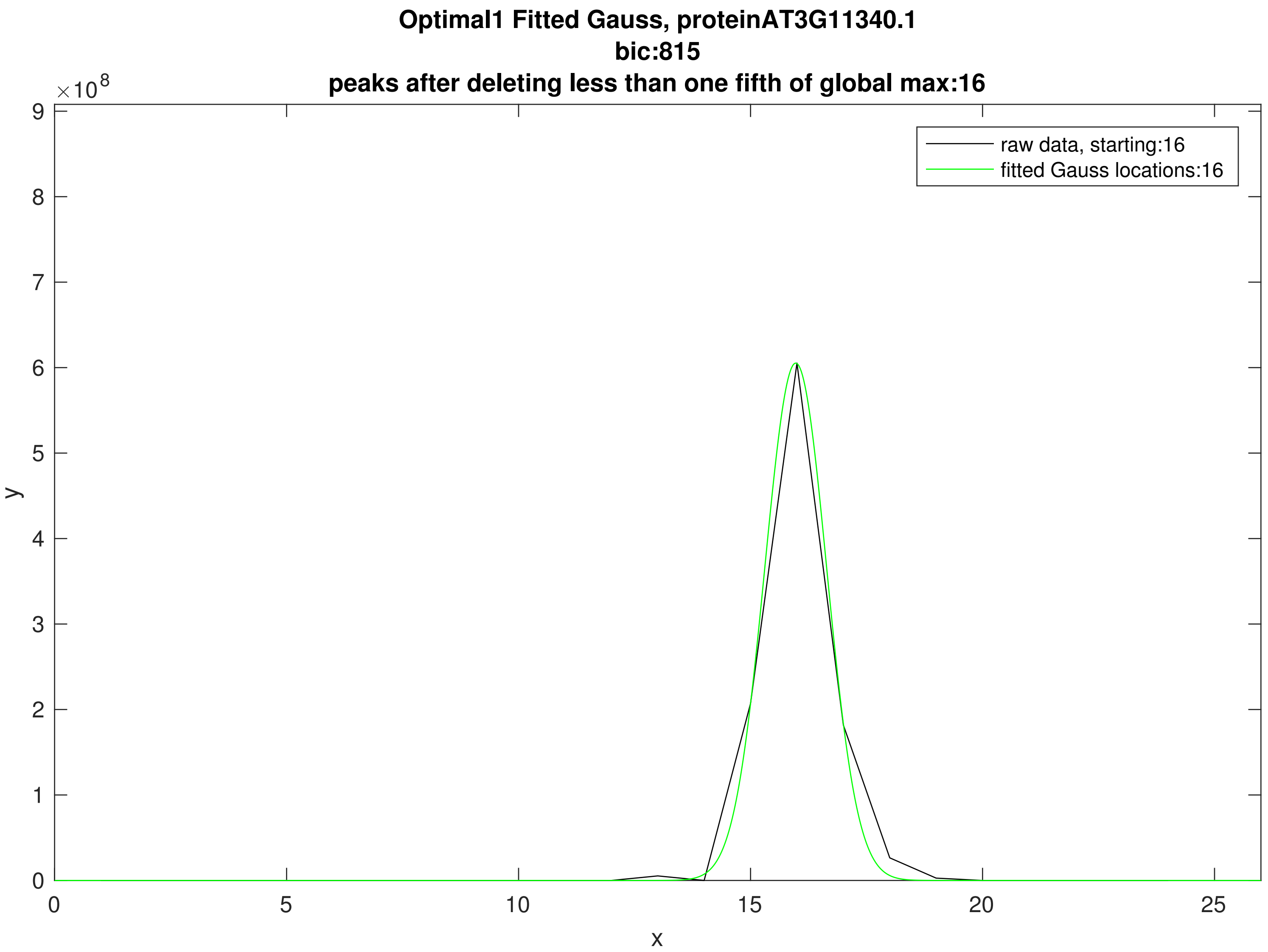

Optimal1 Fitted Gauss, proteinAT3G11340.1
bic:815
peaks after deleting less than one fifth of global max:16
×10^8
9
8
7
6
5
4
3
2
1
0
y
0
5
10
15
20
25
x
raw data, starting:16
fitted Gauss locations:16

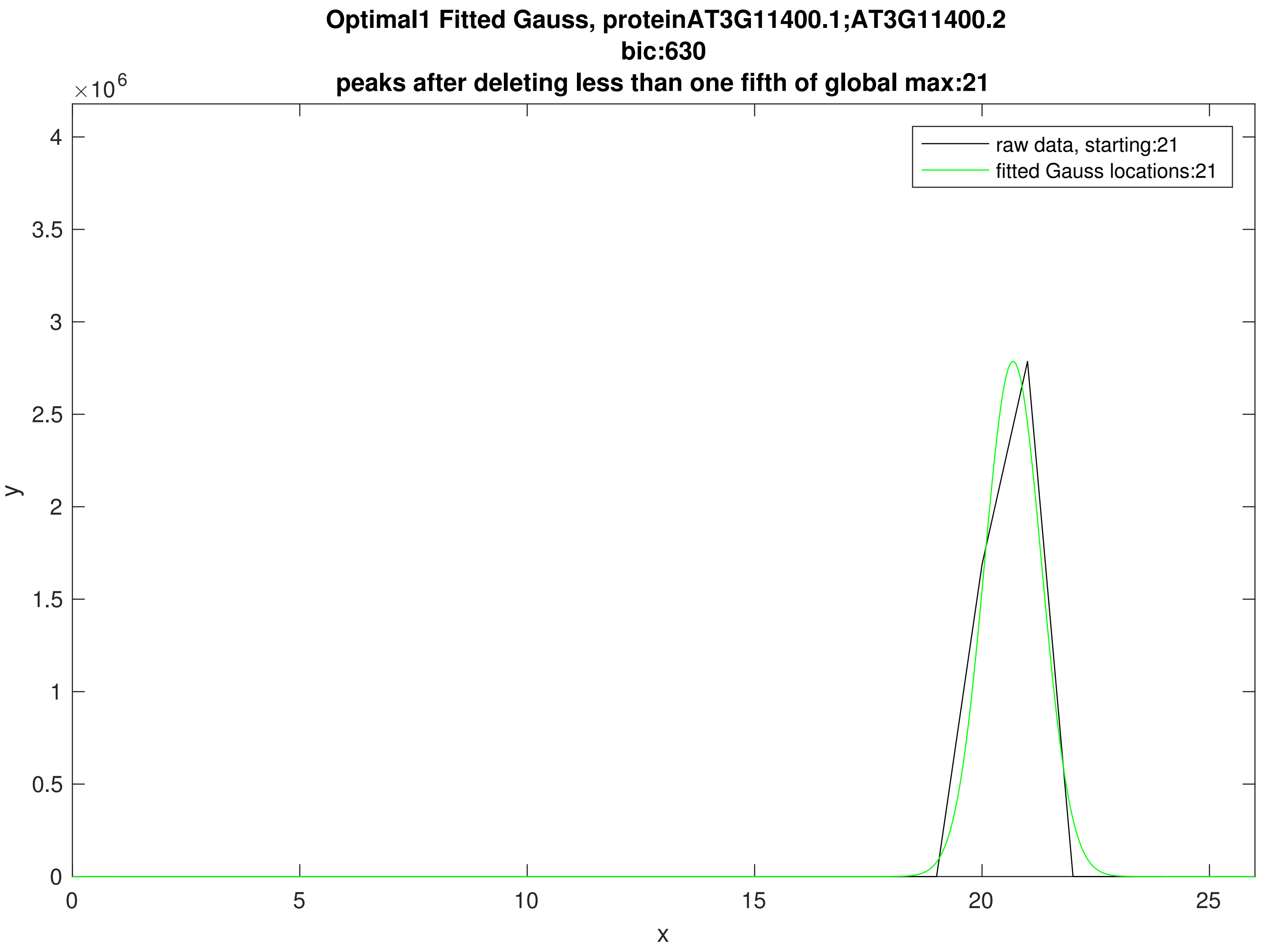
Optimal1 Fitted Gauss, proteinAT3G11400.1;AT3G11400.2
bic:630
peaks after deleting less than one fifth of global max:21
×10^6
raw data, starting:21
fitted Gauss locations:21
y
x
0
0.5
1
1.5
2
2.5
3
3.5
4
5
10
15
20
25

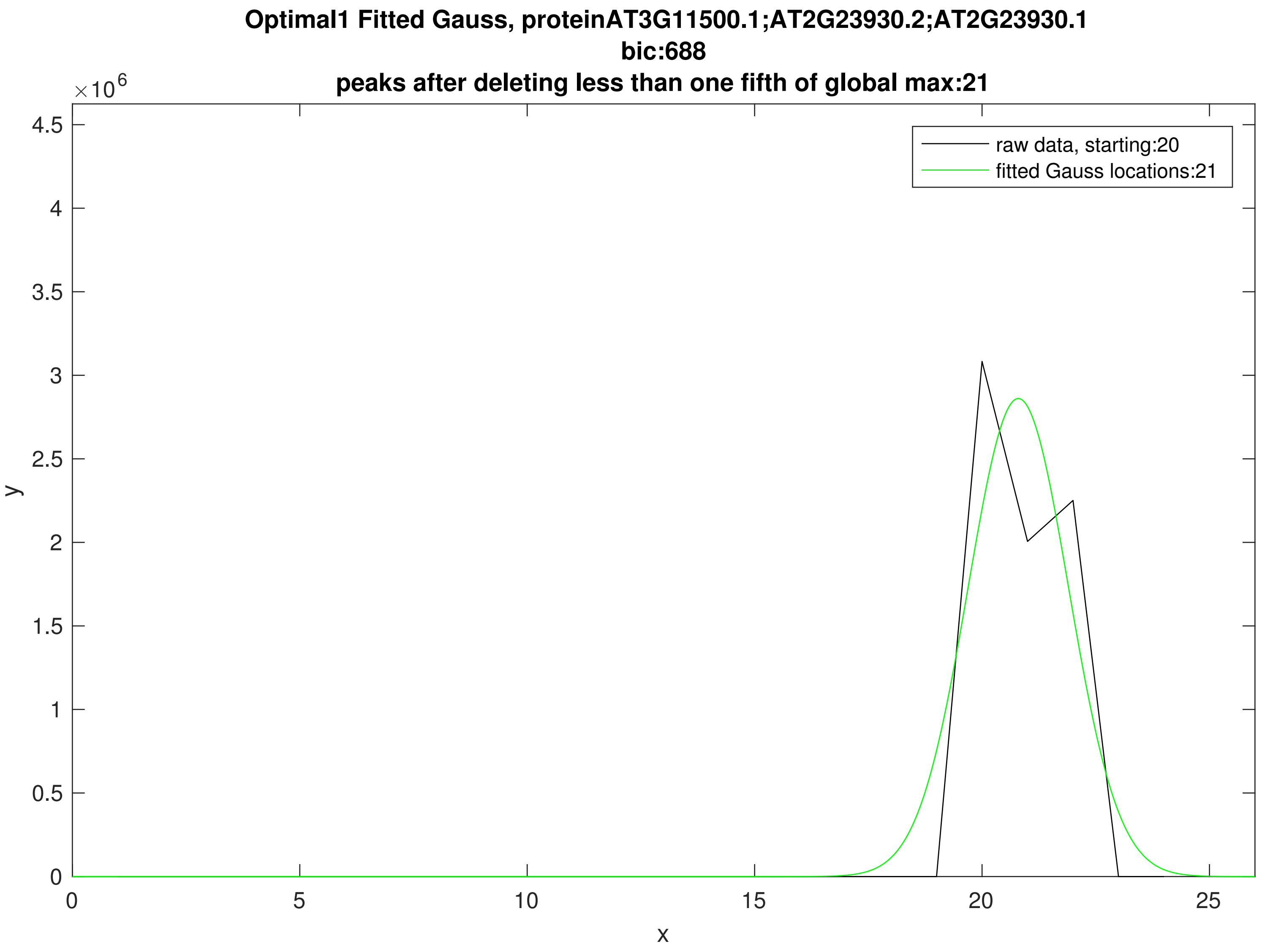

Optimal1 Fitted Gauss, proteinAT3G11500.1;AT2G23930.2;AT2G23930.1
bic:688
peaks after deleting less than one fifth of global max:21
×10^6
4.5
4
3.5
3
2.5
2
1.5
1
0.5
0
y
0
5
10
15
20
25
x
raw data, starting:20
fitted Gauss locations:21

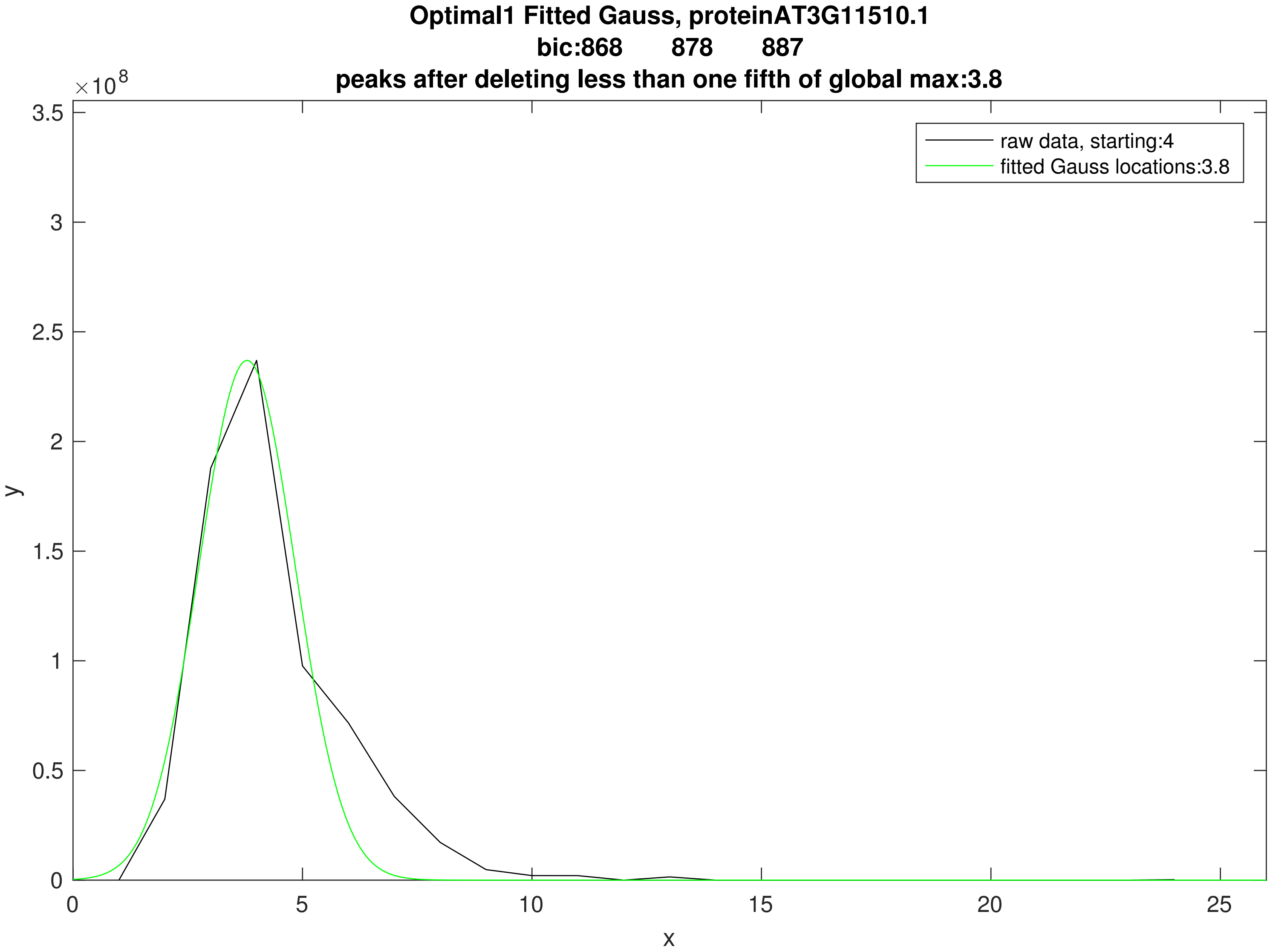
Optimal1 Fitted Gauss, proteinAT3G11510.1
bic:868 878 887
peaks after deleting less than one fifth of global max:3.8
raw data, starting:4
fitted Gauss locations:3.8
×10^8
3.5
3
2.5
2
1.5
1
0.5
0
y
0
5
10
15
20
25
x

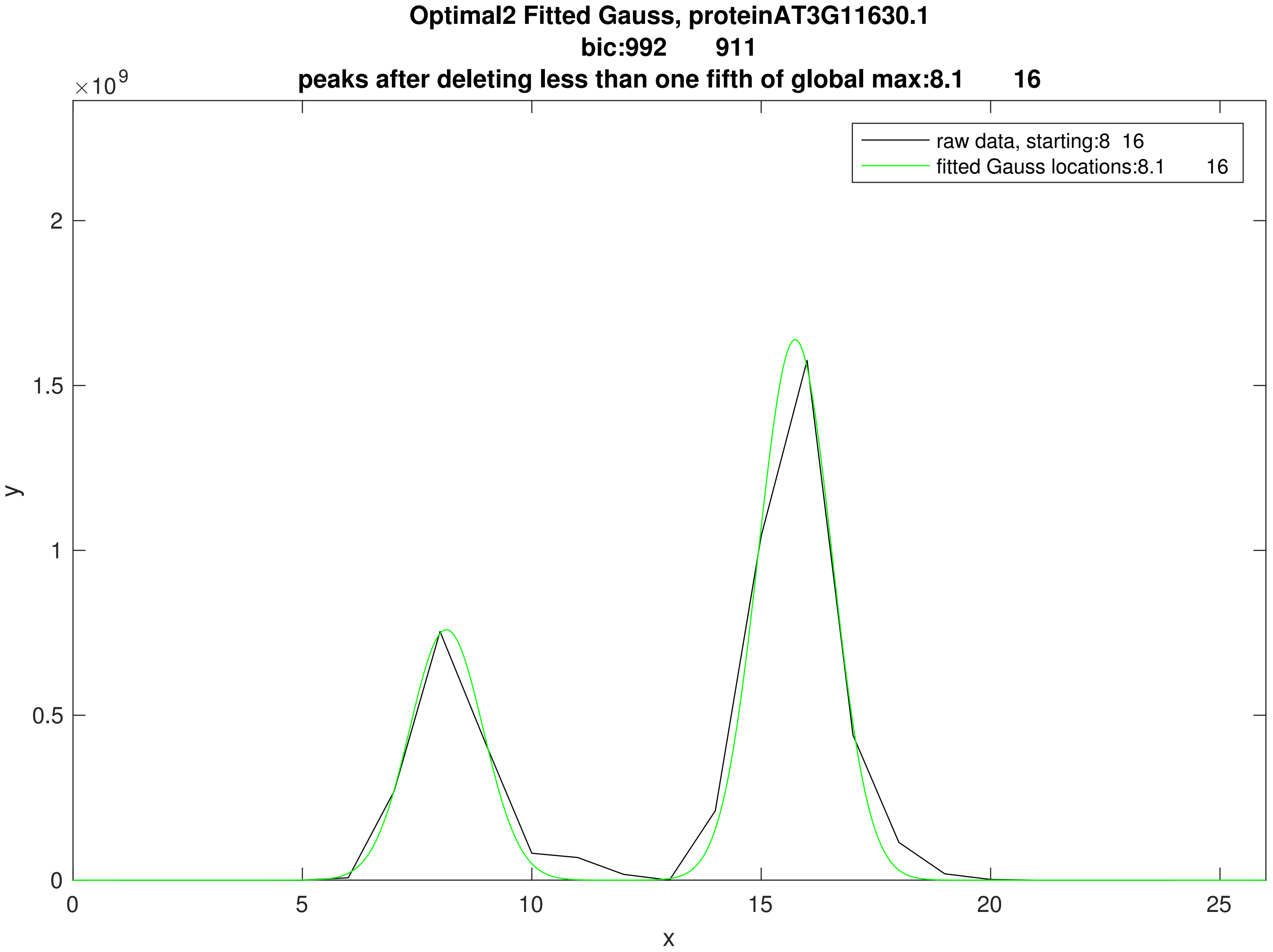
Optimal2 Fitted Gauss, proteinAT3G11630.1
bic:992 911
peaks after deleting less than one fifth of global max:8.1 16
×10^9
raw data, starting:8 16
fitted Gauss locations:8.1 16
y
x
0
0.5
1
1.5
2
5
10
15
20
25

**Optimal1 Fitted Gauss, proteinAT3G11700.1**
**bic:797**
**peaks after deleting less than one fifth of global max:5.1**

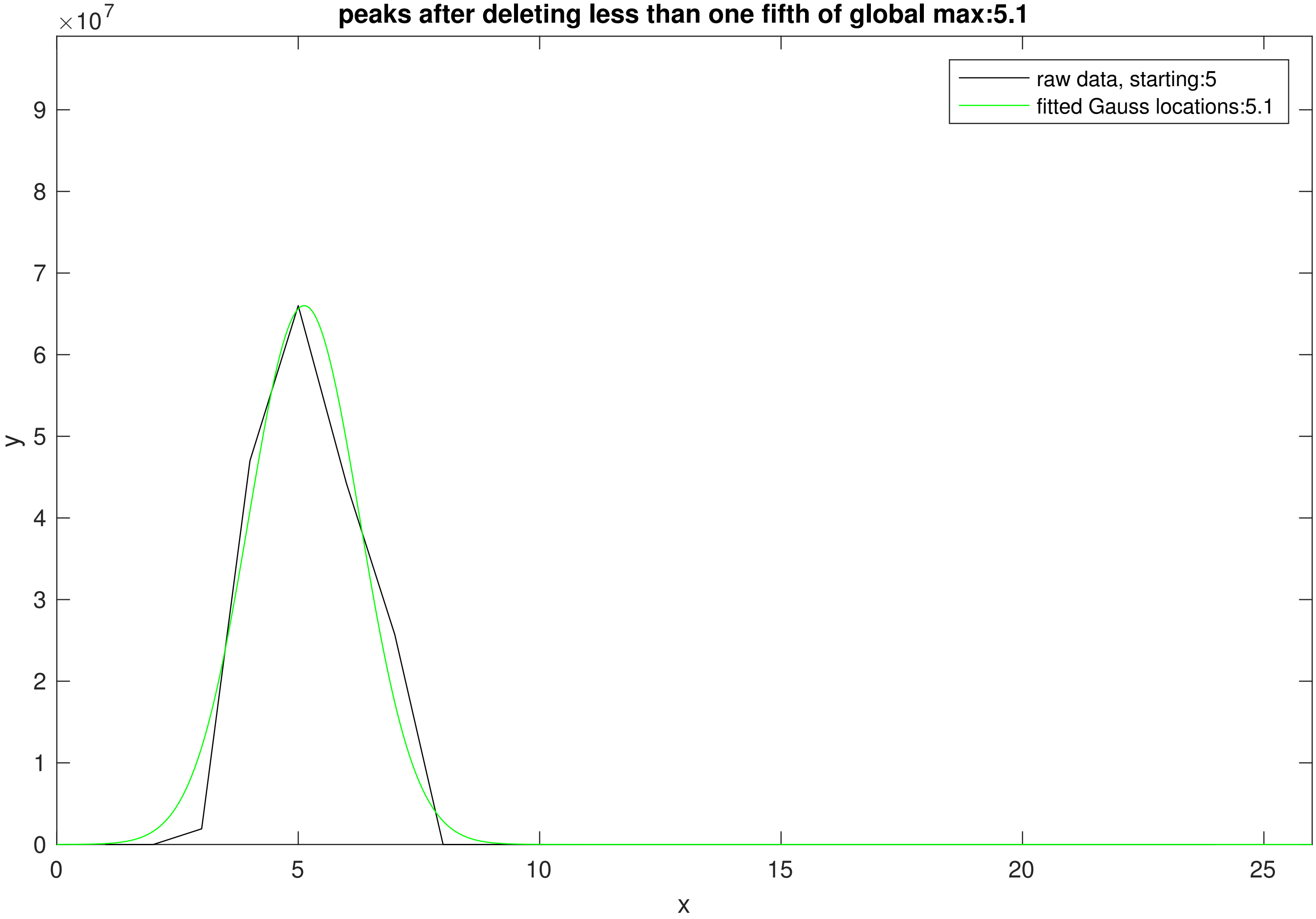

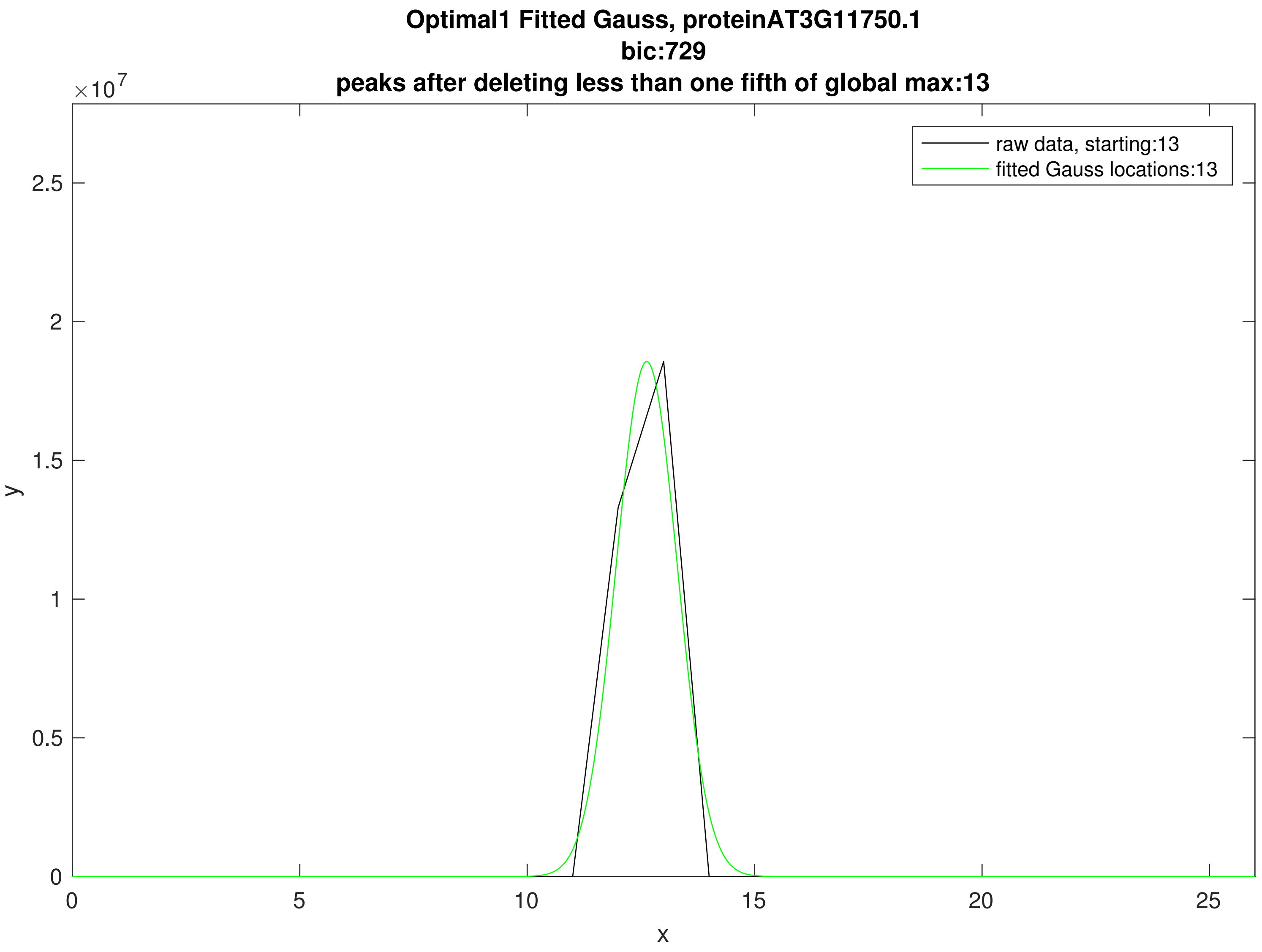
Optimal1 Fitted Gauss, proteinAT3G11750.1
bic:729
peaks after deleting less than one fifth of global max:13
×10^7
raw data, starting:13
fitted Gauss locations:13
2.5
2
1.5
1
0.5
0
y
0
5
10
15
20
25
x

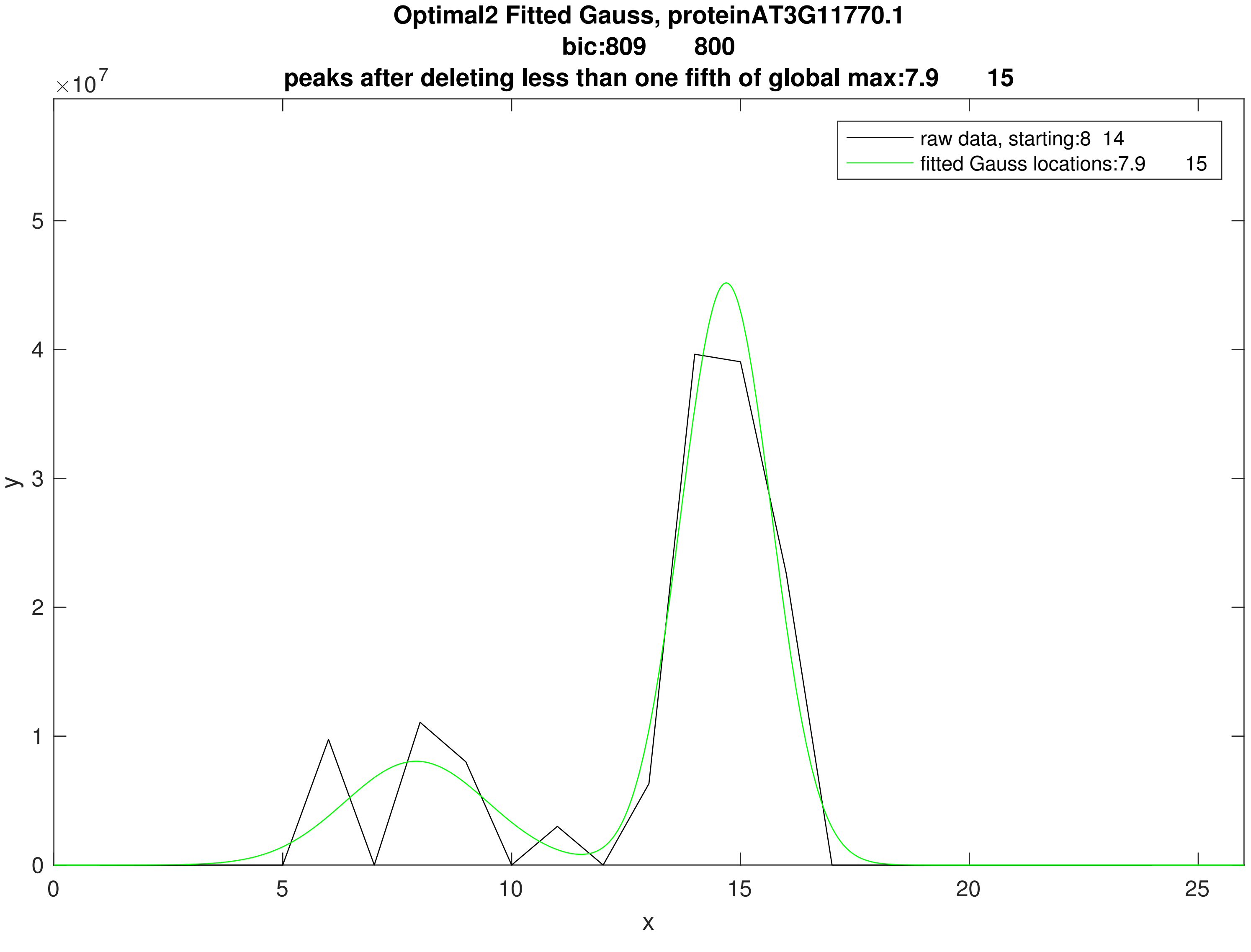

Optimal2 Fitted Gauss, proteinAT3G11770.1
bic:809 800
peaks after deleting less than one fifth of global max:7.9 15
×10^7
raw data, starting:8 14
fitted Gauss locations:7.9 15
5
4
3
2
1
0
y
0
5
10
15
20
25
x

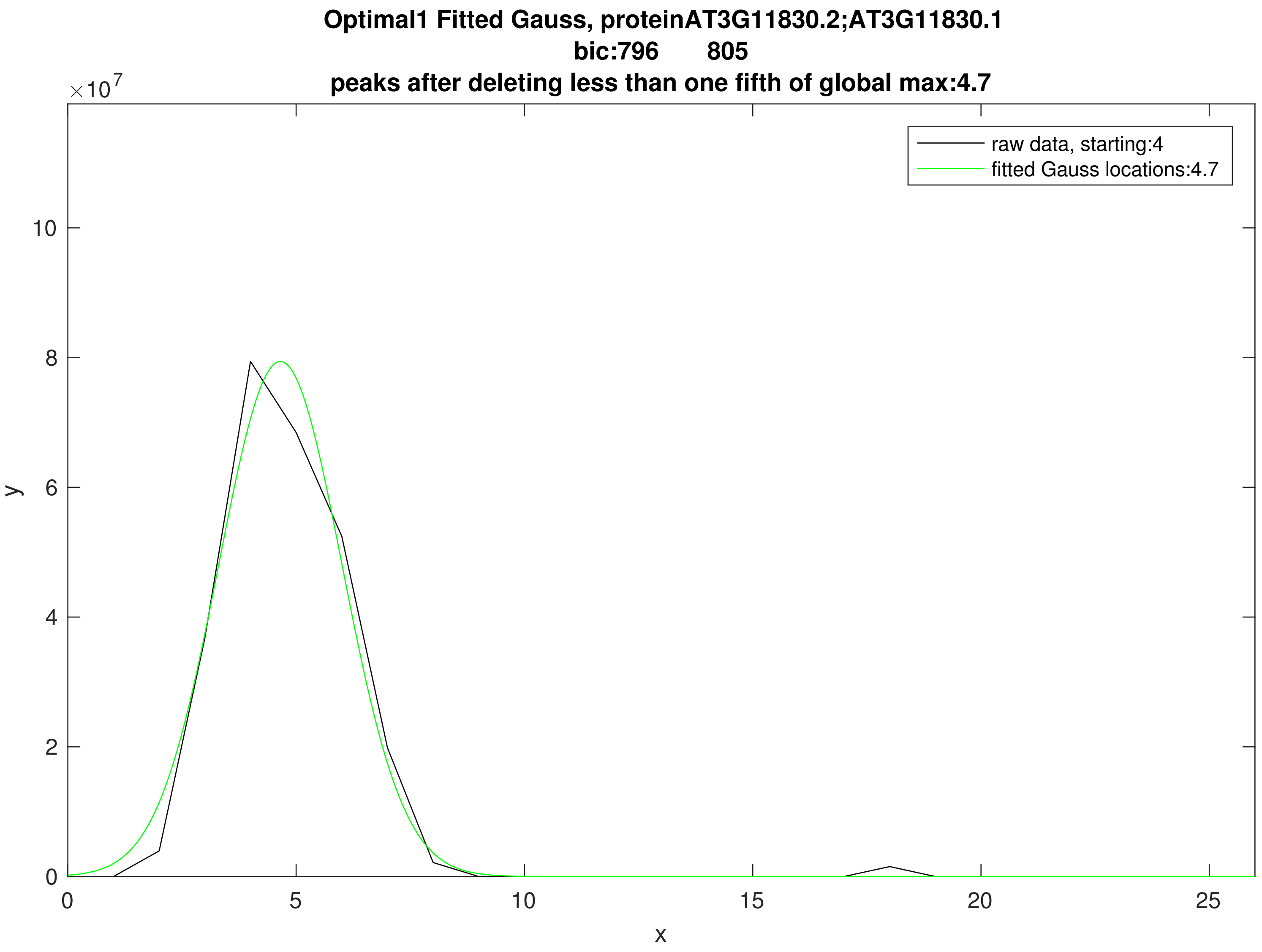
Optimal1 Fitted Gauss, proteinAT3G11830.2;AT3G11830.1
bic:796 805
peaks after deleting less than one fifth of global max:4.7
×10^7
raw data, starting:4
fitted Gauss locations:4.7
10
8
6
4
2
0
y
0
5
10
15
20
25
x

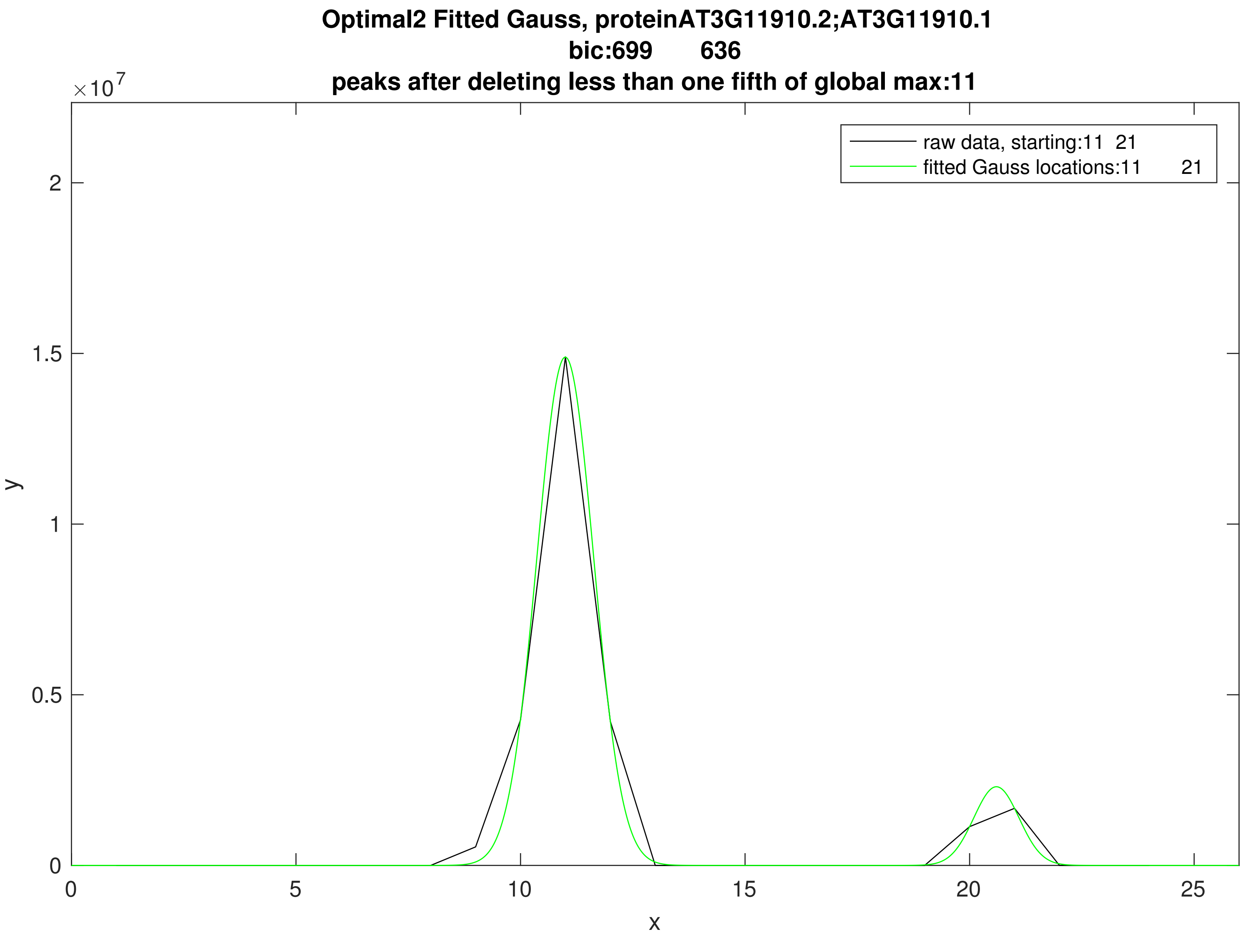

Optimal2 Fitted Gauss, proteinAT3G11910.2;AT3G11910.1
bic:699 636
peaks after deleting less than one fifth of global max:11
×10^7
raw data, starting:11 21
fitted Gauss locations:11 21
y
x
0
0.5
1
1.5
2
5
10
15
20
25

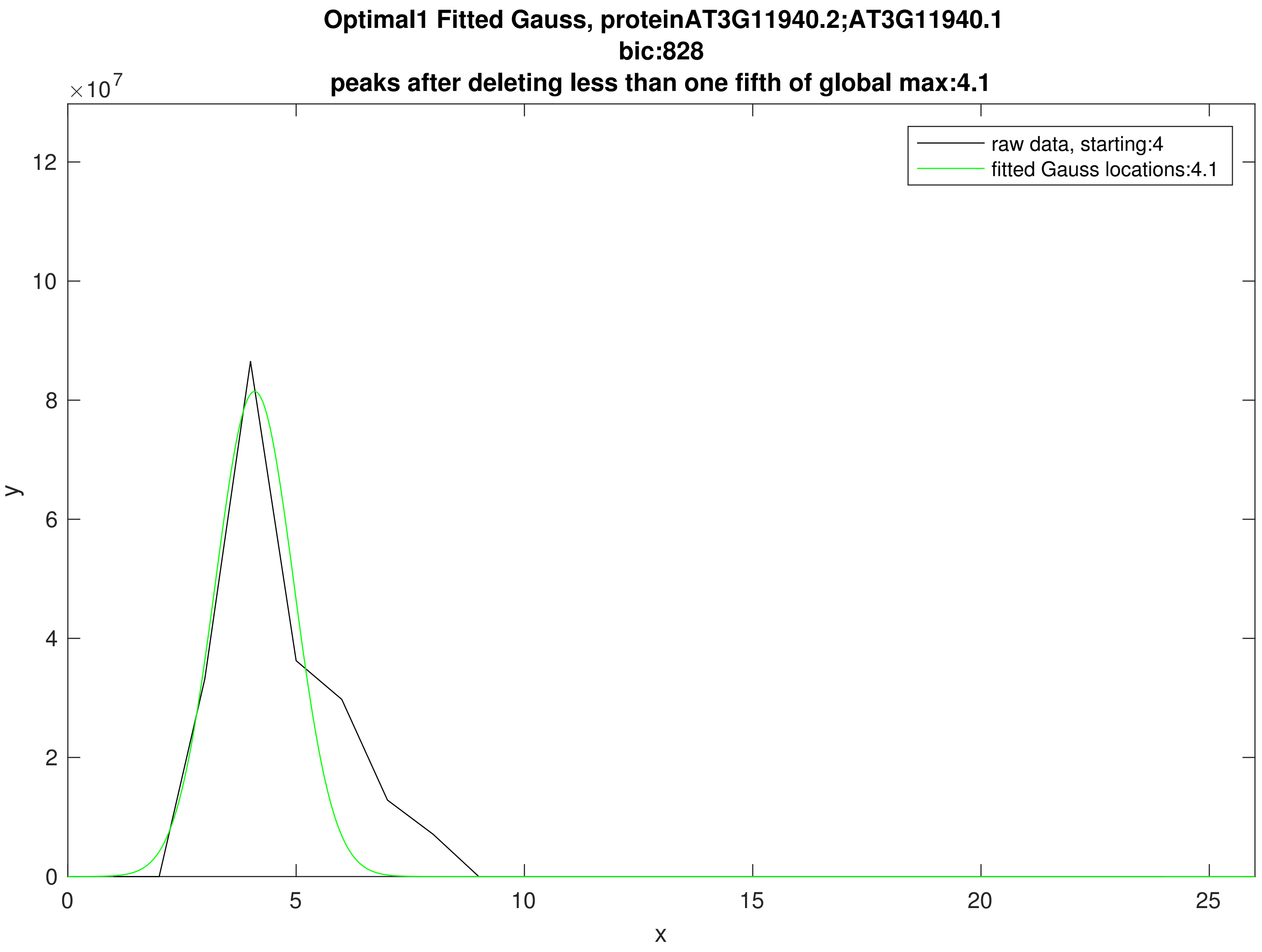

Optimal1 Fitted Gauss, proteinAT3G11940.2;AT3G11940.1
bic:828
peaks after deleting less than one fifth of global max:4.1
×10^7
raw data, starting:4
fitted Gauss locations:4.1
y
x
0
2
4
6
8
10
12
0
5
10
15
20
25

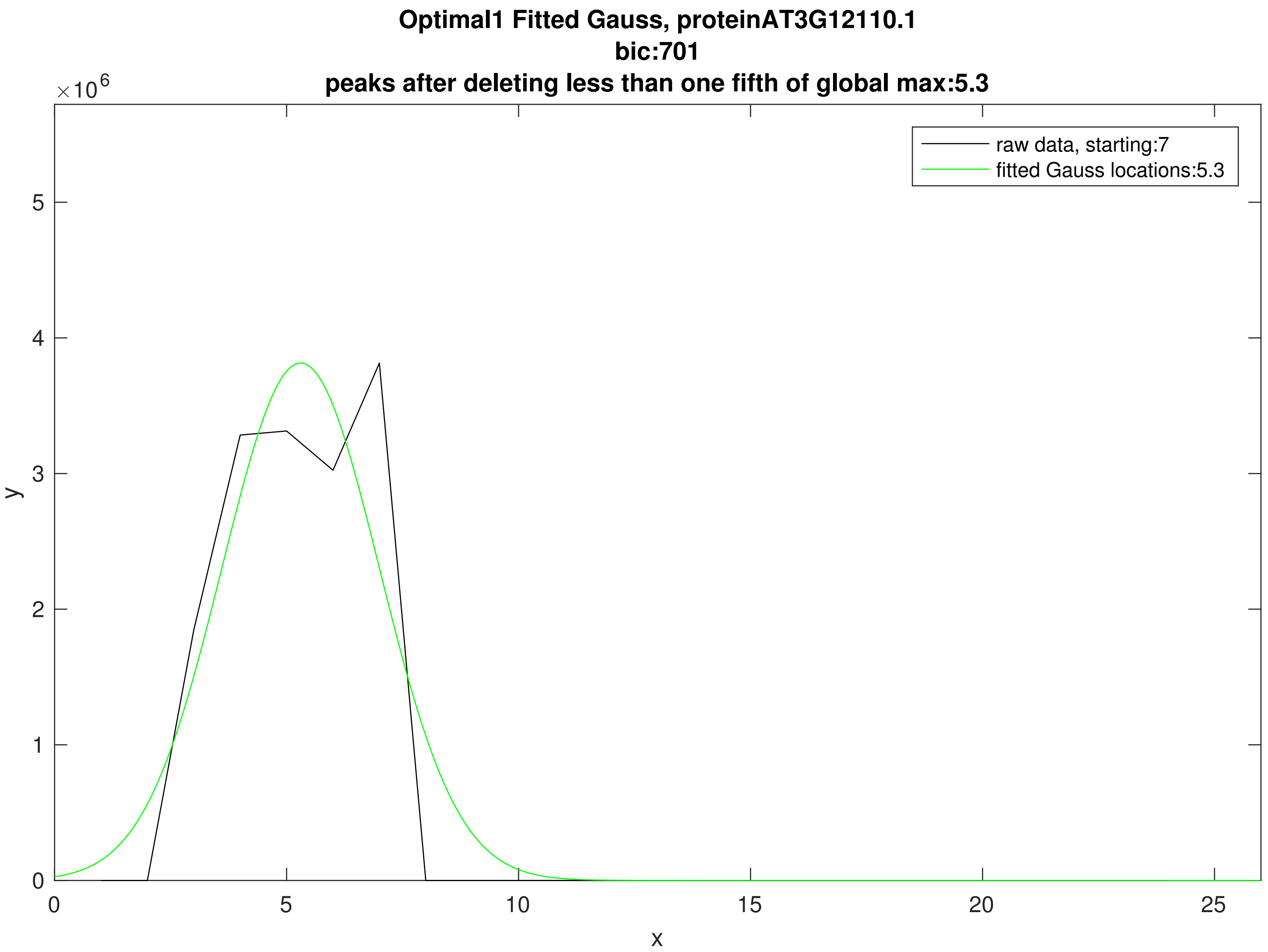
Optimal1 Fitted Gauss, proteinAT3G12110.1
bic:701
peaks after deleting less than one fifth of global max:5.3
×10^6
raw data, starting:7
fitted Gauss locations:5.3
y
x
0
1
2
3
4
5
10
15
20
25

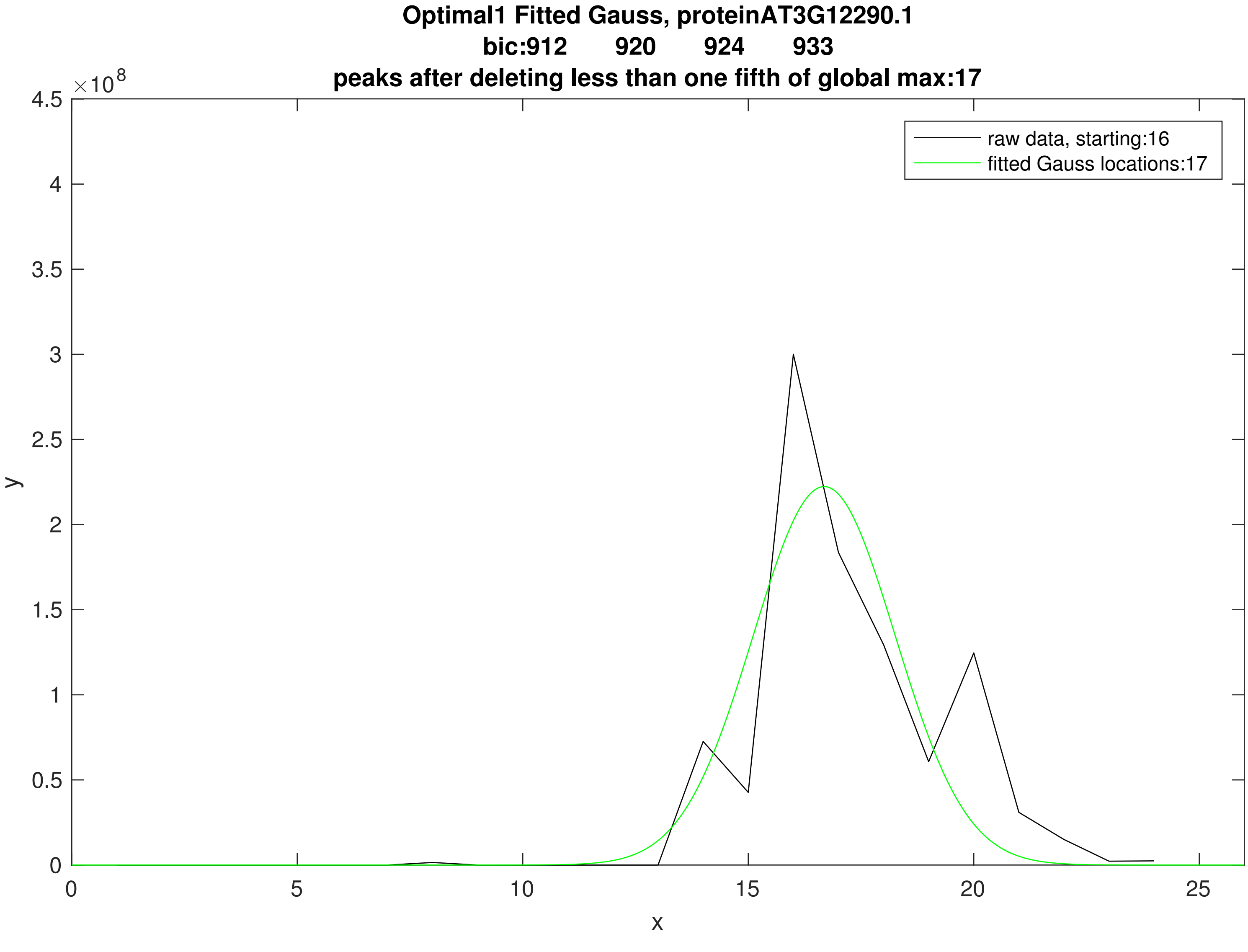

Optimal1 Fitted Gauss, proteinAT3G12290.1
bic:912 920 924 933
peaks after deleting less than one fifth of global max:17
×10^8
4.5
4
3.5
3
2.5
2
1.5
1
0.5
0
y
0
5
10
15
20
25
x
raw data, starting:16
fitted Gauss locations:17

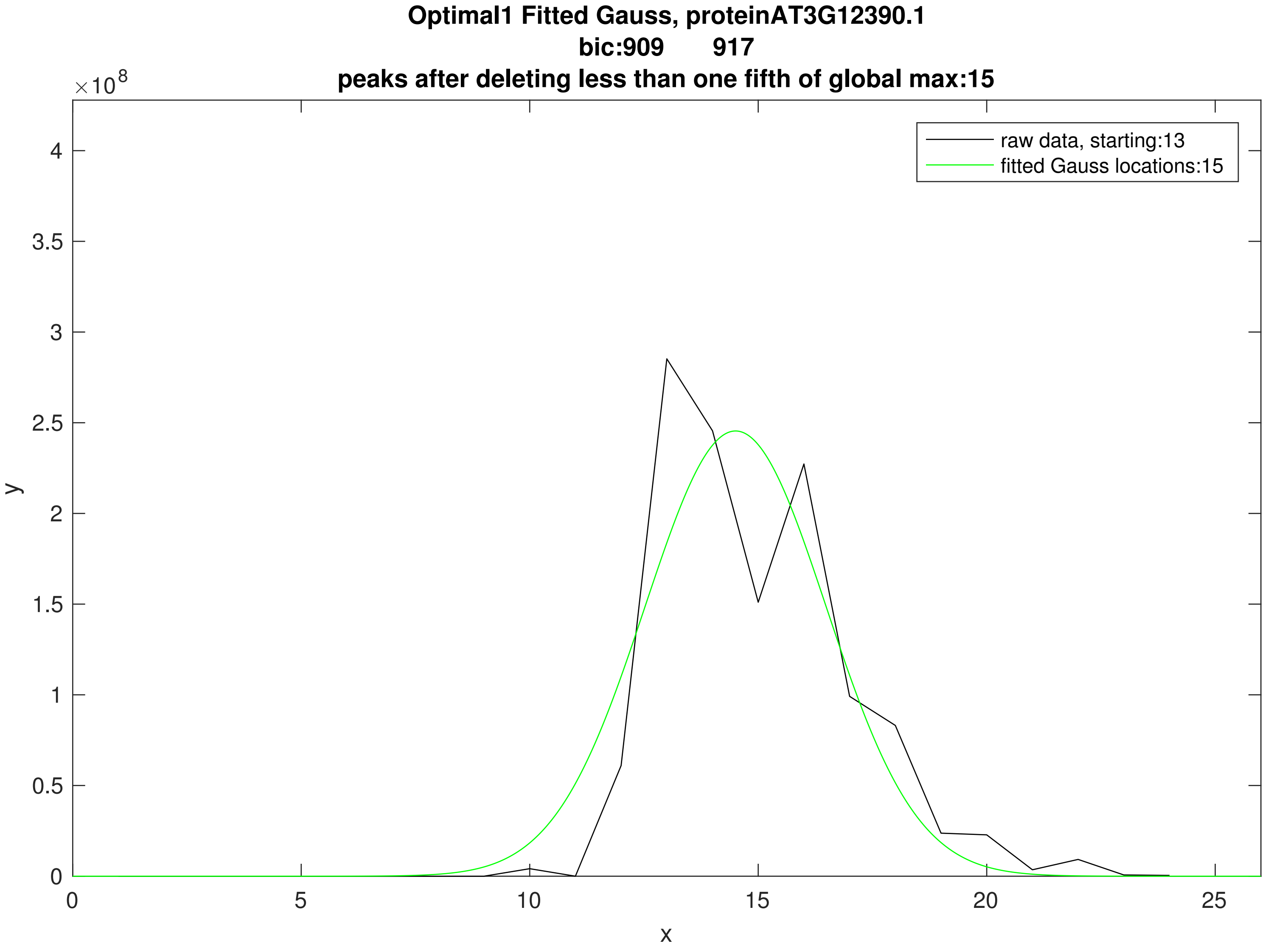

Optimal1 Fitted Gauss, proteinAT3G12390.1
bic:909 917
peaks after deleting less than one fifth of global max:15
×10^8
raw data, starting:13
fitted Gauss locations:15
y
x
0
0.5
1
1.5
2
2.5
3
3.5
4
5
10
15
20
25

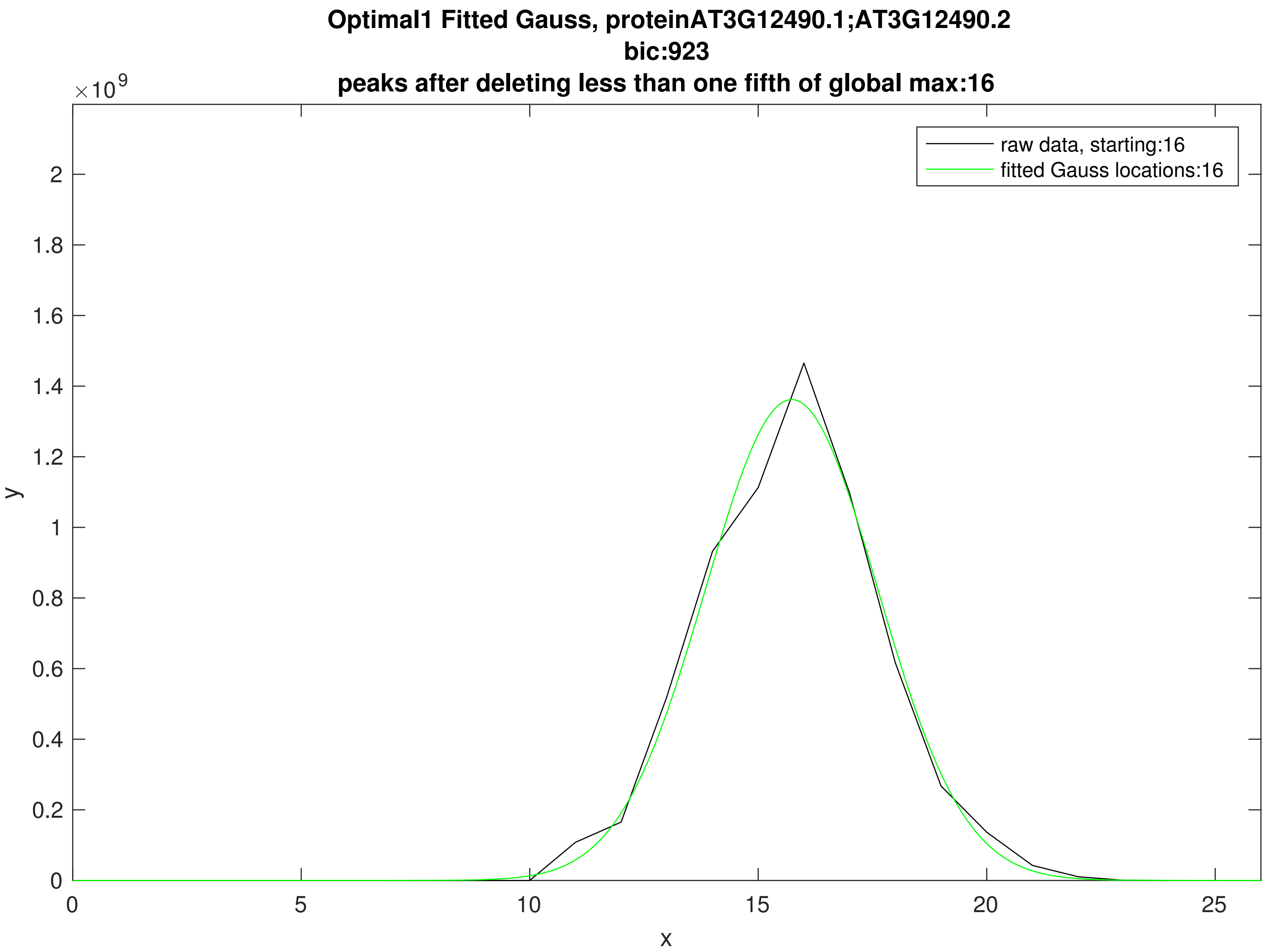
Optimal1 Fitted Gauss, proteinAT3G12490.1;AT3G12490.2
bic:923
peaks after deleting less than one fifth of global max:16
×10^9
raw data, starting:16
fitted Gauss locations:16
y
x
0
0.2
0.4
0.6
0.8
1
1.2
1.4
1.6
1.8
2
0
5
10
15
20
25

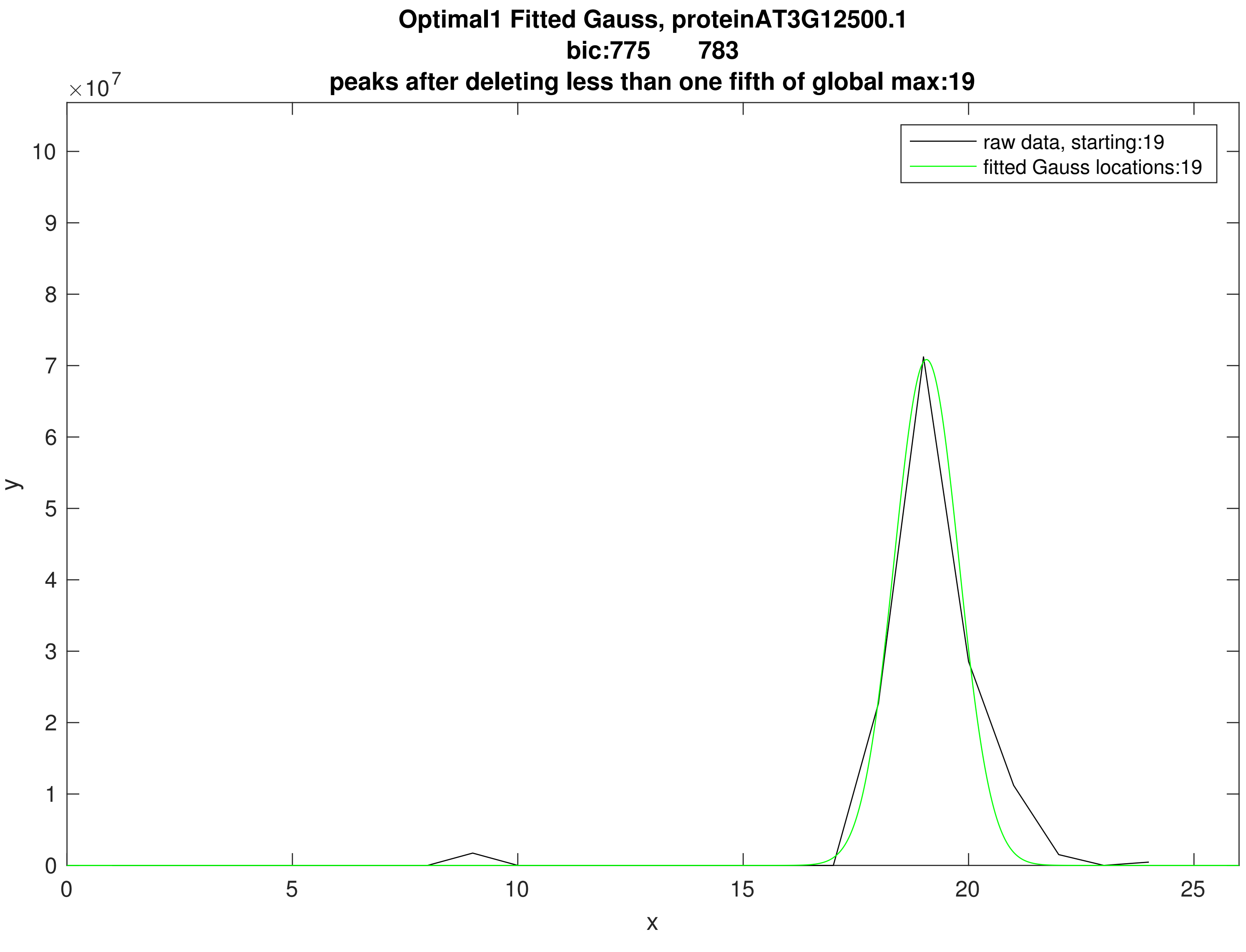

Optimal1 Fitted Gauss, proteinAT3G12500.1
bic:775 783
peaks after deleting less than one fifth of global max:19
×10^7
10
9
8
7
6
5
4
3
2
1
0
y
0
5
10
15
20
25
x
raw data, starting:19
fitted Gauss locations:19

**Optimal1 Fitted Gauss, proteinAT3G12580.1**
**bic:863 868 879**
**peaks after deleting less than one fifth of global max:13**

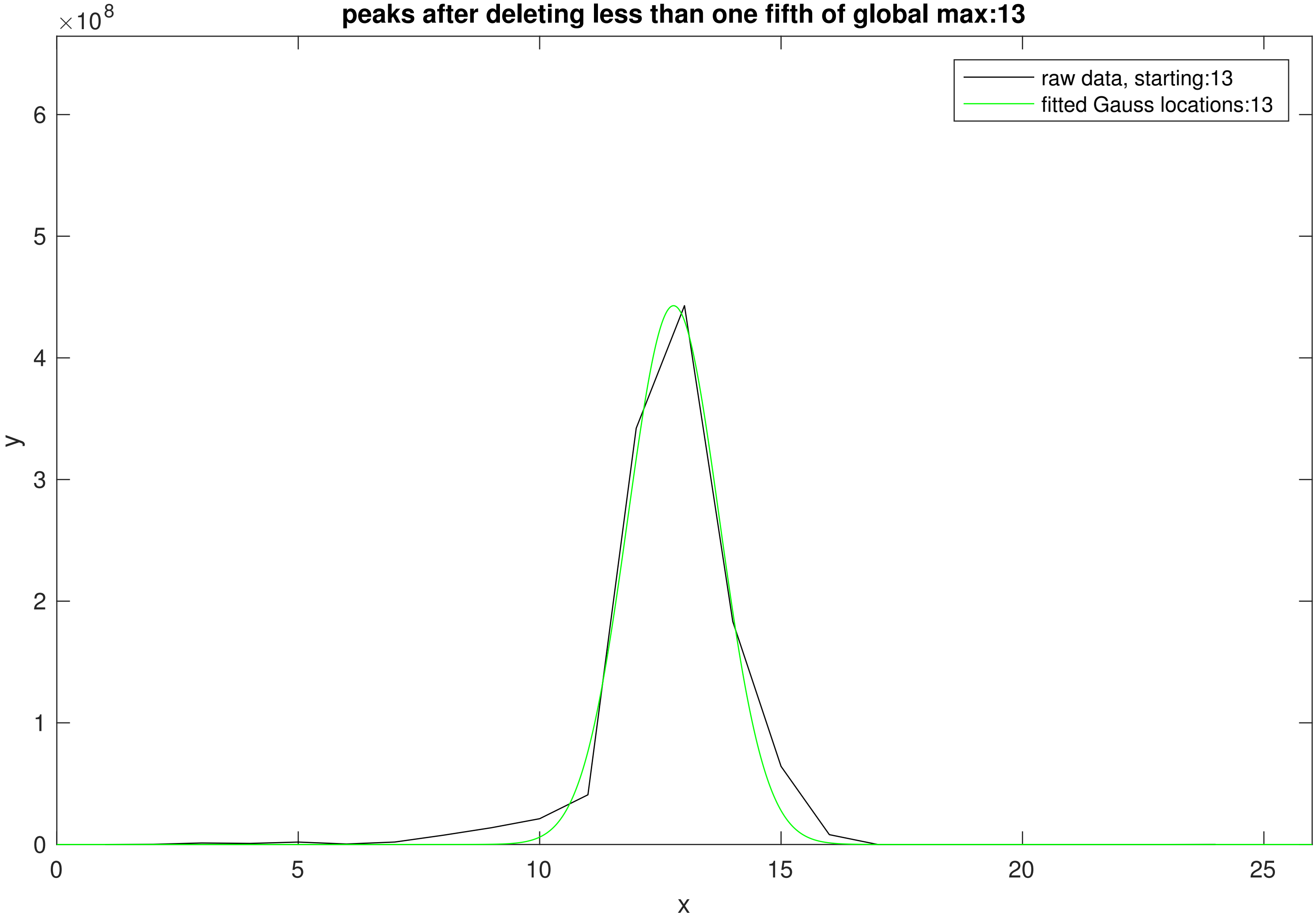

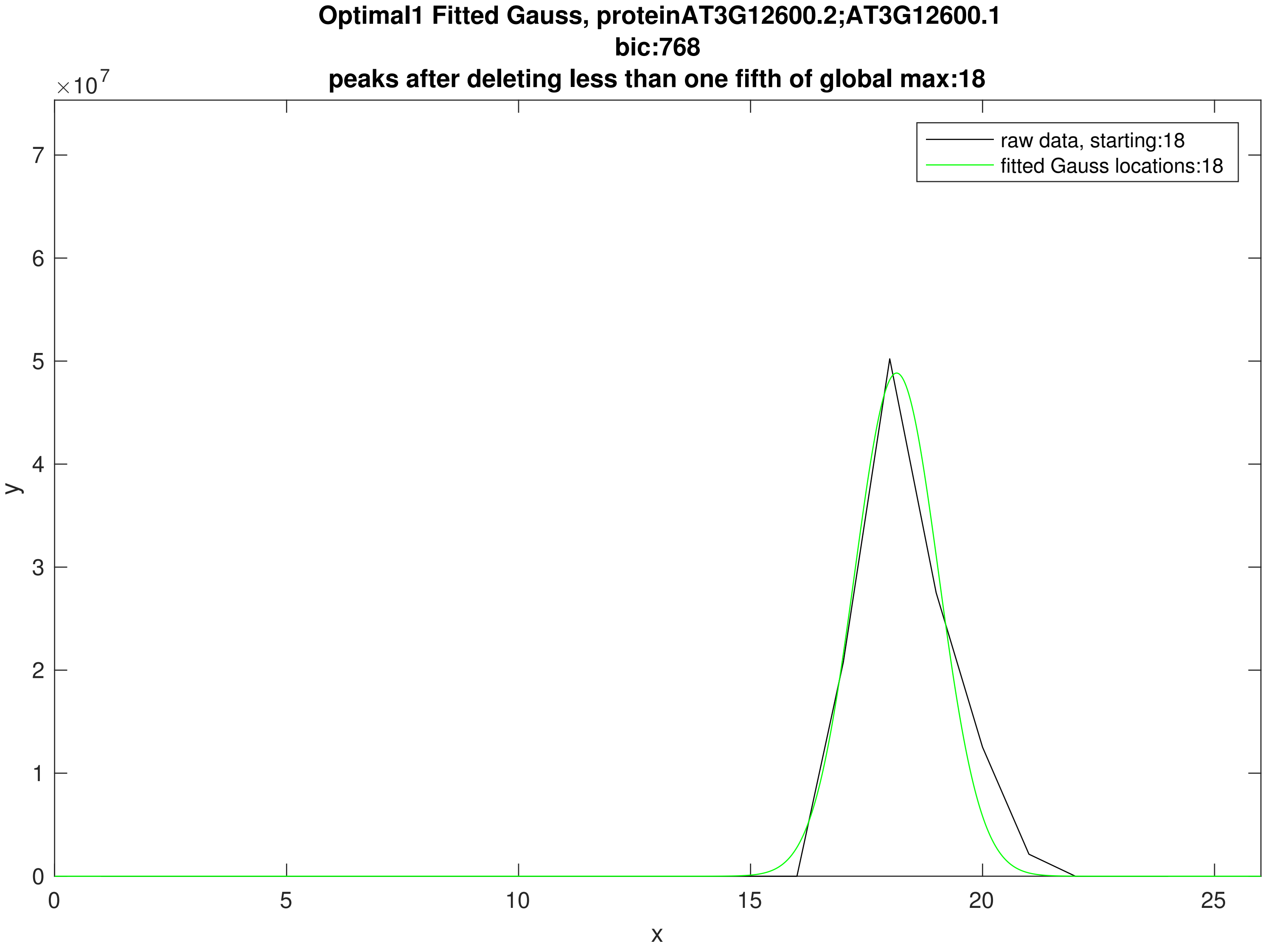

Optimal1 Fitted Gauss, proteinAT3G12600.2;AT3G12600.1
bic:768
peaks after deleting less than one fifth of global max:18
×10^7
raw data, starting:18
fitted Gauss locations:18
y
x
0
1
2
3
4
5
6
7
0
5
10
15
20
25

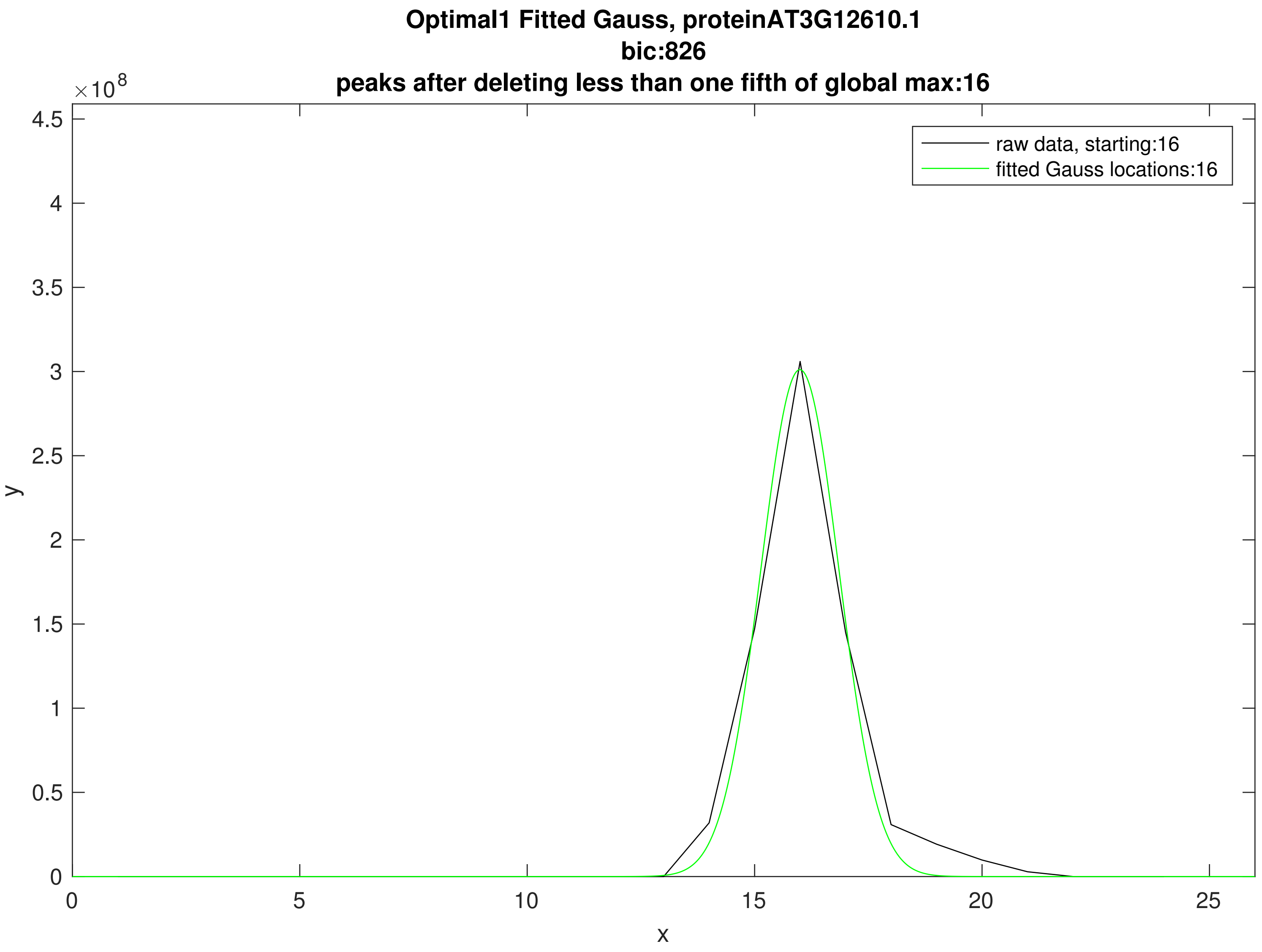
Optimal1 Fitted Gauss, proteinAT3G12610.1
bic:826
peaks after deleting less than one fifth of global max:16
×10^8
4.5
4
3.5
3
2.5
2
1.5
1
0.5
0
y
0
5
10
15
20
25
x
raw data, starting:16
fitted Gauss locations:16

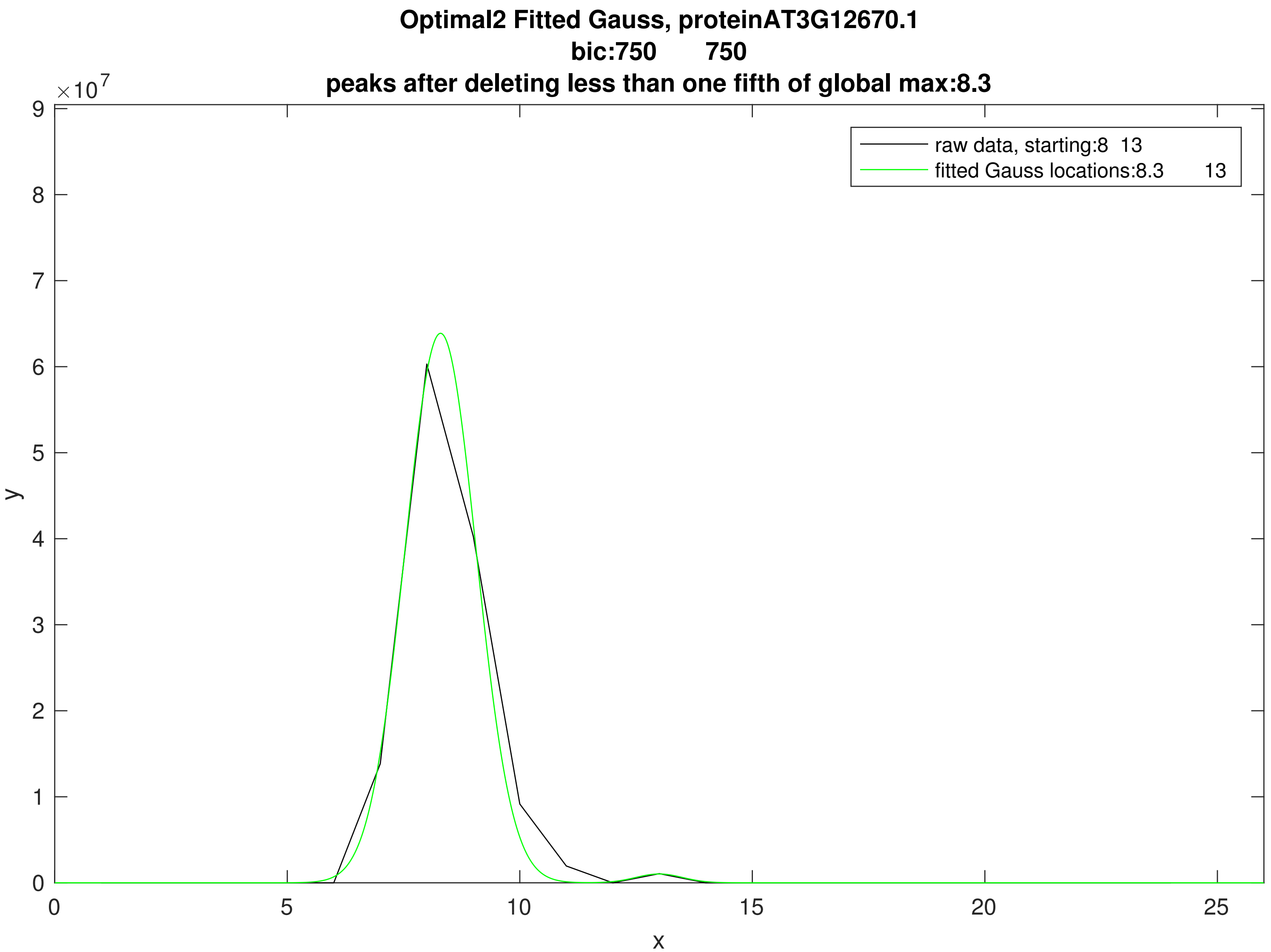

Optimal2 Fitted Gauss, proteinAT3G12670.1
bic:750 750
peaks after deleting less than one fifth of global max:8.3
×10^7
raw data, starting:8 13
fitted Gauss locations:8.3 13
y
x
0
1
2
3
4
5
6
7
8
9
0
5
10
15
20
25

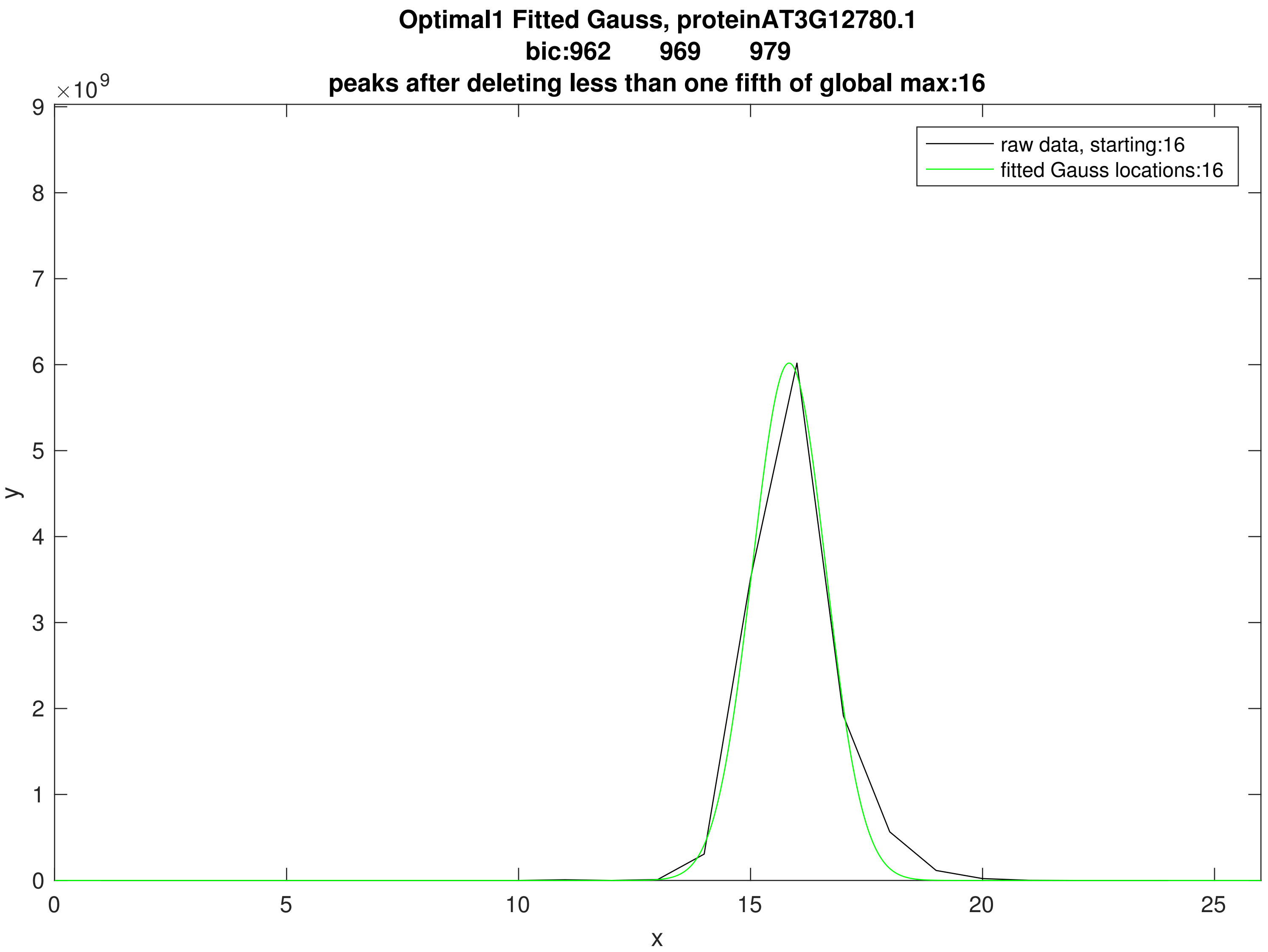

Optimal1 Fitted Gauss, proteinAT3G12780.1
bic:962 969 979
peaks after deleting less than one fifth of global max:16
×10^9
raw data, starting:16
fitted Gauss locations:16
y
x
0
1
2
3
4
5
6
7
8
9
0
5
10
15
20
25

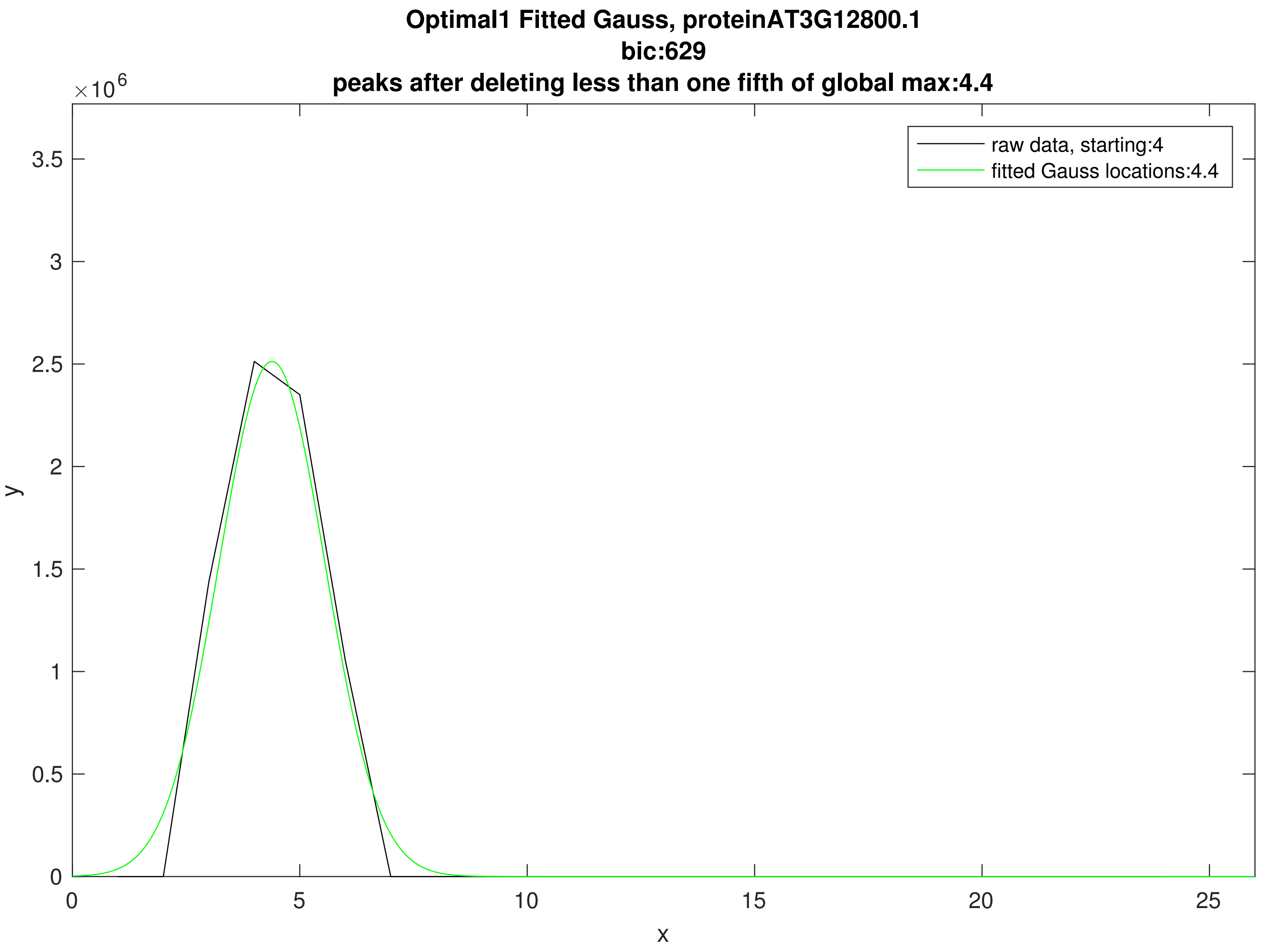

Optimal1 Fitted Gauss, proteinAT3G12800.1
bic:629
peaks after deleting less than one fifth of global max:4.4
×10^6
raw data, starting:4
fitted Gauss locations:4.4
3.5
3
2.5
2
1.5
1
0.5
0
y
0
5
10
15
20
25
x

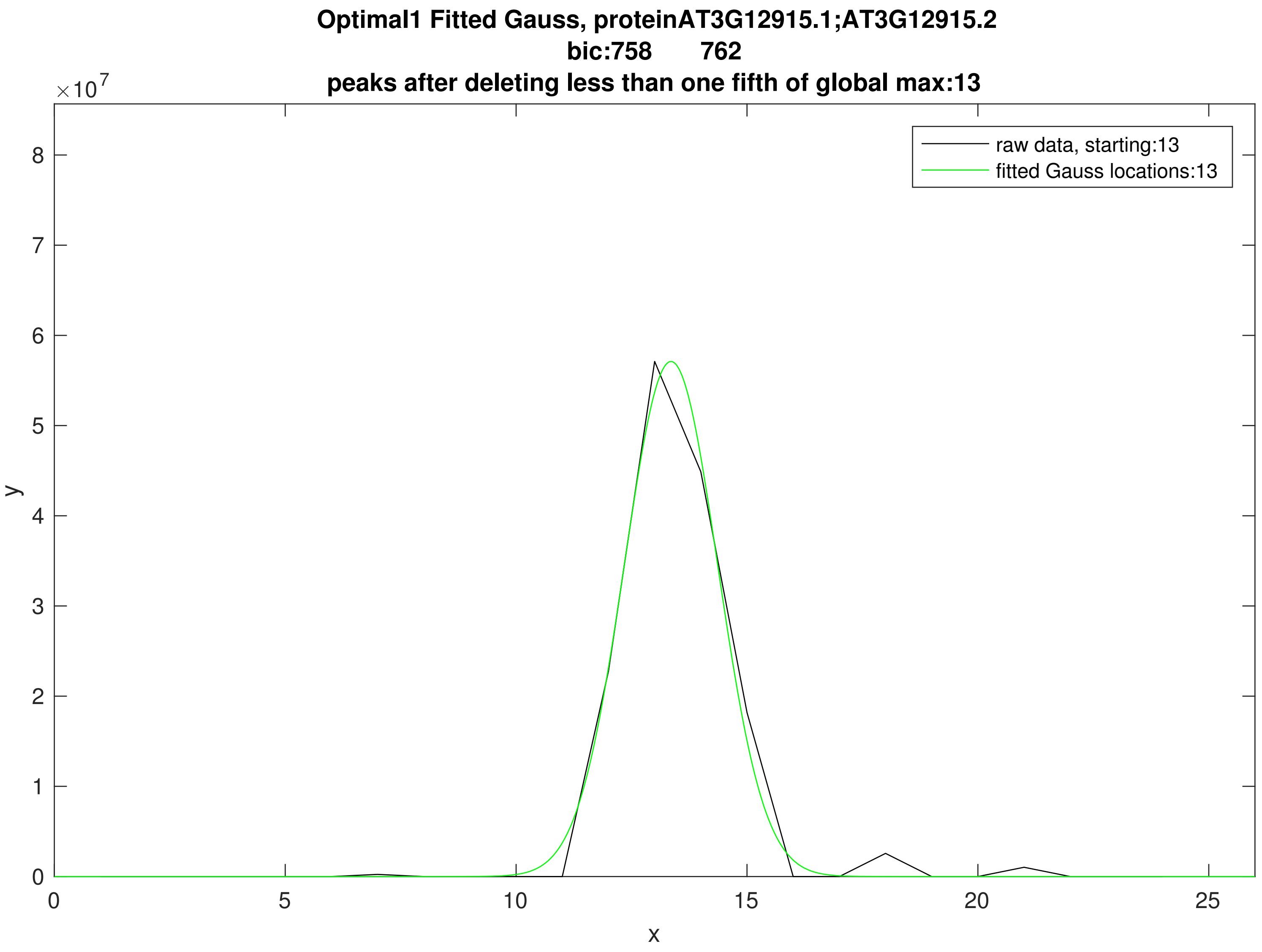
Optimal1 Fitted Gauss, proteinAT3G12915.1;AT3G12915.2
bic:758 762
peaks after deleting less than one fifth of global max:13
×10^7
raw data, starting:13
fitted Gauss locations:13
y
x
0
1
2
3
4
5
6
7
8
0
5
10
15
20
25

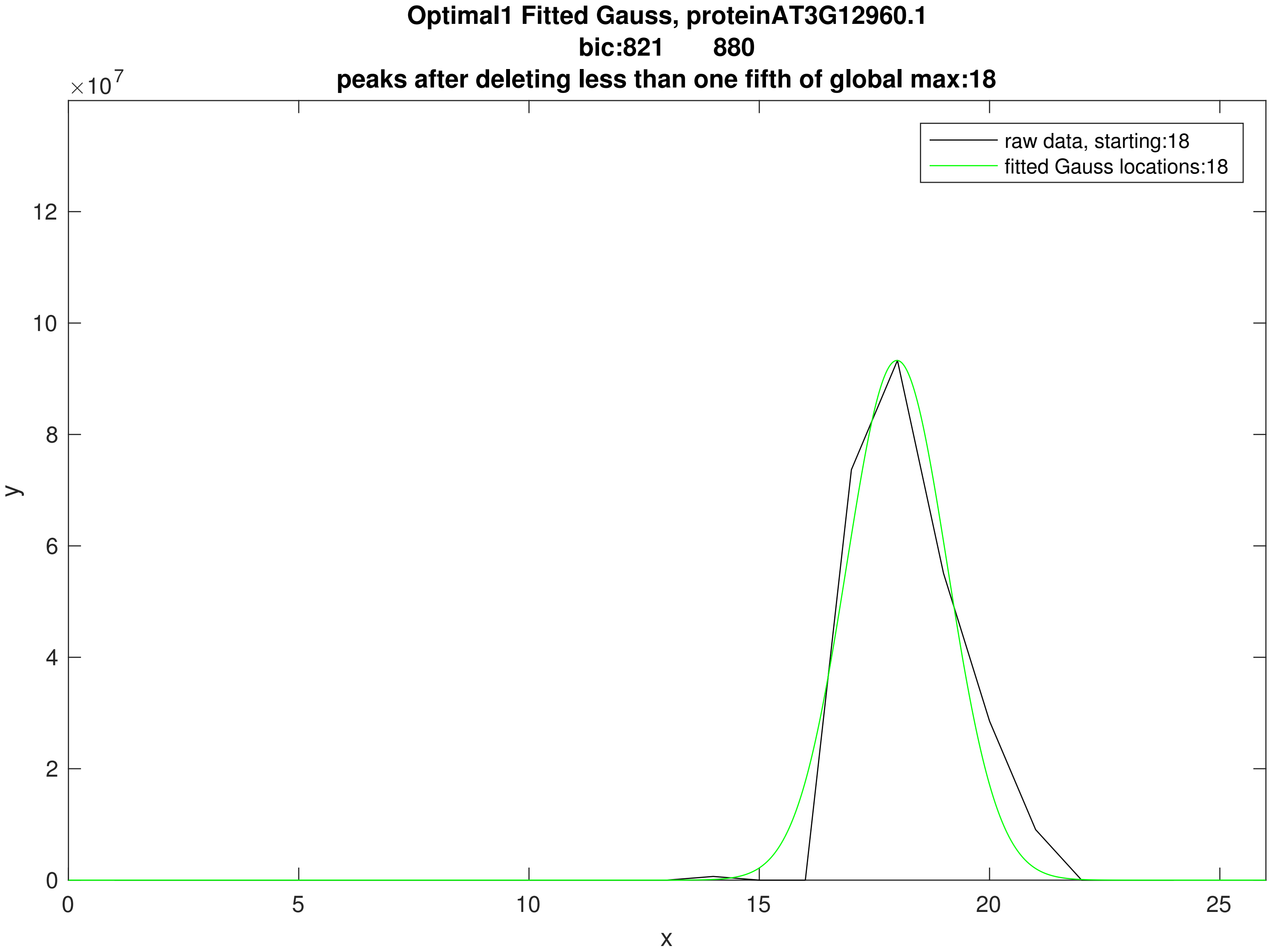
Optimal1 Fitted Gauss, proteinAT3G12960.1
bic:821 880
peaks after deleting less than one fifth of global max:18
×10^7
raw data, starting:18
fitted Gauss locations:18
12
10
8
6
4
2
0
y
0
5
10
15
20
25
x

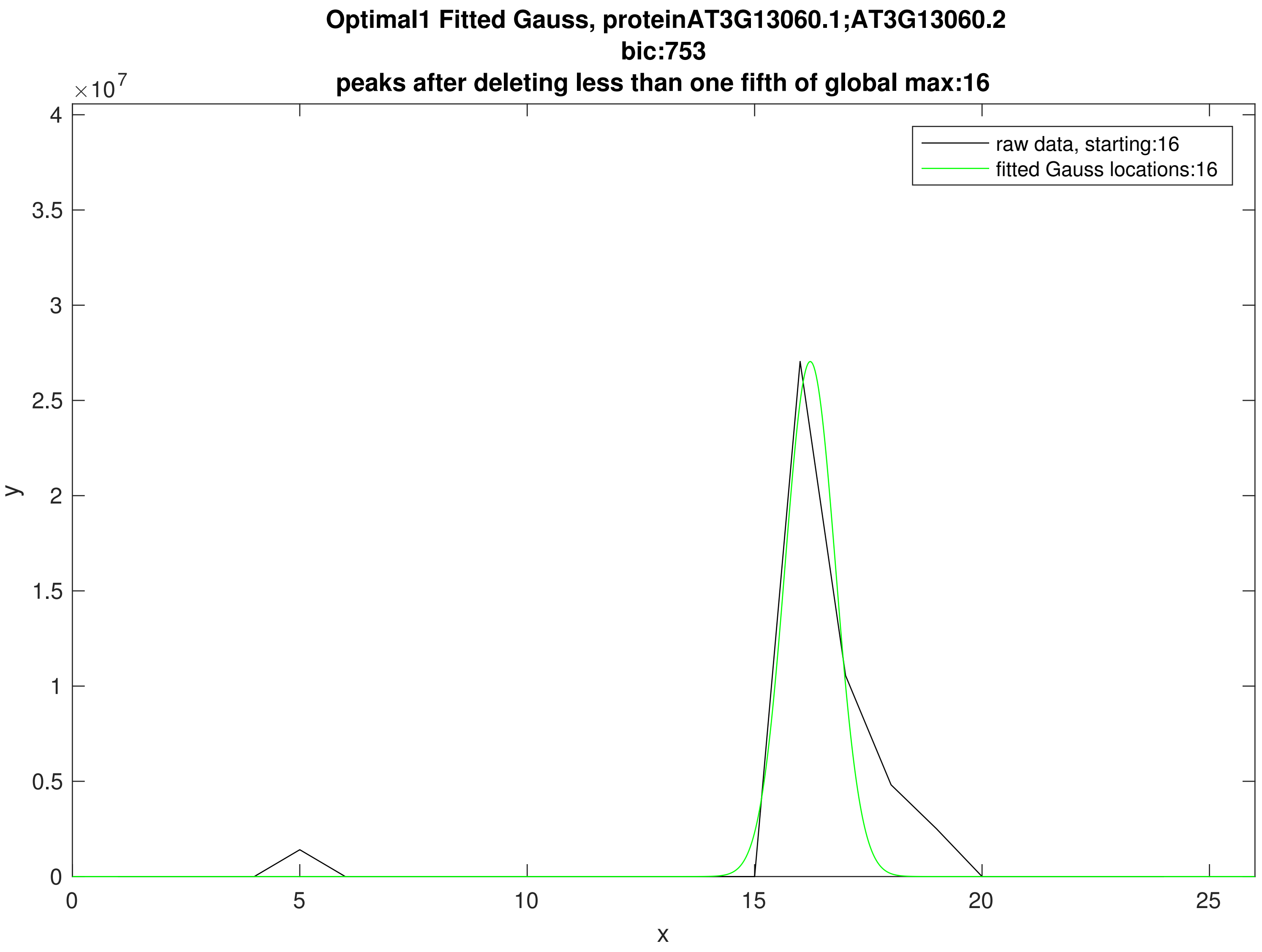

Optimal1 Fitted Gauss, proteinAT3G13060.1;AT3G13060.2
bic:753
peaks after deleting less than one fifth of global max:16
×10^7
4
3.5
3
2.5
2
1.5
1
0.5
0
y
0
5
10
15
20
25
x
raw data, starting:16
fitted Gauss locations:16

#### Optimal2 Fitted Gauss, proteinAT3G13235.3;AT3G13235.2;AT3G13235.1
#### bic:753 752
#### peaks after deleting less than one fifth of global max:13

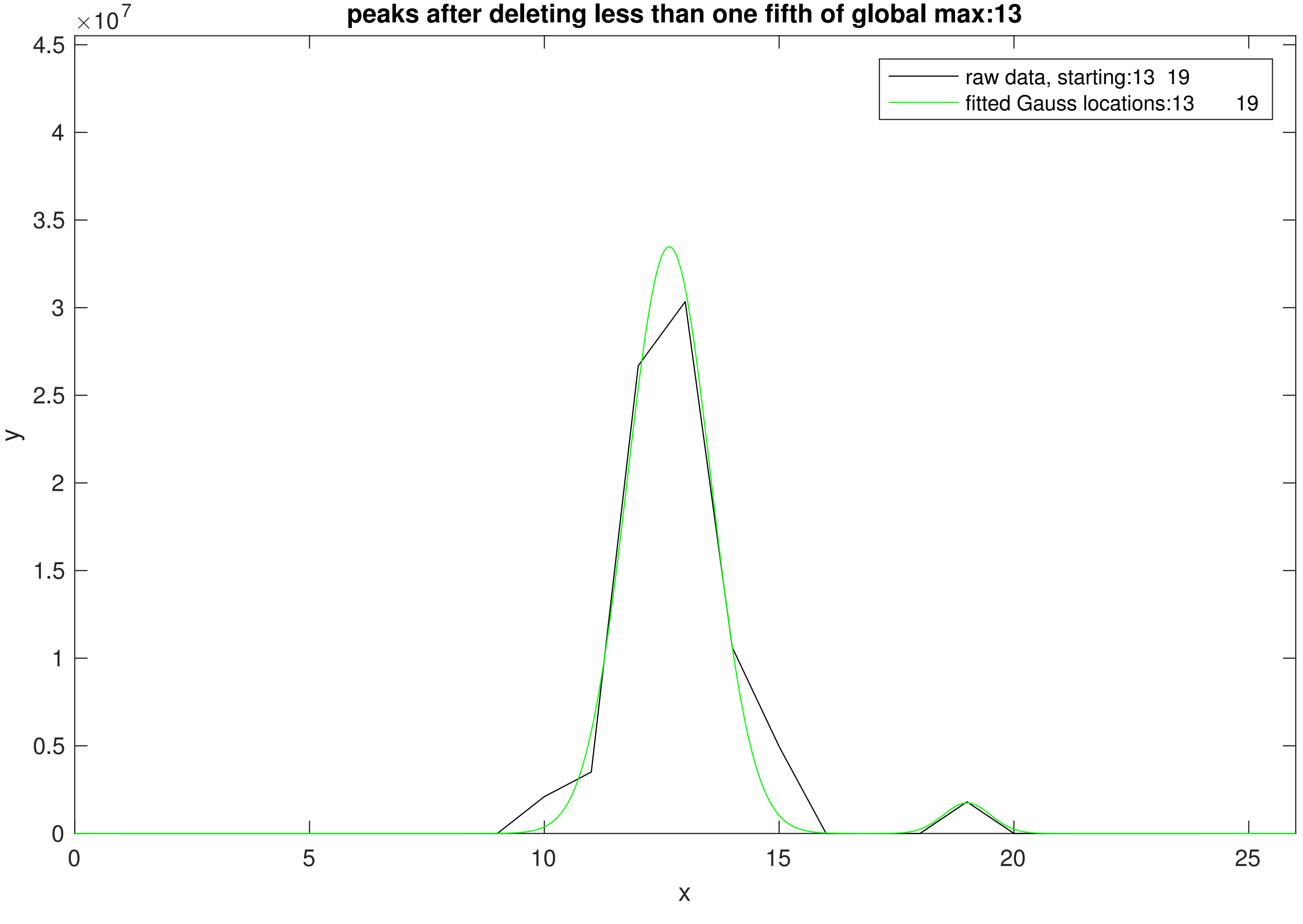

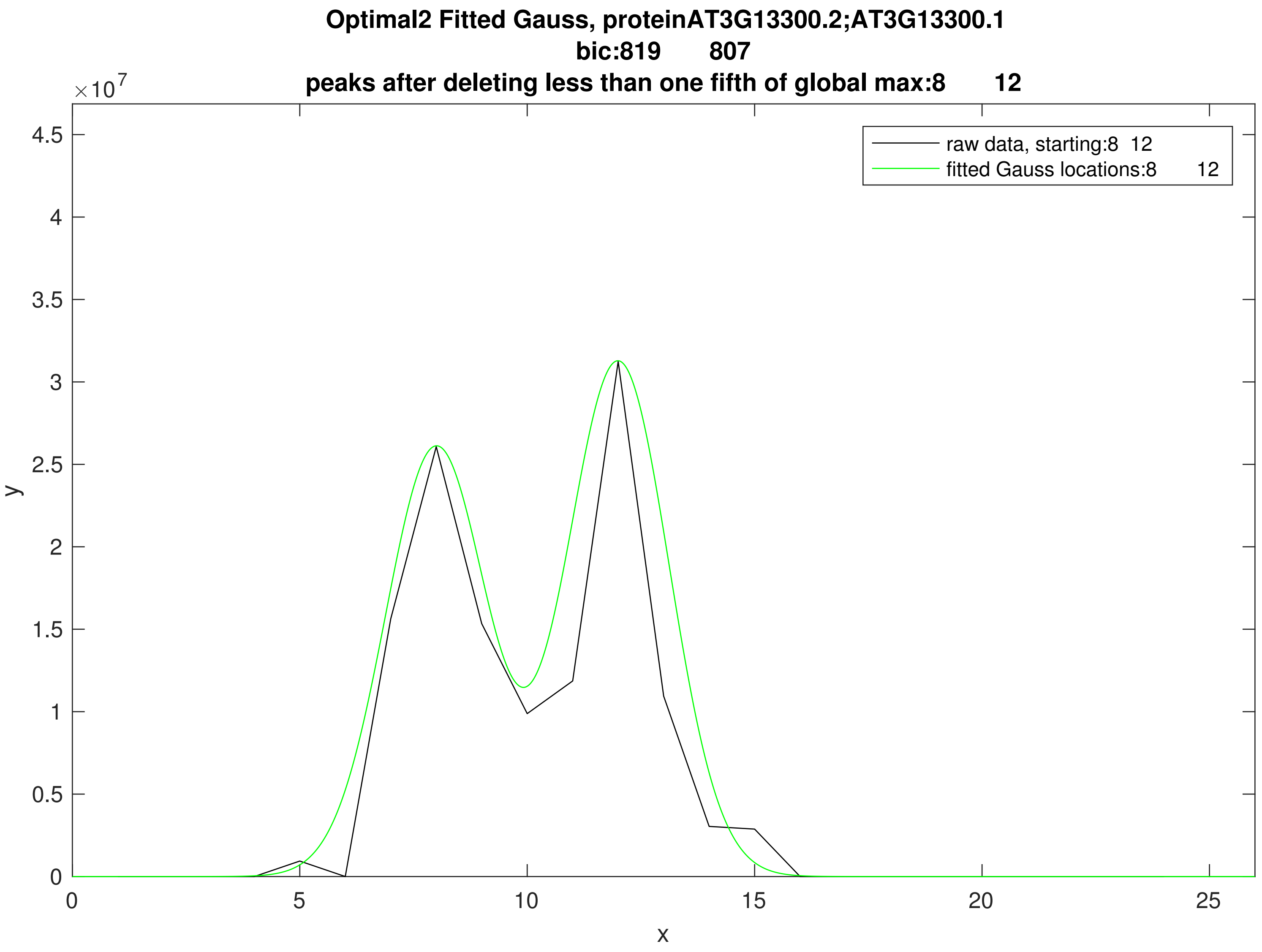
Optimal2 Fitted Gauss, proteinAT3G13300.2;AT3G13300.1
bic:819 807
peaks after deleting less than one fifth of global max:8 12
×10^7
raw data, starting:8 12
fitted Gauss locations:8 12
y
x
0
0.5
1
1.5
2
2.5
3
3.5
4
4.5
0
5
10
15
20
25

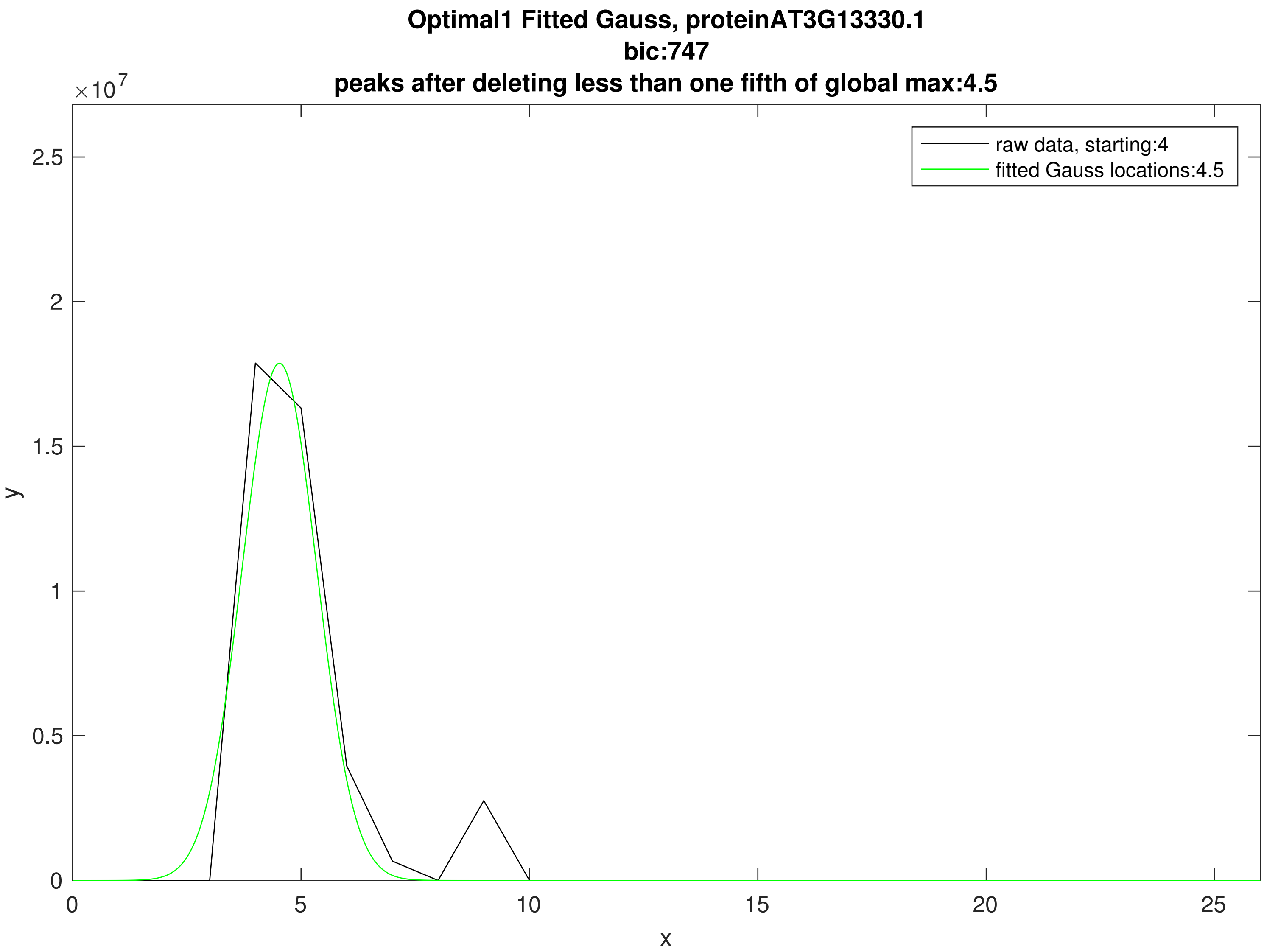
Optimal1 Fitted Gauss, proteinAT3G13330.1
bic:747
peaks after deleting less than one fifth of global max:4.5
×10^7
2.5
2
1.5
1
0.5
0
y
0
5
10
15
20
25
x
raw data, starting:4
fitted Gauss locations:4.5

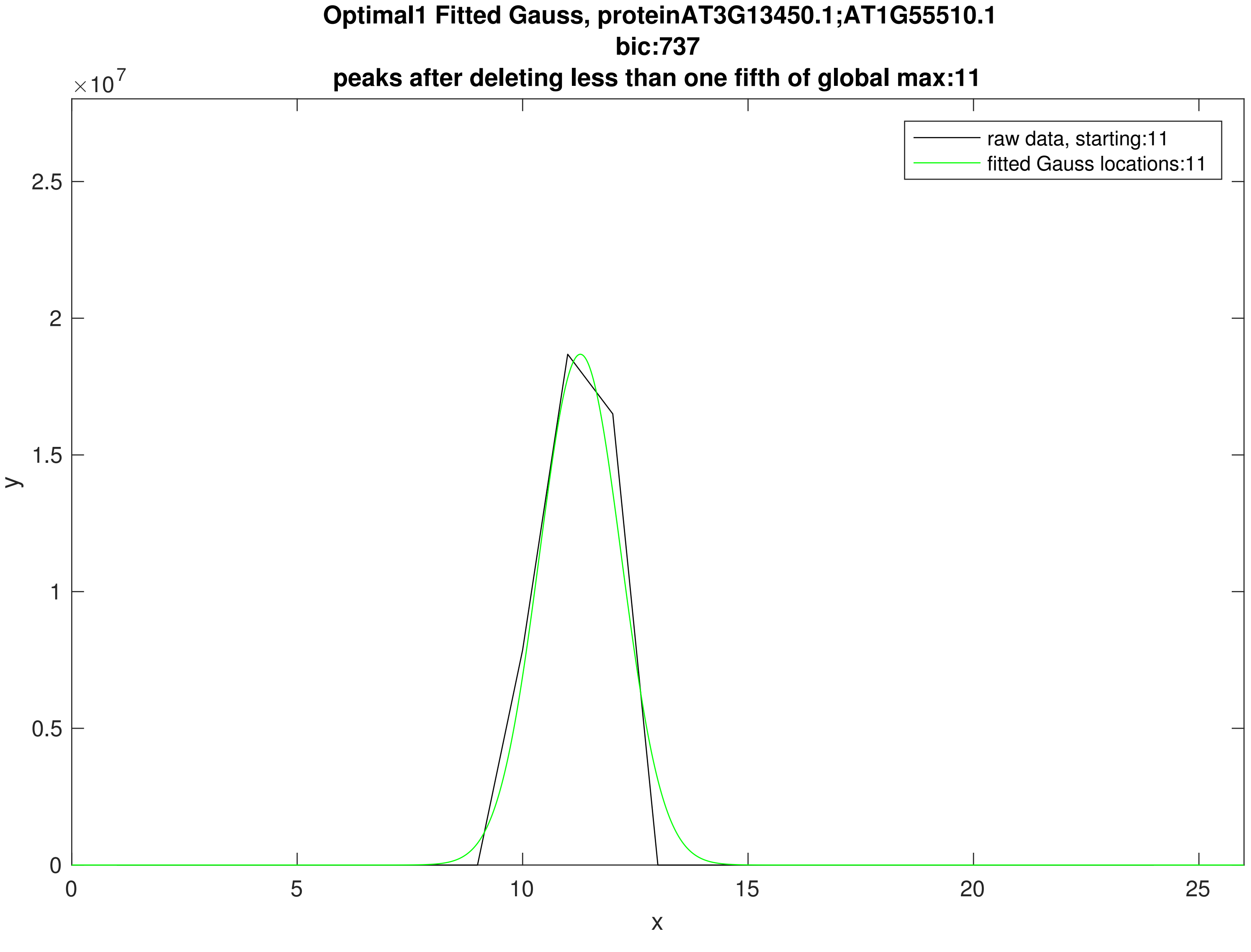
Optimal1 Fitted Gauss, proteinAT3G13450.1;AT1G55510.1
bic:737
peaks after deleting less than one fifth of global max:11
×10^7
raw data, starting:11
fitted Gauss locations:11
y
x
0
0.5
1
1.5
2
2.5
5
10
15
20
25

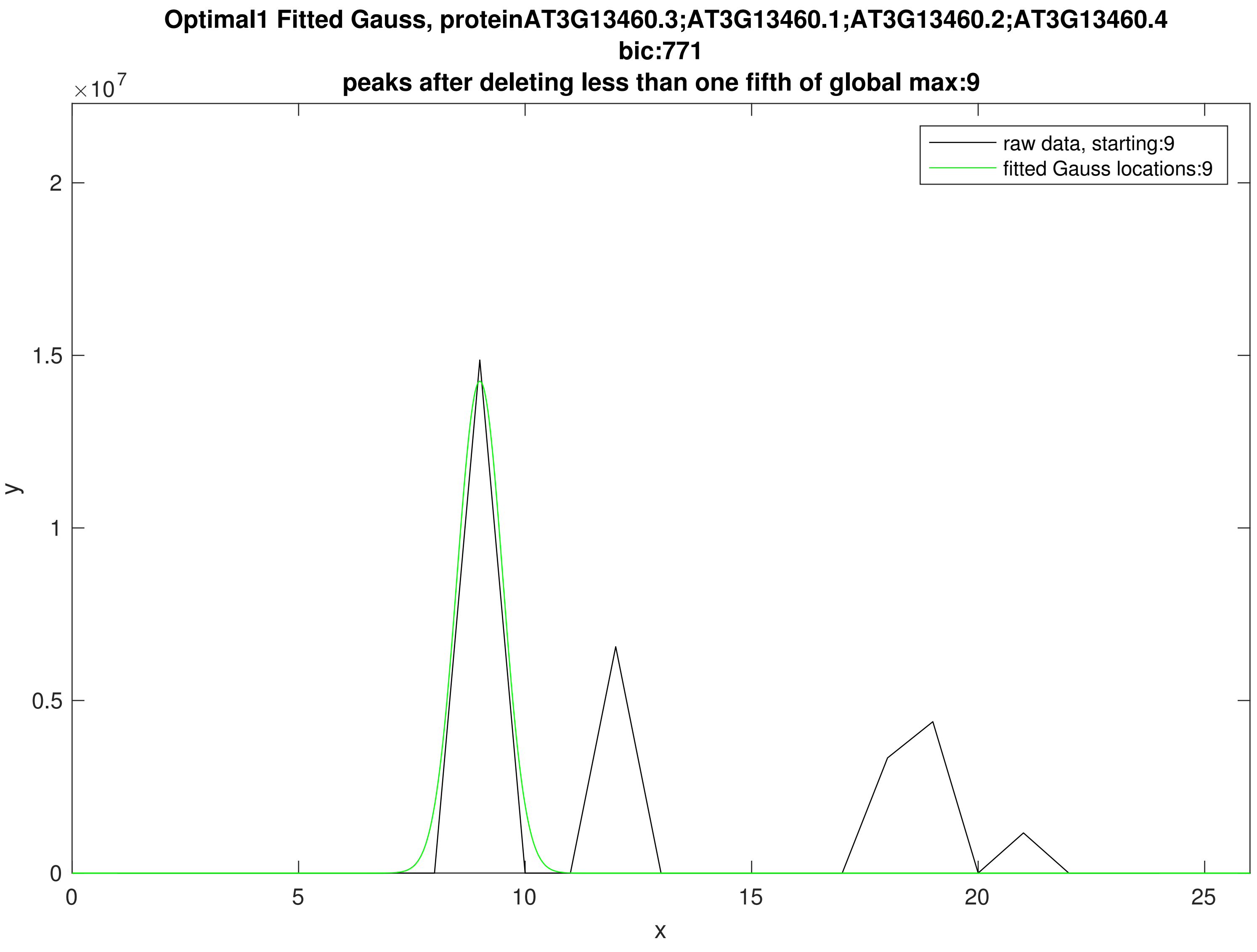

Optimal1 Fitted Gauss, proteinAT3G13460.3;AT3G13460.1;AT3G13460.2;AT3G13460.4
bic:771
peaks after deleting less than one fifth of global max:9
×10^7
2
1.5
1
0.5
0
y
0
5
10
15
20
25
x
raw data, starting:9
fitted Gauss locations:9

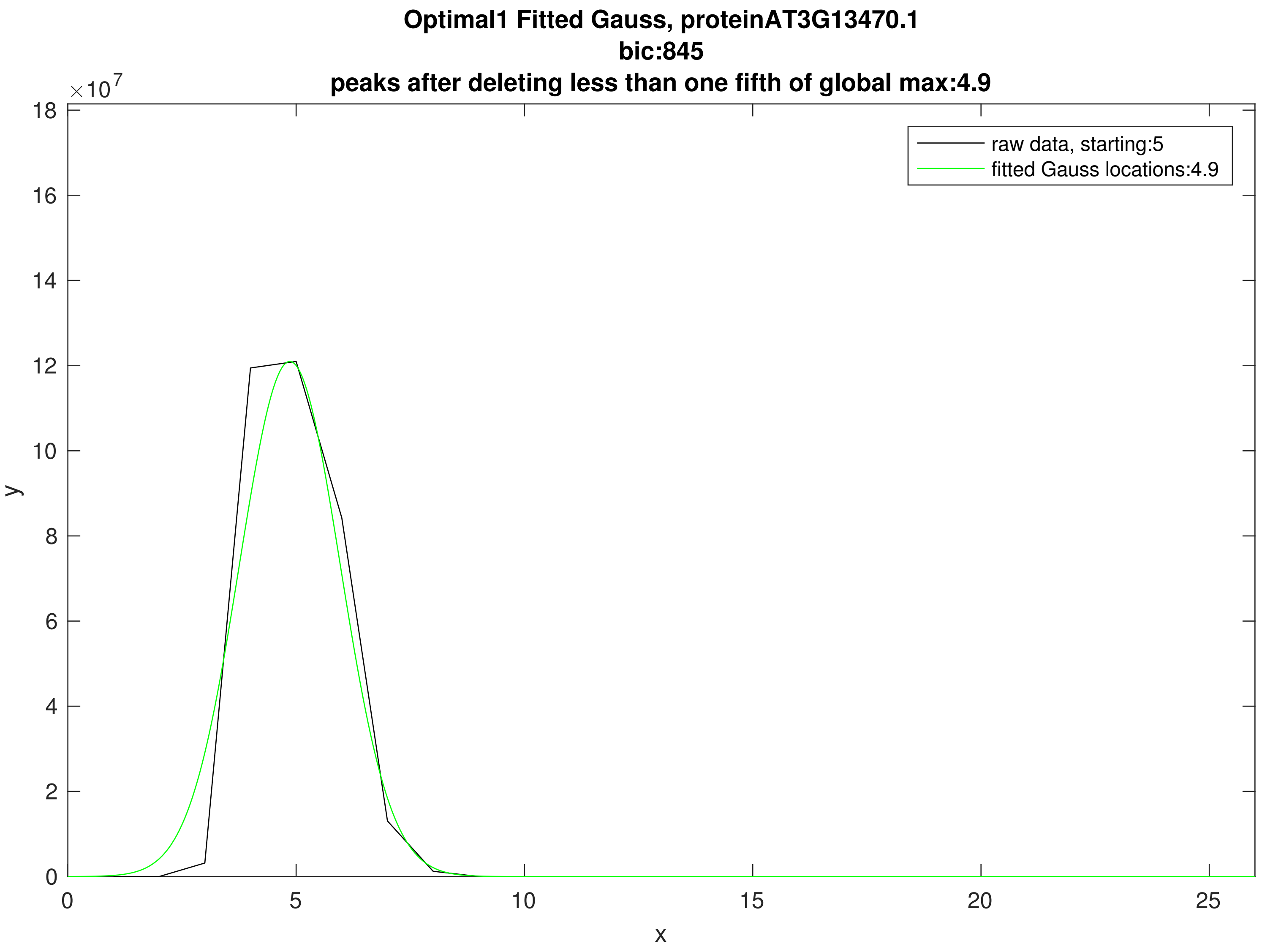
Optimal1 Fitted Gauss, proteinAT3G13470.1
bic:845
peaks after deleting less than one fifth of global max:4.9
×10^7
raw data, starting:5
fitted Gauss locations:4.9
y
x
0
2
4
6
8
10
12
14
16
18
0
5
10
15
20
25

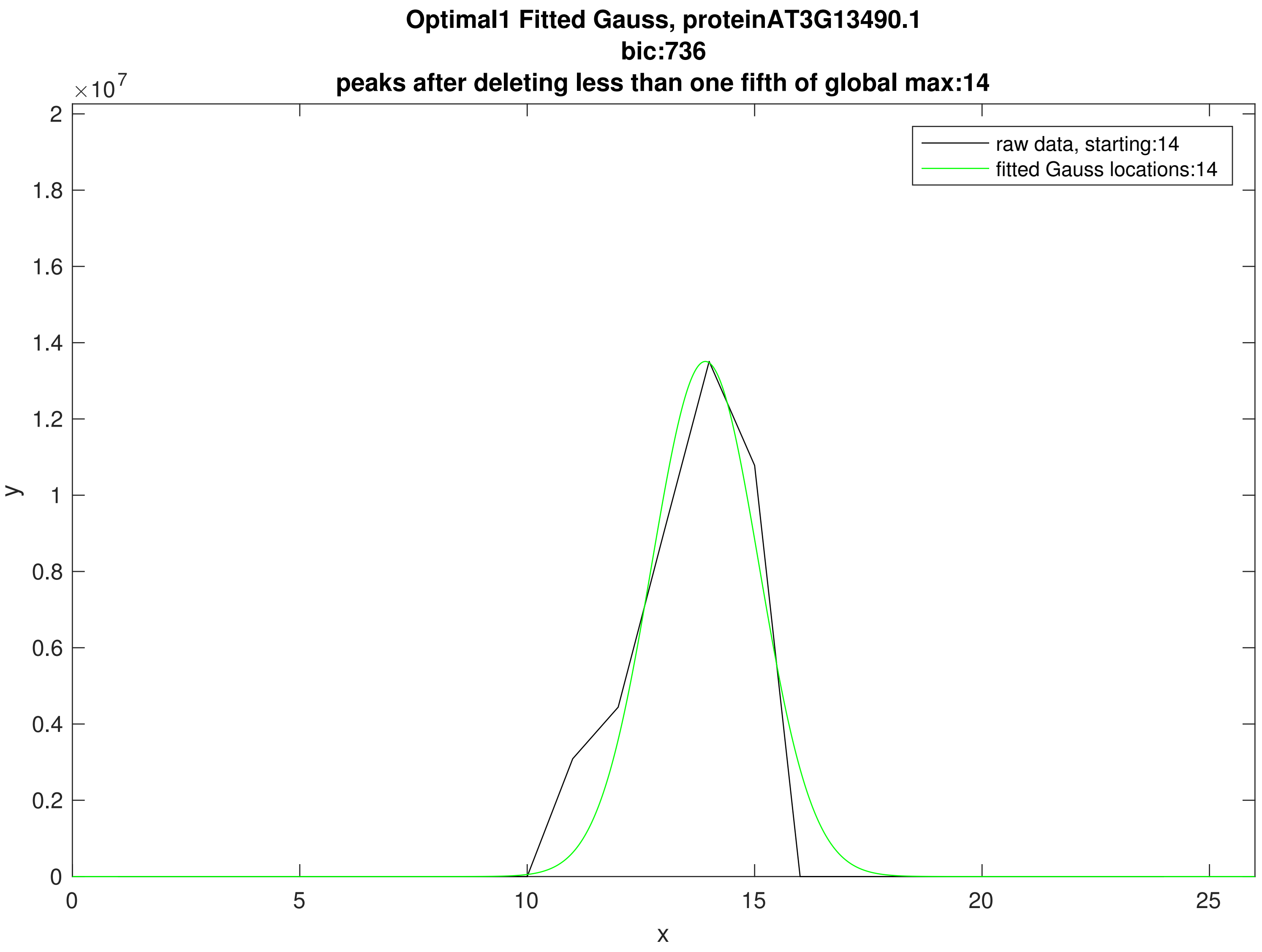
Optimal1 Fitted Gauss, proteinAT3G13490.1
bic:736
peaks after deleting less than one fifth of global max:14
×10^7
2
1.8
1.6
1.4
1.2
1
0.8
0.6
0.4
0.2
0
y
0
5
10
15
20
25
x
raw data, starting:14
fitted Gauss locations:14

Optimal1 Fitted Gauss, proteinAT3G13560.3;AT3G13560.2;AT3G13560.1
bic:739
peaks after deleting less than one fifth of global max:9.8

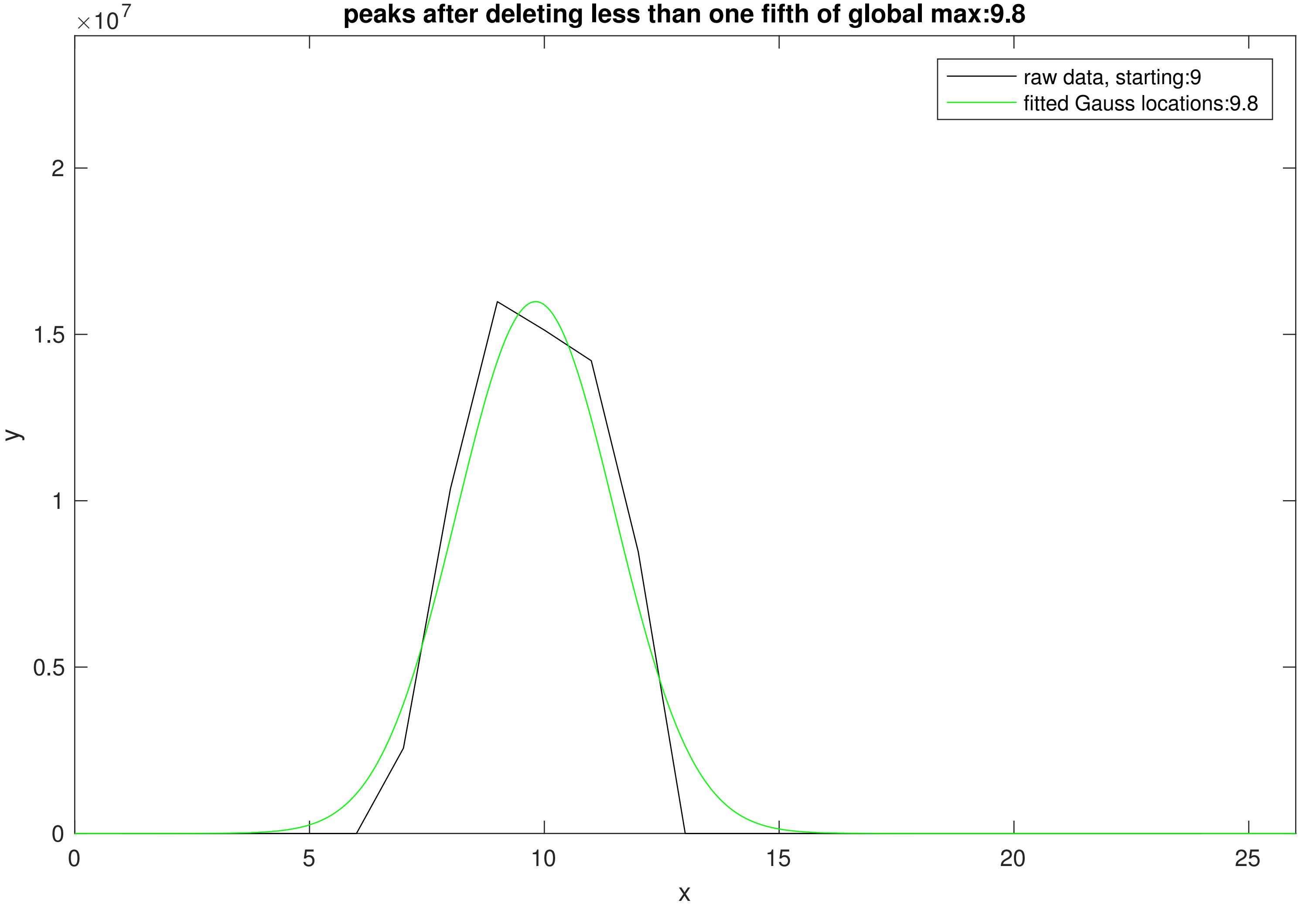

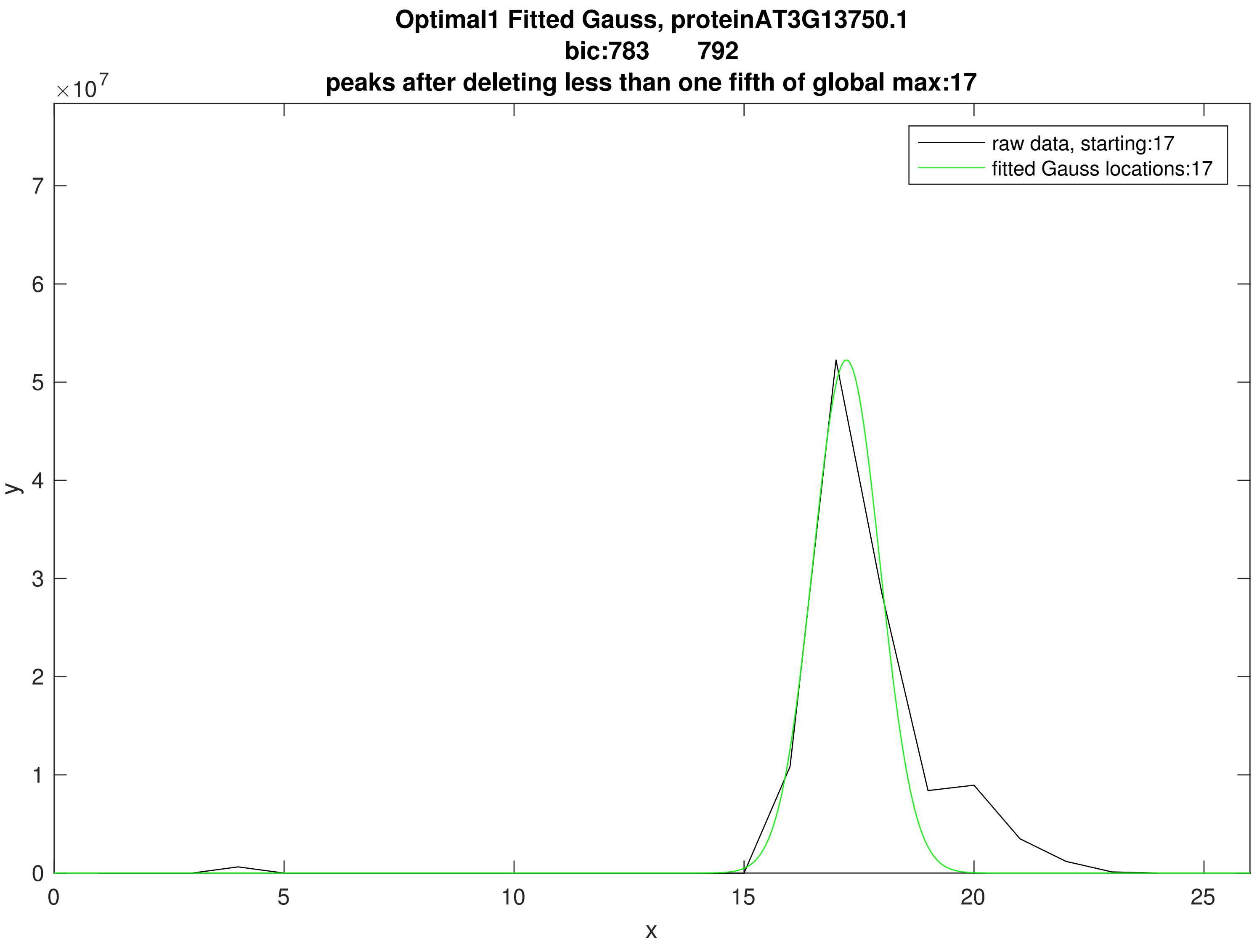
Optimal1 Fitted Gauss, proteinAT3G13750.1
bic:783 792
peaks after deleting less than one fifth of global max:17
×10^7
raw data, starting:17
fitted Gauss locations:17
y
x
0
1
2
3
4
5
6
7
0
5
10
15
20
25

Optimal2 Fitted Gauss, proteinAT3G13860.1
bic:950 914 924
peaks after deleting less than one fifth of global max:5.1 11

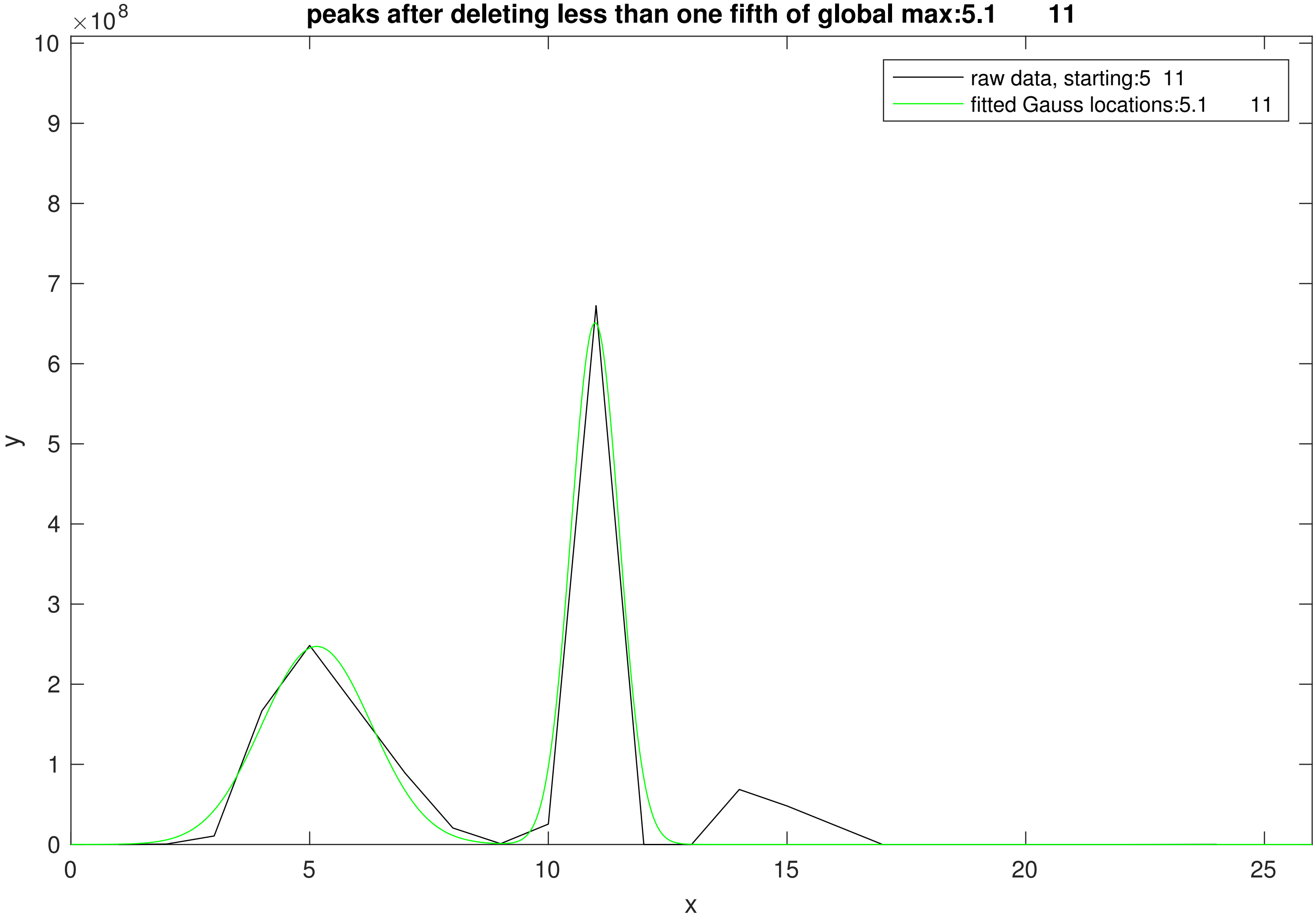

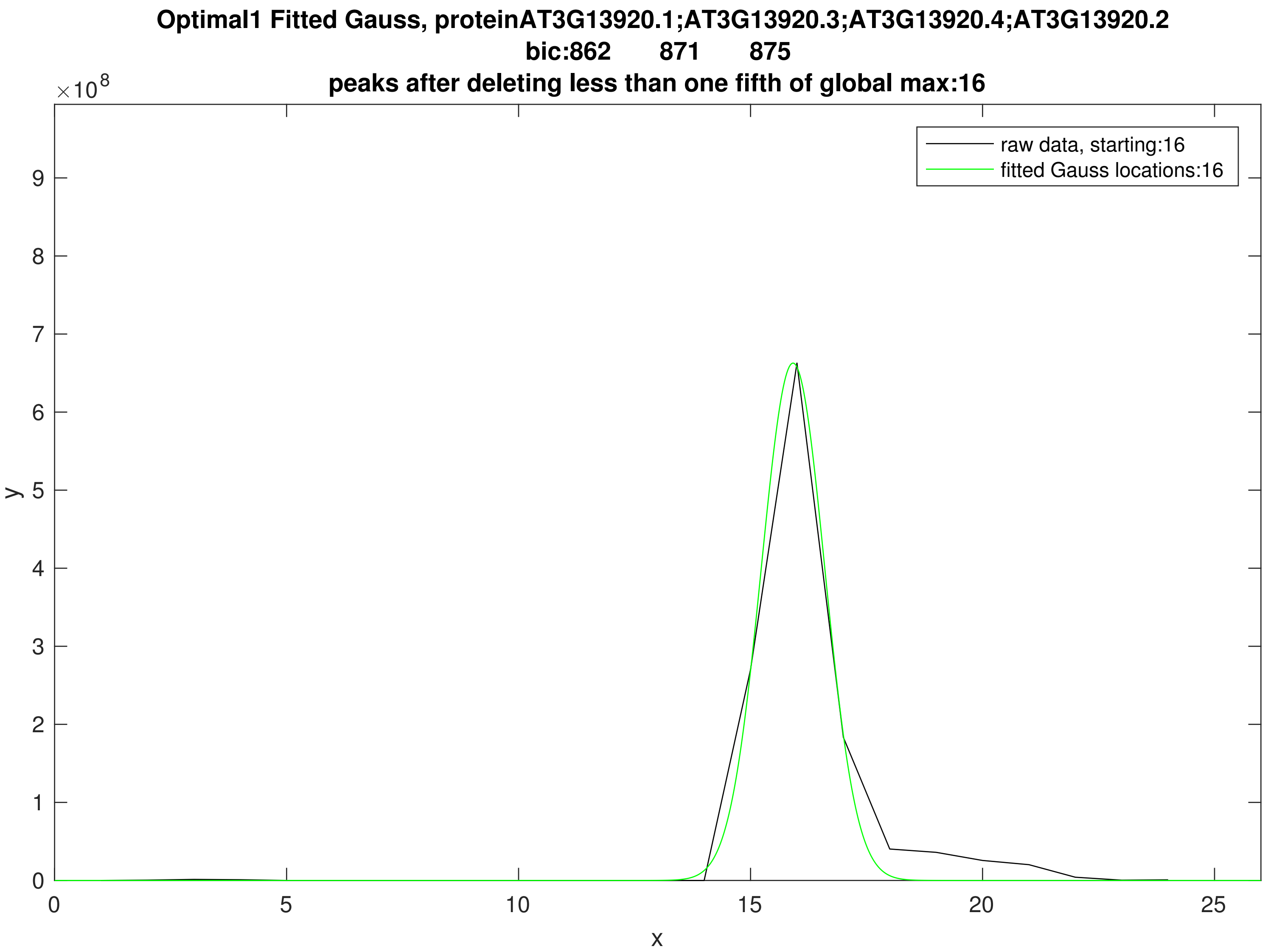

Optimal1 Fitted Gauss, proteinAT3G13920.1;AT3G13920.3;AT3G13920.4;AT3G13920.2
bic:862 871 875
peaks after deleting less than one fifth of global max:16
×10^8
raw data, starting:16
fitted Gauss locations:16
y
x
0
1
2
3
4
5
6
7
8
9
0
5
10
15
20
25

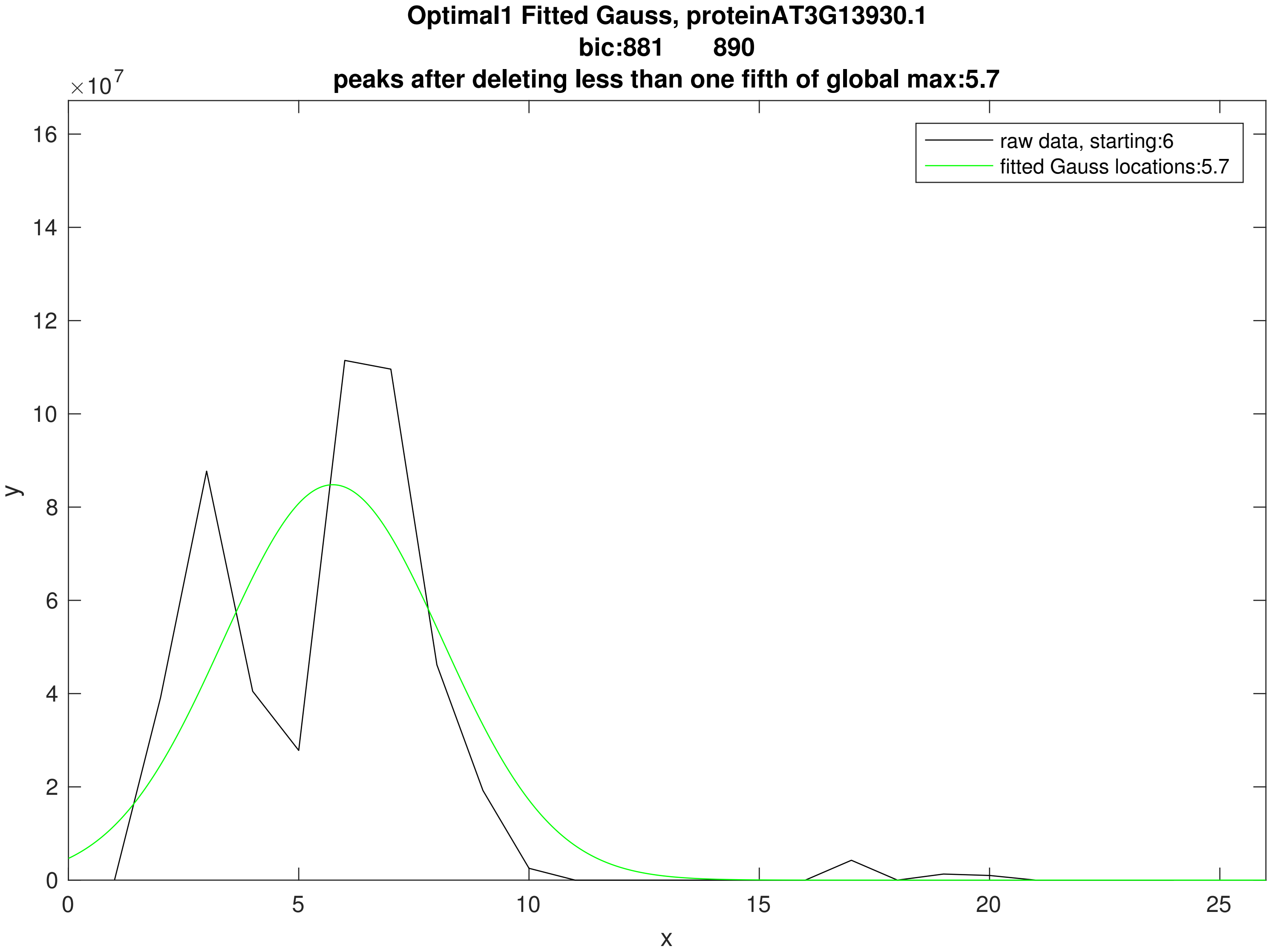
Optimal1 Fitted Gauss, proteinAT3G13930.1
bic:881 890
peaks after deleting less than one fifth of global max:5.7
×10^7
raw data, starting:6
fitted Gauss locations:5.7
y
x
0
2
4
6
8
10
12
14
16
0
5
10
15
20
25

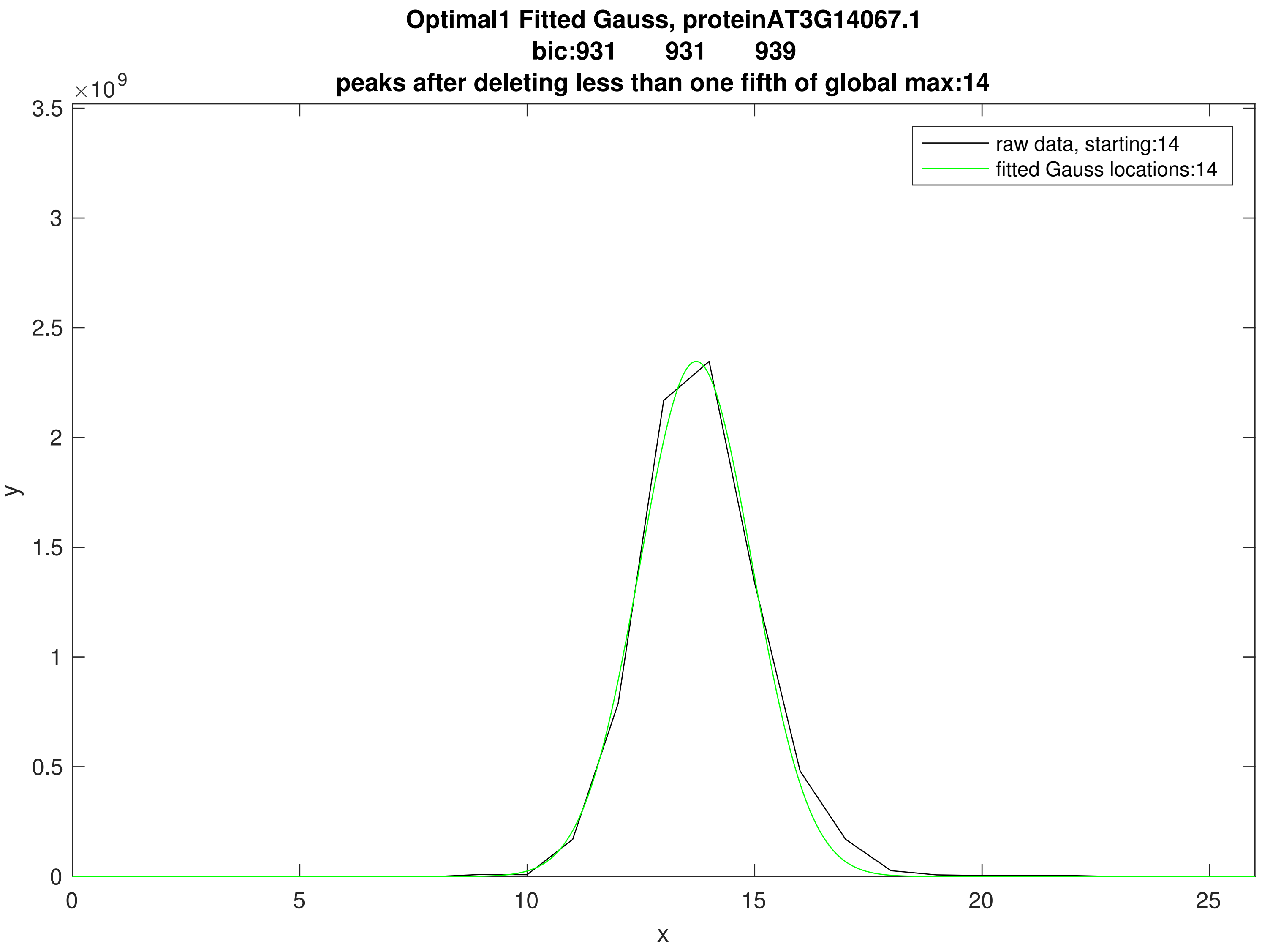

Optimal1 Fitted Gauss, proteinAT3G14067.1
bic:931 931 939
peaks after deleting less than one fifth of global max:14
×10^9
3.5
3
2.5
2
1.5
1
0.5
0
y
0
5
10
15
20
25
x
raw data, starting:14
fitted Gauss locations:14

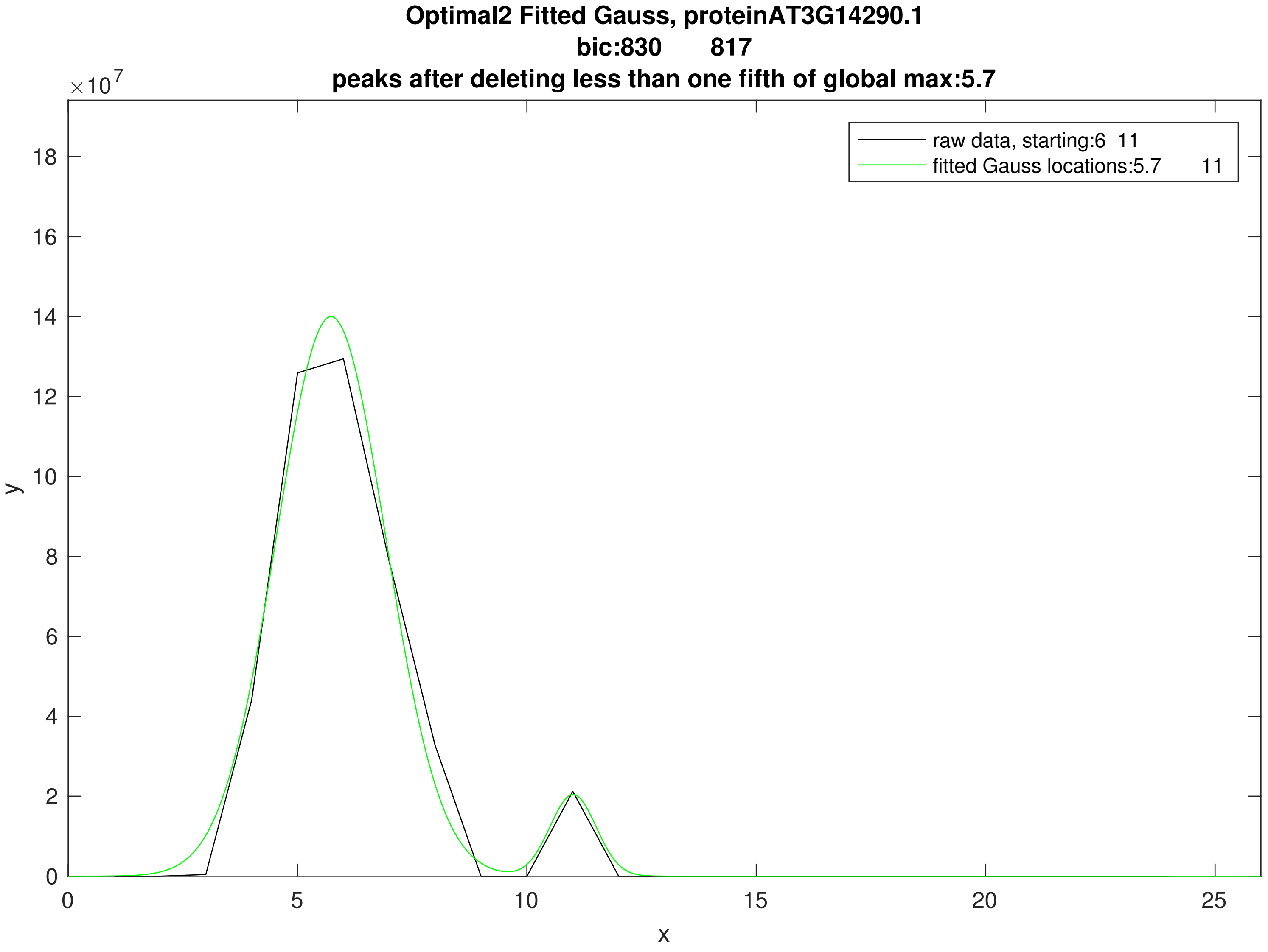

Optimal2 Fitted Gauss, proteinAT3G14290.1
bic:830 817
peaks after deleting less than one fifth of global max:5.7
×10^7
raw data, starting:6 11
fitted Gauss locations:5.7 11
y
x
0
2
4
6
8
10
12
14
16
18
0
5
10
15
20
25

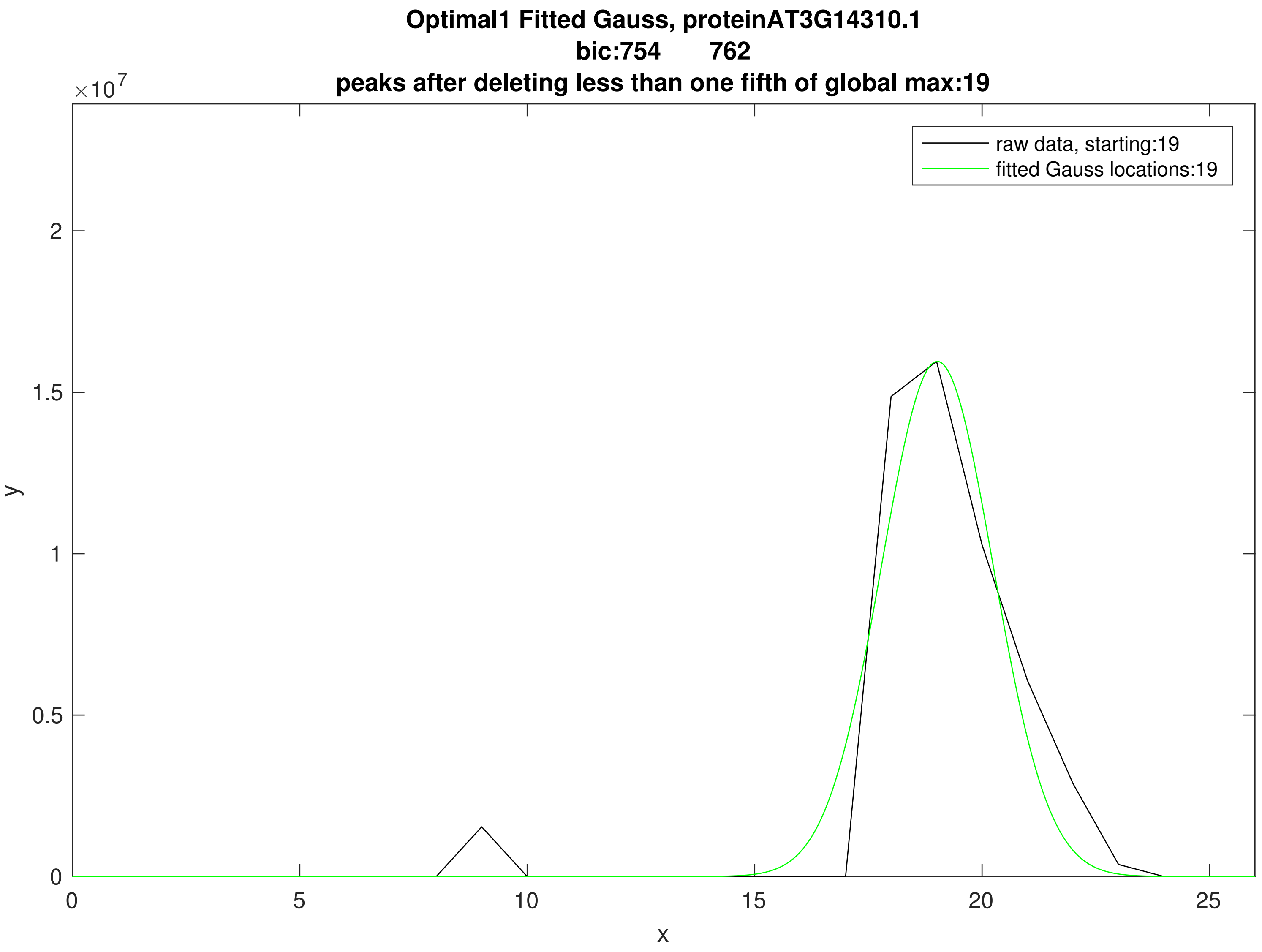
Optimal1 Fitted Gauss, proteinAT3G14310.1
bic:754 762
peaks after deleting less than one fifth of global max:19
×10^7
raw data, starting:19
fitted Gauss locations:19
2
1.5
1
0.5
0
y
0
5
10
15
20
25
x

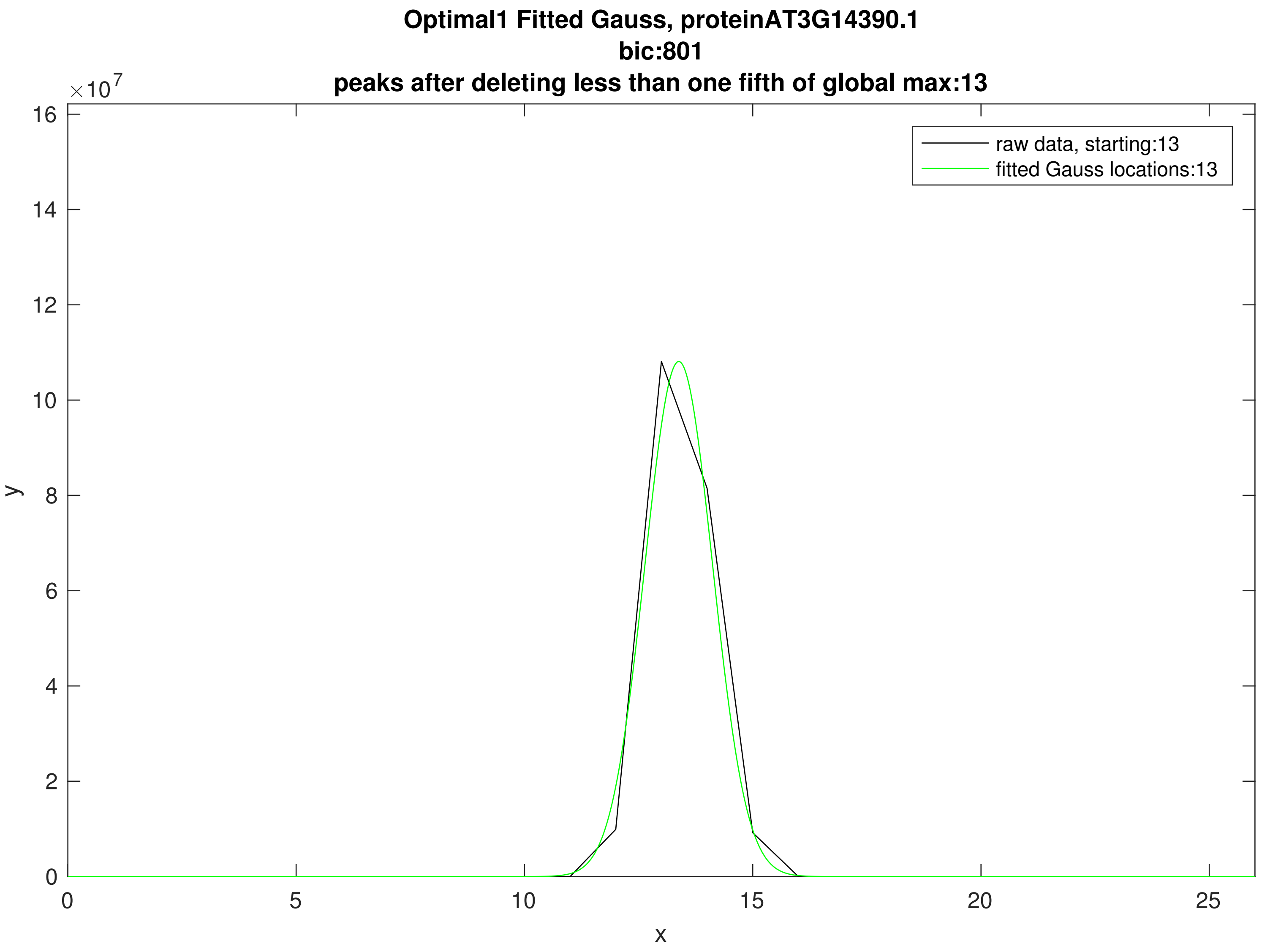

Optimal1 Fitted Gauss, proteinAT3G14390.1
bic:801
peaks after deleting less than one fifth of global max:13
×10^7
16
14
12
10
8
6
4
2
0
y
0
5
10
15
20
25
x
raw data, starting:13
fitted Gauss locations:13

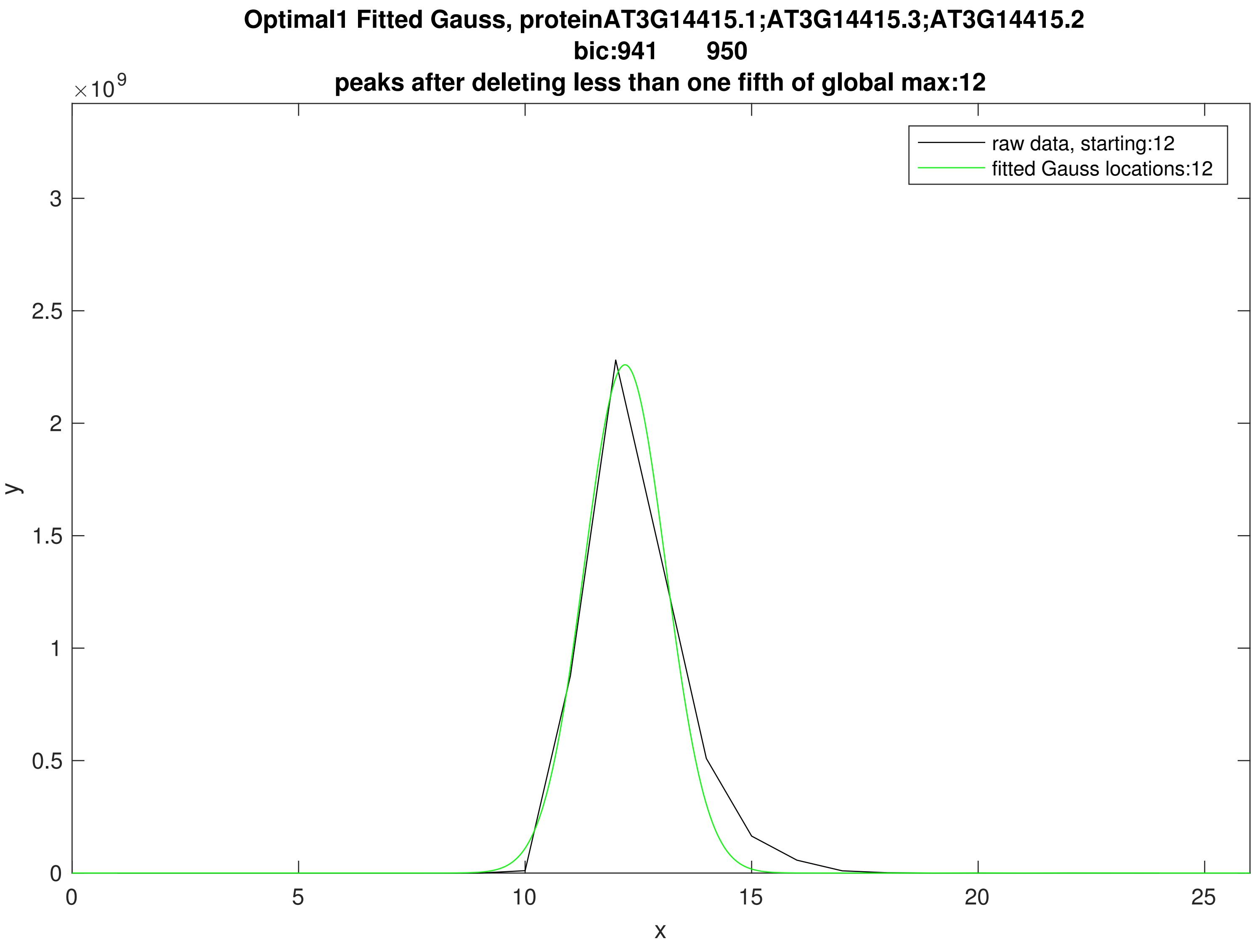

Optimal1 Fitted Gauss, proteinAT3G14415.1;AT3G14415.3;AT3G14415.2
bic:941 950
peaks after deleting less than one fifth of global max:12
×10^9
raw data, starting:12
fitted Gauss locations:12
y
x
0
0.5
1
1.5
2
2.5
3
0
5
10
15
20
25

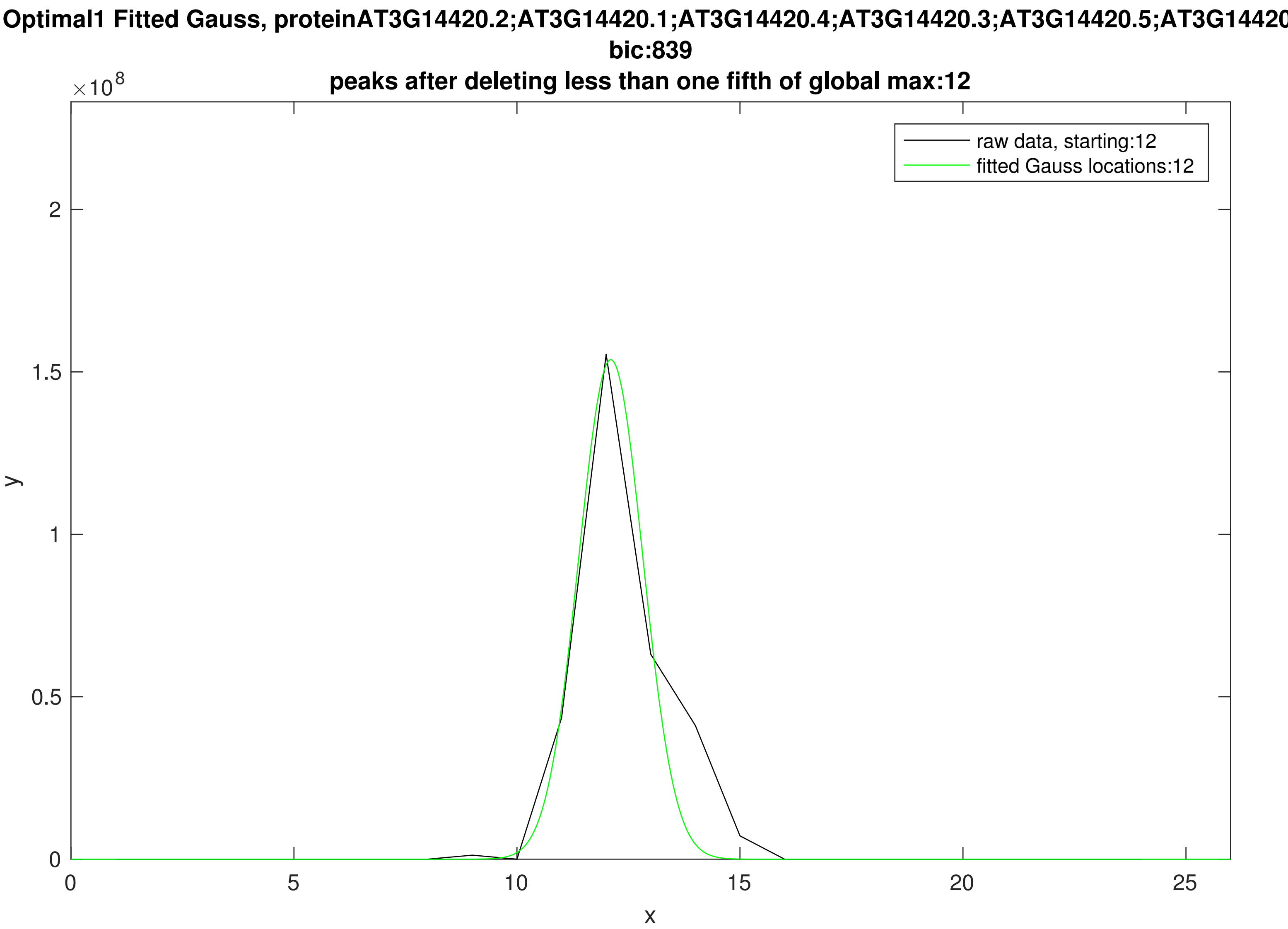
Optimal1 Fitted Gauss, proteinAT3G14420.2;AT3G14420.1;AT3G14420.4;AT3G14420.3;AT3G14420.5;AT3G14420
bic:839
peaks after deleting less than one fifth of global max:12
raw data, starting:12
fitted Gauss locations:12
y
x

**Optimal1 Fitted Gauss, proteinAT3G14600.1**
**bic:560**
**peaks after deleting less than one fifth of global max:2.9**

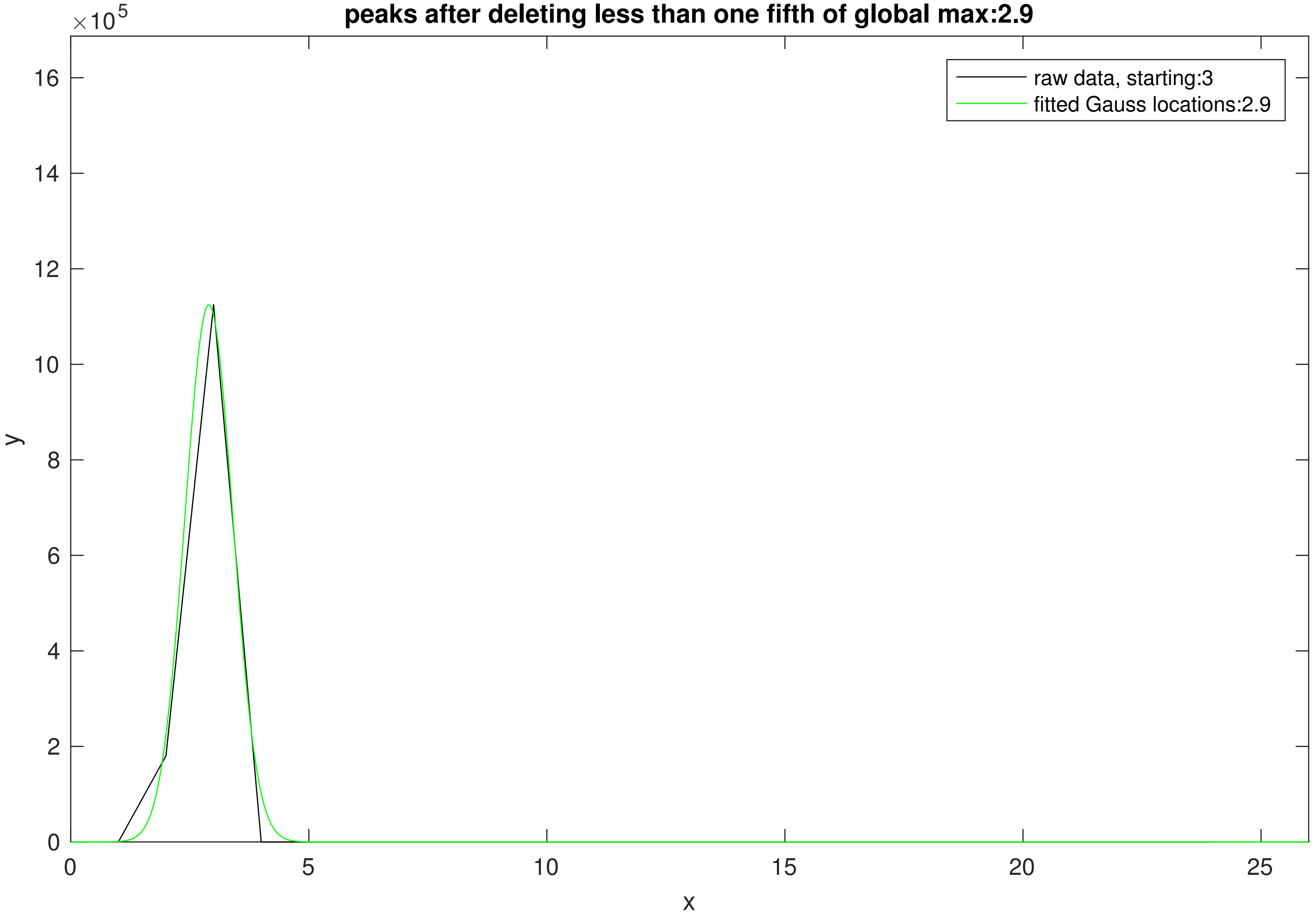

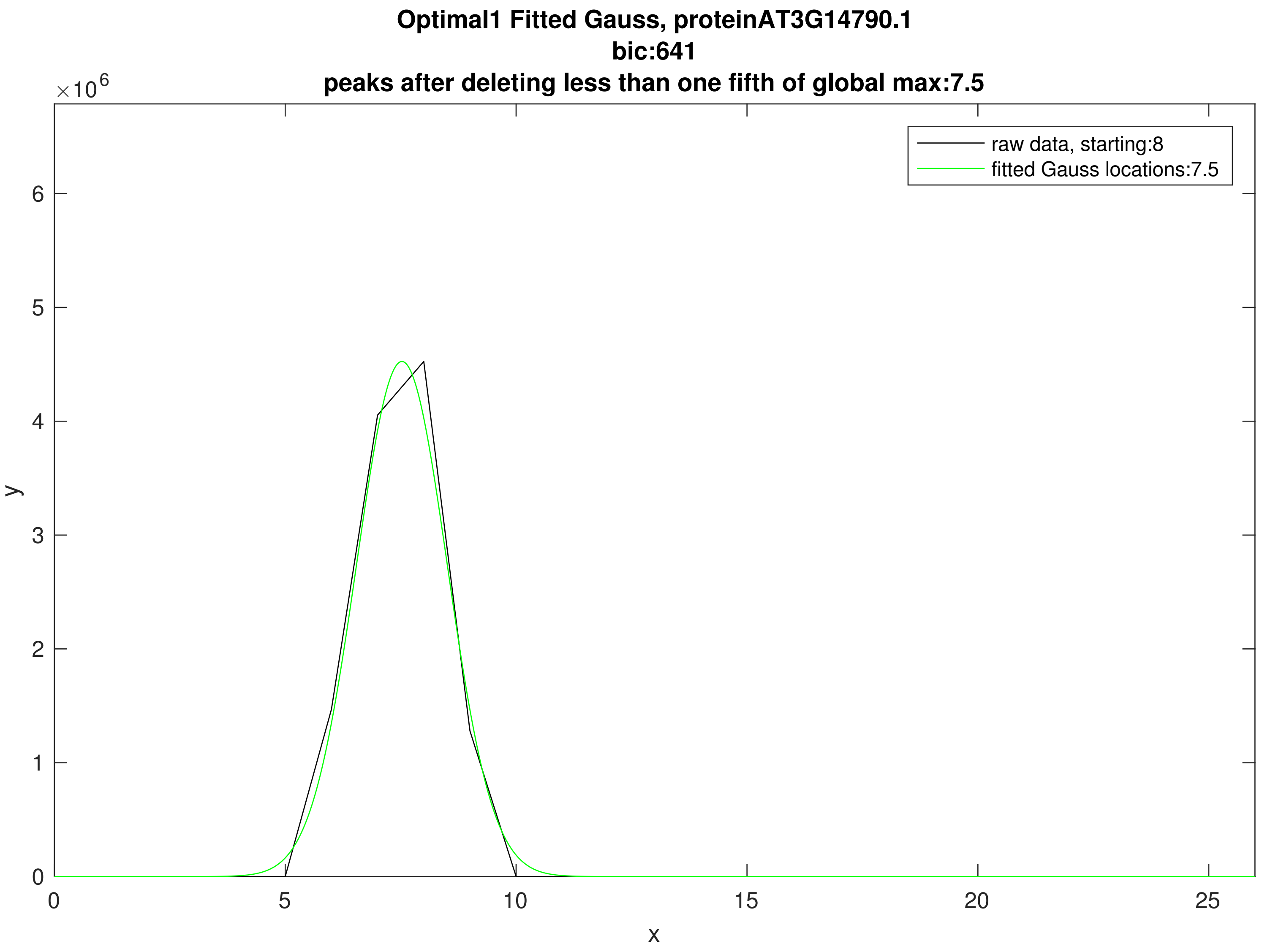

Optimal1 Fitted Gauss, proteinAT3G14790.1
bic:641
peaks after deleting less than one fifth of global max:7.5
×10^6
raw data, starting:8
fitted Gauss locations:7.5
6
5
4
3
2
1
0
y
0
5
10
15
20
25
x

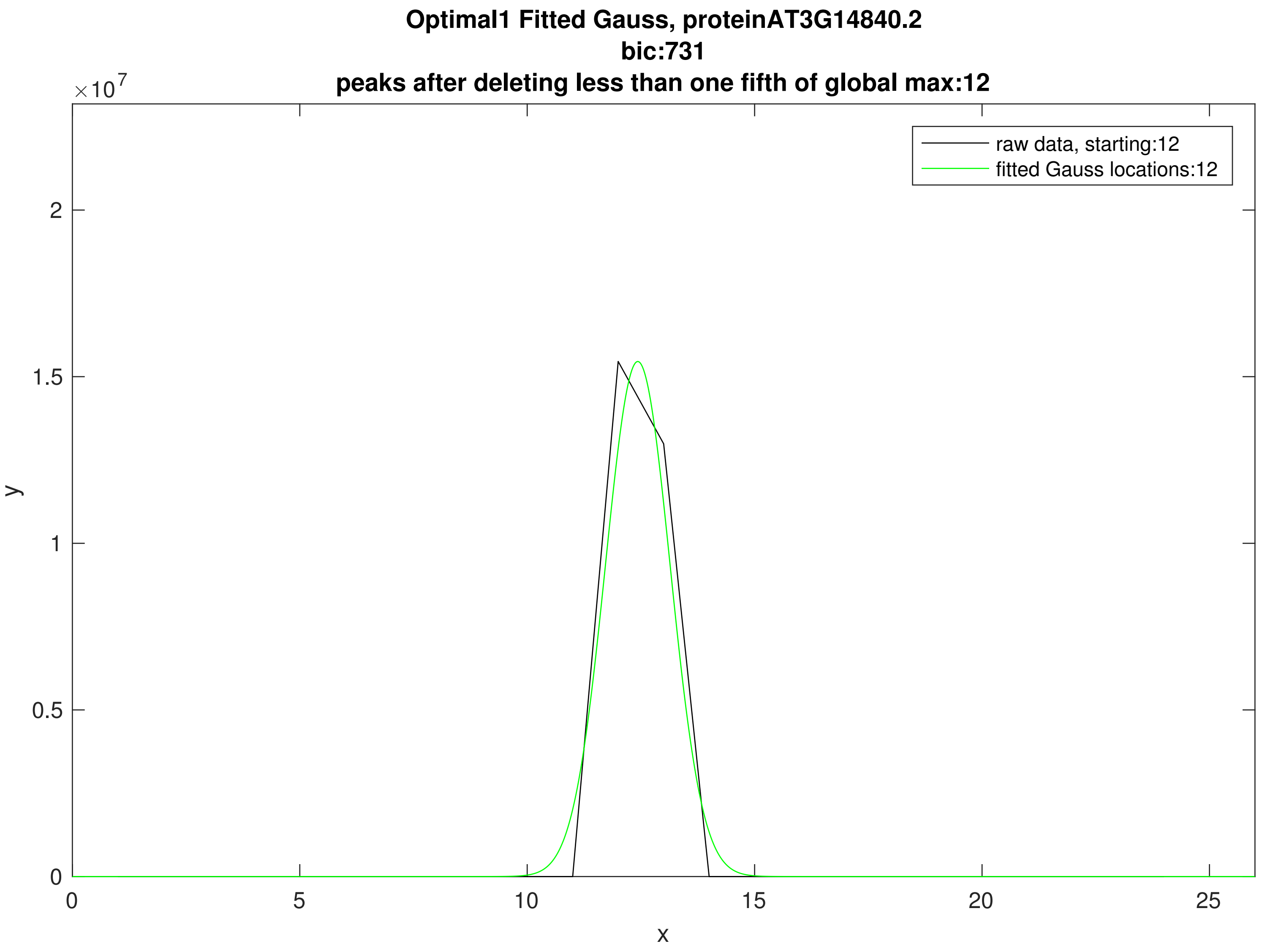
Optimal1 Fitted Gauss, proteinAT3G14840.2
bic:731
peaks after deleting less than one fifth of global max:12
×10^7
raw data, starting:12
fitted Gauss locations:12
2
1.5
1
0.5
0
y
0
5
10
15
20
25
x

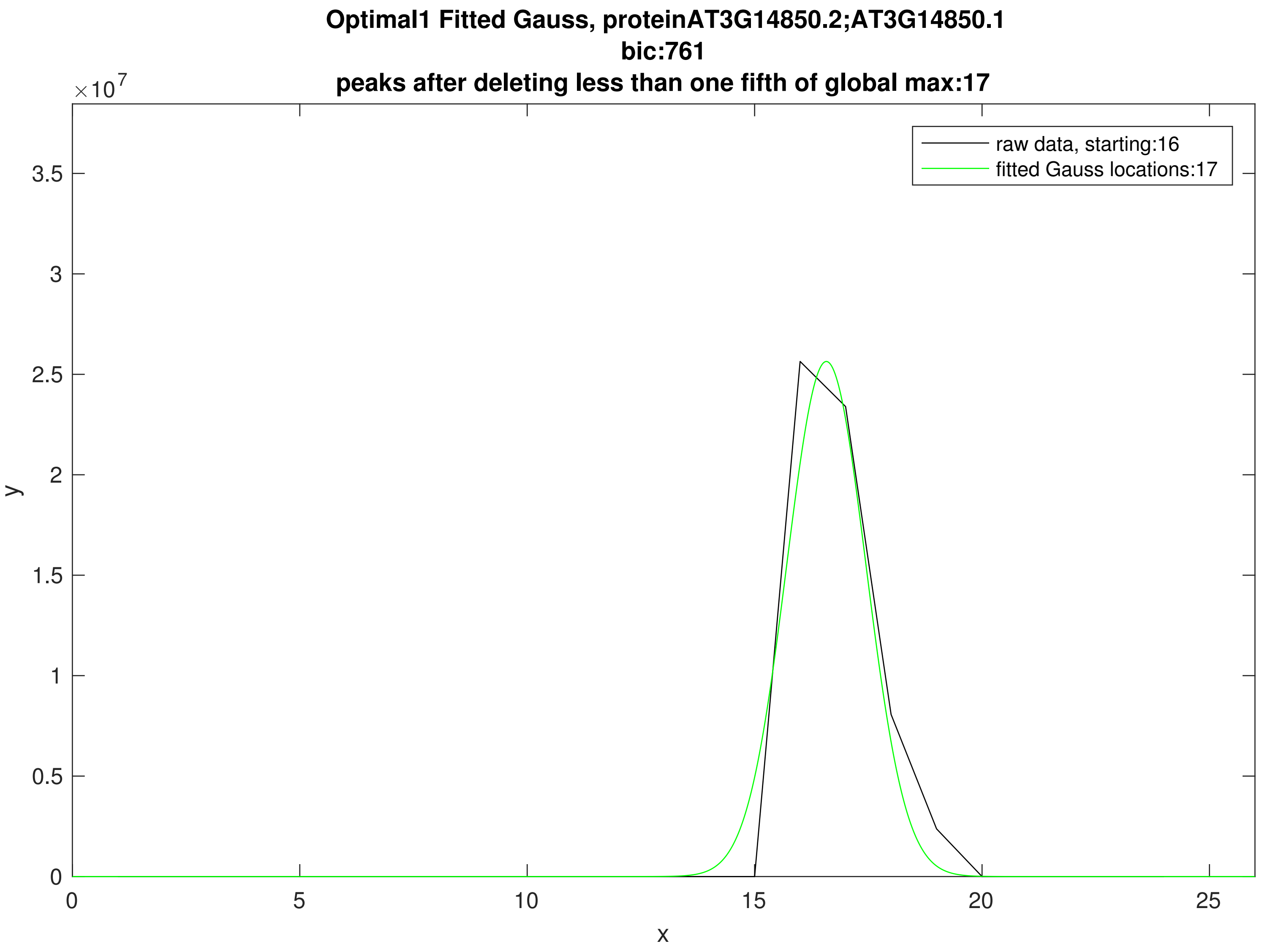
Optimal1 Fitted Gauss, proteinAT3G14850.2;AT3G14850.1
bic:761
peaks after deleting less than one fifth of global max:17
×10^7
raw data, starting:16
fitted Gauss locations:17
3.5
3
2.5
2
1.5
1
0.5
0
y
0
5
10
15
20
25
x

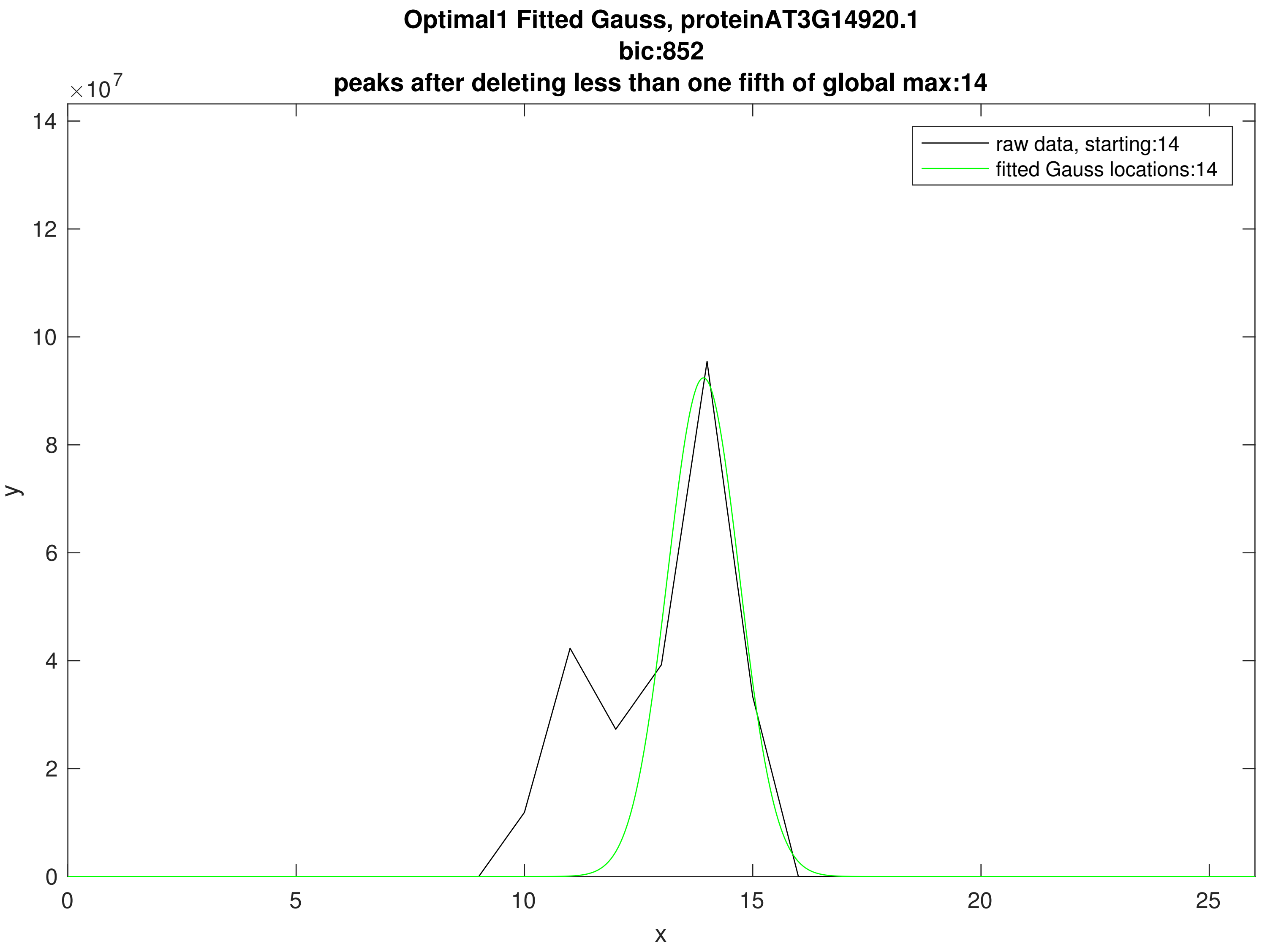

Optimal1 Fitted Gauss, proteinAT3G14920.1
bic:852
peaks after deleting less than one fifth of global max:14
×10^7
14
12
10
8
6
4
2
0
y
0
5
10
15
20
25
x
raw data, starting:14
fitted Gauss locations:14

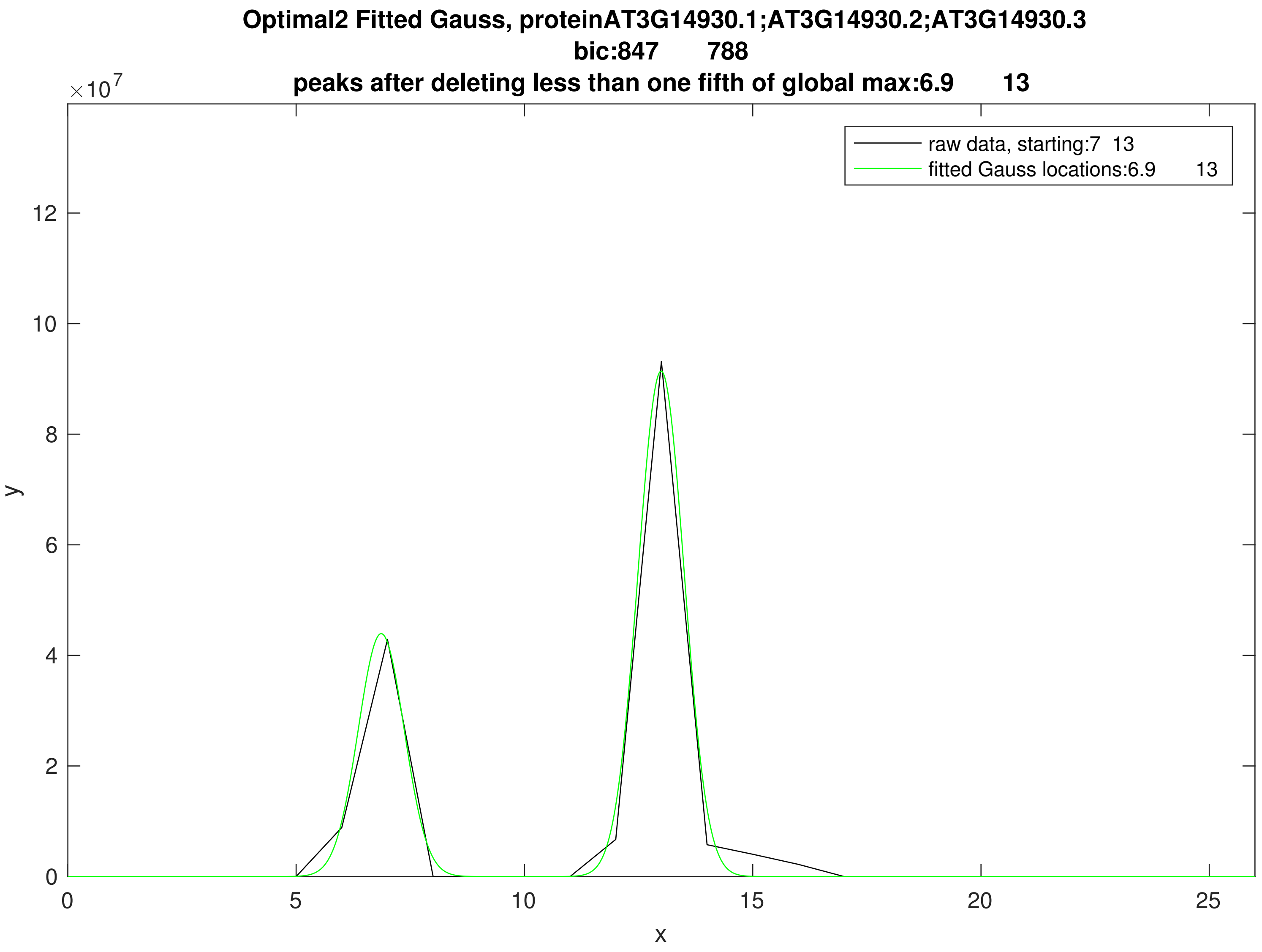

Optimal2 Fitted Gauss, proteinAT3G14930.1;AT3G14930.2;AT3G14930.3
bic:847 788
peaks after deleting less than one fifth of global max:6.9 13
×10^7
raw data, starting:7 13
fitted Gauss locations:6.9 13
y
x

#### Optimal2 Fitted Gauss, proteinAT3G14940.1
#### bic:865 850
#### peaks after deleting less than one fifth of global max:7 11

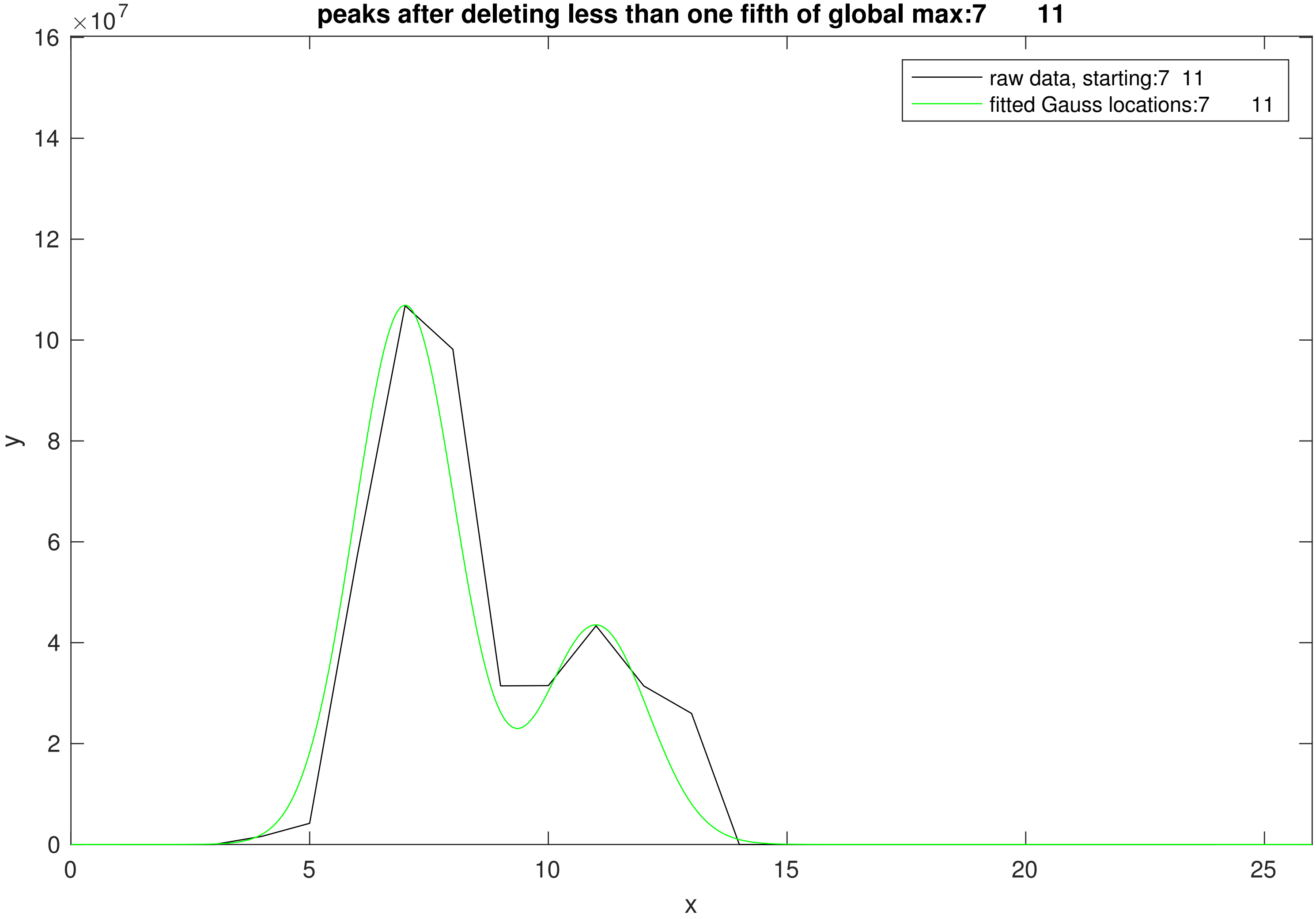

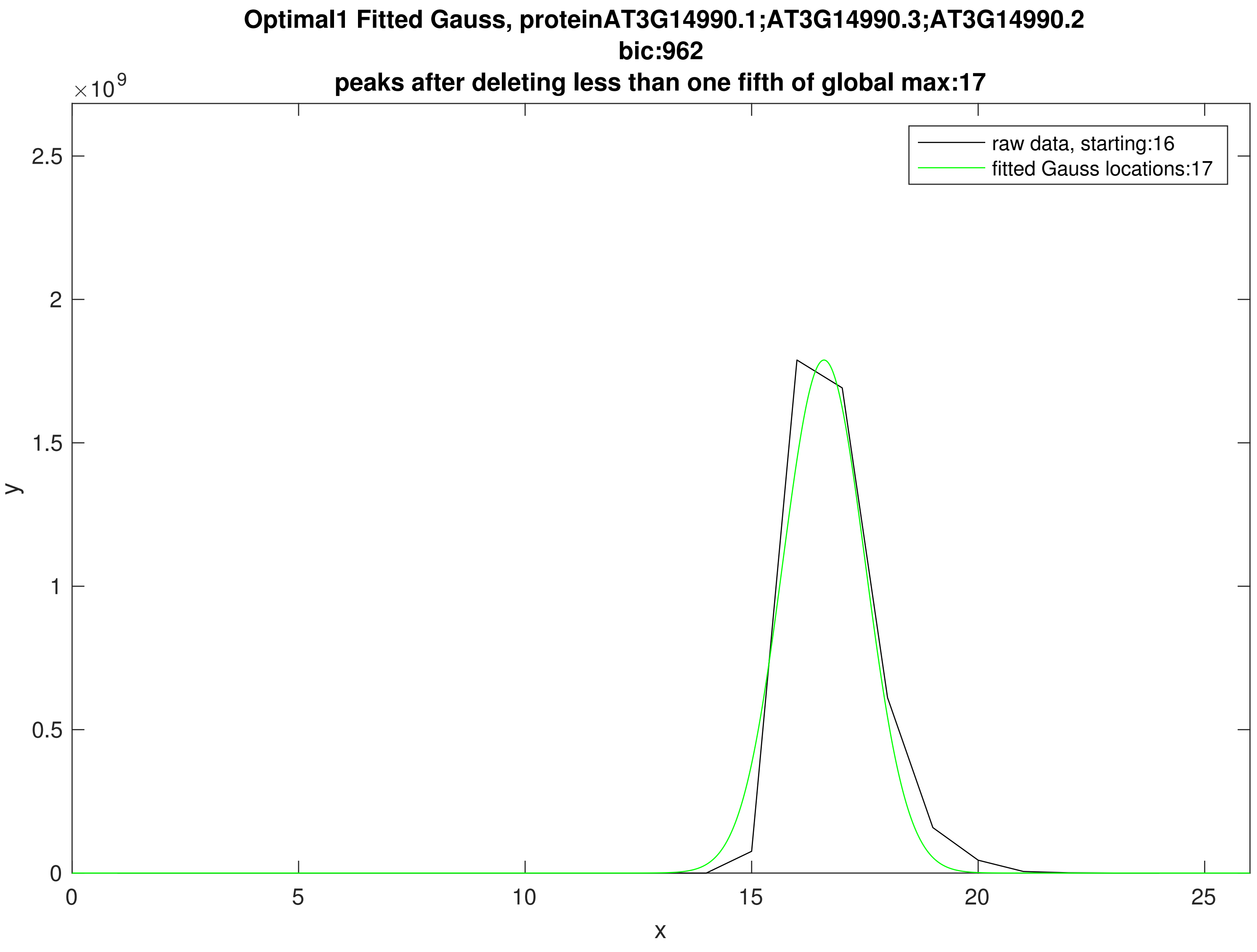

Optimal1 Fitted Gauss, proteinAT3G14990.1;AT3G14990.3;AT3G14990.2
bic:962
peaks after deleting less than one fifth of global max:17
×10^9
2.5
2
1.5
1
0.5
0
y
0
5
10
15
20
25
x
raw data, starting:16
fitted Gauss locations:17

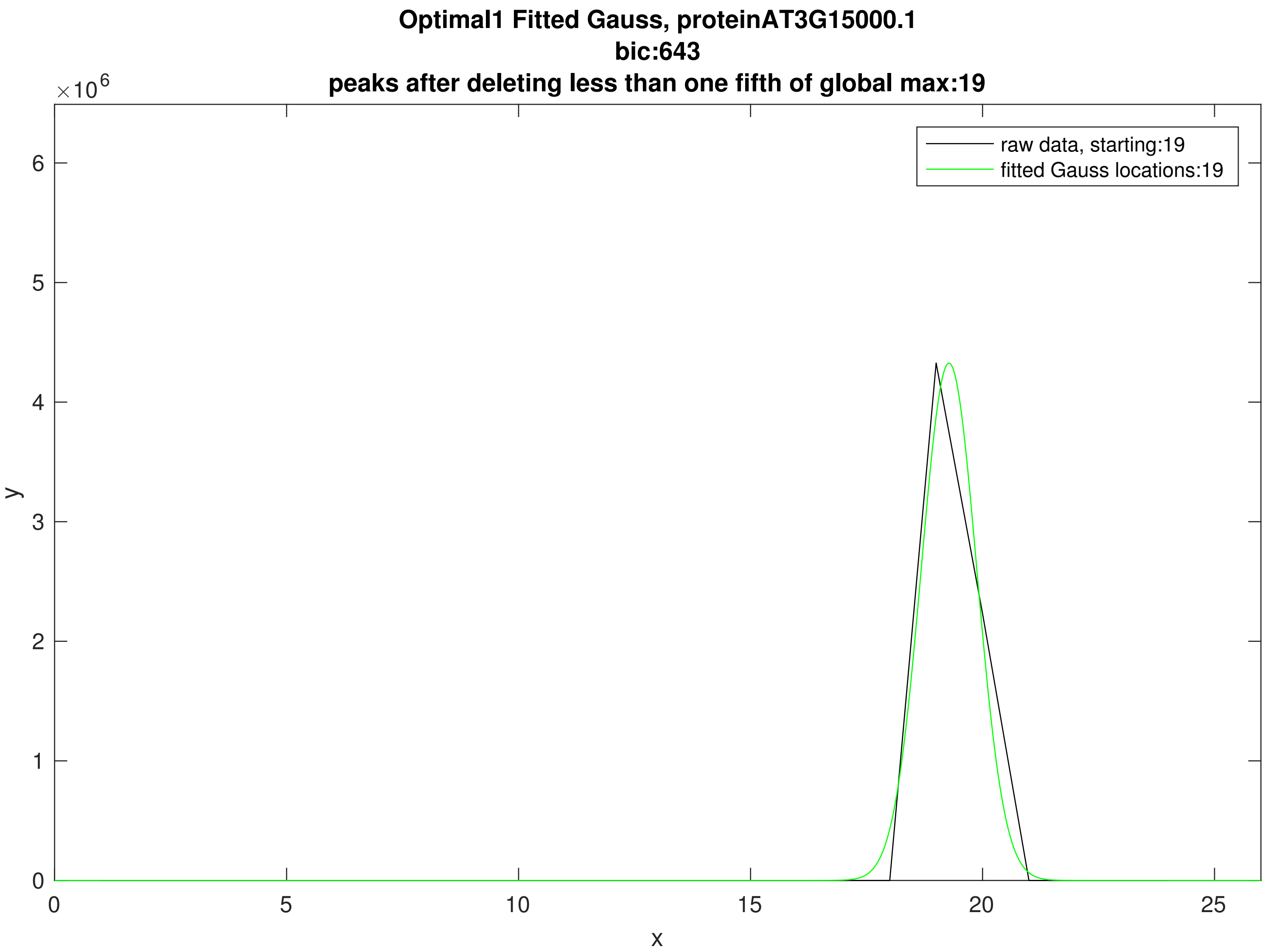

Optimal1 Fitted Gauss, proteinAT3G15000.1
bic:643
peaks after deleting less than one fifth of global max:19
×10^6
raw data, starting:19
fitted Gauss locations:19
y
x
0
1
2
3
4
5
6
0
5
10
15
20
25

Optimal1 Fitted Gauss, proteinAT3G15020.1;AT3G15020.2
bic:871
peaks after deleting less than one fifth of global max:15

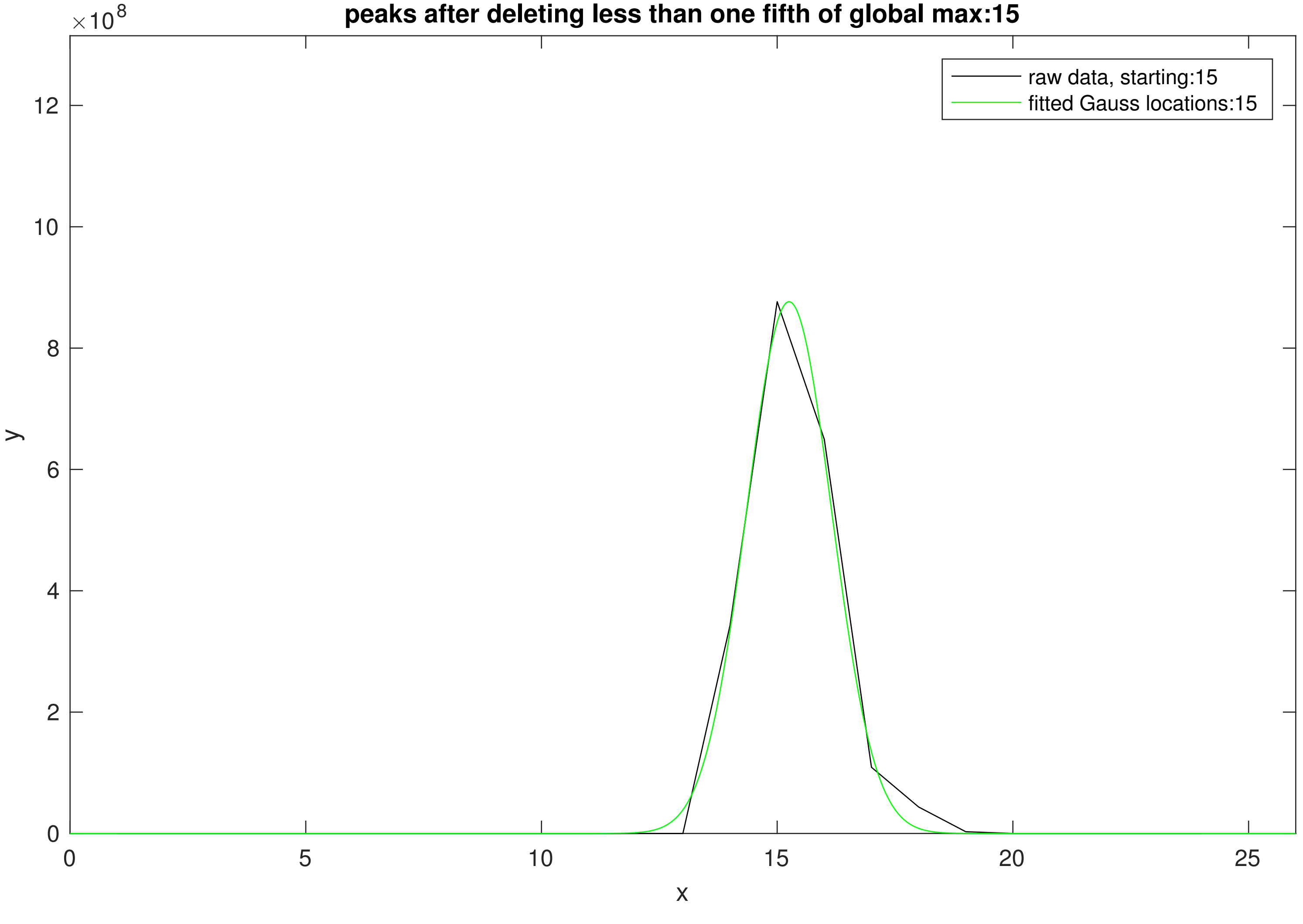

**Optimal1 Fitted Gauss, proteinAT3G15190.1**
**bic:722 778**
**peaks after deleting less than one fifth of global max:5**

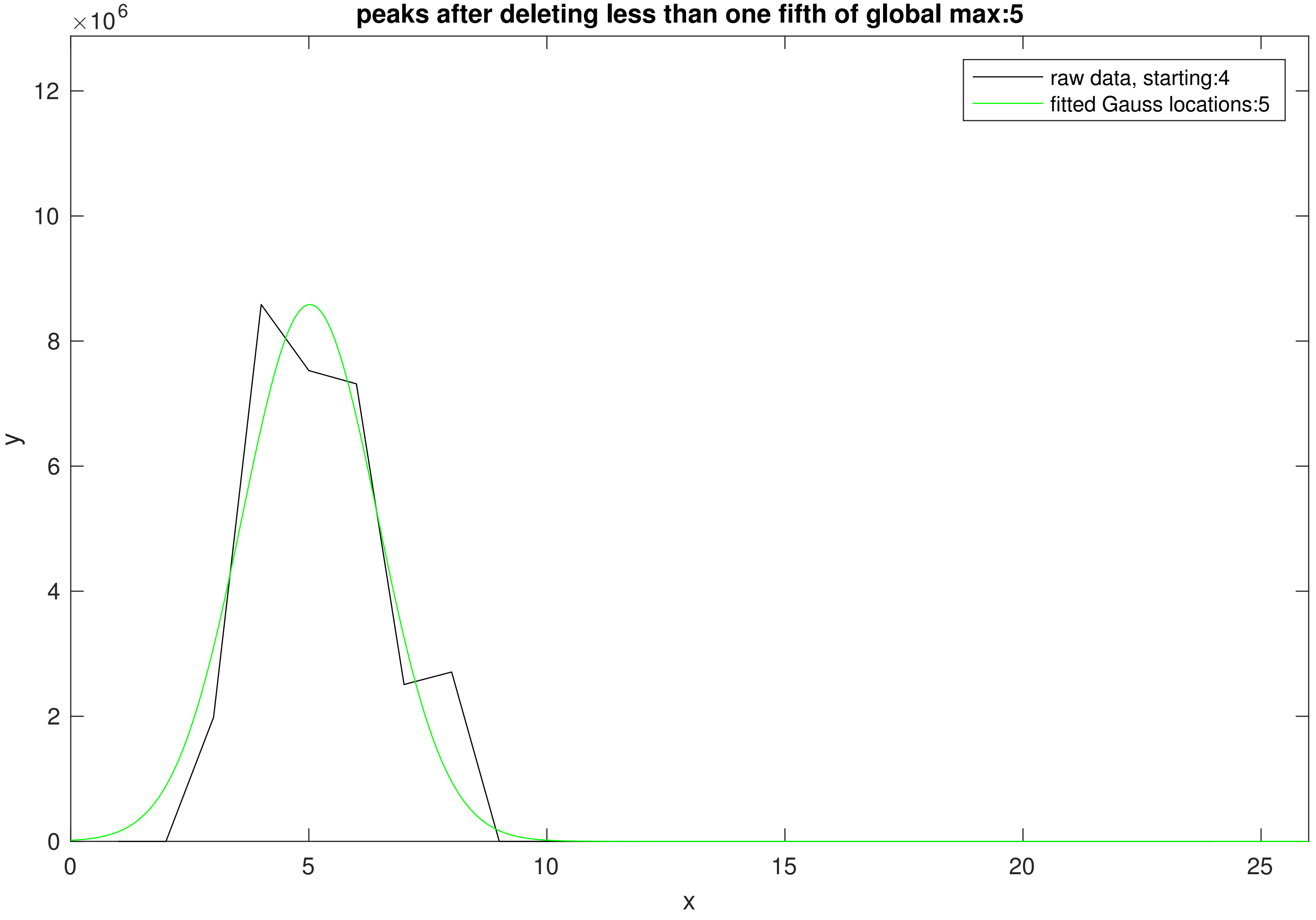

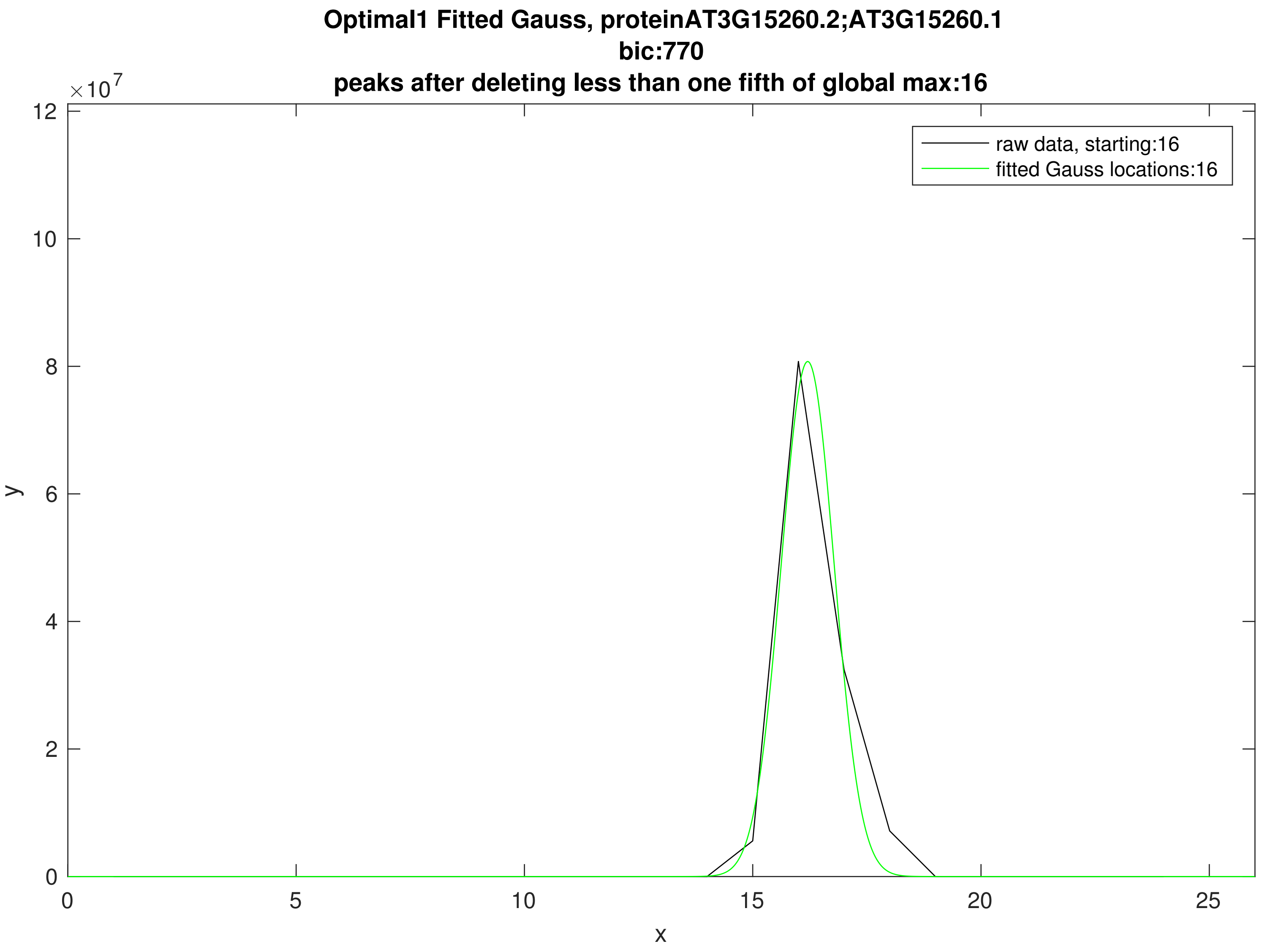
Optimal1 Fitted Gauss, proteinAT3G15260.2;AT3G15260.1
bic:770
peaks after deleting less than one fifth of global max:16
×10^7
raw data, starting:16
fitted Gauss locations:16
y
x
0
2
4
6
8
10
12
0
5
10
15
20
25

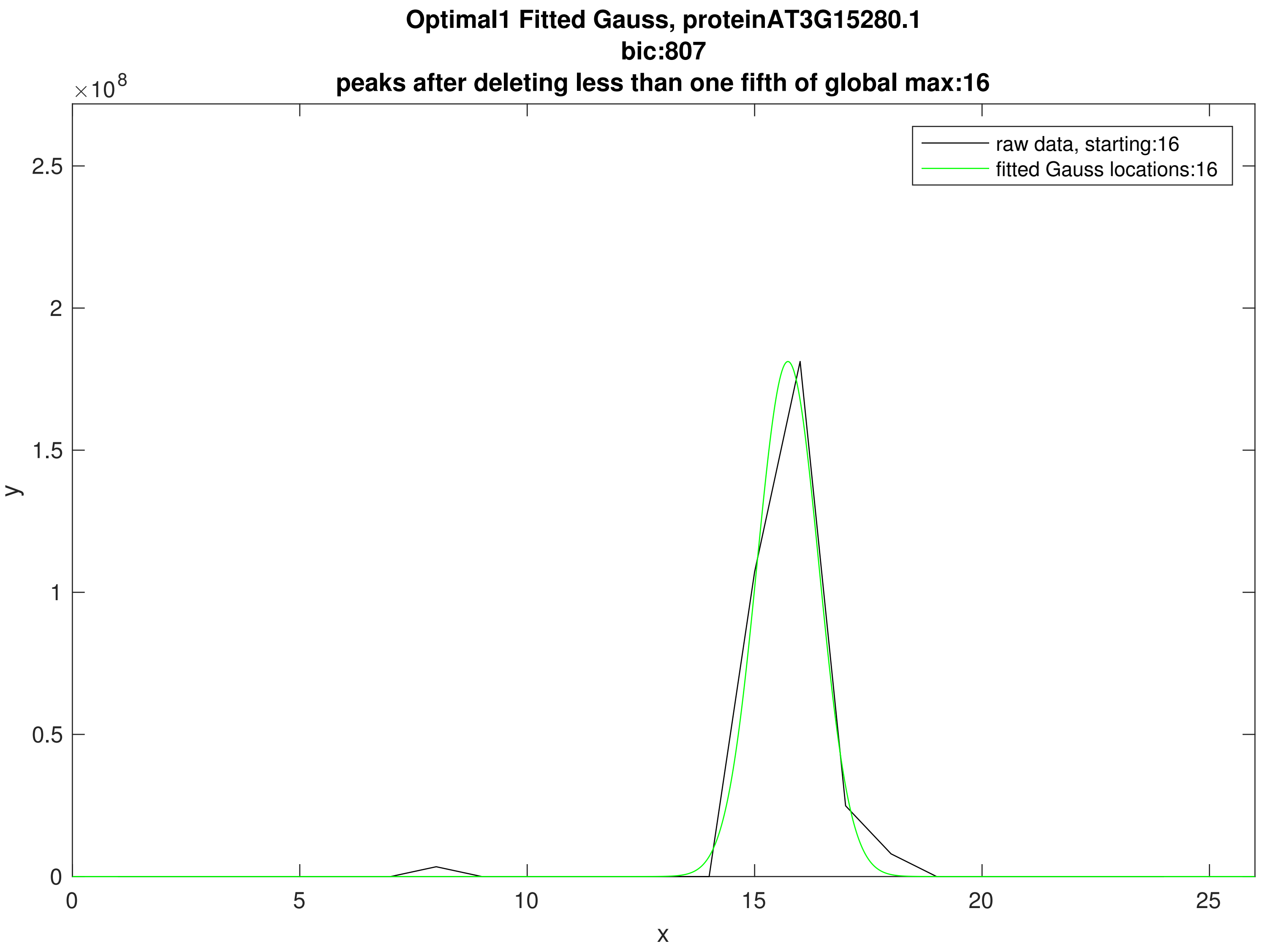

Optimal1 Fitted Gauss, proteinAT3G15280.1
bic:807
peaks after deleting less than one fifth of global max:16
×10^8
2.5
2
1.5
1
0.5
0
y
0
5
10
15
20
25
x
raw data, starting:16
fitted Gauss locations:16

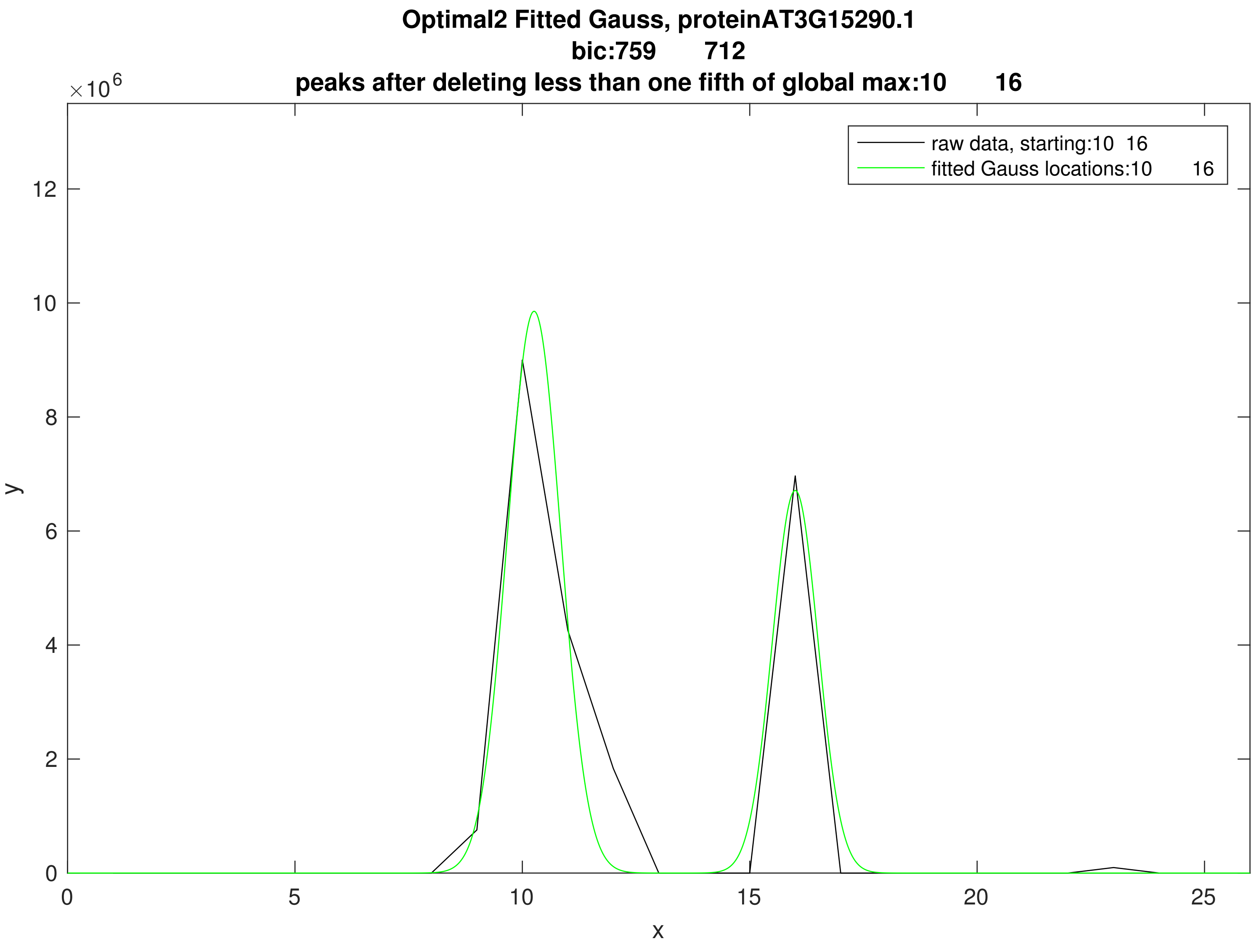
Optimal2 Fitted Gauss, proteinAT3G15290.1
bic:759 712
peaks after deleting less than one fifth of global max:10 16
×10^6
raw data, starting:10 16
fitted Gauss locations:10 16
y
x
0
2
4
6
8
10
12
0
5
10
15
20
25

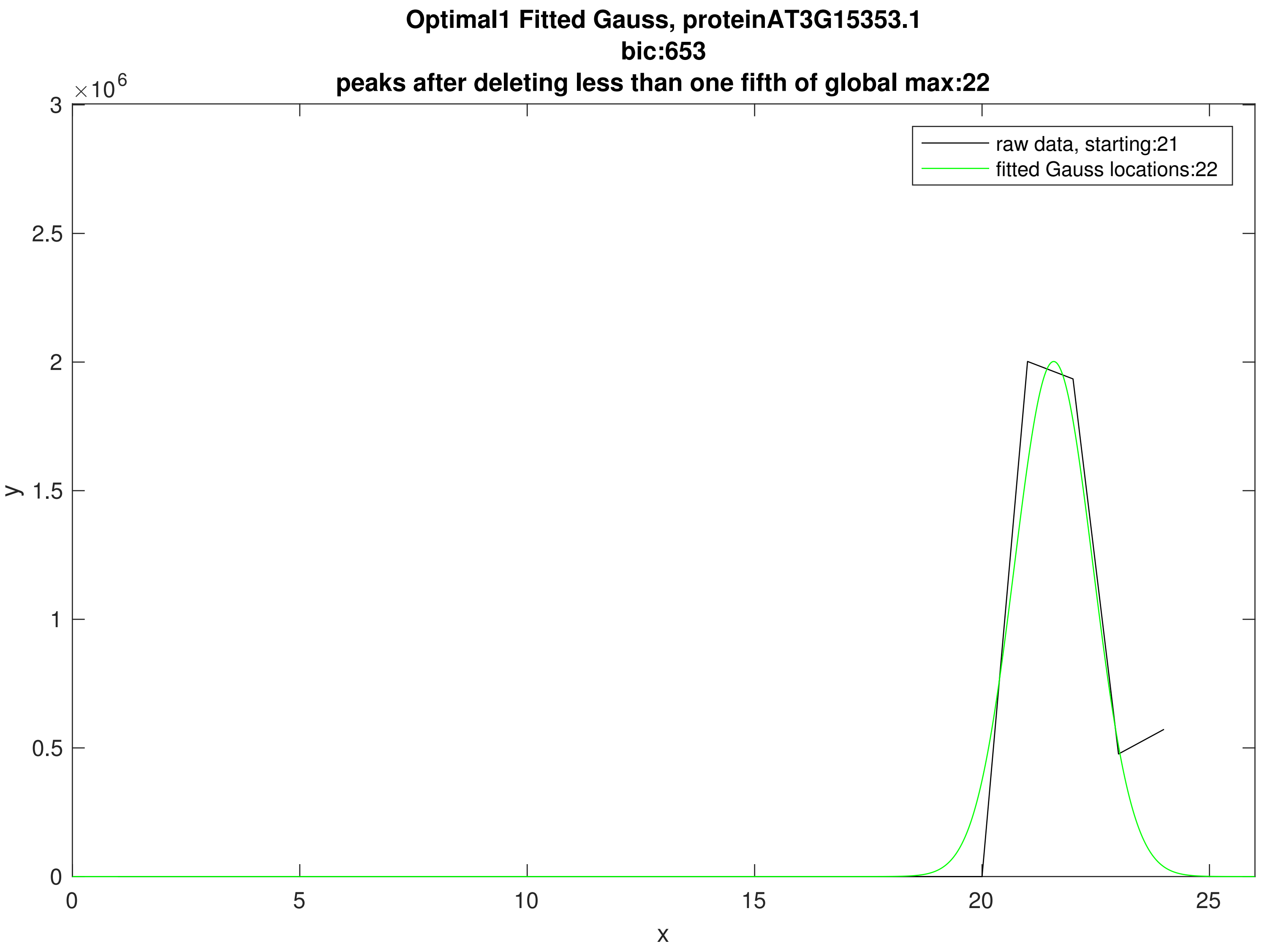

Optimal1 Fitted Gauss, proteinAT3G15353.1
bic:653
peaks after deleting less than one fifth of global max:22
×10^6
3
2.5
2
1.5
1
0.5
0
y
0
5
10
15
20
25
x
raw data, starting:21
fitted Gauss locations:22

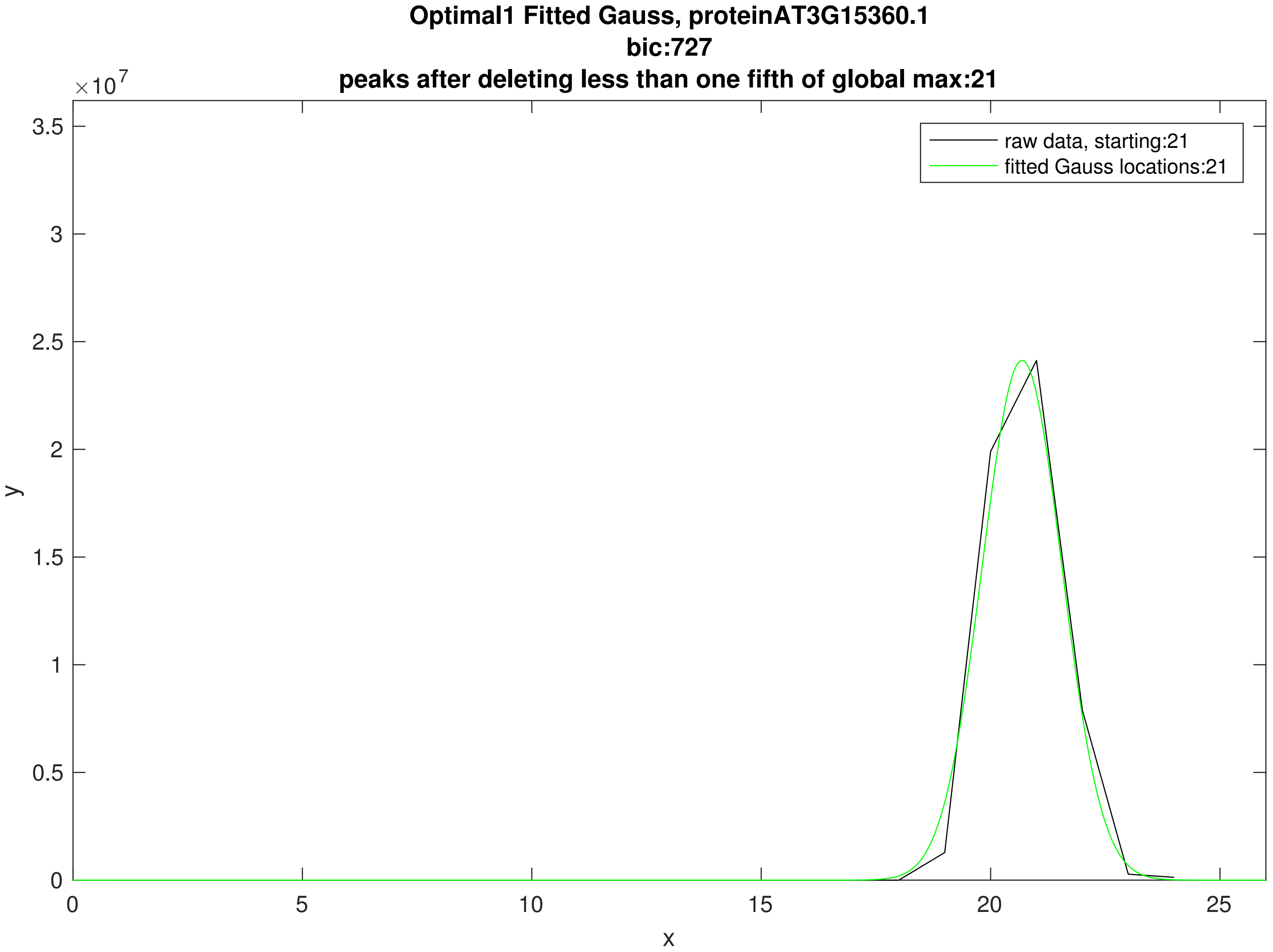

Optimal1 Fitted Gauss, proteinAT3G15360.1
bic:727
peaks after deleting less than one fifth of global max:21
×10^7
3.5
3
2.5
2
1.5
1
0.5
0
y
0
5
10
15
20
25
x
raw data, starting:21
fitted Gauss locations:21

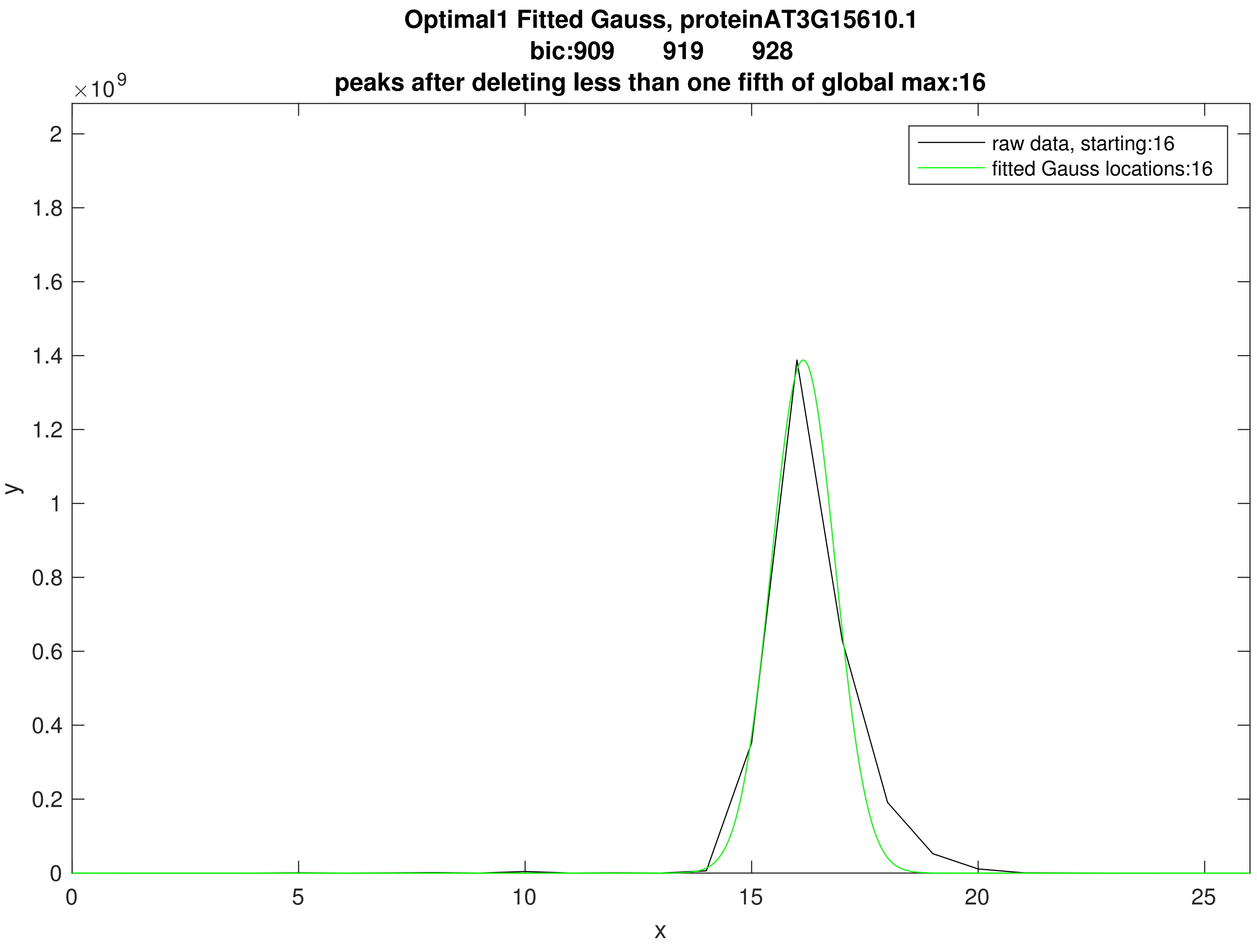

Optimal1 Fitted Gauss, proteinAT3G15610.1
bic:909 919 928
peaks after deleting less than one fifth of global max:16
×10^9
raw data, starting:16
fitted Gauss locations:16
y
x
0
0.2
0.4
0.6
0.8
1
1.2
1.4
1.6
1.8
2
0
5
10
15
20
25

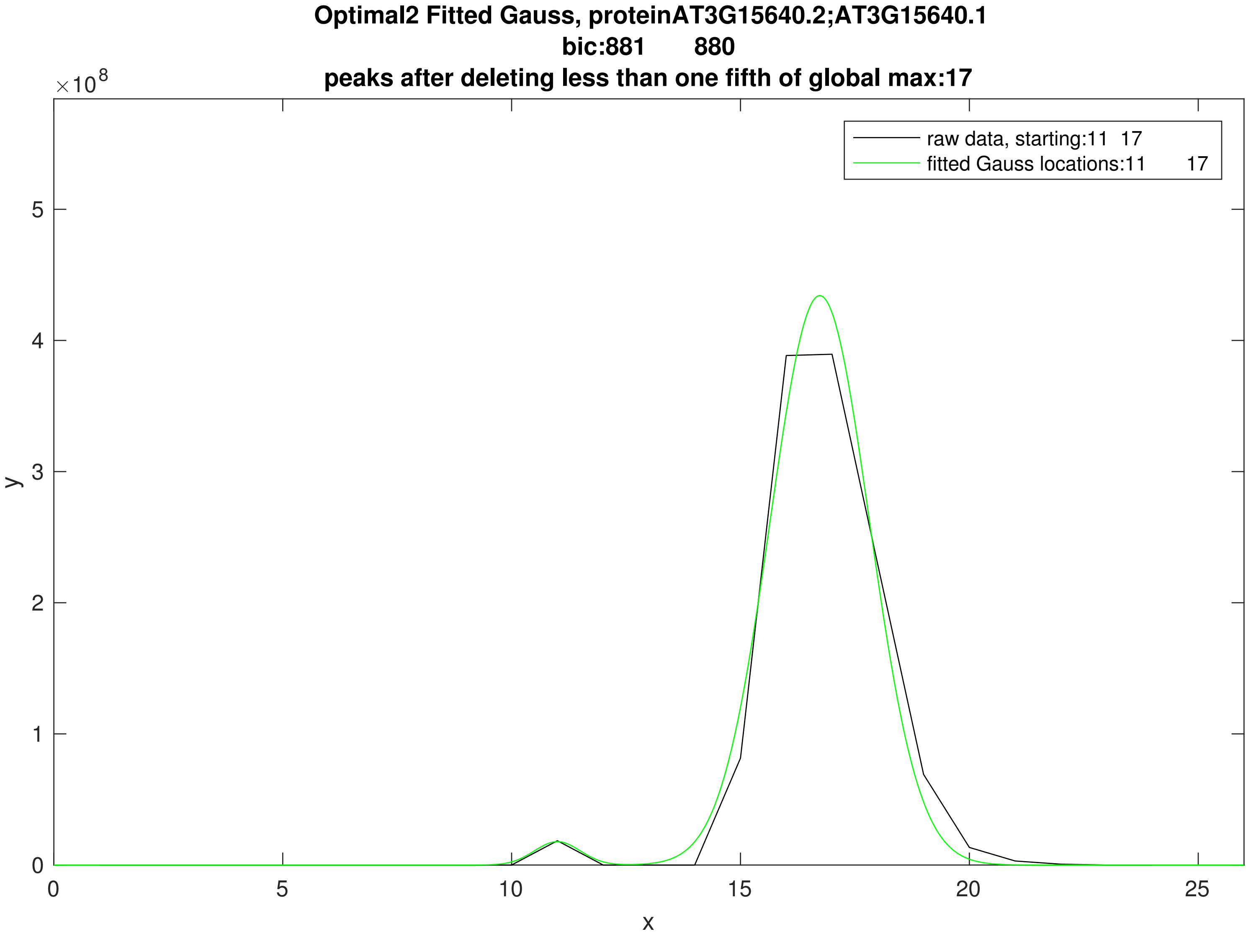

Optimal2 Fitted Gauss, proteinAT3G15640.2;AT3G15640.1
bic:881 880
peaks after deleting less than one fifth of global max:17
×10^8
raw data, starting:11 17
fitted Gauss locations:11 17
y
x
0
1
2
3
4
5
0
5
10
15
20
25

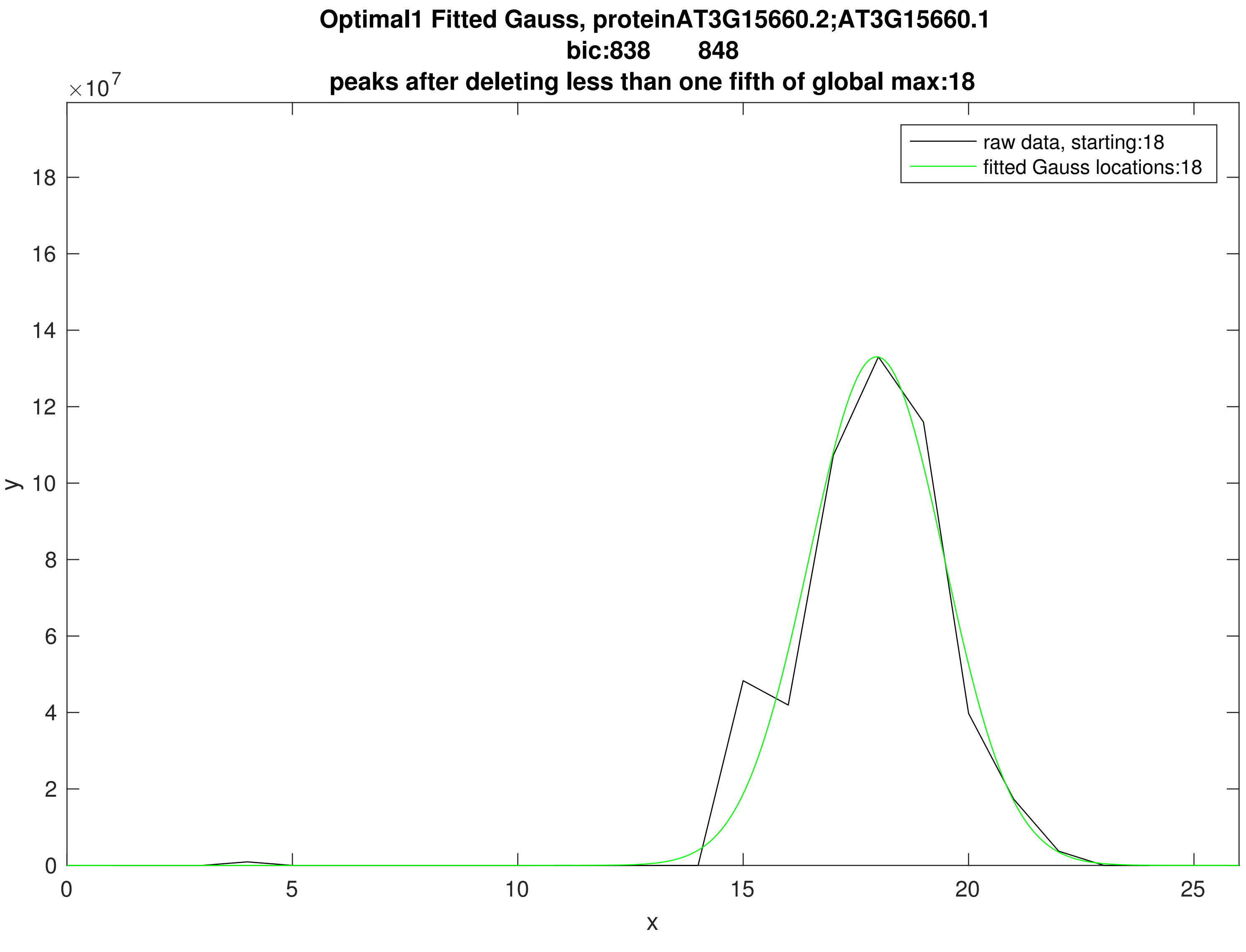
Optimal1 Fitted Gauss, proteinAT3G15660.2;AT3G15660.1
bic:838 848
peaks after deleting less than one fifth of global max:18
×10^7
raw data, starting:18
fitted Gauss locations:18
y
x
0
2
4
6
8
10
12
14
16
18
0
5
10
15
20
25

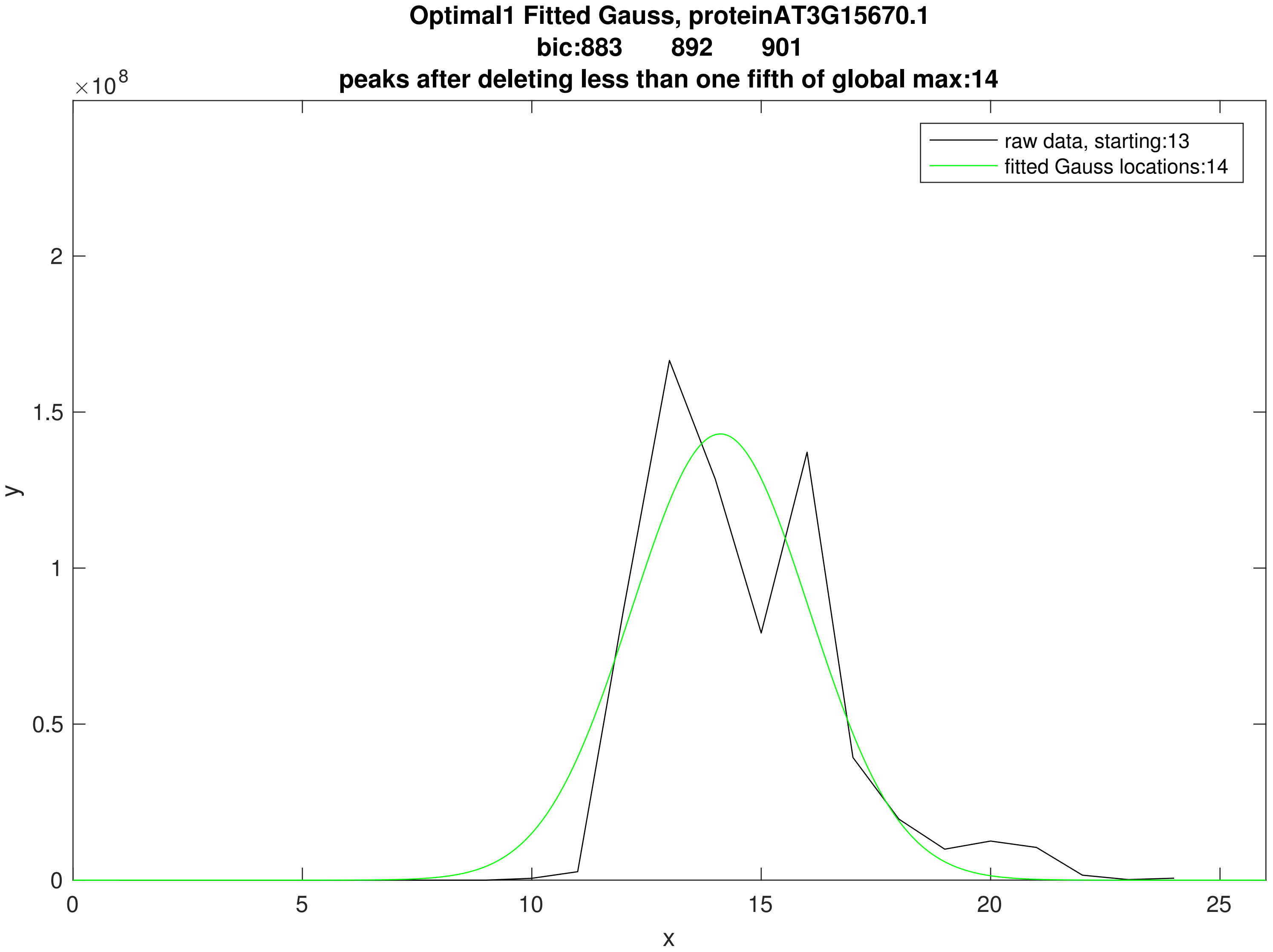
Optimal1 Fitted Gauss, proteinAT3G15670.1
bic:883 892 901
peaks after deleting less than one fifth of global max:14
×10^8
raw data, starting:13
fitted Gauss locations:14
y
x
0
0.5
1
1.5
2
0
5
10
15
20
25

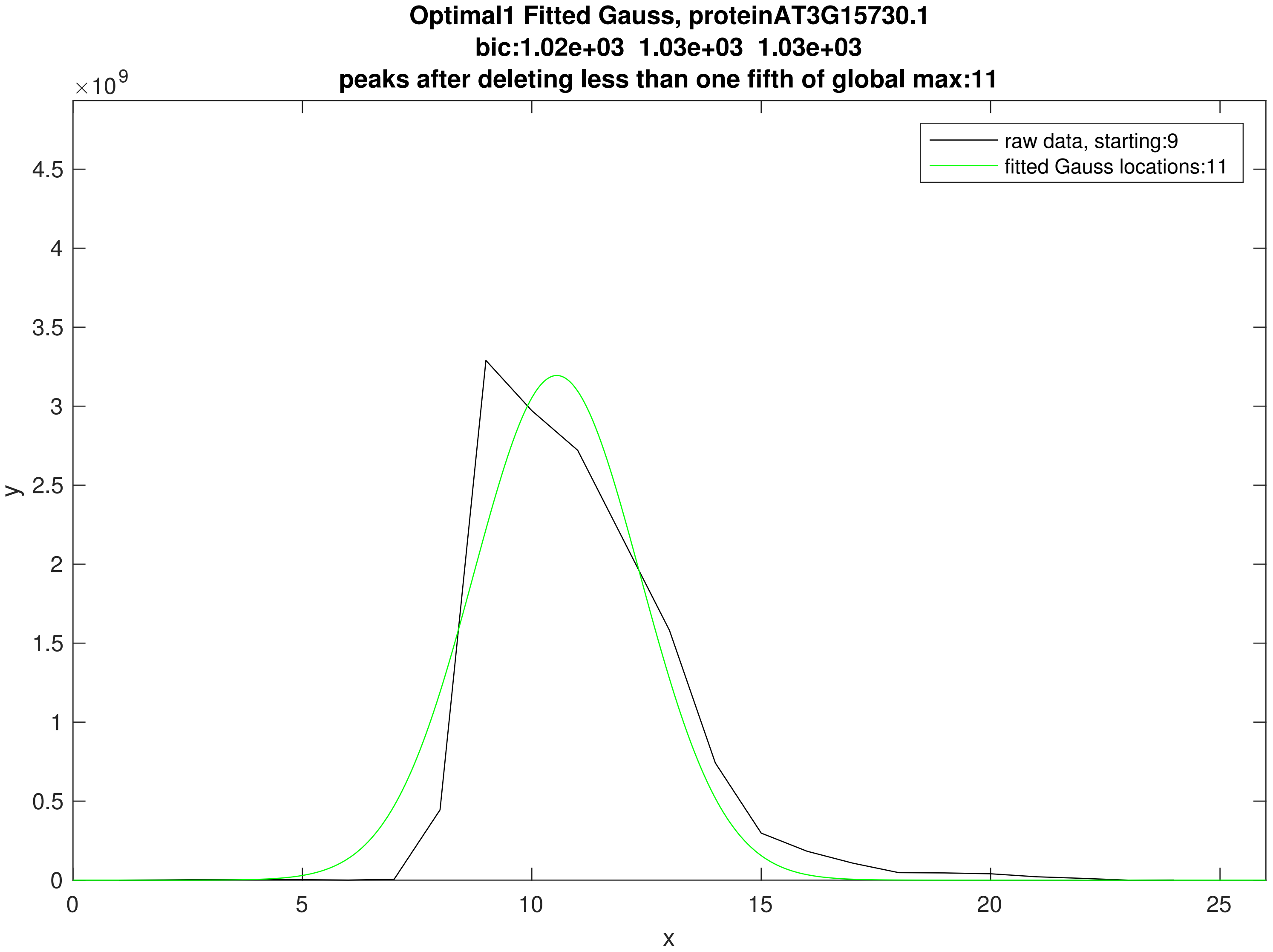
Optimal1 Fitted Gauss, proteinAT3G15730.1
bic:1.02e+03  1.03e+03  1.03e+03
peaks after deleting less than one fifth of global max:11
×10^9
raw data, starting:9
fitted Gauss locations:11
y
x
0
0.5
1
1.5
2
2.5
3
3.5
4
4.5
5
10
15
20
25

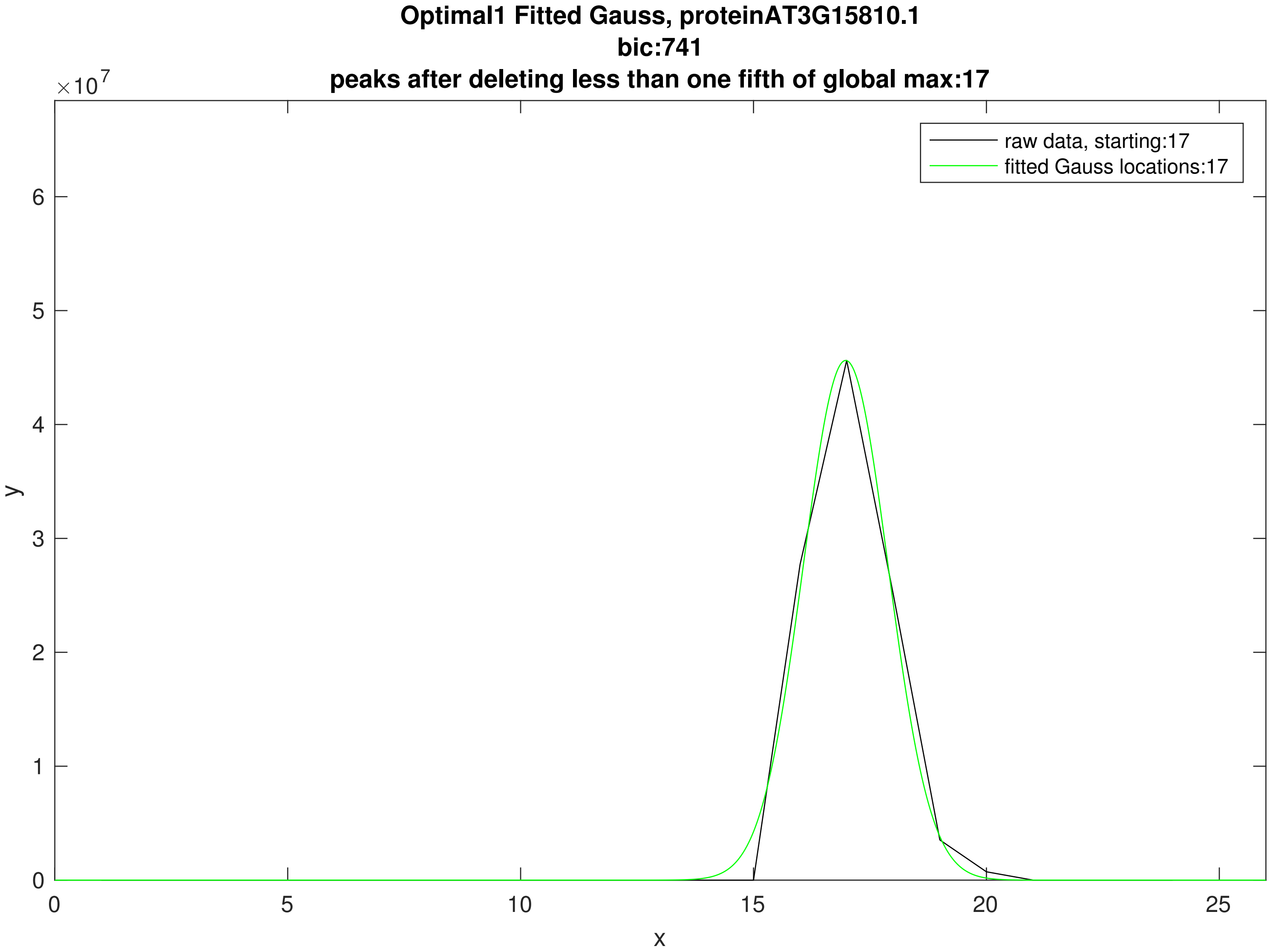
Optimal1 Fitted Gauss, proteinAT3G15810.1
bic:741
peaks after deleting less than one fifth of global max:17
×10^7
raw data, starting:17
fitted Gauss locations:17
y
x
0
1
2
3
4
5
6
0
5
10
15
20
25

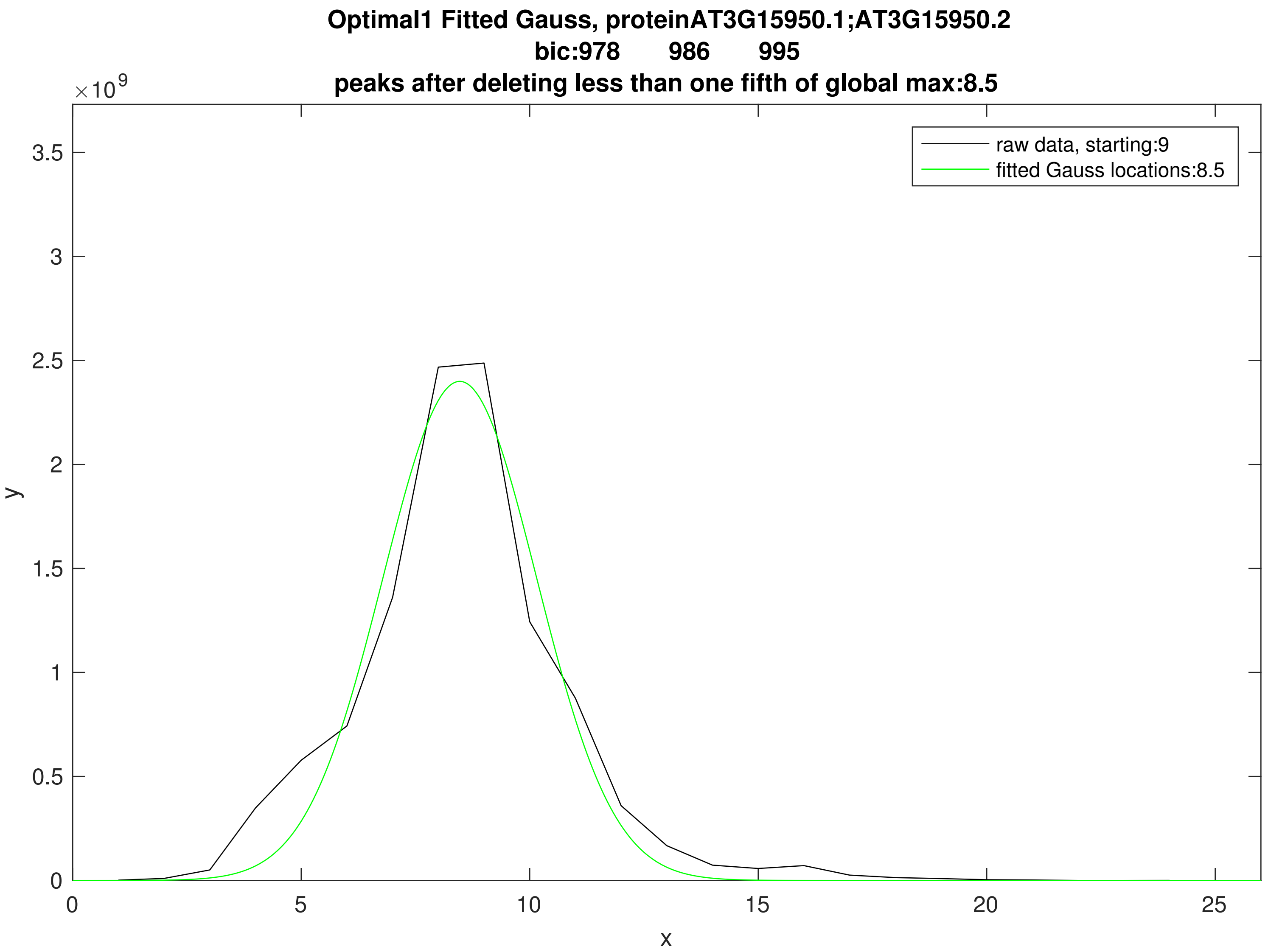

Optimal1 Fitted Gauss, proteinAT3G15950.1;AT3G15950.2
bic:978 986 995
peaks after deleting less than one fifth of global max:8.5
×10^9
raw data, starting:9
fitted Gauss locations:8.5
y
x
0
0.5
1
1.5
2
2.5
3
3.5
5
10
15
20
25

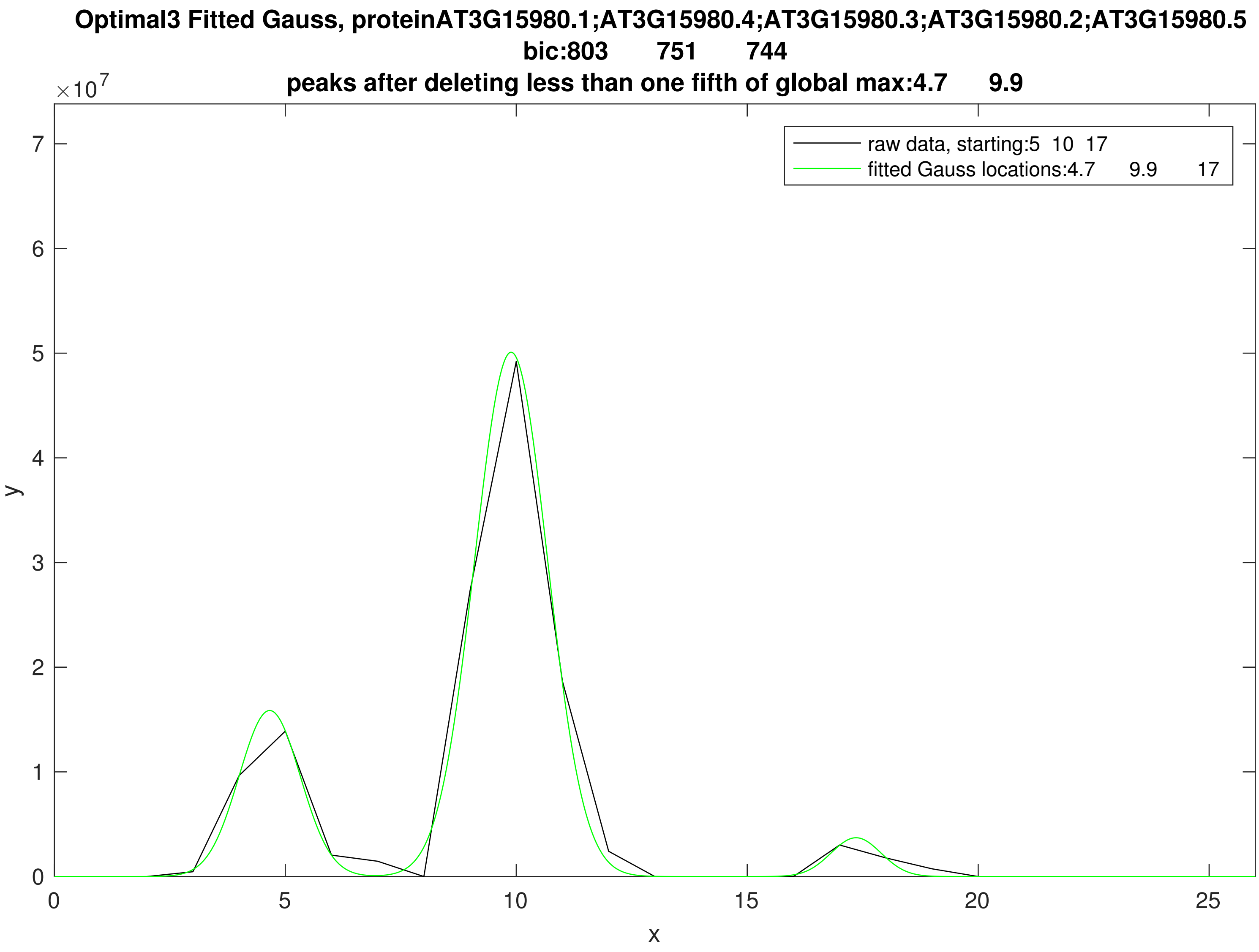

Optimal3 Fitted Gauss, proteinAT3G15980.1;AT3G15980.4;AT3G15980.3;AT3G15980.2;AT3G15980.5
bic:803 751 744
peaks after deleting less than one fifth of global max:4.7 9.9
×10^7
raw data, starting:5 10 17
fitted Gauss locations:4.7 9.9 17
y
x
0
1
2
3
4
5
6
7
0
5
10
15
20
25

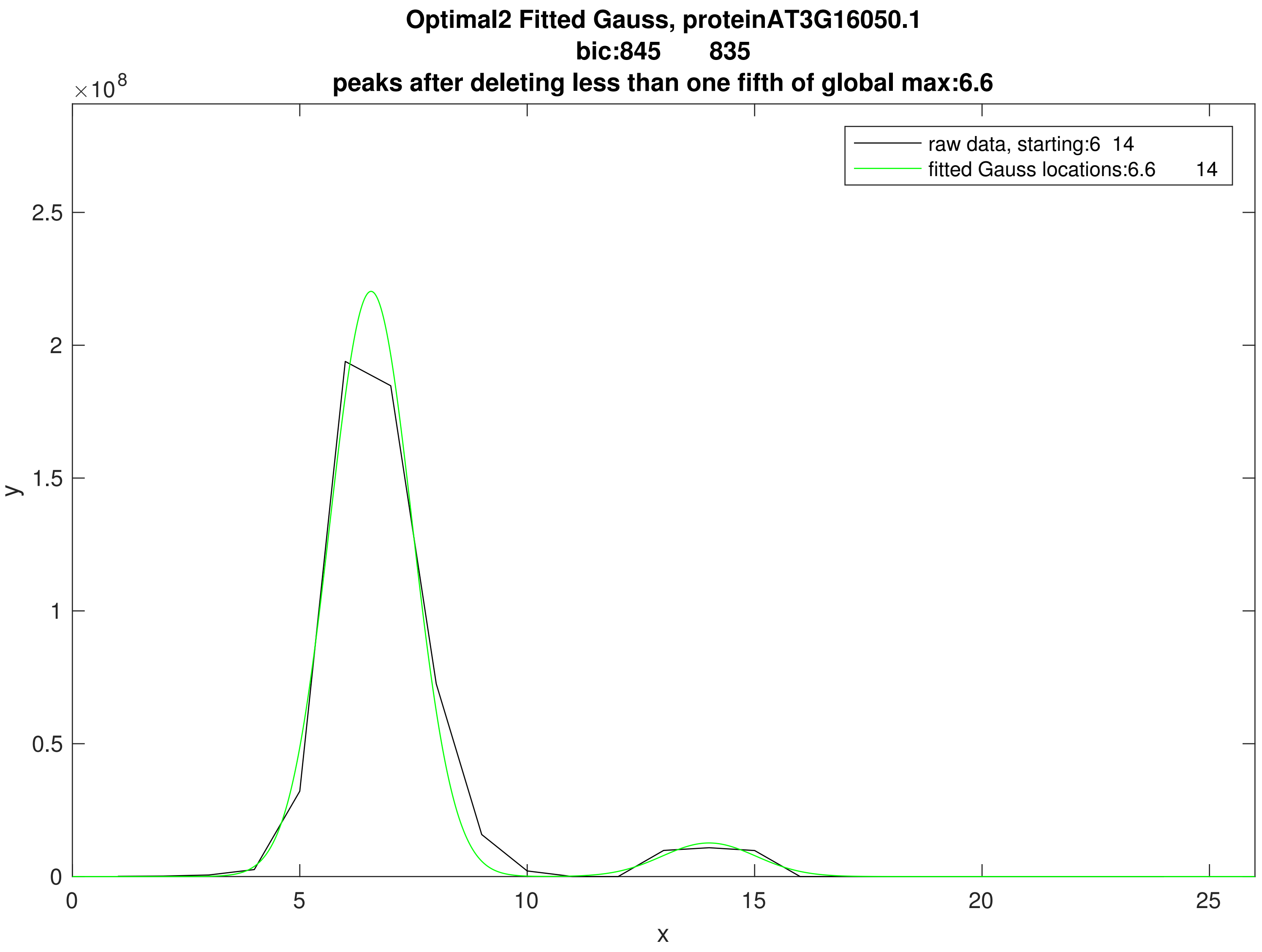

Optimal2 Fitted Gauss, proteinAT3G16050.1
bic:845 835
peaks after deleting less than one fifth of global max:6.6
×10^8
raw data, starting:6 14
fitted Gauss locations:6.6 14
y
x
0
0.5
1
1.5
2
2.5
5
10
15
20
25

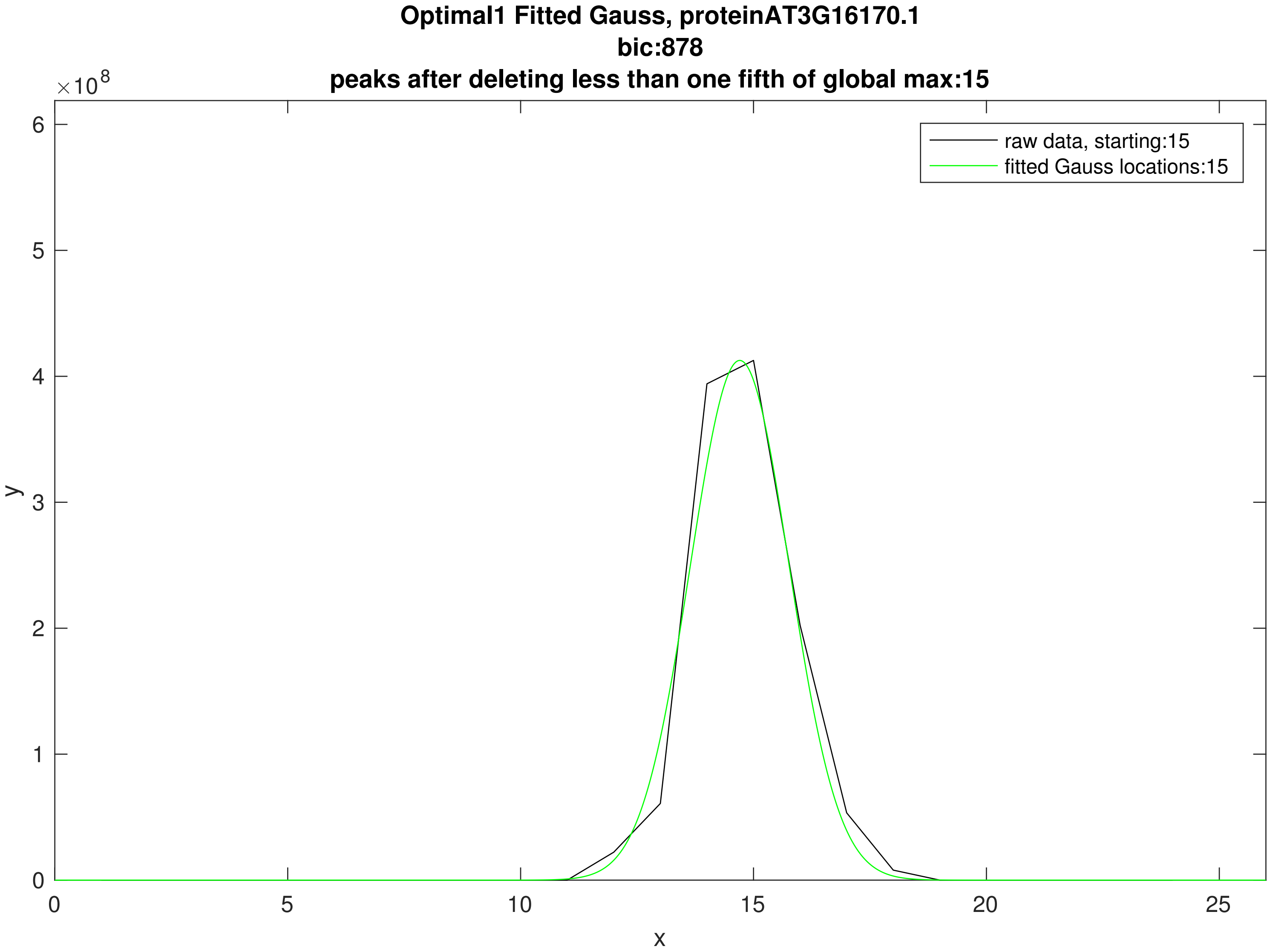

Optimal1 Fitted Gauss, proteinAT3G16170.1
bic:878
peaks after deleting less than one fifth of global max:15
×10^8
raw data, starting:15
fitted Gauss locations:15
y
x
0
1
2
3
4
5
6
0
5
10
15
20
25

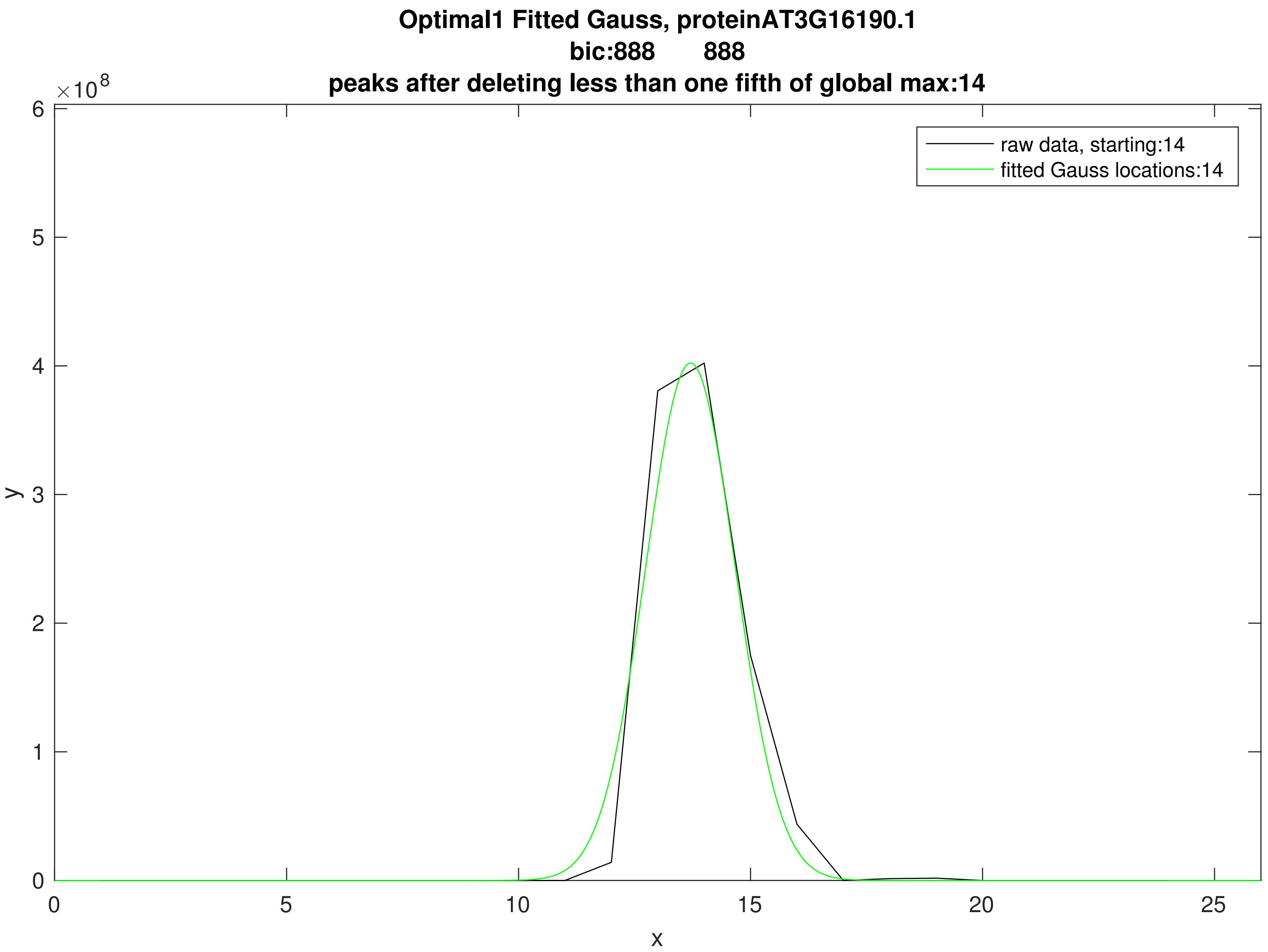
Optimal1 Fitted Gauss, proteinAT3G16190.1
bic:888 888
peaks after deleting less than one fifth of global max:14
×10^8
6
5
4
3
2
1
0
y
0
5
10
15
20
25
x
raw data, starting:14
fitted Gauss locations:14

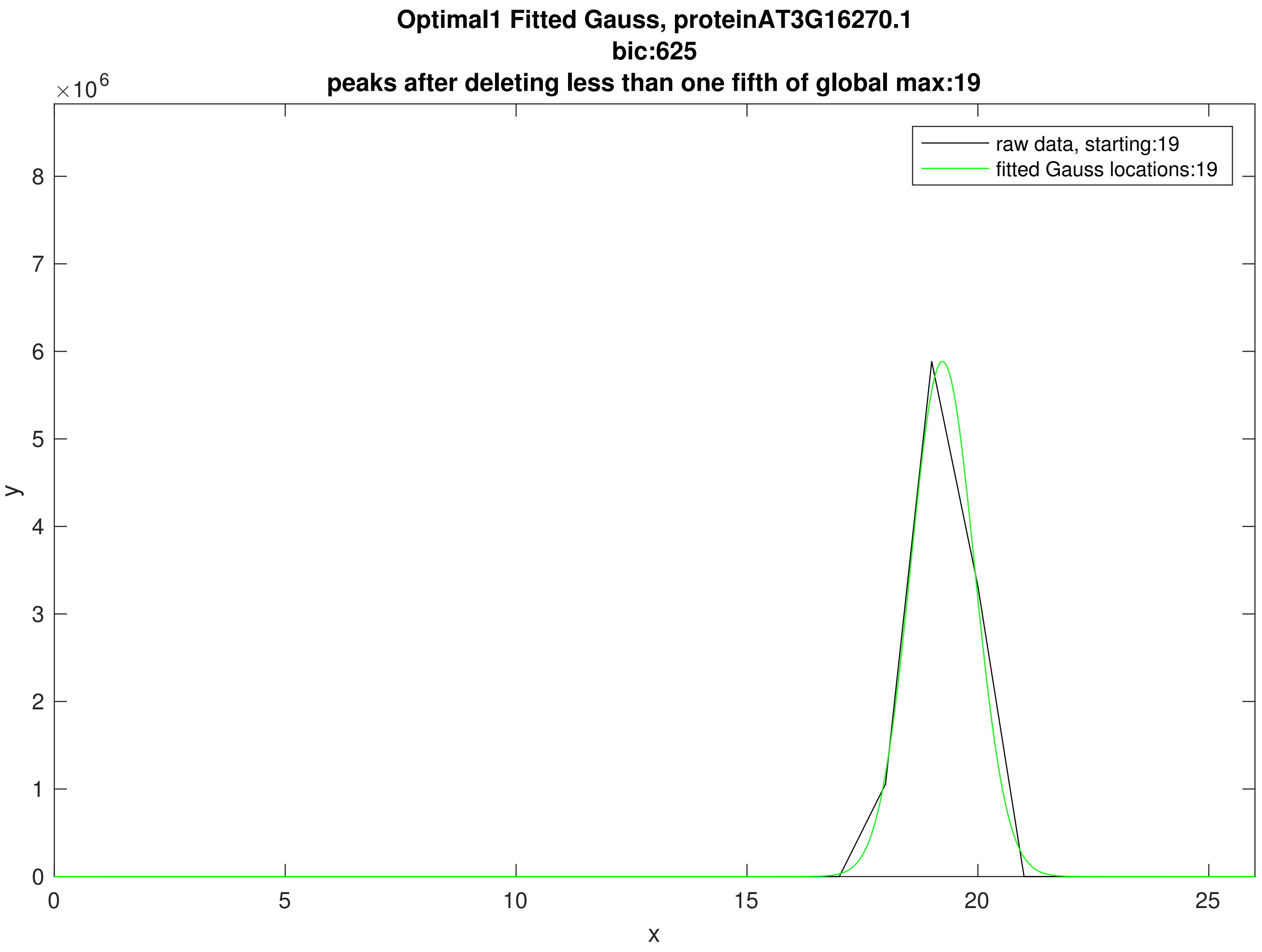
Optimal1 Fitted Gauss, proteinAT3G16270.1
bic:625
peaks after deleting less than one fifth of global max:19
×10^6
raw data, starting:19
fitted Gauss locations:19
y
x
0
1
2
3
4
5
6
7
8
0
5
10
15
20
25

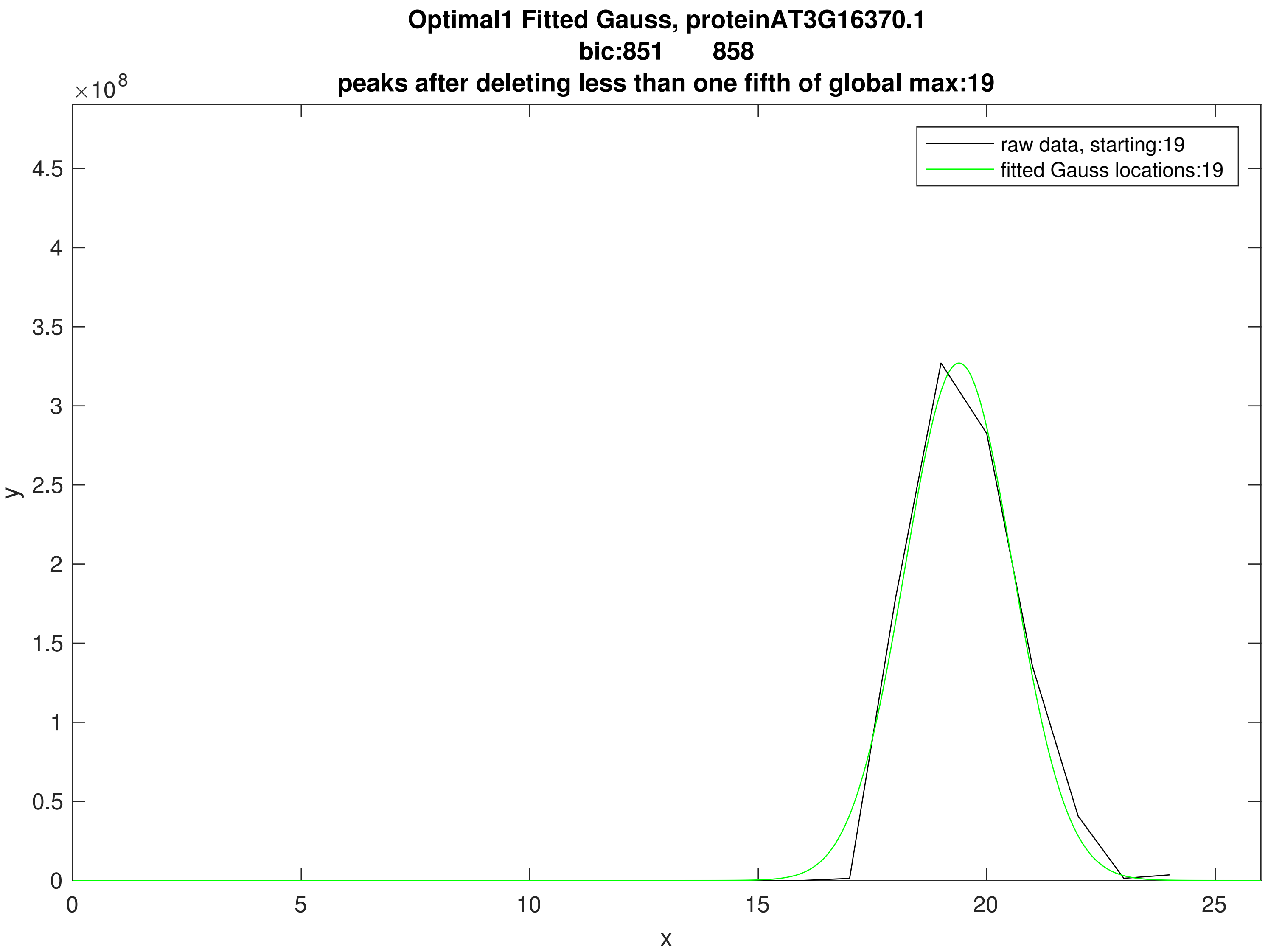
Optimal1 Fitted Gauss, proteinAT3G16370.1
bic:851 858
peaks after deleting less than one fifth of global max:19
×10^8
raw data, starting:19
fitted Gauss locations:19
4.5
4
3.5
3
2.5
2
1.5
1
0.5
0
y
0
5
10
15
20
25
x

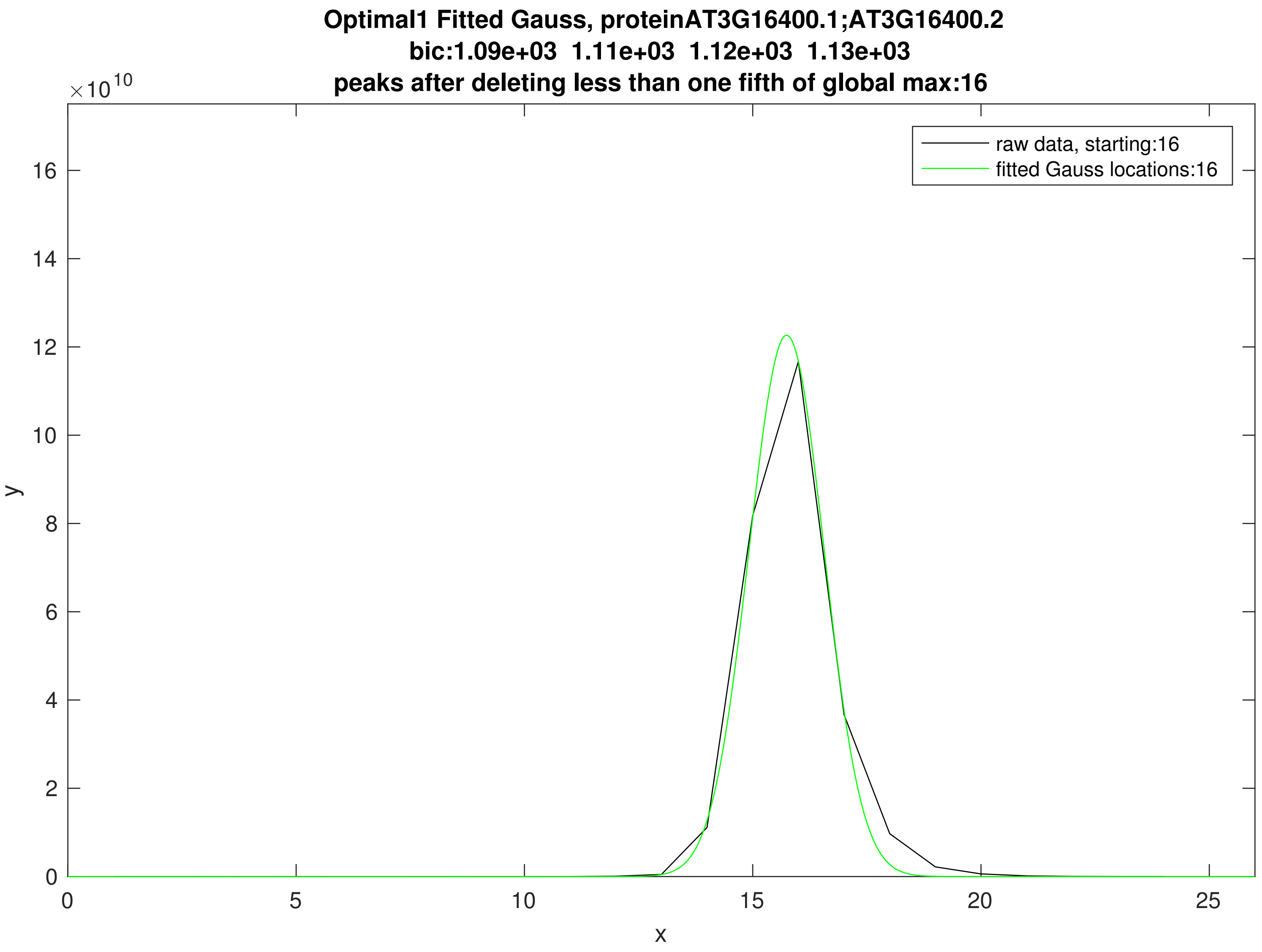
Optimal1 Fitted Gauss, proteinAT3G16400.1;AT3G16400.2
bic:1.09e+03 1.11e+03 1.12e+03 1.13e+03
peaks after deleting less than one fifth of global max:16
×10^10
raw data, starting:16
fitted Gauss locations:16
y
x
0
2
4
6
8
10
12
14
16
0
5
10
15
20
25

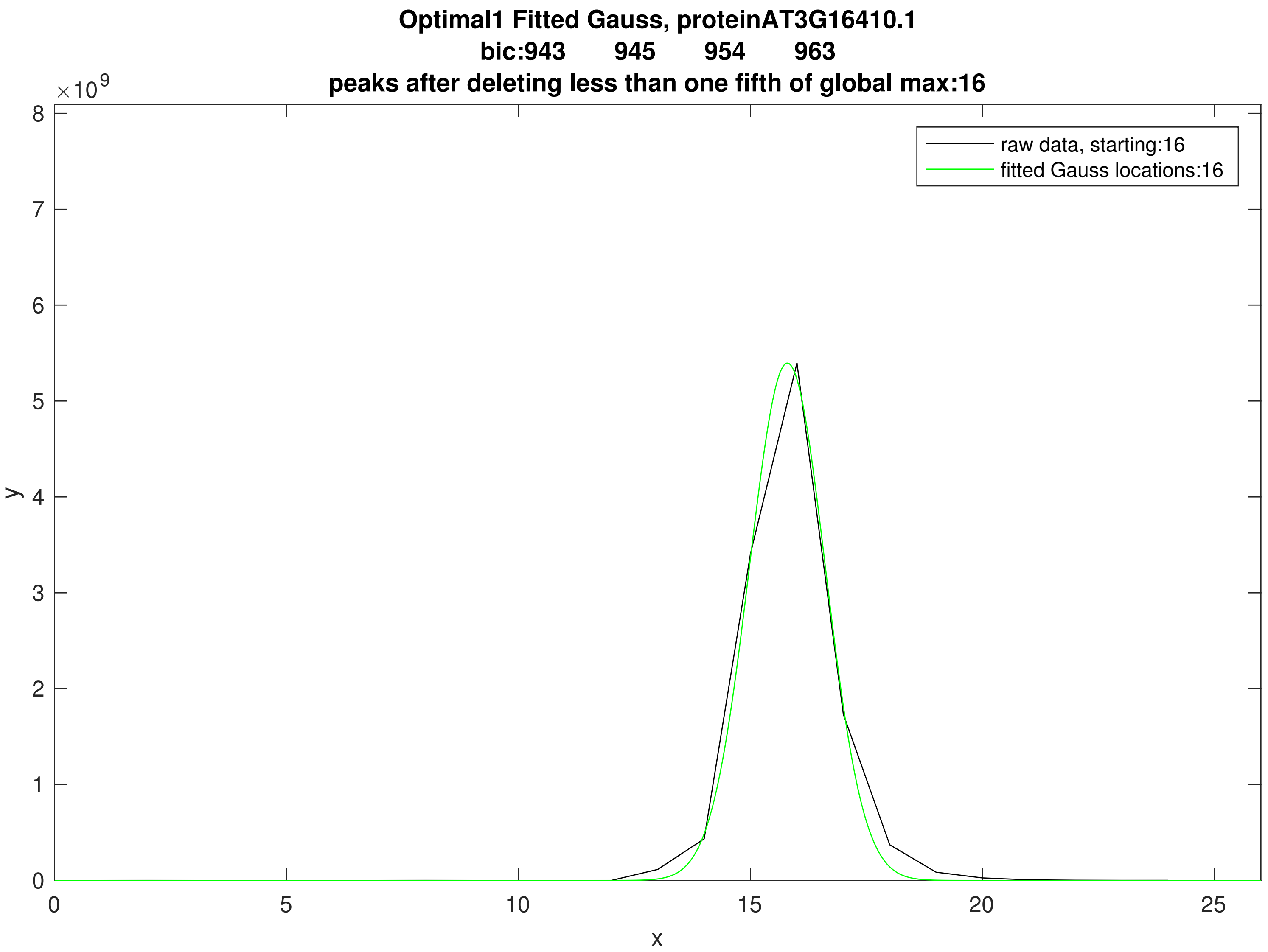
Optimal1 Fitted Gauss, proteinAT3G16410.1
bic:943 945 954 963
peaks after deleting less than one fifth of global max:16
×10^9
raw data, starting:16
fitted Gauss locations:16
y
x
0
1
2
3
4
5
6
7
8
0
5
10
15
20
25

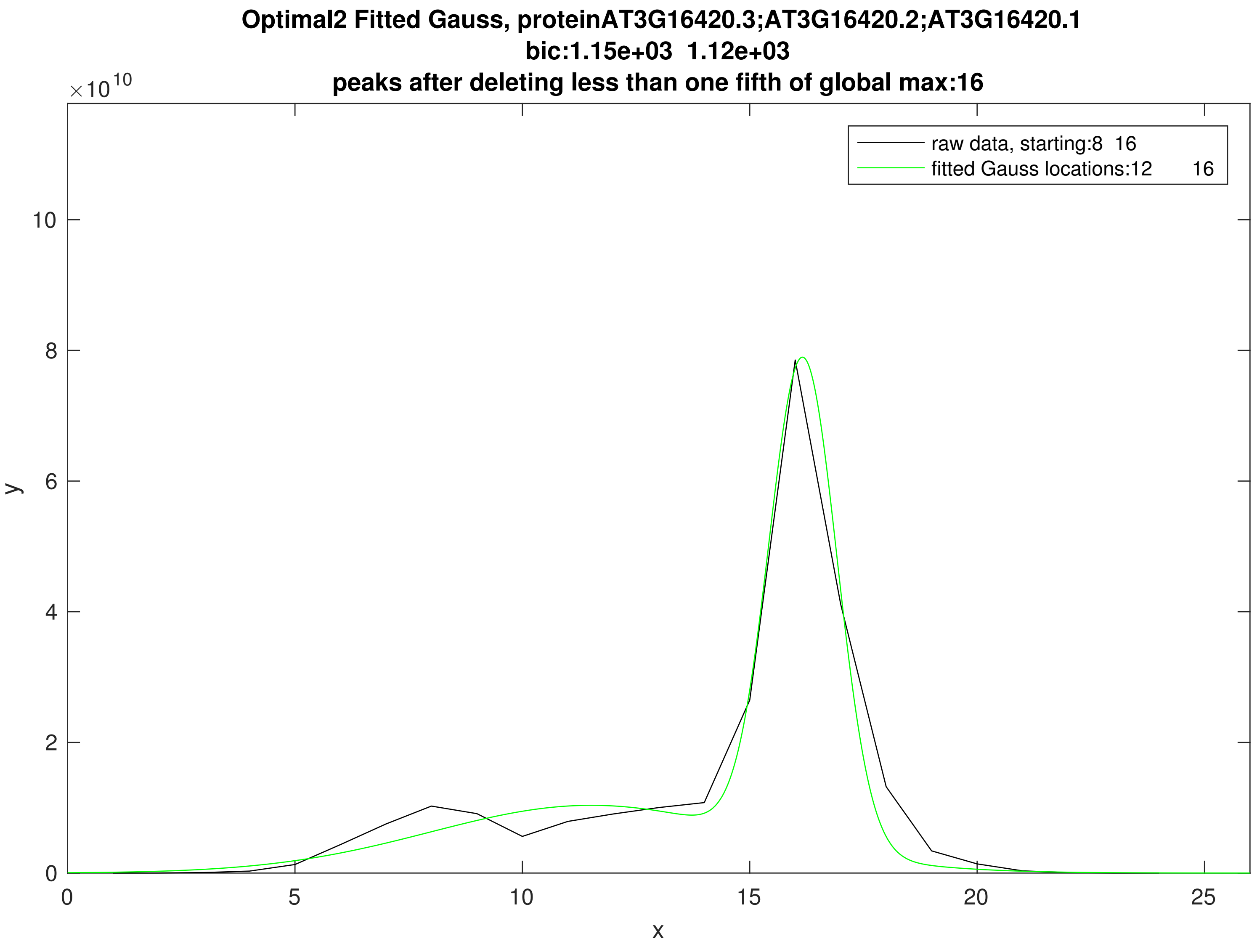
Optimal2 Fitted Gauss, proteinAT3G16420.3;AT3G16420.2;AT3G16420.1
bic:1.15e+03 1.12e+03
peaks after deleting less than one fifth of global max:16
×10^10
raw data, starting:8 16
fitted Gauss locations:12 16
y
x
0
2
4
6
8
10
0
5
10
15
20
25

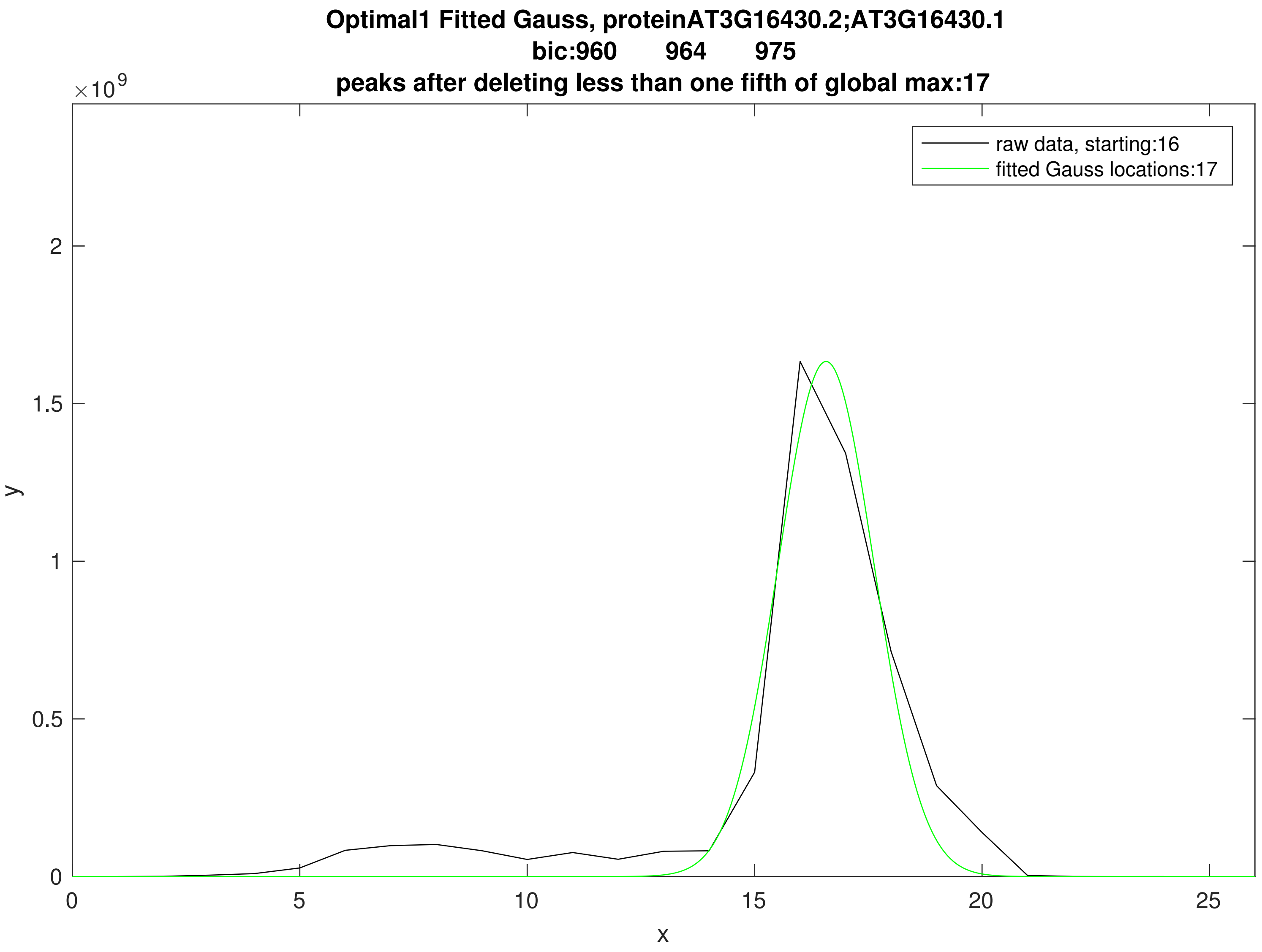
Optimal1 Fitted Gauss, proteinAT3G16430.2;AT3G16430.1
bic:960 964 975
peaks after deleting less than one fifth of global max:17
×10^9
raw data, starting:16
fitted Gauss locations:17
y
x
0
0.5
1
1.5
2
5
10
15
20
25

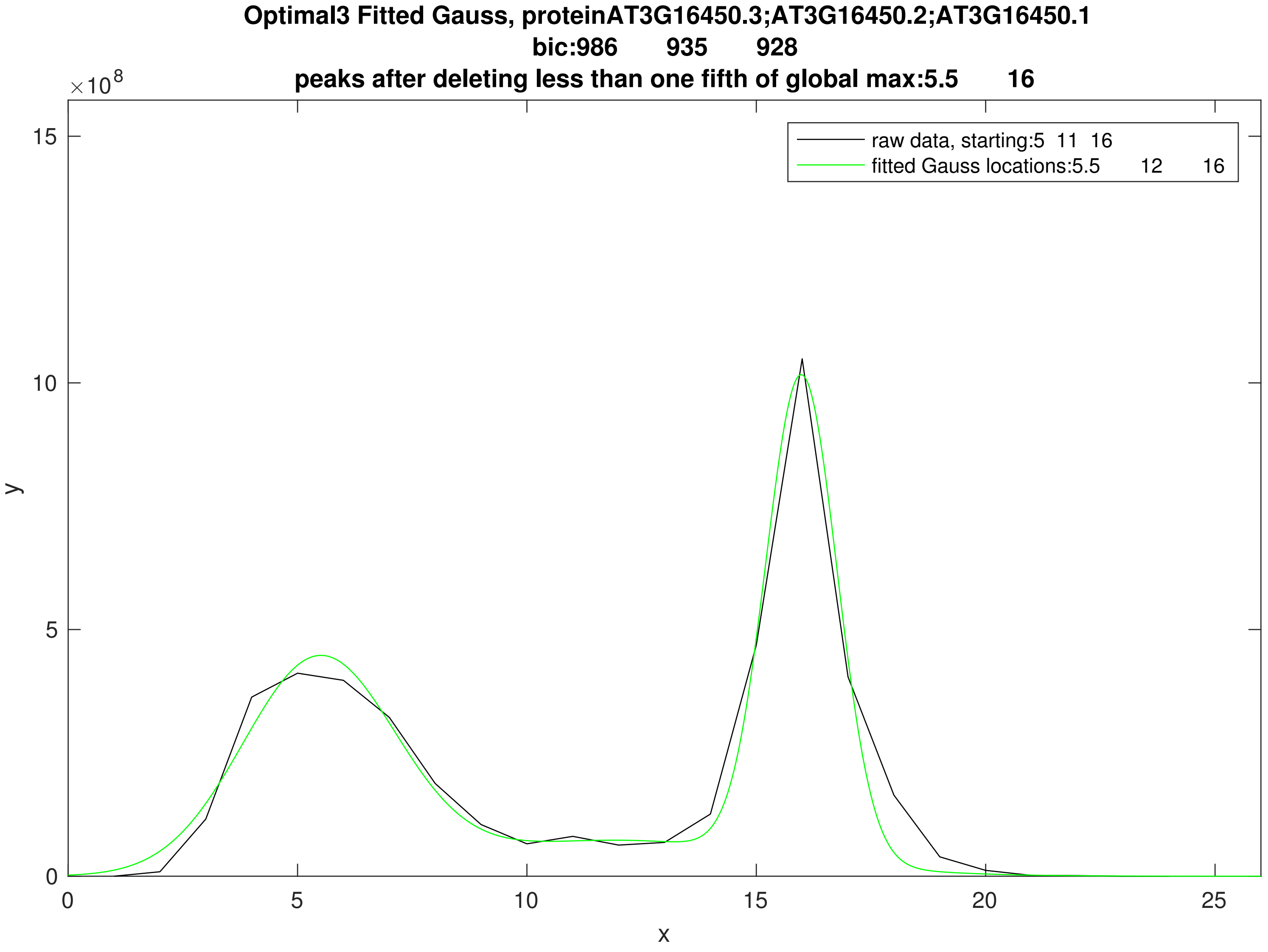
Optimal3 Fitted Gauss, proteinAT3G16450.3;AT3G16450.2;AT3G16450.1
bic:986 935 928
peaks after deleting less than one fifth of global max:5.5 16
×10^8
15
10
5
0
y
0
5
10
15
20
25
x
raw data, starting:5 11 16
fitted Gauss locations:5.5 12 16

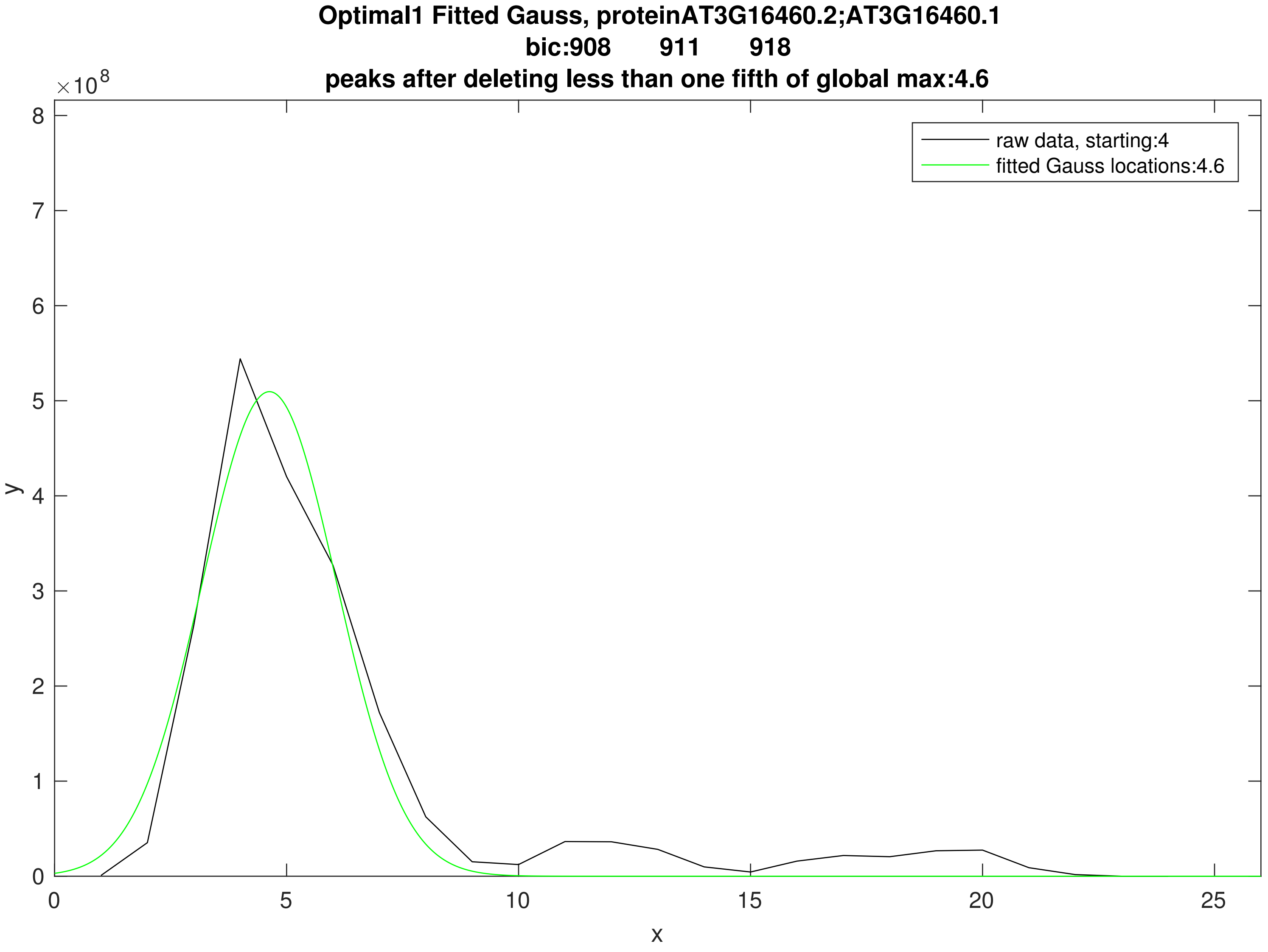
Optimal1 Fitted Gauss, proteinAT3G16460.2;AT3G16460.1
bic:908 911 918
peaks after deleting less than one fifth of global max:4.6
×10^8
raw data, starting:4
fitted Gauss locations:4.6
y
x
0
1
2
3
4
5
6
7
8
10
15
20
25

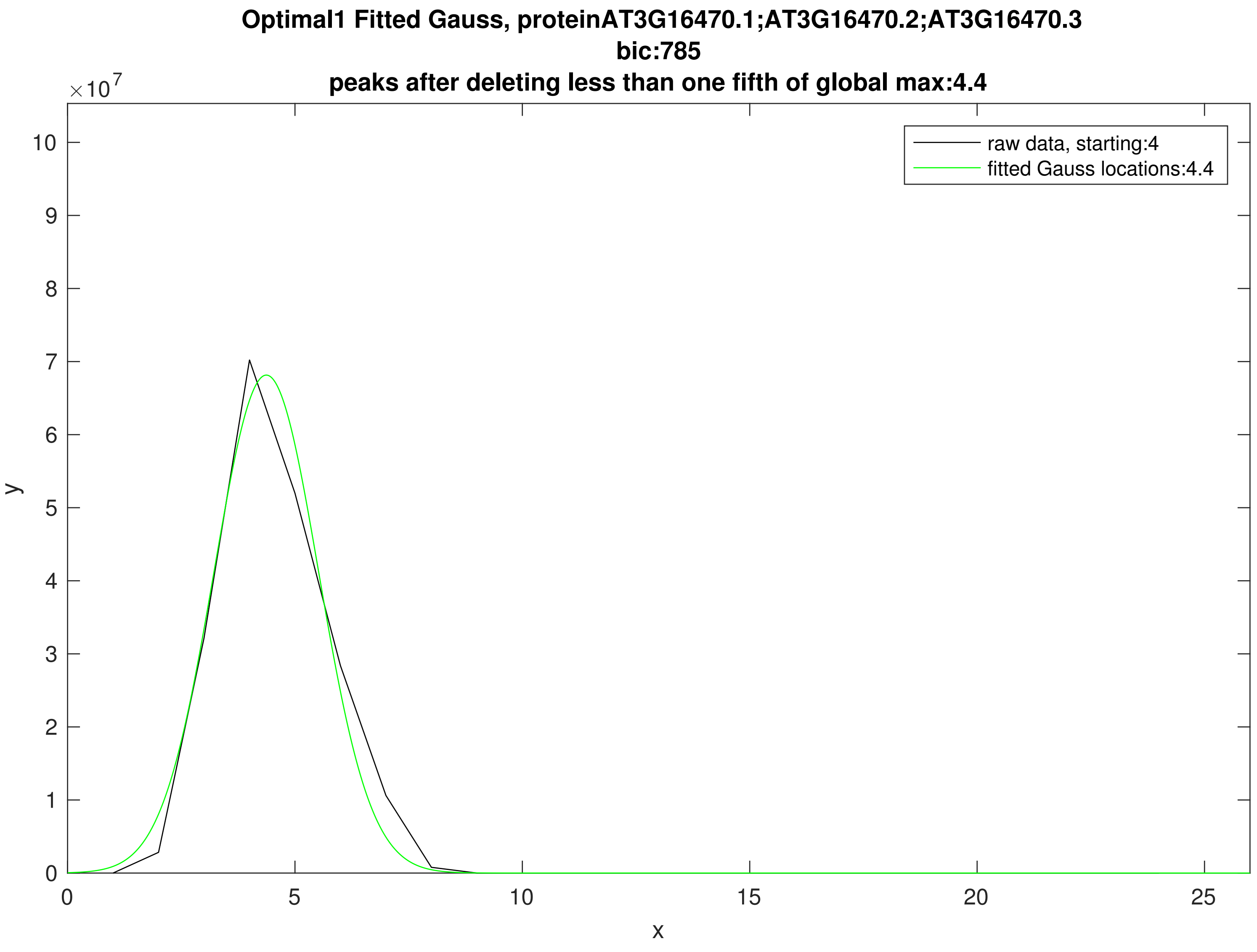

Optimal1 Fitted Gauss, proteinAT3G16470.1;AT3G16470.2;AT3G16470.3
bic:785
peaks after deleting less than one fifth of global max:4.4
×10^7
raw data, starting:4
fitted Gauss locations:4.4
y
x
0
1
2
3
4
5
6
7
8
9
10
0
5
10
15
20
25

**Optimal1 Fitted Gauss, proteinAT3G16640.1**
**bic:965 973**
**peaks after deleting less than one fifth of global max:17**

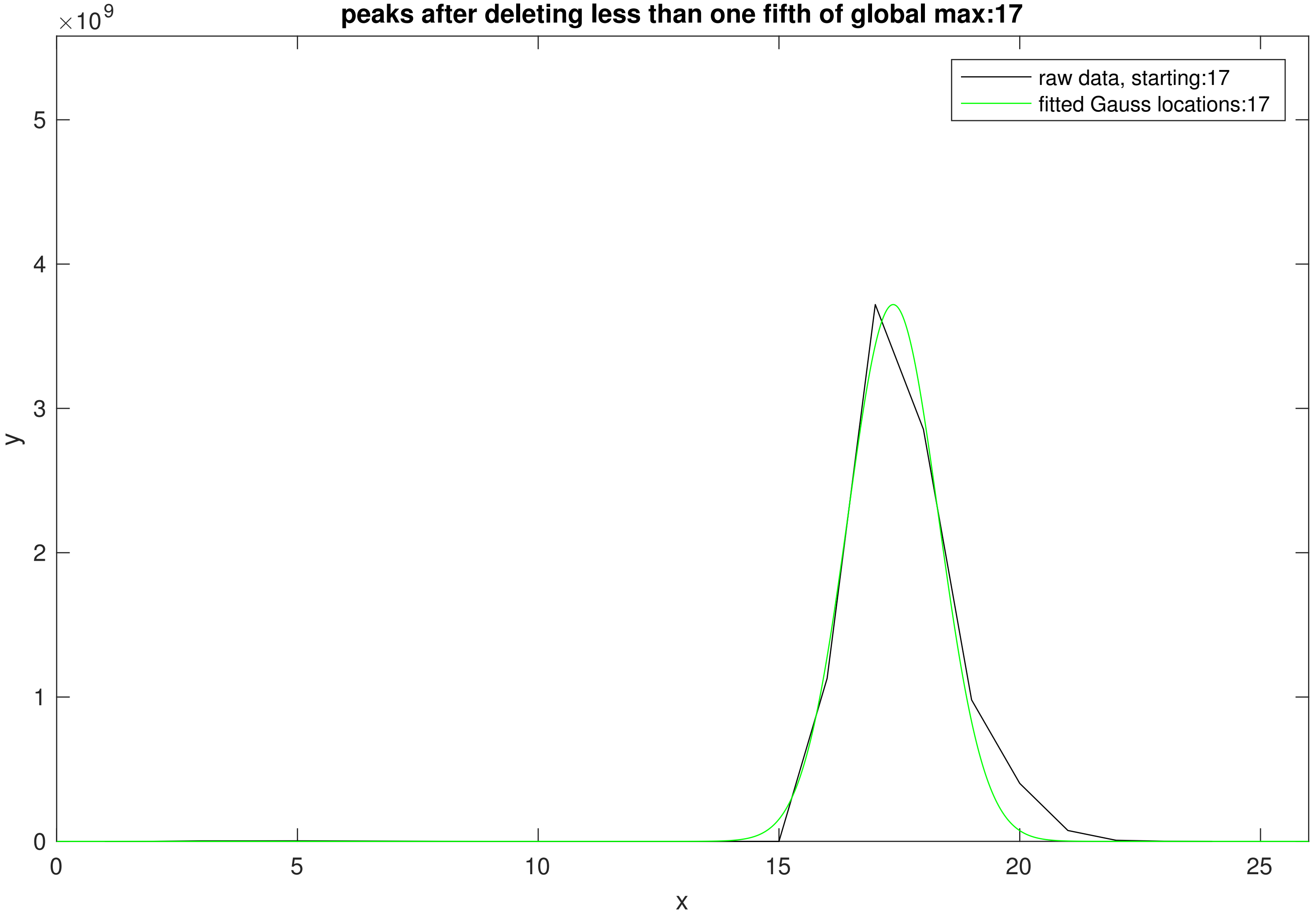

**Optimal1 Fitted Gauss, proteinAT3G16760.1**
**bic:672**
**peaks after deleting less than one fifth of global max:18**

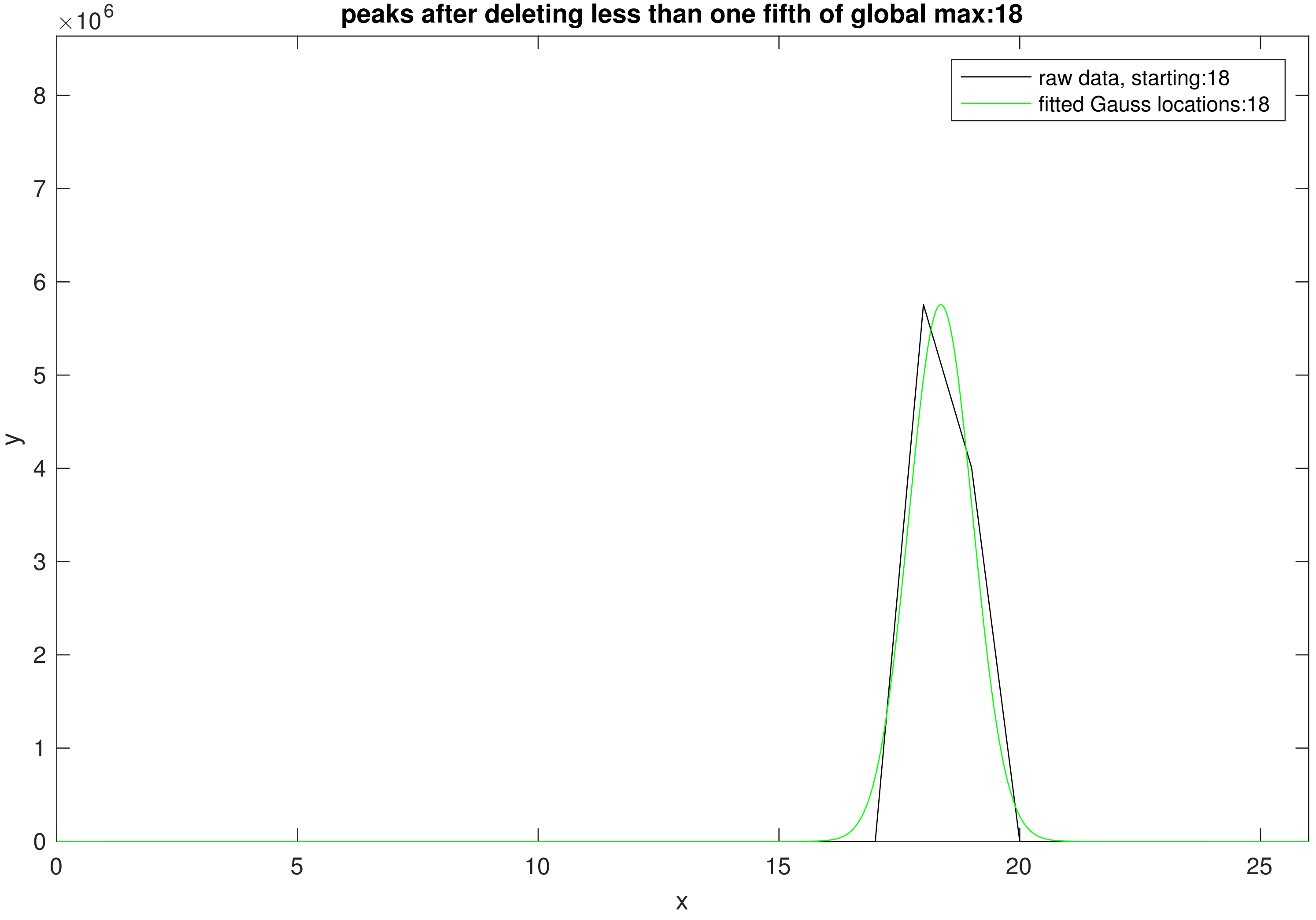

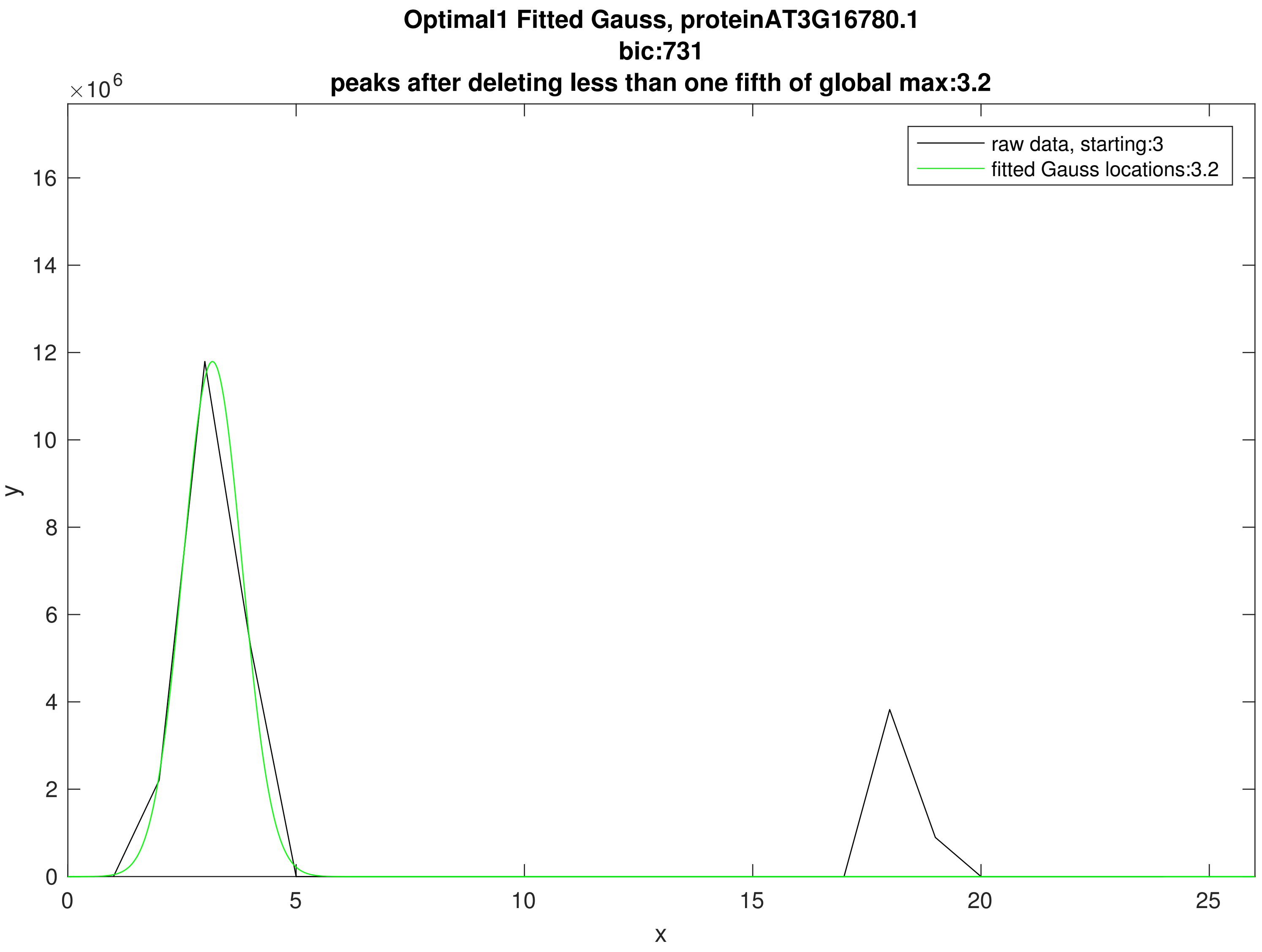

Optimal1 Fitted Gauss, proteinAT3G16780.1
bic:731
peaks after deleting less than one fifth of global max:3.2
×10^6
raw data, starting:3
fitted Gauss locations:3.2
y
x
0
2
4
6
8
10
12
14
16
0
5
10
15
20
25

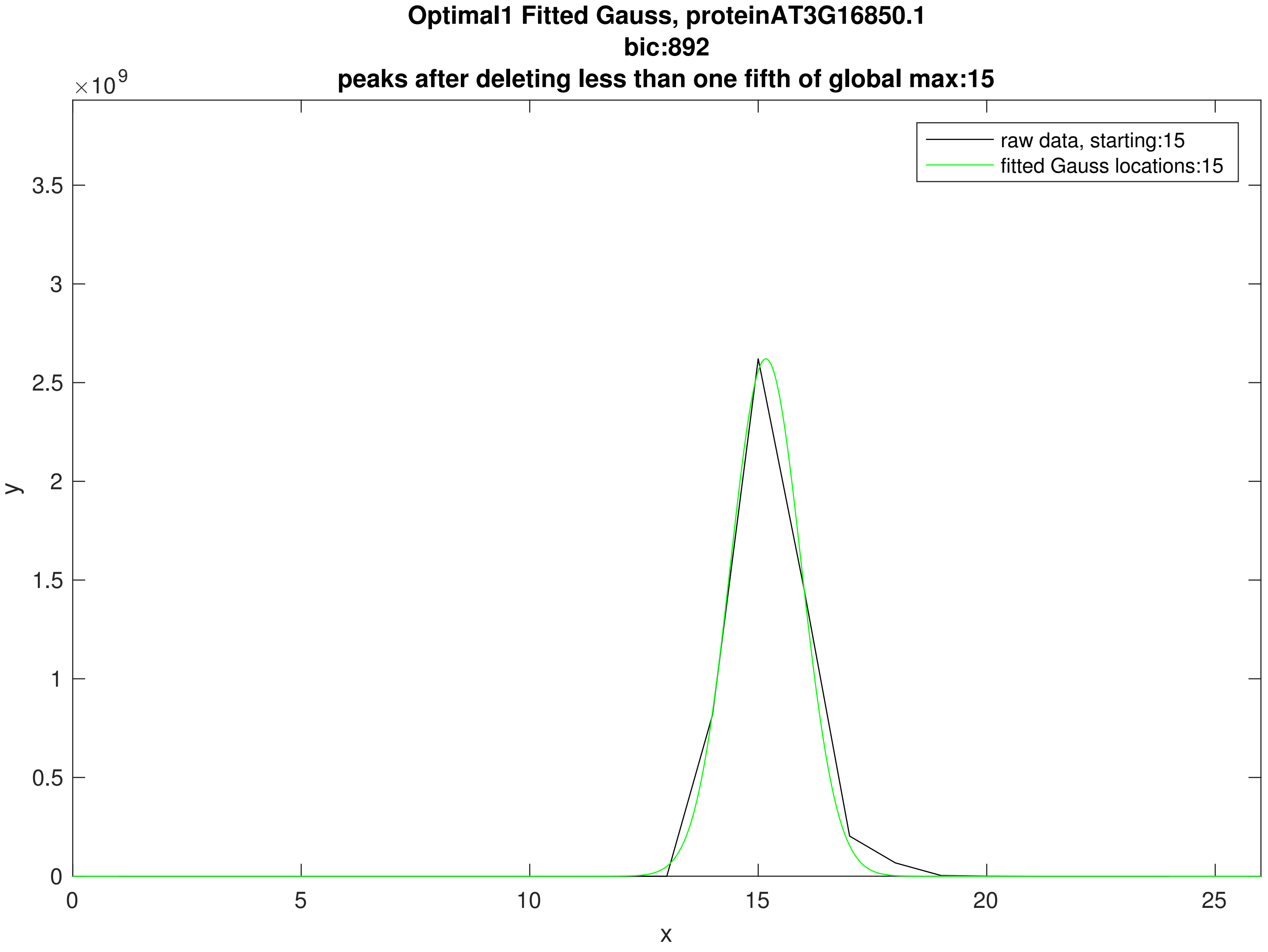
Optimal1 Fitted Gauss, proteinAT3G16850.1
bic:892
peaks after deleting less than one fifth of global max:15
×10^9
raw data, starting:15
fitted Gauss locations:15
3.5
3
2.5
2
1.5
1
0.5
0
y
0
5
10
15
20
25
x

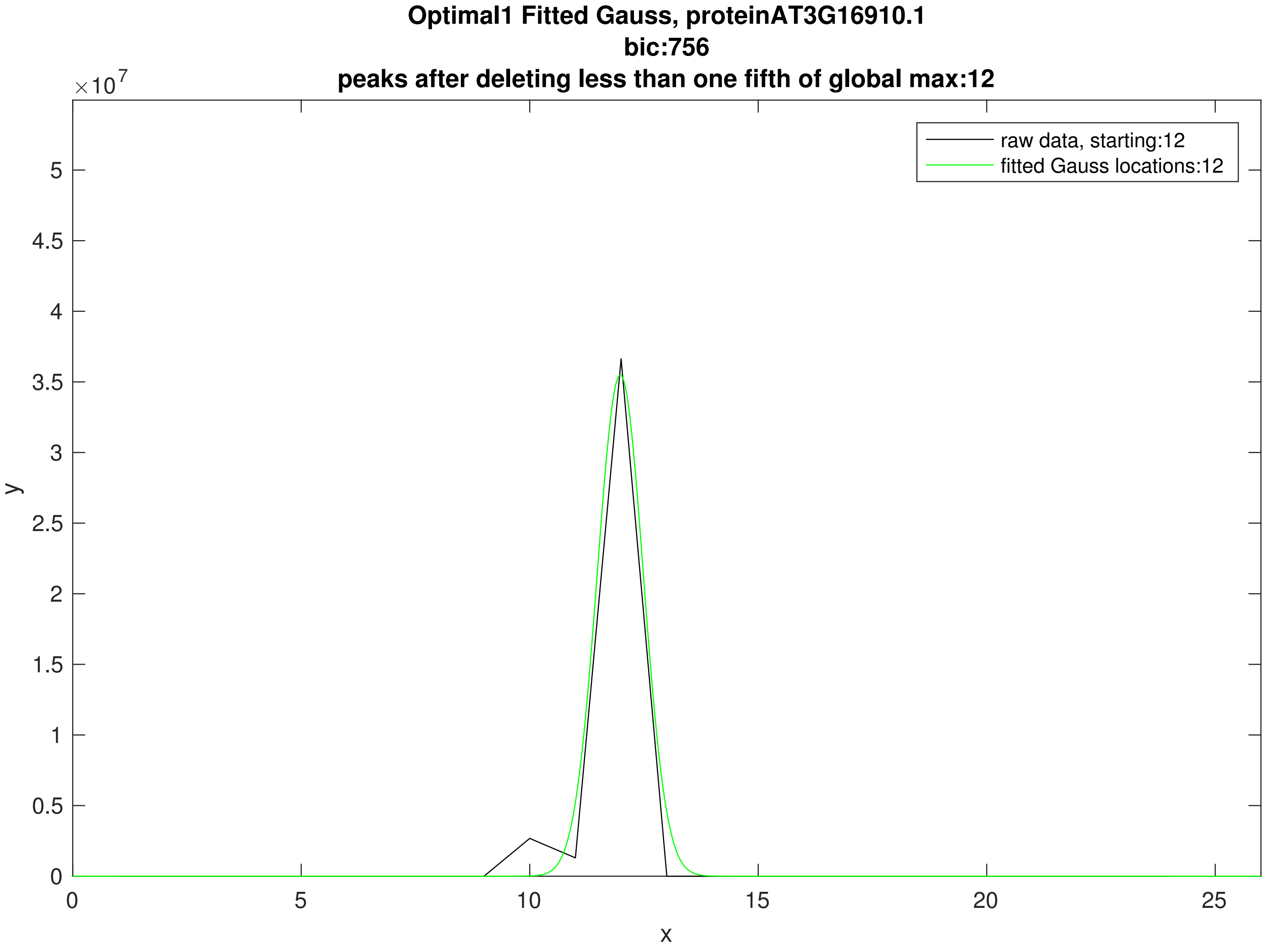

Optimal1 Fitted Gauss, proteinAT3G16910.1
bic:756
peaks after deleting less than one fifth of global max:12
×10^7
raw data, starting:12
fitted Gauss locations:12
5
4.5
4
3.5
3
2.5
2
1.5
1
0.5
0
y
0
5
10
15
20
25
x

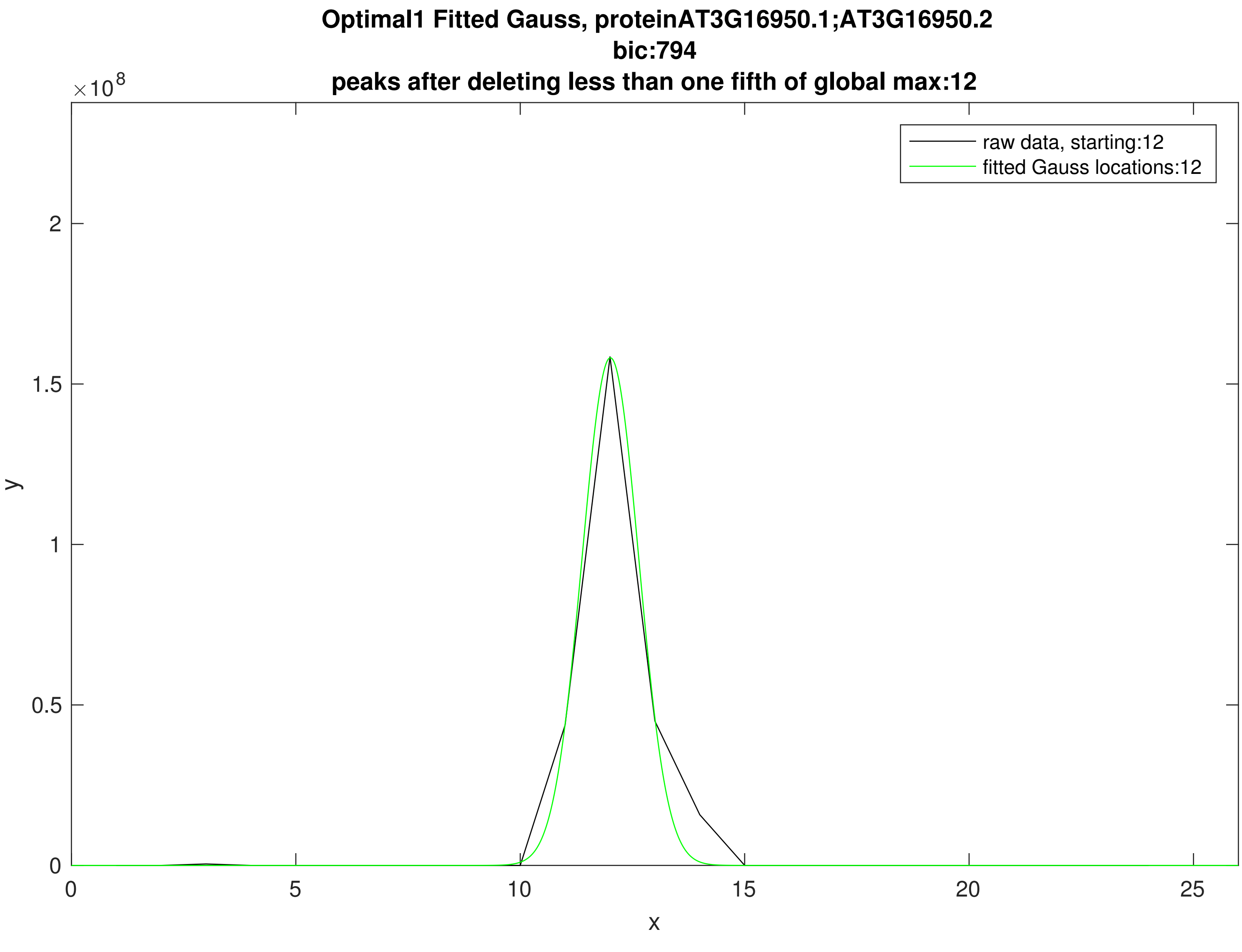
Optimal1 Fitted Gauss, proteinAT3G16950.1;AT3G16950.2
bic:794
peaks after deleting less than one fifth of global max:12
×10^8
raw data, starting:12
fitted Gauss locations:12
y
x
0
0.5
1
1.5
2
5
10
15
20
25

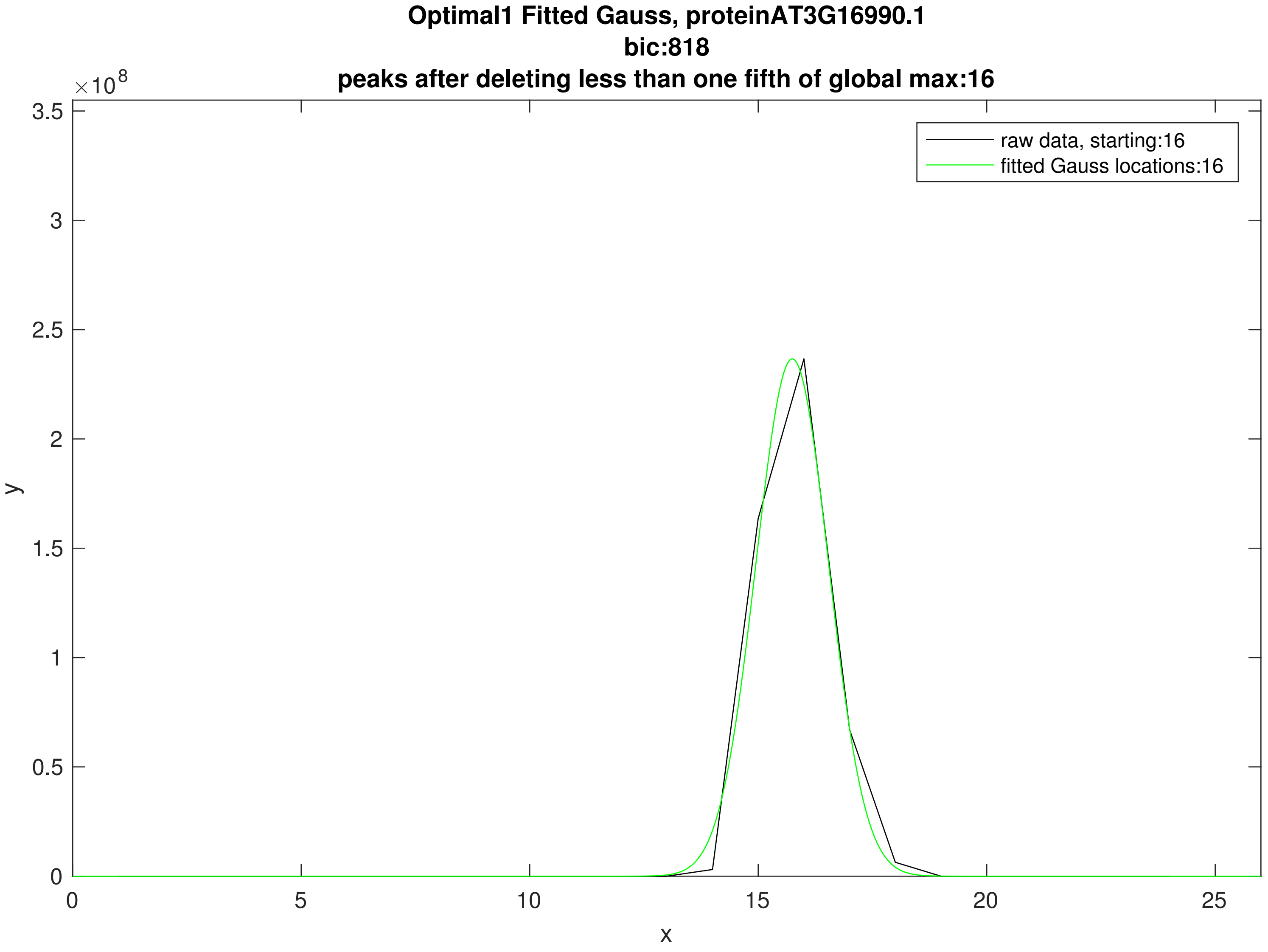

Optimal1 Fitted Gauss, proteinAT3G16990.1
bic:818
peaks after deleting less than one fifth of global max:16
×10^8
3.5
3
2.5
2
1.5
1
0.5
0
y
0
5
10
15
20
25
x
raw data, starting:16
fitted Gauss locations:16

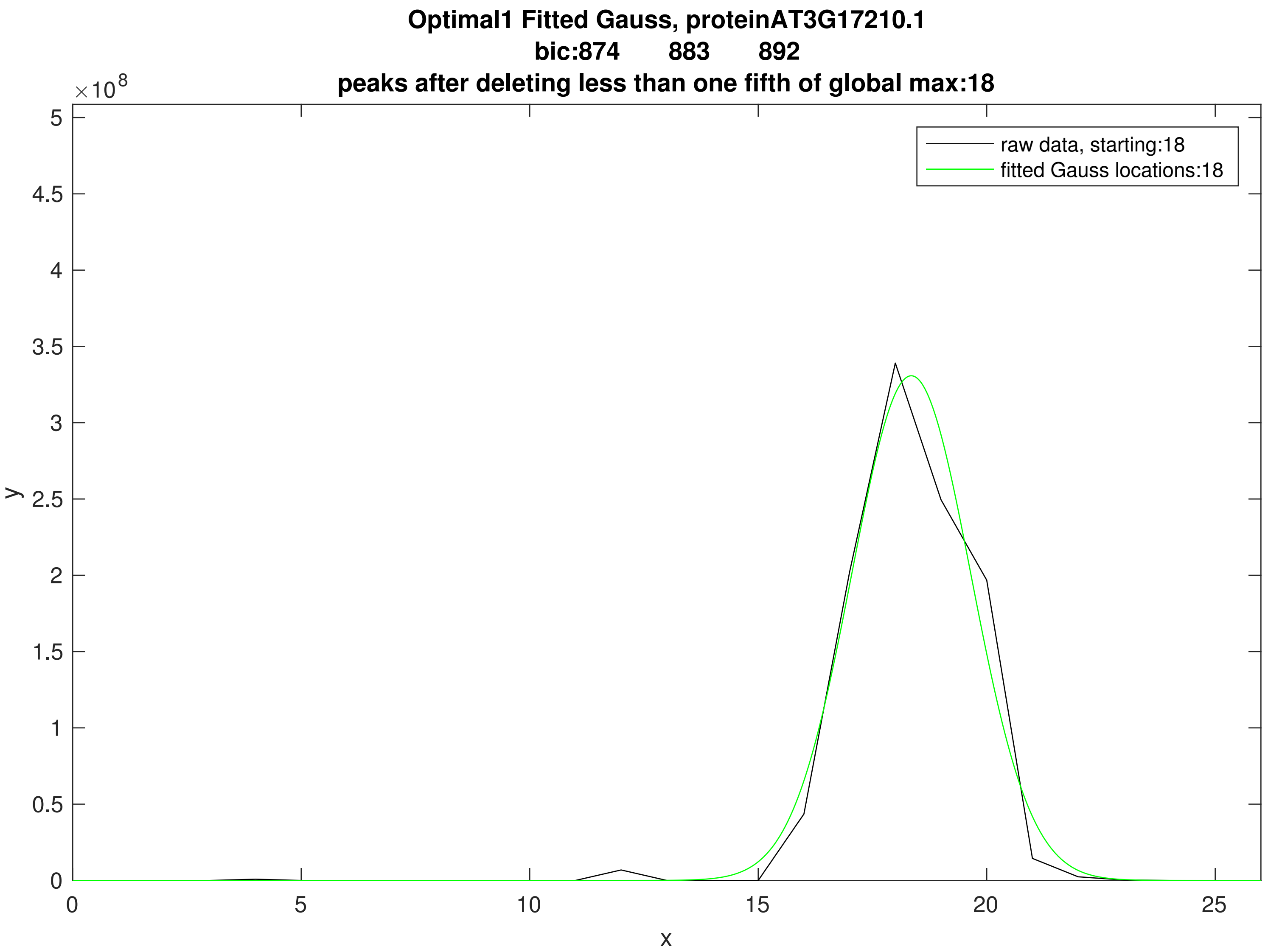

Optimal1 Fitted Gauss, proteinAT3G17210.1
bic:874 883 892
peaks after deleting less than one fifth of global max:18
×10^8
raw data, starting:18
fitted Gauss locations:18
y
x
0
0.5
1
1.5
2
2.5
3
3.5
4
4.5
5
0
5
10
15
20
25

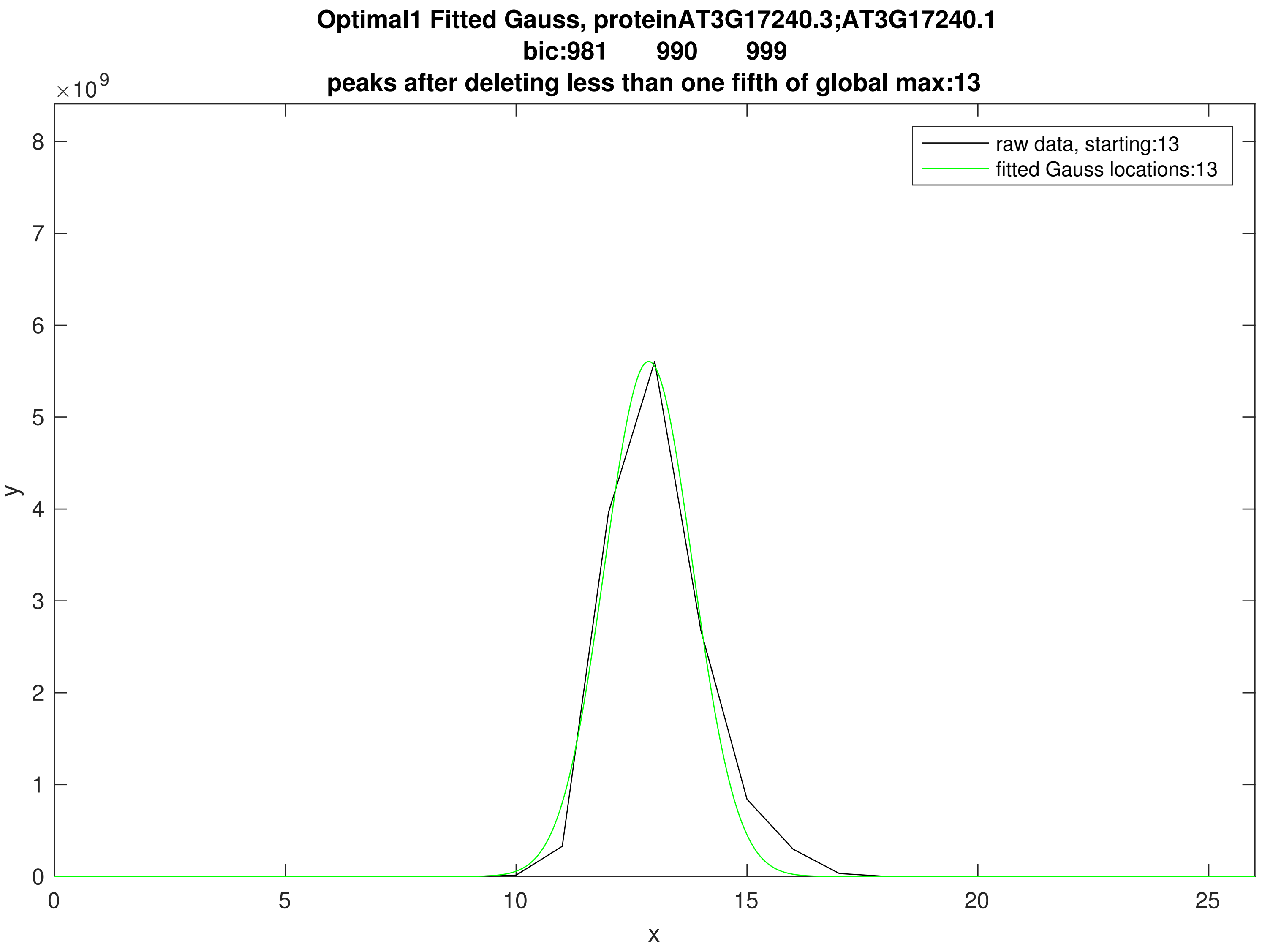
Optimal1 Fitted Gauss, proteinAT3G17240.3;AT3G17240.1
bic:981 990 999
peaks after deleting less than one fifth of global max:13
×10^9
raw data, starting:13
fitted Gauss locations:13
y
x

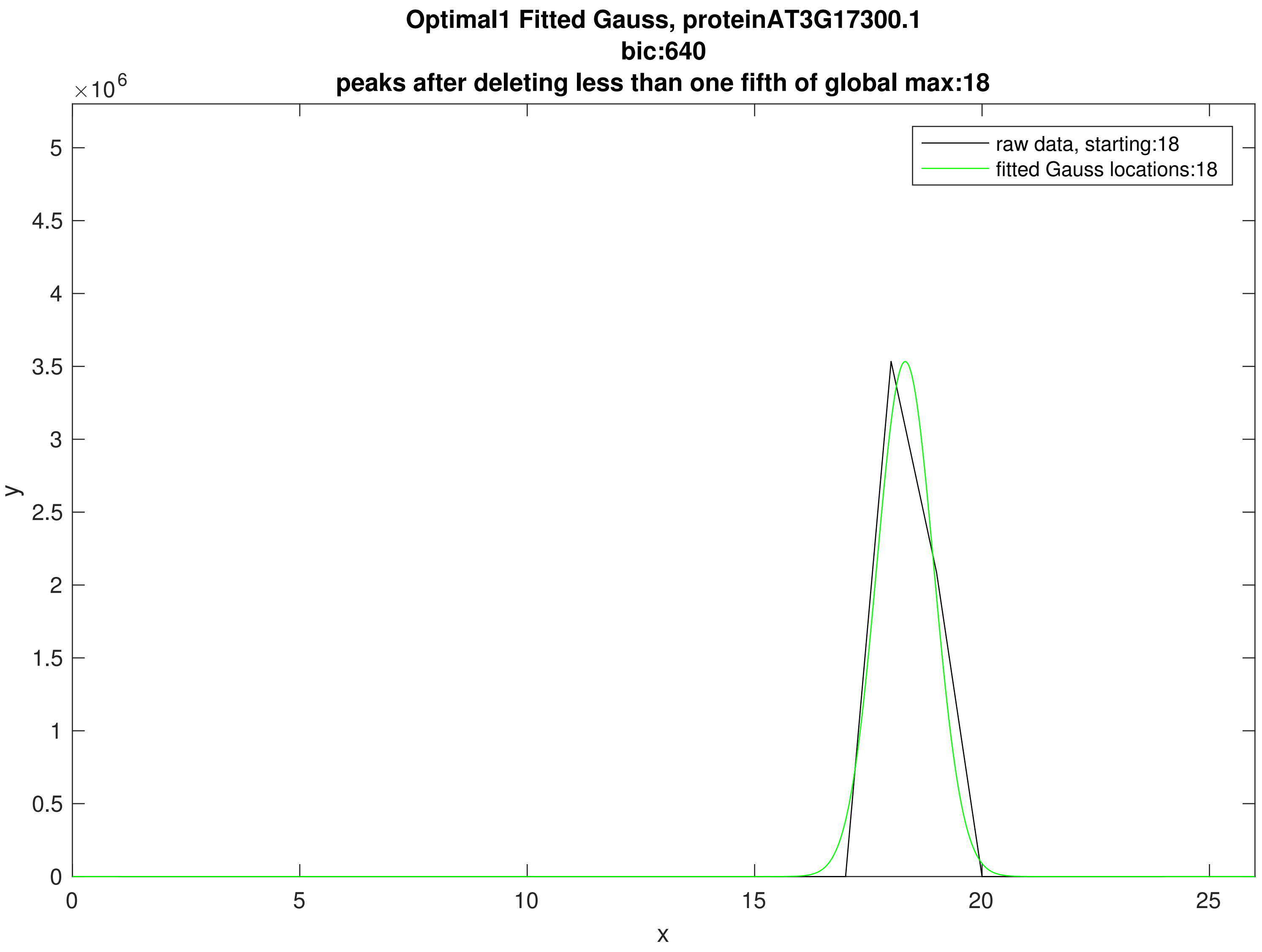
Optimal1 Fitted Gauss, proteinAT3G17300.1
bic:640
peaks after deleting less than one fifth of global max:18
×10^6
raw data, starting:18
fitted Gauss locations:18
y
x
0
0.5
1
1.5
2
2.5
3
3.5
4
4.5
5
0
5
10
15
20
25

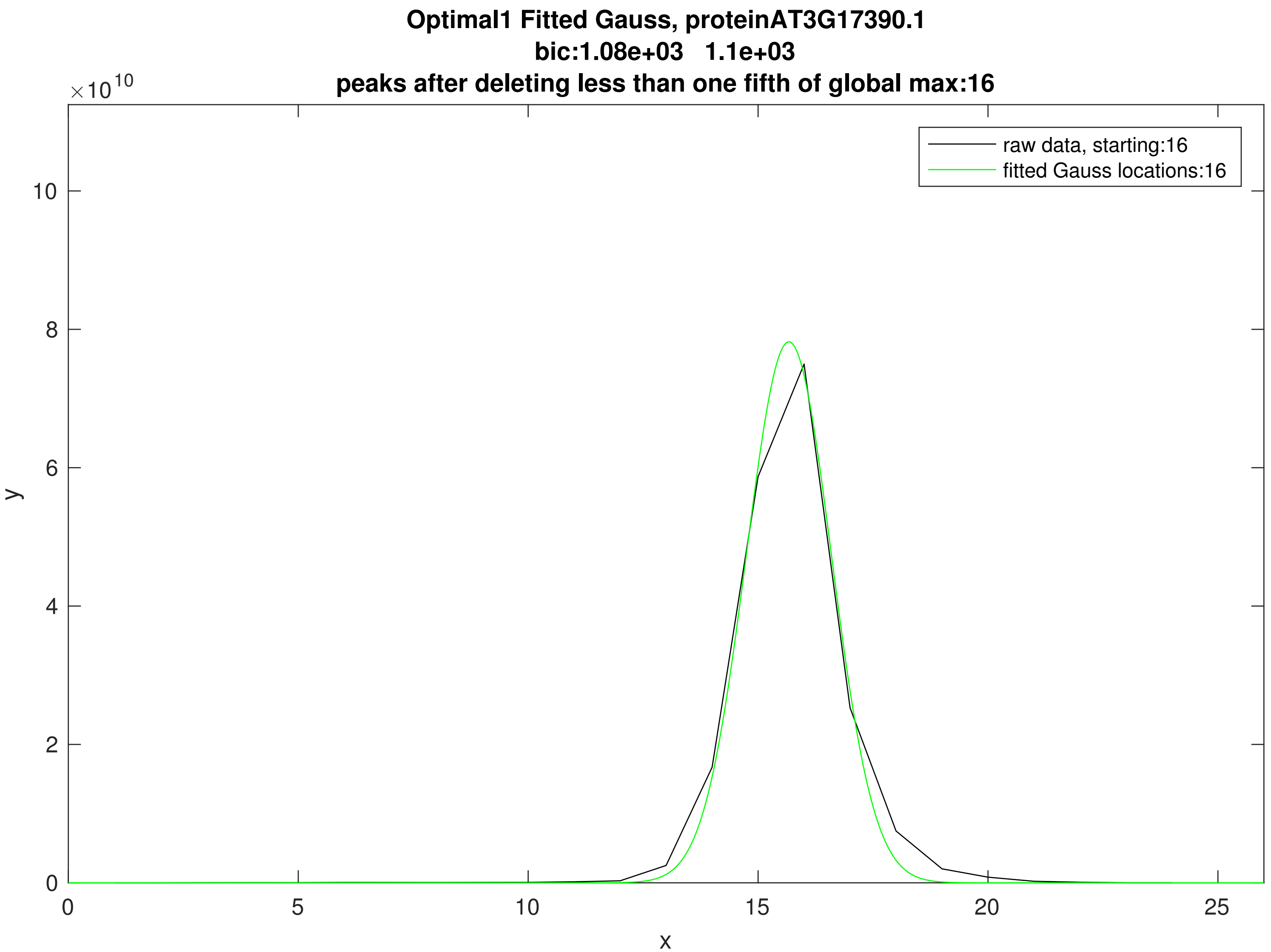
Optimal1 Fitted Gauss, proteinAT3G17390.1
bic:1.08e+03 1.1e+03
peaks after deleting less than one fifth of global max:16
×10^10
raw data, starting:16
fitted Gauss locations:16
y
x
0
2
4
6
8
10
5
15
20
25

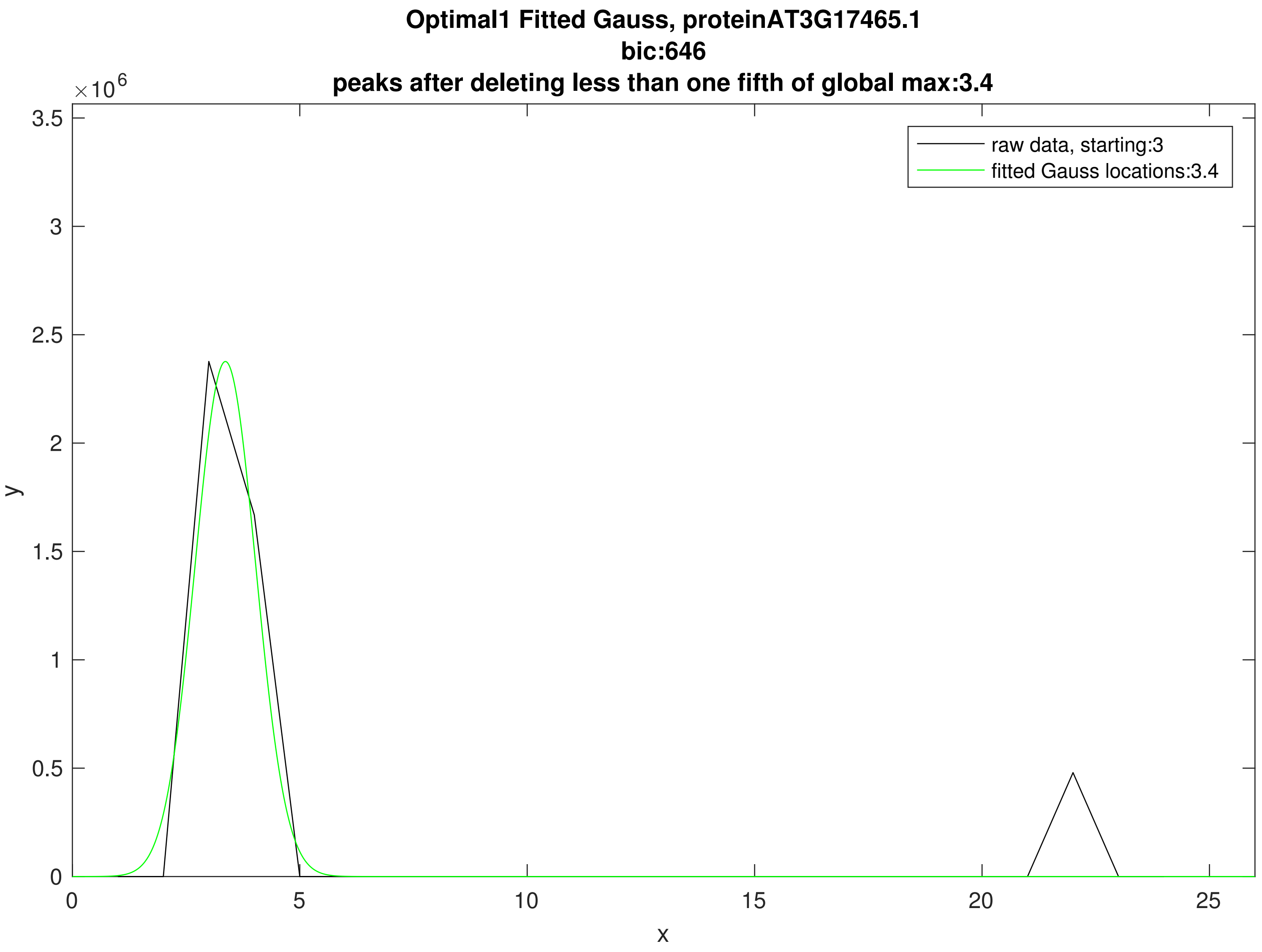
Optimal1 Fitted Gauss, proteinAT3G17465.1
bic:646
peaks after deleting less than one fifth of global max:3.4
×10^6
3.5
3
2.5
2
1.5
1
0.5
0
y
0
5
10
15
20
25
x
raw data, starting:3
fitted Gauss locations:3.4

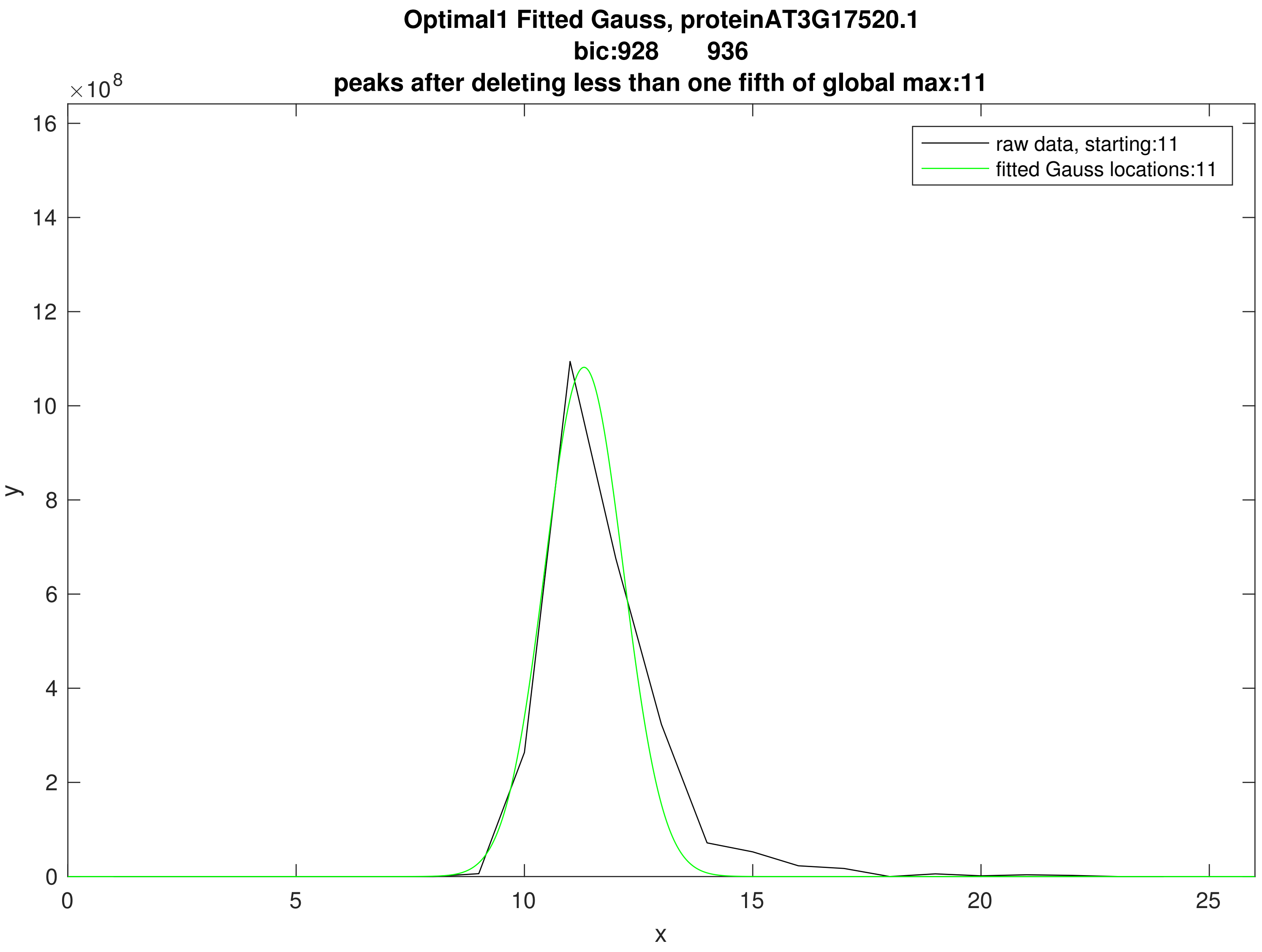
Optimal1 Fitted Gauss, proteinAT3G17520.1
bic:928 936
peaks after deleting less than one fifth of global max:11
×10^8
16
14
12
10
8
6
4
2
0
y
raw data, starting:11
fitted Gauss locations:11
0
5
10
15
20
25
x

Optimal1 Fitted Gauss, proteinAT3G17770.1
bic:724
peaks after deleting less than one fifth of global max:11

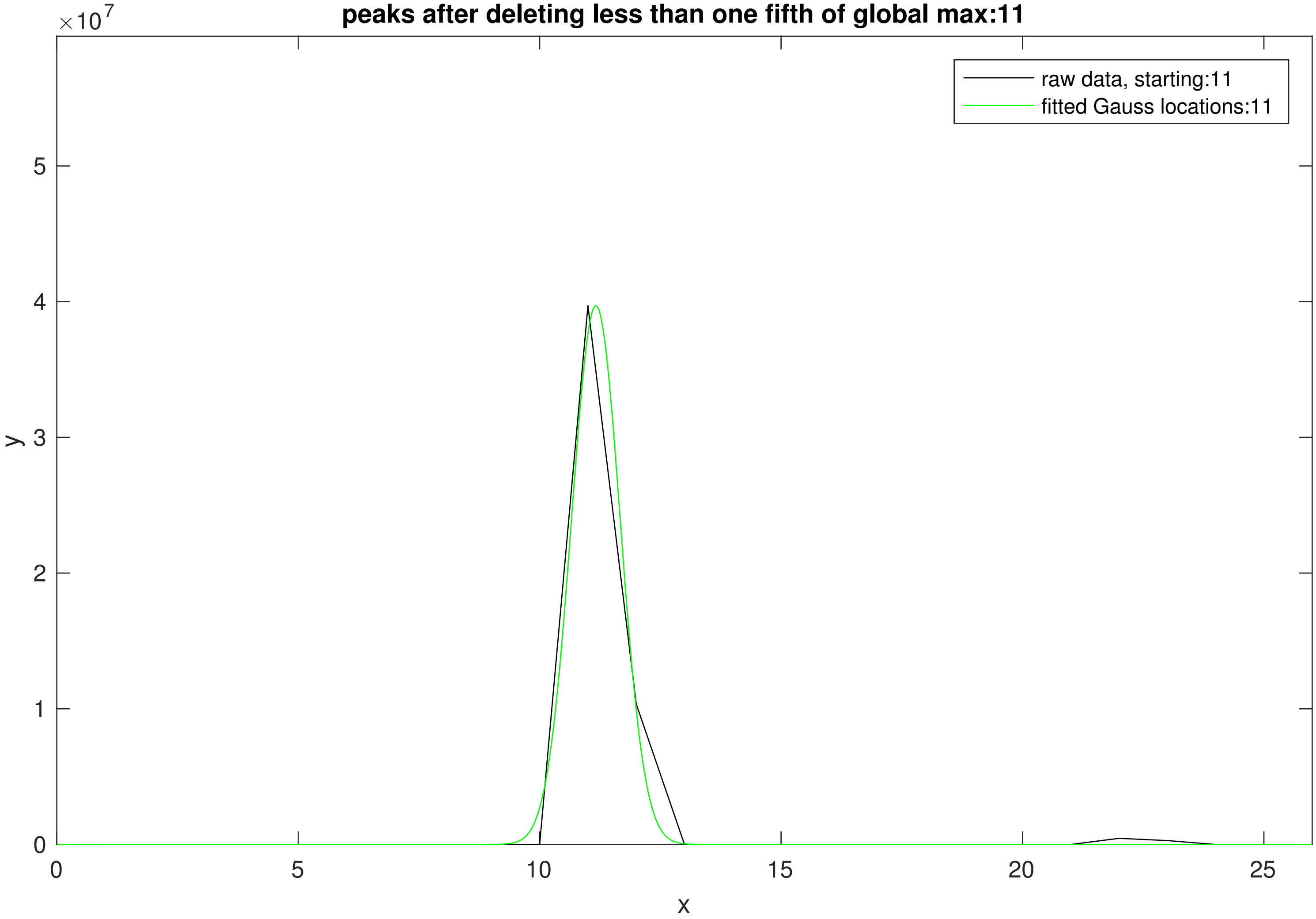

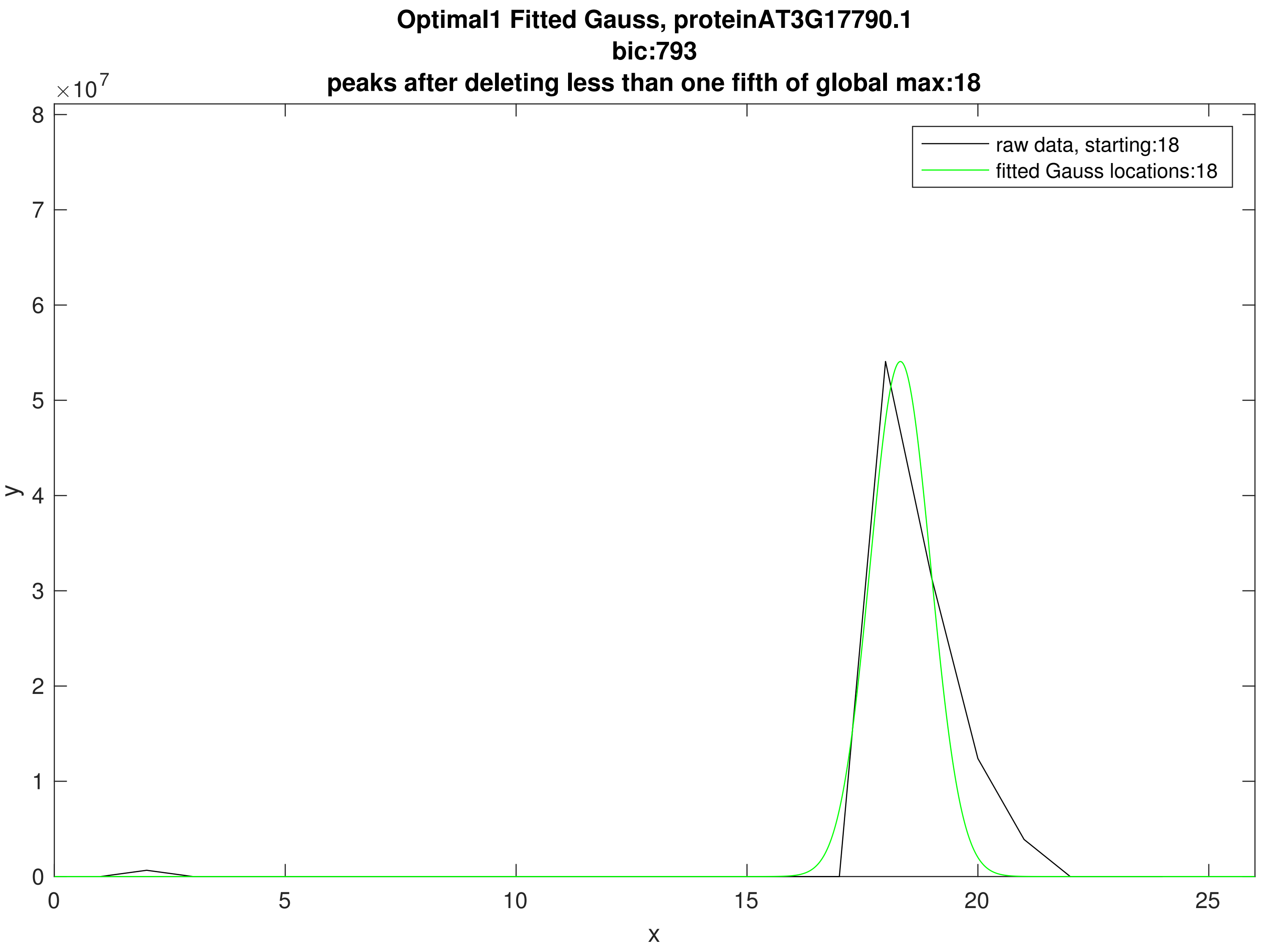

Optimal1 Fitted Gauss, proteinAT3G17790.1
bic:793
peaks after deleting less than one fifth of global max:18
×10^7
raw data, starting:18
fitted Gauss locations:18
y
x
0
1
2
3
4
5
6
7
8
0
5
10
15
20
25

Optimal1 Fitted Gauss, proteinAT3G17810.1
bic:990 1e+03 1.01e+03
peaks after deleting less than one fifth of global max:14

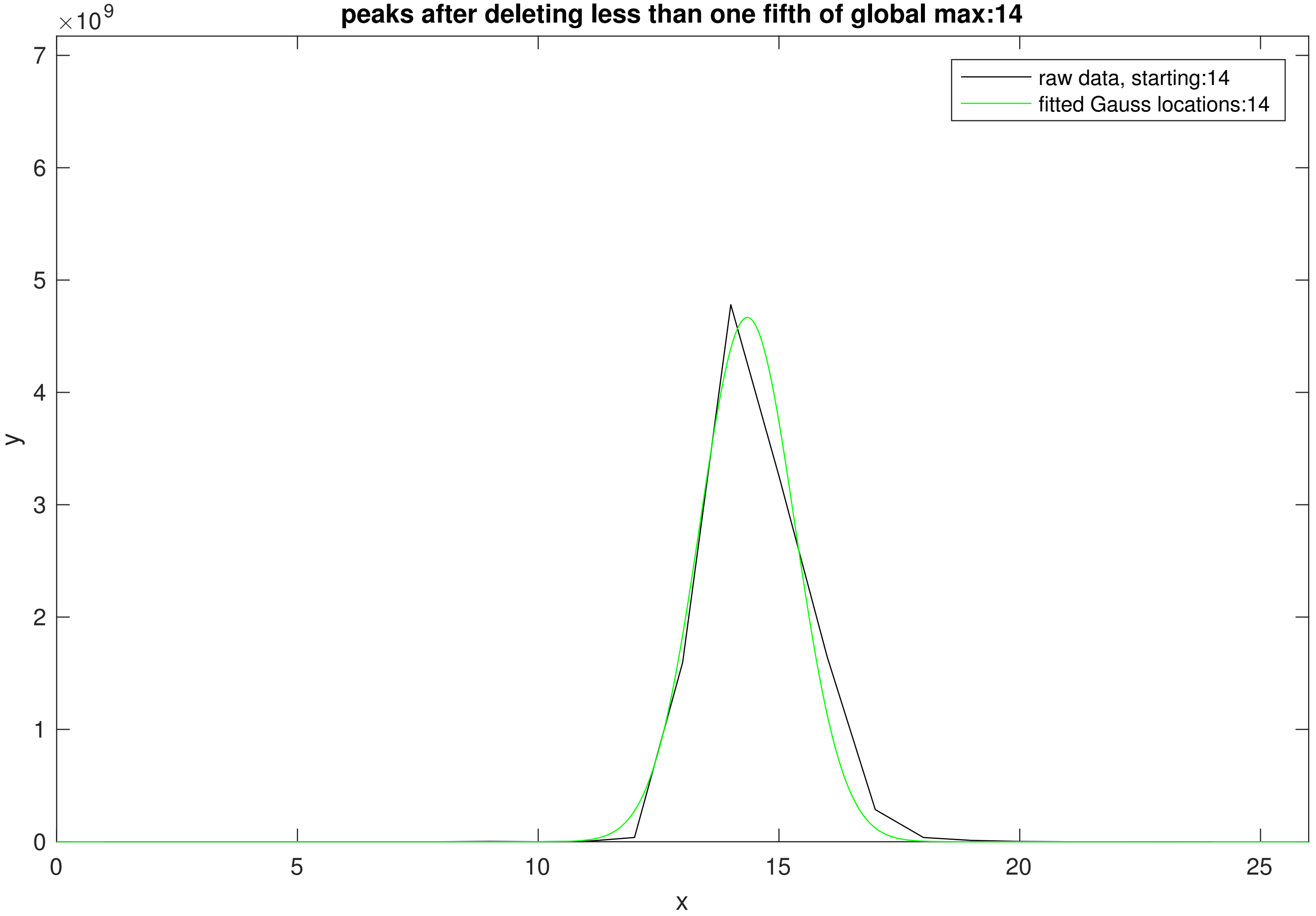

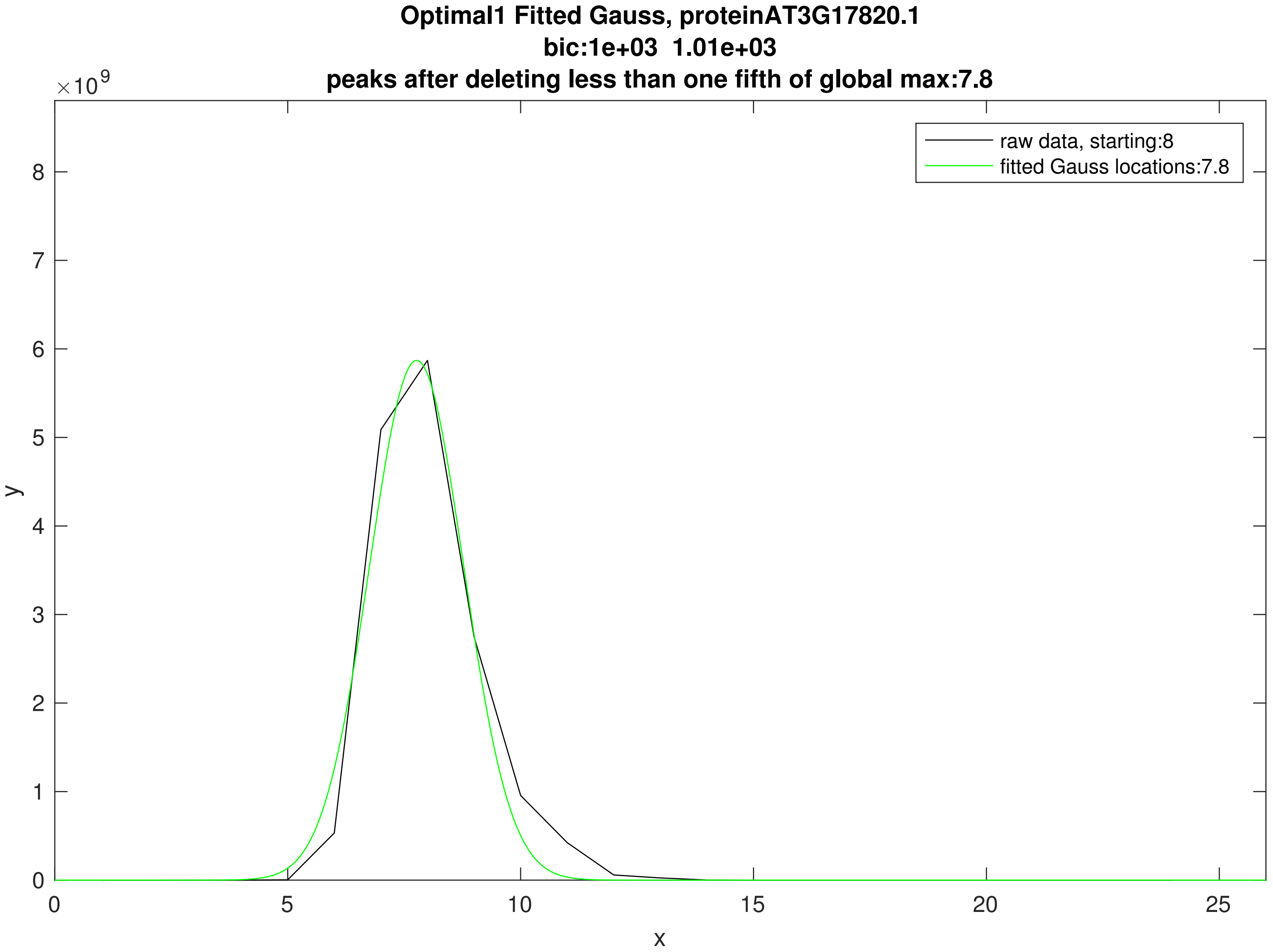
Optimal1 Fitted Gauss, proteinAT3G17820.1
bic:1e+03 1.01e+03
peaks after deleting less than one fifth of global max:7.8
×10^9
raw data, starting:8
fitted Gauss locations:7.8
y
x
0
1
2
3
4
5
6
7
8
0
5
10
15
20
25

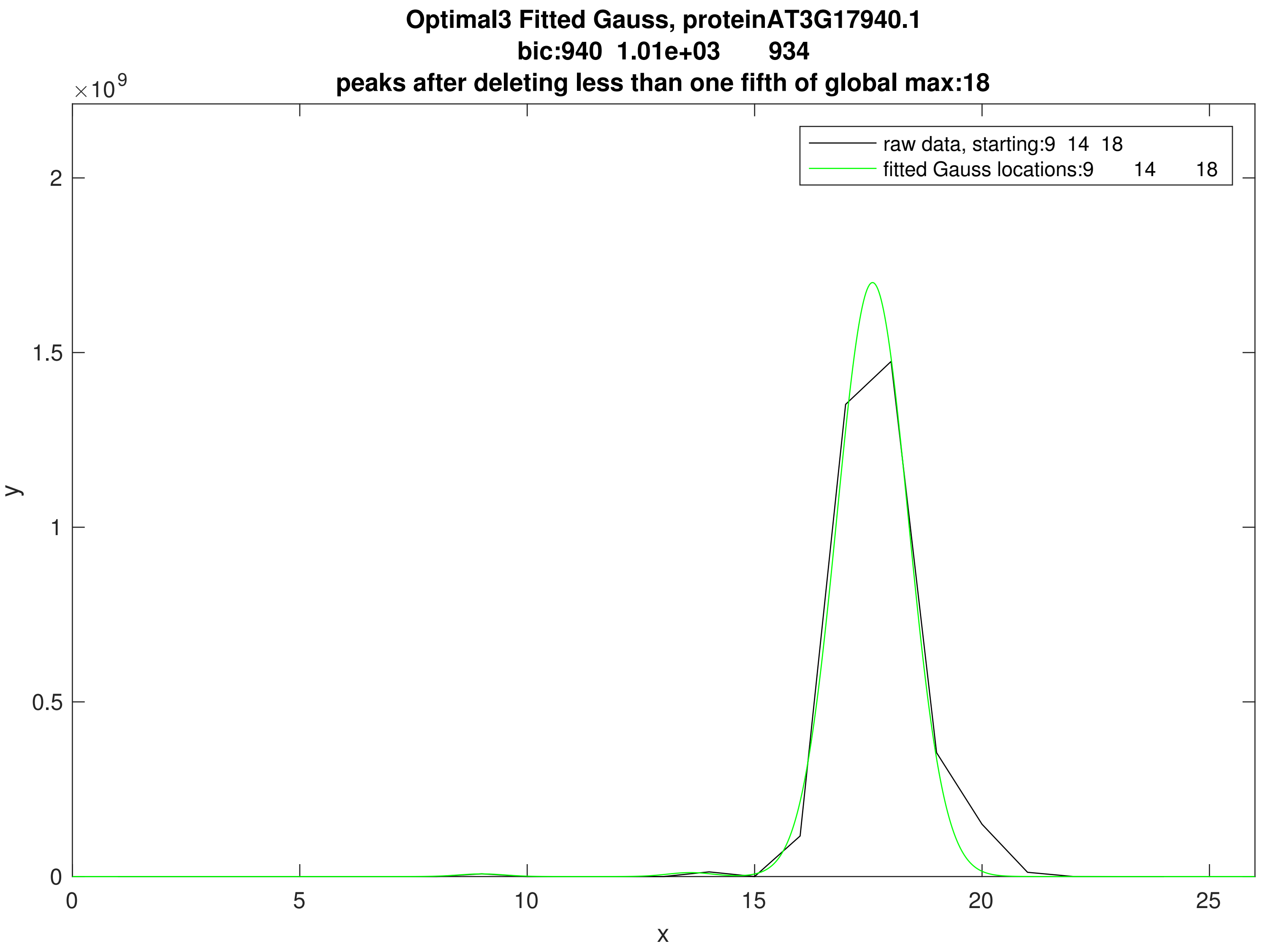
Optimal3 Fitted Gauss, proteinAT3G17940.1
bic:940 1.01e+03 934
peaks after deleting less than one fifth of global max:18
×10^9
raw data, starting:9 14 18
fitted Gauss locations:9 14 18
y
x
0
0.5
1
1.5
2
5
10
15
20
25

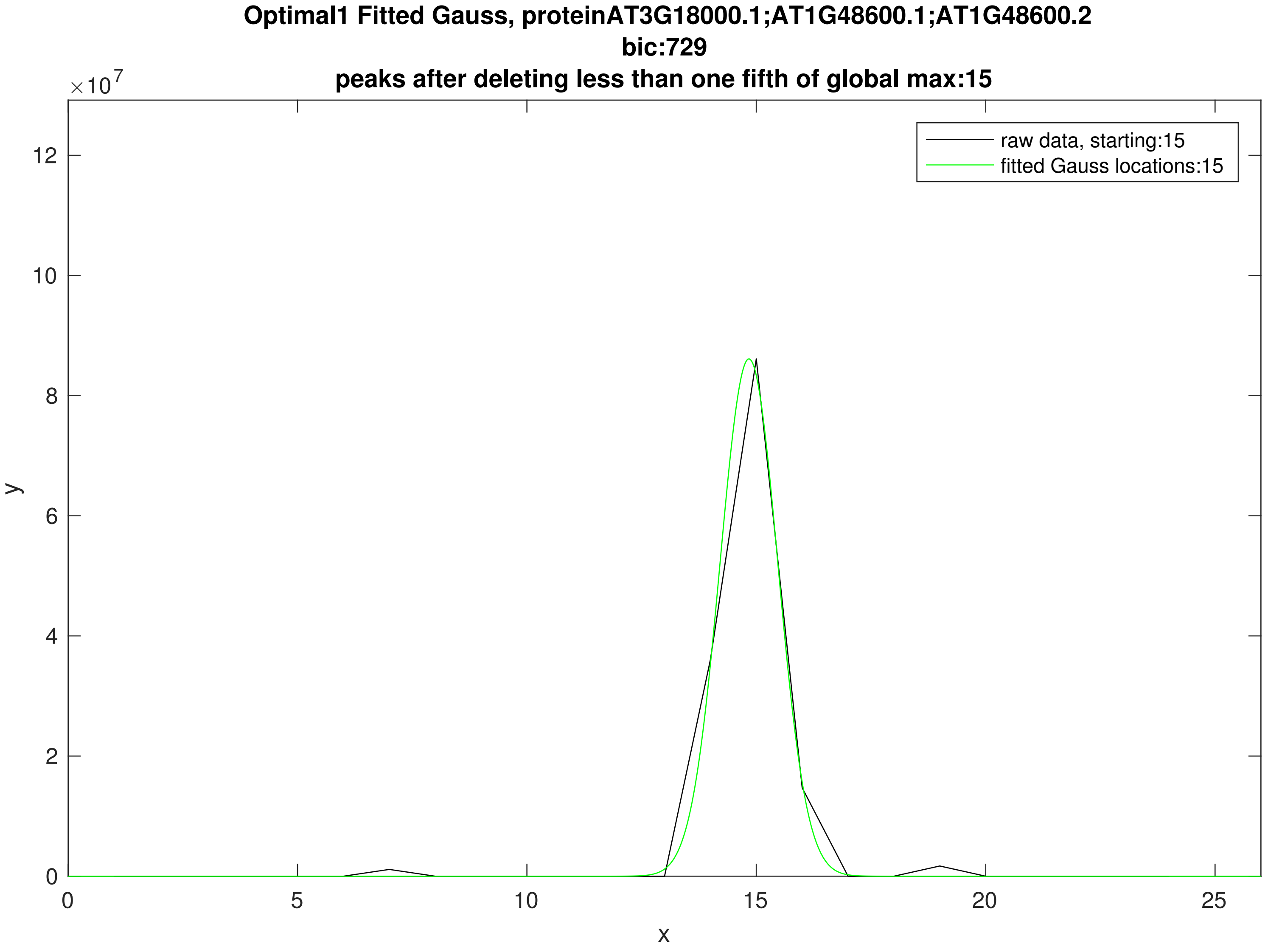

Optimal1 Fitted Gauss, proteinAT3G18000.1;AT1G48600.1;AT1G48600.2
bic:729
peaks after deleting less than one fifth of global max:15
×10^7
raw data, starting:15
fitted Gauss locations:15
12
10
8
6
4
2
0
y
0
5
10
15
20
25
x

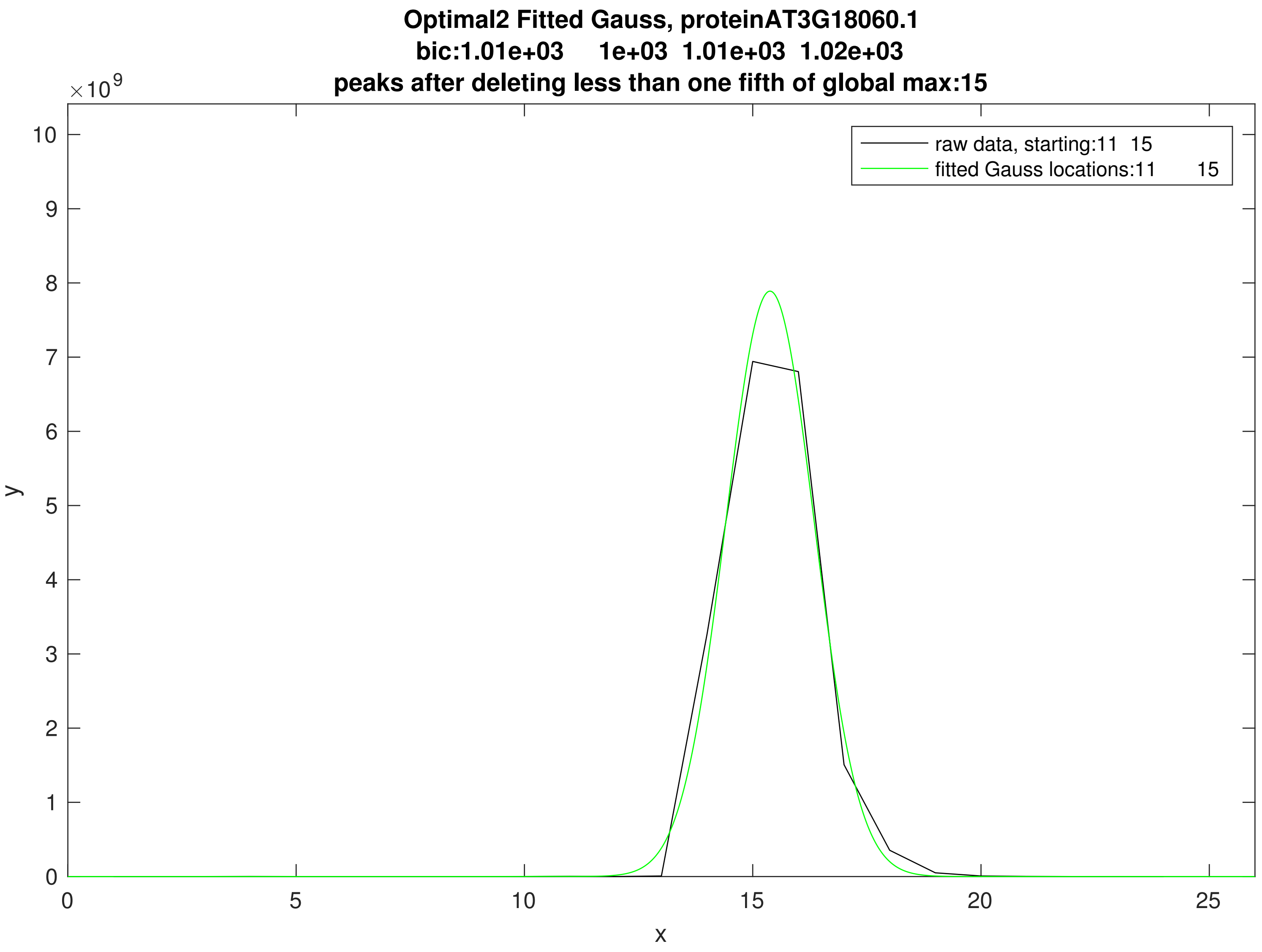

Optimal2 Fitted Gauss, proteinAT3G18060.1
bic:1.01e+03 1e+03 1.01e+03 1.02e+03
peaks after deleting less than one fifth of global max:15
×10^9
raw data, starting:11 15
fitted Gauss locations:11 15
y
x
0
1
2
3
4
5
6
7
8
9
10
0
5
10
15
20
25

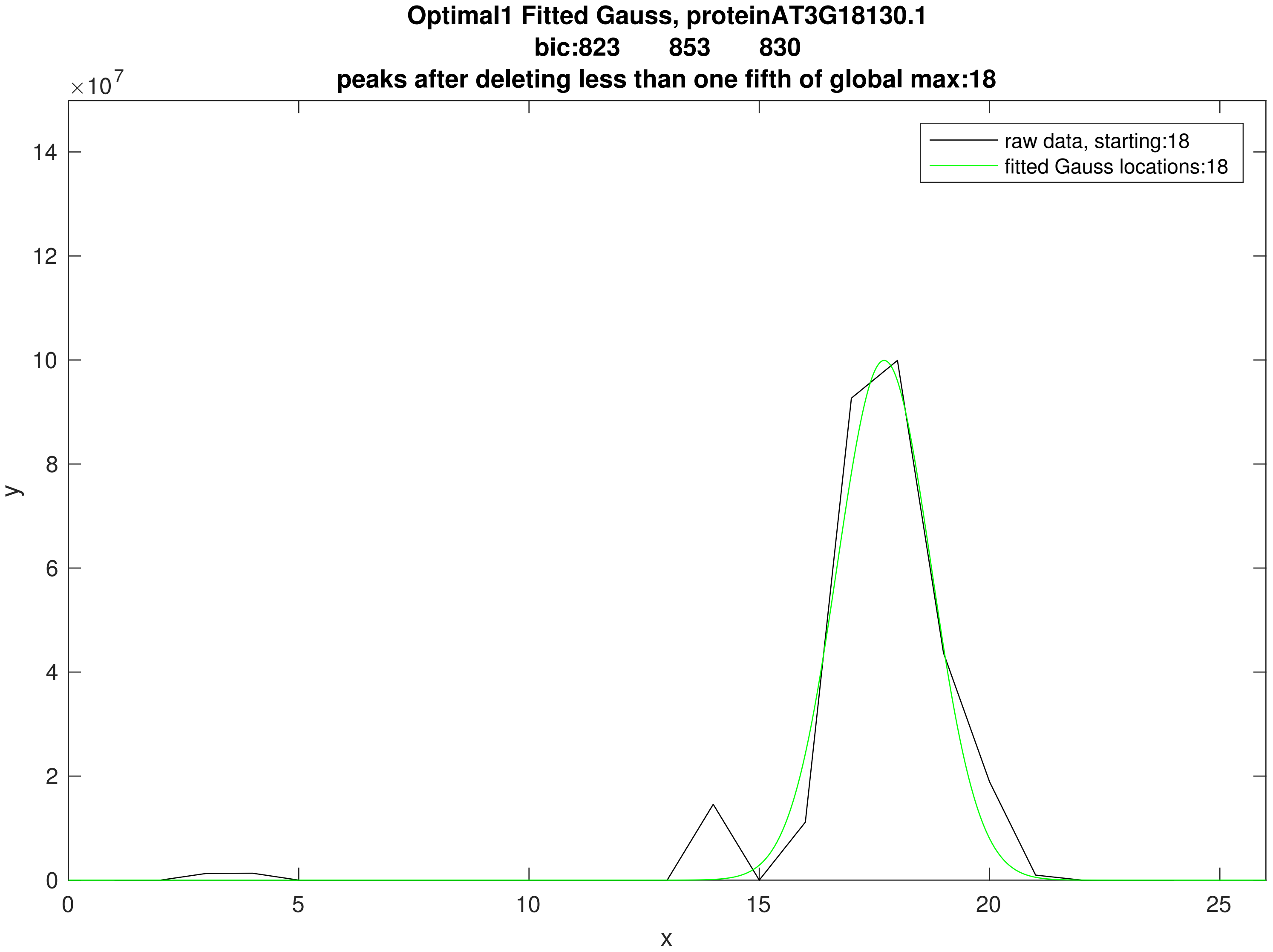

Optimal1 Fitted Gauss, proteinAT3G18130.1
bic:823 853 830
peaks after deleting less than one fifth of global max:18
×10^7
14
12
10
8
6
4
2
0
y
0
5
10
15
20
25
x
raw data, starting:18
fitted Gauss locations:18

**Optimal2 Fitted Gauss, proteinAT3G18140.1;AT3G18140.2**
**bic:717 697**
**peaks after deleting less than one fifth of global max:19**

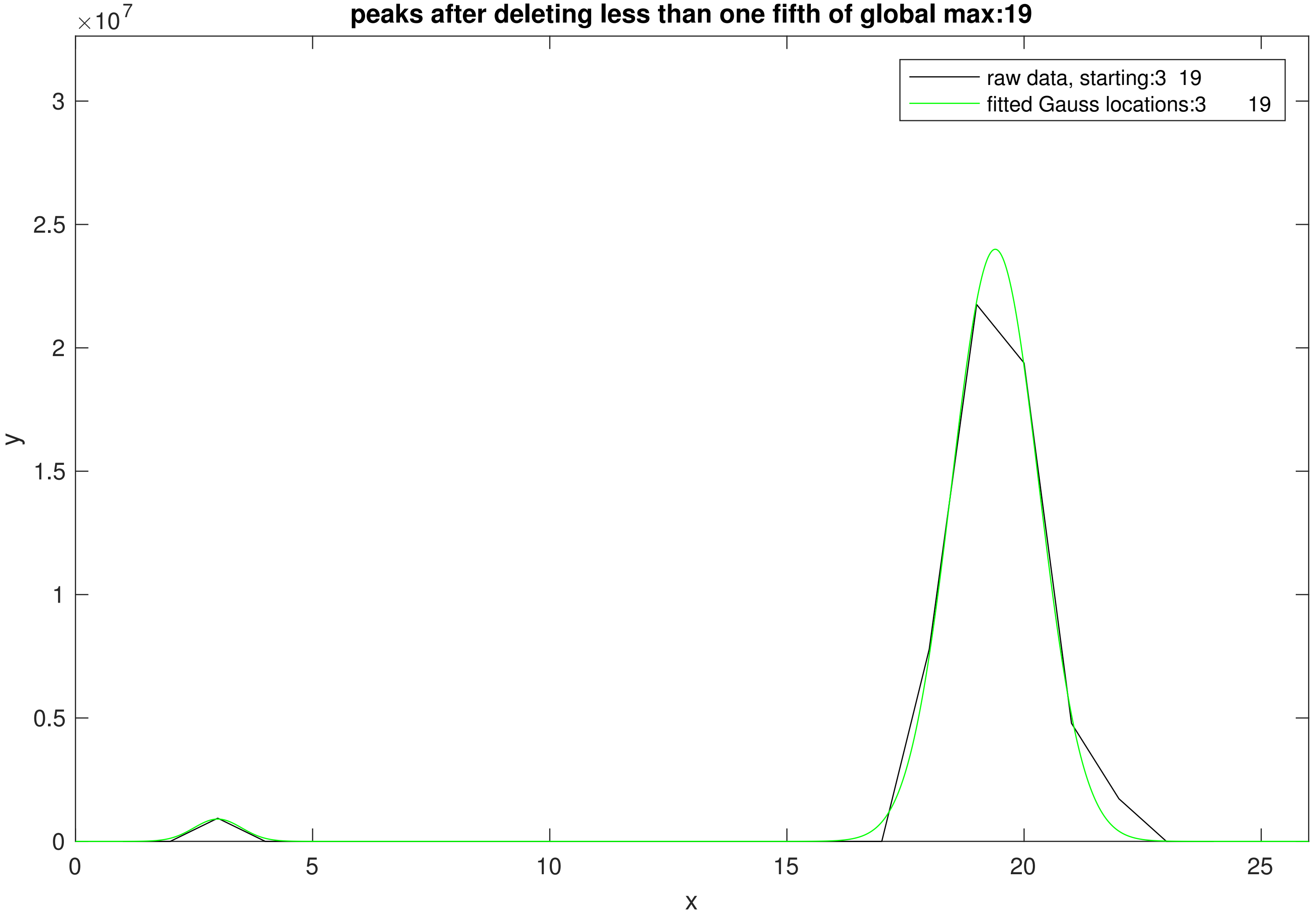

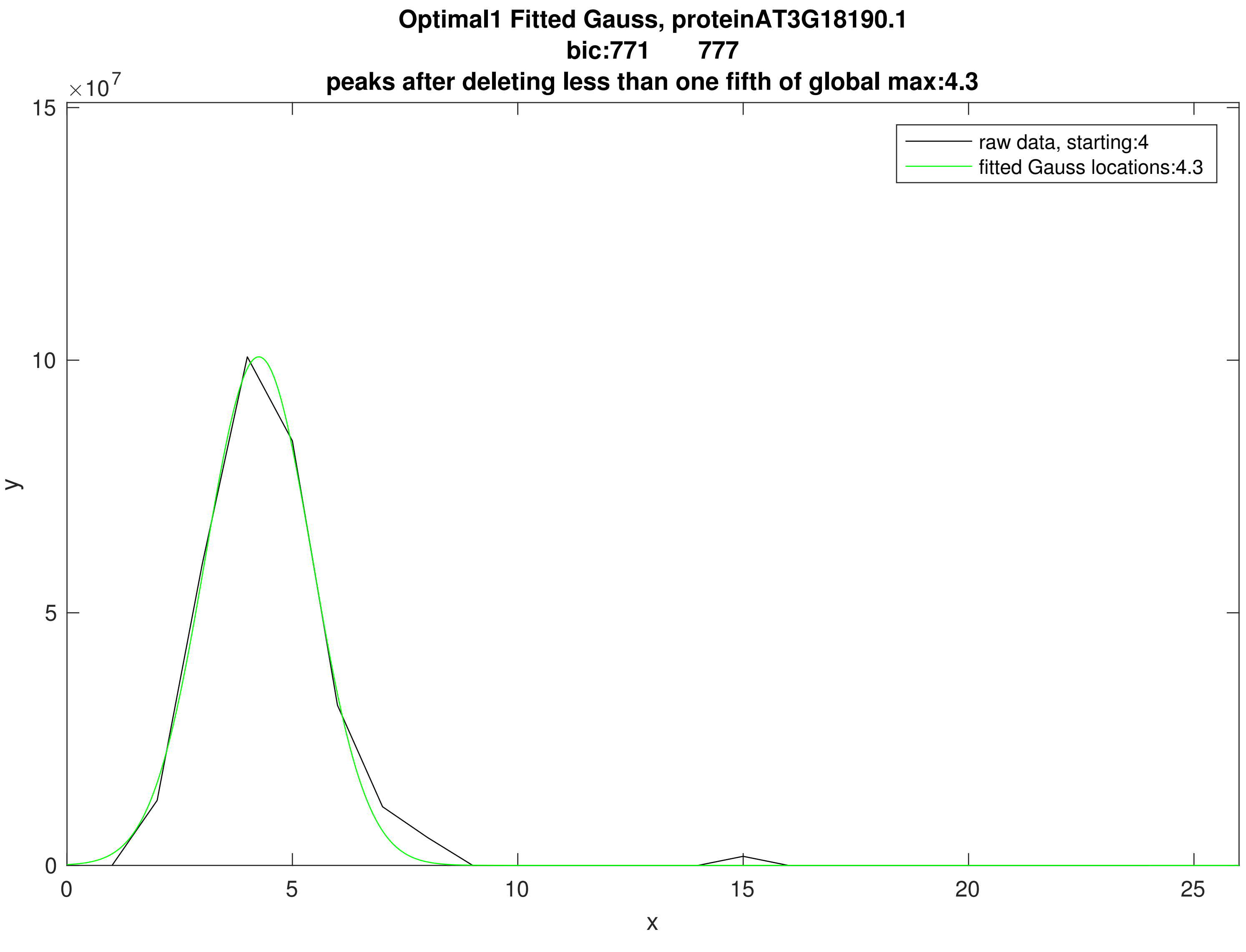
Optimal1 Fitted Gauss, proteinAT3G18190.1
bic:771 777
peaks after deleting less than one fifth of global max:4.3
×10^7
15
10
5
0
y
0
5
10
15
20
25
x
raw data, starting:4
fitted Gauss locations:4.3

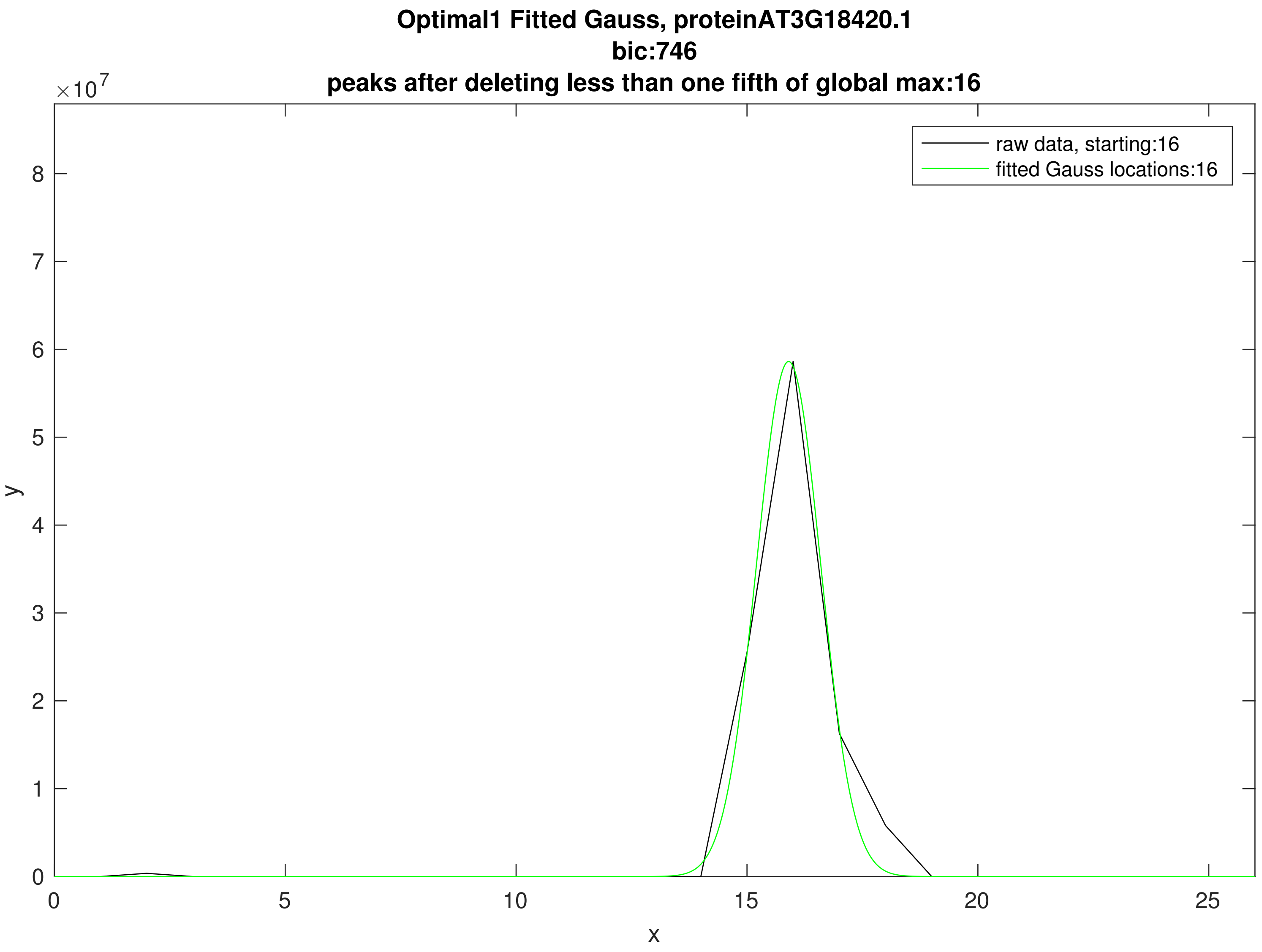

Optimal1 Fitted Gauss, proteinAT3G18420.1
bic:746
peaks after deleting less than one fifth of global max:16
×10^7
raw data, starting:16
fitted Gauss locations:16
y
x
0
1
2
3
4
5
6
7
8
0
5
10
15
20
25

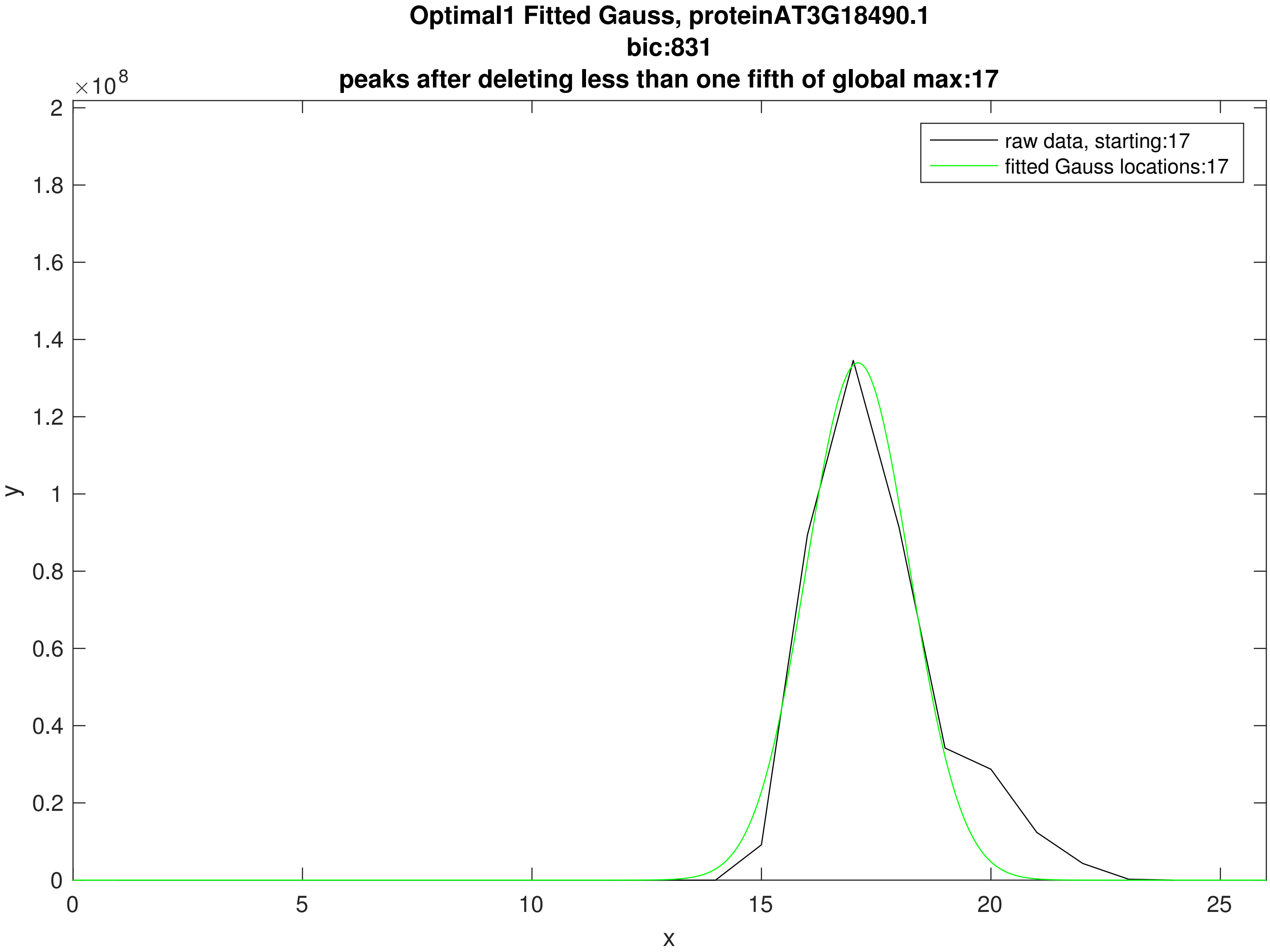
Optimal1 Fitted Gauss, proteinAT3G18490.1
bic:831
peaks after deleting less than one fifth of global max:17
×10^8
2
1.8
1.6
1.4
1.2
1
0.8
0.6
0.4
0.2
0
y
0
5
10
15
20
25
x
raw data, starting:17
fitted Gauss locations:17

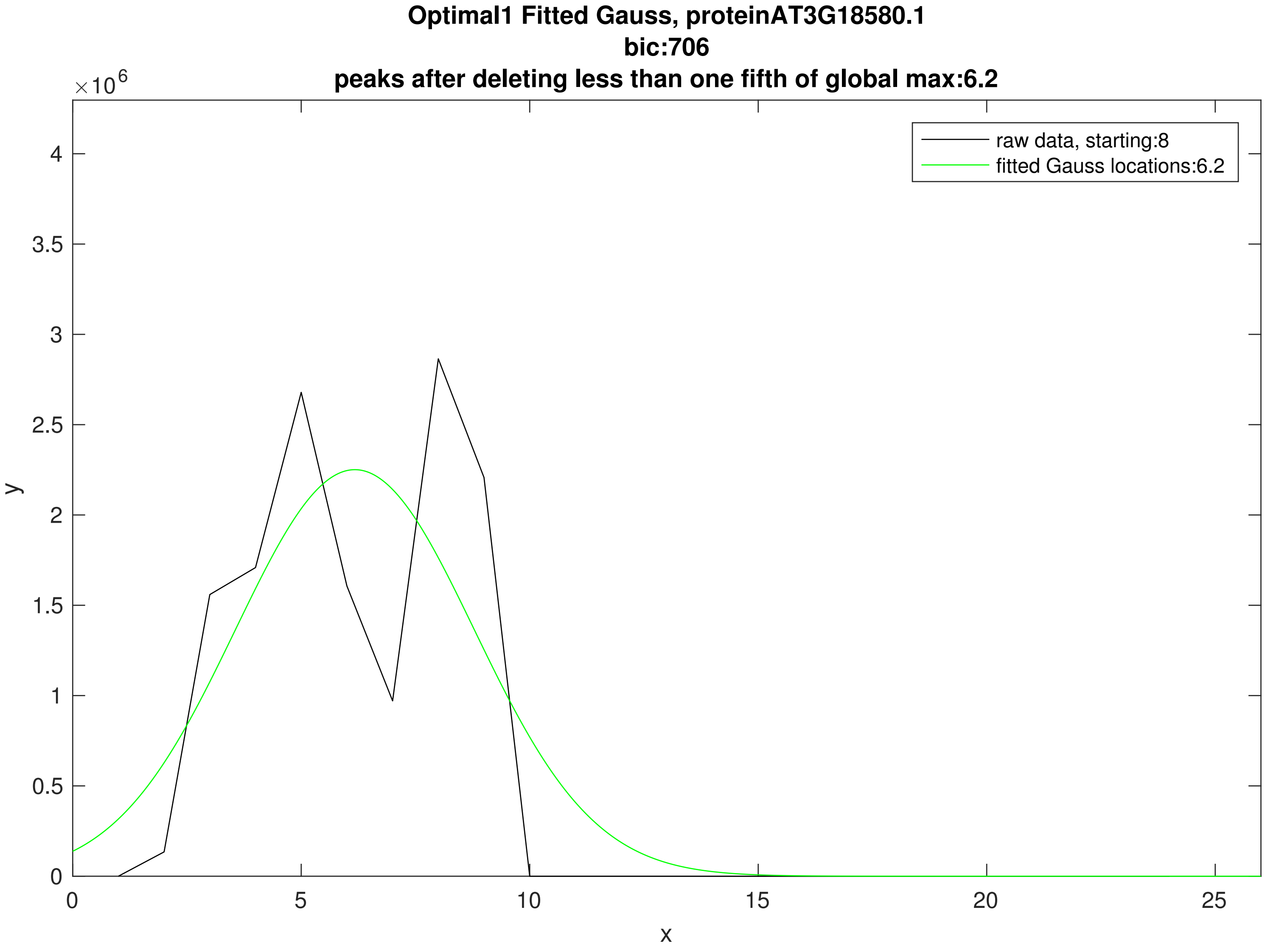

Optimal1 Fitted Gauss, proteinAT3G18580.1
bic:706
peaks after deleting less than one fifth of global max:6.2
×10^6
raw data, starting:8
fitted Gauss locations:6.2
y
x
0
0.5
1
1.5
2
2.5
3
3.5
4
5
10
15
20
25

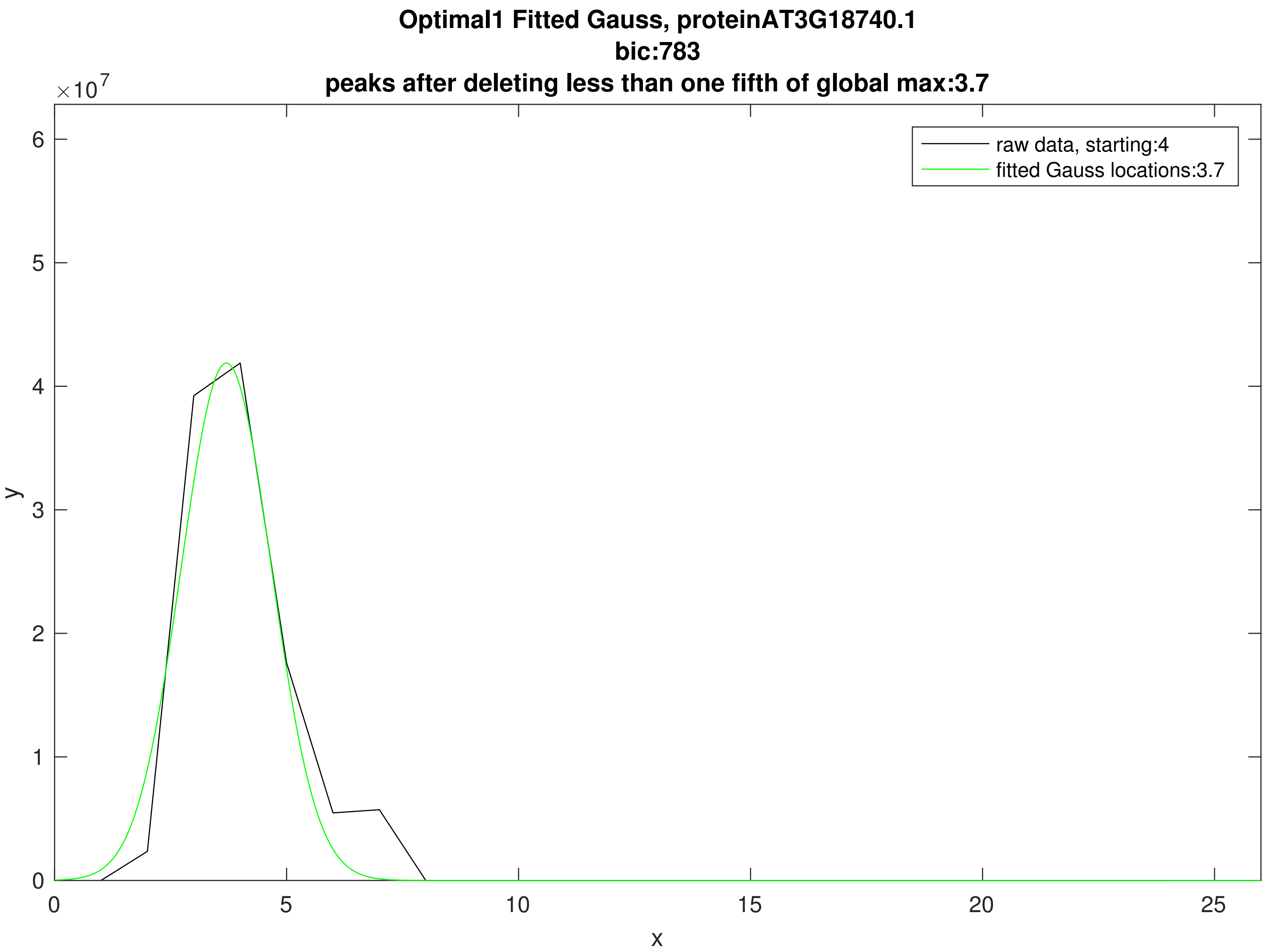
Optimal1 Fitted Gauss, proteinAT3G18740.1
bic:783
peaks after deleting less than one fifth of global max:3.7
×10^7
6
5
4
3
2
1
0
y
0
5
10
15
20
25
x
raw data, starting:4
fitted Gauss locations:3.7

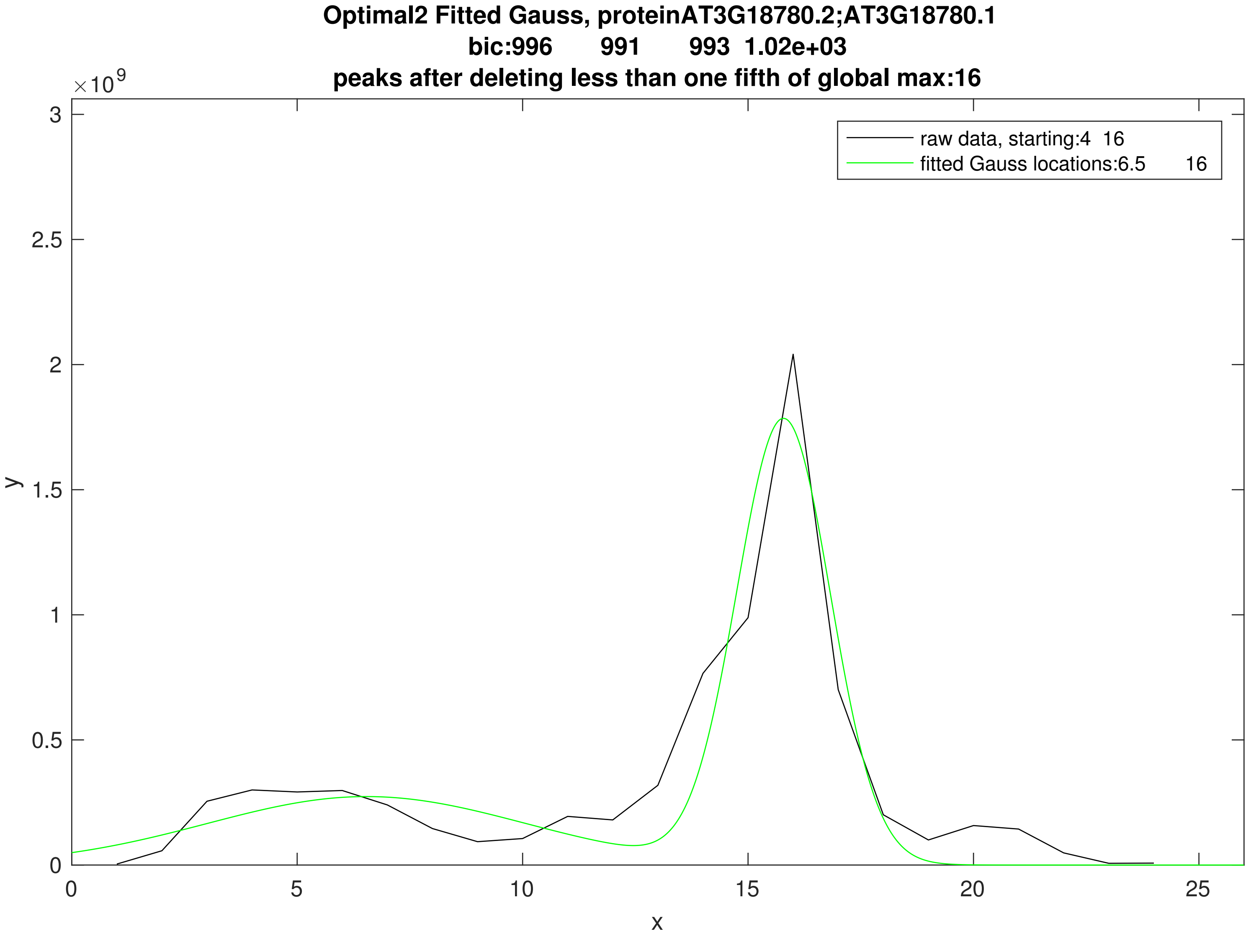
Optimal2 Fitted Gauss, proteinAT3G18780.2;AT3G18780.1
bic:996 991 993 1.02e+03
peaks after deleting less than one fifth of global max:16
×10^9
3
2.5
2
1.5
1
0.5
0
y
0
5
10
15
20
25
x
raw data, starting:4 16
fitted Gauss locations:6.5 16

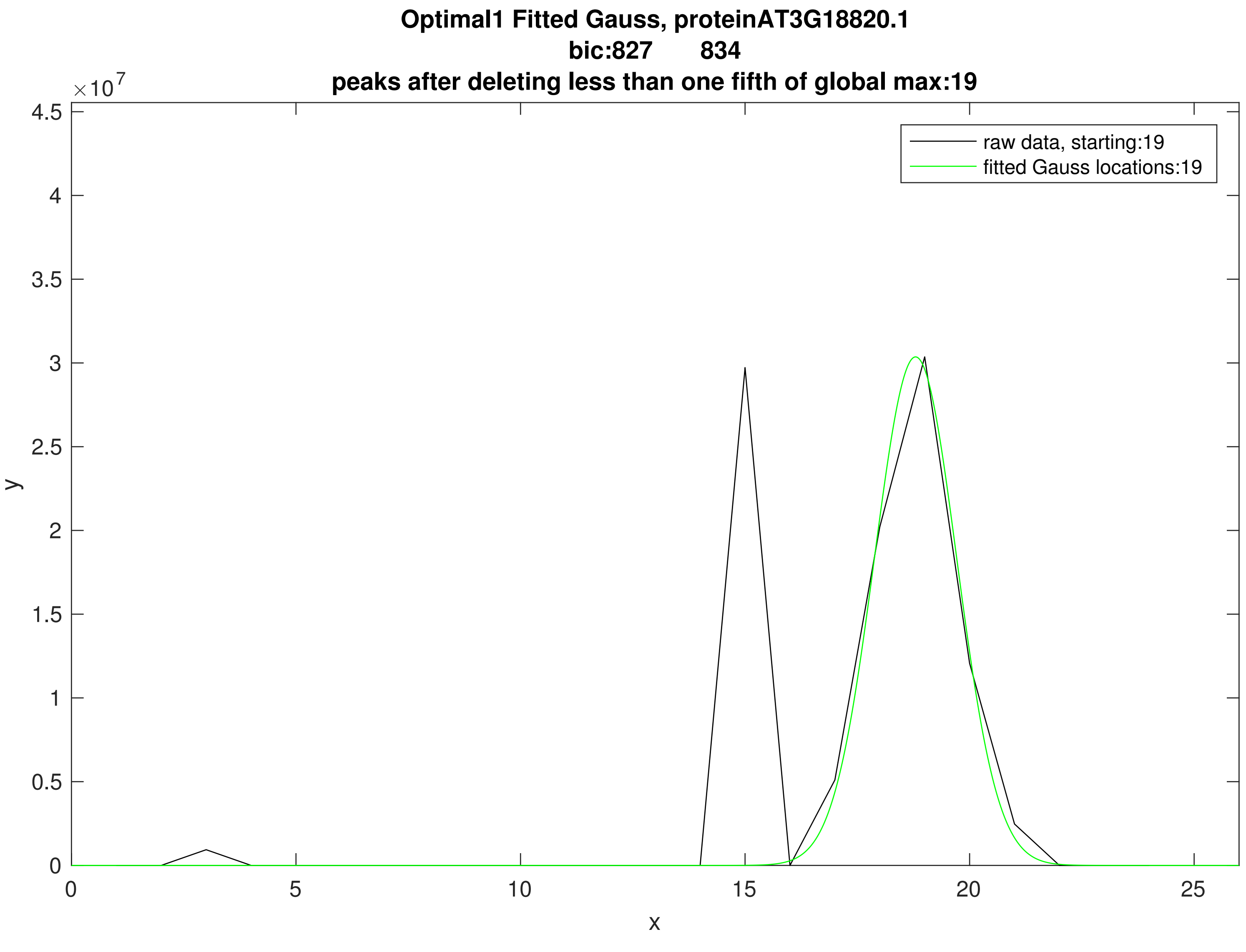
Optimal1 Fitted Gauss, proteinAT3G18820.1
bic:827 834
peaks after deleting less than one fifth of global max:19
×10^7
4.5
4
3.5
3
2.5
2
1.5
1
0.5
0
y
0
5
10
15
20
25
x
raw data, starting:19
fitted Gauss locations:19

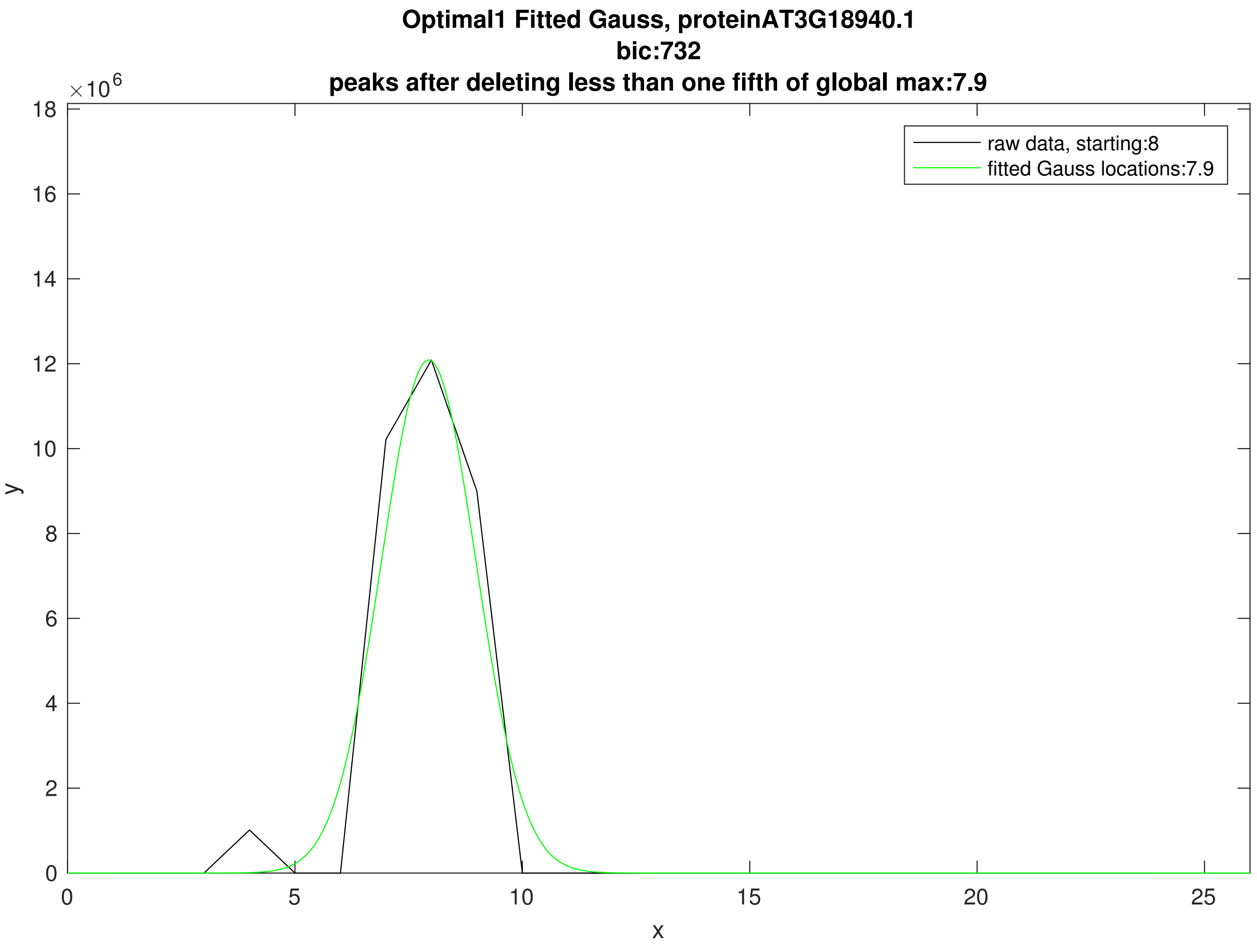
Optimal1 Fitted Gauss, proteinAT3G18940.1
bic:732
peaks after deleting less than one fifth of global max:7.9
×10^6
raw data, starting:8
fitted Gauss locations:7.9
y
x
0
2
4
6
8
10
12
14
16
18
0
5
10
15
20
25

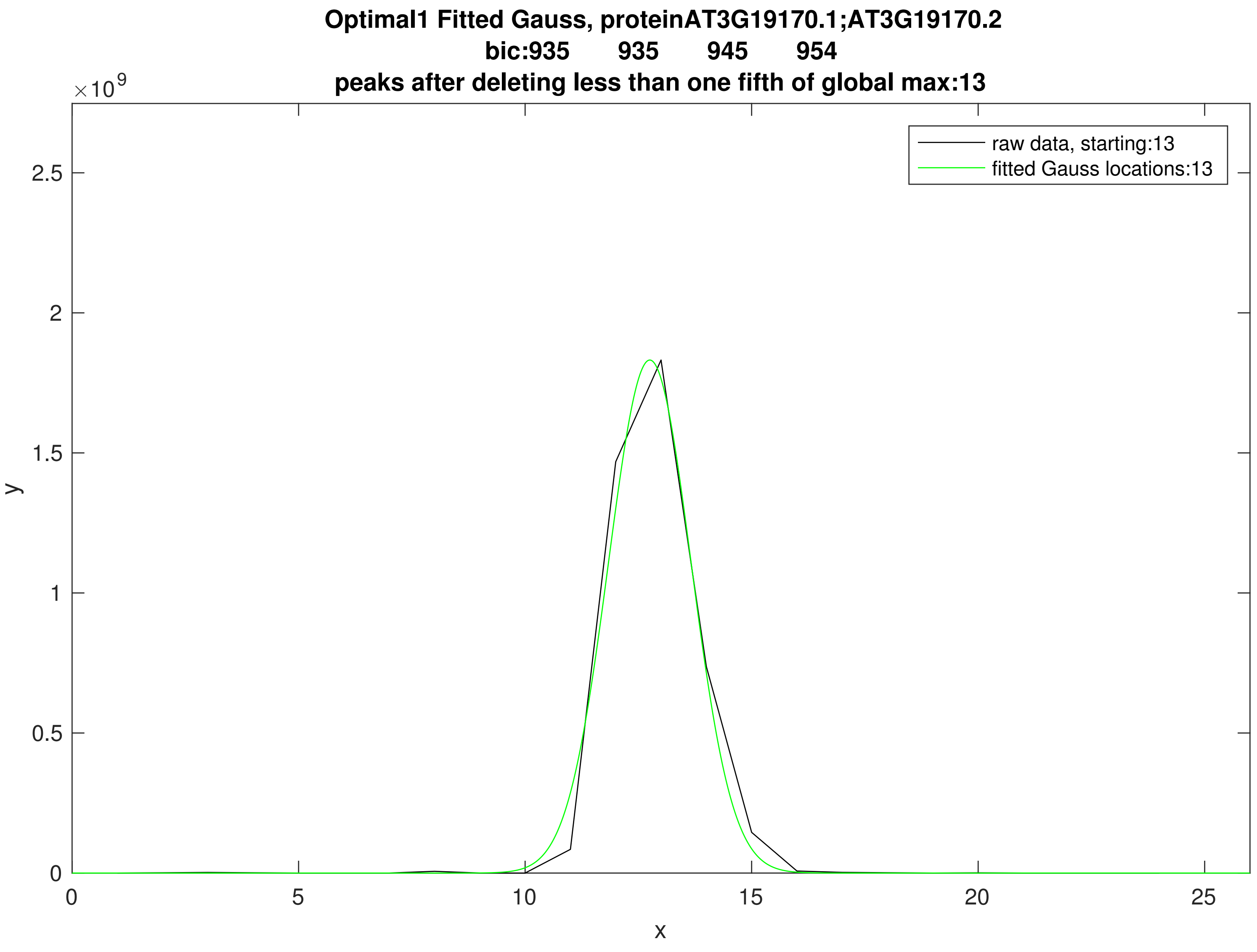

Optimal1 Fitted Gauss, proteinAT3G19170.1;AT3G19170.2
bic:935 935 945 954
peaks after deleting less than one fifth of global max:13
×10^9
2.5
2
1.5
1
0.5
0
y
0
5
10
15
20
25
x
raw data, starting:13
fitted Gauss locations:13

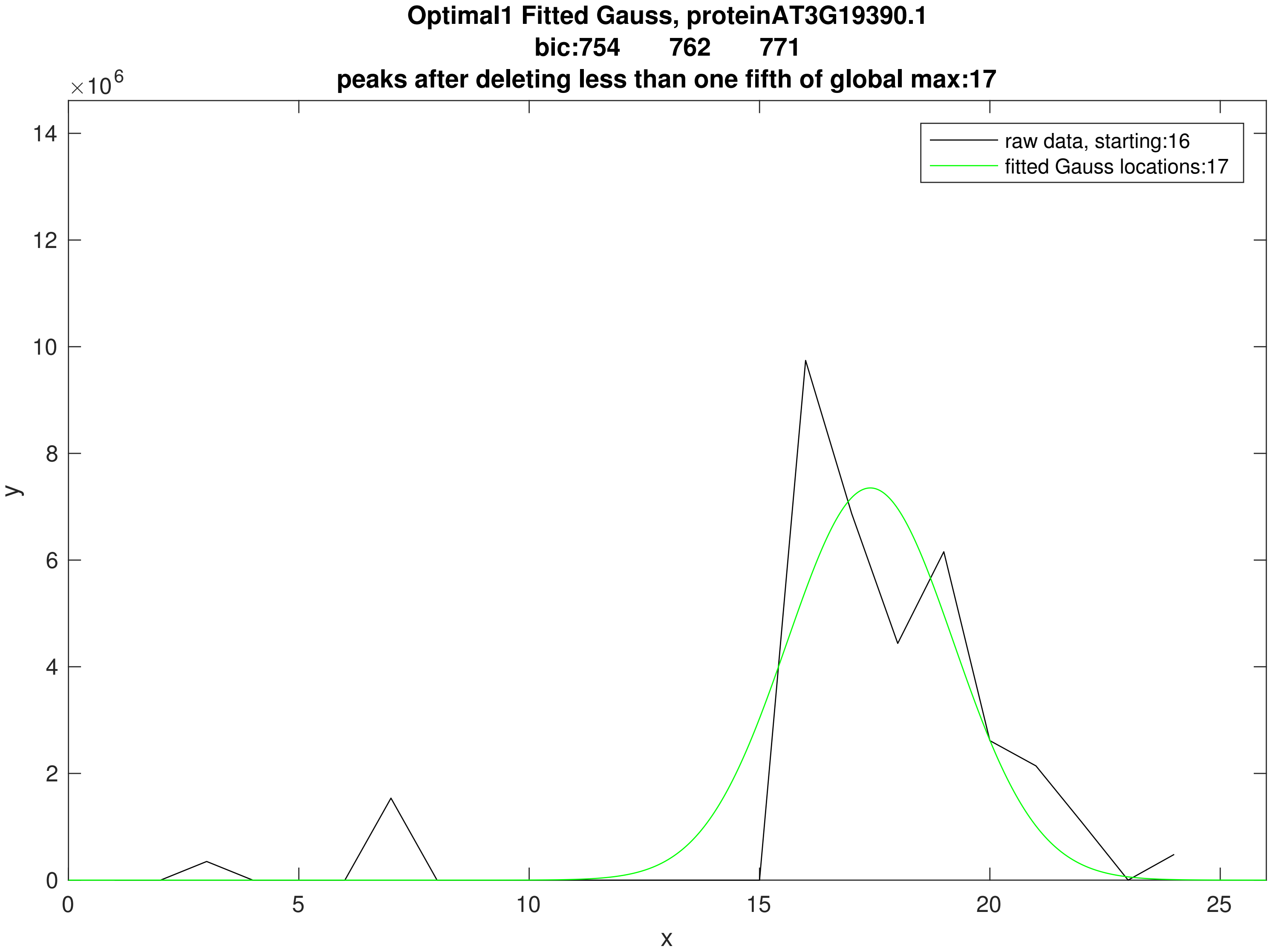
Optimal1 Fitted Gauss, proteinAT3G19390.1
bic:754 762 771
peaks after deleting less than one fifth of global max:17
×10^6
14
12
10
8
6
4
2
0
y
0
5
10
15
20
25
x
raw data, starting:16
fitted Gauss locations:17

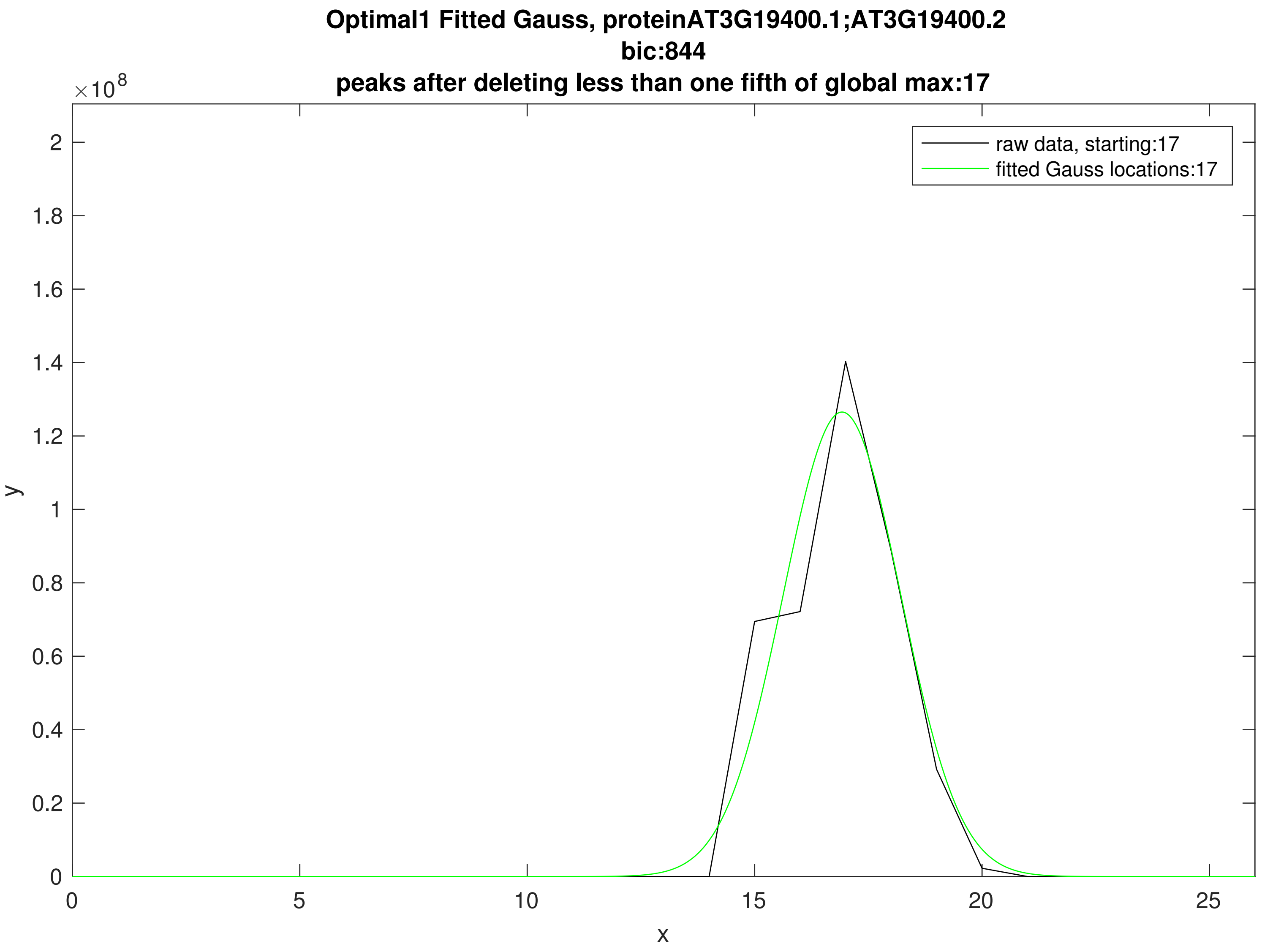
Optimal1 Fitted Gauss, proteinAT3G19400.1;AT3G19400.2
bic:844
peaks after deleting less than one fifth of global max:17
×10^8
raw data, starting:17
fitted Gauss locations:17
y
x
0
0.2
0.4
0.6
0.8
1
1.2
1.4
1.6
1.8
2
0
5
10
15
20
25

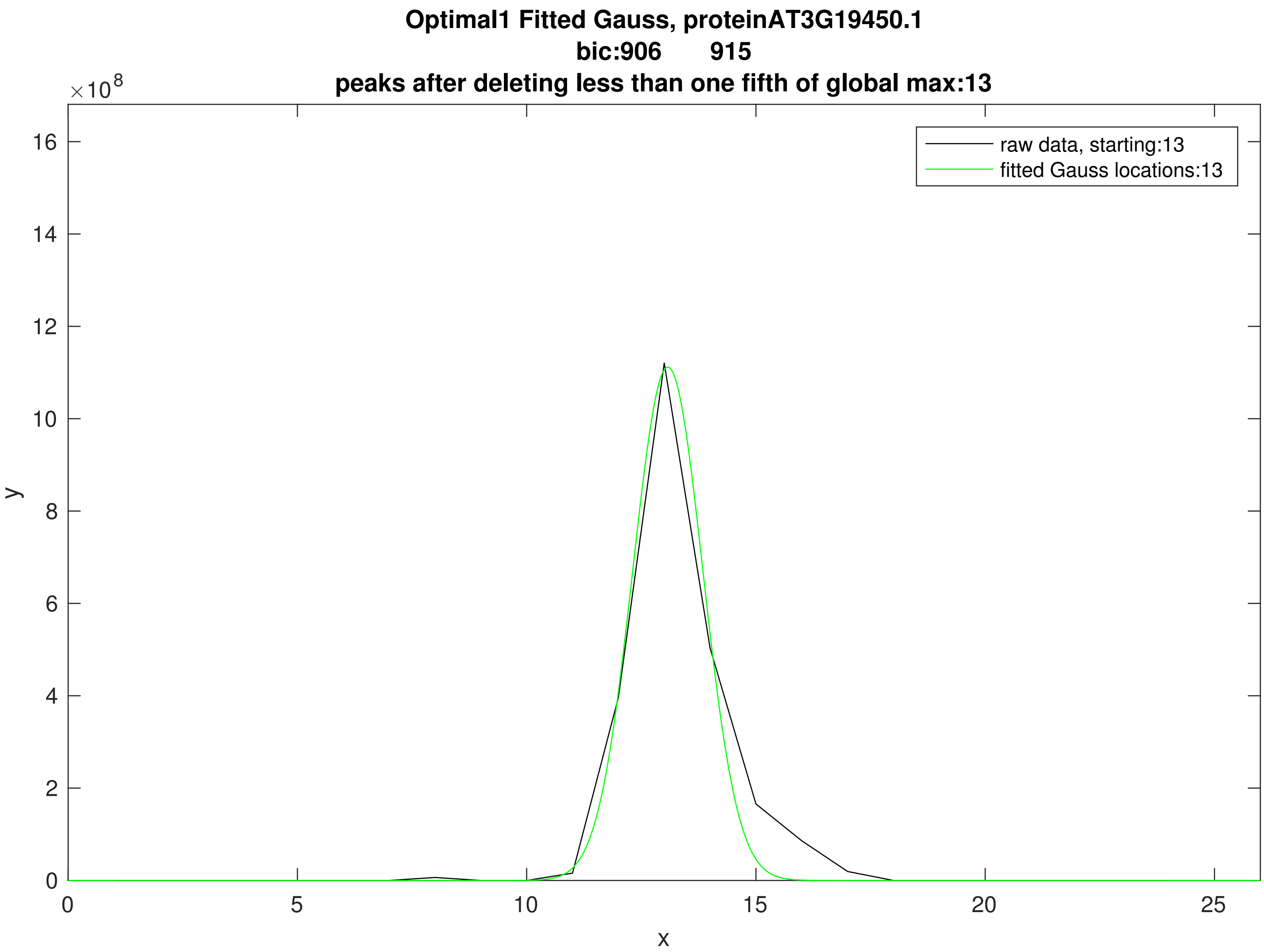

Optimal1 Fitted Gauss, proteinAT3G19450.1
bic:906 915
peaks after deleting less than one fifth of global max:13
×10^8
16
14
12
10
8
6
4
2
0
y
0
5
10
15
20
25
x
raw data, starting:13
fitted Gauss locations:13

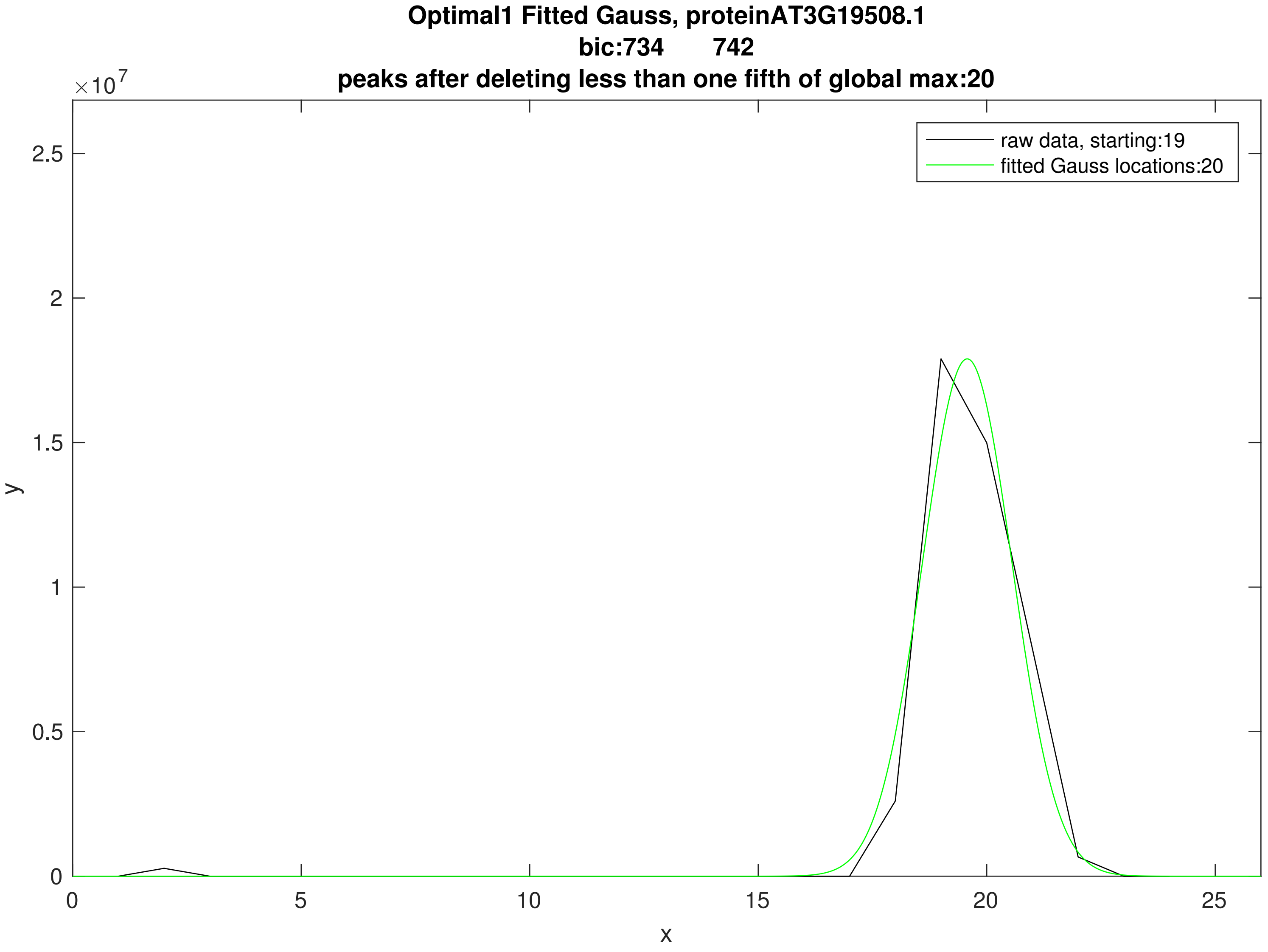

Optimal1 Fitted Gauss, proteinAT3G19508.1
bic:734 742
peaks after deleting less than one fifth of global max:20
×10^7
raw data, starting:19
fitted Gauss locations:20
y
x
0
0.5
1
1.5
2
2.5
0
5
10
15
20
25

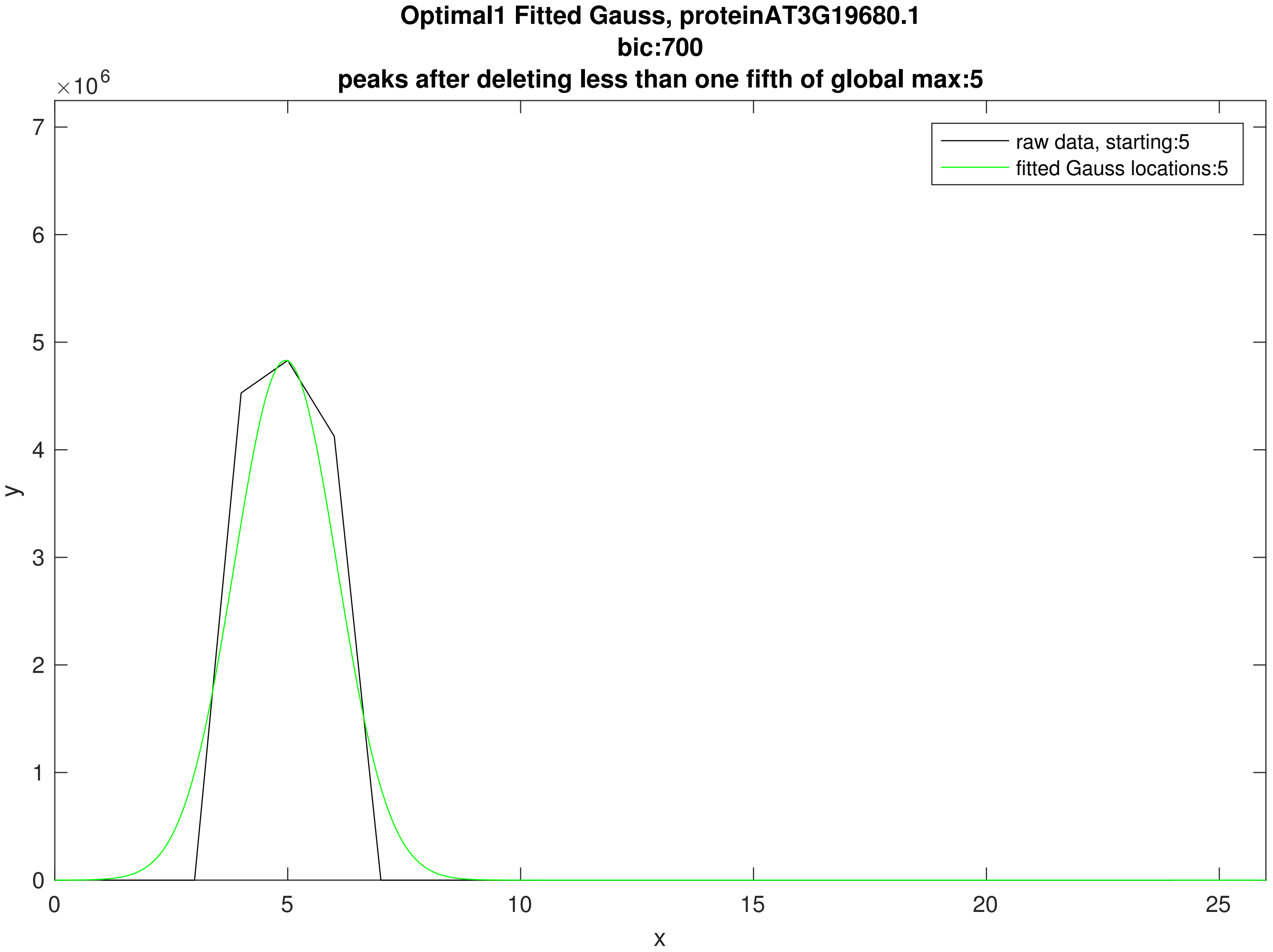
Optimal1 Fitted Gauss, proteinAT3G19680.1
bic:700
peaks after deleting less than one fifth of global max:5
×10^6
raw data, starting:5
fitted Gauss locations:5
y
x
0
1
2
3
4
5
6
7
0
5
10
15
20
25

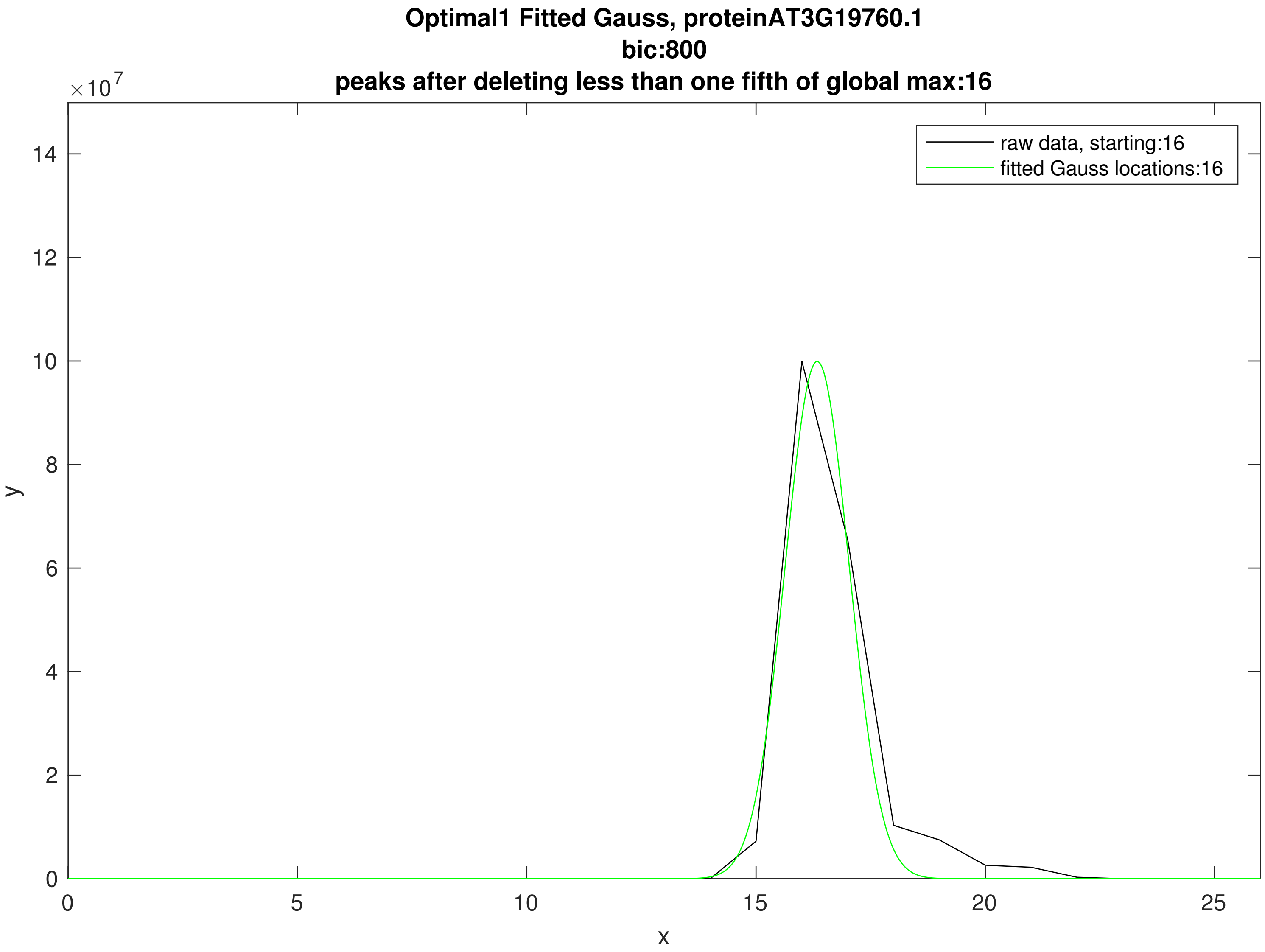

Optimal1 Fitted Gauss, proteinAT3G19760.1
bic:800
peaks after deleting less than one fifth of global max:16
×10^7
raw data, starting:16
fitted Gauss locations:16
y
x
0
2
4
6
8
10
12
14
5
15
20
25

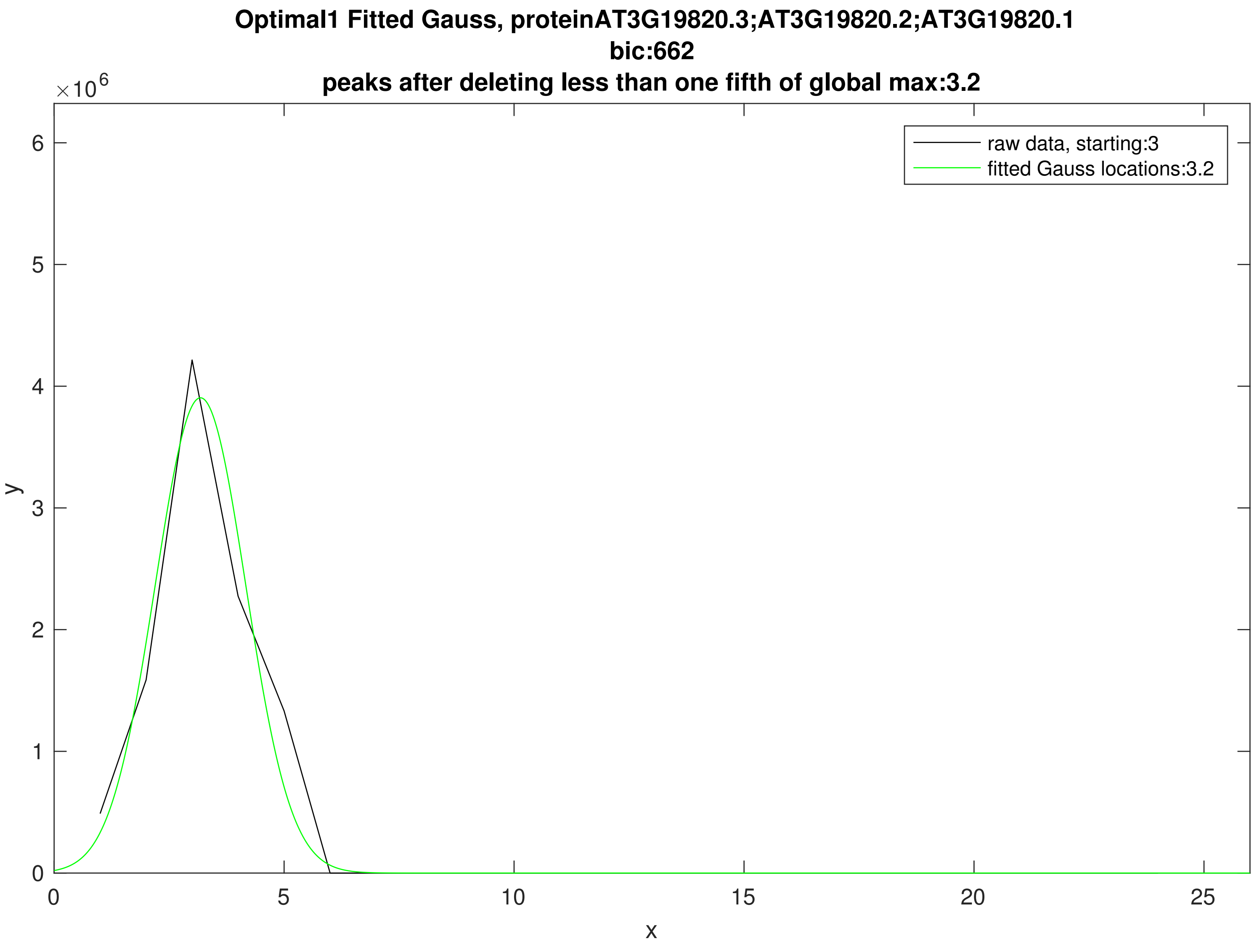
Optimal1 Fitted Gauss, proteinAT3G19820.3;AT3G19820.2;AT3G19820.1
bic:662
peaks after deleting less than one fifth of global max:3.2
×10^6
raw data, starting:3
fitted Gauss locations:3.2
y
x
0
1
2
3
4
5
6
0
5
10
15
20
25

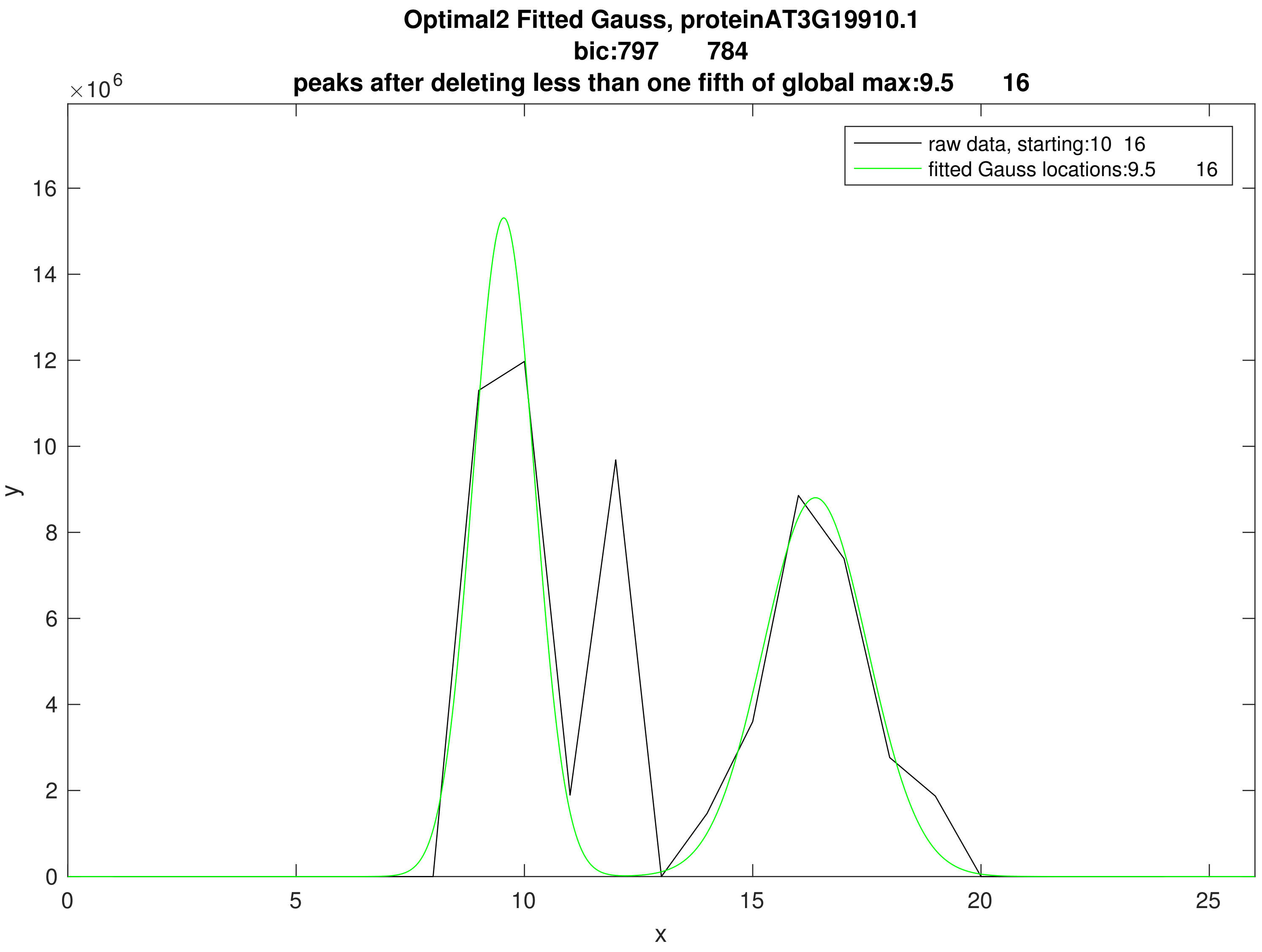
Optimal2 Fitted Gauss, proteinAT3G19910.1
bic:797 784
peaks after deleting less than one fifth of global max:9.5 16
×10^6
raw data, starting:10 16
fitted Gauss locations:9.5 16
y
x
0
2
4
6
8
10
12
14
16
0
5
10
15
20
25

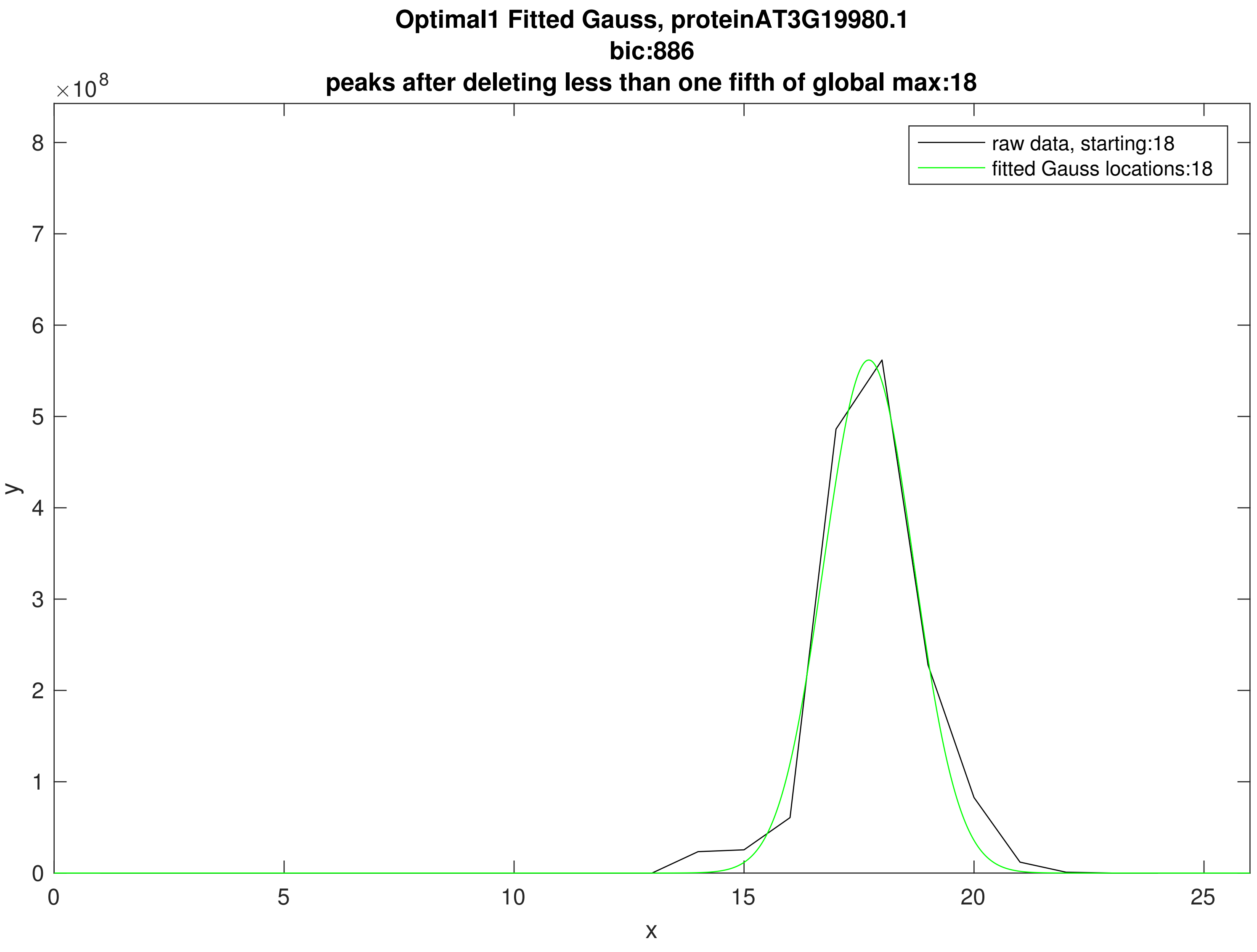
Optimal1 Fitted Gauss, proteinAT3G19980.1
bic:886
peaks after deleting less than one fifth of global max:18
×10^8
raw data, starting:18
fitted Gauss locations:18
y
x
0
1
2
3
4
5
6
7
8
0
5
10
15
20
25

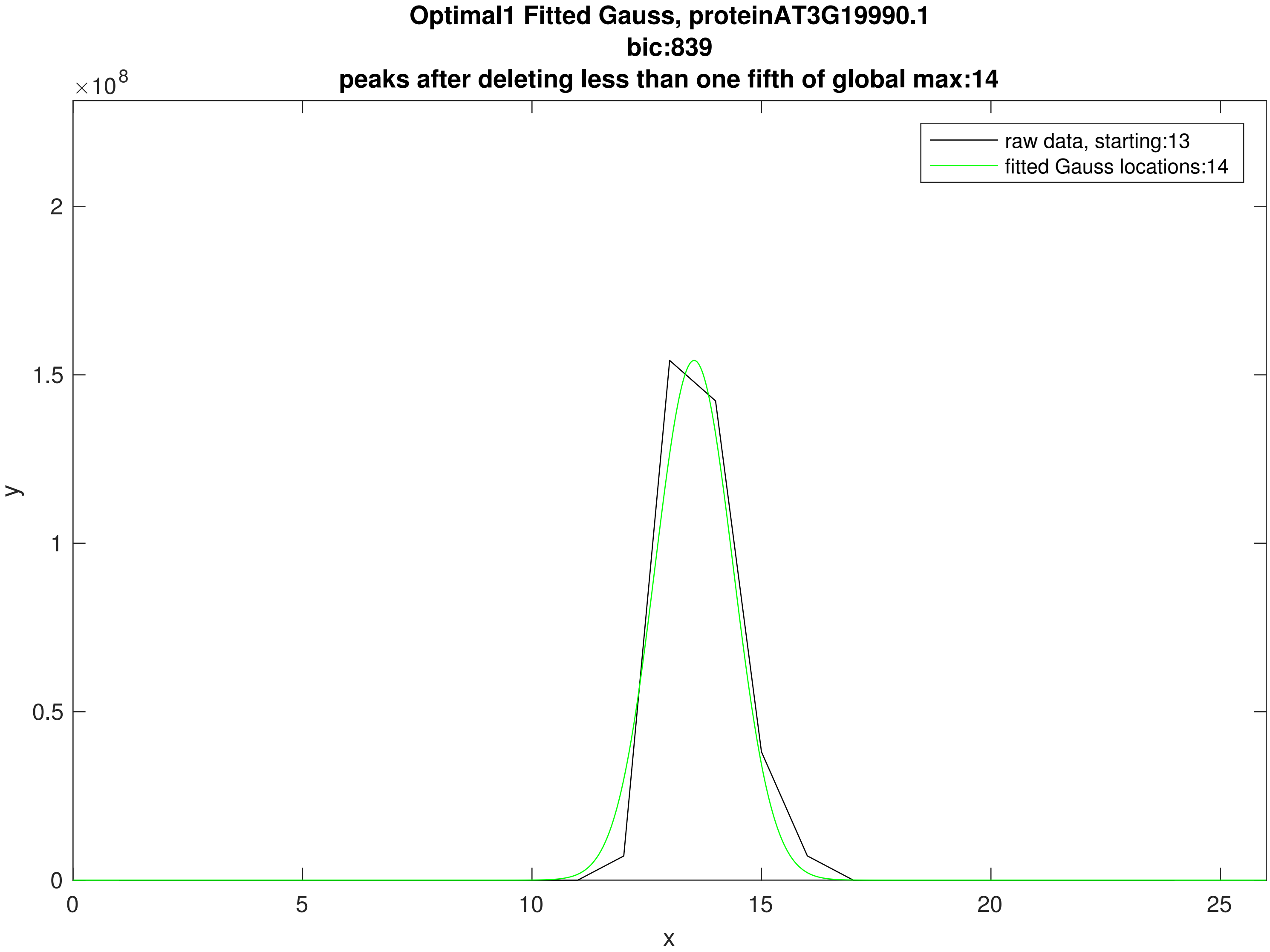

Optimal1 Fitted Gauss, proteinAT3G19990.1
bic:839
peaks after deleting less than one fifth of global max:14
×10^8
raw data, starting:13
fitted Gauss locations:14
y
x
0
0.5
1
1.5
2
0
5
10
15
20
25

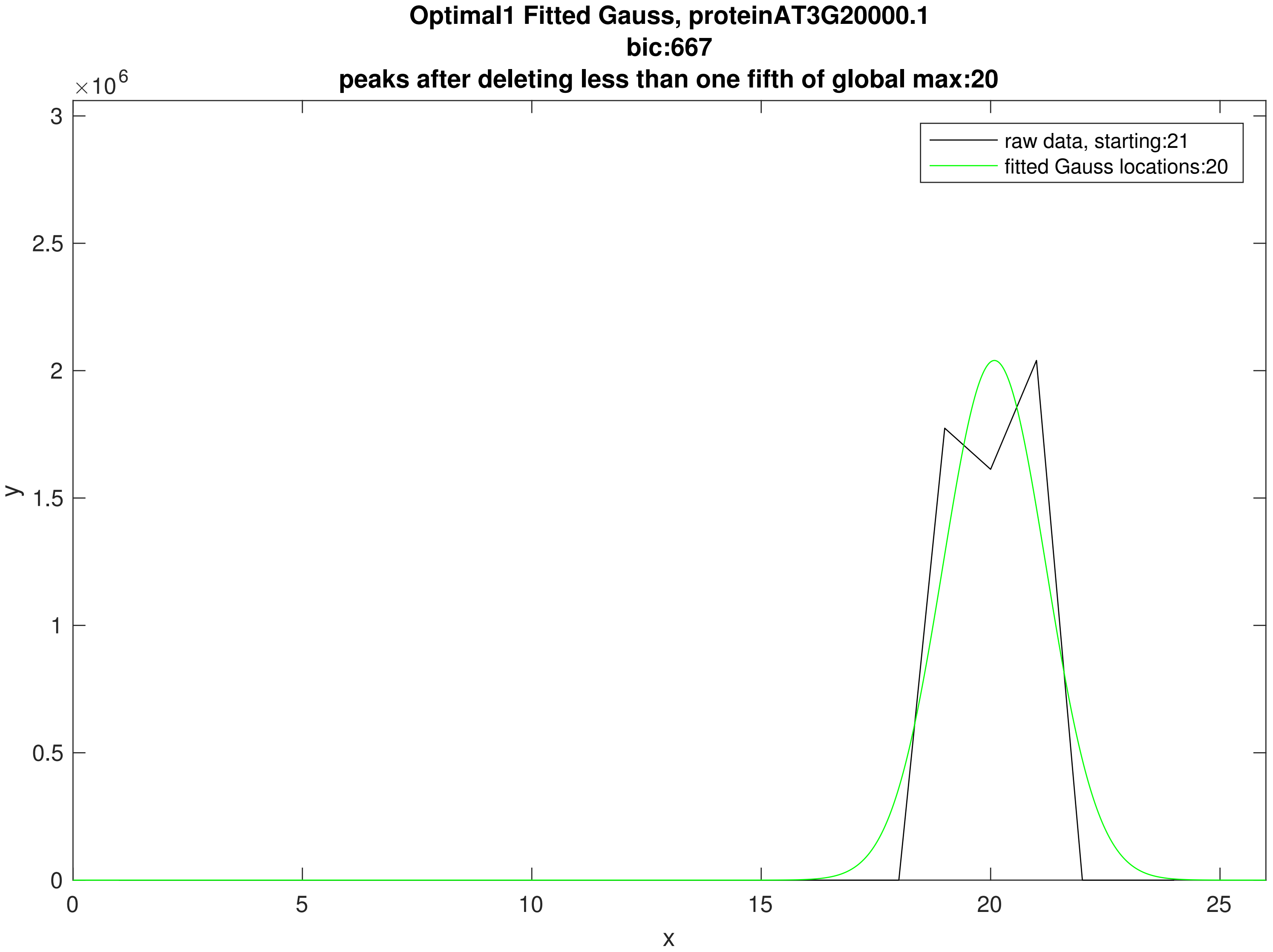
Optimal1 Fitted Gauss, proteinAT3G20000.1
bic:667
peaks after deleting less than one fifth of global max:20
×10^6
3
2.5
2
1.5
1
0.5
0
y
0
5
10
15
20
25
x
raw data, starting:21
fitted Gauss locations:20

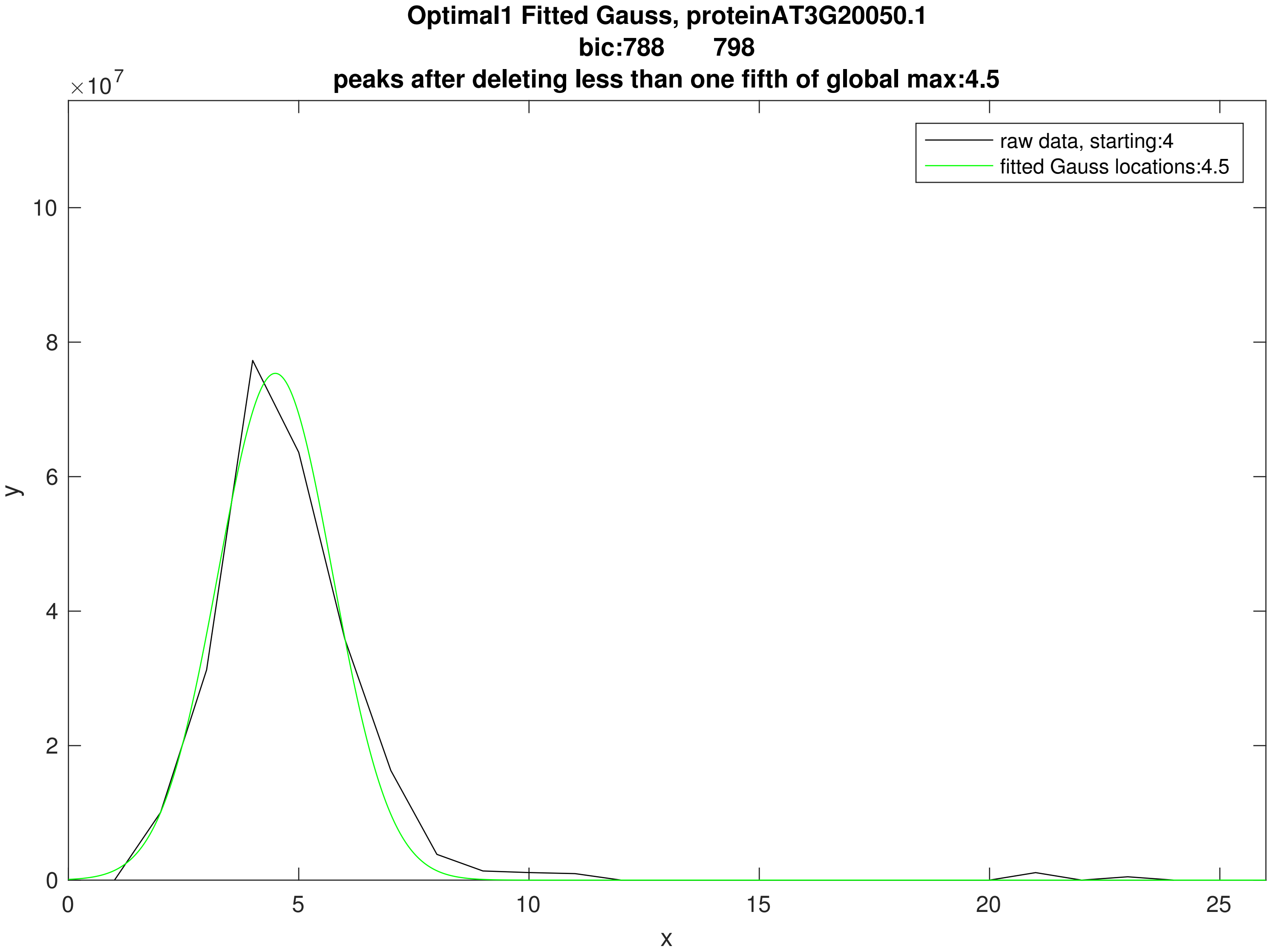

Optimal1 Fitted Gauss, proteinAT3G20050.1
bic:788 798
peaks after deleting less than one fifth of global max:4.5
×10^7
raw data, starting:4
fitted Gauss locations:4.5
y
x
0
2
4
6
8
10
0
5
10
15
20
25

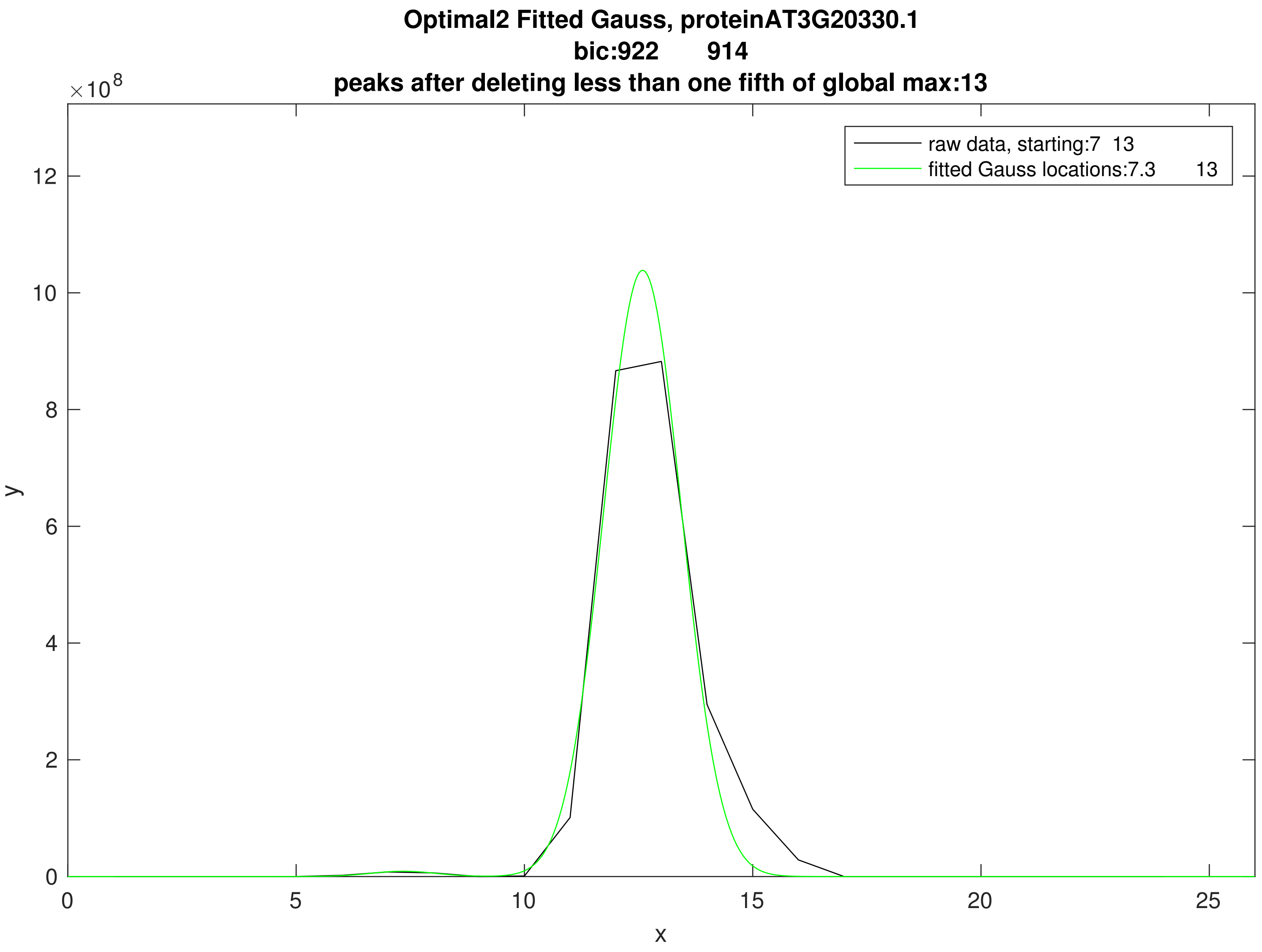
Optimal2 Fitted Gauss, proteinAT3G20330.1
bic:922 914
peaks after deleting less than one fifth of global max:13
×10^8
raw data, starting:7 13
fitted Gauss locations:7.3 13
y
x
0
2
4
6
8
10
12
0
5
10
15
20
25

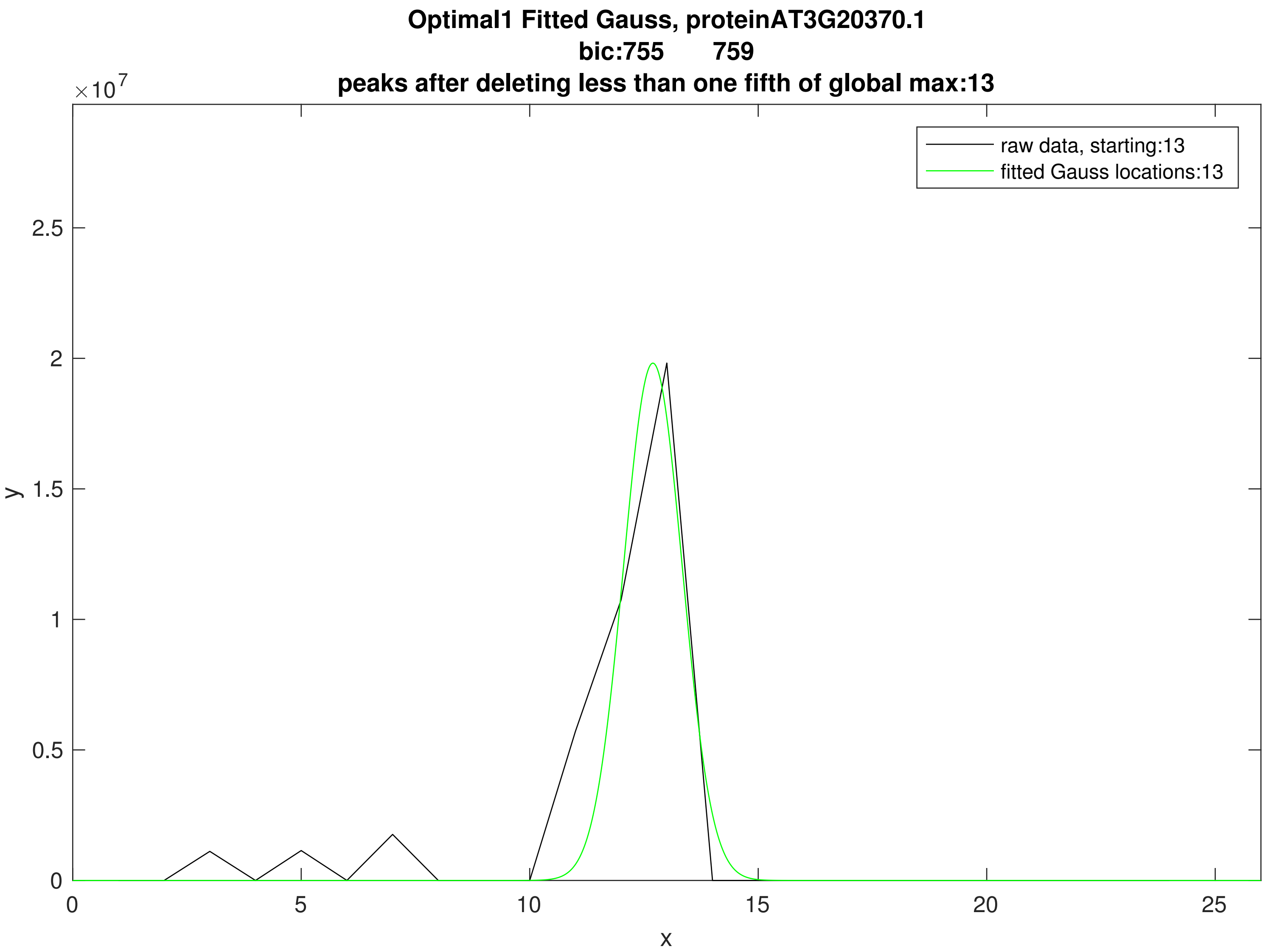

Optimal1 Fitted Gauss, proteinAT3G20370.1
bic:755 759
peaks after deleting less than one fifth of global max:13
×10^7
raw data, starting:13
fitted Gauss locations:13
y
x
0
0.5
1
1.5
2
2.5
0
5
10
15
20
25

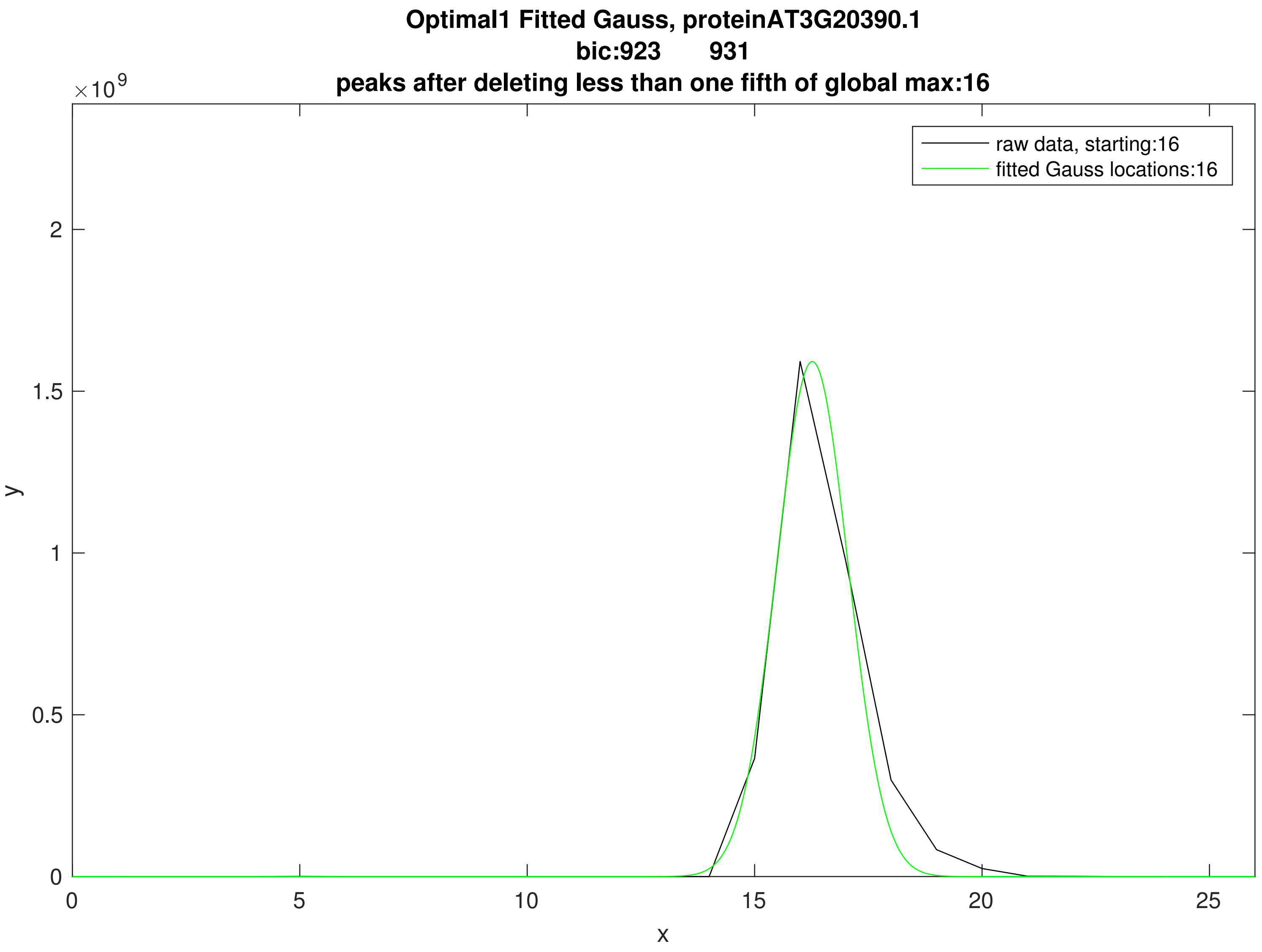
Optimal1 Fitted Gauss, proteinAT3G20390.1
bic:923 931
peaks after deleting less than one fifth of global max:16
×10^9
raw data, starting:16
fitted Gauss locations:16
y
x
0
0.5
1
1.5
2
5
10
15
20
25

#### Optimal1 Fitted Gauss, proteinAT3G20570.1
#### bic:723
#### peaks after deleting less than one fifth of global max:13

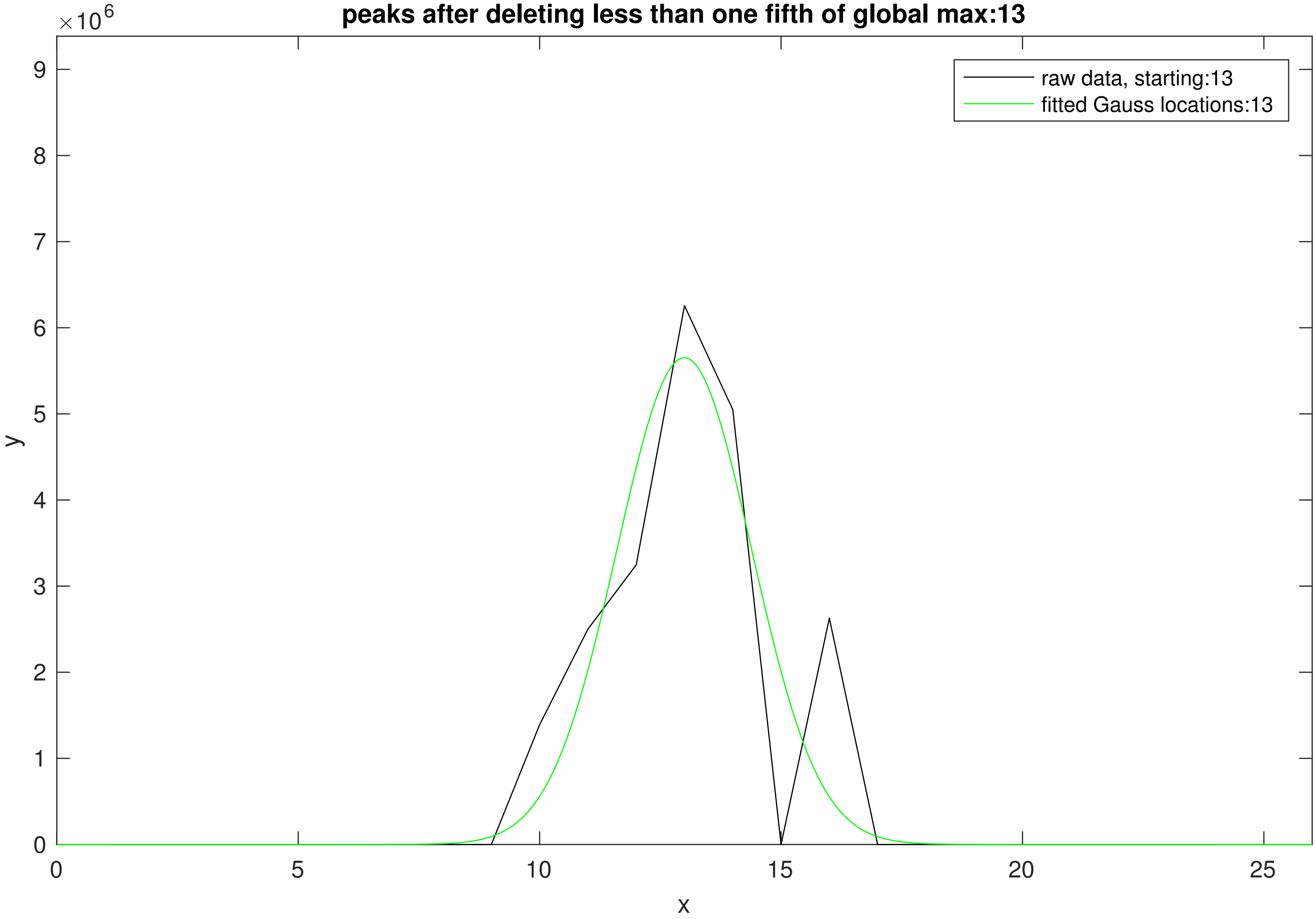

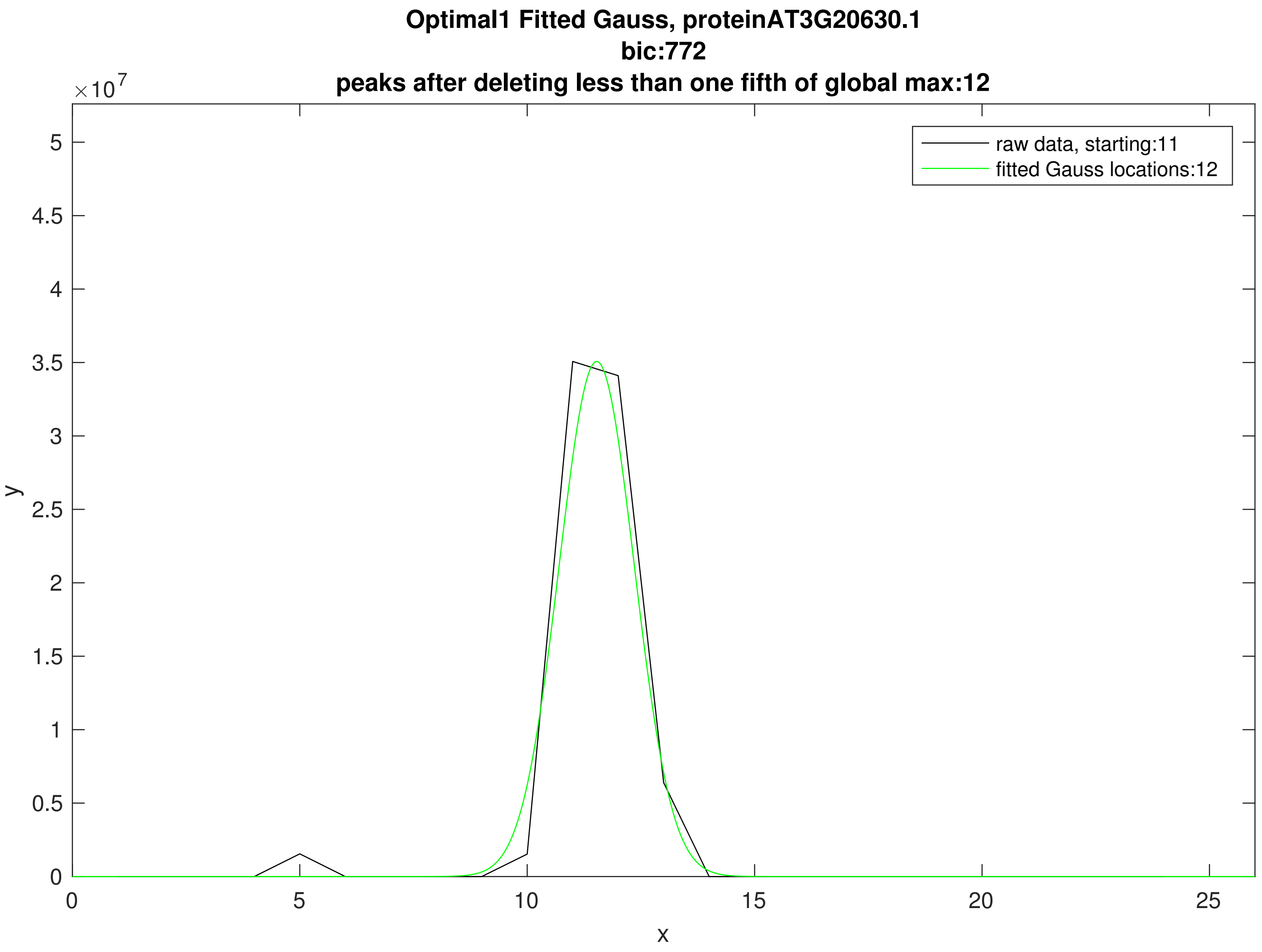
Optimal1 Fitted Gauss, proteinAT3G20630.1
bic:772
peaks after deleting less than one fifth of global max:12
×10^7
raw data, starting:11
fitted Gauss locations:12
y
x
0
0.5
1
1.5
2
2.5
3
3.5
4
4.5
5
0
5
10
15
20
25

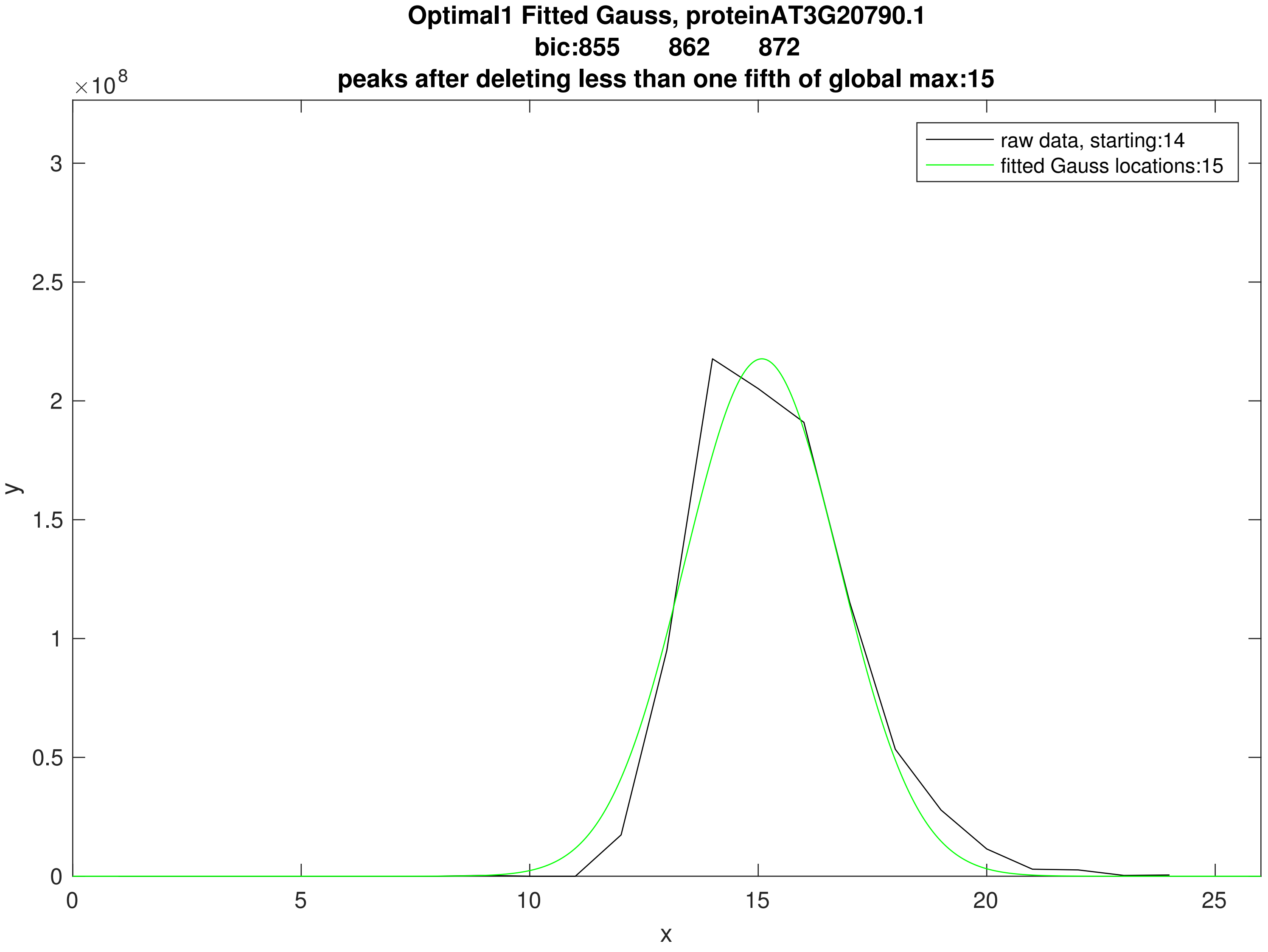

Optimal1 Fitted Gauss, proteinAT3G20790.1
bic:855 862 872
peaks after deleting less than one fifth of global max:15
×10^8
3
2.5
2
1.5
1
0.5
0
y
0
5
10
15
20
25
x
raw data, starting:14
fitted Gauss locations:15

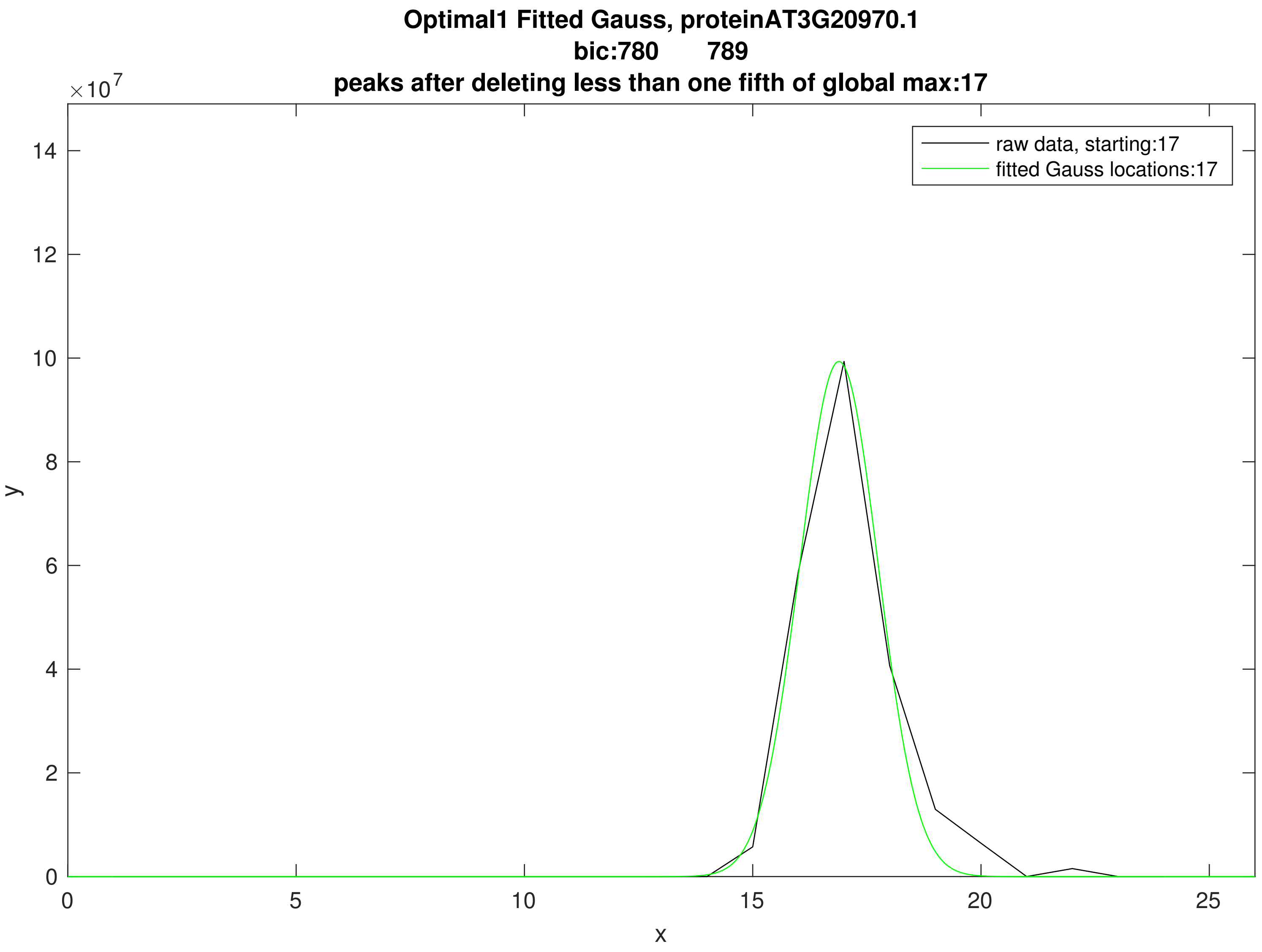
Optimal1 Fitted Gauss, proteinAT3G20970.1
bic:780 789
peaks after deleting less than one fifth of global max:17
×10^7
raw data, starting:17
fitted Gauss locations:17
y
x
0
2
4
6
8
10
12
14
0
5
10
15
20
25

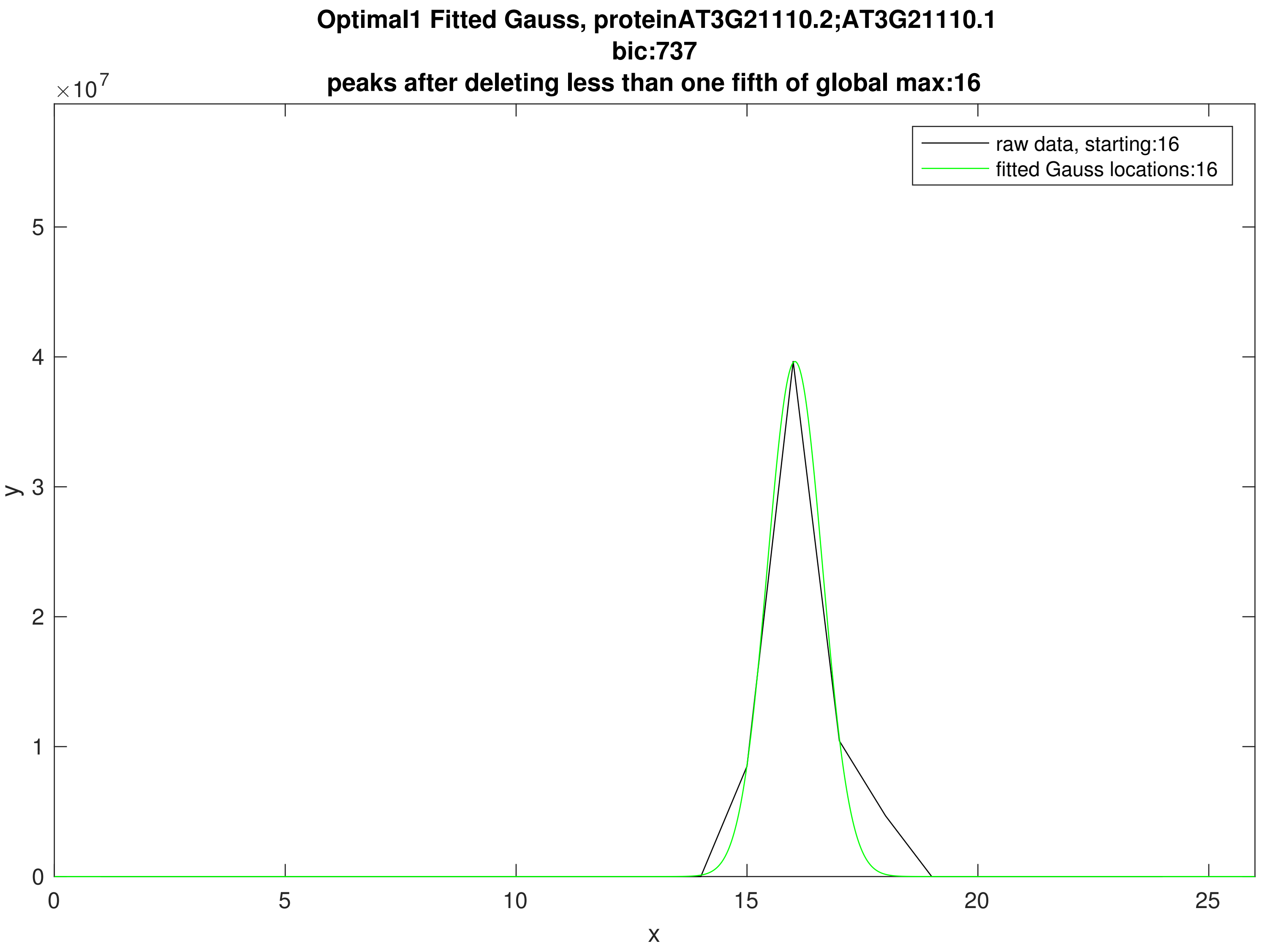

Optimal1 Fitted Gauss, proteinAT3G21110.2;AT3G21110.1
bic:737
peaks after deleting less than one fifth of global max:16
×10^7
raw data, starting:16
fitted Gauss locations:16
y
x
0
1
2
3
4
5
0
5
10
15
20
25

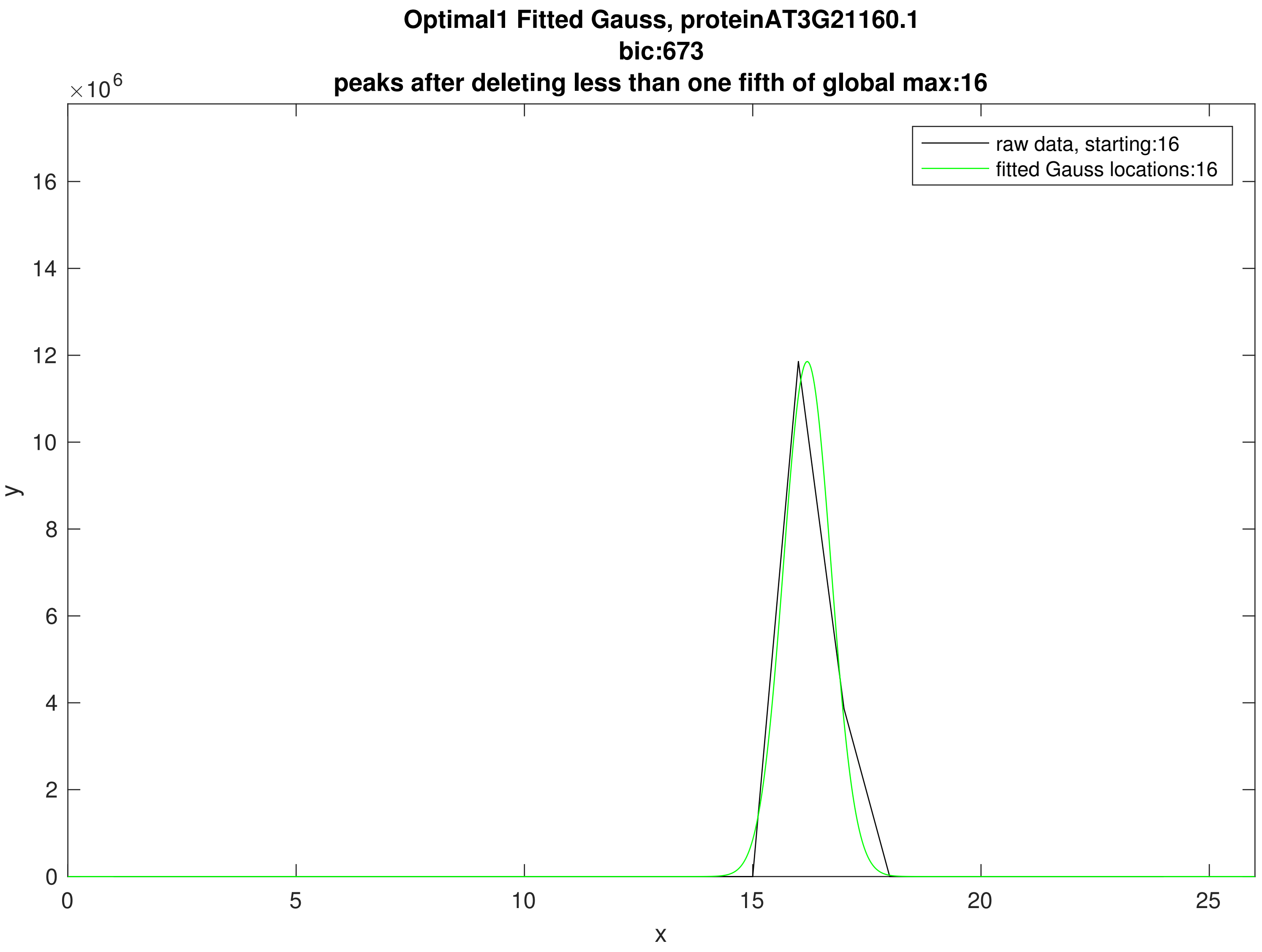

Optimal1 Fitted Gauss, proteinAT3G21160.1
bic:673
peaks after deleting less than one fifth of global max:16
×10^6
raw data, starting:16
fitted Gauss locations:16
16
14
12
10
8
6
4
2
0
y
0
5
10
15
20
25
x

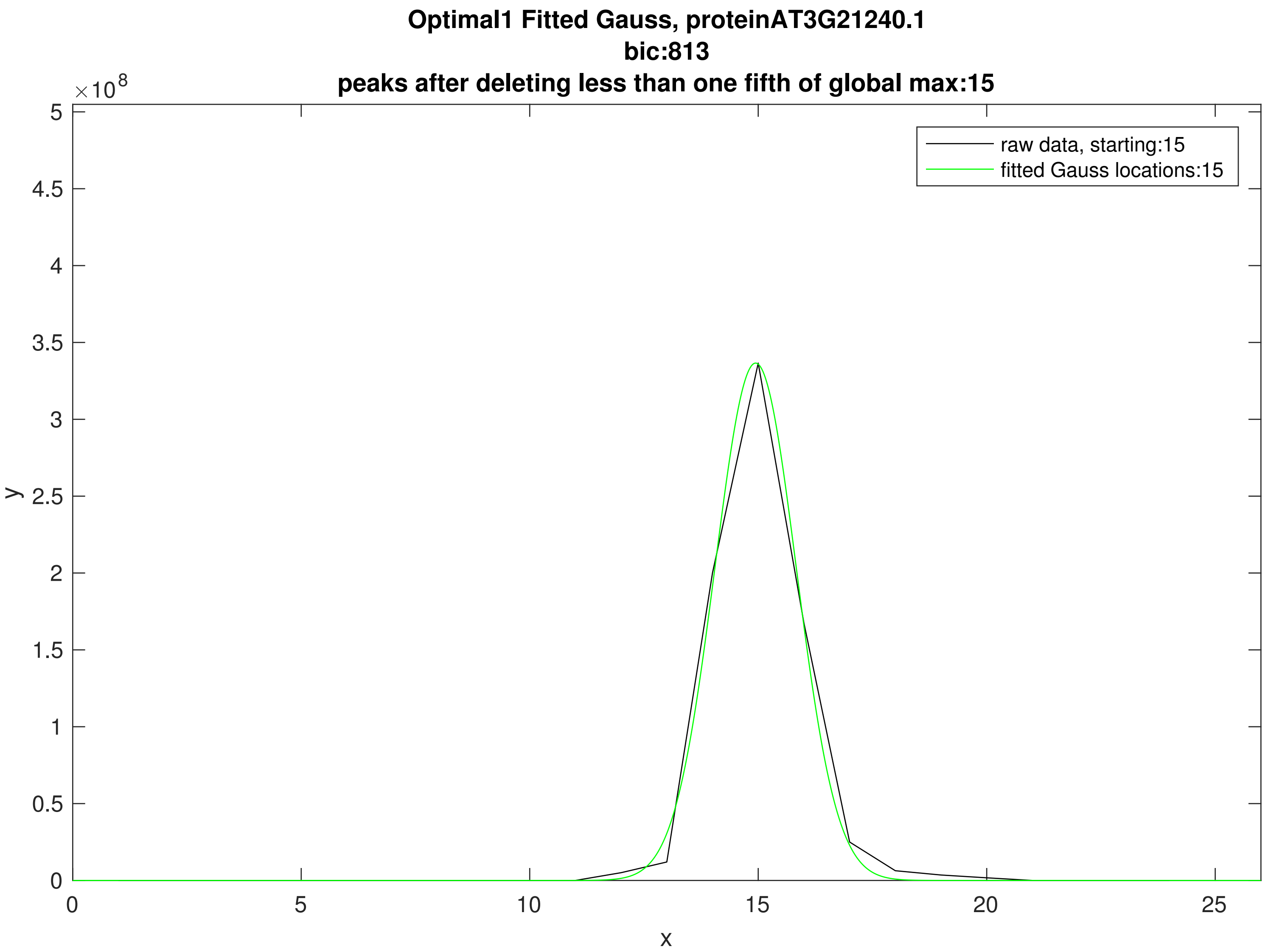

Optimal1 Fitted Gauss, proteinAT3G21240.1
bic:813
peaks after deleting less than one fifth of global max:15
×10^8
5
4.5
4
3.5
3
2.5
2
1.5
1
0.5
0
y
0
5
10
15
20
25
x
raw data, starting:15
fitted Gauss locations:15

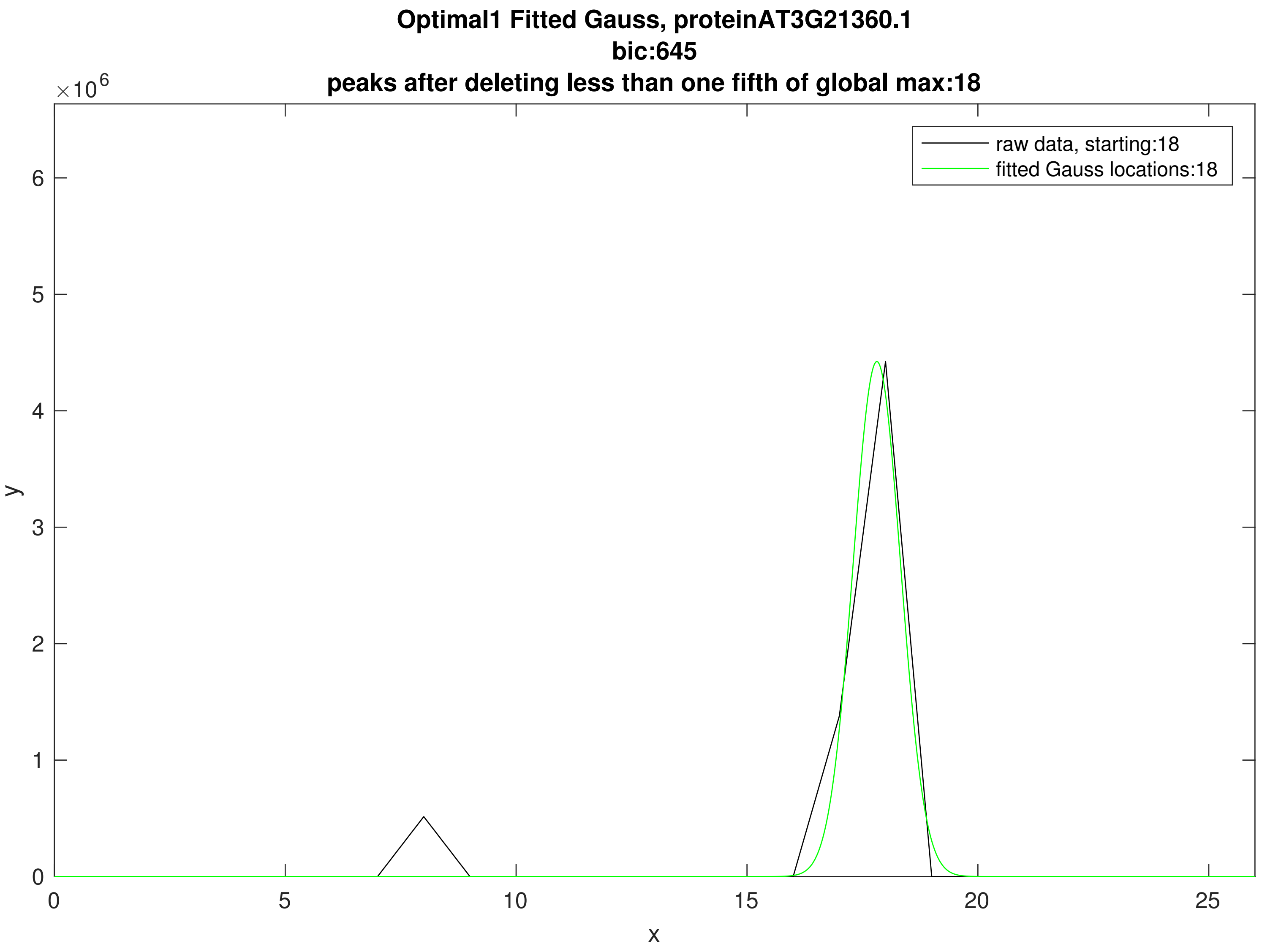

Optimal1 Fitted Gauss, proteinAT3G21360.1
bic:645
peaks after deleting less than one fifth of global max:18
×10^6
raw data, starting:18
fitted Gauss locations:18
y
x
0
1
2
3
4
5
6
0
5
10
15
20
25

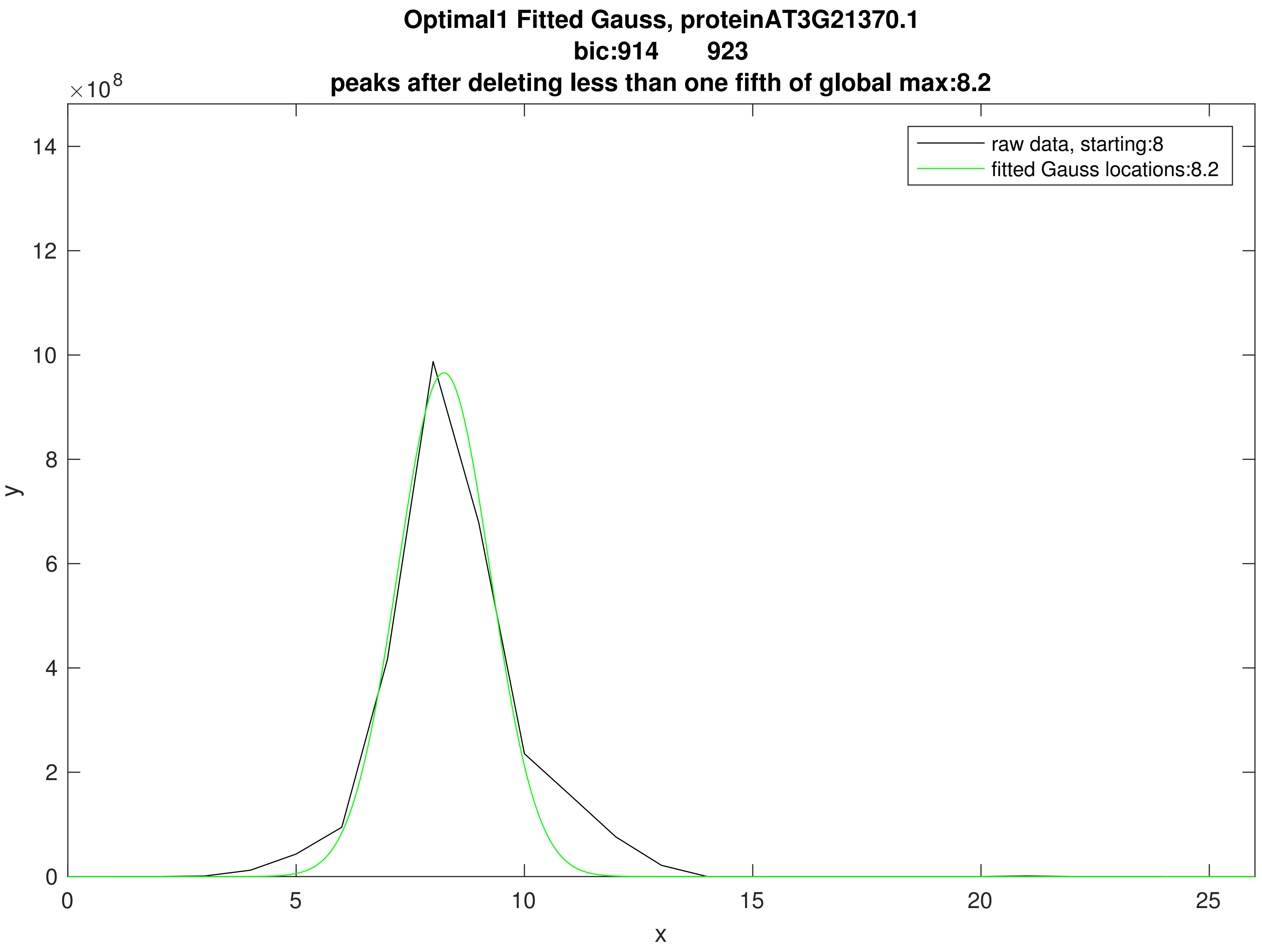
Optimal1 Fitted Gauss, proteinAT3G21370.1
bic:914 923
peaks after deleting less than one fifth of global max:8.2
×10^8
raw data, starting:8
fitted Gauss locations:8.2
y
x
0
2
4
6
8
10
12
14
0
5
10
15
20
25

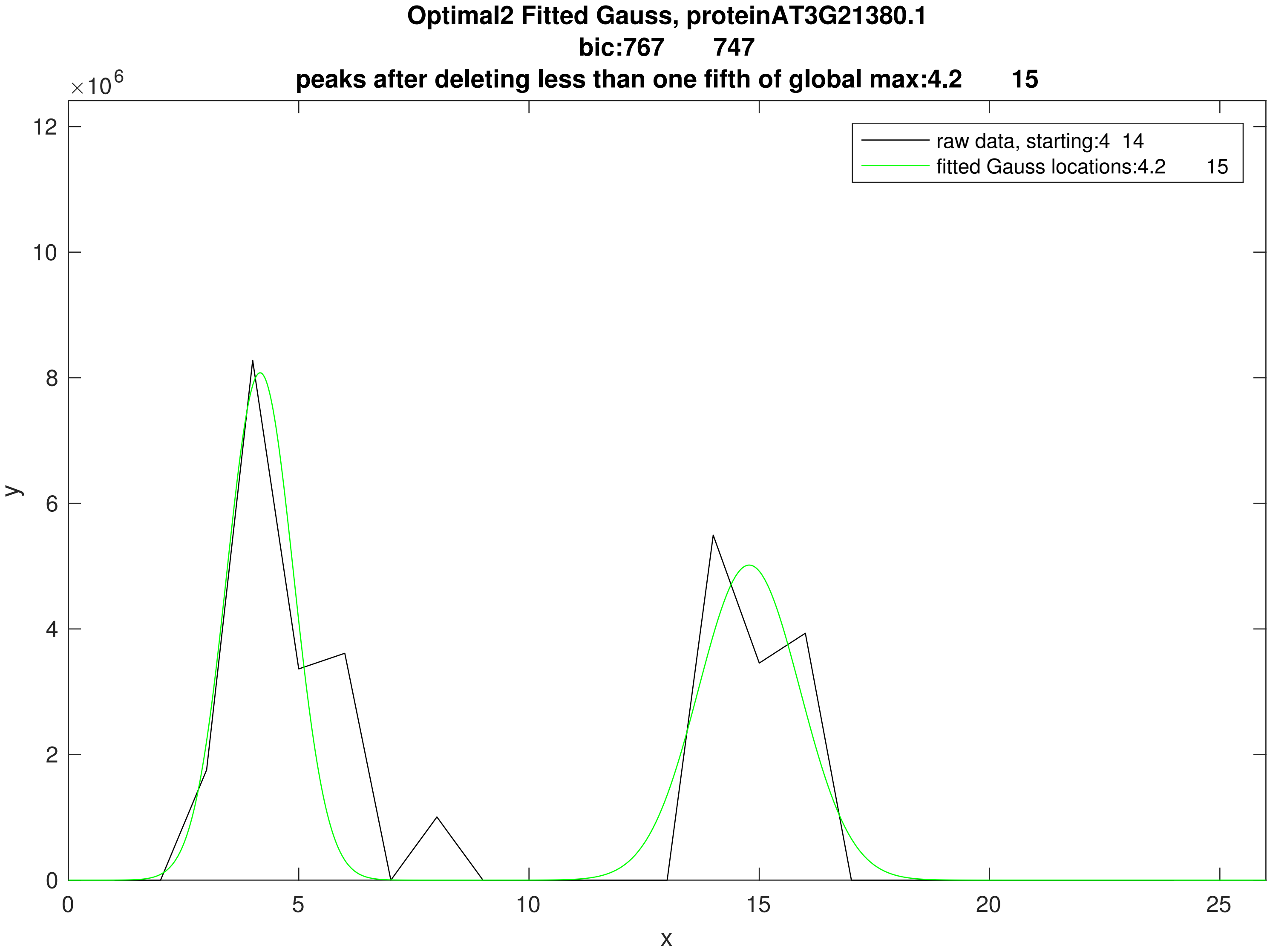

Optimal2 Fitted Gauss, proteinAT3G21380.1
bic:767 747
peaks after deleting less than one fifth of global max:4.2 15
×10^6
raw data, starting:4 14
fitted Gauss locations:4.2 15
y
x
0
2
4
6
8
10
12
0
5
10
15
20
25

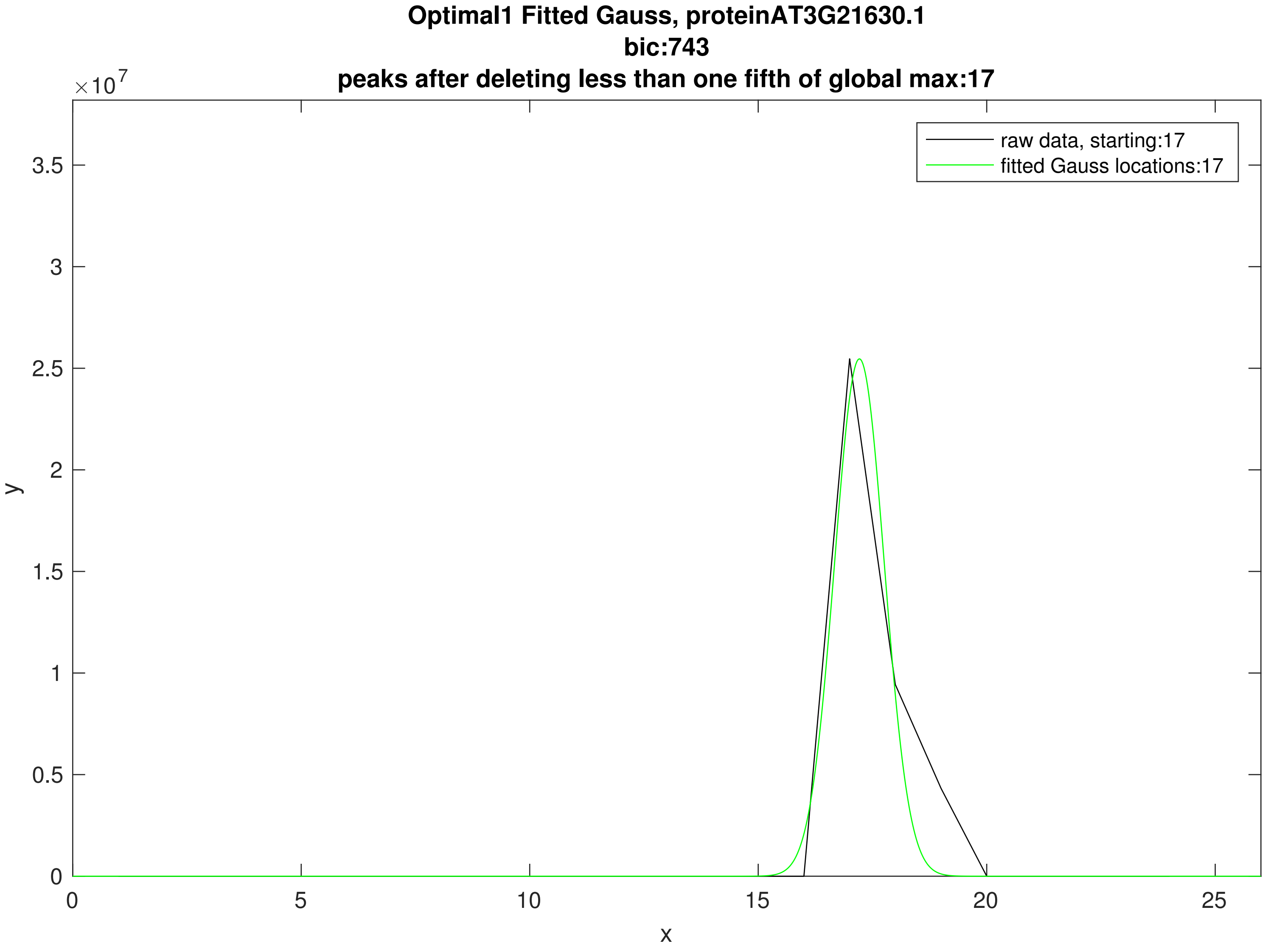

Optimal1 Fitted Gauss, proteinAT3G21630.1
bic:743
peaks after deleting less than one fifth of global max:17
×10^7
raw data, starting:17
fitted Gauss locations:17
3.5
3
2.5
2
1.5
1
0.5
0
y
0
5
10
15
20
25
x

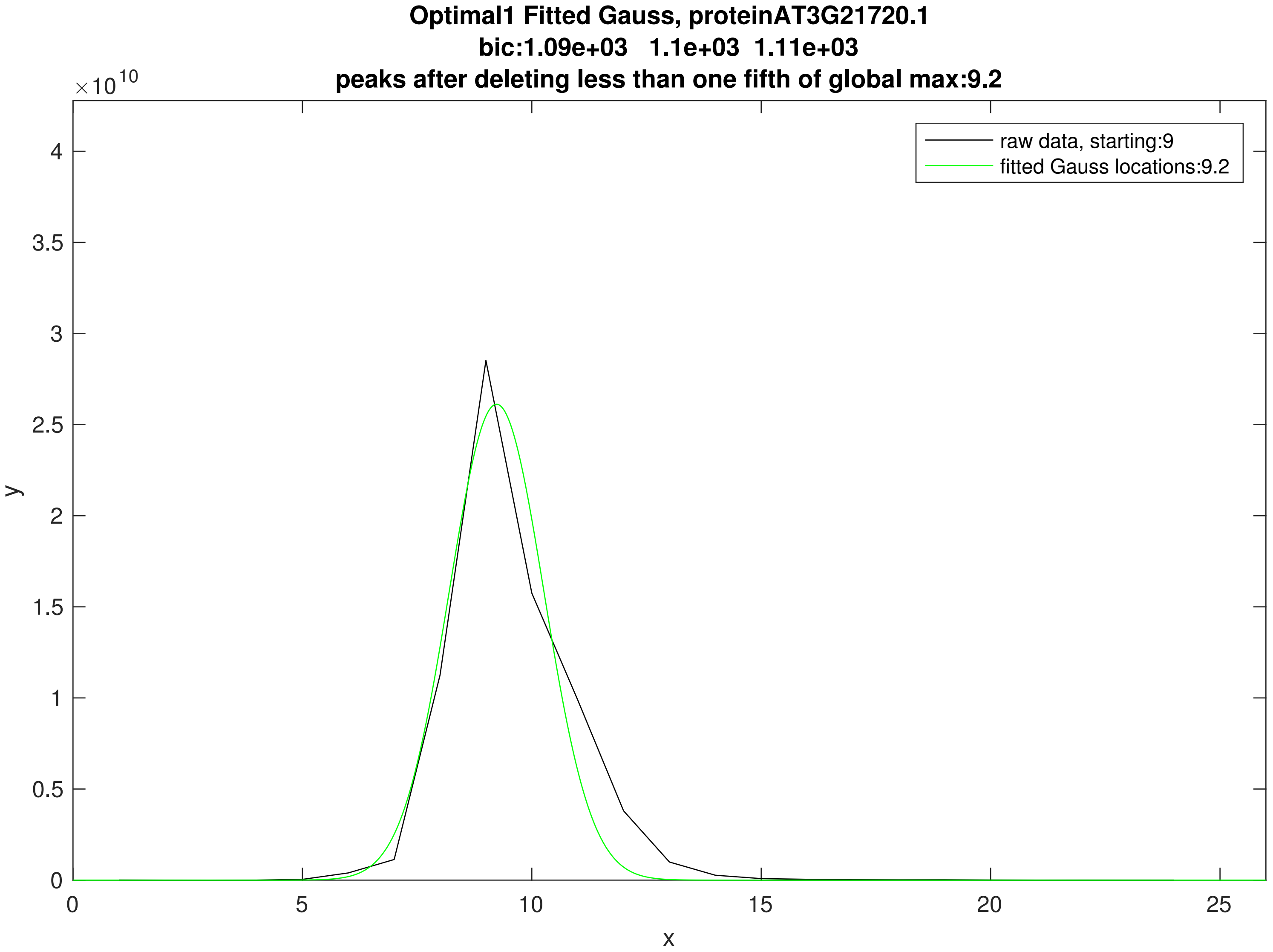
Optimal1 Fitted Gauss, proteinAT3G21720.1
bic:1.09e+03 1.1e+03 1.11e+03
peaks after deleting less than one fifth of global max:9.2
×10^10
raw data, starting:9
fitted Gauss locations:9.2
y
x
0
0.5
1
1.5
2
2.5
3
3.5
4
0
5
10
15
20
25

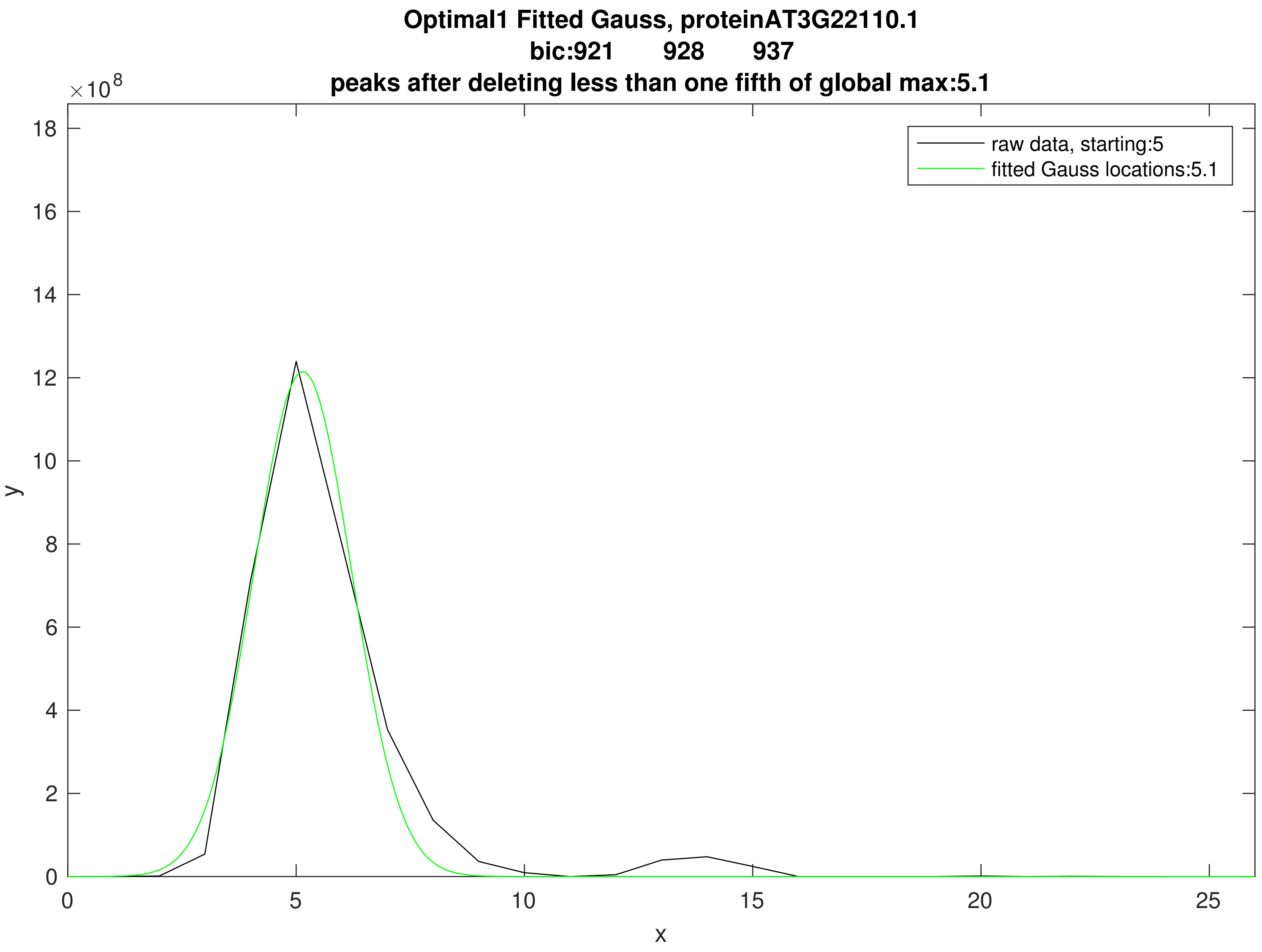

Optimal1 Fitted Gauss, proteinAT3G22110.1
bic:921 928 937
peaks after deleting less than one fifth of global max:5.1
×10^8
18
16
14
12
10
8
6
4
2
0
y
0
5
10
15
20
25
x
raw data, starting:5
fitted Gauss locations:5.1

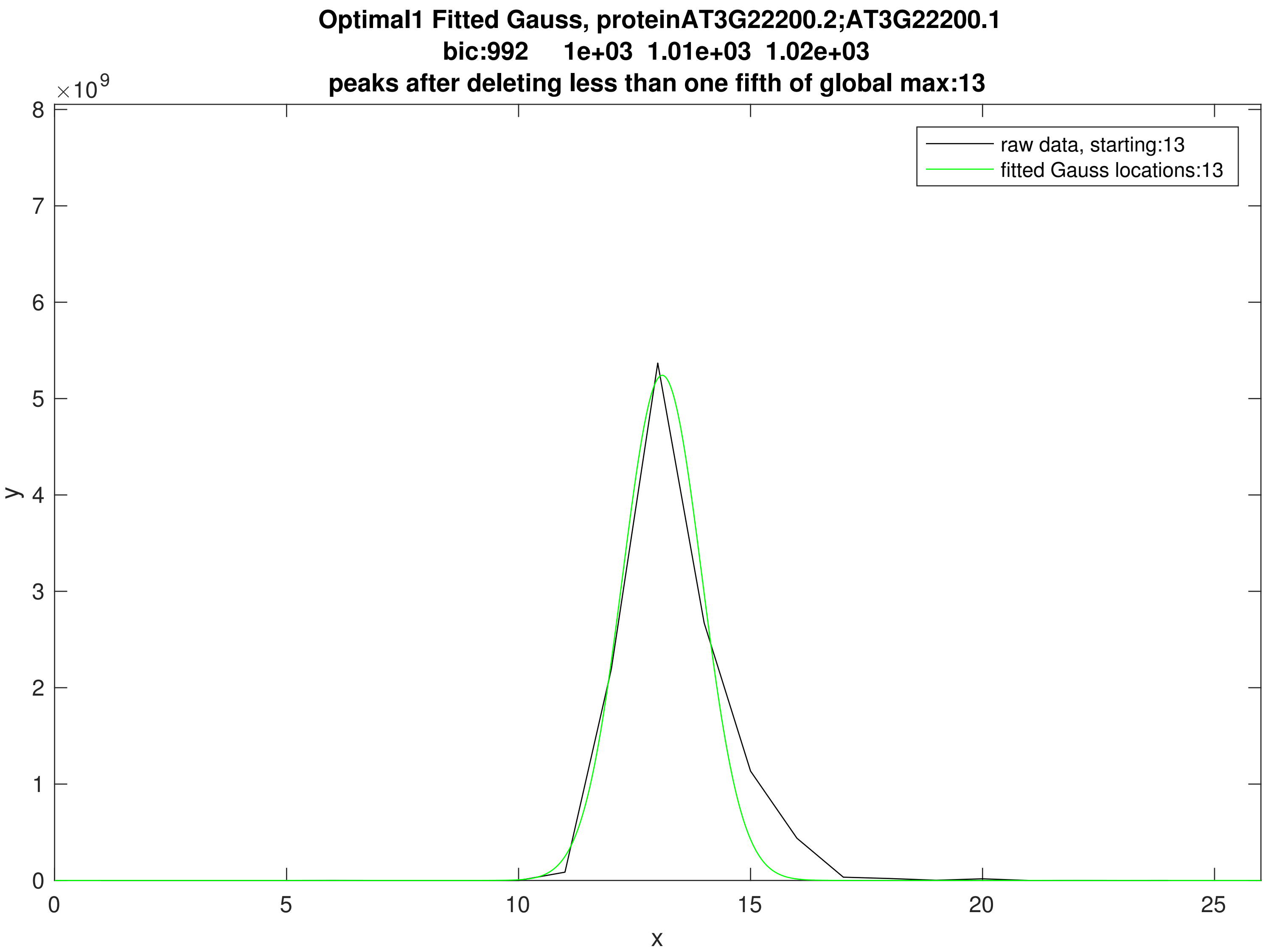

Optimal1 Fitted Gauss, proteinAT3G22200.2;AT3G22200.1
bic:992 1e+03 1.01e+03 1.02e+03
peaks after deleting less than one fifth of global max:13
raw data, starting:13
fitted Gauss locations:13
y
x
×10^9
0
1
2
3
4
5
6
7
8
0
5
10
15
20
25

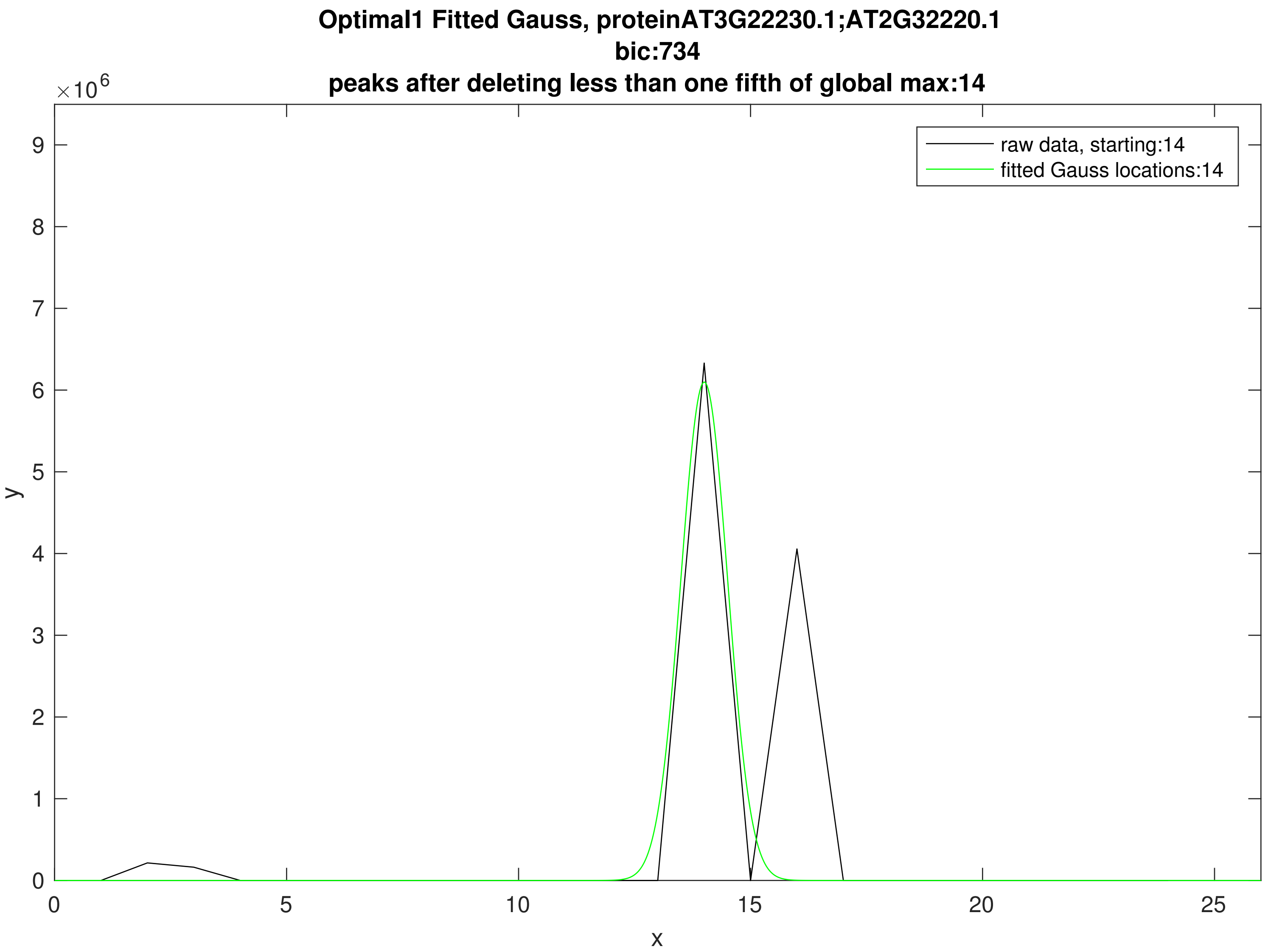

Optimal1 Fitted Gauss, proteinAT3G22230.1;AT2G32220.1
bic:734
peaks after deleting less than one fifth of global max:14
×10^6
raw data, starting:14
fitted Gauss locations:14
y
x
0
1
2
3
4
5
6
7
8
9
0
5
10
15
20
25

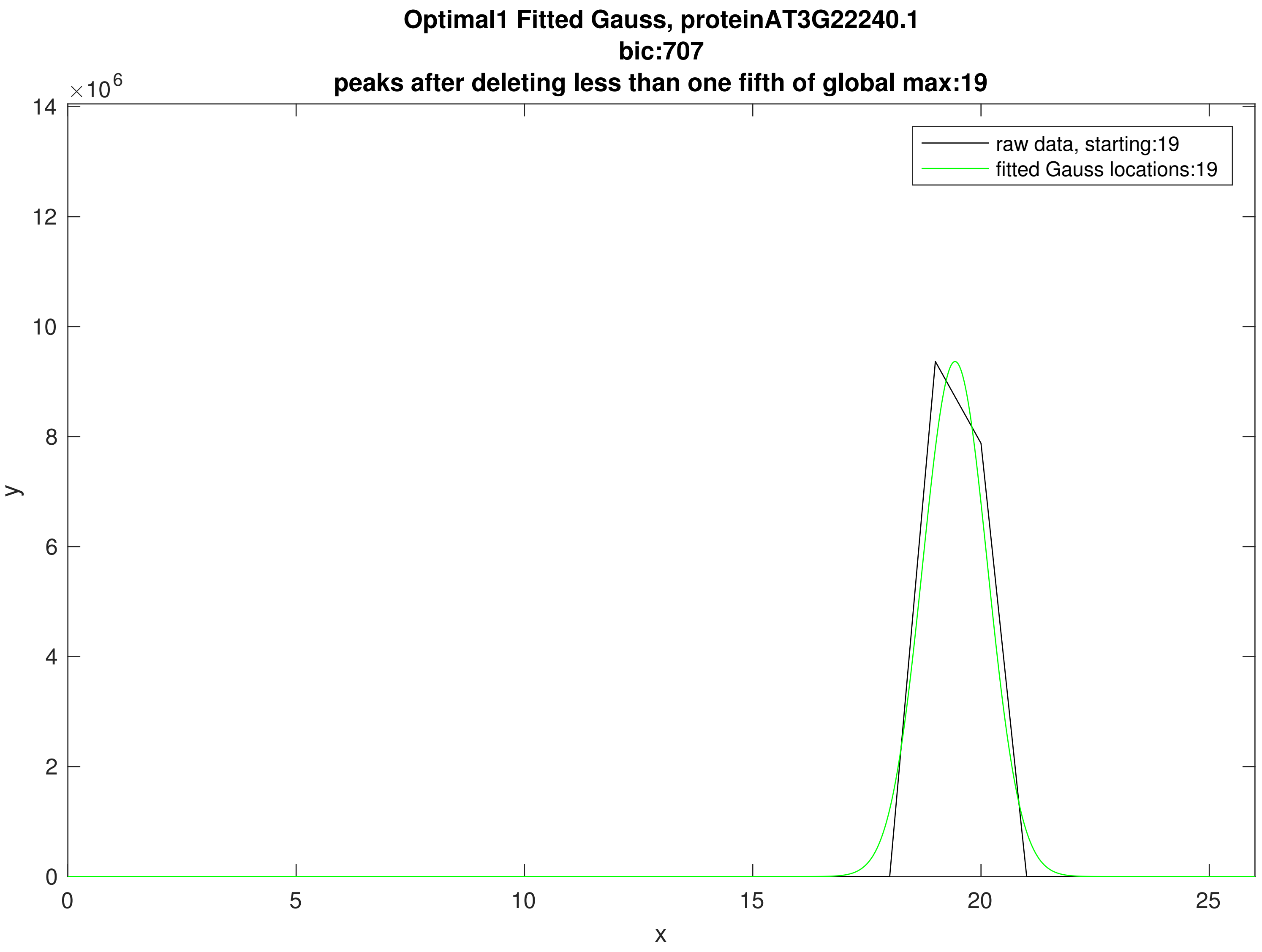

Optimal1 Fitted Gauss, proteinAT3G22240.1
bic:707
peaks after deleting less than one fifth of global max:19
×10^6
14
12
10
8
6
4
2
0
y
0
5
10
15
20
25
x
raw data, starting:19
fitted Gauss locations:19

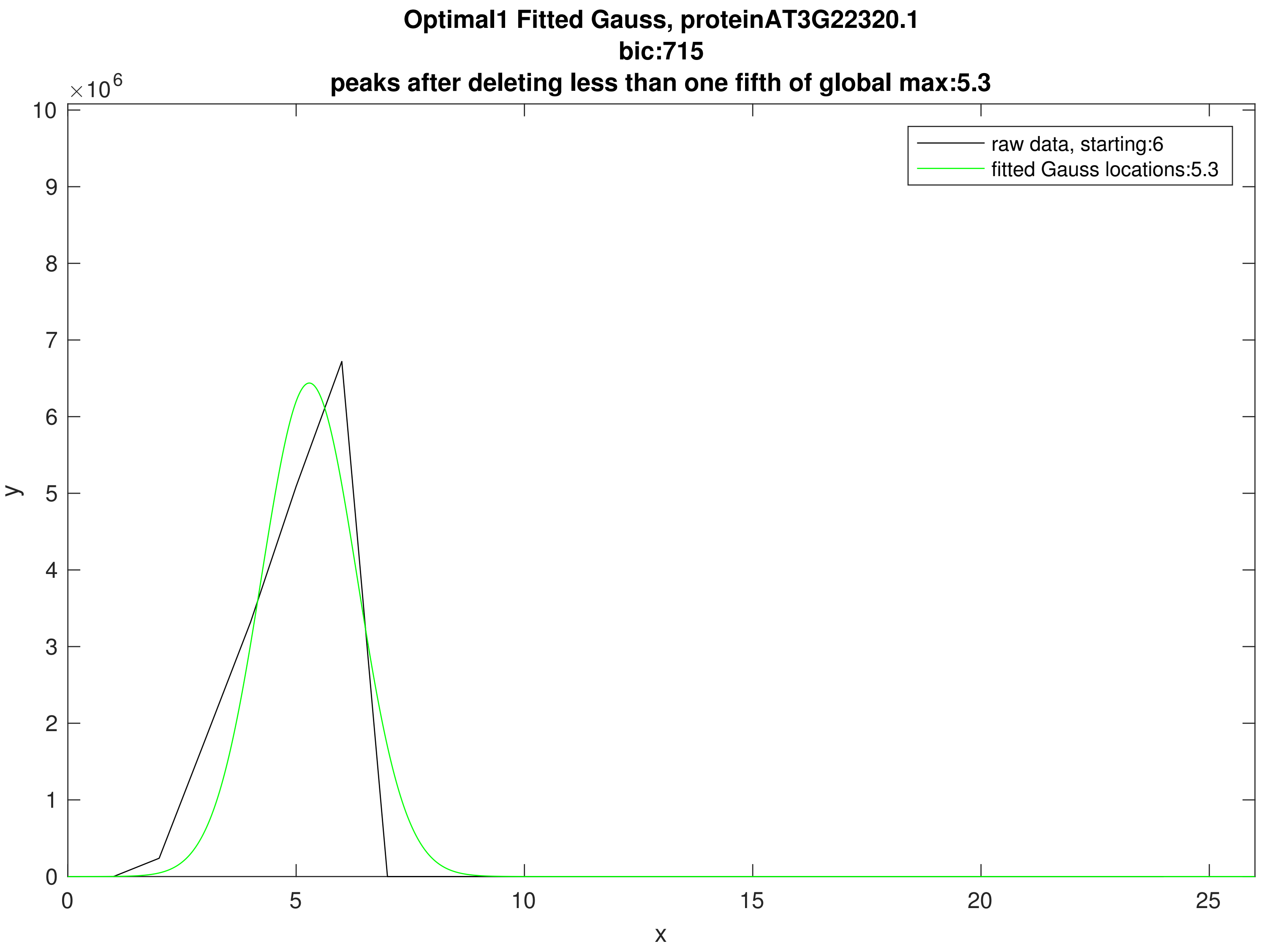
Optimal1 Fitted Gauss, proteinAT3G22320.1
bic:715
peaks after deleting less than one fifth of global max:5.3
×10^6
raw data, starting:6
fitted Gauss locations:5.3
y
x

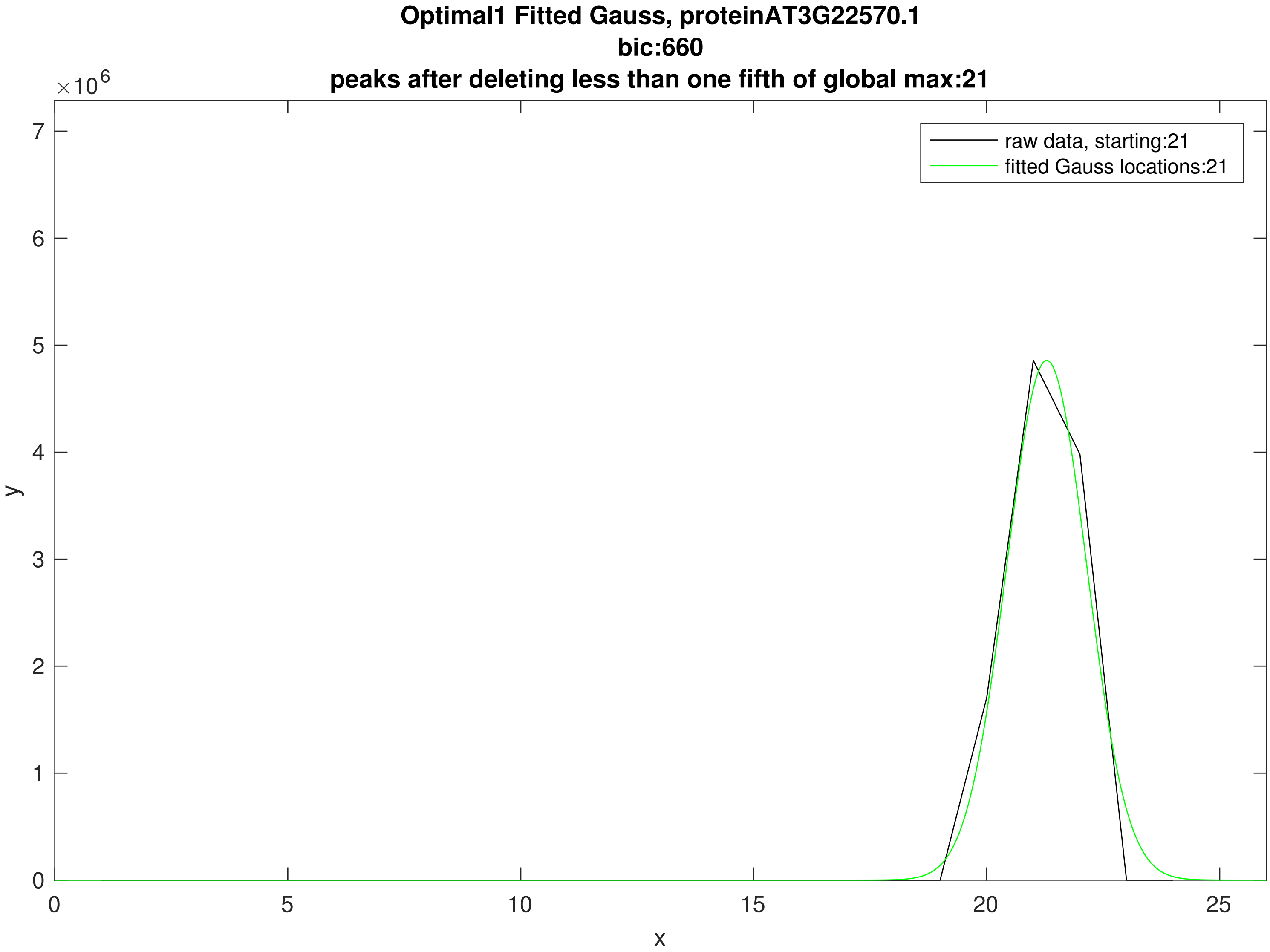

Optimal1 Fitted Gauss, proteinAT3G22570.1
bic:660
peaks after deleting less than one fifth of global max:21
×10^6
7
6
5
4
3
2
1
0
y
0
5
10
15
20
25
x
raw data, starting:21
fitted Gauss locations:21

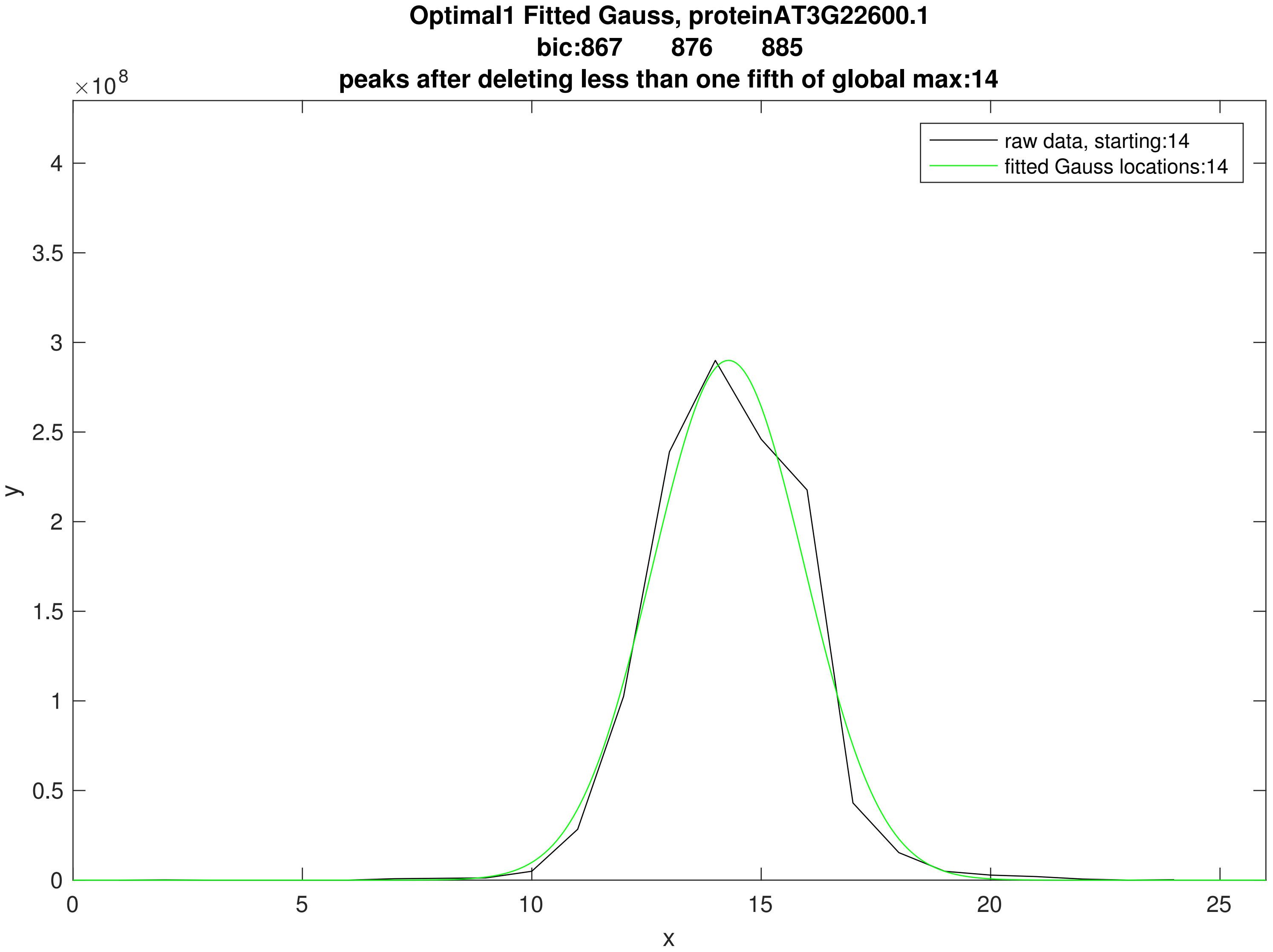

Optimal1 Fitted Gauss, proteinAT3G22600.1
bic:867 876 885
peaks after deleting less than one fifth of global max:14
×10^8
raw data, starting:14
fitted Gauss locations:14
y
x
0
0.5
1
1.5
2
2.5
3
3.5
4
0
5
10
15
20
25

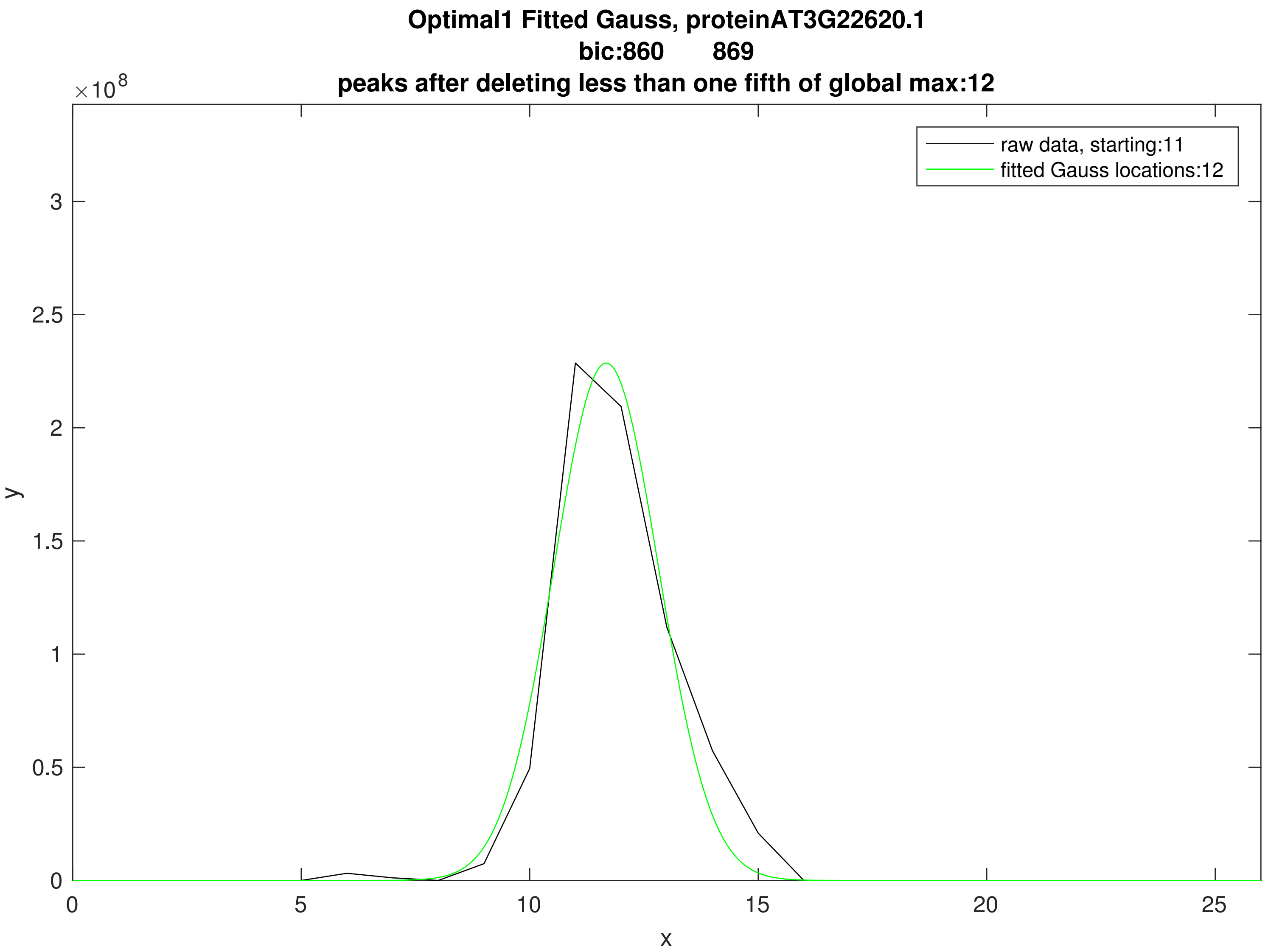

Optimal1 Fitted Gauss, proteinAT3G22620.1
bic:860 869
peaks after deleting less than one fifth of global max:12
×10^8
raw data, starting:11
fitted Gauss locations:12
3
2.5
2
1.5
1
0.5
0
y
0
5
10
15
20
25
x

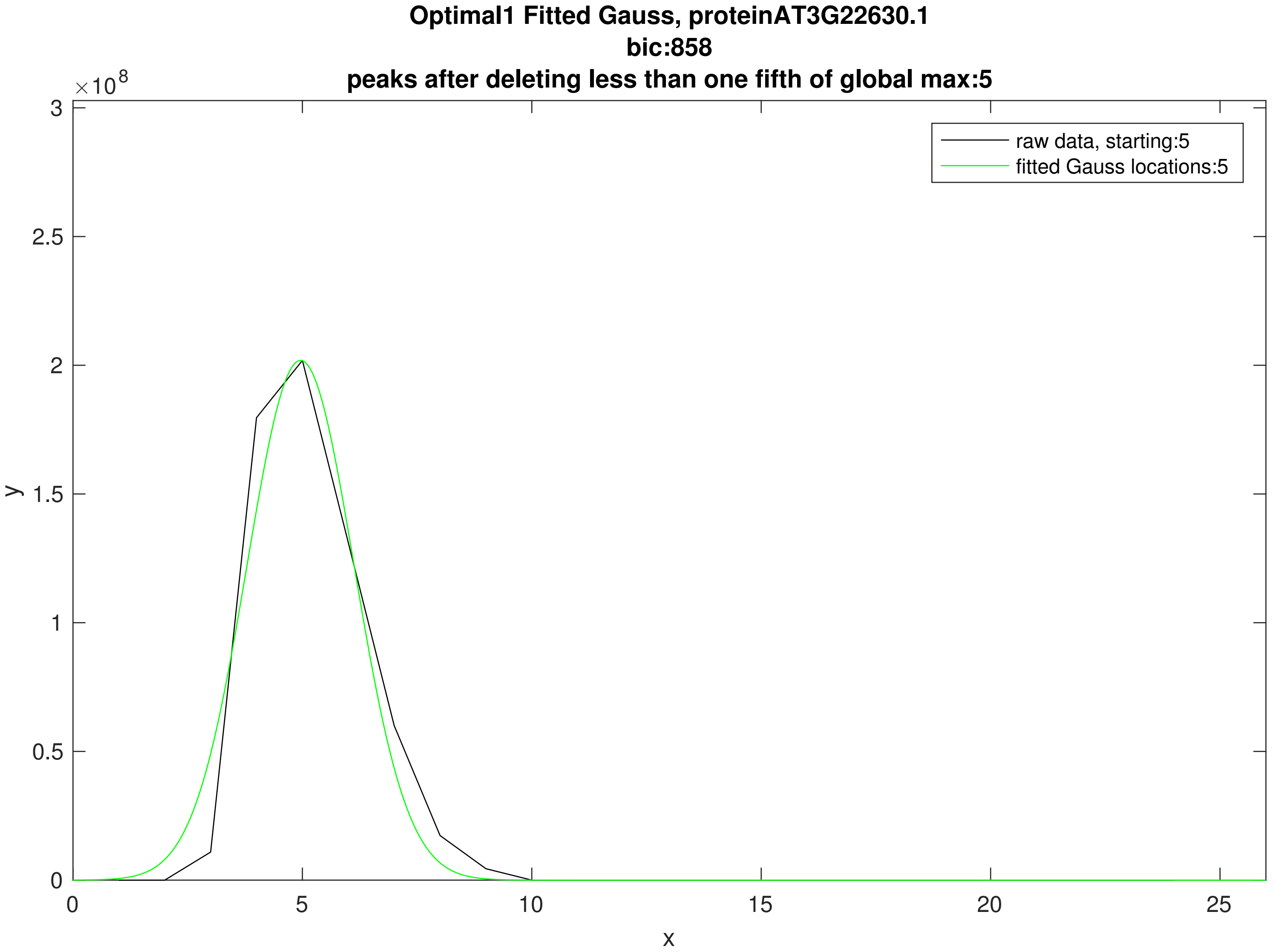

Optimal1 Fitted Gauss, proteinAT3G22630.1
bic:858
peaks after deleting less than one fifth of global max:5
×10^8
raw data, starting:5
fitted Gauss locations:5
y
x
0
0.5
1
1.5
2
2.5
3
0
5
10
15
20
25

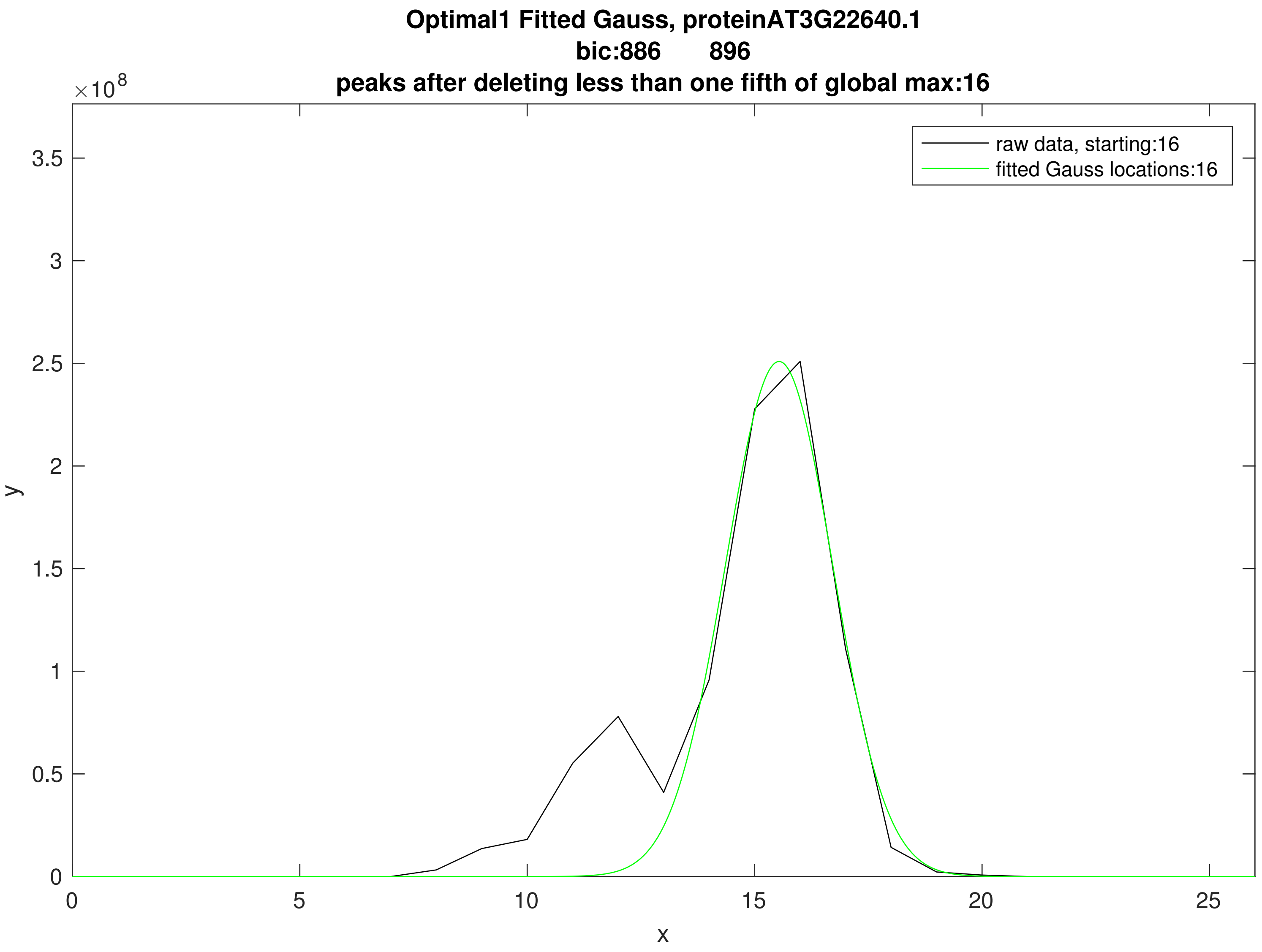
Optimal1 Fitted Gauss, proteinAT3G22640.1
bic:886 896
peaks after deleting less than one fifth of global max:16
×10^8
3.5
3
2.5
2
1.5
1
0.5
0
y
0
5
10
15
20
25
x
raw data, starting:16
fitted Gauss locations:16

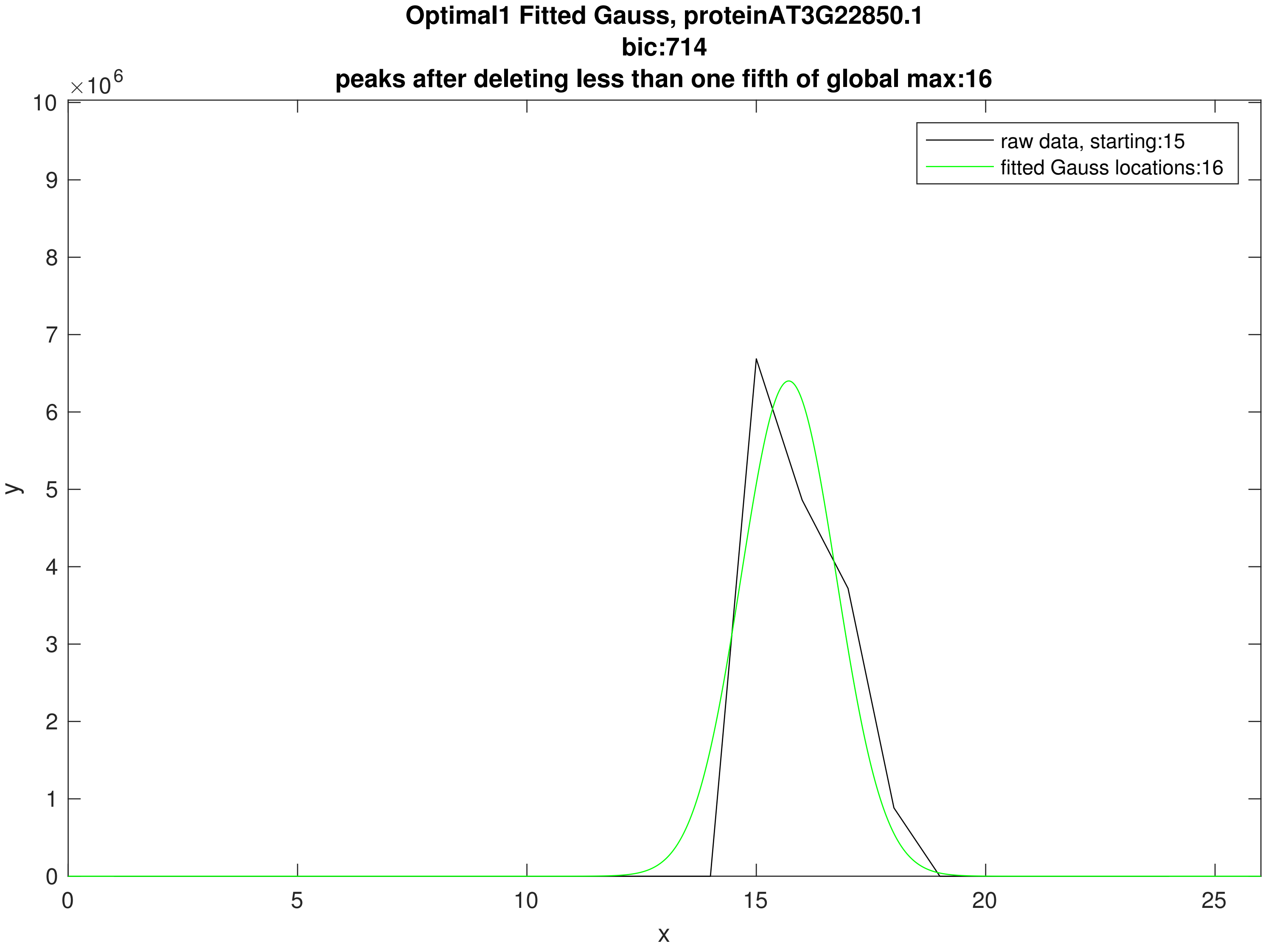

Optimal1 Fitted Gauss, proteinAT3G22850.1
bic:714
peaks after deleting less than one fifth of global max:16
×10^6
raw data, starting:15
fitted Gauss locations:16
y
x
0
1
2
3
4
5
6
7
8
9
10
0
5
10
15
20
25

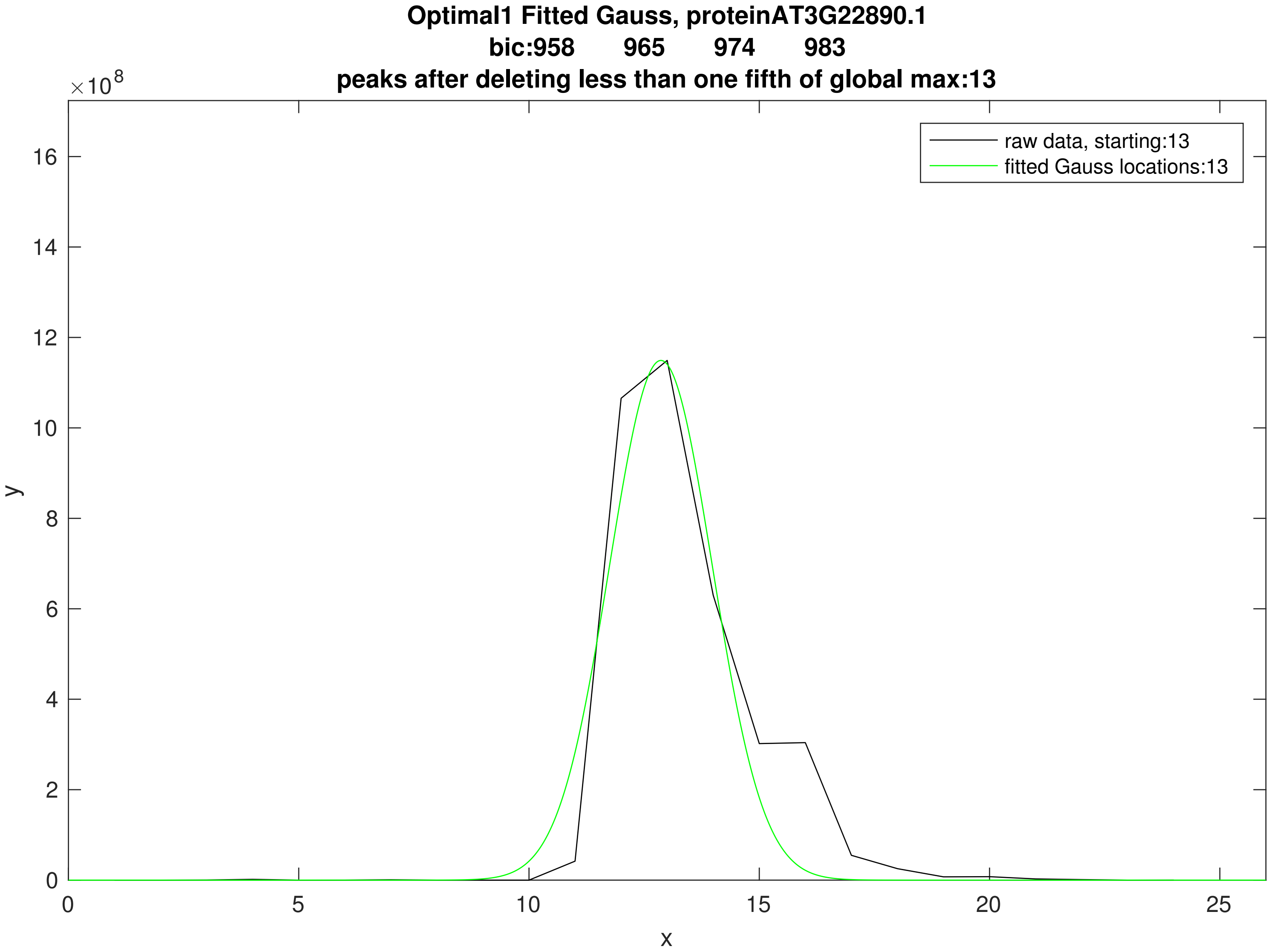
Optimal1 Fitted Gauss, proteinAT3G22890.1
bic:958 965 974 983
peaks after deleting less than one fifth of global max:13
×10^8
16
14
12
10
8
6
4
2
0
y
0
5
10
15
20
25
x
raw data, starting:13
fitted Gauss locations:13

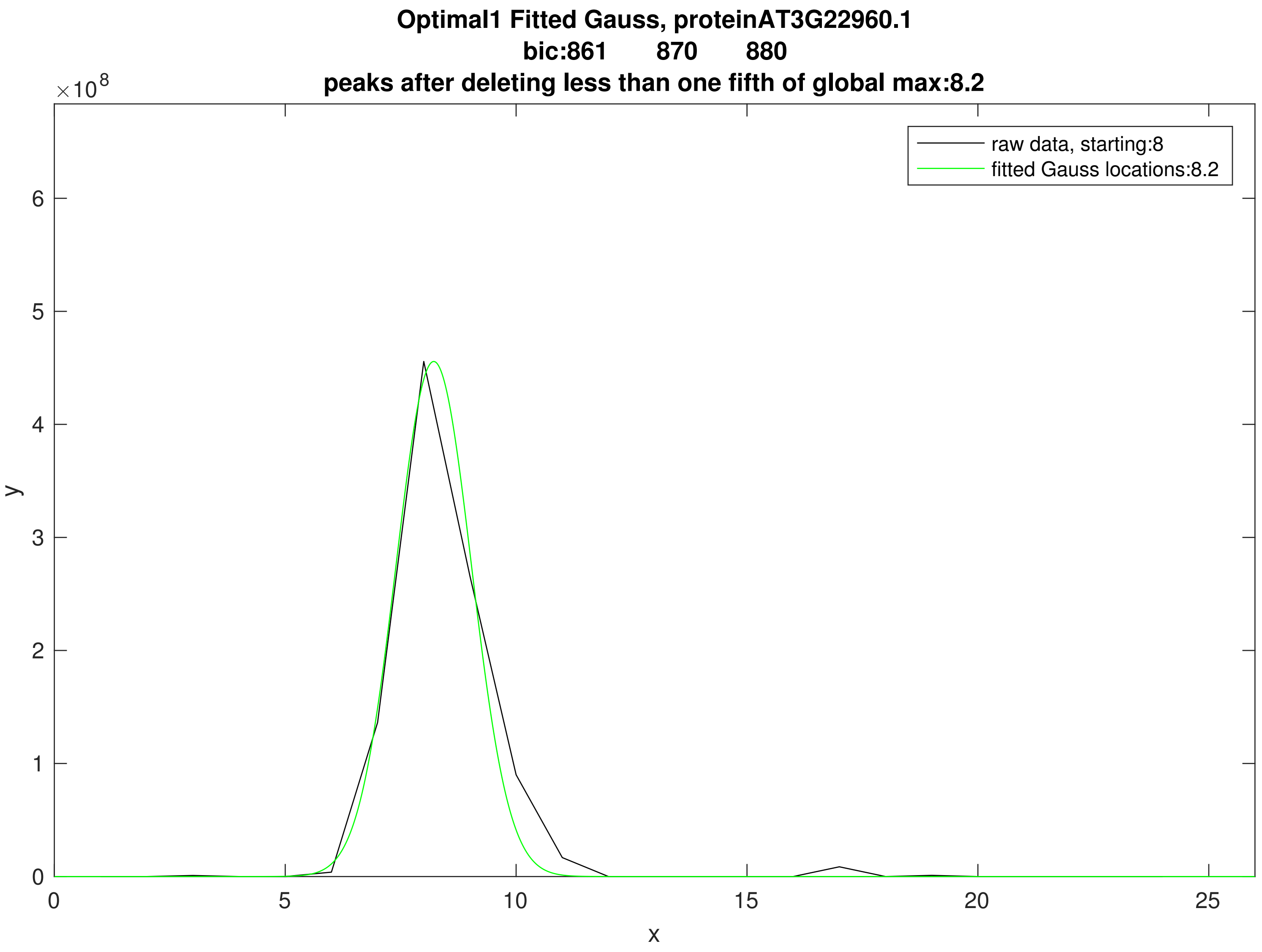
Optimal1 Fitted Gauss, proteinAT3G22960.1
bic:861 870 880
peaks after deleting less than one fifth of global max:8.2
×10^8
raw data, starting:8
fitted Gauss locations:8.2
y
x
0
1
2
3
4
5
6
0
5
10
15
20
25

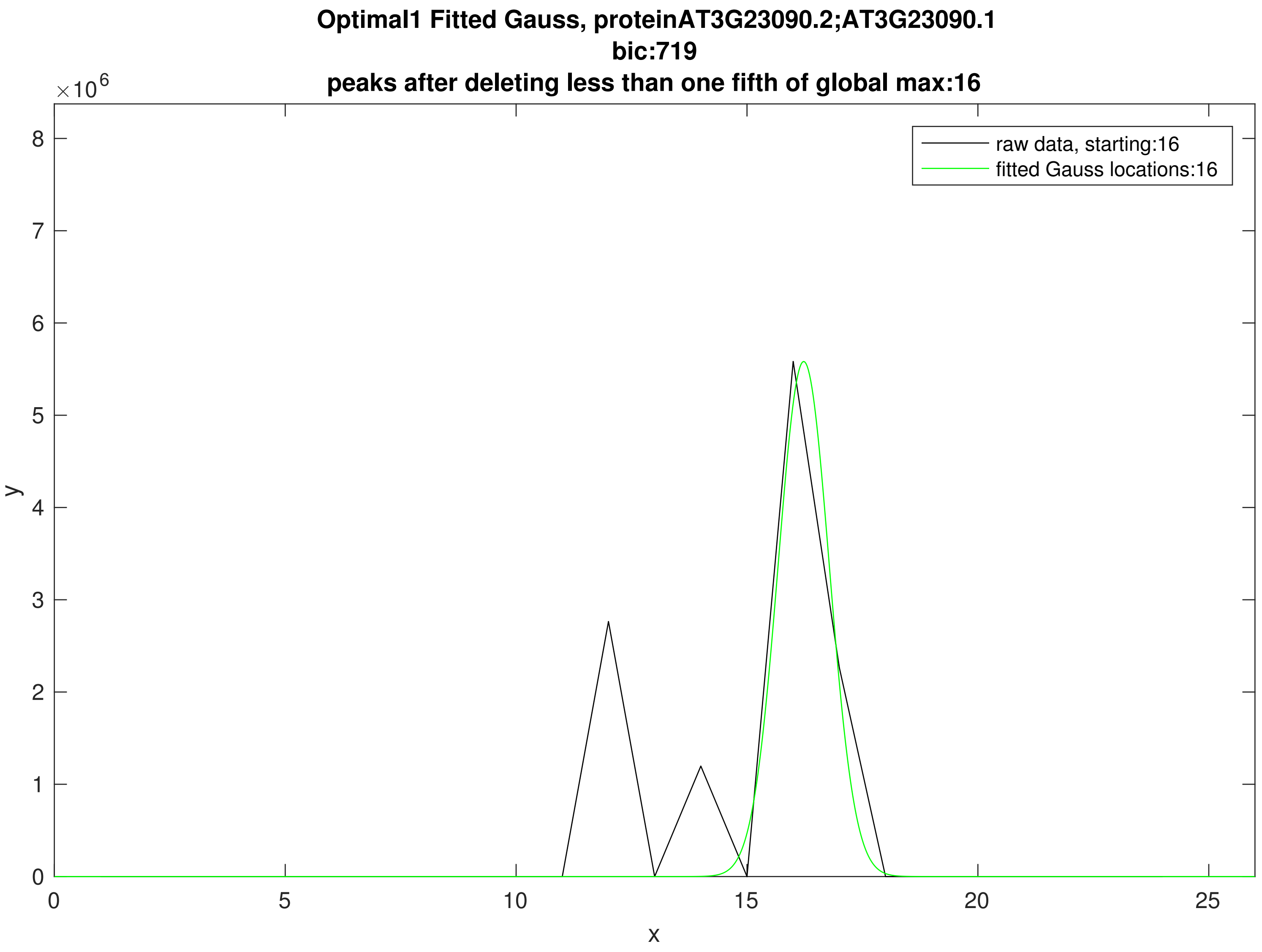
Optimal1 Fitted Gauss, proteinAT3G23090.2;AT3G23090.1
bic:719
peaks after deleting less than one fifth of global max:16
×10^6
8
7
6
5
4
3
2
1
0
y
0
5
10
15
20
25
x
raw data, starting:16
fitted Gauss locations:16

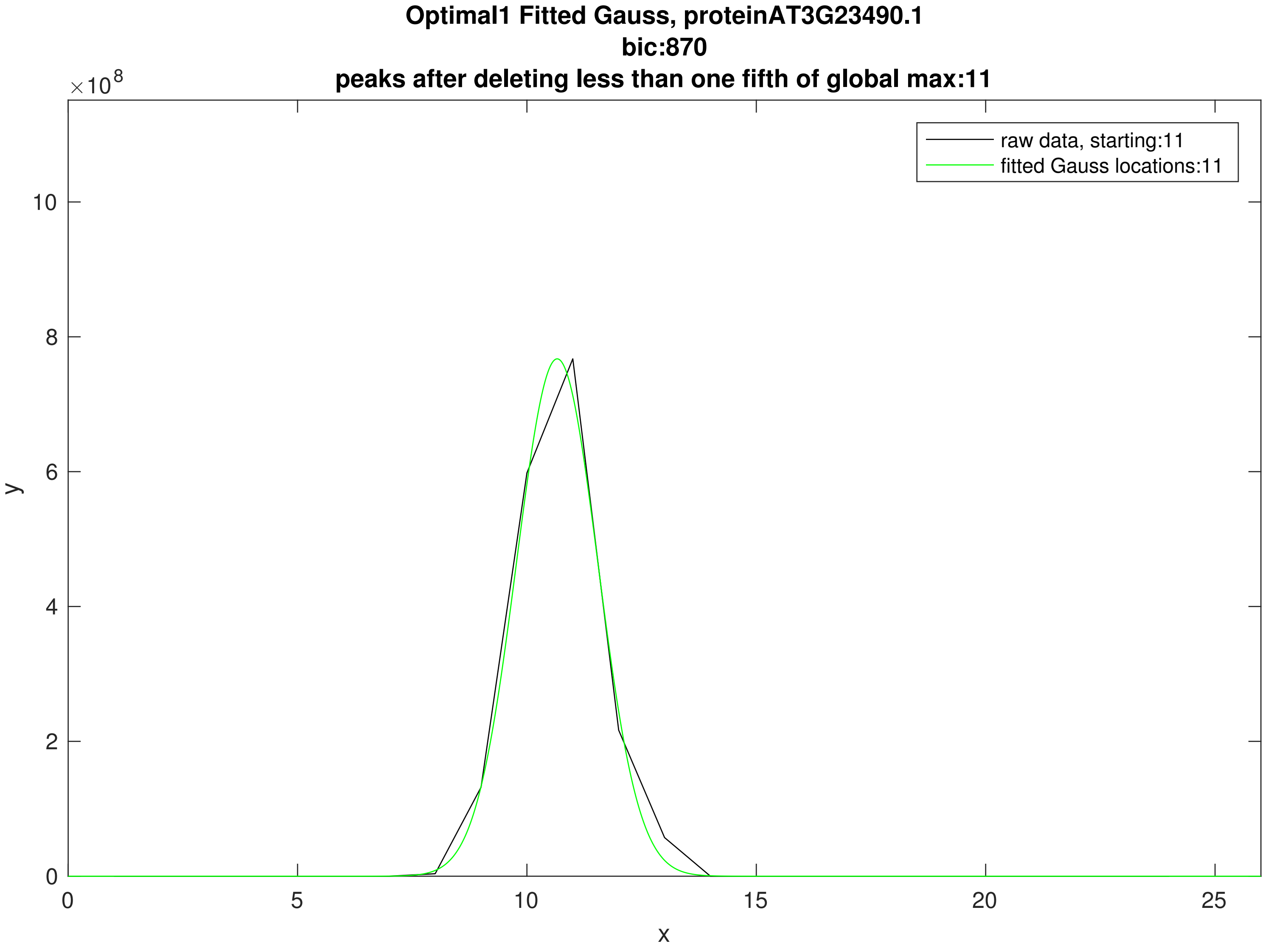

Optimal1 Fitted Gauss, proteinAT3G23490.1
bic:870
peaks after deleting less than one fifth of global max:11
×10^8
raw data, starting:11
fitted Gauss locations:11
y
x
0
2
4
6
8
10
0
5
10
15
20
25

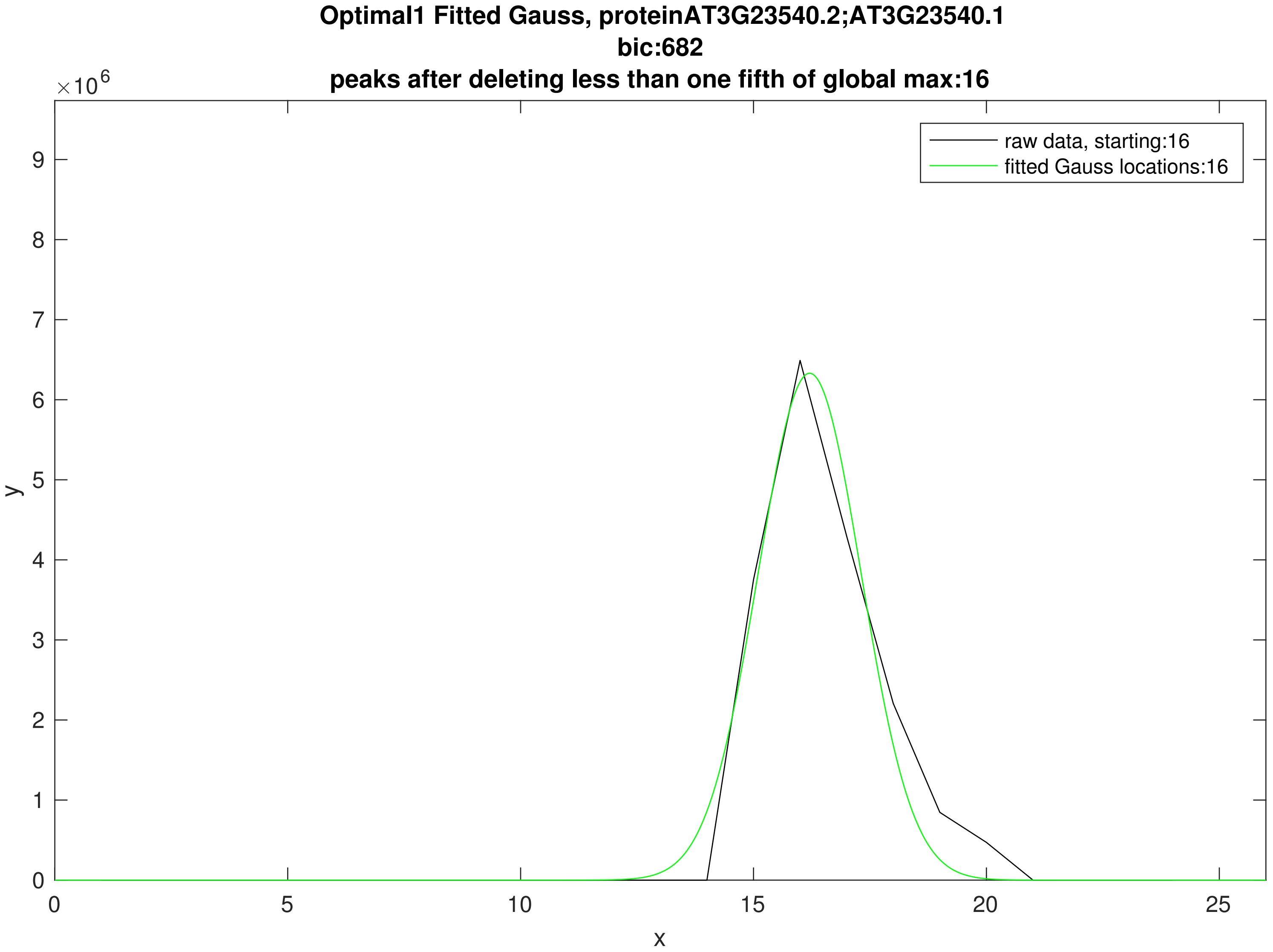
Optimal1 Fitted Gauss, proteinAT3G23540.2;AT3G23540.1
bic:682
peaks after deleting less than one fifth of global max:16
×10^6
raw data, starting:16
fitted Gauss locations:16
y
x
0
1
2
3
4
5
6
7
8
9
0
5
10
15
20
25

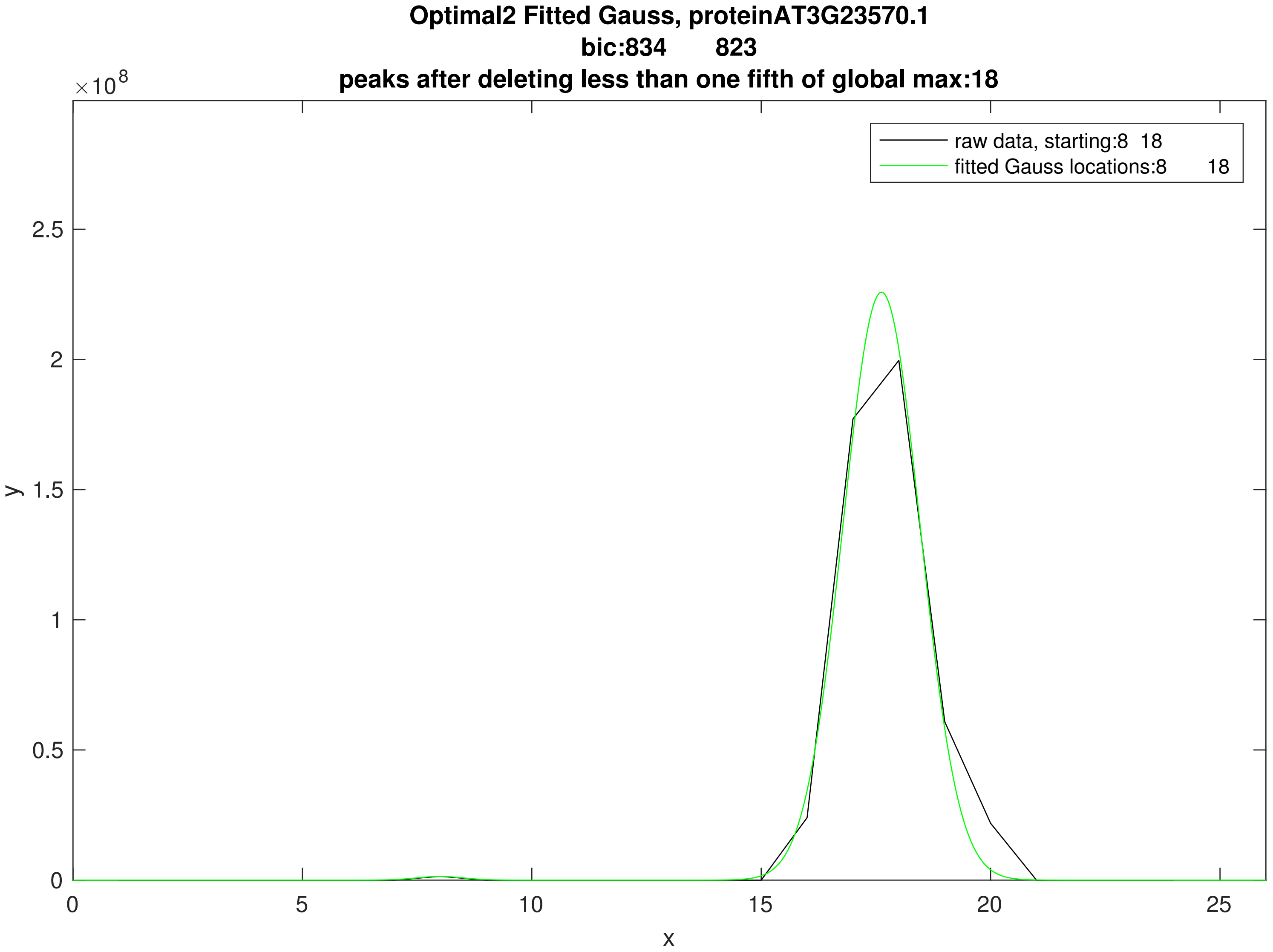

Optimal2 Fitted Gauss, proteinAT3G23570.1
bic:834 823
peaks after deleting less than one fifth of global max:18
×10^8
raw data, starting:8 18
fitted Gauss locations:8 18
2.5
2
1.5
1
0.5
0
y
0
5
10
15
20
25
x

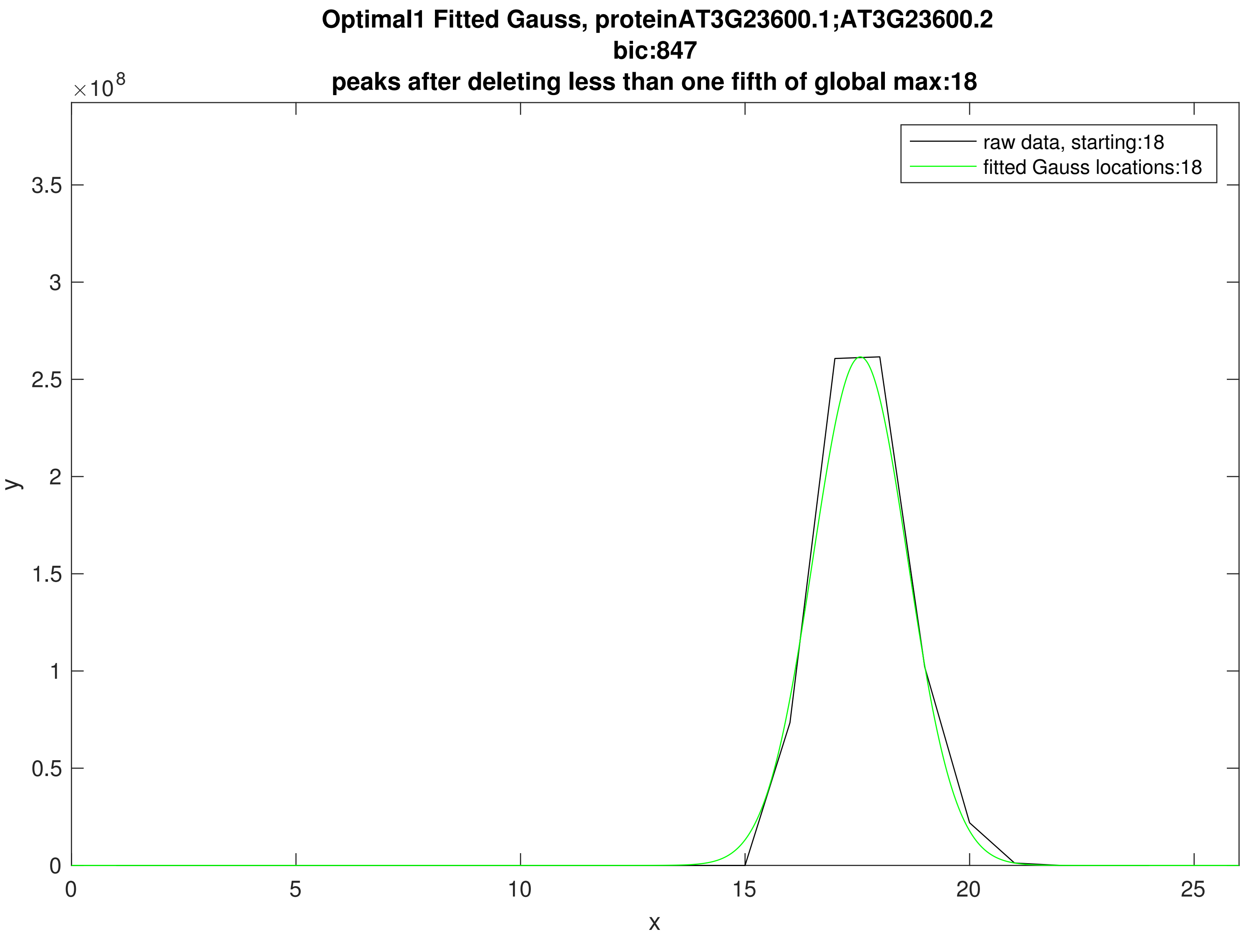
Optimal1 Fitted Gauss, proteinAT3G23600.1;AT3G23600.2
bic:847
peaks after deleting less than one fifth of global max:18
×10^8
raw data, starting:18
fitted Gauss locations:18
y
x
0
0.5
1
1.5
2
2.5
3
3.5
0
5
10
15
20
25

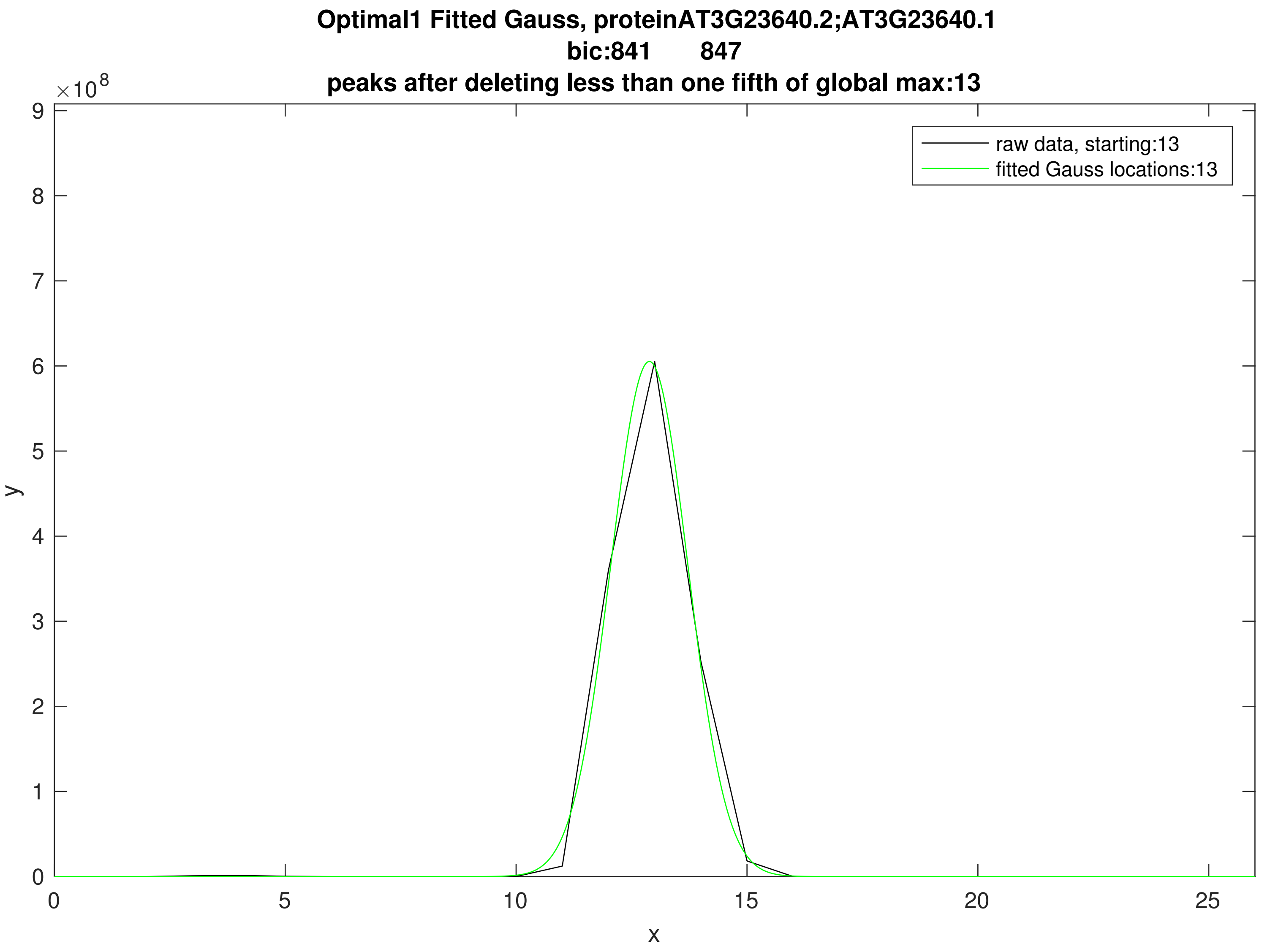
Optimal1 Fitted Gauss, proteinAT3G23640.2;AT3G23640.1
bic:841 847
peaks after deleting less than one fifth of global max:13
×10^8
9
8
7
6
5
4
3
2
1
0
y
0
5
10
15
20
25
x
raw data, starting:13
fitted Gauss locations:13

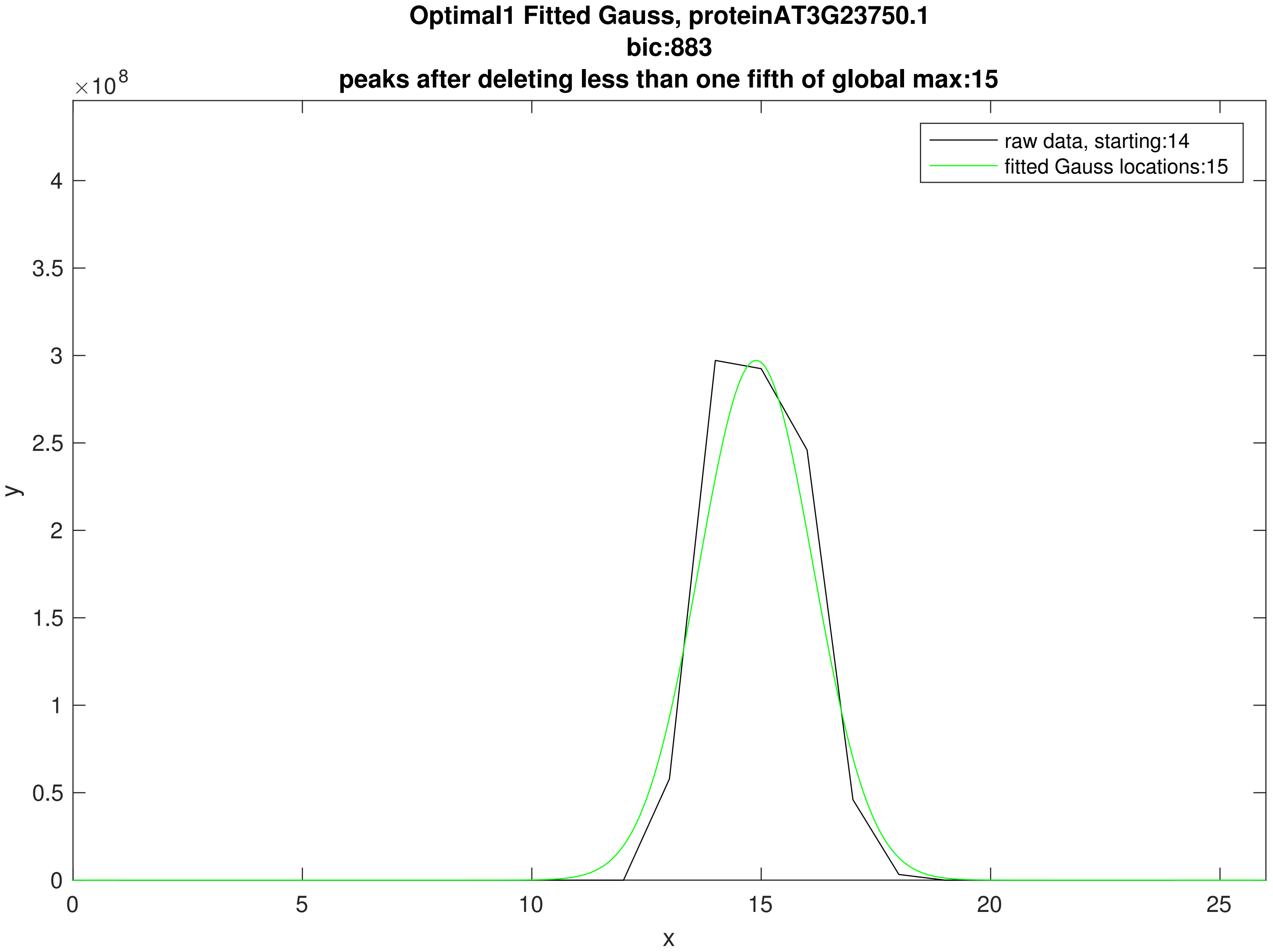
Optimal1 Fitted Gauss, proteinAT3G23750.1
bic:883
peaks after deleting less than one fifth of global max:15
×10^8
raw data, starting:14
fitted Gauss locations:15
y
x
0
0.5
1
1.5
2
2.5
3
3.5
4
0
5
10
15
20
25

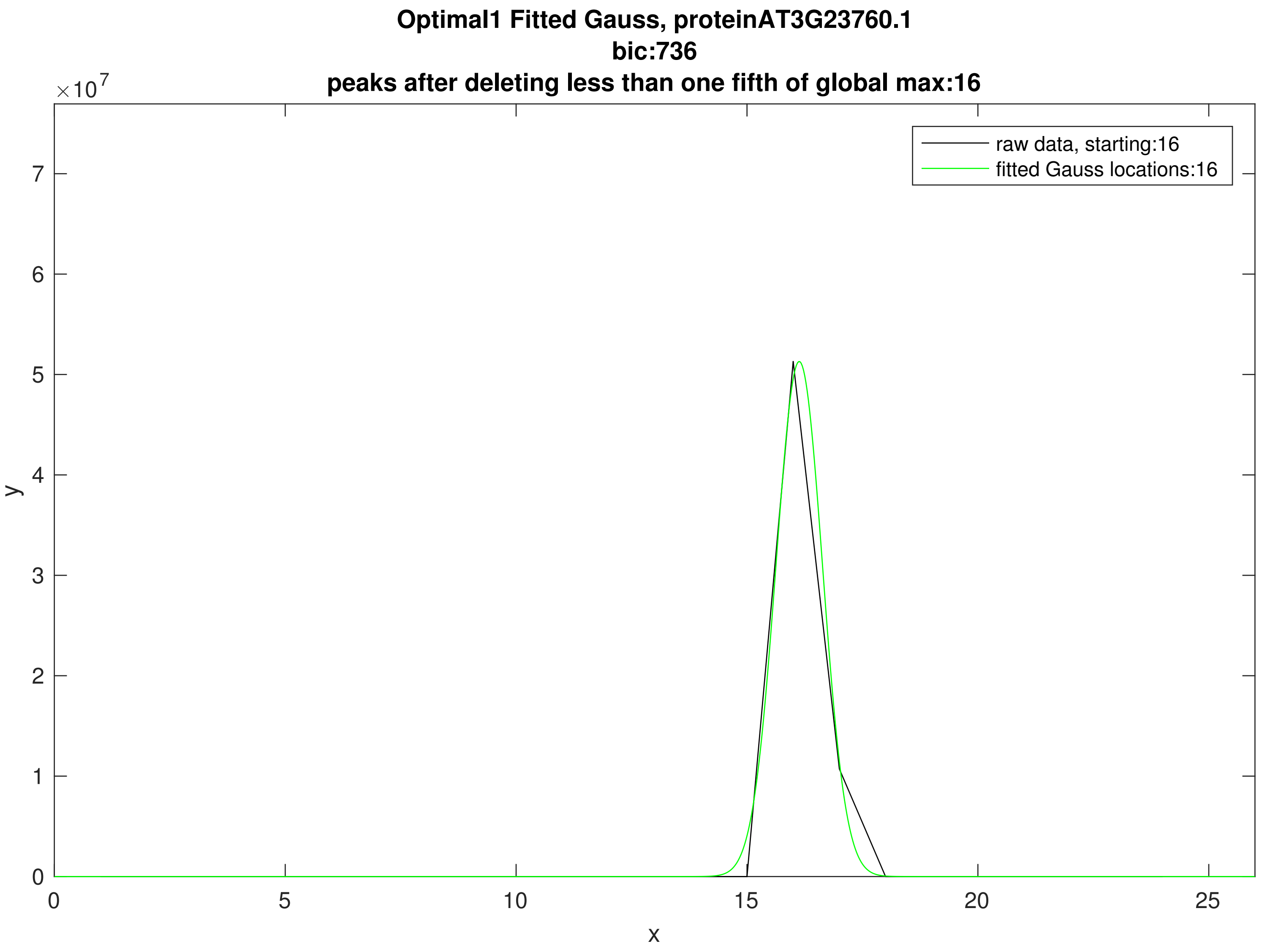
Optimal1 Fitted Gauss, proteinAT3G23760.1
bic:736
peaks after deleting less than one fifth of global max:16
×10^7
raw data, starting:16
fitted Gauss locations:16
y
x
0
1
2
3
4
5
6
7
0
5
10
15
20
25

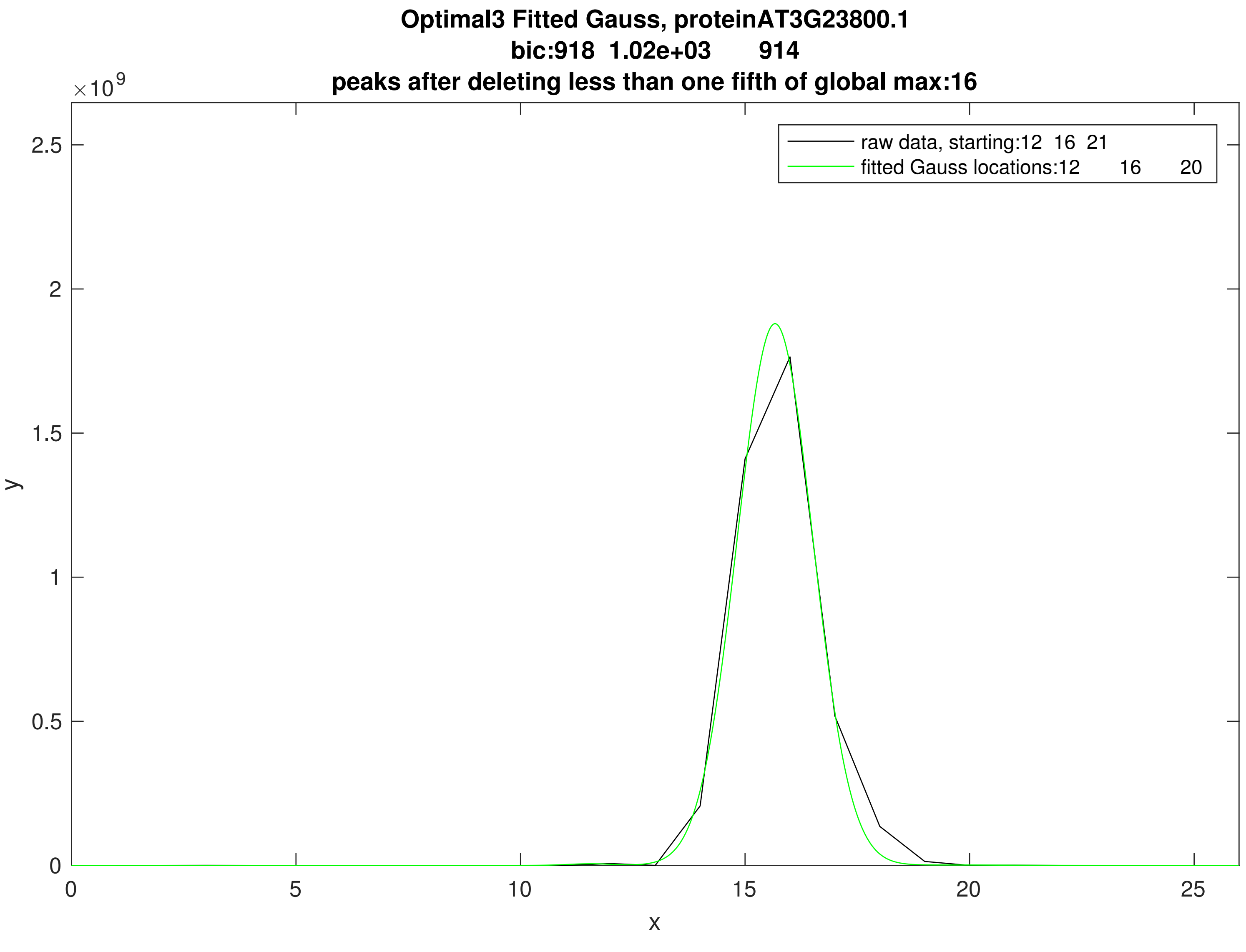
Optimal3 Fitted Gauss, proteinAT3G23800.1
bic:918 1.02e+03 914
peaks after deleting less than one fifth of global max:16
×10^9
raw data, starting:12 16 21
fitted Gauss locations:12 16 20
y
x
0
0.5
1
1.5
2
2.5
0
5
10
15
20
25

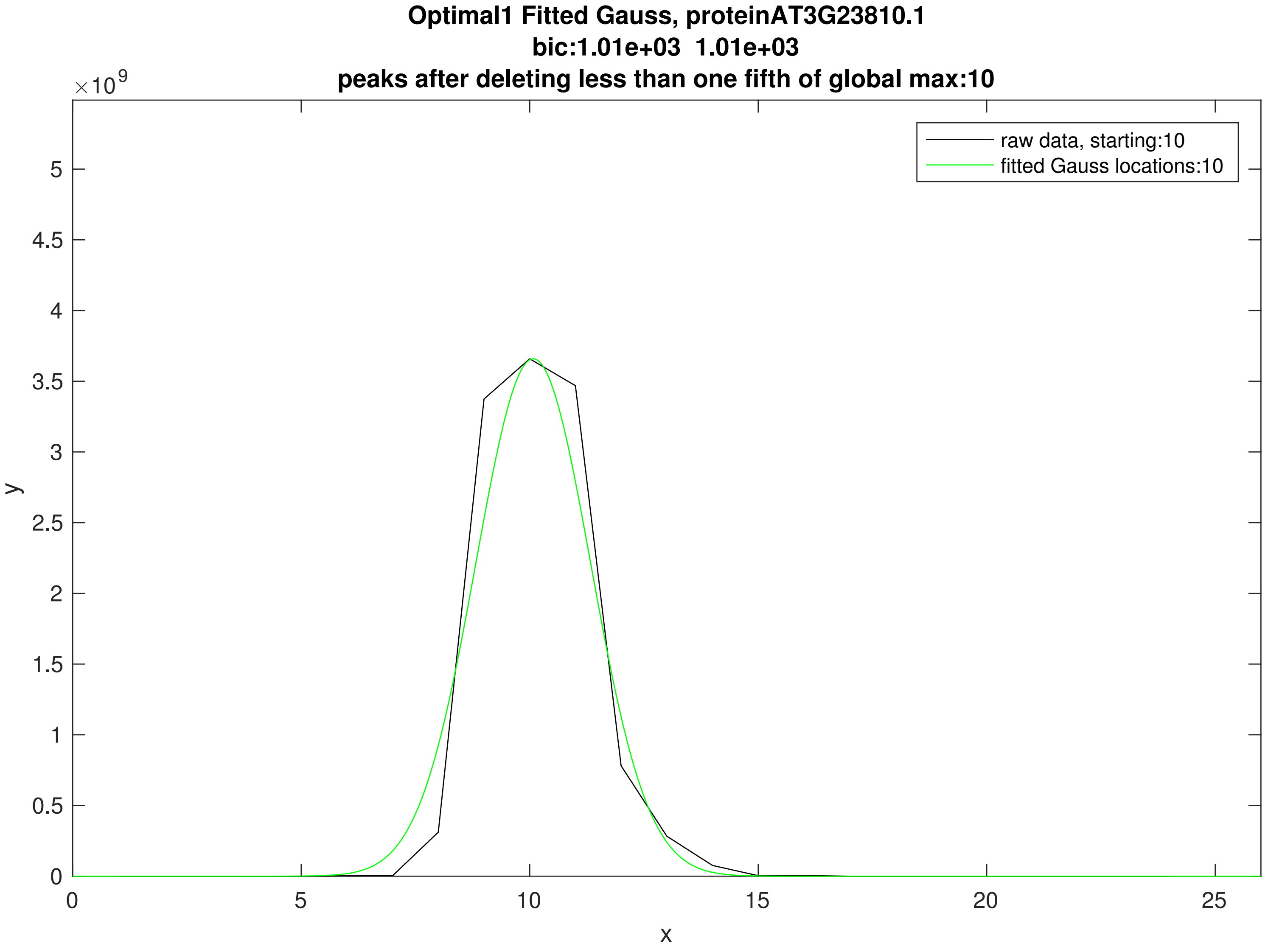
Optimal1 Fitted Gauss, proteinAT3G23810.1
bic:1.01e+03  1.01e+03
peaks after deleting less than one fifth of global max:10
×10^9
raw data, starting:10
fitted Gauss locations:10
y
x
0
0.5
1
1.5
2
2.5
3
3.5
4
4.5
5
0
5
10
15
20
25

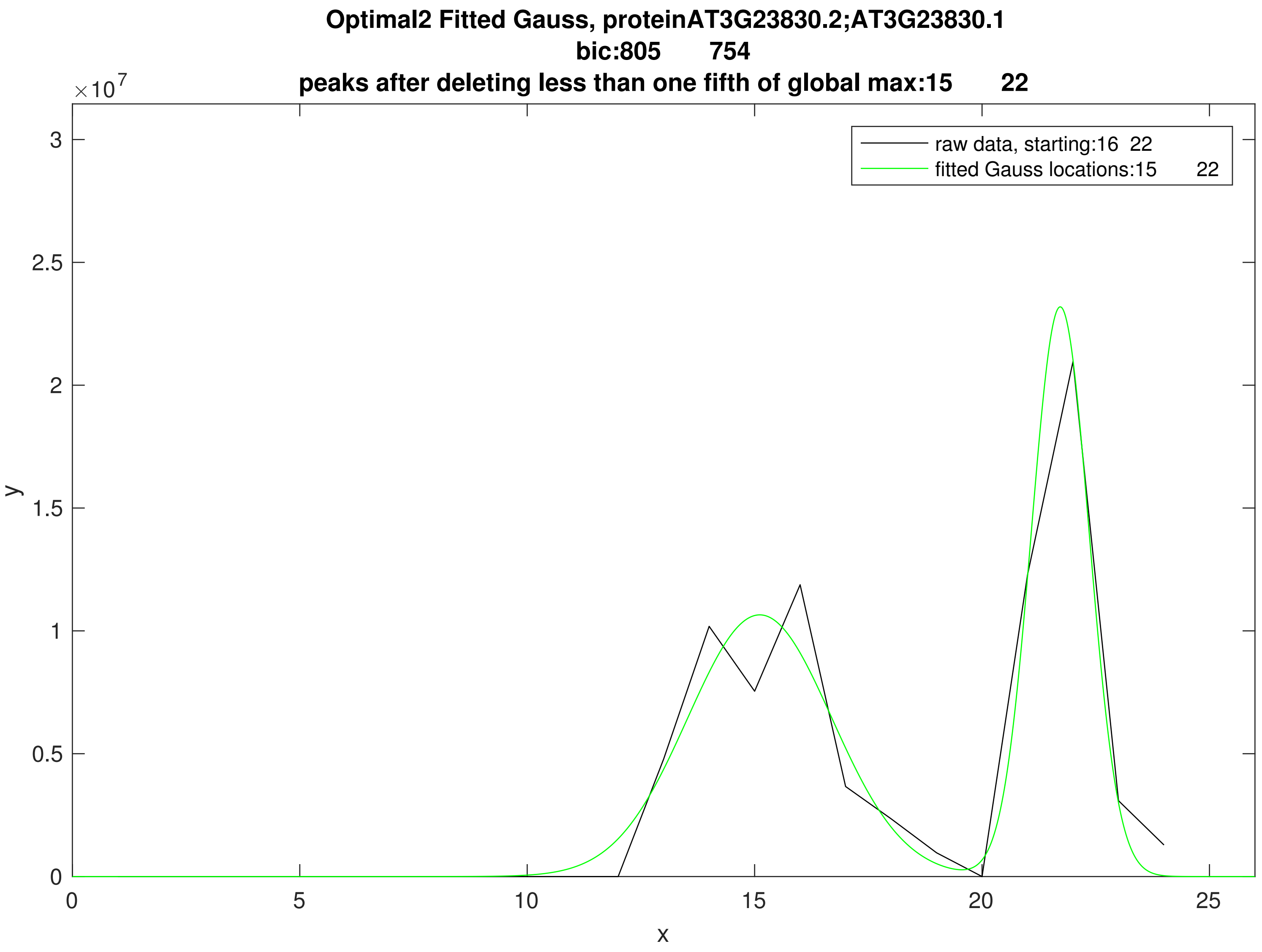
Optimal2 Fitted Gauss, proteinAT3G23830.2;AT3G23830.1
bic:805 754
peaks after deleting less than one fifth of global max:15 22
×10^7
raw data, starting:16 22
fitted Gauss locations:15 22
y
x
0
0.5
1
1.5
2
2.5
3
0
5
10
15
20
25

**Optimal1 Fitted Gauss, proteinAT3G23940.2;AT3G23940.1**
**bic:949 959**
**peaks after deleting less than one fifth of global max:13**

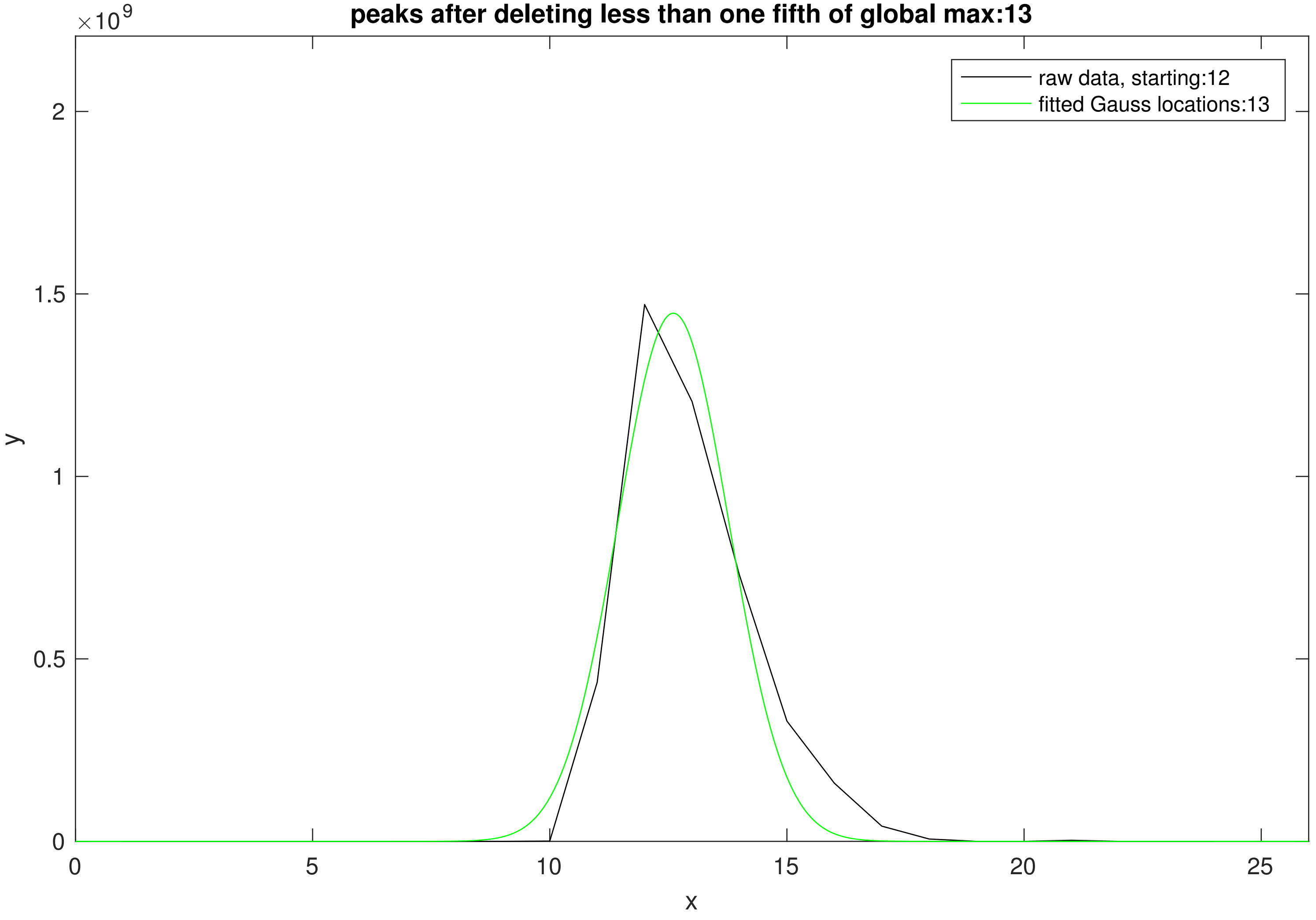

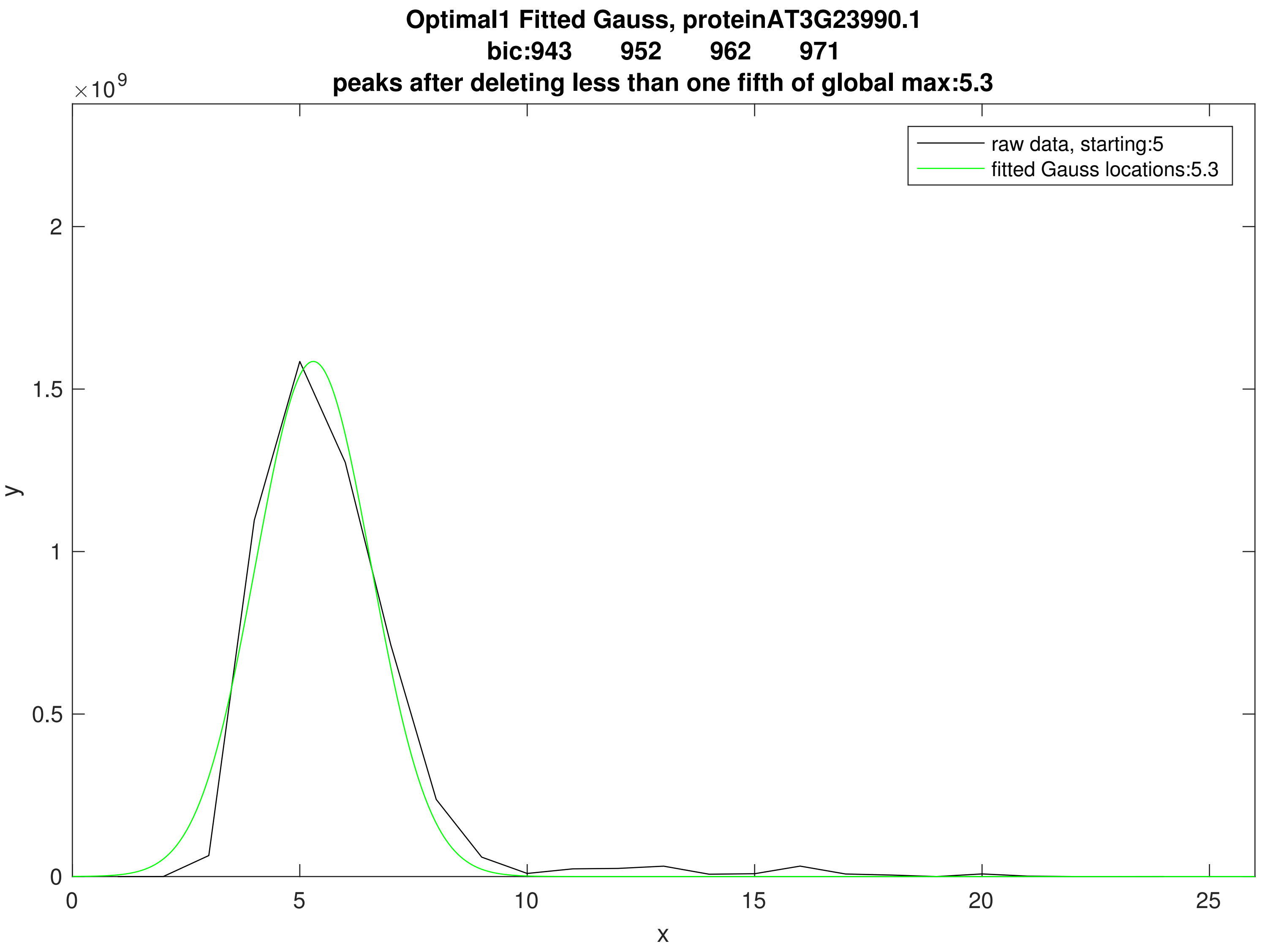

Optimal1 Fitted Gauss, proteinAT3G23990.1
bic:943 952 962 971
peaks after deleting less than one fifth of global max:5.3
×10^9
raw data, starting:5
fitted Gauss locations:5.3
y
x
0
0.5
1
1.5
2
0
5
10
15
20
25

Optimal1 Fitted Gauss, proteinAT3G24020.1;AT4G13580.1
bic:744
peaks after deleting less than one fifth of global max:18

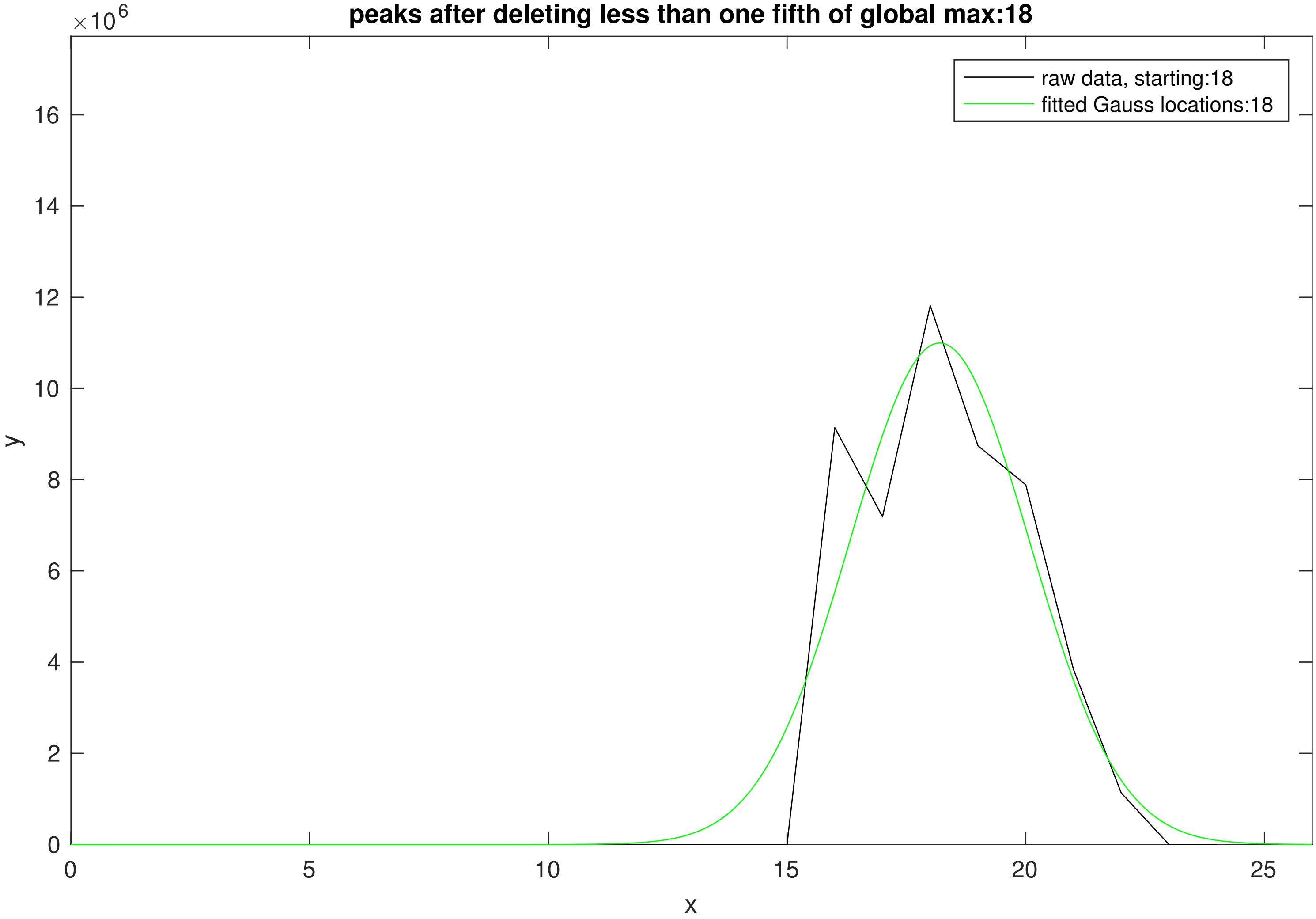

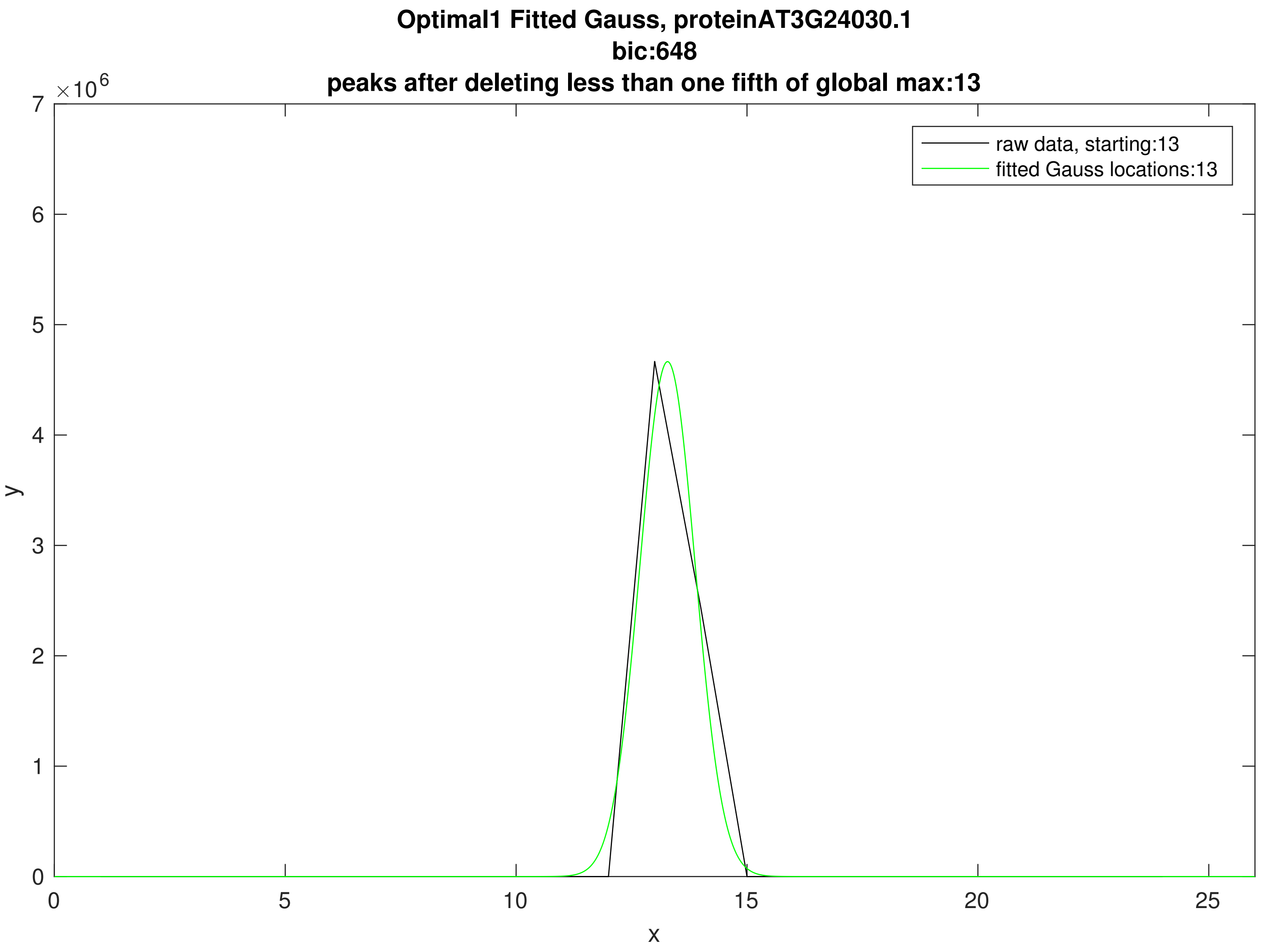
Optimal1 Fitted Gauss, proteinAT3G24030.1
bic:648
peaks after deleting less than one fifth of global max:13
×10^6
7
6
5
4
3
2
1
0
y
0
5
10
15
20
25
x
raw data, starting:13
fitted Gauss locations:13

Optimal1 Fitted Gauss, proteinAT3G24170.3;AT3G24170.2;AT3G24170.1
bic:990 994 1e+03
peaks after deleting less than one fifth of global max:12

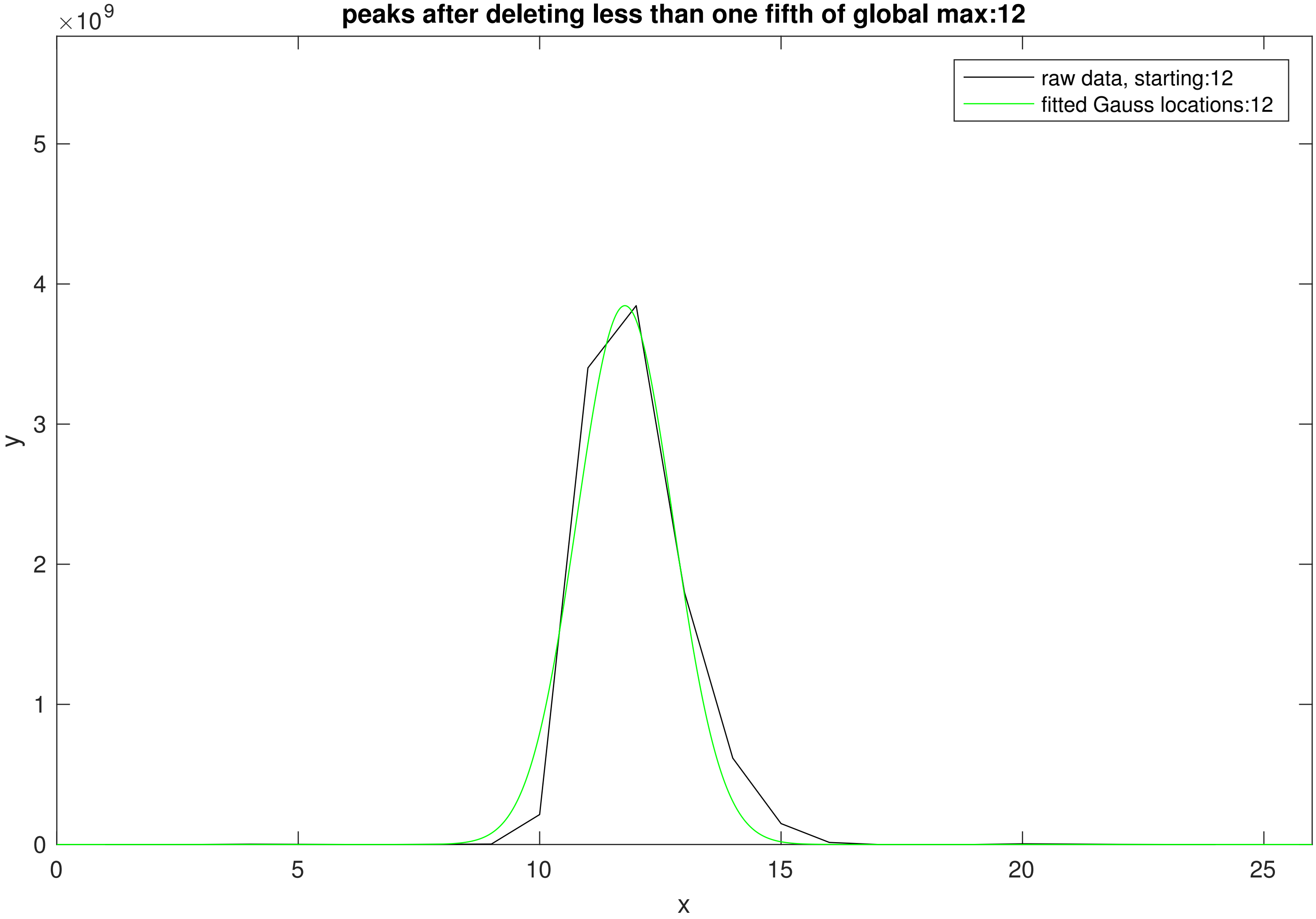

Optimal1 Fitted Gauss, proteinAT3G24420.1
bic:770
peaks after deleting less than one fifth of global max:18

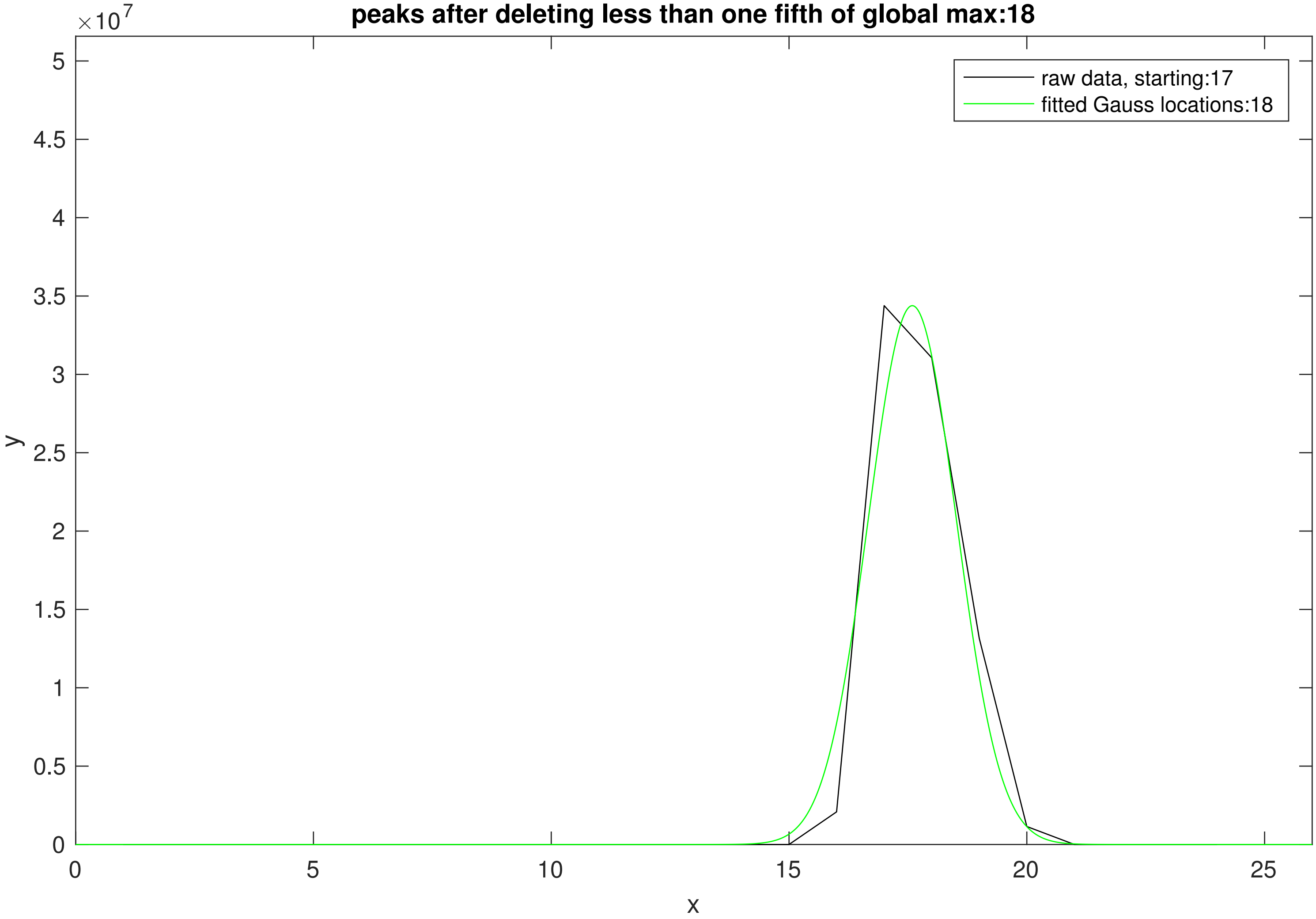

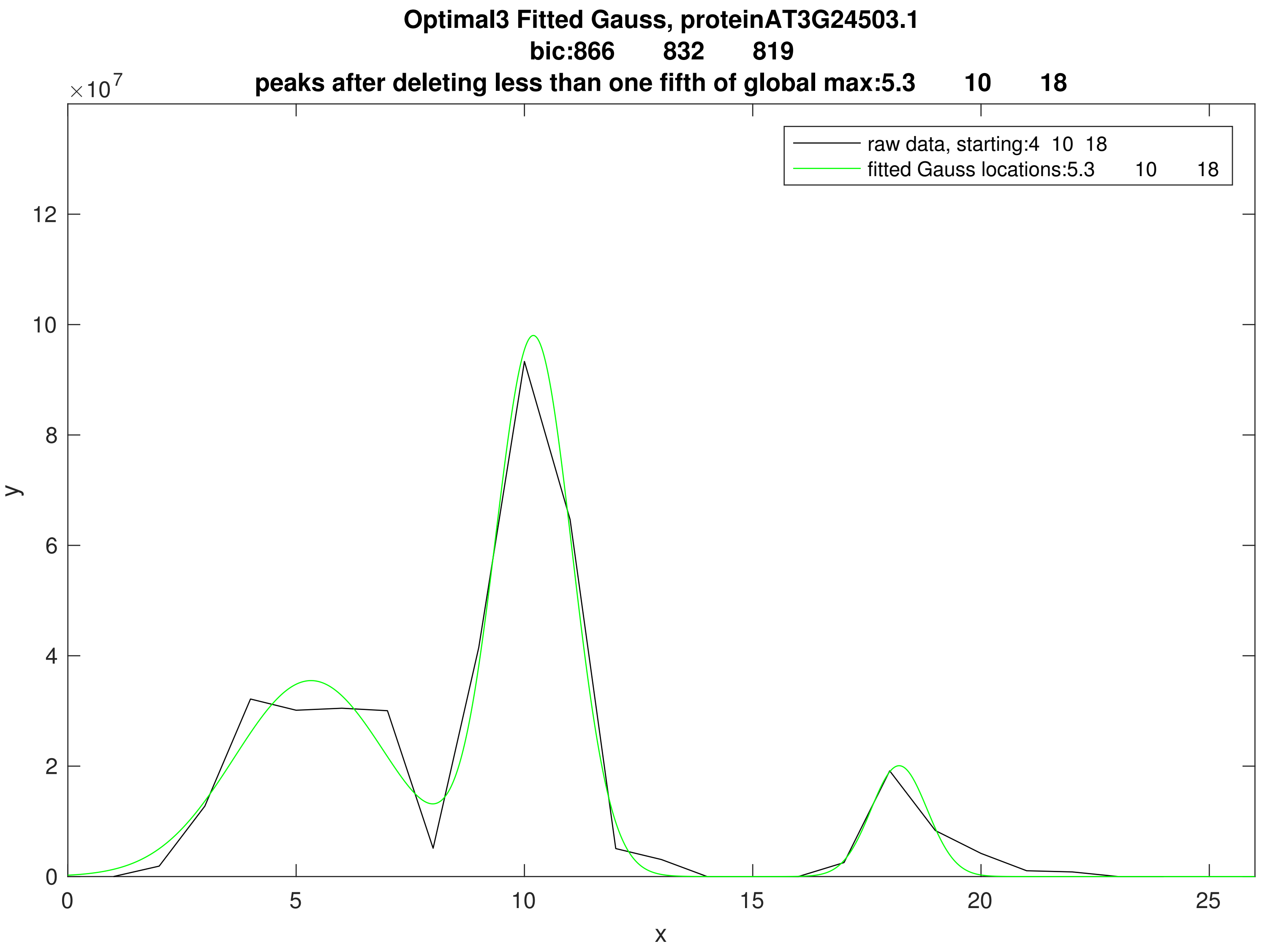
Optimal3 Fitted Gauss, proteinAT3G24503.1
bic:866 832 819
peaks after deleting less than one fifth of global max:5.3 10 18
×10^7
raw data, starting:4 10 18
fitted Gauss locations:5.3 10 18
y
x
0
2
4
6
8
10
12
0
5
10
15
20
25

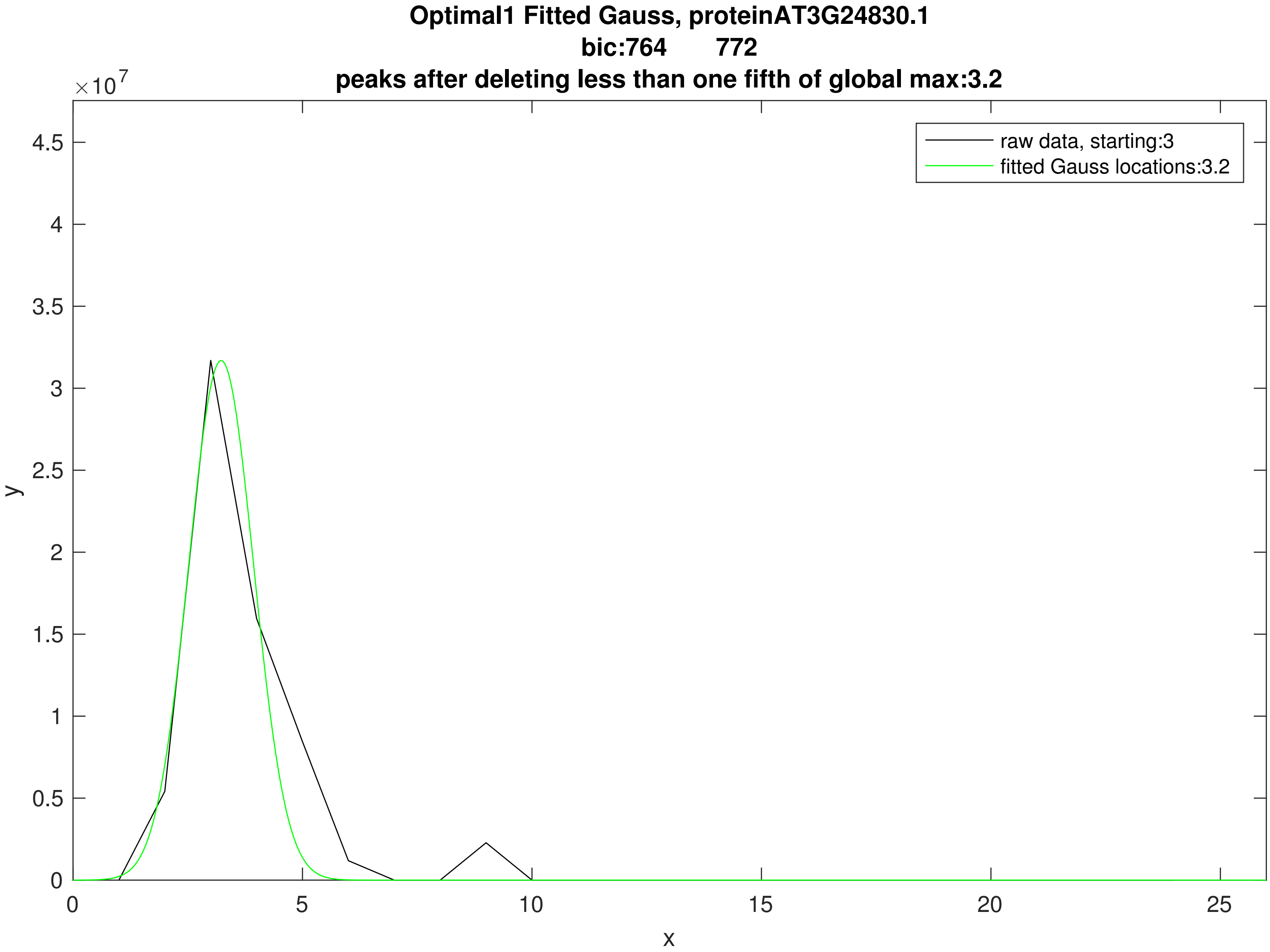
Optimal1 Fitted Gauss, proteinAT3G24830.1
bic:764 772
peaks after deleting less than one fifth of global max:3.2
×10^7
4.5
4
3.5
3
2.5
2
1.5
1
0.5
0
y
0
5
10
15
20
25
x
raw data, starting:3
fitted Gauss locations:3.2

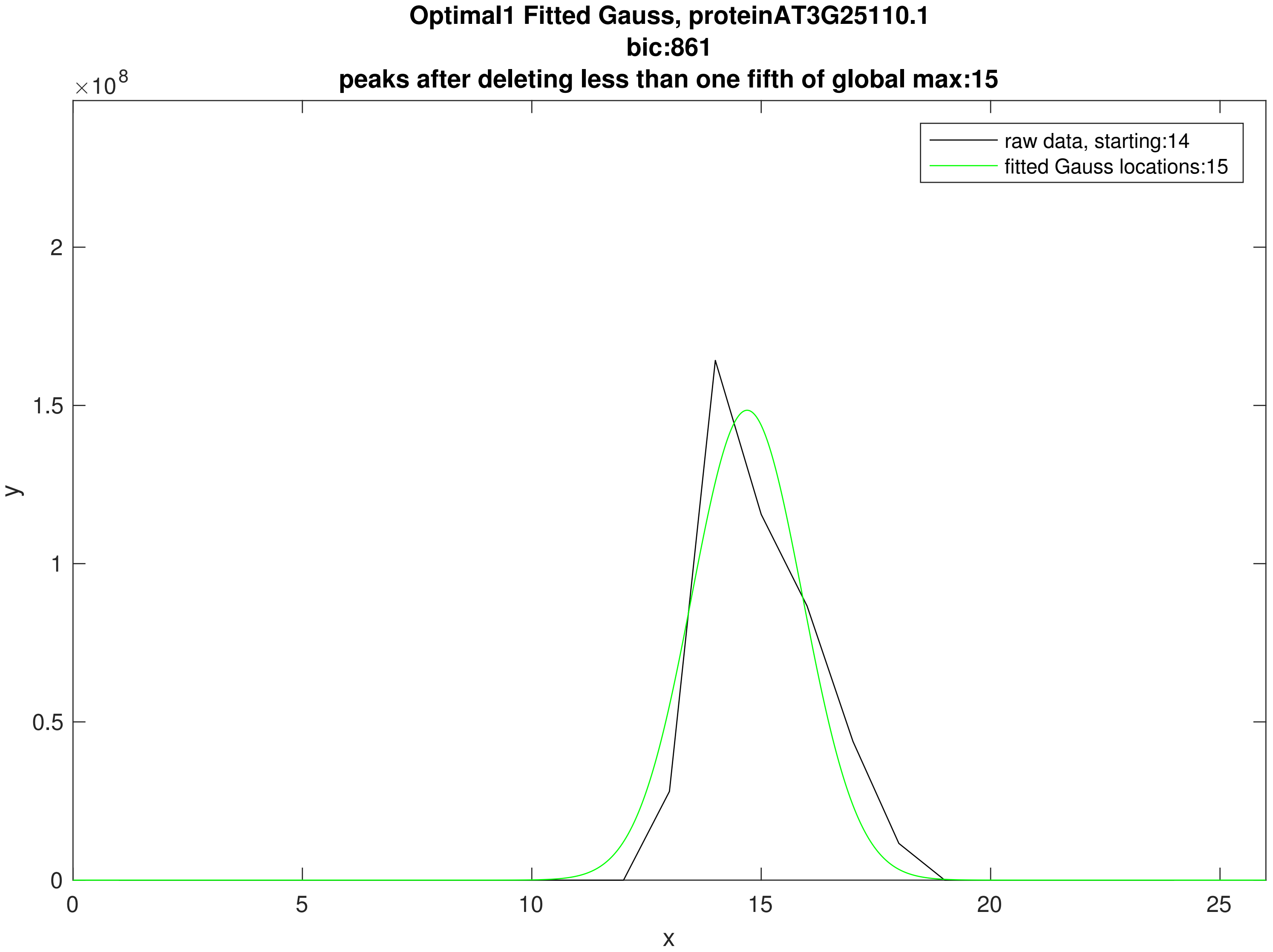
Optimal1 Fitted Gauss, proteinAT3G25110.1
bic:861
peaks after deleting less than one fifth of global max:15
×10^8
raw data, starting:14
fitted Gauss locations:15
y
x
0
0.5
1
1.5
2
0
5
10
15
20
25

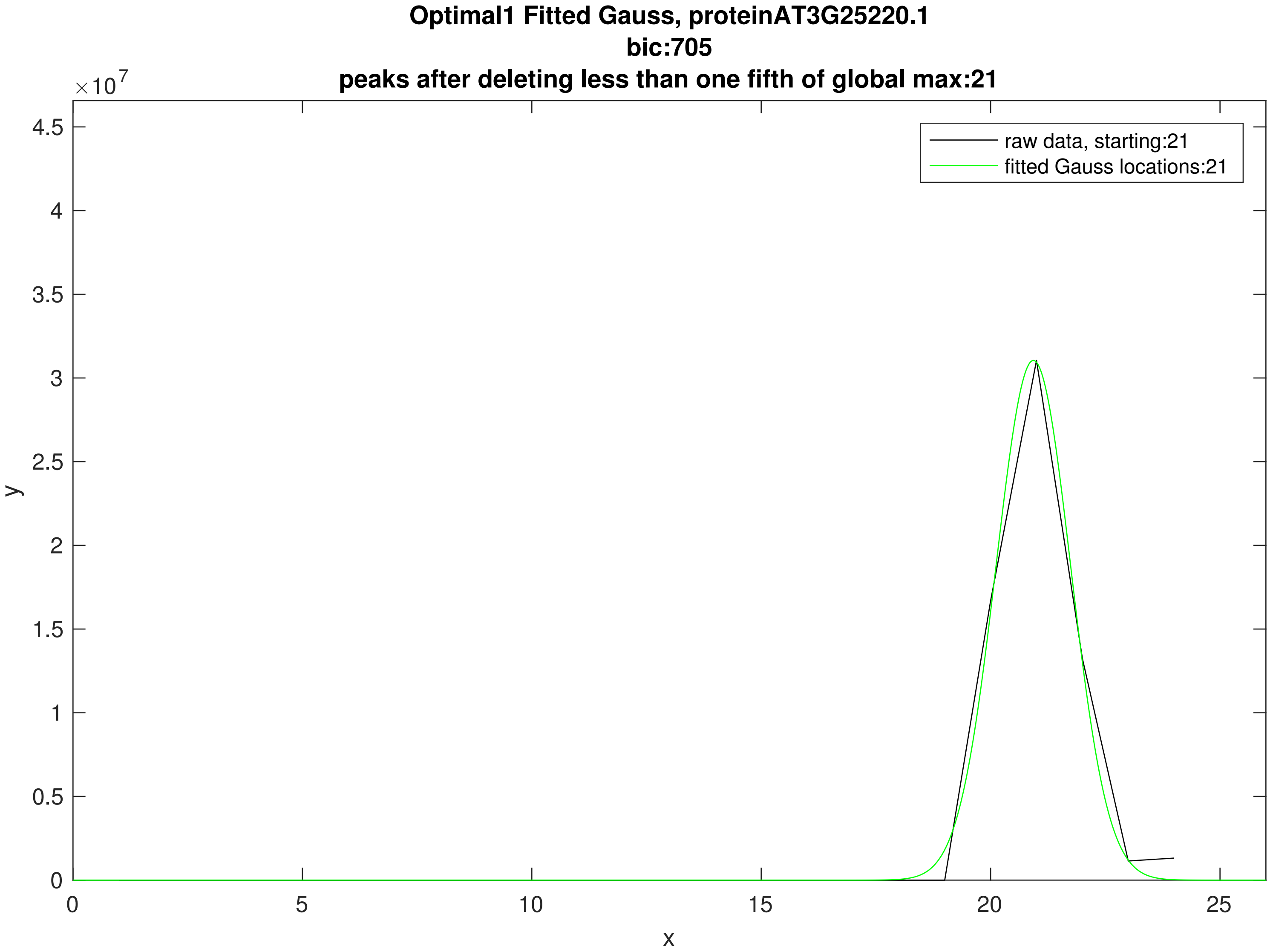

Optimal1 Fitted Gauss, proteinAT3G25220.1
bic:705
peaks after deleting less than one fifth of global max:21
×10^7
4.5
4
3.5
3
2.5
2
1.5
1
0.5
0
y
0
5
10
15
20
25
x
raw data, starting:21
fitted Gauss locations:21

Optimal1 Fitted Gauss, proteinAT3G25230.1;AT3G25230.2
bic:865
peaks after deleting less than one fifth of global max:13

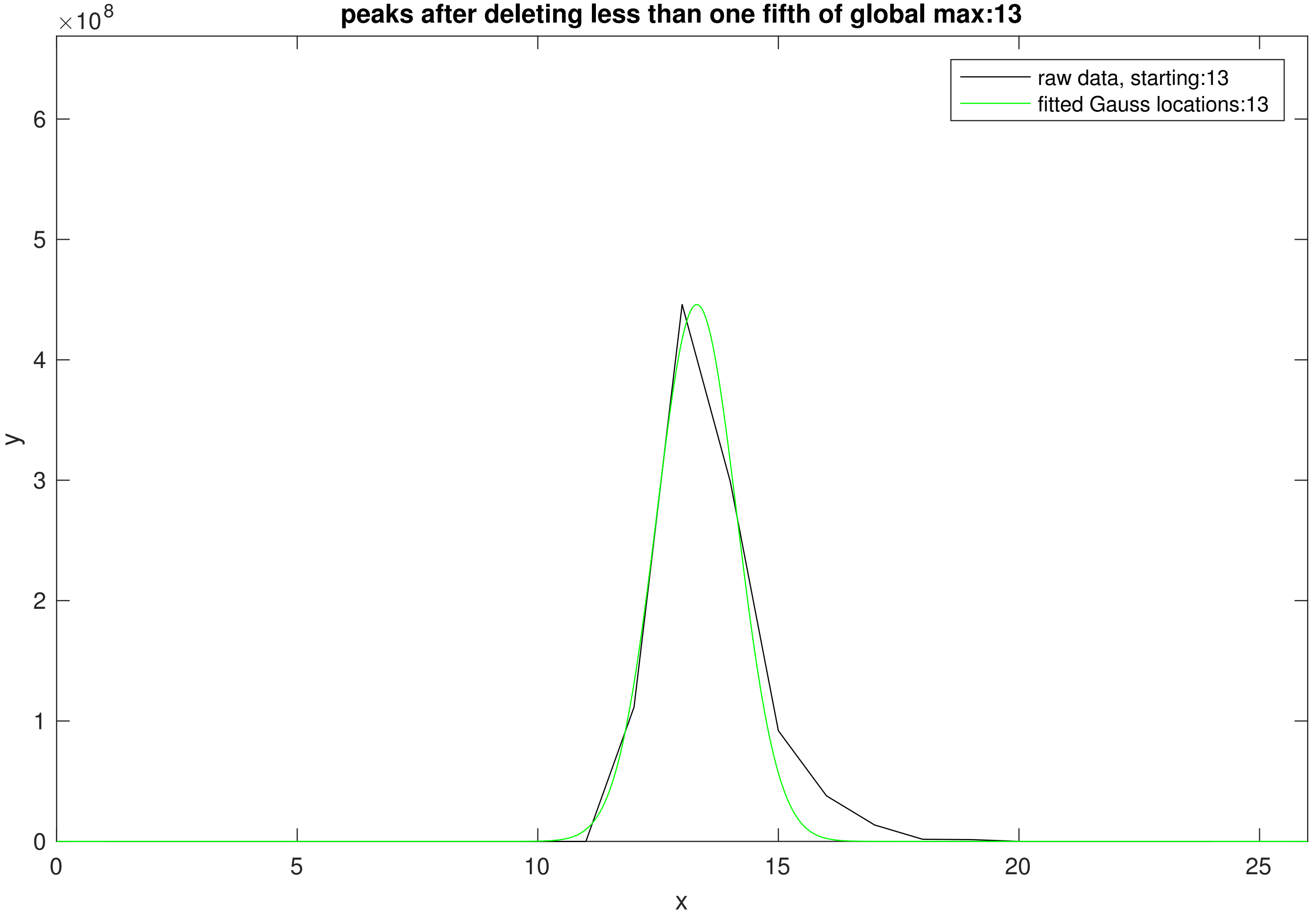

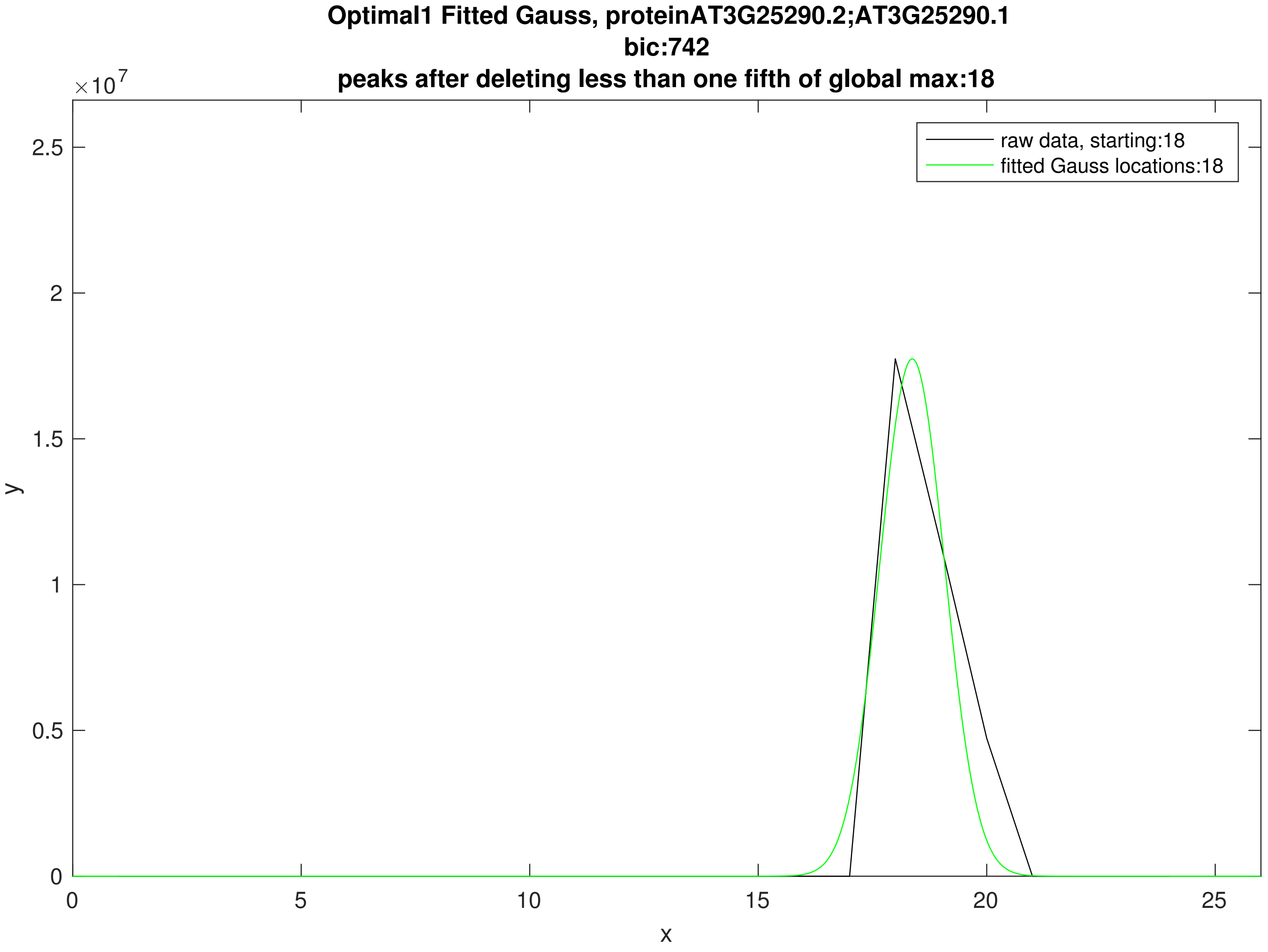

Optimal1 Fitted Gauss, proteinAT3G25290.2;AT3G25290.1
bic:742
peaks after deleting less than one fifth of global max:18
×10^7
2.5
2
1.5
1
0.5
0
y
0
5
10
15
20
25
x
raw data, starting:18
fitted Gauss locations:18

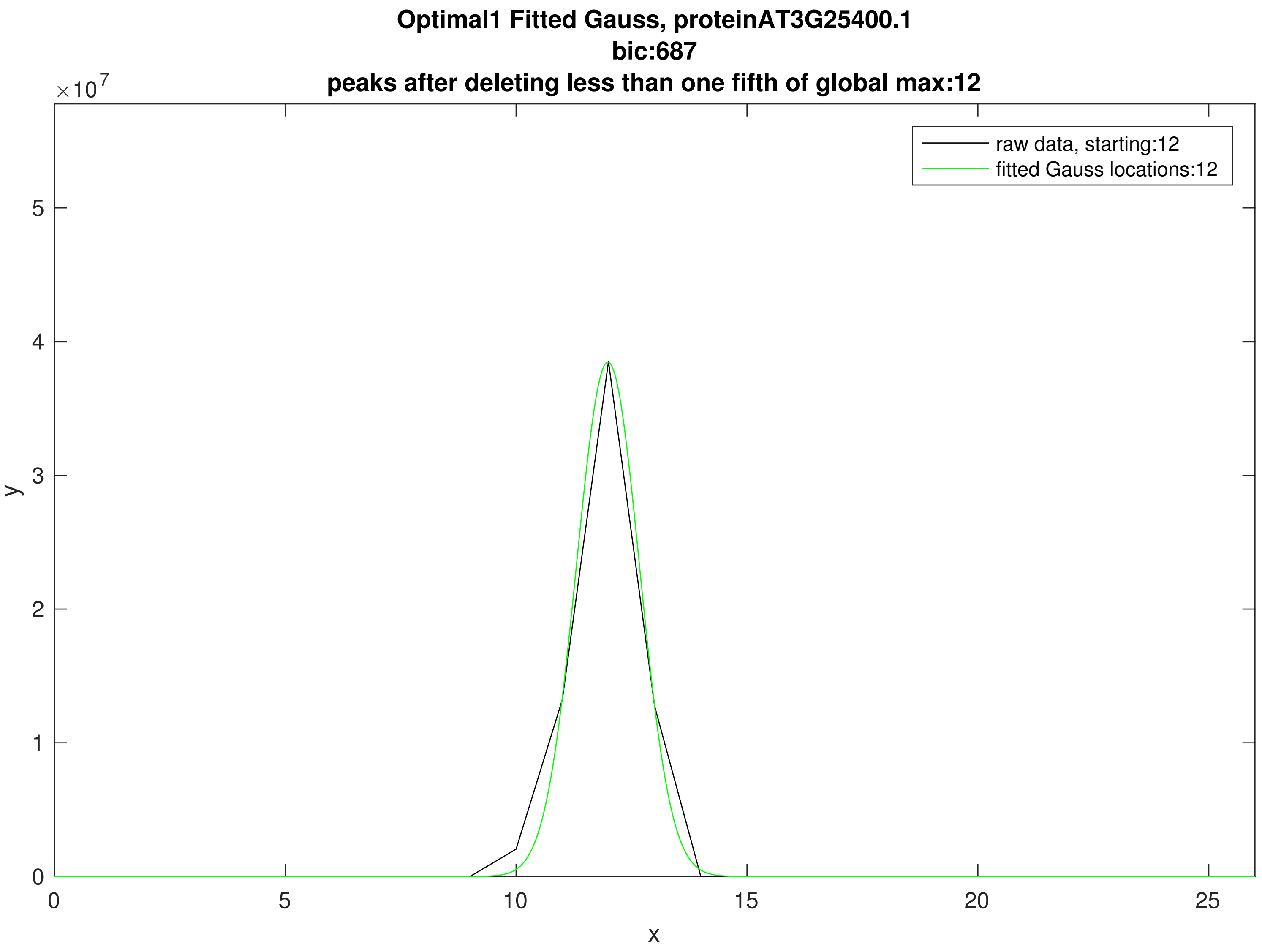
Optimal1 Fitted Gauss, proteinAT3G25400.1
bic:687
peaks after deleting less than one fifth of global max:12
×10^7
raw data, starting:12
fitted Gauss locations:12
y
x
0
1
2
3
4
5
0
5
10
15
20
25

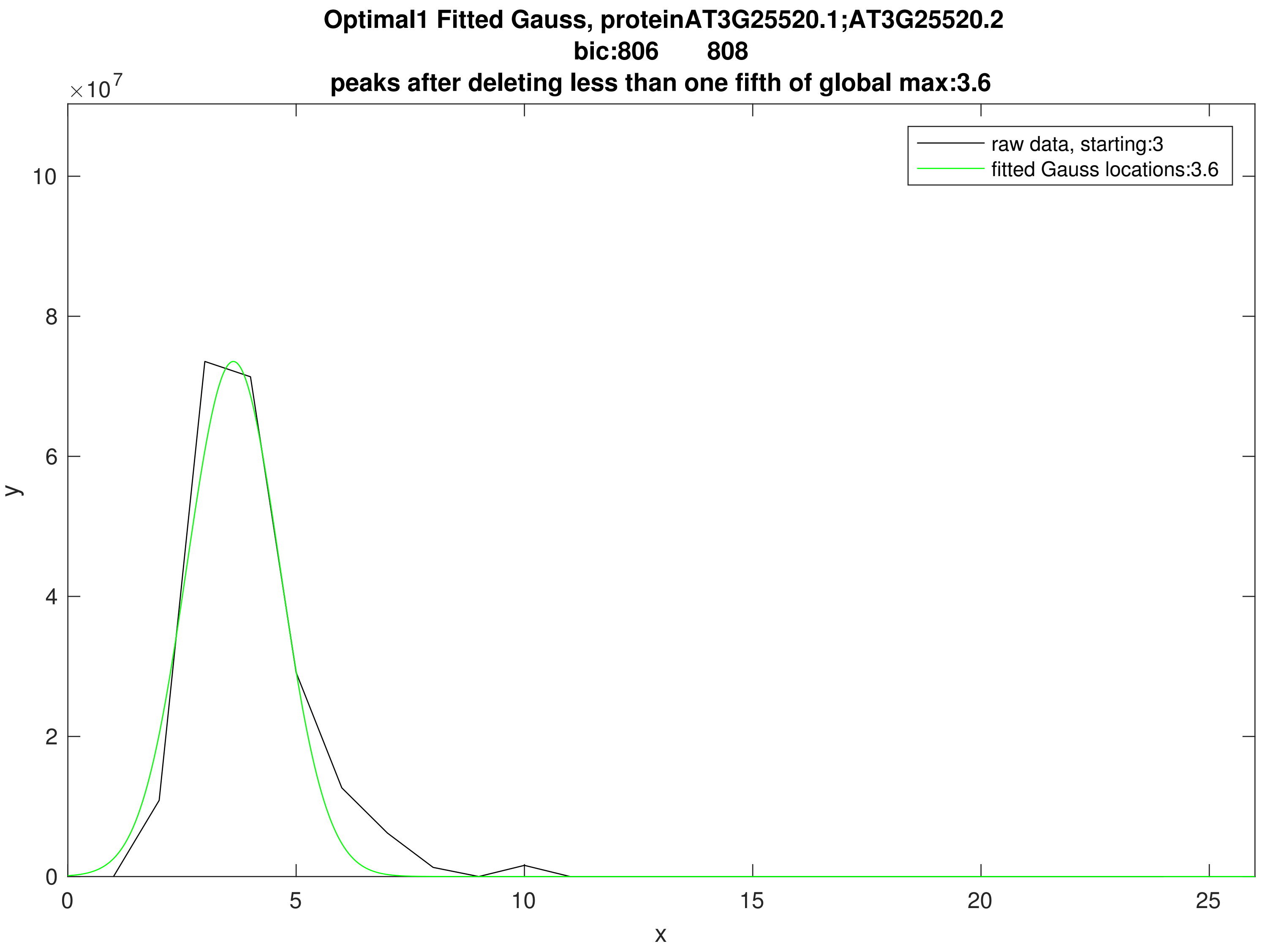

Optimal1 Fitted Gauss, proteinAT3G25520.1;AT3G25520.2
bic:806 808
peaks after deleting less than one fifth of global max:3.6
×10^7
raw data, starting:3
fitted Gauss locations:3.6
y
x
0
2
4
6
8
10
0
5
10
15
20
25

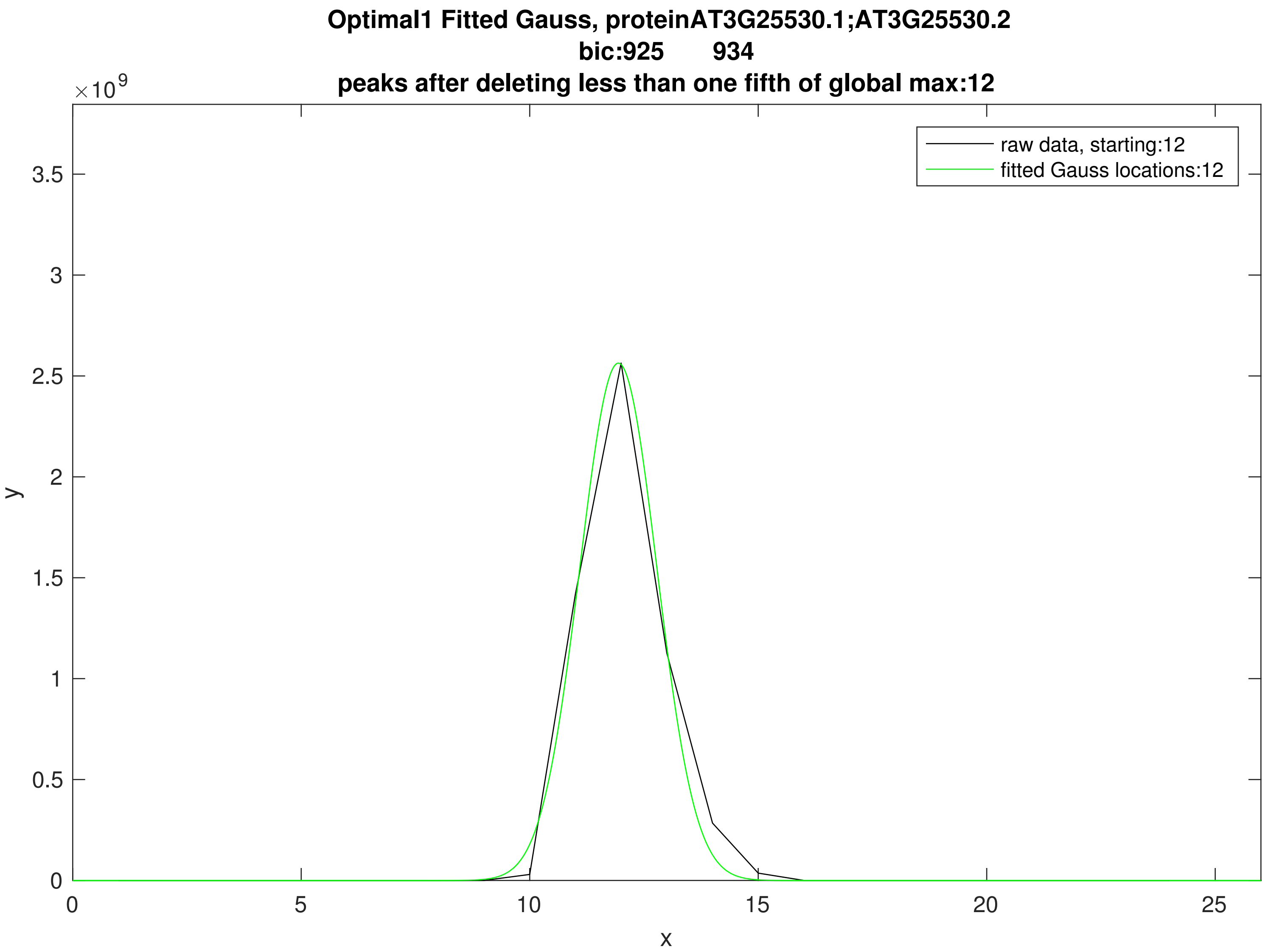

Optimal1 Fitted Gauss, proteinAT3G25530.1;AT3G25530.2
bic:925 934
peaks after deleting less than one fifth of global max:12
×10^9
3.5
3
2.5
2
1.5
1
0.5
0
y
0
5
10
15
20
25
x
raw data, starting:12
fitted Gauss locations:12

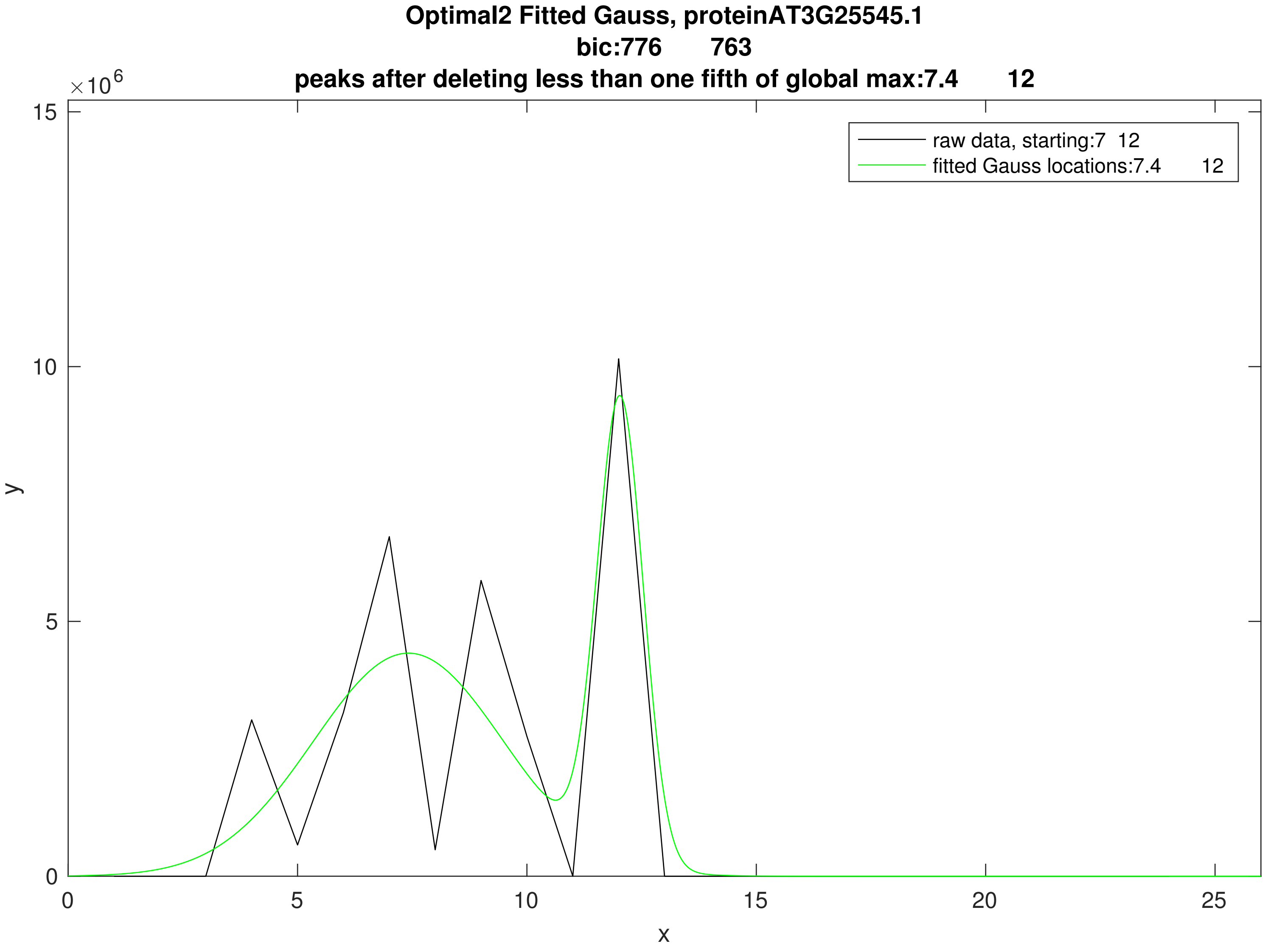

Optimal2 Fitted Gauss, proteinAT3G25545.1
bic:776 763
peaks after deleting less than one fifth of global max:7.4 12
×10^6
15
10
5
0
y
0
5
10
15
20
25
x
raw data, starting:7 12
fitted Gauss locations:7.4 12

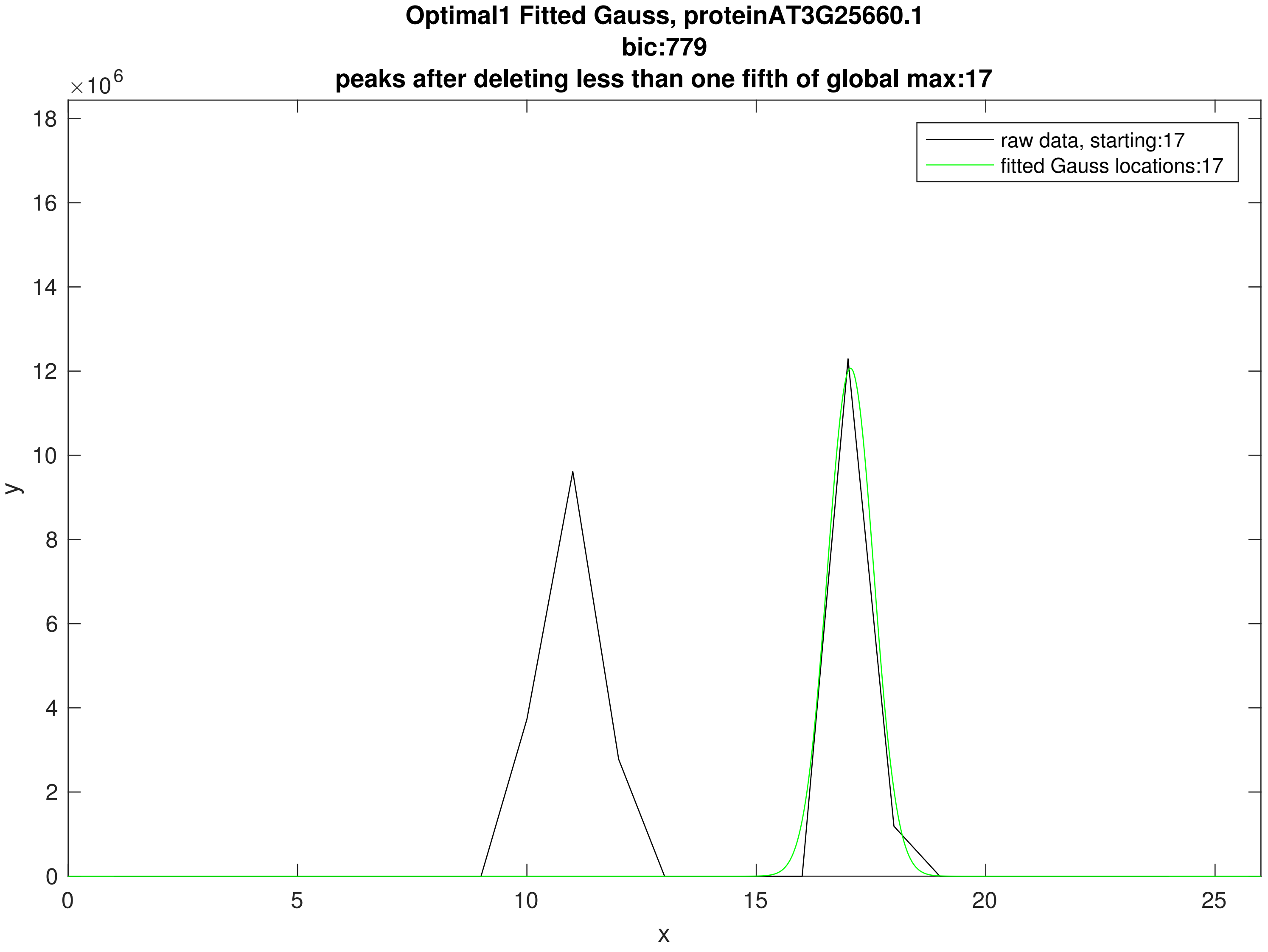
Optimal1 Fitted Gauss, proteinAT3G25660.1
bic:779
peaks after deleting less than one fifth of global max:17
×10^6
raw data, starting:17
fitted Gauss locations:17
y
x
0
2
4
6
8
10
12
14
16
18
0
5
10
15
20
25

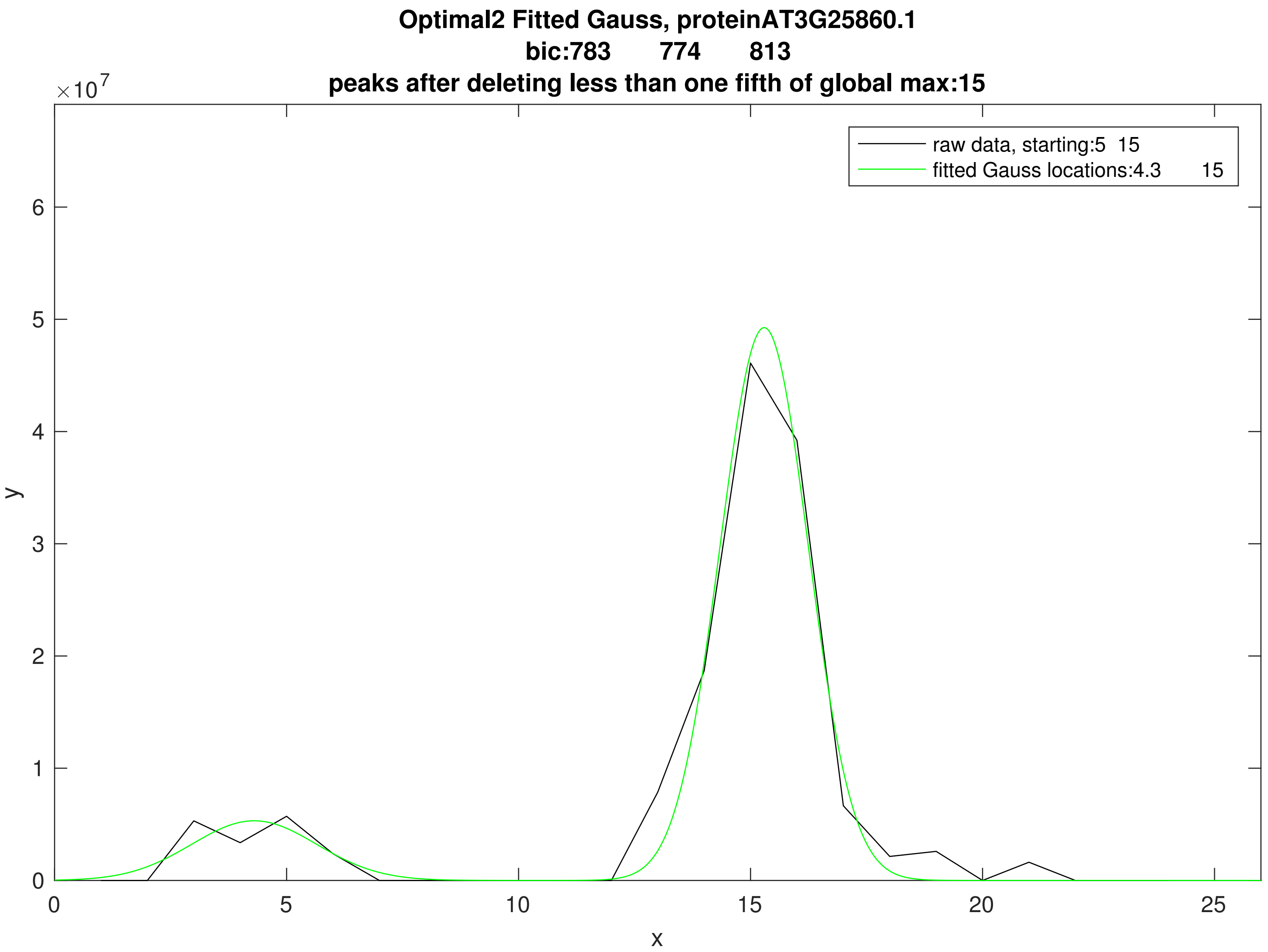
Optimal2 Fitted Gauss, proteinAT3G25860.1
bic:783 774 813
peaks after deleting less than one fifth of global max:15
×10^7
raw data, starting:5 15
fitted Gauss locations:4.3 15
6
5
4
3
2
1
0
y
0
5
10
15
20
25
x

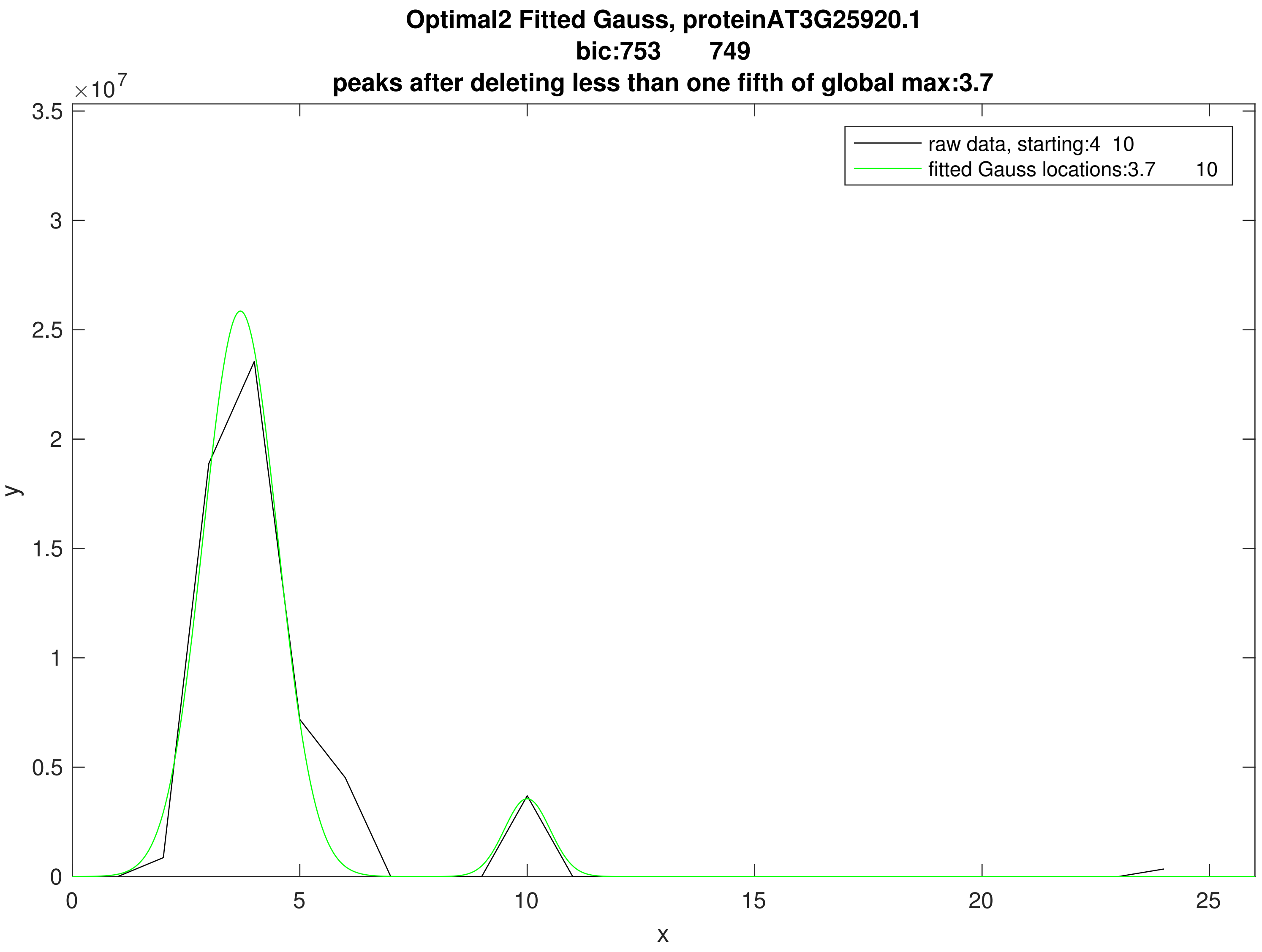
Optimal2 Fitted Gauss, proteinAT3G25920.1
bic:753 749
peaks after deleting less than one fifth of global max:3.7
×10^7
raw data, starting:4 10
fitted Gauss locations:3.7 10
y
x
0
0.5
1
1.5
2
2.5
3
3.5
5
10
15
20
25

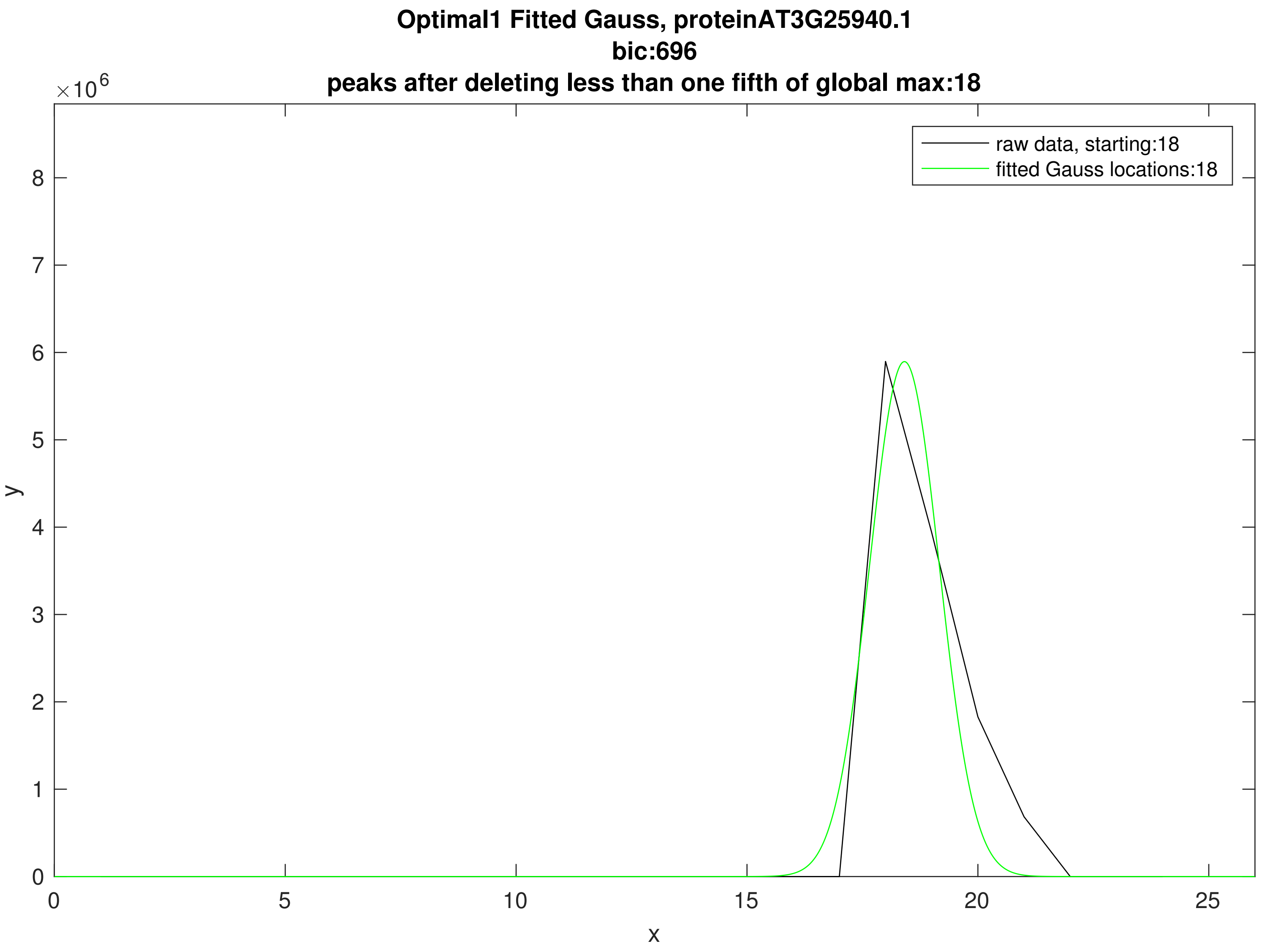
Optimal1 Fitted Gauss, proteinAT3G25940.1
bic:696
peaks after deleting less than one fifth of global max:18
×10^6
raw data, starting:18
fitted Gauss locations:18
y
x
0
1
2
3
4
5
6
7
8
0
5
10
15
20
25

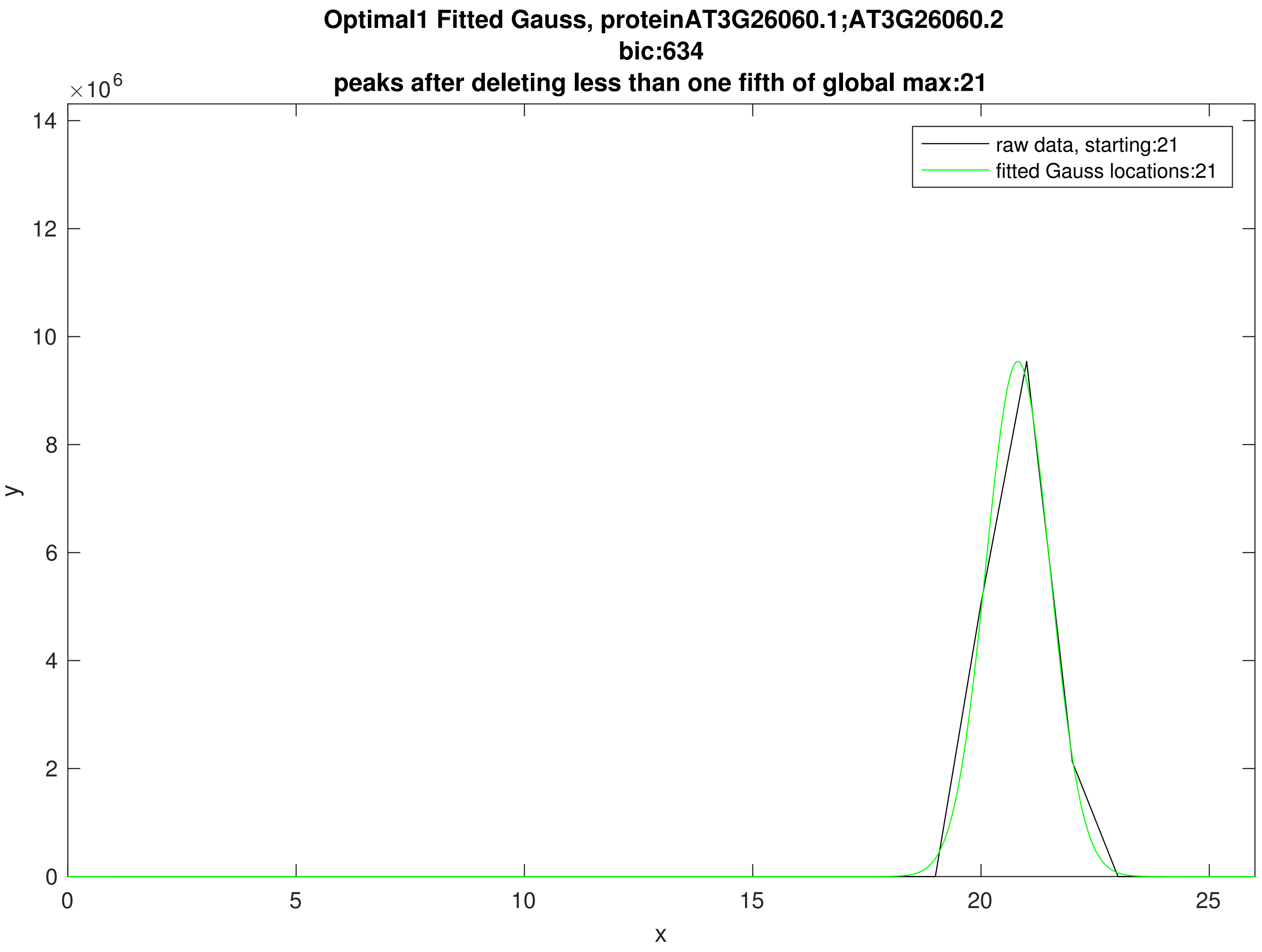
Optimal1 Fitted Gauss, proteinAT3G26060.1;AT3G26060.2
bic:634
peaks after deleting less than one fifth of global max:21
×10^6
14
12
10
8
6
4
2
0
y
0
5
10
15
20
25
x
raw data, starting:21
fitted Gauss locations:21

#### Optimal1 Fitted Gauss, proteinAT3G26340.1
#### bic:776
#### peaks after deleting less than one fifth of global max:5.2

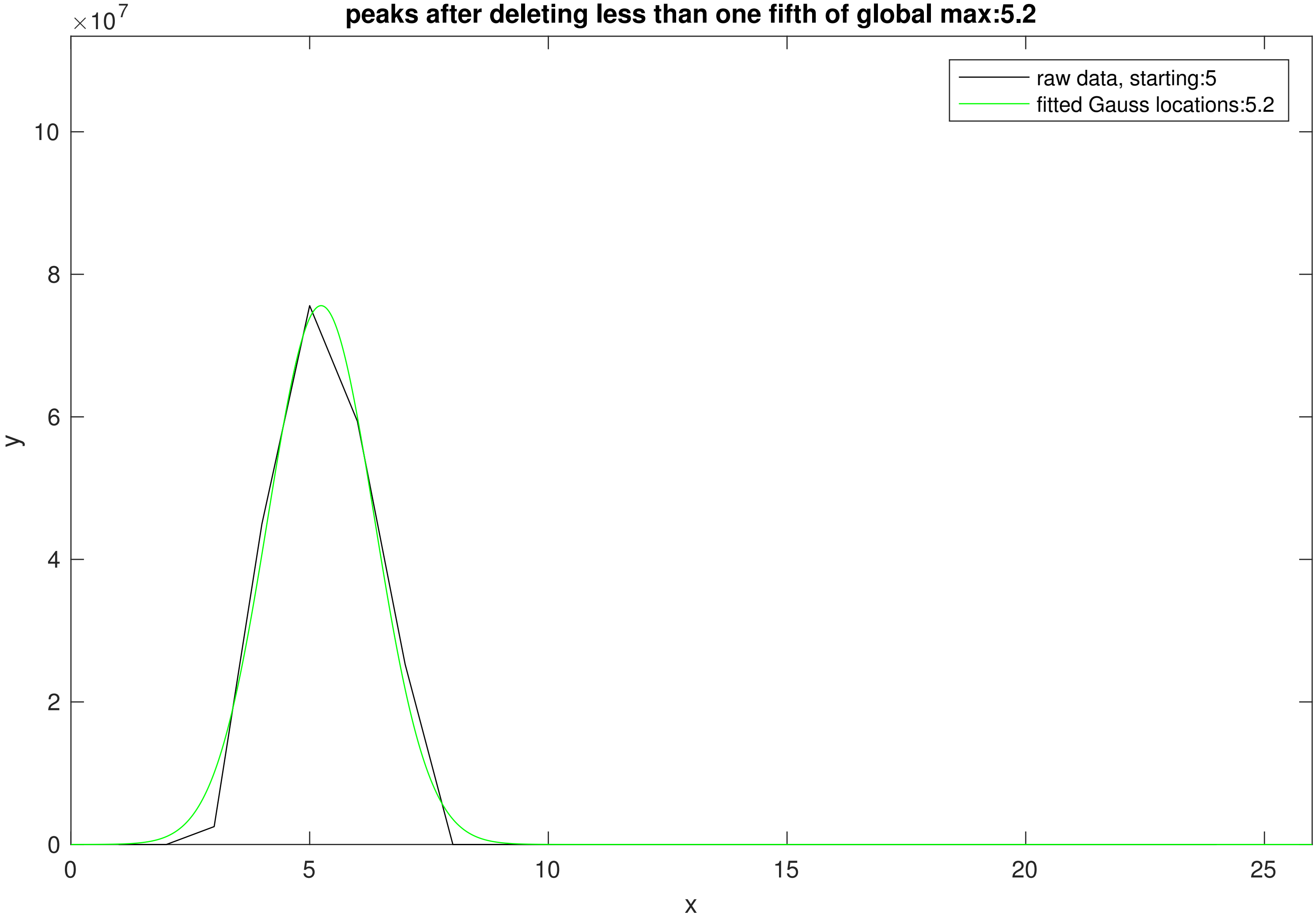

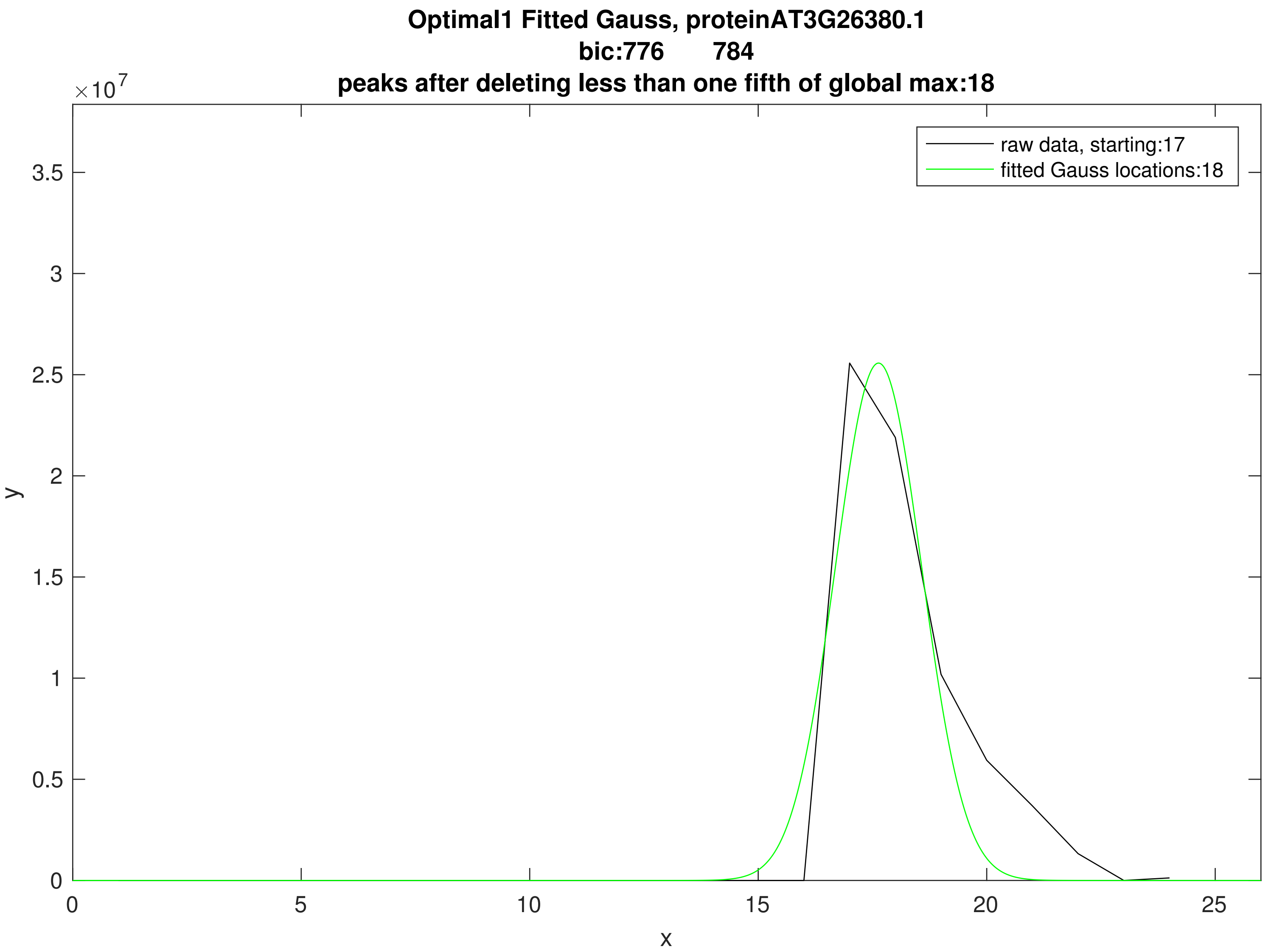
Optimal1 Fitted Gauss, proteinAT3G26380.1
bic:776 784
peaks after deleting less than one fifth of global max:18
×10^7
raw data, starting:17
fitted Gauss locations:18
3.5
3
2.5
2
1.5
1
0.5
0
y
0
5
10
15
20
25
x

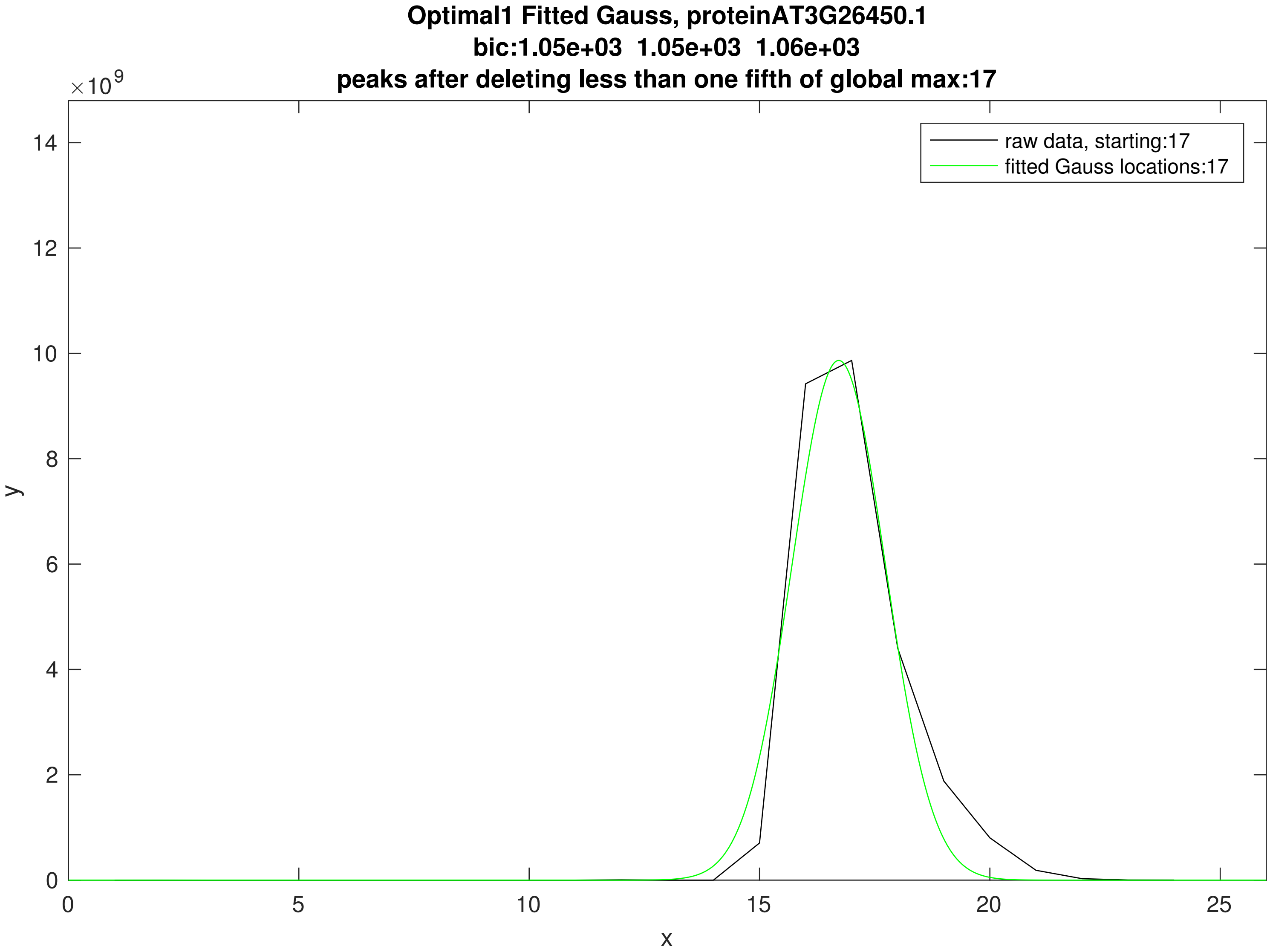

Optimal1 Fitted Gauss, proteinAT3G26450.1
bic:1.05e+03  1.05e+03  1.06e+03
peaks after deleting less than one fifth of global max:17
×10^9
14
12
10
8
6
4
2
0
y
0
5
10
15
20
25
x
raw data, starting:17
fitted Gauss locations:17

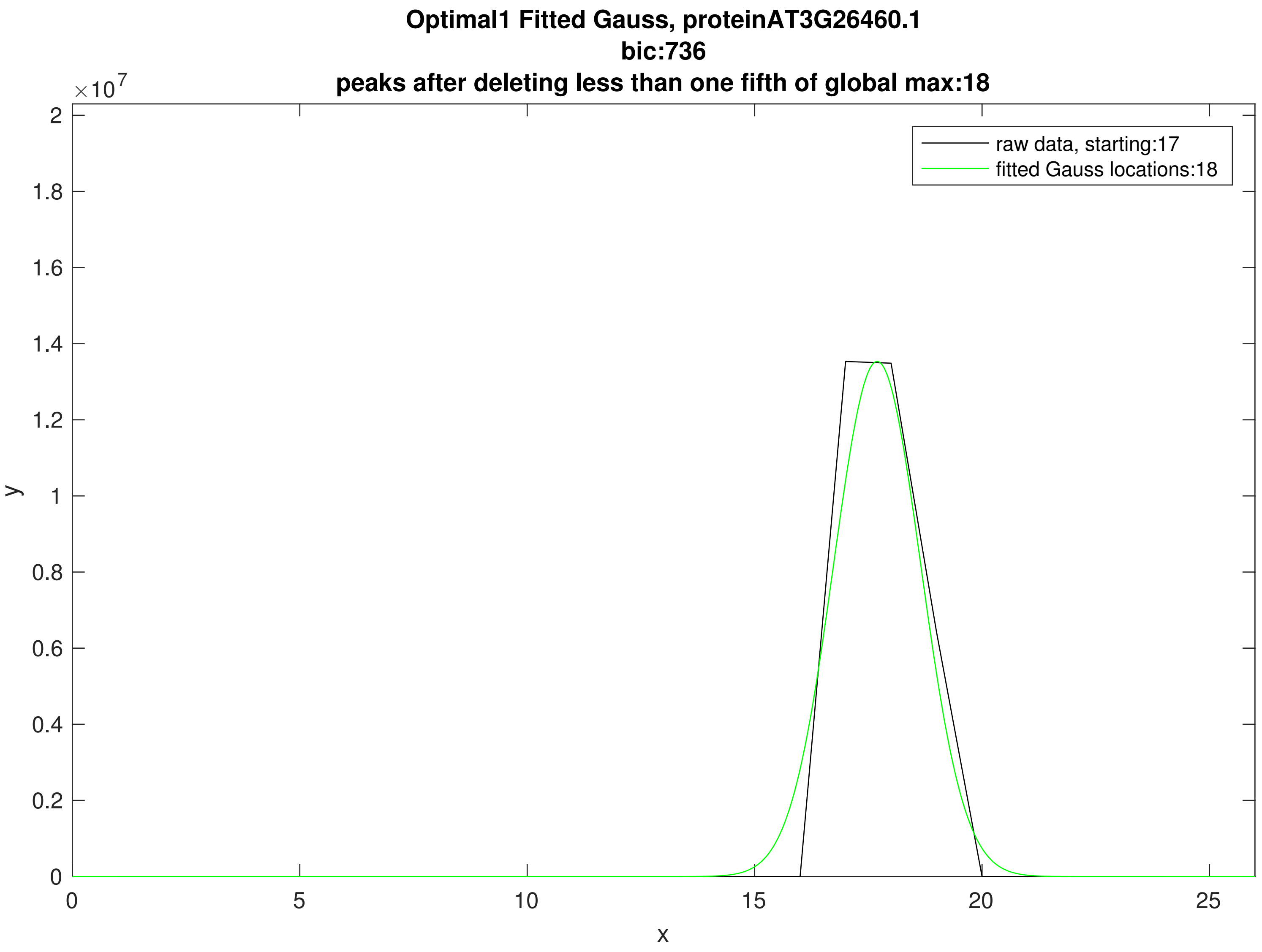

Optimal1 Fitted Gauss, proteinAT3G26460.1
bic:736
peaks after deleting less than one fifth of global max:18
×10^7
2
1.8
1.6
1.4
1.2
1
0.8
0.6
0.4
0.2
0
y
0
5
10
15
20
25
x
raw data, starting:17
fitted Gauss locations:18

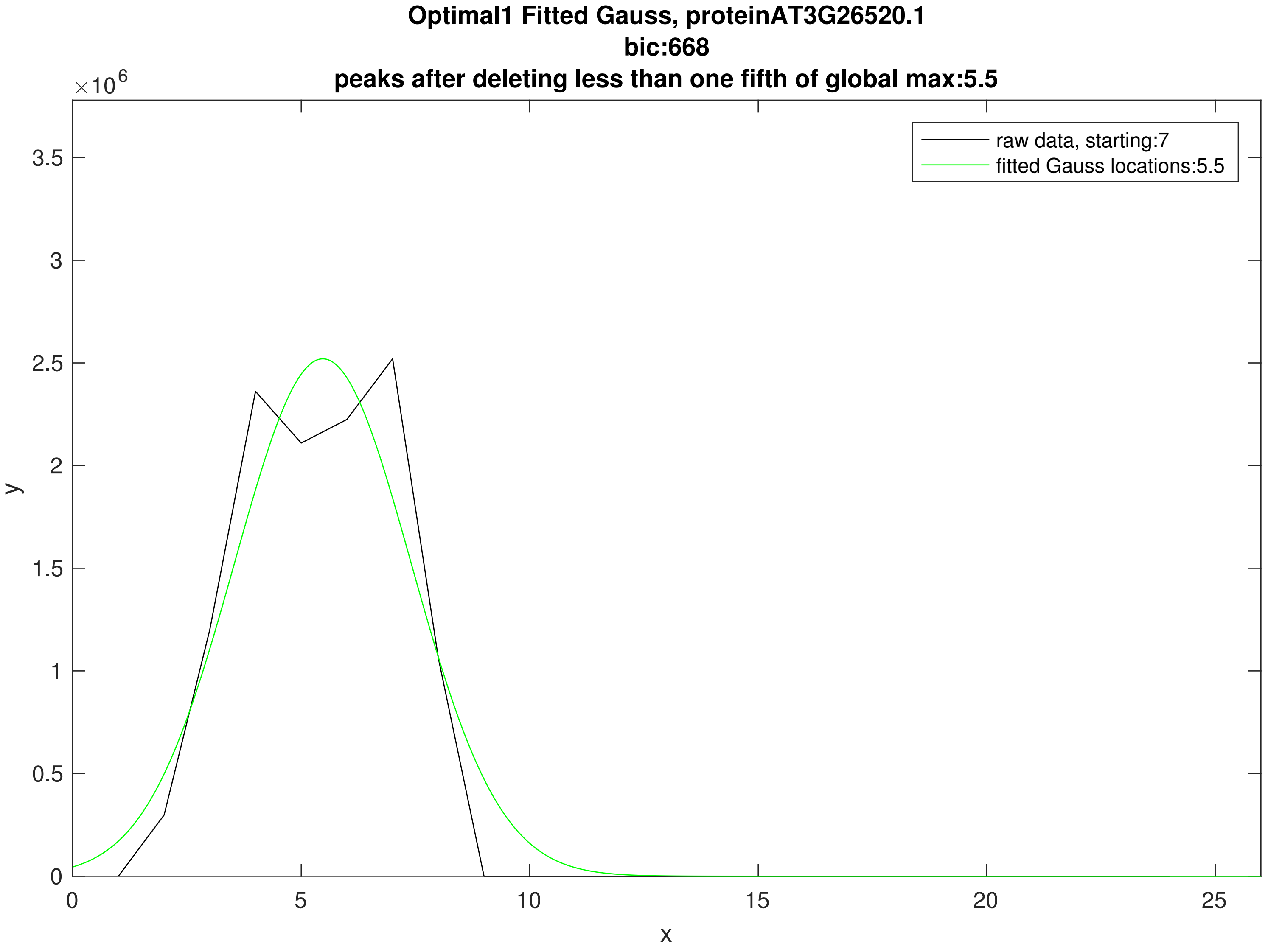
Optimal1 Fitted Gauss, proteinAT3G26520.1
bic:668
peaks after deleting less than one fifth of global max:5.5
×10^6
raw data, starting:7
fitted Gauss locations:5.5
y
x
0
0.5
1
1.5
2
2.5
3
3.5
0
5
10
15
20
25

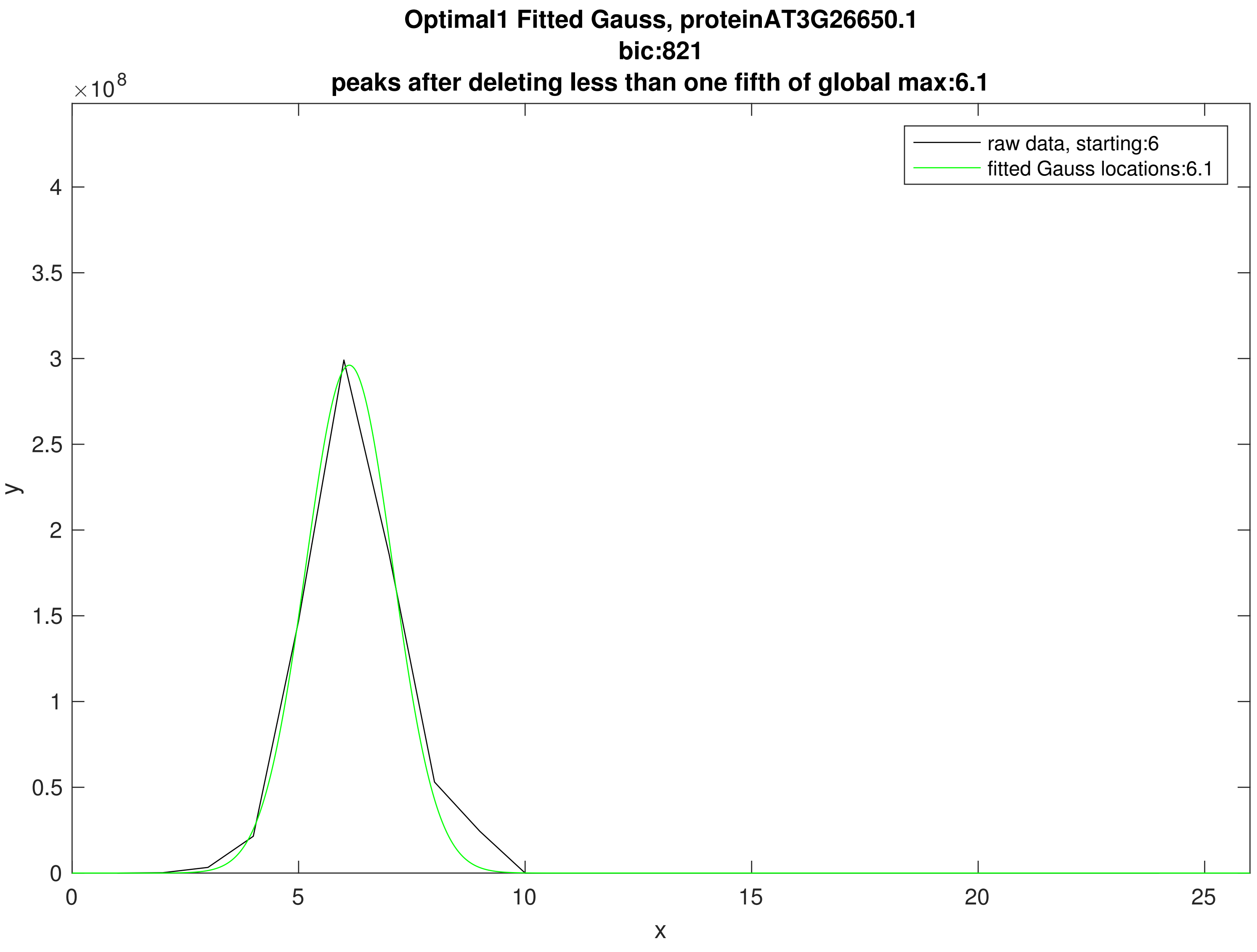
Optimal1 Fitted Gauss, proteinAT3G26650.1
bic:821
peaks after deleting less than one fifth of global max:6.1
raw data, starting:6
fitted Gauss locations:6.1
×10^8
y
x
0
0.5
1
1.5
2
2.5
3
3.5
4
5
10
15
20
25

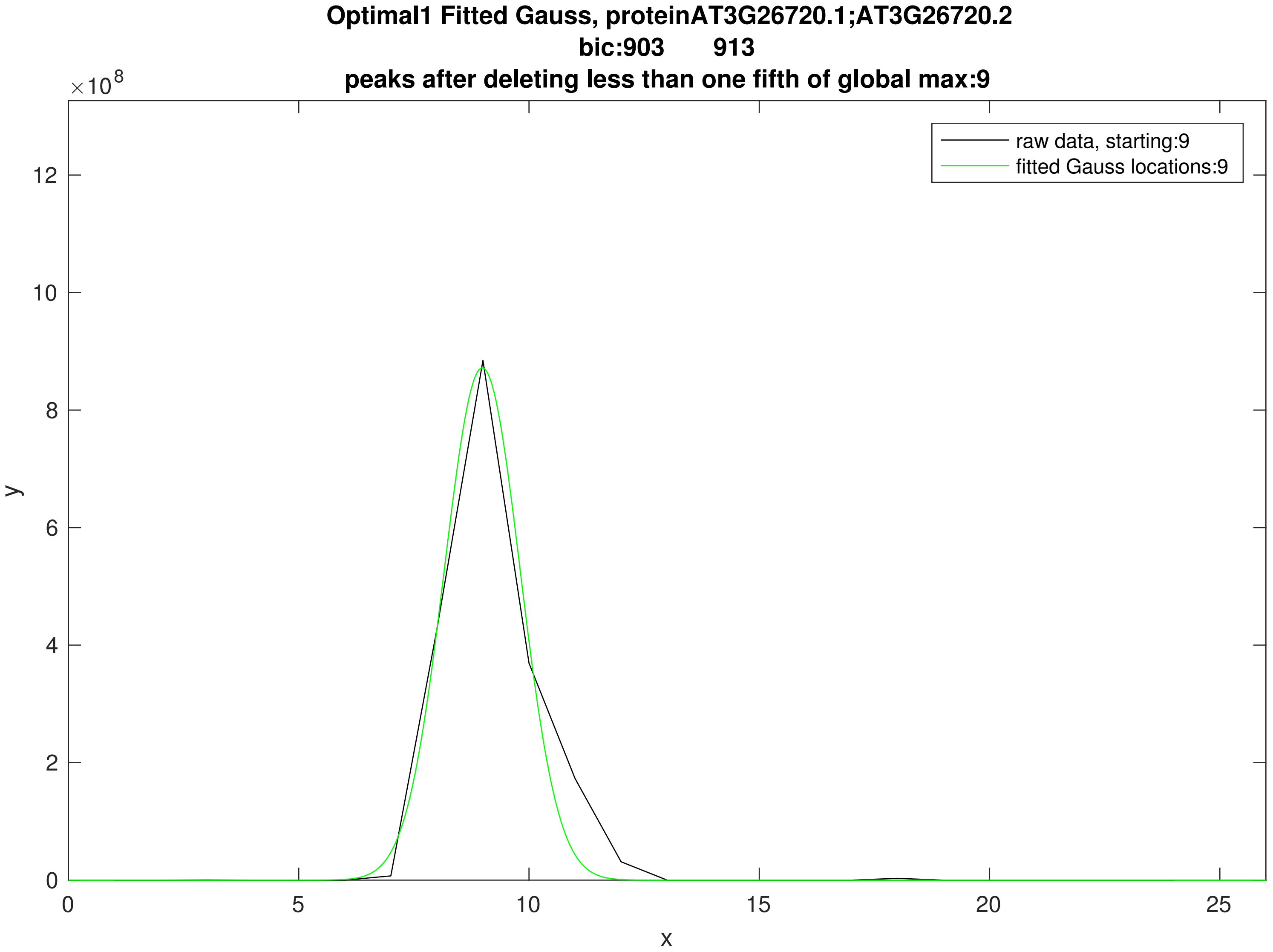

Optimal1 Fitted Gauss, proteinAT3G26720.1;AT3G26720.2
bic:903 913
peaks after deleting less than one fifth of global max:9
×10^8
raw data, starting:9
fitted Gauss locations:9
y
x
0
2
4
6
8
10
12
5
15
20
25

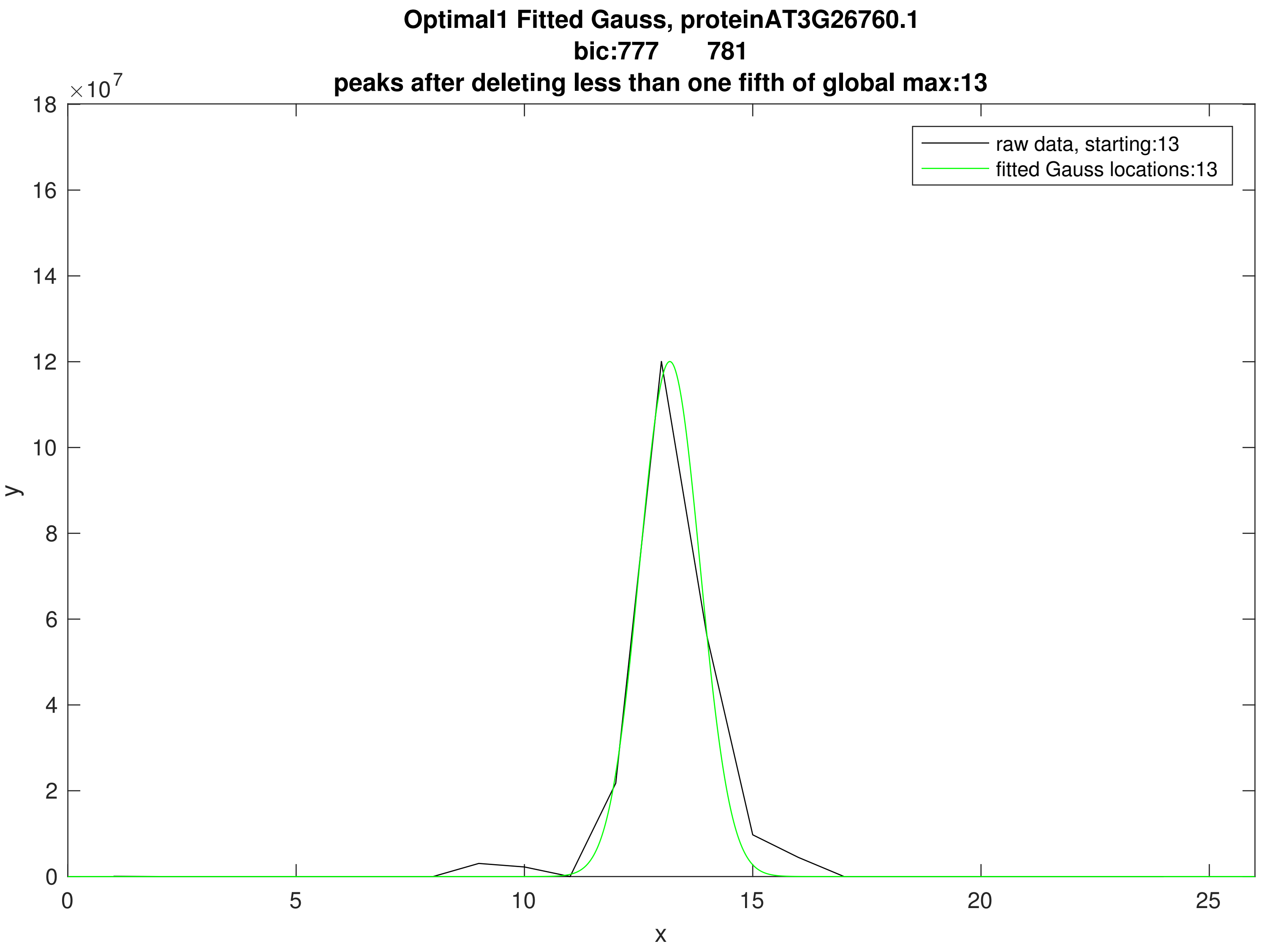
Optimal1 Fitted Gauss, proteinAT3G26760.1
bic:777 781
peaks after deleting less than one fifth of global max:13
×10^7
18
16
14
12
10
8
6
4
2
0
y
0
5
10
15
20
25
x
raw data, starting:13
fitted Gauss locations:13

#### Optimal1 Fitted Gauss, proteinAT3G26770.1
#### bic:806
#### peaks after deleting less than one fifth of global max:12

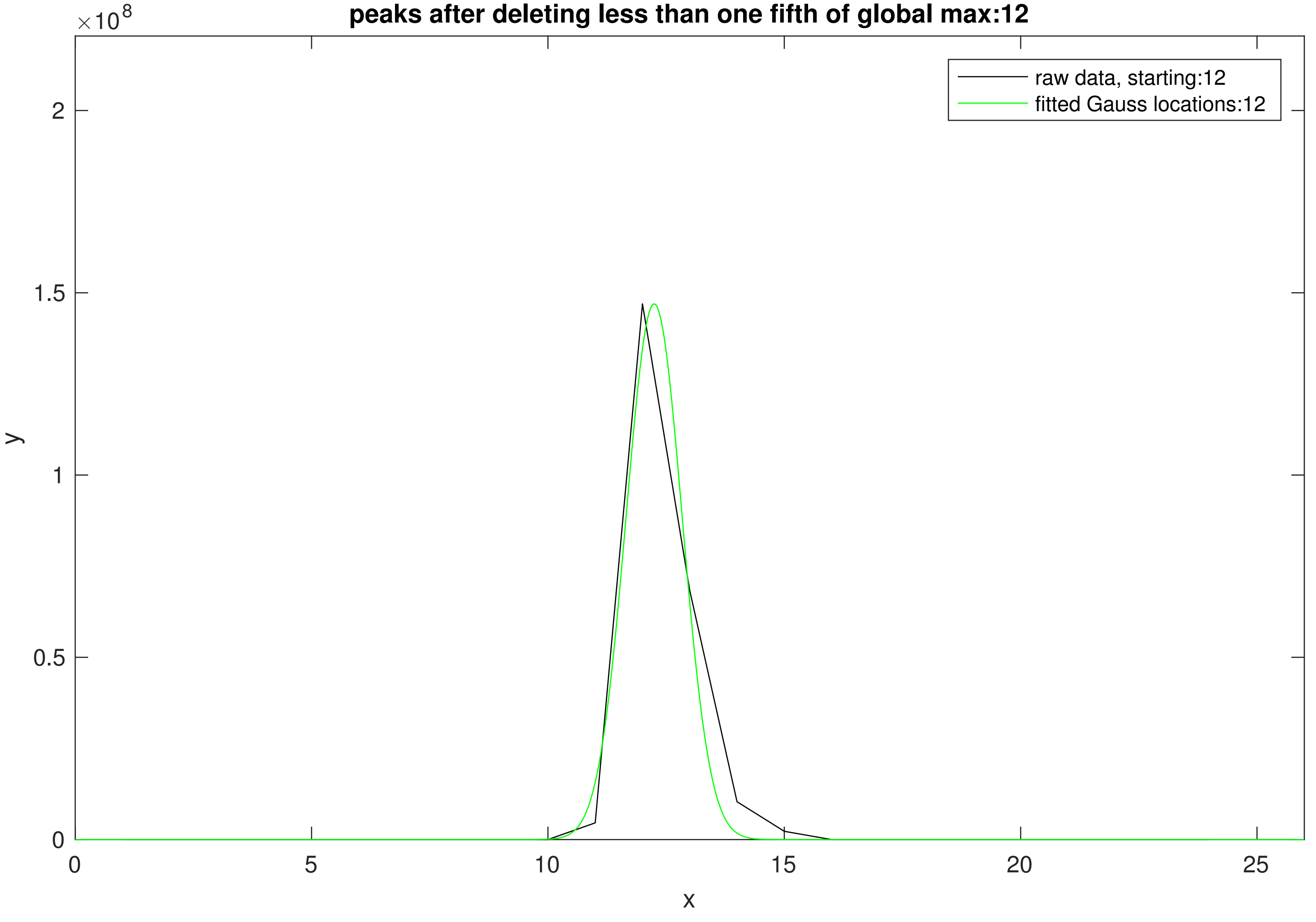

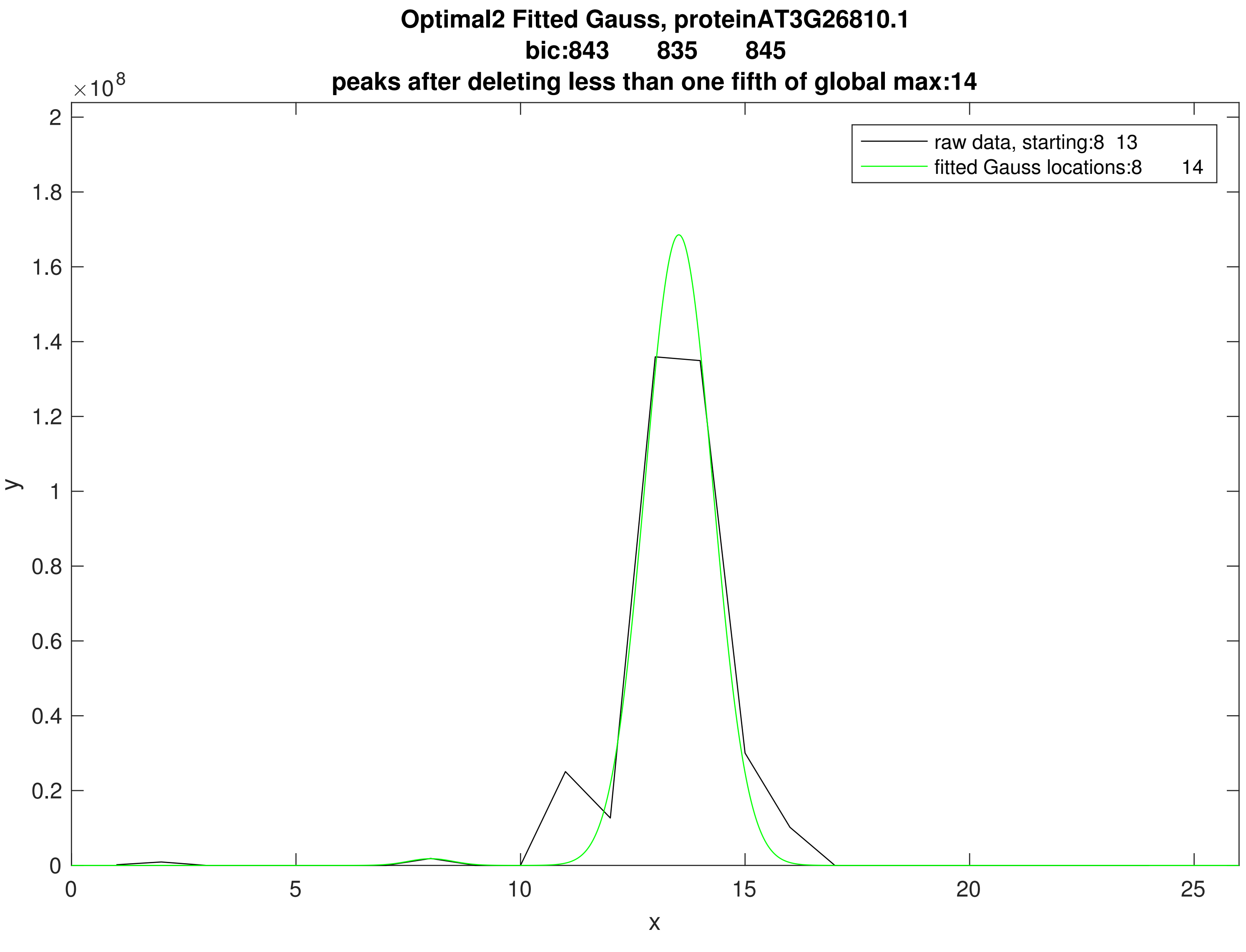
Optimal2 Fitted Gauss, proteinAT3G26810.1
bic:843 835 845
peaks after deleting less than one fifth of global max:14
×10^8
2
1.8
1.6
1.4
1.2
1
0.8
0.6
0.4
0.2
0
y
0
5
10
15
20
25
x
raw data, starting:8 13
fitted Gauss locations:8 14

Optimal1 Fitted Gauss, proteinAT3G27200.1
bic:539
peaks after deleting less than one fifth of global max:18

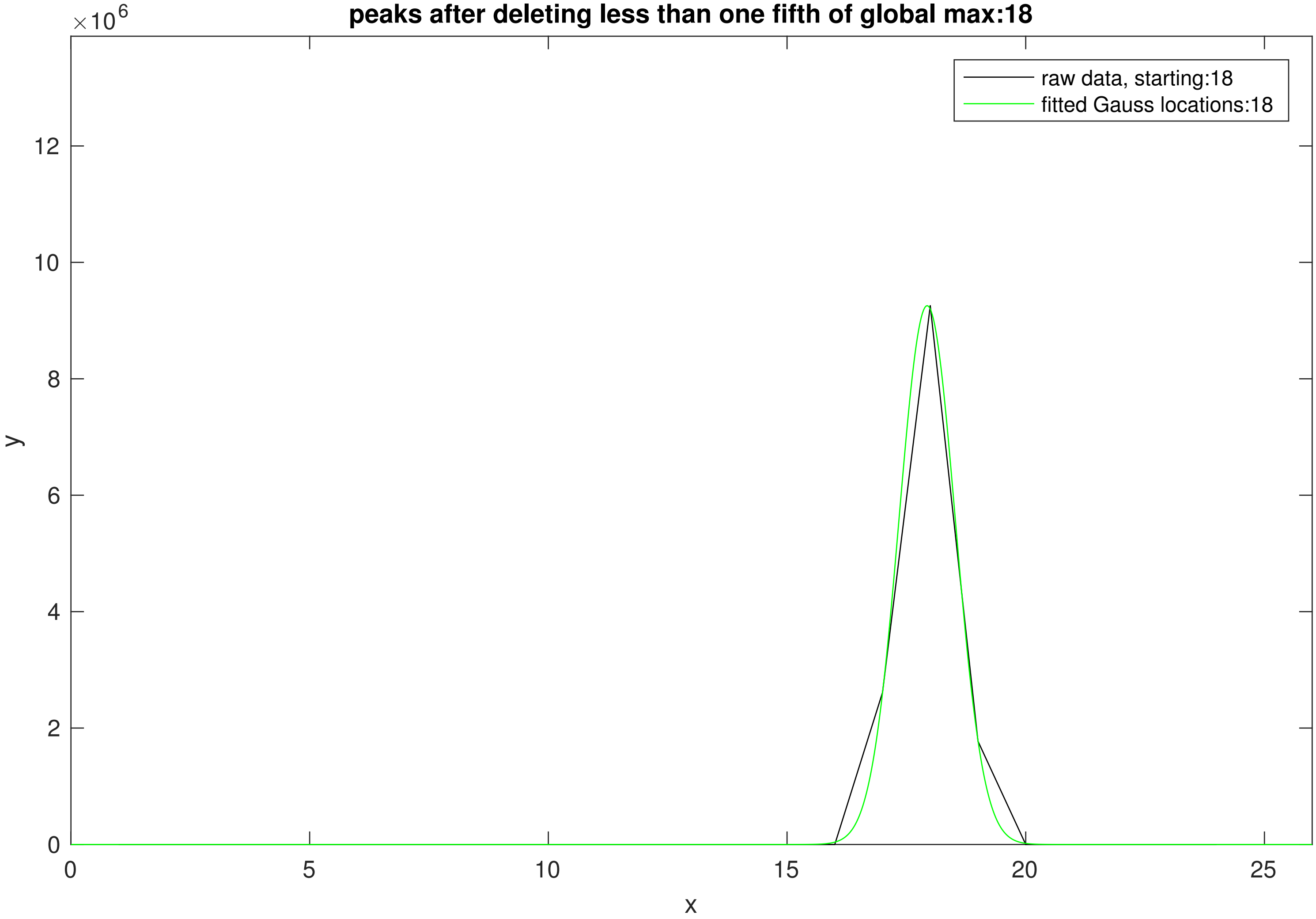

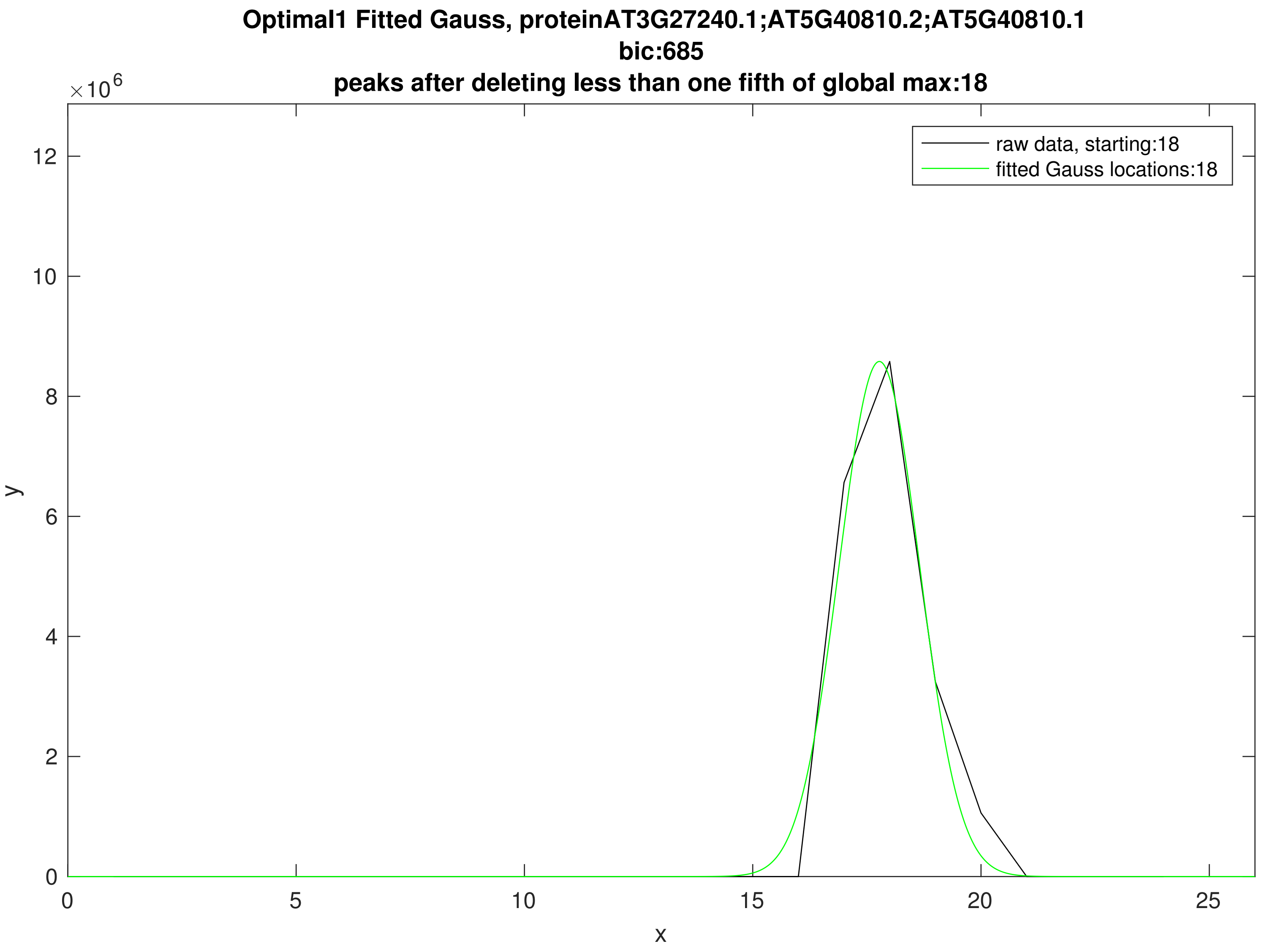
Optimal1 Fitted Gauss, proteinAT3G27240.1;AT5G40810.2;AT5G40810.1
bic:685
peaks after deleting less than one fifth of global max:18
×10^6
raw data, starting:18
fitted Gauss locations:18
12
10
8
6
4
2
0
y
0
5
10
15
20
25
x

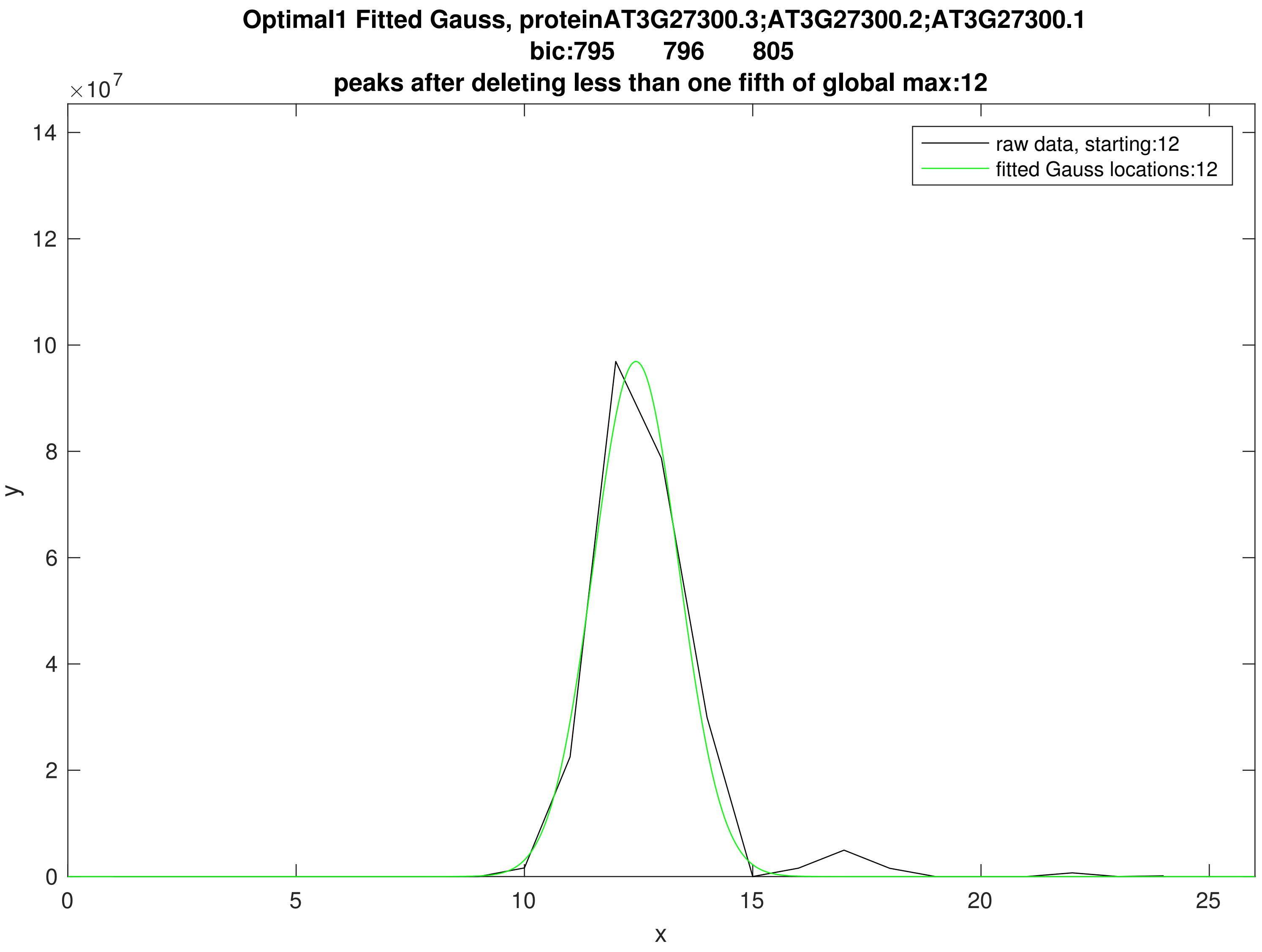

Optimal1 Fitted Gauss, proteinAT3G27300.3;AT3G27300.2;AT3G27300.1
bic:795 796 805
peaks after deleting less than one fifth of global max:12
×10^7
raw data, starting:12
fitted Gauss locations:12
y
x
0
2
4
6
8
10
12
14
0
5
10
15
20
25

Optimal1 Fitted Gauss, proteinAT3G27380.2;AT3G27380.1
bic:854
peaks after deleting less than one fifth of global max:14

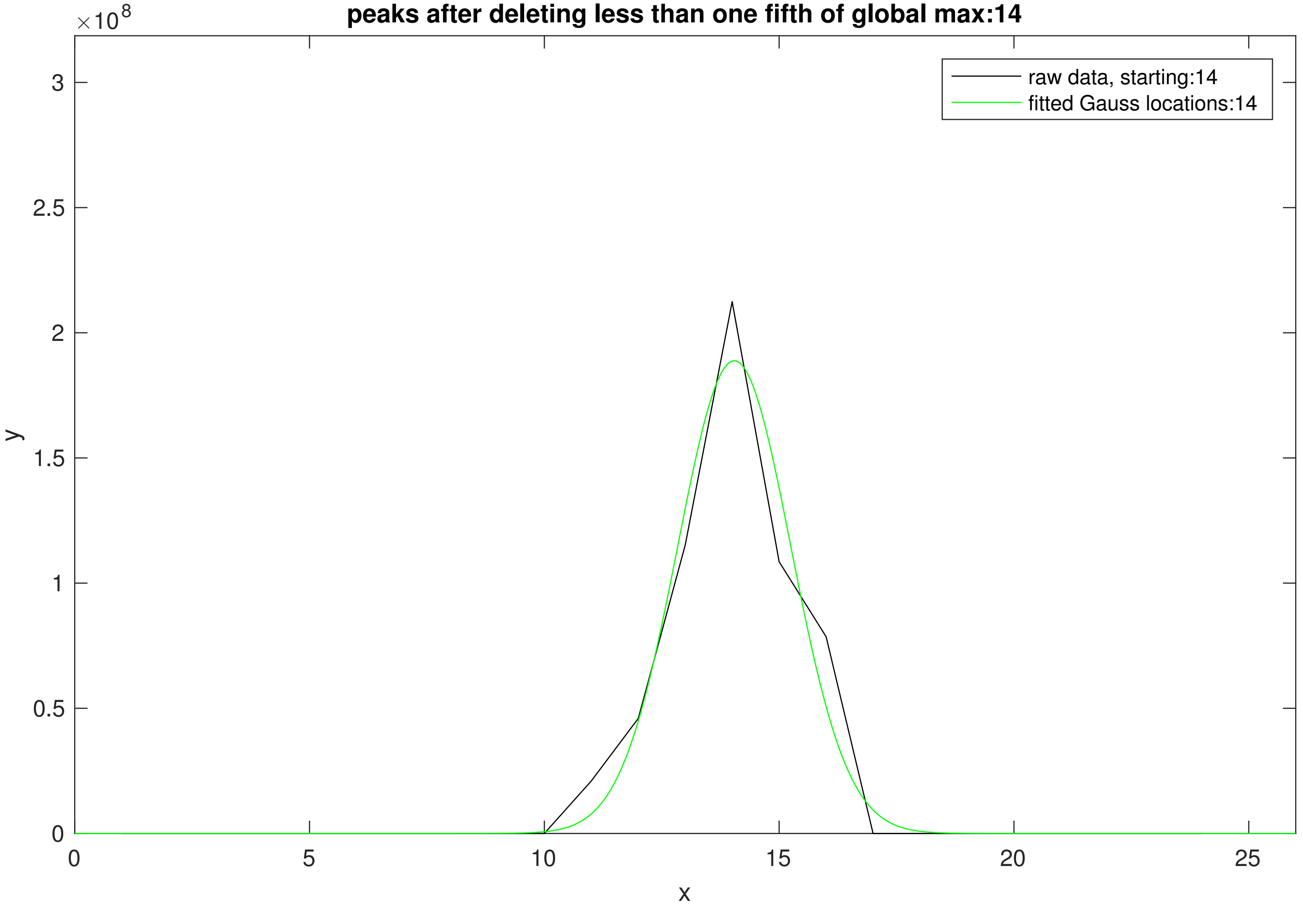

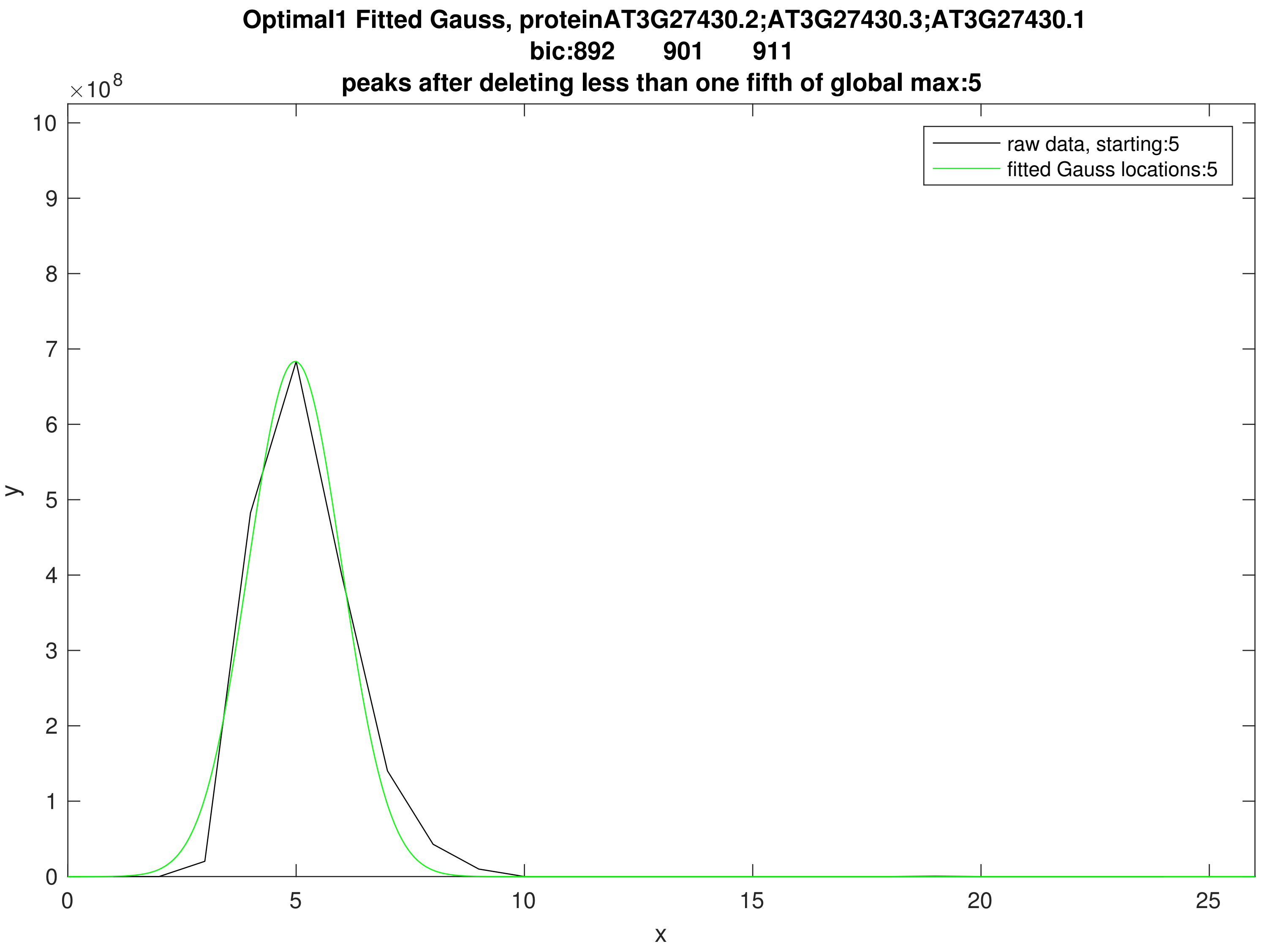
Optimal1 Fitted Gauss, proteinAT3G27430.2;AT3G27430.3;AT3G27430.1
bic:892 901 911
peaks after deleting less than one fifth of global max:5
×10^8
raw data, starting:5
fitted Gauss locations:5
y
x
0
1
2
3
4
5
6
7
8
9
10
0
5
10
15
20
25

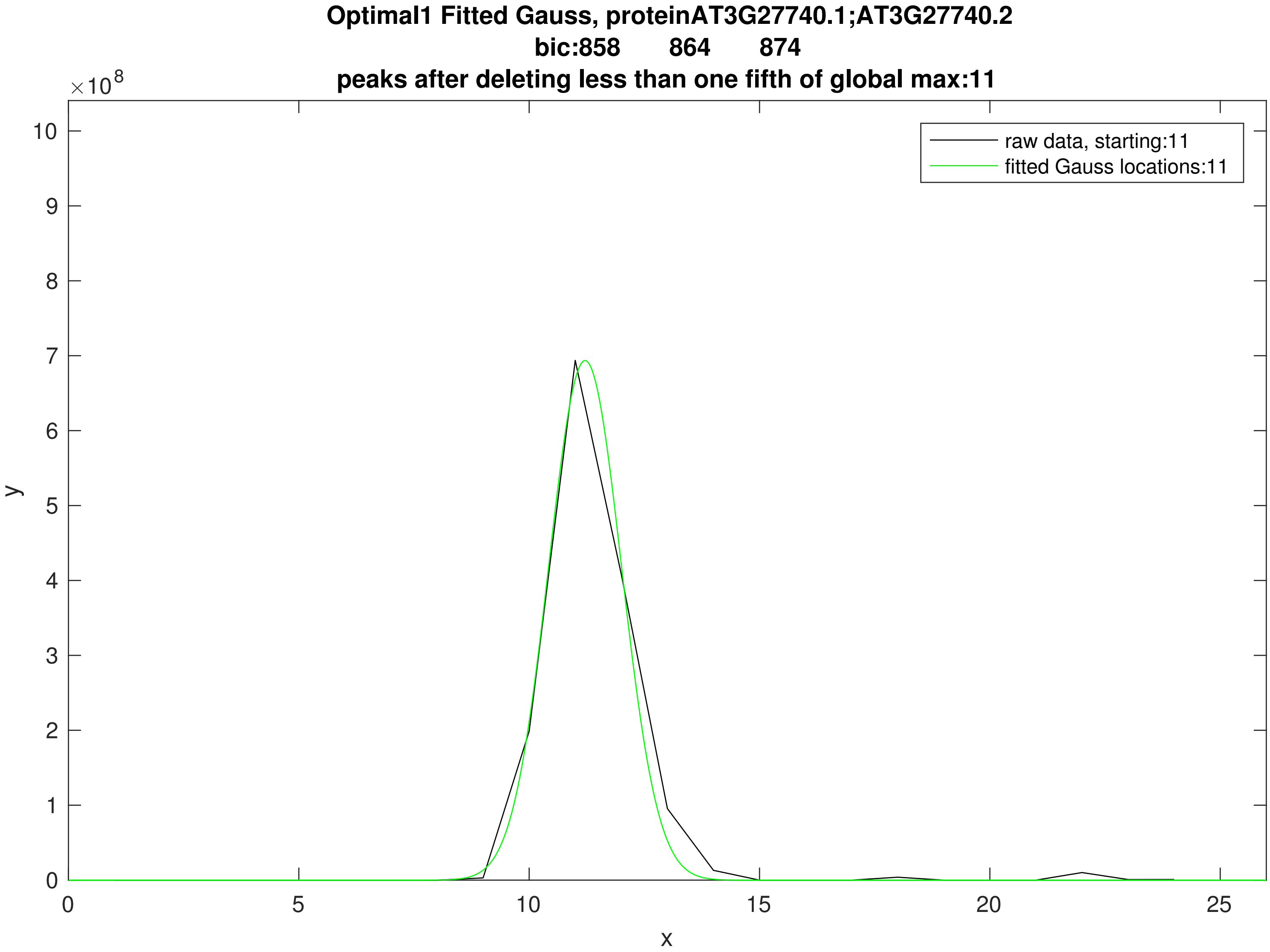

Optimal1 Fitted Gauss, proteinAT3G27740.1;AT3G27740.2
bic:858 864 874
peaks after deleting less than one fifth of global max:11
raw data, starting:11
fitted Gauss locations:11
y
x

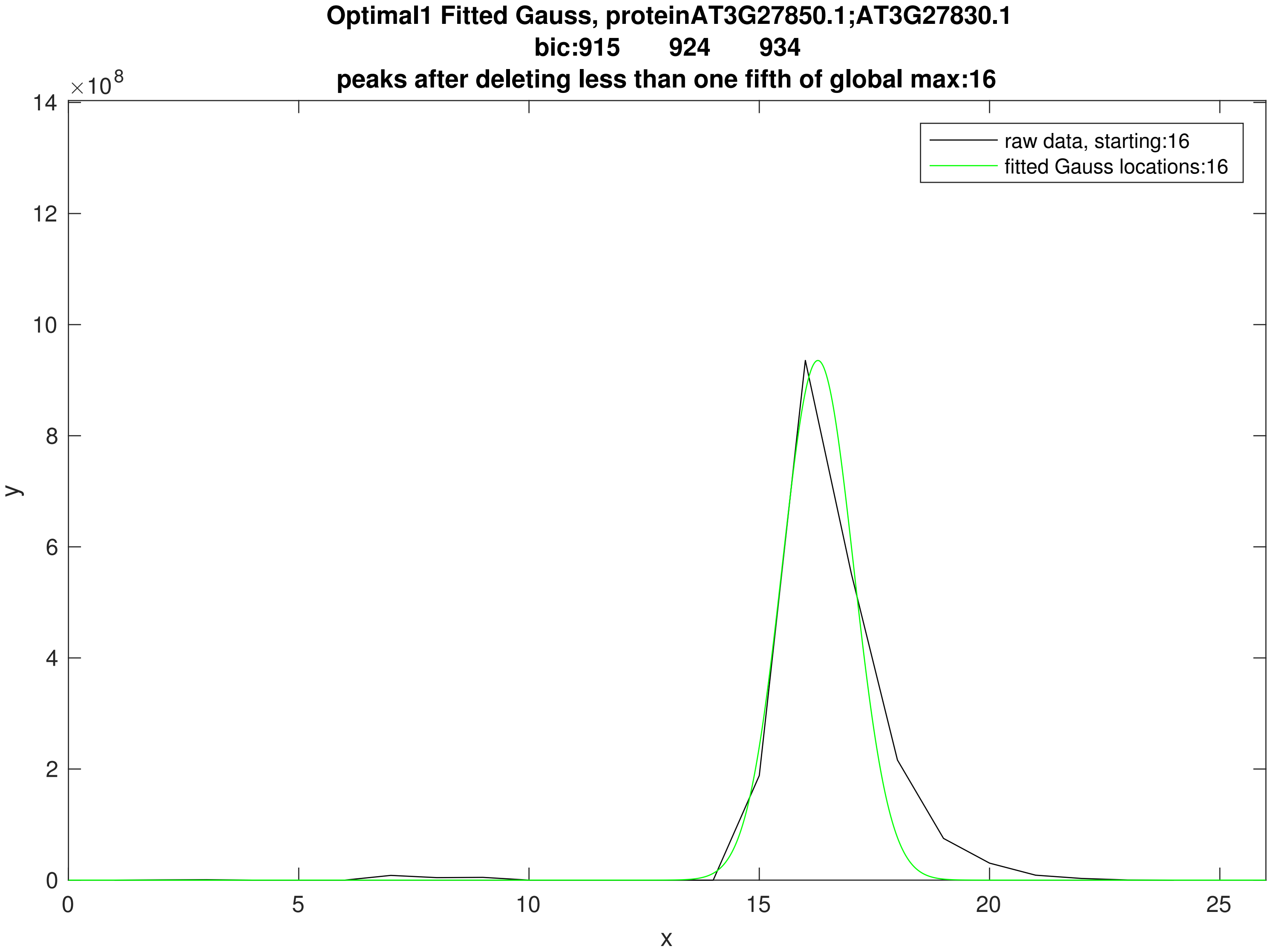

Optimal1 Fitted Gauss, proteinAT3G27850.1;AT3G27830.1
bic:915 924 934
peaks after deleting less than one fifth of global max:16
×10^8
14
12
10
8
6
4
2
0
y
0
5
10
15
20
25
x
raw data, starting:16
fitted Gauss locations:16

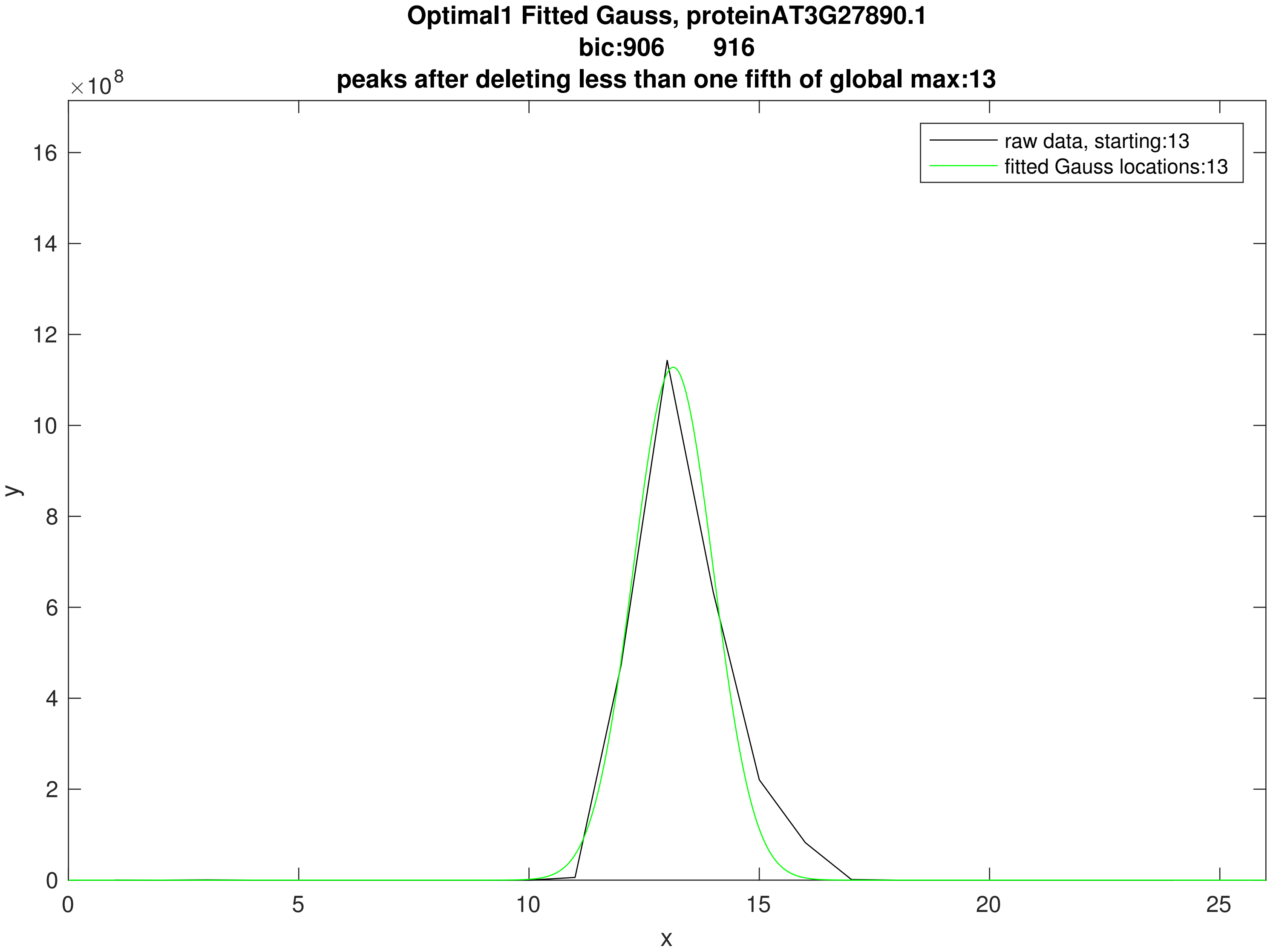
Optimal1 Fitted Gauss, proteinAT3G27890.1
bic:906 916
peaks after deleting less than one fifth of global max:13
×10^8
raw data, starting:13
fitted Gauss locations:13
y
x
0
2
4
6
8
10
12
14
16
0
5
10
15
20
25

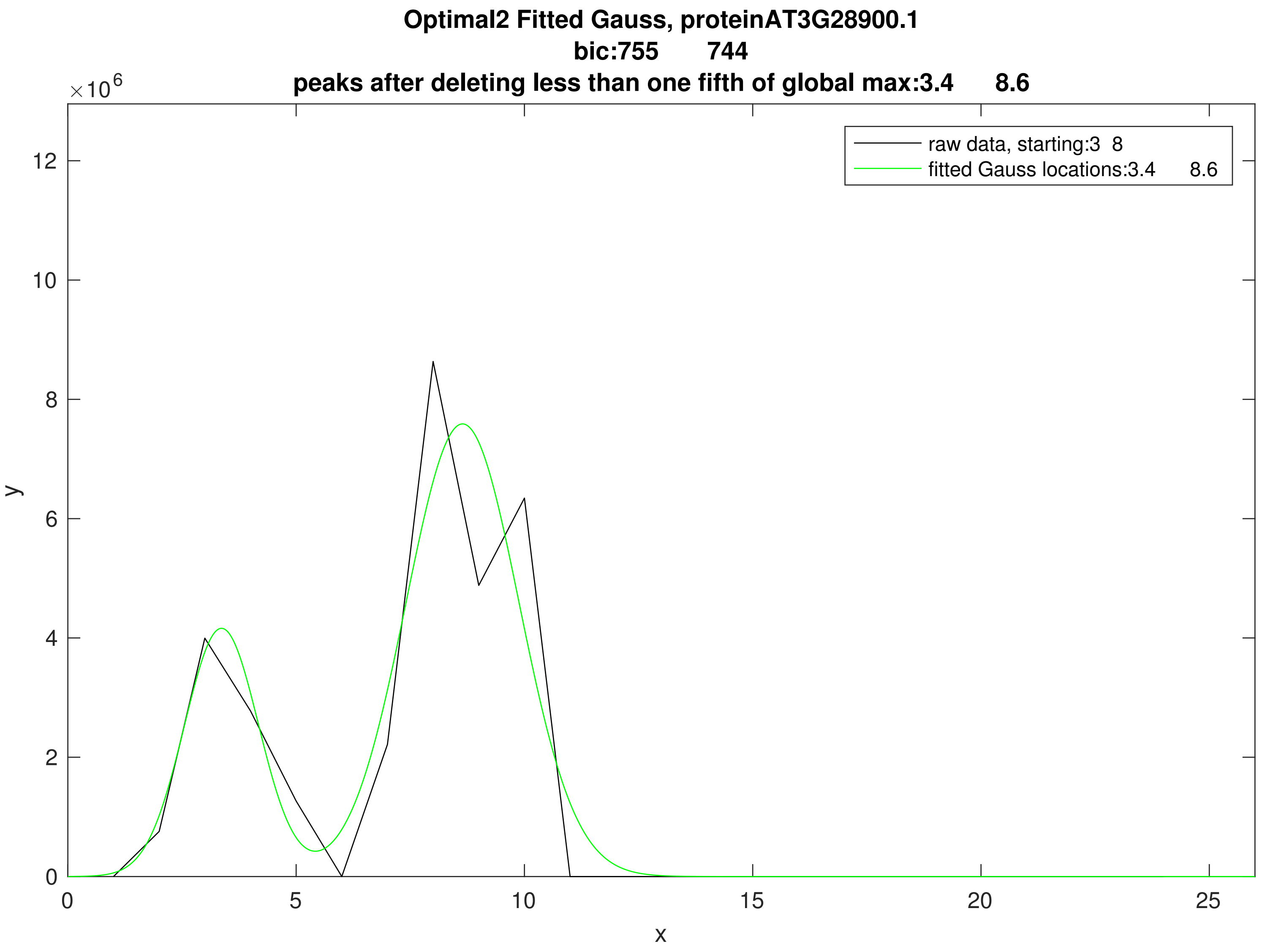
Optimal2 Fitted Gauss, proteinAT3G28900.1
bic:755 744
peaks after deleting less than one fifth of global max:3.4 8.6
×10^6
raw data, starting:3 8
fitted Gauss locations:3.4 8.6
y
x
0
2
4
6
8
10
12
0
5
10
15
20
25

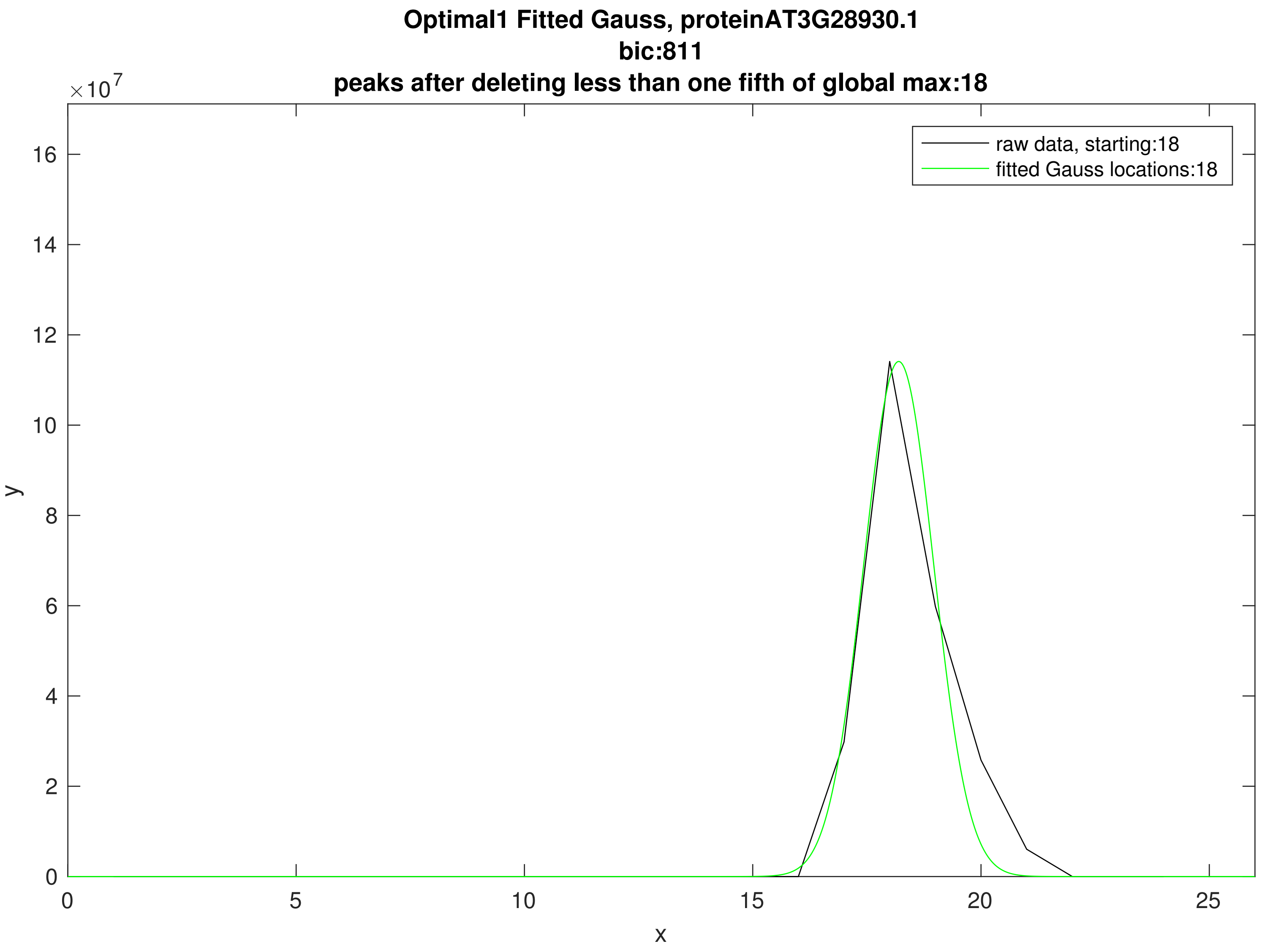
Optimal1 Fitted Gauss, proteinAT3G28930.1
bic:811
peaks after deleting less than one fifth of global max:18
×10^7
raw data, starting:18
fitted Gauss locations:18
y
x
0
2
4
6
8
10
12
14
16
0
5
10
15
20
25

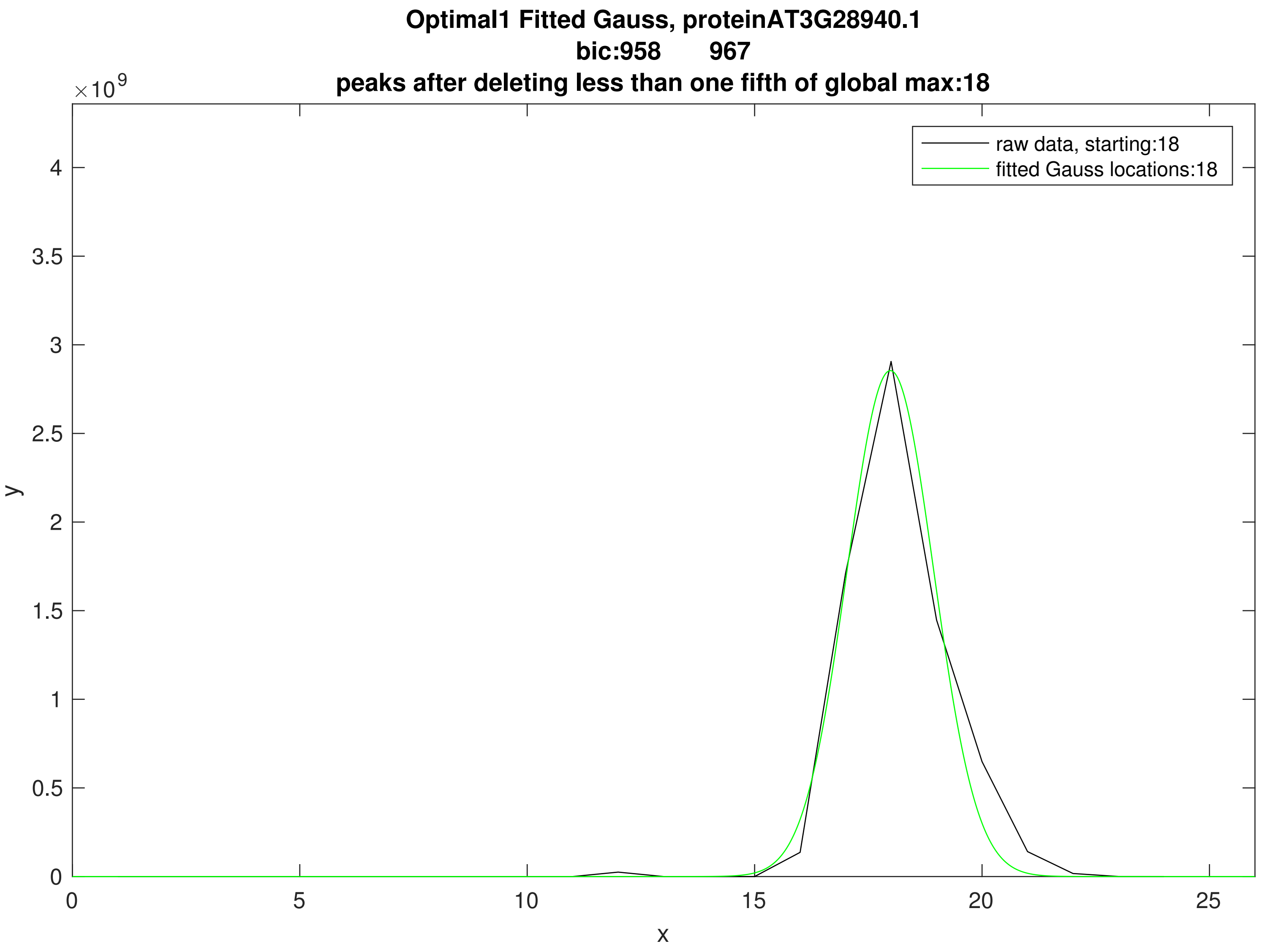
Optimal1 Fitted Gauss, proteinAT3G28940.1
bic:958 967
peaks after deleting less than one fifth of global max:18
×10^9
raw data, starting:18
fitted Gauss locations:18
y
x
0
0.5
1
1.5
2
2.5
3
3.5
4
5
10
15
20
25

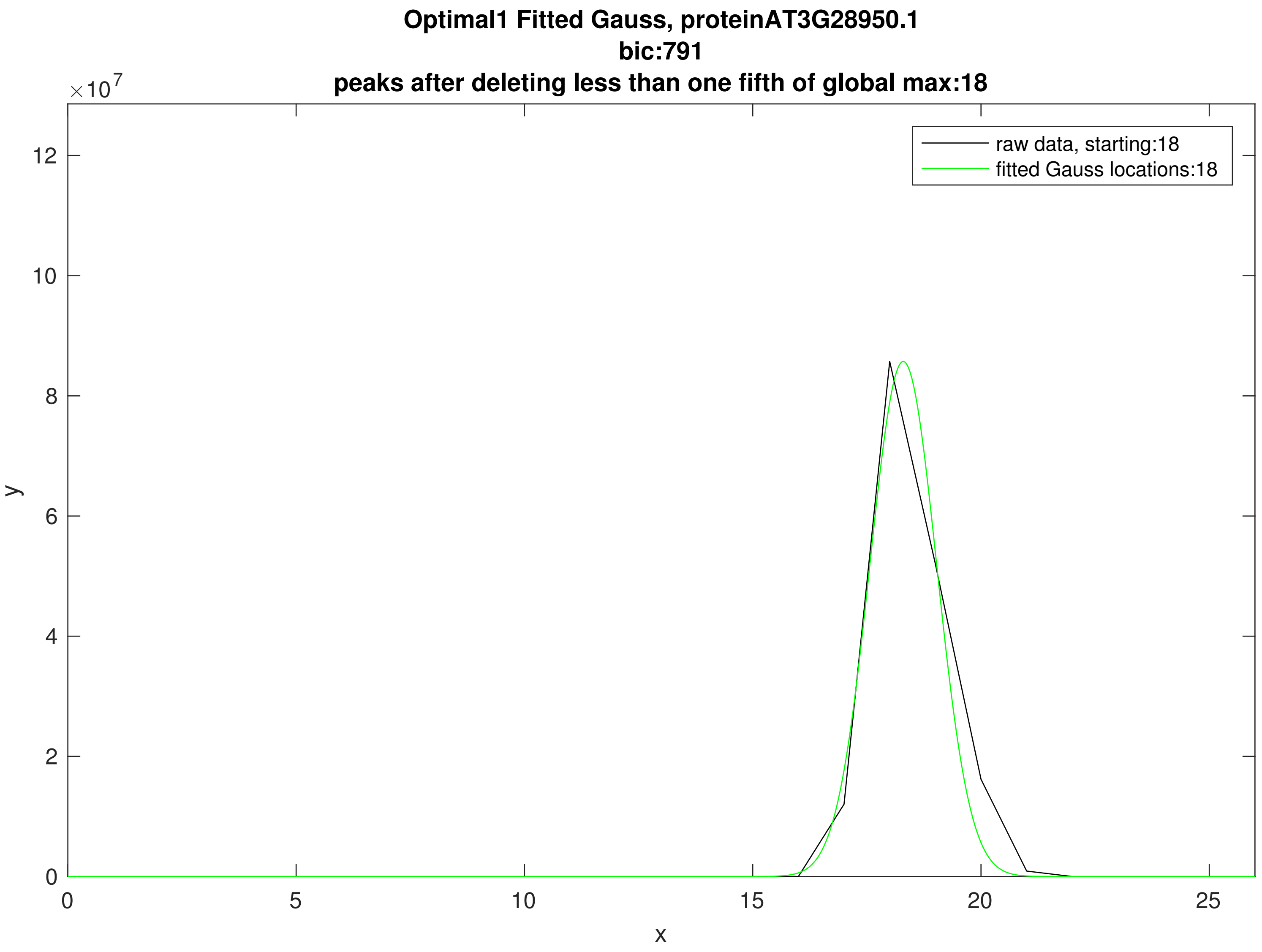
Optimal1 Fitted Gauss, proteinAT3G28950.1
bic:791
peaks after deleting less than one fifth of global max:18
×10^7
raw data, starting:18
fitted Gauss locations:18
y
x
0
2
4
6
8
10
12
0
5
10
15
20
25

Optimal1 Fitted Gauss, proteinAT3G29010.1
bic:567
peaks after deleting less than one fifth of global max:18

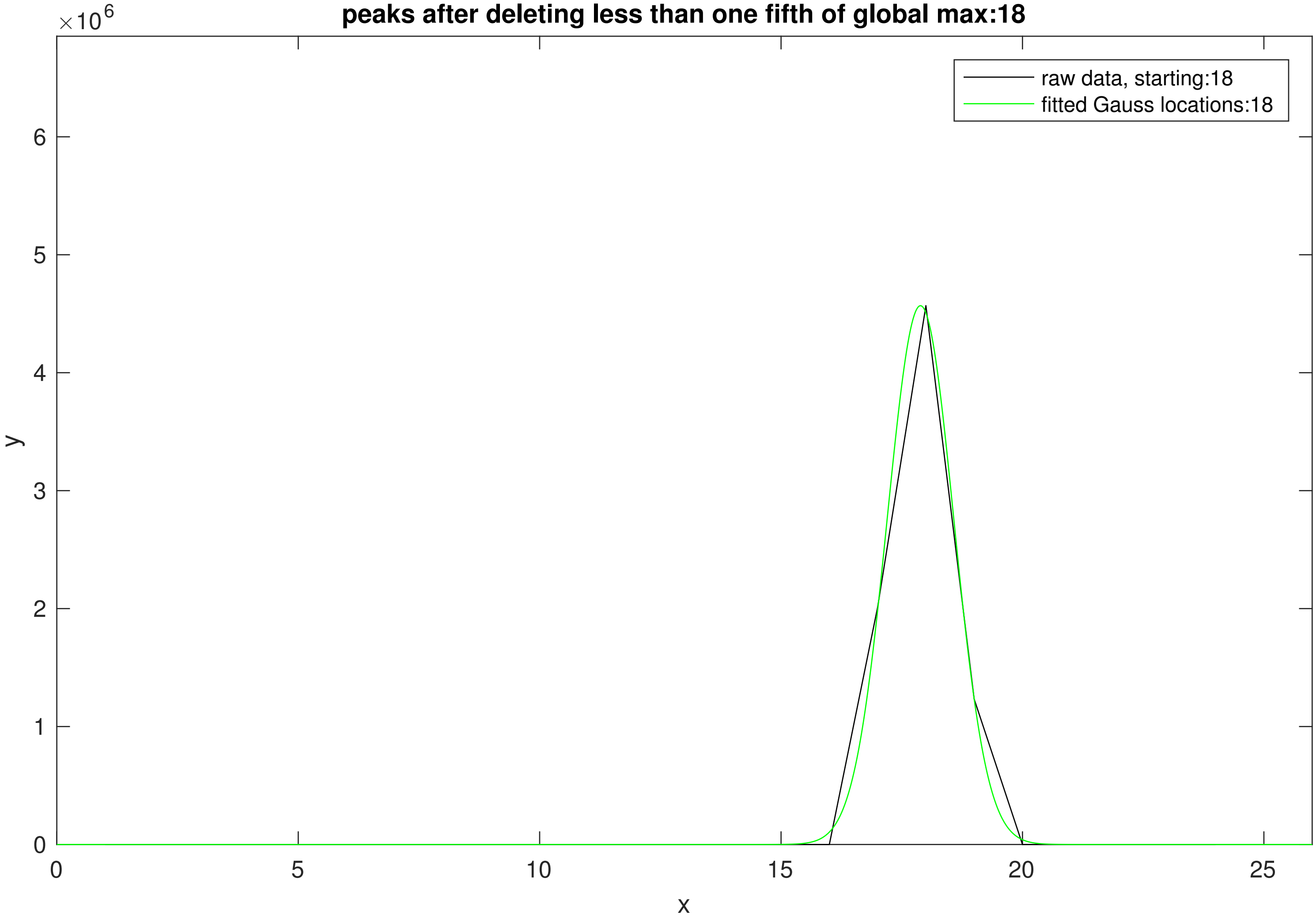

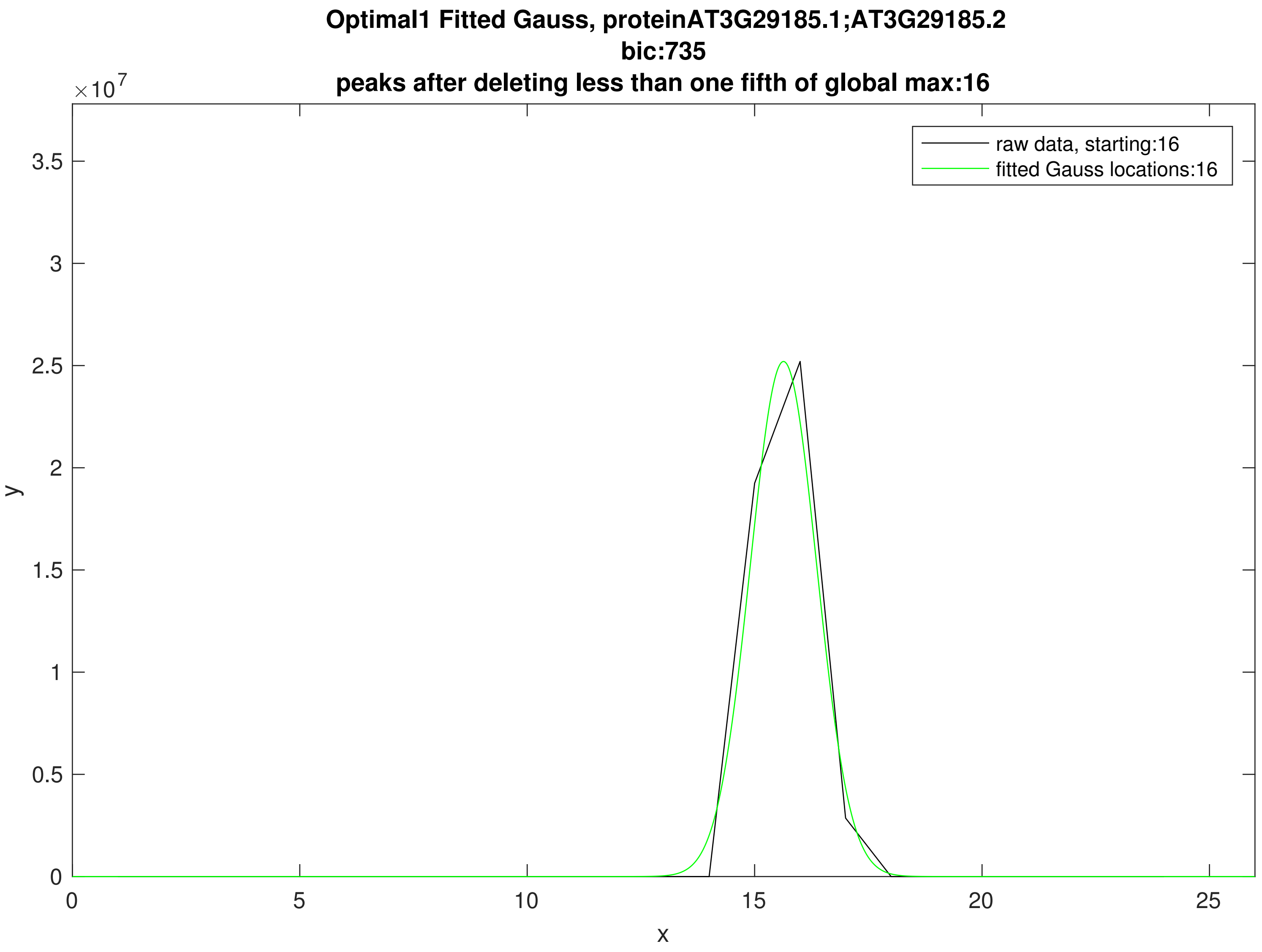

Optimal1 Fitted Gauss, proteinAT3G29185.1;AT3G29185.2
bic:735
peaks after deleting less than one fifth of global max:16
×10^7
3.5
3
2.5
2
1.5
1
0.5
0
y
0
5
10
15
20
25
x
raw data, starting:16
fitted Gauss locations:16

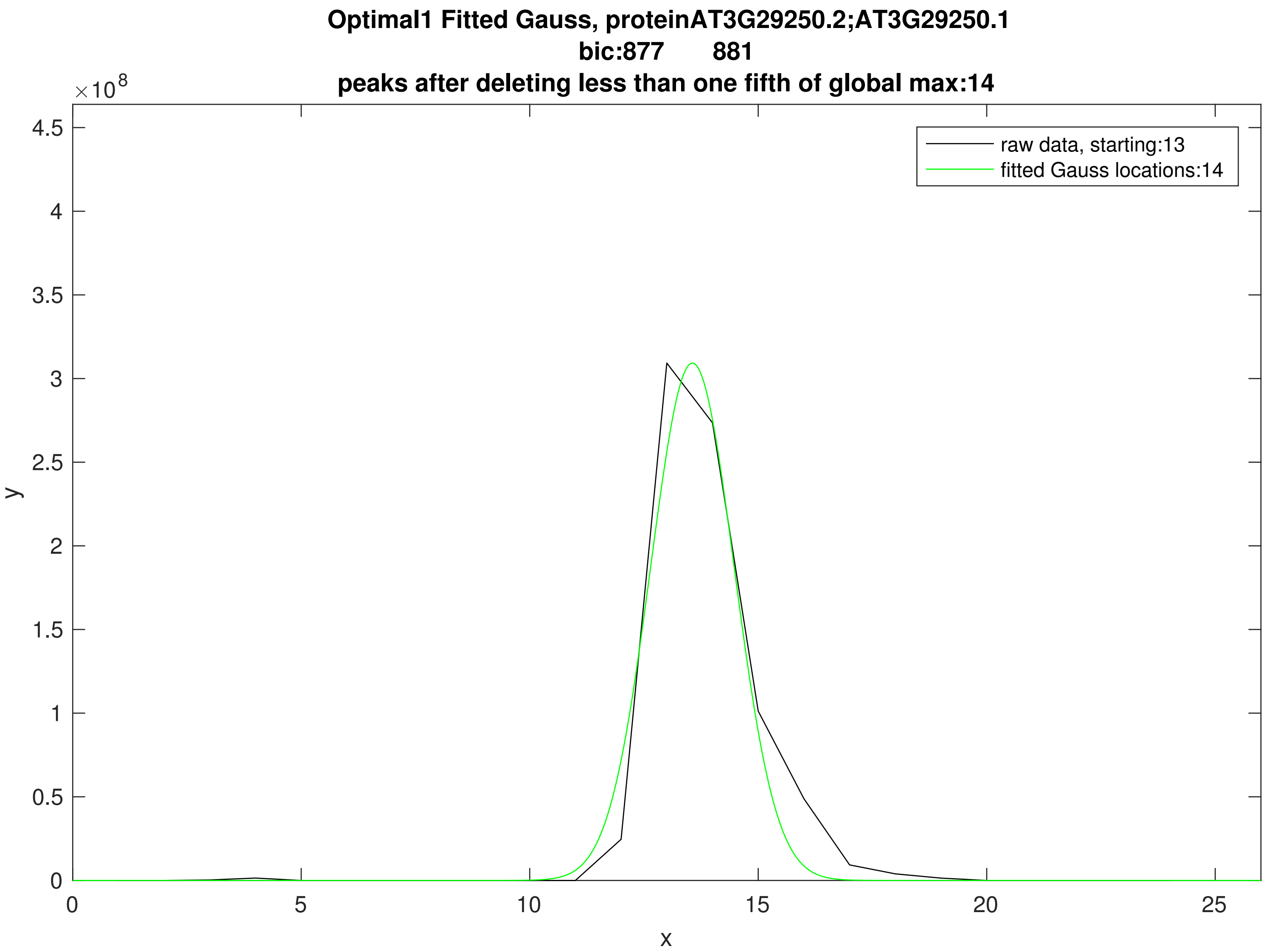
Optimal1 Fitted Gauss, proteinAT3G29250.2;AT3G29250.1
bic:877 881
peaks after deleting less than one fifth of global max:14
×10^8
raw data, starting:13
fitted Gauss locations:14
y
x
0
0.5
1
1.5
2
2.5
3
3.5
4
4.5
5
10
15
20
25

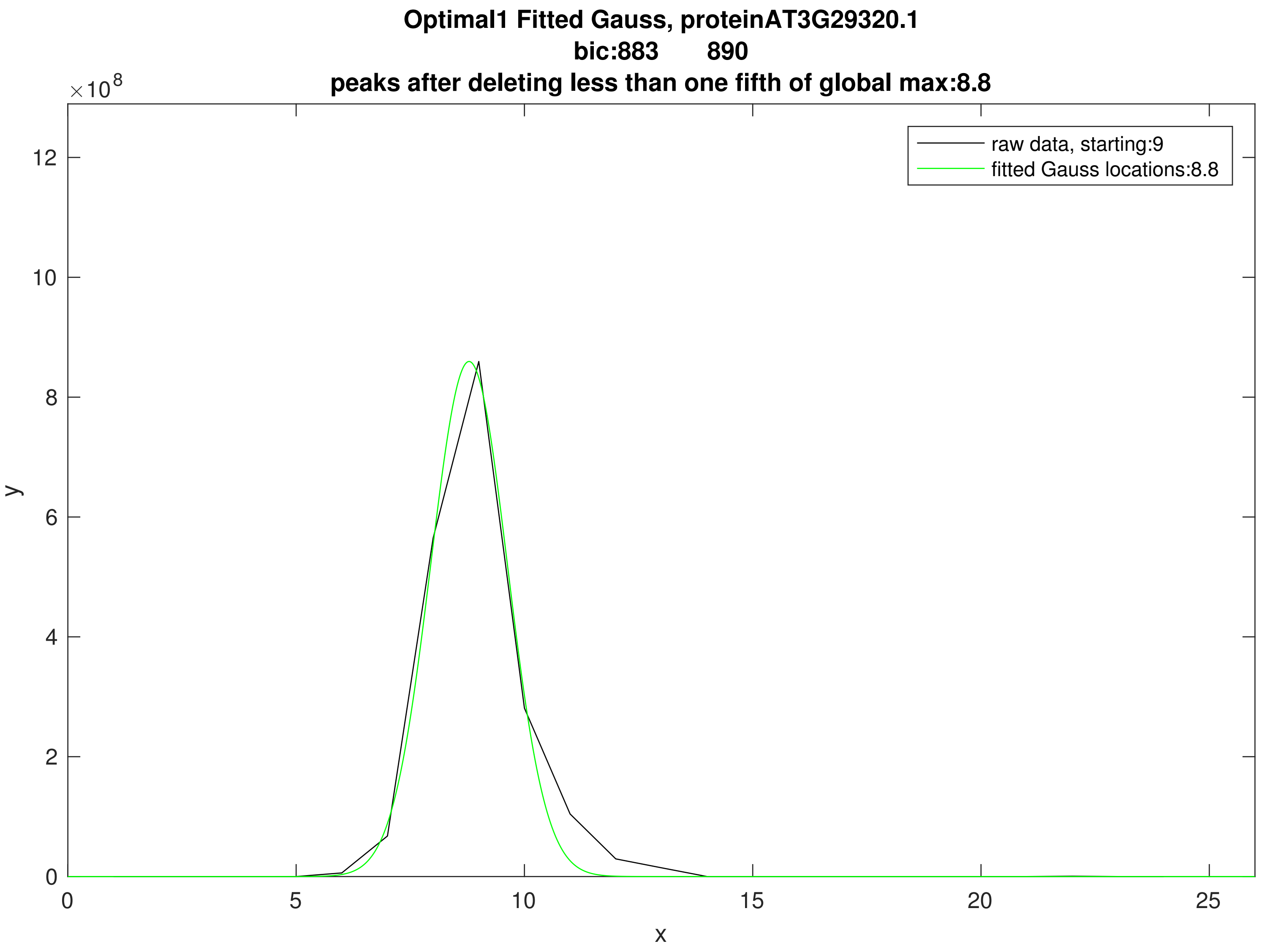
Optimal1 Fitted Gauss, proteinAT3G29320.1
bic:883 890
peaks after deleting less than one fifth of global max:8.8
×10^8
raw data, starting:9
fitted Gauss locations:8.8
y
x
0
2
4
6
8
10
12
0
5
10
15
20
25

**Optimal1 Fitted Gauss, proteinAT3G29350.1;AT3G29350.2**
**bic:684**
**peaks after deleting less than one fifth of global max:19**

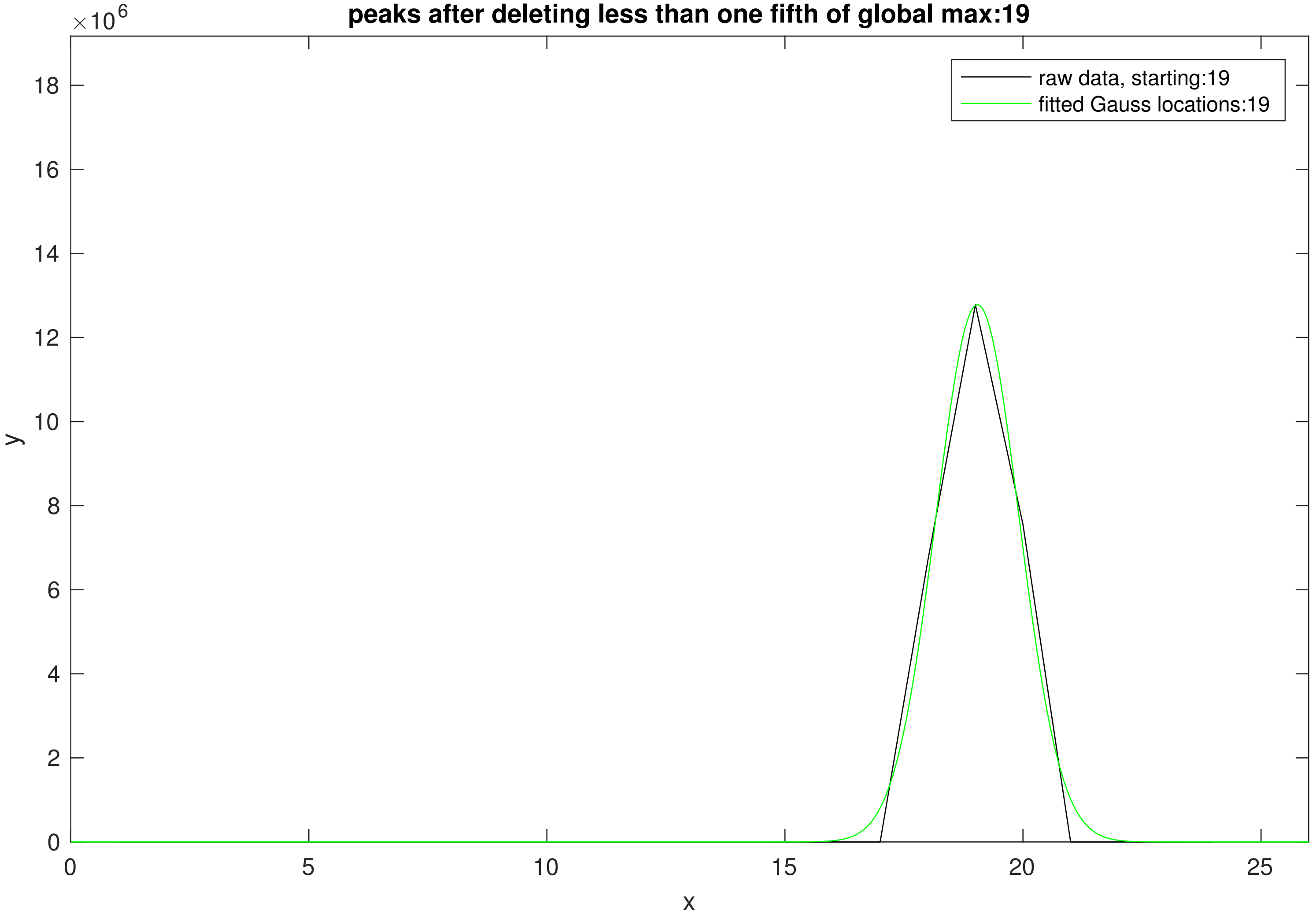

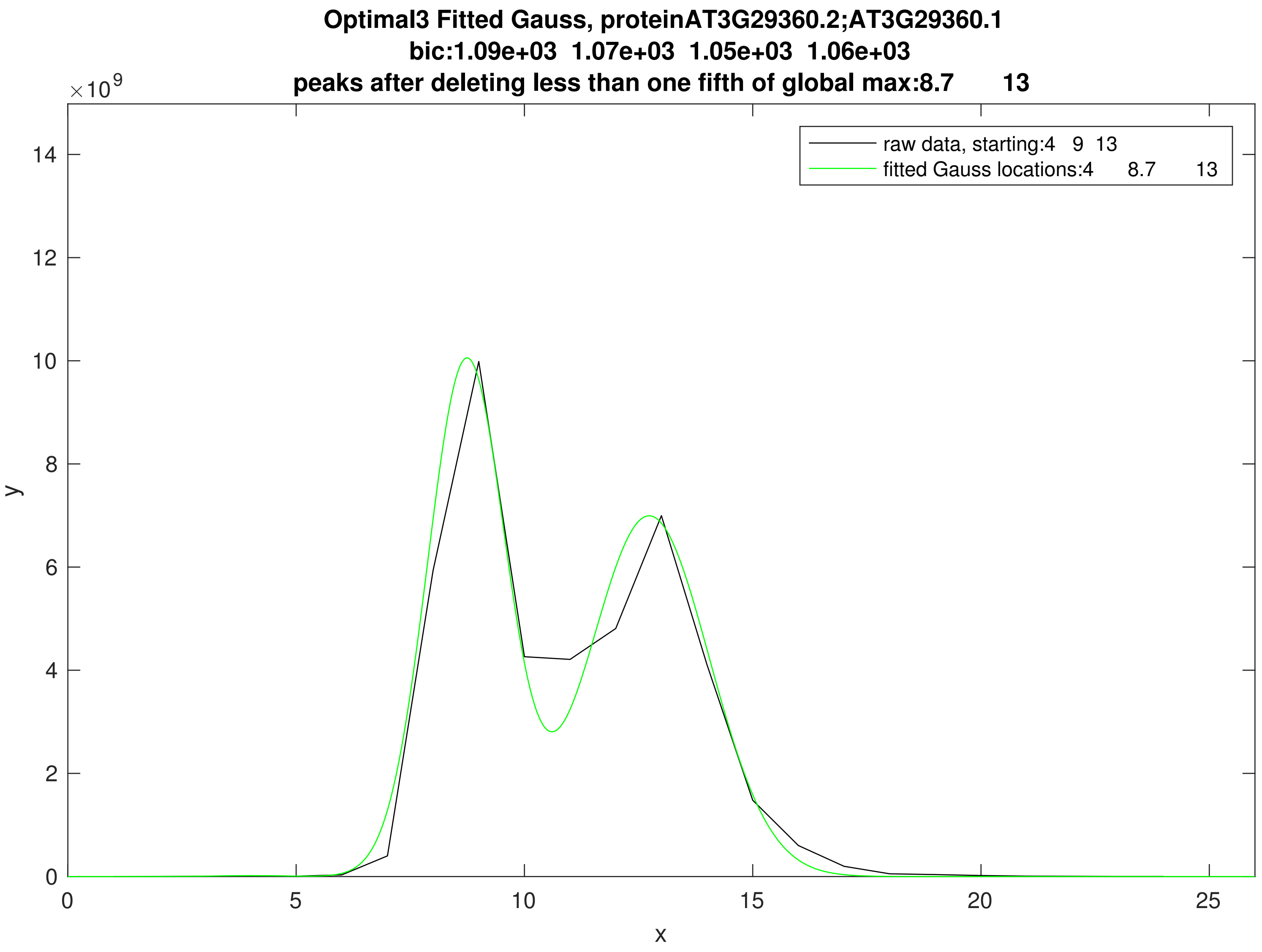
Optimal3 Fitted Gauss, proteinAT3G29360.2;AT3G29360.1
bic:1.09e+03 1.07e+03 1.05e+03 1.06e+03
peaks after deleting less than one fifth of global max:8.7 13
raw data, starting:4 9 13
fitted Gauss locations:4 8.7 13
$\times 10^9$
y
x

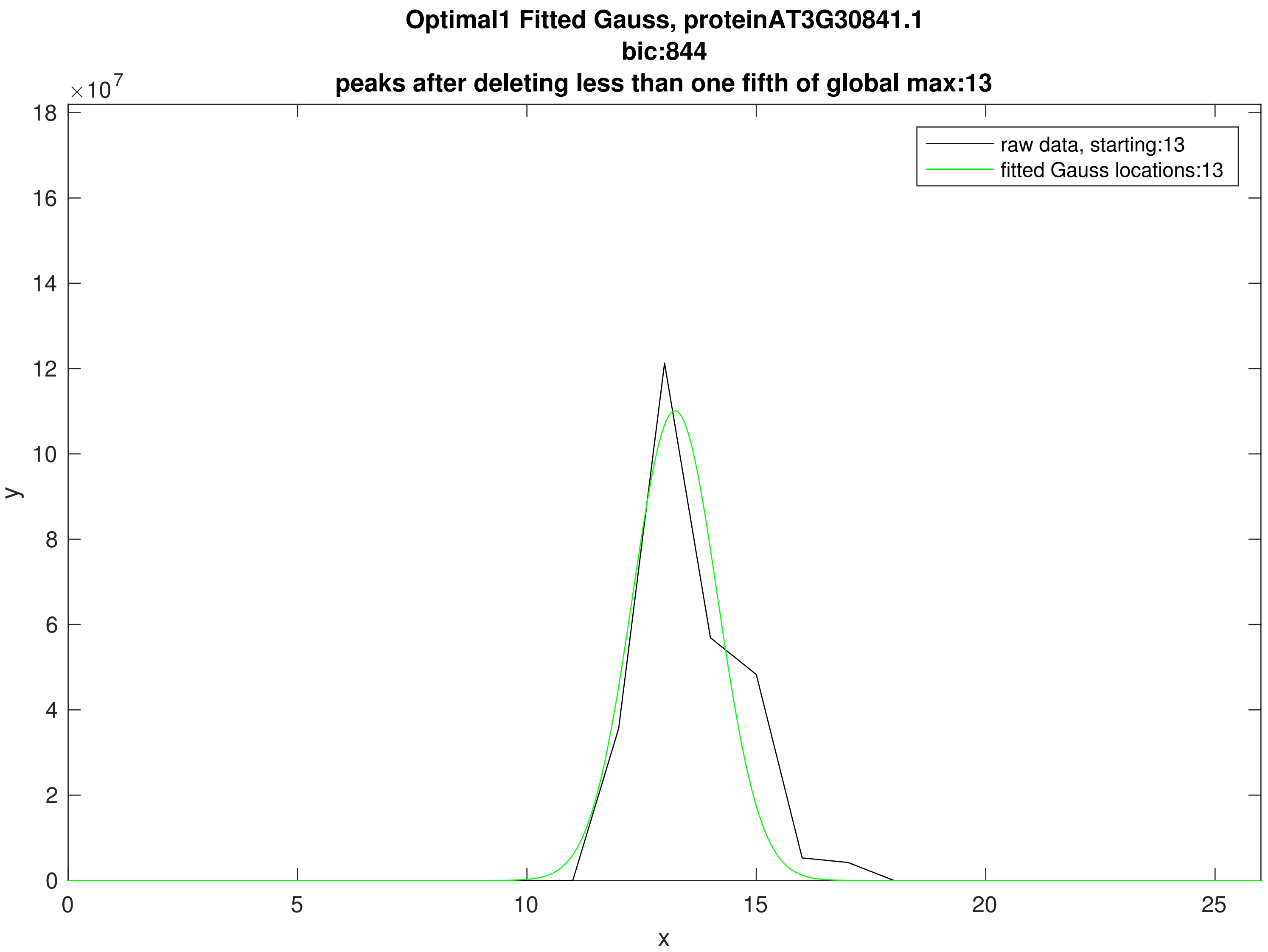
Optimal1 Fitted Gauss, proteinAT3G30841.1
bic:844
peaks after deleting less than one fifth of global max:13
×10^7
raw data, starting:13
fitted Gauss locations:13
y
x
0
2
4
6
8
10
12
14
16
18
0
5
10
15
20
25

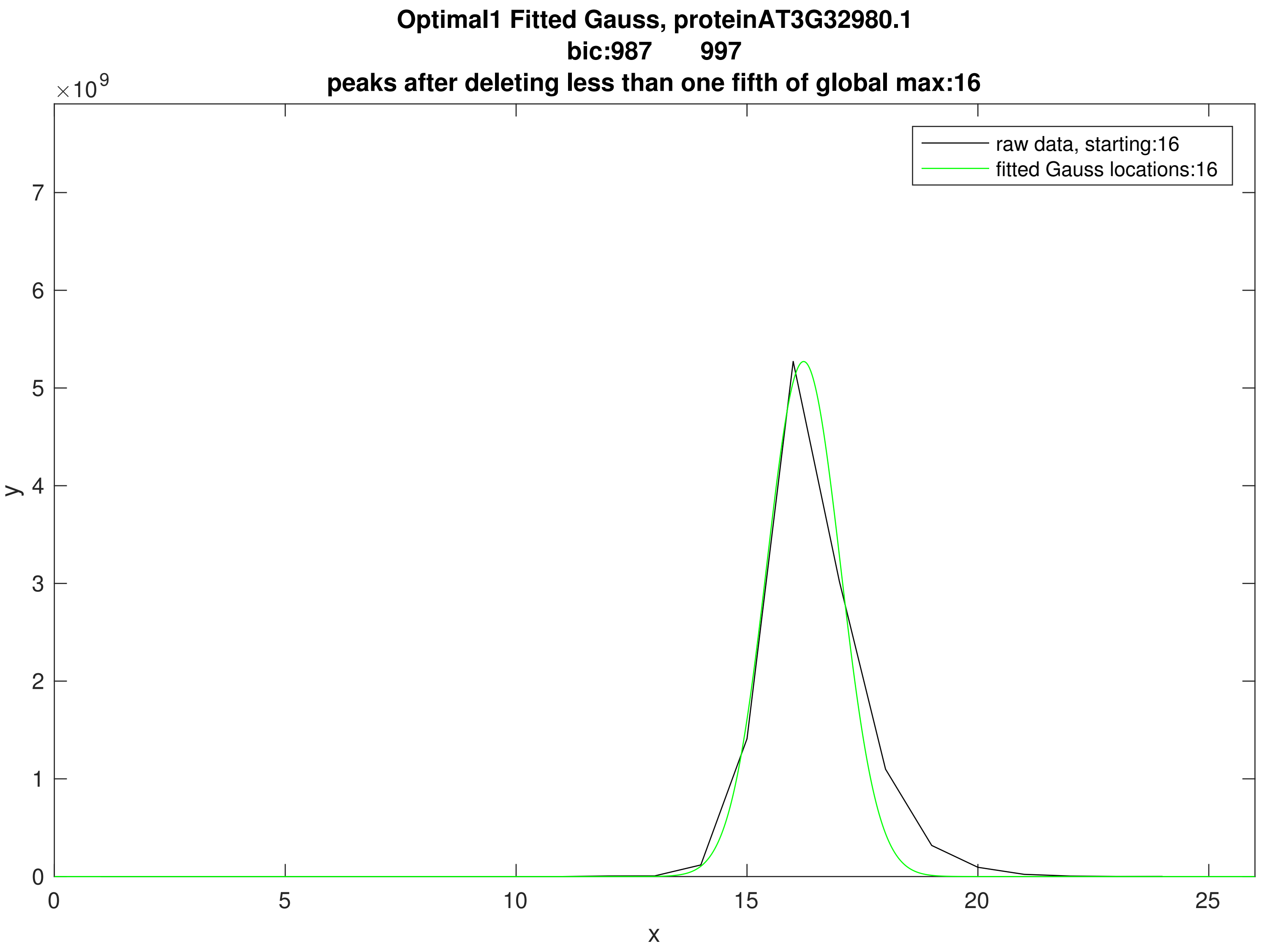
Optimal1 Fitted Gauss, proteinAT3G32980.1
bic:987 997
peaks after deleting less than one fifth of global max:16
×10^9
raw data, starting:16
fitted Gauss locations:16
y
x
0
1
2
3
4
5
6
7
0
5
10
15
20
25

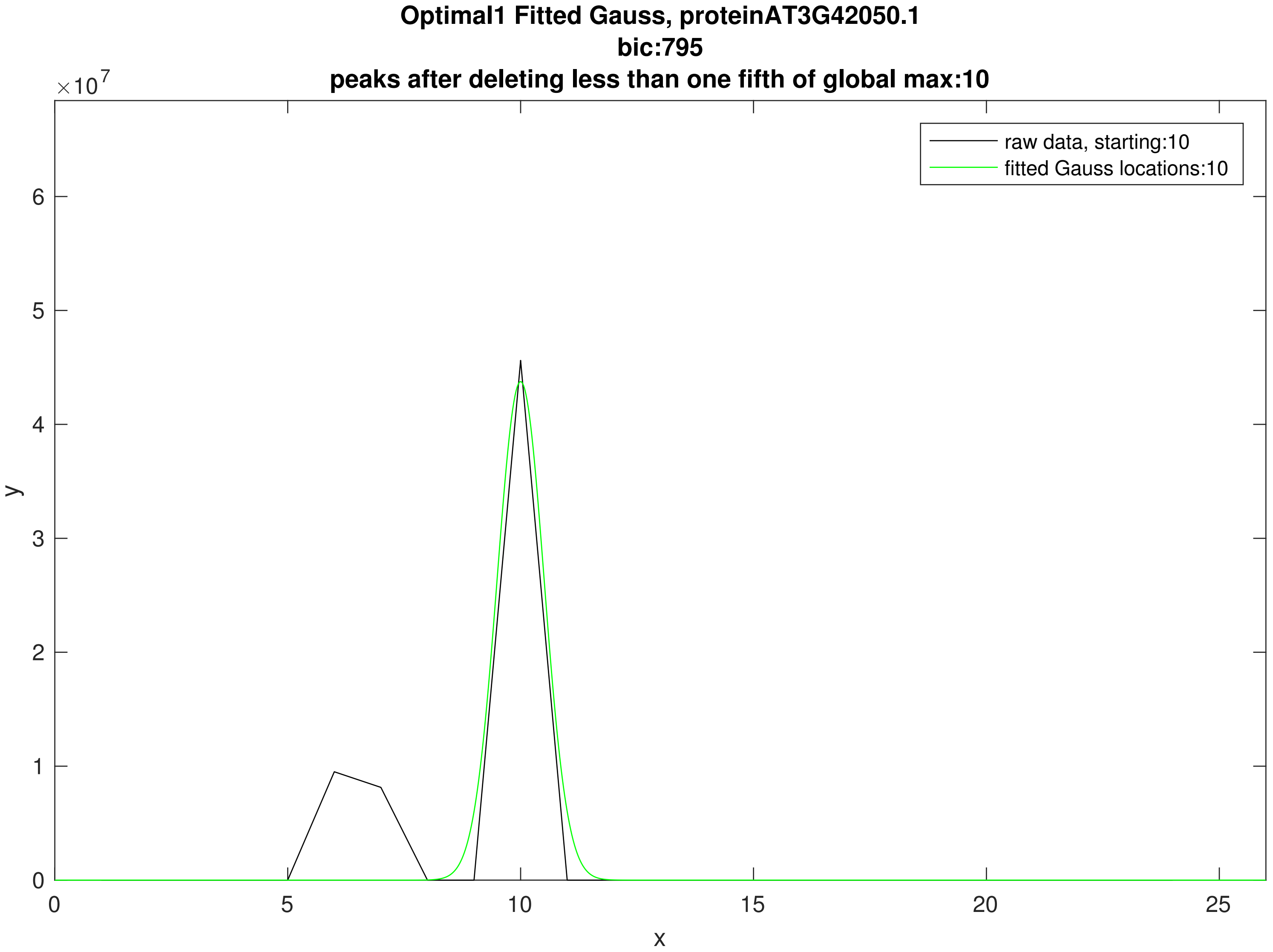

Optimal1 Fitted Gauss, proteinAT3G42050.1
bic:795
peaks after deleting less than one fifth of global max:10
×10^7
raw data, starting:10
fitted Gauss locations:10
y
x
0
1
2
3
4
5
6
0
5
10
15
20
25

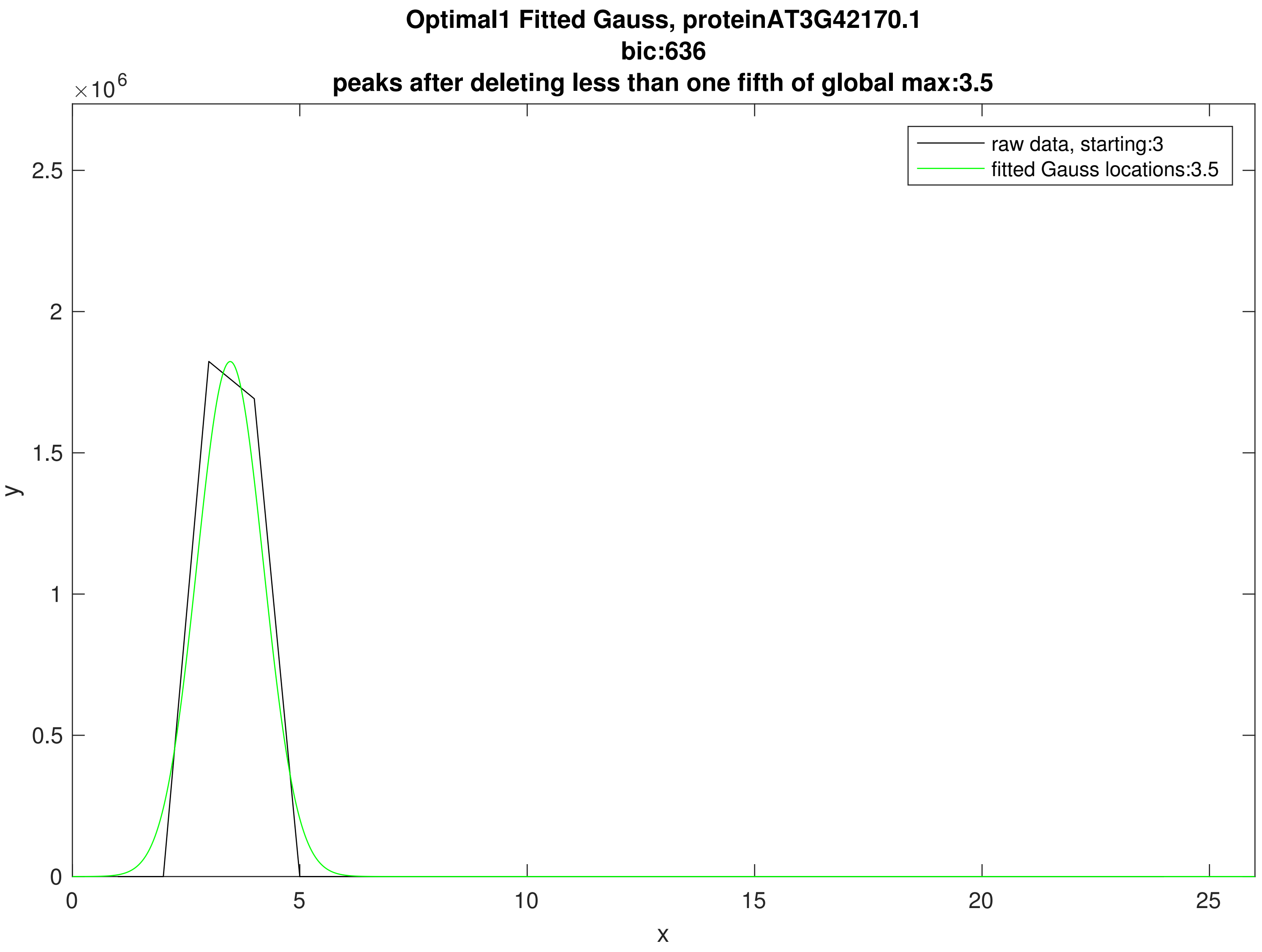
Optimal1 Fitted Gauss, proteinAT3G42170.1
bic:636
peaks after deleting less than one fifth of global max:3.5
×10^6
raw data, starting:3
fitted Gauss locations:3.5
2.5
2
1.5
1
0.5
0
y
0
5
10
15
20
25
x

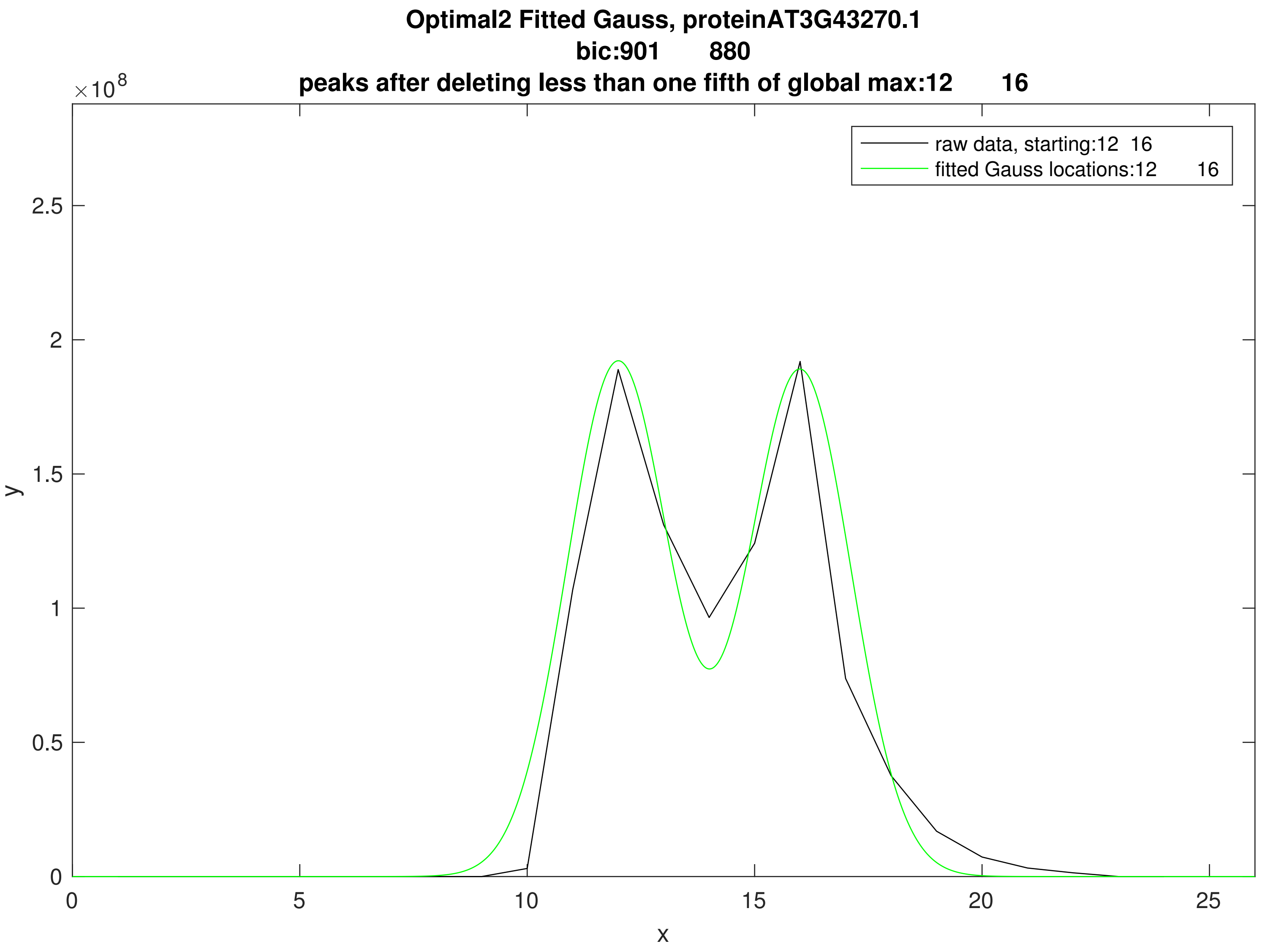

Optimal2 Fitted Gauss, proteinAT3G43270.1
bic:901 880
peaks after deleting less than one fifth of global max:12 16
×10^8
raw data, starting:12 16
fitted Gauss locations:12 16
2.5
2
1.5
1
0.5
0
y
0
5
10
15
20
25
x

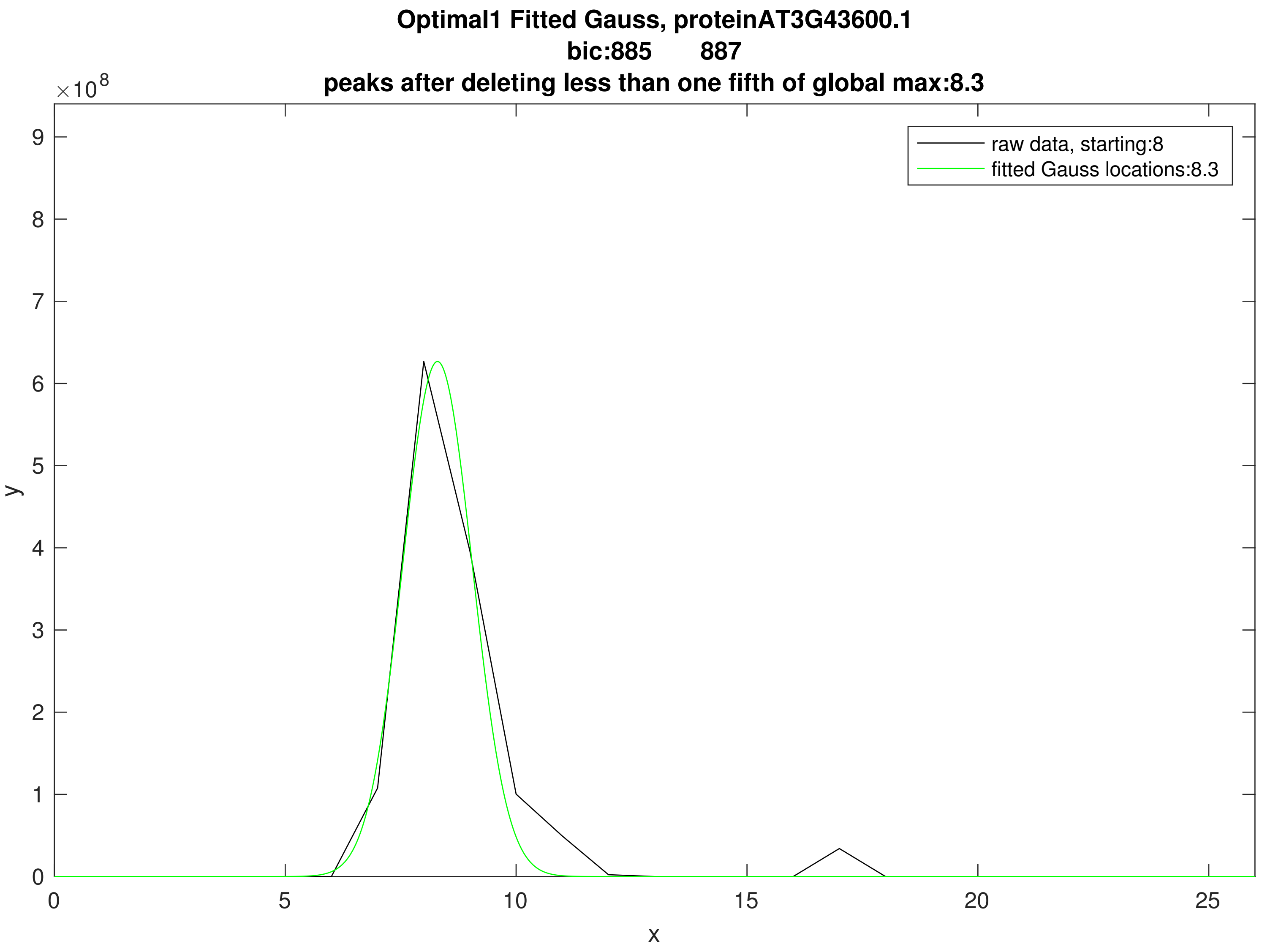
Optimal1 Fitted Gauss, proteinAT3G43600.1
bic:885 887
peaks after deleting less than one fifth of global max:8.3
×10^8
raw data, starting:8
fitted Gauss locations:8.3
y
x
0
1
2
3
4
5
6
7
8
9
0
5
10
15
20
25

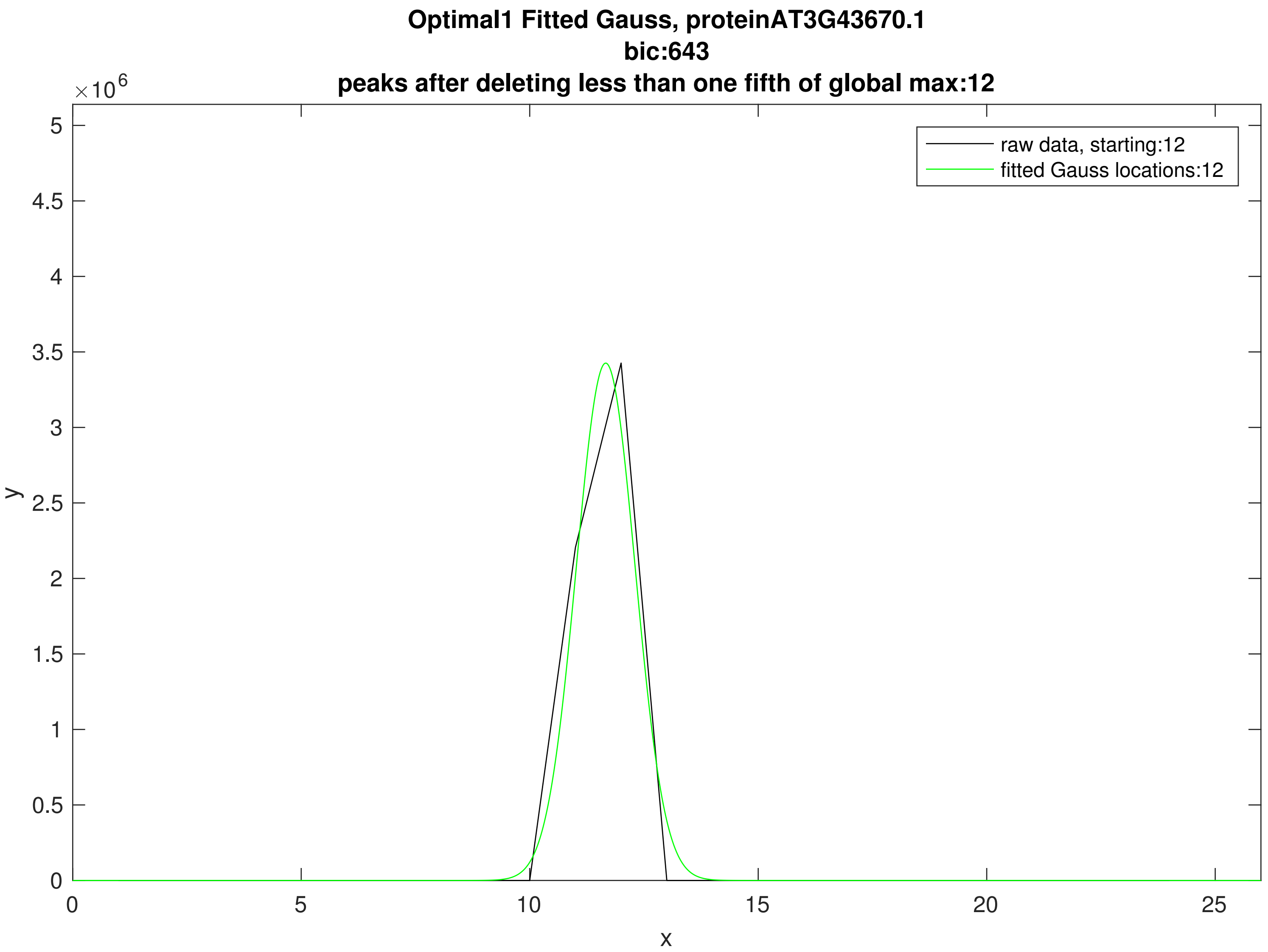
Optimal1 Fitted Gauss, proteinAT3G43670.1
bic:643
peaks after deleting less than one fifth of global max:12
×10^6
raw data, starting:12
fitted Gauss locations:12
y
x
0
0.5
1
1.5
2
2.5
3
3.5
4
4.5
5
0
5
10
15
20
25

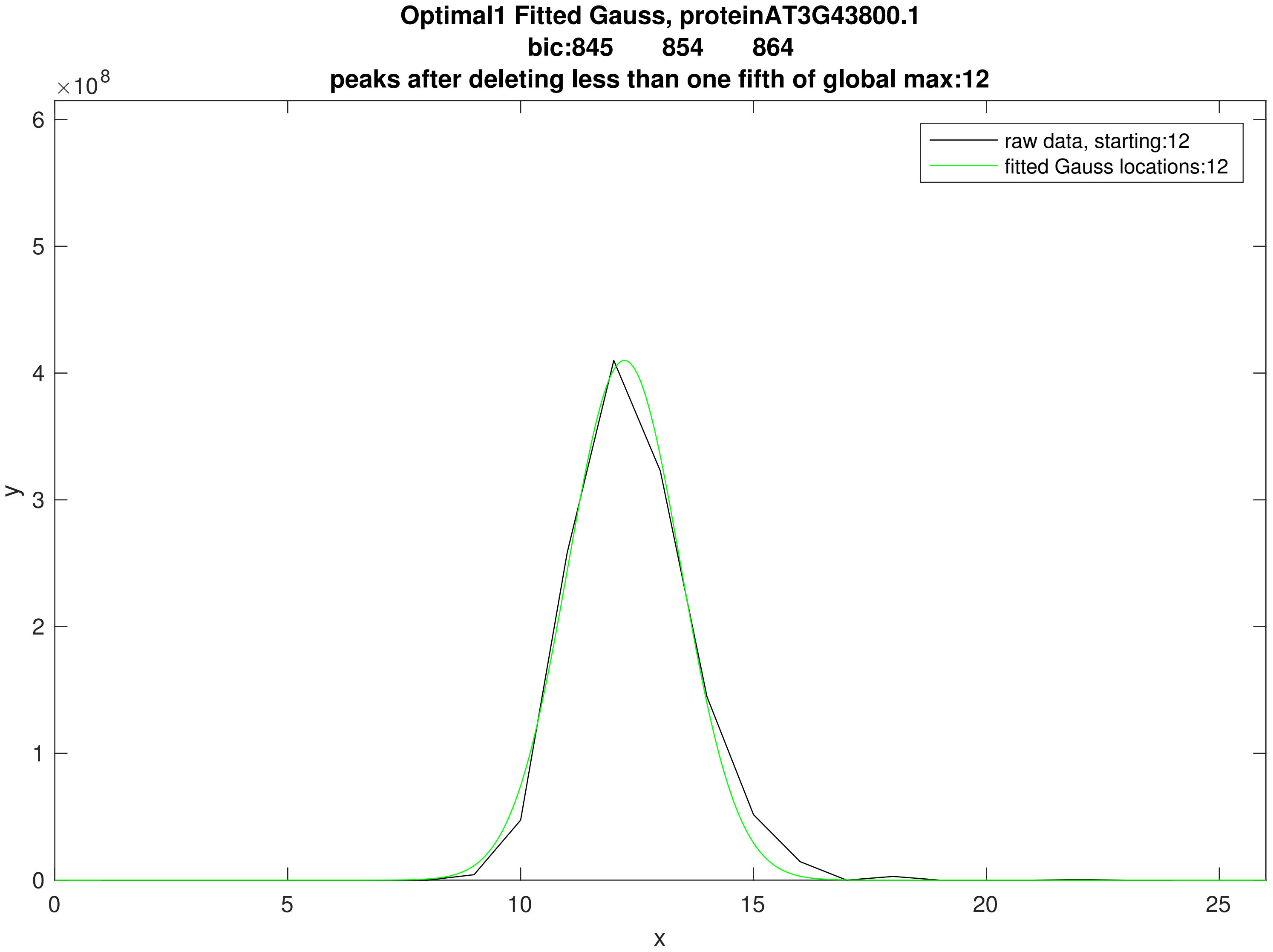

Optimal1 Fitted Gauss, proteinAT3G43800.1
bic:845 854 864
peaks after deleting less than one fifth of global max:12
×10^8
raw data, starting:12
fitted Gauss locations:12
y
x
0
1
2
3
4
5
6
0
5
10
15
20
25

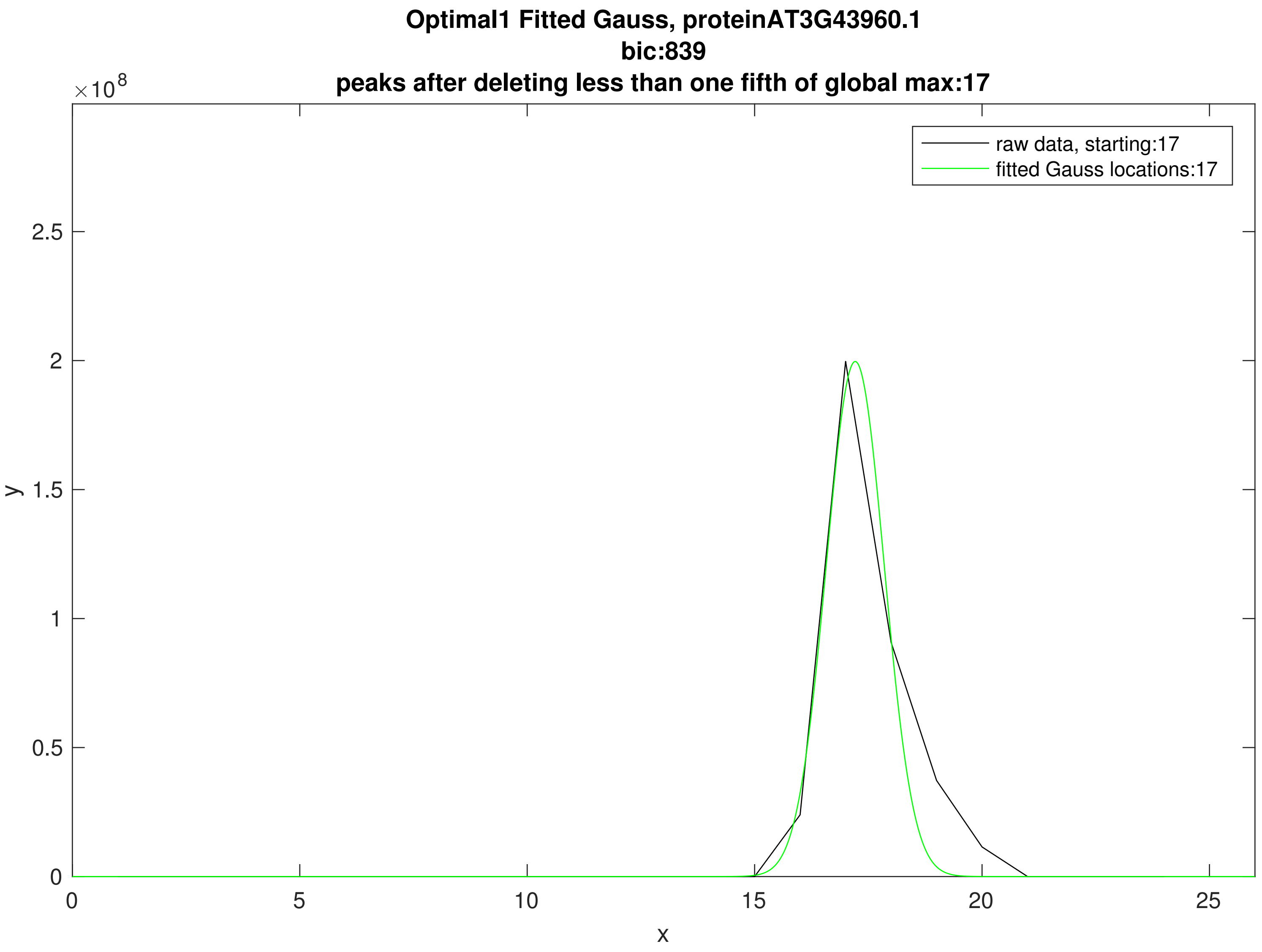

Optimal1 Fitted Gauss, proteinAT3G43960.1
bic:839
peaks after deleting less than one fifth of global max:17
×10^8
raw data, starting:17
fitted Gauss locations:17
y
x
0
0.5
1
1.5
2
2.5
5
10
15
20
25

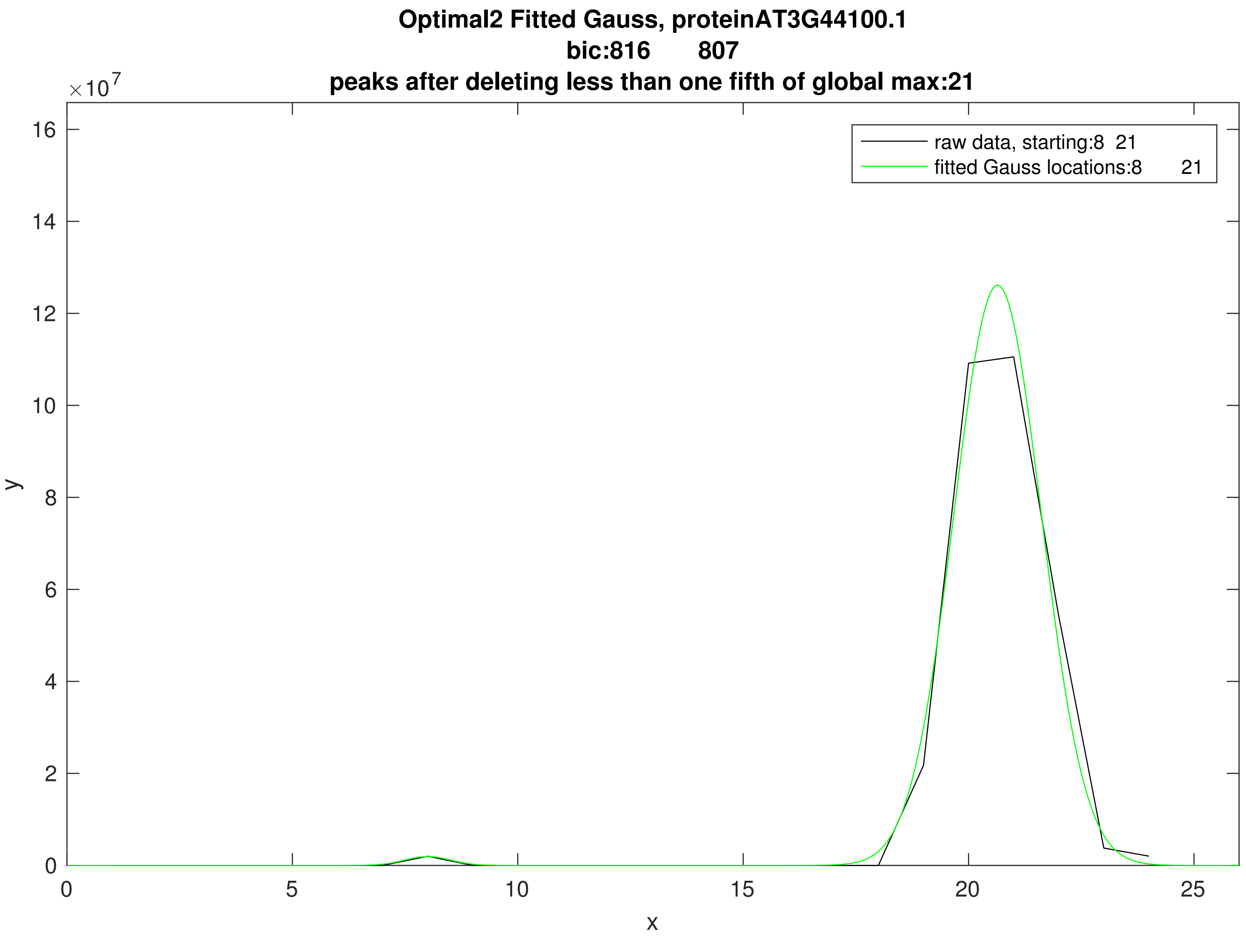
Optimal2 Fitted Gauss, proteinAT3G44100.1
bic:816 807
peaks after deleting less than one fifth of global max:21
×10^7
16
14
12
10
8
6
4
2
0
y
0
5
10
15
20
25
x
raw data, starting:8 21
fitted Gauss locations:8 21

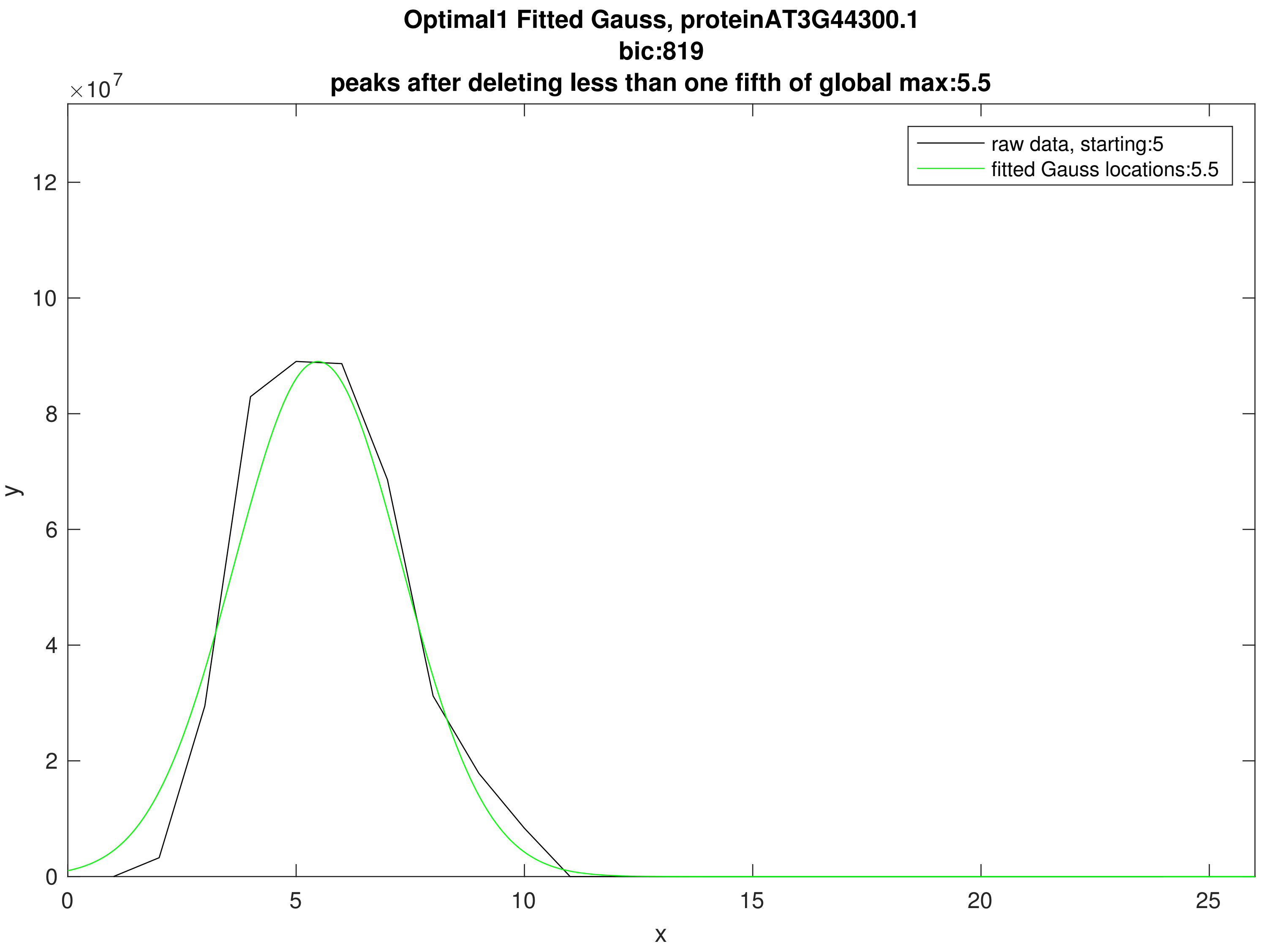
Optimal1 Fitted Gauss, proteinAT3G44300.1
bic:819
peaks after deleting less than one fifth of global max:5.5
×10^7
raw data, starting:5
fitted Gauss locations:5.5
12
10
8
6
4
2
0
y
0
5
10
15
20
25
x

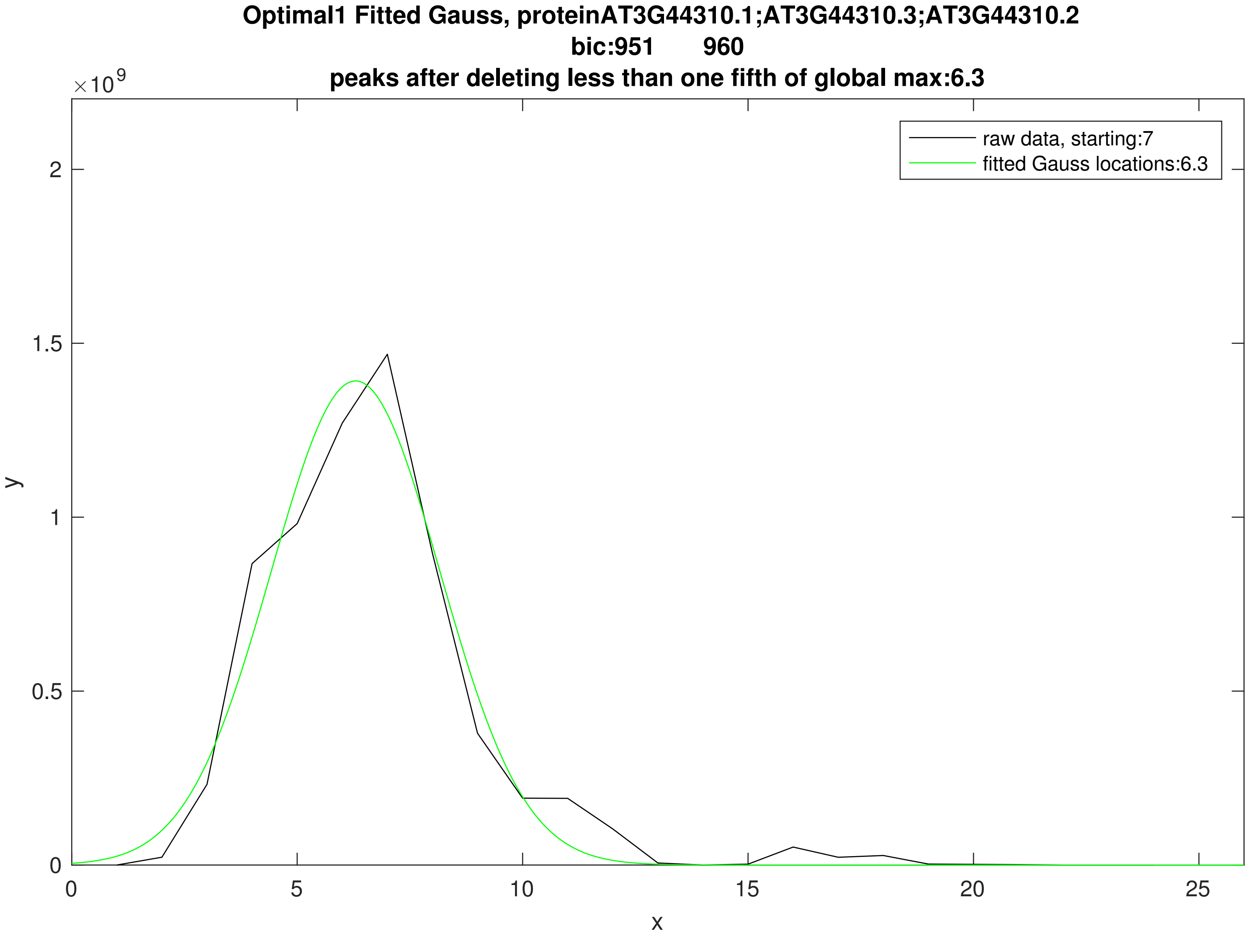

Optimal1 Fitted Gauss, proteinAT3G44310.1;AT3G44310.3;AT3G44310.2
bic:951 960
peaks after deleting less than one fifth of global max:6.3
×10^9
raw data, starting:7
fitted Gauss locations:6.3
y
x
0
0.5
1
1.5
2
5
10
15
20
25

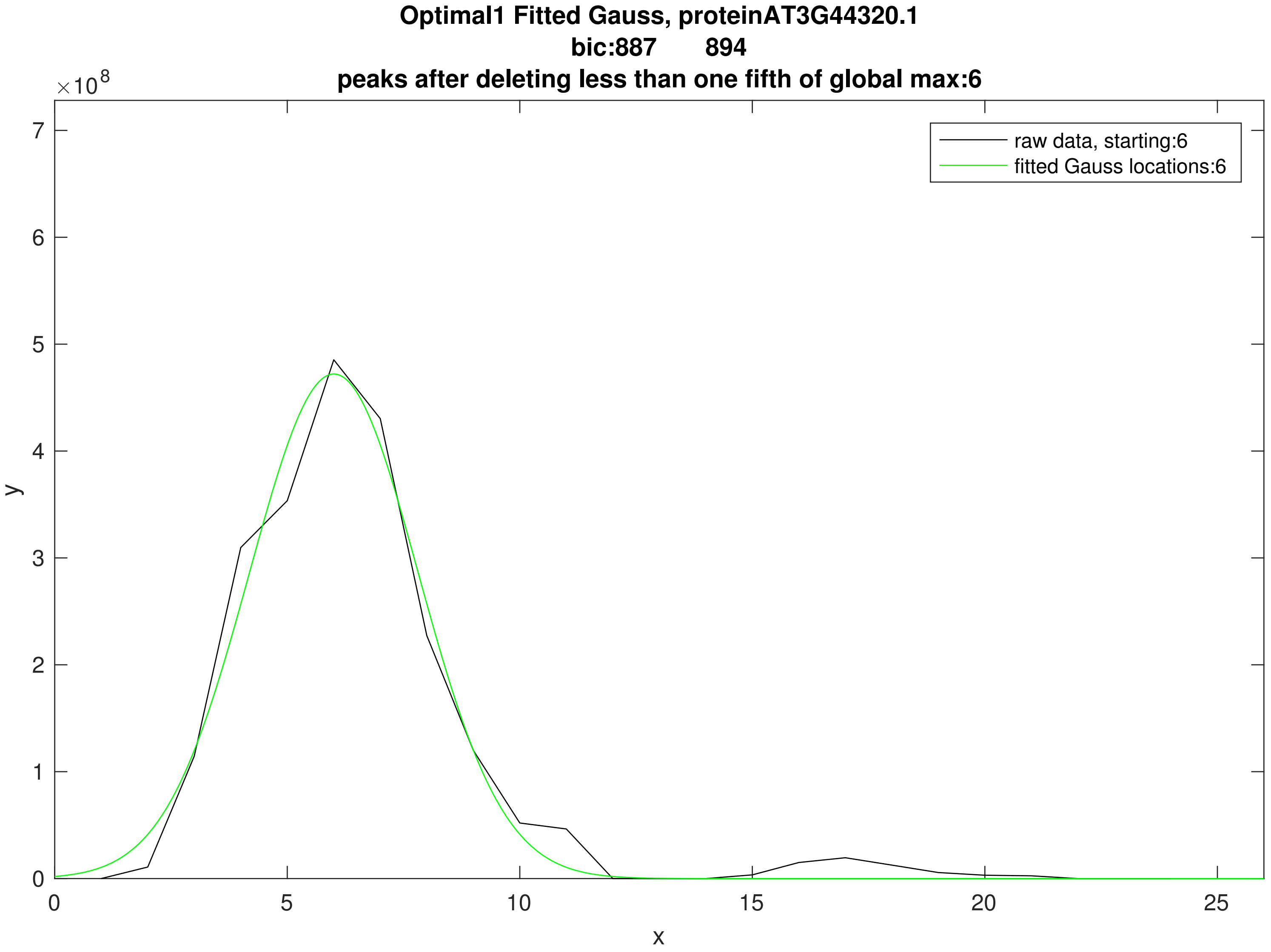
Optimal1 Fitted Gauss, proteinAT3G44320.1
bic:887 894
peaks after deleting less than one fifth of global max:6
×10^8
raw data, starting:6
fitted Gauss locations:6
y
x
0
1
2
3
4
5
6
7
0
5
10
15
20
25

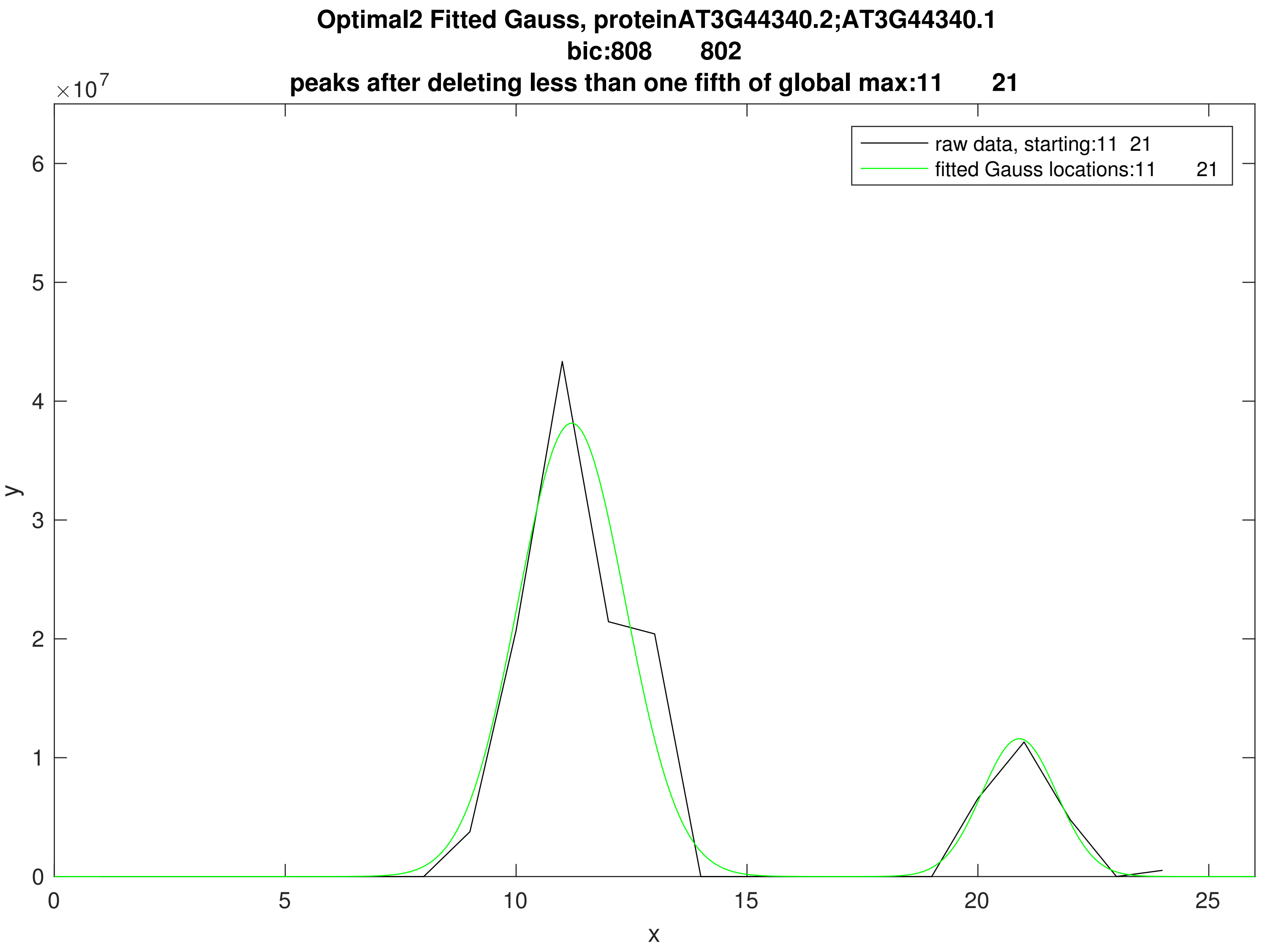
Optimal2 Fitted Gauss, proteinAT3G44340.2;AT3G44340.1
bic:808 802
peaks after deleting less than one fifth of global max:11 21
×10^7
raw data, starting:11 21
fitted Gauss locations:11 21
y
x
0
1
2
3
4
5
6
0
5
10
15
20
25

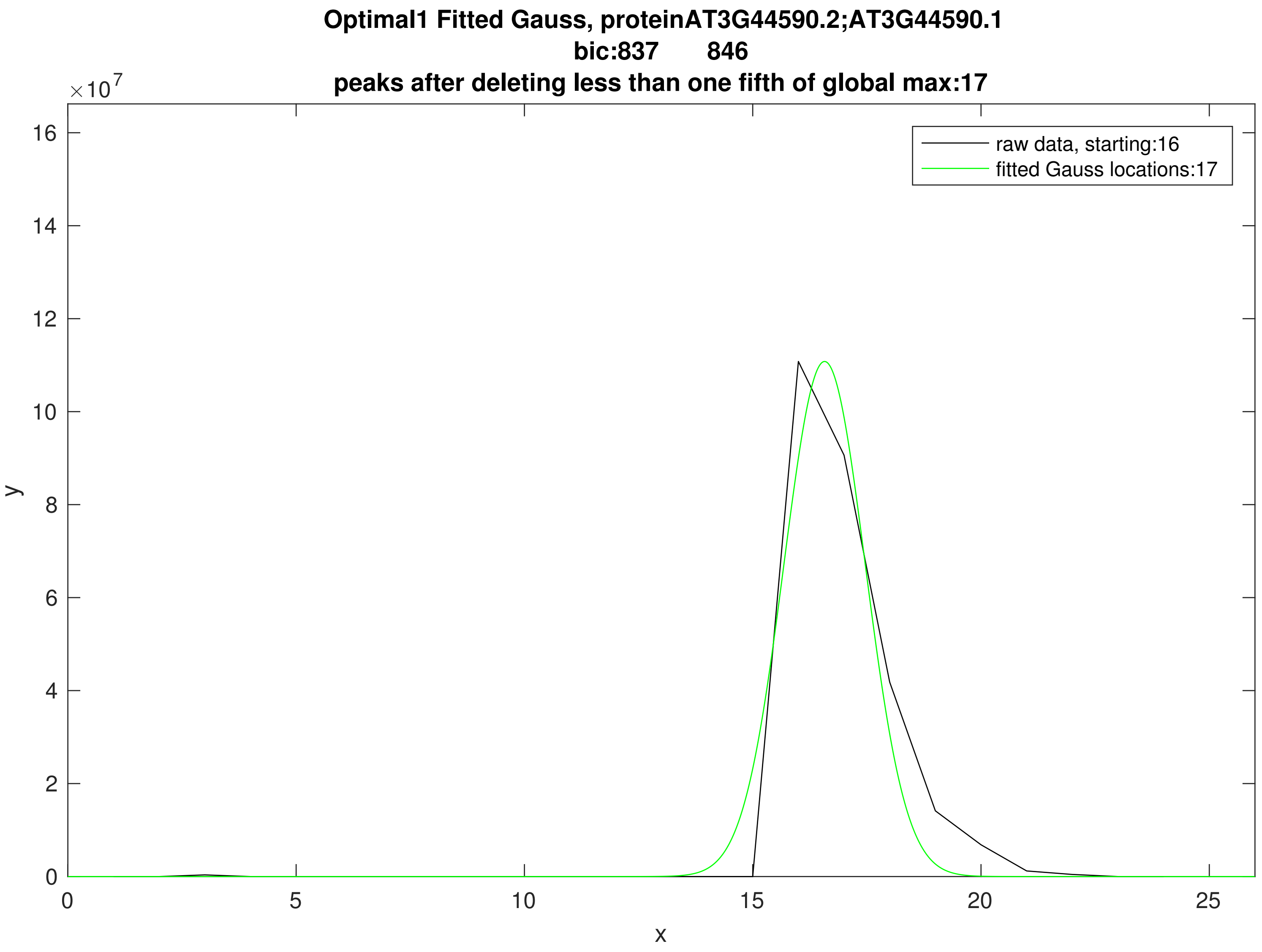
Optimal1 Fitted Gauss, proteinAT3G44590.2;AT3G44590.1
bic:837 846
peaks after deleting less than one fifth of global max:17
×10^7
16
14
12
10
8
6
4
2
0
y
0
5
10
15
20
25
x
raw data, starting:16
fitted Gauss locations:17

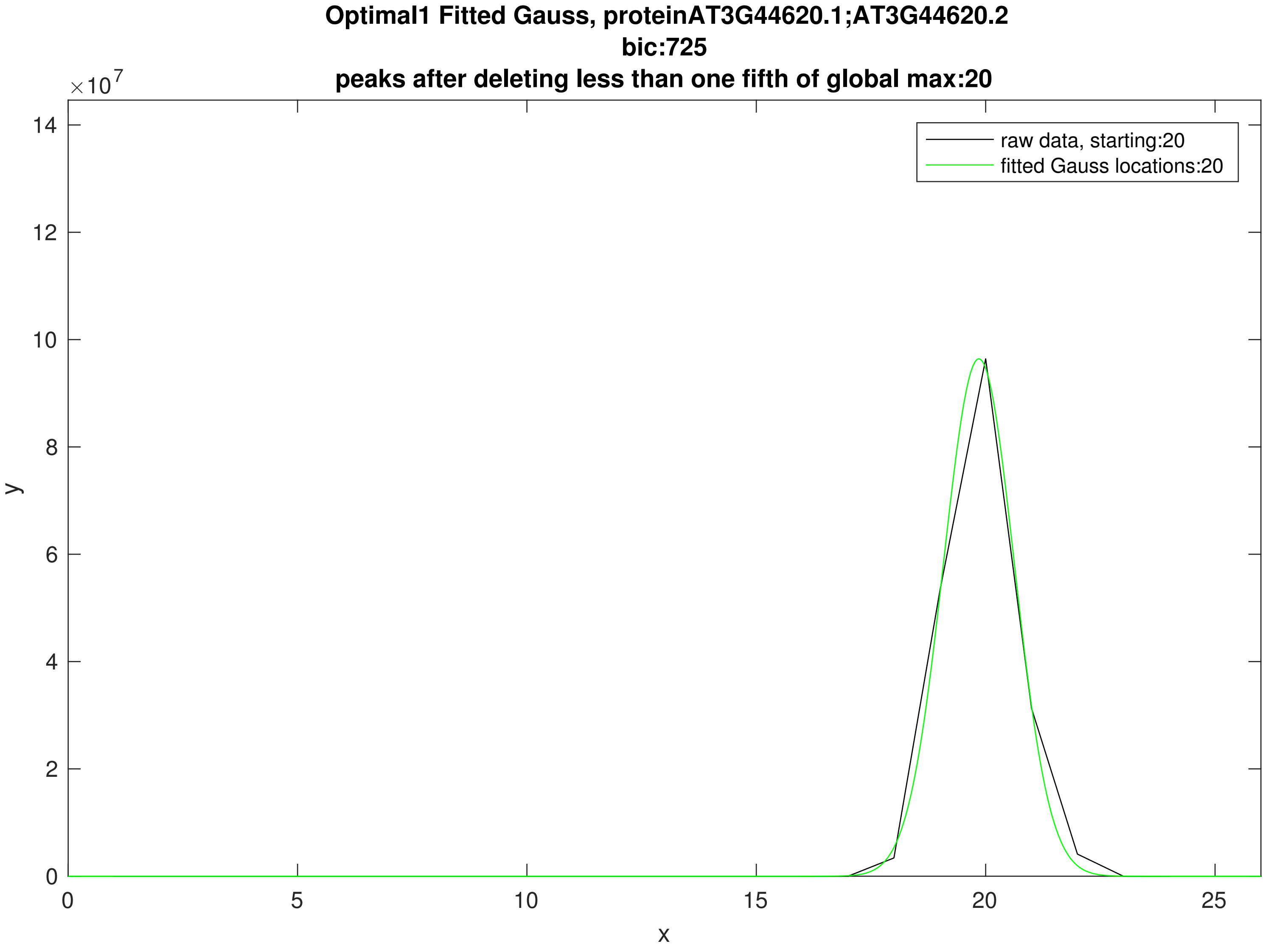

Optimal1 Fitted Gauss, proteinAT3G44620.1;AT3G44620.2
bic:725
peaks after deleting less than one fifth of global max:20
×10^7
raw data, starting:20
fitted Gauss locations:20
y
x
0
2
4
6
8
10
12
14
0
5
10
15
20
25

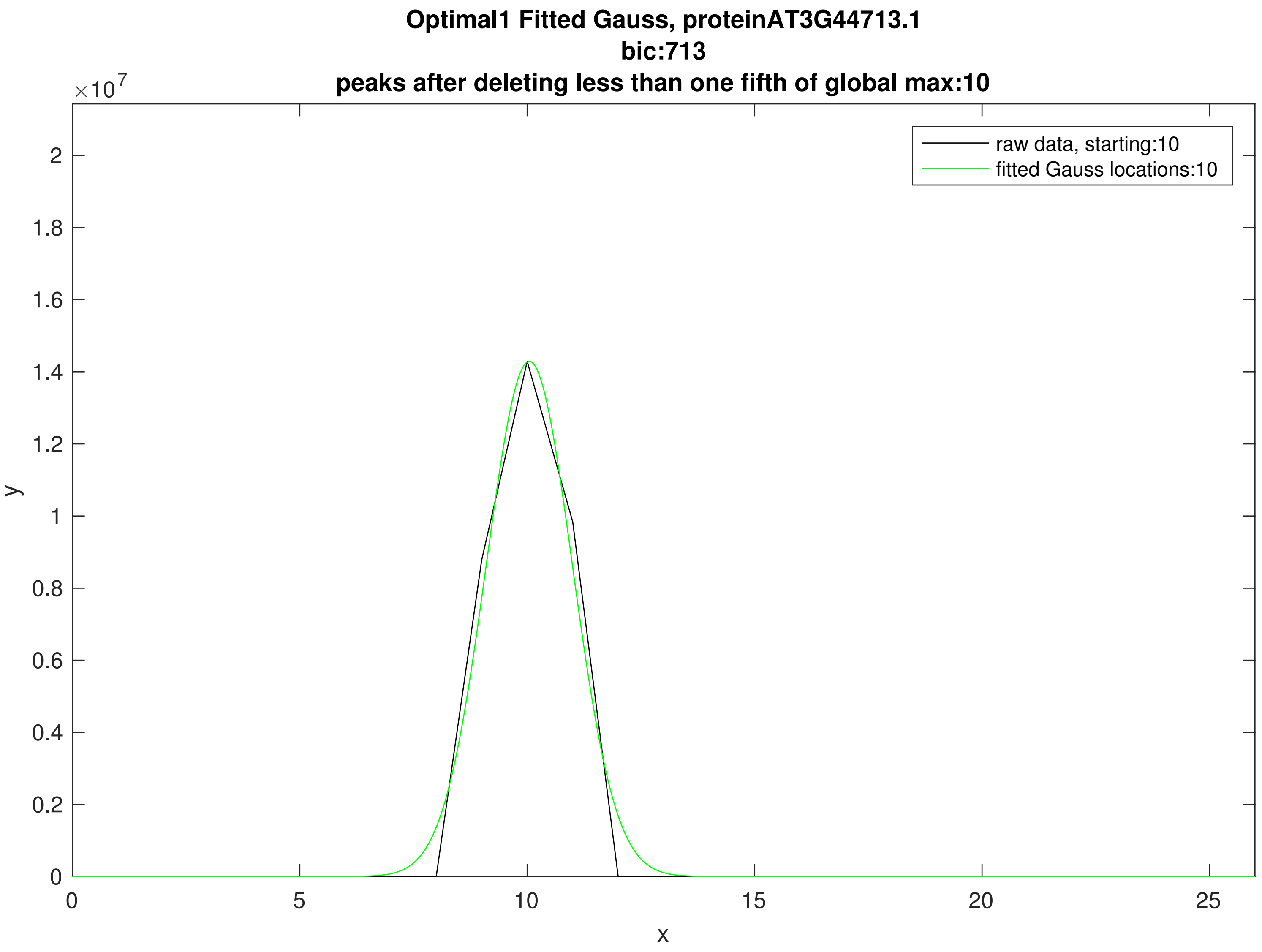
Optimal1 Fitted Gauss, proteinAT3G44713.1
bic:713
peaks after deleting less than one fifth of global max:10
×10^7
raw data, starting:10
fitted Gauss locations:10
y
x
0
5
10
15
20
25
0
0.2
0.4
0.6
0.8
1
1.2
1.4
1.6
1.8
2

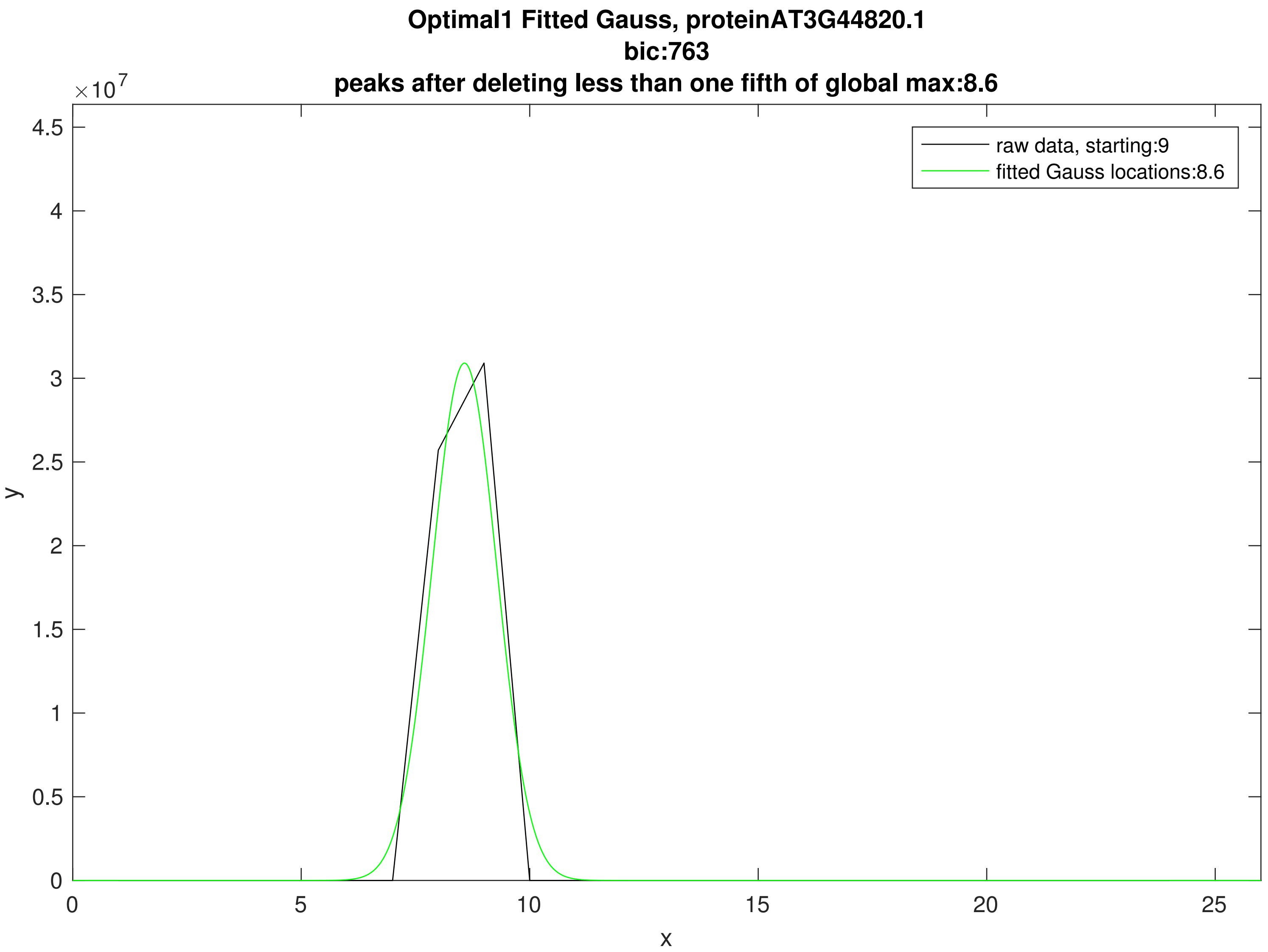

Optimal1 Fitted Gauss, proteinAT3G44820.1
bic:763
peaks after deleting less than one fifth of global max:8.6
×10^7
4.5
4
3.5
3
2.5
2
1.5
1
0.5
0
y
0
5
10
15
20
25
x
raw data, starting:9
fitted Gauss locations:8.6

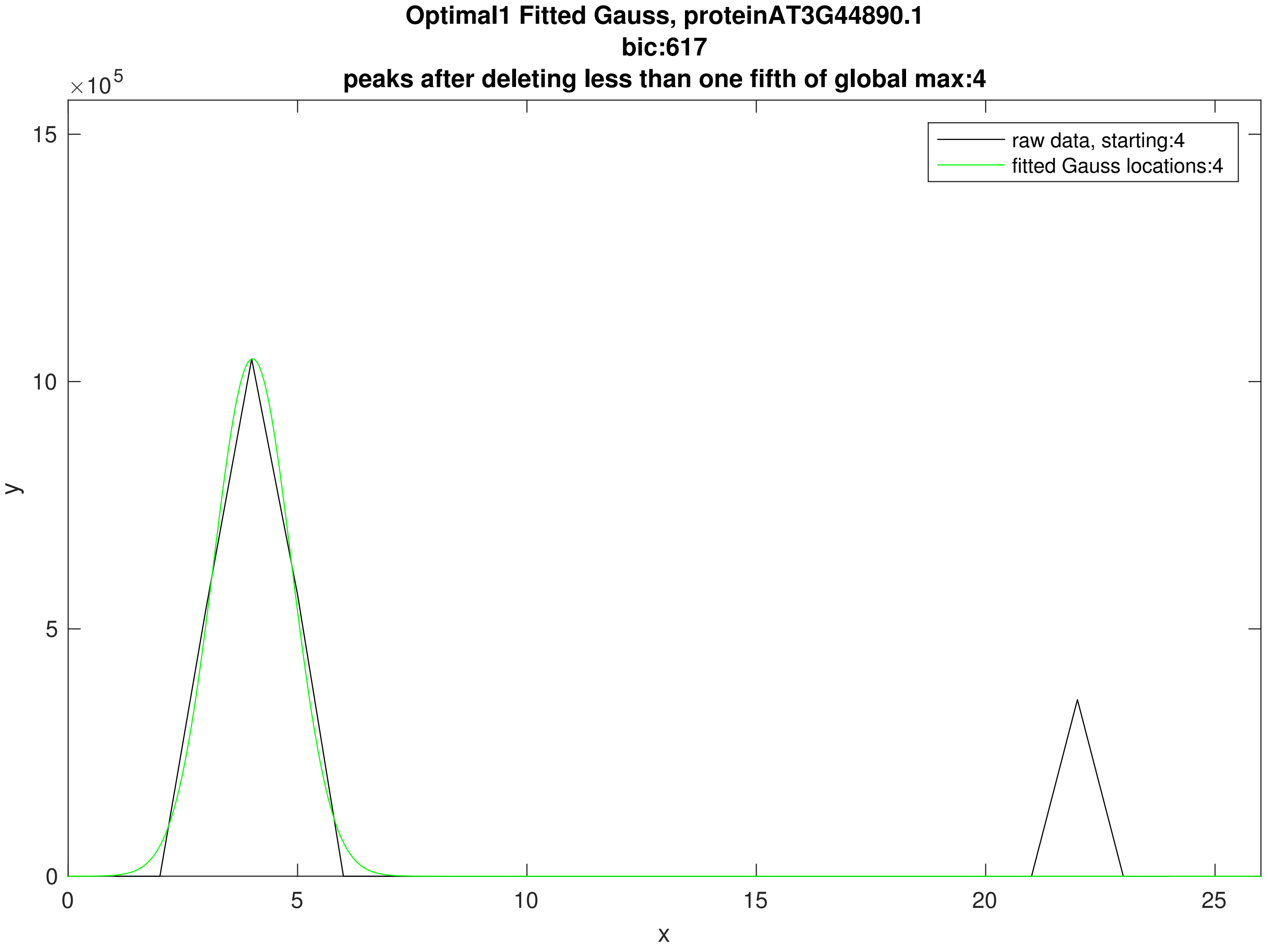
Optimal1 Fitted Gauss, proteinAT3G44890.1
bic:617
peaks after deleting less than one fifth of global max:4
×10^5
raw data, starting:4
fitted Gauss locations:4
15
10
5
0
y
0
5
10
15
20
25
x

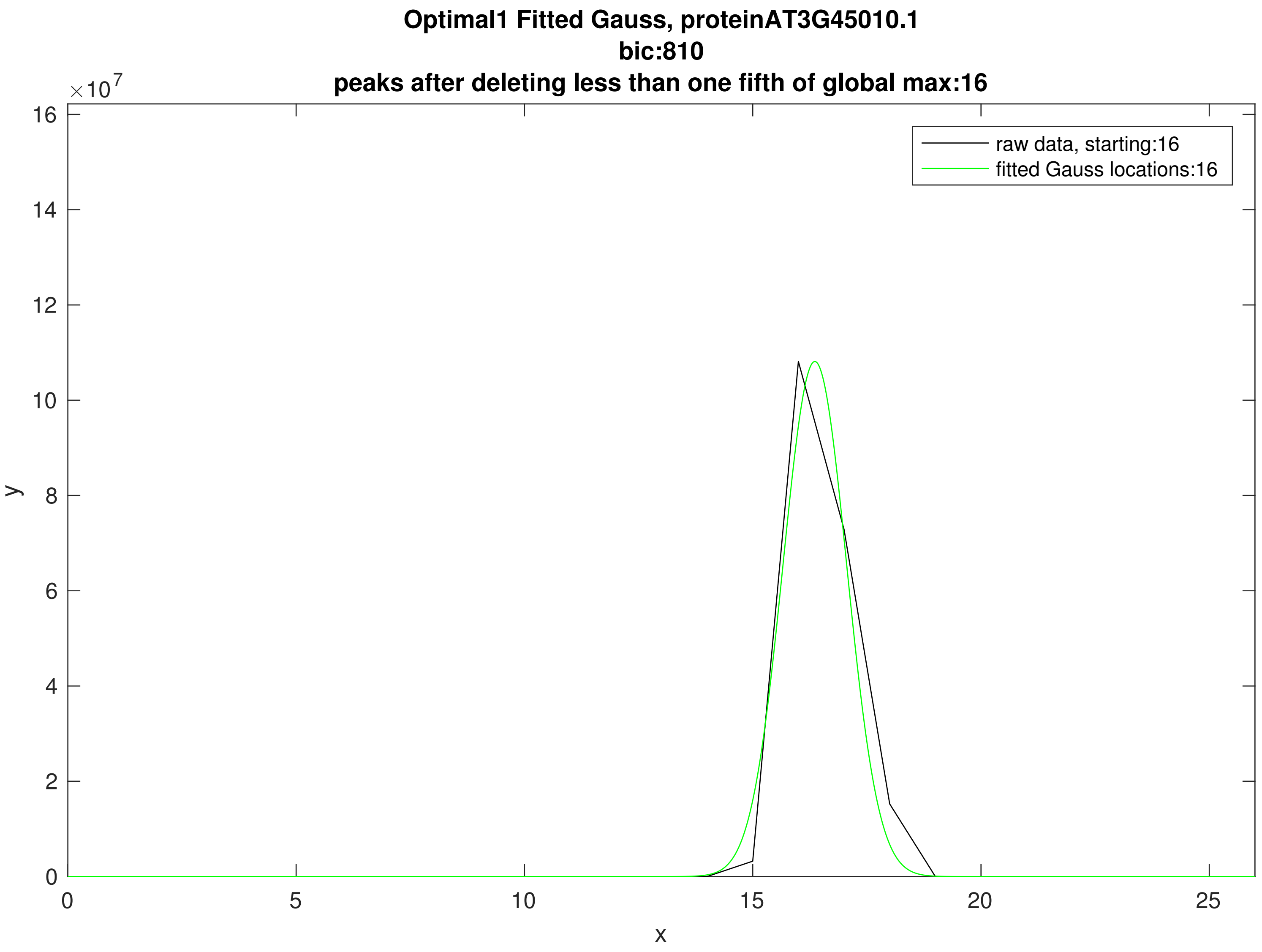
Optimal1 Fitted Gauss, proteinAT3G45010.1
bic:810
peaks after deleting less than one fifth of global max:16
×10^7
16
14
12
10
8
6
4
2
0
y
0
5
10
15
20
25
x
raw data, starting:16
fitted Gauss locations:16

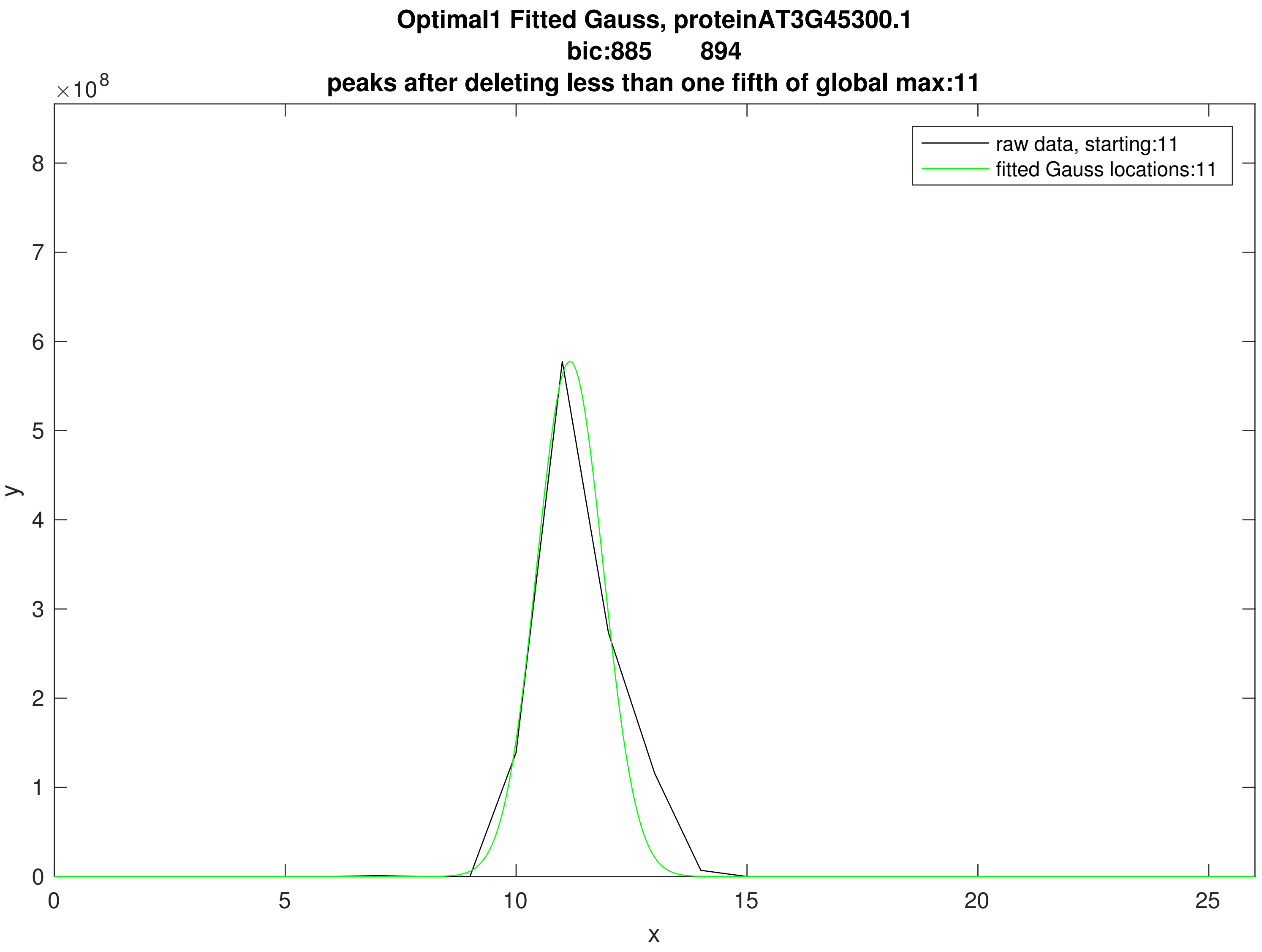
Optimal1 Fitted Gauss, proteinAT3G45300.1
bic:885 894
peaks after deleting less than one fifth of global max:11
×10^8
raw data, starting:11
fitted Gauss locations:11
y
x
0
1
2
3
4
5
6
7
8
0
5
10
15
20
25

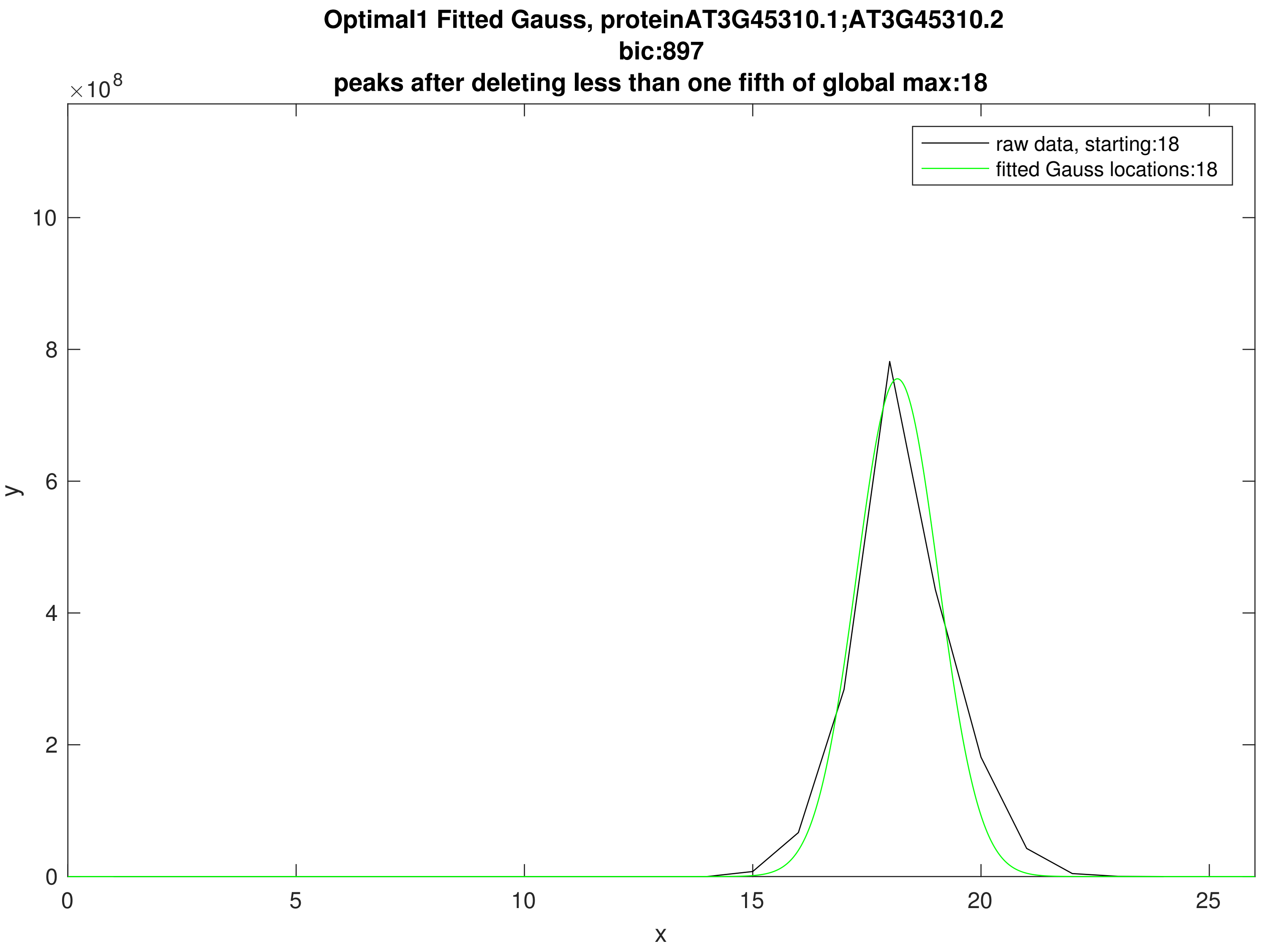

Optimal1 Fitted Gauss, proteinAT3G45310.1;AT3G45310.2
bic:897
peaks after deleting less than one fifth of global max:18
×10^8
raw data, starting:18
fitted Gauss locations:18
y
x
0
2
4
6
8
10
0
5
10
15
20
25

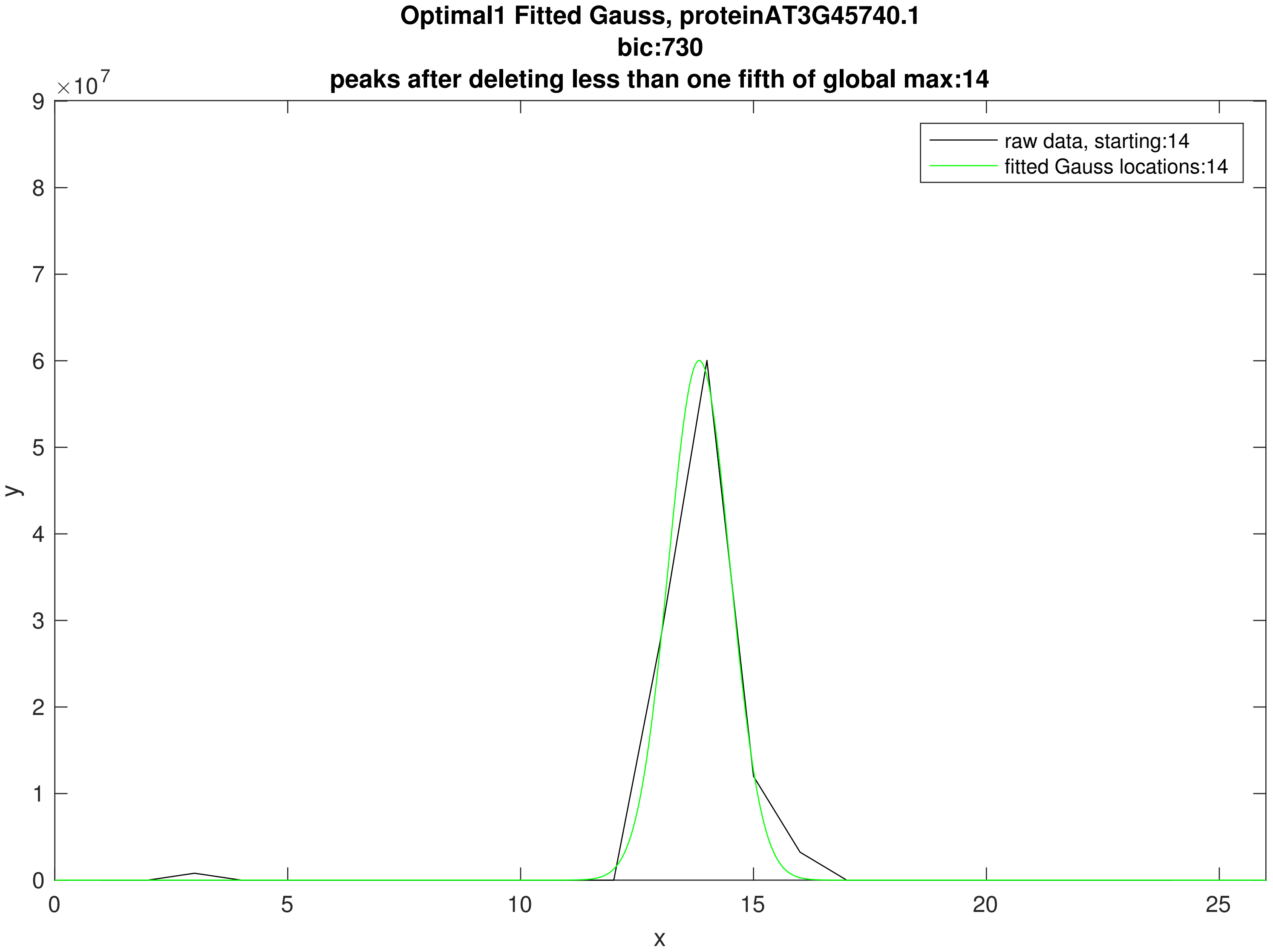
Optimal1 Fitted Gauss, proteinAT3G45740.1
bic:730
peaks after deleting less than one fifth of global max:14
×10^7
y
x
raw data, starting:14
fitted Gauss locations:14
0
1
2
3
4
5
6
7
8
9
0
5
10
15
20
25

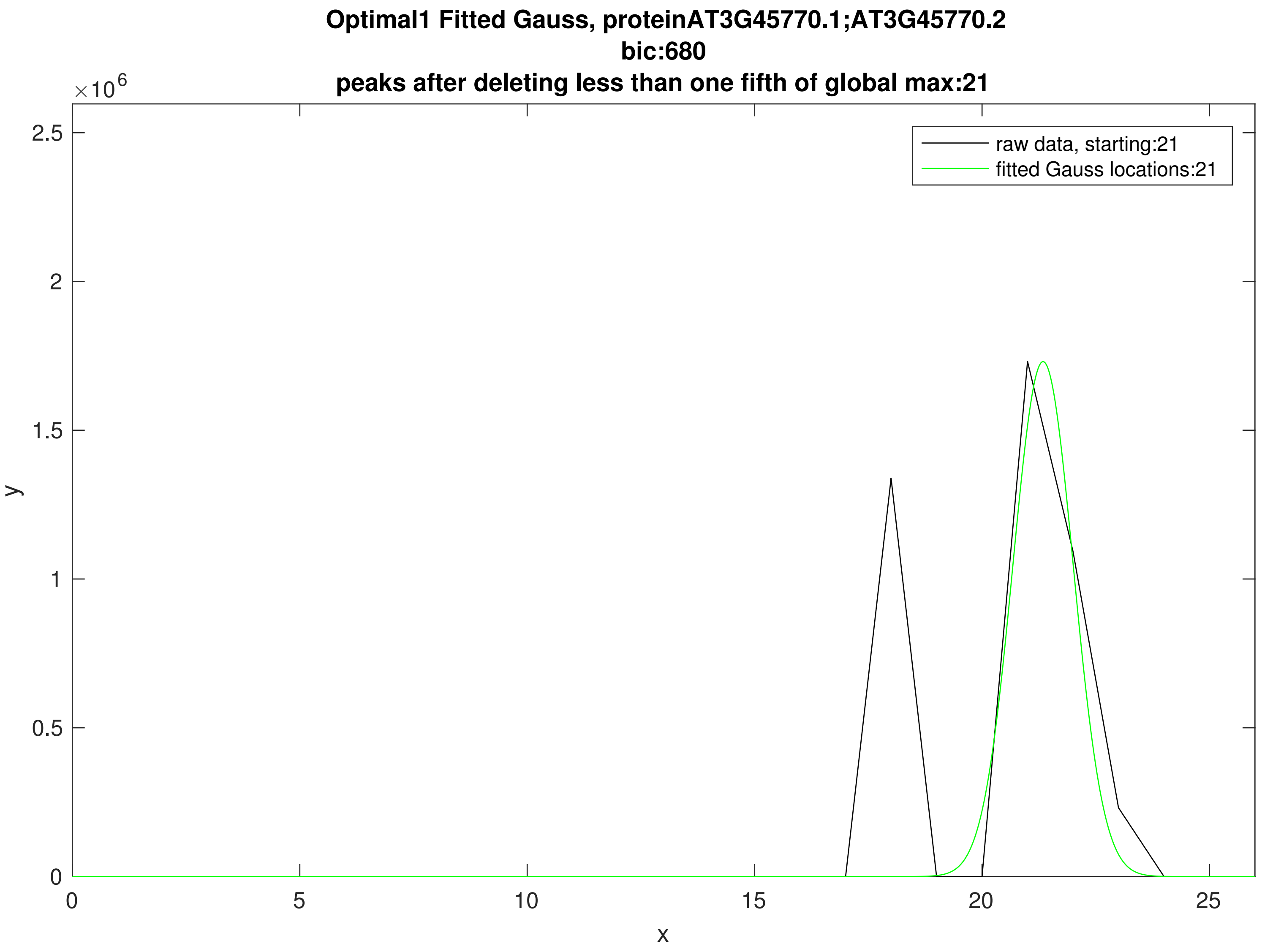
Optimal1 Fitted Gauss, proteinAT3G45770.1;AT3G45770.2
bic:680
peaks after deleting less than one fifth of global max:21
×10^6
2.5
2
1.5
1
0.5
0
y
0
5
10
15
20
25
x
raw data, starting:21
fitted Gauss locations:21

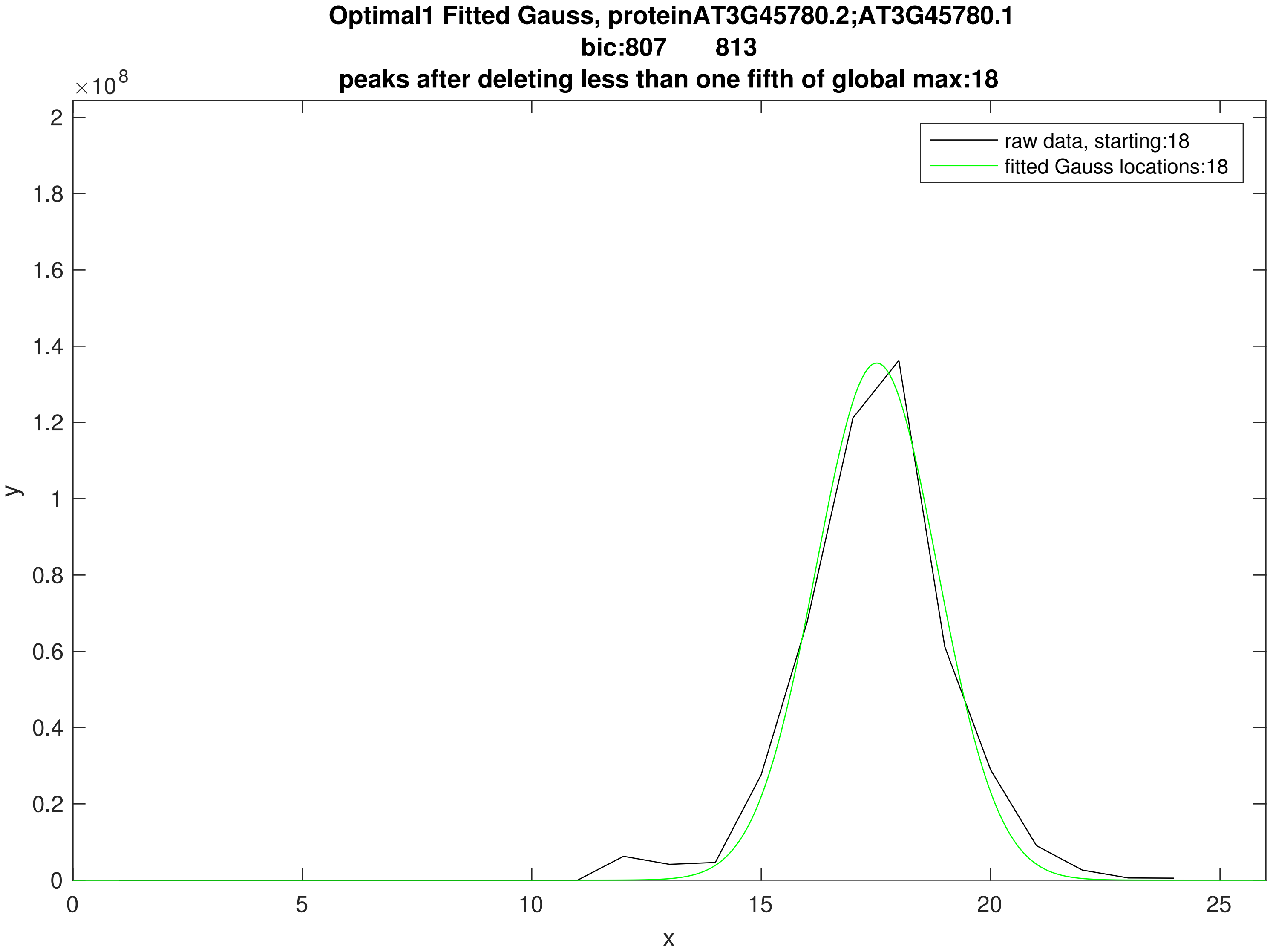

Optimal1 Fitted Gauss, proteinAT3G45780.2;AT3G45780.1
bic:807 813
peaks after deleting less than one fifth of global max:18
×10^8
raw data, starting:18
fitted Gauss locations:18
y
x
0
0.2
0.4
0.6
0.8
1
1.2
1.4
1.6
1.8
2
5
10
15
20
25

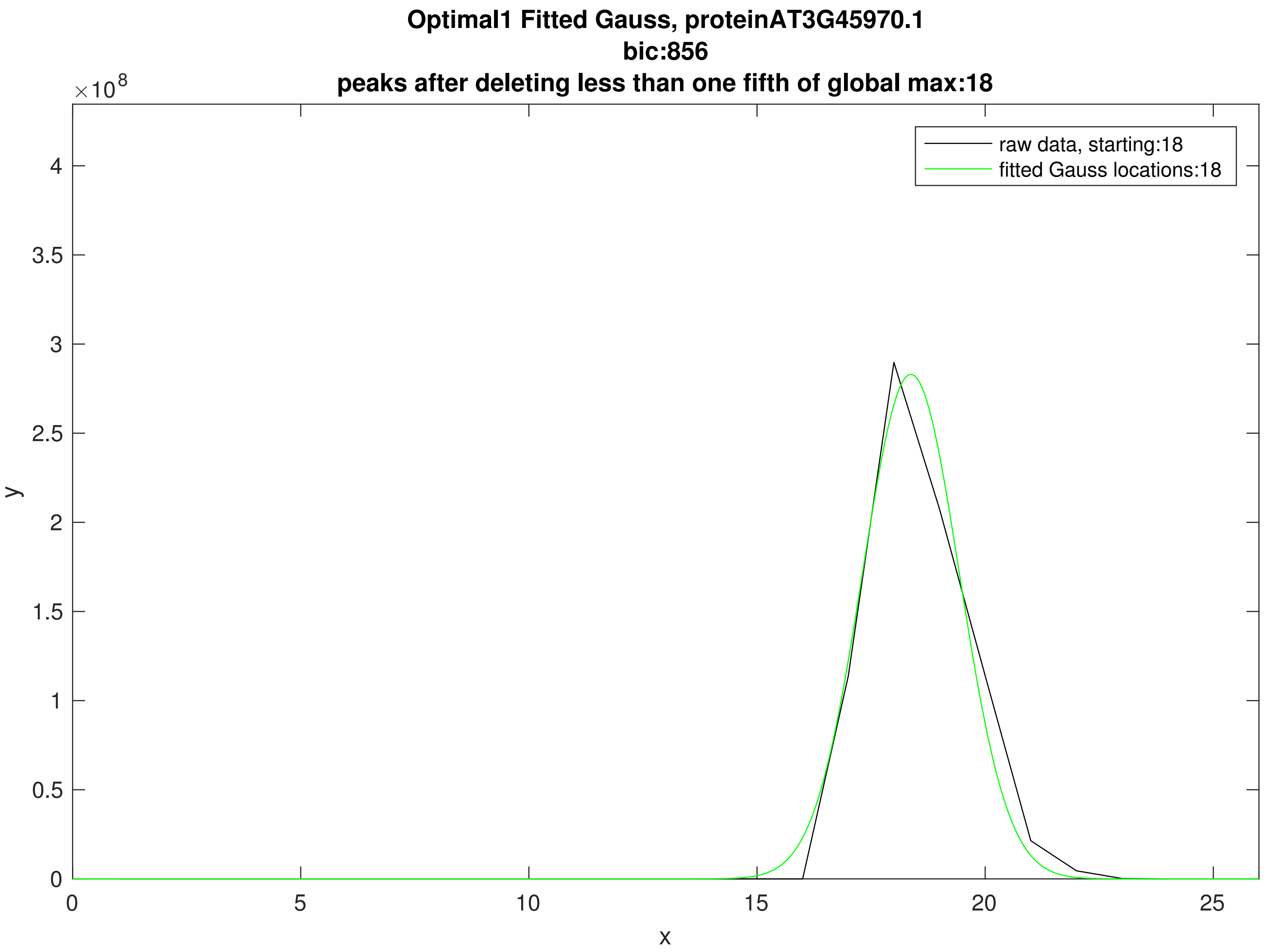
Optimal1 Fitted Gauss, proteinAT3G45970.1
bic:856
peaks after deleting less than one fifth of global max:18
×10^8
4
3.5
3
2.5
2
1.5
1
0.5
0
y
0
5
10
15
20
25
x
raw data, starting:18
fitted Gauss locations:18

**Optimal2 Fitted Gauss, proteinAT3G46000.1**
**bic:869 815**
**peaks after deleting less than one fifth of global max:13 19**

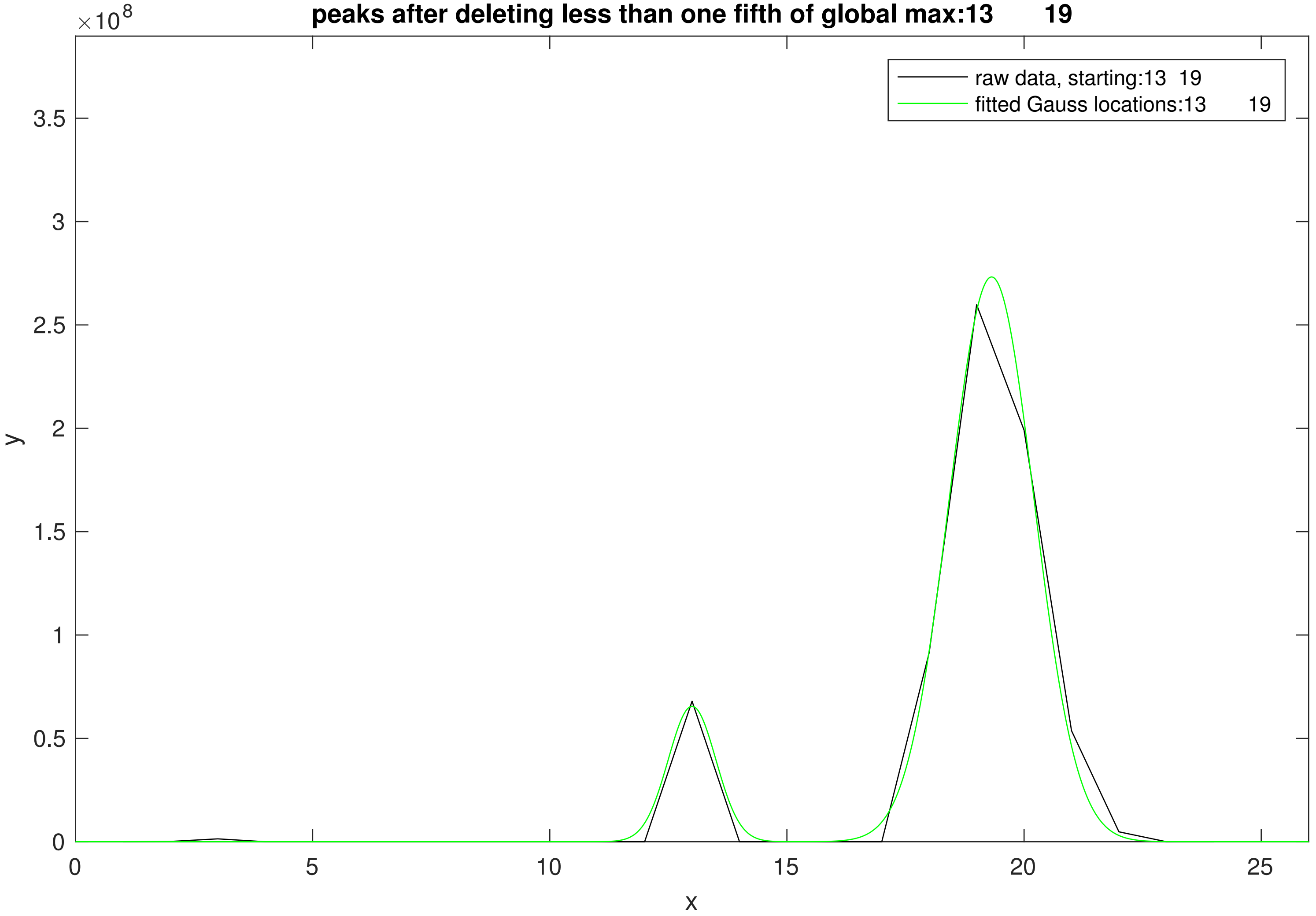

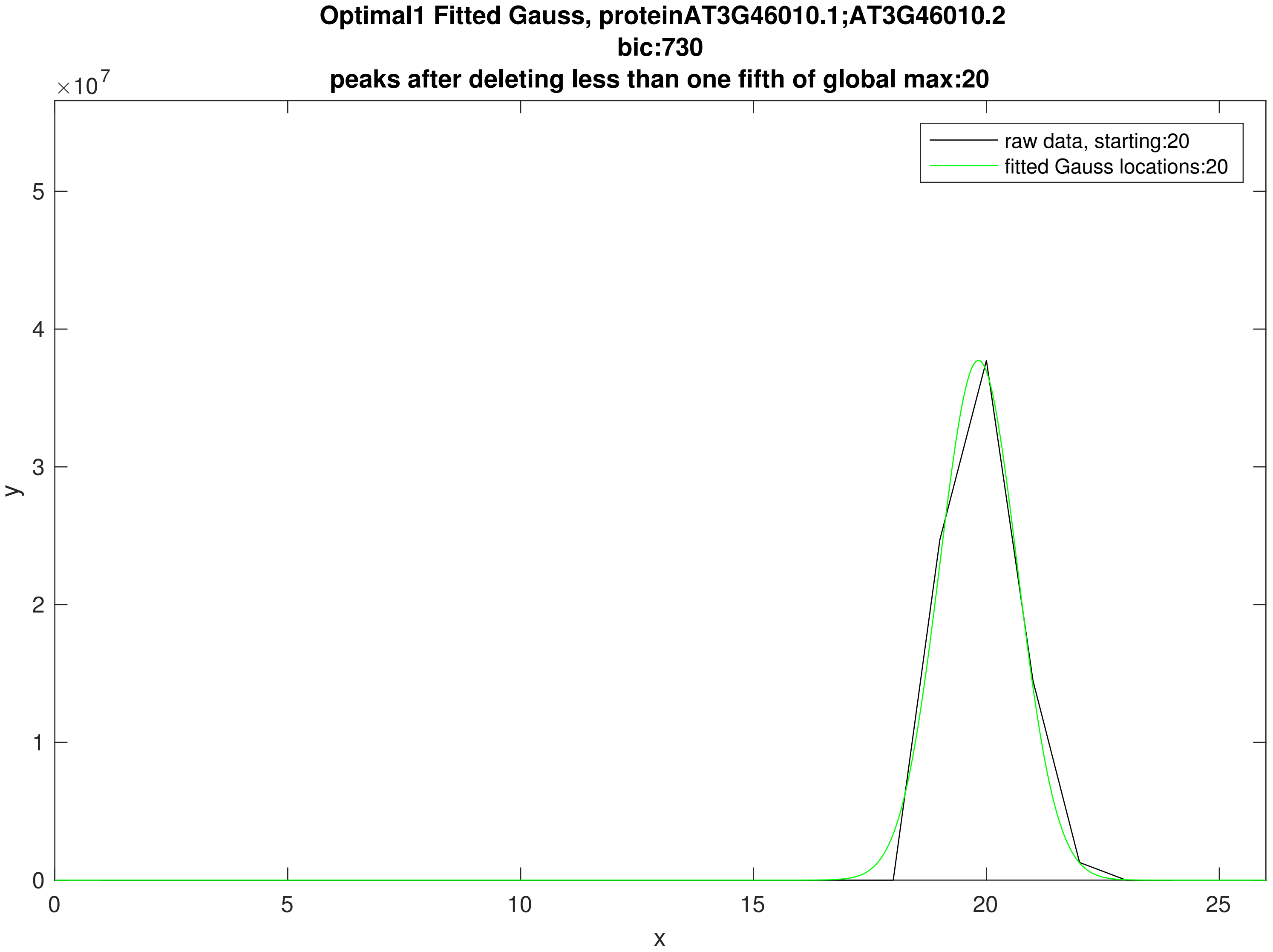
Optimal1 Fitted Gauss, proteinAT3G46010.1;AT3G46010.2
bic:730
peaks after deleting less than one fifth of global max:20
×10^7
raw data, starting:20
fitted Gauss locations:20
y
x
0
1
2
3
4
5
0
5
10
15
20
25

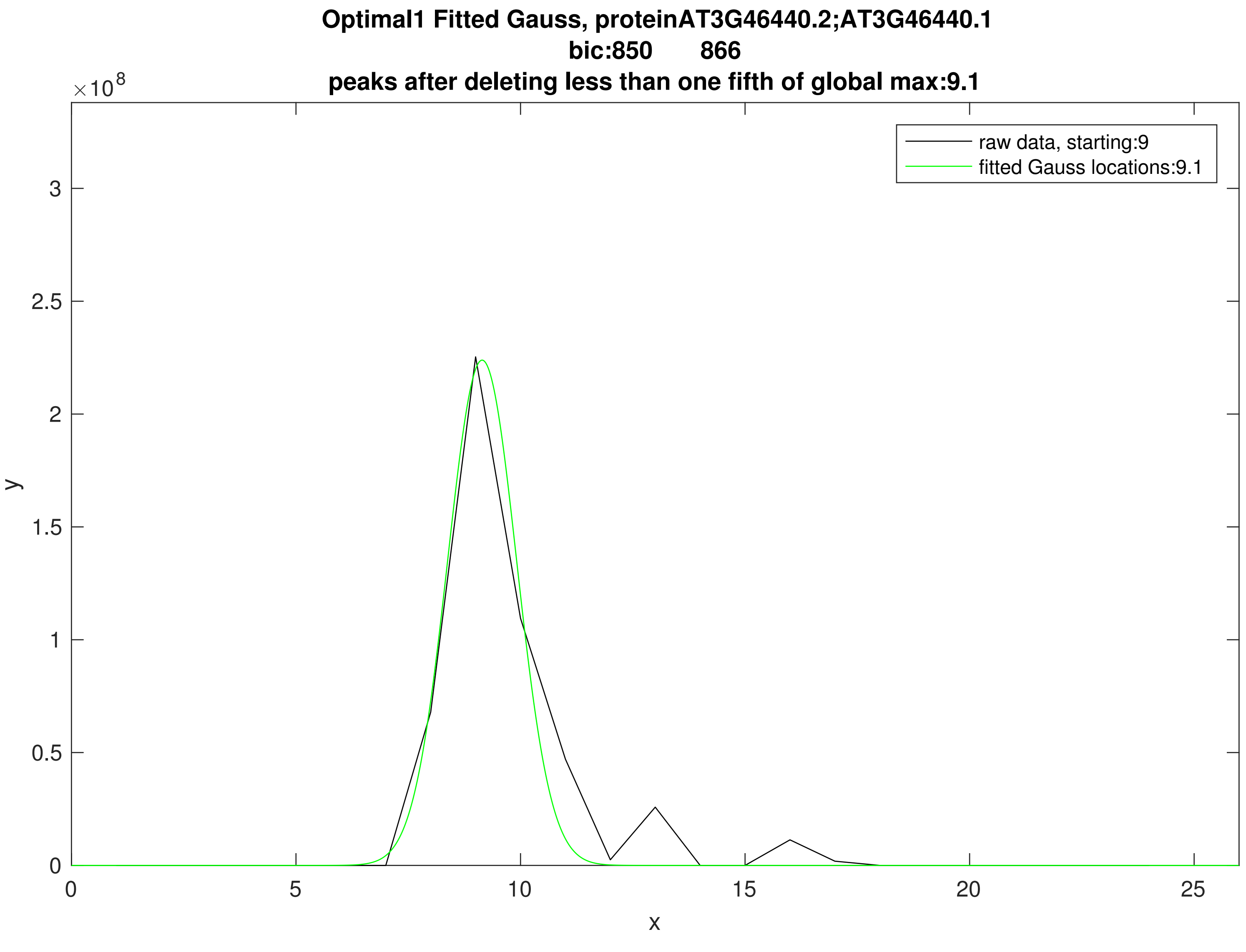
Optimal1 Fitted Gauss, proteinAT3G46440.2;AT3G46440.1
bic:850 866
peaks after deleting less than one fifth of global max:9.1
×10^8
3
2.5
2
1.5
1
0.5
0
y
0
5
10
15
20
25
x
raw data, starting:9
fitted Gauss locations:9.1

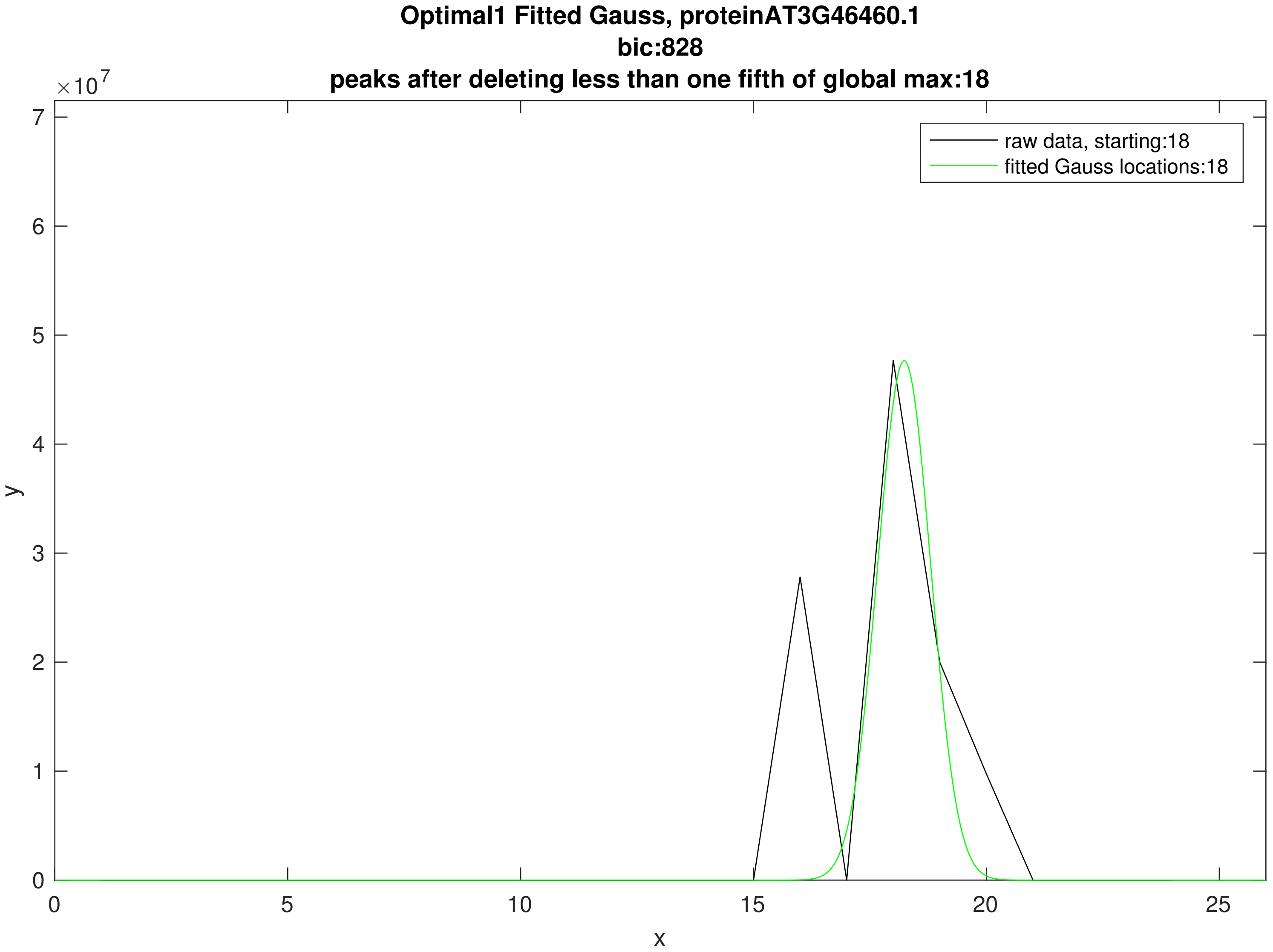
Optimal1 Fitted Gauss, proteinAT3G46460.1
bic:828
peaks after deleting less than one fifth of global max:18
×10^7
7
6
5
4
3
2
1
0
y
0
5
10
15
20
25
x
raw data, starting:18
fitted Gauss locations:18

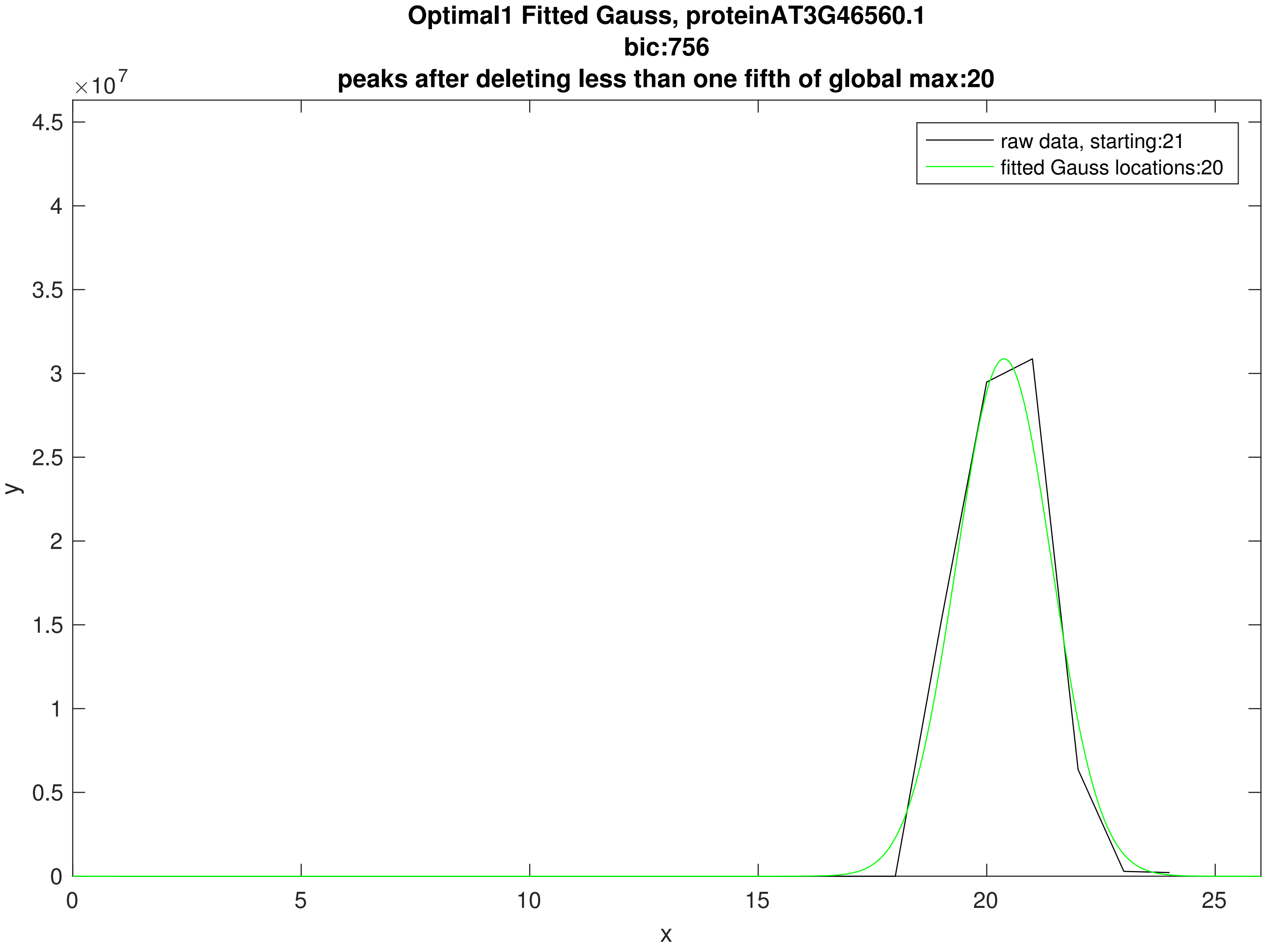

Optimal1 Fitted Gauss, proteinAT3G46560.1
bic:756
peaks after deleting less than one fifth of global max:20
×10^7
4.5
4
3.5
3
2.5
2
1.5
1
0.5
0
y
0
5
10
15
20
25
x
raw data, starting:21
fitted Gauss locations:20

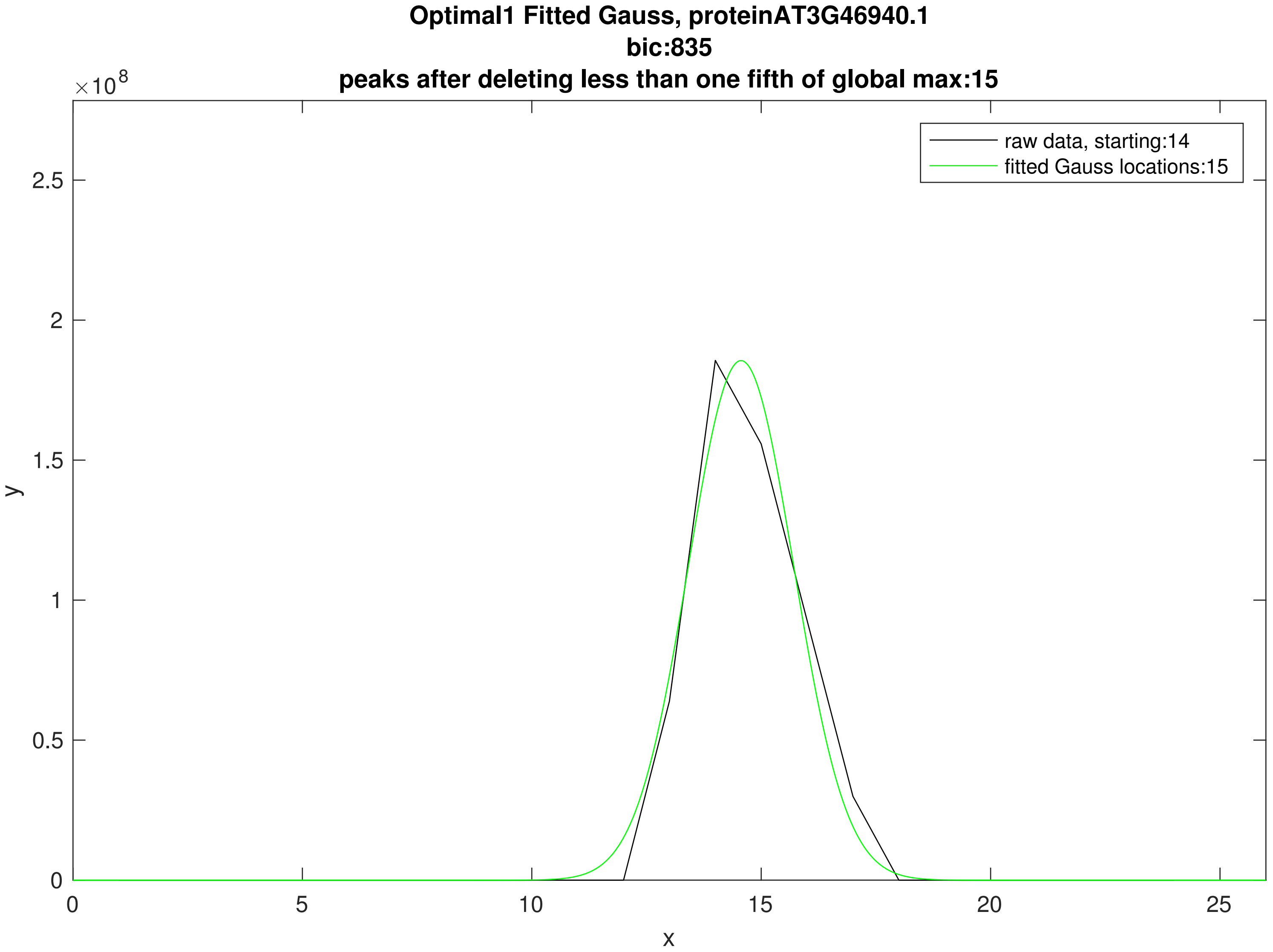

Optimal1 Fitted Gauss, proteinAT3G46940.1
bic:835
peaks after deleting less than one fifth of global max:15
×10^8
raw data, starting:14
fitted Gauss locations:15
2.5
2
1.5
1
0.5
0
y
0
5
10
15
20
25
x

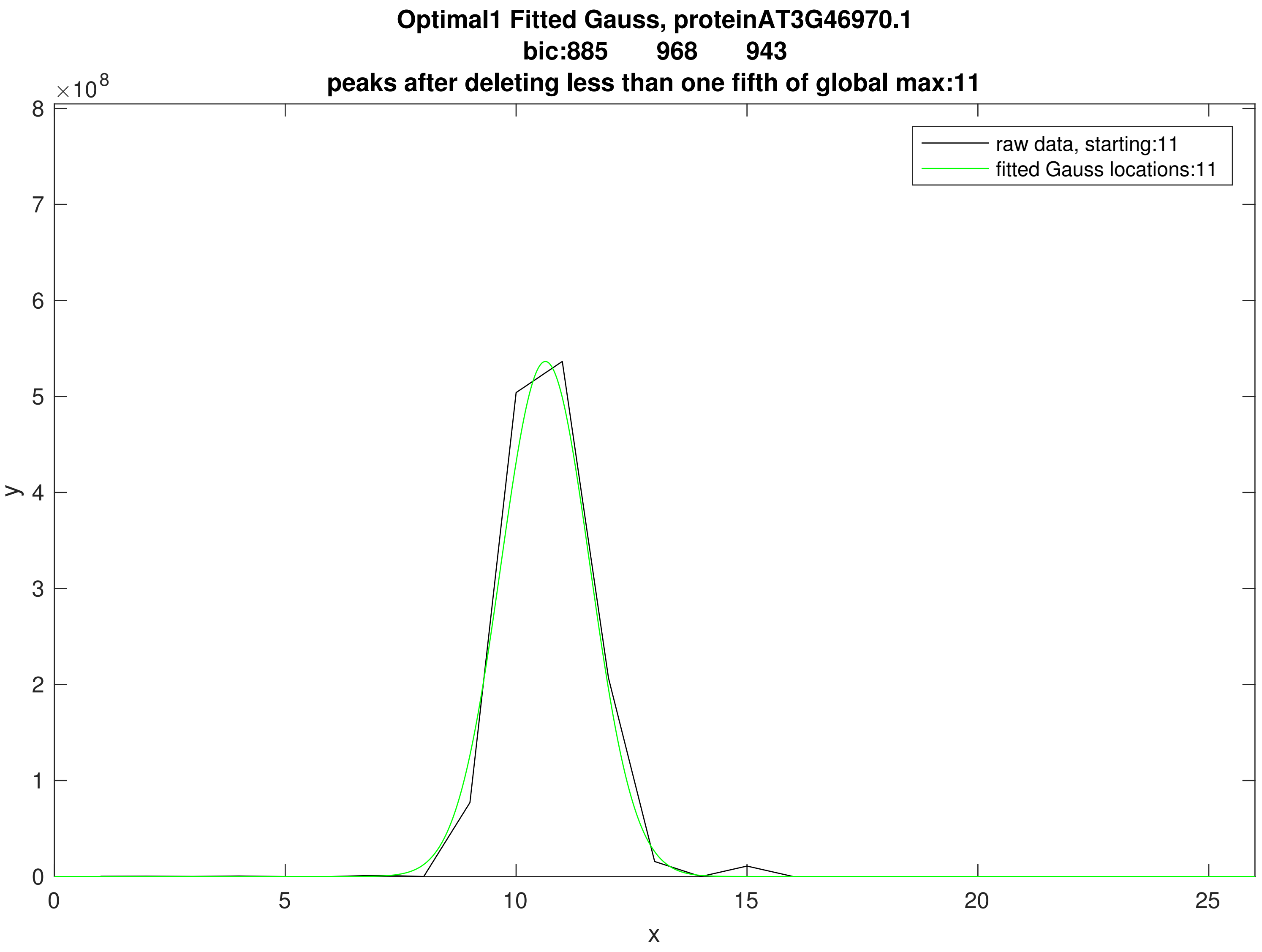
Optimal1 Fitted Gauss, proteinAT3G46970.1
bic:885 968 943
peaks after deleting less than one fifth of global max:11
×10^8
raw data, starting:11
fitted Gauss locations:11
y
x
0
1
2
3
4
5
6
7
8
0
5
10
15
20
25

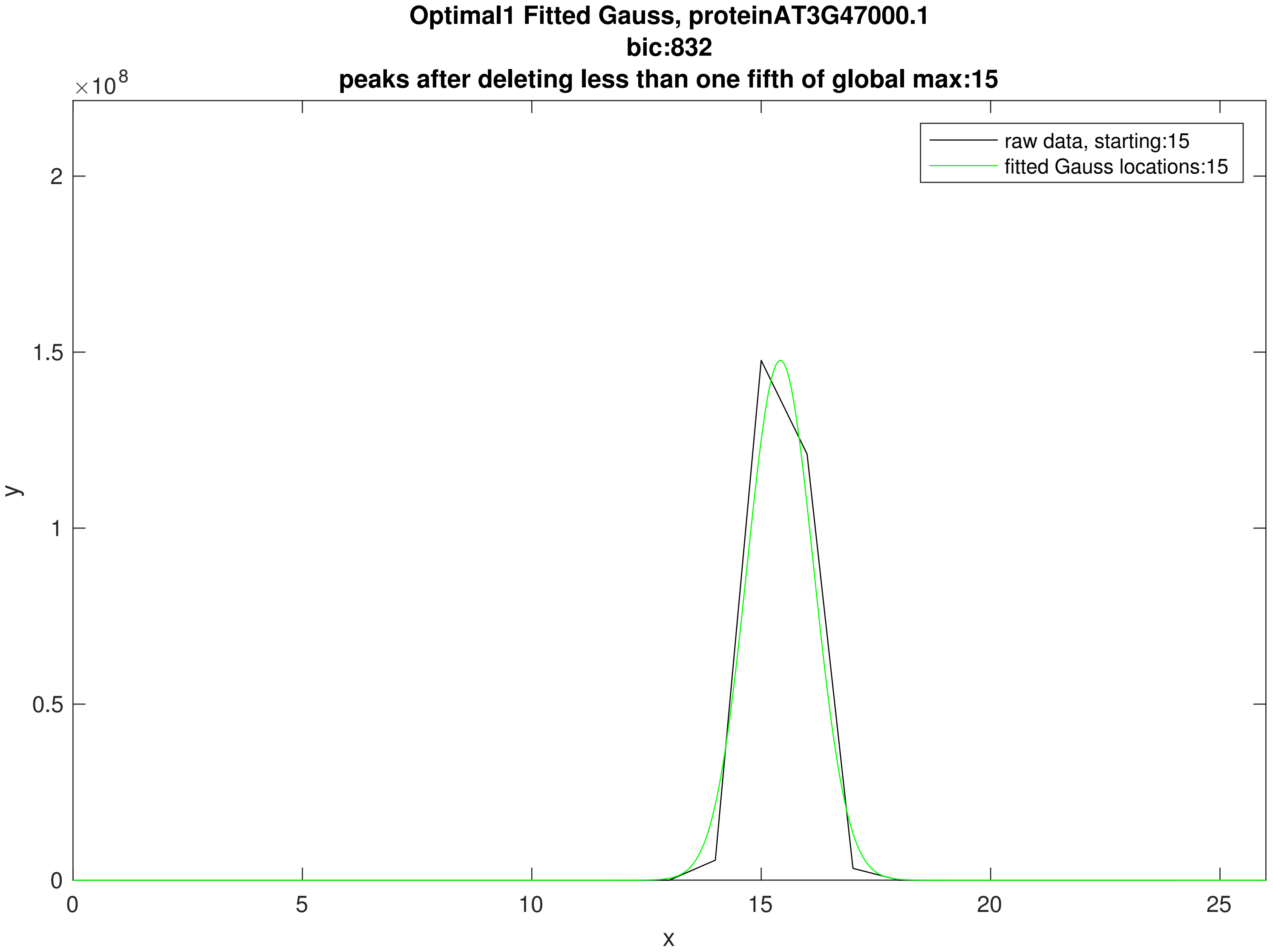

Optimal1 Fitted Gauss, proteinAT3G47000.1
bic:832
peaks after deleting less than one fifth of global max:15
×10^8
raw data, starting:15
fitted Gauss locations:15
y
x
2
1.5
1
0.5
0
0
5
10
15
20
25

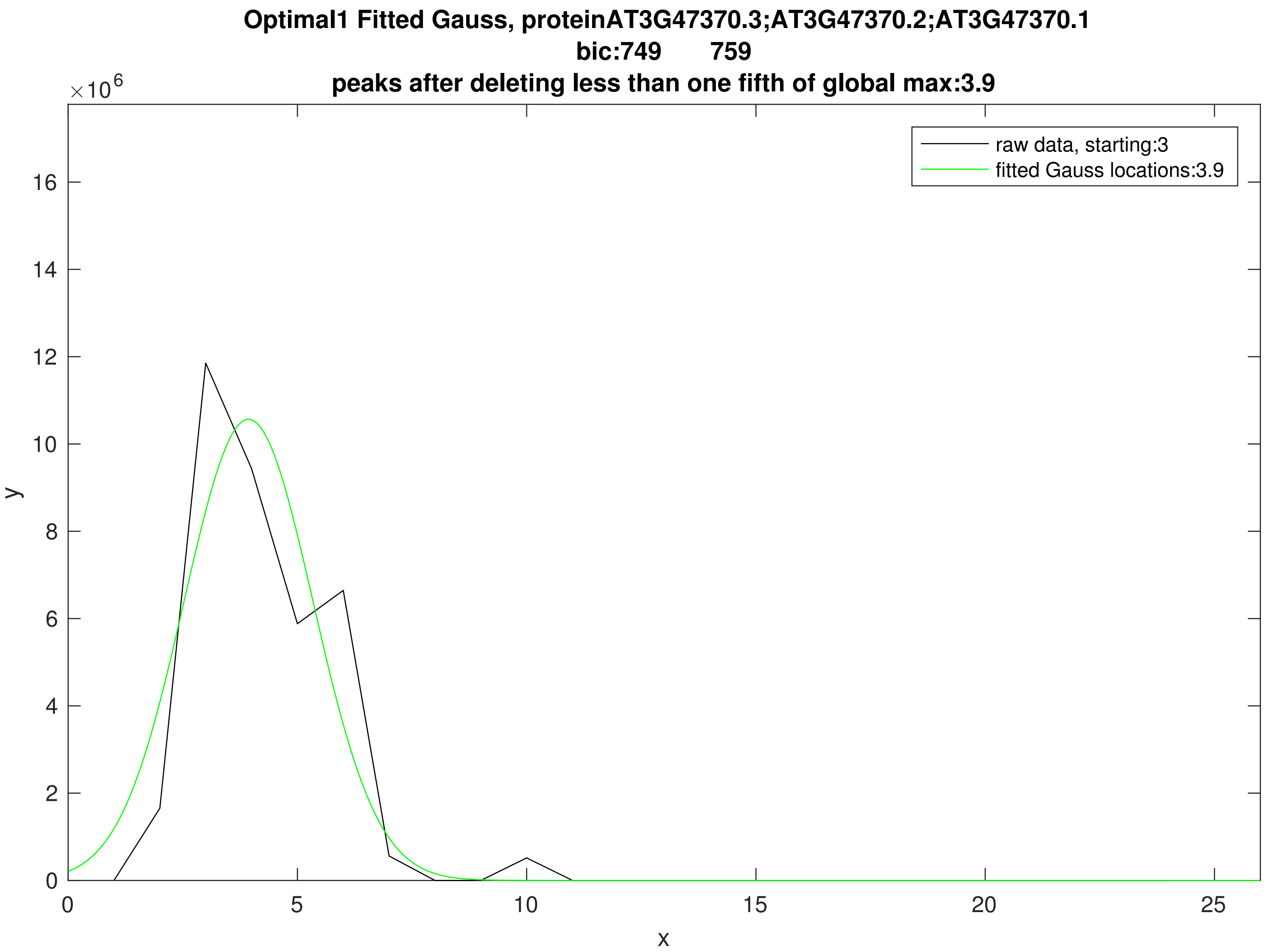

Optimal1 Fitted Gauss, proteinAT3G47370.3;AT3G47370.2;AT3G47370.1
bic:749 759
peaks after deleting less than one fifth of global max:3.9
×10^6
raw data, starting:3
fitted Gauss locations:3.9
y
x
0
2
4
6
8
10
12
14
16
0
5
10
15
20
25

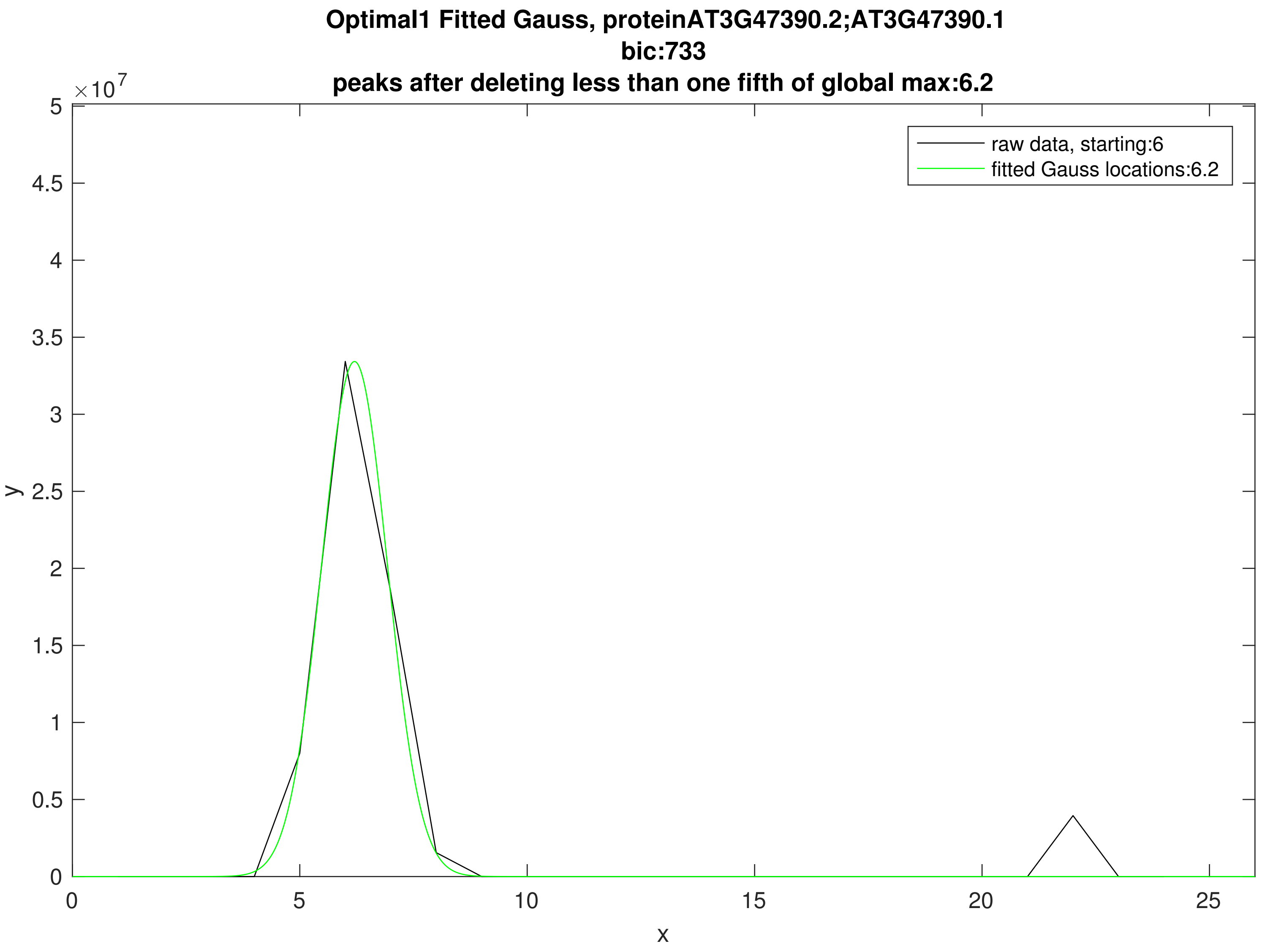
Optimal1 Fitted Gauss, proteinAT3G47390.2;AT3G47390.1
bic:733
peaks after deleting less than one fifth of global max:6.2
×10^7
raw data, starting:6
fitted Gauss locations:6.2
y
x
0
0.5
1
1.5
2
2.5
3
3.5
4
4.5
5
0
5
10
15
20
25

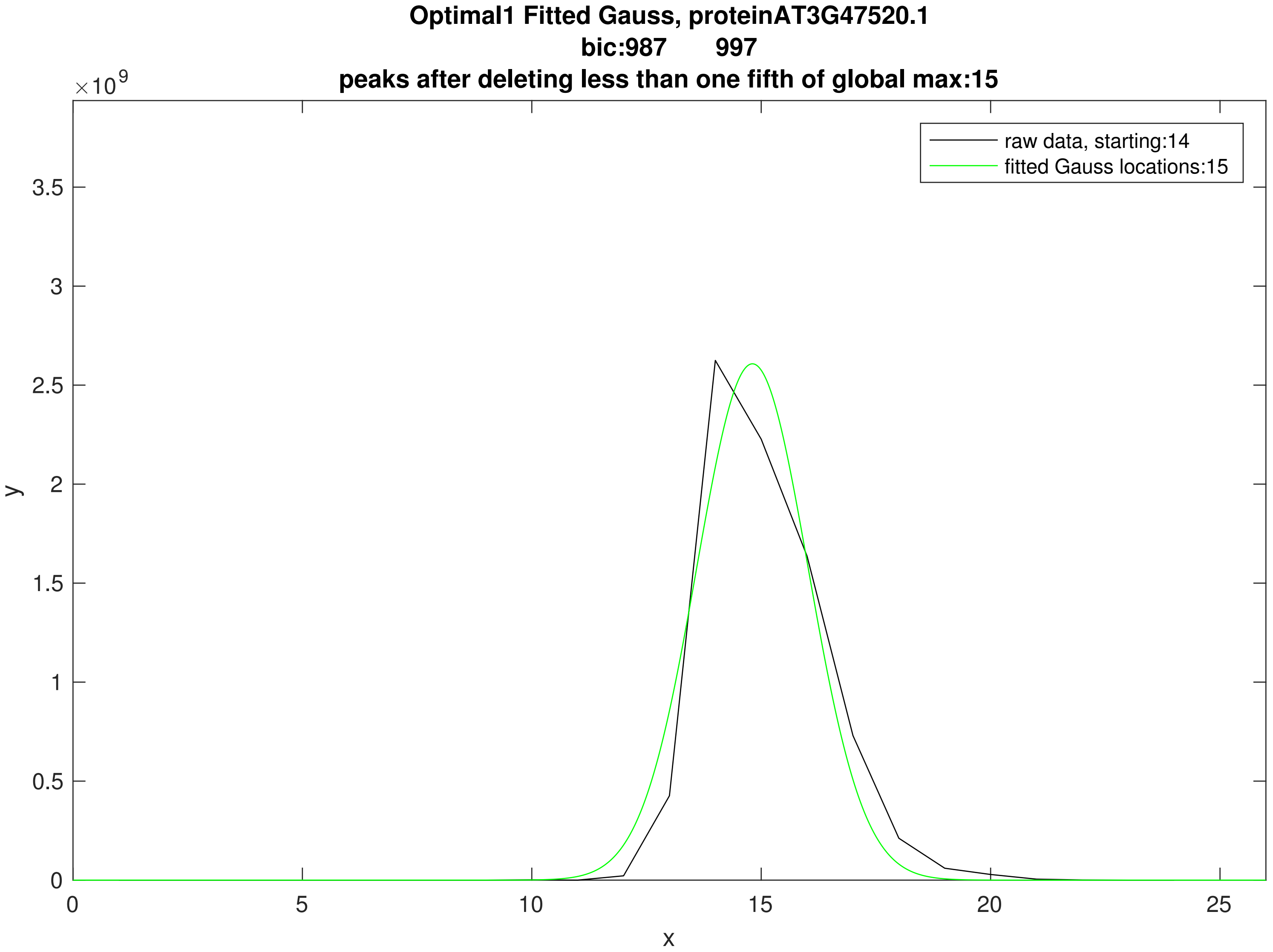
Optimal1 Fitted Gauss, proteinAT3G47520.1
bic:987 997
peaks after deleting less than one fifth of global max:15
×10^9
raw data, starting:14
fitted Gauss locations:15
3.5
3
2.5
2
1.5
1
0.5
0
y
0
5
10
15
20
25
x

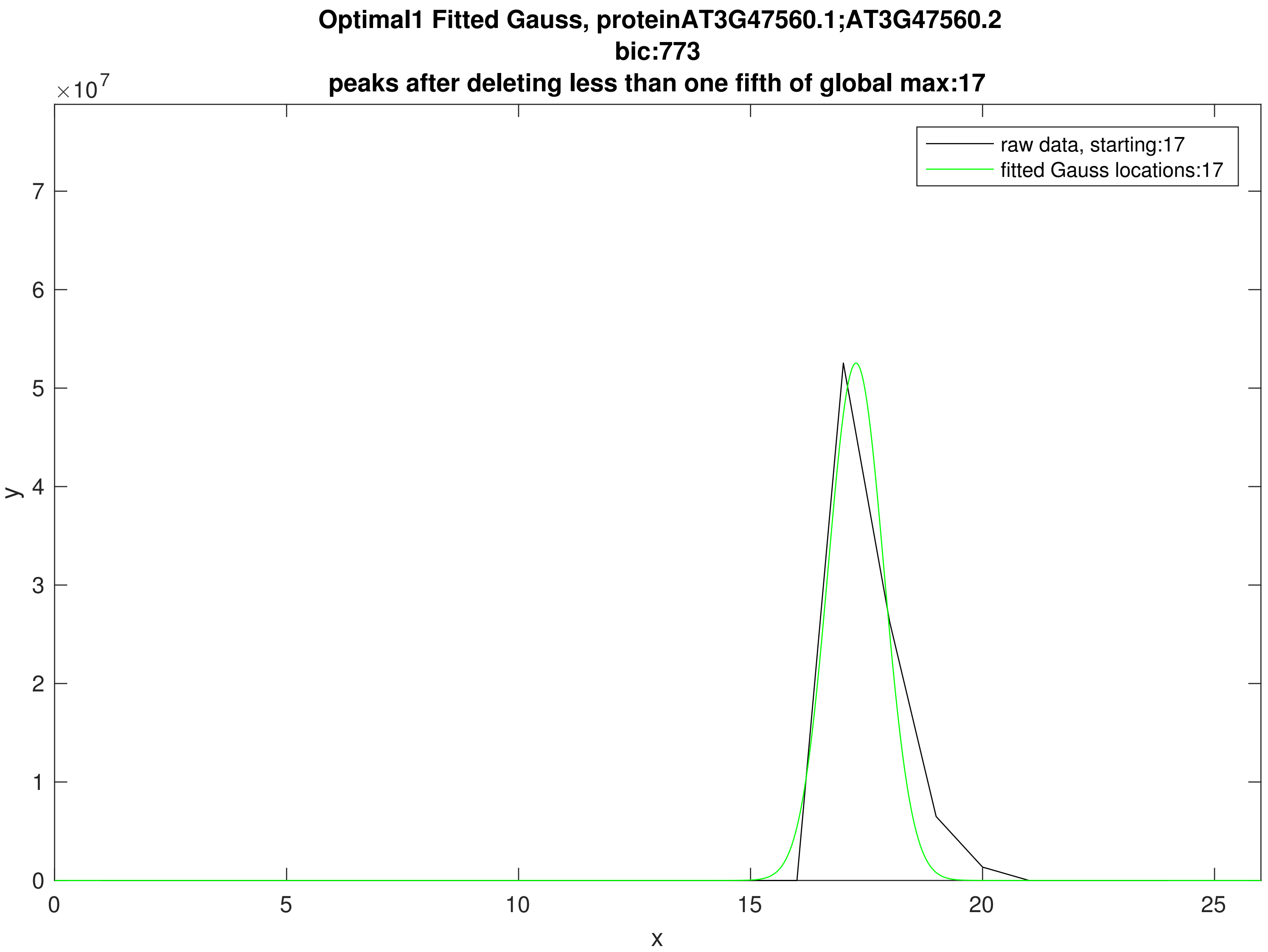

Optimal1 Fitted Gauss, proteinAT3G47560.1;AT3G47560.2
bic:773
peaks after deleting less than one fifth of global max:17
×10^7
raw data, starting:17
fitted Gauss locations:17
y
x
0
1
2
3
4
5
6
7
0
5
10
15
20
25

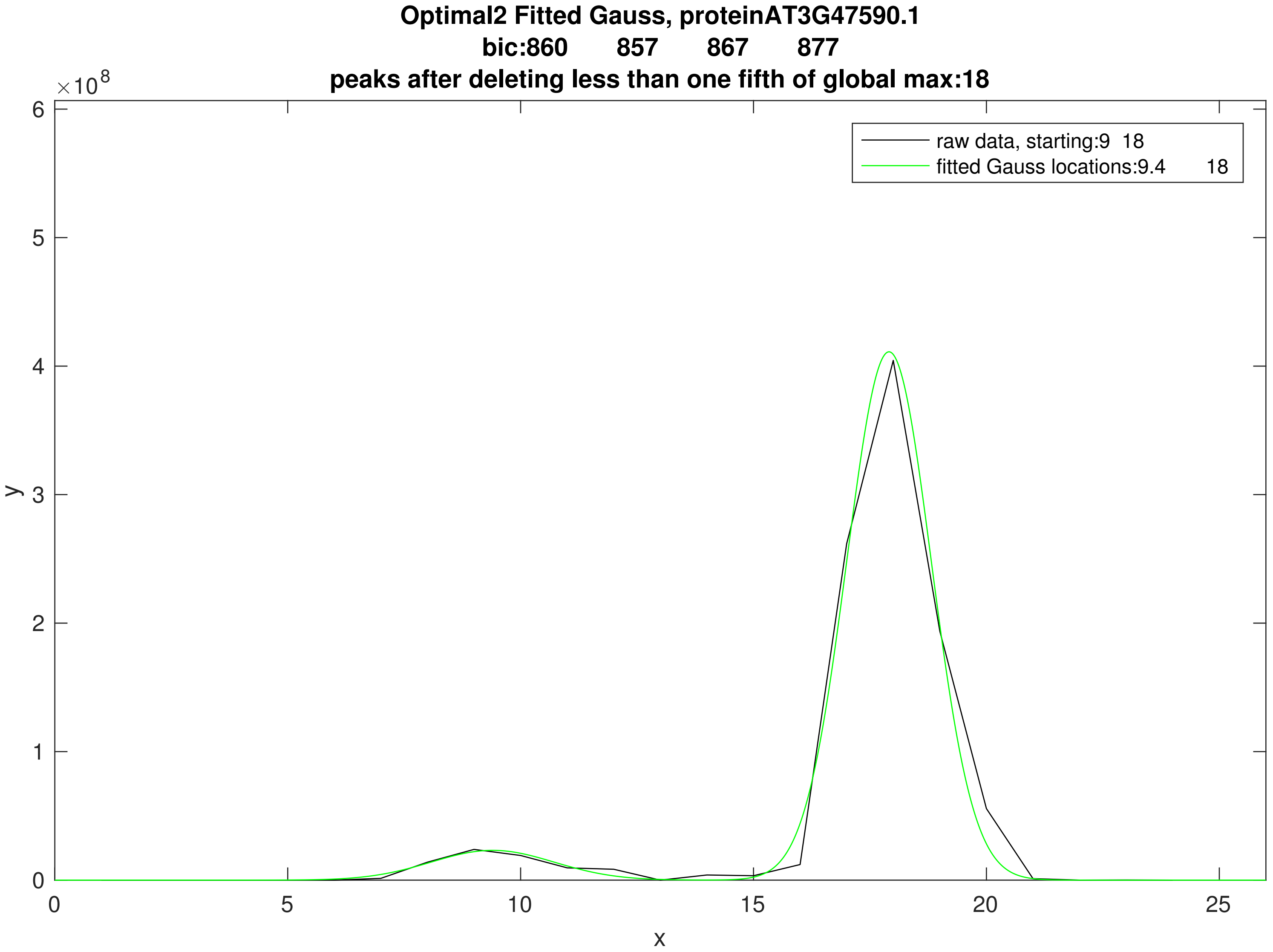

Optimal2 Fitted Gauss, proteinAT3G47590.1
bic:860 857 867 877
peaks after deleting less than one fifth of global max:18
×10^8
raw data, starting:9 18
fitted Gauss locations:9.4 18
y
x
0
1
2
3
4
5
6
0
5
10
15
20
25

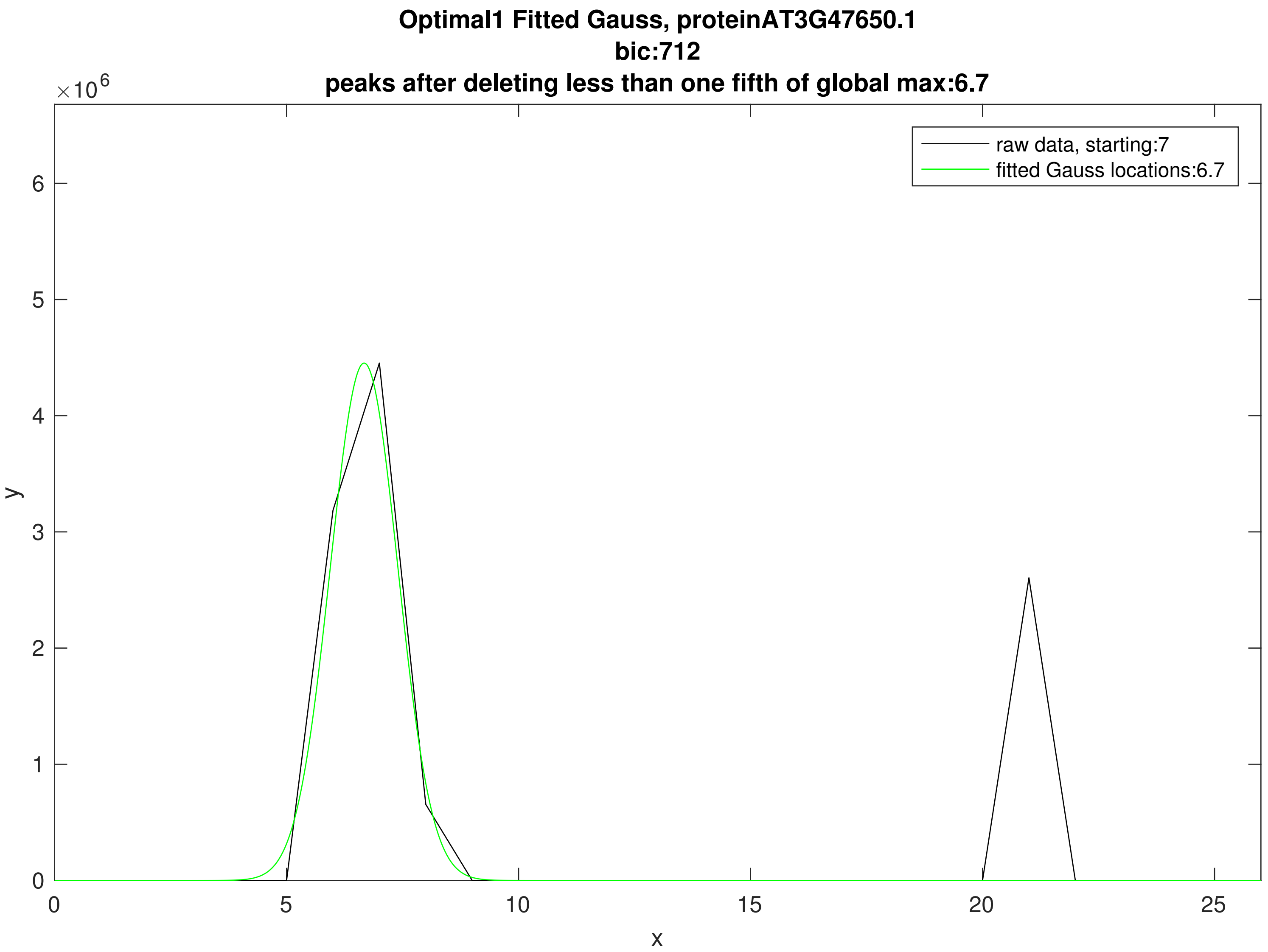
Optimal1 Fitted Gauss, proteinAT3G47650.1
bic:712
peaks after deleting less than one fifth of global max:6.7
×10^6
raw data, starting:7
fitted Gauss locations:6.7
y
x
0
1
2
3
4
5
6
0
5
10
15
20
25

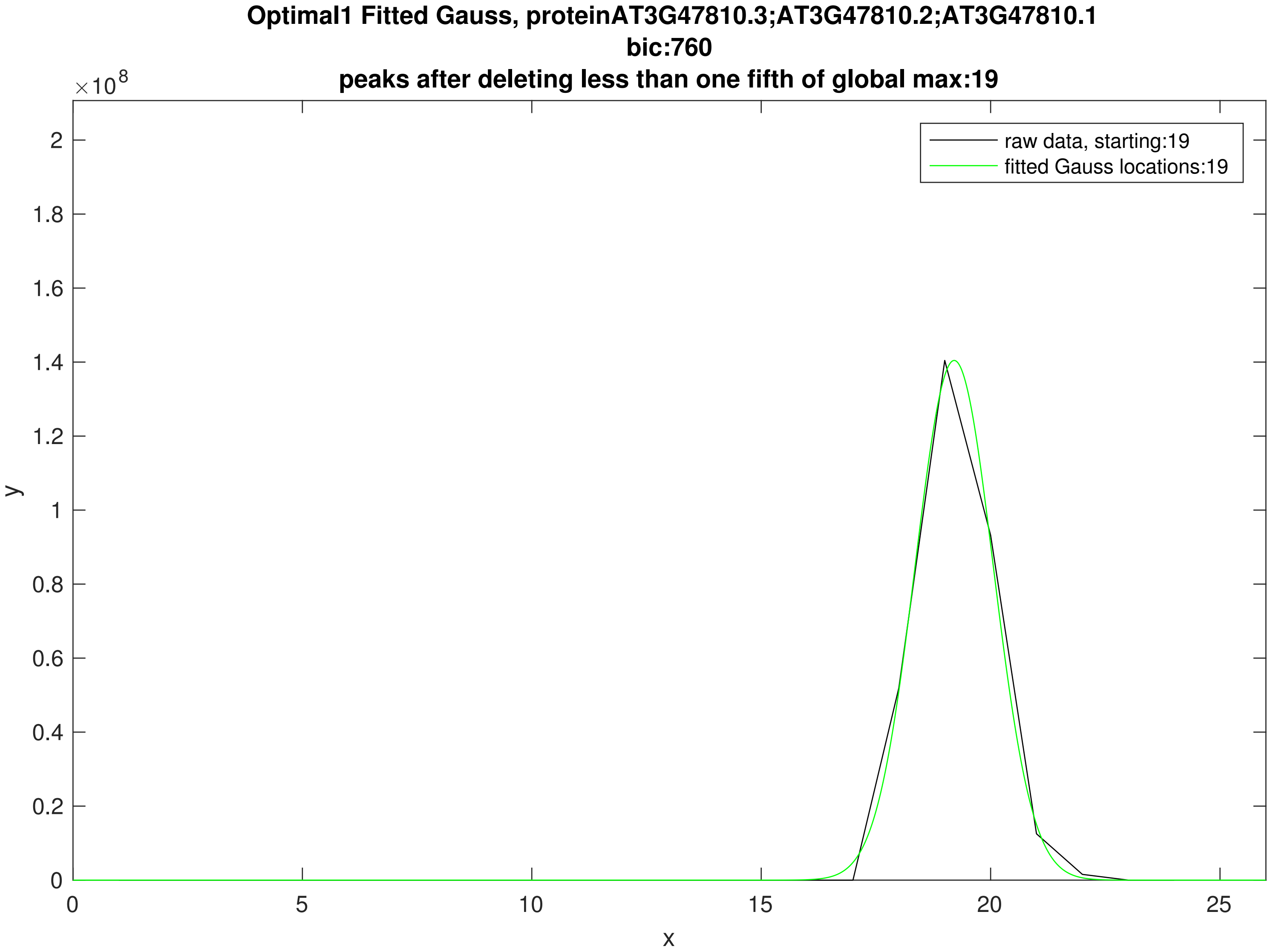

Optimal1 Fitted Gauss, proteinAT3G47810.3;AT3G47810.2;AT3G47810.1
bic:760
peaks after deleting less than one fifth of global max:19
×10^8
2
1.8
1.6
1.4
1.2
1
0.8
0.6
0.4
0.2
0
y
0
5
10
15
20
25
x
raw data, starting:19
fitted Gauss locations:19

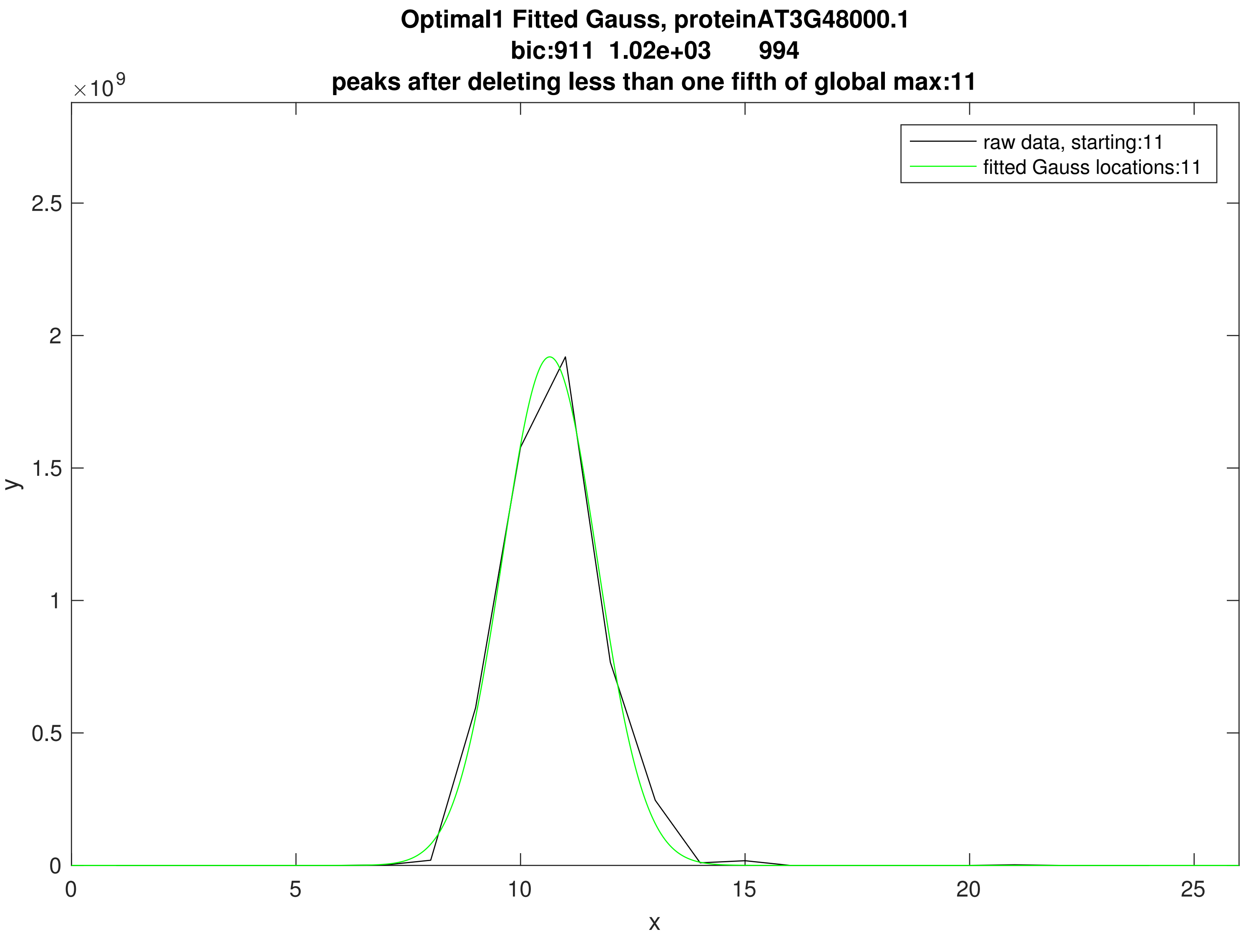
Optimal1 Fitted Gauss, proteinAT3G48000.1
bic:911 1.02e+03 994
peaks after deleting less than one fifth of global max:11
×10^9
raw data, starting:11
fitted Gauss locations:11
2.5
2
1.5
1
0.5
0
y
0
5
10
15
20
25
x

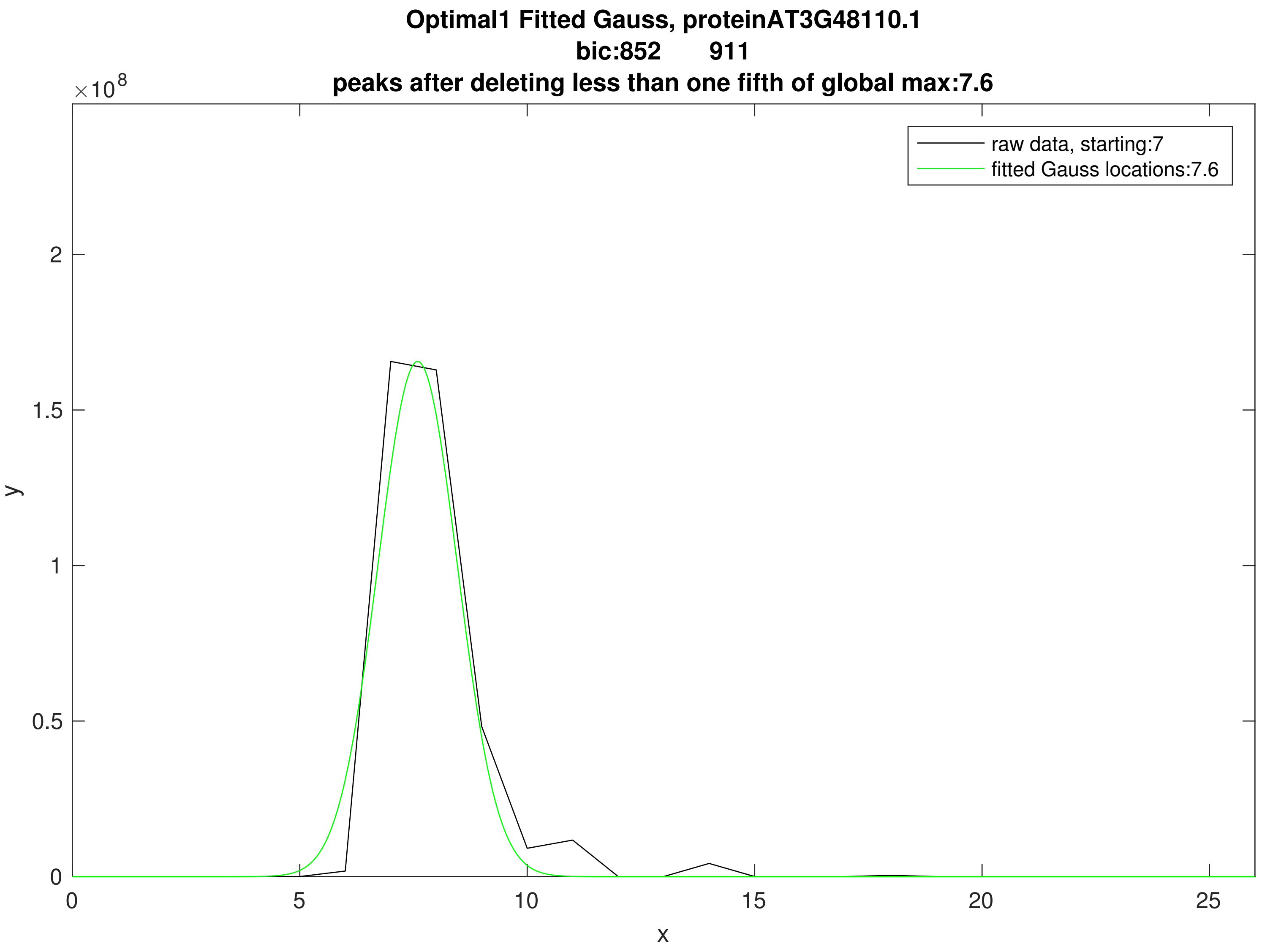
Optimal1 Fitted Gauss, proteinAT3G48110.1
bic:852 911
peaks after deleting less than one fifth of global max:7.6
×10^8
raw data, starting:7
fitted Gauss locations:7.6
y
x
0
0.5
1
1.5
2
0
5
10
15
20
25

Optimal1 Fitted Gauss, proteinAT3G48170.1
bic:932
peaks after deleting less than one fifth of global max:14

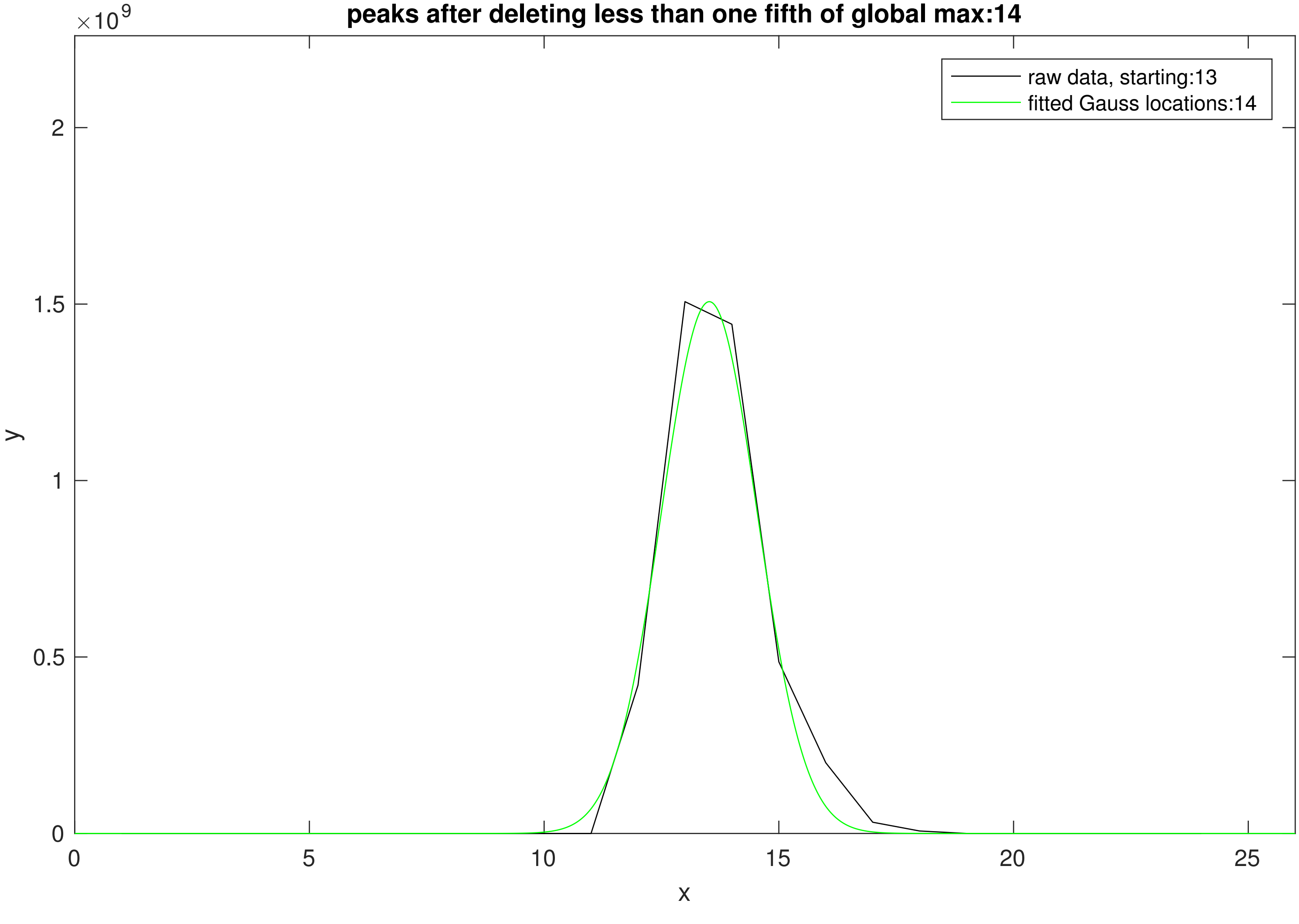

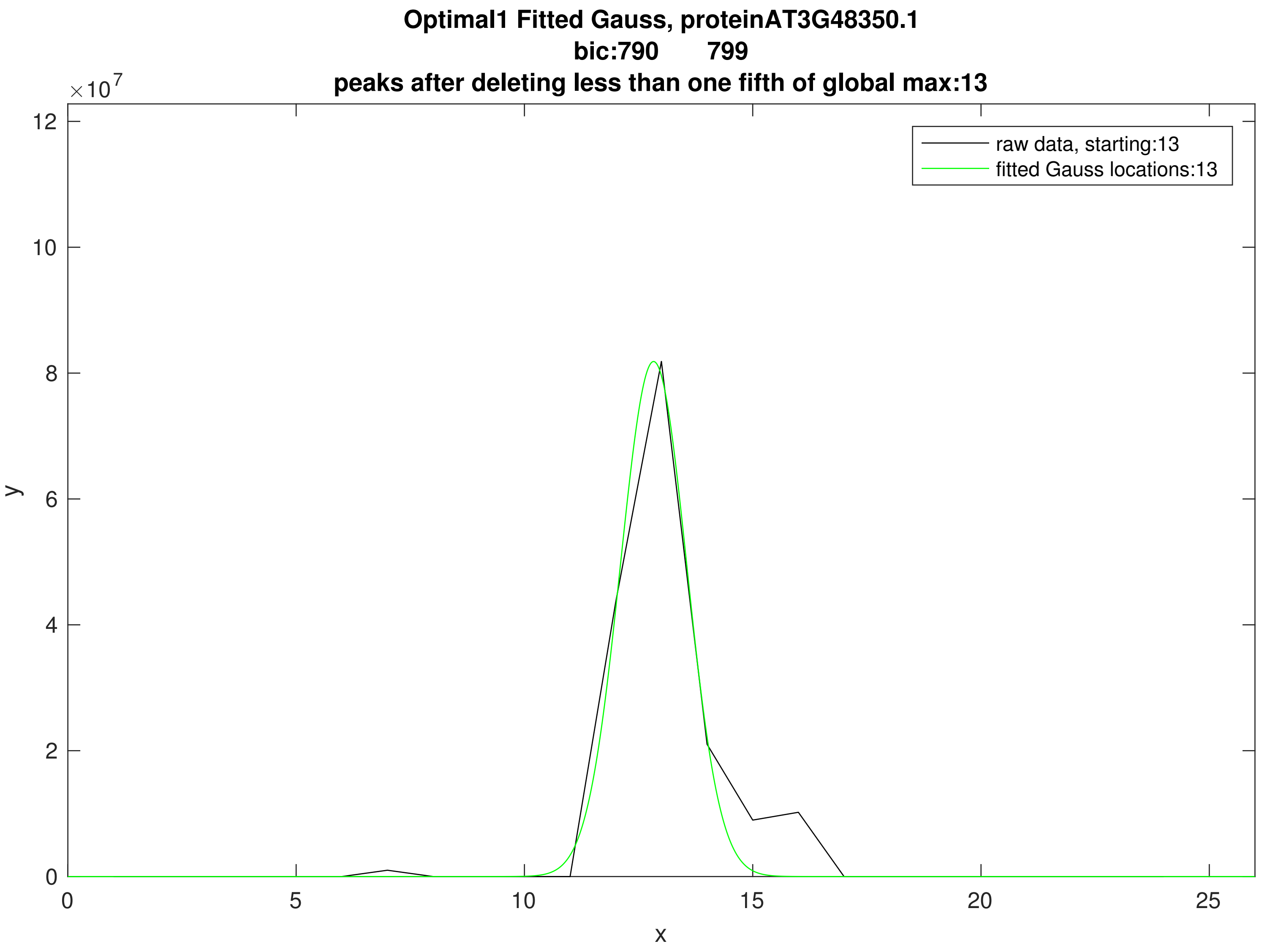
Optimal1 Fitted Gauss, proteinAT3G48350.1
bic:790 799
peaks after deleting less than one fifth of global max:13
×10^7
12
10
8
6
4
2
0
y
0
5
10
15
20
25
x
raw data, starting:13
fitted Gauss locations:13

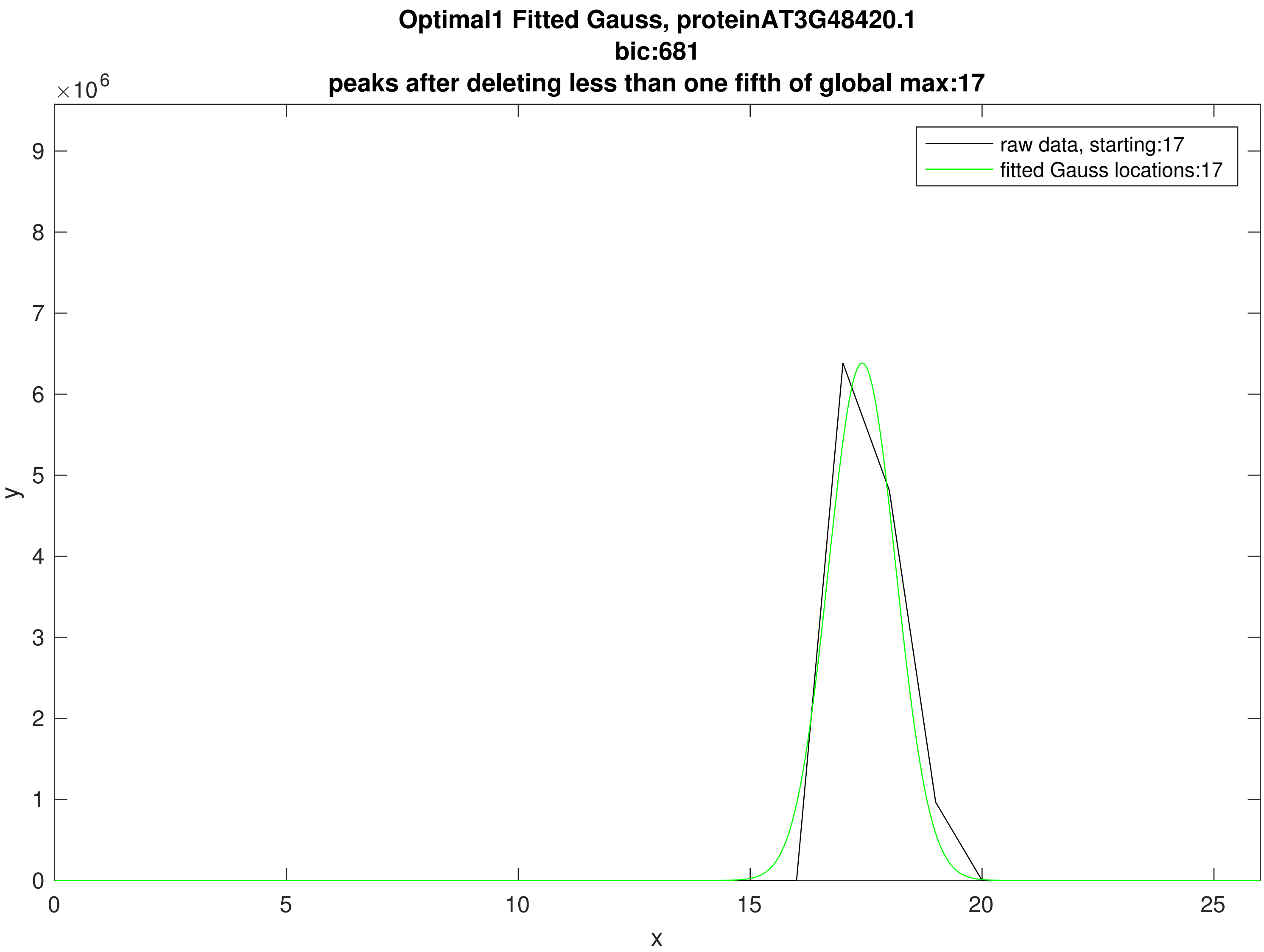

Optimal1 Fitted Gauss, proteinAT3G48420.1
bic:681
peaks after deleting less than one fifth of global max:17
×10^6
9
8
7
6
5
4
3
2
1
0
y
0
5
10
15
20
25
x
raw data, starting:17
fitted Gauss locations:17

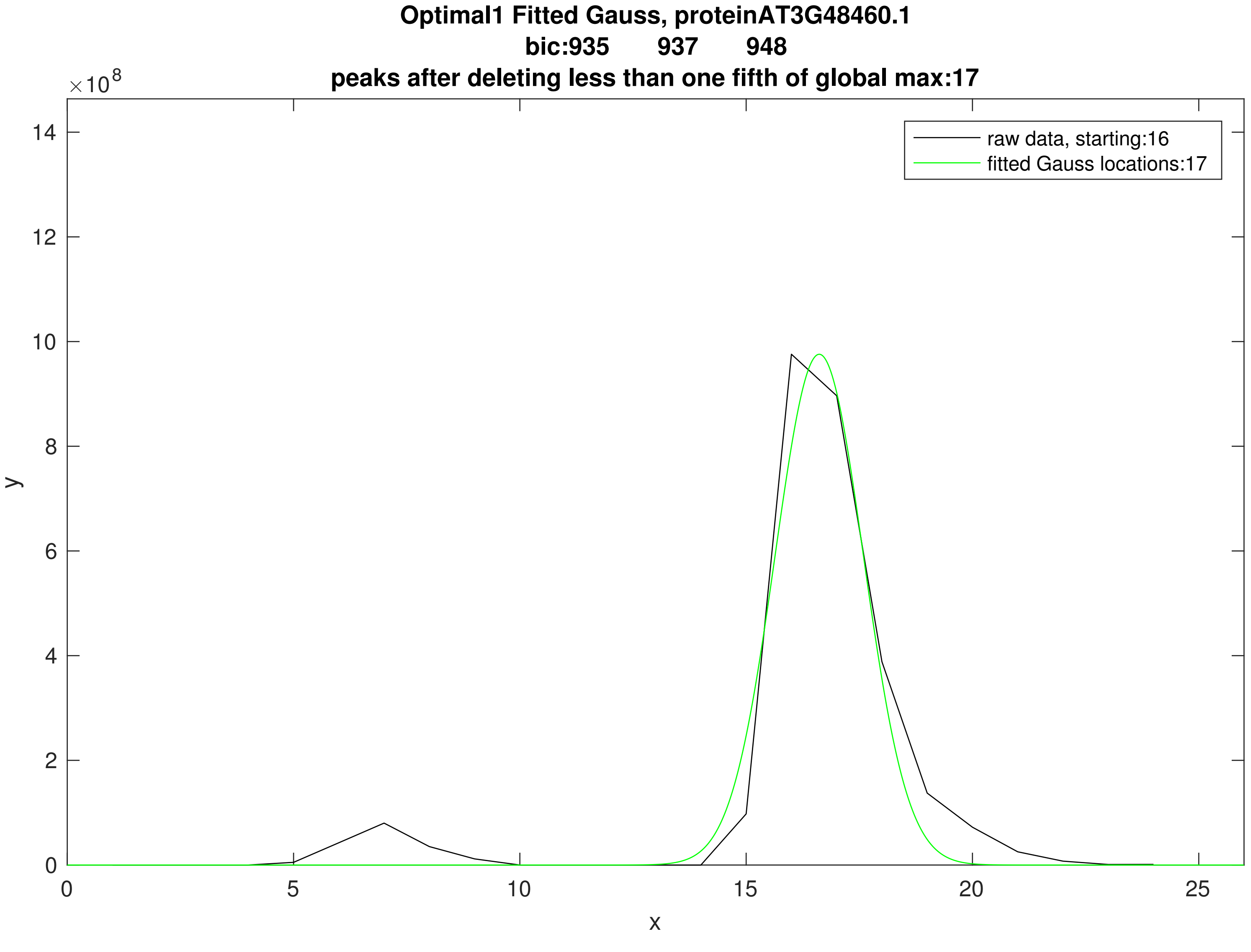

Optimal1 Fitted Gauss, proteinAT3G48460.1
bic:935 937 948
peaks after deleting less than one fifth of global max:17
×10^8
14
12
10
8
6
4
2
0
y
0
5
10
15
20
25
x
raw data, starting:16
fitted Gauss locations:17

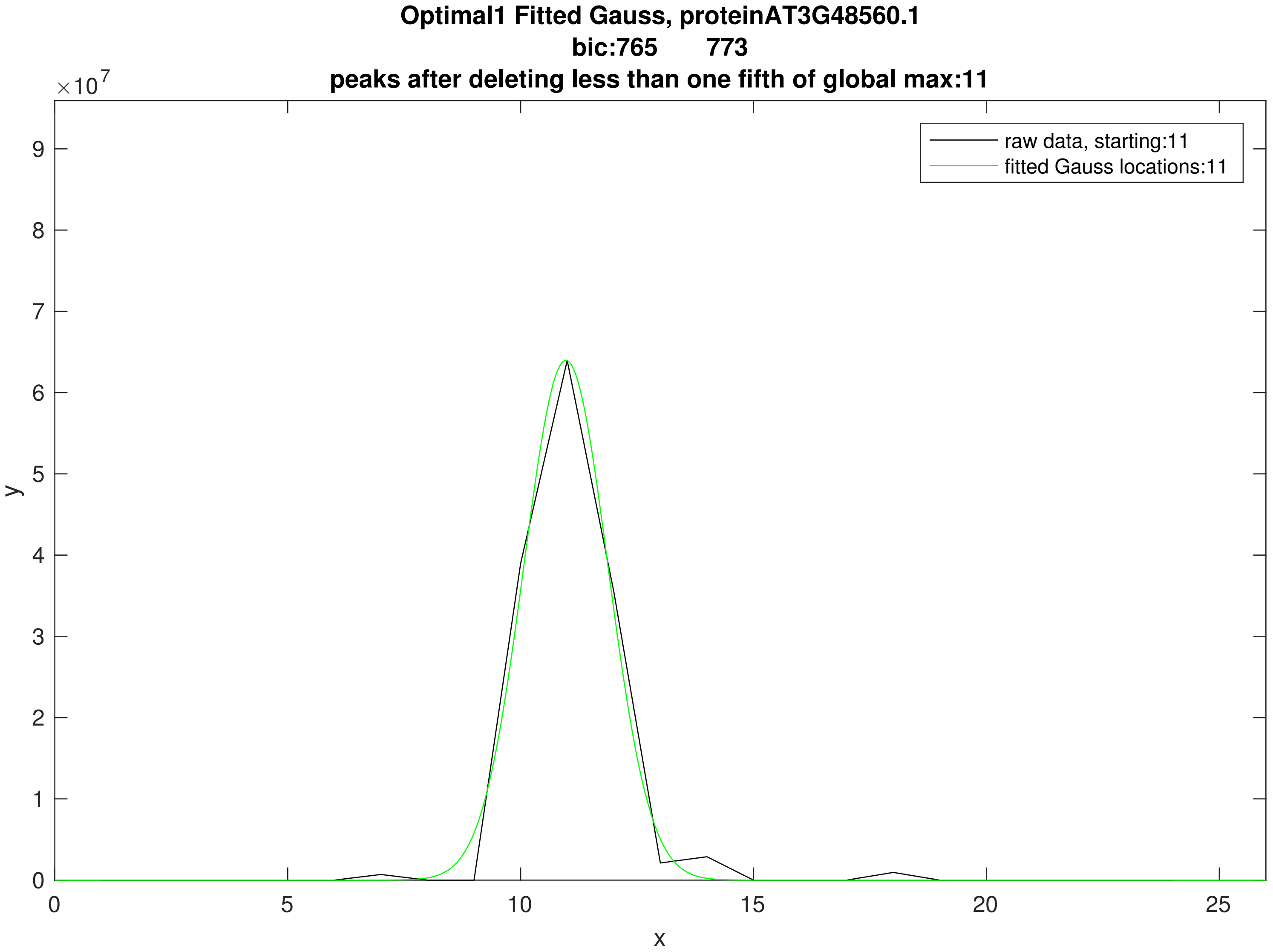
Optimal1 Fitted Gauss, proteinAT3G48560.1
bic:765 773
peaks after deleting less than one fifth of global max:11
×10^7
raw data, starting:11
fitted Gauss locations:11
y
x
0
1
2
3
4
5
6
7
8
9
0
5
10
15
20
25

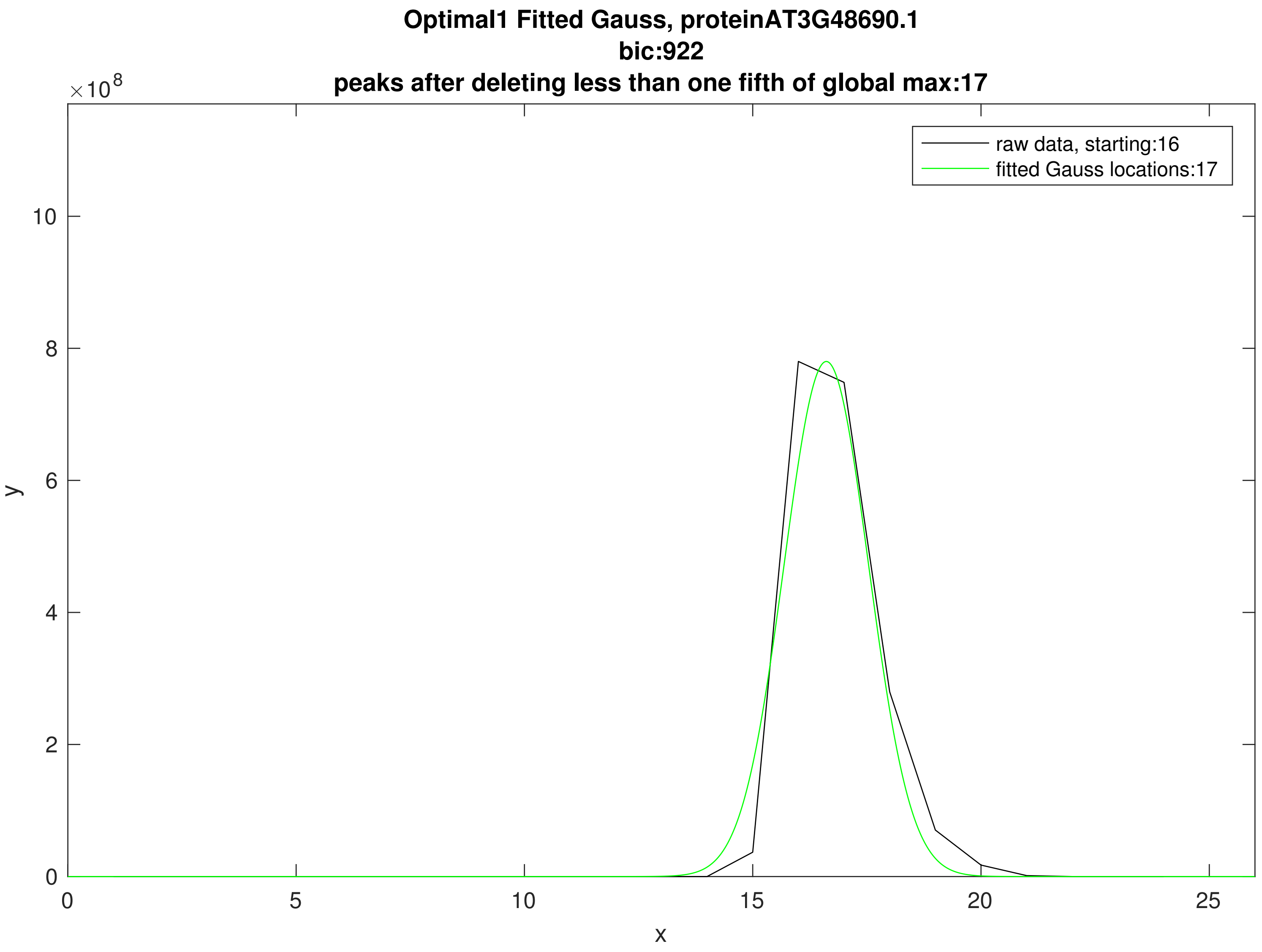
Optimal1 Fitted Gauss, proteinAT3G48690.1
bic:922
peaks after deleting less than one fifth of global max:17
×10^8
raw data, starting:16
fitted Gauss locations:17
10
8
6
4
2
0
y
0
5
10
15
20
25
x

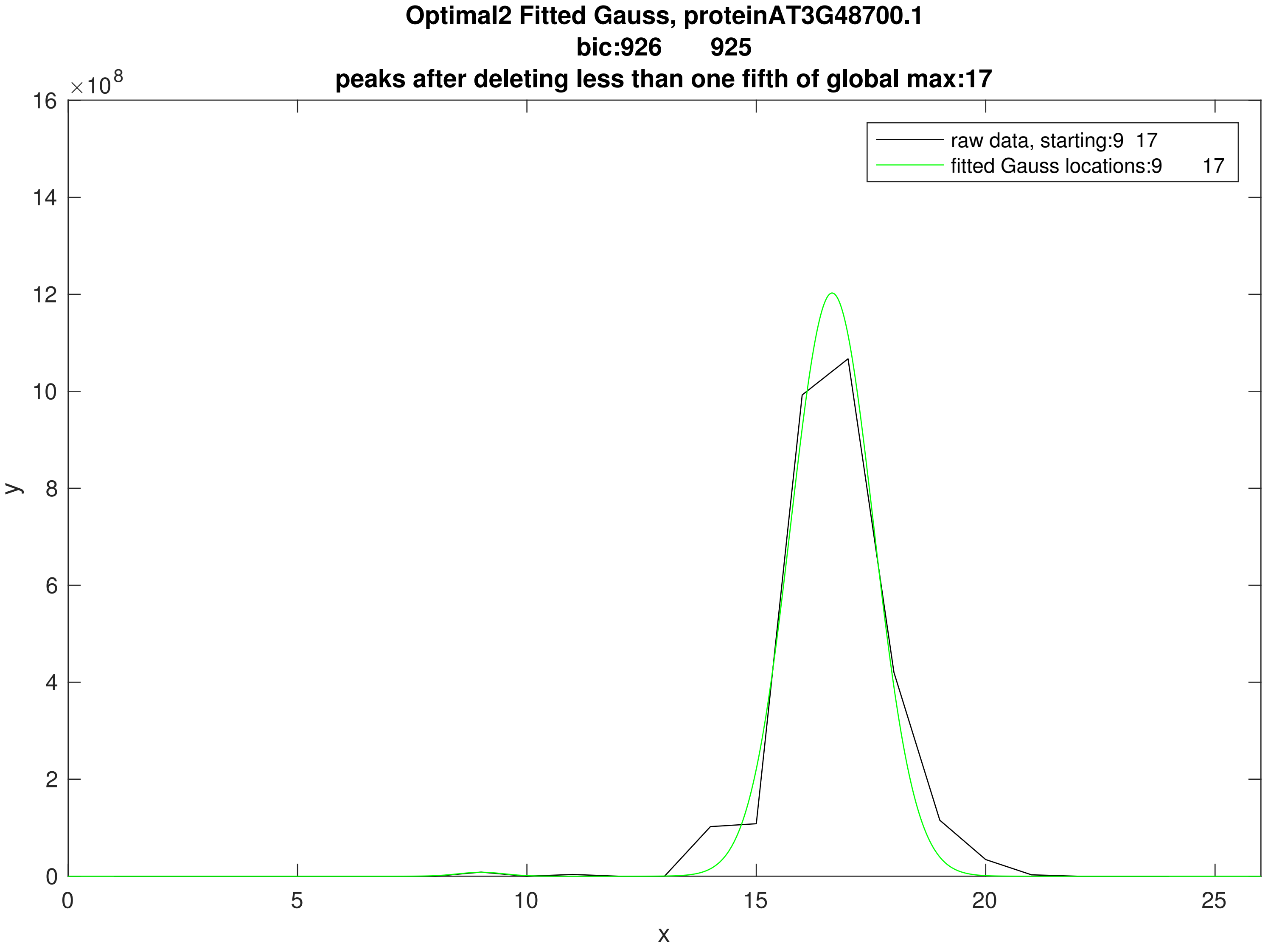

Optimal2 Fitted Gauss, proteinAT3G48700.1
bic:926 925
peaks after deleting less than one fifth of global max:17
×10^8
raw data, starting:9 17
fitted Gauss locations:9 17
y
x
0
2
4
6
8
10
12
14
16
0
5
10
15
20
25

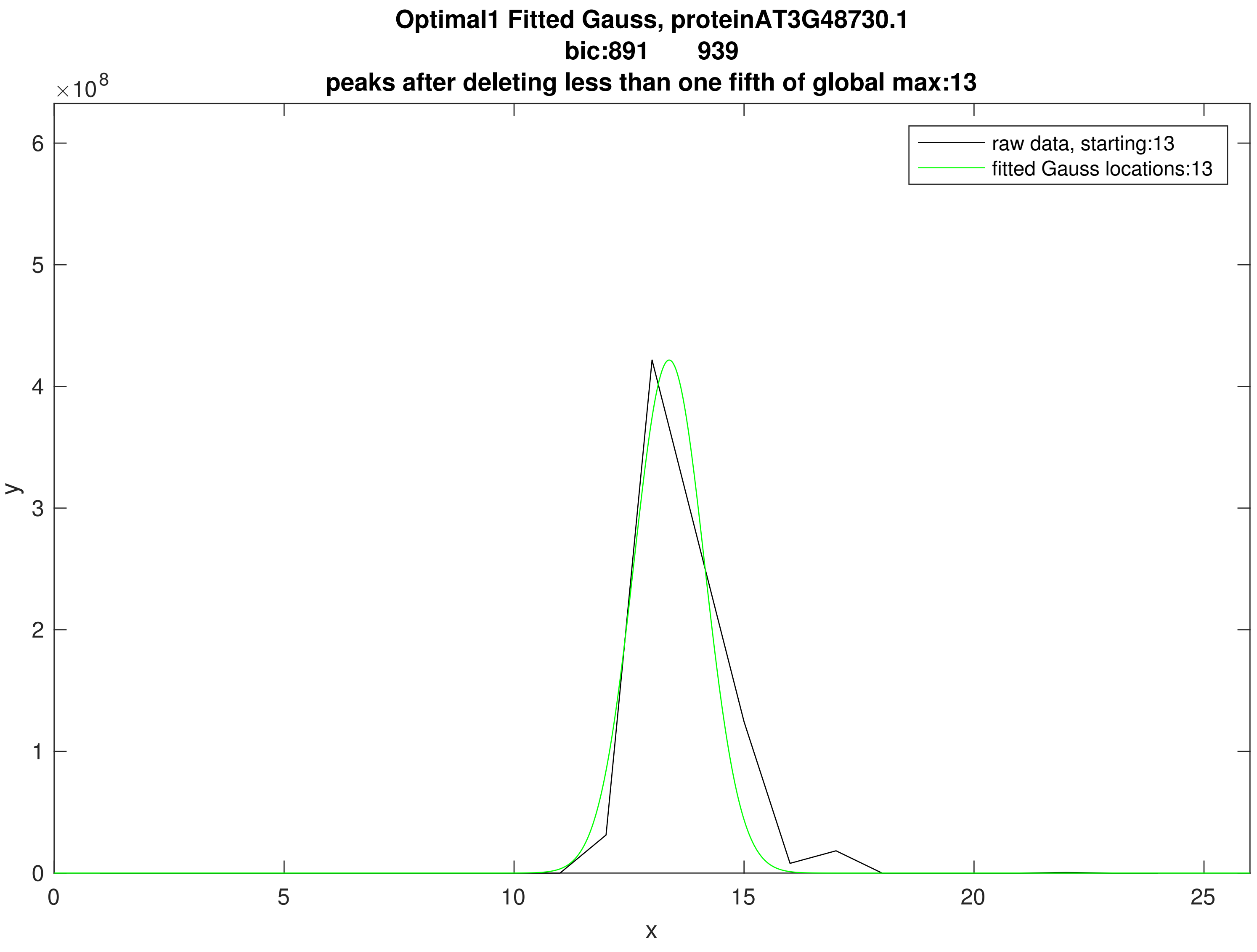
Optimal1 Fitted Gauss, proteinAT3G48730.1
bic:891 939
peaks after deleting less than one fifth of global max:13
×10^8
raw data, starting:13
fitted Gauss locations:13
y
x
0
1
2
3
4
5
6
0
5
10
15
20
25

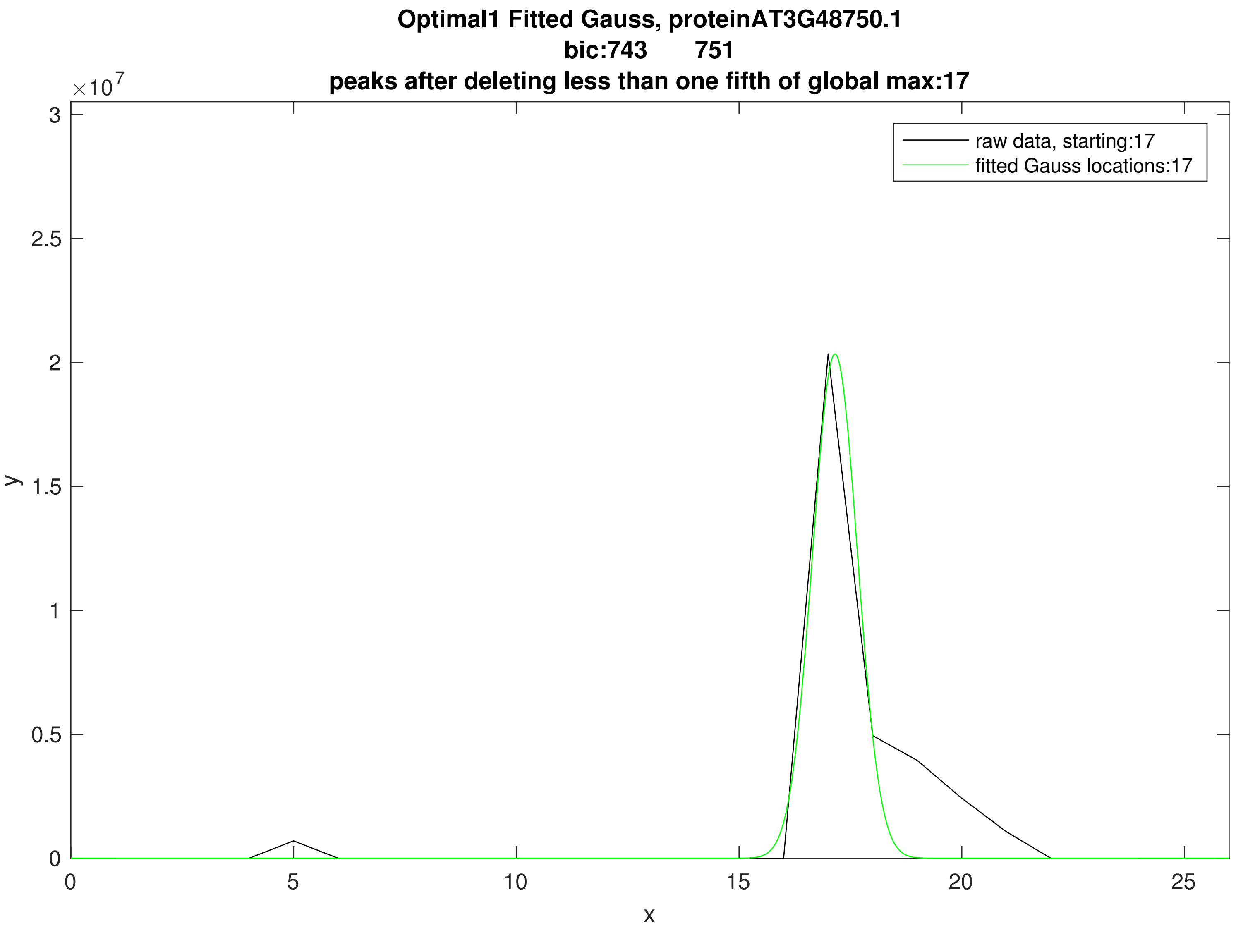

Optimal1 Fitted Gauss, proteinAT3G48750.1
bic:743 751
peaks after deleting less than one fifth of global max:17
×10^7
3
2.5
2
1.5
1
0.5
0
y
0
5
10
15
20
25
x
raw data, starting:17
fitted Gauss locations:17

Optimal1 Fitted Gauss, proteinAT3G48930.1
bic:827 837
peaks after deleting less than one fifth of global max:4.2

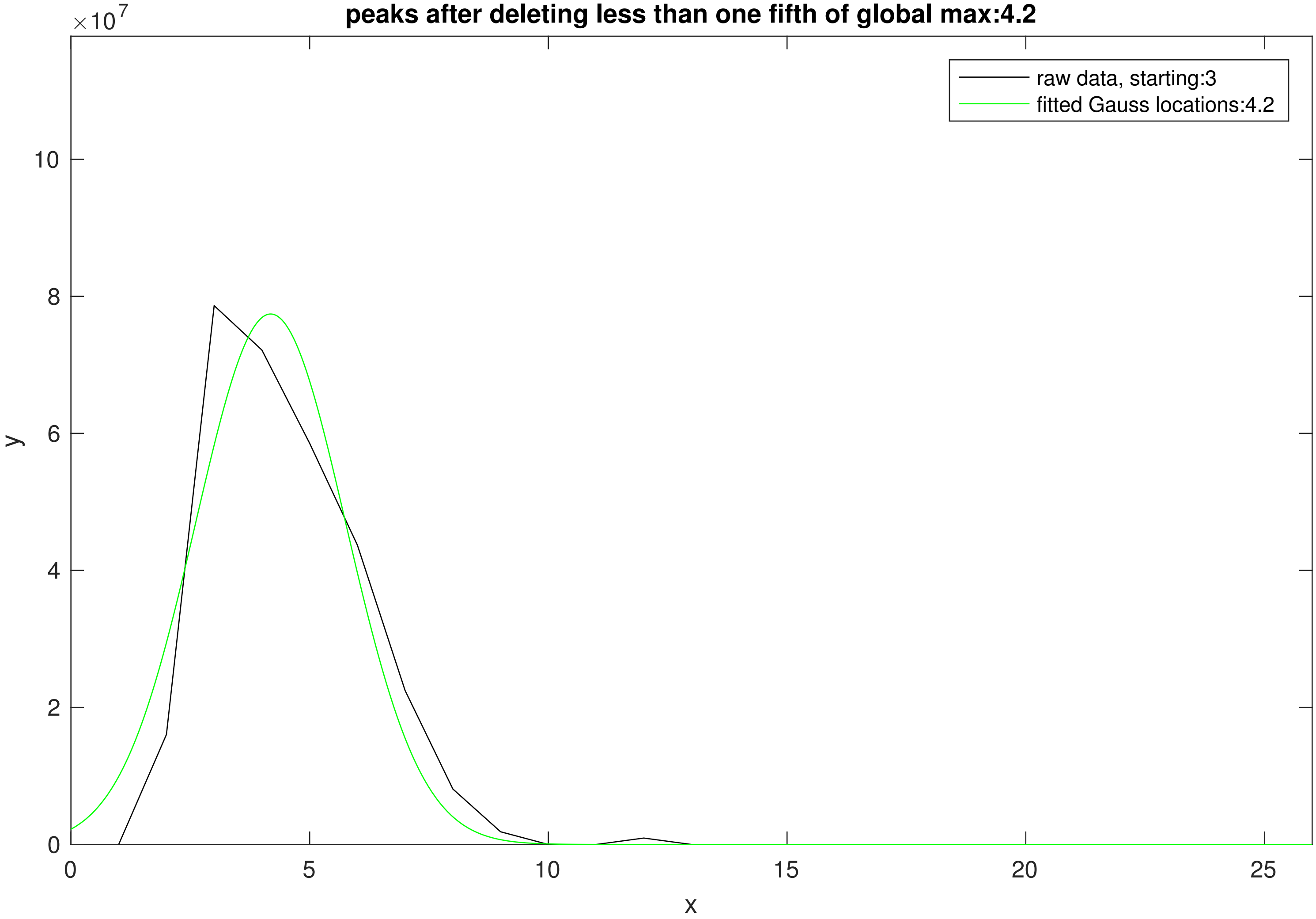

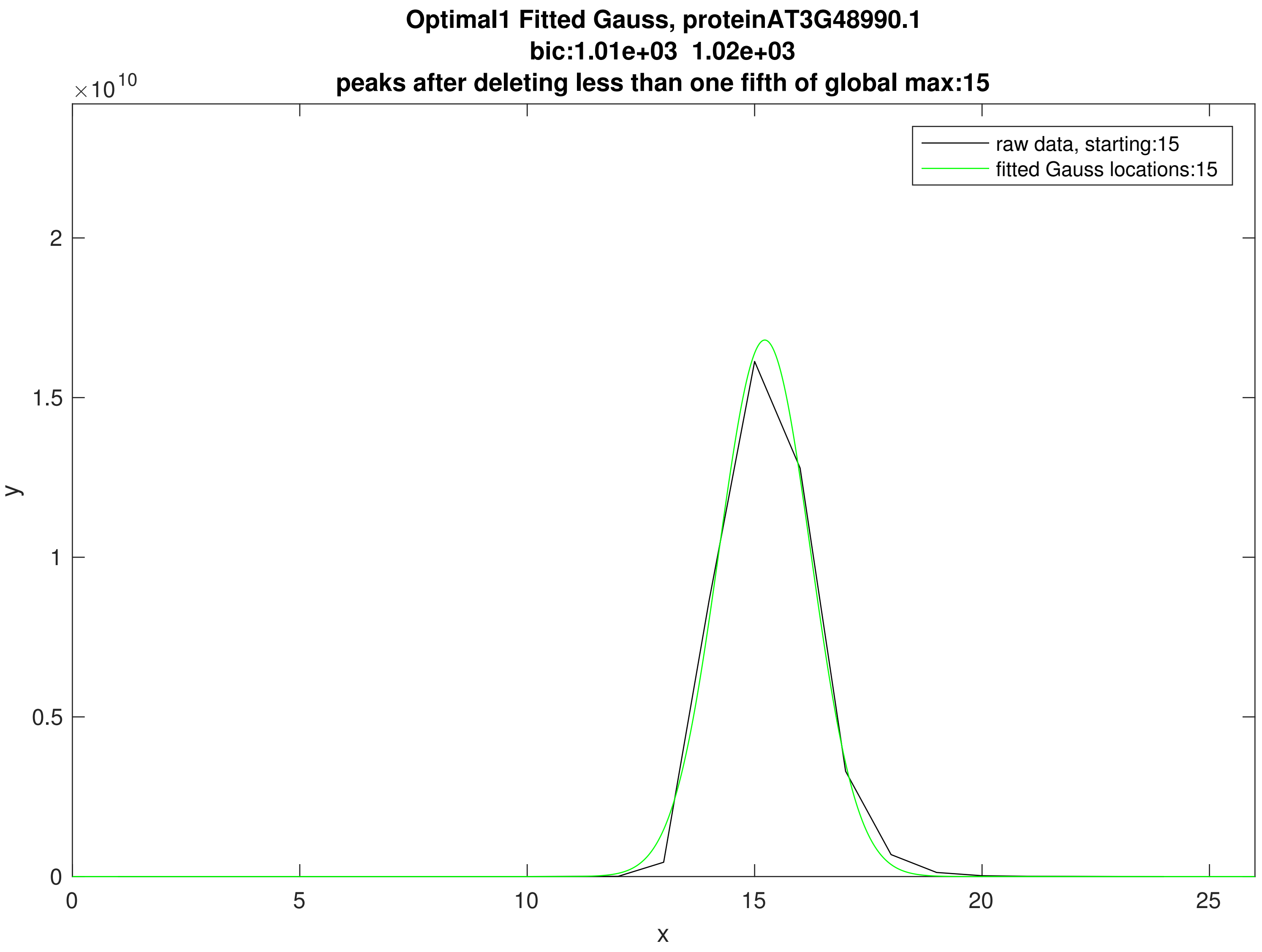
Optimal1 Fitted Gauss, proteinAT3G48990.1
bic:1.01e+03  1.02e+03
peaks after deleting less than one fifth of global max:15
×10^10
raw data, starting:15
fitted Gauss locations:15
2
1.5
1
0.5
0
y
0
5
10
15
20
25
x

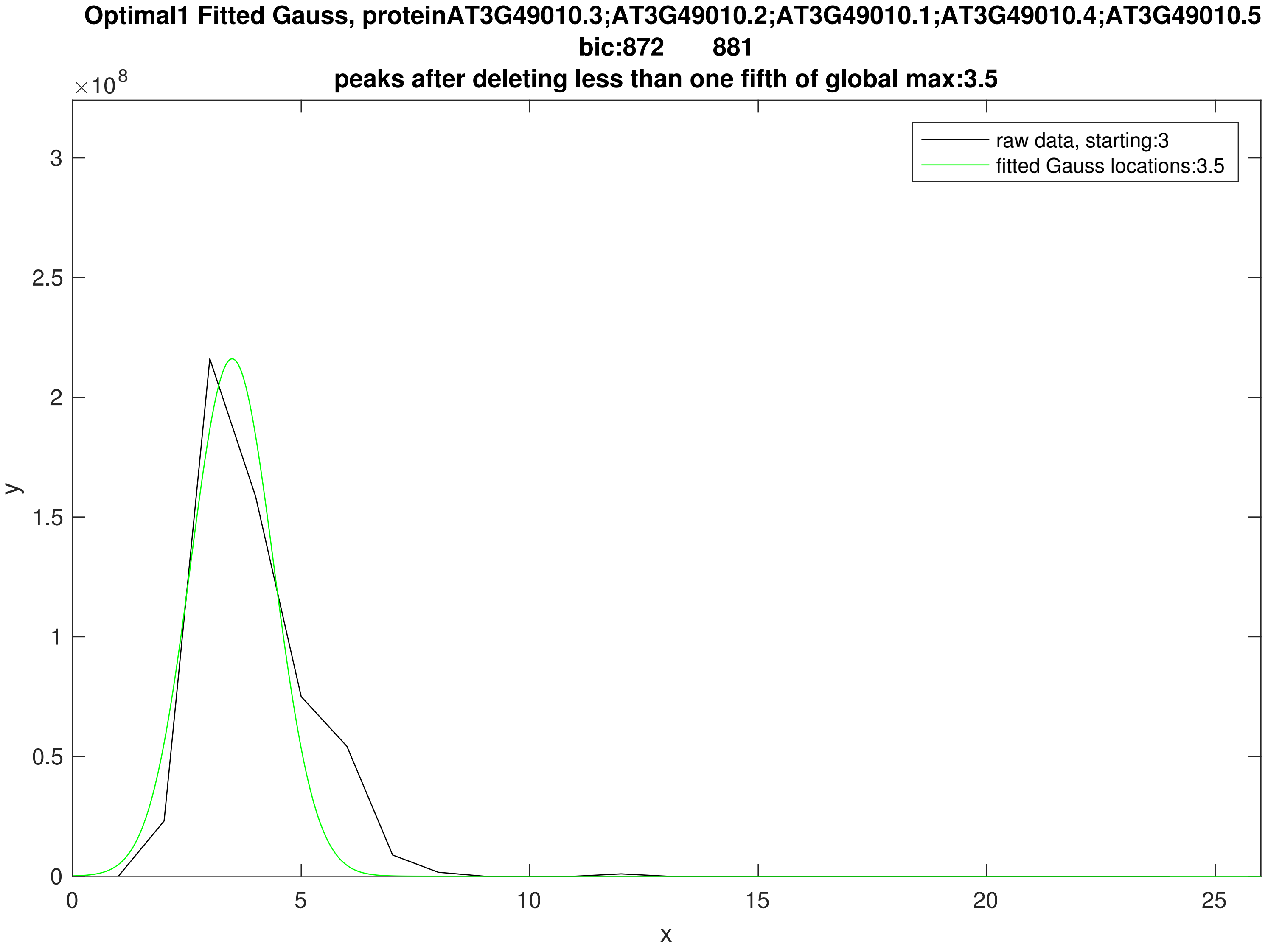

Optimal1 Fitted Gauss, proteinAT3G49010.3;AT3G49010.2;AT3G49010.1;AT3G49010.4;AT3G49010.5
bic:872 881
peaks after deleting less than one fifth of global max:3.5
×10^8
raw data, starting:3
fitted Gauss locations:3.5
y
x
0
0.5
1
1.5
2
2.5
3
5
10
15
20
25

**Optimal1 Fitted Gauss, proteinAT3G49070.1**
**bic:684**
**peaks after deleting less than one fifth of global max:13**

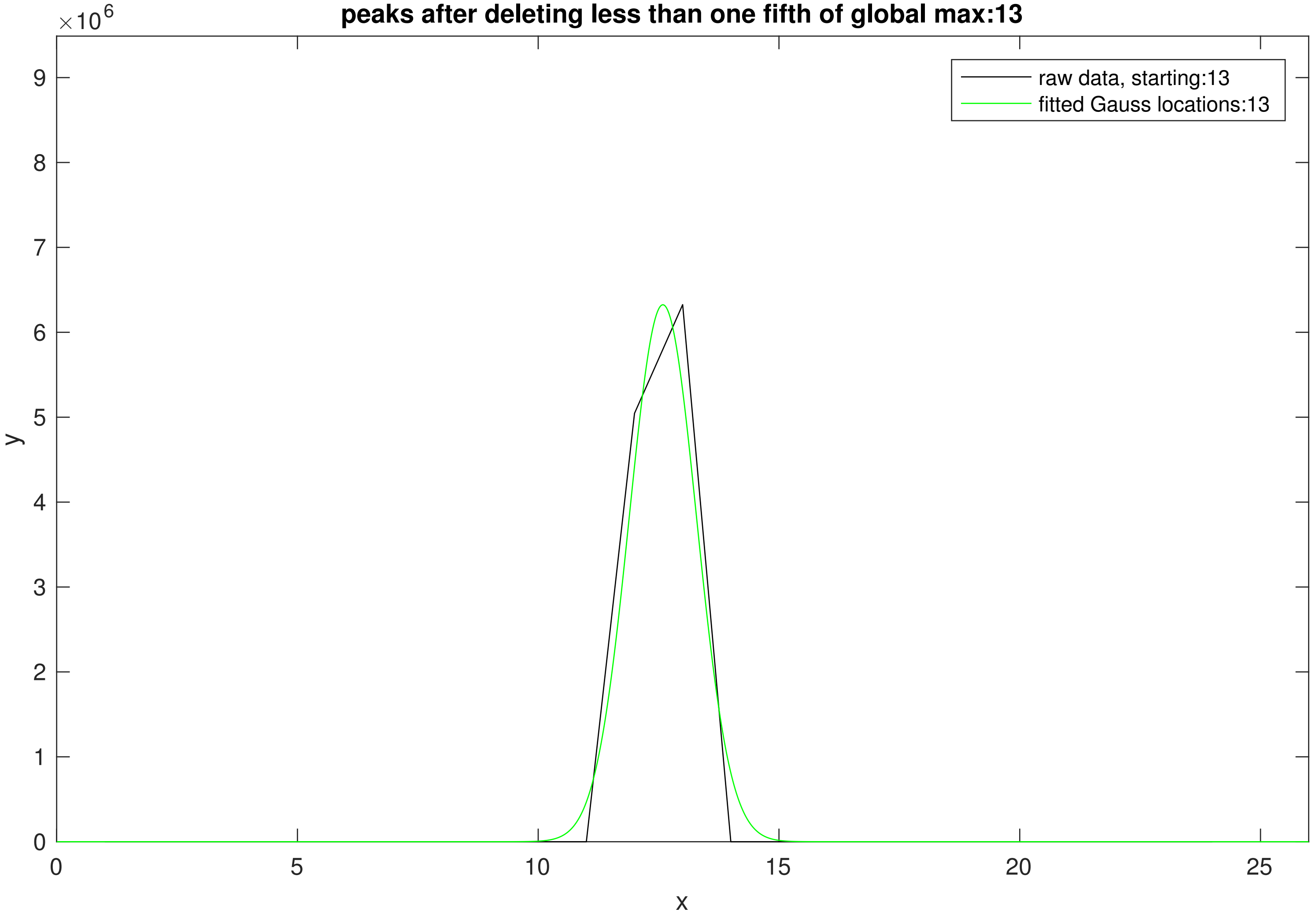

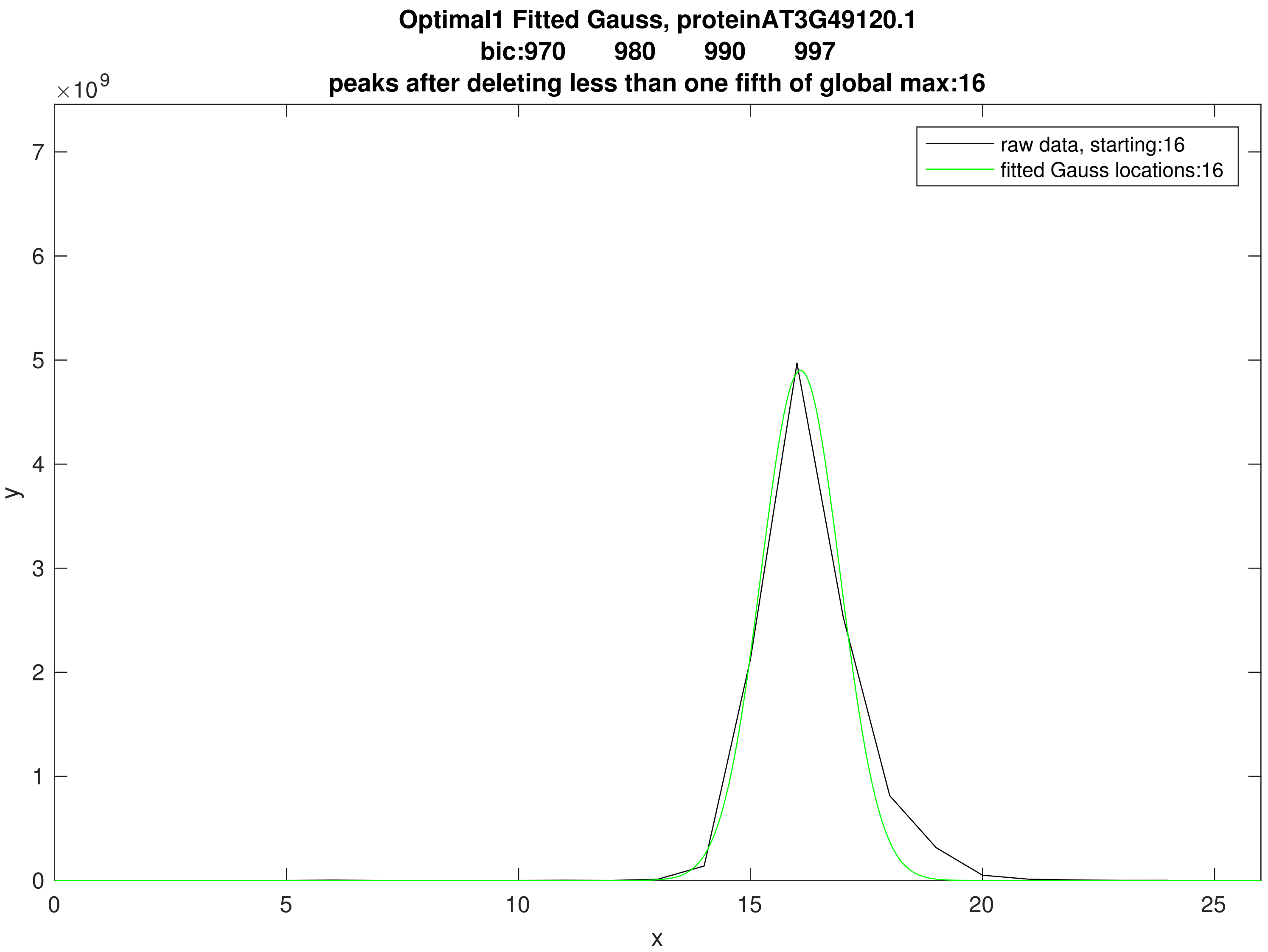
Optimal1 Fitted Gauss, proteinAT3G49120.1
bic:970 980 990 997
peaks after deleting less than one fifth of global max:16
×10^9
raw data, starting:16
fitted Gauss locations:16
y
x
0
1
2
3
4
5
6
7
0
5
10
15
20
25

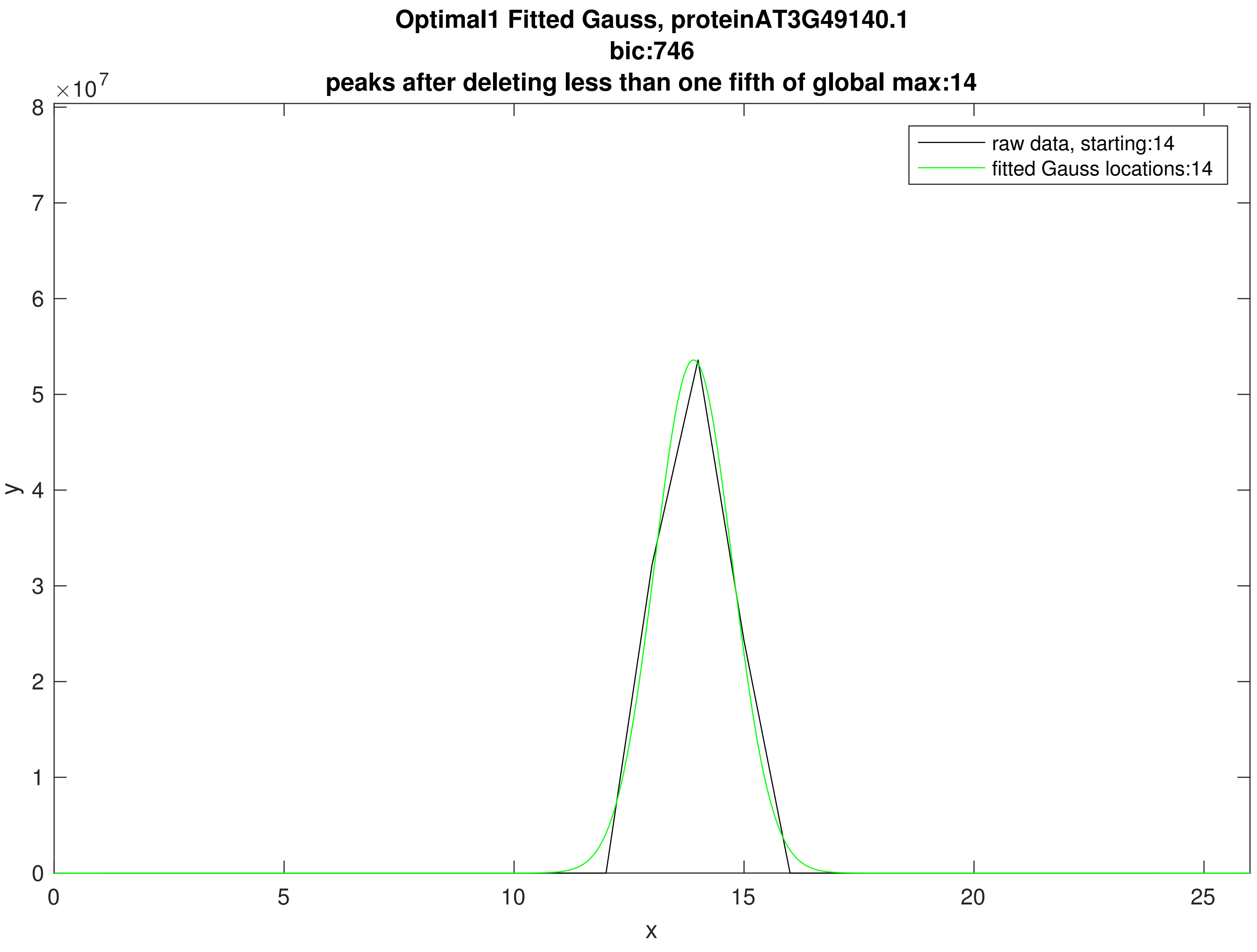

Optimal1 Fitted Gauss, proteinAT3G49140.1
bic:746
peaks after deleting less than one fifth of global max:14
×10^7
raw data, starting:14
fitted Gauss locations:14
y
x
0
1
2
3
4
5
6
7
8
0
5
10
15
20
25

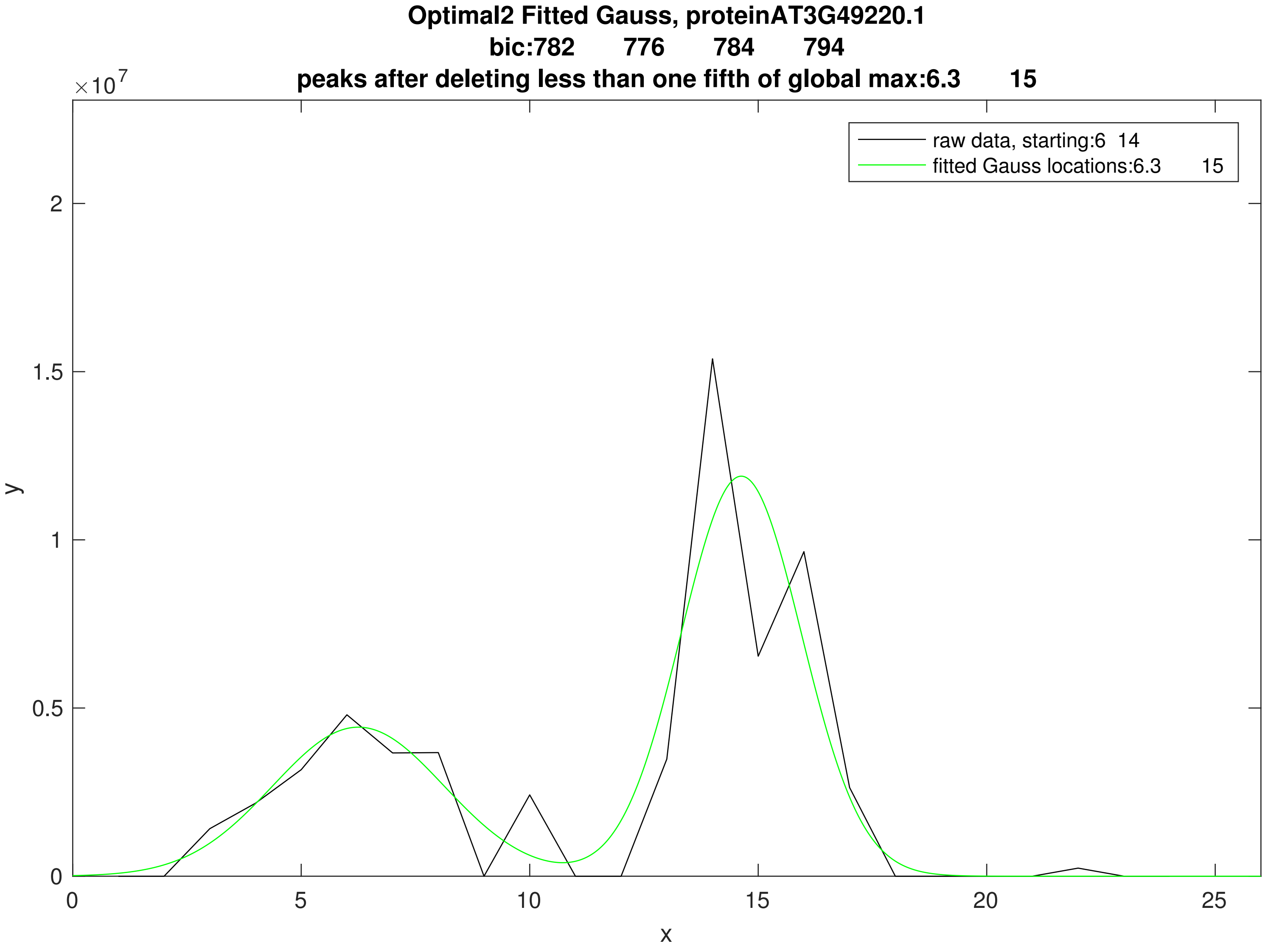
Optimal2 Fitted Gauss, proteinAT3G49220.1
bic:782 776 784 794
peaks after deleting less than one fifth of global max:6.3 15
×10^7
raw data, starting:6 14
fitted Gauss locations:6.3 15
2
1.5
1
0.5
0
0
5
10
15
20
25
y
x

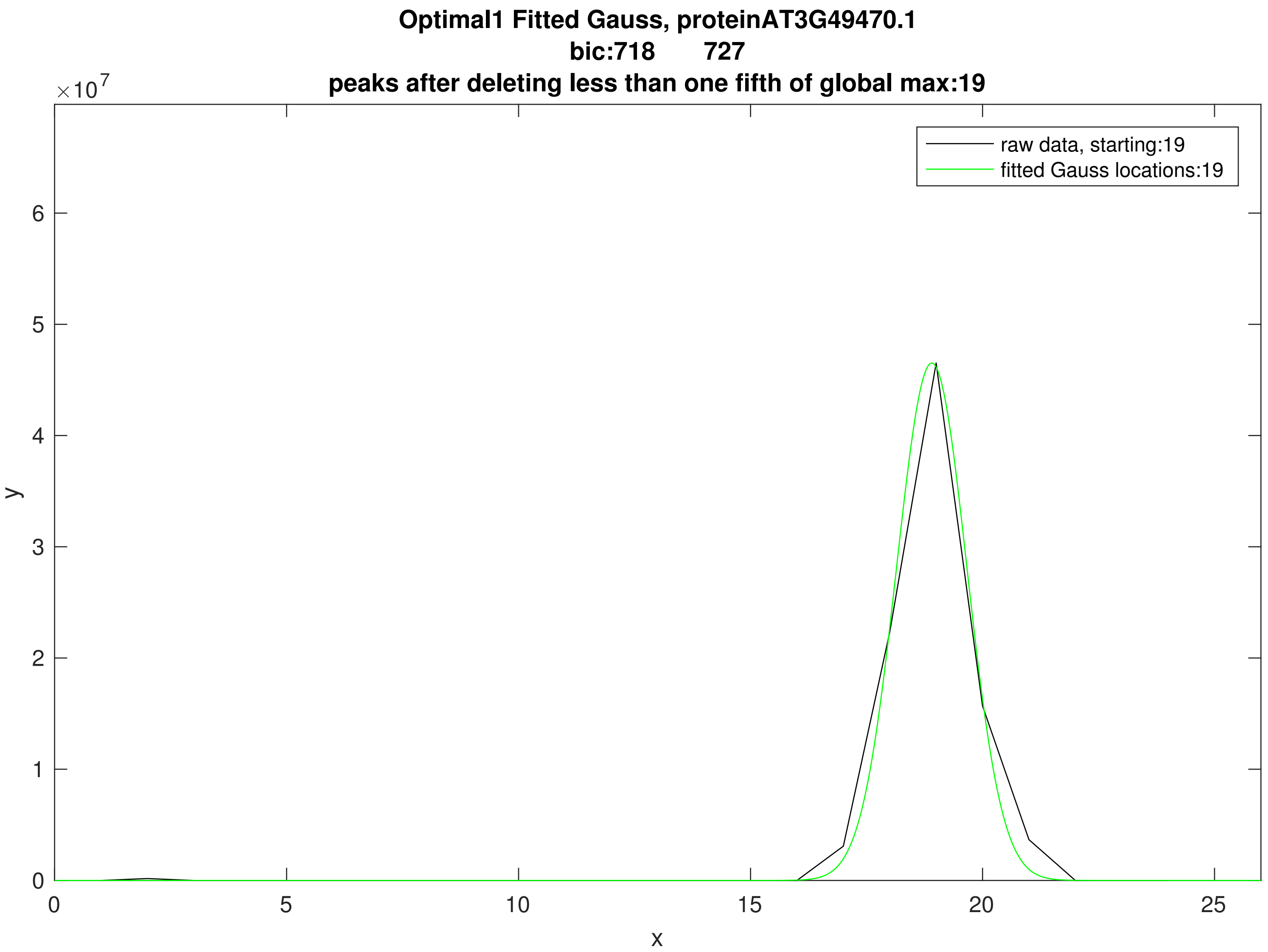
Optimal1 Fitted Gauss, proteinAT3G49470.1
bic:718 727
peaks after deleting less than one fifth of global max:19
×10^7
raw data, starting:19
fitted Gauss locations:19
y
x
0
1
2
3
4
5
6
0
5
10
15
20
25

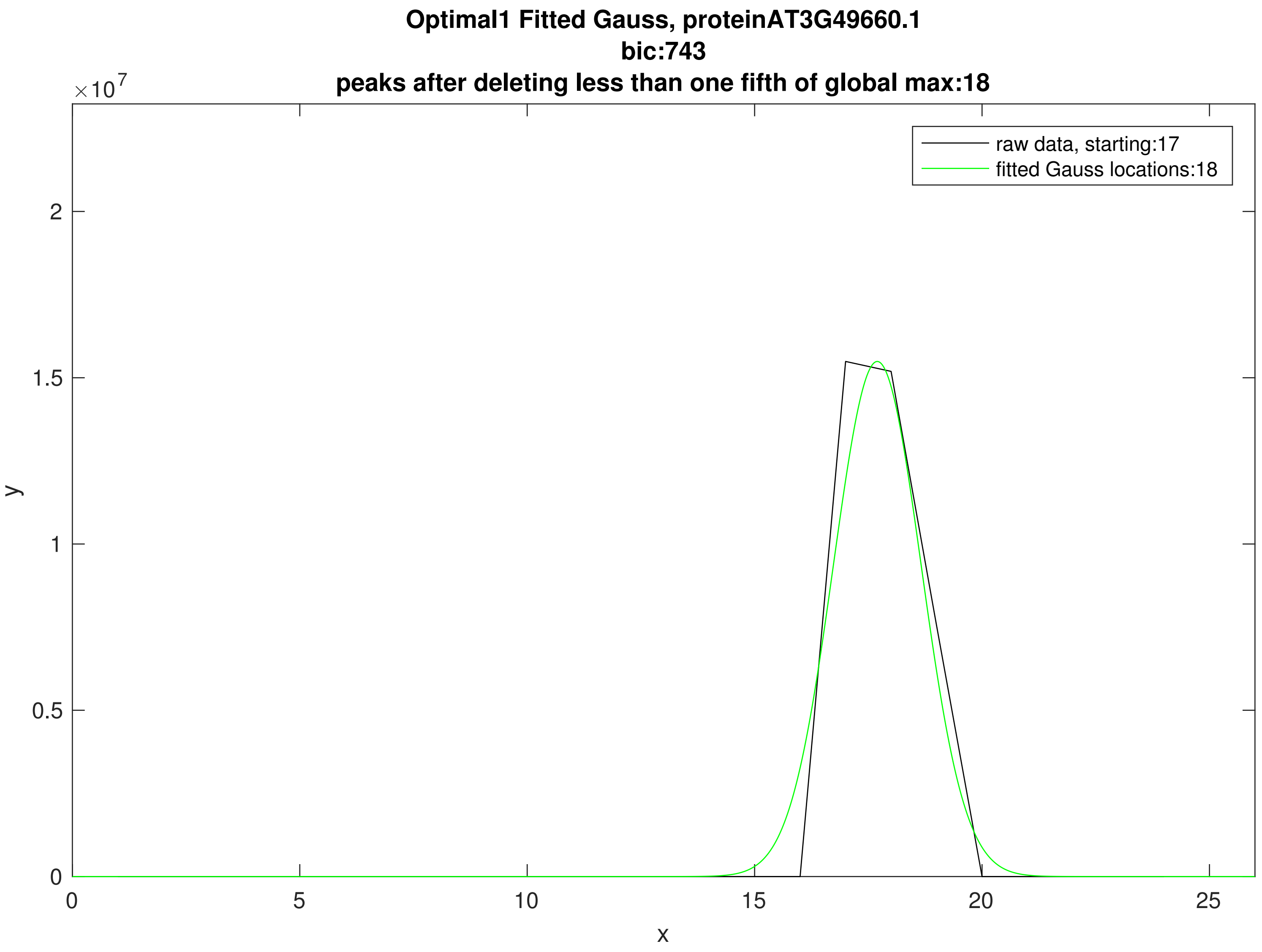
Optimal1 Fitted Gauss, proteinAT3G49660.1
bic:743
peaks after deleting less than one fifth of global max:18
×10^7
raw data, starting:17
fitted Gauss locations:18
y
x
0
0.5
1
1.5
2
0
5
10
15
20
25

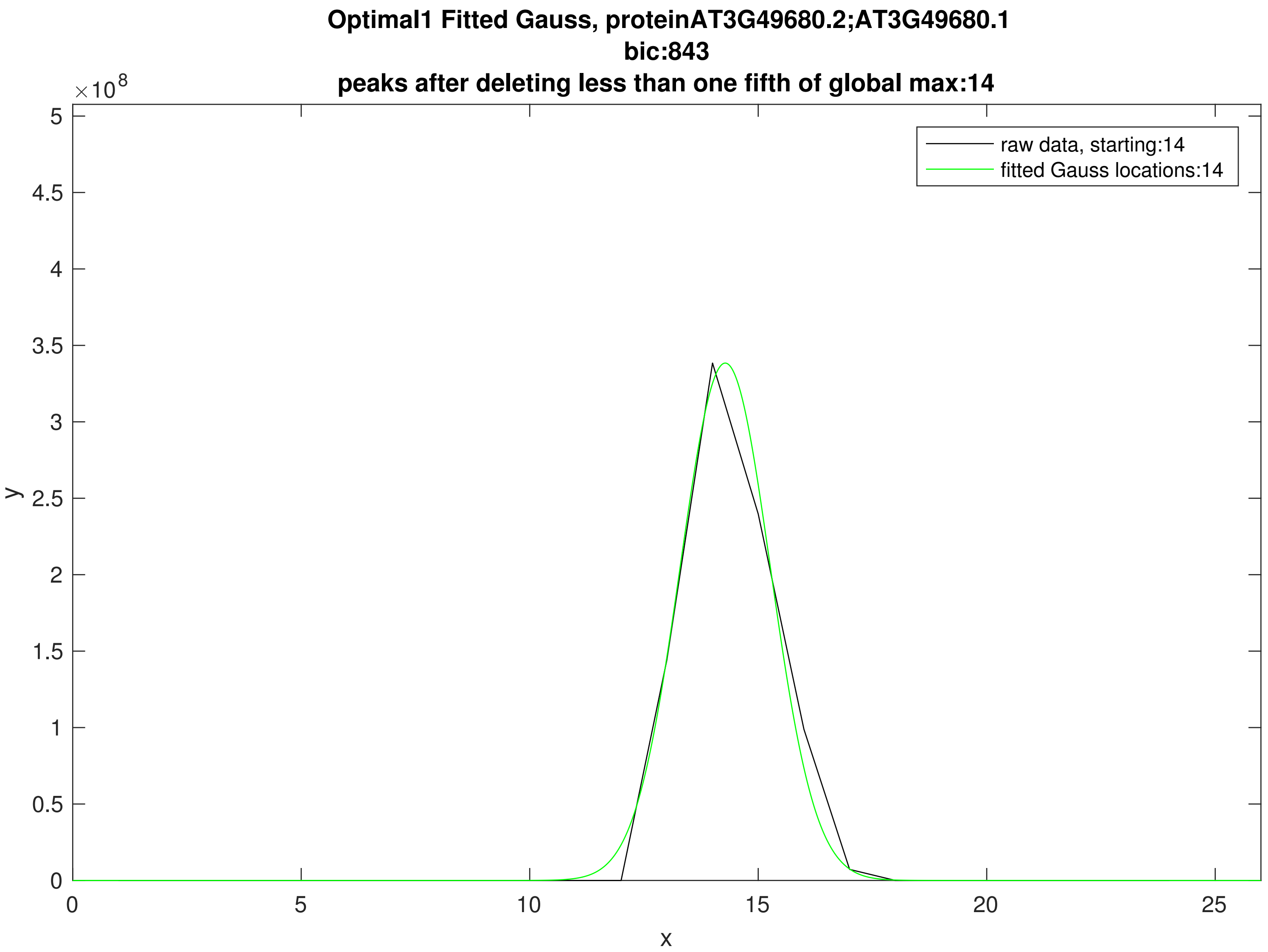
Optimal1 Fitted Gauss, proteinAT3G49680.2;AT3G49680.1
bic:843
peaks after deleting less than one fifth of global max:14
×10^8
raw data, starting:14
fitted Gauss locations:14
y
x
0
0.5
1
1.5
2
2.5
3
3.5
4
4.5
5
0
5
10
15
20
25

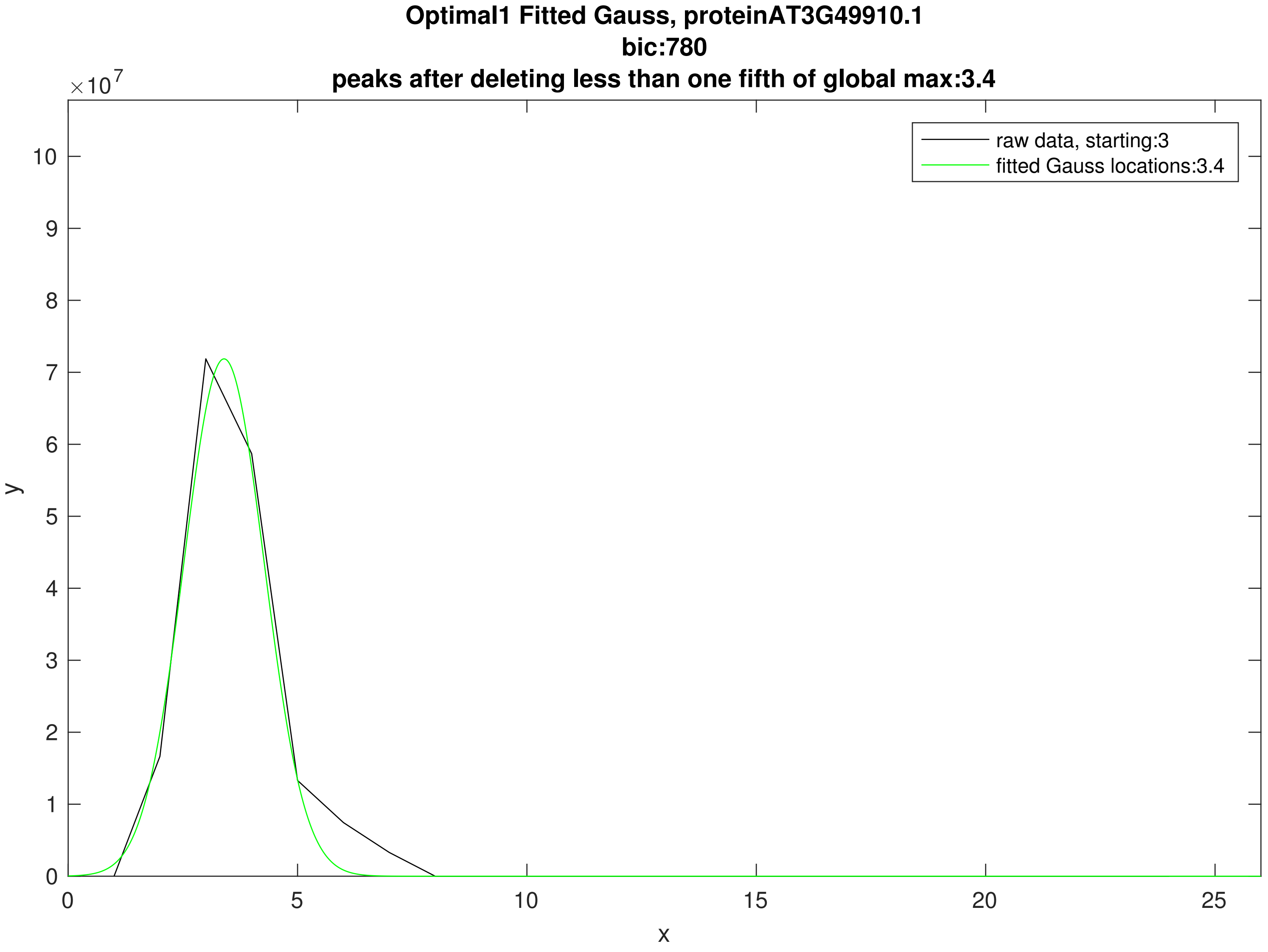
Optimal1 Fitted Gauss, proteinAT3G49910.1
bic:780
peaks after deleting less than one fifth of global max:3.4
×10^7
raw data, starting:3
fitted Gauss locations:3.4
y
x
0
1
2
3
4
5
6
7
8
9
10
0
5
10
15
20
25

Optimal1 Fitted Gauss, proteinAT3G50210.3;AT3G50210.1;AT3G50210.2
bic:748
peaks after deleting less than one fifth of global max:17

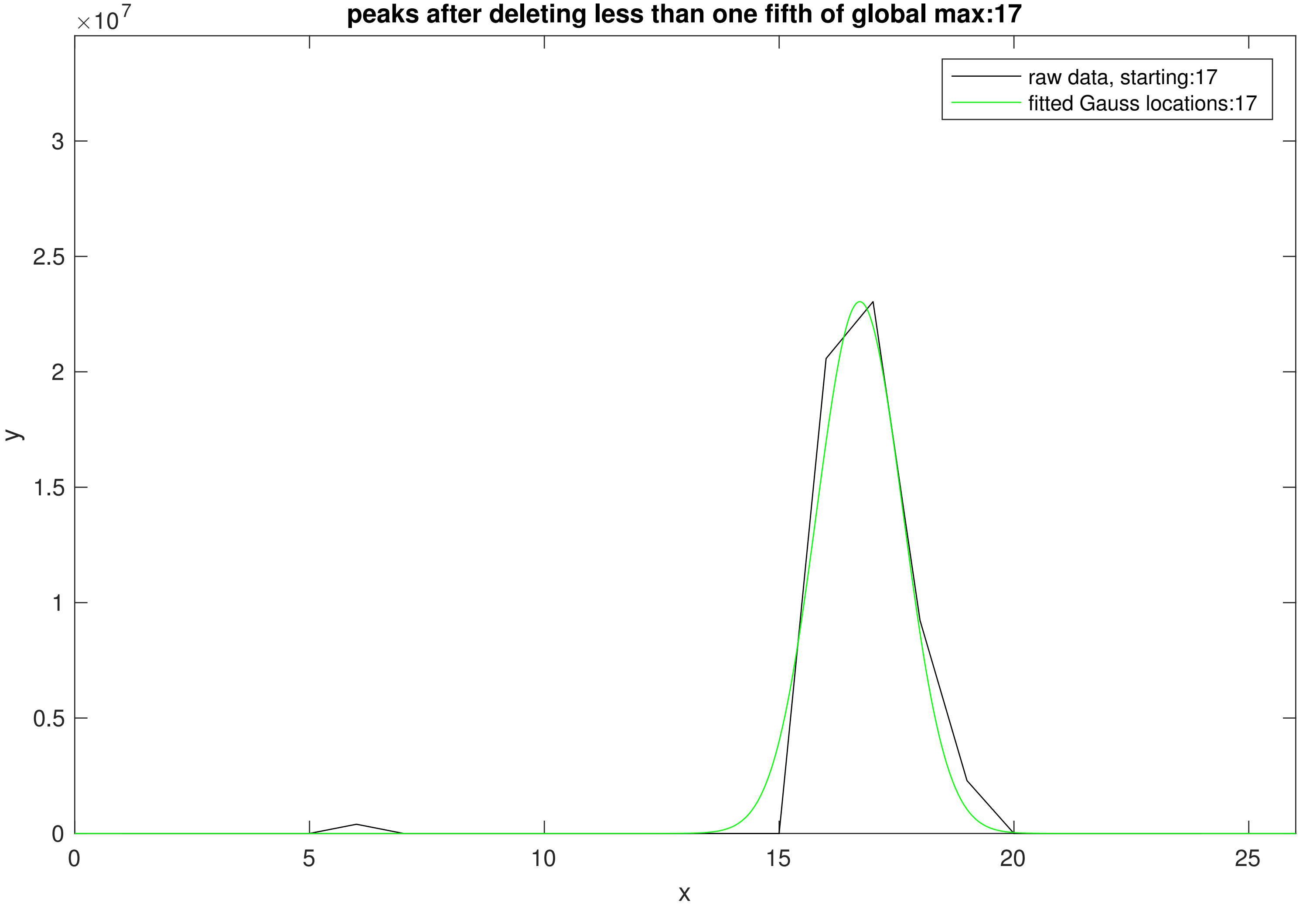

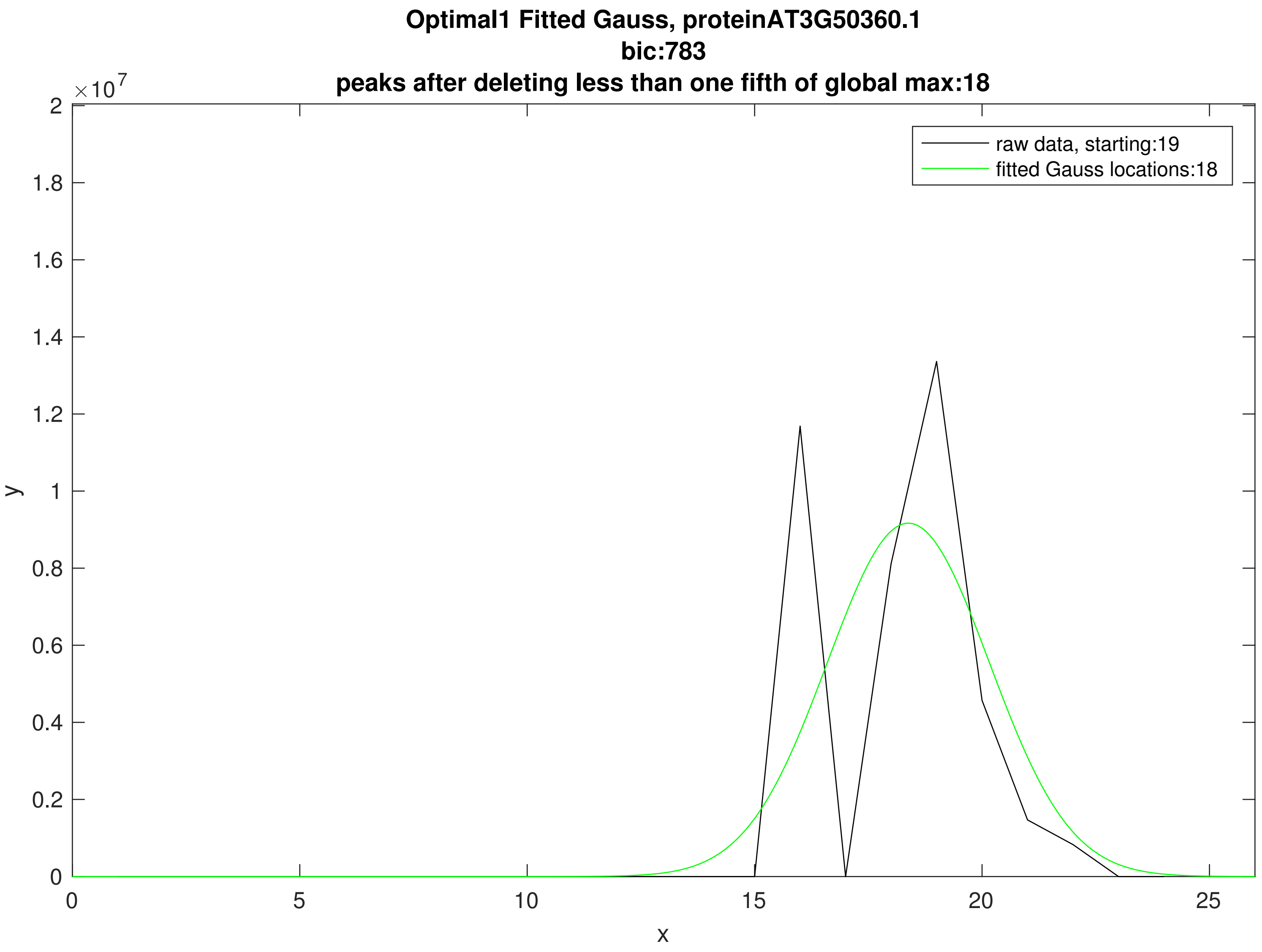
Optimal1 Fitted Gauss, proteinAT3G50360.1
bic:783
peaks after deleting less than one fifth of global max:18
raw data, starting:19
fitted Gauss locations:18
×10^7
y
x
0
0.2
0.4
0.6
0.8
1
1.2
1.4
1.6
1.8
2
0
5
10
15
20
25

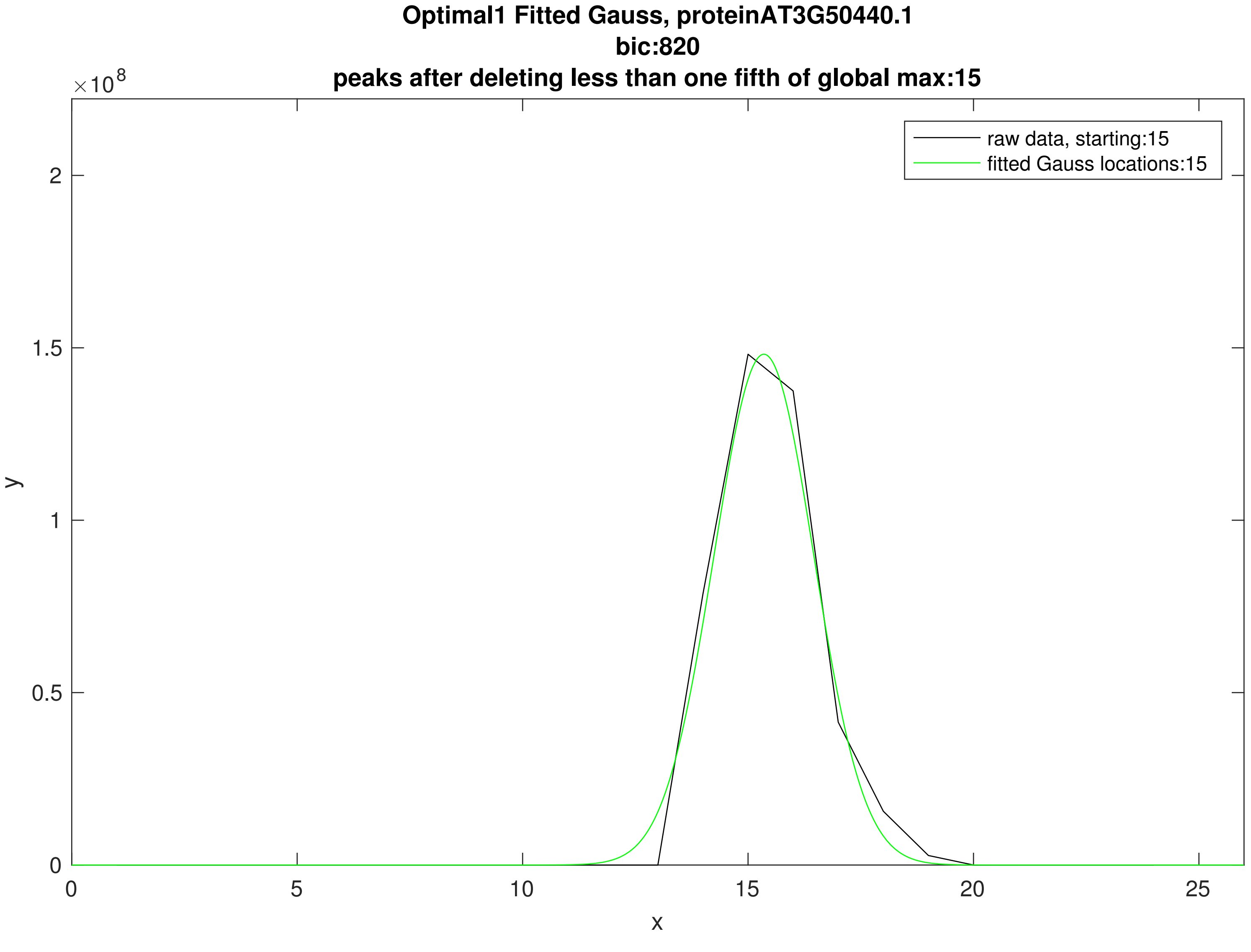
Optimal1 Fitted Gauss, proteinAT3G50440.1
bic:820
peaks after deleting less than one fifth of global max:15
×10^8
raw data, starting:15
fitted Gauss locations:15
y
x
0
0.5
1
1.5
2
0
5
10
15
20
25

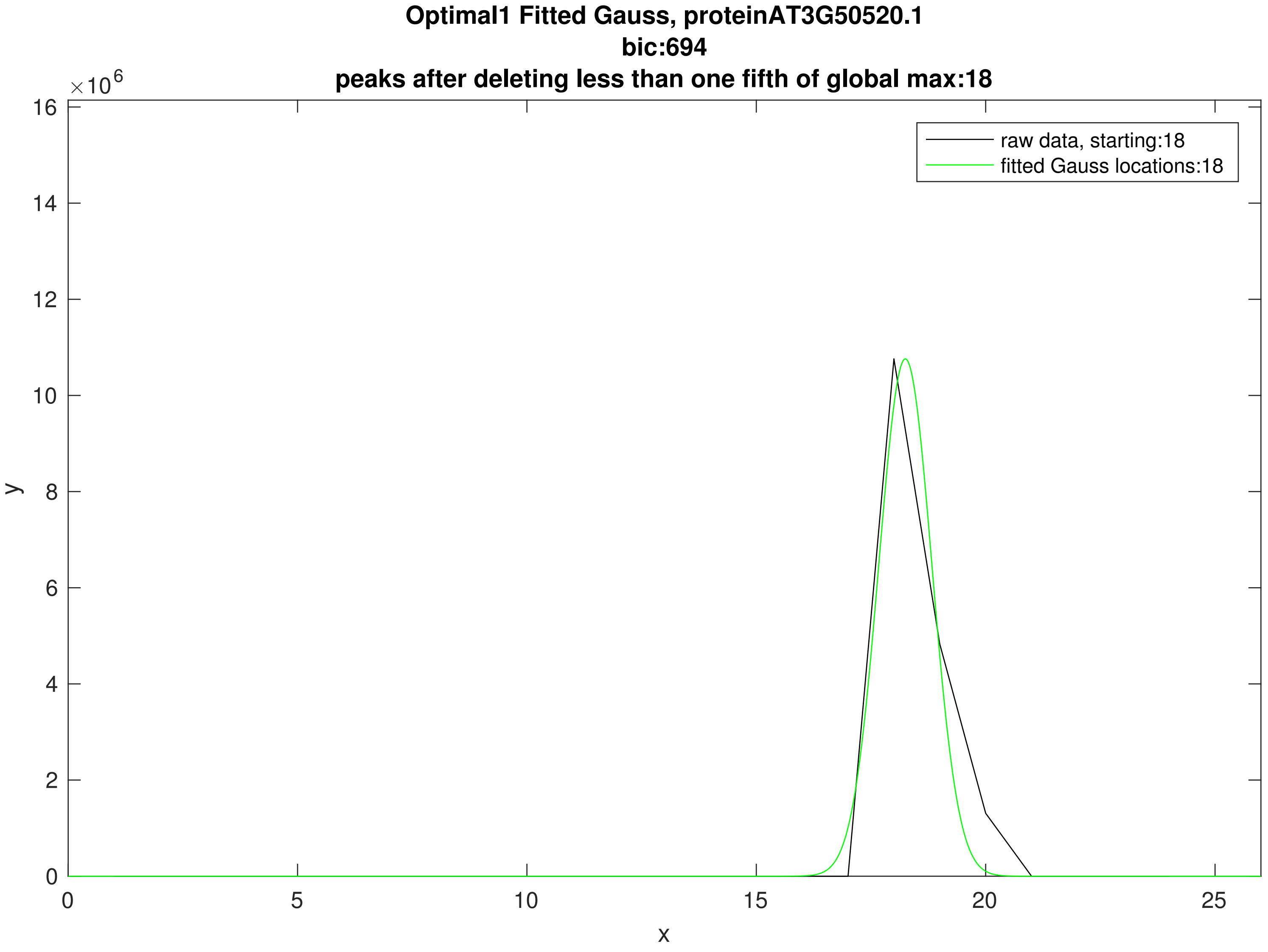
Optimal1 Fitted Gauss, proteinAT3G50520.1
bic:694
peaks after deleting less than one fifth of global max:18
×10^6
raw data, starting:18
fitted Gauss locations:18
y
x
0
2
4
6
8
10
12
14
16
0
5
10
15
20
25

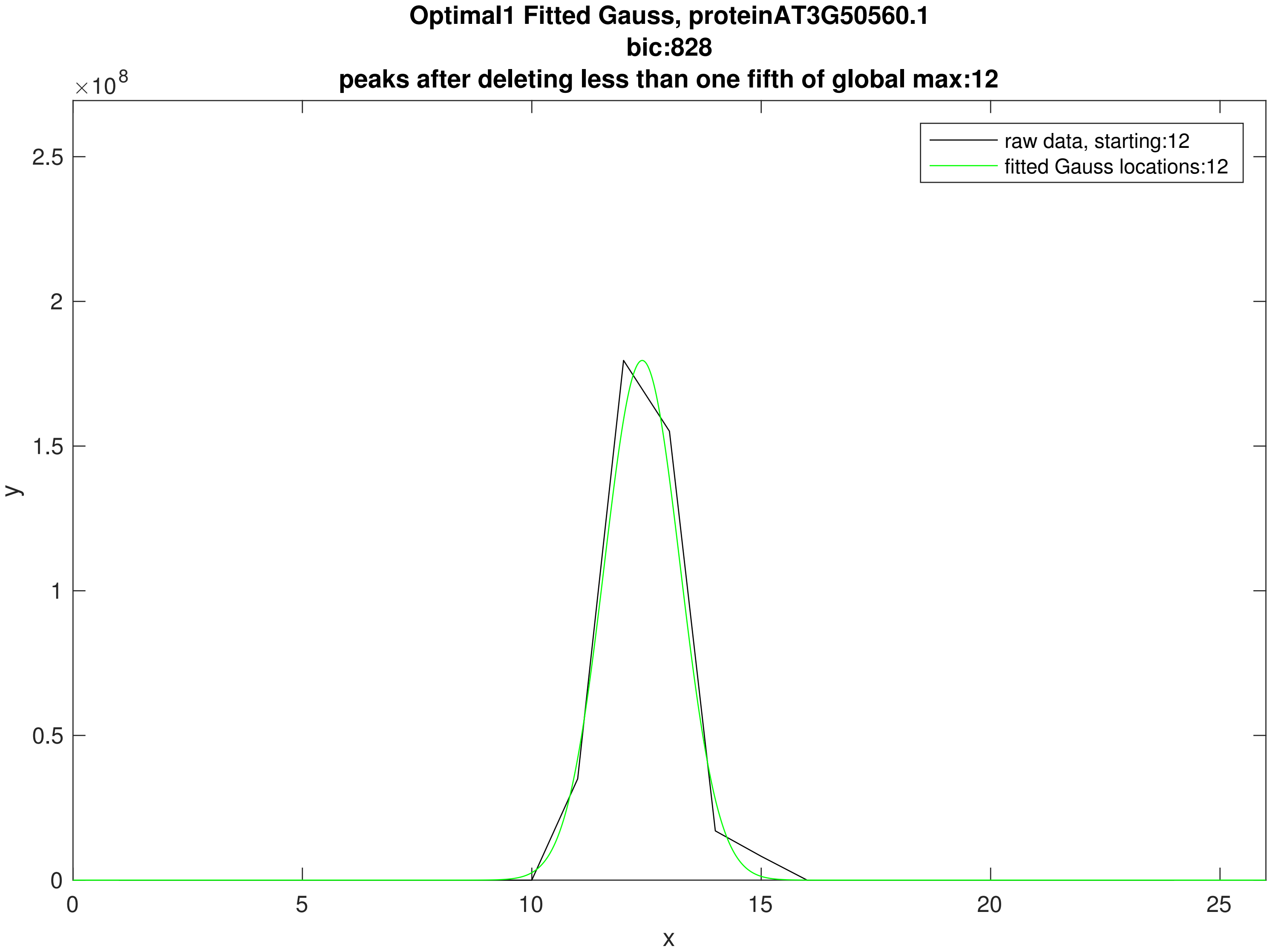

Optimal1 Fitted Gauss, proteinAT3G50560.1
bic:828
peaks after deleting less than one fifth of global max:12
×10^8
raw data, starting:12
fitted Gauss locations:12
y
x

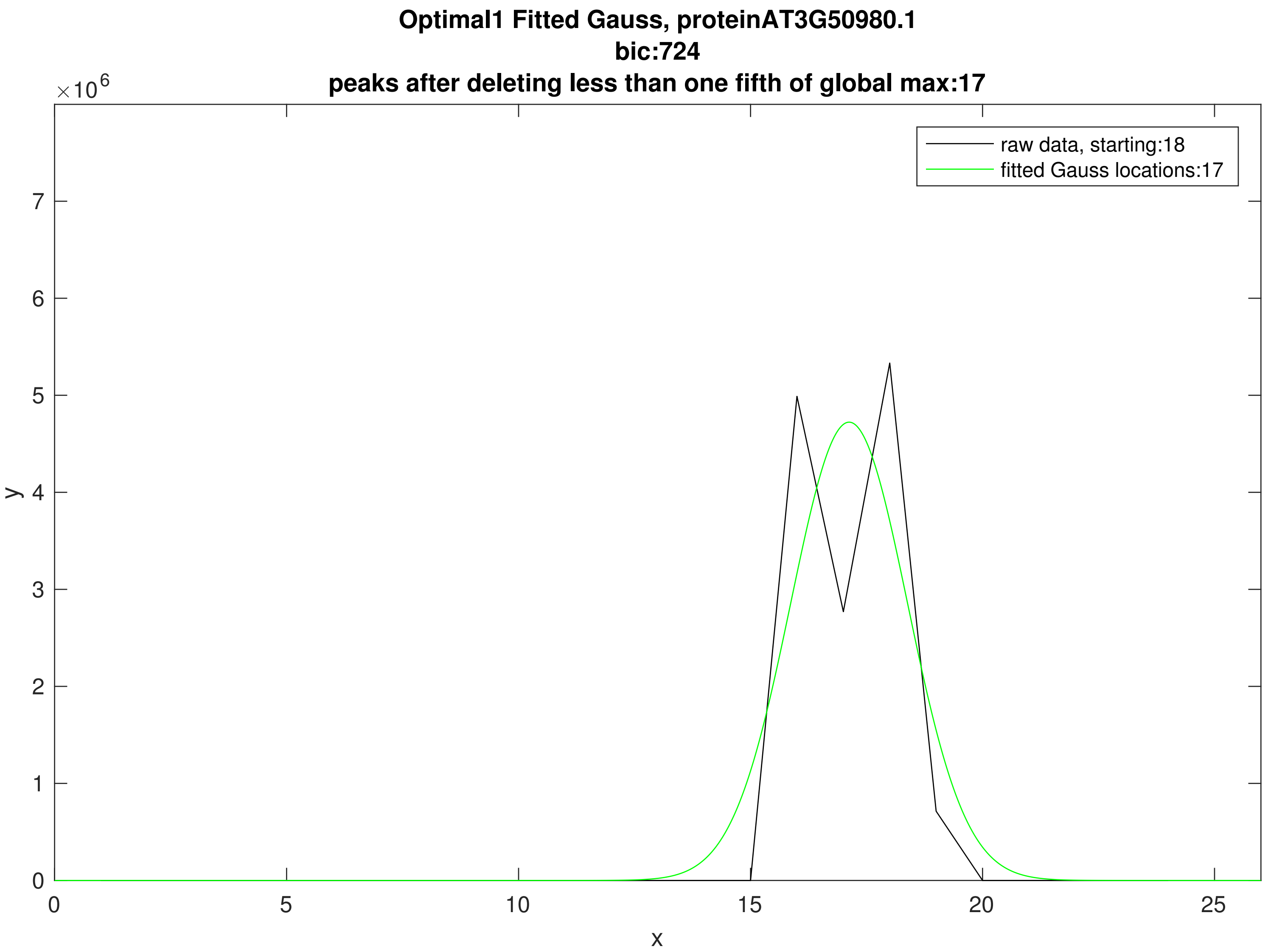
Optimal1 Fitted Gauss, proteinAT3G50980.1
bic:724
peaks after deleting less than one fifth of global max:17
×10^6
raw data, starting:18
fitted Gauss locations:17
y
x
0
1
2
3
4
5
6
7
0
5
10
15
20
25

#### Optimal1 Fitted Gauss, proteinAT3G51000.1
#### bic:823
#### peaks after deleting less than one fifth of global max:17

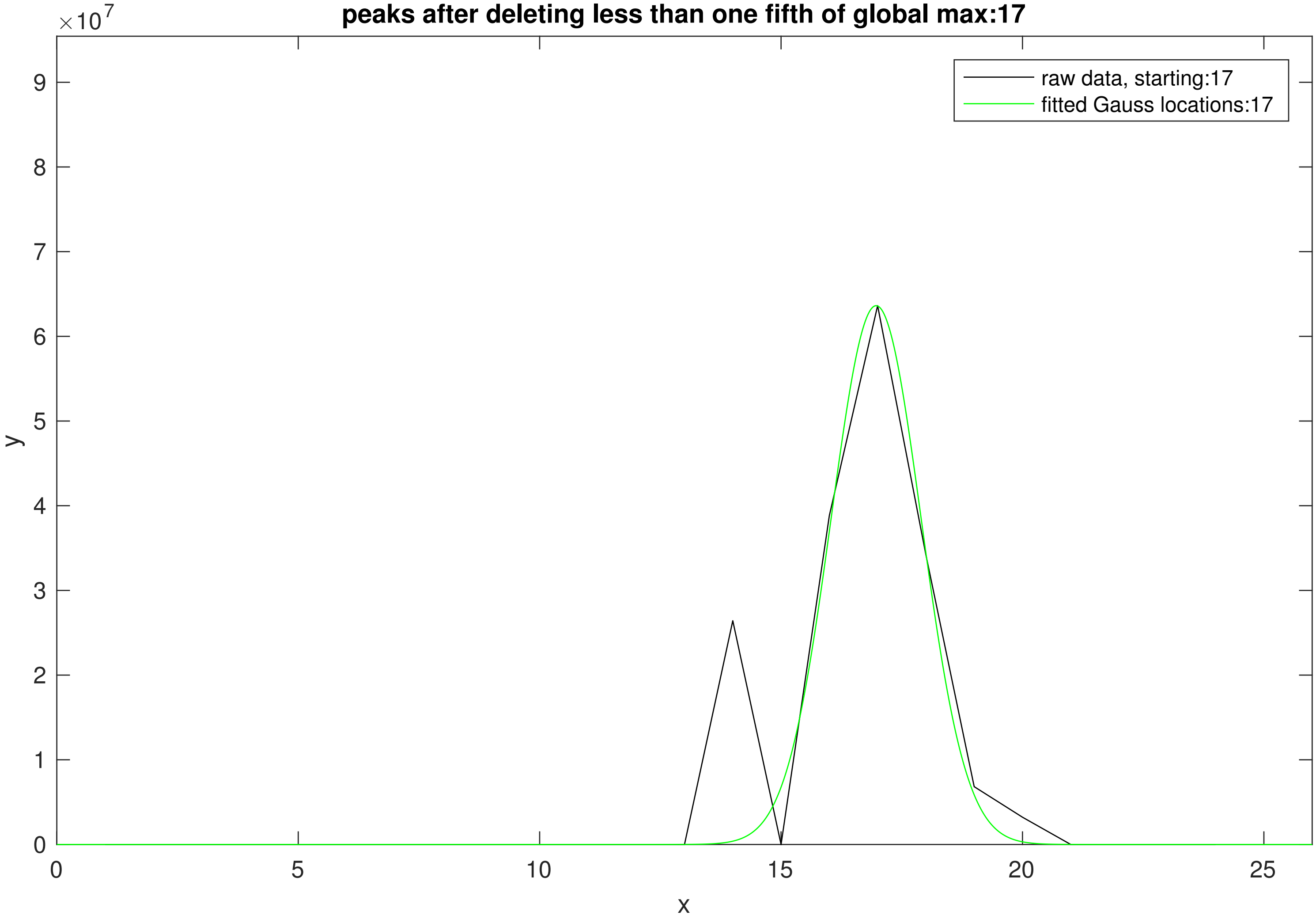

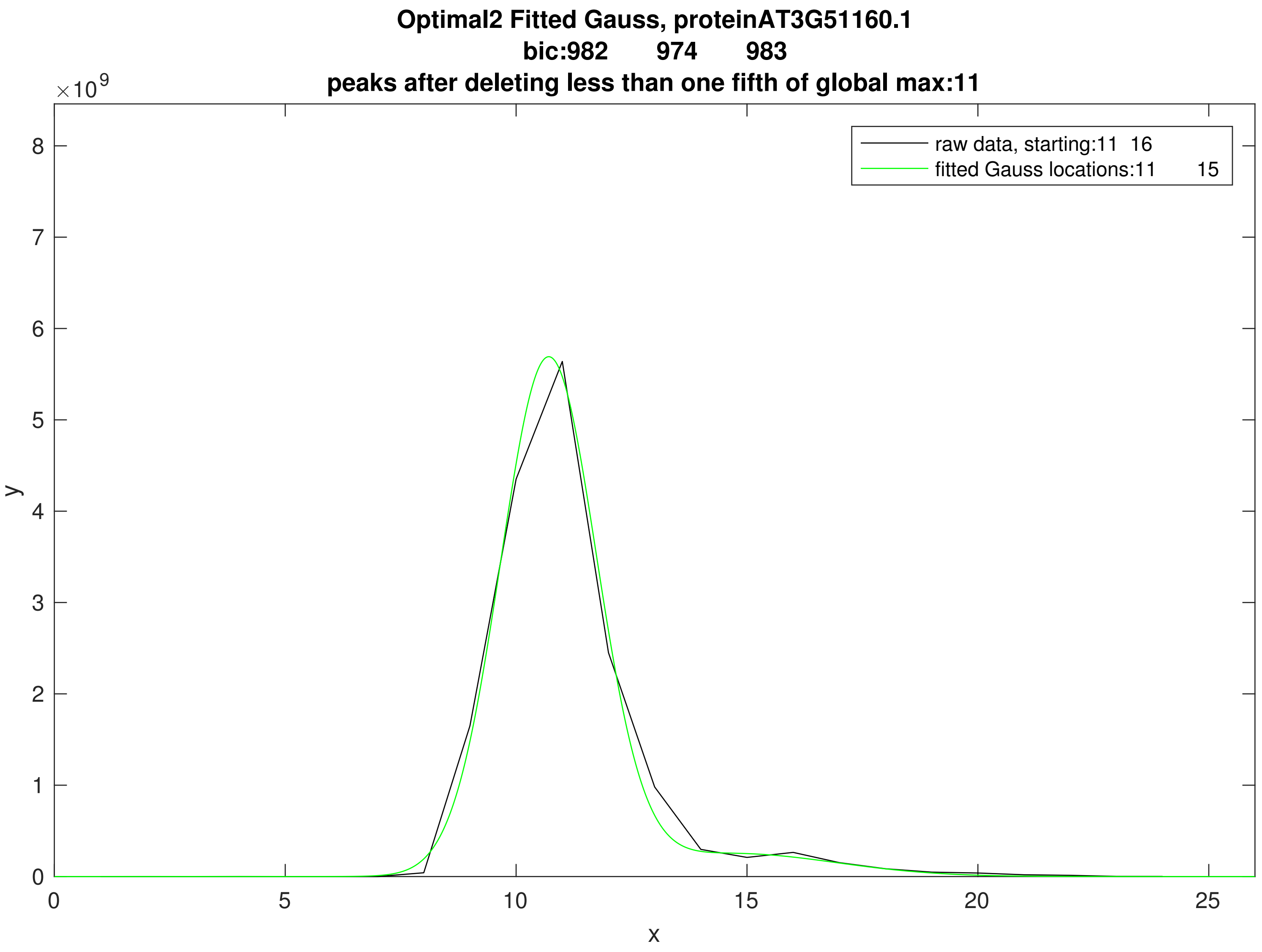

Optimal2 Fitted Gauss, proteinAT3G51160.1
bic:982 974 983
peaks after deleting less than one fifth of global max:11
×10^9
raw data, starting:11 16
fitted Gauss locations:11 15
y
x
0
1
2
3
4
5
6
7
8
0
5
10
15
20
25

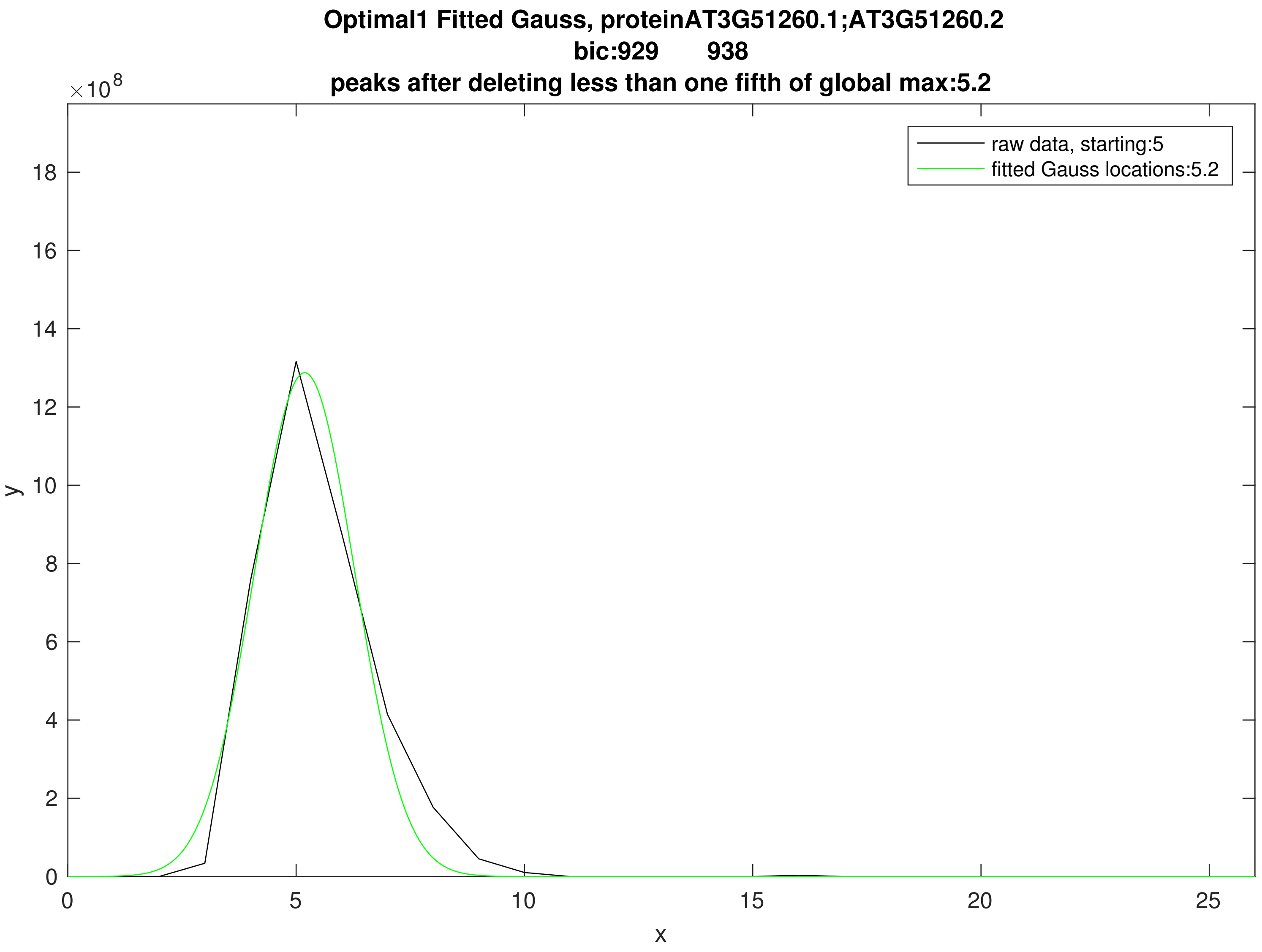
Optimal1 Fitted Gauss, proteinAT3G51260.1;AT3G51260.2
bic:929 938
peaks after deleting less than one fifth of global max:5.2
×10^8
y
x
raw data, starting:5
fitted Gauss locations:5.2
0
2
4
6
8
10
12
14
16
18
0
5
10
15
20
25

Optimal1 Fitted Gauss, proteinAT3G51480.1
bic:702
peaks after deleting less than one fifth of global max:12

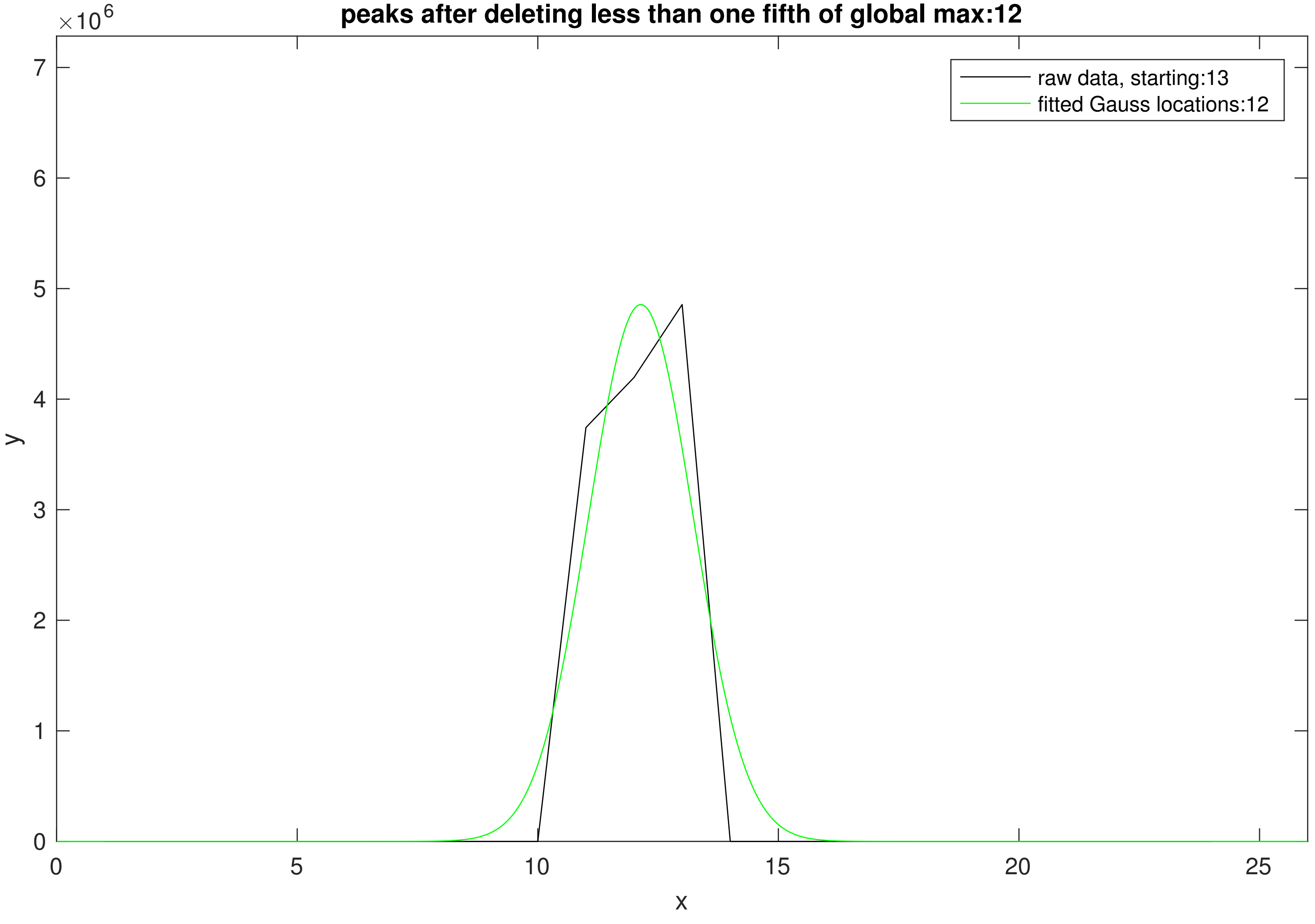

#### Optimal1 Fitted Gauss, proteinAT3G51550.1
#### bic:866 872
#### peaks after deleting less than one fifth of global max:14

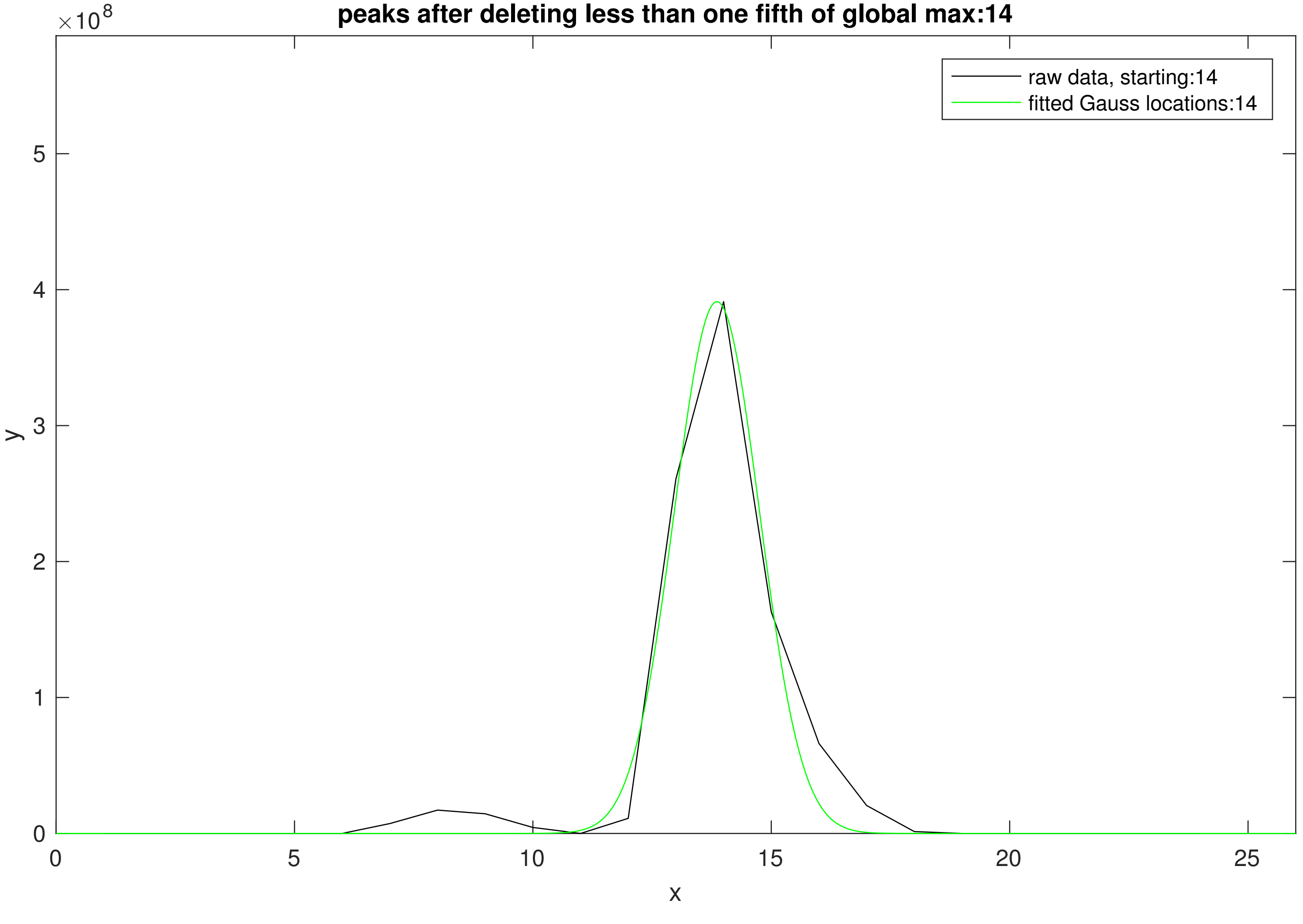

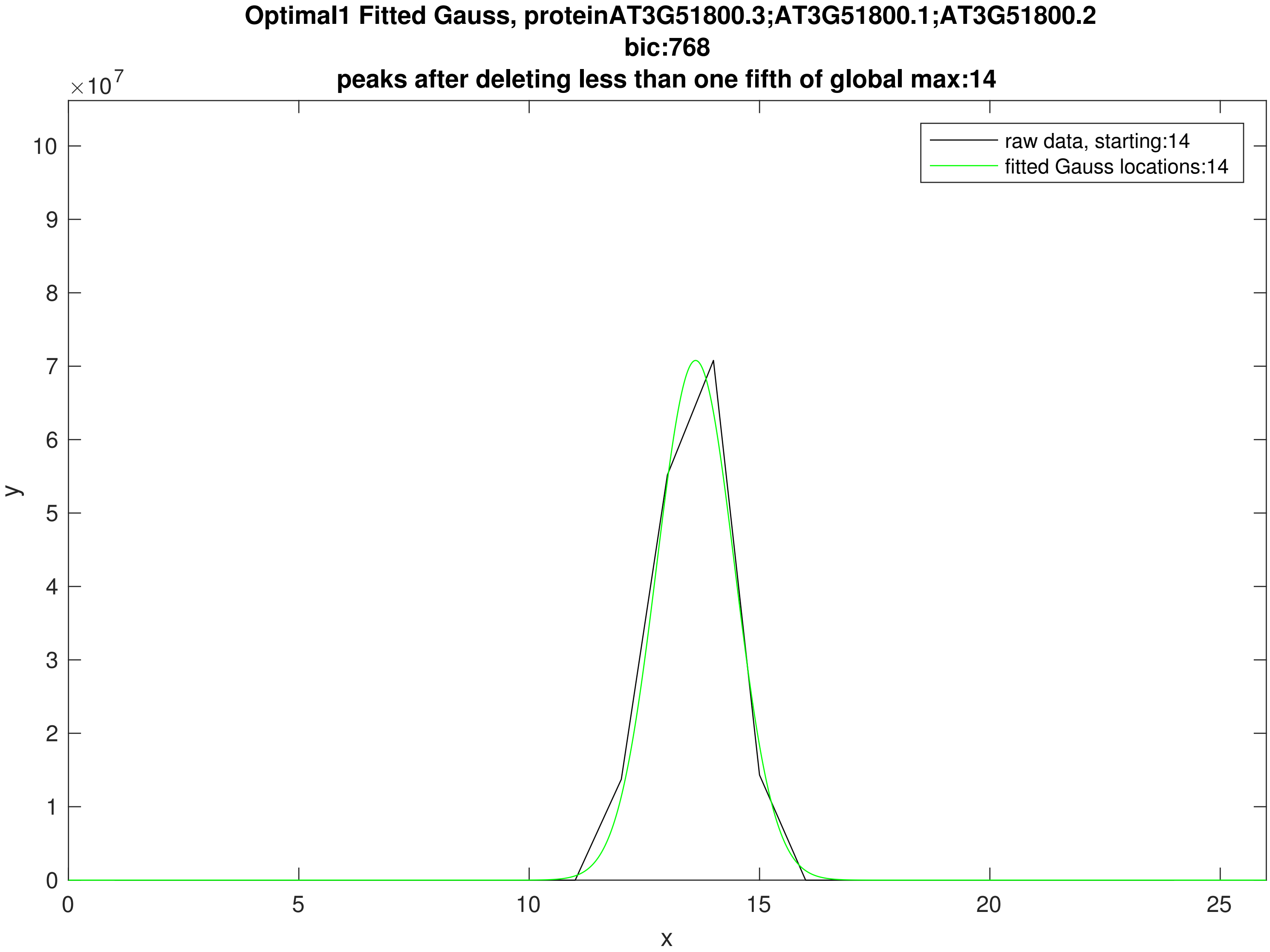

Optimal1 Fitted Gauss, proteinAT3G51800.3;AT3G51800.1;AT3G51800.2
bic:768
peaks after deleting less than one fifth of global max:14
×10^7
10
9
8
7
6
5
4
3
2
1
0
y
0
5
10
15
20
25
x
raw data, starting:14
fitted Gauss locations:14

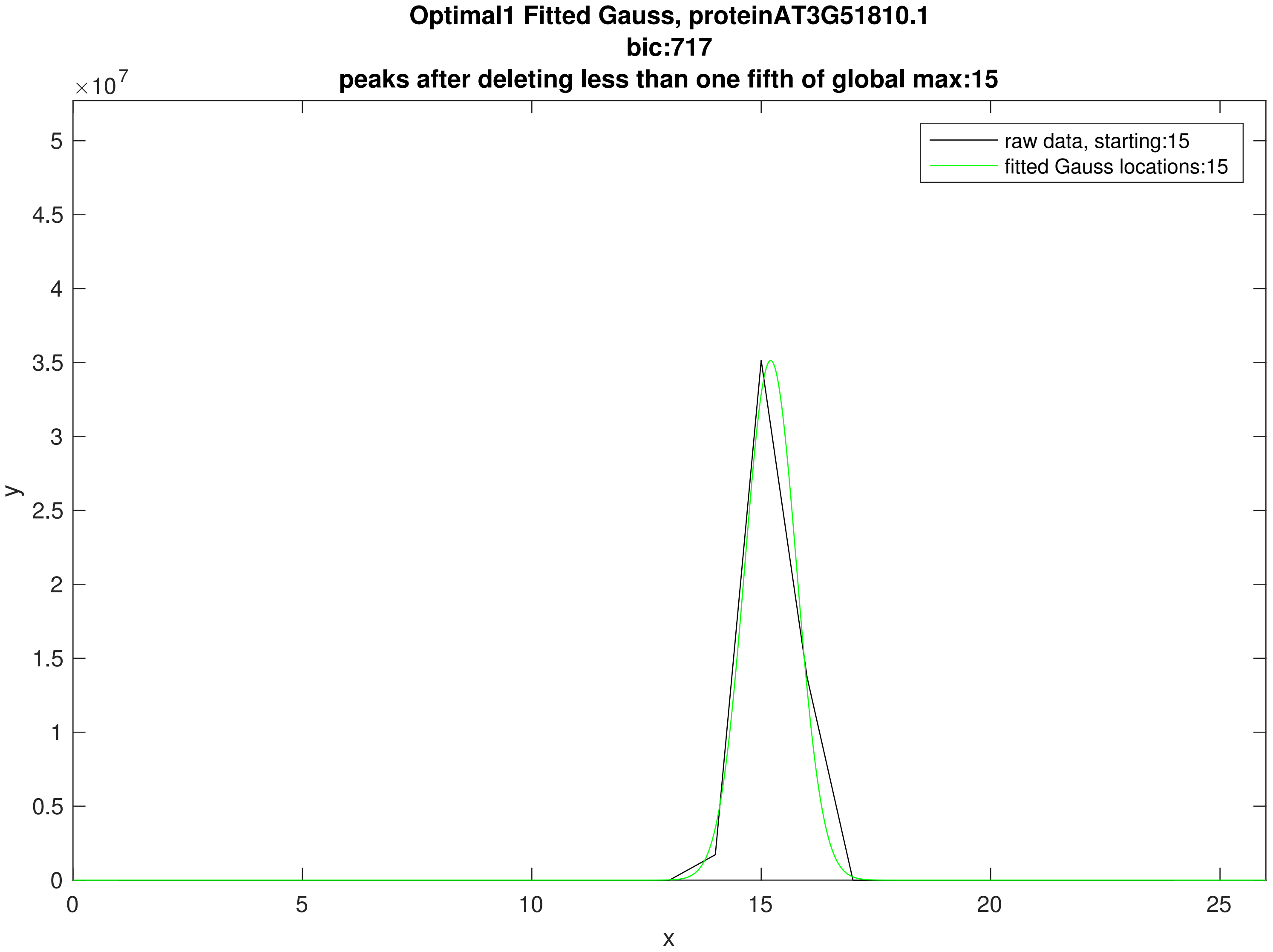

Optimal1 Fitted Gauss, proteinAT3G51810.1
bic:717
peaks after deleting less than one fifth of global max:15
×10^7
raw data, starting:15
fitted Gauss locations:15
y
x
0
0.5
1
1.5
2
2.5
3
3.5
4
4.5
5
0
5
10
15
20
25

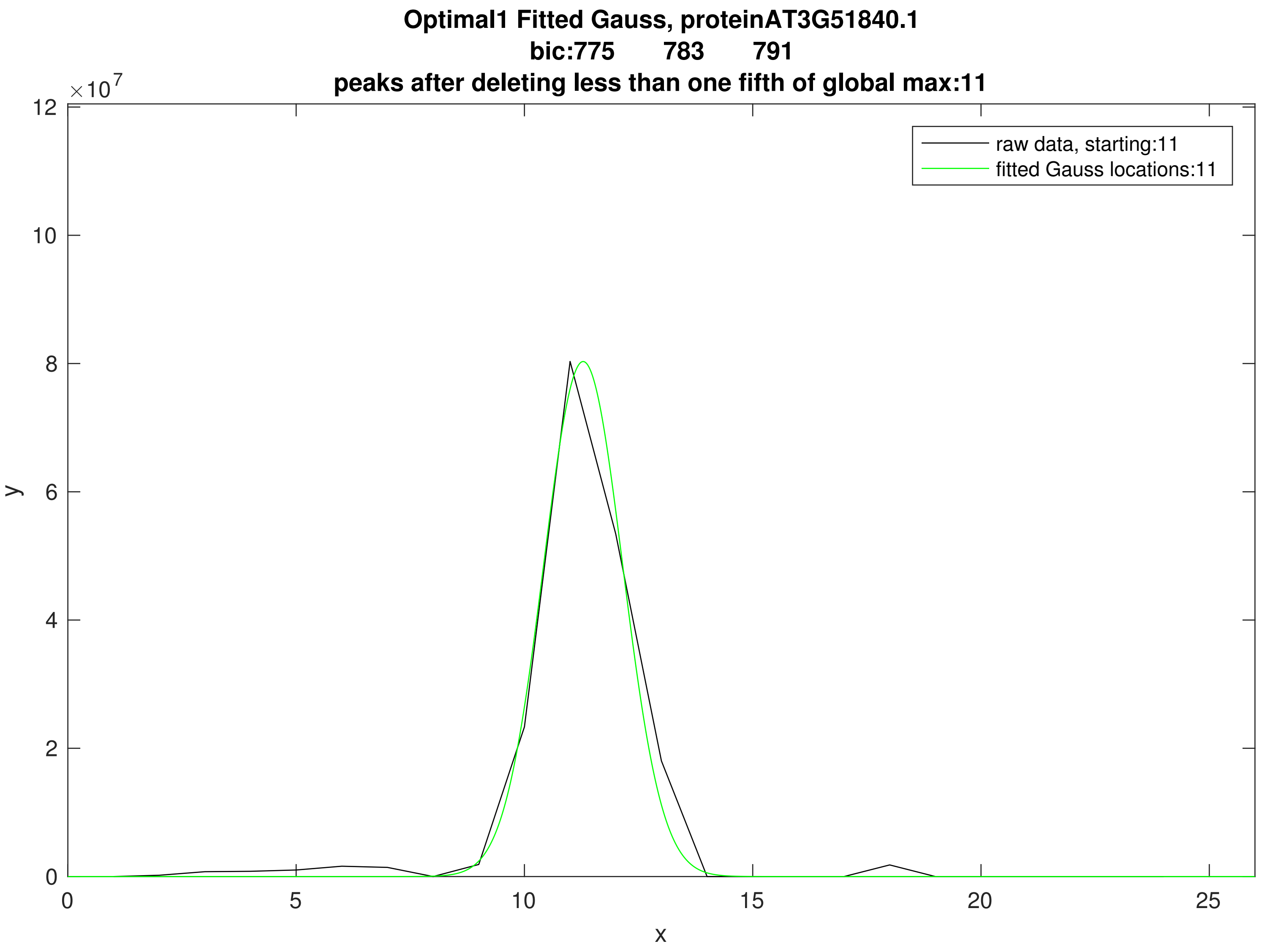
Optimal1 Fitted Gauss, proteinAT3G51840.1
bic:775 783 791
peaks after deleting less than one fifth of global max:11
×10^7
raw data, starting:11
fitted Gauss locations:11
12
10
8
6
4
2
0
y
0
5
10
15
20
25
x

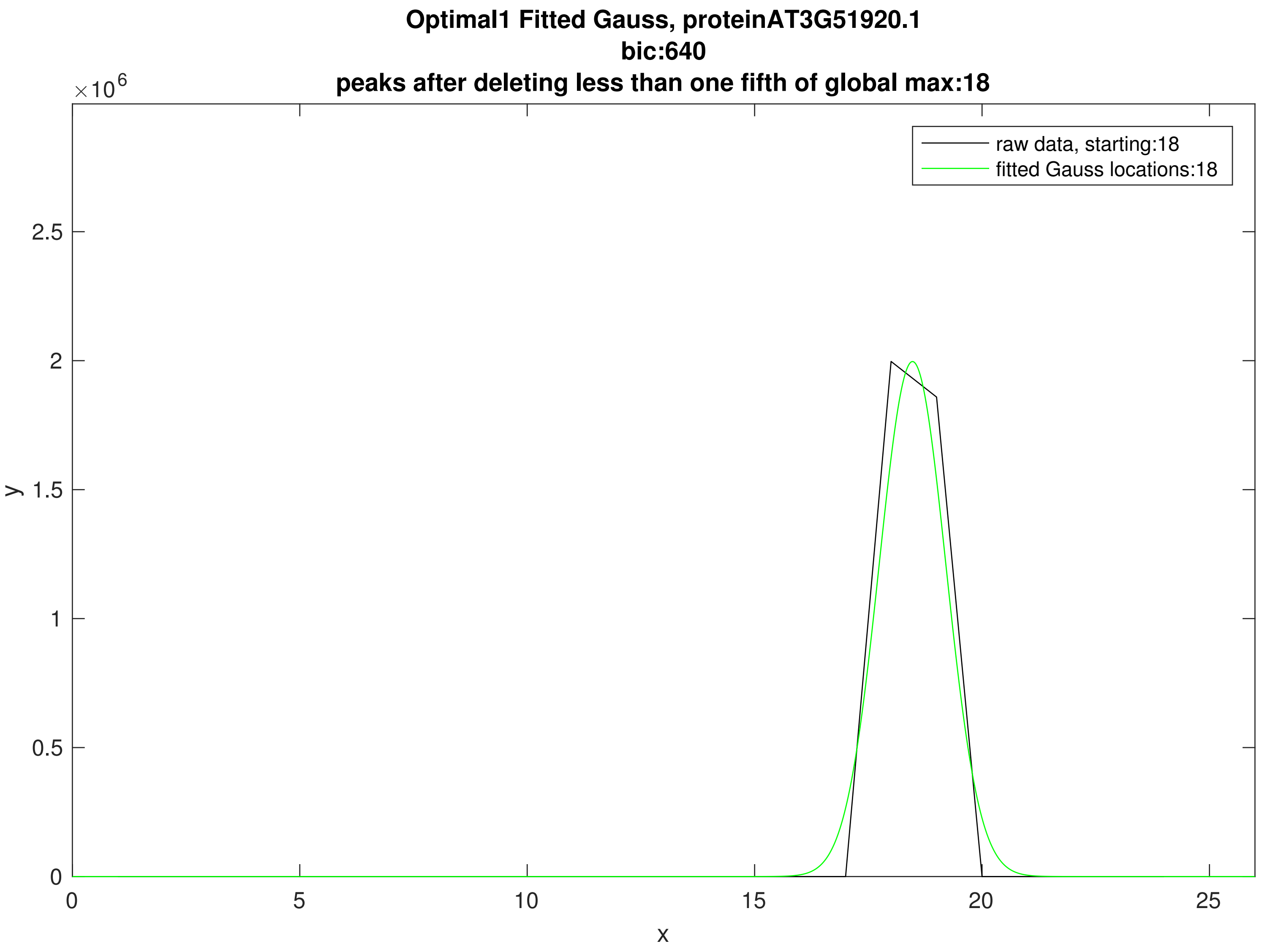

Optimal1 Fitted Gauss, proteinAT3G51920.1
bic:640
peaks after deleting less than one fifth of global max:18
×10^6
raw data, starting:18
fitted Gauss locations:18
2.5
2
1.5
1
0.5
0
y
0
5
10
15
20
25
x

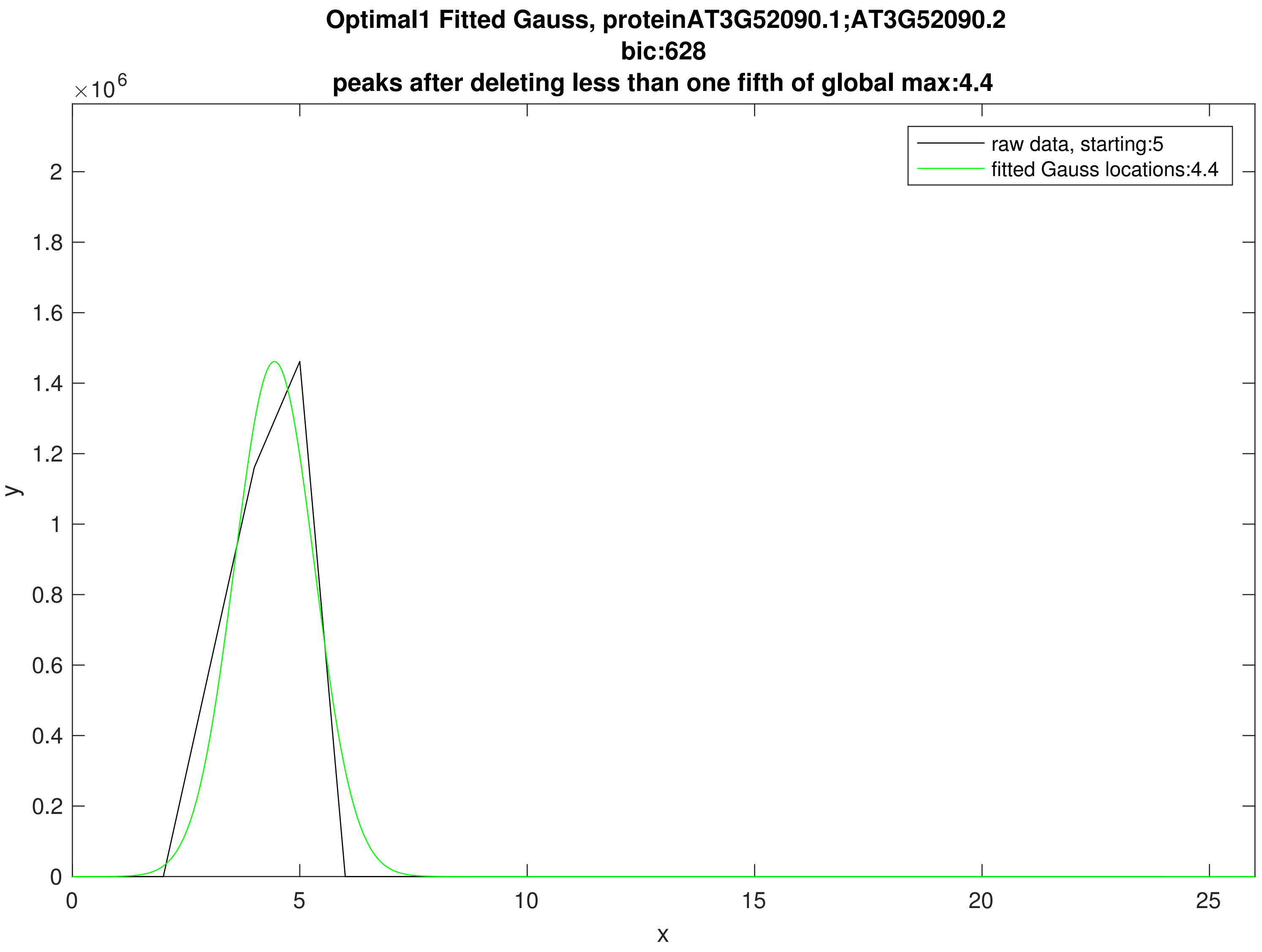

Optimal1 Fitted Gauss, proteinAT3G52090.1;AT3G52090.2
bic:628
peaks after deleting less than one fifth of global max:4.4
×10^6
raw data, starting:5
fitted Gauss locations:4.4
y
x
0
0.2
0.4
0.6
0.8
1
1.2
1.4
1.6
1.8
2
5
10
15
20
25

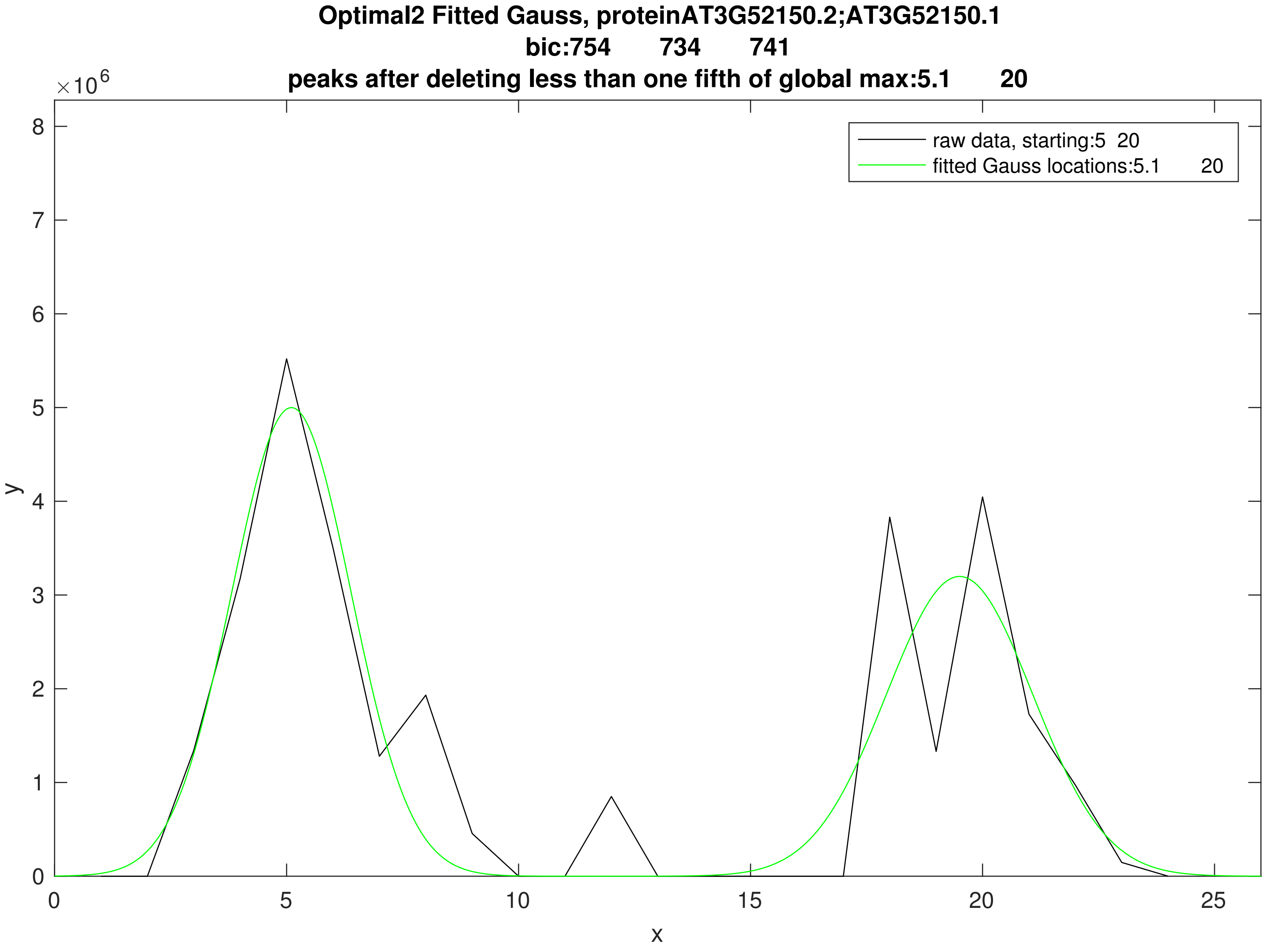

Optimal2 Fitted Gauss, proteinAT3G52150.2;AT3G52150.1
bic:754 734 741
peaks after deleting less than one fifth of global max:5.1 20
×10^6
raw data, starting:5 20
fitted Gauss locations:5.1 20
y
x
0
1
2
3
4
5
6
7
8
0
5
10
15
20
25

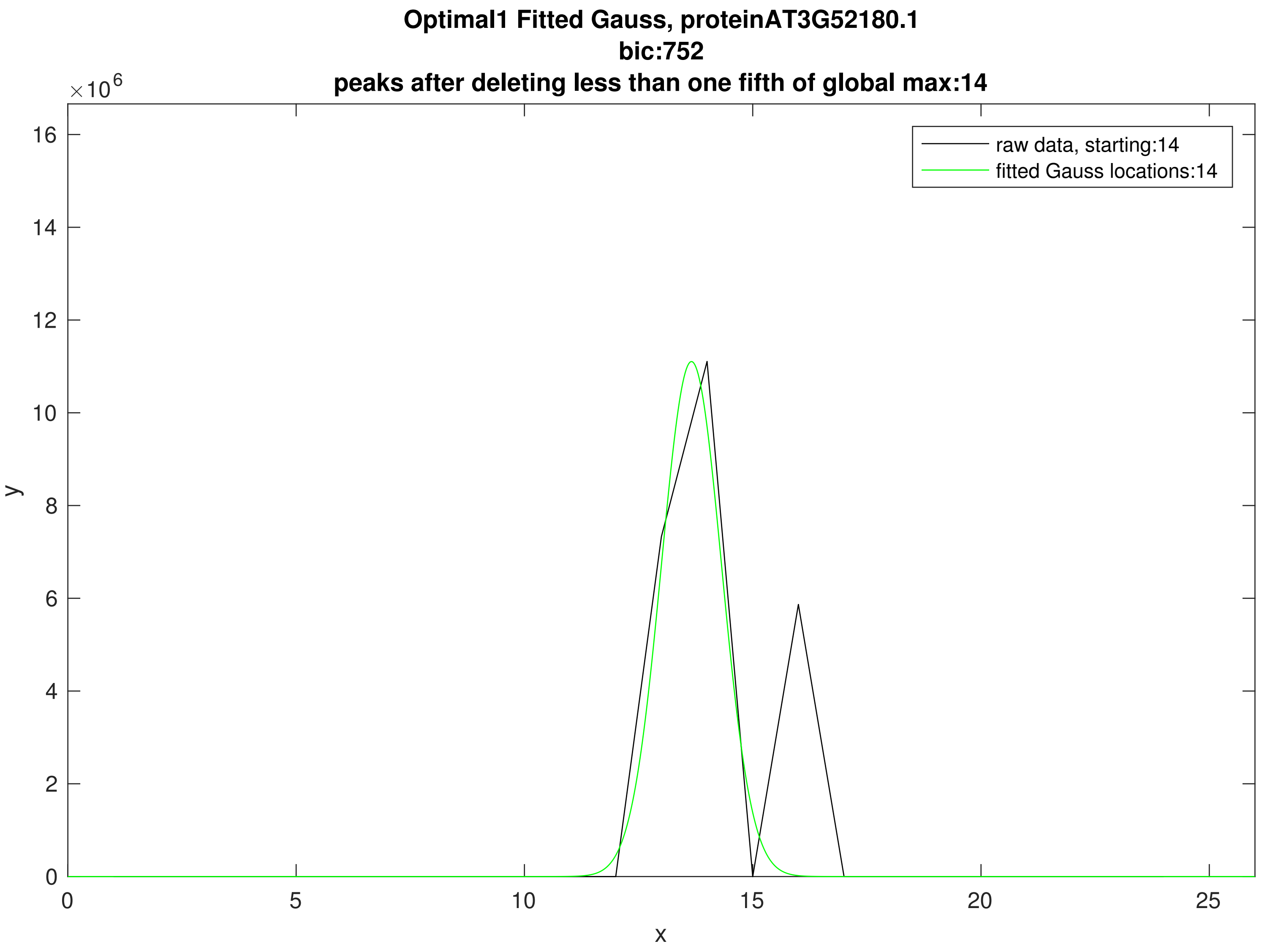
Optimal1 Fitted Gauss, proteinAT3G52180.1
bic:752
peaks after deleting less than one fifth of global max:14
×10^6
16
14
12
10
8
6
4
2
0
y
0
5
10
15
20
25
x
raw data, starting:14
fitted Gauss locations:14

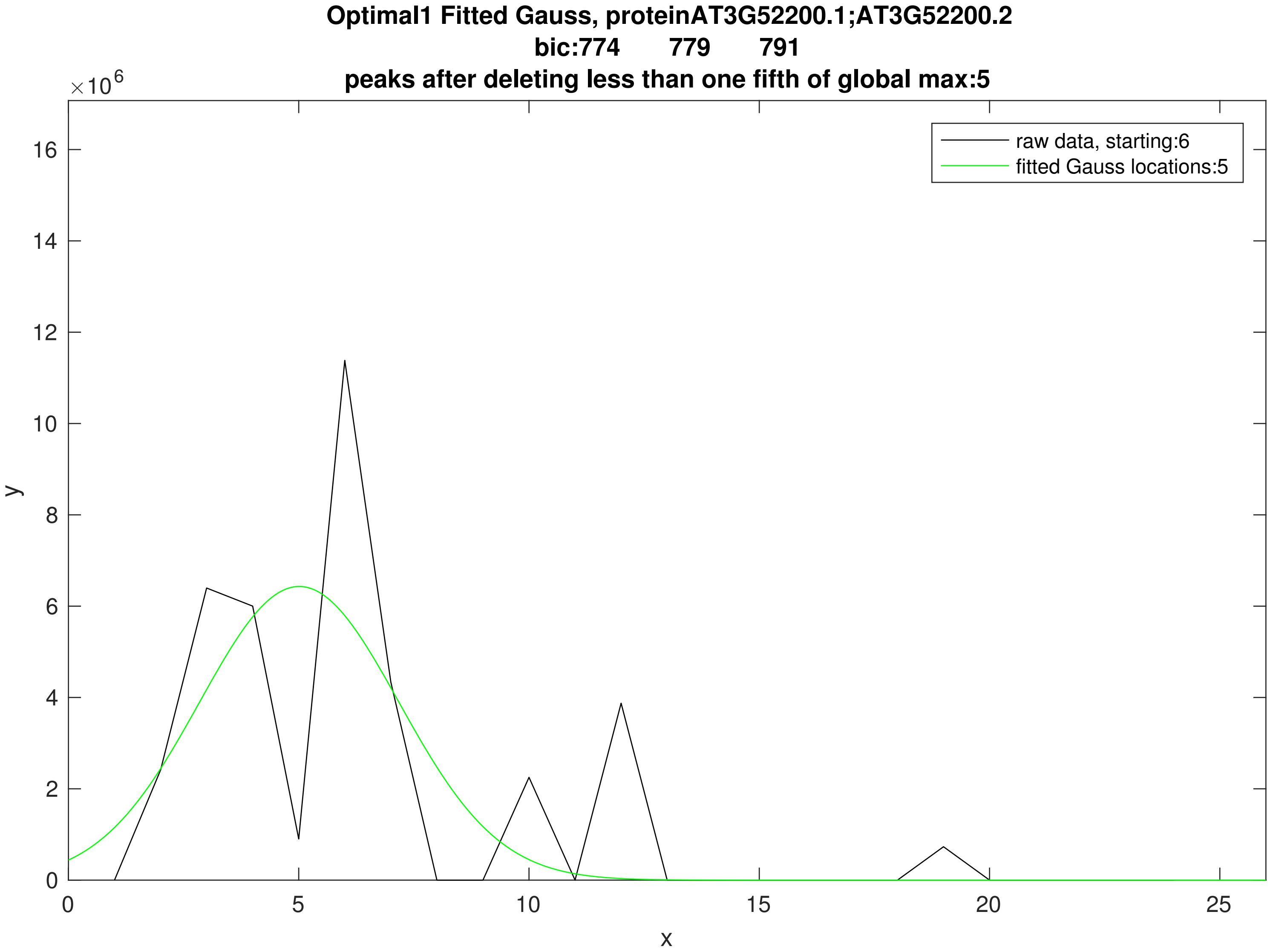

Optimal1 Fitted Gauss, proteinAT3G52200.1;AT3G52200.2
bic:774 779 791
peaks after deleting less than one fifth of global max:5
×10^6
raw data, starting:6
fitted Gauss locations:5
y
x
0
2
4
6
8
10
12
14
16
0
5
10
15
20
25

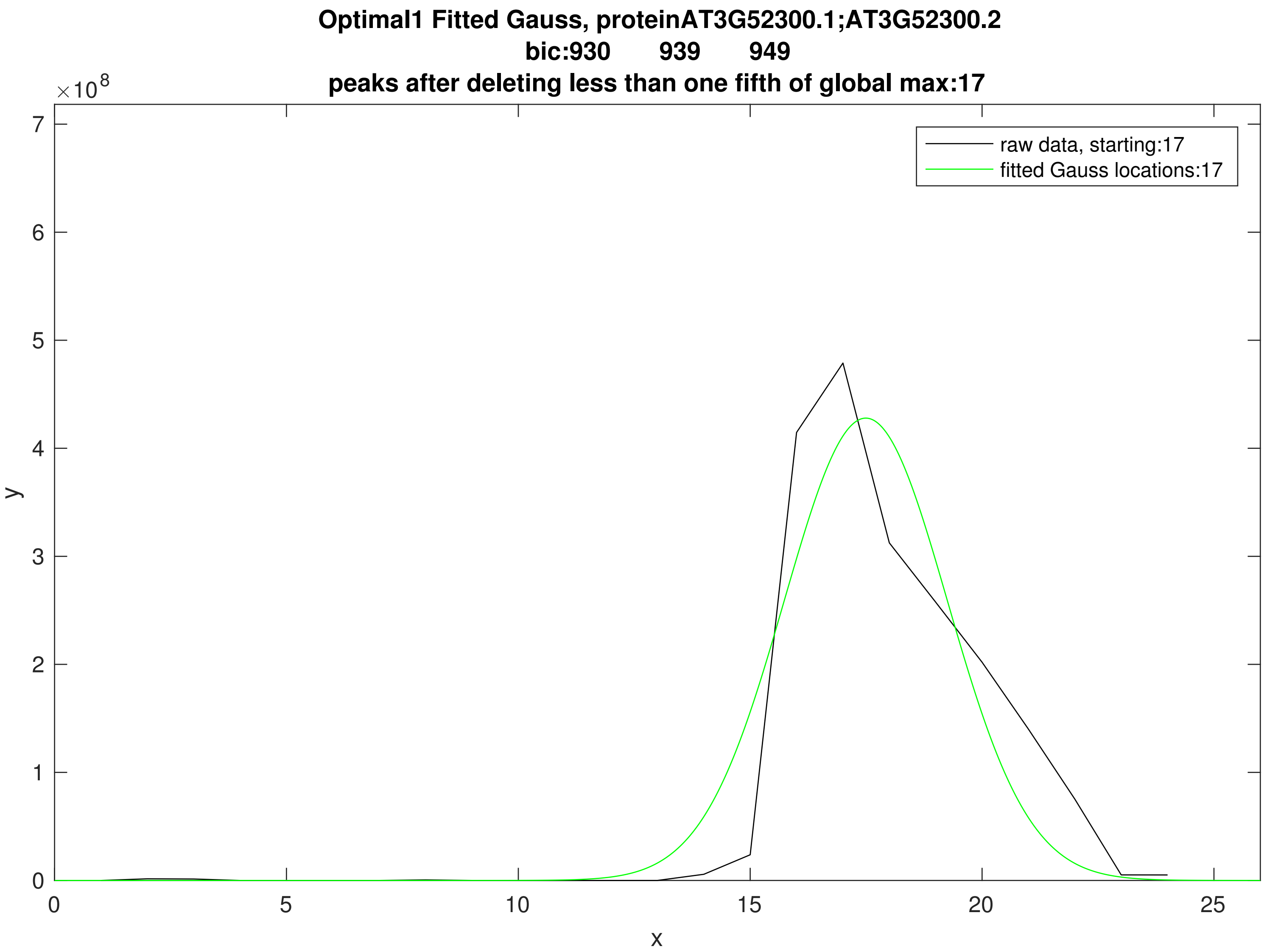
Optimal1 Fitted Gauss, proteinAT3G52300.1;AT3G52300.2
bic:930 939 949
peaks after deleting less than one fifth of global max:17
×10^8
raw data, starting:17
fitted Gauss locations:17
y
x
0
1
2
3
4
5
6
7
10
15
20
25

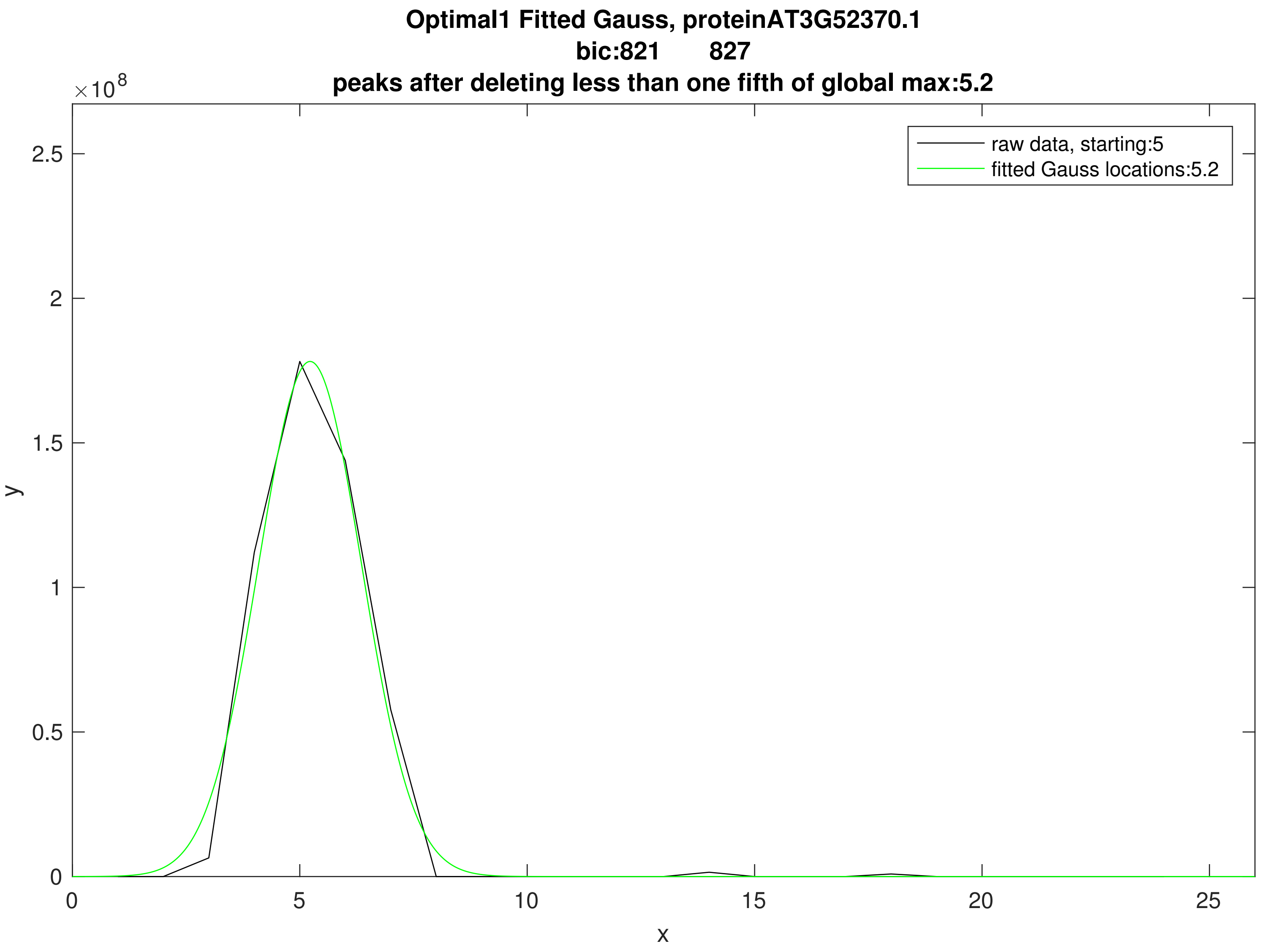
Optimal1 Fitted Gauss, proteinAT3G52370.1
bic:821 827
peaks after deleting less than one fifth of global max:5.2
×10^8
2.5
2
1.5
1
0.5
0
y
0
5
10
15
20
25
x
raw data, starting:5
fitted Gauss locations:5.2

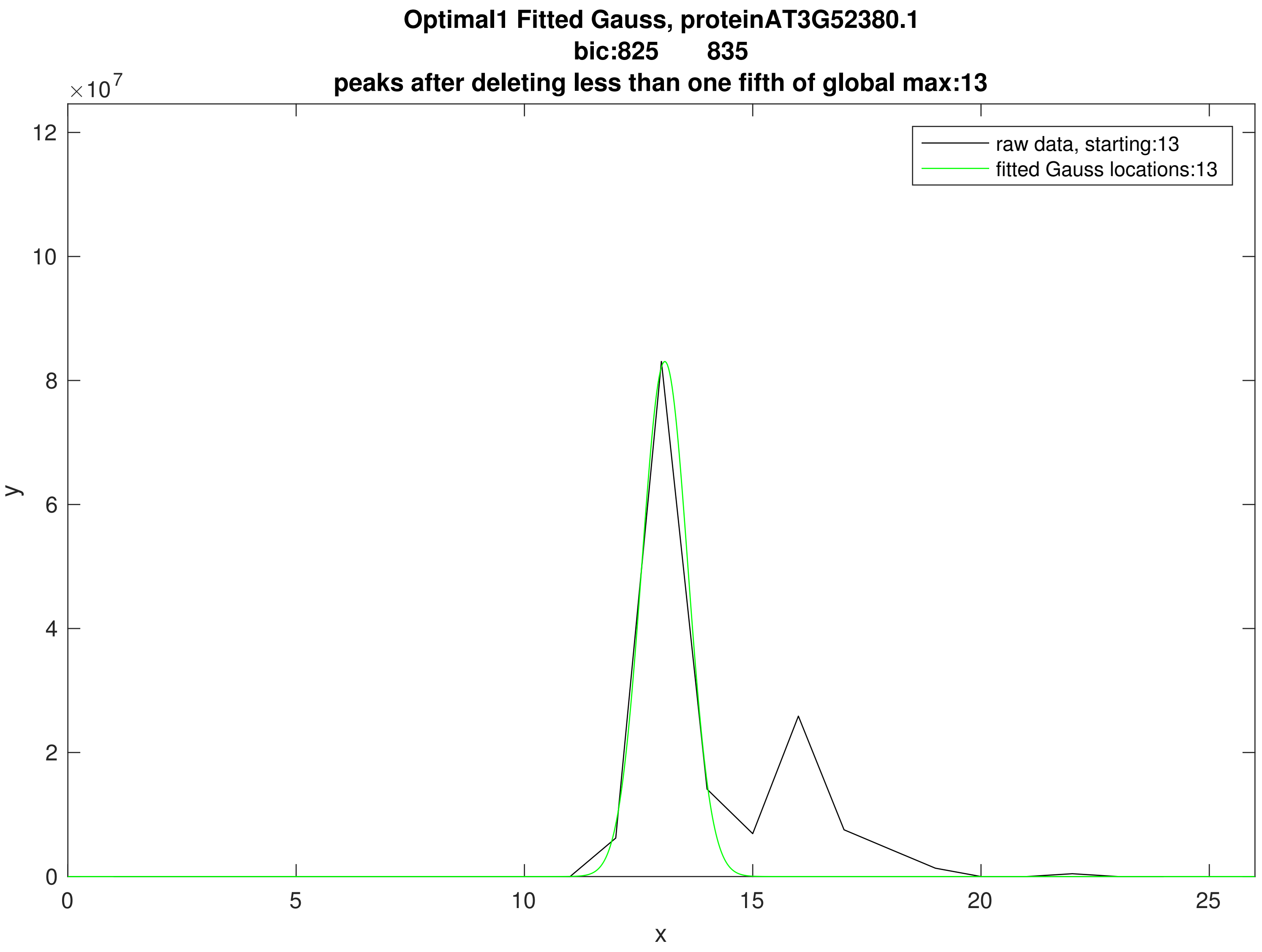
Optimal1 Fitted Gauss, proteinAT3G52380.1
bic:825 835
peaks after deleting less than one fifth of global max:13
×10^7
raw data, starting:13
fitted Gauss locations:13
y
x
0
2
4
6
8
10
12
0
5
10
15
20
25

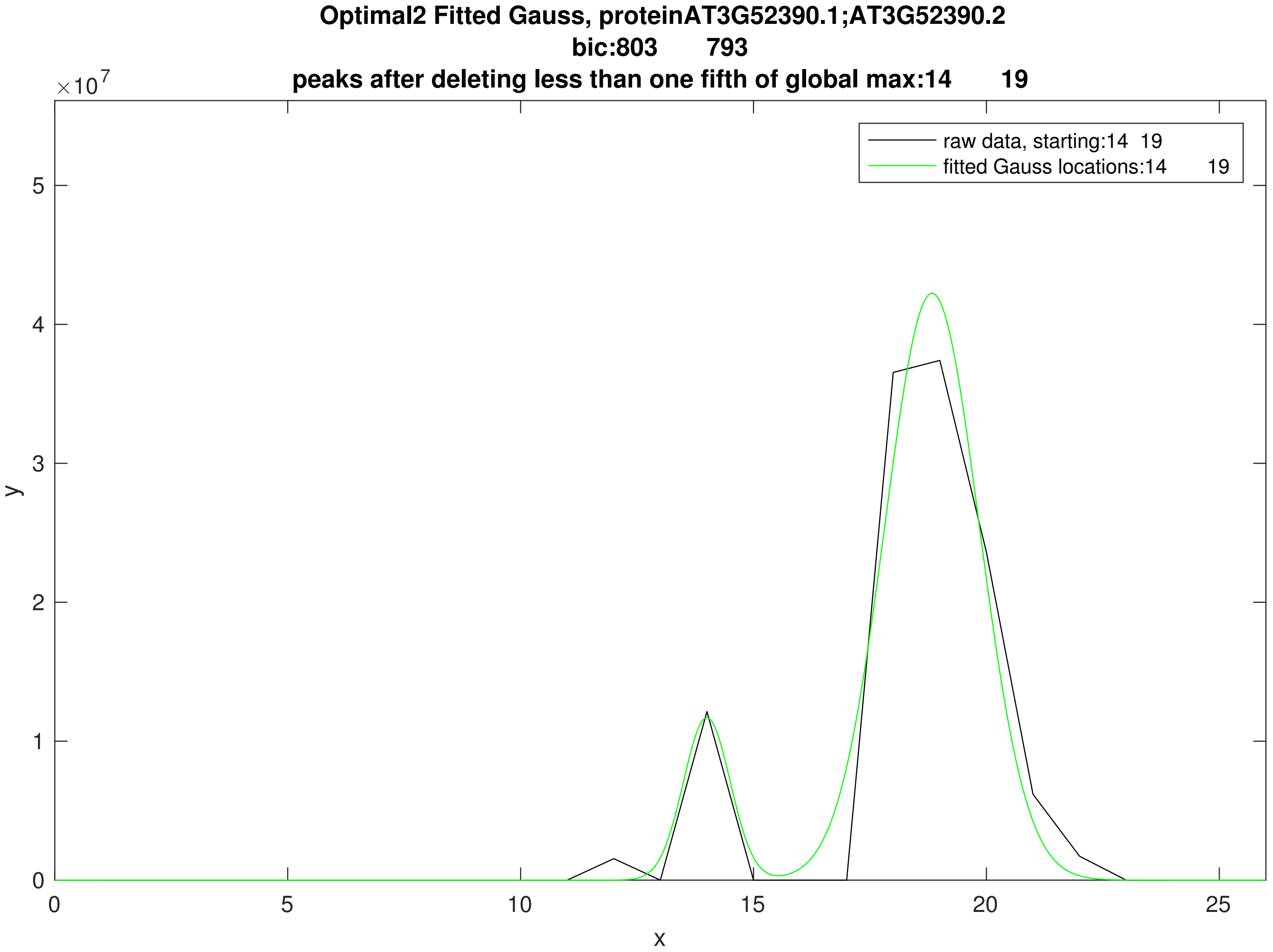

Optimal2 Fitted Gauss, proteinAT3G52390.1;AT3G52390.2
bic:803 793
peaks after deleting less than one fifth of global max:14 19
×10^7
raw data, starting:14 19
fitted Gauss locations:14 19
y
x
0
1
2
3
4
5
0
5
10
15
20
25

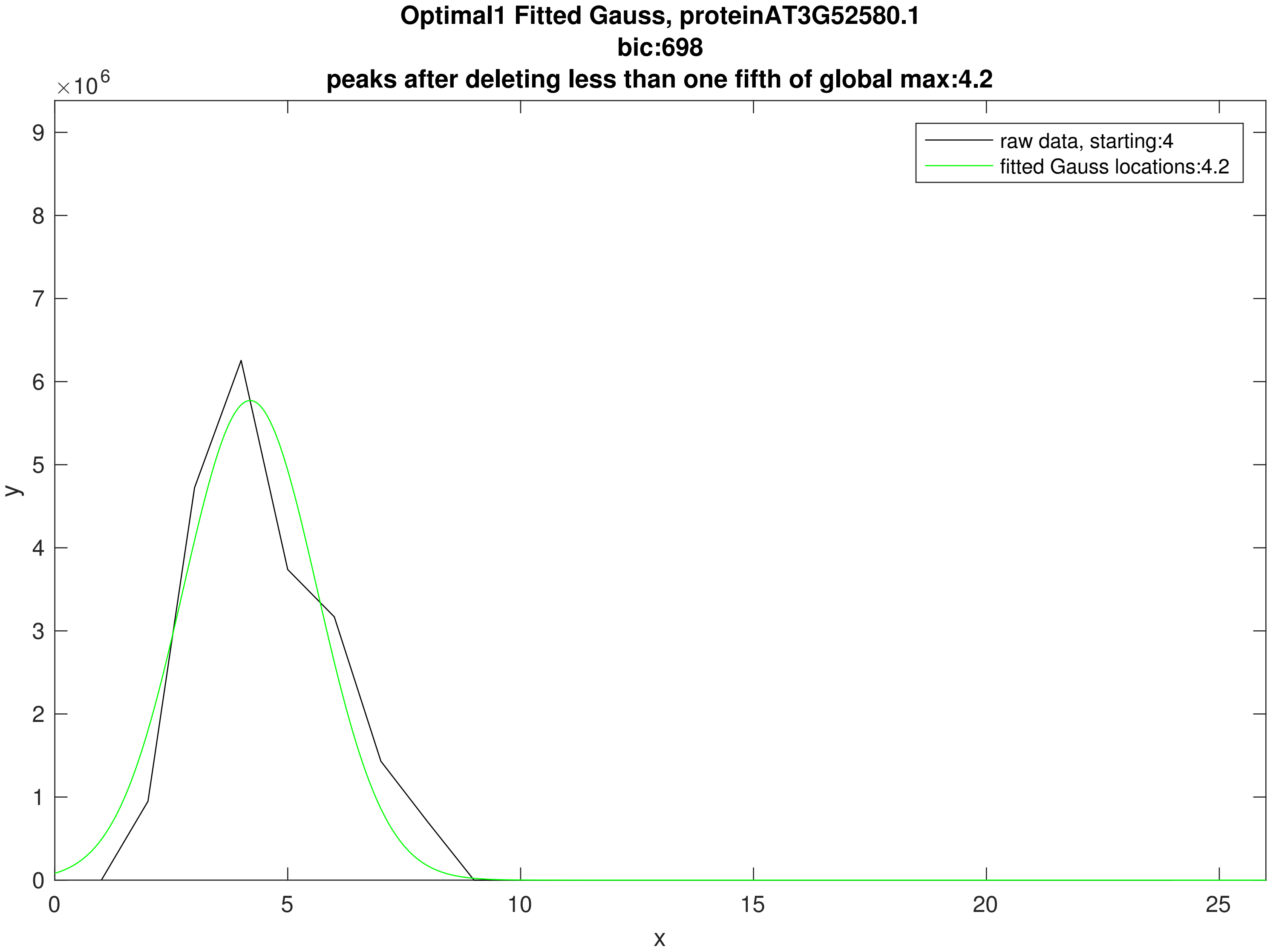
Optimal1 Fitted Gauss, proteinAT3G52580.1
bic:698
peaks after deleting less than one fifth of global max:4.2
×10^6
raw data, starting:4
fitted Gauss locations:4.2
y
x
0
1
2
3
4
5
6
7
8
9
0
5
10
15
20
25

Optimal1 Fitted Gauss, proteinAT3G52610.1
bic:714
peaks after deleting less than one fifth of global max:17

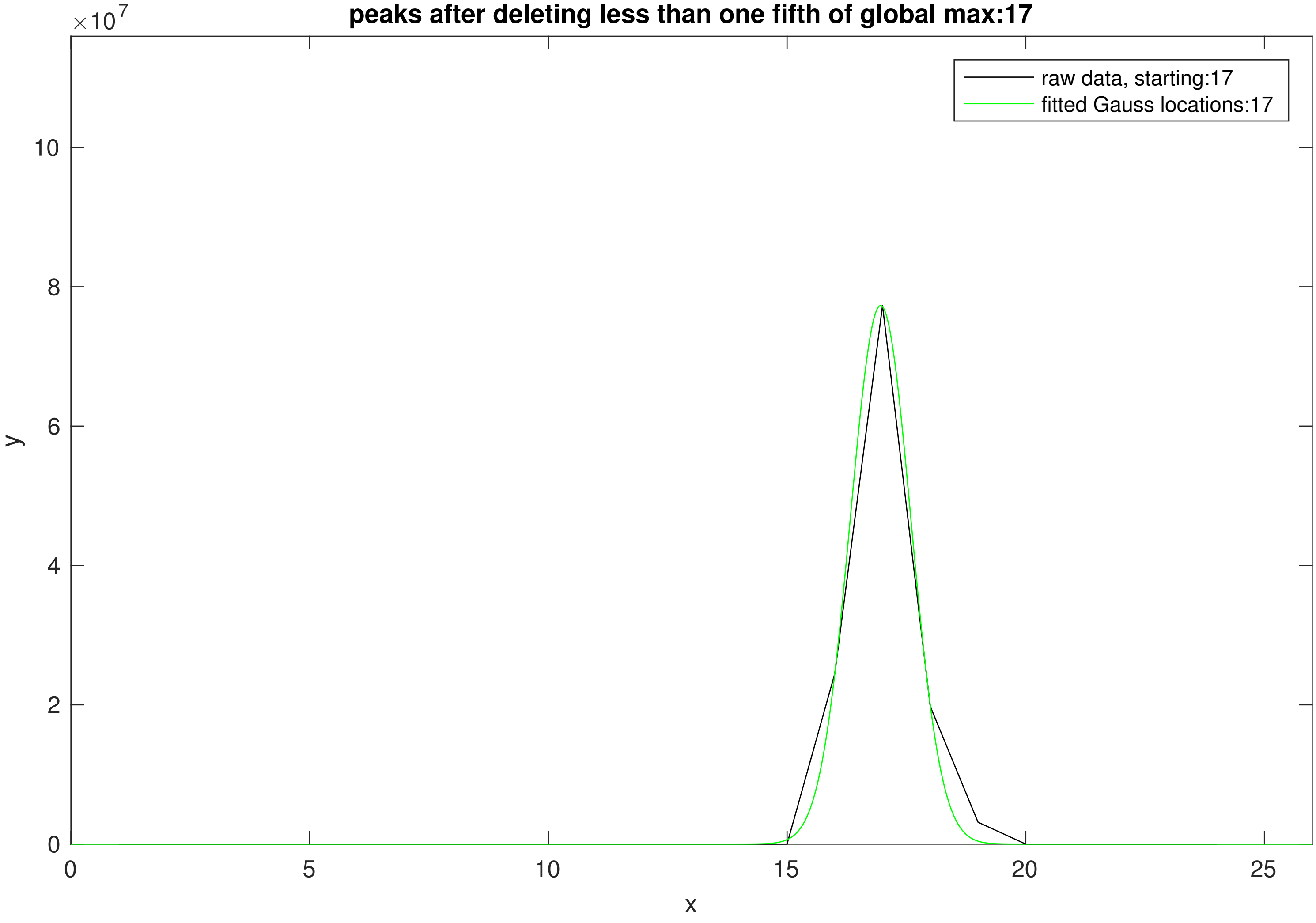

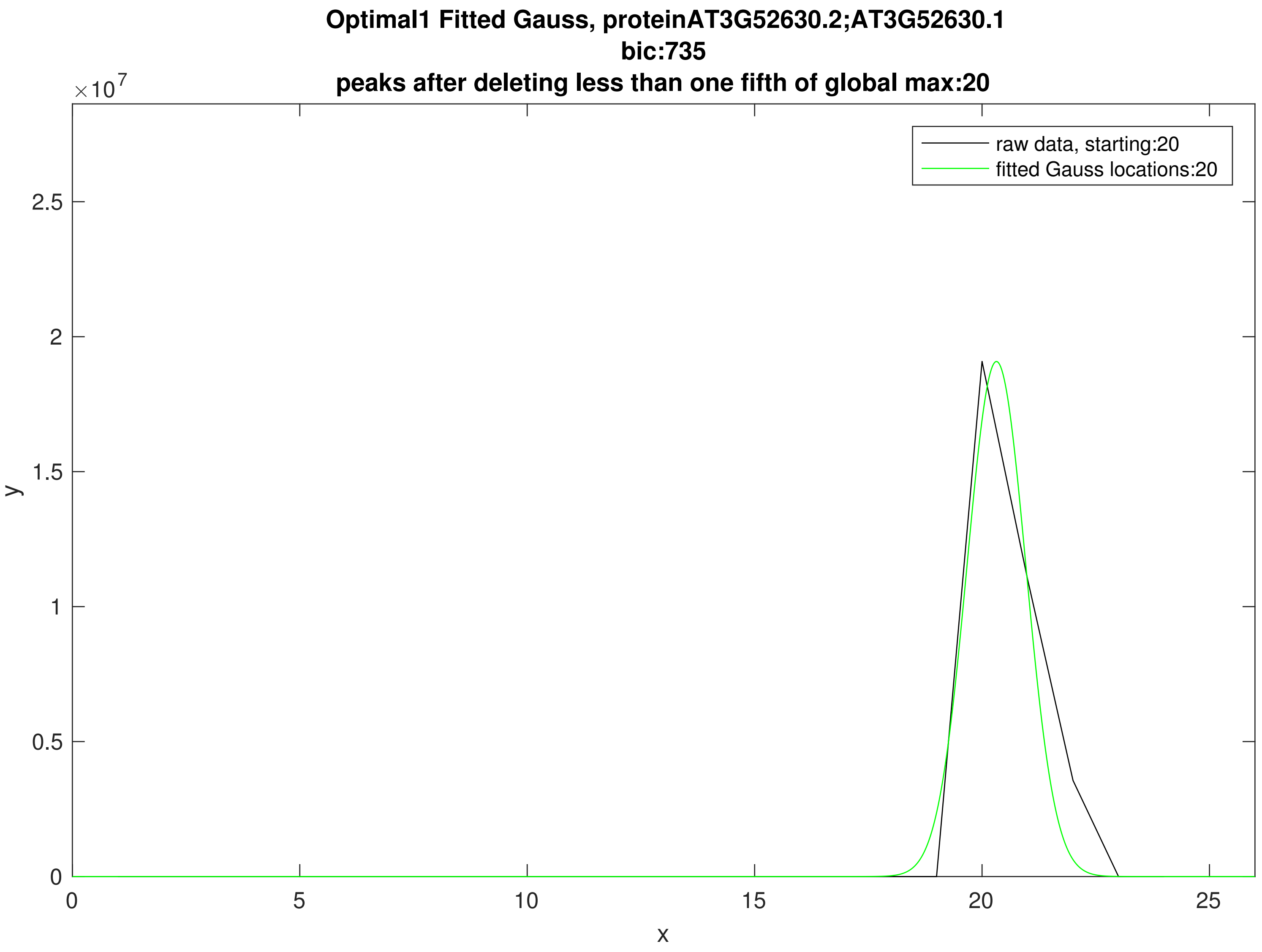
Optimal1 Fitted Gauss, proteinAT3G52630.2;AT3G52630.1
bic:735
peaks after deleting less than one fifth of global max:20
×10^7
2.5
2
1.5
1
0.5
0
y
0
5
10
15
20
25
x
raw data, starting:20
fitted Gauss locations:20

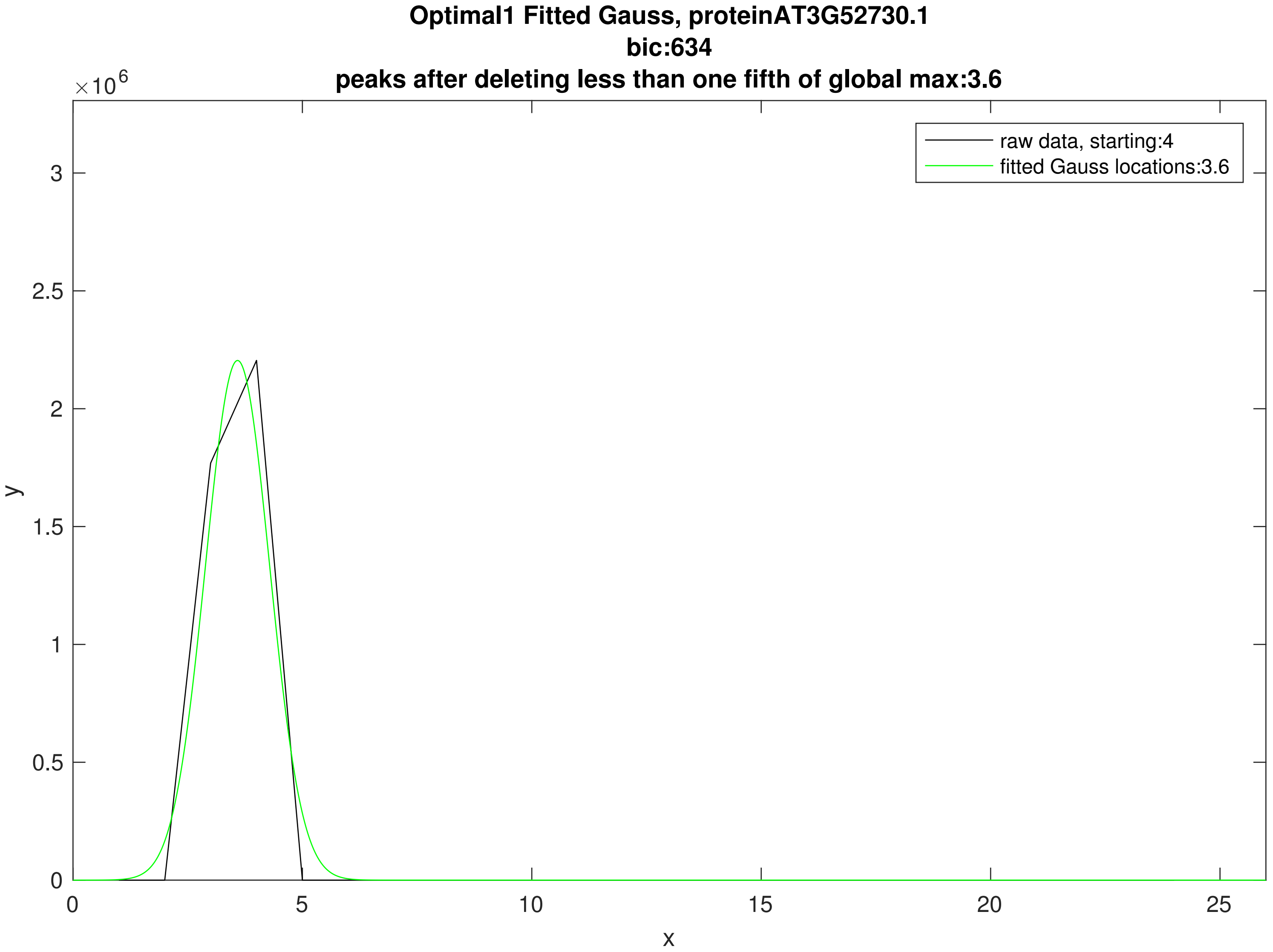
Optimal1 Fitted Gauss, proteinAT3G52730.1
bic:634
peaks after deleting less than one fifth of global max:3.6
×10^6
raw data, starting:4
fitted Gauss locations:3.6
y
x
0
0.5
1
1.5
2
2.5
3
0
5
10
15
20
25

Optimal1 Fitted Gauss, proteinAT3G52840.1
bic:733
peaks after deleting less than one fifth of global max:16

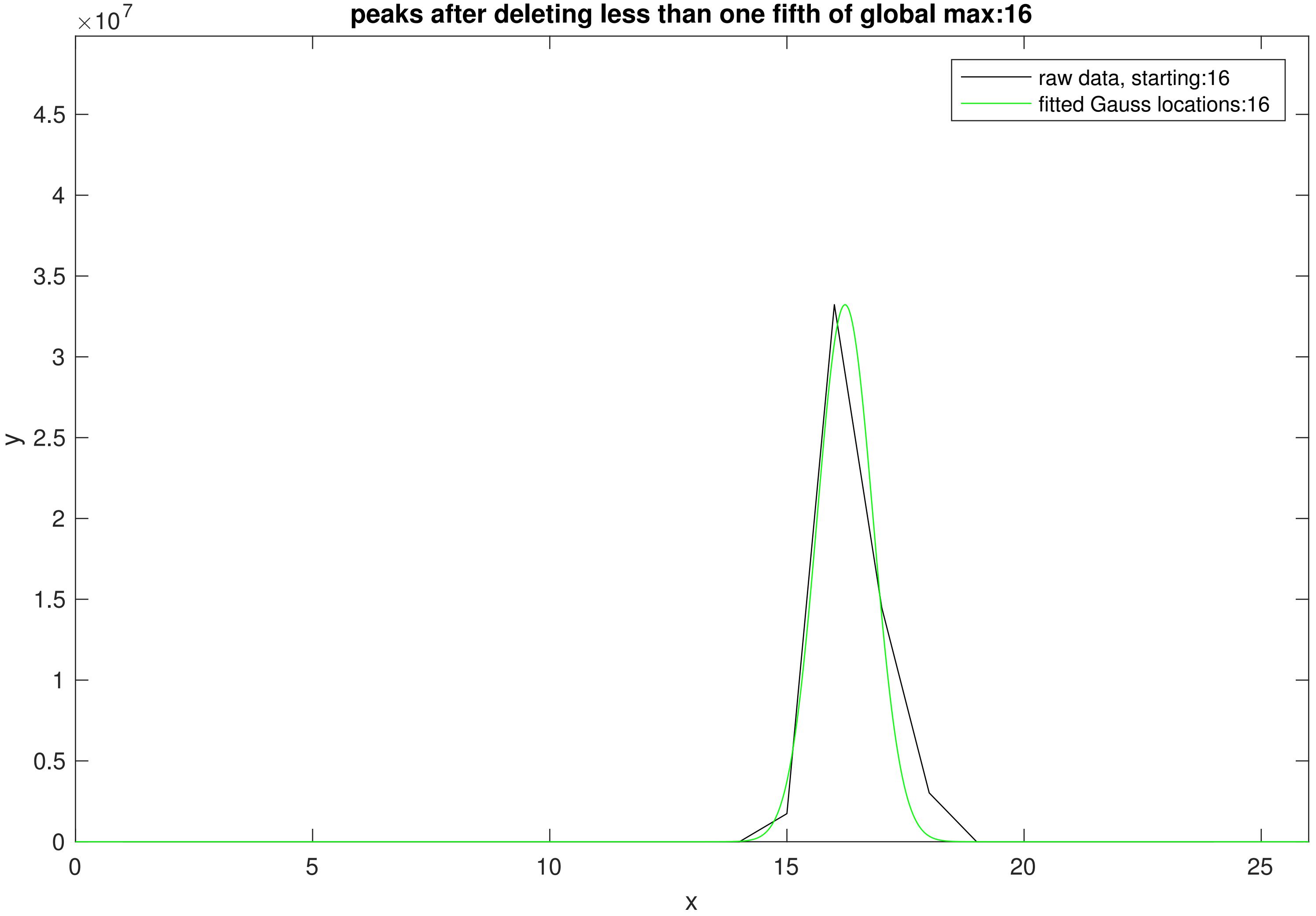

#### Optimal1 Fitted Gauss, proteinAT3G52850.1
#### bic:611
#### peaks after deleting less than one fifth of global max:20

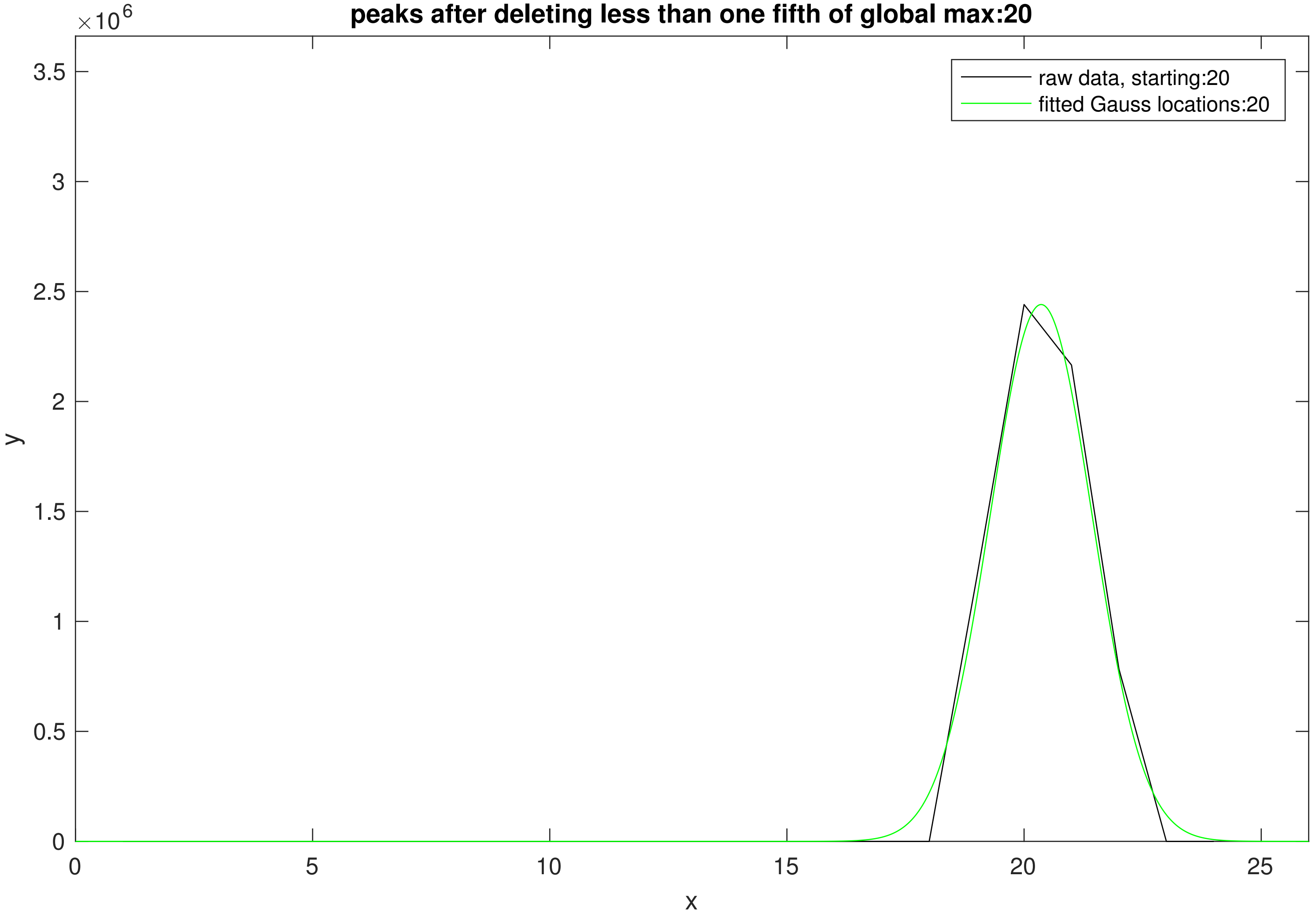

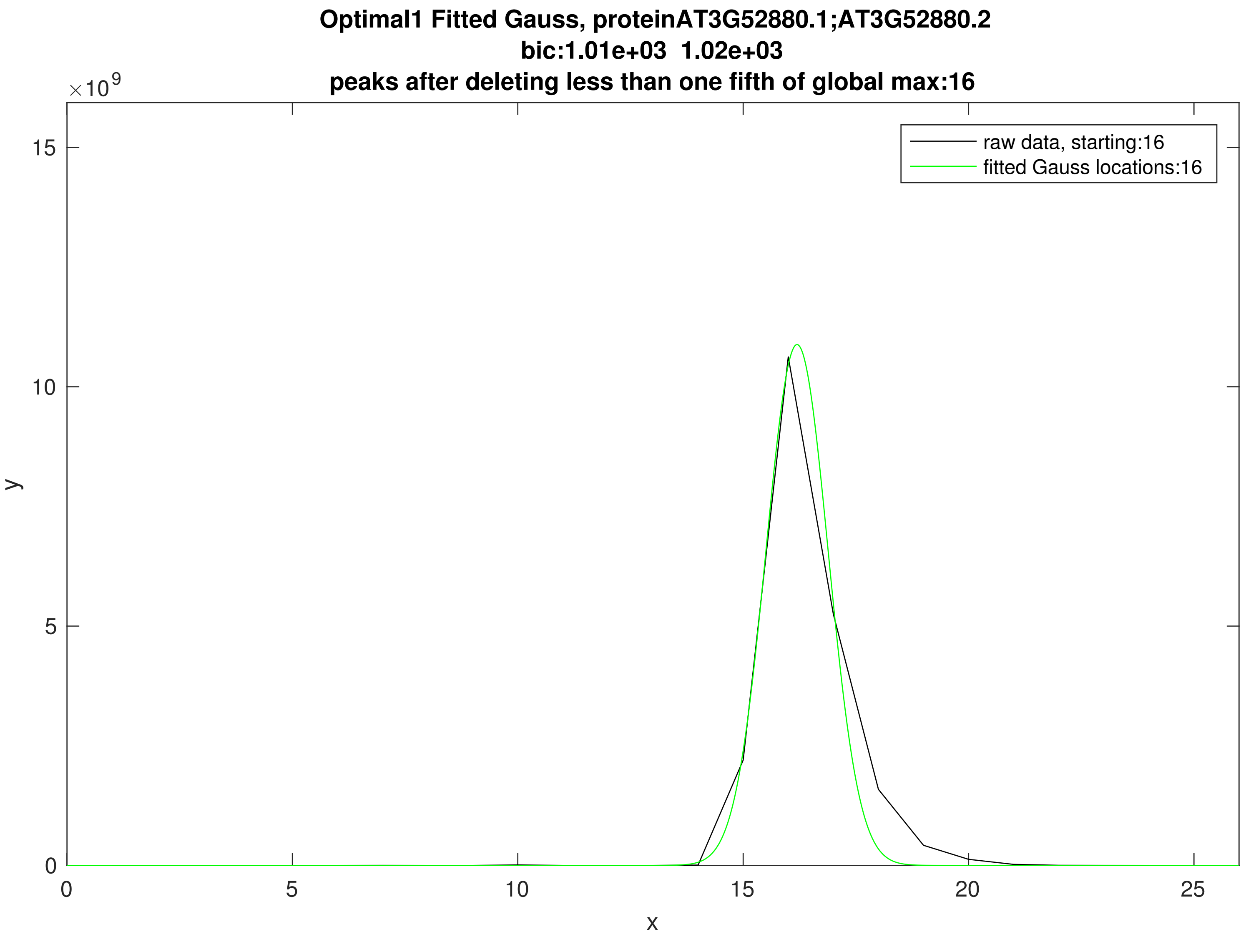
Optimal1 Fitted Gauss, proteinAT3G52880.1;AT3G52880.2
bic:1.01e+03  1.02e+03
peaks after deleting less than one fifth of global max:16
×10^9
raw data, starting:16
fitted Gauss locations:16
y
x
0
5
10
15
20
25

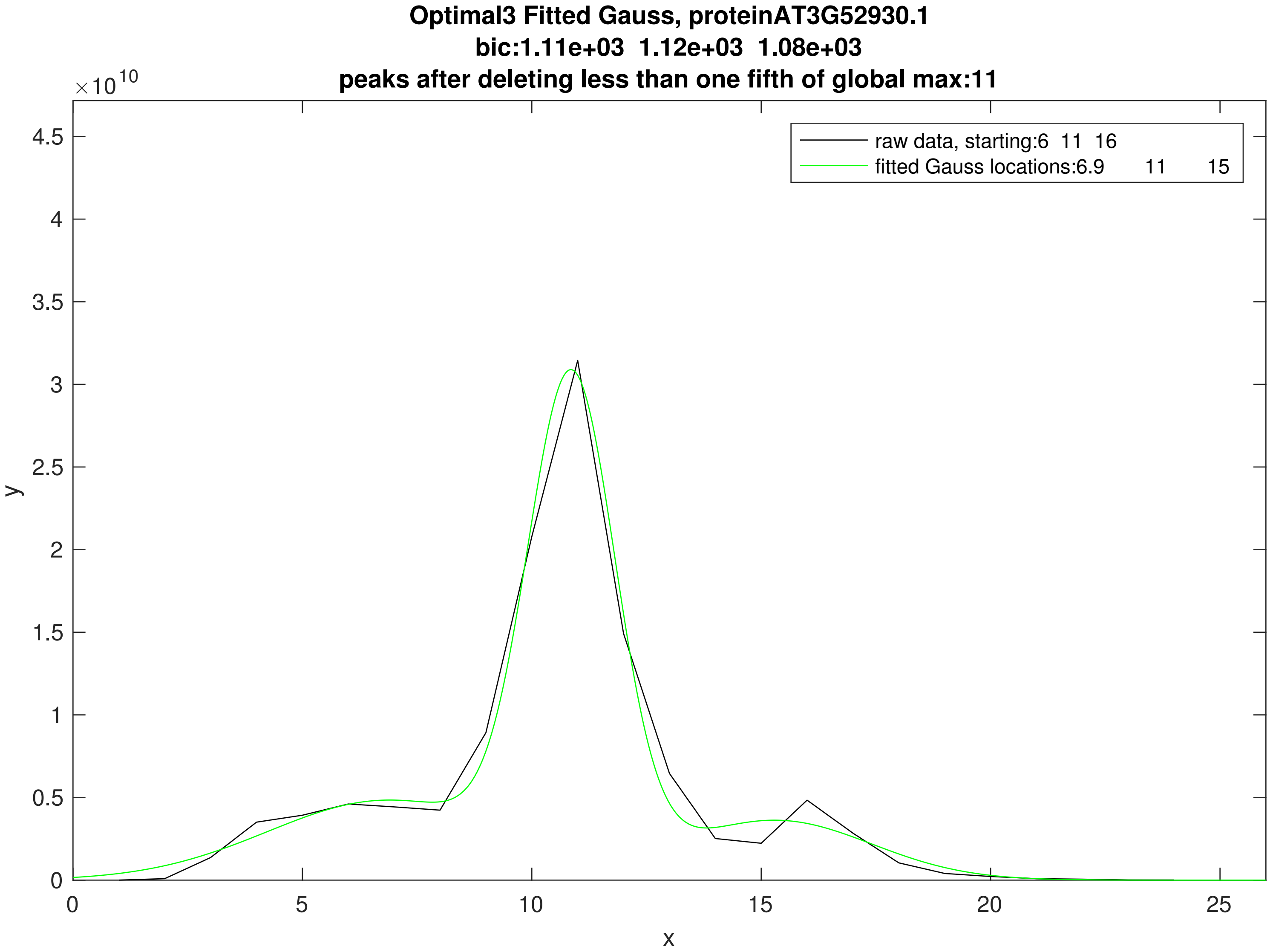

Optimal3 Fitted Gauss, proteinAT3G52930.1
bic:1.11e+03 1.12e+03 1.08e+03
peaks after deleting less than one fifth of global max:11
×10^10
raw data, starting:6 11 16
fitted Gauss locations:6.9 11 15
y
x
0
0.5
1
1.5
2
2.5
3
3.5
4
4.5
0
5
10
15
20
25

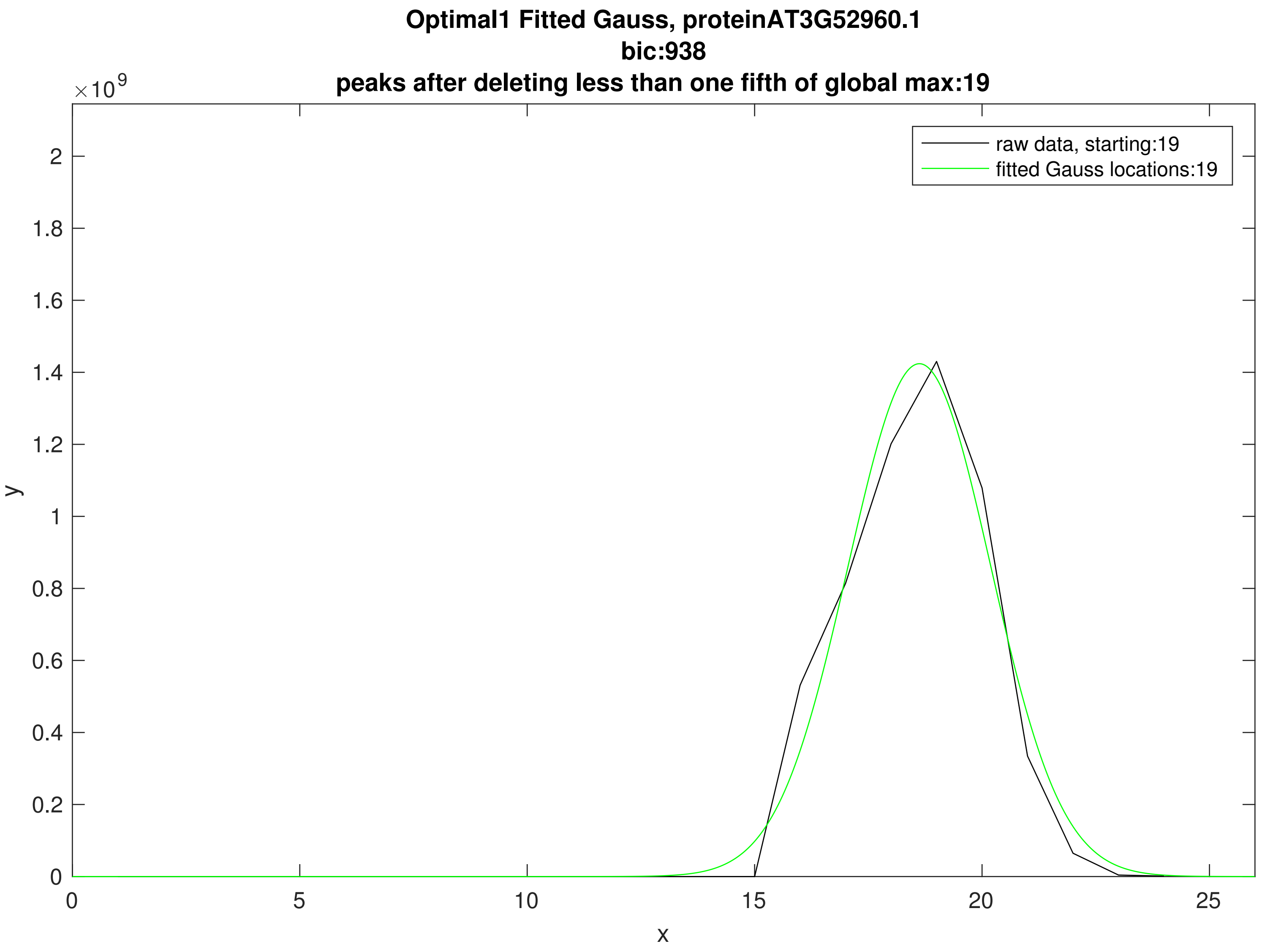
Optimal1 Fitted Gauss, proteinAT3G52960.1
bic:938
peaks after deleting less than one fifth of global max:19
×10^9
raw data, starting:19
fitted Gauss locations:19
y
x
0
0.2
0.4
0.6
0.8
1
1.2
1.4
1.6
1.8
2
0
5
10
15
20
25

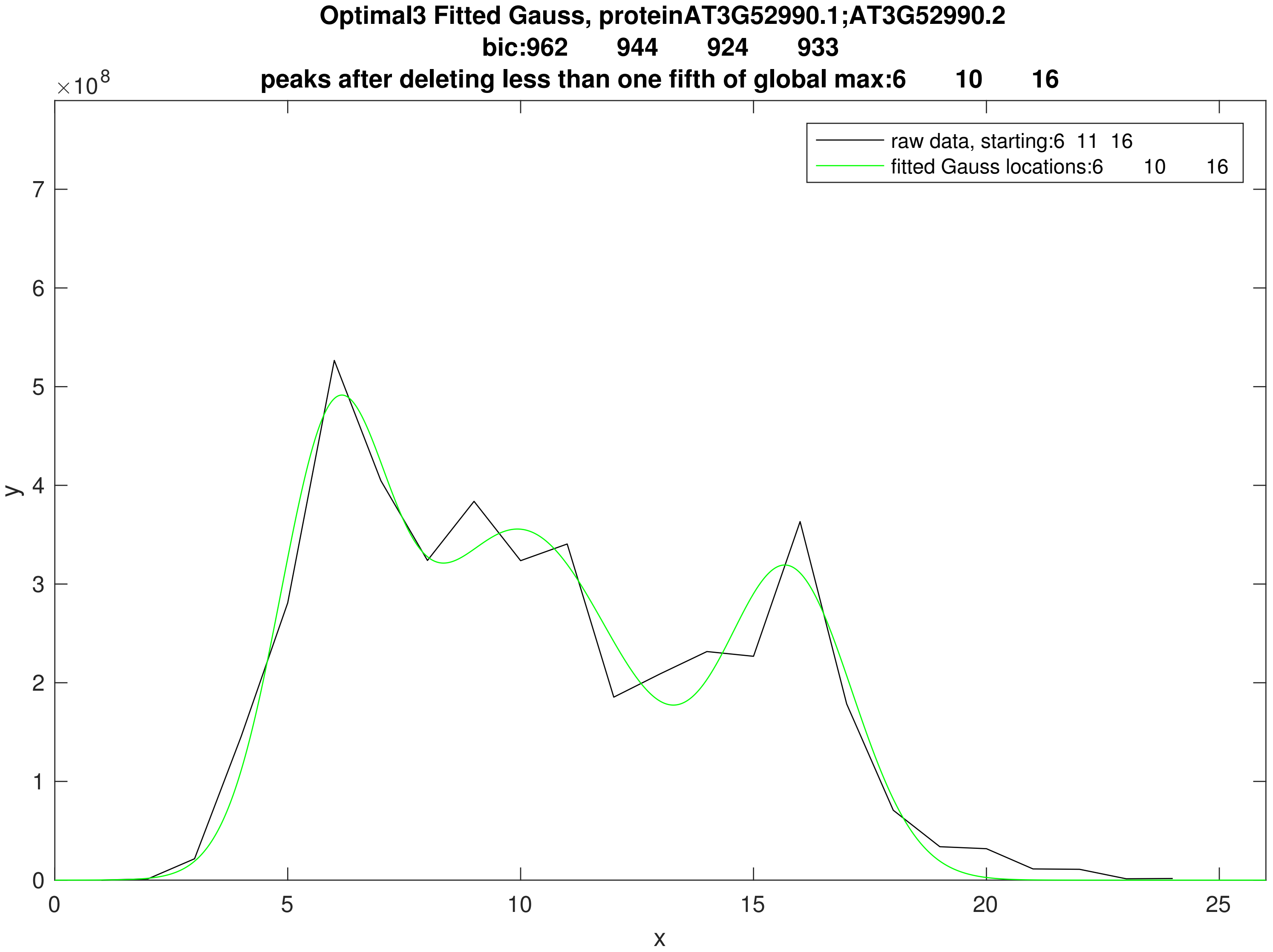

Optimal3 Fitted Gauss, proteinAT3G52990.1;AT3G52990.2
bic:962 944 924 933
peaks after deleting less than one fifth of global max:6 10 16
×10^8
raw data, starting:6 11 16
fitted Gauss locations:6 10 16
y
x
0
1
2
3
4
5
6
7
0
5
10
15
20
25

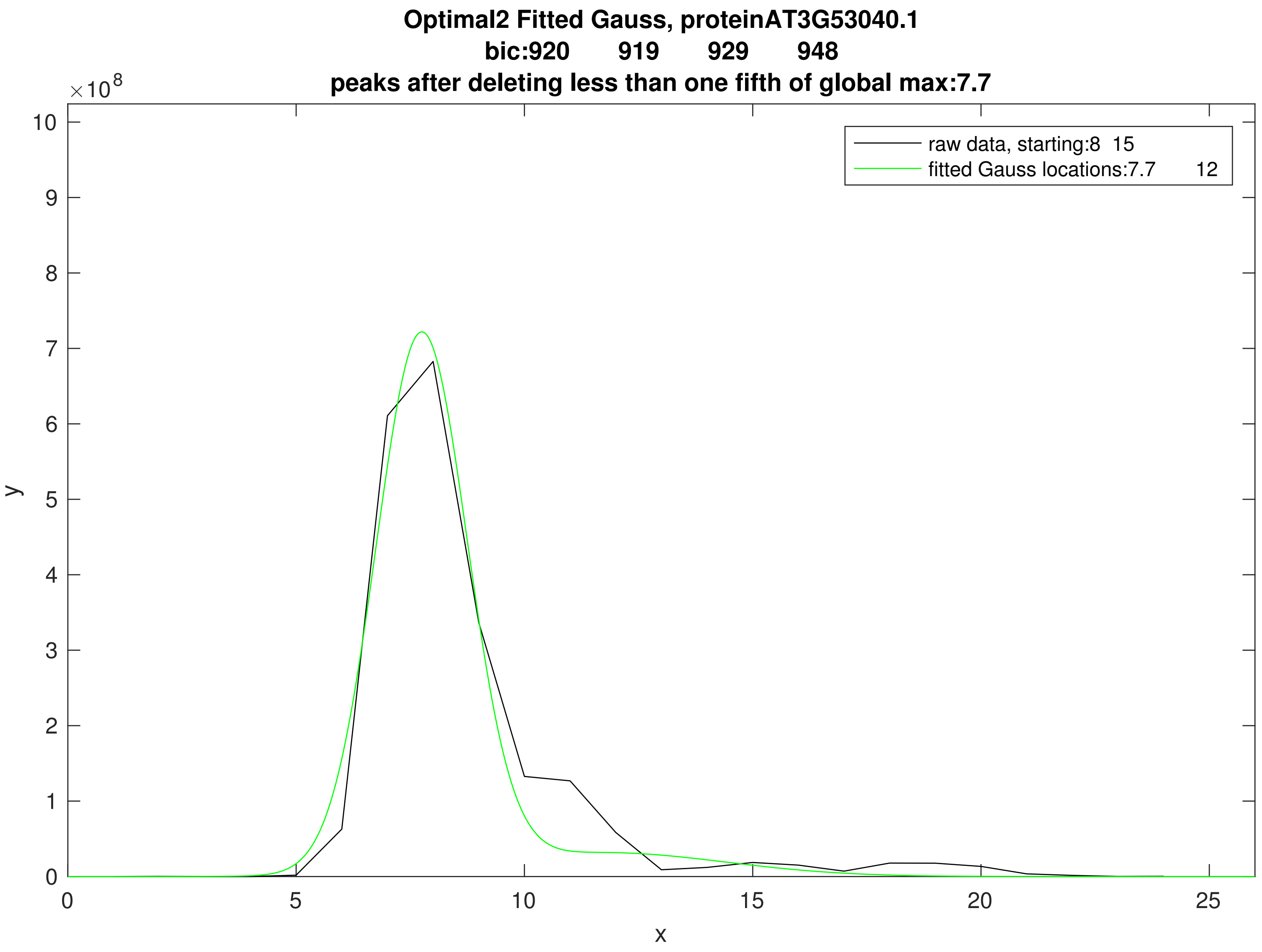
Optimal2 Fitted Gauss, proteinAT3G53040.1
bic:920 919 929 948
peaks after deleting less than one fifth of global max:7.7
×10^8
raw data, starting:8 15
fitted Gauss locations:7.7 12
y
x
0
1
2
3
4
5
6
7
8
9
10
0
5
10
15
20
25

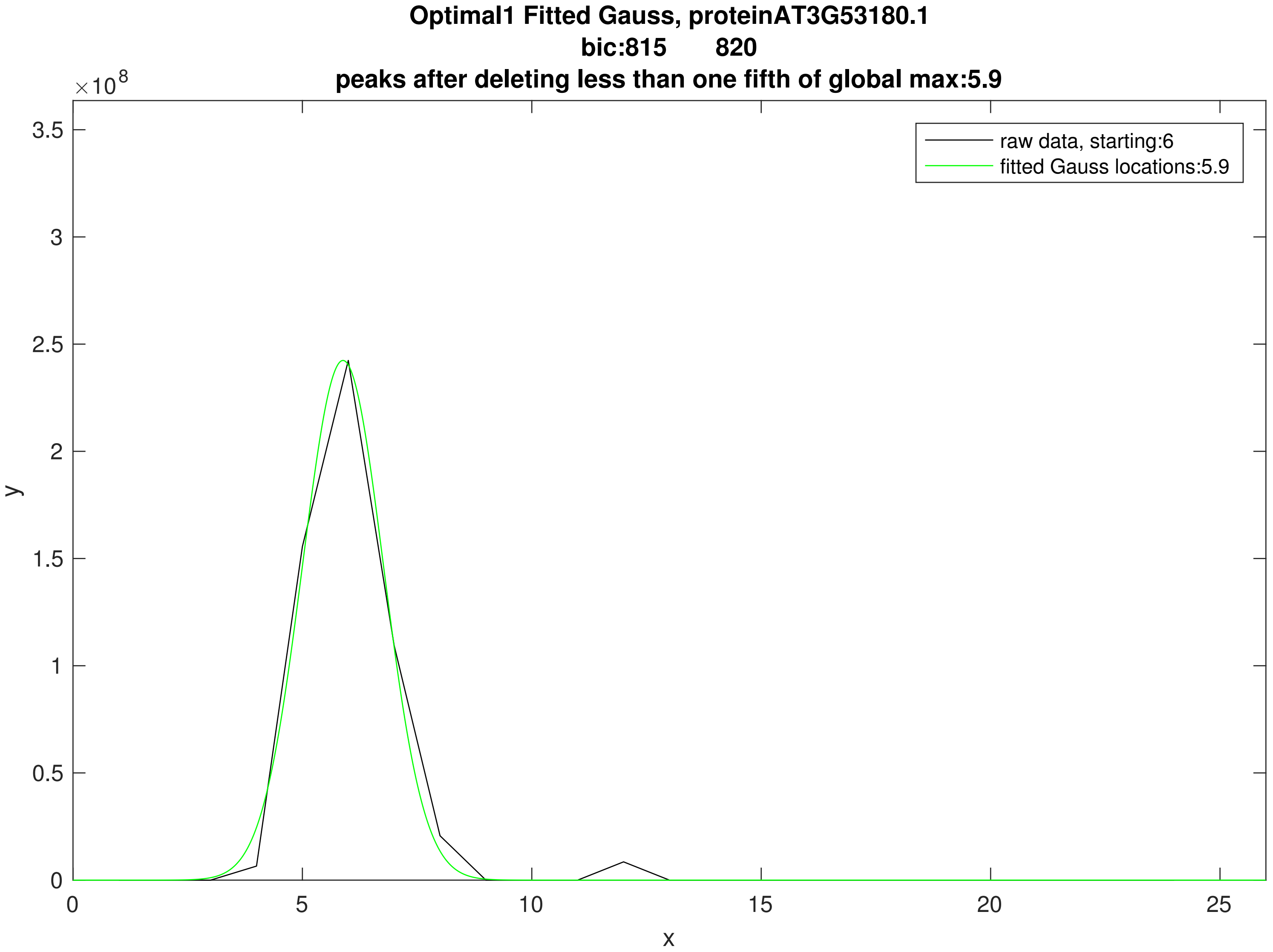

Optimal1 Fitted Gauss, proteinAT3G53180.1
bic:815 820
peaks after deleting less than one fifth of global max:5.9
×10^8
3.5
3
2.5
2
1.5
1
0.5
0
y
0
5
10
15
20
25
x
raw data, starting:6
fitted Gauss locations:5.9

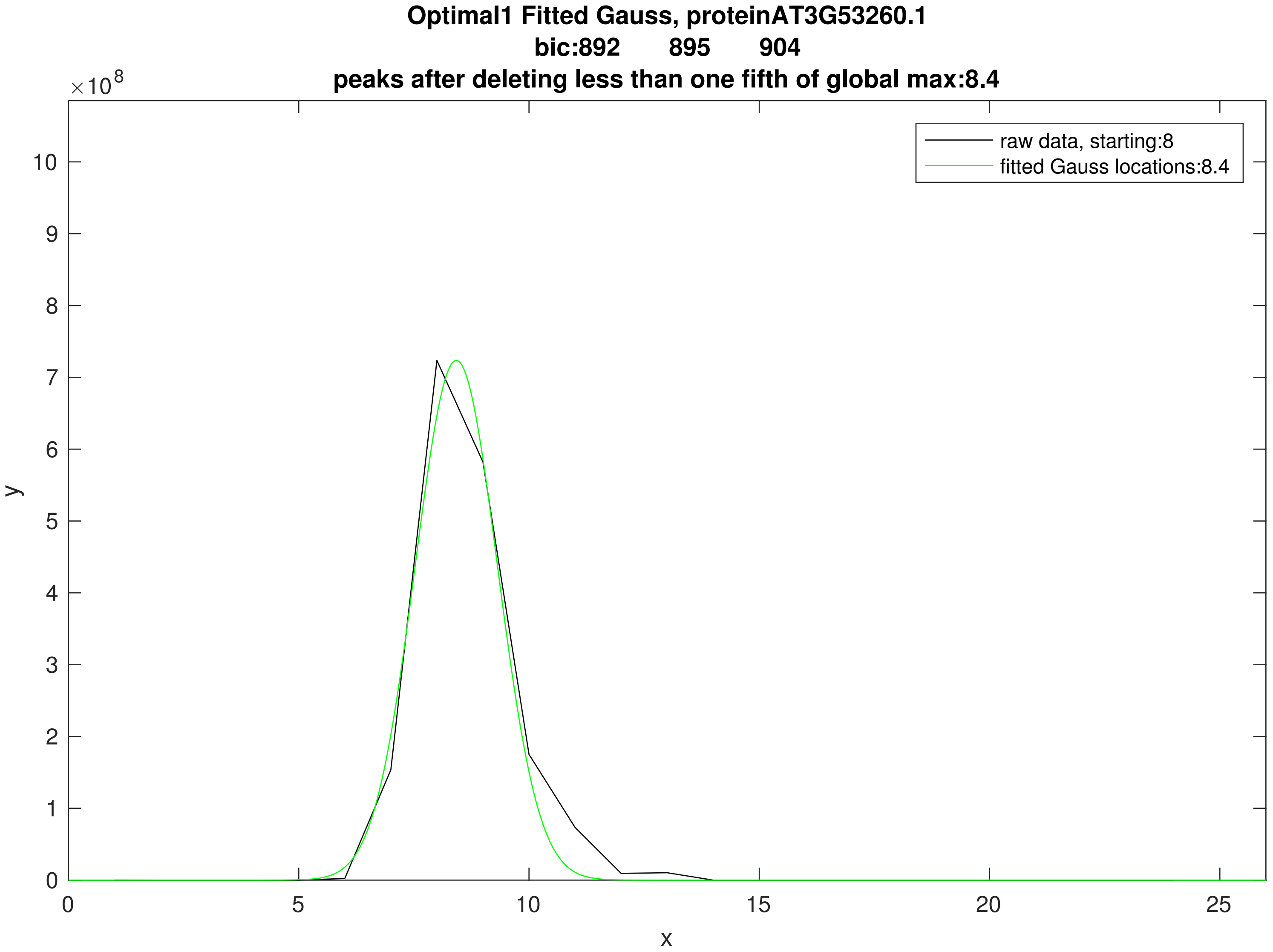

Optimal1 Fitted Gauss, proteinAT3G53260.1
bic:892 895 904
peaks after deleting less than one fifth of global max:8.4
×10^8
raw data, starting:8
fitted Gauss locations:8.4
y
x
0
1
2
3
4
5
6
7
8
9
10
0
5
10
15
20
25

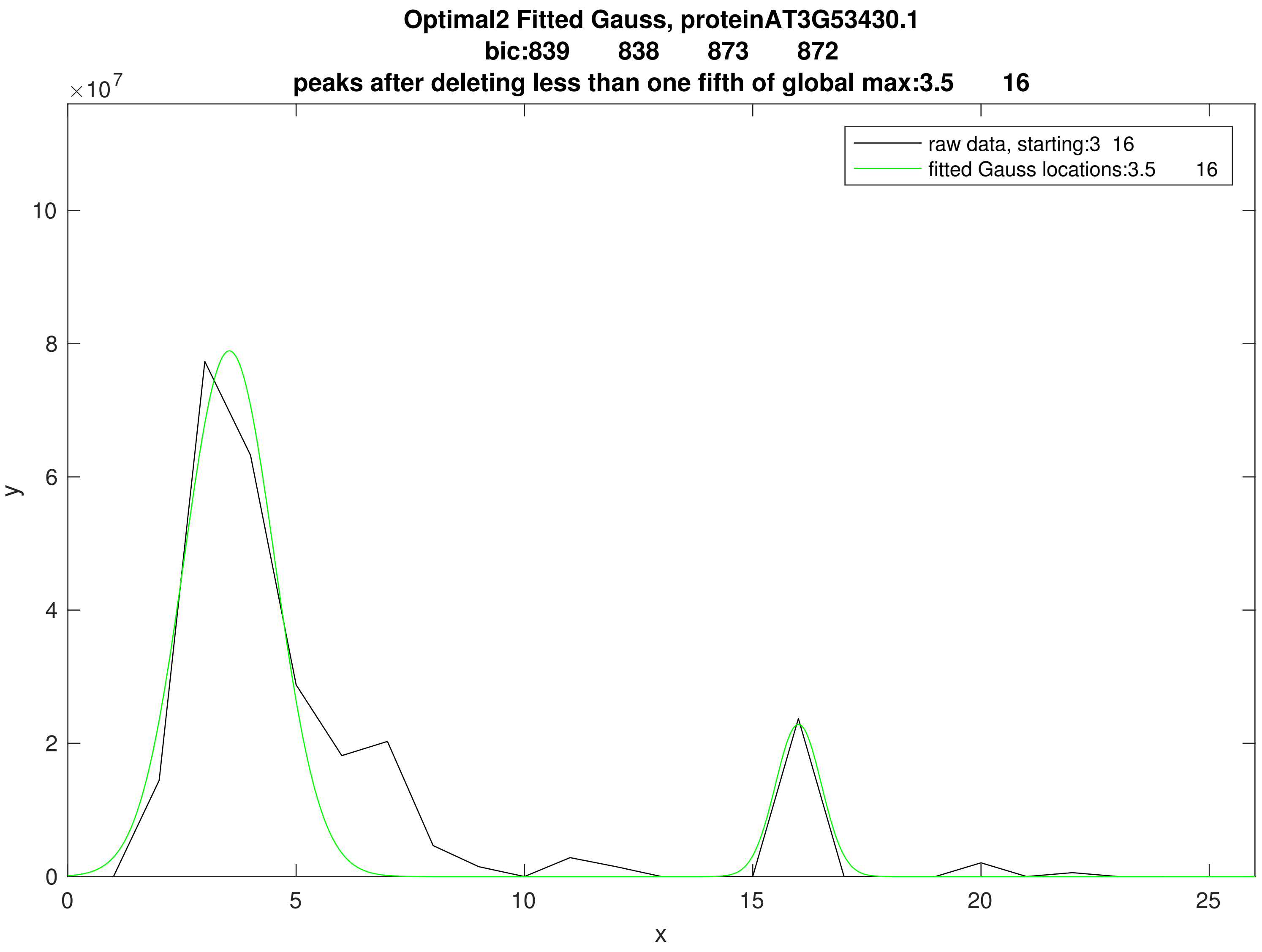
Optimal2 Fitted Gauss, proteinAT3G53430.1
bic:839 838 873 872
peaks after deleting less than one fifth of global max:3.5 16
×10^7
raw data, starting:3 16
fitted Gauss locations:3.5 16
y
x
0
2
4
6
8
10
0
5
10
15
20
25

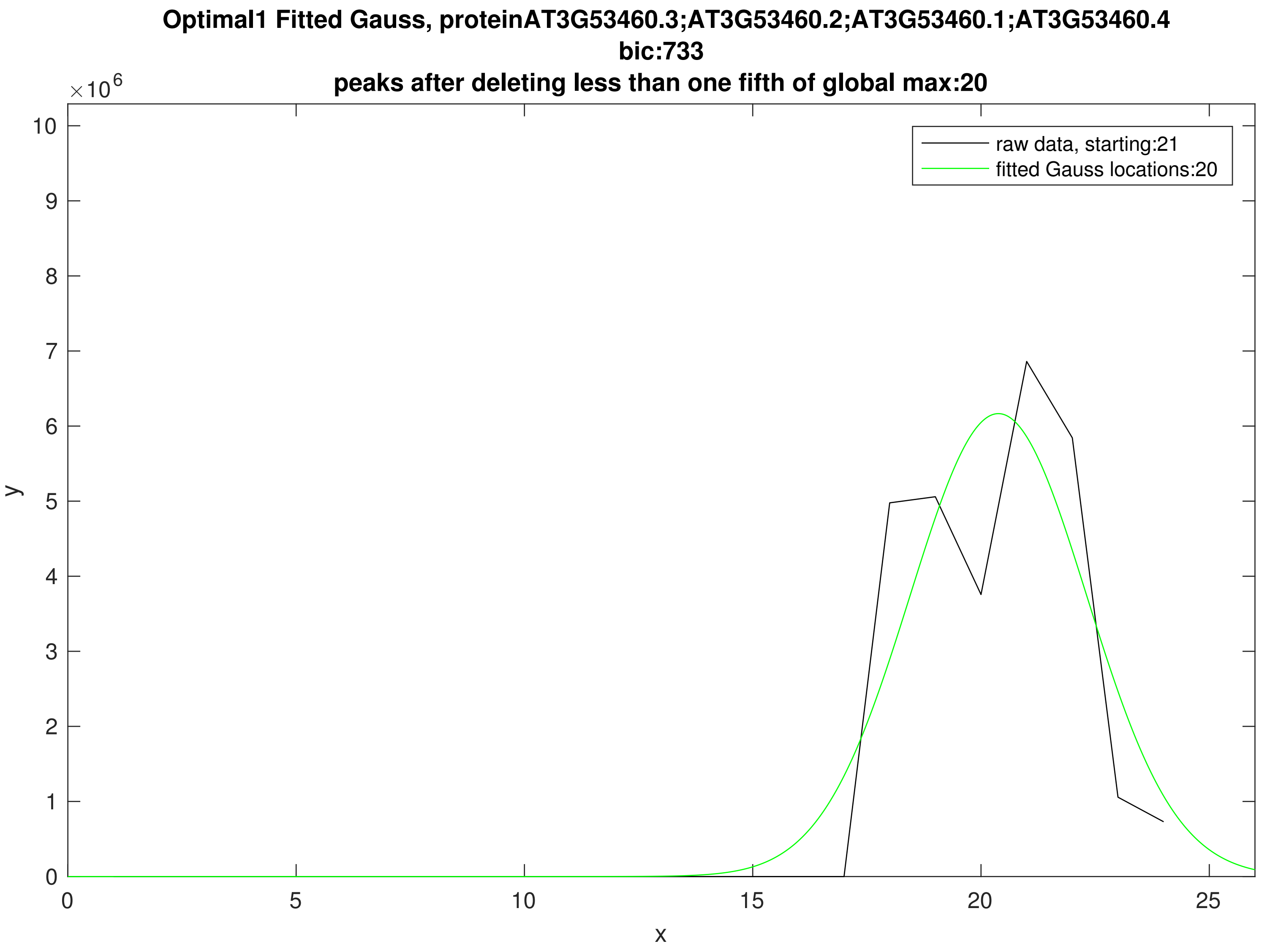
Optimal1 Fitted Gauss, proteinAT3G53460.3;AT3G53460.2;AT3G53460.1;AT3G53460.4
bic:733
peaks after deleting less than one fifth of global max:20
×10^6
raw data, starting:21
fitted Gauss locations:20
y
x
0
1
2
3
4
5
6
7
8
9
10
0
5
10
15
20
25

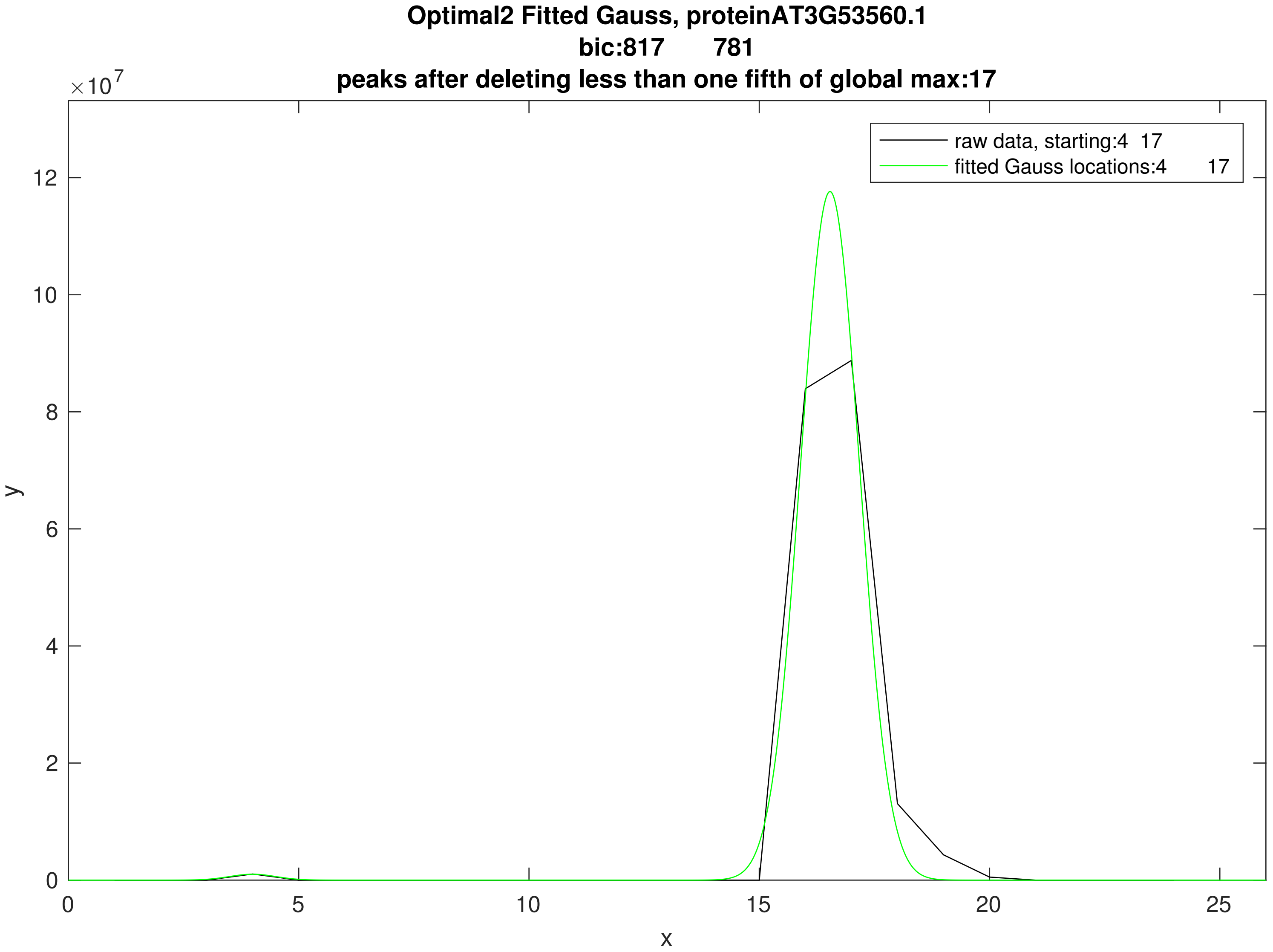

Optimal2 Fitted Gauss, proteinAT3G53560.1
bic:817 781
peaks after deleting less than one fifth of global max:17
×10^7
raw data, starting:4 17
fitted Gauss locations:4 17
12
10
8
6
4
2
0
y
0
5
10
15
20
25
x

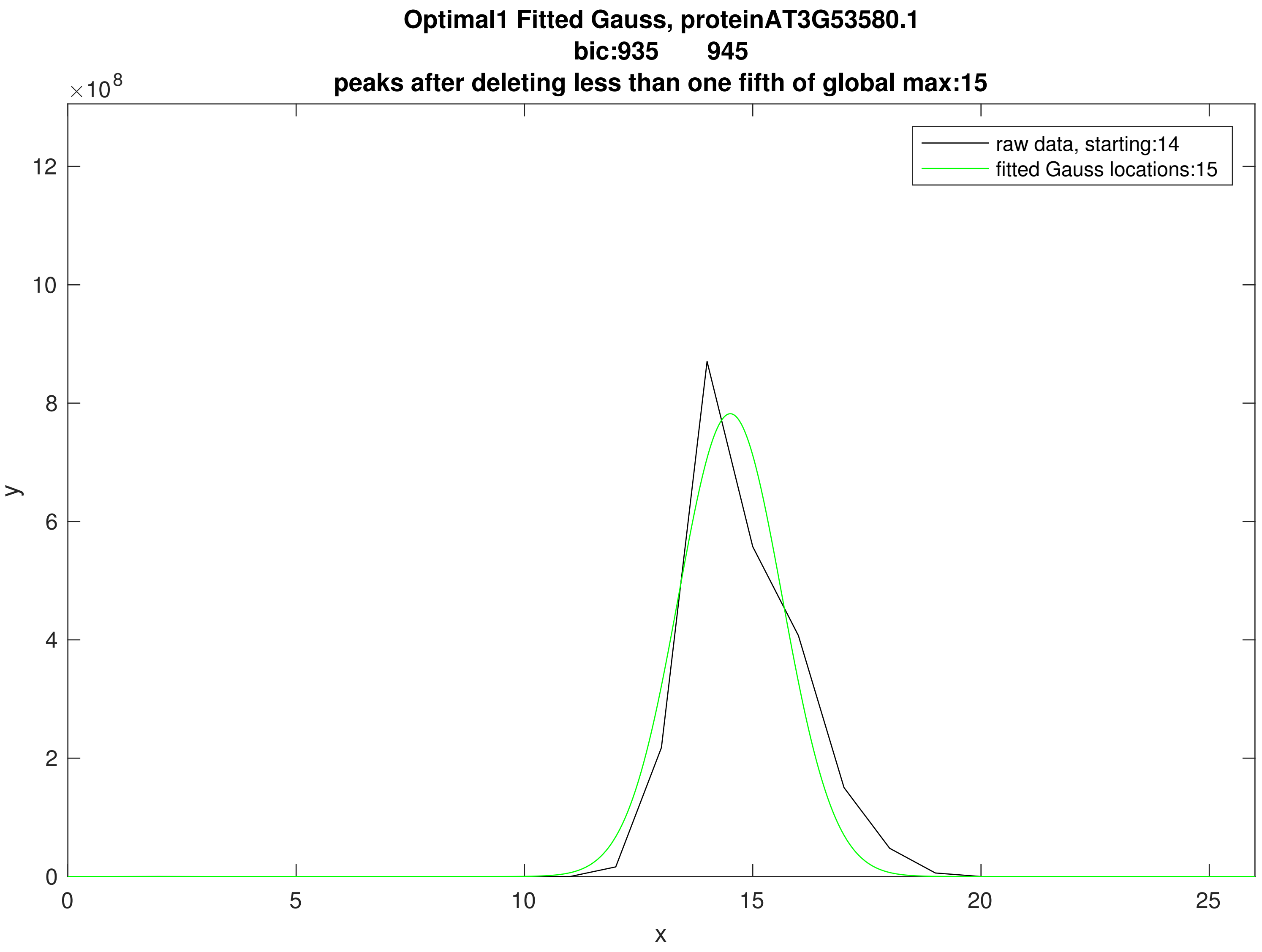
Optimal1 Fitted Gauss, proteinAT3G53580.1
bic:935 945
peaks after deleting less than one fifth of global max:15
×10^8
raw data, starting:14
fitted Gauss locations:15
y
x
0
2
4
6
8
10
12
0
5
10
15
20
25

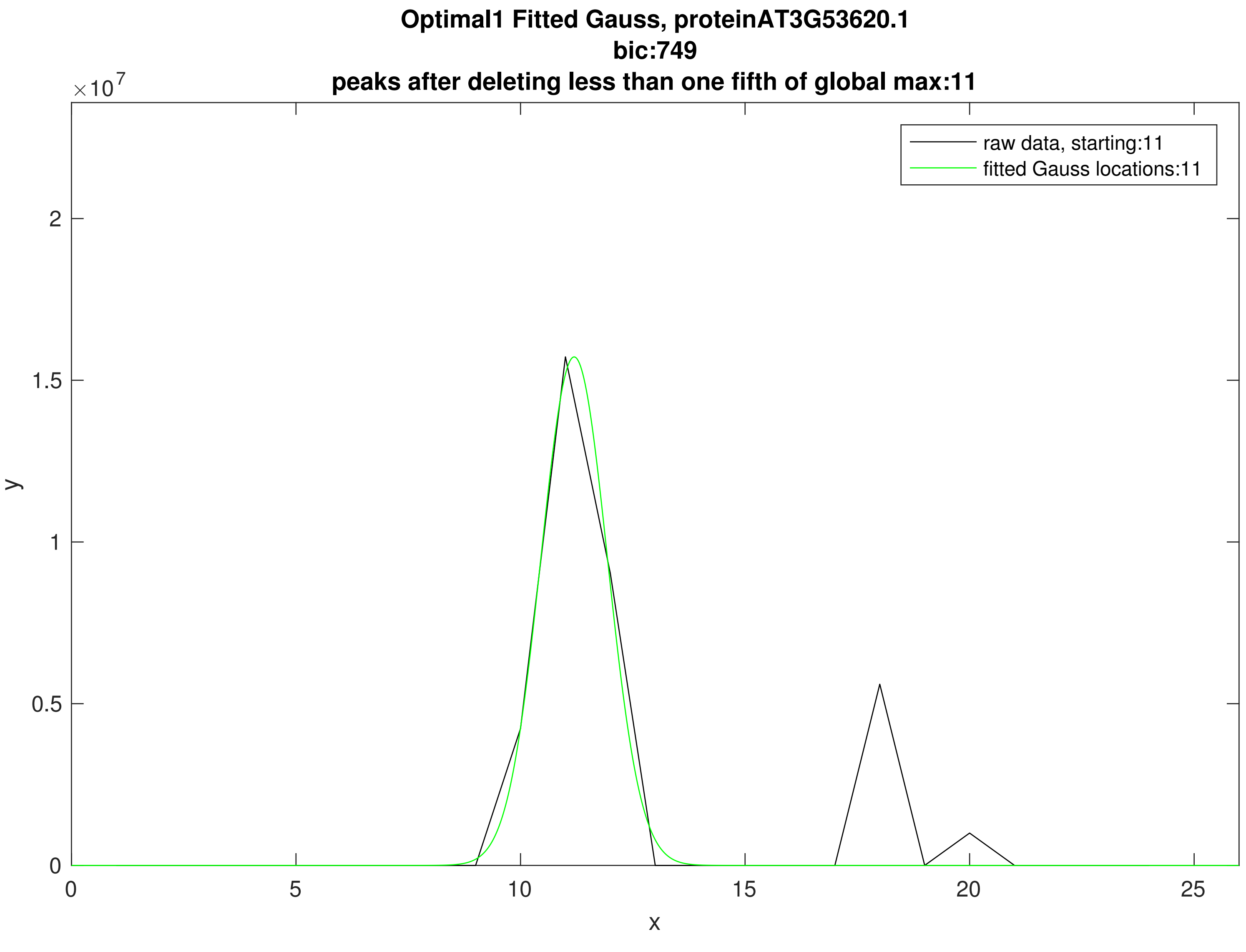
Optimal1 Fitted Gauss, proteinAT3G53620.1
bic:749
peaks after deleting less than one fifth of global max:11
×10^7
raw data, starting:11
fitted Gauss locations:11
y
x
0
0.5
1
1.5
2
0
5
10
15
20
25

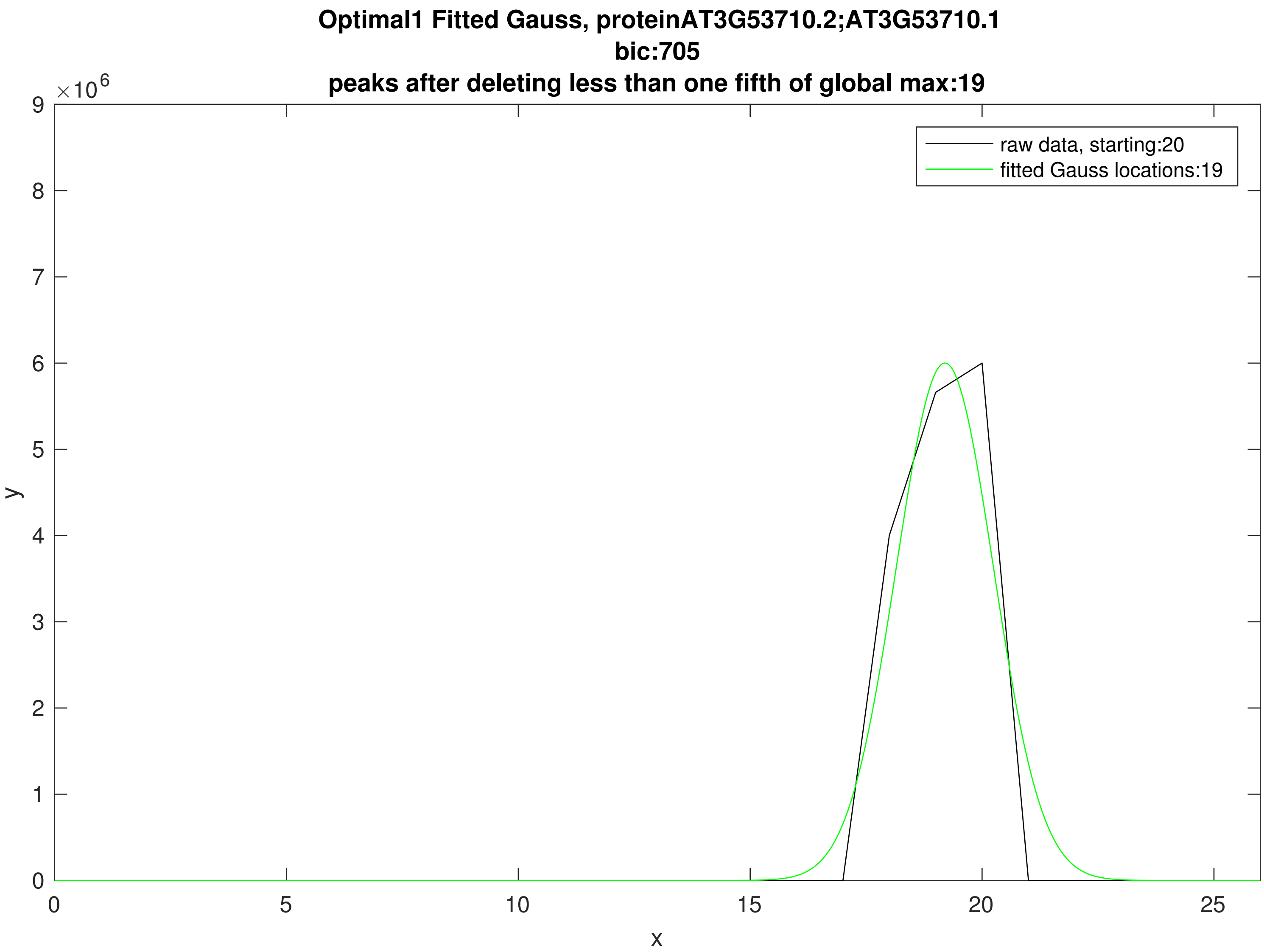
Optimal1 Fitted Gauss, proteinAT3G53710.2;AT3G53710.1
bic:705
peaks after deleting less than one fifth of global max:19
×10^6
raw data, starting:20
fitted Gauss locations:19
y
x

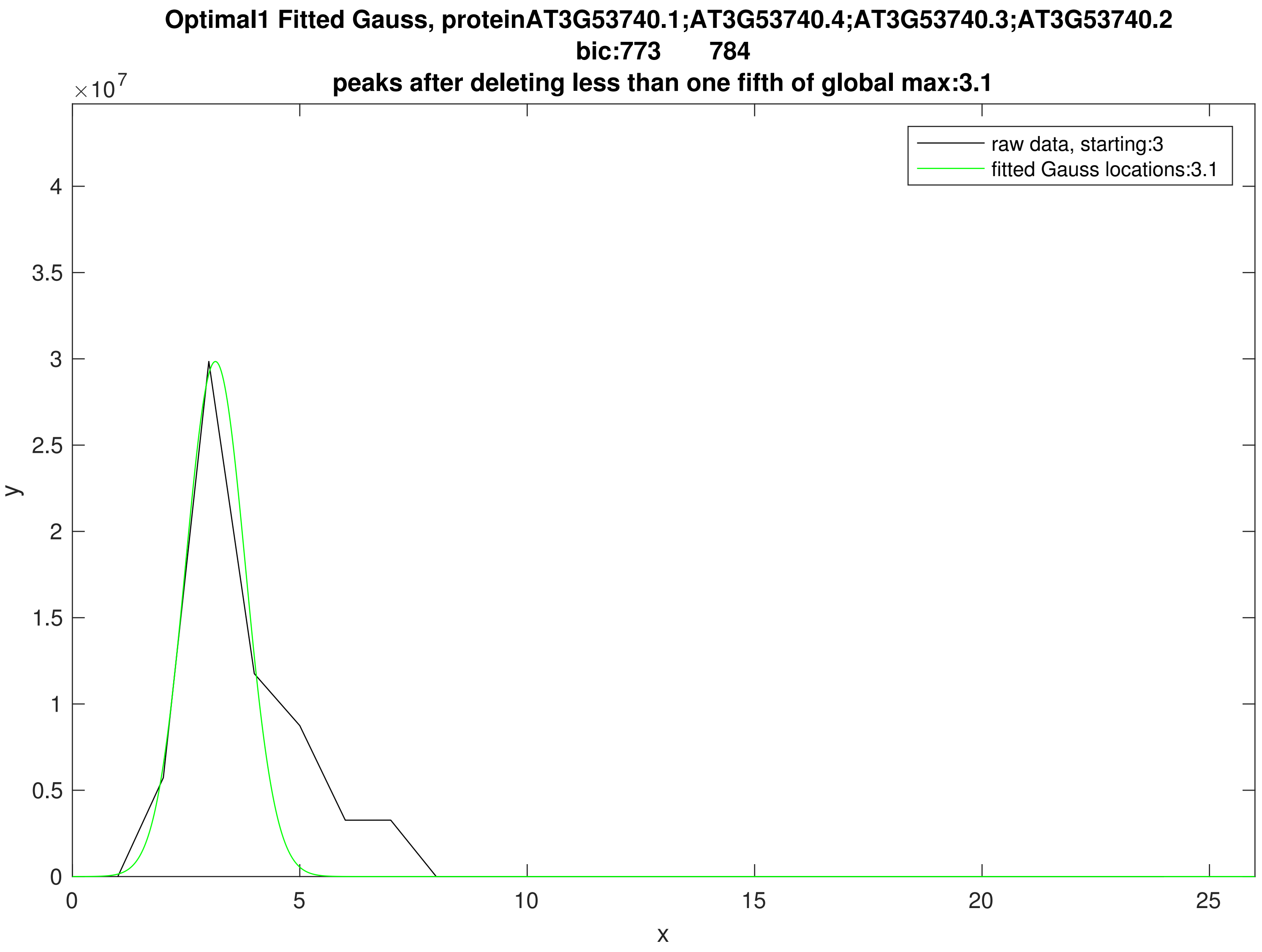

Optimal1 Fitted Gauss, proteinAT3G53740.1;AT3G53740.4;AT3G53740.3;AT3G53740.2
bic:773 784
peaks after deleting less than one fifth of global max:3.1
×10^7
raw data, starting:3
fitted Gauss locations:3.1
y
x
0
0.5
1
1.5
2
2.5
3
3.5
4
5
10
15
20
25

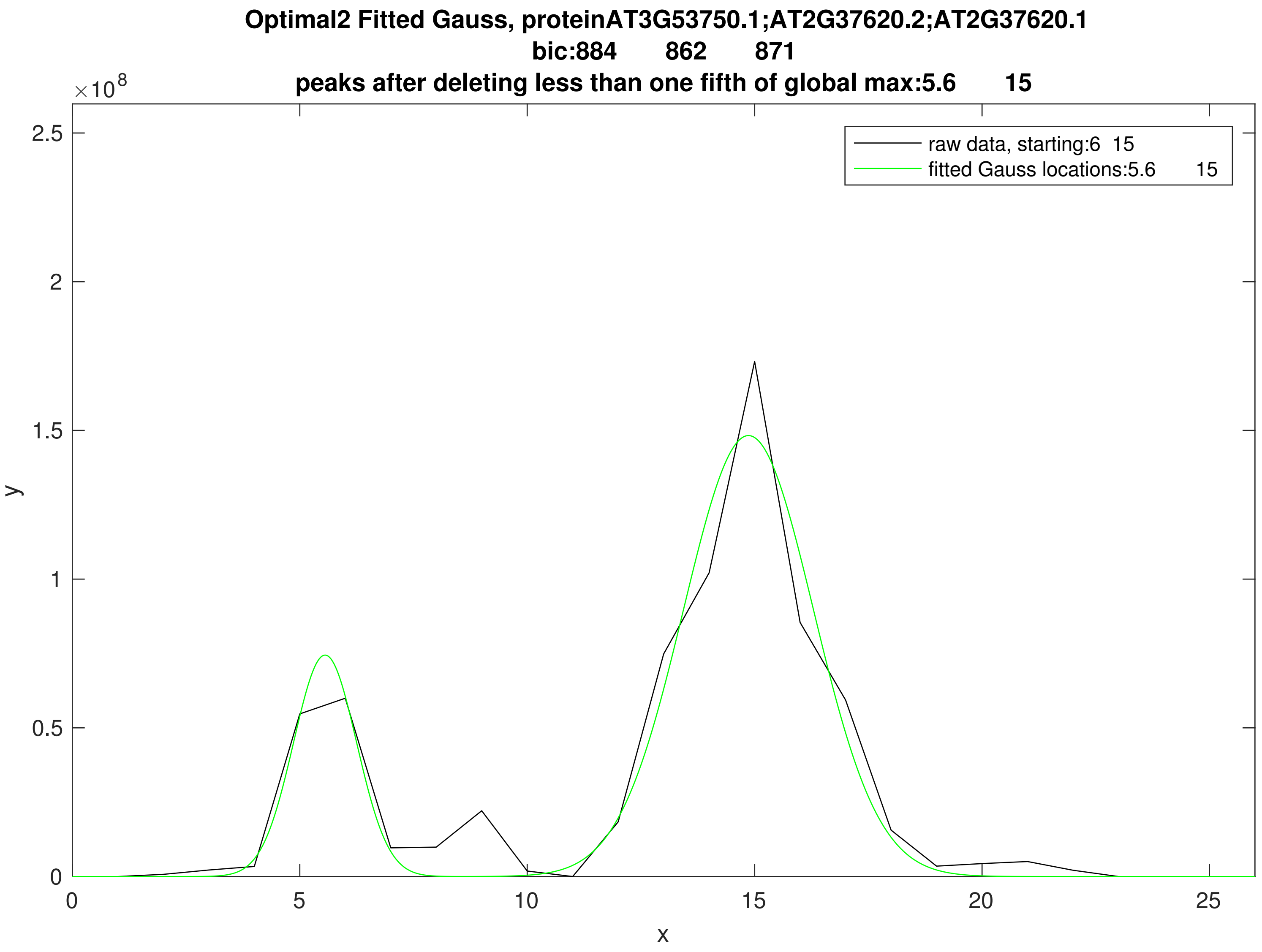

Optimal2 Fitted Gauss, proteinAT3G53750.1;AT2G37620.2;AT2G37620.1
bic:884 862 871
peaks after deleting less than one fifth of global max:5.6 15
×10^8
raw data, starting:6 15
fitted Gauss locations:5.6 15
y
x
0
0.5
1
1.5
2
2.5
0
5
10
15
20
25

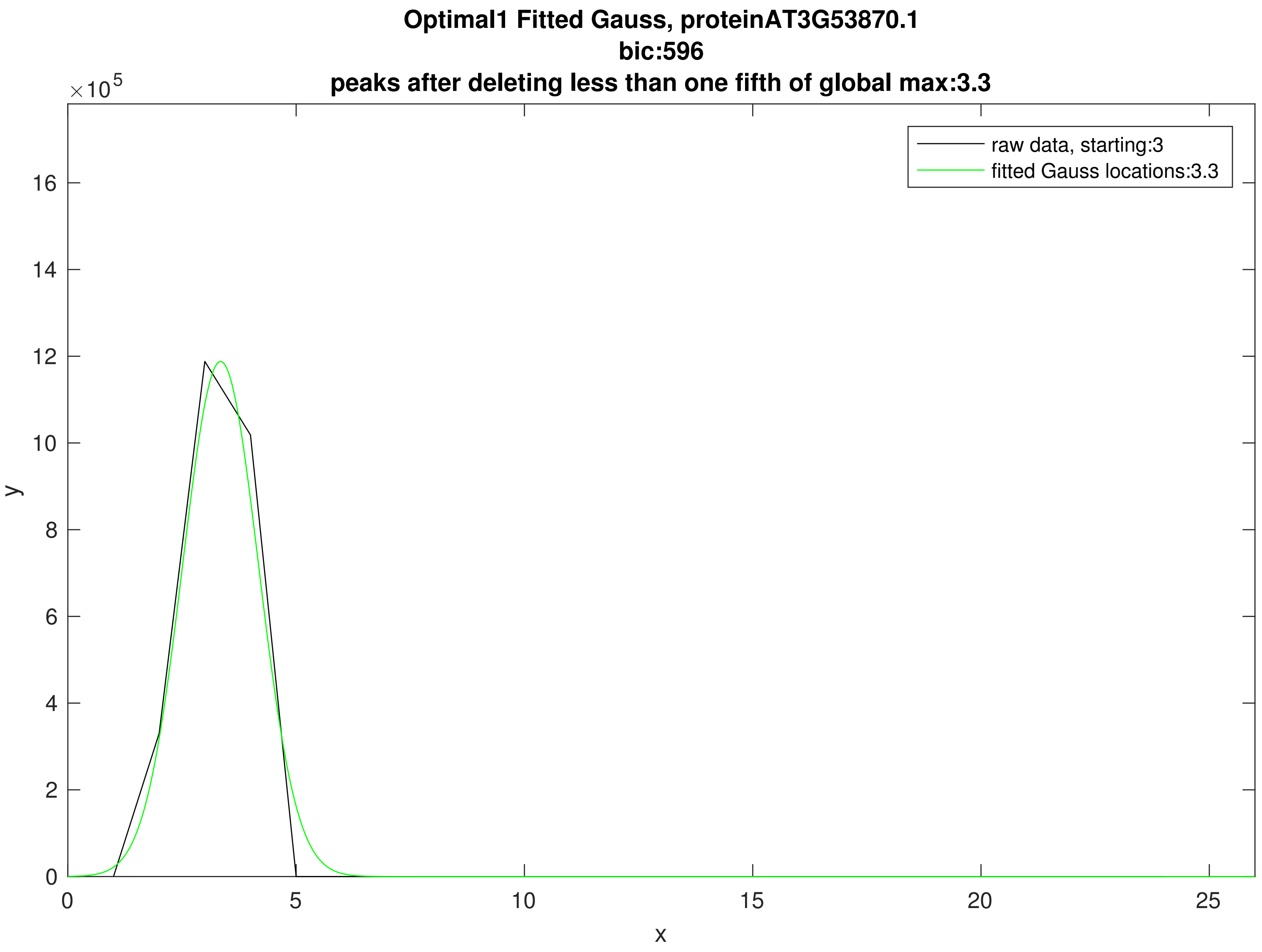
Optimal1 Fitted Gauss, proteinAT3G53870.1
bic:596
peaks after deleting less than one fifth of global max:3.3
×10^5
raw data, starting:3
fitted Gauss locations:3.3
y
x
0
2
4
6
8
10
12
14
16
5
15
20
25

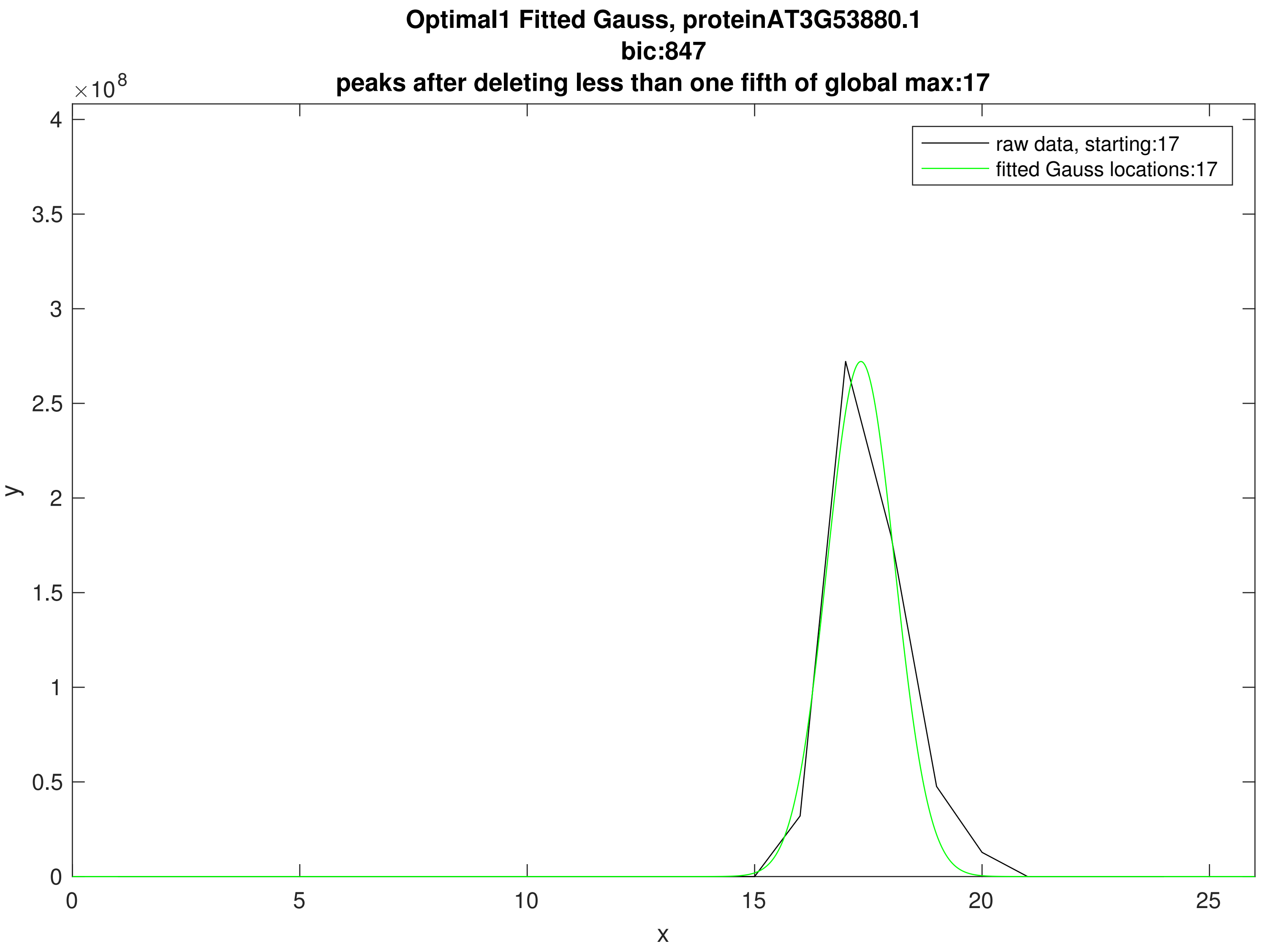

Optimal1 Fitted Gauss, proteinAT3G53880.1
bic:847
peaks after deleting less than one fifth of global max:17
×10^8
raw data, starting:17
fitted Gauss locations:17
y
x
0
0.5
1
1.5
2
2.5
3
3.5
4
0
5
10
15
20
25

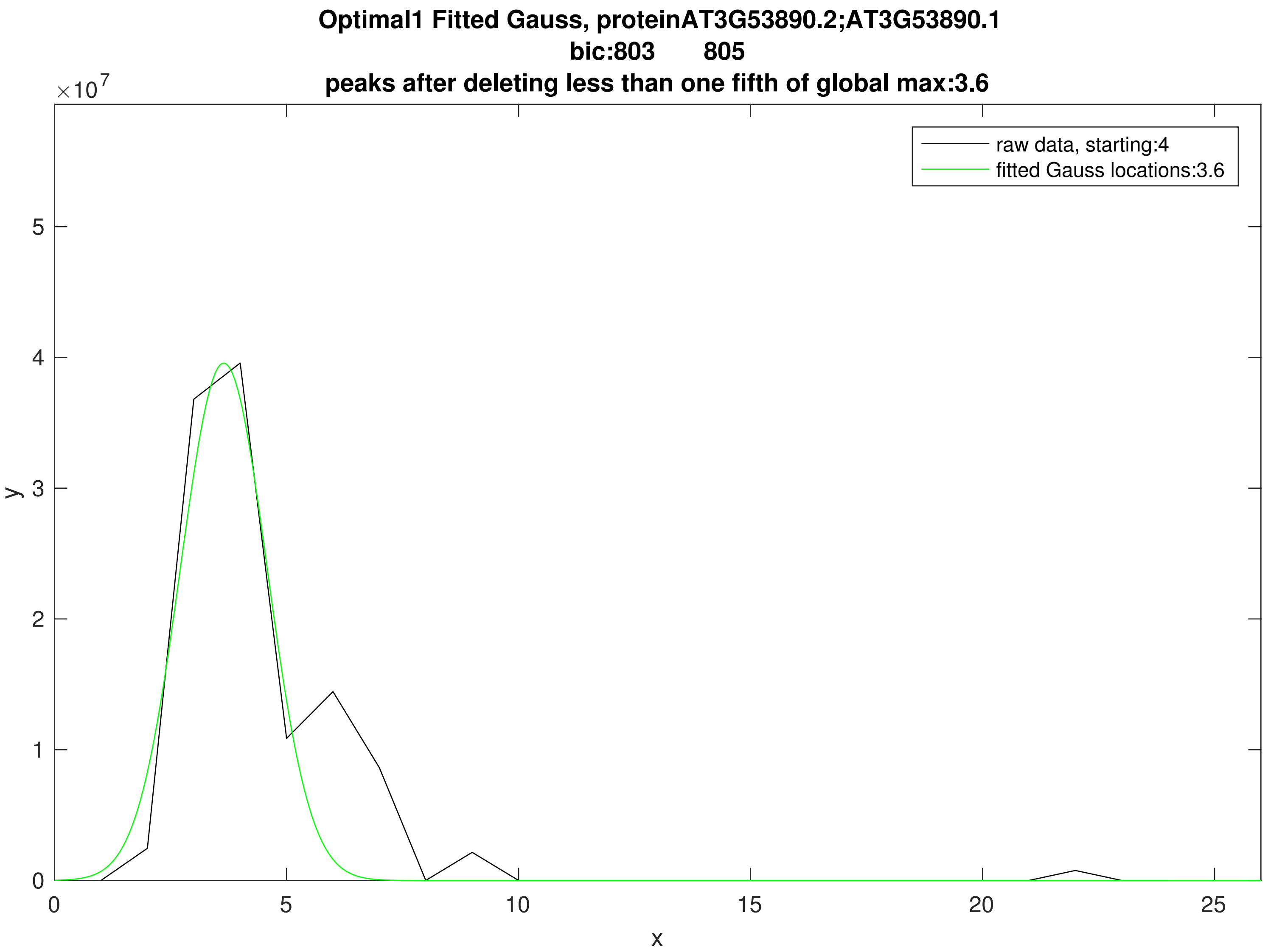
Optimal1 Fitted Gauss, proteinAT3G53890.2;AT3G53890.1
bic:803 805
peaks after deleting less than one fifth of global max:3.6
×10^7
raw data, starting:4
fitted Gauss locations:3.6
y
x
0
1
2
3
4
5
0
5
10
15
20
25

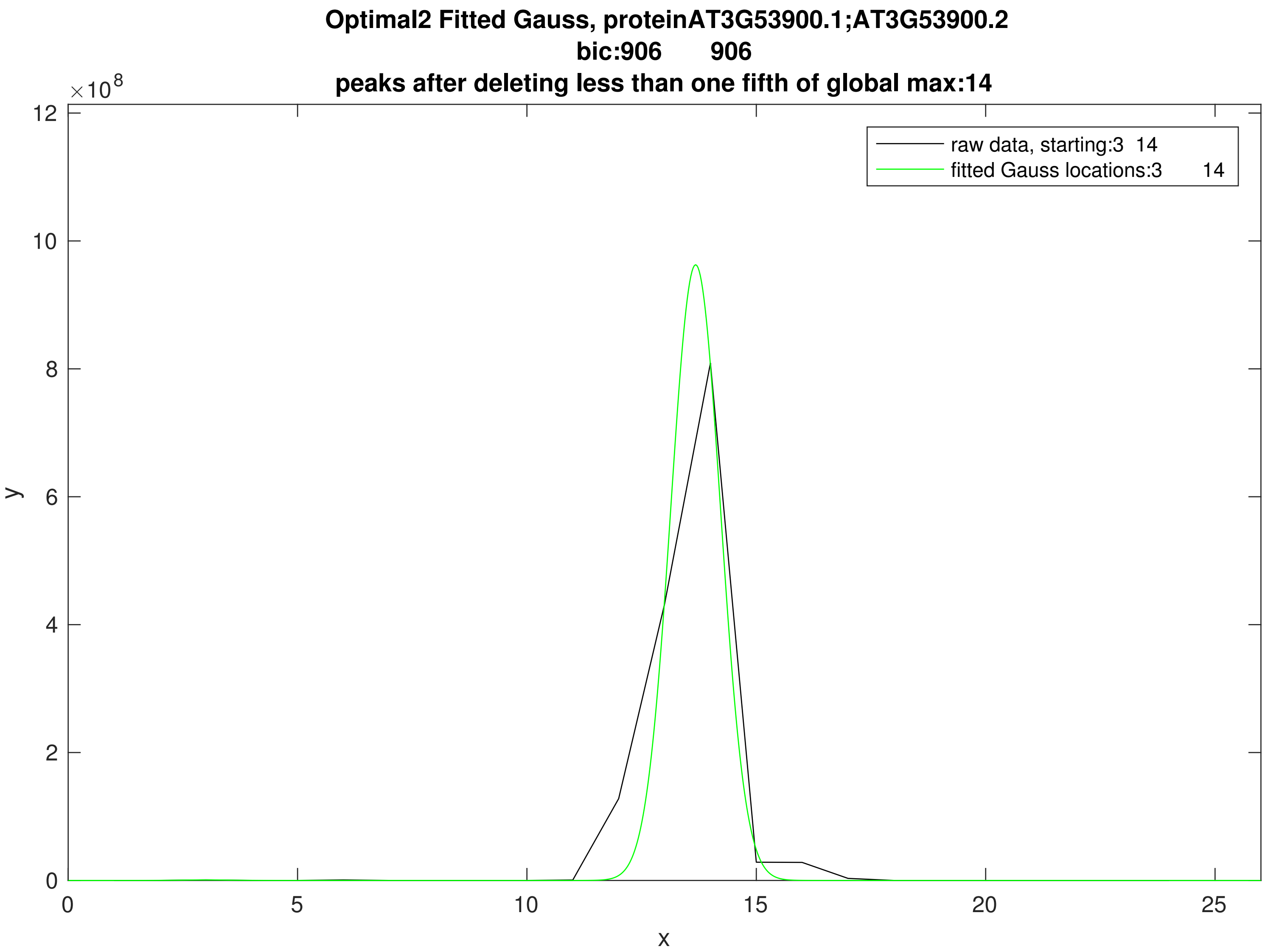

Optimal2 Fitted Gauss, proteinAT3G53900.1;AT3G53900.2
bic:906 906
peaks after deleting less than one fifth of global max:14
×10^8
raw data, starting:3 14
fitted Gauss locations:3 14
y
x
0
2
4
6
8
10
12
5
15
20
25

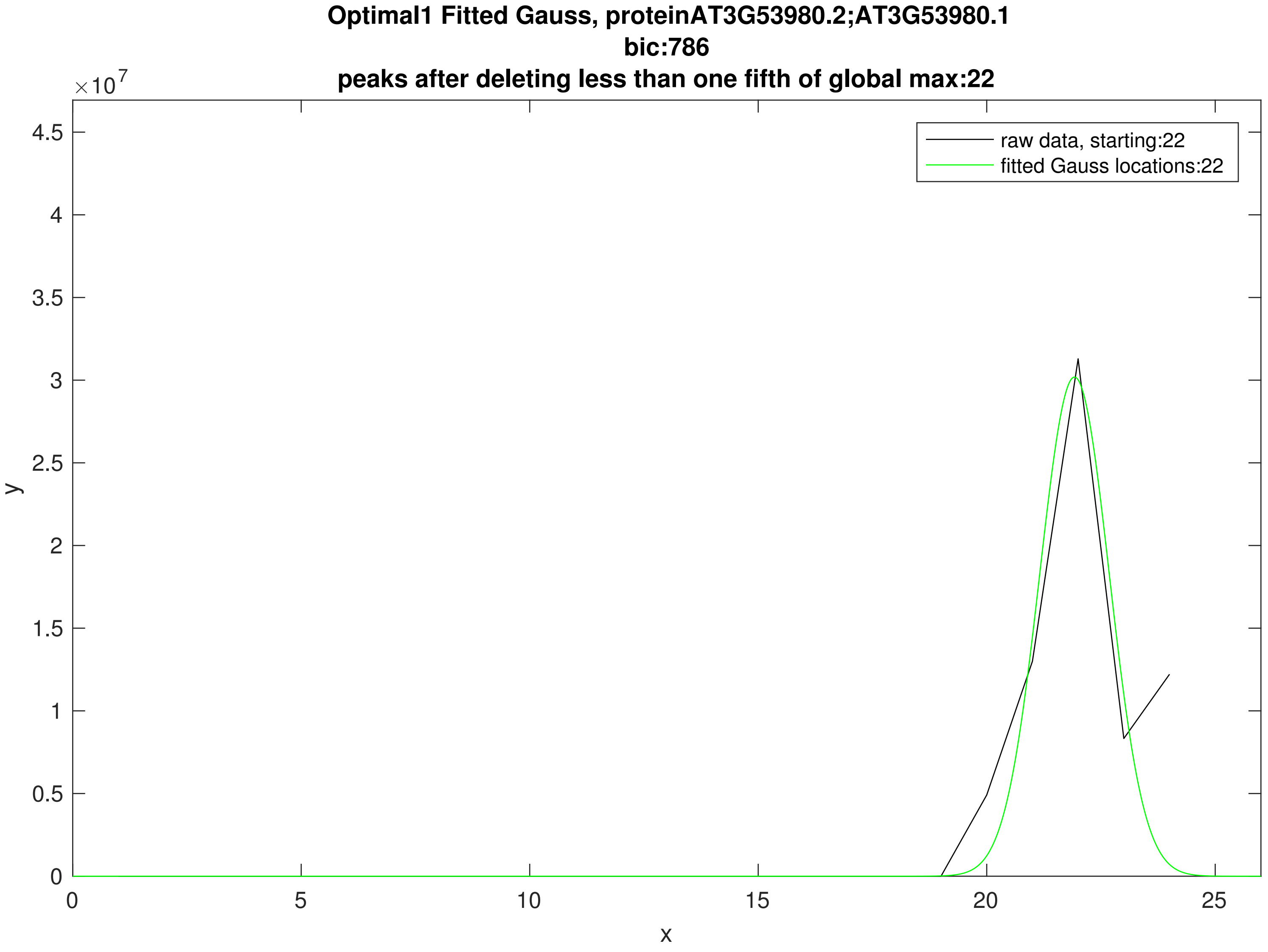

Optimal1 Fitted Gauss, proteinAT3G53980.2;AT3G53980.1
bic:786
peaks after deleting less than one fifth of global max:22
×10^7
raw data, starting:22
fitted Gauss locations:22
4.5
4
3.5
3
2.5
2
1.5
1
0.5
0
y
0
5
10
15
20
25
x

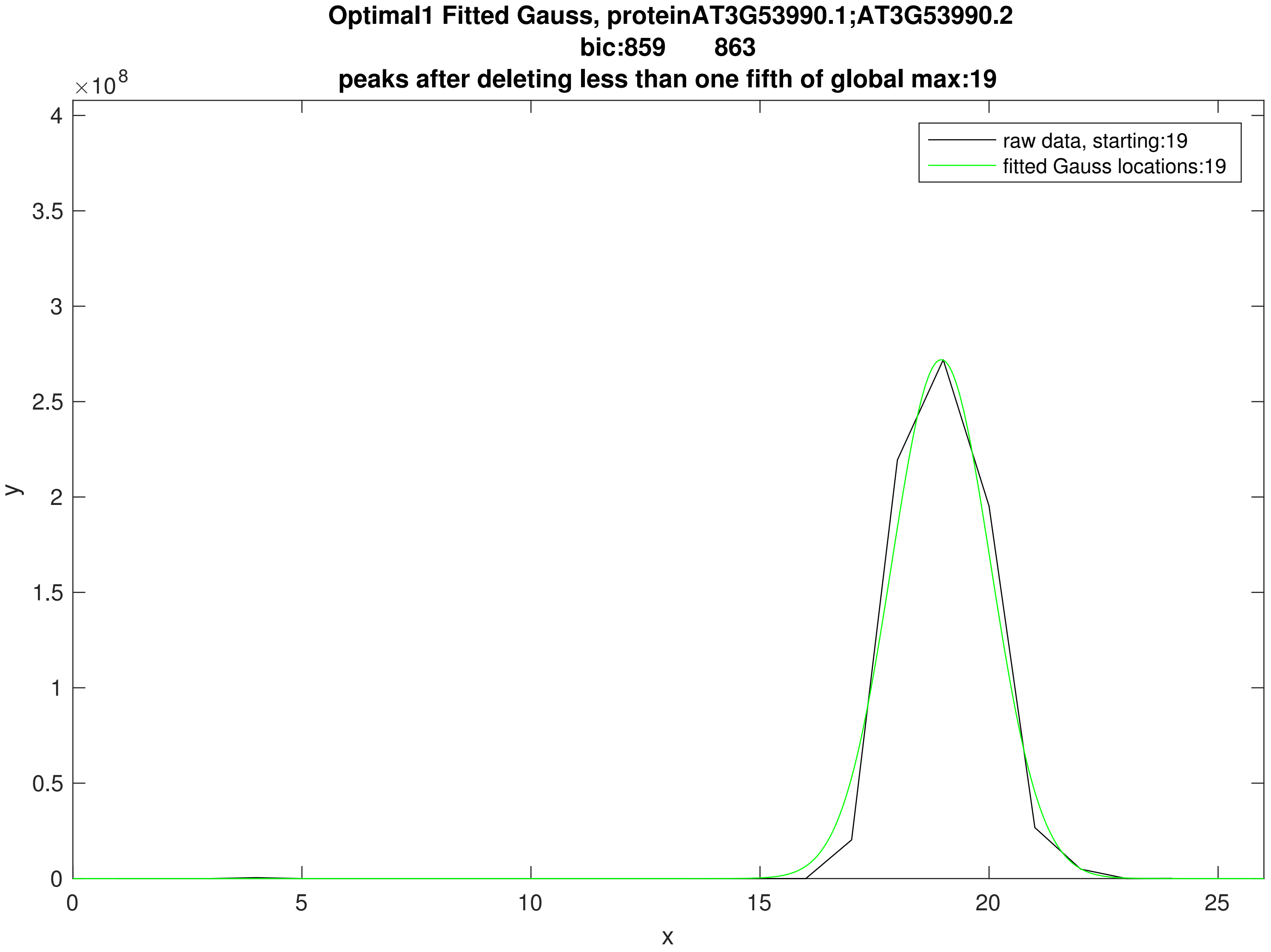

Optimal1 Fitted Gauss, proteinAT3G53990.1;AT3G53990.2
bic:859 863
peaks after deleting less than one fifth of global max:19
×10^8
4
3.5
3
2.5
2
1.5
1
0.5
0
y
0
5
10
15
20
25
x
raw data, starting:19
fitted Gauss locations:19

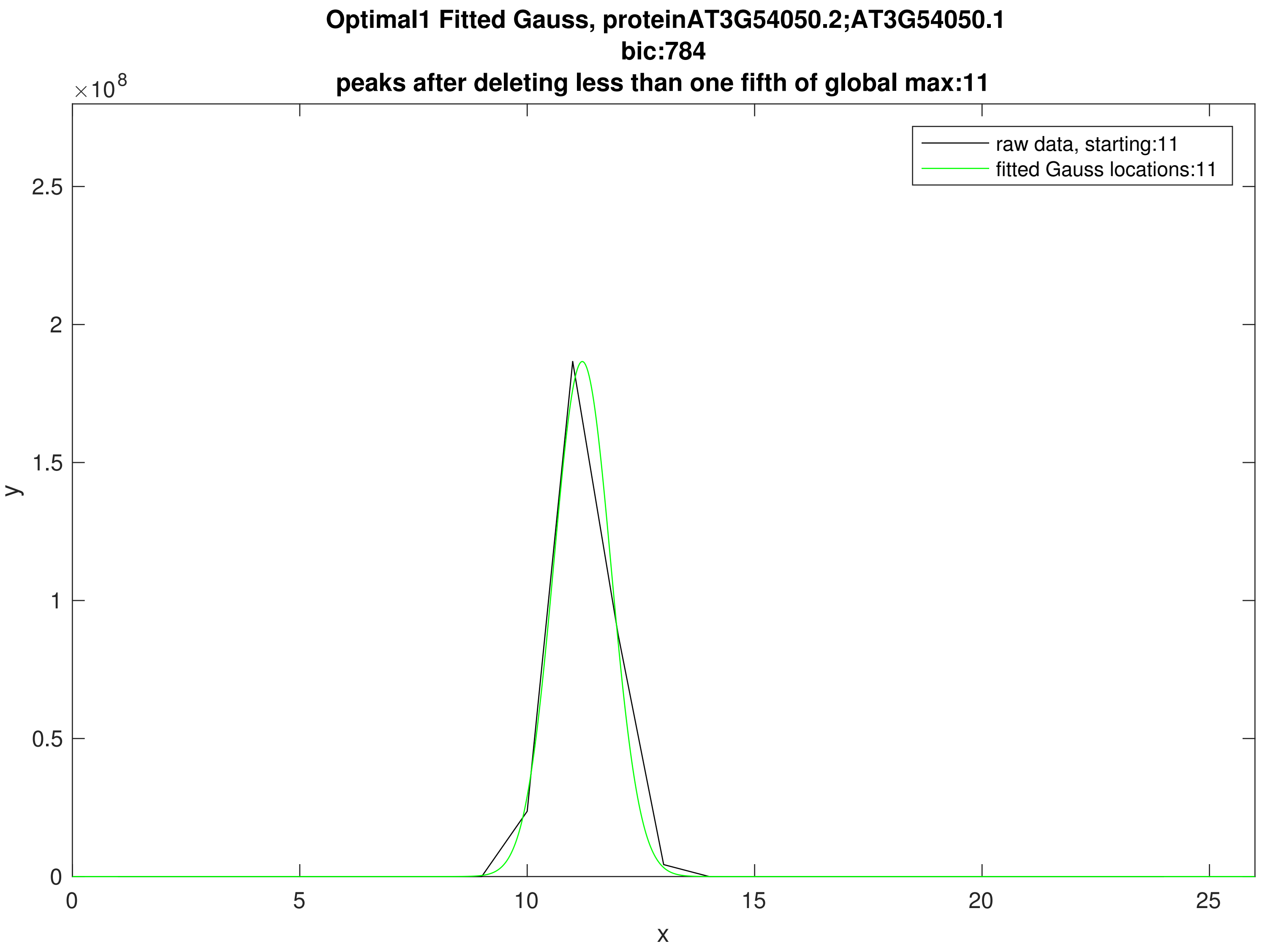

Optimal1 Fitted Gauss, proteinAT3G54050.2;AT3G54050.1
bic:784
peaks after deleting less than one fifth of global max:11
×10^8
2.5
2
1.5
1
0.5
0
y
0
5
10
15
20
25
x
raw data, starting:11
fitted Gauss locations:11

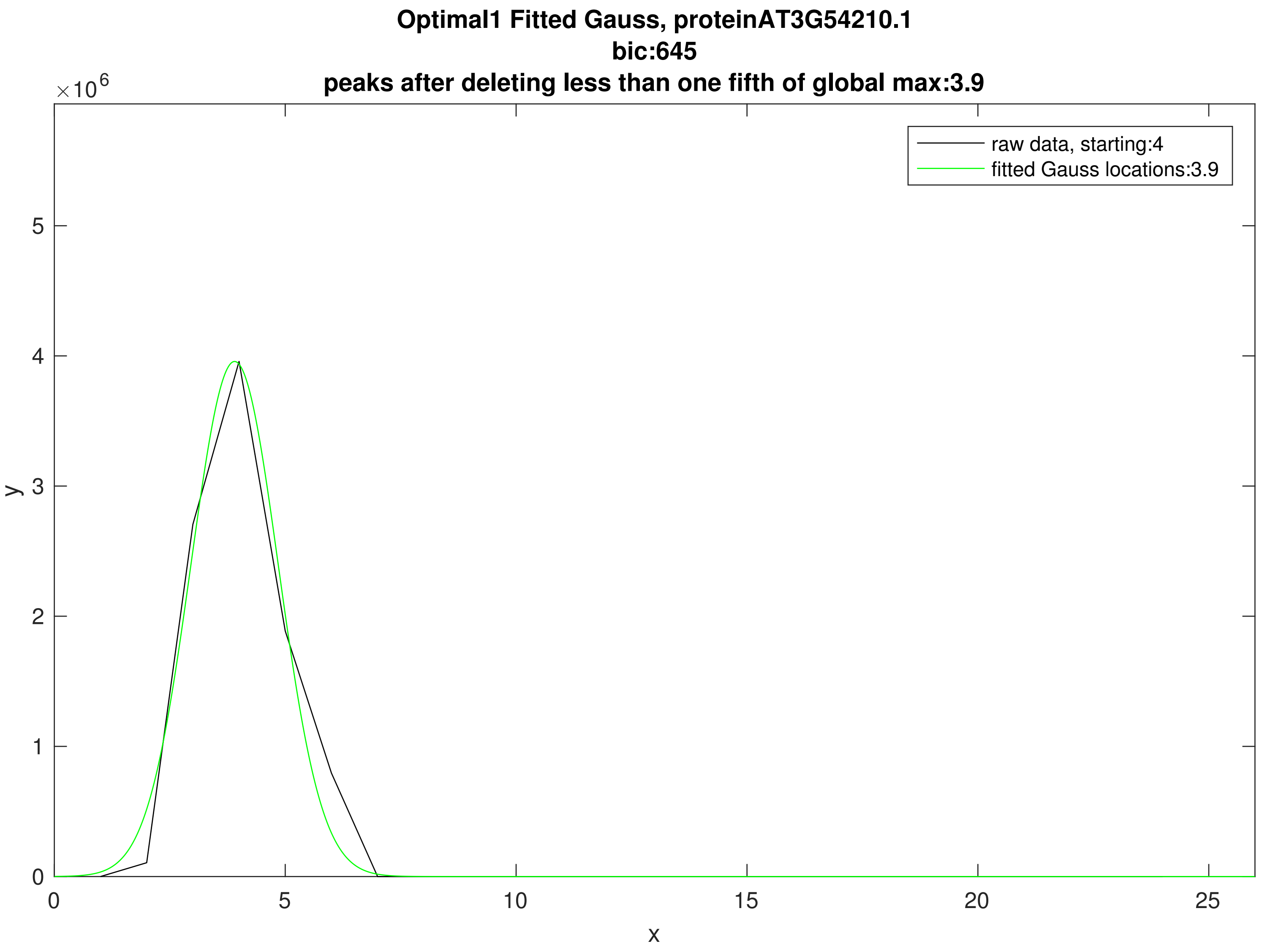
Optimal1 Fitted Gauss, proteinAT3G54210.1
bic:645
peaks after deleting less than one fifth of global max:3.9
×10^6
raw data, starting:4
fitted Gauss locations:3.9
y
x
0
1
2
3
4
5
10
15
20
25

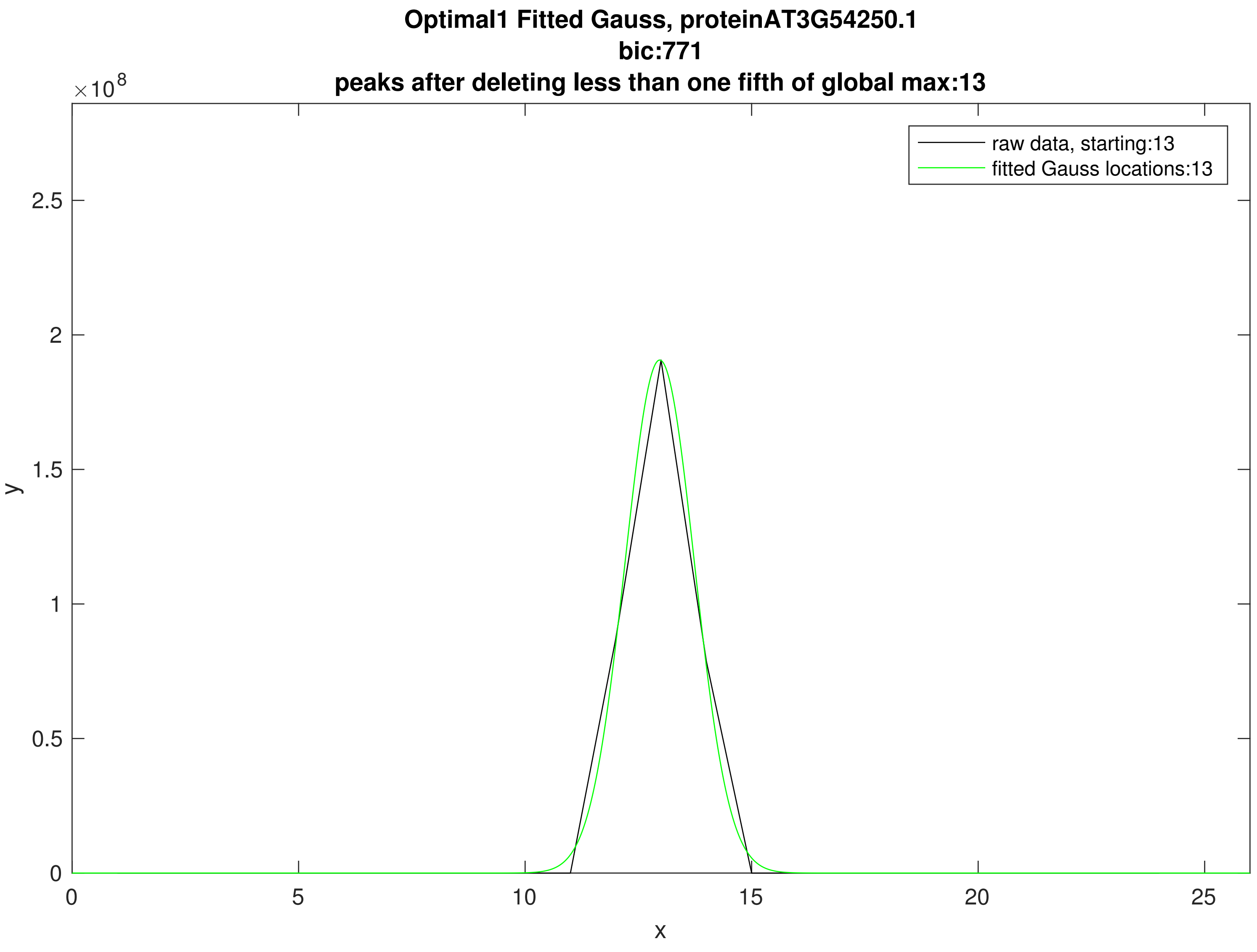
Optimal1 Fitted Gauss, proteinAT3G54250.1
bic:771
peaks after deleting less than one fifth of global max:13
×10^8
raw data, starting:13
fitted Gauss locations:13
y
x
0
0.5
1
1.5
2
2.5
0
5
10
15
20
25

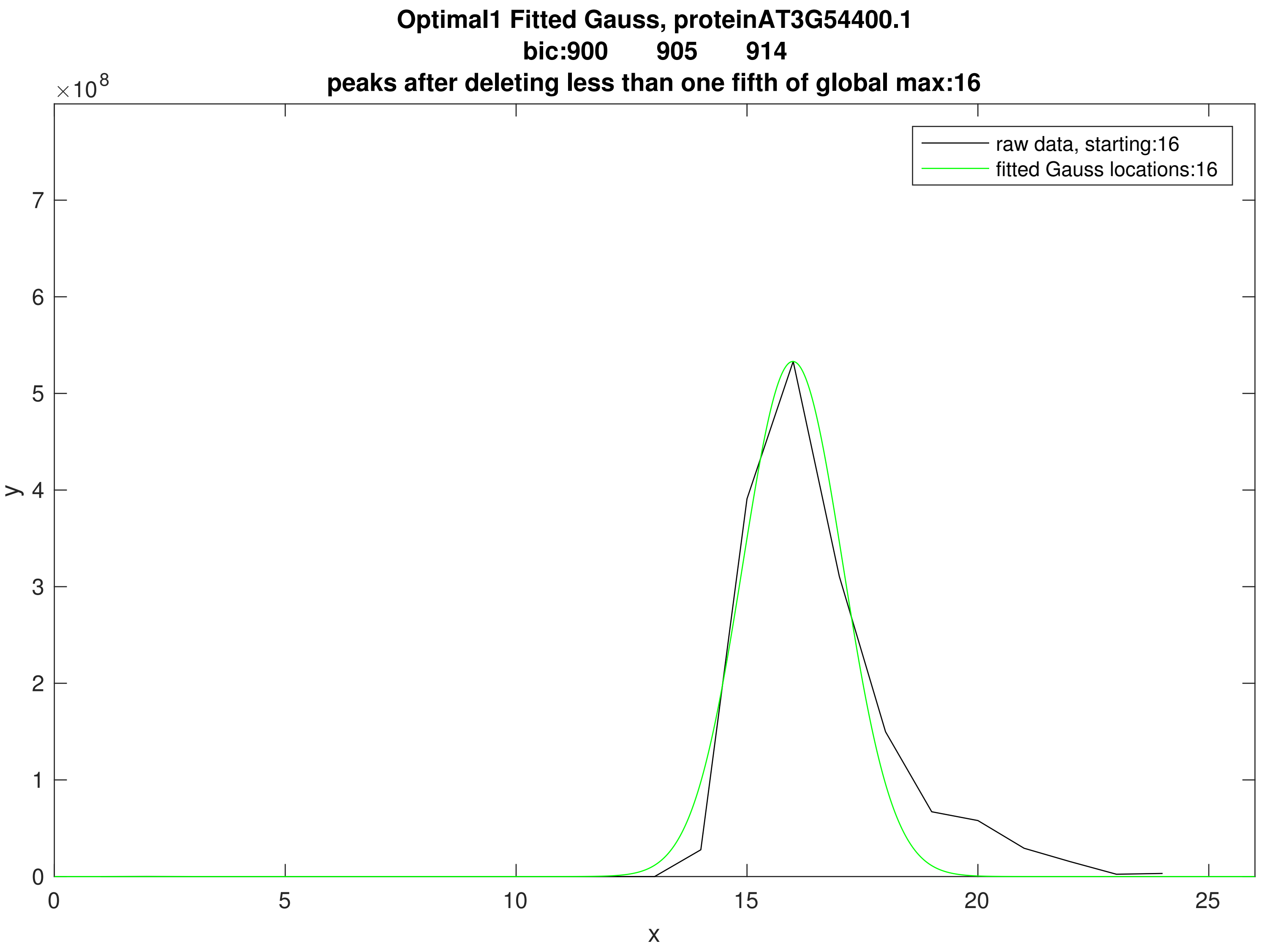

Optimal1 Fitted Gauss, proteinAT3G54400.1
bic:900 905 914
peaks after deleting less than one fifth of global max:16
×10^8
raw data, starting:16
fitted Gauss locations:16
y
x
0
1
2
3
4
5
6
7
0
5
10
15
20
25

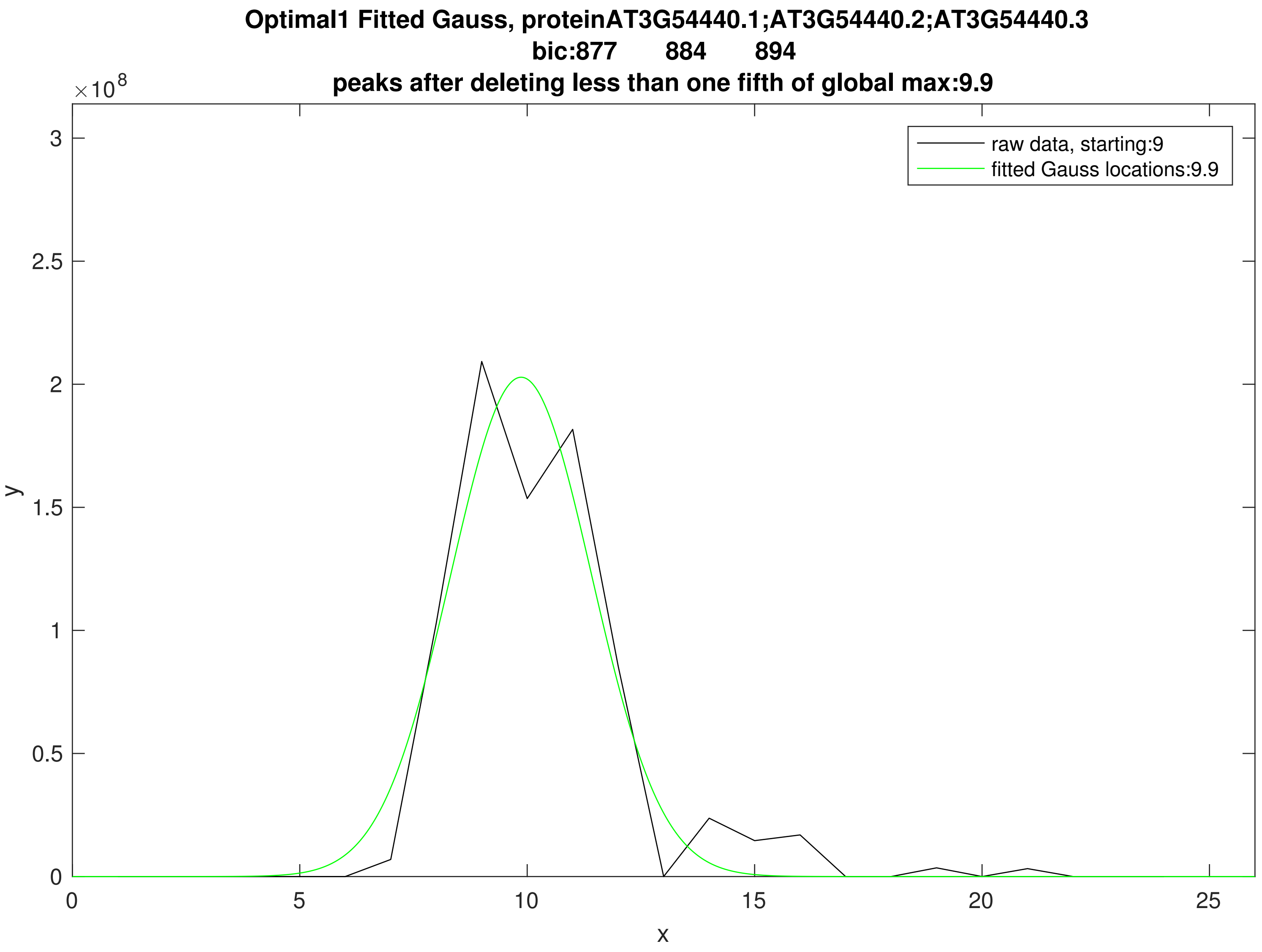
Optimal1 Fitted Gauss, proteinAT3G54440.1;AT3G54440.2;AT3G54440.3
bic:877 884 894
peaks after deleting less than one fifth of global max:9.9
×10^8
3
2.5
2
1.5
1
0.5
0
y
0
5
10
15
20
25
x
raw data, starting:9
fitted Gauss locations:9.9

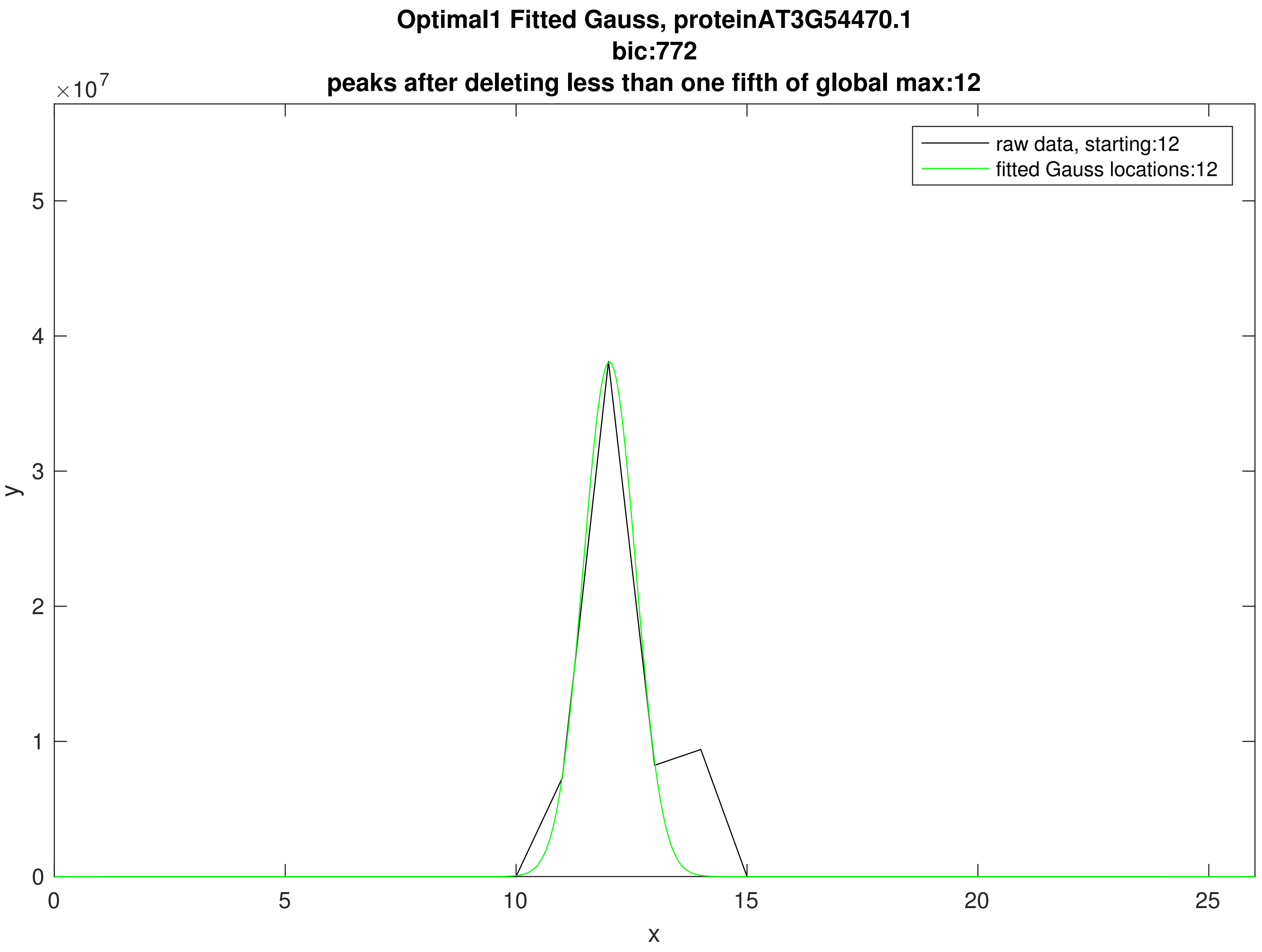

Optimal1 Fitted Gauss, proteinAT3G54470.1
bic:772
peaks after deleting less than one fifth of global max:12
×10^7
raw data, starting:12
fitted Gauss locations:12
y
x
0
1
2
3
4
5
0
5
10
15
20
25

#### Optimal1 Fitted Gauss, proteinAT3G54600.1
#### bic:705
#### peaks after deleting less than one fifth of global max:13

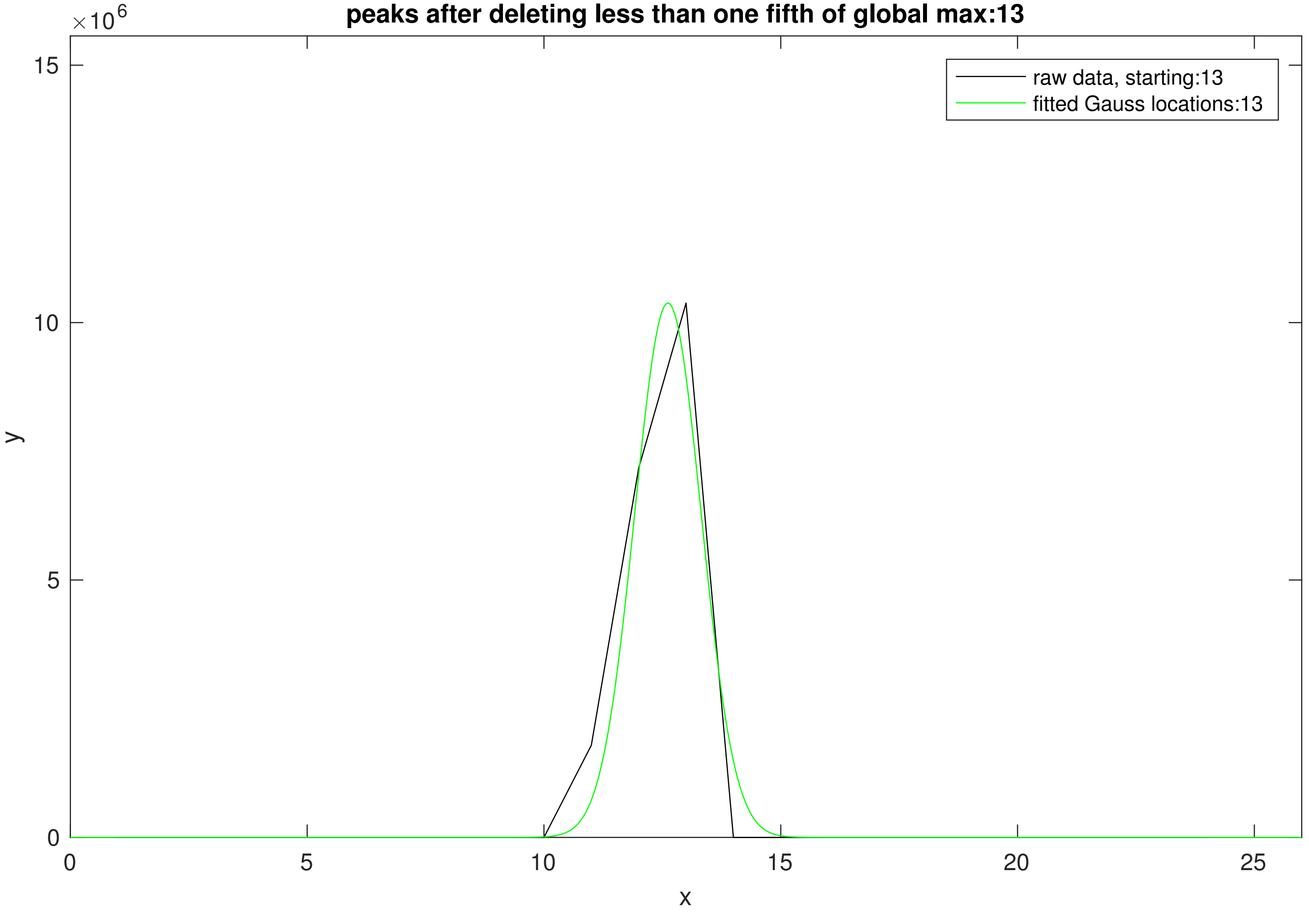

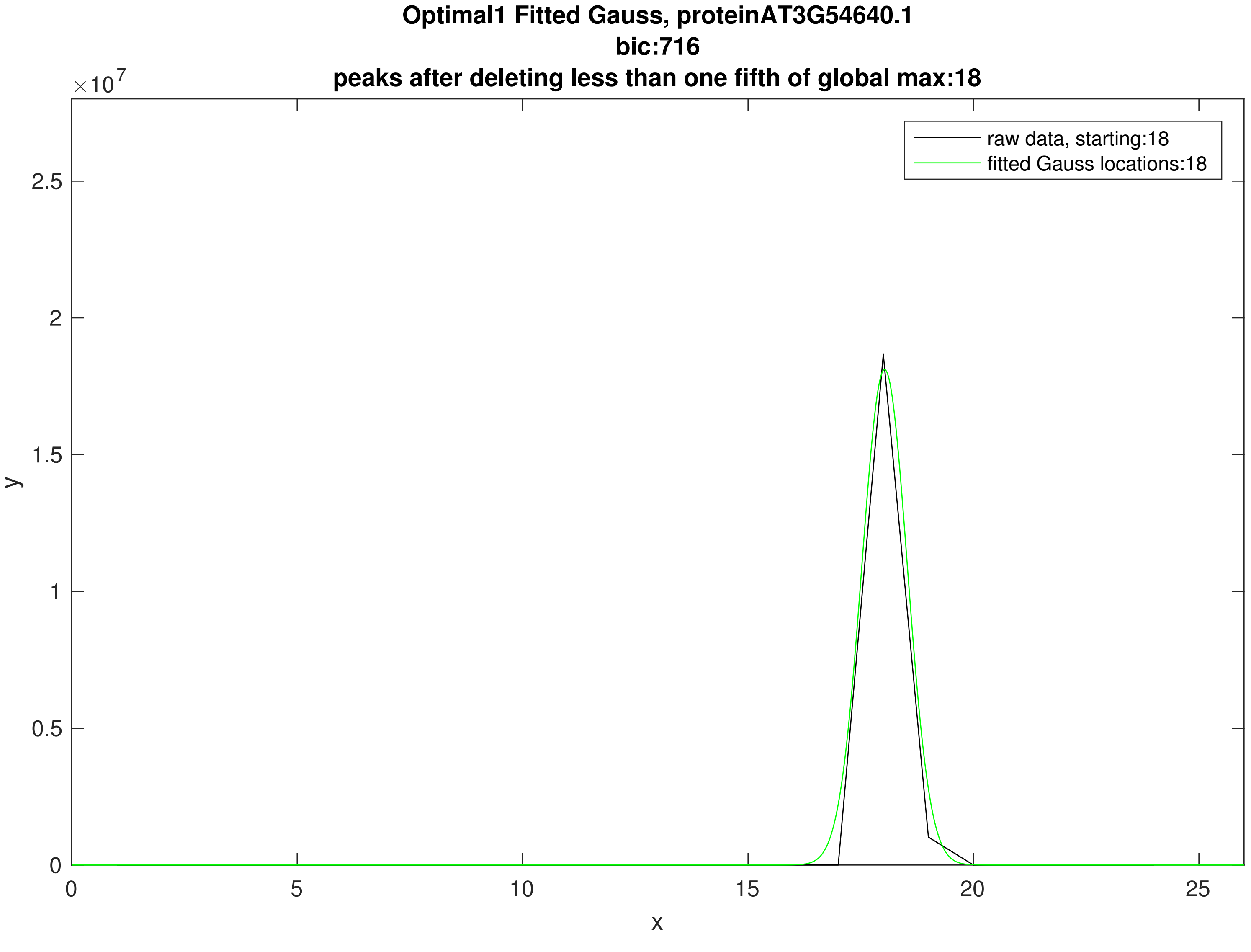
Optimal1 Fitted Gauss, proteinAT3G54640.1
bic:716
peaks after deleting less than one fifth of global max:18
×10^7
2.5
2
1.5
1
0.5
0
y
0
5
10
15
20
25
x
raw data, starting:18
fitted Gauss locations:18

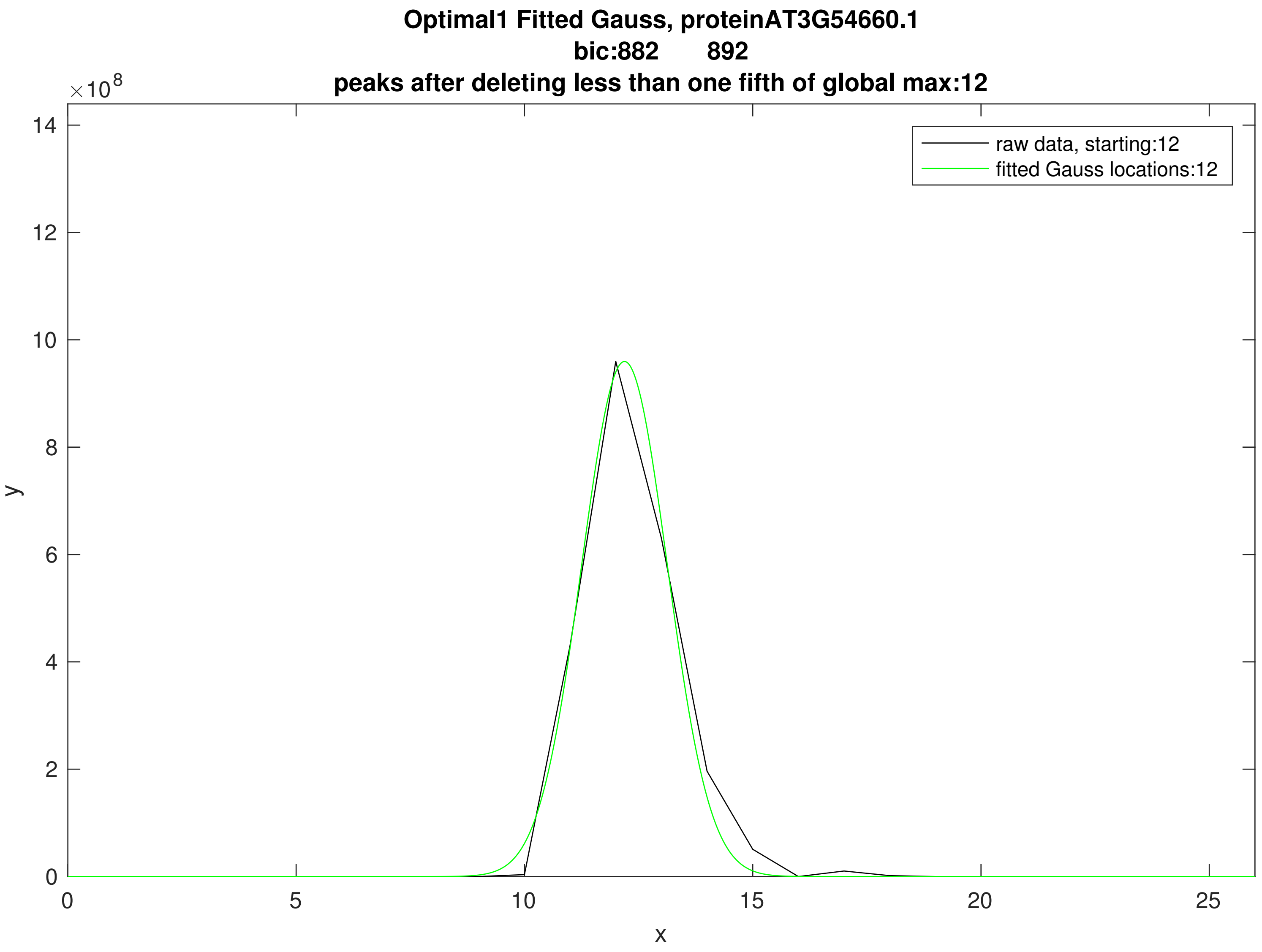
Optimal1 Fitted Gauss, proteinAT3G54660.1
bic:882 892
peaks after deleting less than one fifth of global max:12
×10^8
14
12
10
8
6
4
2
0
y
0
5
10
15
20
25
x
raw data, starting:12
fitted Gauss locations:12

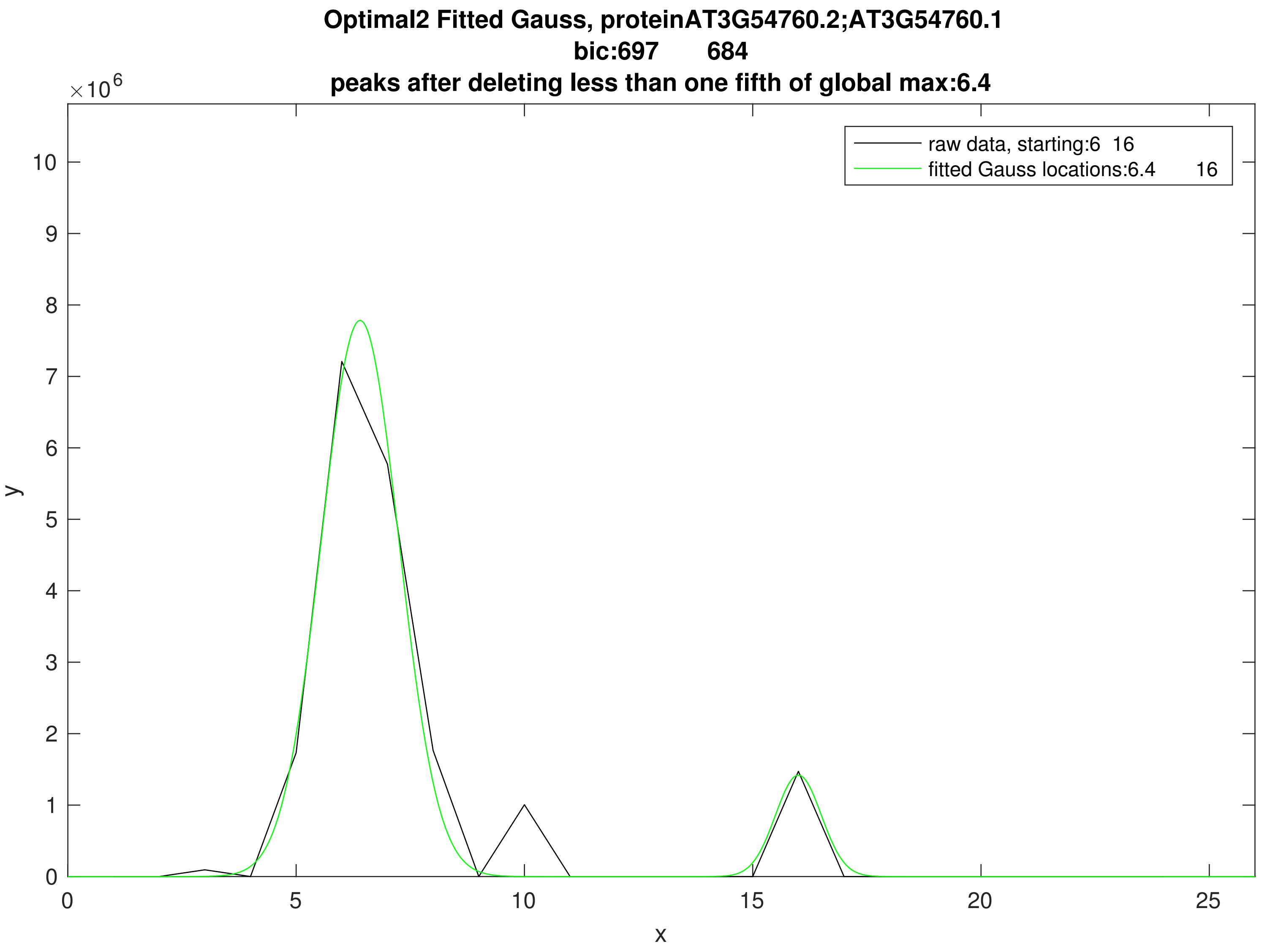

Optimal2 Fitted Gauss, proteinAT3G54760.2;AT3G54760.1
bic:697 684
peaks after deleting less than one fifth of global max:6.4
×10^6
raw data, starting:6 16
fitted Gauss locations:6.4 16
y
x
0
1
2
3
4
5
6
7
8
9
10
0
5
10
15
20
25

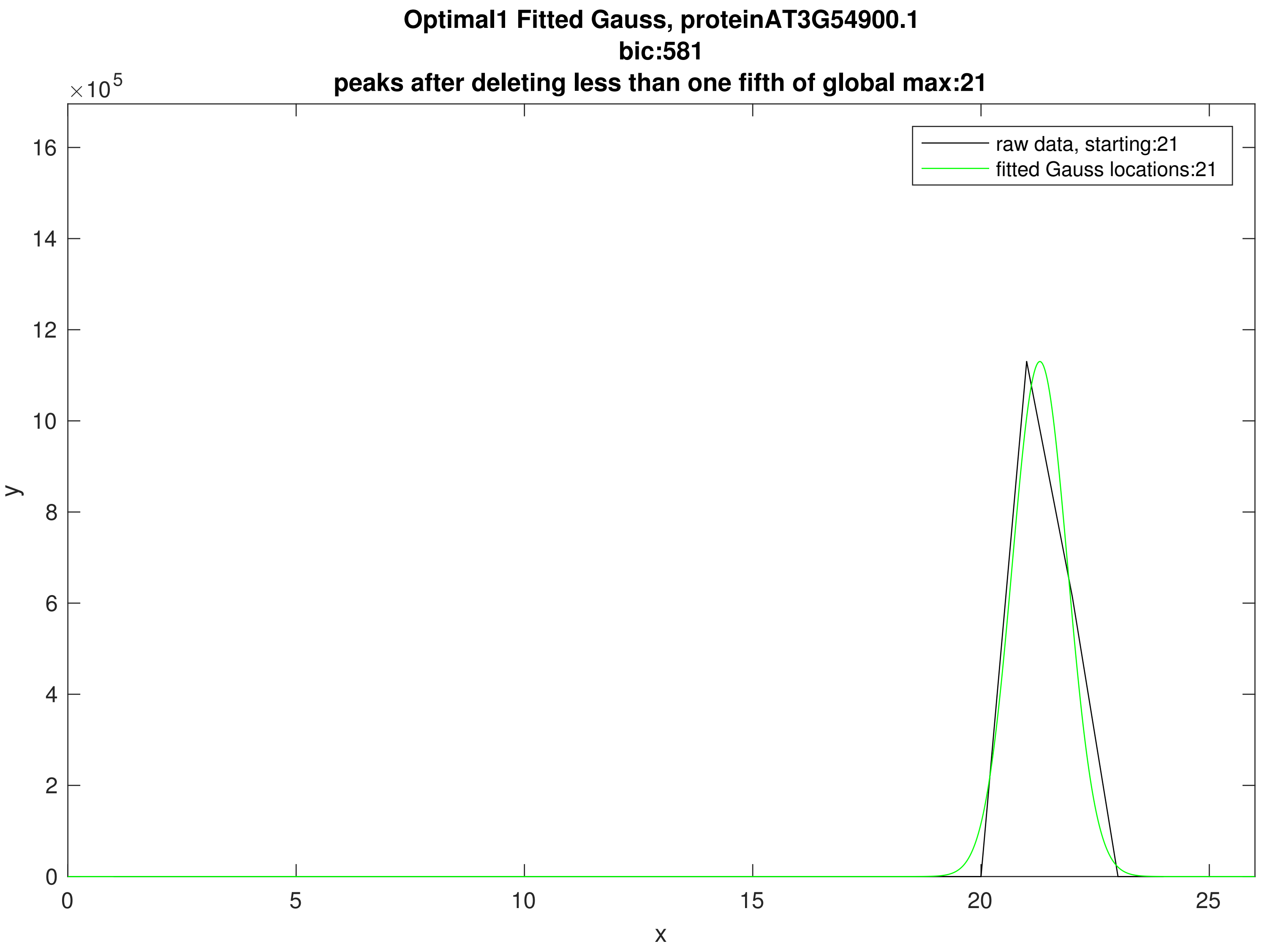

Optimal1 Fitted Gauss, proteinAT3G54900.1
bic:581
peaks after deleting less than one fifth of global max:21
×10^5
16
14
12
10
8
6
4
2
0
y
0
5
10
15
20
25
x
raw data, starting:21
fitted Gauss locations:21

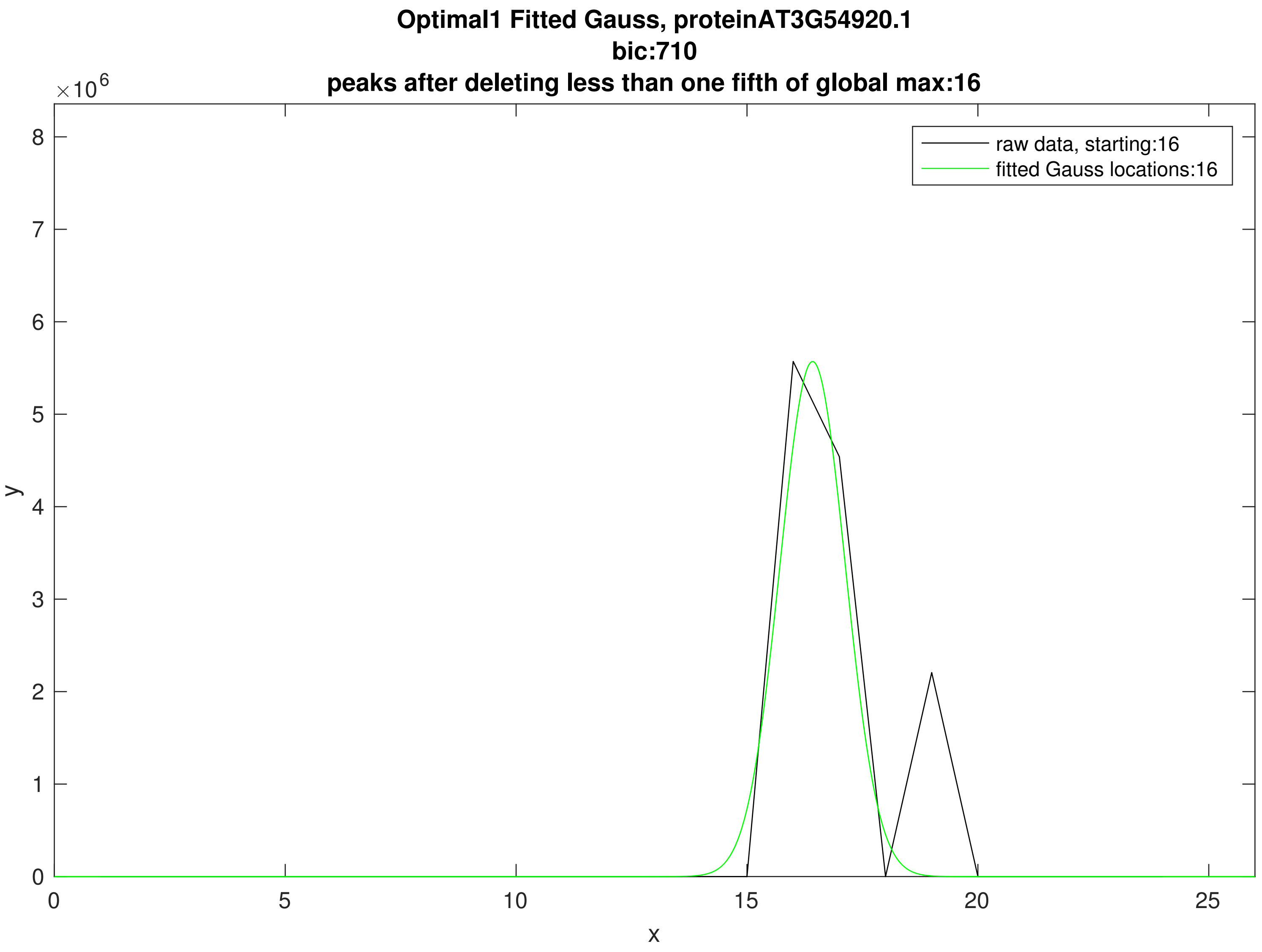
Optimal1 Fitted Gauss, proteinAT3G54920.1
bic:710
peaks after deleting less than one fifth of global max:16
×10^6
raw data, starting:16
fitted Gauss locations:16
y
x
0
1
2
3
4
5
6
7
8
0
5
10
15
20
25

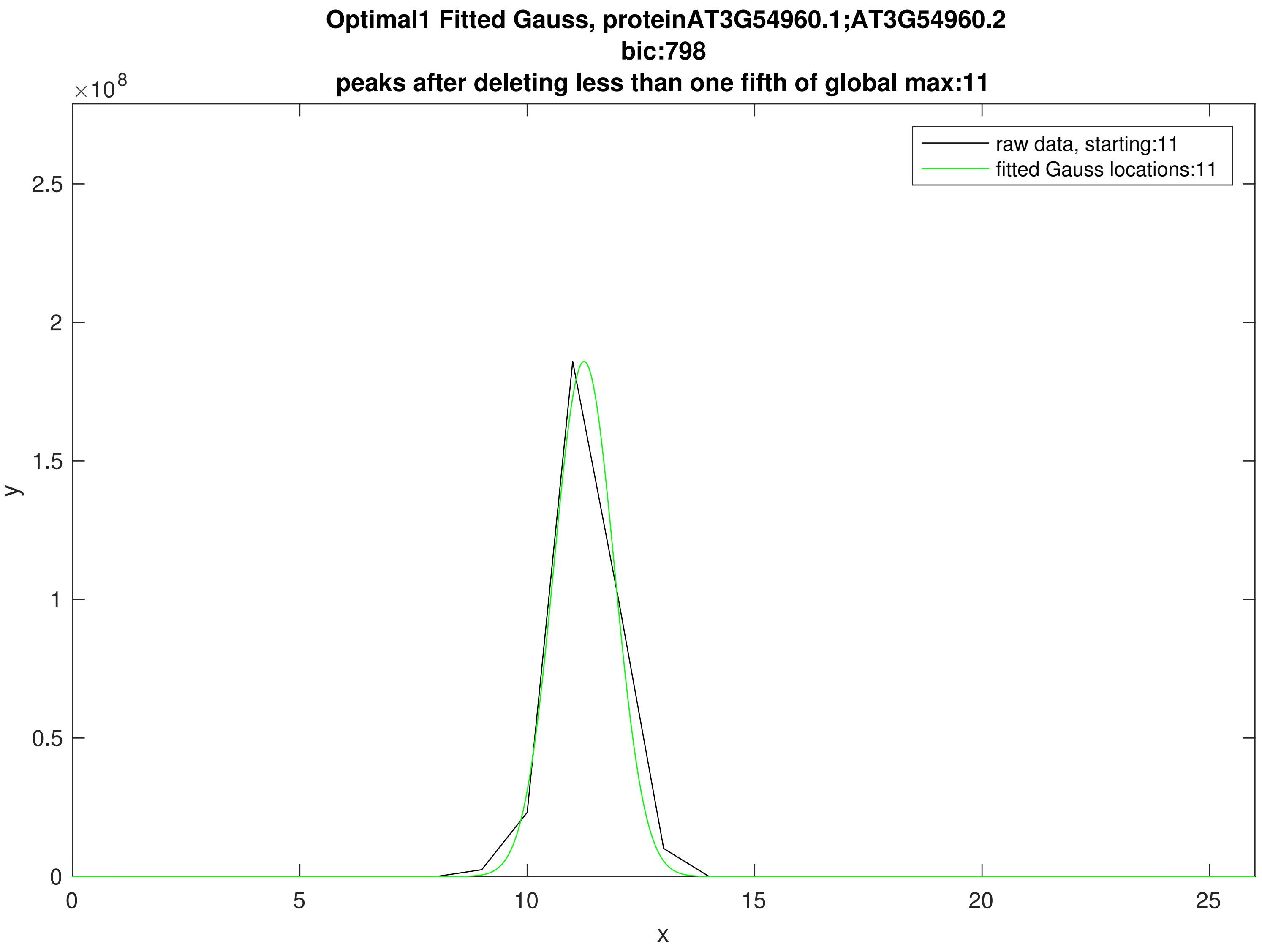

Optimal1 Fitted Gauss, proteinAT3G54960.1;AT3G54960.2
bic:798
peaks after deleting less than one fifth of global max:11
×10^8
raw data, starting:11
fitted Gauss locations:11
y
x
0
0.5
1
1.5
2
2.5
0
5
10
15
20
25

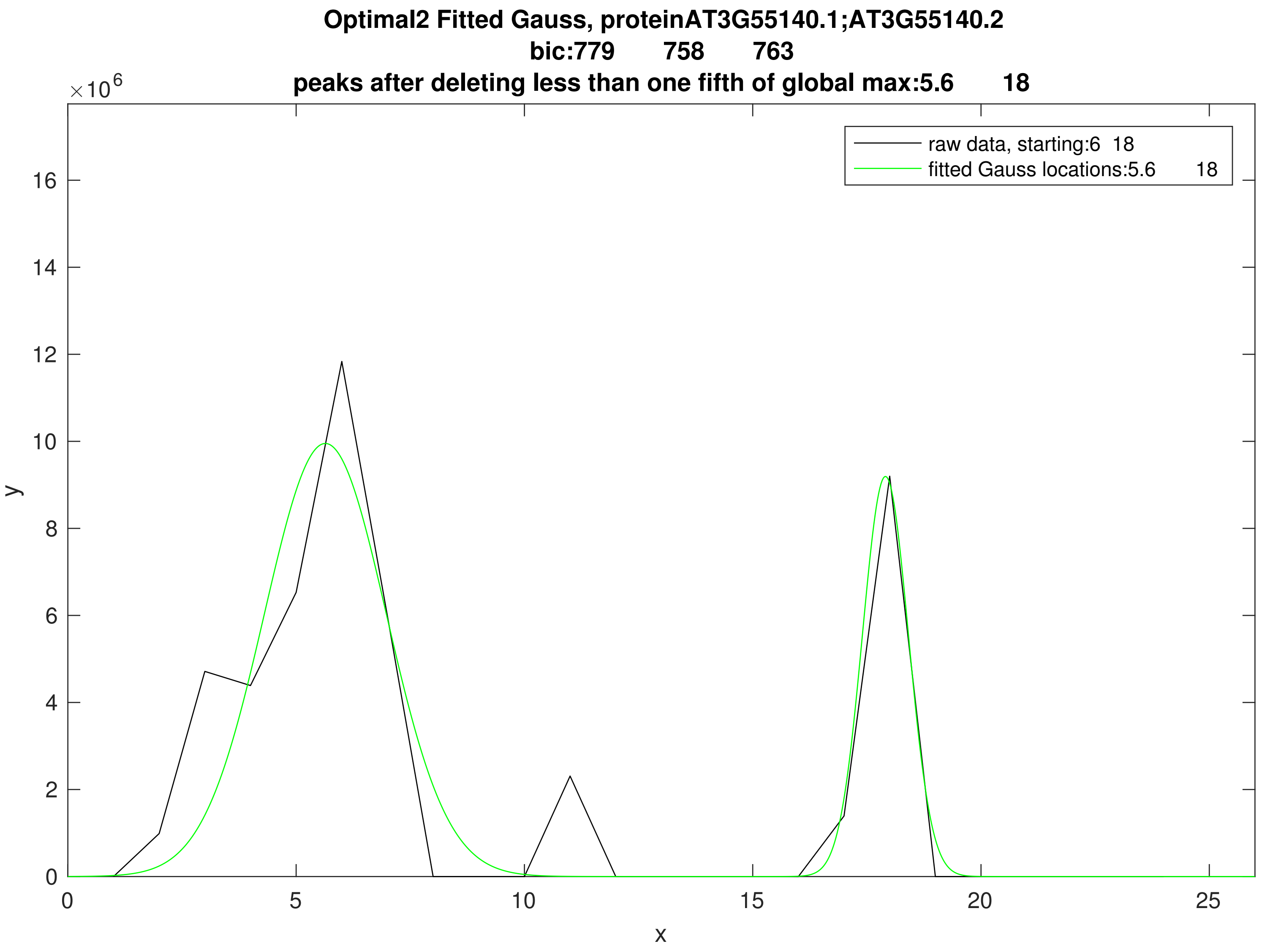

Optimal2 Fitted Gauss, proteinAT3G55140.1;AT3G55140.2
bic:779 758 763
peaks after deleting less than one fifth of global max:5.6 18
×10^6
raw data, starting:6 18
fitted Gauss locations:5.6 18
y
x
0
2
4
6
8
10
12
14
16
0
5
10
15
20
25

Optimal1 Fitted Gauss, proteinAT3G55220.1;AT3G55200.1
bic:881 886
peaks after deleting less than one fifth of global max:11

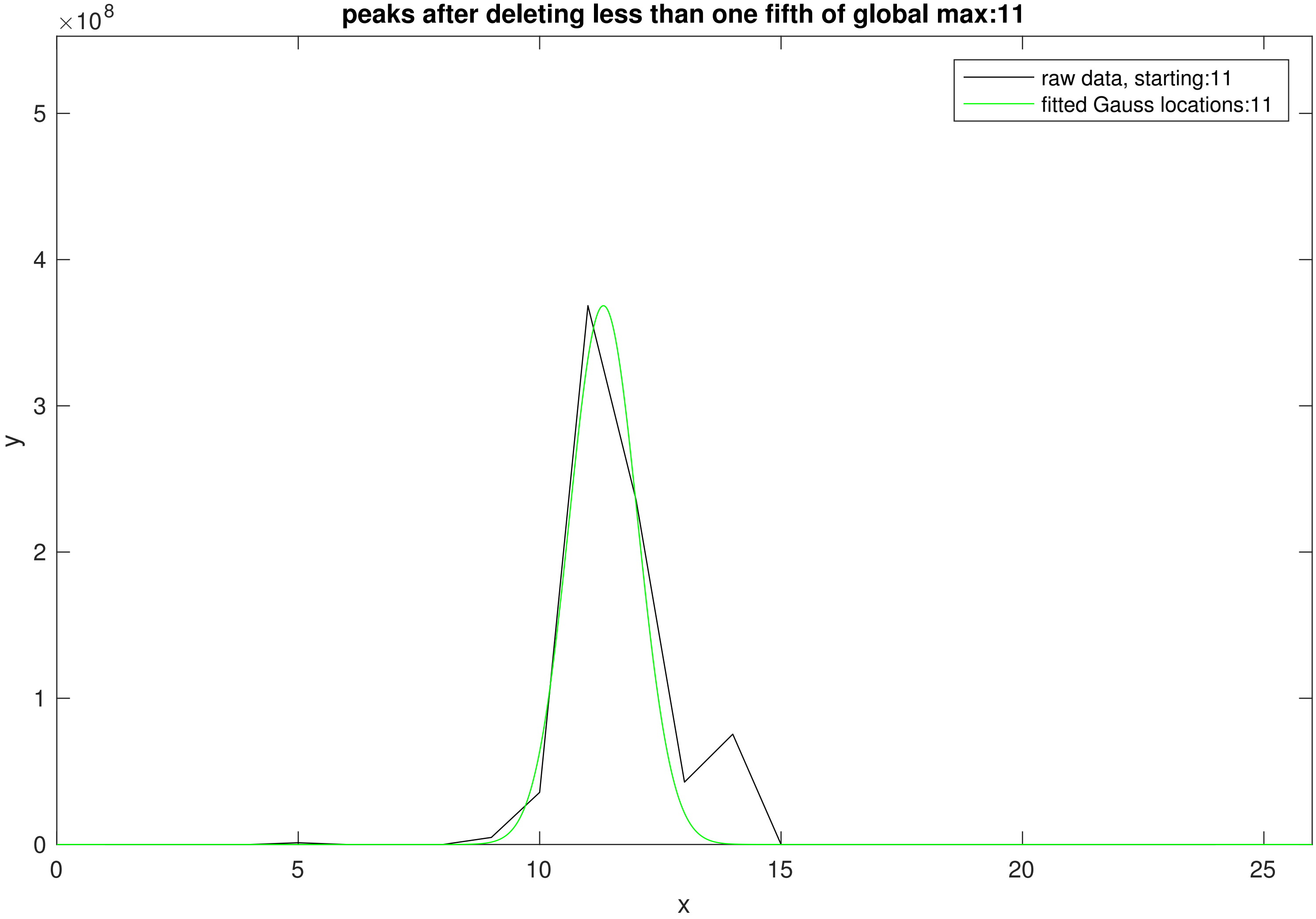

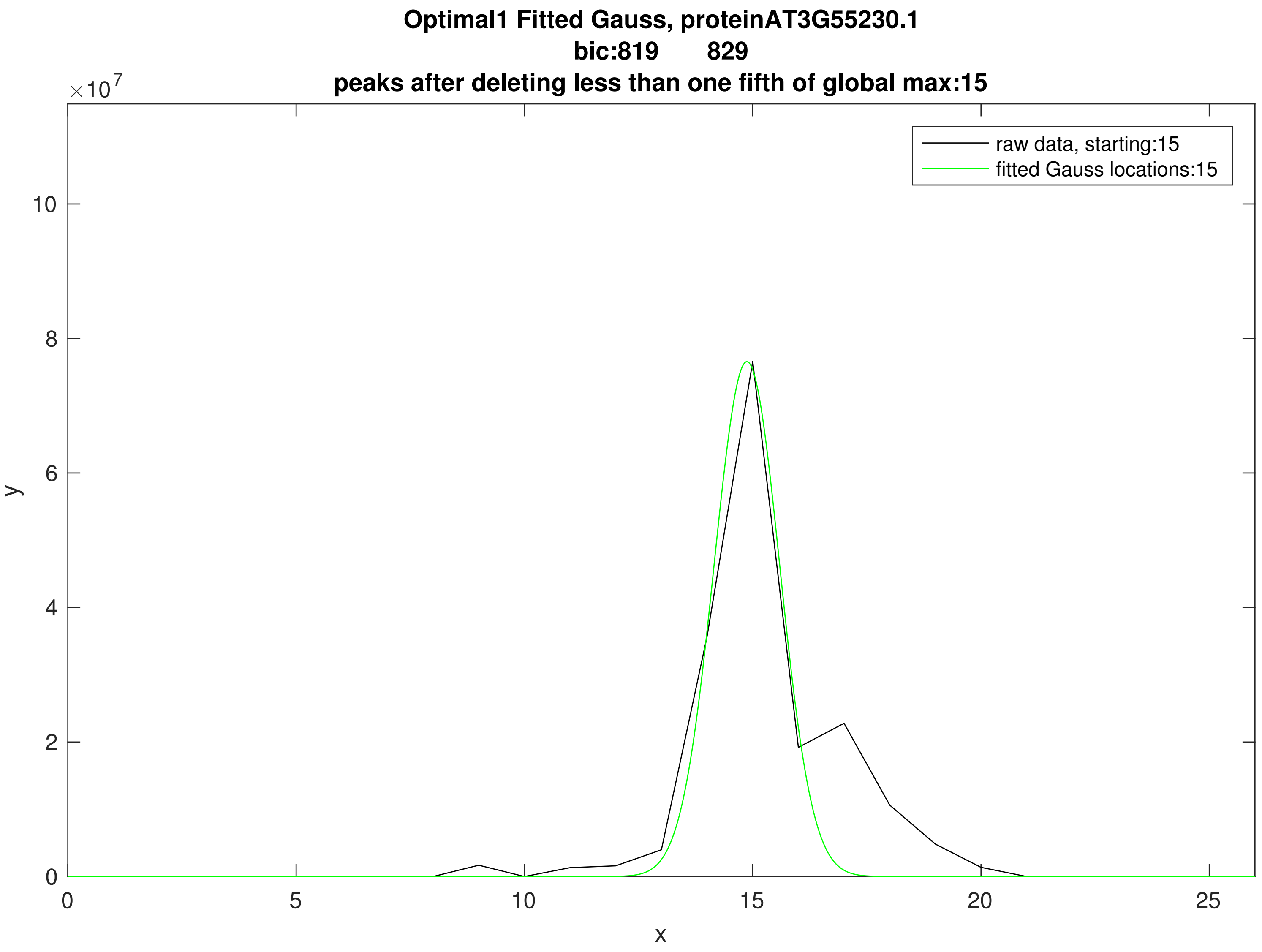
Optimal1 Fitted Gauss, proteinAT3G55230.1
bic:819 829
peaks after deleting less than one fifth of global max:15
×10^7
raw data, starting:15
fitted Gauss locations:15
10
8
6
4
2
0
y
0
5
10
15
20
25
x

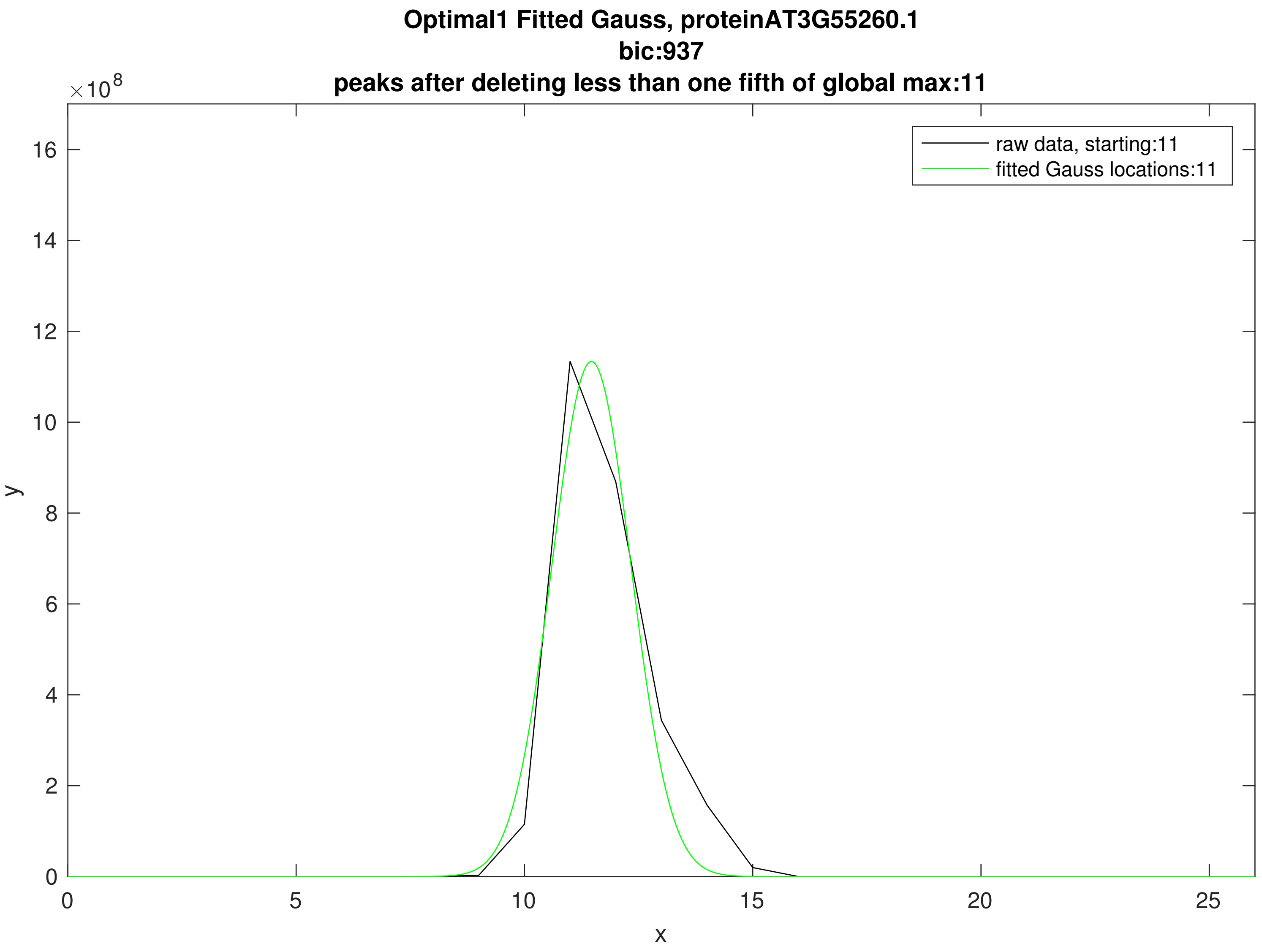

Optimal1 Fitted Gauss, proteinAT3G55260.1
bic:937
peaks after deleting less than one fifth of global max:11
×10^8
16
14
12
10
8
6
4
2
0
y
0
5
10
15
20
25
x
raw data, starting:11
fitted Gauss locations:11

**Optimal1 Fitted Gauss, proteinAT3G55280.3;AT3G55280.2;AT3G55280.1;AT2G39460.2;AT2G39460.1**
**bic:814 817**
**peaks after deleting less than one fifth of global max:3.7**

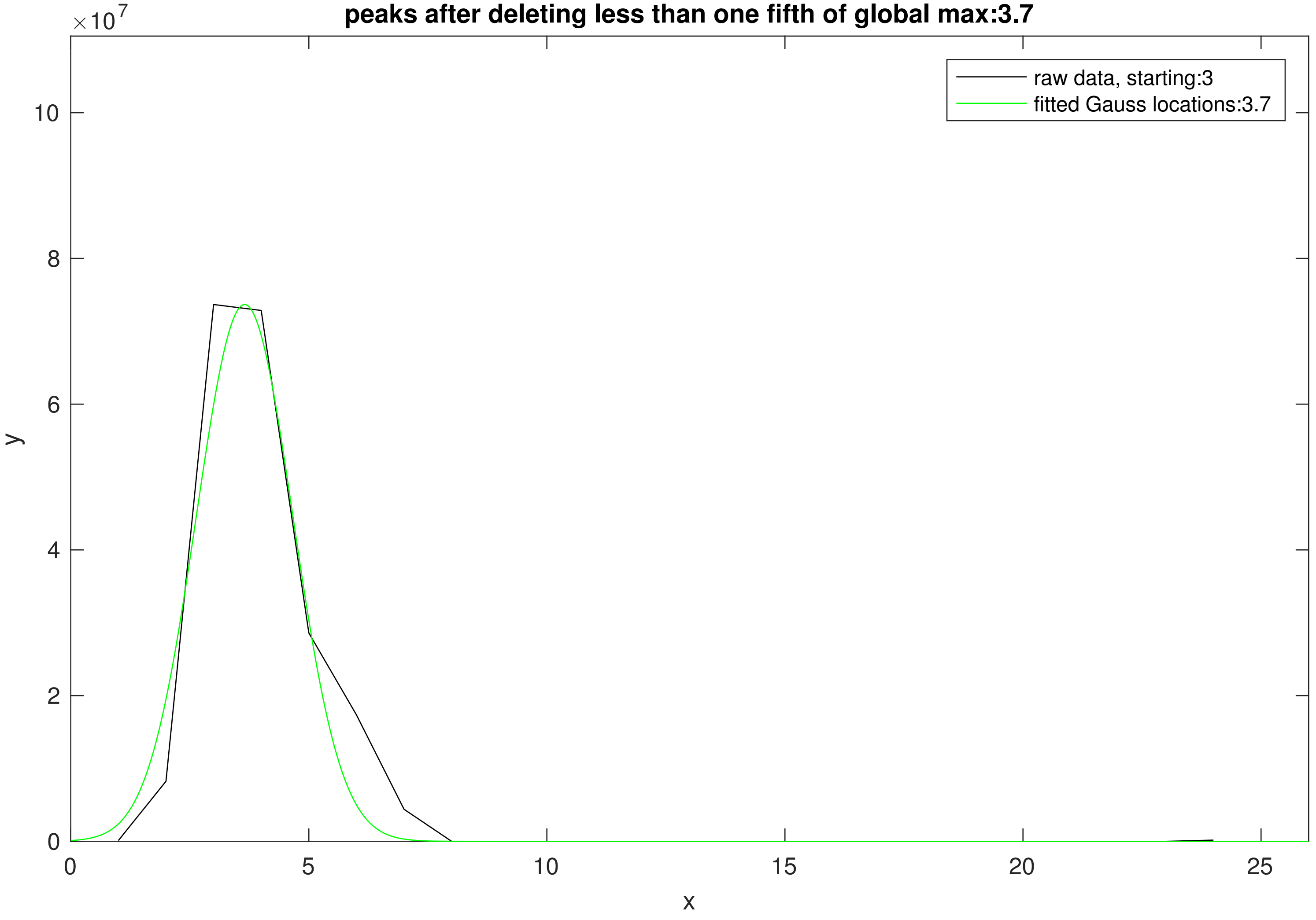

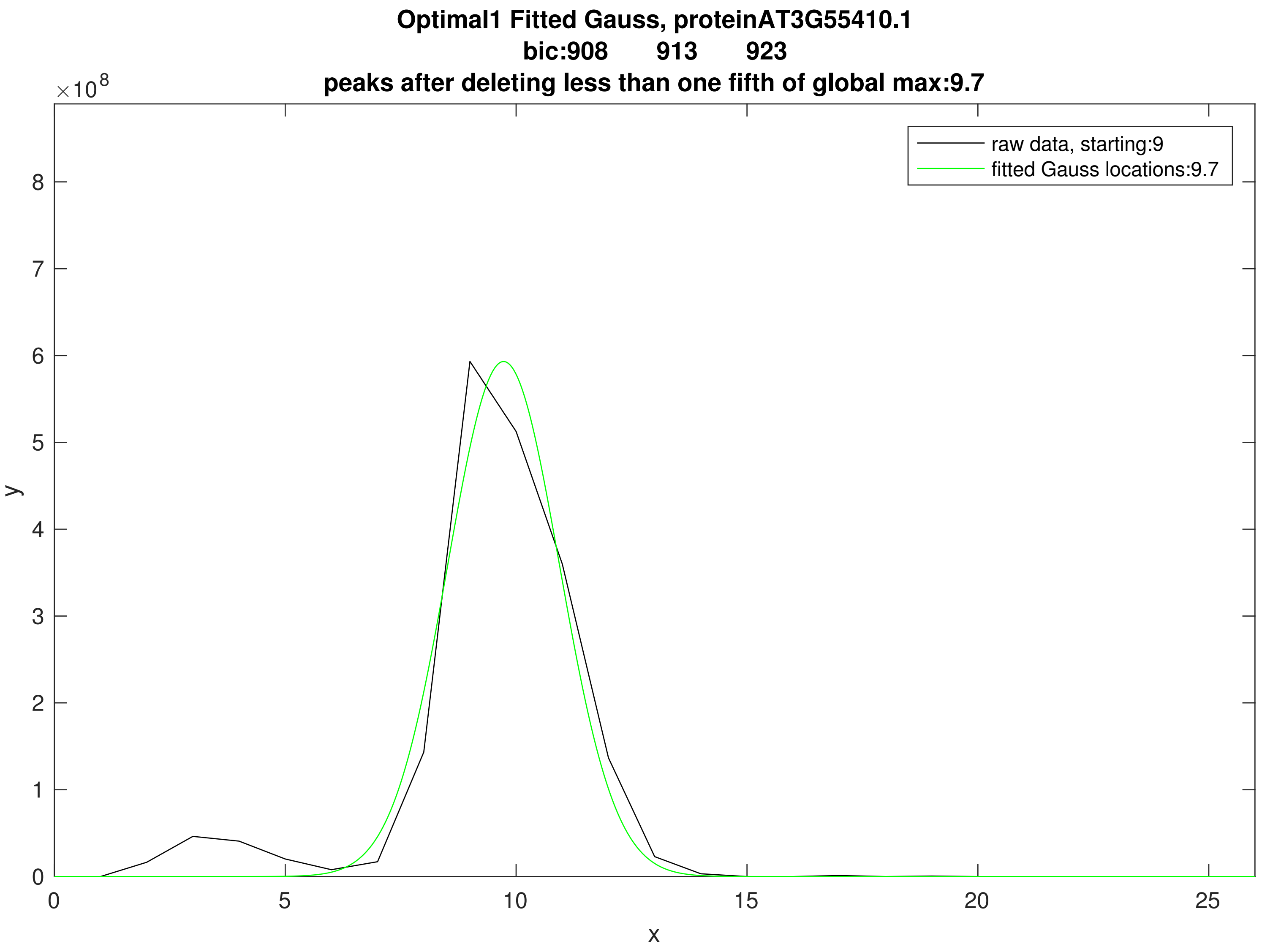
Optimal1 Fitted Gauss, proteinAT3G55410.1
bic:908 913 923
peaks after deleting less than one fifth of global max:9.7
×10^8
raw data, starting:9
fitted Gauss locations:9.7
y
x
0
1
2
3
4
5
6
7
8
0
5
10
15
20
25

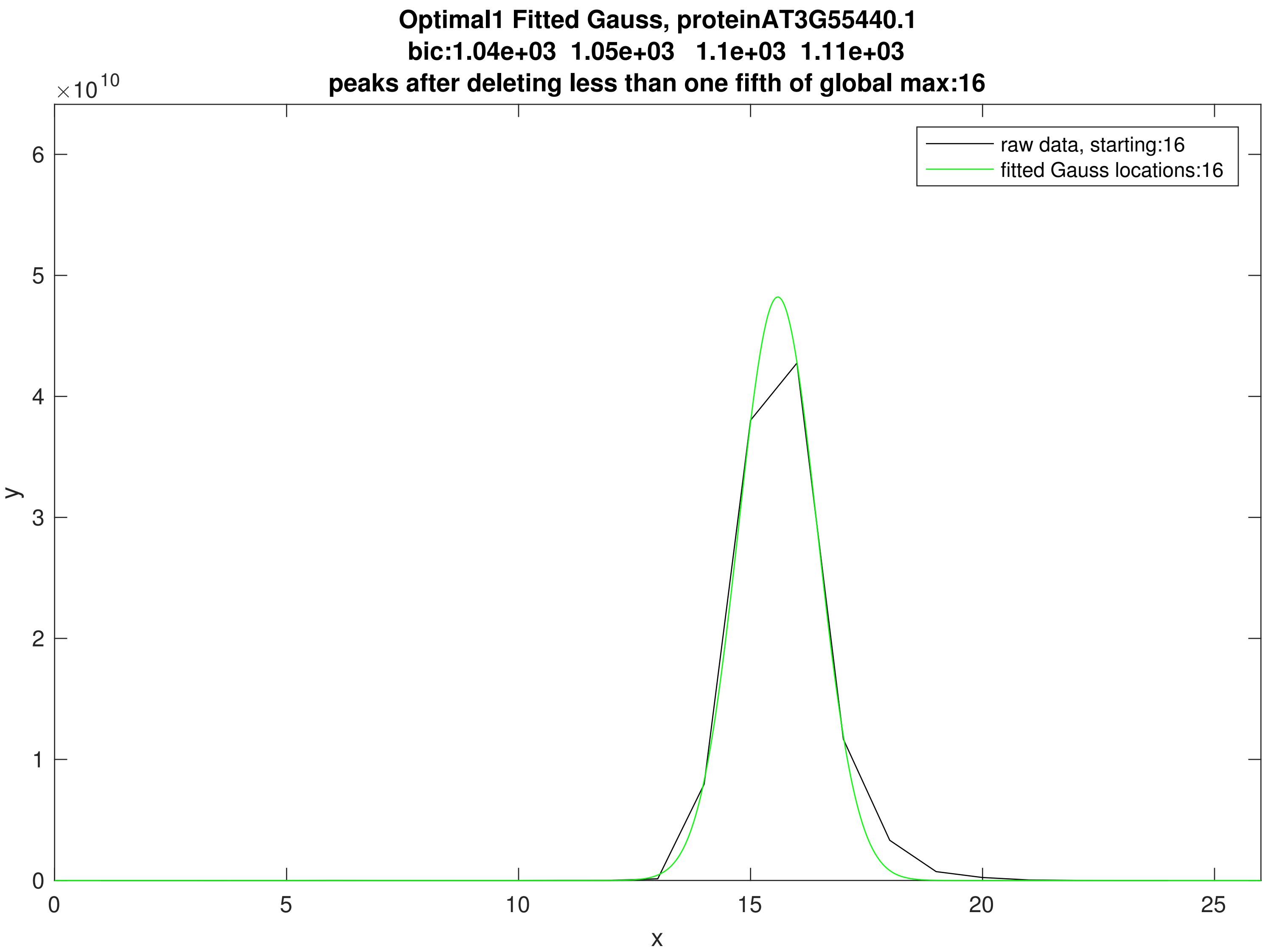
Optimal1 Fitted Gauss, proteinAT3G55440.1
bic:1.04e+03 1.05e+03 1.1e+03 1.11e+03
peaks after deleting less than one fifth of global max:16
×10^10
raw data, starting:16
fitted Gauss locations:16
y
x
0
1
2
3
4
5
6
0
5
10
15
20
25

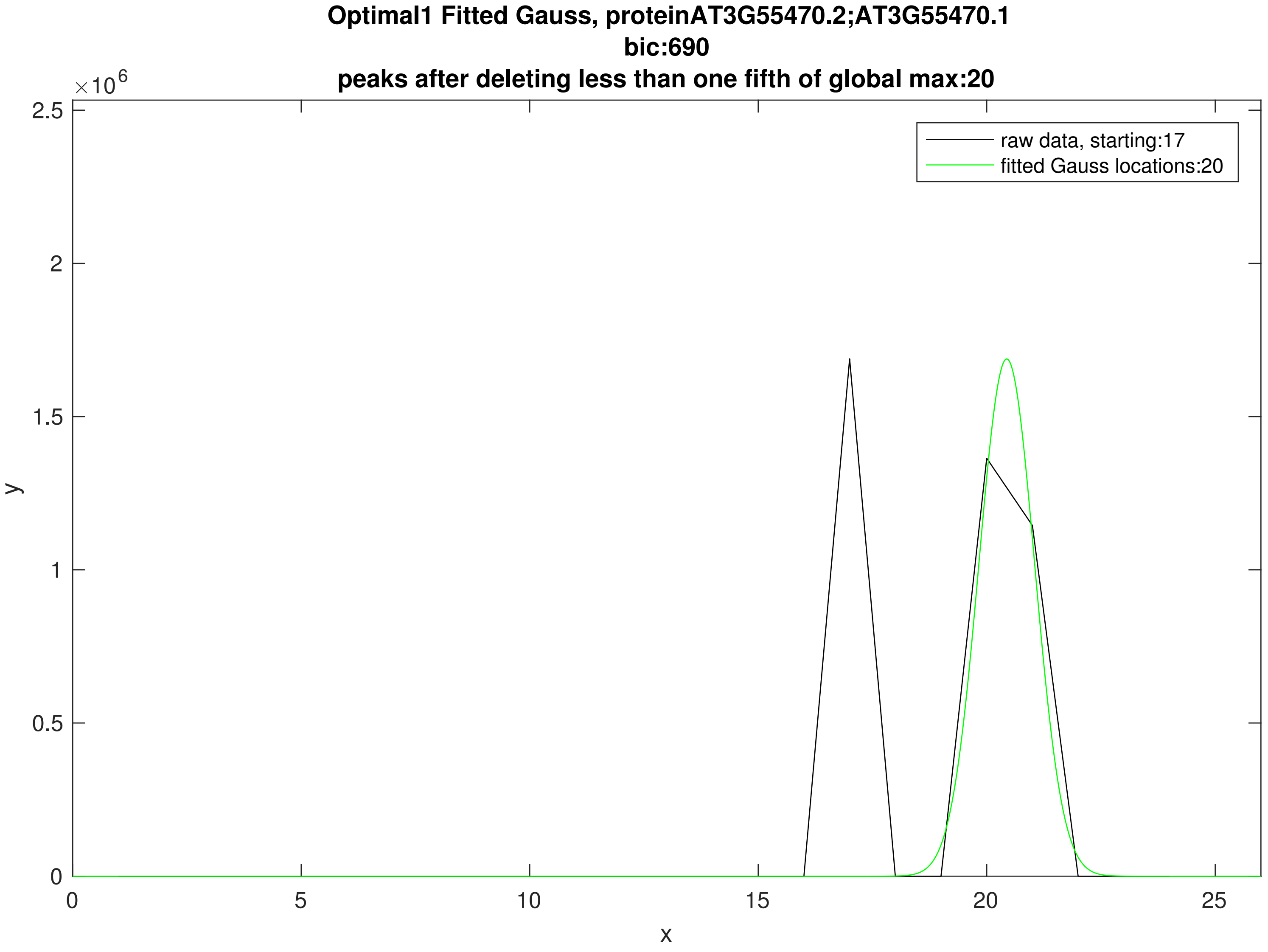
Optimal1 Fitted Gauss, proteinAT3G55470.2;AT3G55470.1
bic:690
peaks after deleting less than one fifth of global max:20
×10^6
2.5
2
1.5
1
0.5
0
y
0
5
10
15
20
25
x
raw data, starting:17
fitted Gauss locations:20

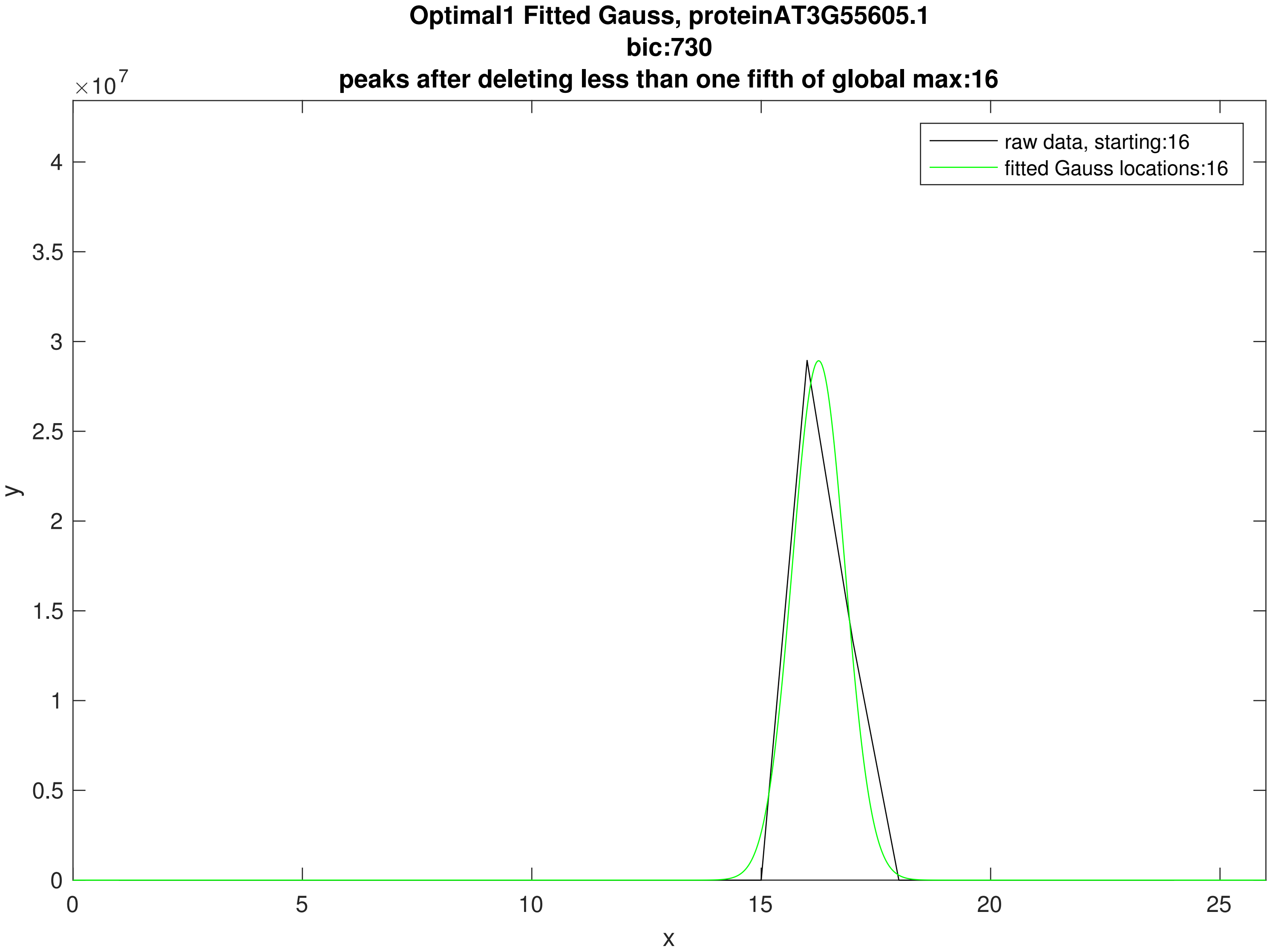
Optimal1 Fitted Gauss, proteinAT3G55605.1
bic:730
peaks after deleting less than one fifth of global max:16
×10^7
raw data, starting:16
fitted Gauss locations:16
4
3.5
3
2.5
2
1.5
1
0.5
0
y
0
5
10
15
20
25
x

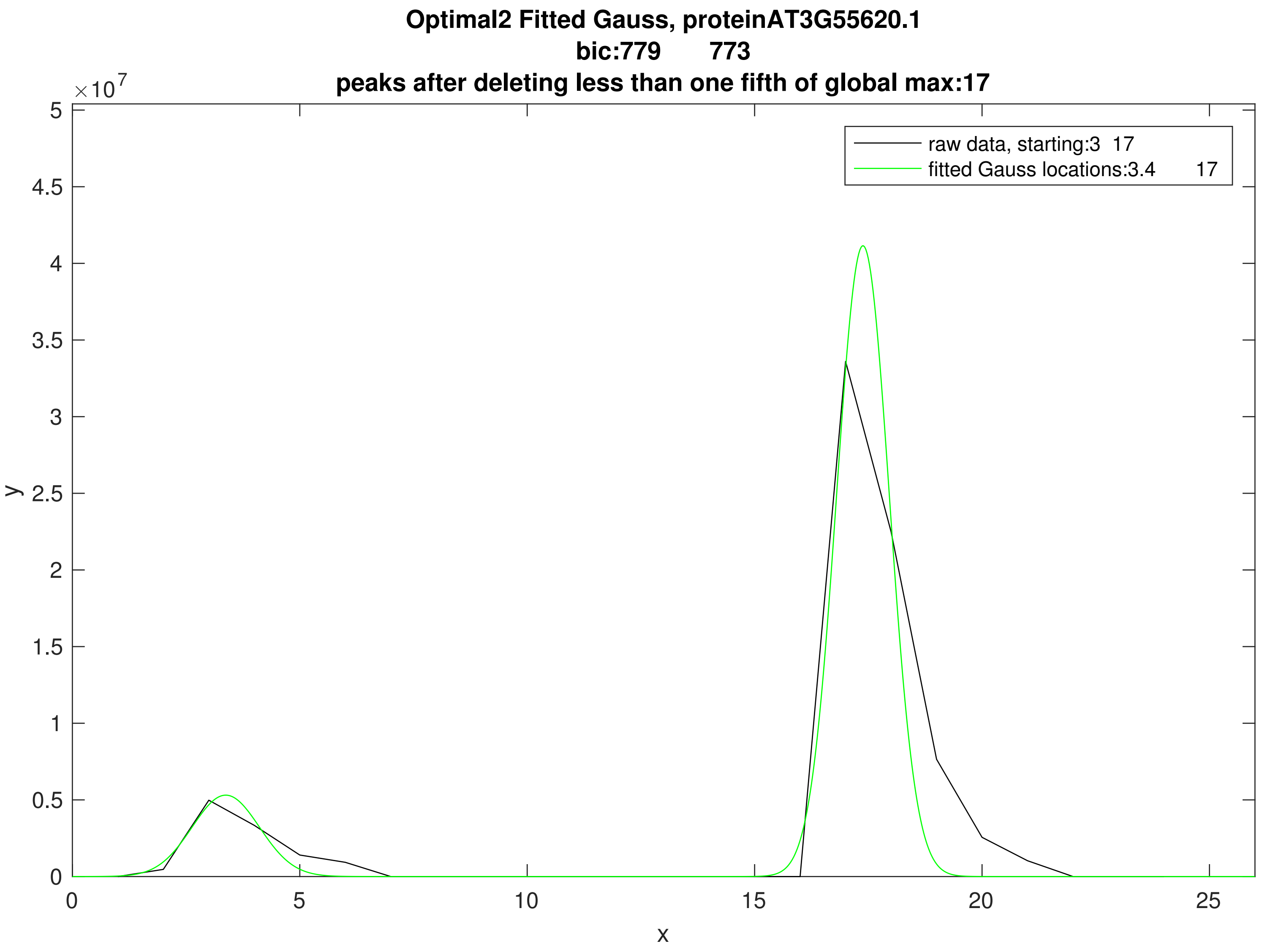
Optimal2 Fitted Gauss, proteinAT3G55620.1
bic:779 773
peaks after deleting less than one fifth of global max:17
×10^7
raw data, starting:3 17
fitted Gauss locations:3.4 17
y
x
0
0.5
1
1.5
2
2.5
3
3.5
4
4.5
5
0
5
10
15
20
25

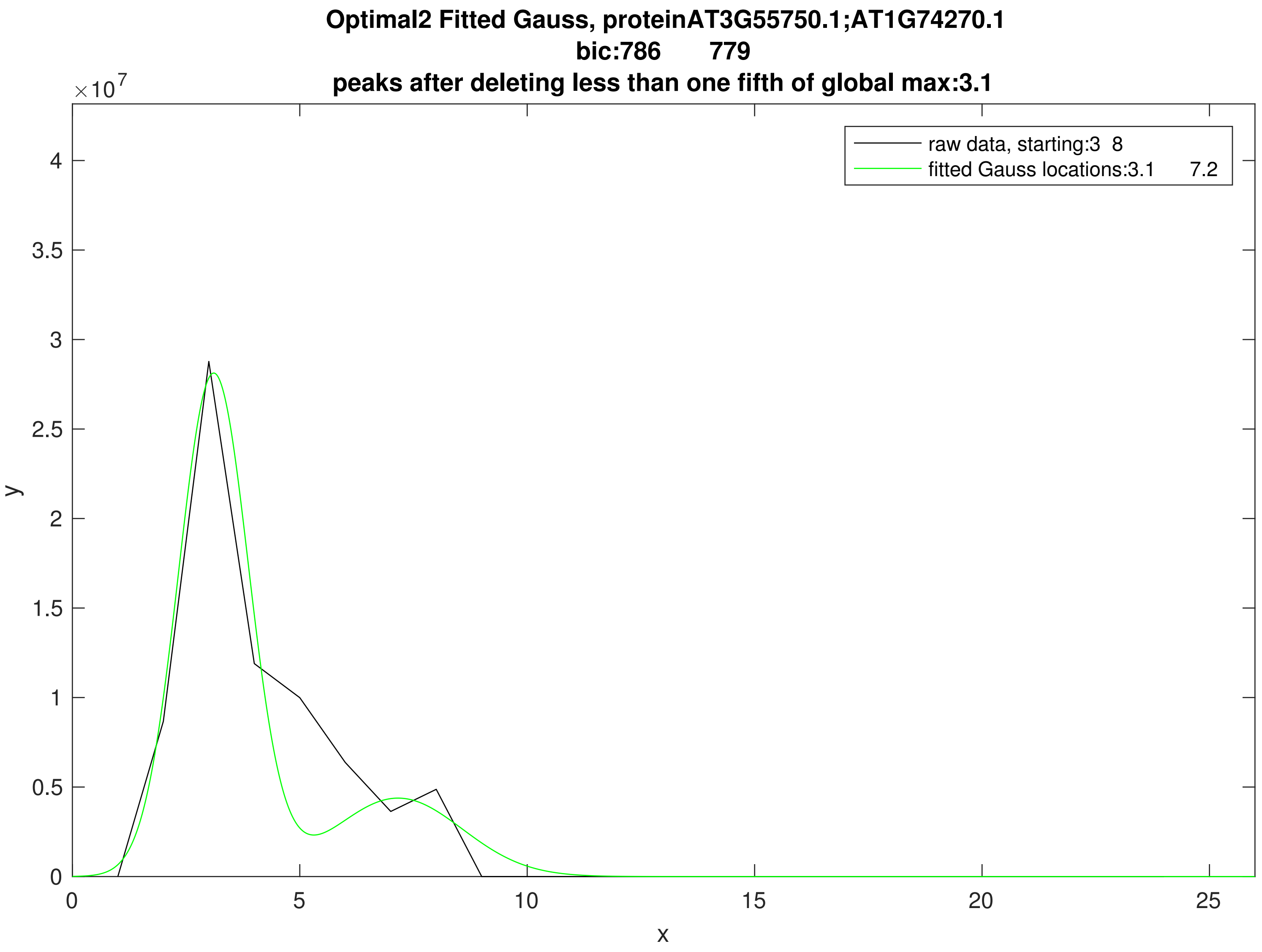

Optimal2 Fitted Gauss, proteinAT3G55750.1;AT1G74270.1
bic:786 779
peaks after deleting less than one fifth of global max:3.1
×10^7
raw data, starting:3 8
fitted Gauss locations:3.1 7.2
y
x
0
0.5
1
1.5
2
2.5
3
3.5
4
5
10
15
20
25

**Optimal1 Fitted Gauss, proteinAT3G55800.1**
**bic:865 875**
**peaks after deleting less than one fifth of global max:14**

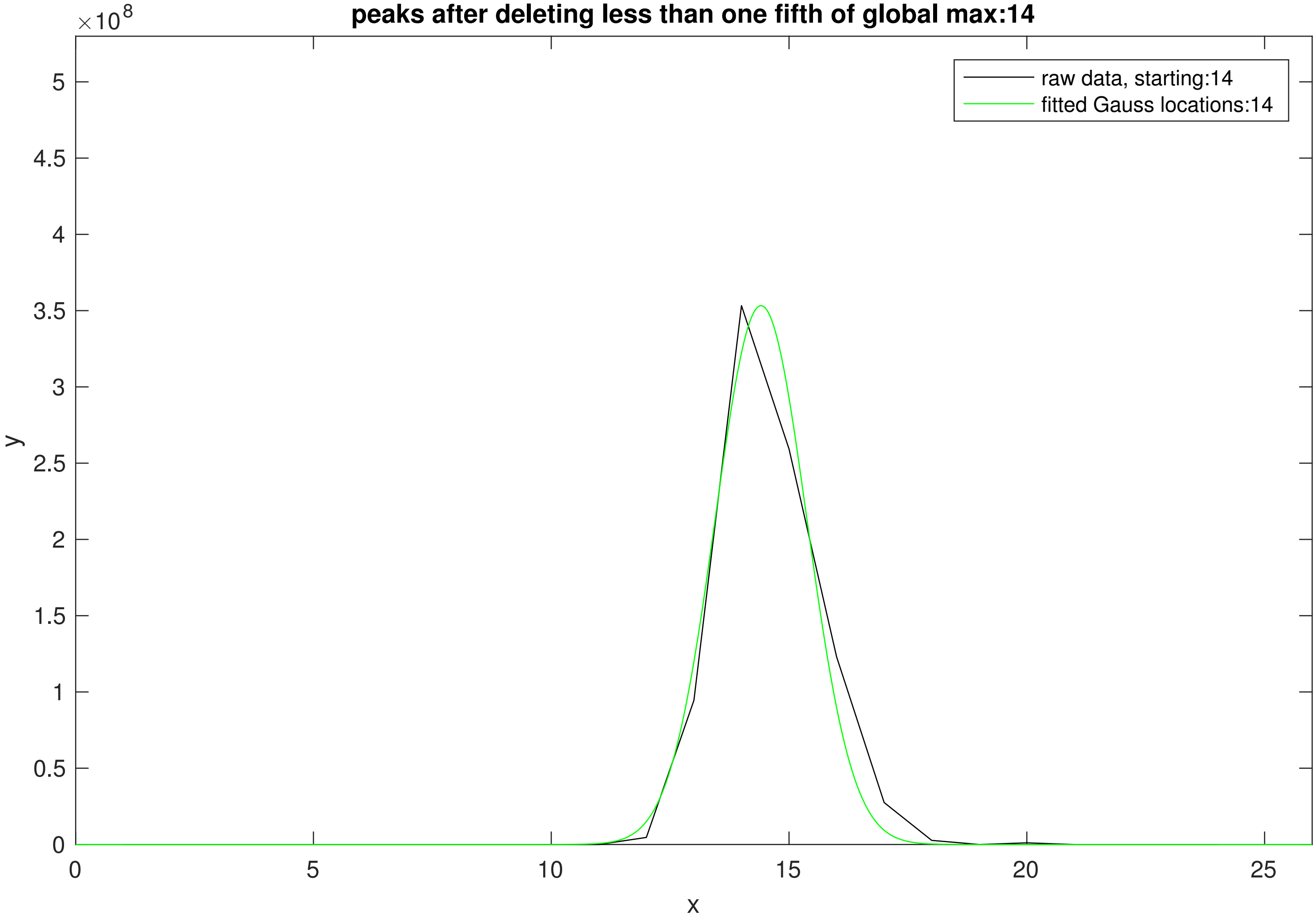

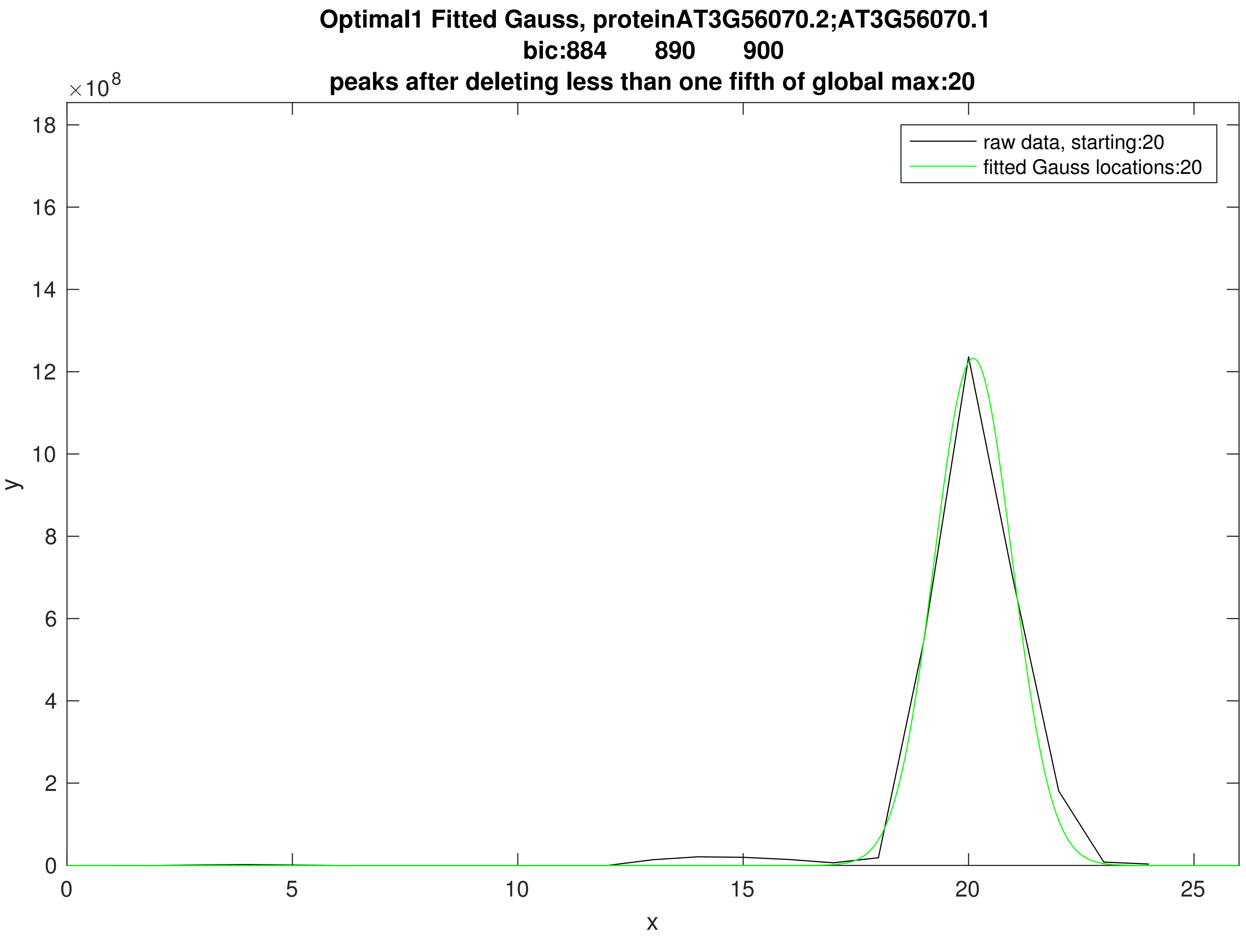
Optimal1 Fitted Gauss, proteinAT3G56070.2;AT3G56070.1
bic:884 890 900
peaks after deleting less than one fifth of global max:20
×10^8
raw data, starting:20
fitted Gauss locations:20
y
x
0
2
4
6
8
10
12
14
16
18
0
5
10
15
20
25

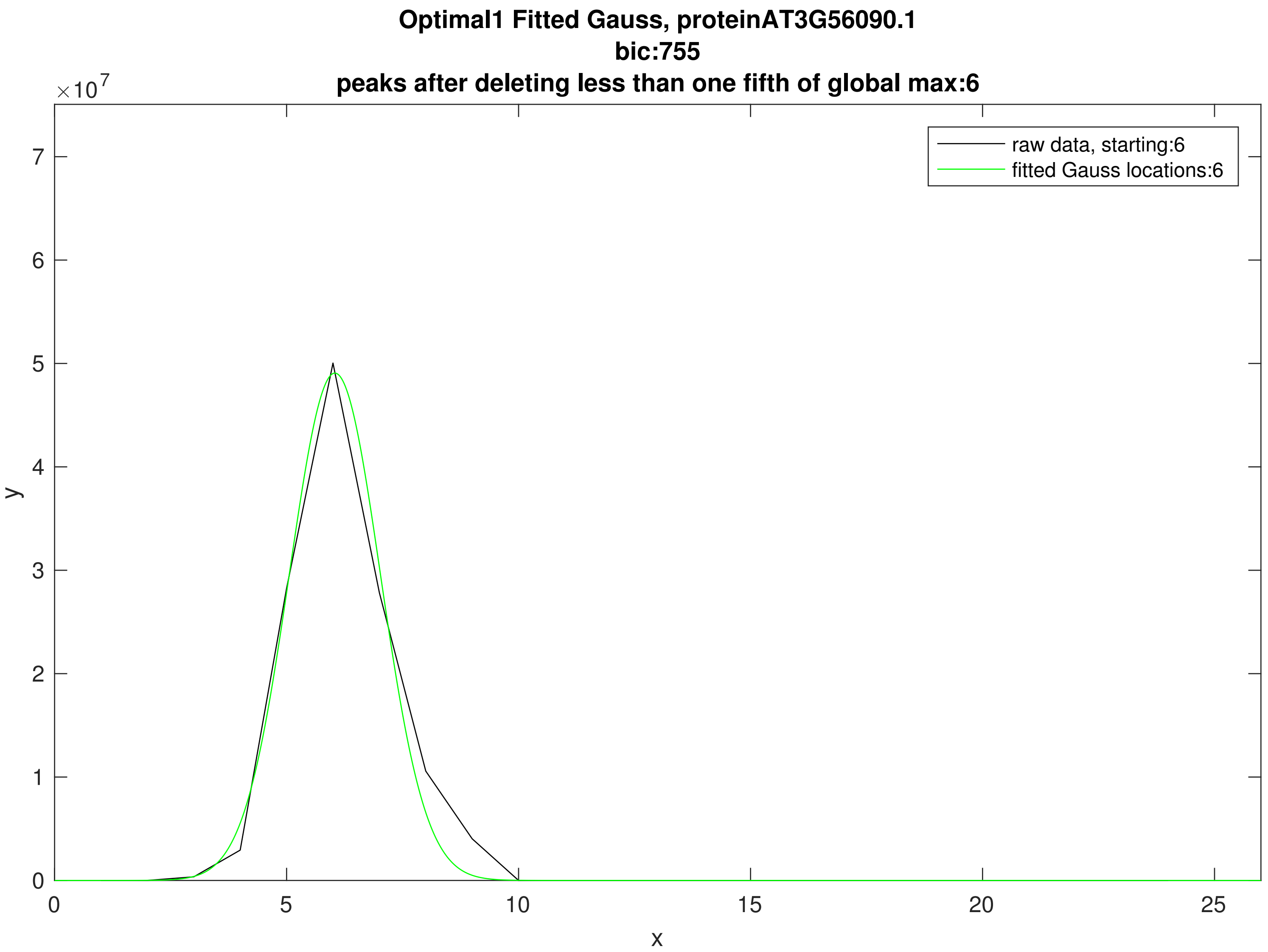
Optimal1 Fitted Gauss, proteinAT3G56090.1
bic:755
peaks after deleting less than one fifth of global max:6
$\times 10^7$
raw data, starting:6
fitted Gauss locations:6
y
x
0
1
2
3
4
5
6
7
0
5
10
15
20
25

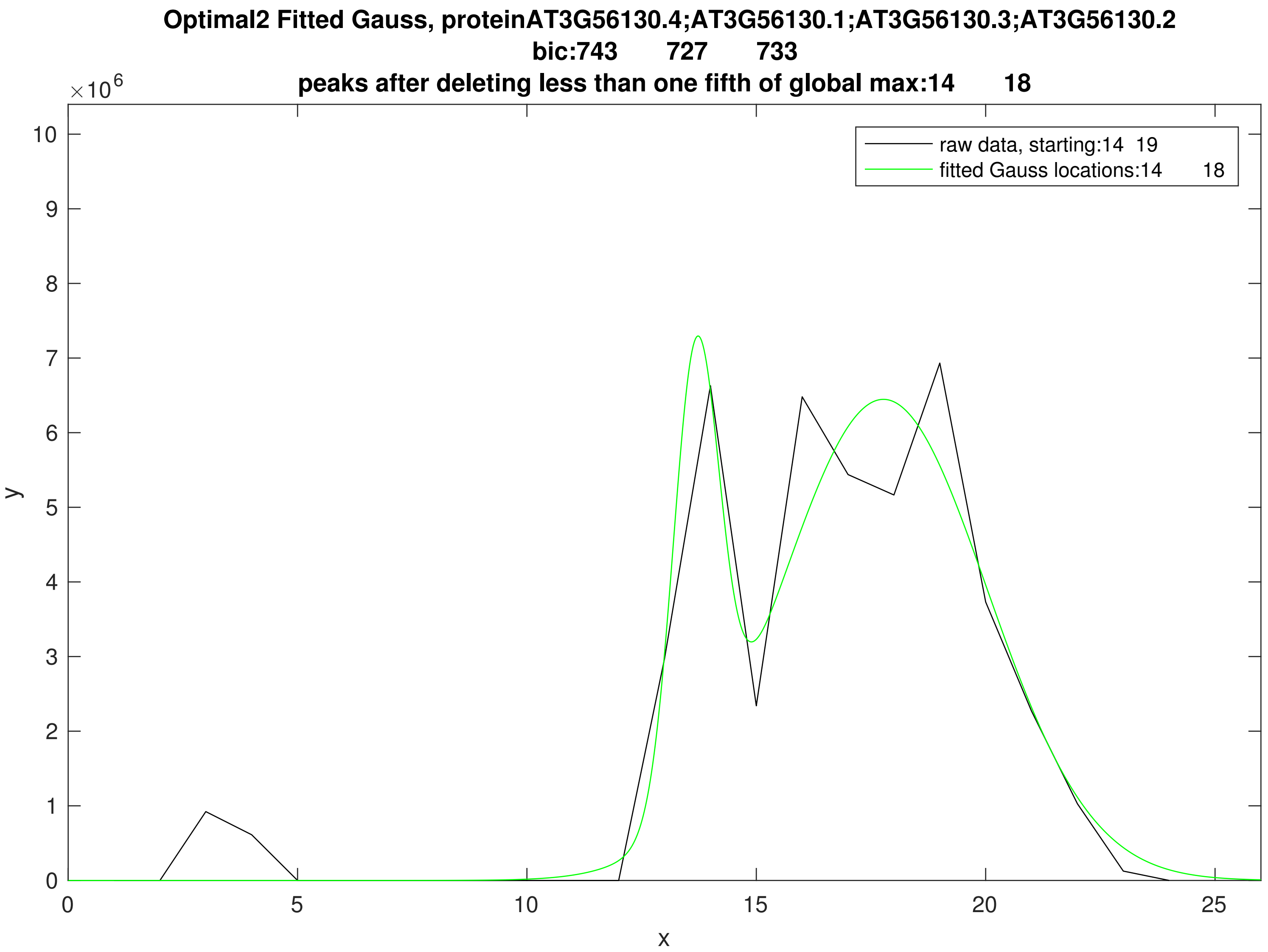
Optimal2 Fitted Gauss, proteinAT3G56130.4;AT3G56130.1;AT3G56130.3;AT3G56130.2
bic:743 727 733
peaks after deleting less than one fifth of global max:14 18
×10^6
raw data, starting:14 19
fitted Gauss locations:14 18
y
x
0
1
2
3
4
5
6
7
8
9
10
0
5
10
15
20
25

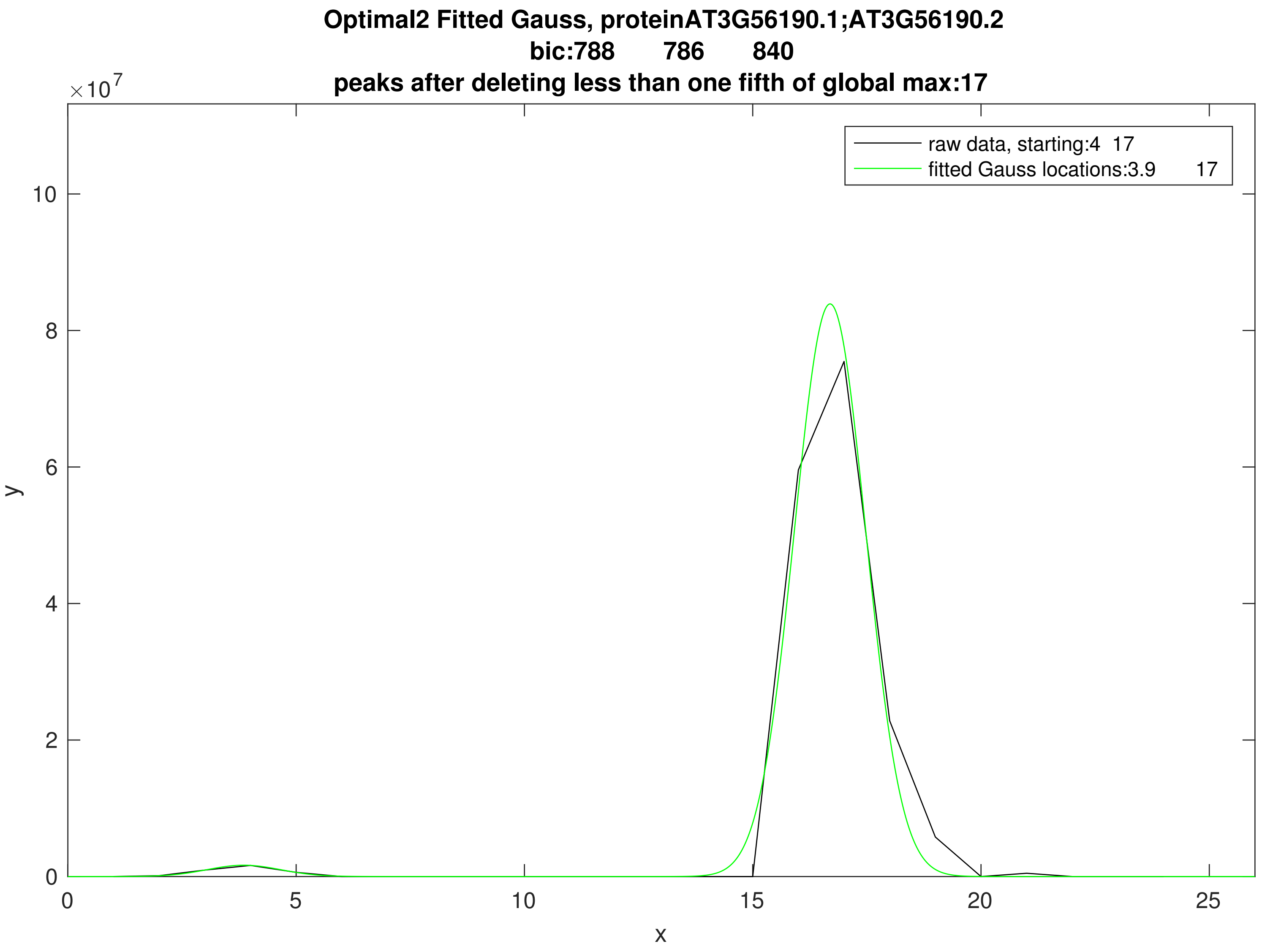
Optimal2 Fitted Gauss, proteinAT3G56190.1;AT3G56190.2
bic:788 786 840
peaks after deleting less than one fifth of global max:17
×10^7
raw data, starting:4 17
fitted Gauss locations:3.9 17
10
8
6
4
2
0
y
0
5
10
15
20
25
x

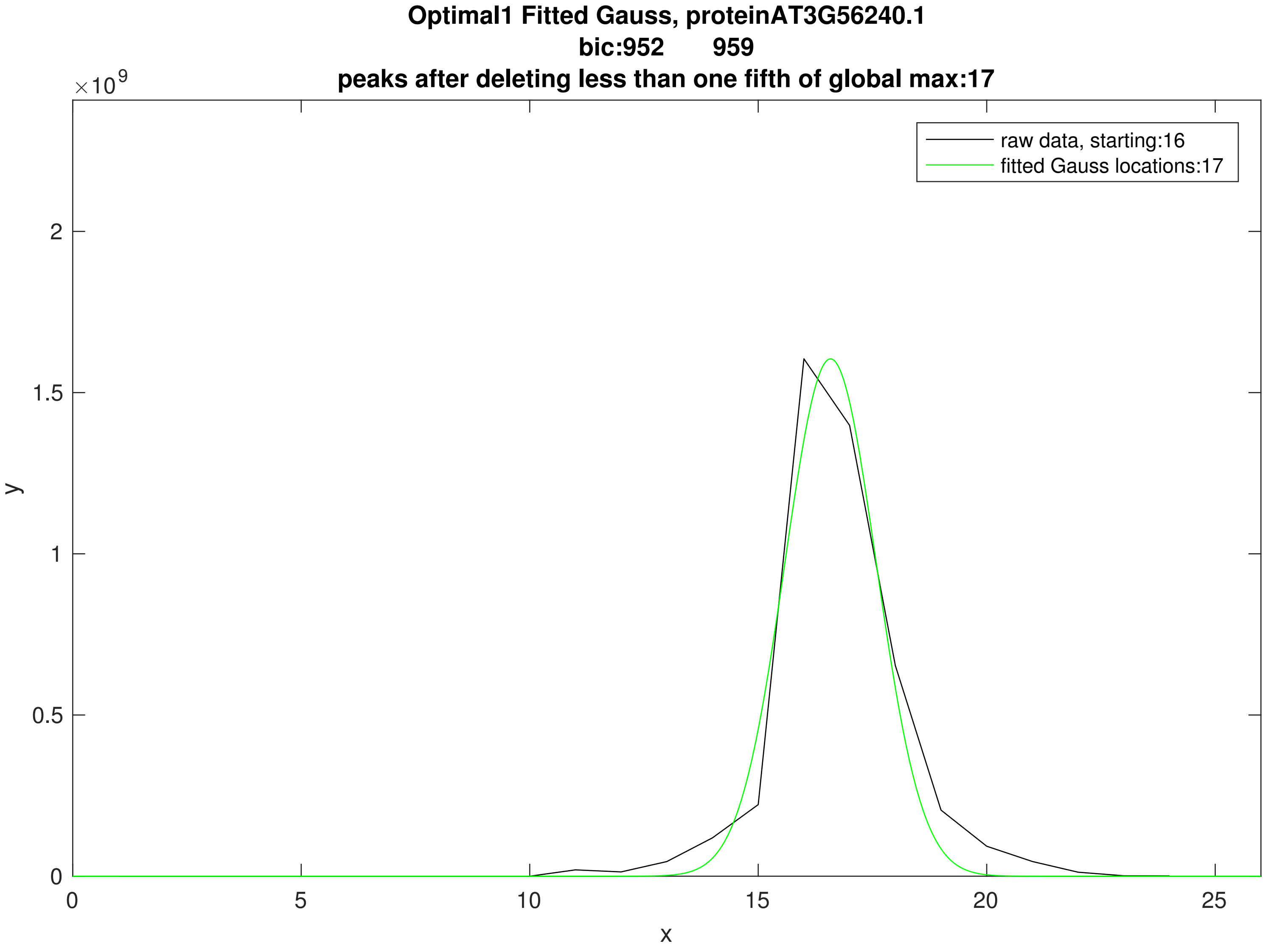
Optimal1 Fitted Gauss, proteinAT3G56240.1
bic:952 959
peaks after deleting less than one fifth of global max:17
×10^9
raw data, starting:16
fitted Gauss locations:17
y
x
0
0.5
1
1.5
2
0
5
10
15
20
25

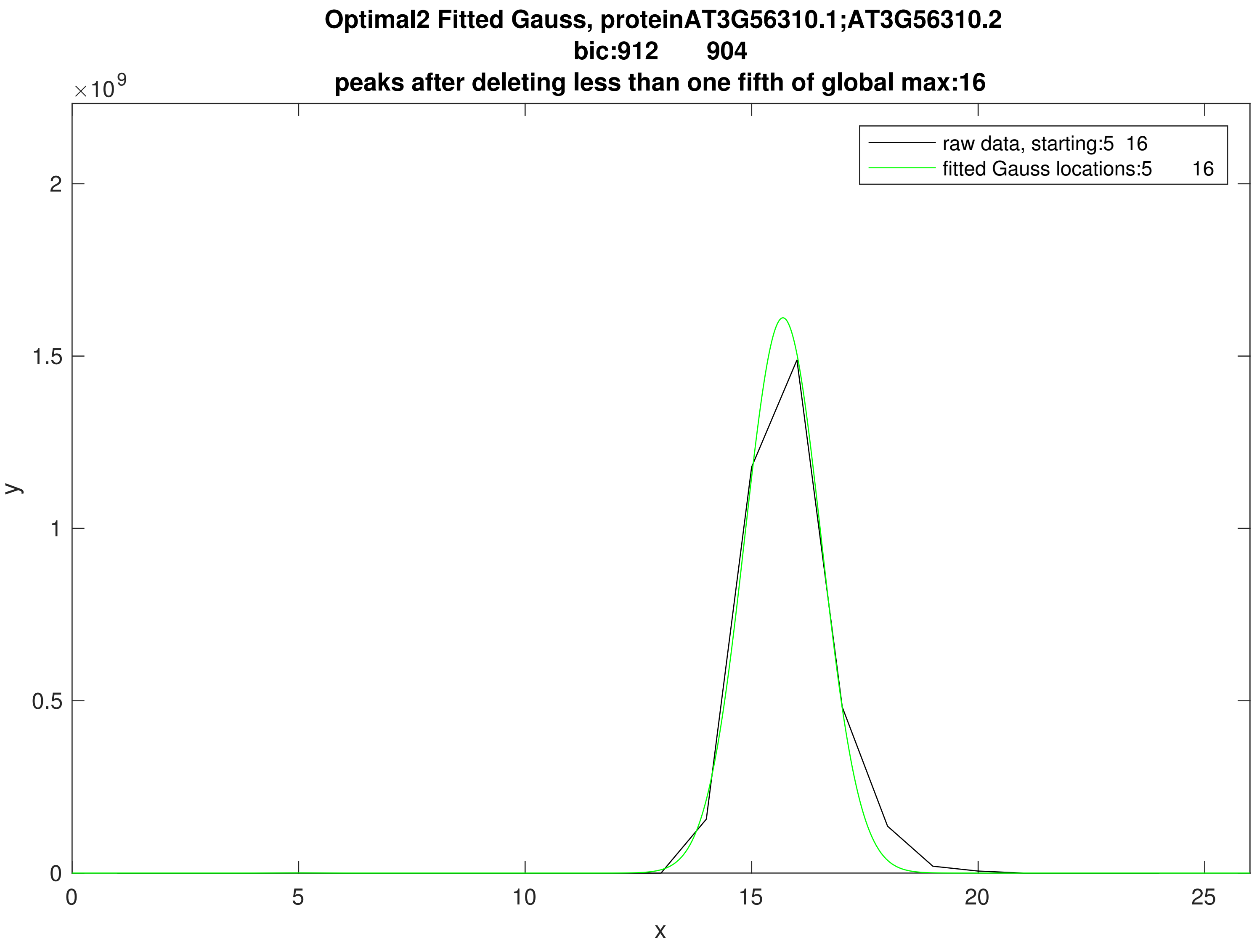

Optimal2 Fitted Gauss, proteinAT3G56310.1;AT3G56310.2
bic:912 904
peaks after deleting less than one fifth of global max:16
×10^9
raw data, starting:5 16
fitted Gauss locations:5 16
y
x
0
0.5
1
1.5
2
0
5
10
15
20
25

Optimal1 Fitted Gauss, proteinAT3G56340.1
bic:691
peaks after deleting less than one fifth of global max:2.8

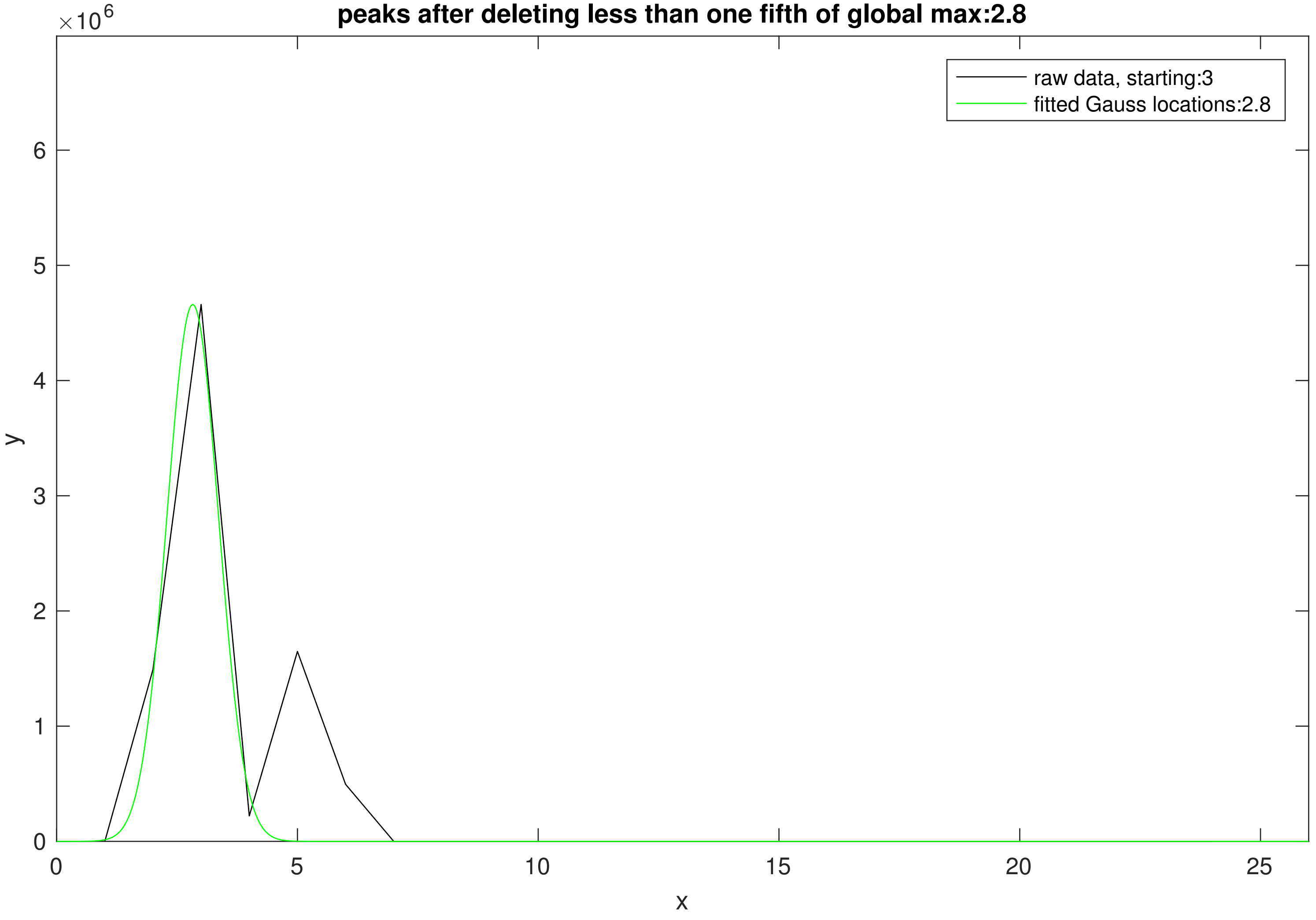

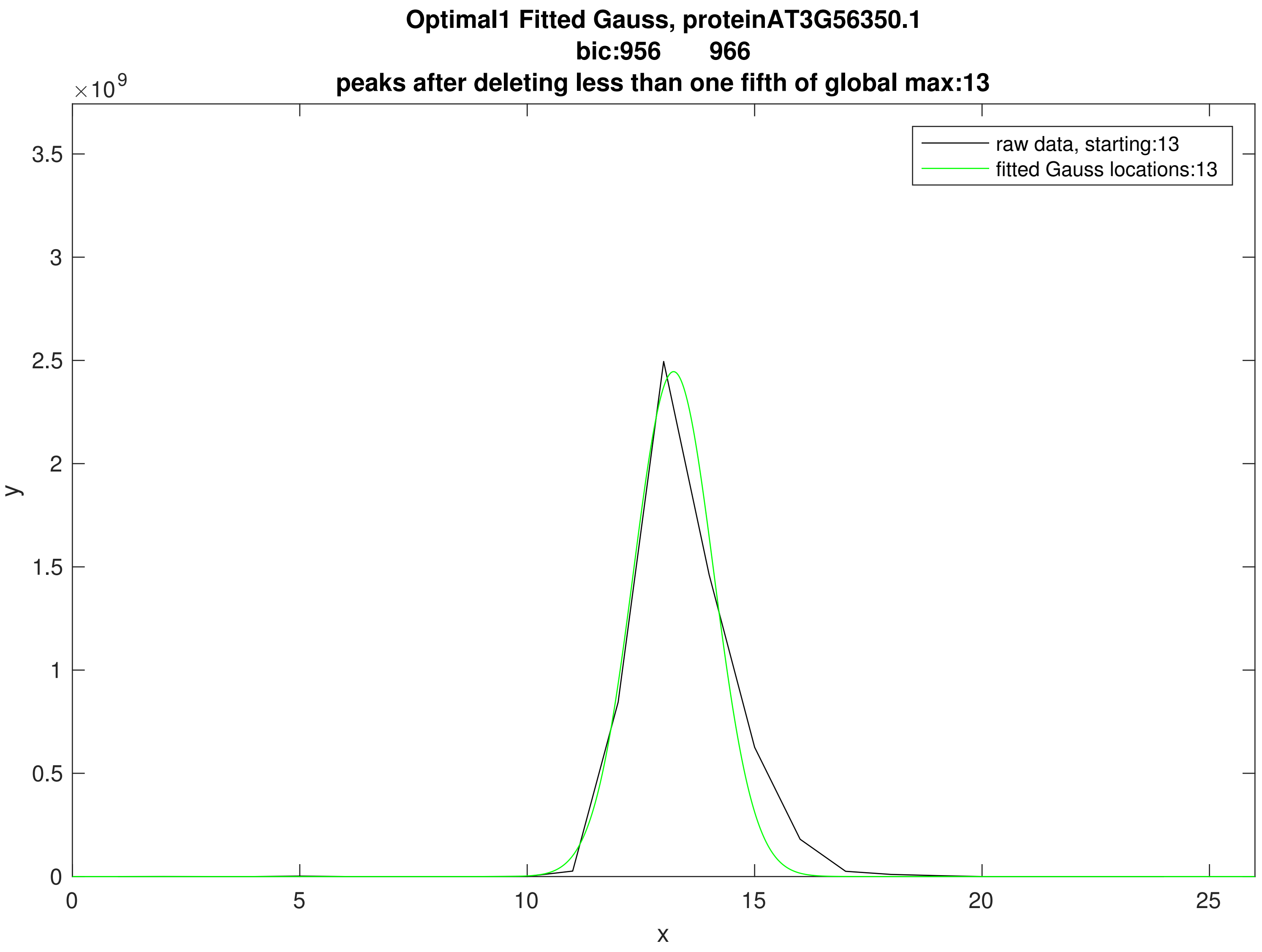

Optimal1 Fitted Gauss, proteinAT3G56350.1
bic:956 966
peaks after deleting less than one fifth of global max:13
×10^9
3.5
3
2.5
2
1.5
1
0.5
0
y
0
5
10
15
20
25
x
raw data, starting:13
fitted Gauss locations:13

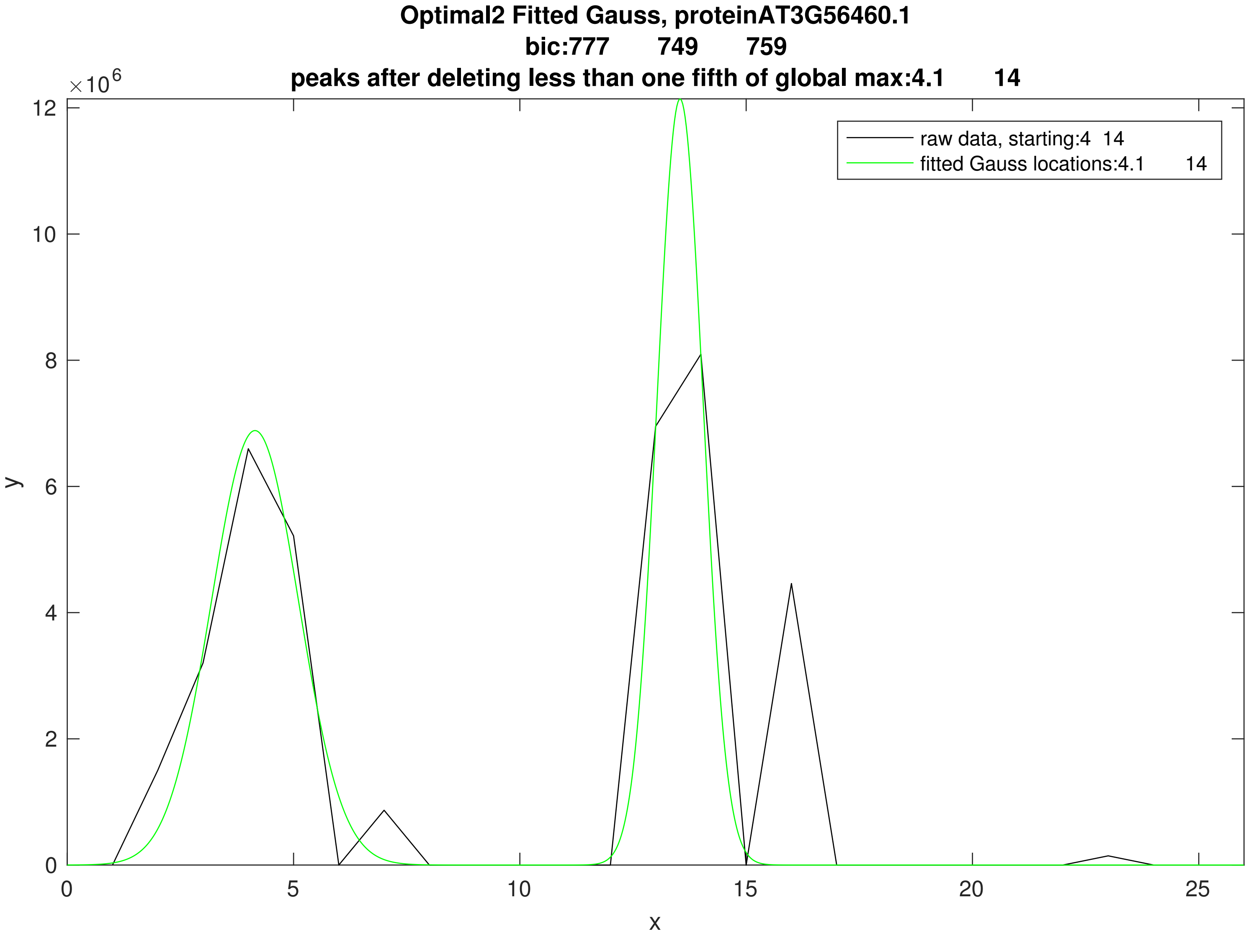

Optimal2 Fitted Gauss, proteinAT3G56460.1
bic:777 749 759
peaks after deleting less than one fifth of global max:4.1 14
×10^6
raw data, starting:4 14
fitted Gauss locations:4.1 14
y
x
0
2
4
6
8
10
12
0
5
10
15
20
25

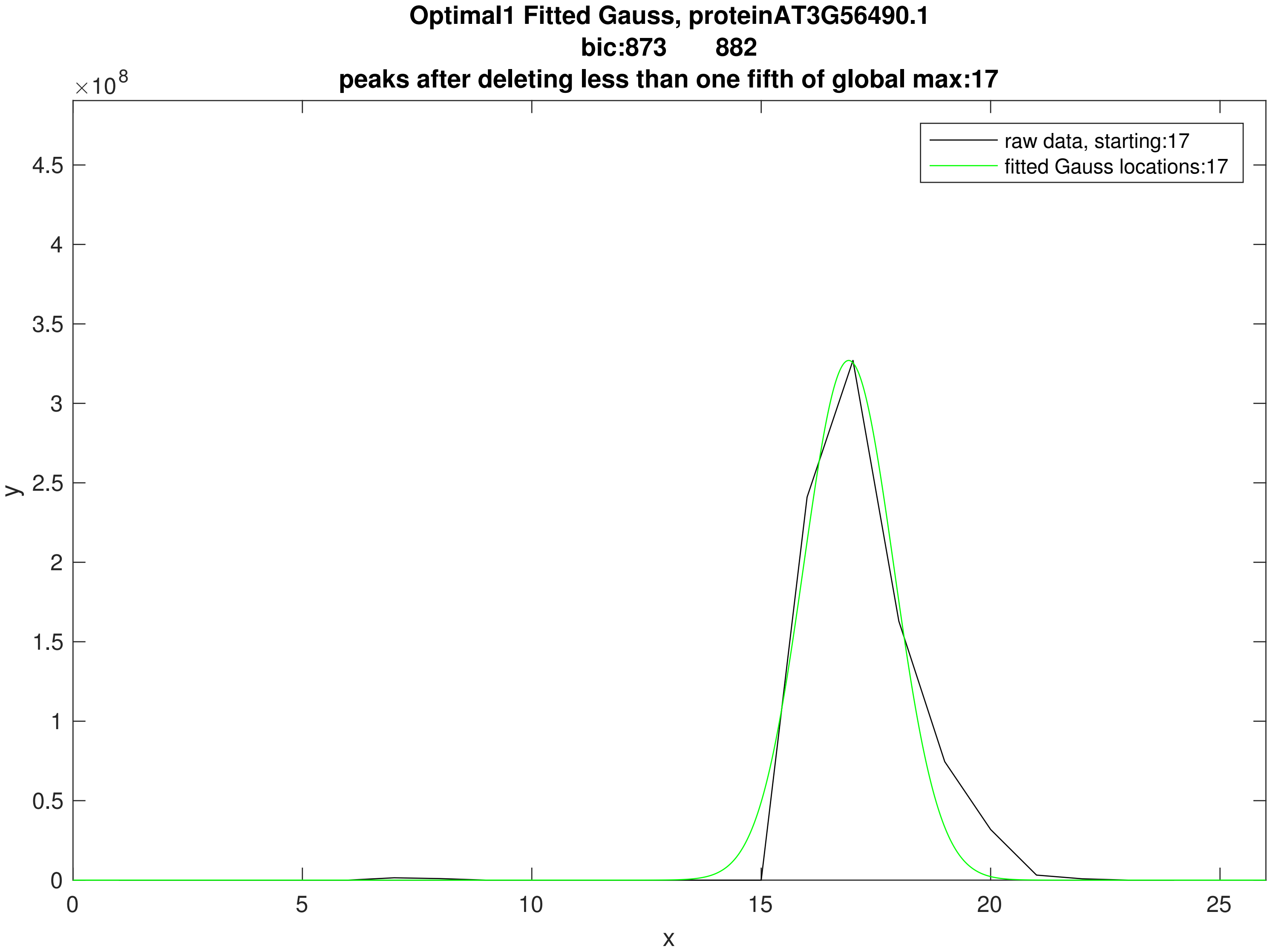
Optimal1 Fitted Gauss, proteinAT3G56490.1
bic:873 882
peaks after deleting less than one fifth of global max:17
×10^8
raw data, starting:17
fitted Gauss locations:17
y
x
0
0.5
1
1.5
2
2.5
3
3.5
4
4.5
0
5
10
15
20
25

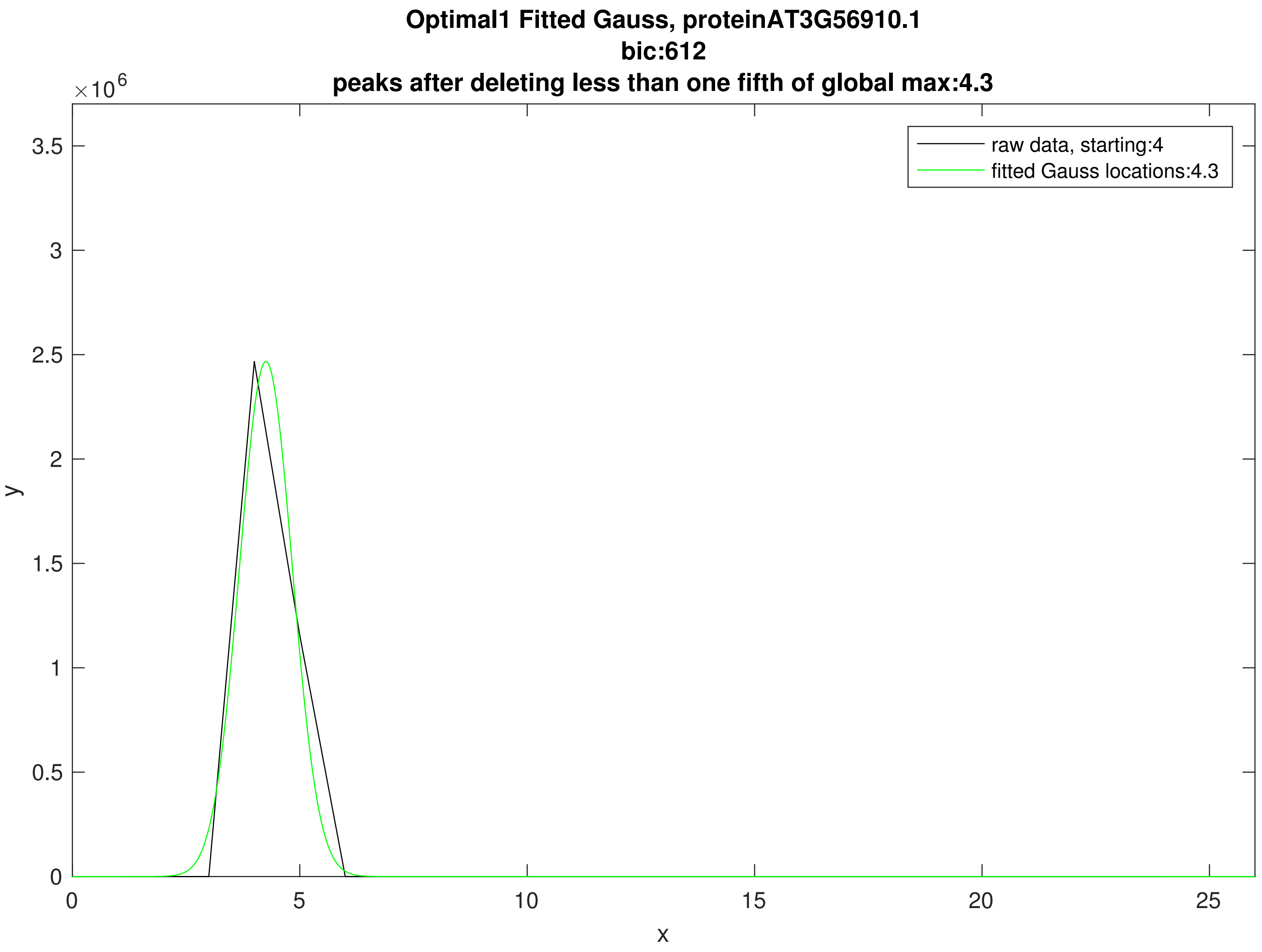

Optimal1 Fitted Gauss, proteinAT3G56910.1
bic:612
peaks after deleting less than one fifth of global max:4.3
×10^6
raw data, starting:4
fitted Gauss locations:4.3
y
x
0
0.5
1
1.5
2
2.5
3
3.5
0
5
10
15
20
25

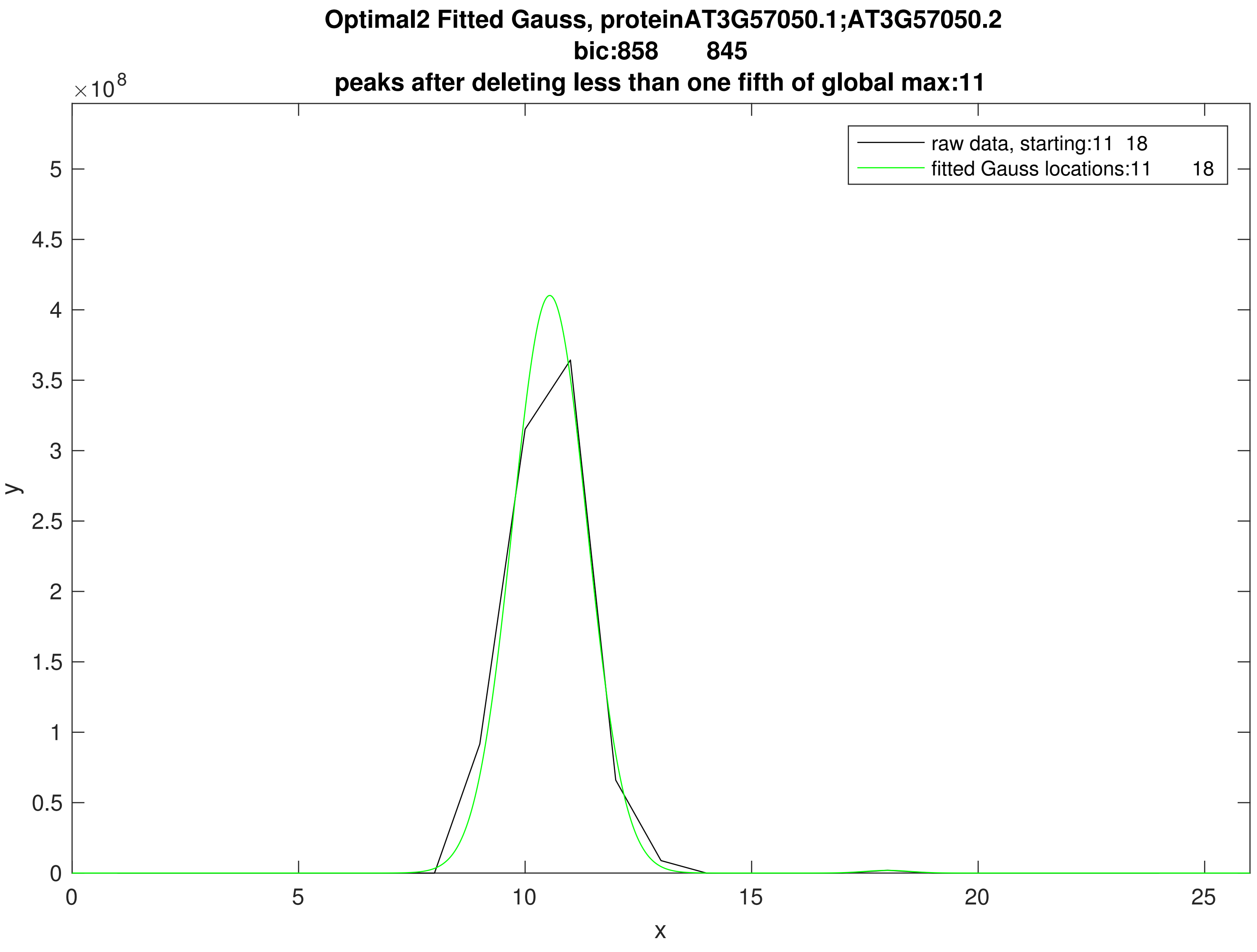
Optimal2 Fitted Gauss, proteinAT3G57050.1;AT3G57050.2
bic:858 845
peaks after deleting less than one fifth of global max:11
raw data, starting:11 18
fitted Gauss locations:11 18
×10^8
y
x
0
0.5
1
1.5
2
2.5
3
3.5
4
4.5
5
0
5
10
15
20
25

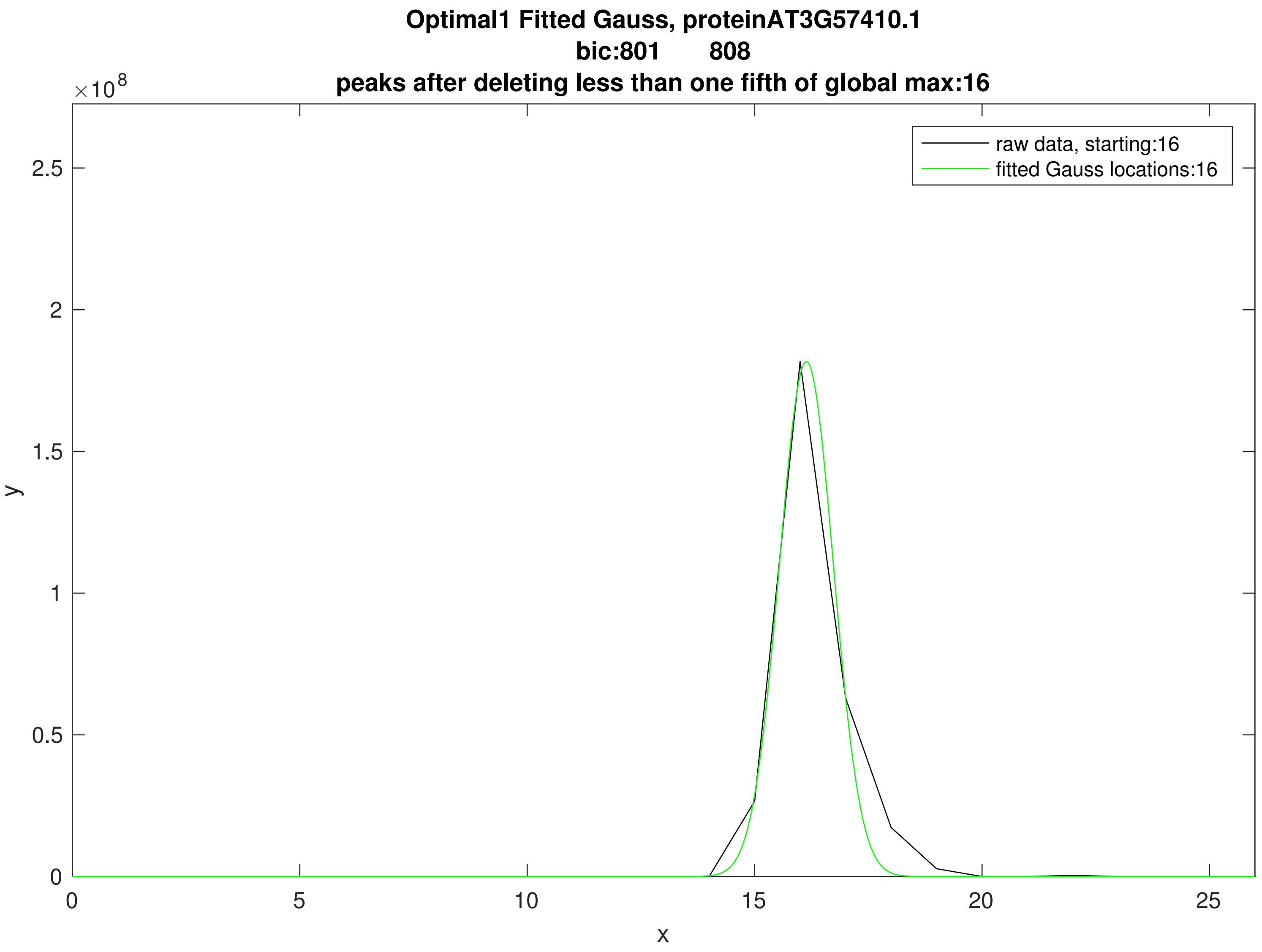

Optimal1 Fitted Gauss, proteinAT3G57410.1
bic:801 808
peaks after deleting less than one fifth of global max:16
×10^8
2.5
2
1.5
1
0.5
0
y
0
5
10
15
20
25
x
raw data, starting:16
fitted Gauss locations:16

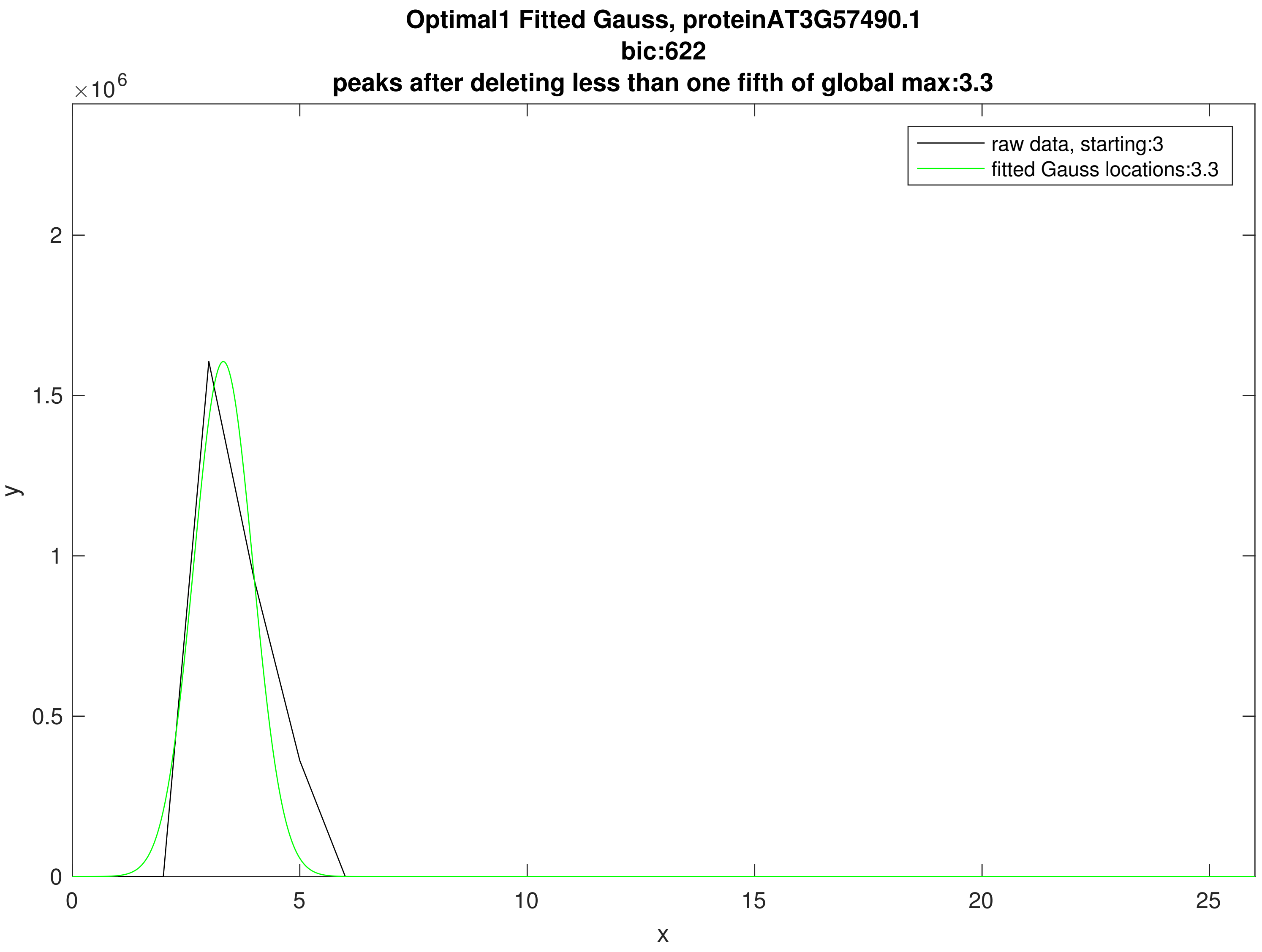
Optimal1 Fitted Gauss, proteinAT3G57490.1
bic:622
peaks after deleting less than one fifth of global max:3.3
×10^6
raw data, starting:3
fitted Gauss locations:3.3
2
1.5
1
0.5
0
y
0
5
10
15
20
25
x

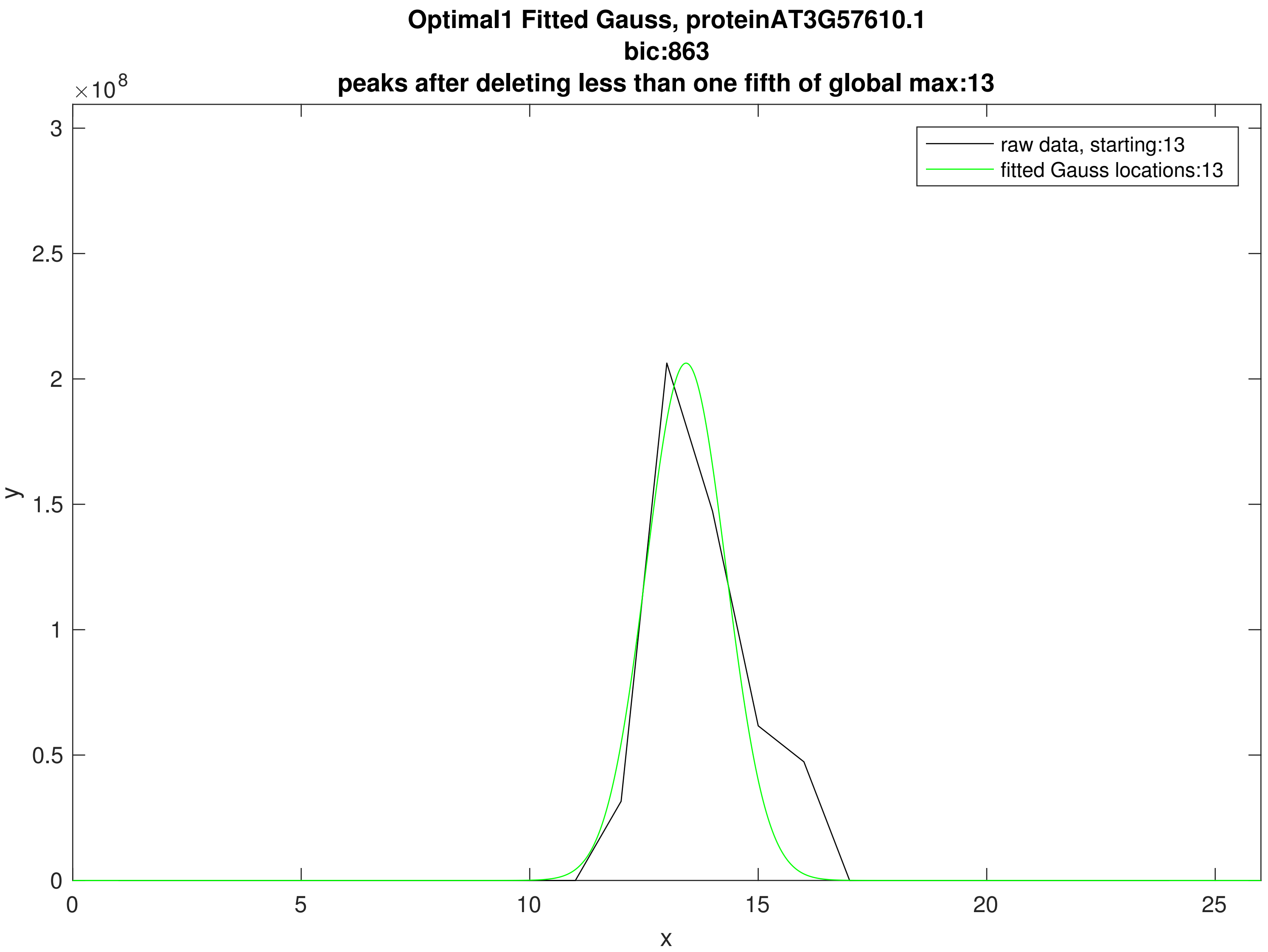

Optimal1 Fitted Gauss, proteinAT3G57610.1
bic:863
peaks after deleting less than one fifth of global max:13
×10^8
3
2.5
2
1.5
1
0.5
0
y
0
5
10
15
20
25
x
raw data, starting:13
fitted Gauss locations:13

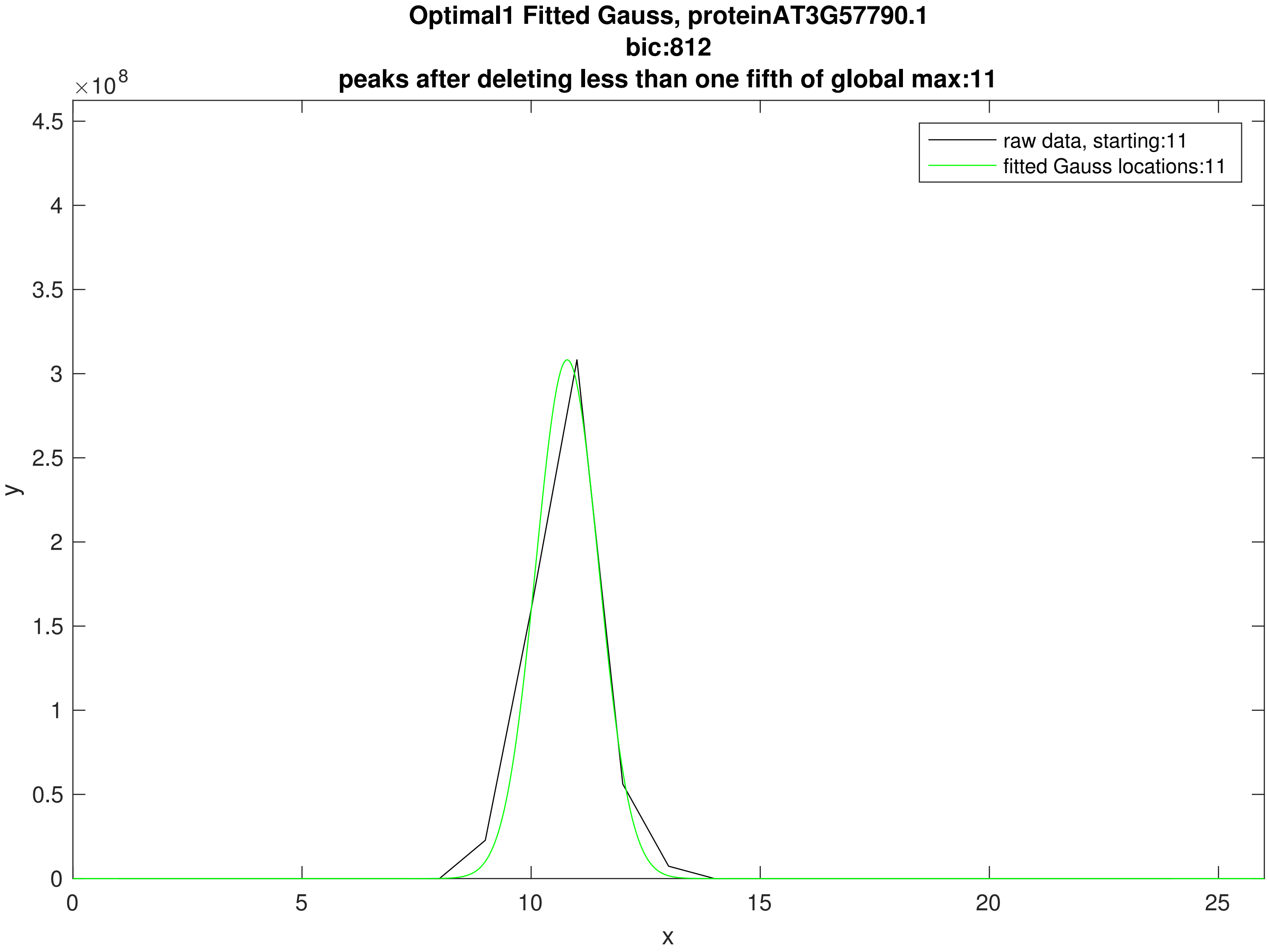

Optimal1 Fitted Gauss, proteinAT3G57790.1
bic:812
peaks after deleting less than one fifth of global max:11
×10^8
4.5
4
3.5
3
2.5
2
1.5
1
0.5
0
y
0
5
10
15
20
25
x
raw data, starting:11
fitted Gauss locations:11

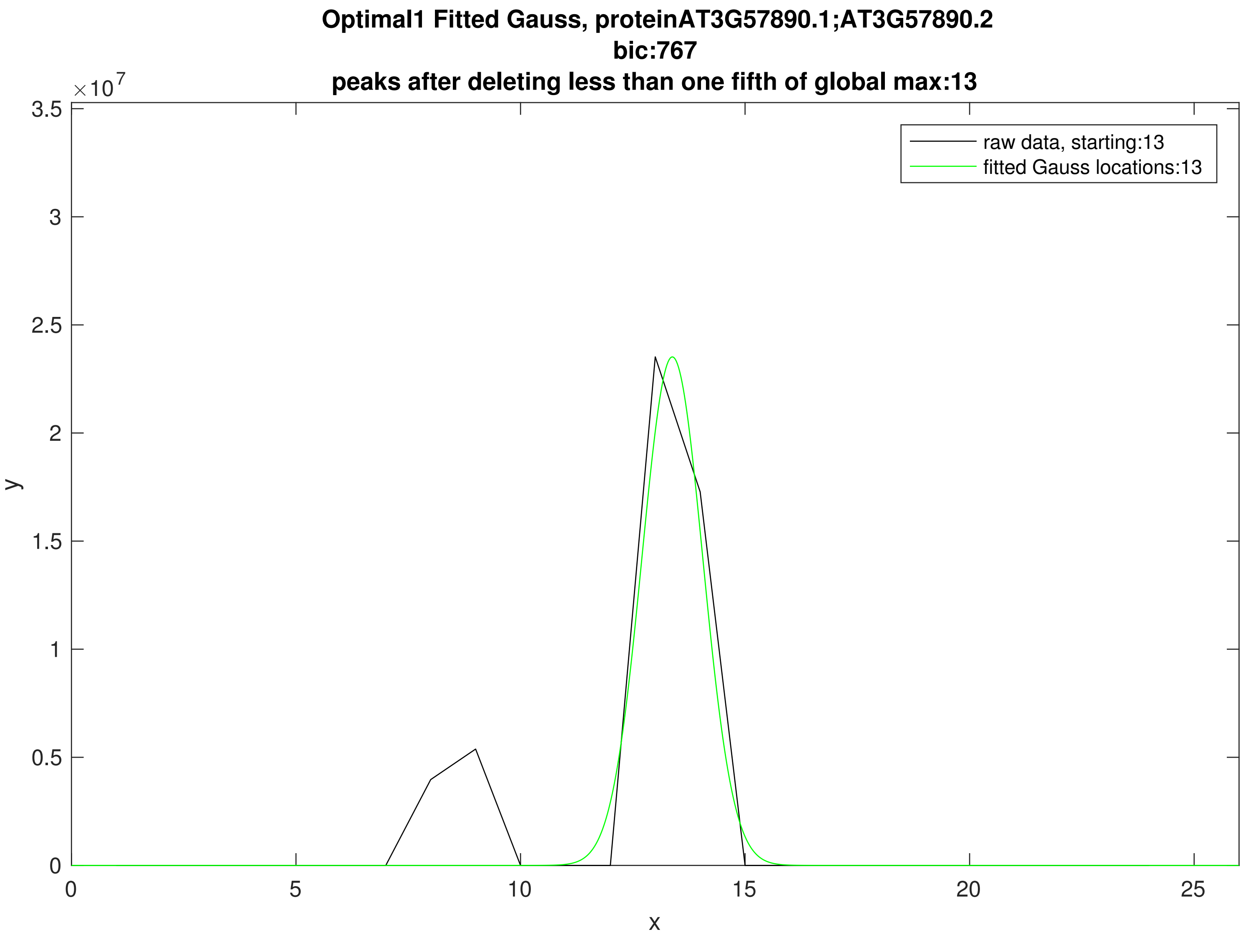

Optimal1 Fitted Gauss, proteinAT3G57890.1;AT3G57890.2
bic:767
peaks after deleting less than one fifth of global max:13
×10^7
3.5
3
2.5
2
1.5
1
0.5
0
y
0
5
10
15
20
25
x
raw data, starting:13
fitted Gauss locations:13

Optimal2 Fitted Gauss, proteinAT3G58140.1
bic:861 847
peaks after deleting less than one fifth of global max:16

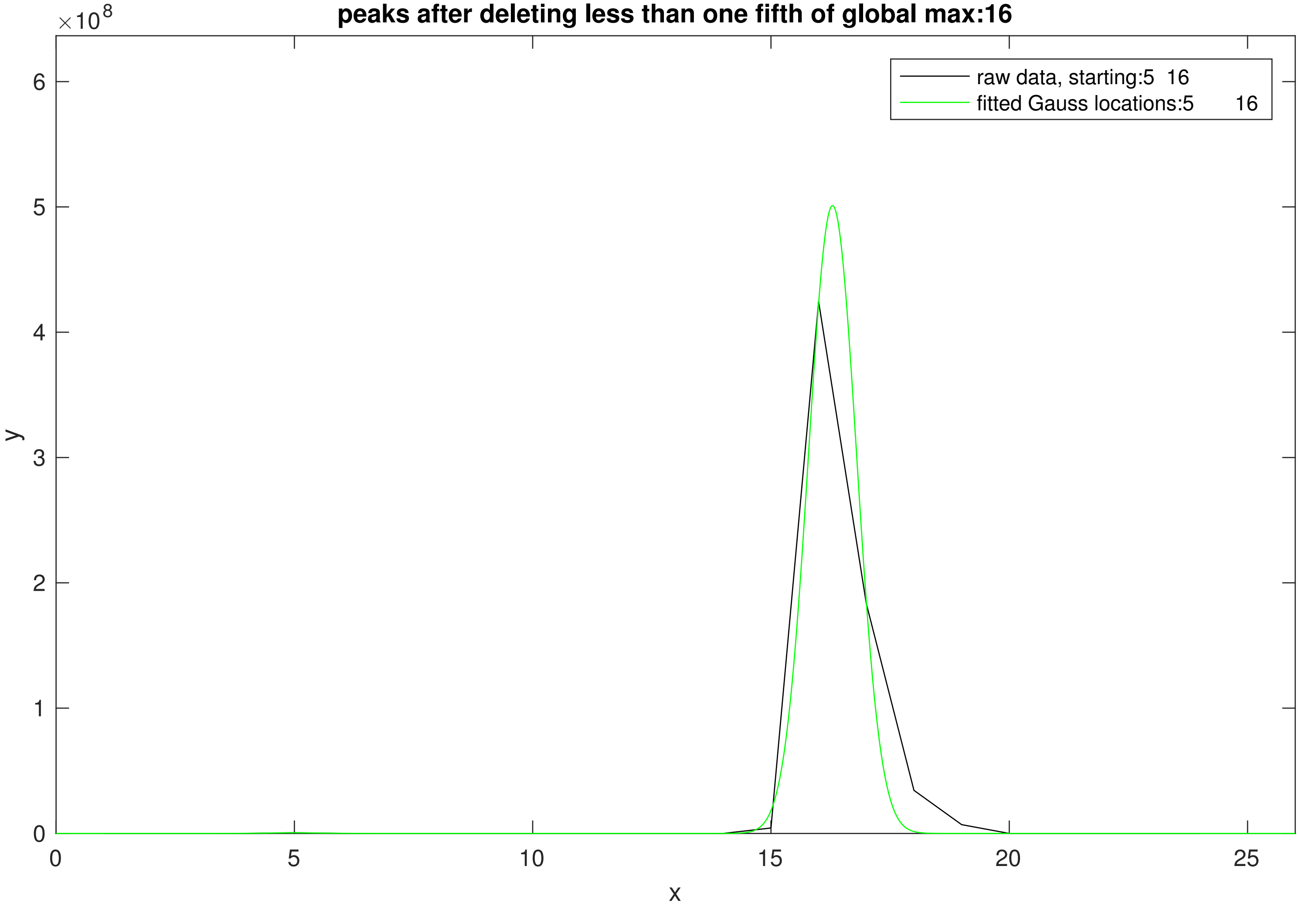

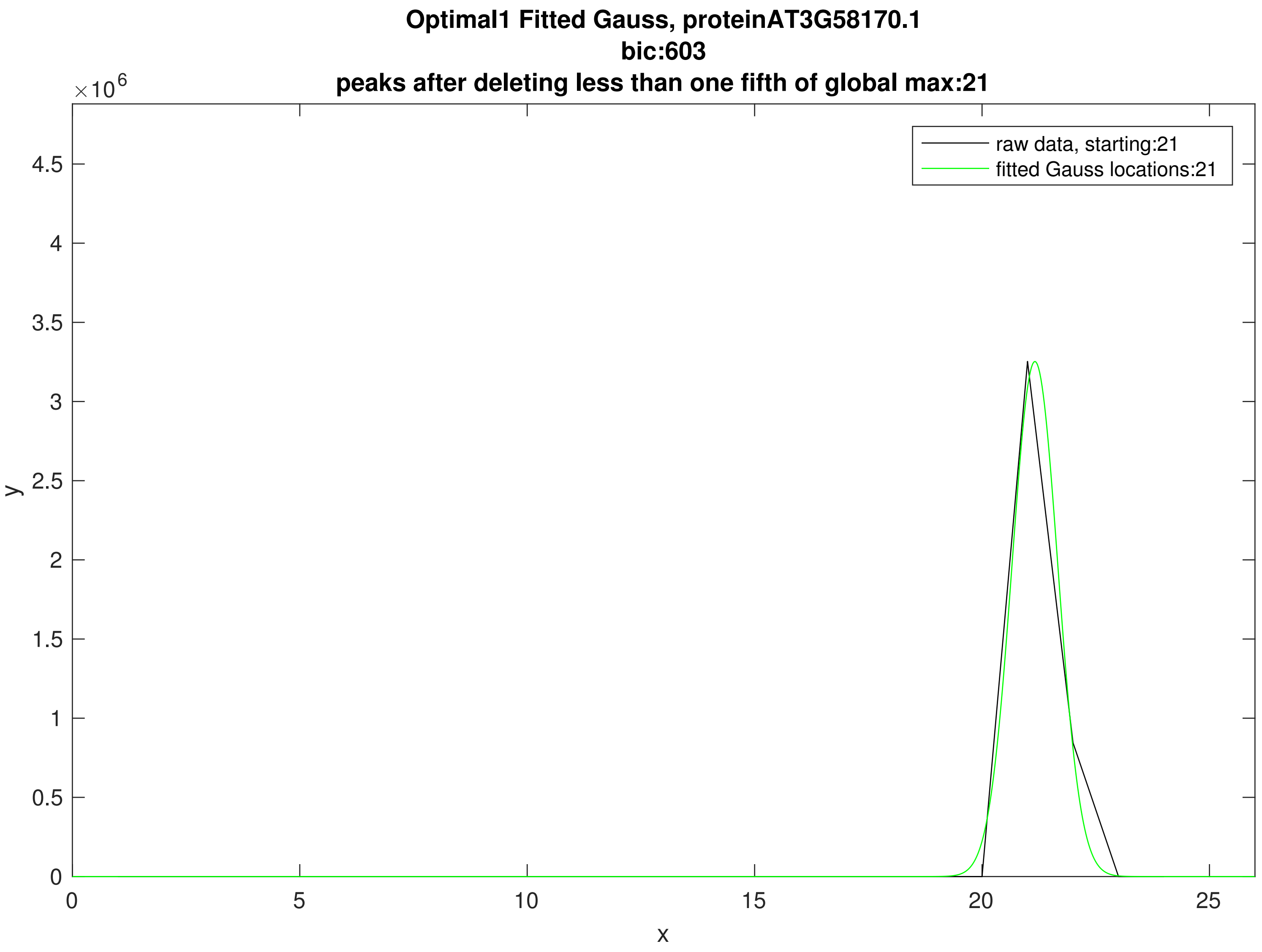
Optimal1 Fitted Gauss, proteinAT3G58170.1
bic:603
peaks after deleting less than one fifth of global max:21
×10^6
raw data, starting:21
fitted Gauss locations:21
4.5
4
3.5
3
2.5
2
1.5
1
0.5
0
y
0
5
10
15
20
25
x

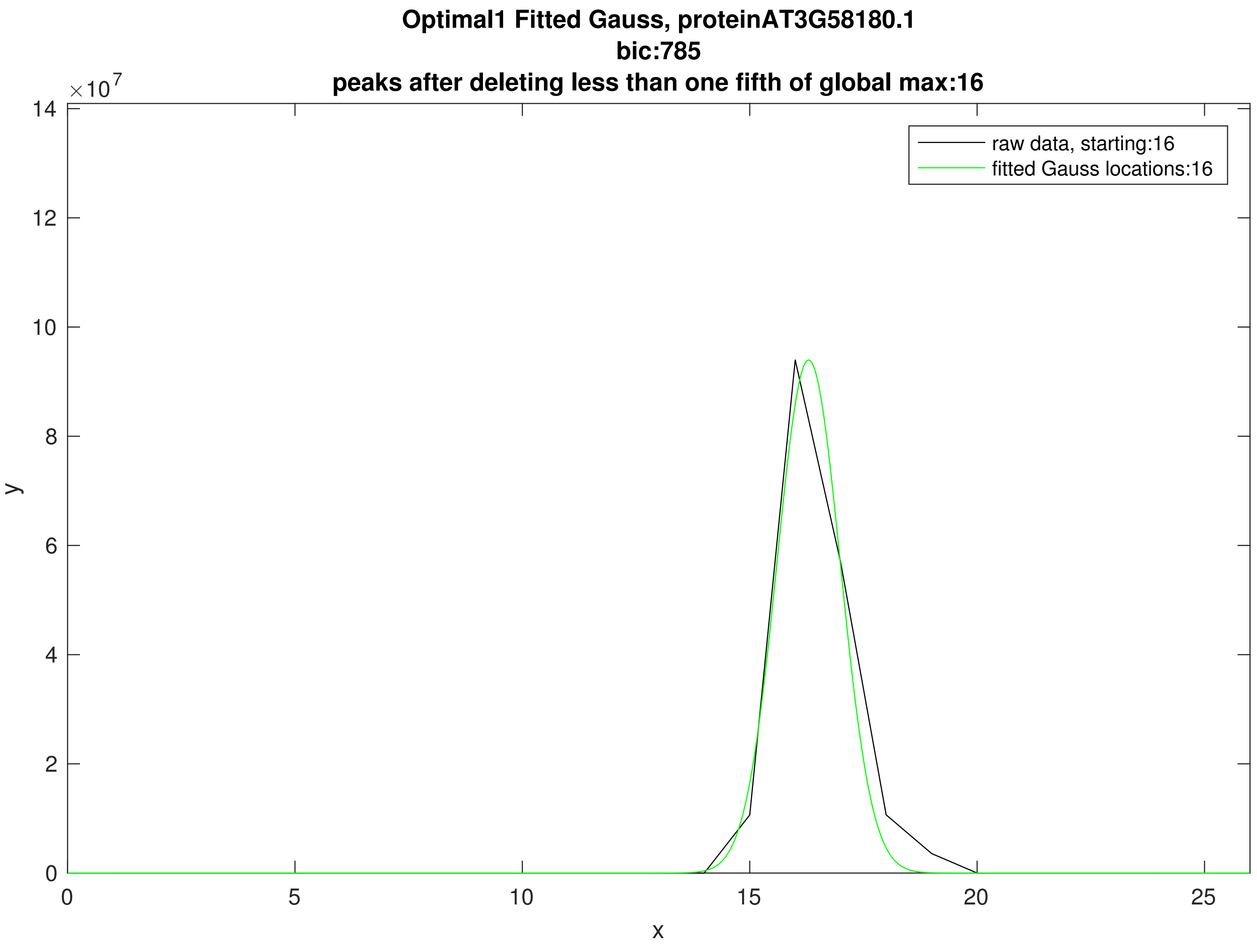
Optimal1 Fitted Gauss, proteinAT3G58180.1
bic:785
peaks after deleting less than one fifth of global max:16
×10^7
raw data, starting:16
fitted Gauss locations:16
y
x
0
2
4
6
8
10
12
14
0
5
10
15
20
25

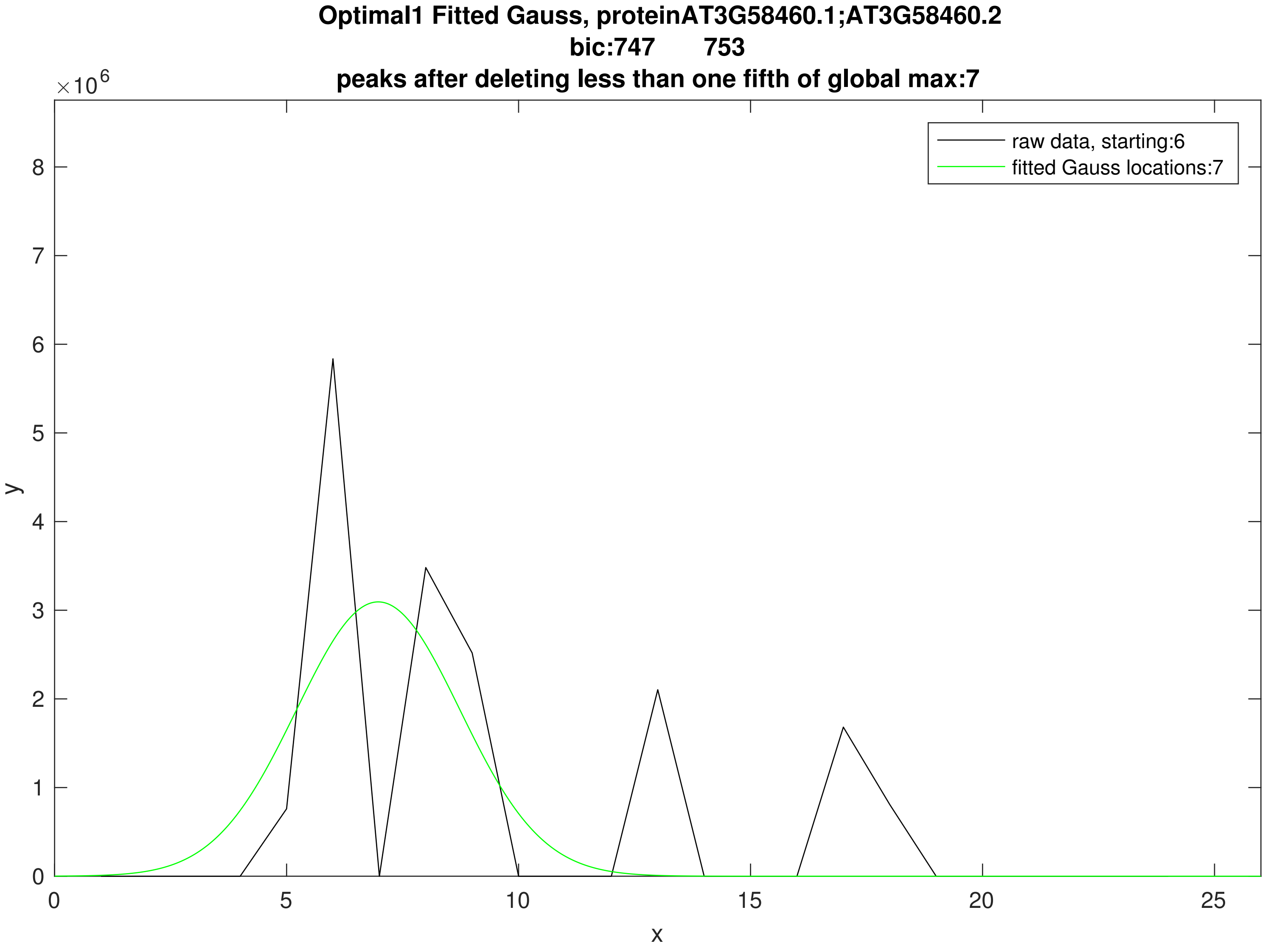

Optimal1 Fitted Gauss, proteinAT3G58460.1;AT3G58460.2
bic:747 753
peaks after deleting less than one fifth of global max:7
×10^6
raw data, starting:6
fitted Gauss locations:7
y
x
0
1
2
3
4
5
6
7
8
0
5
10
15
20
25

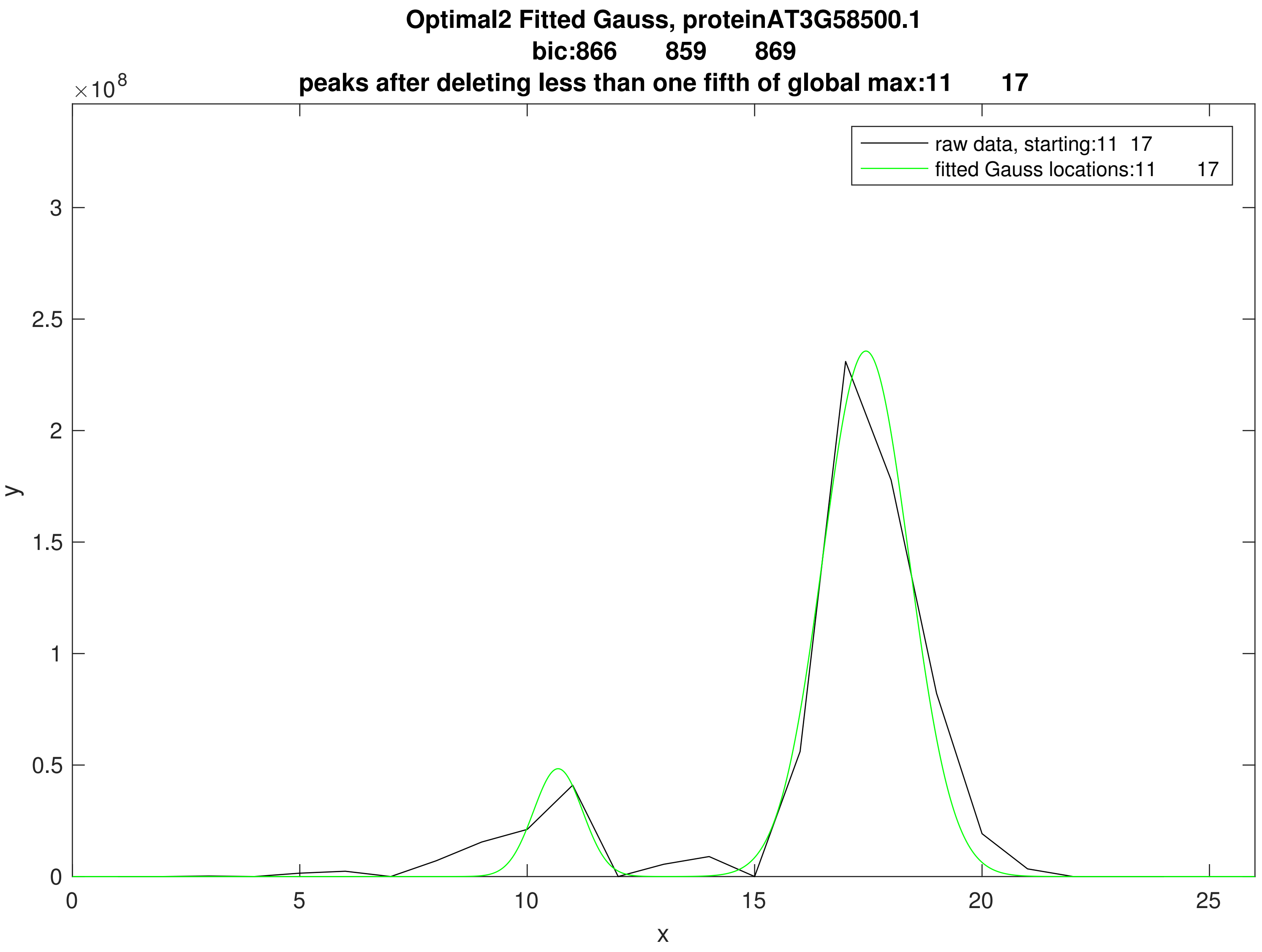
Optimal2 Fitted Gauss, proteinAT3G58500.1
bic:866 859 869
peaks after deleting less than one fifth of global max:11 17
×10^8
raw data, starting:11 17
fitted Gauss locations:11 17
y
x
0
0.5
1
1.5
2
2.5
3
0
5
10
15
20
25

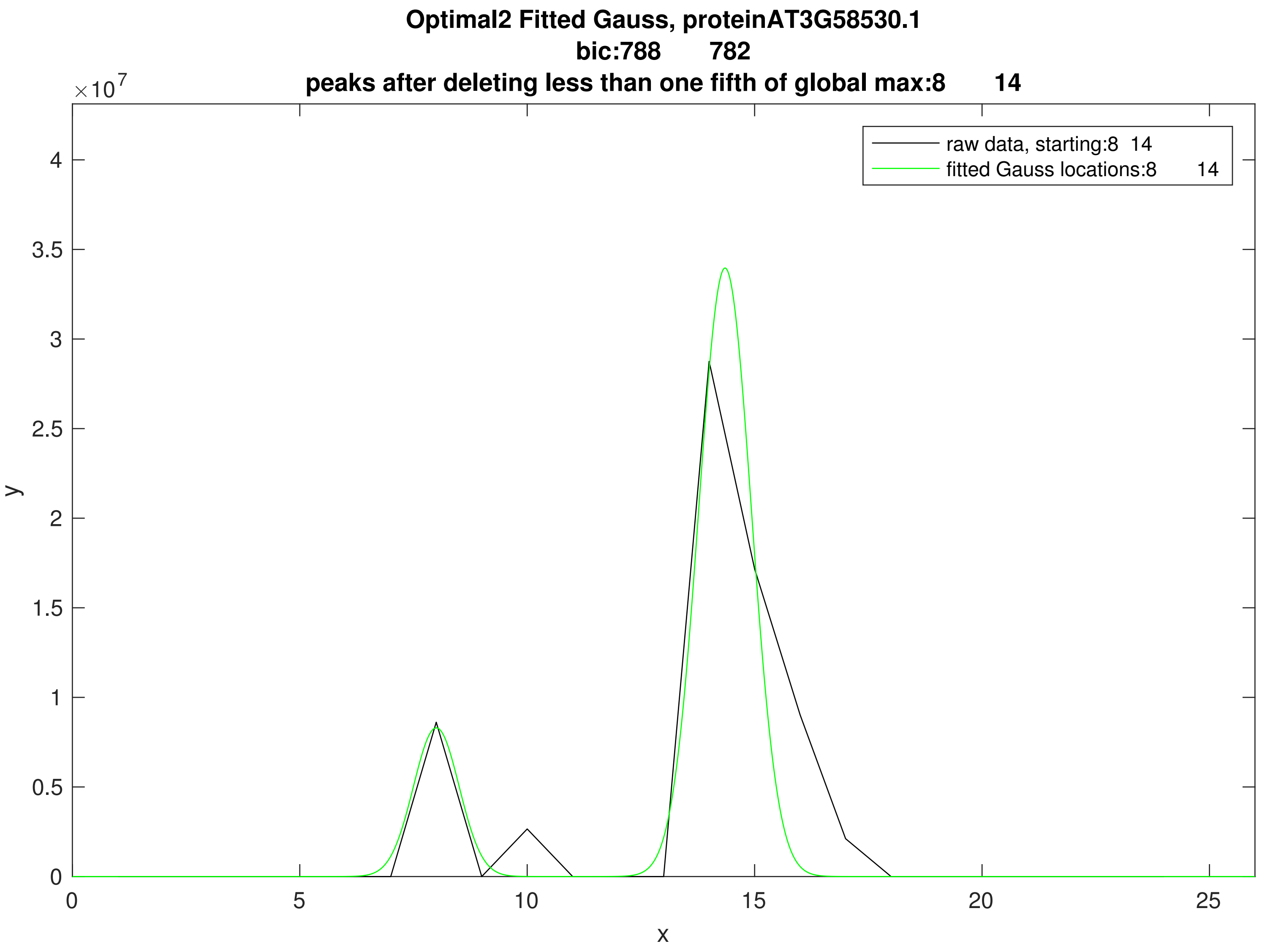

Optimal2 Fitted Gauss, proteinAT3G58530.1
bic:788 782
peaks after deleting less than one fifth of global max:8 14
×10^7
raw data, starting:8 14
fitted Gauss locations:8 14
y
x
0
0.5
1
1.5
2
2.5
3
3.5
4
0
5
10
15
20
25

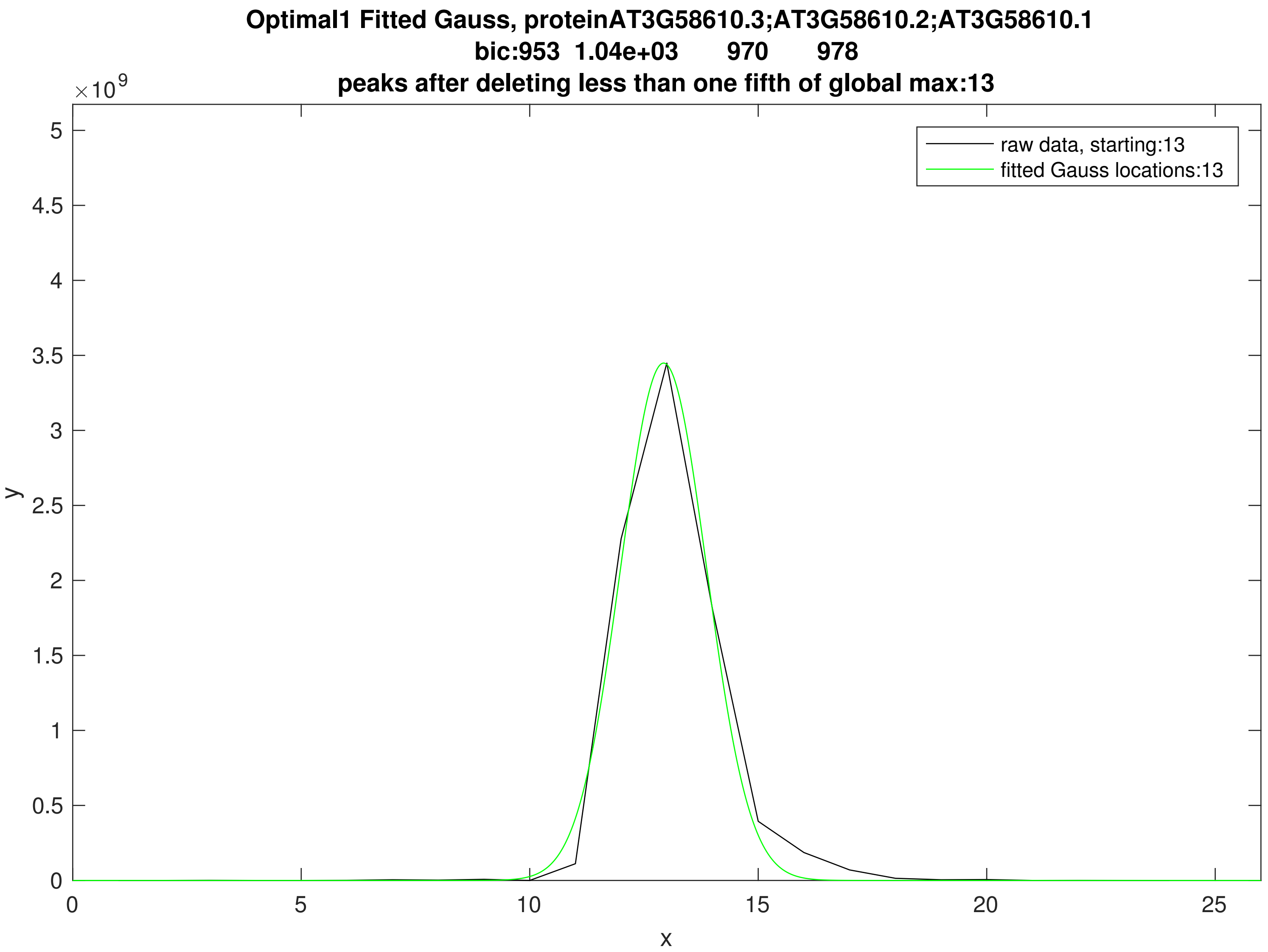

Optimal1 Fitted Gauss, proteinAT3G58610.3;AT3G58610.2;AT3G58610.1
bic:953 1.04e+03 970 978
peaks after deleting less than one fifth of global max:13
×10^9
raw data, starting:13
fitted Gauss locations:13
y
x
0
0.5
1
1.5
2
2.5
3
3.5
4
4.5
5
0
5
10
15
20
25

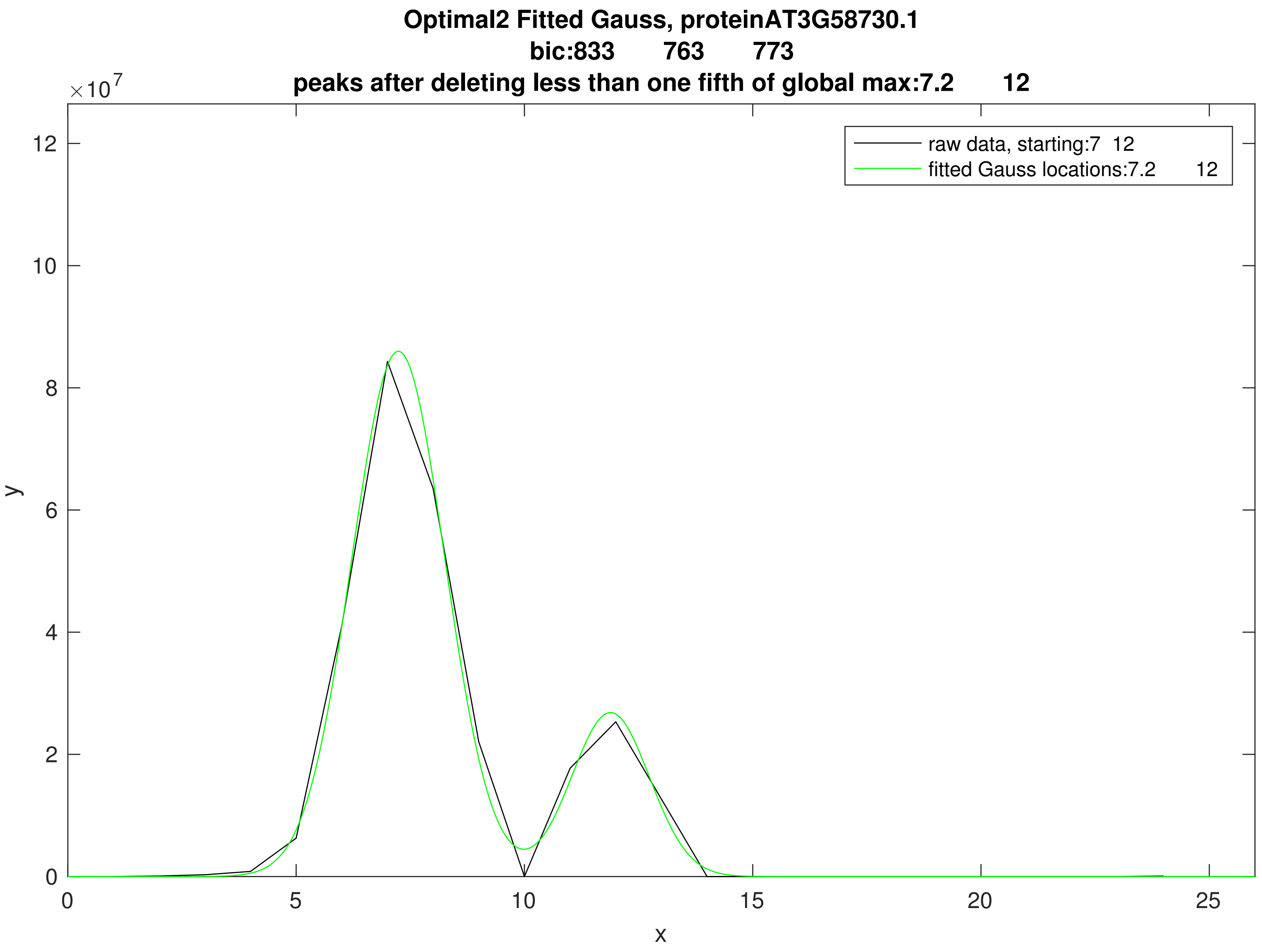

Optimal2 Fitted Gauss, proteinAT3G58730.1
bic:833 763 773
peaks after deleting less than one fifth of global max:7.2 12
×10^7
raw data, starting:7 12
fitted Gauss locations:7.2 12
y
x
0
2
4
6
8
10
12
0
5
10
15
20
25

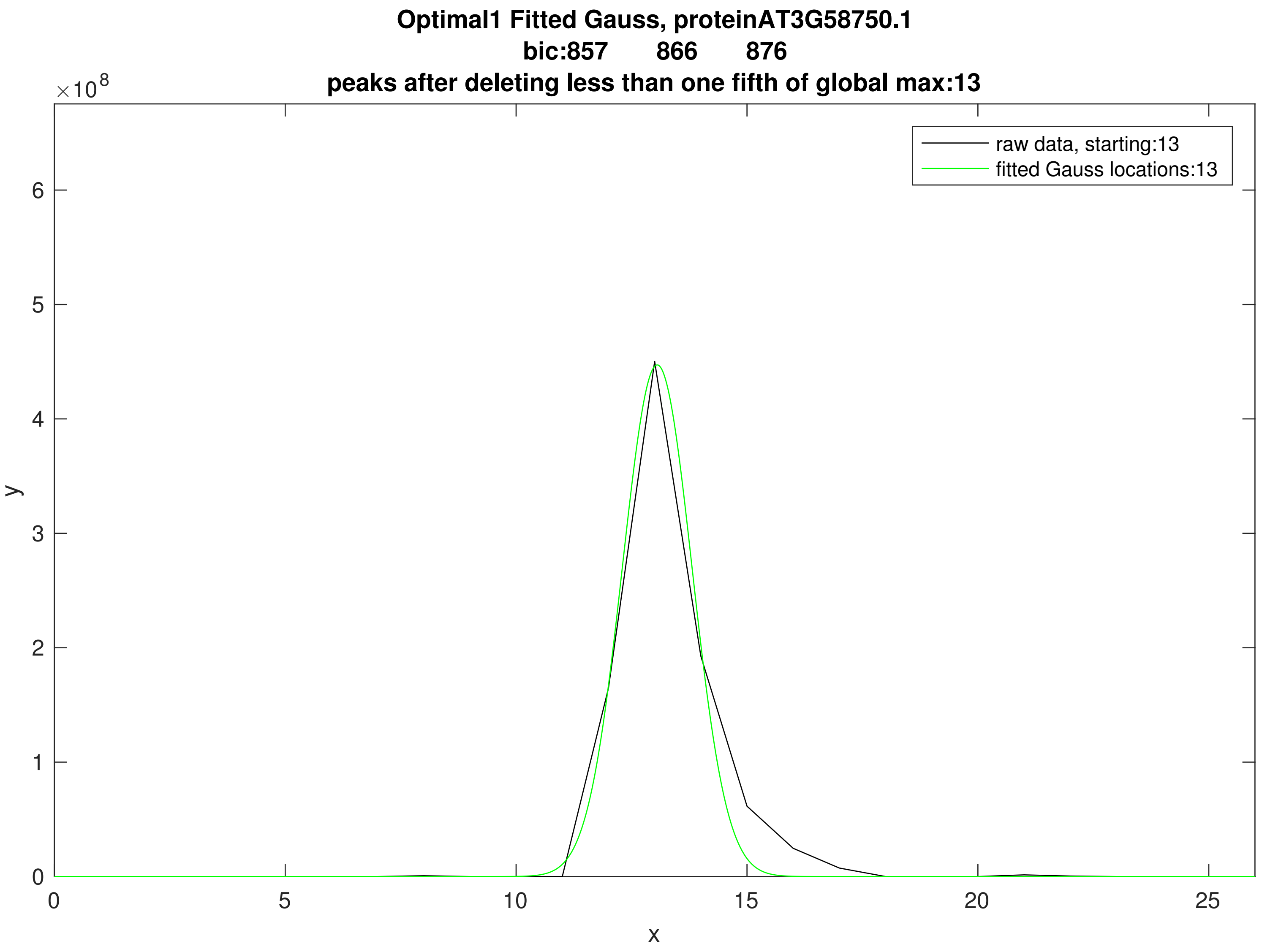

Optimal1 Fitted Gauss, proteinAT3G58750.1
bic:857 866 876
peaks after deleting less than one fifth of global max:13
×10^8
raw data, starting:13
fitted Gauss locations:13
y
x
0
1
2
3
4
5
6
0
5
10
15
20
25

Optimal1 Fitted Gauss, proteinAT3G58840.2;AT3G58840.1
bic:771 774
peaks after deleting less than one fifth of global max:12

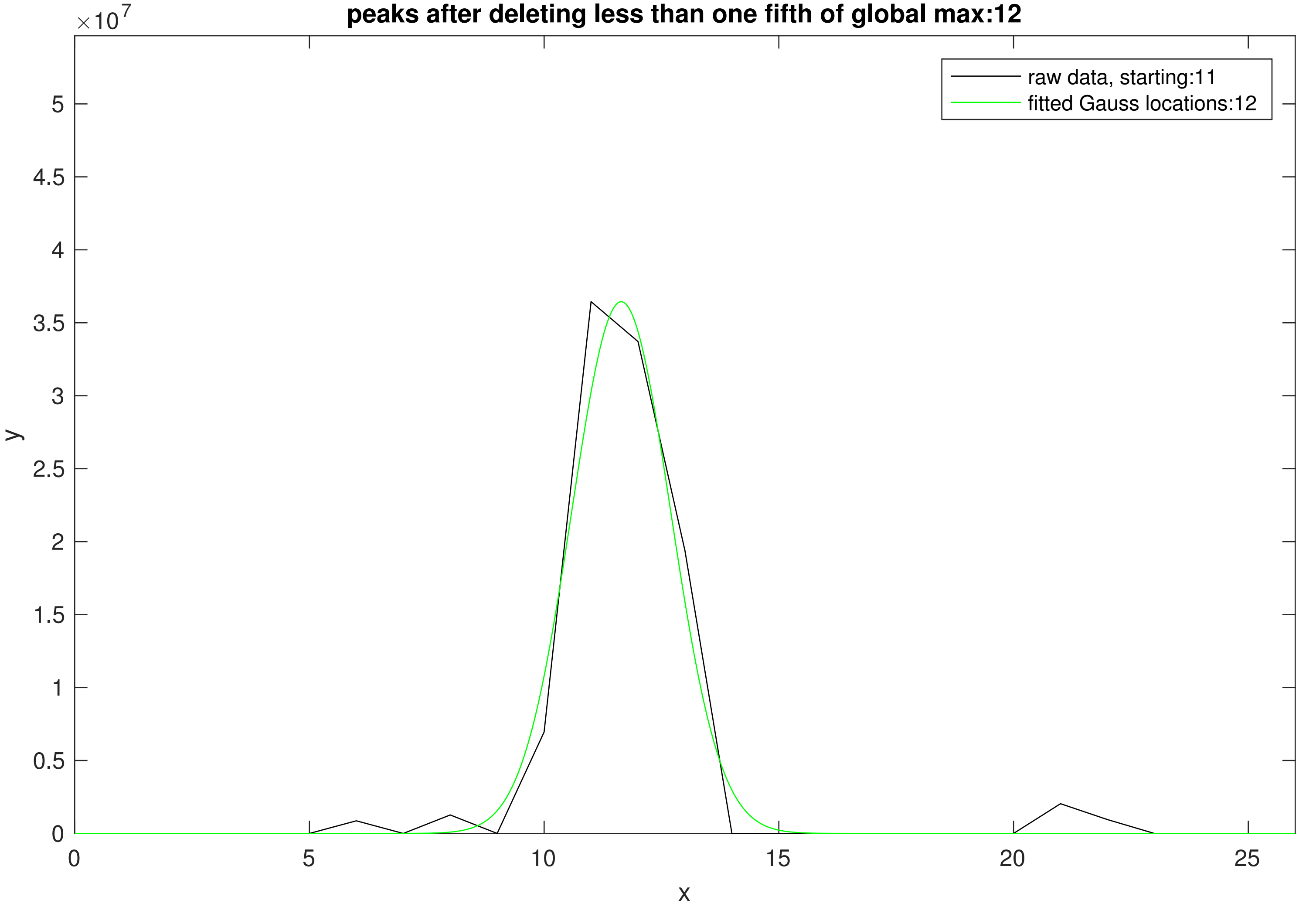

Optimal1 Fitted Gauss, proteinAT3G58990.1
bic:836
peaks after deleting less than one fifth of global max:14

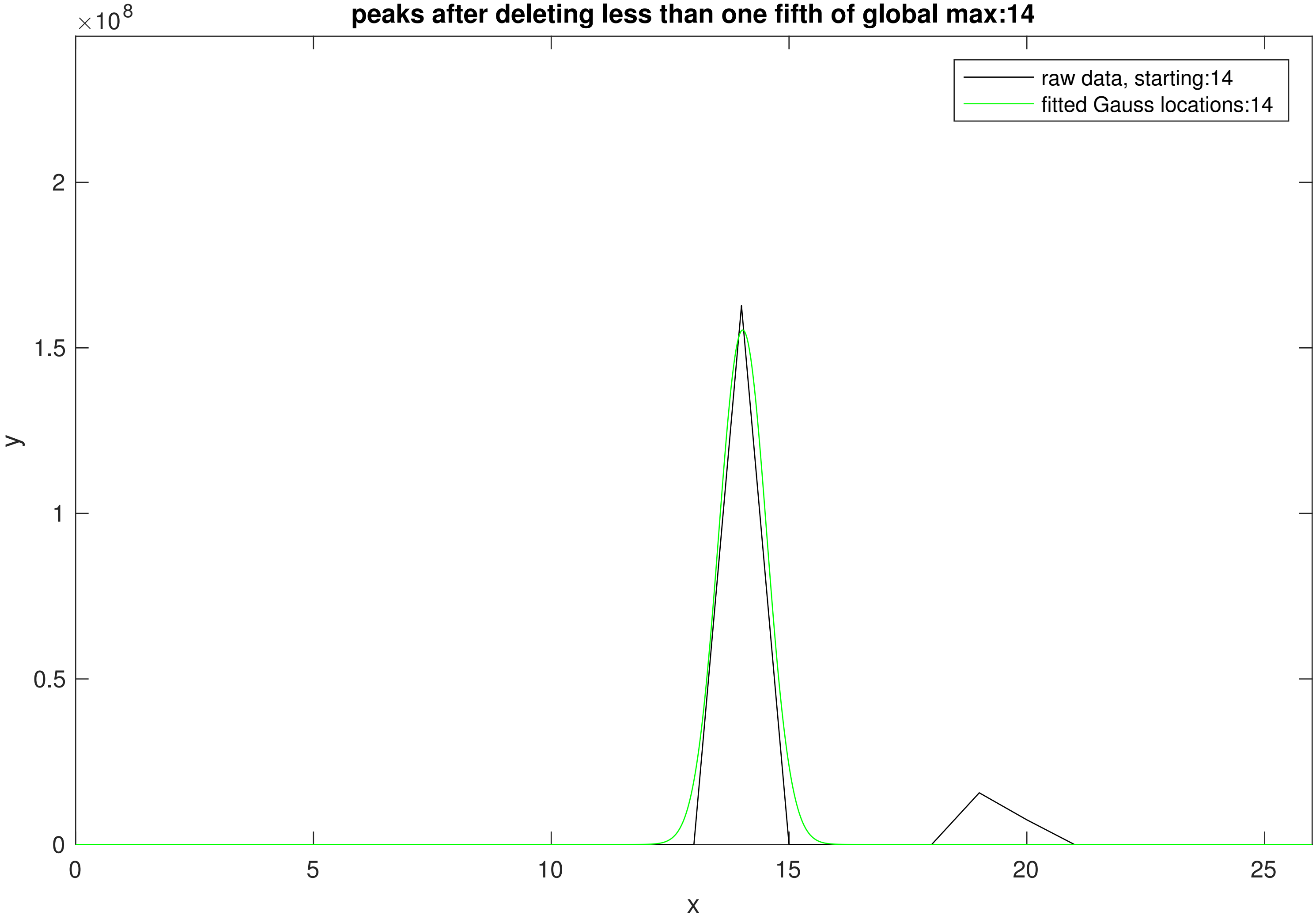

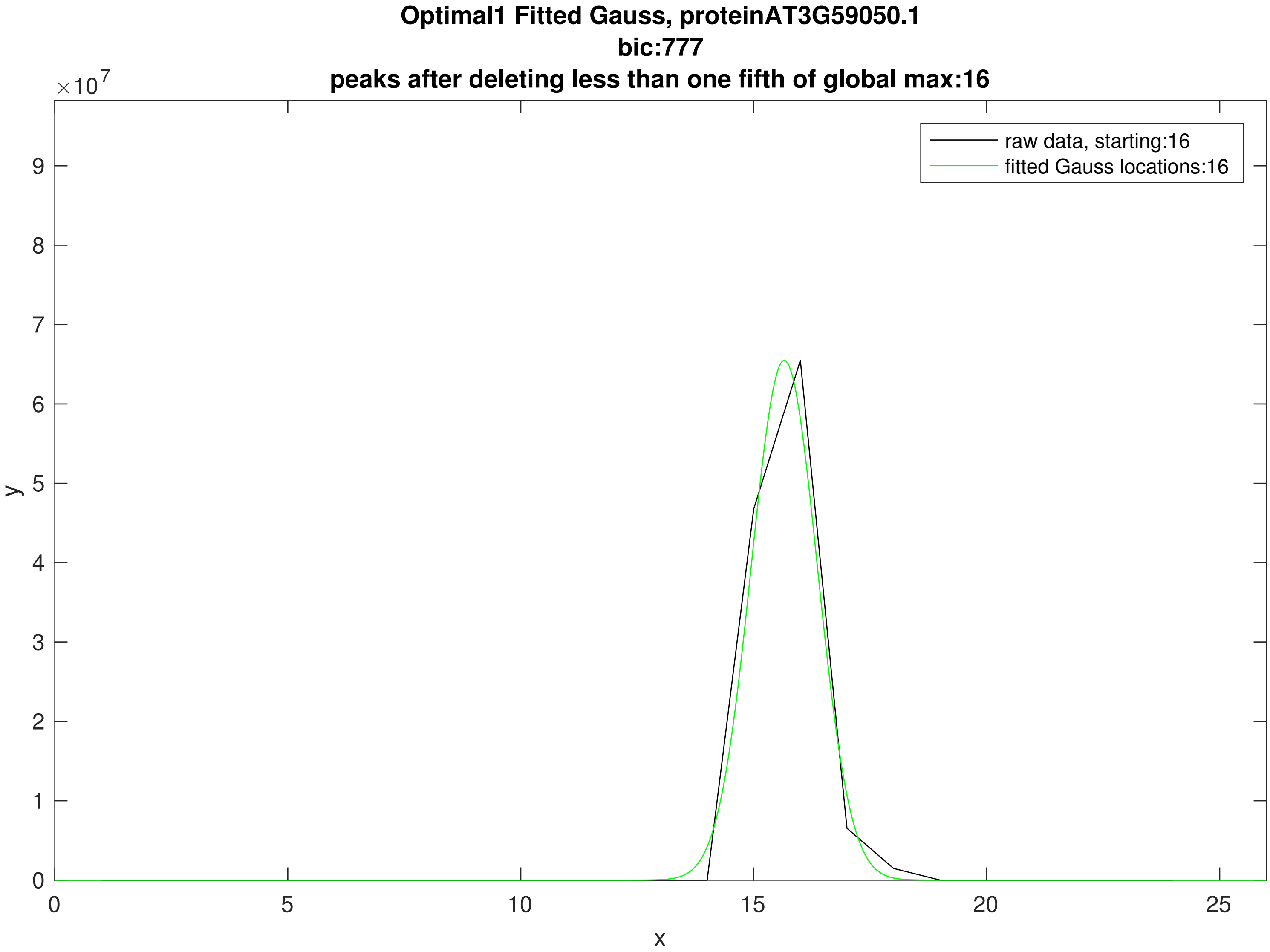
Optimal1 Fitted Gauss, proteinAT3G59050.1
bic:777
peaks after deleting less than one fifth of global max:16
×10^7
raw data, starting:16
fitted Gauss locations:16
y
x
0
1
2
3
4
5
6
7
8
9
0
5
10
15
20
25

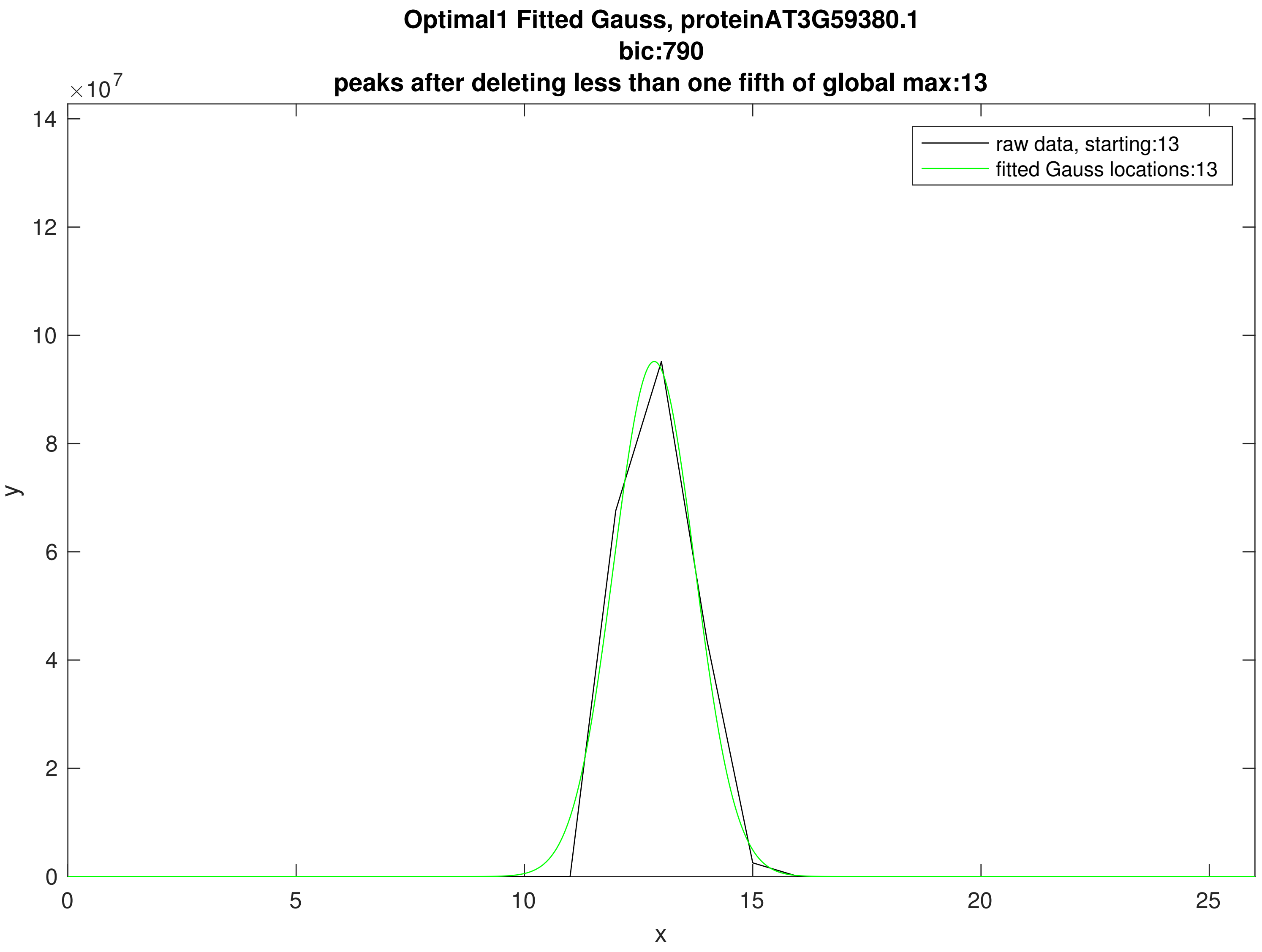

Optimal1 Fitted Gauss, proteinAT3G59380.1
bic:790
peaks after deleting less than one fifth of global max:13
×10^7
14
12
10
8
6
4
2
0
y
0
5
10
15
20
25
x
raw data, starting:13
fitted Gauss locations:13

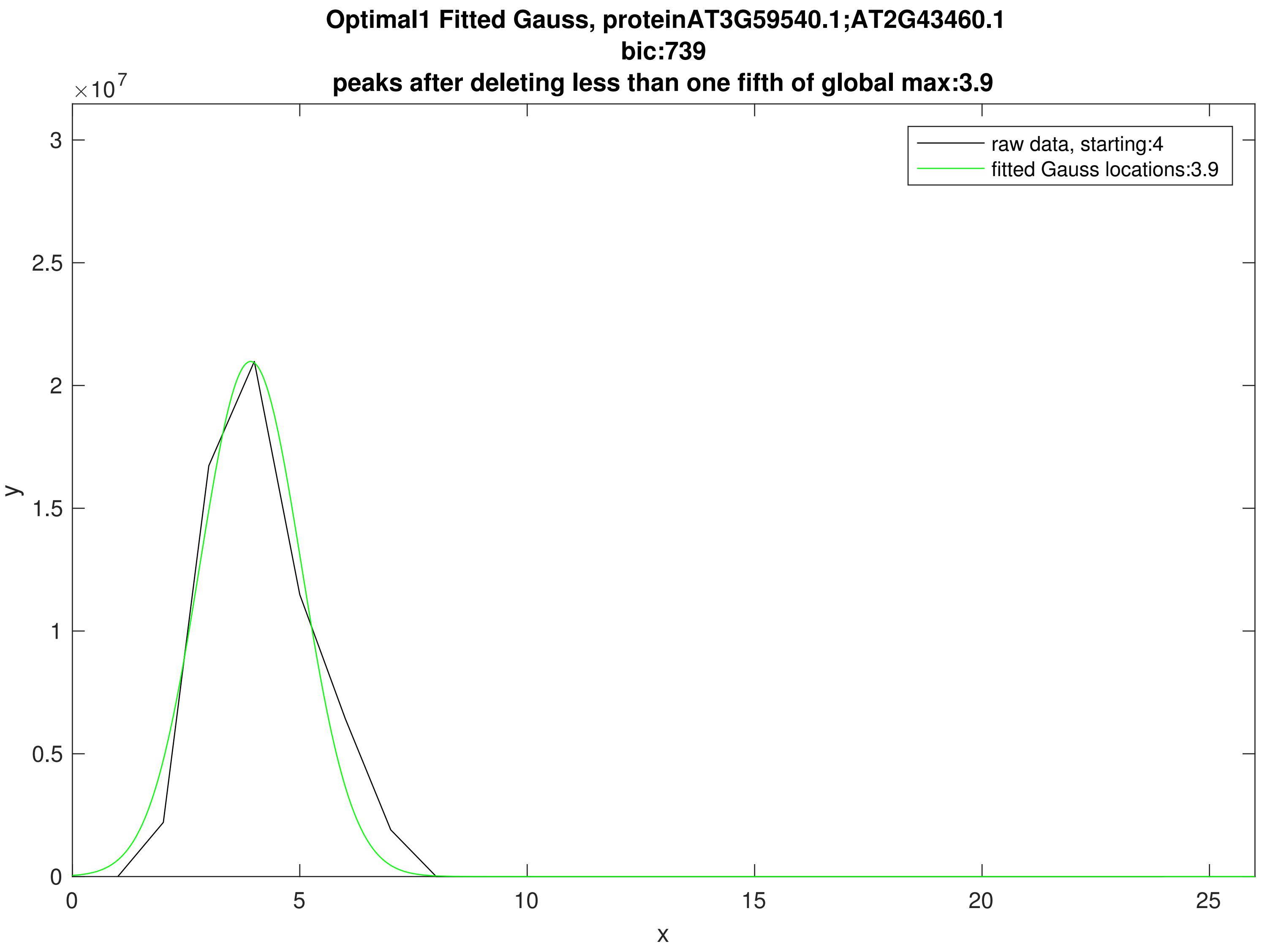
Optimal1 Fitted Gauss, proteinAT3G59540.1;AT2G43460.1
bic:739
peaks after deleting less than one fifth of global max:3.9
×10^7
3
2.5
2
1.5
1
0.5
0
y
0
5
10
15
20
25
x
raw data, starting:4
fitted Gauss locations:3.9

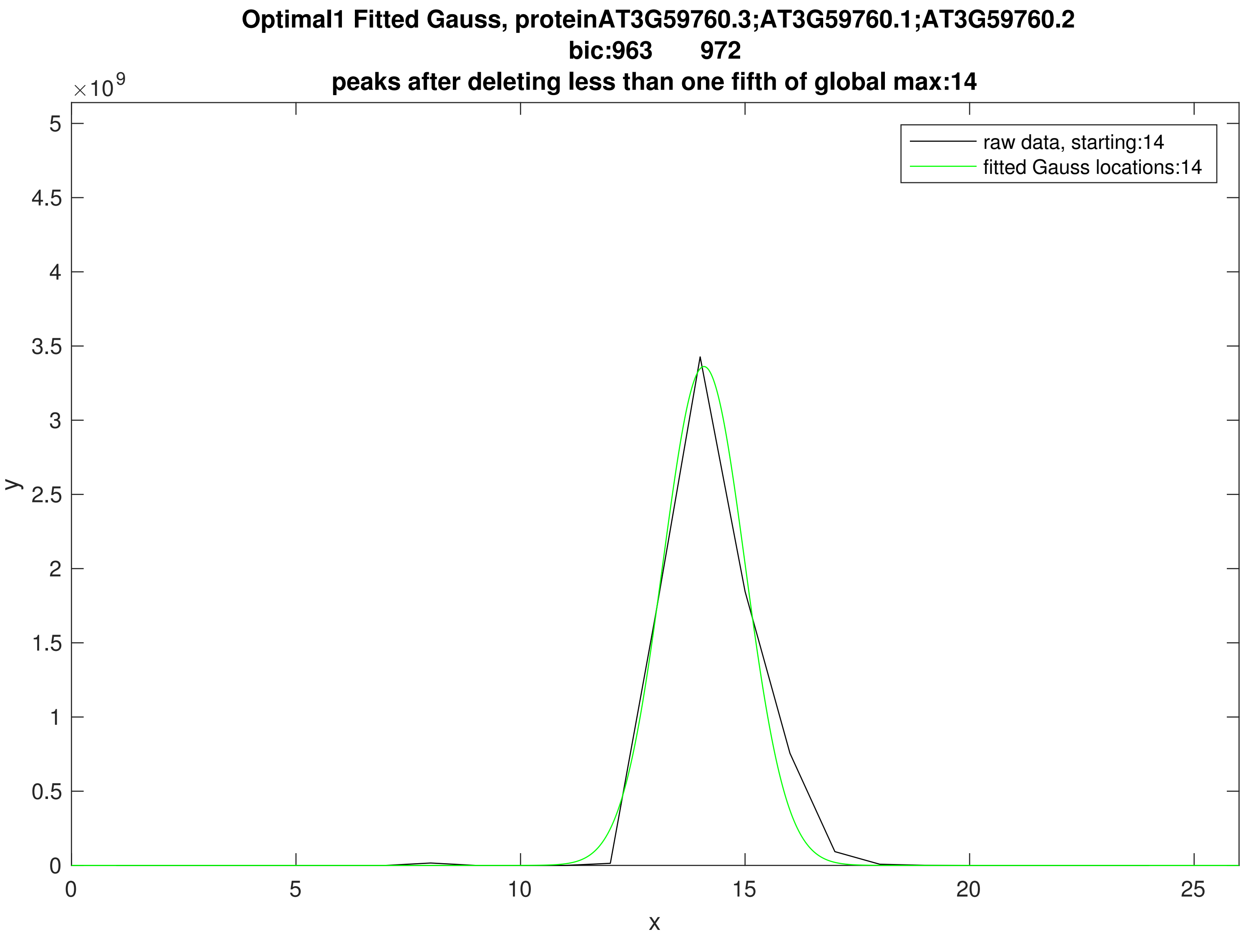

Optimal1 Fitted Gauss, proteinAT3G59760.3;AT3G59760.1;AT3G59760.2
bic:963 972
peaks after deleting less than one fifth of global max:14
×10^9
5
4.5
4
3.5
3
2.5
2
1.5
1
0.5
0
y
0
5
10
15
20
25
x
raw data, starting:14
fitted Gauss locations:14

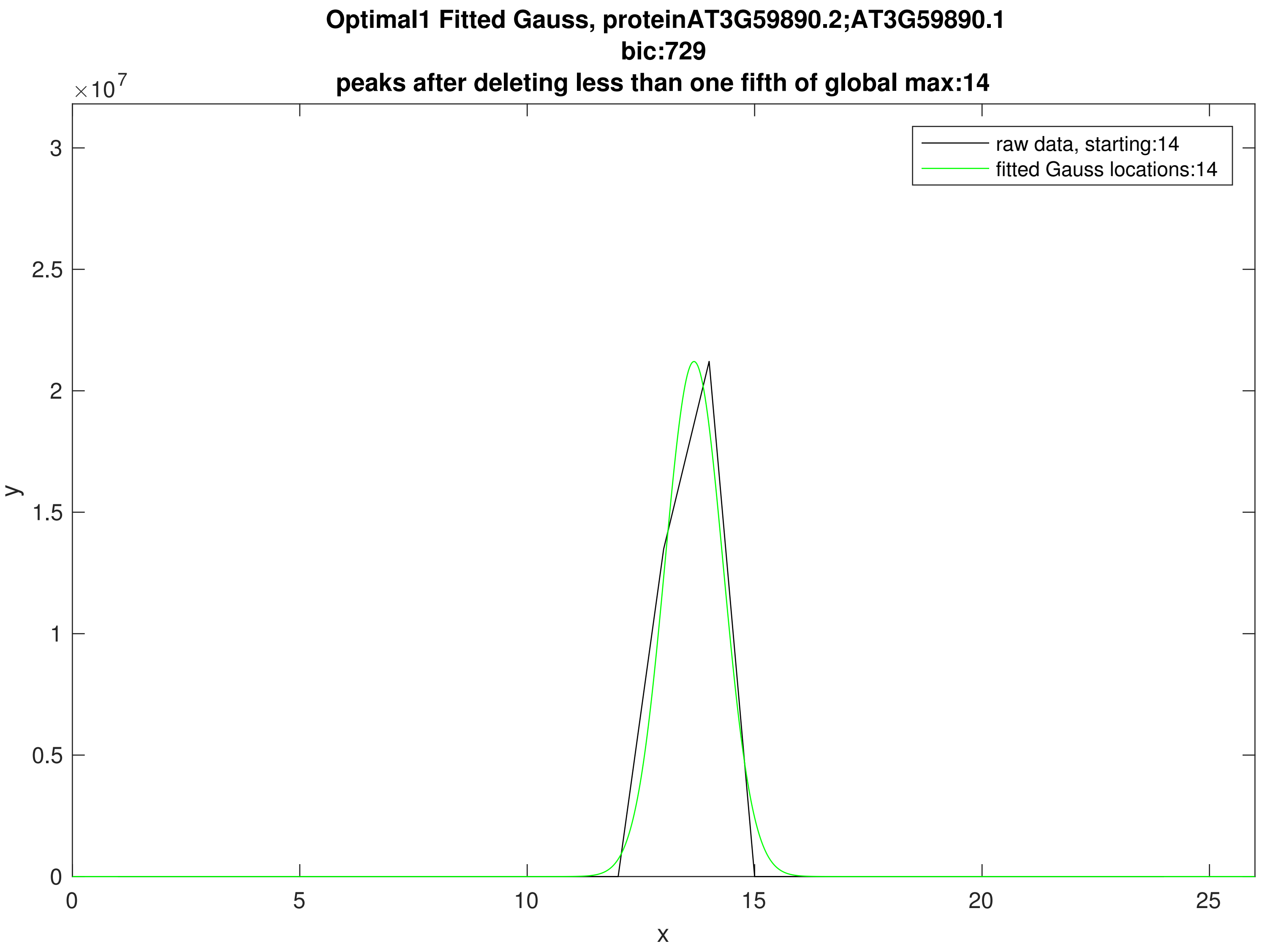

Optimal1 Fitted Gauss, proteinAT3G59890.2;AT3G59890.1
bic:729
peaks after deleting less than one fifth of global max:14
×10^7
3
2.5
2
1.5
1
0.5
0
y
0
5
10
15
20
25
x
raw data, starting:14
fitted Gauss locations:14

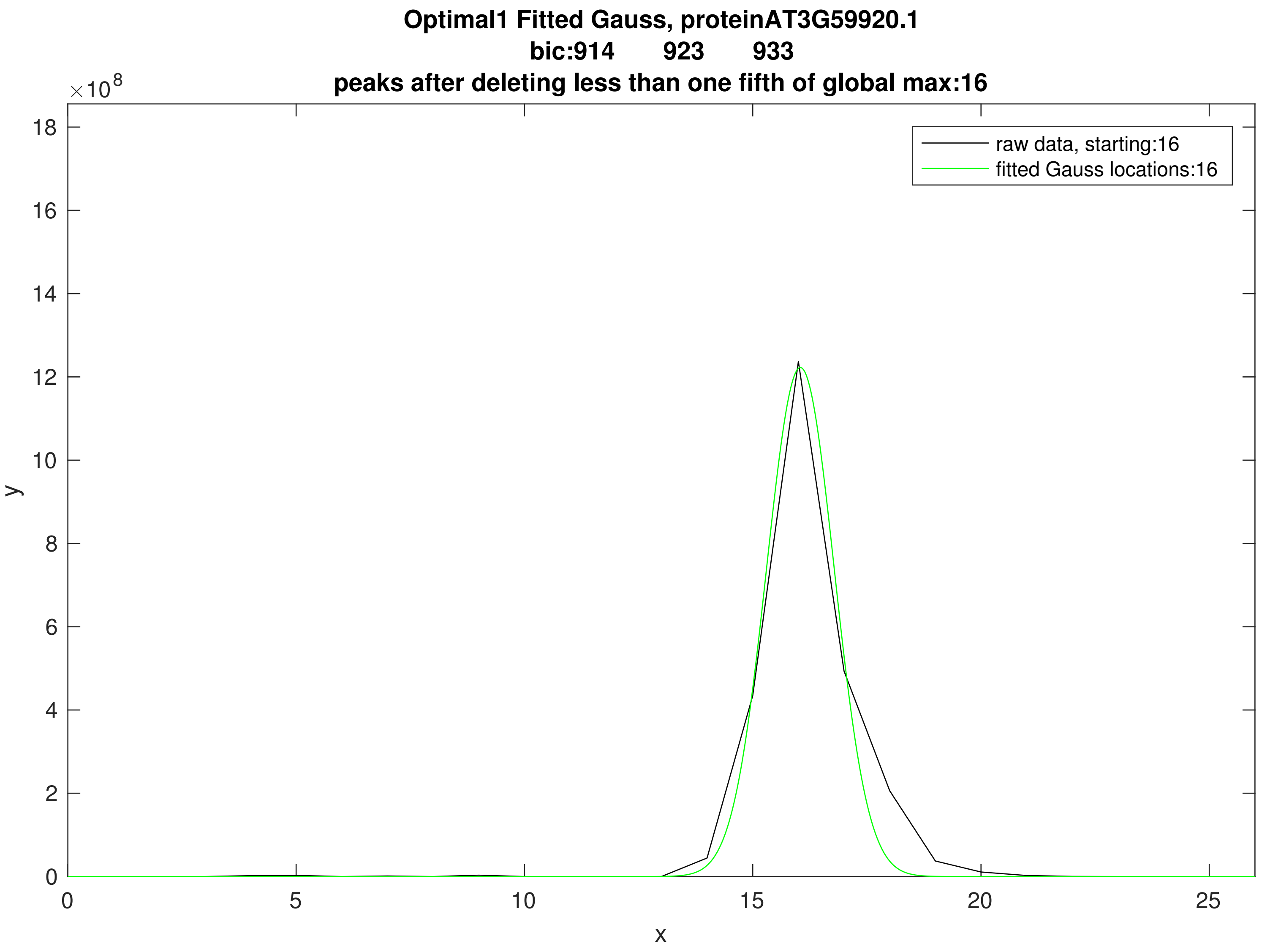

Optimal1 Fitted Gauss, proteinAT3G59920.1
bic:914 923 933
peaks after deleting less than one fifth of global max:16
×10^8
raw data, starting:16
fitted Gauss locations:16
y
x
0
2
4
6
8
10
12
14
16
18
0
5
10
15
20
25

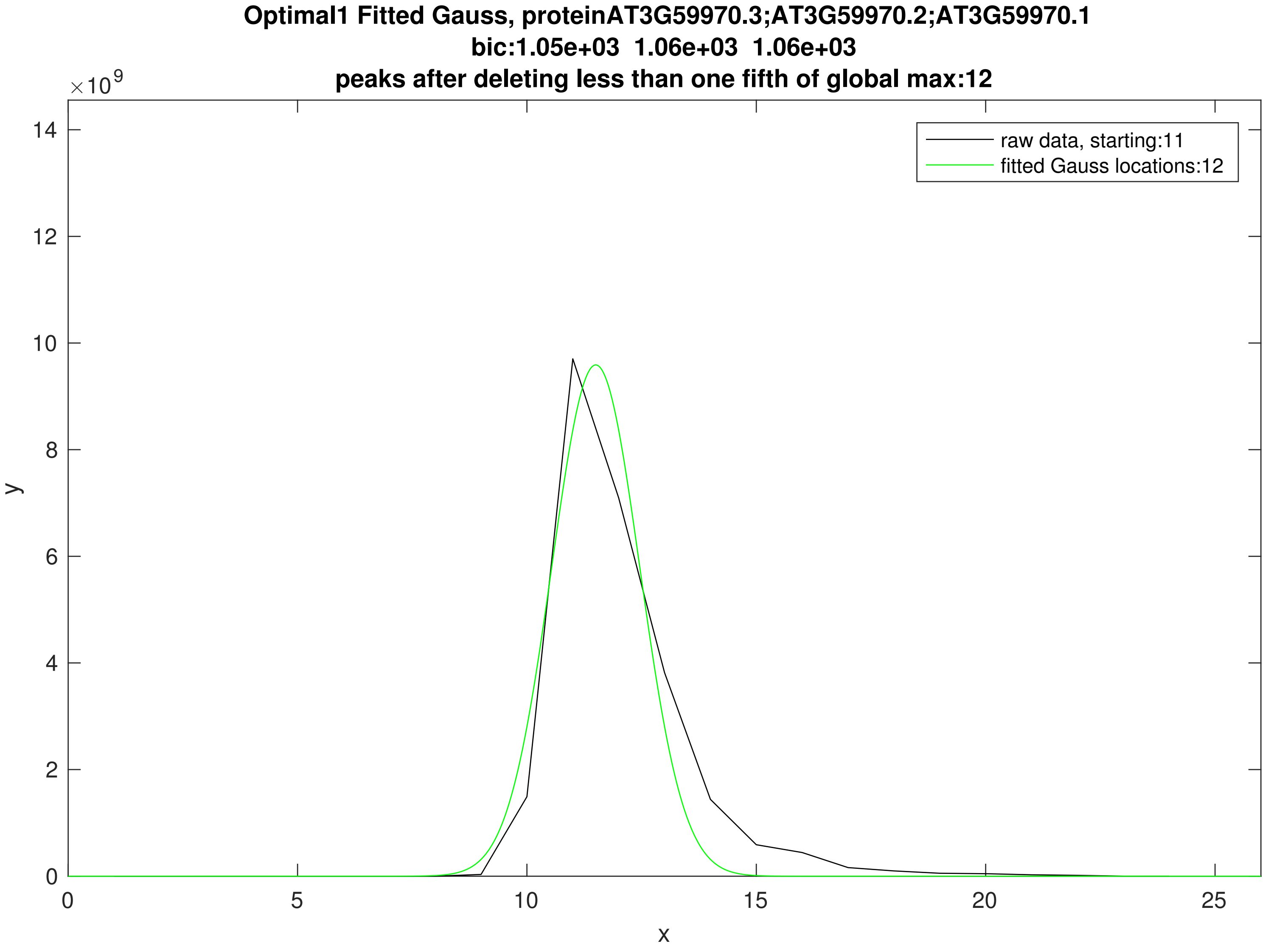
Optimal1 Fitted Gauss, proteinAT3G59970.3;AT3G59970.2;AT3G59970.1
bic:1.05e+03  1.06e+03  1.06e+03
peaks after deleting less than one fifth of global max:12
×10^9
raw data, starting:11
fitted Gauss locations:12
y
x
0
2
4
6
8
10
12
14
0
5
10
15
20
25

**Optimal1 Fitted Gauss, proteinAT3G59980.1**
**bic:776**
**peaks after deleting less than one fifth of global max:17**

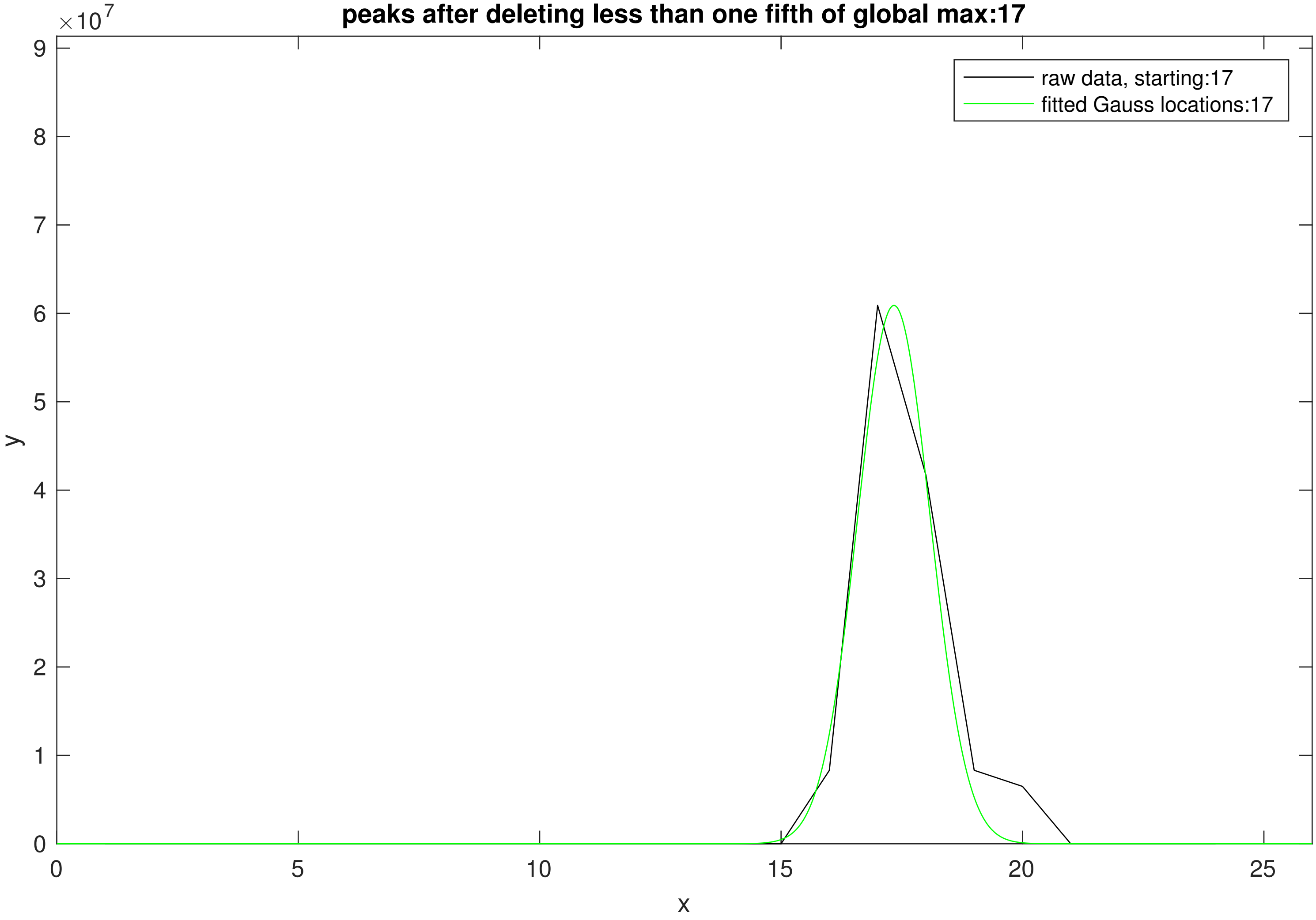

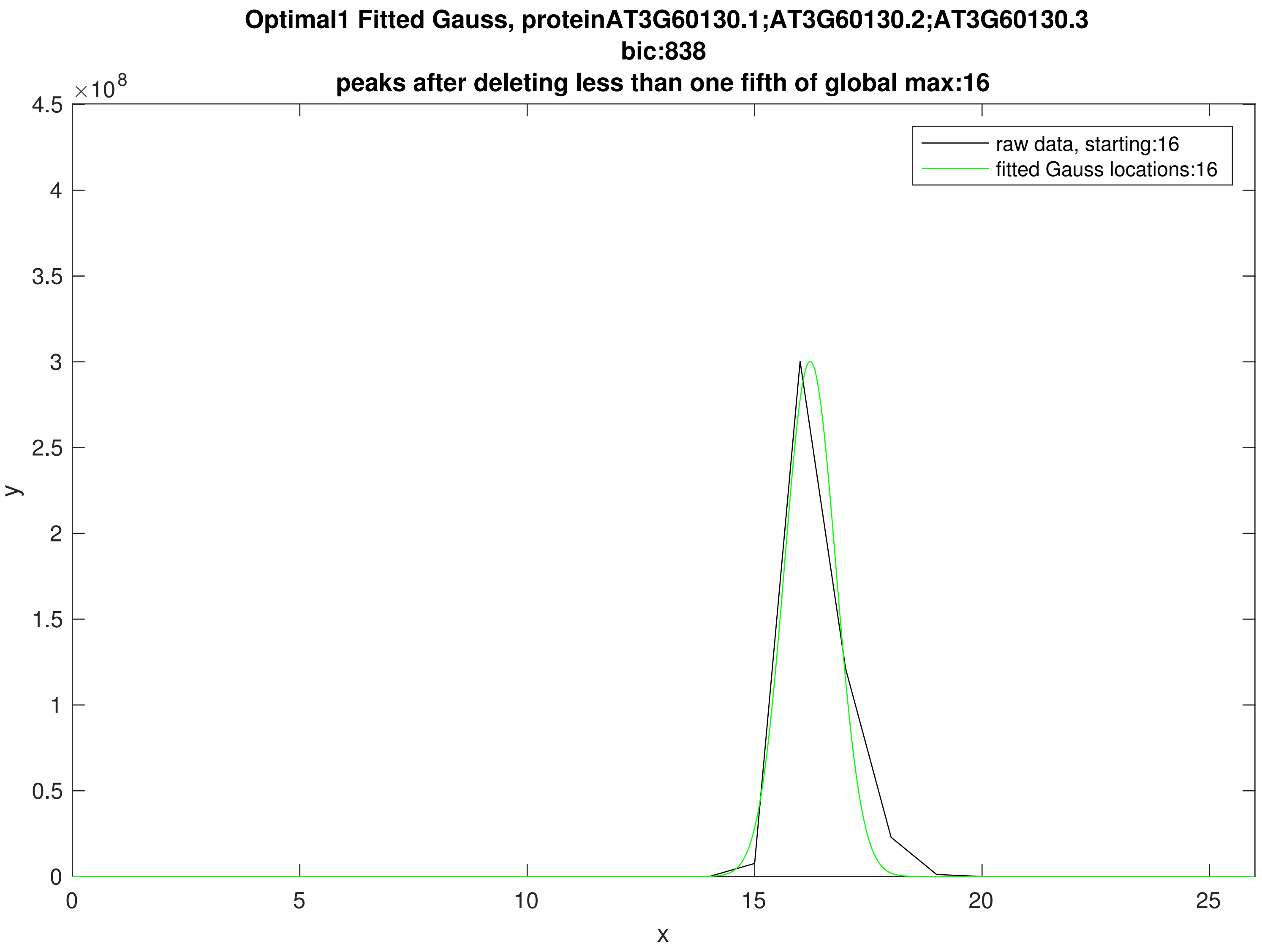

Optimal1 Fitted Gauss, proteinAT3G60130.1;AT3G60130.2;AT3G60130.3
bic:838
peaks after deleting less than one fifth of global max:16
×10^8
4.5
4
3.5
3
2.5
2
1.5
1
0.5
0
y
0
5
10
15
20
25
x
raw data, starting:16
fitted Gauss locations:16

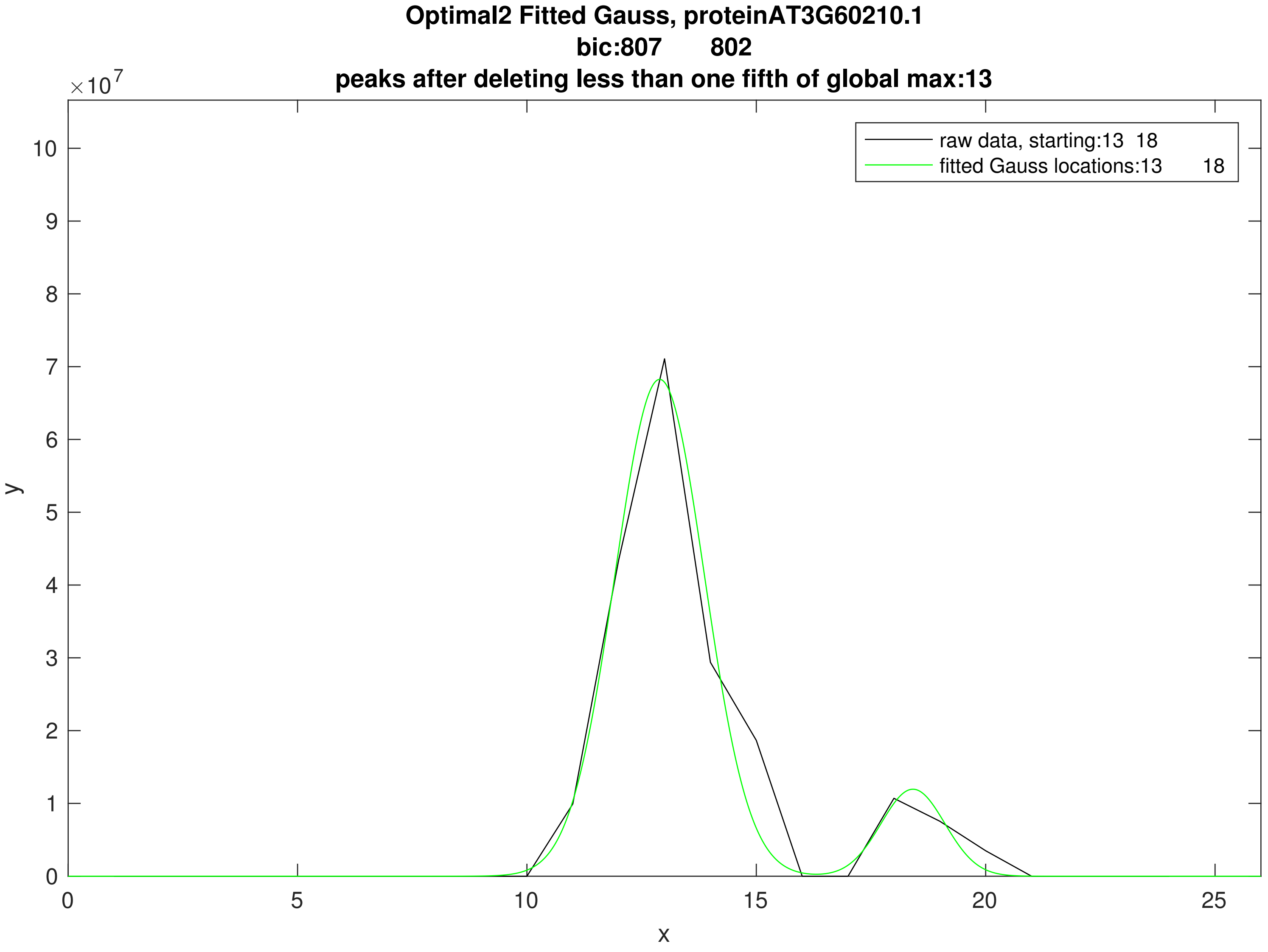
Optimal2 Fitted Gauss, proteinAT3G60210.1
bic:807 802
peaks after deleting less than one fifth of global max:13
×10^7
raw data, starting:13 18
fitted Gauss locations:13 18
y
x
0
1
2
3
4
5
6
7
8
9
10
0
5
10
15
20
25

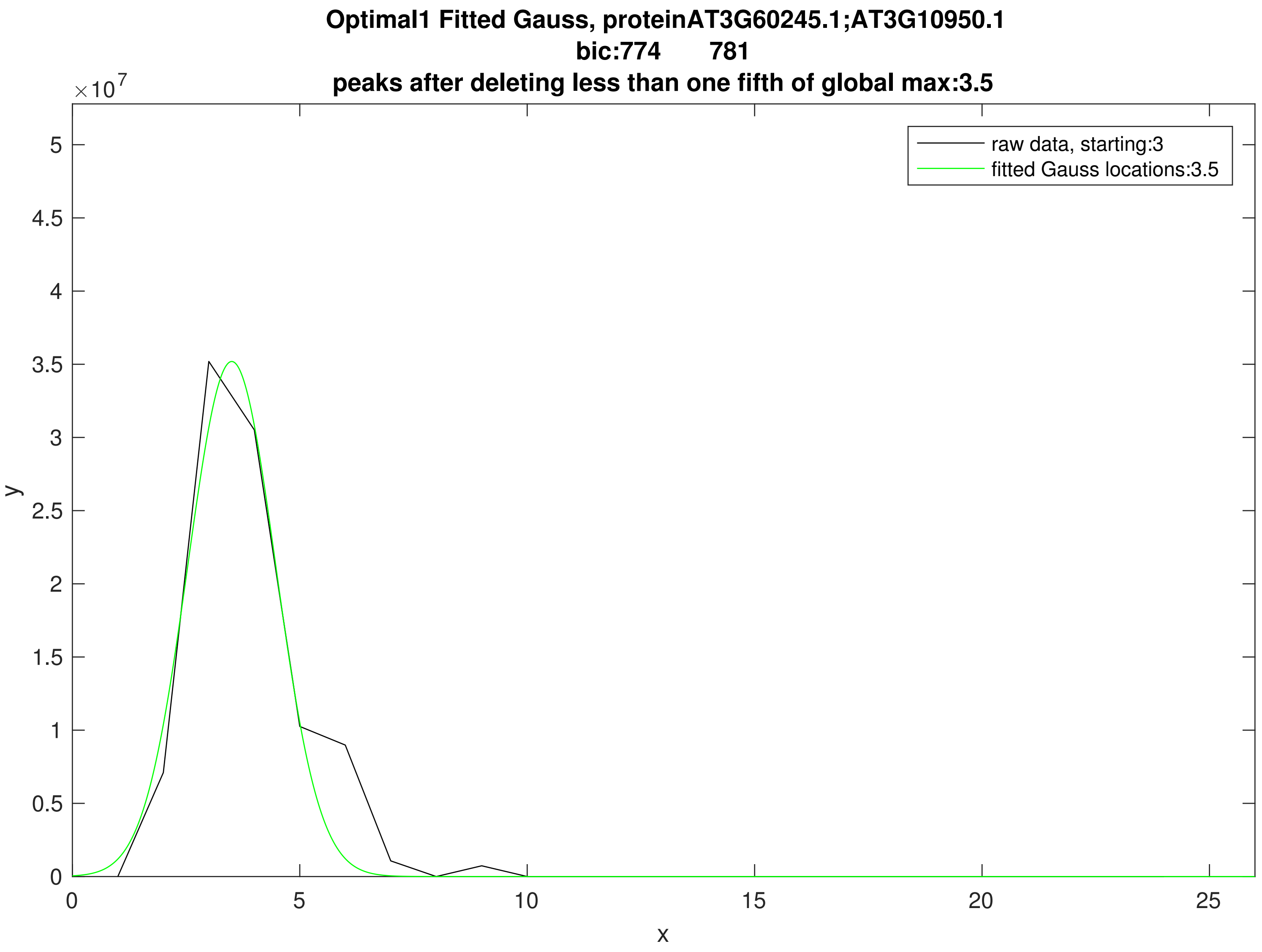
Optimal1 Fitted Gauss, proteinAT3G60245.1;AT3G10950.1
bic:774 781
peaks after deleting less than one fifth of global max:3.5
×10^7
raw data, starting:3
fitted Gauss locations:3.5
y
x
0
0.5
1
1.5
2
2.5
3
3.5
4
4.5
5
0
5
10
15
20
25

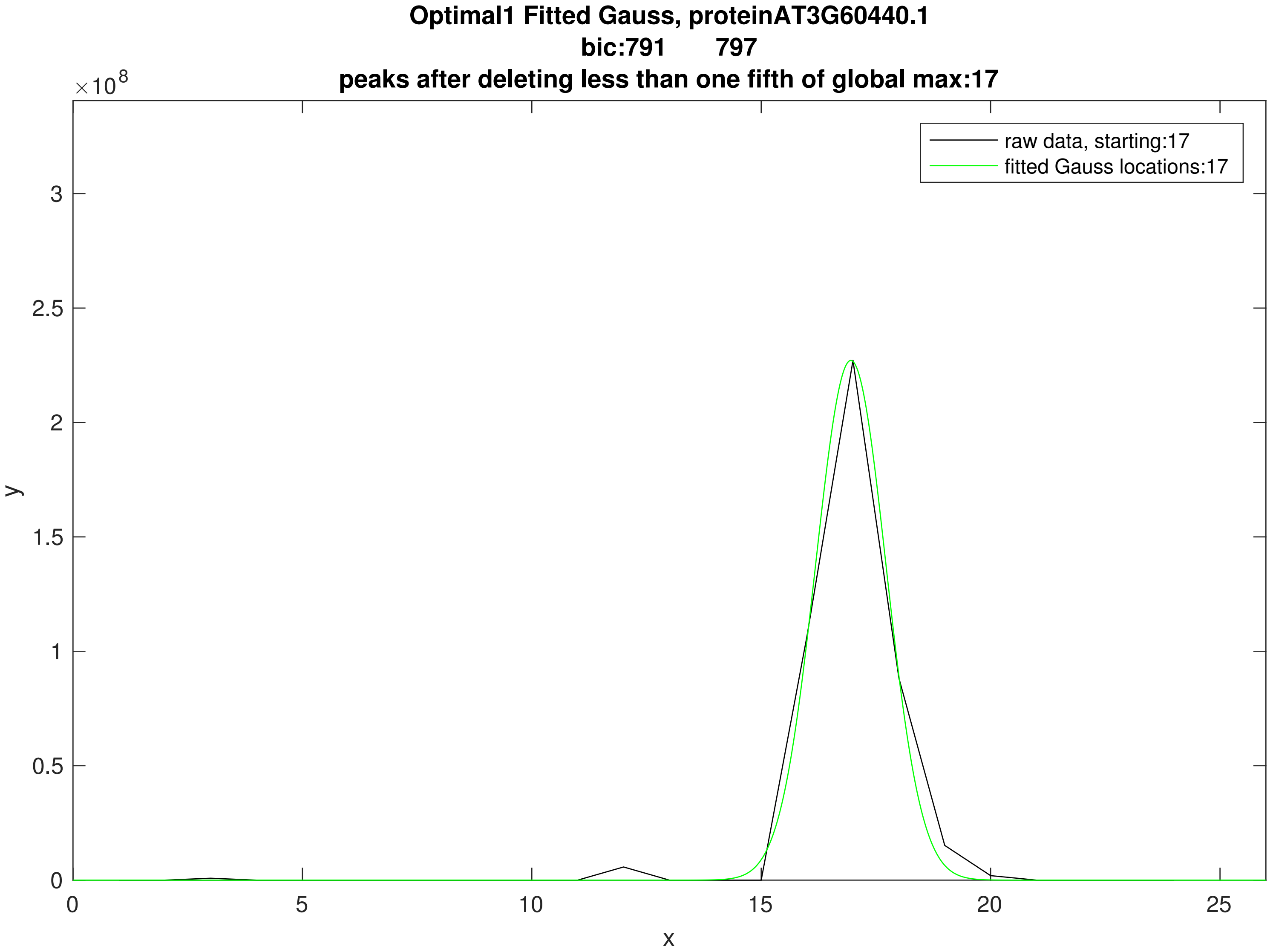

Optimal1 Fitted Gauss, proteinAT3G60440.1
bic:791 797
peaks after deleting less than one fifth of global max:17
×10^8
raw data, starting:17
fitted Gauss locations:17
3
2.5
2
1.5
1
0.5
0
y
0
5
10
15
20
25
x

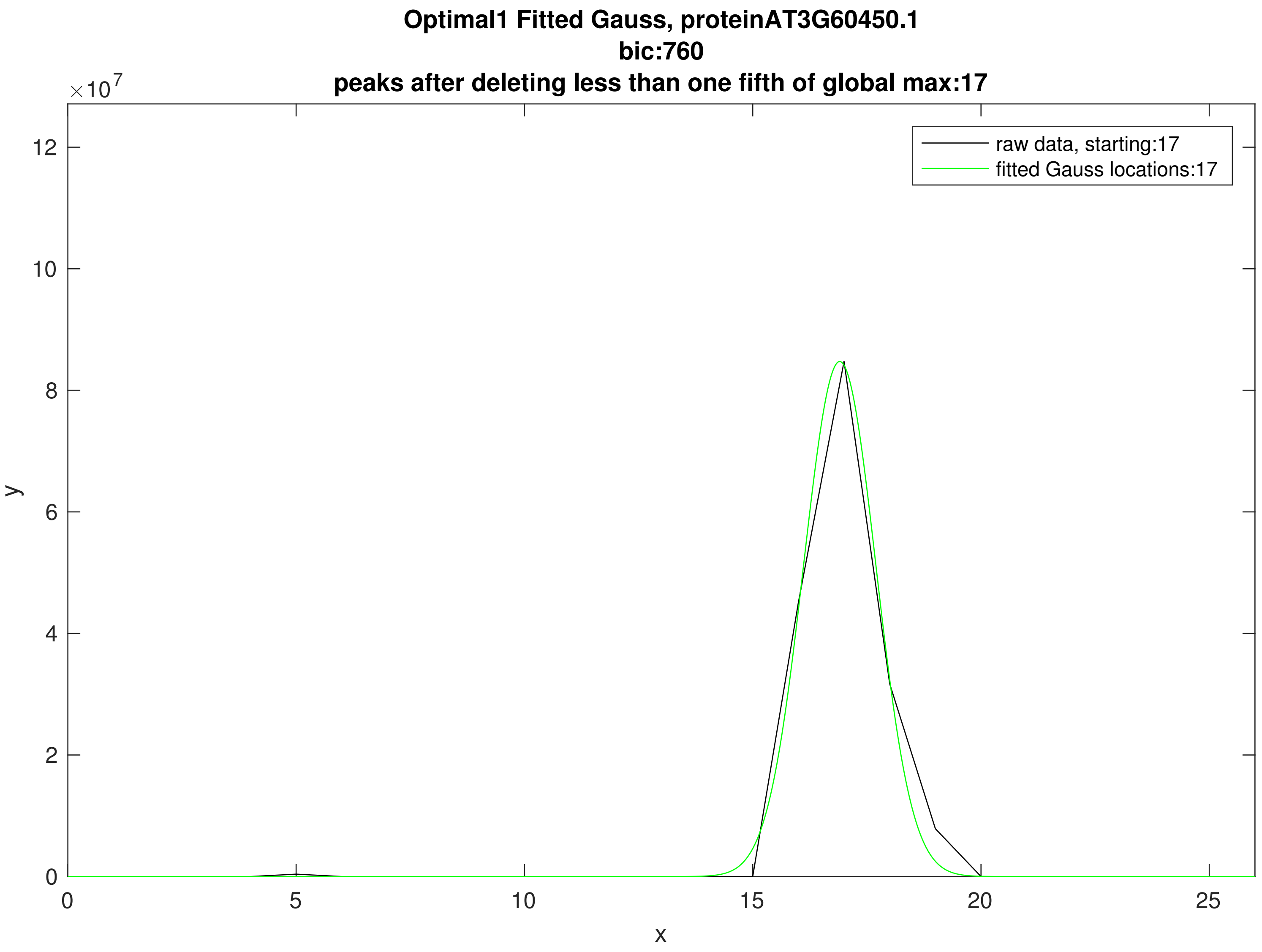

Optimal1 Fitted Gauss, proteinAT3G60450.1
bic:760
peaks after deleting less than one fifth of global max:17
×10^7
12
10
8
6
4
2
0
y
0
5
10
15
20
25
x
raw data, starting:17
fitted Gauss locations:17

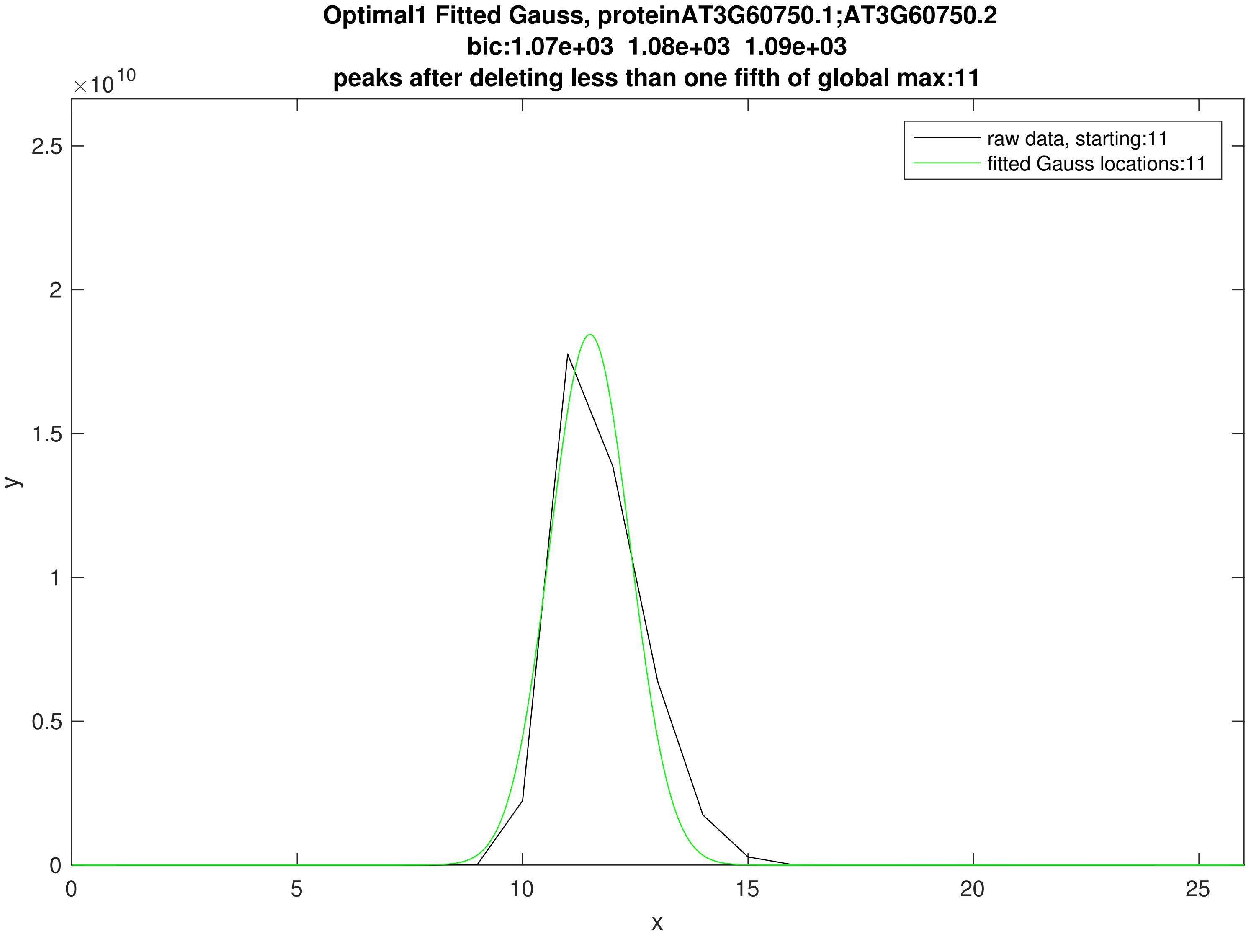

Optimal1 Fitted Gauss, proteinAT3G60750.1;AT3G60750.2
bic:1.07e+03 1.08e+03 1.09e+03
peaks after deleting less than one fifth of global max:11
×10^10
raw data, starting:11
fitted Gauss locations:11
y
x
0
0.5
1
1.5
2
2.5
5
10
15
20
25

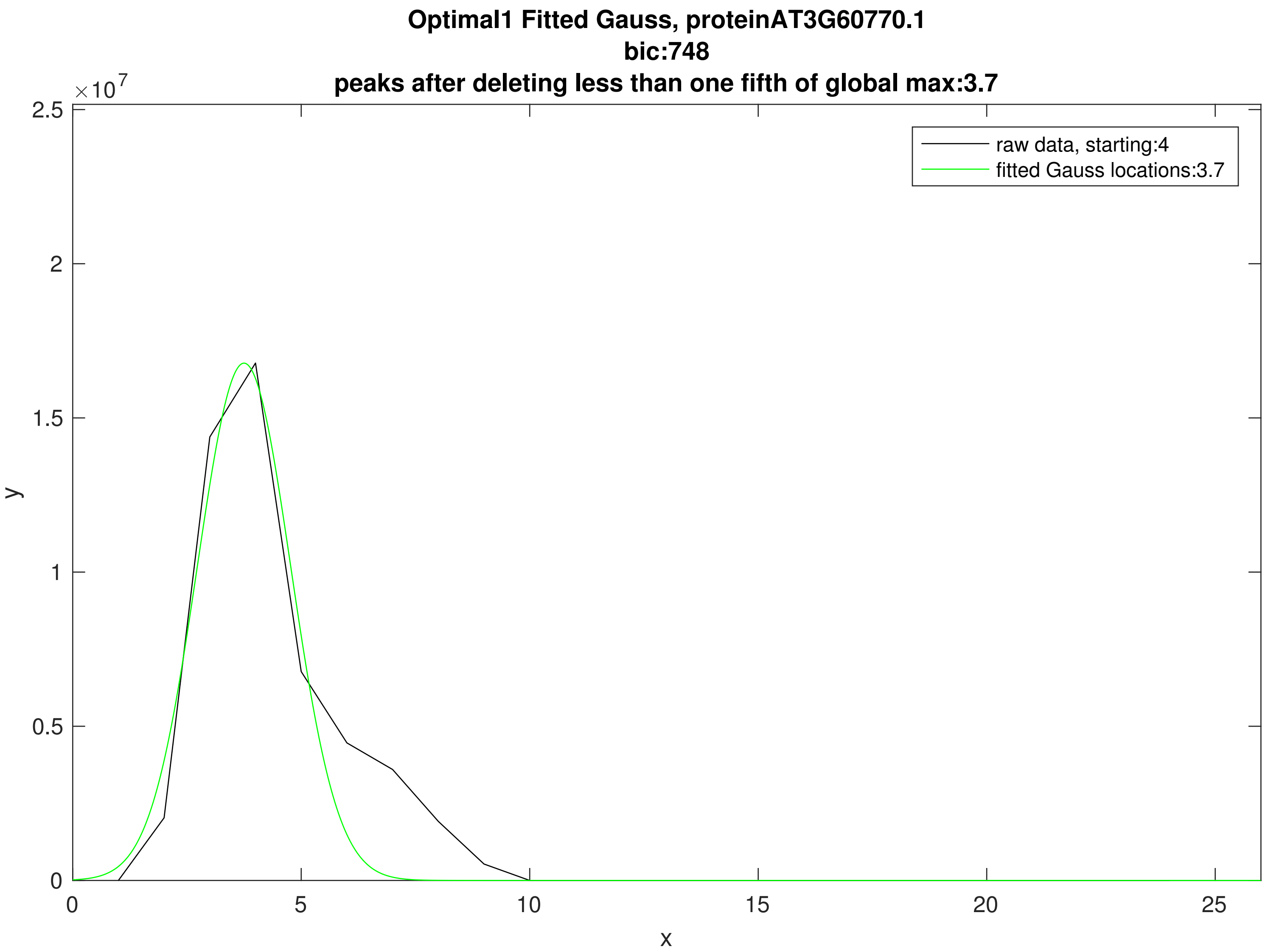
Optimal1 Fitted Gauss, proteinAT3G60770.1
bic:748
peaks after deleting less than one fifth of global max:3.7
×10^7
2.5
2
1.5
1
0.5
0
y
0
5
10
15
20
25
x
raw data, starting:4
fitted Gauss locations:3.7

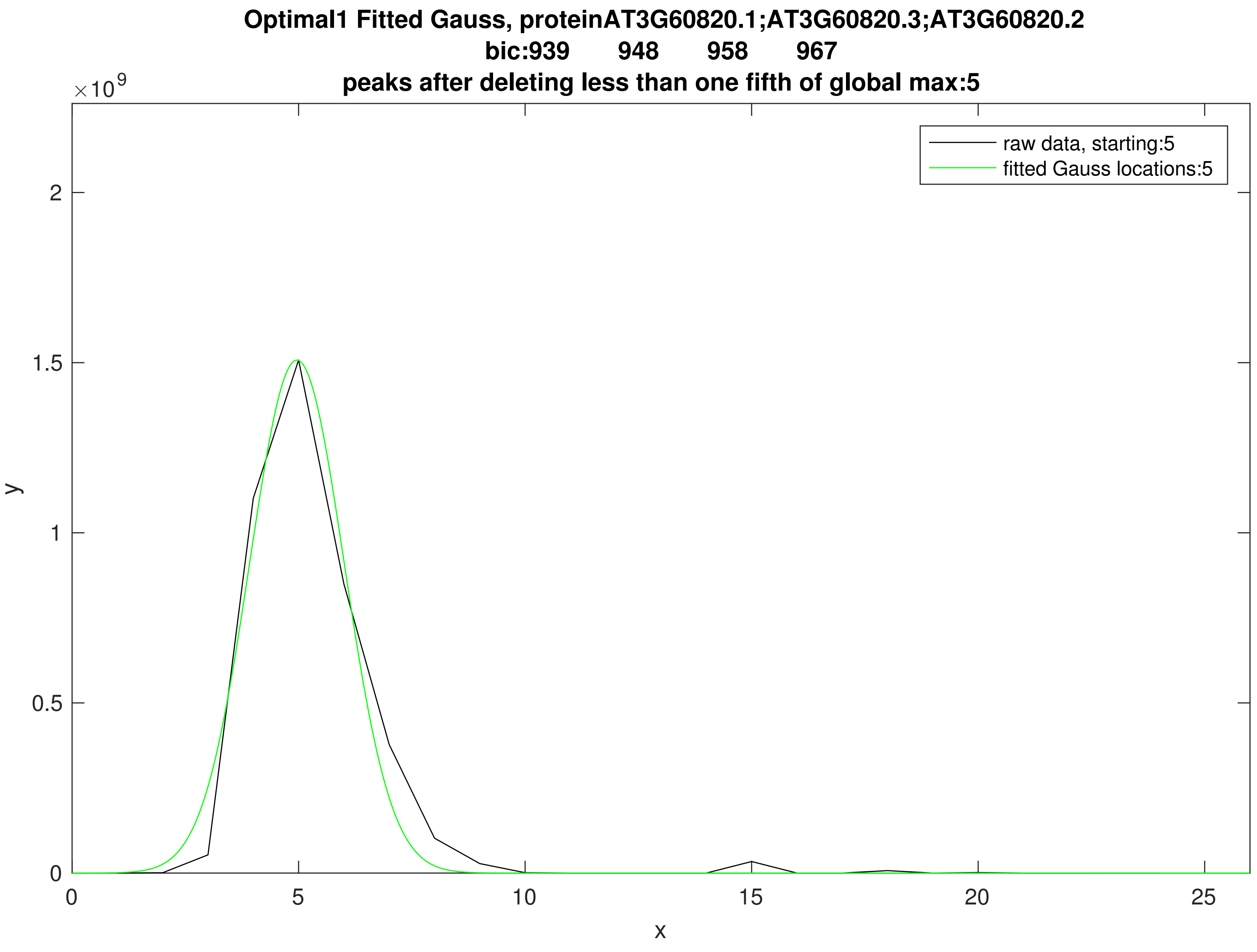
Optimal1 Fitted Gauss, proteinAT3G60820.1;AT3G60820.3;AT3G60820.2
bic:939 948 958 967
peaks after deleting less than one fifth of global max:5
×10^9
raw data, starting:5
fitted Gauss locations:5
y
x
0
0.5
1
1.5
2
5
10
15
20
25

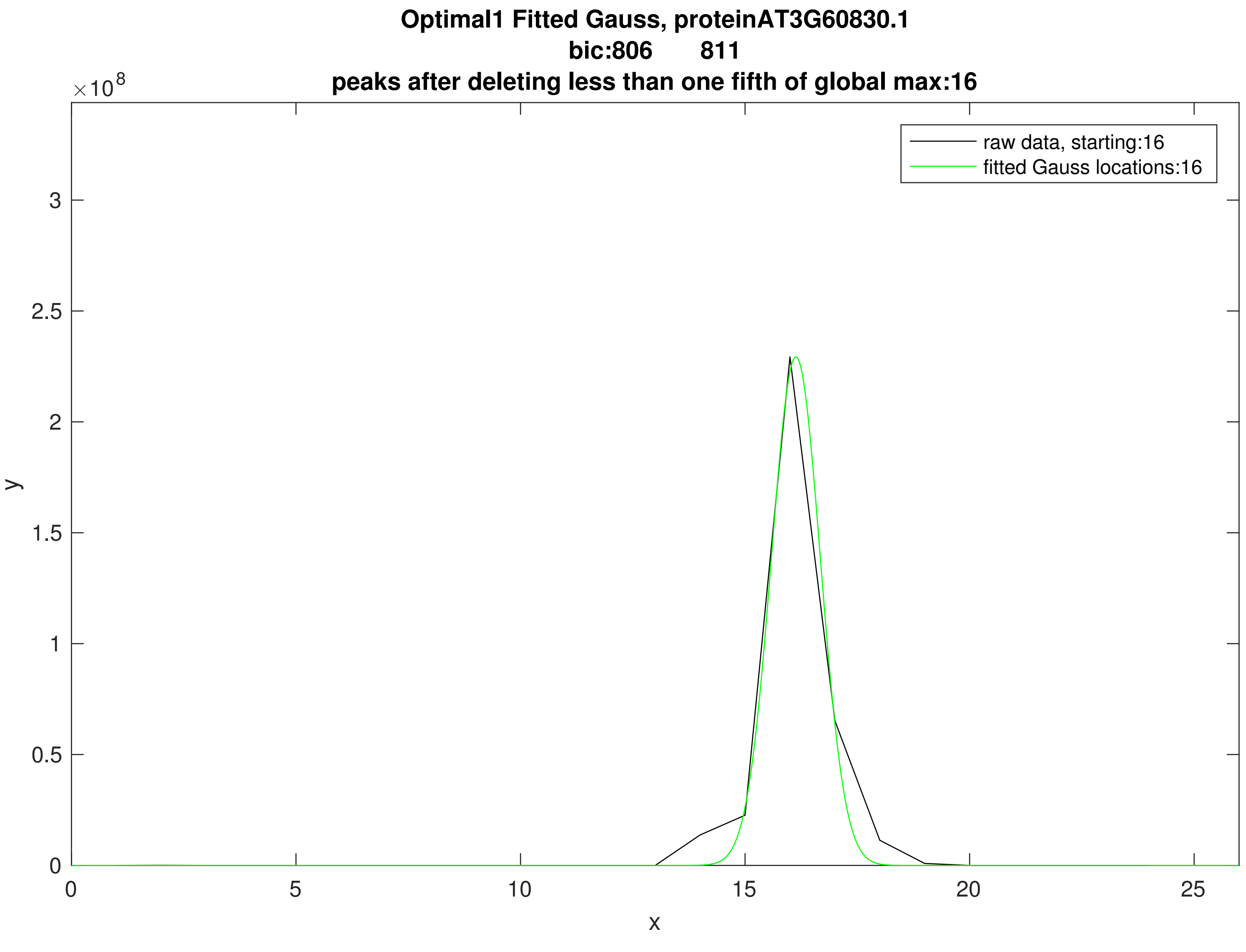

Optimal1 Fitted Gauss, proteinAT3G60830.1
bic:806 811
peaks after deleting less than one fifth of global max:16
×10^8
raw data, starting:16
fitted Gauss locations:16
y
x
0
0.5
1
1.5
2
2.5
3
0
5
10
15
20
25

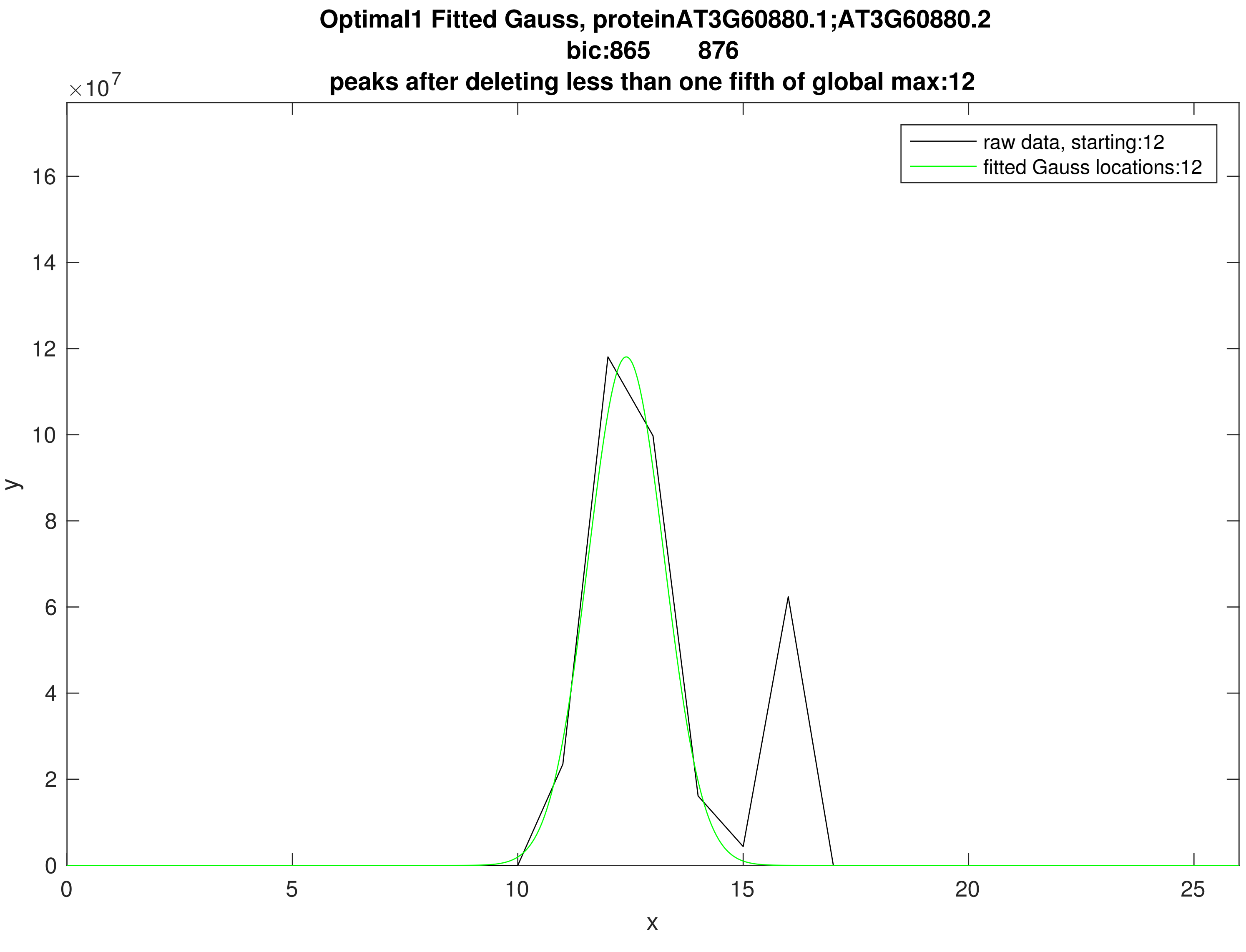
Optimal1 Fitted Gauss, proteinAT3G60880.1;AT3G60880.2
bic:865 876
peaks after deleting less than one fifth of global max:12
×10^7
raw data, starting:12
fitted Gauss locations:12
y
x
0
2
4
6
8
10
12
14
16
0
5
10
15
20
25

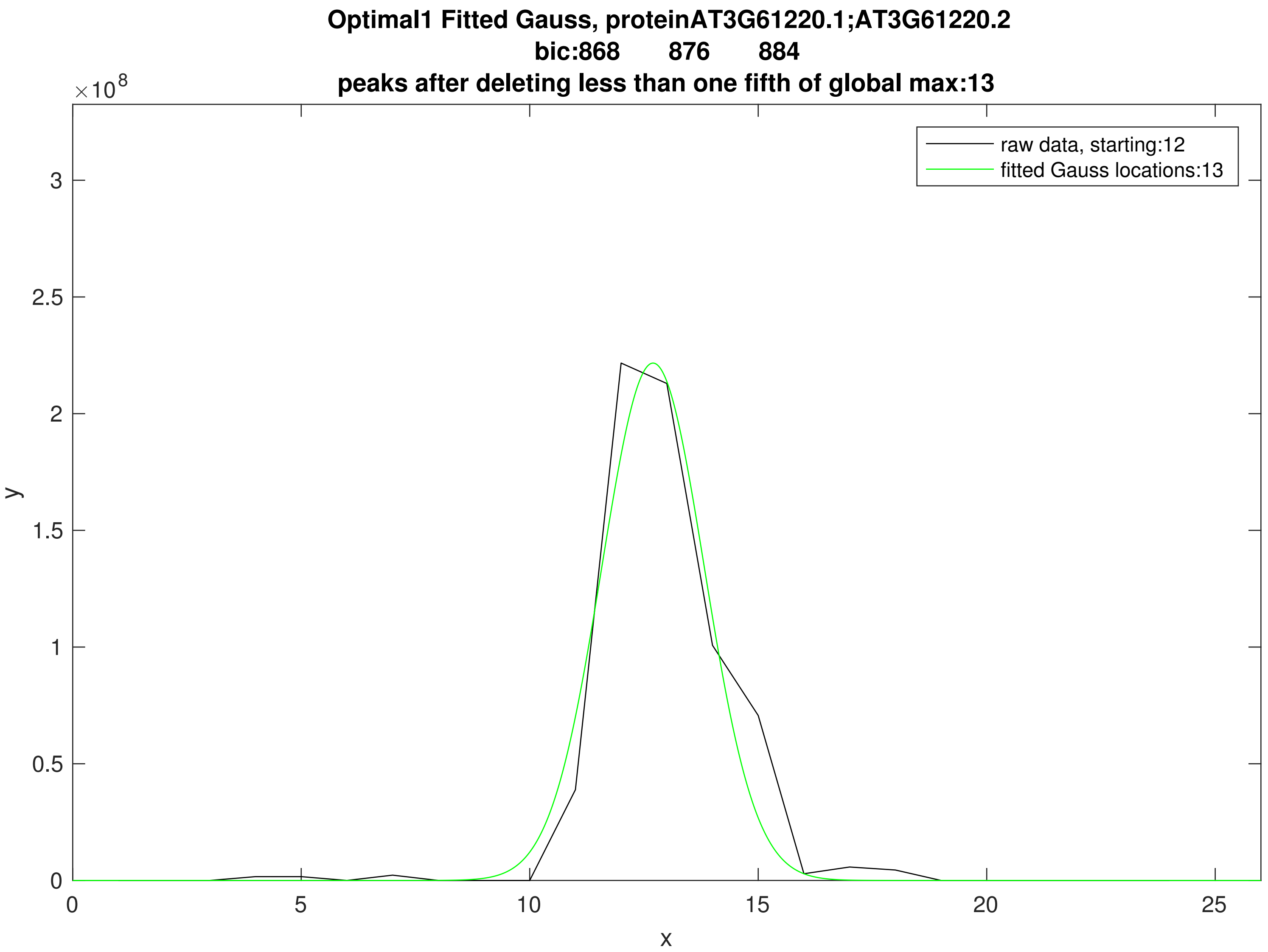

Optimal1 Fitted Gauss, proteinAT3G61220.1;AT3G61220.2
bic:868 876 884
peaks after deleting less than one fifth of global max:13
×10^8
3
2.5
2
1.5
1
0.5
0
y
0
5
10
15
20
25
x
raw data, starting:12
fitted Gauss locations:13

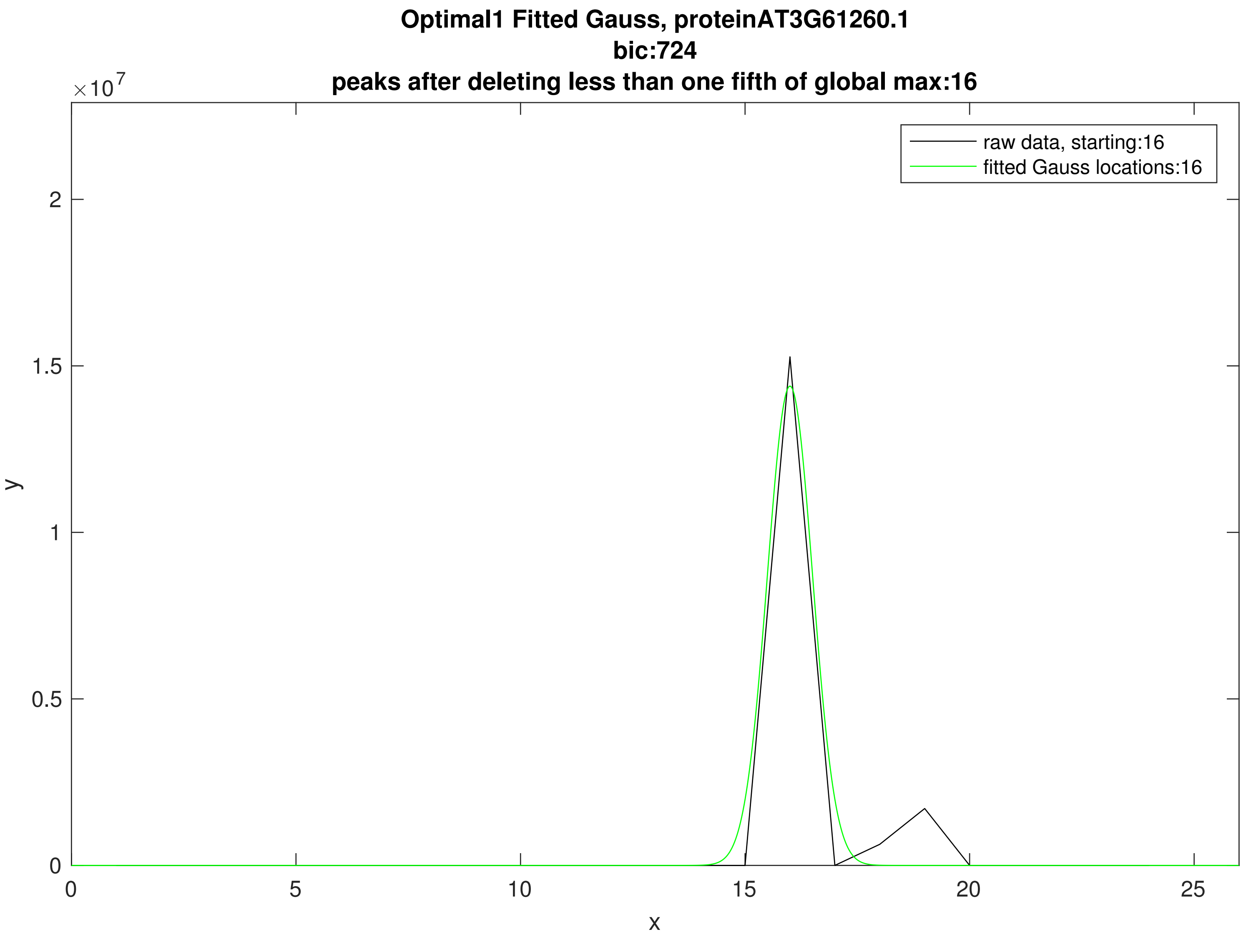
Optimal1 Fitted Gauss, proteinAT3G61260.1
bic:724
peaks after deleting less than one fifth of global max:16
×10^7
raw data, starting:16
fitted Gauss locations:16
y
x
0
0.5
1
1.5
2
0
5
10
15
20
25

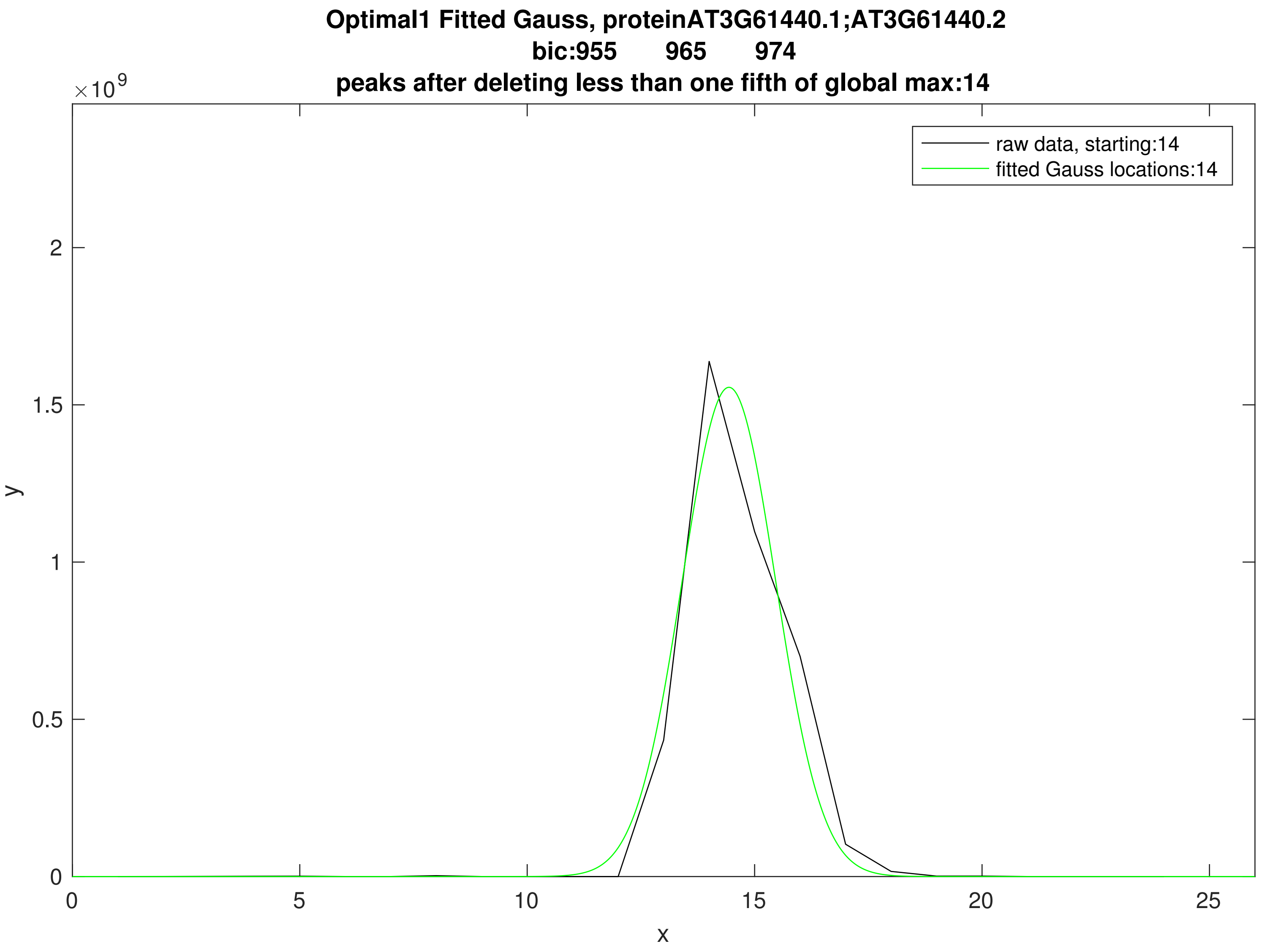

Optimal1 Fitted Gauss, proteinAT3G61440.1;AT3G61440.2
bic:955 965 974
peaks after deleting less than one fifth of global max:14
×10^9
raw data, starting:14
fitted Gauss locations:14
y
x
0
0.5
1
1.5
2
0
5
10
15
20
25

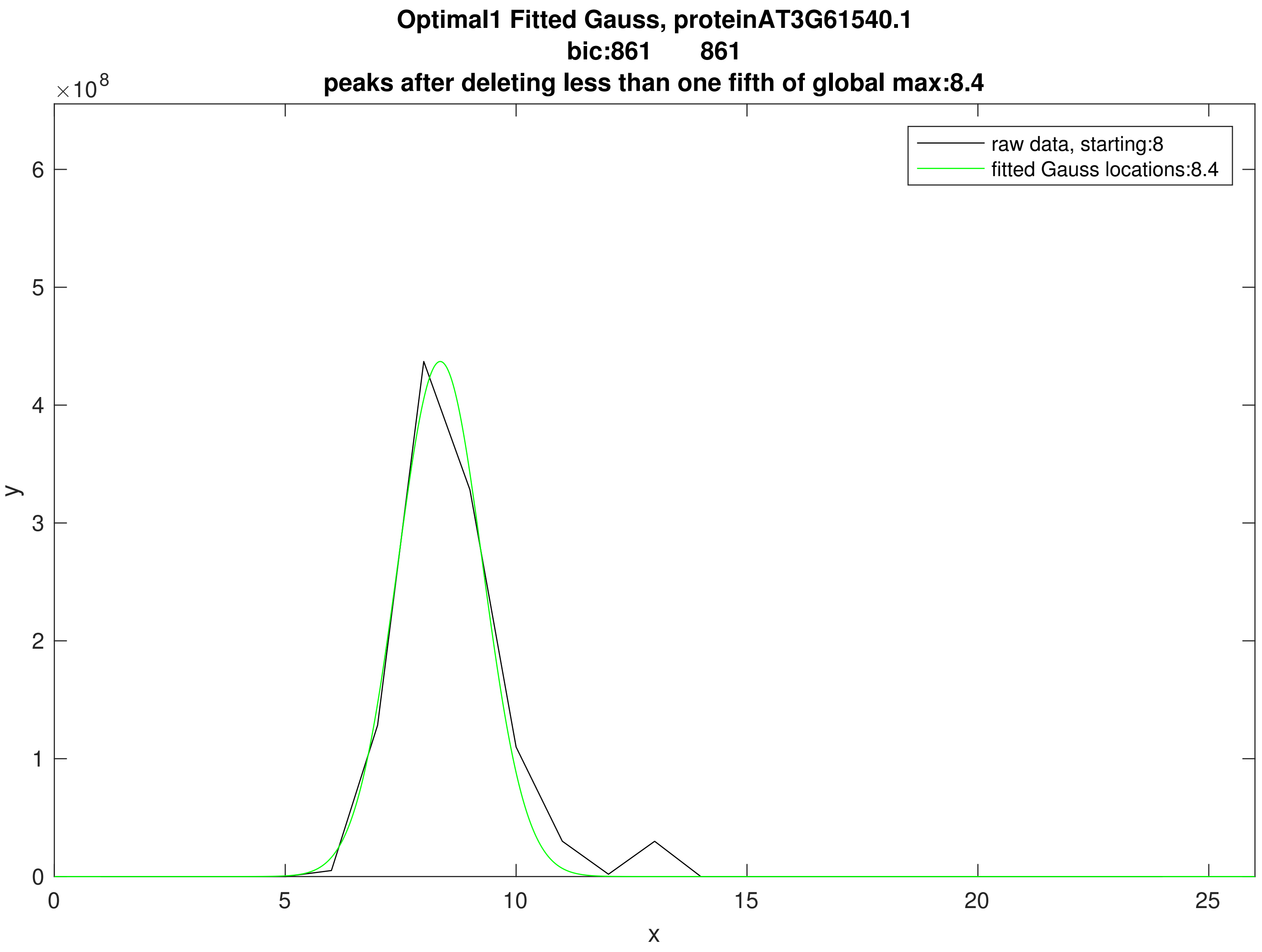
Optimal1 Fitted Gauss, proteinAT3G61540.1
bic:861 861
peaks after deleting less than one fifth of global max:8.4
×10^8
raw data, starting:8
fitted Gauss locations:8.4
y
x
0
1
2
3
4
5
6
0
5
10
15
20
25

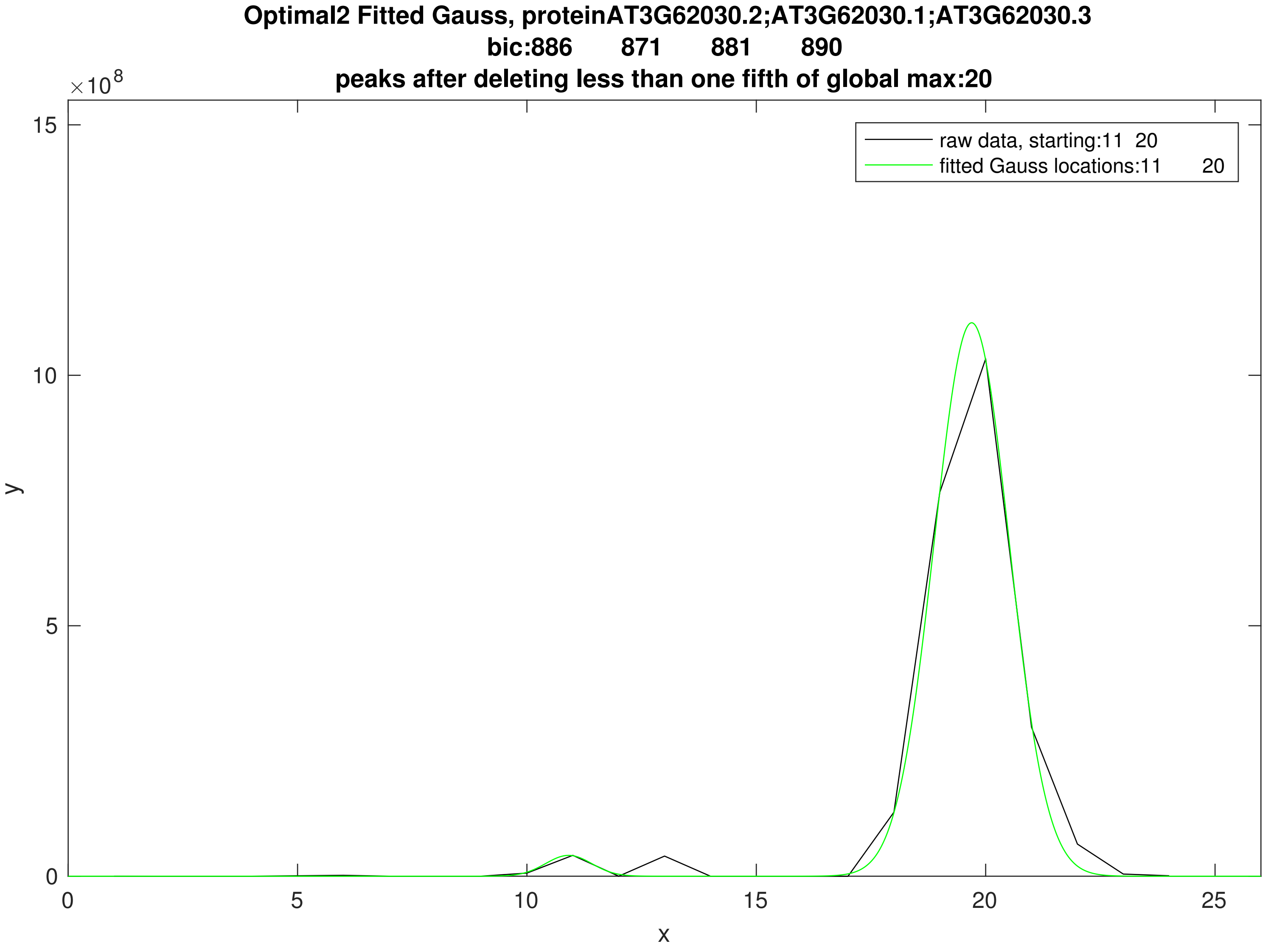

Optimal2 Fitted Gauss, proteinAT3G62030.2;AT3G62030.1;AT3G62030.3
bic:886 871 881 890
peaks after deleting less than one fifth of global max:20
×10^8
raw data, starting:11 20
fitted Gauss locations:11 20
y
x

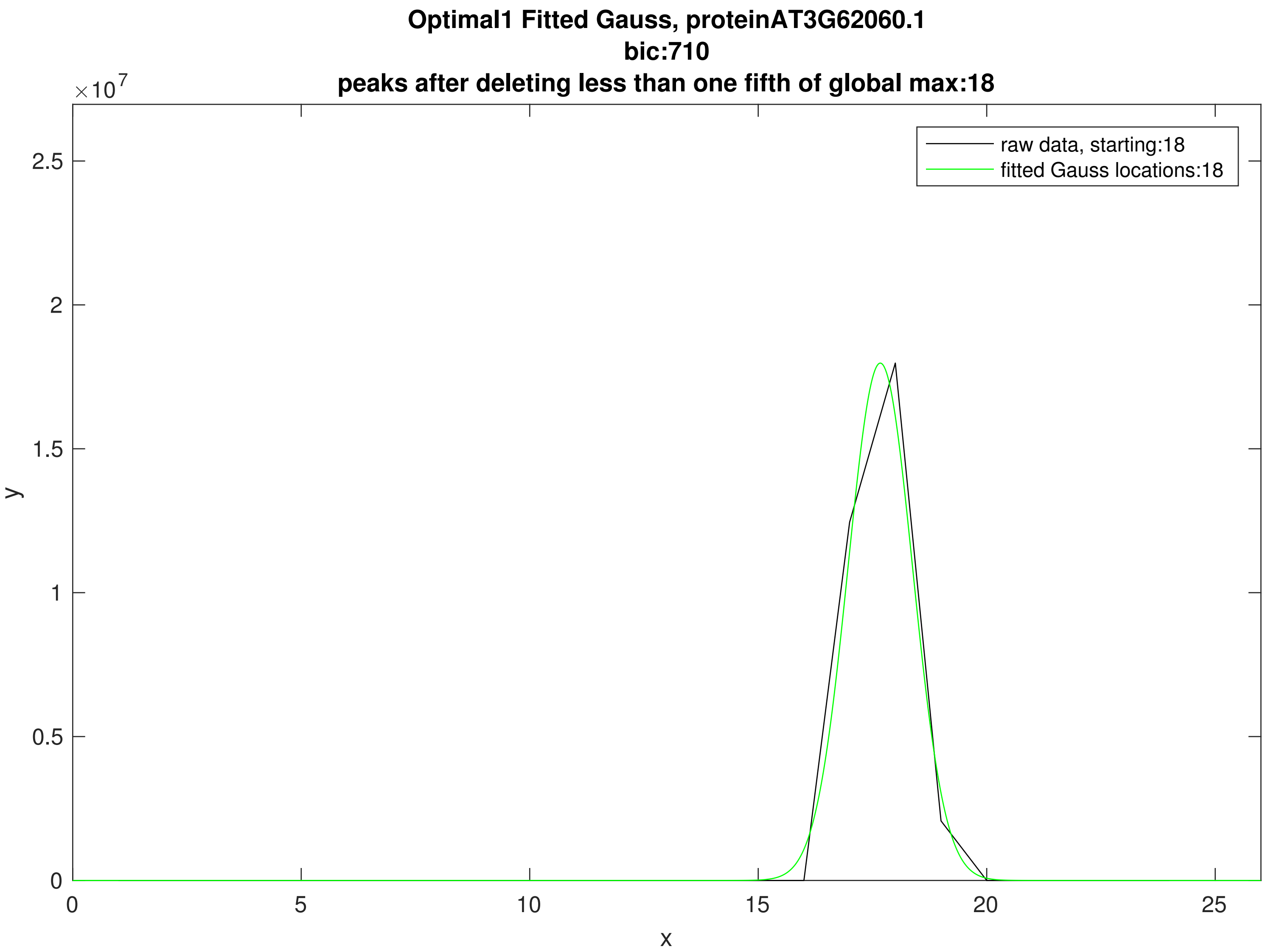
Optimal1 Fitted Gauss, proteinAT3G62060.1
bic:710
peaks after deleting less than one fifth of global max:18
×10^7
raw data, starting:18
fitted Gauss locations:18
y
x
0
0.5
1
1.5
2
2.5
0
5
10
15
20
25

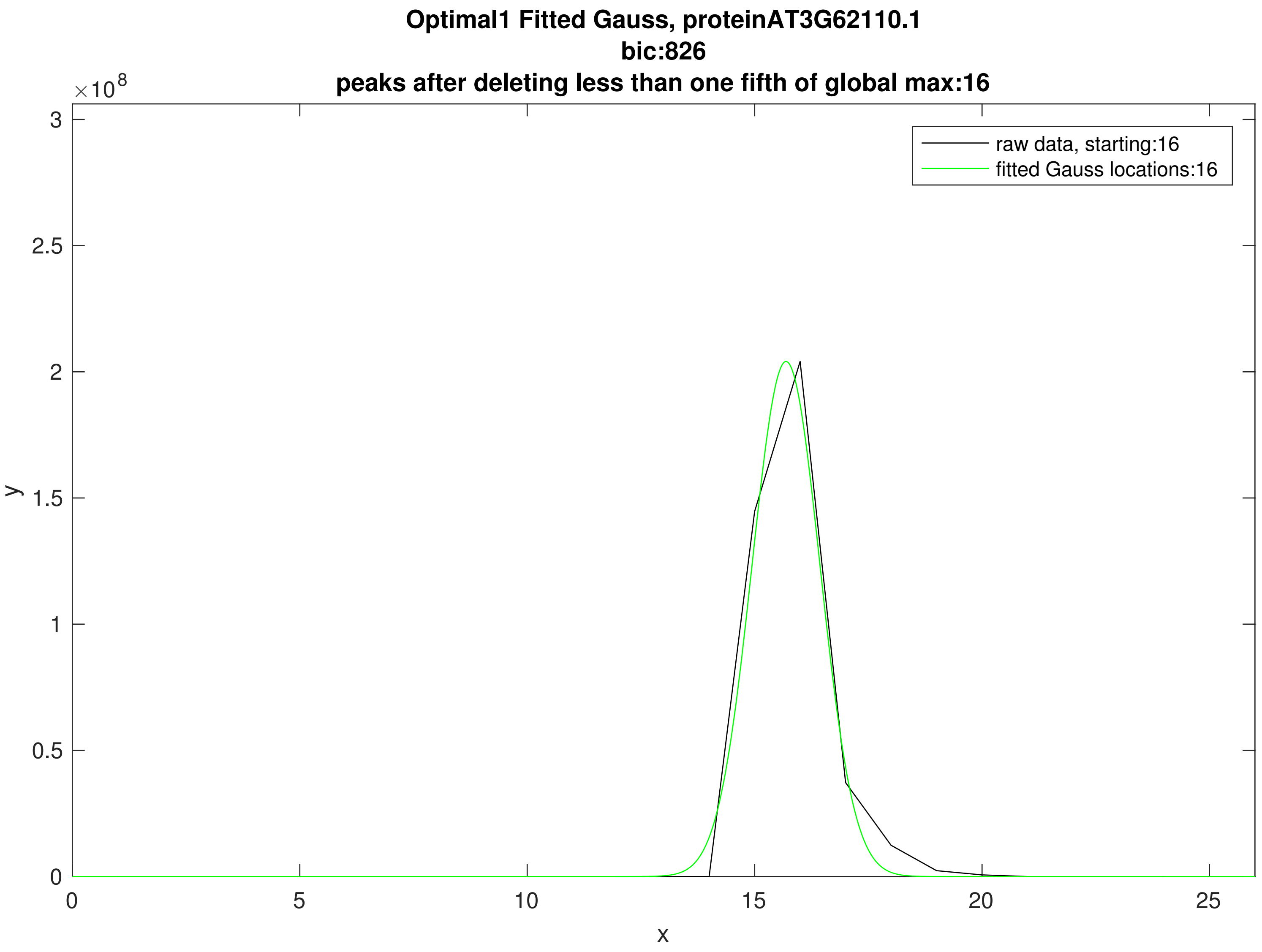

Optimal1 Fitted Gauss, proteinAT3G62110.1
bic:826
peaks after deleting less than one fifth of global max:16
×10^8
raw data, starting:16
fitted Gauss locations:16
y
x

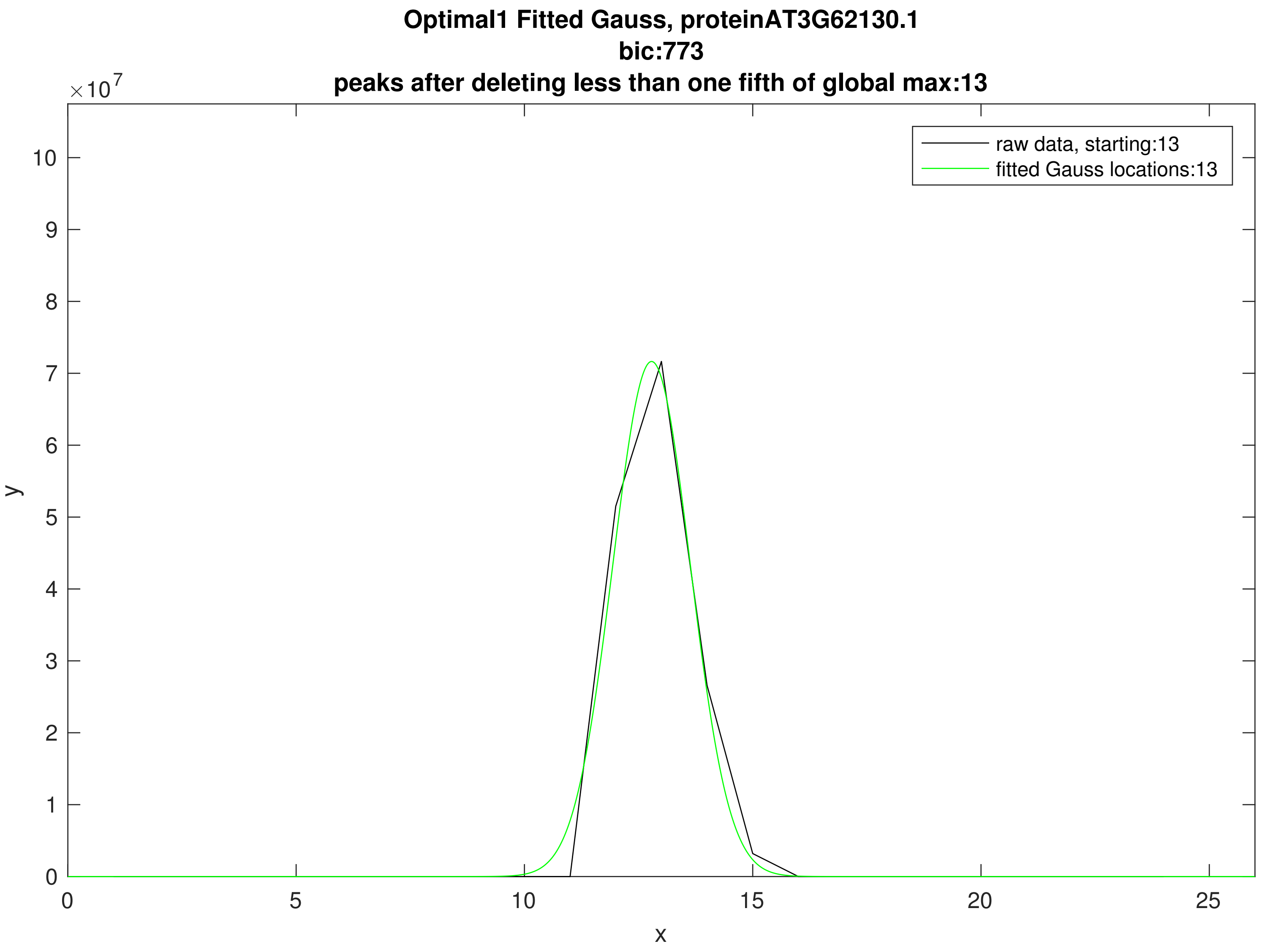

Optimal1 Fitted Gauss, proteinAT3G62130.1
bic:773
peaks after deleting less than one fifth of global max:13
×10^7
raw data, starting:13
fitted Gauss locations:13
y
x
0
1
2
3
4
5
6
7
8
9
10
0
5
10
15
20
25

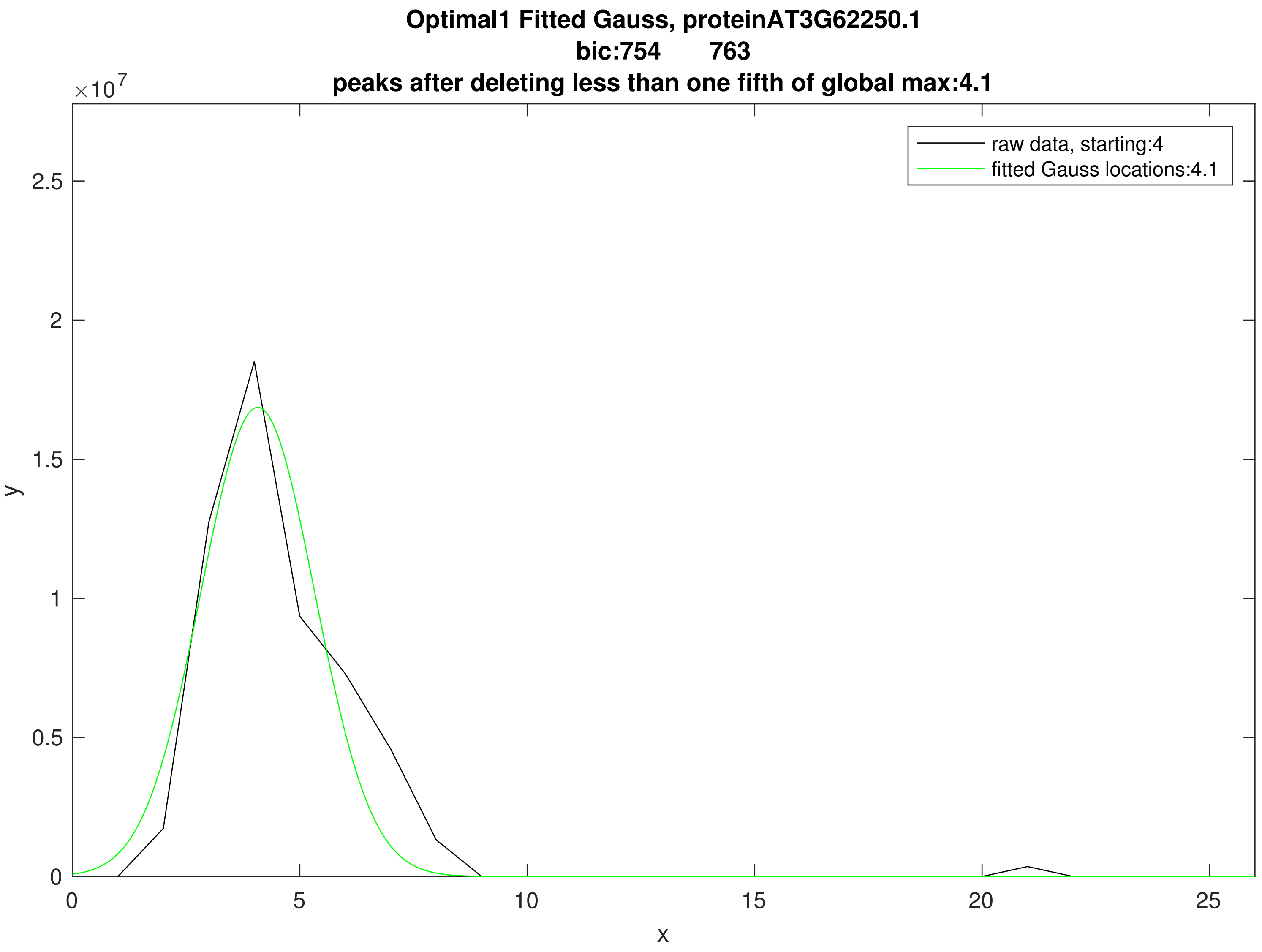
Optimal1 Fitted Gauss, proteinAT3G62250.1
bic:754 763
peaks after deleting less than one fifth of global max:4.1
×10^7
raw data, starting:4
fitted Gauss locations:4.1
y
x
0
0.5
1
1.5
2
2.5
0
5
10
15
20
25

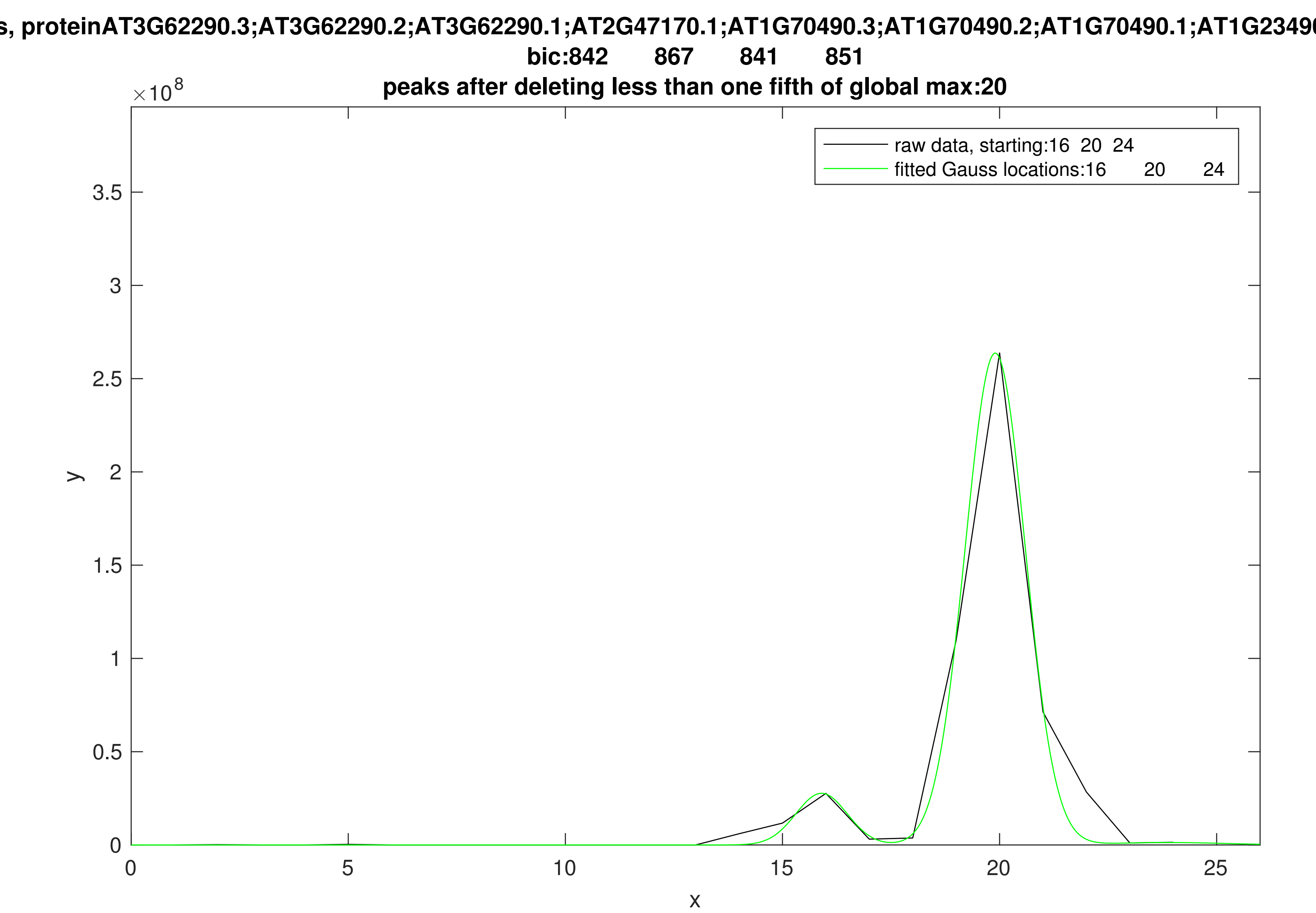

s, proteinAT3G62290.3;AT3G62290.2;AT3G62290.1;AT2G47170.1;AT1G70490.3;AT1G70490.2;AT1G70490.1;AT1G2349
bic:842 867 841 851
peaks after deleting less than one fifth of global max:20
×10^8
raw data, starting:16 20 24
fitted Gauss locations:16 20 24
y
x

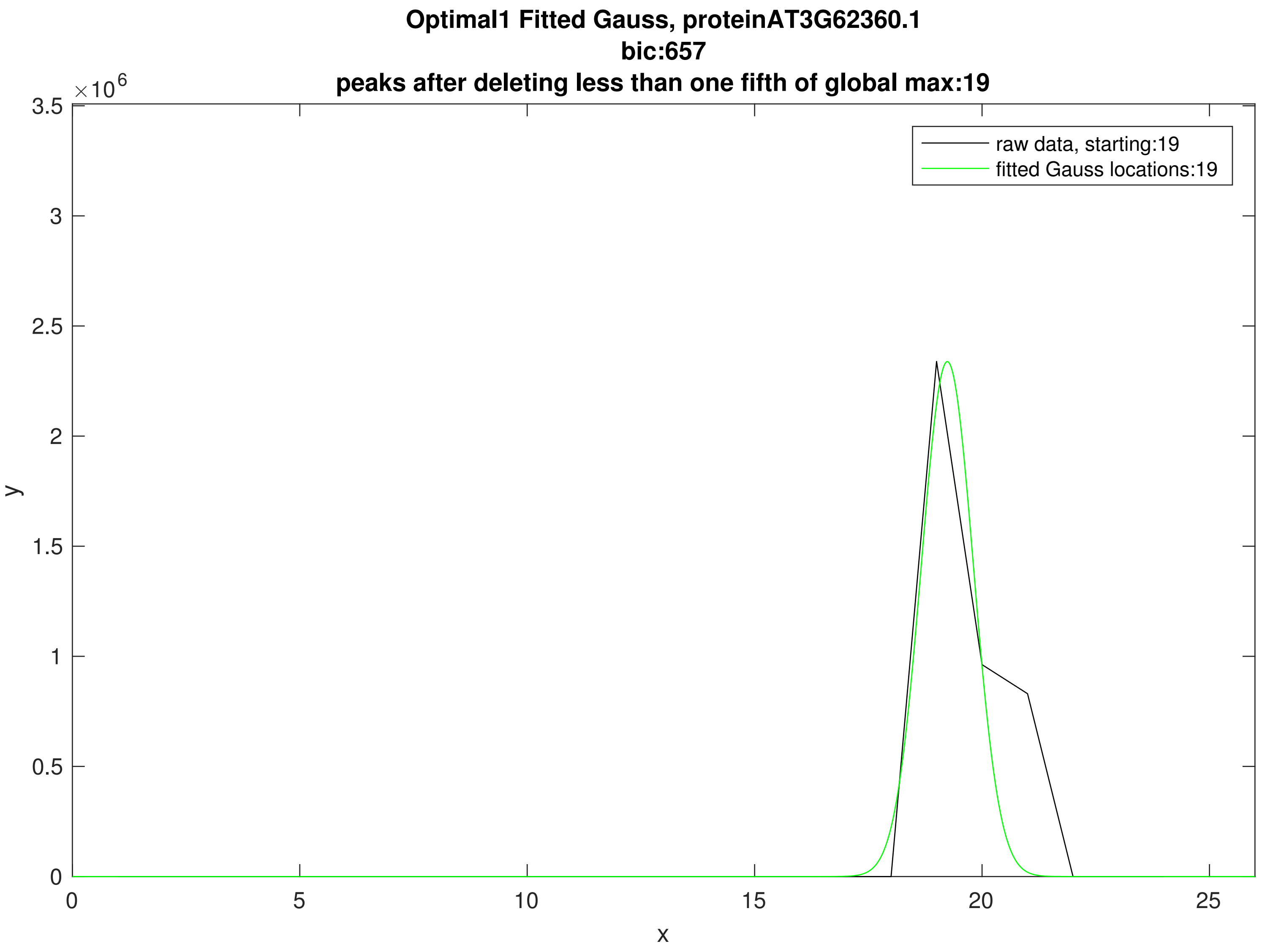
Optimal1 Fitted Gauss, proteinAT3G62360.1
bic:657
peaks after deleting less than one fifth of global max:19
×10^6
3.5
3
2.5
2
1.5
1
0.5
0
y
0
5
10
15
20
25
x
raw data, starting:19
fitted Gauss locations:19

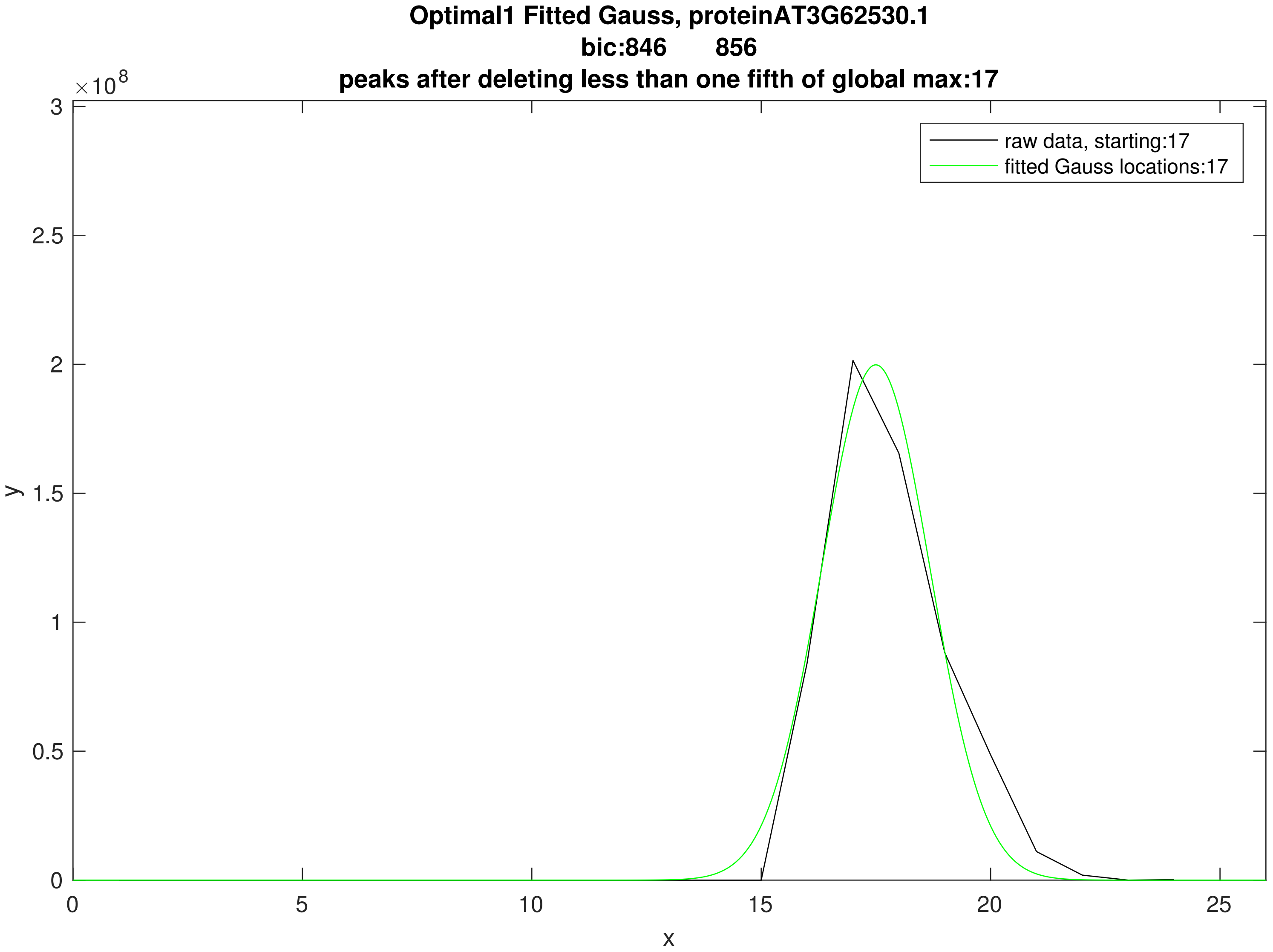
Optimal1 Fitted Gauss, proteinAT3G62530.1
bic:846 856
peaks after deleting less than one fifth of global max:17
×10^8
raw data, starting:17
fitted Gauss locations:17
y
x
0
0.5
1
1.5
2
2.5
3
0
5
10
15
20
25

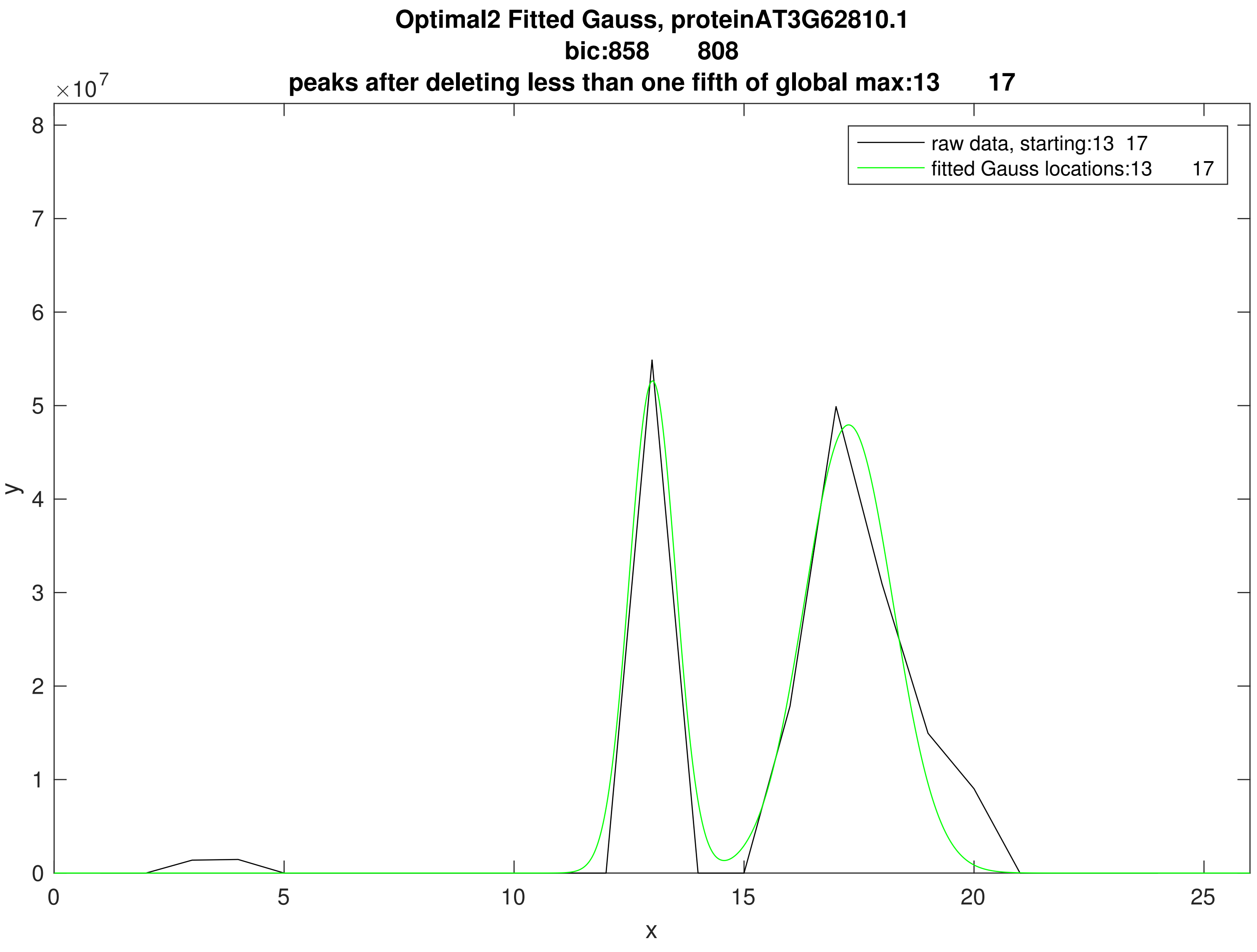

Optimal2 Fitted Gauss, proteinAT3G62810.1
bic:858 808
peaks after deleting less than one fifth of global max:13 17
×10^7
raw data, starting:13 17
fitted Gauss locations:13 17
y
x
0
1
2
3
4
5
6
7
8
0
5
10
15
20
25

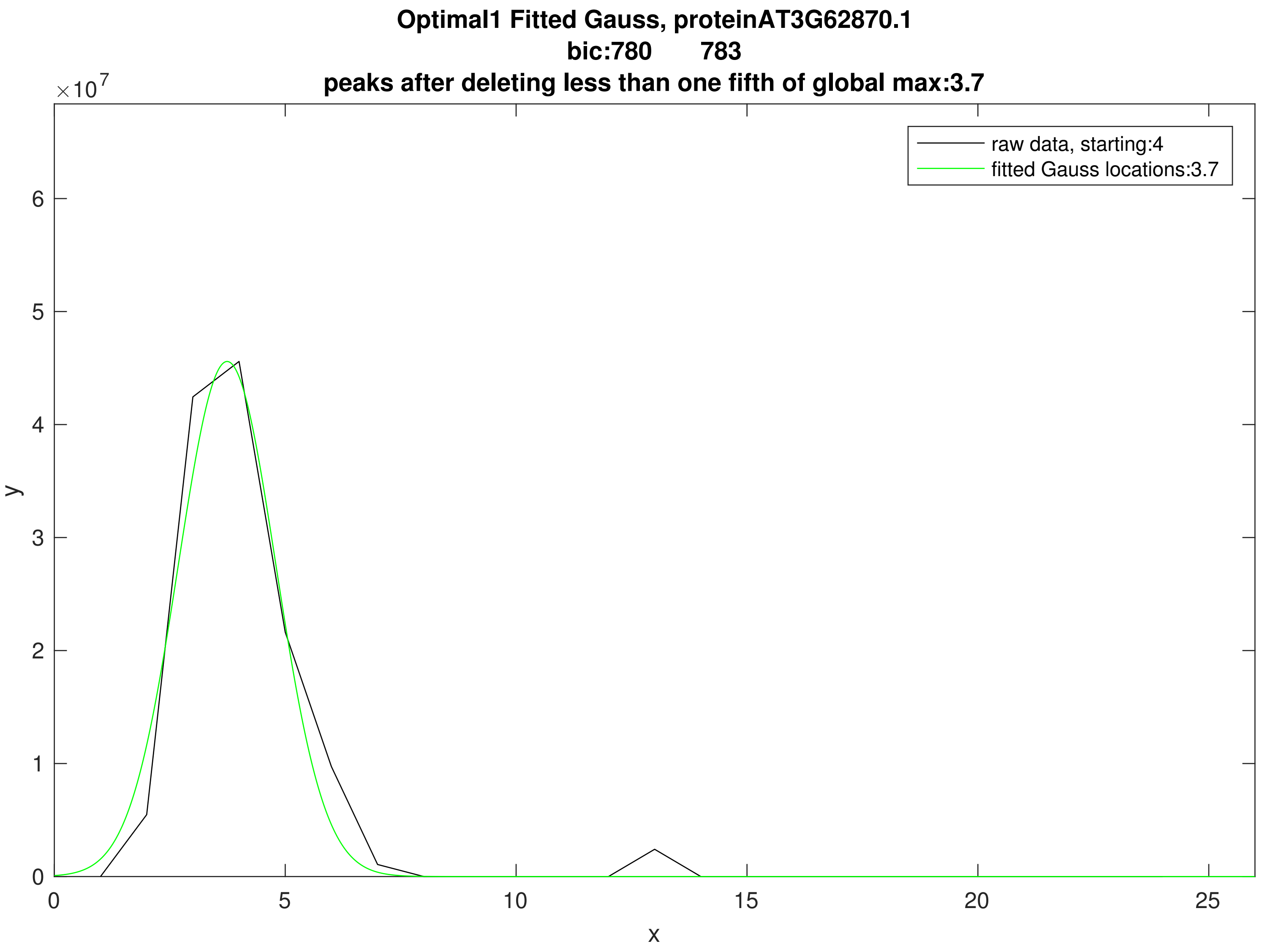
Optimal1 Fitted Gauss, proteinAT3G62870.1
bic:780 783
peaks after deleting less than one fifth of global max:3.7
×10^7
raw data, starting:4
fitted Gauss locations:3.7
y
x
0
1
2
3
4
5
6
0
5
10
15
20
25

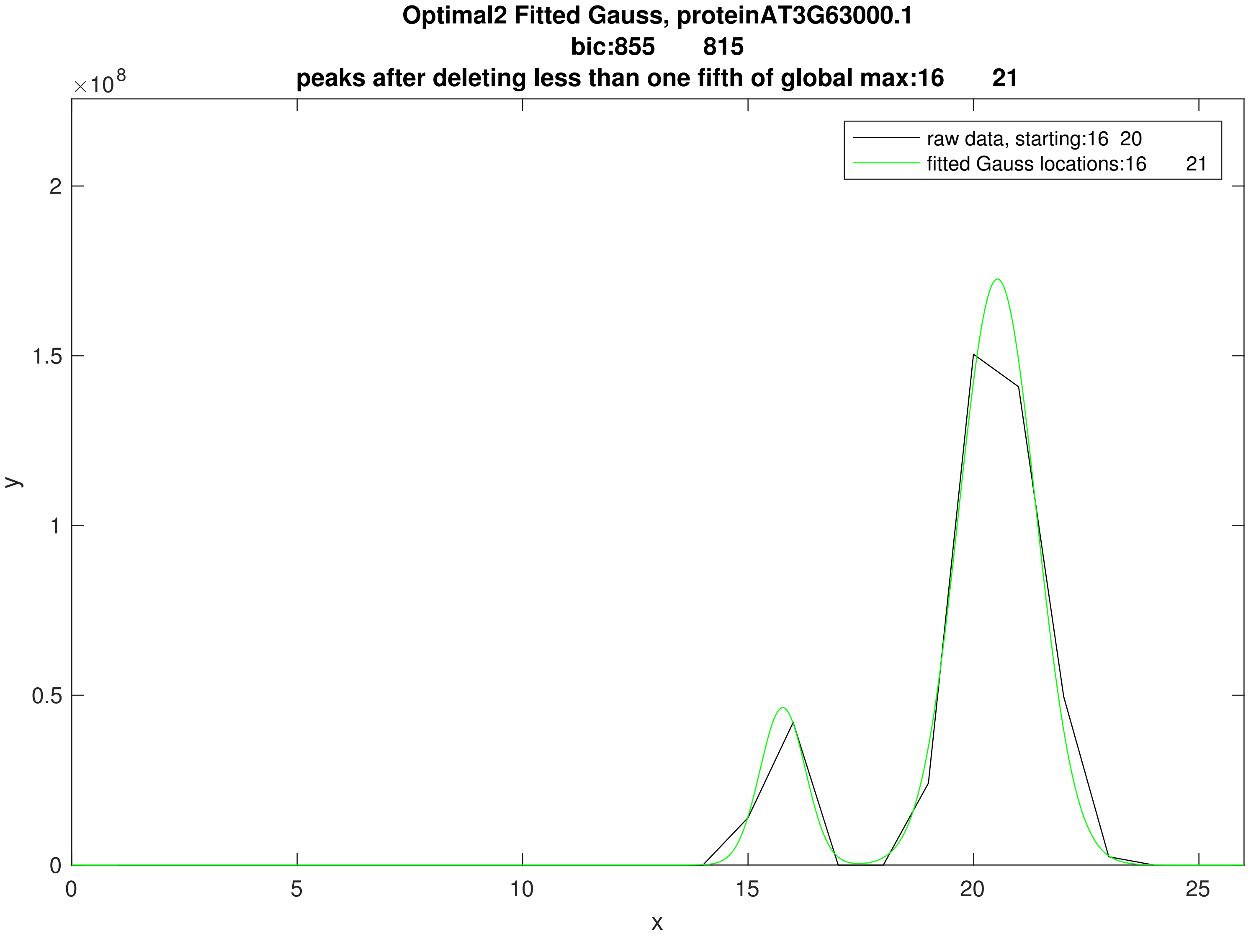

Optimal2 Fitted Gauss, proteinAT3G63000.1
bic:855 815
peaks after deleting less than one fifth of global max:16 21
×10^8
raw data, starting:16 20
fitted Gauss locations:16 21
2
1.5
1
0.5
0
y
0
5
10
15
20
25
x

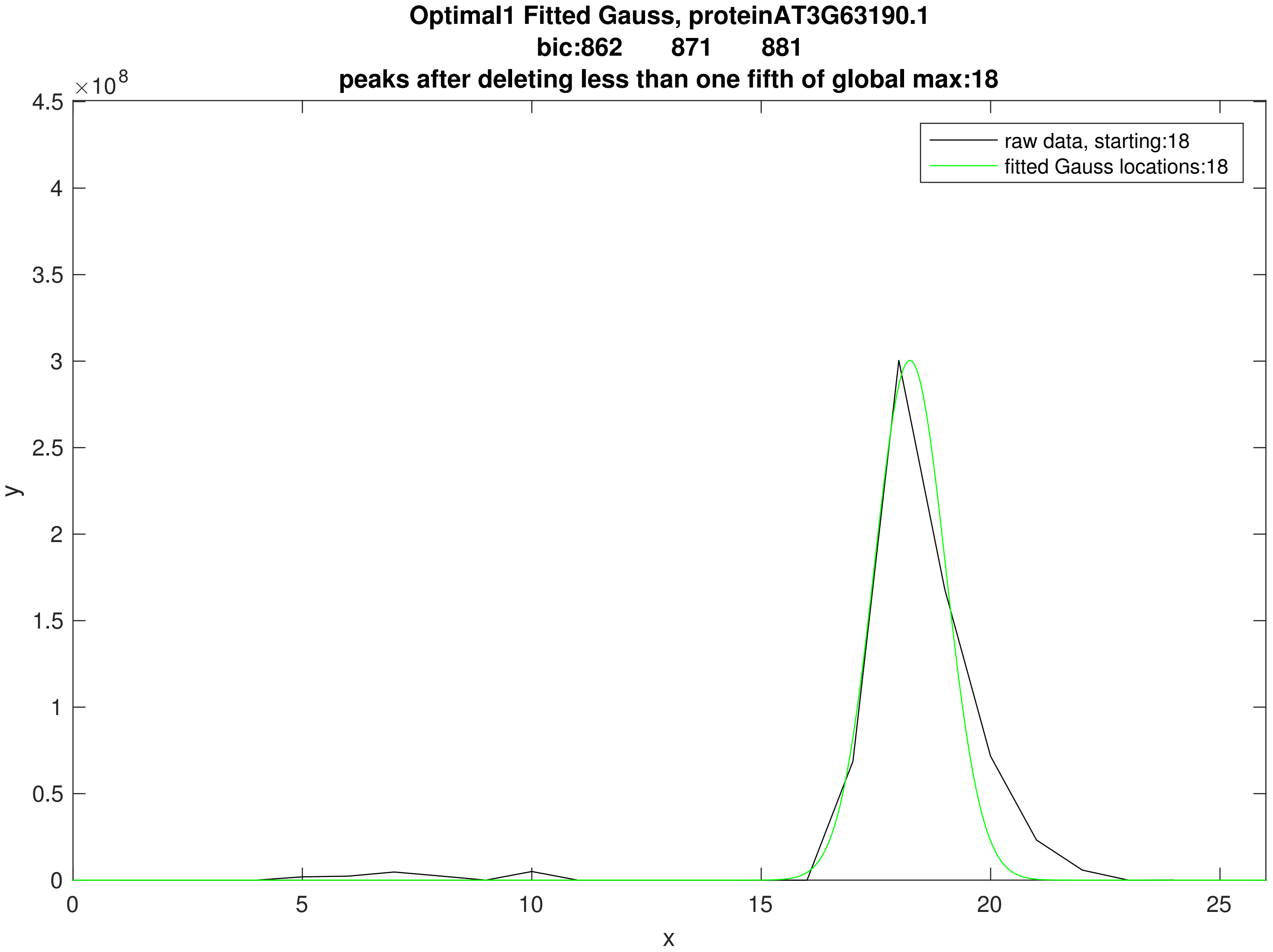
Optimal1 Fitted Gauss, proteinAT3G63190.1
bic:862 871 881
peaks after deleting less than one fifth of global max:18
×10^8
4.5
4
3.5
3
2.5
2
1.5
1
0.5
0
y
0
5
10
15
20
25
x
raw data, starting:18
fitted Gauss locations:18

**Optimal1 Fitted Gauss, proteinAT3G63250.1;AT3G63250.2**
**bic:817**
**peaks after deleting less than one fifth of global max:17**

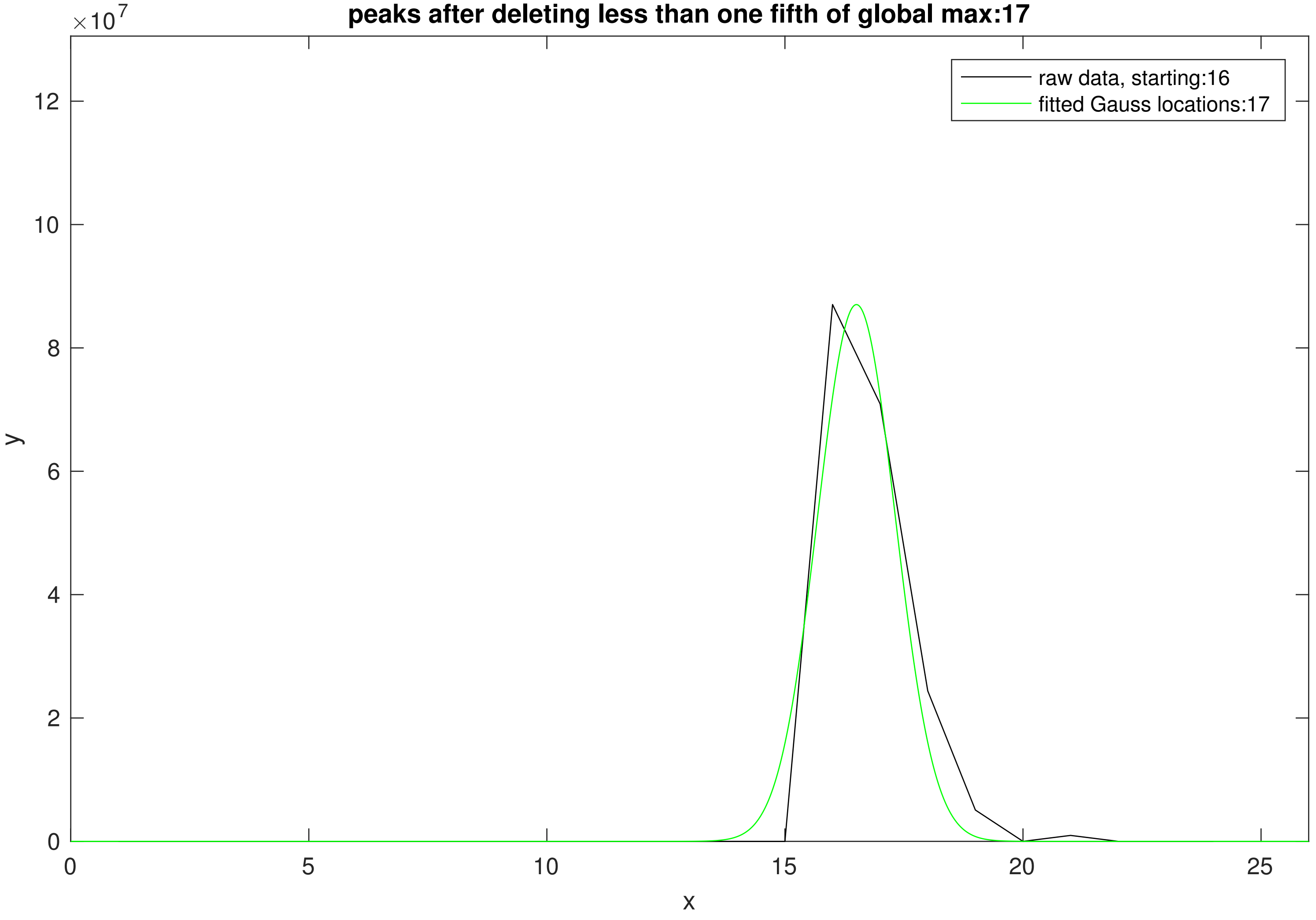

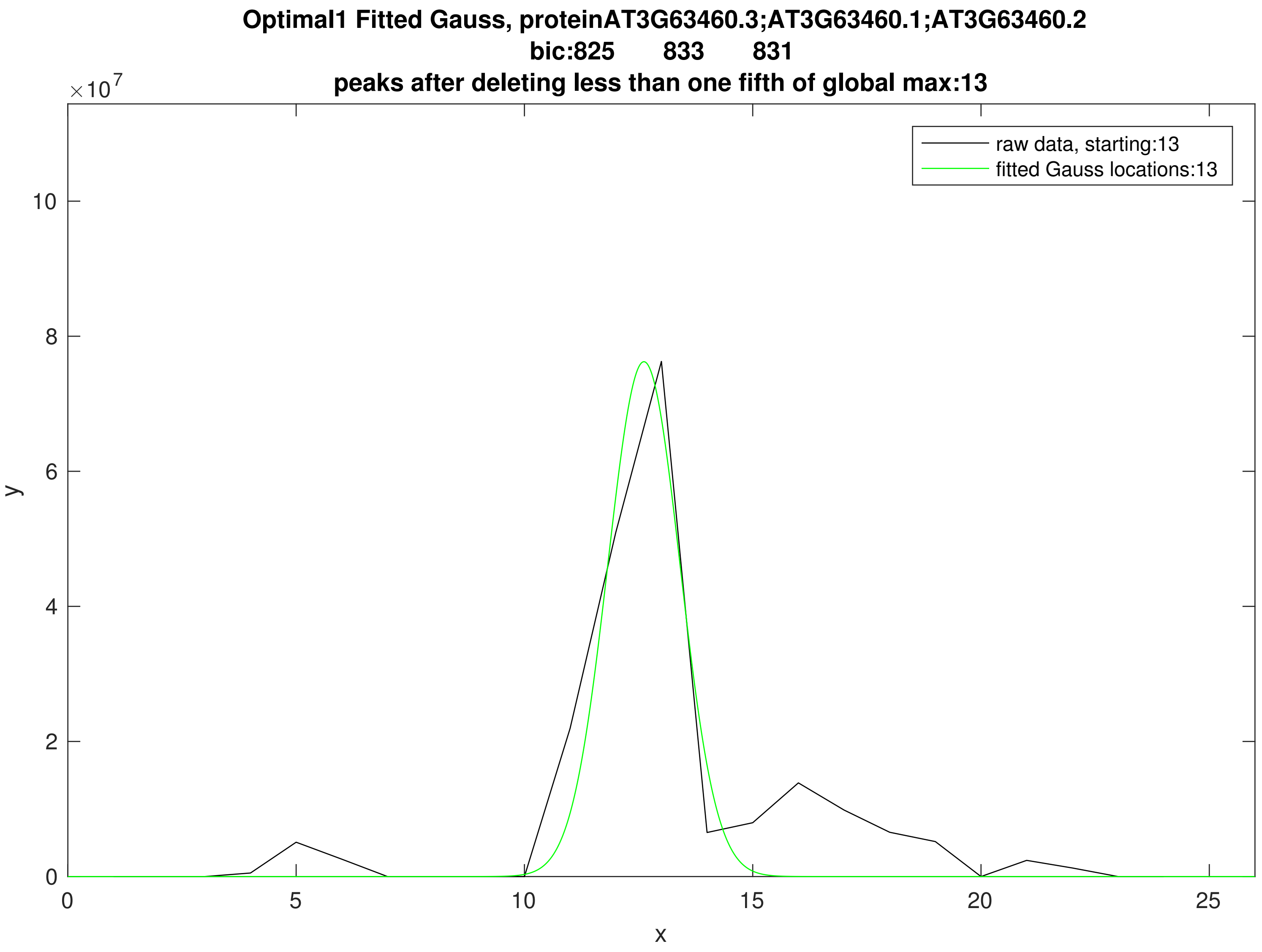

Optimal1 Fitted Gauss, proteinAT3G63460.3;AT3G63460.1;AT3G63460.2
bic:825 833 831
peaks after deleting less than one fifth of global max:13
×10^7
raw data, starting:13
fitted Gauss locations:13
y
x
0
2
4
6
8
10
0
5
10
15
20
25

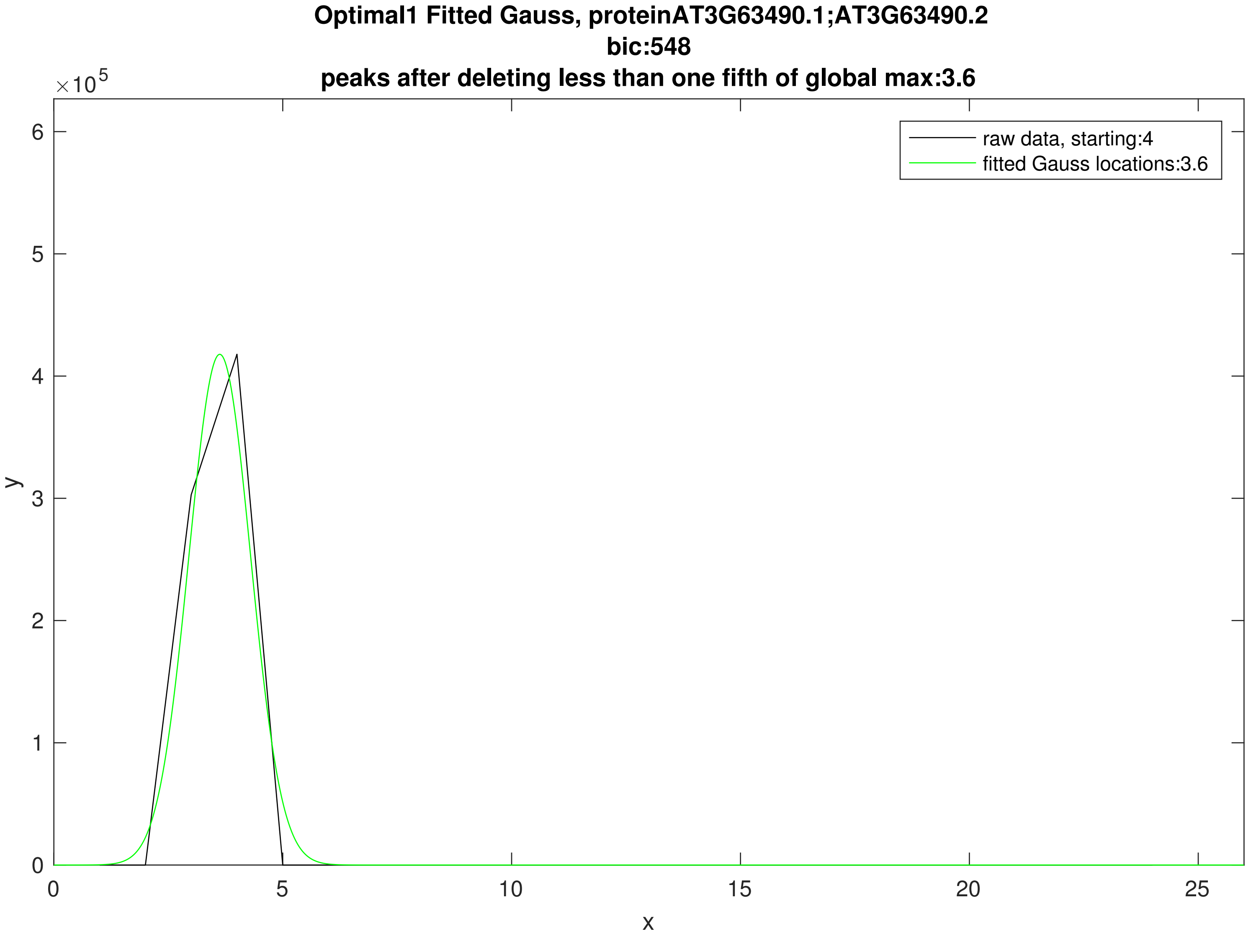

Optimal1 Fitted Gauss, proteinAT3G63490.1;AT3G63490.2
bic:548
peaks after deleting less than one fifth of global max:3.6
×10^5
raw data, starting:4
fitted Gauss locations:3.6
y
x
0
1
2
3
4
5
6
0
5
10
15
20
25

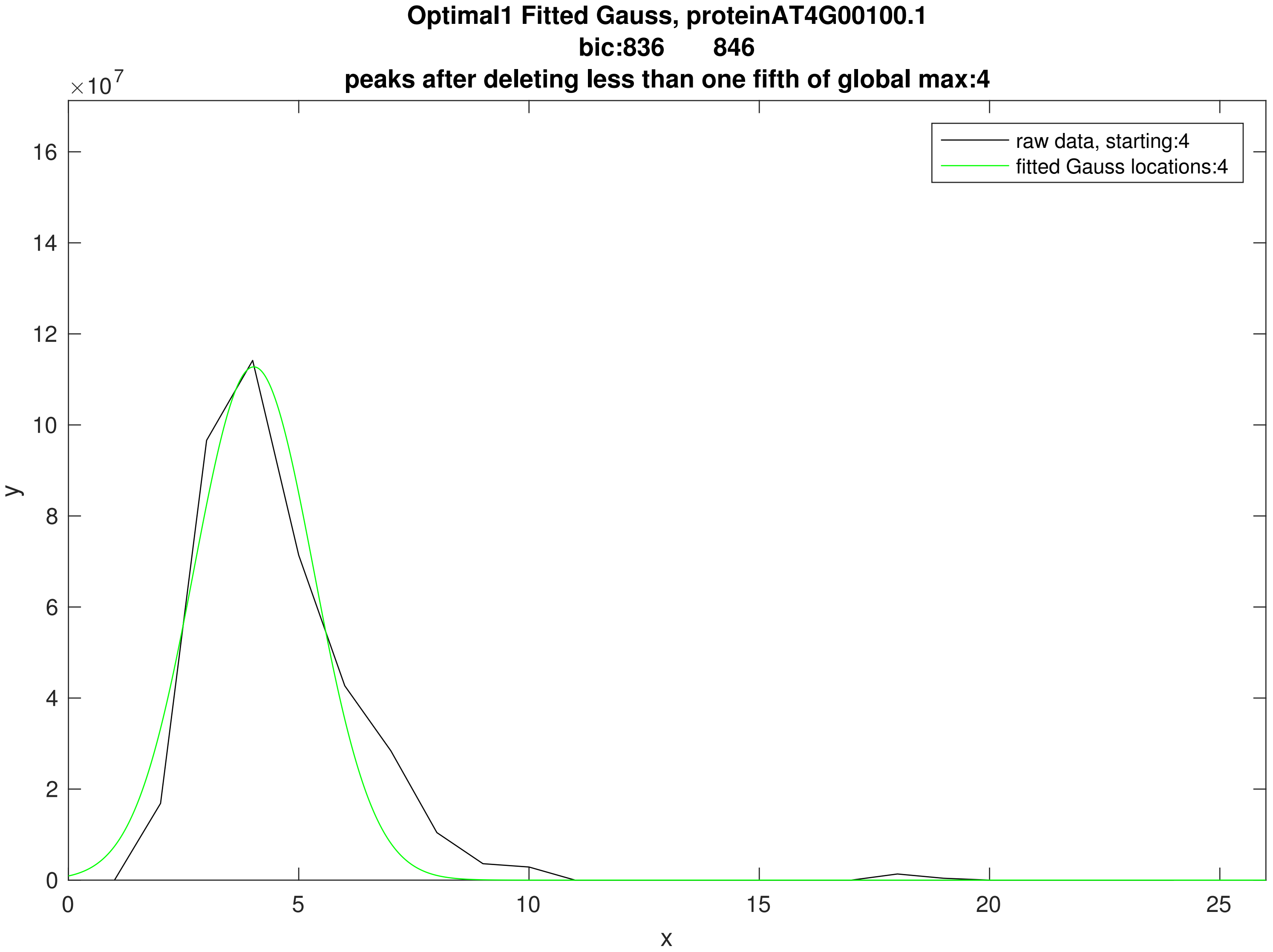
Optimal1 Fitted Gauss, proteinAT4G00100.1
bic:836 846
peaks after deleting less than one fifth of global max:4
×10^7
raw data, starting:4
fitted Gauss locations:4
y
x
0
2
4
6
8
10
12
14
16
0
5
10
15
20
25

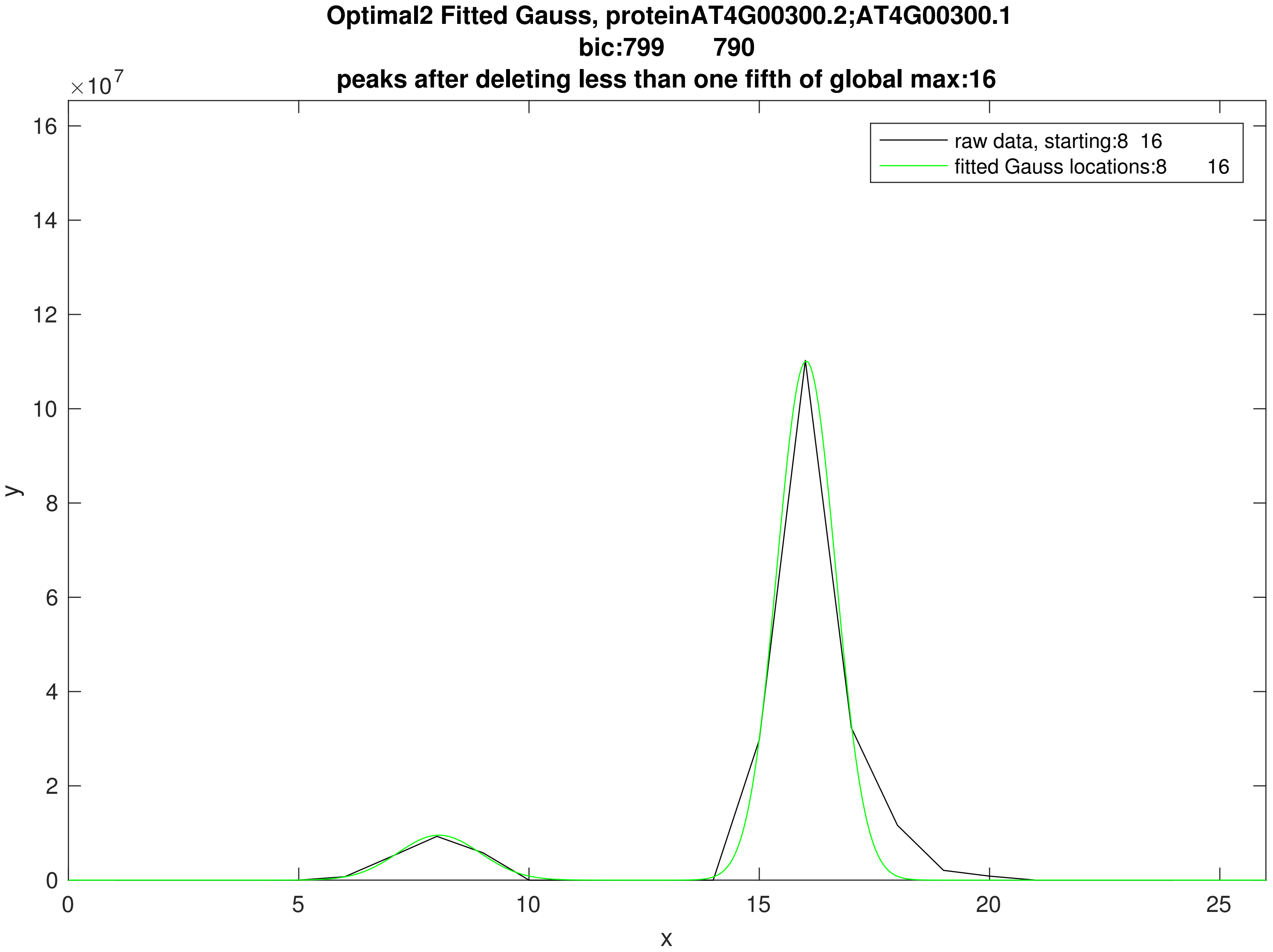

Optimal2 Fitted Gauss, proteinAT4G00300.2;AT4G00300.1
bic:799 790
peaks after deleting less than one fifth of global max:16
×10^7
raw data, starting:8 16
fitted Gauss locations:8 16
y
x
0
2
4
6
8
10
12
14
16
0
5
10
15
20
25

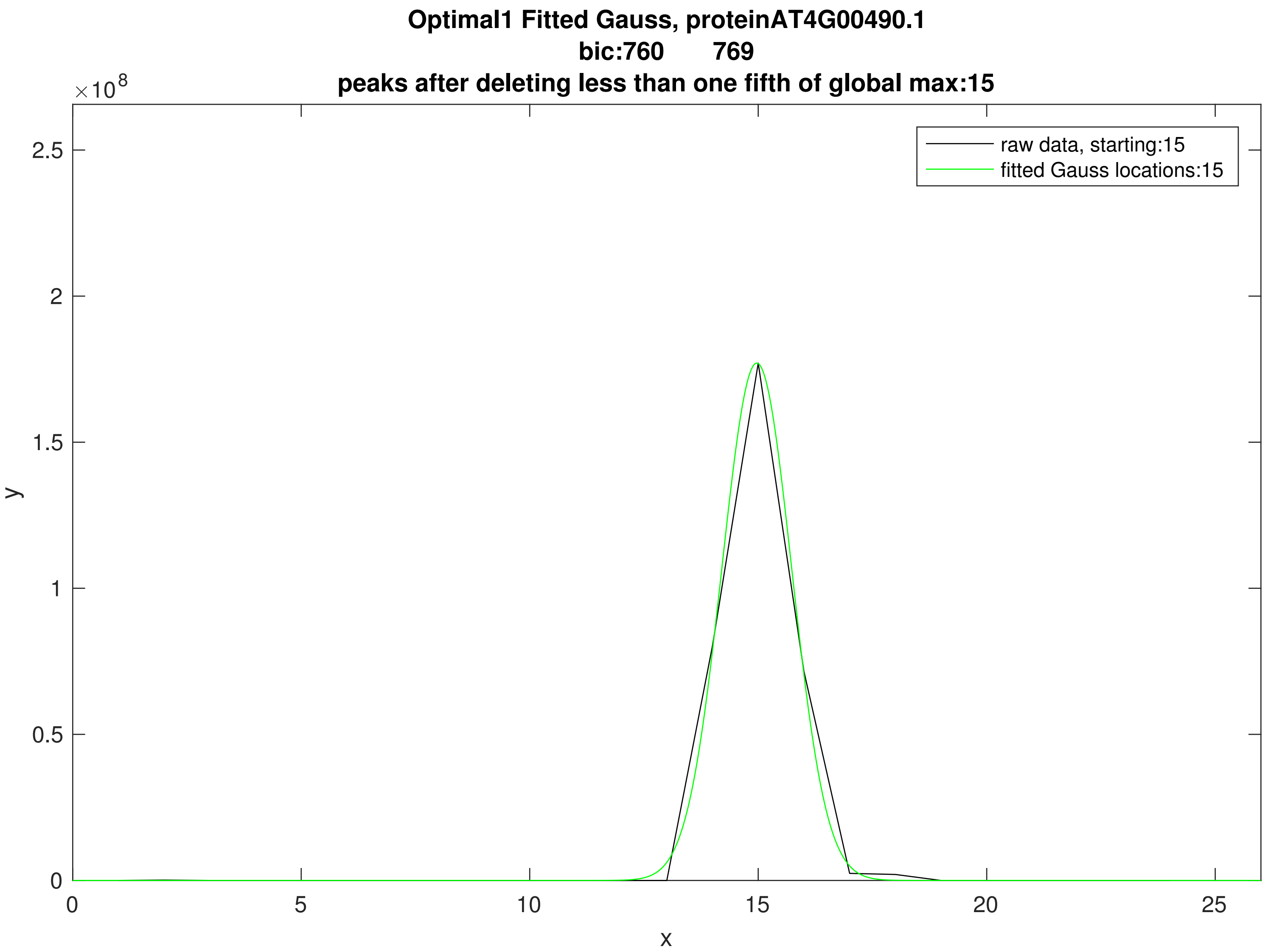
Optimal1 Fitted Gauss, proteinAT4G00490.1
bic:760 769
peaks after deleting less than one fifth of global max:15
×10^8
2.5
2
1.5
1
0.5
0
y
0
5
10
15
20
25
x
raw data, starting:15
fitted Gauss locations:15

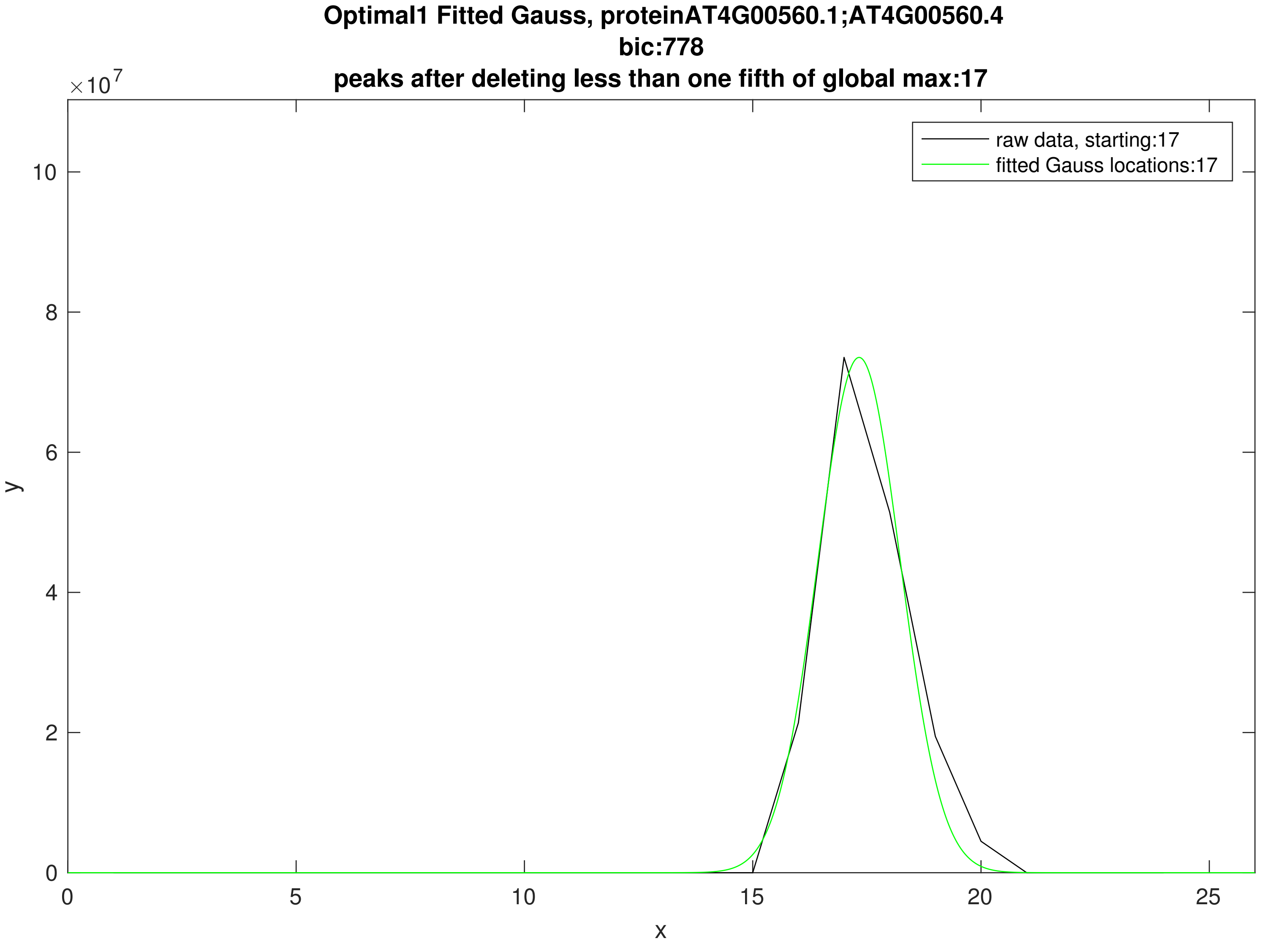
Optimal1 Fitted Gauss, proteinAT4G00560.1;AT4G00560.4
bic:778
peaks after deleting less than one fifth of global max:17
×10^7
raw data, starting:17
fitted Gauss locations:17
y
x
0
2
4
6
8
10
0
5
10
15
20
25

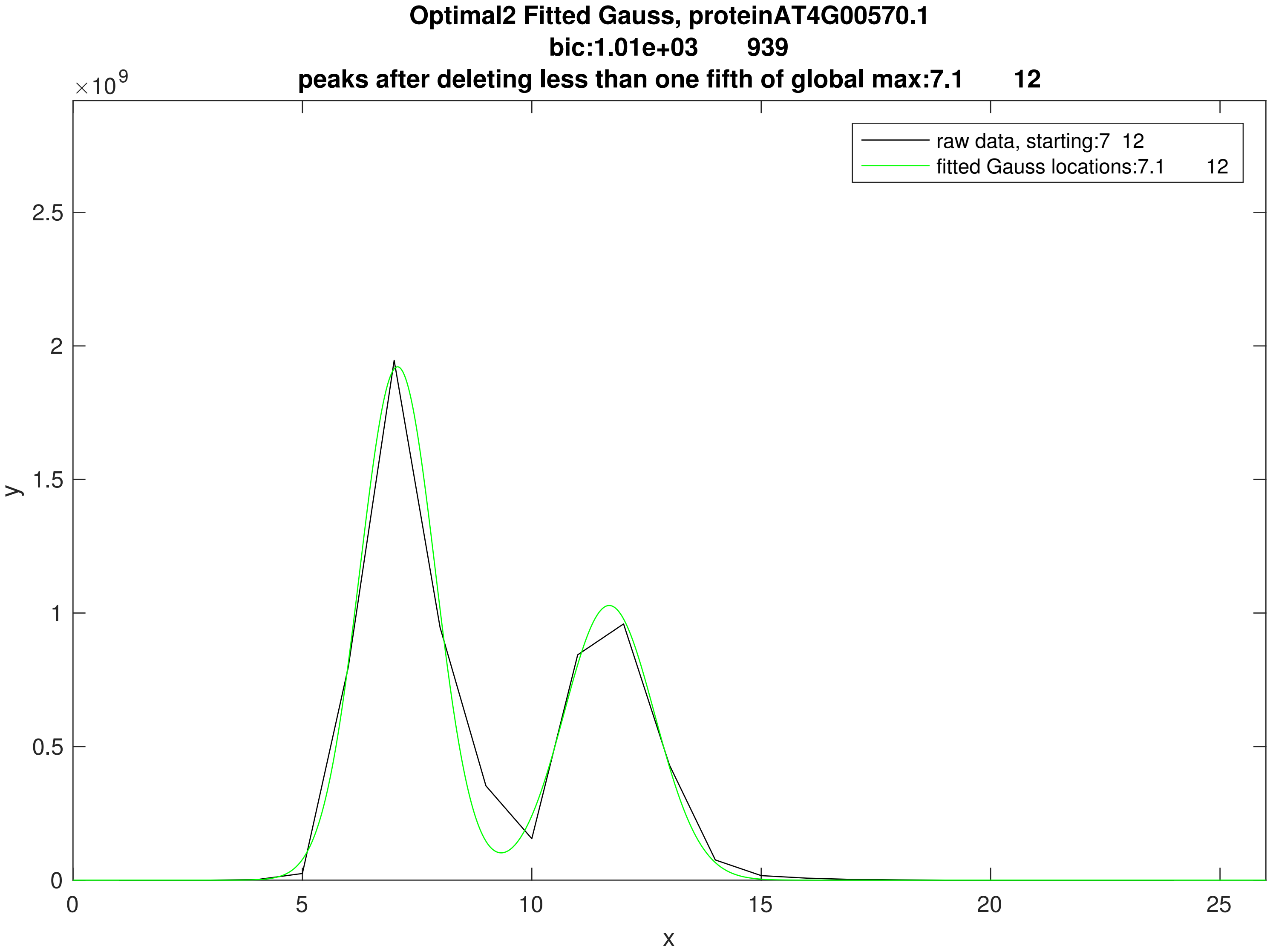
Optimal2 Fitted Gauss, proteinAT4G00570.1
bic:1.01e+03 939
peaks after deleting less than one fifth of global max:7.1 12
×10^9
raw data, starting:7 12
fitted Gauss locations:7.1 12
y
x
0
0.5
1
1.5
2
2.5
0
5
10
15
20
25

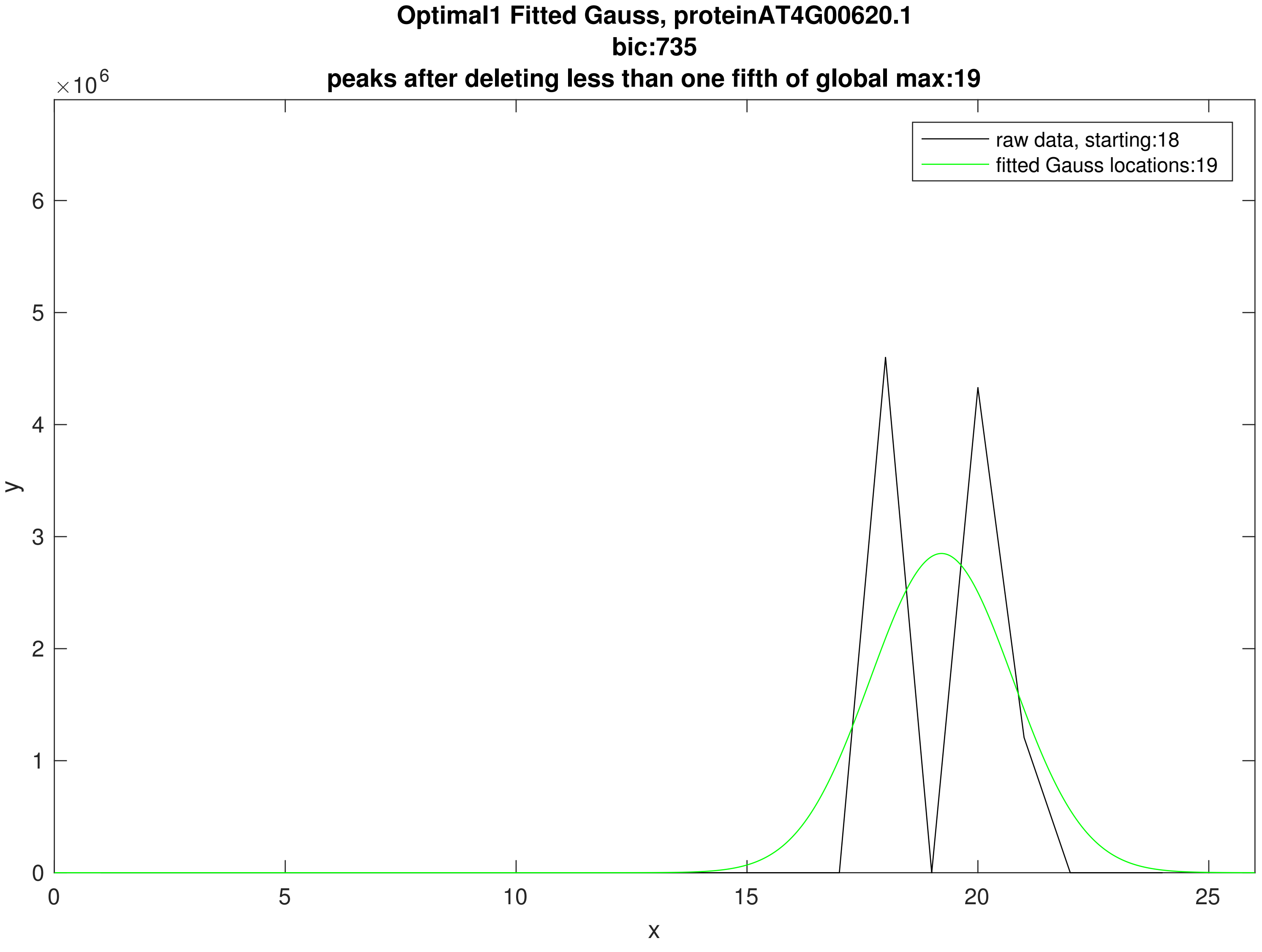
Optimal1 Fitted Gauss, proteinAT4G00620.1
bic:735
peaks after deleting less than one fifth of global max:19
×10^6
raw data, starting:18
fitted Gauss locations:19
y
x
0
1
2
3
4
5
6
0
5
10
15
20
25

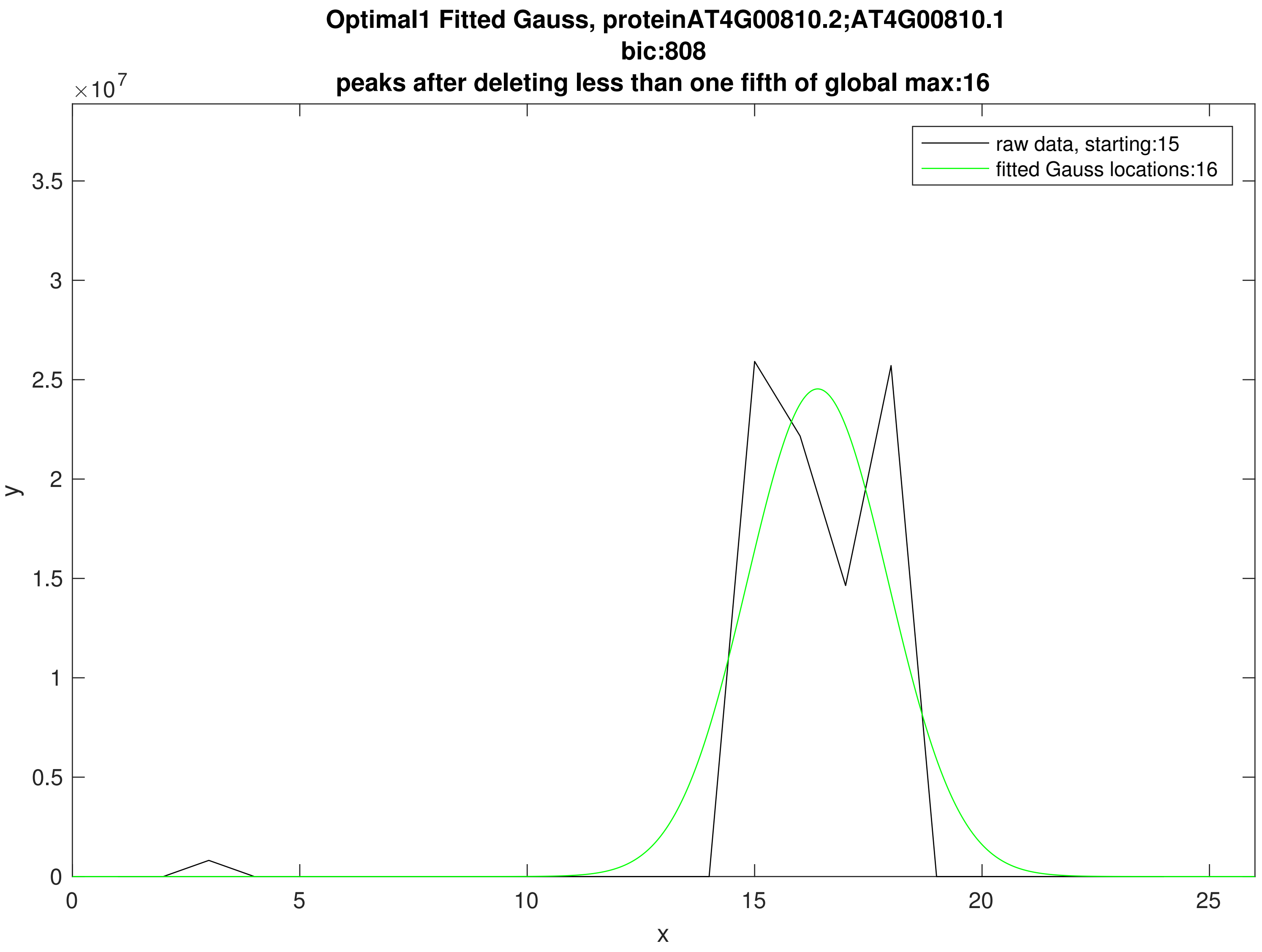
Optimal1 Fitted Gauss, proteinAT4G00810.2;AT4G00810.1
bic:808
peaks after deleting less than one fifth of global max:16
×10^7
raw data, starting:15
fitted Gauss locations:16
y
x
0
0.5
1
1.5
2
2.5
3
3.5
0
5
10
15
20
25

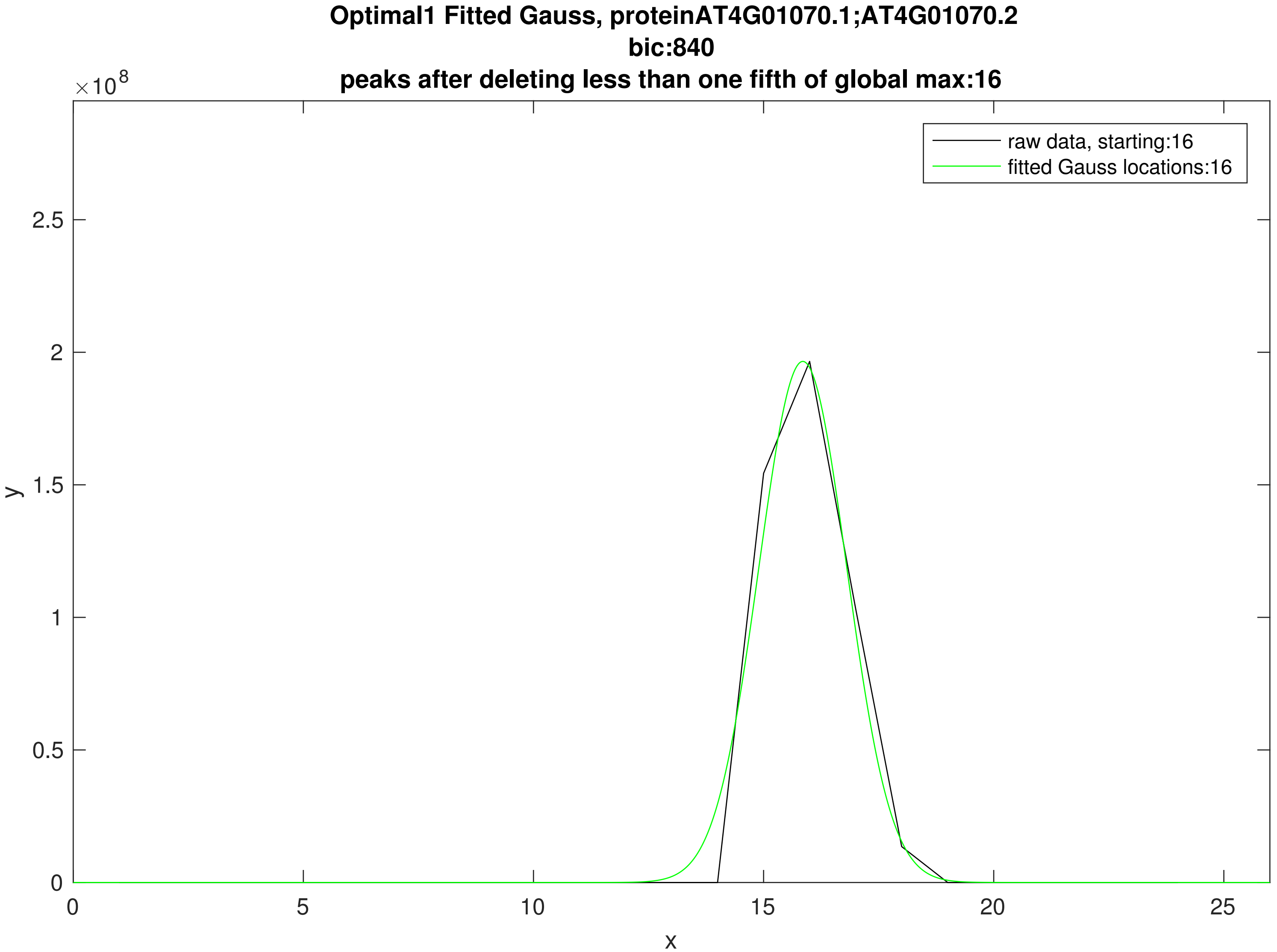
Optimal1 Fitted Gauss, proteinAT4G01070.1;AT4G01070.2
bic:840
peaks after deleting less than one fifth of global max:16
×10^8
raw data, starting:16
fitted Gauss locations:16
y
x
0
0.5
1
1.5
2
2.5
0
5
10
15
20
25

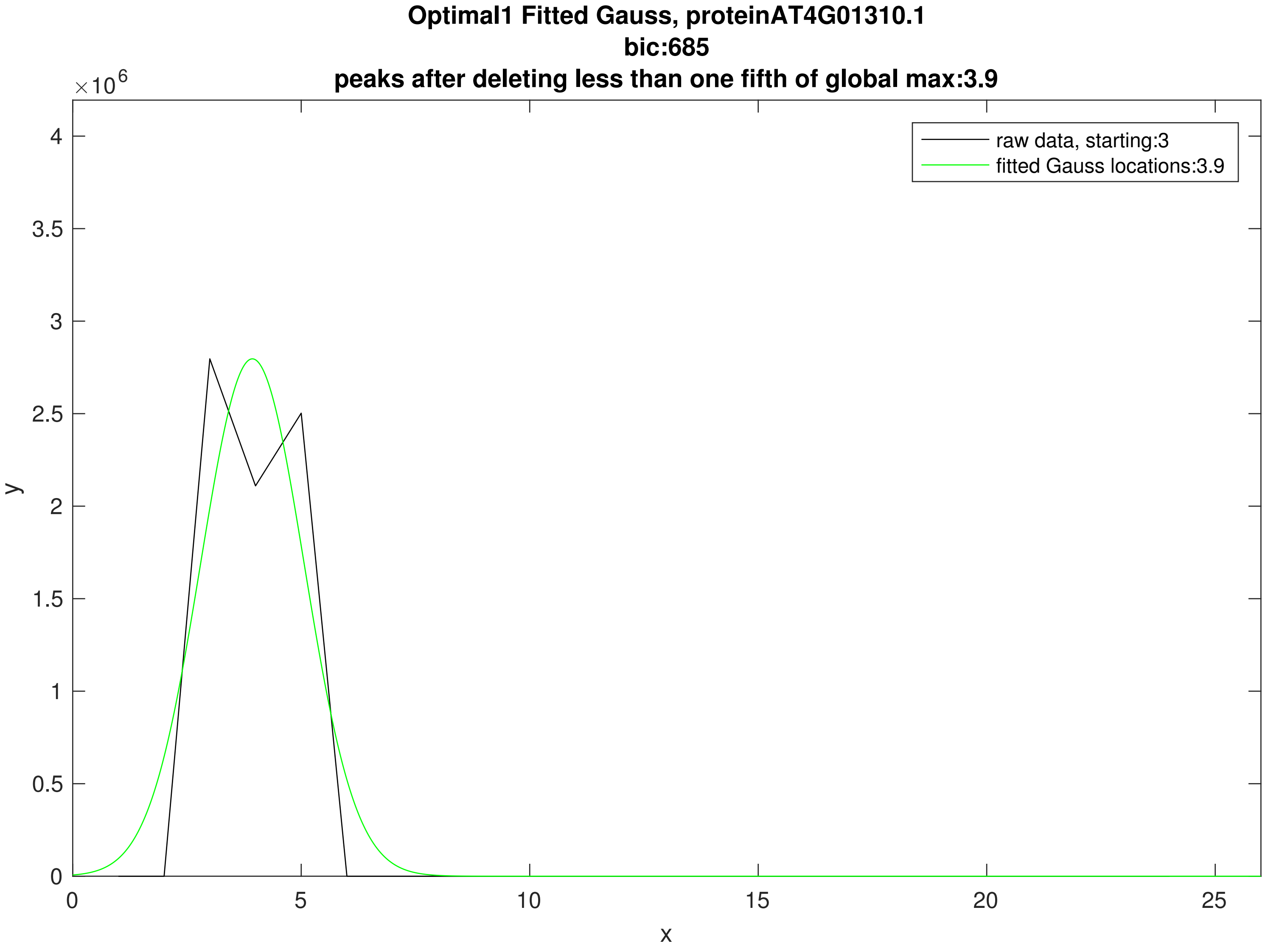
Optimal1 Fitted Gauss, proteinAT4G01310.1
bic:685
peaks after deleting less than one fifth of global max:3.9
×10^6
raw data, starting:3
fitted Gauss locations:3.9
y
x
0
0.5
1
1.5
2
2.5
3
3.5
4
0
5
10
15
20
25

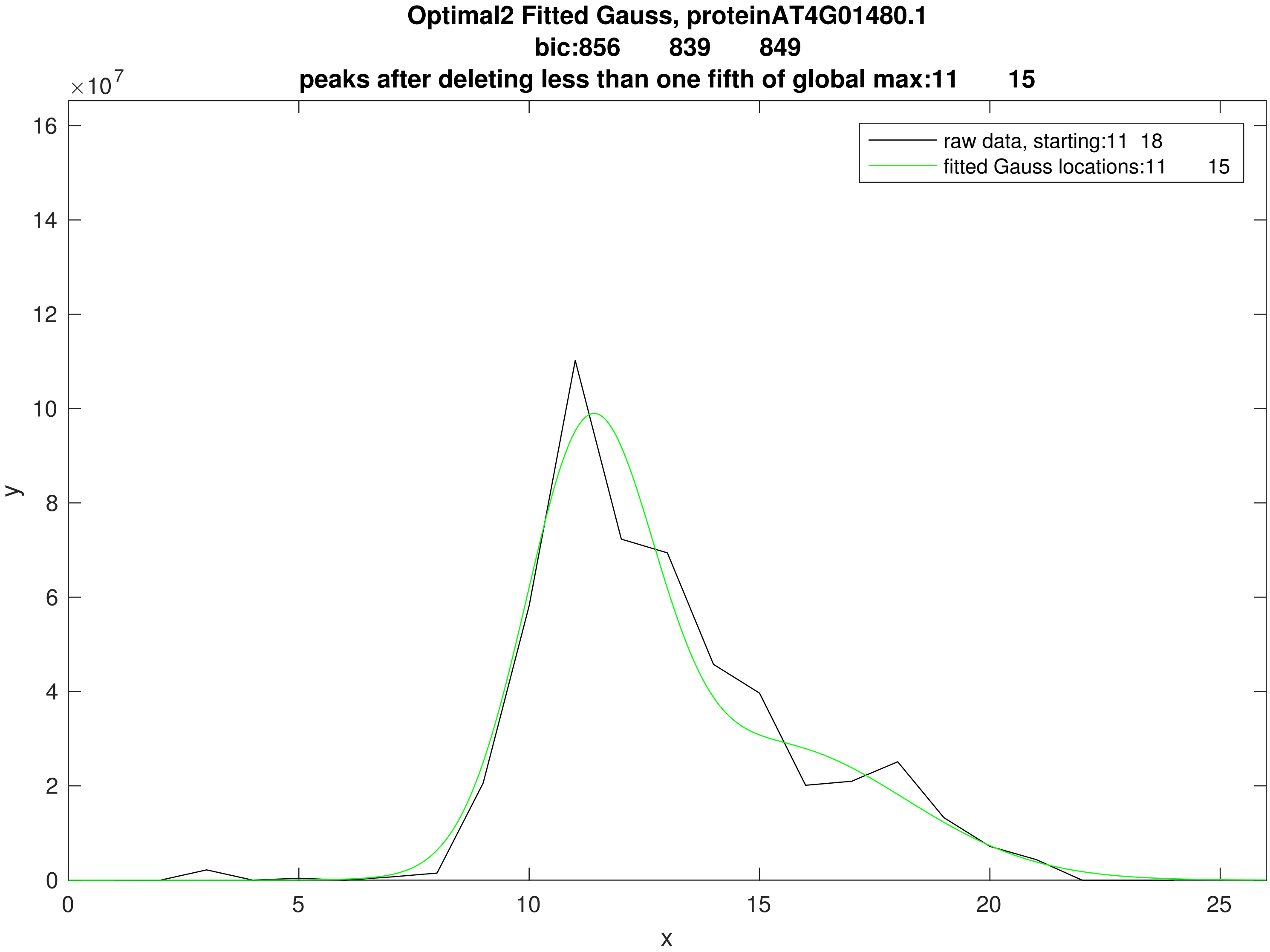

Optimal2 Fitted Gauss, proteinAT4G01480.1
bic:856 839 849
peaks after deleting less than one fifth of global max:11 15
×10^7
raw data, starting:11 18
fitted Gauss locations:11 15
y
x
0
2
4
6
8
10
12
14
16
0
5
10
15
20
25

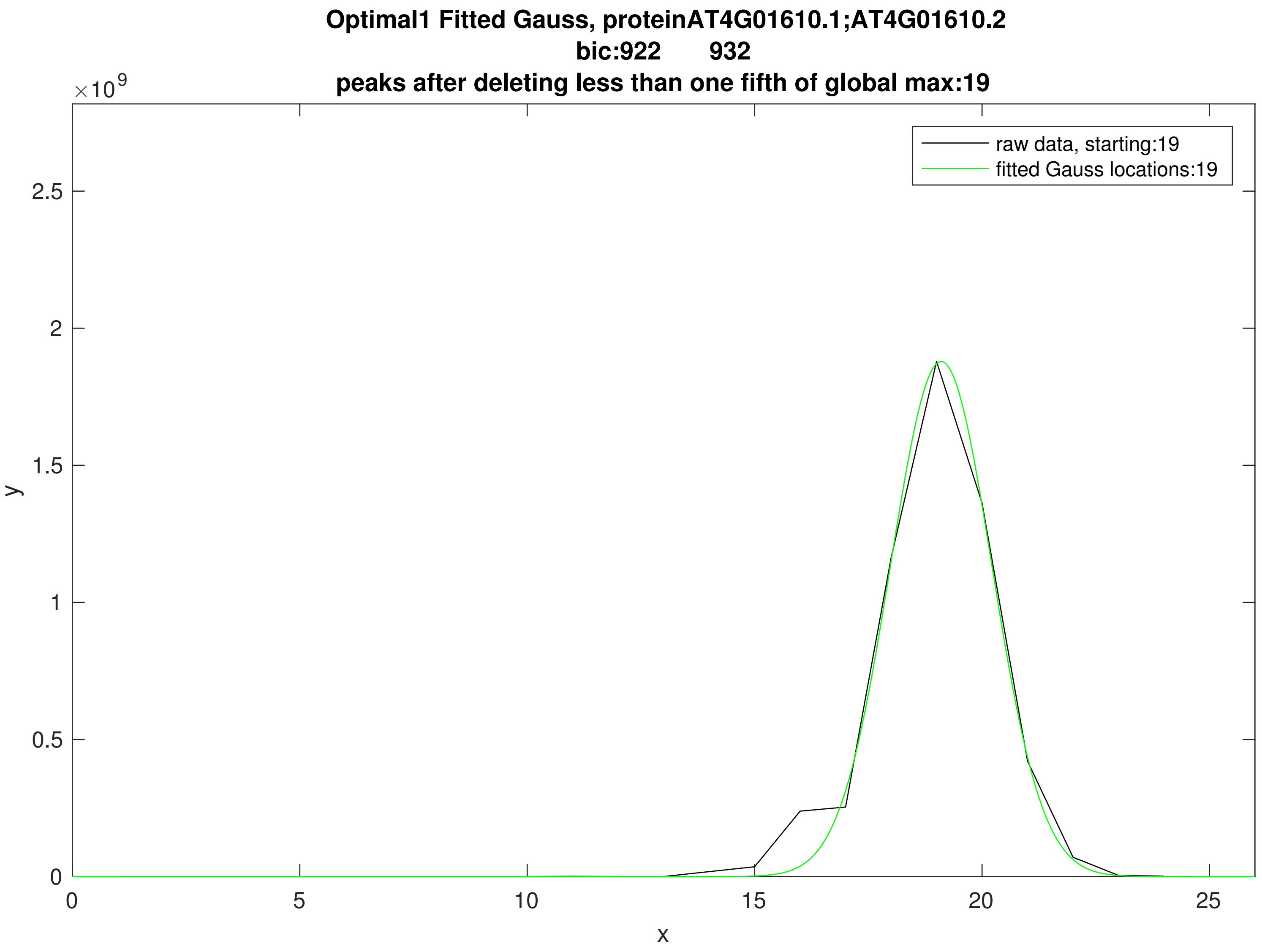
Optimal1 Fitted Gauss, proteinAT4G01610.1;AT4G01610.2
bic:922 932
peaks after deleting less than one fifth of global max:19
×10^9
raw data, starting:19
fitted Gauss locations:19
2.5
2
1.5
1
0.5
0
y
0
5
10
15
20
25
x

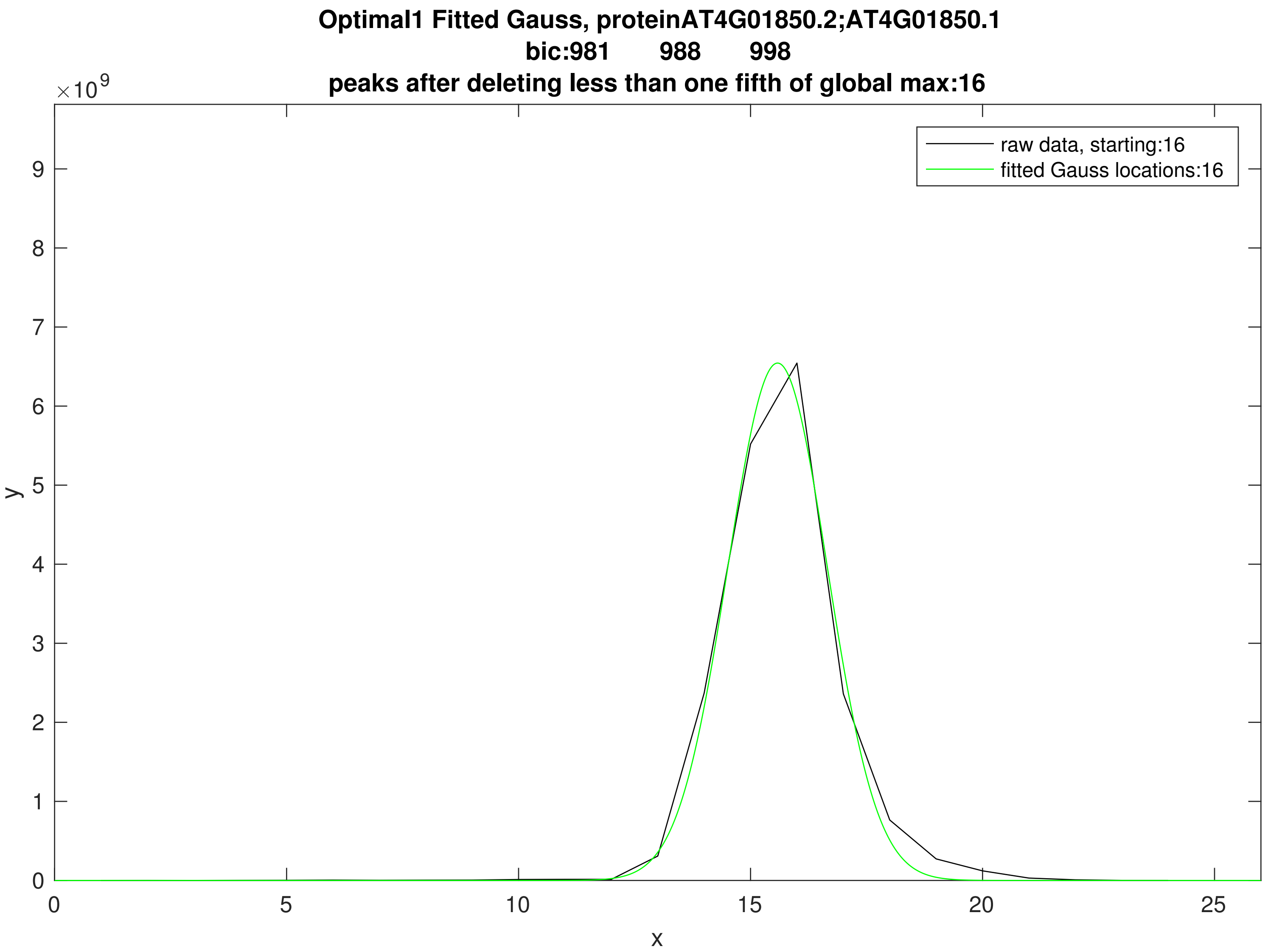
Optimal1 Fitted Gauss, proteinAT4G01850.2;AT4G01850.1
bic:981 988 998
peaks after deleting less than one fifth of global max:16
×10^9
raw data, starting:16
fitted Gauss locations:16
y
x
0
1
2
3
4
5
6
7
8
9
0
5
10
15
20
25

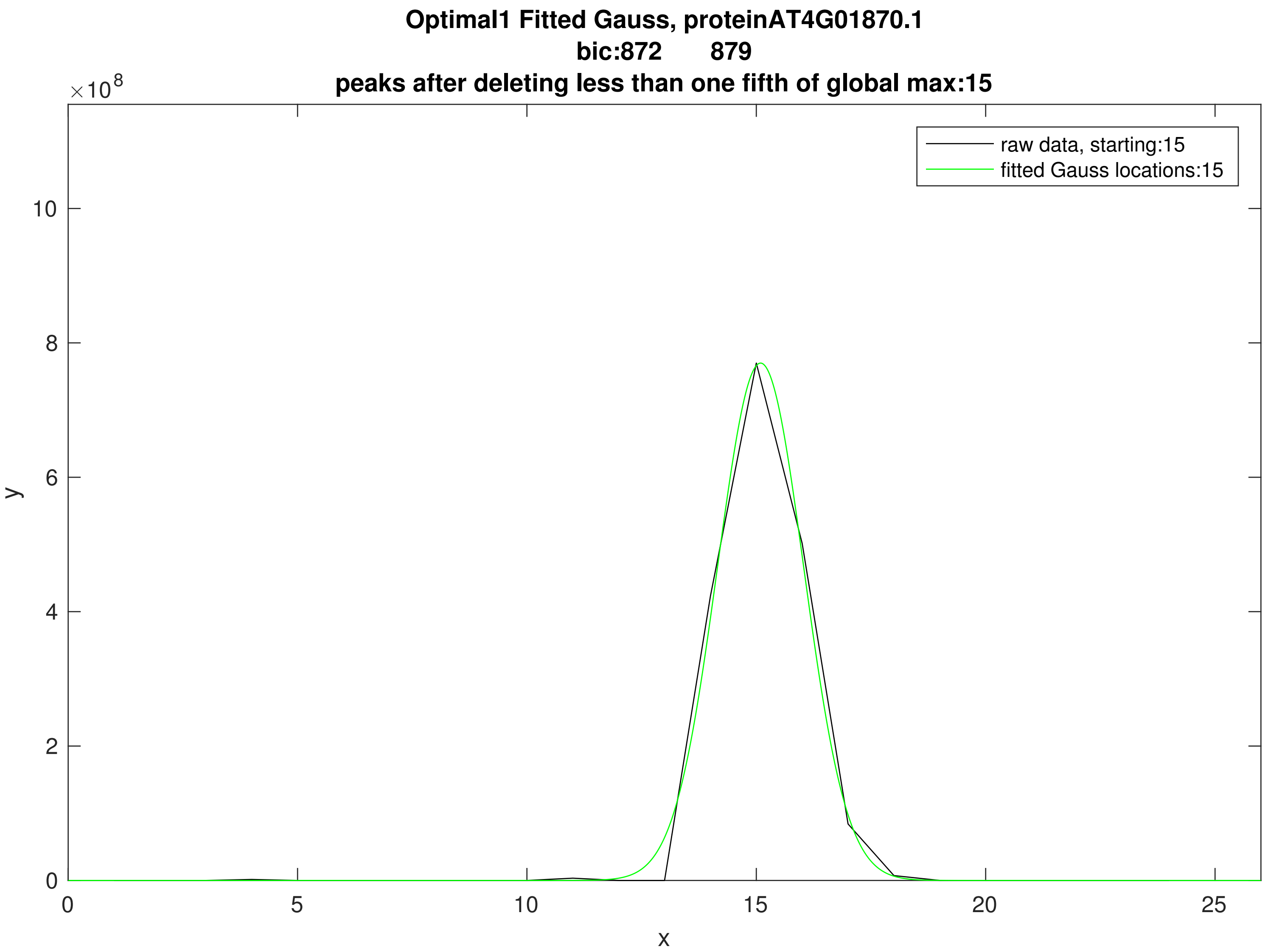

Optimal1 Fitted Gauss, proteinAT4G01870.1
bic:872 879
peaks after deleting less than one fifth of global max:15
×10^8
raw data, starting:15
fitted Gauss locations:15
y
x
0
2
4
6
8
10
0
5
10
15
20
25

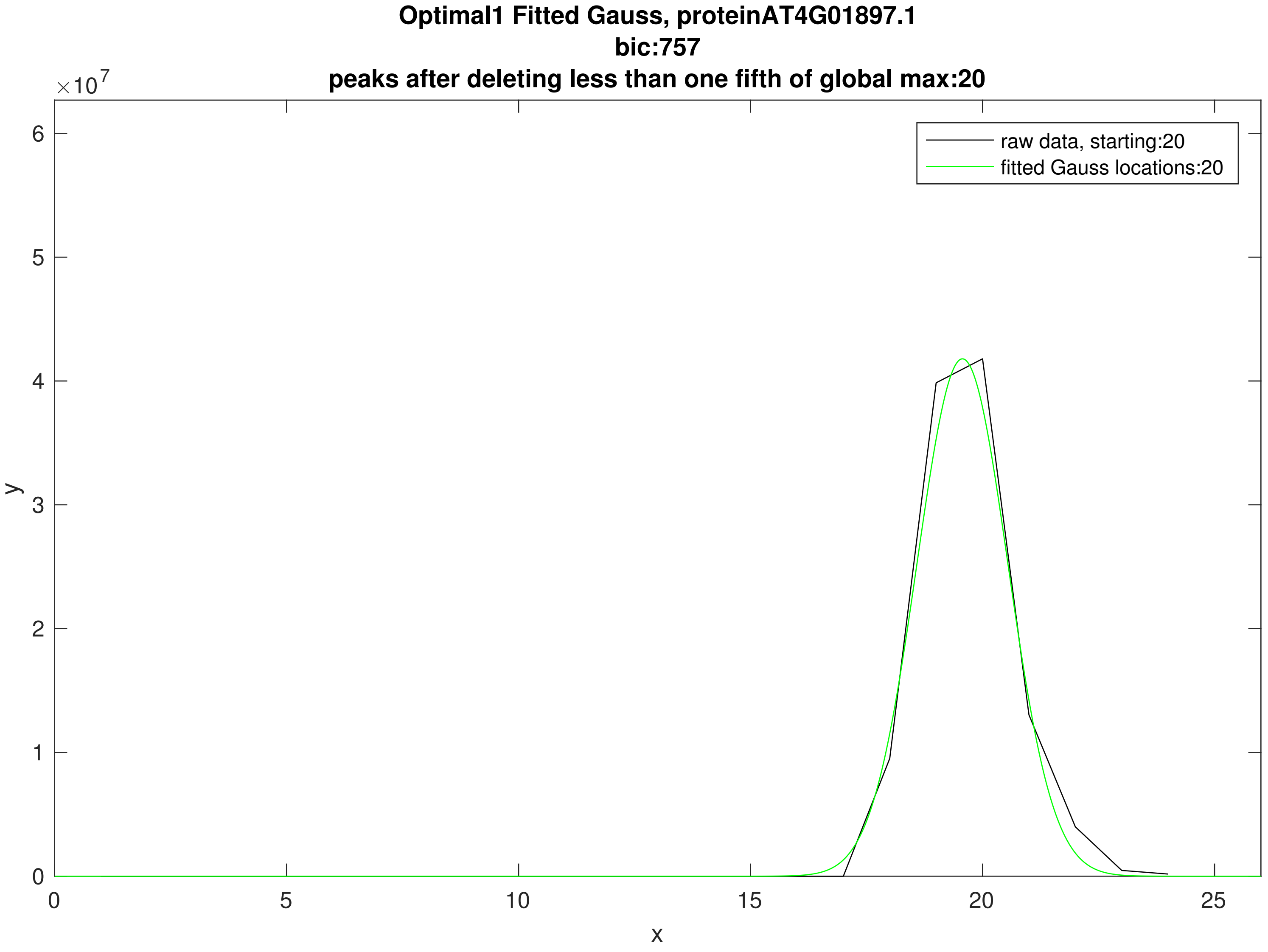

Optimal1 Fitted Gauss, proteinAT4G01897.1
bic:757
peaks after deleting less than one fifth of global max:20
×10^7
raw data, starting:20
fitted Gauss locations:20
y
x
0
1
2
3
4
5
6
0
5
10
15
20
25

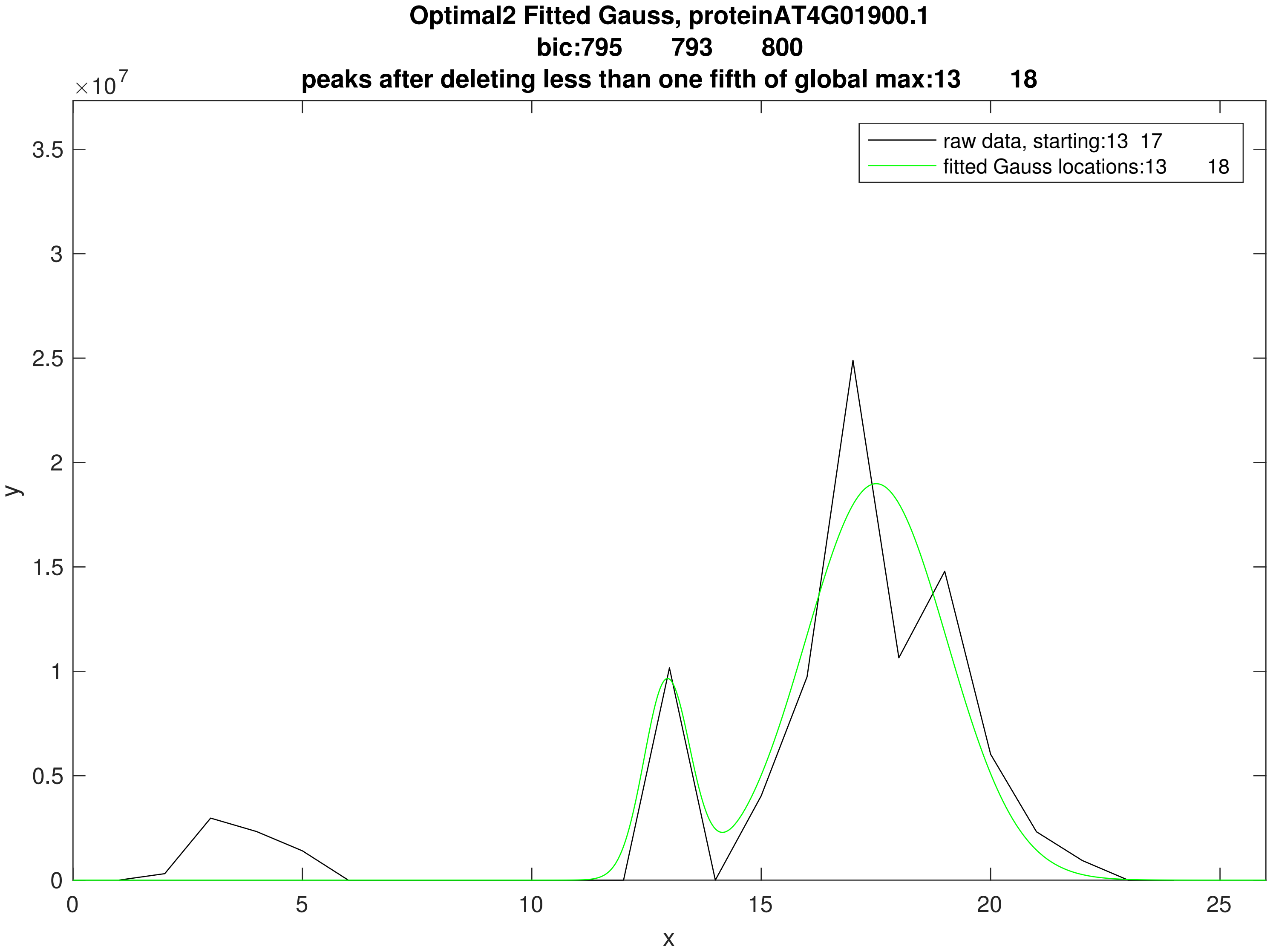
Optimal2 Fitted Gauss, proteinAT4G01900.1
bic:795 793 800
peaks after deleting less than one fifth of global max:13 18
×10^7
raw data, starting:13 17
fitted Gauss locations:13 18
y
x
0
0.5
1
1.5
2
2.5
3
3.5
0
5
10
15
20
25

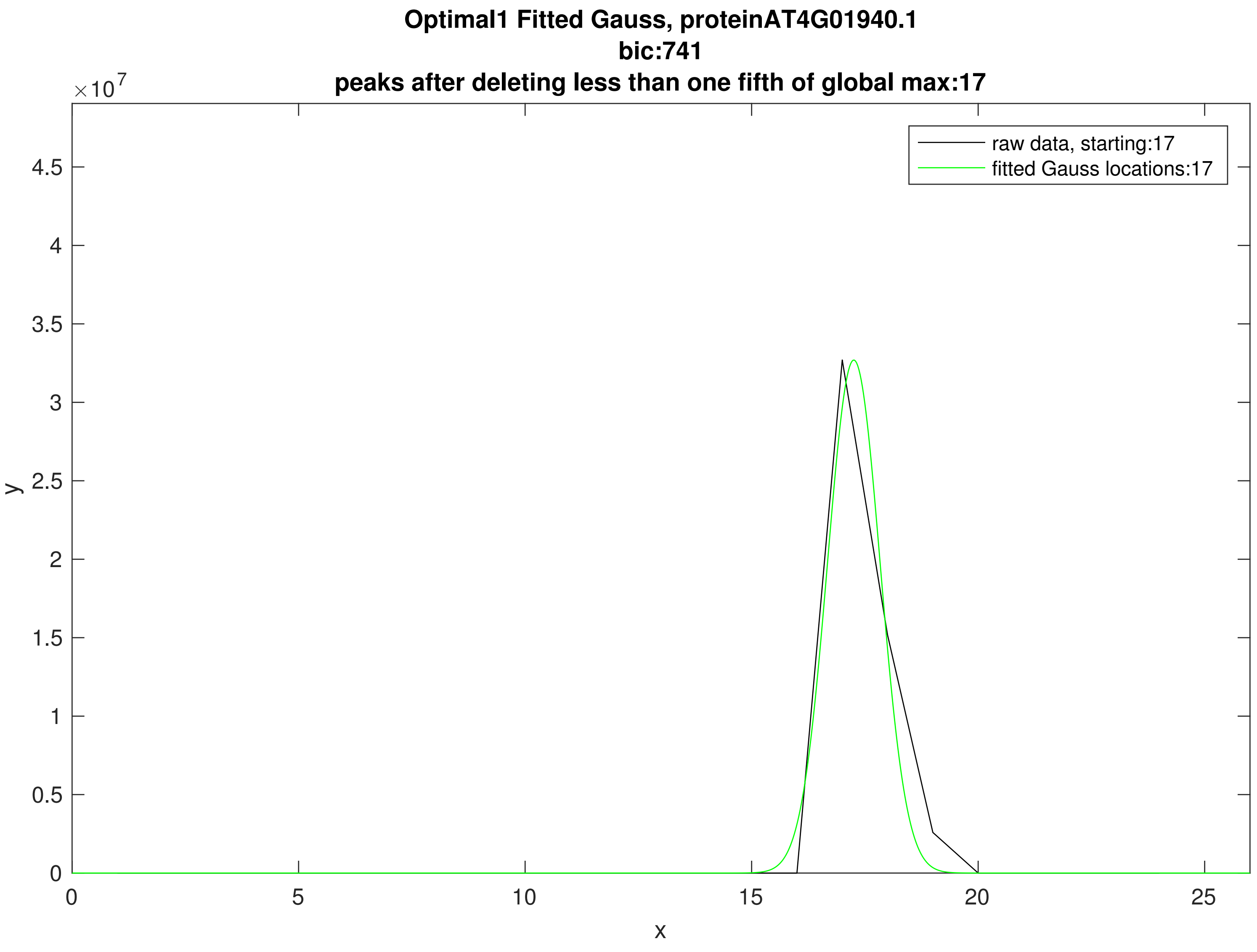

Optimal1 Fitted Gauss, proteinAT4G01940.1
bic:741
peaks after deleting less than one fifth of global max:17
raw data, starting:17
fitted Gauss locations:17
×10^7
y
x

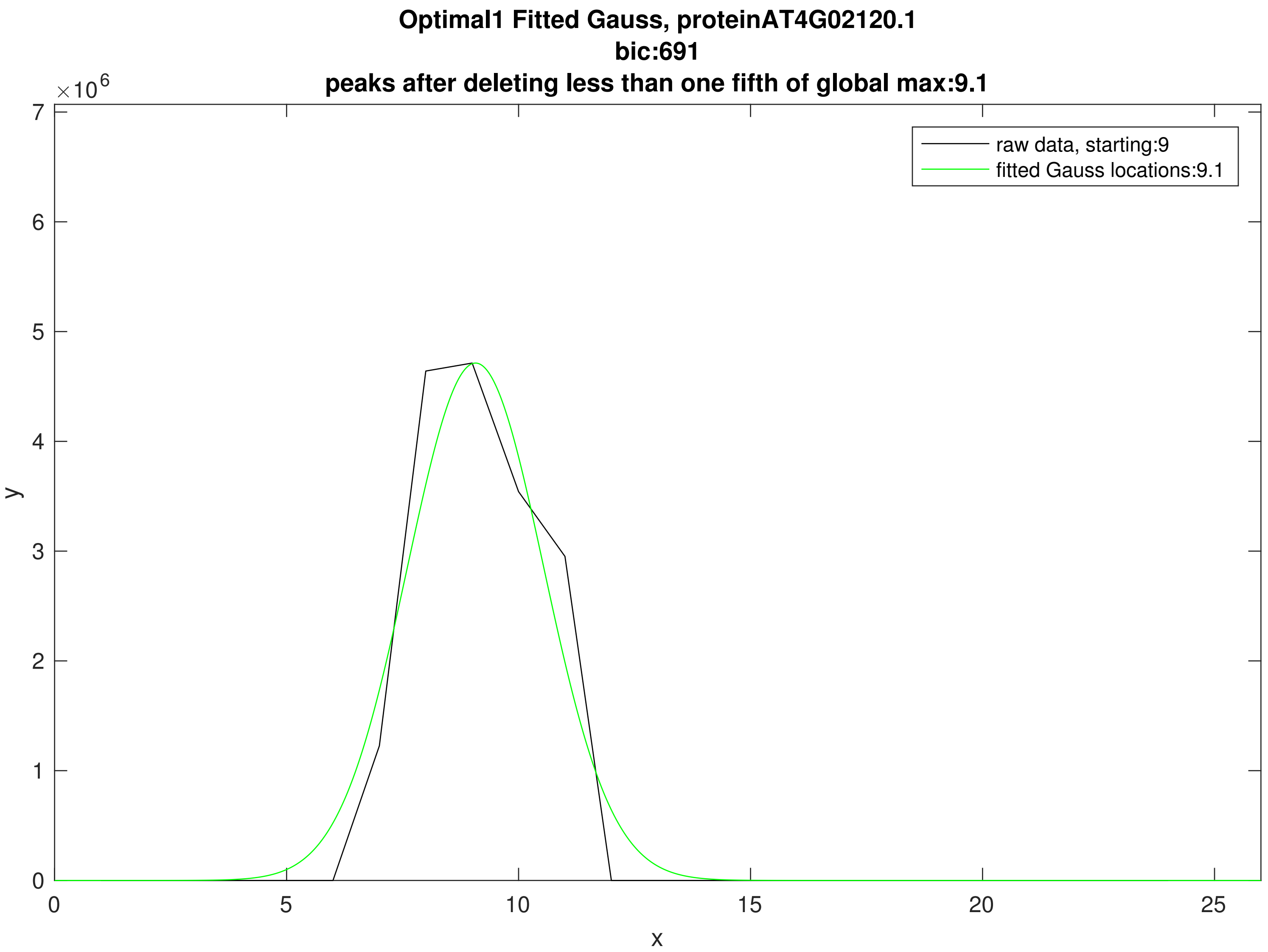
Optimal1 Fitted Gauss, proteinAT4G02120.1
bic:691
peaks after deleting less than one fifth of global max:9.1
$\times10^6$
7
6
5
4
3
2
1
0
y
0
5
10
15
20
25
x
raw data, starting:9
fitted Gauss locations:9.1

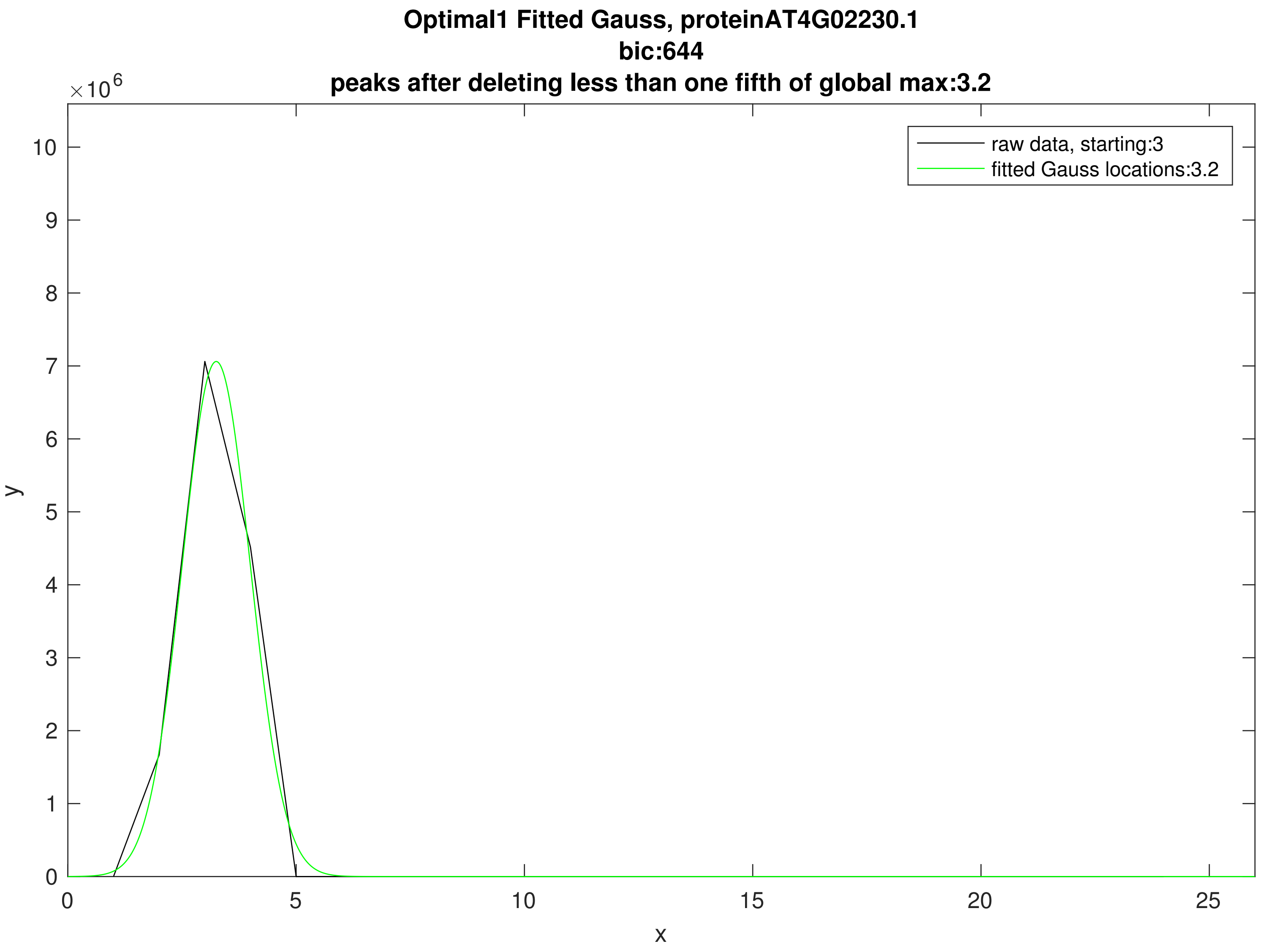

Optimal1 Fitted Gauss, proteinAT4G02230.1
bic:644
peaks after deleting less than one fifth of global max:3.2
×10^6
raw data, starting:3
fitted Gauss locations:3.2
y
x
0
1
2
3
4
5
6
7
8
9
10
0
5
10
15
20
25

#### Optimal1 Fitted Gauss, proteinAT4G02280.1
#### bic:842
#### peaks after deleting less than one fifth of global max:7.3

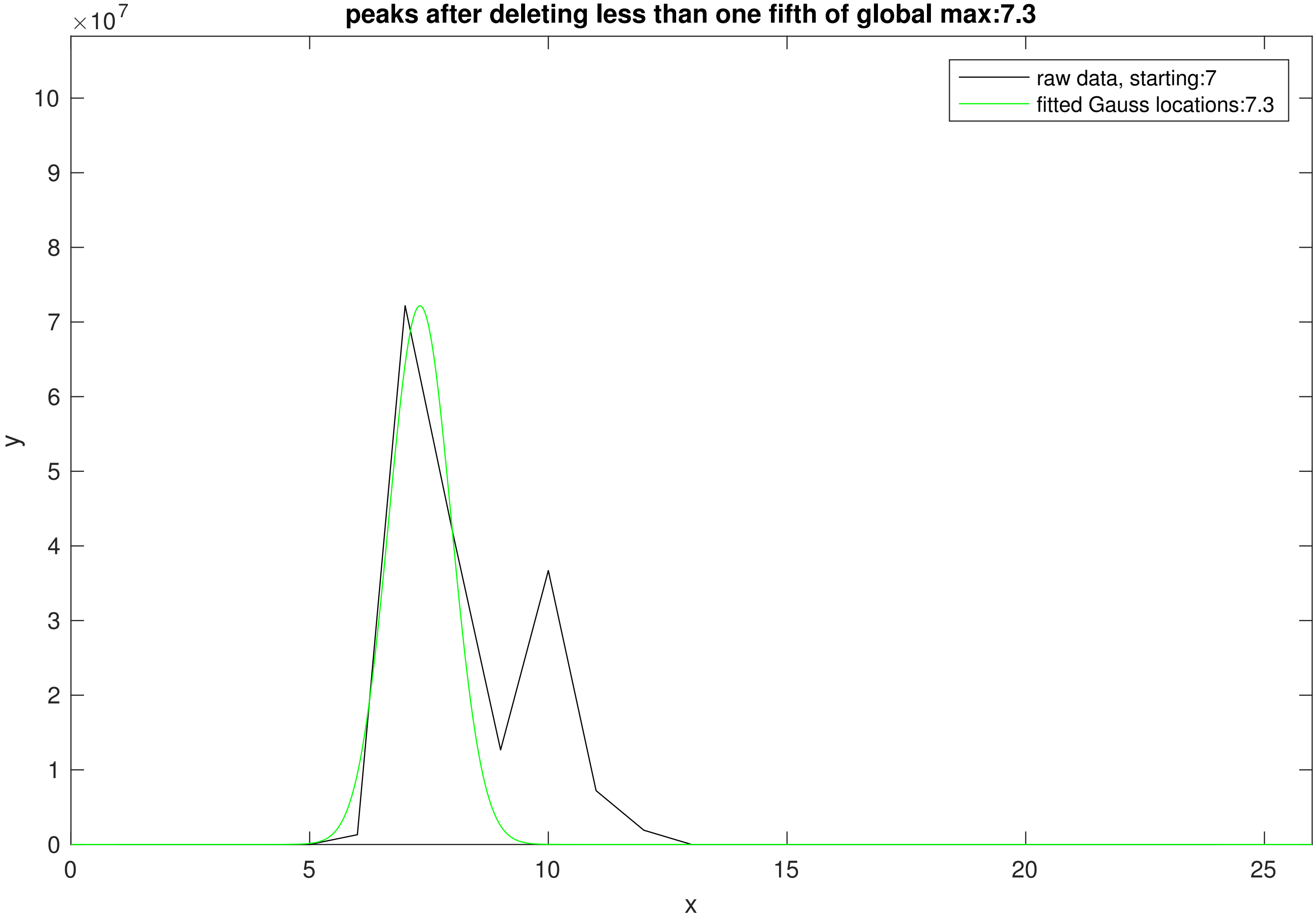

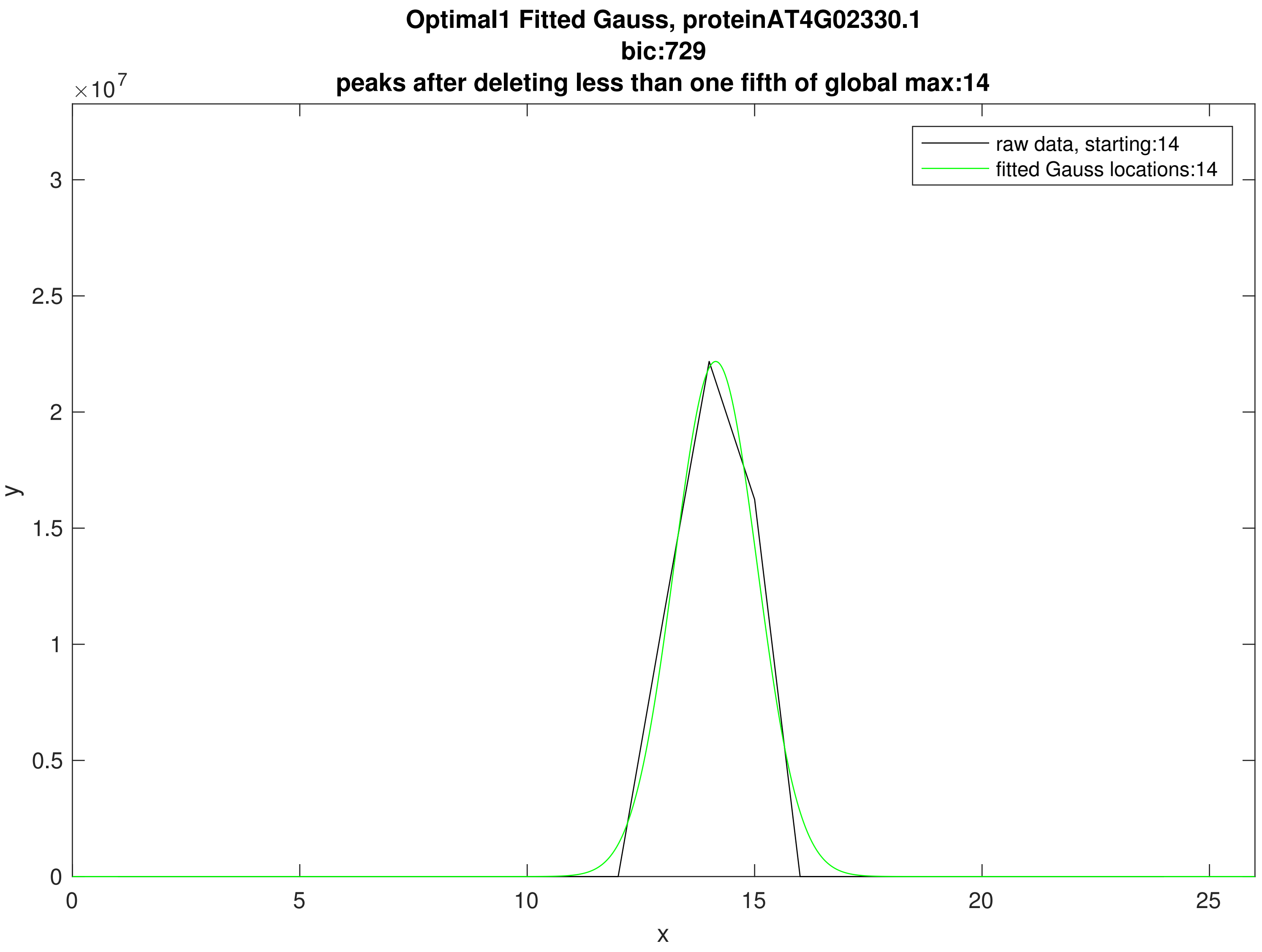

Optimal1 Fitted Gauss, proteinAT4G02330.1
bic:729
peaks after deleting less than one fifth of global max:14
×10^7
raw data, starting:14
fitted Gauss locations:14
y
x
0
0.5
1
1.5
2
2.5
3
5
10
15
20
25

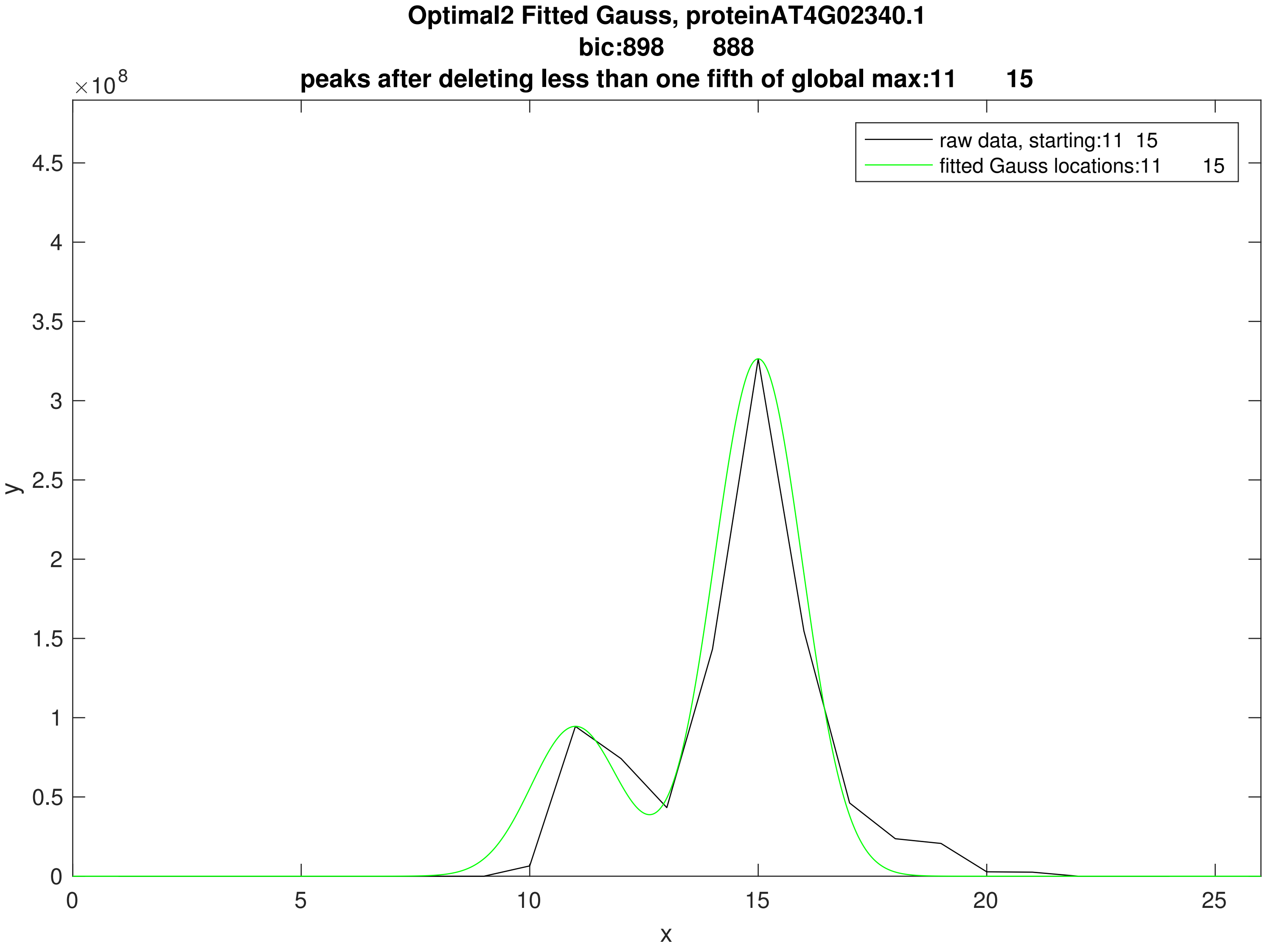
Optimal2 Fitted Gauss, proteinAT4G02340.1
bic:898 888
peaks after deleting less than one fifth of global max:11 15
×10^8
raw data, starting:11 15
fitted Gauss locations:11 15
y
x
0
0.5
1
1.5
2
2.5
3
3.5
4
4.5
5
10
15
20
25

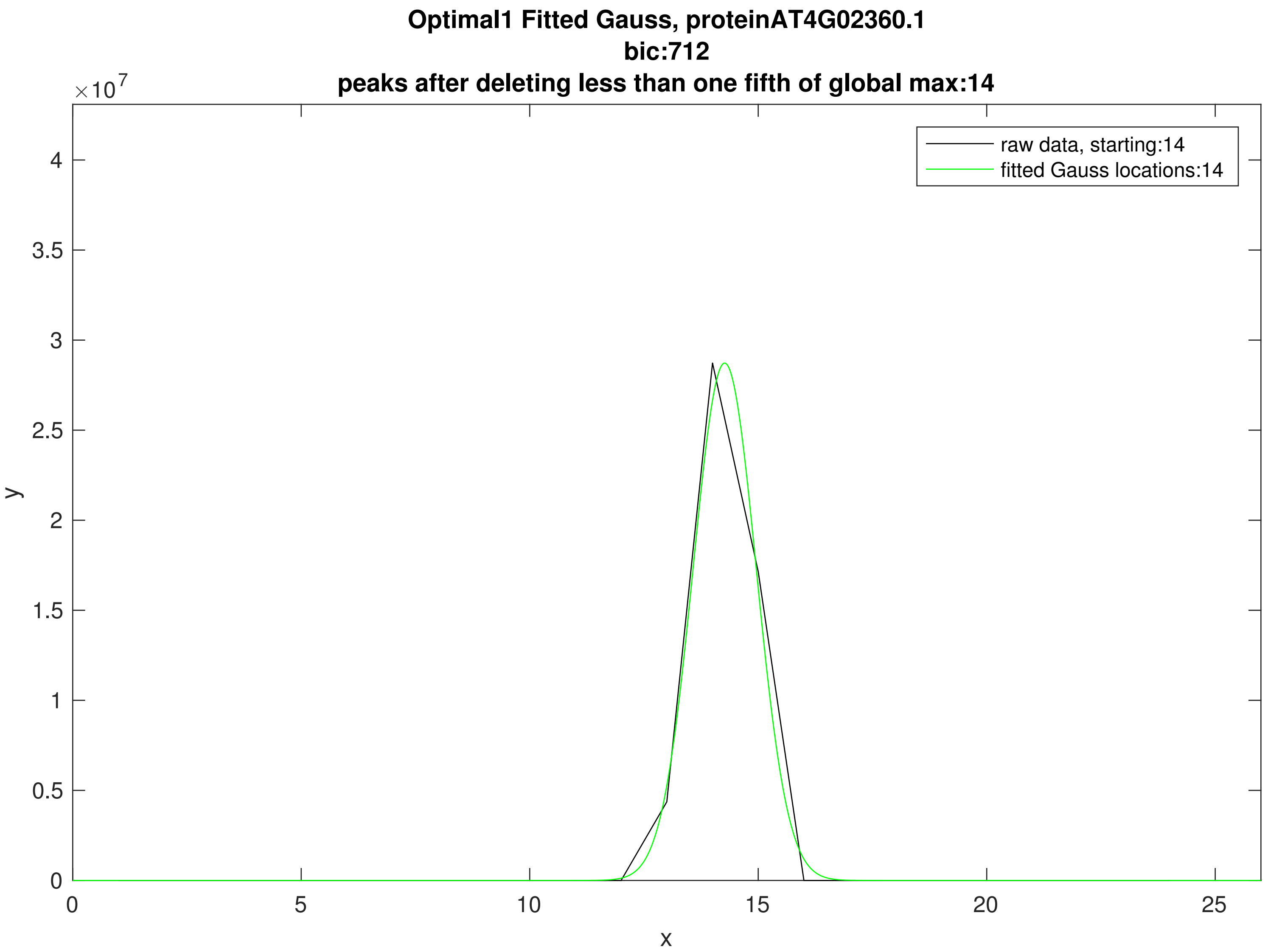

Optimal1 Fitted Gauss, proteinAT4G02360.1
bic:712
peaks after deleting less than one fifth of global max:14
×10^7
raw data, starting:14
fitted Gauss locations:14
y
x
0
0.5
1
1.5
2
2.5
3
3.5
4
0
5
10
15
20
25

Optimal1 Fitted Gauss, proteinAT4G02380.2;AT4G02380.1
bic:746
peaks after deleting less than one fifth of global max:21

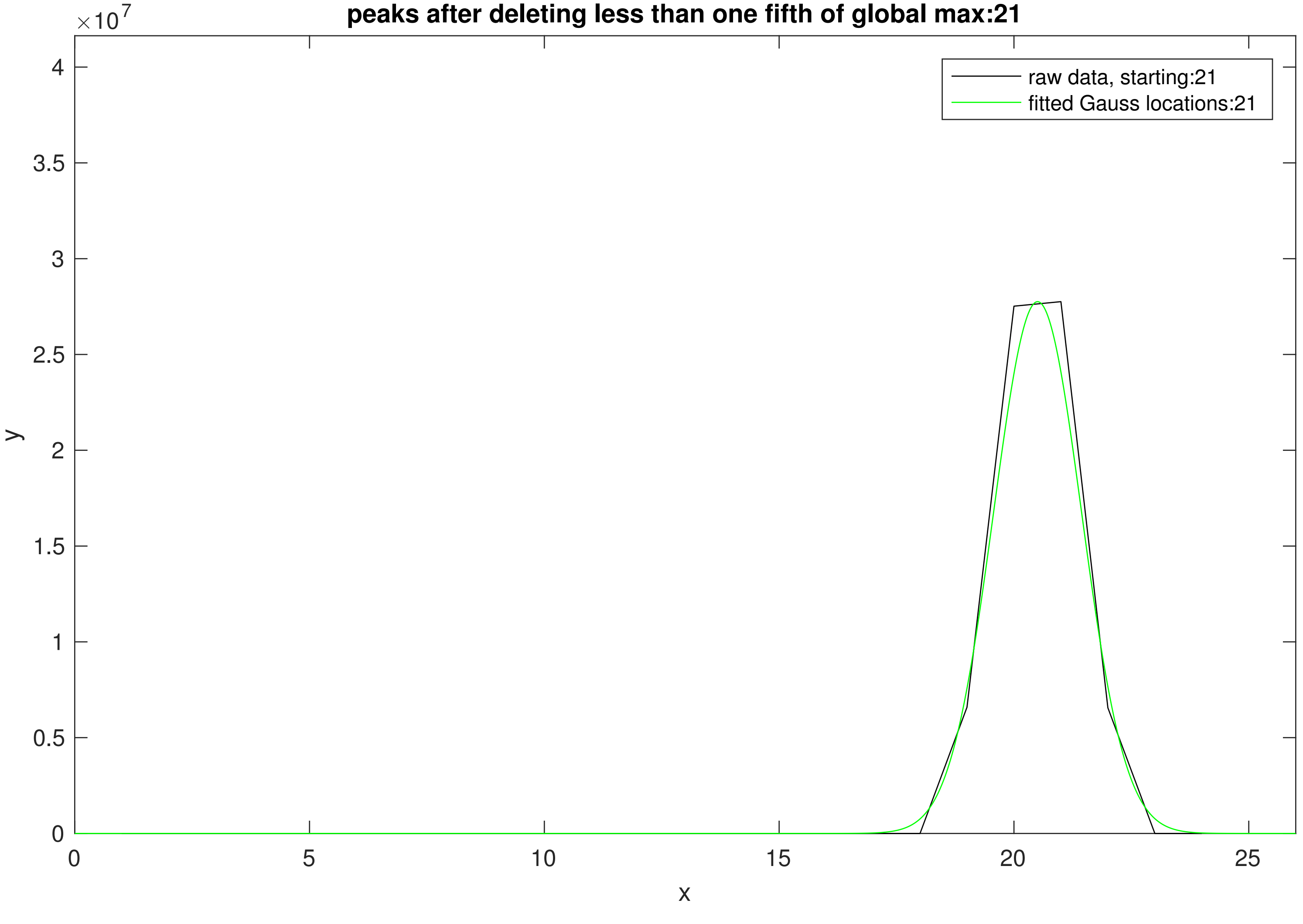

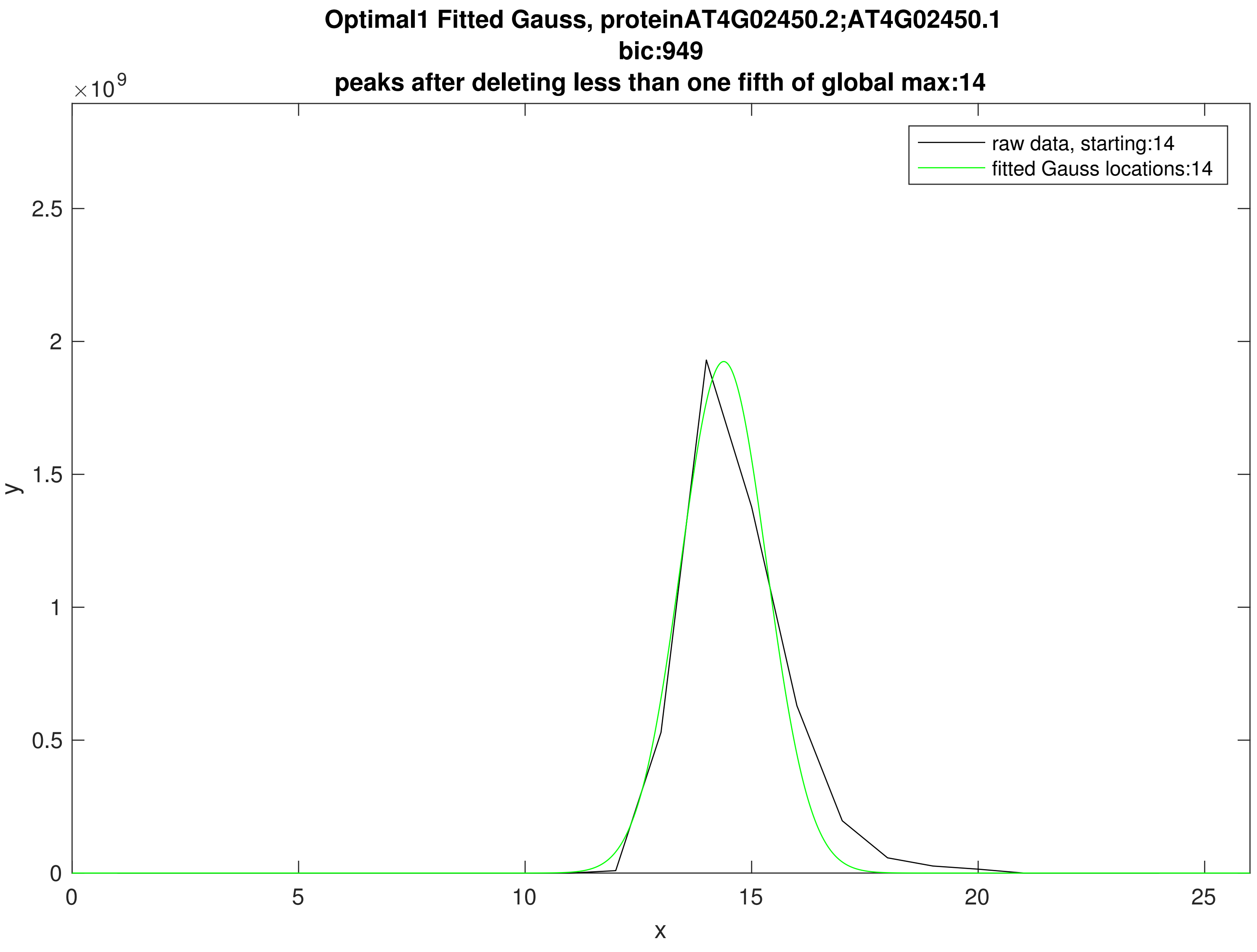

Optimal1 Fitted Gauss, proteinAT4G02450.2;AT4G02450.1
bic:949
peaks after deleting less than one fifth of global max:14
×10^9
raw data, starting:14
fitted Gauss locations:14
2.5
2
1.5
1
0.5
0
y
0
5
10
15
20
25
x

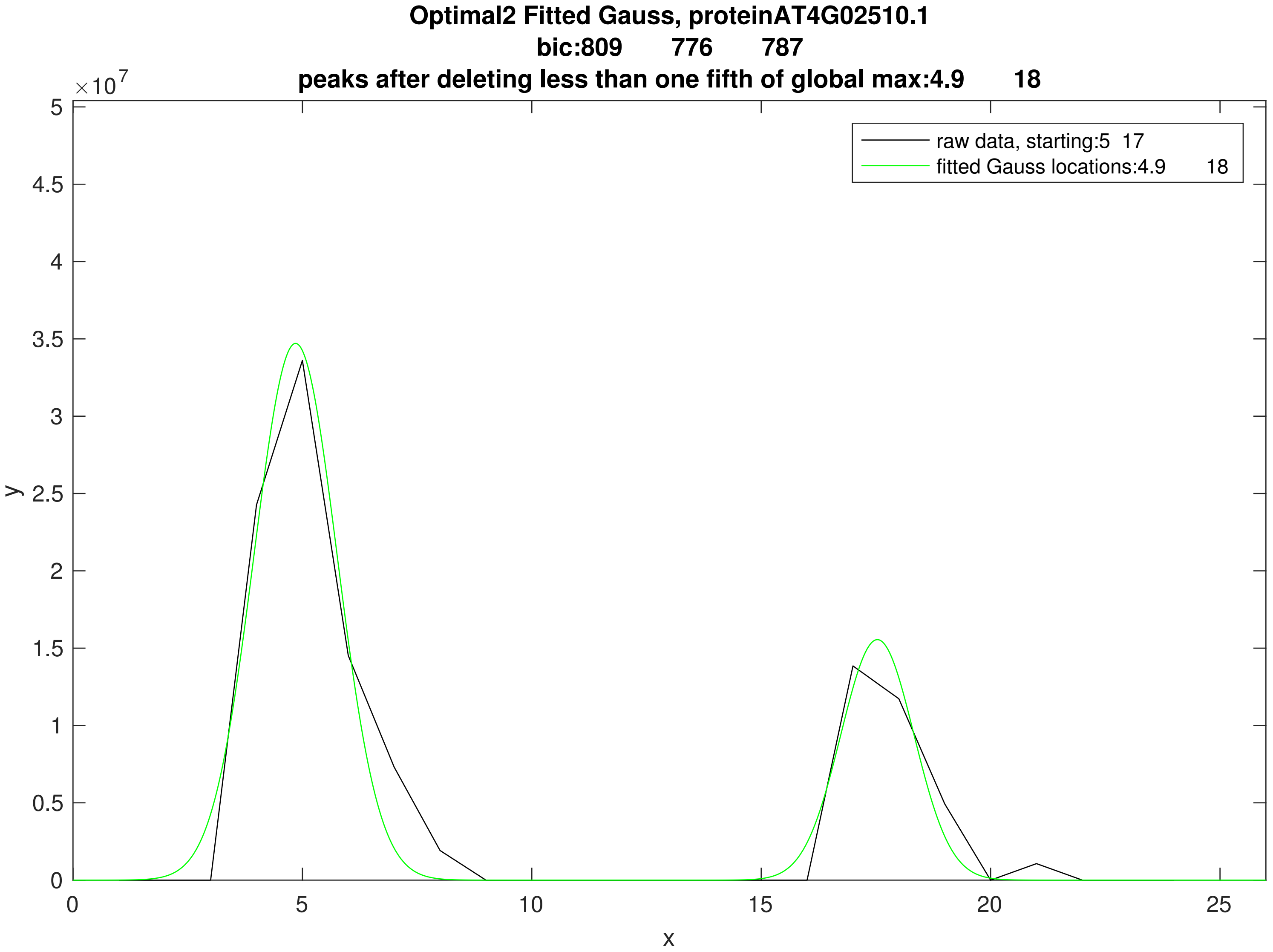
Optimal2 Fitted Gauss, proteinAT4G02510.1
bic:809 776 787
peaks after deleting less than one fifth of global max:4.9 18
×10^7
raw data, starting:5 17
fitted Gauss locations:4.9 18
y
x
0
0.5
1
1.5
2
2.5
3
3.5
4
4.5
5
0
5
10
15
20
25

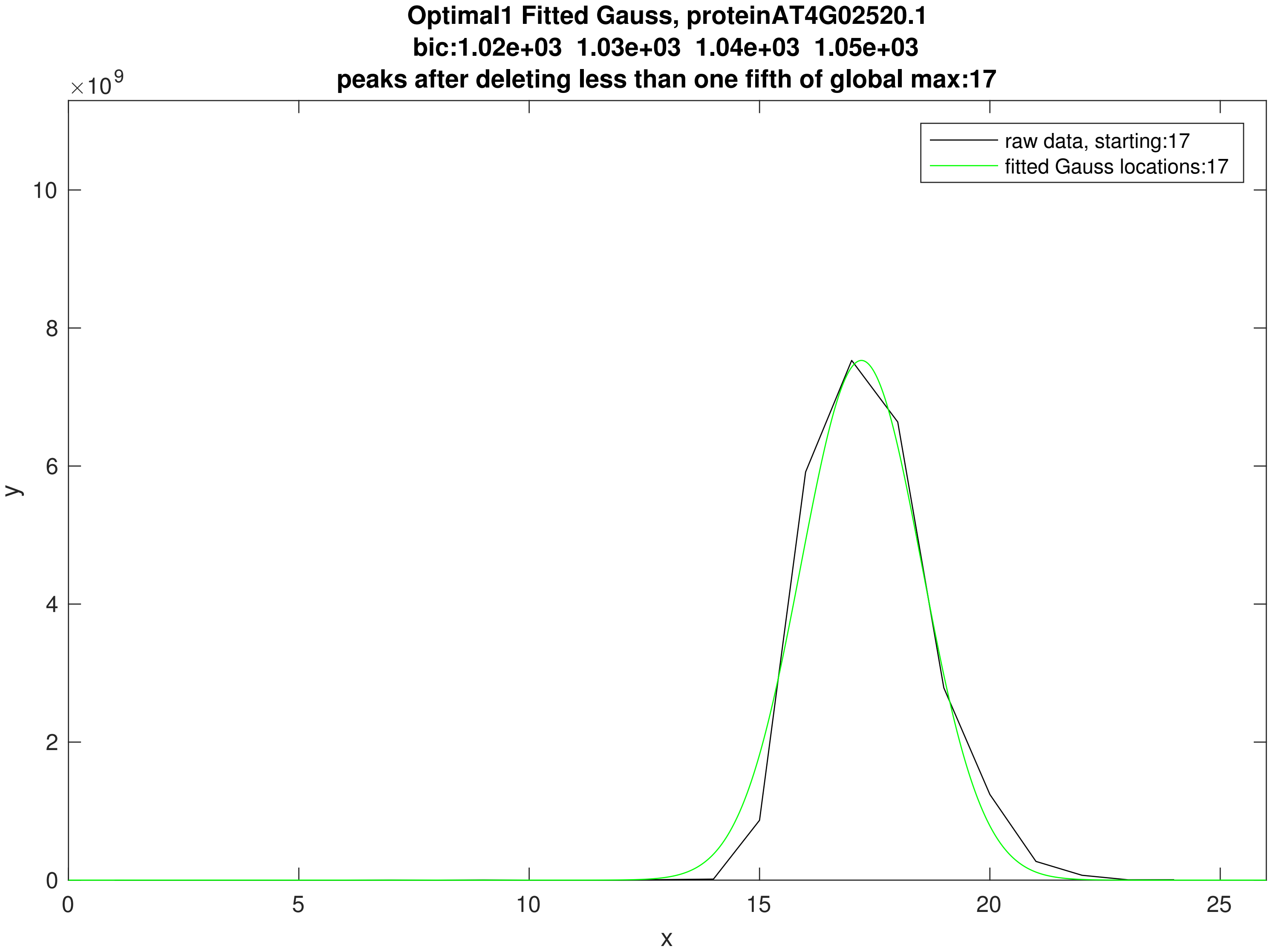
Optimal1 Fitted Gauss, proteinAT4G02520.1
bic:1.02e+03  1.03e+03  1.04e+03  1.05e+03
peaks after deleting less than one fifth of global max:17
×10^9
raw data, starting:17
fitted Gauss locations:17
10
8
6
4
2
0
y
0
5
10
15
20
25
x

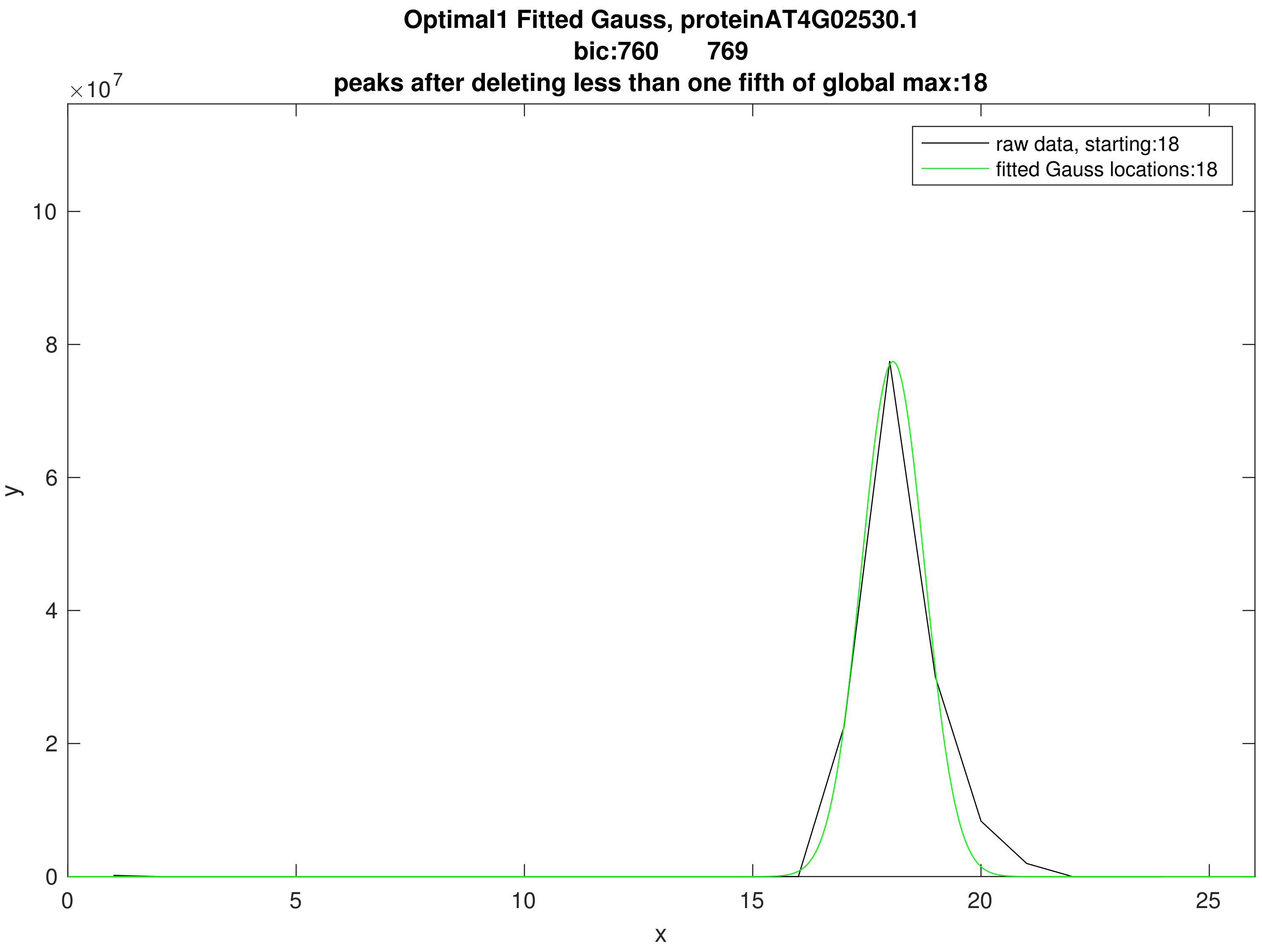

Optimal1 Fitted Gauss, proteinAT4G02530.1
bic:760 769
peaks after deleting less than one fifth of global max:18
×10^7
raw data, starting:18
fitted Gauss locations:18
y
x
0
2
4
6
8
10
5
15
20
25

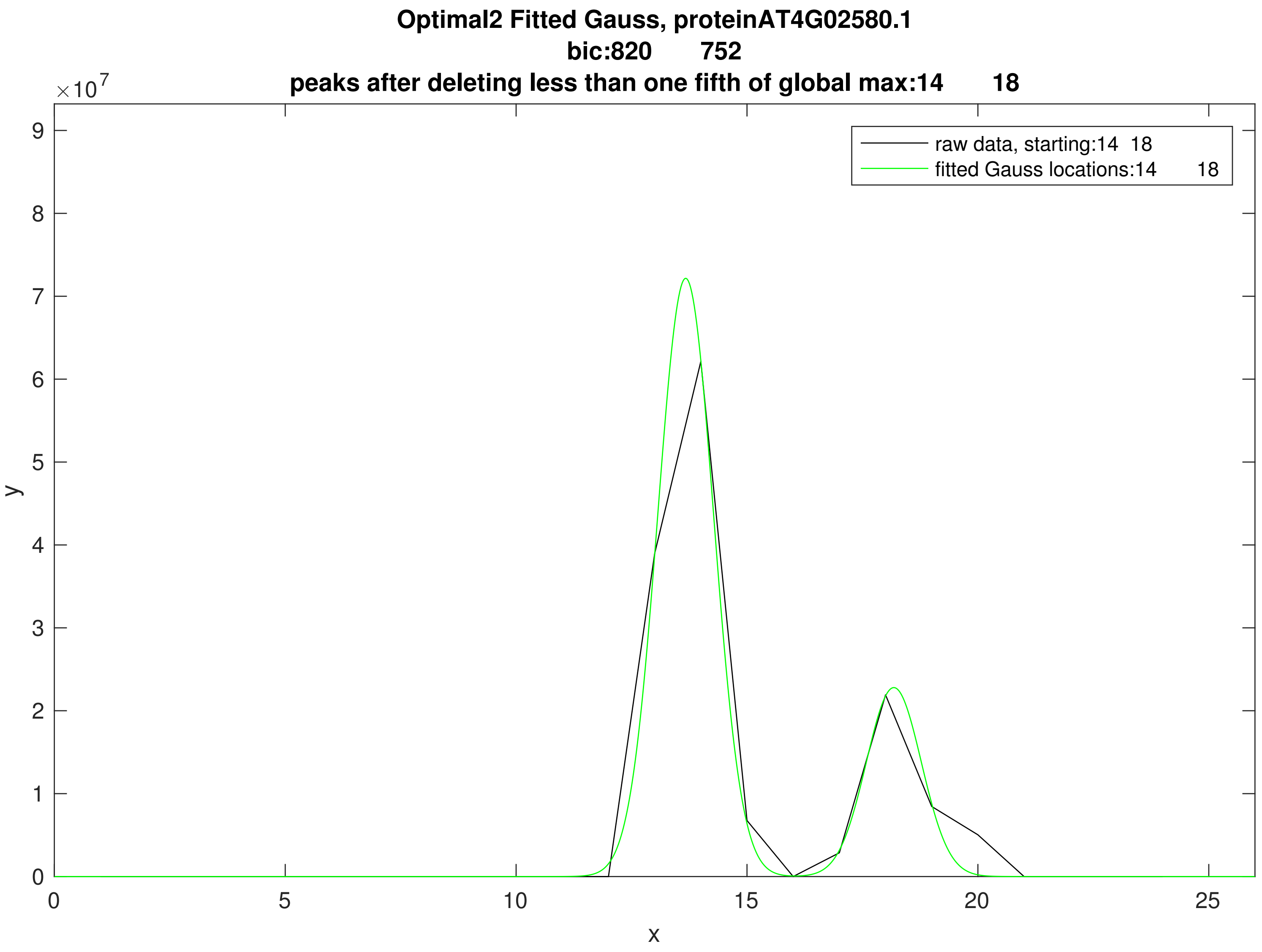

Optimal2 Fitted Gauss, proteinAT4G02580.1
bic:820 752
peaks after deleting less than one fifth of global max:14 18
×10^7
raw data, starting:14 18
fitted Gauss locations:14 18
y
x
0
1
2
3
4
5
6
7
8
9
0
5
10
15
20
25

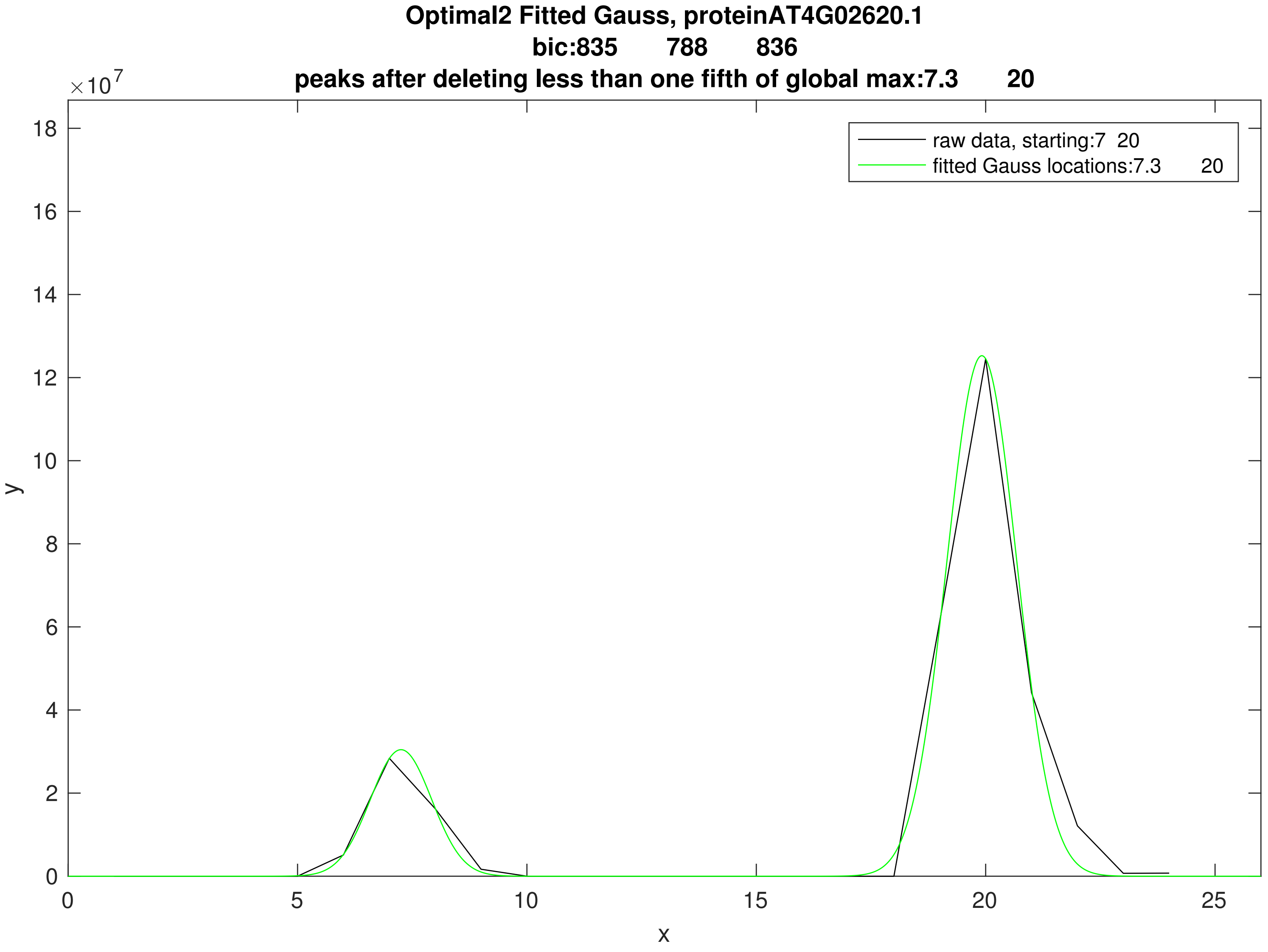

Optimal2 Fitted Gauss, proteinAT4G02620.1
bic:835 788 836
peaks after deleting less than one fifth of global max:7.3 20
×10^7
raw data, starting:7 20
fitted Gauss locations:7.3 20
y
x
0
2
4
6
8
10
12
14
16
18
5
15
20
25

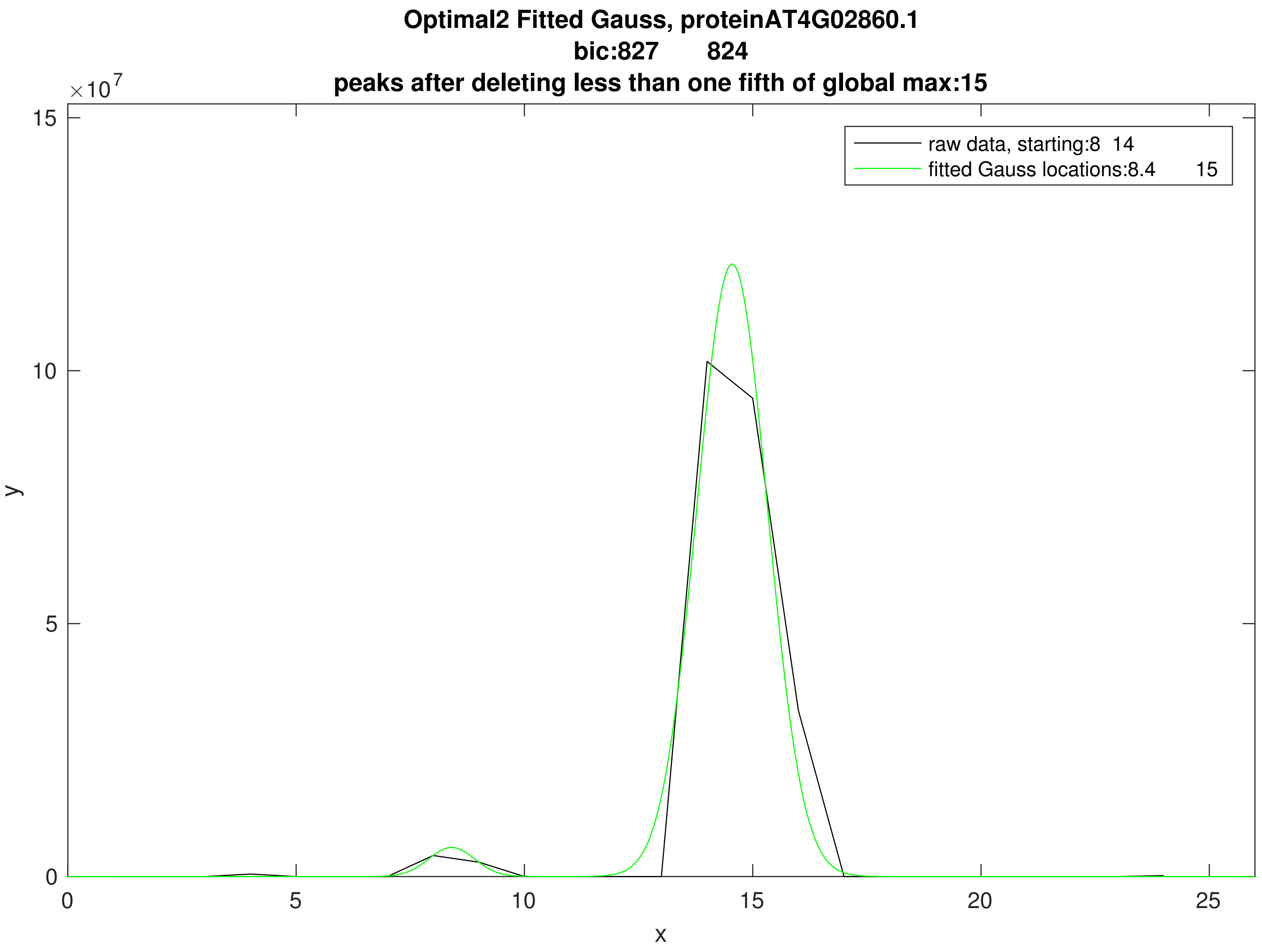

Optimal2 Fitted Gauss, proteinAT4G02860.1
bic:827 824
peaks after deleting less than one fifth of global max:15
×10^7
raw data, starting:8 14
fitted Gauss locations:8.4 15
y
x
0
5
10
15
20
25

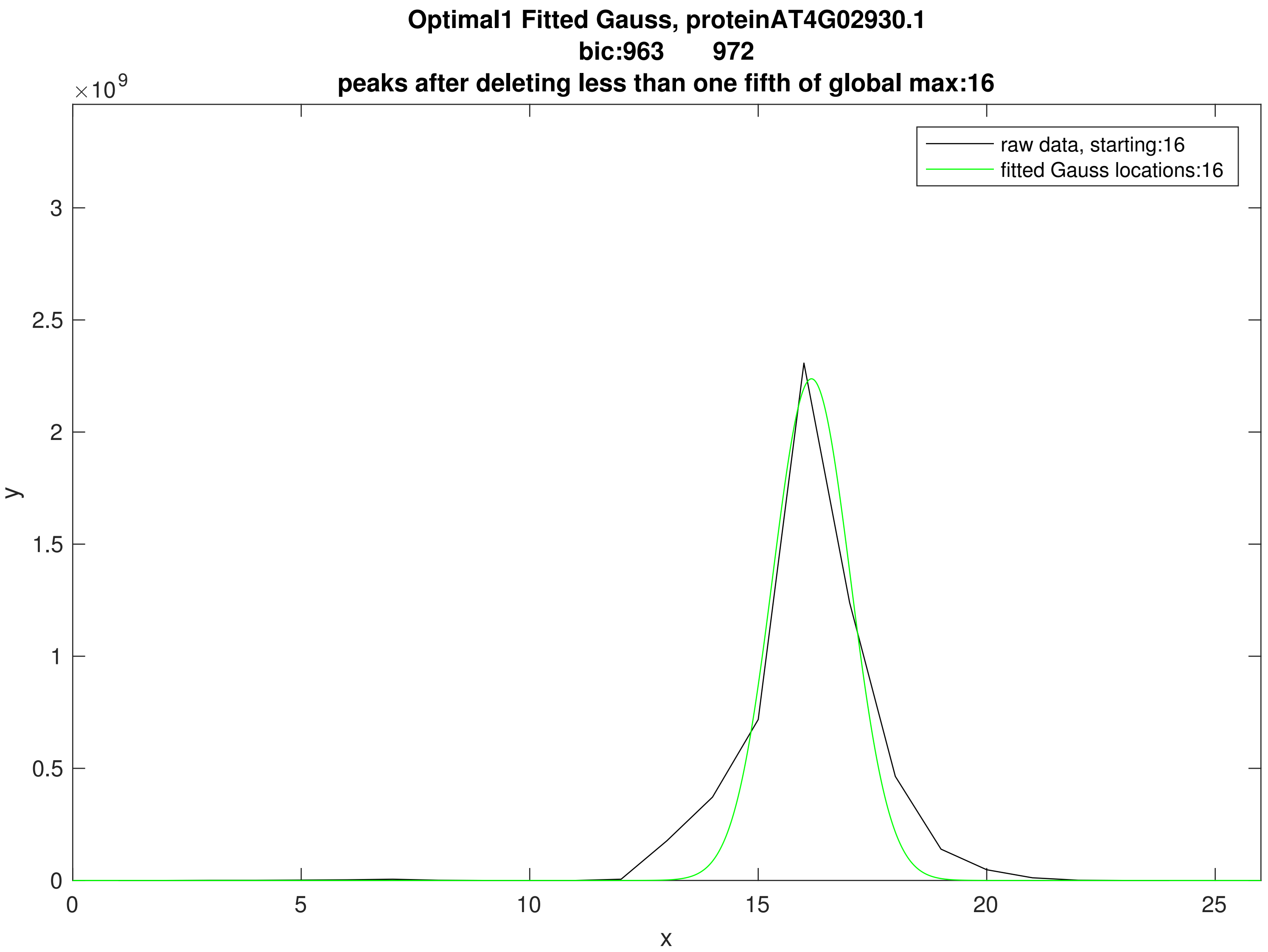
Optimal1 Fitted Gauss, proteinAT4G02930.1
bic:963 972
peaks after deleting less than one fifth of global max:16
×10^9
raw data, starting:16
fitted Gauss locations:16
y
x
0
0.5
1
1.5
2
2.5
3
0
5
10
15
20
25

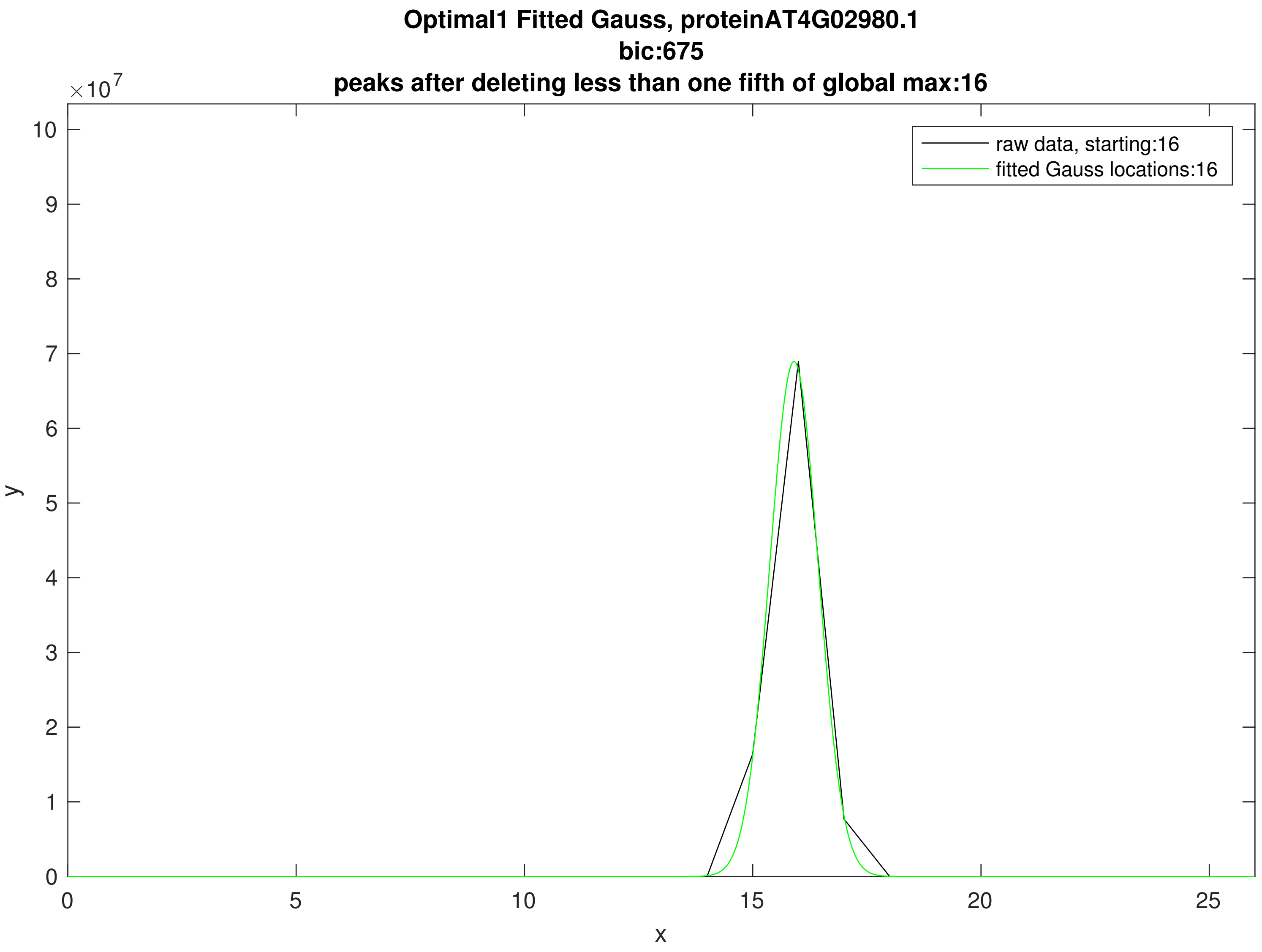
Optimal1 Fitted Gauss, proteinAT4G02980.1
bic:675
peaks after deleting less than one fifth of global max:16
×10^7
raw data, starting:16
fitted Gauss locations:16
y
x
0
1
2
3
4
5
6
7
8
9
10
0
5
10
15
20
25

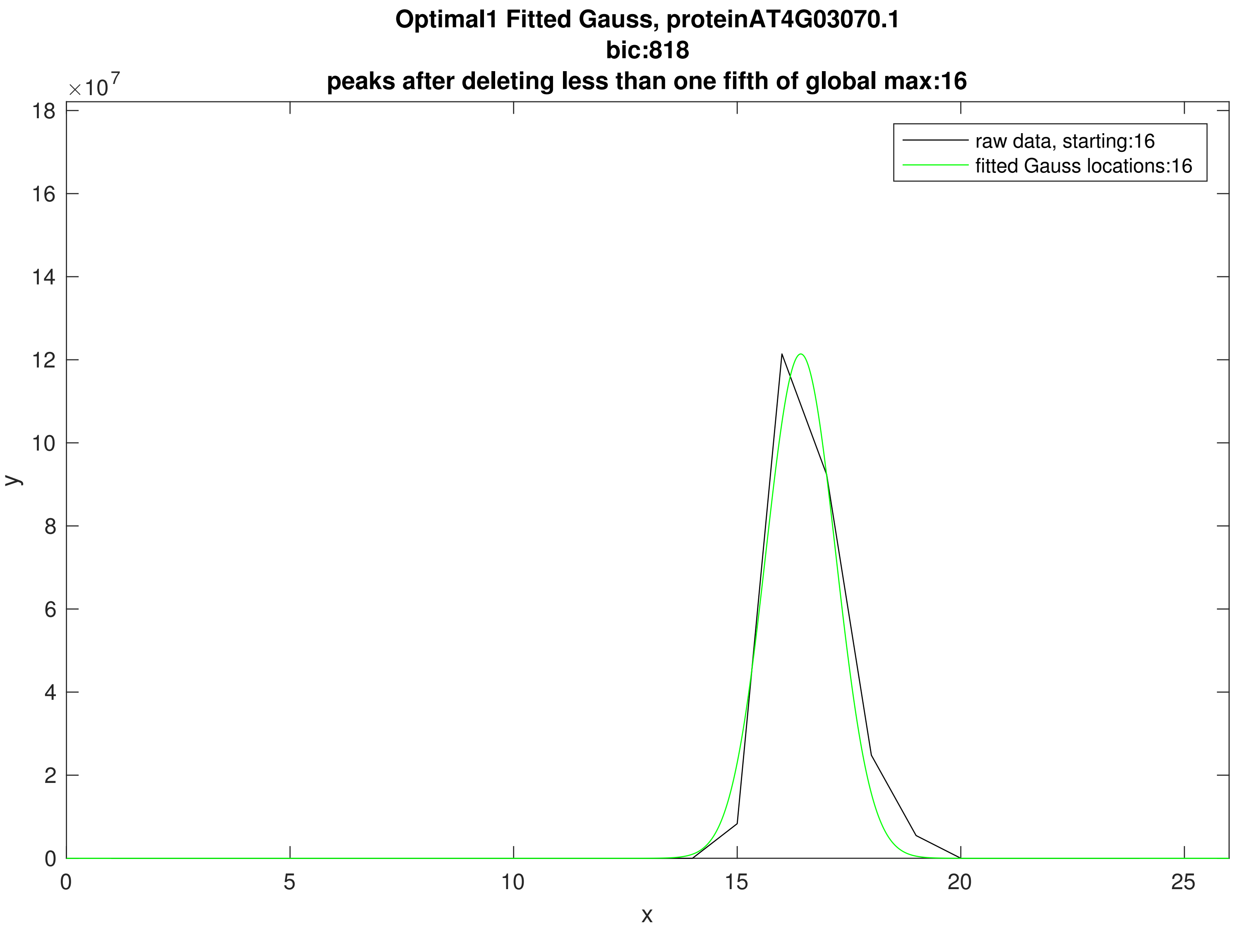
Optimal1 Fitted Gauss, proteinAT4G03070.1
bic:818
peaks after deleting less than one fifth of global max:16
×10^7
raw data, starting:16
fitted Gauss locations:16
y
x
0
2
4
6
8
10
12
14
16
18
0
5
10
15
20
25

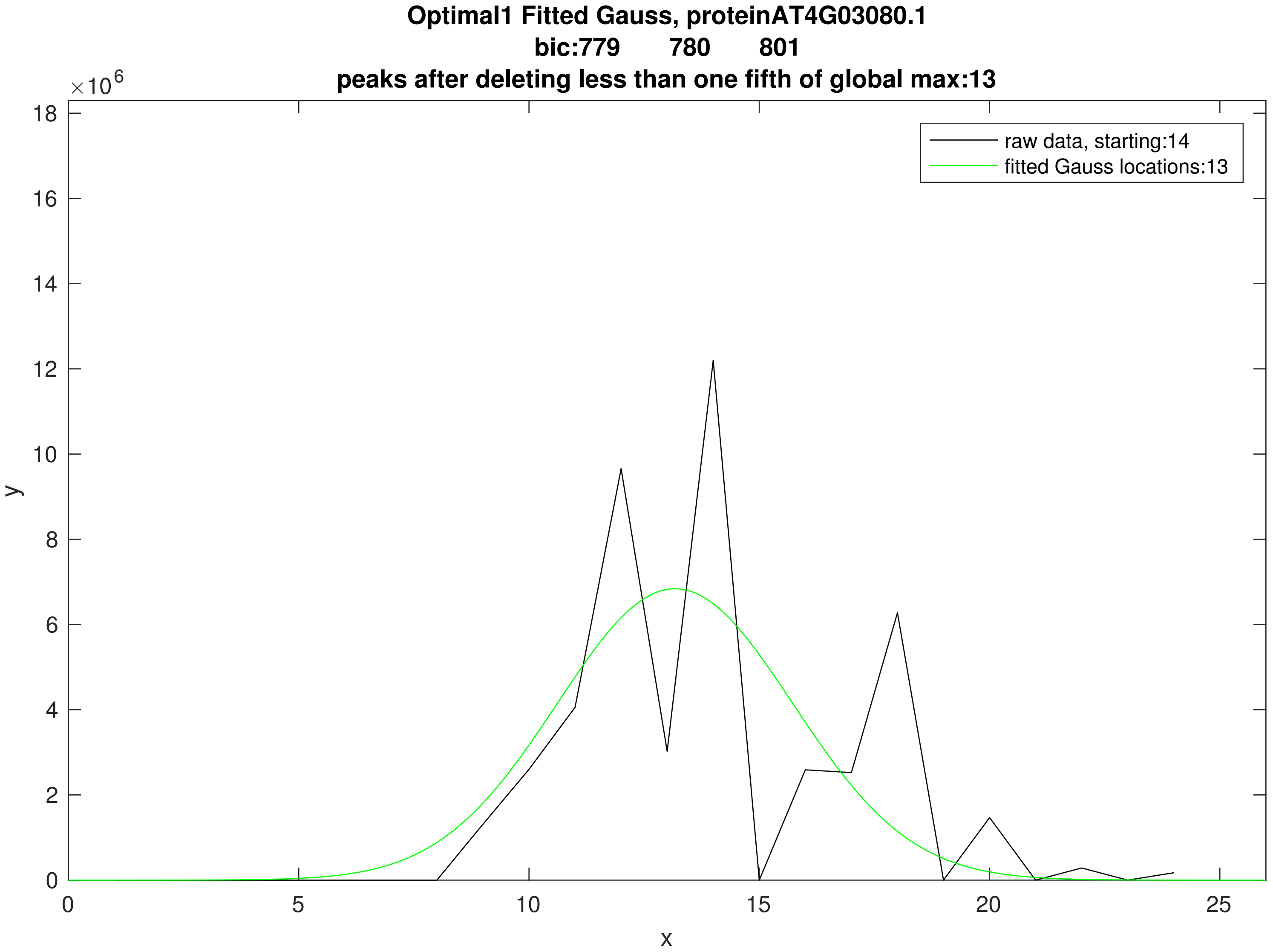

Optimal1 Fitted Gauss, proteinAT4G03080.1
bic:779 780 801
peaks after deleting less than one fifth of global max:13
×10^6
raw data, starting:14
fitted Gauss locations:13
y
x
0
2
4
6
8
10
12
14
16
18
0
5
10
15
20
25

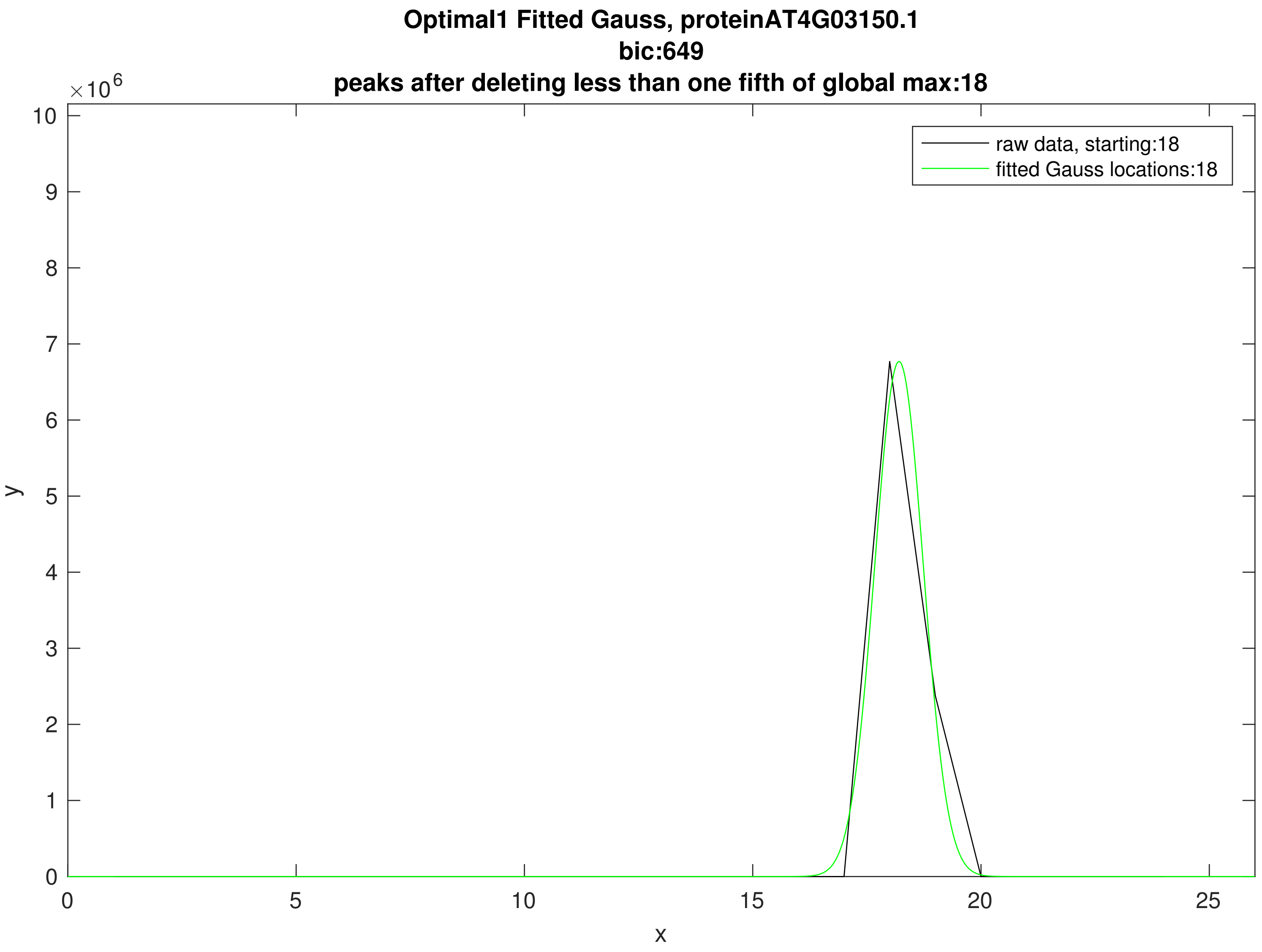

Optimal1 Fitted Gauss, proteinAT4G03150.1
bic:649
peaks after deleting less than one fifth of global max:18
×10^6
10
9
8
7
6
5
4
3
2
1
0
y
0
5
10
15
20
25
x
raw data, starting:18
fitted Gauss locations:18

**Optimal1 Fitted Gauss, proteinAT4G03190.1**
**bic:780**
**peaks after deleting less than one fifth of global max:13**

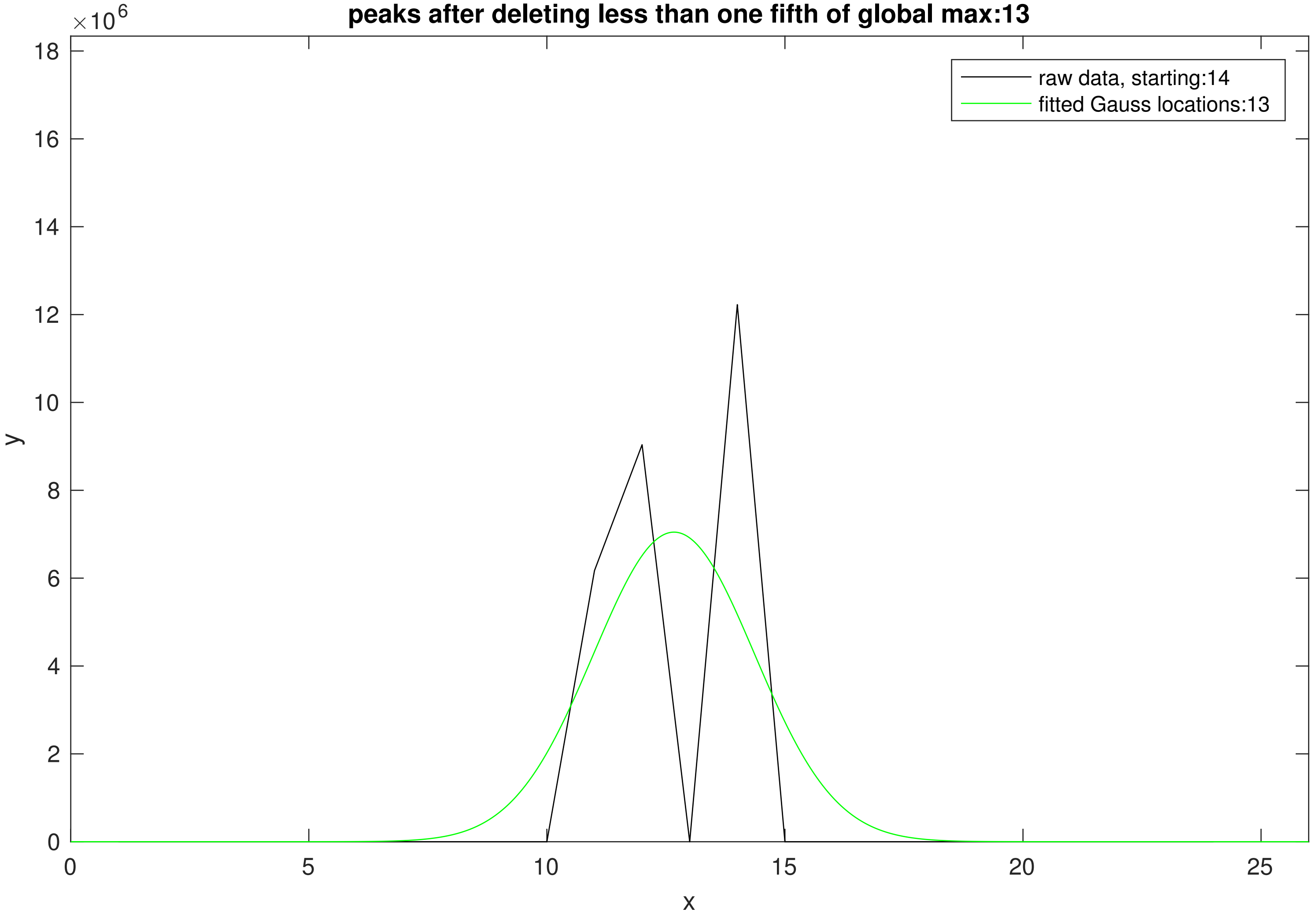

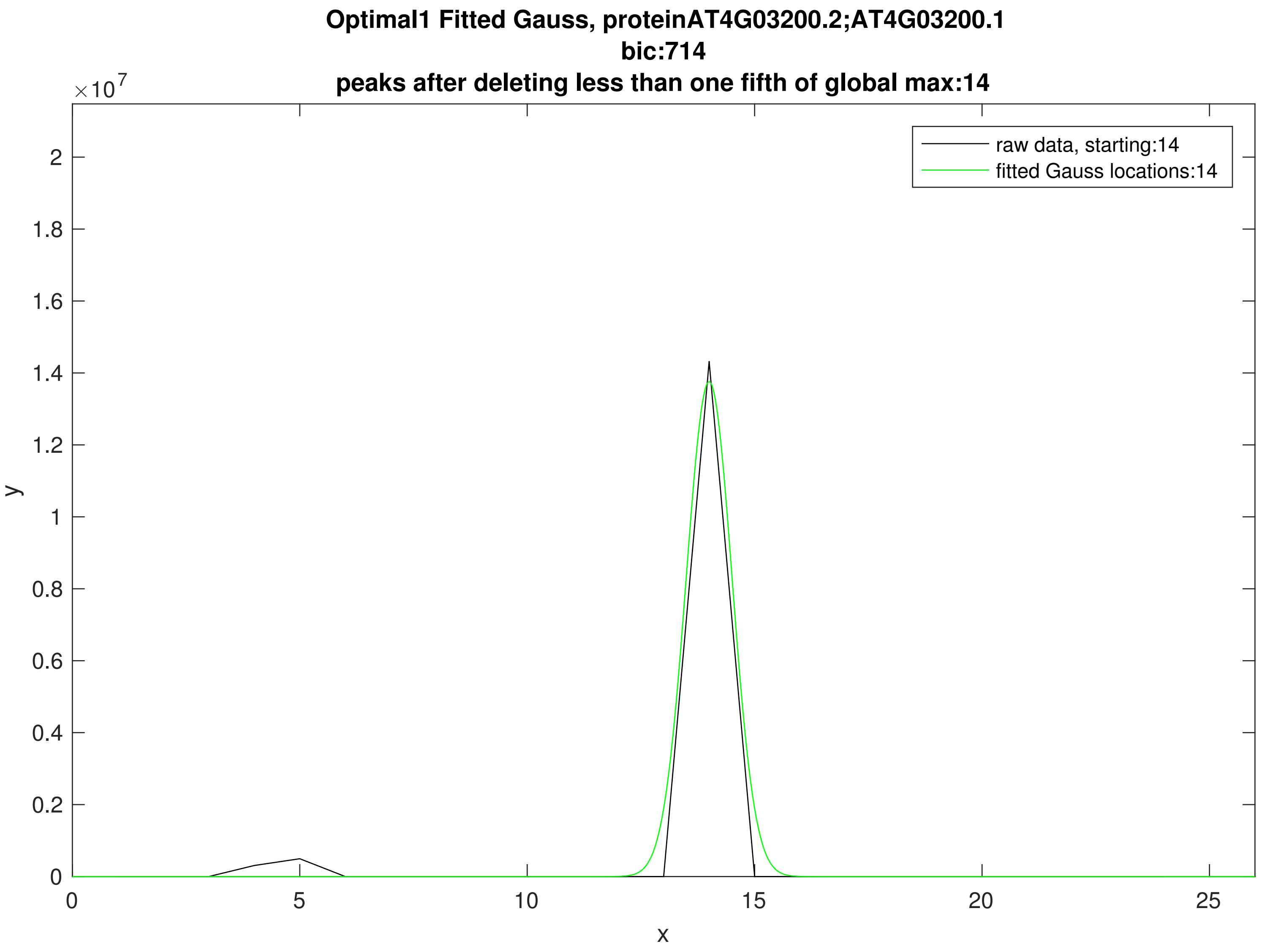

Optimal1 Fitted Gauss, proteinAT4G03200.2;AT4G03200.1
bic:714
peaks after deleting less than one fifth of global max:14
raw data, starting:14
fitted Gauss locations:14
y
x

Optimal1 Fitted Gauss, proteinAT4G03240.1
bic:756
peaks after deleting less than one fifth of global max:19

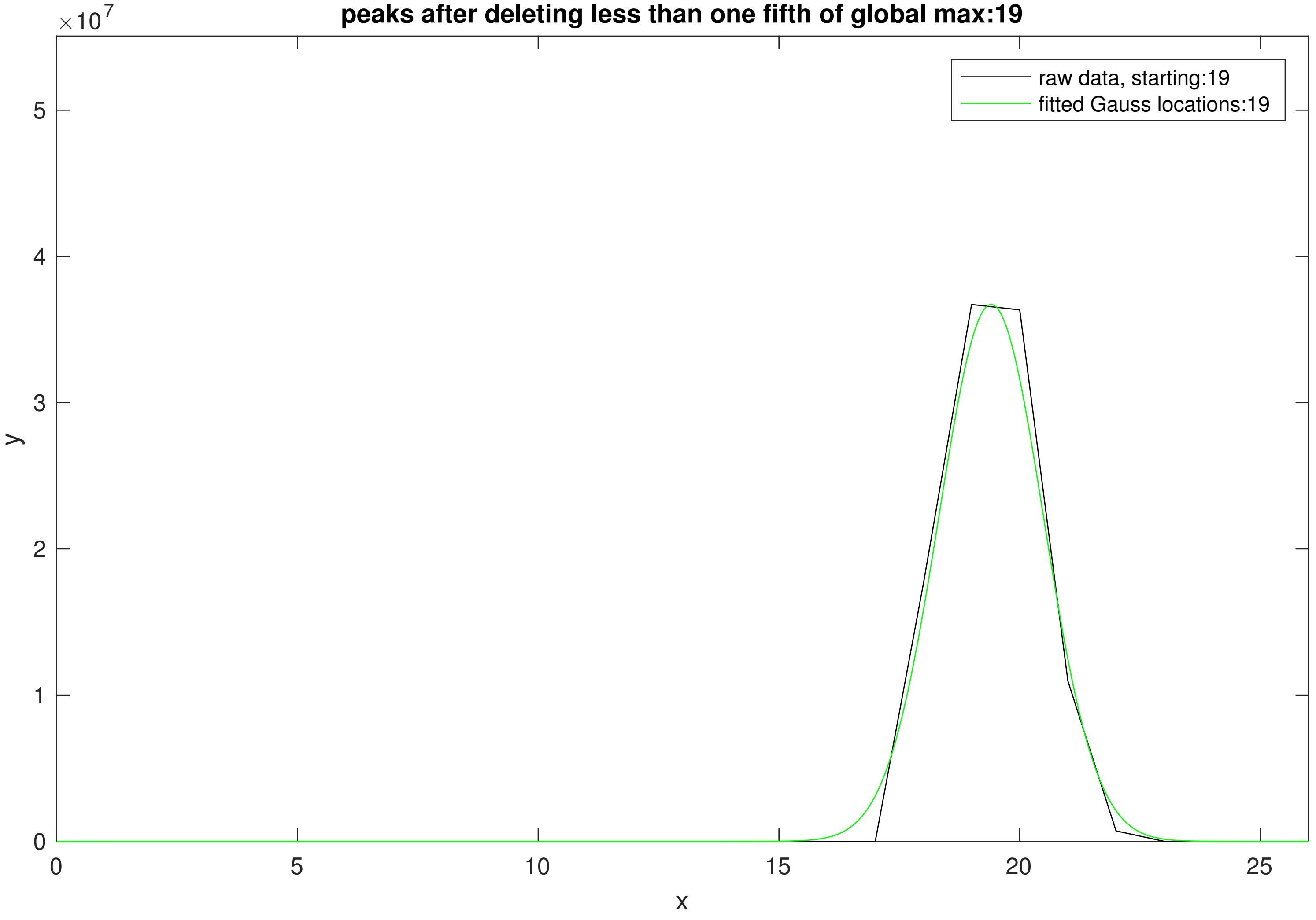

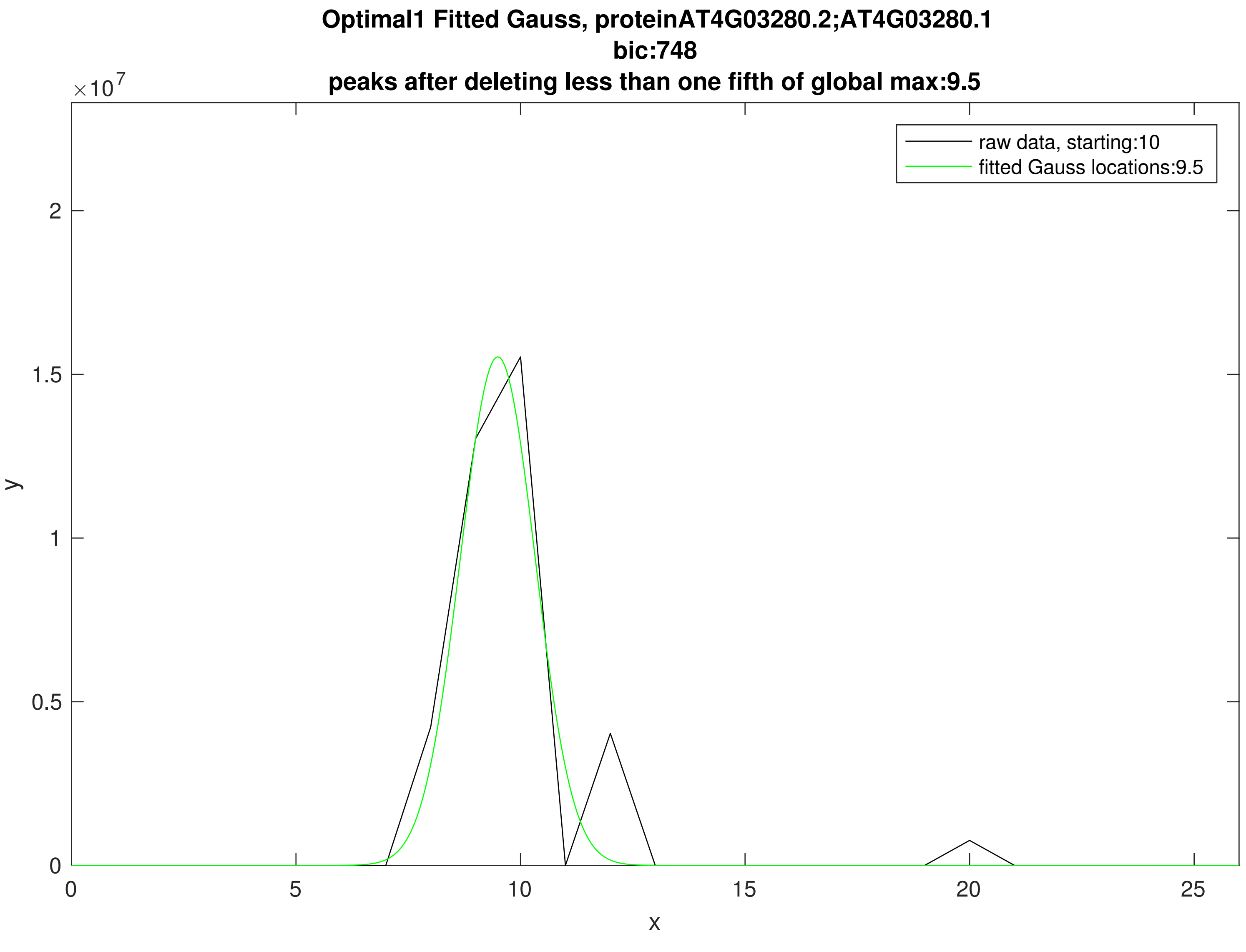
Optimal1 Fitted Gauss, proteinAT4G03280.2;AT4G03280.1
bic:748
peaks after deleting less than one fifth of global max:9.5
×10^7
raw data, starting:10
fitted Gauss locations:9.5
2
1.5
1
0.5
0
y
0
5
10
15
20
25
x

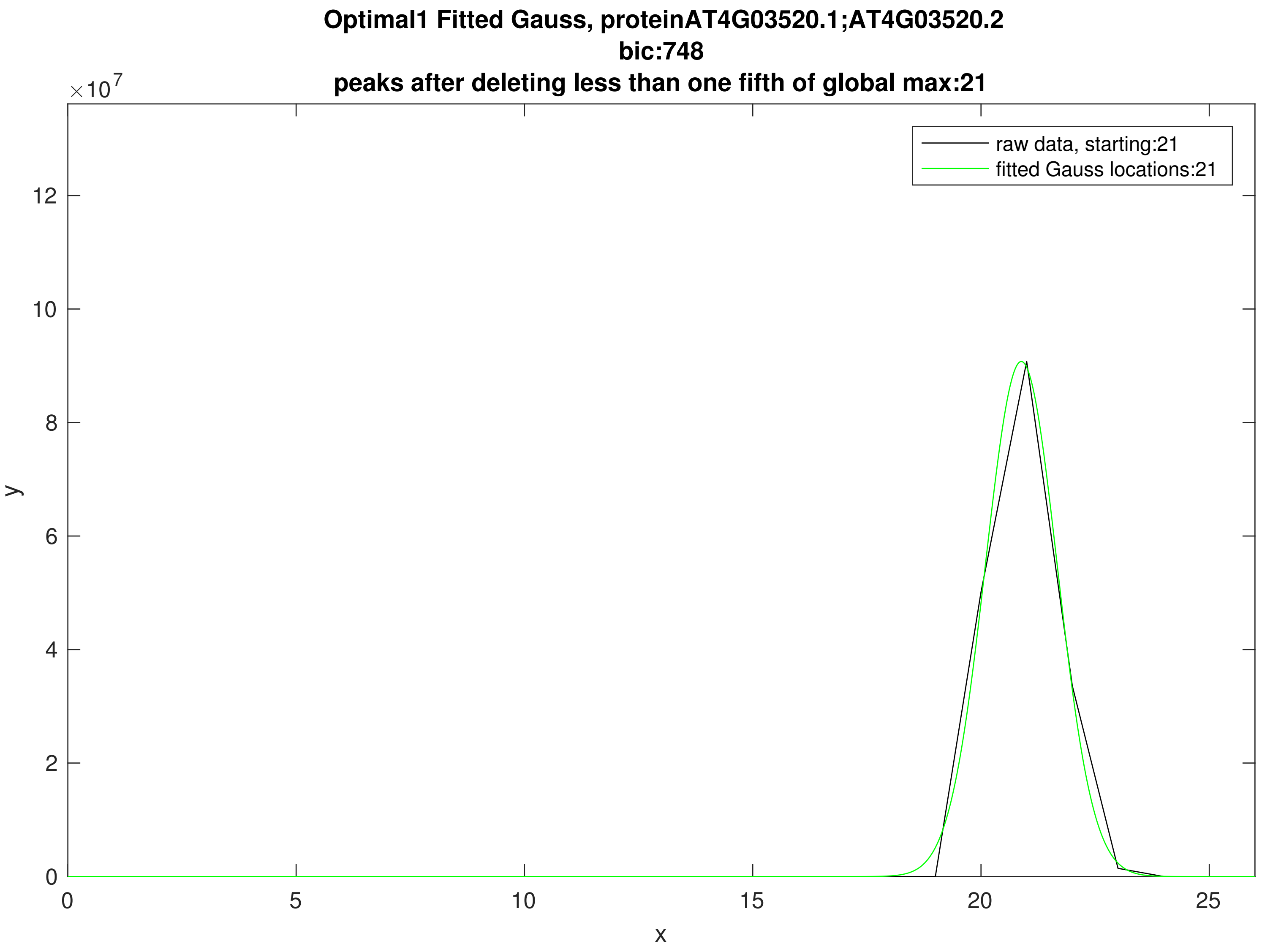
Optimal1 Fitted Gauss, proteinAT4G03520.1;AT4G03520.2
bic:748
peaks after deleting less than one fifth of global max:21
raw data, starting:21
fitted Gauss locations:21
y
x

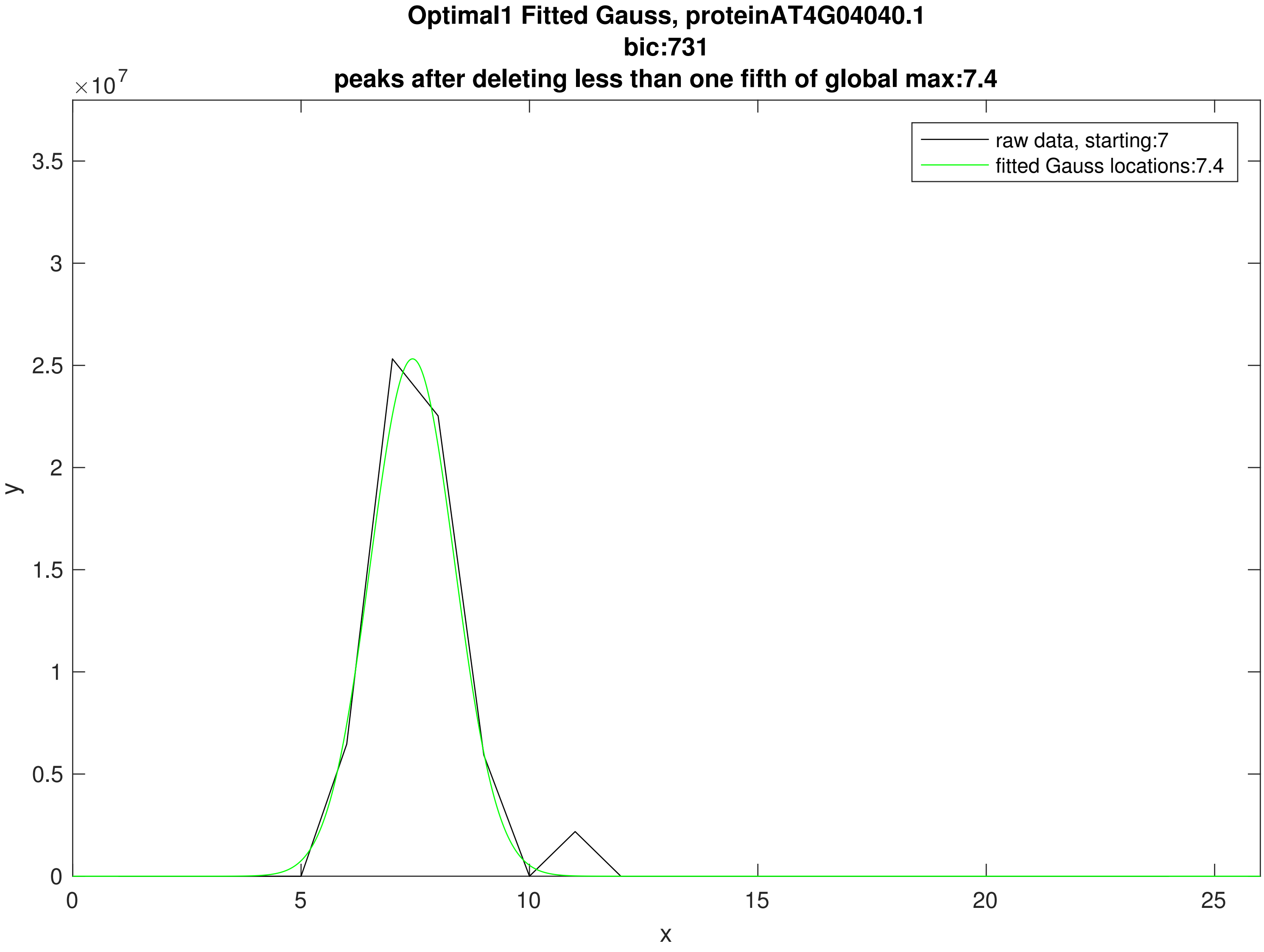
Optimal1 Fitted Gauss, proteinAT4G04040.1
bic:731
peaks after deleting less than one fifth of global max:7.4
×10^7
3.5
3
2.5
2
1.5
1
0.5
0
y
0
5
10
15
20
25
x
raw data, starting:7
fitted Gauss locations:7.4

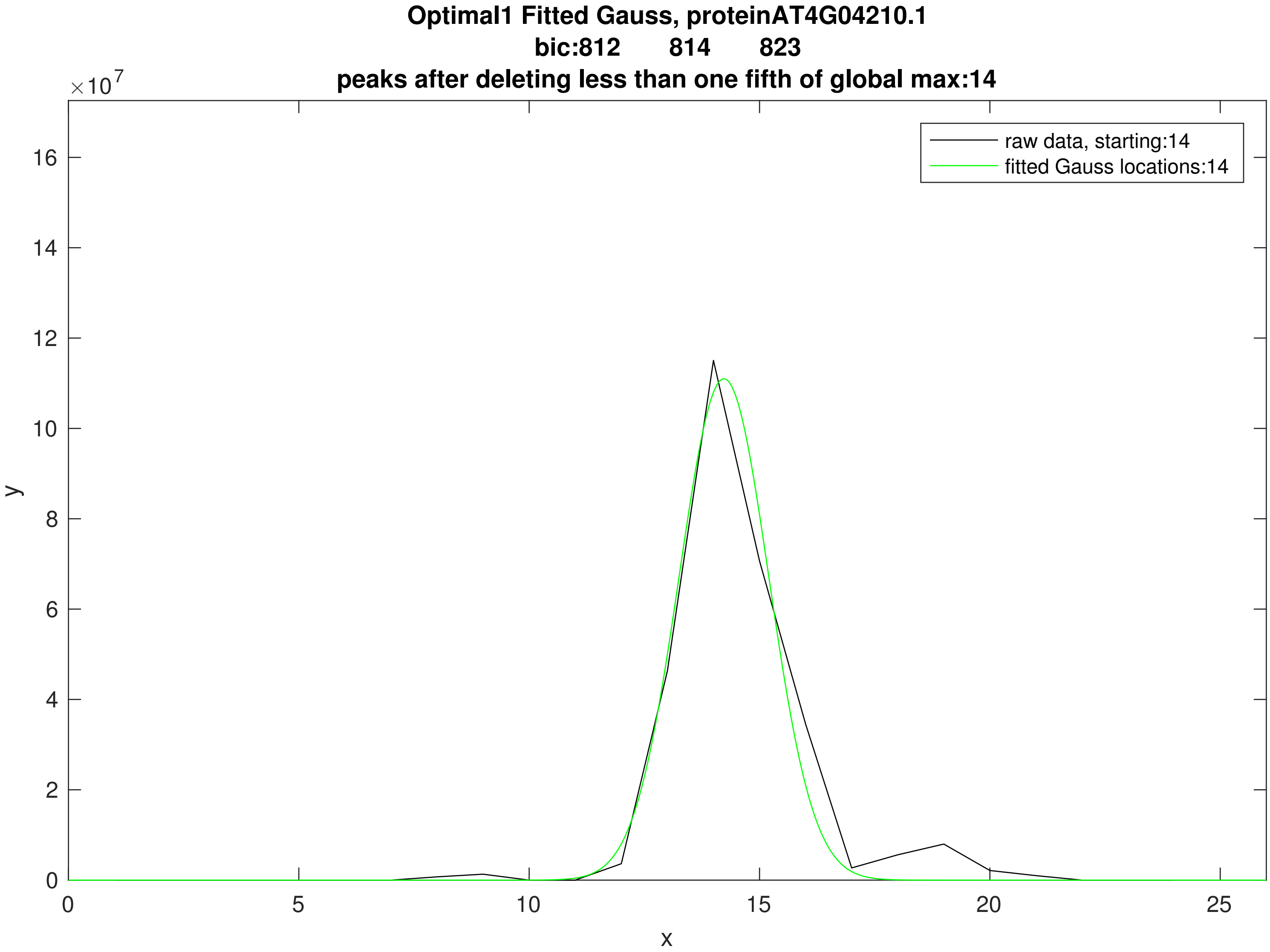
Optimal1 Fitted Gauss, proteinAT4G04210.1
bic:812 814 823
peaks after deleting less than one fifth of global max:14
×10^7
raw data, starting:14
fitted Gauss locations:14
y
x
0
2
4
6
8
10
12
14
16
0
5
10
15
20
25

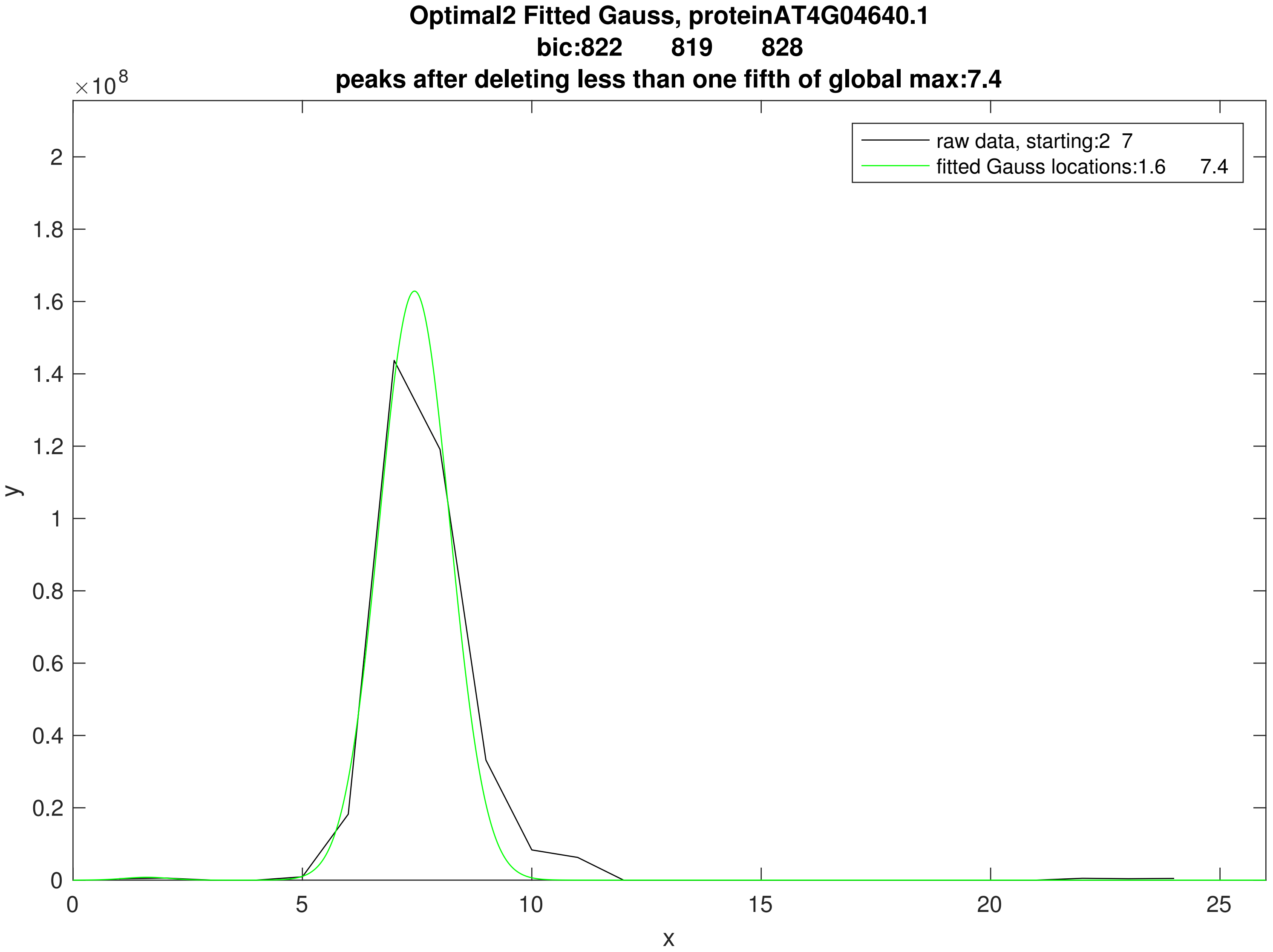
Optimal2 Fitted Gauss, proteinAT4G04640.1
bic:822 819 828
peaks after deleting less than one fifth of global max:7.4
×10^8
raw data, starting:2 7
fitted Gauss locations:1.6 7.4
y
x
0
0.2
0.4
0.6
0.8
1
1.2
1.4
1.6
1.8
2
5
10
15
20
25

#### Optimal1 Fitted Gauss, proteinAT4G04800.1
#### bic:691
#### peaks after deleting less than one fifth of global max:19

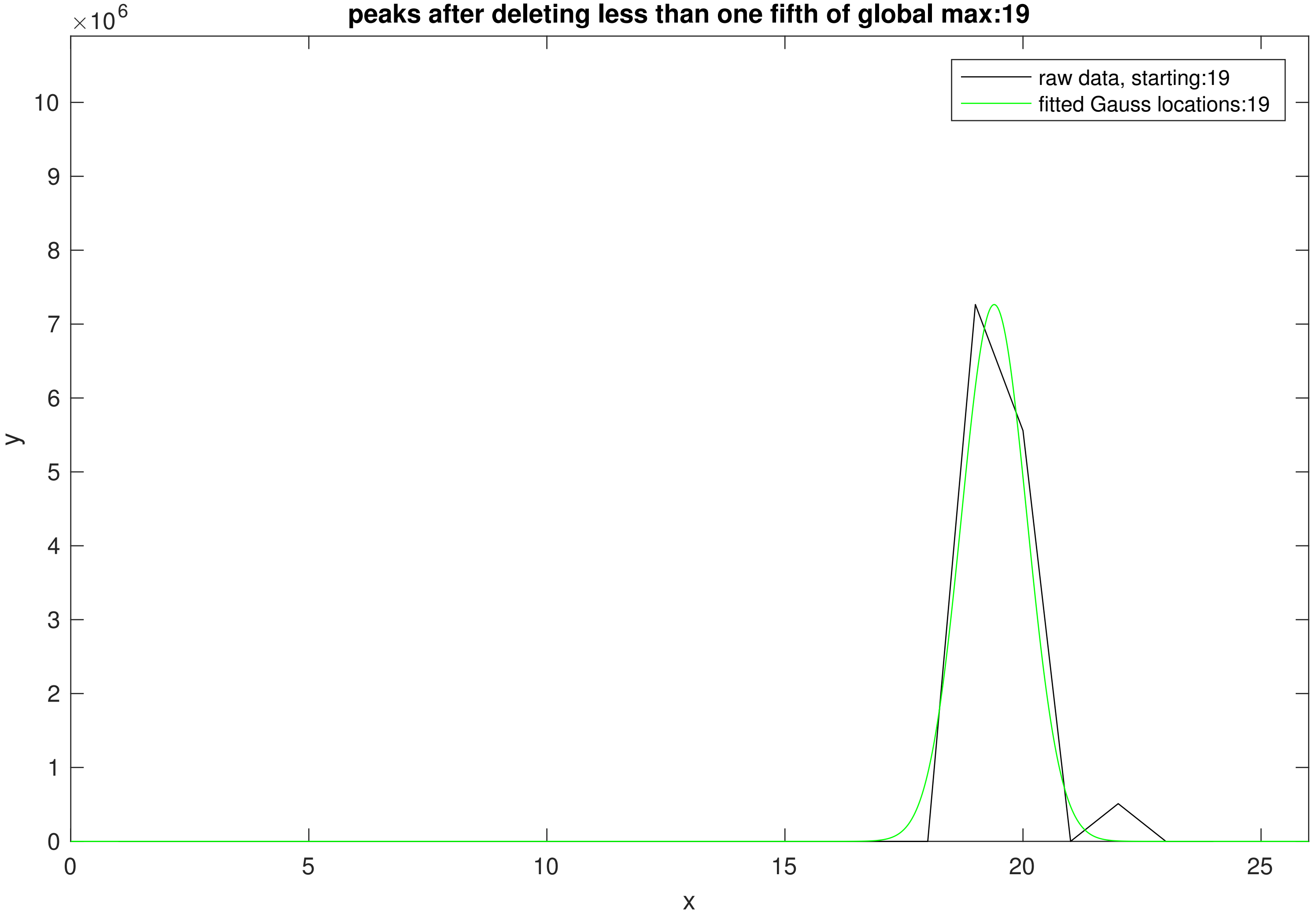

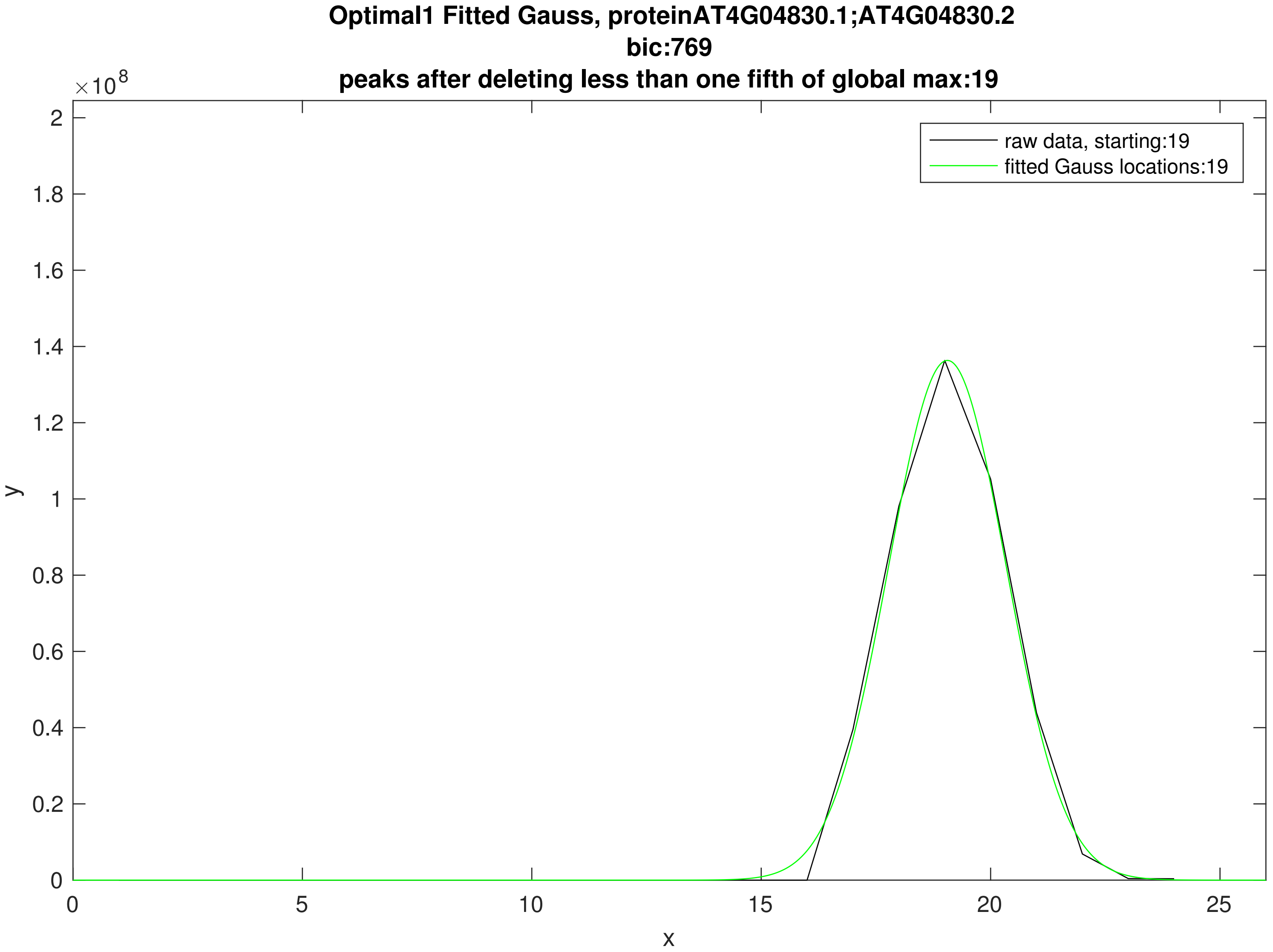
Optimal1 Fitted Gauss, proteinAT4G04830.1;AT4G04830.2
bic:769
peaks after deleting less than one fifth of global max:19
×10^8
raw data, starting:19
fitted Gauss locations:19
y
x

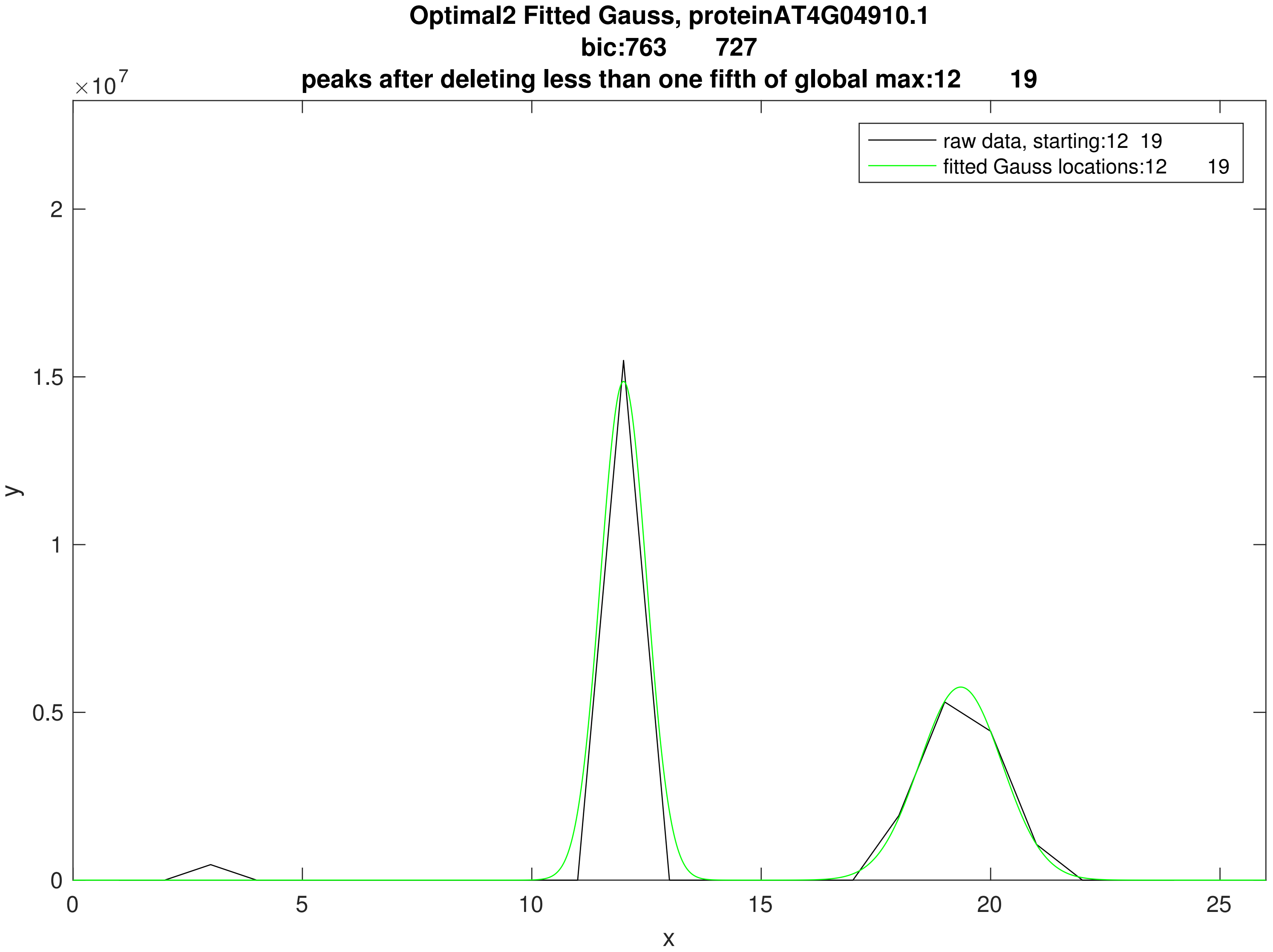

Optimal2 Fitted Gauss, proteinAT4G04910.1
bic:763 727
peaks after deleting less than one fifth of global max:12 19
×10^7
raw data, starting:12 19
fitted Gauss locations:12 19
y
x
0
0.5
1
1.5
2
0
5
10
15
20
25

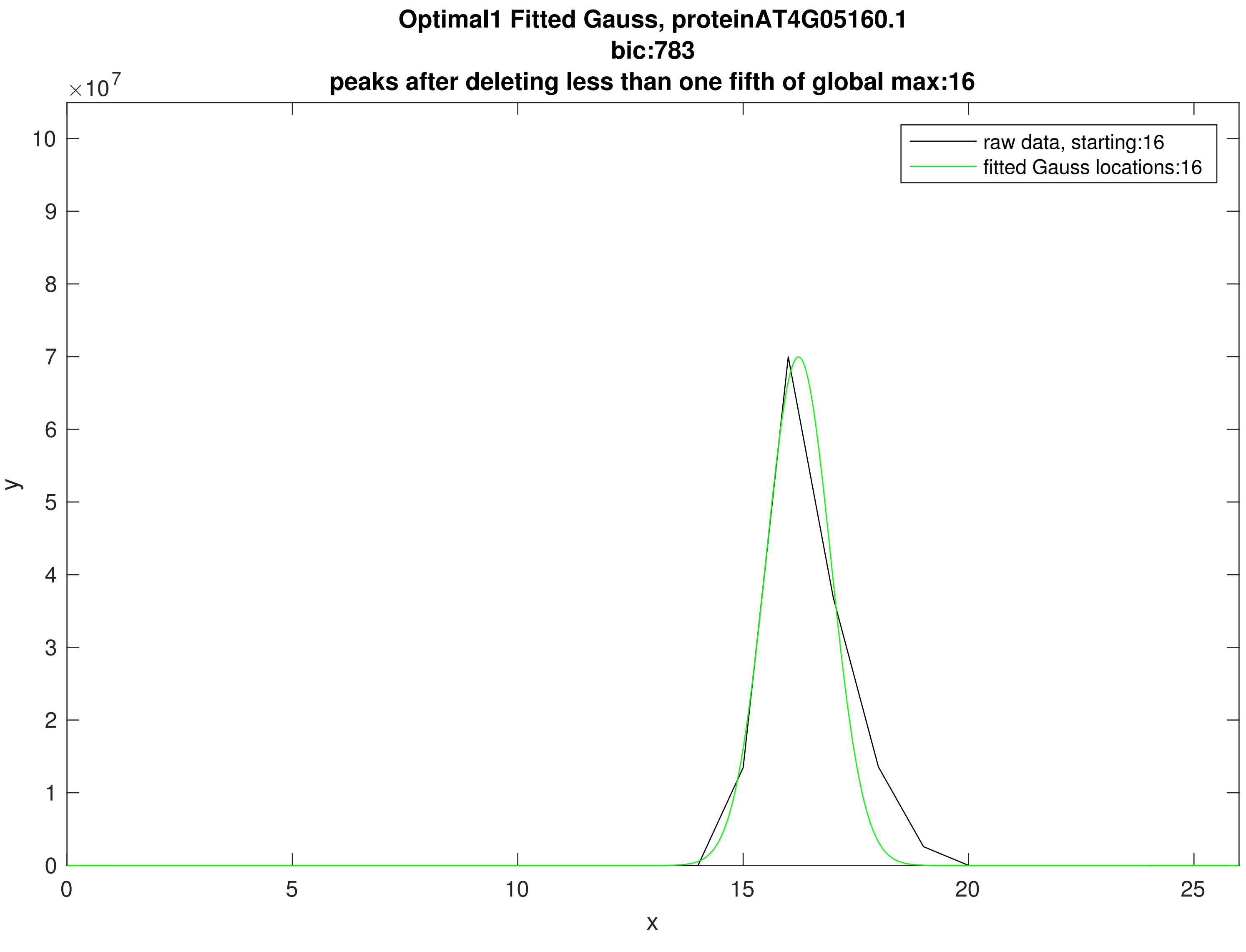
Optimal1 Fitted Gauss, proteinAT4G05160.1
bic:783
peaks after deleting less than one fifth of global max:16
×10^7
10
9
8
7
6
5
4
3
2
1
0
y
0
5
10
15
20
25
x
raw data, starting:16
fitted Gauss locations:16

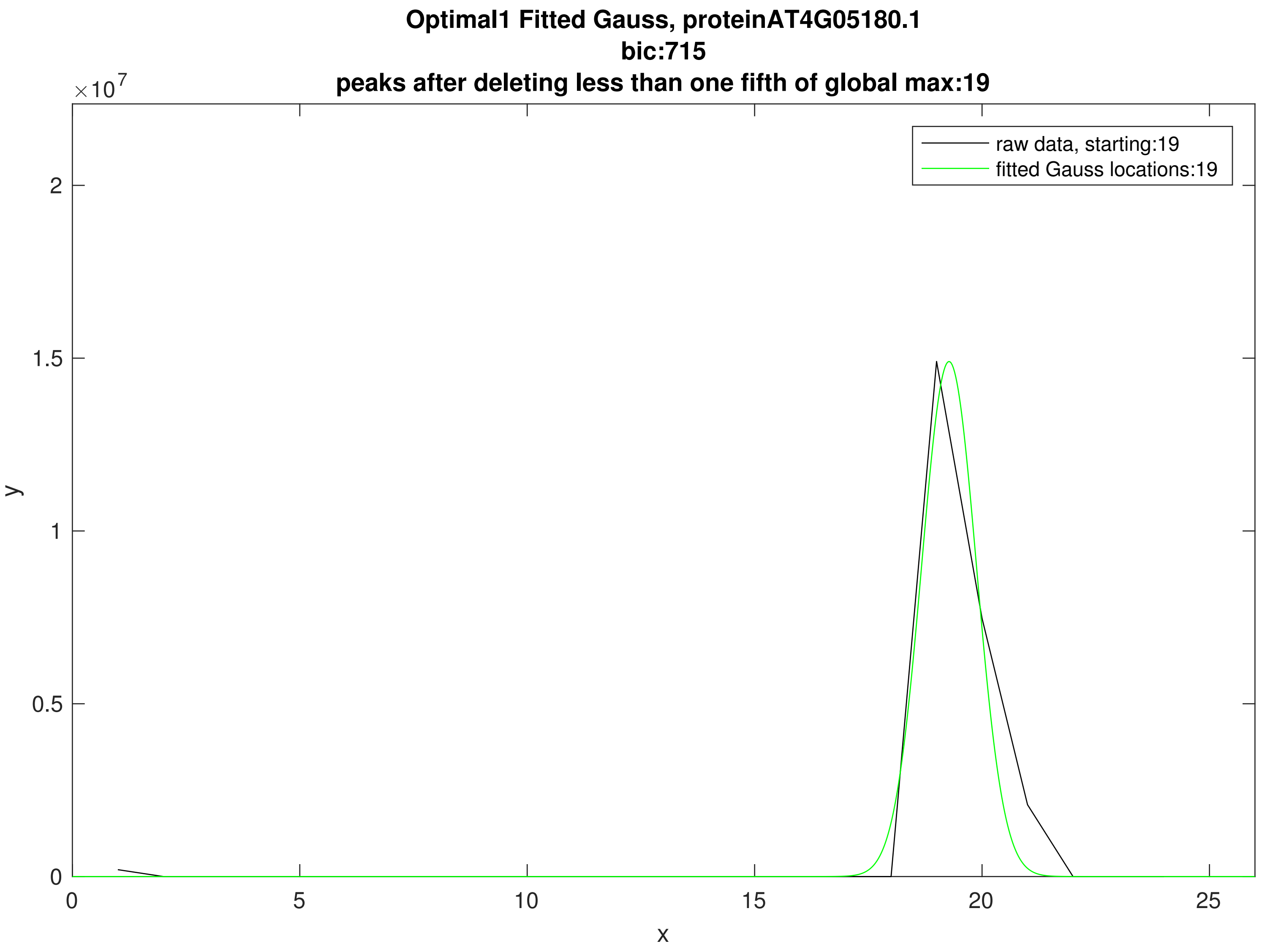

Optimal1 Fitted Gauss, proteinAT4G05180.1
bic:715
peaks after deleting less than one fifth of global max:19
×10^7
raw data, starting:19
fitted Gauss locations:19
2
1.5
1
0.5
0
y
0
5
10
15
20
25
x

Optimal1 Fitted Gauss, proteinAT4G05390.1;AT4G05390.2
bic:962 968
peaks after deleting less than one fifth of global max:16

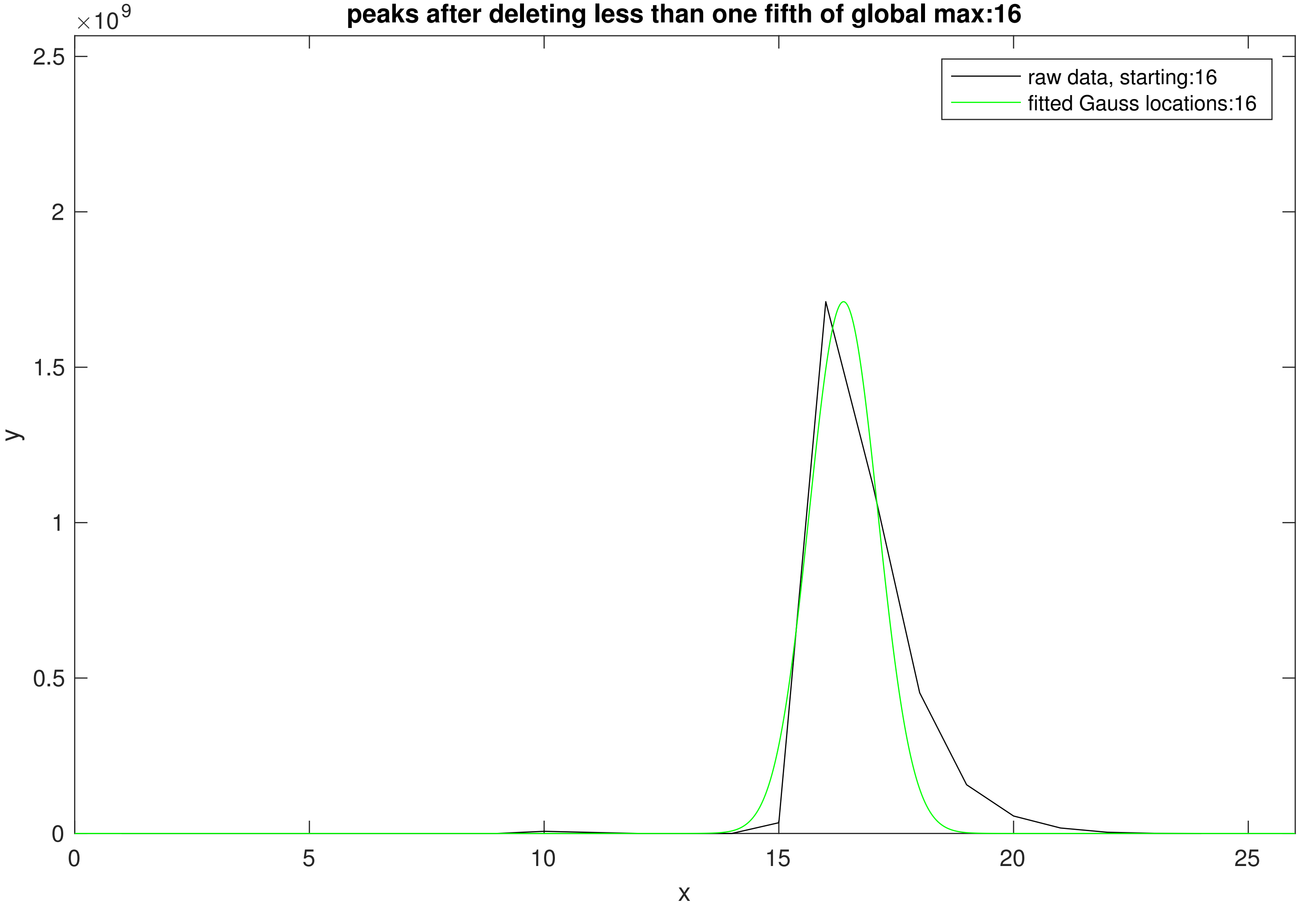

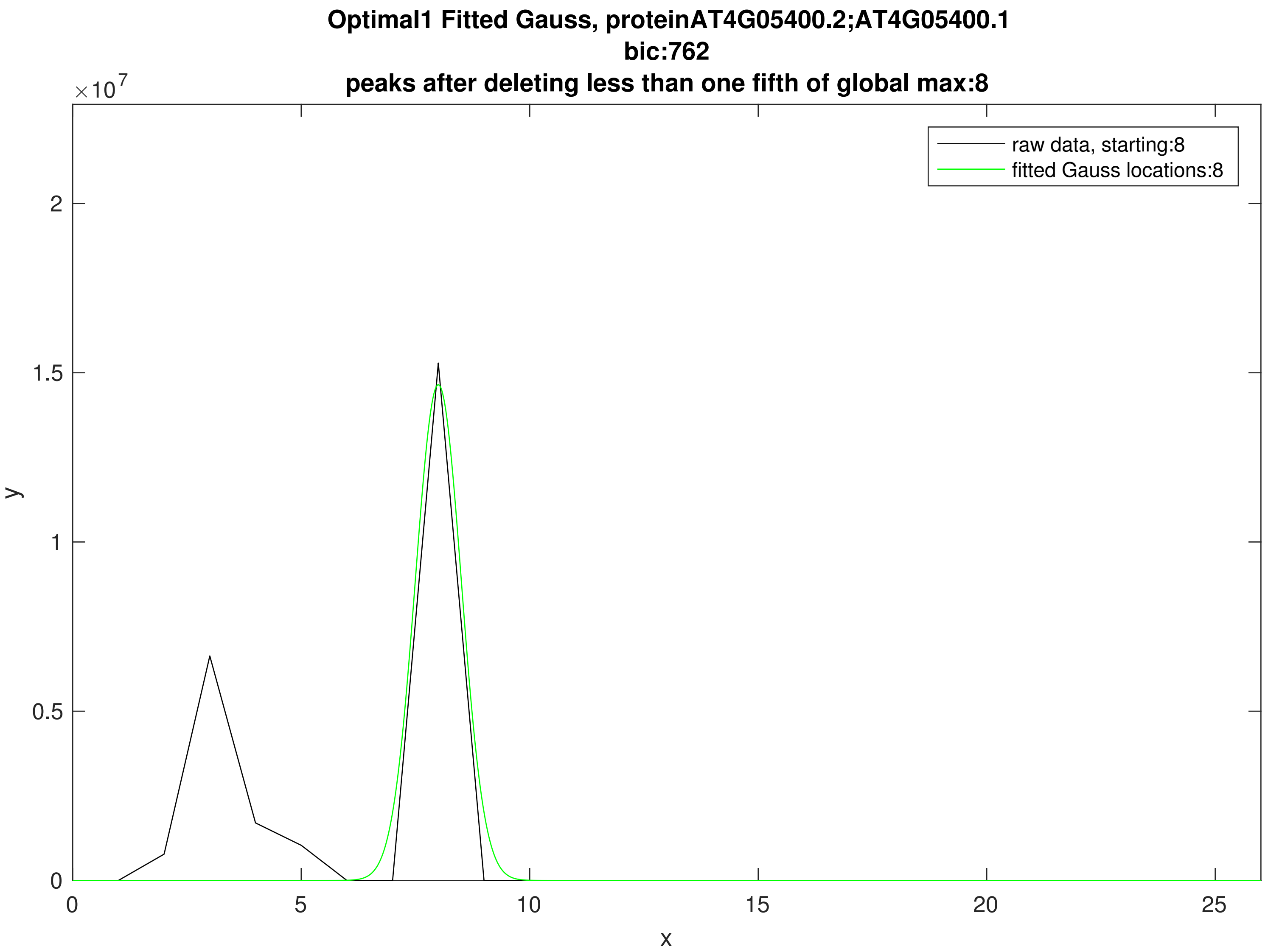
Optimal1 Fitted Gauss, proteinAT4G05400.2;AT4G05400.1
bic:762
peaks after deleting less than one fifth of global max:8
×10^7
raw data, starting:8
fitted Gauss locations:8
y
x
0
0.5
1
1.5
2
0
5
10
15
20
25

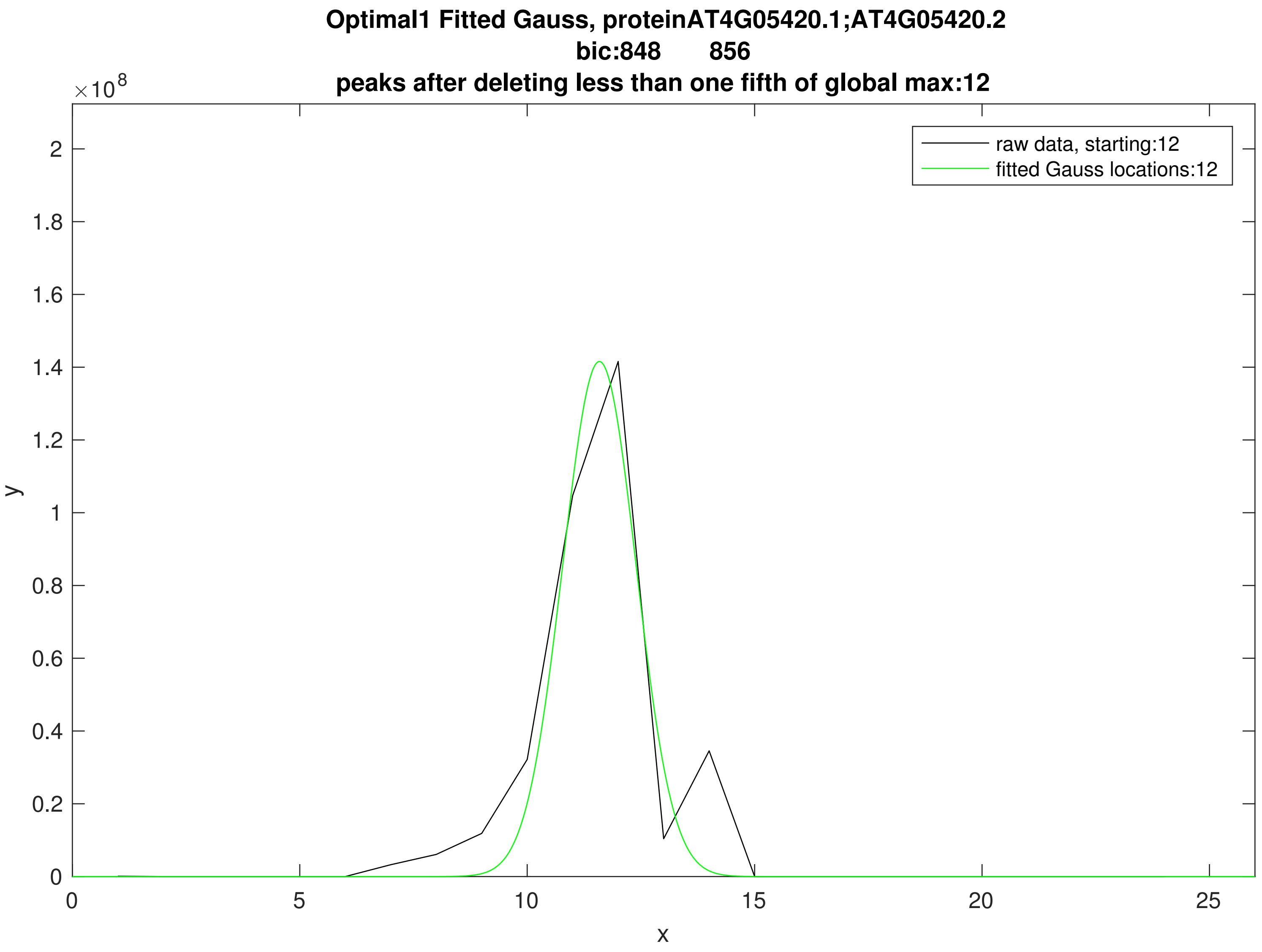

Optimal1 Fitted Gauss, proteinAT4G05420.1;AT4G05420.2
bic:848 856
peaks after deleting less than one fifth of global max:12
×10^8
raw data, starting:12
fitted Gauss locations:12
y
x
0
0.2
0.4
0.6
0.8
1
1.2
1.4
1.6
1.8
2
0
5
10
15
20
25

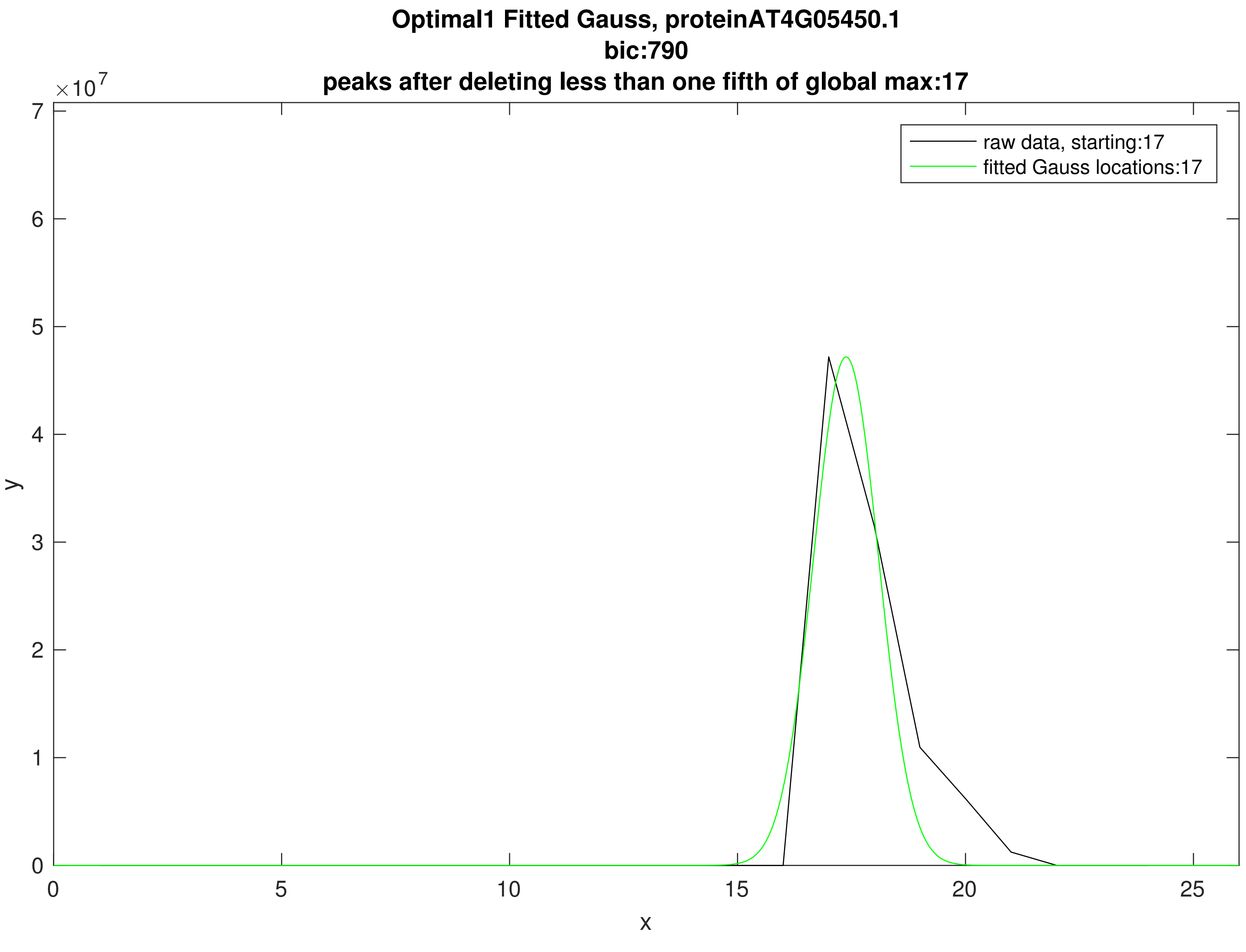

Optimal1 Fitted Gauss, proteinAT4G05450.1
bic:790
peaks after deleting less than one fifth of global max:17
×10^7
7
6
5
4
3
2
1
0
y
0
5
10
15
20
25
x
raw data, starting:17
fitted Gauss locations:17

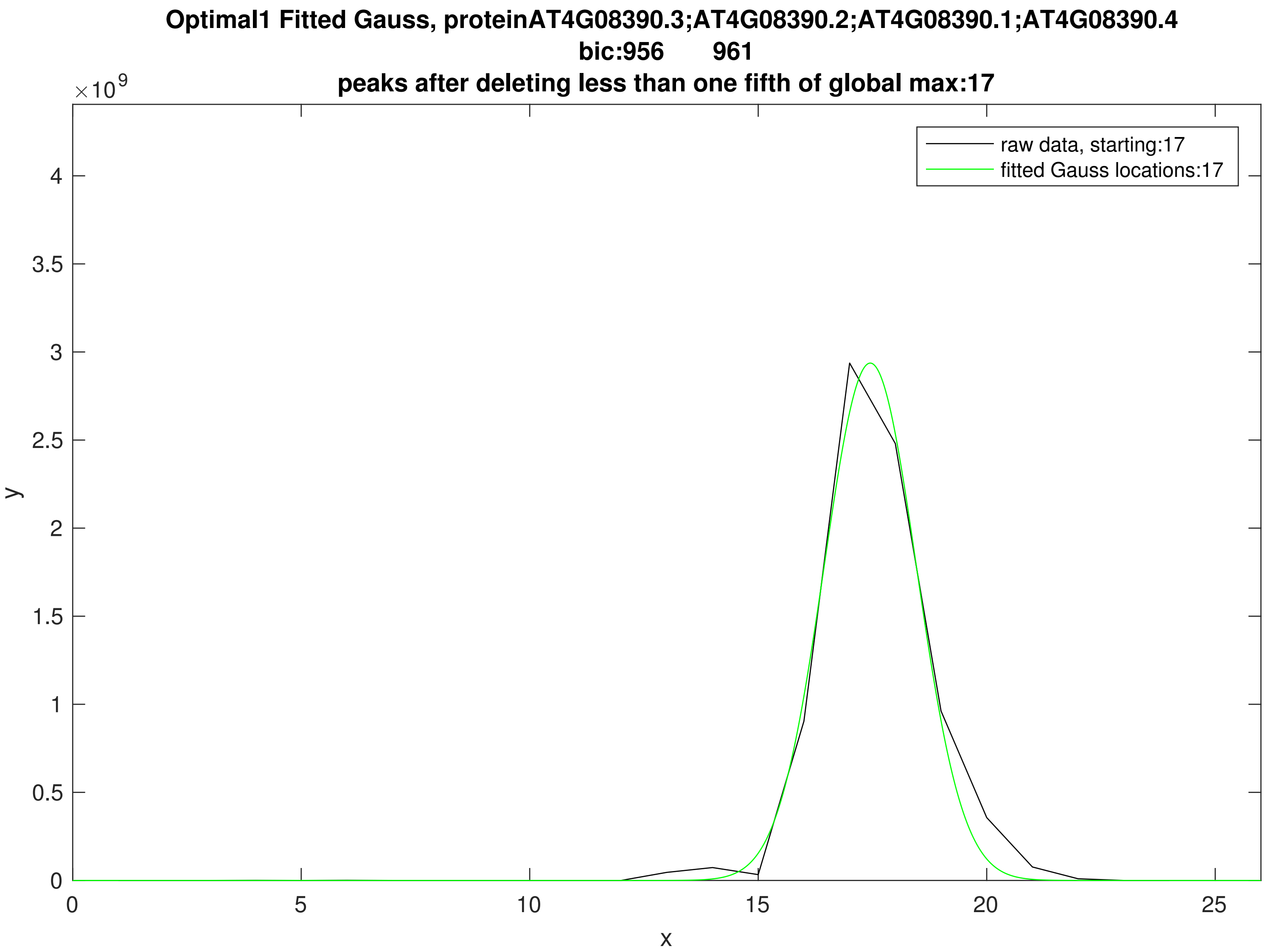
Optimal1 Fitted Gauss, proteinAT4G08390.3;AT4G08390.2;AT4G08390.1;AT4G08390.4
bic:956 961
peaks after deleting less than one fifth of global max:17
×10^9
raw data, starting:17
fitted Gauss locations:17
y
x
0
0.5
1
1.5
2
2.5
3
3.5
4
5
10
15
20
25

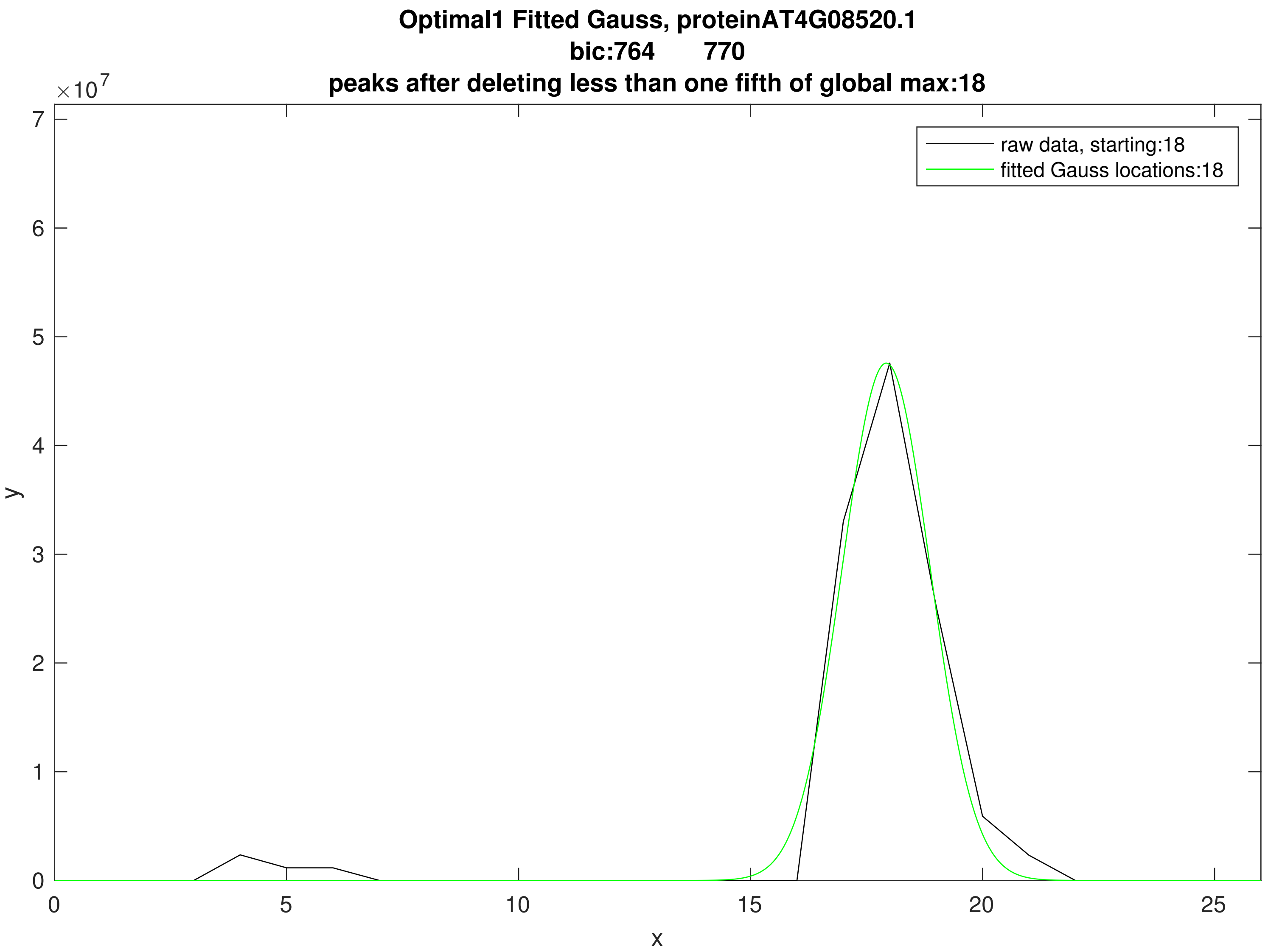
Optimal1 Fitted Gauss, proteinAT4G08520.1
bic:764 770
peaks after deleting less than one fifth of global max:18
×10^7
raw data, starting:18
fitted Gauss locations:18
y
x
0
1
2
3
4
5
6
7
0
5
10
15
20
25

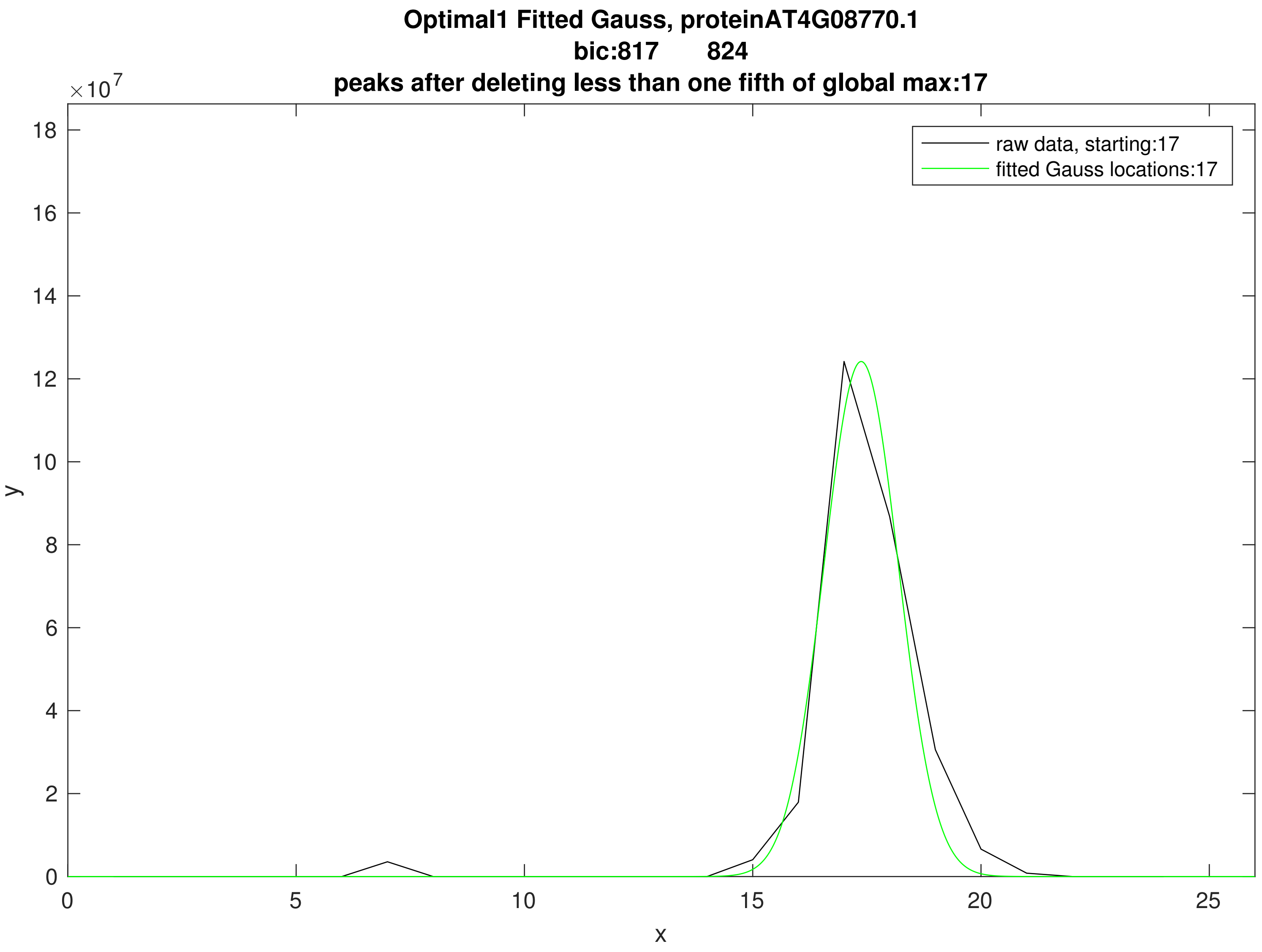
Optimal1 Fitted Gauss, proteinAT4G08770.1
bic:817 824
peaks after deleting less than one fifth of global max:17
×10^7
raw data, starting:17
fitted Gauss locations:17
y
x
0
2
4
6
8
10
12
14
16
18
0
5
10
15
20
25

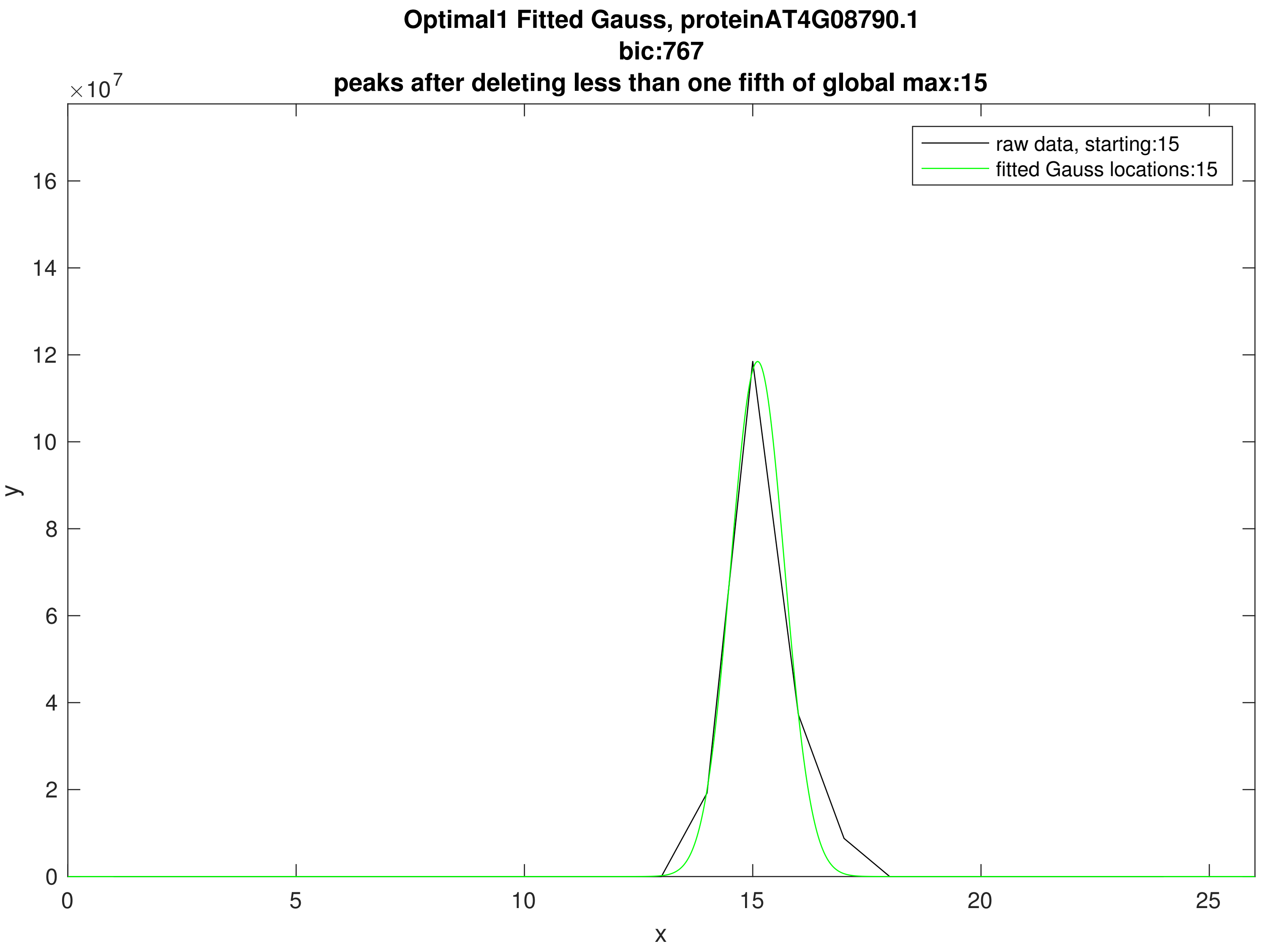
Optimal1 Fitted Gauss, proteinAT4G08790.1
bic:767
peaks after deleting less than one fifth of global max:15
×10^7
raw data, starting:15
fitted Gauss locations:15
y
x
0
2
4
6
8
10
12
14
16
0
5
10
15
20
25

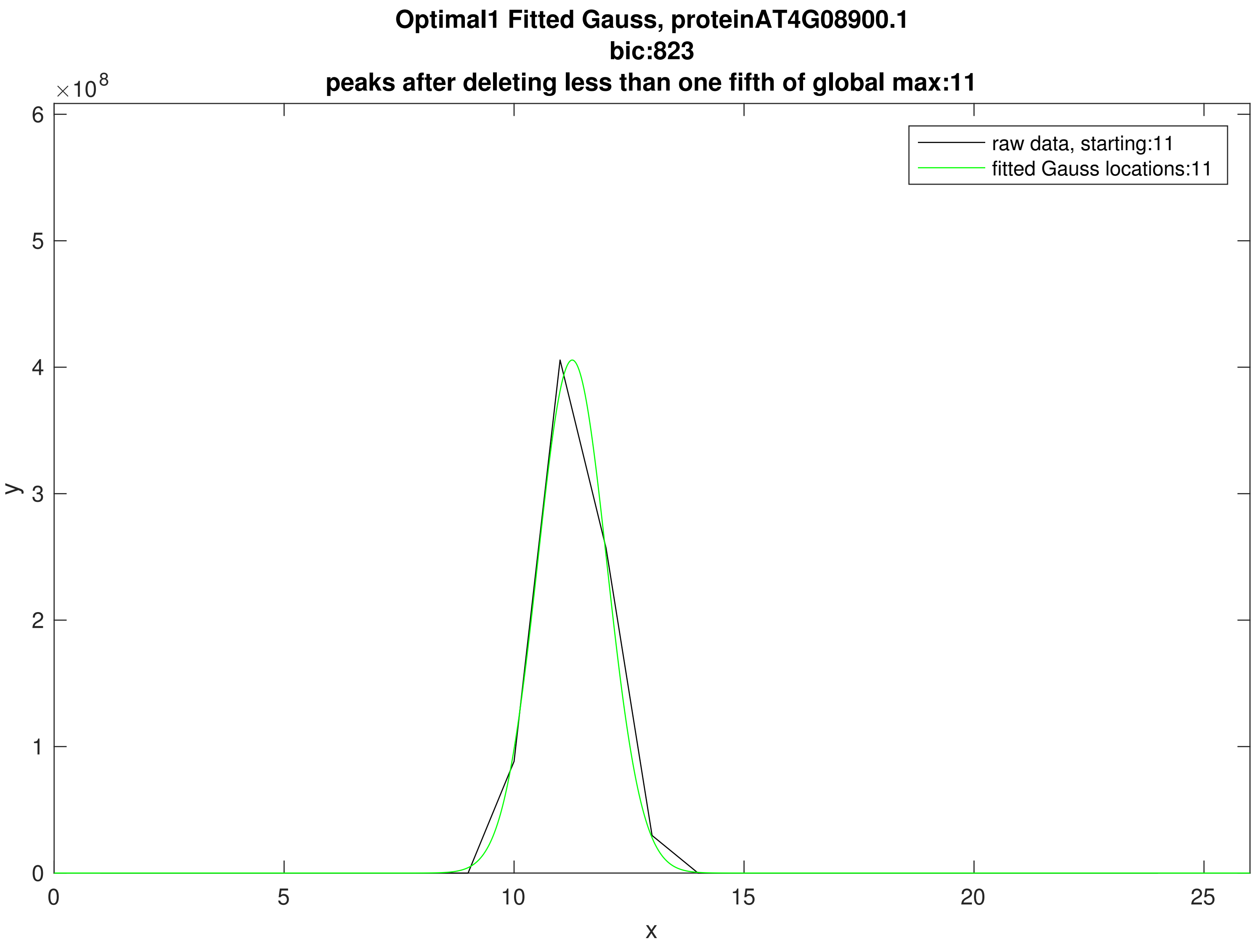

Optimal1 Fitted Gauss, proteinAT4G08900.1
bic:823
peaks after deleting less than one fifth of global max:11
×10^8
6
5
4
3
2
1
0
y
0
5
10
15
20
25
x
raw data, starting:11
fitted Gauss locations:11

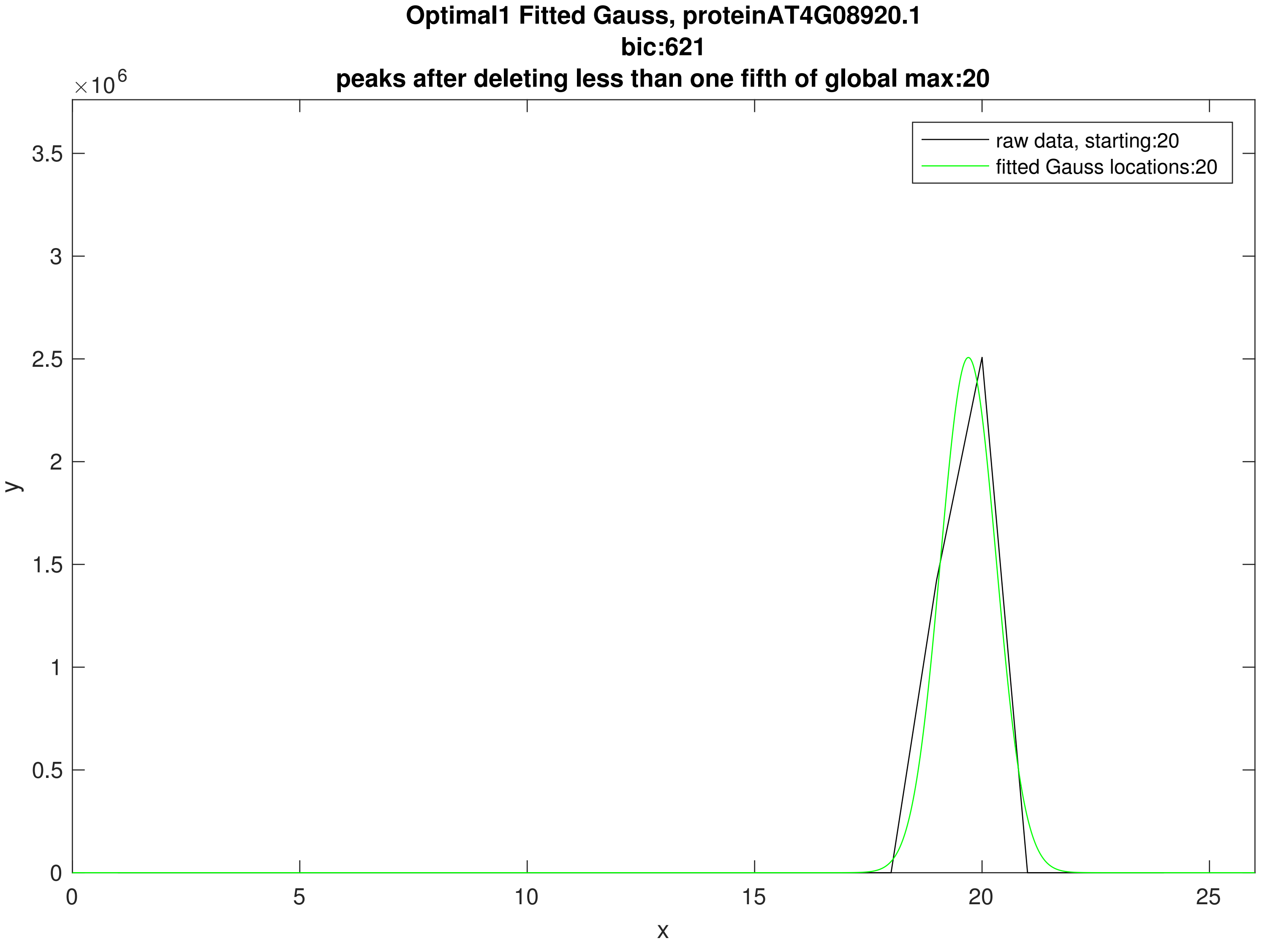
Optimal1 Fitted Gauss, proteinAT4G08920.1
bic:621
peaks after deleting less than one fifth of global max:20
×10^6
raw data, starting:20
fitted Gauss locations:20
3.5
3
2.5
2
1.5
1
0.5
0
y
0
5
10
15
20
25
x

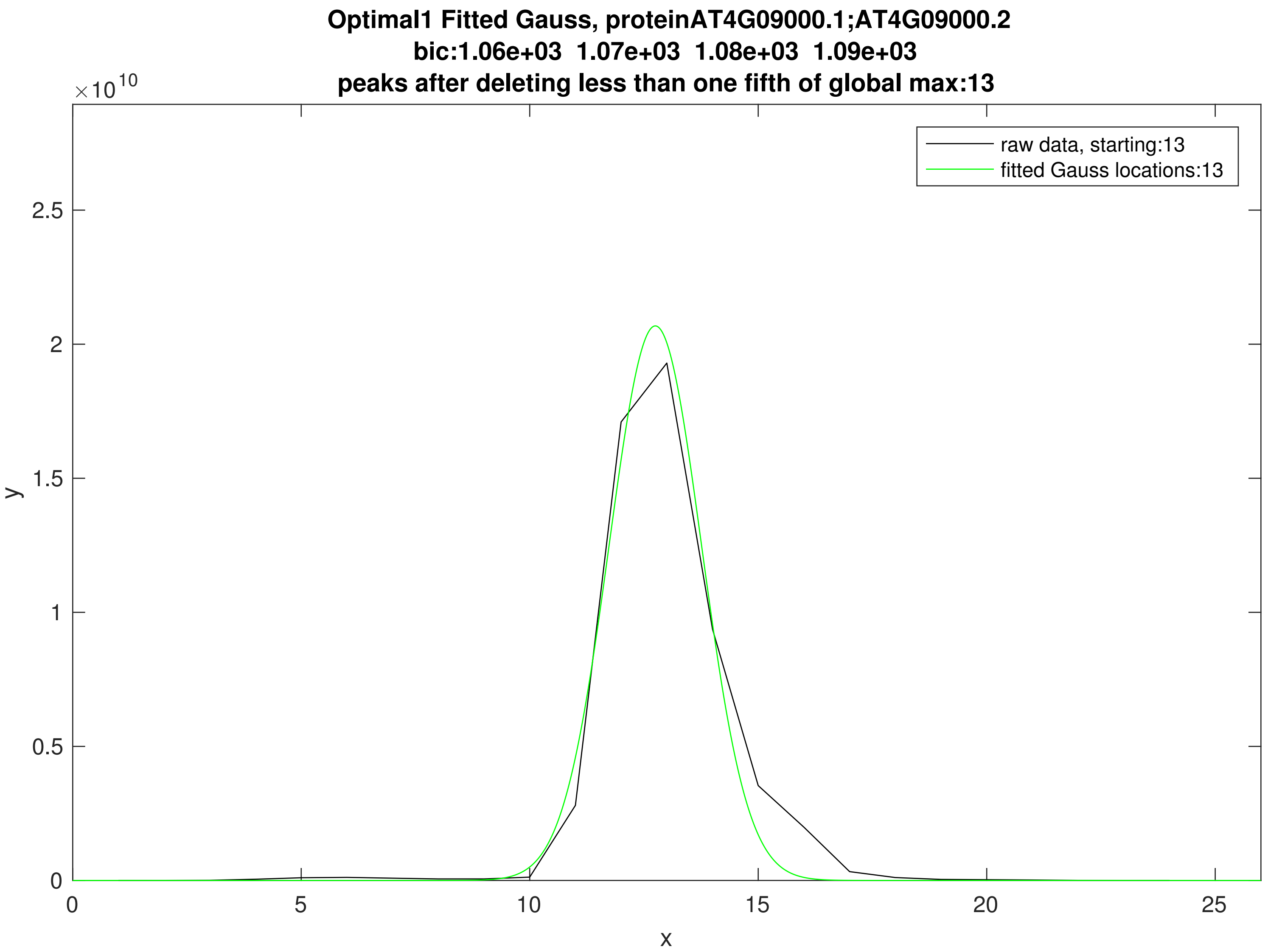
Optimal1 Fitted Gauss, proteinAT4G09000.1;AT4G09000.2
bic:1.06e+03  1.07e+03  1.08e+03  1.09e+03
peaks after deleting less than one fifth of global max:13
×10^10
raw data, starting:13
fitted Gauss locations:13
y
x
0
0.5
1
1.5
2
2.5
5
10
15
20
25

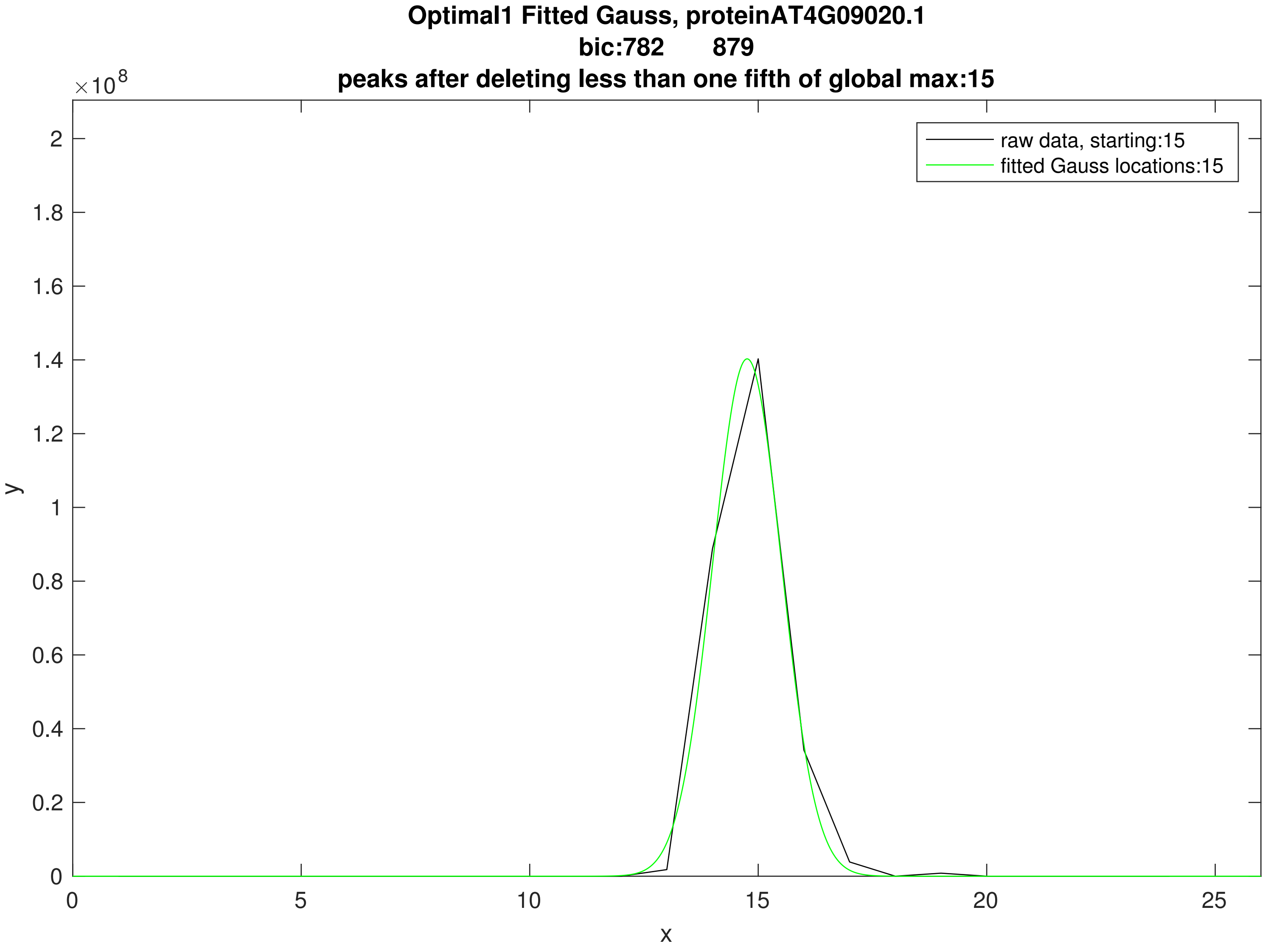

Optimal1 Fitted Gauss, proteinAT4G09020.1
bic:782 879
peaks after deleting less than one fifth of global max:15
×10^8
raw data, starting:15
fitted Gauss locations:15
y
x
0
0.2
0.4
0.6
0.8
1
1.2
1.4
1.6
1.8
2
5
10
15
20
25

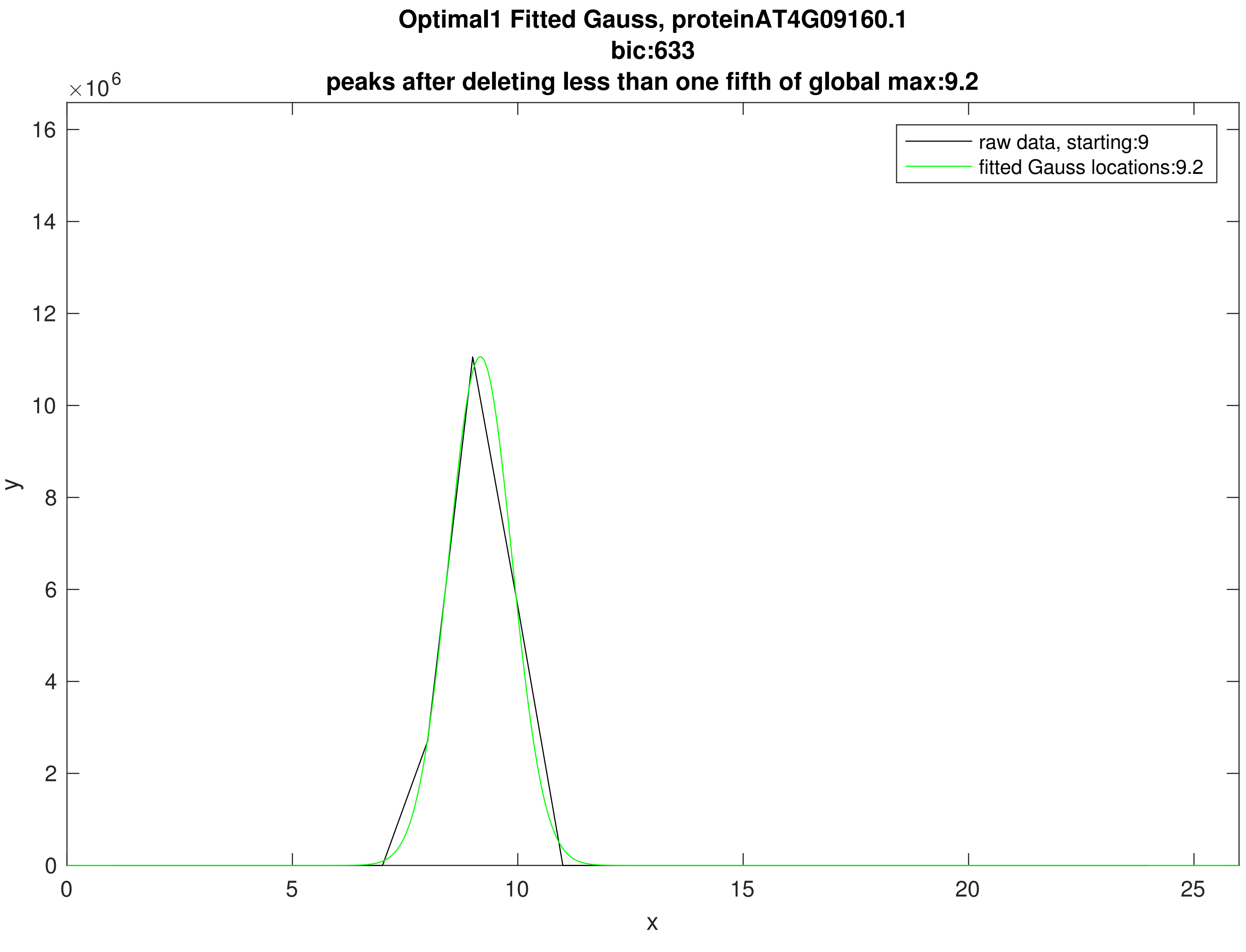

Optimal1 Fitted Gauss, proteinAT4G09160.1
bic:633
peaks after deleting less than one fifth of global max:9.2
×10^6
raw data, starting:9
fitted Gauss locations:9.2
y
x
0
2
4
6
8
10
12
14
16
0
5
10
15
20
25

#### Optimal2 Fitted Gauss, proteinAT4G09320.1
#### bic:992 969 978 988
#### peaks after deleting less than one fifth of global max:16

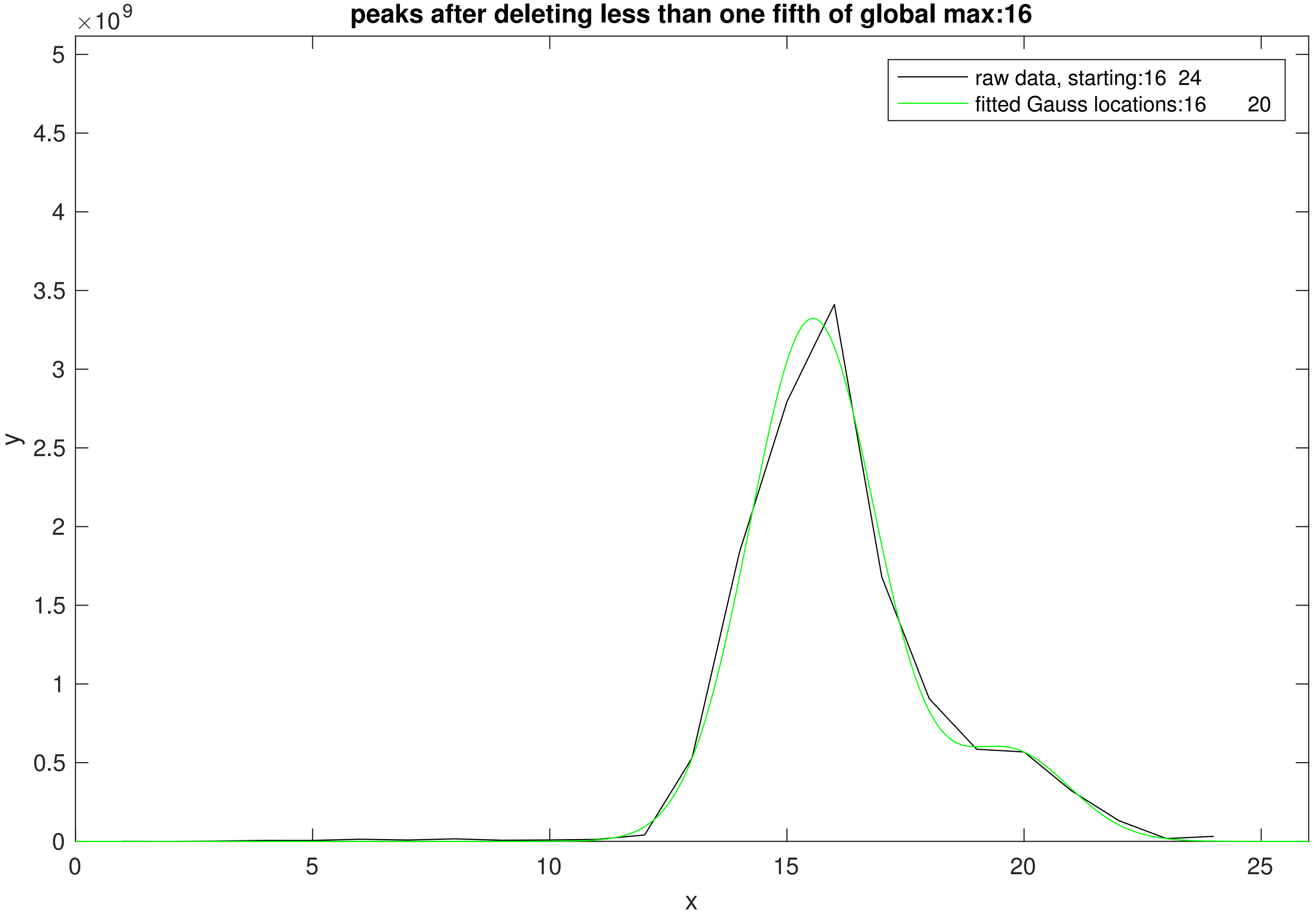

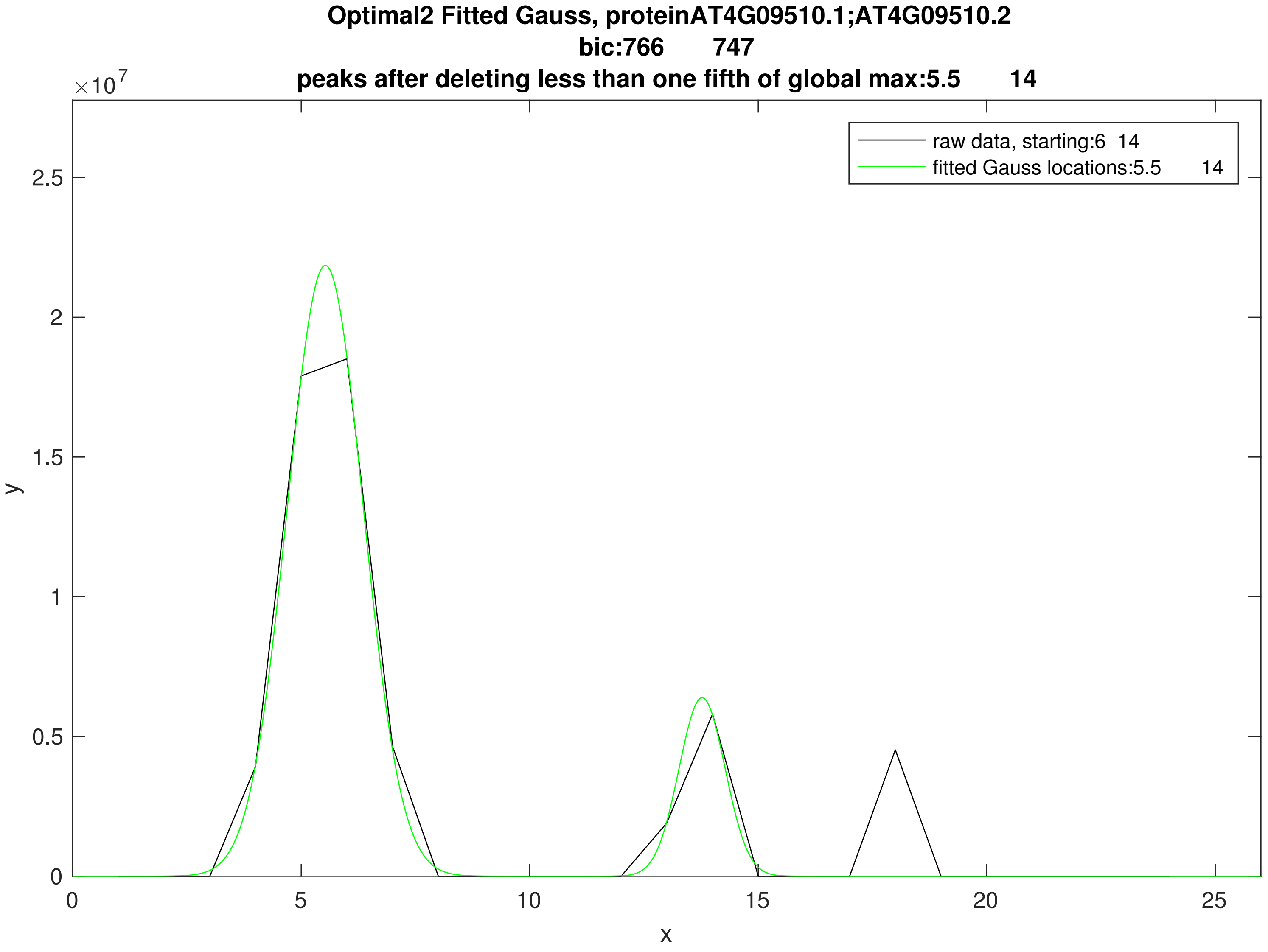
Optimal2 Fitted Gauss, proteinAT4G09510.1;AT4G09510.2
bic:766 747
peaks after deleting less than one fifth of global max:5.5 14
×10^7
raw data, starting:6 14
fitted Gauss locations:5.5 14
y
x
0
0.5
1
1.5
2
2.5
5
10
15
20
25

**Optimal1 Fitted Gauss, proteinAT4G09520.1**
**bic:863**
**peaks after deleting less than one fifth of global max:13**

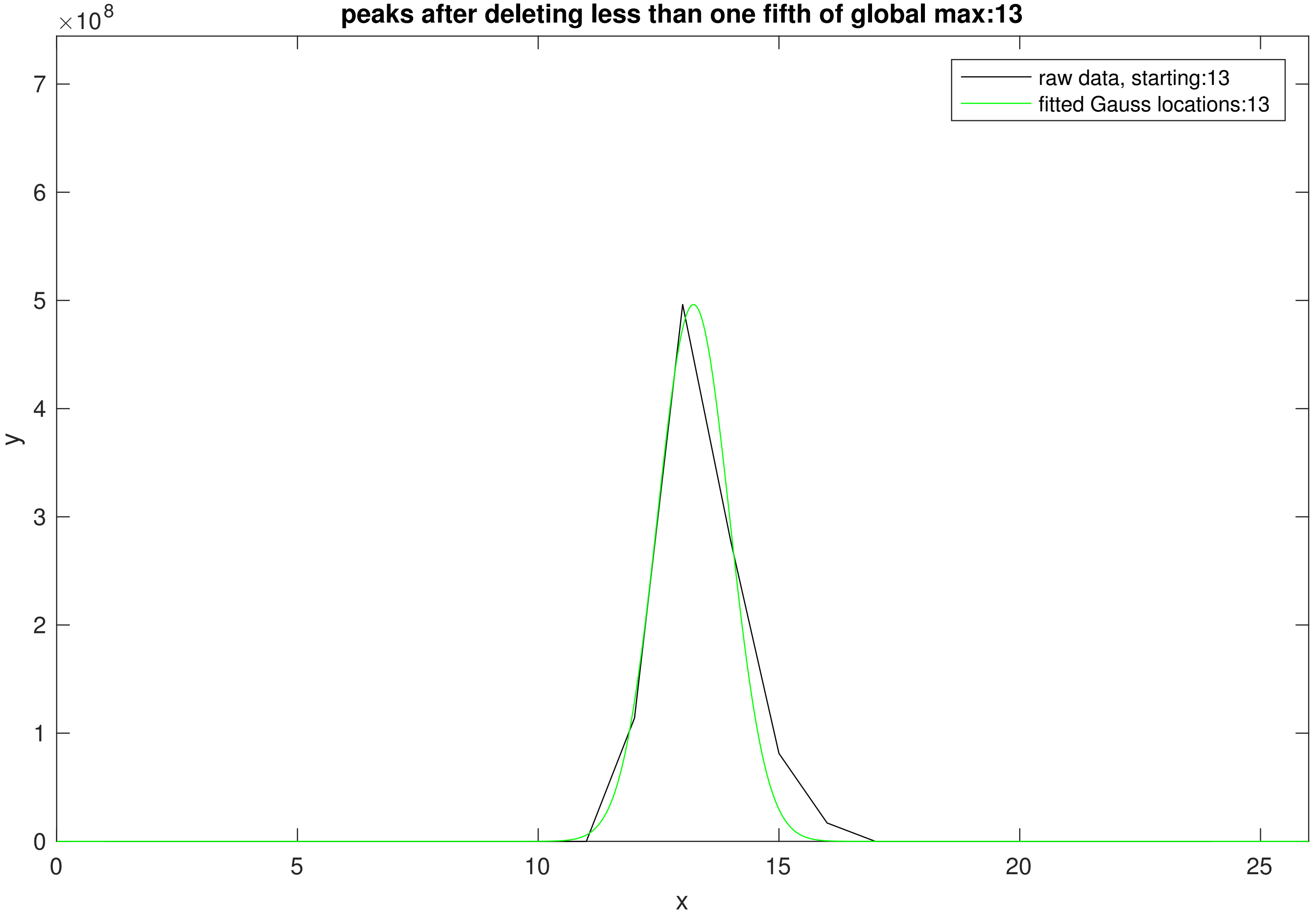

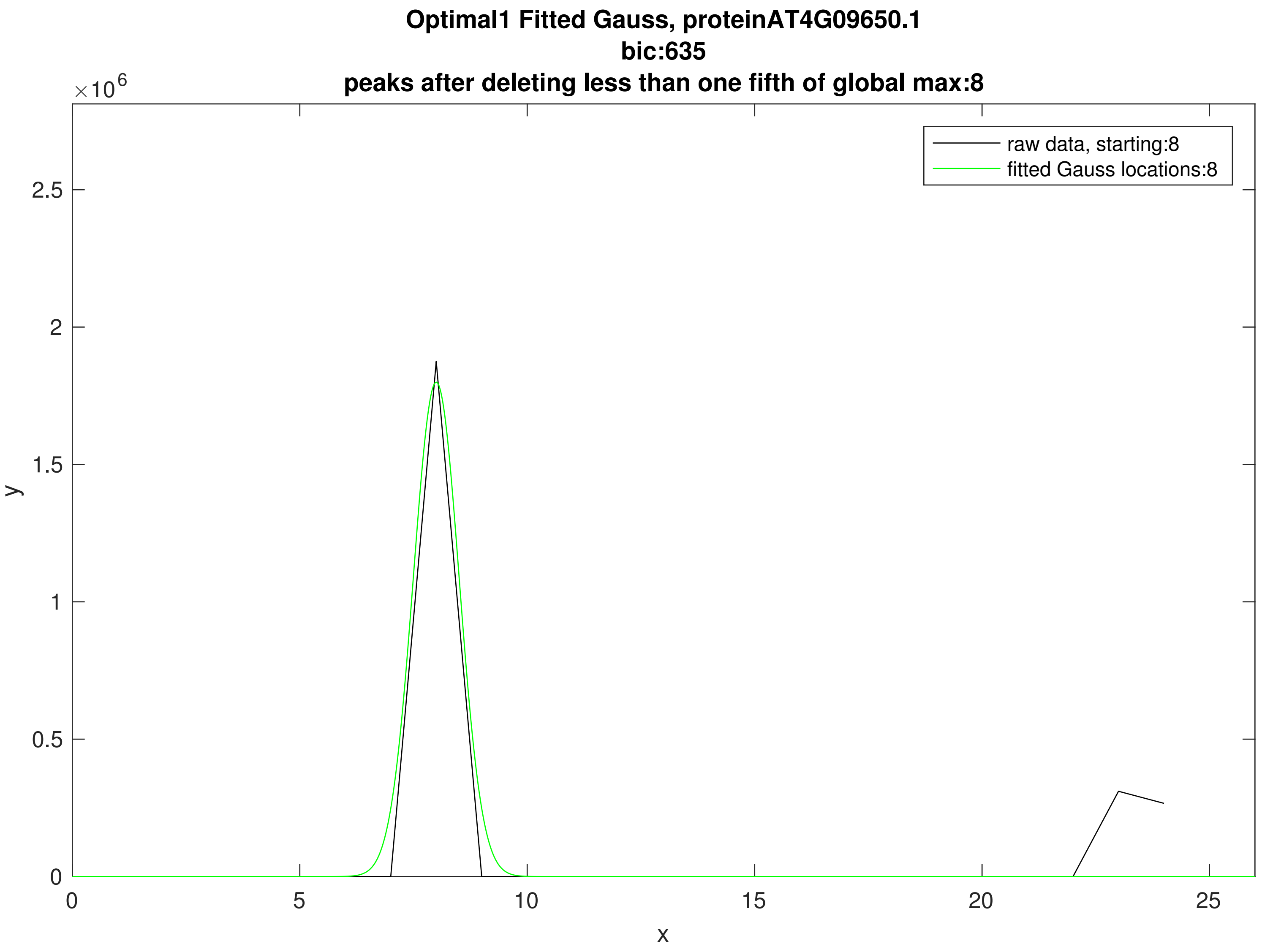

Optimal1 Fitted Gauss, proteinAT4G09650.1
bic:635
peaks after deleting less than one fifth of global max:8
×10^6
raw data, starting:8
fitted Gauss locations:8
2.5
2
1.5
1
0.5
0
y
0
5
10
15
20
25
x

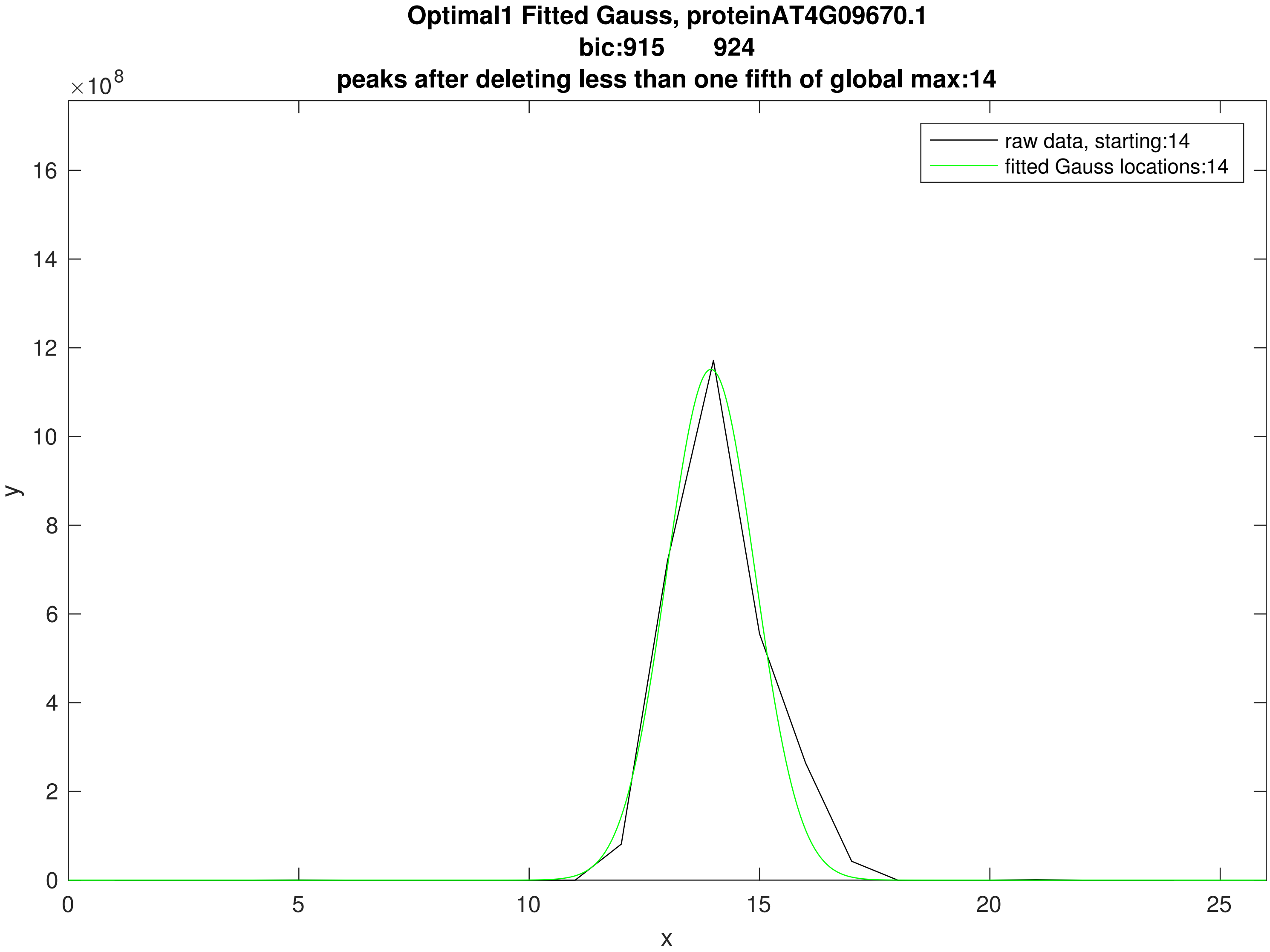

Optimal1 Fitted Gauss, proteinAT4G09670.1
bic:915 924
peaks after deleting less than one fifth of global max:14
×10^8
raw data, starting:14
fitted Gauss locations:14
16
14
12
10
8
6
4
2
0
y
0
5
10
15
20
25
x

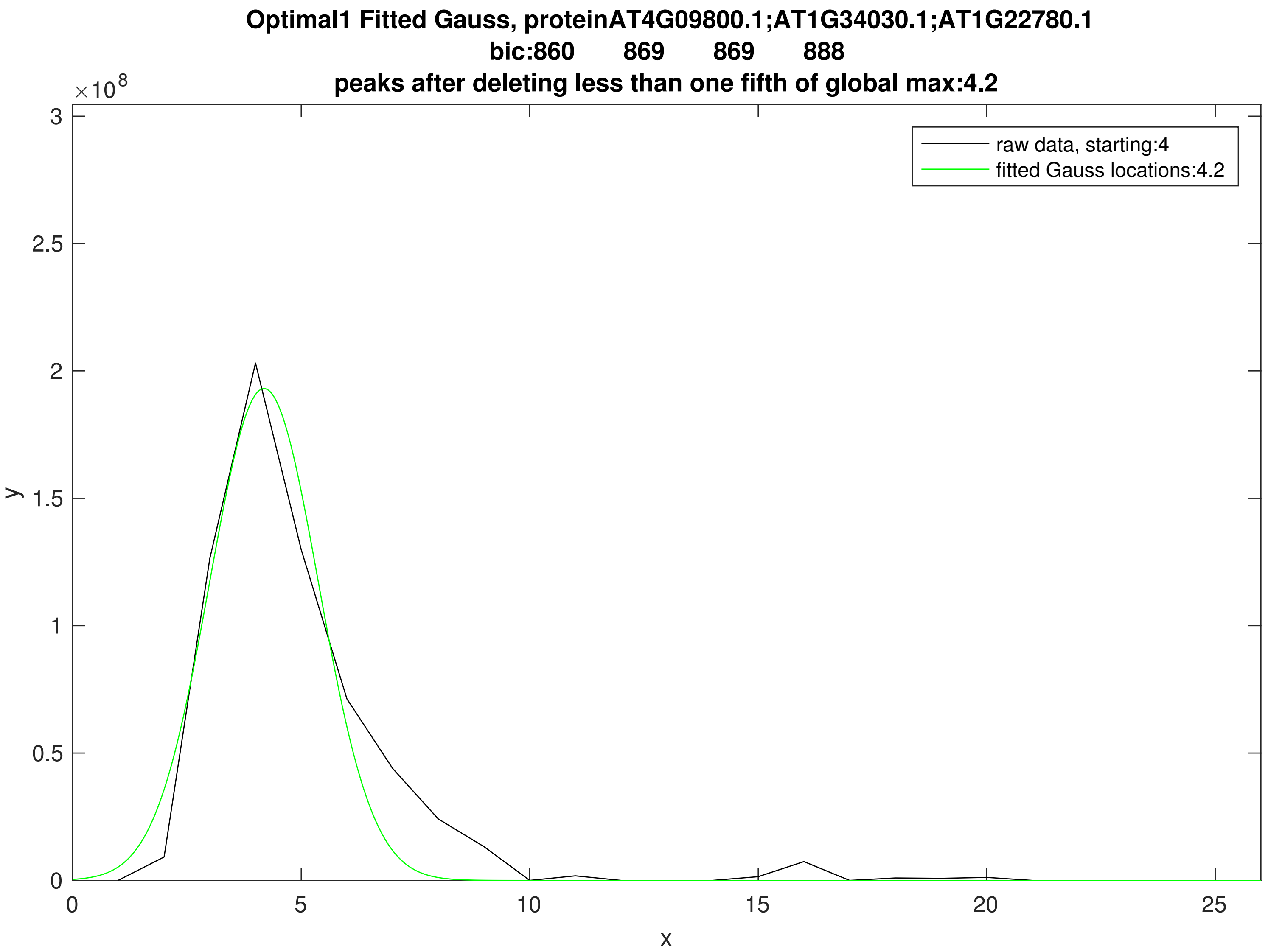
Optimal1 Fitted Gauss, proteinAT4G09800.1;AT1G34030.1;AT1G22780.1
bic:860 869 869 888
peaks after deleting less than one fifth of global max:4.2
×10^8
raw data, starting:4
fitted Gauss locations:4.2
y
x

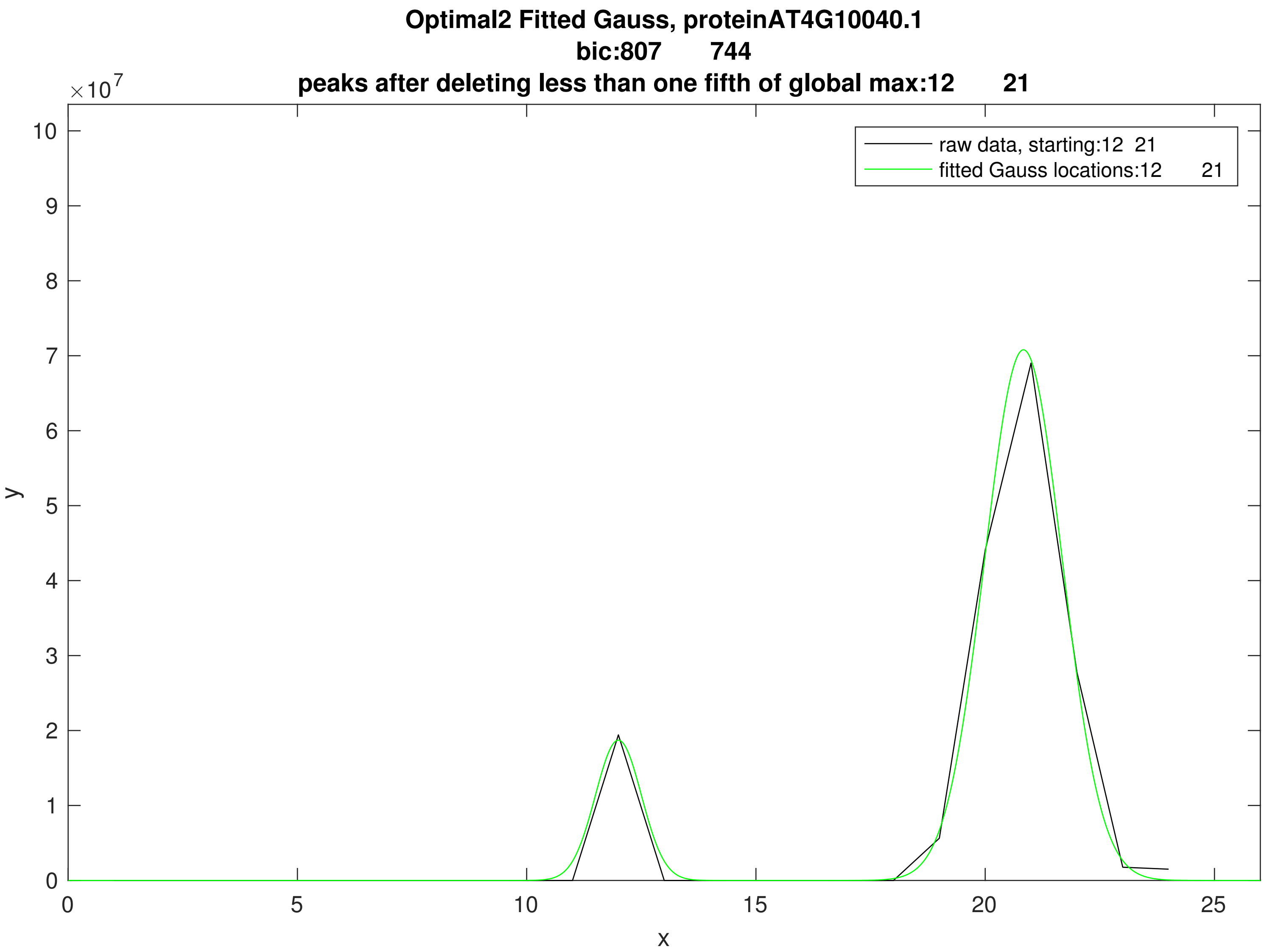
Optimal2 Fitted Gauss, proteinAT4G10040.1
bic:807 744
peaks after deleting less than one fifth of global max:12 21
raw data, starting:12 21
fitted Gauss locations:12 21
×10^7
y
x
0
1
2
3
4
5
6
7
8
9
10
0
5
10
15
20
25

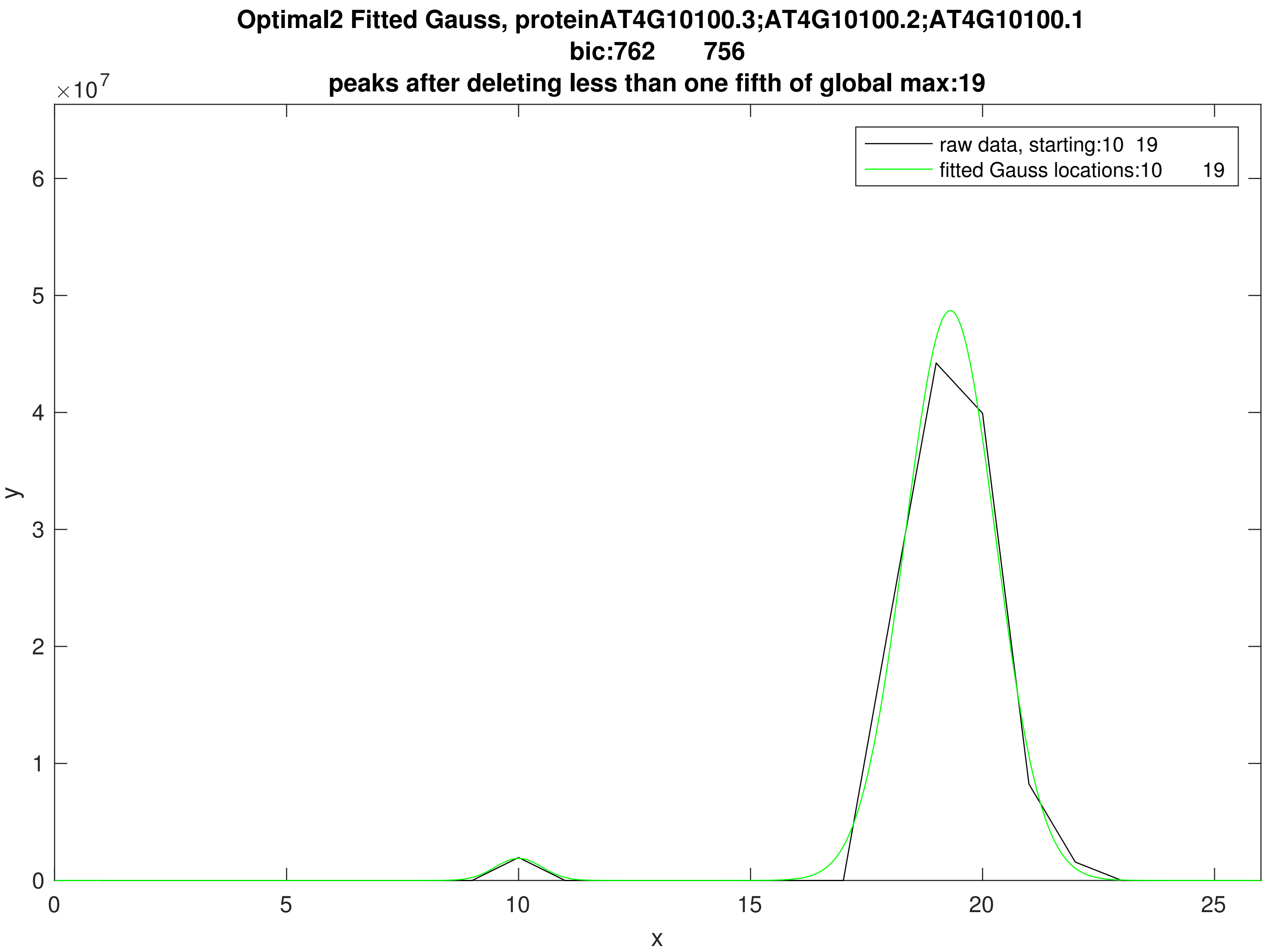
Optimal2 Fitted Gauss, proteinAT4G10100.3;AT4G10100.2;AT4G10100.1
bic:762 756
peaks after deleting less than one fifth of global max:19
×10^7
raw data, starting:10 19
fitted Gauss locations:10 19
0
1
2
3
4
5
6
y
0
5
10
15
20
25
x

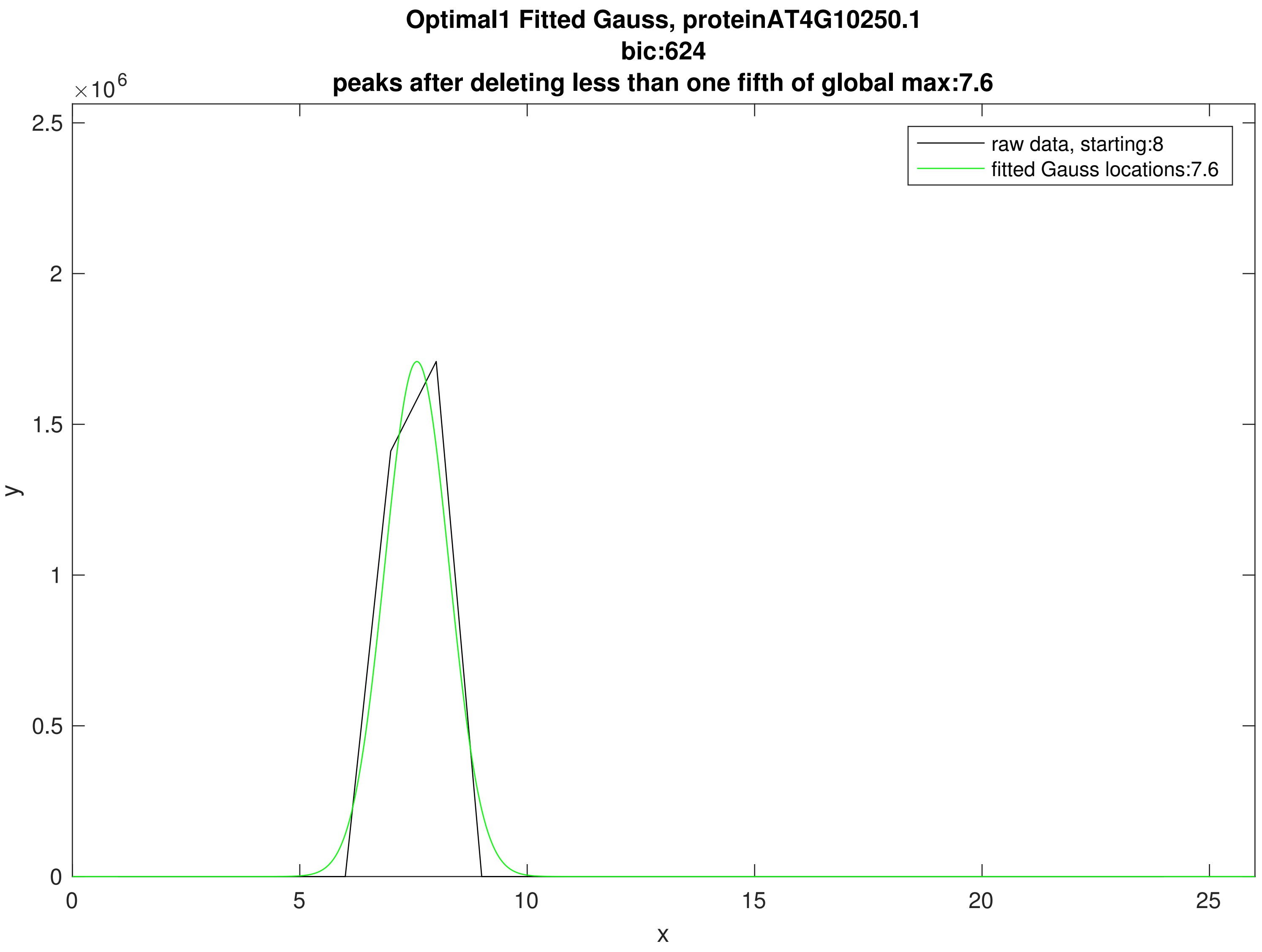
Optimal1 Fitted Gauss, proteinAT4G10250.1
bic:624
peaks after deleting less than one fifth of global max:7.6
×10^6
2.5
2
1.5
1
0.5
0
y
0
5
10
15
20
25
x
raw data, starting:8
fitted Gauss locations:7.6

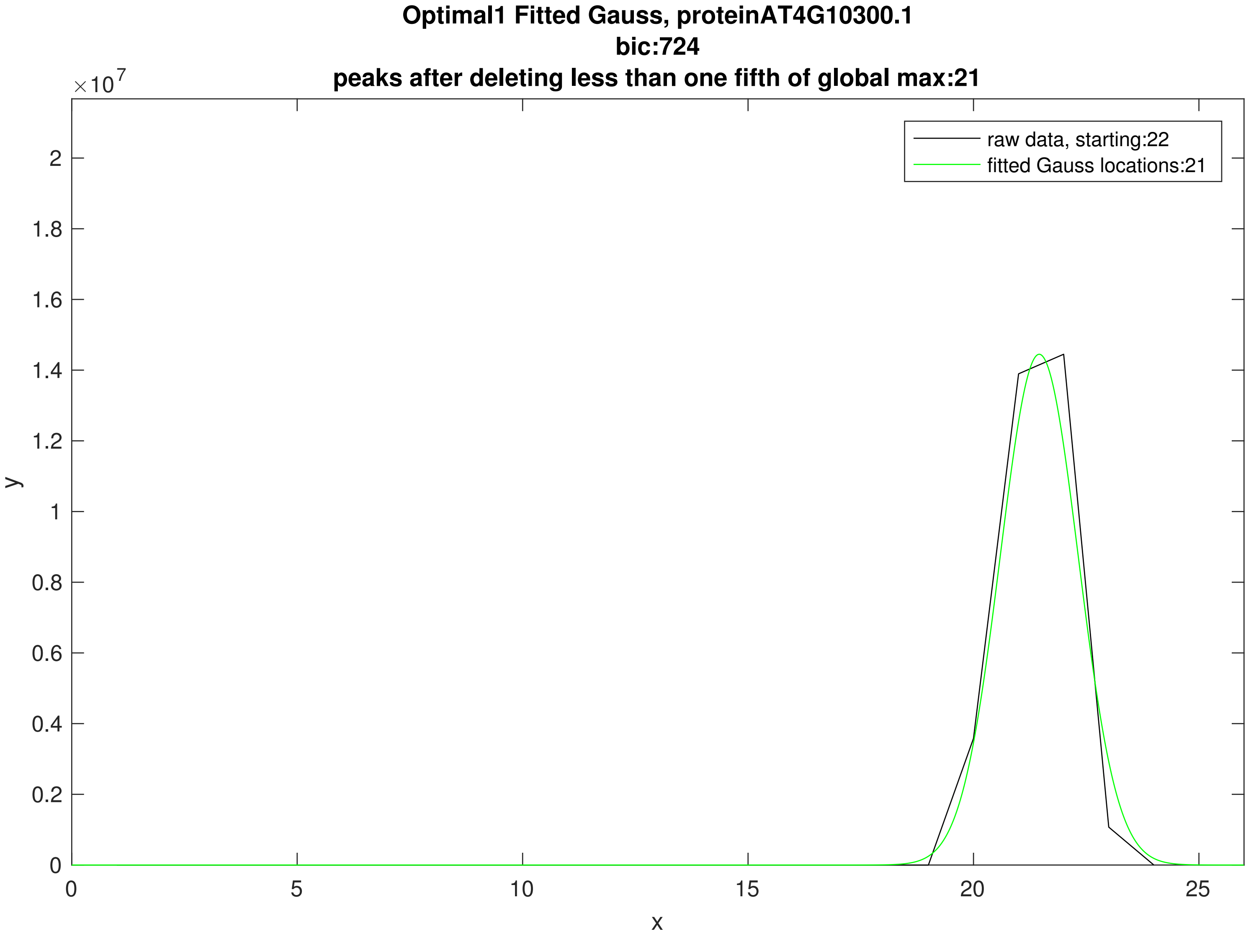
Optimal1 Fitted Gauss, proteinAT4G10300.1
bic:724
peaks after deleting less than one fifth of global max:21
×10^7
raw data, starting:22
fitted Gauss locations:21
y
x
0
0.2
0.4
0.6
0.8
1
1.2
1.4
1.6
1.8
2
0
5
10
15
20
25

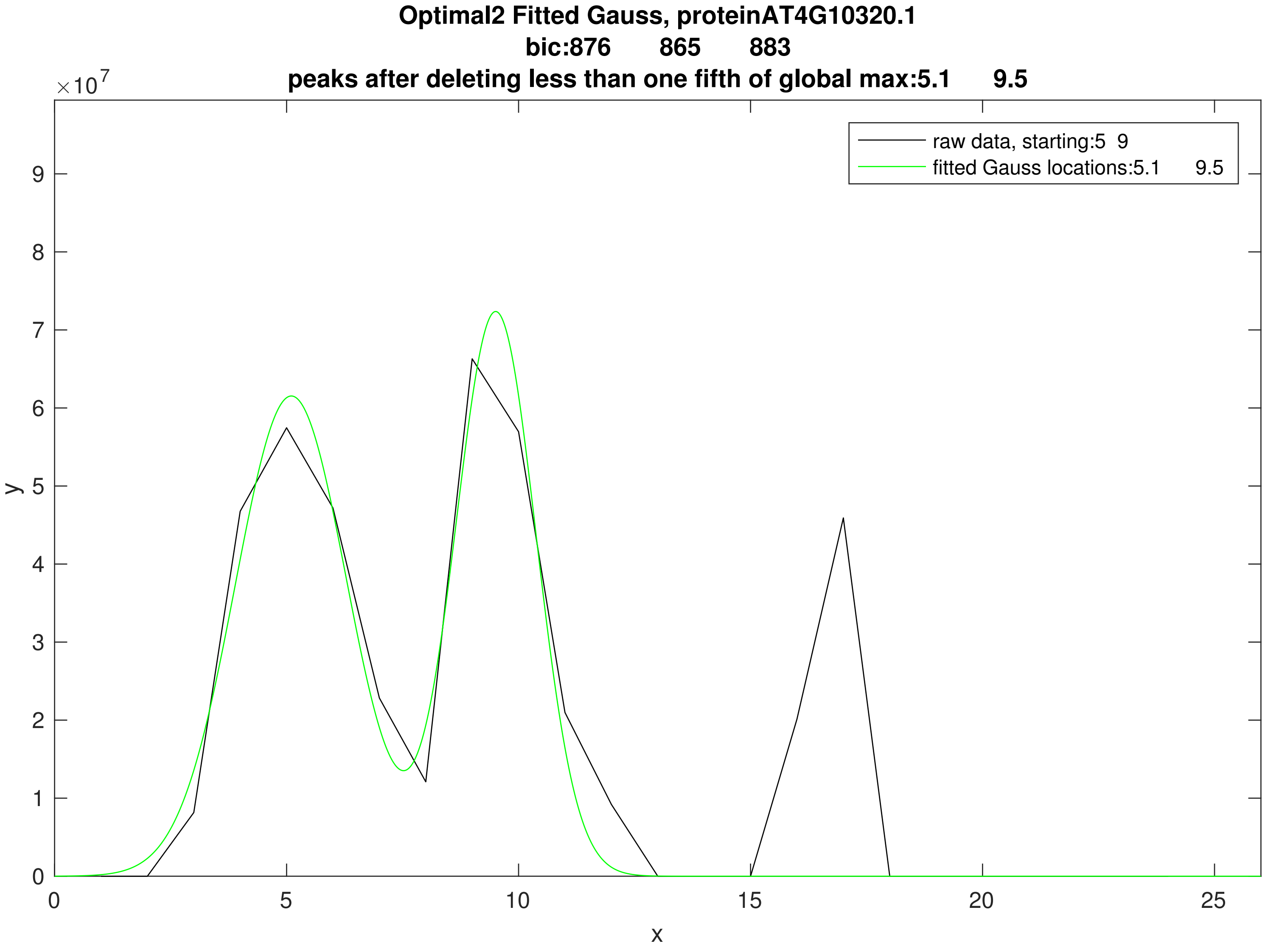

Optimal2 Fitted Gauss, proteinAT4G10320.1
bic:876 865 883
peaks after deleting less than one fifth of global max:5.1 9.5
×10^7
raw data, starting:5 9
fitted Gauss locations:5.1 9.5
y
x
0
1
2
3
4
5
6
7
8
9
0
5
10
15
20
25

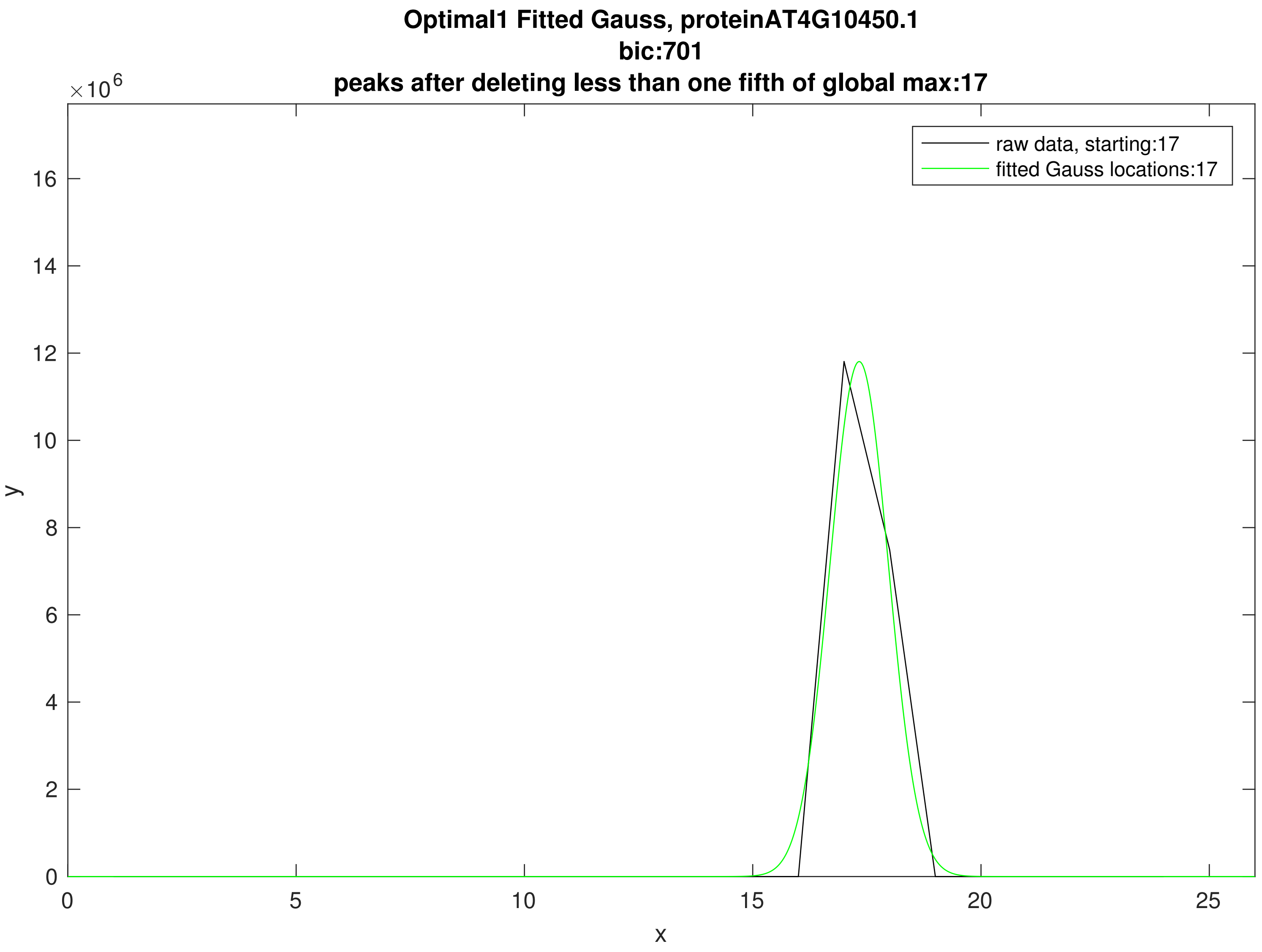
Optimal1 Fitted Gauss, proteinAT4G10450.1
bic:701
peaks after deleting less than one fifth of global max:17
×10^6
raw data, starting:17
fitted Gauss locations:17
y
x
0
2
4
6
8
10
12
14
16
0
5
10
15
20
25

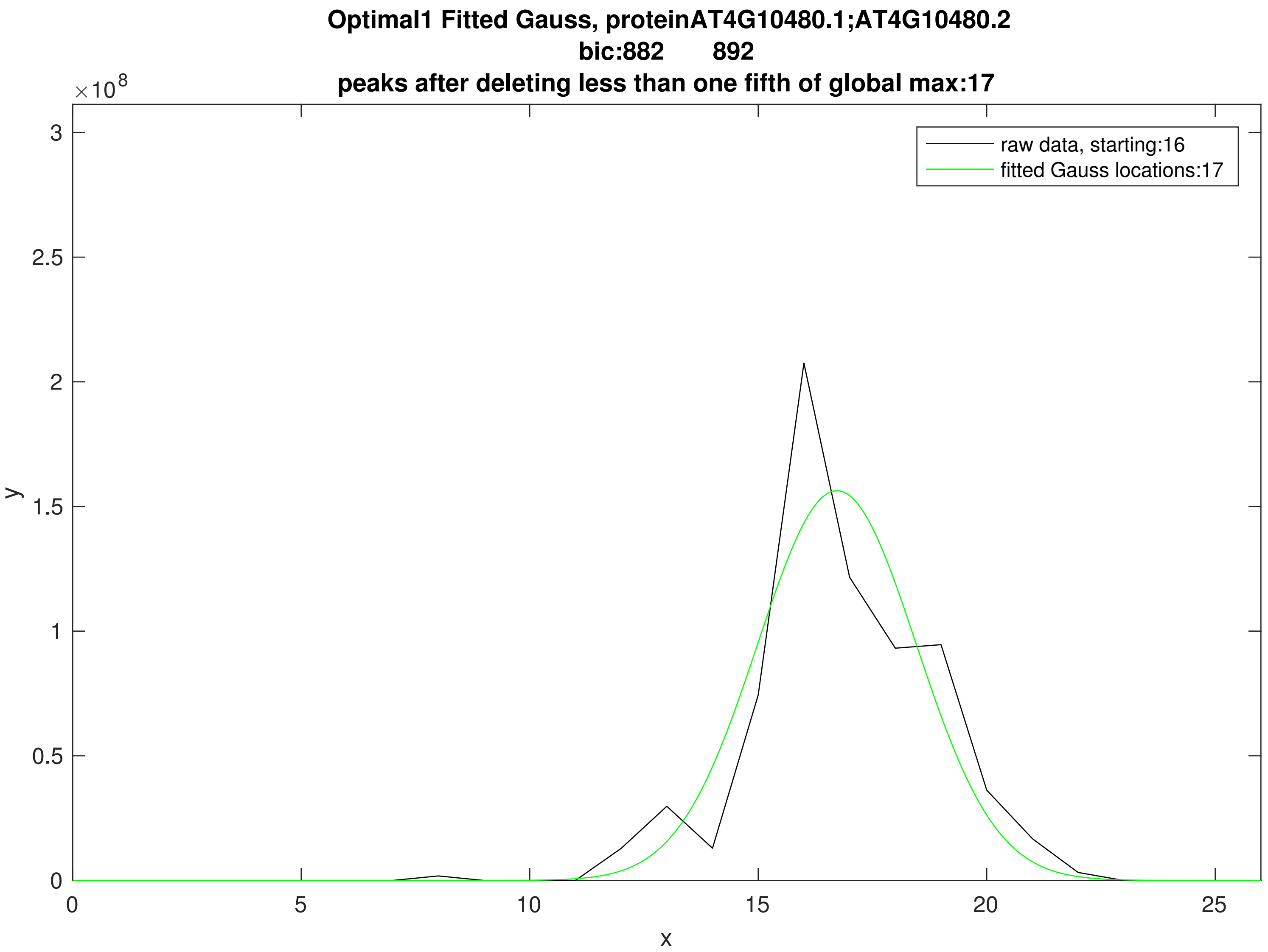

Optimal1 Fitted Gauss, proteinAT4G10480.1;AT4G10480.2
bic:882 892
peaks after deleting less than one fifth of global max:17
×10^8
3
2.5
2
1.5
1
0.5
0
y
0
5
10
15
20
25
x
raw data, starting:16
fitted Gauss locations:17

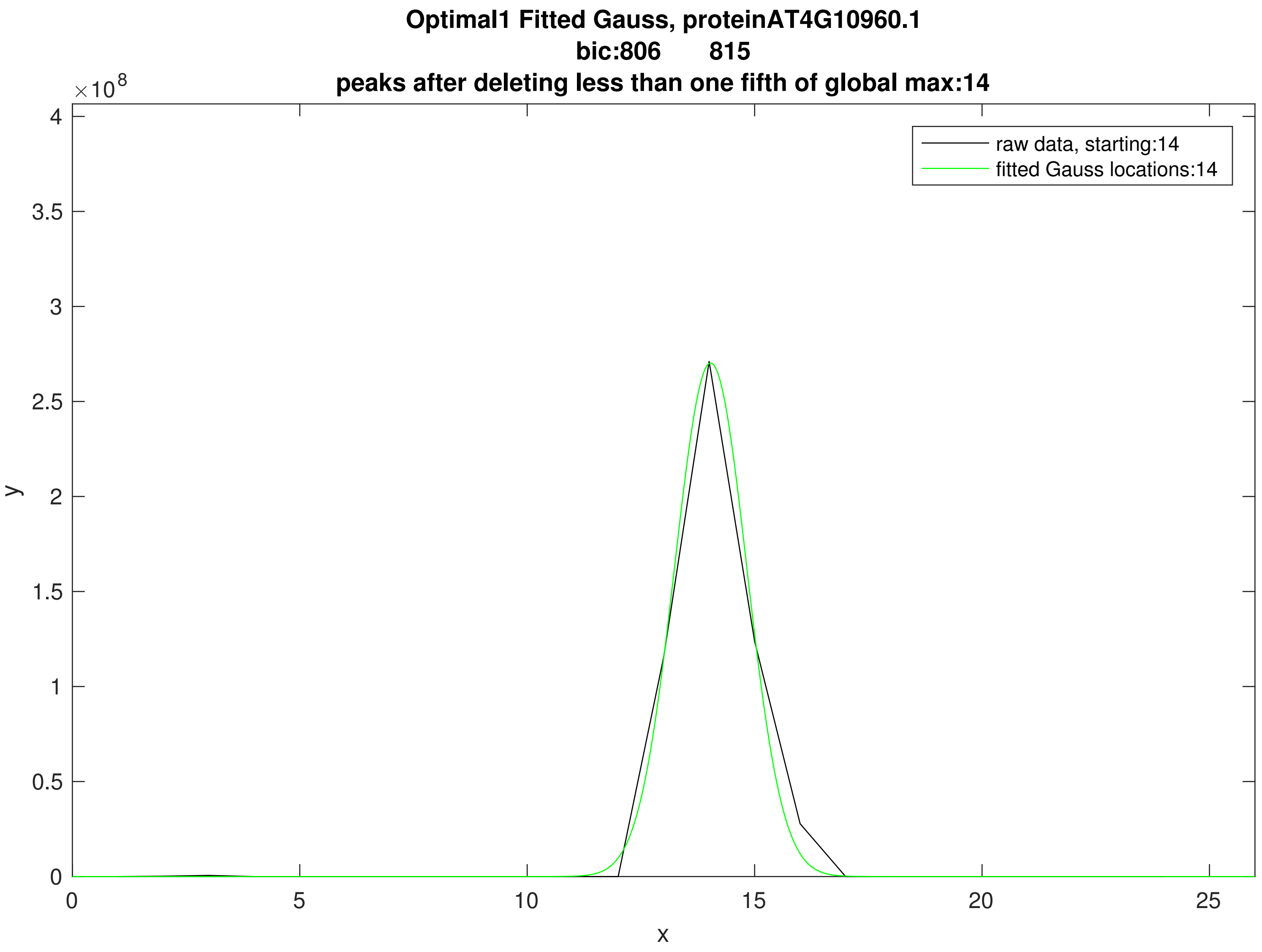

Optimal1 Fitted Gauss, proteinAT4G10960.1
bic:806 815
peaks after deleting less than one fifth of global max:14
×10^8
4
3.5
3
2.5
2
1.5
1
0.5
0
y
0
5
10
15
20
25
x
raw data, starting:14
fitted Gauss locations:14

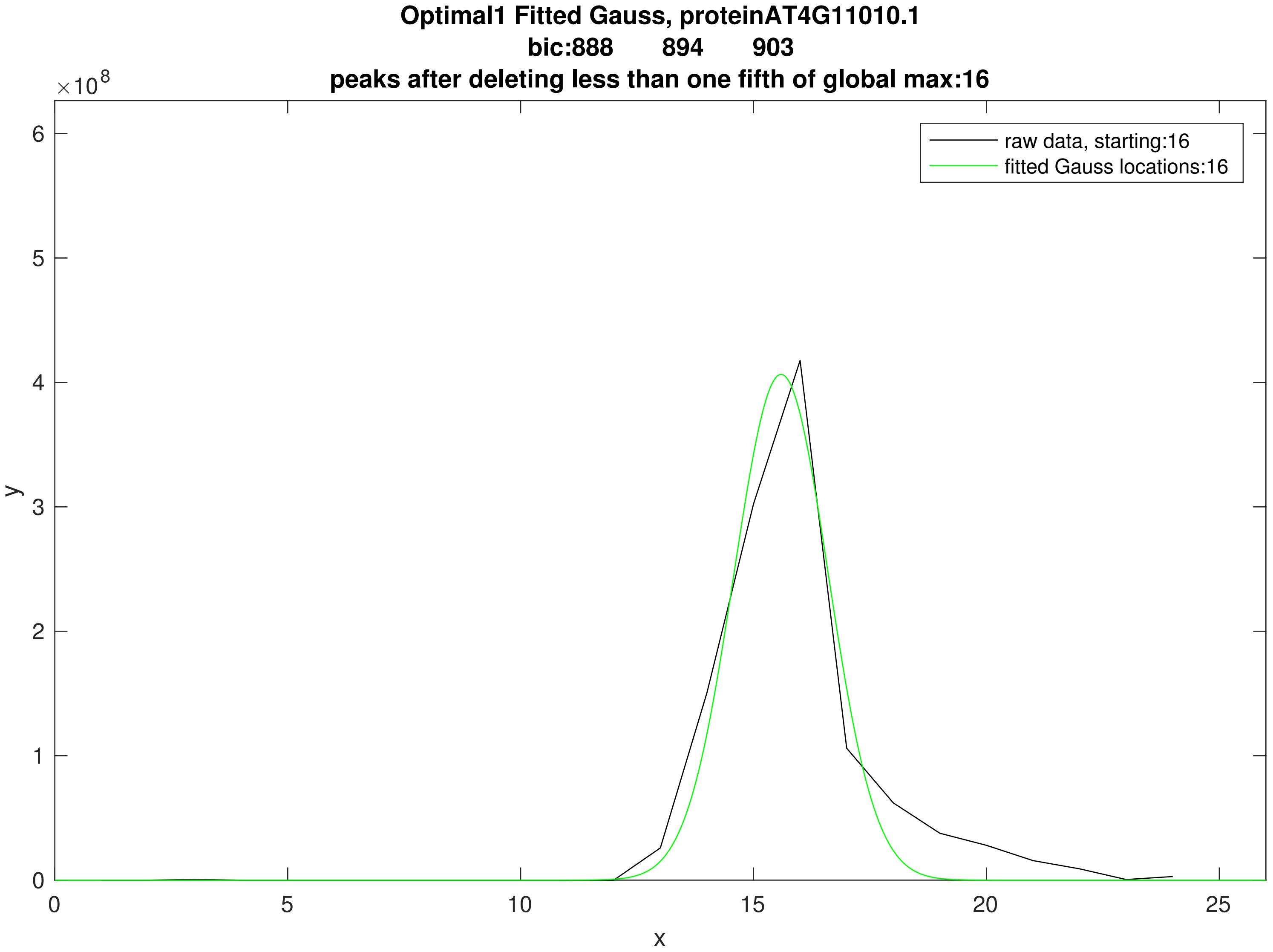
Optimal1 Fitted Gauss, proteinAT4G11010.1
bic:888 894 903
peaks after deleting less than one fifth of global max:16
×10^8
6
5
4
3
2
1
0
y
0
5
10
15
20
25
x
raw data, starting:16
fitted Gauss locations:16

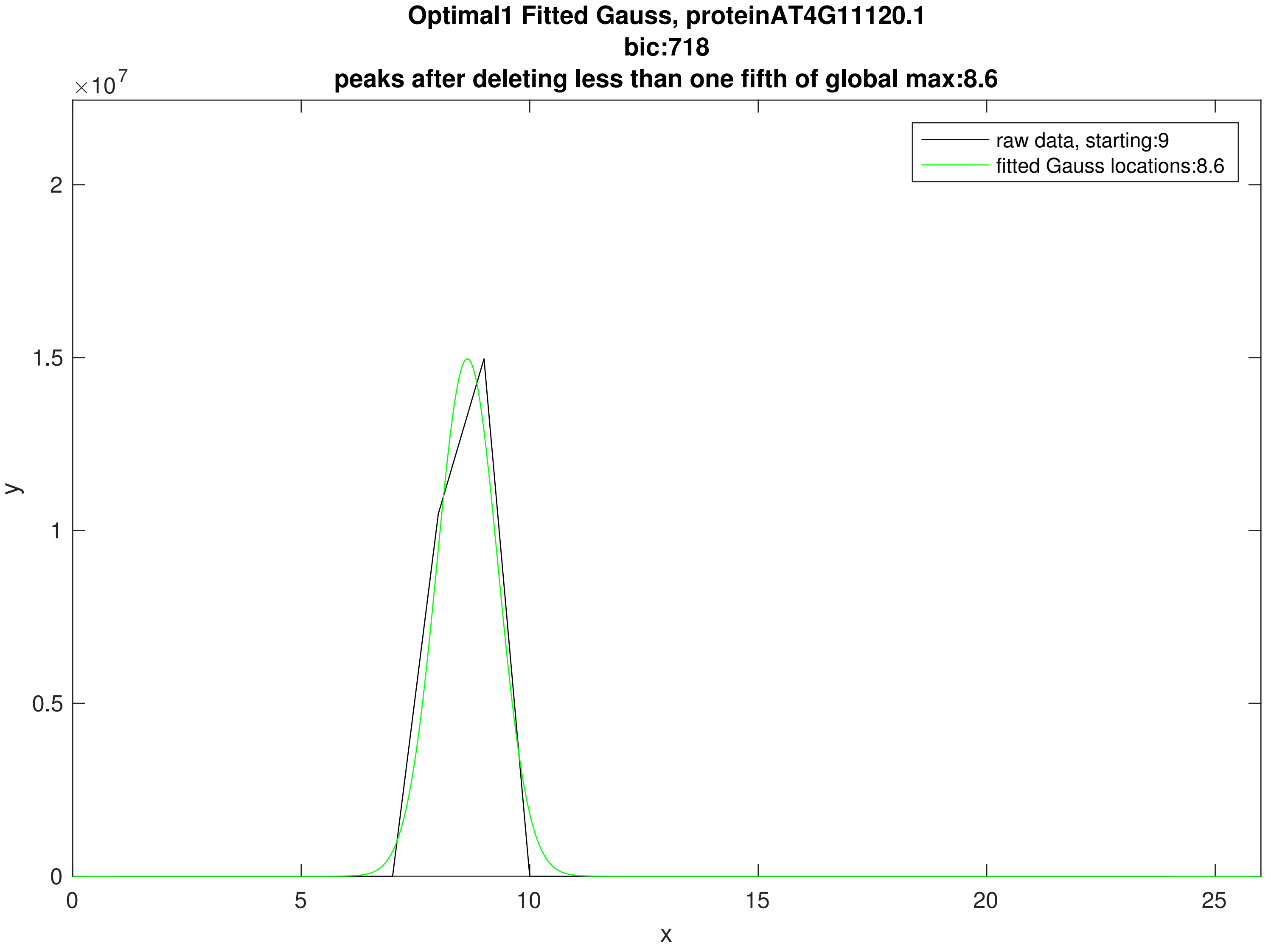

Optimal1 Fitted Gauss, proteinAT4G11120.1
bic:718
peaks after deleting less than one fifth of global max:8.6
×10^7
2
1.5
1
0.5
0
y
0
5
10
15
20
25
x
raw data, starting:9
fitted Gauss locations:8.6

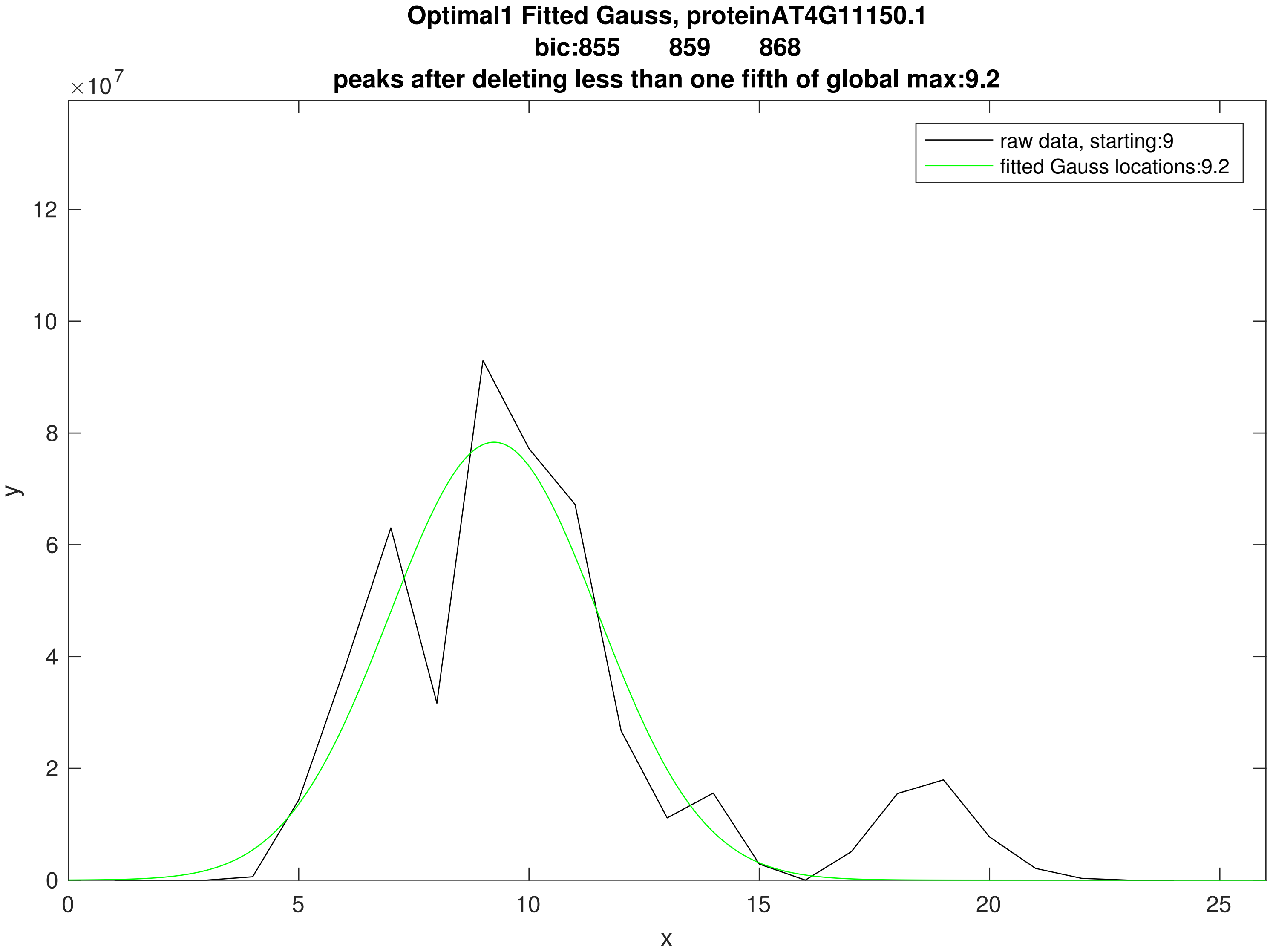
Optimal1 Fitted Gauss, proteinAT4G11150.1
bic:855 859 868
peaks after deleting less than one fifth of global max:9.2
×10^7
raw data, starting:9
fitted Gauss locations:9.2
y
x
0
2
4
6
8
10
12
5
15
20
25

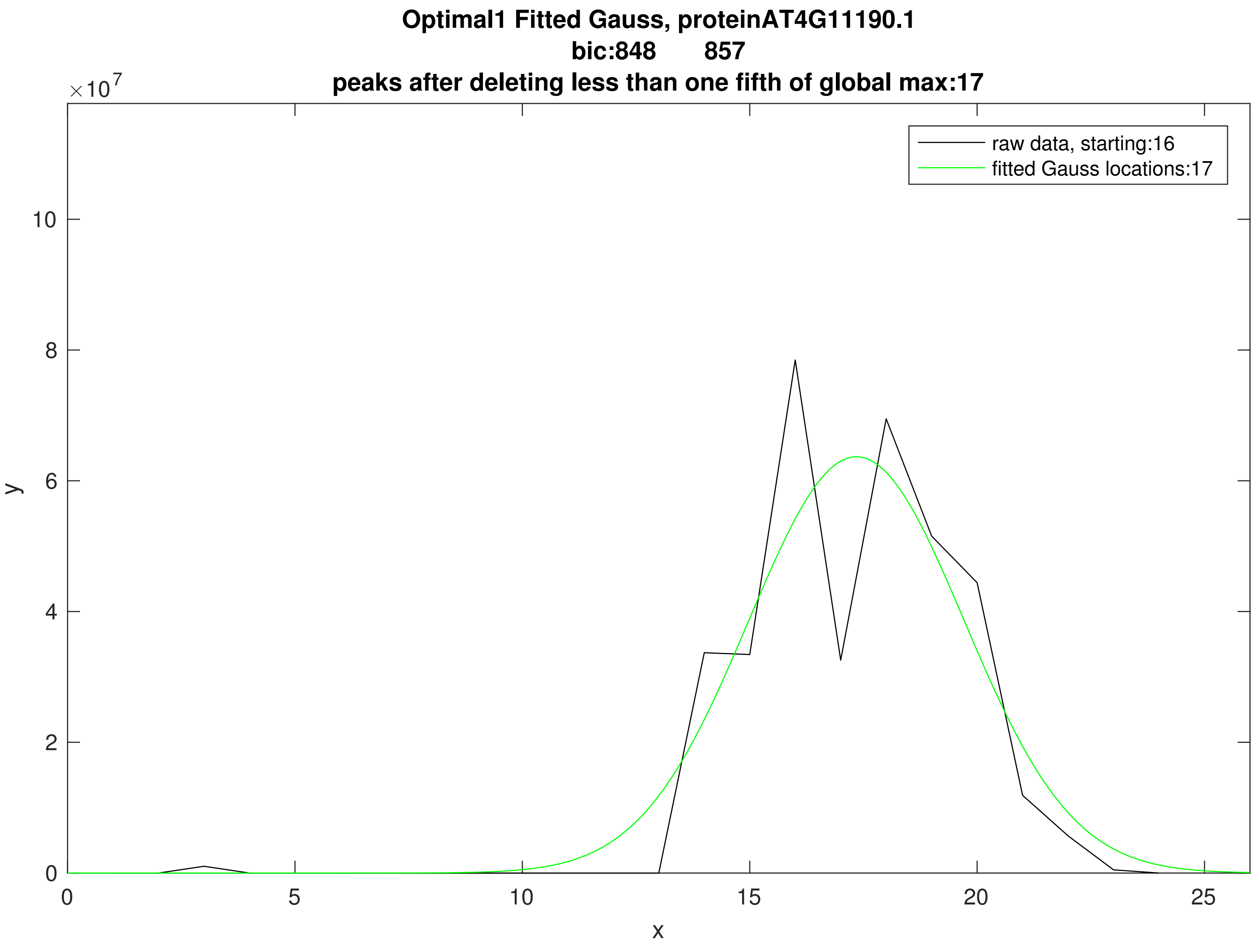

Optimal1 Fitted Gauss, proteinAT4G11190.1
bic:848 857
peaks after deleting less than one fifth of global max:17
×10^7
raw data, starting:16
fitted Gauss locations:17
10
8
6
4
2
0
y
0
5
10
15
20
25
x

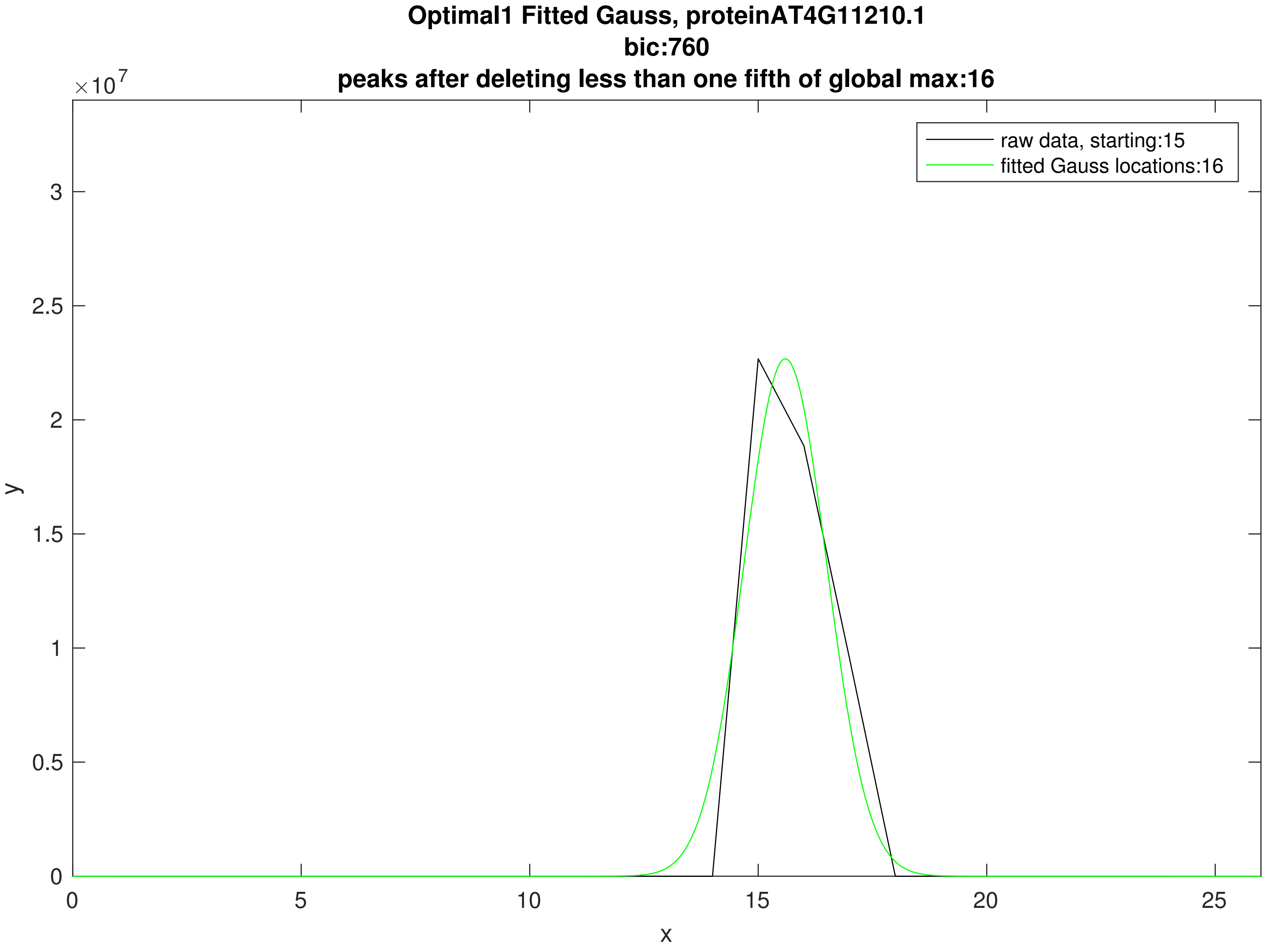
Optimal1 Fitted Gauss, proteinAT4G11210.1
bic:760
peaks after deleting less than one fifth of global max:16
×10^7
raw data, starting:15
fitted Gauss locations:16
y
x
0
0.5
1
1.5
2
2.5
3
5
10
15
20
25

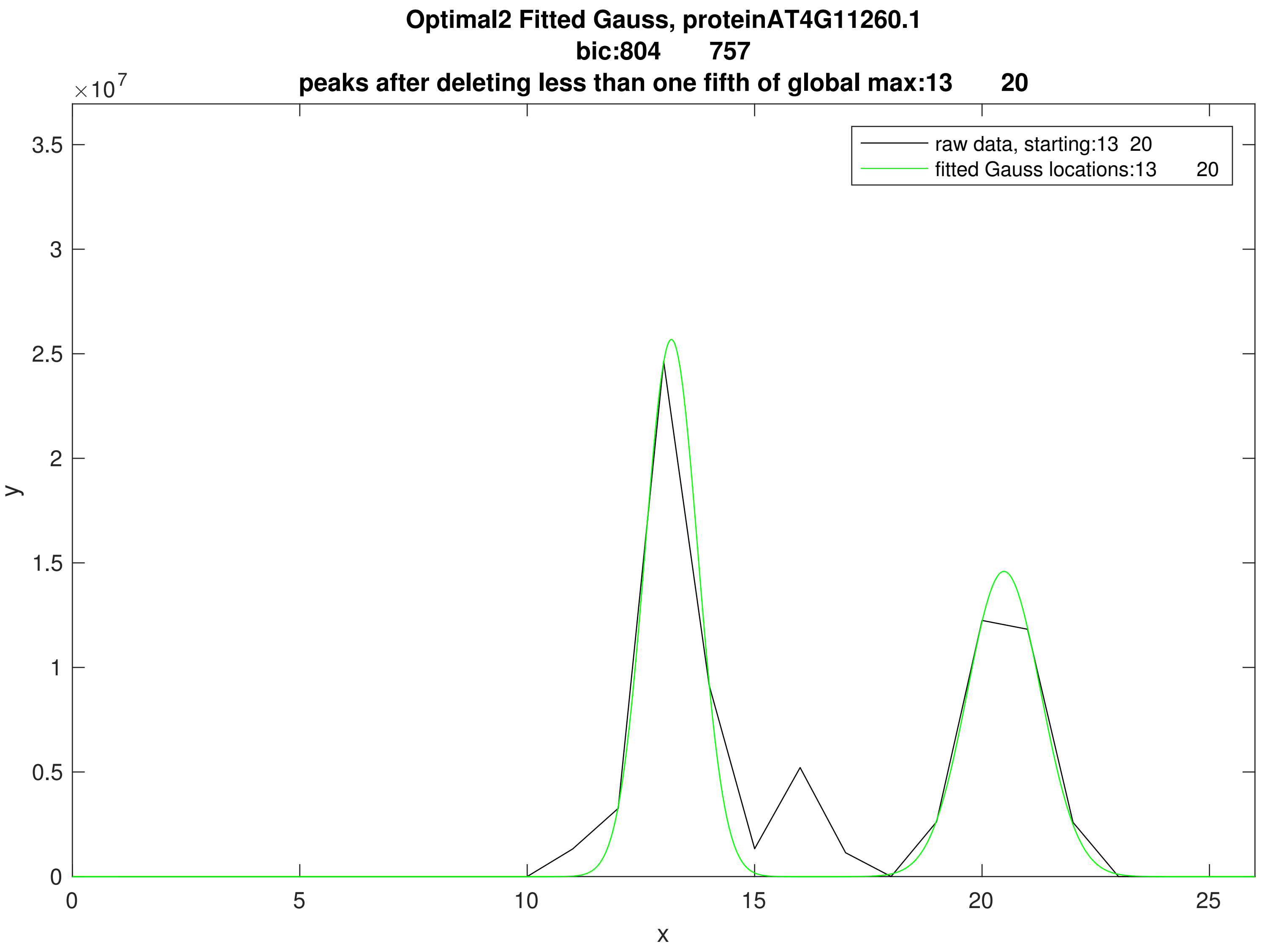
Optimal2 Fitted Gauss, proteinAT4G11260.1
bic:804 757
peaks after deleting less than one fifth of global max:13 20
×10^7
raw data, starting:13 20
fitted Gauss locations:13 20
3.5
3
2.5
2
1.5
1
0.5
0
y
0
5
10
15
20
25
x

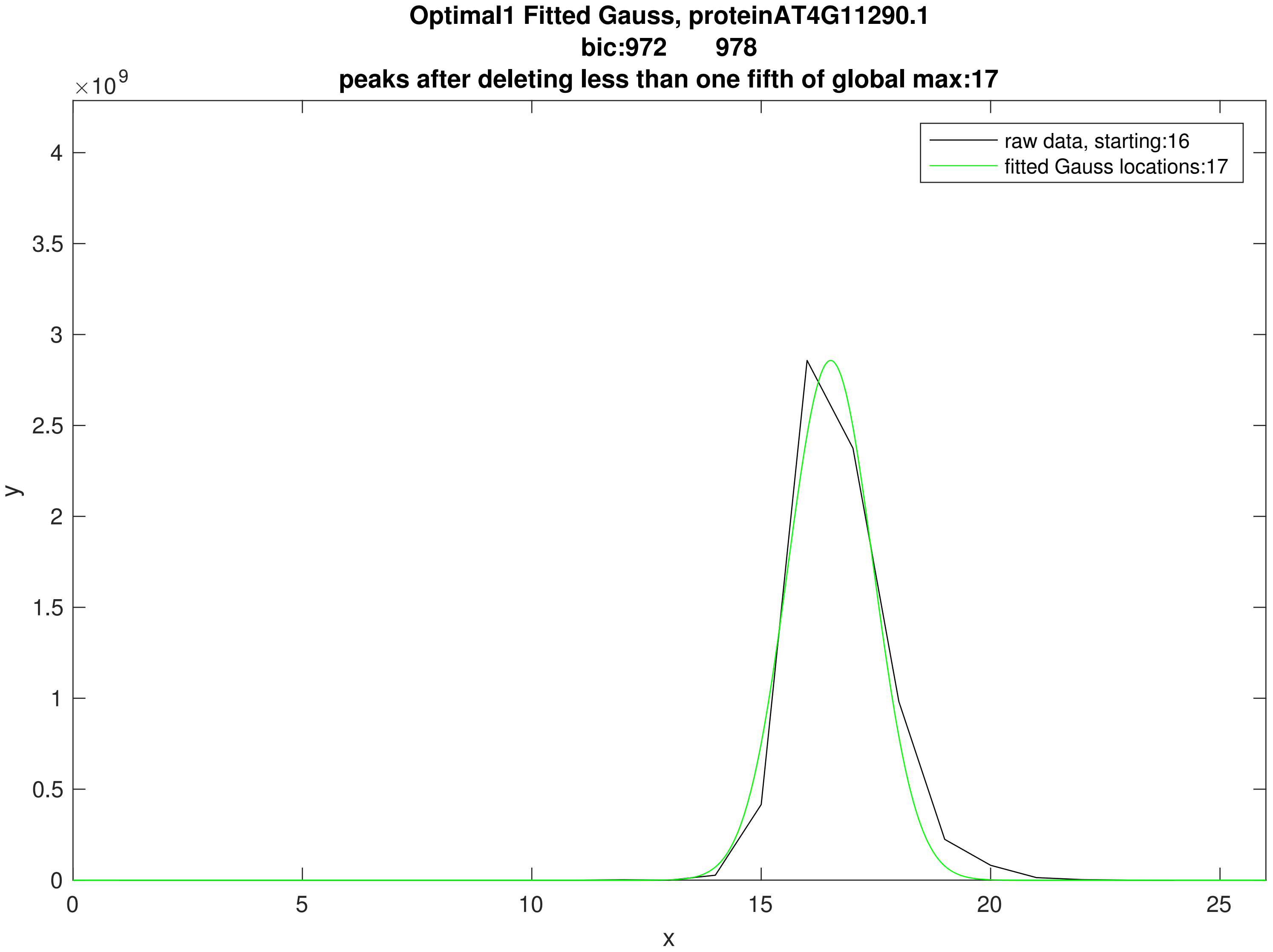
Optimal1 Fitted Gauss, proteinAT4G11290.1
bic:972 978
peaks after deleting less than one fifth of global max:17
×10^9
raw data, starting:16
fitted Gauss locations:17
y
x
0
0.5
1
1.5
2
2.5
3
3.5
4
0
5
10
15
20
25

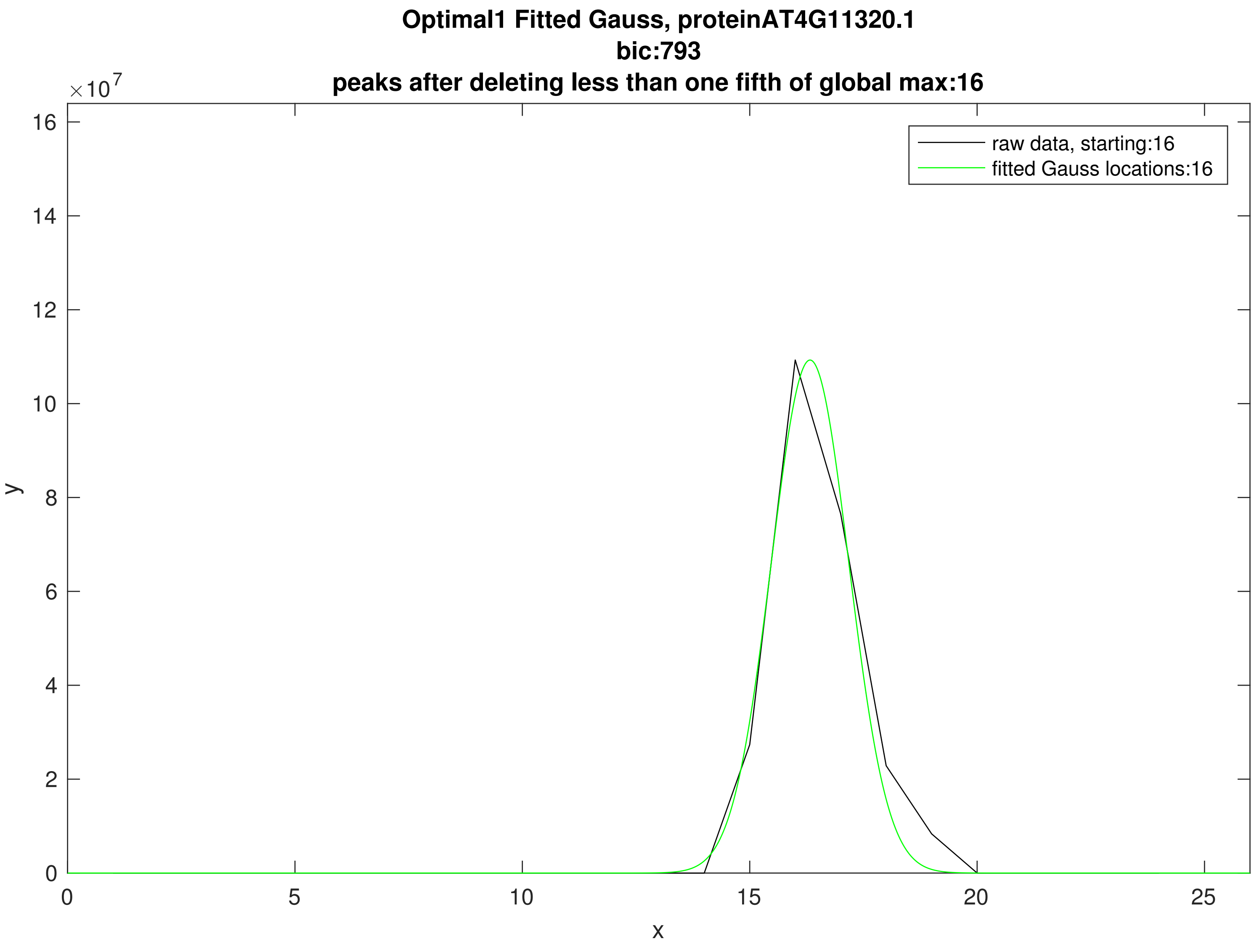

Optimal1 Fitted Gauss, proteinAT4G11320.1
bic:793
peaks after deleting less than one fifth of global max:16
×10^7
16
14
12
10
8
6
4
2
0
y
0
5
10
15
20
25
x
raw data, starting:16
fitted Gauss locations:16

#### Optimal1 Fitted Gauss, proteinAT4G11600.1
#### bic:842 851
#### peaks after deleting less than one fifth of global max:19

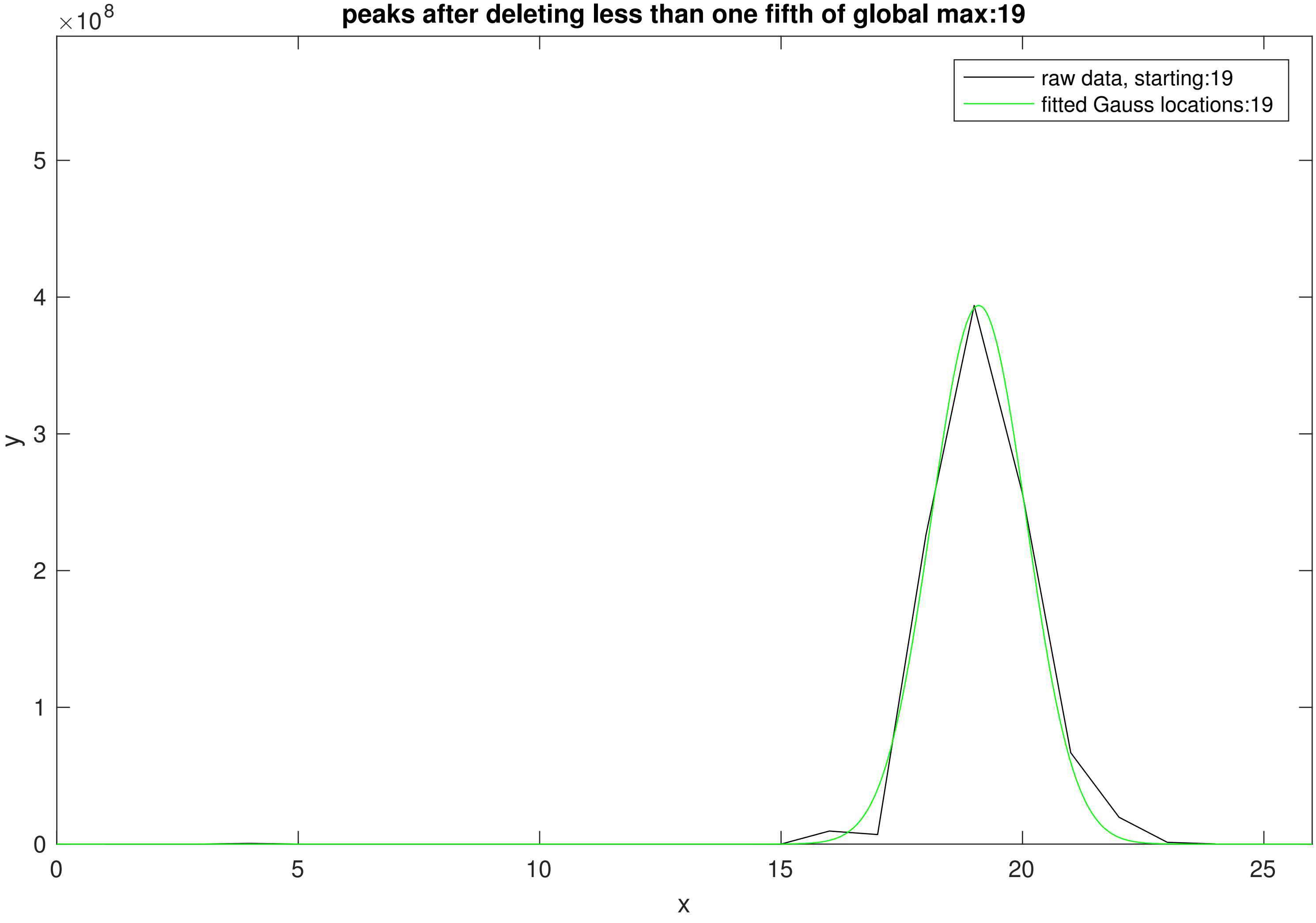

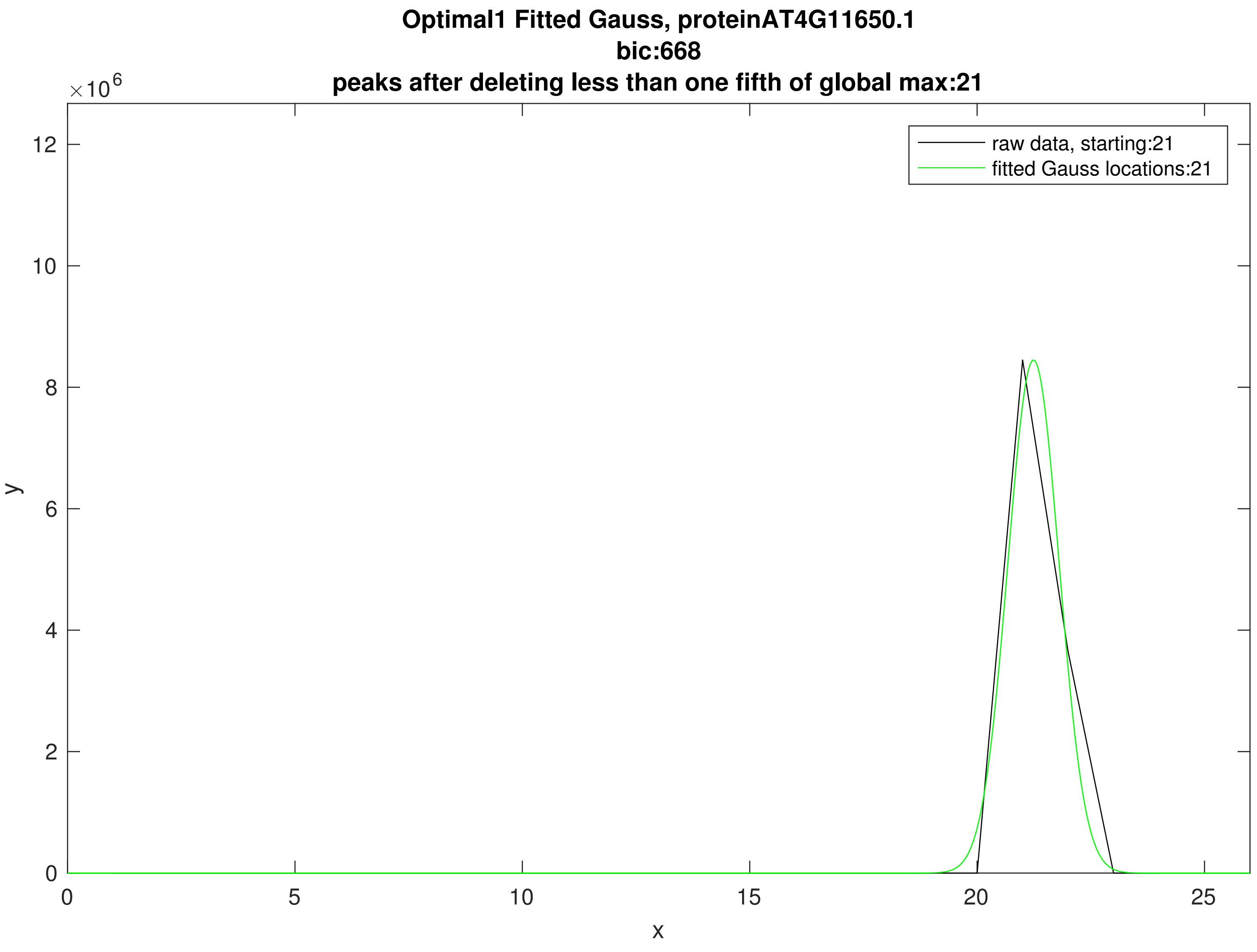
Optimal1 Fitted Gauss, proteinAT4G11650.1
bic:668
peaks after deleting less than one fifth of global max:21
×10^6
raw data, starting:21
fitted Gauss locations:21
y
x
0
2
4
6
8
10
12
0
5
10
15
20
25

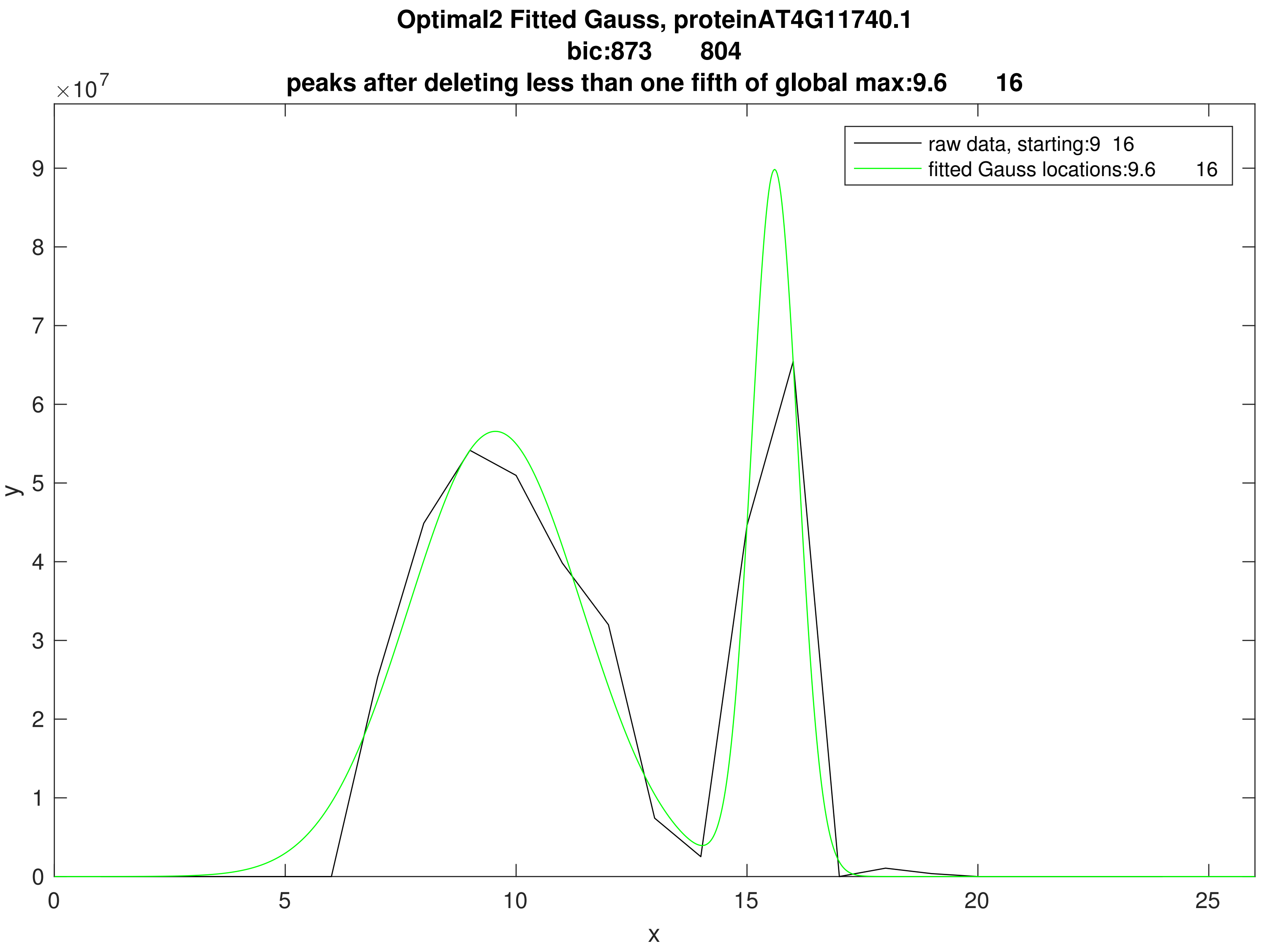

Optimal2 Fitted Gauss, proteinAT4G11740.1
bic:873 804
peaks after deleting less than one fifth of global max:9.6 16
×10^7
raw data, starting:9 16
fitted Gauss locations:9.6 16
y
x
0
1
2
3
4
5
6
7
8
9
0
5
10
15
20
25

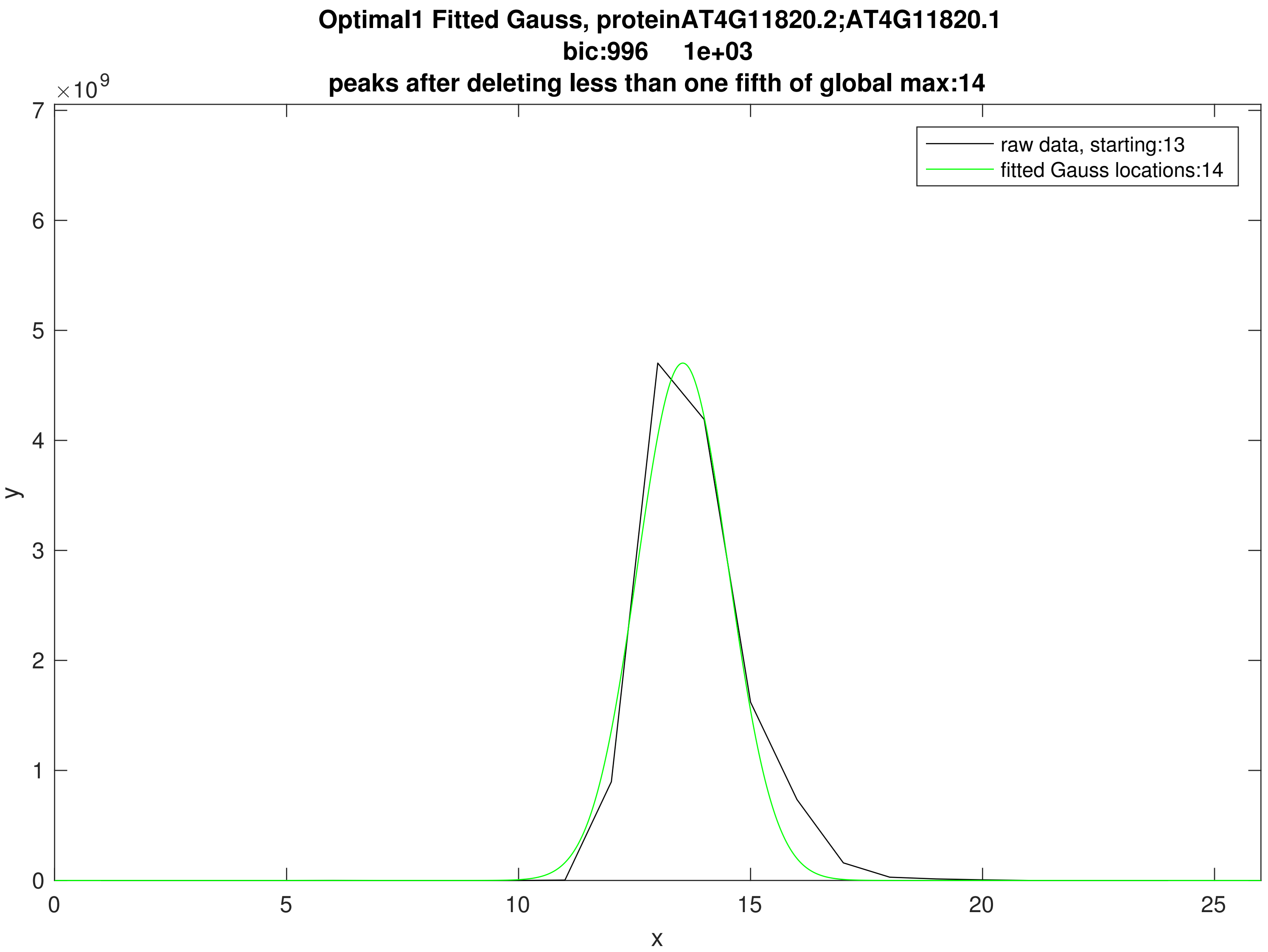

Optimal1 Fitted Gauss, proteinAT4G11820.2;AT4G11820.1
bic:996 1e+03
peaks after deleting less than one fifth of global max:14
×10^9
raw data, starting:13
fitted Gauss locations:14
y
x
0
1
2
3
4
5
6
7
0
5
10
15
20
25

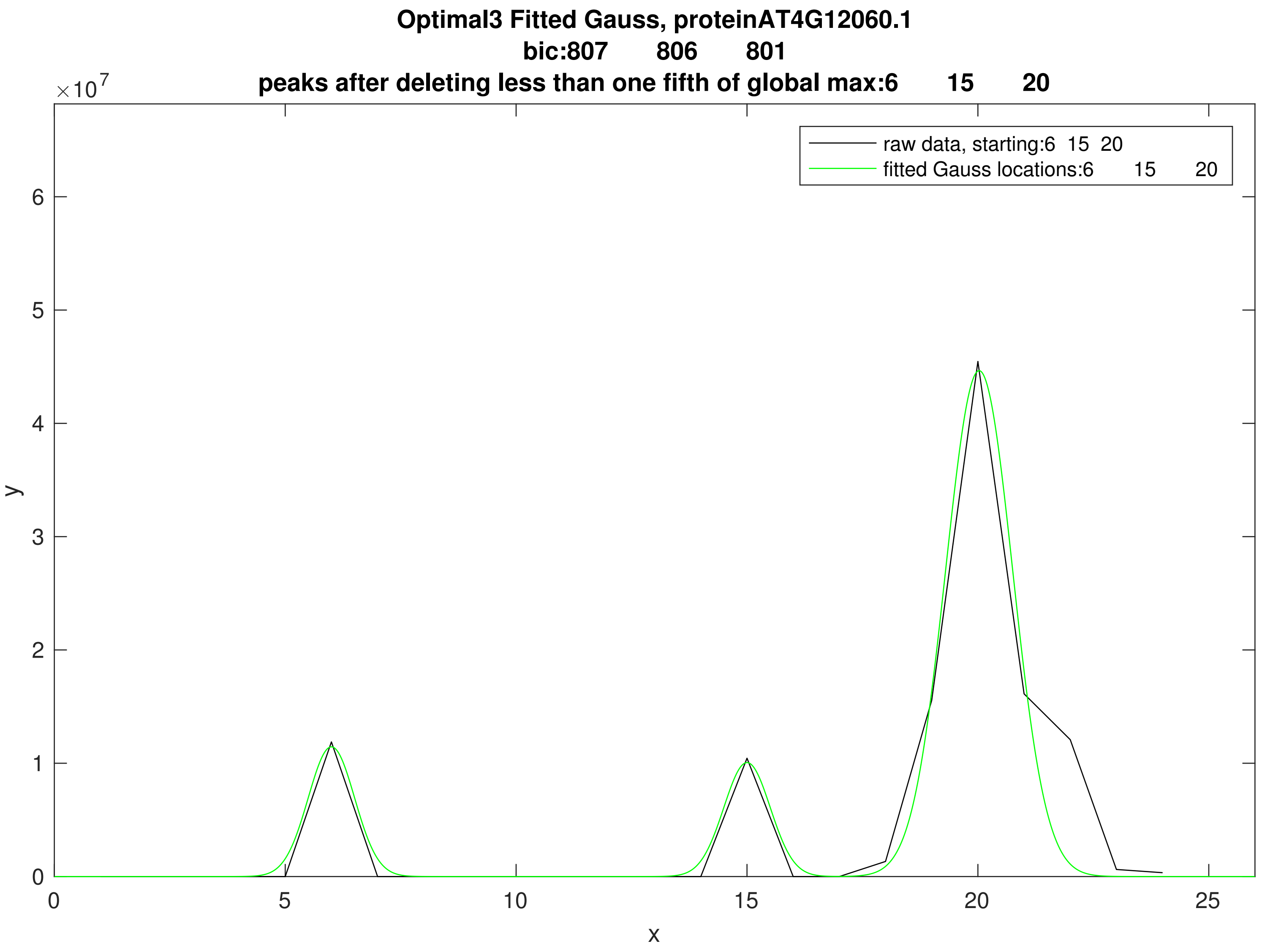
Optimal3 Fitted Gauss, proteinAT4G12060.1
bic:807 806 801
peaks after deleting less than one fifth of global max:6 15 20
×10^7
raw data, starting:6 15 20
fitted Gauss locations:6 15 20
y
x
0
1
2
3
4
5
6
0
5
10
15
20
25

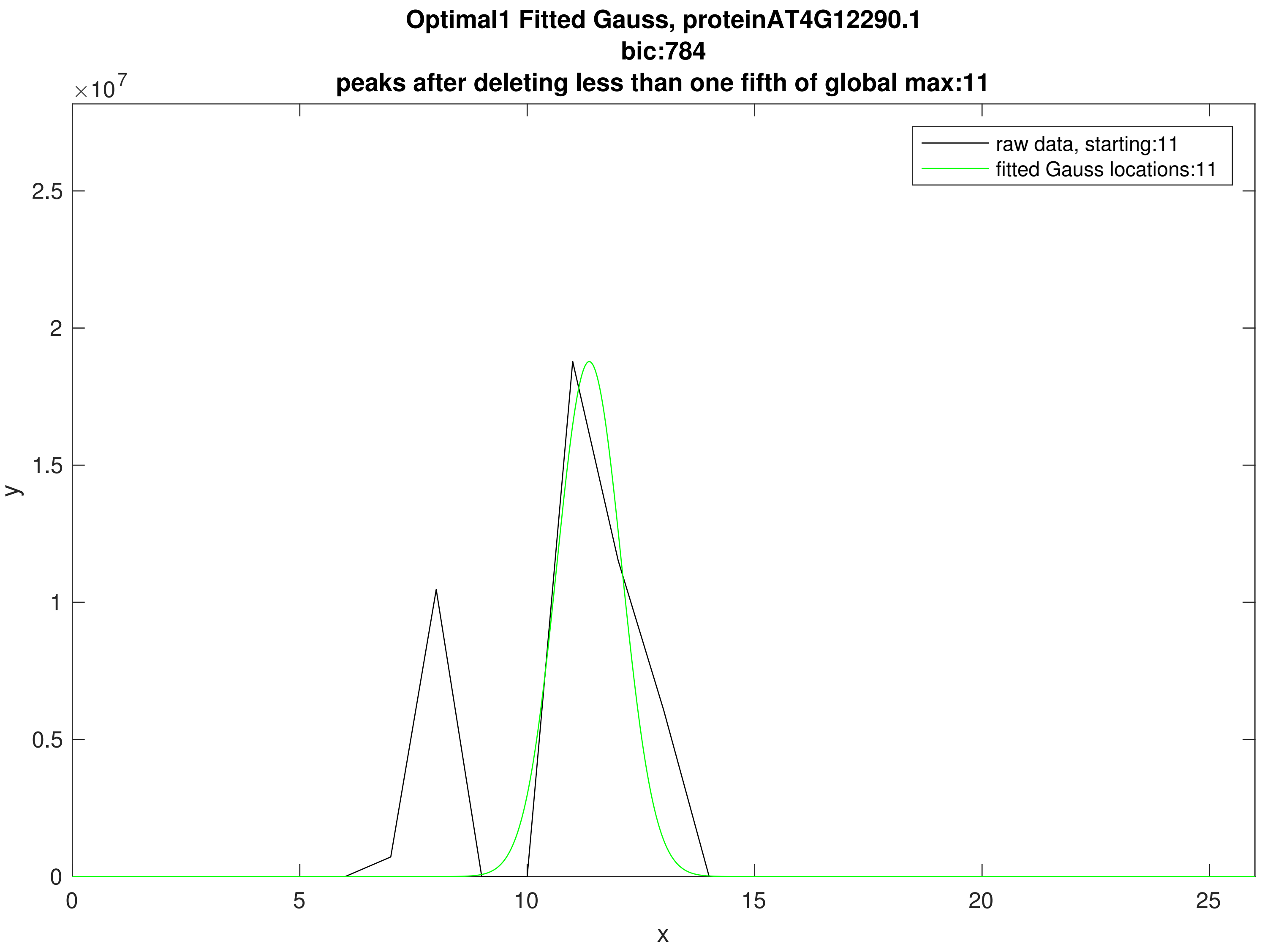
Optimal1 Fitted Gauss, proteinAT4G12290.1
bic:784
peaks after deleting less than one fifth of global max:11
×10^7
raw data, starting:11
fitted Gauss locations:11
2.5
2
1.5
1
0.5
0
y
0
5
10
15
20
25
x

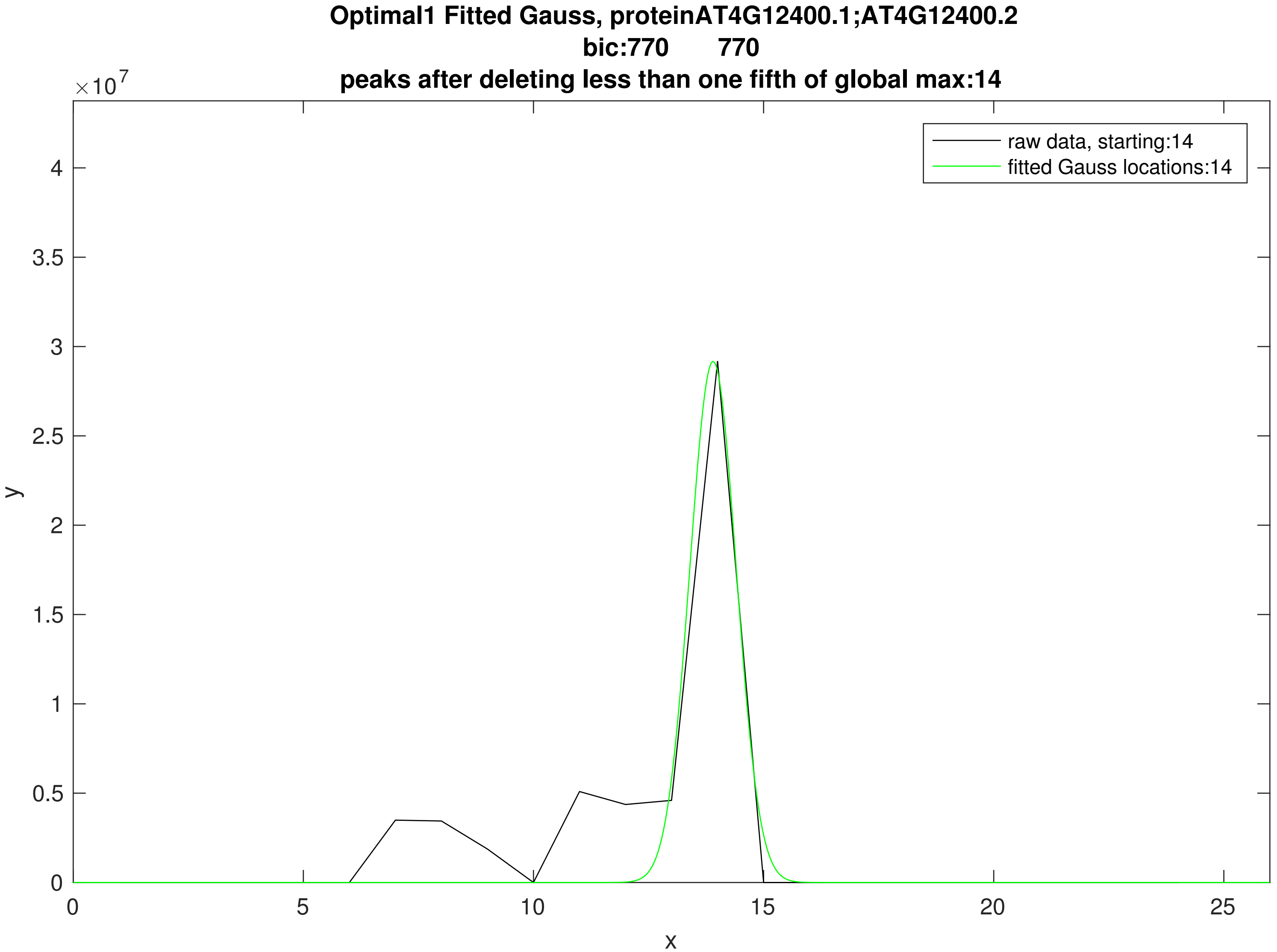
Optimal1 Fitted Gauss, proteinAT4G12400.1;AT4G12400.2
bic:770 770
peaks after deleting less than one fifth of global max:14
×10^7
raw data, starting:14
fitted Gauss locations:14
4
3.5
3
2.5
2
1.5
1
0.5
0
y
0
5
10
15
20
25
x

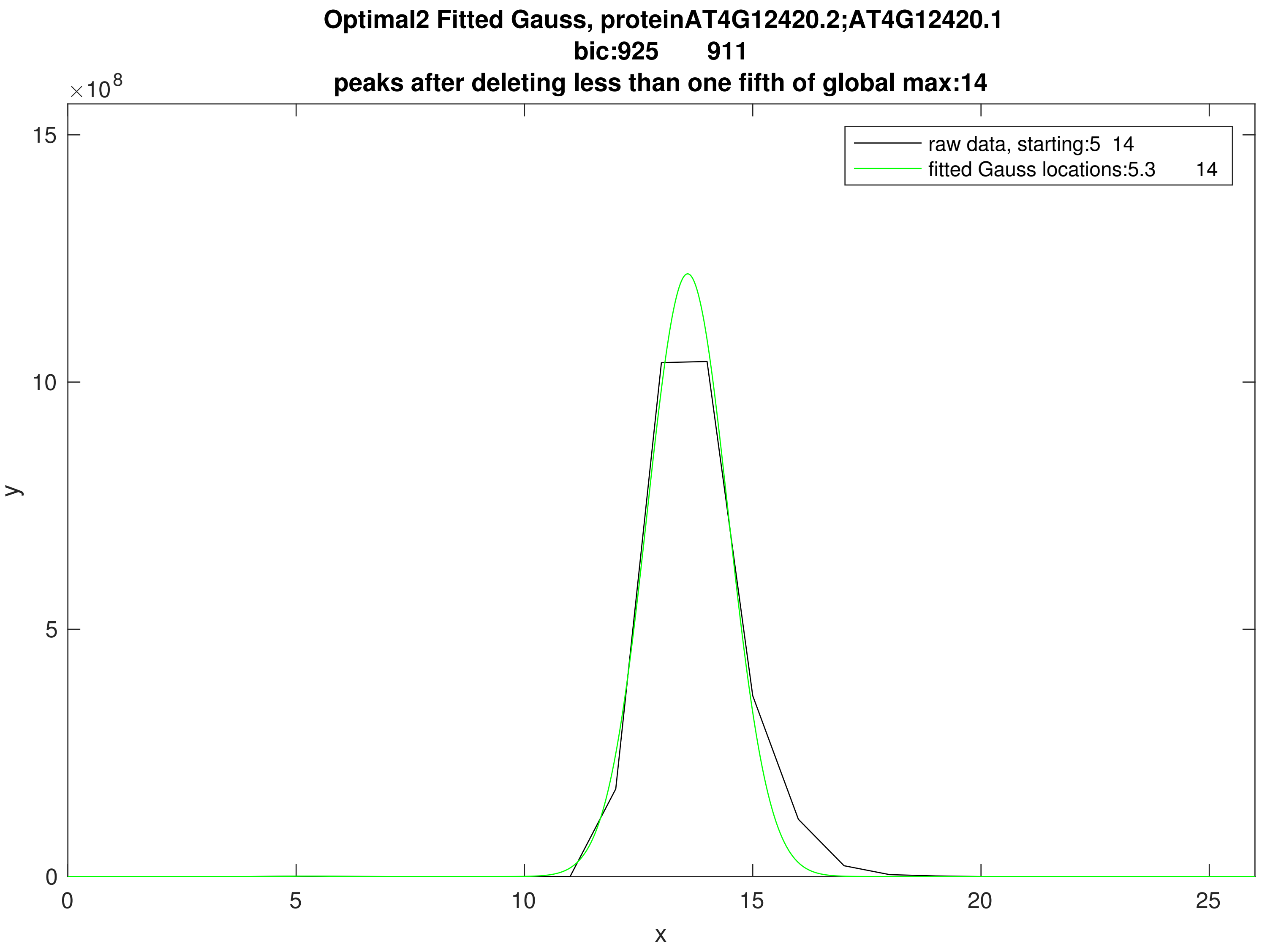
Optimal2 Fitted Gauss, proteinAT4G12420.2;AT4G12420.1
bic:925 911
peaks after deleting less than one fifth of global max:14
×10^8
raw data, starting:5 14
fitted Gauss locations:5.3 14
y
x
0
5
10
15
20
25

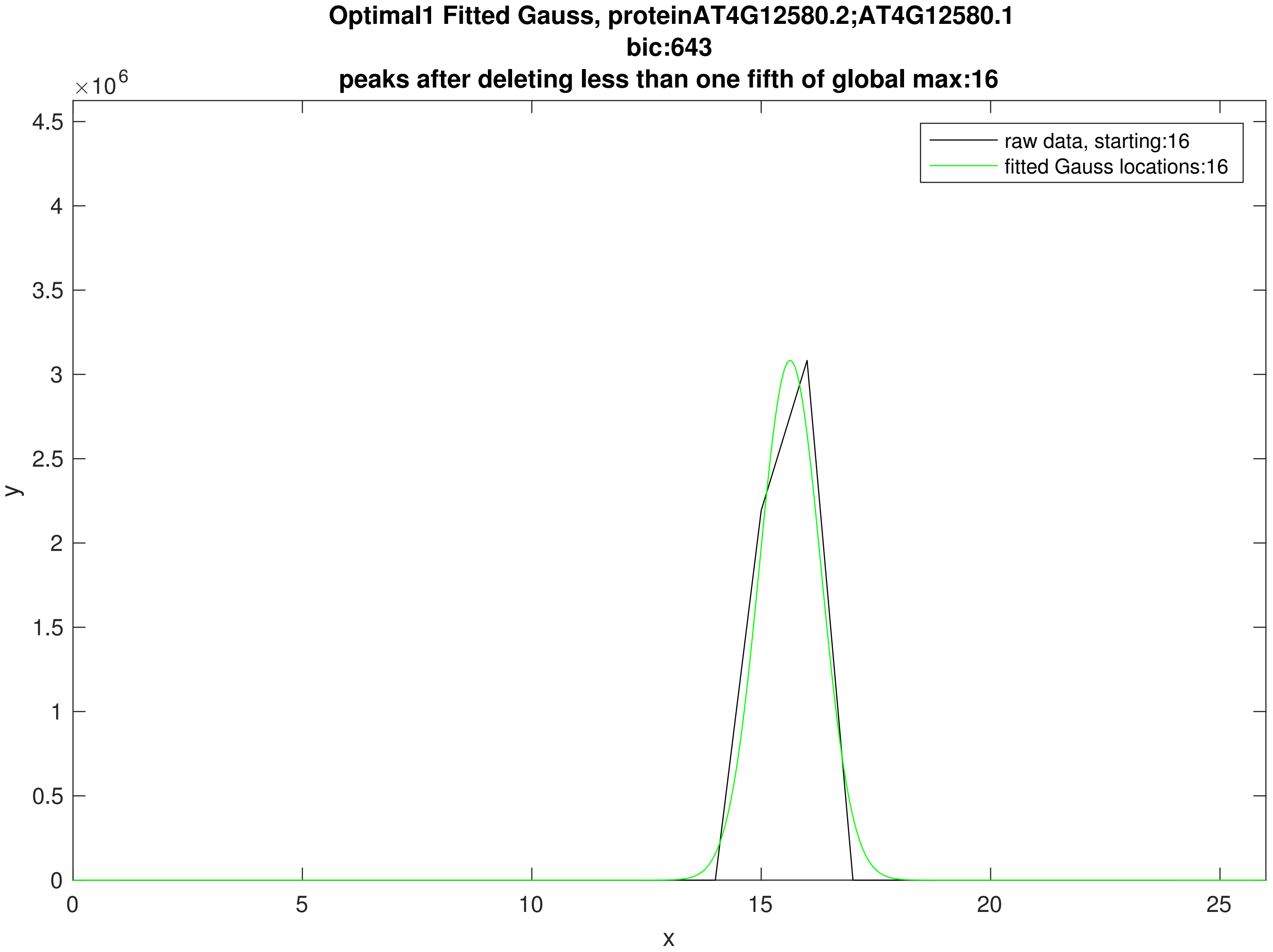
Optimal1 Fitted Gauss, proteinAT4G12580.2;AT4G12580.1
bic:643
peaks after deleting less than one fifth of global max:16
×10^6
4.5
4
3.5
3
2.5
2
1.5
1
0.5
0
y
0
5
10
15
20
25
x
raw data, starting:16
fitted Gauss locations:16

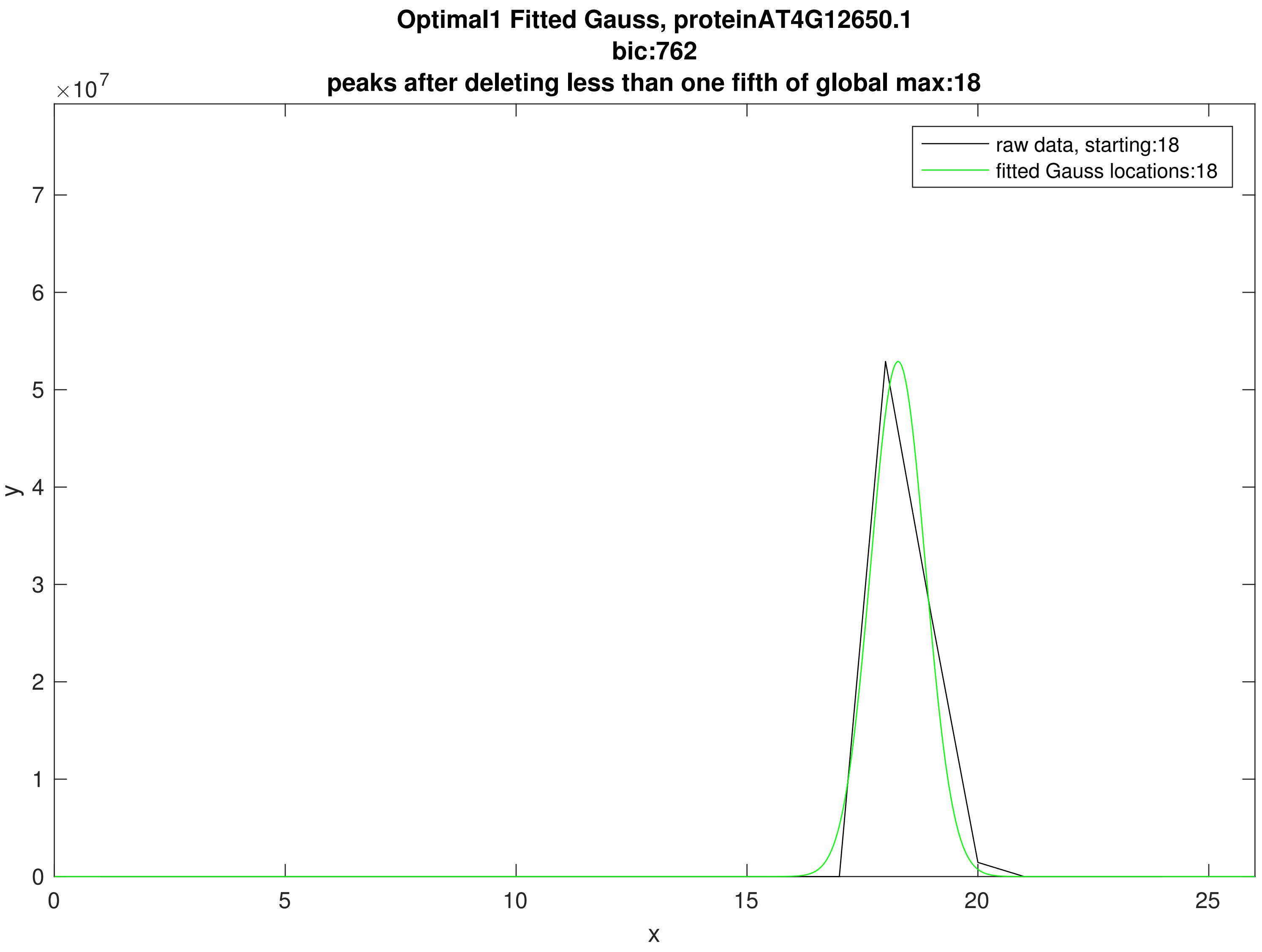
Optimal1 Fitted Gauss, proteinAT4G12650.1
bic:762
peaks after deleting less than one fifth of global max:18
×10^7
raw data, starting:18
fitted Gauss locations:18
y
x
0
1
2
3
4
5
6
7
0
5
10
15
20
25

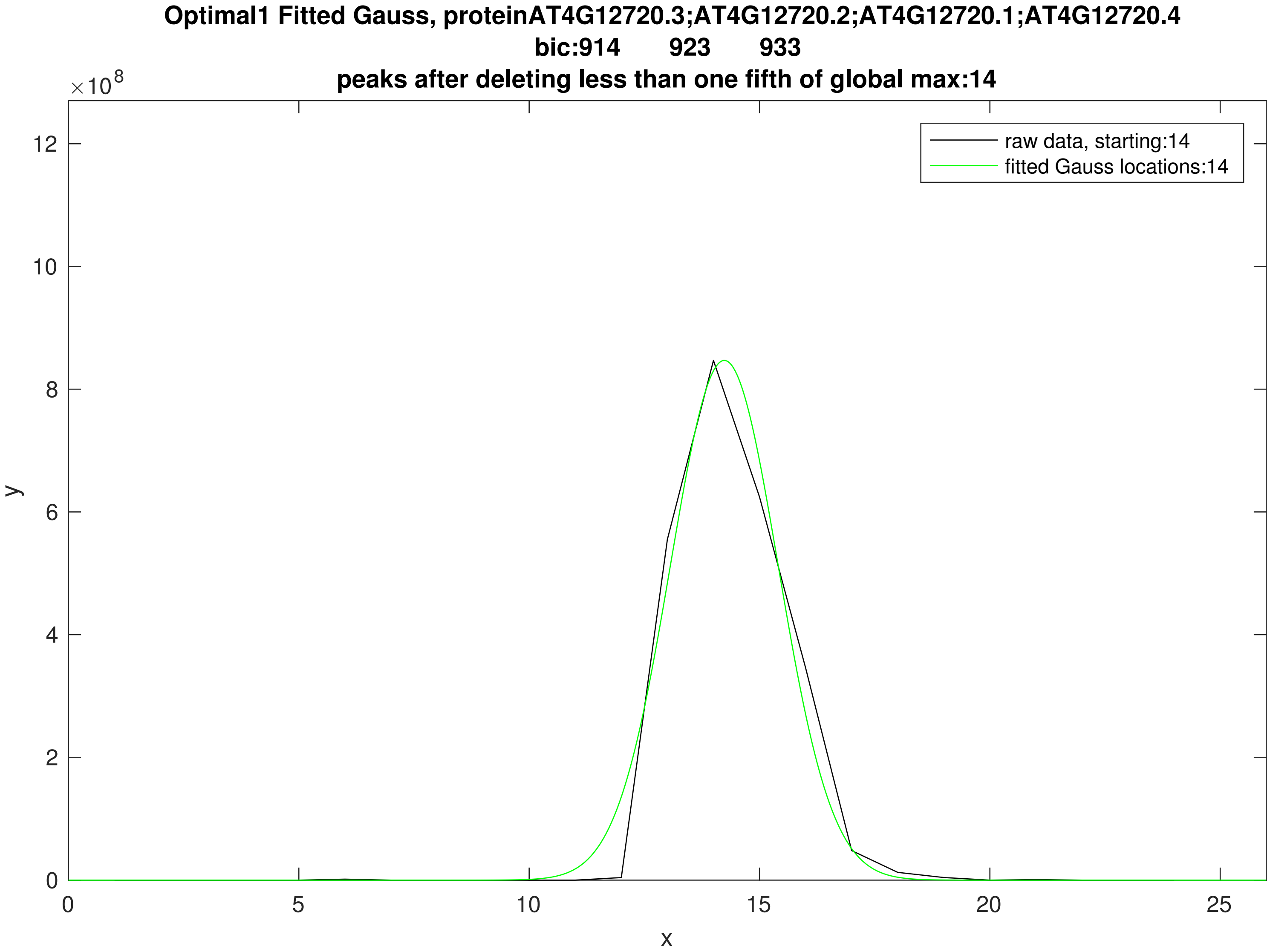
Optimal1 Fitted Gauss, proteinAT4G12720.3;AT4G12720.2;AT4G12720.1;AT4G12720.4
bic:914 923 933
peaks after deleting less than one fifth of global max:14
×10^8
raw data, starting:14
fitted Gauss locations:14
y
x
0
2
4
6
8
10
12
0
5
10
15
20
25

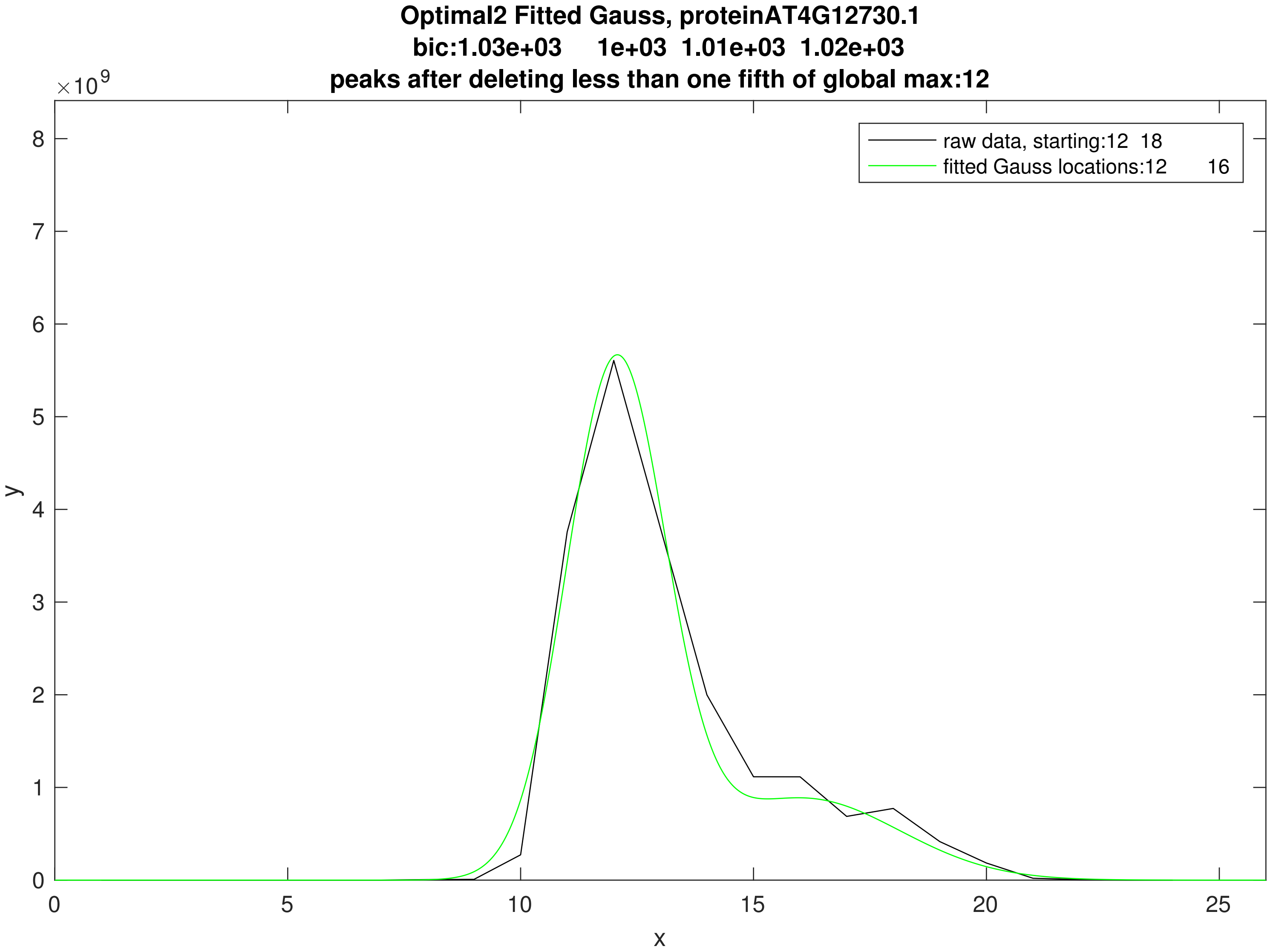
Optimal2 Fitted Gauss, proteinAT4G12730.1
bic:1.03e+03 1e+03 1.01e+03 1.02e+03
peaks after deleting less than one fifth of global max:12
×10^9
raw data, starting:12 18
fitted Gauss locations:12 16
y
x
0
1
2
3
4
5
6
7
8
0
5
10
15
20
25

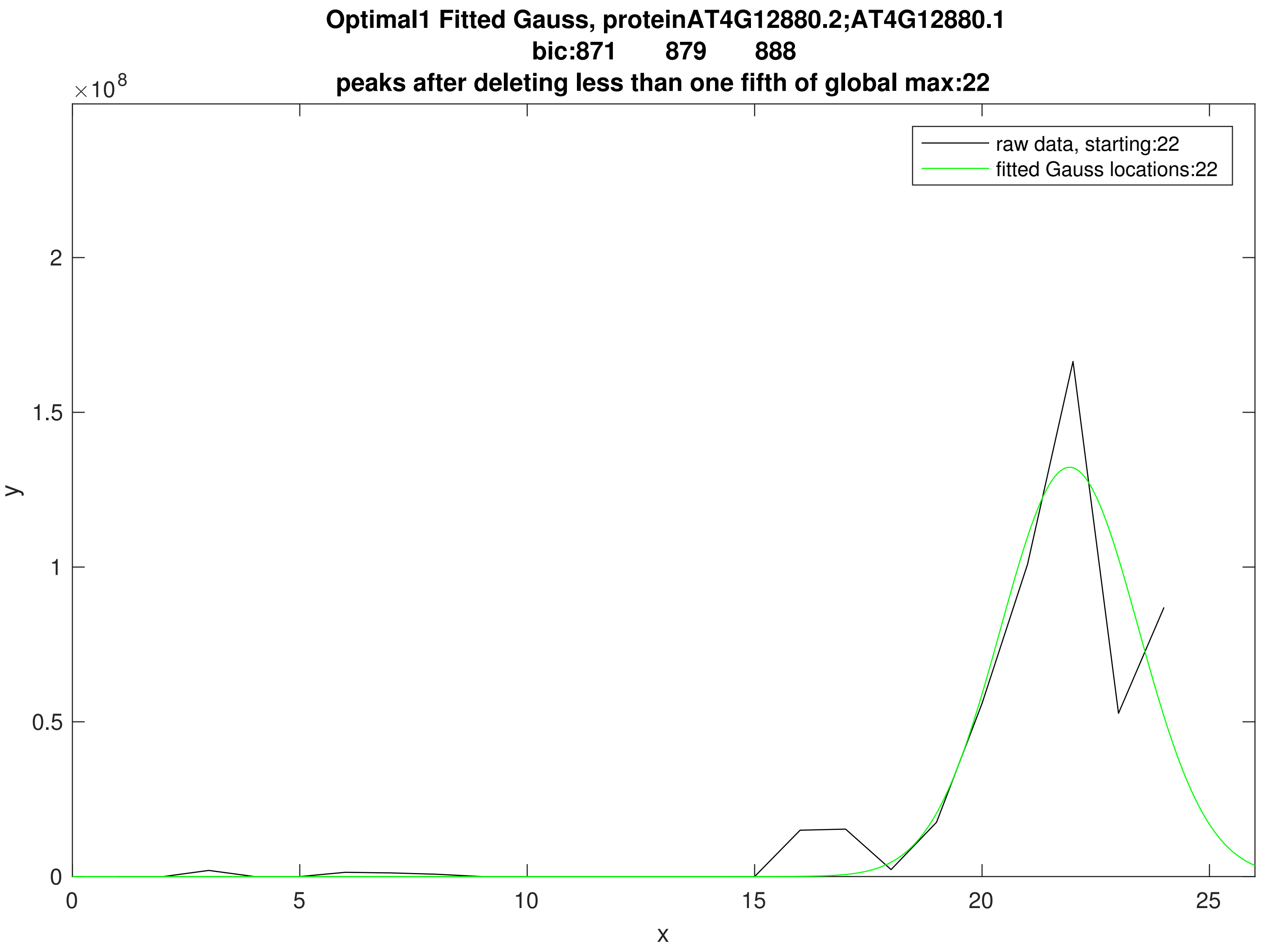
Optimal1 Fitted Gauss, proteinAT4G12880.2;AT4G12880.1
bic:871 879 888
peaks after deleting less than one fifth of global max:22
×10^8
raw data, starting:22
fitted Gauss locations:22
2
1.5
1
0.5
0
y
0
5
10
15
20
25
x

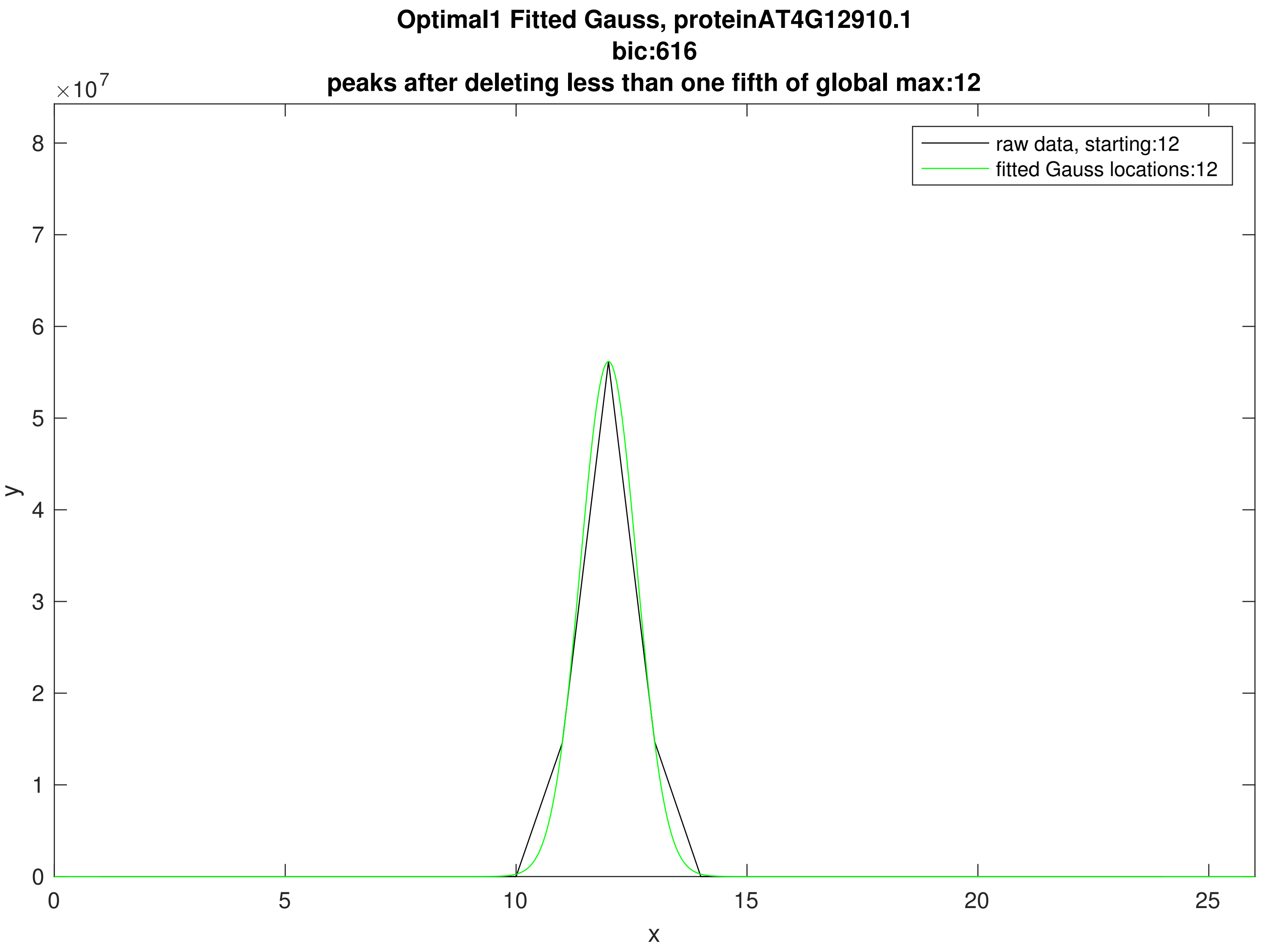

Optimal1 Fitted Gauss, proteinAT4G12910.1
bic:616
peaks after deleting less than one fifth of global max:12
×10^7
raw data, starting:12
fitted Gauss locations:12
y
x
0
1
2
3
4
5
6
7
8
0
5
10
15
20
25

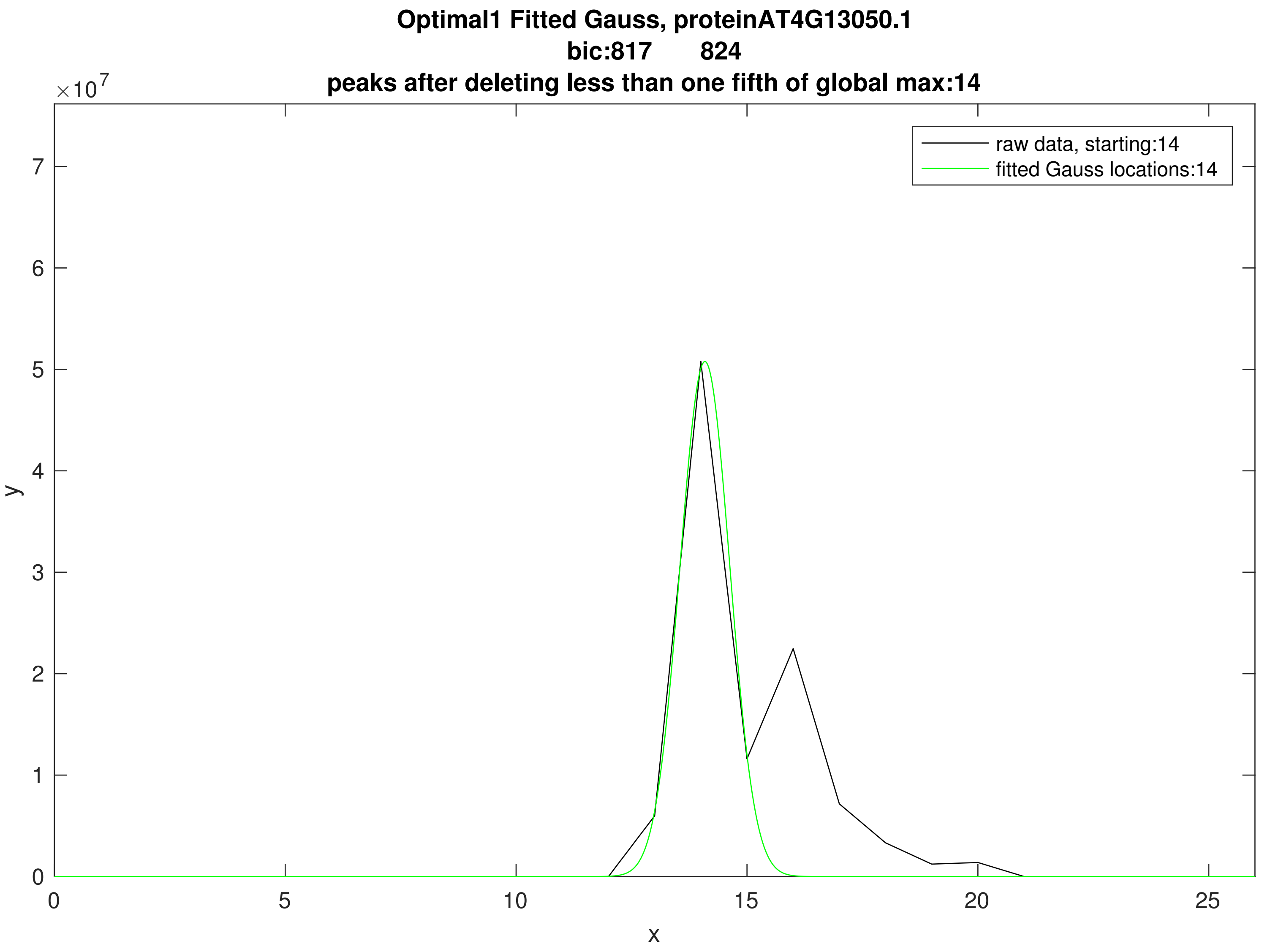
Optimal1 Fitted Gauss, proteinAT4G13050.1
bic:817 824
peaks after deleting less than one fifth of global max:14
×10^7
raw data, starting:14
fitted Gauss locations:14
y
x
0
1
2
3
4
5
6
7
0
5
10
15
20
25

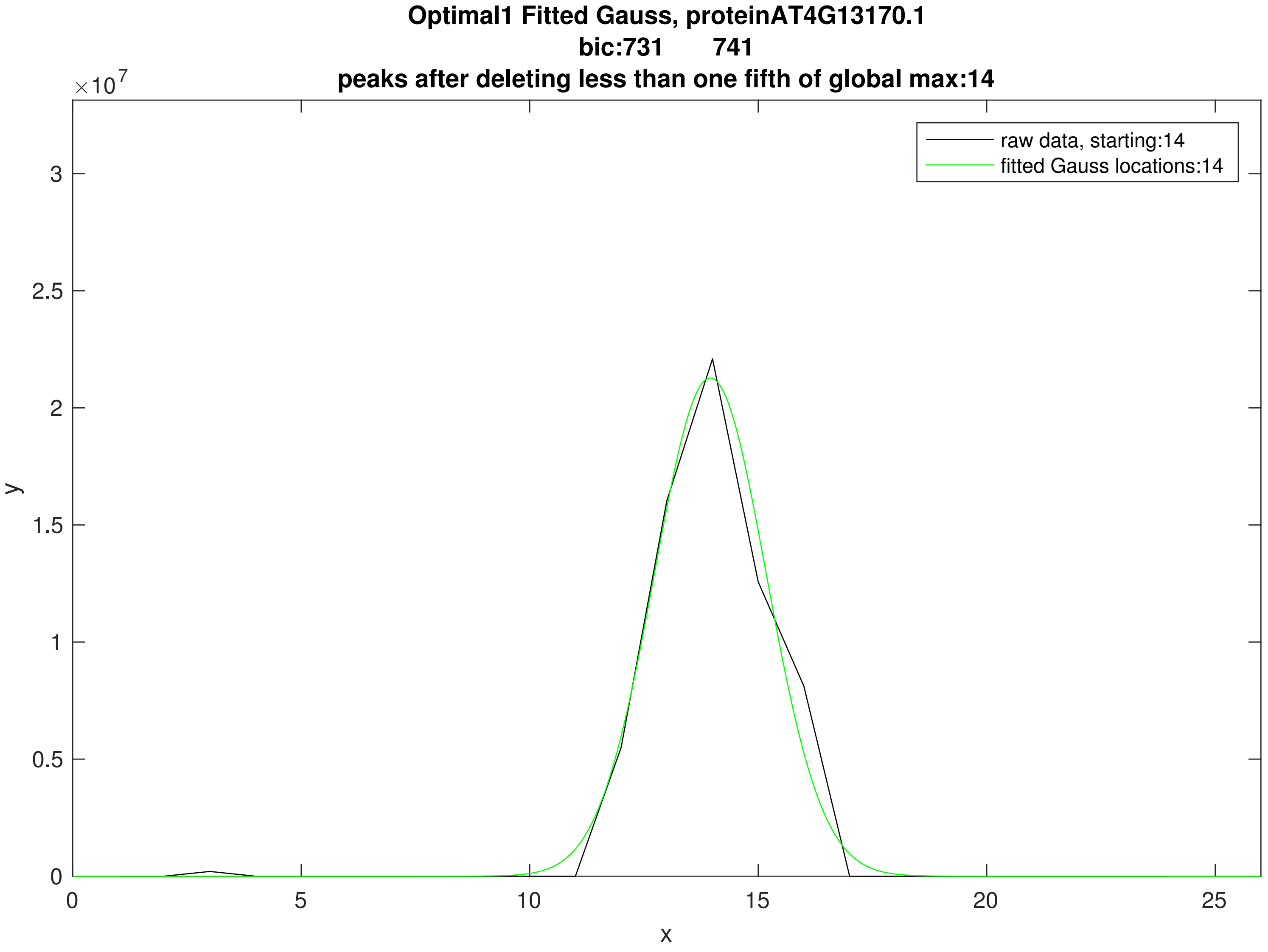
Optimal1 Fitted Gauss, proteinAT4G13170.1
bic:731 741
peaks after deleting less than one fifth of global max:14
×10^7
3
2.5
2
1.5
1
0.5
0
y
0
5
10
15
20
25
x
raw data, starting:14
fitted Gauss locations:14

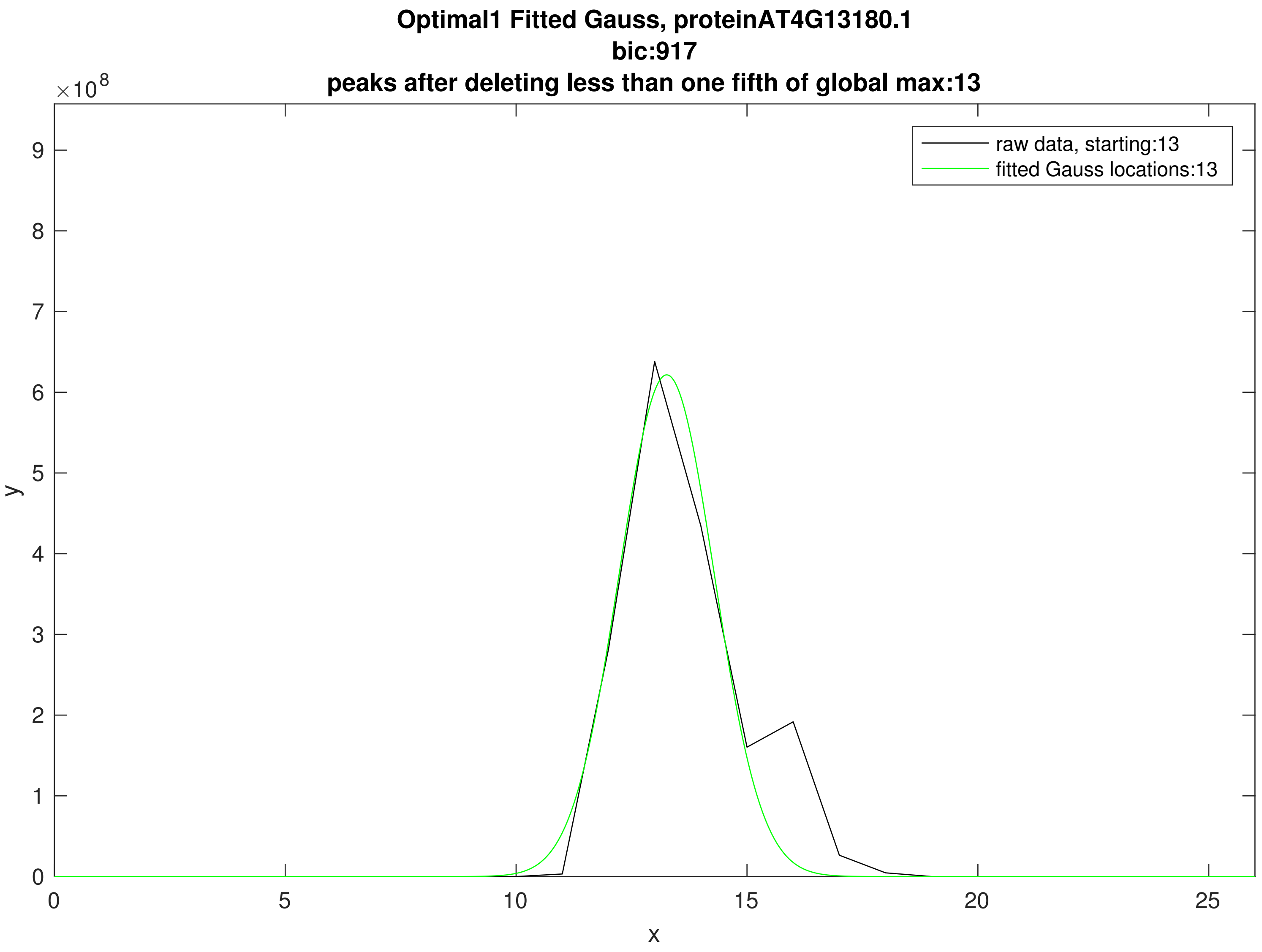

Optimal1 Fitted Gauss, proteinAT4G13180.1
bic:917
peaks after deleting less than one fifth of global max:13
×10^8
raw data, starting:13
fitted Gauss locations:13
y
x
0
1
2
3
4
5
6
7
8
9
0
5
10
15
20
25

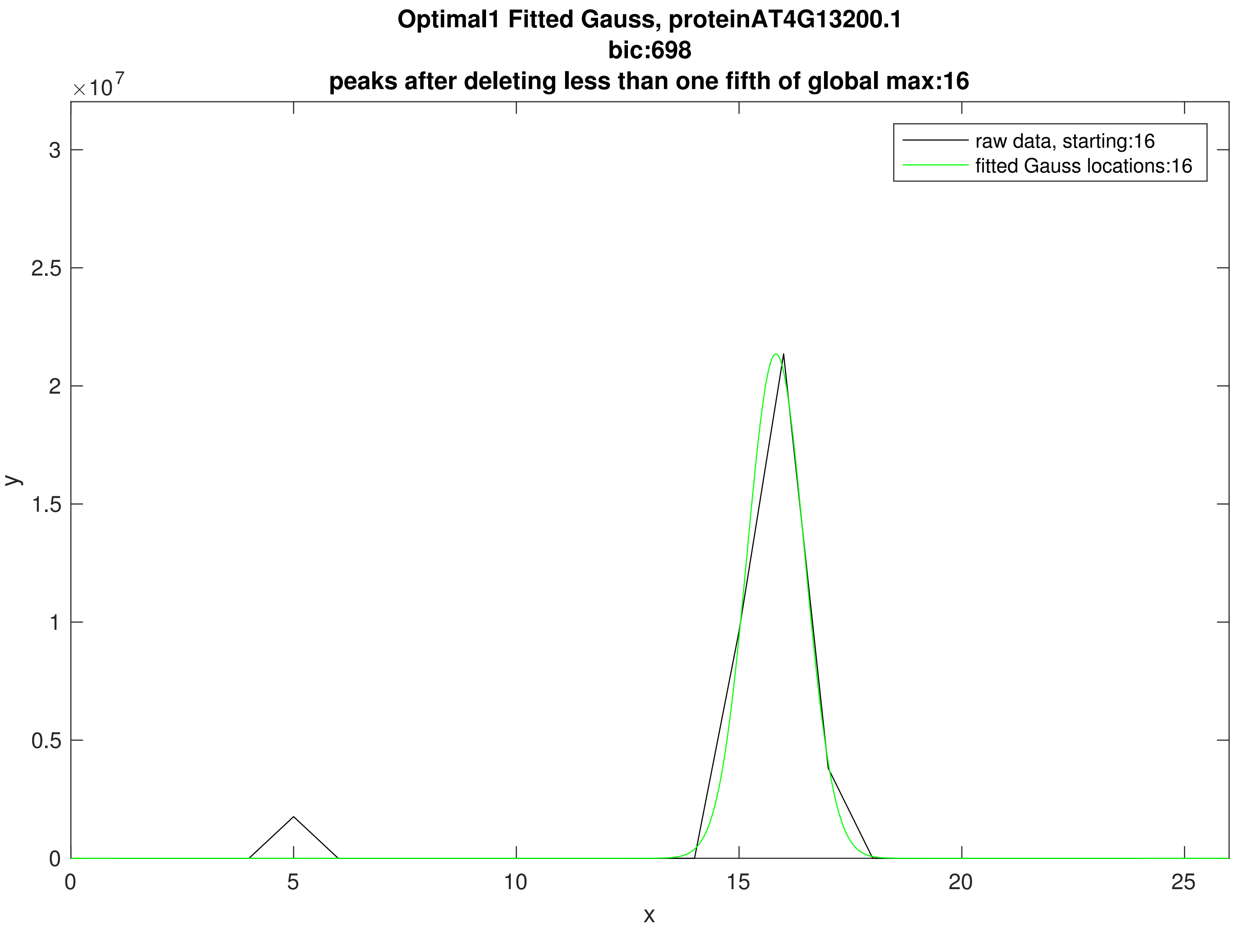
Optimal1 Fitted Gauss, proteinAT4G13200.1
bic:698
peaks after deleting less than one fifth of global max:16
×10^7
raw data, starting:16
fitted Gauss locations:16
y
x
0
0.5
1
1.5
2
2.5
3
0
5
10
15
20
25

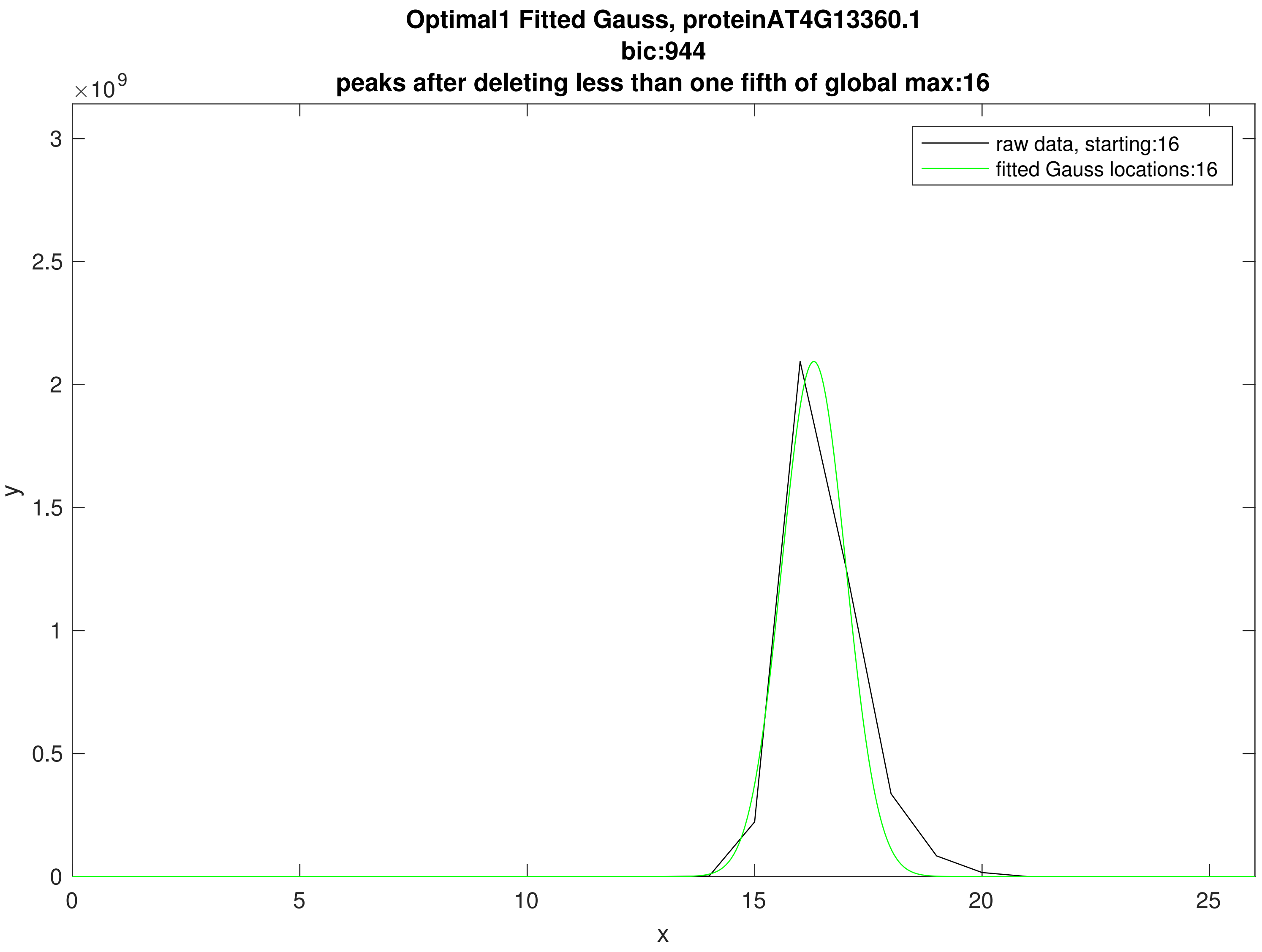
Optimal1 Fitted Gauss, proteinAT4G13360.1
bic:944
peaks after deleting less than one fifth of global max:16
×10^9
raw data, starting:16
fitted Gauss locations:16
y
x
0
0.5
1
1.5
2
2.5
3
0
5
10
15
20
25

**Optimal1 Fitted Gauss, proteinAT4G13430.1**
**bic:965 972 982 991**
**peaks after deleting less than one fifth of global max:15**

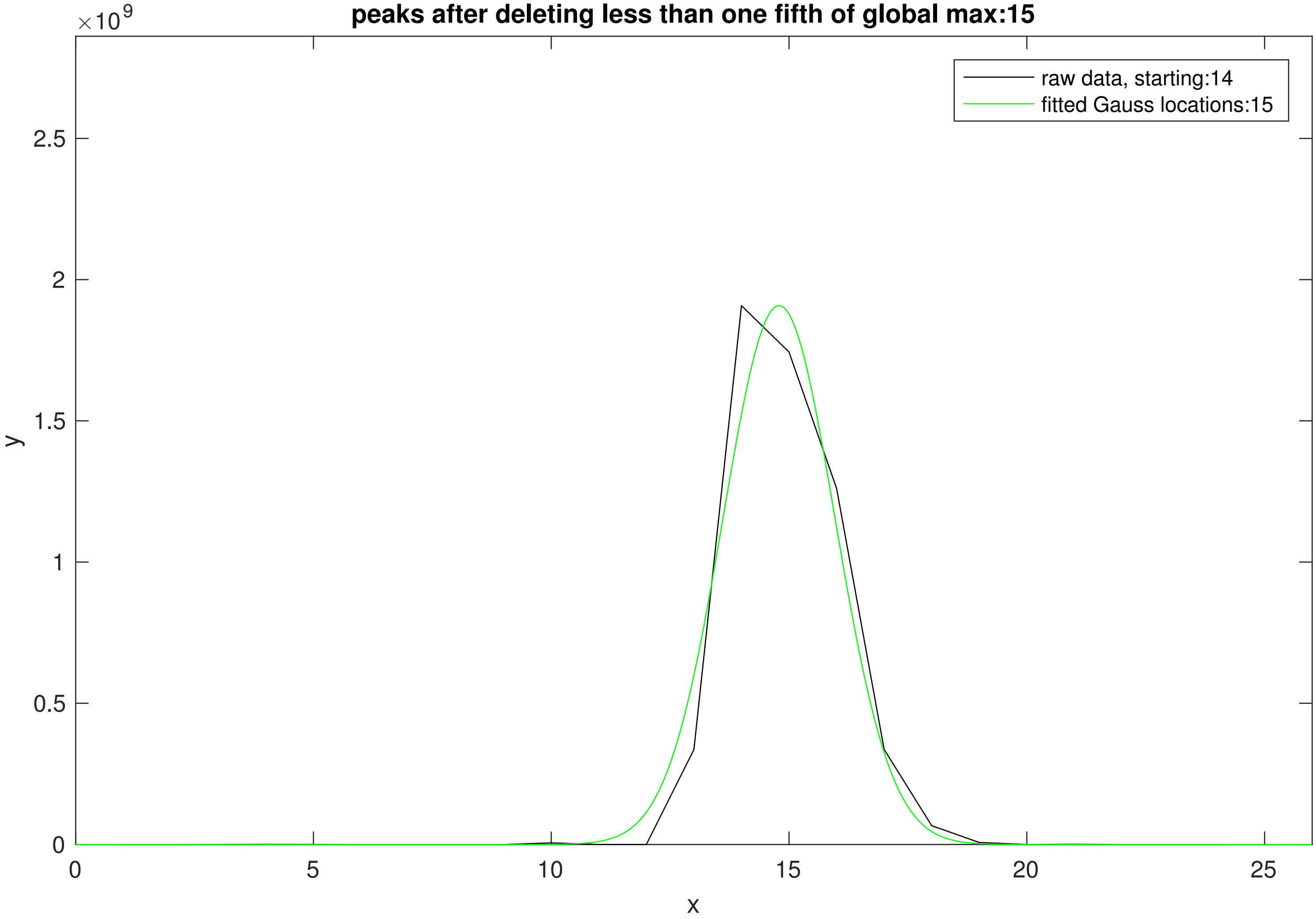

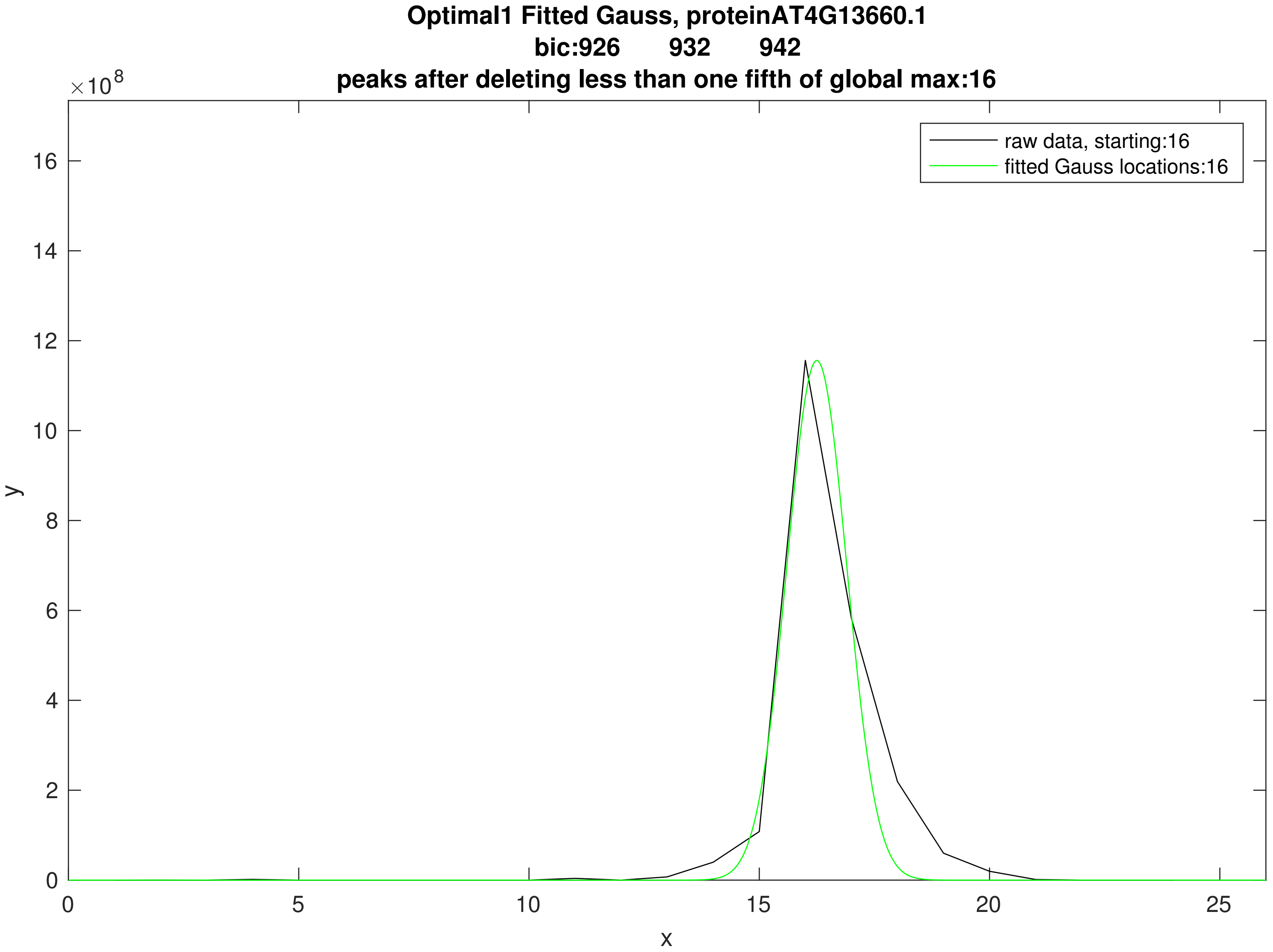

Optimal1 Fitted Gauss, proteinAT4G13660.1
bic:926 932 942
peaks after deleting less than one fifth of global max:16
×10^8
16
14
12
10
8
6
4
2
0
y
0
5
10
15
20
25
x
raw data, starting:16
fitted Gauss locations:16

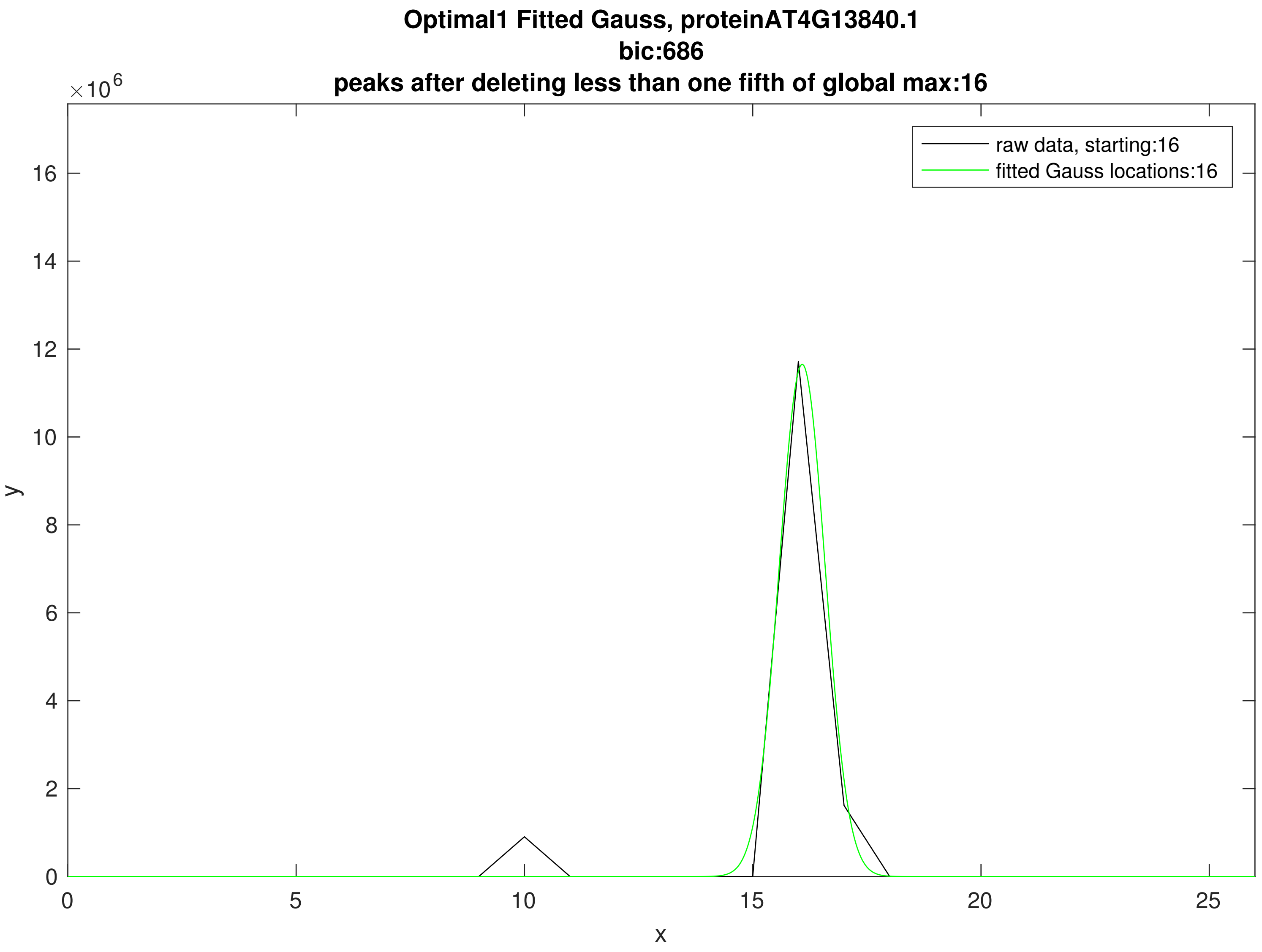

Optimal1 Fitted Gauss, proteinAT4G13840.1
bic:686
peaks after deleting less than one fifth of global max:16
×10^6
raw data, starting:16
fitted Gauss locations:16
y
x
0
2
4
6
8
10
12
14
16
0
5
10
15
20
25

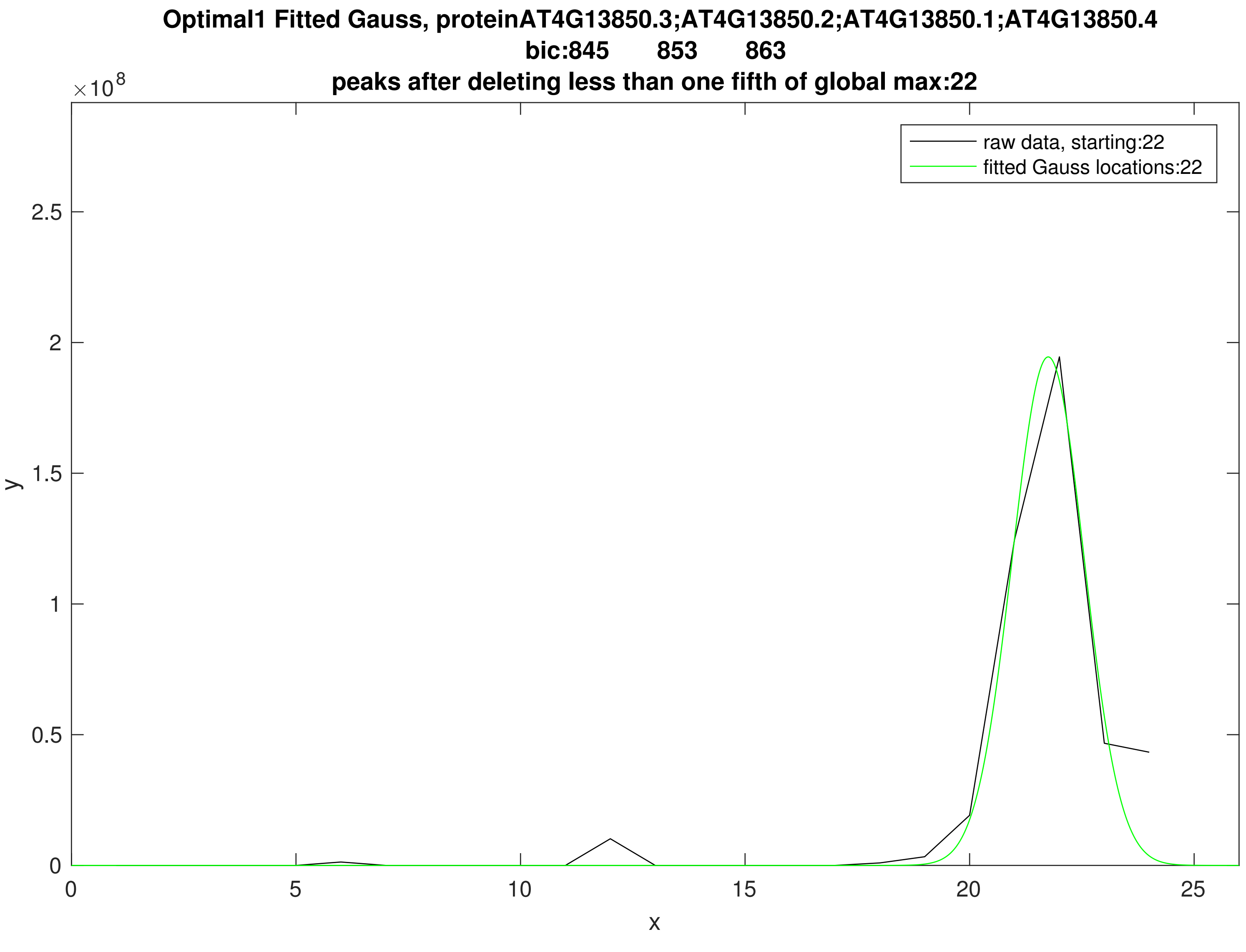
Optimal1 Fitted Gauss, proteinAT4G13850.3;AT4G13850.2;AT4G13850.1;AT4G13850.4
bic:845 853 863
peaks after deleting less than one fifth of global max:22
×10^8
raw data, starting:22
fitted Gauss locations:22
y
x
0
0.5
1
1.5
2
2.5
5
10
15
20
25

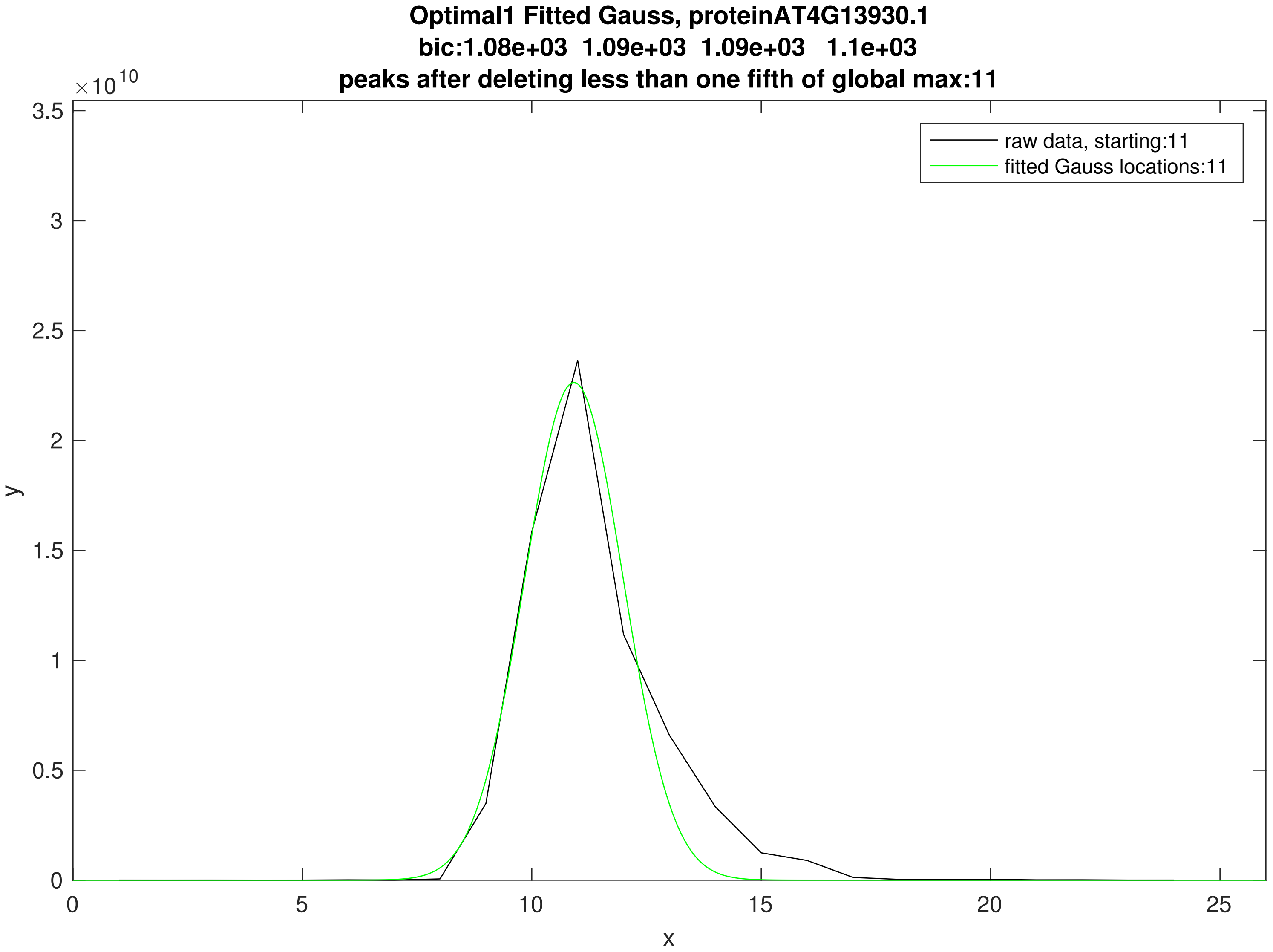
Optimal1 Fitted Gauss, proteinAT4G13930.1
bic:1.08e+03  1.09e+03  1.09e+03   1.1e+03
peaks after deleting less than one fifth of global max:11
×10^10
3.5
3
2.5
2
1.5
1
0.5
0
y
0
5
10
15
20
25
x
raw data, starting:11
fitted Gauss locations:11

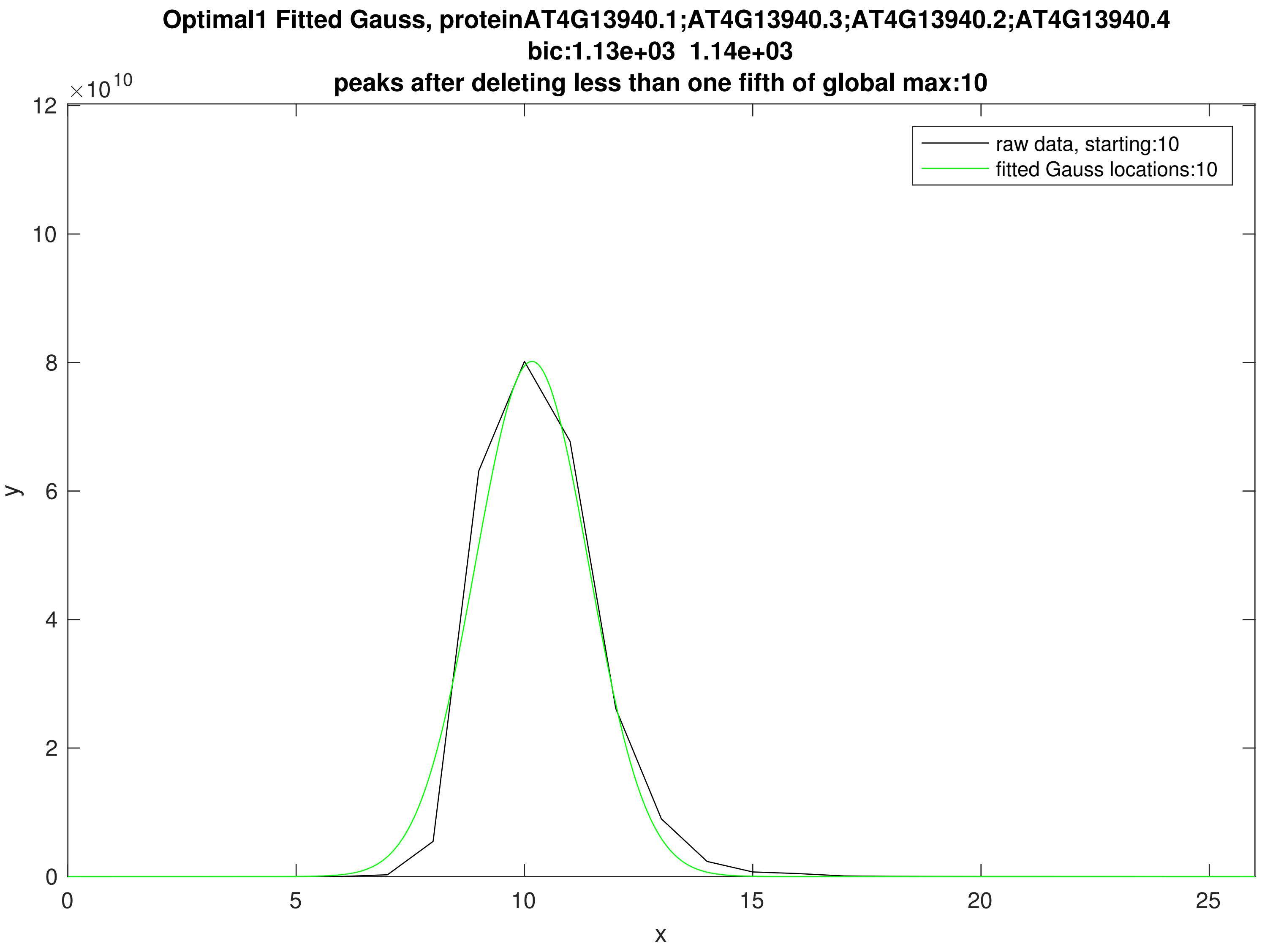
Optimal1 Fitted Gauss, proteinAT4G13940.1;AT4G13940.3;AT4G13940.2;AT4G13940.4
bic:1.13e+03  1.14e+03
peaks after deleting less than one fifth of global max:10
×10^10
12
10
8
6
4
2
0
y
0
5
10
15
20
25
x
raw data, starting:10
fitted Gauss locations:10

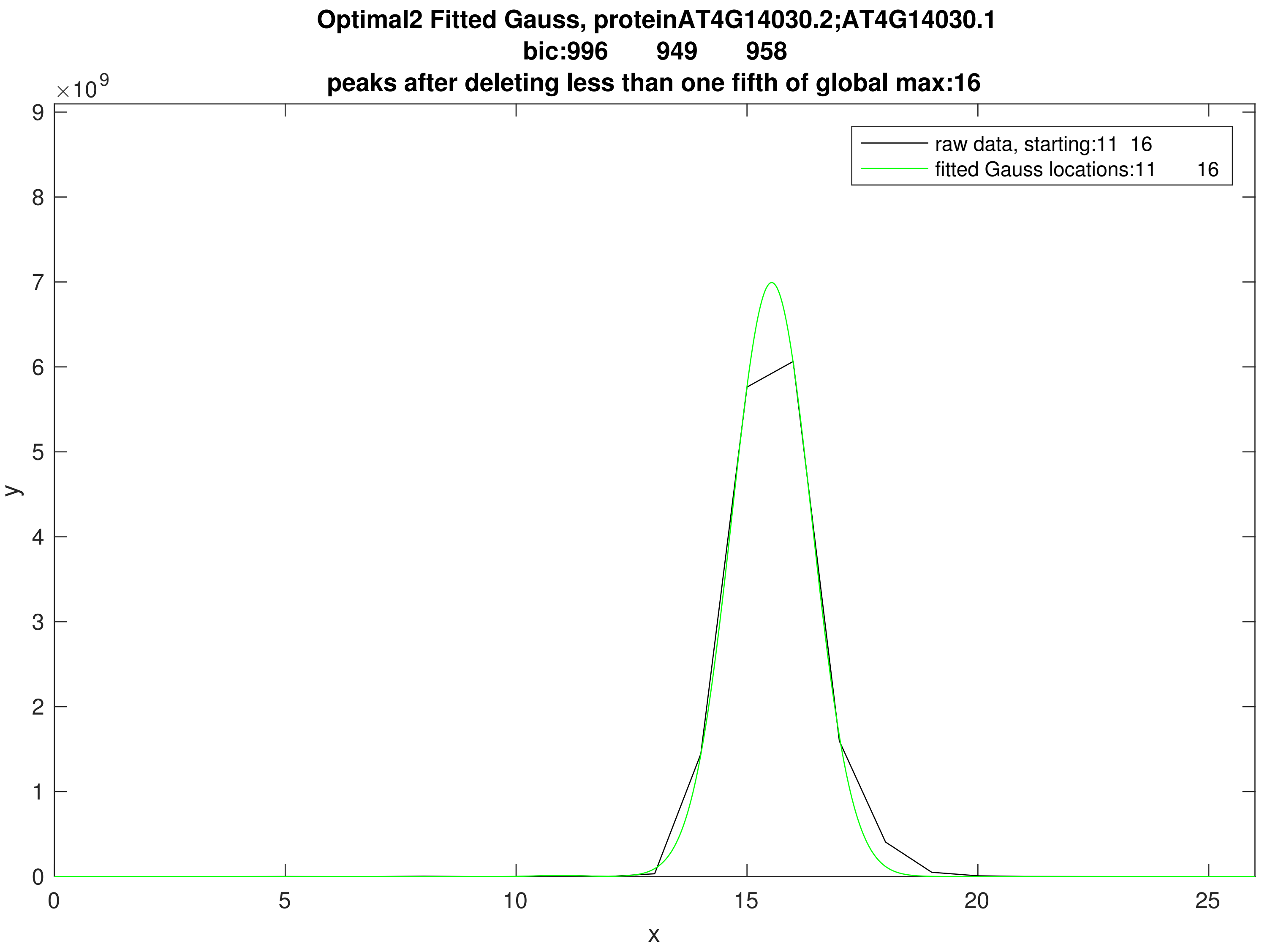
Optimal2 Fitted Gauss, proteinAT4G14030.2;AT4G14030.1
bic:996 949 958
peaks after deleting less than one fifth of global max:16
×10^9
9
8
7
6
5
4
3
2
1
0
y
0
5
10
15
20
25
x
raw data, starting:11 16
fitted Gauss locations:11 16

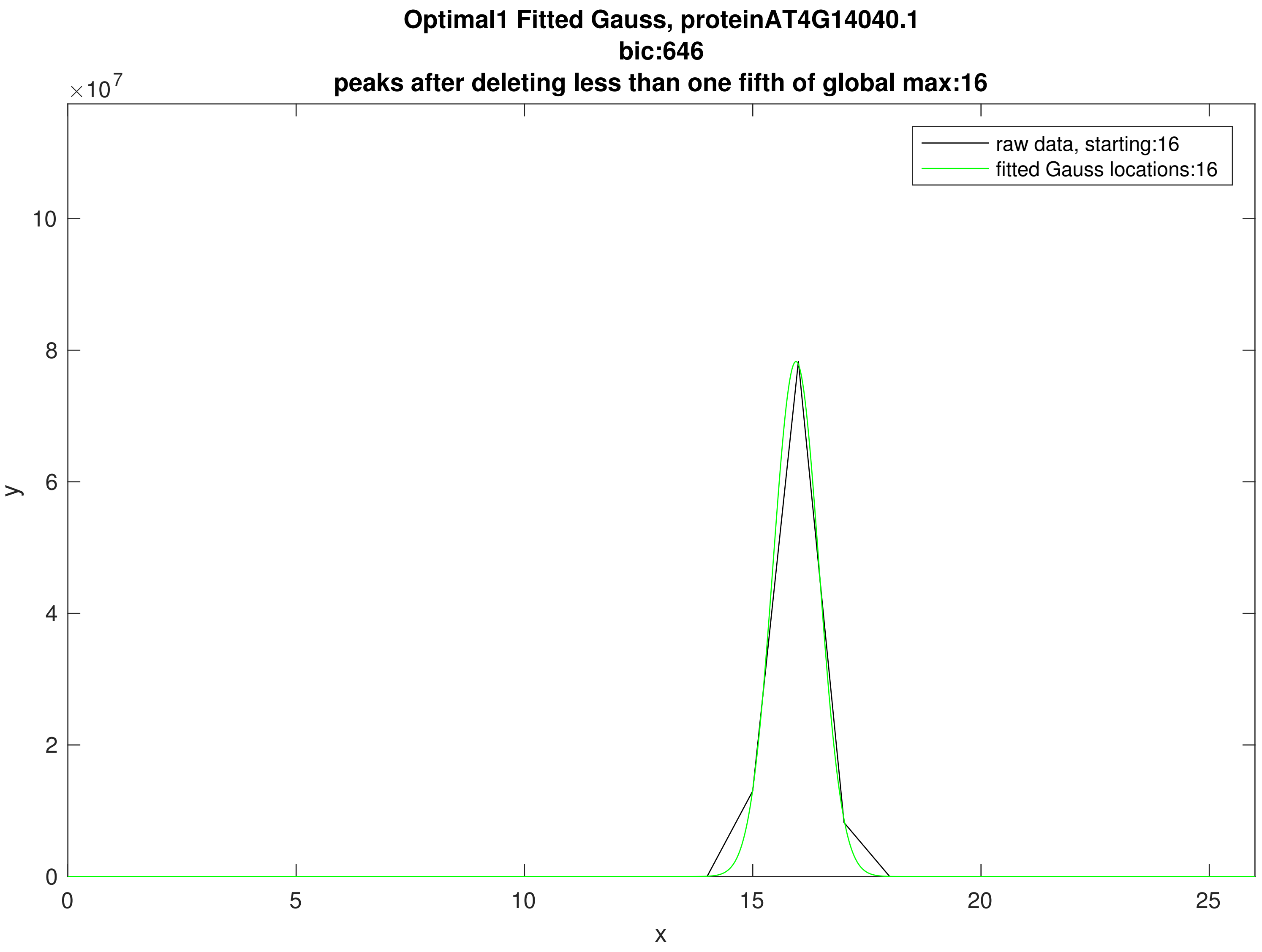

Optimal1 Fitted Gauss, proteinAT4G14040.1
bic:646
peaks after deleting less than one fifth of global max:16
×10^7
raw data, starting:16
fitted Gauss locations:16
y
x
0
2
4
6
8
10
0
5
10
15
20
25

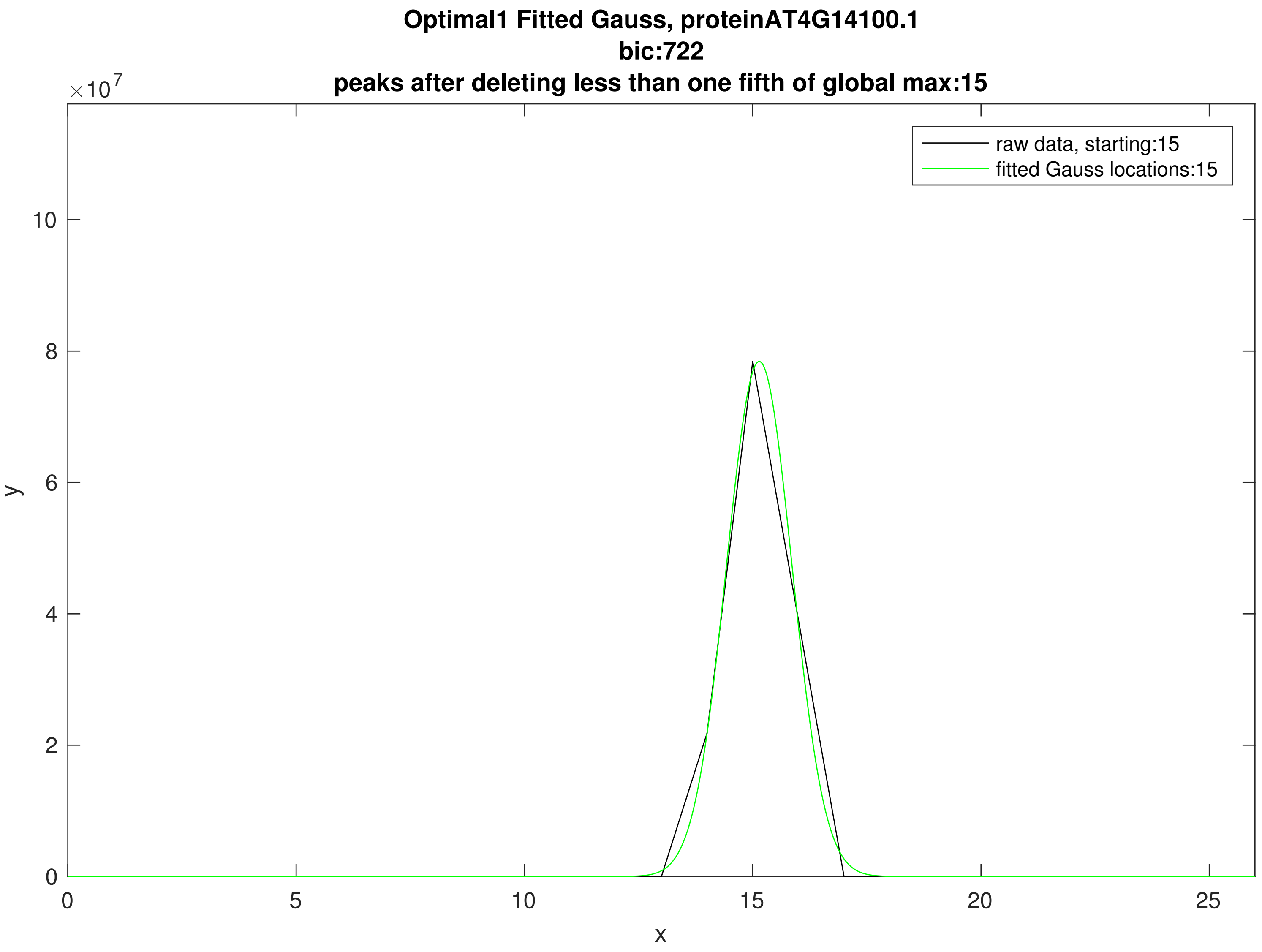
Optimal1 Fitted Gauss, proteinAT4G14100.1
bic:722
peaks after deleting less than one fifth of global max:15
×10^7
raw data, starting:15
fitted Gauss locations:15
10
8
6
4
2
0
y
0
5
10
15
20
25
x

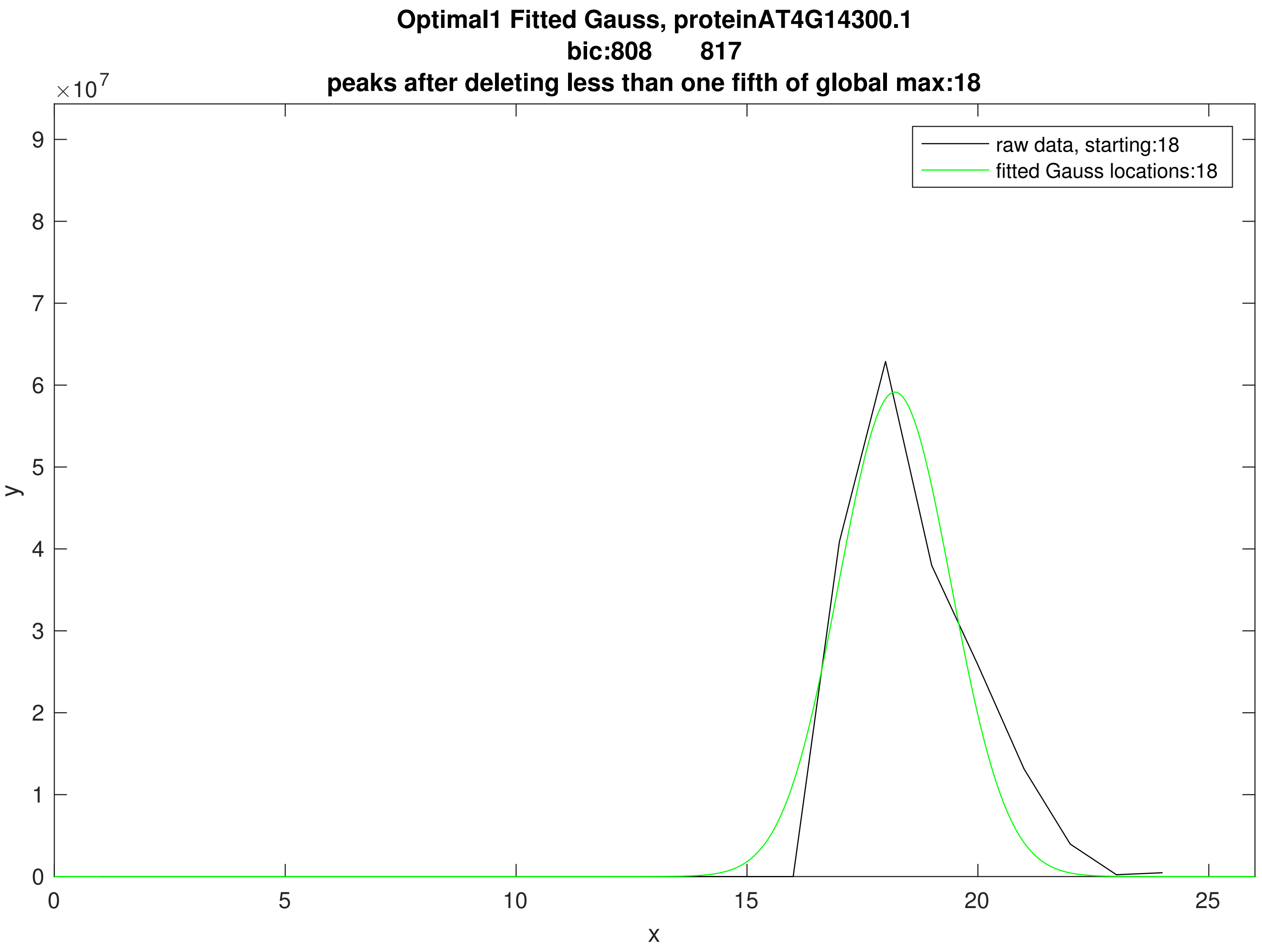

Optimal1 Fitted Gauss, proteinAT4G14300.1
bic:808 817
peaks after deleting less than one fifth of global max:18
×10^7
9
8
7
6
5
4
3
2
1
0
y
0
5
10
15
20
25
x
raw data, starting:18
fitted Gauss locations:18

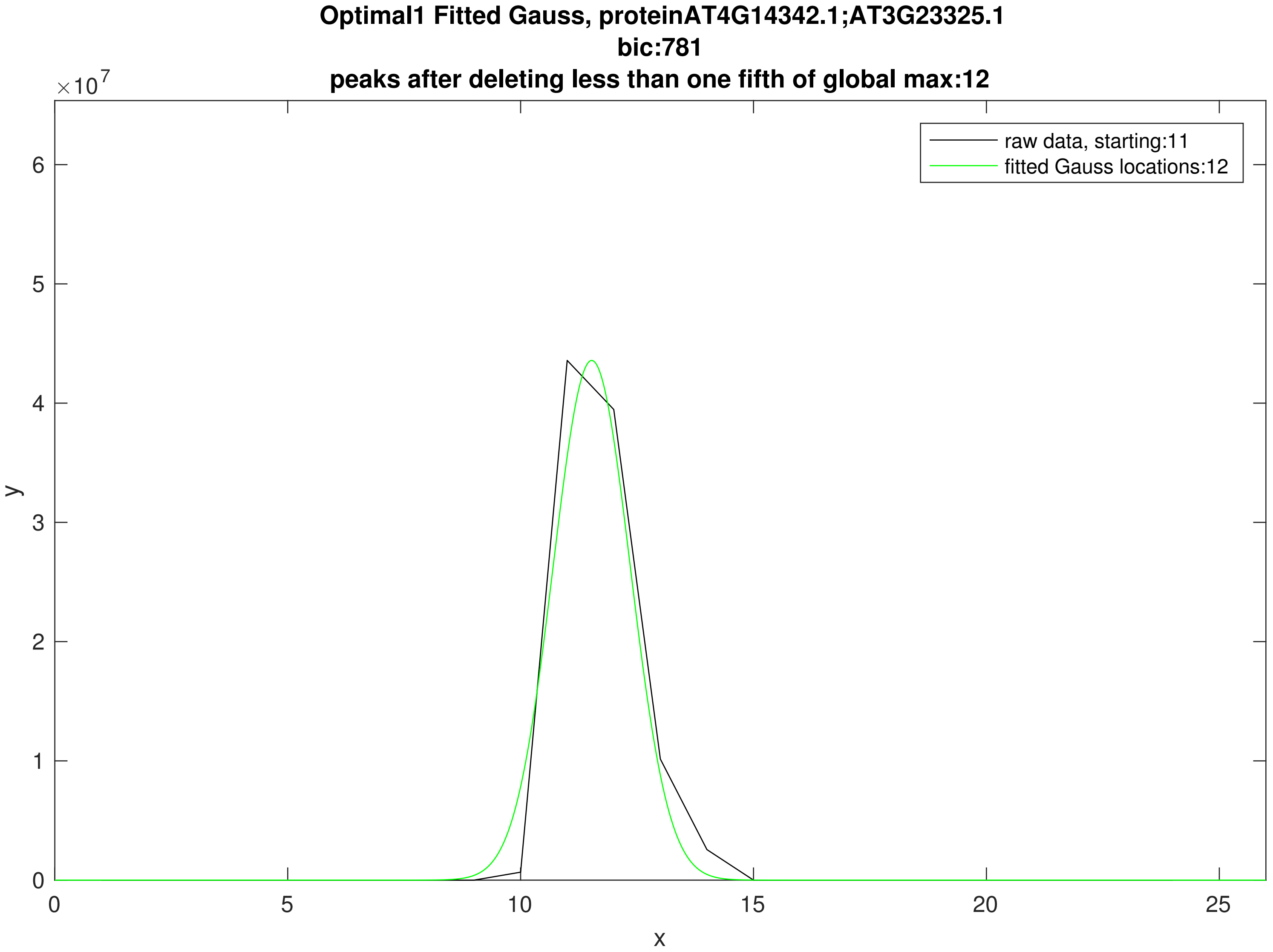

Optimal1 Fitted Gauss, proteinAT4G14342.1;AT3G23325.1
bic:781
peaks after deleting less than one fifth of global max:12
×10^7
raw data, starting:11
fitted Gauss locations:12
y
x
0
1
2
3
4
5
6
0
5
10
15
20
25

Optimal1 Fitted Gauss, proteinAT4G14440.1
bic:772
peaks after deleting less than one fifth of global max:16

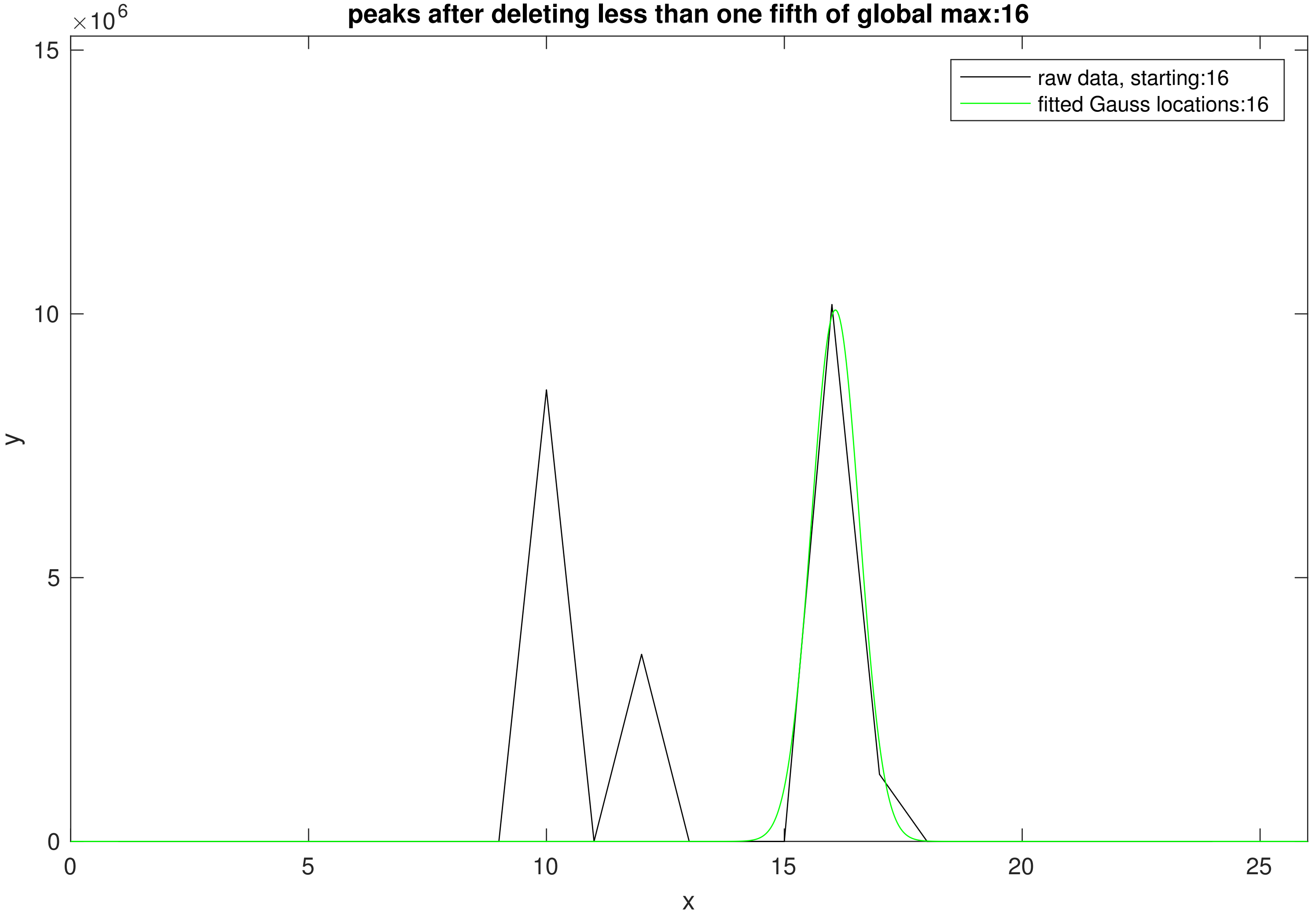

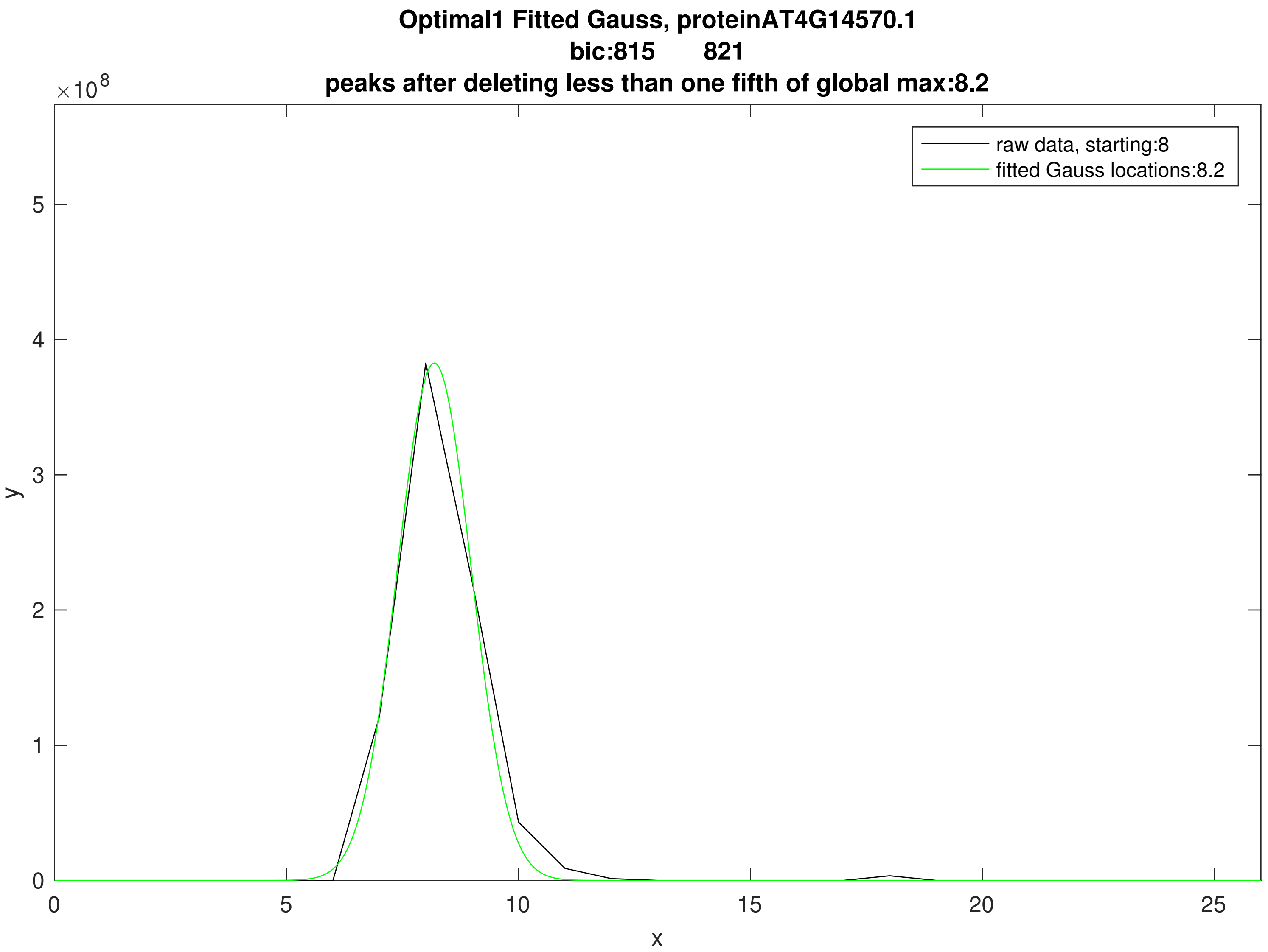

Optimal1 Fitted Gauss, proteinAT4G14570.1
bic:815 821
peaks after deleting less than one fifth of global max:8.2
×10^8
raw data, starting:8
fitted Gauss locations:8.2
y
x
0
1
2
3
4
5
10
15
20
25

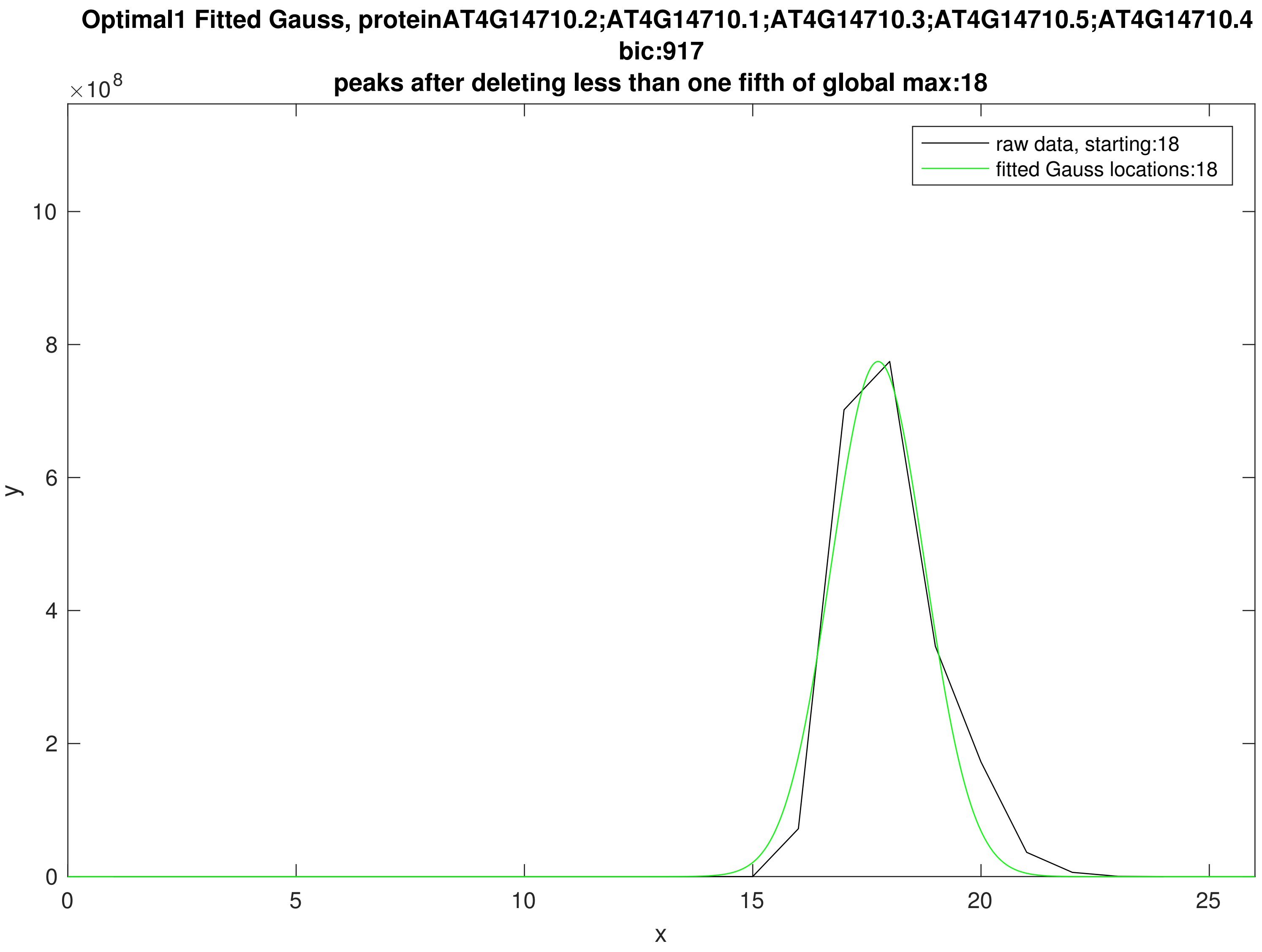
Optimal1 Fitted Gauss, proteinAT4G14710.2;AT4G14710.1;AT4G14710.3;AT4G14710.5;AT4G14710.4
bic:917
peaks after deleting less than one fifth of global max:18
×10^8
raw data, starting:18
fitted Gauss locations:18
y
x
0
2
4
6
8
10
0
5
10
15
20
25

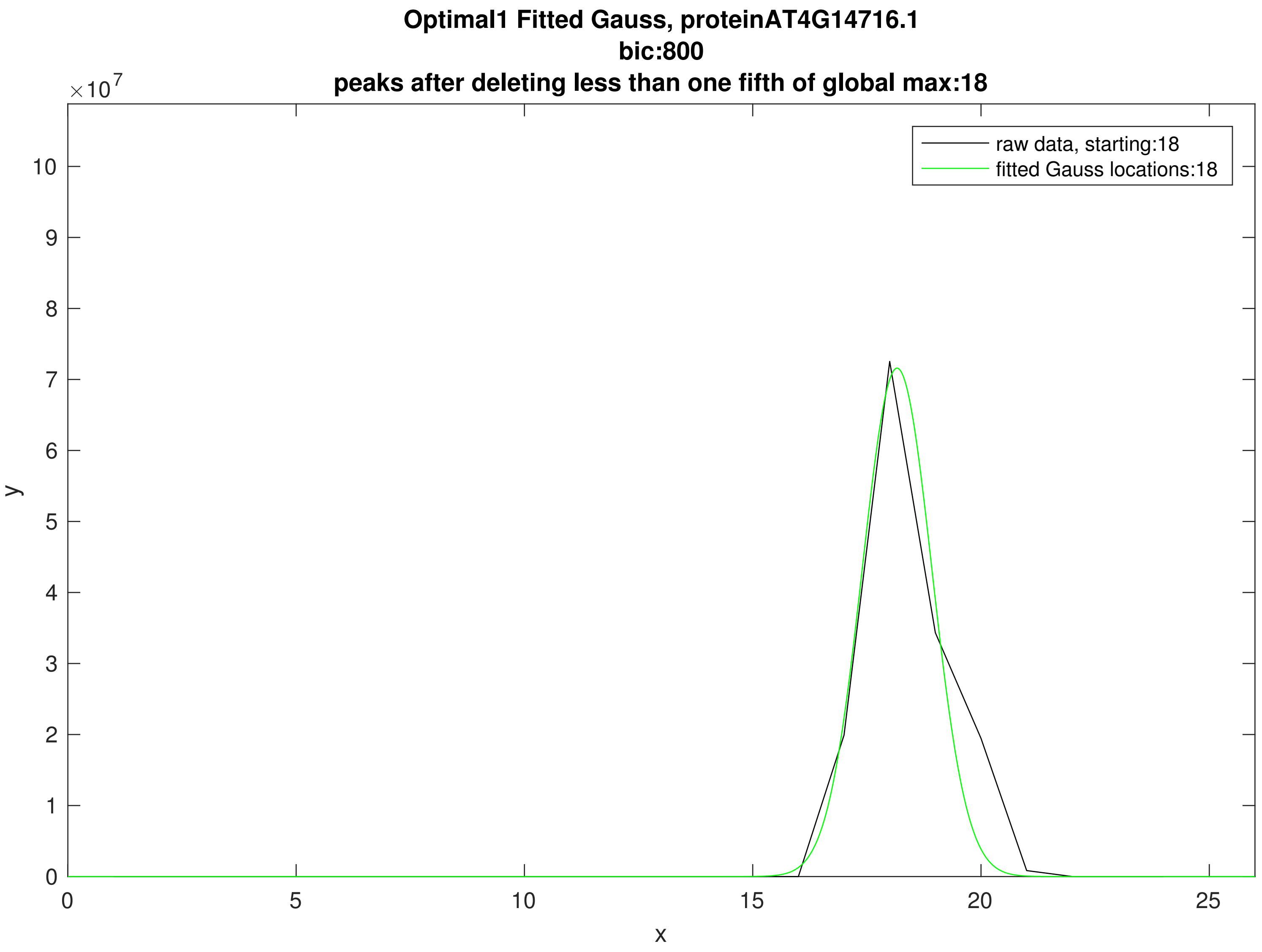
Optimal1 Fitted Gauss, proteinAT4G14716.1
bic:800
peaks after deleting less than one fifth of global max:18
×10^7
raw data, starting:18
fitted Gauss locations:18
y
x
0
1
2
3
4
5
6
7
8
9
10
0
5
10
15
20
25

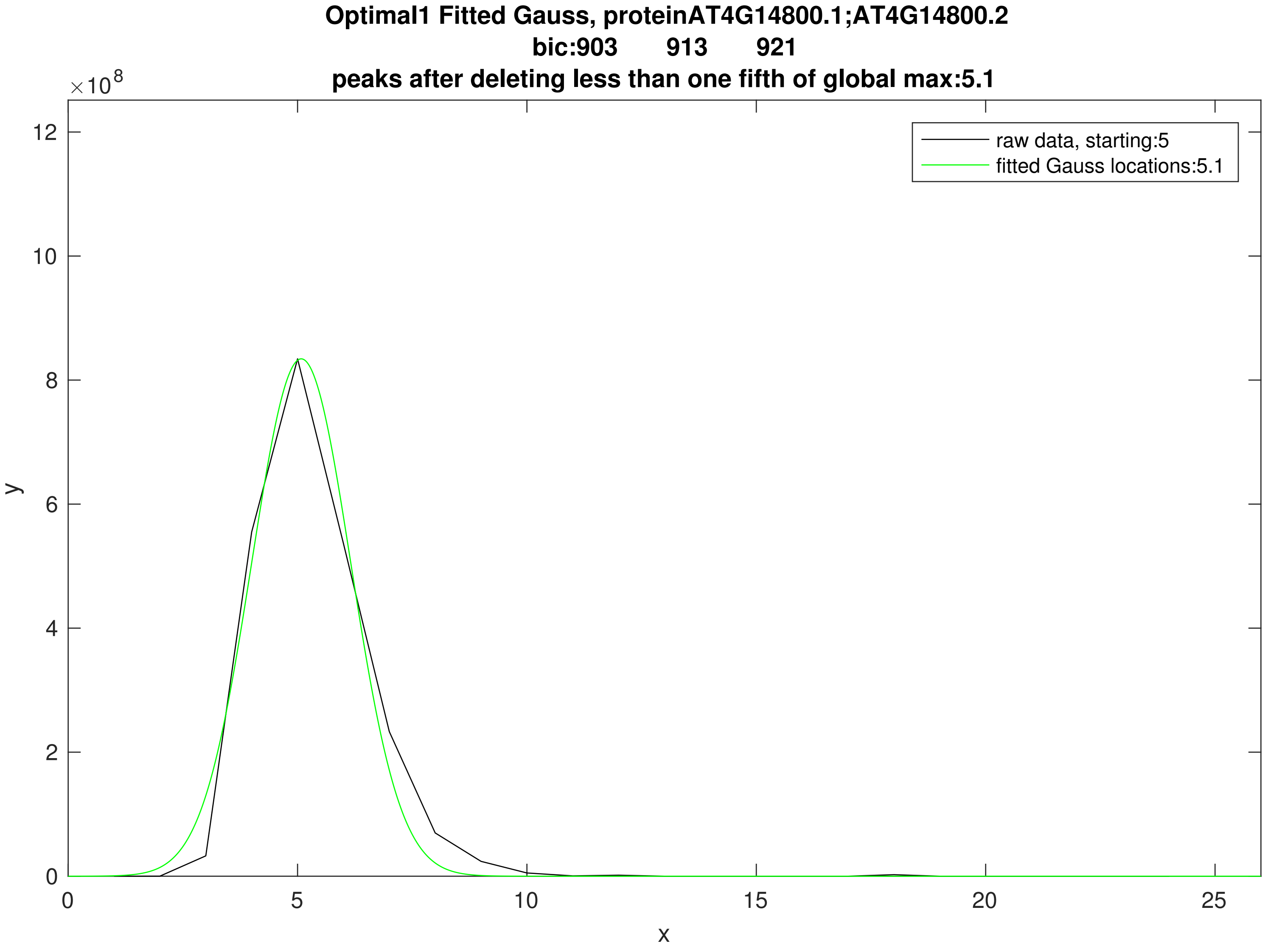

Optimal1 Fitted Gauss, proteinAT4G14800.1;AT4G14800.2
bic:903 913 921
peaks after deleting less than one fifth of global max:5.1
×10^8
12
10
8
6
4
2
0
y
0
5
10
15
20
25
x
raw data, starting:5
fitted Gauss locations:5.1

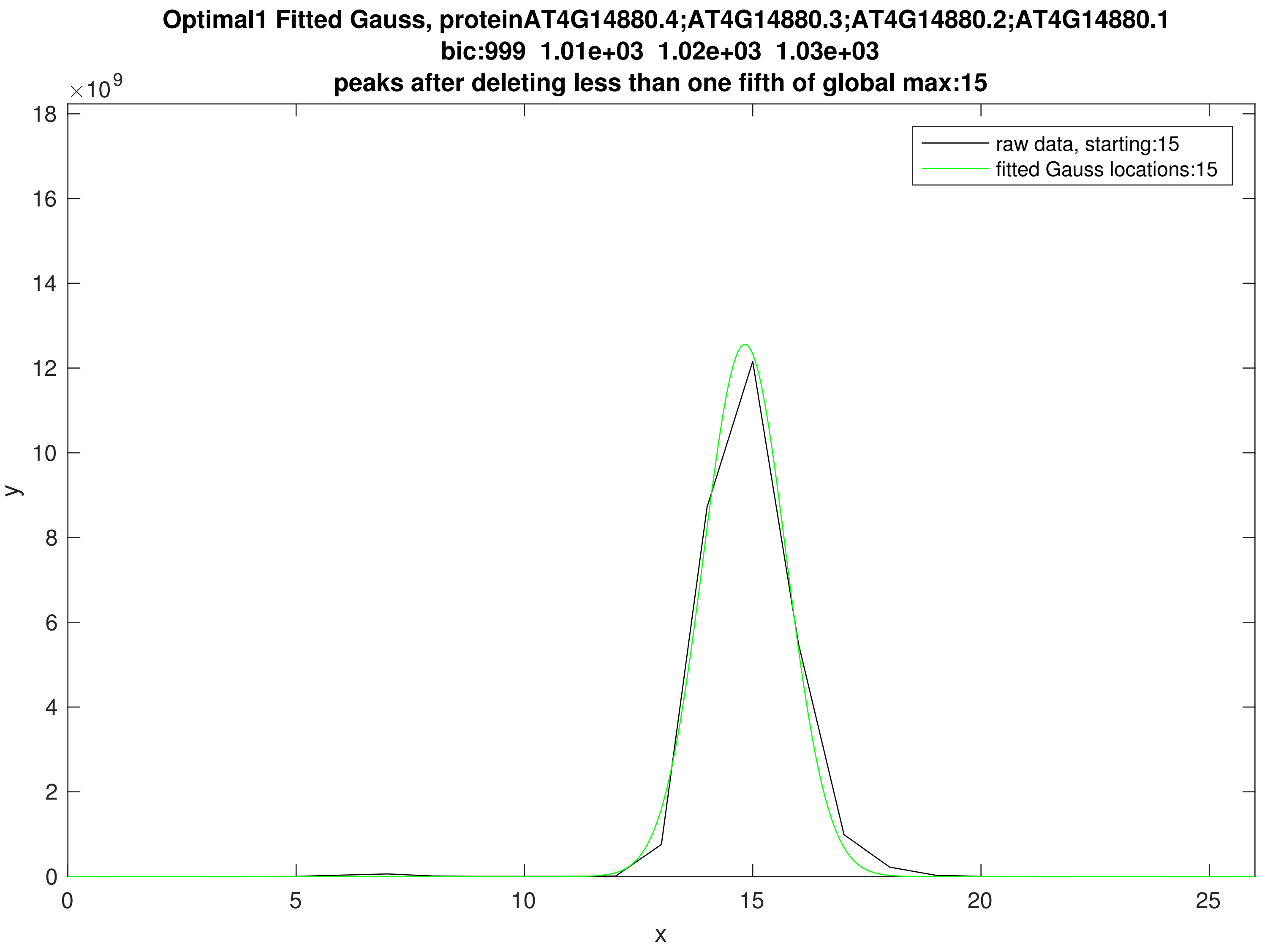

Optimal1 Fitted Gauss, proteinAT4G14880.4;AT4G14880.3;AT4G14880.2;AT4G14880.1
bic:999 1.01e+03 1.02e+03 1.03e+03
peaks after deleting less than one fifth of global max:15
×10^9
18
16
14
12
10
8
6
4
2
0
y
0
5
10
15
20
25
x
raw data, starting:15
fitted Gauss locations:15

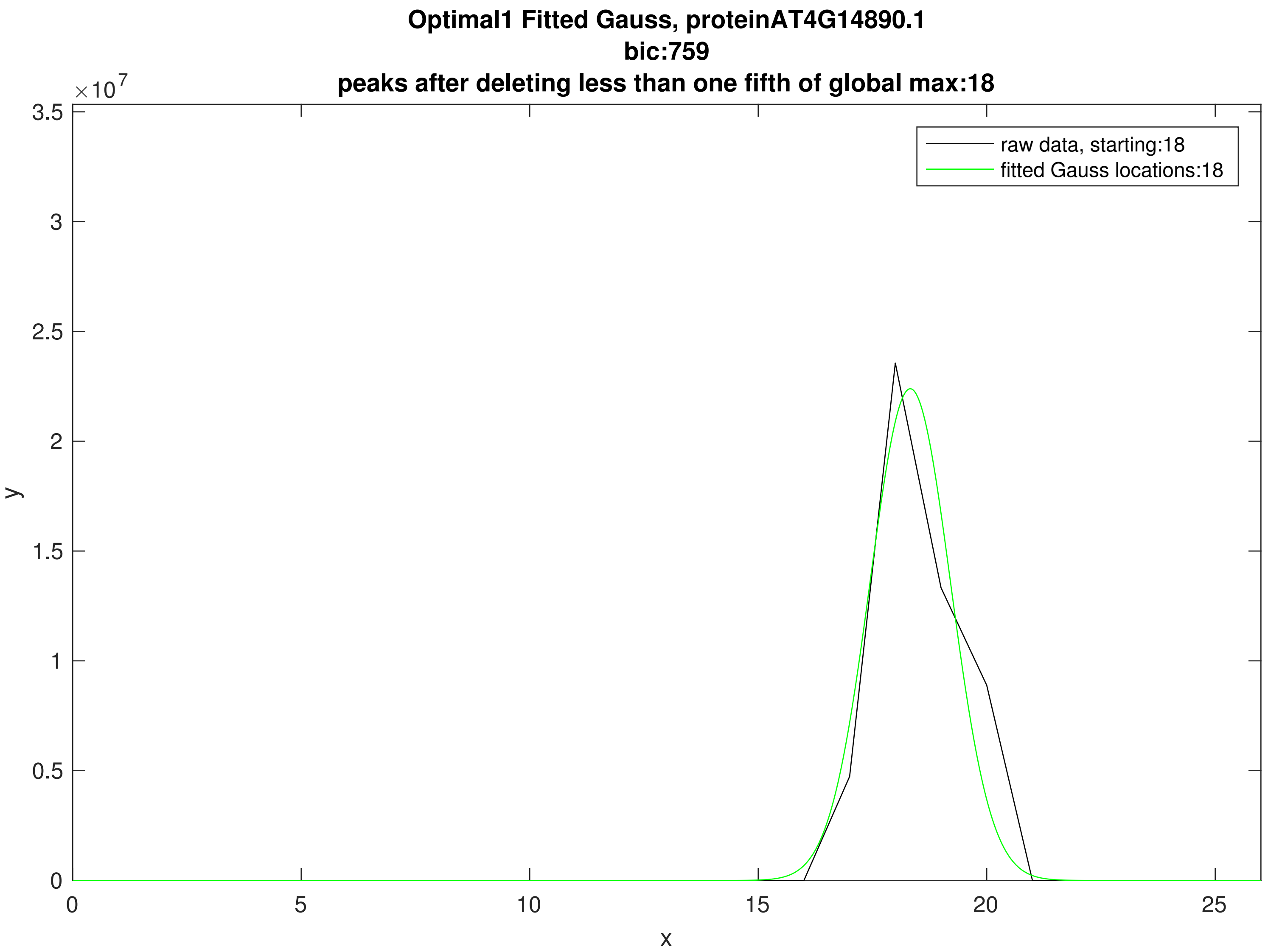

Optimal1 Fitted Gauss, proteinAT4G14890.1
bic:759
peaks after deleting less than one fifth of global max:18
×10^7
3.5
3
2.5
2
1.5
1
0.5
0
y
0
5
10
15
20
25
x
raw data, starting:18
fitted Gauss locations:18

**Optimal1 Fitted Gauss, proteinAT4G14930.1**
**bic:833**
**peaks after deleting less than one fifth of global max:13**

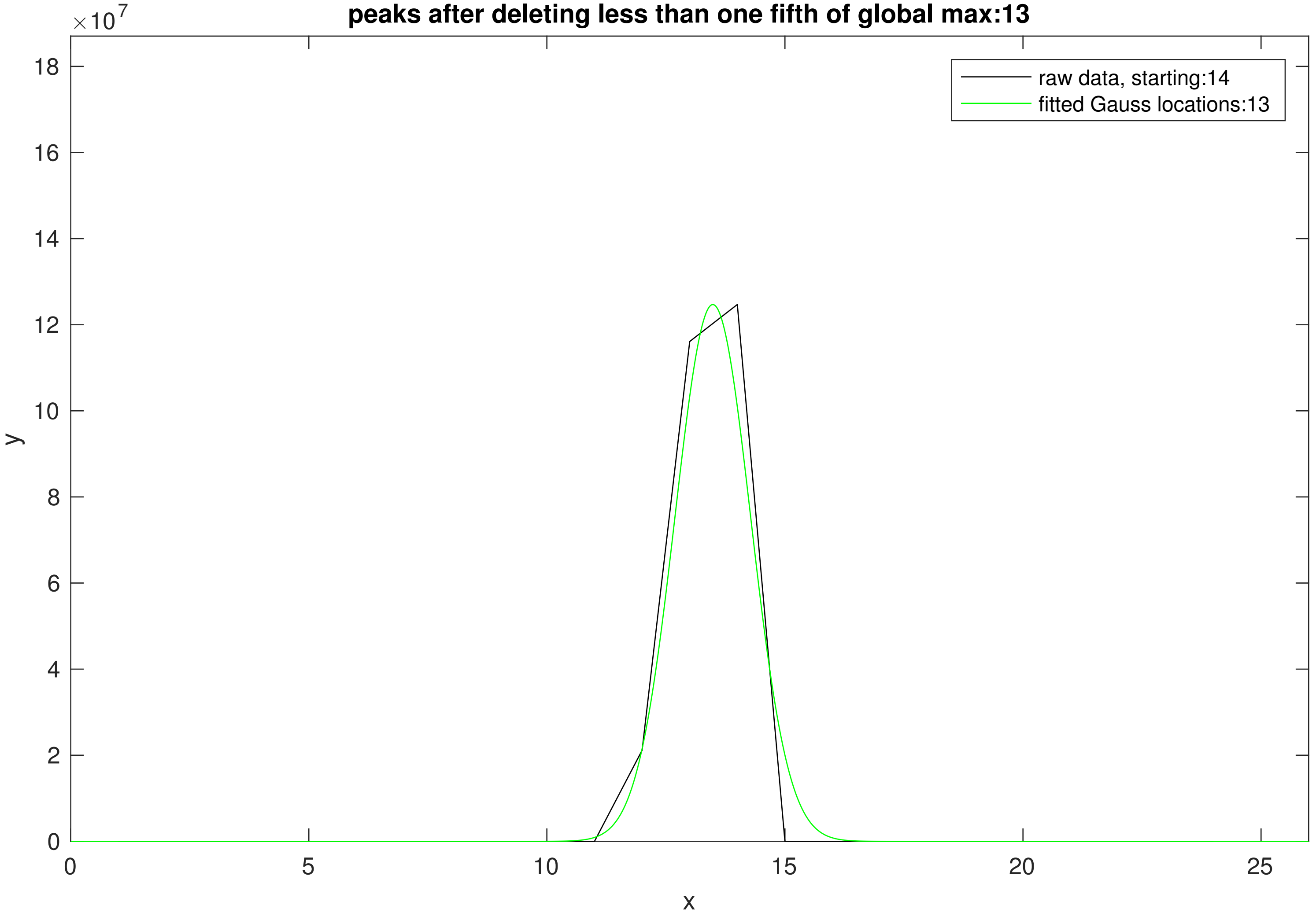

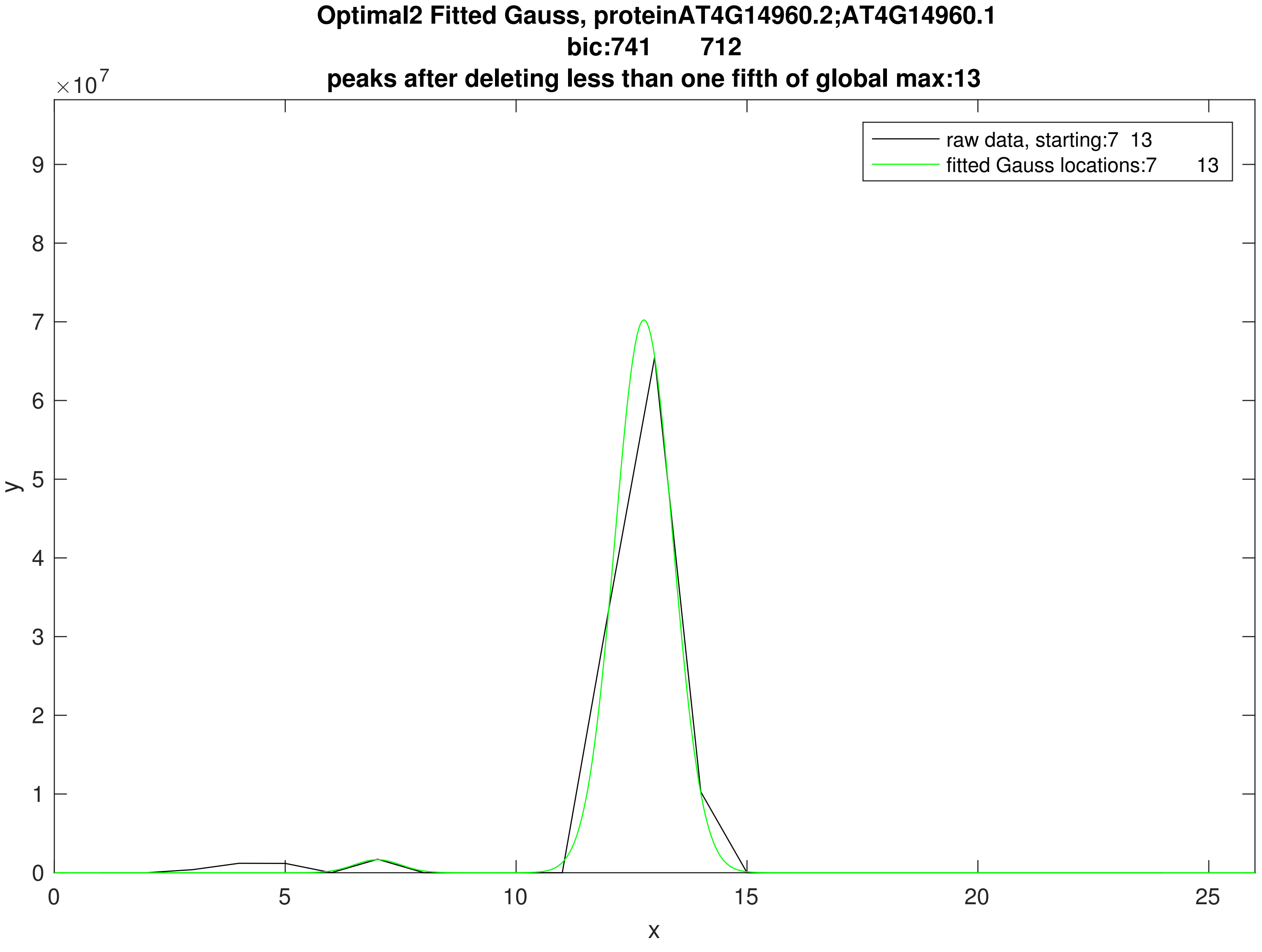
Optimal2 Fitted Gauss, proteinAT4G14960.2;AT4G14960.1
bic:741 712
peaks after deleting less than one fifth of global max:13
×10^7
raw data, starting:7 13
fitted Gauss locations:7 13
y
x
0
1
2
3
4
5
6
7
8
9
0
5
10
15
20
25

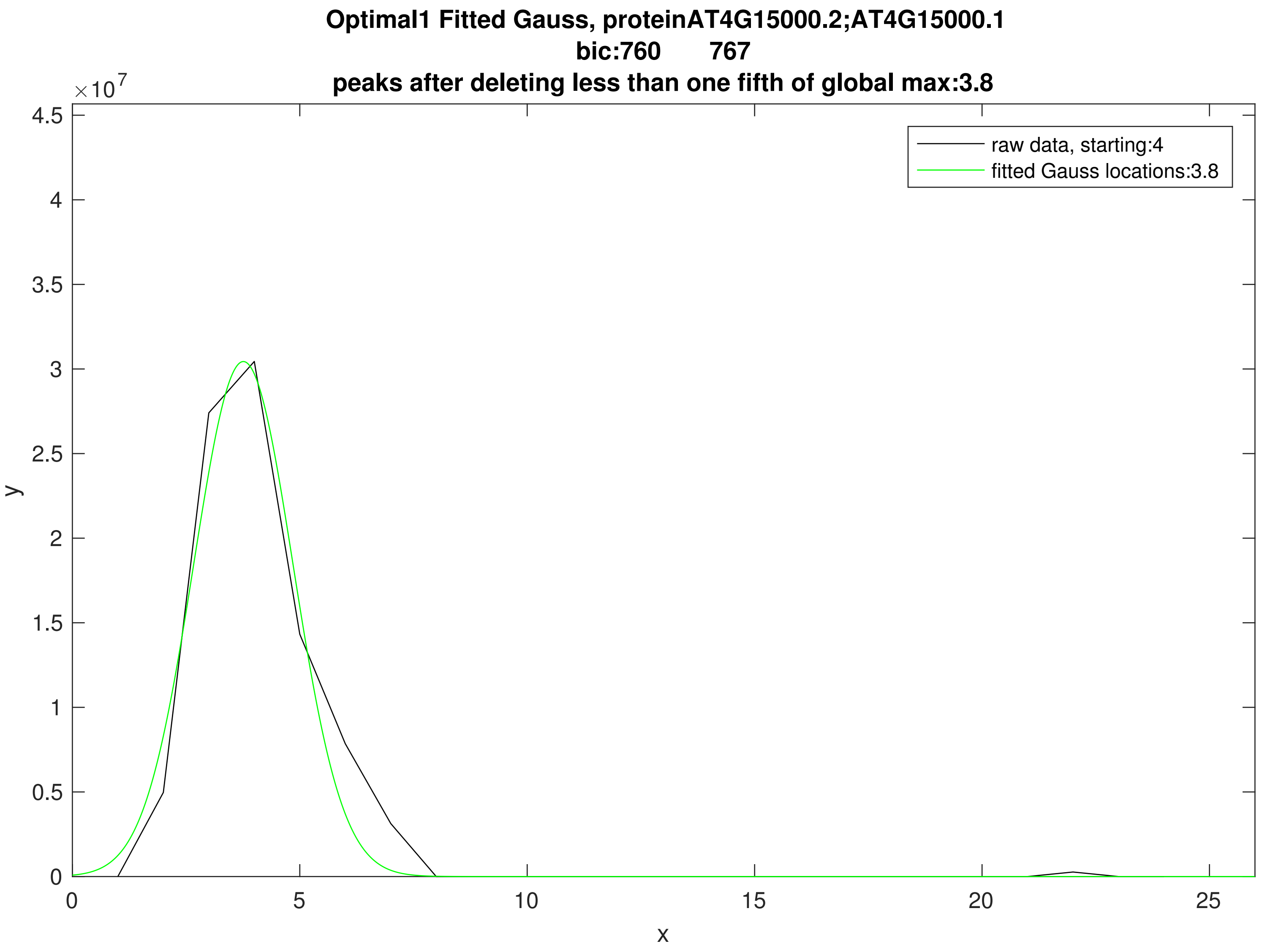

Optimal1 Fitted Gauss, proteinAT4G15000.2;AT4G15000.1
bic:760 767
peaks after deleting less than one fifth of global max:3.8
×10^7
4.5
4
3.5
3
2.5
2
1.5
1
0.5
0
y
0
5
10
15
20
25
x
raw data, starting:4
fitted Gauss locations:3.8

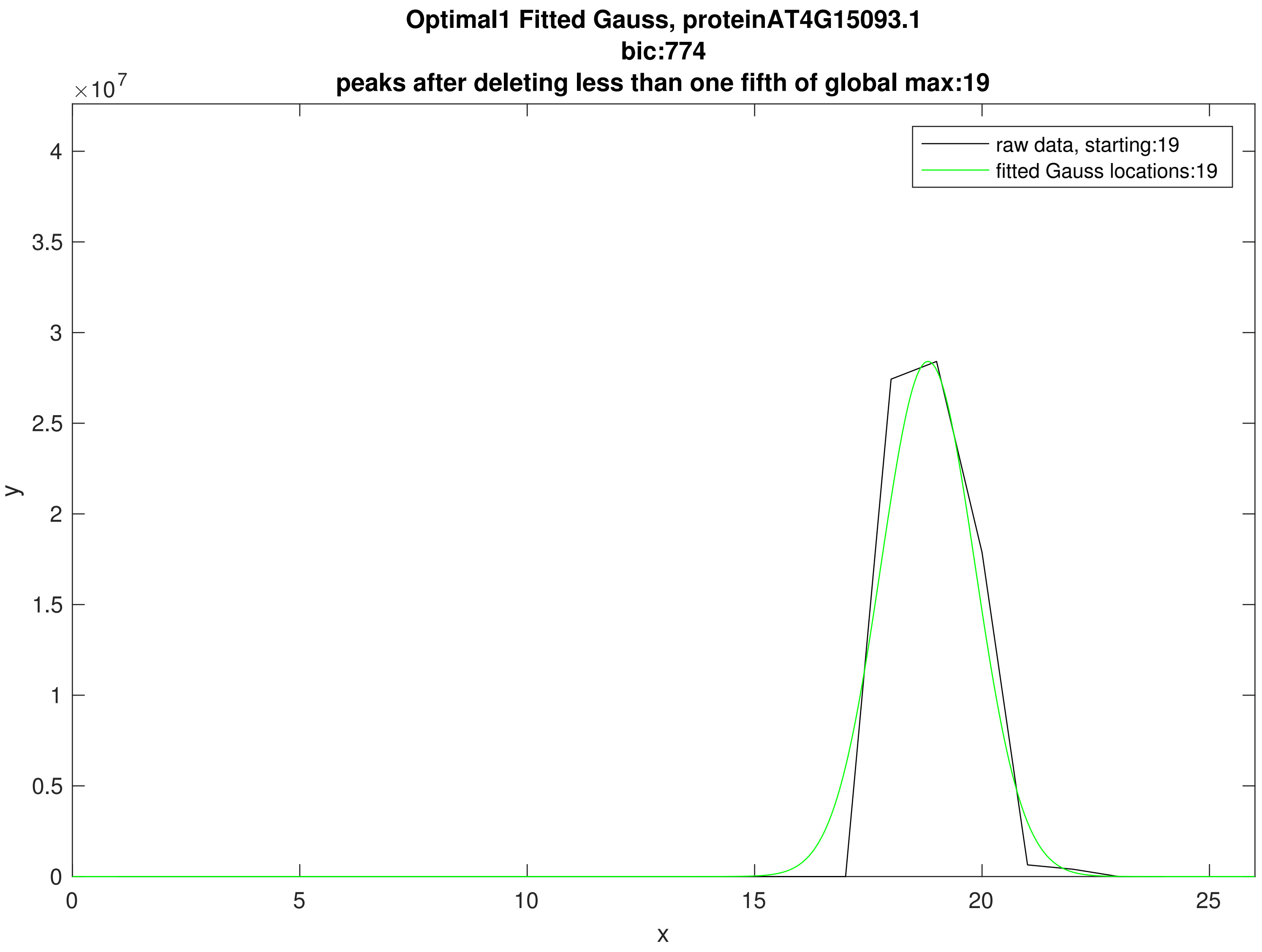
Optimal1 Fitted Gauss, proteinAT4G15093.1
bic:774
peaks after deleting less than one fifth of global max:19
×10^7
raw data, starting:19
fitted Gauss locations:19
y
x
0
0.5
1
1.5
2
2.5
3
3.5
4
5
10
15
20
25

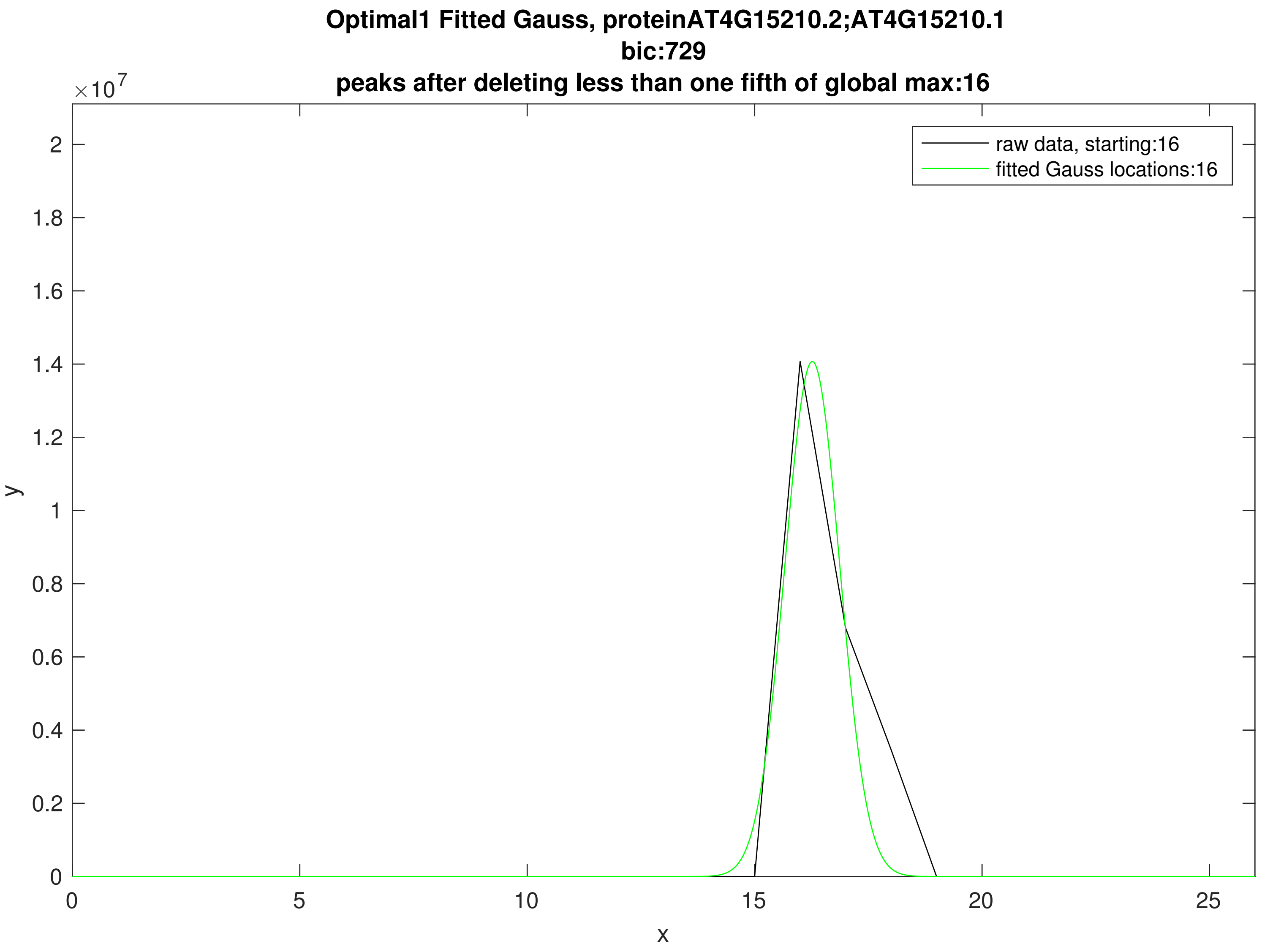

Optimal1 Fitted Gauss, proteinAT4G15210.2;AT4G15210.1
bic:729
peaks after deleting less than one fifth of global max:16
×10^7
2
1.8
1.6
1.4
1.2
1
0.8
0.6
0.4
0.2
0
y
0
5
10
15
20
25
x
raw data, starting:16
fitted Gauss locations:16

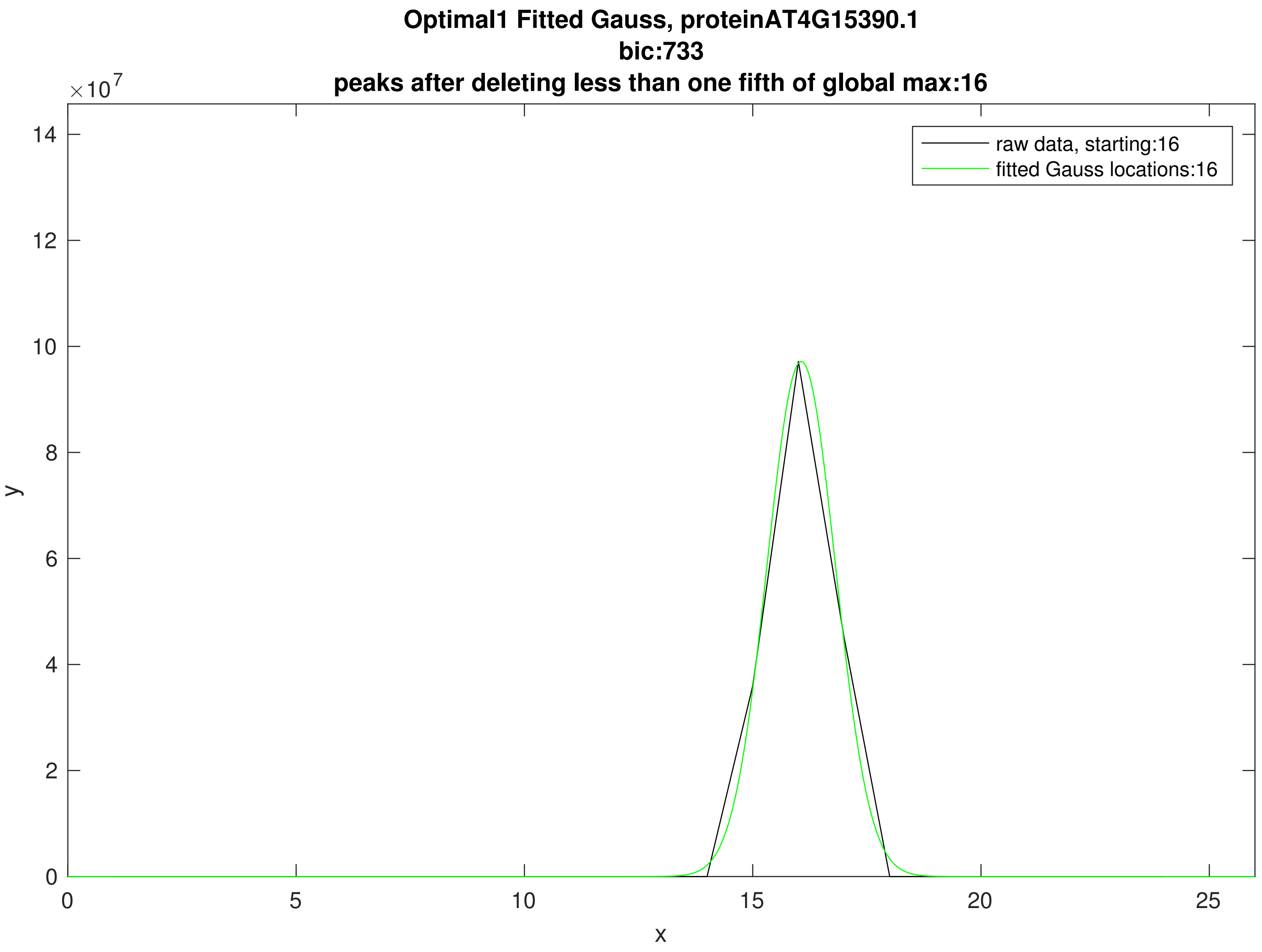
Optimal1 Fitted Gauss, proteinAT4G15390.1
bic:733
peaks after deleting less than one fifth of global max:16
×10^7
raw data, starting:16
fitted Gauss locations:16
y
x
0
2
4
6
8
10
12
14
0
5
10
15
20
25

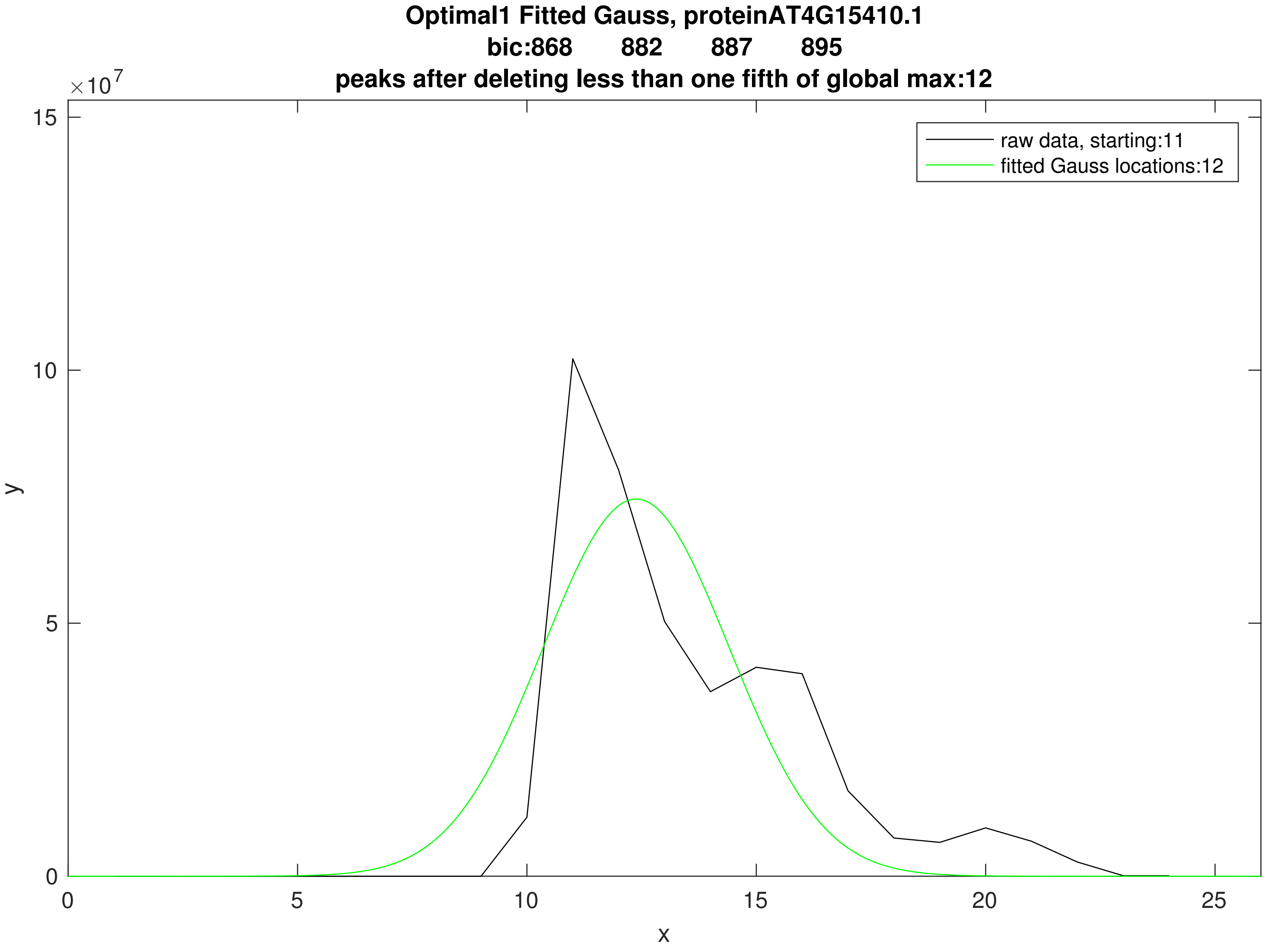

Optimal1 Fitted Gauss, proteinAT4G15410.1
bic:868 882 887 895
peaks after deleting less than one fifth of global max:12
×10^7
15
10
5
0
0
5
10
15
20
25
x
y
raw data, starting:11
fitted Gauss locations:12

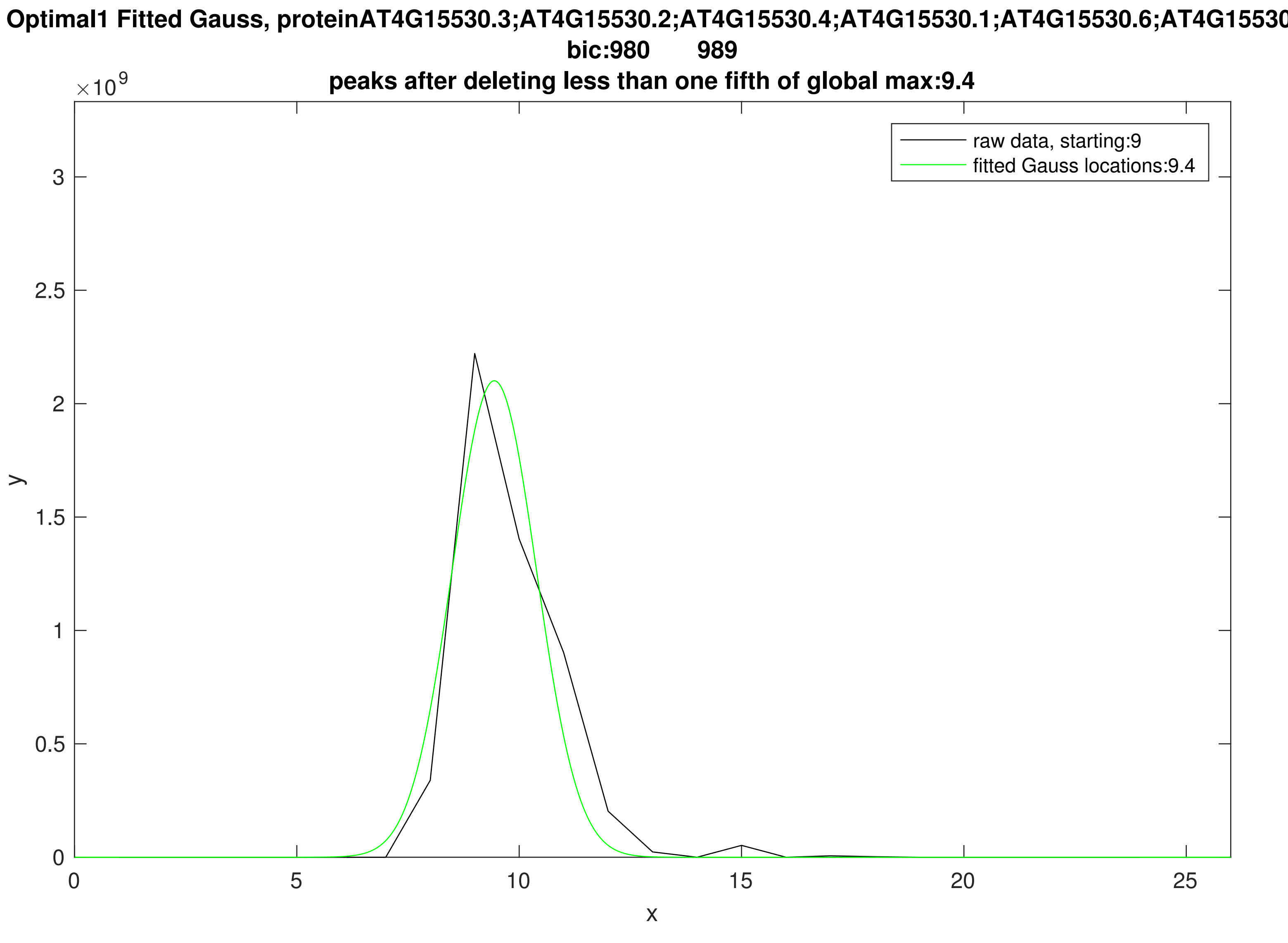

Optimal1 Fitted Gauss, proteinAT4G15530.3;AT4G15530.2;AT4G15530.4;AT4G15530.1;AT4G15530.6;AT4G15530
bic:980 989
peaks after deleting less than one fifth of global max:9.4
×10^9
raw data, starting:9
fitted Gauss locations:9.4
y
x
0
0.5
1
1.5
2
2.5
3
0
5
10
15
20
25

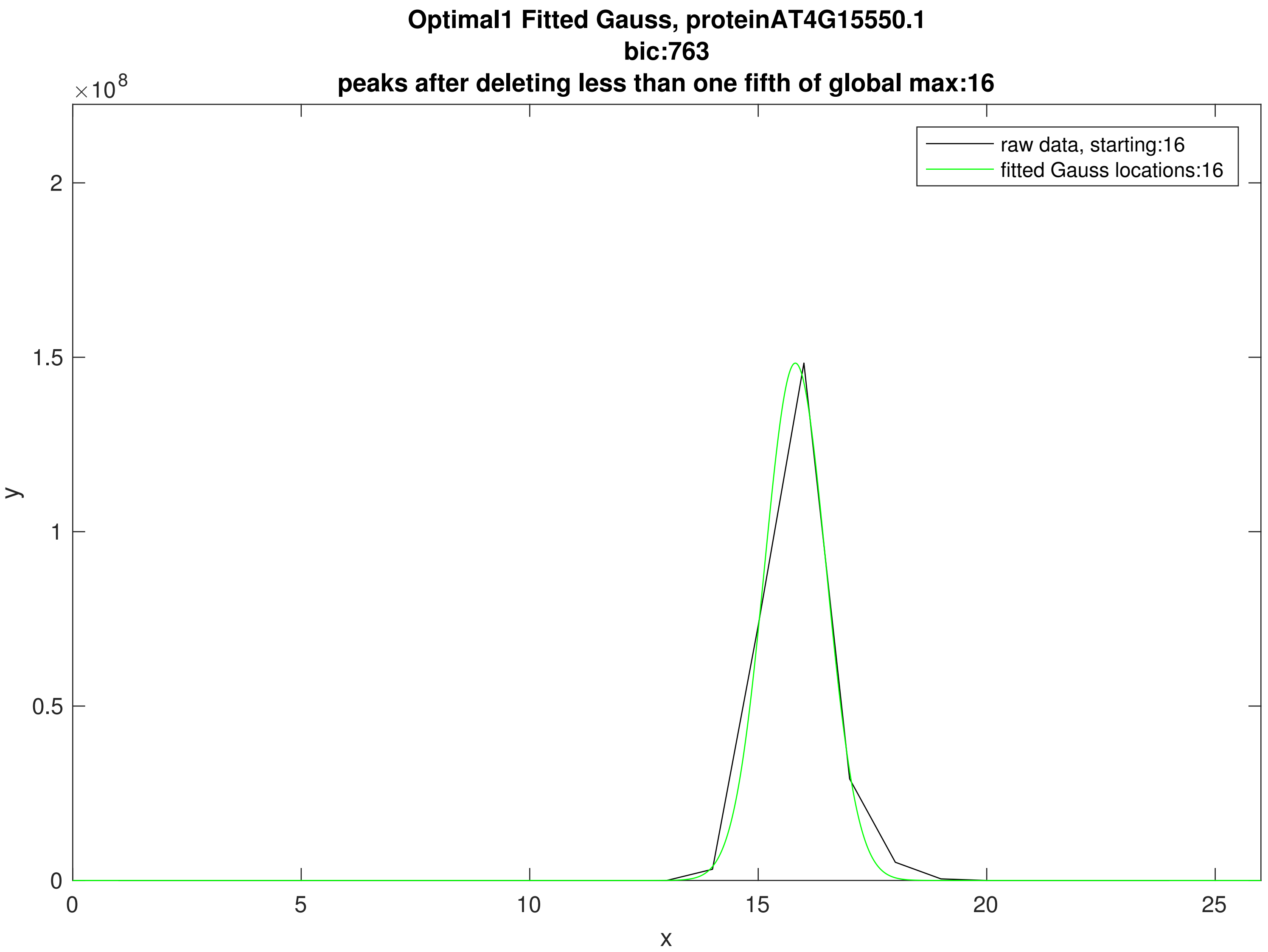

Optimal1 Fitted Gauss, proteinAT4G15550.1
bic:763
peaks after deleting less than one fifth of global max:16
×10^8
raw data, starting:16
fitted Gauss locations:16
y
x
0
0.5
1
1.5
2
5
10
15
20
25

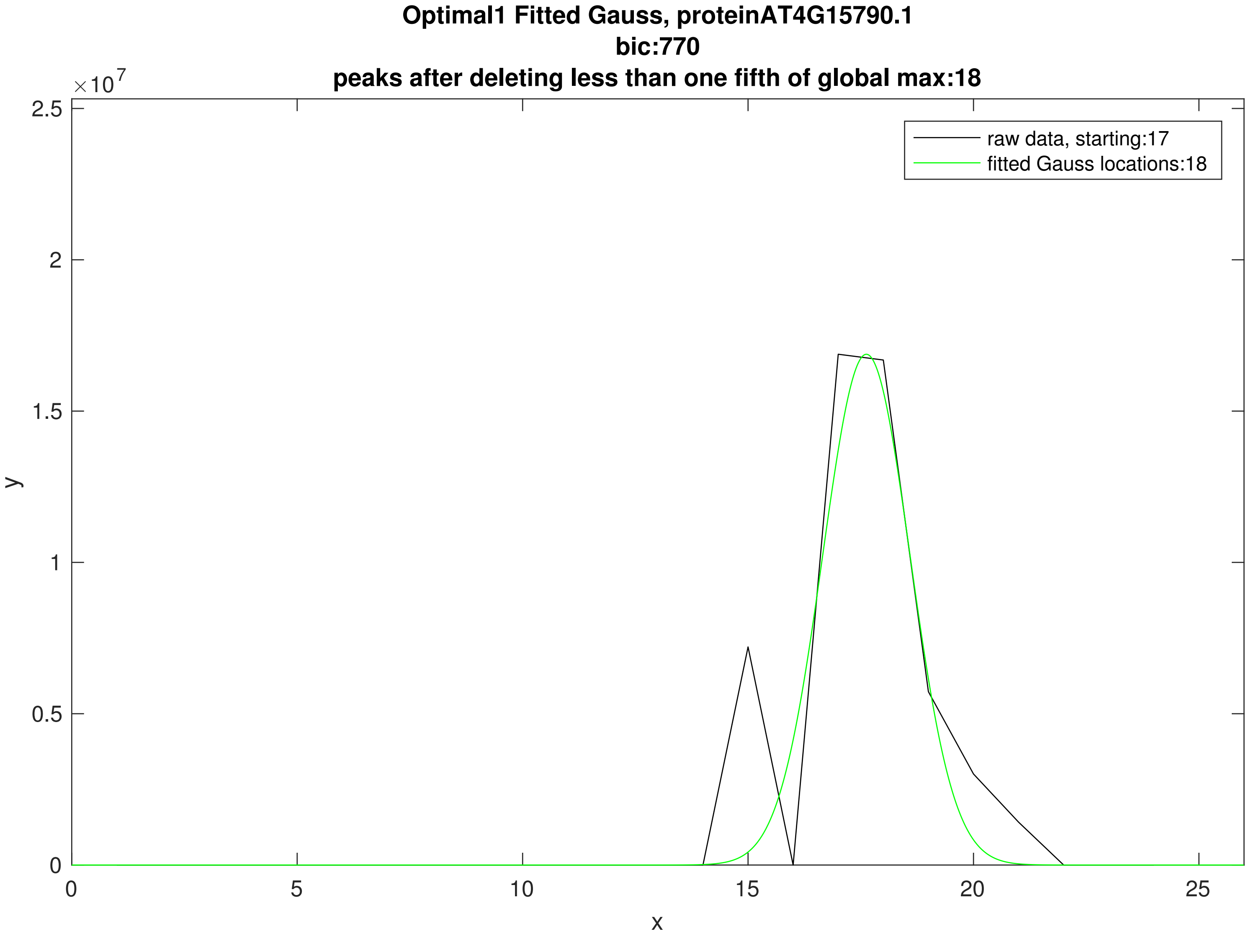

Optimal1 Fitted Gauss, proteinAT4G15790.1
bic:770
peaks after deleting less than one fifth of global max:18
×10^7
2.5
2
1.5
1
0.5
0
y
0
5
10
15
20
25
x
raw data, starting:17
fitted Gauss locations:18

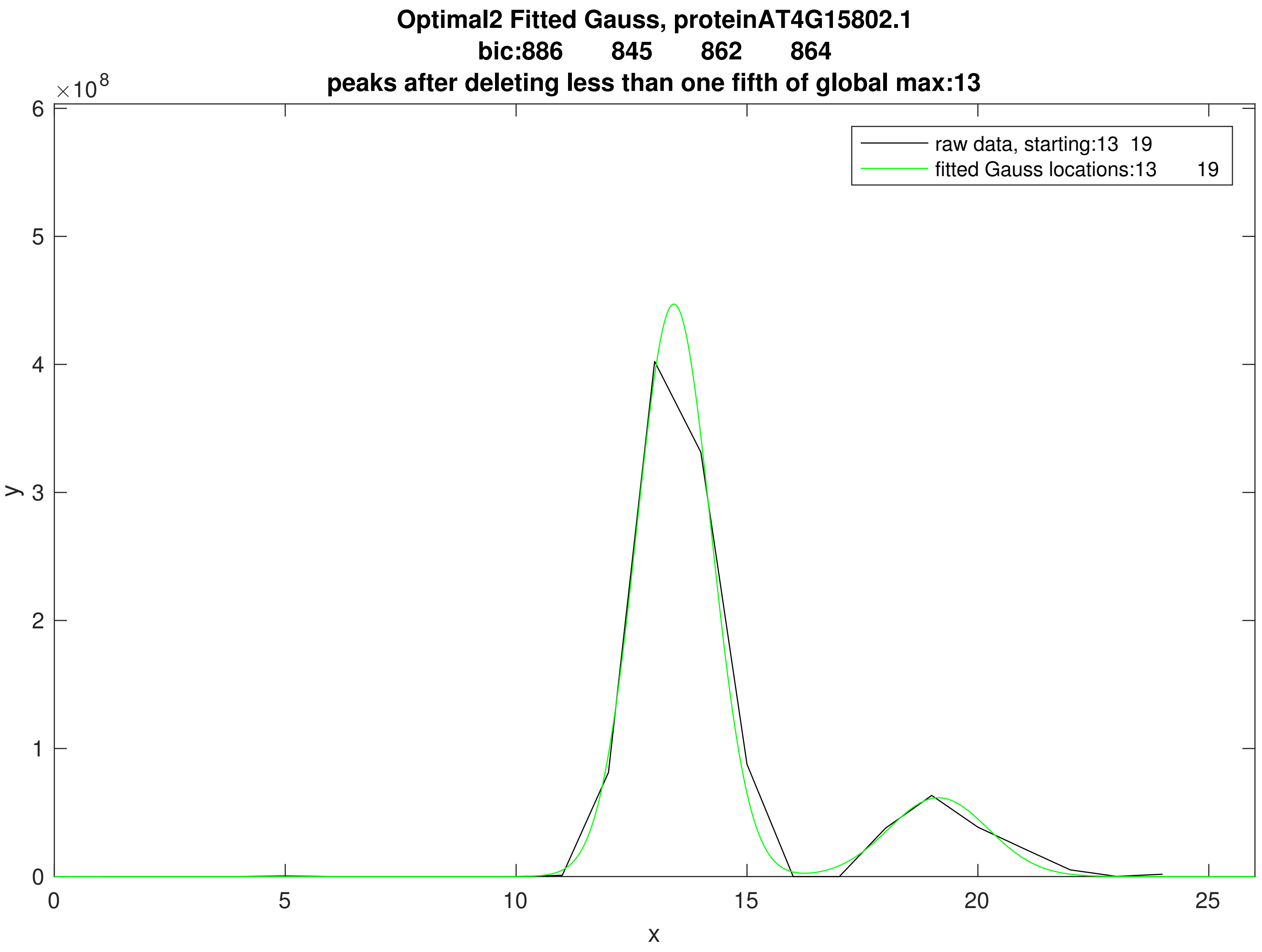

Optimal2 Fitted Gauss, proteinAT4G15802.1
bic:886 845 862 864
peaks after deleting less than one fifth of global max:13
×10^8
raw data, starting:13 19
fitted Gauss locations:13 19
y
x

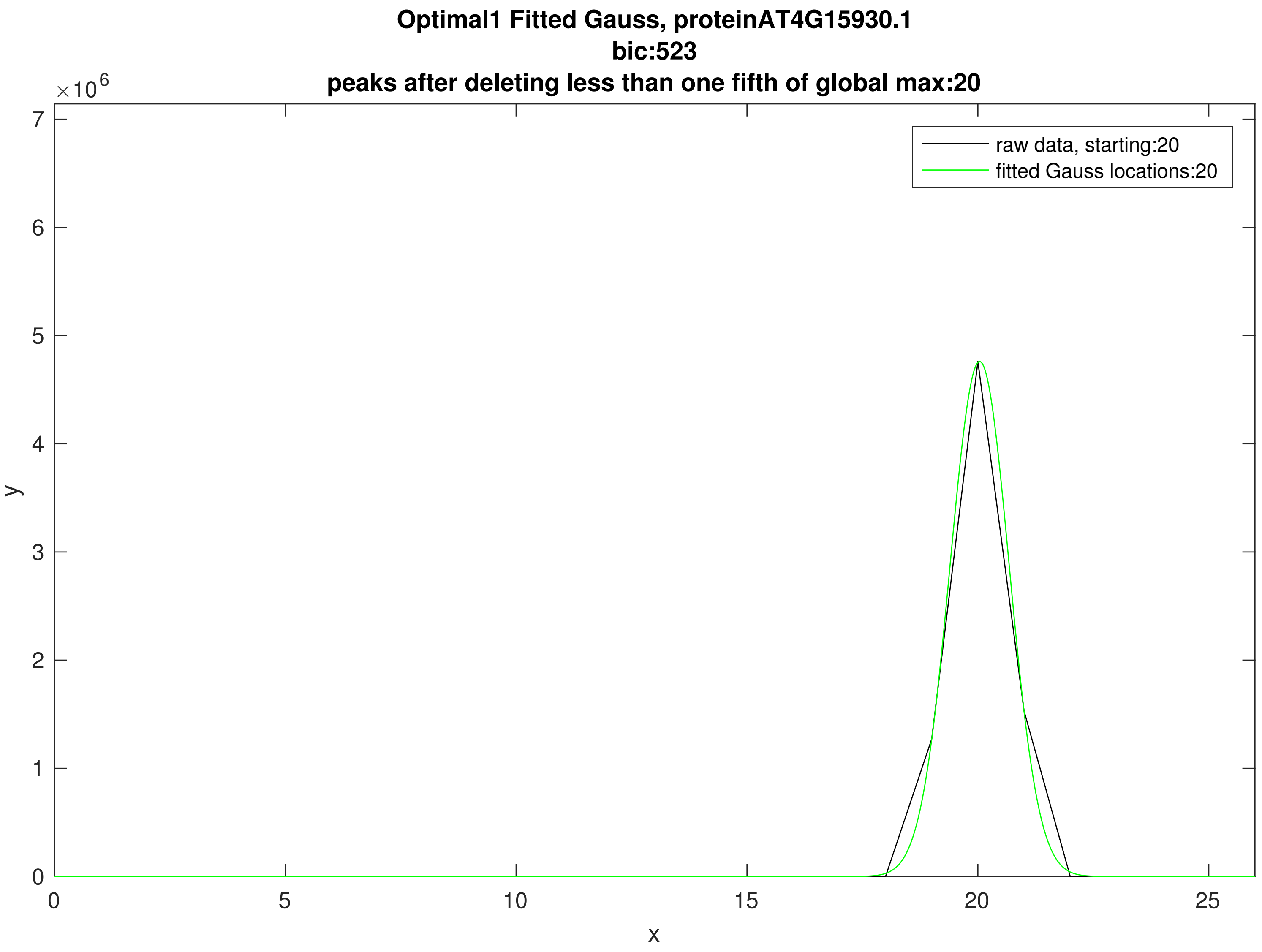
Optimal1 Fitted Gauss, proteinAT4G15930.1
bic:523
peaks after deleting less than one fifth of global max:20
×10^6
raw data, starting:20
fitted Gauss locations:20
y
x
0
1
2
3
4
5
6
7
0
5
10
15
20
25

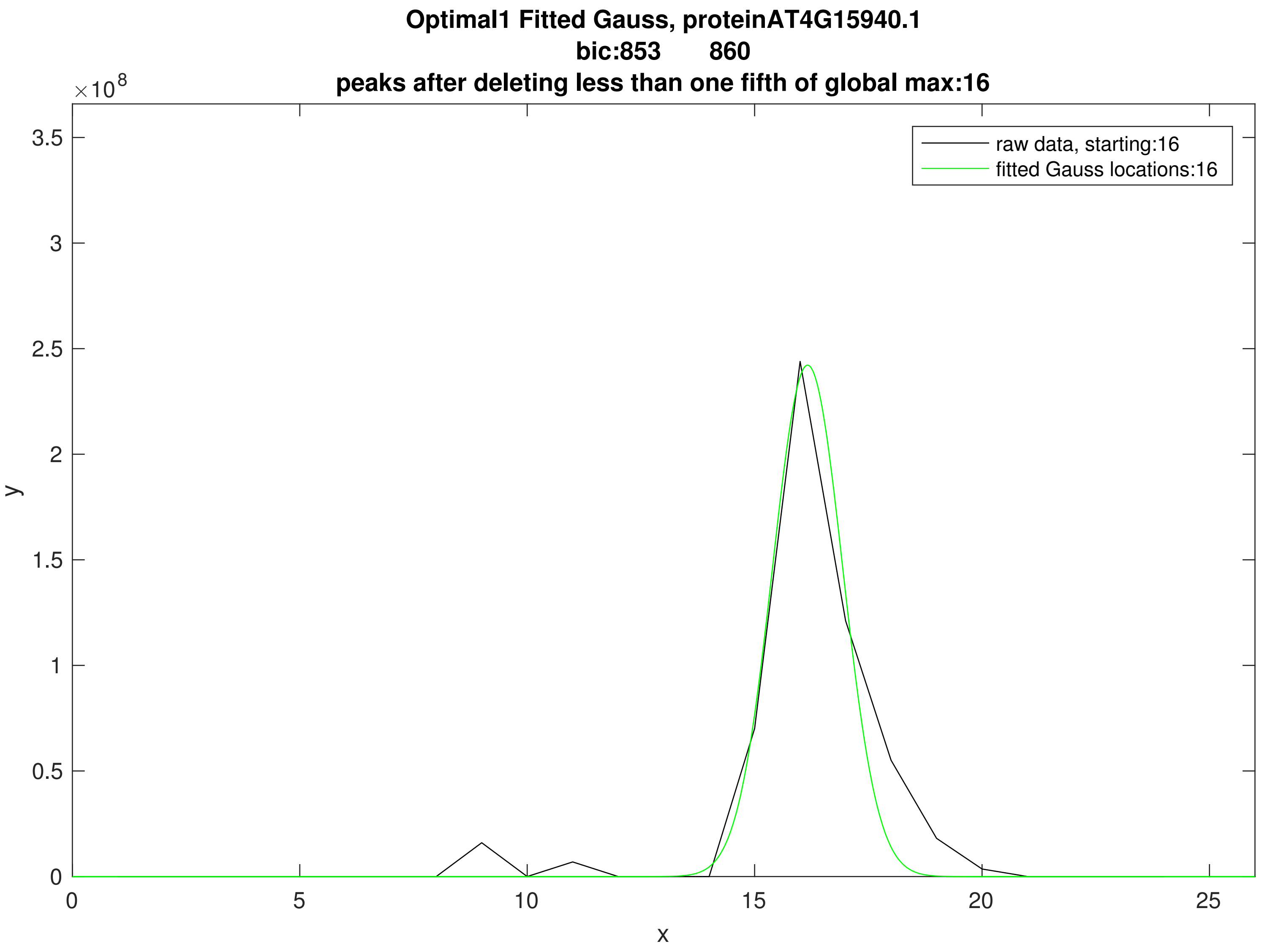
Optimal1 Fitted Gauss, proteinAT4G15940.1
bic:853 860
peaks after deleting less than one fifth of global max:16
×10^8
raw data, starting:16
fitted Gauss locations:16
y
x
0
0.5
1
1.5
2
2.5
3
3.5
0
5
10
15
20
25

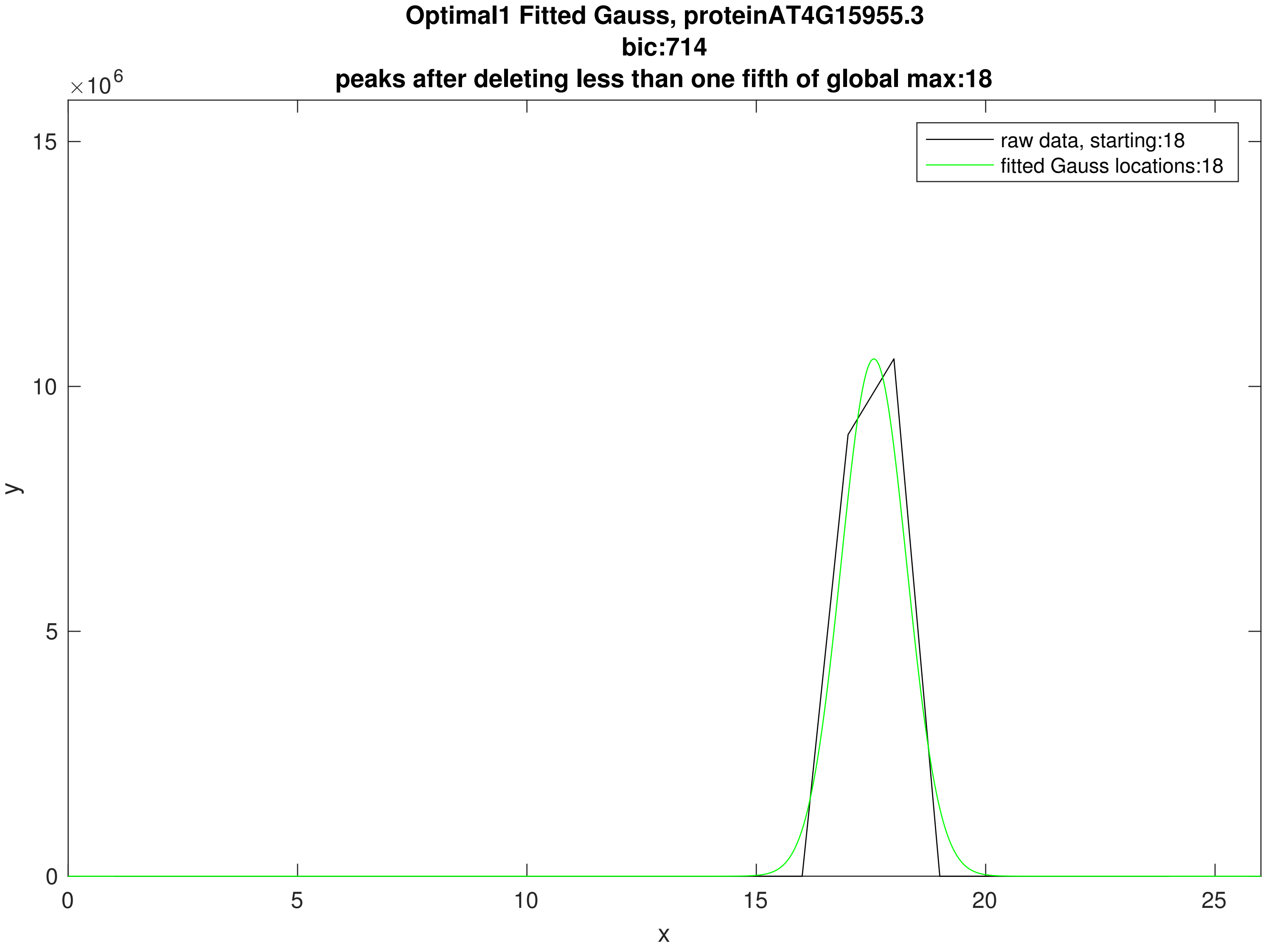

Optimal1 Fitted Gauss, proteinAT4G15955.3
bic:714
peaks after deleting less than one fifth of global max:18
×10^6
15
10
5
0
y
0
5
10
15
20
25
x
raw data, starting:18
fitted Gauss locations:18

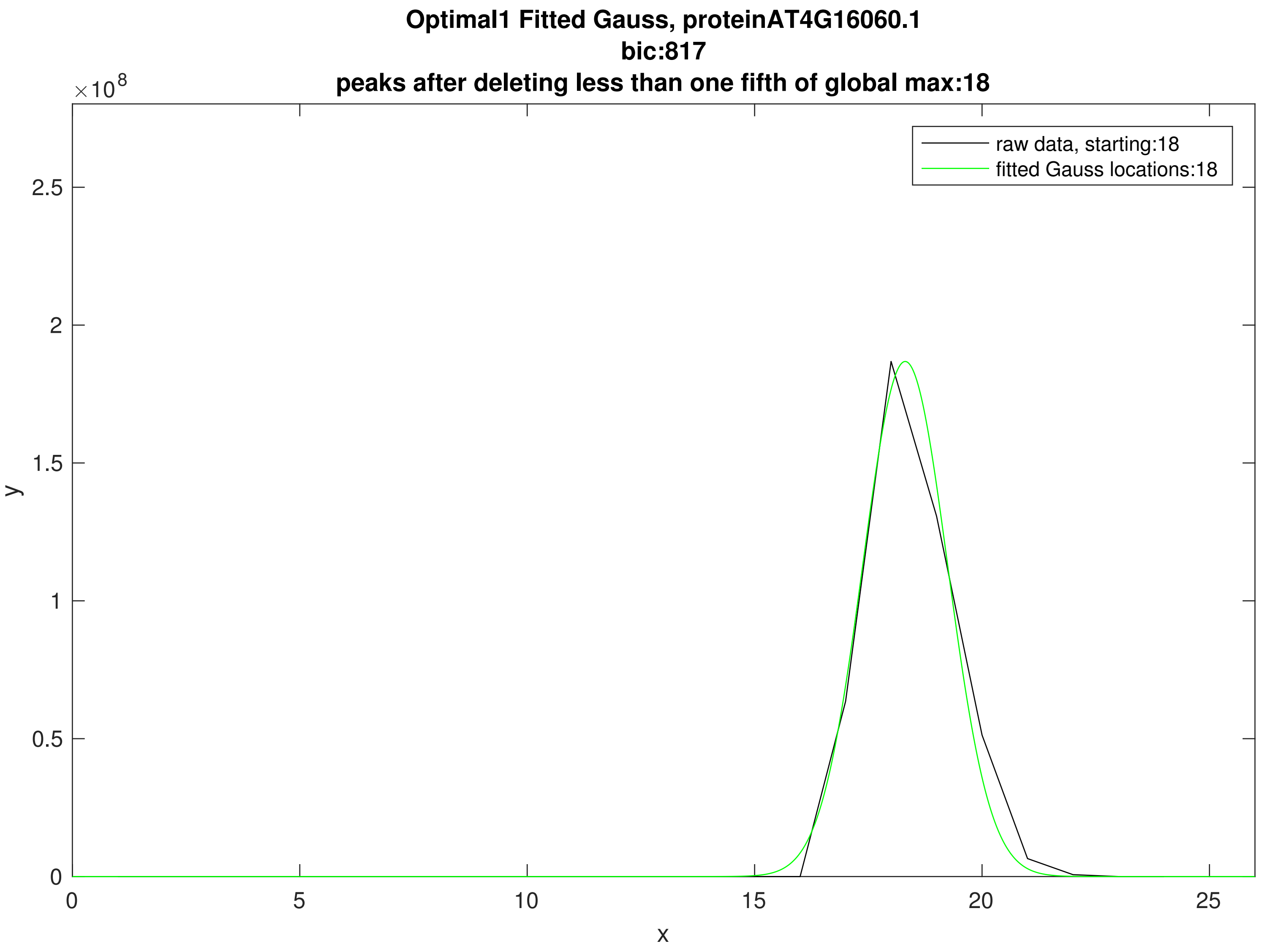
Optimal1 Fitted Gauss, proteinAT4G16060.1
bic:817
peaks after deleting less than one fifth of global max:18
×10^8
raw data, starting:18
fitted Gauss locations:18
2.5
2
1.5
1
0.5
0
y
0
5
10
15
20
25
x

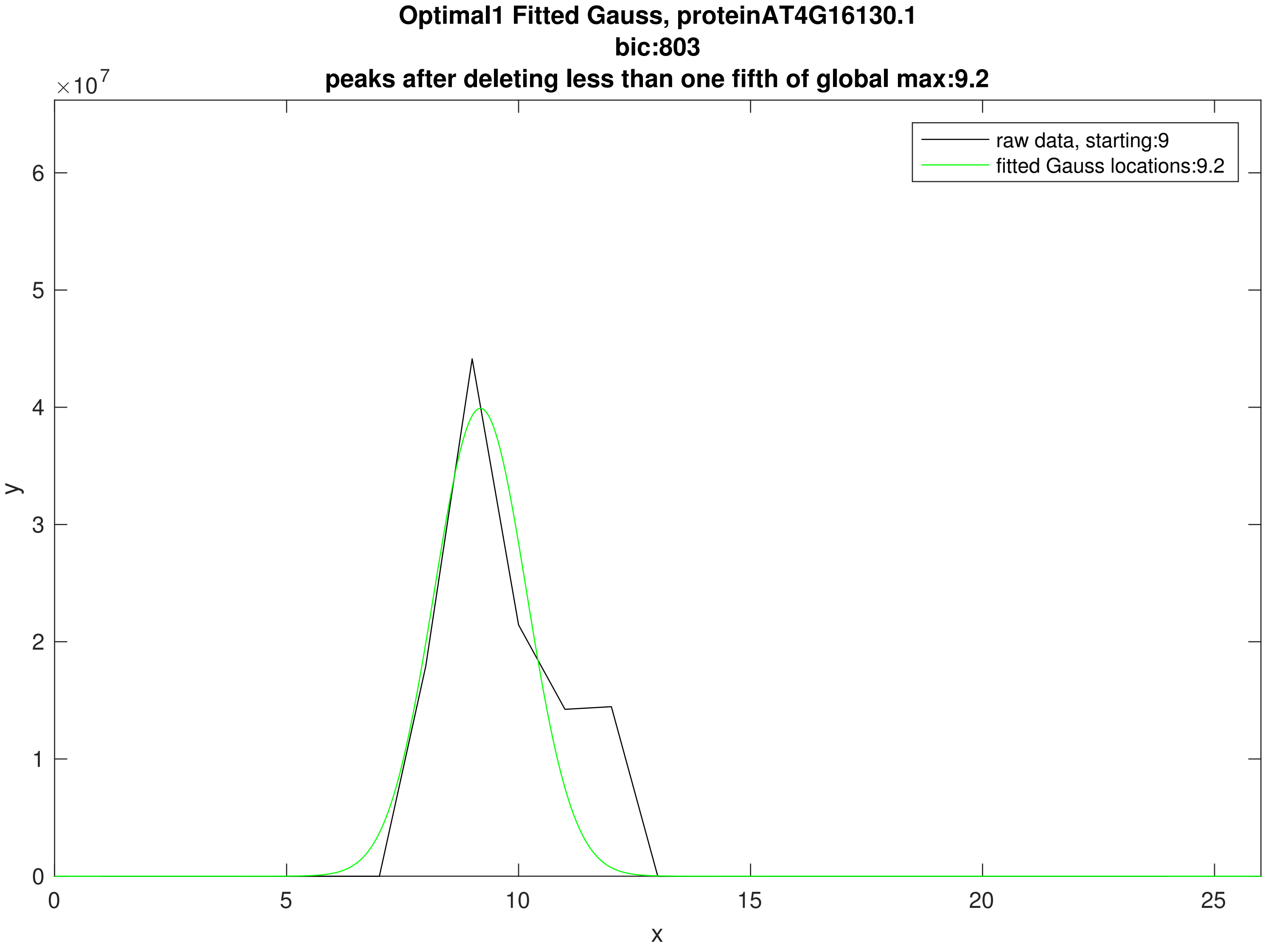
Optimal1 Fitted Gauss, proteinAT4G16130.1
bic:803
peaks after deleting less than one fifth of global max:9.2
×10^7
raw data, starting:9
fitted Gauss locations:9.2
y
x
0
1
2
3
4
5
6
0
5
10
15
20
25

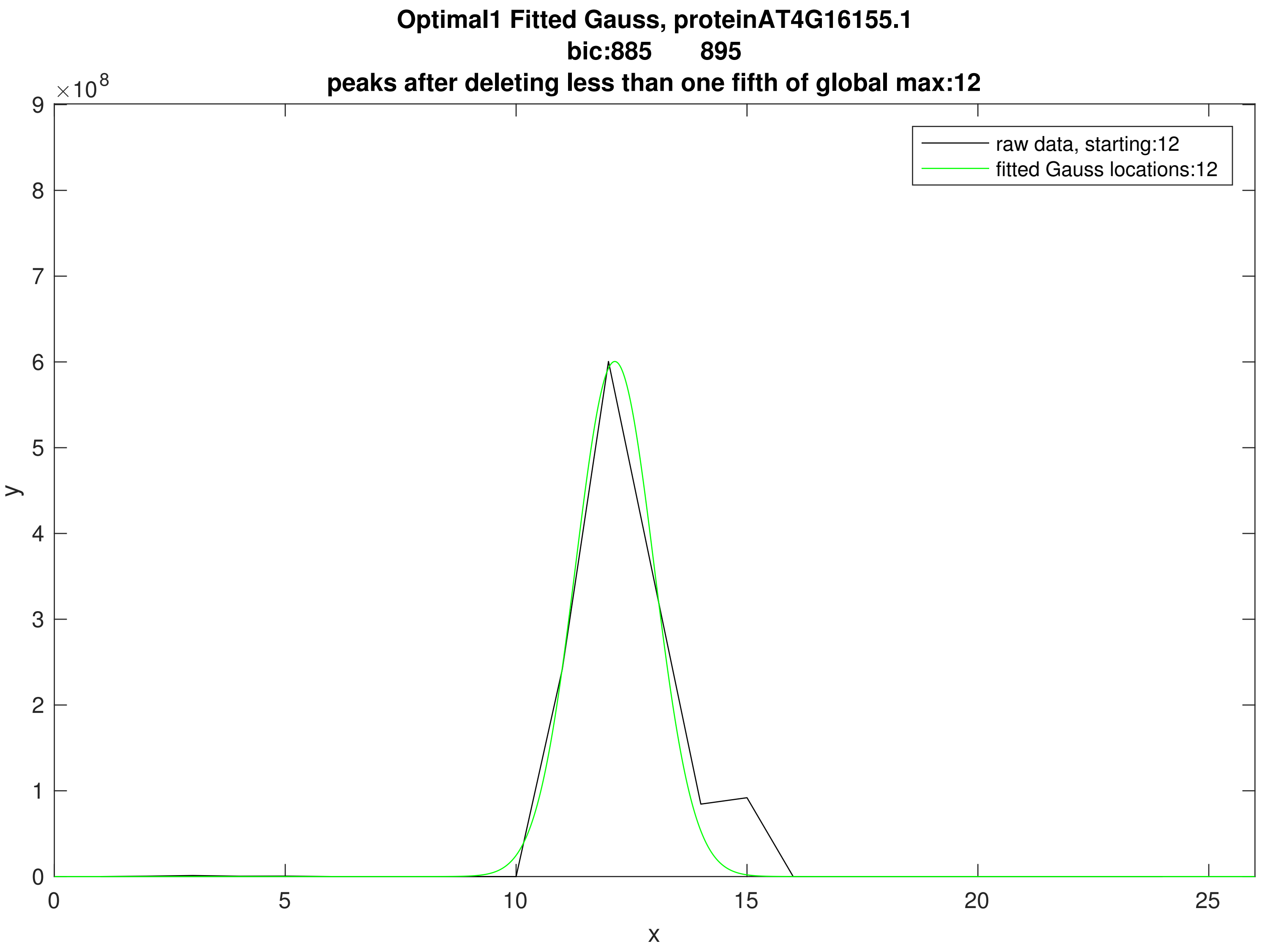
Optimal1 Fitted Gauss, proteinAT4G16155.1
bic:885 895
peaks after deleting less than one fifth of global max:12
×10^8
9
8
7
6
5
4
3
2
1
0
y
0
5
10
15
20
25
x
raw data, starting:12
fitted Gauss locations:12

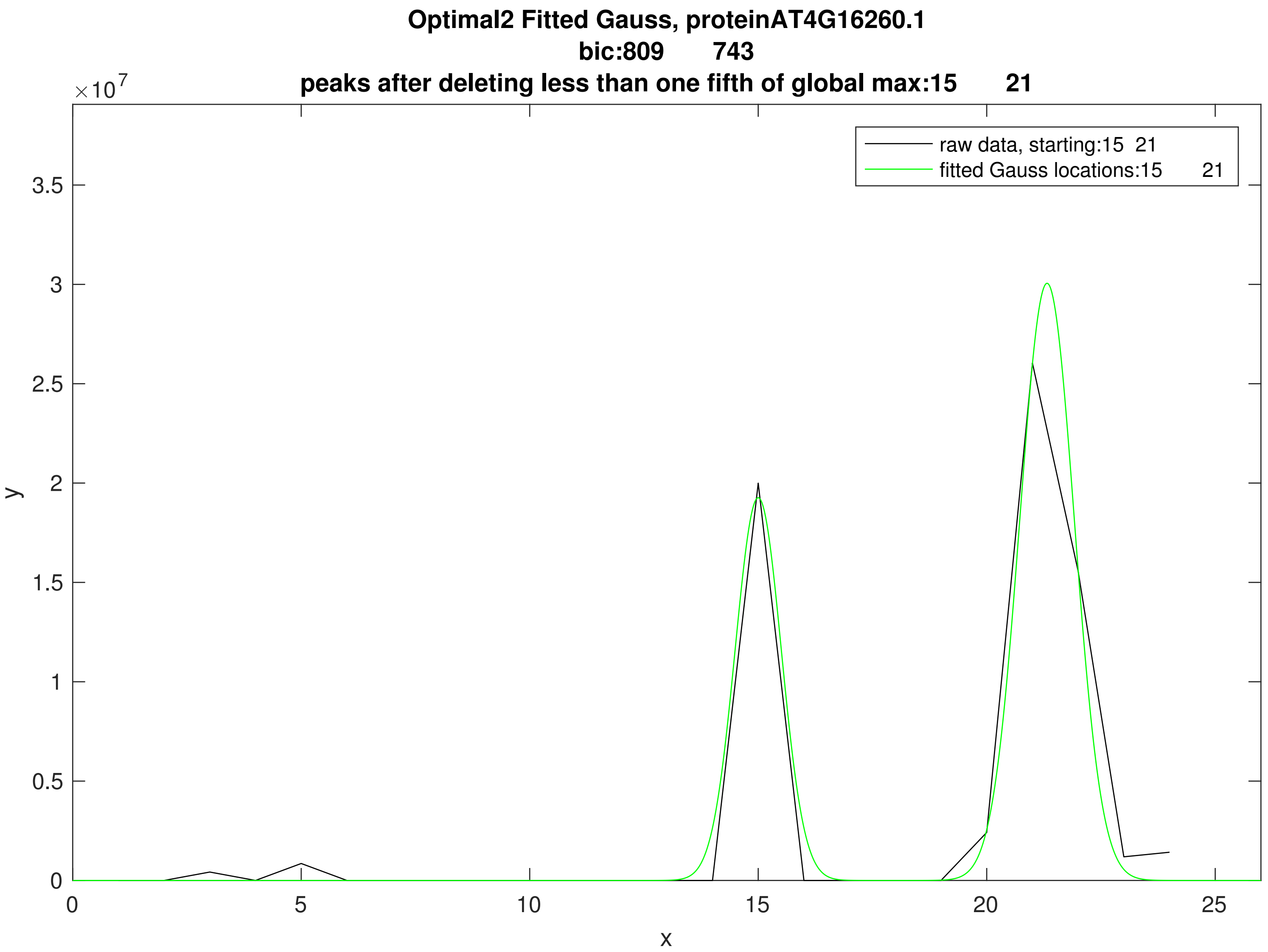
Optimal2 Fitted Gauss, proteinAT4G16260.1
bic:809 743
peaks after deleting less than one fifth of global max:15 21
×10^7
raw data, starting:15 21
fitted Gauss locations:15 21
3.5
3
2.5
2
1.5
1
0.5
0
y
0
5
10
15
20
25
x

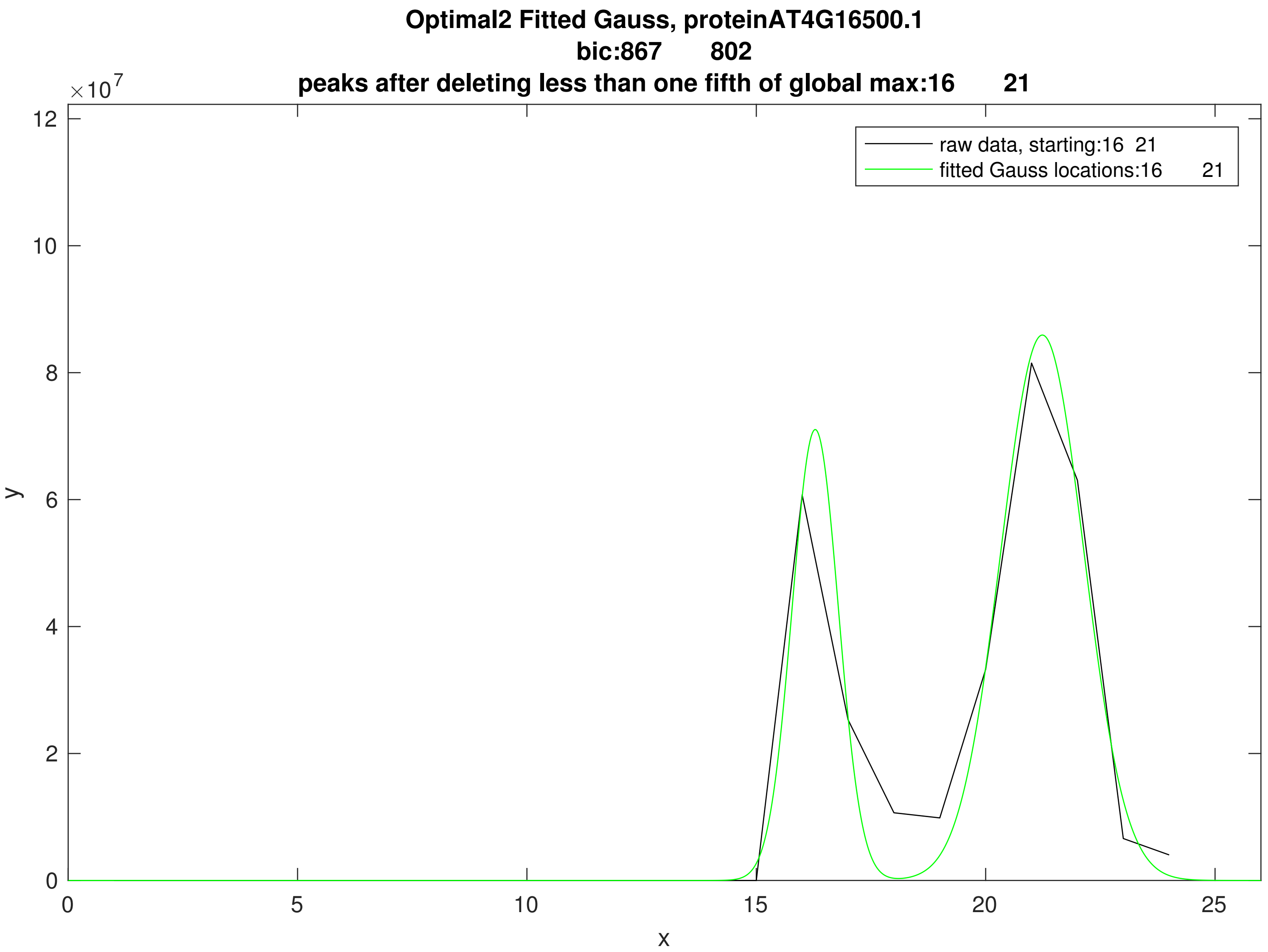

Optimal2 Fitted Gauss, proteinAT4G16500.1
bic:867 802
peaks after deleting less than one fifth of global max:16 21
×10^7
raw data, starting:16 21
fitted Gauss locations:16 21
y
x
0
2
4
6
8
10
12
0
5
10
15
20
25

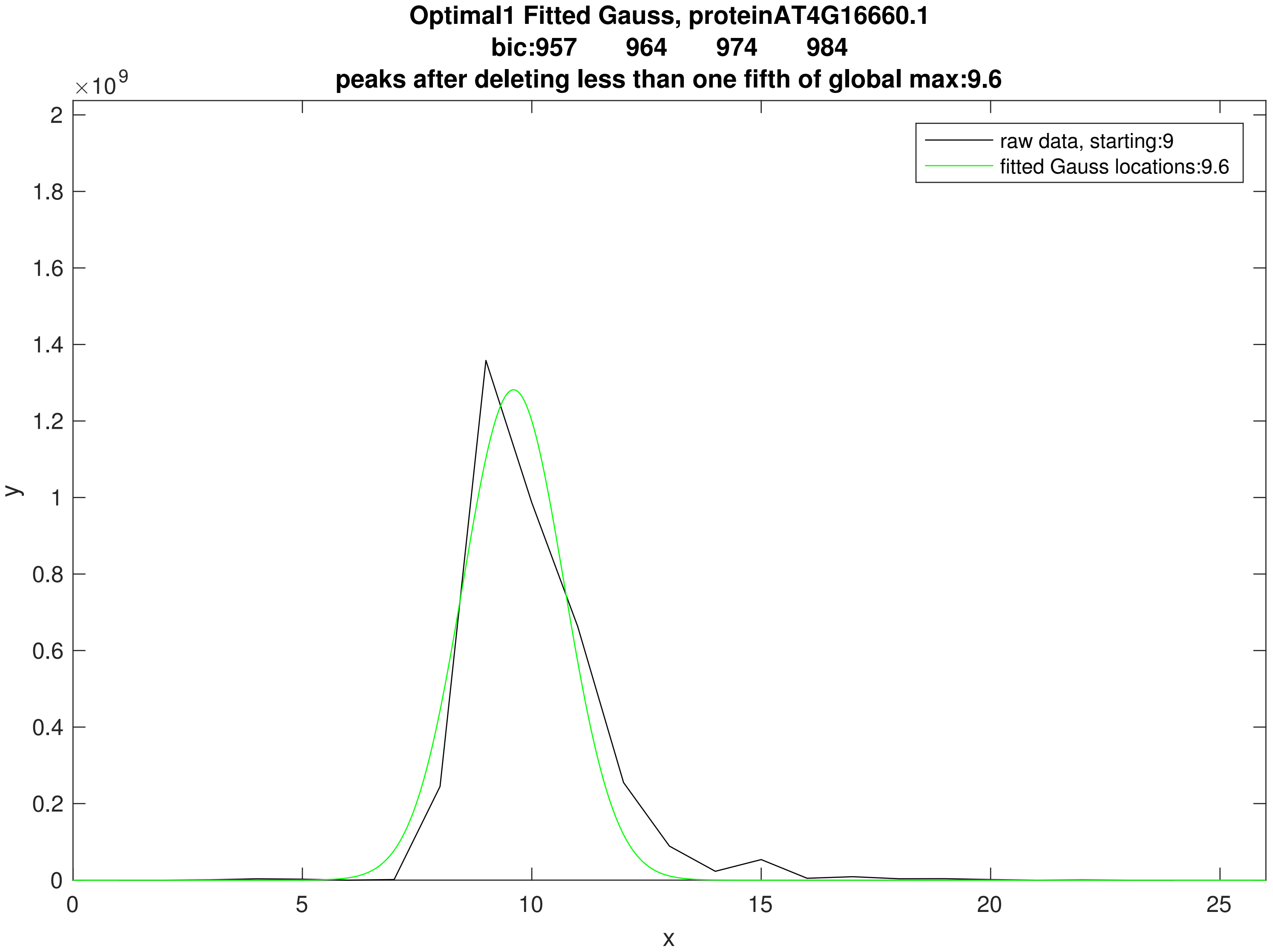
Optimal1 Fitted Gauss, proteinAT4G16660.1
bic:957 964 974 984
peaks after deleting less than one fifth of global max:9.6
×10^9
raw data, starting:9
fitted Gauss locations:9.6
y
x
0
0.2
0.4
0.6
0.8
1
1.2
1.4
1.6
1.8
2
0
5
10
15
20
25

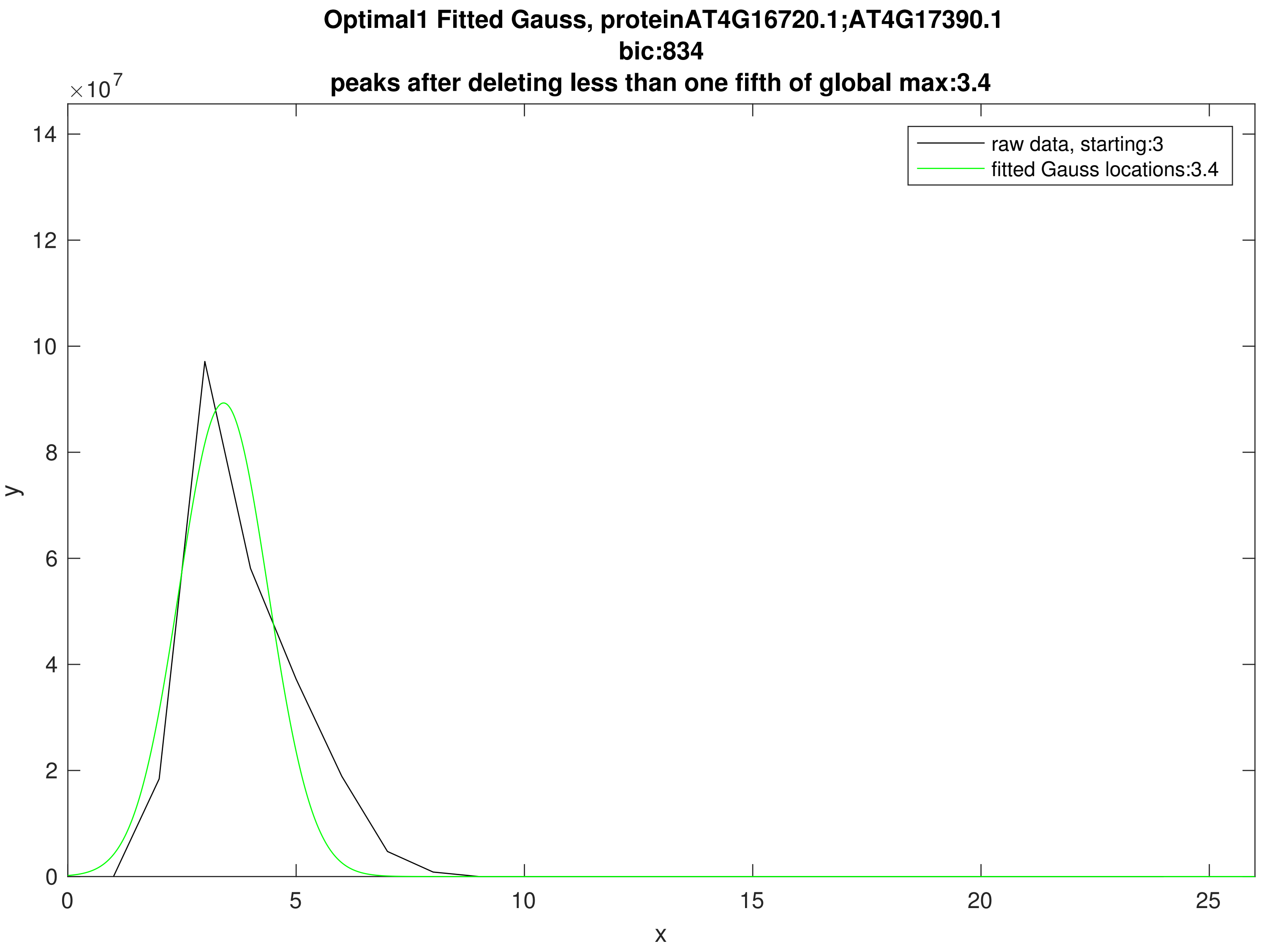
Optimal1 Fitted Gauss, proteinAT4G16720.1;AT4G17390.1
bic:834
peaks after deleting less than one fifth of global max:3.4
×10^7
raw data, starting:3
fitted Gauss locations:3.4
y
x
0
2
4
6
8
10
12
14
5
15
20
25

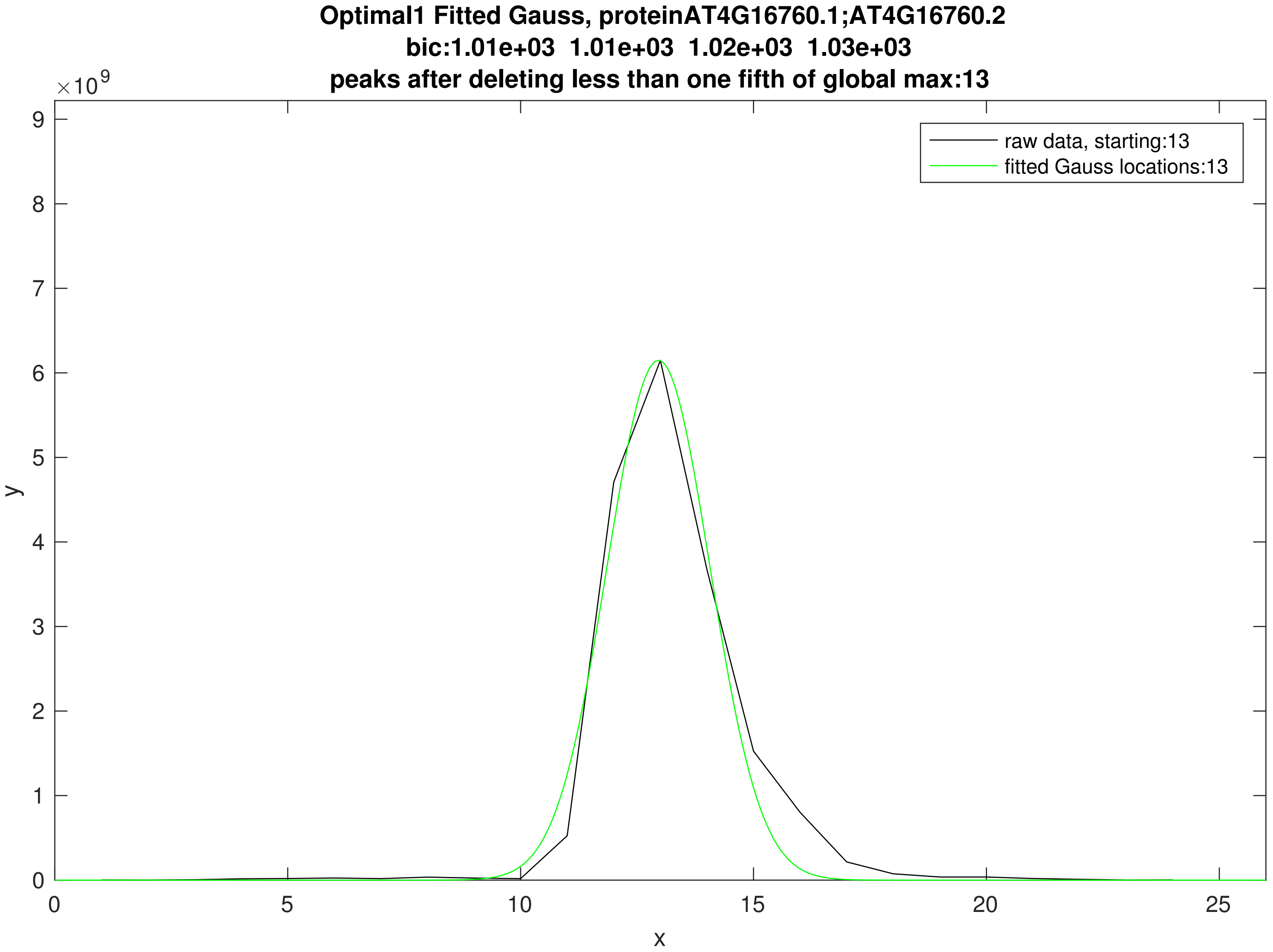
Optimal1 Fitted Gauss, proteinAT4G16760.1;AT4G16760.2
bic:1.01e+03 1.01e+03 1.02e+03 1.03e+03
peaks after deleting less than one fifth of global max:13
×10^9
raw data, starting:13
fitted Gauss locations:13
y
x
0
1
2
3
4
5
6
7
8
9
0
5
10
15
20
25

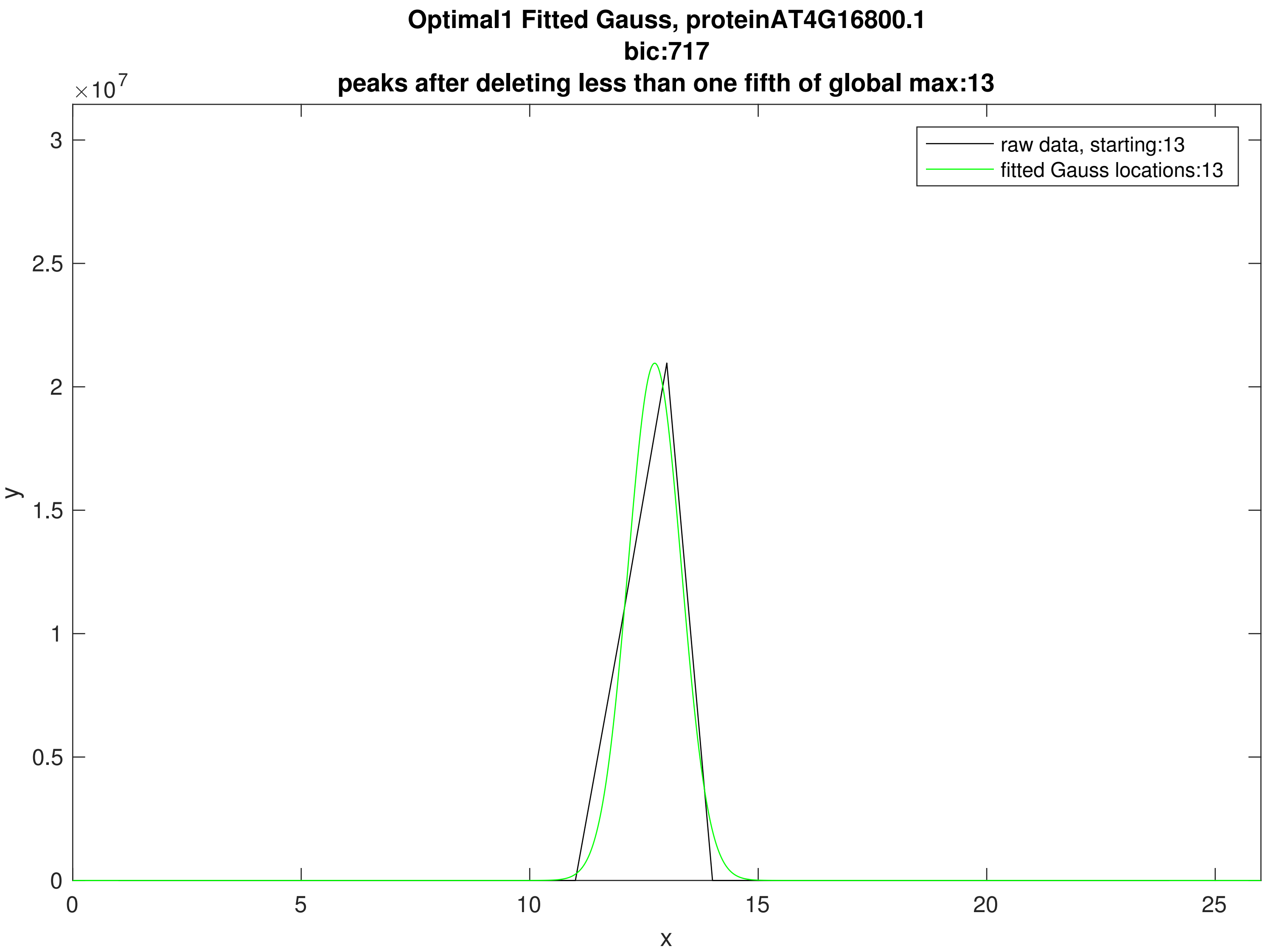
Optimal1 Fitted Gauss, proteinAT4G16800.1
bic:717
peaks after deleting less than one fifth of global max:13
×10^7
3
2.5
2
1.5
1
0.5
0
y
0
5
10
15
20
25
x
raw data, starting:13
fitted Gauss locations:13

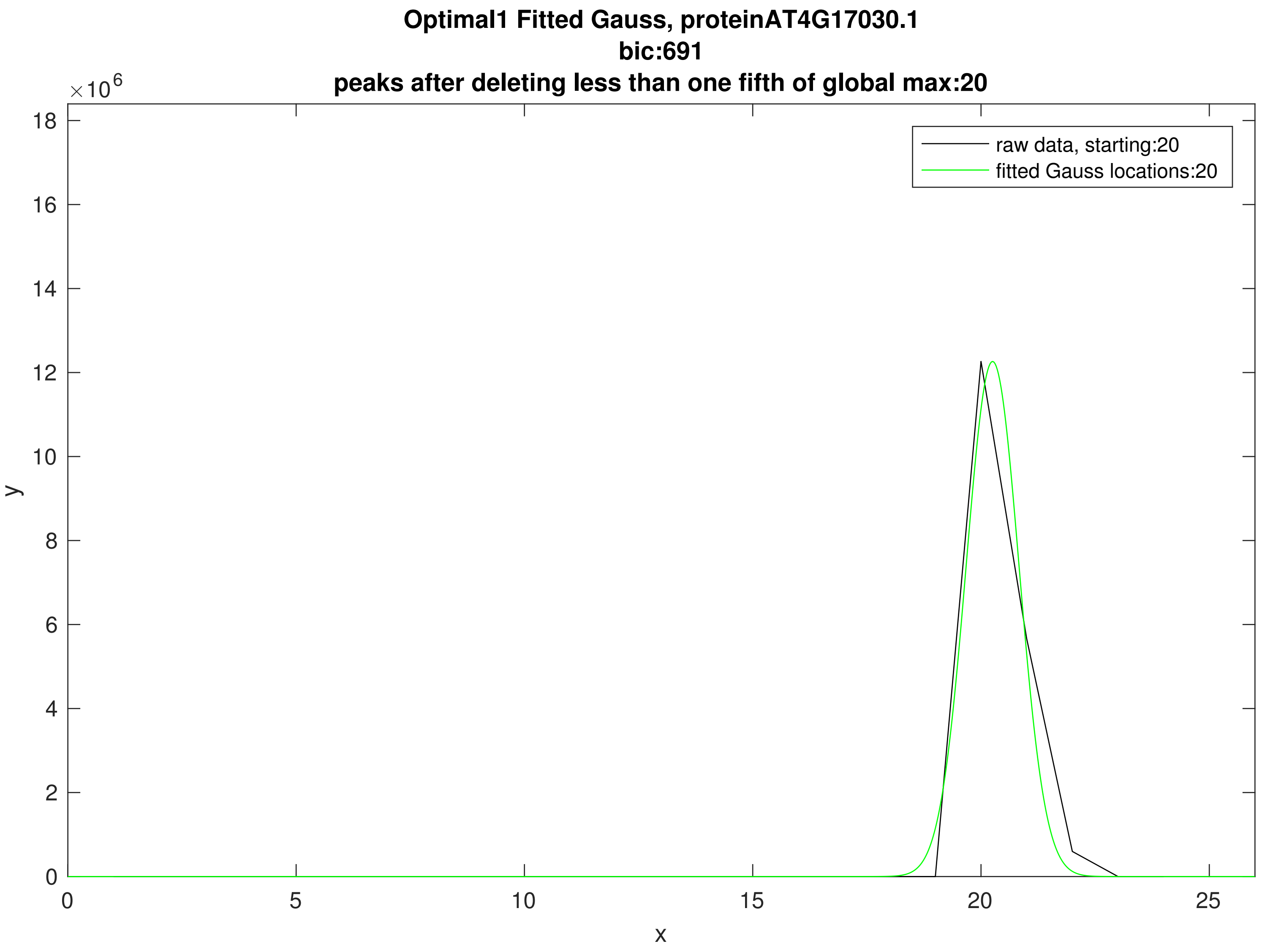
Optimal1 Fitted Gauss, proteinAT4G17030.1
bic:691
peaks after deleting less than one fifth of global max:20
×10^6
raw data, starting:20
fitted Gauss locations:20
y
x
0
2
4
6
8
10
12
14
16
18
0
5
10
15
20
25

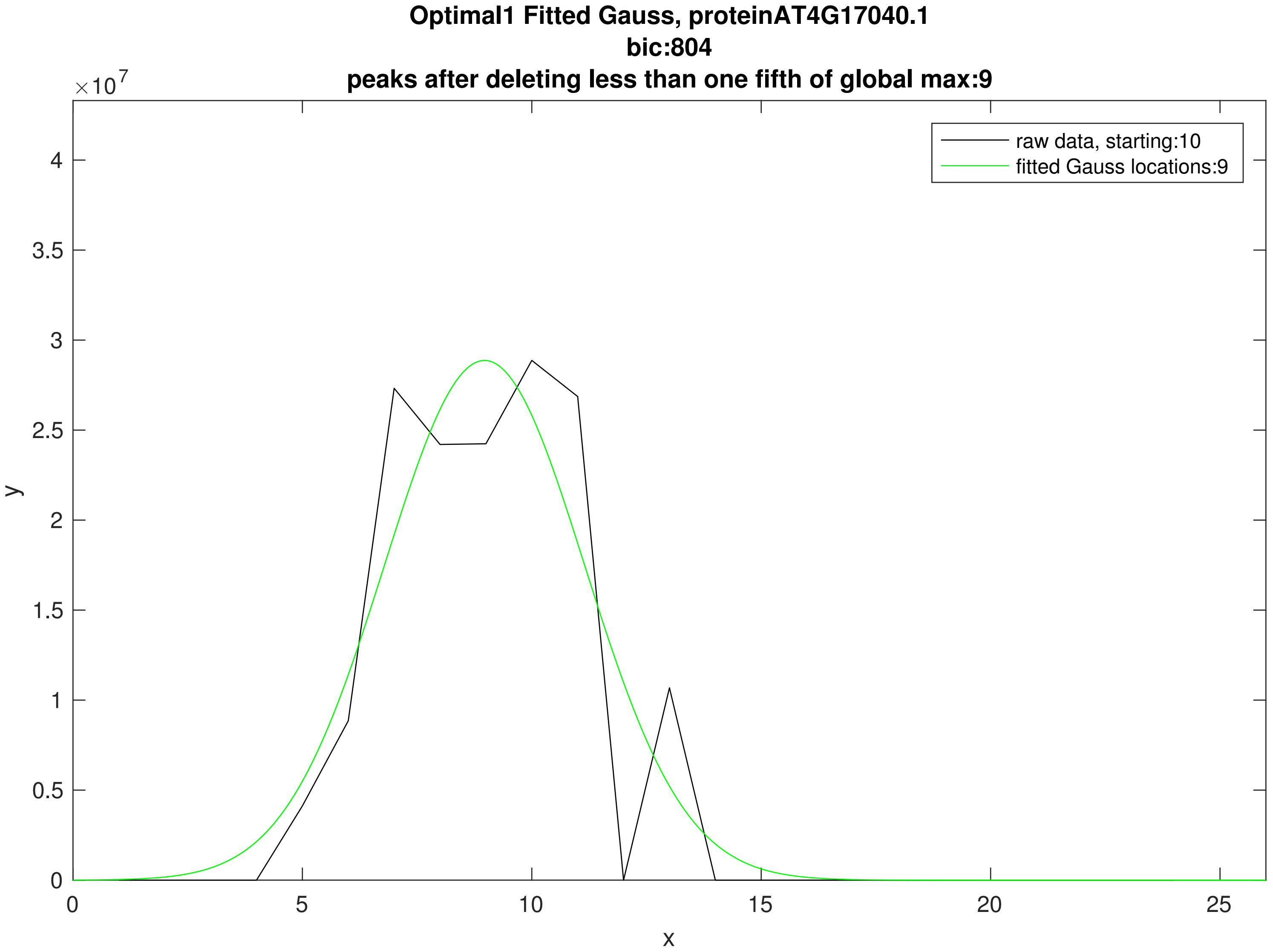
Optimal1 Fitted Gauss, proteinAT4G17040.1
bic:804
peaks after deleting less than one fifth of global max:9
×10^7
raw data, starting:10
fitted Gauss locations:9
y
x
0
0.5
1
1.5
2
2.5
3
3.5
4
5
10
15
20
25

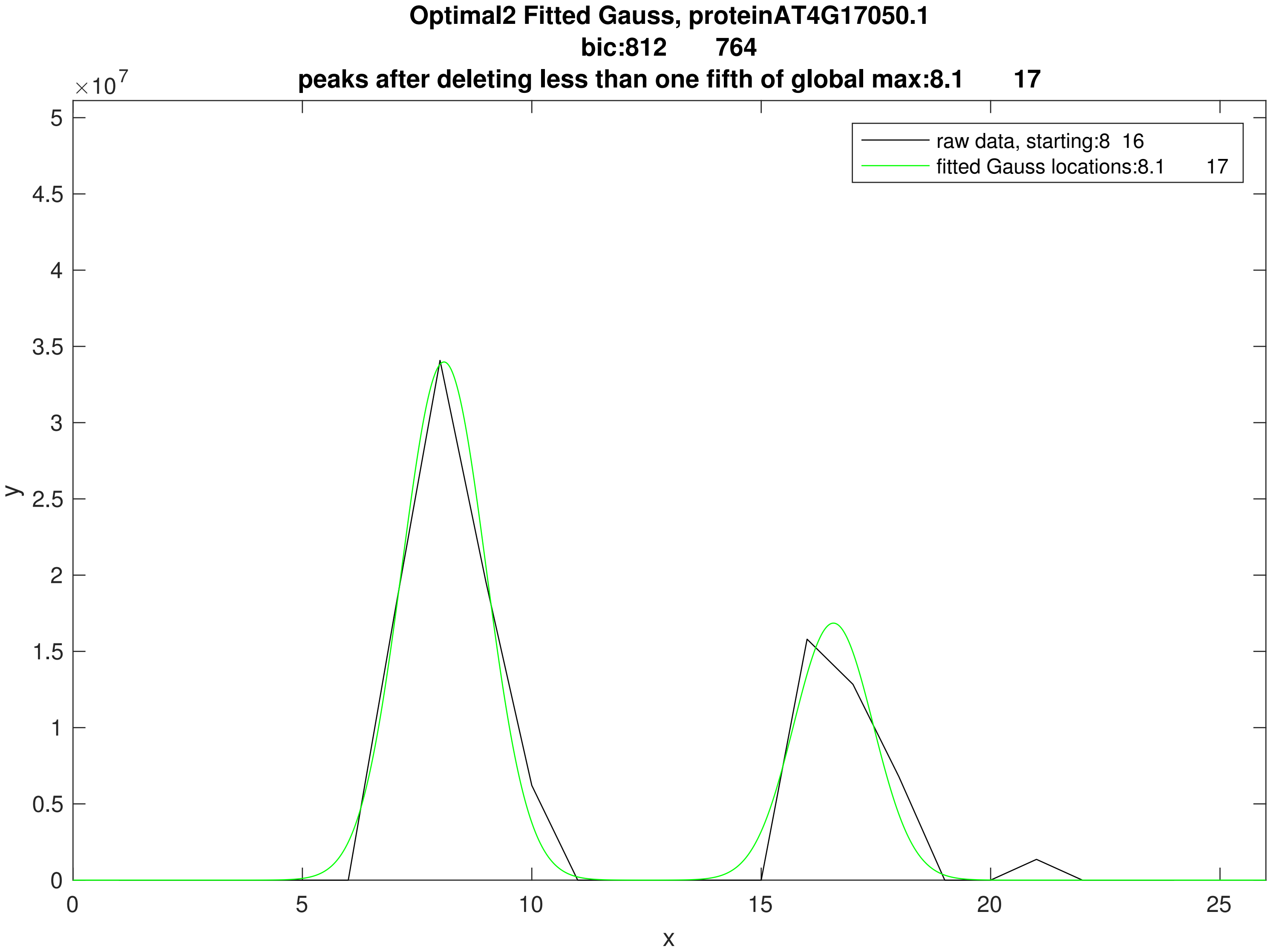
Optimal2 Fitted Gauss, proteinAT4G17050.1
bic:812 764
peaks after deleting less than one fifth of global max:8.1 17
raw data, starting:8 16
fitted Gauss locations:8.1 17
$\times 10^7$
y
x
0
0.5
1
1.5
2
2.5
3
3.5
4
4.5
5
0
5
10
15
20
25

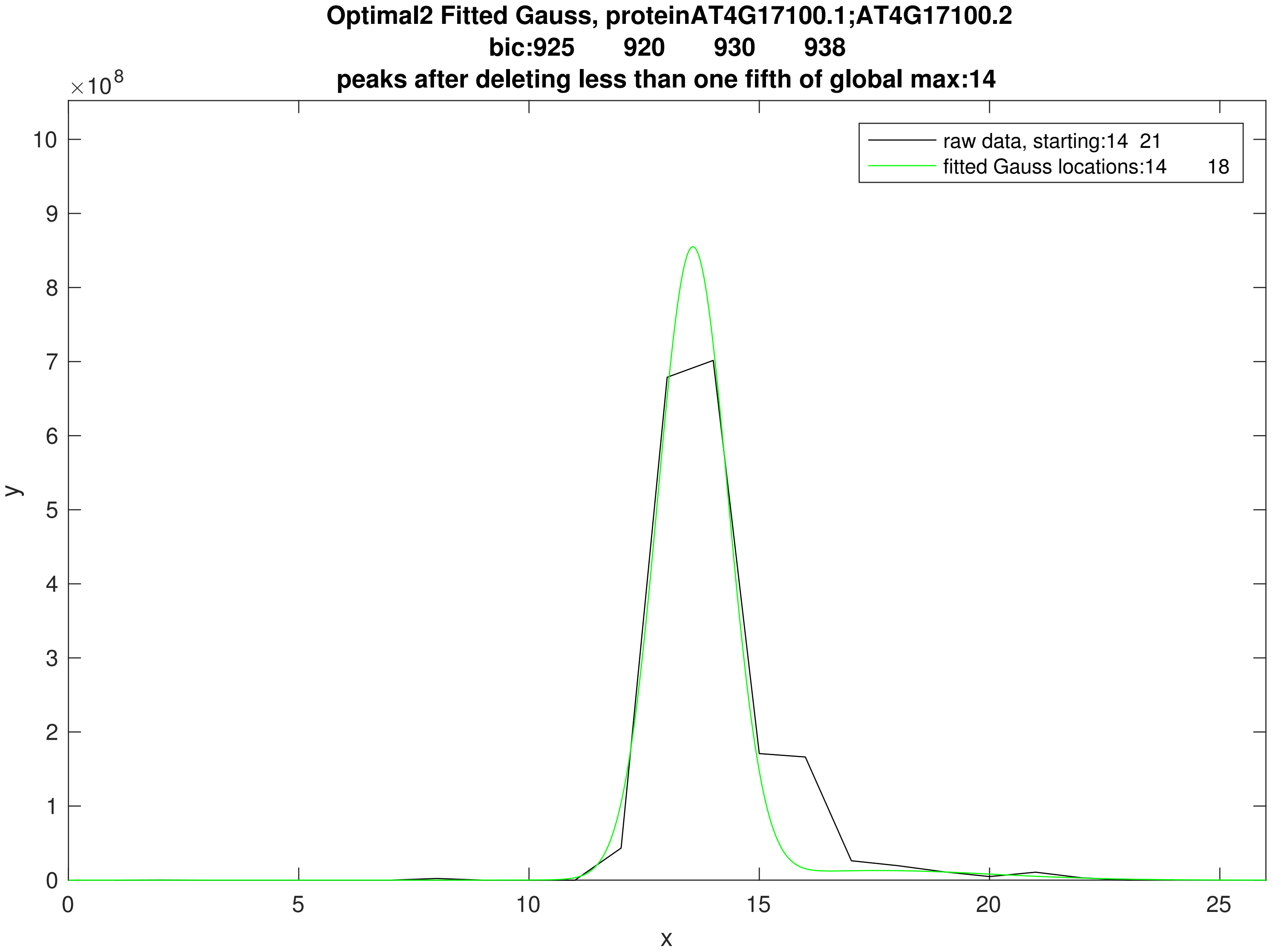

Optimal2 Fitted Gauss, proteinAT4G17100.1;AT4G17100.2
bic:925 920 930 938
peaks after deleting less than one fifth of global max:14
×10^8
raw data, starting:14 21
fitted Gauss locations:14 18
y
x
0
5
10
15
20
25
1
2
3
4
6
7
8
9

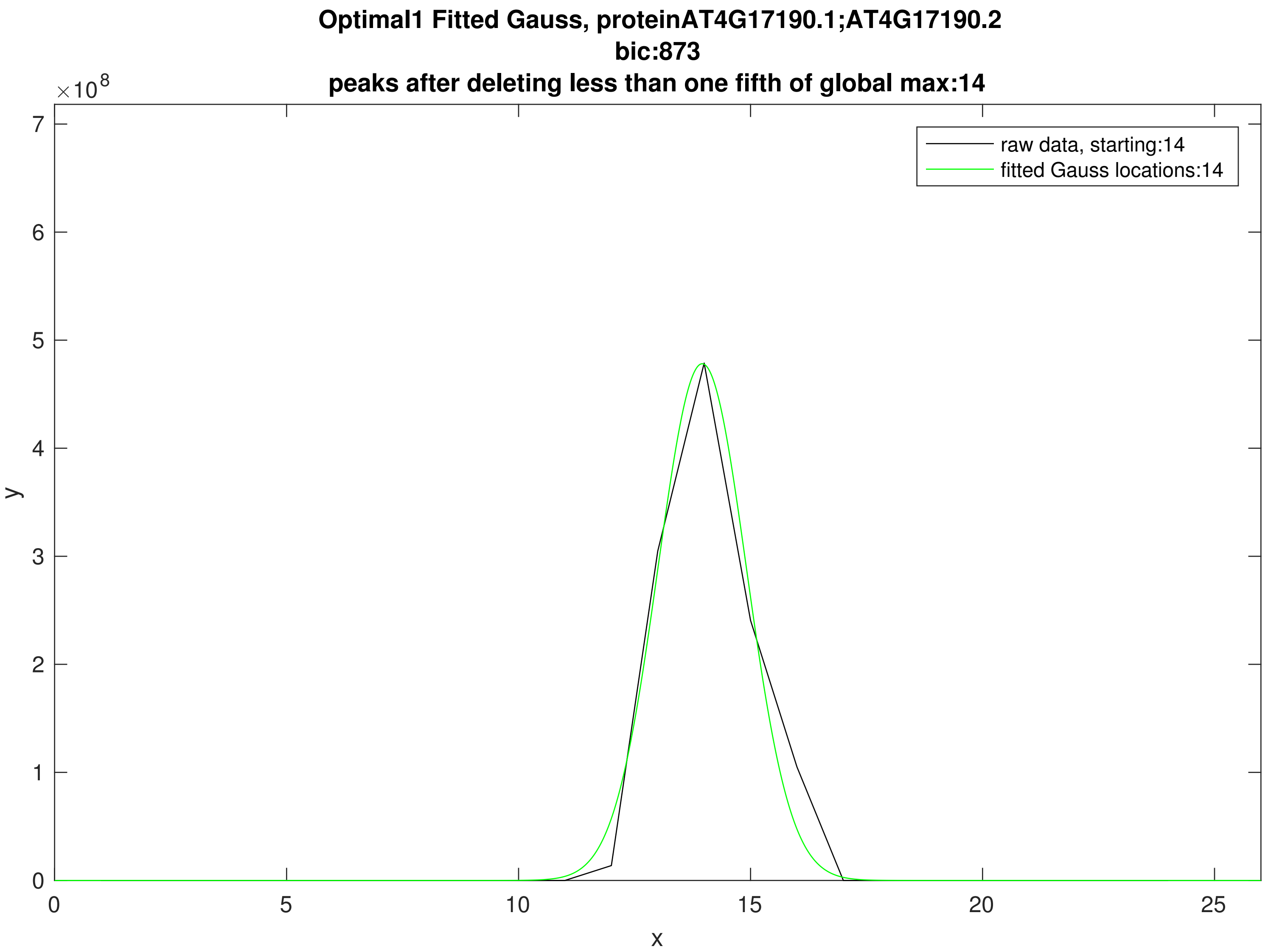

Optimal1 Fitted Gauss, proteinAT4G17190.1;AT4G17190.2
bic:873
peaks after deleting less than one fifth of global max:14
×10^8
raw data, starting:14
fitted Gauss locations:14
y
x
0
1
2
3
4
5
6
7
0
5
10
15
20
25

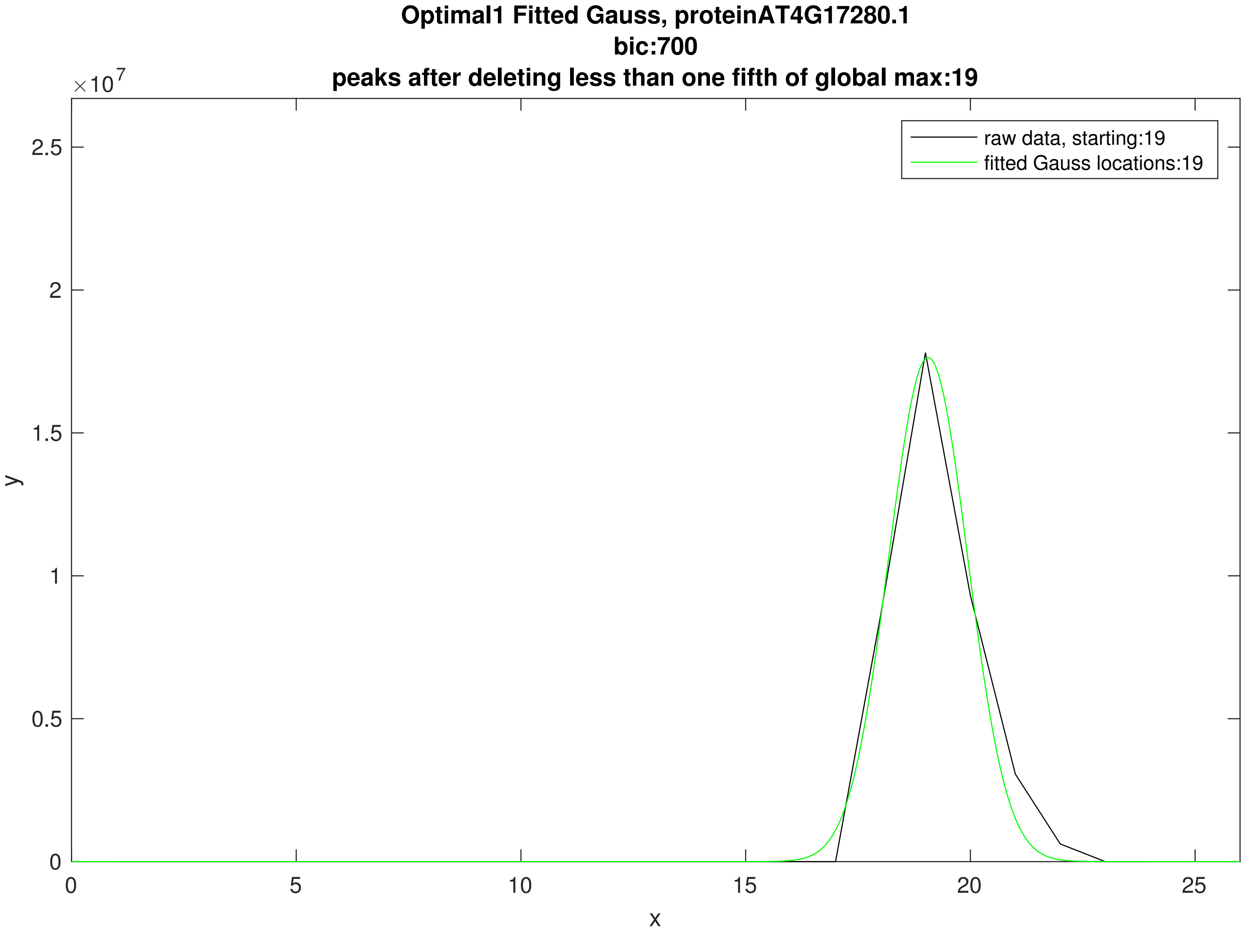
Optimal1 Fitted Gauss, proteinAT4G17280.1
bic:700
peaks after deleting less than one fifth of global max:19
×10^7
2.5
2
1.5
1
0.5
0
y
0
5
10
15
20
25
x
raw data, starting:19
fitted Gauss locations:19

Optimal1 Fitted Gauss, proteinAT4G17300.1
bic:817 869
peaks after deleting less than one fifth of global max:12

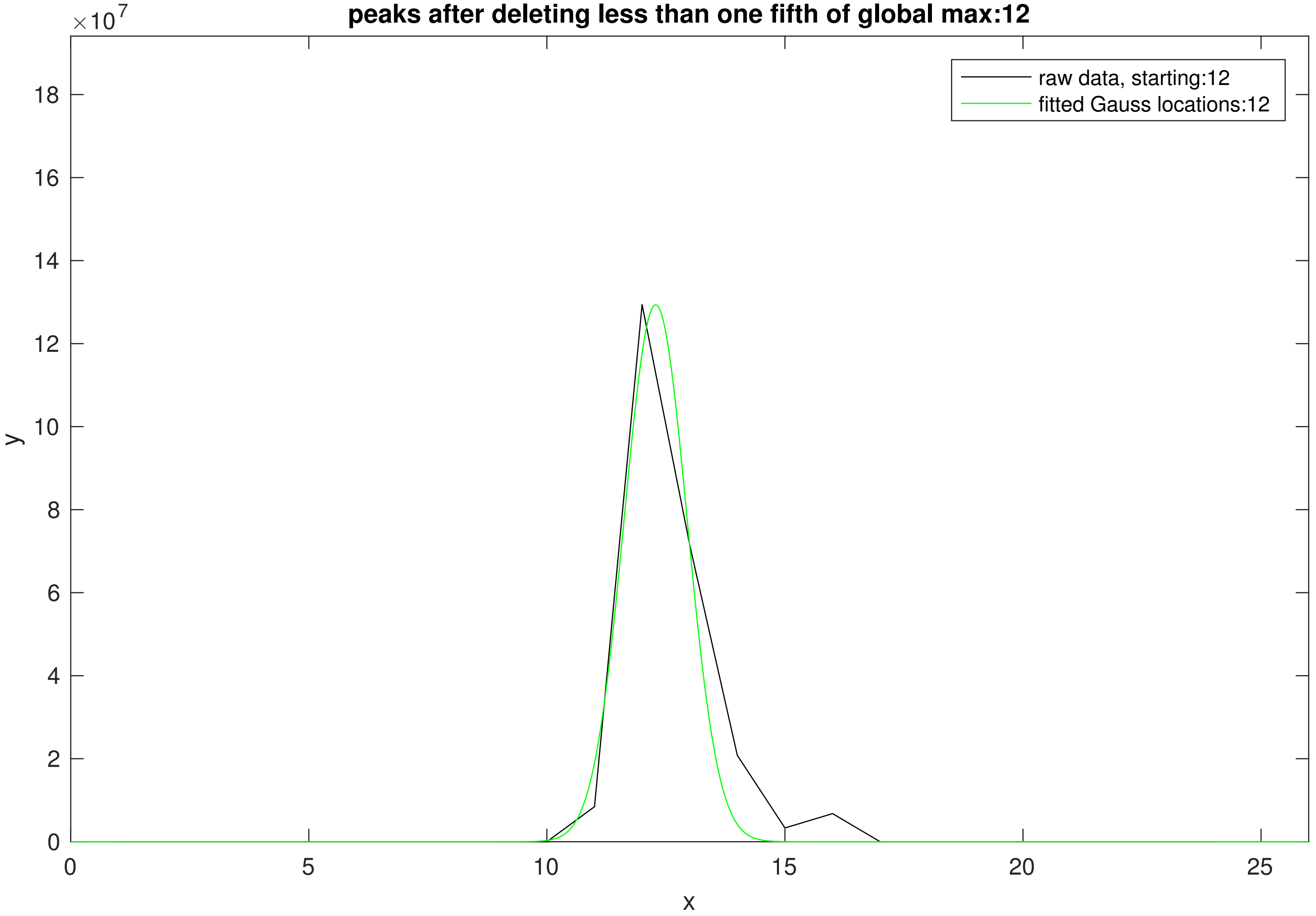

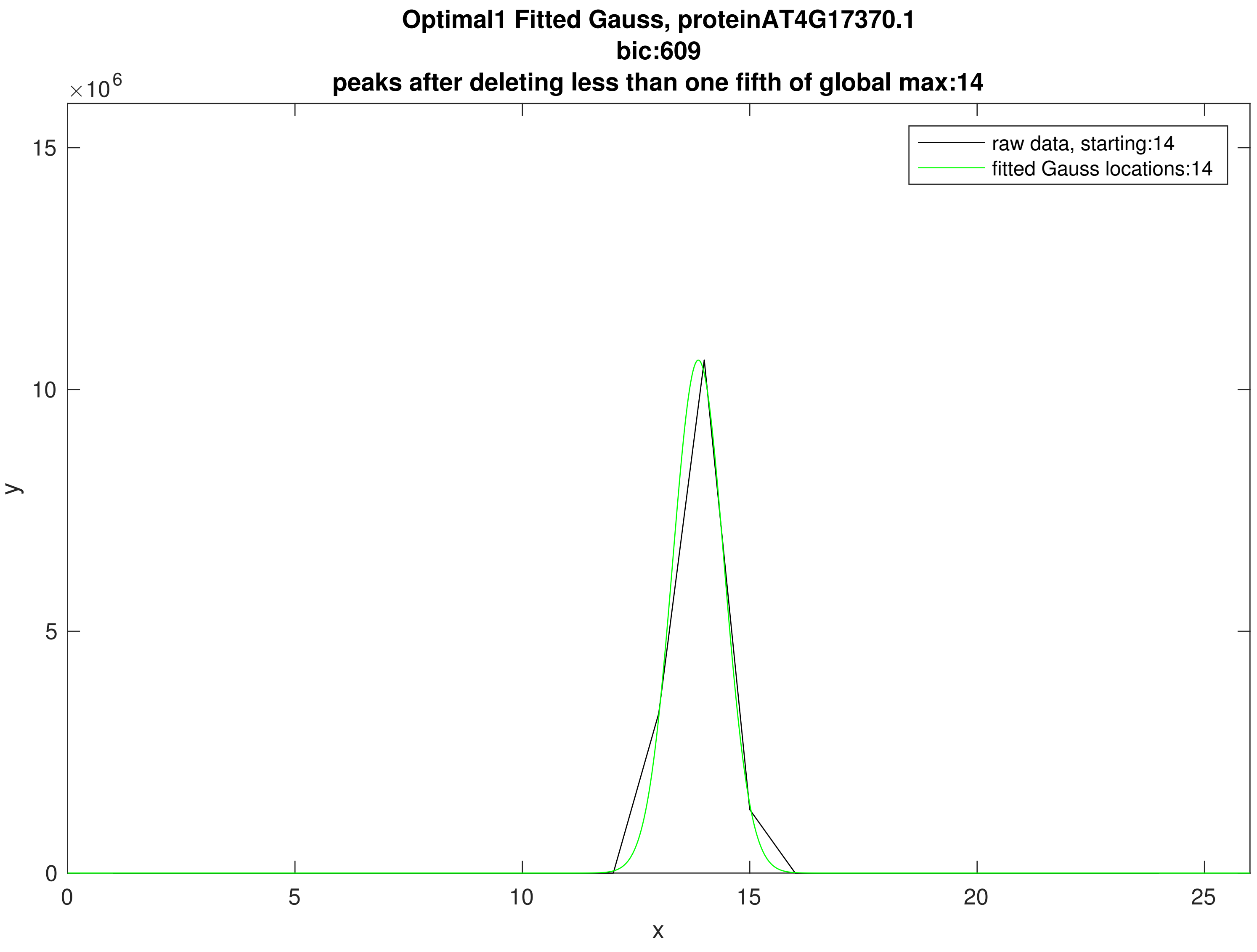
Optimal1 Fitted Gauss, proteinAT4G17370.1
bic:609
peaks after deleting less than one fifth of global max:14
×10^6
15
10
5
0
y
0
5
10
15
20
25
x
raw data, starting:14
fitted Gauss locations:14

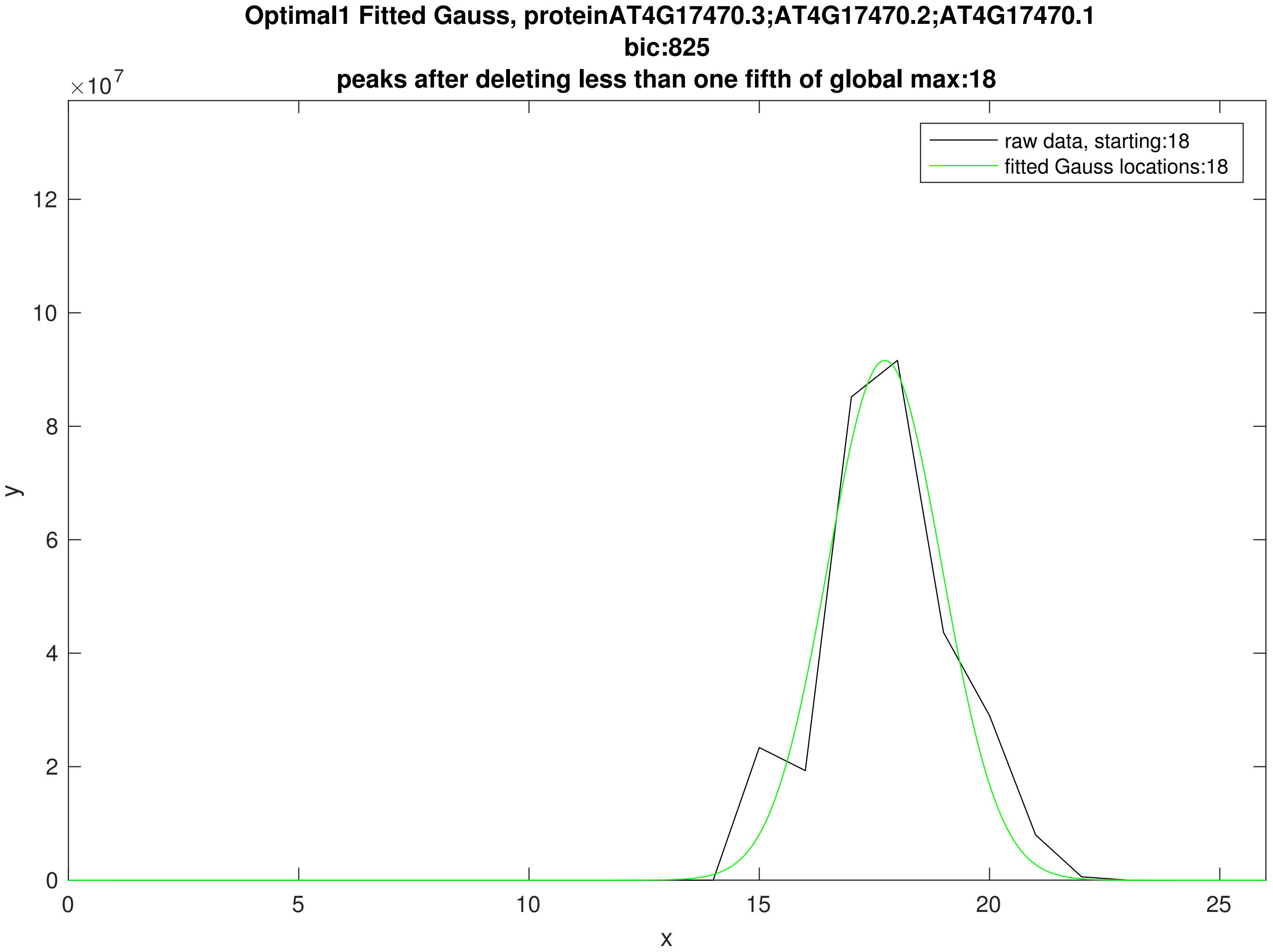
Optimal1 Fitted Gauss, proteinAT4G17470.3;AT4G17470.2;AT4G17470.1
bic:825
peaks after deleting less than one fifth of global max:18
×10^7
raw data, starting:18
fitted Gauss locations:18
y
x
0
2
4
6
8
10
12
0
5
10
15
20
25

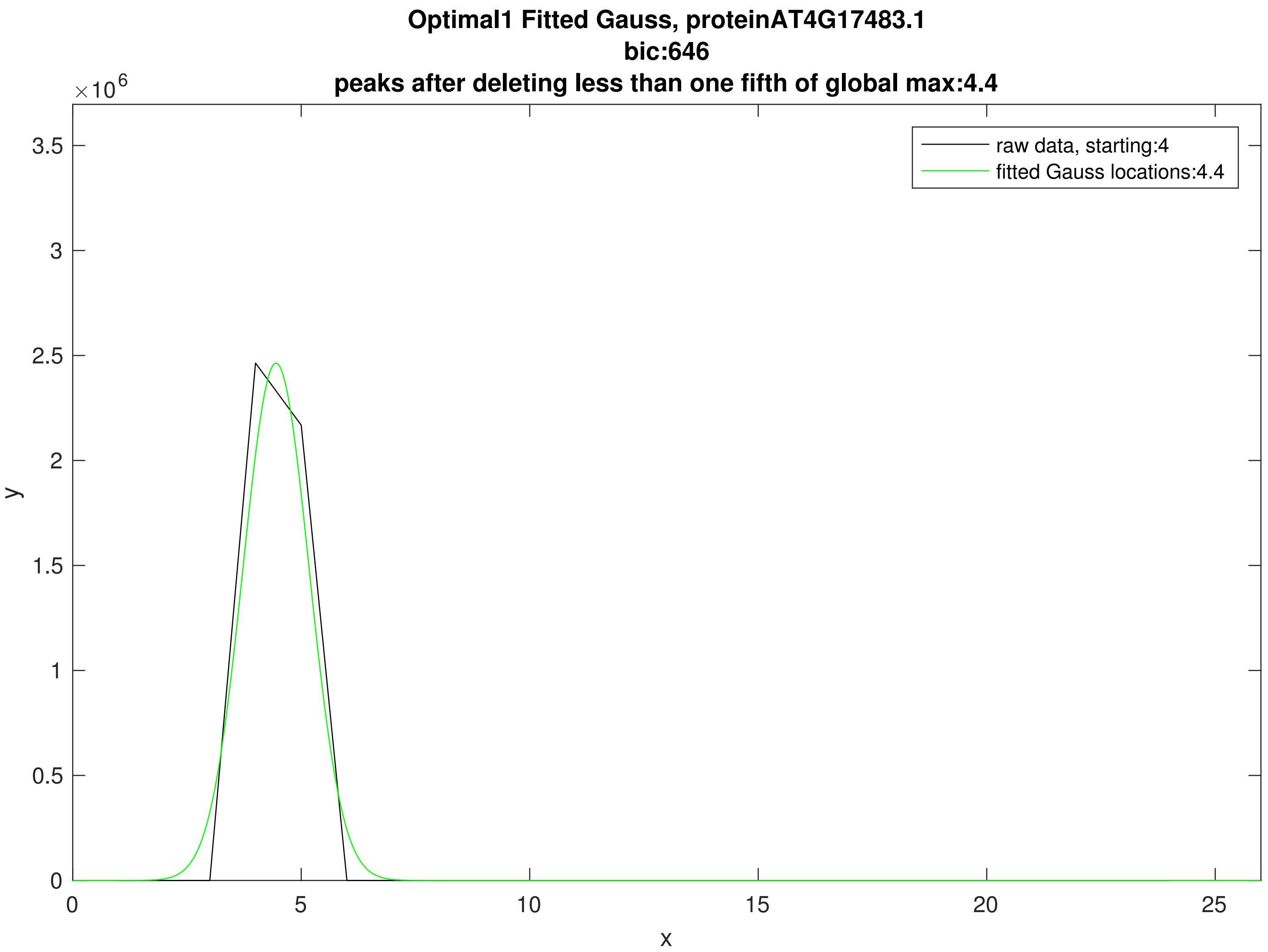
Optimal1 Fitted Gauss, proteinAT4G17483.1
bic:646
peaks after deleting less than one fifth of global max:4.4
×10^6
3.5
3
2.5
2
1.5
1
0.5
0
y
0
5
10
15
20
25
x
raw data, starting:4
fitted Gauss locations:4.4

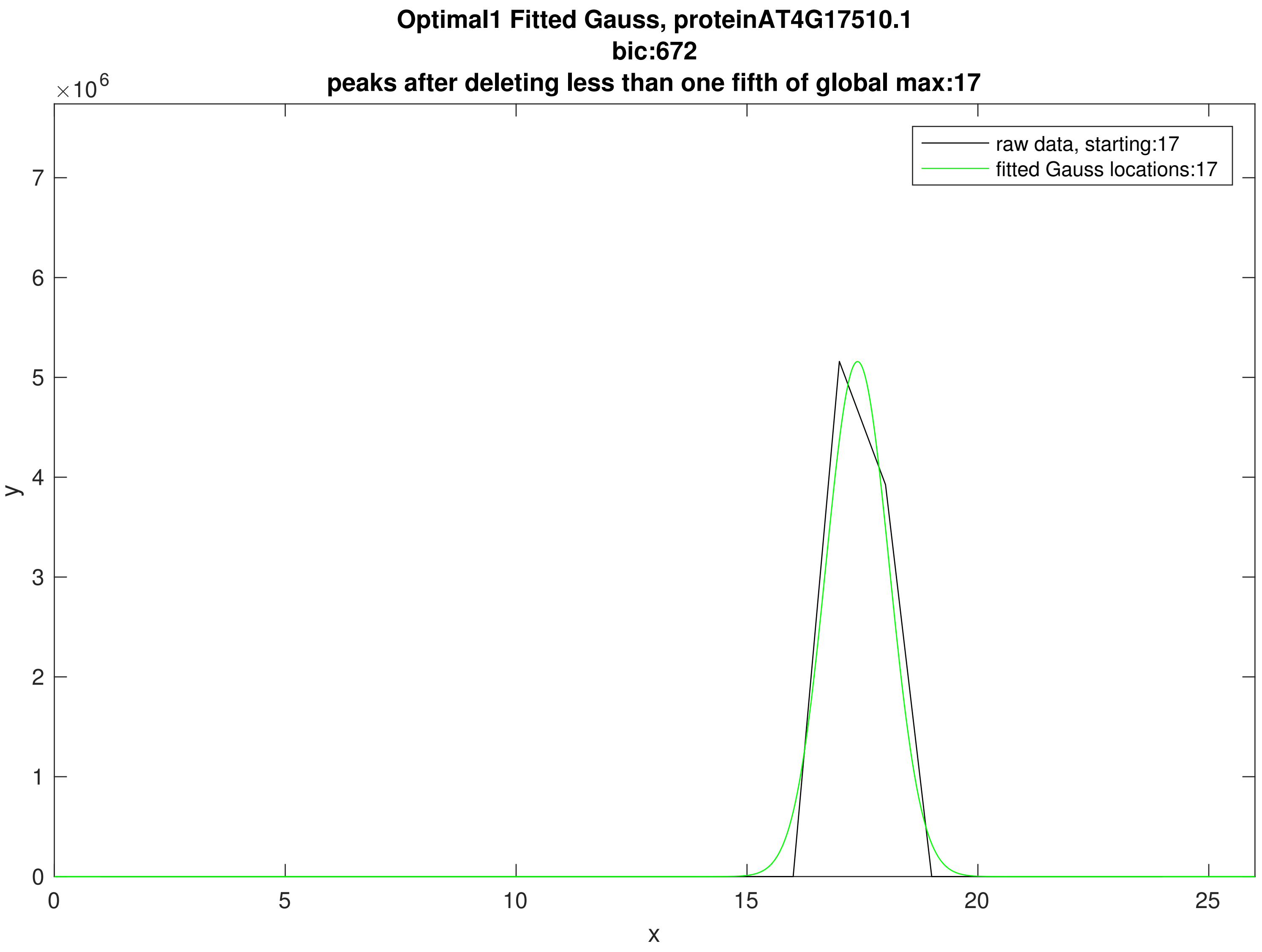

Optimal1 Fitted Gauss, proteinAT4G17510.1
bic:672
peaks after deleting less than one fifth of global max:17
×10^6
raw data, starting:17
fitted Gauss locations:17
y
x
0
1
2
3
4
5
6
7
0
5
10
15
20
25

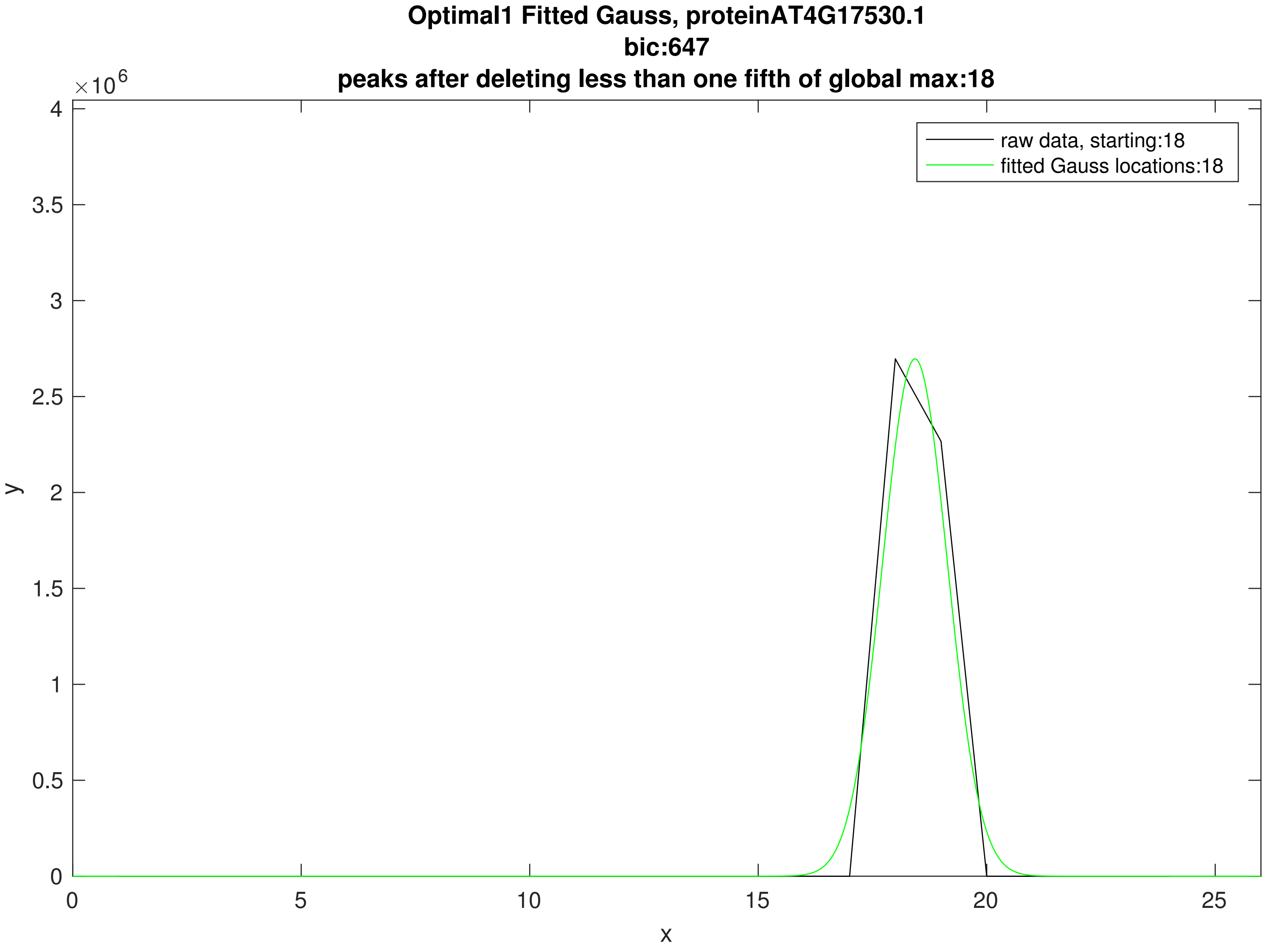
Optimal1 Fitted Gauss, proteinAT4G17530.1
bic:647
peaks after deleting less than one fifth of global max:18
×10^6
4
3.5
3
2.5
2
1.5
1
0.5
0
y
0
5
10
15
20
25
x
raw data, starting:18
fitted Gauss locations:18

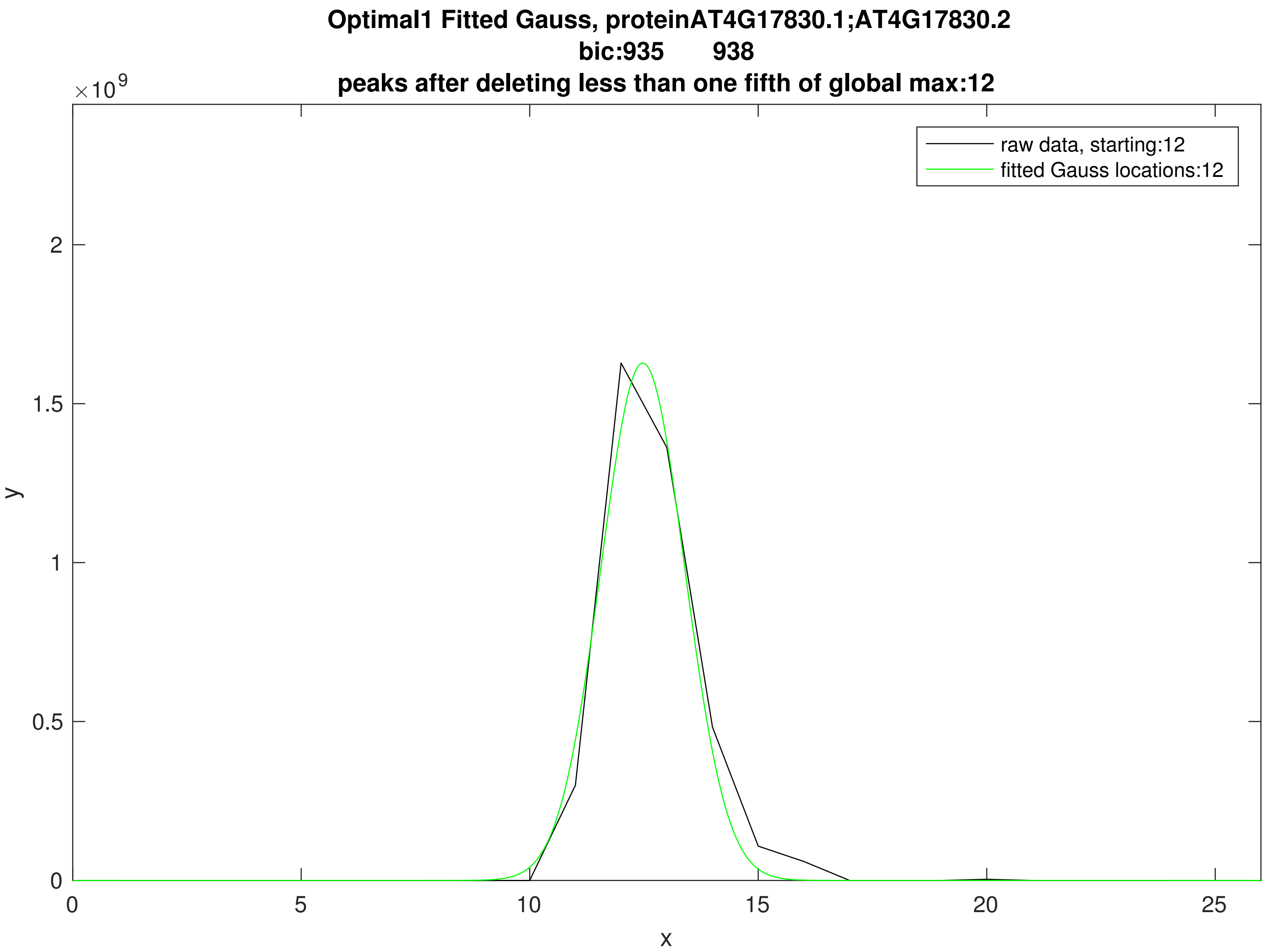
Optimal1 Fitted Gauss, proteinAT4G17830.1;AT4G17830.2
bic:935 938
peaks after deleting less than one fifth of global max:12
×10^9
raw data, starting:12
fitted Gauss locations:12
y
x
0
0.5
1
1.5
2
0
5
10
15
20
25

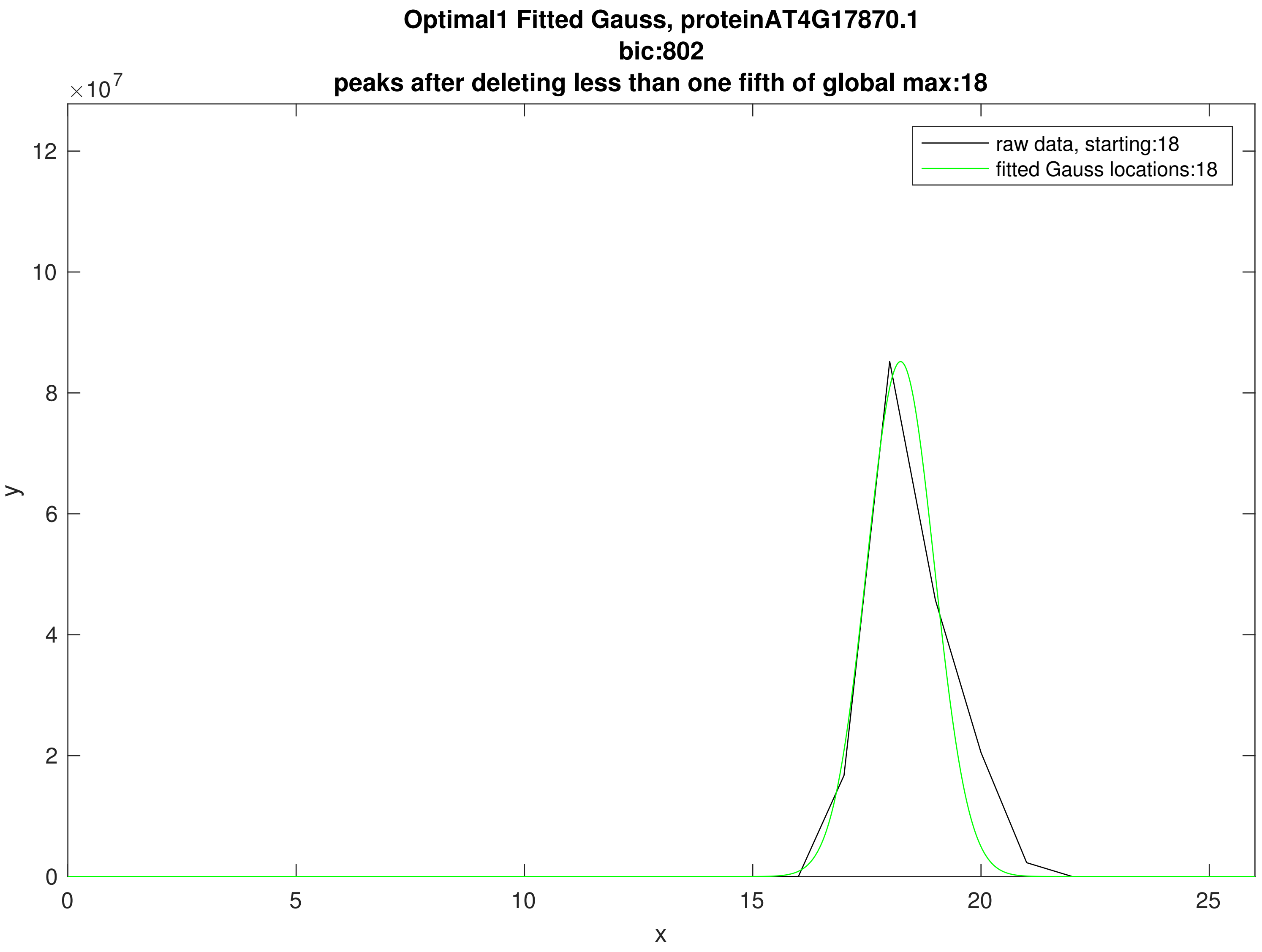

Optimal1 Fitted Gauss, proteinAT4G17870.1
bic:802
peaks after deleting less than one fifth of global max:18
×10^7
12
10
8
6
4
2
0
y
0
5
10
15
20
25
x
raw data, starting:18
fitted Gauss locations:18

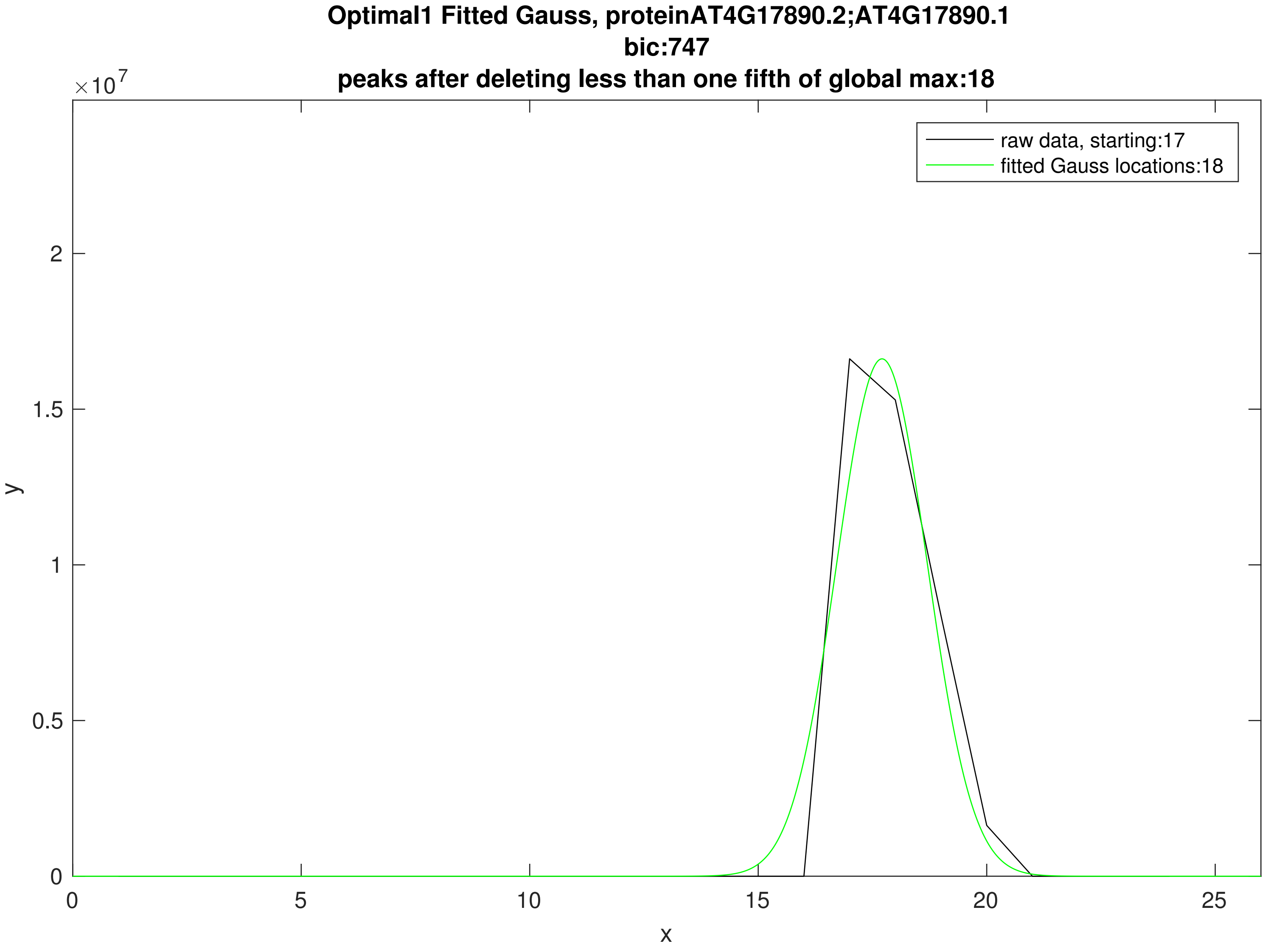
Optimal1 Fitted Gauss, proteinAT4G17890.2;AT4G17890.1
bic:747
peaks after deleting less than one fifth of global max:18
×10^7
raw data, starting:17
fitted Gauss locations:18
2
1.5
1
0.5
0
y
0
5
10
15
20
25
x

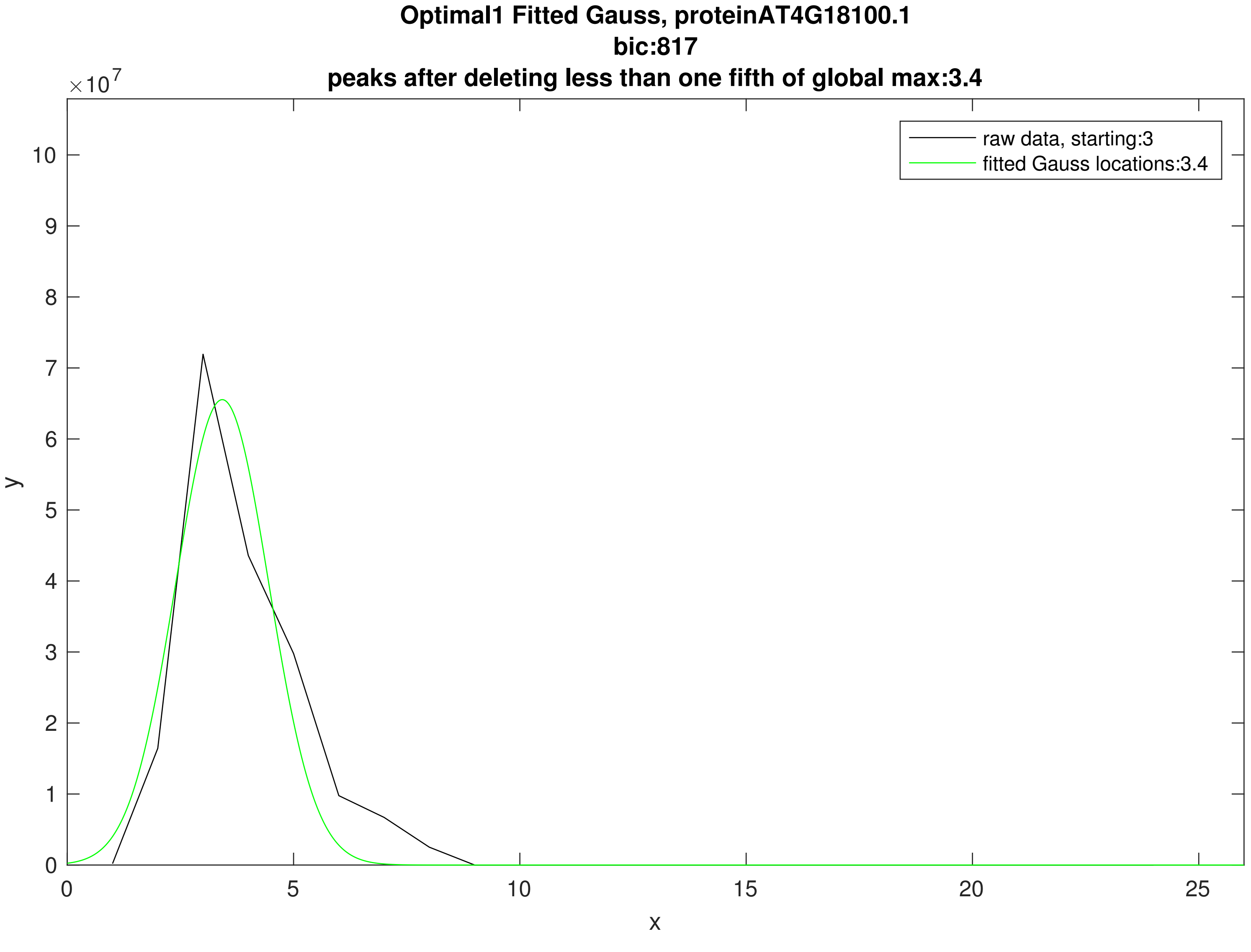

Optimal1 Fitted Gauss, proteinAT4G18100.1
bic:817
peaks after deleting less than one fifth of global max:3.4
×10^7
raw data, starting:3
fitted Gauss locations:3.4
y
x
0
1
2
3
4
5
6
7
8
9
10
0
5
10
15
20
25

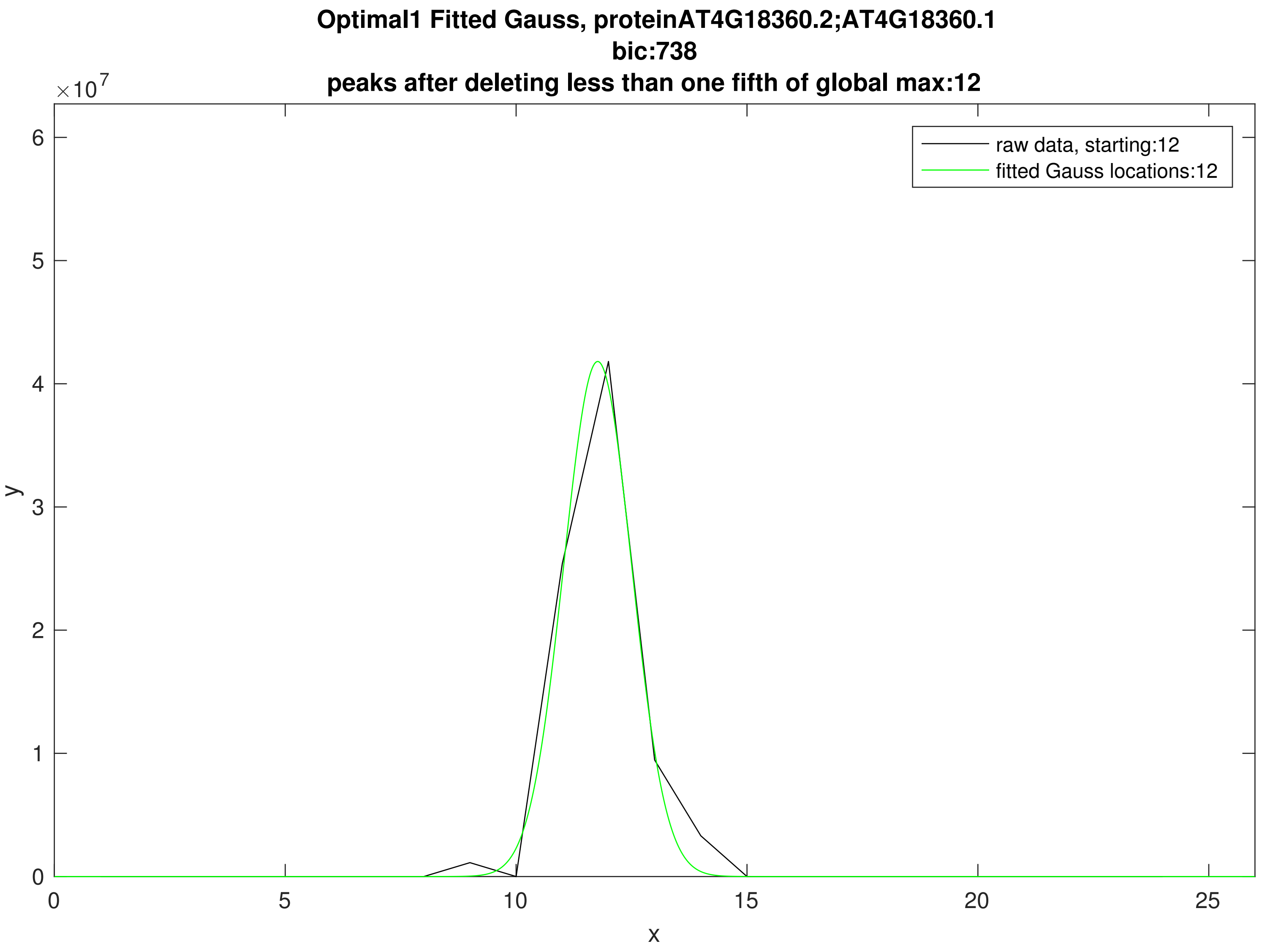
Optimal1 Fitted Gauss, proteinAT4G18360.2;AT4G18360.1
bic:738
peaks after deleting less than one fifth of global max:12
×10^7
raw data, starting:12
fitted Gauss locations:12
y
x
0
1
2
3
4
5
6
0
5
10
15
20
25

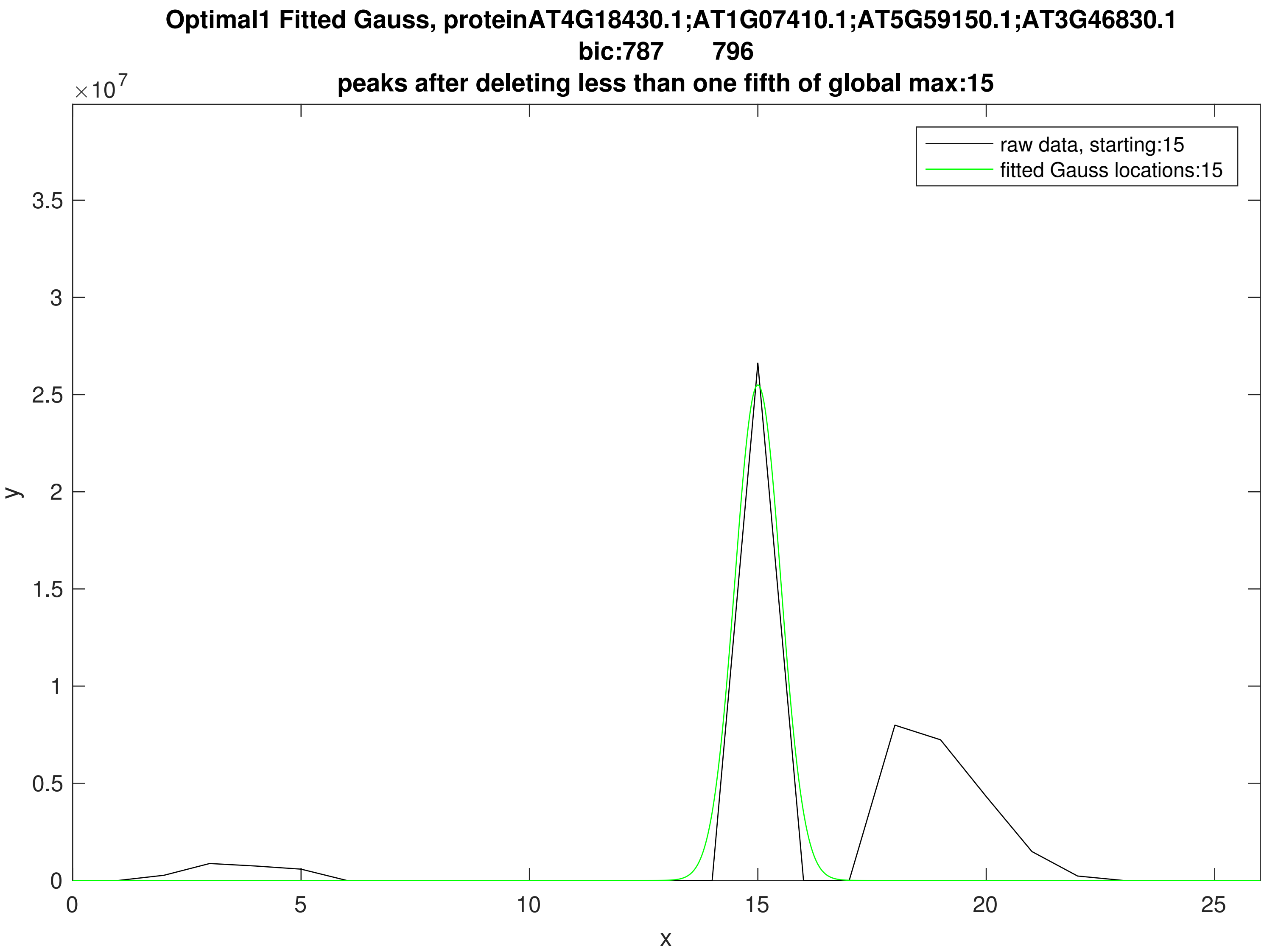
Optimal1 Fitted Gauss, proteinAT4G18430.1;AT1G07410.1;AT5G59150.1;AT3G46830.1
bic:787 796
peaks after deleting less than one fifth of global max:15
×10^7
raw data, starting:15
fitted Gauss locations:15
3.5
3
2.5
2
1.5
1
0.5
0
y
0
5
10
15
20
25
x

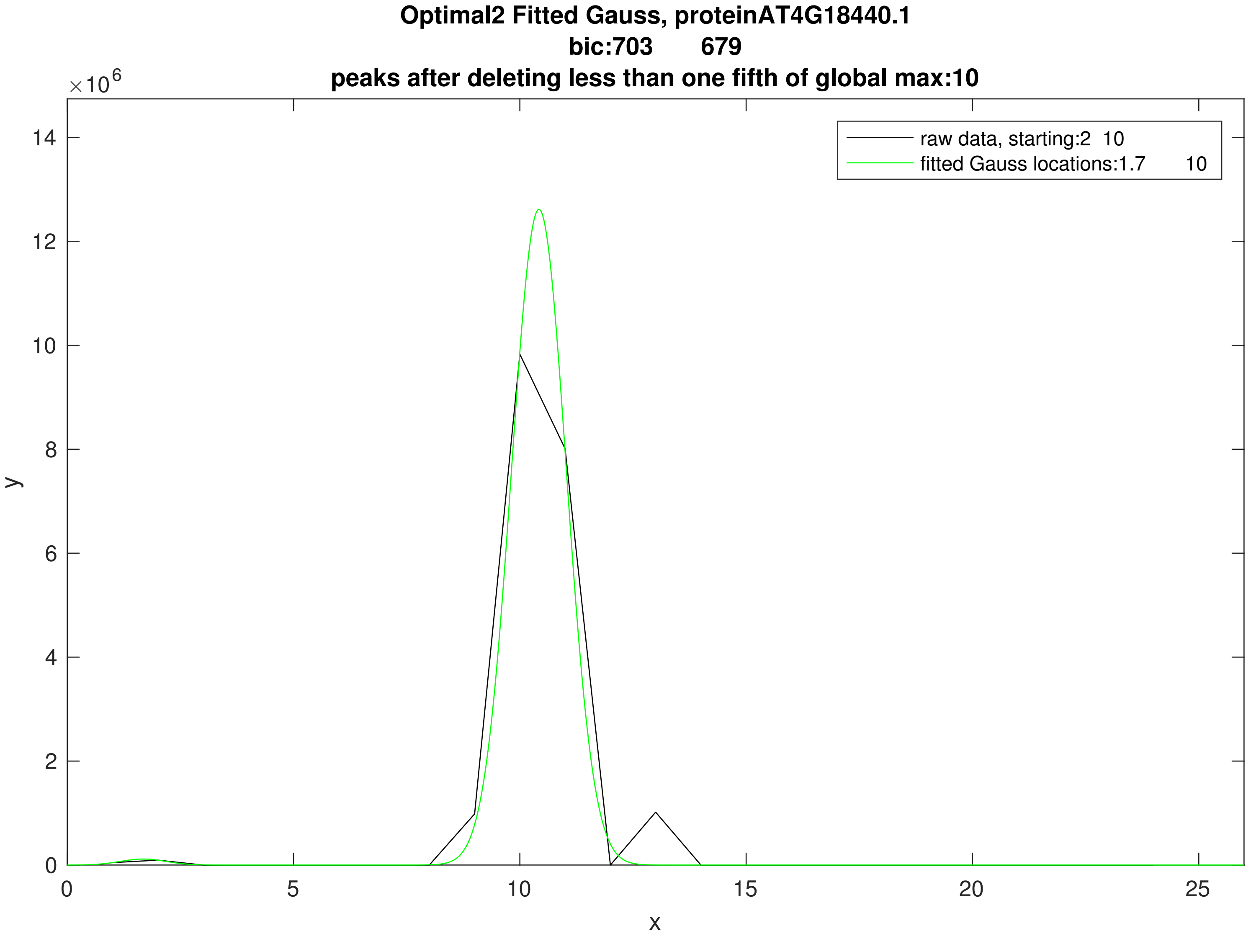

Optimal2 Fitted Gauss, proteinAT4G18440.1
bic:703 679
peaks after deleting less than one fifth of global max:10
×10^6
14
12
10
8
6
4
2
0
y
0
5
10
15
20
25
x
raw data, starting:2 10
fitted Gauss locations:1.7 10

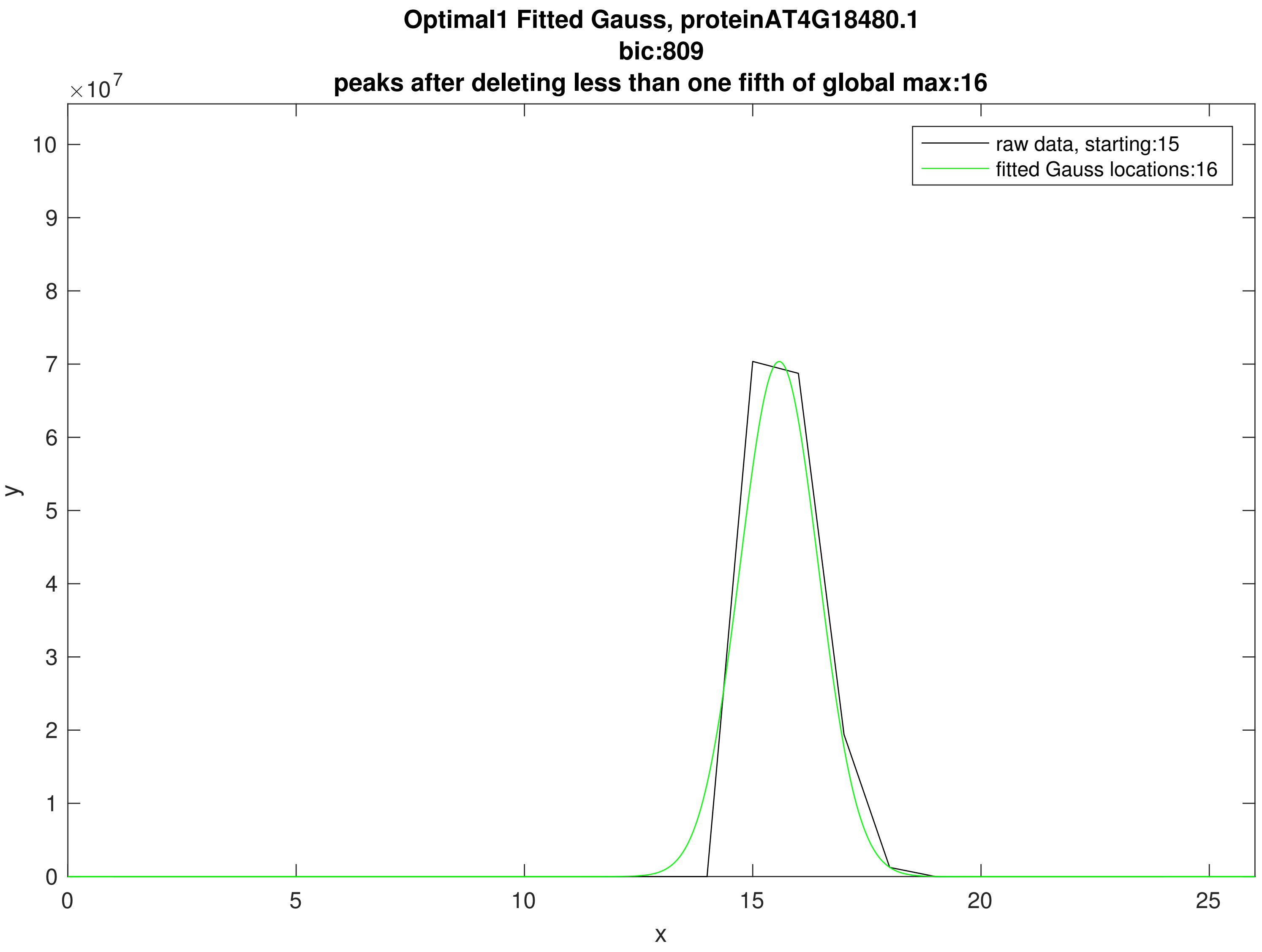
Optimal1 Fitted Gauss, proteinAT4G18480.1
bic:809
peaks after deleting less than one fifth of global max:16
×10^7
raw data, starting:15
fitted Gauss locations:16
y
x
0
1
2
3
4
5
6
7
8
9
10
0
5
10
15
20
25

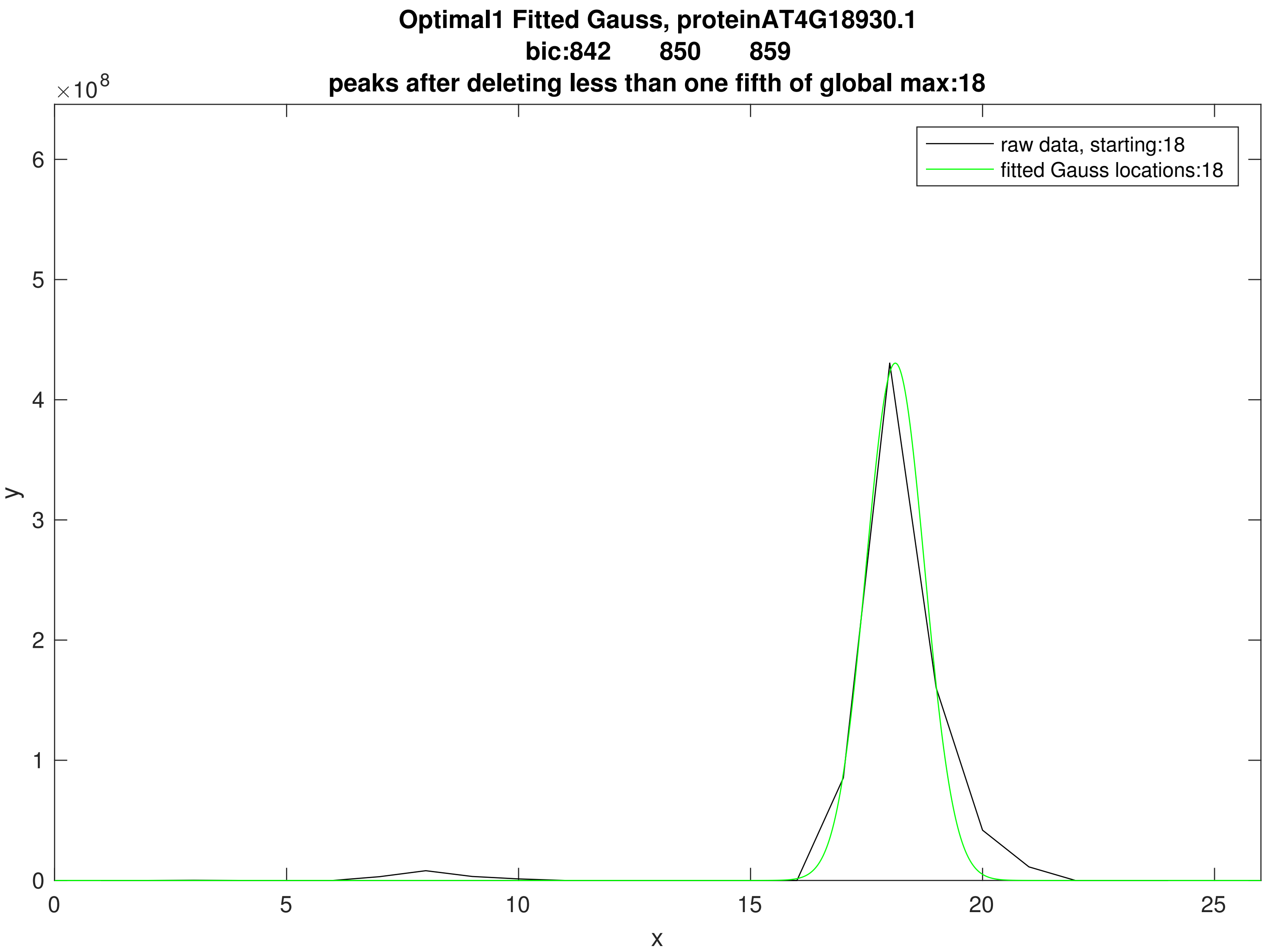
Optimal1 Fitted Gauss, proteinAT4G18930.1
bic:842 850 859
peaks after deleting less than one fifth of global max:18
×10^8
raw data, starting:18
fitted Gauss locations:18
y
x

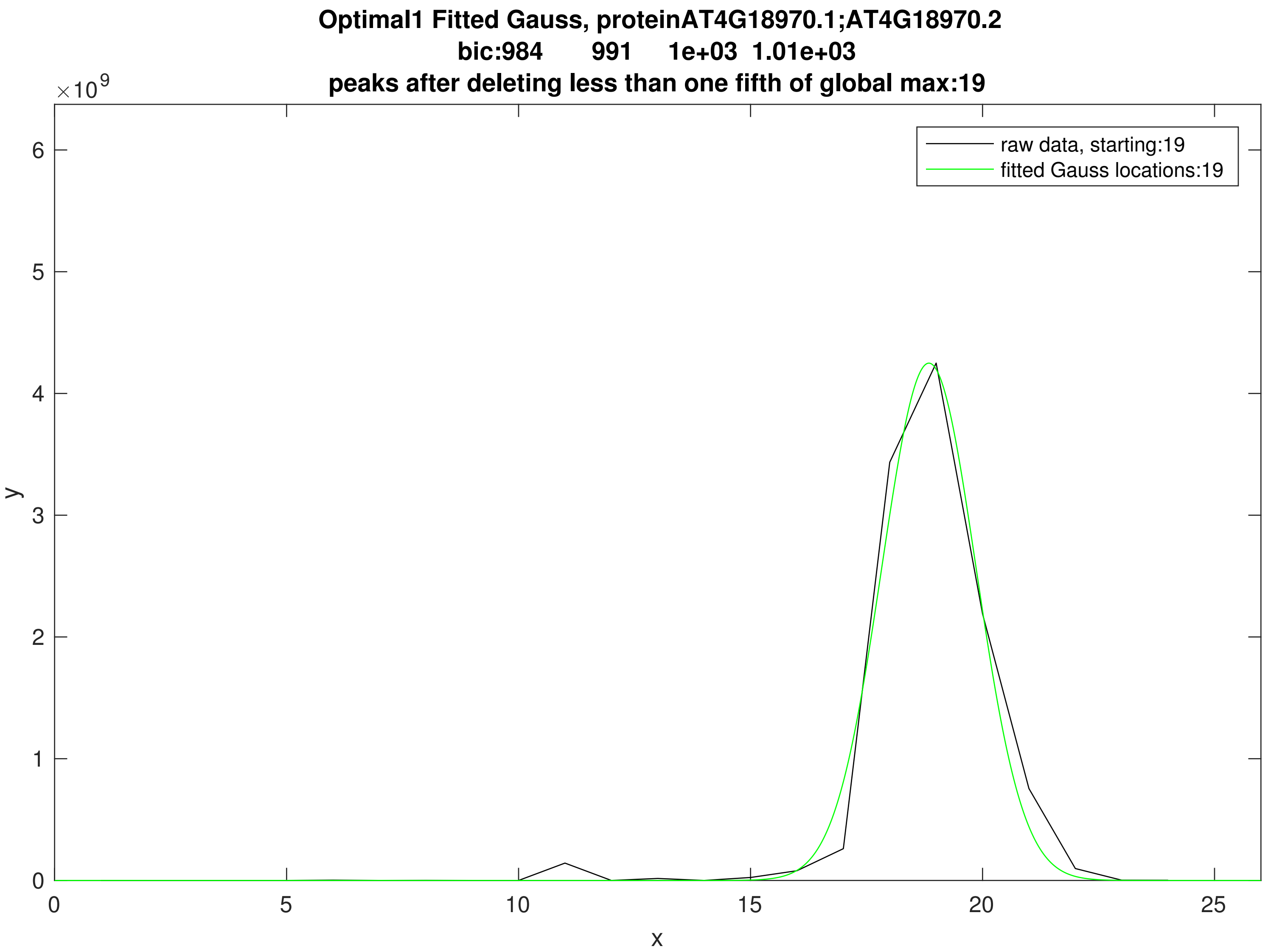
Optimal1 Fitted Gauss, proteinAT4G18970.1;AT4G18970.2
bic:984 991 1e+03 1.01e+03
peaks after deleting less than one fifth of global max:19
×10^9
raw data, starting:19
fitted Gauss locations:19
y
x
0
1
2
3
4
5
6
0
5
10
15
20
25

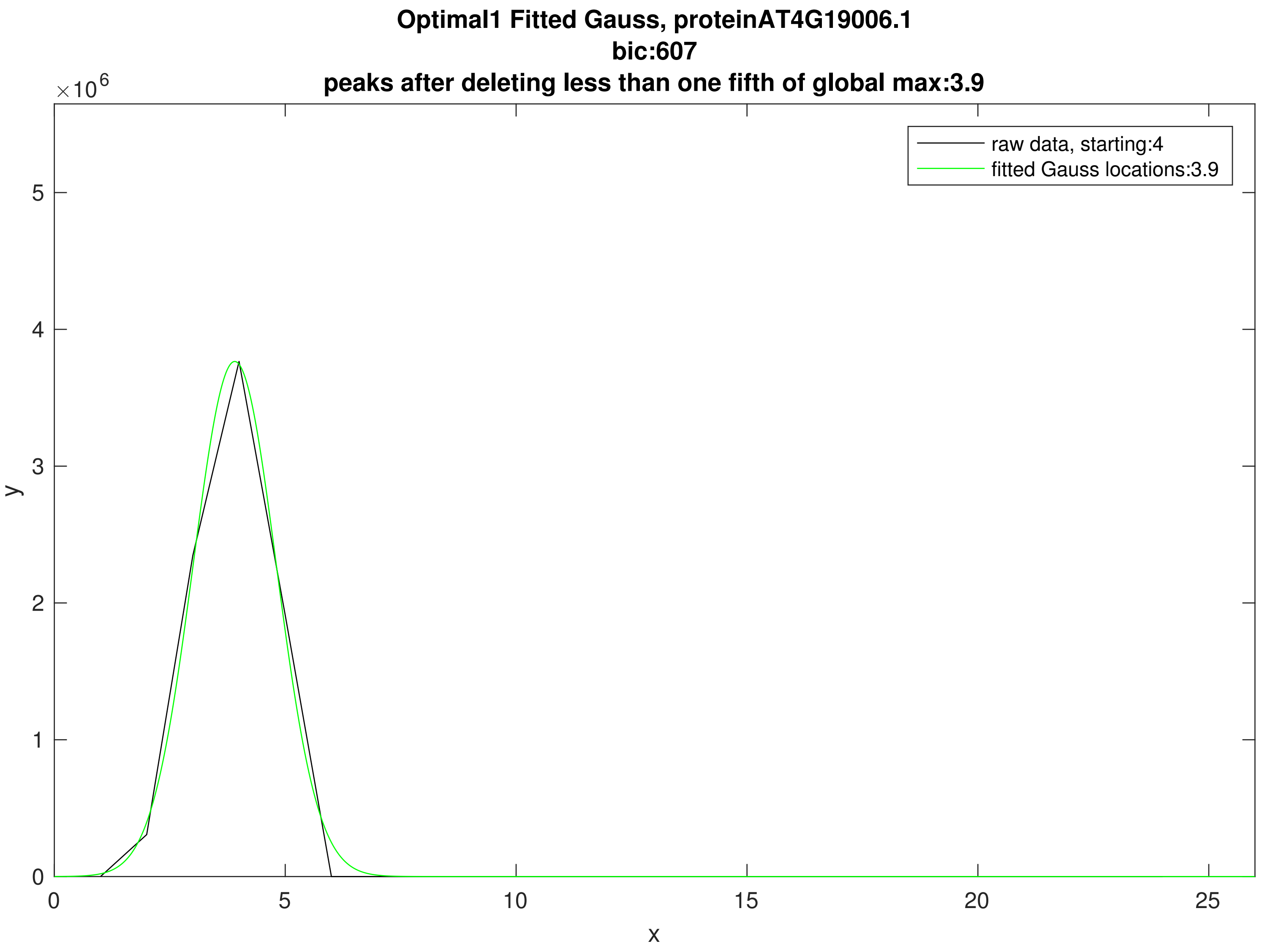

Optimal1 Fitted Gauss, proteinAT4G19006.1
bic:607
peaks after deleting less than one fifth of global max:3.9
×10^6
raw data, starting:4
fitted Gauss locations:3.9
5
4
3
2
1
0
y
0
5
10
15
20
25
x

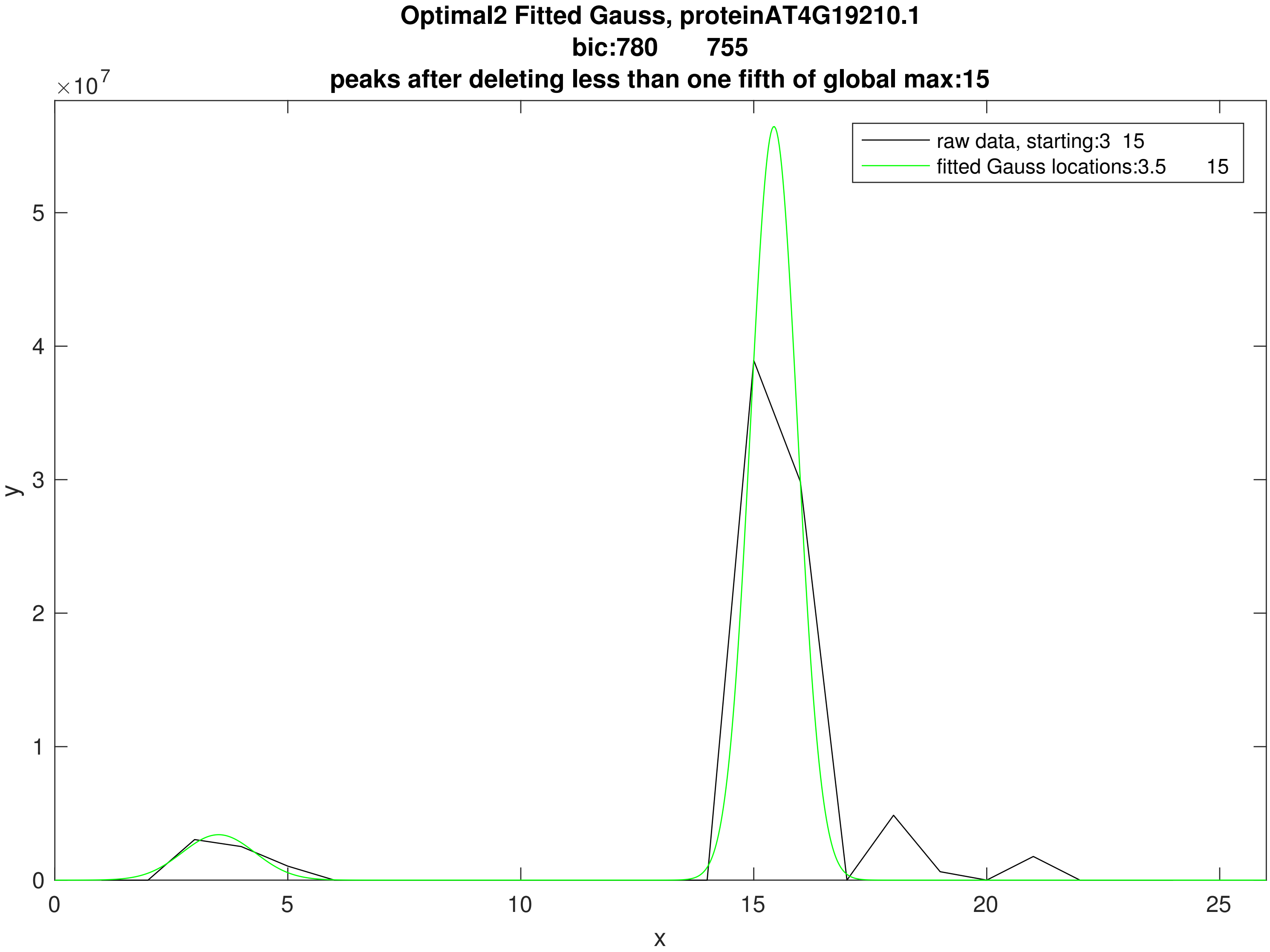
Optimal2 Fitted Gauss, proteinAT4G19210.1
bic:780 755
peaks after deleting less than one fifth of global max:15
×10^7
raw data, starting:3 15
fitted Gauss locations:3.5 15
y
x
0
1
2
3
4
5
0
5
10
15
20
25

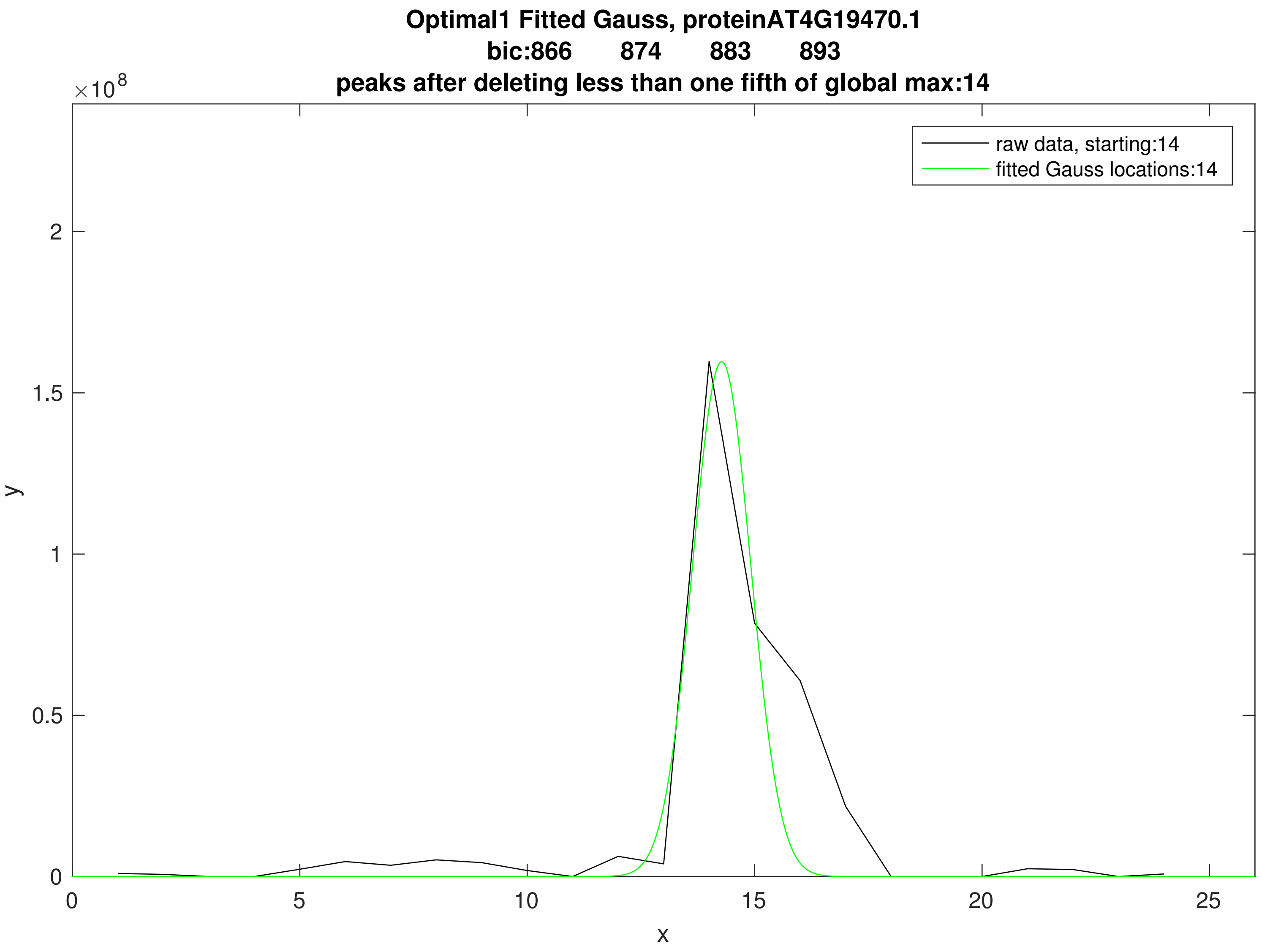
Optimal1 Fitted Gauss, proteinAT4G19470.1
bic:866 874 883 893
peaks after deleting less than one fifth of global max:14
×10^8
raw data, starting:14
fitted Gauss locations:14
y
x
0
0.5
1
1.5
2
0
5
10
15
20
25

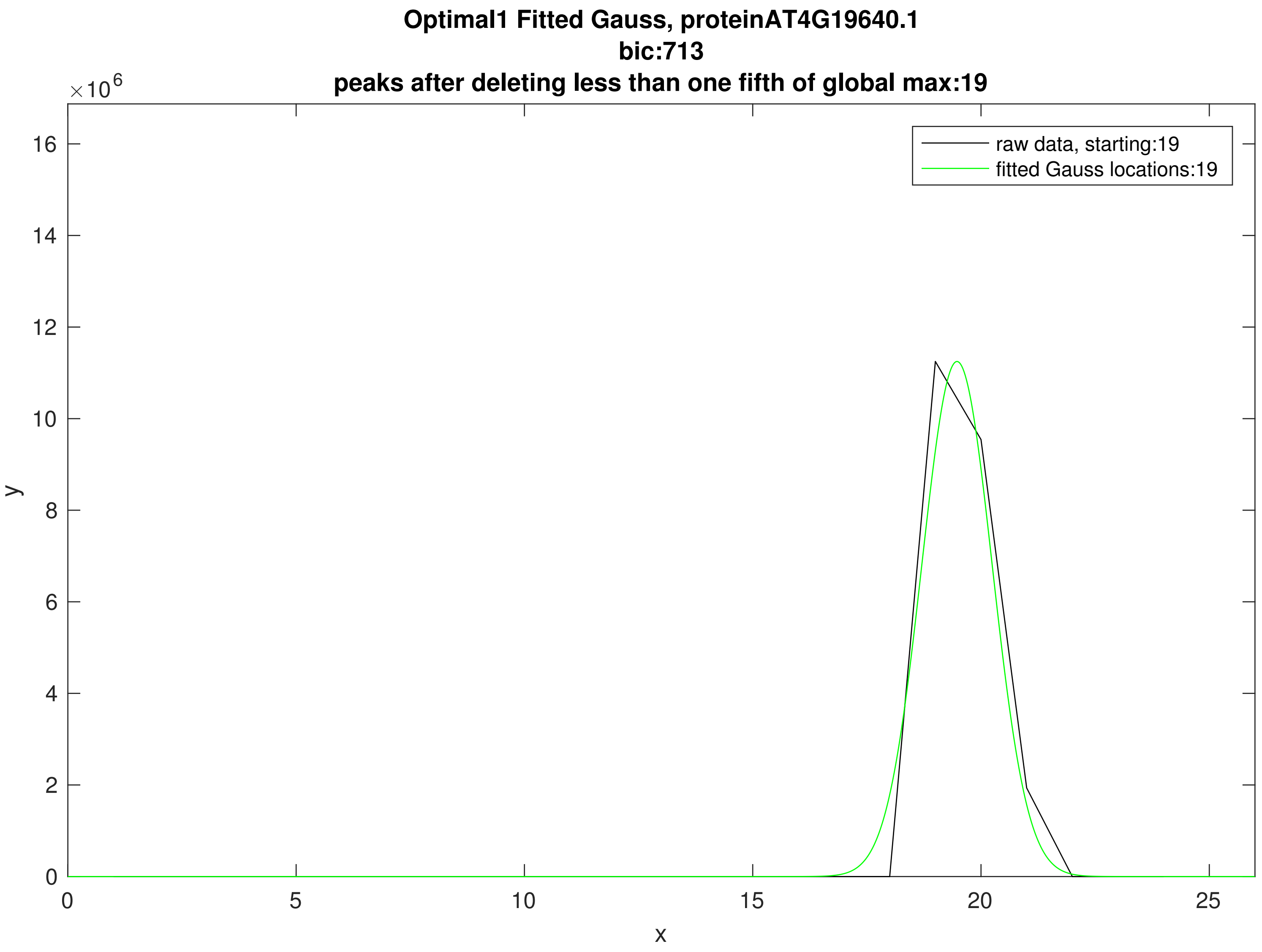
Optimal1 Fitted Gauss, proteinAT4G19640.1
bic:713
peaks after deleting less than one fifth of global max:19
×10^6
raw data, starting:19
fitted Gauss locations:19
16
14
12
10
8
6
4
2
0
y
0
5
10
15
20
25
x

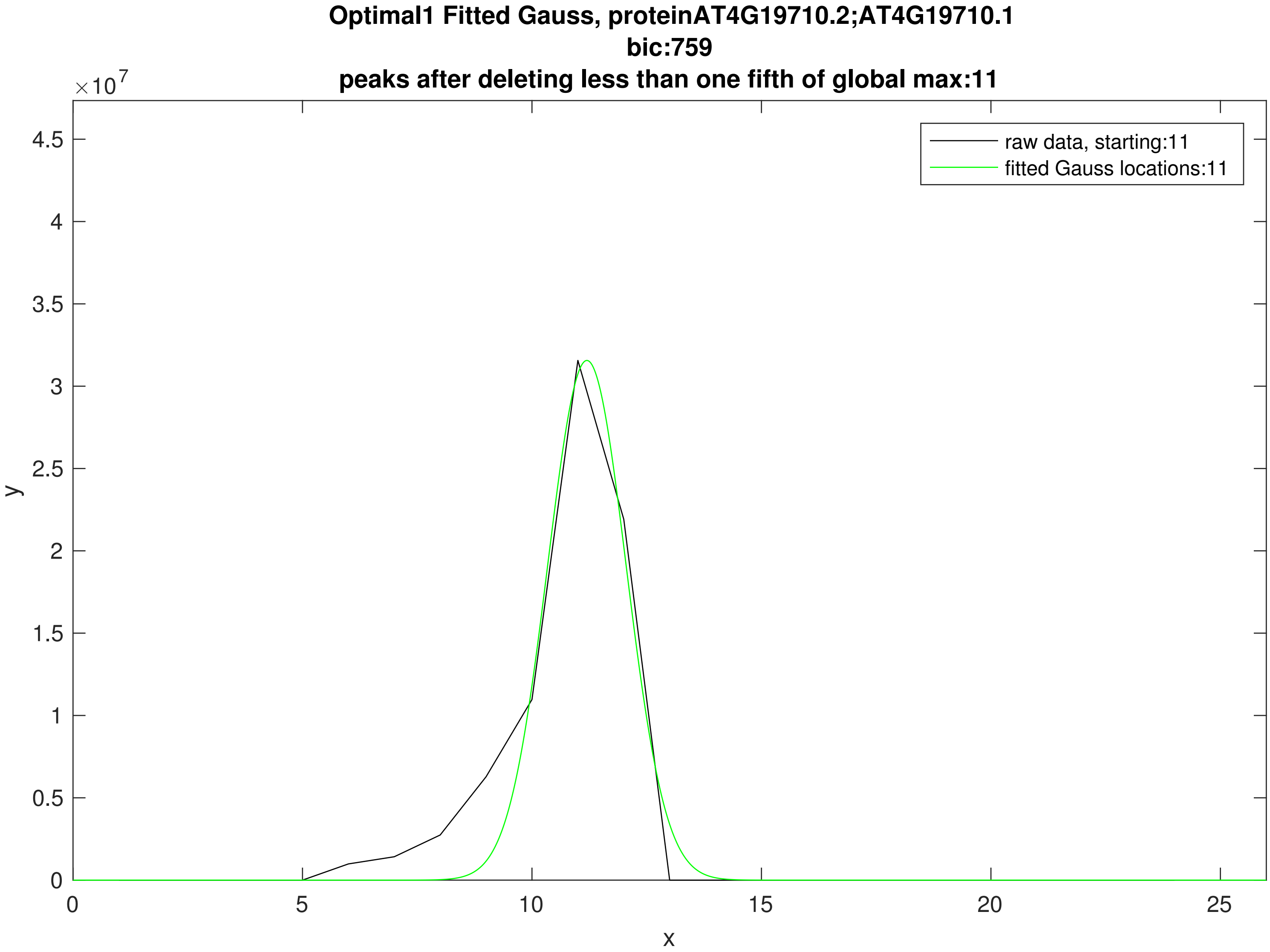
Optimal1 Fitted Gauss, proteinAT4G19710.2;AT4G19710.1
bic:759
peaks after deleting less than one fifth of global max:11
×10^7
raw data, starting:11
fitted Gauss locations:11
4.5
4
3.5
3
2.5
2
1.5
1
0.5
0
y
0
5
10
15
20
25
x

**Optimal1 Fitted Gauss, proteinAT4G19810.1**
**bic:650**
**peaks after deleting less than one fifth of global max:19**

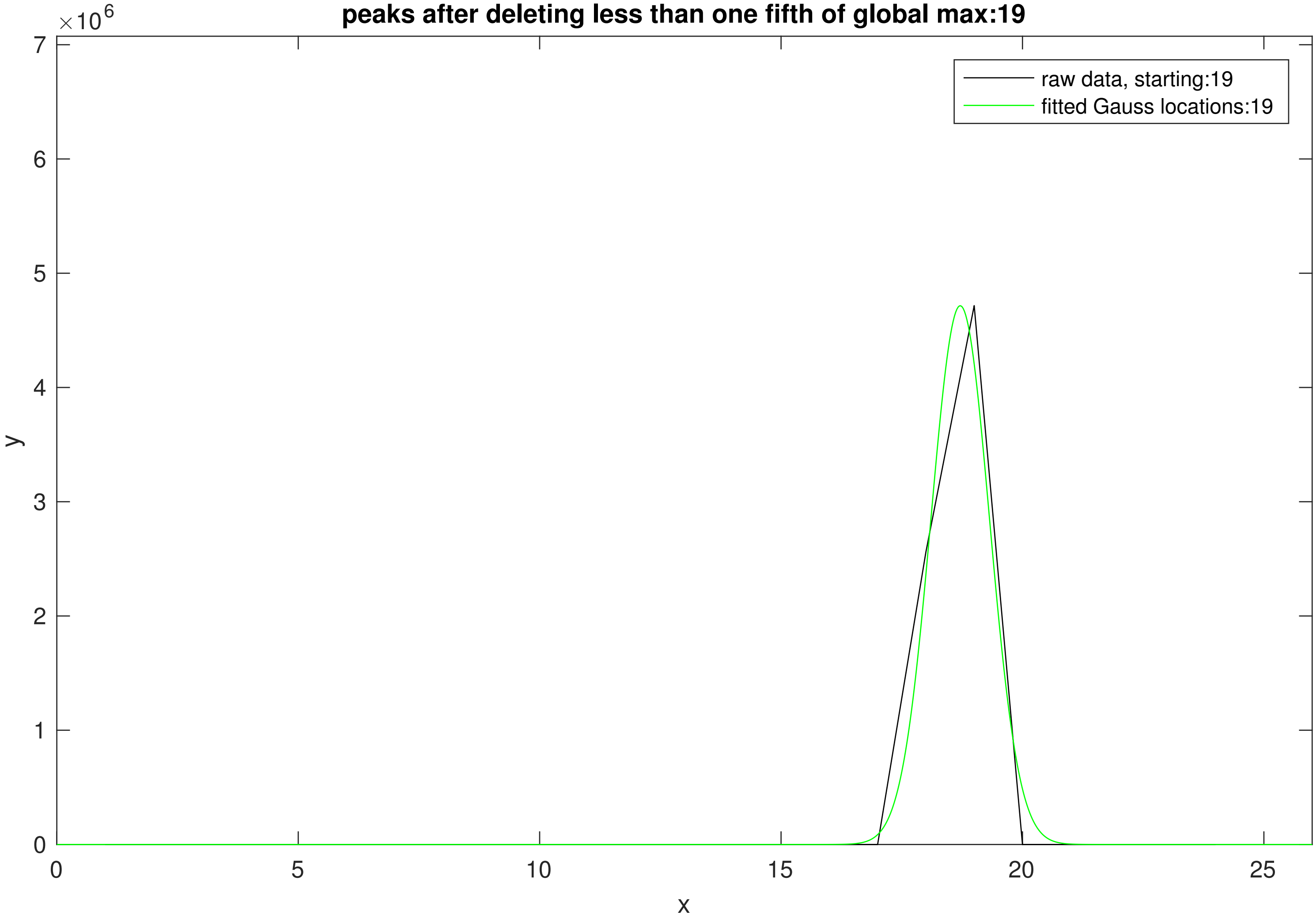

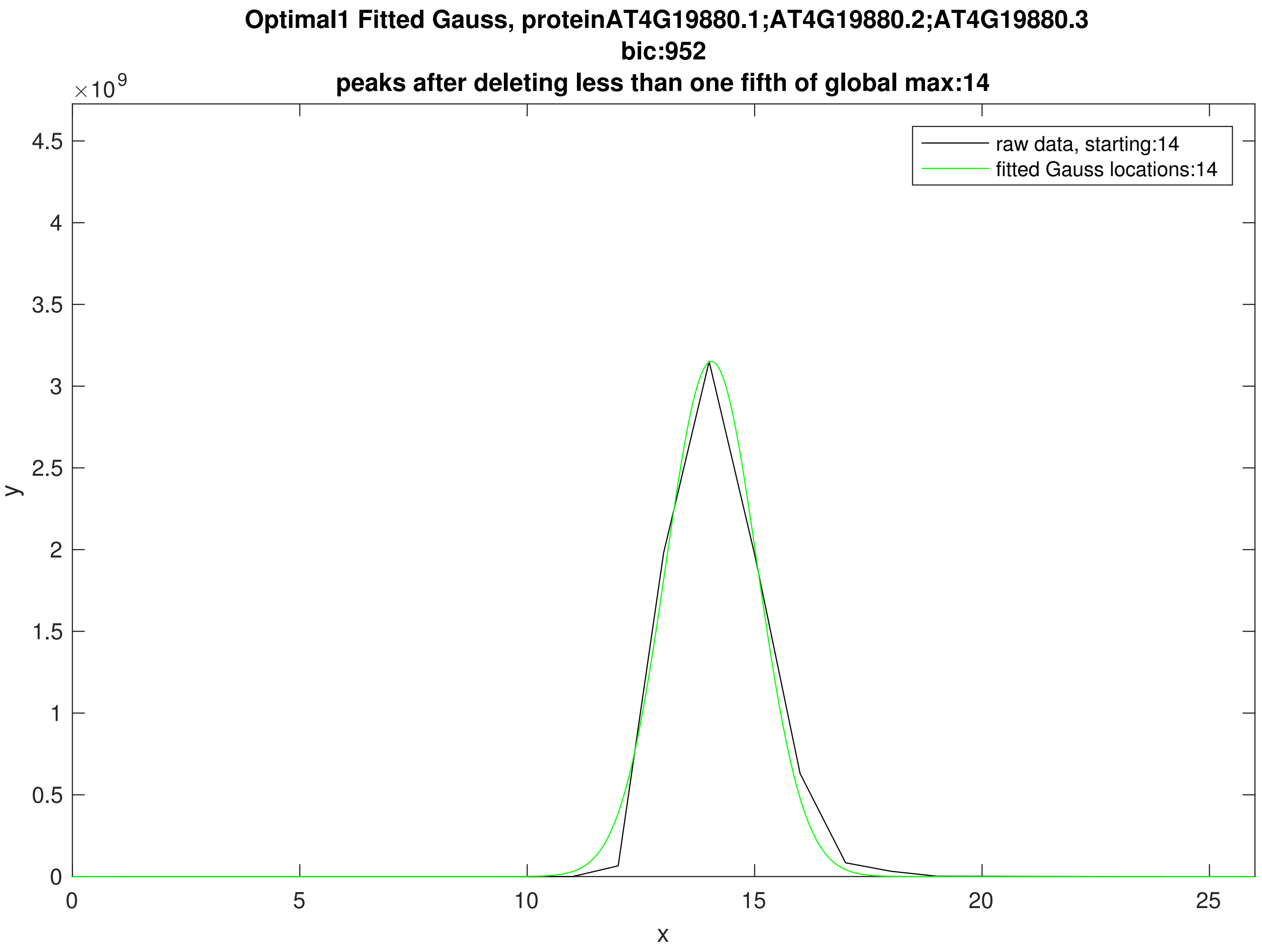
Optimal1 Fitted Gauss, proteinAT4G19880.1;AT4G19880.2;AT4G19880.3
bic:952
peaks after deleting less than one fifth of global max:14
×10^9
raw data, starting:14
fitted Gauss locations:14
y
x
0
0.5
1
1.5
2
2.5
3
3.5
4
4.5
0
5
10
15
20
25

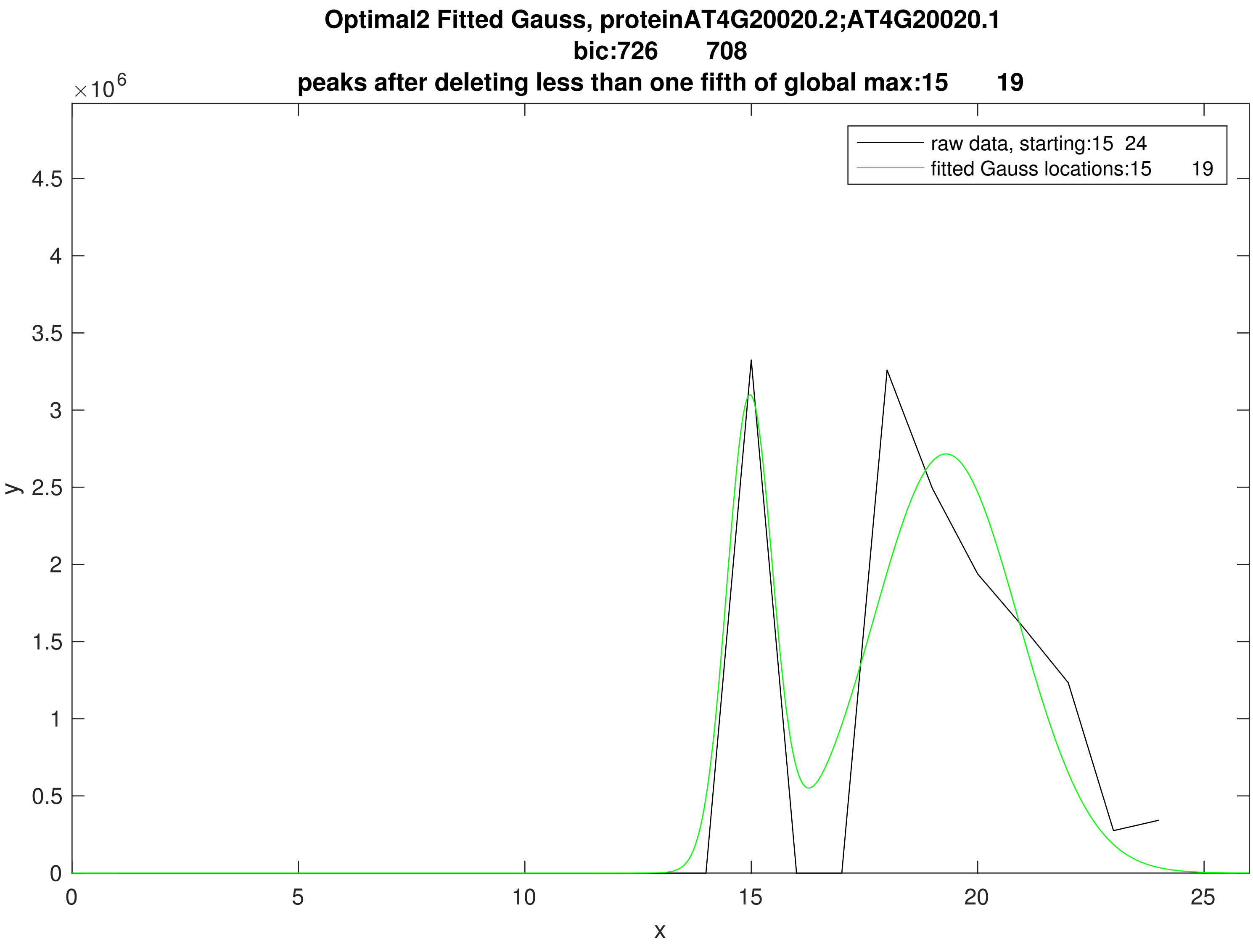

Optimal2 Fitted Gauss, proteinAT4G20020.2;AT4G20020.1
bic:726 708
peaks after deleting less than one fifth of global max:15 19
×10^6
raw data, starting:15 24
fitted Gauss locations:15 19
y
x
0
0.5
1
1.5
2
2.5
3
3.5
4
4.5
5
10
15
20
25

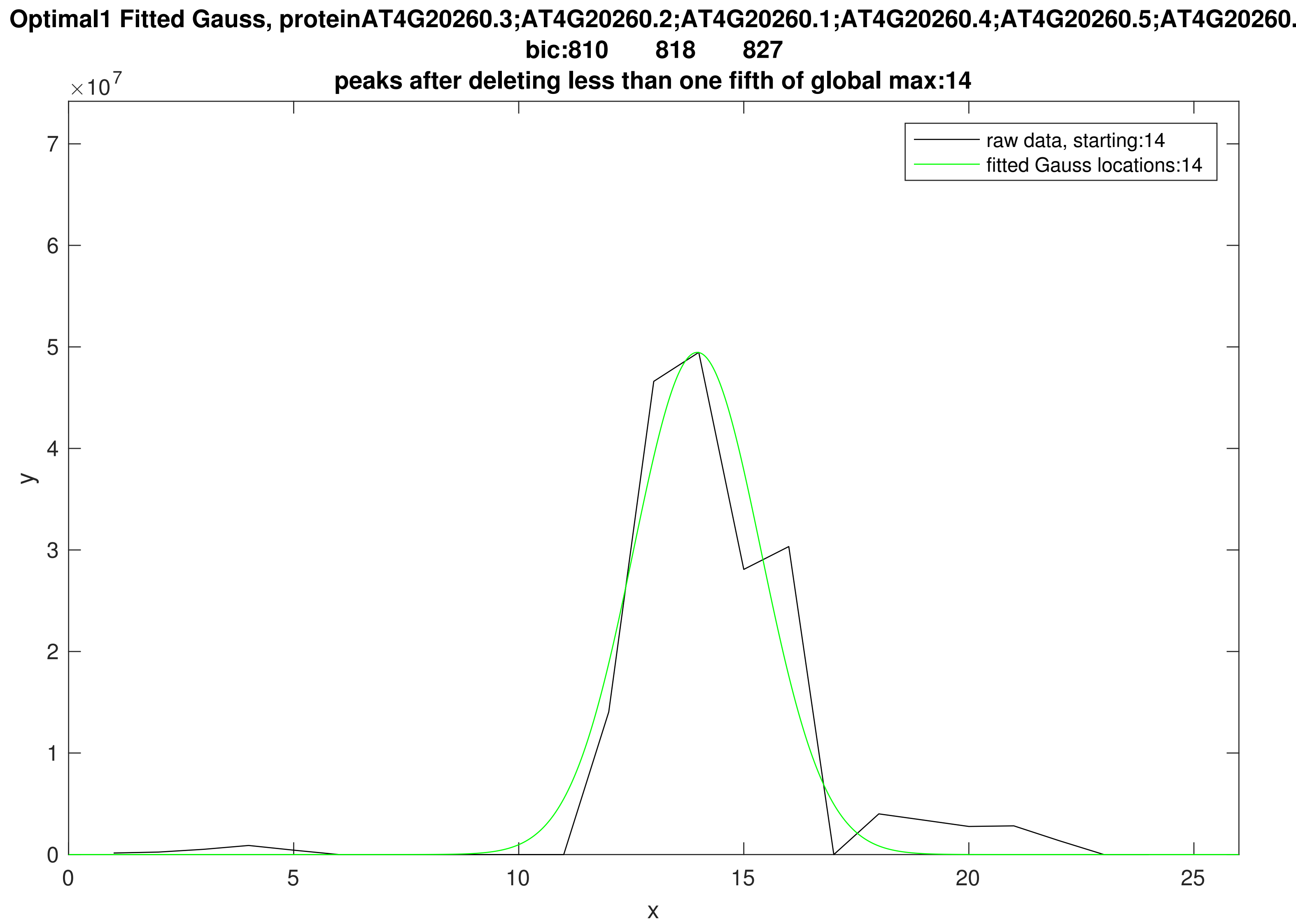
Optimal1 Fitted Gauss, proteinAT4G20260.3;AT4G20260.2;AT4G20260.1;AT4G20260.4;AT4G20260.5;AT4G20260.
bic:810 818 827
peaks after deleting less than one fifth of global max:14
×10^7
raw data, starting:14
fitted Gauss locations:14
y
x
0
1
2
3
4
5
6
7
10
15
20
25

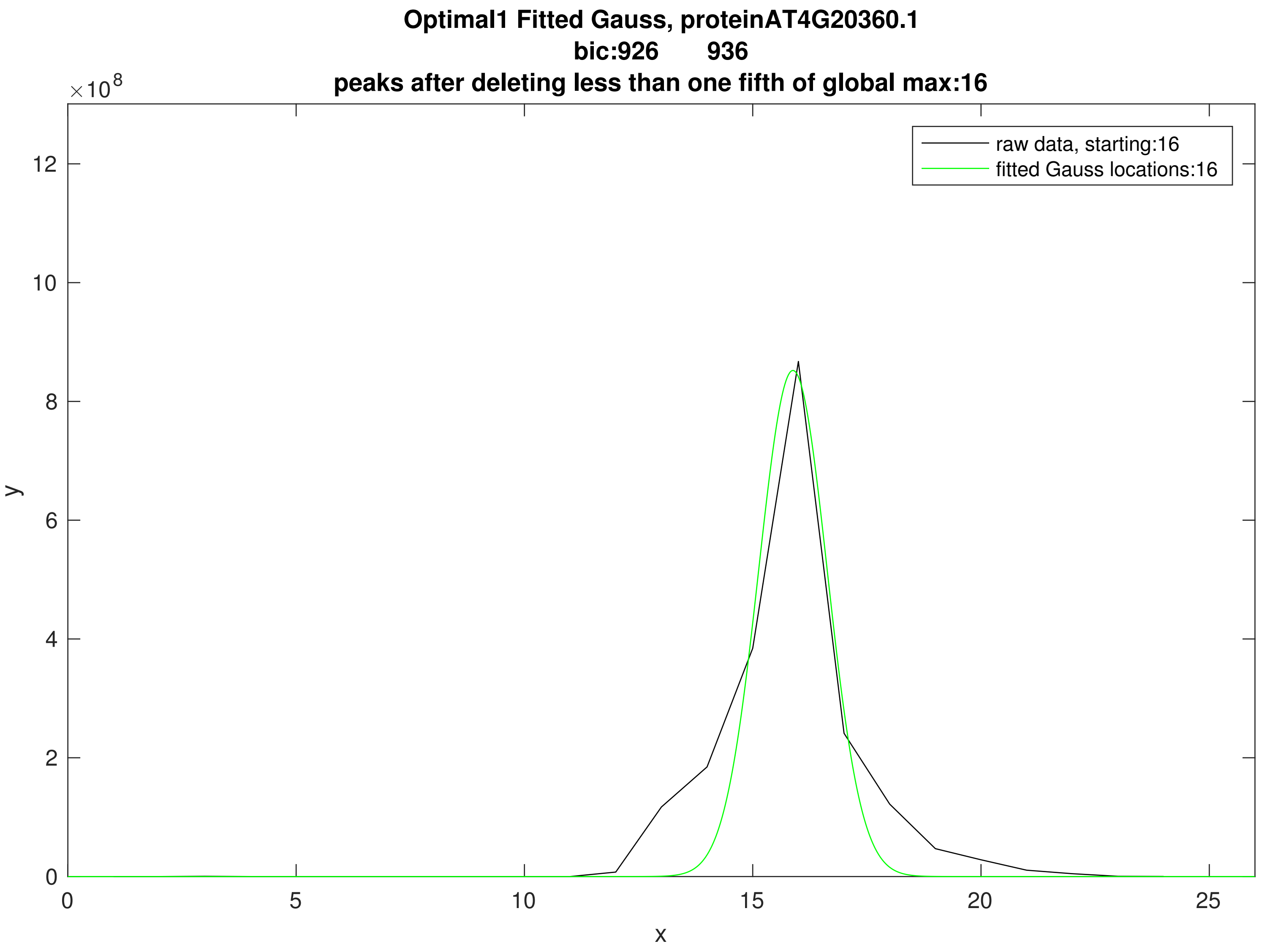
Optimal1 Fitted Gauss, proteinAT4G20360.1
bic:926 936
peaks after deleting less than one fifth of global max:16
×10^8
raw data, starting:16
fitted Gauss locations:16
y
x
0
2
4
6
8
10
12
0
5
10
15
20
25

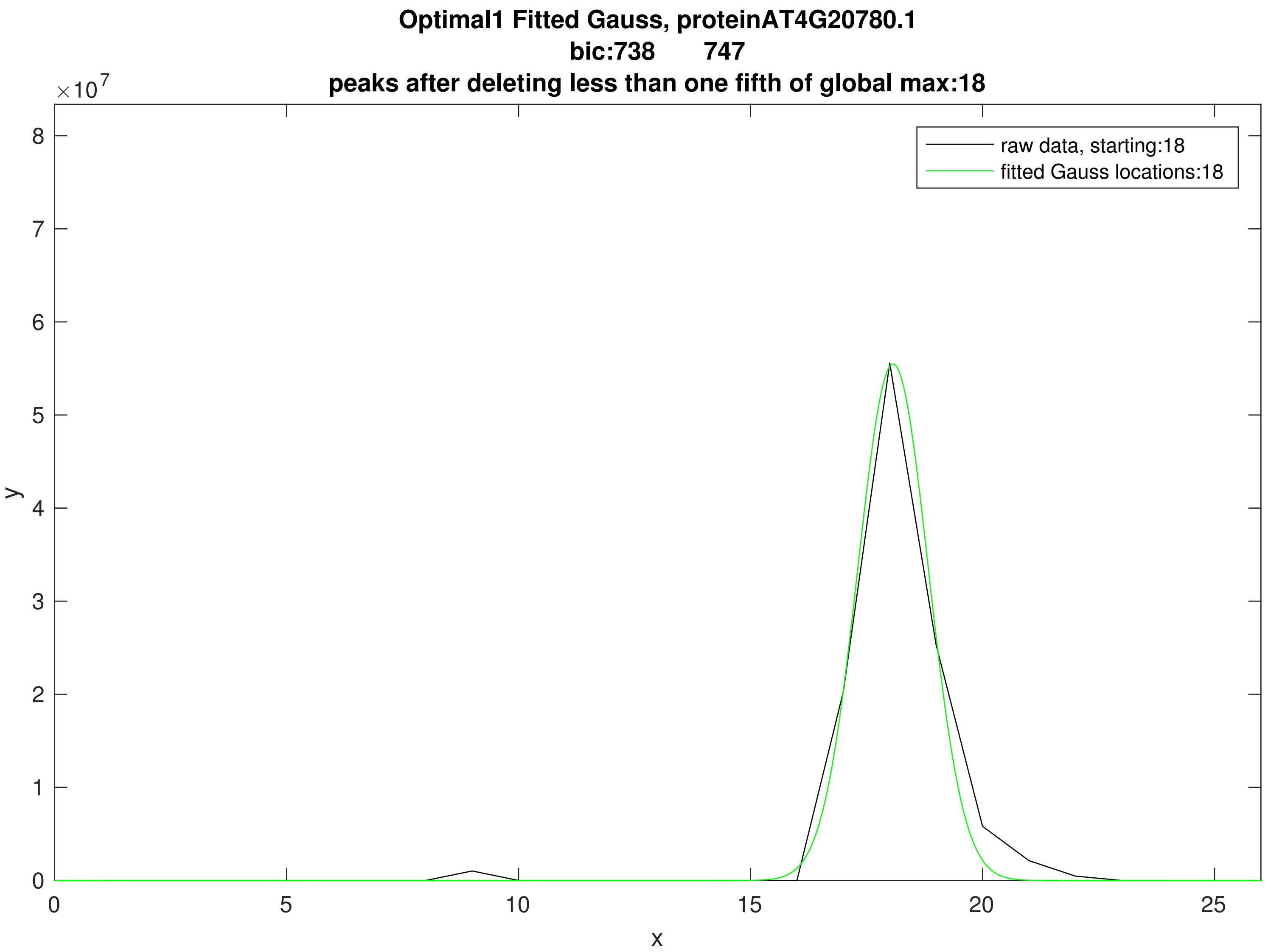
Optimal1 Fitted Gauss, proteinAT4G20780.1
bic:738 747
peaks after deleting less than one fifth of global max:18
×10^7
raw data, starting:18
fitted Gauss locations:18
y
x
0
1
2
3
4
5
6
7
8
0
5
10
15
20
25

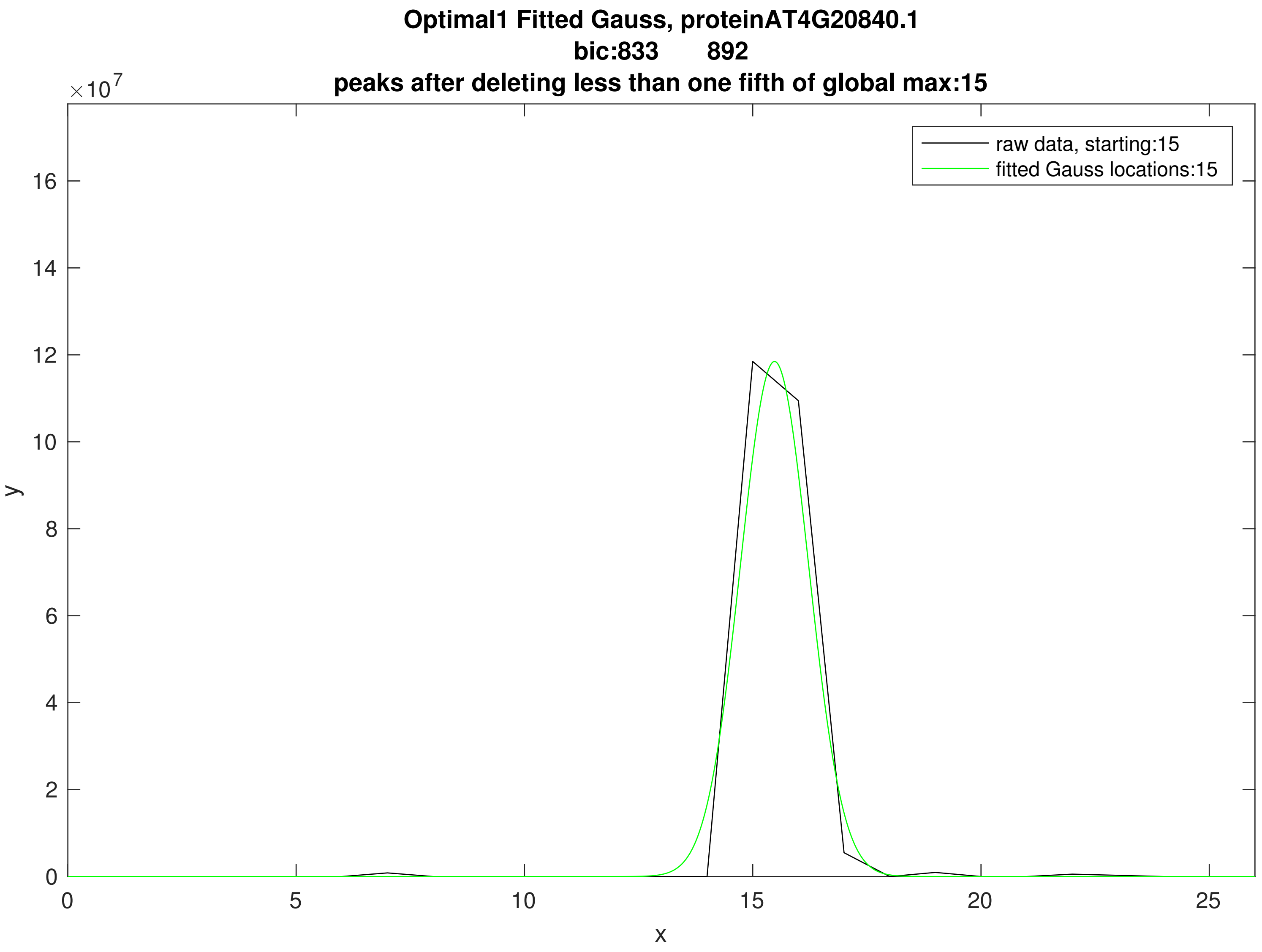

Optimal1 Fitted Gauss, proteinAT4G20840.1
bic:833 892
peaks after deleting less than one fifth of global max:15
$\times 10^7$
16
14
12
10
8
6
4
2
0
y
0
5
10
15
20
25
x
raw data, starting:15
fitted Gauss locations:15

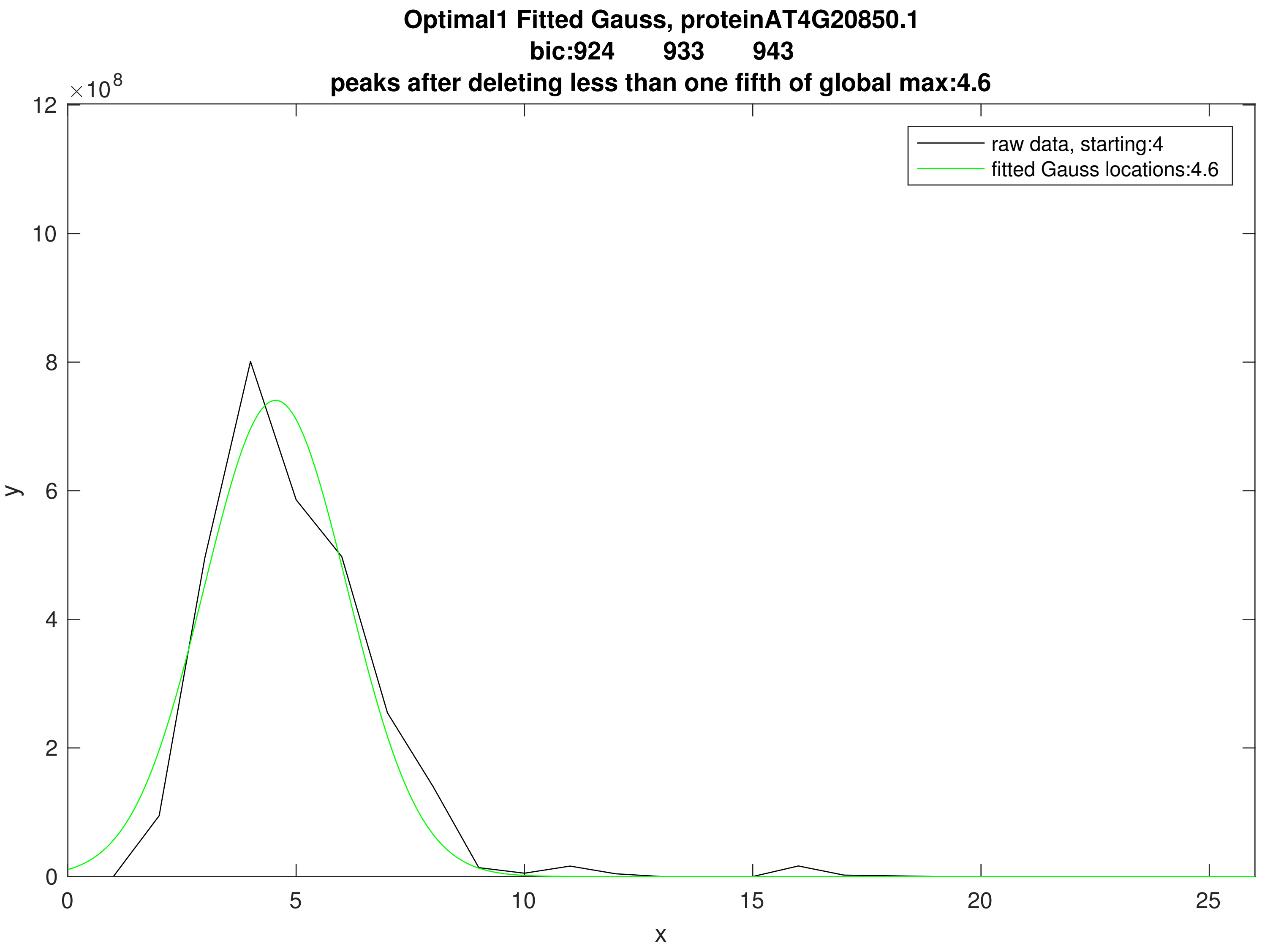
Optimal1 Fitted Gauss, proteinAT4G20850.1
bic:924 933 943
peaks after deleting less than one fifth of global max:4.6
×10^8
12
10
8
6
4
2
0
y
0
5
10
15
20
25
x
raw data, starting:4
fitted Gauss locations:4.6

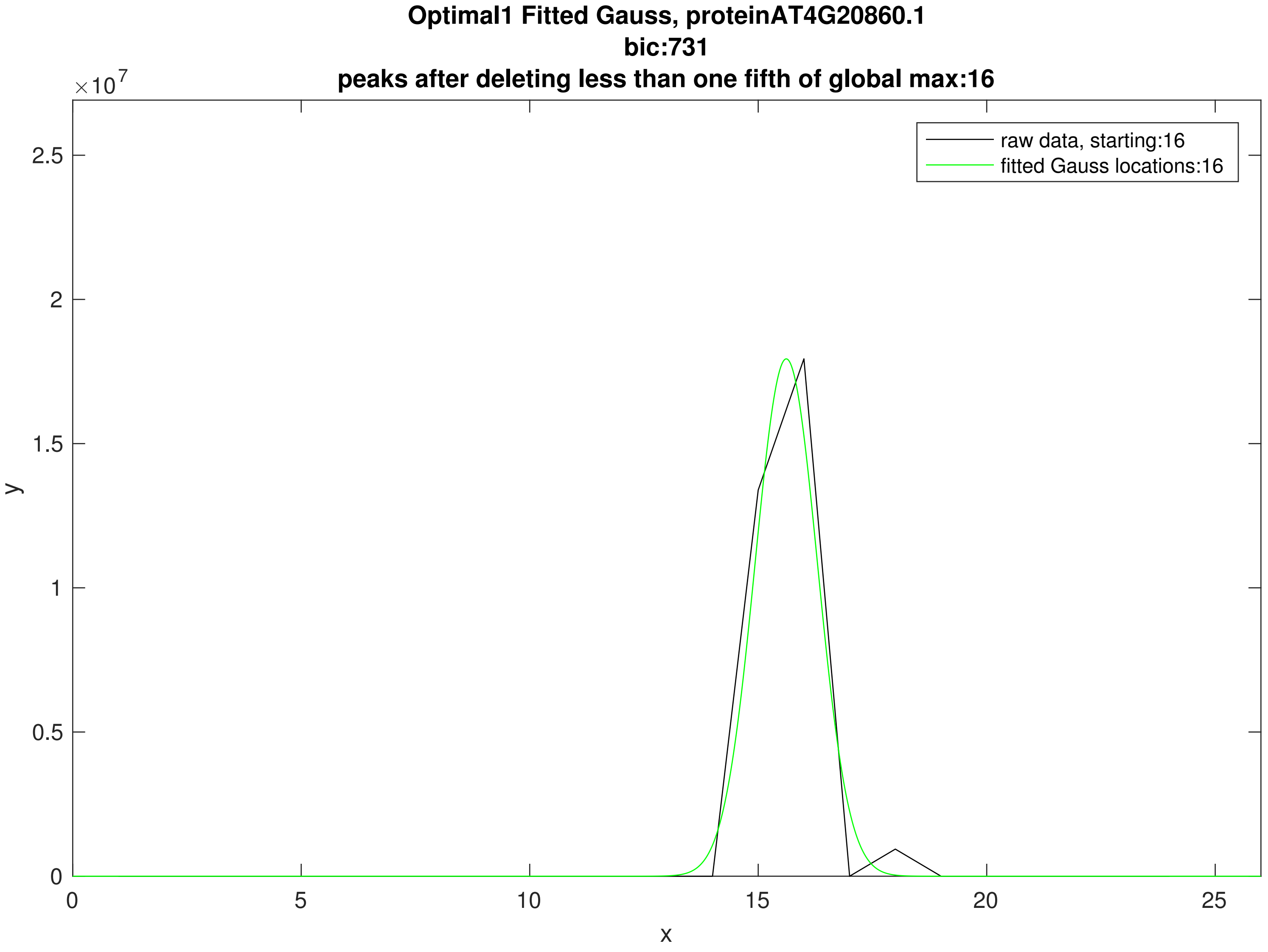
Optimal1 Fitted Gauss, proteinAT4G20860.1
bic:731
peaks after deleting less than one fifth of global max:16
raw data, starting:16
fitted Gauss locations:16
y
x

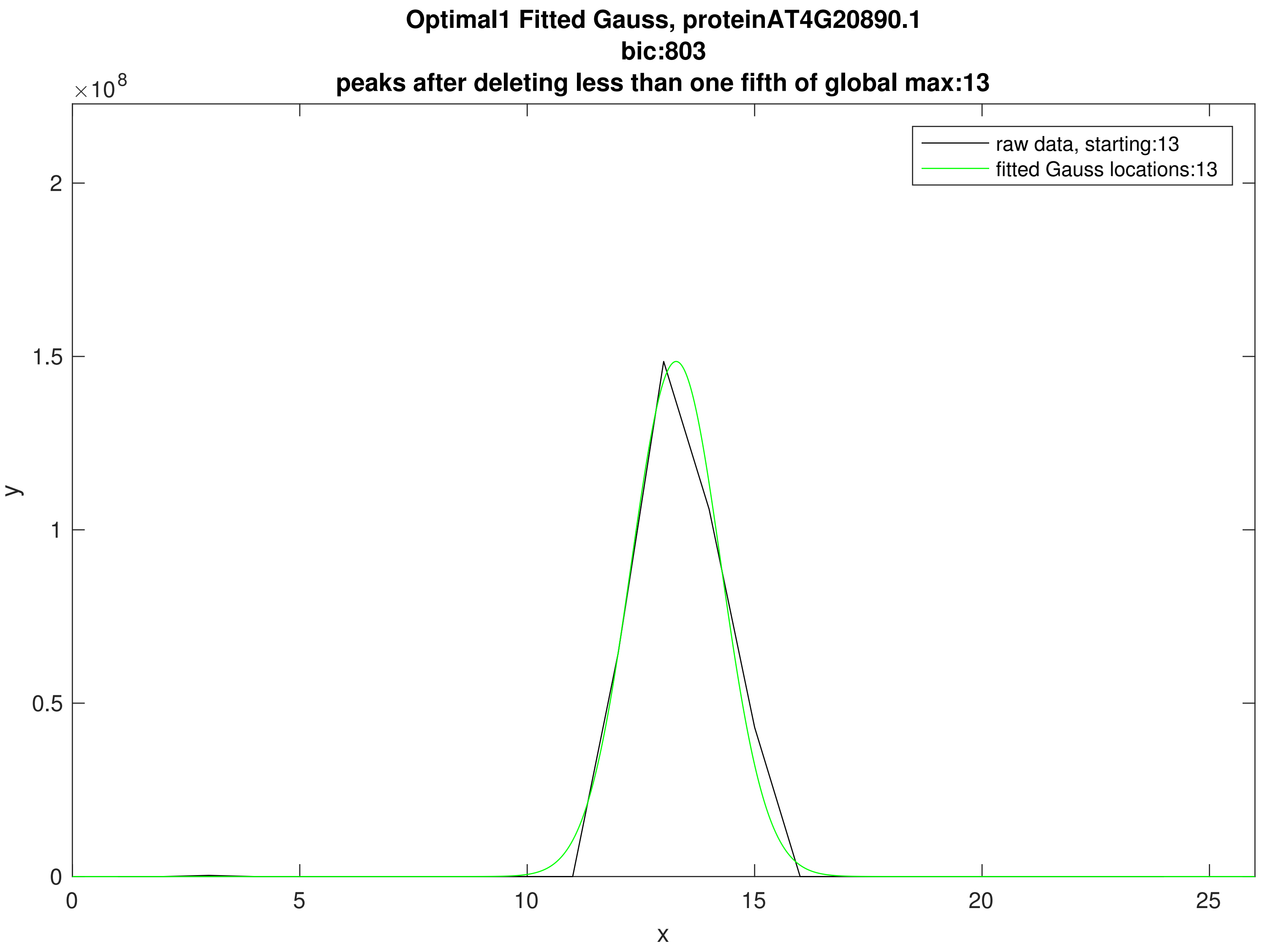
Optimal1 Fitted Gauss, proteinAT4G20890.1
bic:803
peaks after deleting less than one fifth of global max:13
×10^8
raw data, starting:13
fitted Gauss locations:13
y
x
0
0.5
1
1.5
2
0
5
10
15
20
25

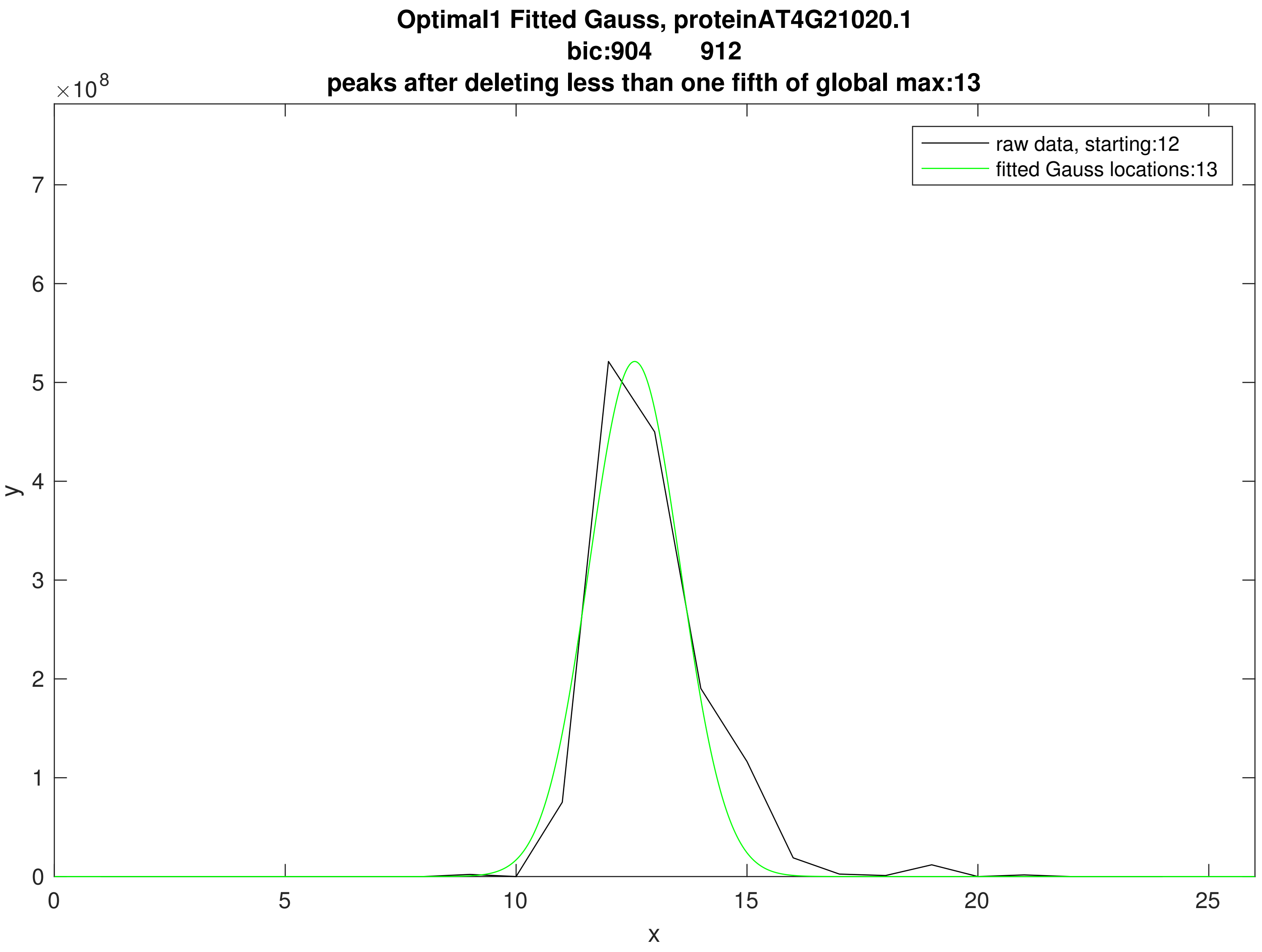
Optimal1 Fitted Gauss, proteinAT4G21020.1
bic:904 912
peaks after deleting less than one fifth of global max:13
×10^8
raw data, starting:12
fitted Gauss locations:13
y
x
0
1
2
3
4
5
6
7
0
5
10
15
20
25

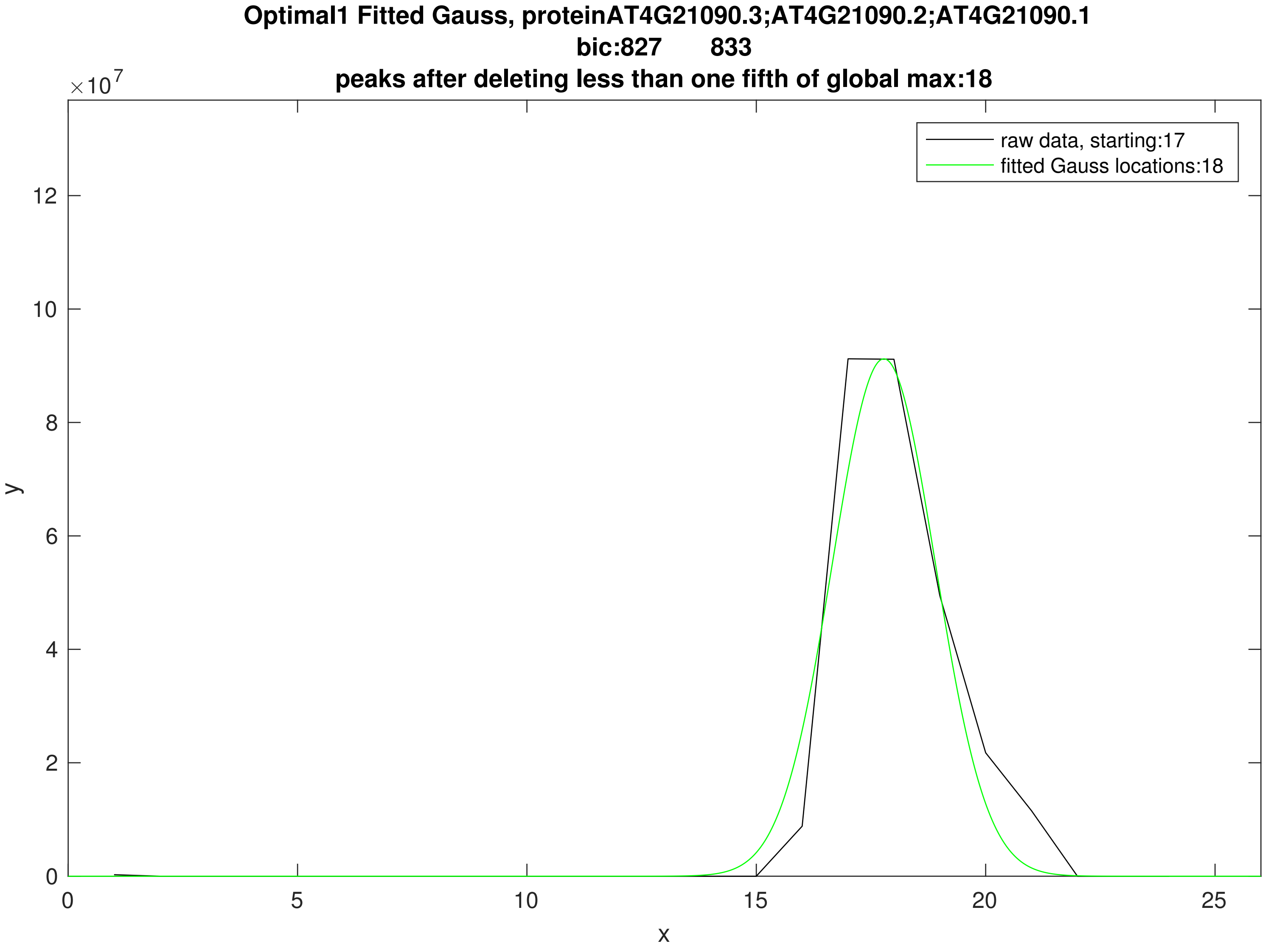

Optimal1 Fitted Gauss, proteinAT4G21090.3;AT4G21090.2;AT4G21090.1
bic:827 833
peaks after deleting less than one fifth of global max:18
×10^7
raw data, starting:17
fitted Gauss locations:18
y
x
0
2
4
6
8
10
12
0
5
10
15
20
25

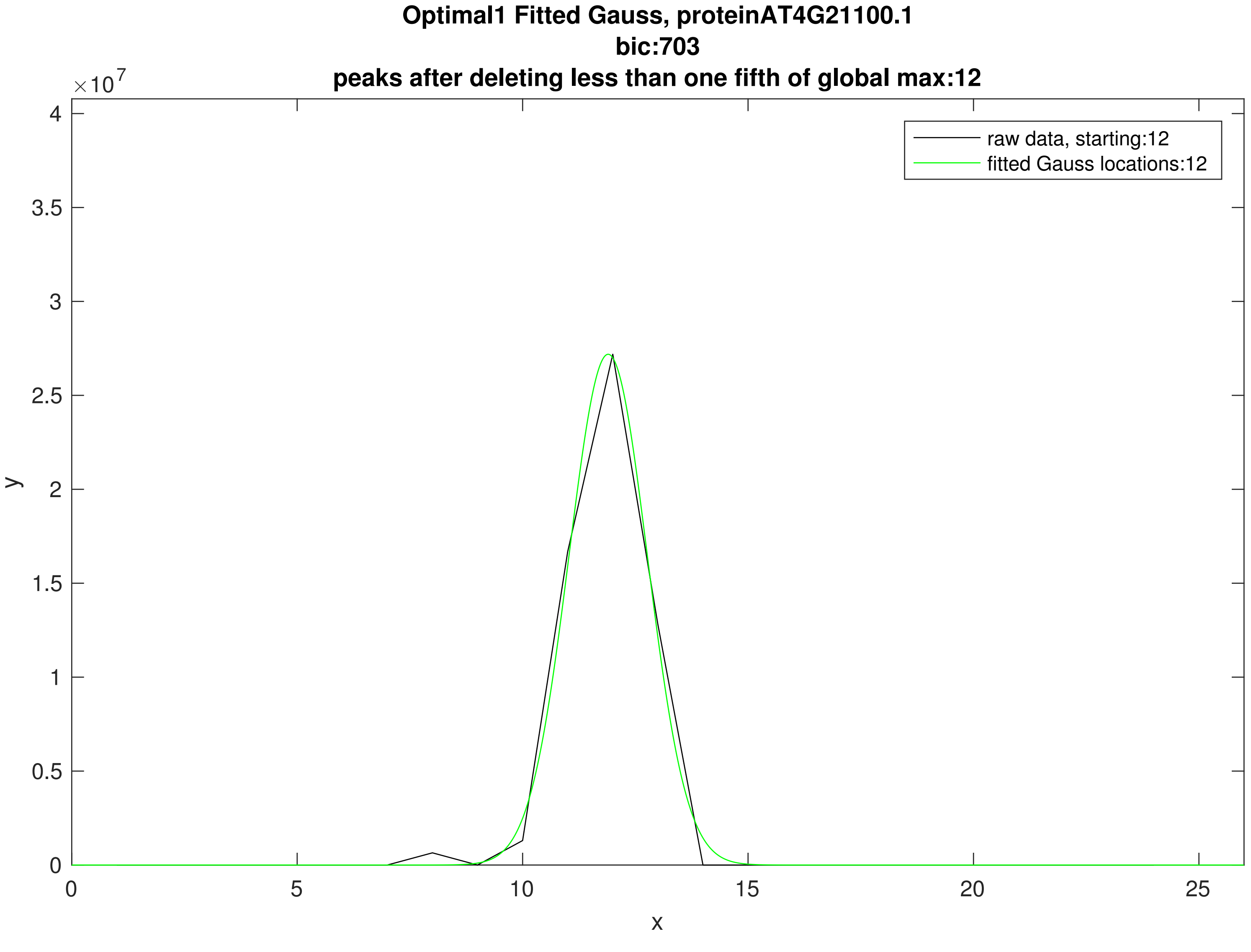
Optimal1 Fitted Gauss, proteinAT4G21100.1
bic:703
peaks after deleting less than one fifth of global max:12
×10^7
4
3.5
3
2.5
2
1.5
1
0.5
0
y
0
5
10
15
20
25
x
raw data, starting:12
fitted Gauss locations:12

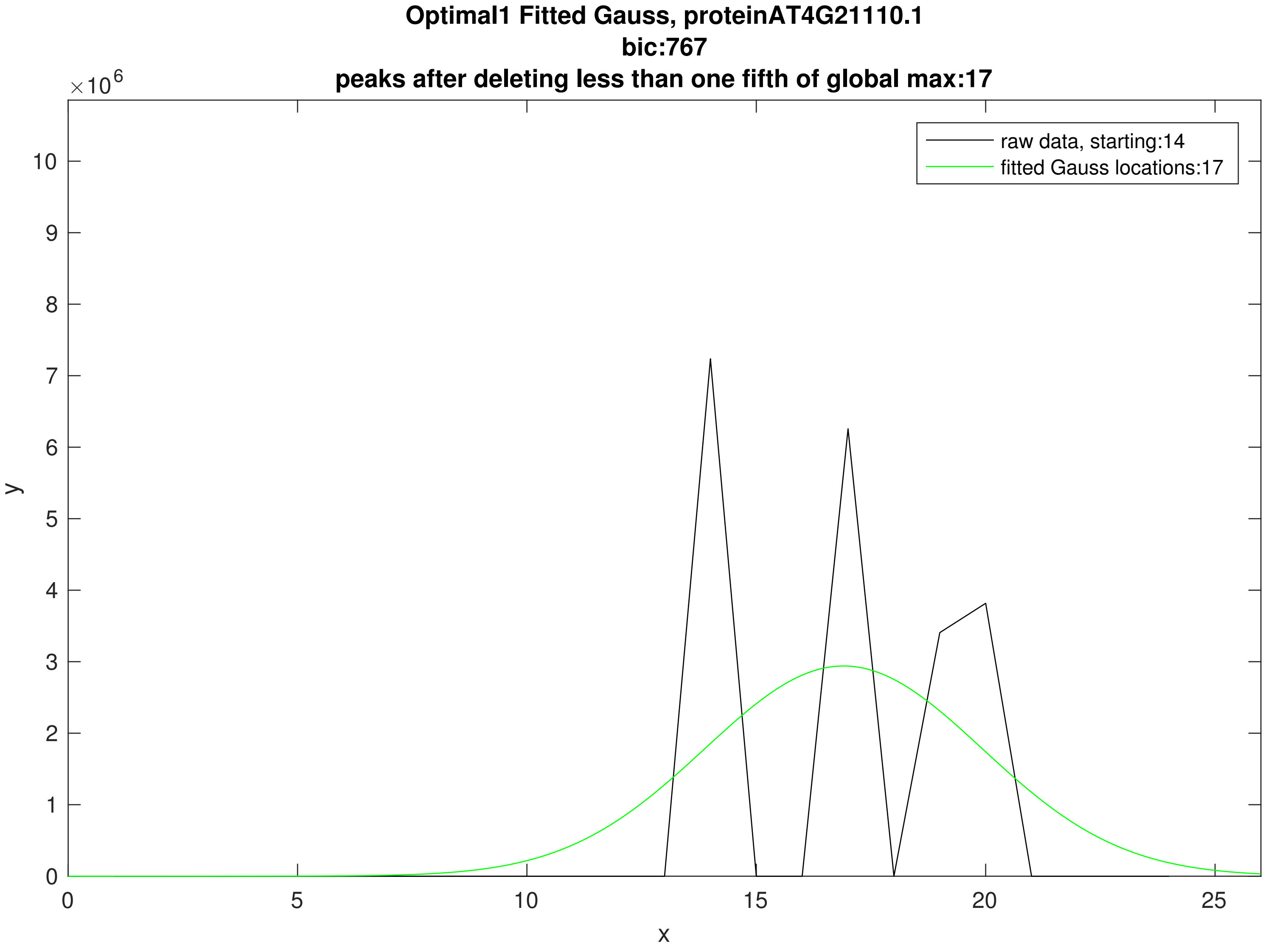
Optimal1 Fitted Gauss, proteinAT4G21110.1
bic:767
peaks after deleting less than one fifth of global max:17
×10^6
raw data, starting:14
fitted Gauss locations:17
y
x
0
1
2
3
4
5
6
7
8
9
10
0
5
10
15
20
25

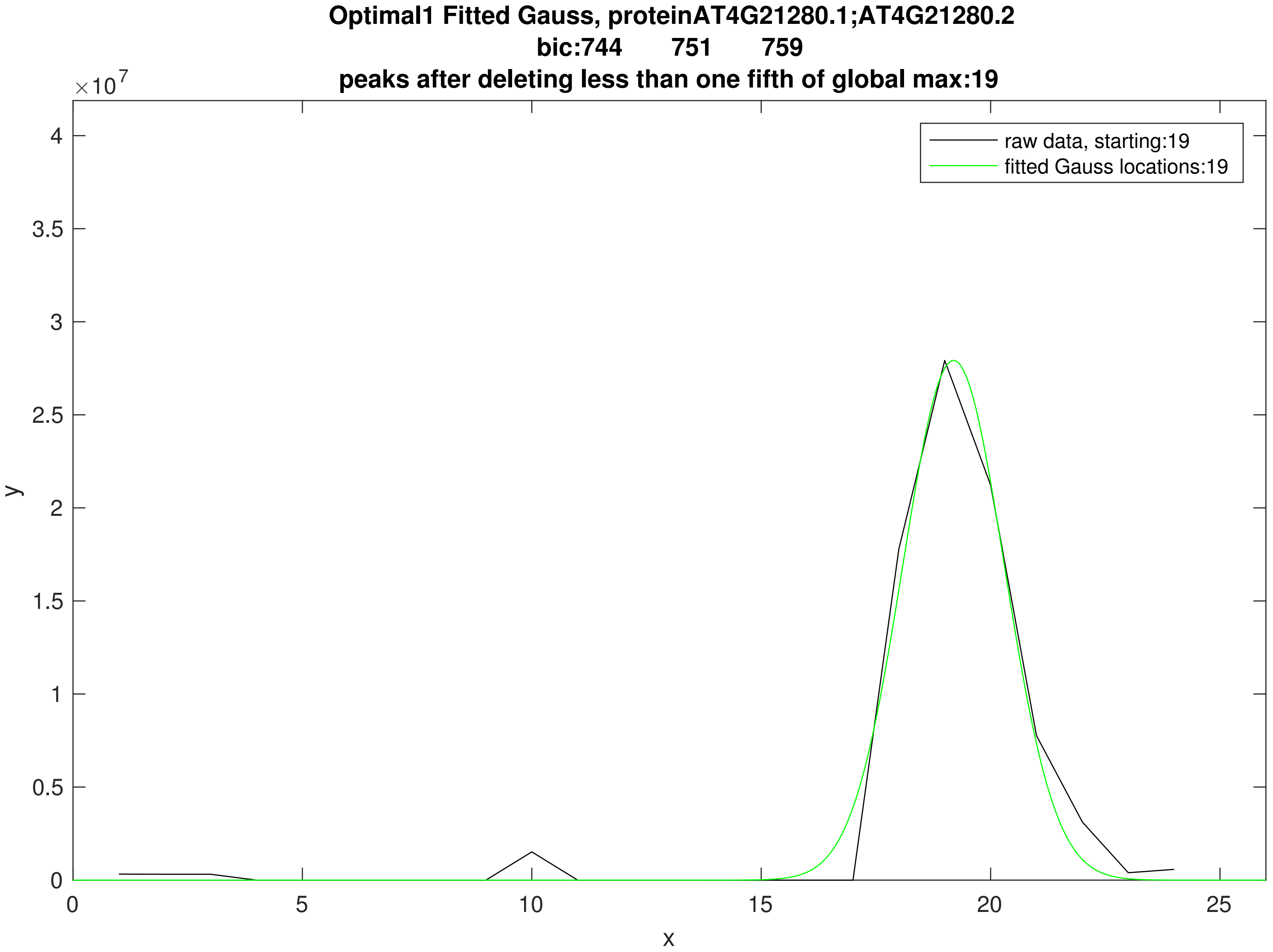
Optimal1 Fitted Gauss, proteinAT4G21280.1;AT4G21280.2
bic:744 751 759
peaks after deleting less than one fifth of global max:19
×10^7
raw data, starting:19
fitted Gauss locations:19
y
x
0
0.5
1
1.5
2
2.5
3
3.5
4
0
5
10
15
20
25

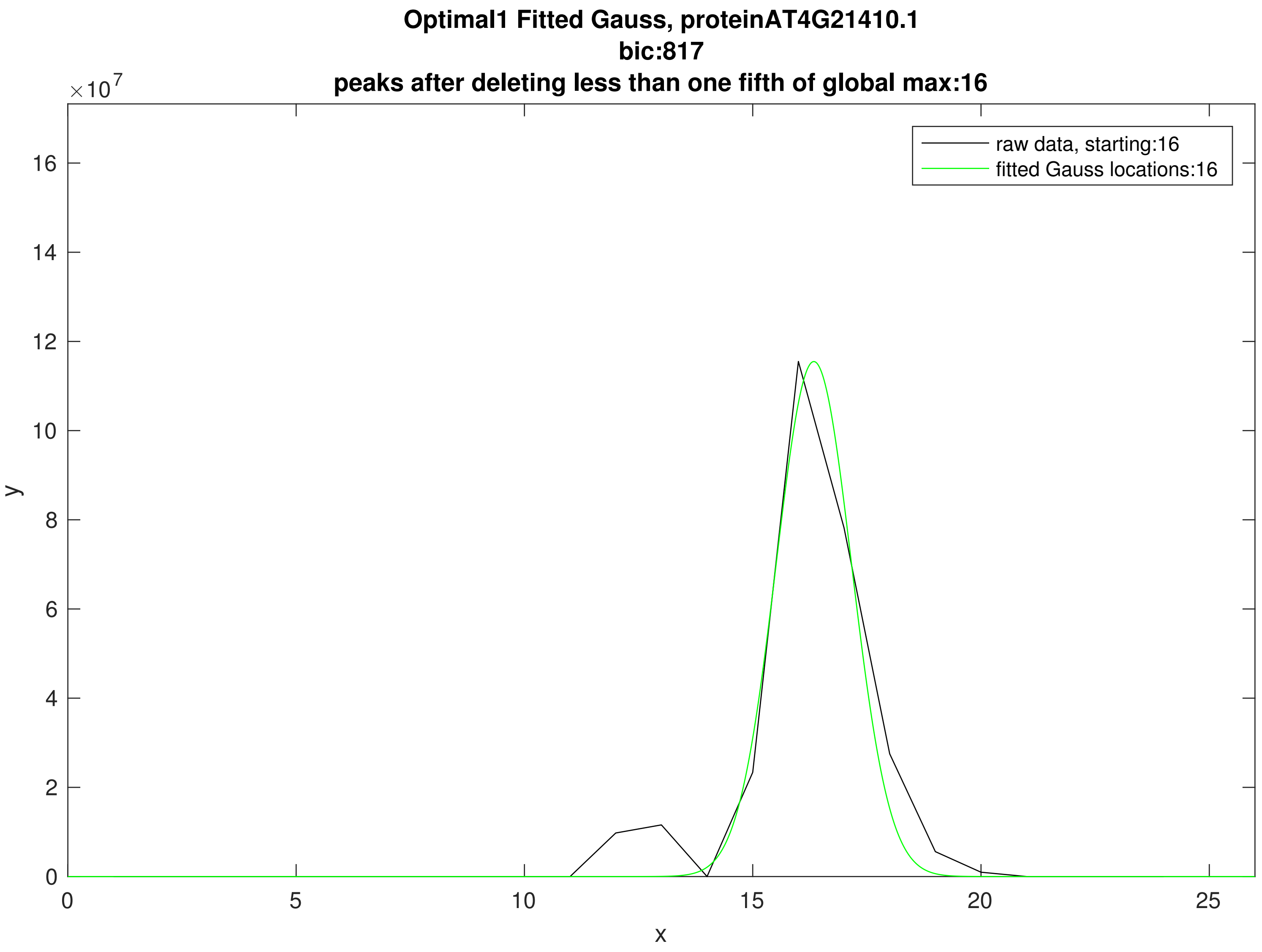
Optimal1 Fitted Gauss, proteinAT4G21410.1
bic:817
peaks after deleting less than one fifth of global max:16
×10^7
raw data, starting:16
fitted Gauss locations:16
y
x
0
2
4
6
8
10
12
14
16
0
5
10
15
20
25

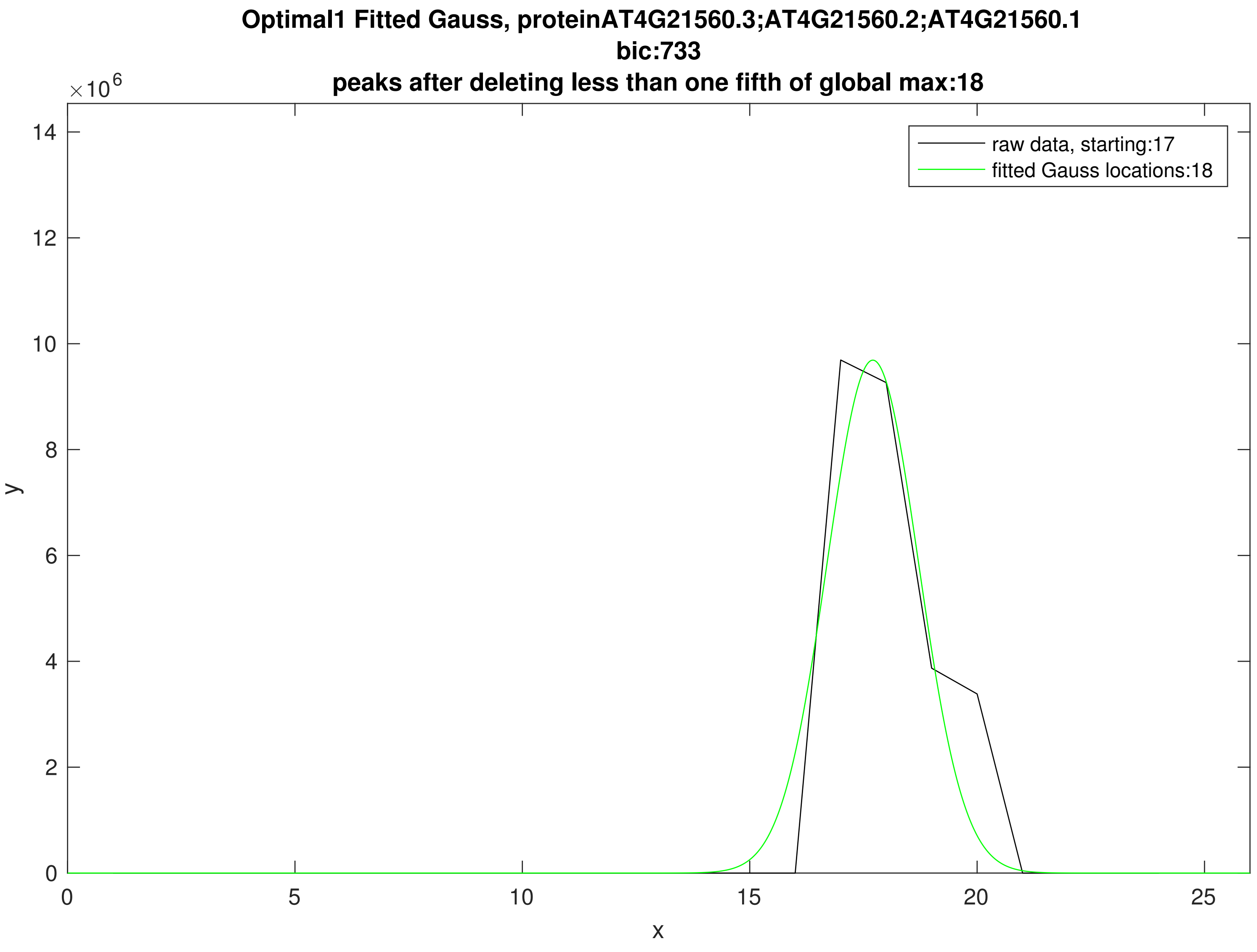
Optimal1 Fitted Gauss, proteinAT4G21560.3;AT4G21560.2;AT4G21560.1
bic:733
peaks after deleting less than one fifth of global max:18
×10^6
raw data, starting:17
fitted Gauss locations:18
y
x
0
2
4
6
8
10
12
14
5
15
20
25

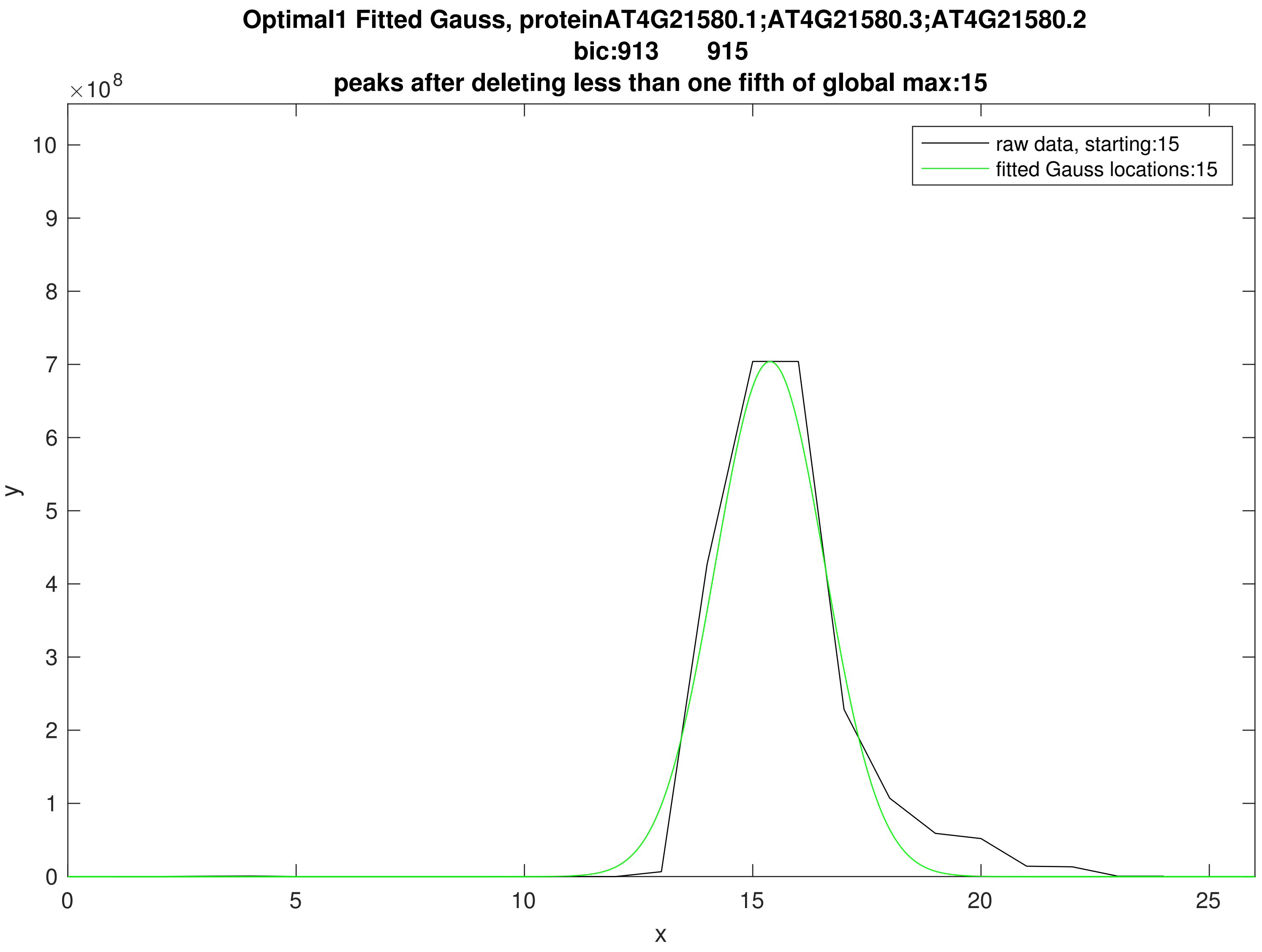
Optimal1 Fitted Gauss, proteinAT4G21580.1;AT4G21580.3;AT4G21580.2
bic:913 915
peaks after deleting less than one fifth of global max:15
raw data, starting:15
fitted Gauss locations:15
$\times 10^8$
y
x
0
5
10
15
20
25
1
2
3
4
6
7
8
9

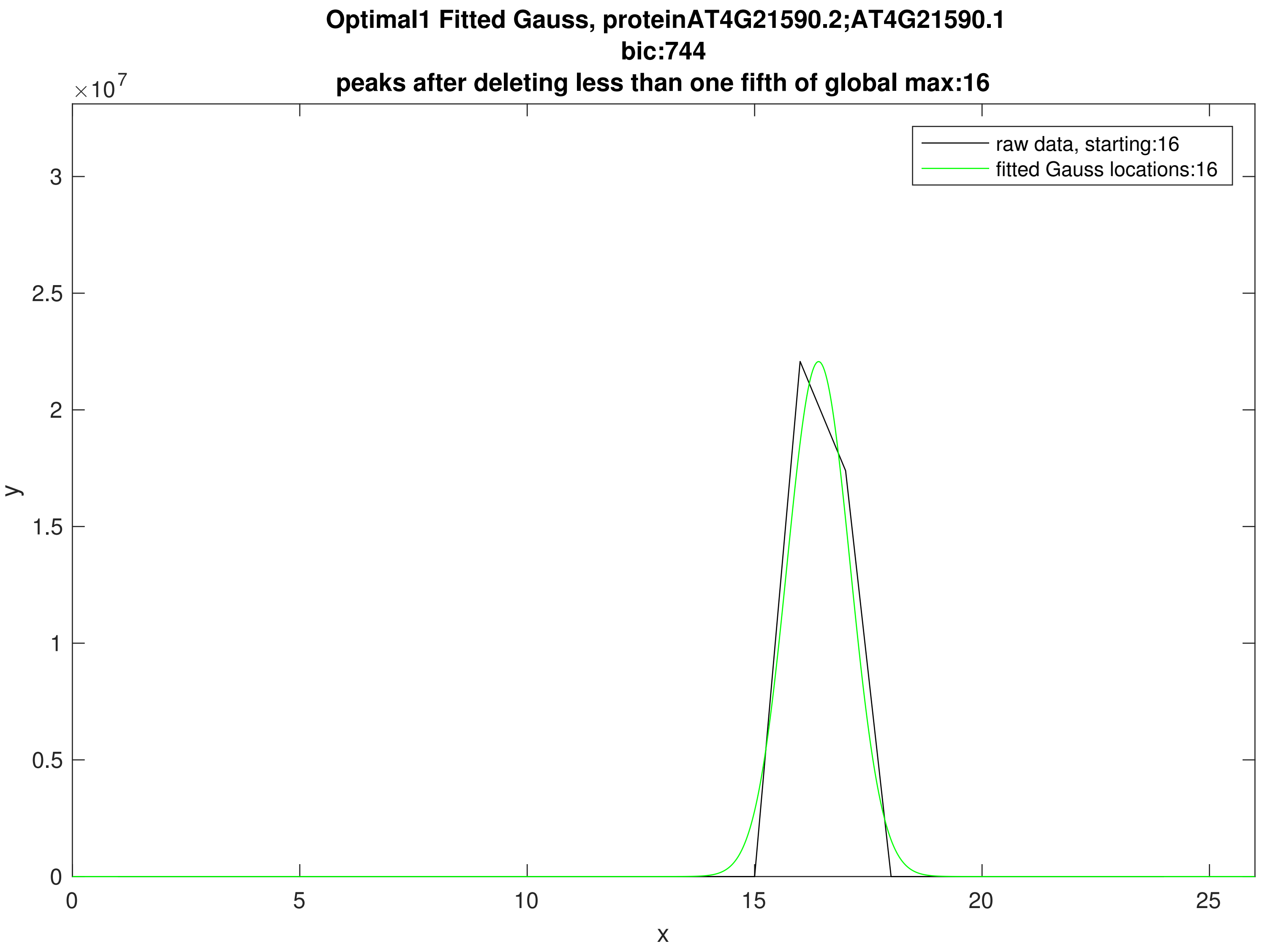

Optimal1 Fitted Gauss, proteinAT4G21590.2;AT4G21590.1
bic:744
peaks after deleting less than one fifth of global max:16
×10^7
raw data, starting:16
fitted Gauss locations:16
y
x
0
0.5
1
1.5
2
2.5
3
0
5
10
15
20
25

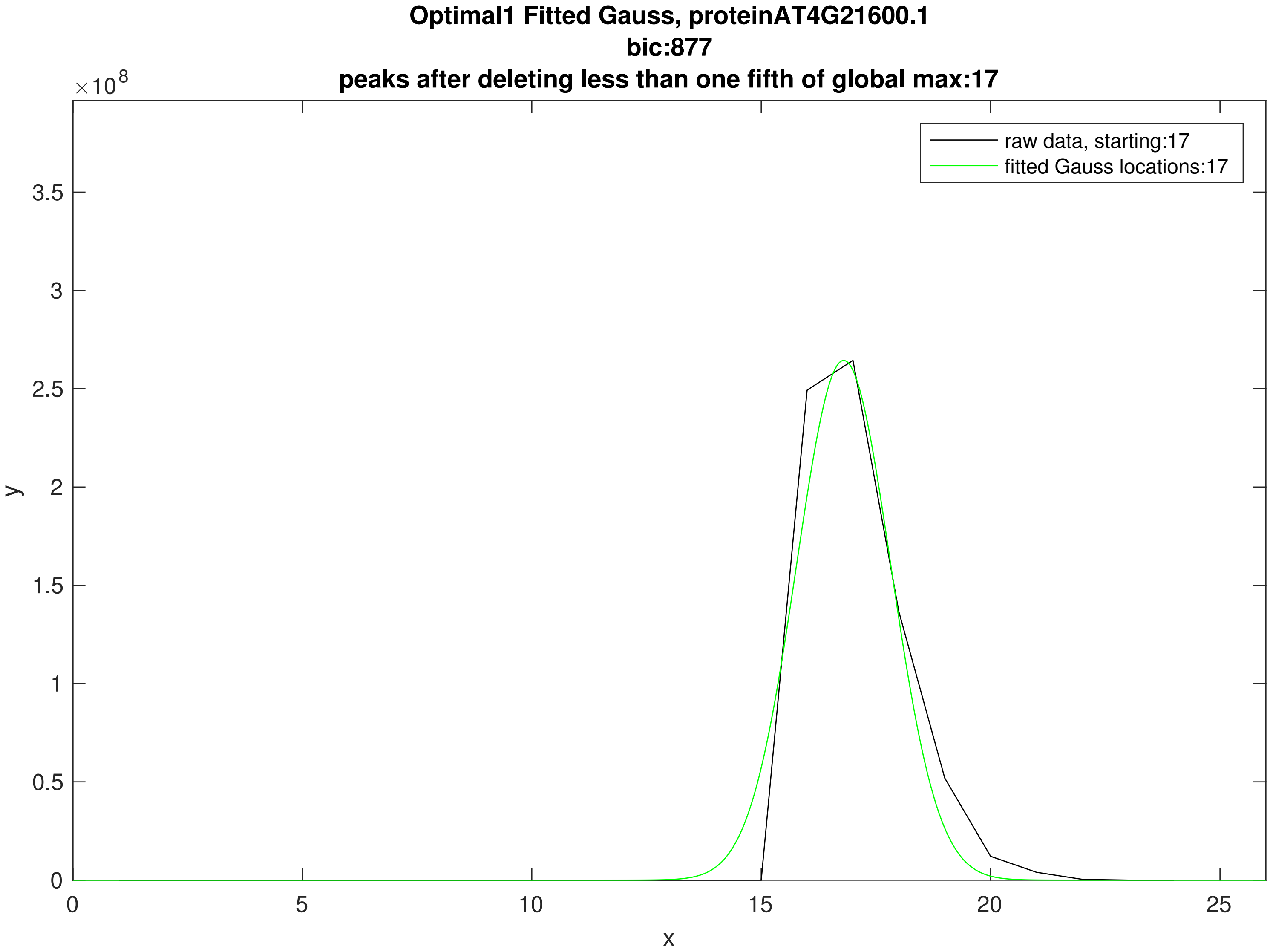
Optimal1 Fitted Gauss, proteinAT4G21600.1
bic:877
peaks after deleting less than one fifth of global max:17
×10^8
raw data, starting:17
fitted Gauss locations:17
3.5
3
2.5
2
1.5
1
0.5
0
y
0
5
10
15
20
25
x

#### Optimal2 Fitted Gauss, proteinAT4G21710.1
#### bic:741 712
#### peaks after deleting less than one fifth of global max:4.8 12

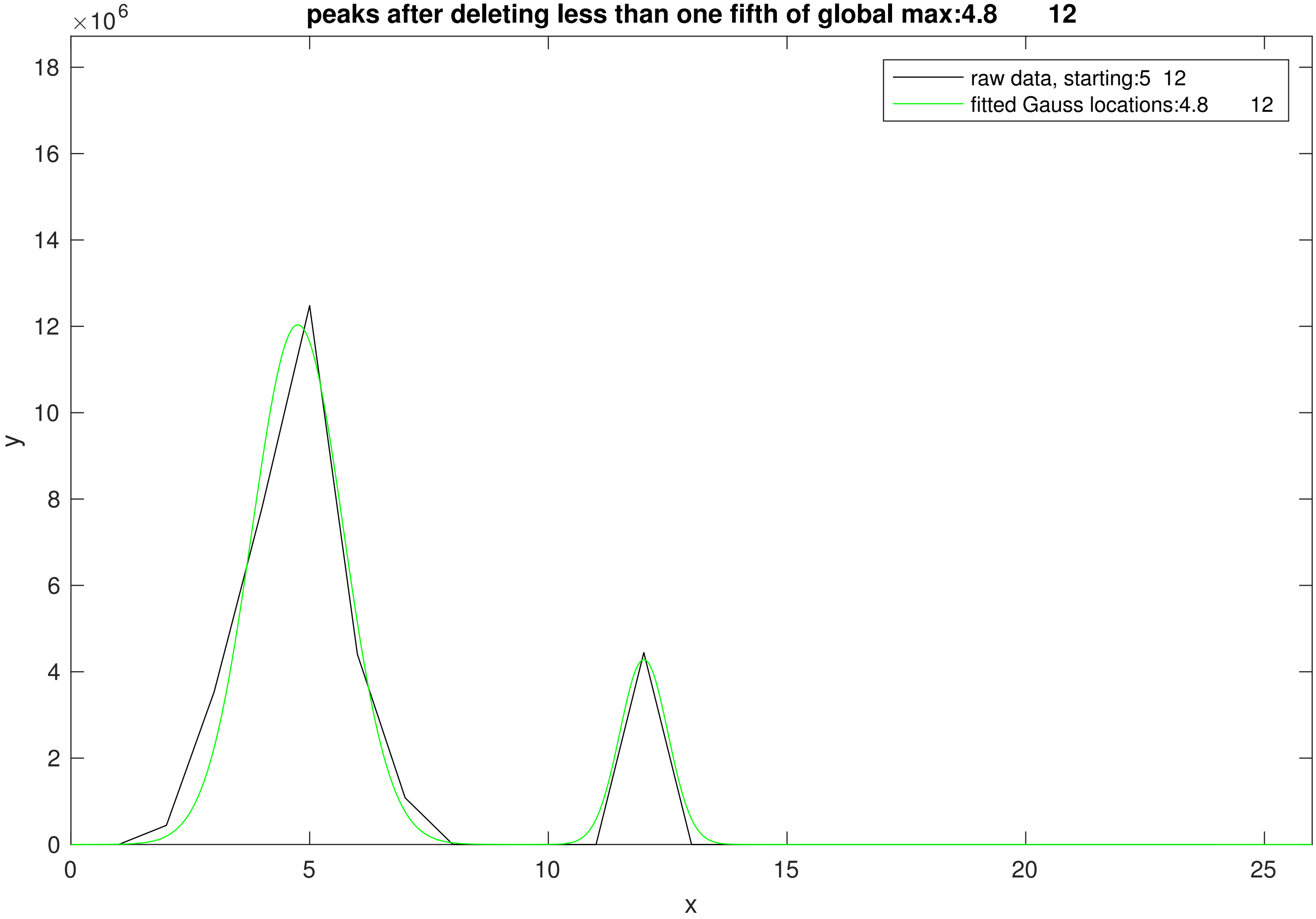

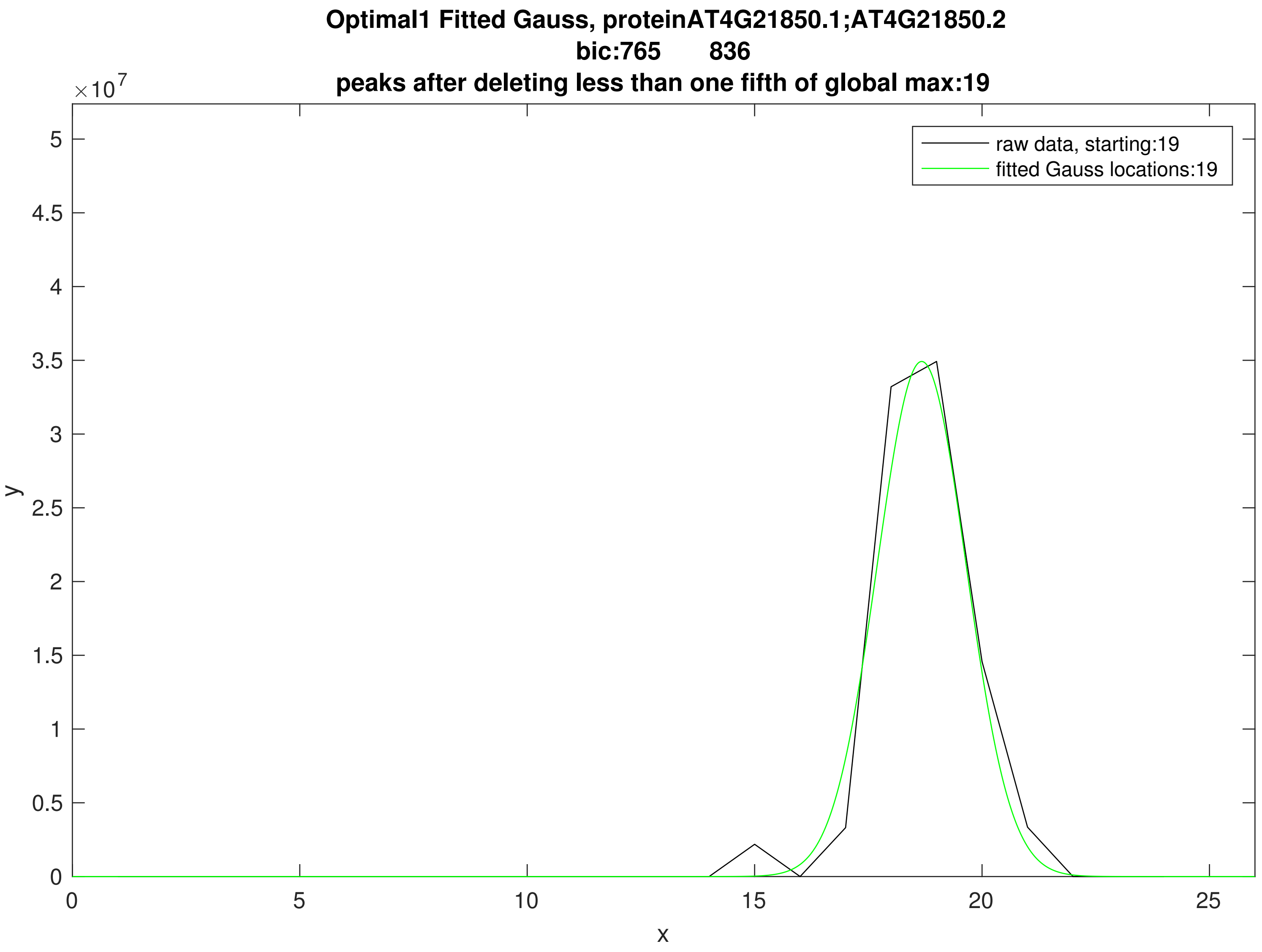

Optimal1 Fitted Gauss, proteinAT4G21850.1;AT4G21850.2
bic:765 836
peaks after deleting less than one fifth of global max:19
×10^7
5
4.5
4
3.5
3
2.5
2
1.5
1
0.5
0
y
0
5
10
15
20
25
x
raw data, starting:19
fitted Gauss locations:19

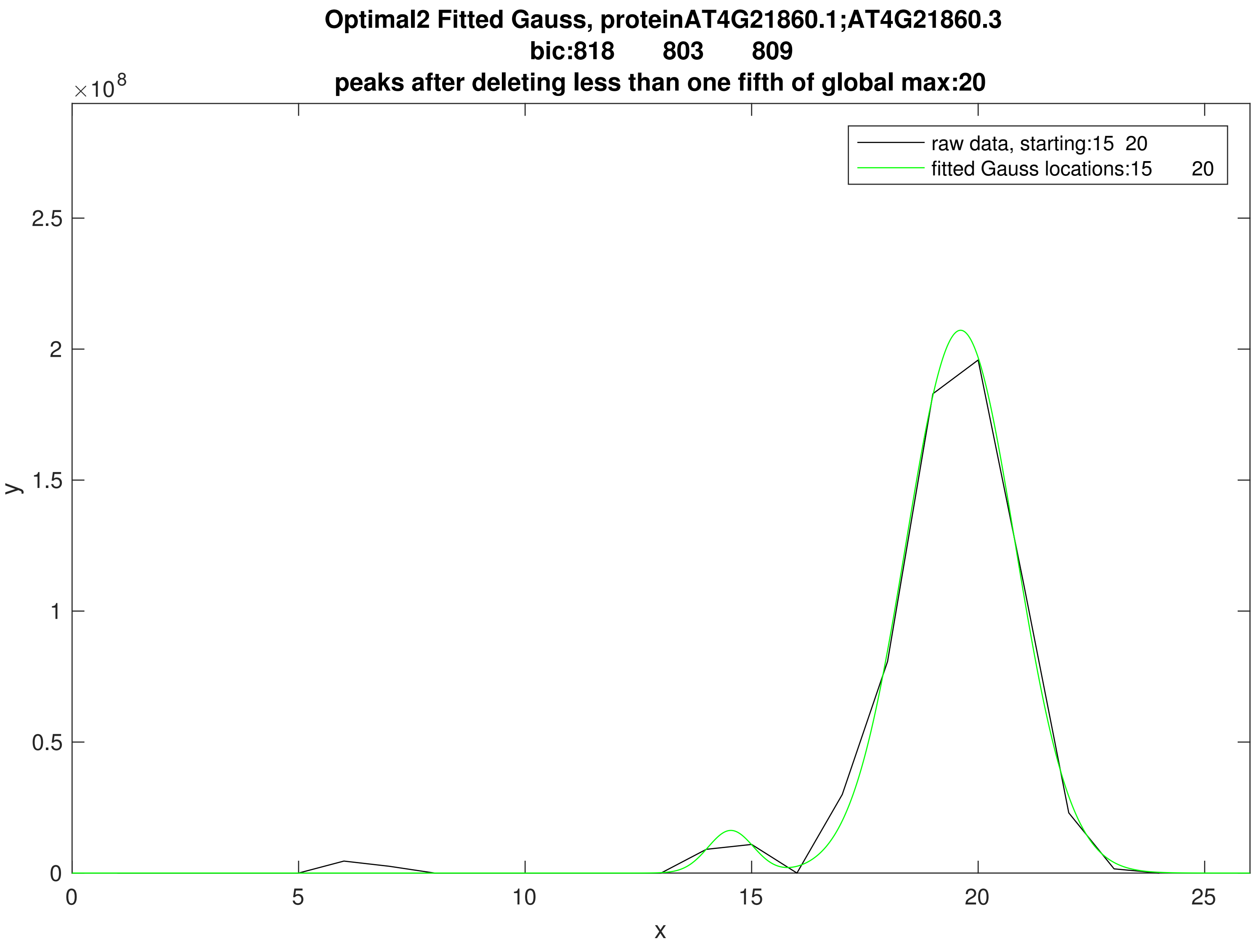

Optimal2 Fitted Gauss, proteinAT4G21860.1;AT4G21860.3
bic:818 803 809
peaks after deleting less than one fifth of global max:20
×10^8
raw data, starting:15 20
fitted Gauss locations:15 20
y
x
0
0.5
1
1.5
2
2.5
0
5
10
15
20
25

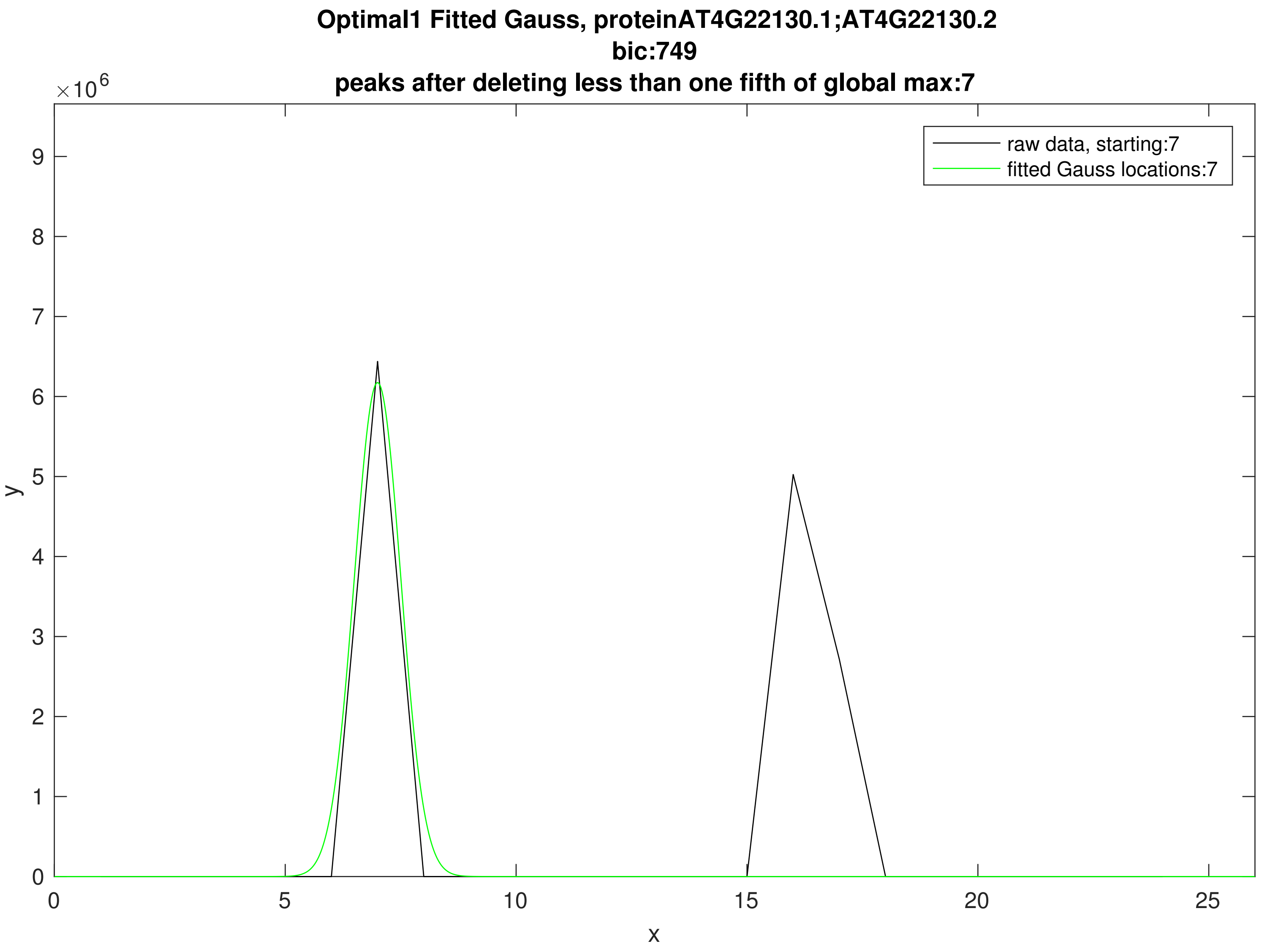
Optimal1 Fitted Gauss, proteinAT4G22130.1;AT4G22130.2
bic:749
peaks after deleting less than one fifth of global max:7
×10^6
raw data, starting:7
fitted Gauss locations:7
y
x
0
1
2
3
4
5
6
7
8
9
0
5
10
15
20
25

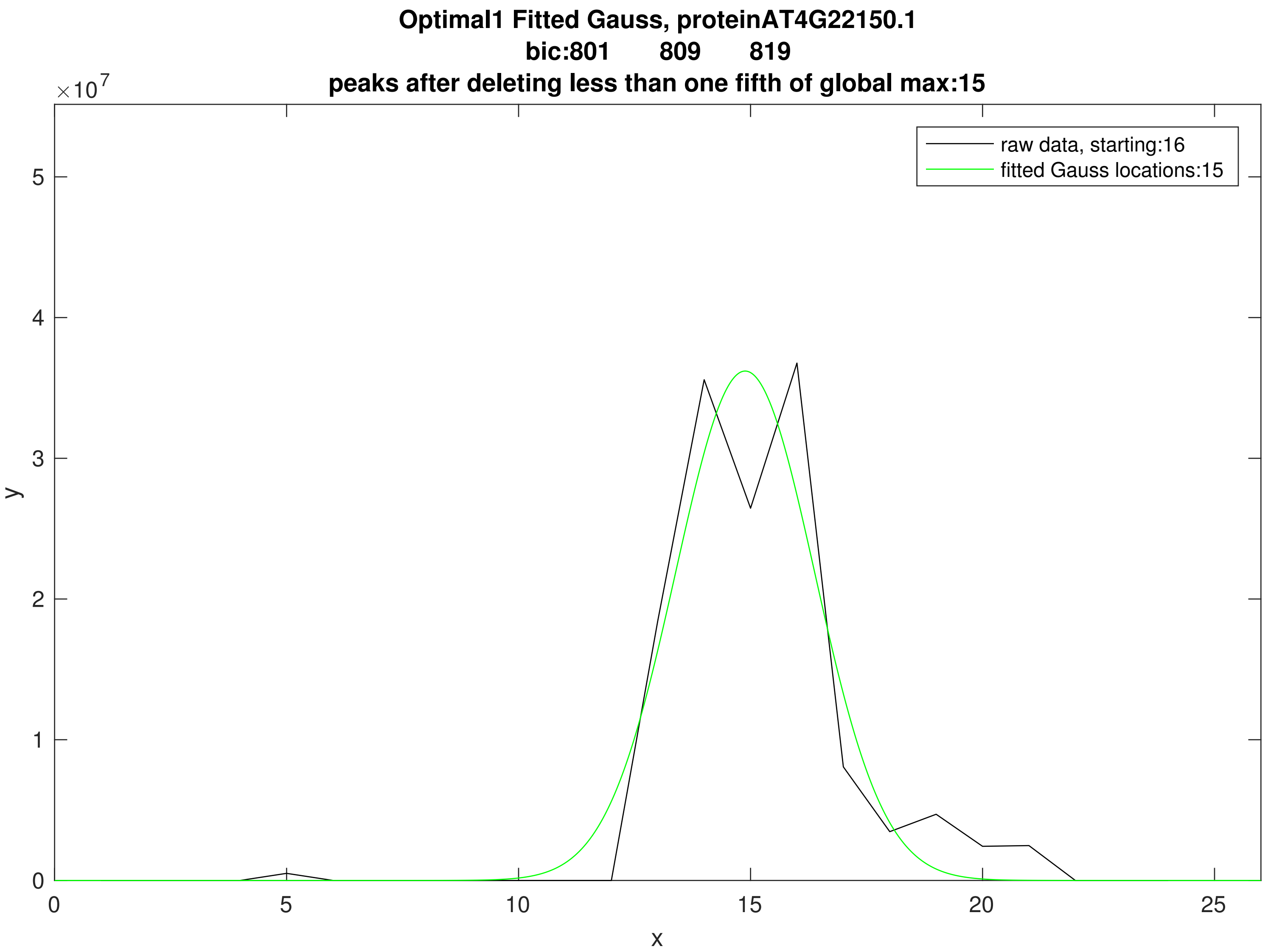

Optimal1 Fitted Gauss, proteinAT4G22150.1
bic:801 809 819
peaks after deleting less than one fifth of global max:15
×10^7
raw data, starting:16
fitted Gauss locations:15
y
x
0
1
2
3
4
5
10
15
20
25

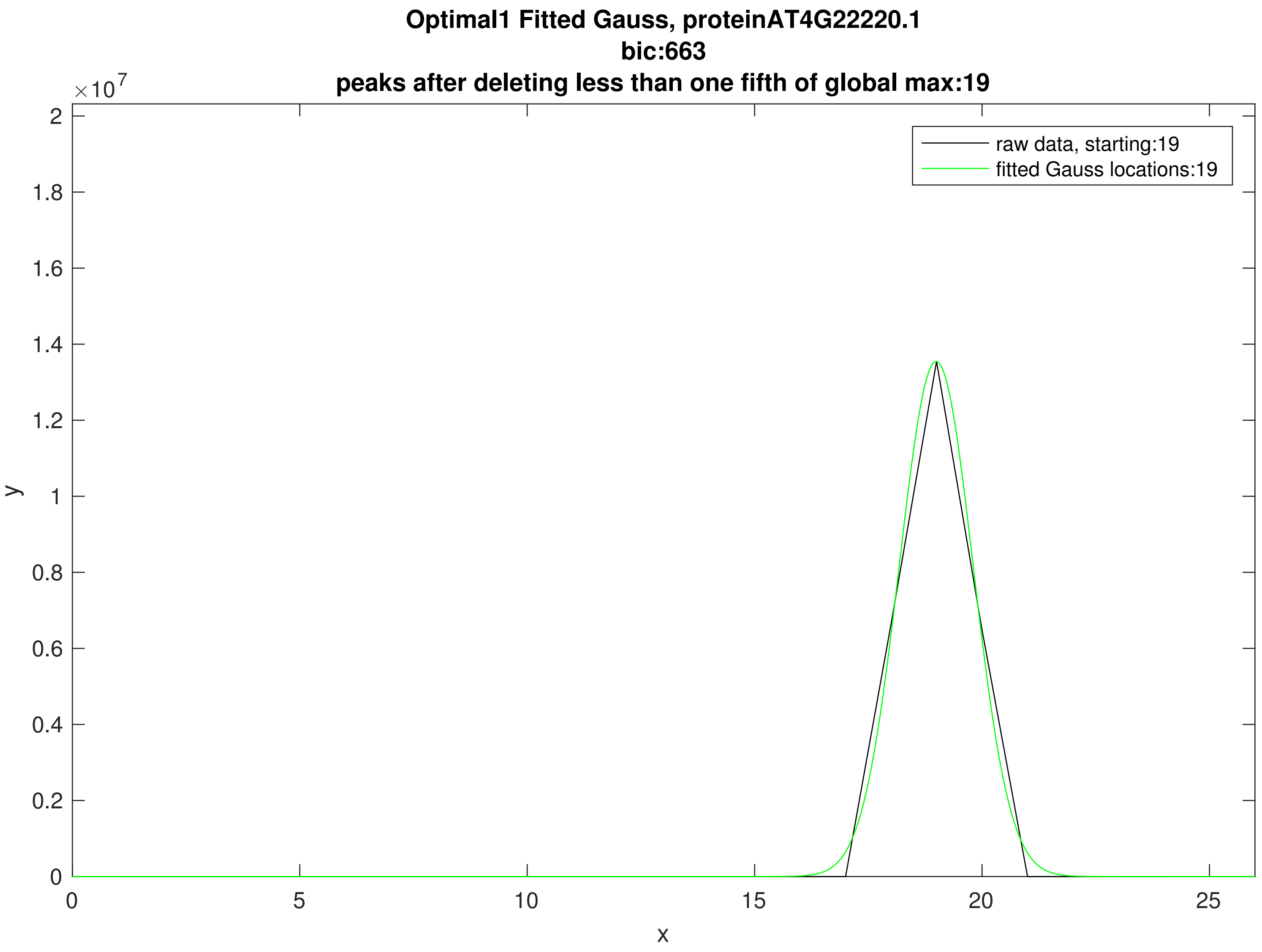
Optimal1 Fitted Gauss, proteinAT4G22220.1
bic:663
peaks after deleting less than one fifth of global max:19
×10^7
2
1.8
1.6
1.4
1.2
1
0.8
0.6
0.4
0.2
0
y
0
5
10
15
20
25
x
raw data, starting:19
fitted Gauss locations:19

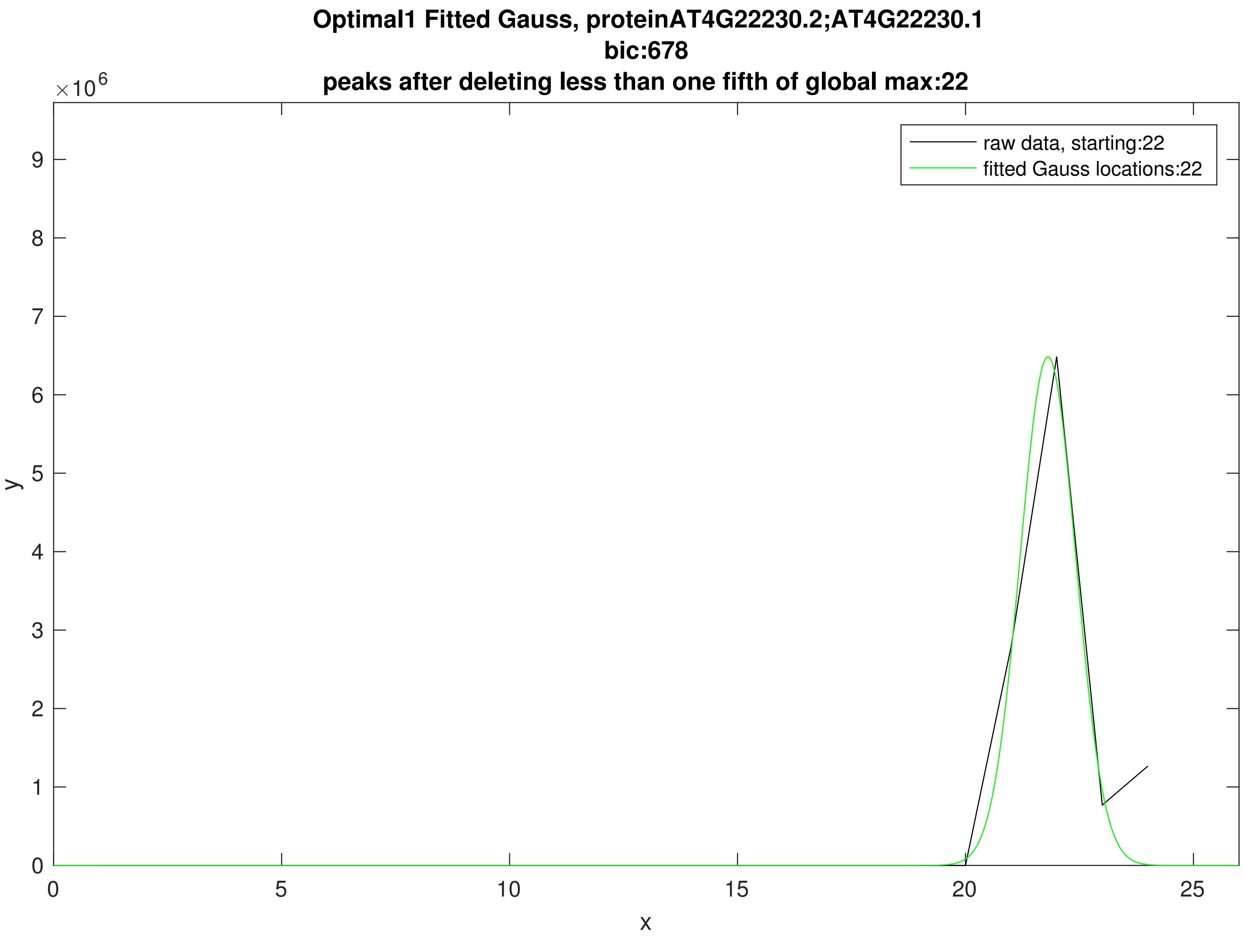

Optimal1 Fitted Gauss, proteinAT4G22230.2;AT4G22230.1
bic:678
peaks after deleting less than one fifth of global max:22
×10^6
raw data, starting:22
fitted Gauss locations:22
y
x
0
1
2
3
4
5
6
7
8
9
0
5
10
15
20
25

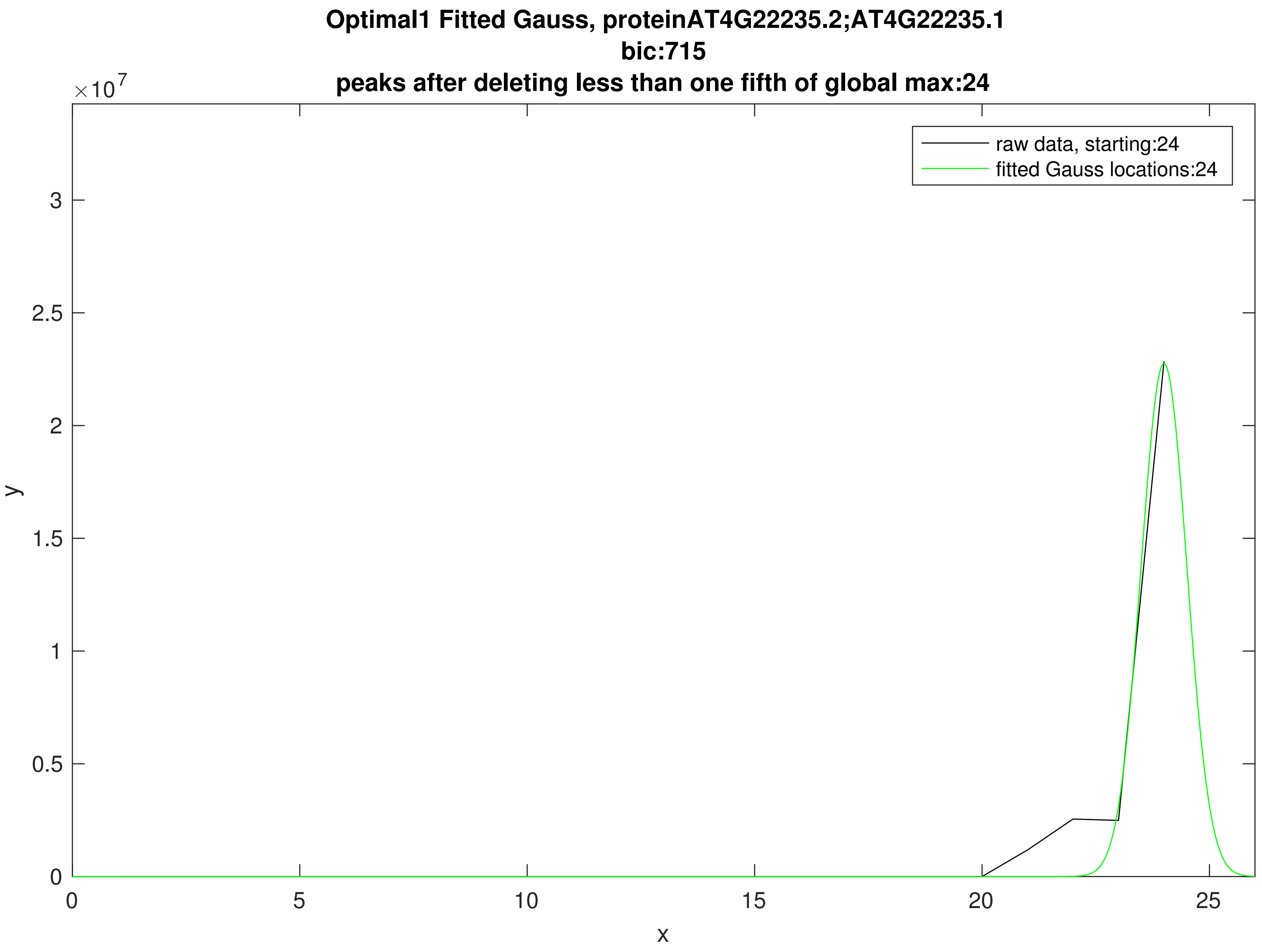
Optimal1 Fitted Gauss, proteinAT4G22235.2;AT4G22235.1
bic:715
peaks after deleting less than one fifth of global max:24
×10^7
raw data, starting:24
fitted Gauss locations:24
y
x
0
0.5
1
1.5
2
2.5
3
0
5
10
15
20
25

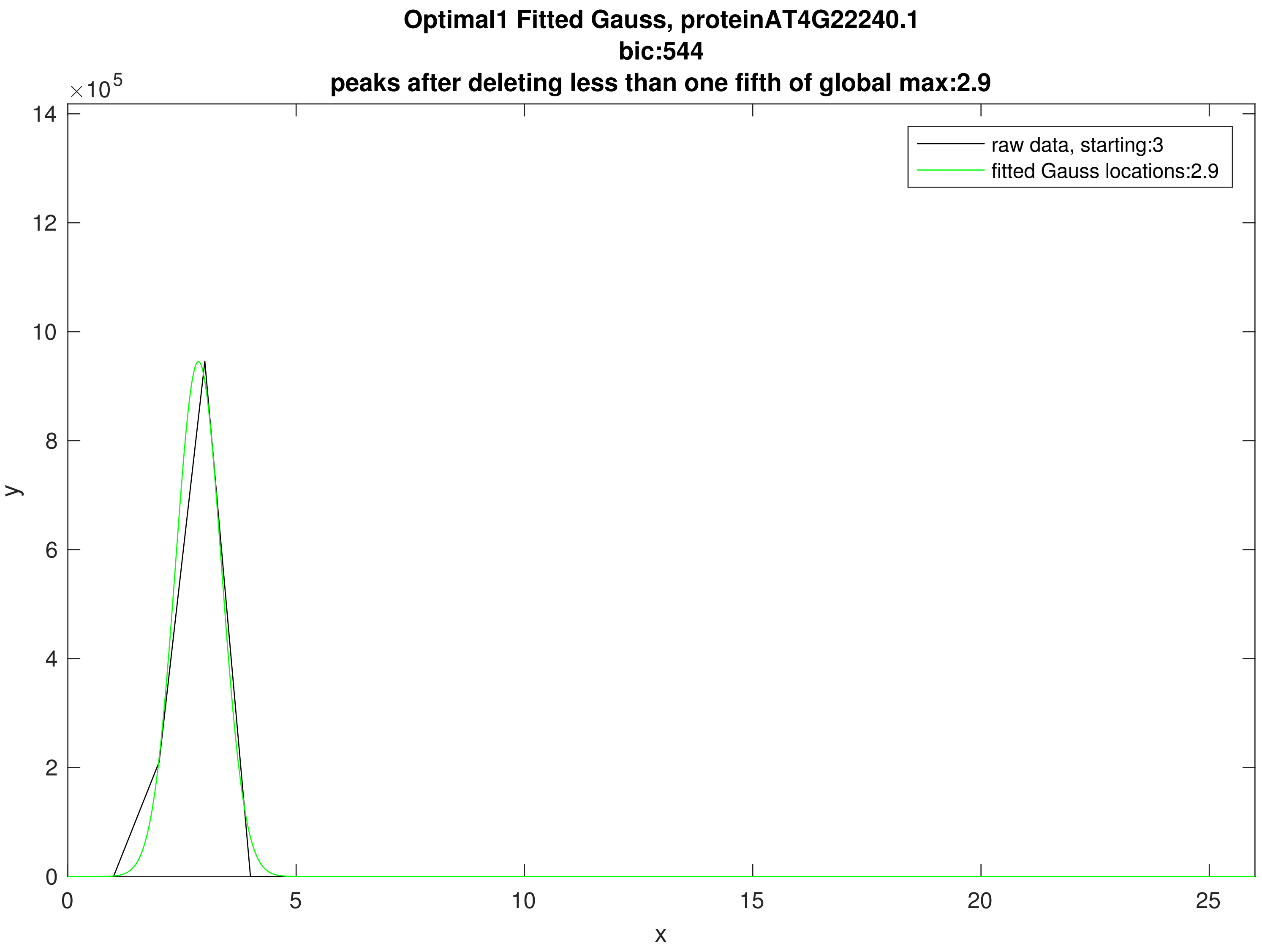

Optimal1 Fitted Gauss, proteinAT4G22240.1
bic:544
peaks after deleting less than one fifth of global max:2.9
×10^5
14
12
10
8
6
4
2
0
y
0
5
10
15
20
25
x
raw data, starting:3
fitted Gauss locations:2.9

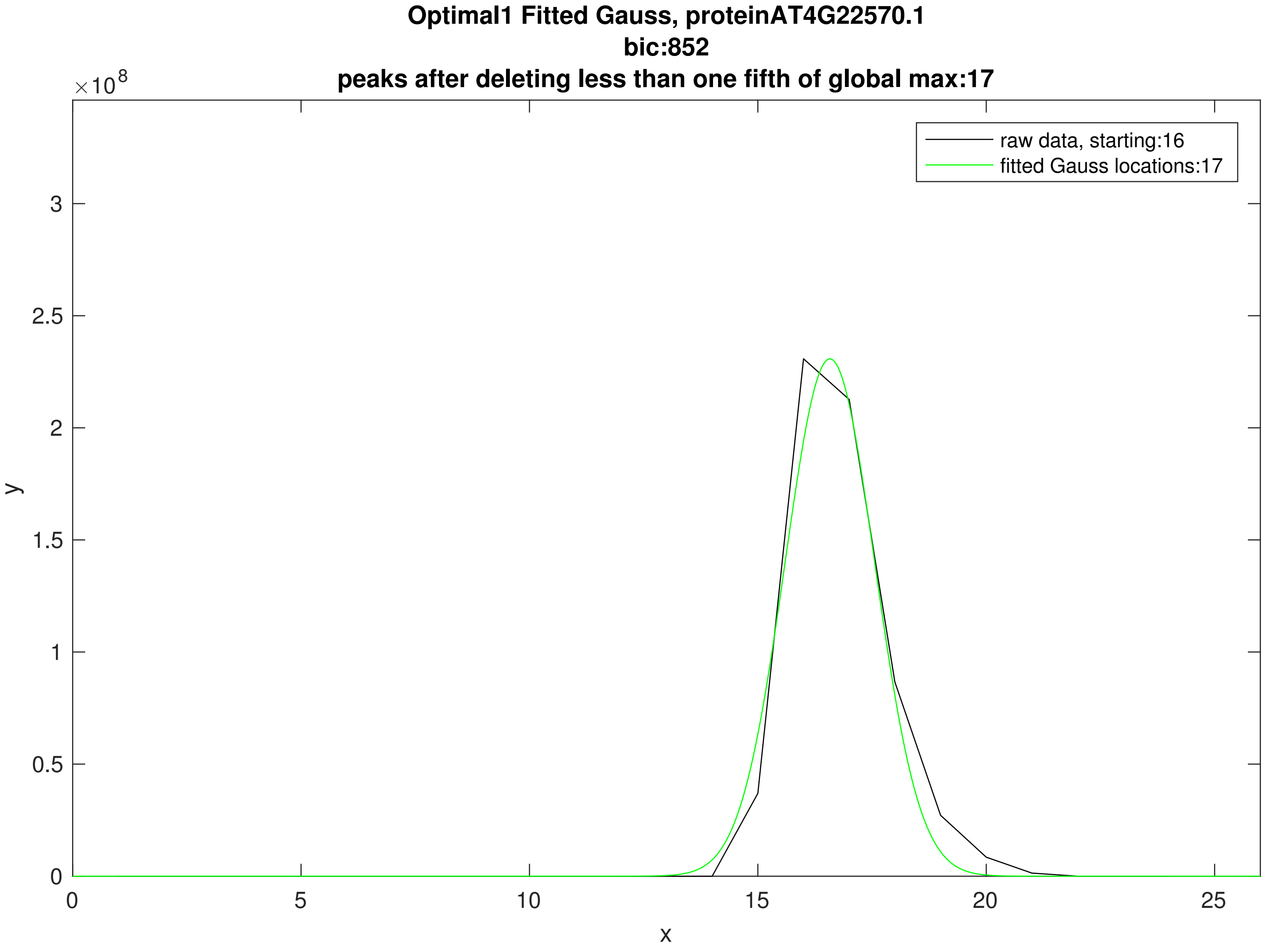

Optimal1 Fitted Gauss, proteinAT4G22570.1
bic:852
peaks after deleting less than one fifth of global max:17
×10^8
raw data, starting:16
fitted Gauss locations:17
y
x
0
0.5
1
1.5
2
2.5
3
0
5
10
15
20
25

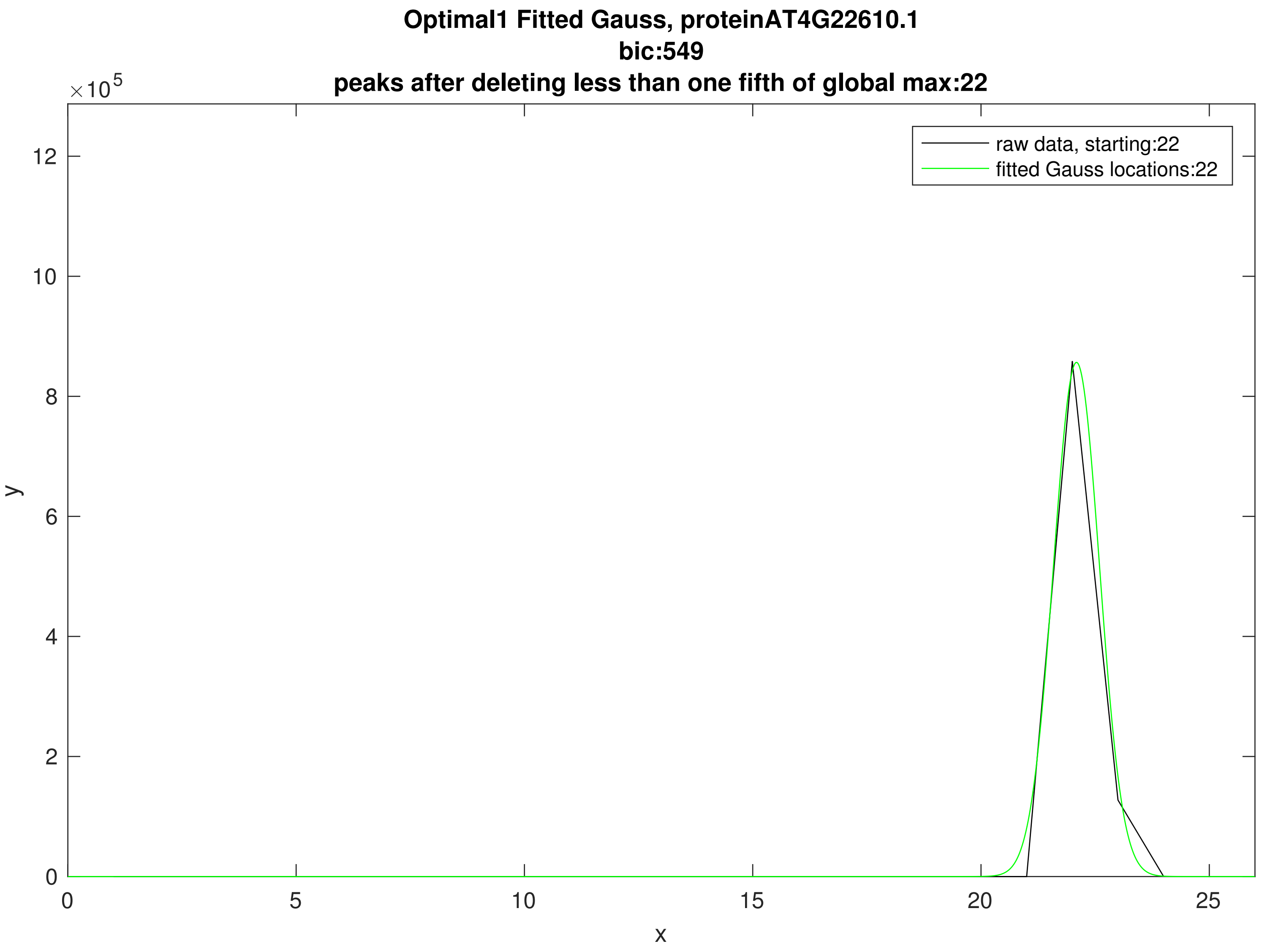

Optimal1 Fitted Gauss, proteinAT4G22610.1
bic:549
peaks after deleting less than one fifth of global max:22
×10^5
raw data, starting:22
fitted Gauss locations:22
y
x
0
2
4
6
8
10
12
0
5
10
15
20
25

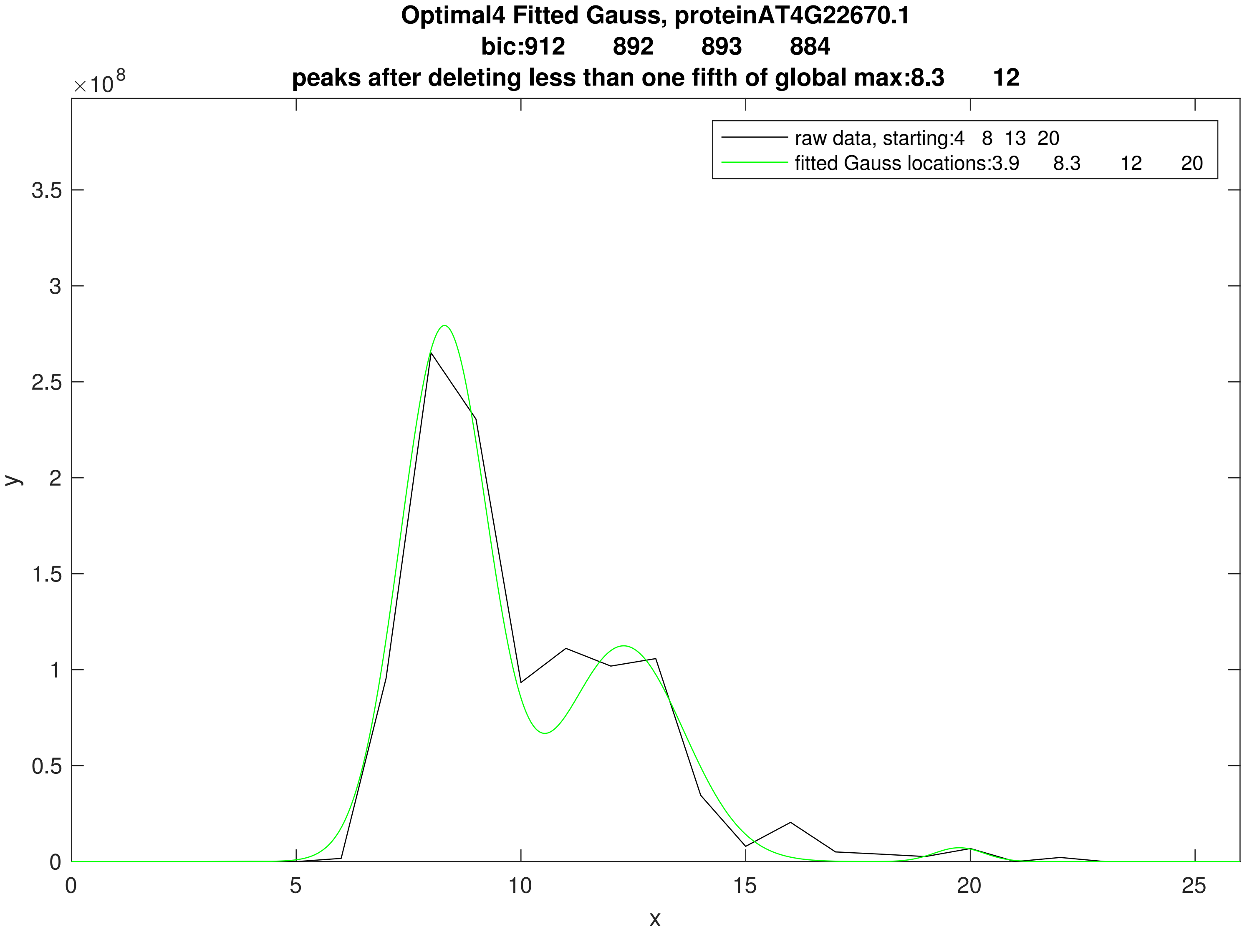

Optimal4 Fitted Gauss, proteinAT4G22670.1
bic:912 892 893 884
peaks after deleting less than one fifth of global max:8.3 12
×10^8
raw data, starting:4 8 13 20
fitted Gauss locations:3.9 8.3 12 20
y
x
0
0.5
1
1.5
2
2.5
3
3.5
5
10
15
20
25

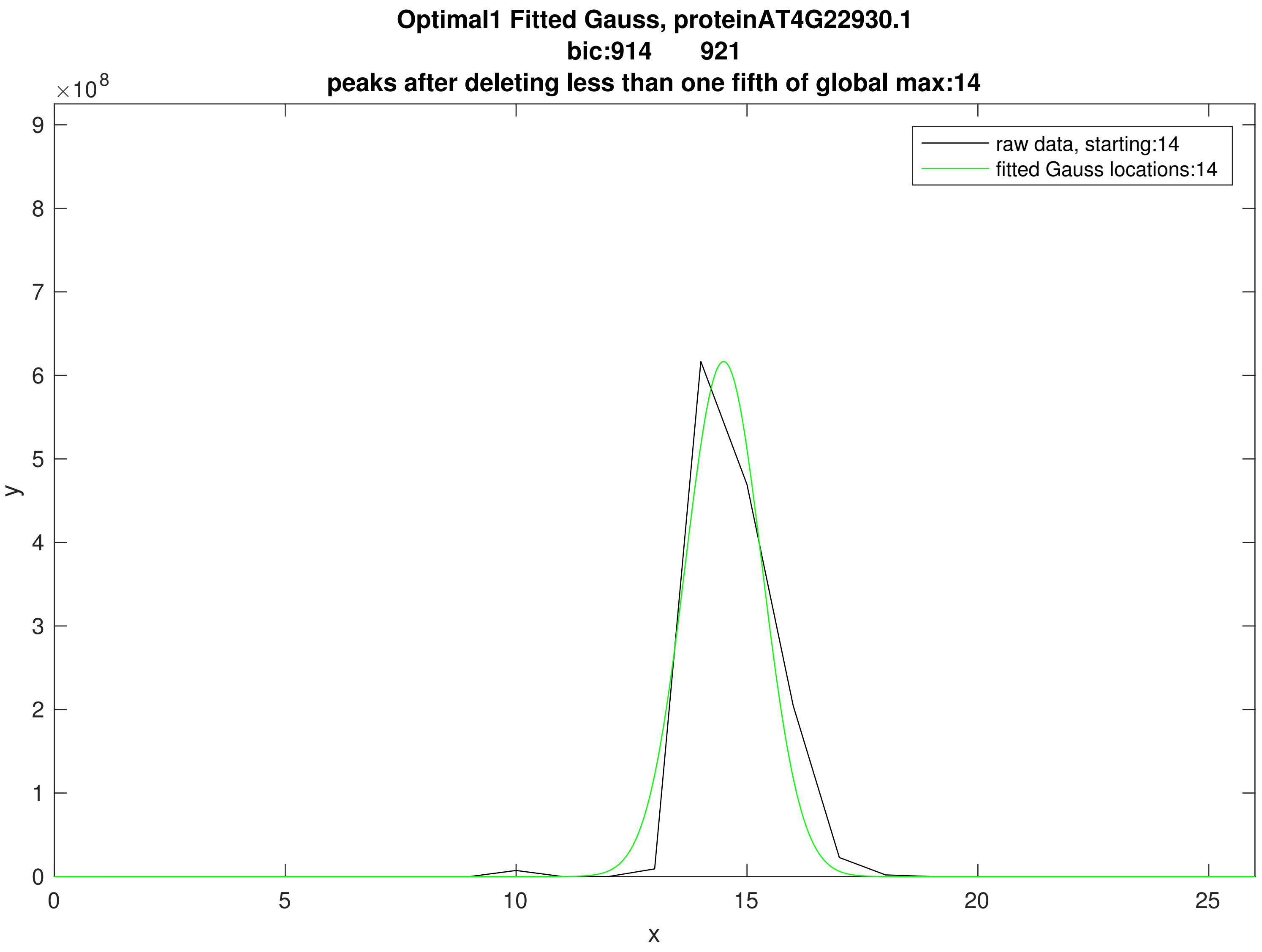
Optimal1 Fitted Gauss, proteinAT4G22930.1
bic:914 921
peaks after deleting less than one fifth of global max:14
×10^8
raw data, starting:14
fitted Gauss locations:14
y
x
0
1
2
3
4
5
6
7
8
9
0
5
10
15
20
25

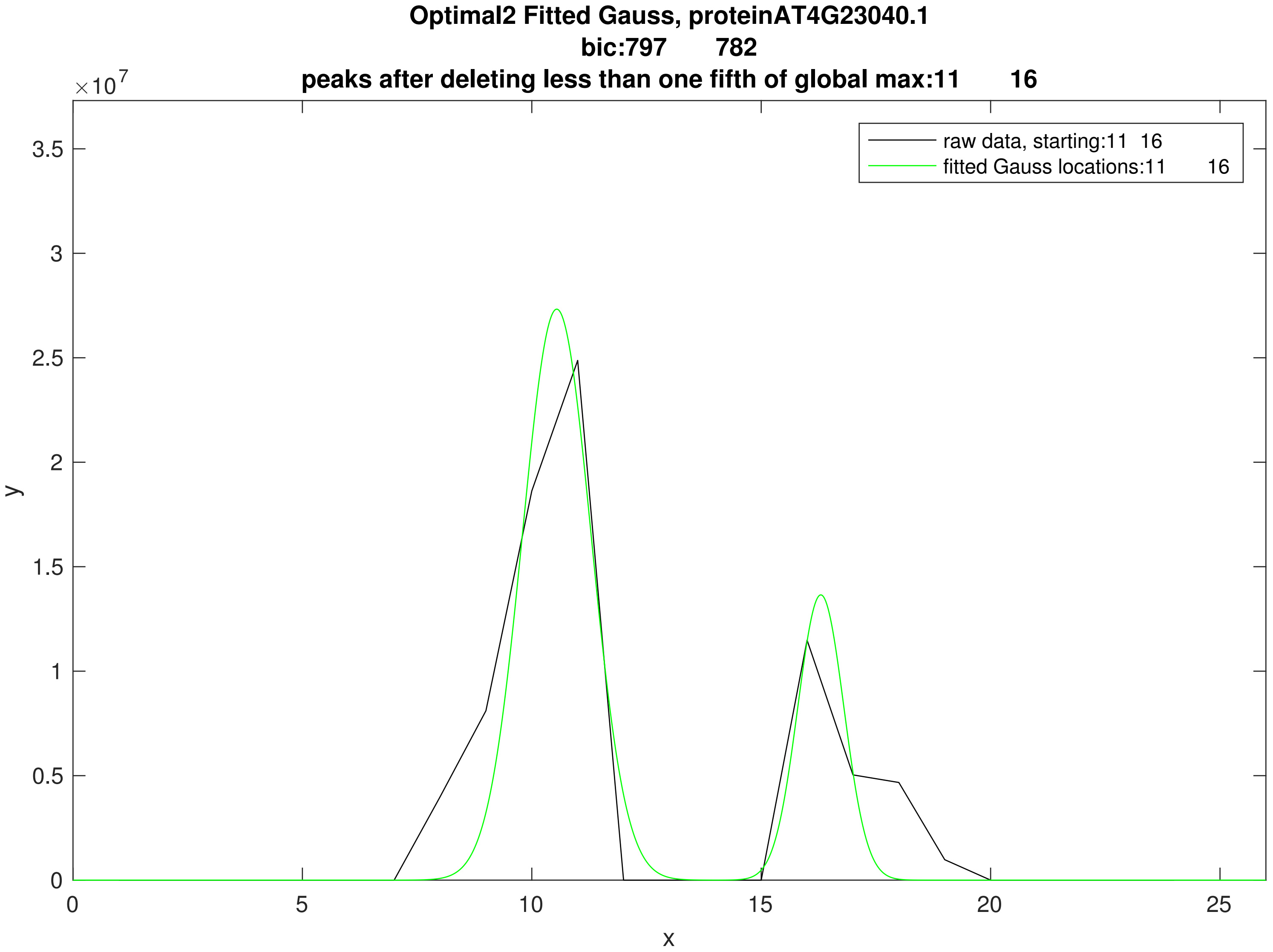

Optimal2 Fitted Gauss, proteinAT4G23040.1
bic:797 782
peaks after deleting less than one fifth of global max:11 16
×10^7
3.5
3
2.5
2
1.5
1
0.5
0
y
0
5
10
15
20
25
x
raw data, starting:11 16
fitted Gauss locations:11 16

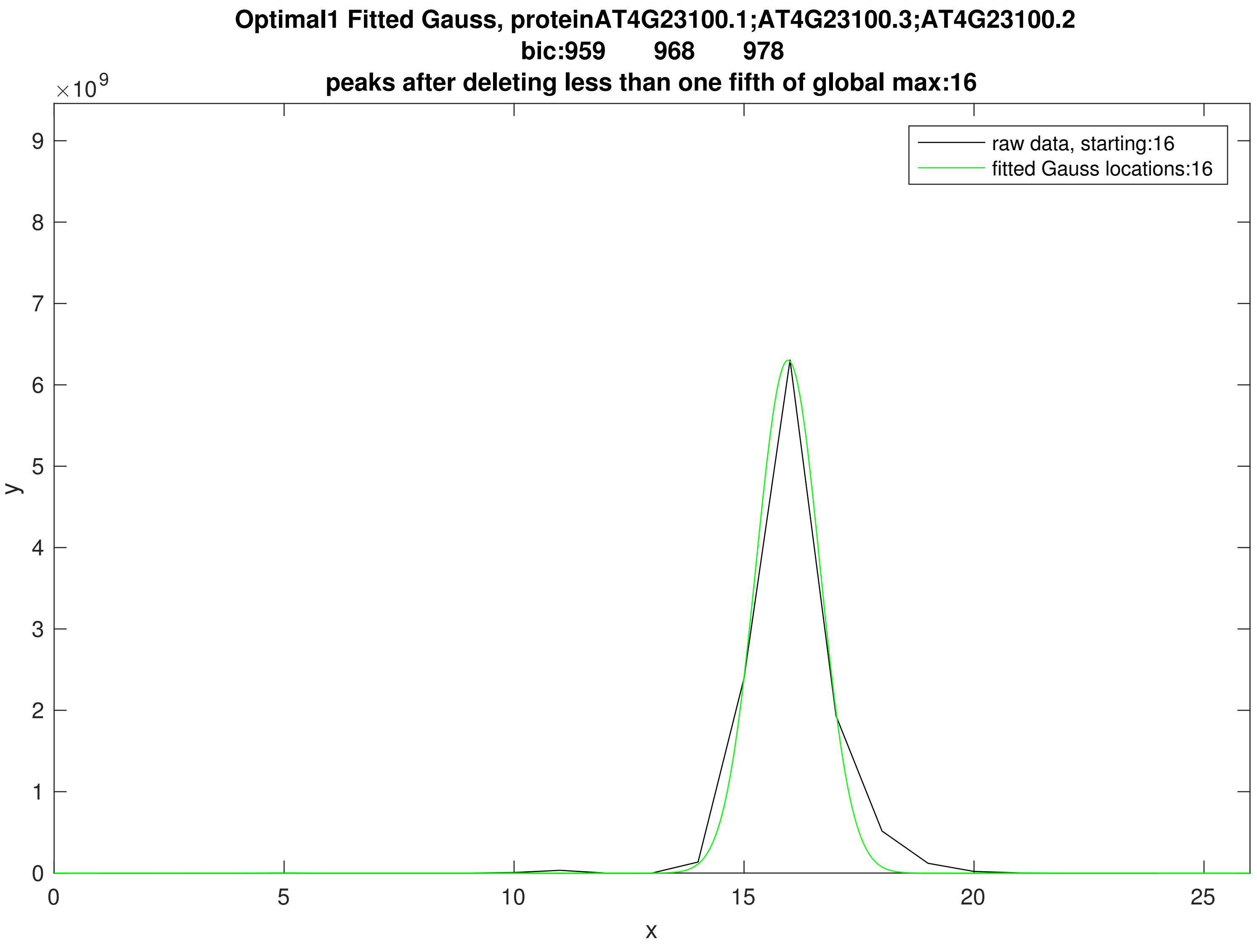

Optimal1 Fitted Gauss, proteinAT4G23100.1;AT4G23100.3;AT4G23100.2
bic:959 968 978
peaks after deleting less than one fifth of global max:16
×10^9
raw data, starting:16
fitted Gauss locations:16
y
x
0
1
2
3
4
5
6
7
8
9
0
5
10
15
20
25

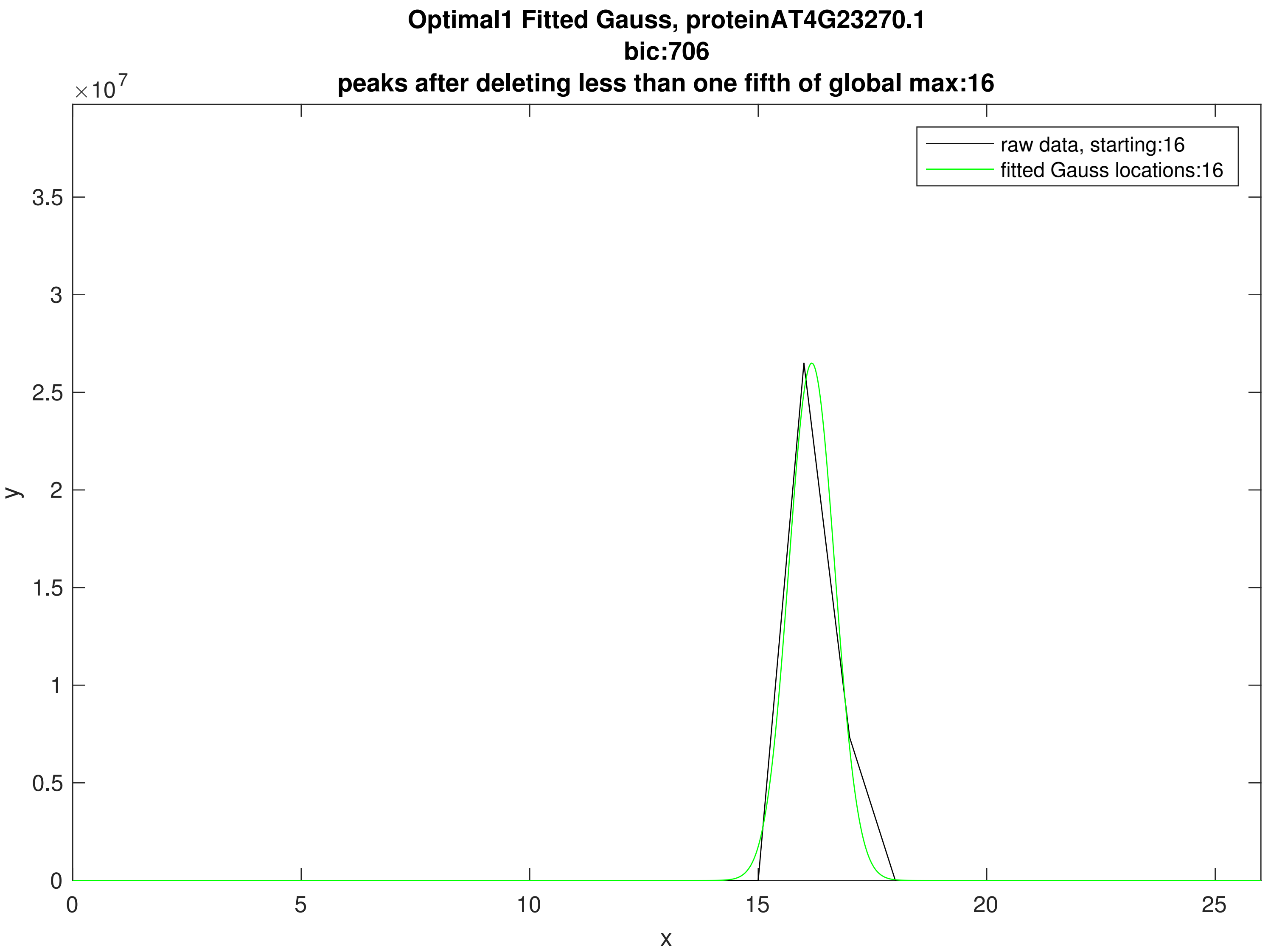
Optimal1 Fitted Gauss, proteinAT4G23270.1
bic:706
peaks after deleting less than one fifth of global max:16
×10^7
raw data, starting:16
fitted Gauss locations:16
y
x
0
0.5
1
1.5
2
2.5
3
3.5
0
5
10
15
20
25

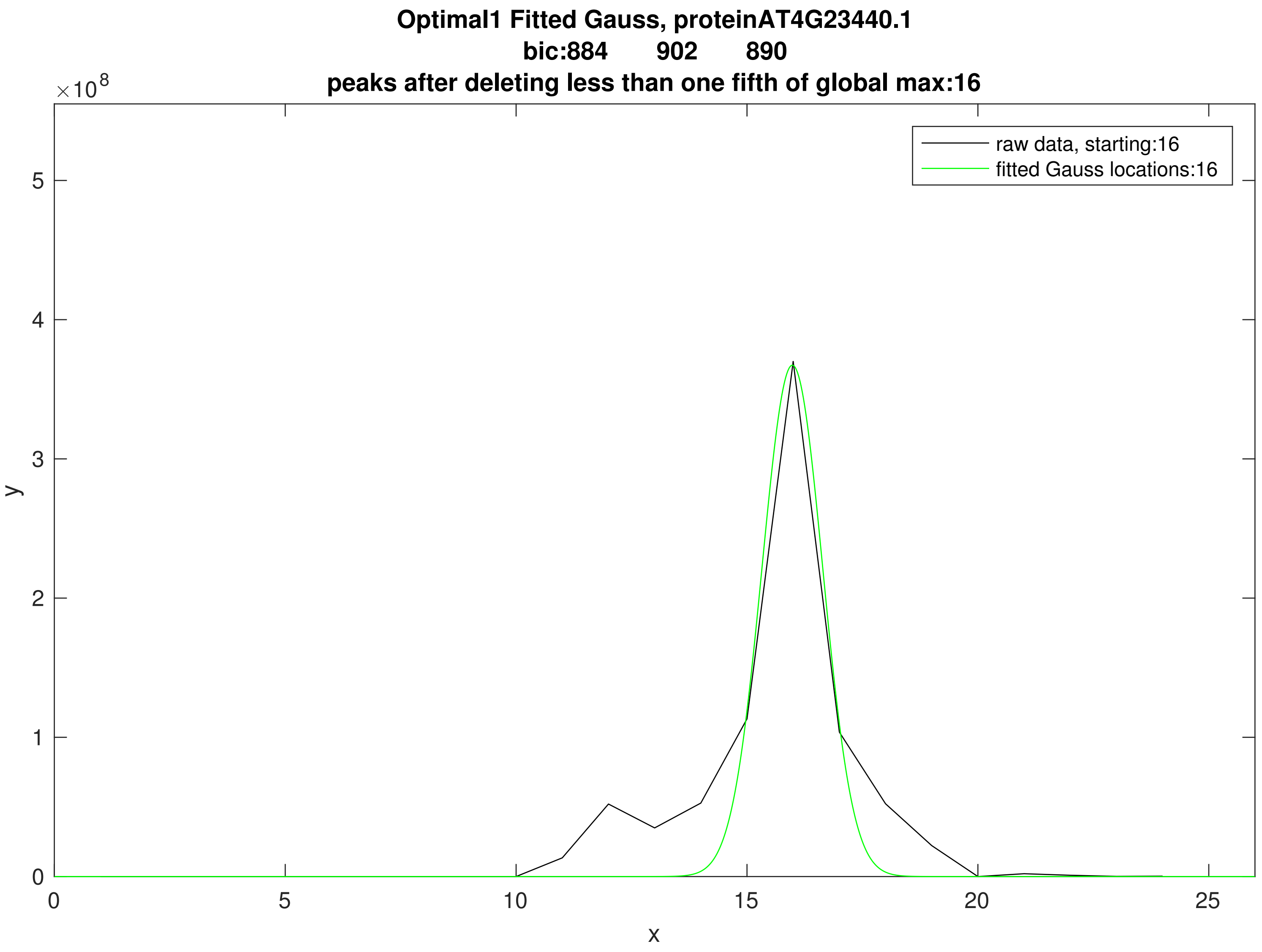

Optimal1 Fitted Gauss, proteinAT4G23440.1
bic:884 902 890
peaks after deleting less than one fifth of global max:16
×10^8
raw data, starting:16
fitted Gauss locations:16
y
x
0
1
2
3
4
5
0
5
10
15
20
25

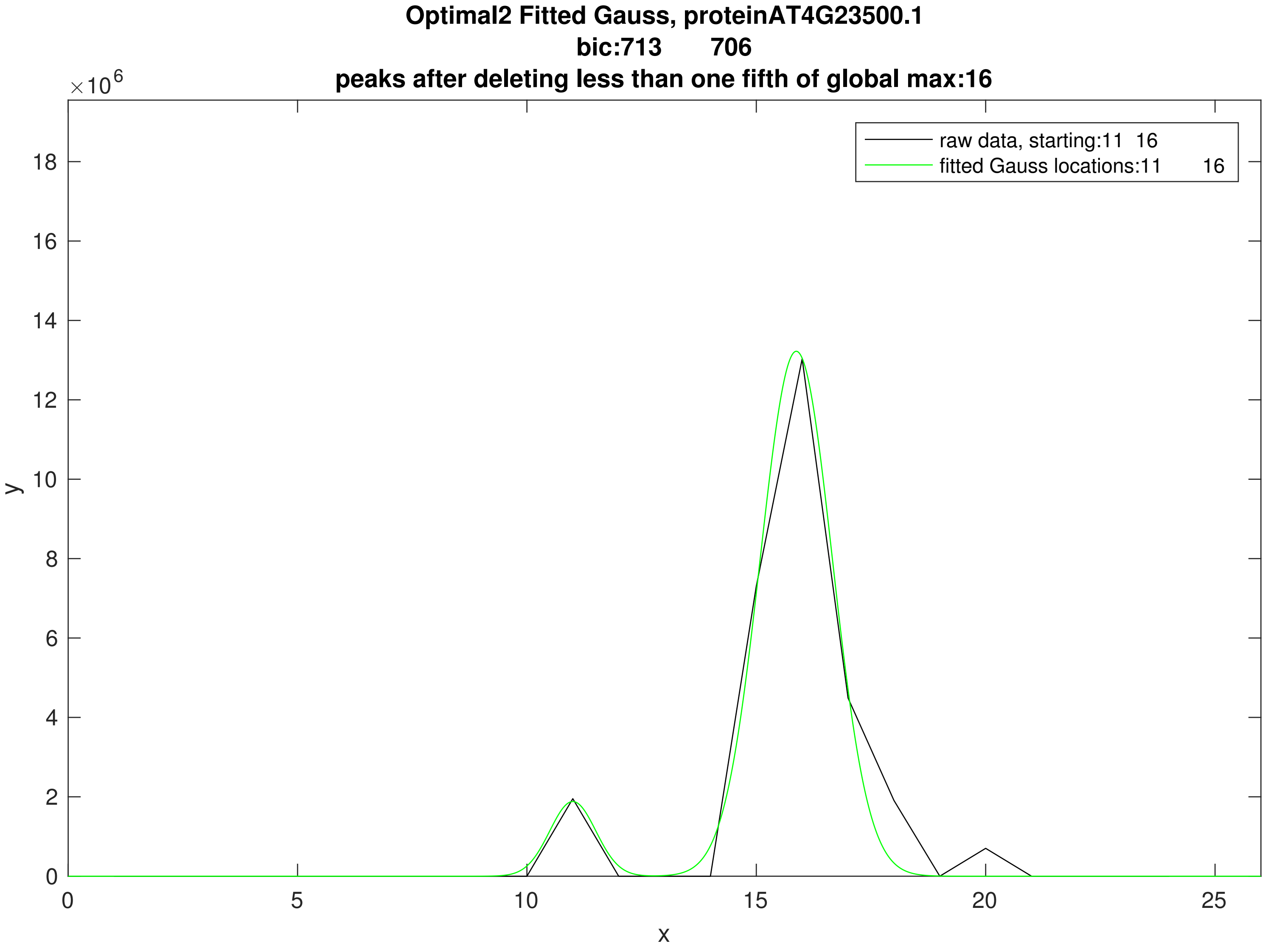
Optimal2 Fitted Gauss, proteinAT4G23500.1
bic:713 706
peaks after deleting less than one fifth of global max:16
×10^6
raw data, starting:11 16
fitted Gauss locations:11 16
y
x
0
2
4
6
8
10
12
14
16
18
5
10
15
20
25

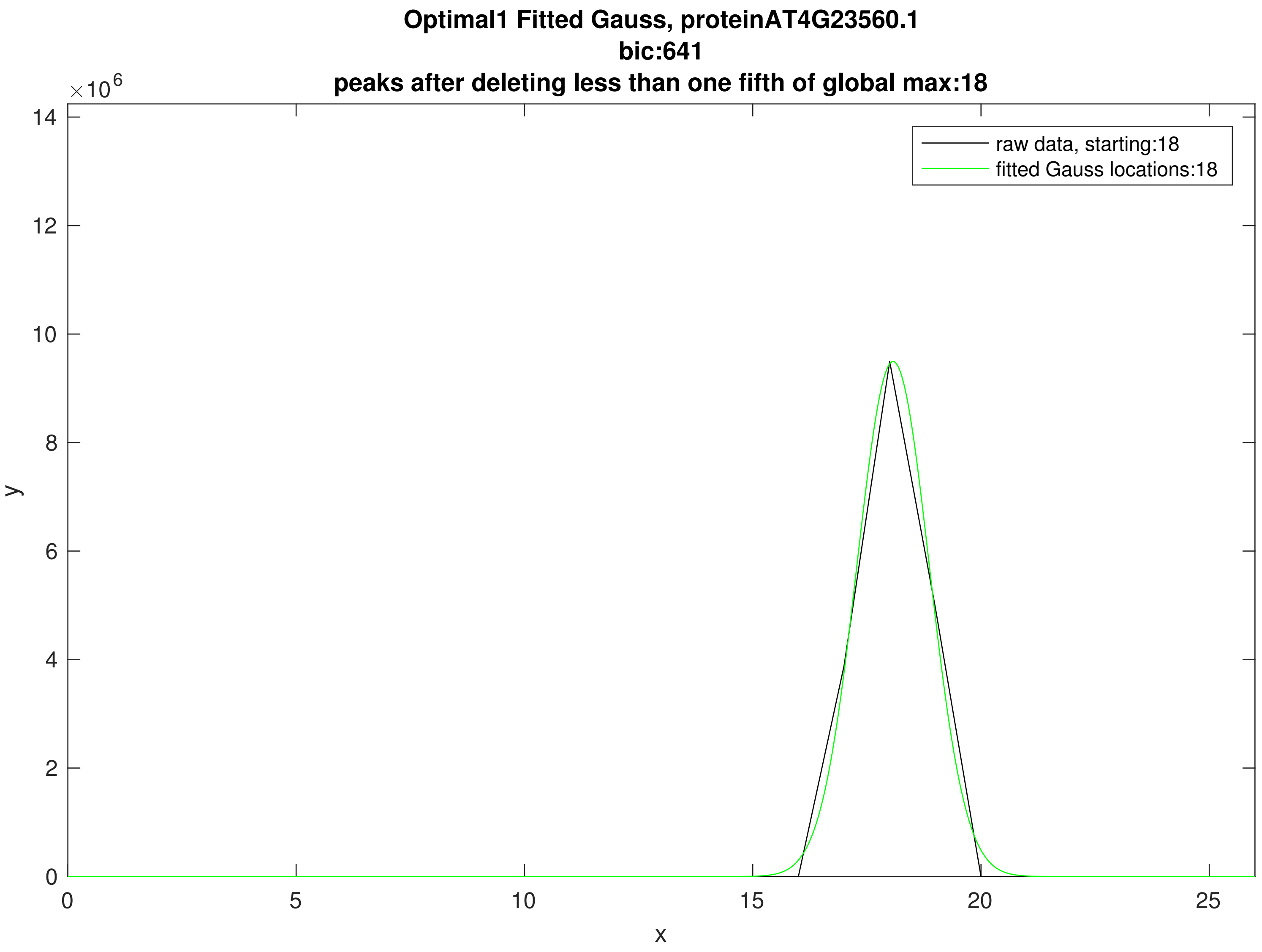

Optimal1 Fitted Gauss, proteinAT4G23560.1
bic:641
peaks after deleting less than one fifth of global max:18
×10^6
raw data, starting:18
fitted Gauss locations:18
y
x
0
2
4
6
8
10
12
14
0
5
10
15
20
25

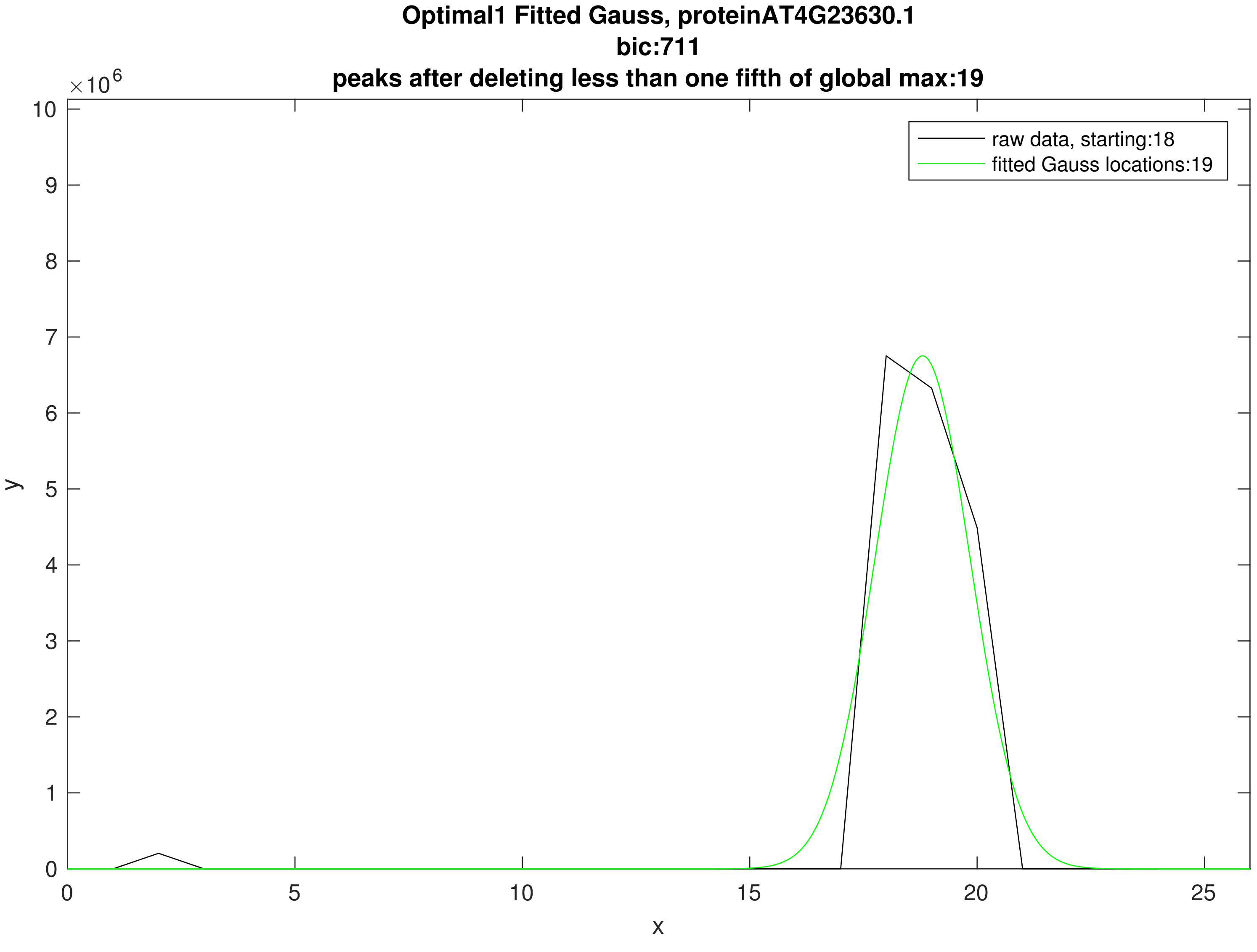
Optimal1 Fitted Gauss, proteinAT4G23630.1
bic:711
peaks after deleting less than one fifth of global max:19
×10^6
raw data, starting:18
fitted Gauss locations:19
y
x
0
1
2
3
4
5
6
7
8
9
10
0
5
10
15
20
25

#### Optimal1 Fitted Gauss, proteinAT4G23650.1
#### bic:718
#### peaks after deleting less than one fifth of global max:15

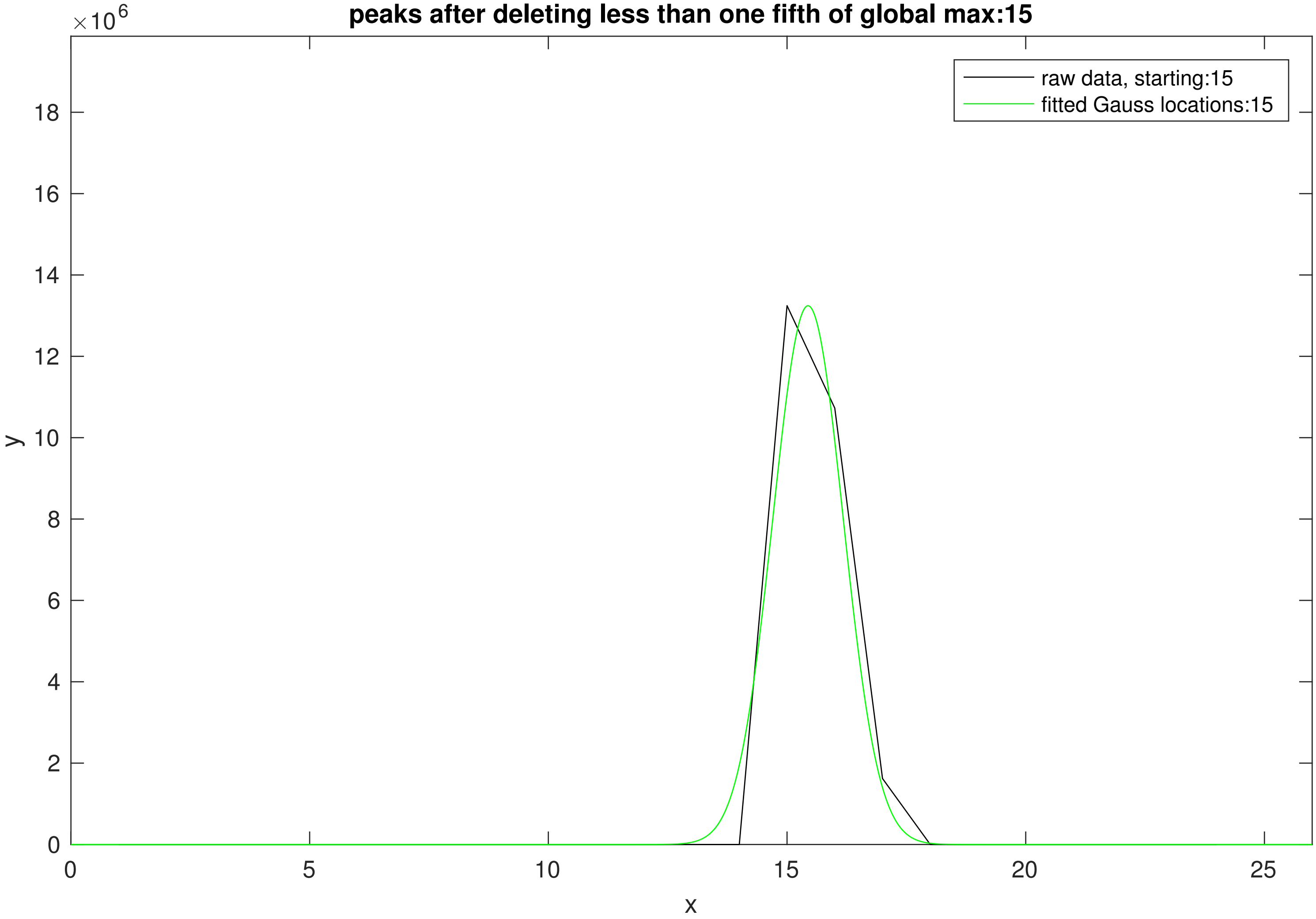

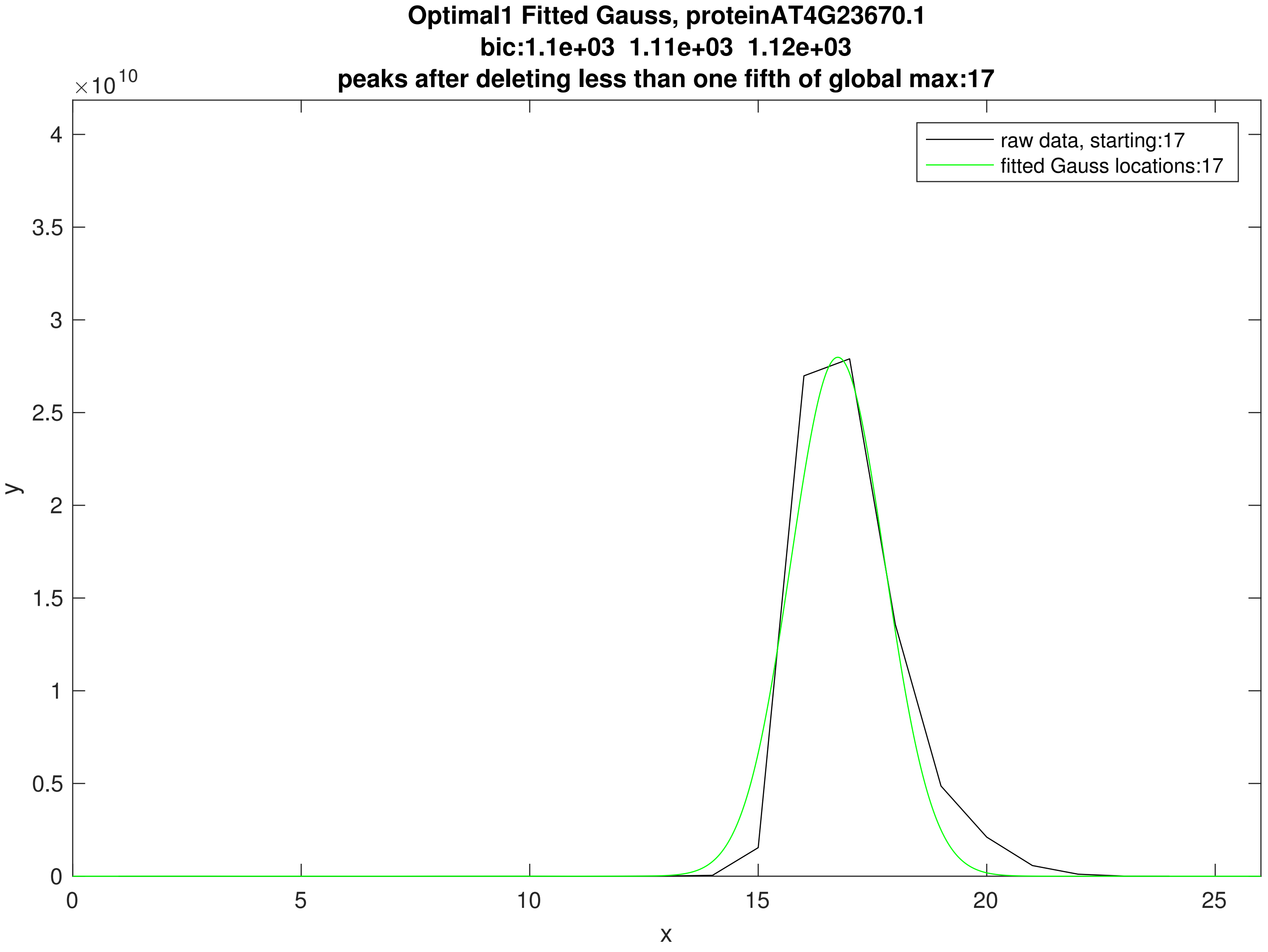
Optimal1 Fitted Gauss, proteinAT4G23670.1
bic:1.1e+03  1.11e+03  1.12e+03
peaks after deleting less than one fifth of global max:17
×10^10
raw data, starting:17
fitted Gauss locations:17
y
x
0
0.5
1
1.5
2
2.5
3
3.5
4
0
5
10
15
20
25

**Optimal1 Fitted Gauss, proteinAT4G23680.1**
**bic:967 976**
**peaks after deleting less than one fifth of global max:17**

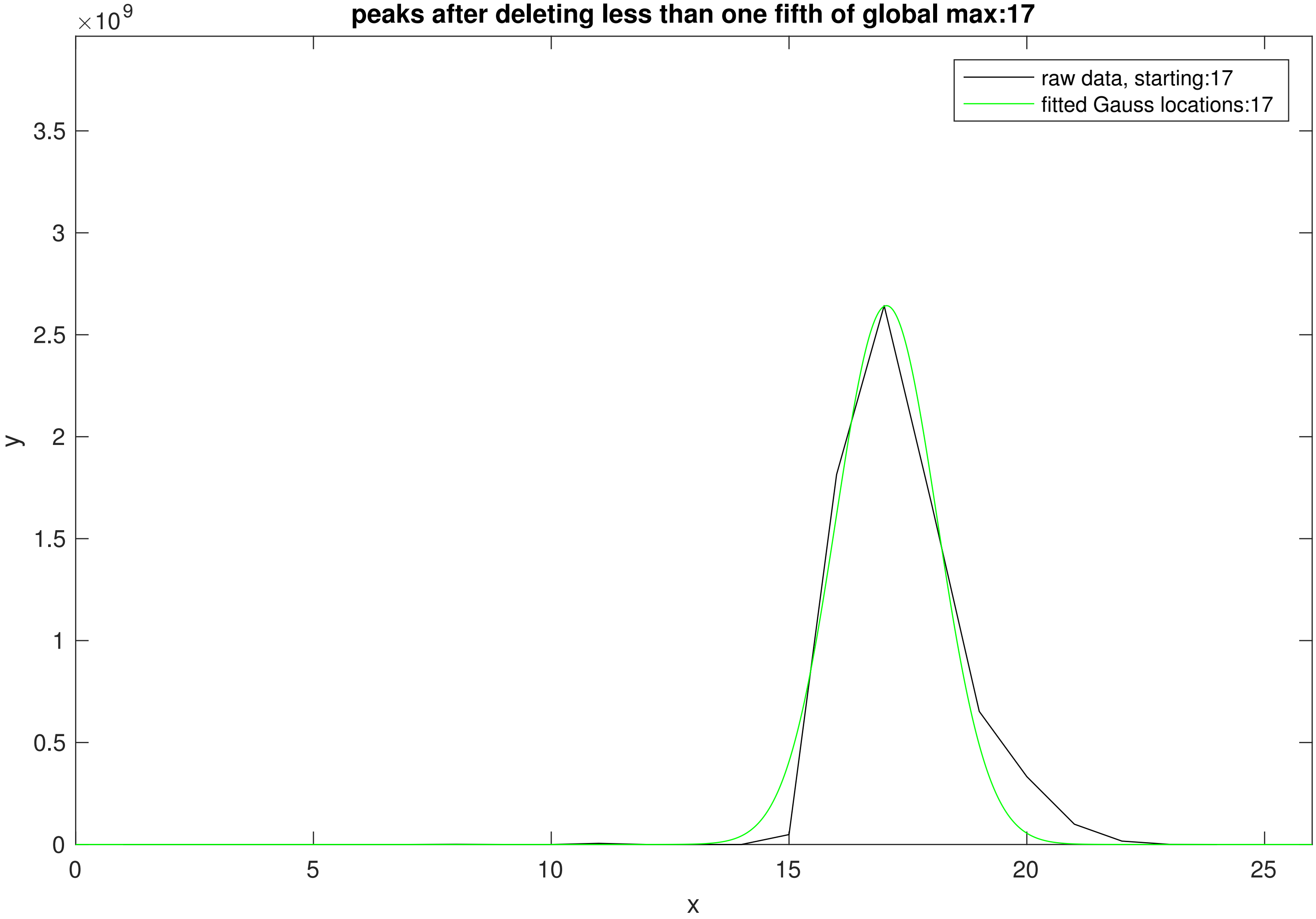

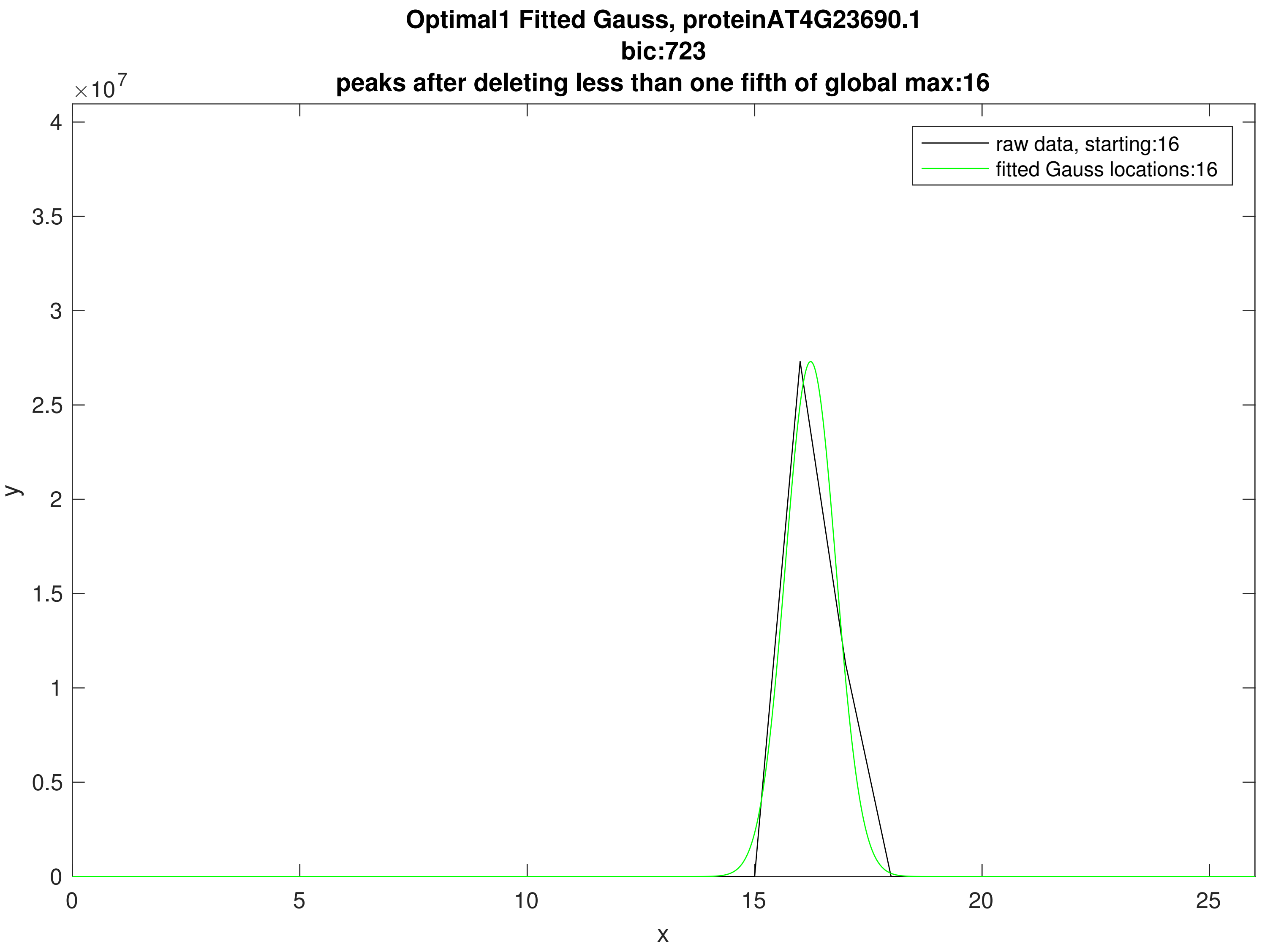

Optimal1 Fitted Gauss, proteinAT4G23690.1
bic:723
peaks after deleting less than one fifth of global max:16
$\times 10^7$
raw data, starting:16
fitted Gauss locations:16
y
x
0
0.5
1
1.5
2
2.5
3
3.5
4
0
5
10
15
20
25

**Optimal1 Fitted Gauss, proteinAT4G23710.1**
**bic:943 953 962**
**peaks after deleting less than one fifth of global max:17**

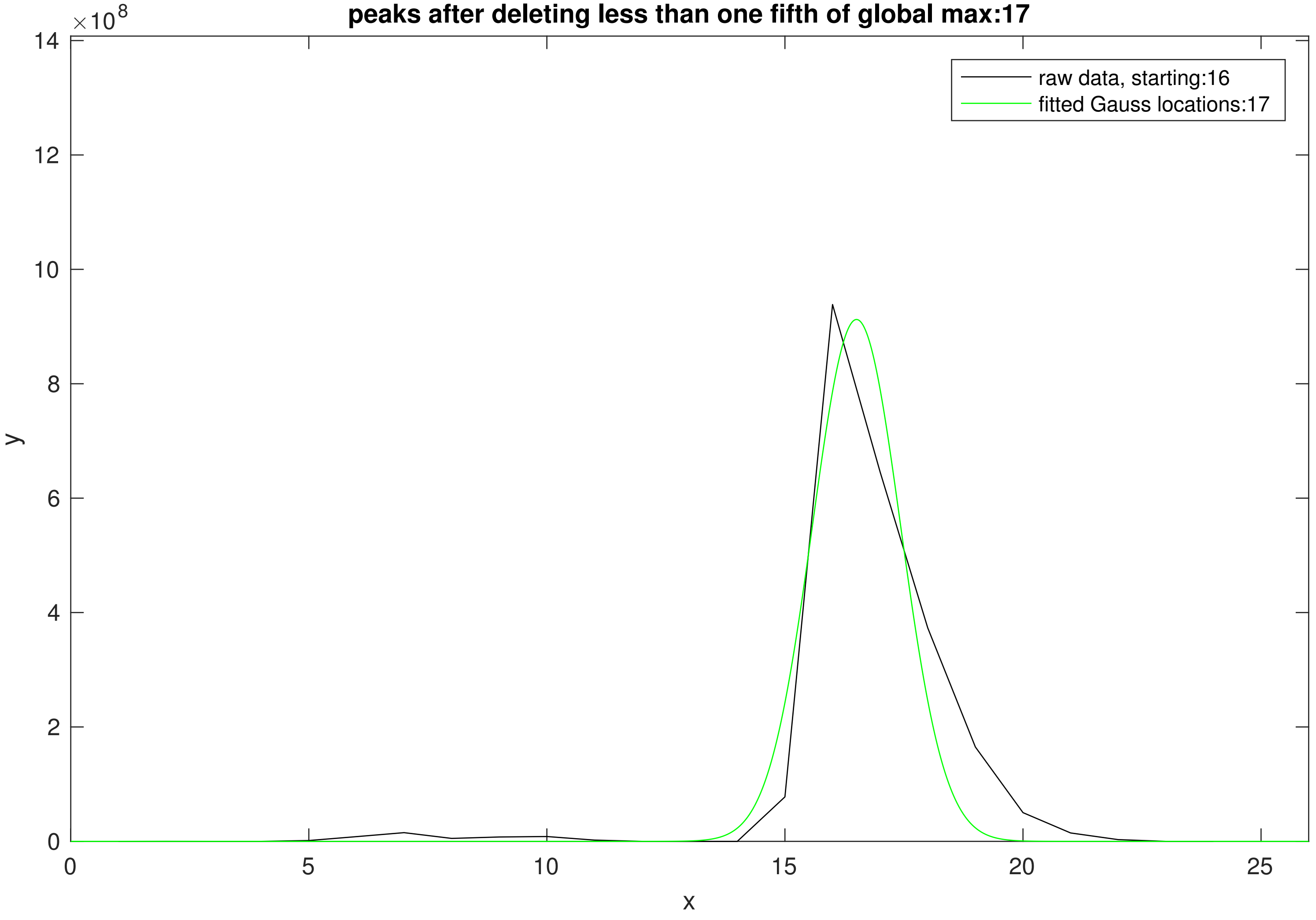

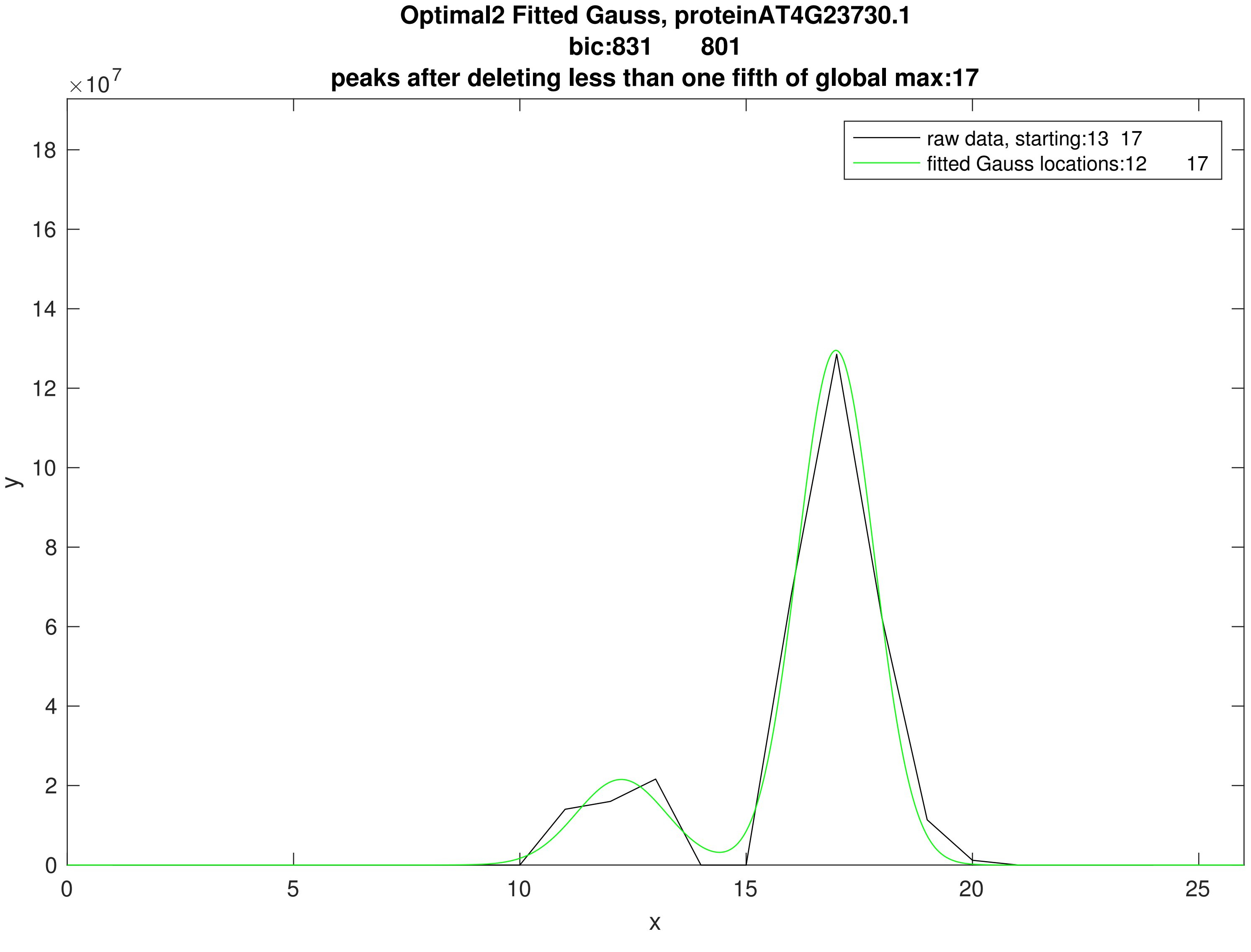
Optimal2 Fitted Gauss, proteinAT4G23730.1
bic:831 801
peaks after deleting less than one fifth of global max:17
×10^7
18
16
14
12
10
8
6
4
2
0
y
0
5
10
15
20
25
x
raw data, starting:13 17
fitted Gauss locations:12 17

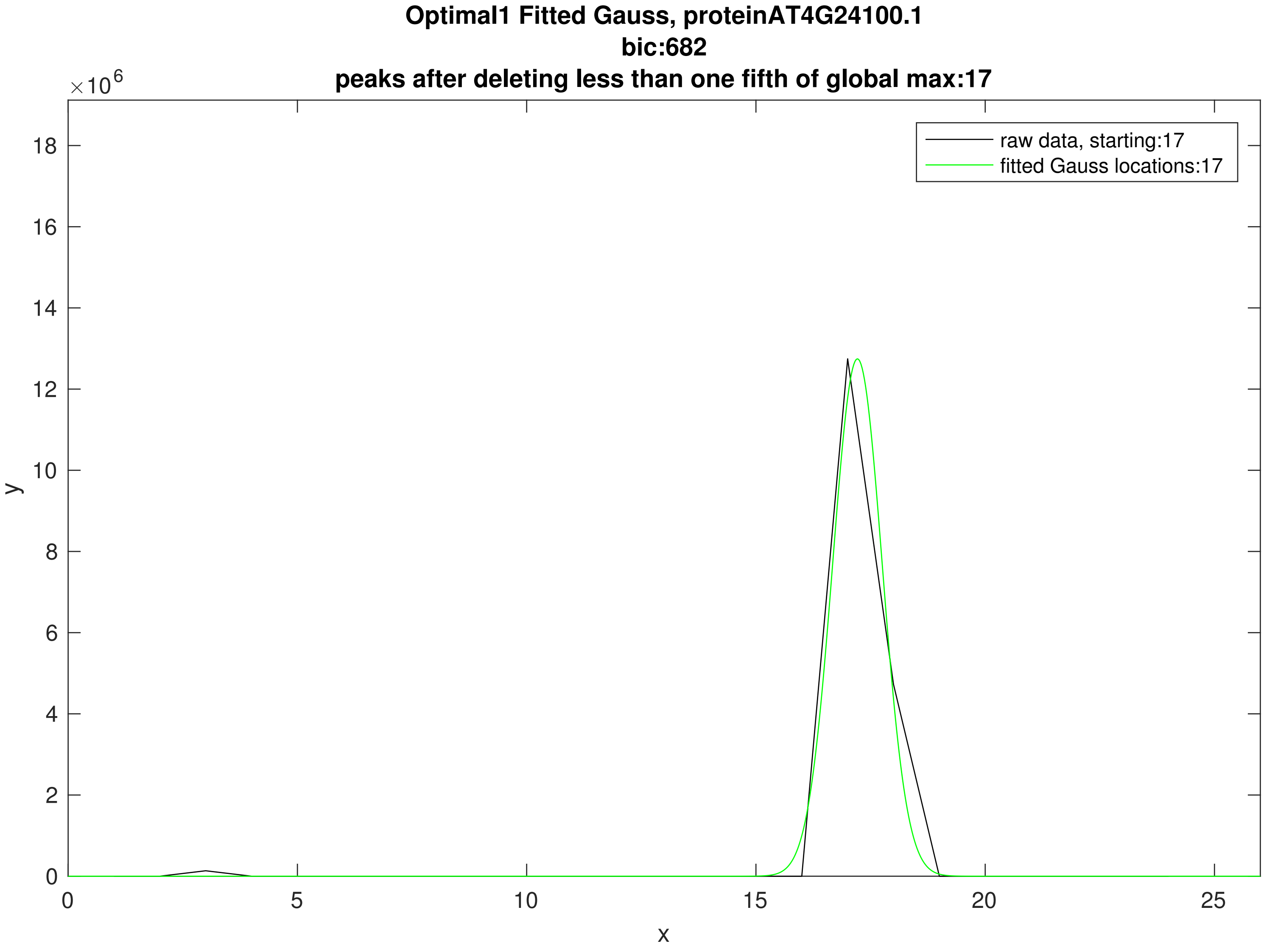

Optimal1 Fitted Gauss, proteinAT4G24100.1
bic:682
peaks after deleting less than one fifth of global max:17
×10^6
raw data, starting:17
fitted Gauss locations:17
y
x
0
2
4
6
8
10
12
14
16
18
5
15
20
25

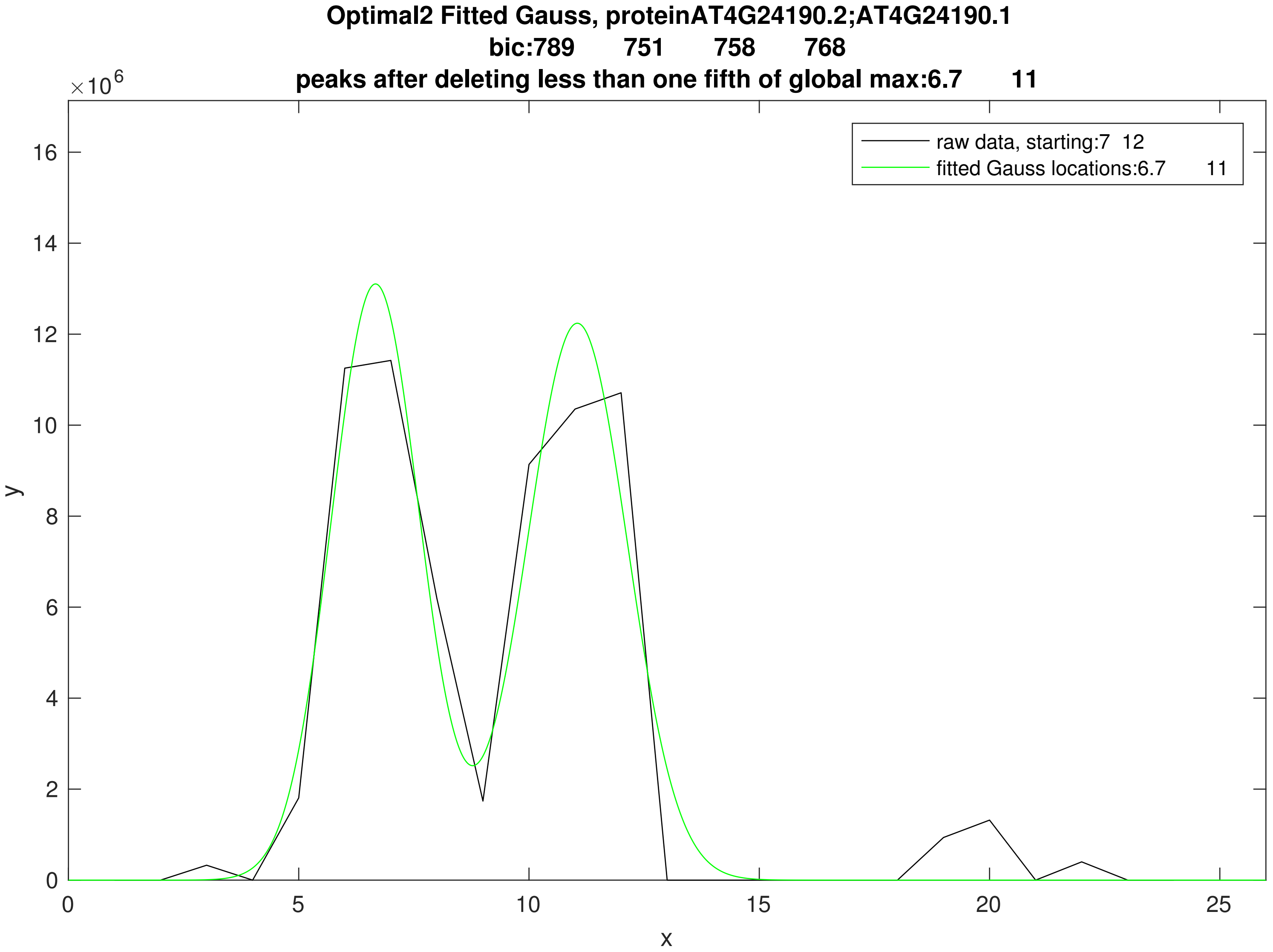
Optimal2 Fitted Gauss, proteinAT4G24190.2;AT4G24190.1
bic:789 751 758 768
peaks after deleting less than one fifth of global max:6.7 11
raw data, starting:7 12
fitted Gauss locations:6.7 11
×10^6
y
x
0
2
4
6
8
10
12
14
16
5
15
20
25

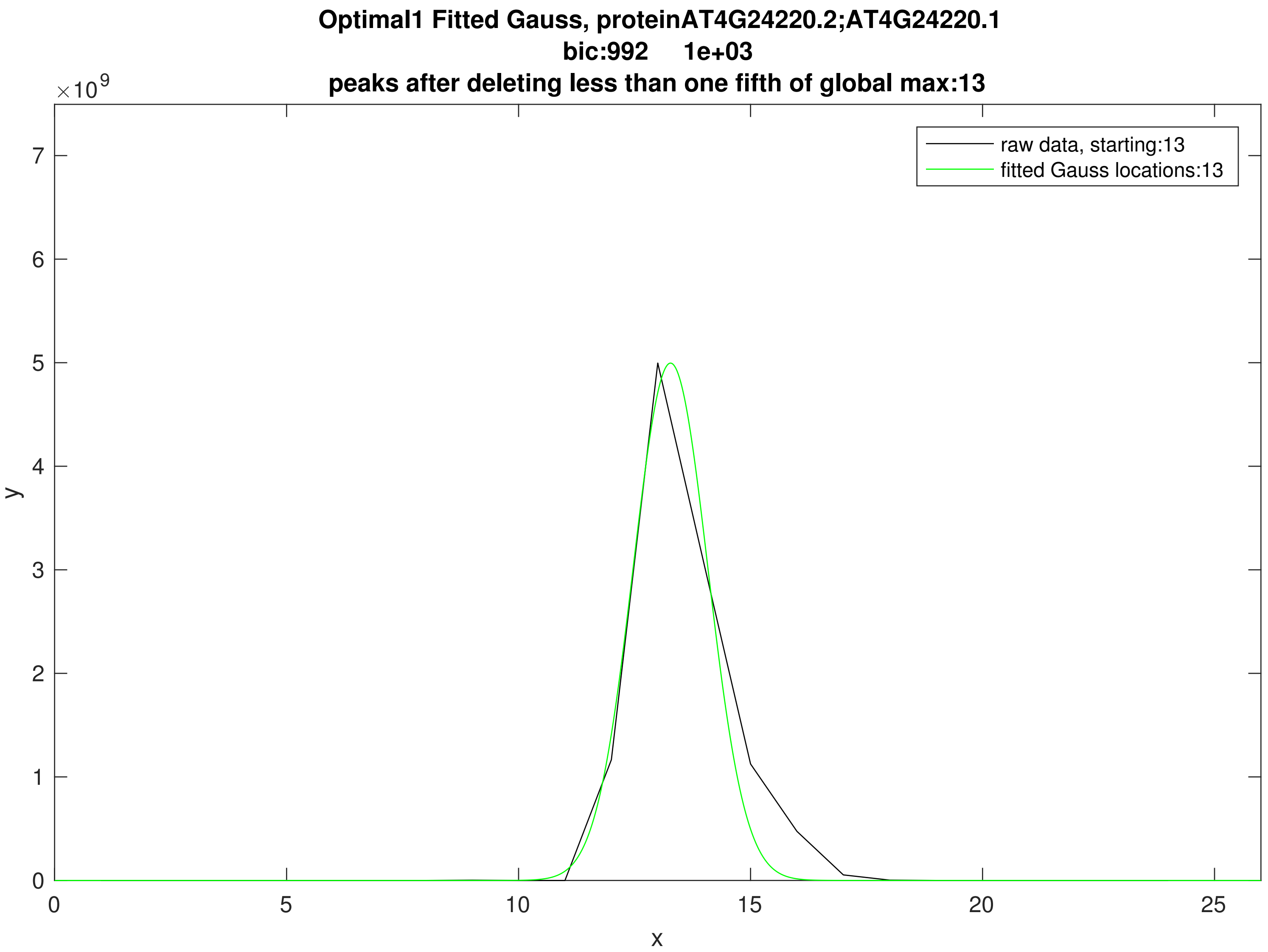

Optimal1 Fitted Gauss, proteinAT4G24220.2;AT4G24220.1
bic:992 1e+03
peaks after deleting less than one fifth of global max:13
×10^9
raw data, starting:13
fitted Gauss locations:13
y
x
0
1
2
3
4
5
6
7
0
5
10
15
20
25

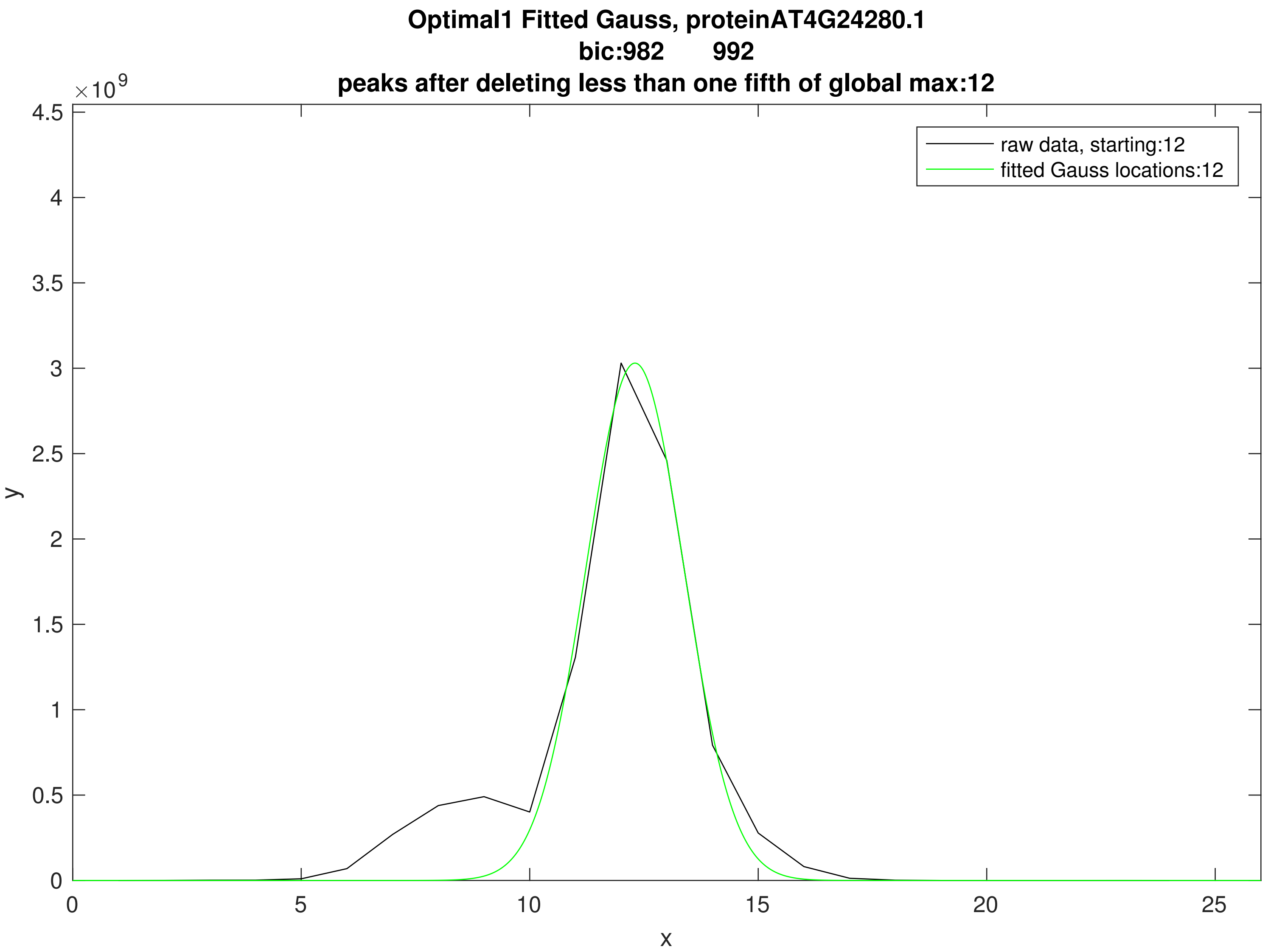

Optimal1 Fitted Gauss, proteinAT4G24280.1
bic:982 992
peaks after deleting less than one fifth of global max:12
×10^9
4.5
4
3.5
3
2.5
2
1.5
1
0.5
0
y
0
5
10
15
20
25
x
raw data, starting:12
fitted Gauss locations:12

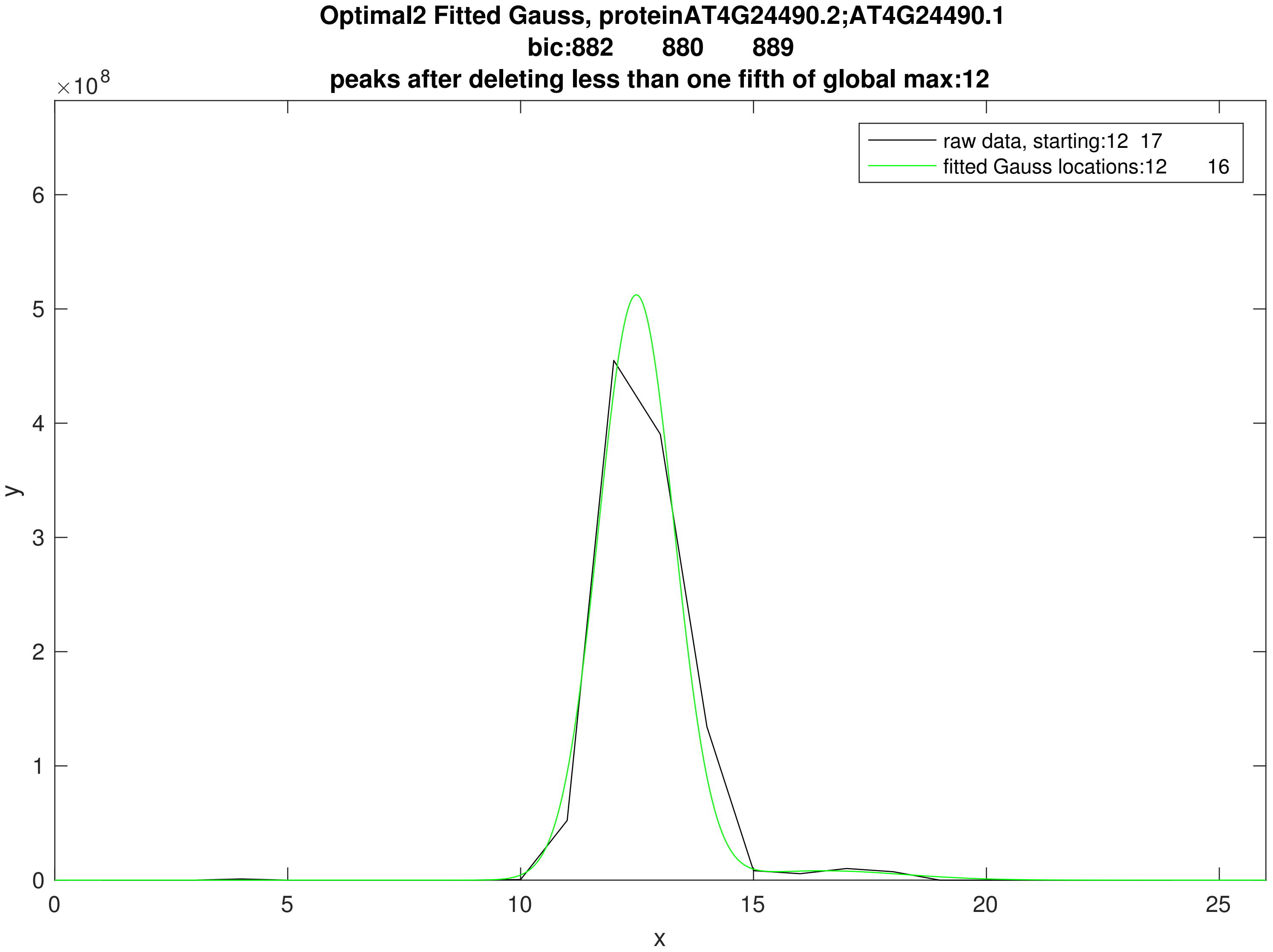
Optimal2 Fitted Gauss, proteinAT4G24490.2;AT4G24490.1
bic:882 880 889
peaks after deleting less than one fifth of global max:12
×10^8
raw data, starting:12 17
fitted Gauss locations:12 16
y
x
0
1
2
3
4
5
6
0
5
10
15
20
25

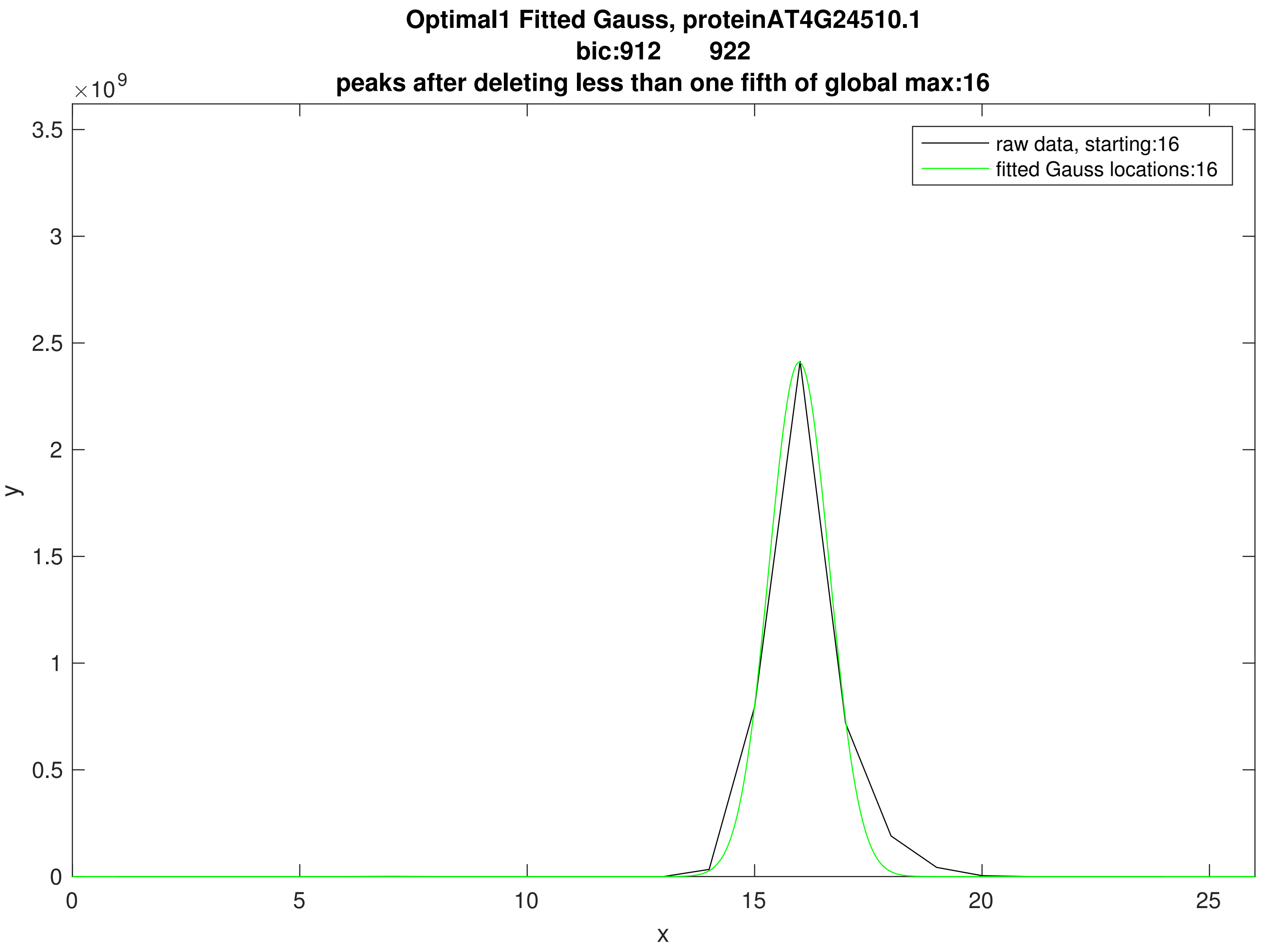
Optimal1 Fitted Gauss, proteinAT4G24510.1
bic:912 922
peaks after deleting less than one fifth of global max:16
×10^9
raw data, starting:16
fitted Gauss locations:16
y
x
0
0.5
1
1.5
2
2.5
3
3.5
0
5
10
15
20
25

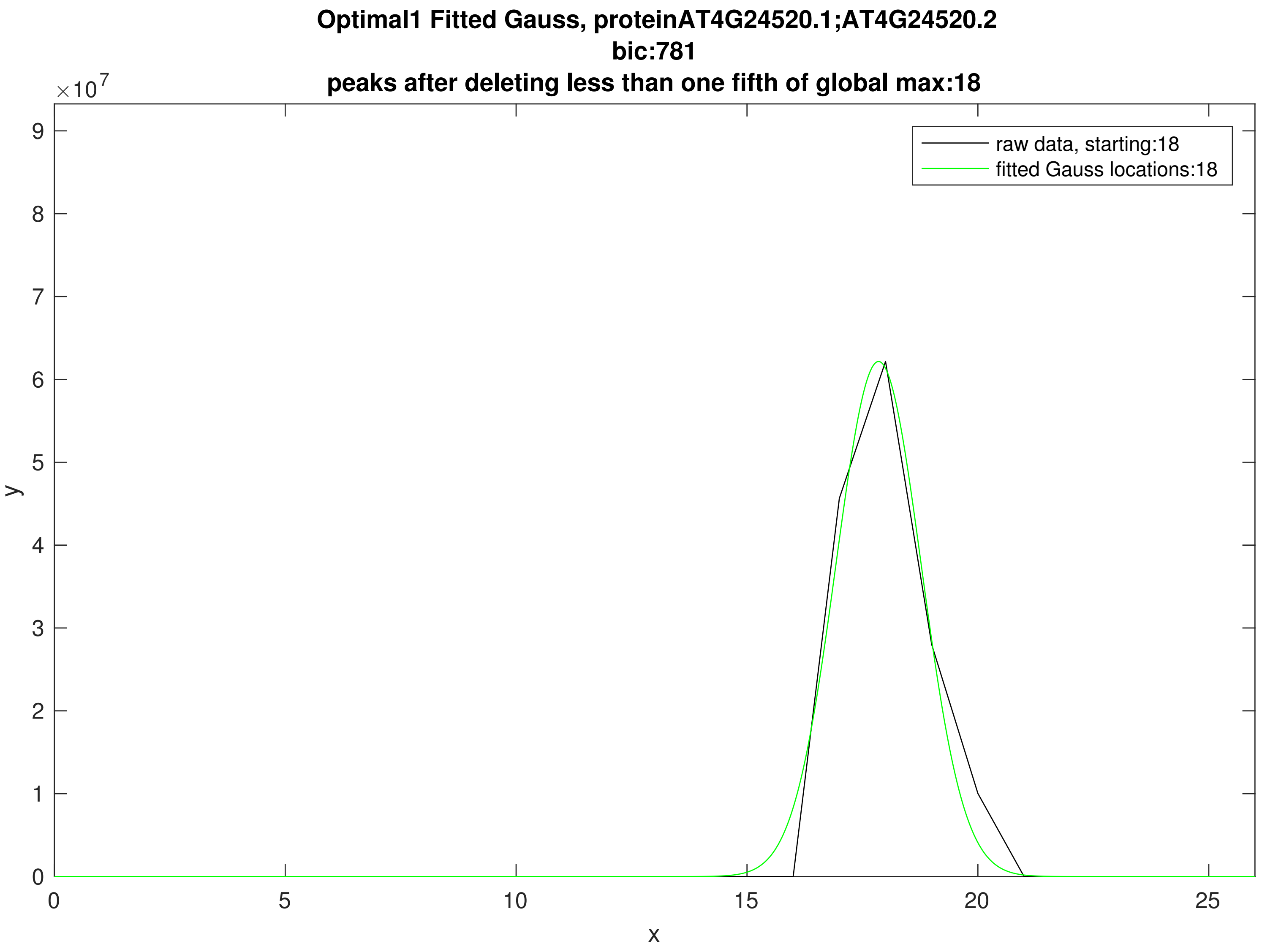
Optimal1 Fitted Gauss, proteinAT4G24520.1;AT4G24520.2
bic:781
peaks after deleting less than one fifth of global max:18
×10^7
9
8
7
6
5
4
3
2
1
0
y
0
5
10
15
20
25
x
raw data, starting:18
fitted Gauss locations:18

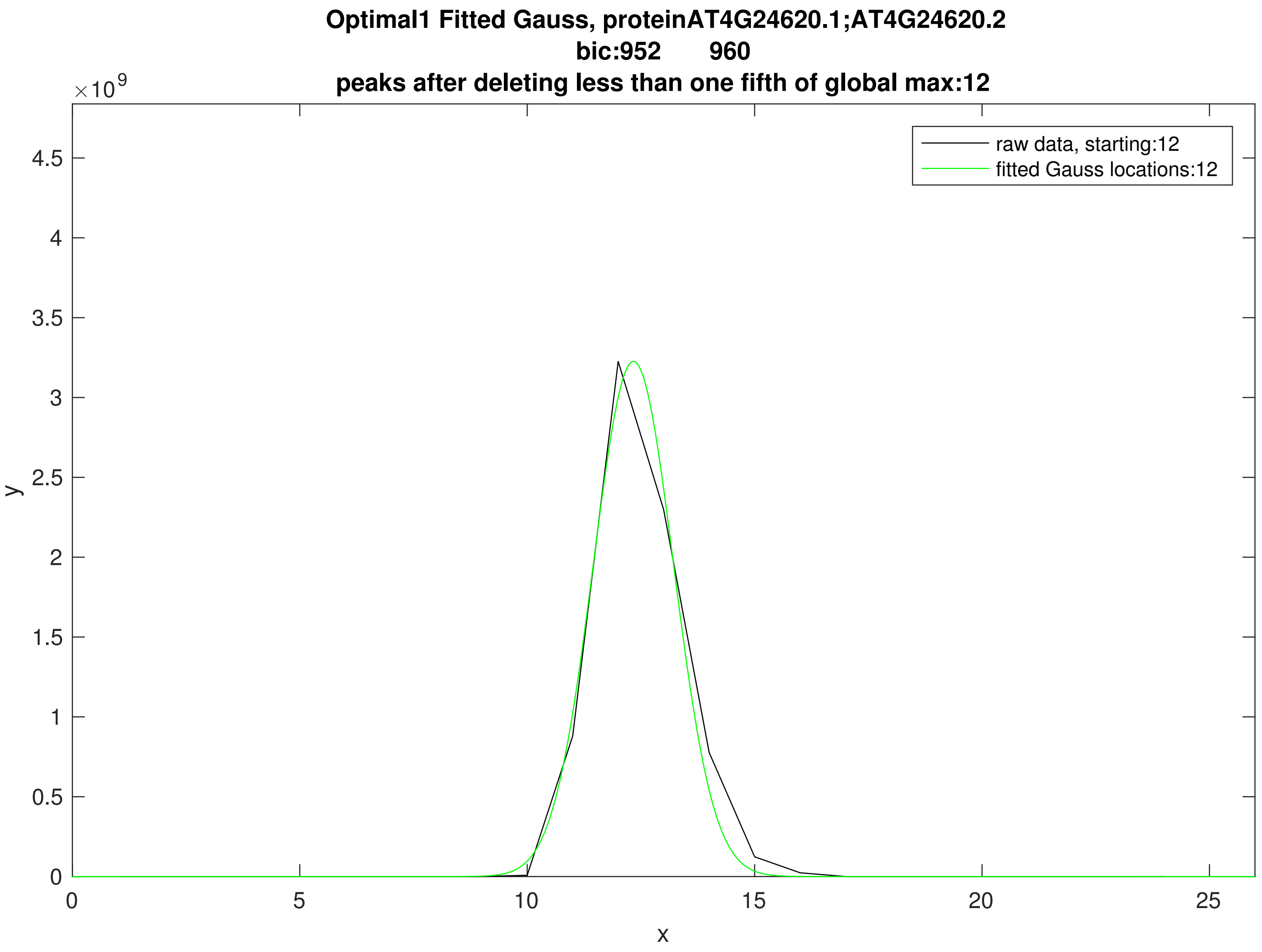
Optimal1 Fitted Gauss, proteinAT4G24620.1;AT4G24620.2
bic:952 960
peaks after deleting less than one fifth of global max:12
×10^9
4.5
4
3.5
3
2.5
2
1.5
1
0.5
0
y
0
5
10
15
20
25
x
raw data, starting:12
fitted Gauss locations:12

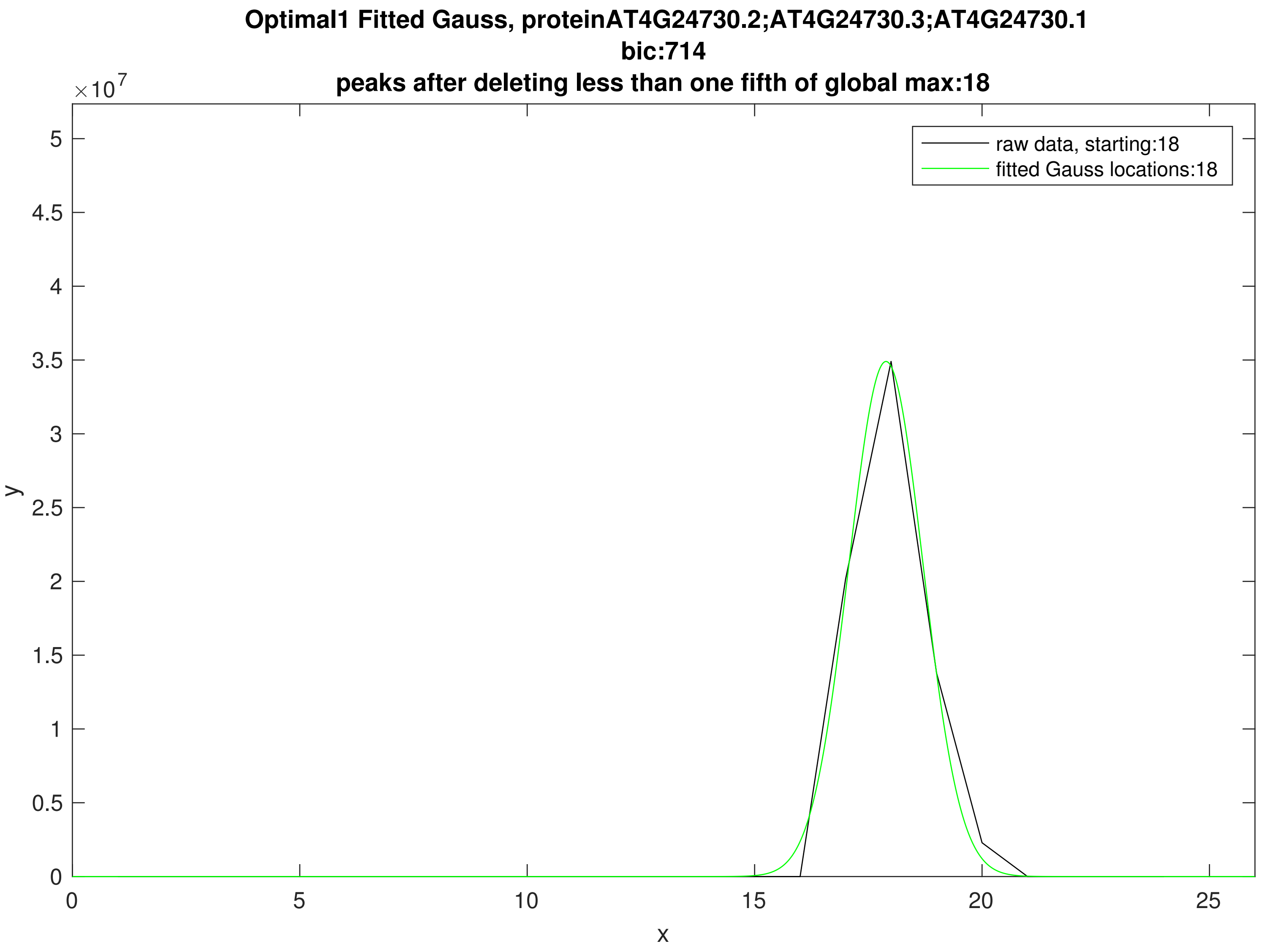
Optimal1 Fitted Gauss, proteinAT4G24730.2;AT4G24730.3;AT4G24730.1
bic:714
peaks after deleting less than one fifth of global max:18
×10^7
raw data, starting:18
fitted Gauss locations:18
y
x
0
0.5
1
1.5
2
2.5
3
3.5
4
4.5
5
0
5
10
15
20
25

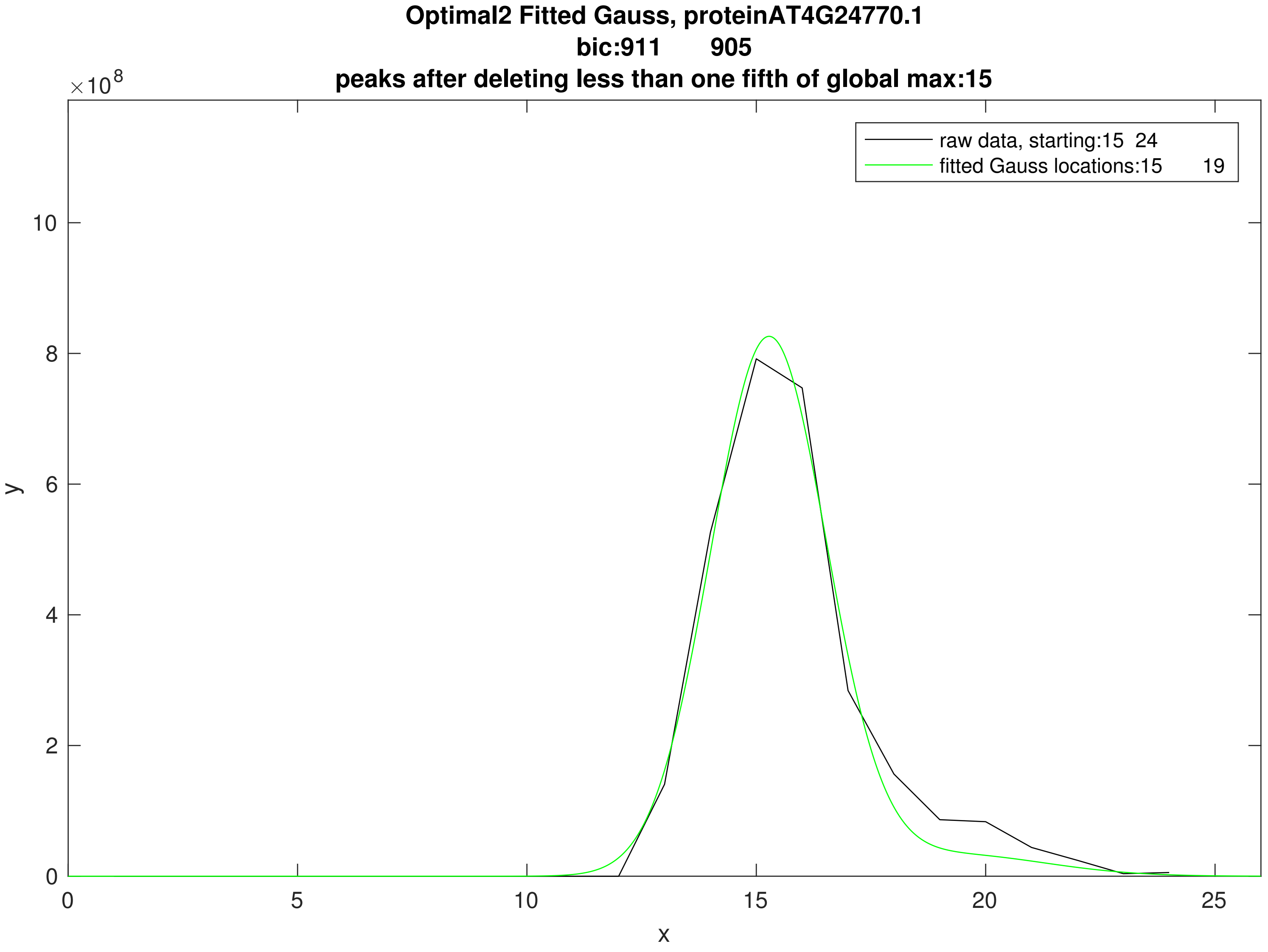
Optimal2 Fitted Gauss, proteinAT4G24770.1
bic:911 905
peaks after deleting less than one fifth of global max:15
×10^8
raw data, starting:15 24
fitted Gauss locations:15 19
y
x
0
2
4
6
8
10
0
5
10
15
20
25

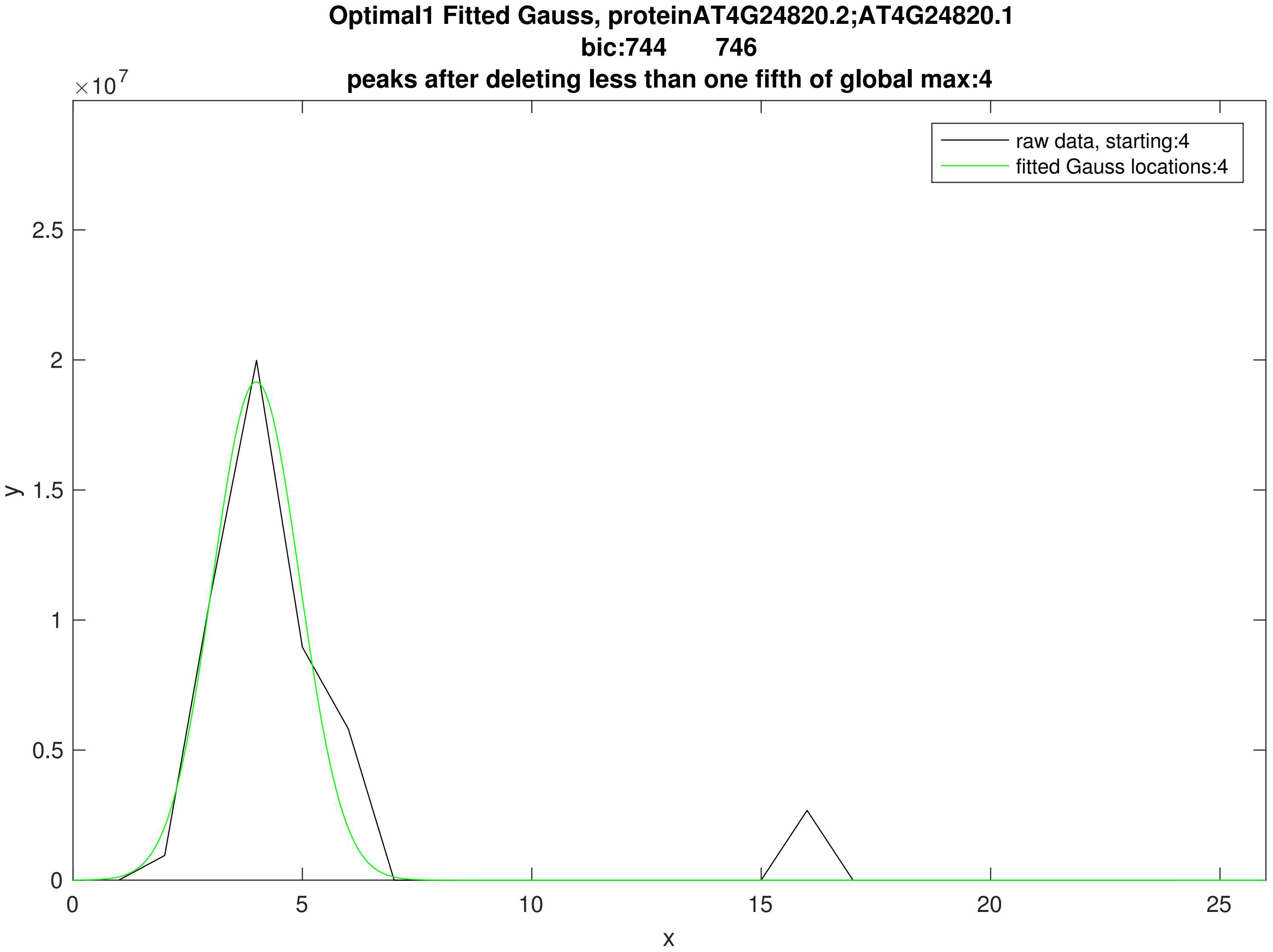

Optimal1 Fitted Gauss, proteinAT4G24820.2;AT4G24820.1
bic:744 746
peaks after deleting less than one fifth of global max:4
×10^7
raw data, starting:4
fitted Gauss locations:4
y
x
0
0.5
1
1.5
2
2.5
0
5
10
15
20
25

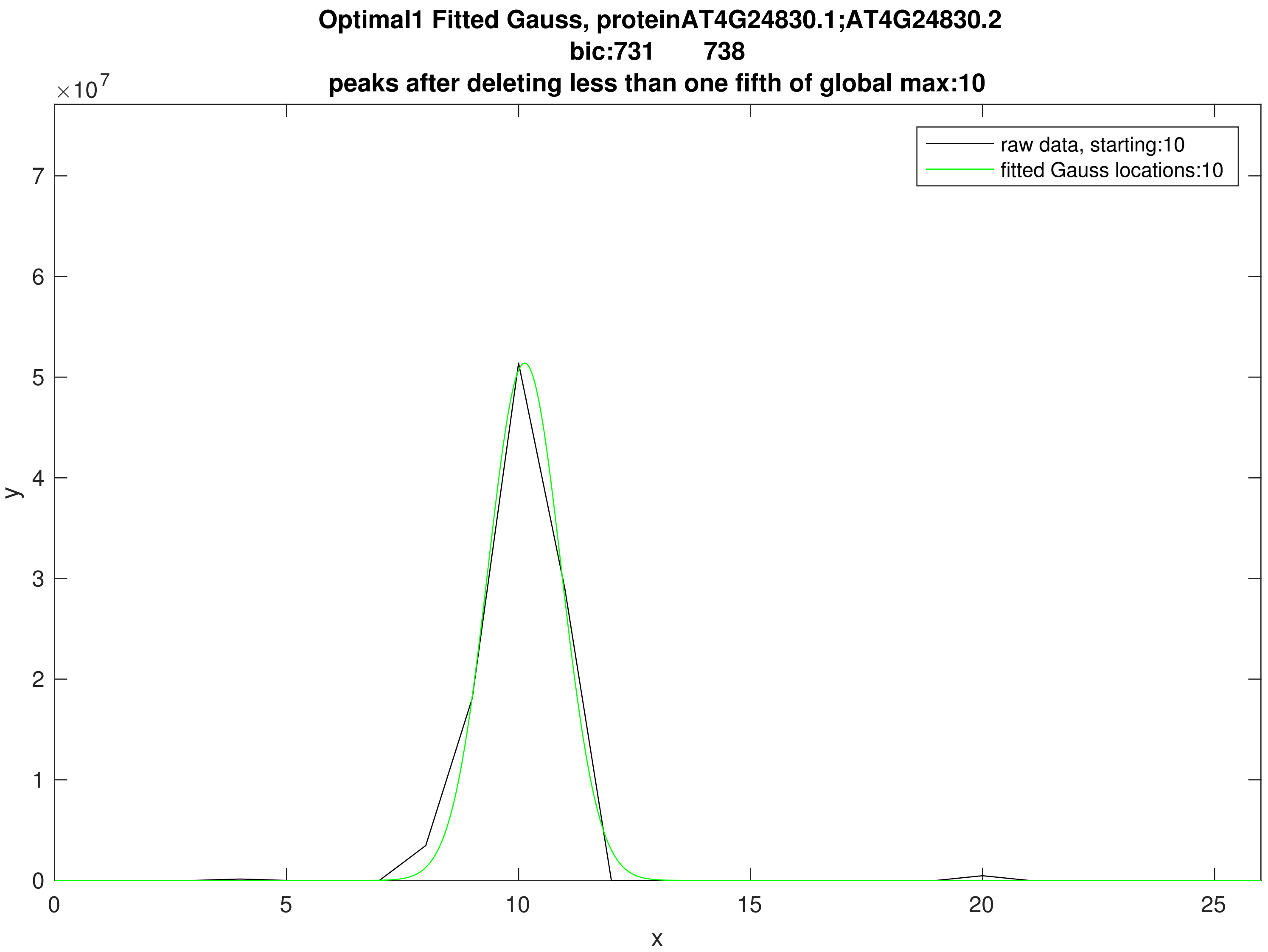

Optimal1 Fitted Gauss, proteinAT4G24830.1;AT4G24830.2
bic:731 738
peaks after deleting less than one fifth of global max:10
×10^7
raw data, starting:10
fitted Gauss locations:10
y
x
0
1
2
3
4
5
6
7
10
15
20
25

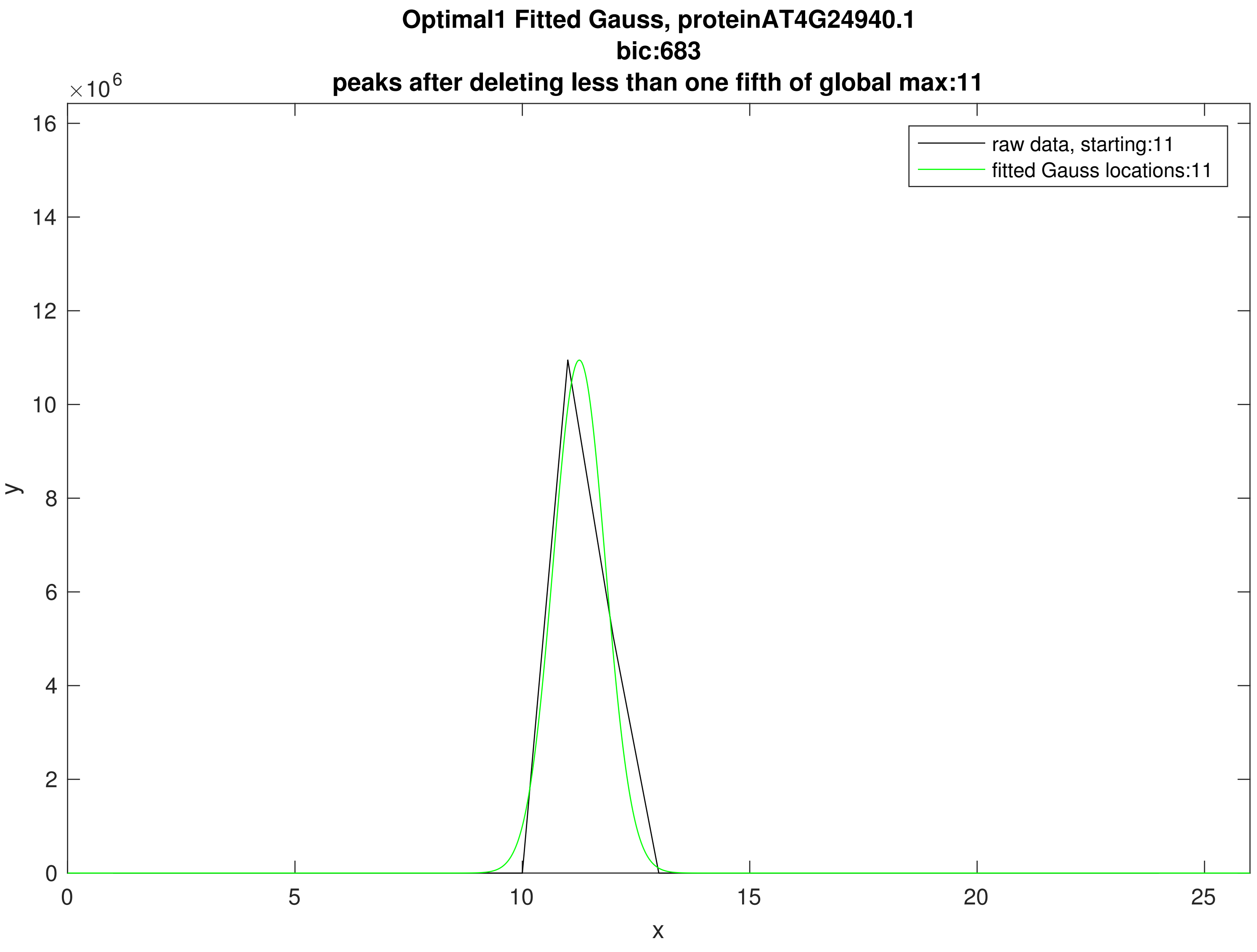

Optimal1 Fitted Gauss, proteinAT4G24940.1
bic:683
peaks after deleting less than one fifth of global max:11
×10^6
16
14
12
10
8
6
4
2
0
y
0
5
10
15
20
25
x
raw data, starting:11
fitted Gauss locations:11

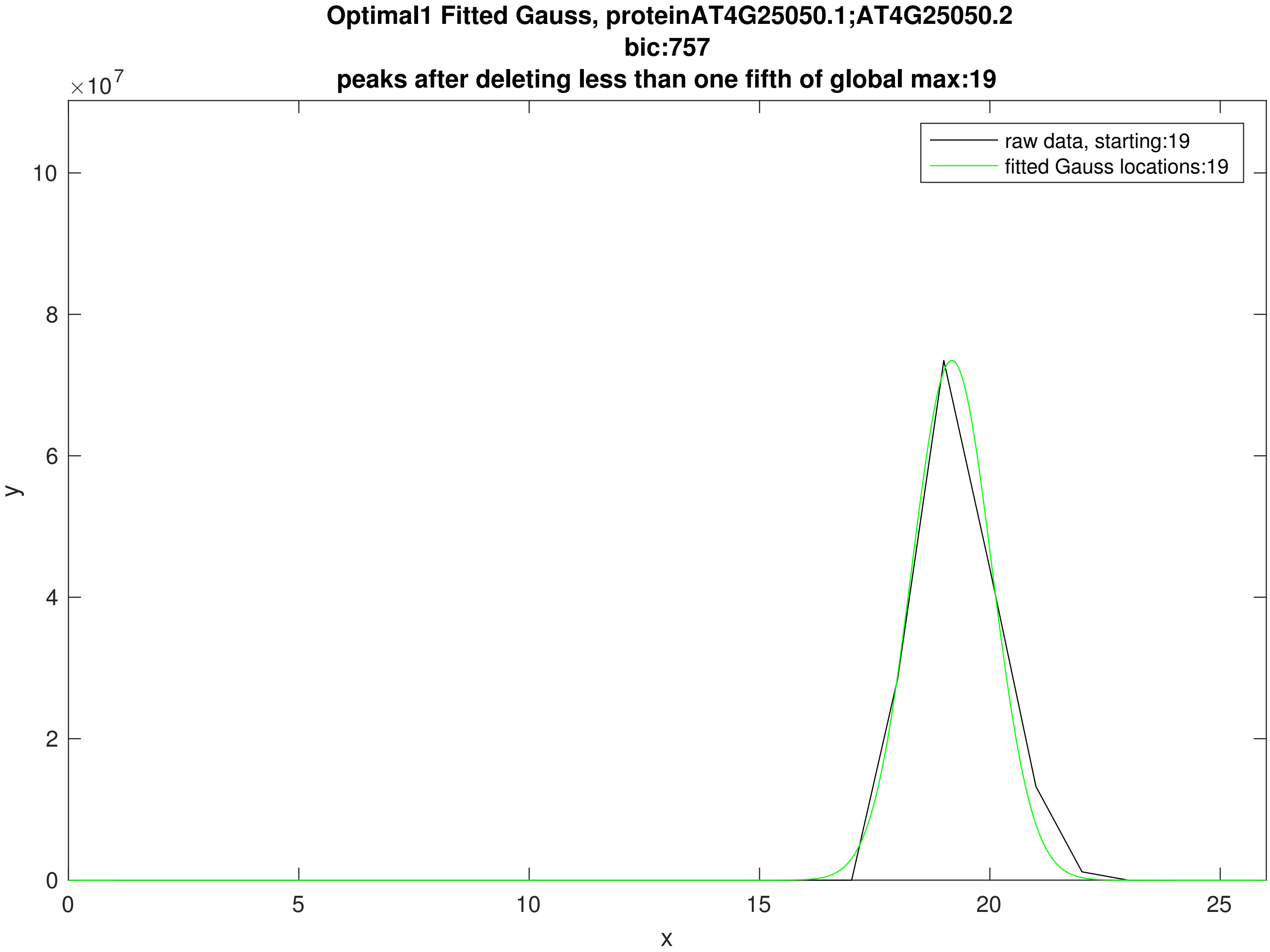

Optimal1 Fitted Gauss, proteinAT4G25050.1;AT4G25050.2
bic:757
peaks after deleting less than one fifth of global max:19
×10^7
raw data, starting:19
fitted Gauss locations:19
y
x
0
2
4
6
8
10
0
5
10
15
20
25

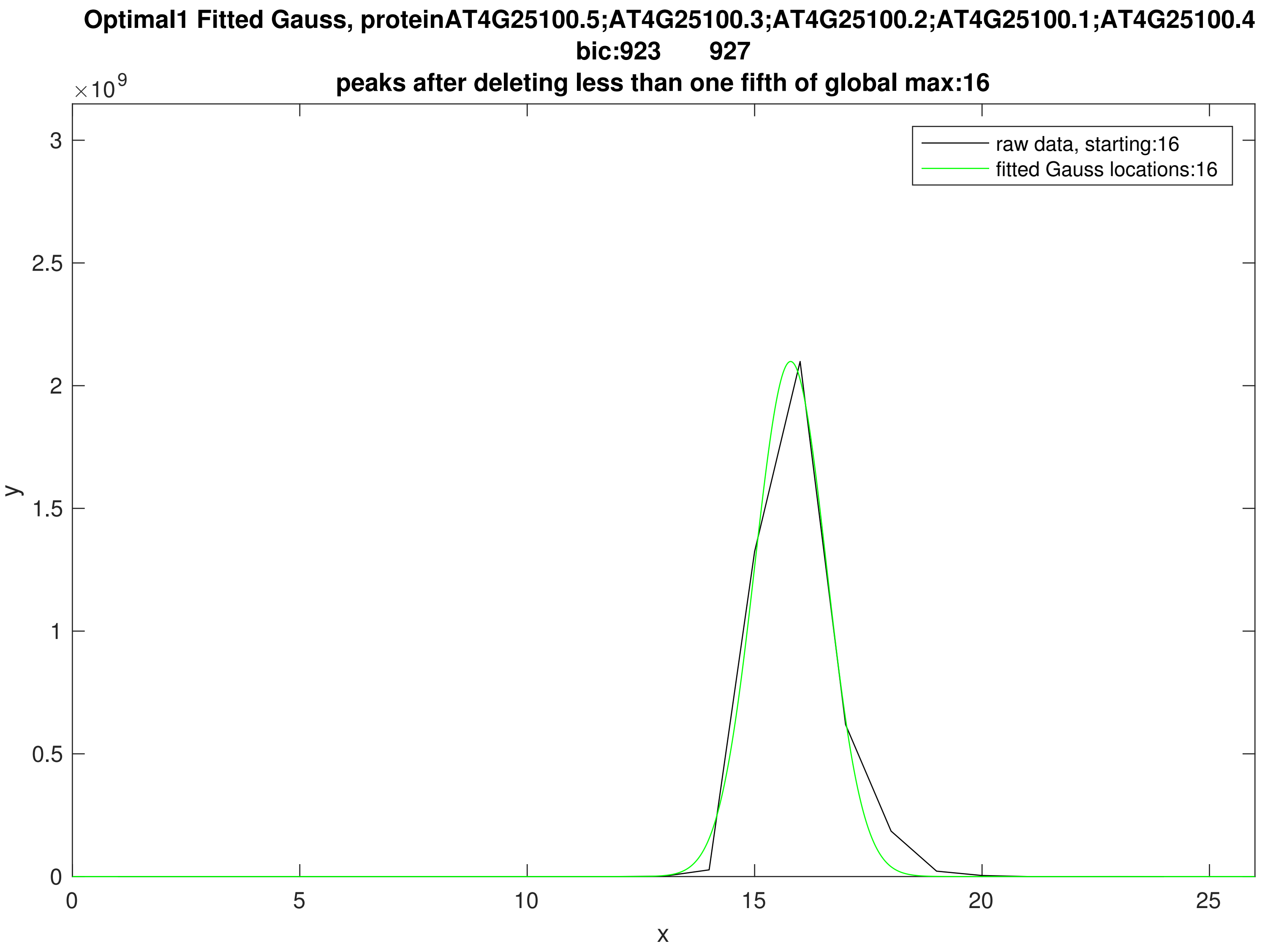
Optimal1 Fitted Gauss, proteinAT4G25100.5;AT4G25100.3;AT4G25100.2;AT4G25100.1;AT4G25100.4
bic:923 927
peaks after deleting less than one fifth of global max:16
×10^9
raw data, starting:16
fitted Gauss locations:16
y
x
0
0.5
1
1.5
2
2.5
3
5
10
15
20
25

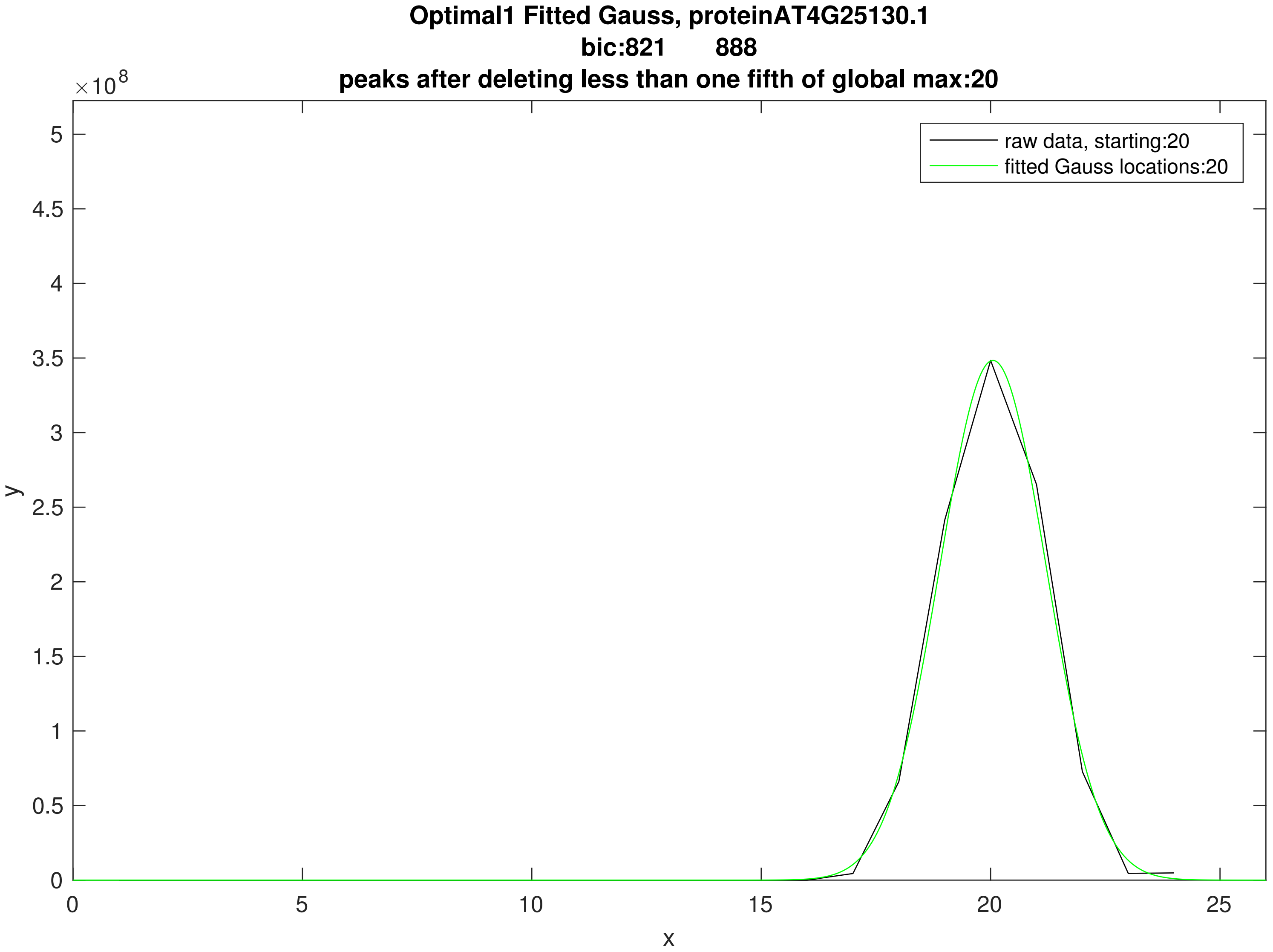
Optimal1 Fitted Gauss, proteinAT4G25130.1
bic:821 888
peaks after deleting less than one fifth of global max:20
×10^8
5
4.5
4
3.5
3
2.5
2
1.5
1
0.5
0
y
0
5
10
15
20
25
x
raw data, starting:20
fitted Gauss locations:20

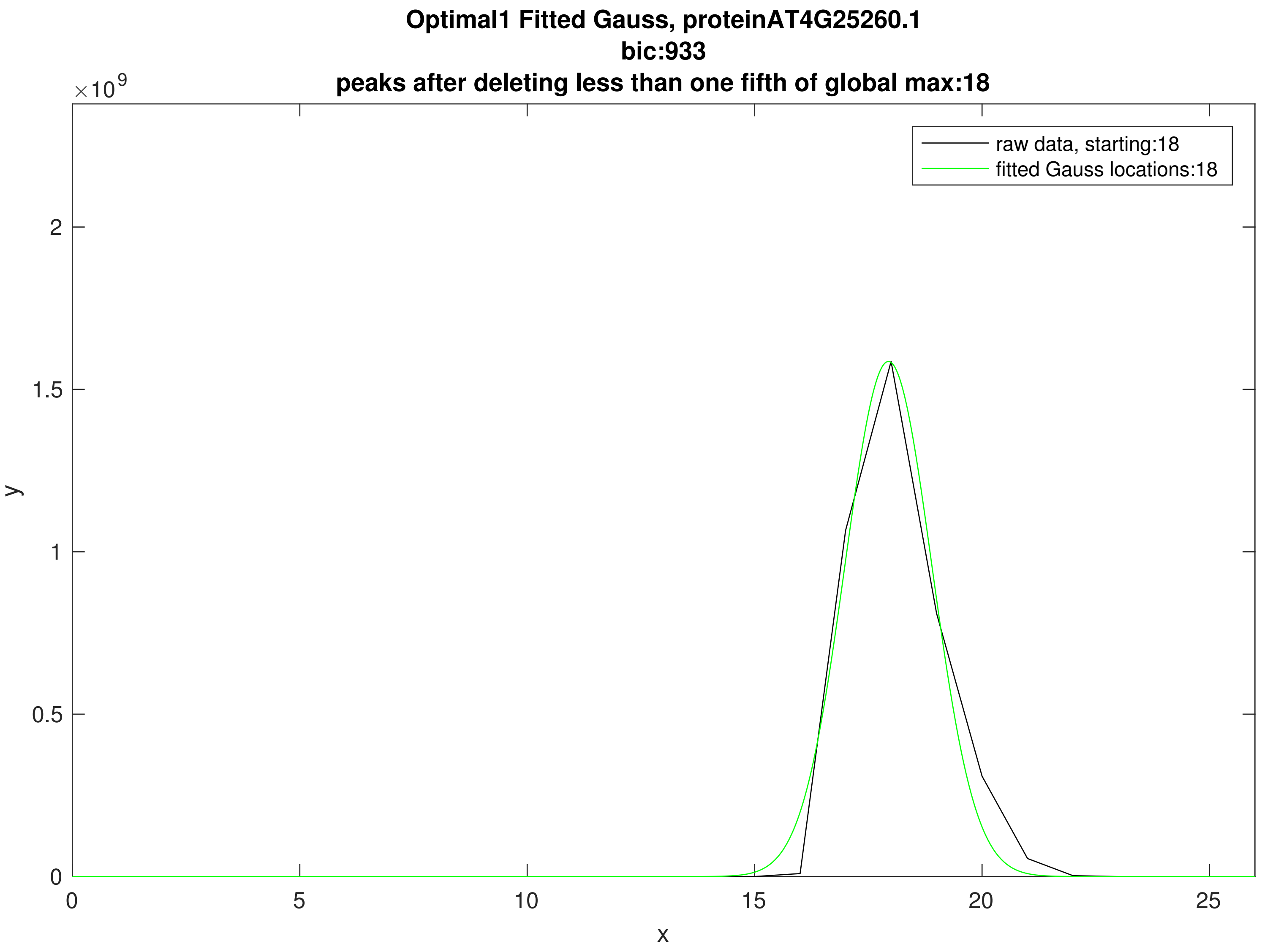
Optimal1 Fitted Gauss, proteinAT4G25260.1
bic:933
peaks after deleting less than one fifth of global max:18
×10^9
raw data, starting:18
fitted Gauss locations:18
y
x
0
0.5
1
1.5
2
0
5
10
15
20
25

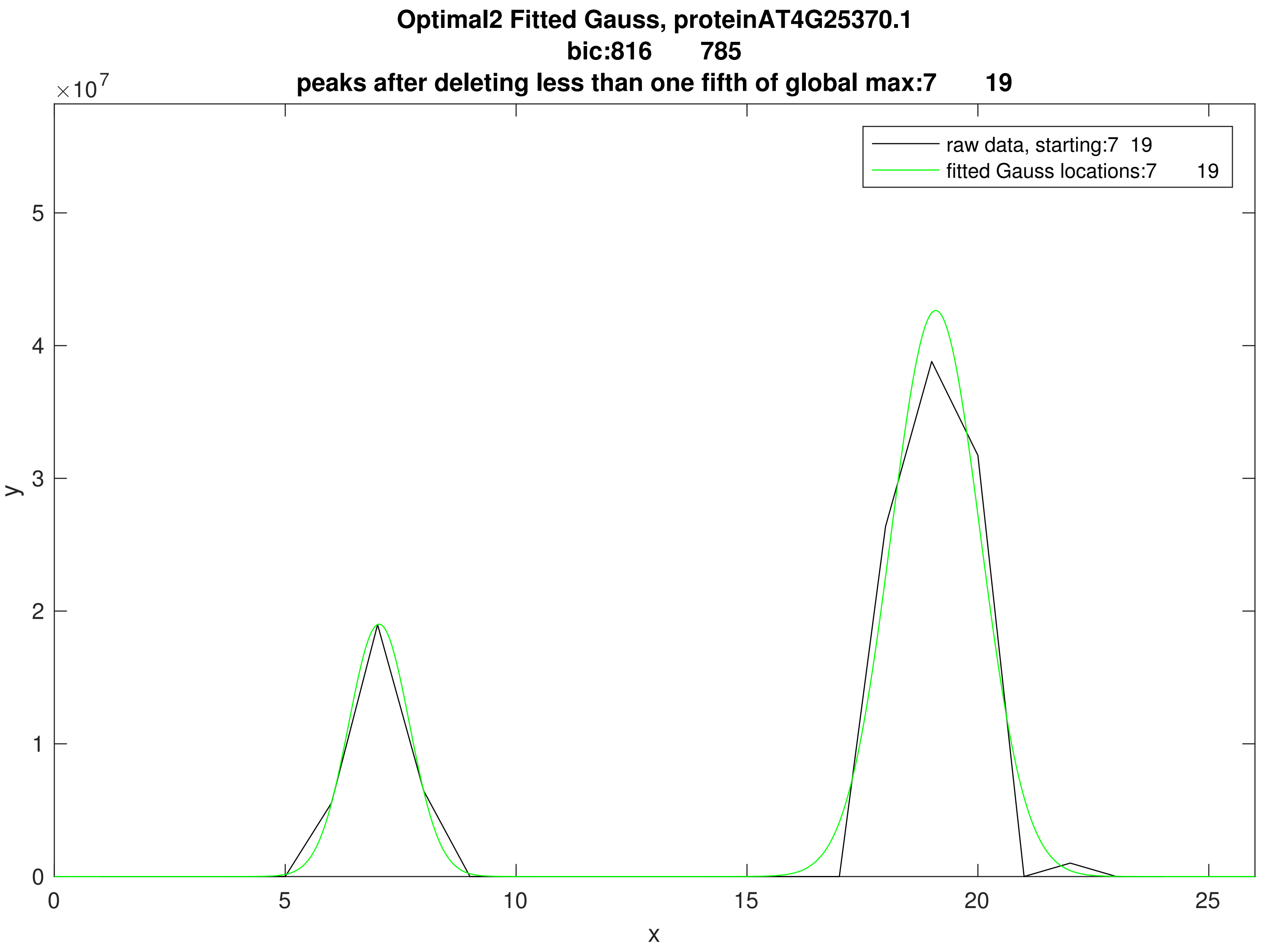
Optimal2 Fitted Gauss, proteinAT4G25370.1
bic:816 785
peaks after deleting less than one fifth of global max:7 19
×10^7
raw data, starting:7 19
fitted Gauss locations:7 19
y
x
0
1
2
3
4
5
0
5
10
15
20
25

**Optimal1 Fitted Gauss, proteinAT4G25810.1**
**bic:767**
**peaks after deleting less than one fifth of global max:19**

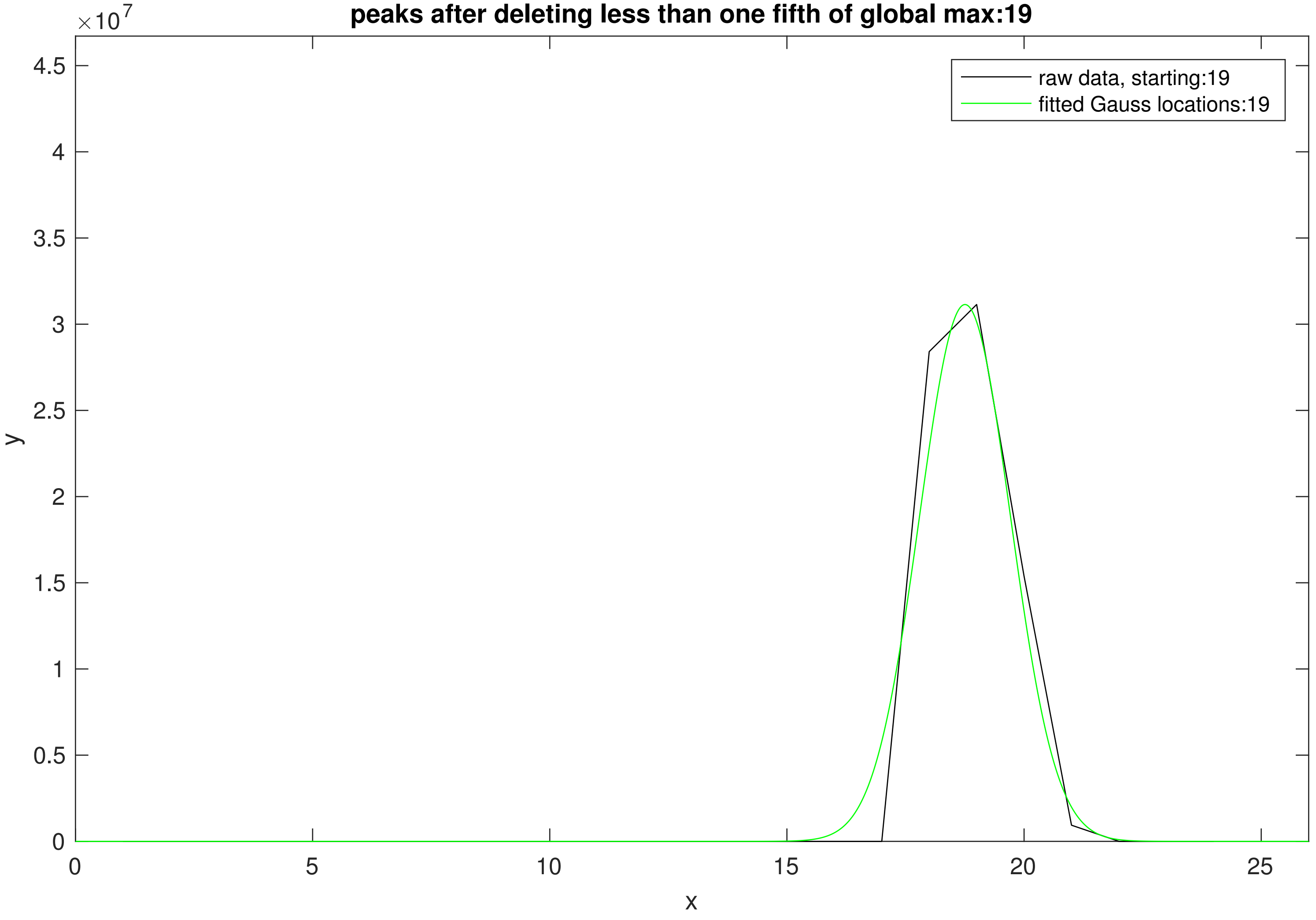

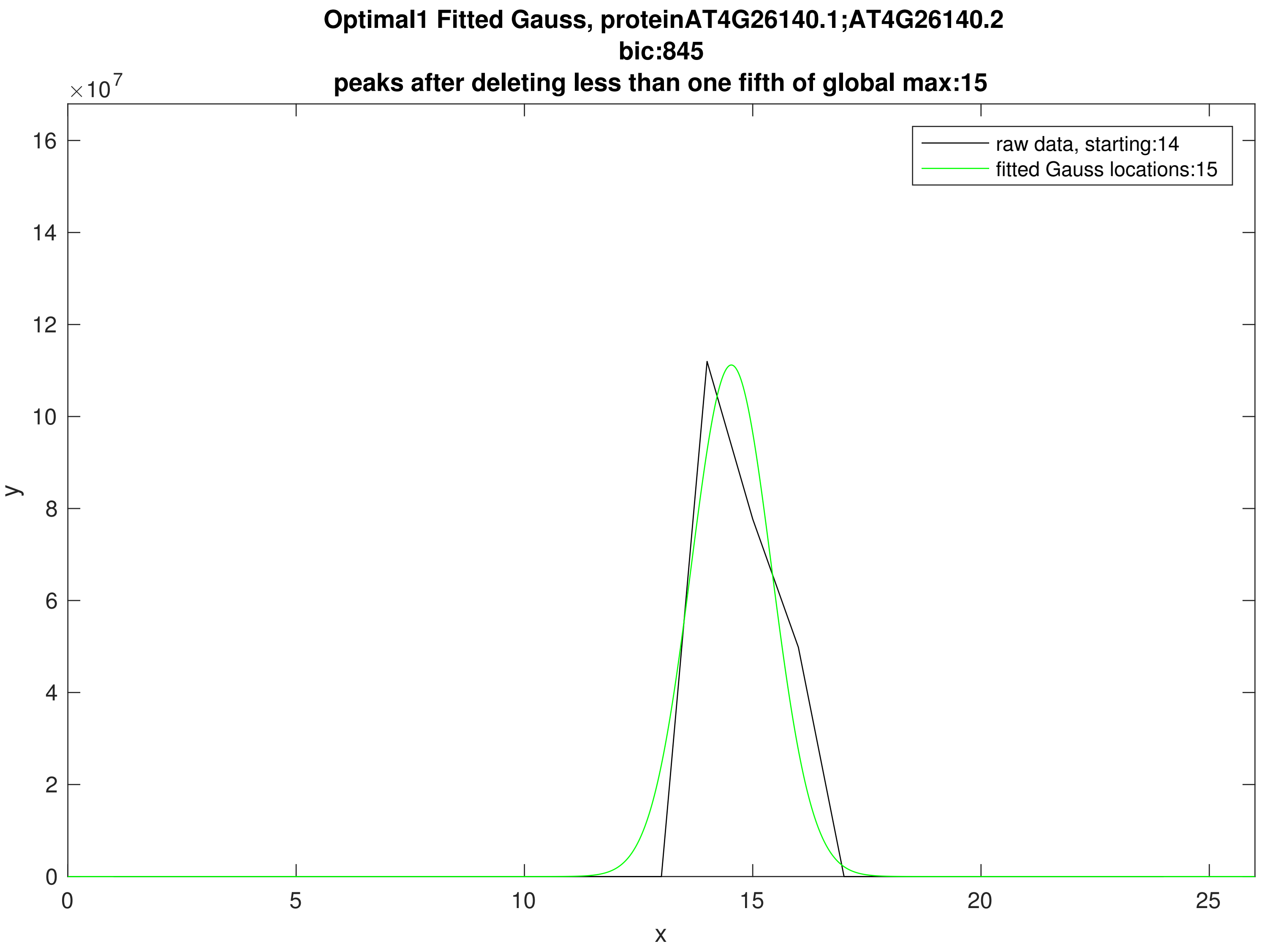

Optimal1 Fitted Gauss, proteinAT4G26140.1;AT4G26140.2
bic:845
peaks after deleting less than one fifth of global max:15
×10^7
raw data, starting:14
fitted Gauss locations:15
y
x
0
2
4
6
8
10
12
14
16
0
5
10
15
20
25

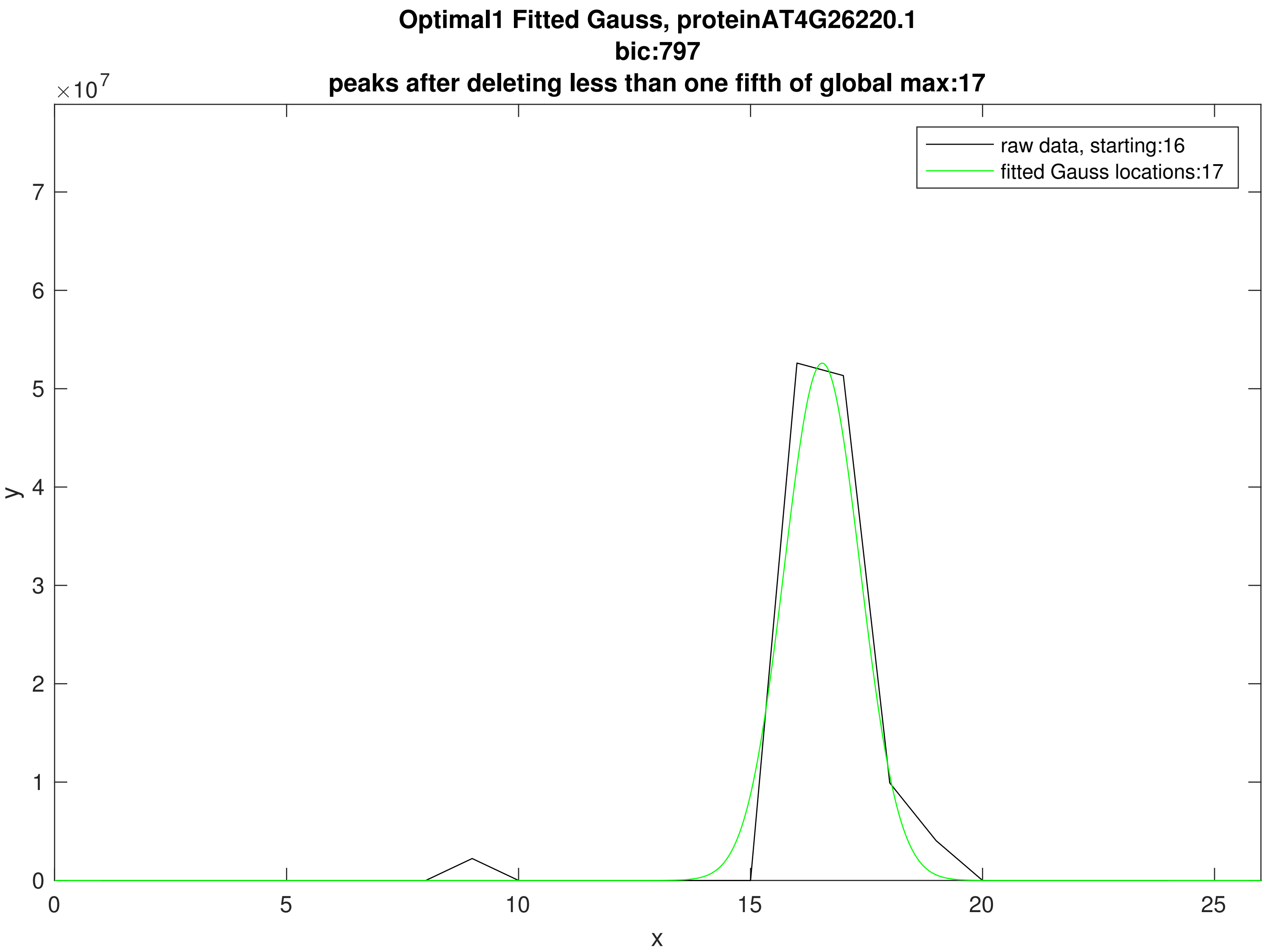
Optimal1 Fitted Gauss, proteinAT4G26220.1
bic:797
peaks after deleting less than one fifth of global max:17
×10^7
raw data, starting:16
fitted Gauss locations:17
y
x
0
1
2
3
4
5
6
7
0
5
10
15
20
25

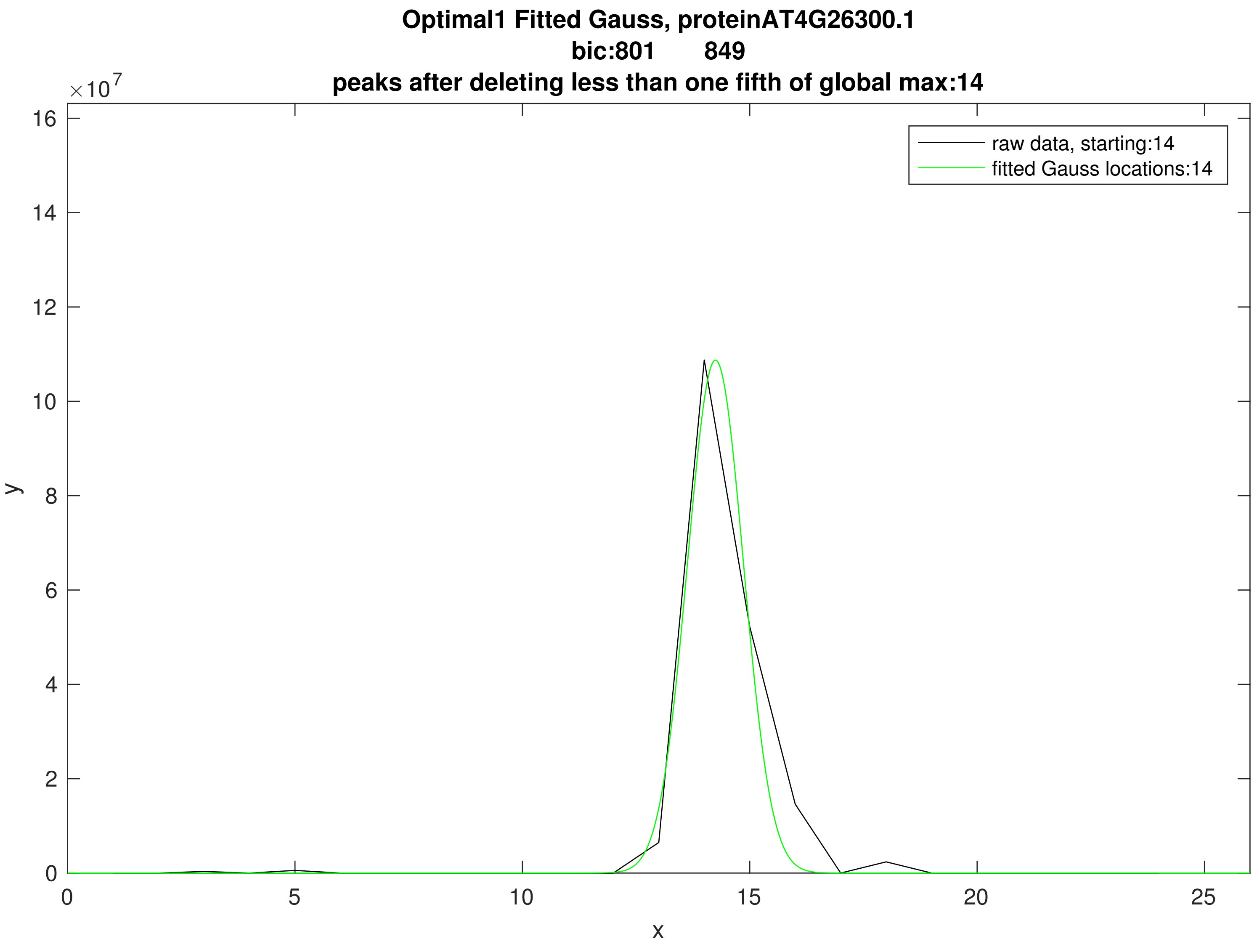
Optimal1 Fitted Gauss, proteinAT4G26300.1
bic:801 849
peaks after deleting less than one fifth of global max:14
×10^7
16
14
12
10
8
6
4
2
0
y
0
5
10
15
20
25
x
raw data, starting:14
fitted Gauss locations:14

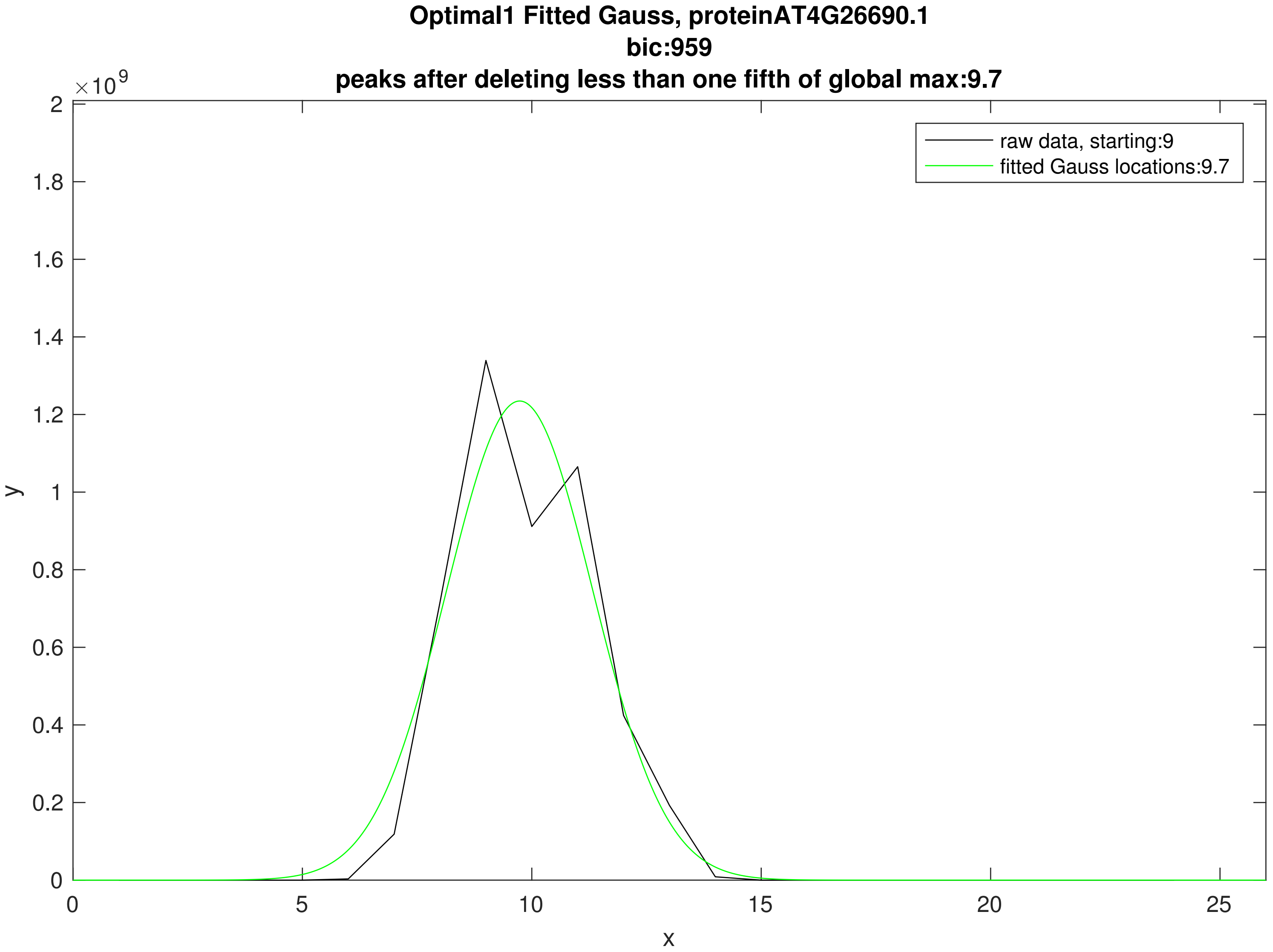

Optimal1 Fitted Gauss, proteinAT4G26690.1
bic:959
peaks after deleting less than one fifth of global max:9.7
×10^9
raw data, starting:9
fitted Gauss locations:9.7
y
x
0
0.2
0.4
0.6
0.8
1
1.2
1.4
1.6
1.8
2
0
5
10
15
20
25

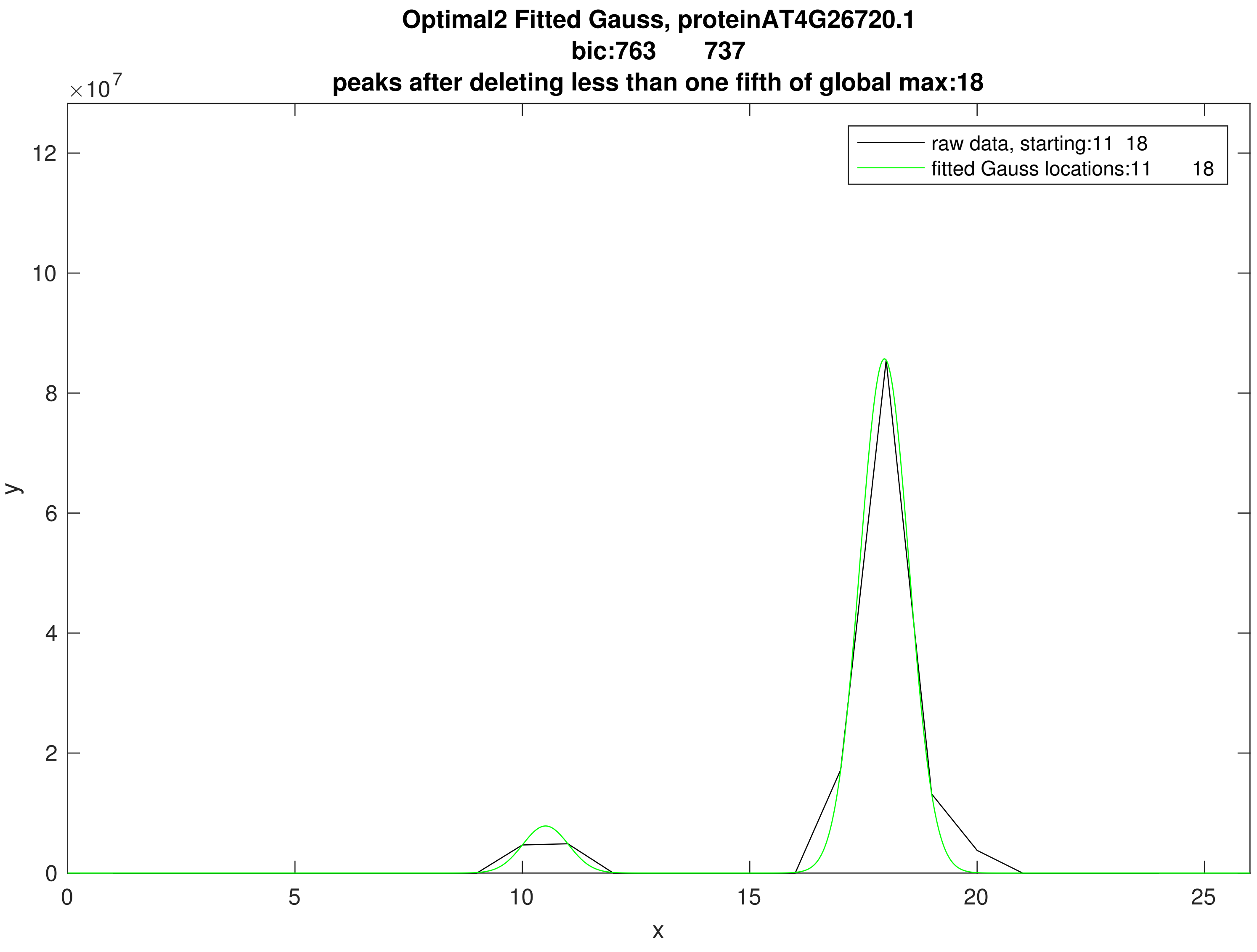

Optimal2 Fitted Gauss, proteinAT4G26720.1
bic:763 737
peaks after deleting less than one fifth of global max:18
raw data, starting:11 18
fitted Gauss locations:11 18
×10^7
y
x
0
2
4
6
8
10
12
0
5
10
15
20
25

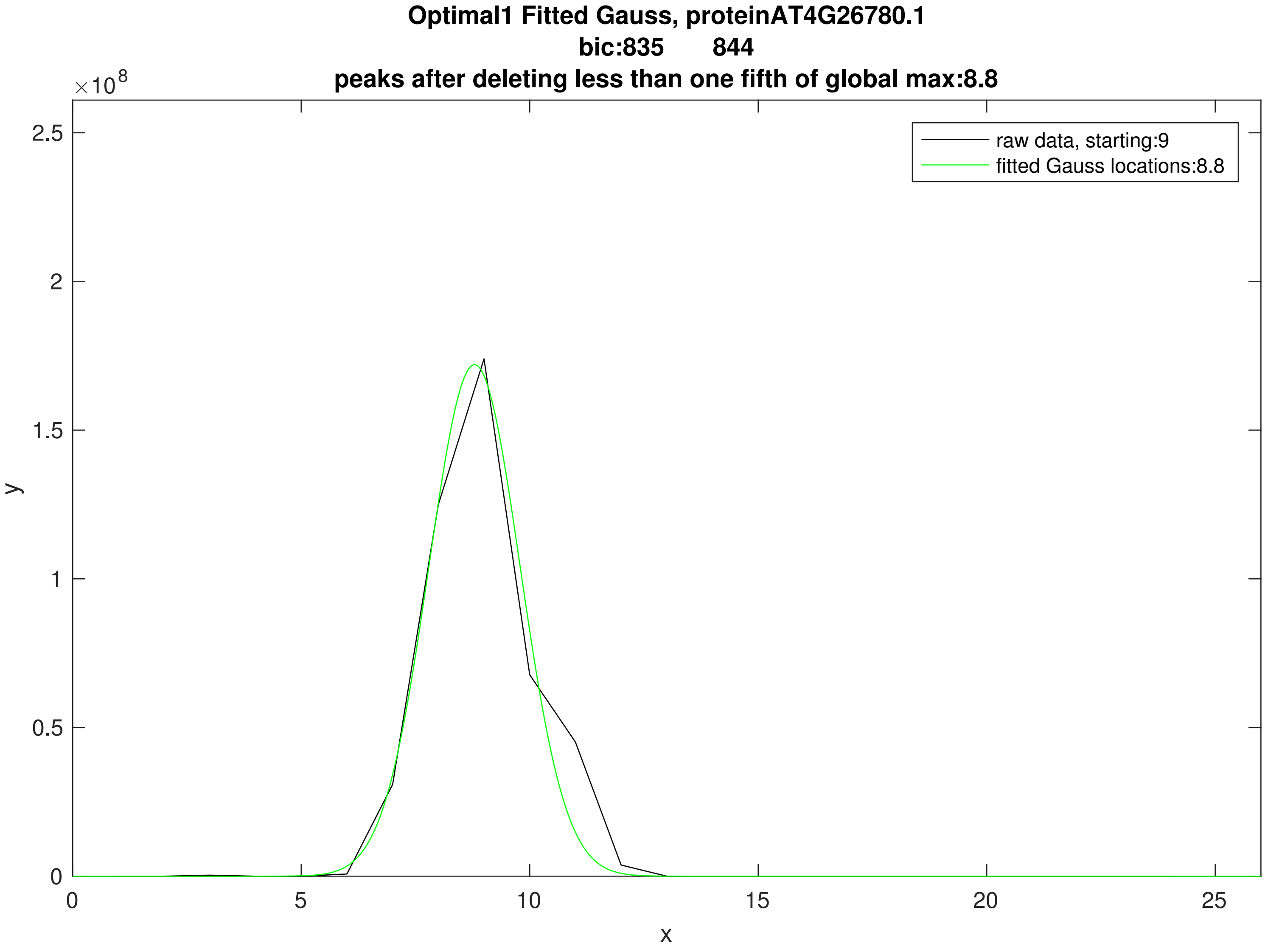

Optimal1 Fitted Gauss, proteinAT4G26780.1
bic:835 844
peaks after deleting less than one fifth of global max:8.8
×10^8
2.5
2
1.5
1
0.5
0
y
0
5
10
15
20
25
x
raw data, starting:9
fitted Gauss locations:8.8

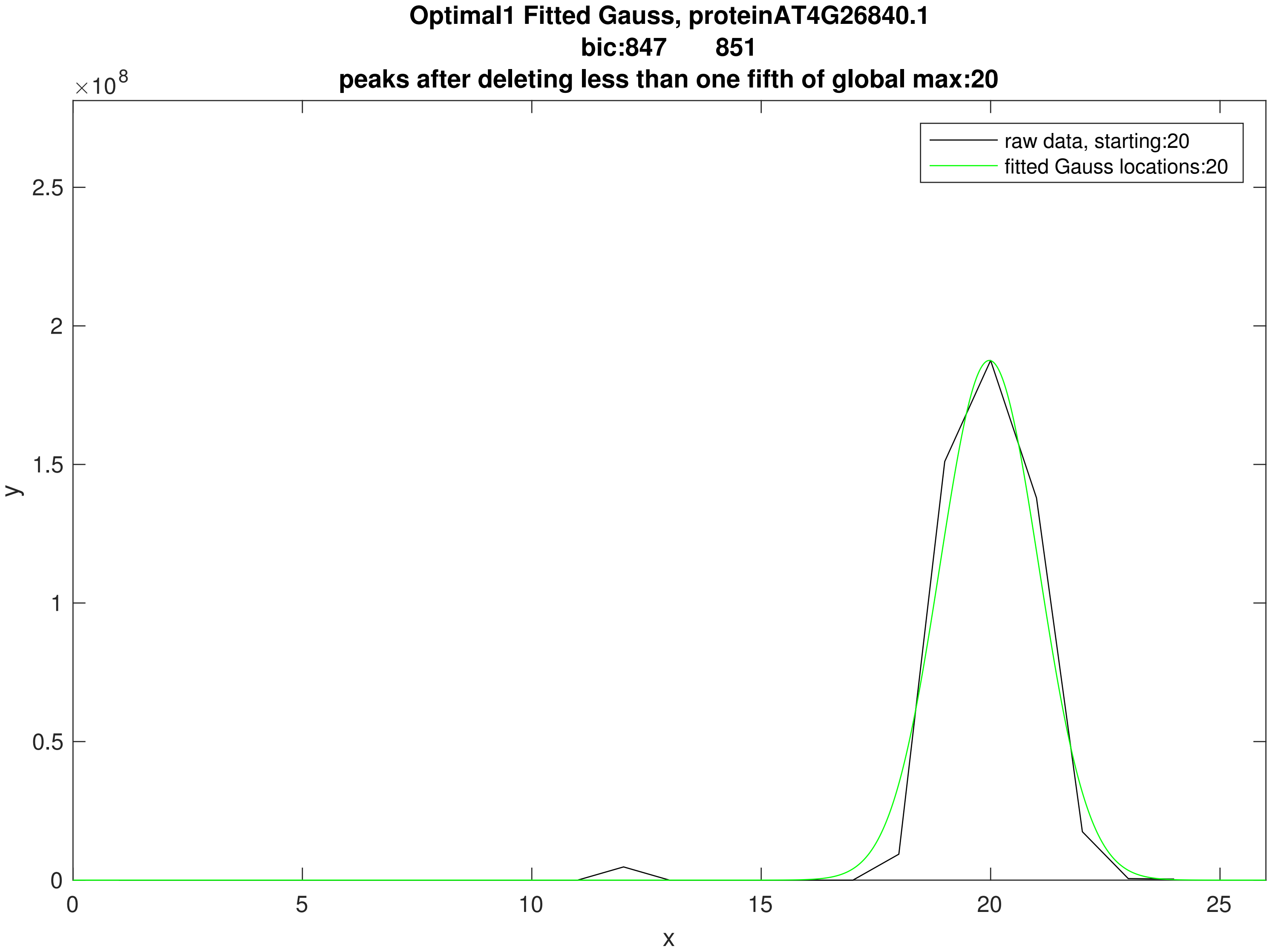
Optimal1 Fitted Gauss, proteinAT4G26840.1
bic:847 851
peaks after deleting less than one fifth of global max:20
×10^8
2.5
2
1.5
1
0.5
0
y
0
5
10
15
20
25
x
raw data, starting:20
fitted Gauss locations:20

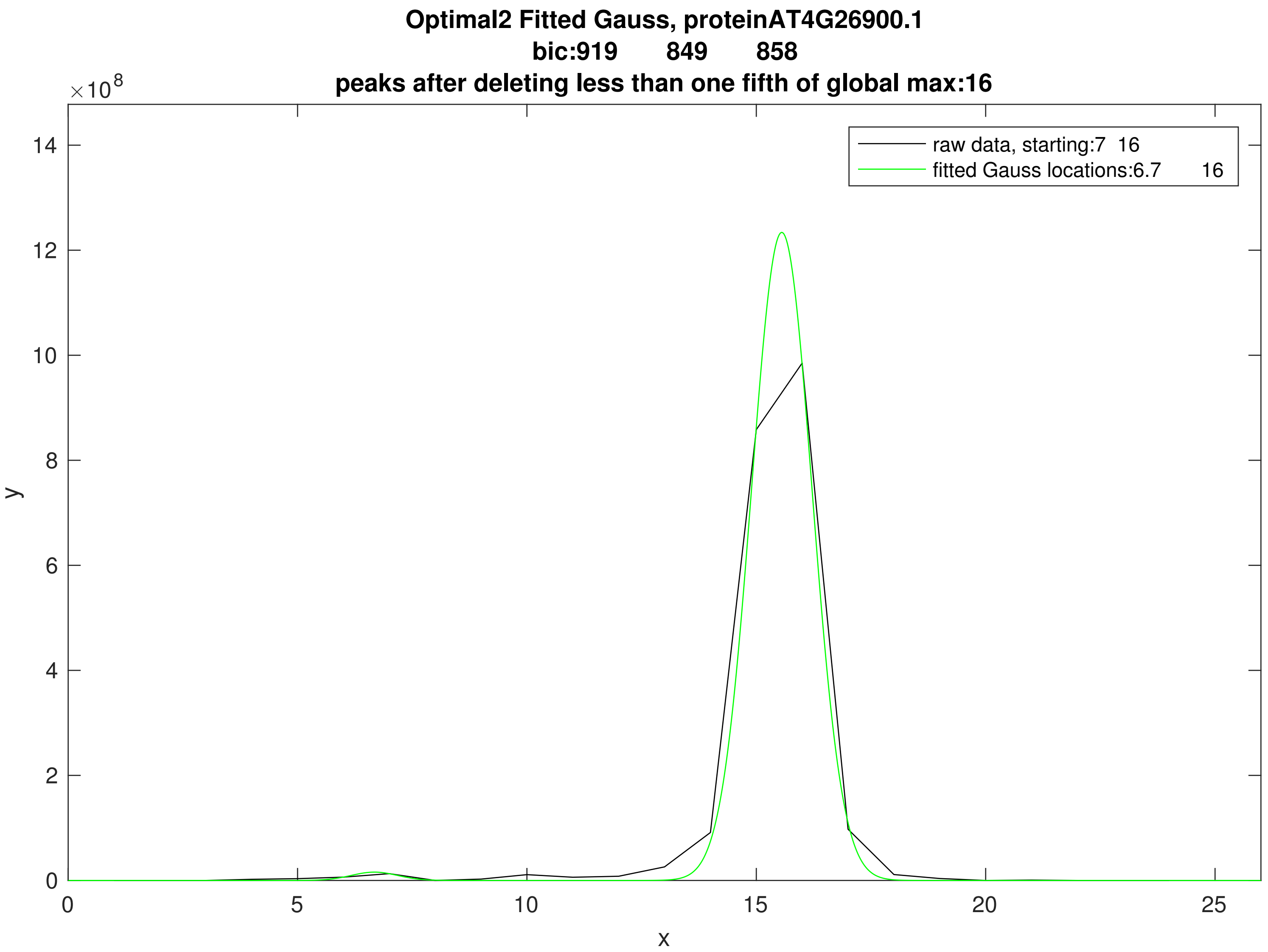

Optimal2 Fitted Gauss, proteinAT4G26900.1
bic:919 849 858
peaks after deleting less than one fifth of global max:16
×10^8
raw data, starting:7 16
fitted Gauss locations:6.7 16
y
x
0
2
4
6
8
10
12
14
0
5
10
15
20
25

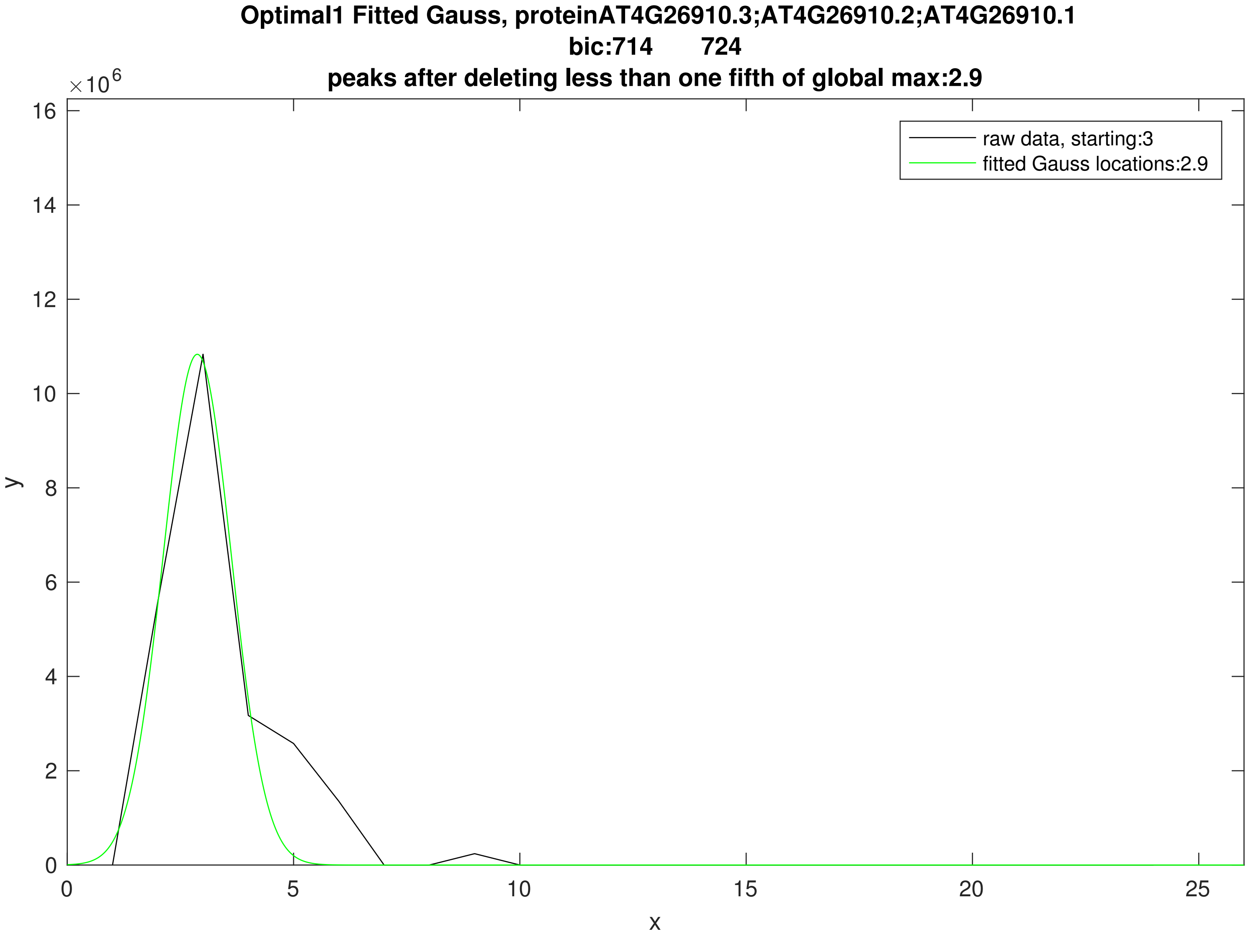

Optimal1 Fitted Gauss, proteinAT4G26910.3;AT4G26910.2;AT4G26910.1
bic:714 724
peaks after deleting less than one fifth of global max:2.9
×10^6
16
14
12
10
8
6
4
2
0
y
0
5
10
15
20
25
x
raw data, starting:3
fitted Gauss locations:2.9

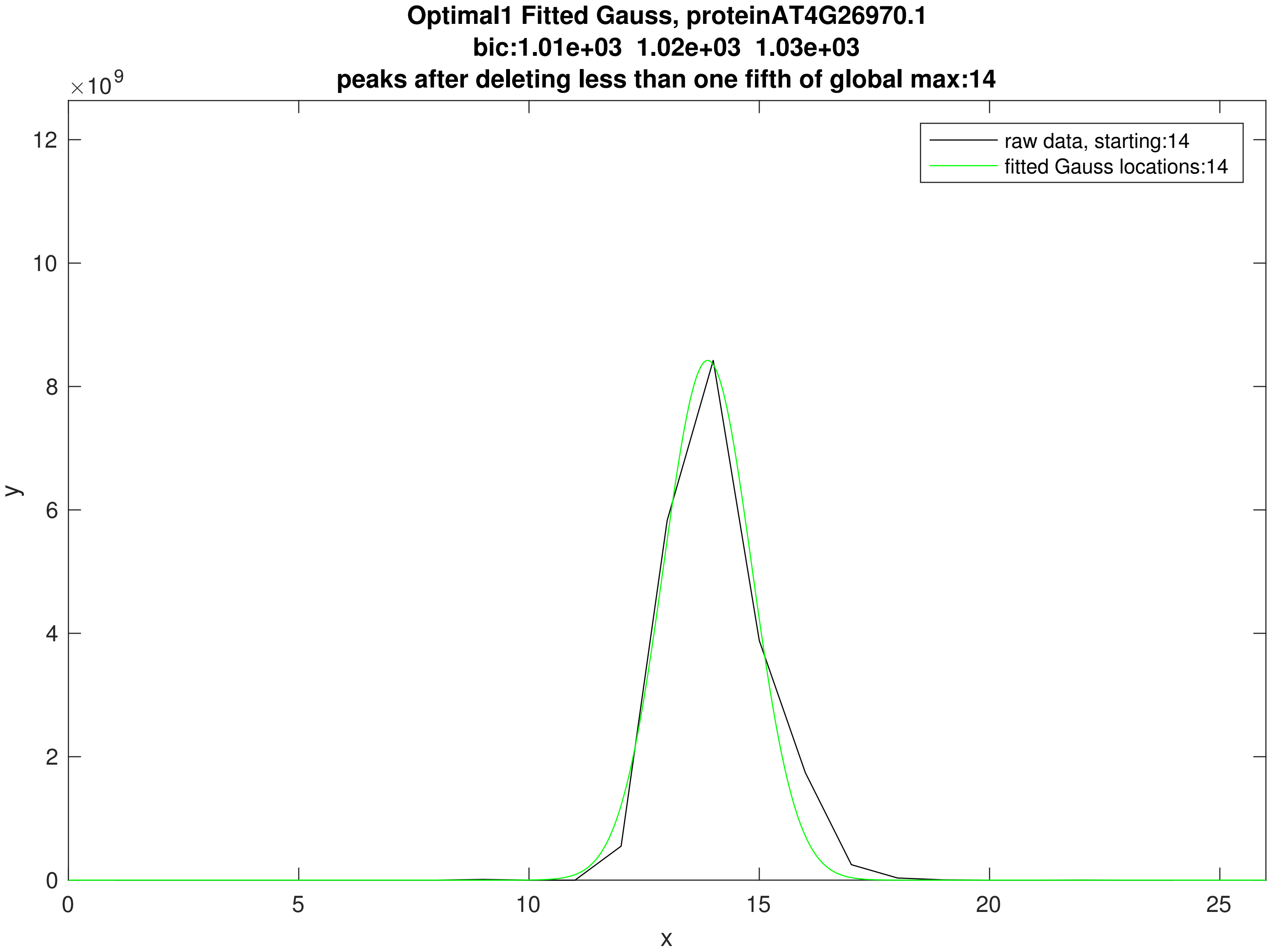

Optimal1 Fitted Gauss, proteinAT4G26970.1
bic:1.01e+03  1.02e+03  1.03e+03
peaks after deleting less than one fifth of global max:14
×10^9
raw data, starting:14
fitted Gauss locations:14
y
x
0
2
4
6
8
10
12
0
5
10
15
20
25

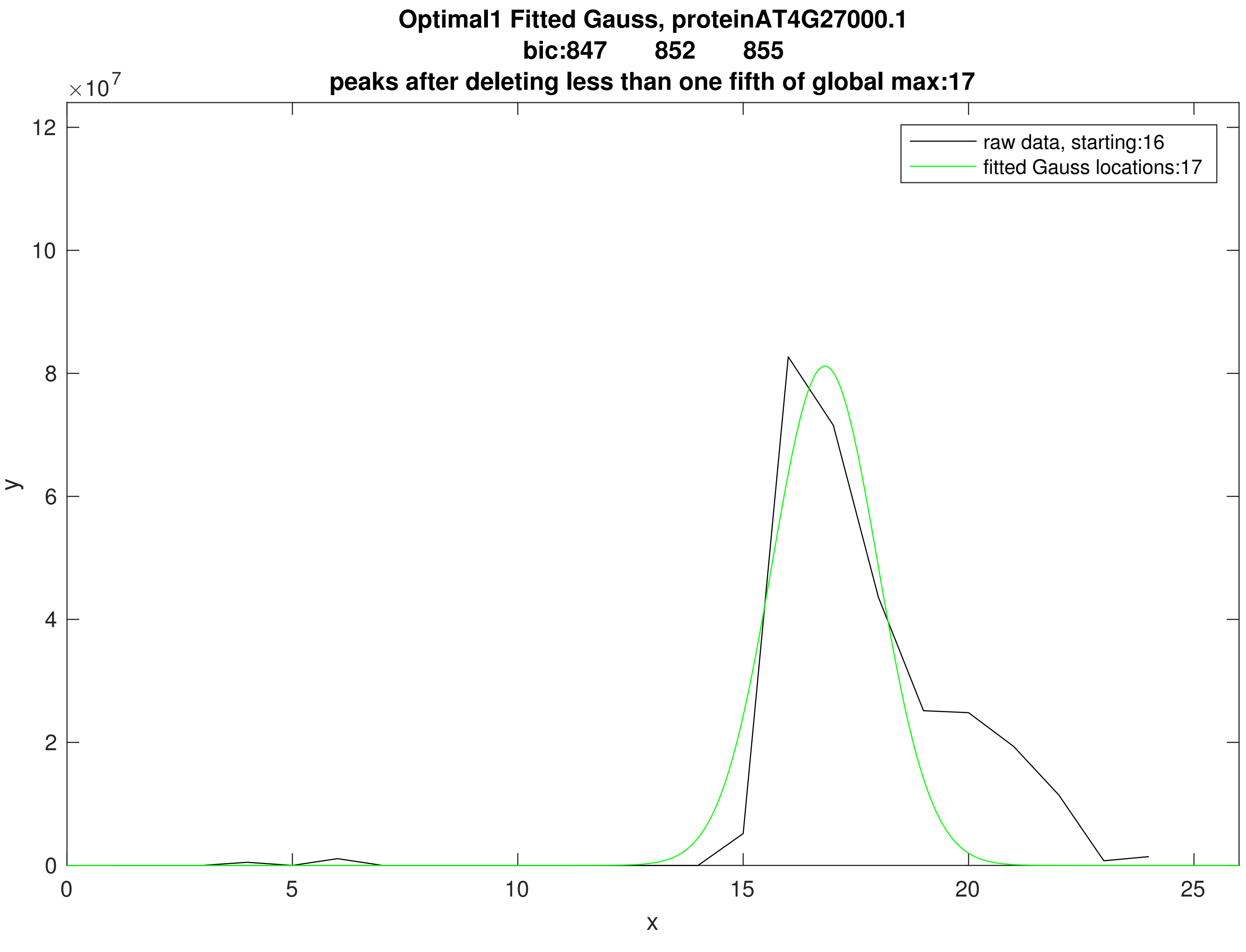

Optimal1 Fitted Gauss, proteinAT4G27000.1
bic:847 852 855
peaks after deleting less than one fifth of global max:17
×10^7
raw data, starting:16
fitted Gauss locations:17
y
x
0
2
4
6
8
10
12
0
5
10
15
20
25

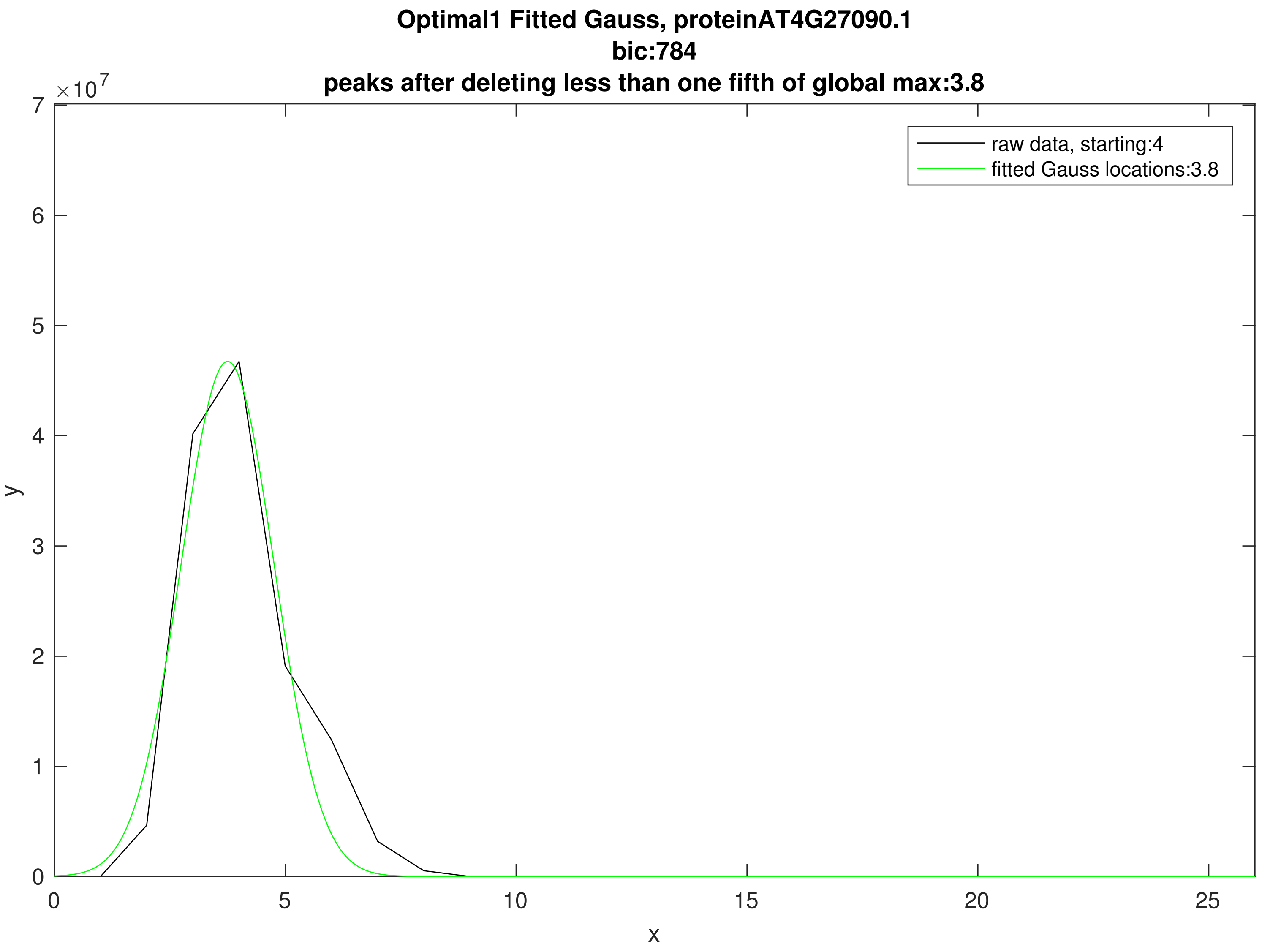
Optimal1 Fitted Gauss, proteinAT4G27090.1
bic:784
peaks after deleting less than one fifth of global max:3.8
×10^7
raw data, starting:4
fitted Gauss locations:3.8
y
x
0
1
2
3
4
5
6
7
0
5
10
15
20
25

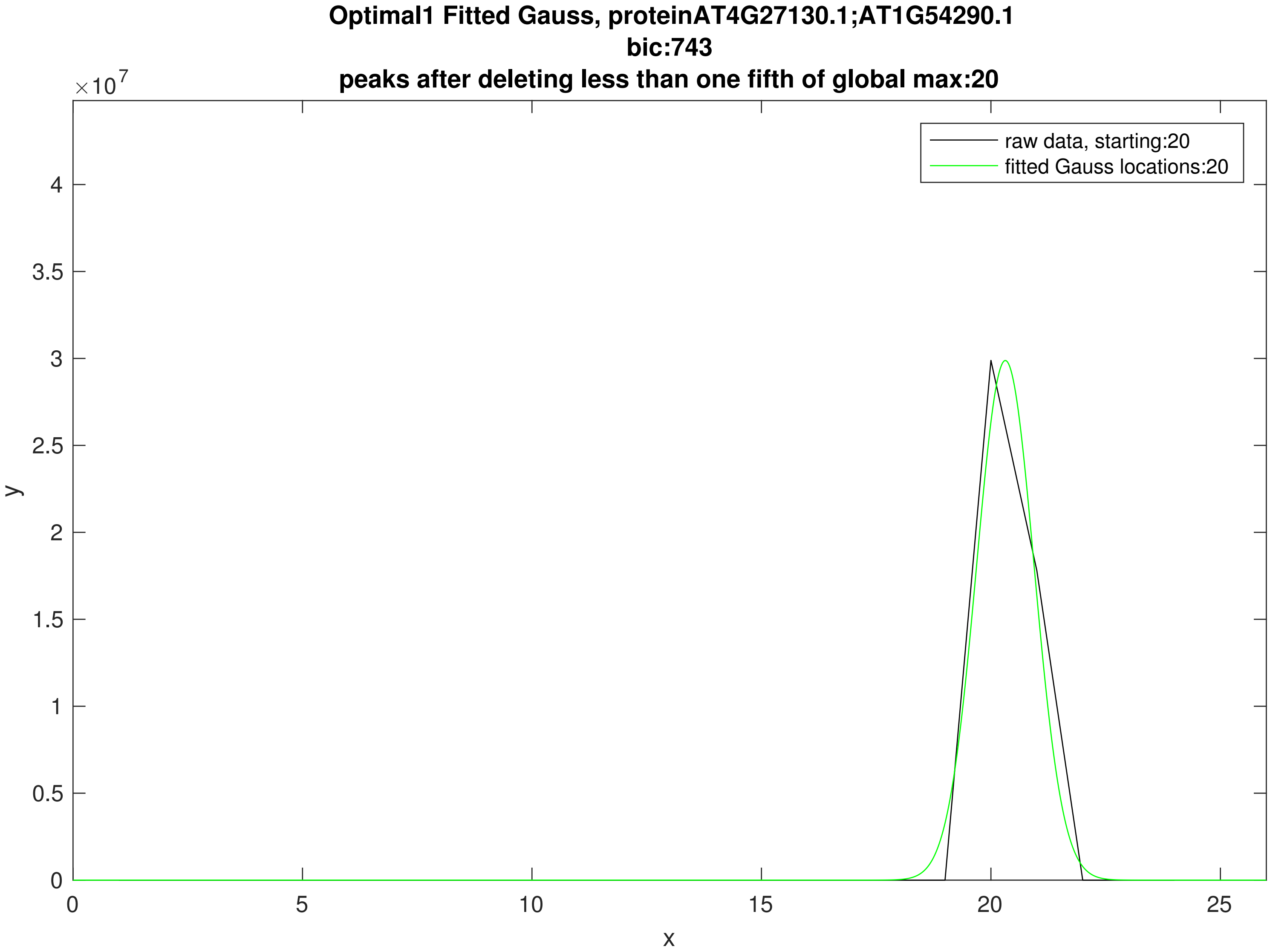
Optimal1 Fitted Gauss, proteinAT4G27130.1;AT1G54290.1
bic:743
peaks after deleting less than one fifth of global max:20
×10^7
raw data, starting:20
fitted Gauss locations:20
4
3.5
3
2.5
2
1.5
1
0.5
0
y
0
5
10
15
20
25
x

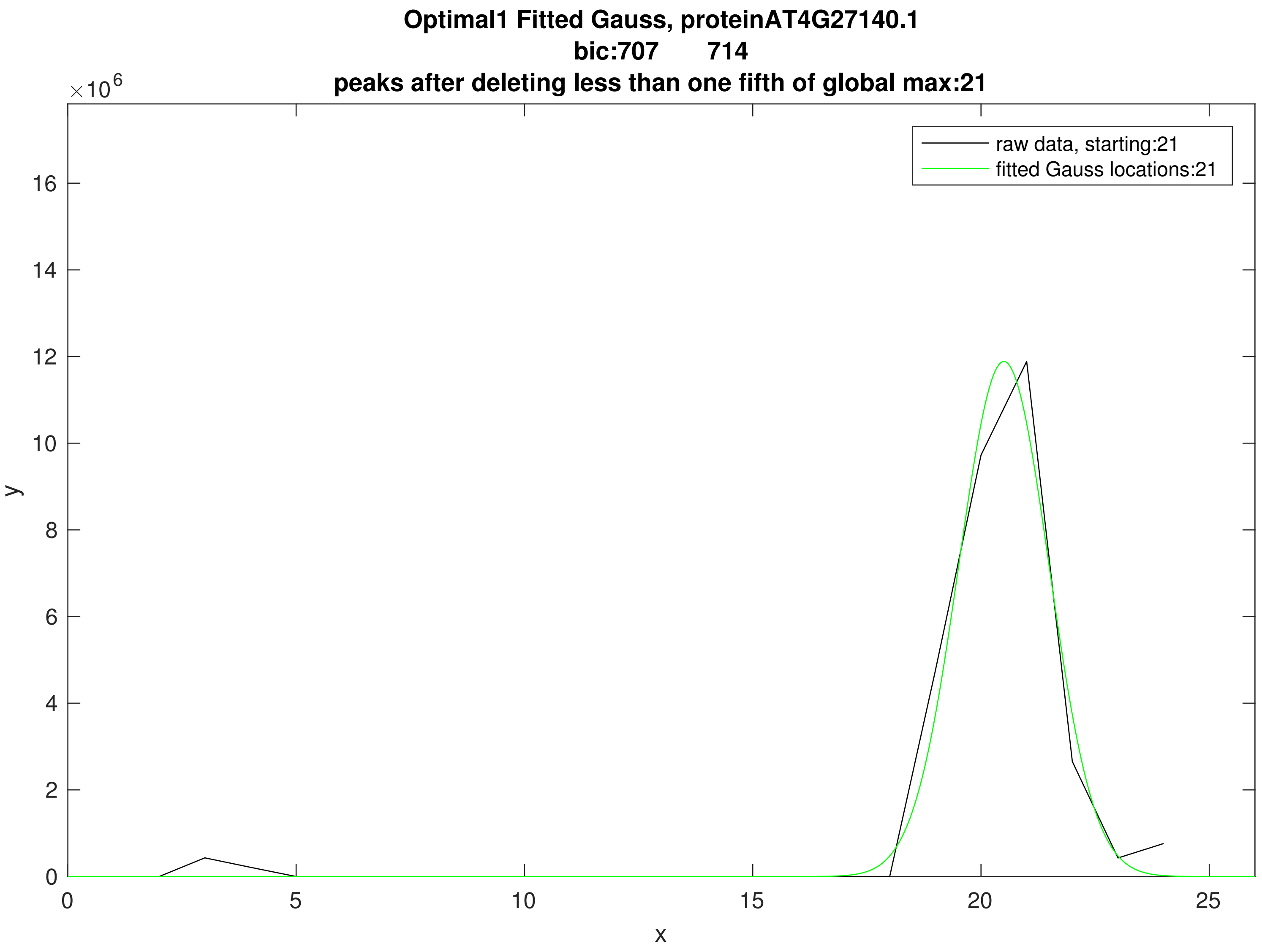

Optimal1 Fitted Gauss, proteinAT4G27140.1
bic:707 714
peaks after deleting less than one fifth of global max:21
×10^6
raw data, starting:21
fitted Gauss locations:21
16
14
12
10
8
6
4
2
0
y
0
5
10
15
20
25
x

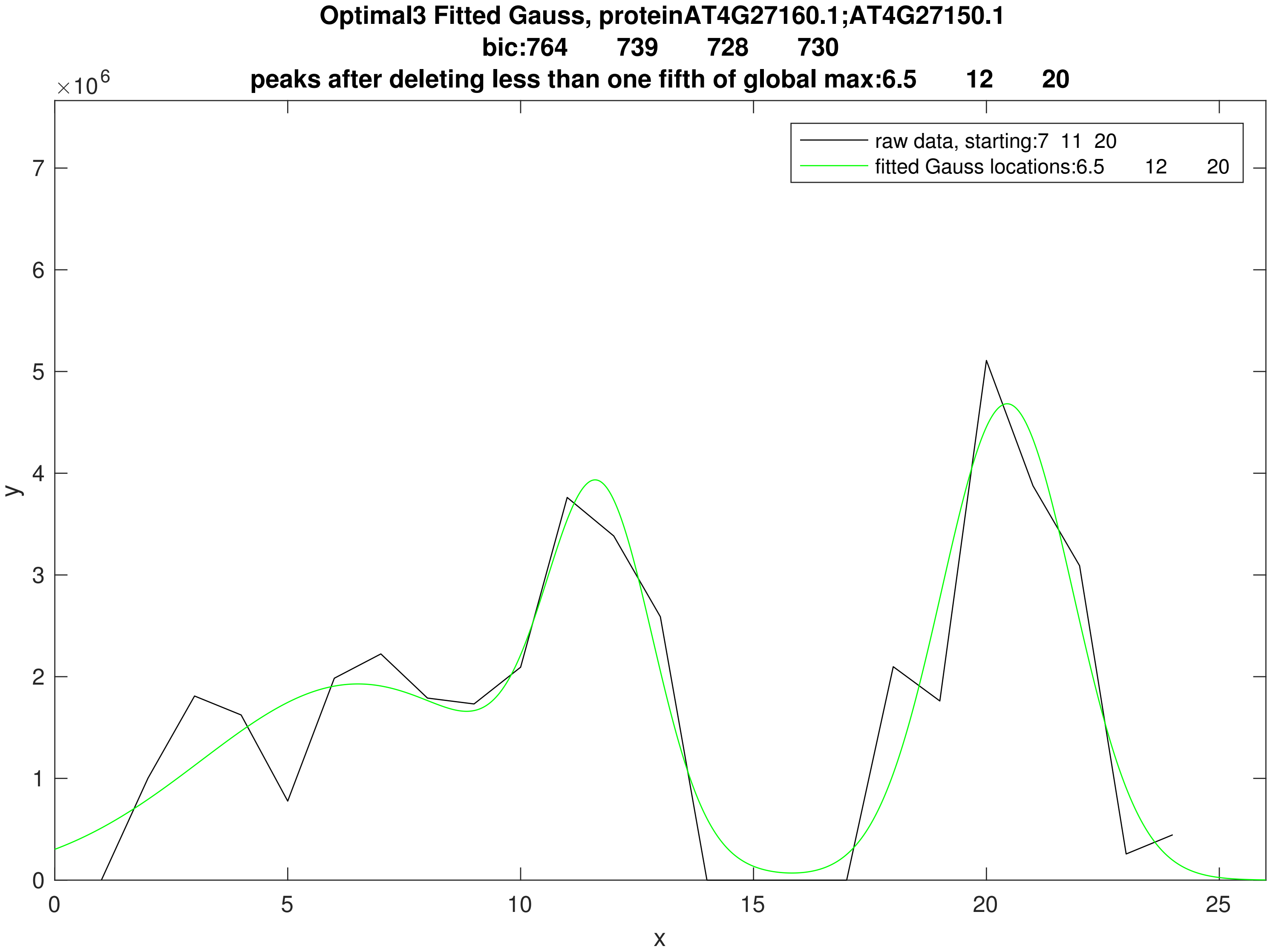

Optimal3 Fitted Gauss, proteinAT4G27160.1;AT4G27150.1
bic:764 739 728 730
peaks after deleting less than one fifth of global max:6.5 12 20
×10^6
raw data, starting:7 11 20
fitted Gauss locations:6.5 12 20
y
x

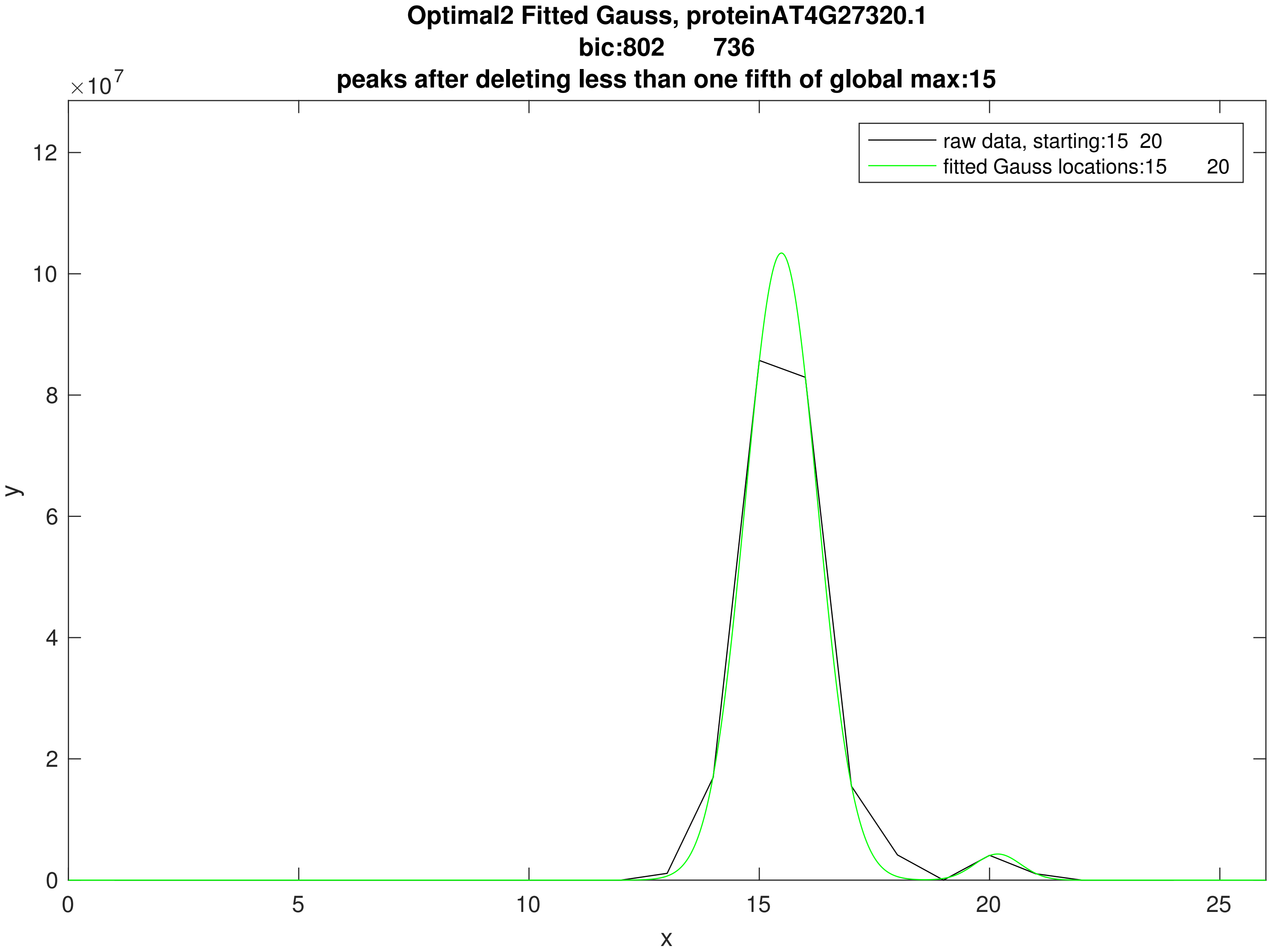

Optimal2 Fitted Gauss, proteinAT4G27320.1
bic:802 736
peaks after deleting less than one fifth of global max:15
×10^7
raw data, starting:15 20
fitted Gauss locations:15 20
y
x
0
2
4
6
8
10
12
0
5
10
15
20
25

**Optimal2 Fitted Gauss, proteinAT4G27440.2;AT4G27440.1**
**bic:713 701**
**peaks after deleting less than one fifth of global max:4.2 21**

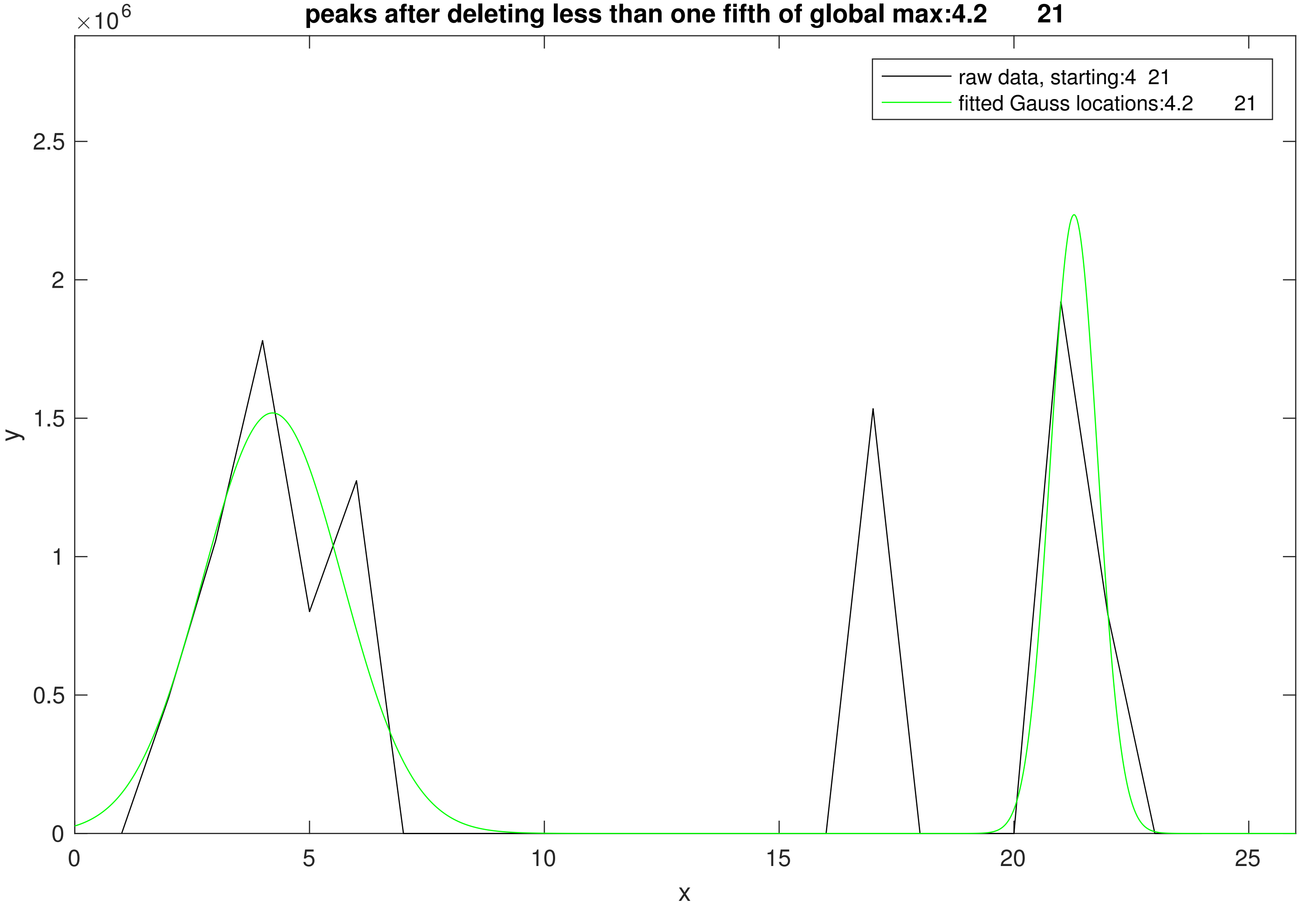

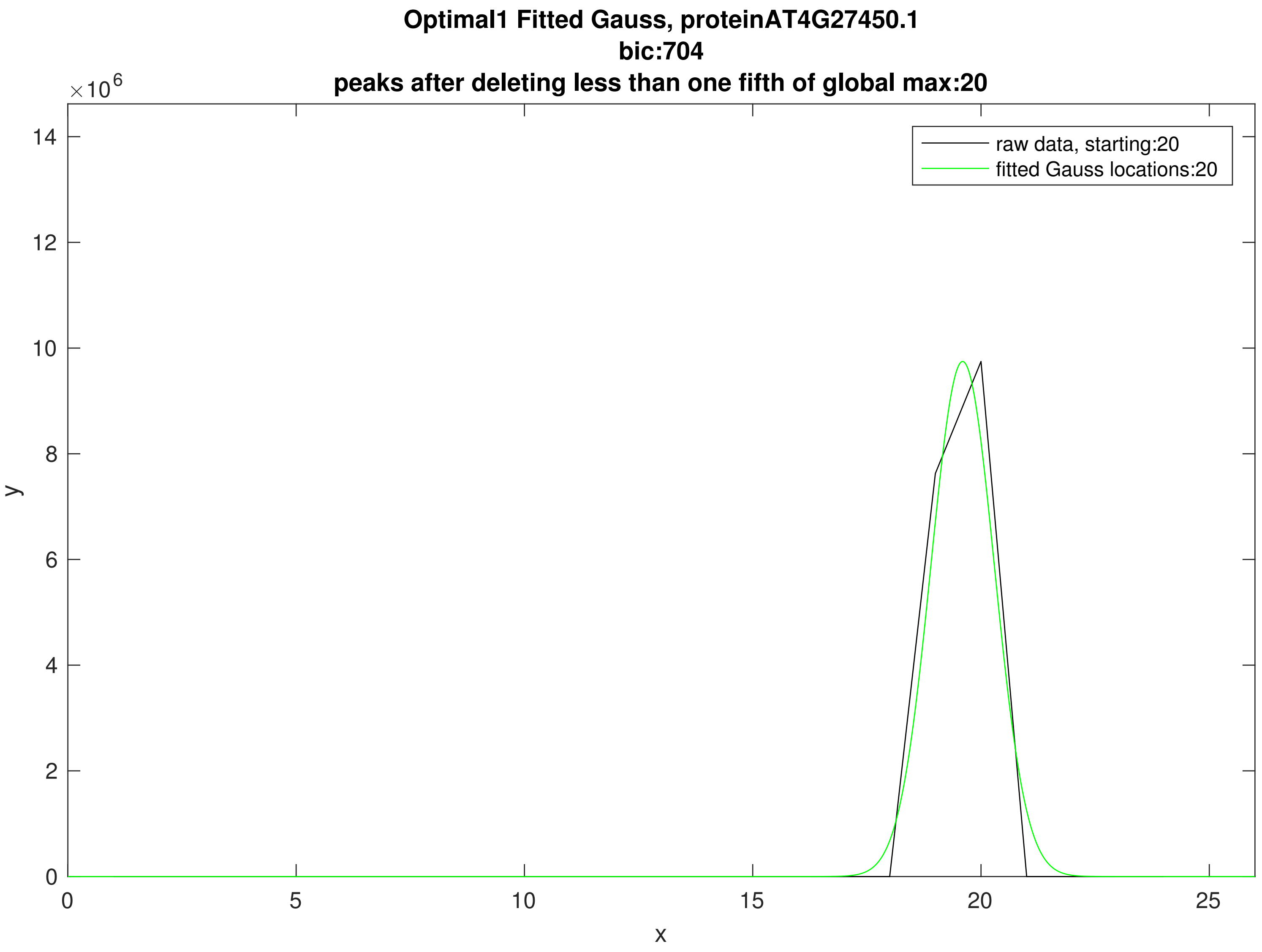

Optimal1 Fitted Gauss, proteinAT4G27450.1
bic:704
peaks after deleting less than one fifth of global max:20
×10^6
raw data, starting:20
fitted Gauss locations:20
14
12
10
8
6
4
2
0
y
0
5
10
15
20
25
x

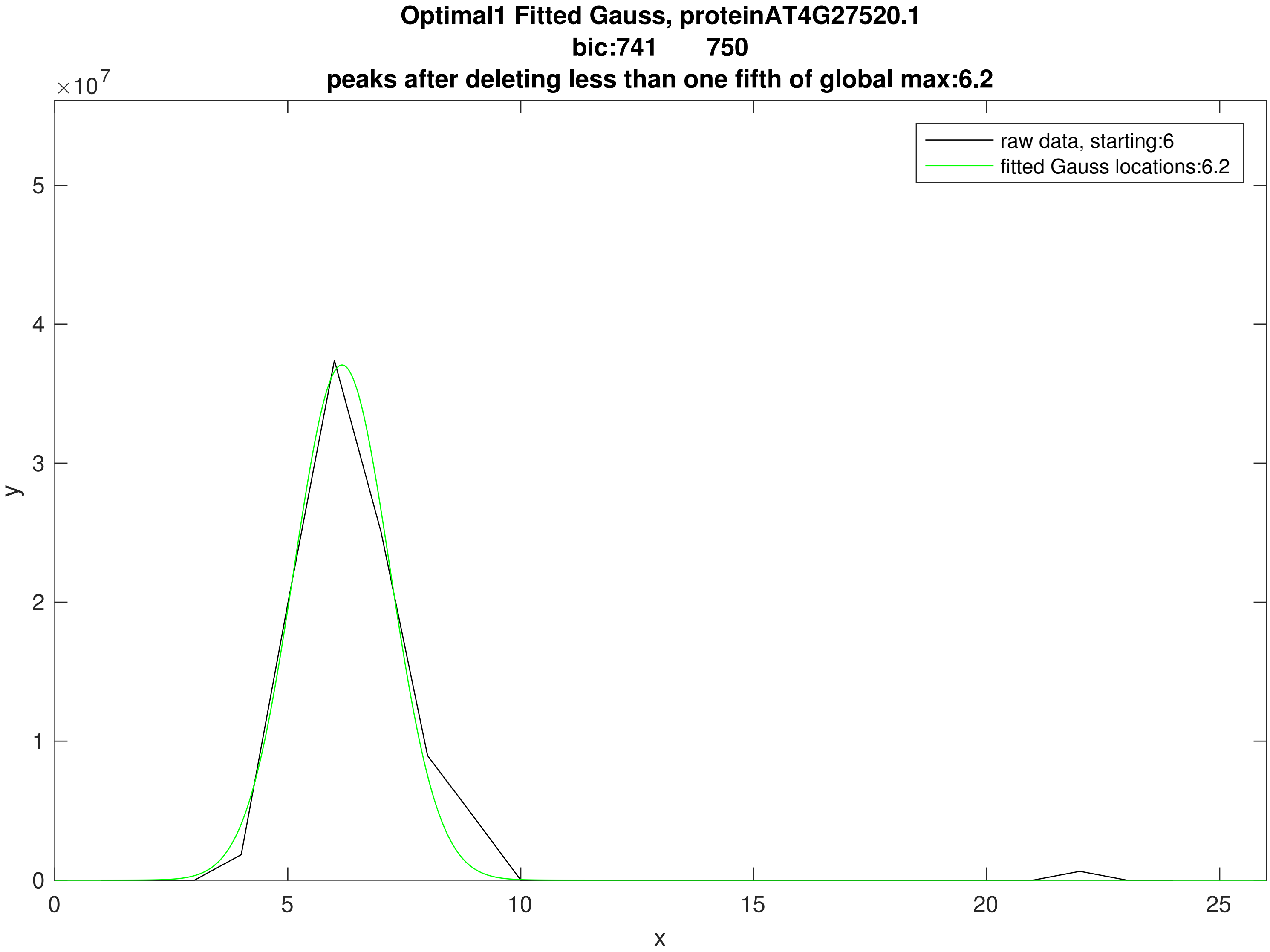

Optimal1 Fitted Gauss, proteinAT4G27520.1
bic:741 750
peaks after deleting less than one fifth of global max:6.2
×10^7
raw data, starting:6
fitted Gauss locations:6.2
0
1
2
3
4
5
y
0
5
10
15
20
25
x

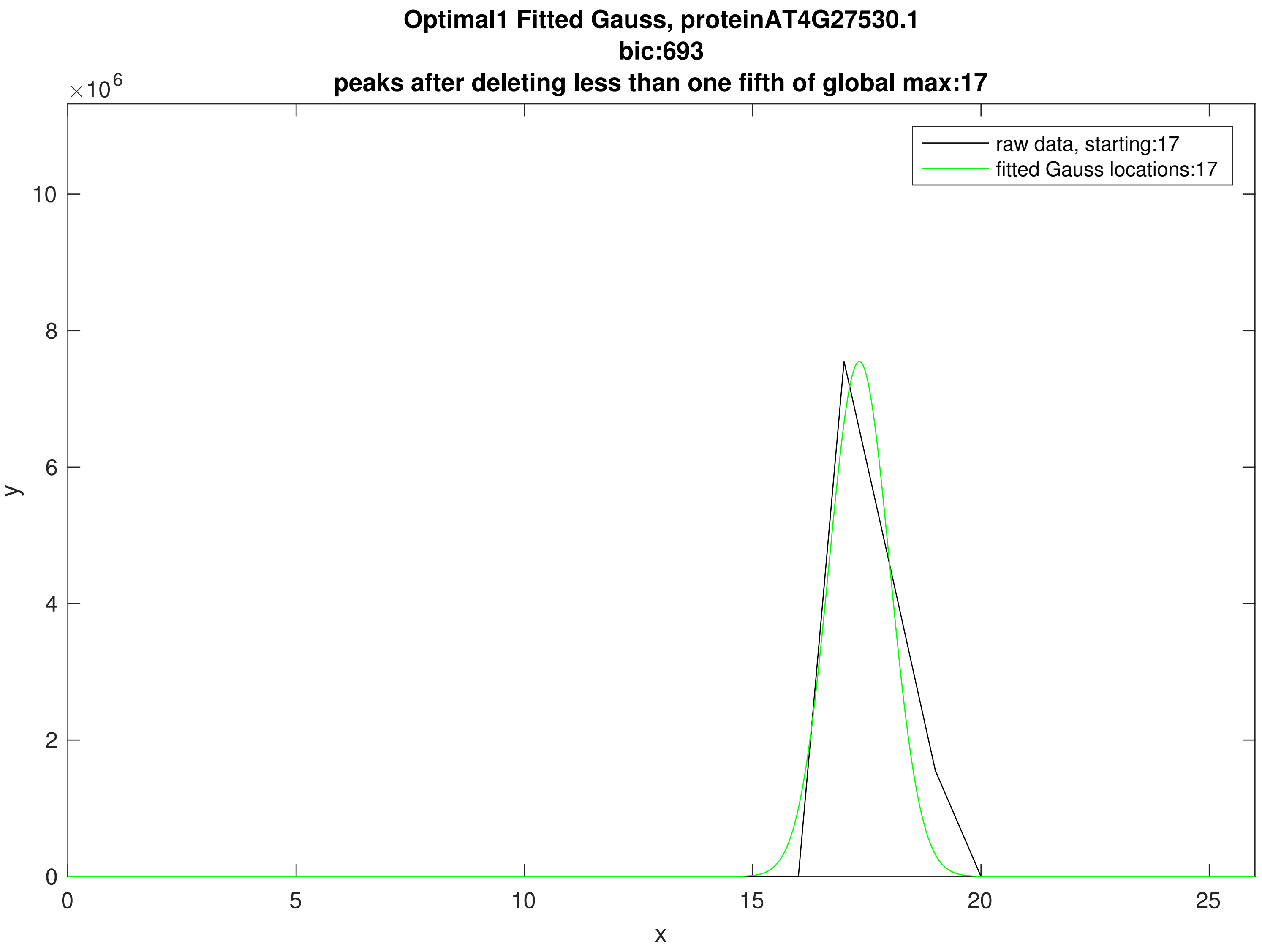

Optimal1 Fitted Gauss, proteinAT4G27530.1
bic:693
peaks after deleting less than one fifth of global max:17
×10^6
raw data, starting:17
fitted Gauss locations:17
10
8
6
4
2
0
y
0
5
10
15
20
25
x

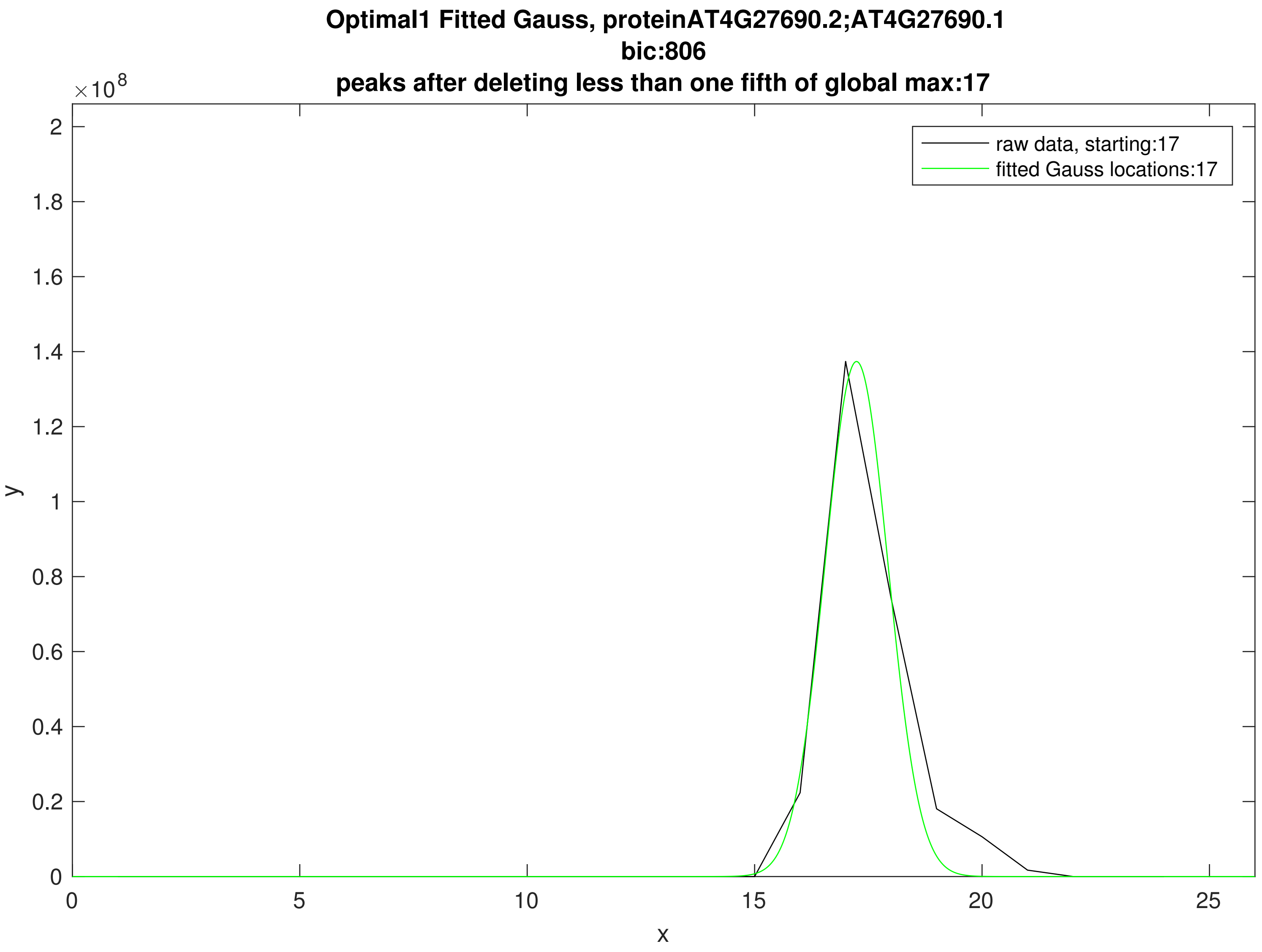

Optimal1 Fitted Gauss, proteinAT4G27690.2;AT4G27690.1
bic:806
peaks after deleting less than one fifth of global max:17
×10^8
raw data, starting:17
fitted Gauss locations:17
y
x
0
0.2
0.4
0.6
0.8
1
1.2
1.4
1.6
1.8
2
0
5
10
15
20
25

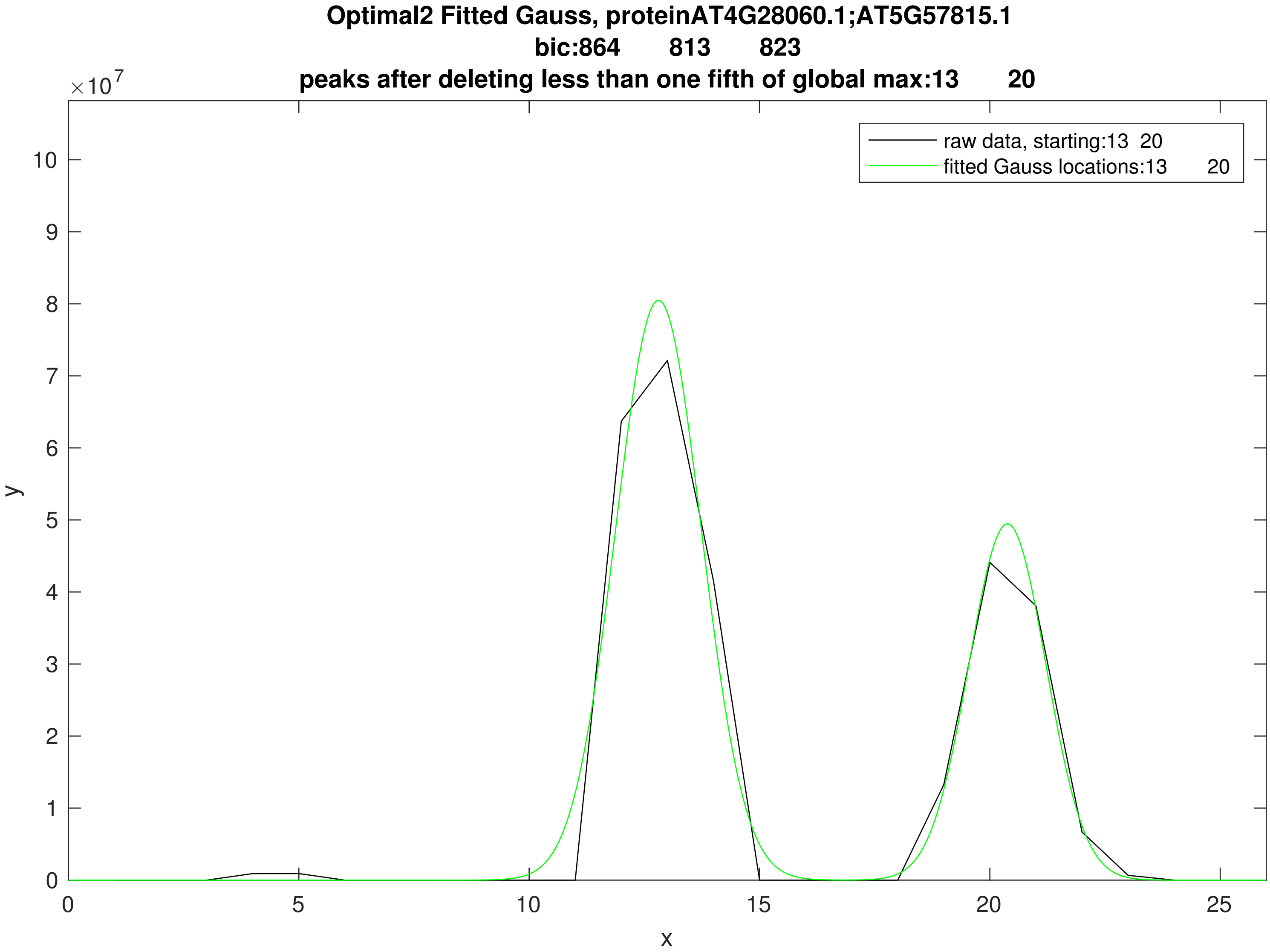
Optimal2 Fitted Gauss, proteinAT4G28060.1;AT5G57815.1
bic:864 813 823
peaks after deleting less than one fifth of global max:13 20
×10^7
raw data, starting:13 20
fitted Gauss locations:13 20
y
x
0
1
2
3
4
5
6
7
8
9
10
0
5
10
15
20
25

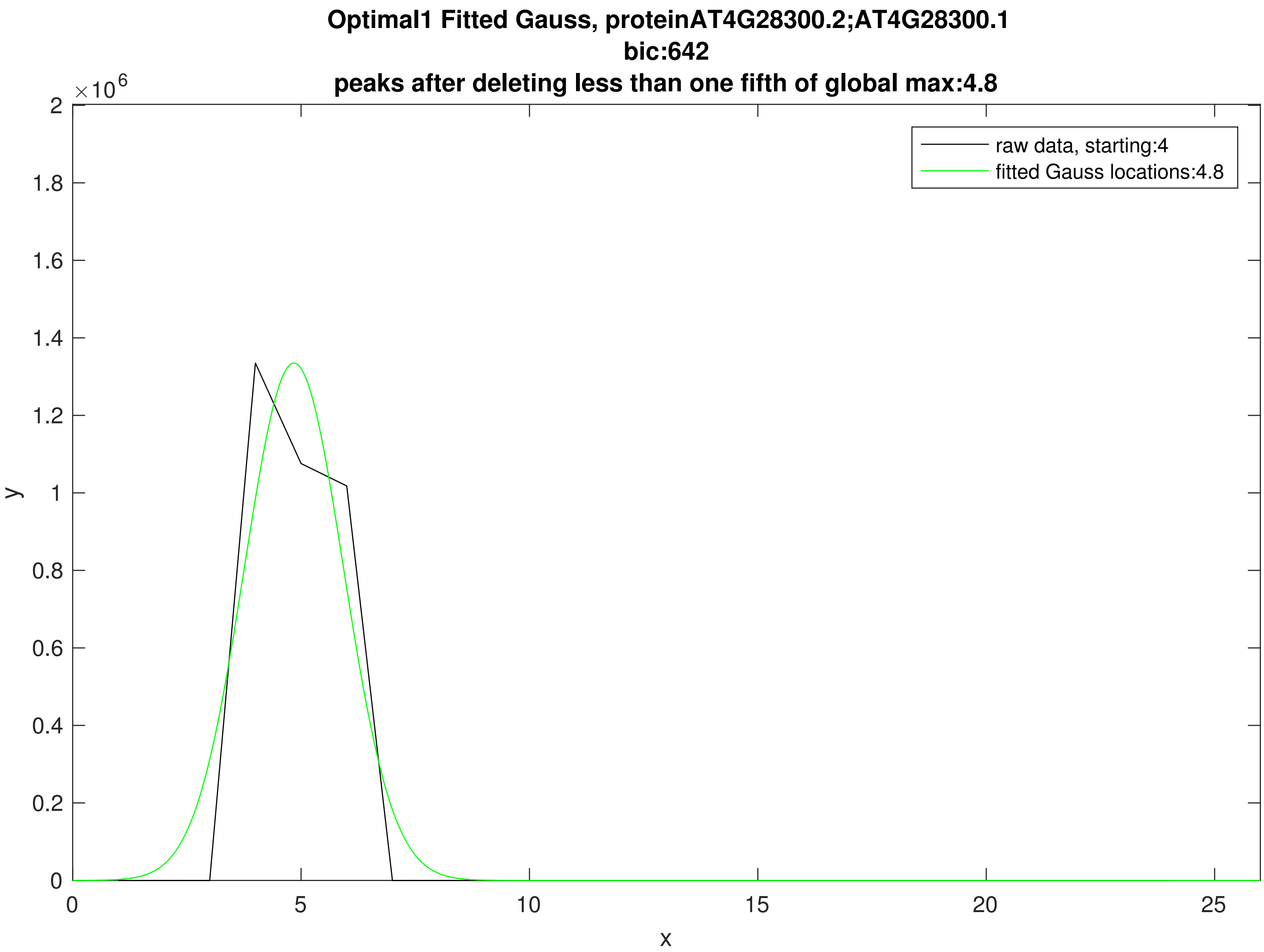
Optimal1 Fitted Gauss, proteinAT4G28300.2;AT4G28300.1
bic:642
peaks after deleting less than one fifth of global max:4.8
×10^6
raw data, starting:4
fitted Gauss locations:4.8
y
x
0
0.2
0.4
0.6
0.8
1
1.2
1.4
1.6
1.8
2
0
5
10
15
20
25

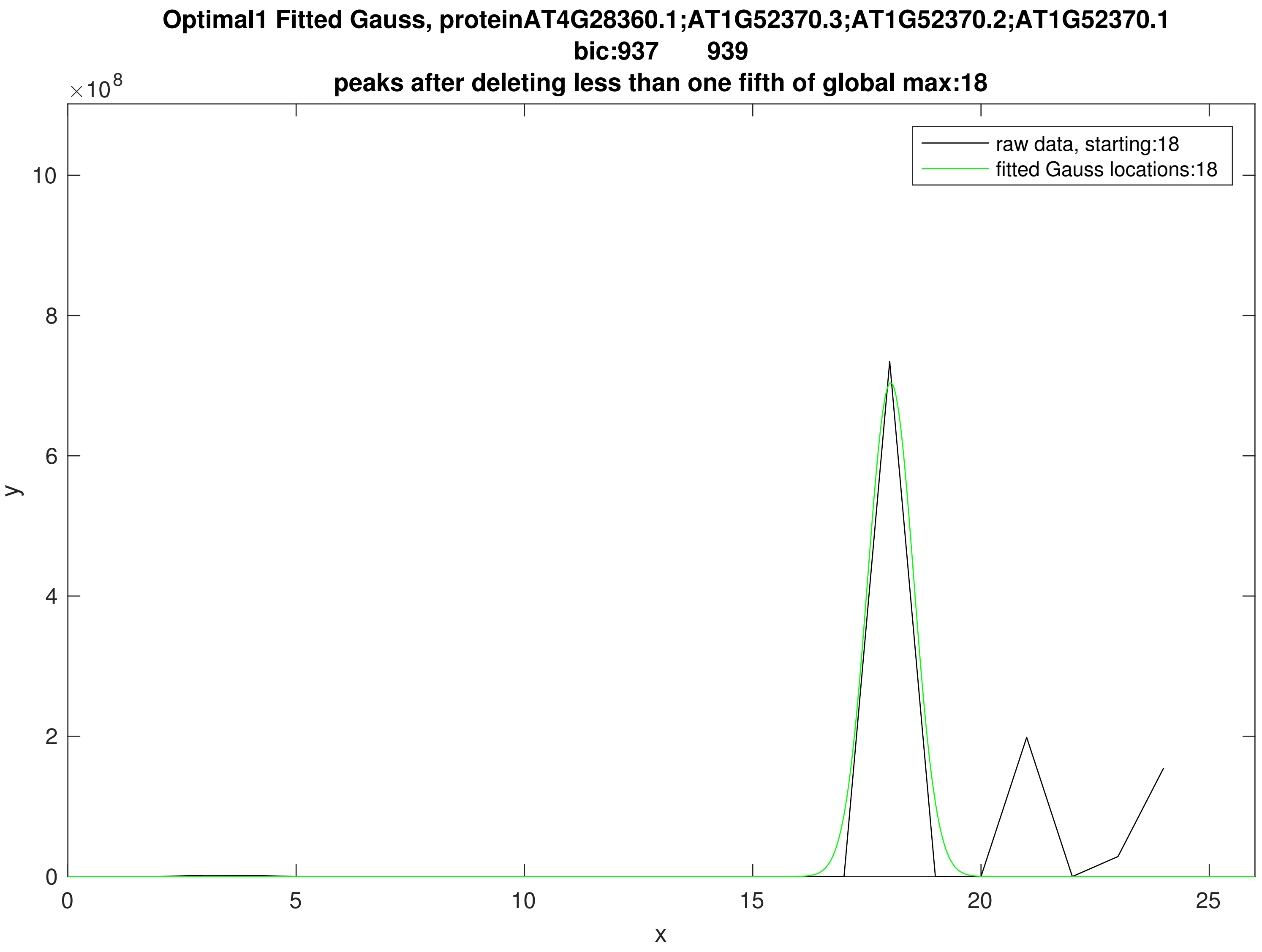

Optimal1 Fitted Gauss, proteinAT4G28360.1;AT1G52370.3;AT1G52370.2;AT1G52370.1
bic:937 939
peaks after deleting less than one fifth of global max:18
×10^8
raw data, starting:18
fitted Gauss locations:18
y
x
0
2
4
6
8
10
0
5
10
15
20
25

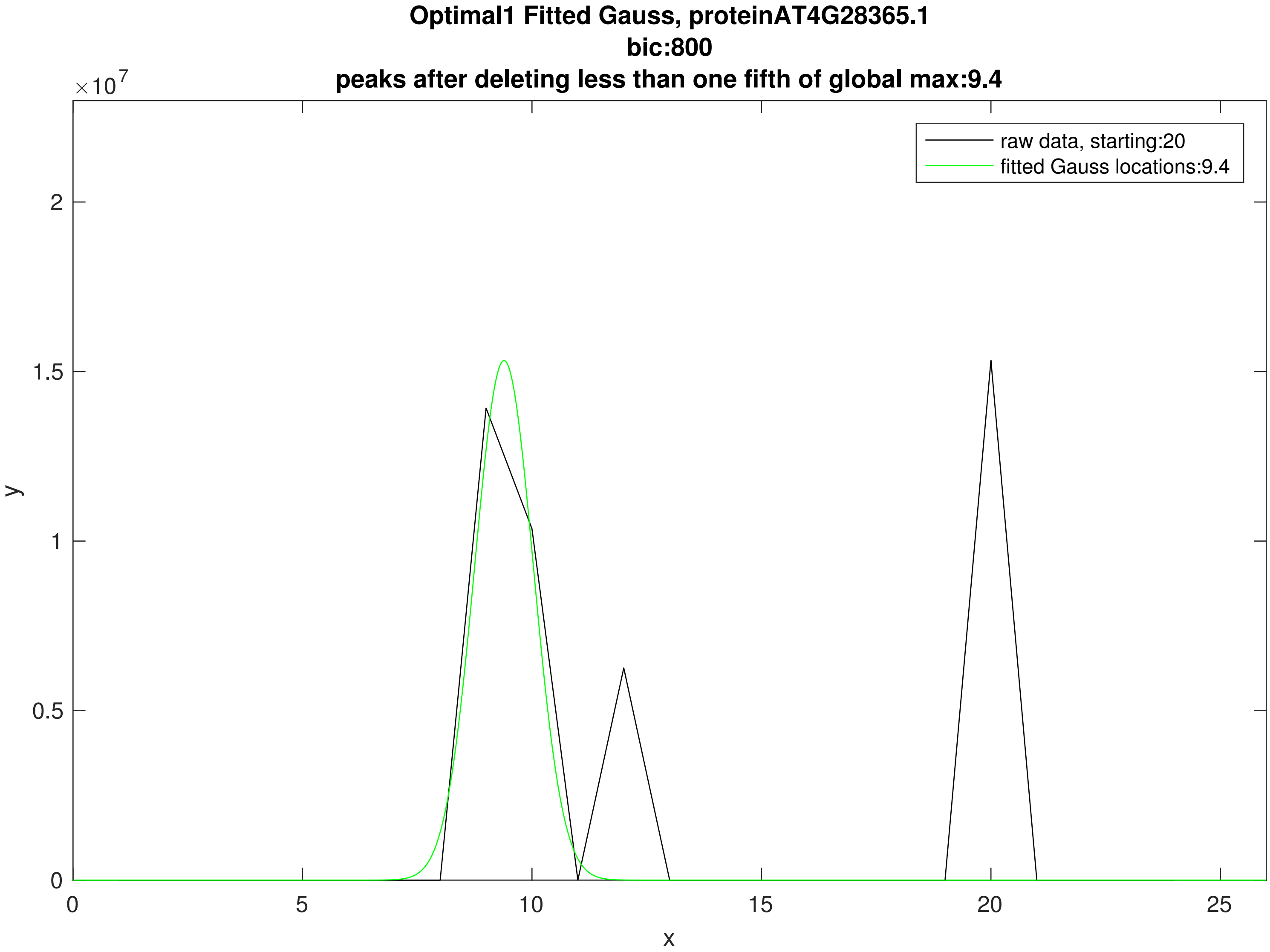
Optimal1 Fitted Gauss, proteinAT4G28365.1
bic:800
peaks after deleting less than one fifth of global max:9.4
×10^7
y
x
0
0.5
1
1.5
2
0
5
10
15
20
25
raw data, starting:20
fitted Gauss locations:9.4

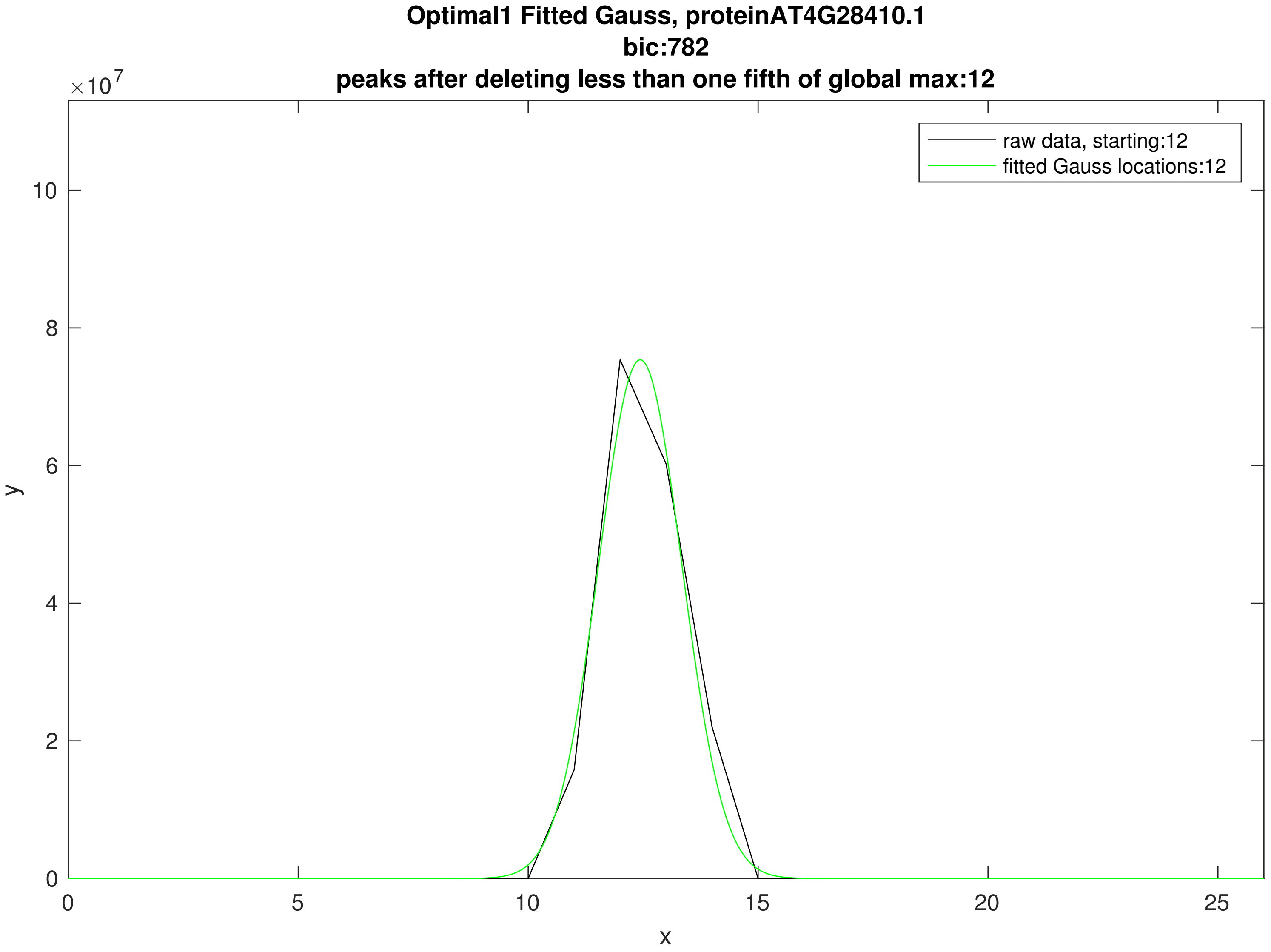

Optimal1 Fitted Gauss, proteinAT4G28410.1
bic:782
peaks after deleting less than one fifth of global max:12
×10^7
raw data, starting:12
fitted Gauss locations:12
10
8
6
4
2
0
y
0
5
10
15
20
25
x

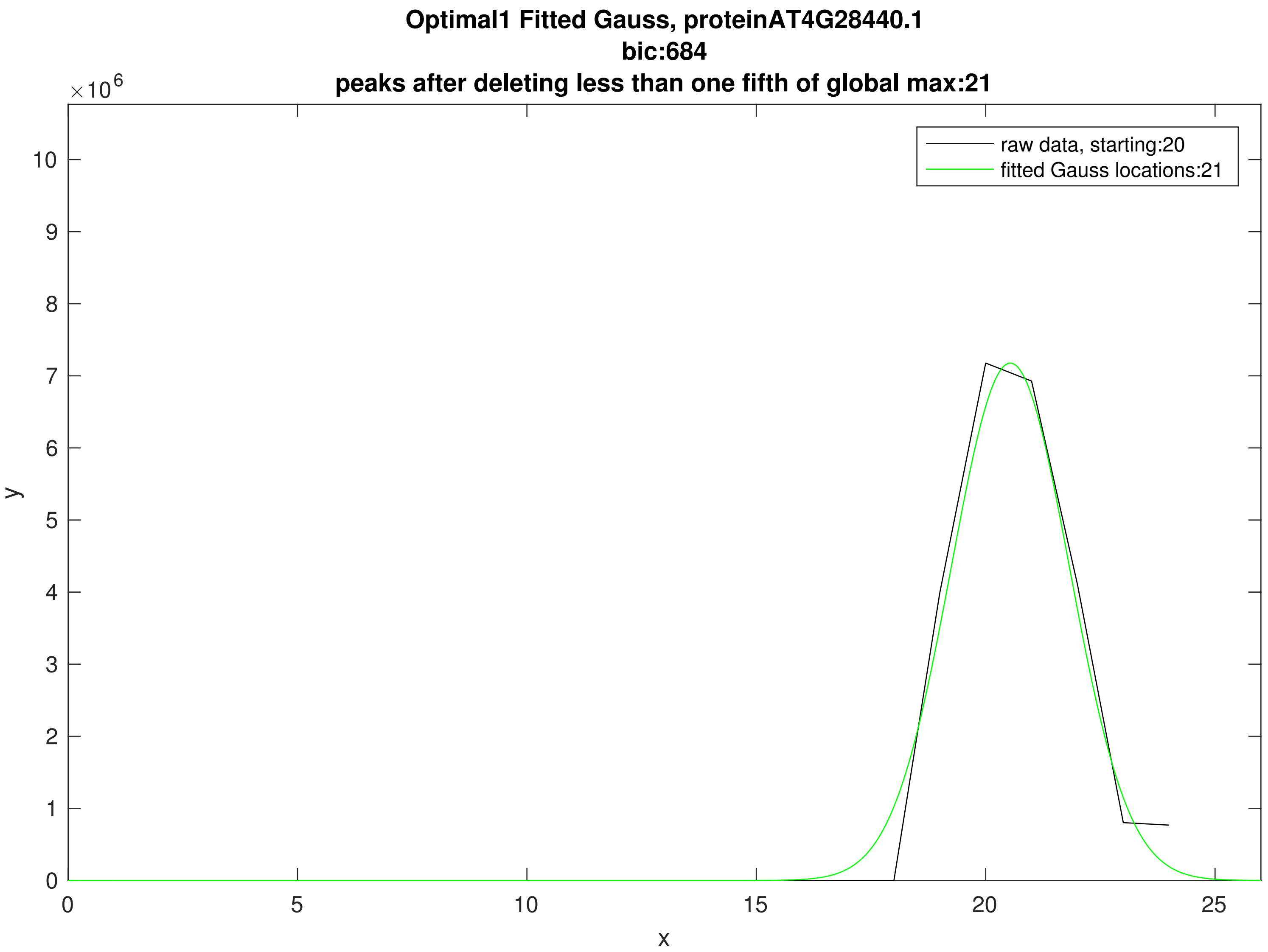

Optimal1 Fitted Gauss, proteinAT4G28440.1
bic:684
peaks after deleting less than one fifth of global max:21
×10^6
raw data, starting:20
fitted Gauss locations:21
y
x
0
1
2
3
4
5
6
7
8
9
10
0
5
10
15
20
25

**Optimal1 Fitted Gauss, proteinAT4G28520.1;AT4G28520.3;AT4G28520.2;AT4G28520.4**
**bic:986 995 992**
**peaks after deleting less than one fifth of global max:11**

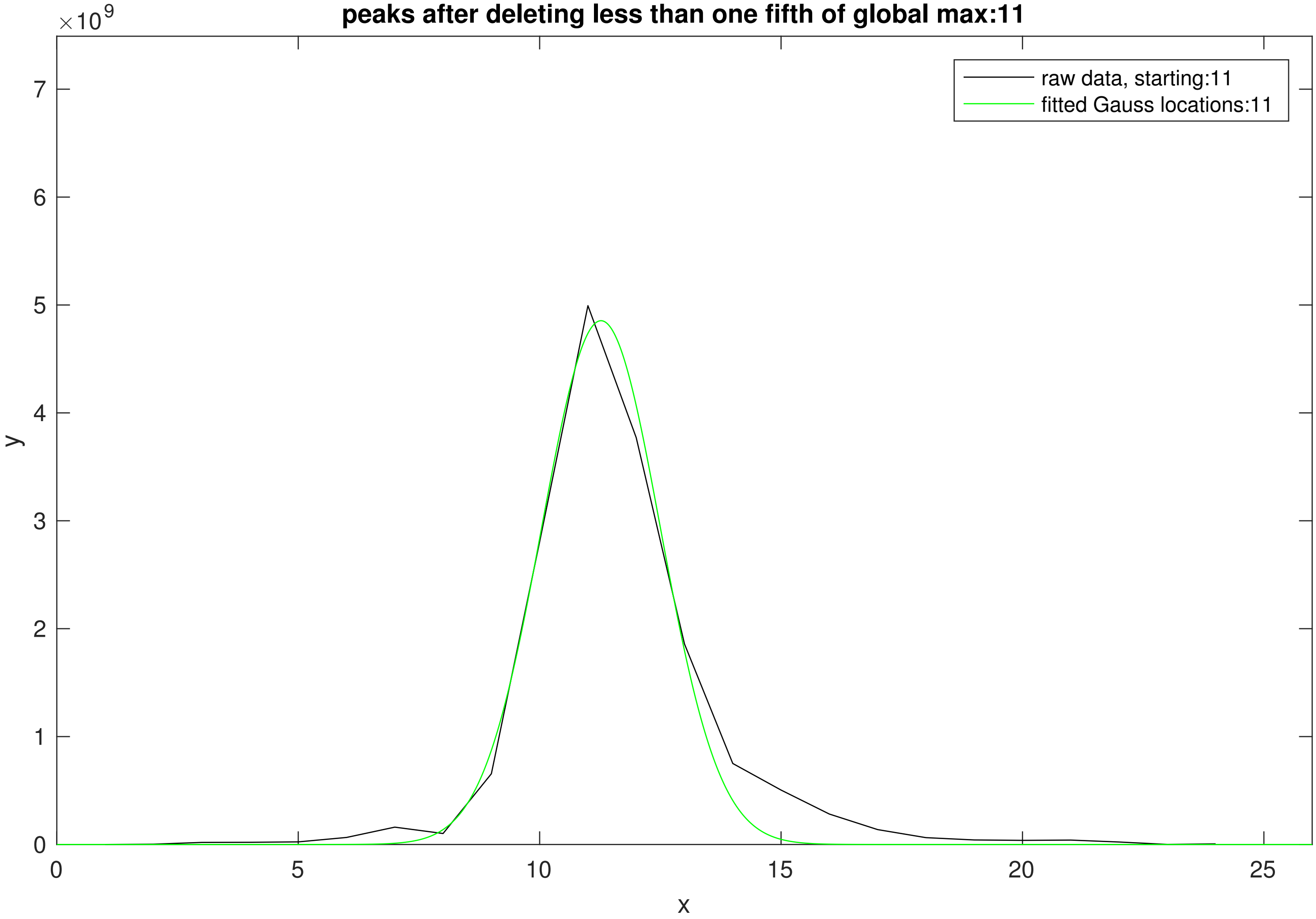

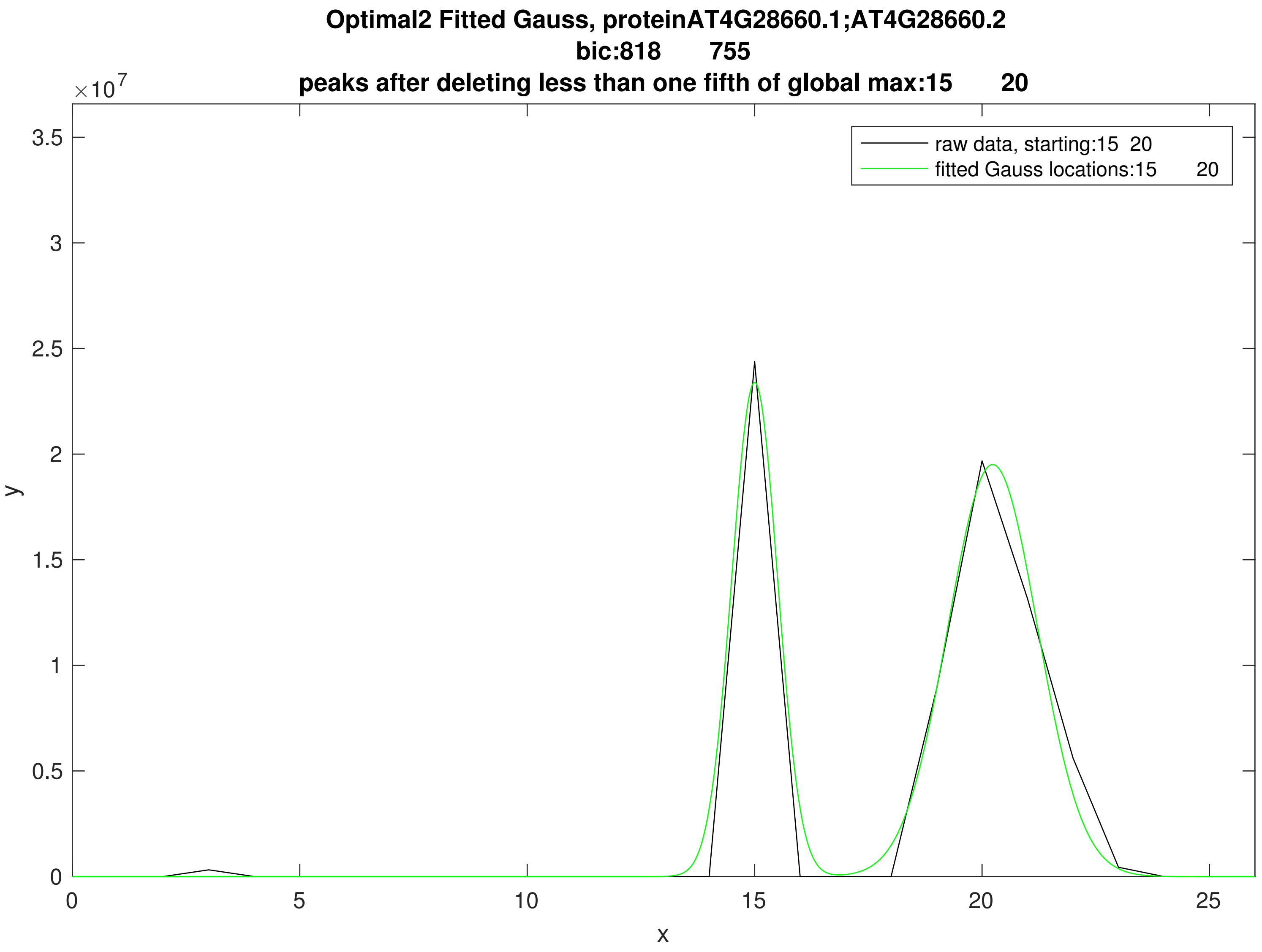

Optimal2 Fitted Gauss, proteinAT4G28660.1;AT4G28660.2
bic:818 755
peaks after deleting less than one fifth of global max:15 20
×10^7
3.5
3
2.5
2
1.5
1
0.5
0
y
0
5
10
15
20
25
x
raw data, starting:15 20
fitted Gauss locations:15 20

**Optimal1 Fitted Gauss, proteinAT4G28750.1**
**bic:628**
**peaks after deleting less than one fifth of global max:20**

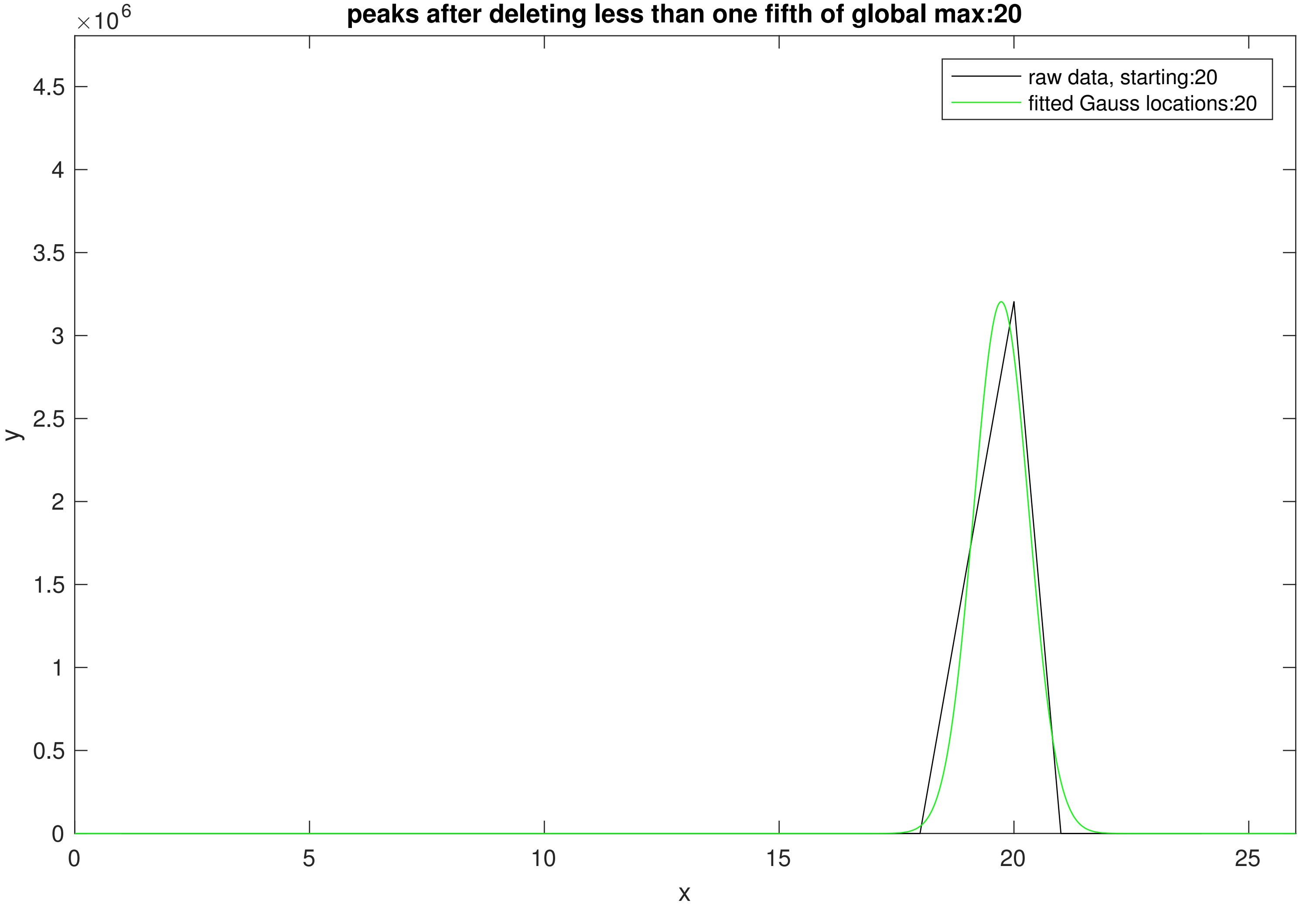

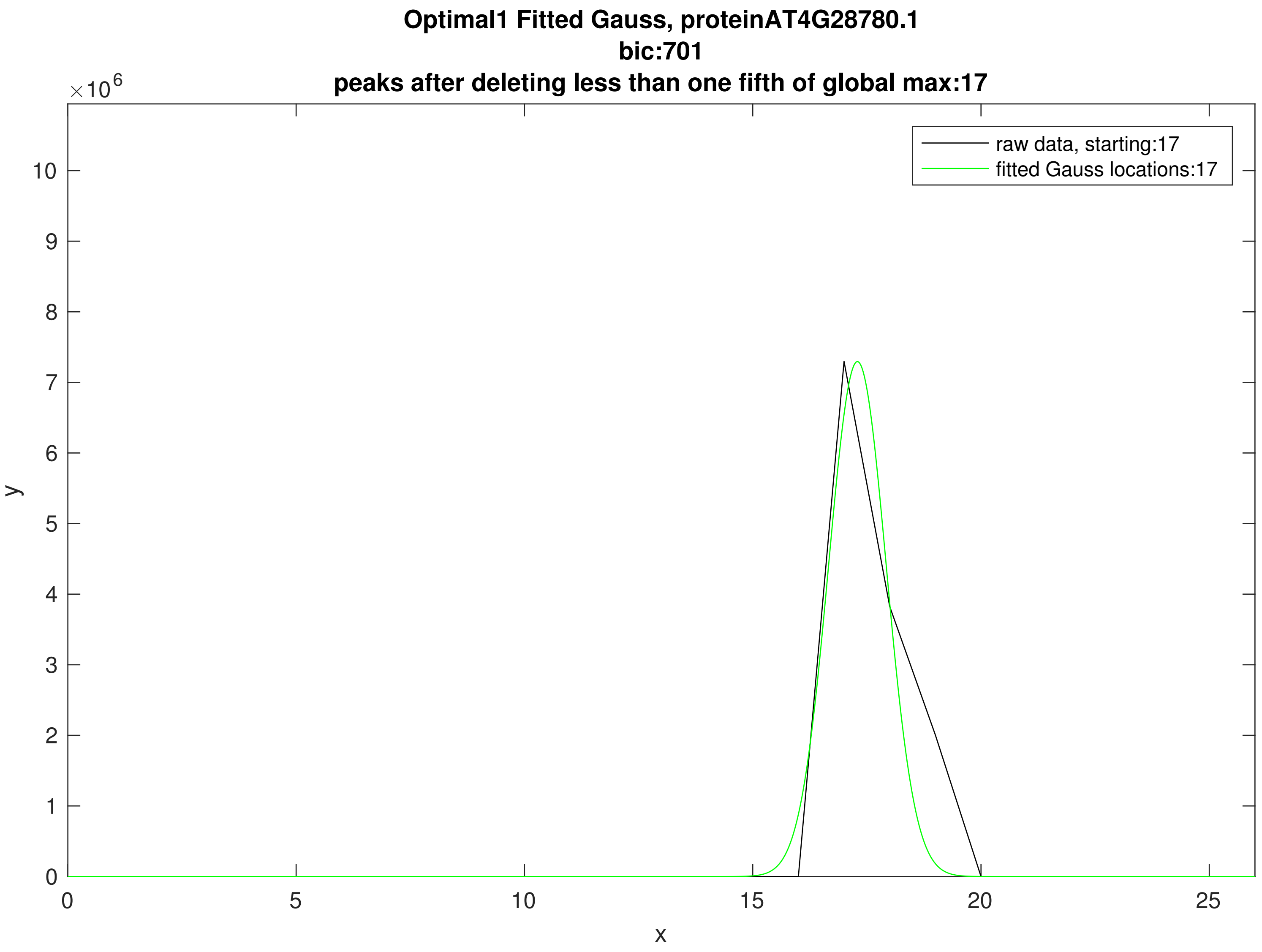

Optimal1 Fitted Gauss, proteinAT4G28780.1
bic:701
peaks after deleting less than one fifth of global max:17
×10^6
raw data, starting:17
fitted Gauss locations:17
y
x
0
1
2
3
4
5
6
7
8
9
10
0
5
10
15
20
25

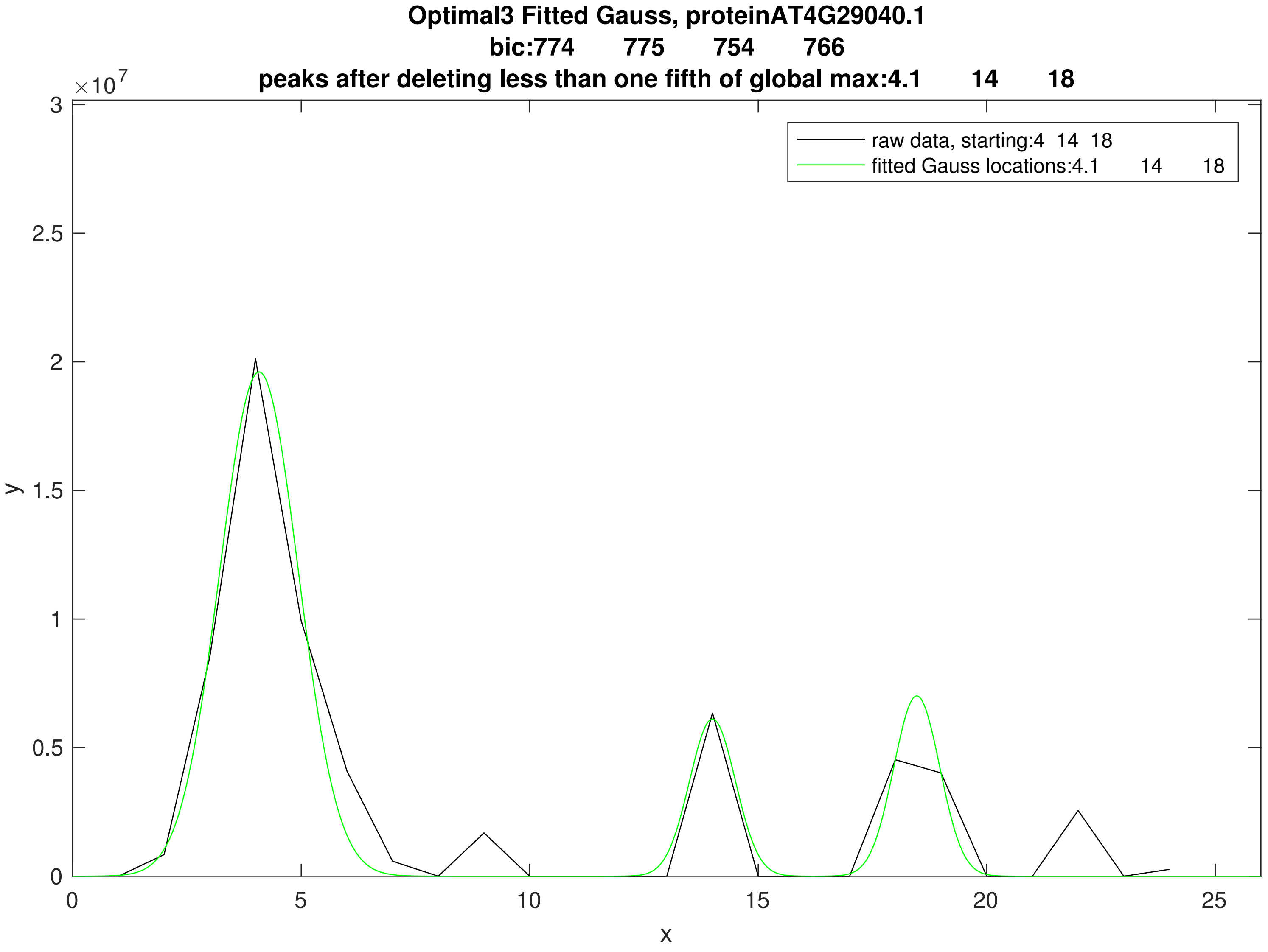

Optimal3 Fitted Gauss, proteinAT4G29040.1
bic:774 775 754 766
peaks after deleting less than one fifth of global max:4.1 14 18
×10^7
raw data, starting:4 14 18
fitted Gauss locations:4.1 14 18
y
x
0
0.5
1
1.5
2
2.5
3
0
5
10
15
20
25

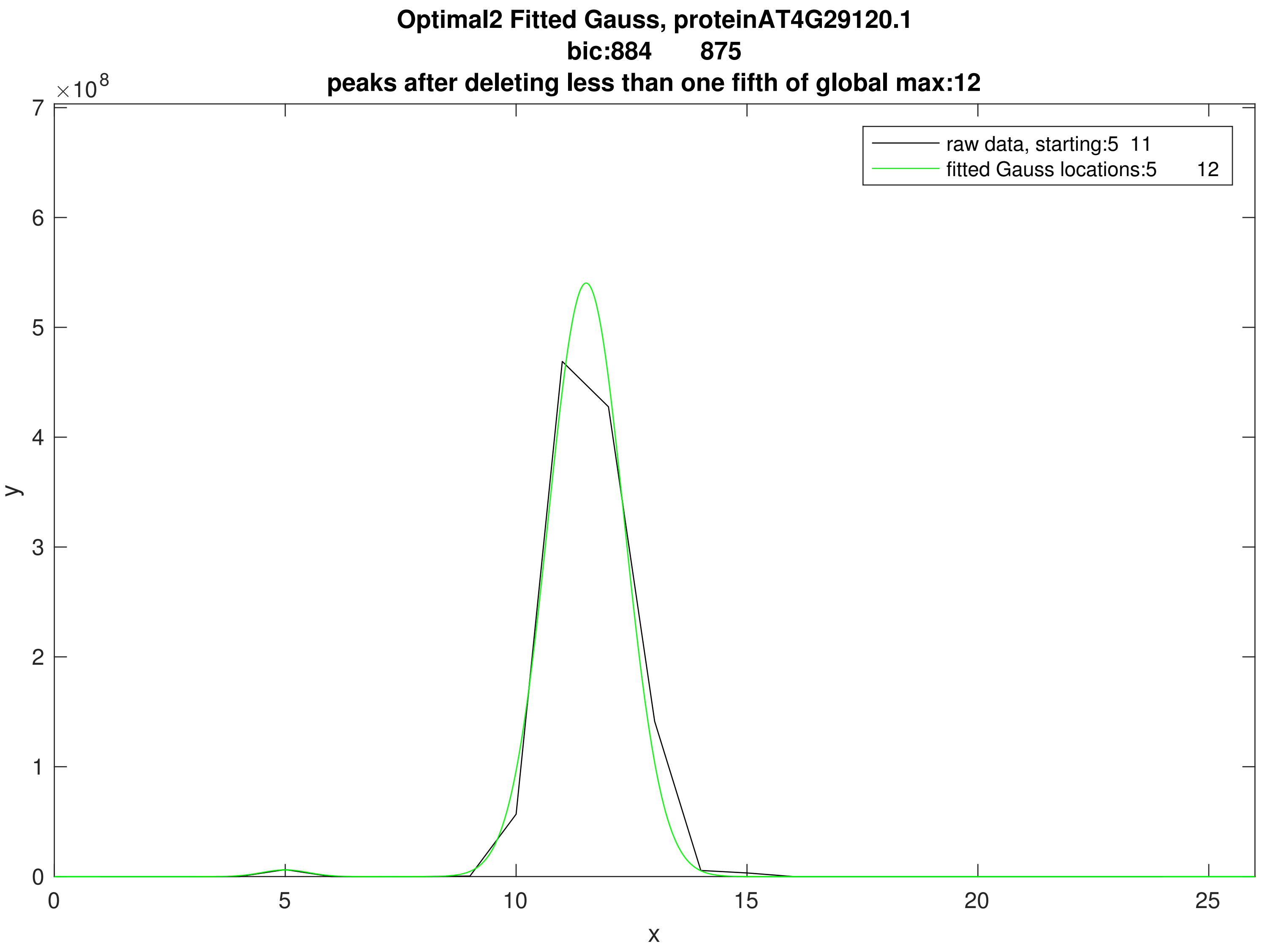

Optimal2 Fitted Gauss, proteinAT4G29120.1
bic:884 875
peaks after deleting less than one fifth of global max:12
×10^8
7
6
5
4
3
2
1
0
y
0
5
10
15
20
25
x
raw data, starting:5 11
fitted Gauss locations:5 12

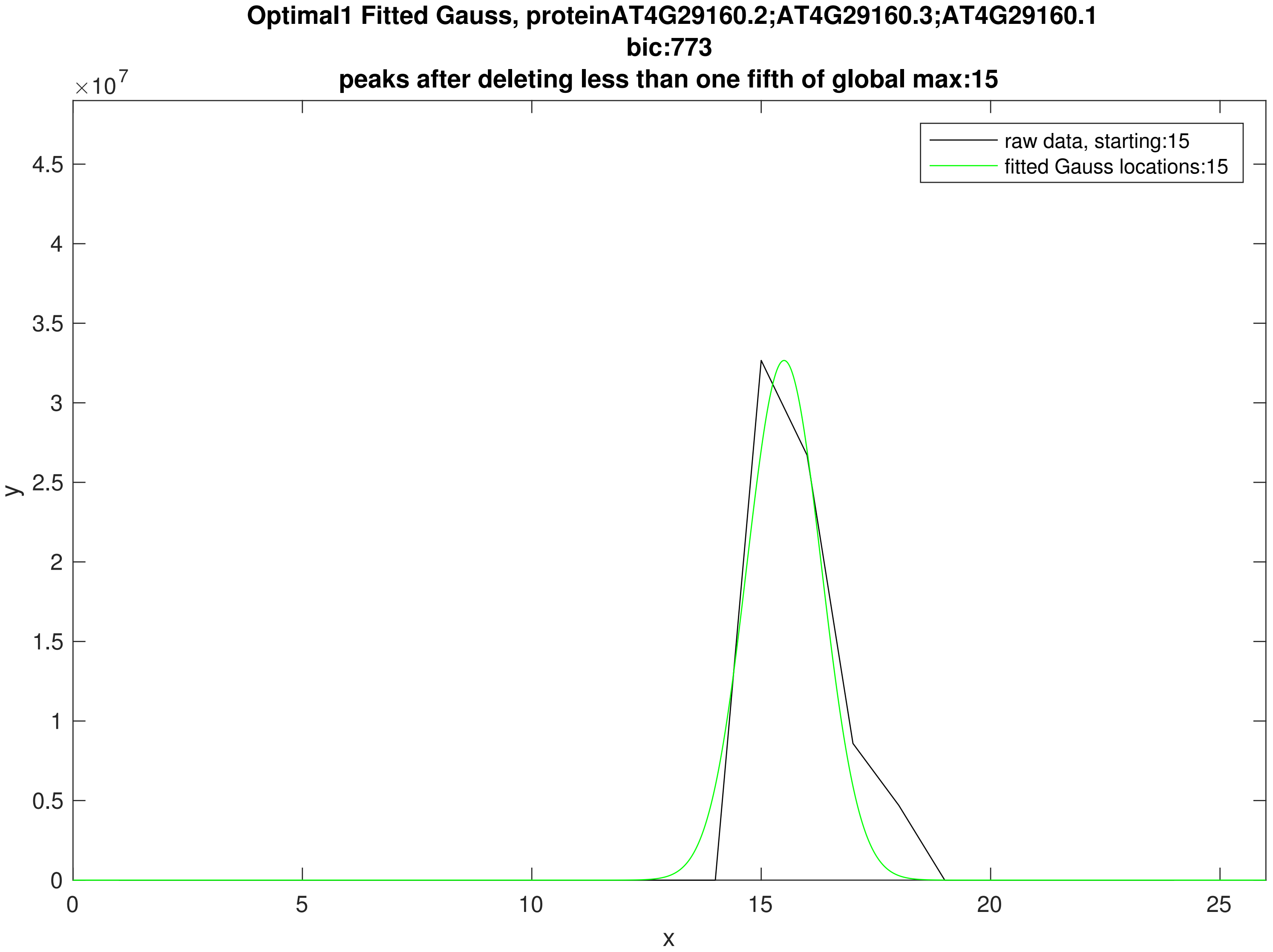
Optimal1 Fitted Gauss, proteinAT4G29160.2;AT4G29160.3;AT4G29160.1
bic:773
peaks after deleting less than one fifth of global max:15
×10^7
4.5
4
3.5
3
2.5
2
1.5
1
0.5
0
y
0
5
10
15
20
25
x
raw data, starting:15
fitted Gauss locations:15

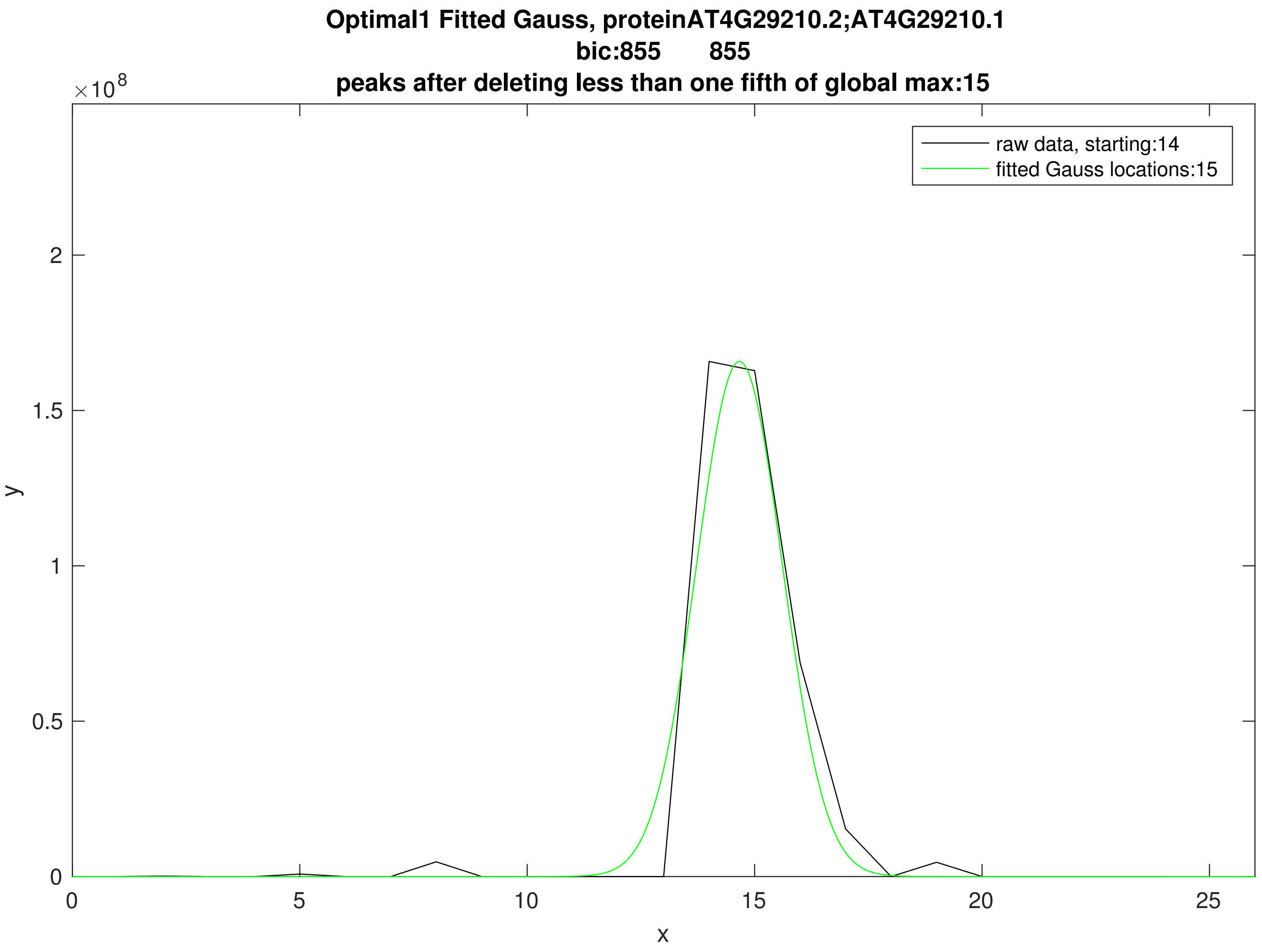

Optimal1 Fitted Gauss, proteinAT4G29210.2;AT4G29210.1
bic:855 855
peaks after deleting less than one fifth of global max:15
×10^8
raw data, starting:14
fitted Gauss locations:15
y
x
0
0.5
1
1.5
2
0
5
10
15
20
25

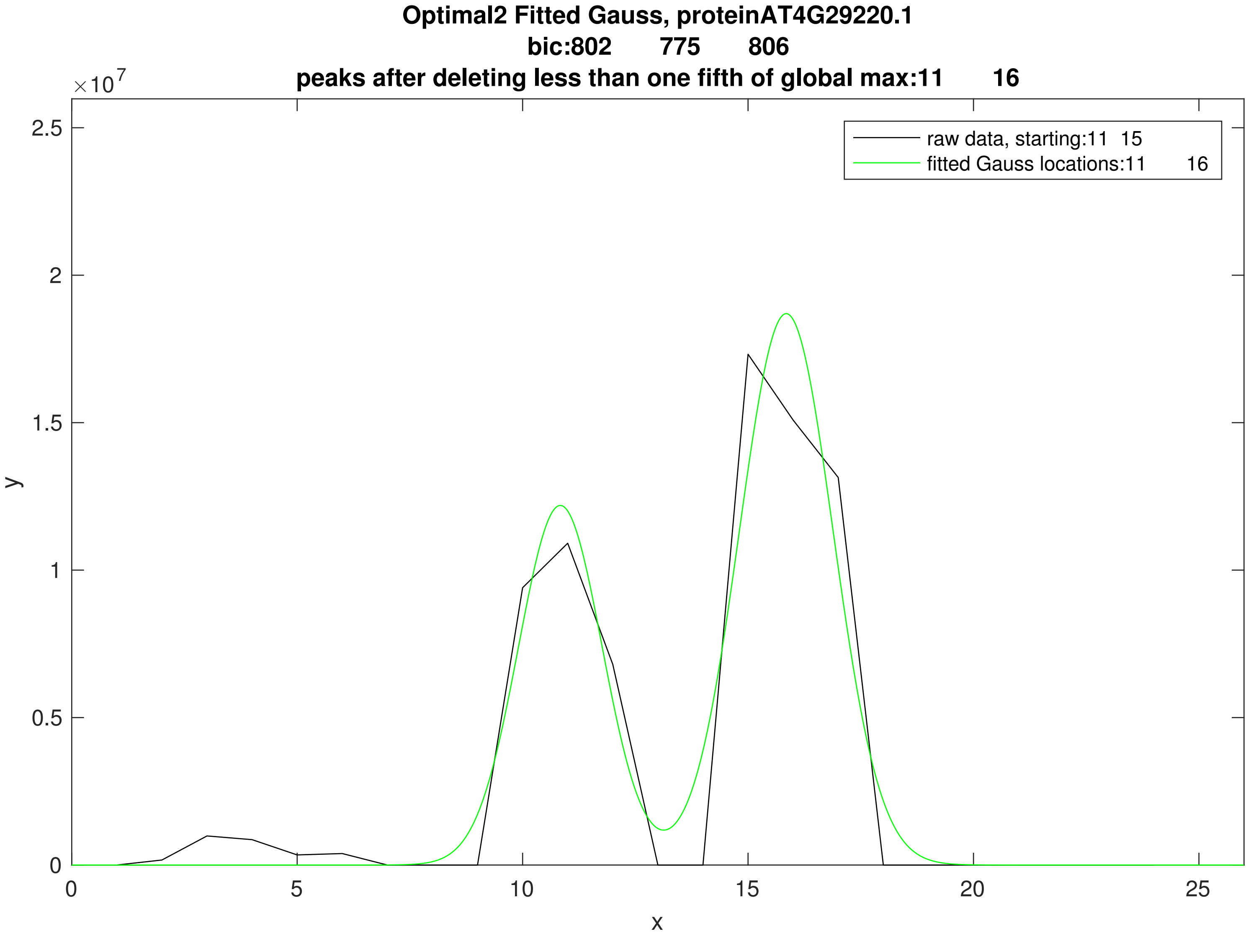

Optimal2 Fitted Gauss, proteinAT4G29220.1
bic:802 775 806
peaks after deleting less than one fifth of global max:11 16
×10^7
raw data, starting:11 15
fitted Gauss locations:11 16
y
x
2.5
2
1.5
1
0.5
0
5
10
15
20
25

**Optimal1 Fitted Gauss, proteinAT4G29260.1**
**bic:878 888**
**peaks after deleting less than one fifth of global max:16**

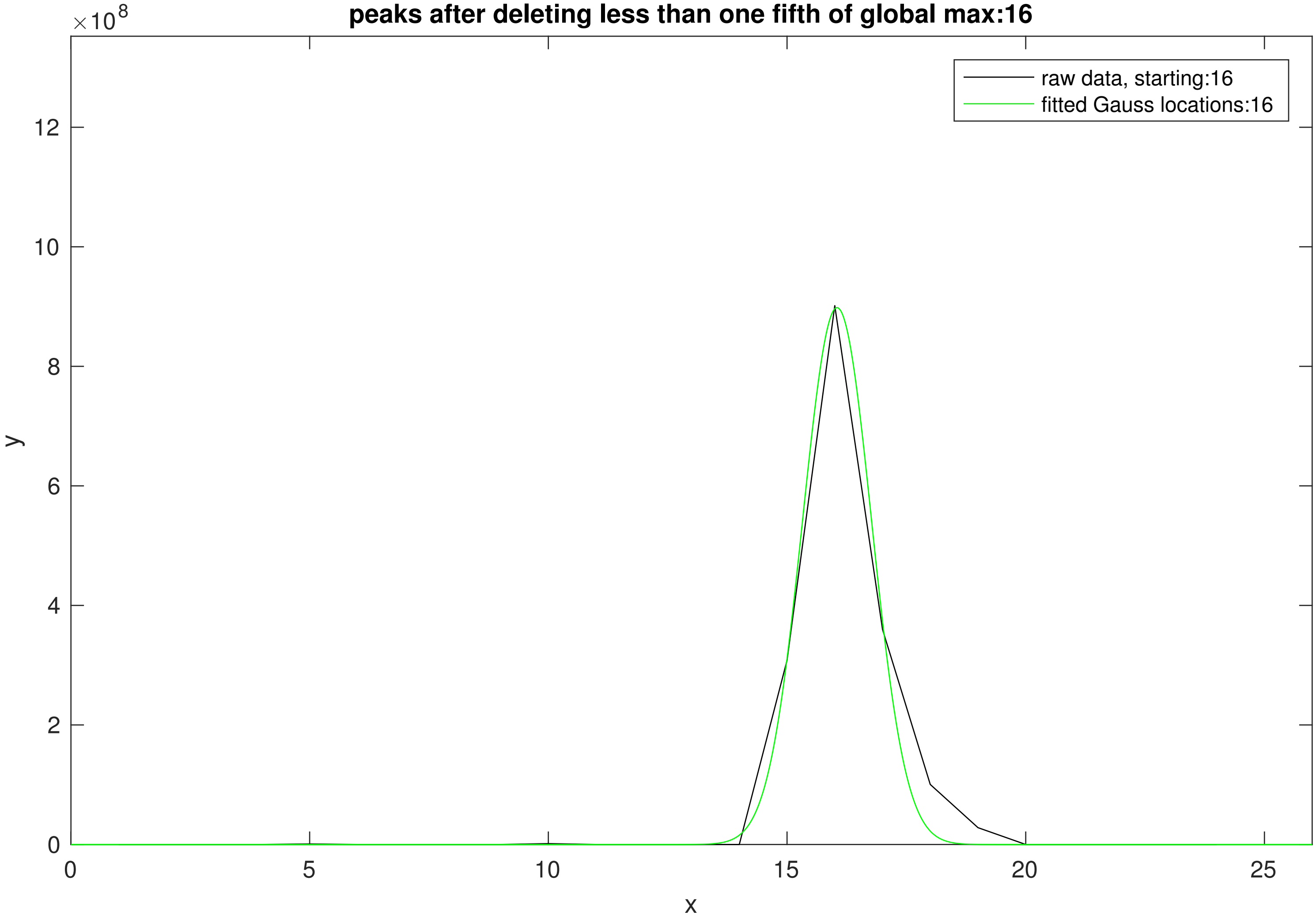

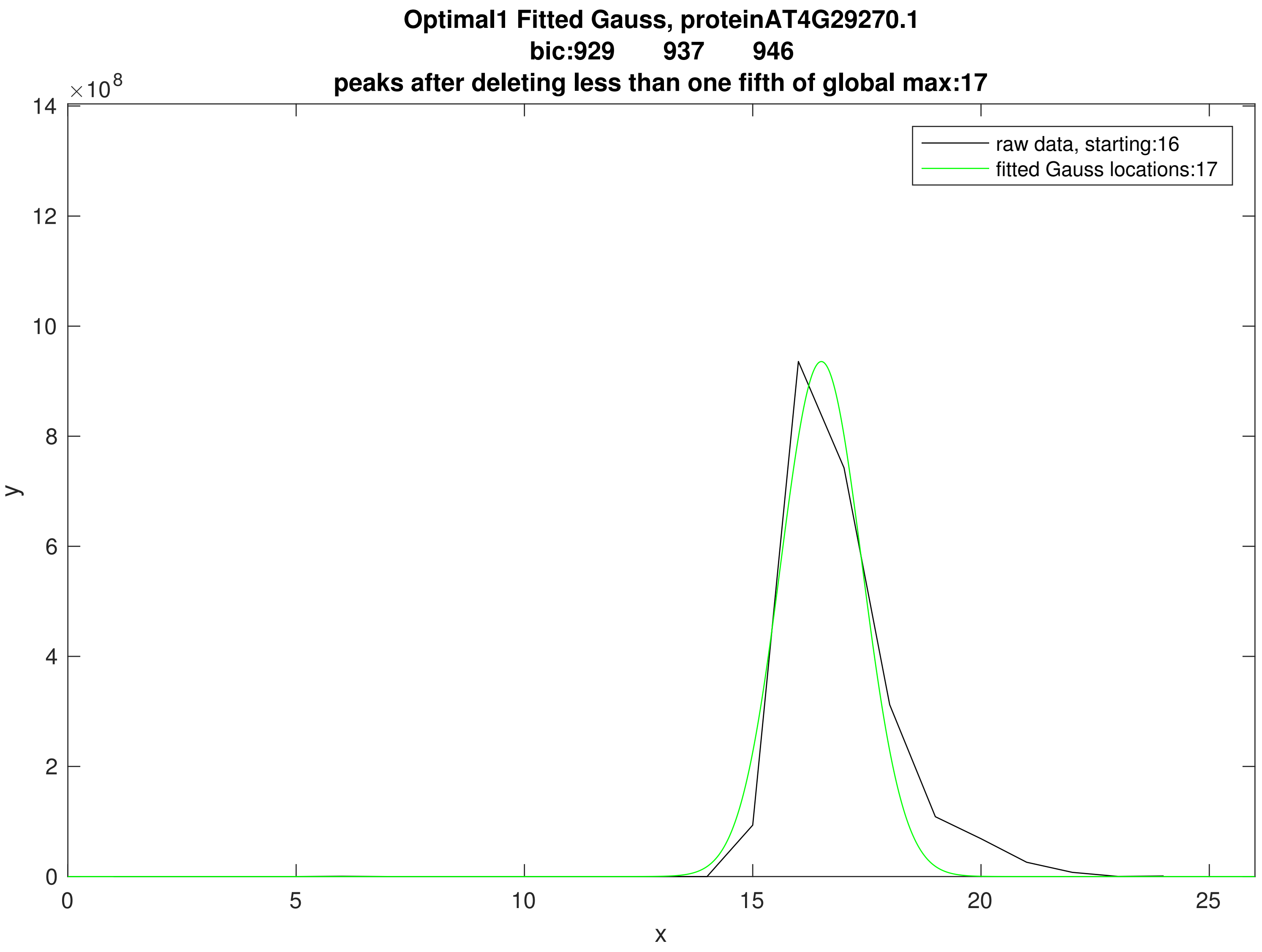
Optimal1 Fitted Gauss, proteinAT4G29270.1
bic:929 937 946
peaks after deleting less than one fifth of global max:17
×10^8
raw data, starting:16
fitted Gauss locations:17
y
x
0
2
4
6
8
10
12
14
0
5
10
15
20
25

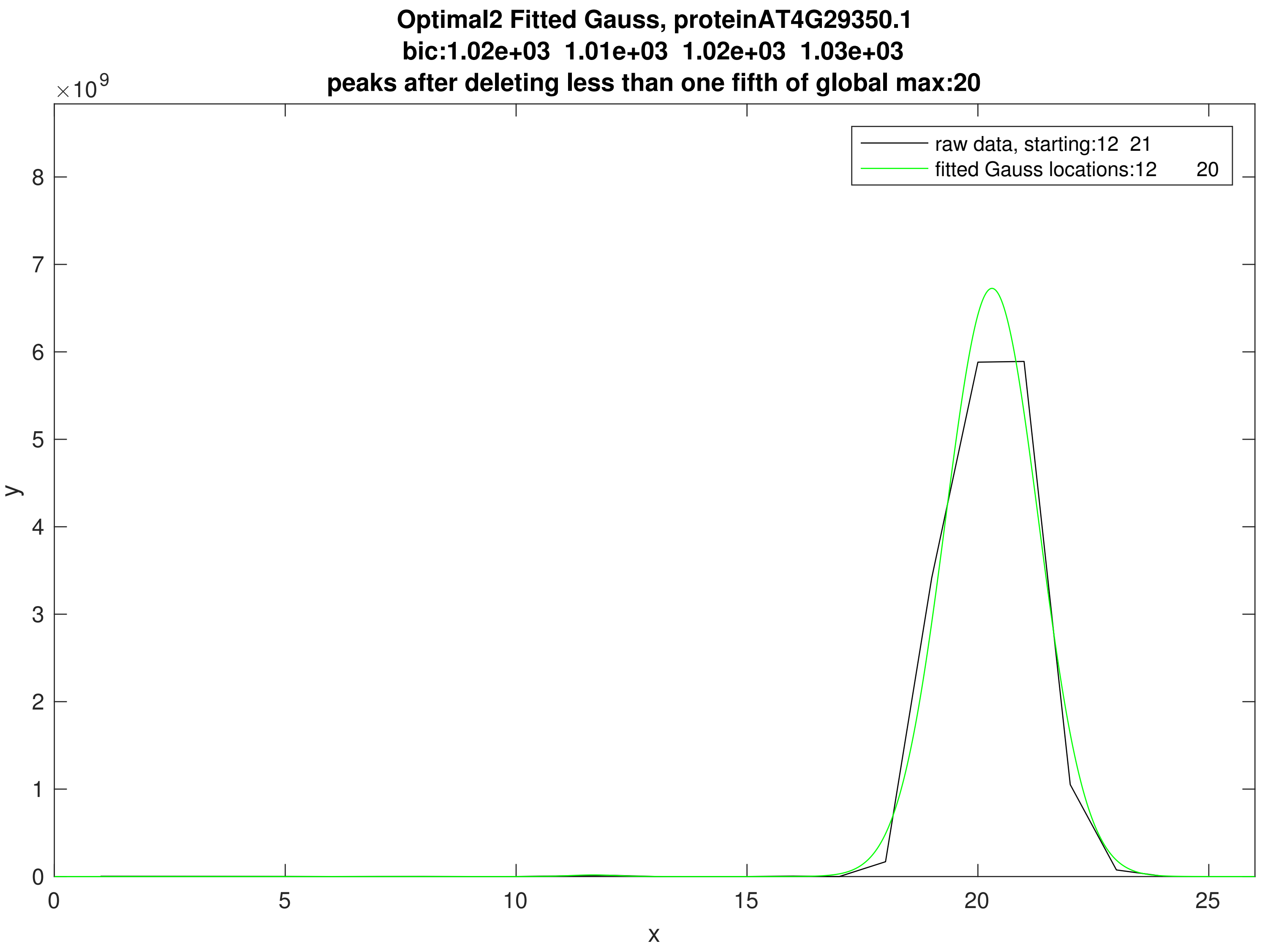
Optimal2 Fitted Gauss, proteinAT4G29350.1
bic:1.02e+03 1.01e+03 1.02e+03 1.03e+03
peaks after deleting less than one fifth of global max:20
×10^9
raw data, starting:12 21
fitted Gauss locations:12 20
y
x
0
1
2
3
4
5
6
7
8
0
5
10
15
20
25

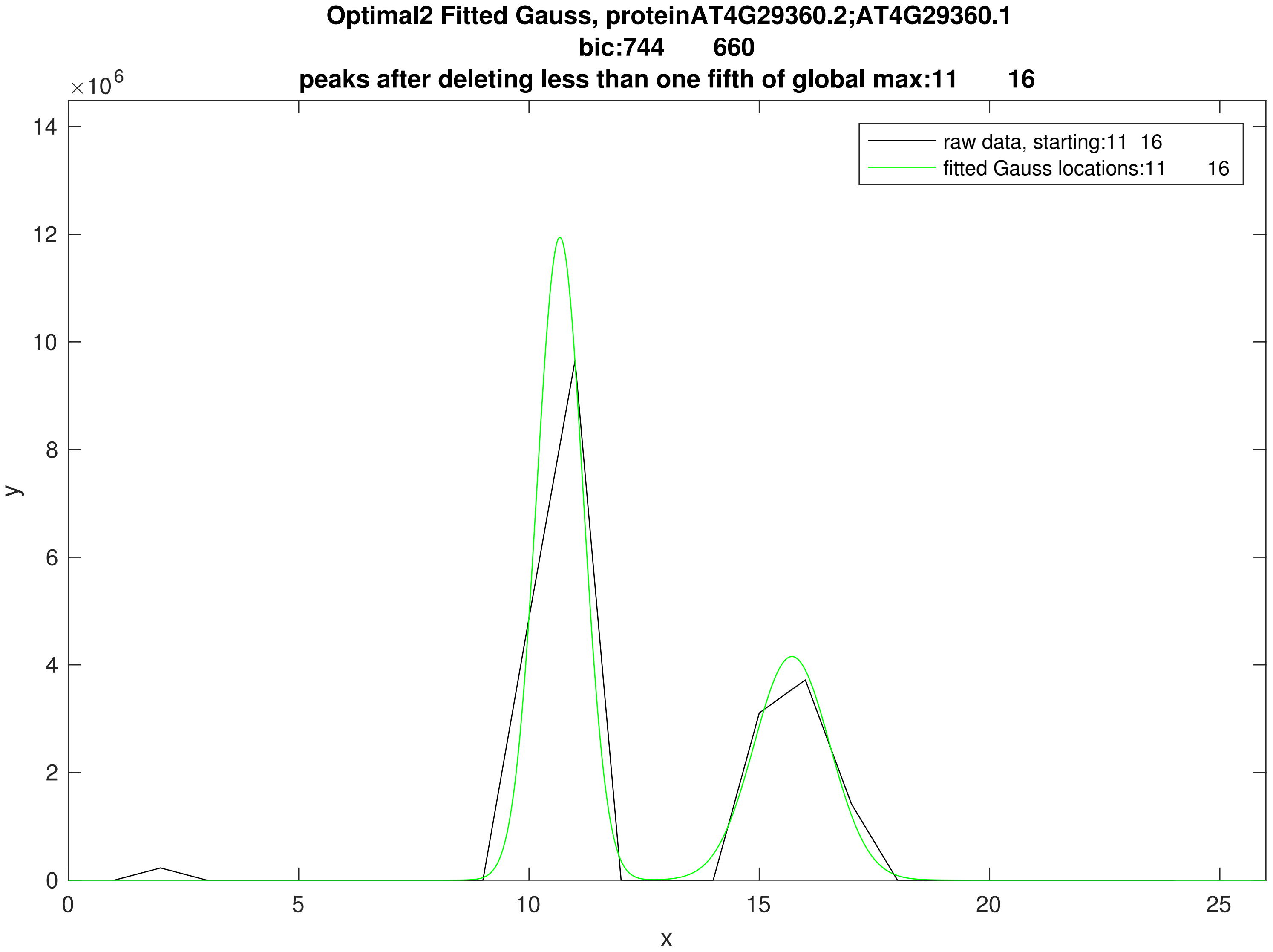
Optimal2 Fitted Gauss, proteinAT4G29360.2;AT4G29360.1
bic:744 660
peaks after deleting less than one fifth of global max:11 16
×10^6
raw data, starting:11 16
fitted Gauss locations:11 16
y
x
0
2
4
6
8
10
12
14
5
15
20
25

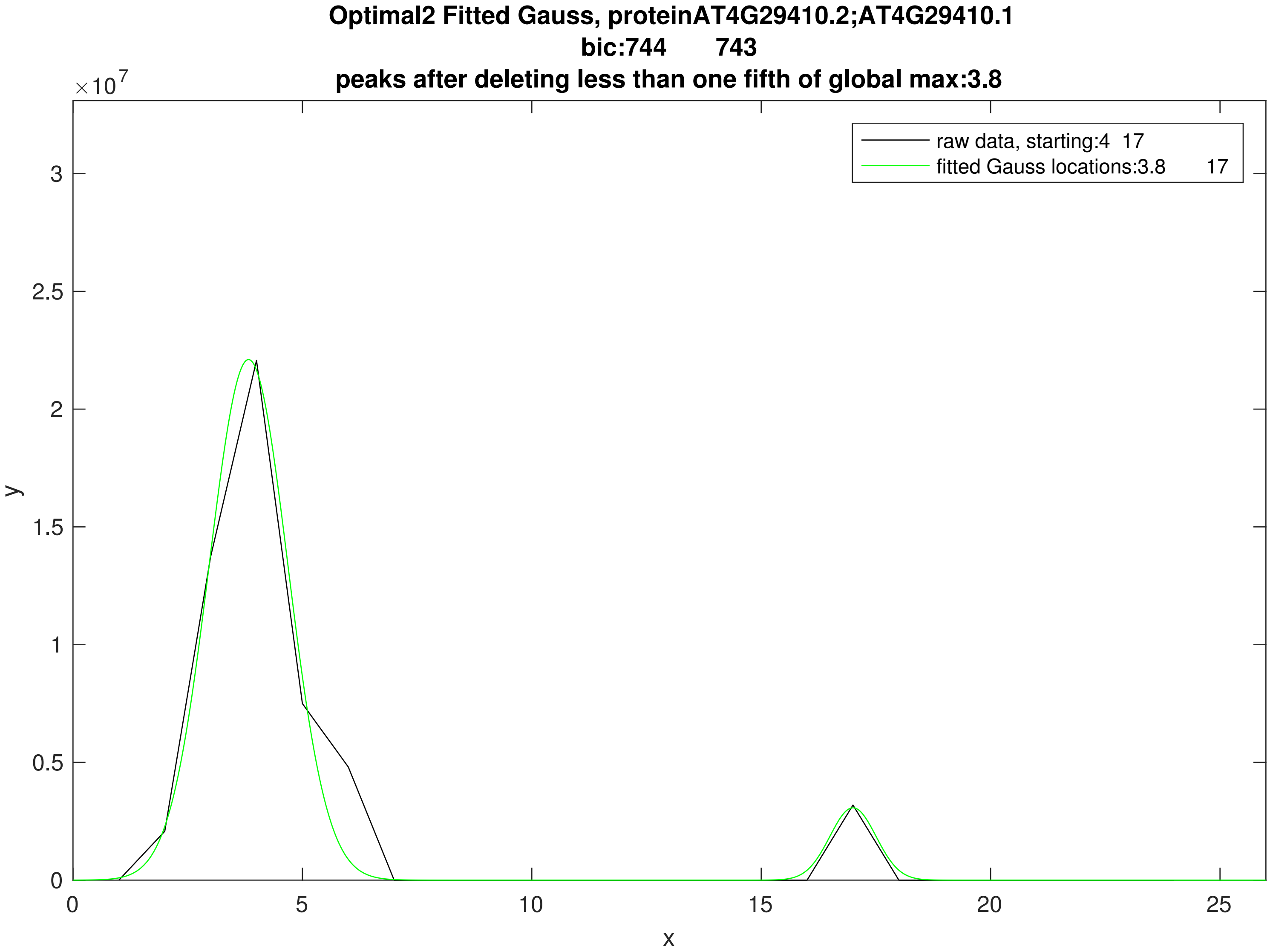
Optimal2 Fitted Gauss, proteinAT4G29410.2;AT4G29410.1
bic:744 743
peaks after deleting less than one fifth of global max:3.8
×10^7
raw data, starting:4 17
fitted Gauss locations:3.8 17
y
x
0
0.5
1
1.5
2
2.5
3
5
10
15
20
25

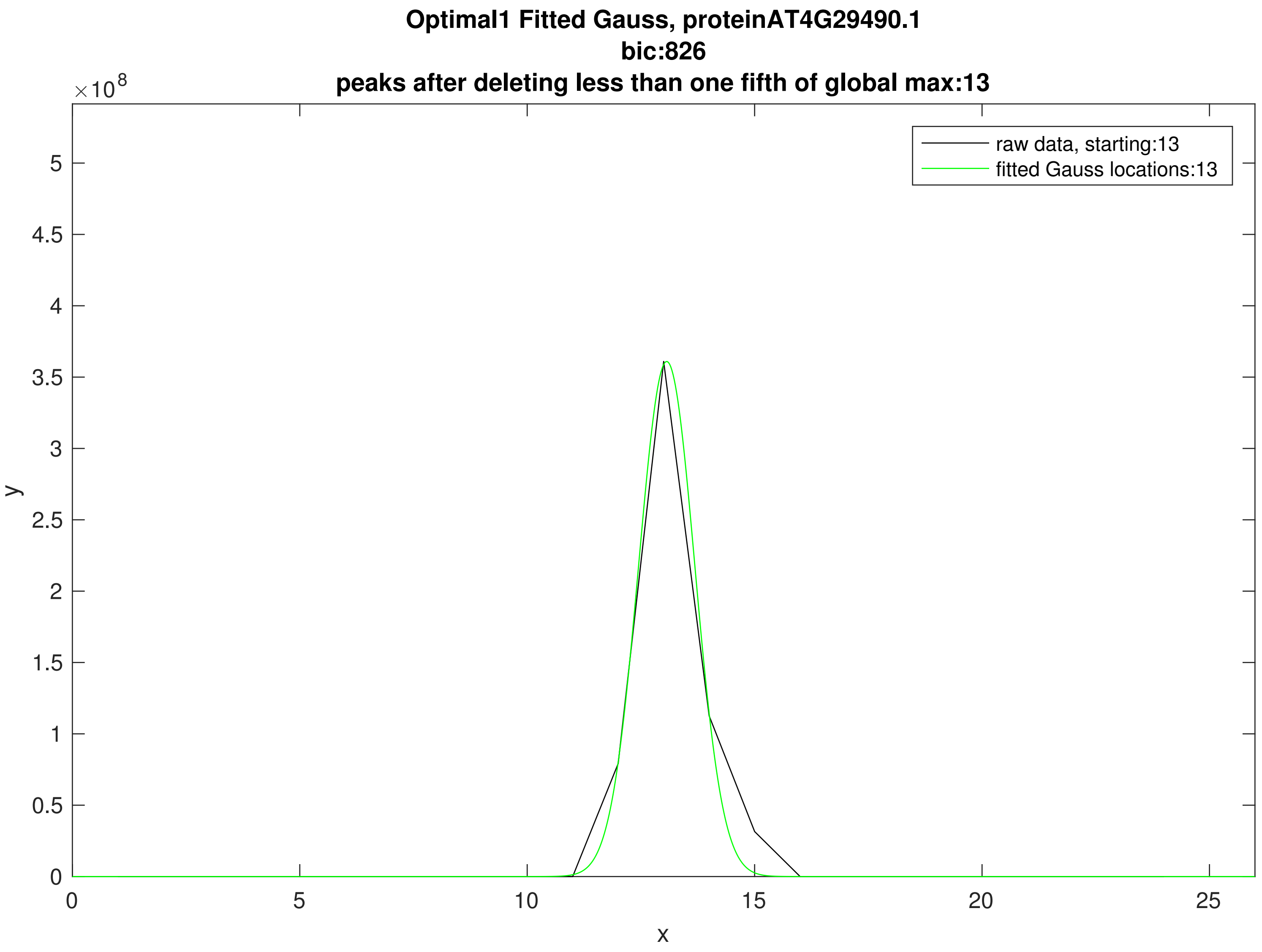

Optimal1 Fitted Gauss, proteinAT4G29490.1
bic:826
peaks after deleting less than one fifth of global max:13
×10^8
raw data, starting:13
fitted Gauss locations:13
y
x
0
0.5
1
1.5
2
2.5
3
3.5
4
4.5
5
0
5
10
15
20
25

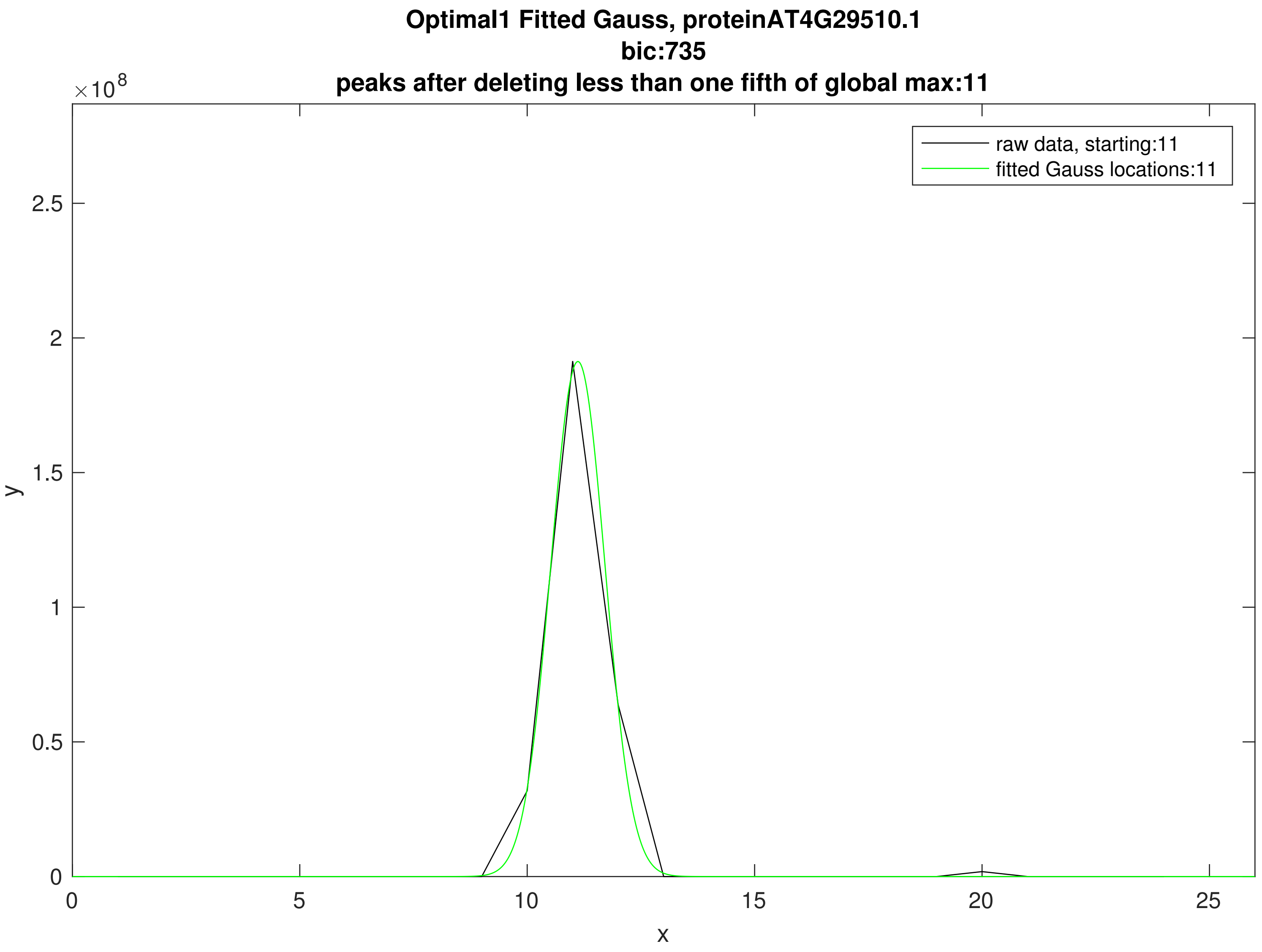
Optimal1 Fitted Gauss, proteinAT4G29510.1
bic:735
peaks after deleting less than one fifth of global max:11
×10^8
raw data, starting:11
fitted Gauss locations:11
2.5
2
1.5
1
0.5
0
y
0
5
10
15
20
25
x

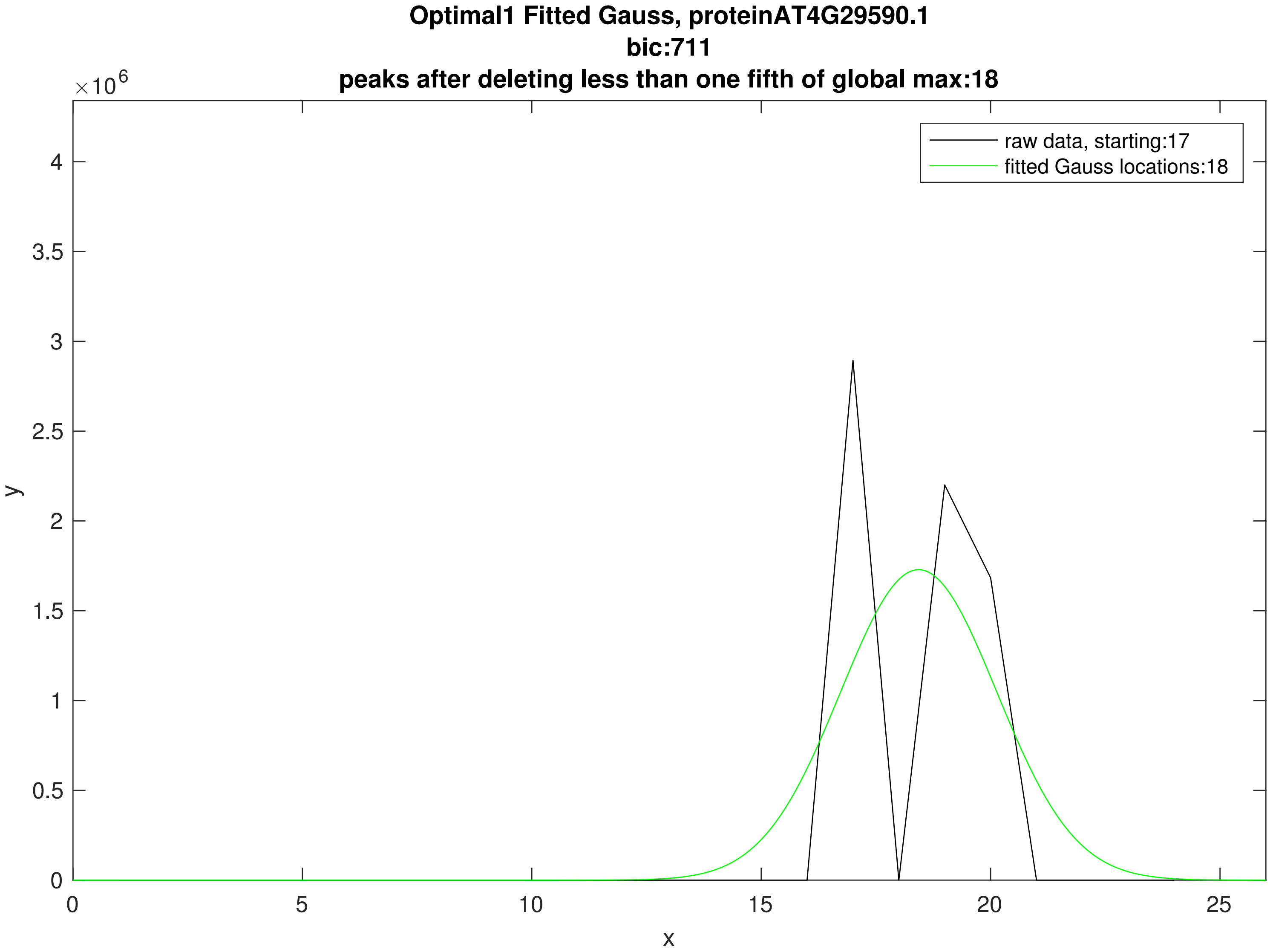
Optimal1 Fitted Gauss, proteinAT4G29590.1
bic:711
peaks after deleting less than one fifth of global max:18
×10^6
raw data, starting:17
fitted Gauss locations:18
y
x
0
0.5
1
1.5
2
2.5
3
3.5
4
0
5
10
15
20
25

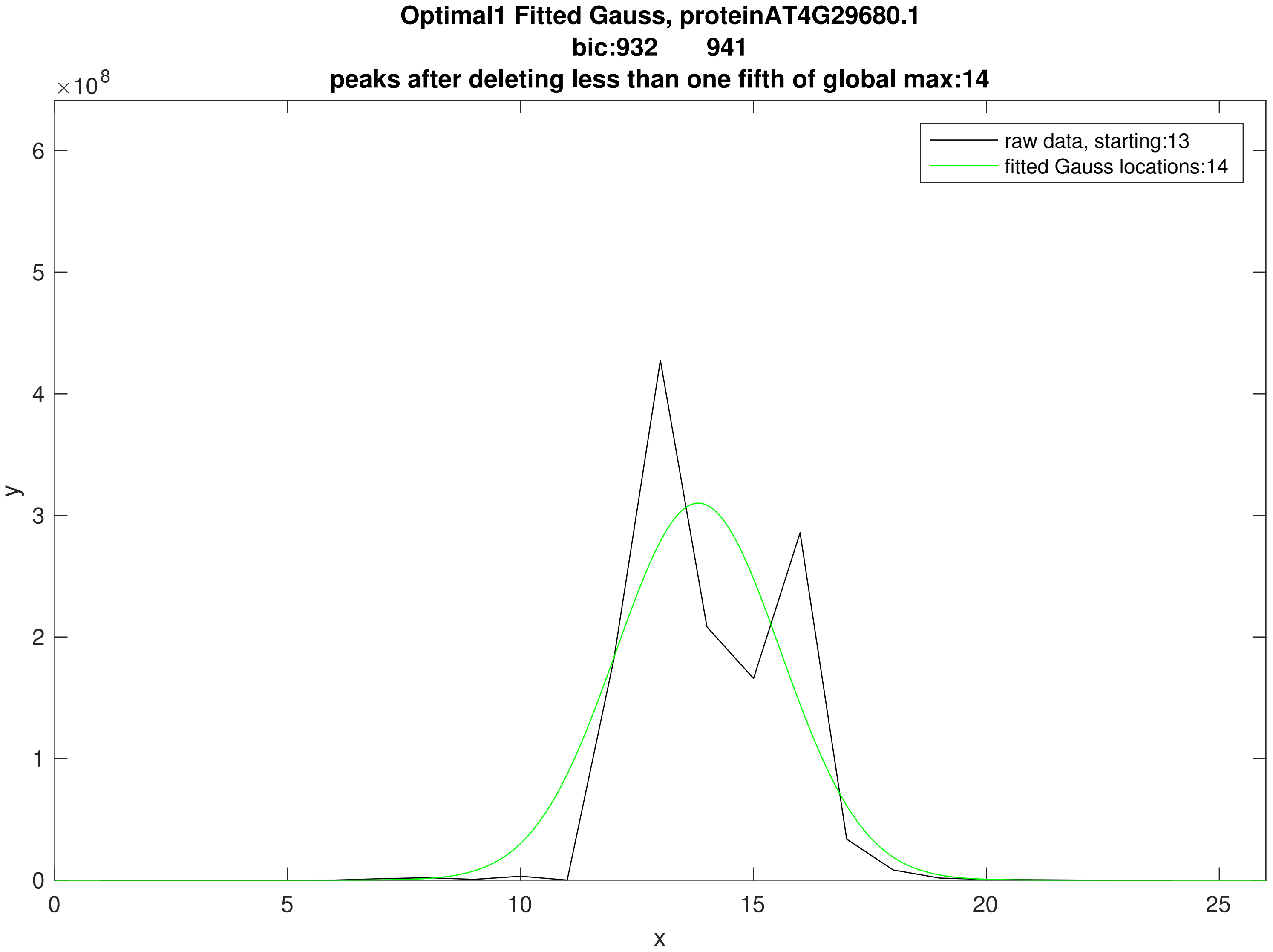
Optimal1 Fitted Gauss, proteinAT4G29680.1
bic:932 941
peaks after deleting less than one fifth of global max:14
×10^8
raw data, starting:13
fitted Gauss locations:14
y
x
0
1
2
3
4
5
6
0
5
10
15
20
25

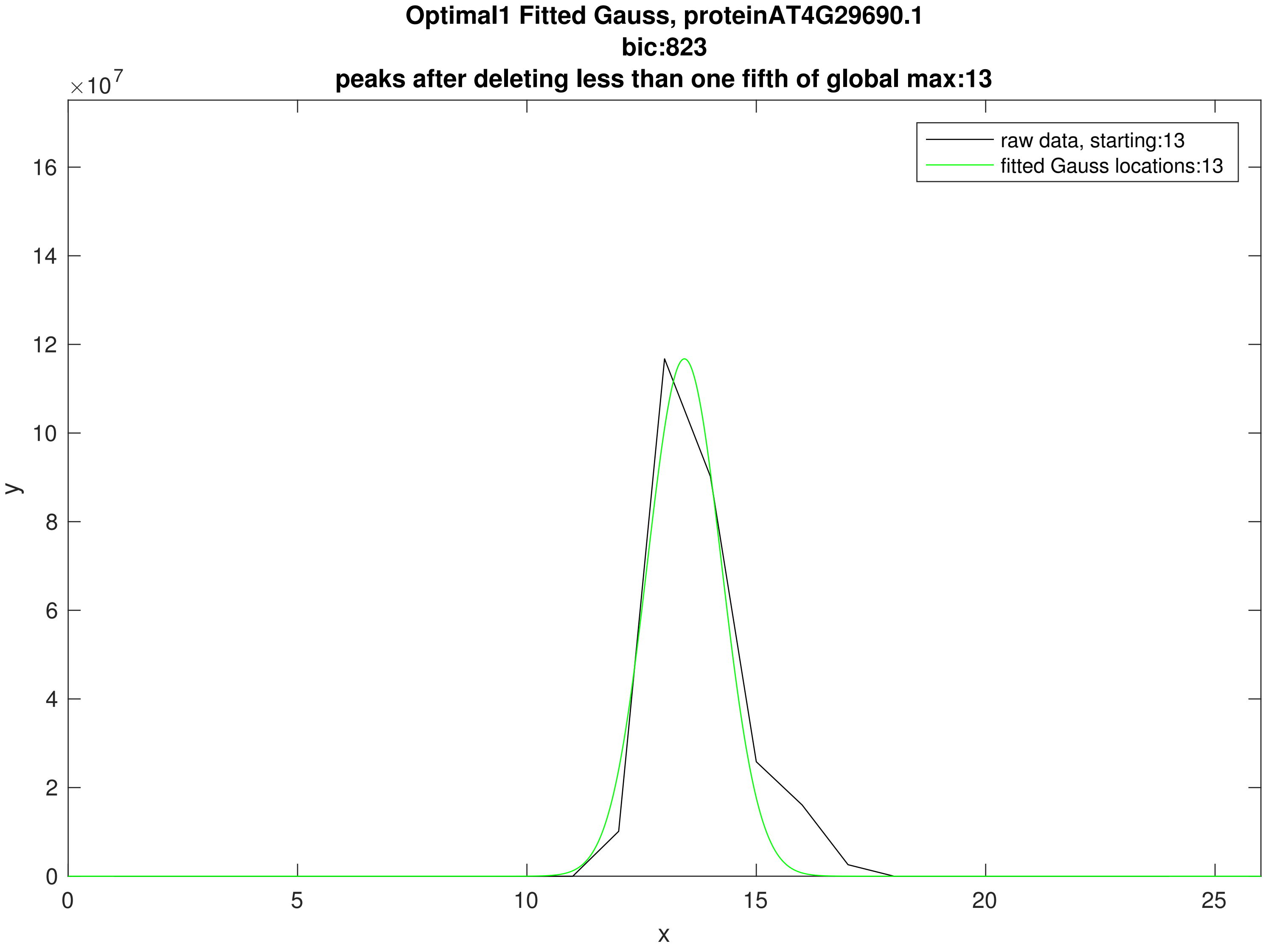
Optimal1 Fitted Gauss, proteinAT4G29690.1
bic:823
peaks after deleting less than one fifth of global max:13
×10^7
raw data, starting:13
fitted Gauss locations:13
y
x
0
2
4
6
8
10
12
14
16
0
5
10
15
20
25

Optimal1 Fitted Gauss, proteinAT4G29700.1
bic:797
peaks after deleting less than one fifth of global max:13

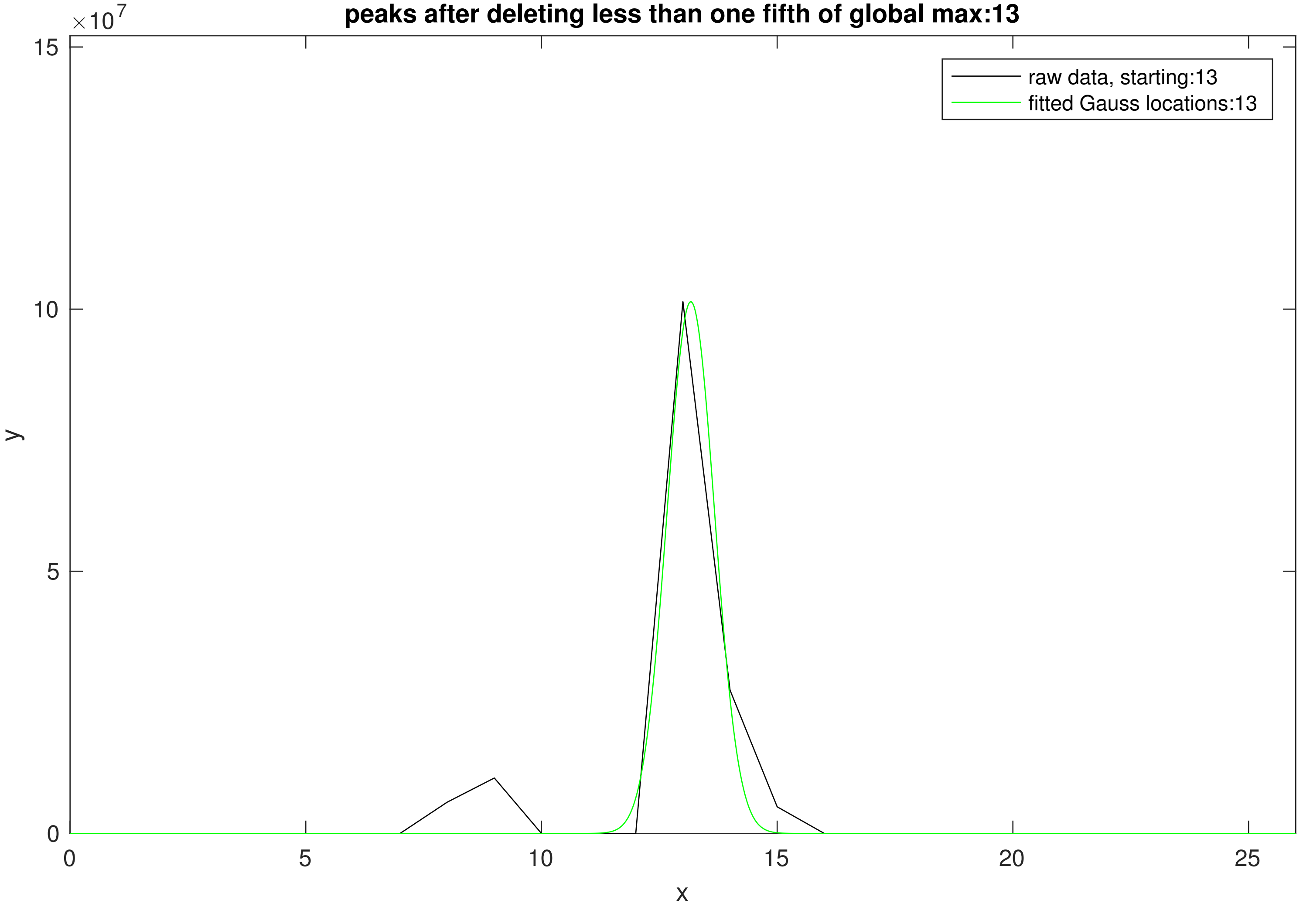

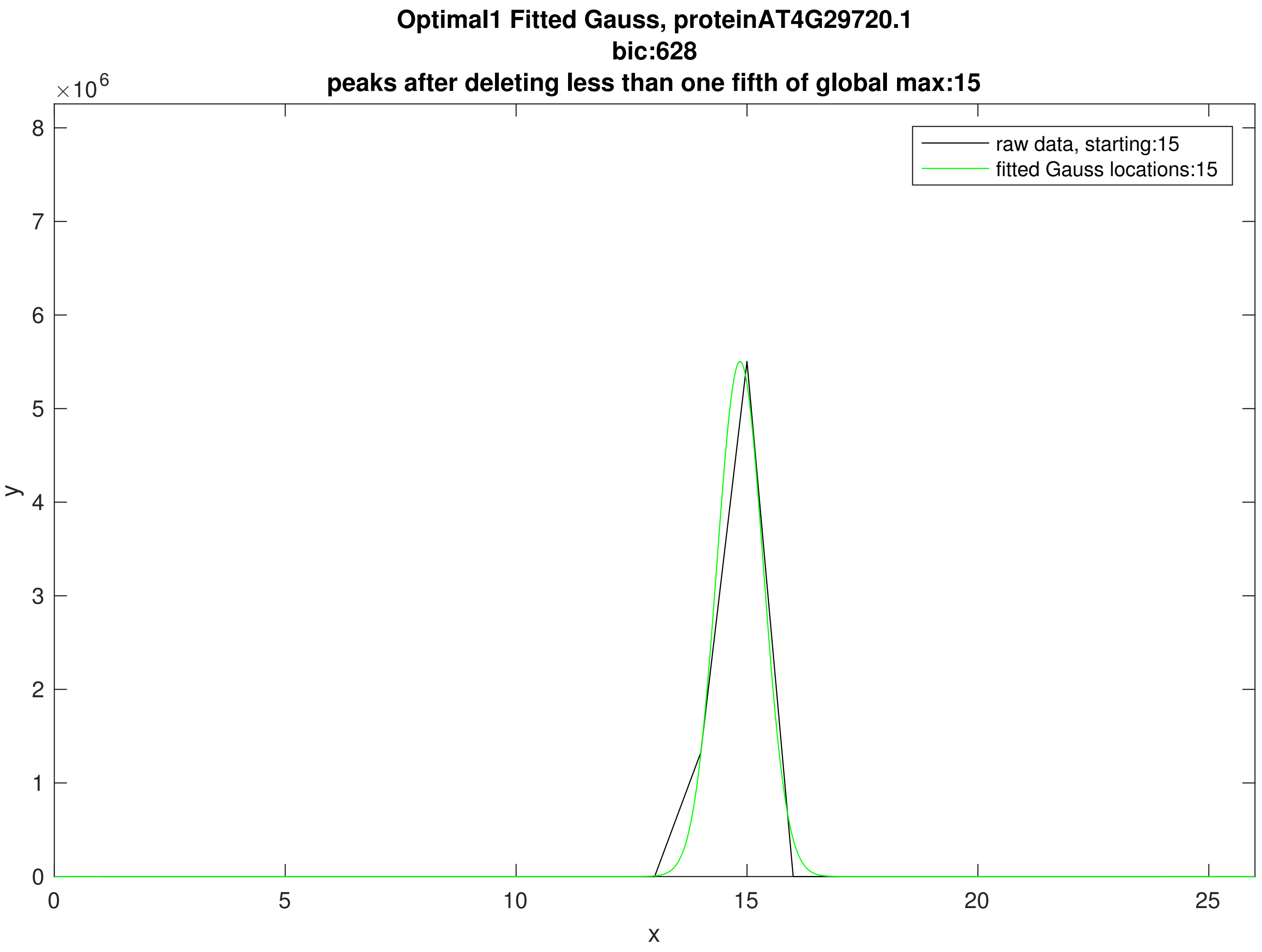
Optimal1 Fitted Gauss, proteinAT4G29720.1
bic:628
peaks after deleting less than one fifth of global max:15
×10^6
raw data, starting:15
fitted Gauss locations:15
y
x
0
1
2
3
4
5
6
7
8
0
5
10
15
20
25

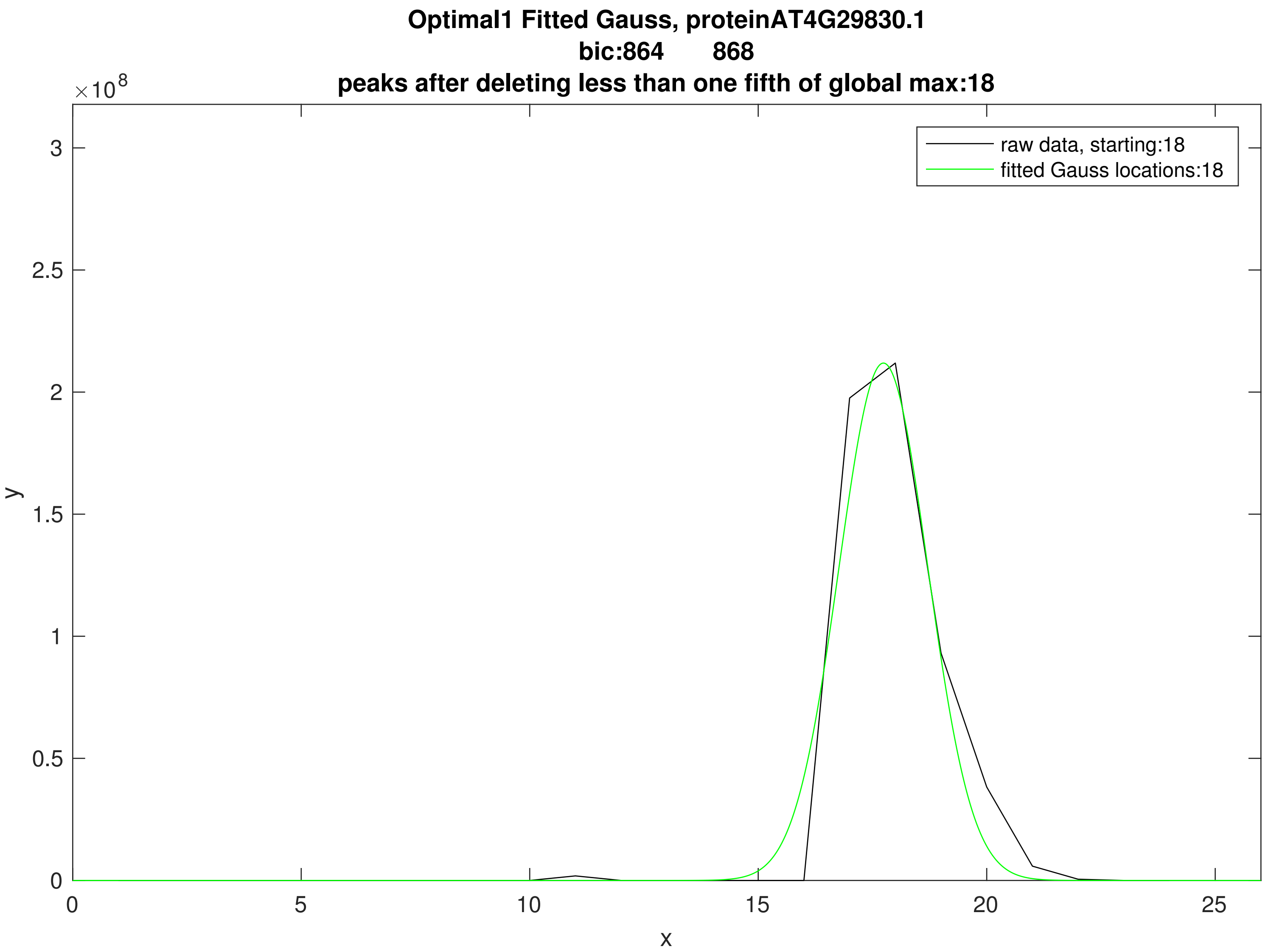
Optimal1 Fitted Gauss, proteinAT4G29830.1
bic:864 868
peaks after deleting less than one fifth of global max:18
×10^8
3
2.5
2
1.5
1
0.5
0
y
0
5
10
15
20
25
x
raw data, starting:18
fitted Gauss locations:18

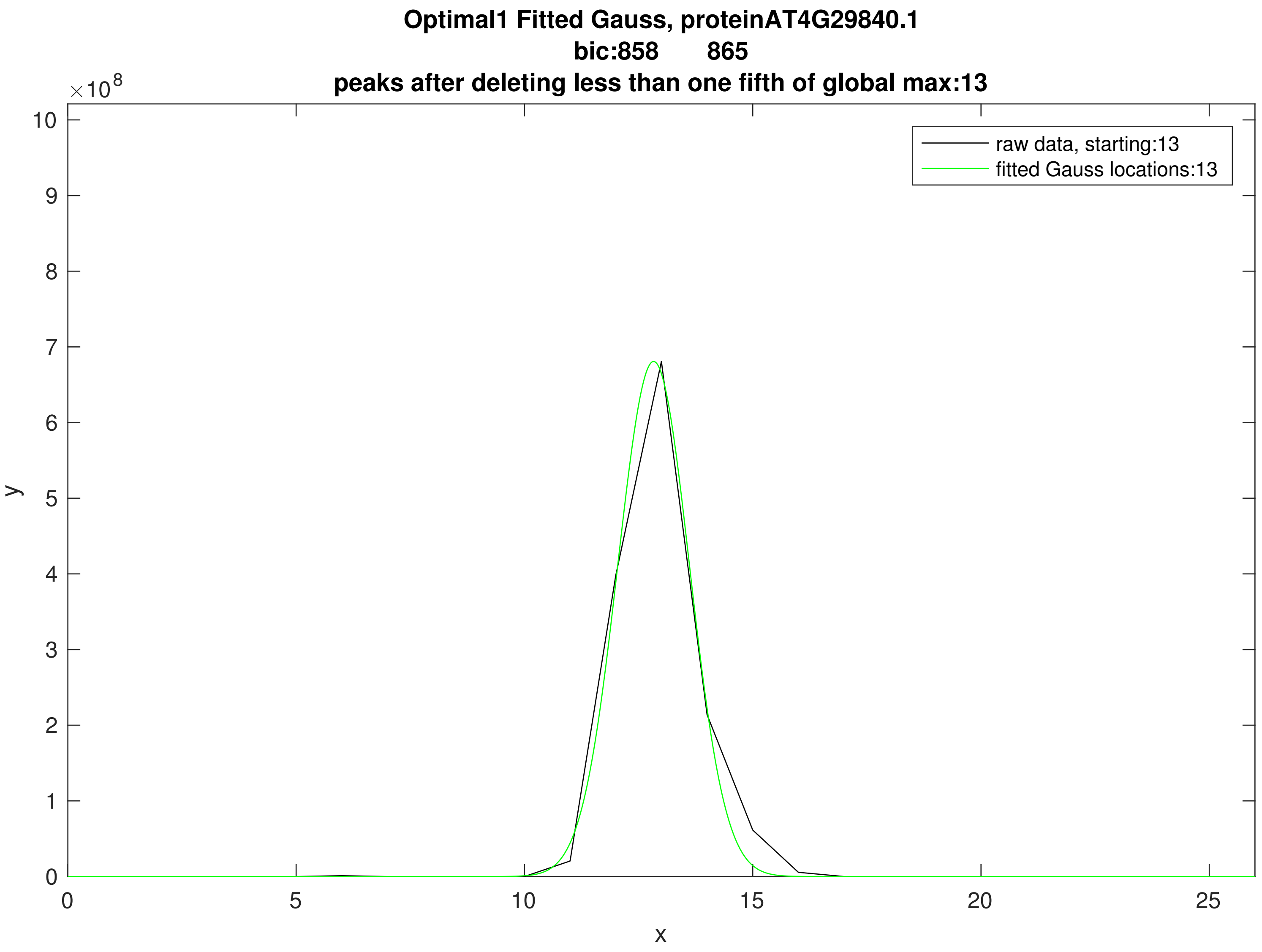

Optimal1 Fitted Gauss, proteinAT4G29840.1
bic:858 865
peaks after deleting less than one fifth of global max:13
×10^8
10
9
8
7
6
5
4
3
2
1
0
y
0
5
10
15
20
25
x
raw data, starting:13
fitted Gauss locations:13

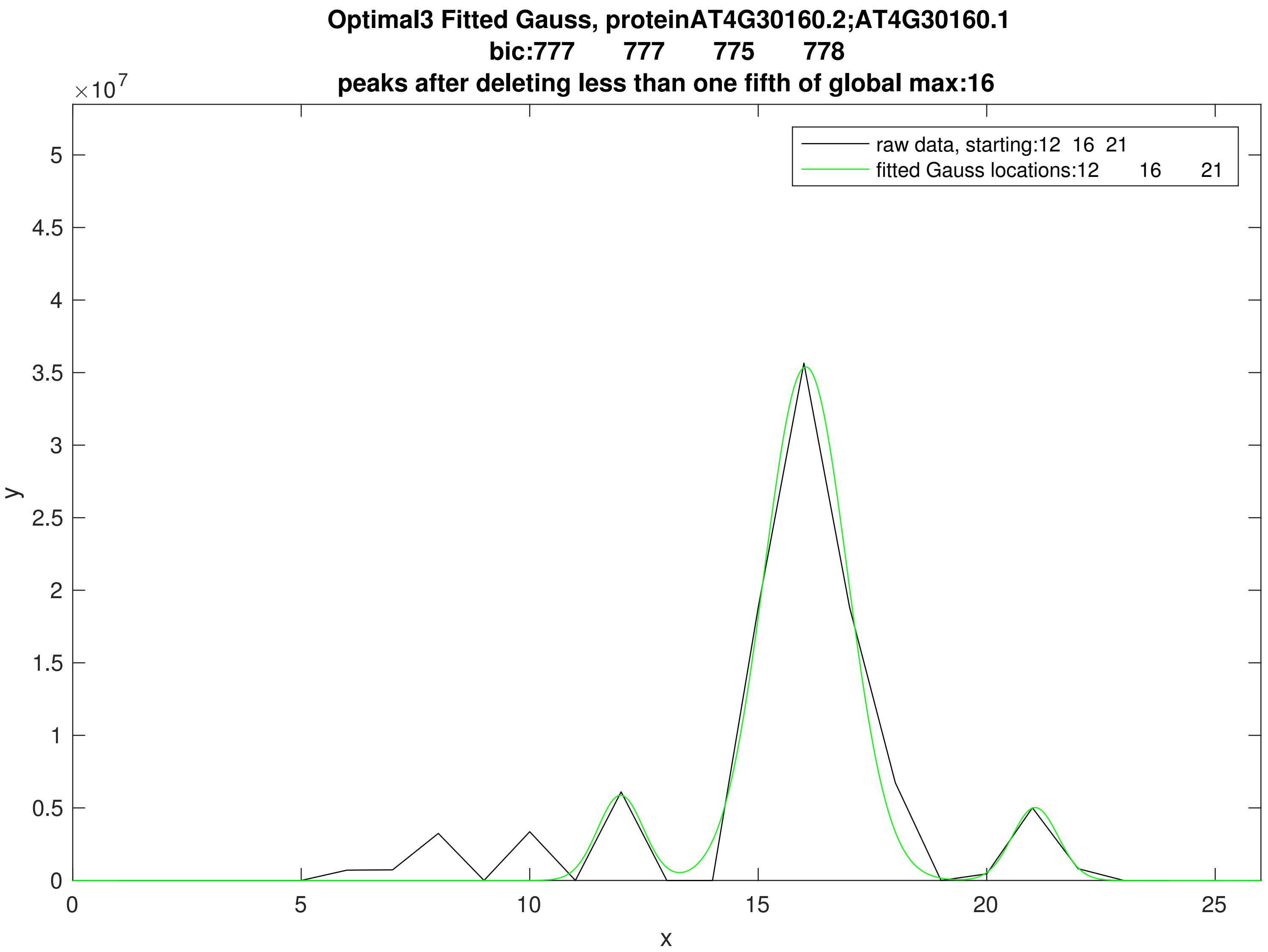
Optimal3 Fitted Gauss, proteinAT4G30160.2;AT4G30160.1
bic:777 777 775 778
peaks after deleting less than one fifth of global max:16
×10^7
raw data, starting:12 16 21
fitted Gauss locations:12 16 21
y
x
0
0.5
1
1.5
2
2.5
3
3.5
4
4.5
5
0
5
10
15
20
25

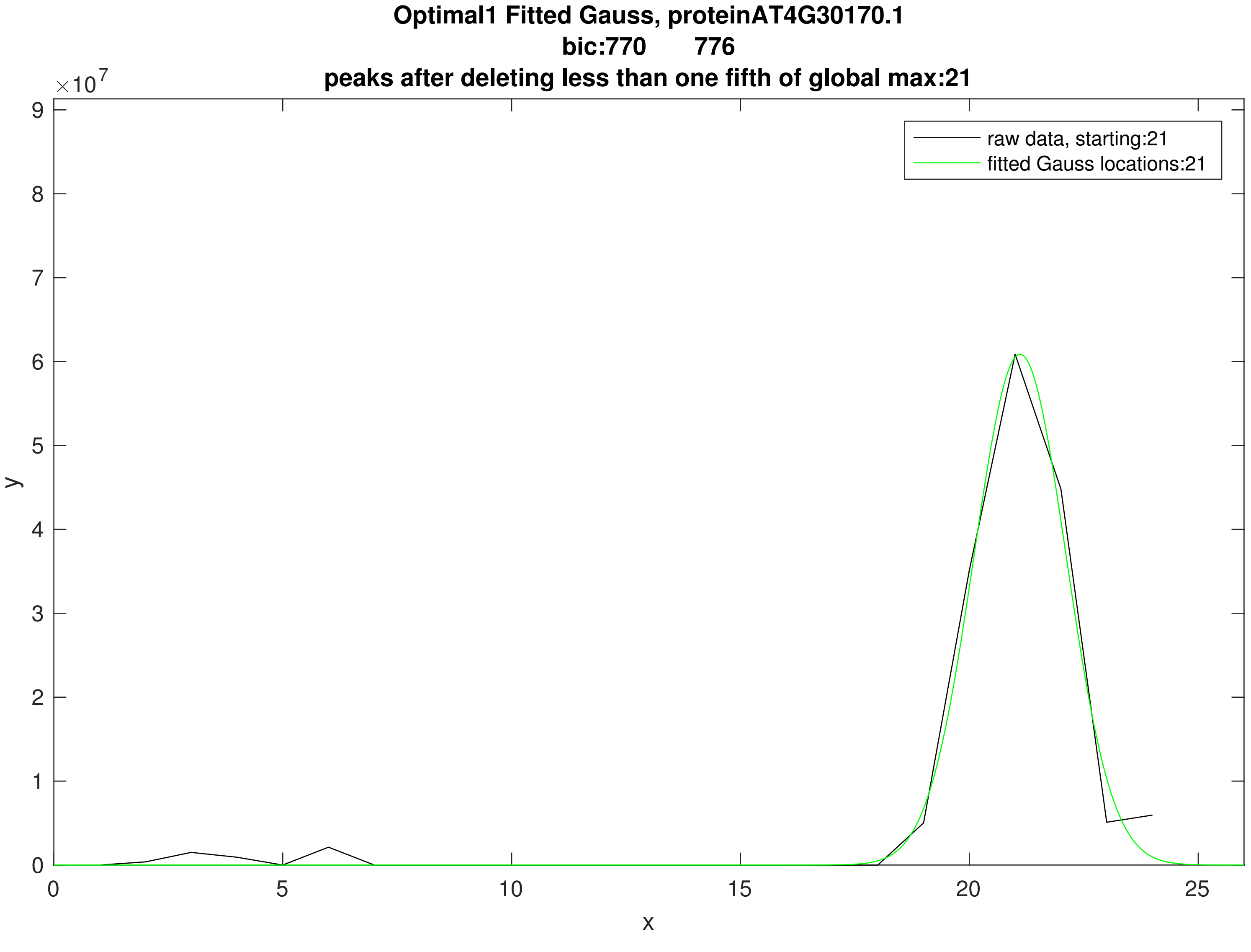
Optimal1 Fitted Gauss, proteinAT4G30170.1
bic:770 776
peaks after deleting less than one fifth of global max:21
×10^7
9
8
7
6
5
4
3
2
1
0
y
0
5
10
15
20
25
x
raw data, starting:21
fitted Gauss locations:21

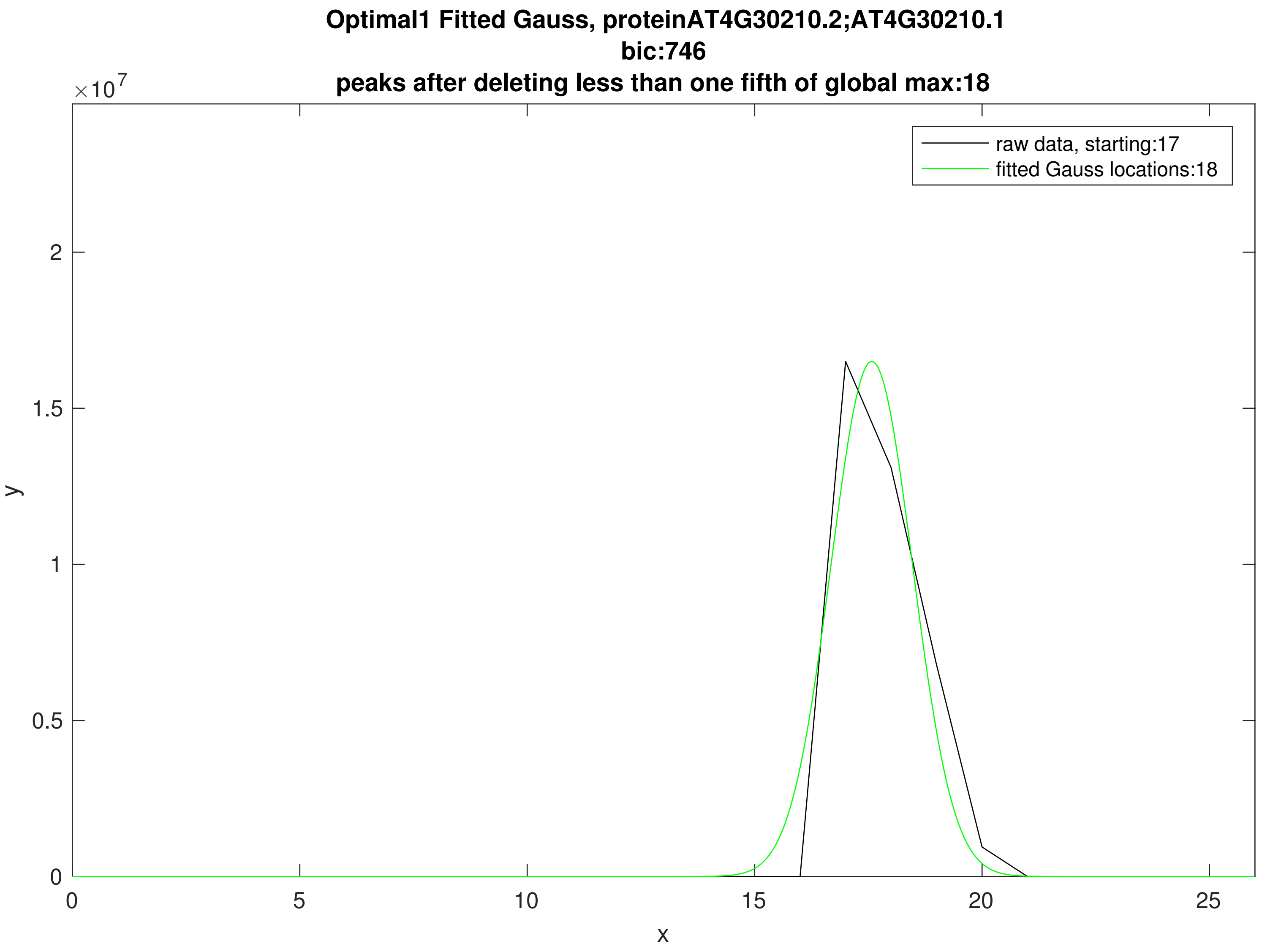

Optimal1 Fitted Gauss, proteinAT4G30210.2;AT4G30210.1
bic:746
peaks after deleting less than one fifth of global max:18
×10^7
raw data, starting:17
fitted Gauss locations:18
y
x
0
0.5
1
1.5
2
0
5
10
15
20
25

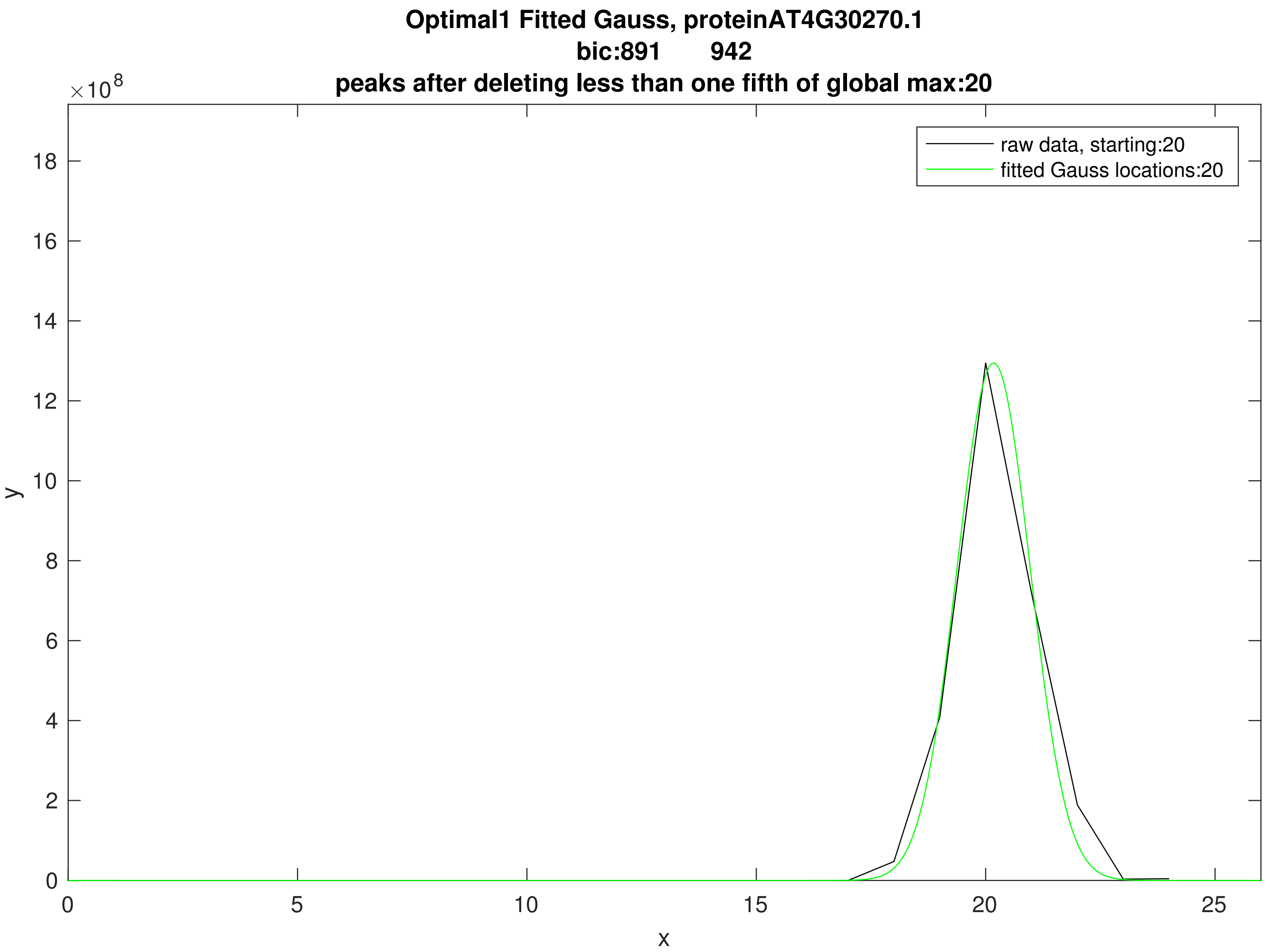

Optimal1 Fitted Gauss, proteinAT4G30270.1
bic:891 942
peaks after deleting less than one fifth of global max:20
×10^8
18
16
14
12
10
8
6
4
2
0
y
0
5
10
15
20
25
x
raw data, starting:20
fitted Gauss locations:20

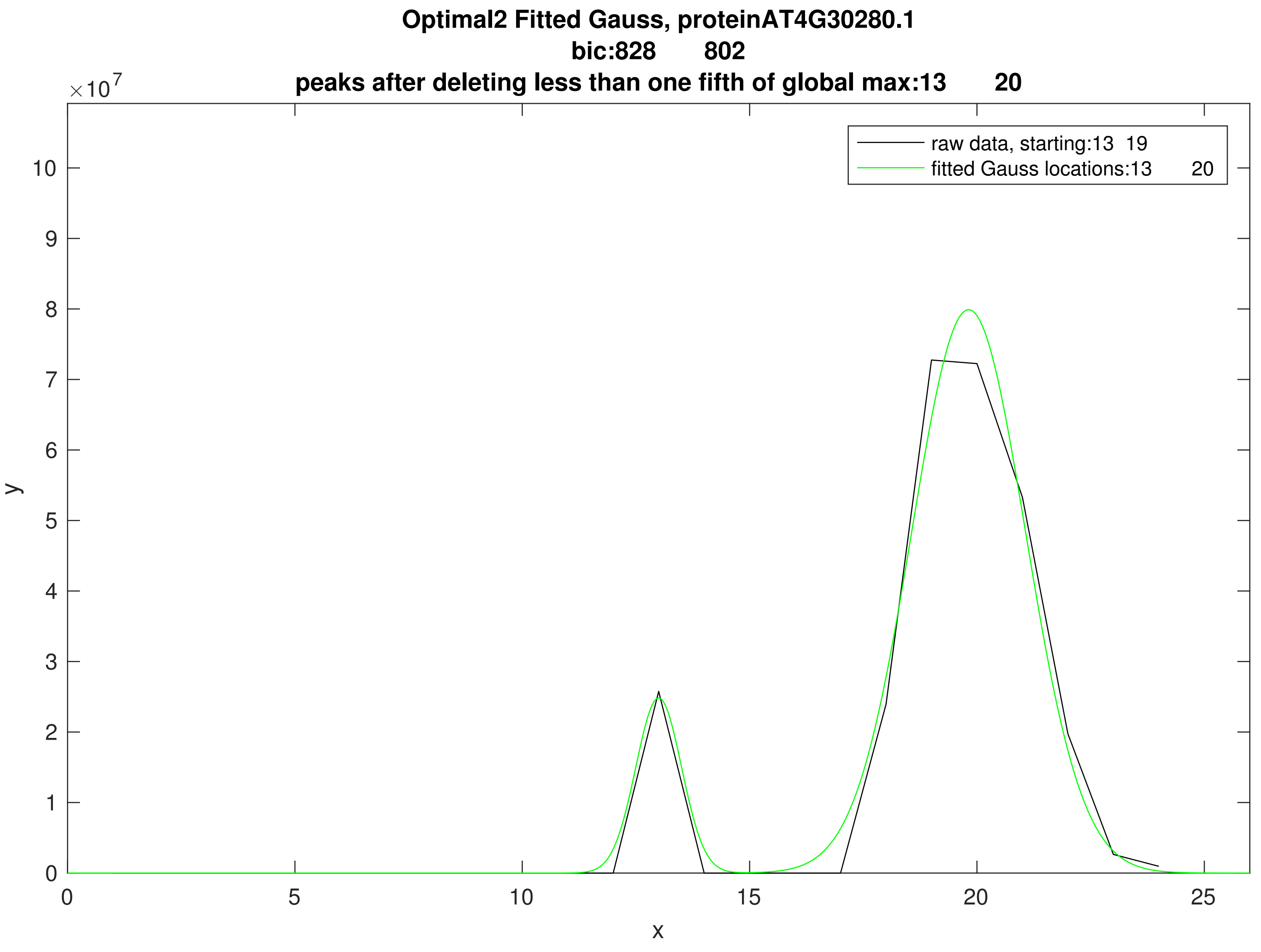
Optimal2 Fitted Gauss, proteinAT4G30280.1
bic:828 802
peaks after deleting less than one fifth of global max:13 20
×10^7
raw data, starting:13 19
fitted Gauss locations:13 20
y
x
0
1
2
3
4
5
6
7
8
9
10
0
5
10
15
20
25

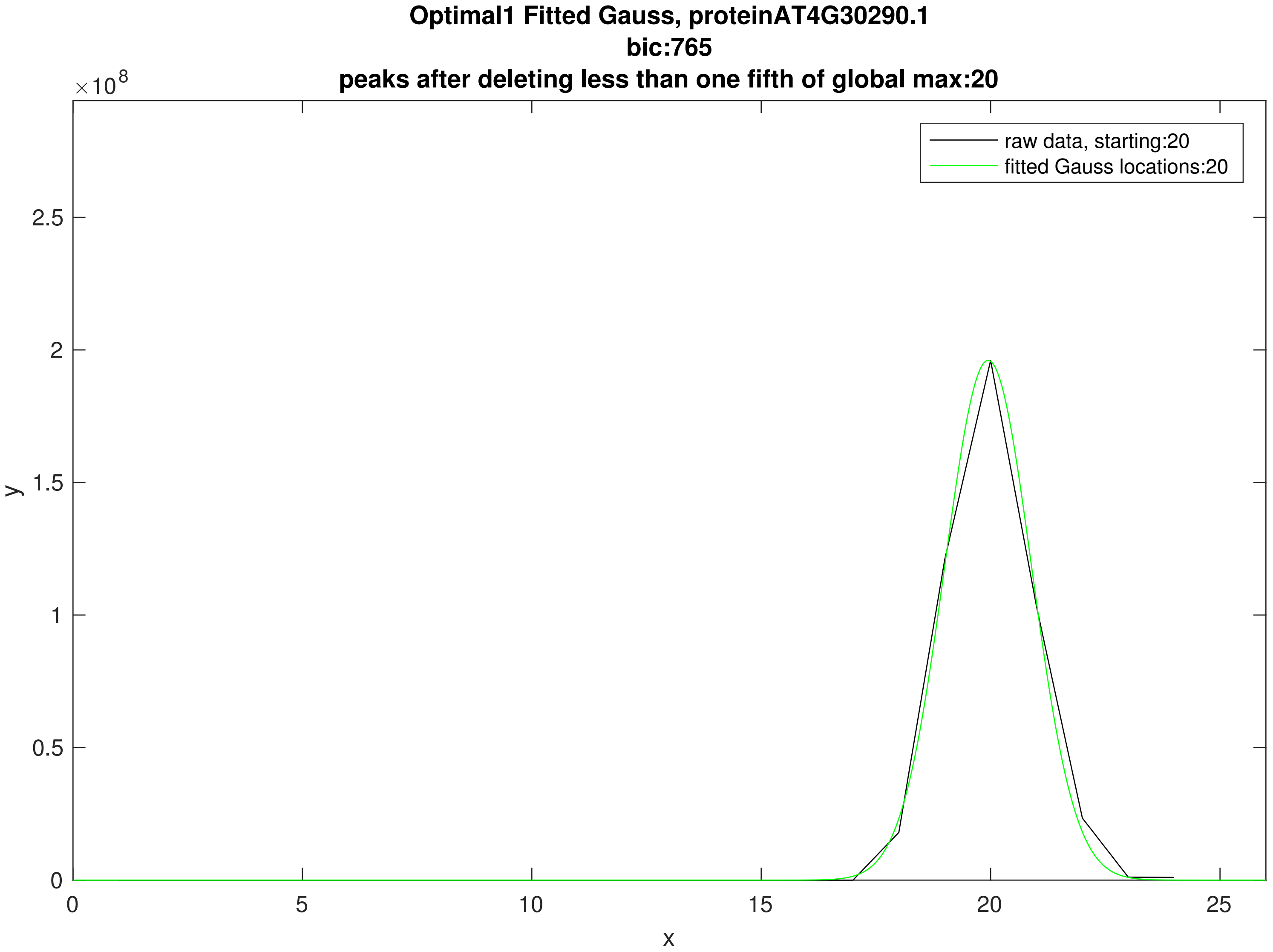
Optimal1 Fitted Gauss, proteinAT4G30290.1
bic:765
peaks after deleting less than one fifth of global max:20
×10^8
raw data, starting:20
fitted Gauss locations:20
y
x
0
0.5
1
1.5
2
2.5
0
5
10
15
20
25

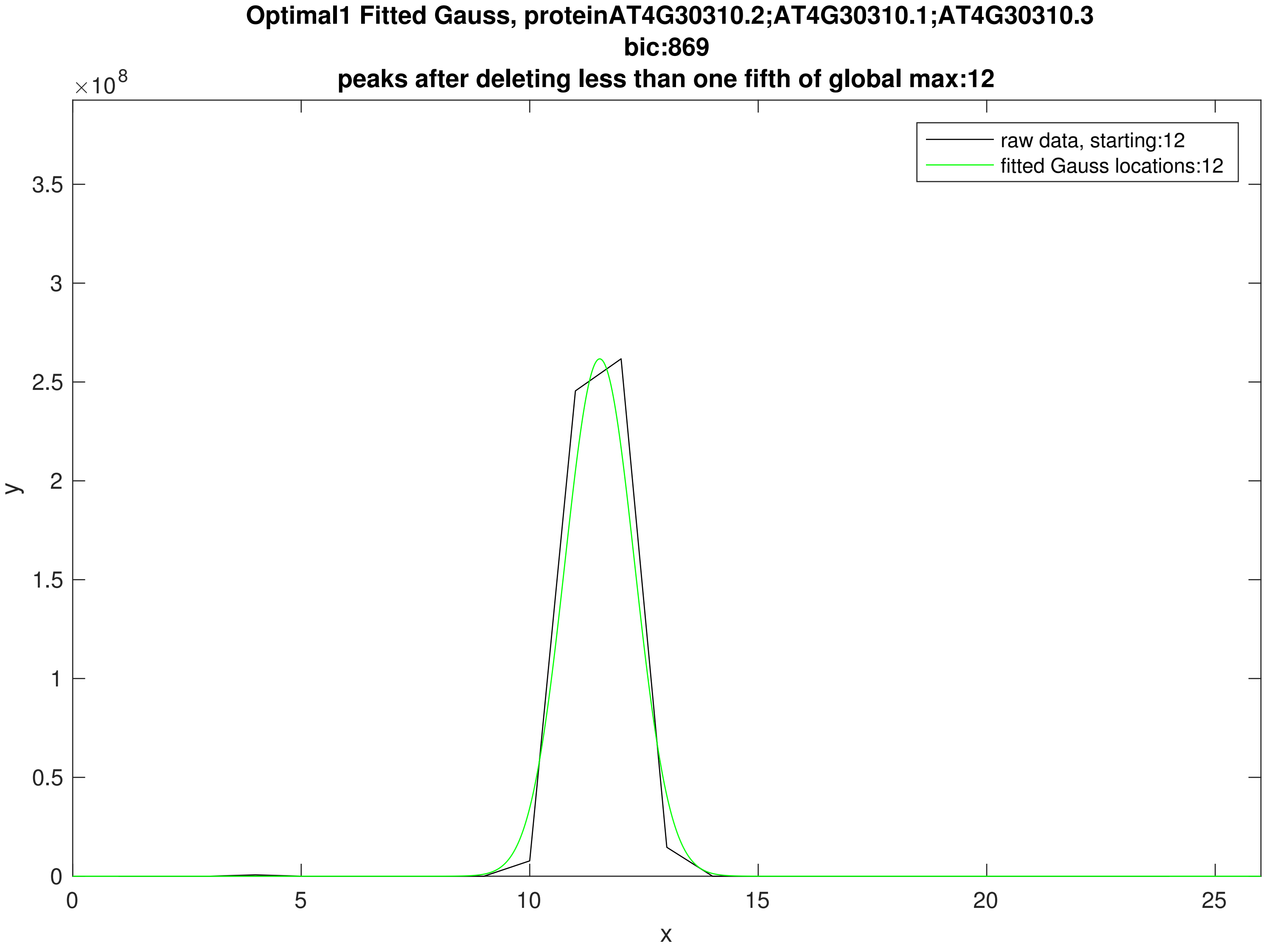
Optimal1 Fitted Gauss, proteinAT4G30310.2;AT4G30310.1;AT4G30310.3
bic:869
peaks after deleting less than one fifth of global max:12
×10^8
raw data, starting:12
fitted Gauss locations:12
y
x
0
0.5
1
1.5
2
2.5
3
3.5
0
5
10
15
20
25

**Optimal1 Fitted Gauss, proteinAT4G30450.1**
**bic:636**
**peaks after deleting less than one fifth of global max:21**

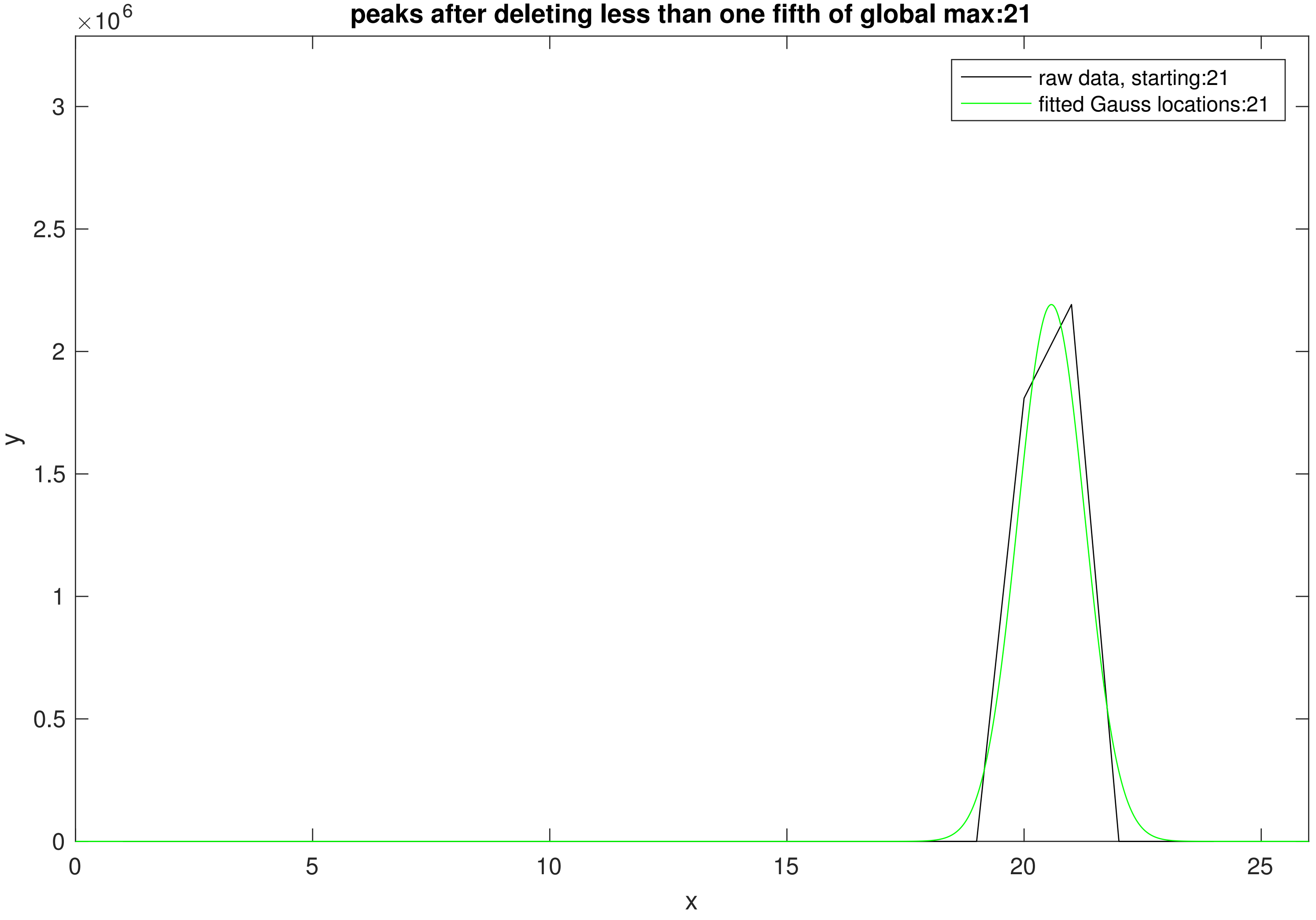

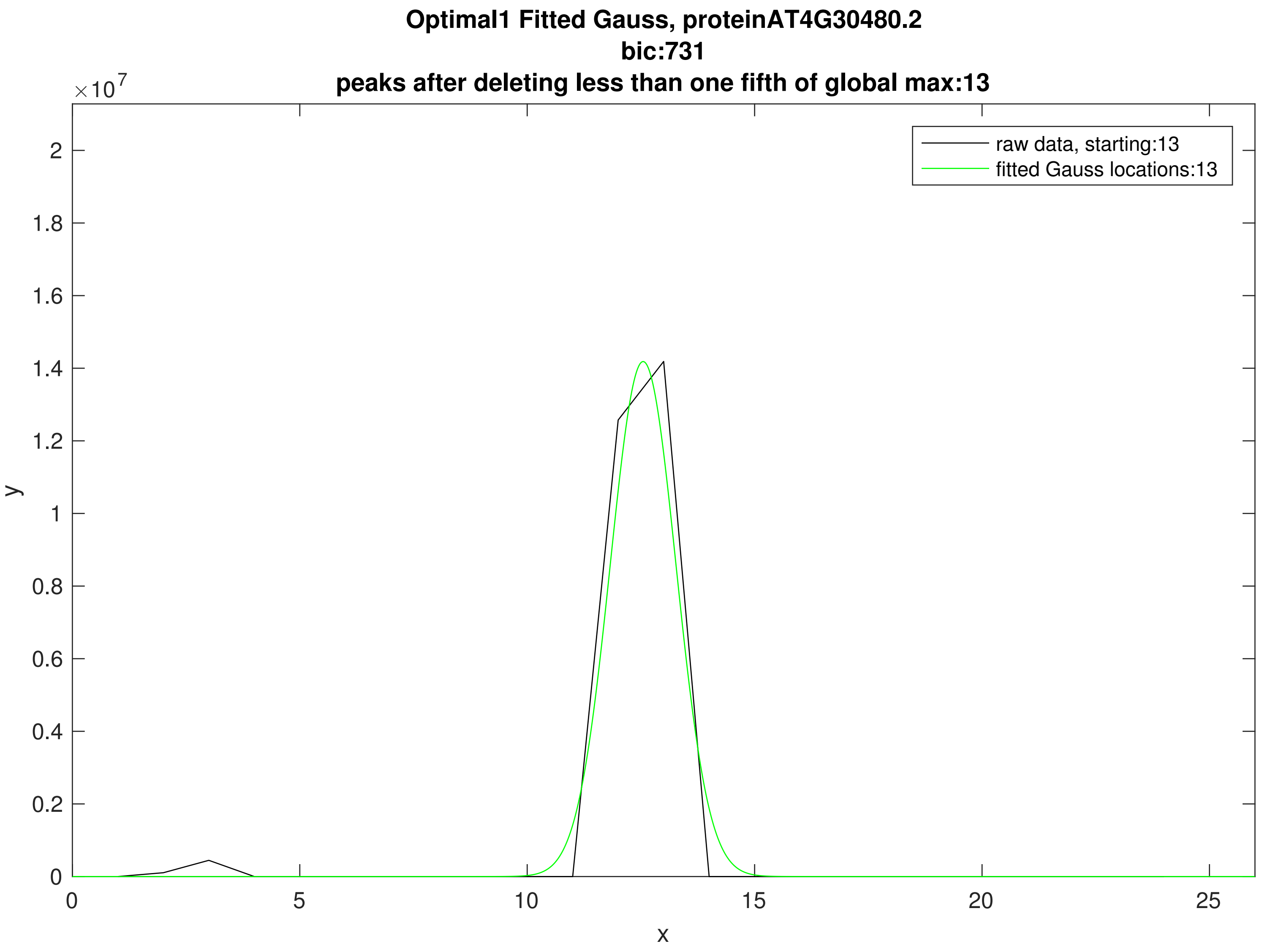
Optimal1 Fitted Gauss, proteinAT4G30480.2
bic:731
peaks after deleting less than one fifth of global max:13
×10^7
raw data, starting:13
fitted Gauss locations:13
y
x
0
0.2
0.4
0.6
0.8
1
1.2
1.4
1.6
1.8
2
5
10
15
20
25

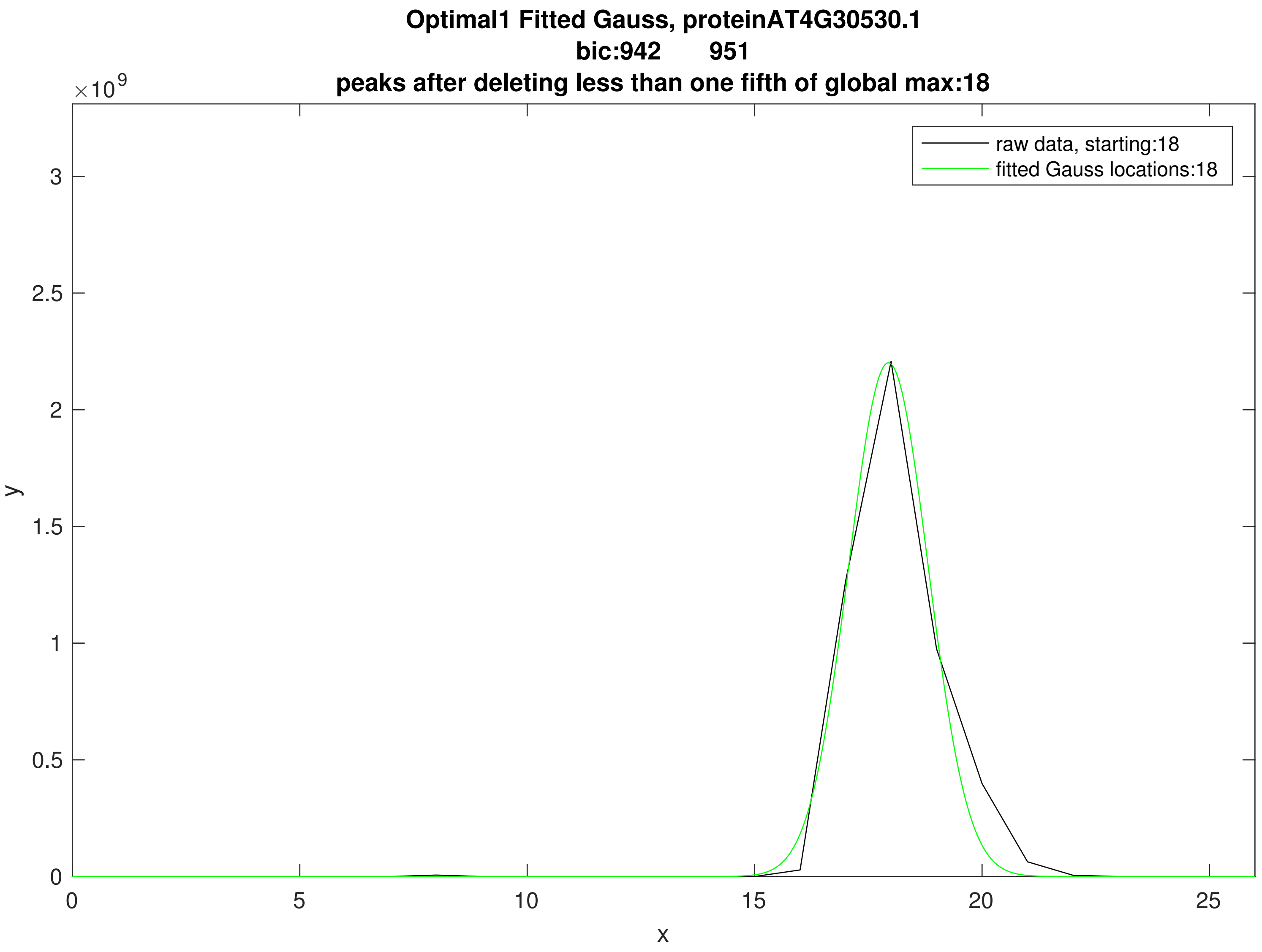

Optimal1 Fitted Gauss, proteinAT4G30530.1
bic:942 951
peaks after deleting less than one fifth of global max:18
×10^9
raw data, starting:18
fitted Gauss locations:18
y
x
0
0.5
1
1.5
2
2.5
3
0
5
10
15
20
25

**Optimal1 Fitted Gauss, proteinAT4G30550.1**
**bic:859 868**
**peaks after deleting less than one fifth of global max:18**

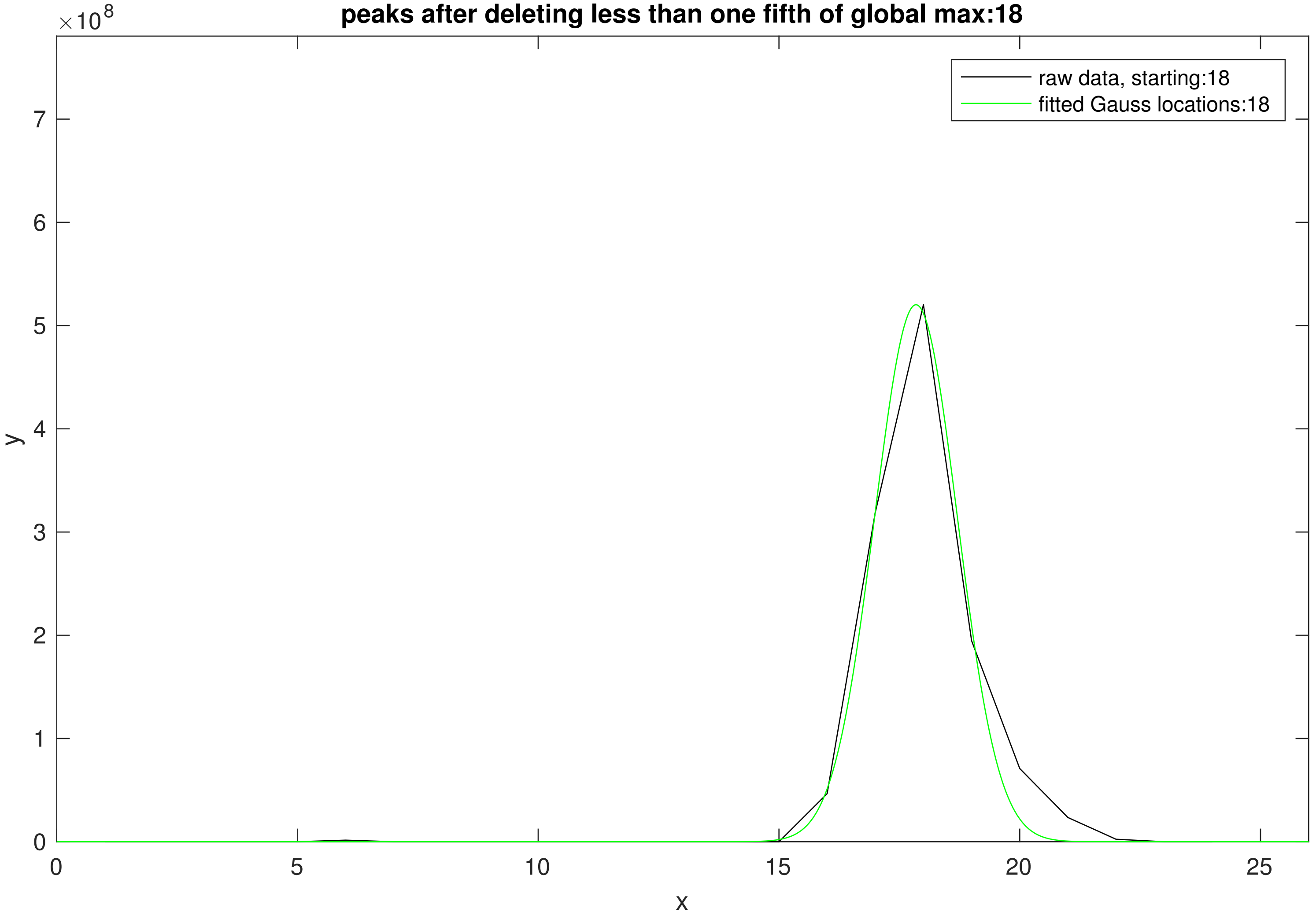

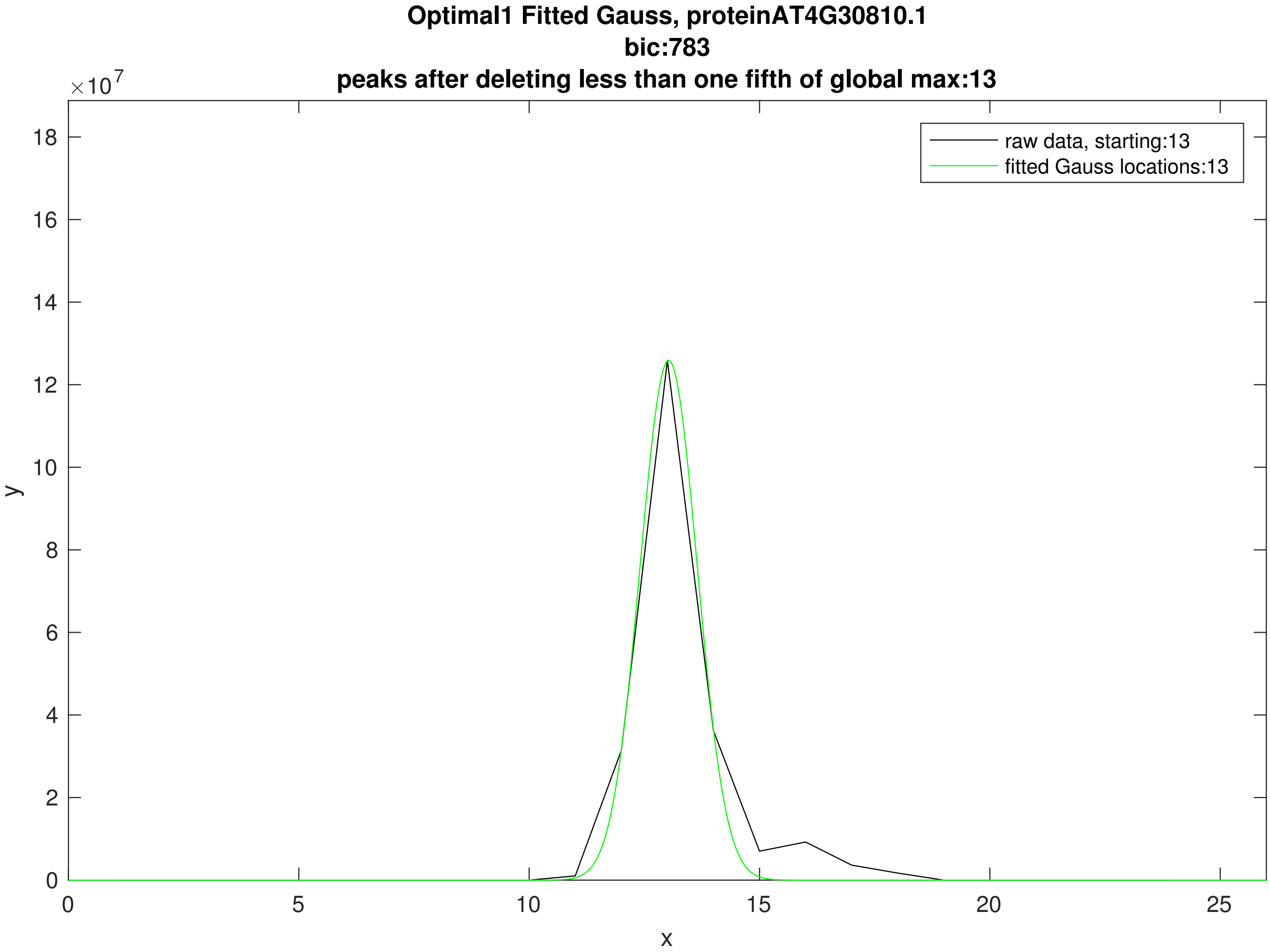
Optimal1 Fitted Gauss, proteinAT4G30810.1
bic:783
peaks after deleting less than one fifth of global max:13
×10^7
raw data, starting:13
fitted Gauss locations:13
y
x
0
2
4
6
8
10
12
14
16
18
0
5
10
15
20
25

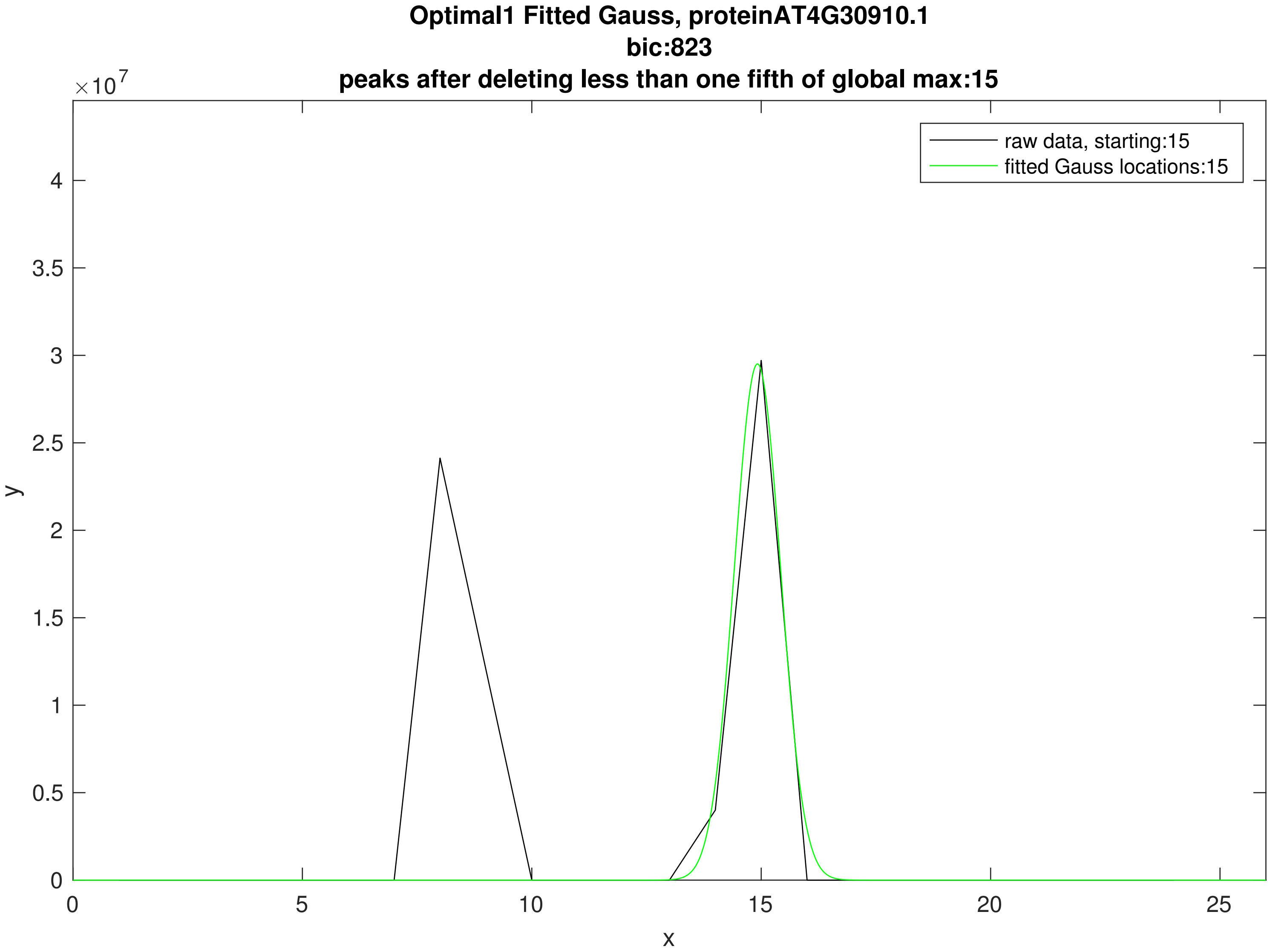
Optimal1 Fitted Gauss, proteinAT4G30910.1
bic:823
peaks after deleting less than one fifth of global max:15
×10^7
raw data, starting:15
fitted Gauss locations:15
y
x
0
0.5
1
1.5
2
2.5
3
3.5
4
0
5
10
15
20
25

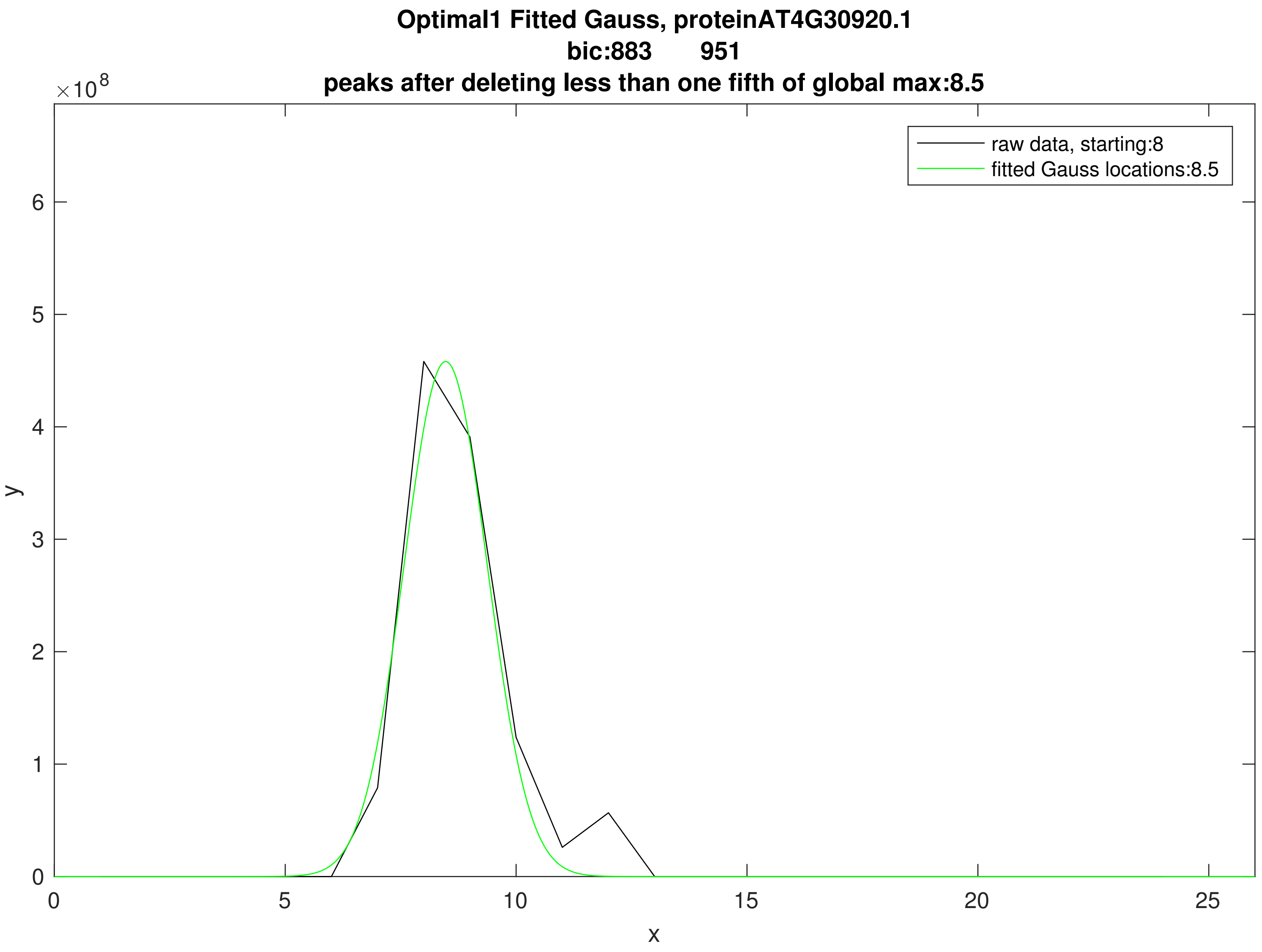

Optimal1 Fitted Gauss, proteinAT4G30920.1
bic:883 951
peaks after deleting less than one fifth of global max:8.5
×10^8
raw data, starting:8
fitted Gauss locations:8.5
0
1
2
3
4
5
6
y
0
5
10
15
20
25
x

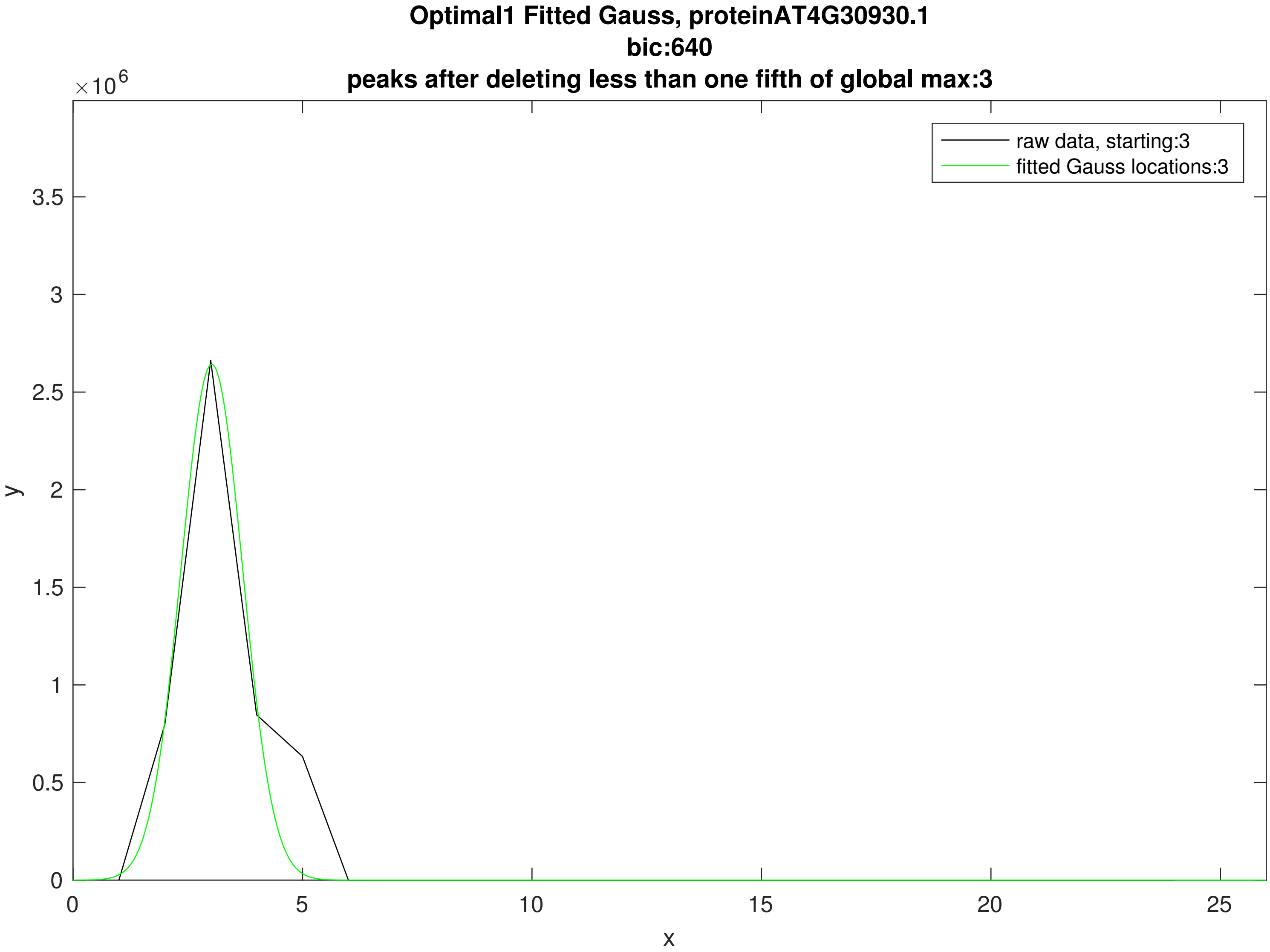
Optimal1 Fitted Gauss, proteinAT4G30930.1
bic:640
peaks after deleting less than one fifth of global max:3
×10^6
raw data, starting:3
fitted Gauss locations:3
y
x
0
0.5
1
1.5
2
2.5
3
3.5
0
5
10
15
20
25

**Optimal1 Fitted Gauss, proteinAT4G30996.1**
**bic:583**
**peaks after deleting less than one fifth of global max:21**

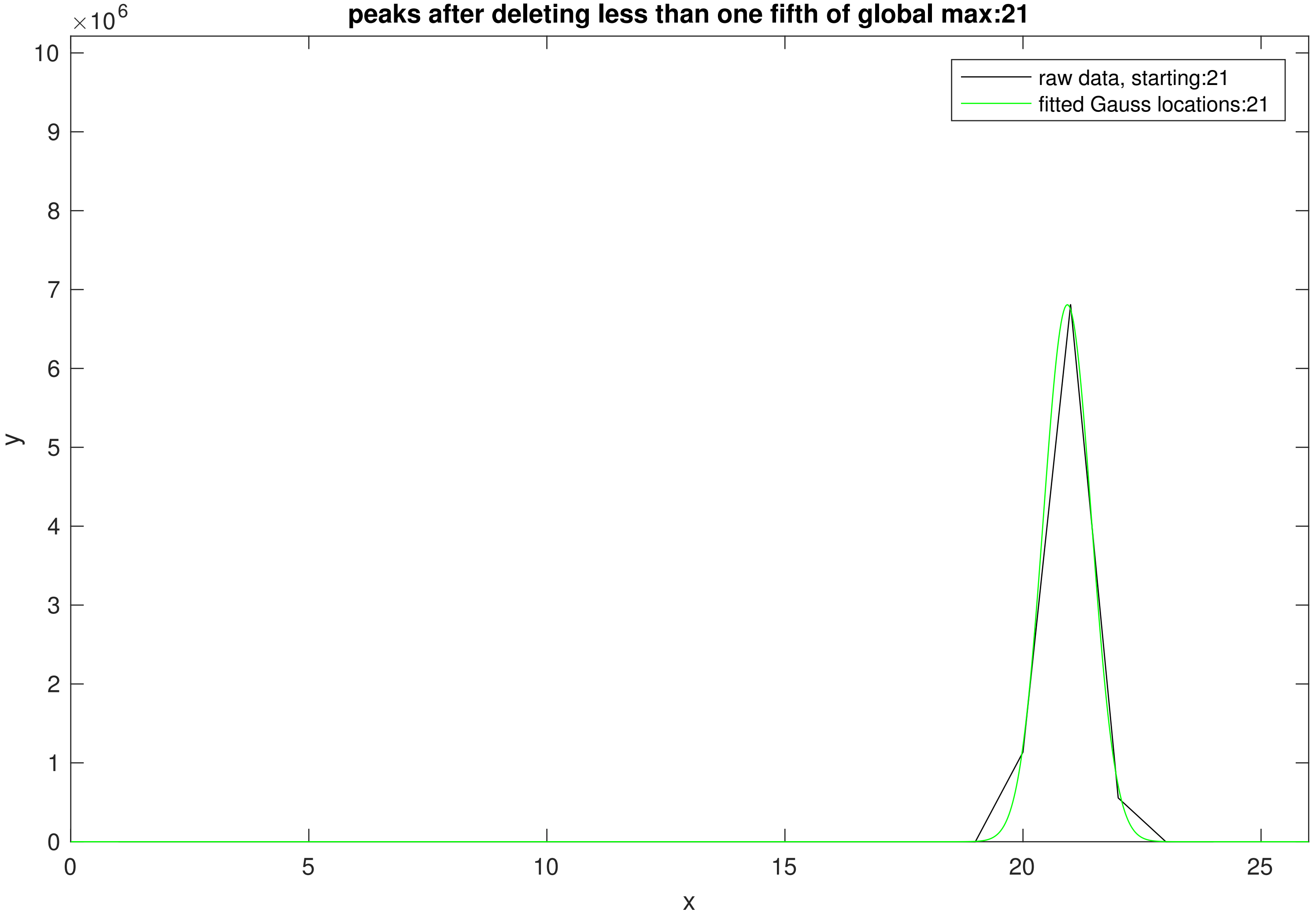

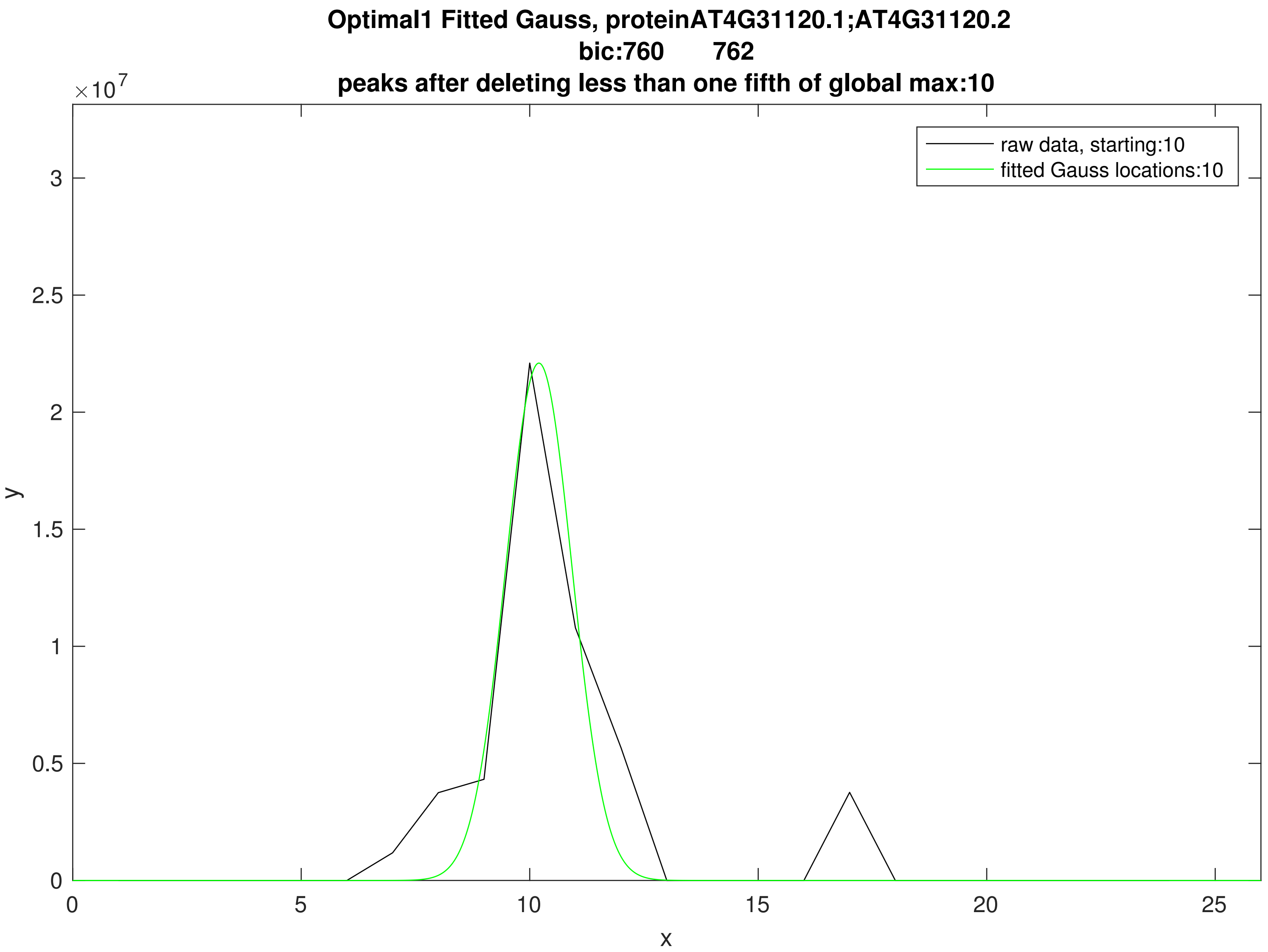
Optimal1 Fitted Gauss, proteinAT4G31120.1;AT4G31120.2
bic:760 762
peaks after deleting less than one fifth of global max:10
×10^7
3
2.5
2
1.5
1
0.5
0
y
0
5
10
15
20
25
x
raw data, starting:10
fitted Gauss locations:10

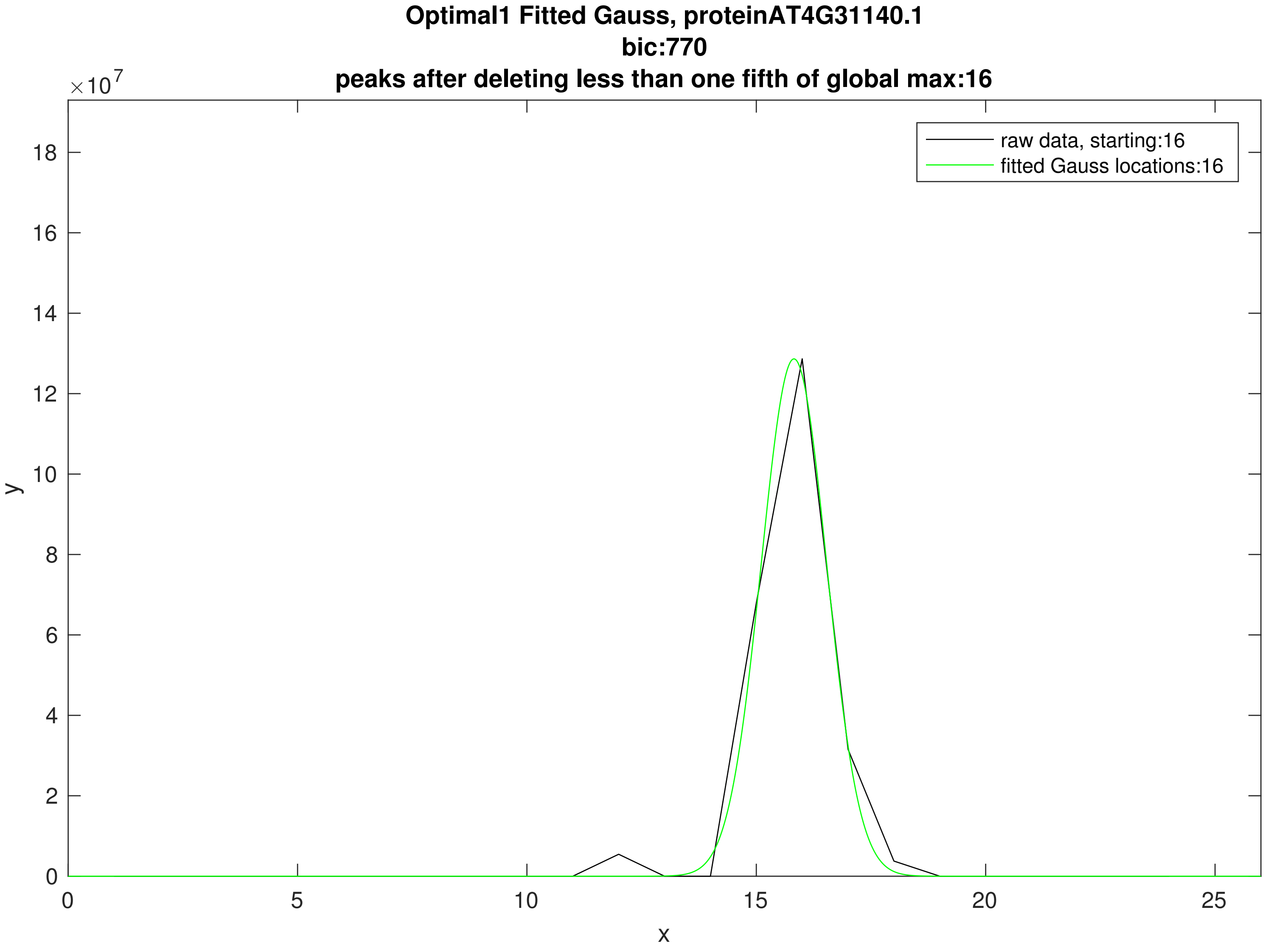
Optimal1 Fitted Gauss, proteinAT4G31140.1
bic:770
peaks after deleting less than one fifth of global max:16
×10^7
raw data, starting:16
fitted Gauss locations:16
y
x
0
2
4
6
8
10
12
14
16
18
0
5
10
15
20
25

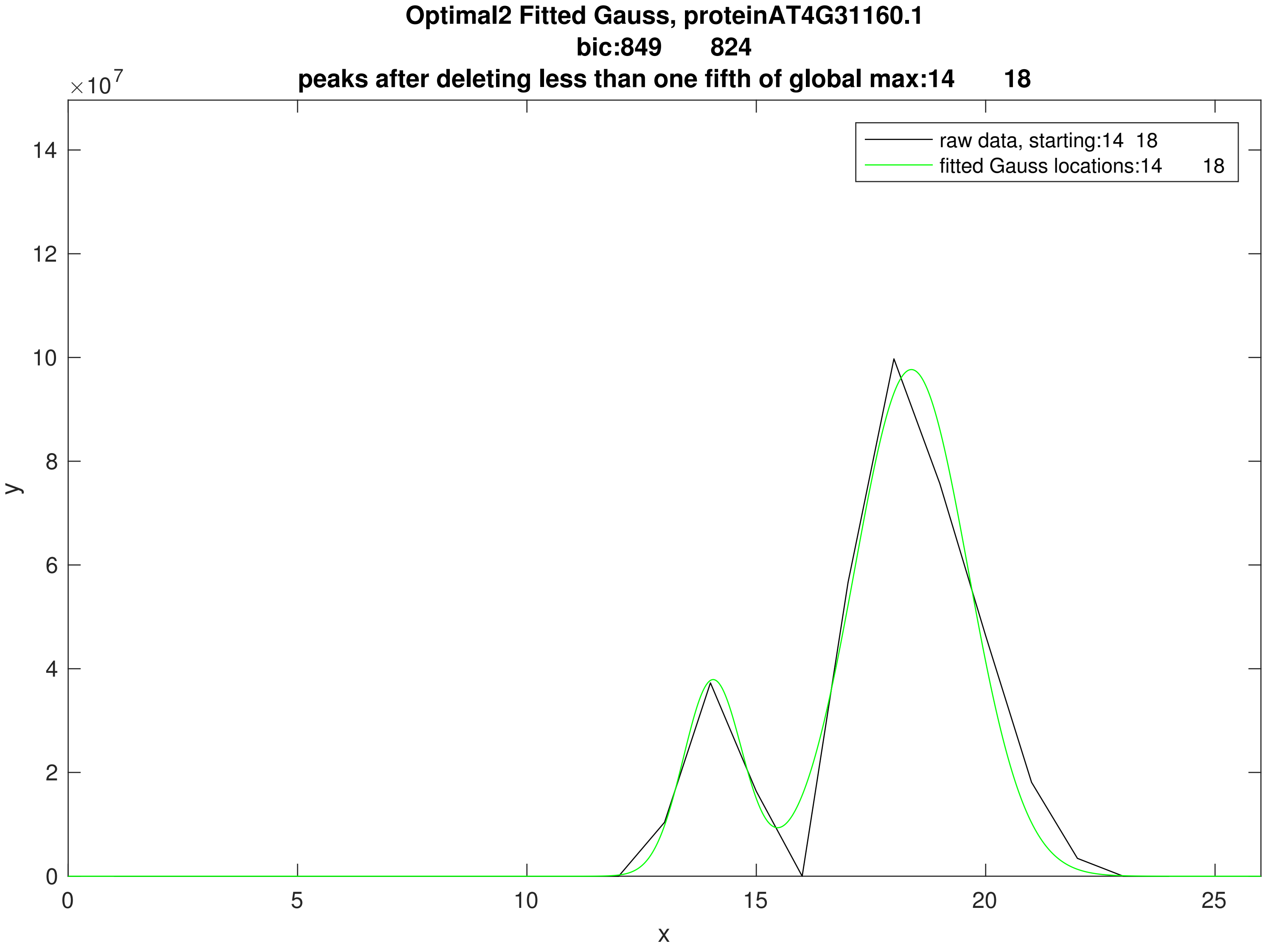
Optimal2 Fitted Gauss, proteinAT4G31160.1
bic:849 824
peaks after deleting less than one fifth of global max:14 18
×10^7
raw data, starting:14 18
fitted Gauss locations:14 18
y
x
0
2
4
6
8
10
12
14
0
5
10
15
20
25

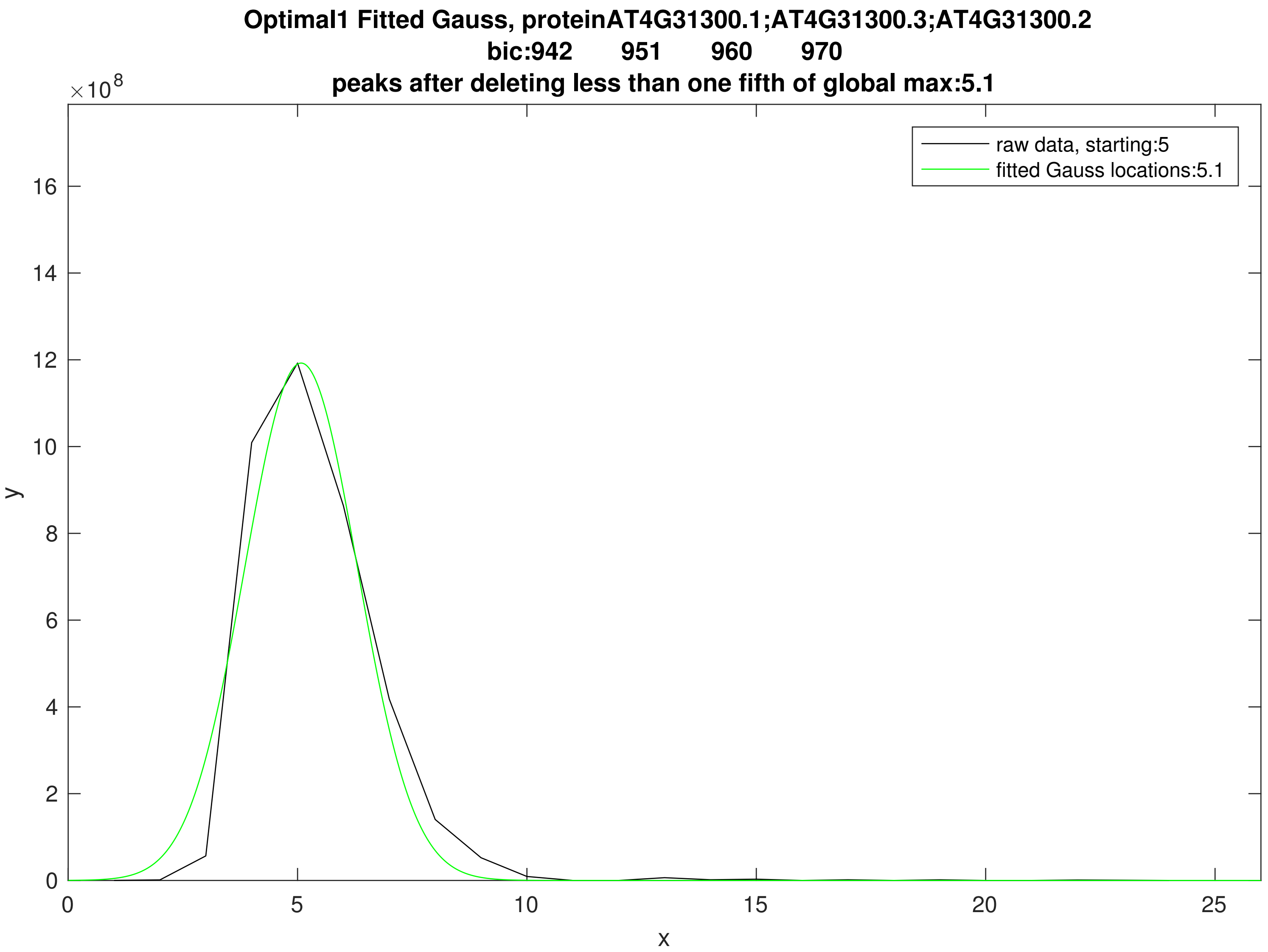

Optimal1 Fitted Gauss, proteinAT4G31300.1;AT4G31300.3;AT4G31300.2
bic:942 951 960 970
peaks after deleting less than one fifth of global max:5.1
×10^8
raw data, starting:5
fitted Gauss locations:5.1
16
14
12
10
8
6
4
2
0
y
0
5
10
15
20
25
x

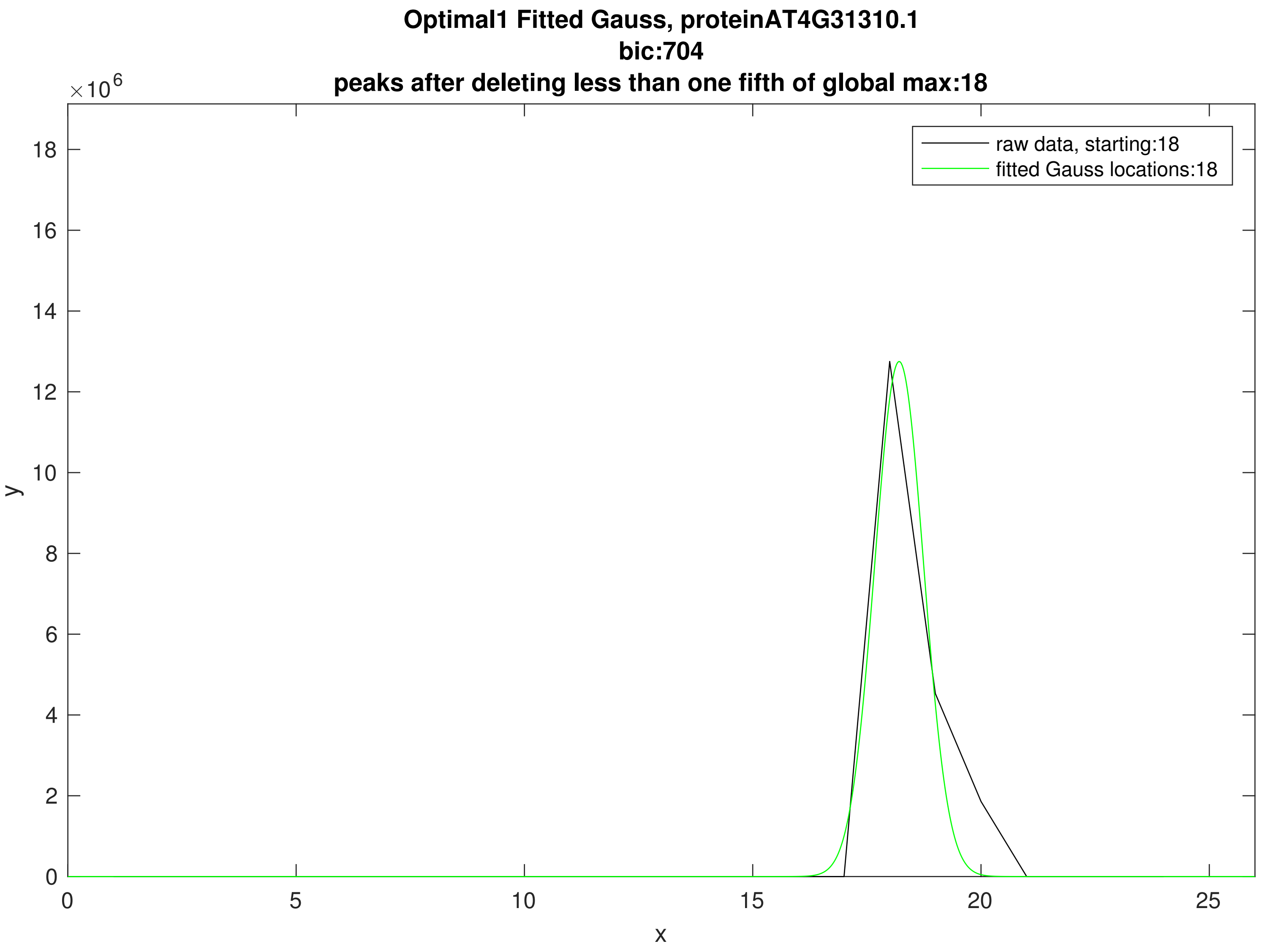

Optimal1 Fitted Gauss, proteinAT4G31310.1
bic:704
peaks after deleting less than one fifth of global max:18
×10^6
raw data, starting:18
fitted Gauss locations:18
y
x
0
2
4
6
8
10
12
14
16
18
0
5
10
15
20
25

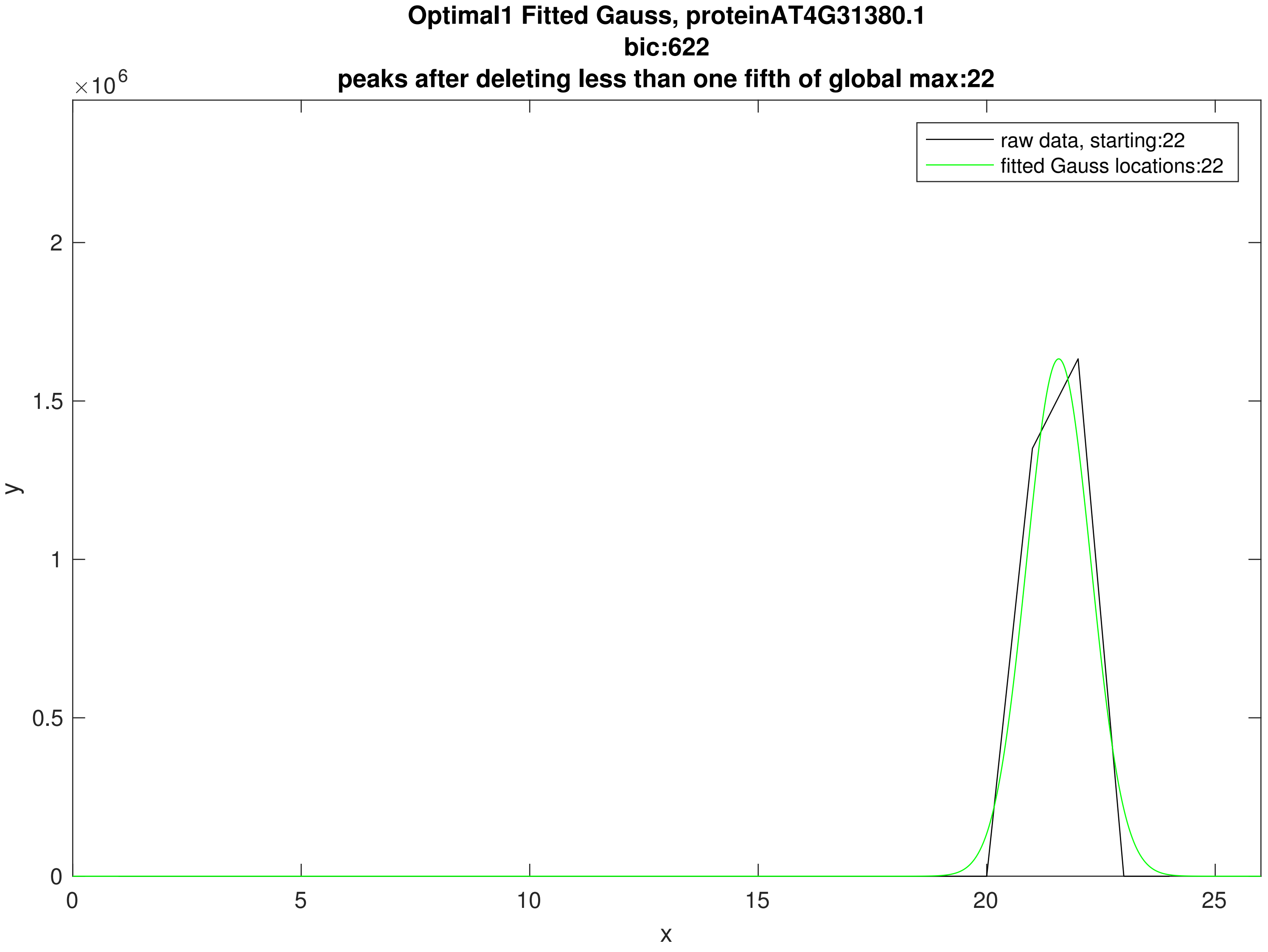

Optimal1 Fitted Gauss, proteinAT4G31380.1
bic:622
peaks after deleting less than one fifth of global max:22
×10^6
raw data, starting:22
fitted Gauss locations:22
y
x
0
0.5
1
1.5
2
0
5
10
15
20
25

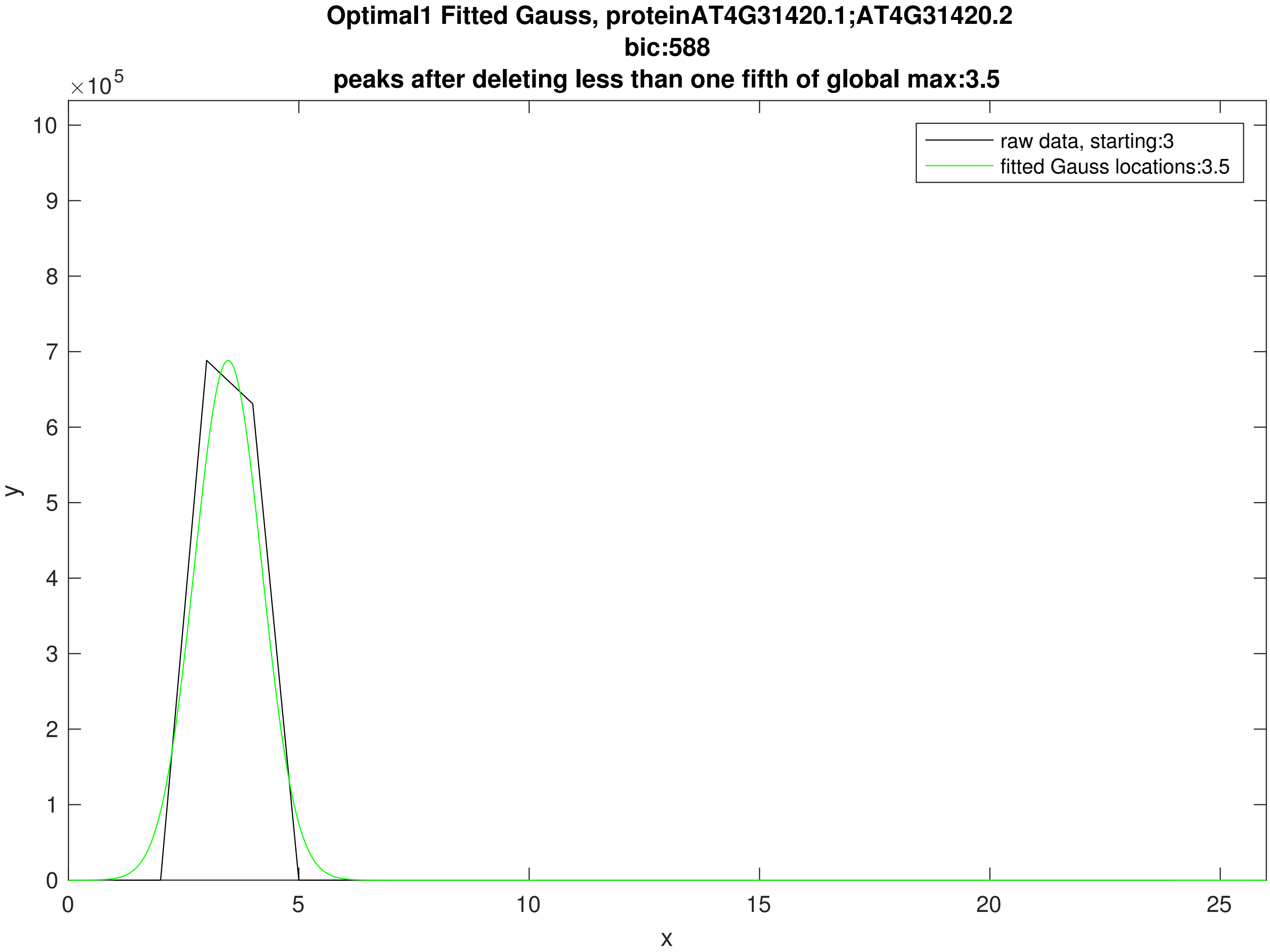
Optimal1 Fitted Gauss, proteinAT4G31420.1;AT4G31420.2
bic:588
peaks after deleting less than one fifth of global max:3.5
×10^5
raw data, starting:3
fitted Gauss locations:3.5
y
x
0
1
2
3
4
5
6
7
8
9
10
0
5
10
15
20
25

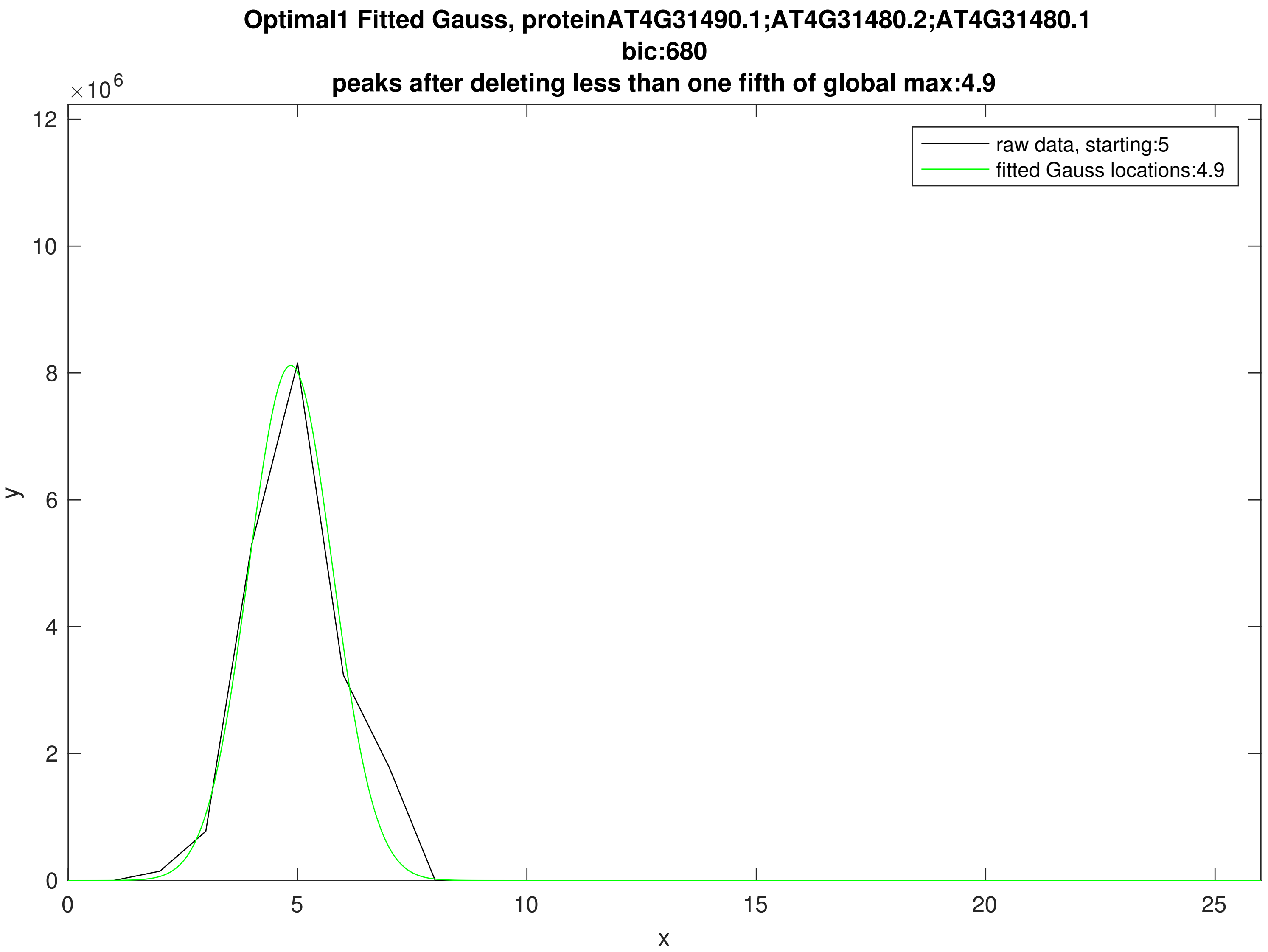
Optimal1 Fitted Gauss, proteinAT4G31490.1;AT4G31480.2;AT4G31480.1
bic:680
peaks after deleting less than one fifth of global max:4.9
×10^6
raw data, starting:5
fitted Gauss locations:4.9
y
x

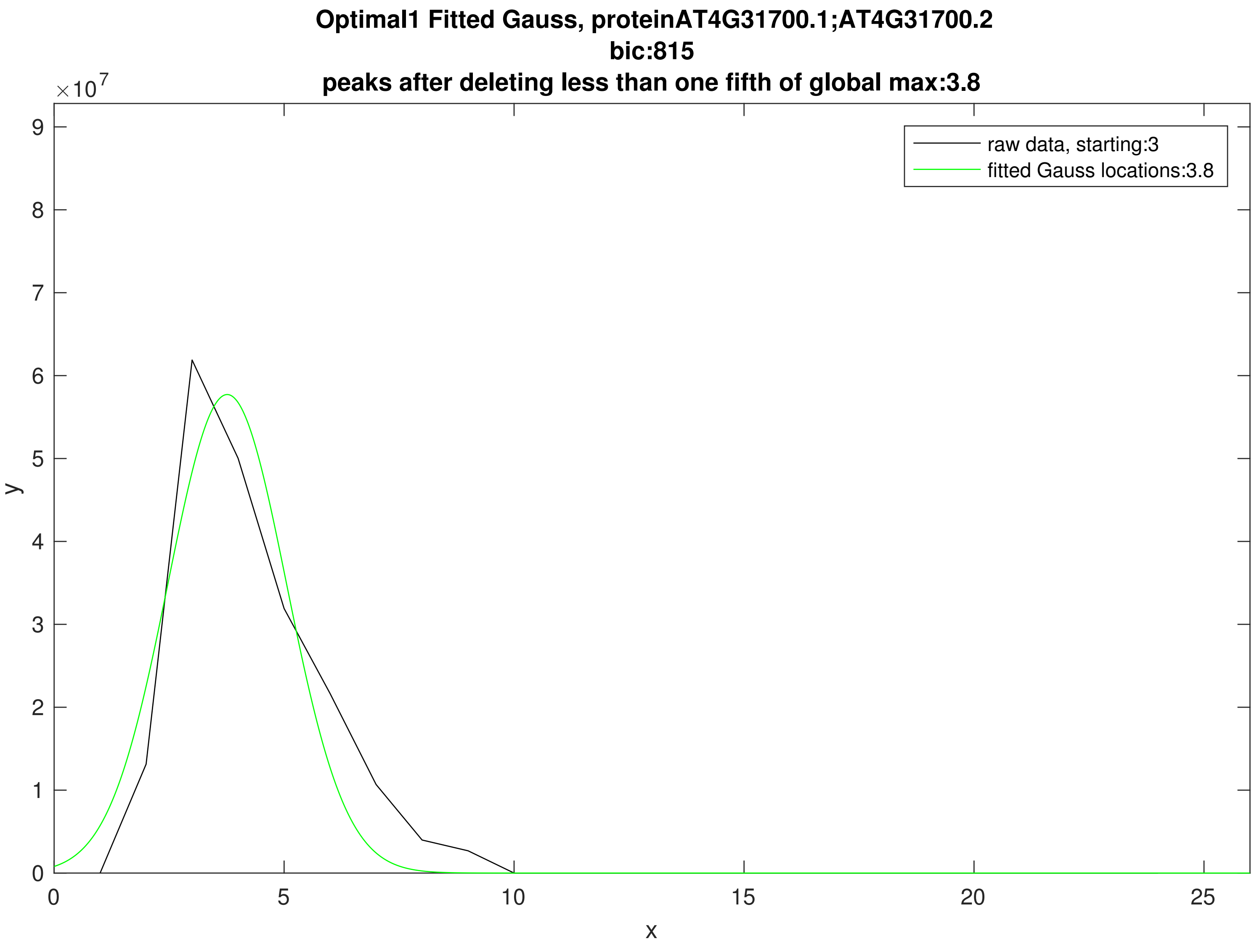
Optimal1 Fitted Gauss, proteinAT4G31700.1;AT4G31700.2
bic:815
peaks after deleting less than one fifth of global max:3.8
×10^7
raw data, starting:3
fitted Gauss locations:3.8
y
x
0
1
2
3
4
5
6
7
8
9
0
5
10
15
20
25

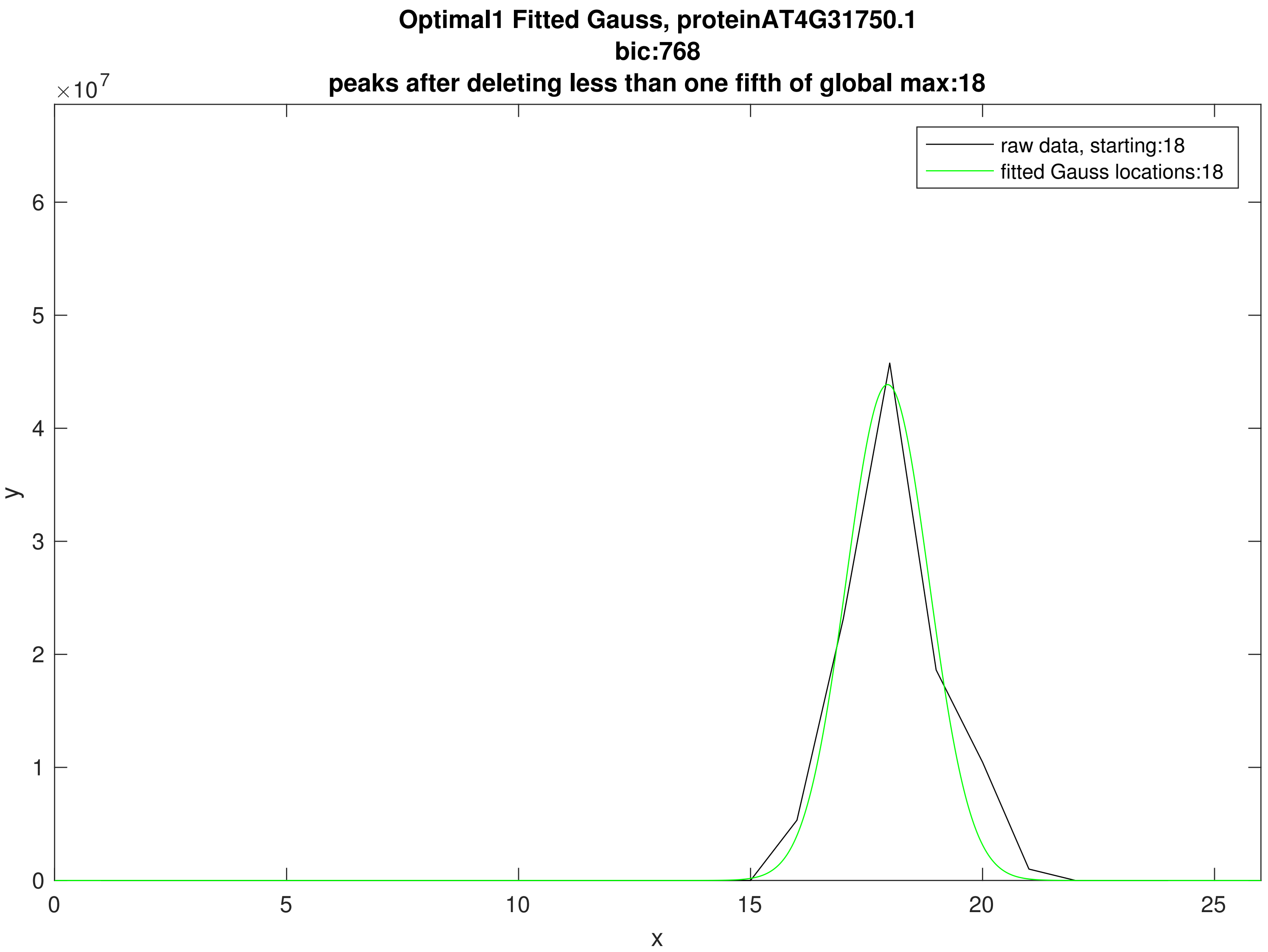
Optimal1 Fitted Gauss, proteinAT4G31750.1
bic:768
peaks after deleting less than one fifth of global max:18
×10^7
raw data, starting:18
fitted Gauss locations:18
y
x
0
1
2
3
4
5
6
0
5
10
15
20
25

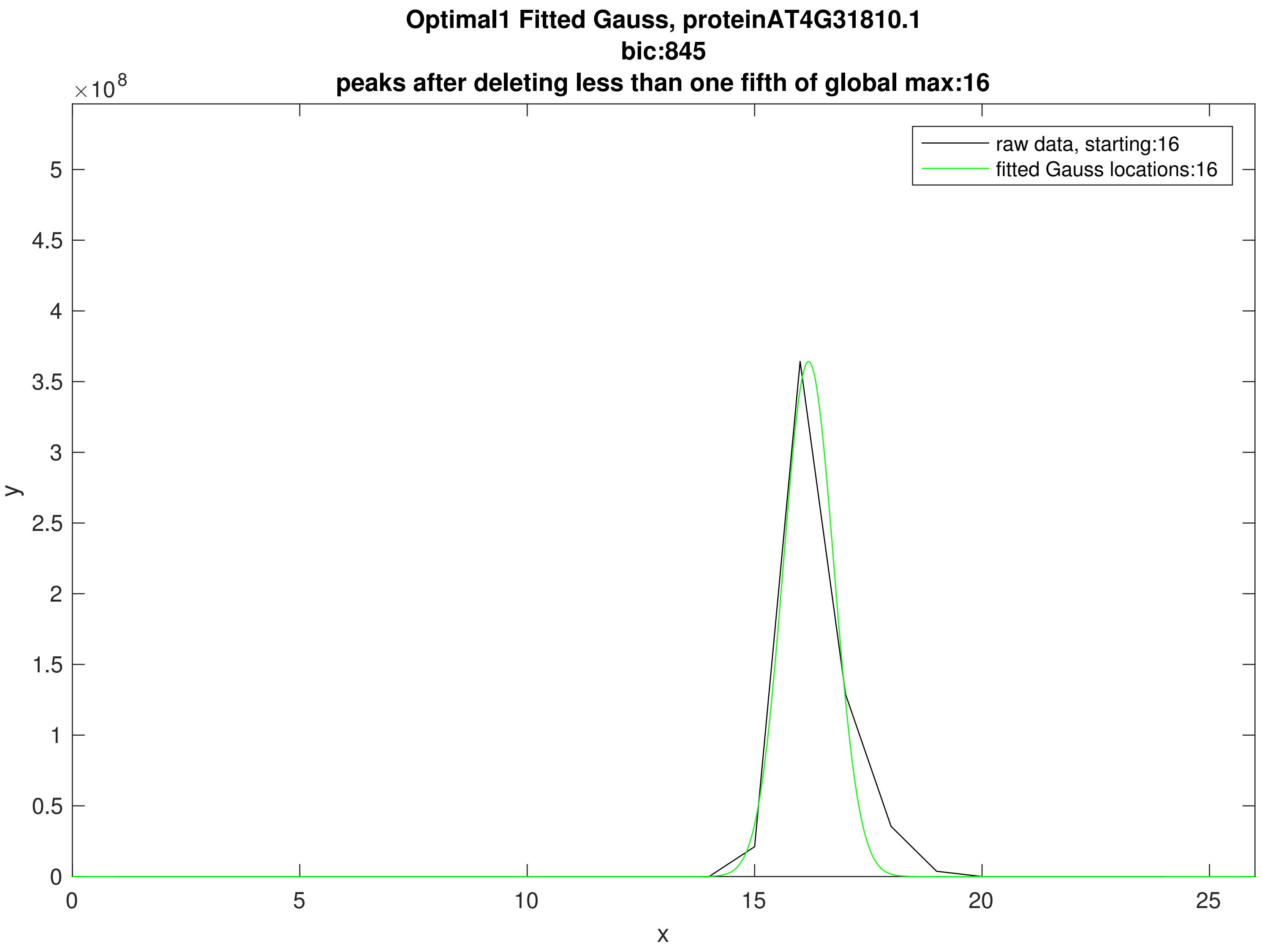
Optimal1 Fitted Gauss, proteinAT4G31810.1
bic:845
peaks after deleting less than one fifth of global max:16
×10^8
raw data, starting:16
fitted Gauss locations:16
5
4.5
4
3.5
3
2.5
2
1.5
1
0.5
0
y
0
5
10
15
20
25
x

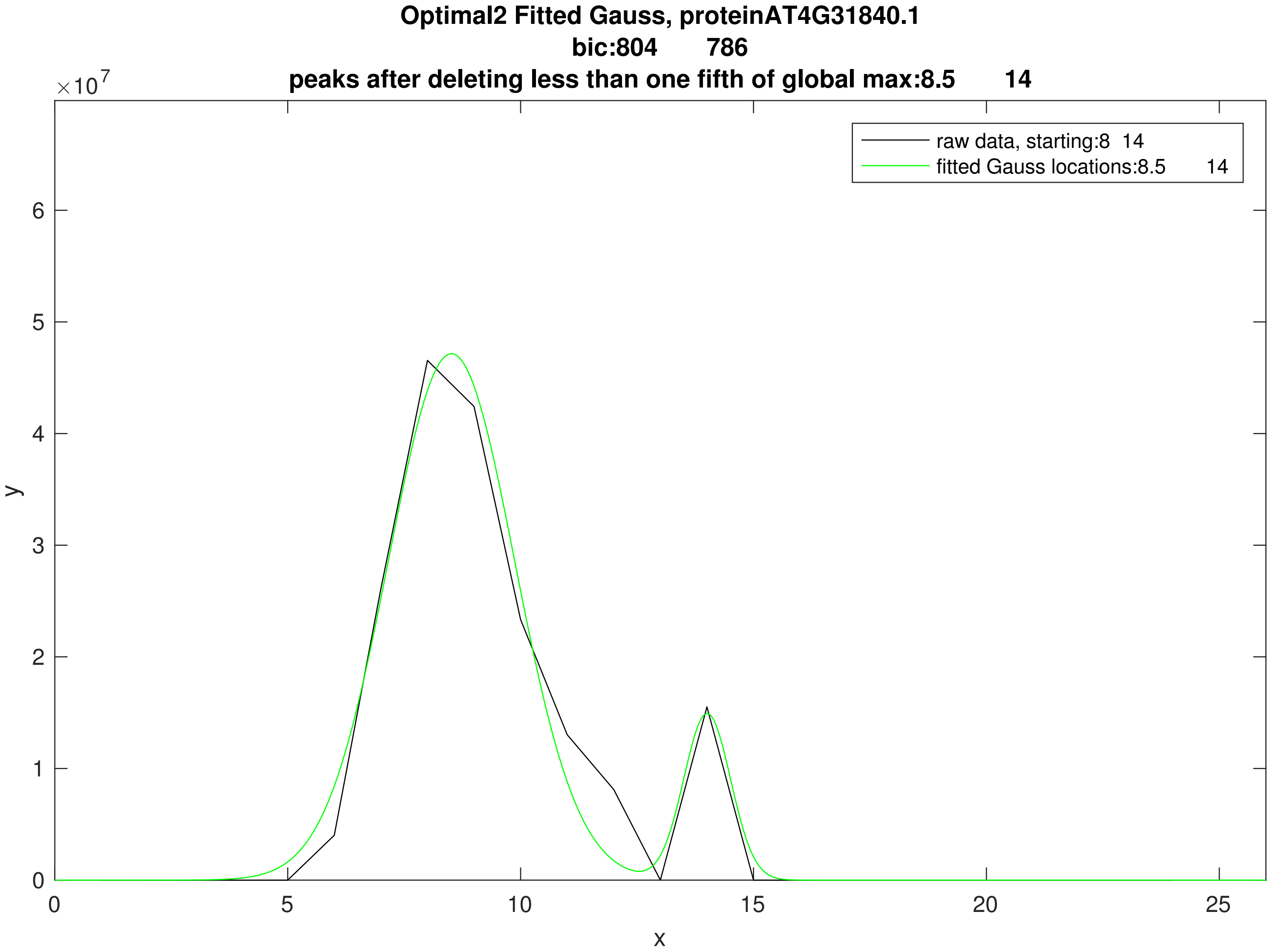
Optimal2 Fitted Gauss, proteinAT4G31840.1
bic:804 786
peaks after deleting less than one fifth of global max:8.5 14
×10^7
raw data, starting:8 14
fitted Gauss locations:8.5 14
y
x
0
1
2
3
4
5
6
0
5
10
15
20
25

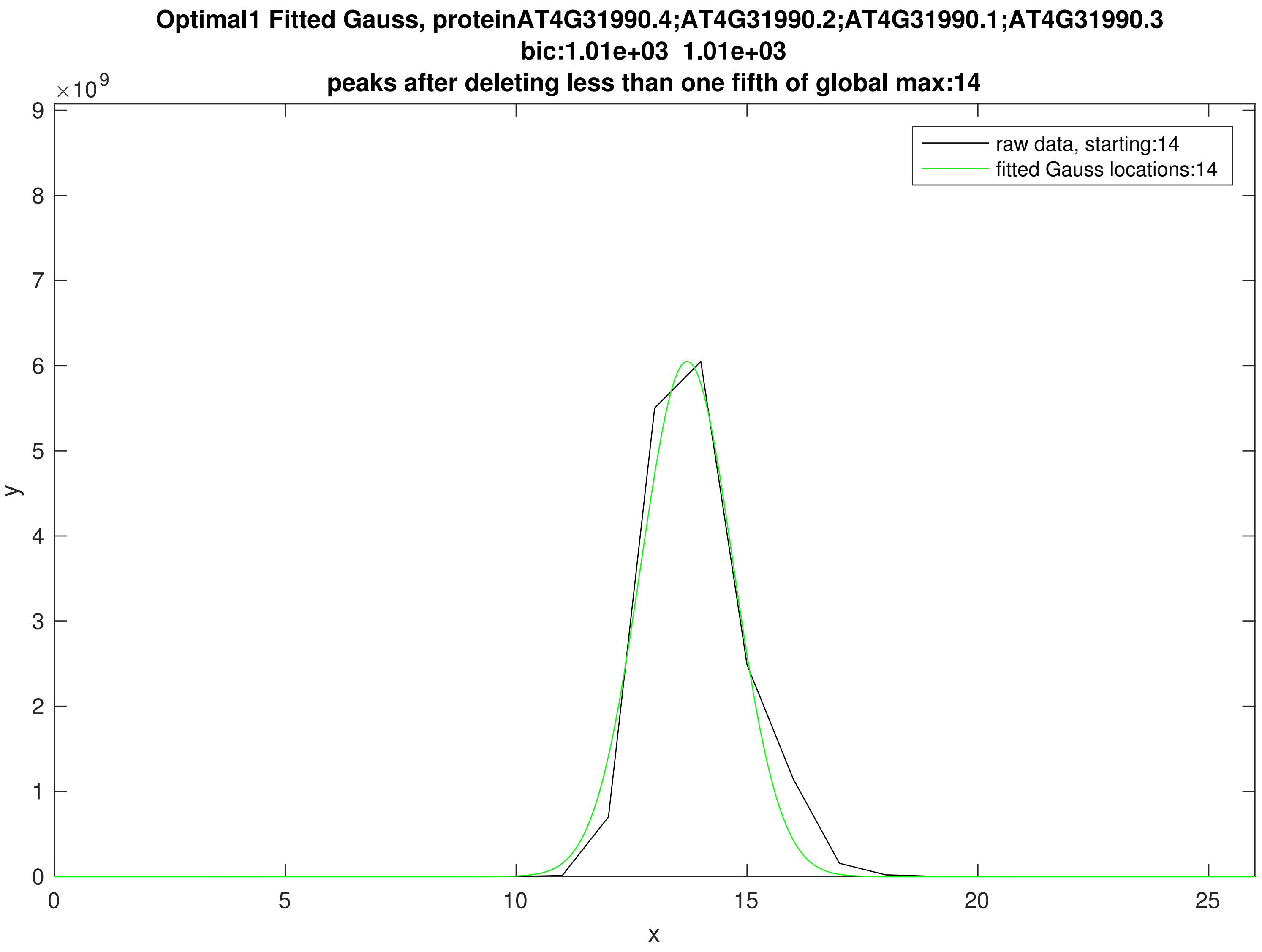

Optimal1 Fitted Gauss, proteinAT4G31990.4;AT4G31990.2;AT4G31990.1;AT4G31990.3
bic:1.01e+03 1.01e+03
peaks after deleting less than one fifth of global max:14
×10^9
9
8
7
6
5
4
3
2
1
0
y
0
5
10
15
20
25
x
raw data, starting:14
fitted Gauss locations:14

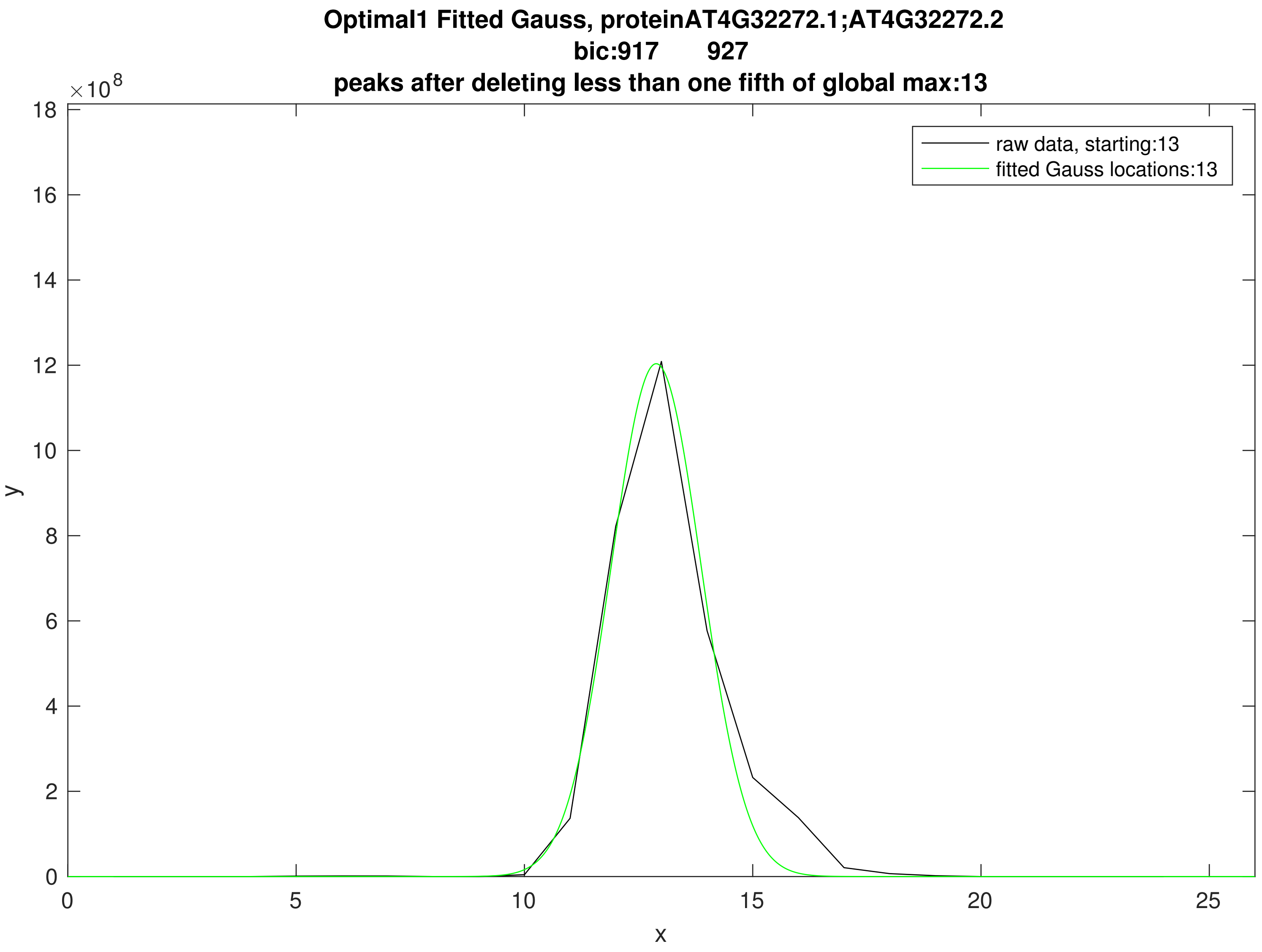
Optimal1 Fitted Gauss, proteinAT4G32272.1;AT4G32272.2
bic:917 927
peaks after deleting less than one fifth of global max:13
×10^8
raw data, starting:13
fitted Gauss locations:13
y
x
0
2
4
6
8
10
12
14
16
18
0
5
10
15
20
25

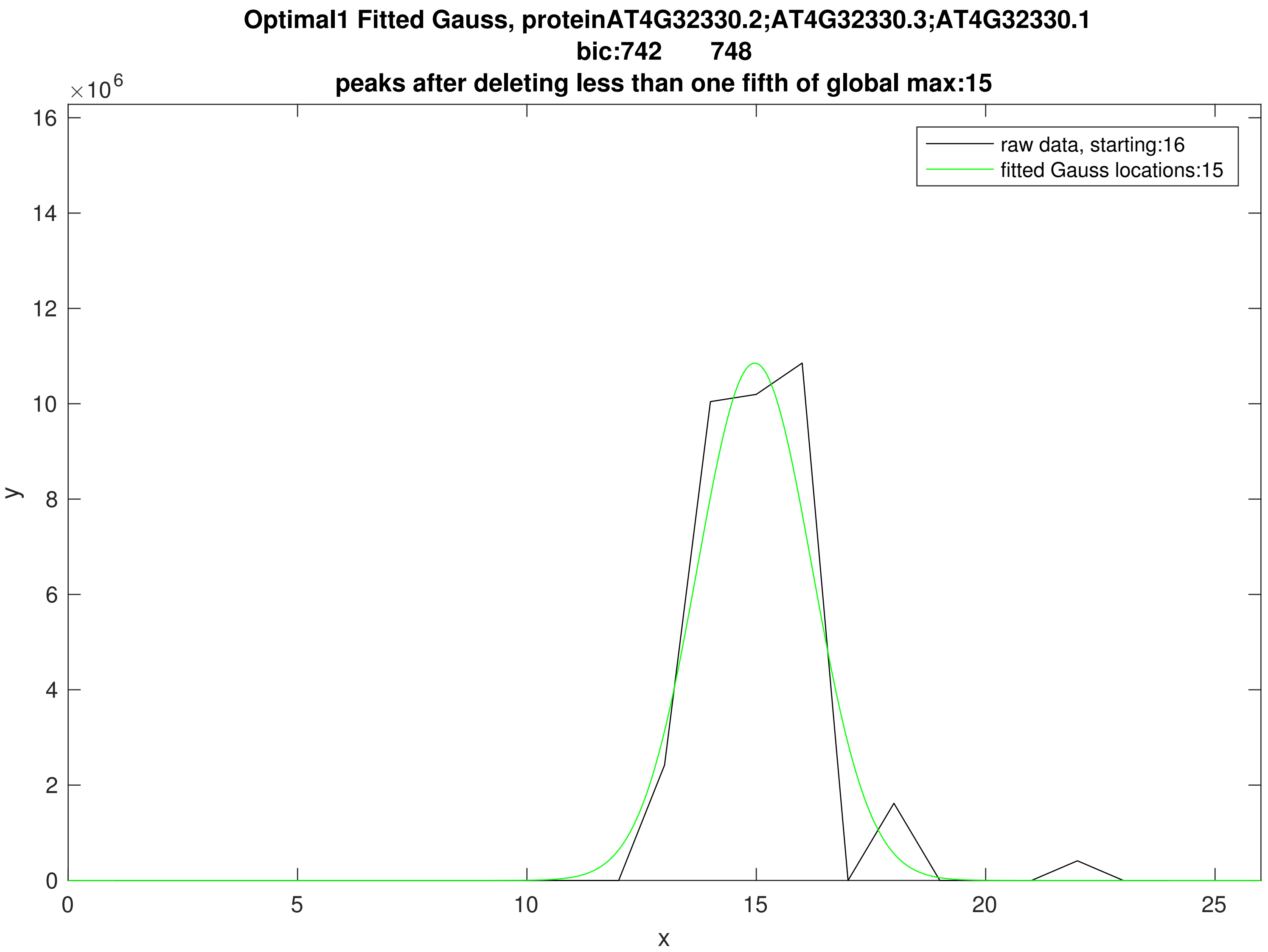
Optimal1 Fitted Gauss, proteinAT4G32330.2;AT4G32330.3;AT4G32330.1
bic:742 748
peaks after deleting less than one fifth of global max:15
×10^6
16
14
12
10
8
6
4
2
0
y
0
5
10
15
20
25
x
raw data, starting:16
fitted Gauss locations:15

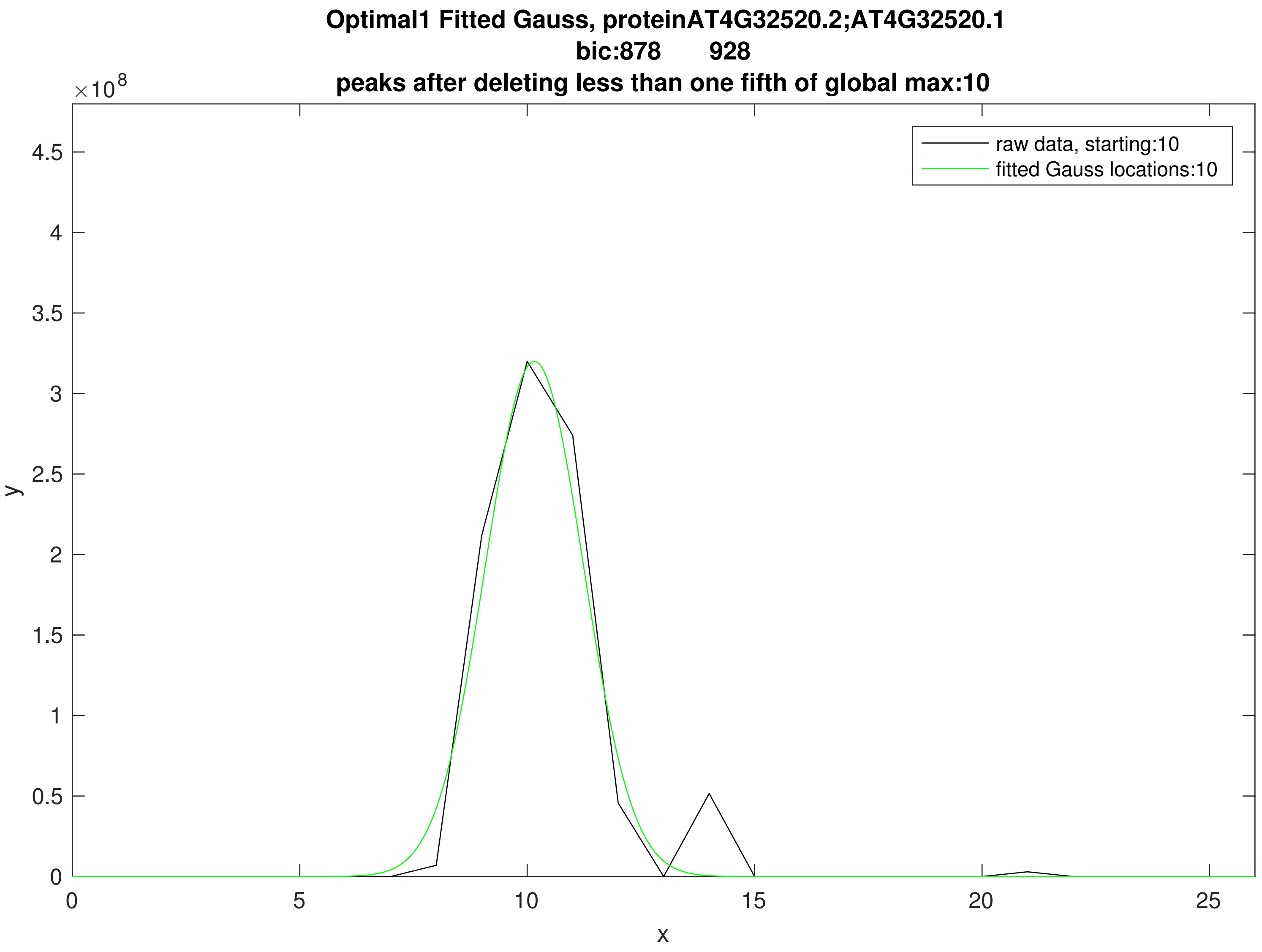

Optimal1 Fitted Gauss, proteinAT4G32520.2;AT4G32520.1
bic:878 928
peaks after deleting less than one fifth of global max:10
×10^8
4.5
4
3.5
3
2.5
2
1.5
1
0.5
0
y
0
5
10
15
20
25
x
raw data, starting:10
fitted Gauss locations:10

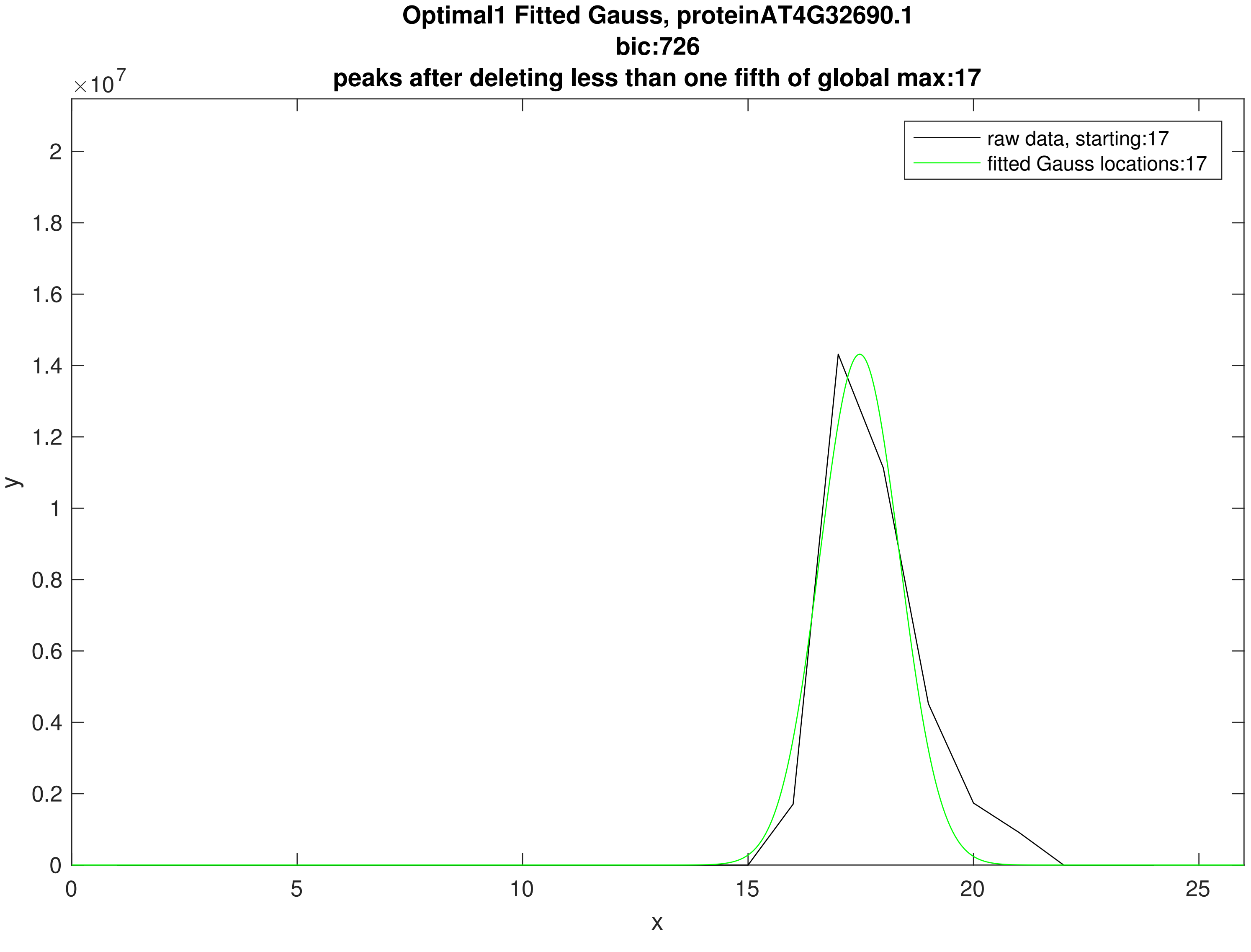
Optimal1 Fitted Gauss, proteinAT4G32690.1
bic:726
peaks after deleting less than one fifth of global max:17
×10^7
2
1.8
1.6
1.4
1.2
1
0.8
0.6
0.4
0.2
0
y
0
5
10
15
20
25
x
raw data, starting:17
fitted Gauss locations:17

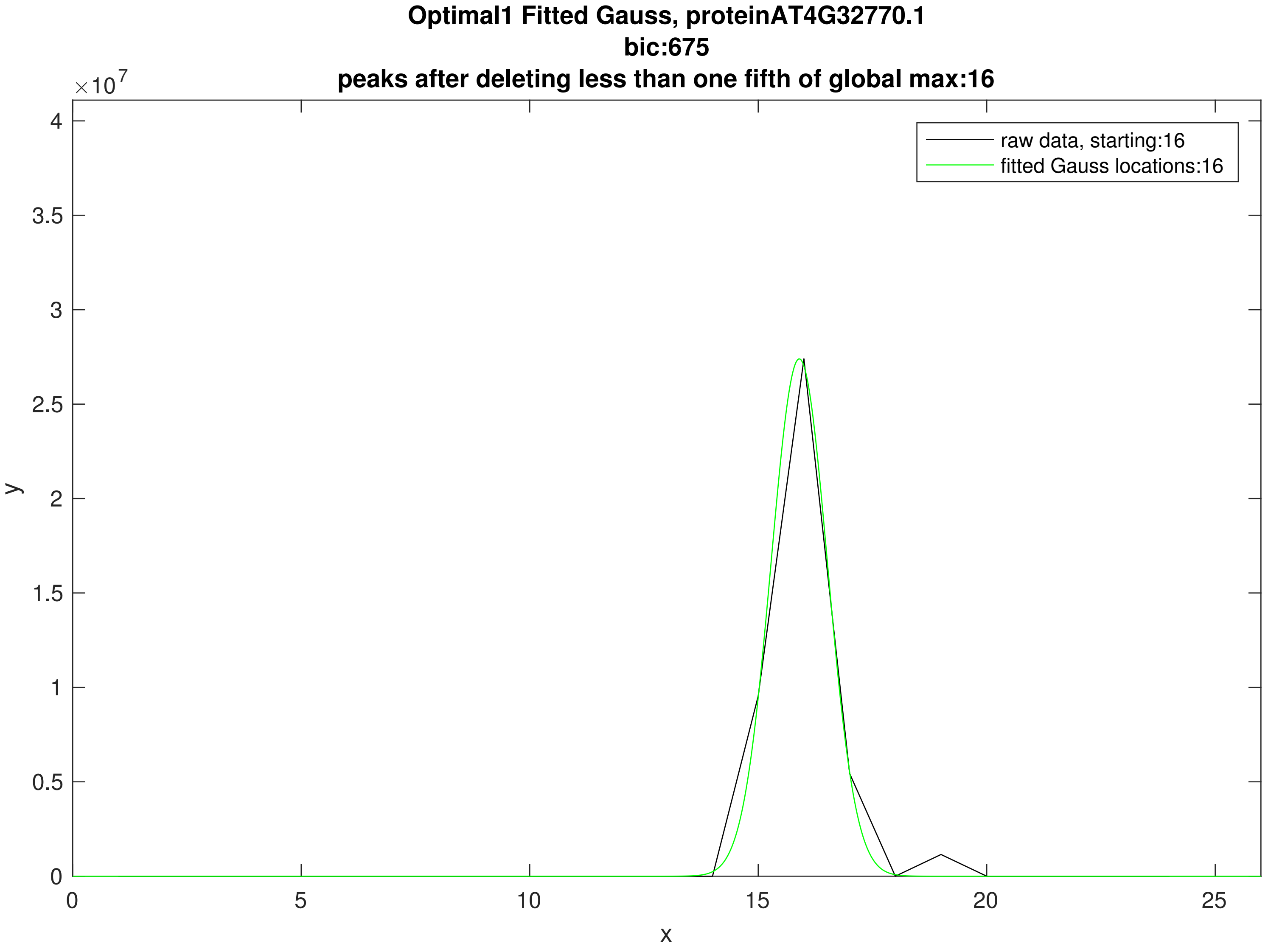

Optimal1 Fitted Gauss, proteinAT4G32770.1
bic:675
peaks after deleting less than one fifth of global max:16
×10^7
4
3.5
3
2.5
2
1.5
1
0.5
0
y
0
5
10
15
20
25
x
raw data, starting:16
fitted Gauss locations:16

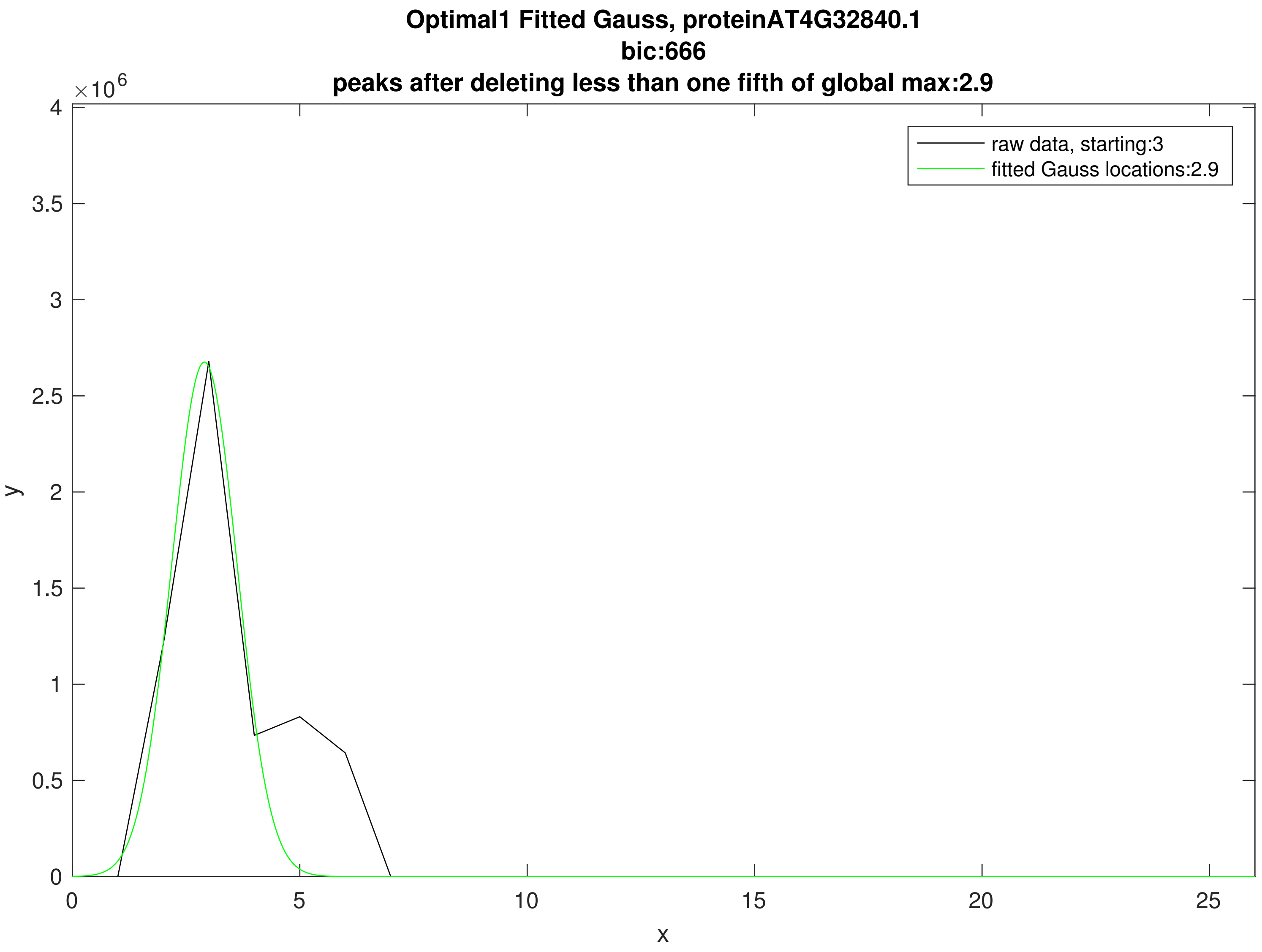
Optimal1 Fitted Gauss, proteinAT4G32840.1
bic:666
peaks after deleting less than one fifth of global max:2.9
×10^6
raw data, starting:3
fitted Gauss locations:2.9
y
x
0
0.5
1
1.5
2
2.5
3
3.5
4
5
10
15
20
25

Optimal1 Fitted Gauss, proteinAT4G32870.1
bic:655
peaks after deleting less than one fifth of global max:19

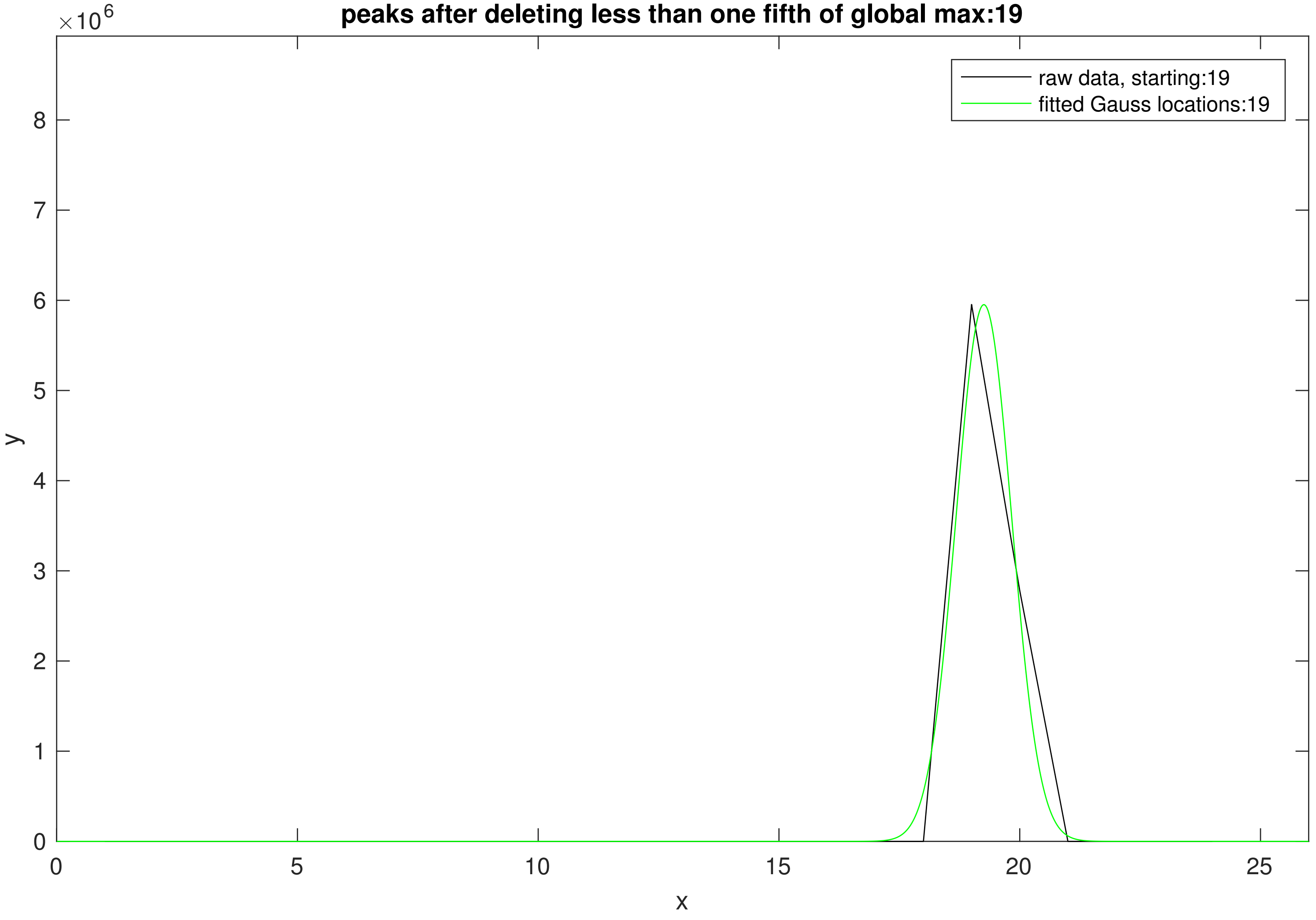

**Optimal1 Fitted Gauss, proteinAT4G32915.1**
**bic:698**
**peaks after deleting less than one fifth of global max:18**

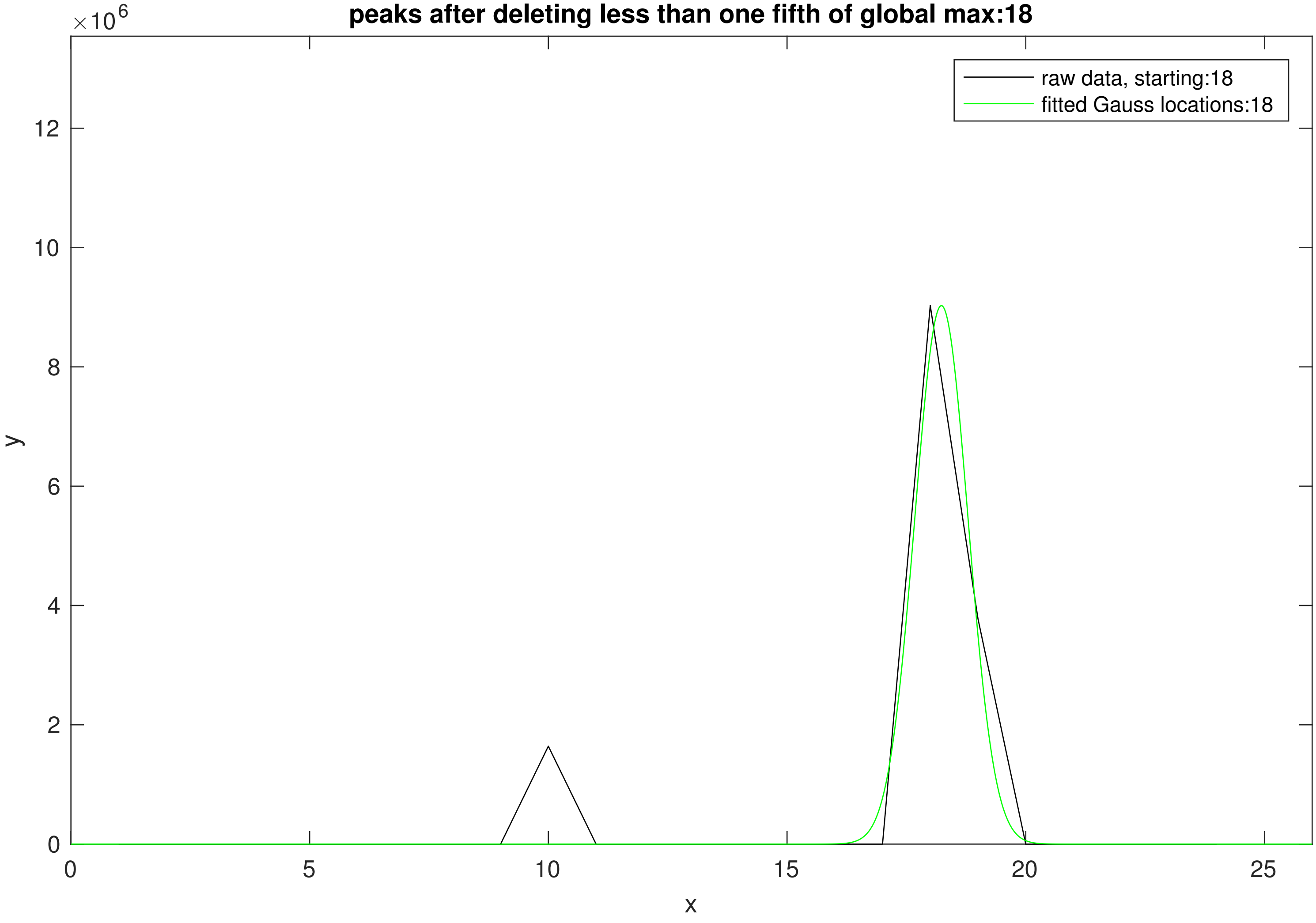

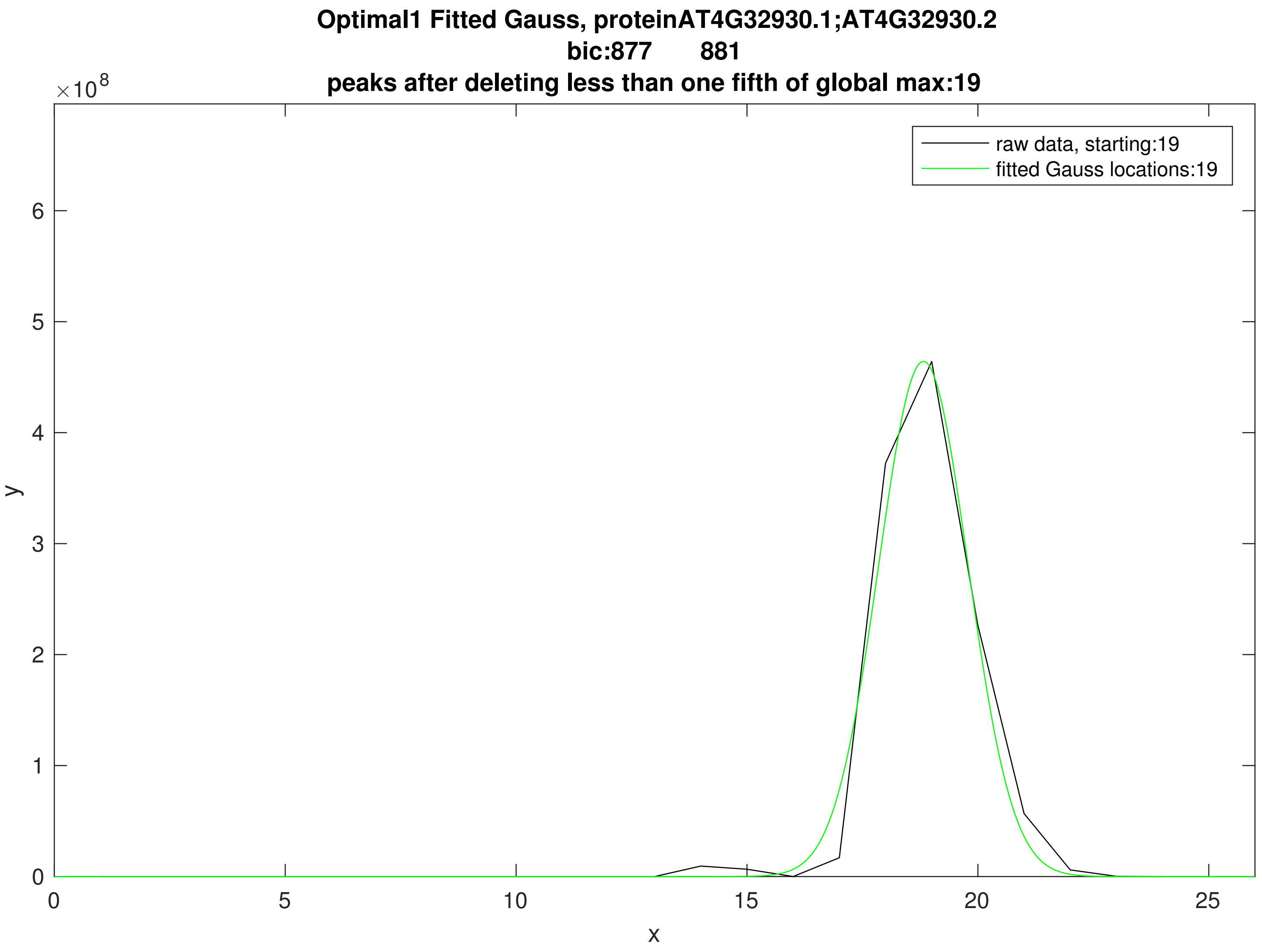
Optimal1 Fitted Gauss, proteinAT4G32930.1;AT4G32930.2
bic:877 881
peaks after deleting less than one fifth of global max:19
×10^8
raw data, starting:19
fitted Gauss locations:19
y
x
0
1
2
3
4
5
6
0
5
10
15
20
25

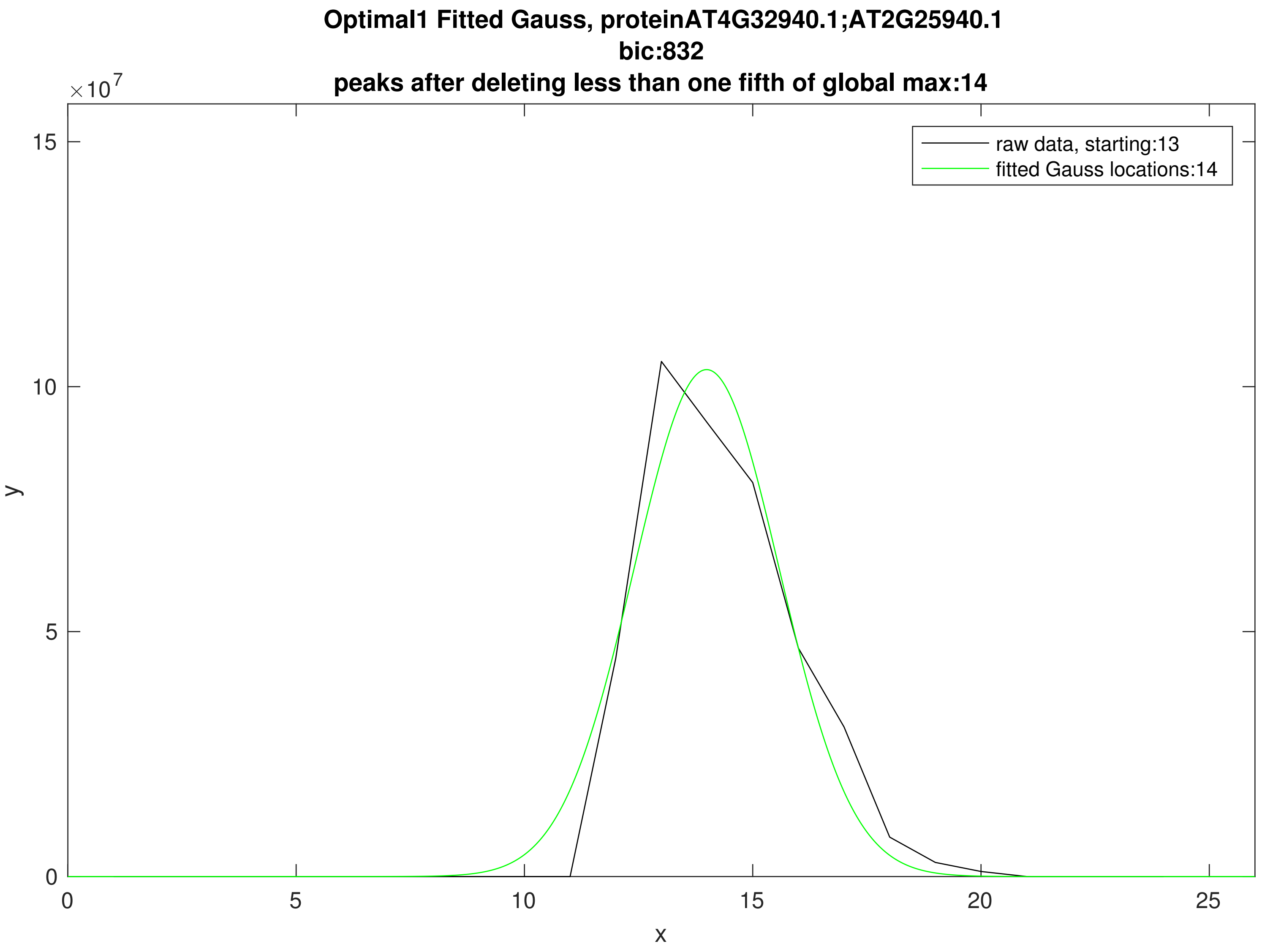

Optimal1 Fitted Gauss, proteinAT4G32940.1;AT2G25940.1
bic:832
peaks after deleting less than one fifth of global max:14
×10^7
raw data, starting:13
fitted Gauss locations:14
15
10
5
0
y
0
5
10
15
20
25
x

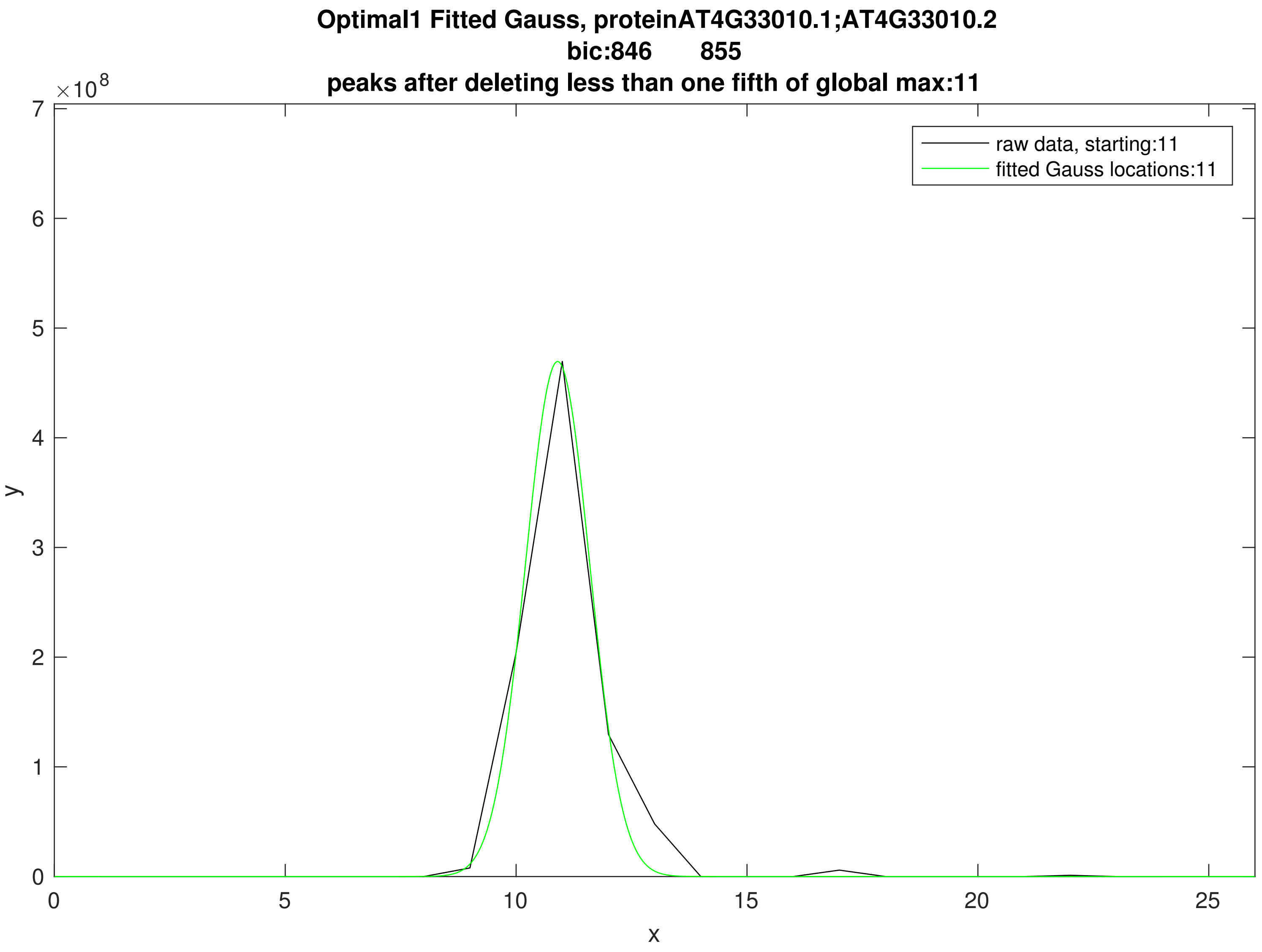
Optimal1 Fitted Gauss, proteinAT4G33010.1;AT4G33010.2
bic:846 855
peaks after deleting less than one fifth of global max:11
×10^8
7
6
5
4
3
2
1
0
y
0
5
10
15
20
25
x
raw data, starting:11
fitted Gauss locations:11

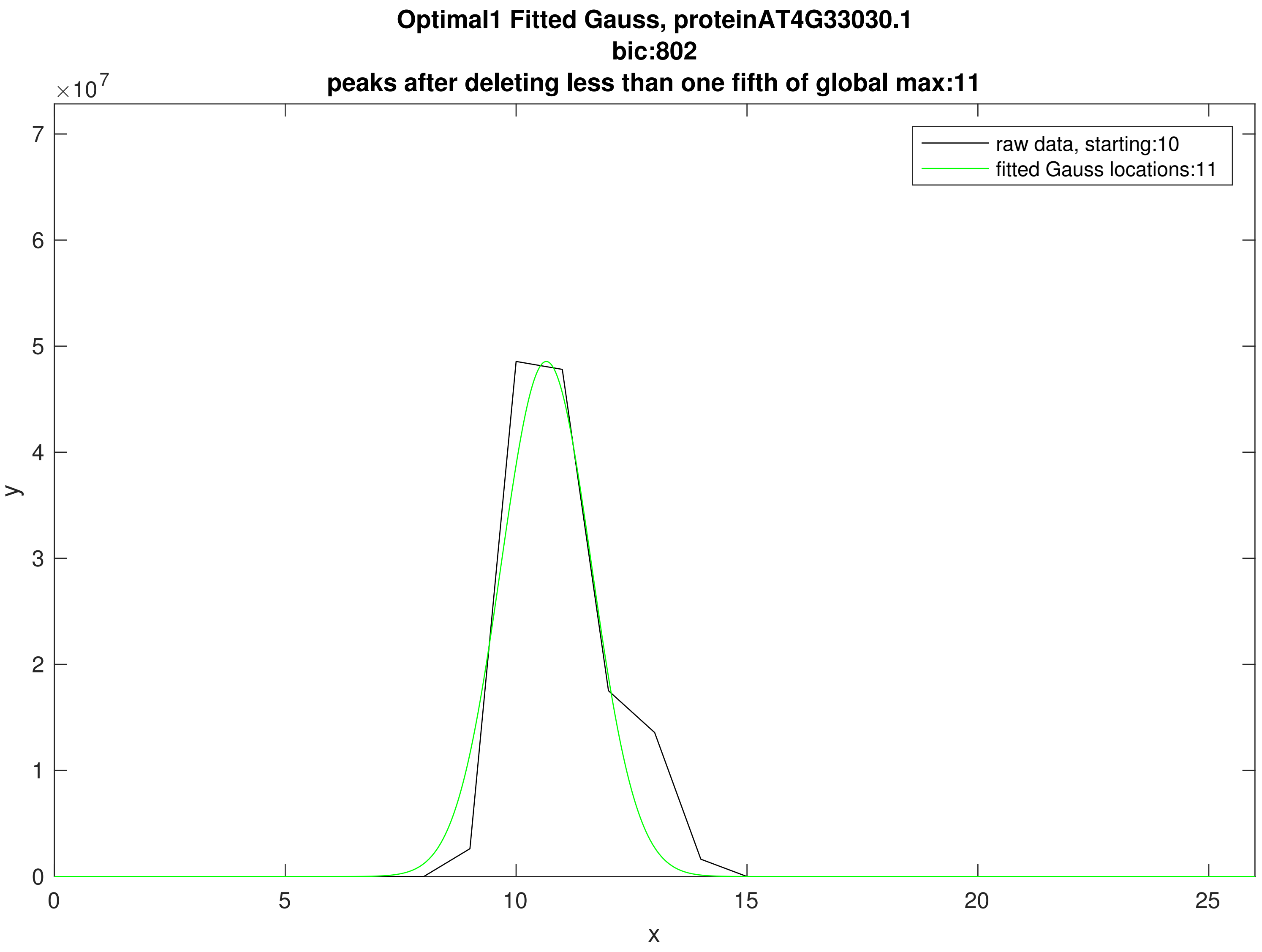
Optimal1 Fitted Gauss, proteinAT4G33030.1
bic:802
peaks after deleting less than one fifth of global max:11
×10^7
raw data, starting:10
fitted Gauss locations:11
y
x
0
1
2
3
4
5
6
7
0
5
10
15
20
25

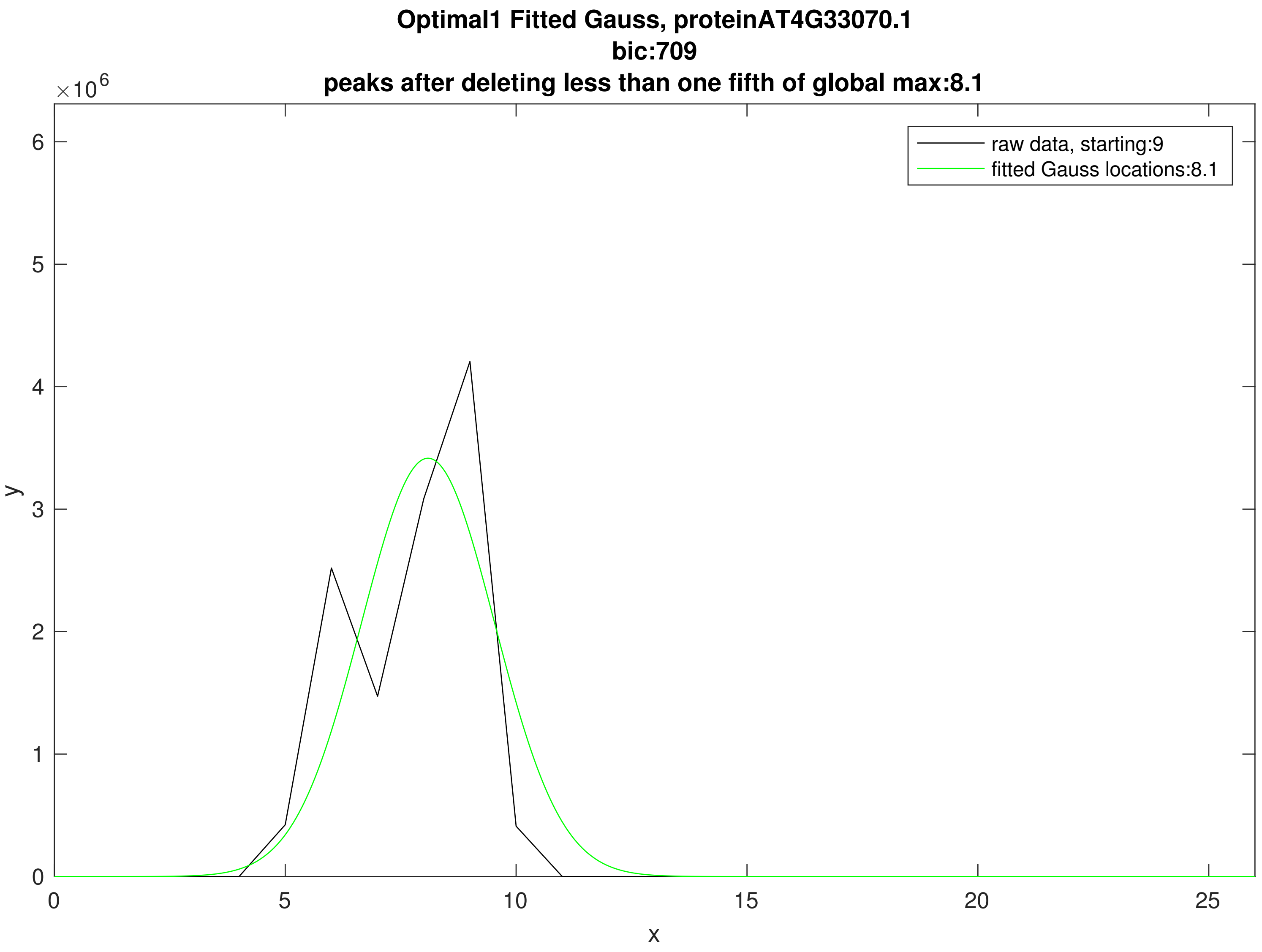

Optimal1 Fitted Gauss, proteinAT4G33070.1
bic:709
peaks after deleting less than one fifth of global max:8.1
×10^6
raw data, starting:9
fitted Gauss locations:8.1
y
x
0
1
2
3
4
5
6
0
5
10
15
20
25

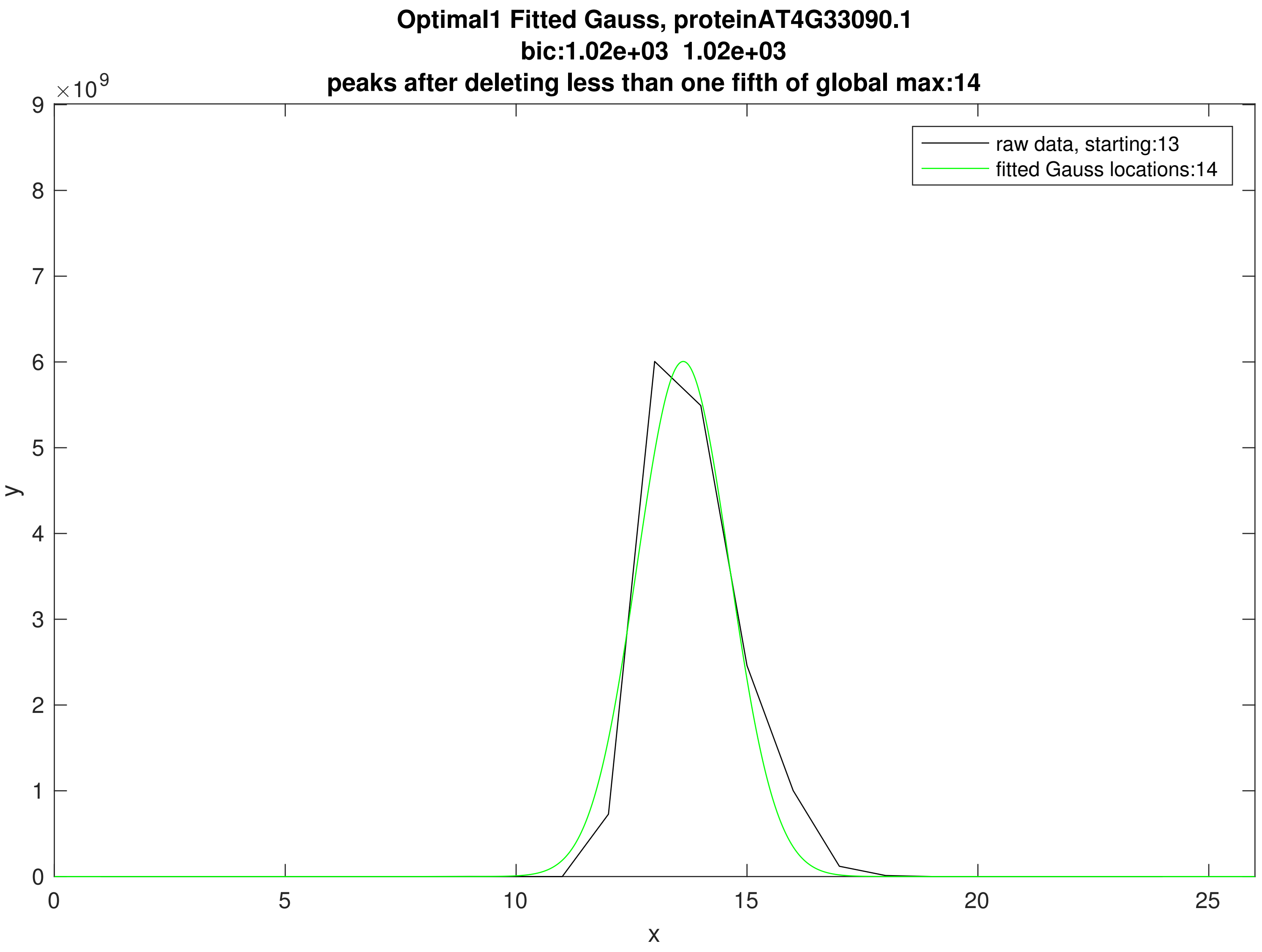

Optimal1 Fitted Gauss, proteinAT4G33090.1
bic:1.02e+03 1.02e+03
peaks after deleting less than one fifth of global max:14
×10^9
raw data, starting:13
fitted Gauss locations:14
y
x
0
1
2
3
4
5
6
7
8
9
0
5
10
15
20
25

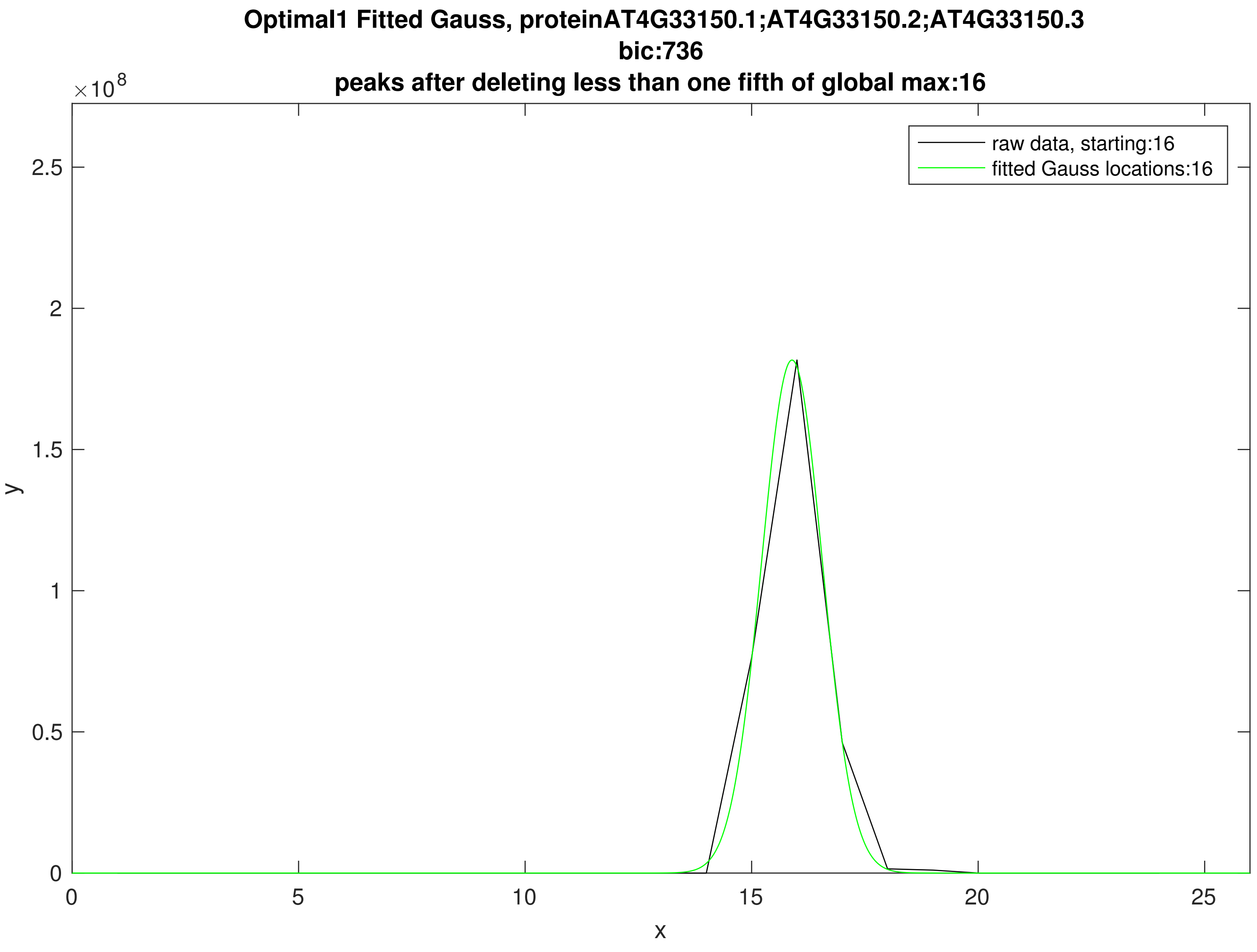

Optimal1 Fitted Gauss, proteinAT4G33150.1;AT4G33150.2;AT4G33150.3
bic:736
peaks after deleting less than one fifth of global max:16
×10^8
2.5
2
1.5
1
0.5
0
y
0
5
10
15
20
25
x
raw data, starting:16
fitted Gauss locations:16

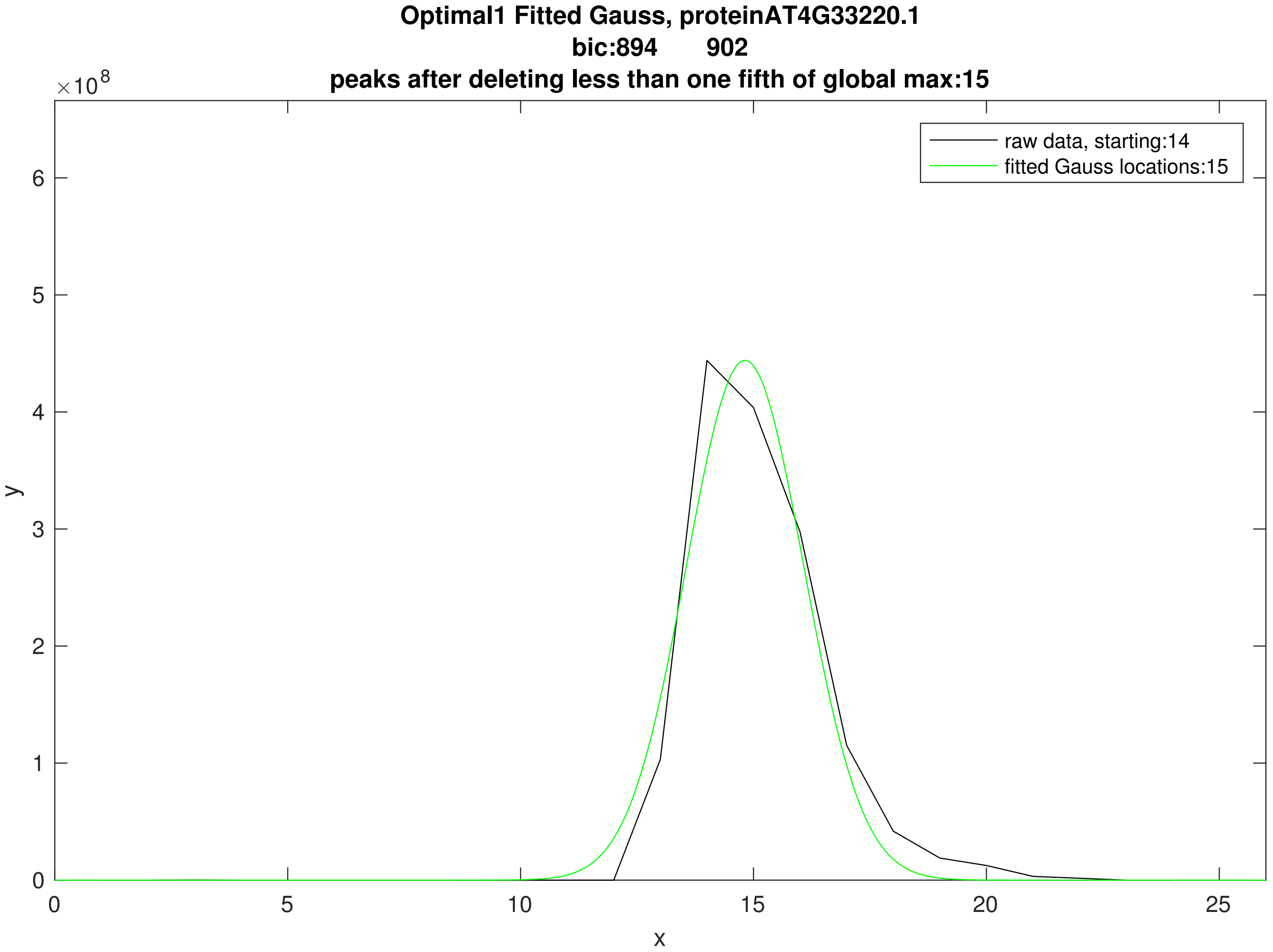
Optimal1 Fitted Gauss, proteinAT4G33220.1
bic:894 902
peaks after deleting less than one fifth of global max:15
×10^8
raw data, starting:14
fitted Gauss locations:15
y
x
0
1
2
3
4
5
6
0
5
10
15
20
25

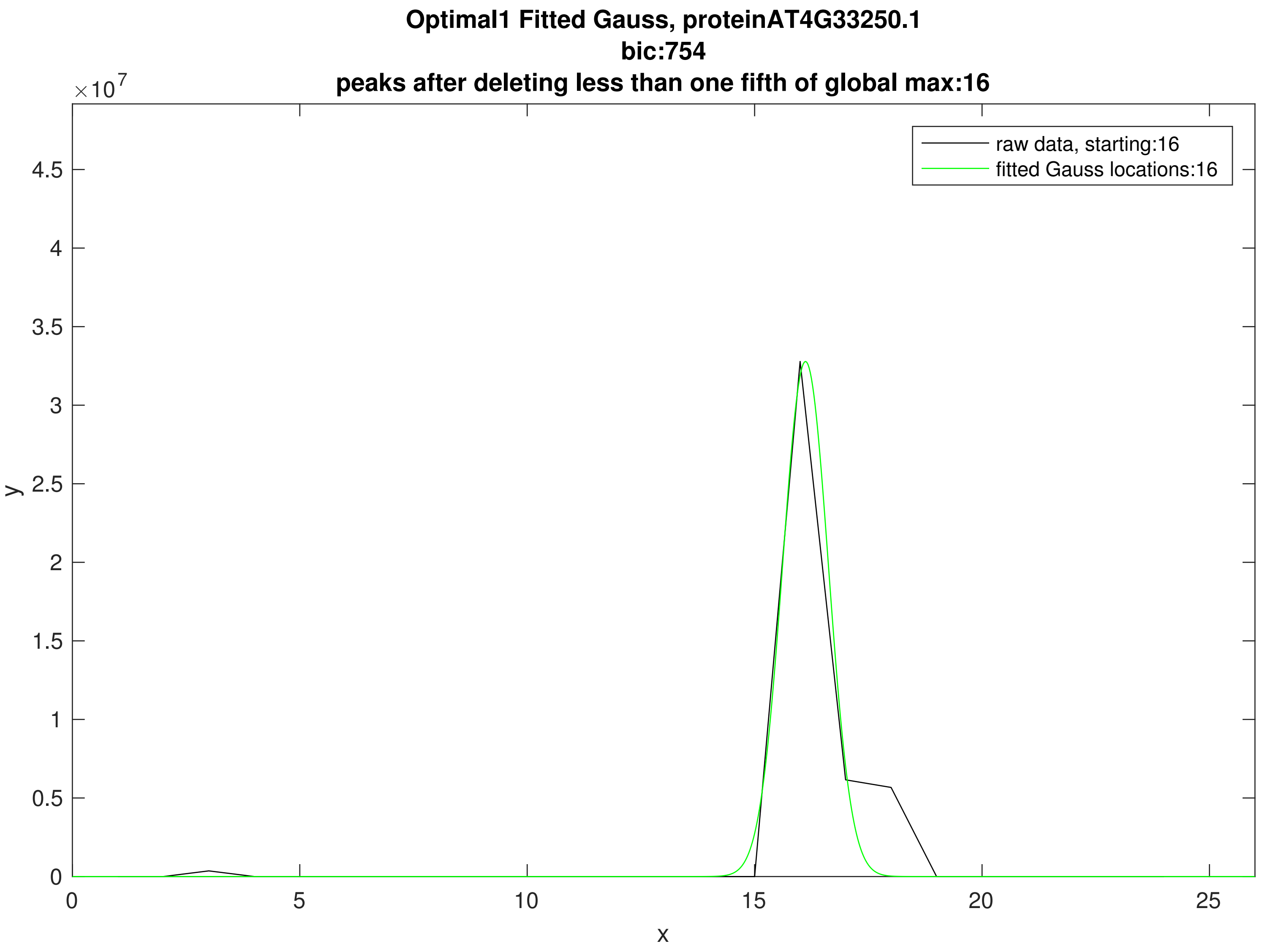
Optimal1 Fitted Gauss, proteinAT4G33250.1
bic:754
peaks after deleting less than one fifth of global max:16
×10^7
raw data, starting:16
fitted Gauss locations:16
y
x
0
0.5
1
1.5
2
2.5
3
3.5
4
4.5
5
10
15
20
25

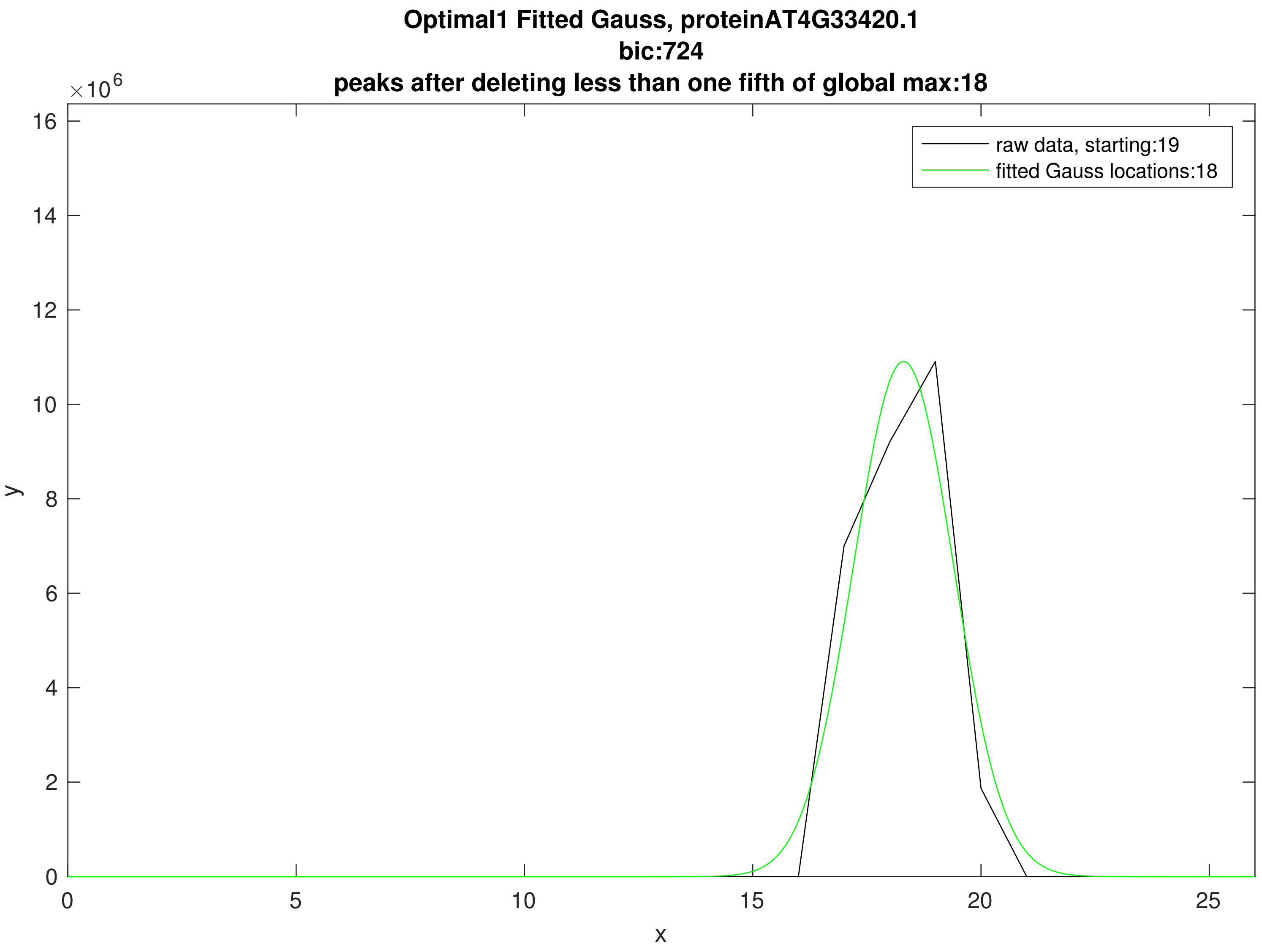

Optimal1 Fitted Gauss, proteinAT4G33420.1
bic:724
peaks after deleting less than one fifth of global max:18
×10^6
16
14
12
10
8
6
4
2
0
y
0
5
10
15
20
25
x
raw data, starting:19
fitted Gauss locations:18

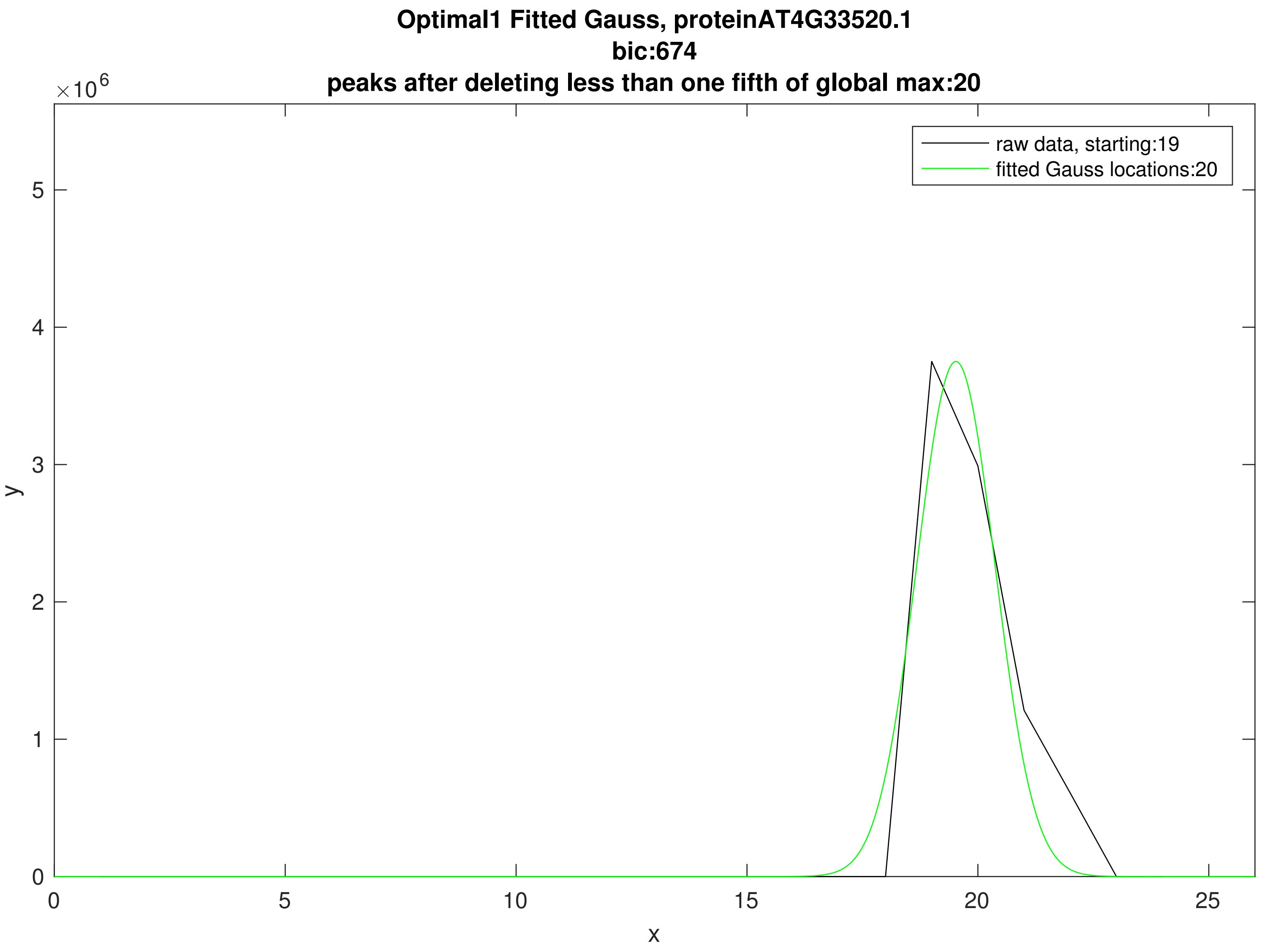
Optimal1 Fitted Gauss, proteinAT4G33520.1
bic:674
peaks after deleting less than one fifth of global max:20
×10^6
raw data, starting:19
fitted Gauss locations:20
y
x
0
1
2
3
4
5
0
5
10
15
20
25

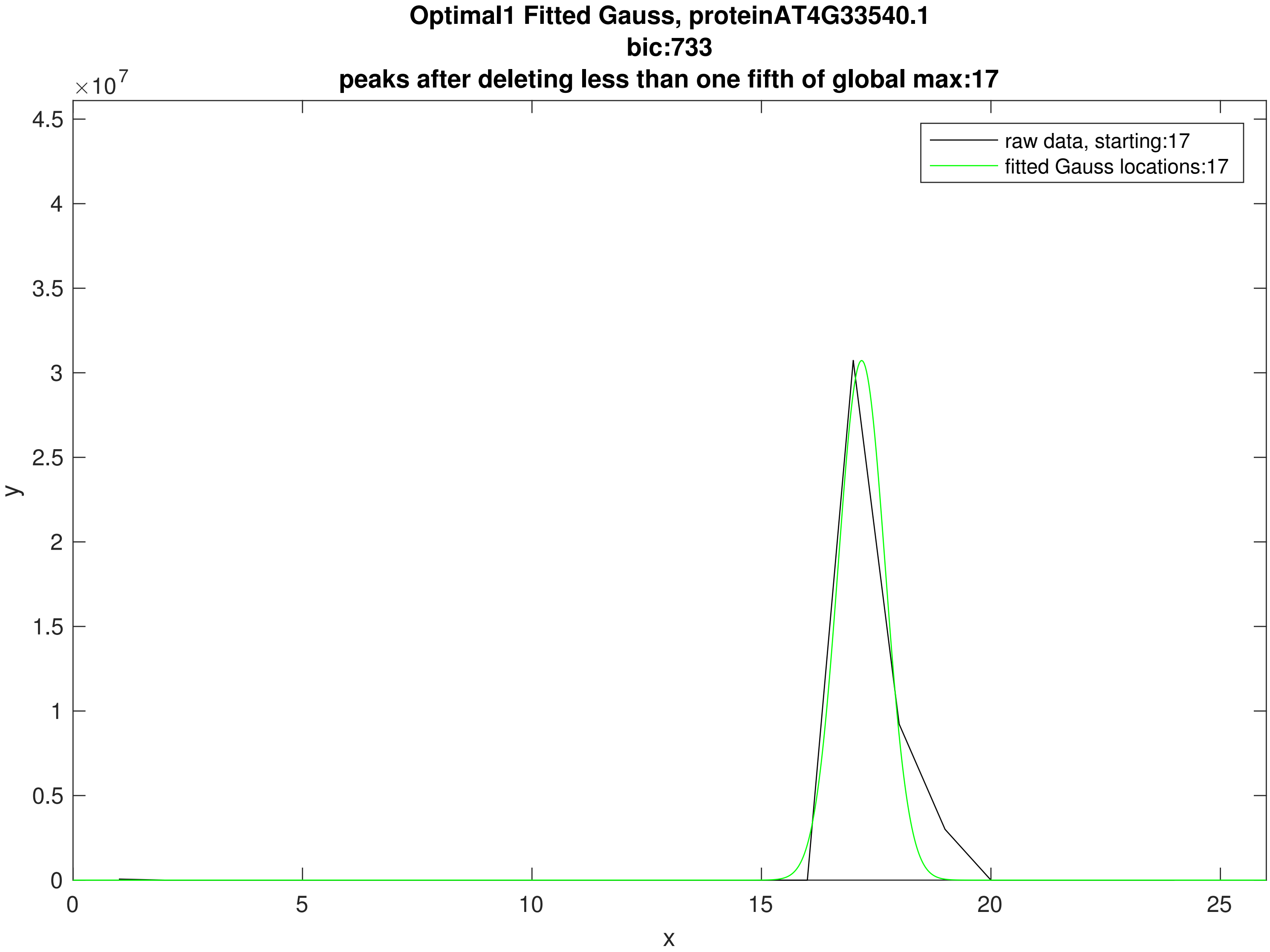

Optimal1 Fitted Gauss, proteinAT4G33540.1
bic:733
peaks after deleting less than one fifth of global max:17
×10^7
4.5
4
3.5
3
2.5
2
1.5
1
0.5
0
y
0
5
10
15
20
25
x
raw data, starting:17
fitted Gauss locations:17

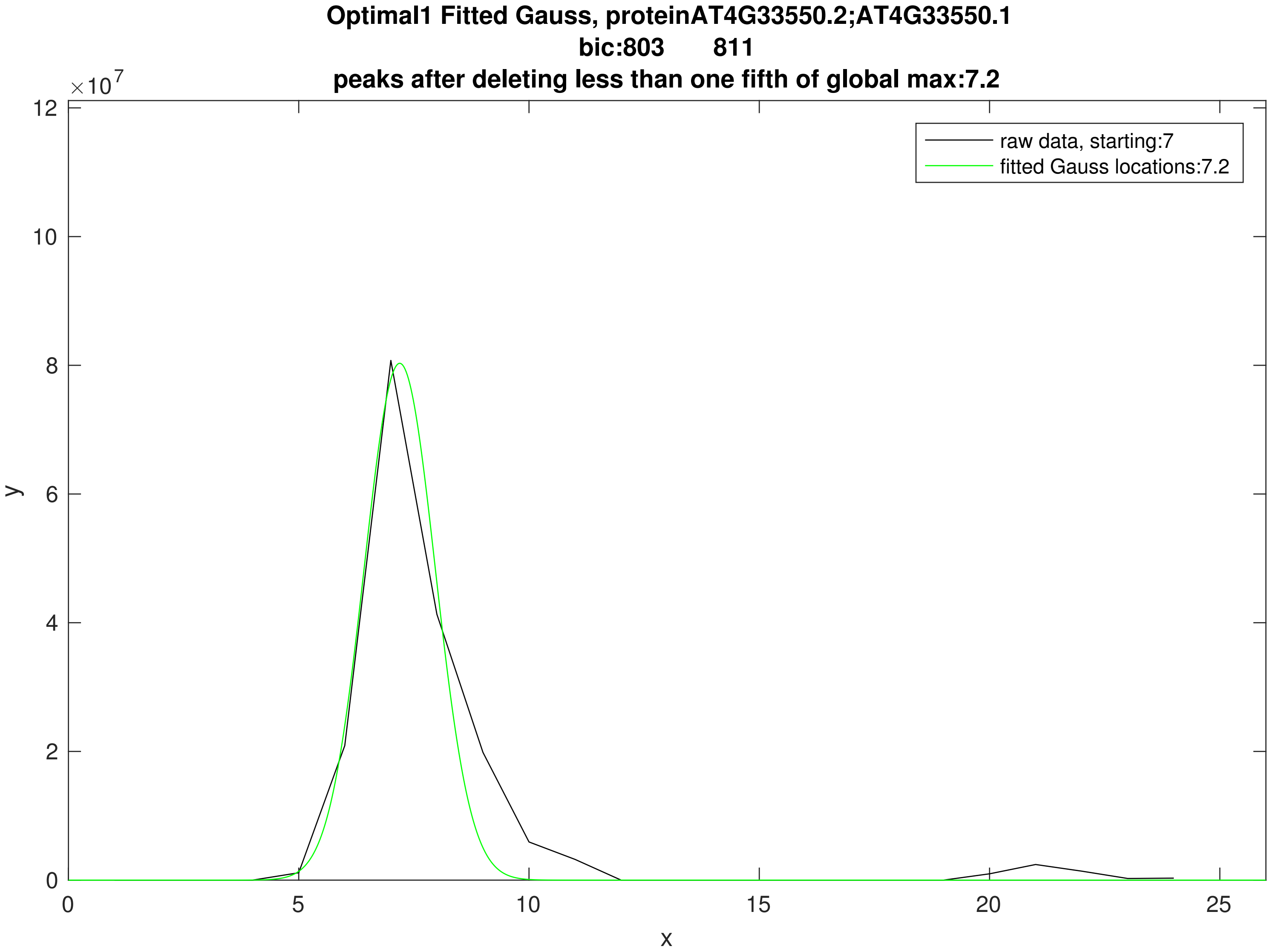
Optimal1 Fitted Gauss, proteinAT4G33550.2;AT4G33550.1
bic:803 811
peaks after deleting less than one fifth of global max:7.2
×10^7
12
10
8
6
4
2
0
y
0
5
10
15
20
25
x
raw data, starting:7
fitted Gauss locations:7.2

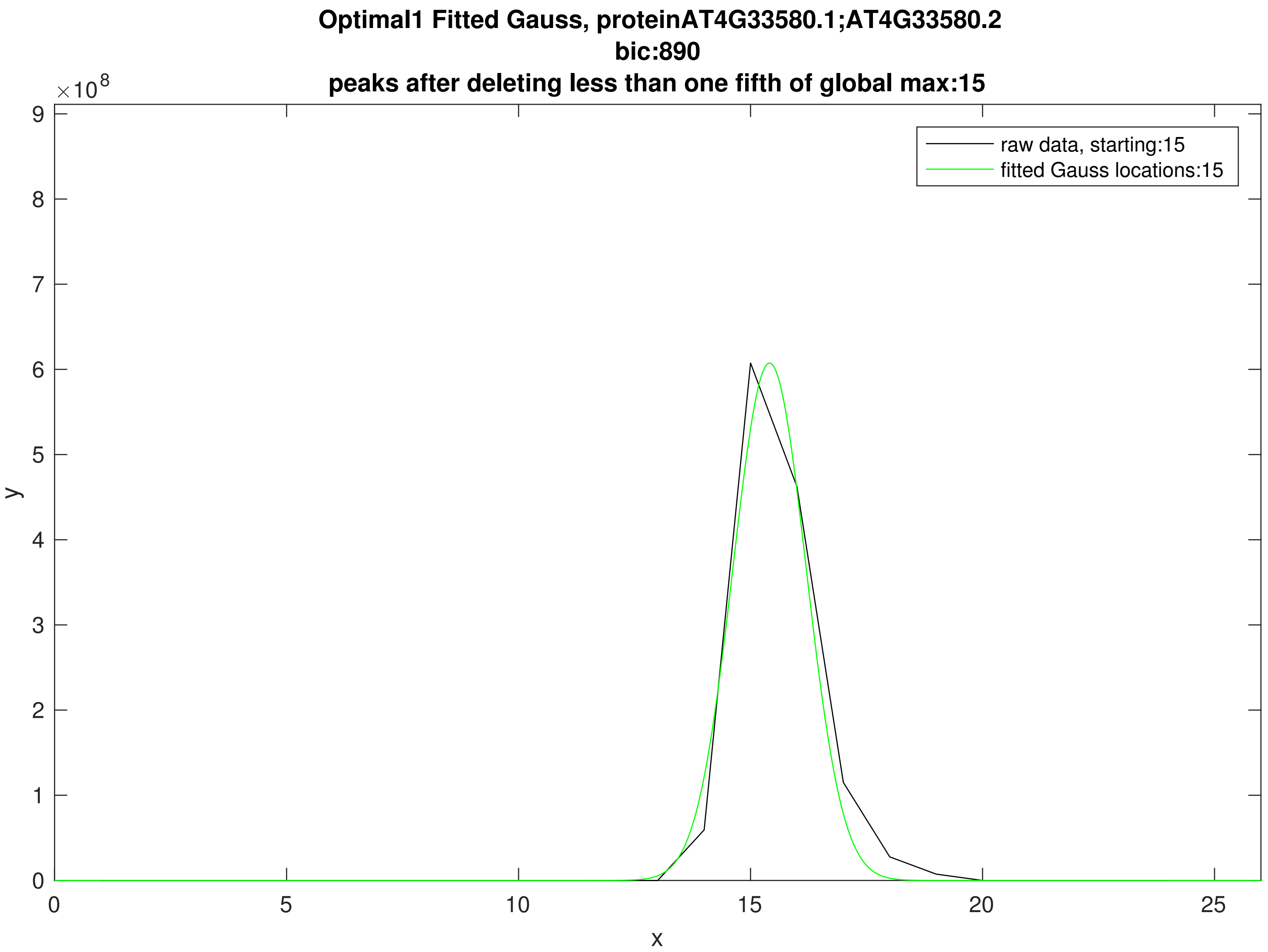

Optimal1 Fitted Gauss, proteinAT4G33580.1;AT4G33580.2
bic:890
peaks after deleting less than one fifth of global max:15
×10^8
9
8
7
6
5
4
3
2
1
0
y
0
5
10
15
20
25
x
raw data, starting:15
fitted Gauss locations:15

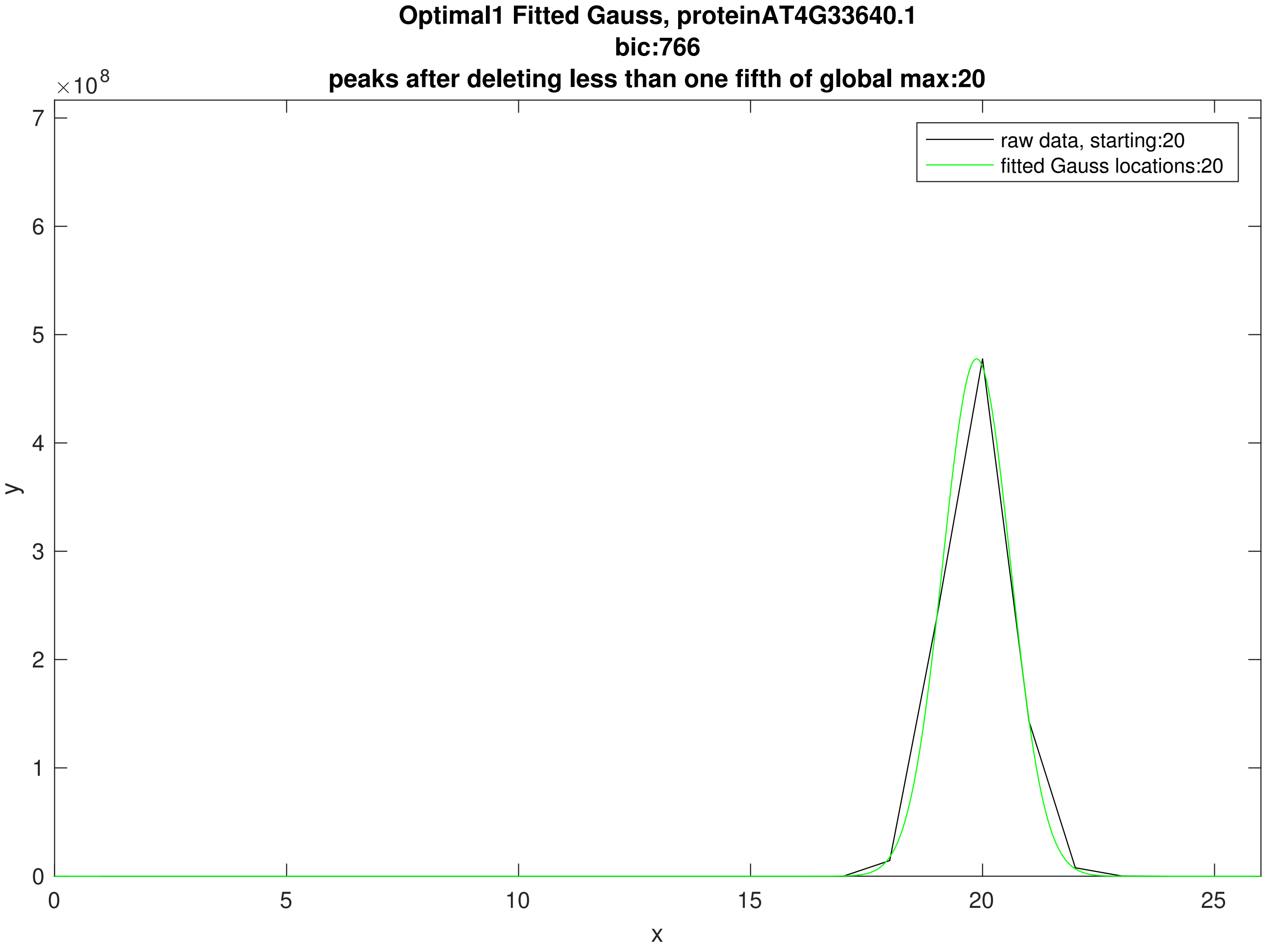

Optimal1 Fitted Gauss, proteinAT4G33640.1
bic:766
peaks after deleting less than one fifth of global max:20
×10^8
raw data, starting:20
fitted Gauss locations:20
y
x
0
1
2
3
4
5
6
7
0
5
10
15
20
25

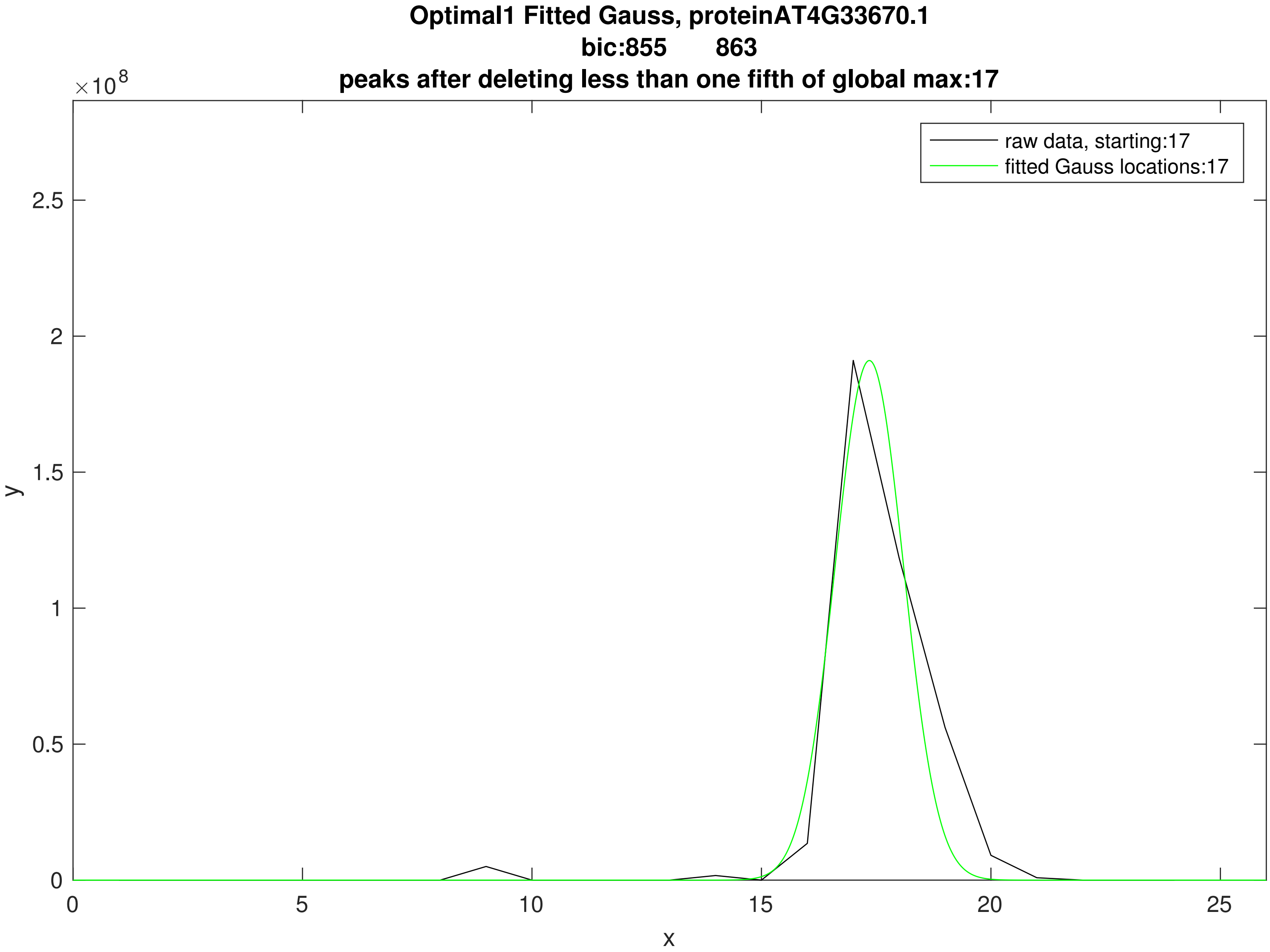
Optimal1 Fitted Gauss, proteinAT4G33670.1
bic:855 863
peaks after deleting less than one fifth of global max:17
×10^8
raw data, starting:17
fitted Gauss locations:17
y
2.5
2
1.5
1
0.5
0
x
0
5
10
15
20
25

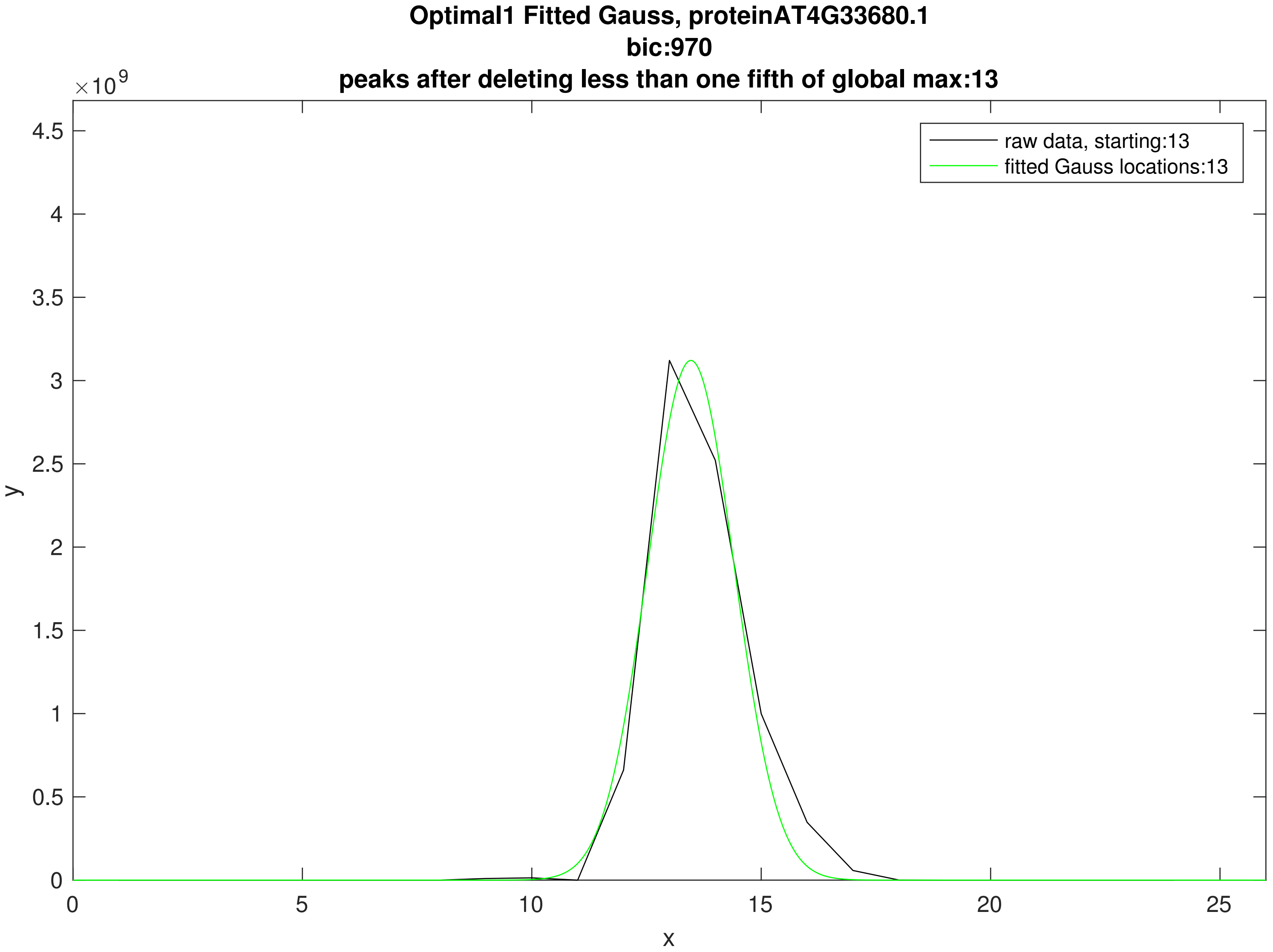

Optimal1 Fitted Gauss, proteinAT4G33680.1
bic:970
peaks after deleting less than one fifth of global max:13
×10^9
4.5
4
3.5
3
2.5
2
1.5
1
0.5
0
y
0
5
10
15
20
25
x
raw data, starting:13
fitted Gauss locations:13

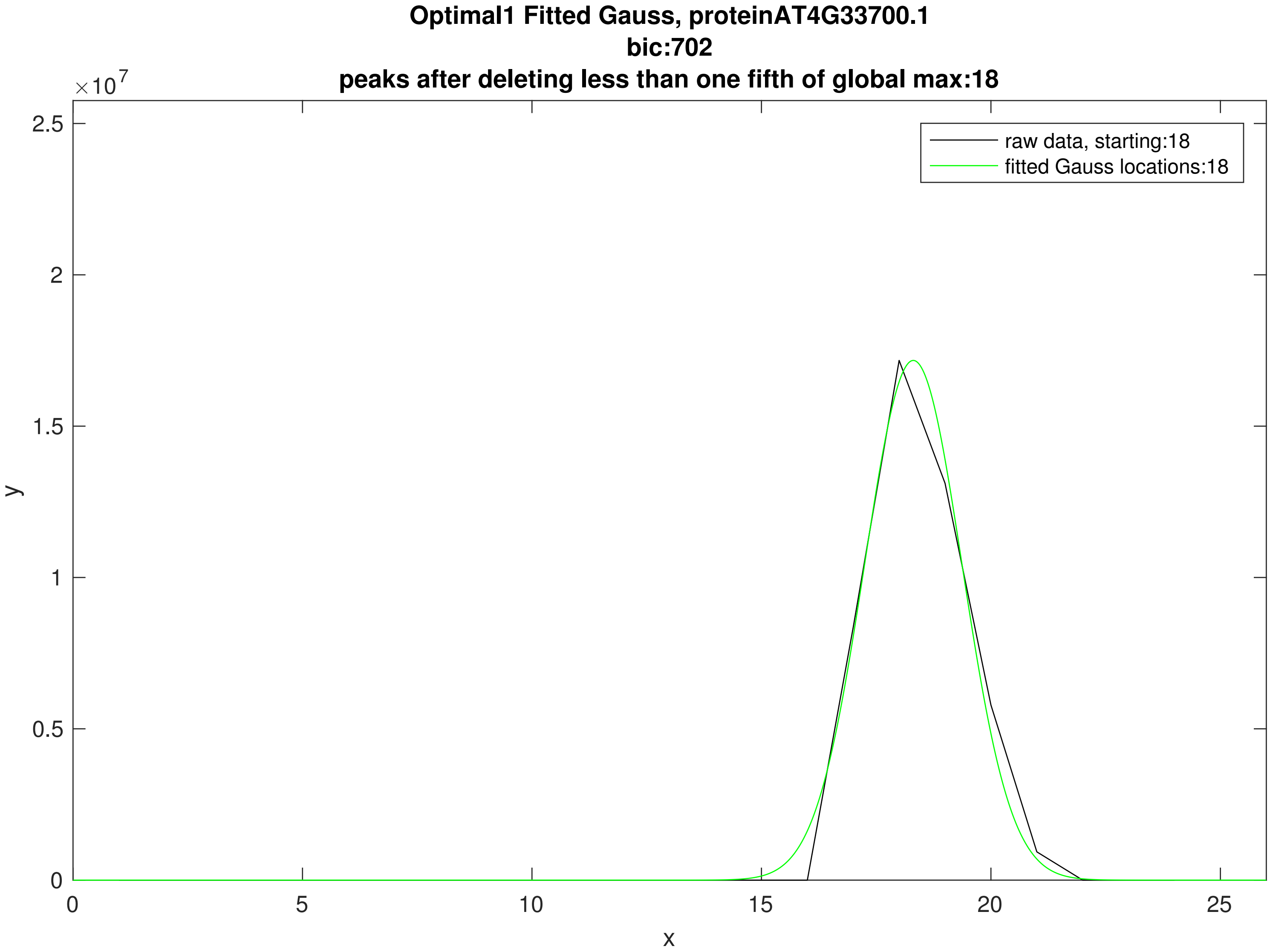

Optimal1 Fitted Gauss, proteinAT4G33700.1
bic:702
peaks after deleting less than one fifth of global max:18
×10^7
2.5
2
1.5
1
0.5
0
y
0
5
10
15
20
25
x
raw data, starting:18
fitted Gauss locations:18

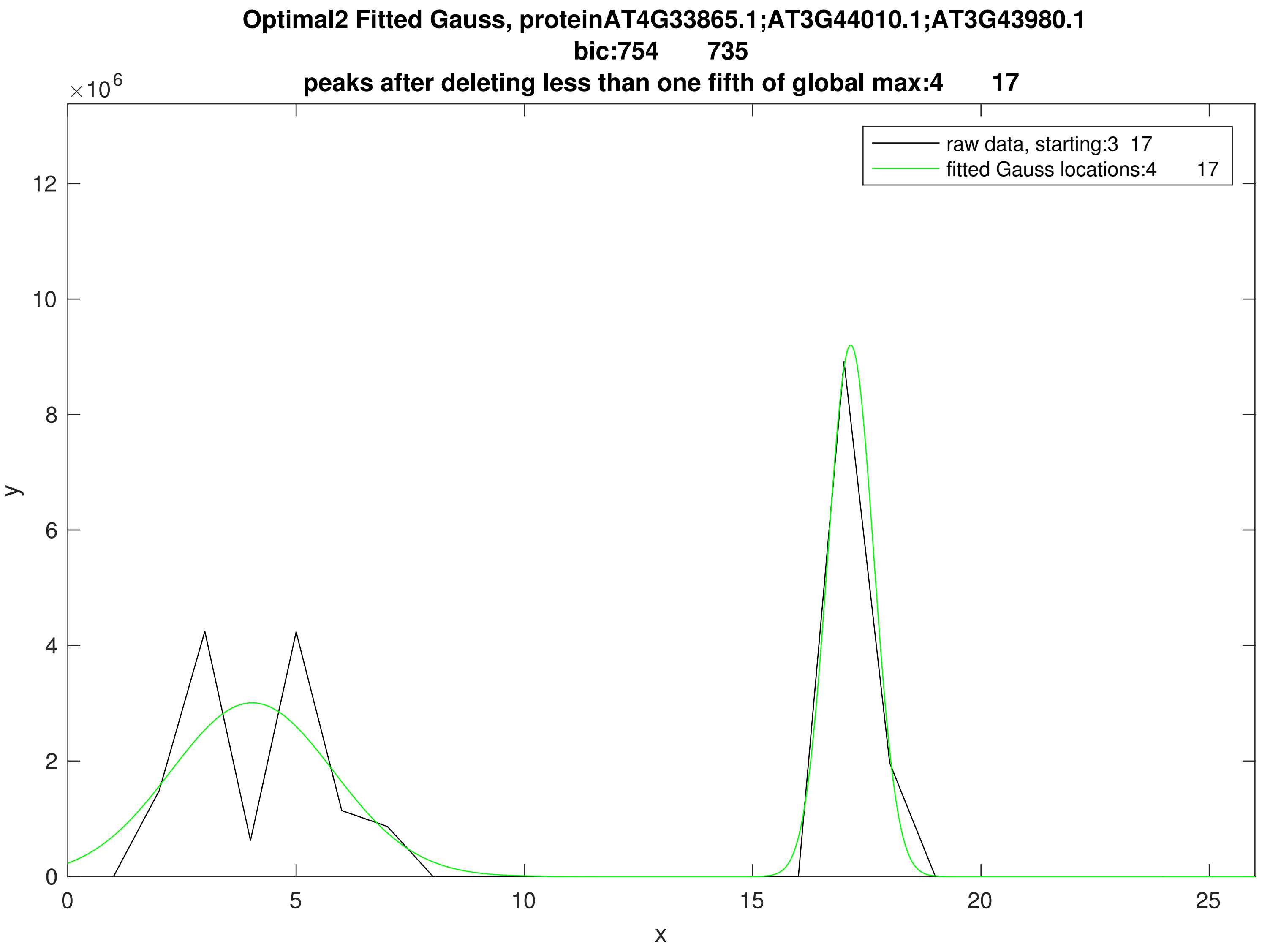

Optimal2 Fitted Gauss, proteinAT4G33865.1;AT3G44010.1;AT3G43980.1
bic:754 735
peaks after deleting less than one fifth of global max:4 17
×10^6
raw data, starting:3 17
fitted Gauss locations:4 17
y
x
0
2
4
6
8
10
12
0
5
10
15
20
25

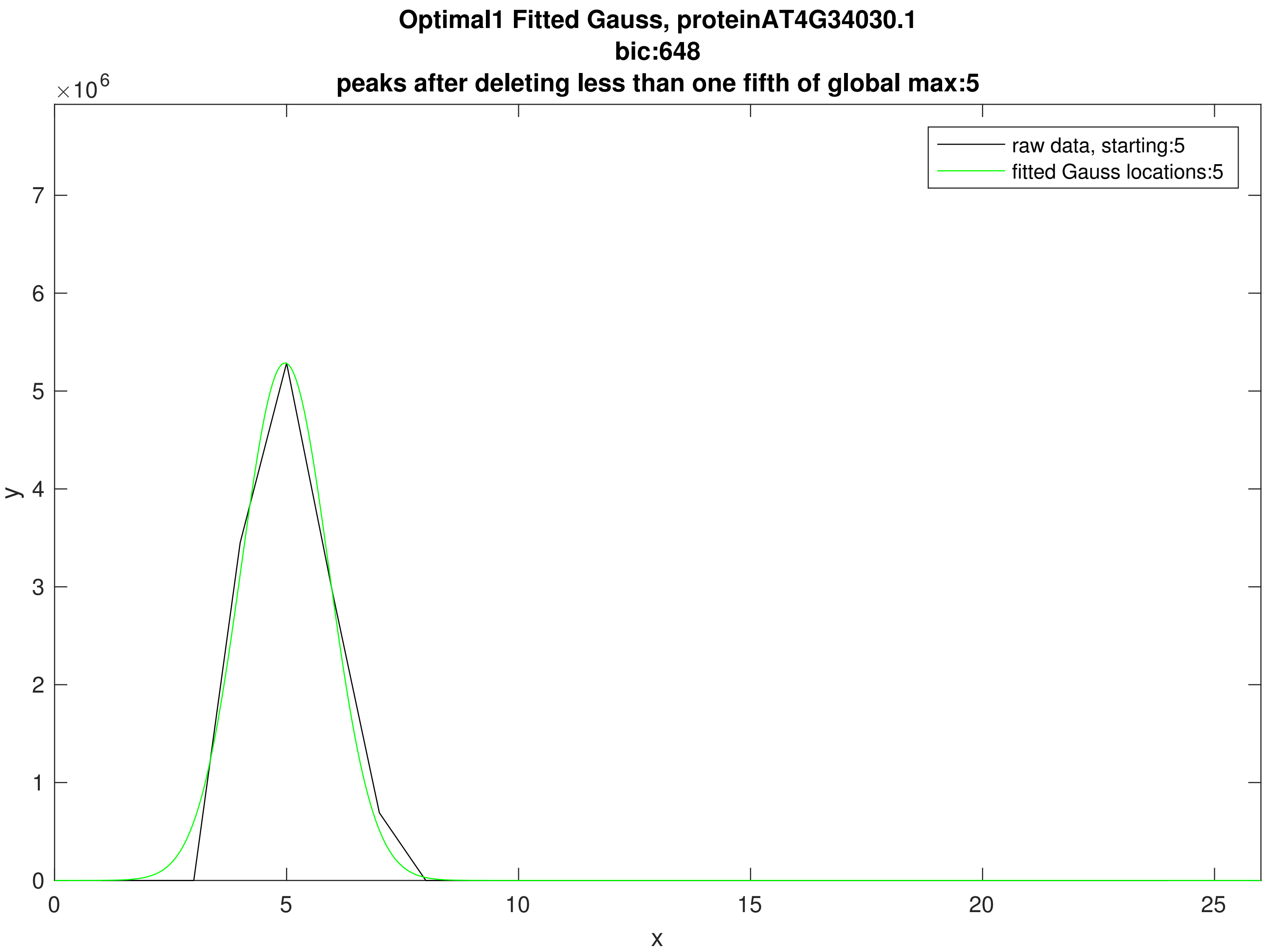

Optimal1 Fitted Gauss, proteinAT4G34030.1
bic:648
peaks after deleting less than one fifth of global max:5
×10^6
raw data, starting:5
fitted Gauss locations:5
y
x
0
1
2
3
4
5
6
7
0
5
10
15
20
25

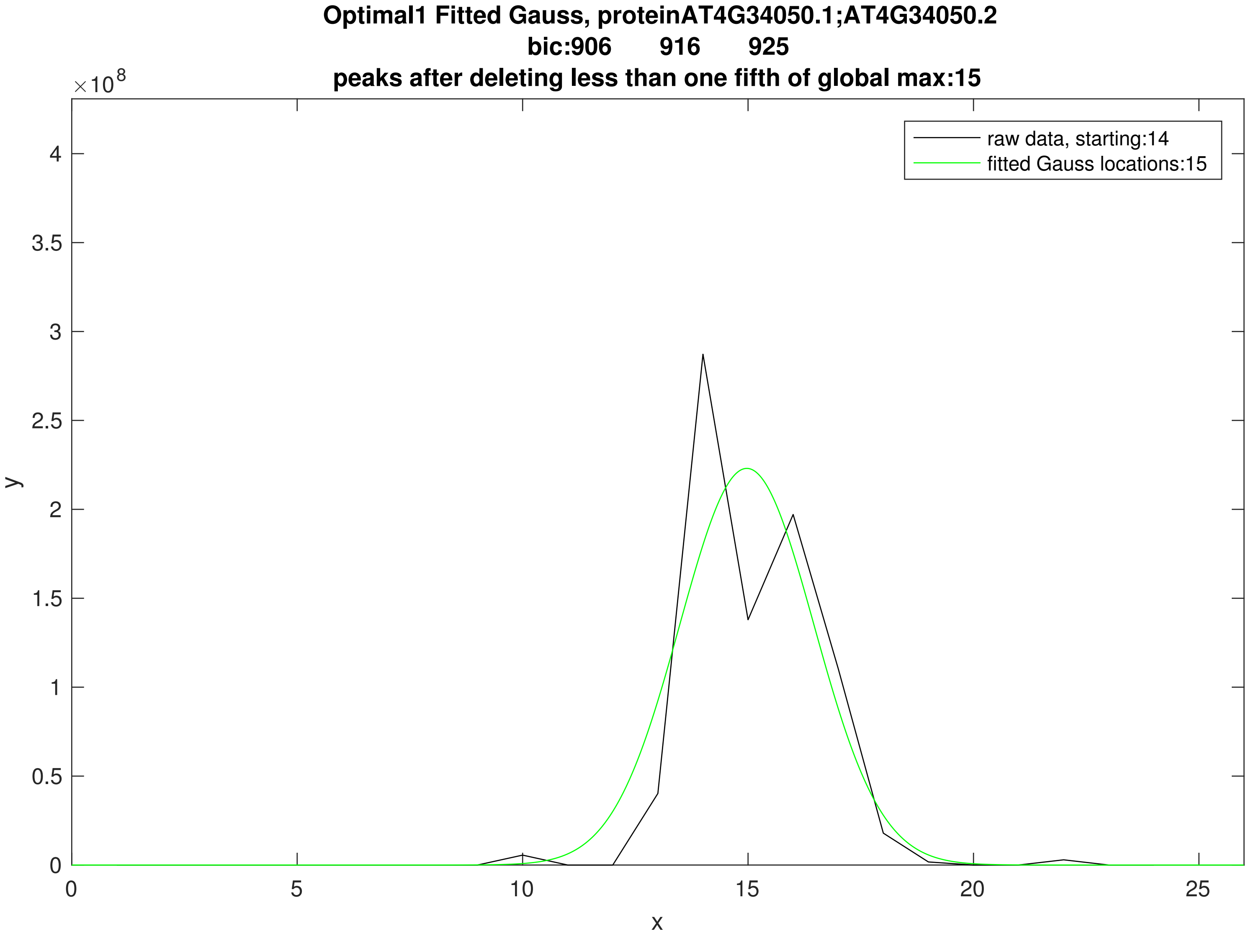

Optimal1 Fitted Gauss, proteinAT4G34050.1;AT4G34050.2
bic:906 916 925
peaks after deleting less than one fifth of global max:15
×10^8
raw data, starting:14
fitted Gauss locations:15
y
x
0
0.5
1
1.5
2
2.5
3
3.5
4
0
5
10
15
20
25

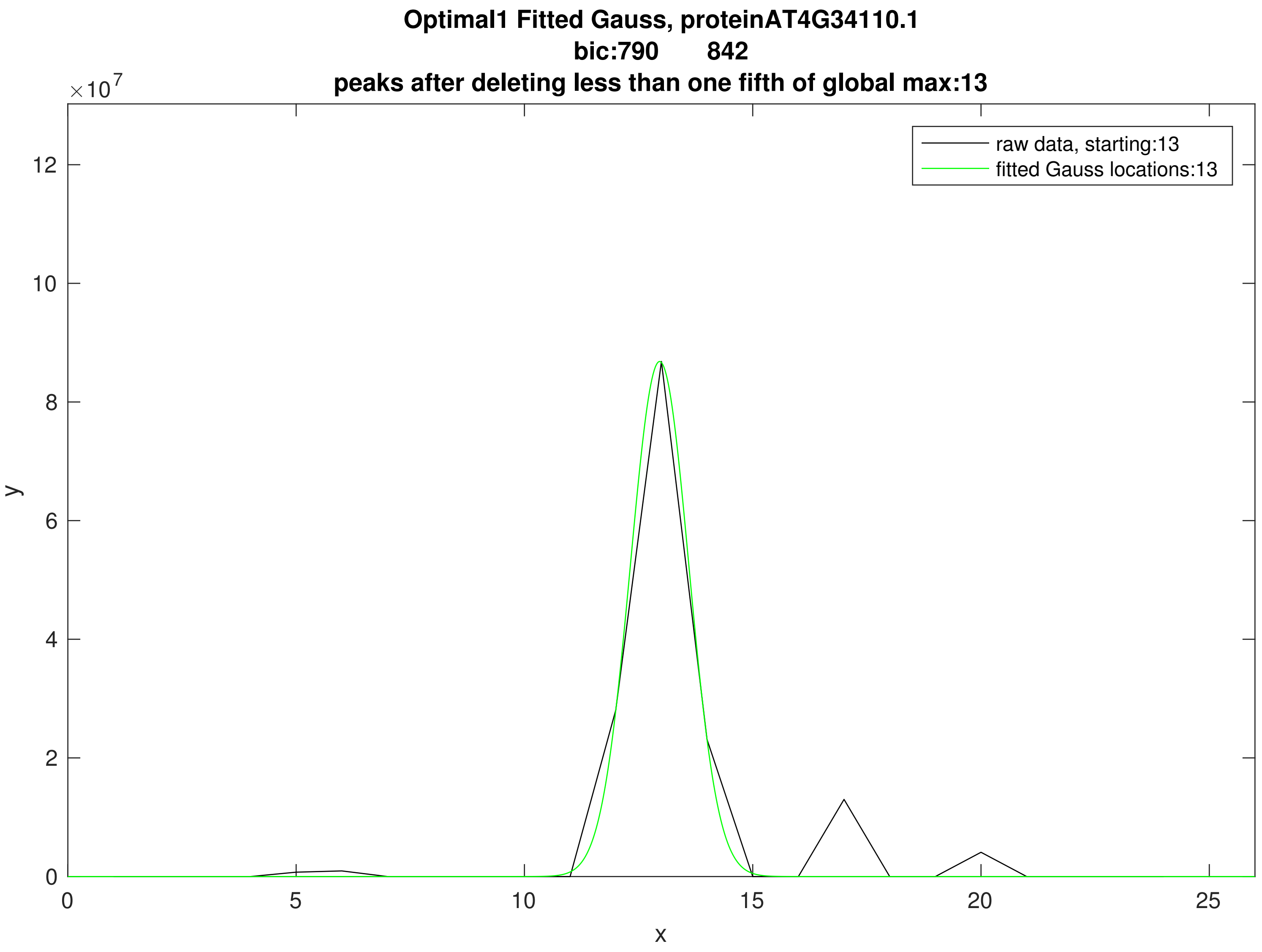

Optimal1 Fitted Gauss, proteinAT4G34110.1
bic:790 842
peaks after deleting less than one fifth of global max:13
×10^7
raw data, starting:13
fitted Gauss locations:13
y
x
0
2
4
6
8
10
12
0
5
10
15
20
25

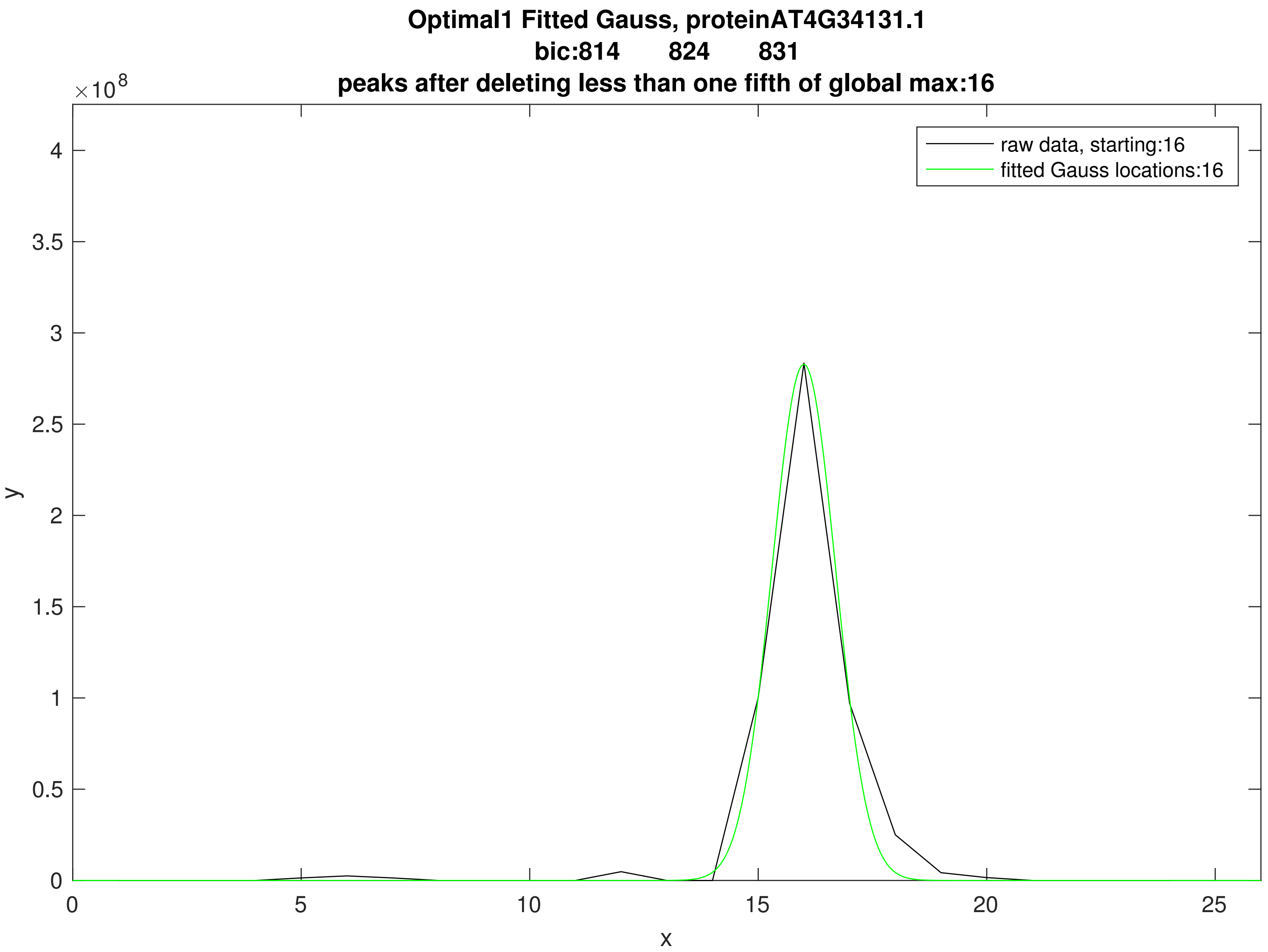
Optimal1 Fitted Gauss, proteinAT4G34131.1
bic:814 824 831
peaks after deleting less than one fifth of global max:16
×10^8
raw data, starting:16
fitted Gauss locations:16
y
x
0
0.5
1
1.5
2
2.5
3
3.5
4
5
10
15
20
25

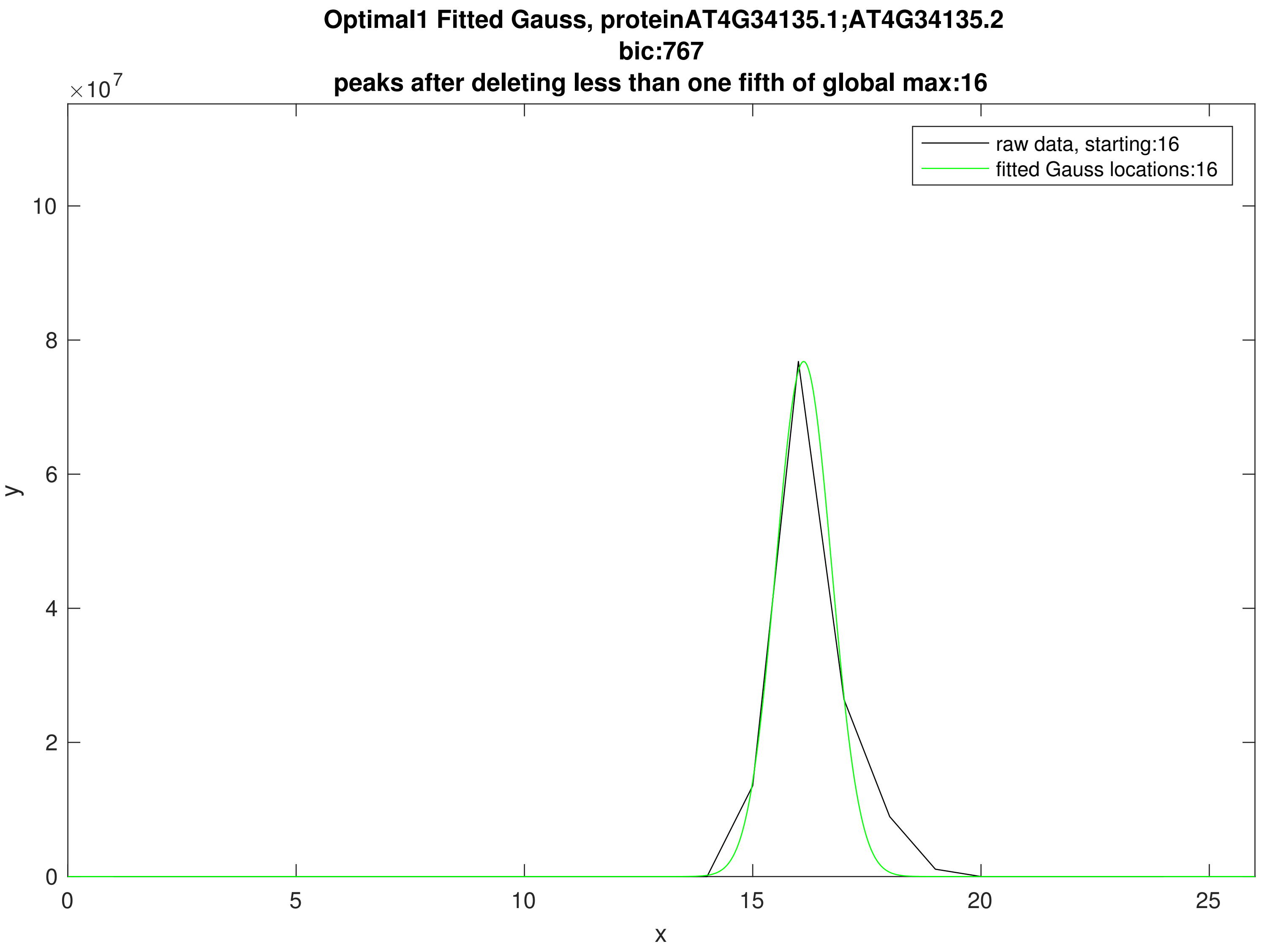

Optimal1 Fitted Gauss, proteinAT4G34135.1;AT4G34135.2
bic:767
peaks after deleting less than one fifth of global max:16
×10^7
raw data, starting:16
fitted Gauss locations:16
y
x
0
2
4
6
8
10
0
5
10
15
20
25

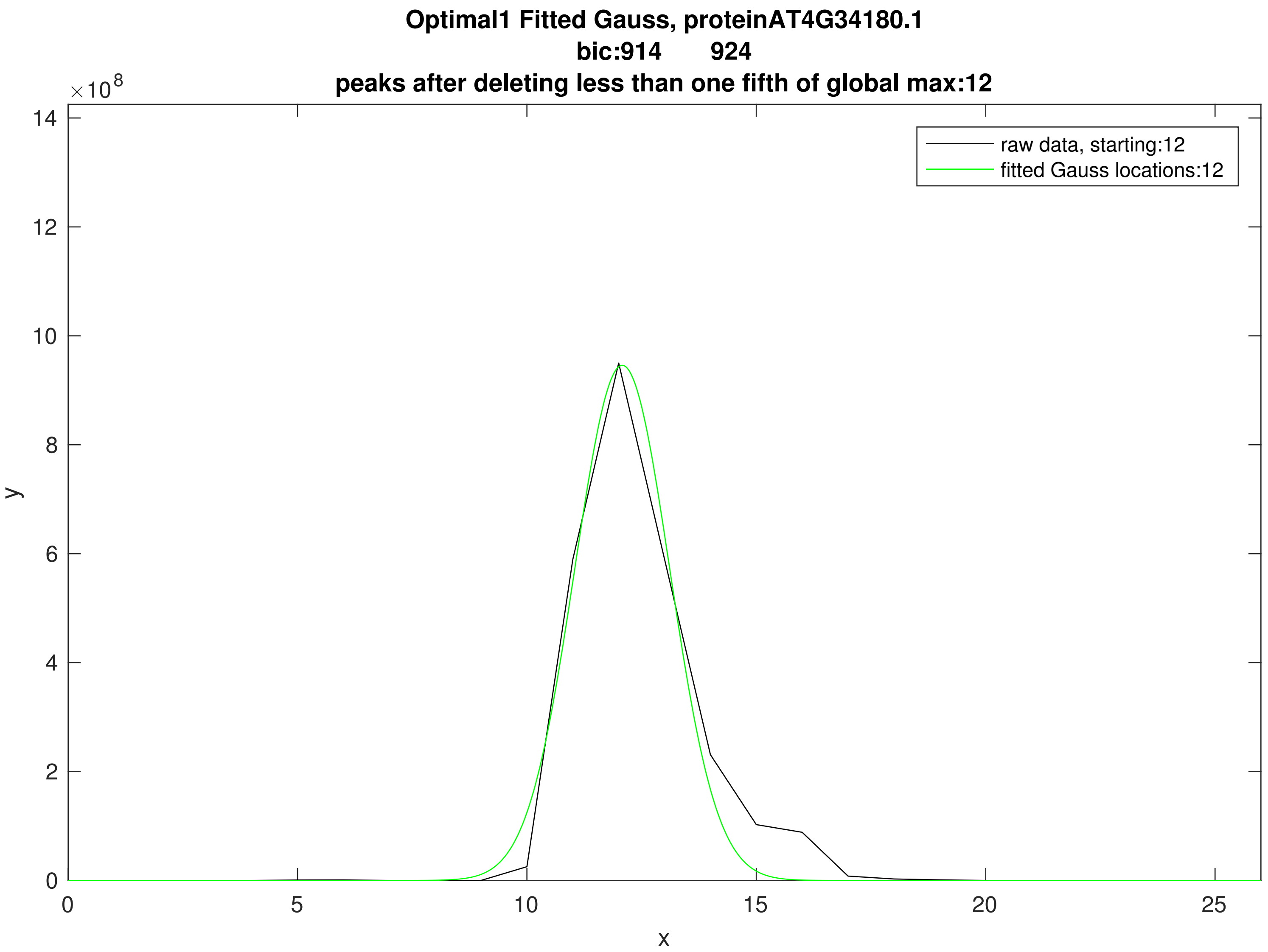
Optimal1 Fitted Gauss, proteinAT4G34180.1
bic:914 924
peaks after deleting less than one fifth of global max:12
×10^8
14
12
10
8
6
4
2
0
y
0
5
10
15
20
25
x
raw data, starting:12
fitted Gauss locations:12

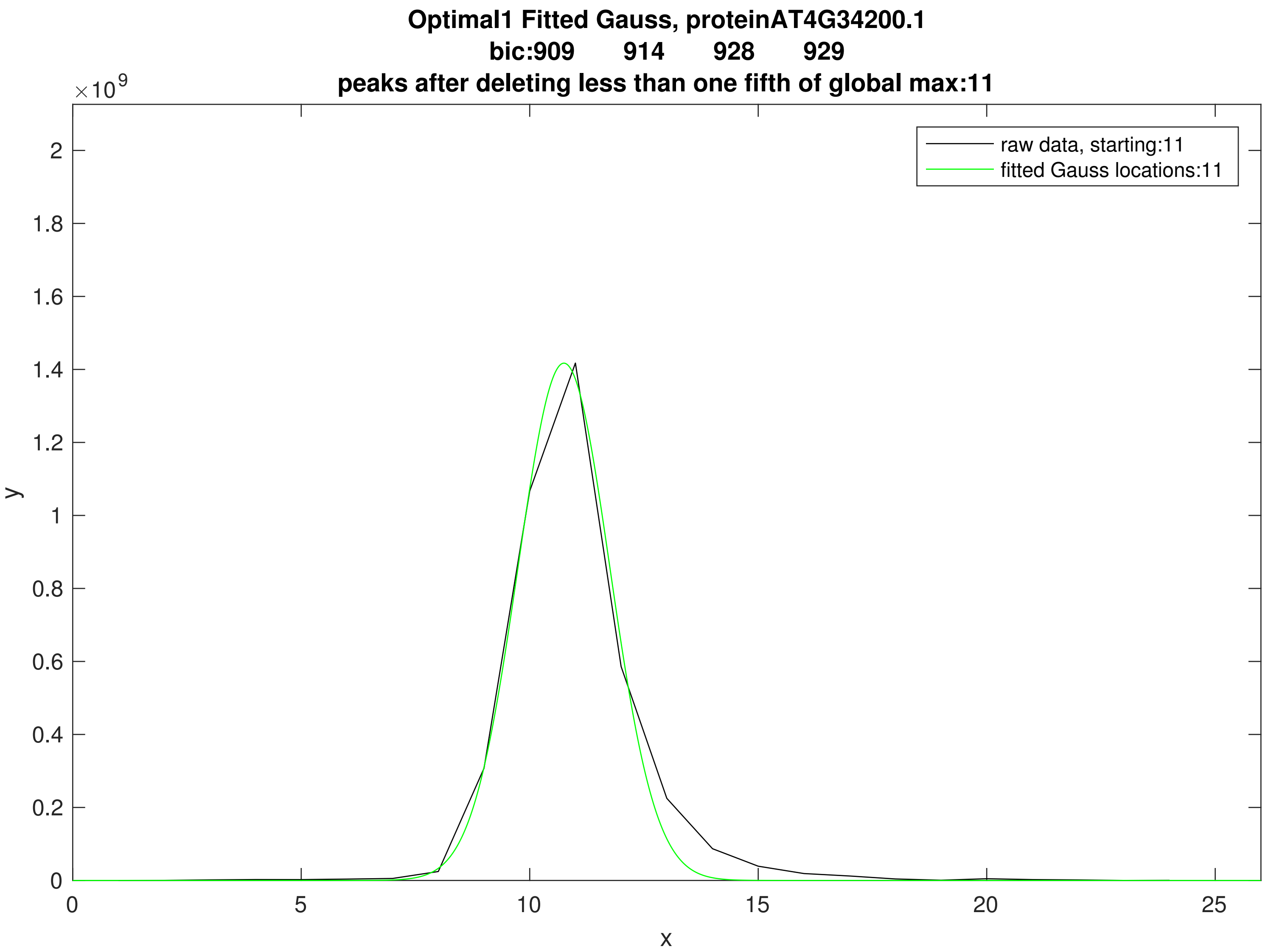

Optimal1 Fitted Gauss, proteinAT4G34200.1
bic:909 914 928 929
peaks after deleting less than one fifth of global max:11
×10^9
raw data, starting:11
fitted Gauss locations:11
y
x
0
0.2
0.4
0.6
0.8
1
1.2
1.4
1.6
1.8
2
0
5
10
15
20
25

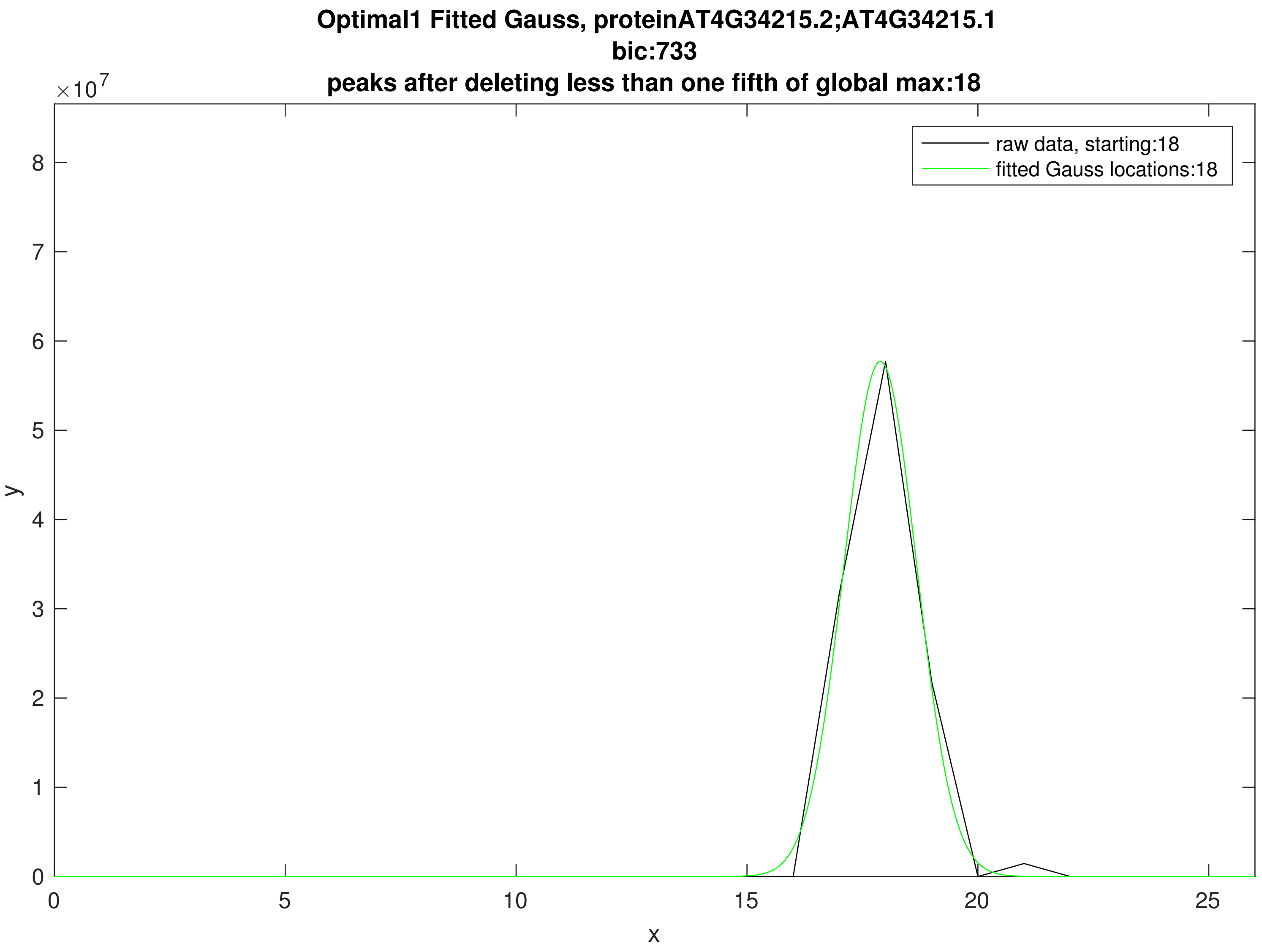
Optimal1 Fitted Gauss, proteinAT4G34215.2;AT4G34215.1
bic:733
peaks after deleting less than one fifth of global max:18
×10^7
raw data, starting:18
fitted Gauss locations:18
y
x
0
1
2
3
4
5
6
7
8
0
5
10
15
20
25

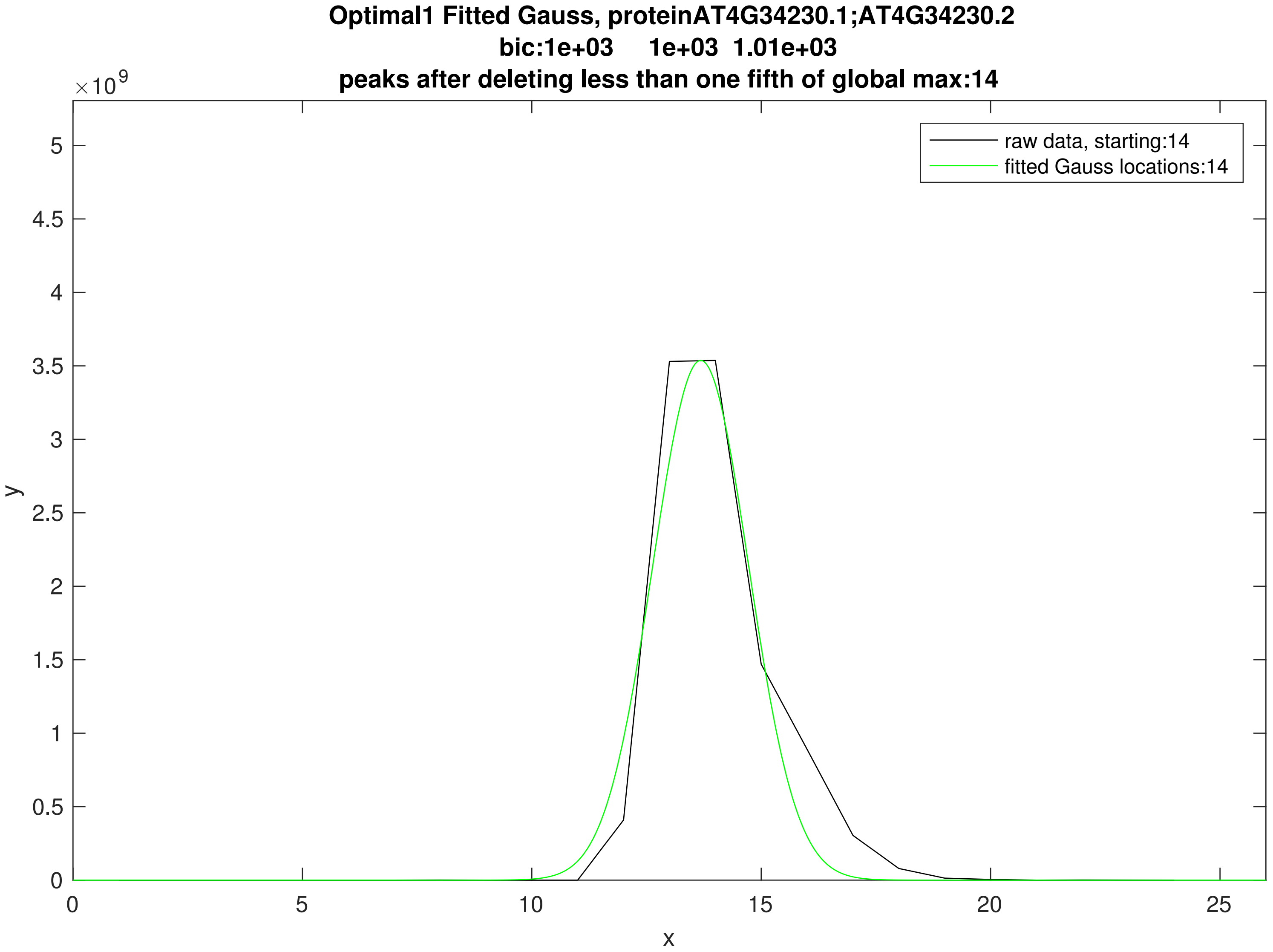
Optimal1 Fitted Gauss, proteinAT4G34230.1;AT4G34230.2
bic:1e+03 1e+03 1.01e+03
peaks after deleting less than one fifth of global max:14
×10^9
raw data, starting:14
fitted Gauss locations:14
y
x
0
0.5
1
1.5
2
2.5
3
3.5
4
4.5
5
0
5
10
15
20
25

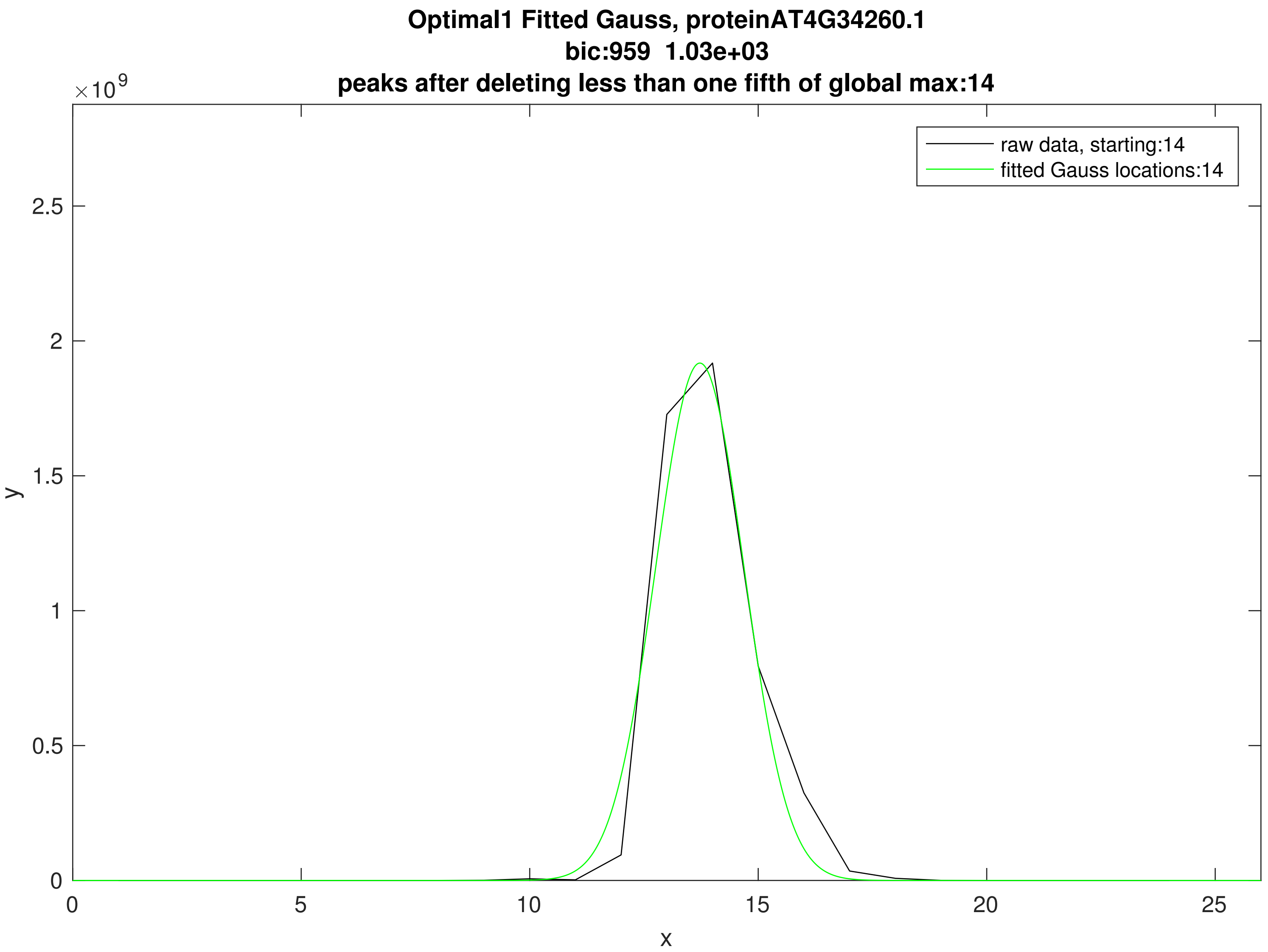

Optimal1 Fitted Gauss, proteinAT4G34260.1
bic:959 1.03e+03
peaks after deleting less than one fifth of global max:14
×10^9
raw data, starting:14
fitted Gauss locations:14
2.5
2
1.5
1
0.5
0
y
0
5
10
15
20
25
x

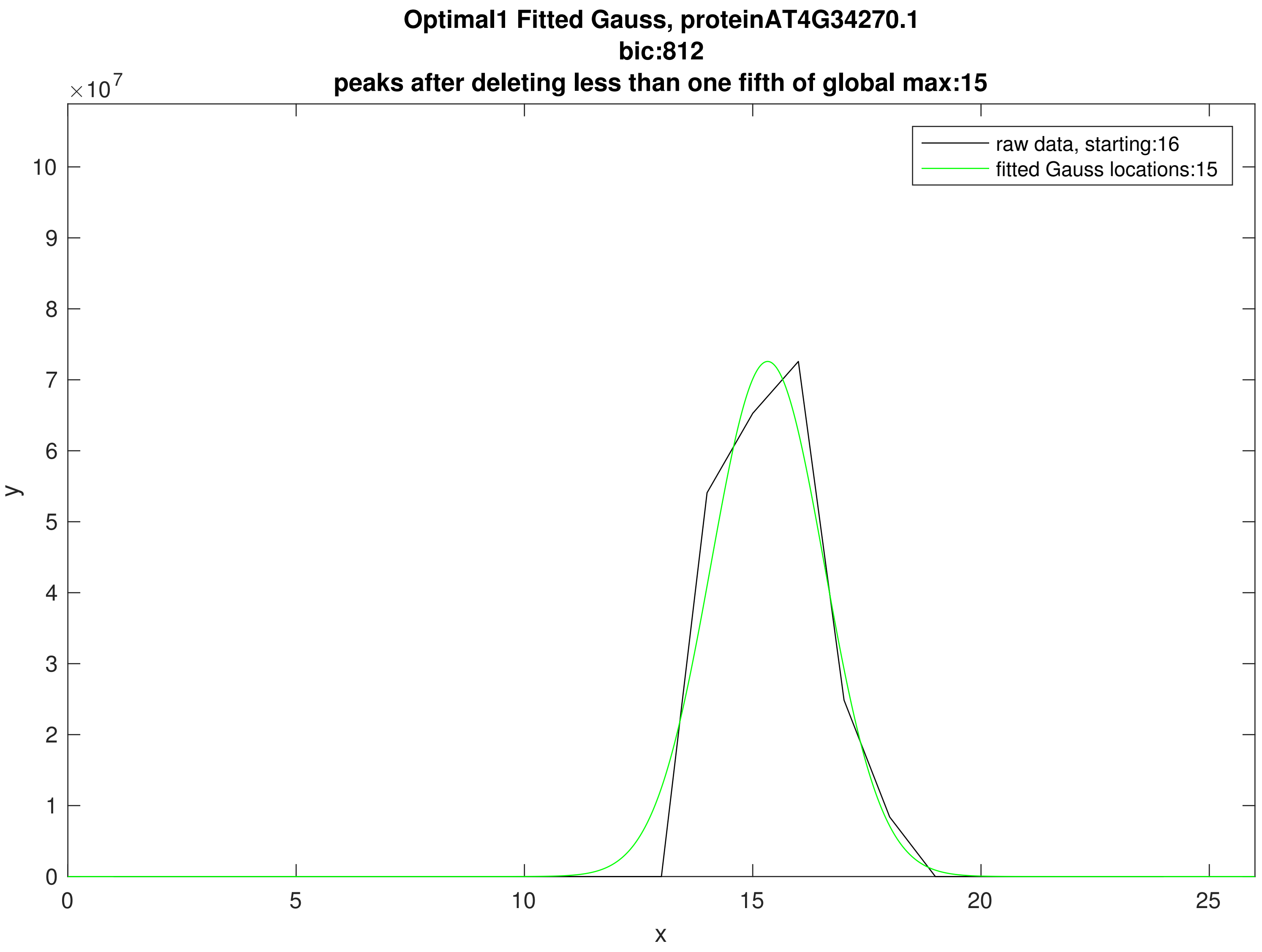

Optimal1 Fitted Gauss, proteinAT4G34270.1
bic:812
peaks after deleting less than one fifth of global max:15
×10^7
raw data, starting:16
fitted Gauss locations:15
y
x
0
1
2
3
4
5
6
7
8
9
10
0
5
10
15
20
25

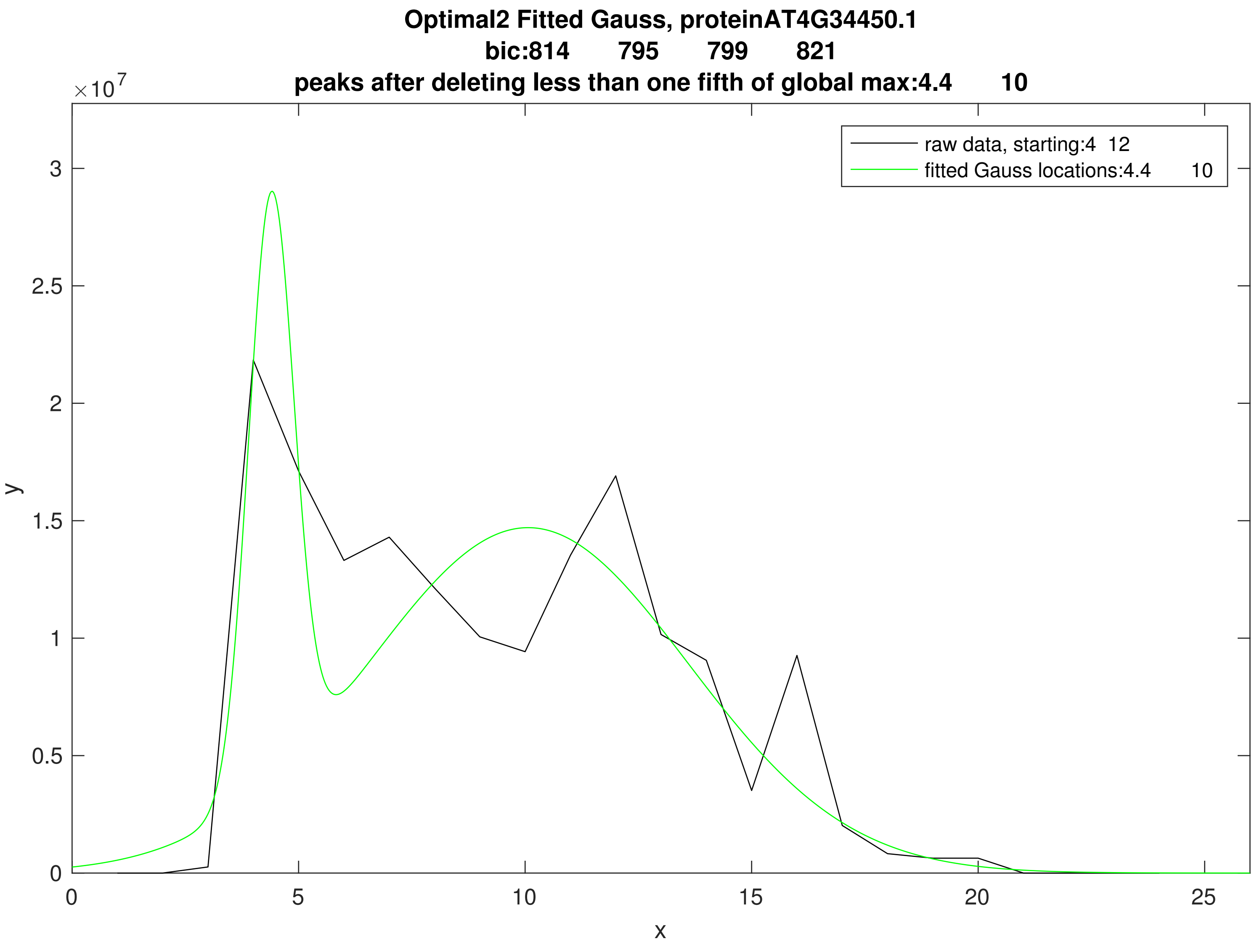

Optimal2 Fitted Gauss, proteinAT4G34450.1
bic:814 795 799 821
peaks after deleting less than one fifth of global max:4.4 10
raw data, starting:4 12
fitted Gauss locations:4.4 10
×10^7
y
x
0
0.5
1
1.5
2
2.5
3
5
10
15
20
25

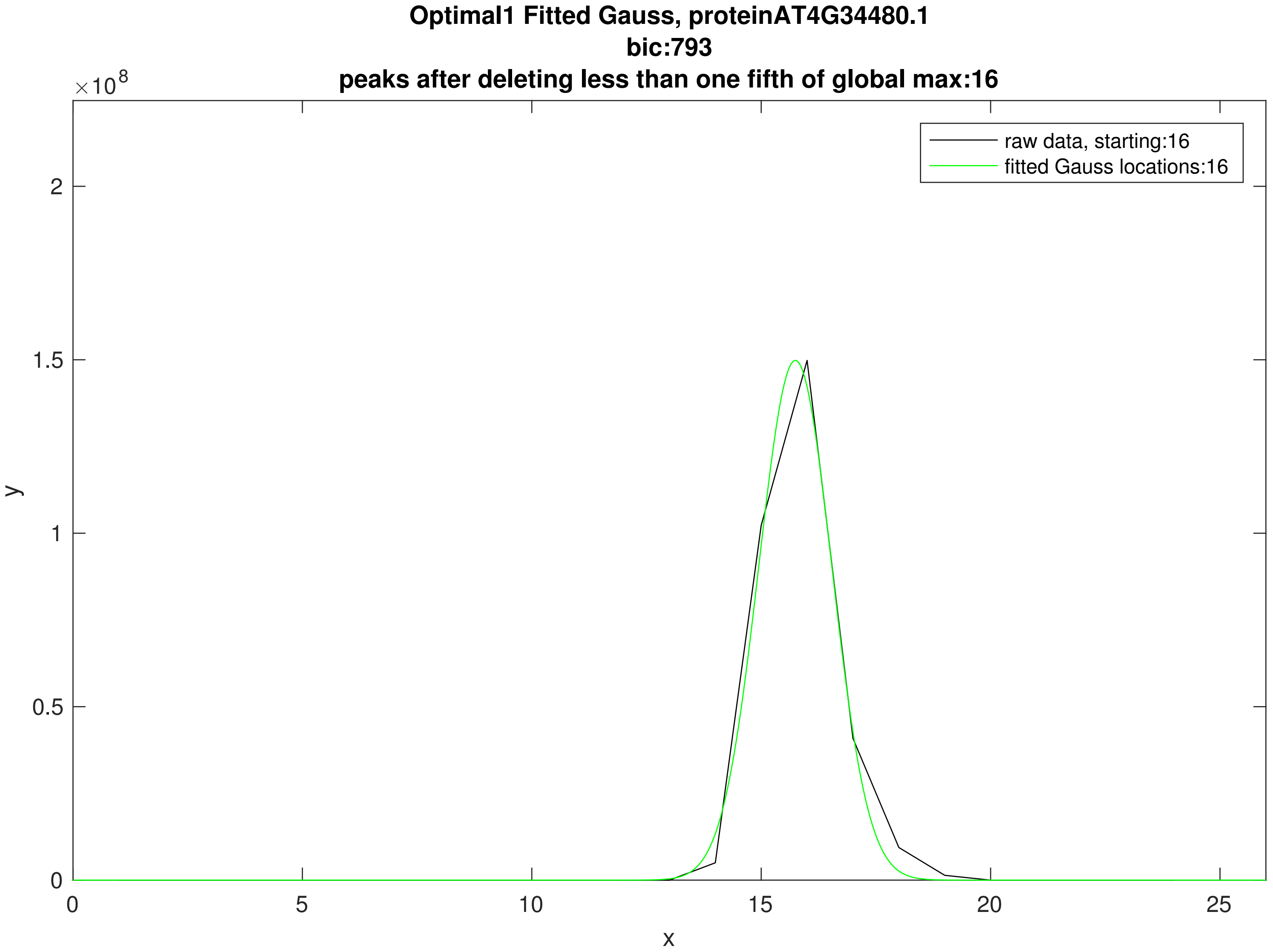
Optimal1 Fitted Gauss, proteinAT4G34480.1
bic:793
peaks after deleting less than one fifth of global max:16
×10^8
raw data, starting:16
fitted Gauss locations:16
y
x
0
0.5
1
1.5
2
0
5
10
15
20
25

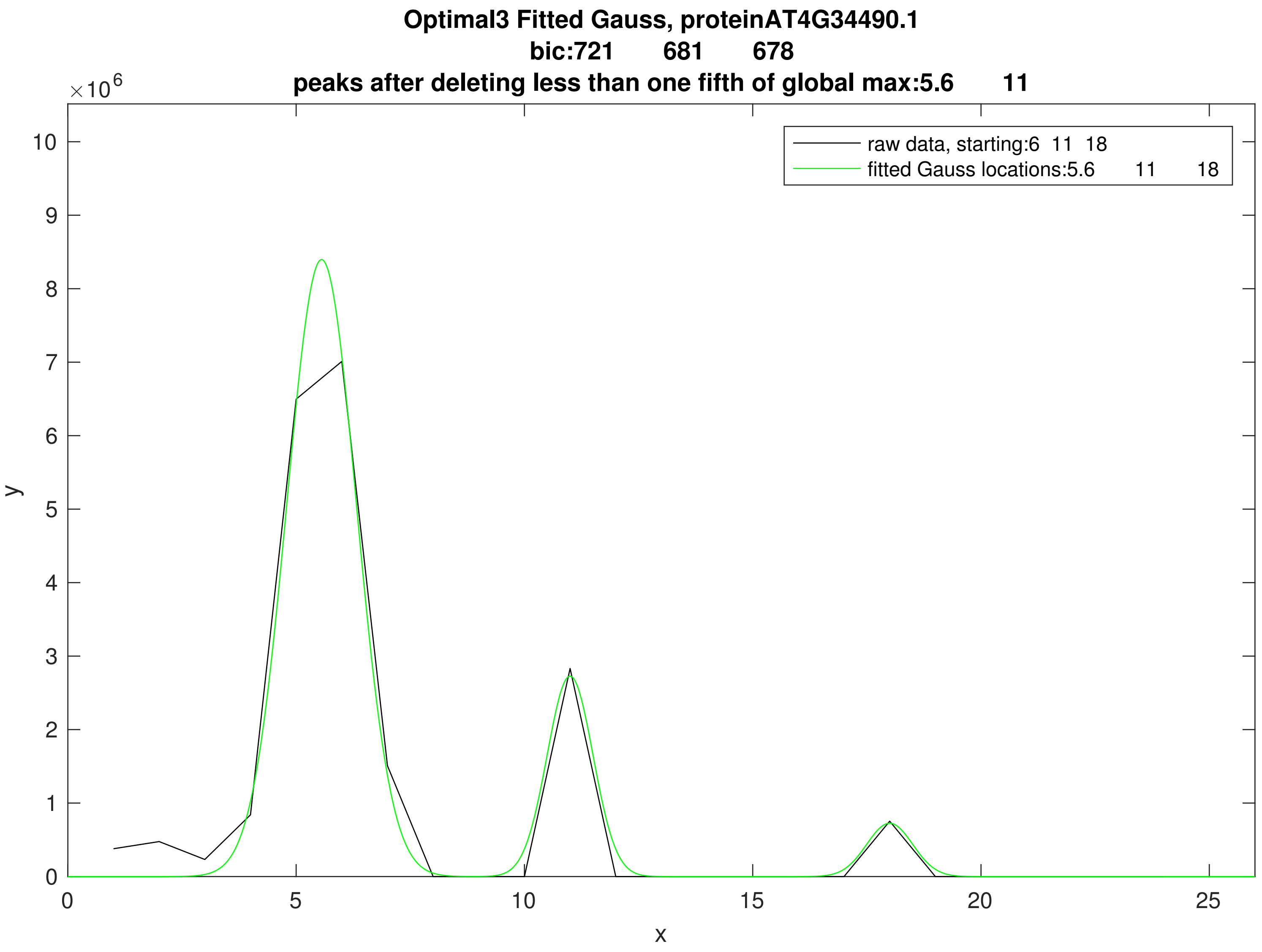
Optimal3 Fitted Gauss, proteinAT4G34490.1
bic:721 681 678
peaks after deleting less than one fifth of global max:5.6 11
×10^6
raw data, starting:6 11 18
fitted Gauss locations:5.6 11 18
y
x
0
1
2
3
4
5
6
7
8
9
10
0
5
10
15
20
25

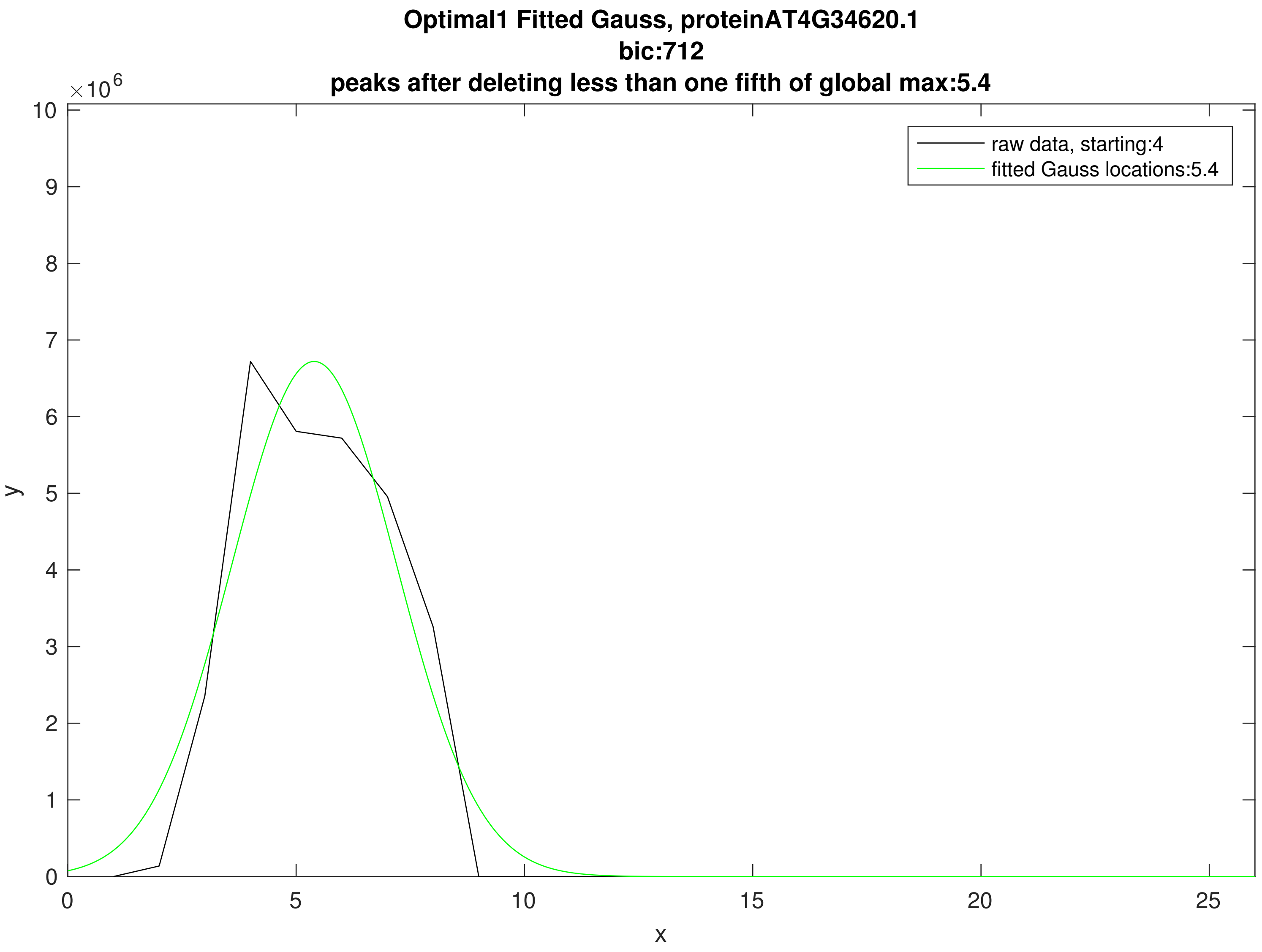
Optimal1 Fitted Gauss, proteinAT4G34620.1
bic:712
peaks after deleting less than one fifth of global max:5.4
×10^6
raw data, starting:4
fitted Gauss locations:5.4
y
x
0
1
2
3
4
5
6
7
8
9
10
0
5
10
15
20
25

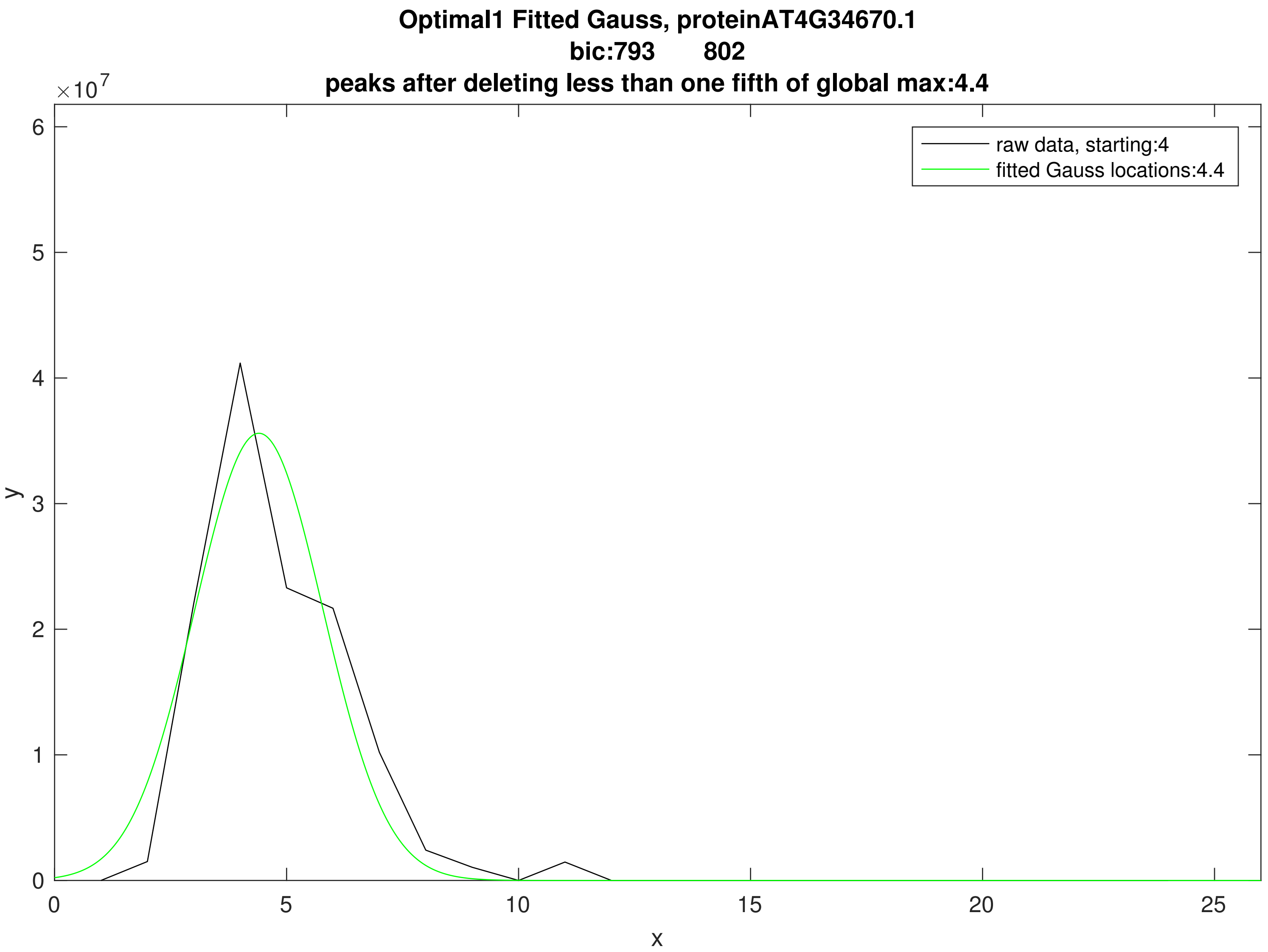
Optimal1 Fitted Gauss, proteinAT4G34670.1
bic:793 802
peaks after deleting less than one fifth of global max:4.4
×10^7
raw data, starting:4
fitted Gauss locations:4.4
y
x
0
1
2
3
4
5
6
0
5
10
15
20
25

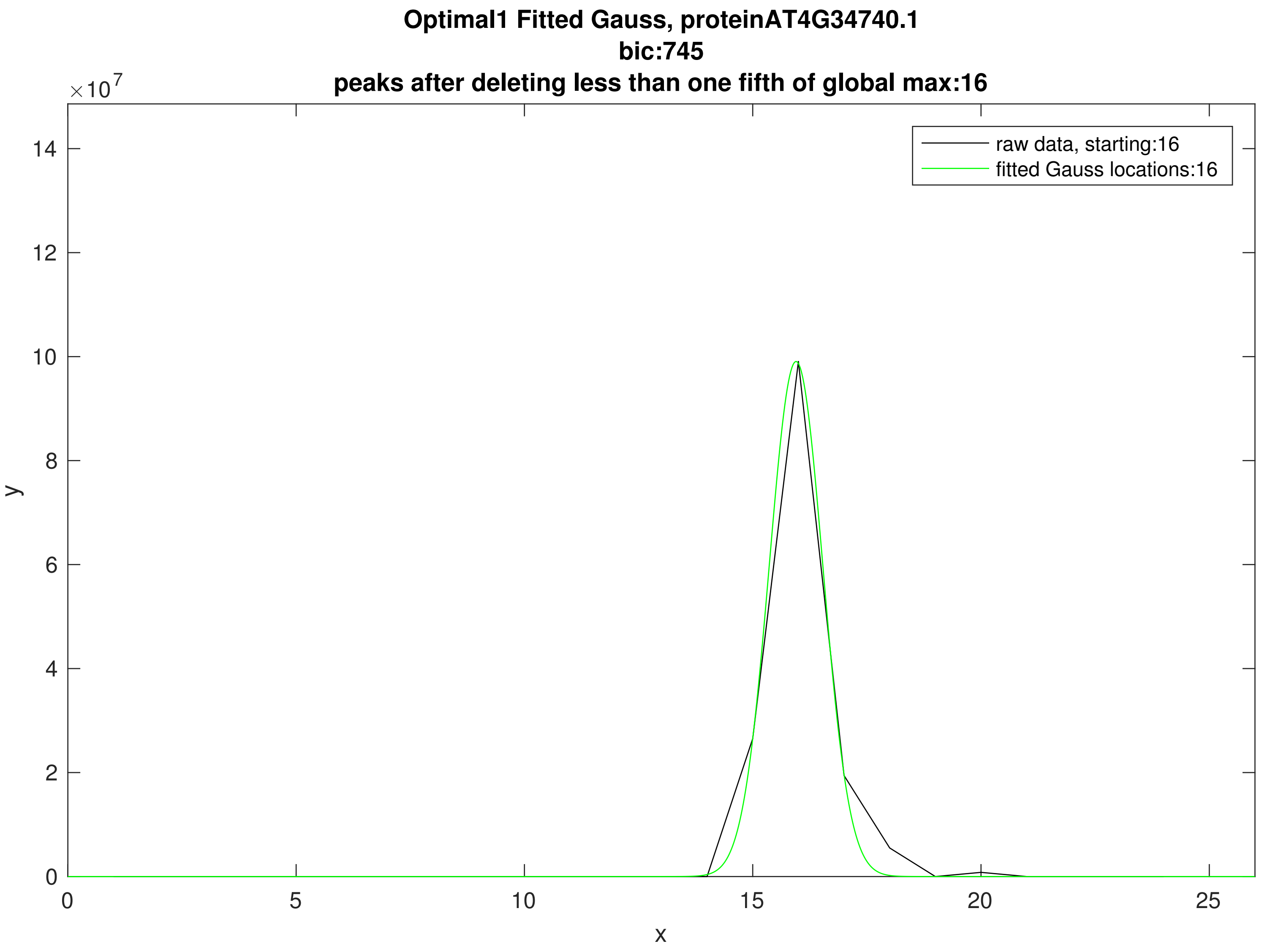

Optimal1 Fitted Gauss, proteinAT4G34740.1
bic:745
peaks after deleting less than one fifth of global max:16
×10^7
raw data, starting:16
fitted Gauss locations:16
y
x
0
2
4
6
8
10
12
14
0
5
10
15
20
25

Optimal2 Fitted Gauss, proteinAT4G34860.2;AT4G34860.1
bic:705 655
peaks after deleting less than one fifth of global max:5.3

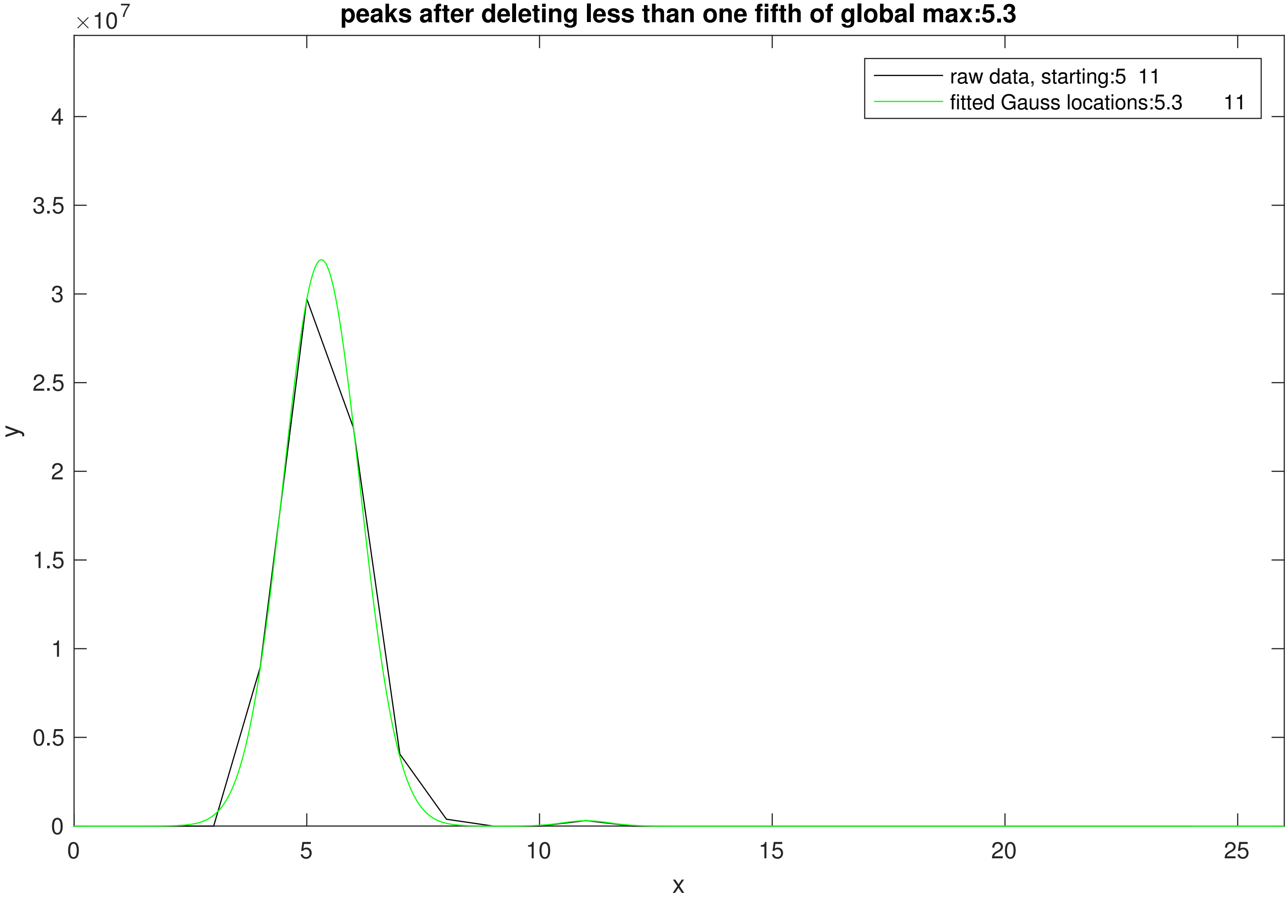

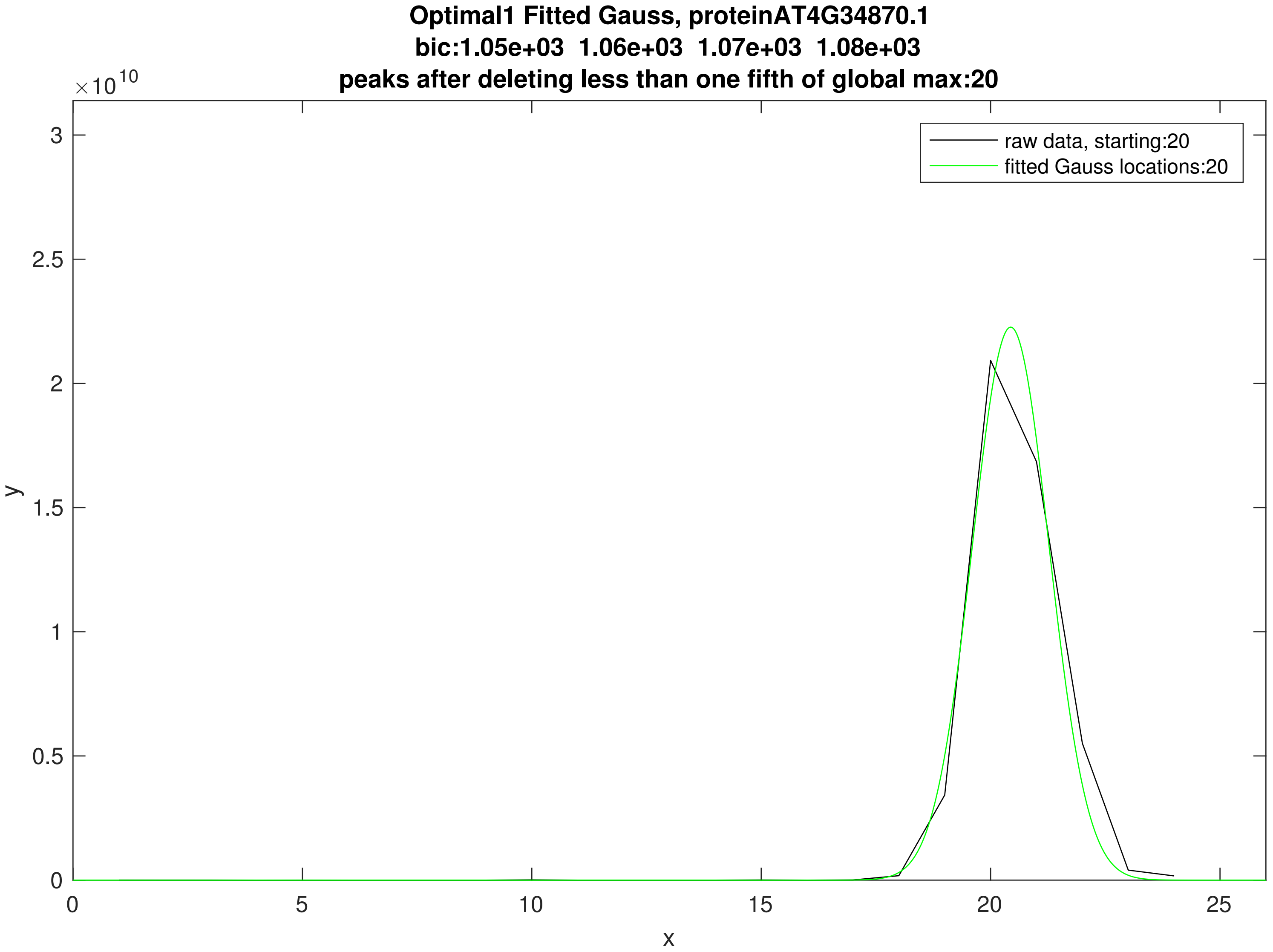
Optimal1 Fitted Gauss, proteinAT4G34870.1
bic:1.05e+03 1.06e+03 1.07e+03 1.08e+03
peaks after deleting less than one fifth of global max:20
×10^10
3
2.5
2
1.5
1
0.5
0
y
0
5
10
15
20
25
x
raw data, starting:20
fitted Gauss locations:20

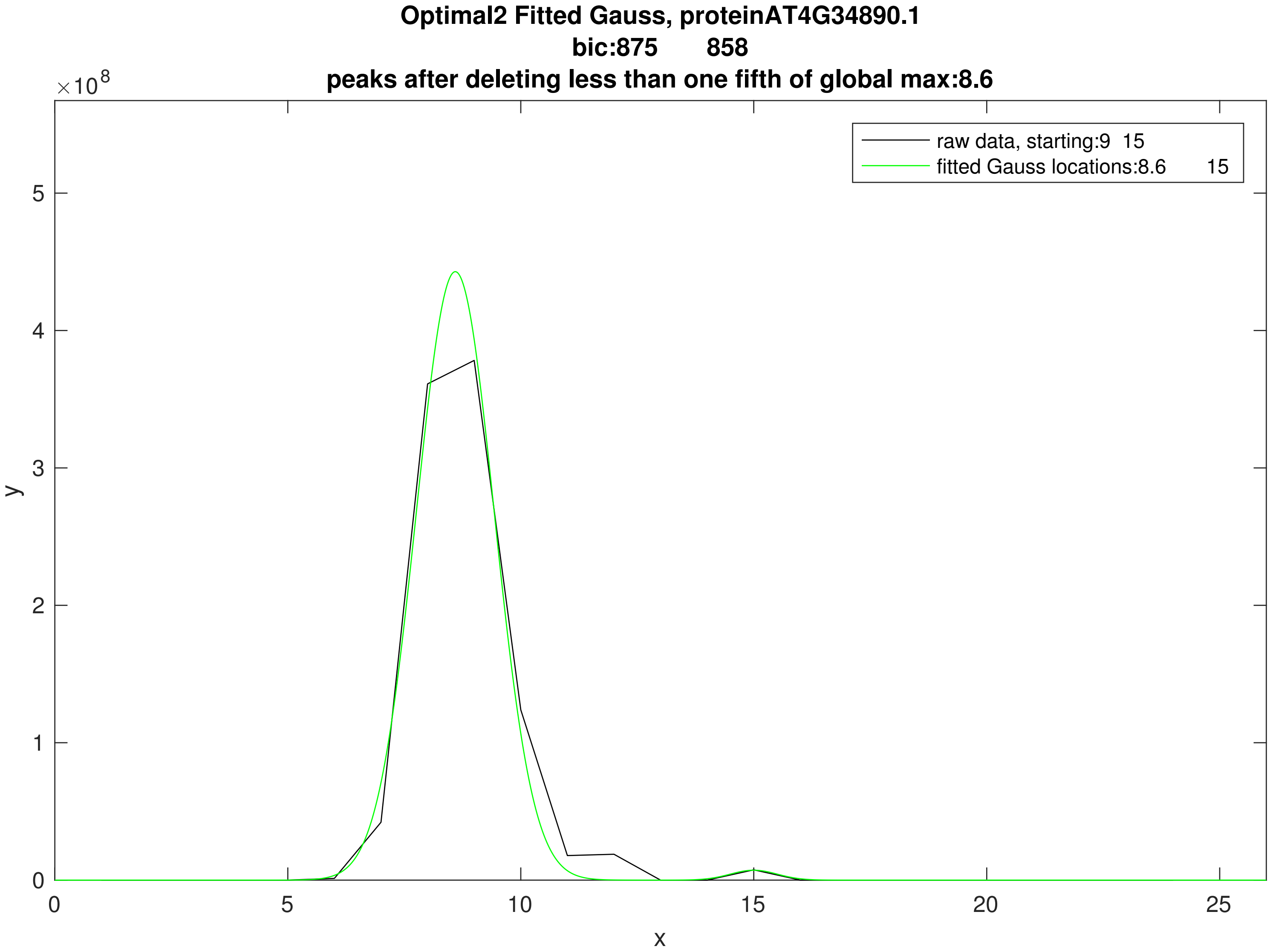

Optimal2 Fitted Gauss, proteinAT4G34890.1
bic:875 858
peaks after deleting less than one fifth of global max:8.6
×10^8
raw data, starting:9 15
fitted Gauss locations:8.6 15
y
x
0
1
2
3
4
5
0
5
10
15
20
25

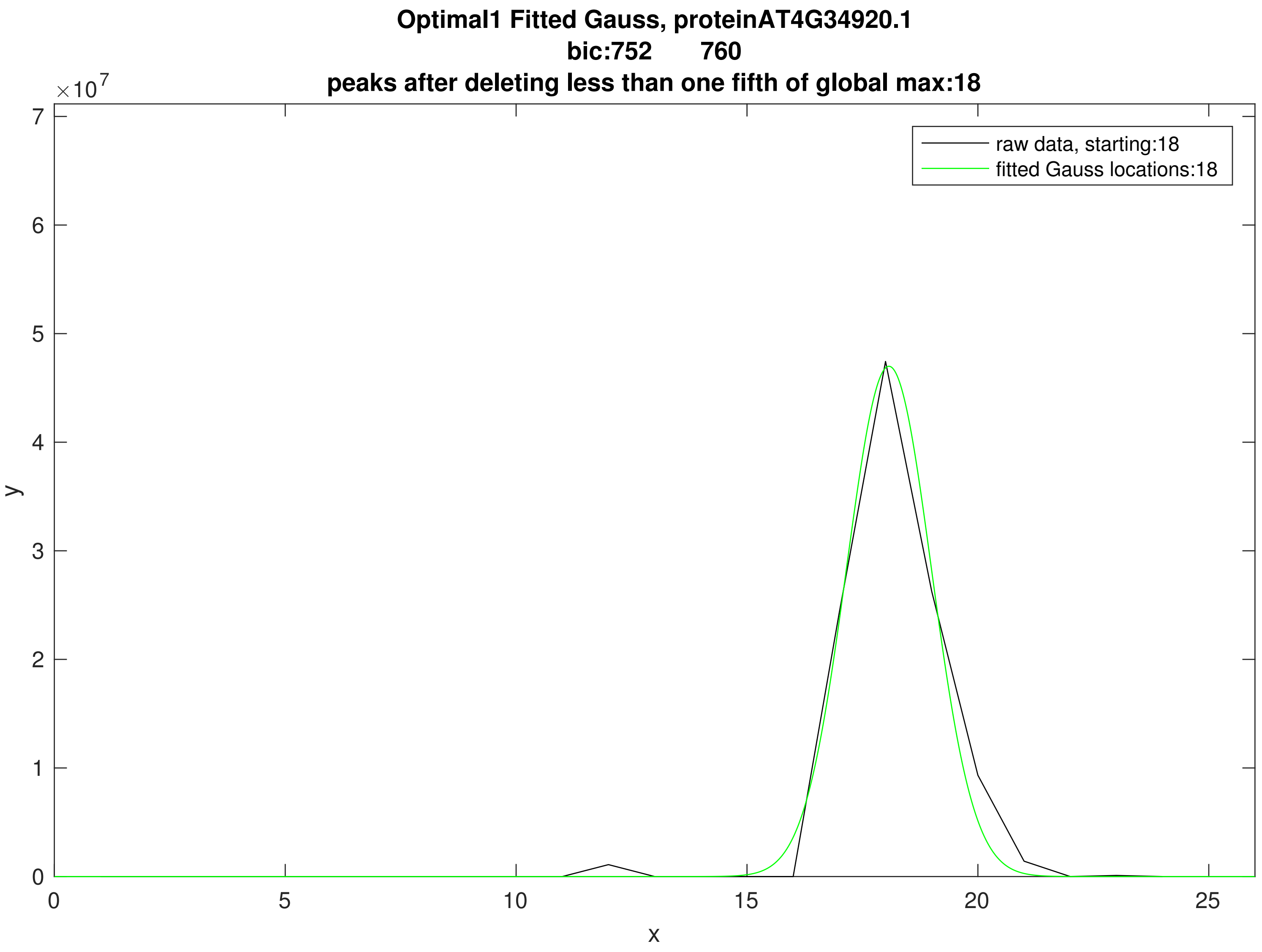
Optimal1 Fitted Gauss, proteinAT4G34920.1
bic:752 760
peaks after deleting less than one fifth of global max:18
×10^7
raw data, starting:18
fitted Gauss locations:18
y
x
0
1
2
3
4
5
6
7
0
5
10
15
20
25

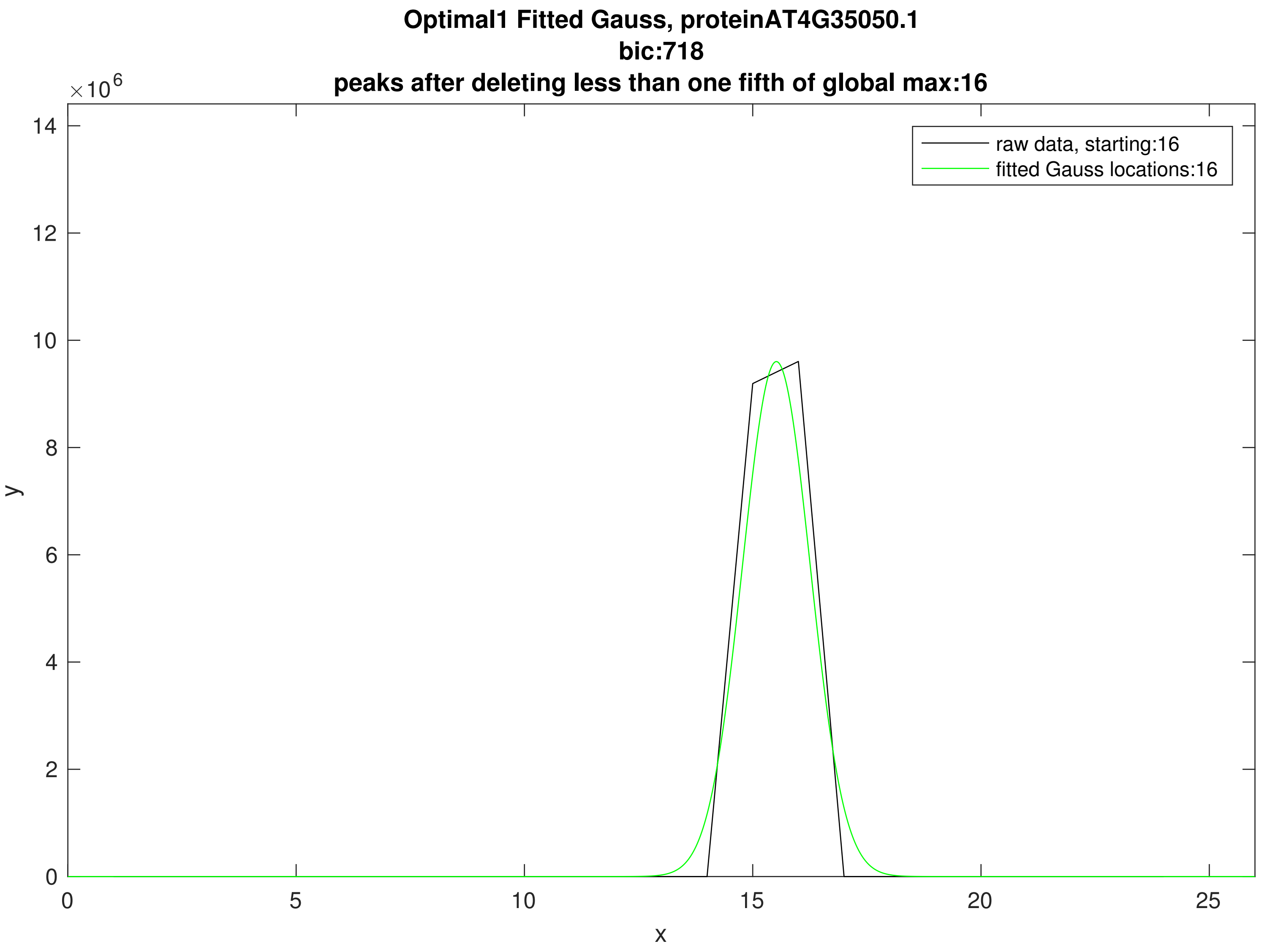
Optimal1 Fitted Gauss, proteinAT4G35050.1
bic:718
peaks after deleting less than one fifth of global max:16
×10^6
raw data, starting:16
fitted Gauss locations:16
y
x
0
2
4
6
8
10
12
14
0
5
10
15
20
25

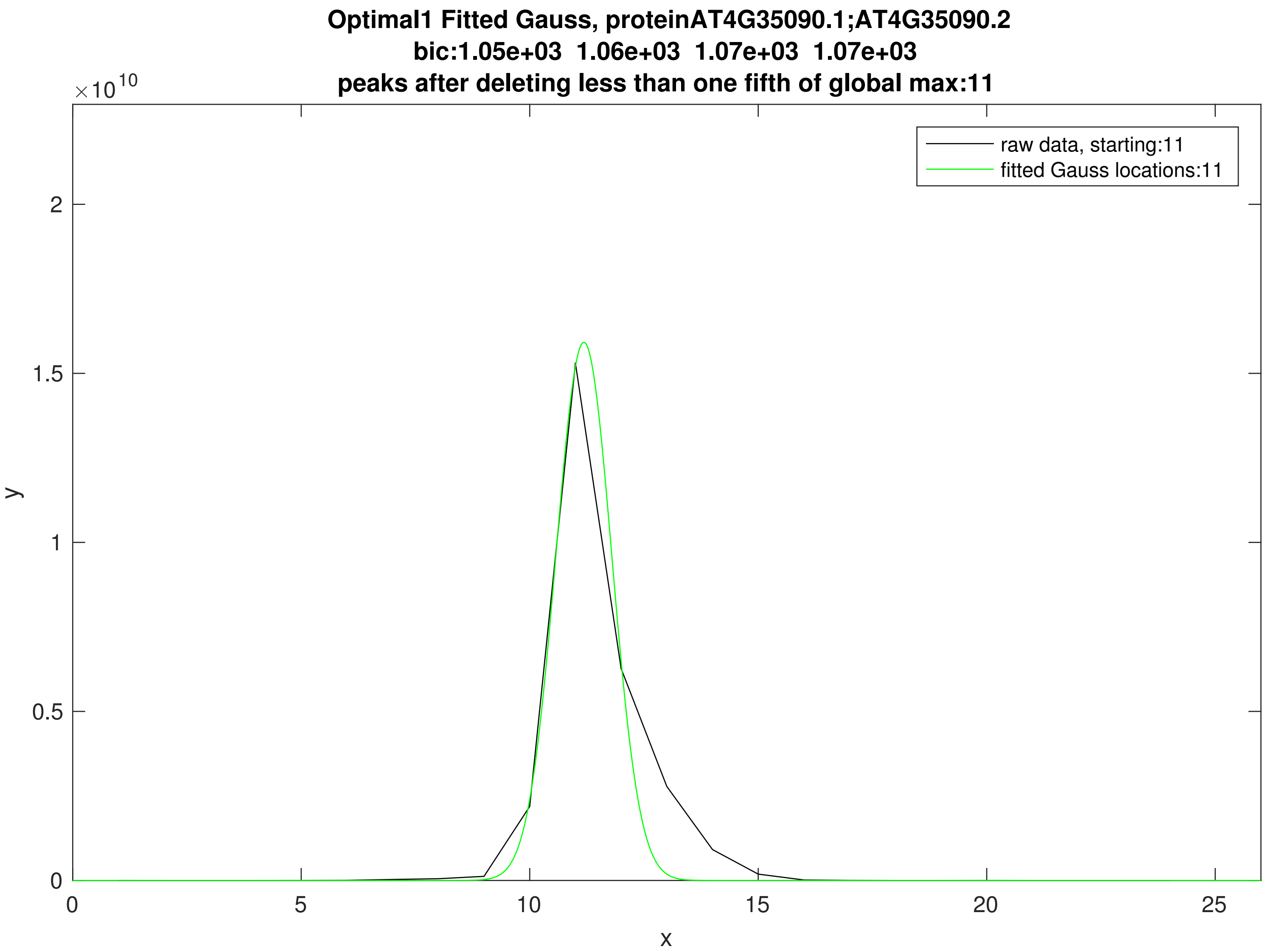
Optimal1 Fitted Gauss, proteinAT4G35090.1;AT4G35090.2
bic:1.05e+03 1.06e+03 1.07e+03 1.07e+03
peaks after deleting less than one fifth of global max:11
×10^10
raw data, starting:11
fitted Gauss locations:11
y
x
0
0.5
1
1.5
2
0
5
10
15
20
25

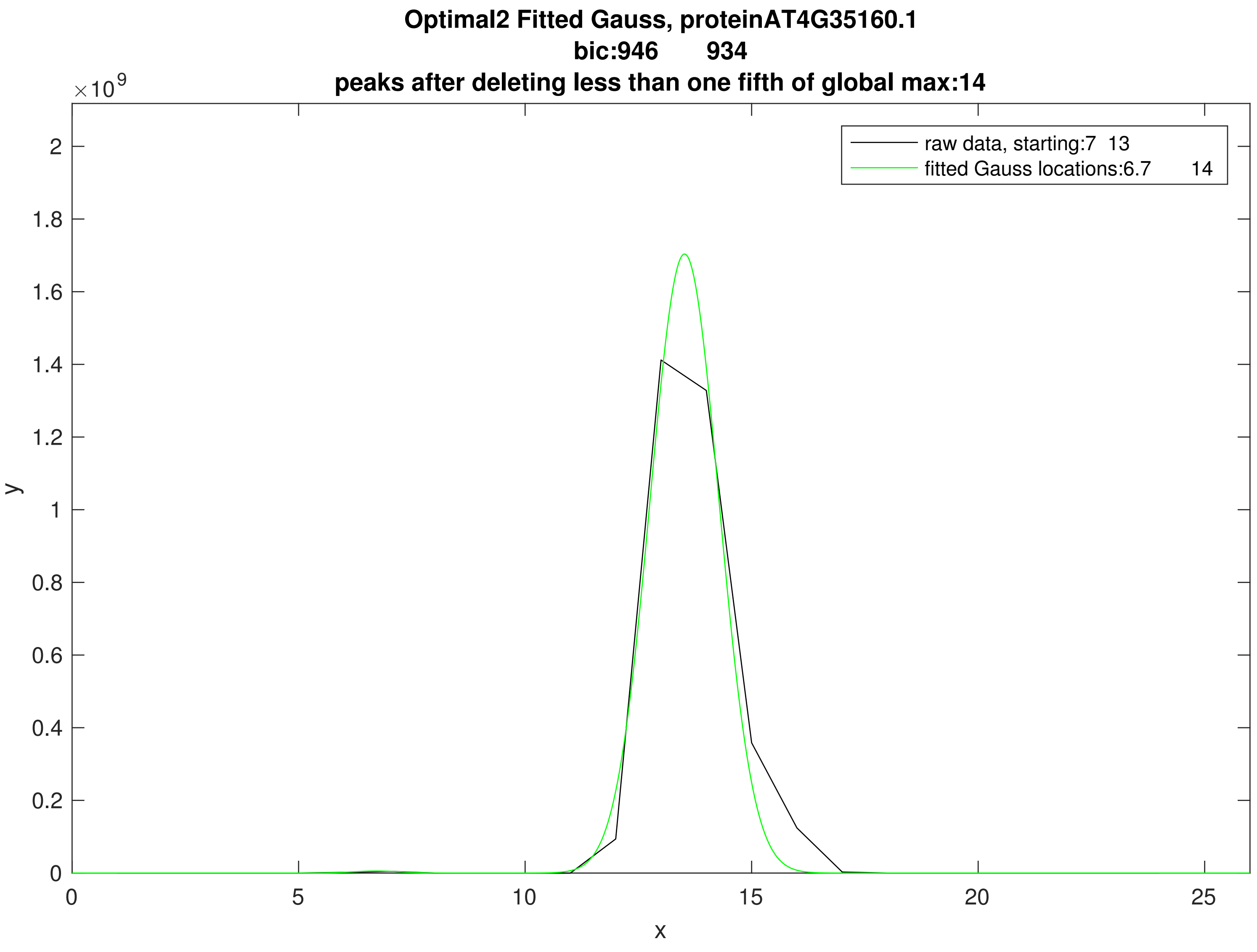

Optimal2 Fitted Gauss, proteinAT4G35160.1
bic:946 934
peaks after deleting less than one fifth of global max:14
×10^9
2
1.8
1.6
1.4
1.2
1
0.8
0.6
0.4
0.2
0
y
0
5
10
15
20
25
x
raw data, starting:7 13
fitted Gauss locations:6.7 14

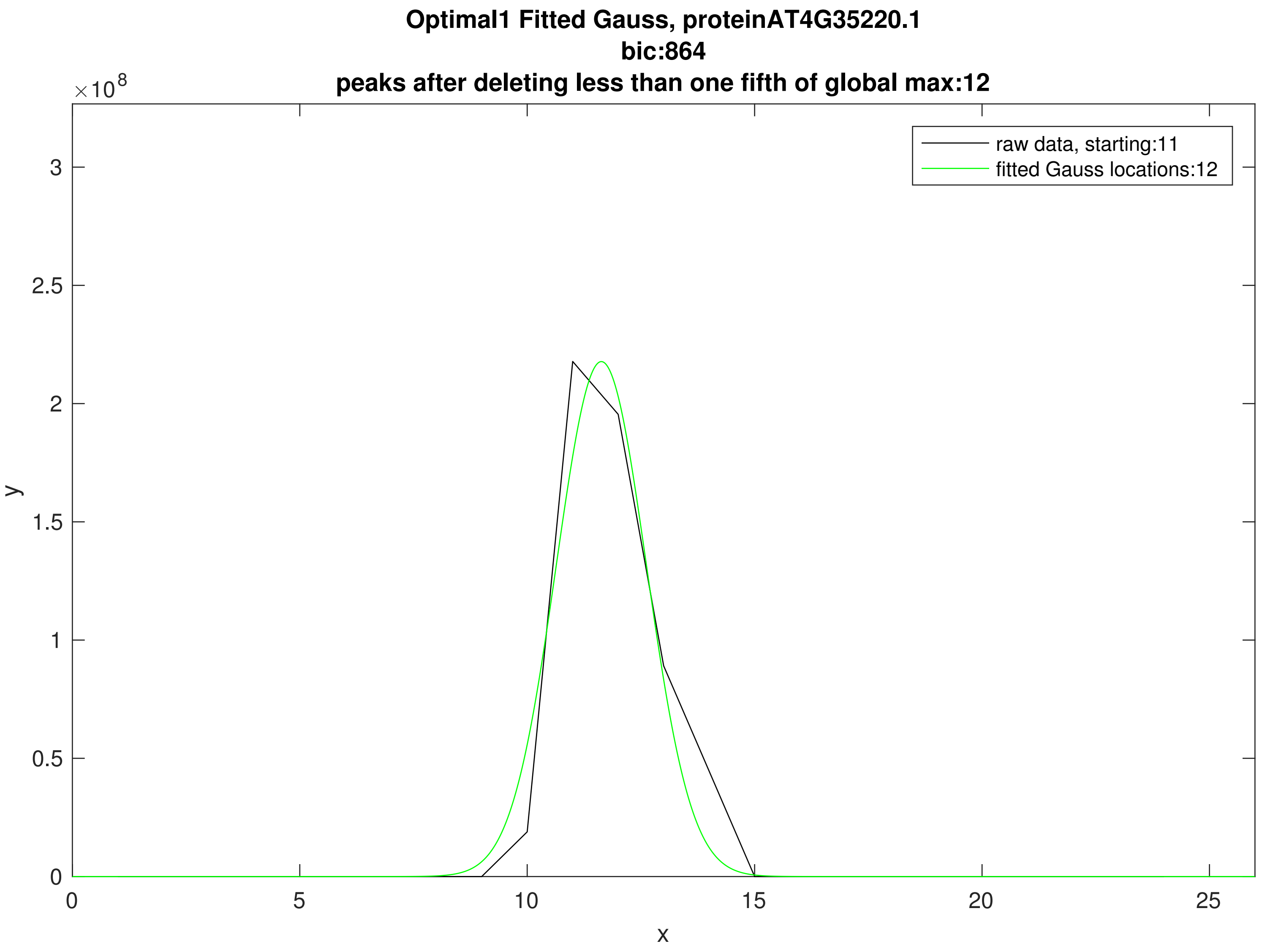
Optimal1 Fitted Gauss, proteinAT4G35220.1
bic:864
peaks after deleting less than one fifth of global max:12
×10^8
raw data, starting:11
fitted Gauss locations:12
0
0.5
1
1.5
2
2.5
3
y
0
5
10
15
20
25
x

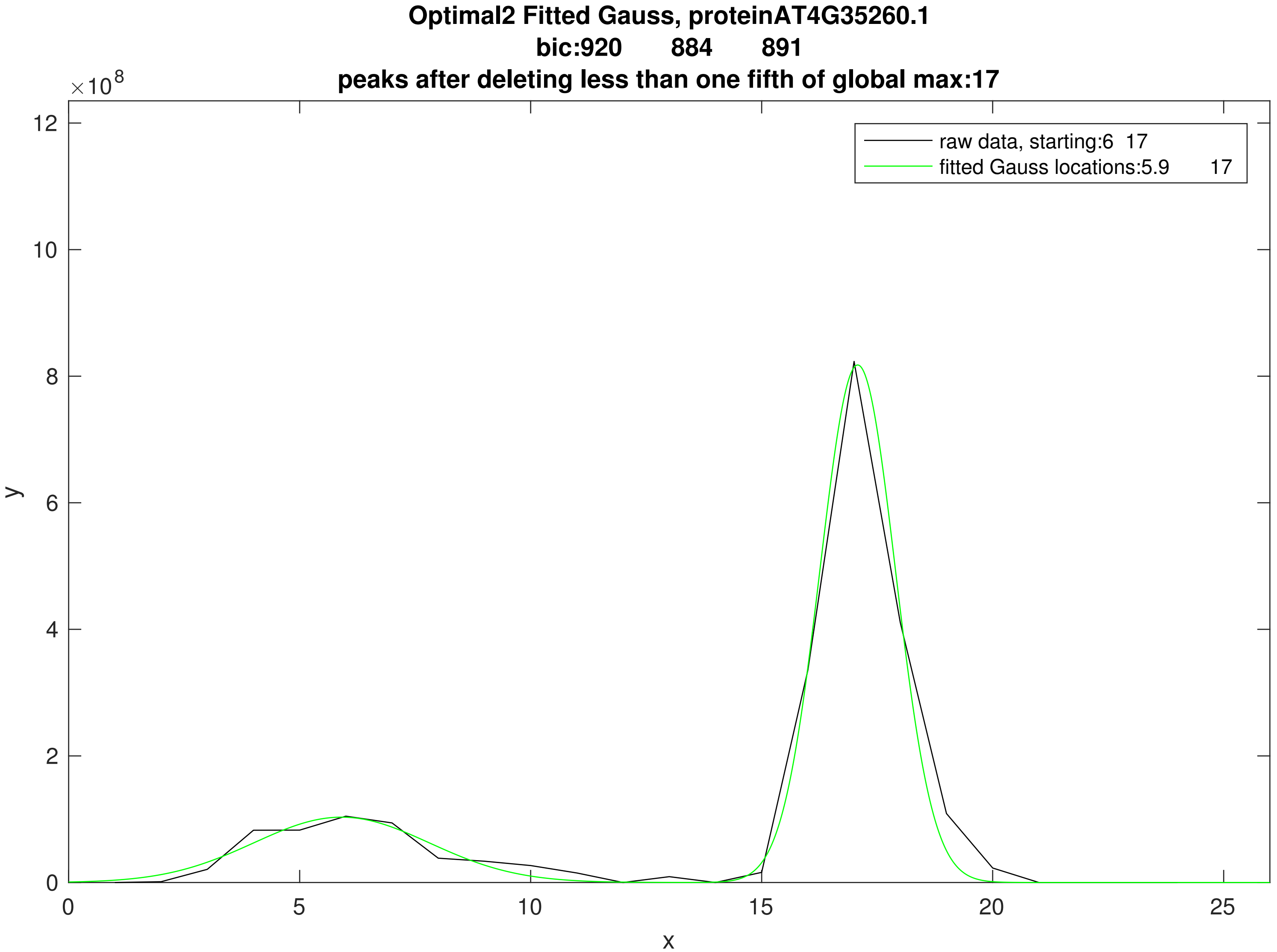

Optimal2 Fitted Gauss, proteinAT4G35260.1
bic:920 884 891
peaks after deleting less than one fifth of global max:17
×10^8
raw data, starting:6 17
fitted Gauss locations:5.9 17
y
x
0
2
4
6
8
10
12
0
5
10
15
20
25

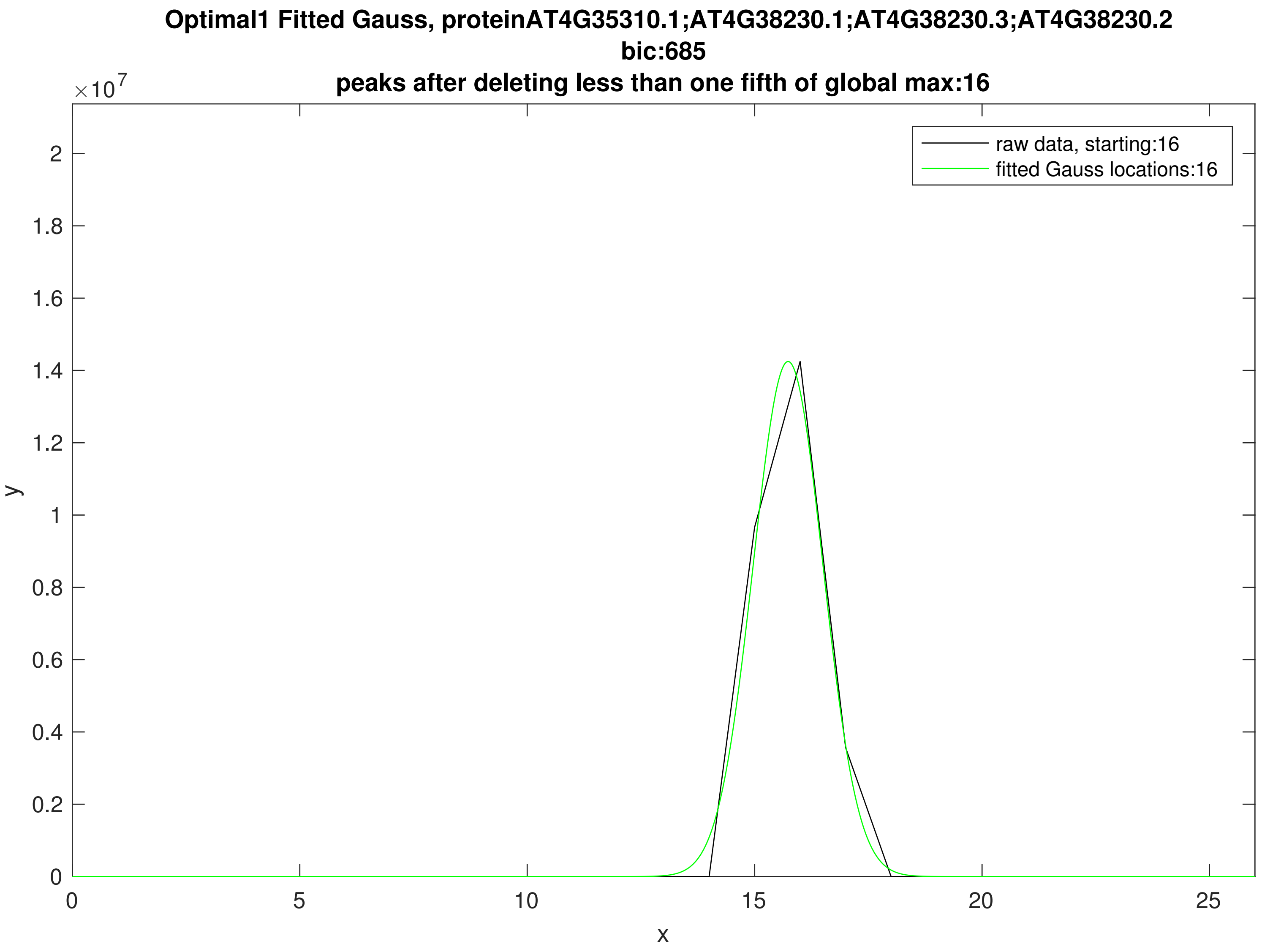

Optimal1 Fitted Gauss, proteinAT4G35310.1;AT4G38230.1;AT4G38230.3;AT4G38230.2
bic:685
peaks after deleting less than one fifth of global max:16
×10^7
2
1.8
1.6
1.4
1.2
1
0.8
0.6
0.4
0.2
0
y
0
5
10
15
20
25
x
raw data, starting:16
fitted Gauss locations:16

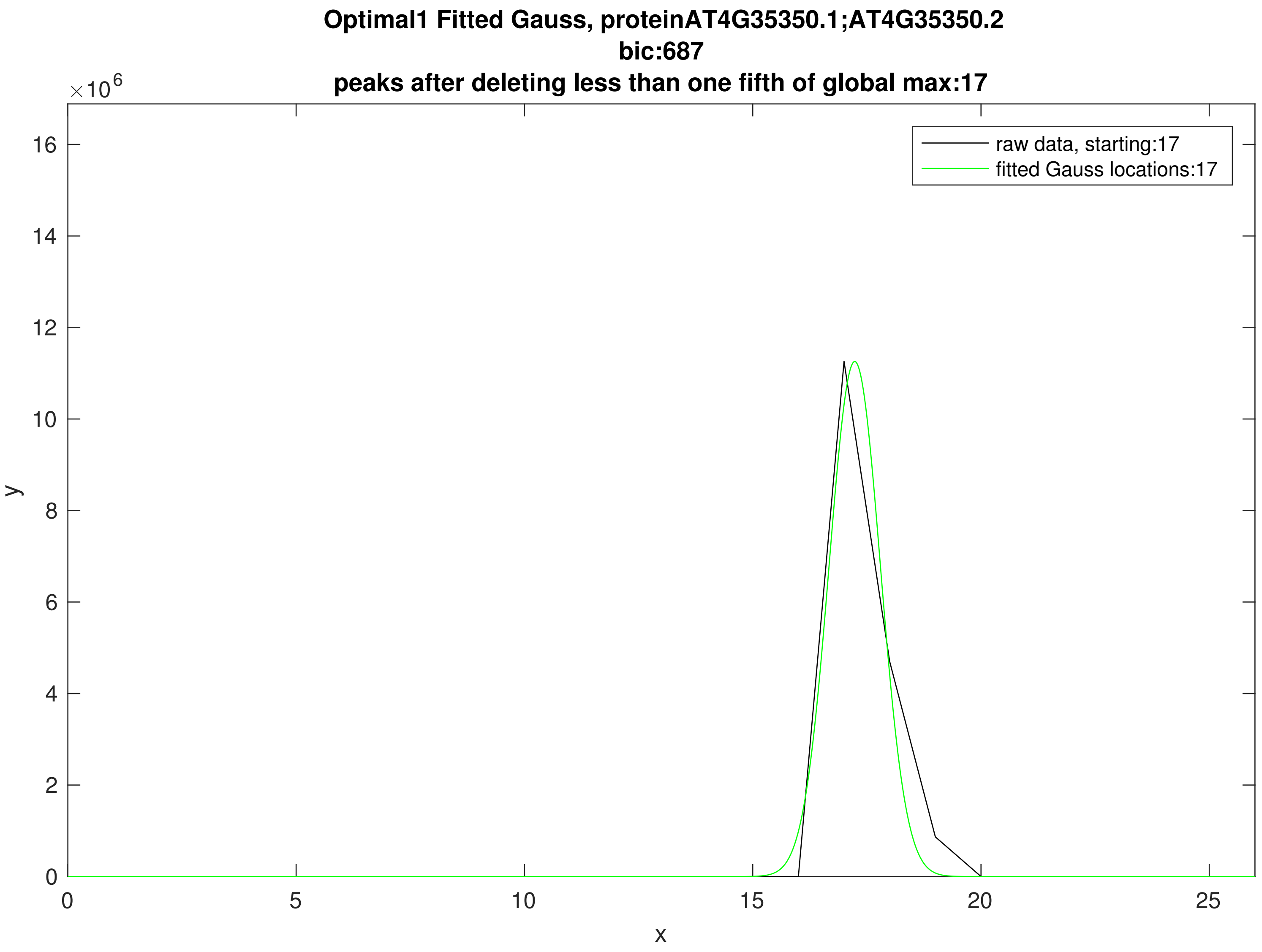
Optimal1 Fitted Gauss, proteinAT4G35350.1;AT4G35350.2
bic:687
peaks after deleting less than one fifth of global max:17
×10^6
raw data, starting:17
fitted Gauss locations:17
y
x
0
2
4
6
8
10
12
14
16
0
5
10
15
20
25

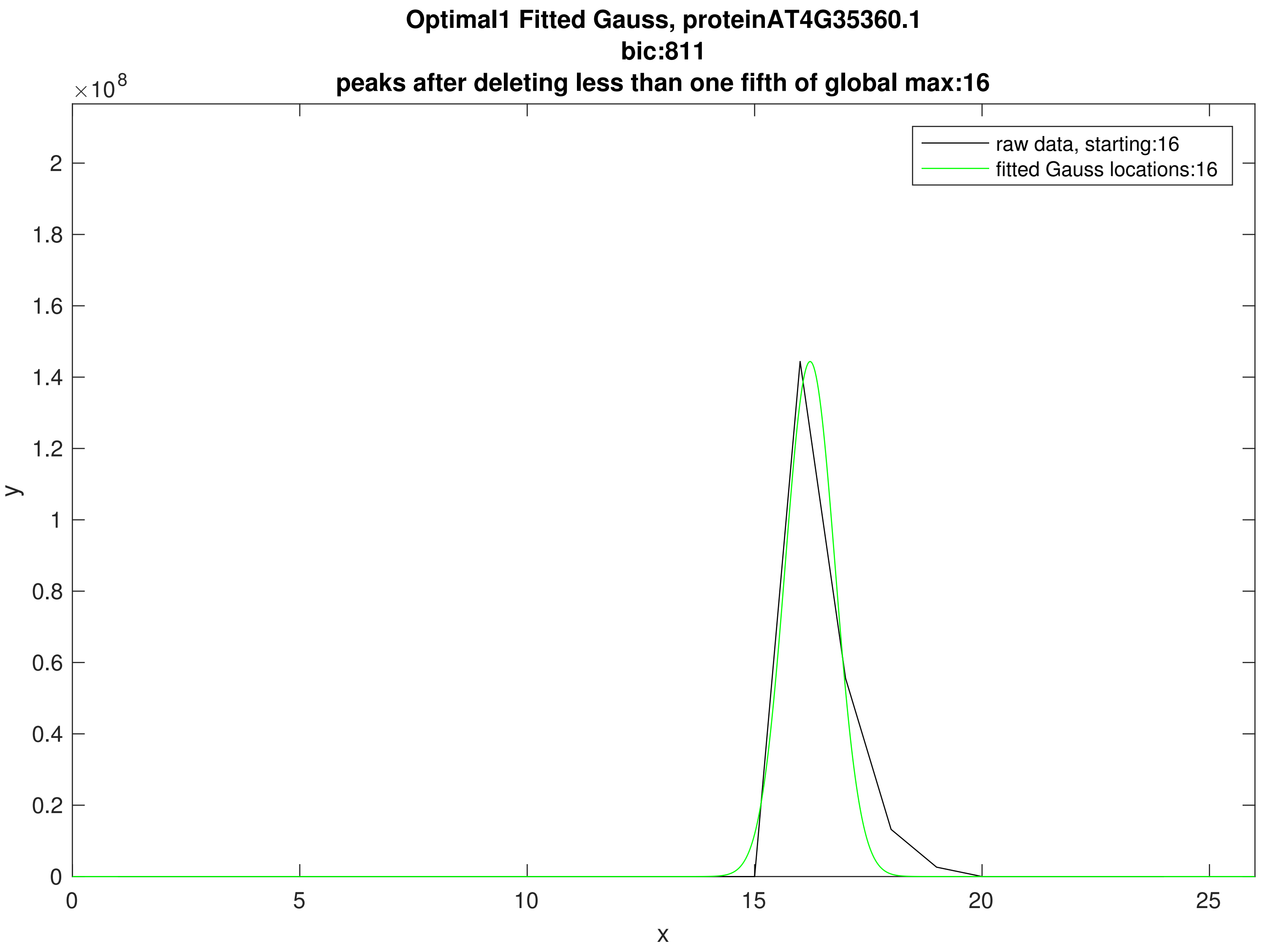
Optimal1 Fitted Gauss, proteinAT4G35360.1
bic:811
peaks after deleting less than one fifth of global max:16
×10^8
2
1.8
1.6
1.4
1.2
1
0.8
0.6
0.4
0.2
0
y
0
5
10
15
20
25
x
raw data, starting:16
fitted Gauss locations:16

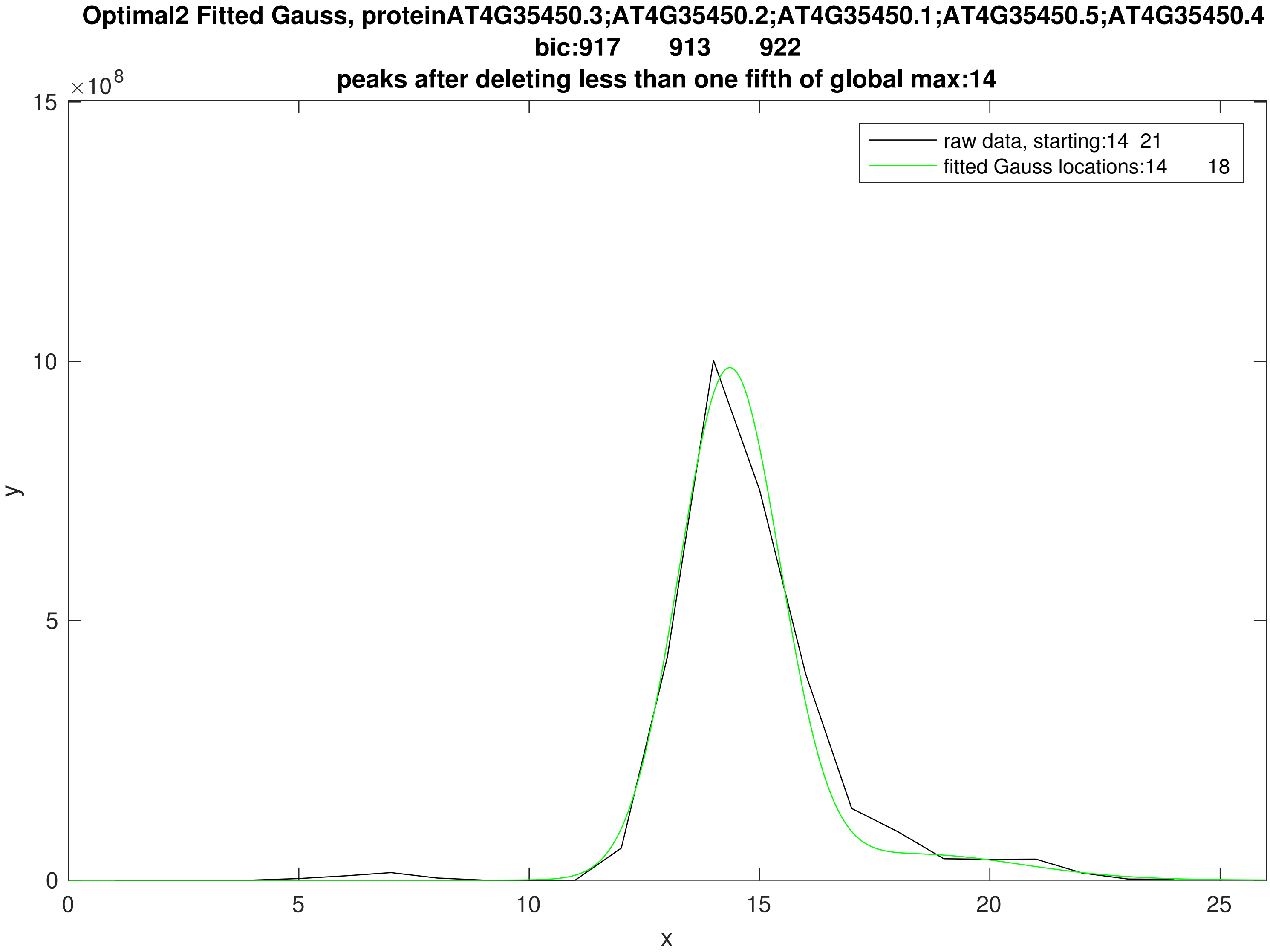
Optimal2 Fitted Gauss, proteinAT4G35450.3;AT4G35450.2;AT4G35450.1;AT4G35450.5;AT4G35450.4
bic:917 913 922
peaks after deleting less than one fifth of global max:14
×10^8
raw data, starting:14 21
fitted Gauss locations:14 18
y
x
0
5
10
15
20
25

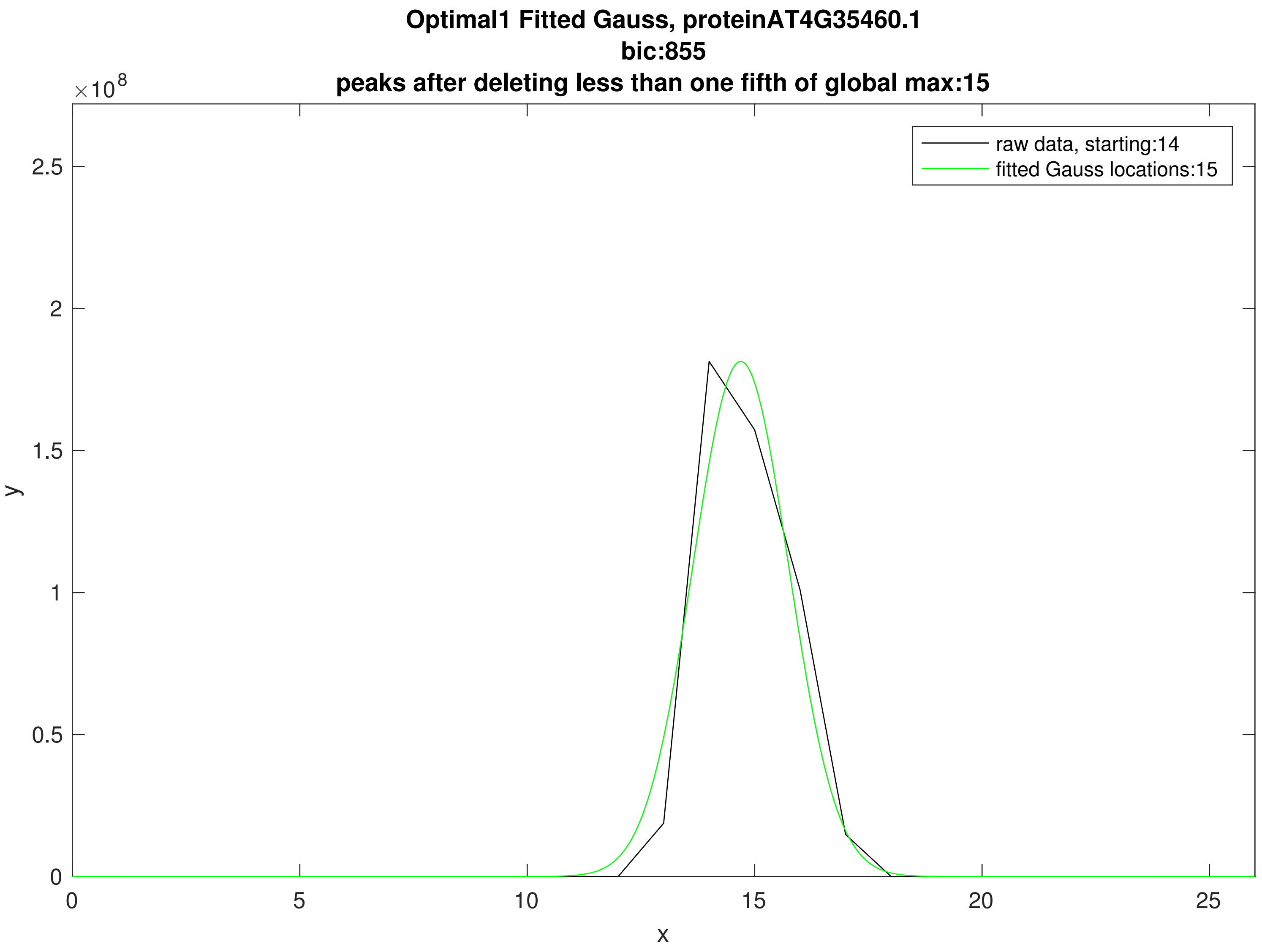
Optimal1 Fitted Gauss, proteinAT4G35460.1
bic:855
peaks after deleting less than one fifth of global max:15
×10^8
2.5
2
1.5
1
0.5
0
y
0
5
10
15
20
25
x
raw data, starting:14
fitted Gauss locations:15

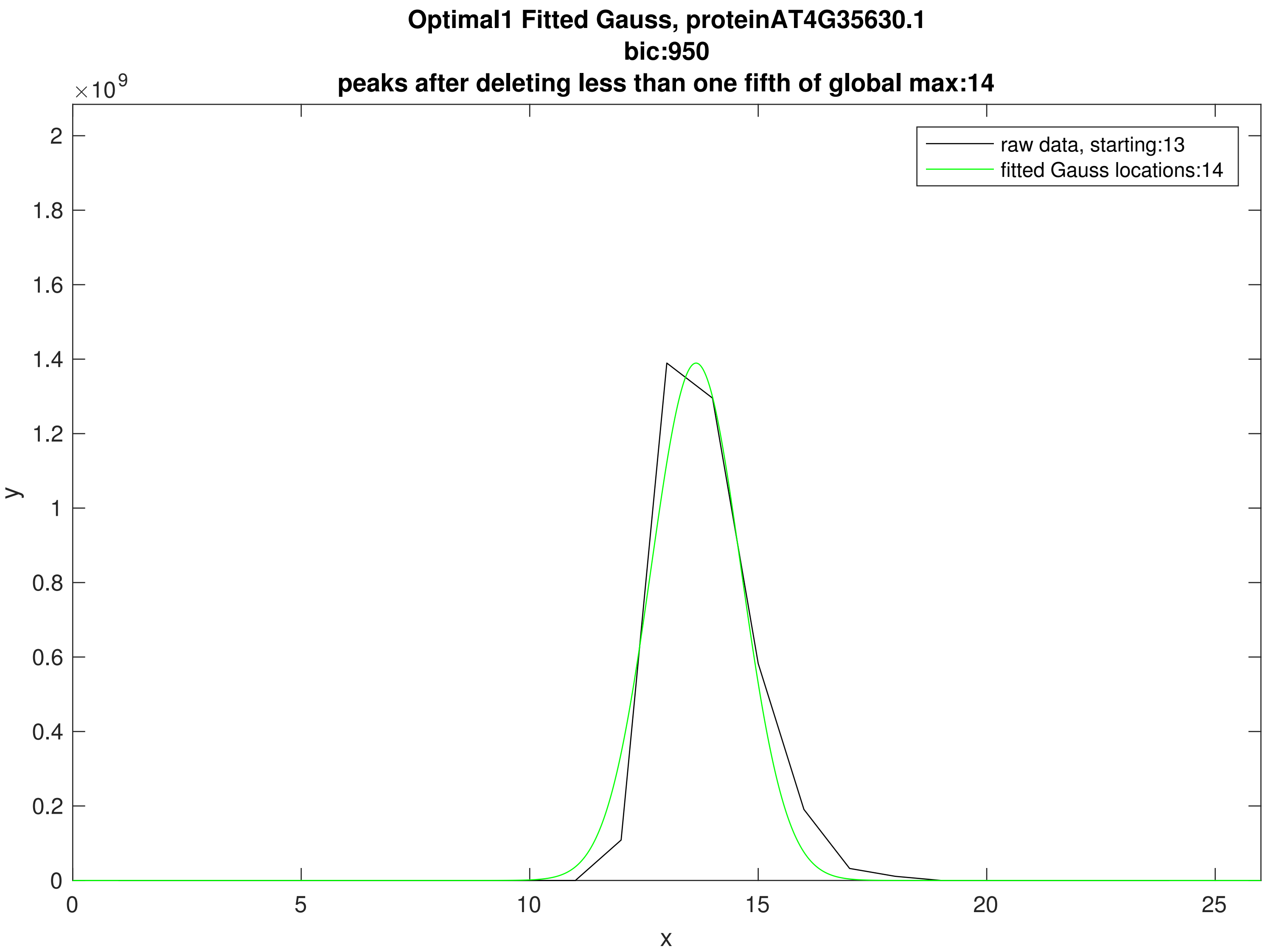
Optimal1 Fitted Gauss, proteinAT4G35630.1
bic:950
peaks after deleting less than one fifth of global max:14
×10^9
raw data, starting:13
fitted Gauss locations:14
y
x
0
0.2
0.4
0.6
0.8
1
1.2
1.4
1.6
1.8
2
0
5
10
15
20
25

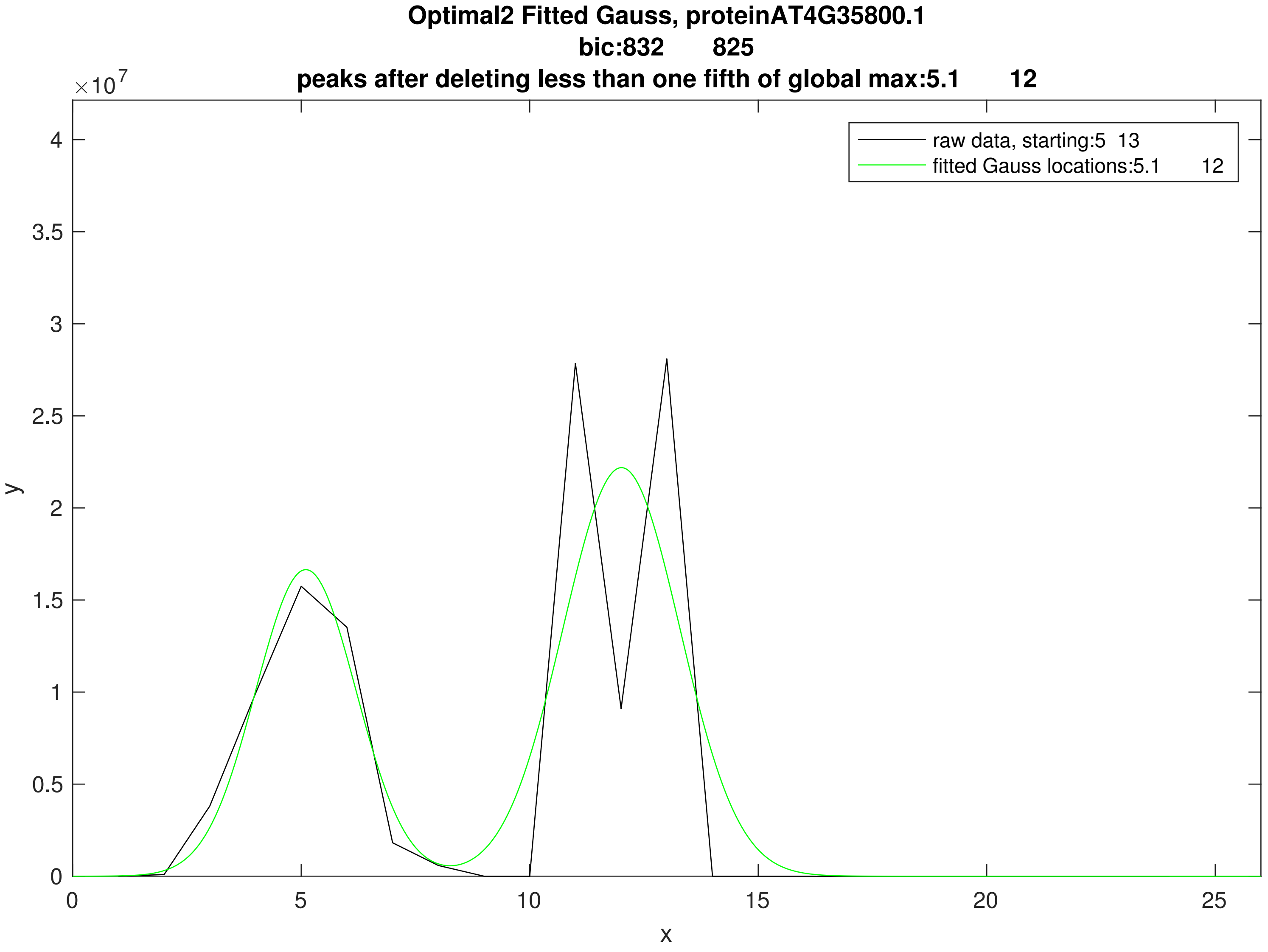
Optimal2 Fitted Gauss, proteinAT4G35800.1
bic:832 825
peaks after deleting less than one fifth of global max:5.1 12
×10^7
raw data, starting:5 13
fitted Gauss locations:5.1 12
y
x
0
0.5
1
1.5
2
2.5
3
3.5
4
5
10
15
20
25

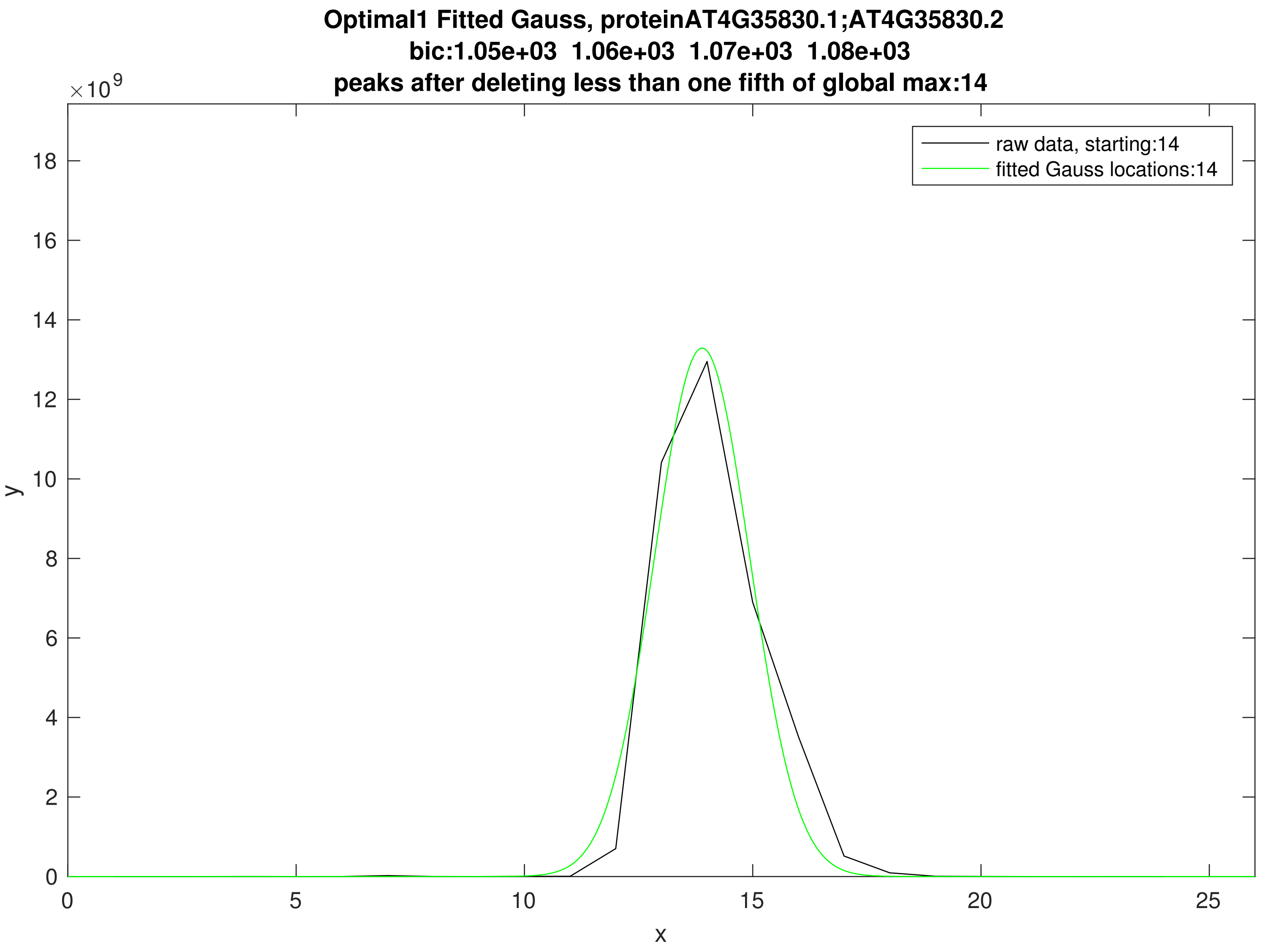

Optimal1 Fitted Gauss, proteinAT4G35830.1;AT4G35830.2
bic:1.05e+03 1.06e+03 1.07e+03 1.08e+03
peaks after deleting less than one fifth of global max:14
×10^9
raw data, starting:14
fitted Gauss locations:14
y
x
0
2
4
6
8
10
12
14
16
18
5
15
20
25

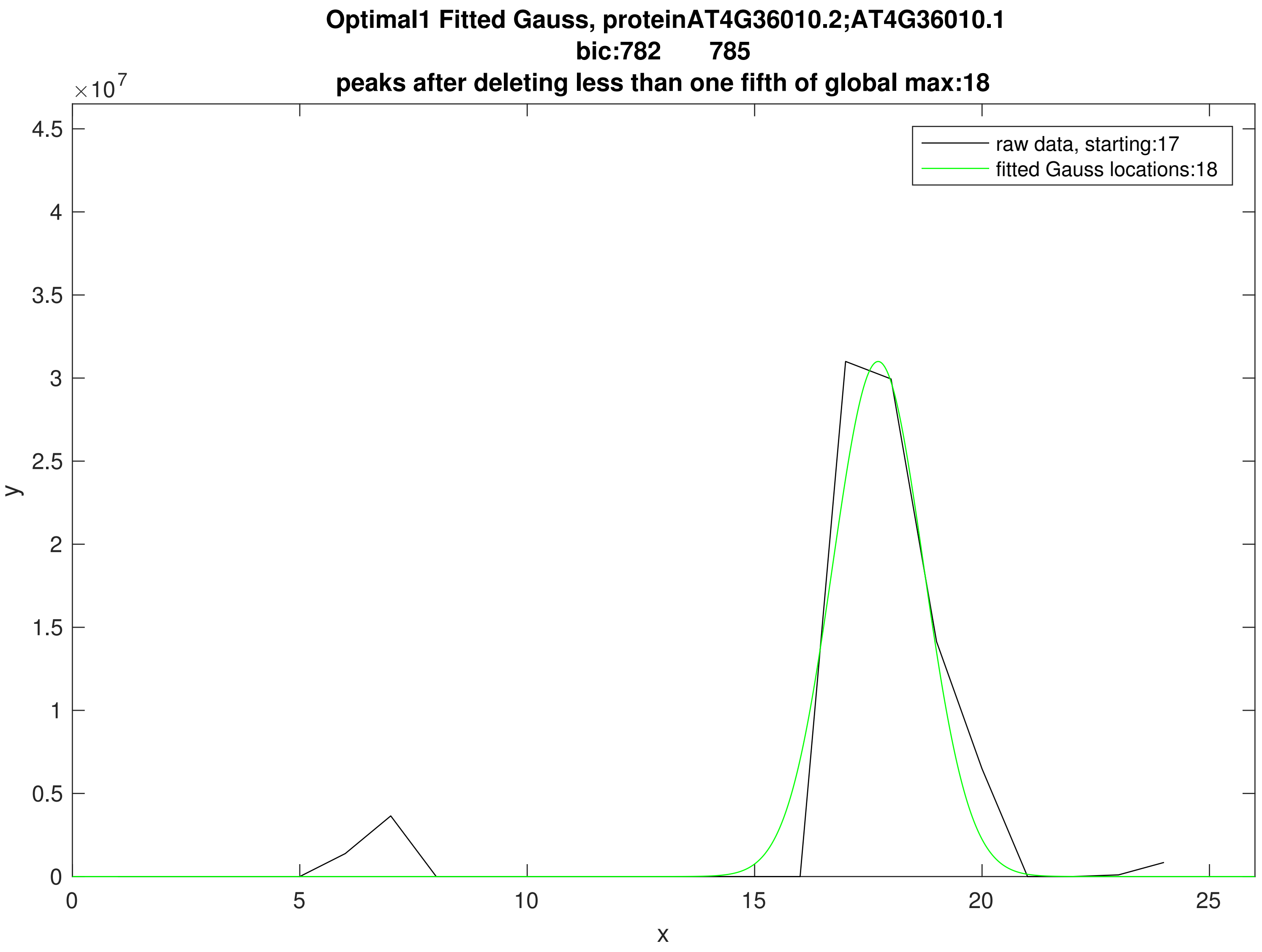
Optimal1 Fitted Gauss, proteinAT4G36010.2;AT4G36010.1
bic:782 785
peaks after deleting less than one fifth of global max:18
×10^7
raw data, starting:17
fitted Gauss locations:18
y
x
0
0.5
1
1.5
2
2.5
3
3.5
4
4.5
5
10
15
20
25

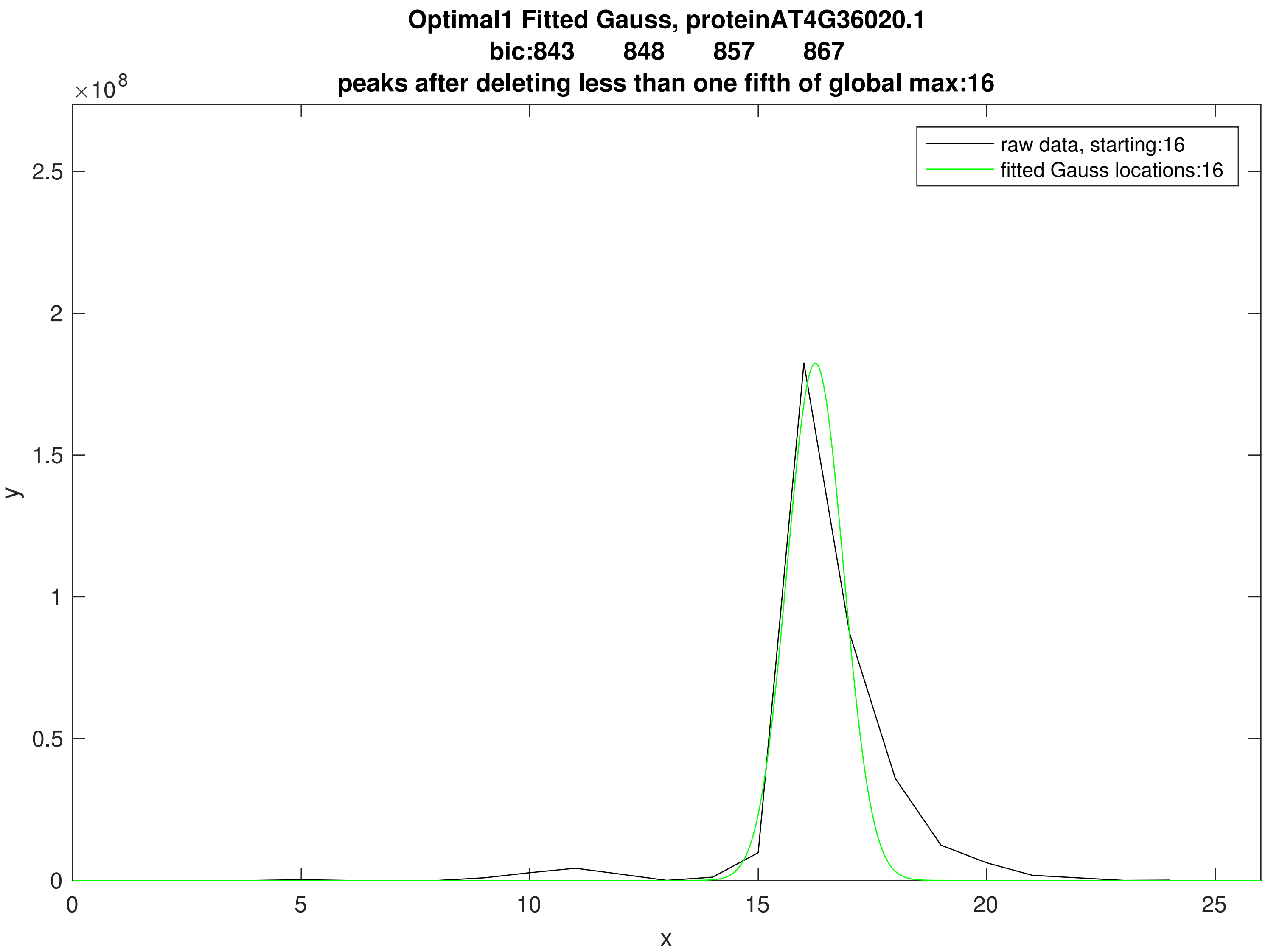
Optimal1 Fitted Gauss, proteinAT4G36020.1
bic:843 848 857 867
peaks after deleting less than one fifth of global max:16
×10^8
raw data, starting:16
fitted Gauss locations:16
y
x
2.5
2
1.5
1
0.5
0
5
10
15
20
25

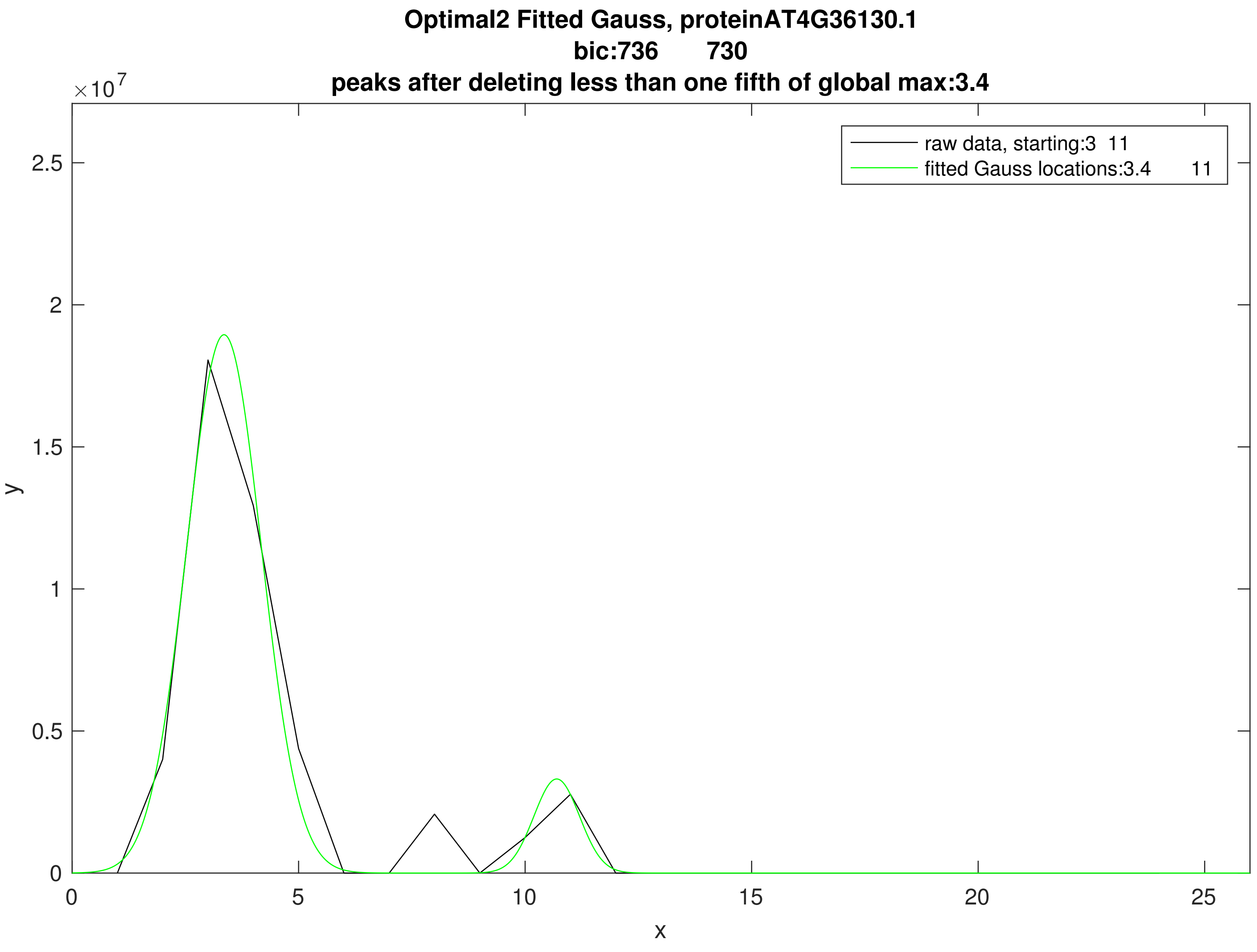
Optimal2 Fitted Gauss, proteinAT4G36130.1
bic:736 730
peaks after deleting less than one fifth of global max:3.4
×10^7
raw data, starting:3 11
fitted Gauss locations:3.4 11
y
x
0
0.5
1
1.5
2
2.5
5
10
15
20
25

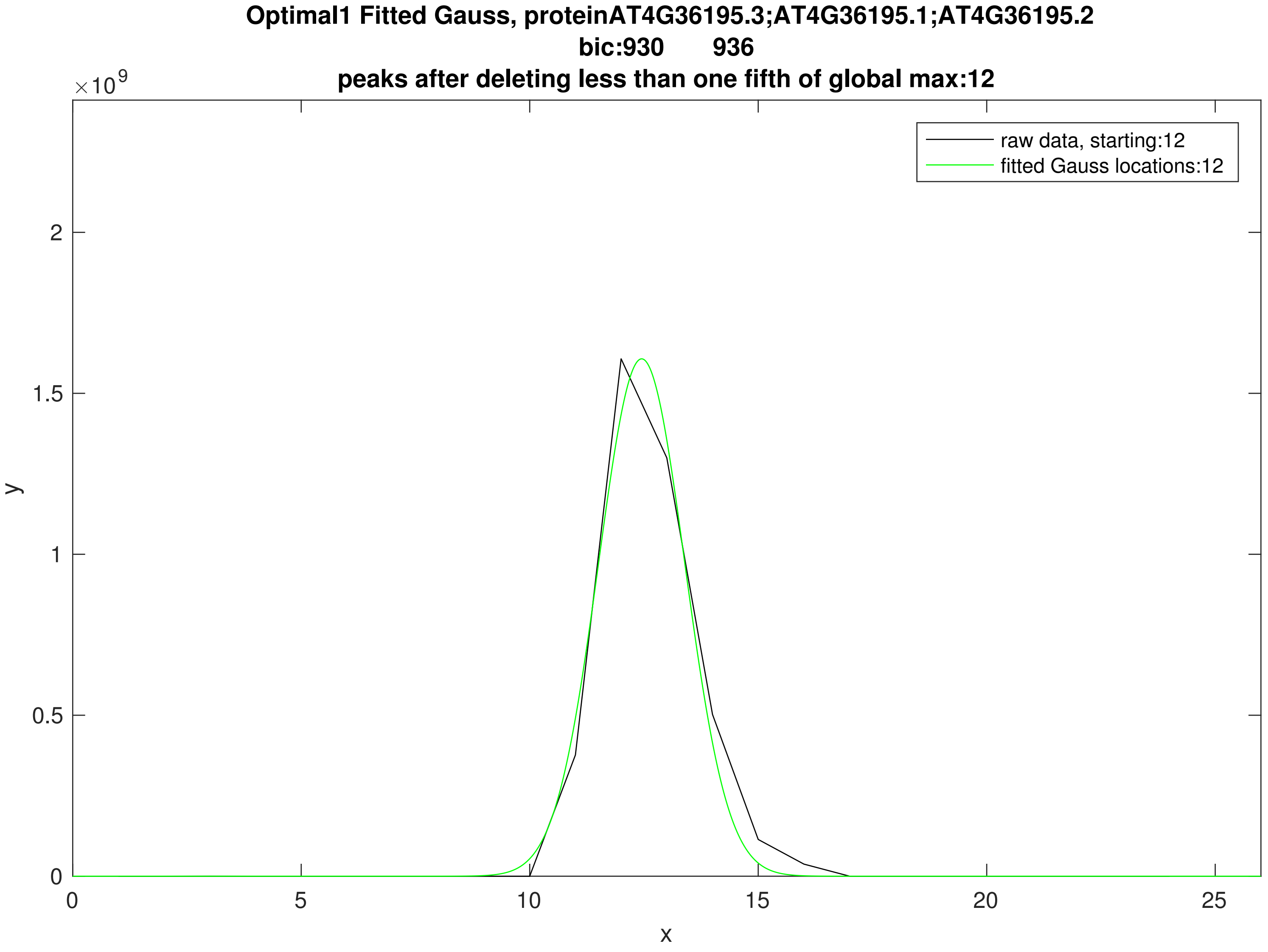

Optimal1 Fitted Gauss, proteinAT4G36195.3;AT4G36195.1;AT4G36195.2
bic:930 936
peaks after deleting less than one fifth of global max:12
×10^9
raw data, starting:12
fitted Gauss locations:12
2
1.5
1
0.5
0
y
0
5
10
15
20
25
x

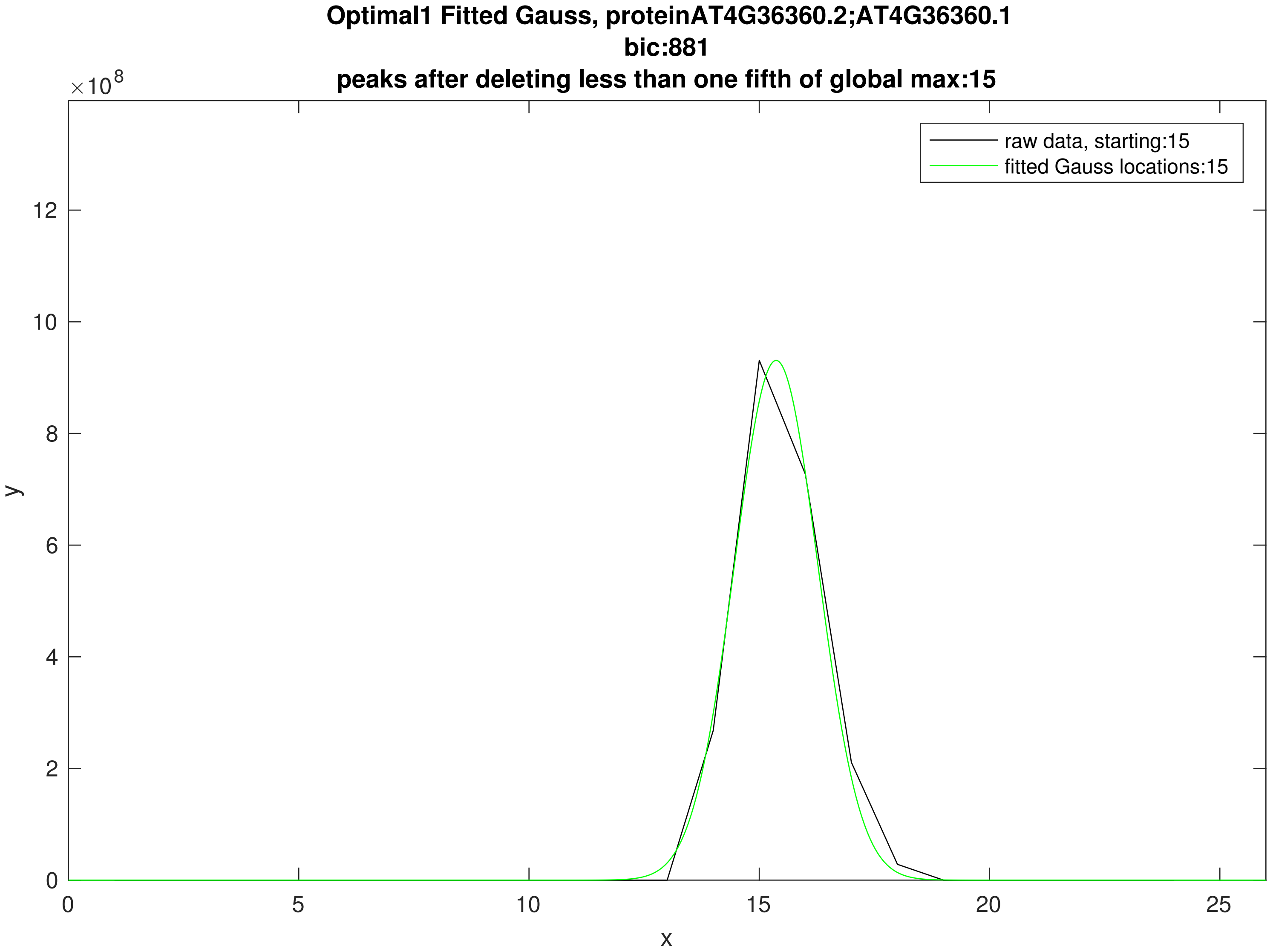
Optimal1 Fitted Gauss, proteinAT4G36360.2;AT4G36360.1
bic:881
peaks after deleting less than one fifth of global max:15
×10^8
raw data, starting:15
fitted Gauss locations:15
12
10
8
6
4
2
0
y
0
5
10
15
20
25
x

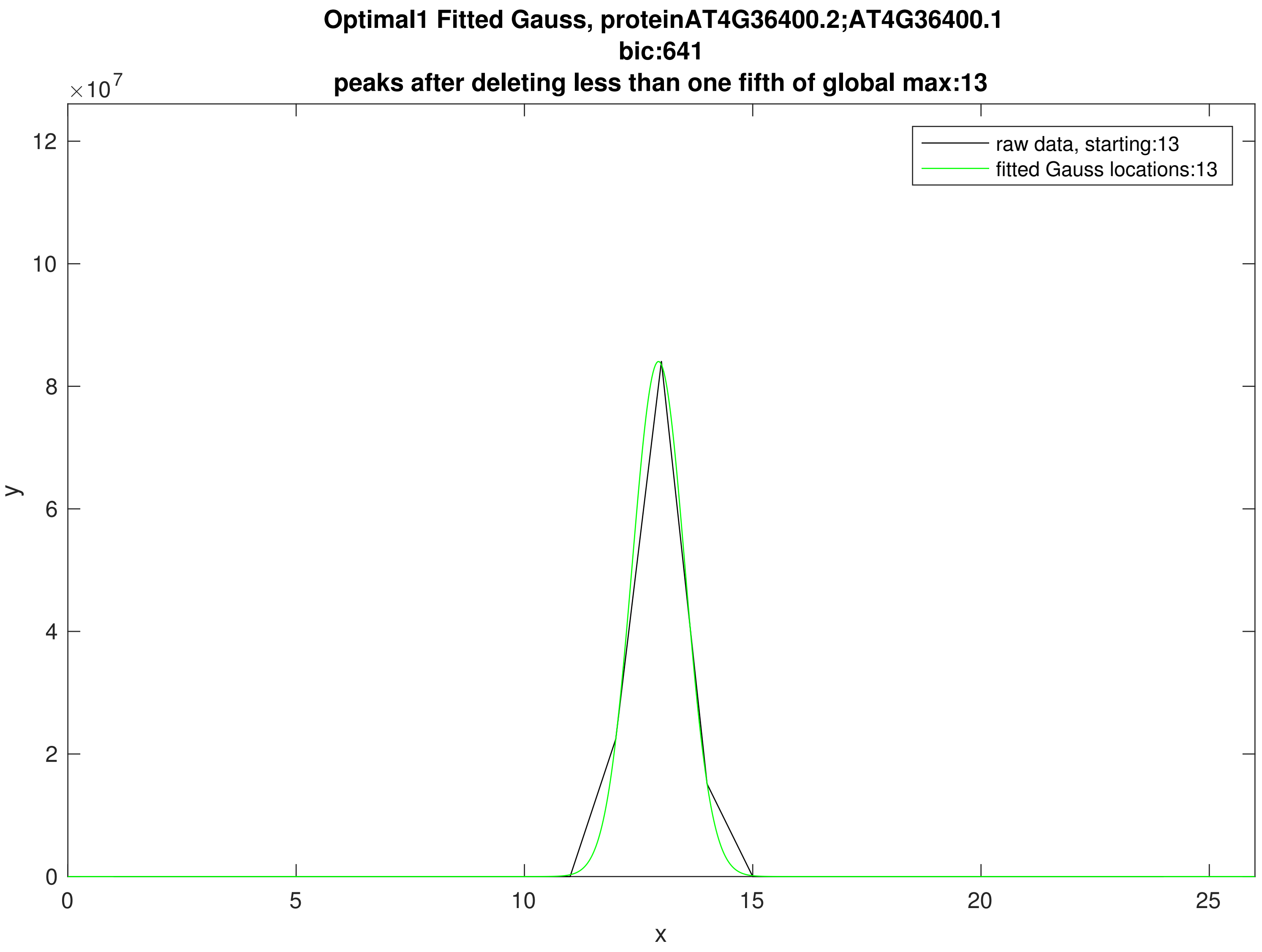
Optimal1 Fitted Gauss, proteinAT4G36400.2;AT4G36400.1
bic:641
peaks after deleting less than one fifth of global max:13
×10^7
12
10
8
6
4
2
0
y
0
5
10
15
20
25
x
raw data, starting:13
fitted Gauss locations:13

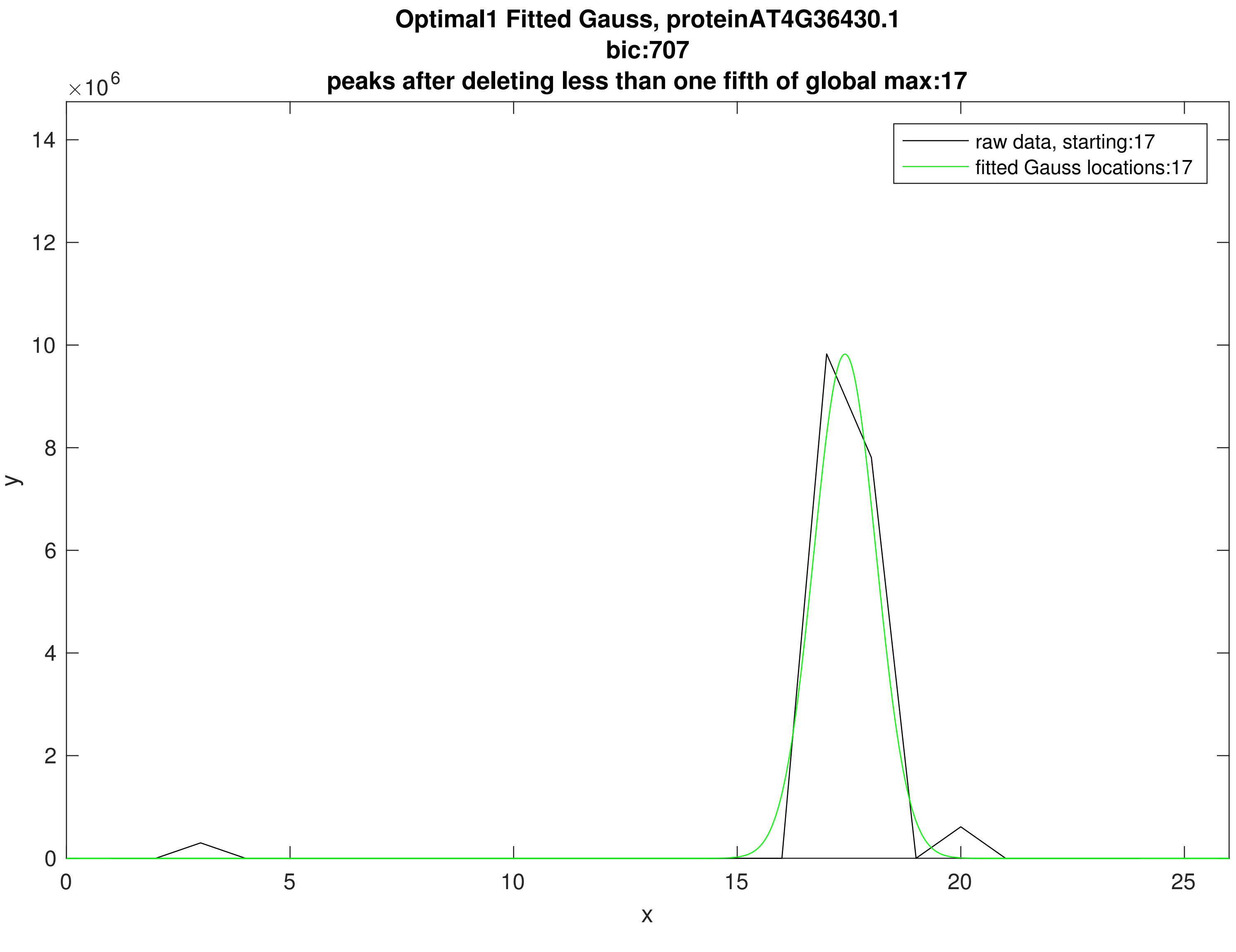
Optimal1 Fitted Gauss, proteinAT4G36430.1
bic:707
peaks after deleting less than one fifth of global max:17
×10^6
raw data, starting:17
fitted Gauss locations:17
y
x
0
2
4
6
8
10
12
14
0
5
10
15
20
25

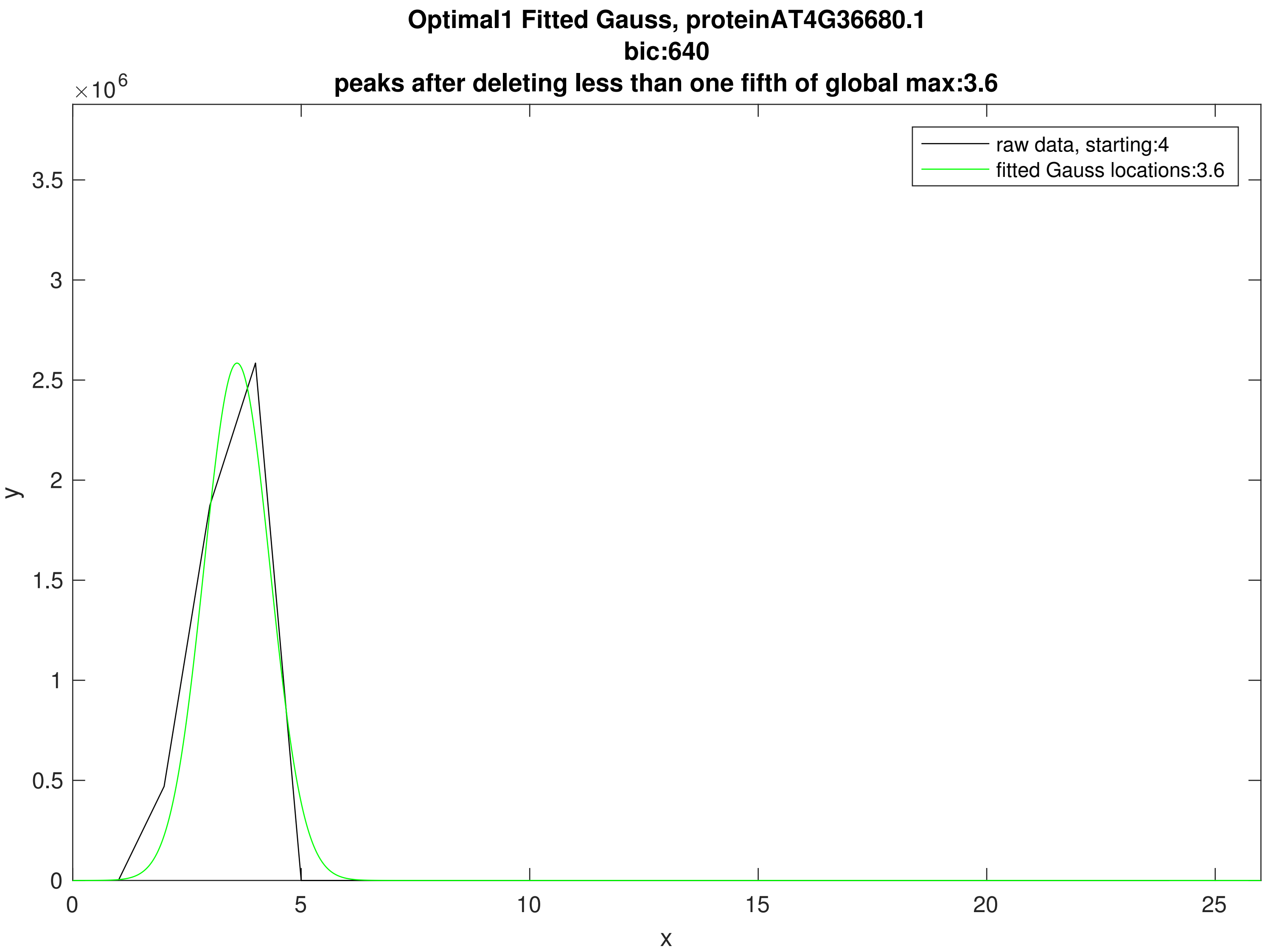
Optimal1 Fitted Gauss, proteinAT4G36680.1
bic:640
peaks after deleting less than one fifth of global max:3.6
×10^6
raw data, starting:4
fitted Gauss locations:3.6
y
x
0
0.5
1
1.5
2
2.5
3
3.5
0
5
10
15
20
25

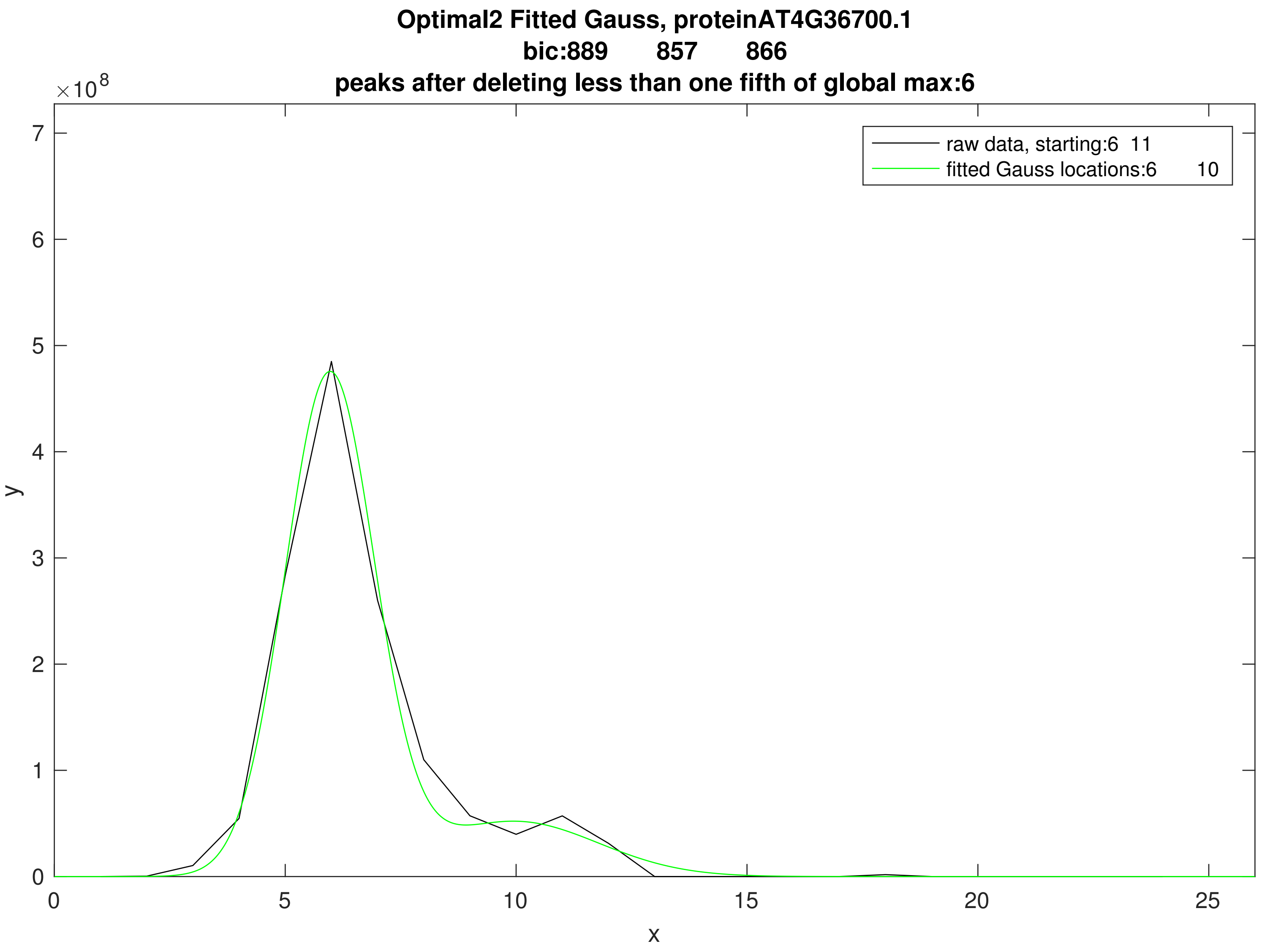

Optimal2 Fitted Gauss, proteinAT4G36700.1
bic:889 857 866
peaks after deleting less than one fifth of global max:6
×10^8
raw data, starting:6 11
fitted Gauss locations:6 10
y
x
0
1
2
3
4
5
6
7
0
5
10
15
20
25

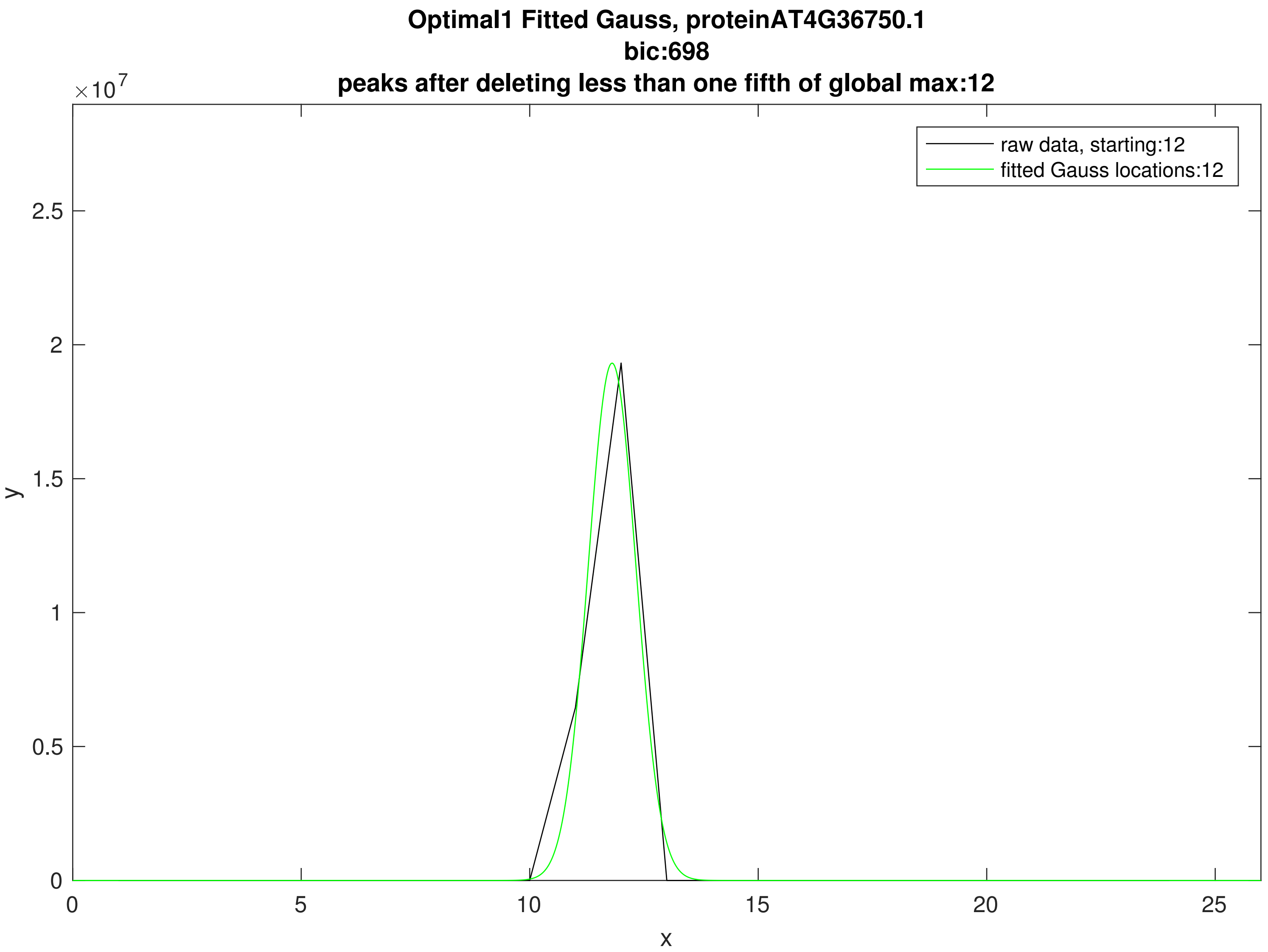
Optimal1 Fitted Gauss, proteinAT4G36750.1
bic:698
peaks after deleting less than one fifth of global max:12
×10^7
raw data, starting:12
fitted Gauss locations:12
0
0.5
1
1.5
2
2.5
y
0
5
10
15
20
25
x

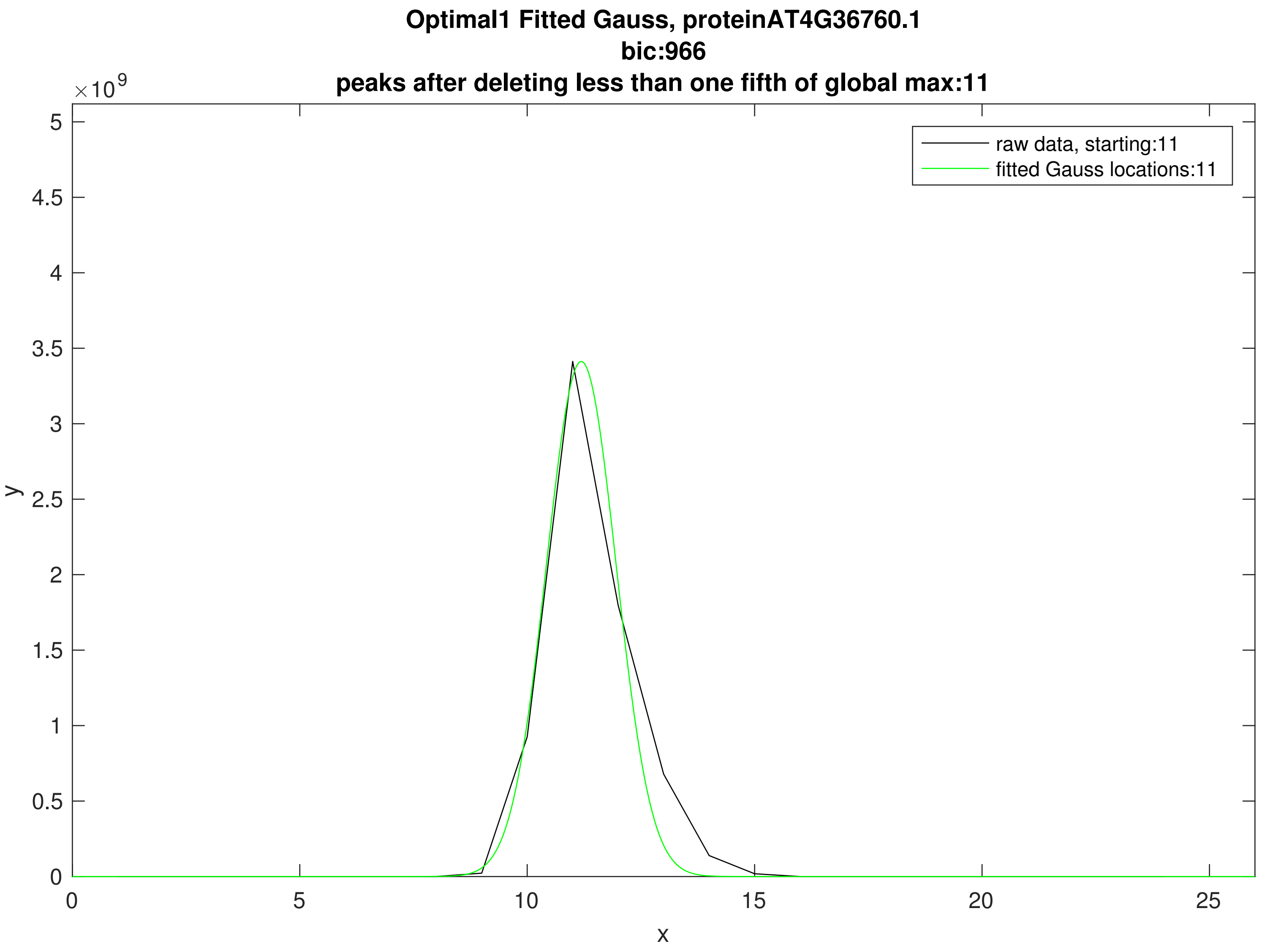
Optimal1 Fitted Gauss, proteinAT4G36760.1
bic:966
peaks after deleting less than one fifth of global max:11
×10^9
raw data, starting:11
fitted Gauss locations:11
y
x
0
0.5
1
1.5
2
2.5
3
3.5
4
4.5
5
0
5
10
15
20
25

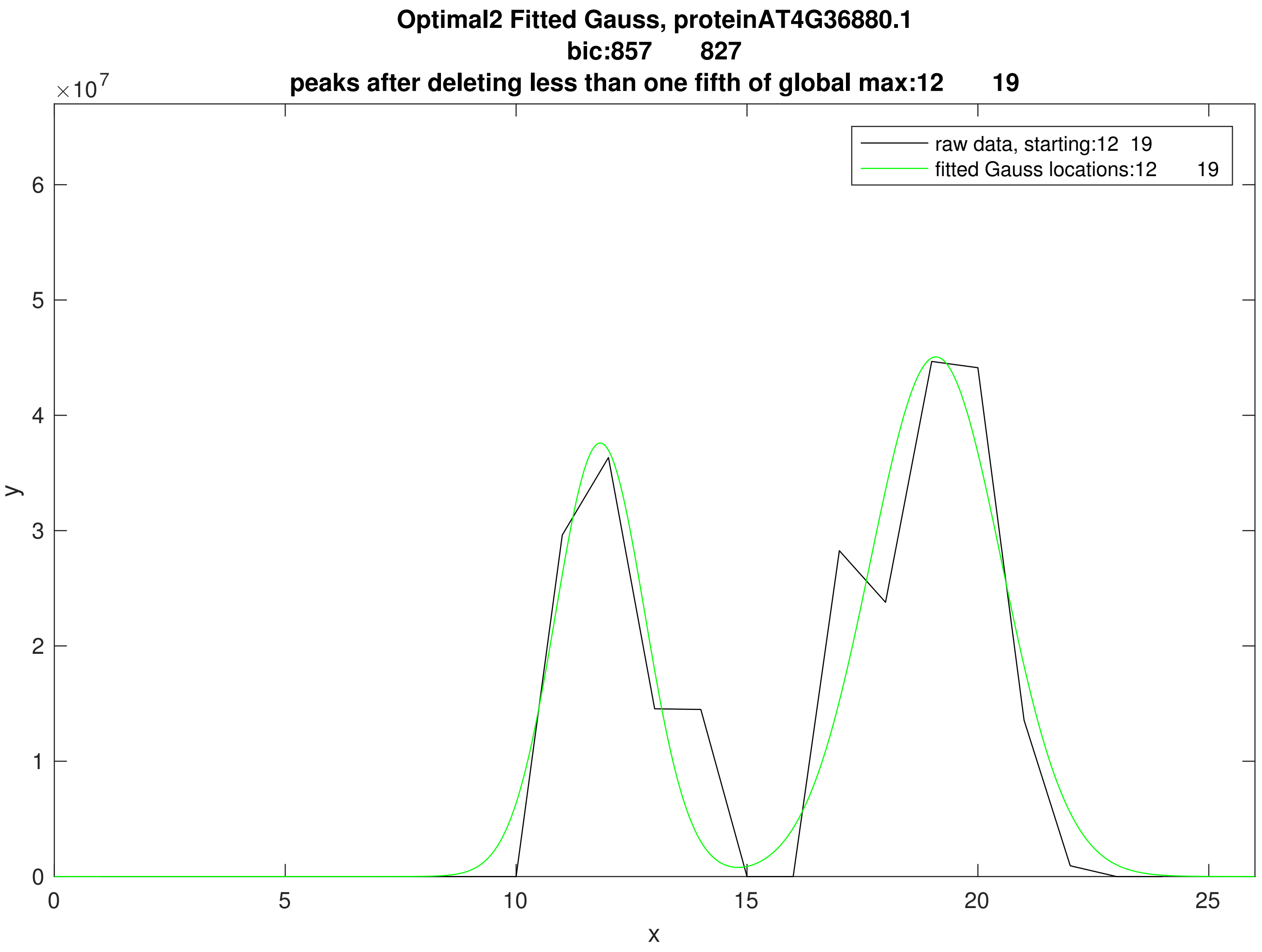

Optimal2 Fitted Gauss, proteinAT4G36880.1
bic:857 827
peaks after deleting less than one fifth of global max:12 19
×10^7
raw data, starting:12 19
fitted Gauss locations:12 19
0
1
2
3
4
5
6
y
0
5
10
15
20
25
x

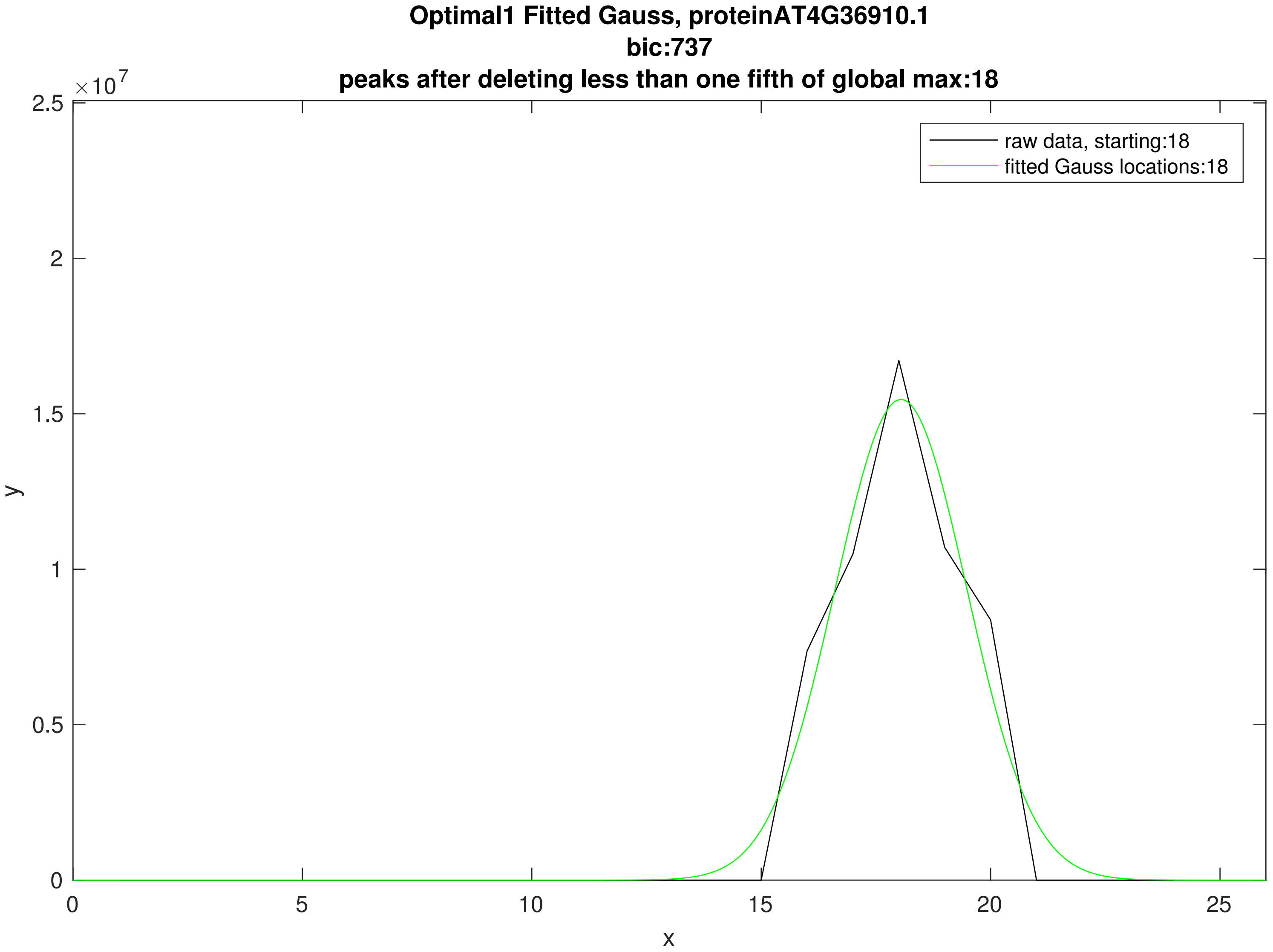

Optimal1 Fitted Gauss, proteinAT4G36910.1
bic:737
peaks after deleting less than one fifth of global max:18
×10^7
2.5
2
1.5
1
0.5
0
y
0
5
10
15
20
25
x
raw data, starting:18
fitted Gauss locations:18

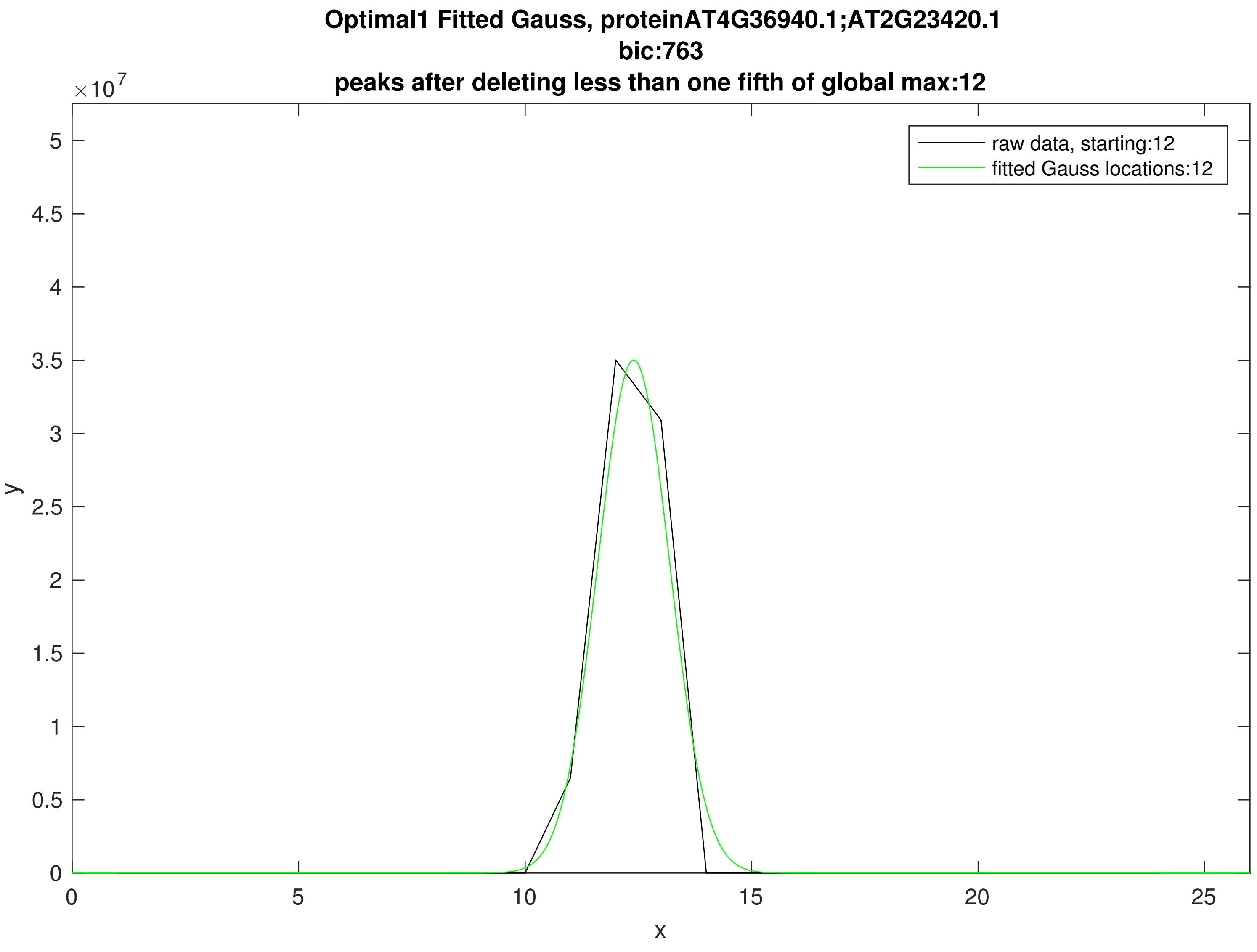
Optimal1 Fitted Gauss, proteinAT4G36940.1;AT2G23420.1
bic:763
peaks after deleting less than one fifth of global max:12
×10^7
5
4.5
4
3.5
3
2.5
2
1.5
1
0.5
0
y
0
5
10
15
20
25
x
raw data, starting:12
fitted Gauss locations:12

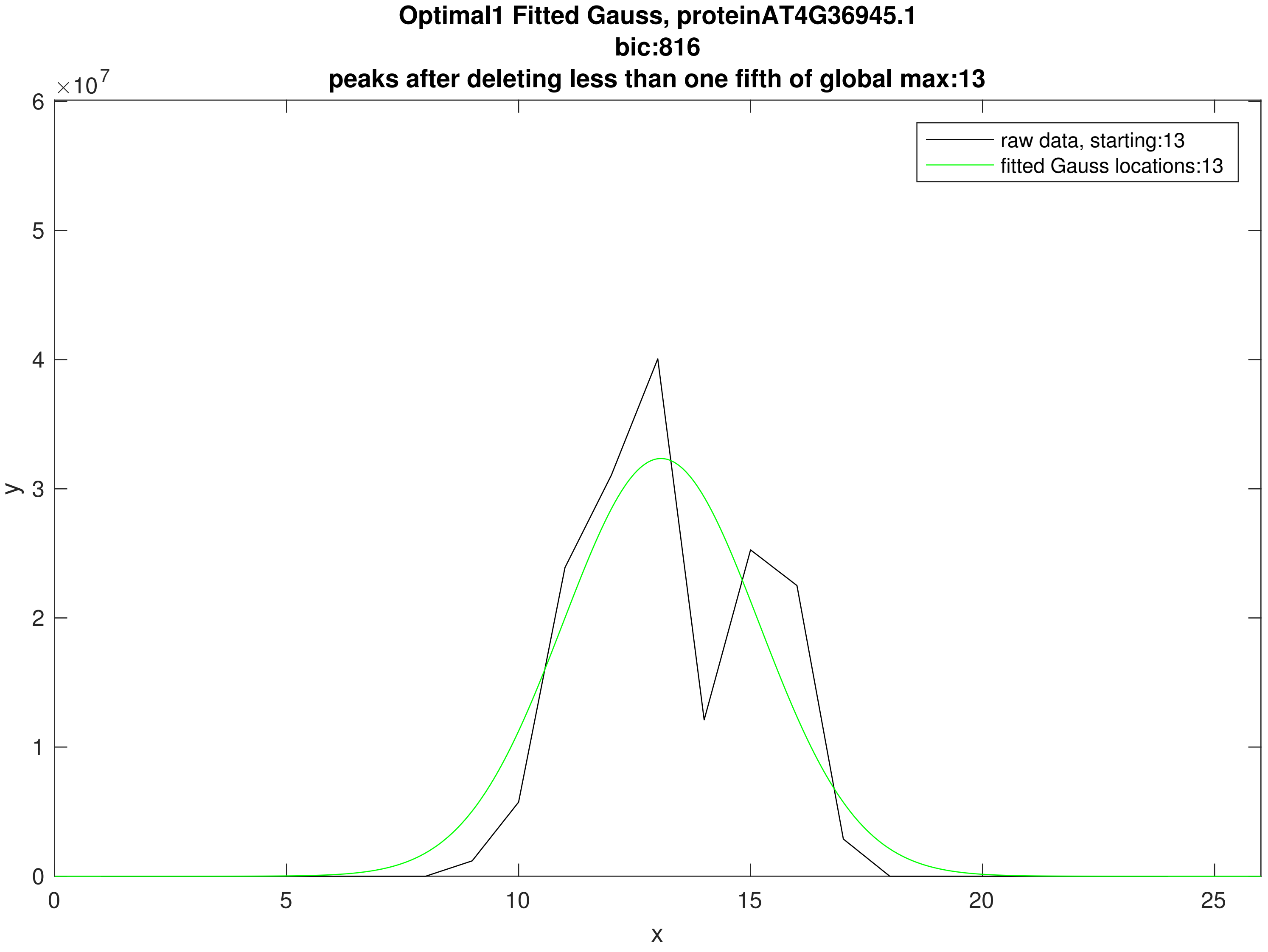
Optimal1 Fitted Gauss, proteinAT4G36945.1
bic:816
peaks after deleting less than one fifth of global max:13
×10^7
6
5
4
3
2
1
0
y
0
5
10
15
20
25
x
raw data, starting:13
fitted Gauss locations:13

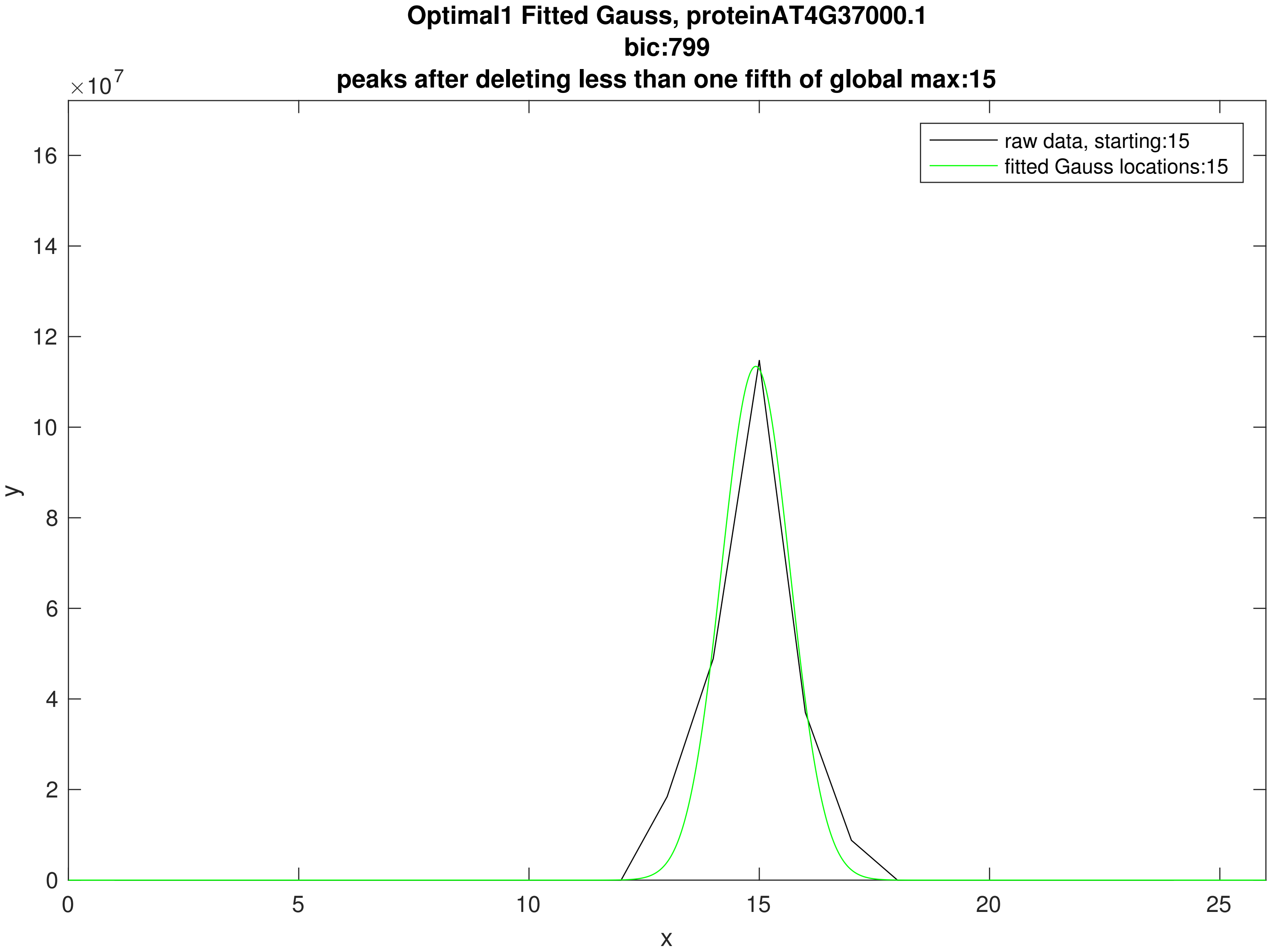
Optimal1 Fitted Gauss, proteinAT4G37000.1
bic:799
peaks after deleting less than one fifth of global max:15
×10^7
raw data, starting:15
fitted Gauss locations:15
y
x
0
2
4
6
8
10
12
14
16
0
5
10
15
20
25

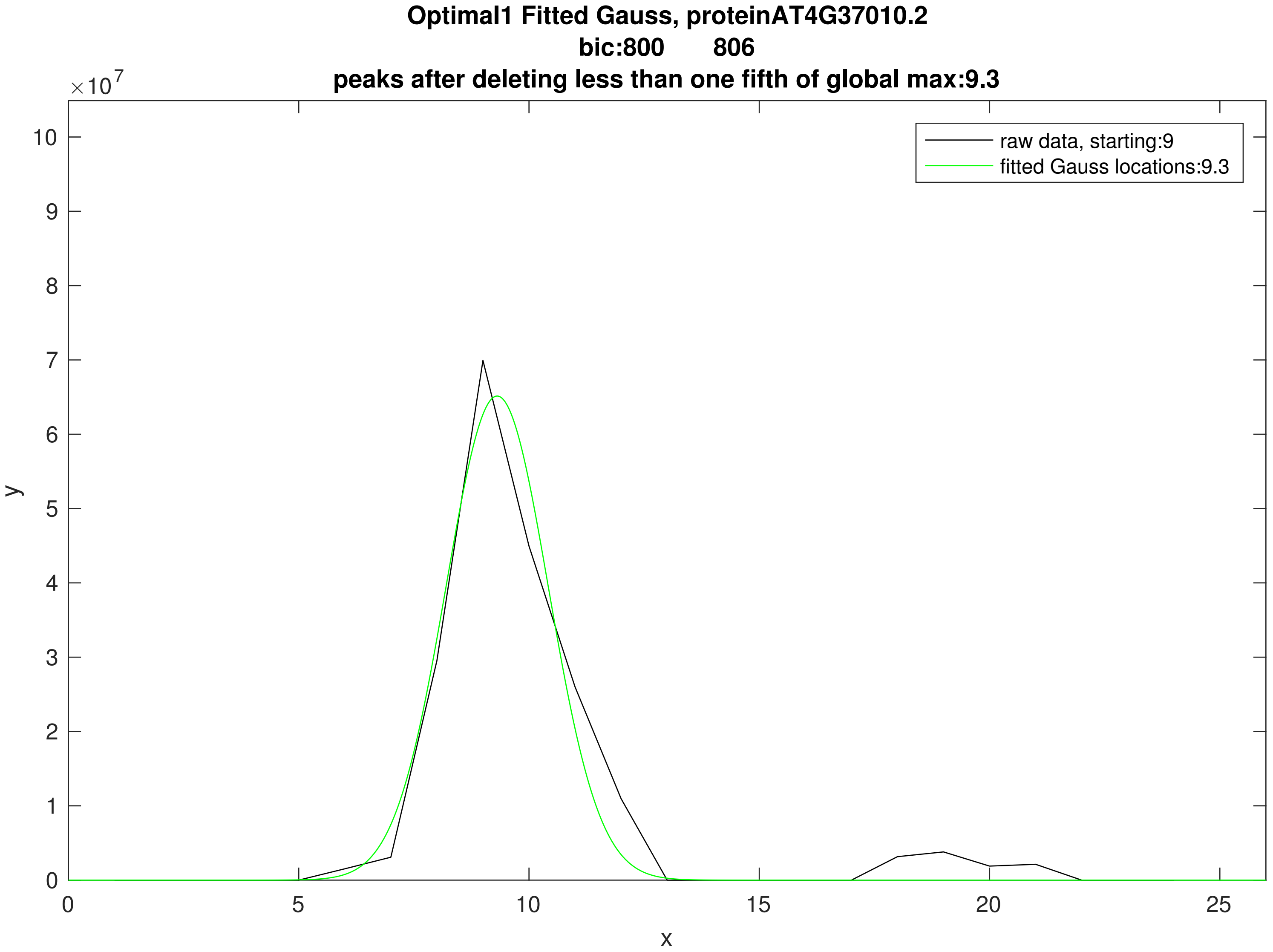
Optimal1 Fitted Gauss, proteinAT4G37010.2
bic:800 806
peaks after deleting less than one fifth of global max:9.3
×10^7
raw data, starting:9
fitted Gauss locations:9.3
y
x
0
1
2
3
4
5
6
7
8
9
10
0
5
10
15
20
25

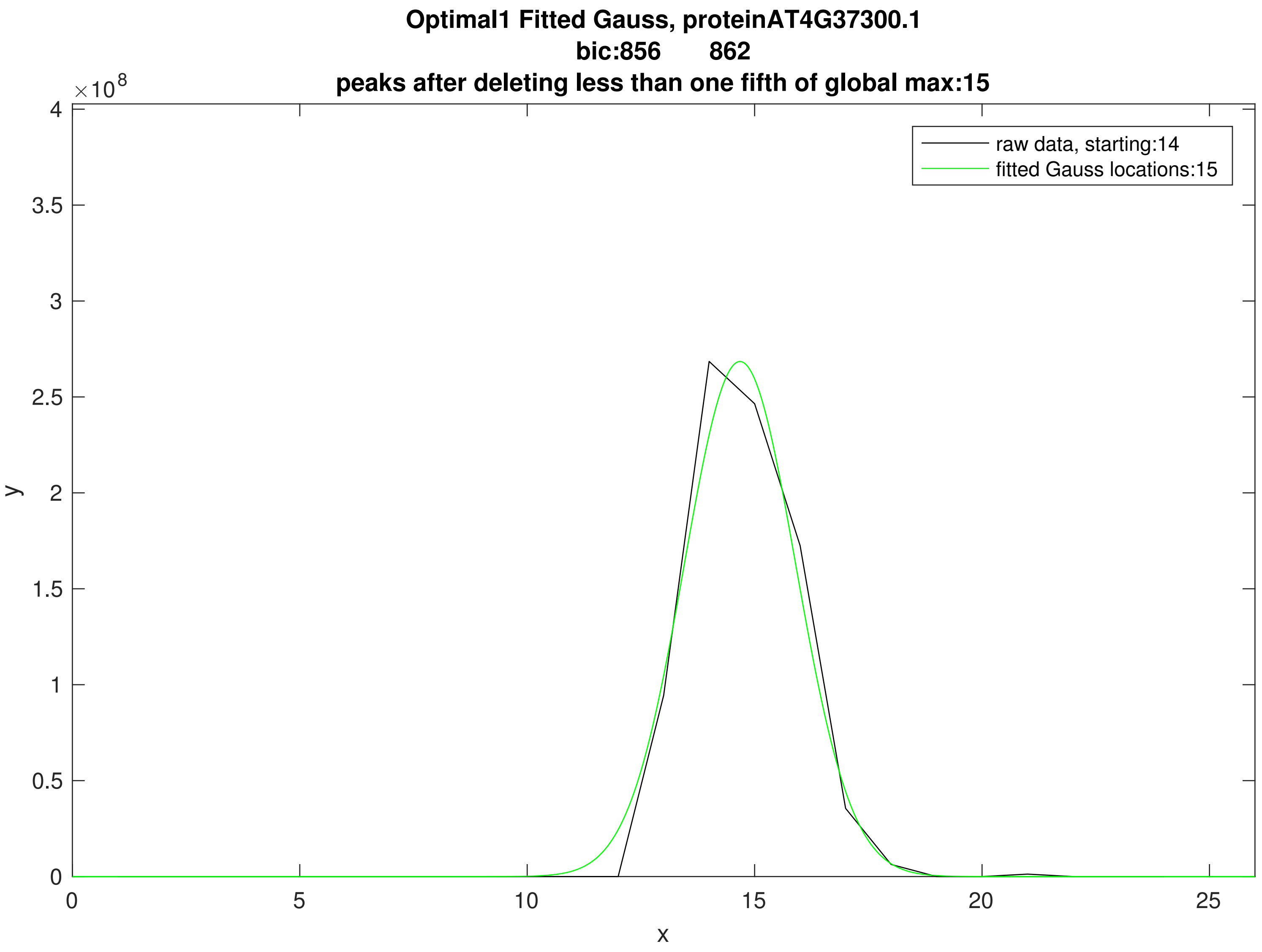
Optimal1 Fitted Gauss, proteinAT4G37300.1
bic:856 862
peaks after deleting less than one fifth of global max:15
×10^8
4
3.5
3
2.5
2
1.5
1
0.5
0
y
0
5
10
15
20
25
x
raw data, starting:14
fitted Gauss locations:15

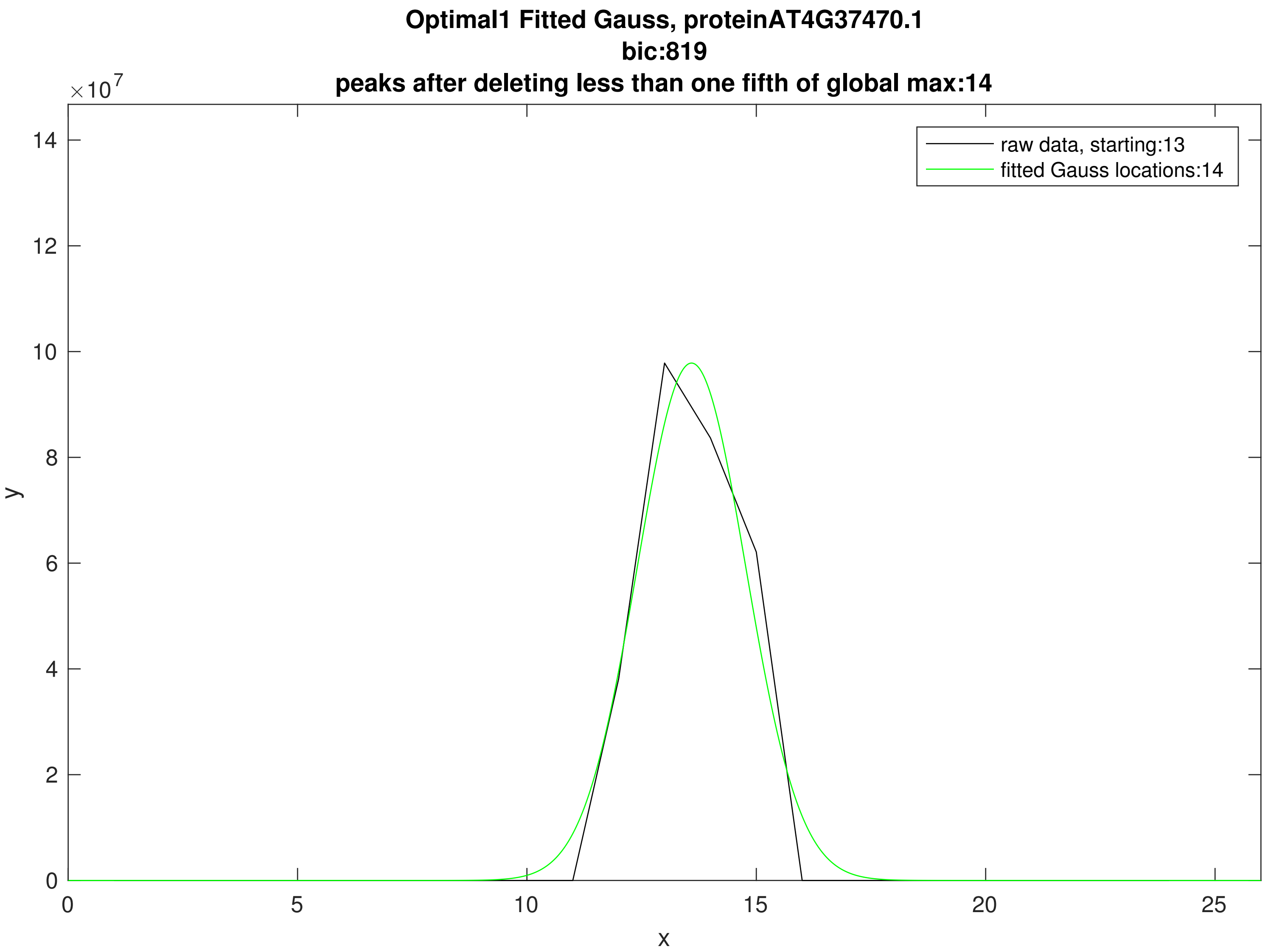
Optimal1 Fitted Gauss, proteinAT4G37470.1
bic:819
peaks after deleting less than one fifth of global max:14
×10^7
14
12
10
8
6
4
2
0
y
0
5
10
15
20
25
x
raw data, starting:13
fitted Gauss locations:14

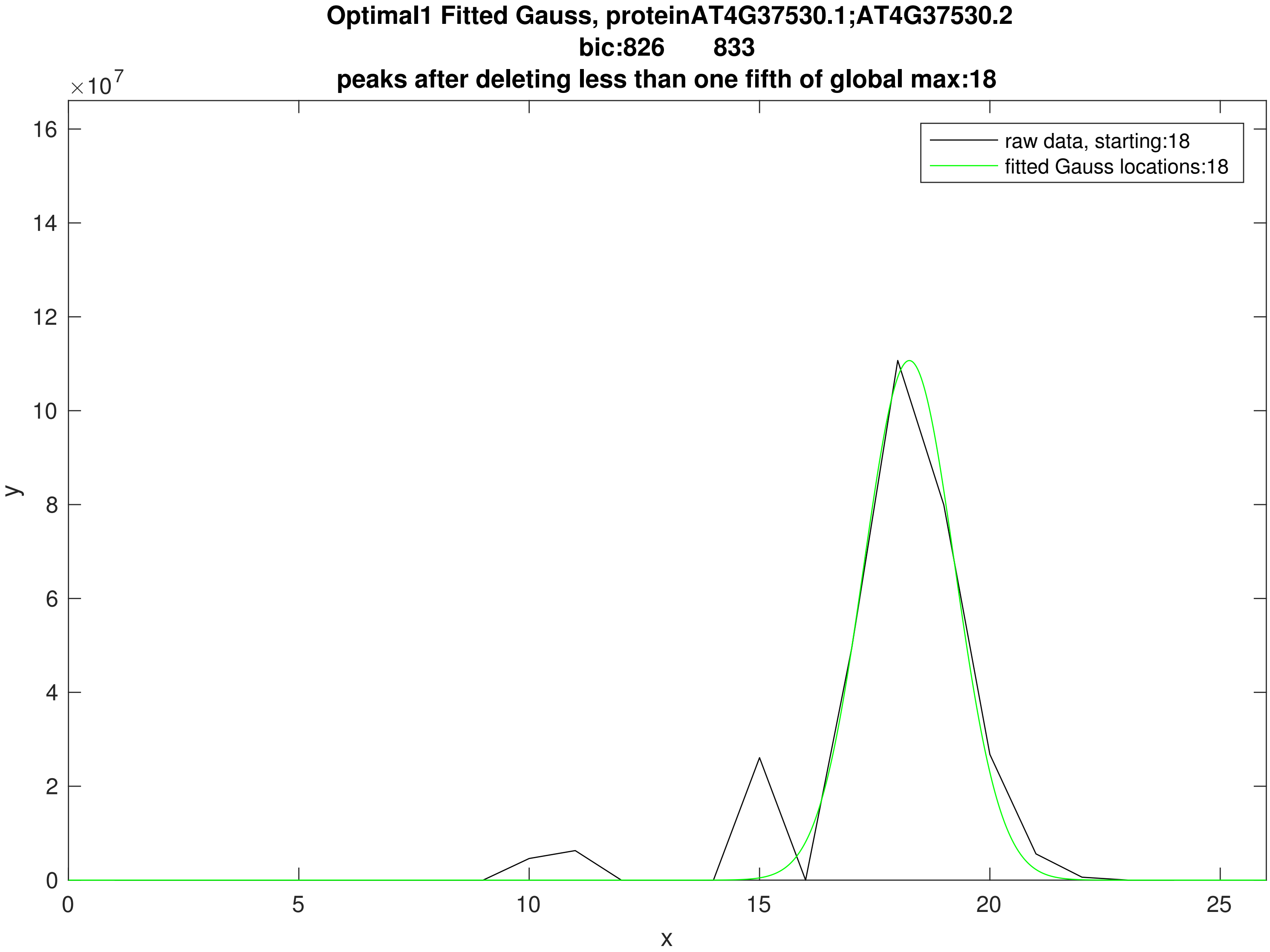

Optimal1 Fitted Gauss, proteinAT4G37530.1;AT4G37530.2
bic:826 833
peaks after deleting less than one fifth of global max:18
×10^7
raw data, starting:18
fitted Gauss locations:18
y
x
0
2
4
6
8
10
12
14
16
0
5
10
15
20
25

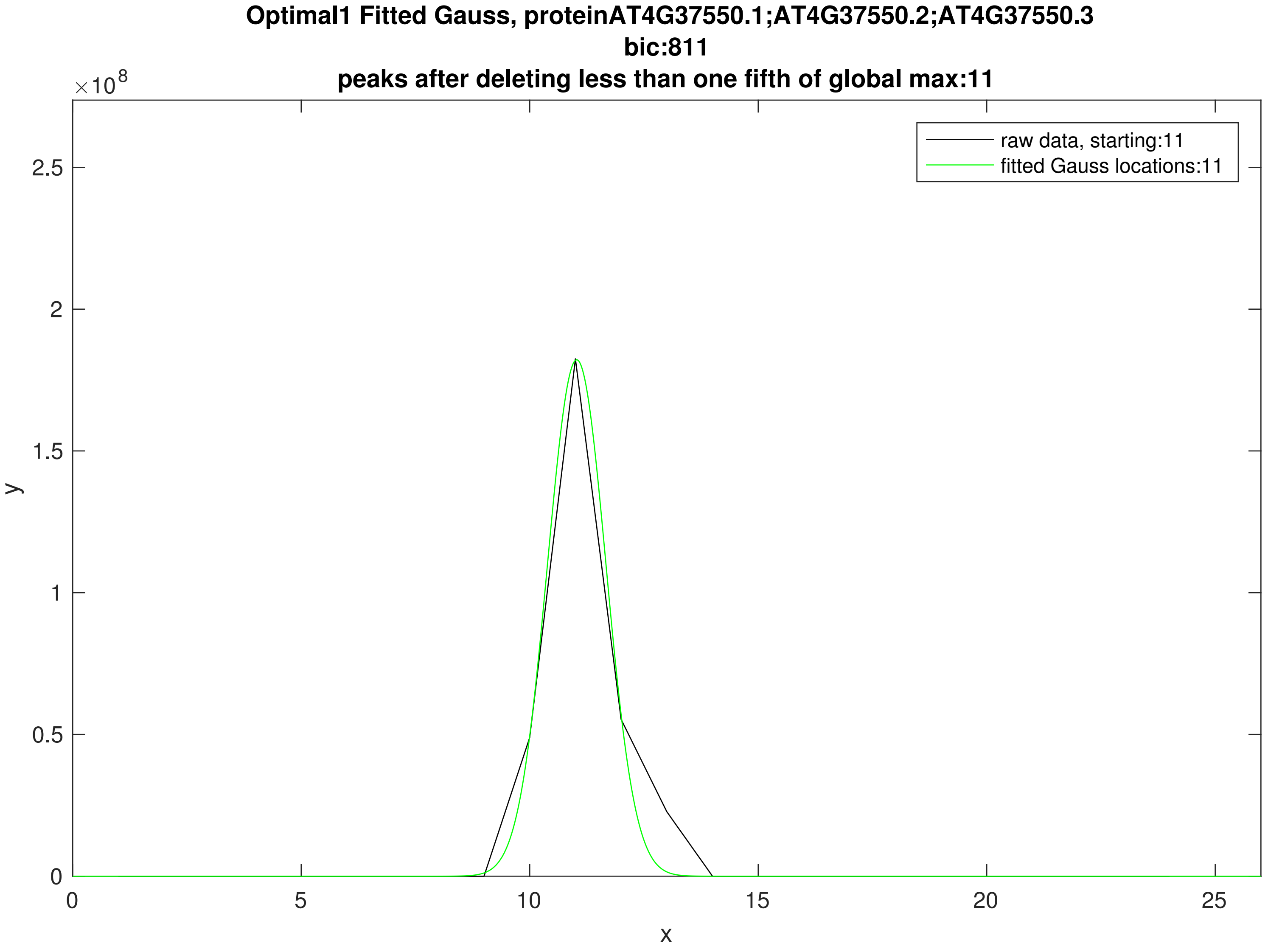
Optimal1 Fitted Gauss, proteinAT4G37550.1;AT4G37550.2;AT4G37550.3
bic:811
peaks after deleting less than one fifth of global max:11
×10^8
2.5
2
1.5
1
0.5
0
y
0
5
10
15
20
25
x
raw data, starting:11
fitted Gauss locations:11

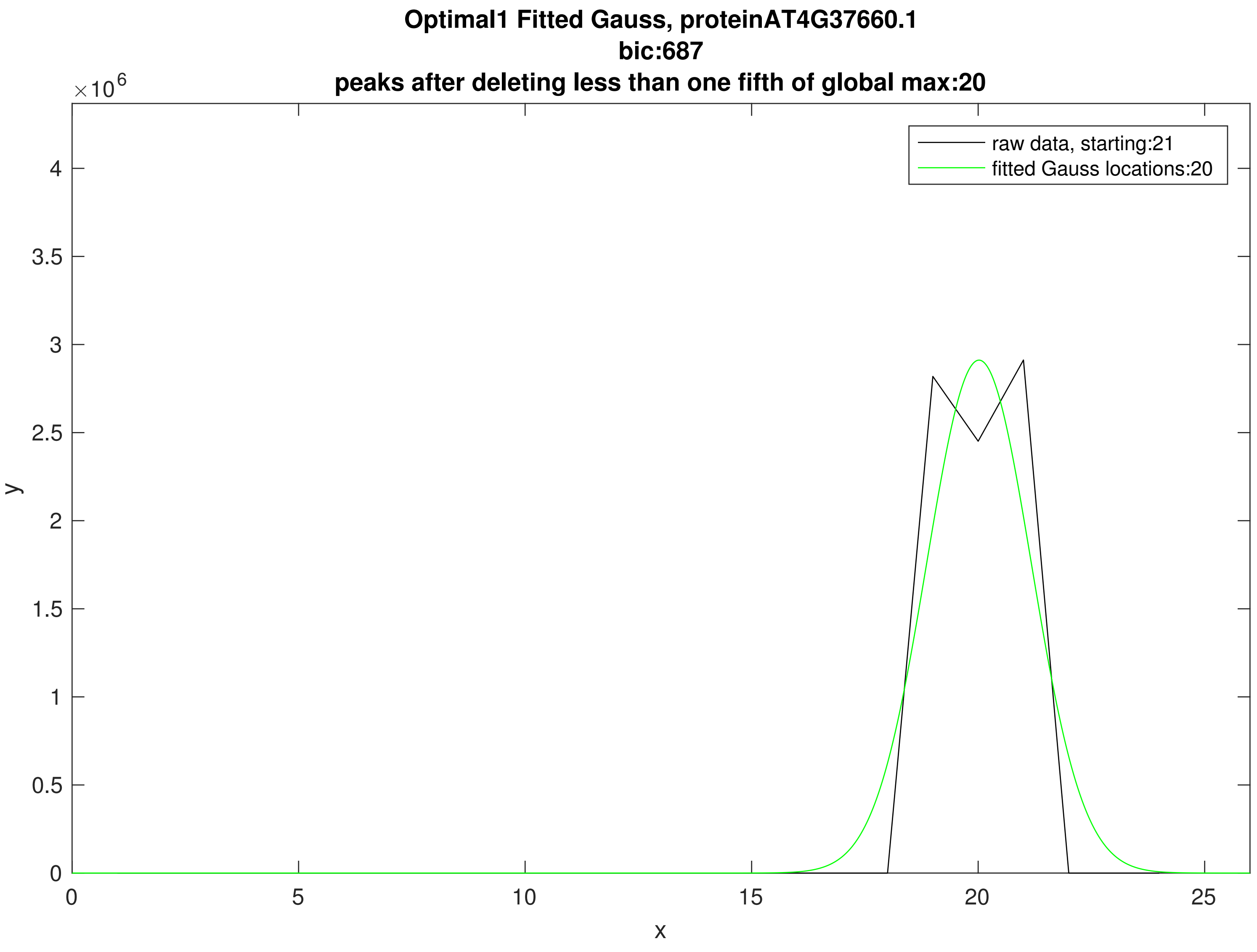

Optimal1 Fitted Gauss, proteinAT4G37660.1
bic:687
peaks after deleting less than one fifth of global max:20
×10^6
raw data, starting:21
fitted Gauss locations:20
4
3.5
3
2.5
2
1.5
1
0.5
0
y
0
5
10
15
20
25
x

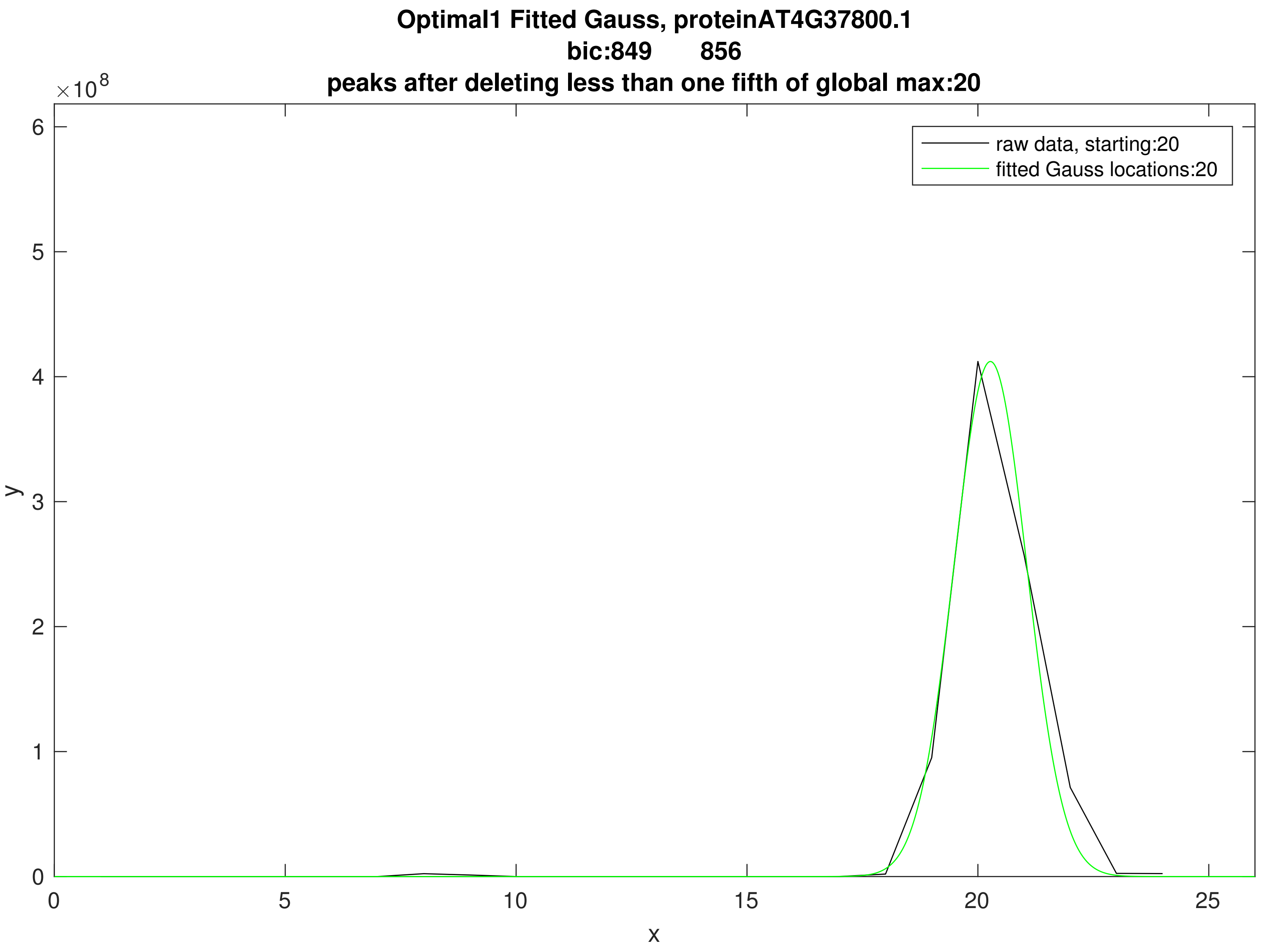
Optimal1 Fitted Gauss, proteinAT4G37800.1
bic:849 856
peaks after deleting less than one fifth of global max:20
×10^8
raw data, starting:20
fitted Gauss locations:20
y
x
0
1
2
3
4
5
6
0
5
10
15
20
25

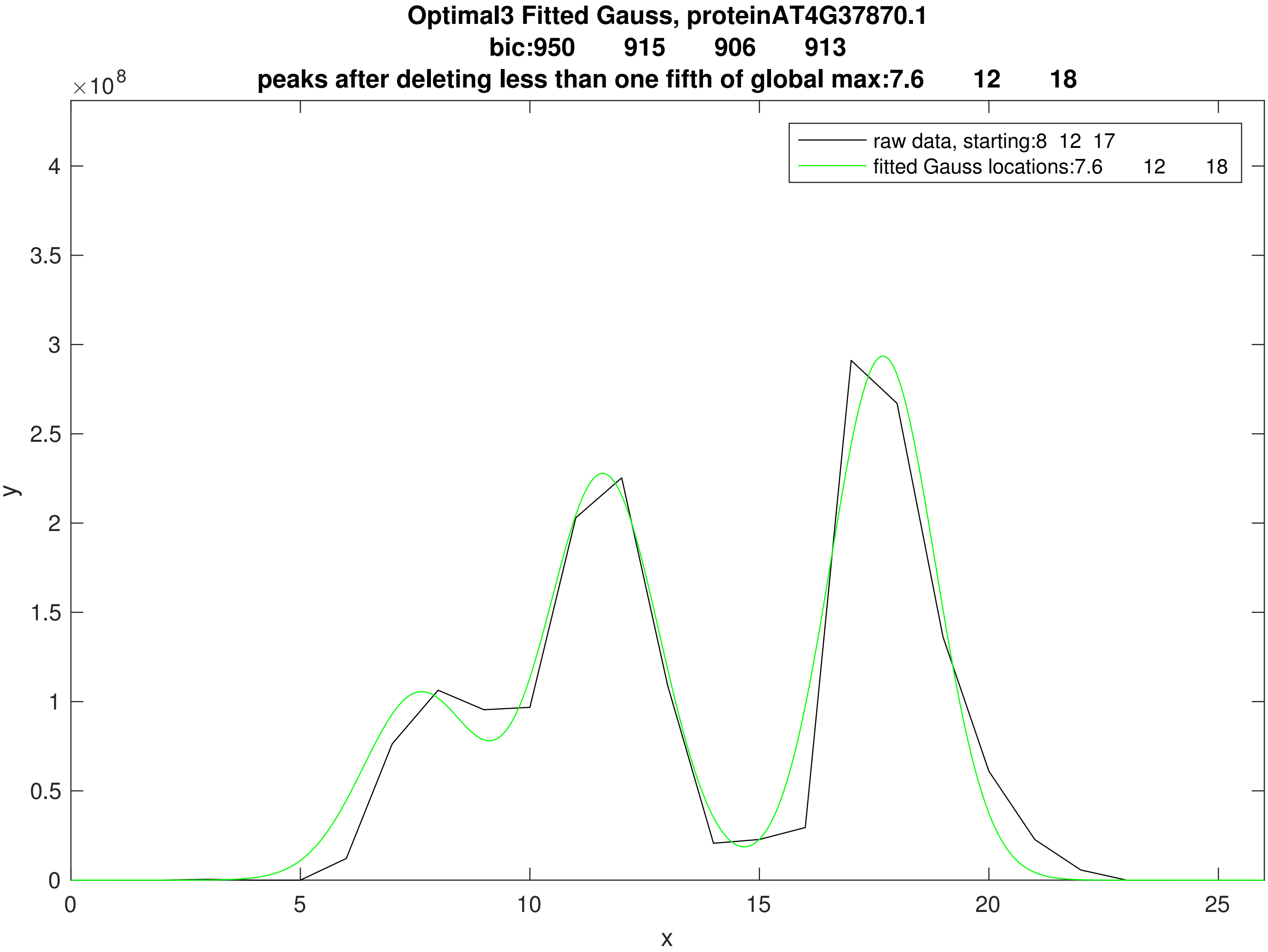
Optimal3 Fitted Gauss, proteinAT4G37870.1
bic:950 915 906 913
peaks after deleting less than one fifth of global max:7.6 12 18
×10^8
raw data, starting:8 12 17
fitted Gauss locations:7.6 12 18
y
x
0
0.5
1
1.5
2
2.5
3
3.5
4
0
5
10
15
20
25

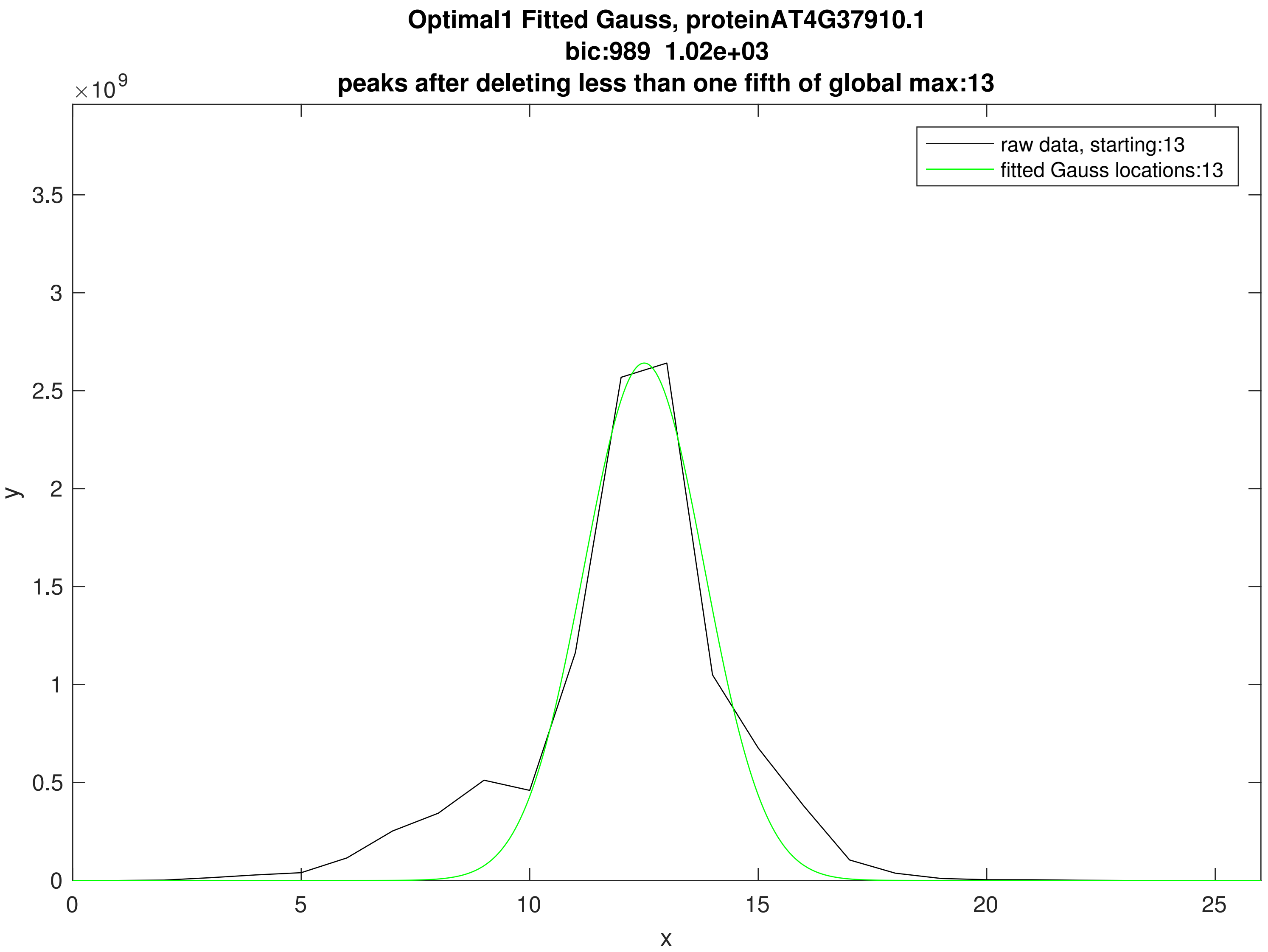
Optimal1 Fitted Gauss, proteinAT4G37910.1
bic:989 1.02e+03
peaks after deleting less than one fifth of global max:13
×10^9
raw data, starting:13
fitted Gauss locations:13
y
x
0
0.5
1
1.5
2
2.5
3
3.5
0
5
10
15
20
25

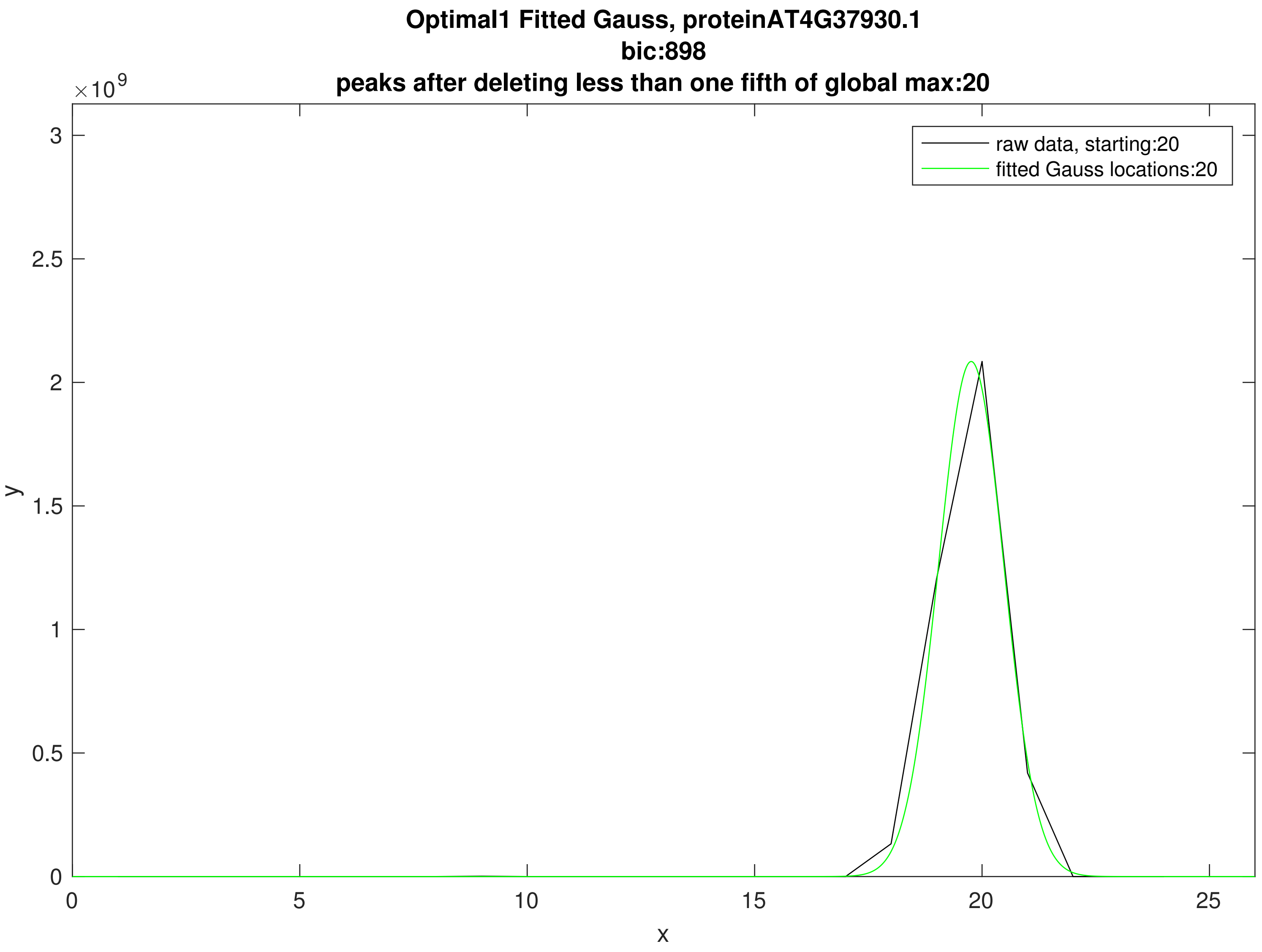

Optimal1 Fitted Gauss, proteinAT4G37930.1
bic:898
peaks after deleting less than one fifth of global max:20
×10^9
3
2.5
2
1.5
1
0.5
0
y
0
5
10
15
20
25
x
raw data, starting:20
fitted Gauss locations:20

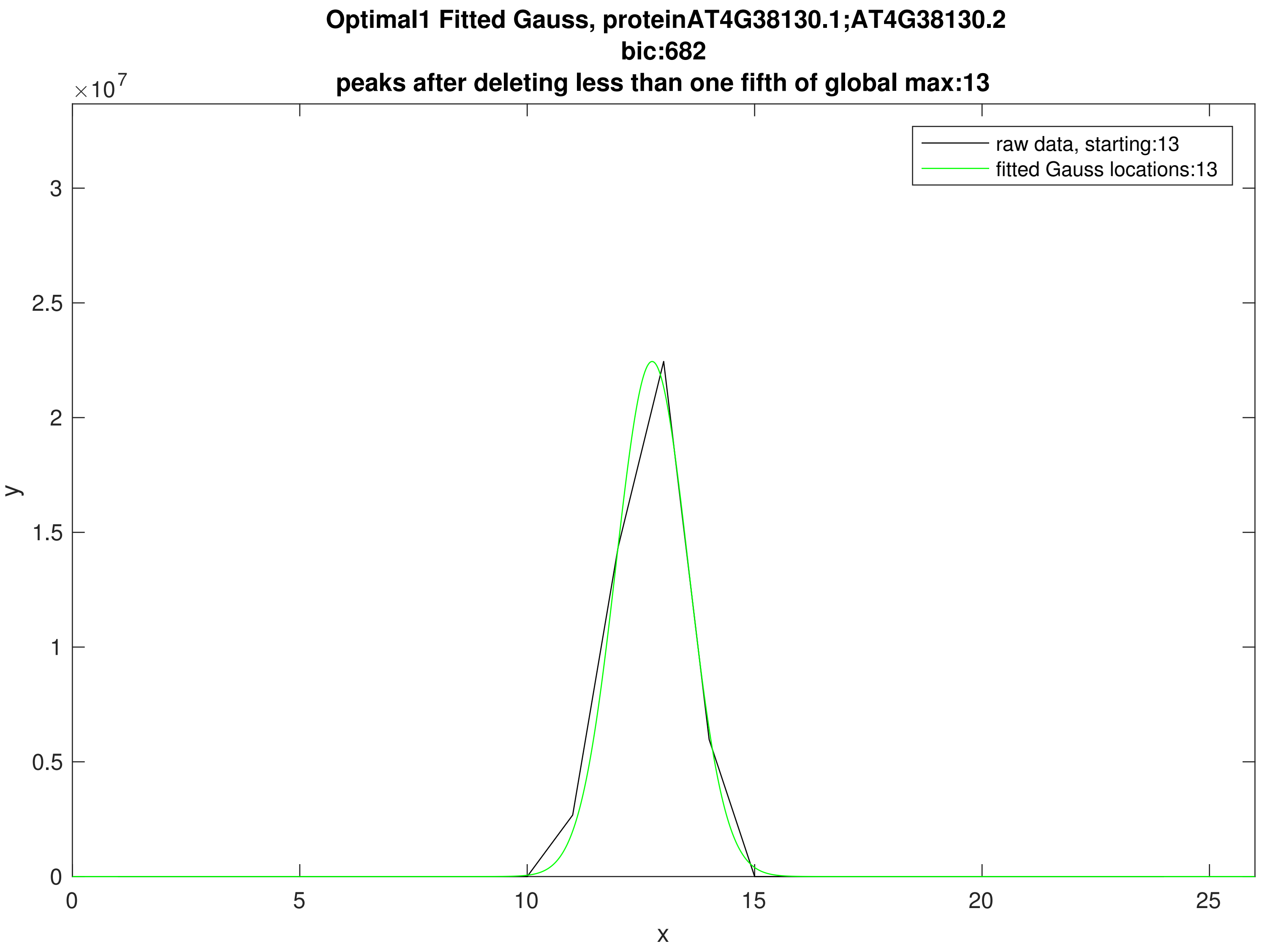

Optimal1 Fitted Gauss, proteinAT4G38130.1;AT4G38130.2
bic:682
peaks after deleting less than one fifth of global max:13
×10^7
3
2.5
2
1.5
1
0.5
0
y
0
5
10
15
20
25
x
raw data, starting:13
fitted Gauss locations:13

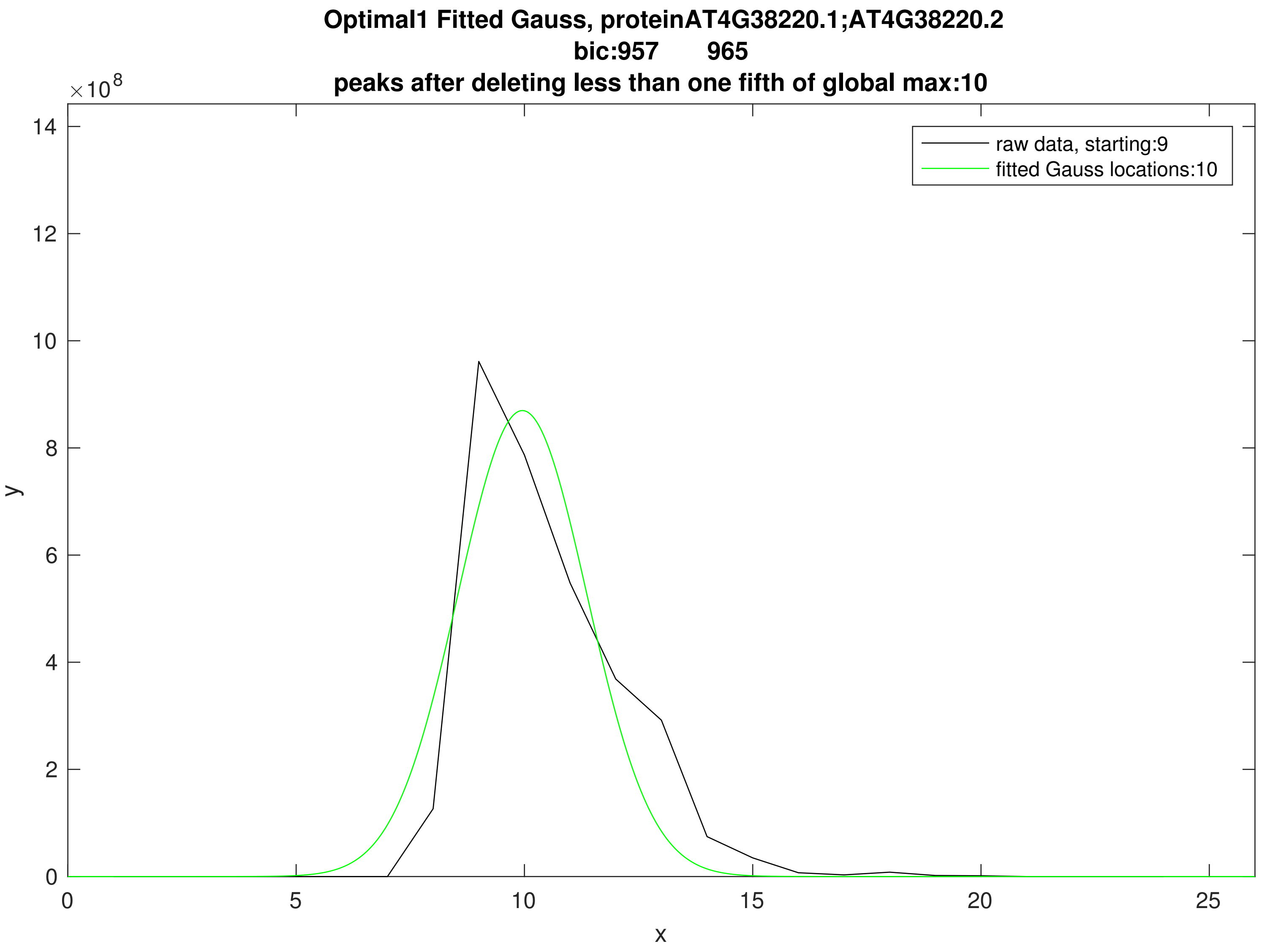

Optimal1 Fitted Gauss, proteinAT4G38220.1;AT4G38220.2
bic:957 965
peaks after deleting less than one fifth of global max:10
×10^8
raw data, starting:9
fitted Gauss locations:10
y
x
0
2
4
6
8
10
12
14
0
5
10
15
20
25

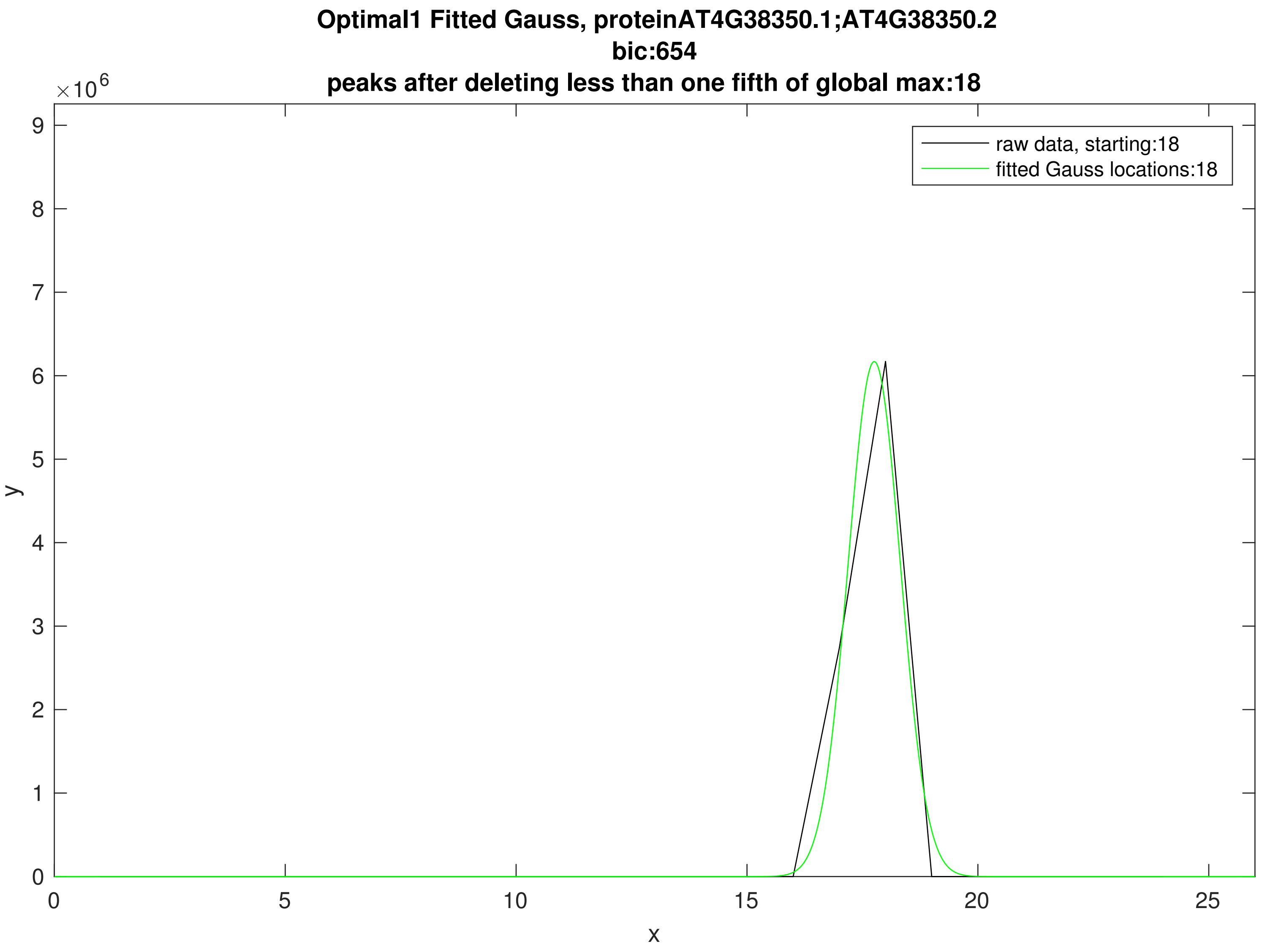
Optimal1 Fitted Gauss, proteinAT4G38350.1;AT4G38350.2
bic:654
peaks after deleting less than one fifth of global max:18
×10^6
9
8
7
6
5
4
3
2
1
0
y
0
5
10
15
20
25
x
raw data, starting:18
fitted Gauss locations:18

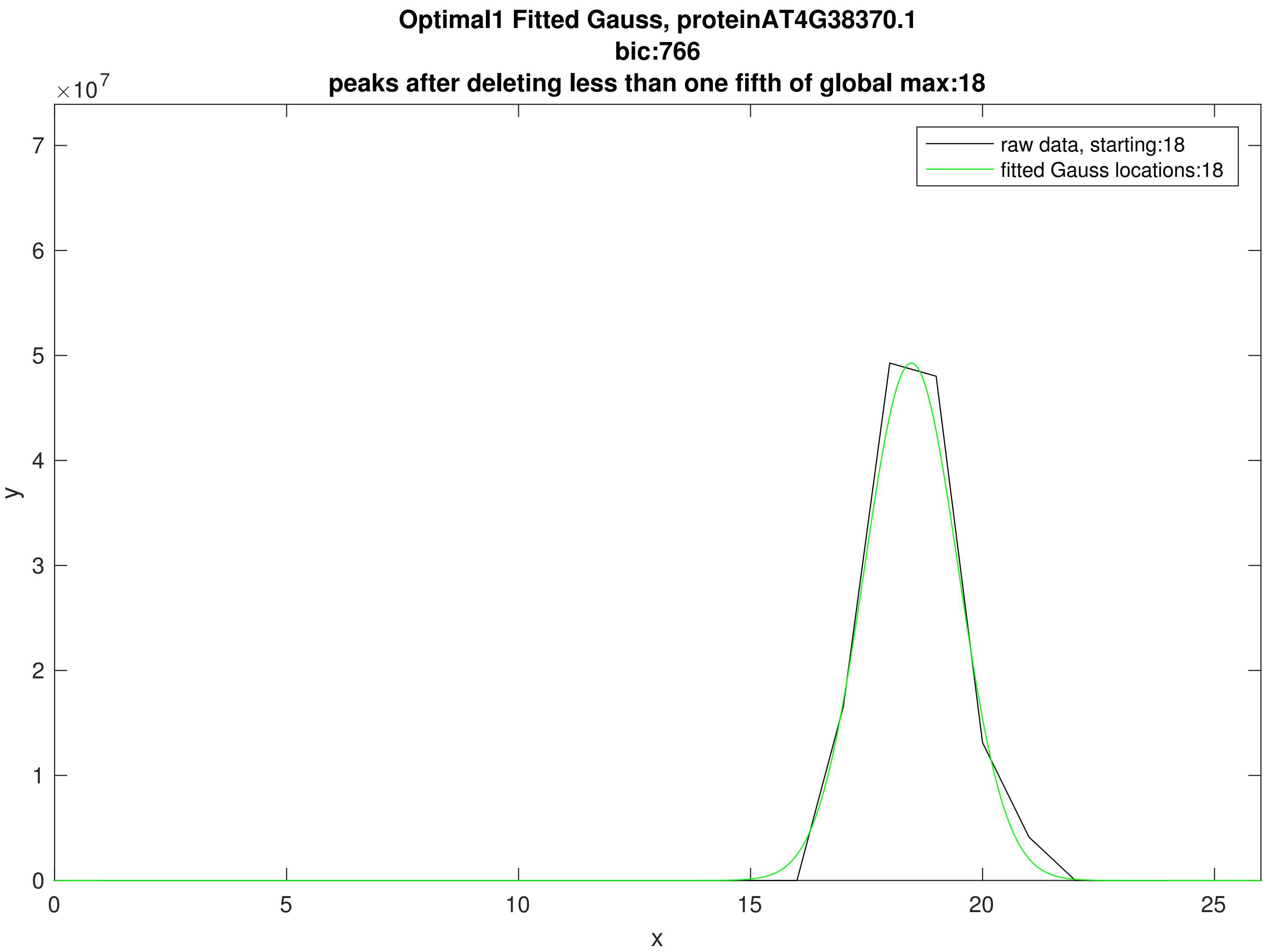

Optimal1 Fitted Gauss, proteinAT4G38370.1
bic:766
peaks after deleting less than one fifth of global max:18
×10^7
7
6
5
4
3
2
1
0
y
0
5
10
15
20
25
x
raw data, starting:18
fitted Gauss locations:18

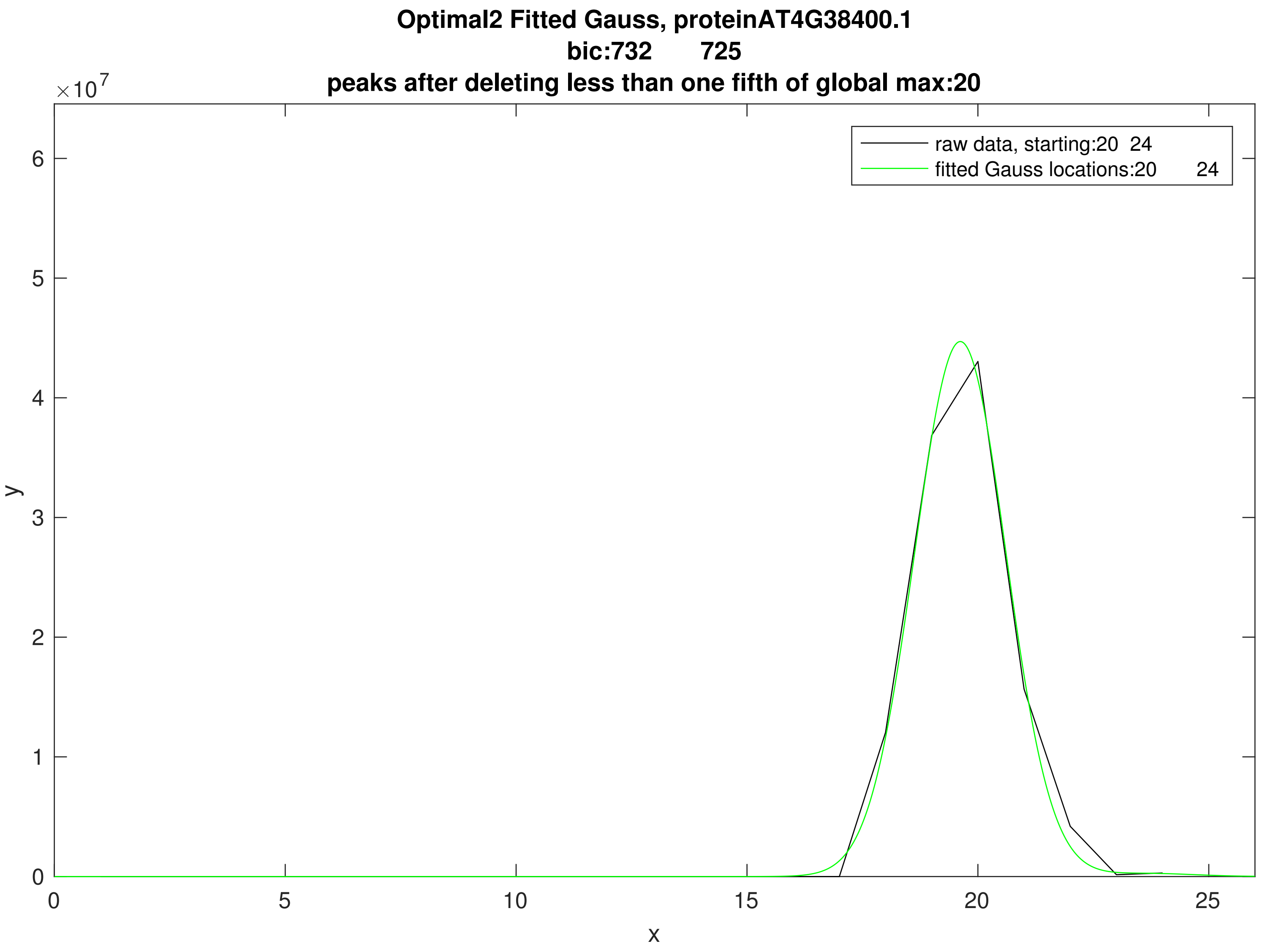
Optimal2 Fitted Gauss, proteinAT4G38400.1
bic:732 725
peaks after deleting less than one fifth of global max:20
×10^7
raw data, starting:20 24
fitted Gauss locations:20 24
y
x
0
1
2
3
4
5
6
0
5
10
15
20
25

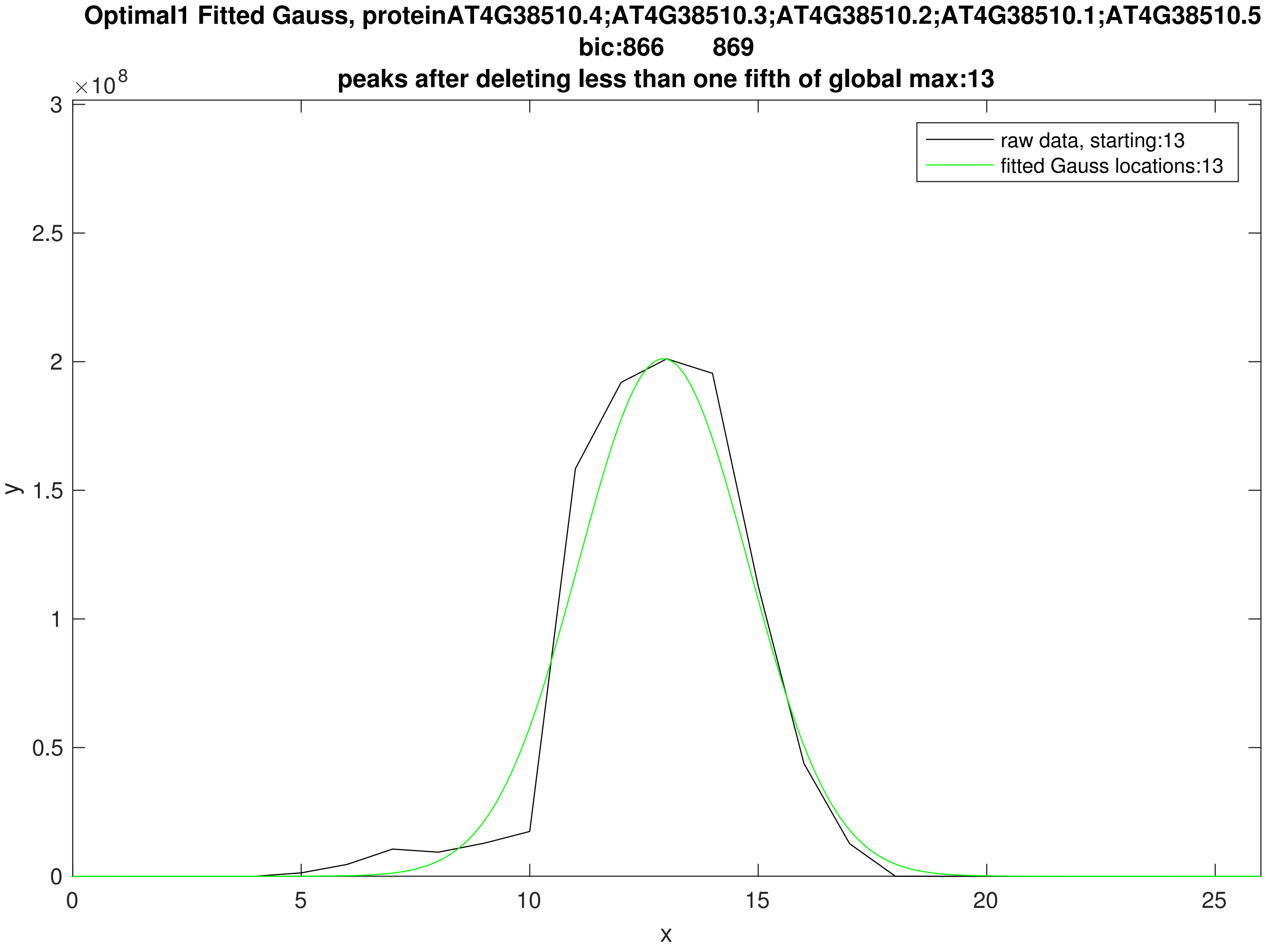
Optimal1 Fitted Gauss, proteinAT4G38510.4;AT4G38510.3;AT4G38510.2;AT4G38510.1;AT4G38510.5
bic:866 869
peaks after deleting less than one fifth of global max:13
×10^8
3
2.5
2
1.5
1
0.5
0
y
raw data, starting:13
fitted Gauss locations:13
0
5
10
15
20
25
x

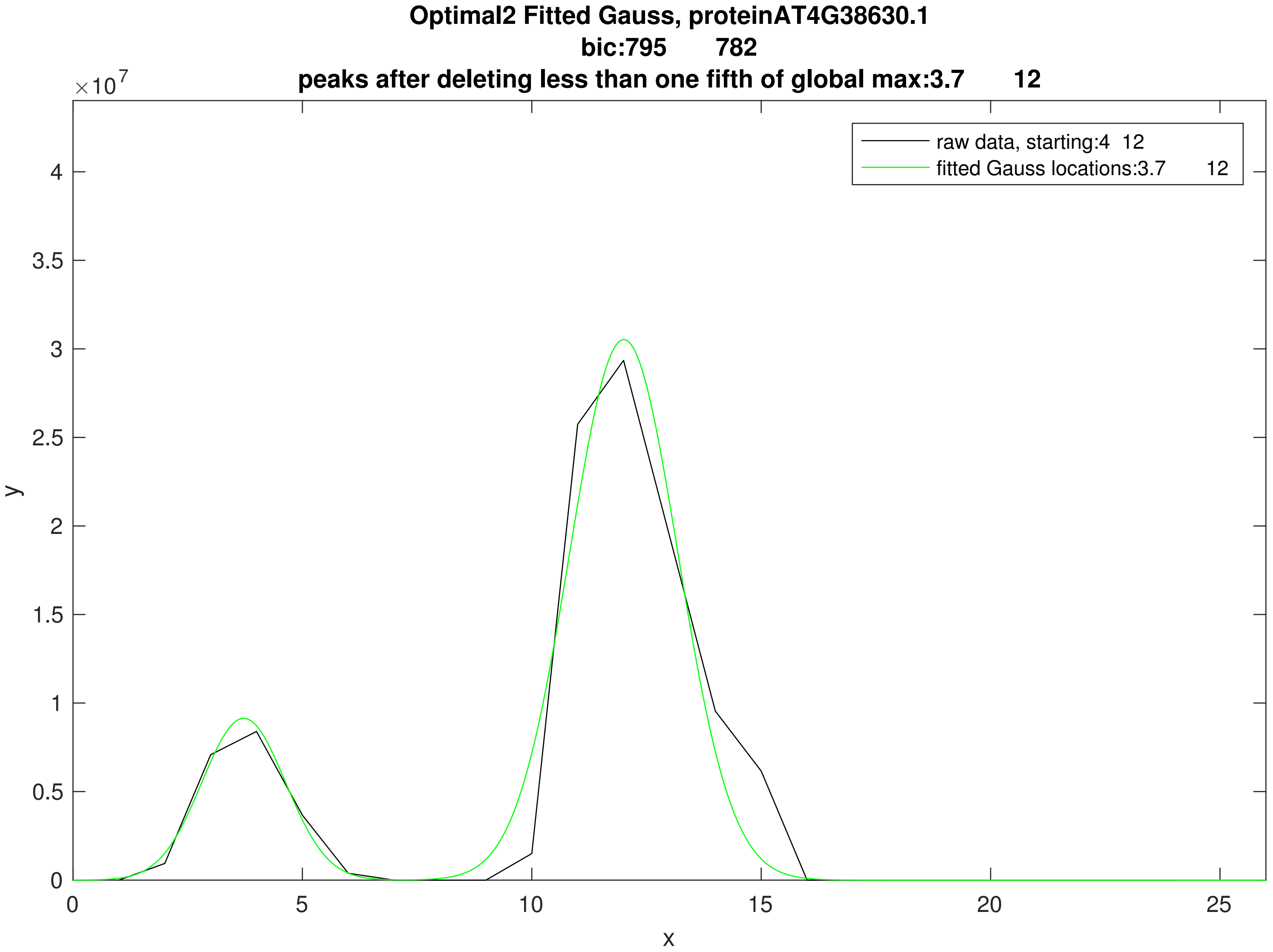
Optimal2 Fitted Gauss, proteinAT4G38630.1
bic:795 782
peaks after deleting less than one fifth of global max:3.7 12
×10^7
raw data, starting:4 12
fitted Gauss locations:3.7 12
y
x
0
0.5
1
1.5
2
2.5
3
3.5
4
5
10
15
20
25

Optimal1 Fitted Gauss, proteinAT4G38680.1
bic:880 883 891
peaks after deleting less than one fifth of global max:19

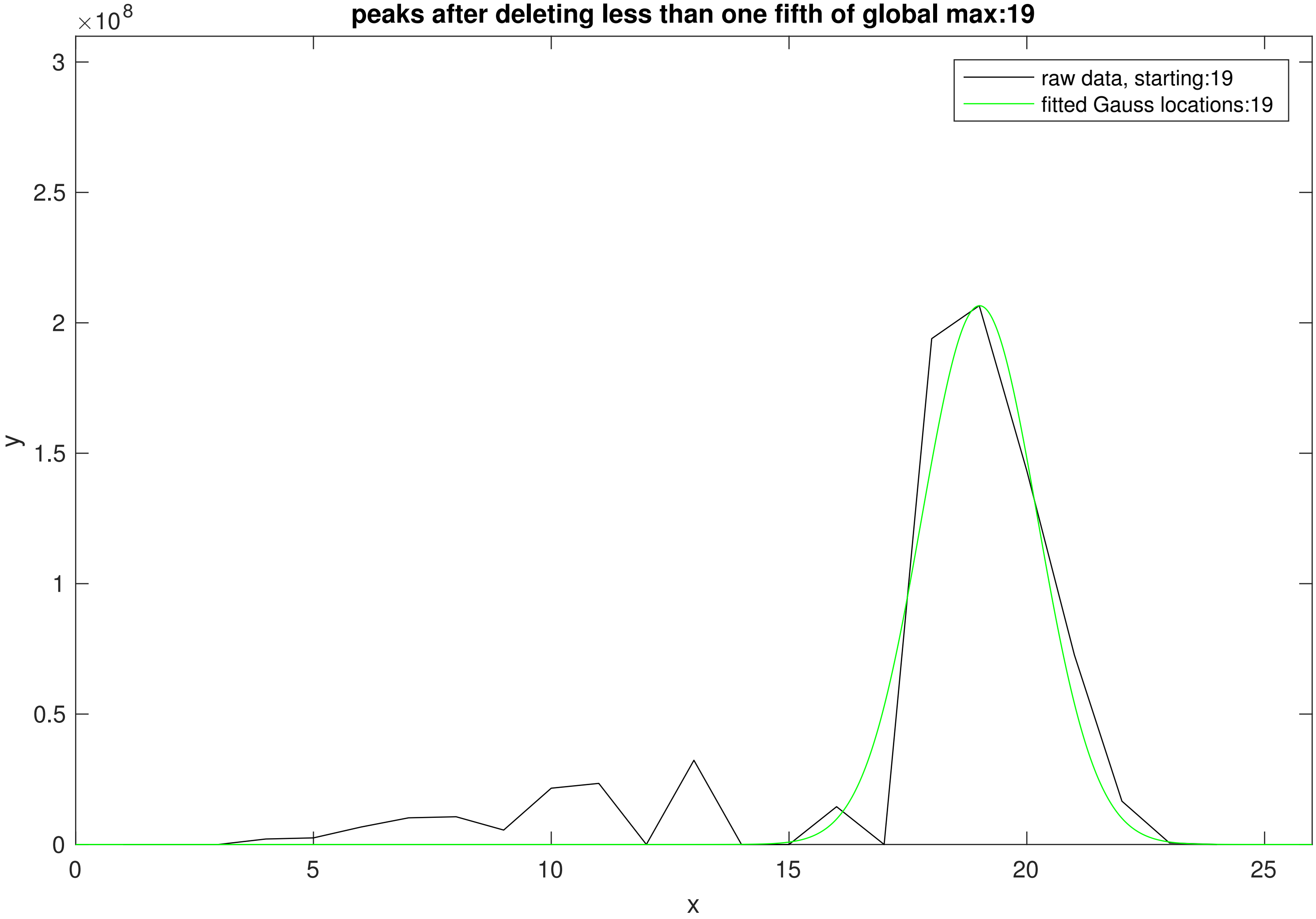

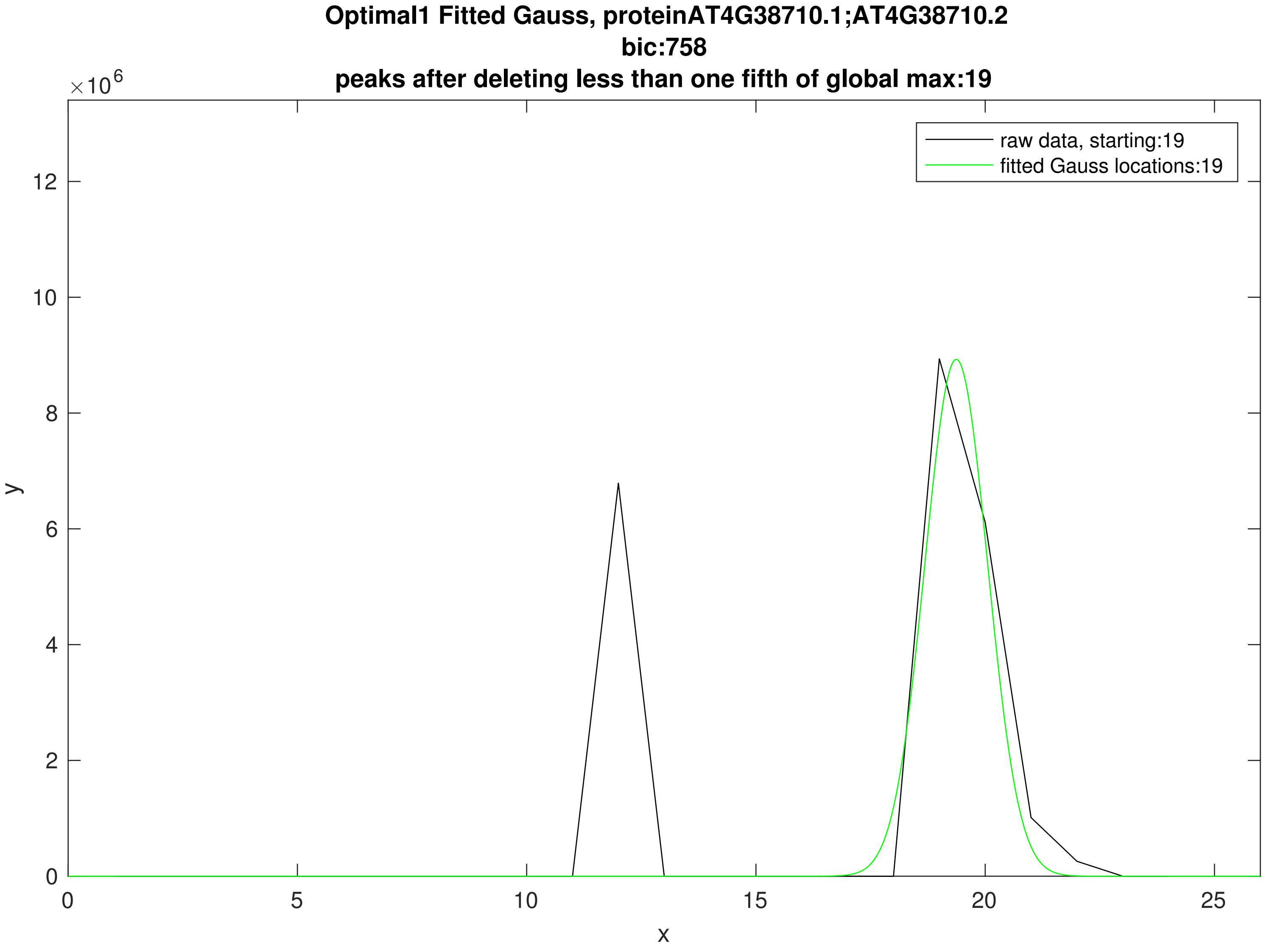

Optimal1 Fitted Gauss, proteinAT4G38710.1;AT4G38710.2
bic:758
peaks after deleting less than one fifth of global max:19
×10^6
raw data, starting:19
fitted Gauss locations:19
y
x
0
2
4
6
8
10
12
0
5
10
15
20
25

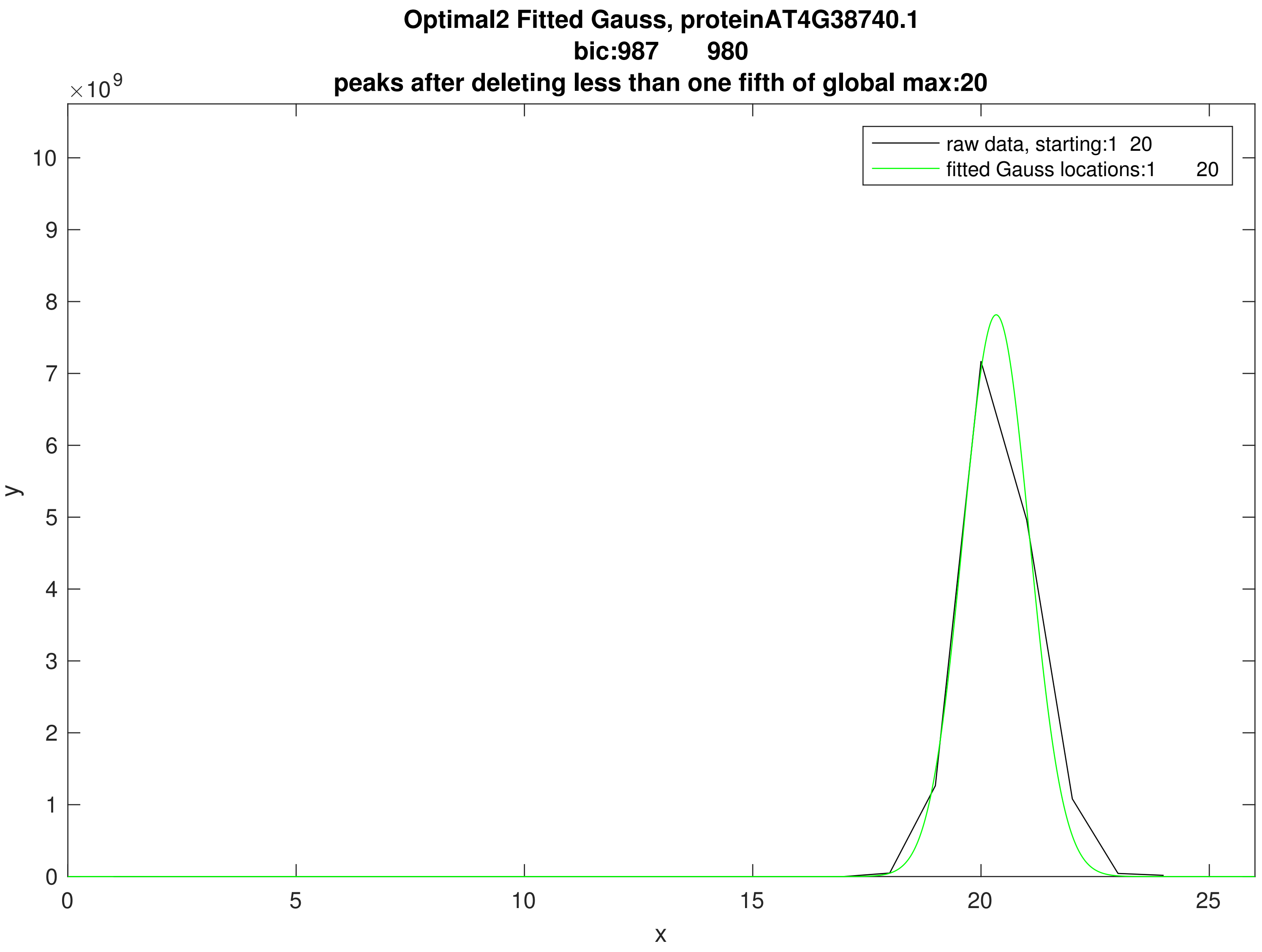

Optimal2 Fitted Gauss, proteinAT4G38740.1
bic:987 980
peaks after deleting less than one fifth of global max:20
×10^9
raw data, starting:1 20
fitted Gauss locations:1 20
y
x
0
1
2
3
4
5
6
7
8
9
10
0
5
10
15
20
25

**Optimal1 Fitted Gauss, proteinAT4G38800.1**
**bic:726**
**peaks after deleting less than one fifth of global max:14**

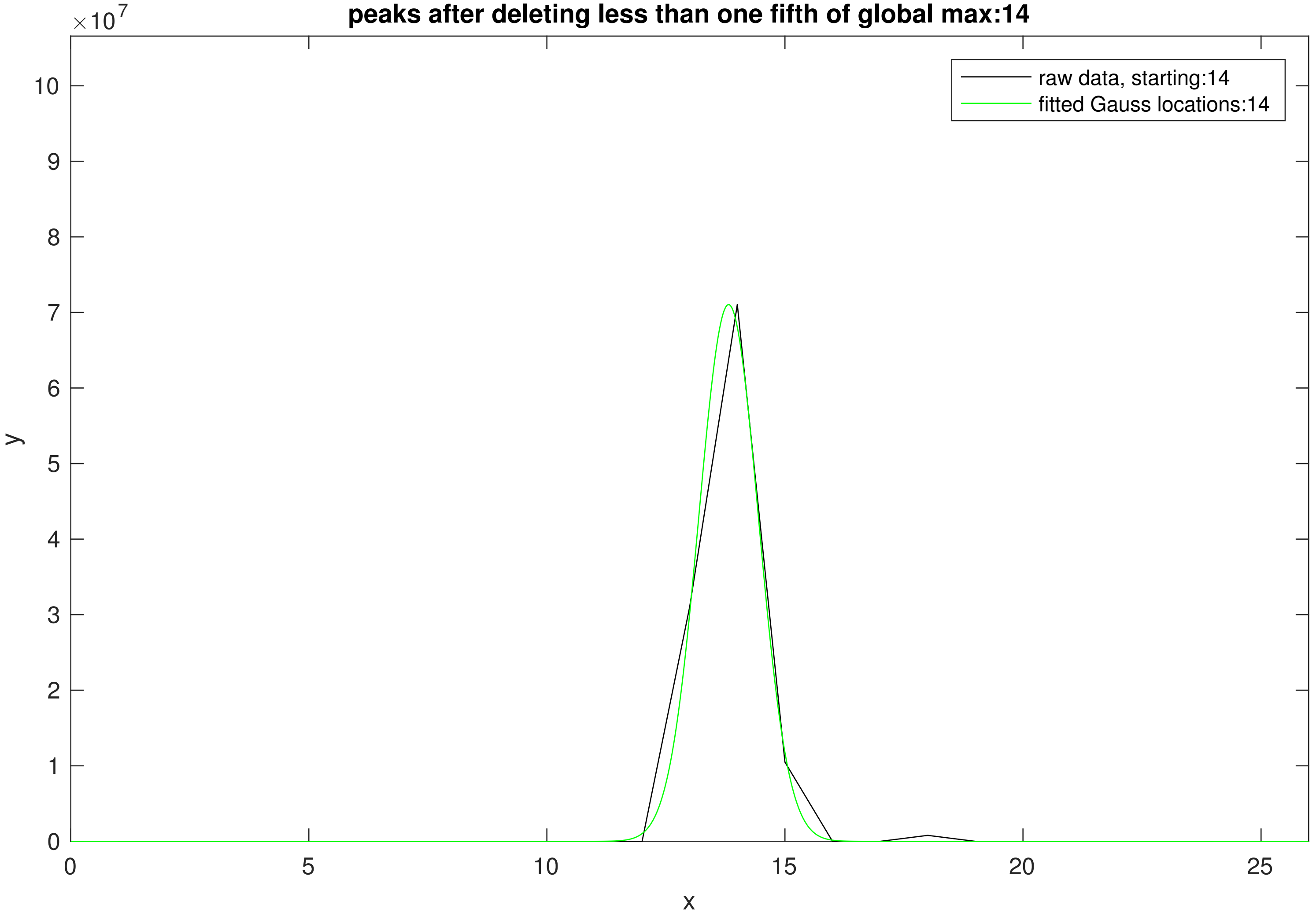

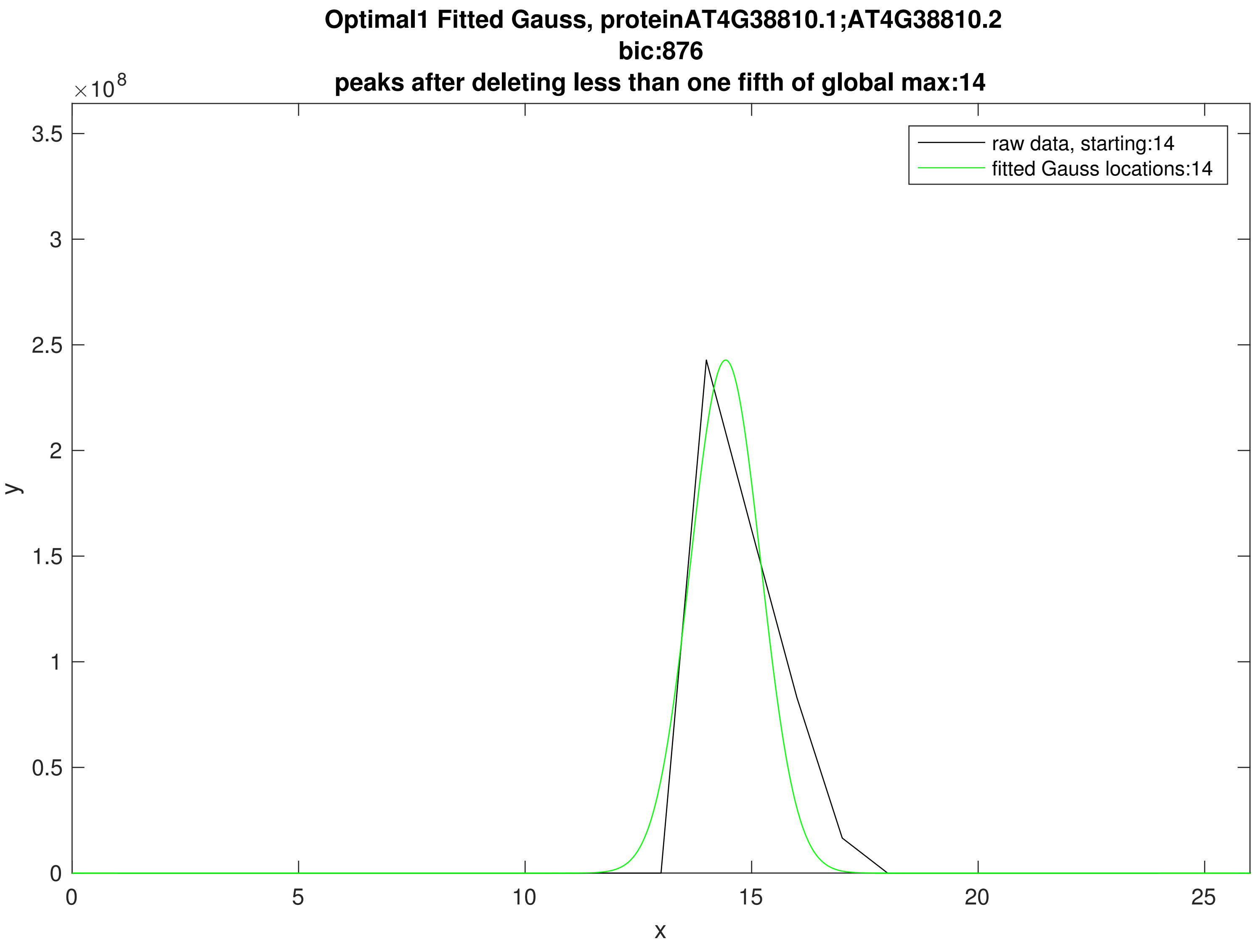
Optimal1 Fitted Gauss, proteinAT4G38810.1;AT4G38810.2
bic:876
peaks after deleting less than one fifth of global max:14
×10^8
3.5
3
2.5
2
1.5
1
0.5
0
y
0
5
10
15
20
25
x
raw data, starting:14
fitted Gauss locations:14

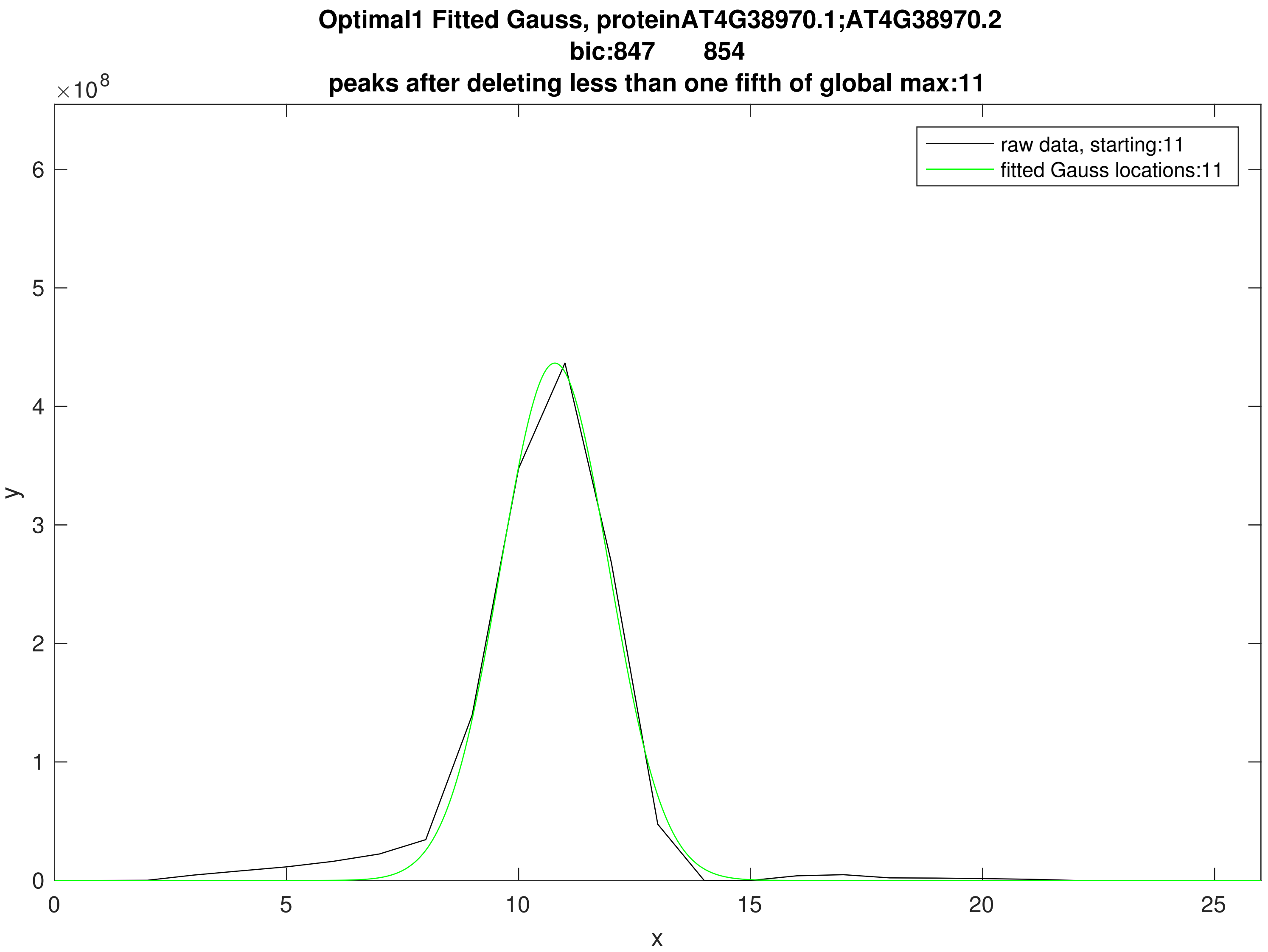

Optimal1 Fitted Gauss, proteinAT4G38970.1;AT4G38970.2
bic:847 854
peaks after deleting less than one fifth of global max:11
×10^8
raw data, starting:11
fitted Gauss locations:11
y
x
0
1
2
3
4
5
6
0
5
10
15
20
25

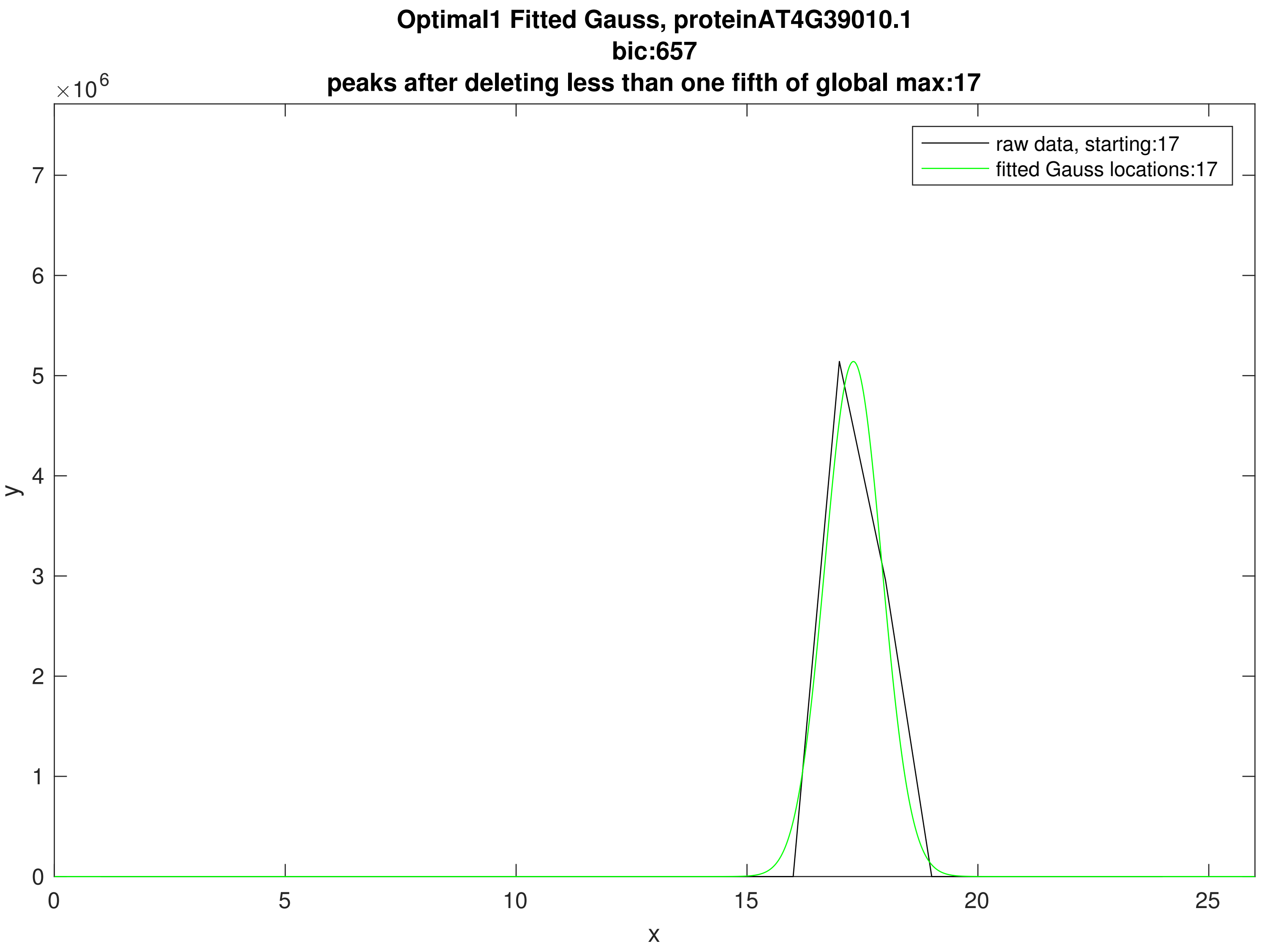
Optimal1 Fitted Gauss, proteinAT4G39010.1
bic:657
peaks after deleting less than one fifth of global max:17
×10^6
raw data, starting:17
fitted Gauss locations:17
y
x
0
1
2
3
4
5
6
7
0
5
10
15
20
25

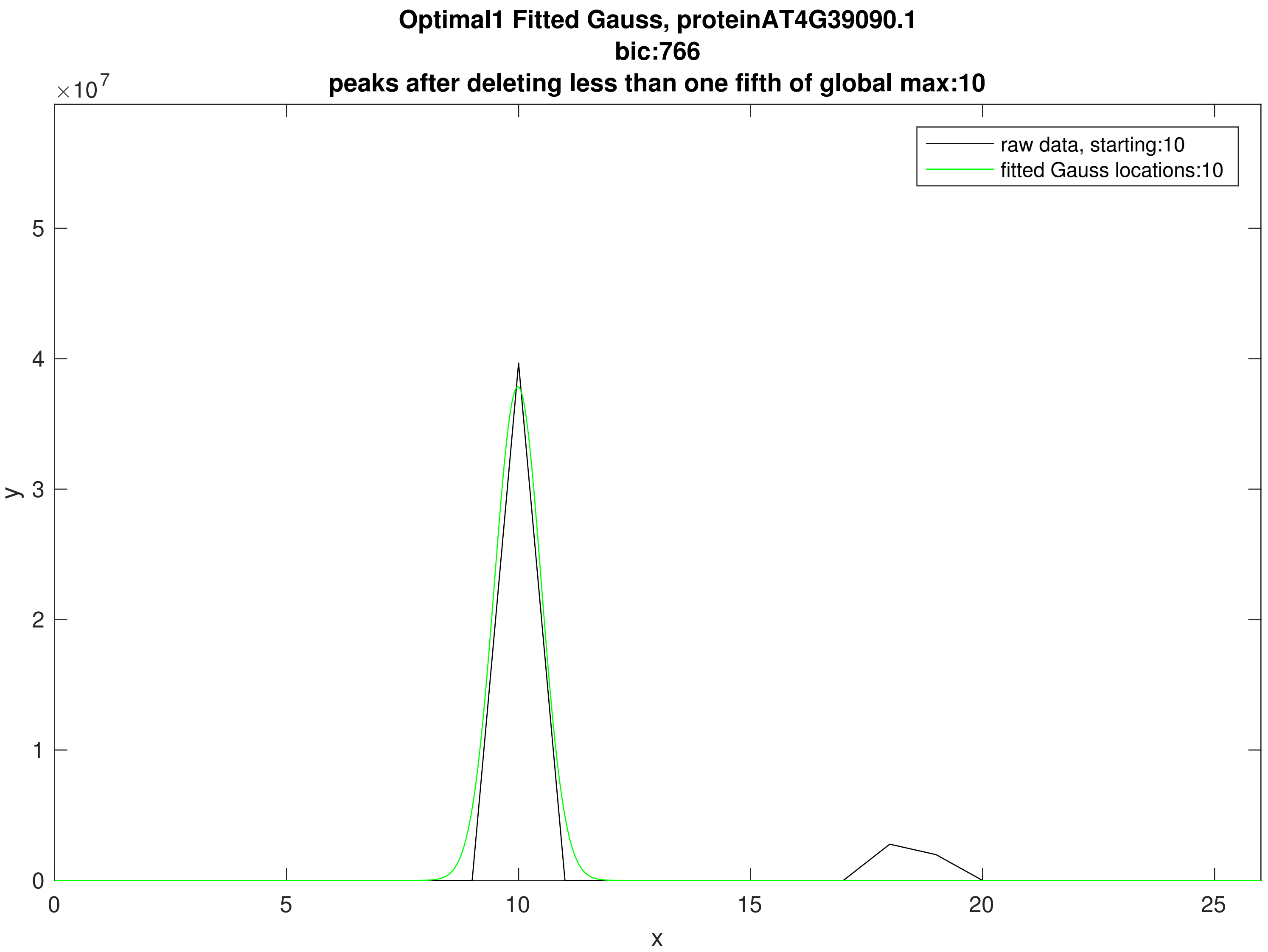
Optimal1 Fitted Gauss, proteinAT4G39090.1
bic:766
peaks after deleting less than one fifth of global max:10
×10^7
raw data, starting:10
fitted Gauss locations:10
y
x
0
1
2
3
4
5
10
15
20
25

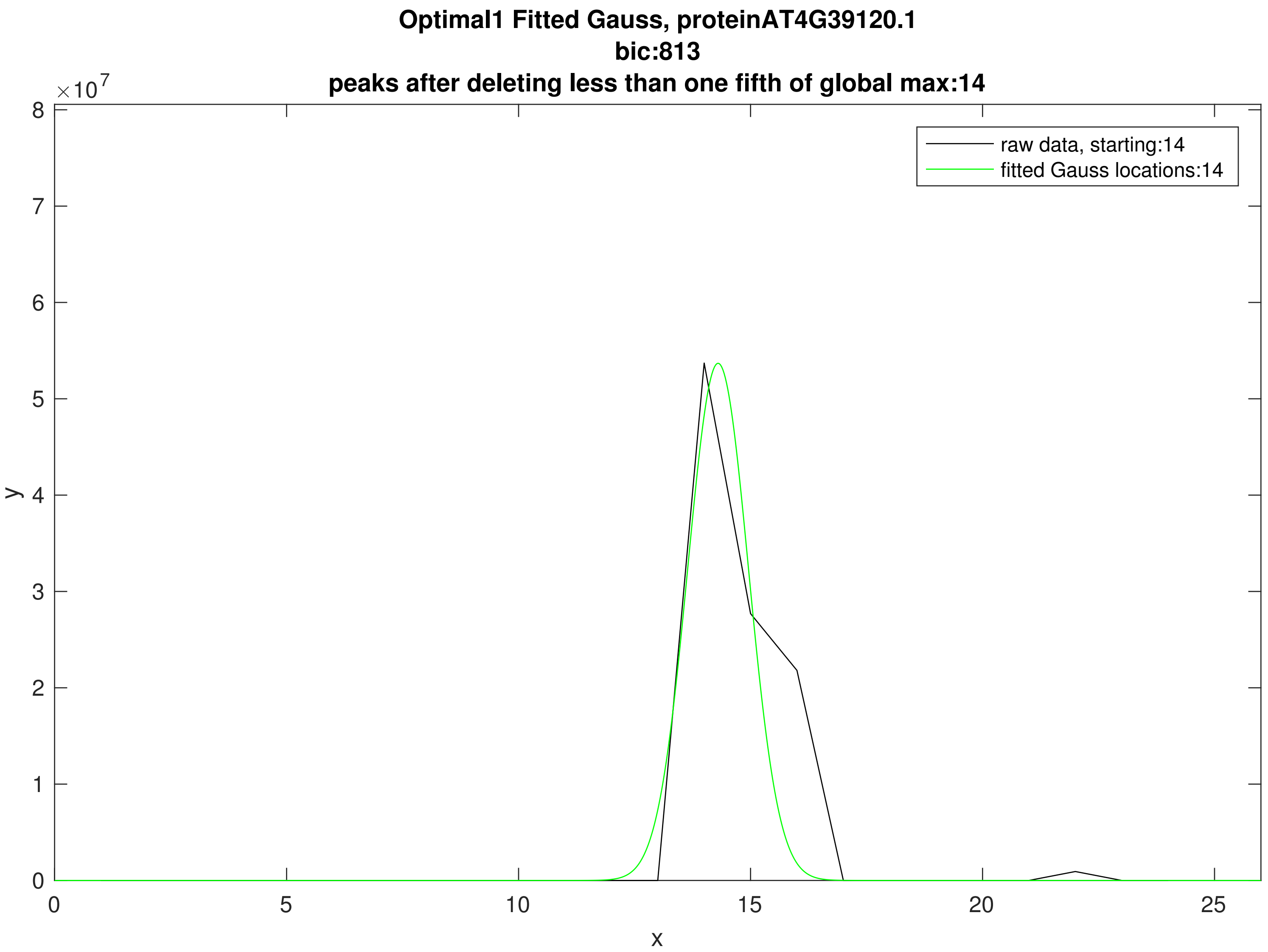

Optimal1 Fitted Gauss, proteinAT4G39120.1
bic:813
peaks after deleting less than one fifth of global max:14
×10^7
raw data, starting:14
fitted Gauss locations:14
y
x
0
1
2
3
4
5
6
7
8
0
5
10
15
20
25

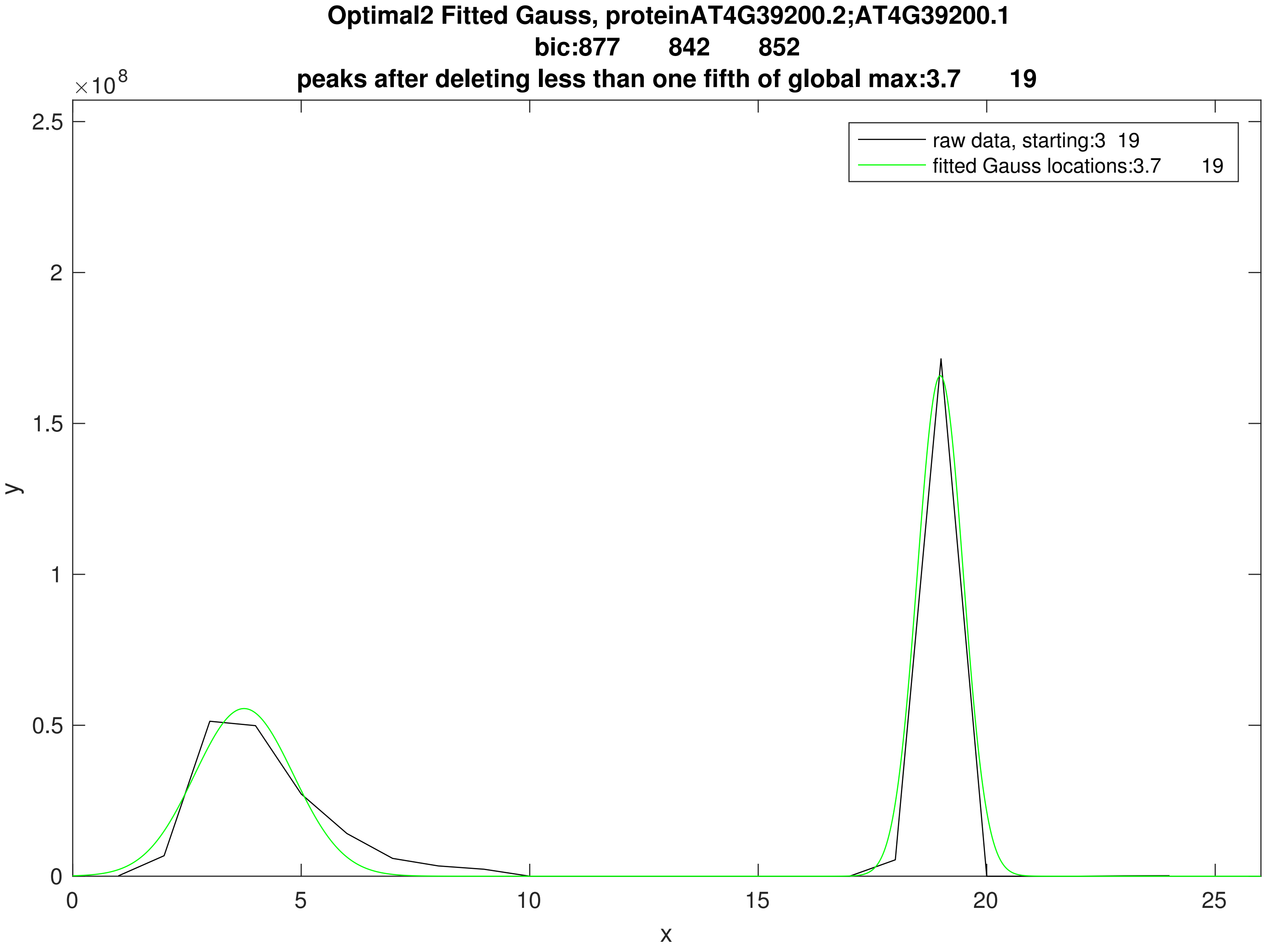
Optimal2 Fitted Gauss, proteinAT4G39200.2;AT4G39200.1
bic:877 842 852
peaks after deleting less than one fifth of global max:3.7 19
×10^8
2.5
2
1.5
1
0.5
0
y
0
5
10
15
20
25
x
raw data, starting:3 19
fitted Gauss locations:3.7 19

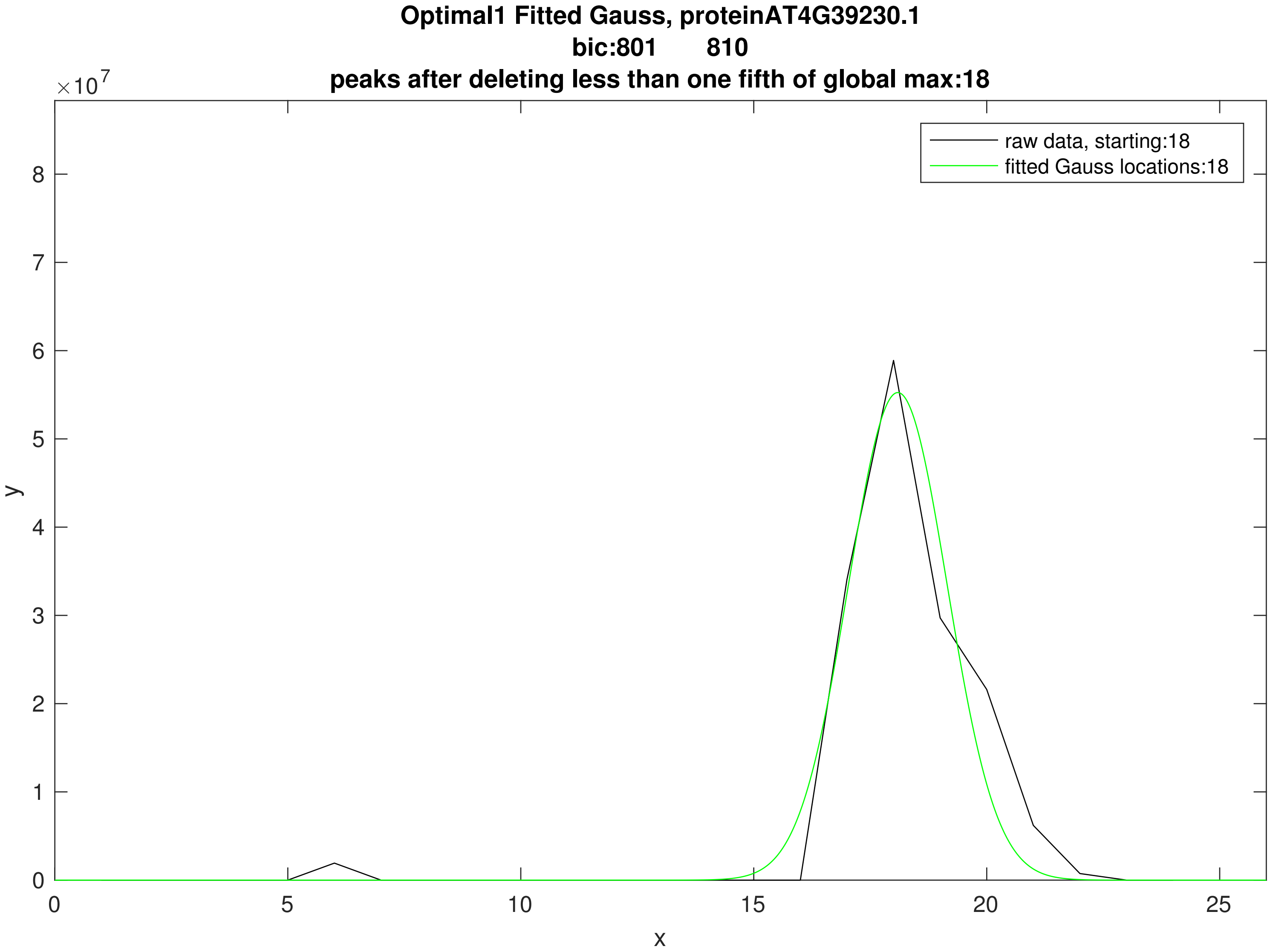
Optimal1 Fitted Gauss, proteinAT4G39230.1
bic:801 810
peaks after deleting less than one fifth of global max:18
×10^7
raw data, starting:18
fitted Gauss locations:18
y
x
0
1
2
3
4
5
6
7
8
0
5
10
15
20
25

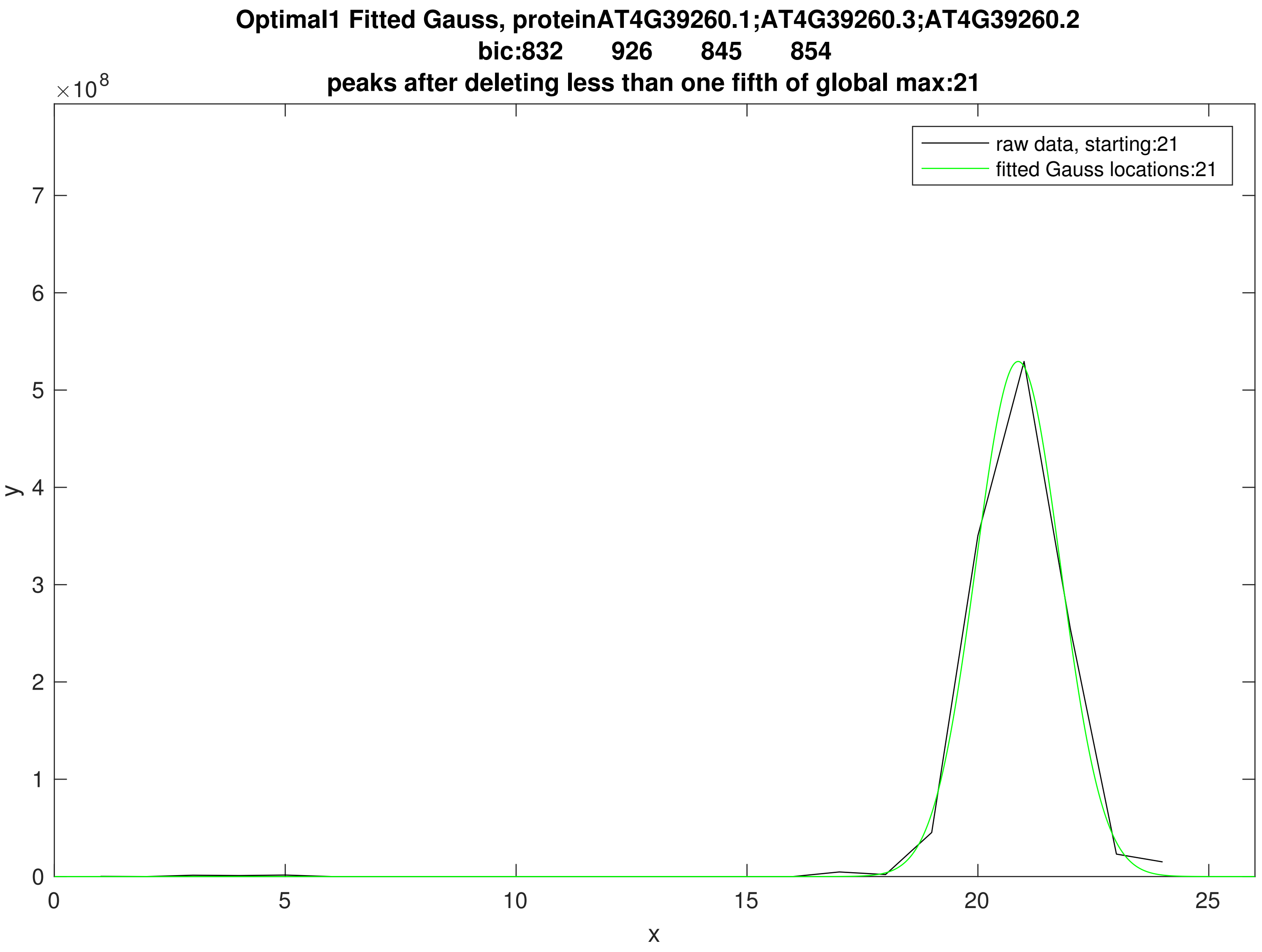

Optimal1 Fitted Gauss, proteinAT4G39260.1;AT4G39260.3;AT4G39260.2
bic:832 926 845 854
peaks after deleting less than one fifth of global max:21
×10^8
raw data, starting:21
fitted Gauss locations:21
y
x
0
1
2
3
4
5
6
7
0
5
10
15
20
25

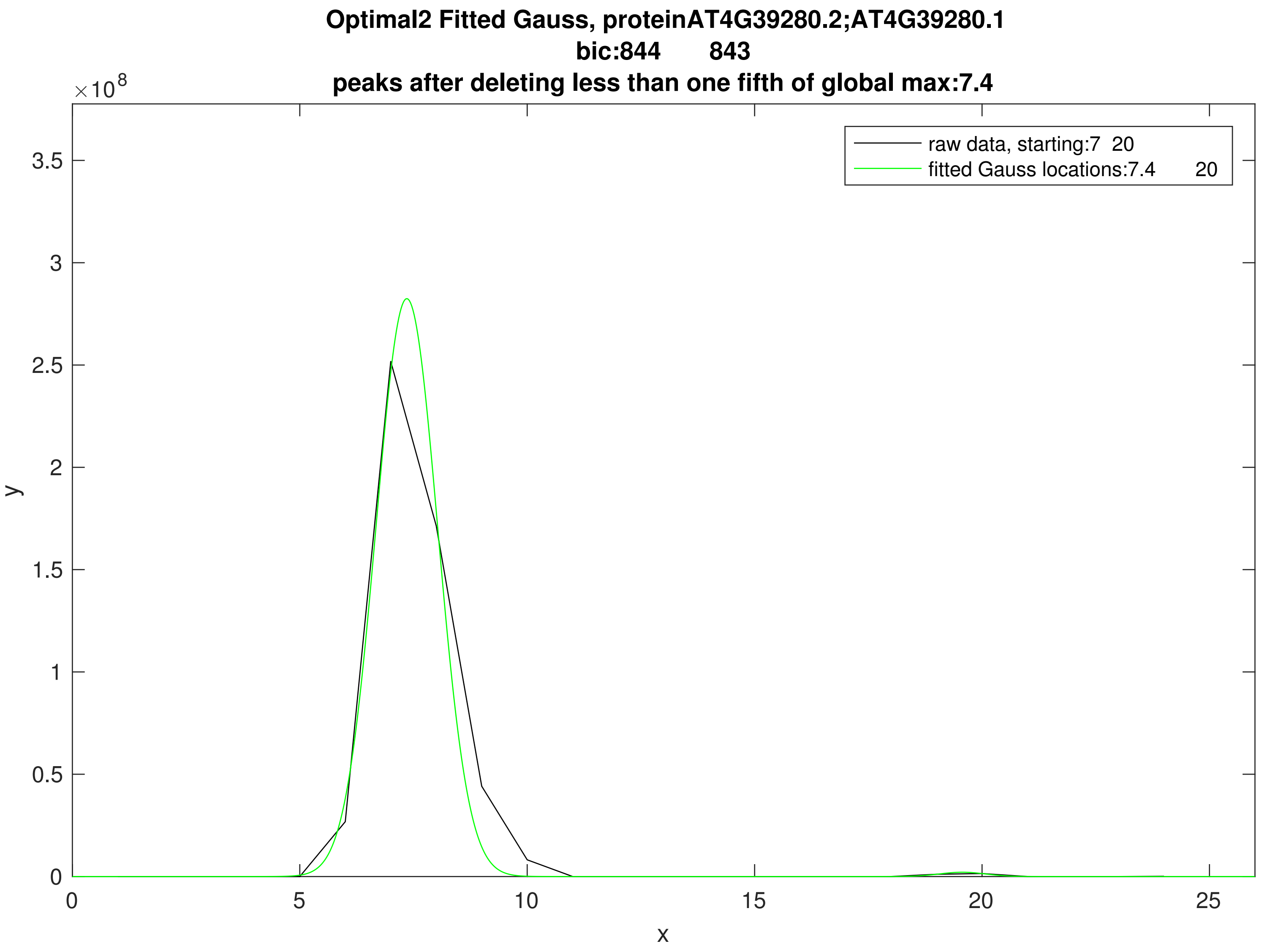

Optimal2 Fitted Gauss, proteinAT4G39280.2;AT4G39280.1
bic:844 843
peaks after deleting less than one fifth of global max:7.4
×10^8
raw data, starting:7 20
fitted Gauss locations:7.4 20
y
x
0
0.5
1
1.5
2
2.5
3
3.5
0
5
10
15
20
25

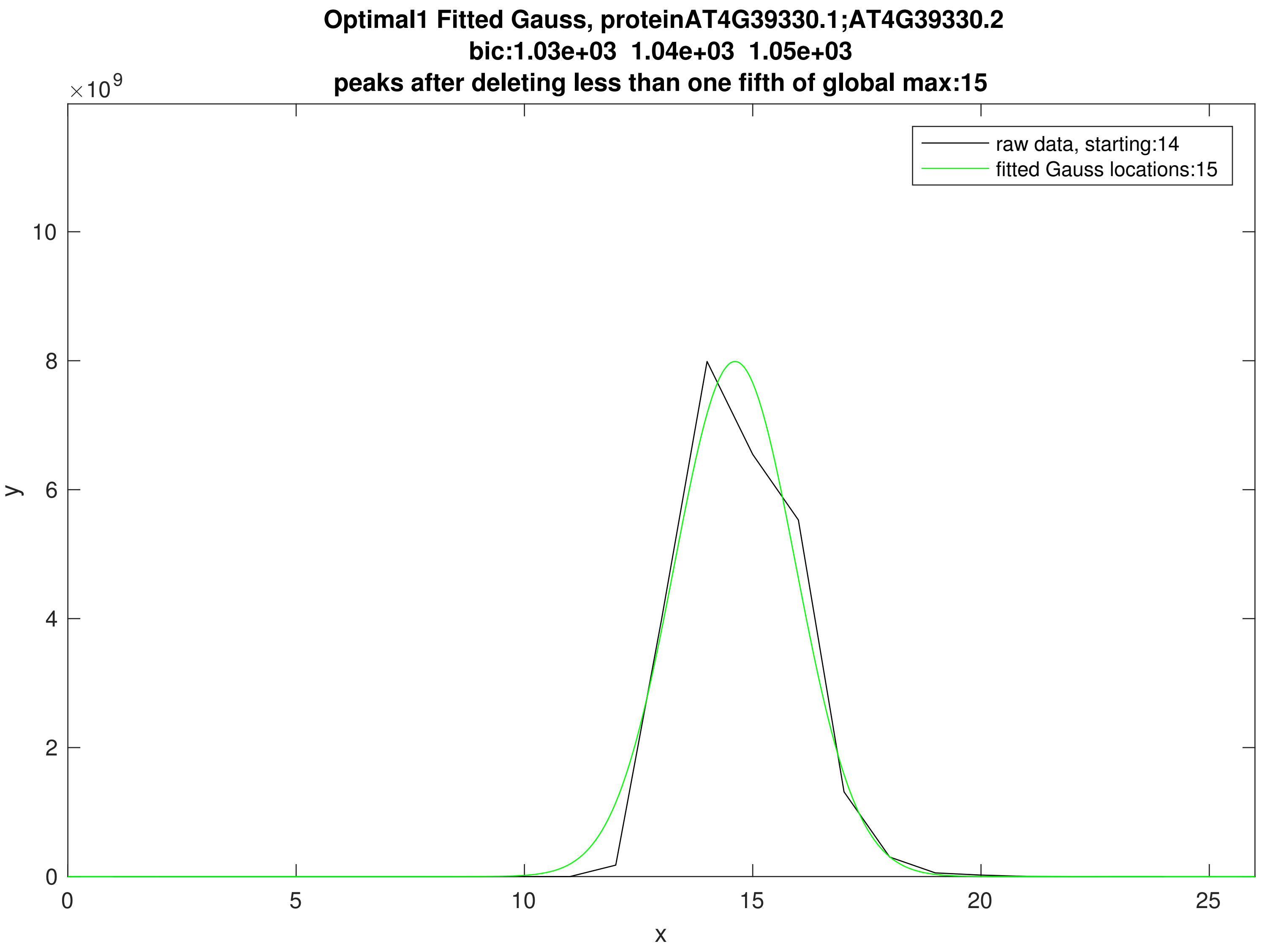

Optimal1 Fitted Gauss, proteinAT4G39330.1;AT4G39330.2
bic:1.03e+03 1.04e+03 1.05e+03
peaks after deleting less than one fifth of global max:15
×10^9
raw data, starting:14
fitted Gauss locations:15
10
8
6
4
2
0
y
0
5
10
15
20
25
x

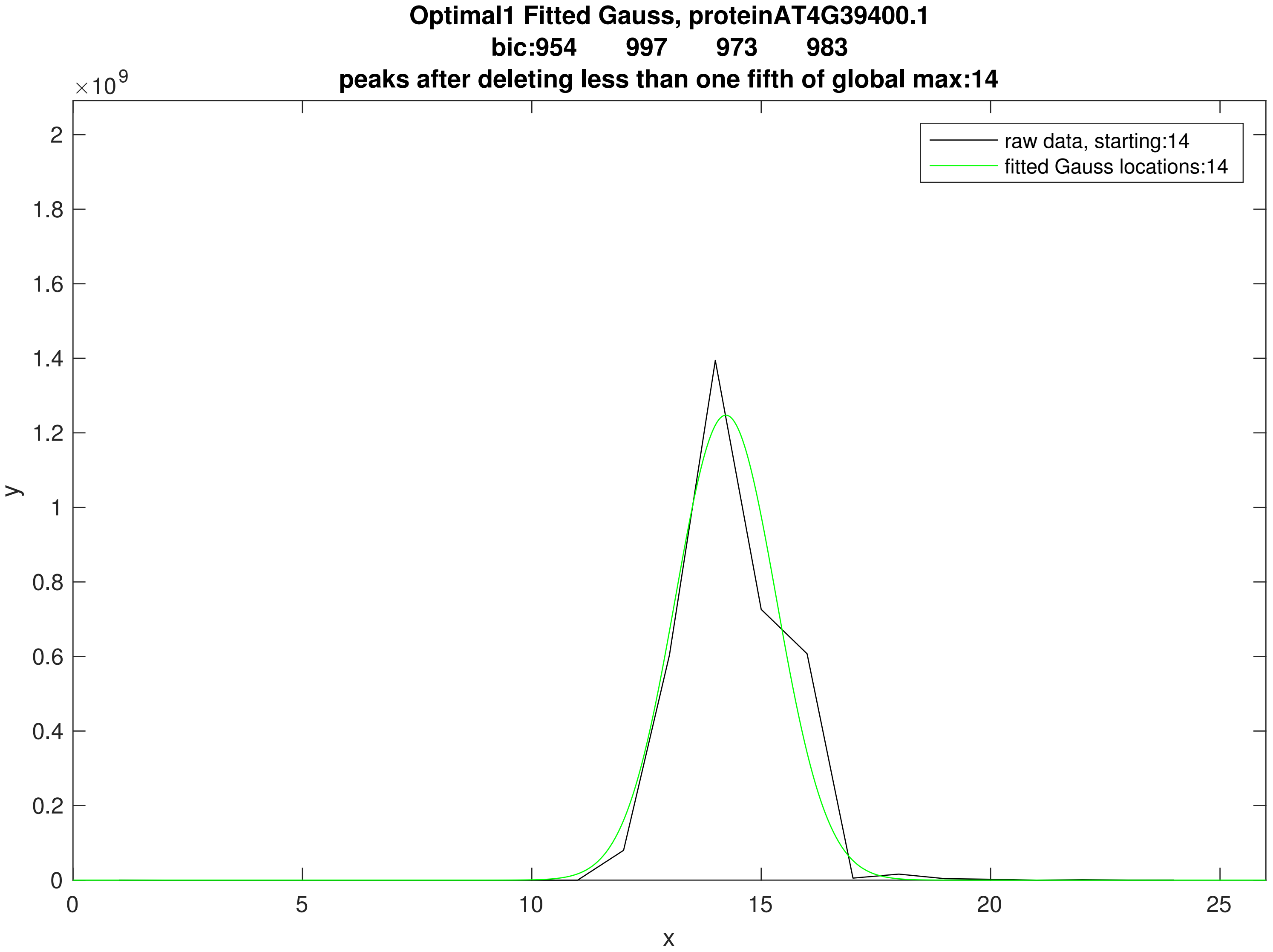
Optimal1 Fitted Gauss, proteinAT4G39400.1
bic:954 997 973 983
peaks after deleting less than one fifth of global max:14
×10^9
2
1.8
1.6
1.4
1.2
1
0.8
0.6
0.4
0.2
0
y
0
5
10
15
20
25
x
raw data, starting:14
fitted Gauss locations:14

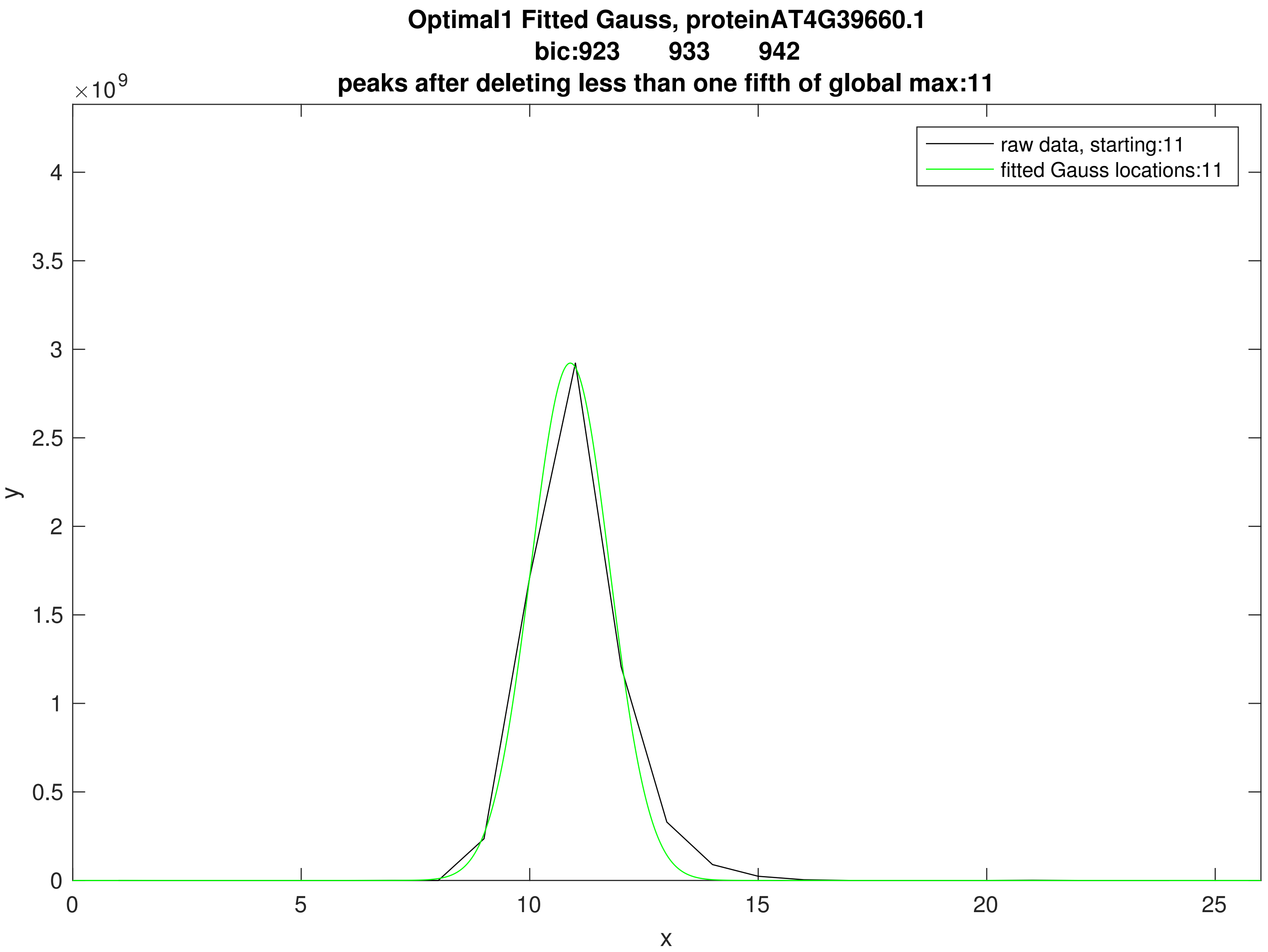

Optimal1 Fitted Gauss, proteinAT4G39660.1
bic:923 933 942
peaks after deleting less than one fifth of global max:11
×10^9
raw data, starting:11
fitted Gauss locations:11
4
3.5
3
2.5
2
1.5
1
0.5
0
y
0
5
10
15
20
25
x

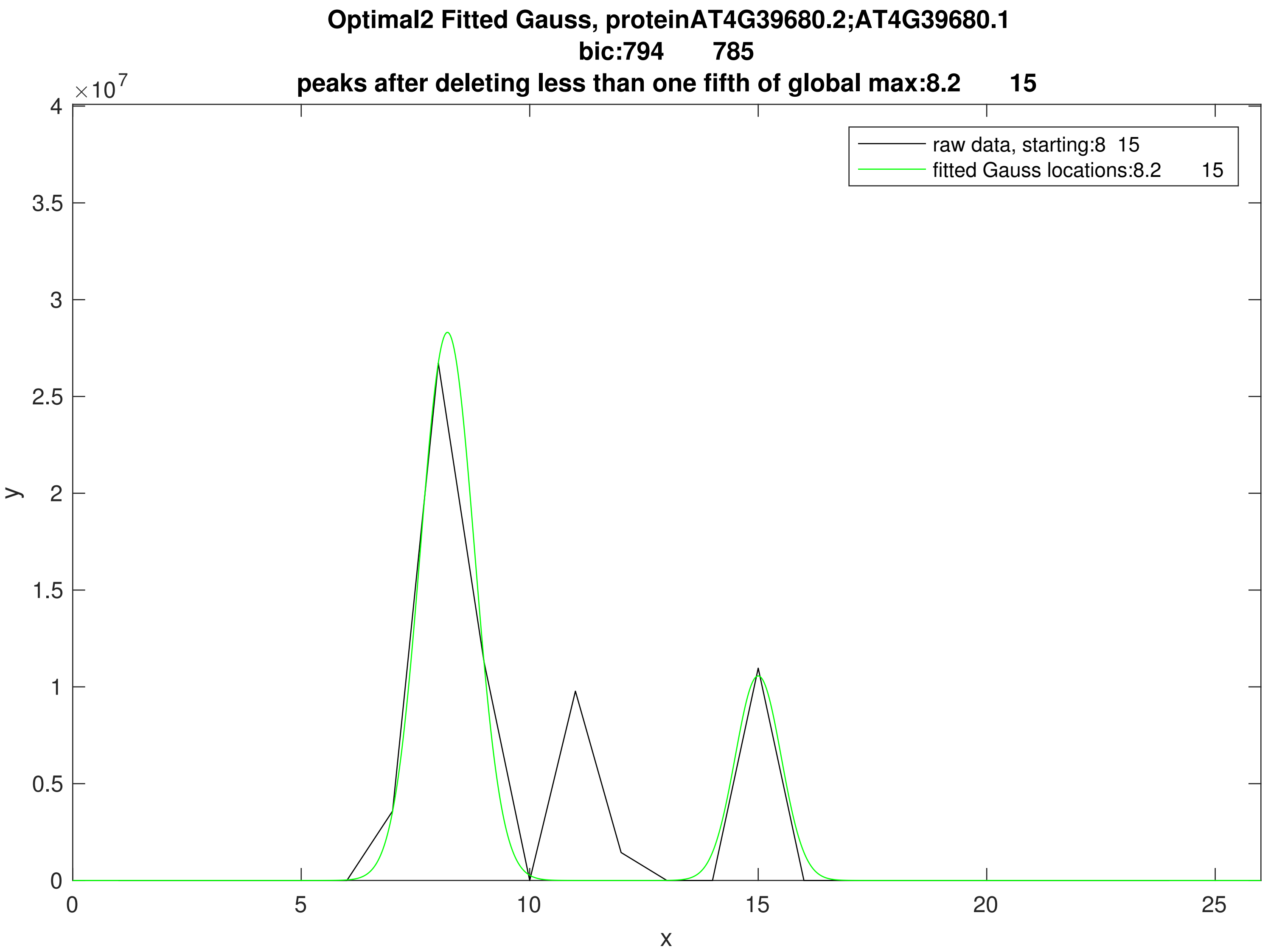
Optimal2 Fitted Gauss, proteinAT4G39680.2;AT4G39680.1
bic:794 785
peaks after deleting less than one fifth of global max:8.2 15
×10^7
raw data, starting:8 15
fitted Gauss locations:8.2 15
y
x

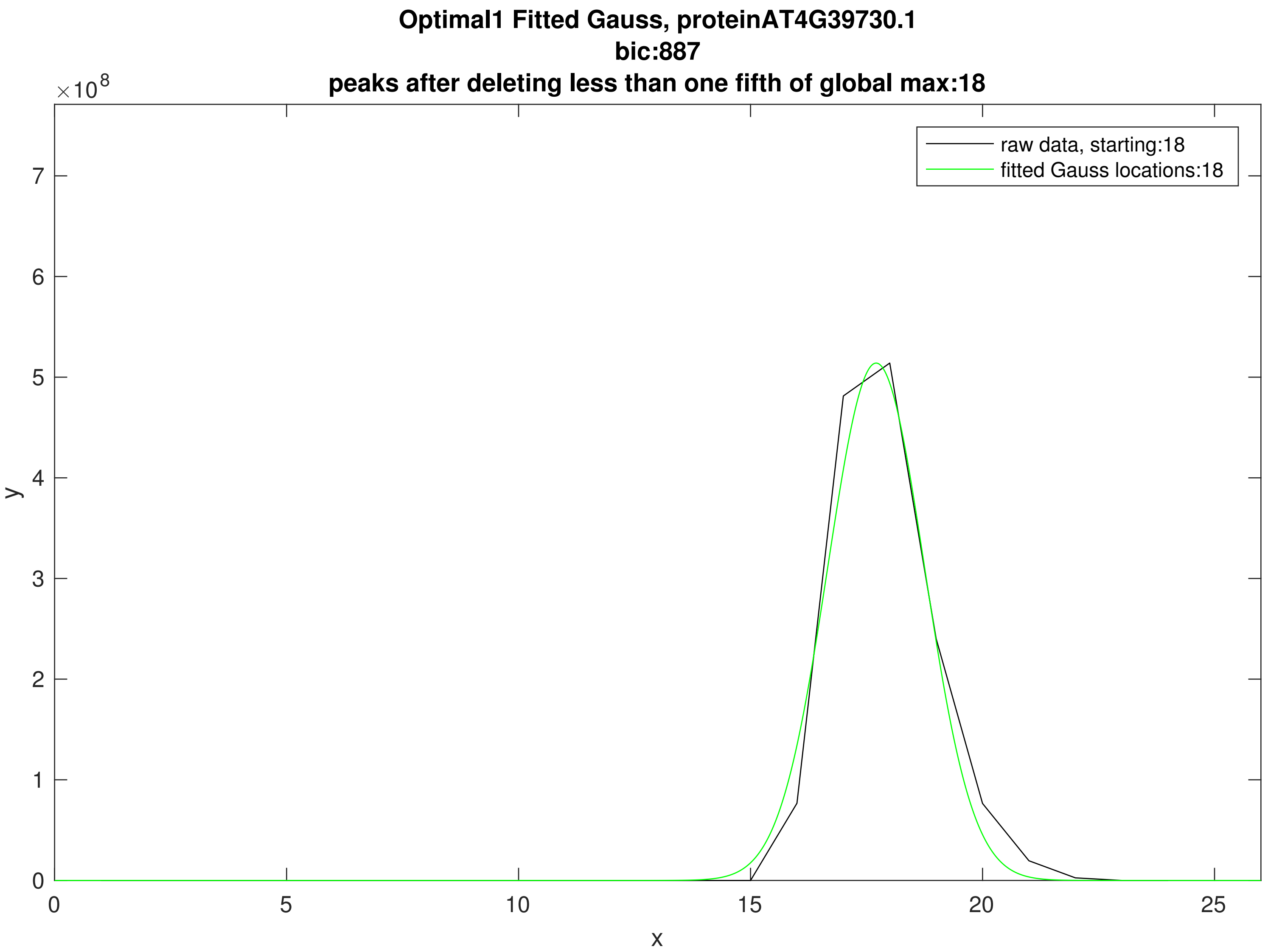
Optimal1 Fitted Gauss, proteinAT4G39730.1
bic:887
peaks after deleting less than one fifth of global max:18
×10^8
raw data, starting:18
fitted Gauss locations:18
y
x
0
1
2
3
4
5
6
7
0
5
10
15
20
25

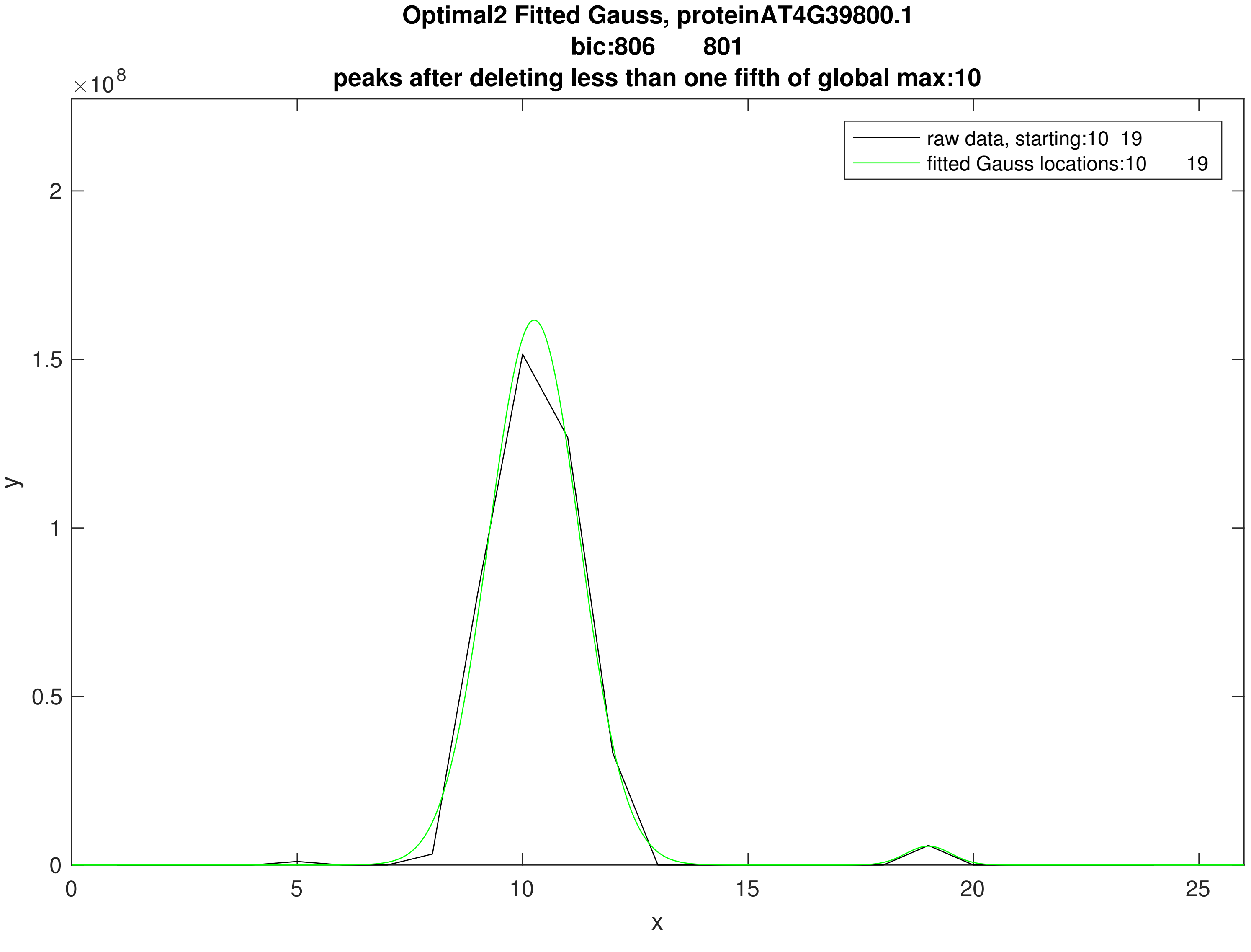
Optimal2 Fitted Gauss, proteinAT4G39800.1
bic:806 801
peaks after deleting less than one fifth of global max:10
raw data, starting:10 19
fitted Gauss locations:10 19
$\times 10^8$
y
x
0
0.5
1
1.5
2
0
5
10
15
20
25

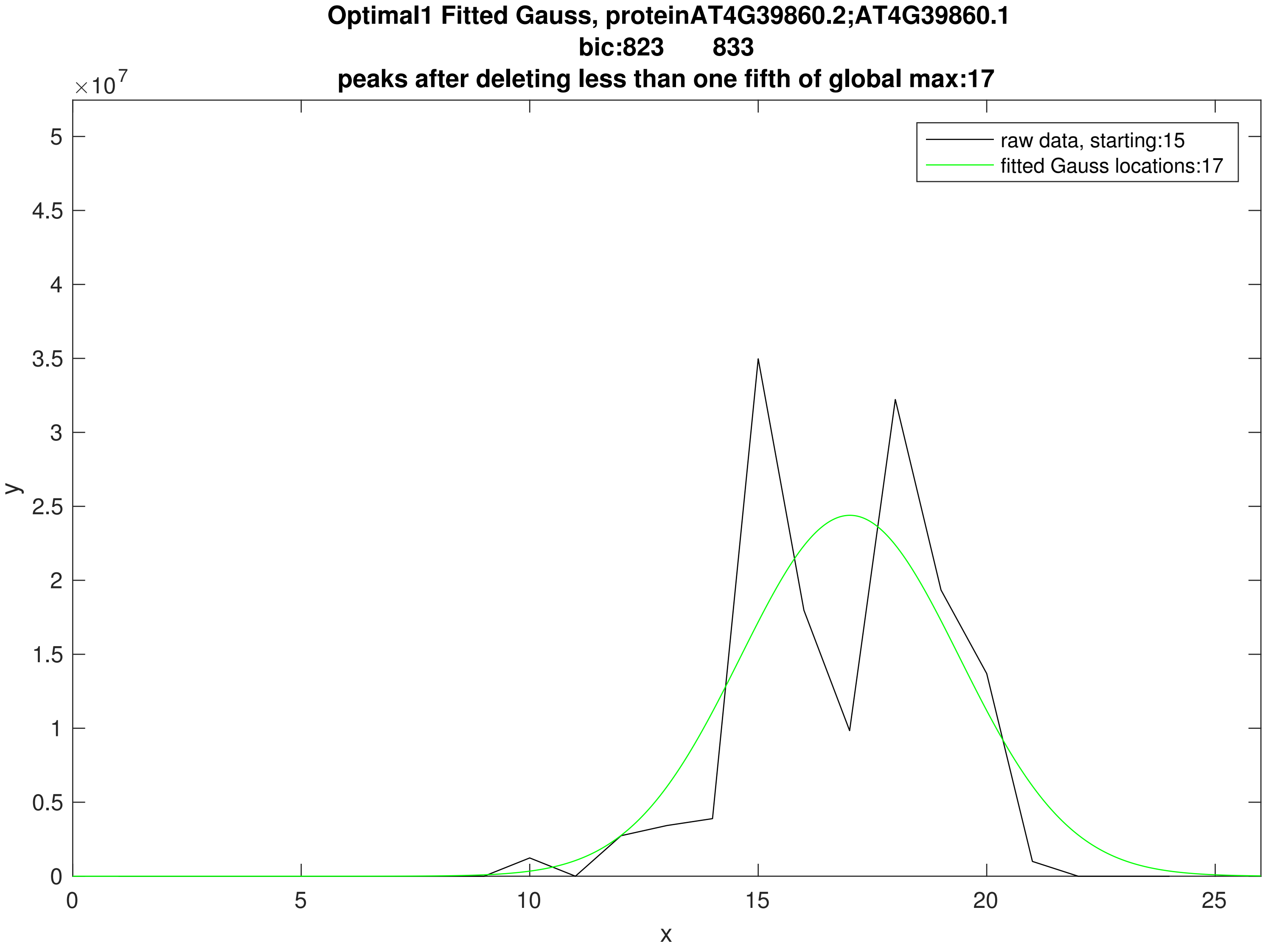
Optimal1 Fitted Gauss, proteinAT4G39860.2;AT4G39860.1
bic:823 833
peaks after deleting less than one fifth of global max:17
×10^7
raw data, starting:15
fitted Gauss locations:17
y
x
0
0.5
1
1.5
2
2.5
3
3.5
4
4.5
5
0
5
10
15
20
25

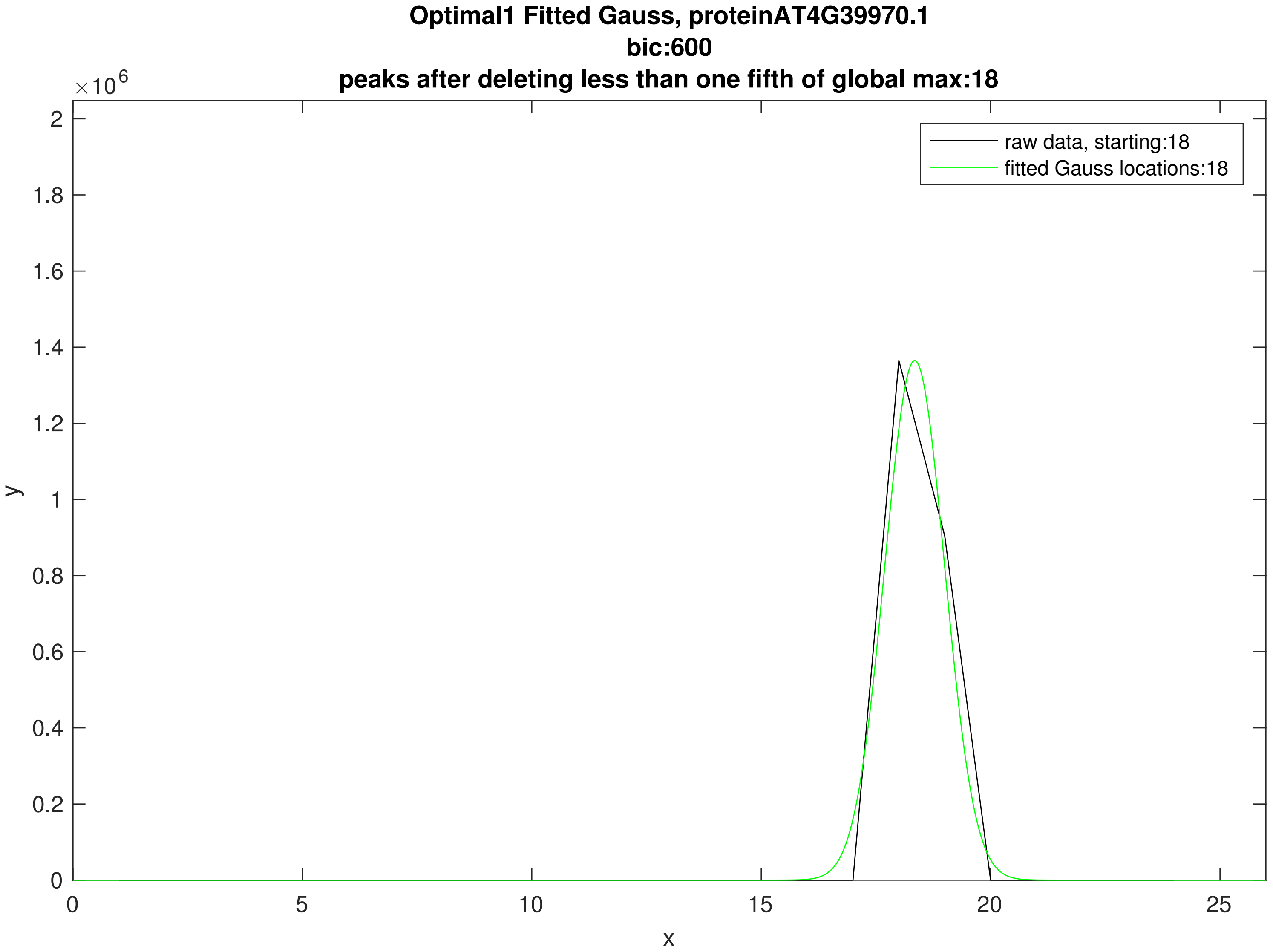

Optimal1 Fitted Gauss, proteinAT4G39970.1
bic:600
peaks after deleting less than one fifth of global max:18
×10^6
2
1.8
1.6
1.4
1.2
1
0.8
0.6
0.4
0.2
0
y
0
5
10
15
20
25
x
raw data, starting:18
fitted Gauss locations:18

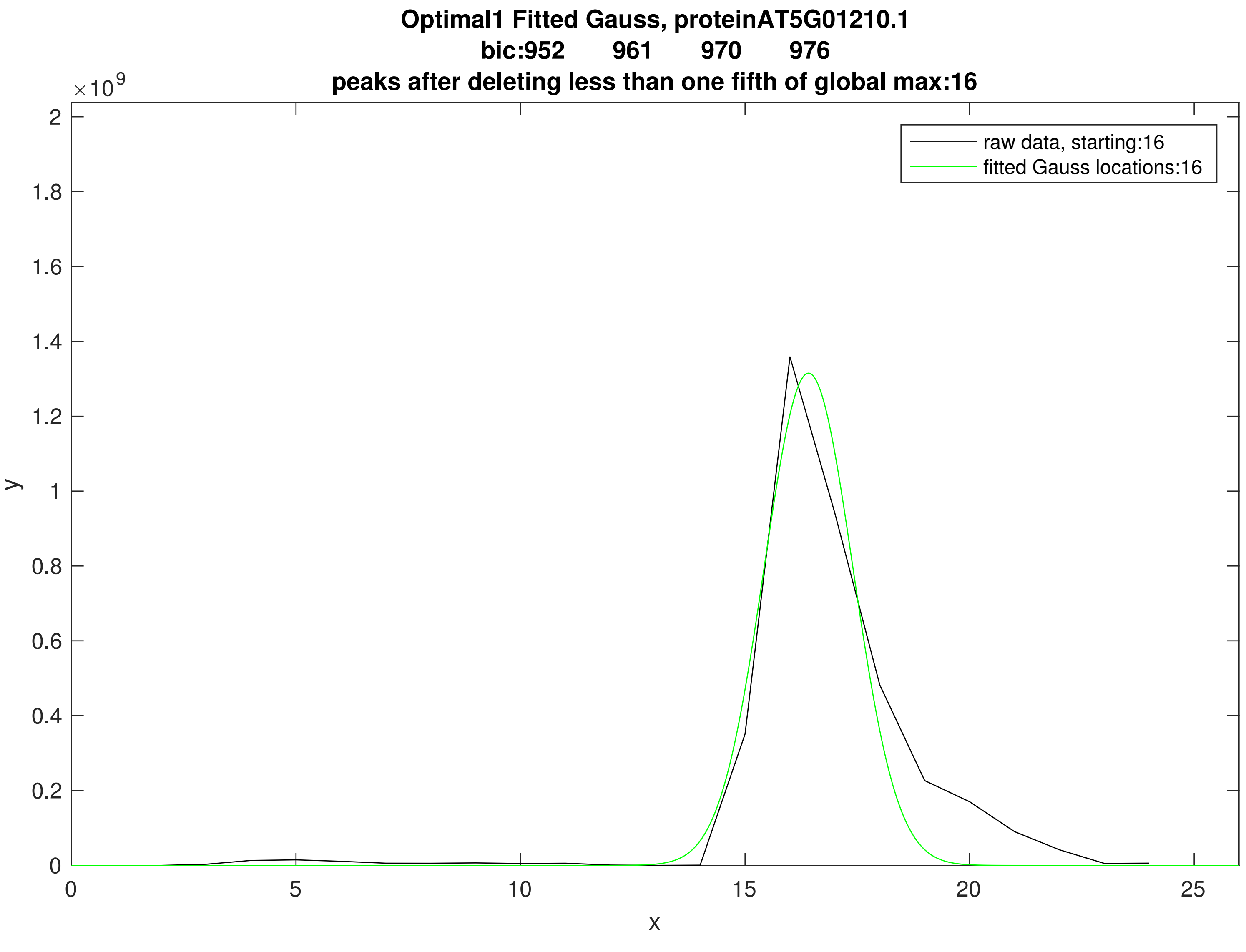
Optimal1 Fitted Gauss, proteinAT5G01210.1
bic:952 961 970 976
peaks after deleting less than one fifth of global max:16
×10^9
2
1.8
1.6
1.4
1.2
1
0.8
0.6
0.4
0.2
0
y
0
5
10
15
20
25
x
raw data, starting:16
fitted Gauss locations:16

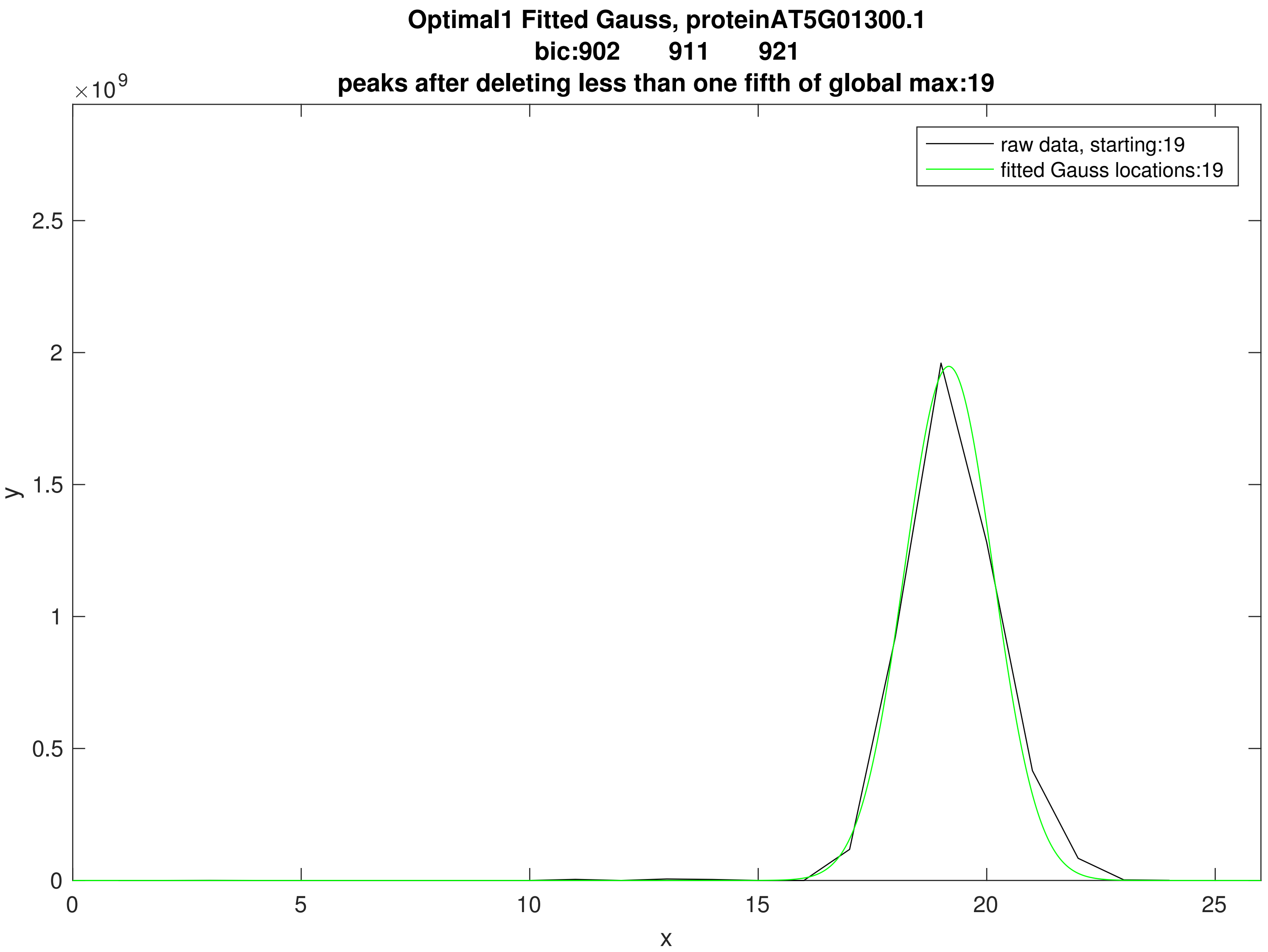
Optimal1 Fitted Gauss, proteinAT5G01300.1
bic:902 911 921
peaks after deleting less than one fifth of global max:19
×10^9
2.5
2
1.5
1
0.5
0
y
0
5
10
15
20
25
x
raw data, starting:19
fitted Gauss locations:19

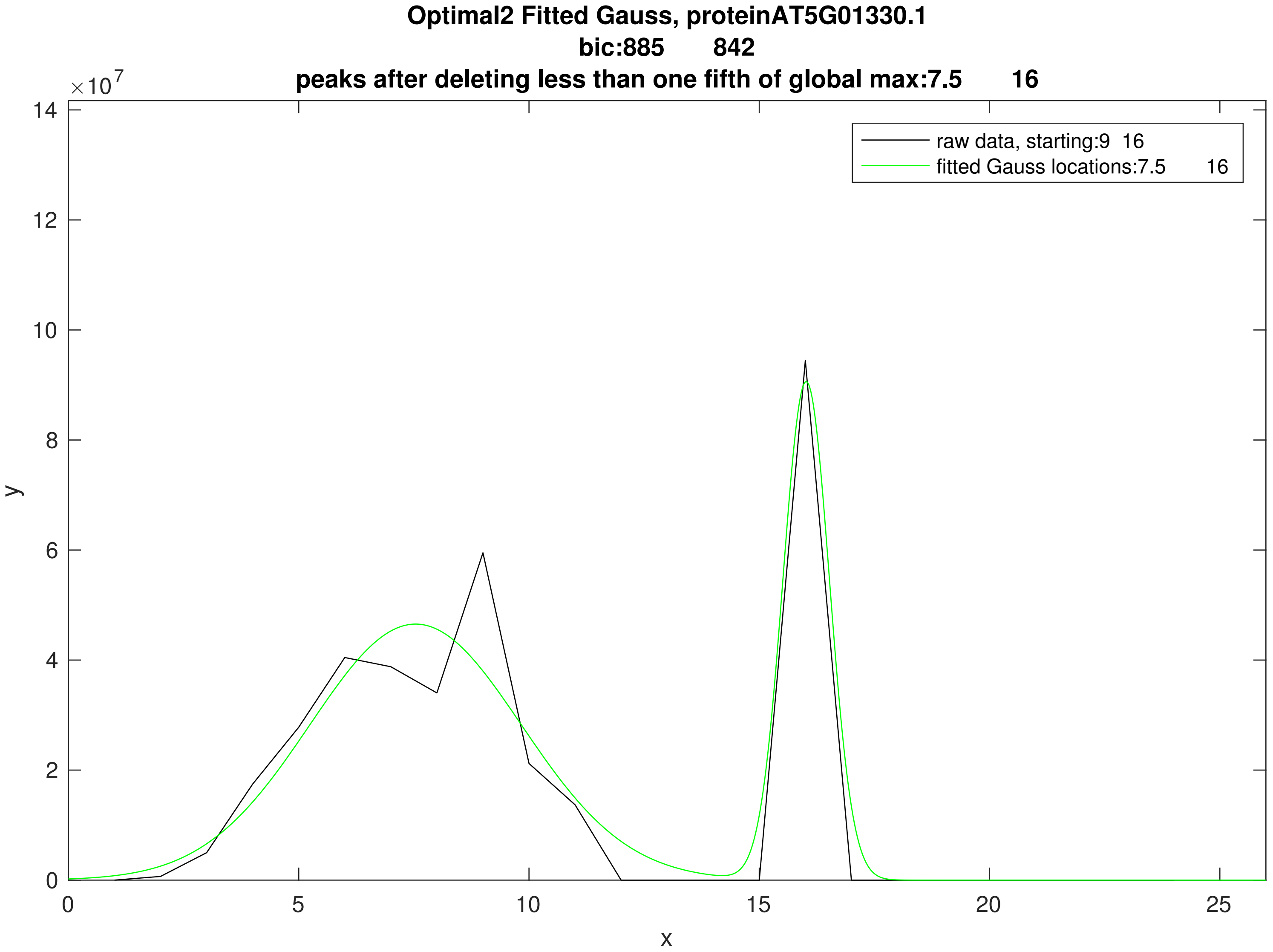
Optimal2 Fitted Gauss, proteinAT5G01330.1
bic:885 842
peaks after deleting less than one fifth of global max:7.5 16
×10^7
raw data, starting:9 16
fitted Gauss locations:7.5 16
y
x
0
2
4
6
8
10
12
14
5
15
20
25

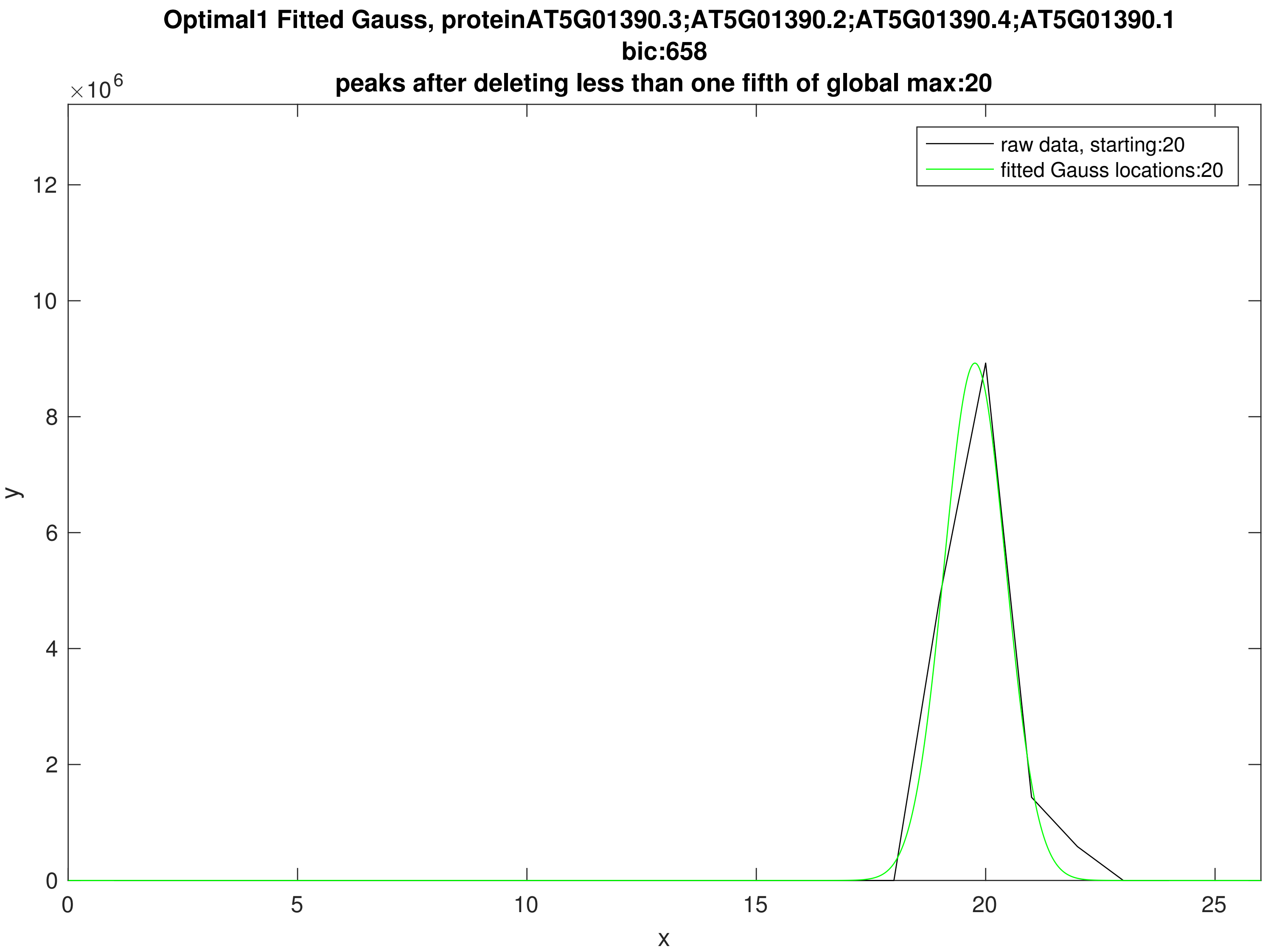
Optimal1 Fitted Gauss, proteinAT5G01390.3;AT5G01390.2;AT5G01390.4;AT5G01390.1
bic:658
peaks after deleting less than one fifth of global max:20
×10^6
raw data, starting:20
fitted Gauss locations:20
y
x
0
2
4
6
8
10
12
0
5
10
15
20
25

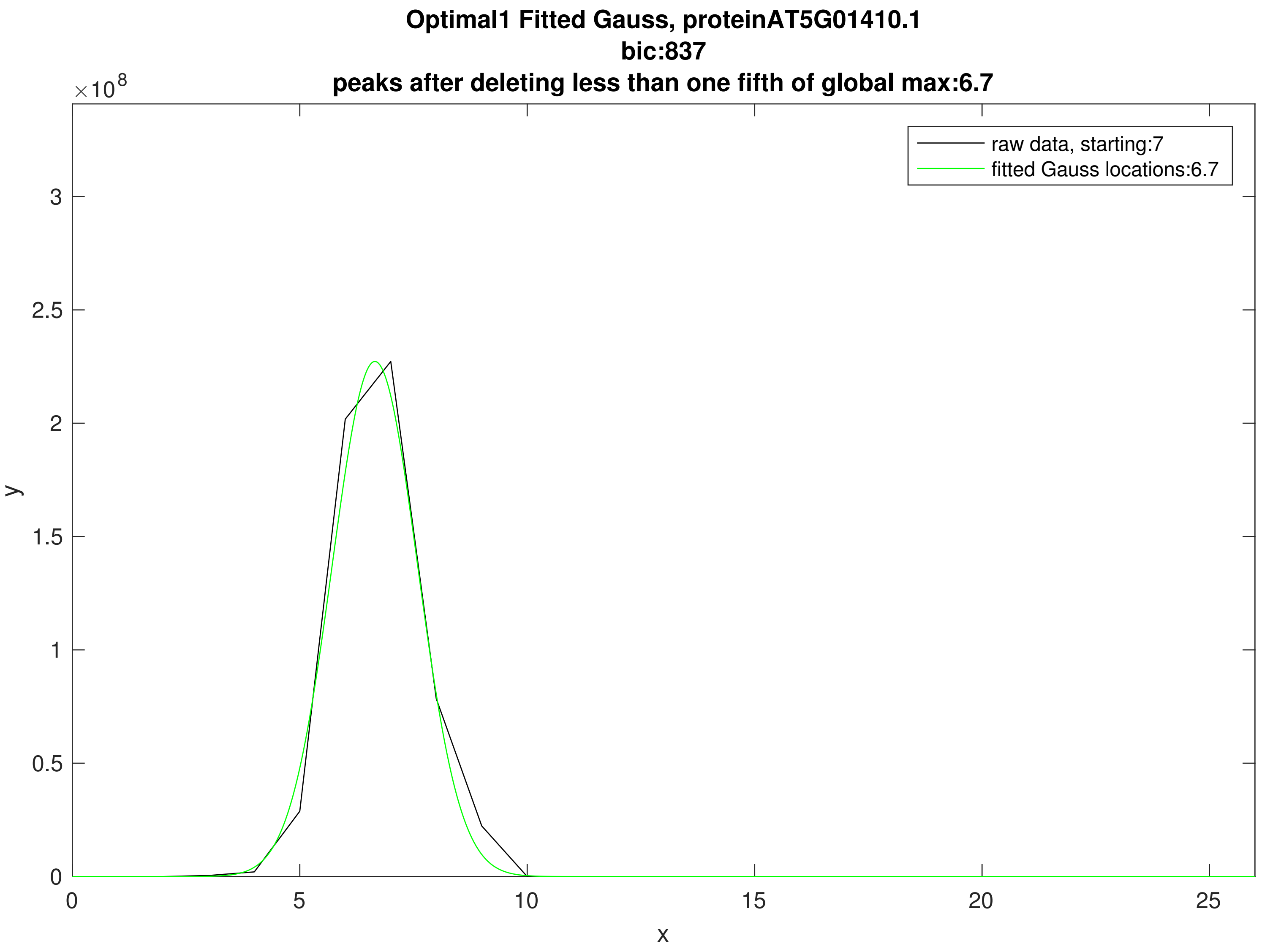
Optimal1 Fitted Gauss, proteinAT5G01410.1
bic:837
peaks after deleting less than one fifth of global max:6.7
×10^8
raw data, starting:7
fitted Gauss locations:6.7
y
x
0
0.5
1
1.5
2
2.5
3
0
5
10
15
20
25

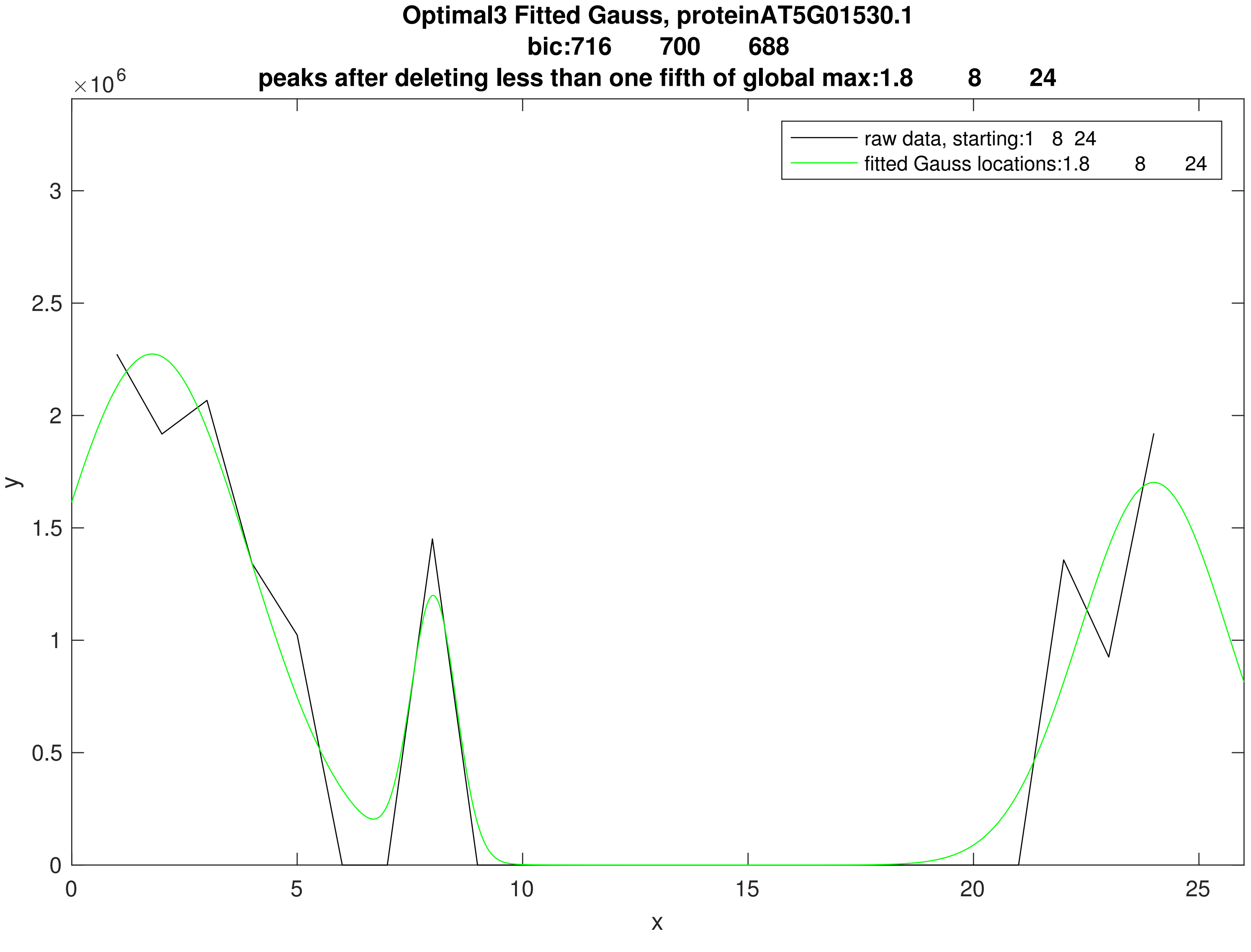

Optimal3 Fitted Gauss, proteinAT5G01530.1
bic:716 700 688
peaks after deleting less than one fifth of global max:1.8 8 24
×10^6
raw data, starting:1 8 24
fitted Gauss locations:1.8 8 24
y
x
0
0.5
1
1.5
2
2.5
3
0
5
10
15
20
25

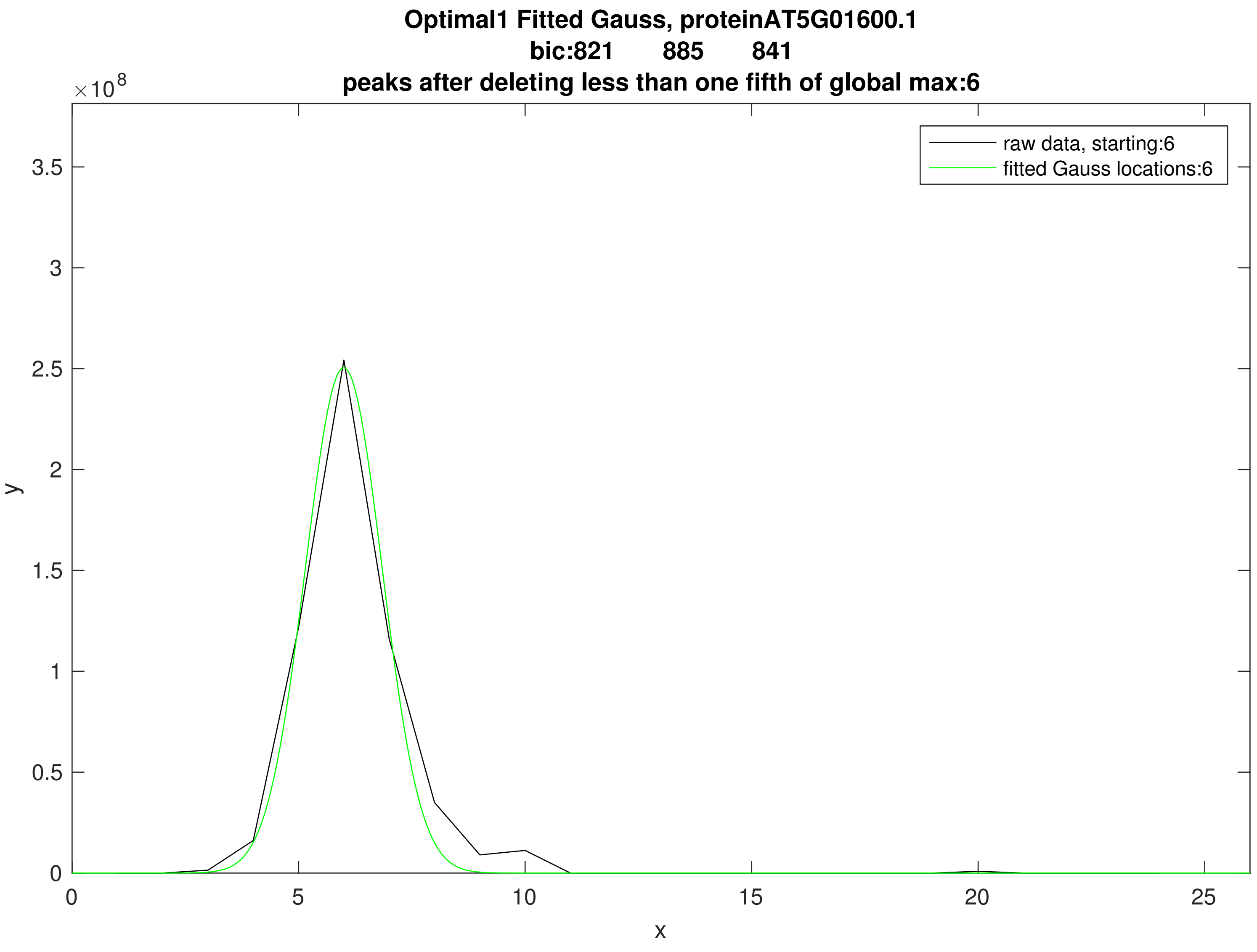

Optimal1 Fitted Gauss, proteinAT5G01600.1
bic:821 885 841
peaks after deleting less than one fifth of global max:6
×10^8
3.5
3
2.5
2
1.5
1
0.5
0
y
0
5
10
15
20
25
x
raw data, starting:6
fitted Gauss locations:6

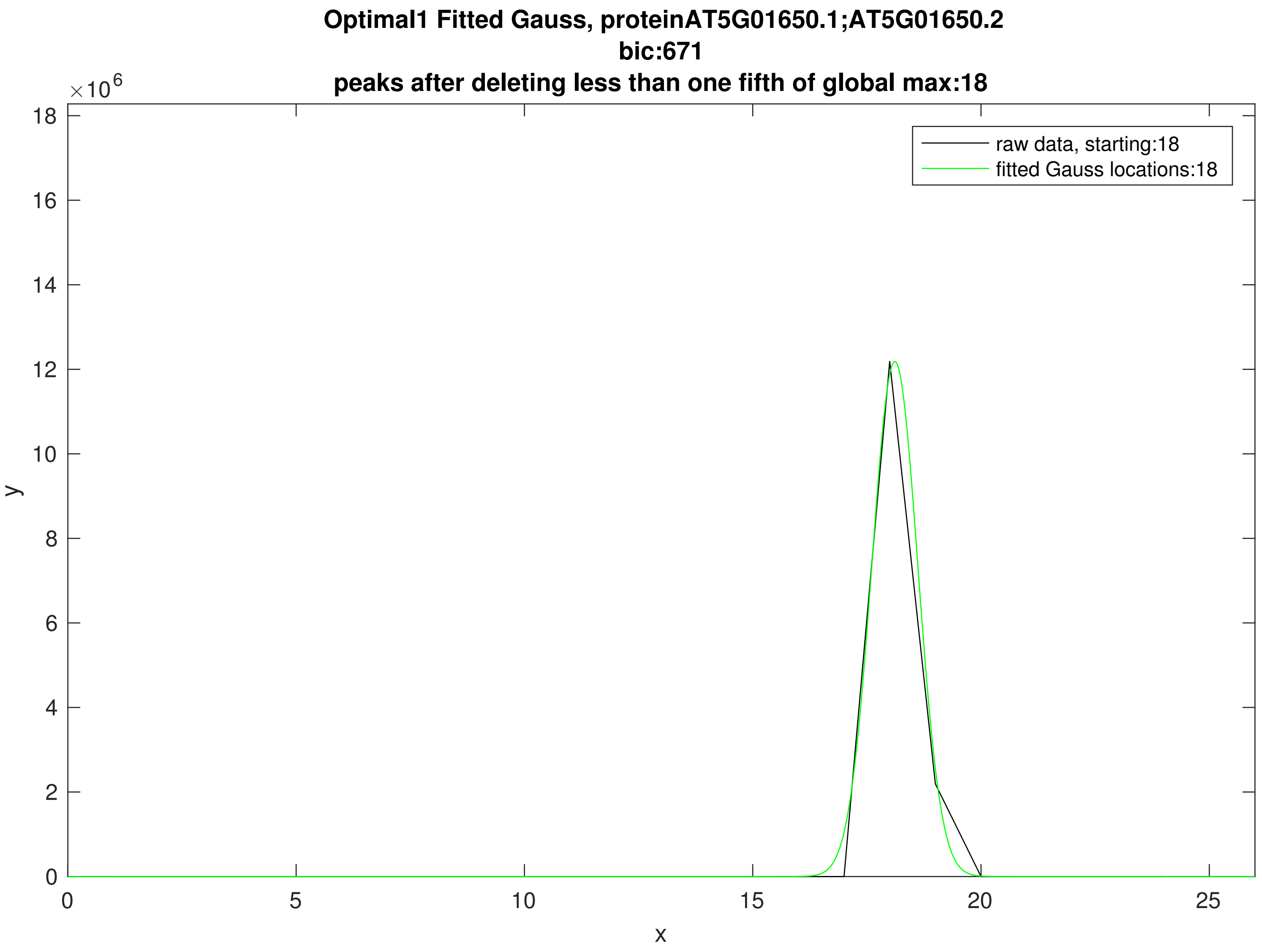
Optimal1 Fitted Gauss, proteinAT5G01650.1;AT5G01650.2
bic:671
peaks after deleting less than one fifth of global max:18
×10^6
raw data, starting:18
fitted Gauss locations:18
y
x
0
2
4
6
8
10
12
14
16
18
0
5
10
15
20
25

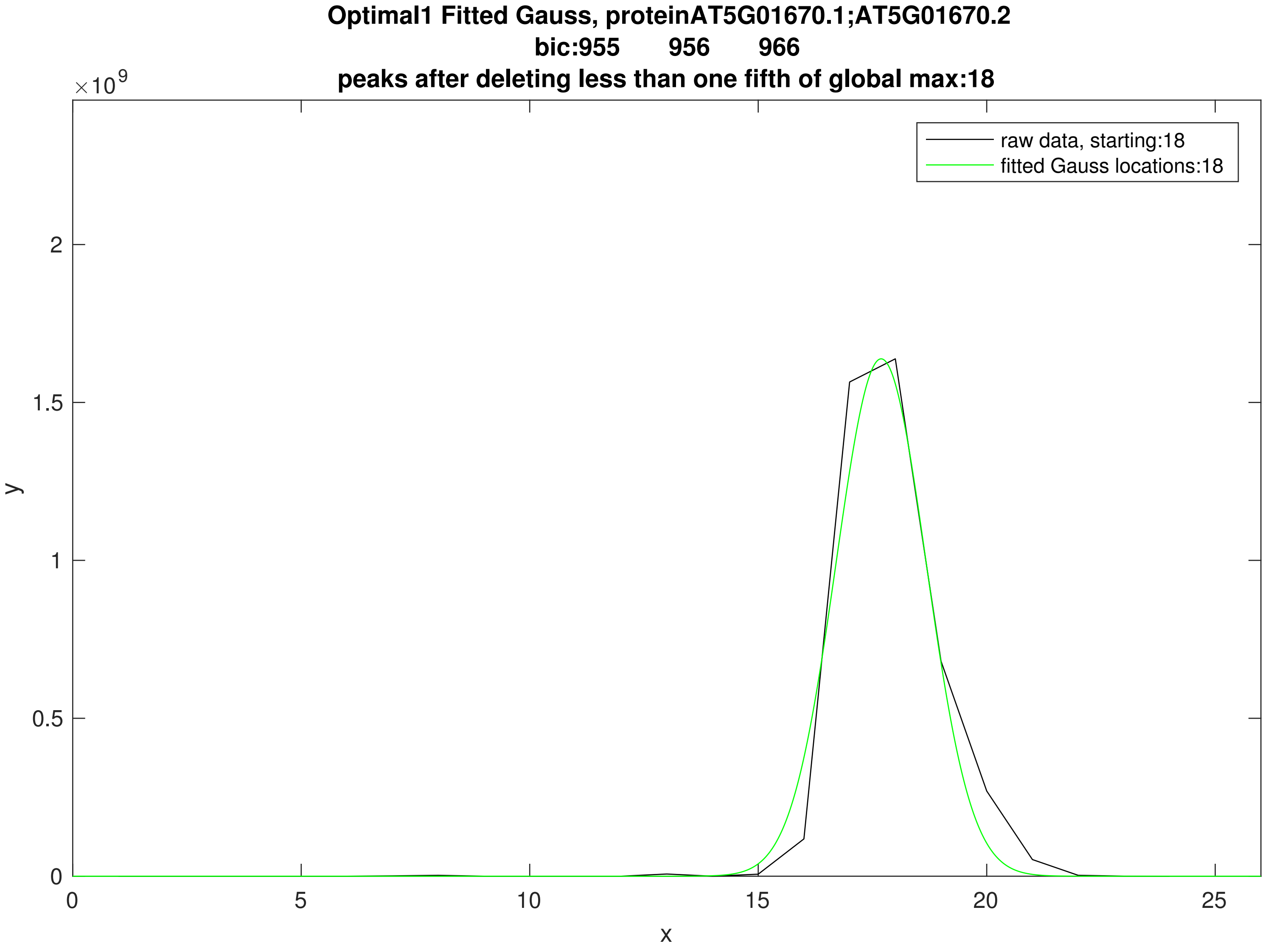
Optimal1 Fitted Gauss, proteinAT5G01670.1;AT5G01670.2
bic:955 956 966
peaks after deleting less than one fifth of global max:18
×10^9
raw data, starting:18
fitted Gauss locations:18
y
x
0
0.5
1
1.5
2
0
5
10
15
20
25

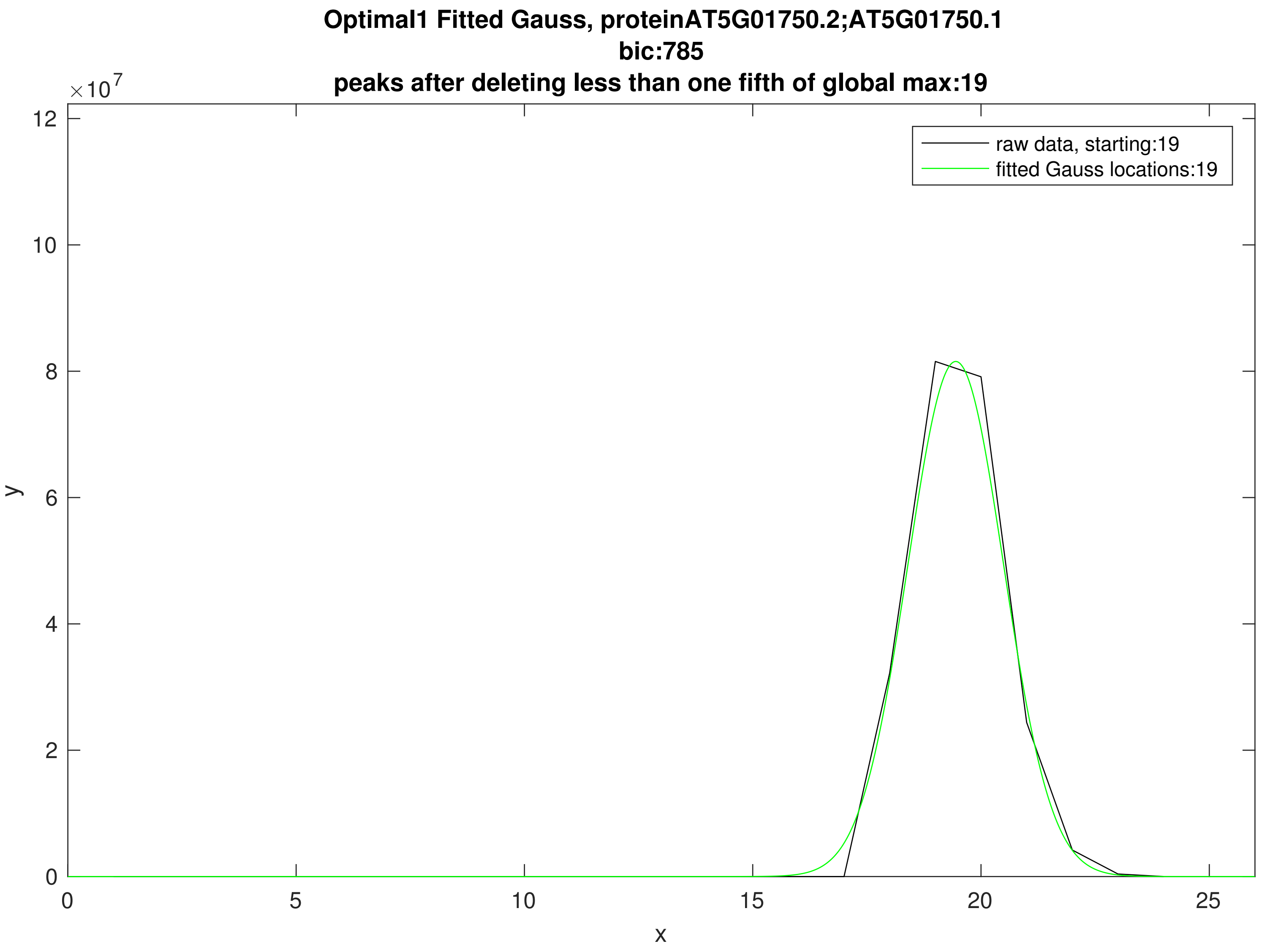
Optimal1 Fitted Gauss, proteinAT5G01750.2;AT5G01750.1
bic:785
peaks after deleting less than one fifth of global max:19
×10^7
12
10
8
6
4
2
0
0
5
10
15
20
25
x
y
raw data, starting:19
fitted Gauss locations:19

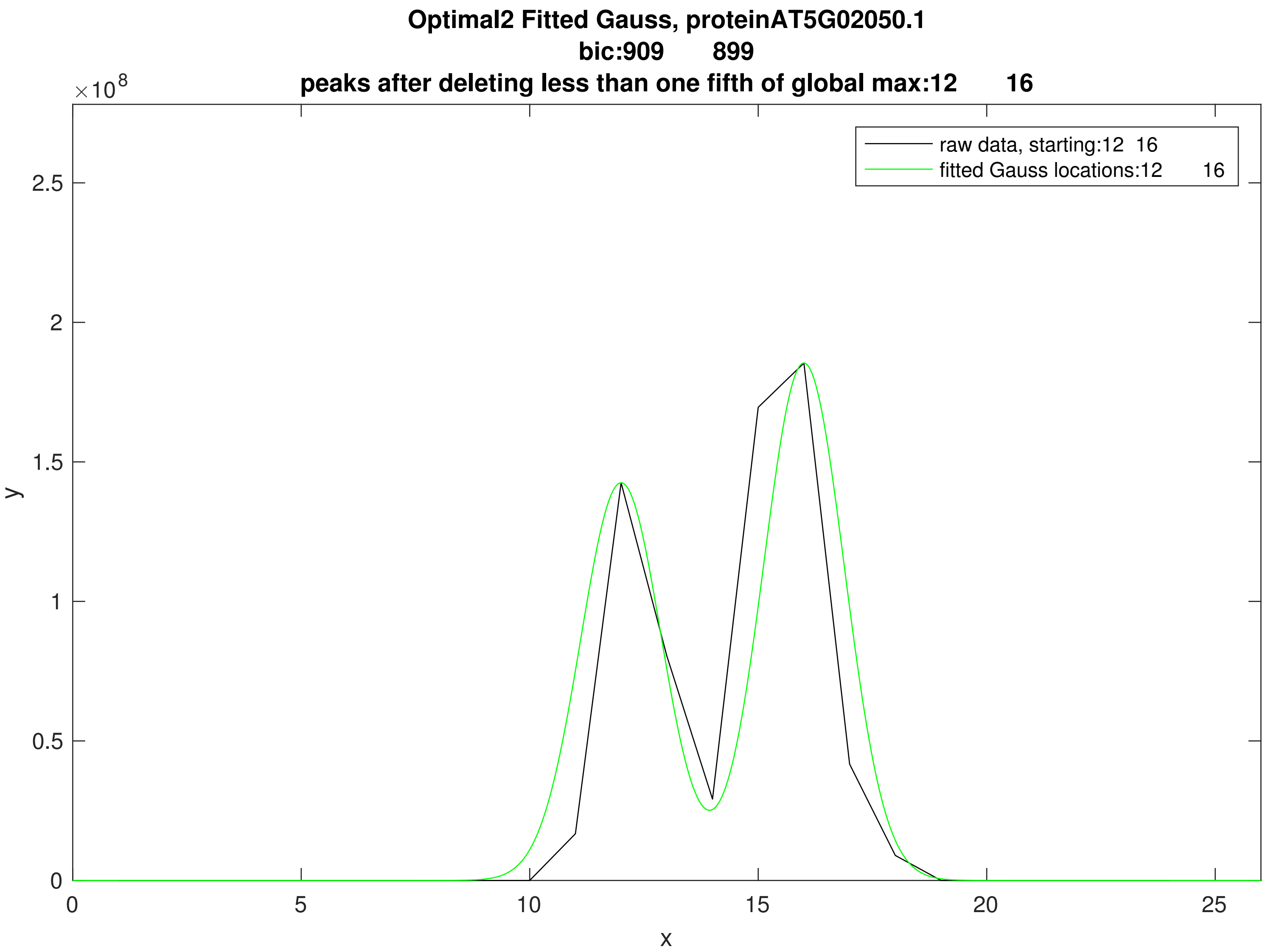
Optimal2 Fitted Gauss, proteinAT5G02050.1
bic:909 899
peaks after deleting less than one fifth of global max:12 16
×10^8
raw data, starting:12 16
fitted Gauss locations:12 16
y
x
0
0.5
1
1.5
2
2.5
5
10
15
20
25

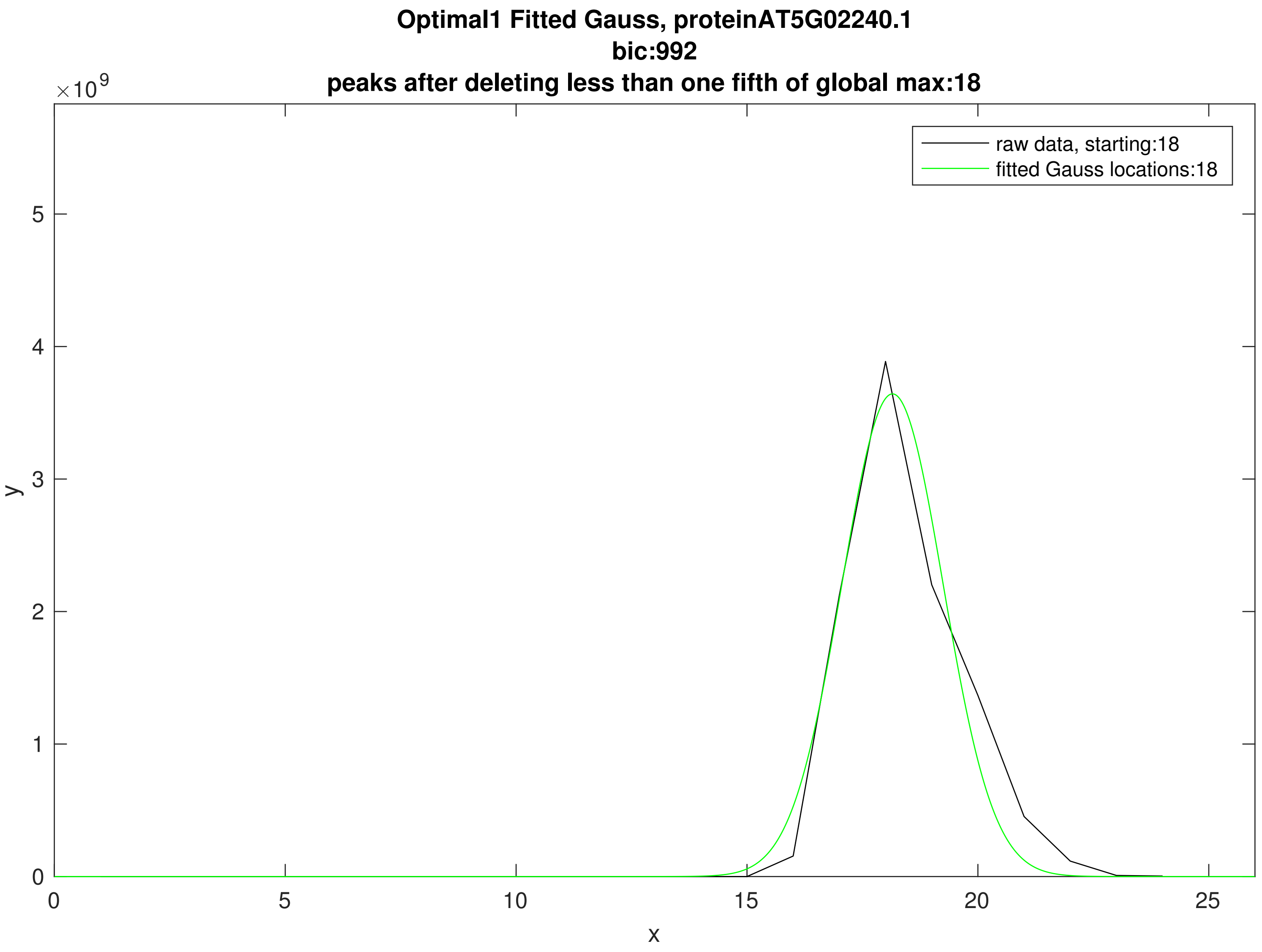

Optimal1 Fitted Gauss, proteinAT5G02240.1
bic:992
peaks after deleting less than one fifth of global max:18
×10^9
raw data, starting:18
fitted Gauss locations:18
y
x
0
1
2
3
4
5
0
5
10
15
20
25

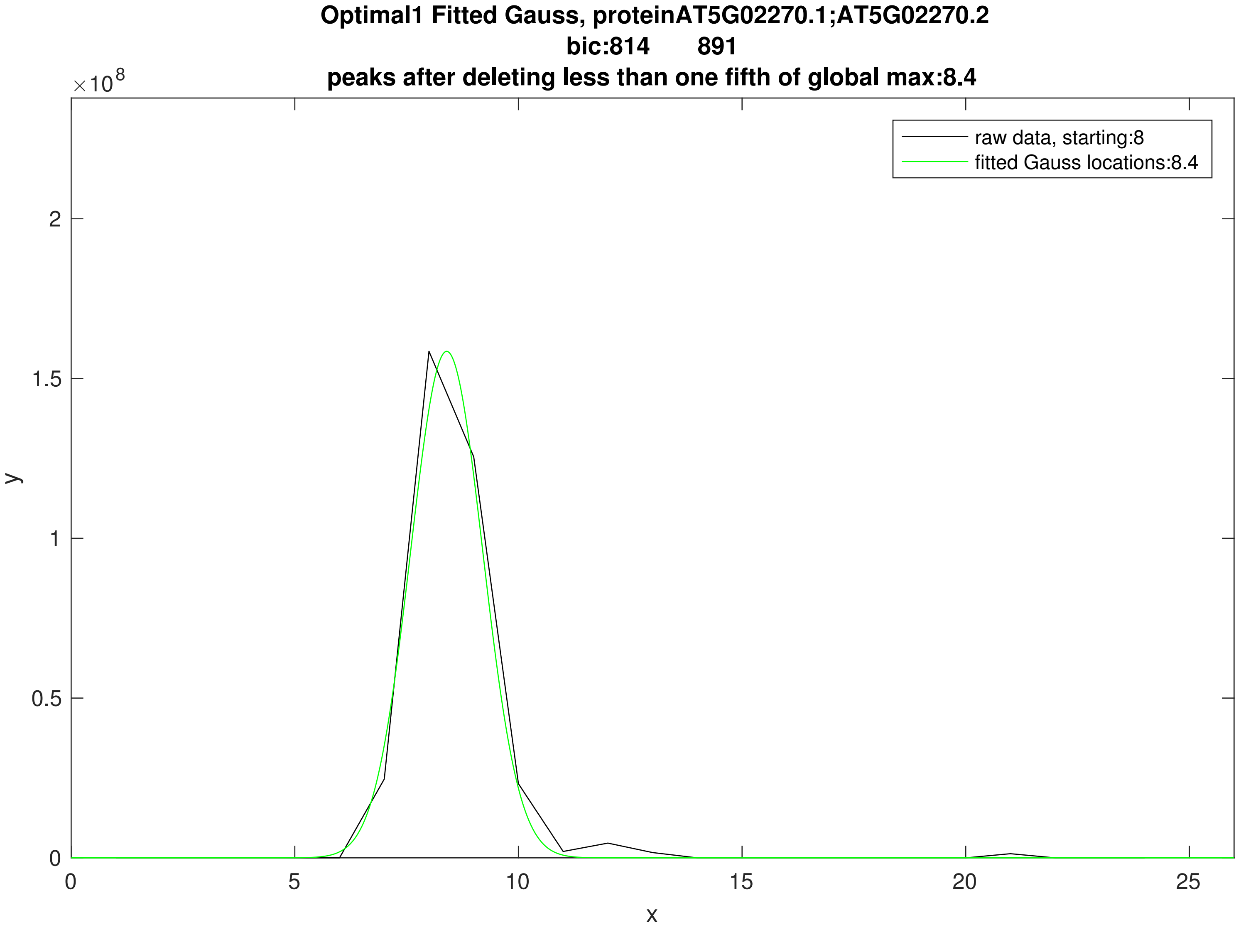
Optimal1 Fitted Gauss, proteinAT5G02270.1;AT5G02270.2
bic:814 891
peaks after deleting less than one fifth of global max:8.4
×10^8
raw data, starting:8
fitted Gauss locations:8.4
y
x
0
0.5
1
1.5
2
0
5
10
15
20
25

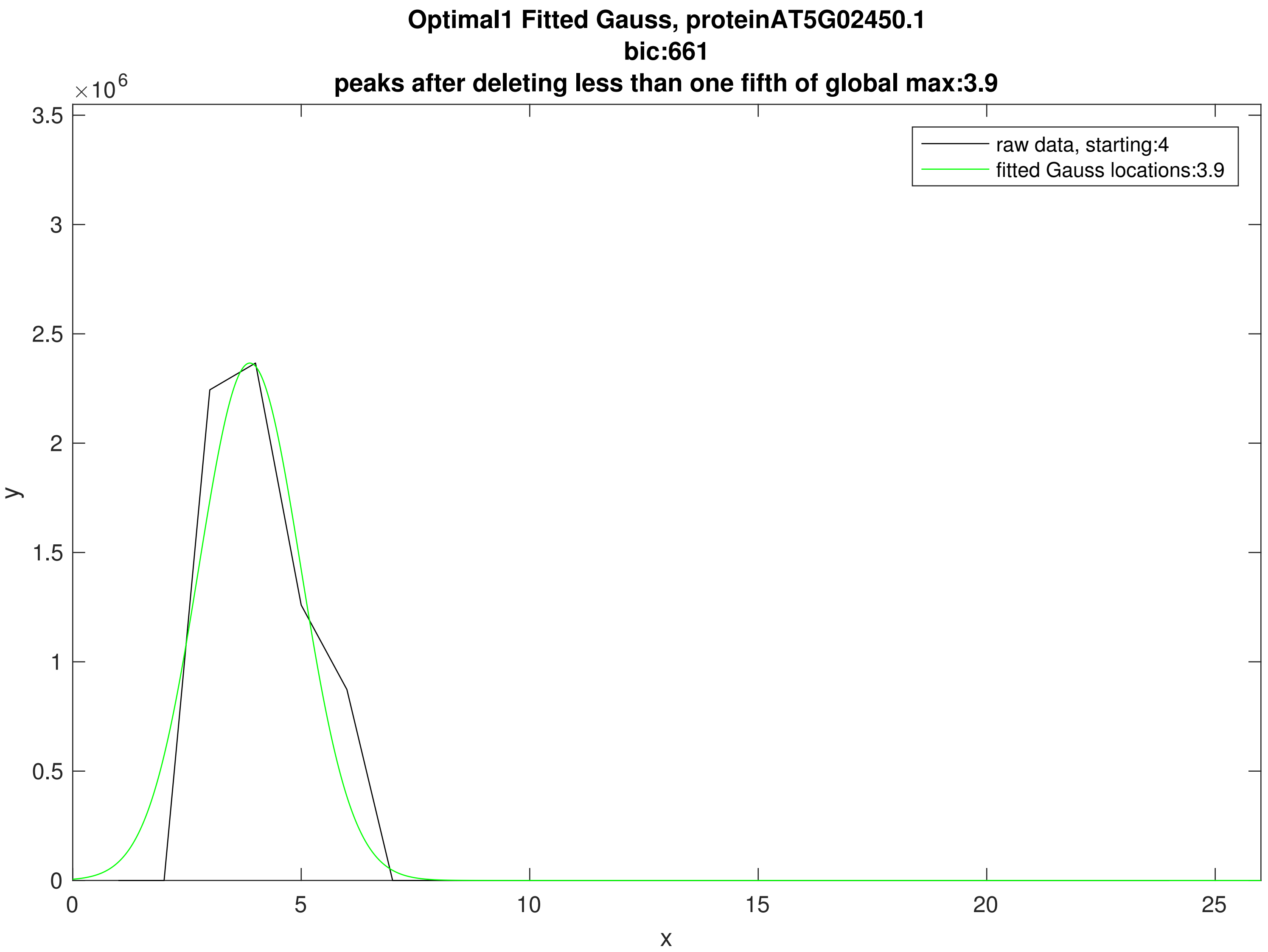
Optimal1 Fitted Gauss, proteinAT5G02450.1
bic:661
peaks after deleting less than one fifth of global max:3.9
×10^6
3.5
3
2.5
2
1.5
1
0.5
0
y
0
5
10
15
20
25
x
raw data, starting:4
fitted Gauss locations:3.9

**Optimal1 Fitted Gauss, proteinAT5G02490.1**
**bic:793       829**
**peaks after deleting less than one fifth of global max:13**

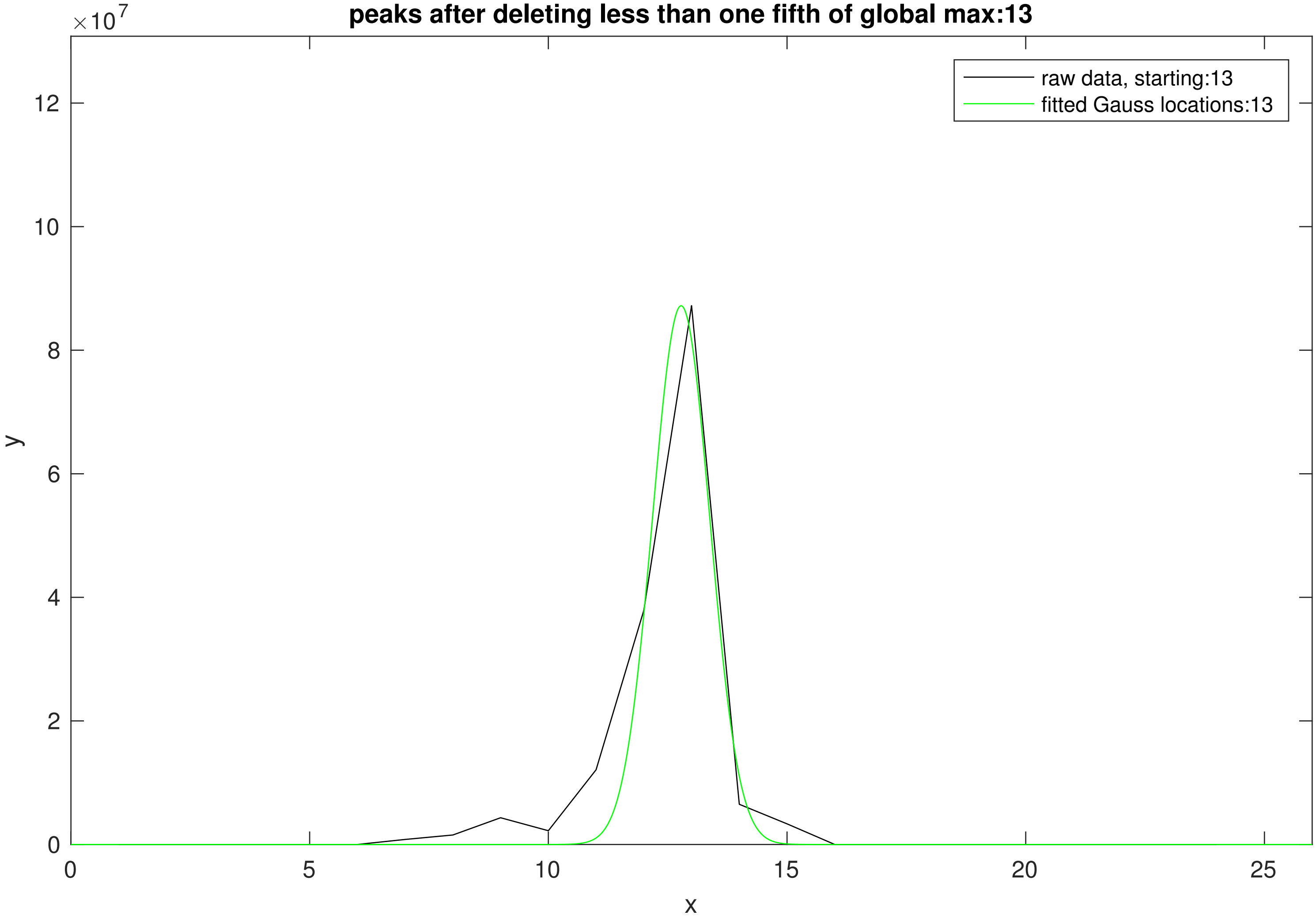

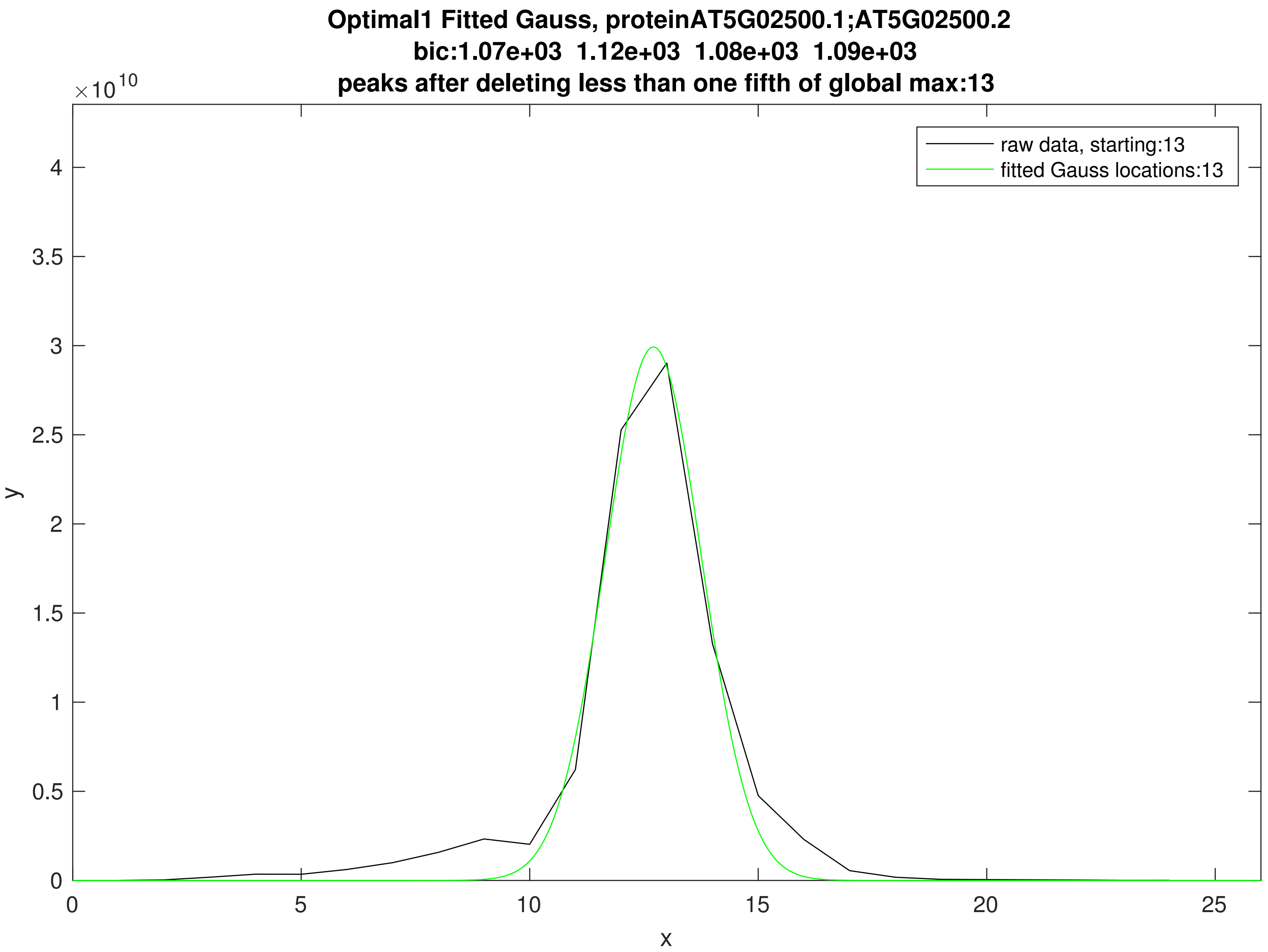
Optimal1 Fitted Gauss, proteinAT5G02500.1;AT5G02500.2
bic:1.07e+03  1.12e+03  1.08e+03  1.09e+03
peaks after deleting less than one fifth of global max:13
×10^10
raw data, starting:13
fitted Gauss locations:13
y
x
0
0.5
1
1.5
2
2.5
3
3.5
4
0
5
10
15
20
25

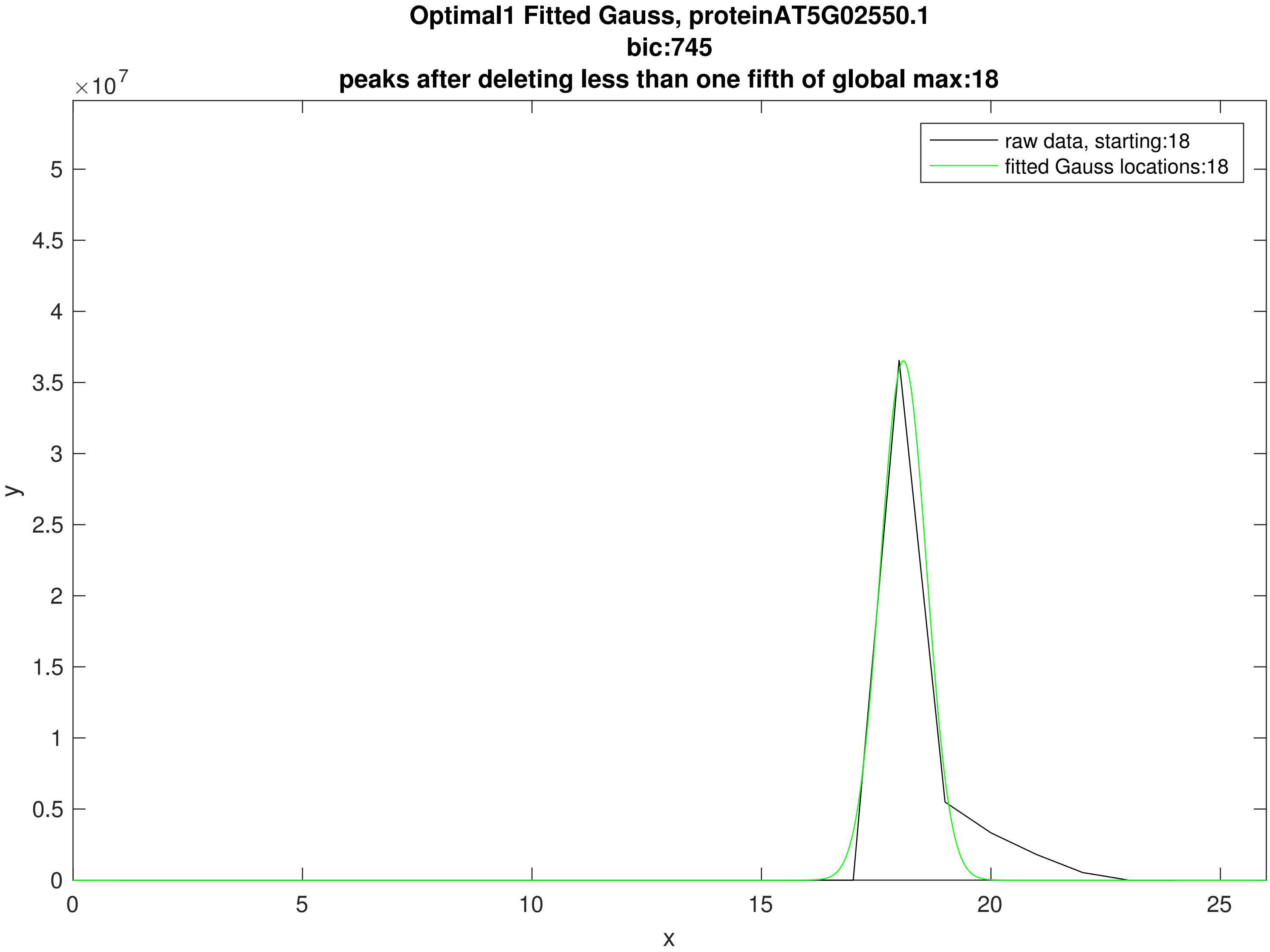
Optimal1 Fitted Gauss, proteinAT5G02550.1
bic:745
peaks after deleting less than one fifth of global max:18
×10^7
5
4.5
4
3.5
3
2.5
2
1.5
1
0.5
0
y
0
5
10
15
20
25
x
raw data, starting:18
fitted Gauss locations:18

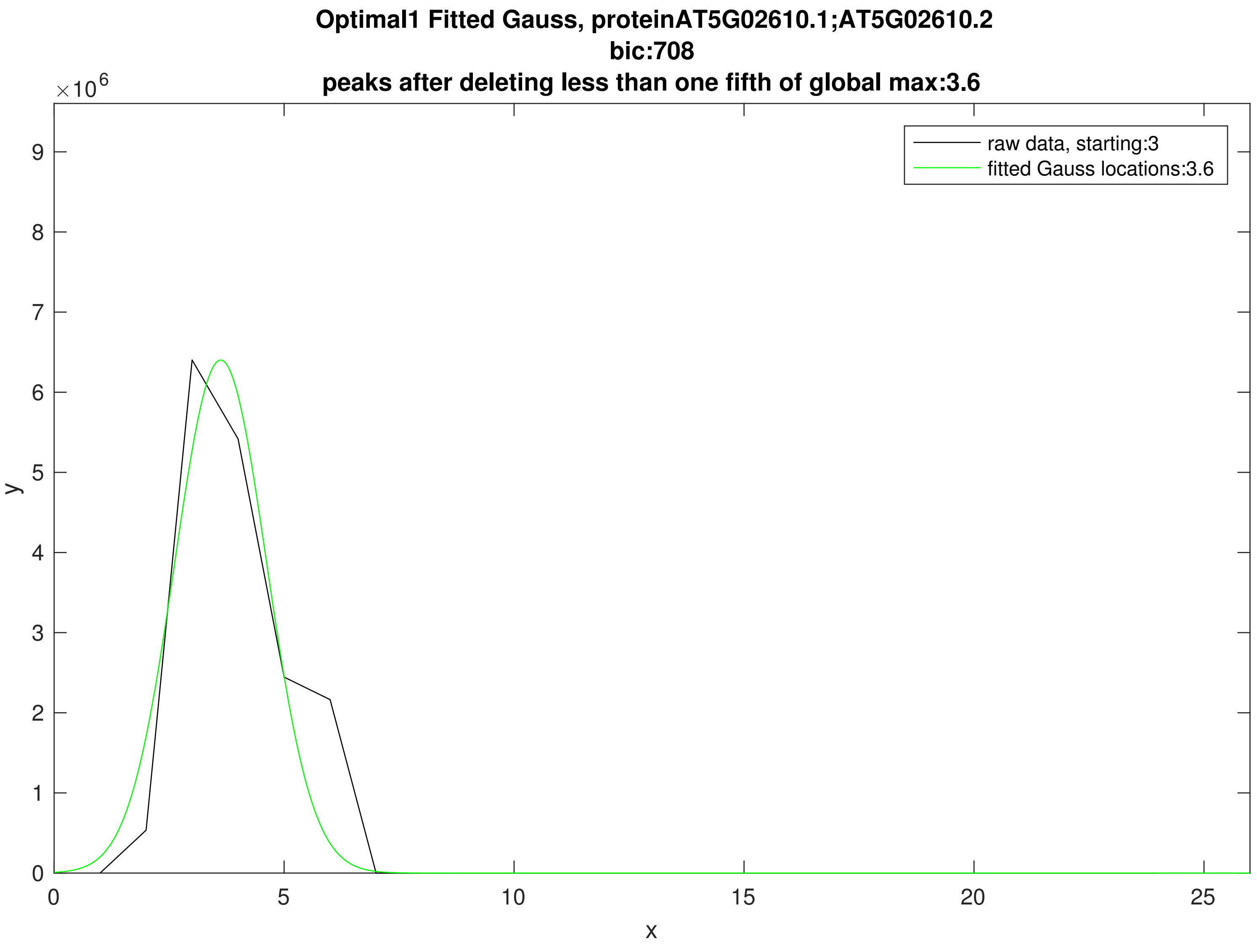
Optimal1 Fitted Gauss, proteinAT5G02610.1;AT5G02610.2
bic:708
peaks after deleting less than one fifth of global max:3.6
×10^6
y
x
raw data, starting:3
fitted Gauss locations:3.6
0
1
2
3
4
5
6
7
8
9
0
5
10
15
20
25

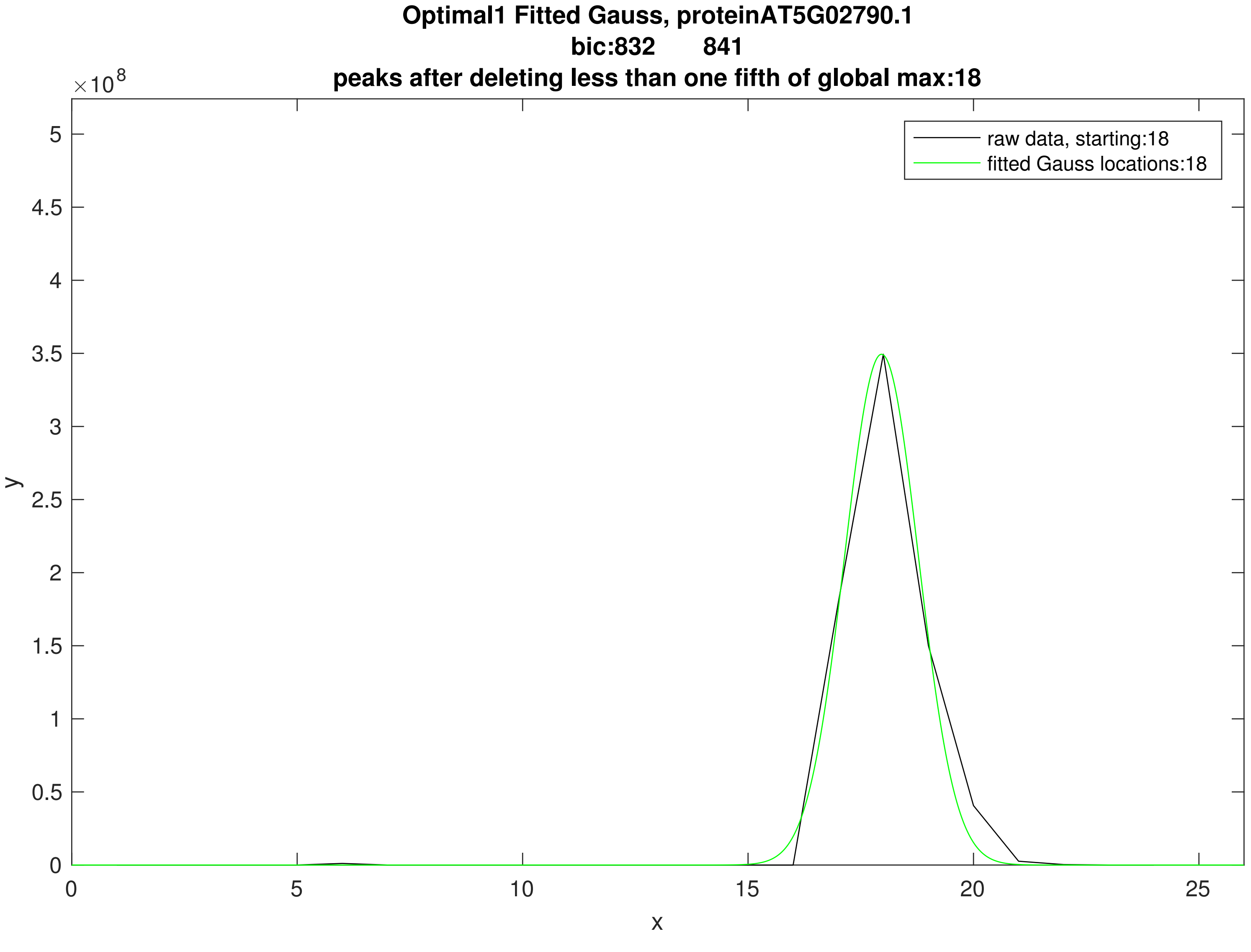
Optimal1 Fitted Gauss, proteinAT5G02790.1
bic:832 841
peaks after deleting less than one fifth of global max:18
×10^8
5
4.5
4
3.5
3
2.5
2
1.5
1
0.5
0
y
0
5
10
15
20
25
x
raw data, starting:18
fitted Gauss locations:18

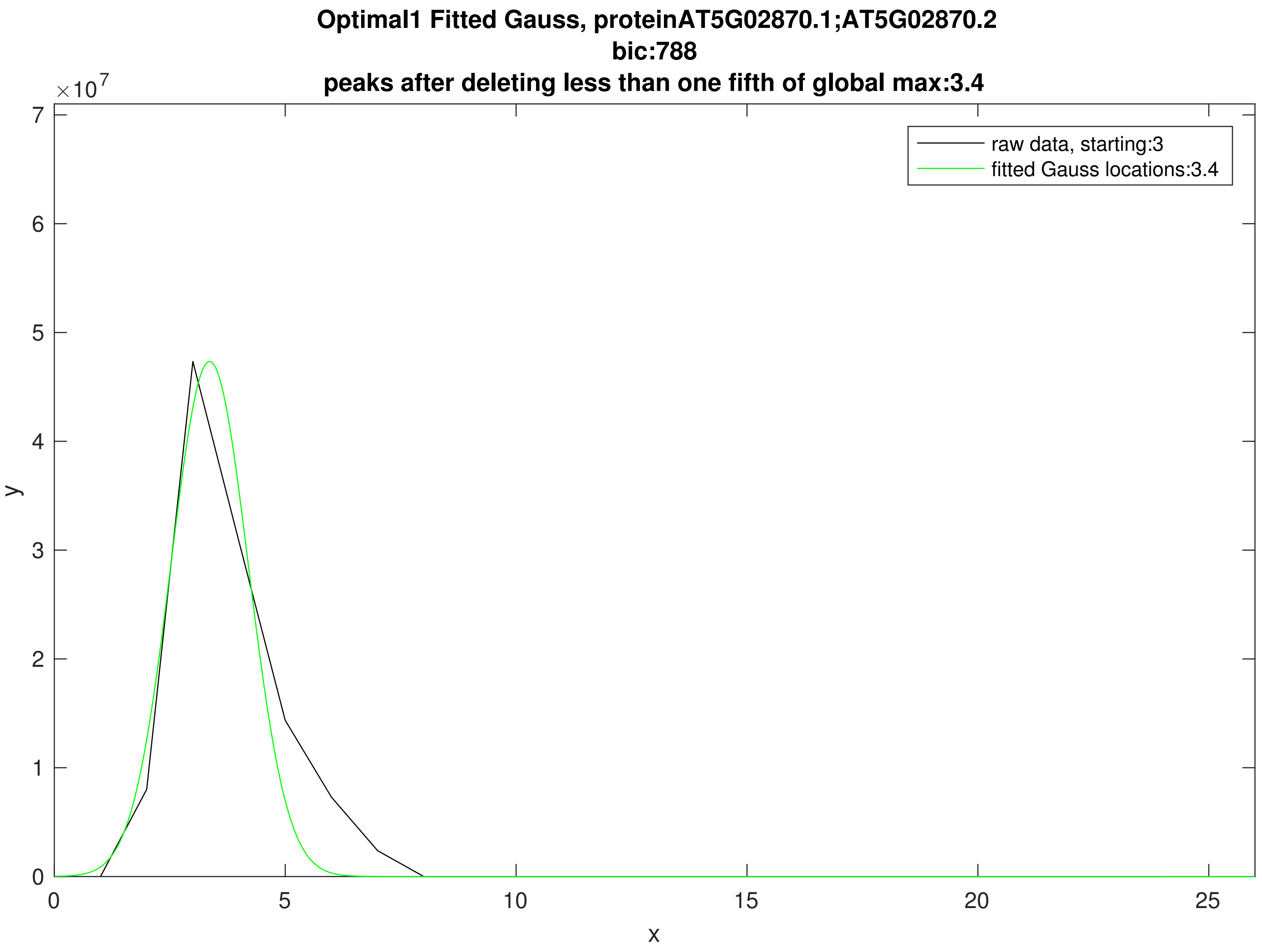
Optimal1 Fitted Gauss, proteinAT5G02870.1;AT5G02870.2
bic:788
peaks after deleting less than one fifth of global max:3.4
×10^7
raw data, starting:3
fitted Gauss locations:3.4
y
x
0
1
2
3
4
5
6
7
0
5
10
15
20
25

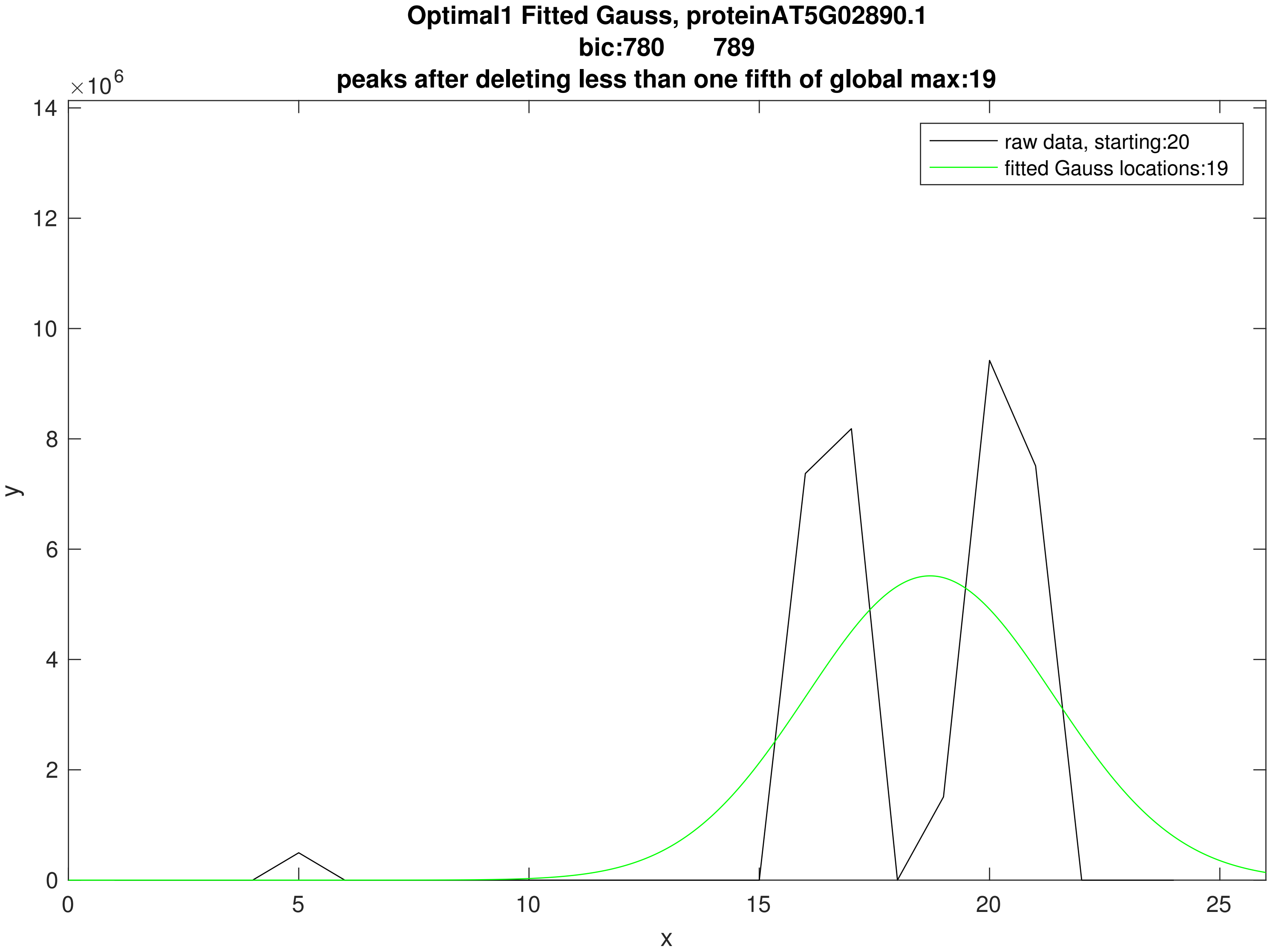
Optimal1 Fitted Gauss, proteinAT5G02890.1
bic:780 789
peaks after deleting less than one fifth of global max:19
×10^6
raw data, starting:20
fitted Gauss locations:19
y
x
0
2
4
6
8
10
12
14
0
5
10
15
20
25

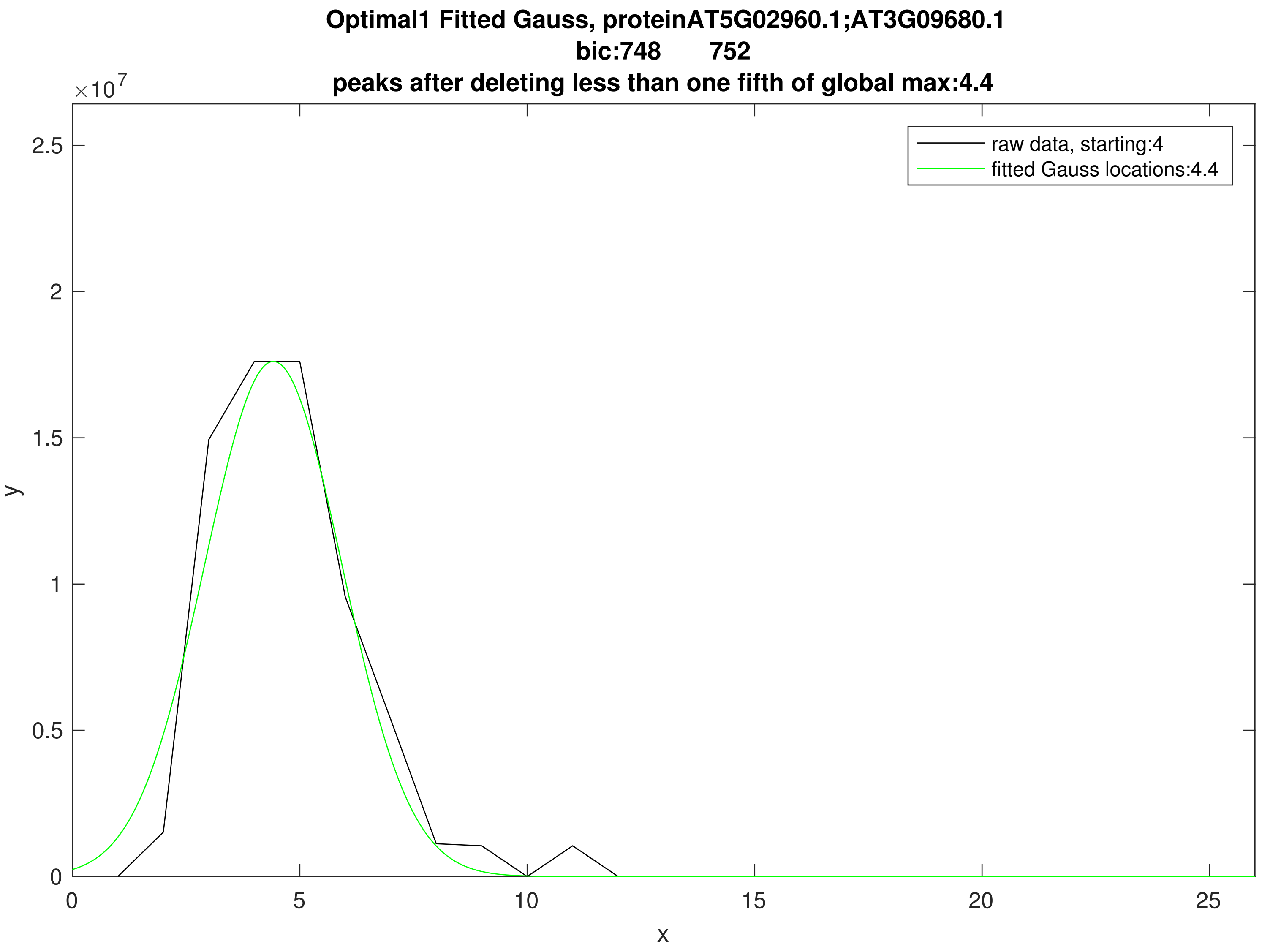

Optimal1 Fitted Gauss, proteinAT5G02960.1;AT3G09680.1
bic:748 752
peaks after deleting less than one fifth of global max:4.4
×10^7
2.5
2
1.5
1
0.5
0
y
0
5
10
15
20
25
x
raw data, starting:4
fitted Gauss locations:4.4

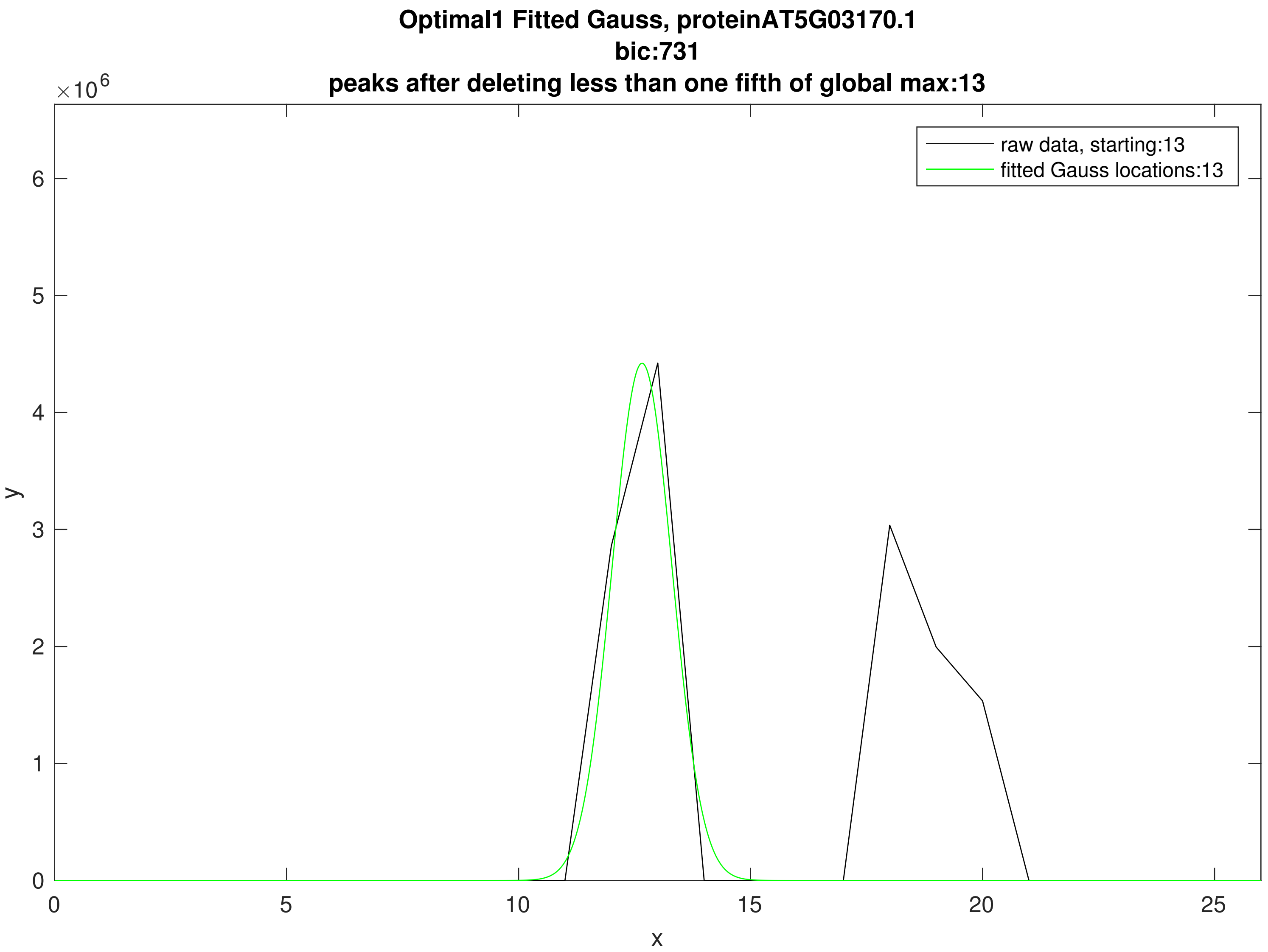

Optimal1 Fitted Gauss, proteinAT5G03170.1
bic:731
peaks after deleting less than one fifth of global max:13
×10^6
raw data, starting:13
fitted Gauss locations:13
y
x
0
1
2
3
4
5
6
0
5
10
15
20
25

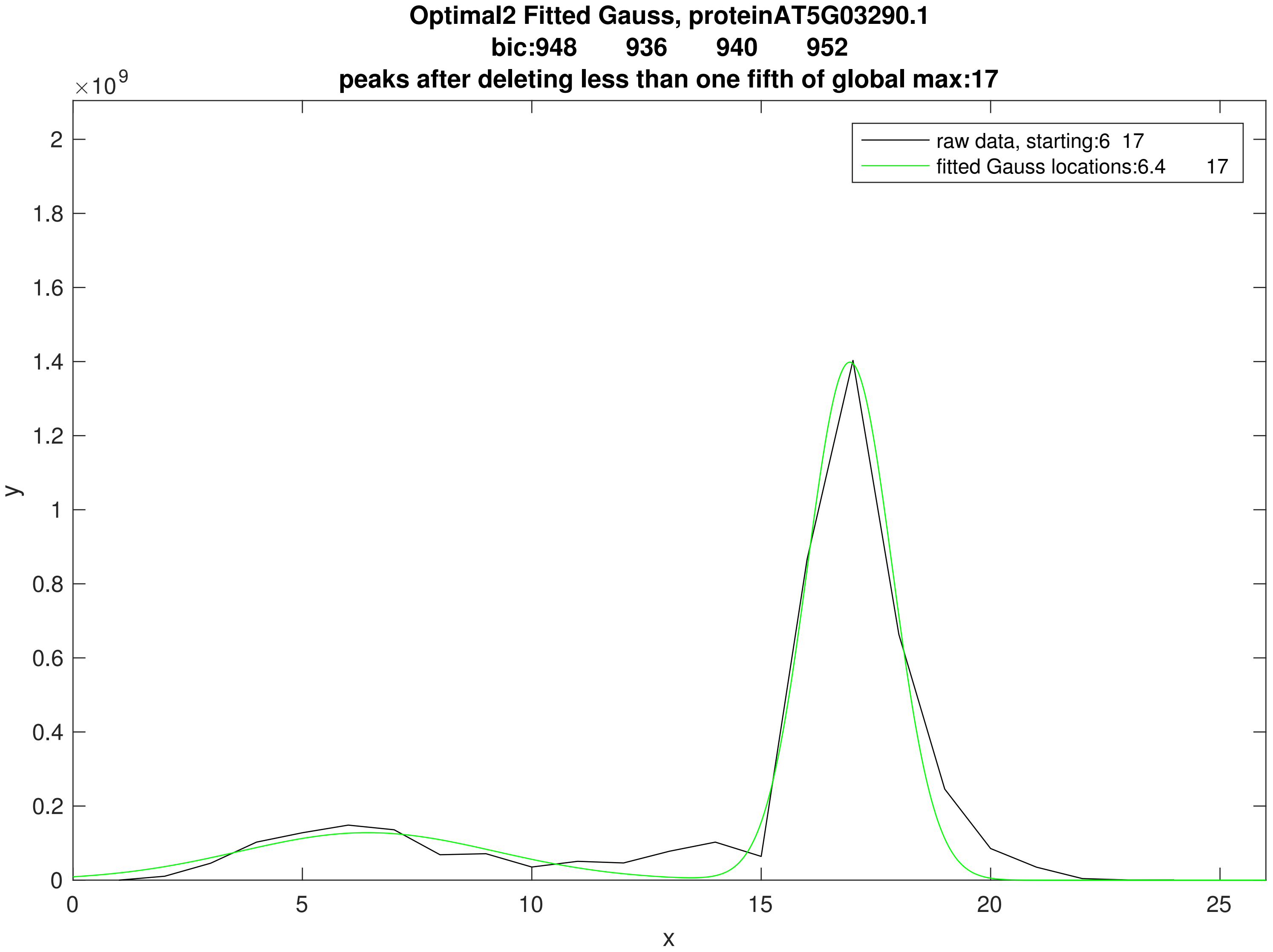

Optimal2 Fitted Gauss, proteinAT5G03290.1
bic:948 936 940 952
peaks after deleting less than one fifth of global max:17
×10^9
raw data, starting:6 17
fitted Gauss locations:6.4 17
y
x
0
0.2
0.4
0.6
0.8
1
1.2
1.4
1.6
1.8
2
0
5
10
15
20
25

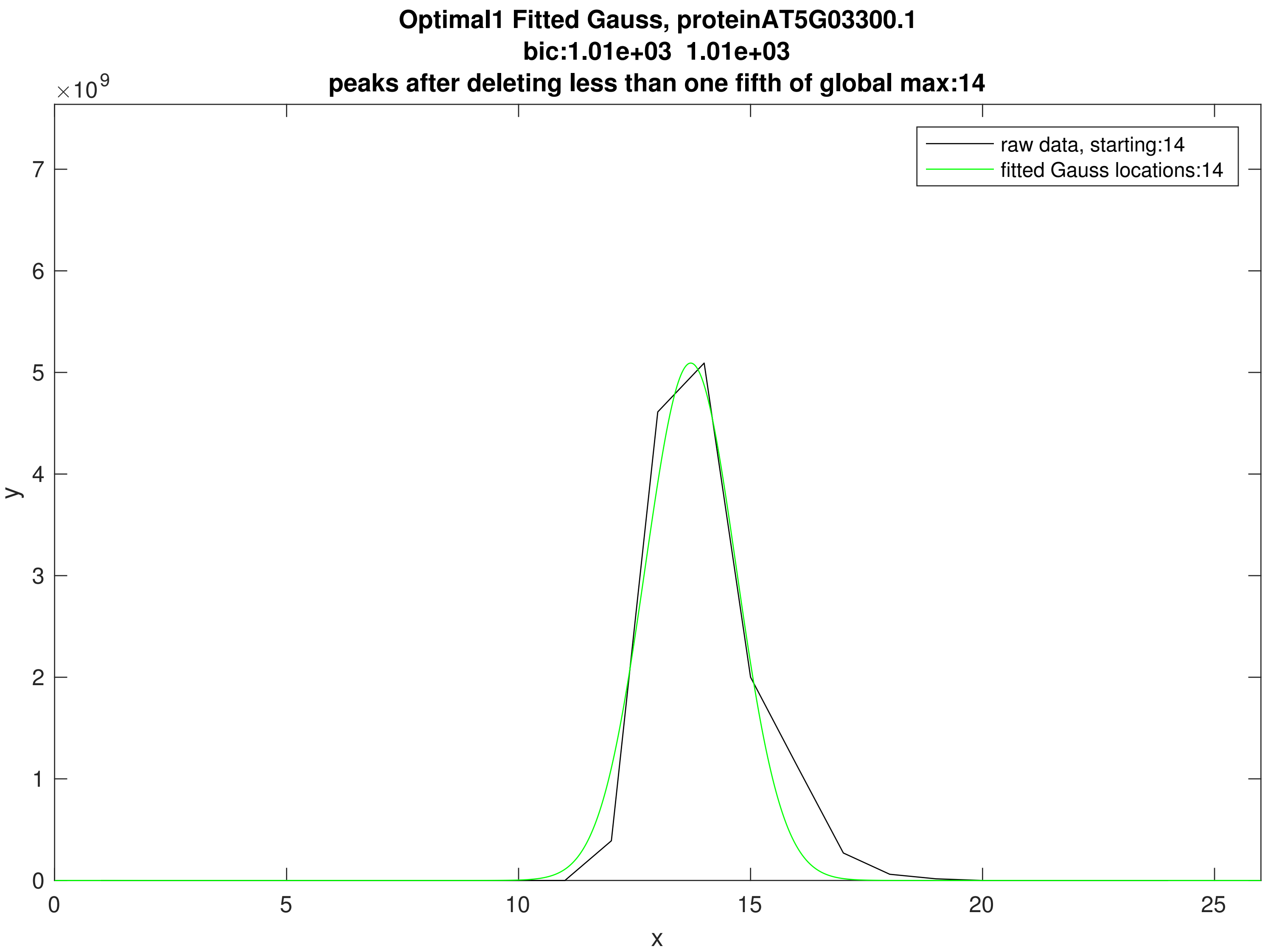
Optimal1 Fitted Gauss, proteinAT5G03300.1
bic:1.01e+03  1.01e+03
peaks after deleting less than one fifth of global max:14
×10^9
raw data, starting:14
fitted Gauss locations:14
y
x
0
1
2
3
4
5
6
7
0
5
10
15
20
25

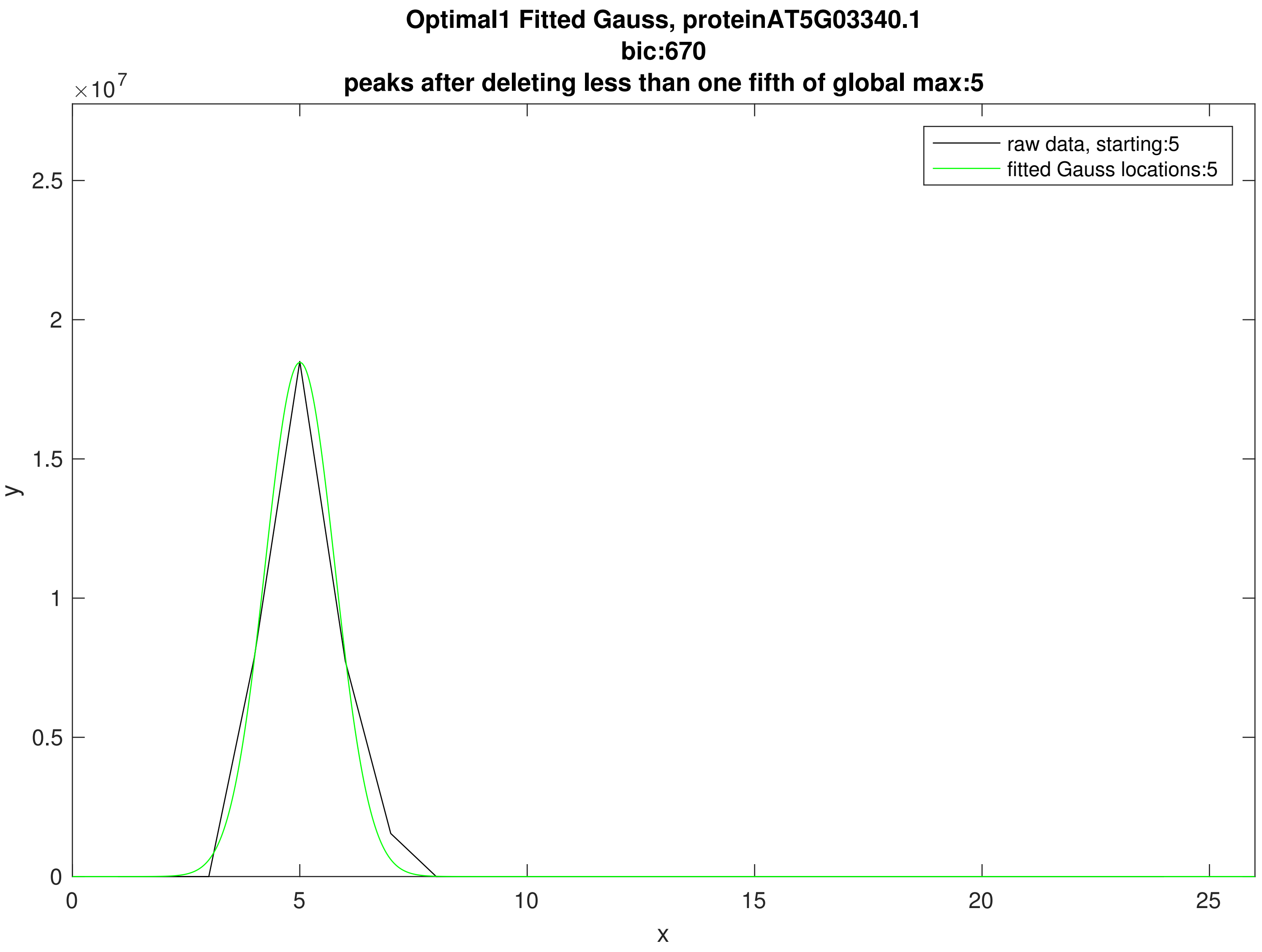
Optimal1 Fitted Gauss, proteinAT5G03340.1
bic:670
peaks after deleting less than one fifth of global max:5
×10^7
2.5
2
1.5
1
0.5
0
y
0
5
10
15
20
25
x
raw data, starting:5
fitted Gauss locations:5

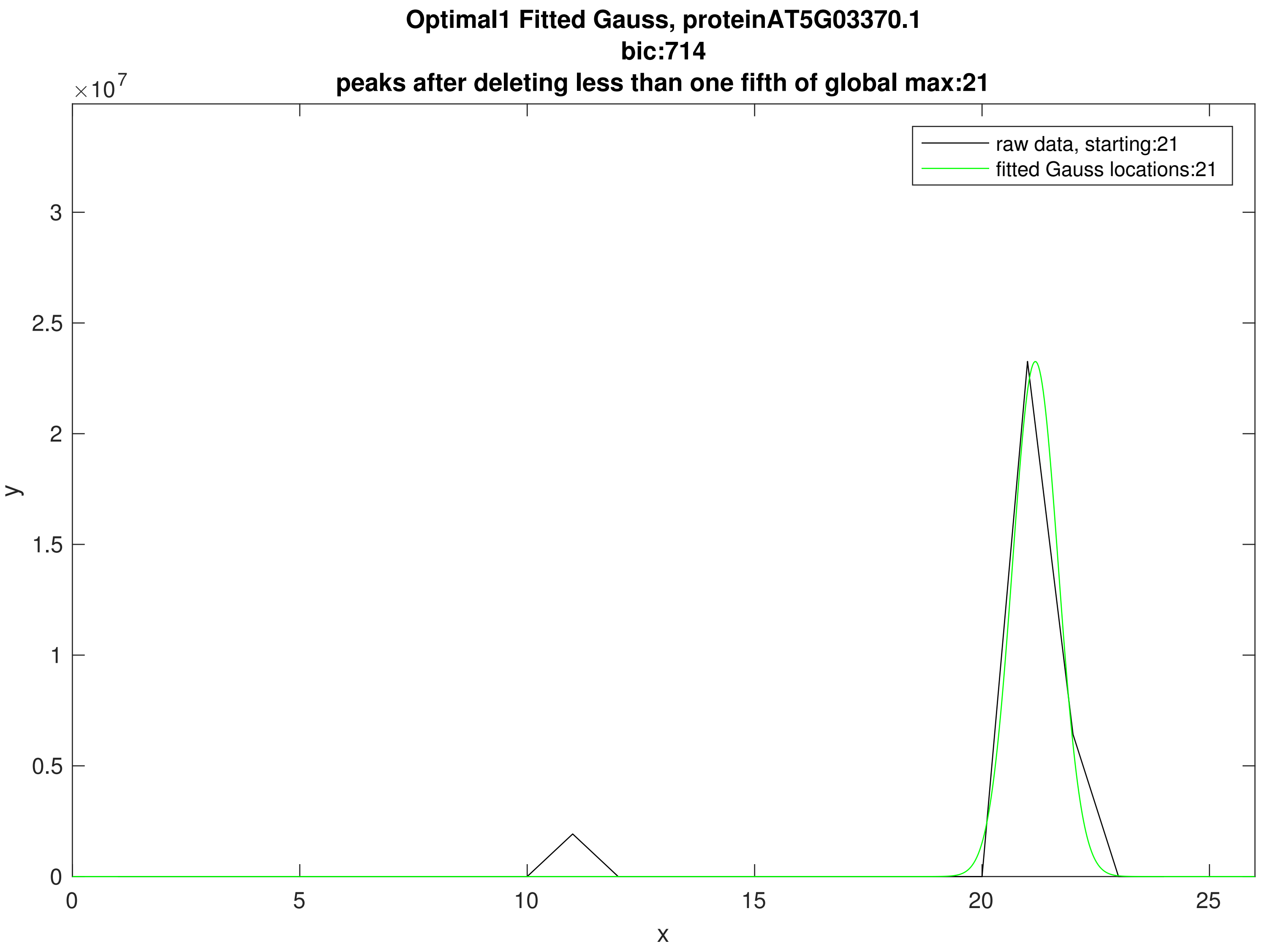
Optimal1 Fitted Gauss, proteinAT5G03370.1
bic:714
peaks after deleting less than one fifth of global max:21
×10^7
raw data, starting:21
fitted Gauss locations:21
y
x
0
0.5
1
1.5
2
2.5
3
0
5
10
15
20
25

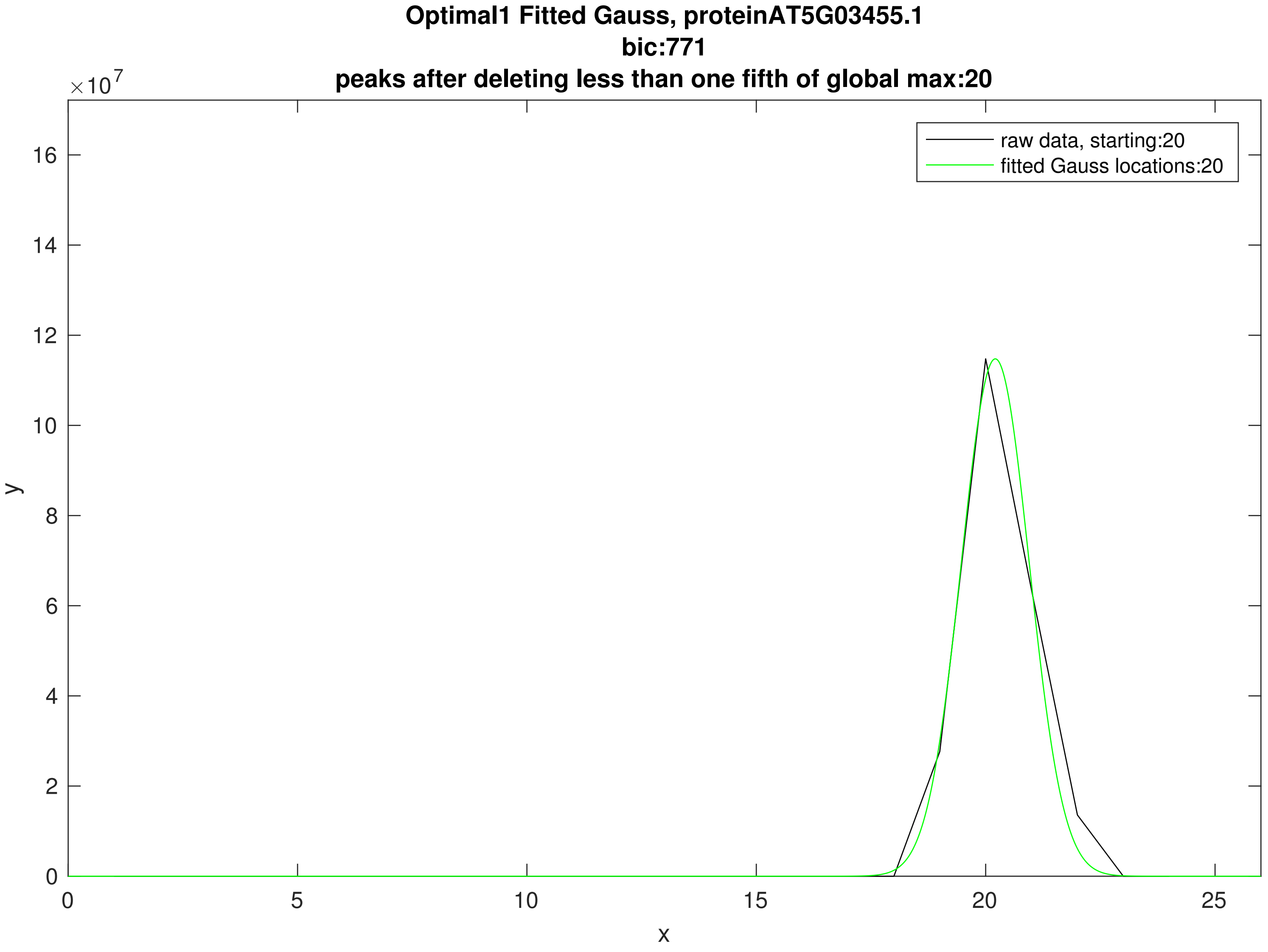

Optimal1 Fitted Gauss, proteinAT5G03455.1
bic:771
peaks after deleting less than one fifth of global max:20
×10^7
16
14
12
10
8
6
4
2
0
y
0
5
10
15
20
25
x
raw data, starting:20
fitted Gauss locations:20

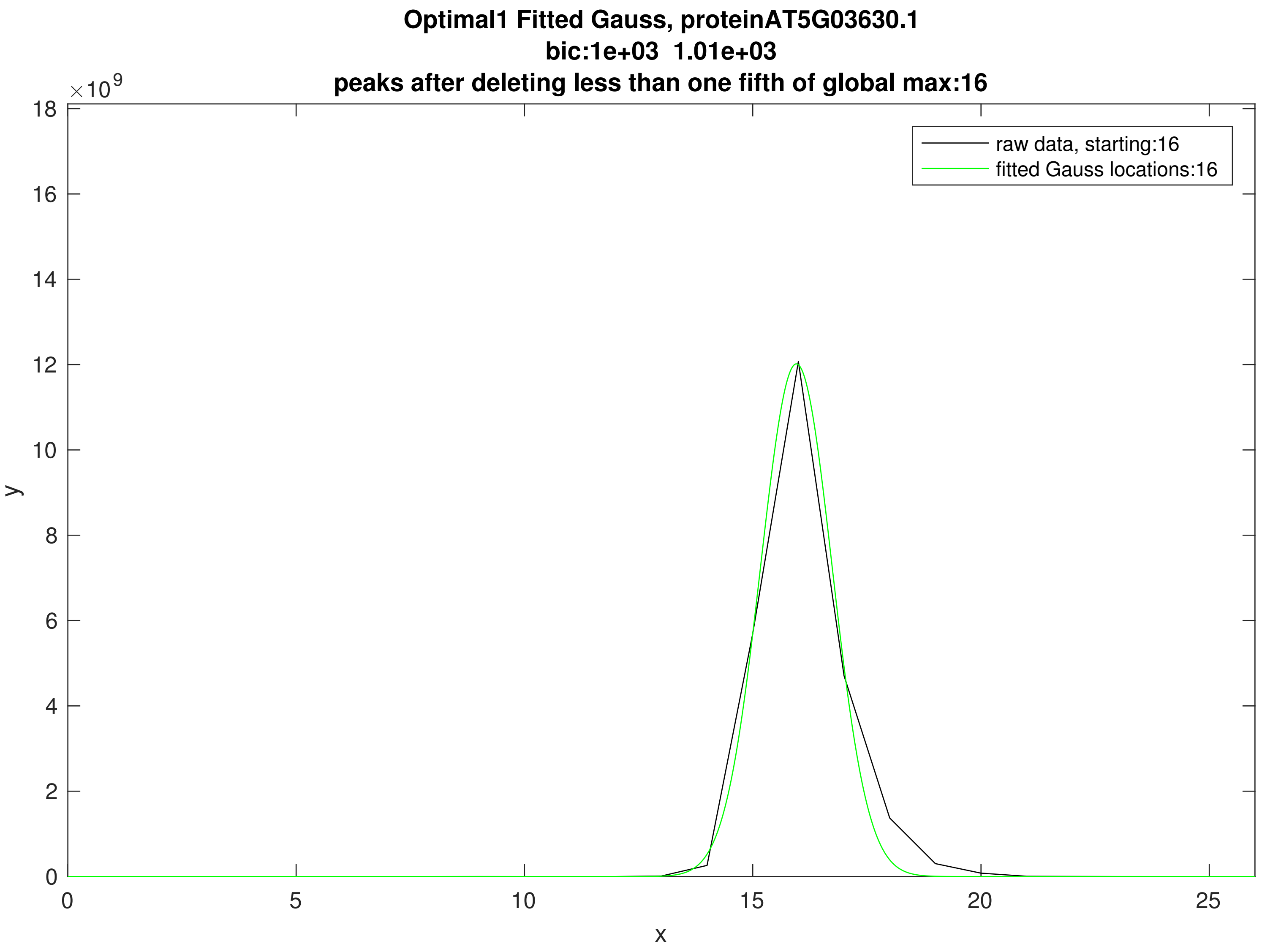
Optimal1 Fitted Gauss, proteinAT5G03630.1
bic:1e+03  1.01e+03
peaks after deleting less than one fifth of global max:16
×10^9
raw data, starting:16
fitted Gauss locations:16
y
x
0
2
4
6
8
10
12
14
16
18
0
5
10
15
20
25

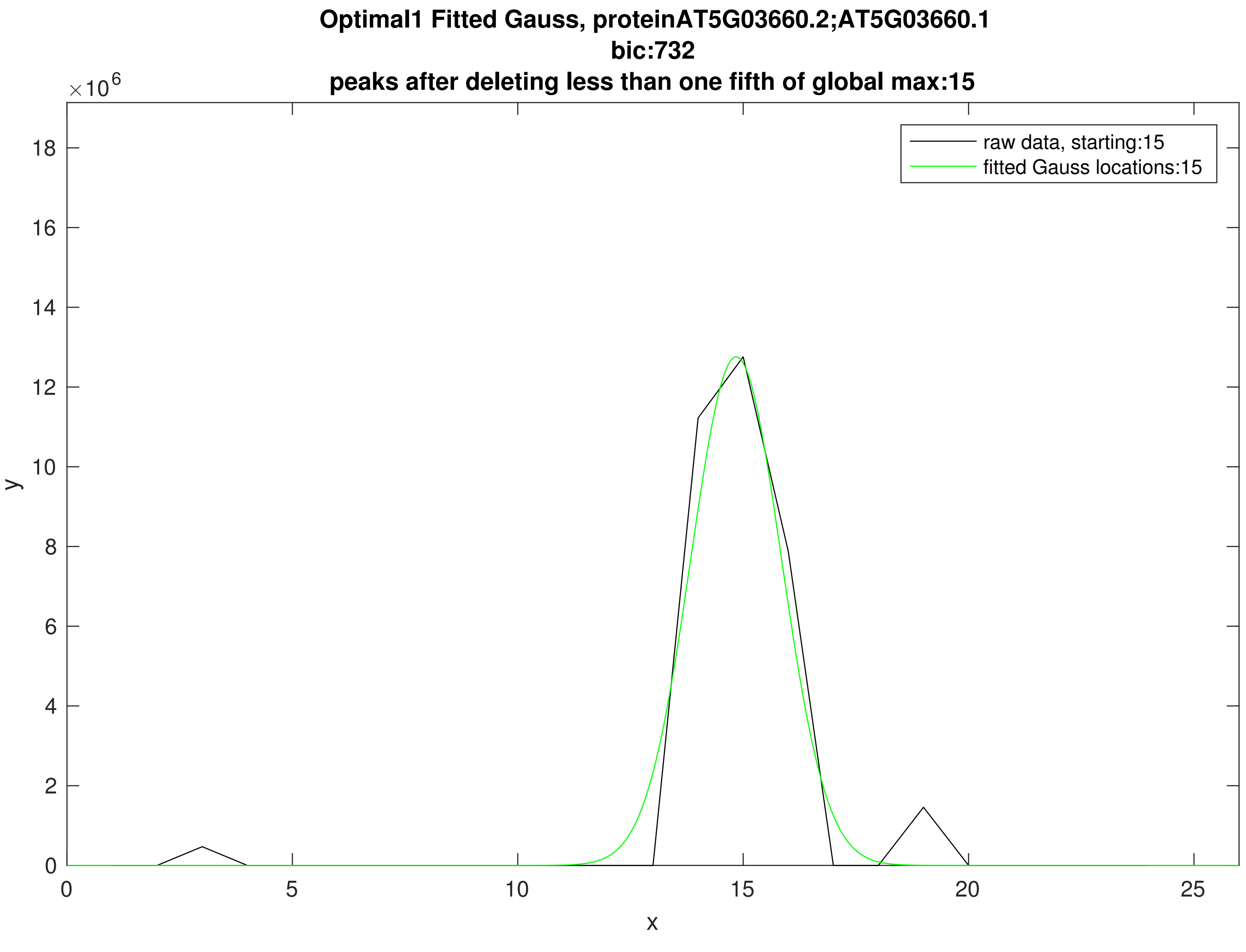

Optimal1 Fitted Gauss, proteinAT5G03660.2;AT5G03660.1
bic:732
peaks after deleting less than one fifth of global max:15
×10^6
18
16
14
12
10
8
6
4
2
0
y
0
5
10
15
20
25
x
raw data, starting:15
fitted Gauss locations:15

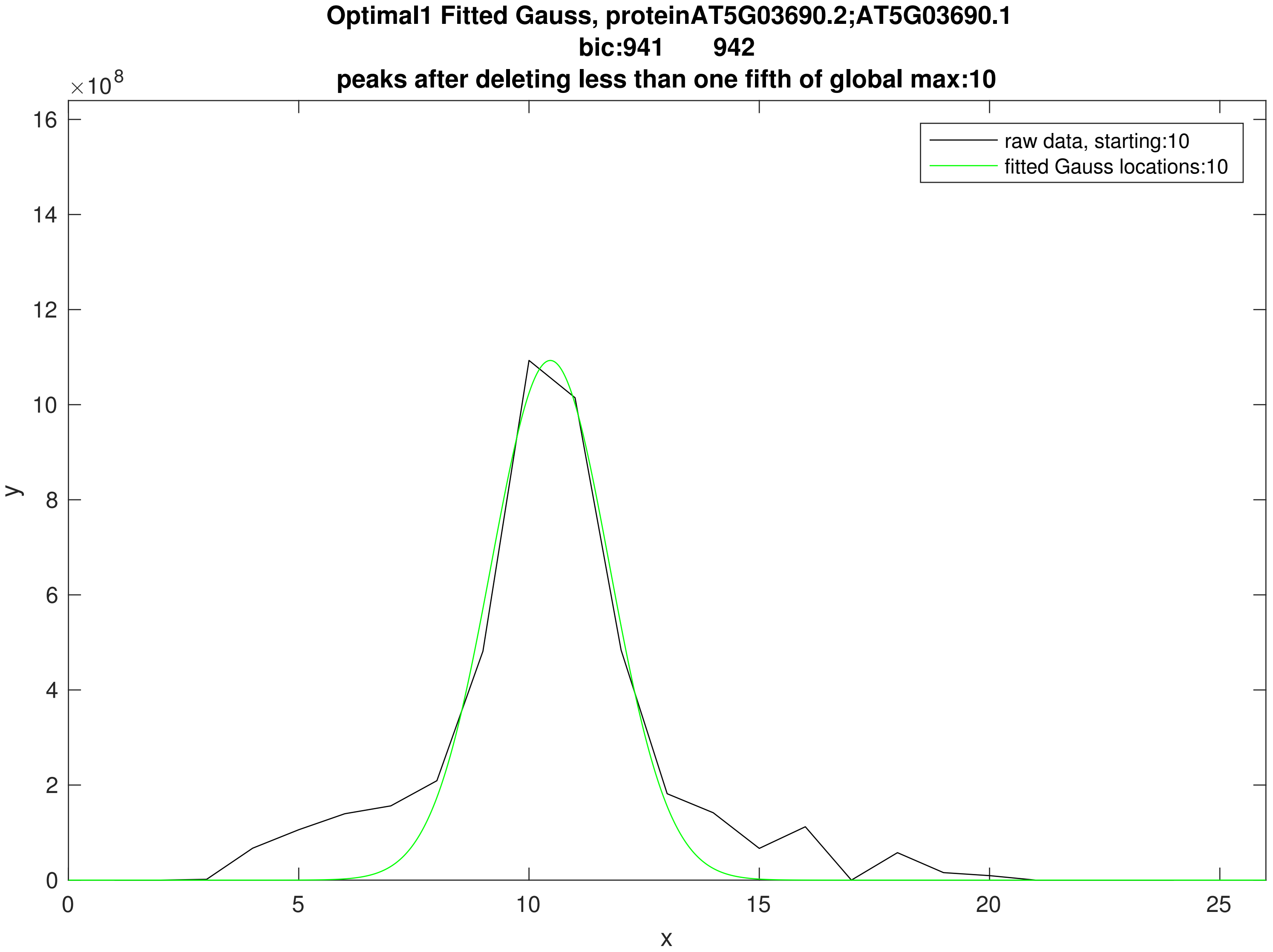

Optimal1 Fitted Gauss, proteinAT5G03690.2;AT5G03690.1
bic:941 942
peaks after deleting less than one fifth of global max:10
×10^8
16
14
12
10
8
6
4
2
0
y
0
5
10
15
20
25
x
raw data, starting:10
fitted Gauss locations:10

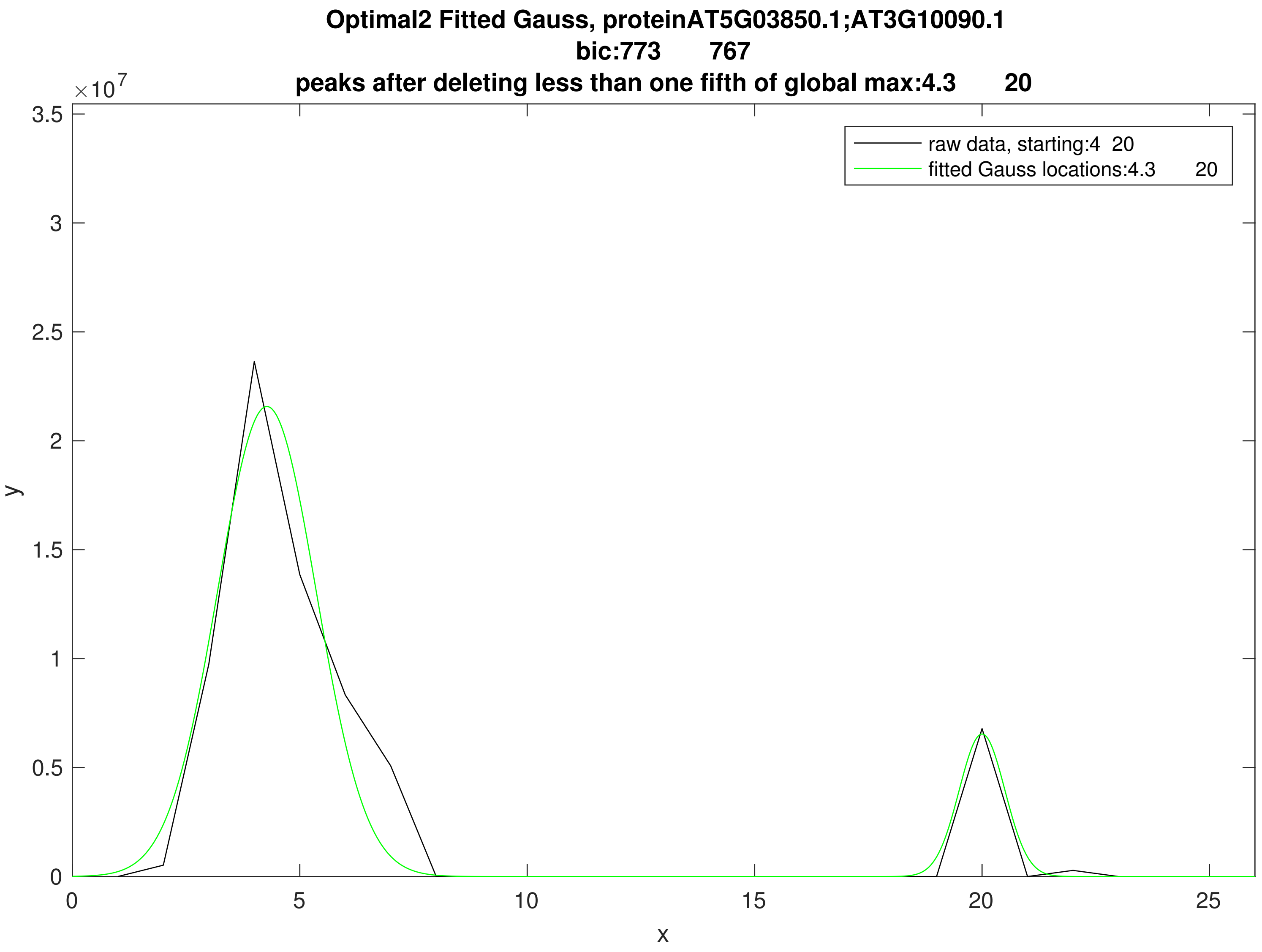
Optimal2 Fitted Gauss, proteinAT5G03850.1;AT3G10090.1
bic:773 767
peaks after deleting less than one fifth of global max:4.3 20
×10^7
3.5
3
2.5
2
1.5
1
0.5
0
y
0
5
10
15
20
25
x
raw data, starting:4 20
fitted Gauss locations:4.3 20

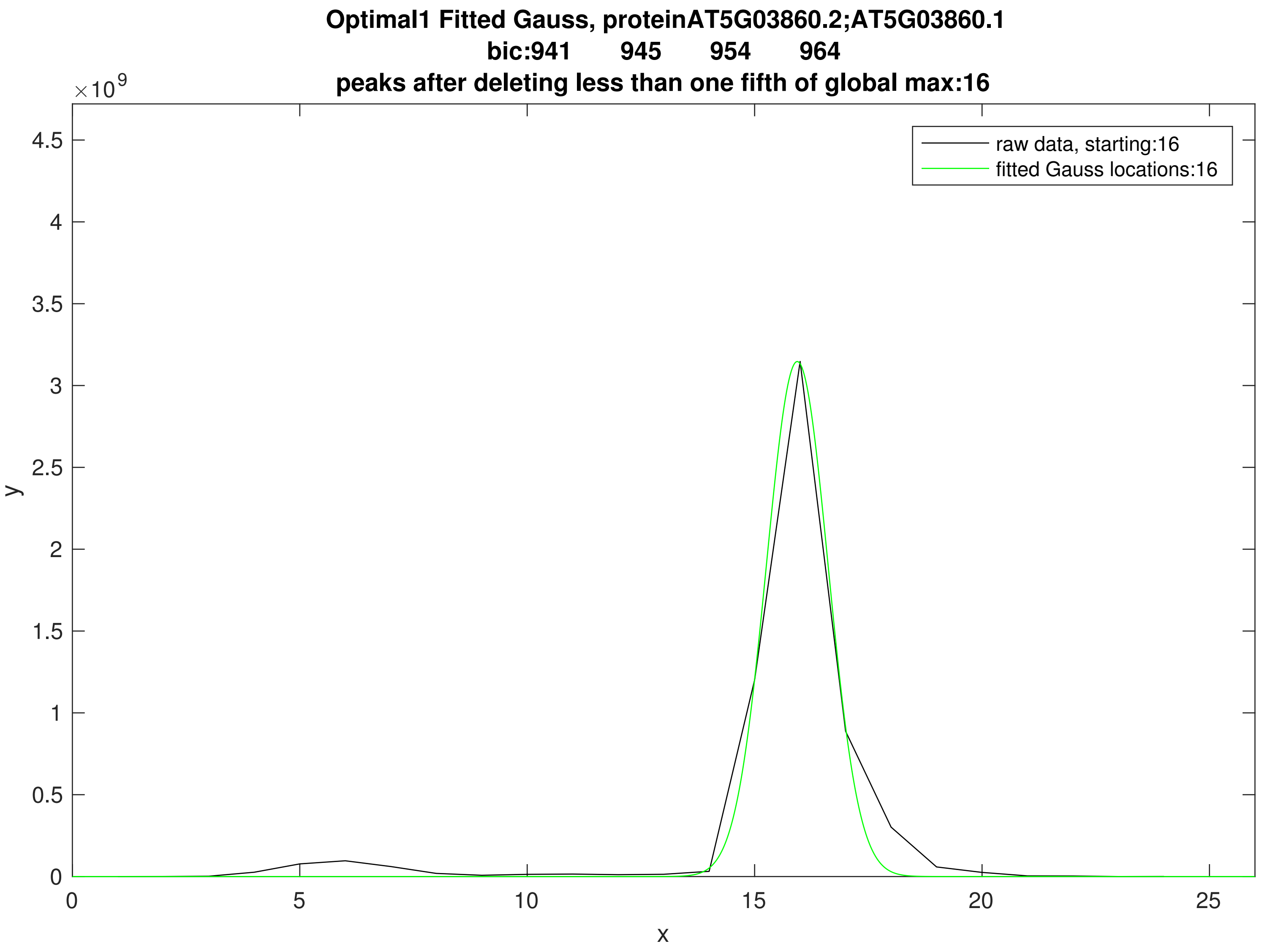

Optimal1 Fitted Gauss, proteinAT5G03860.2;AT5G03860.1
bic:941 945 954 964
peaks after deleting less than one fifth of global max:16
×10^9
4.5
4
3.5
3
2.5
2
1.5
1
0.5
0
y
0
5
10
15
20
25
x
raw data, starting:16
fitted Gauss locations:16

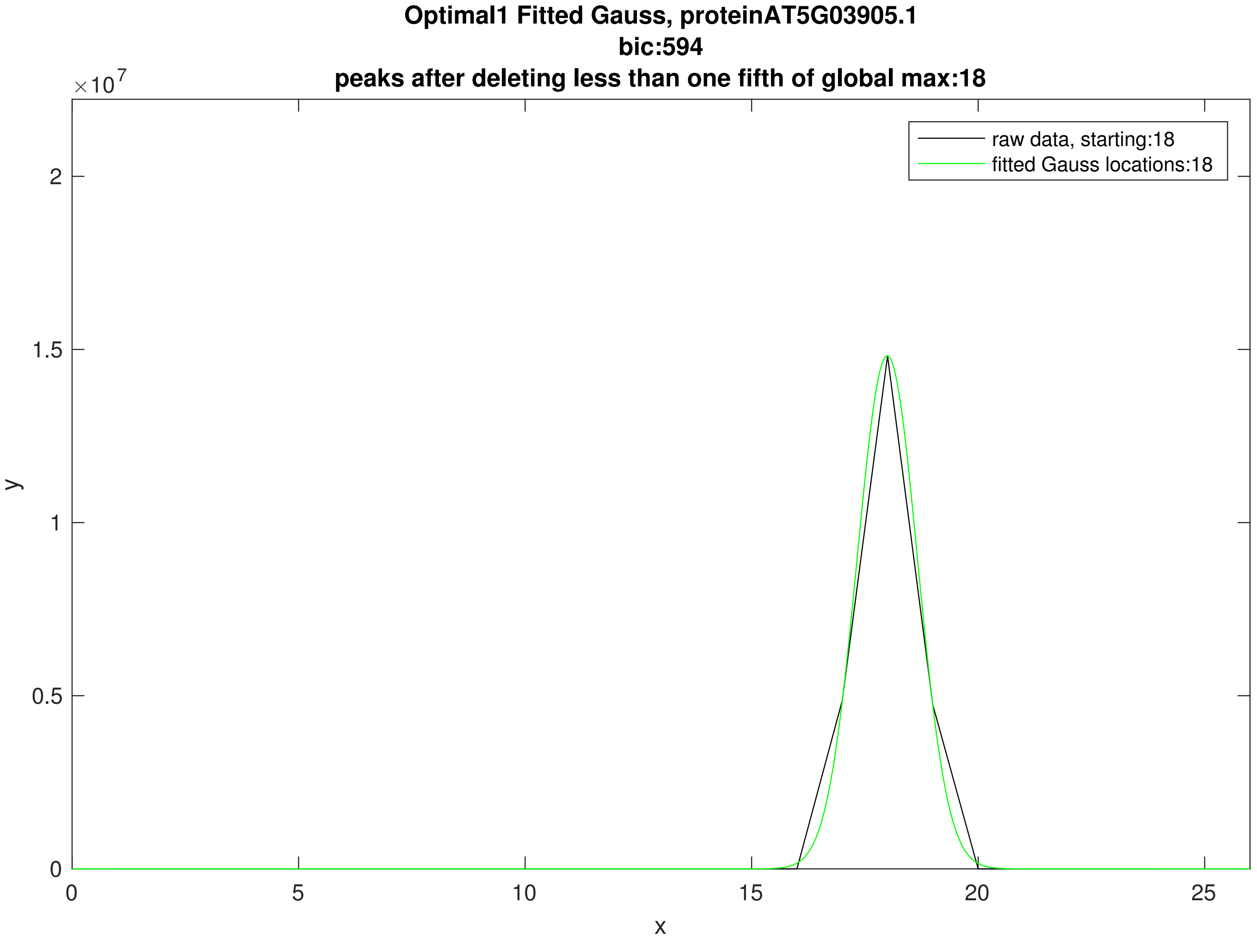
Optimal1 Fitted Gauss, proteinAT5G03905.1
bic:594
peaks after deleting less than one fifth of global max:18
×10^7
raw data, starting:18
fitted Gauss locations:18
2
1.5
1
0.5
0
y
0
5
10
15
20
25
x

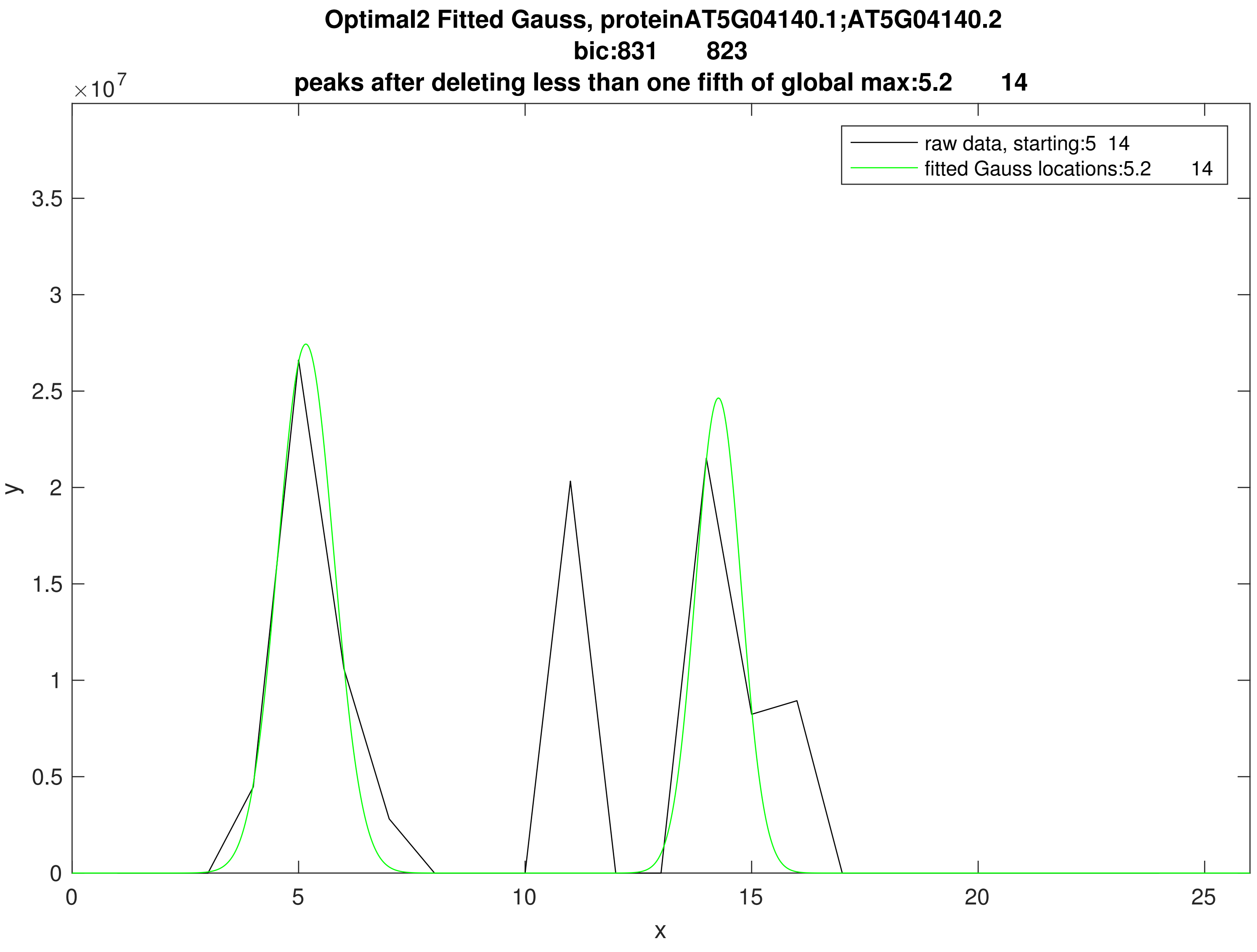

Optimal2 Fitted Gauss, proteinAT5G04140.1;AT5G04140.2
bic:831 823
peaks after deleting less than one fifth of global max:5.2 14
×10^7
raw data, starting:5 14
fitted Gauss locations:5.2 14
3.5
3
2.5
2
1.5
1
0.5
0
y
0
5
10
15
20
25
x

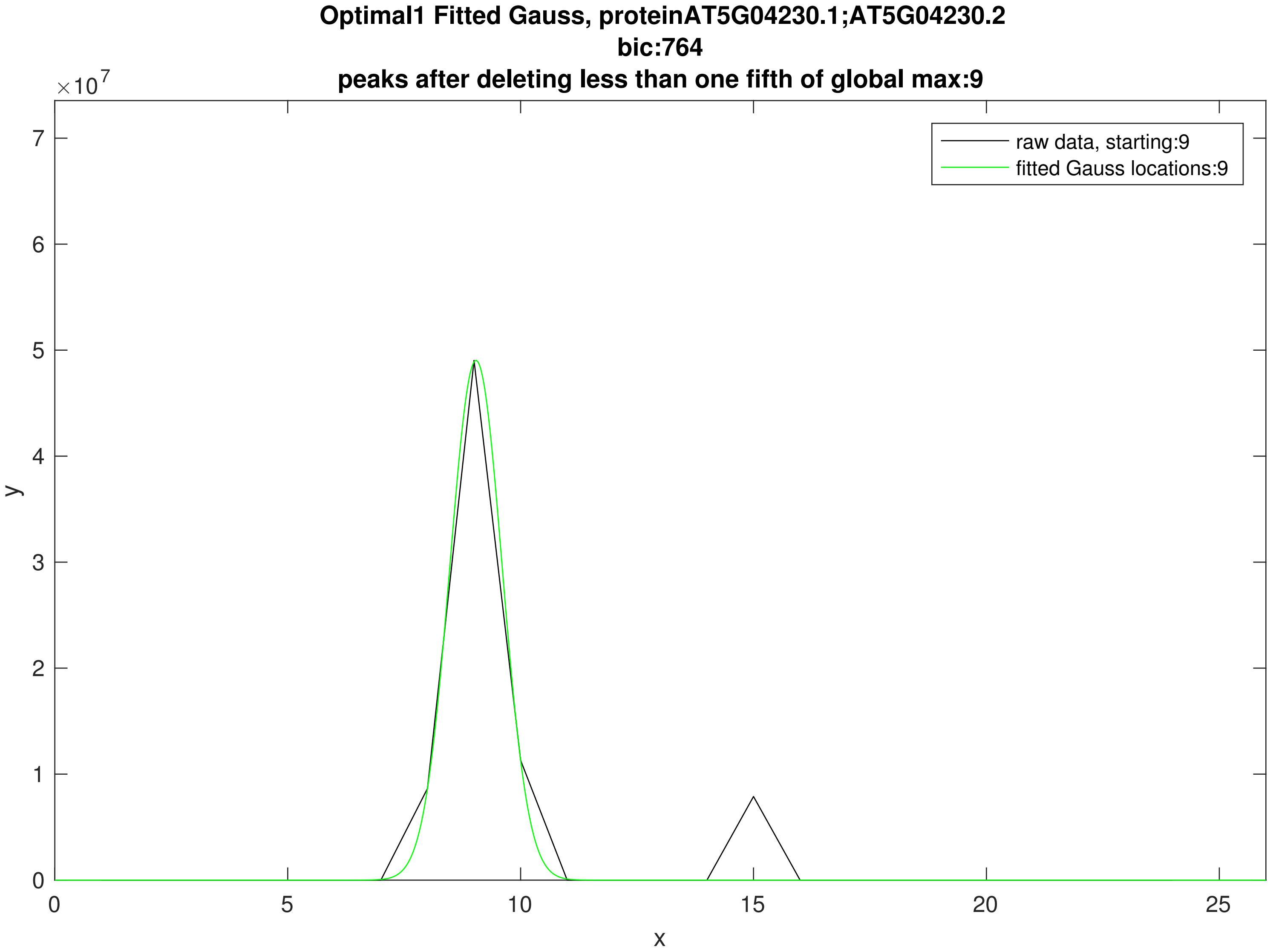
Optimal1 Fitted Gauss, proteinAT5G04230.1;AT5G04230.2
bic:764
peaks after deleting less than one fifth of global max:9
×10^7
raw data, starting:9
fitted Gauss locations:9
y
x
0
1
2
3
4
5
6
7
0
5
10
15
20
25

**Optimal1 Fitted Gauss, proteinAT5G04260.1**
**bic:605**
**peaks after deleting less than one fifth of global max:6**

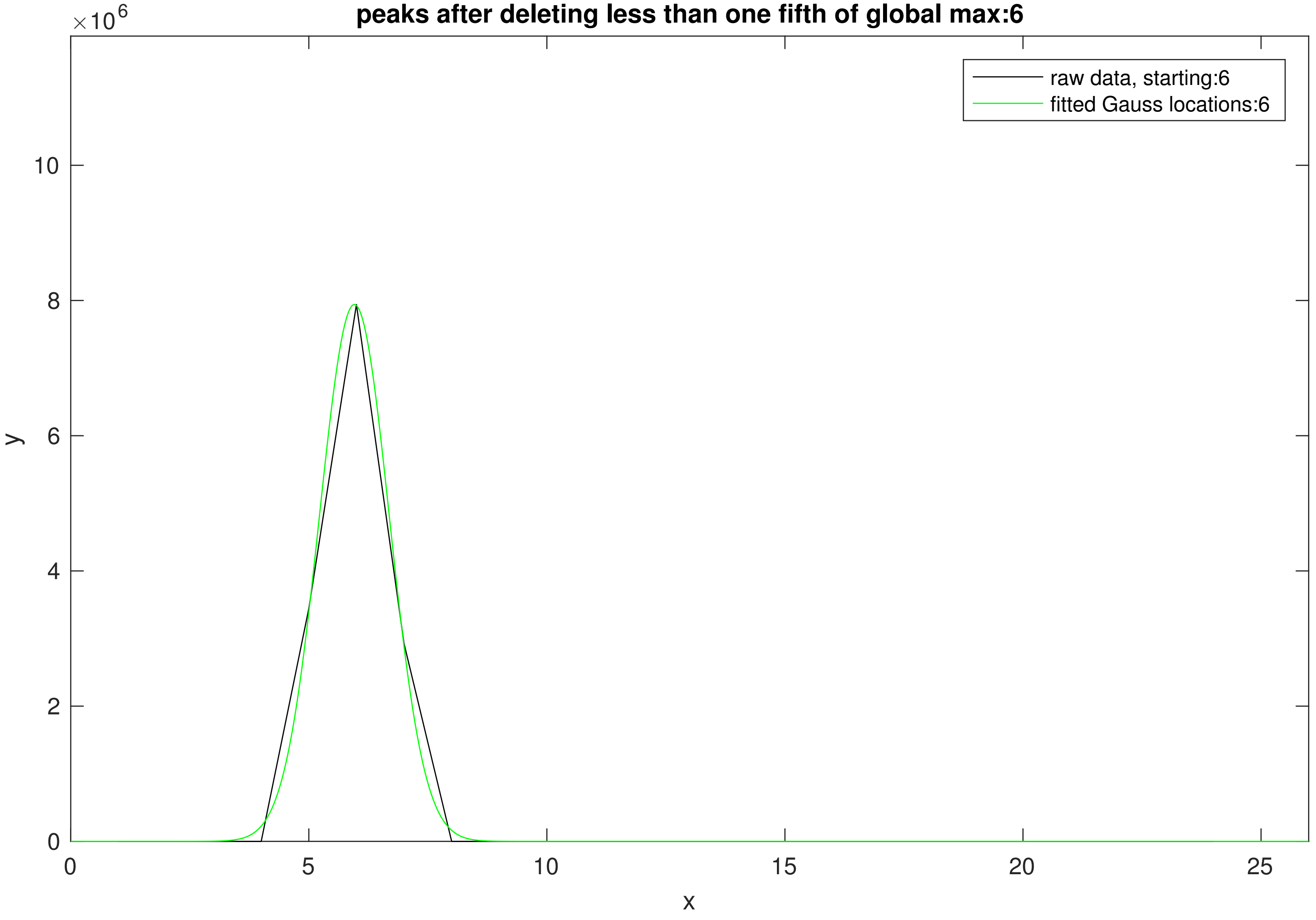

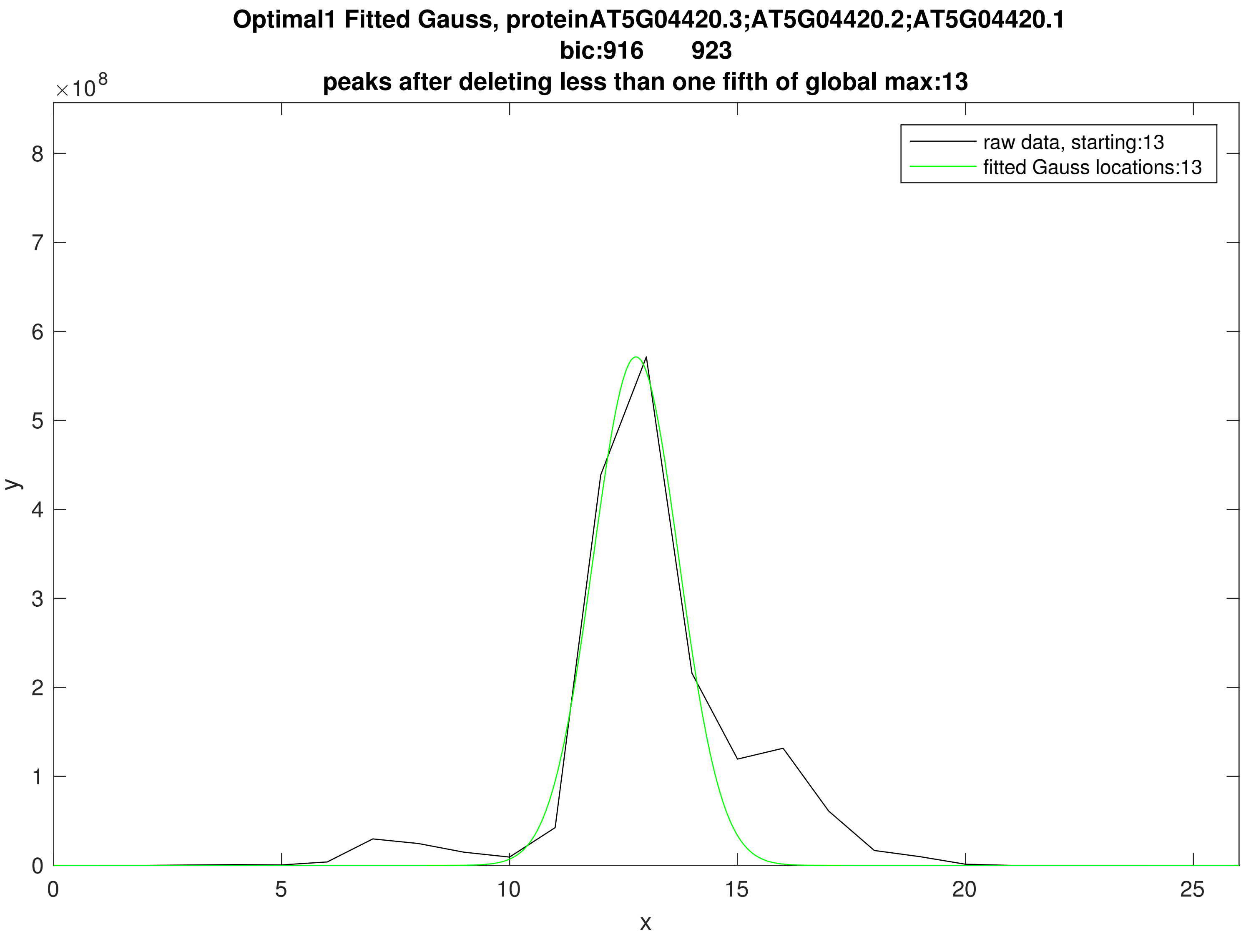

Optimal1 Fitted Gauss, proteinAT5G04420.3;AT5G04420.2;AT5G04420.1
bic:916 923
peaks after deleting less than one fifth of global max:13
×10^8
raw data, starting:13
fitted Gauss locations:13
y
x
0
1
2
3
4
5
6
7
8
0
5
10
15
20
25

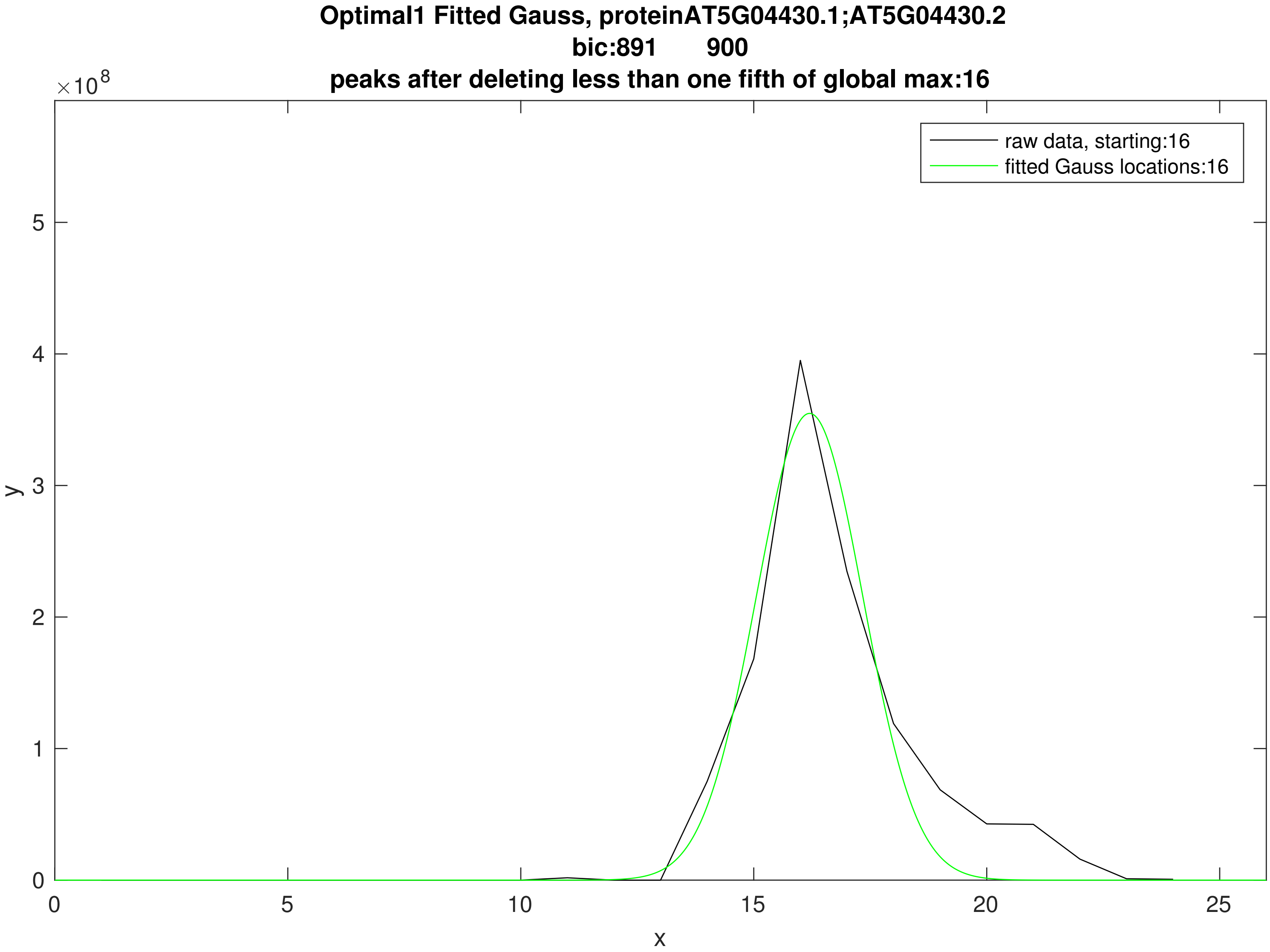

Optimal1 Fitted Gauss, proteinAT5G04430.1;AT5G04430.2
bic:891 900
peaks after deleting less than one fifth of global max:16
×10^8
raw data, starting:16
fitted Gauss locations:16
y
x
0
1
2
3
4
5
0
5
10
15
20
25

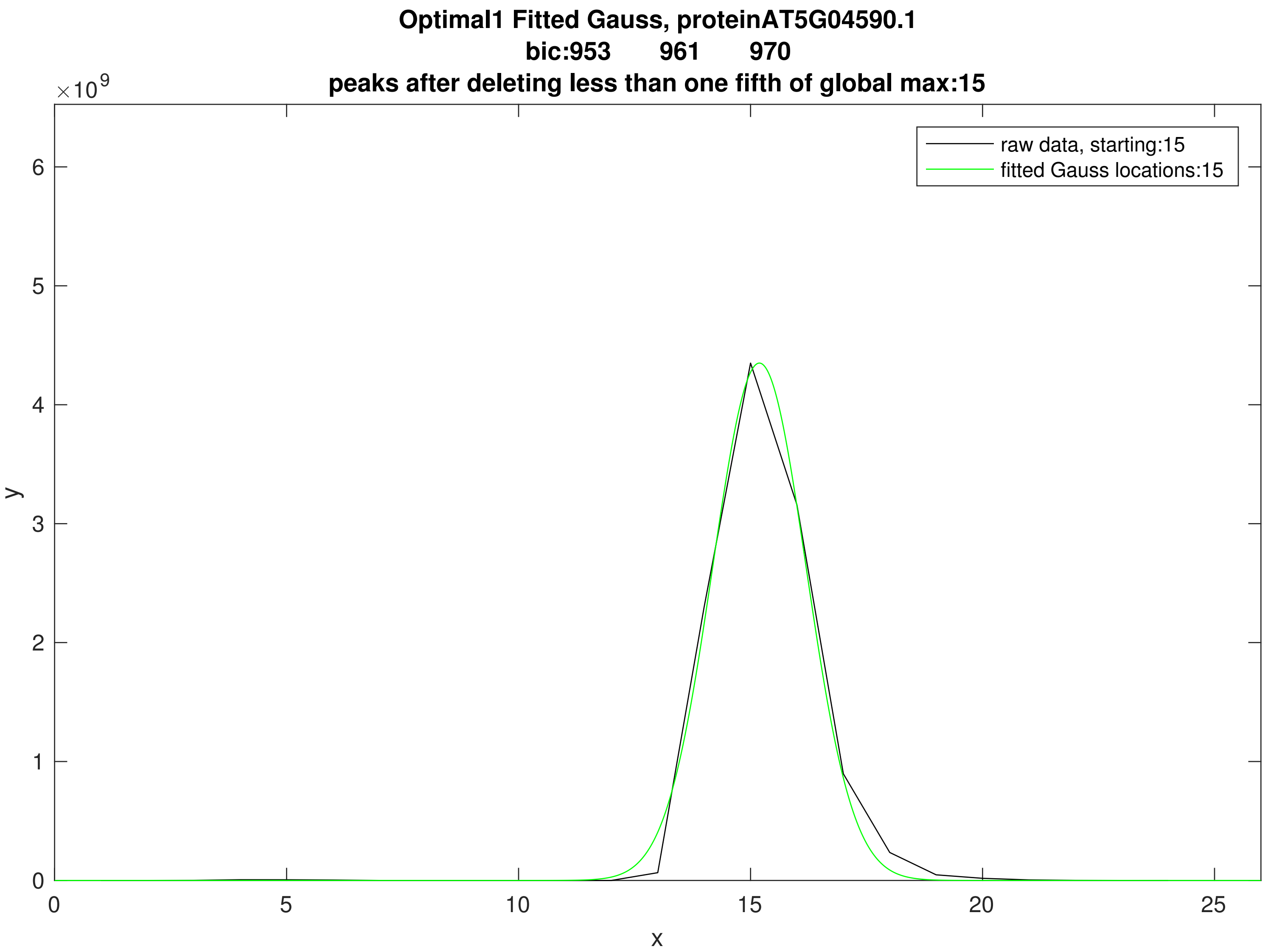

Optimal1 Fitted Gauss, proteinAT5G04590.1
bic:953 961 970
peaks after deleting less than one fifth of global max:15
×10^9
raw data, starting:15
fitted Gauss locations:15
y
x
0
1
2
3
4
5
6
0
5
10
15
20
25

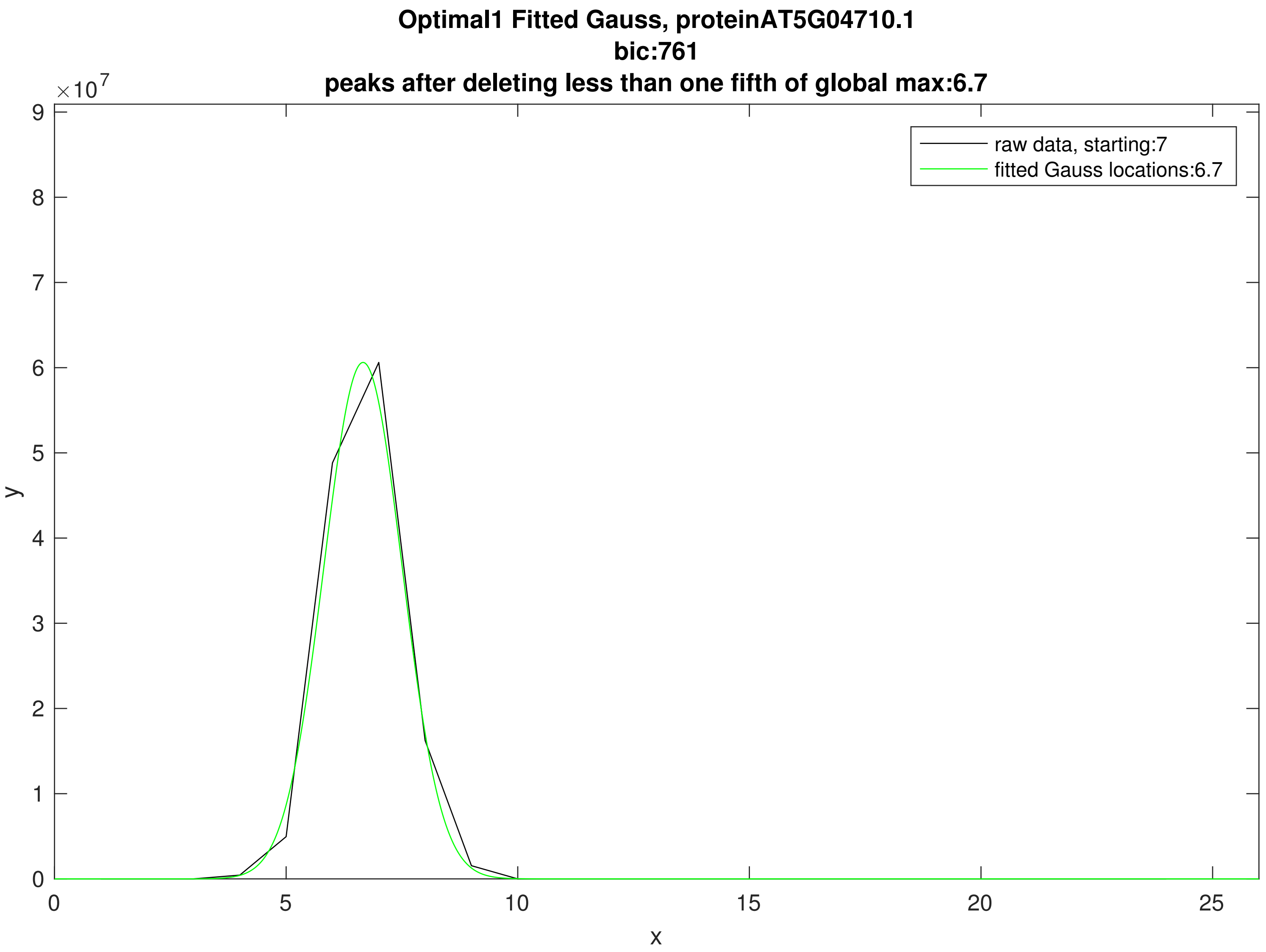
Optimal1 Fitted Gauss, proteinAT5G04710.1
bic:761
peaks after deleting less than one fifth of global max:6.7
×10^7
raw data, starting:7
fitted Gauss locations:6.7
y
x
0
1
2
3
4
5
6
7
8
9
0
5
10
15
20
25

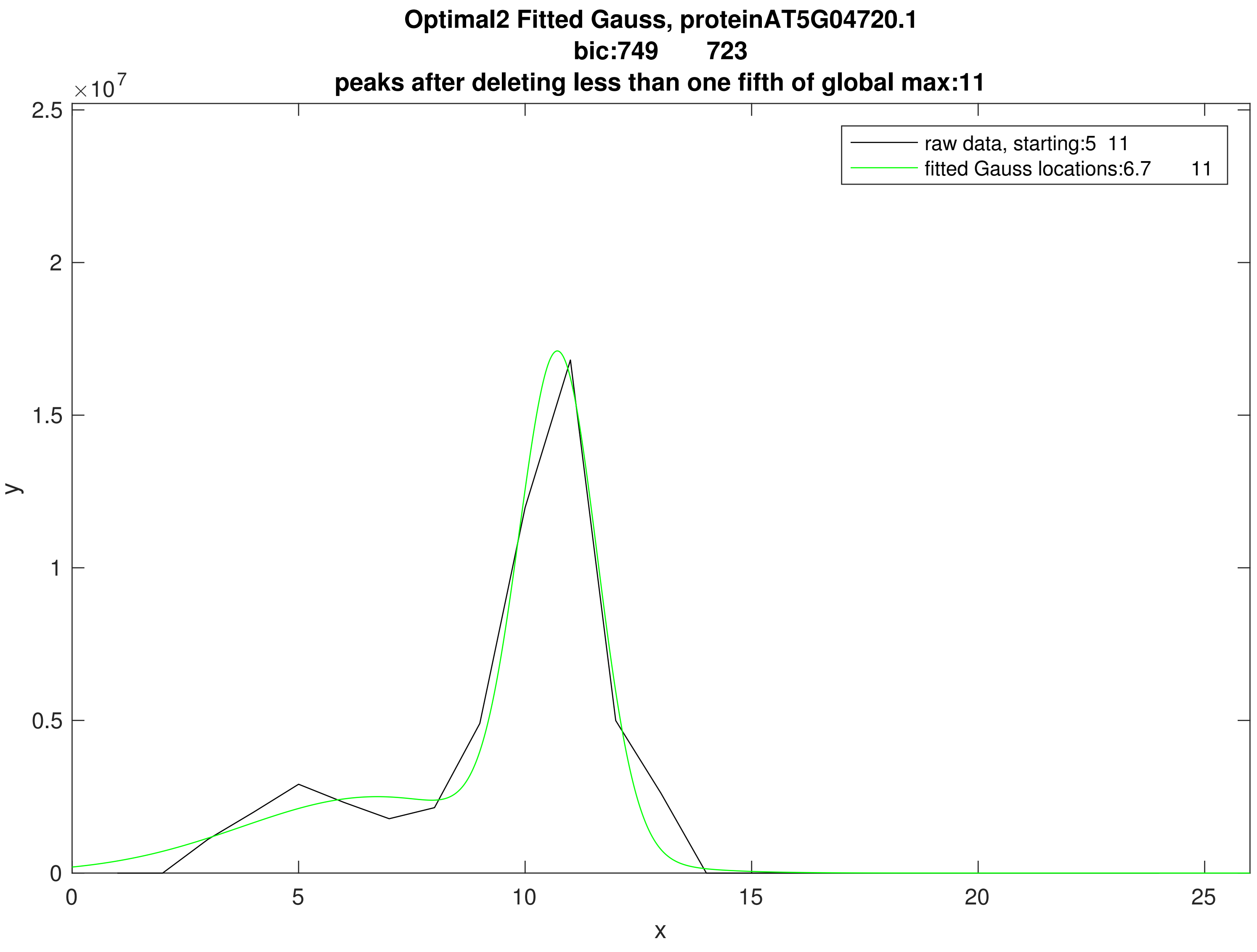

Optimal2 Fitted Gauss, proteinAT5G04720.1
bic:749 723
peaks after deleting less than one fifth of global max:11
×10^7
2.5
2
1.5
1
0.5
0
y
0
5
10
15
20
25
x
raw data, starting:5 11
fitted Gauss locations:6.7 11

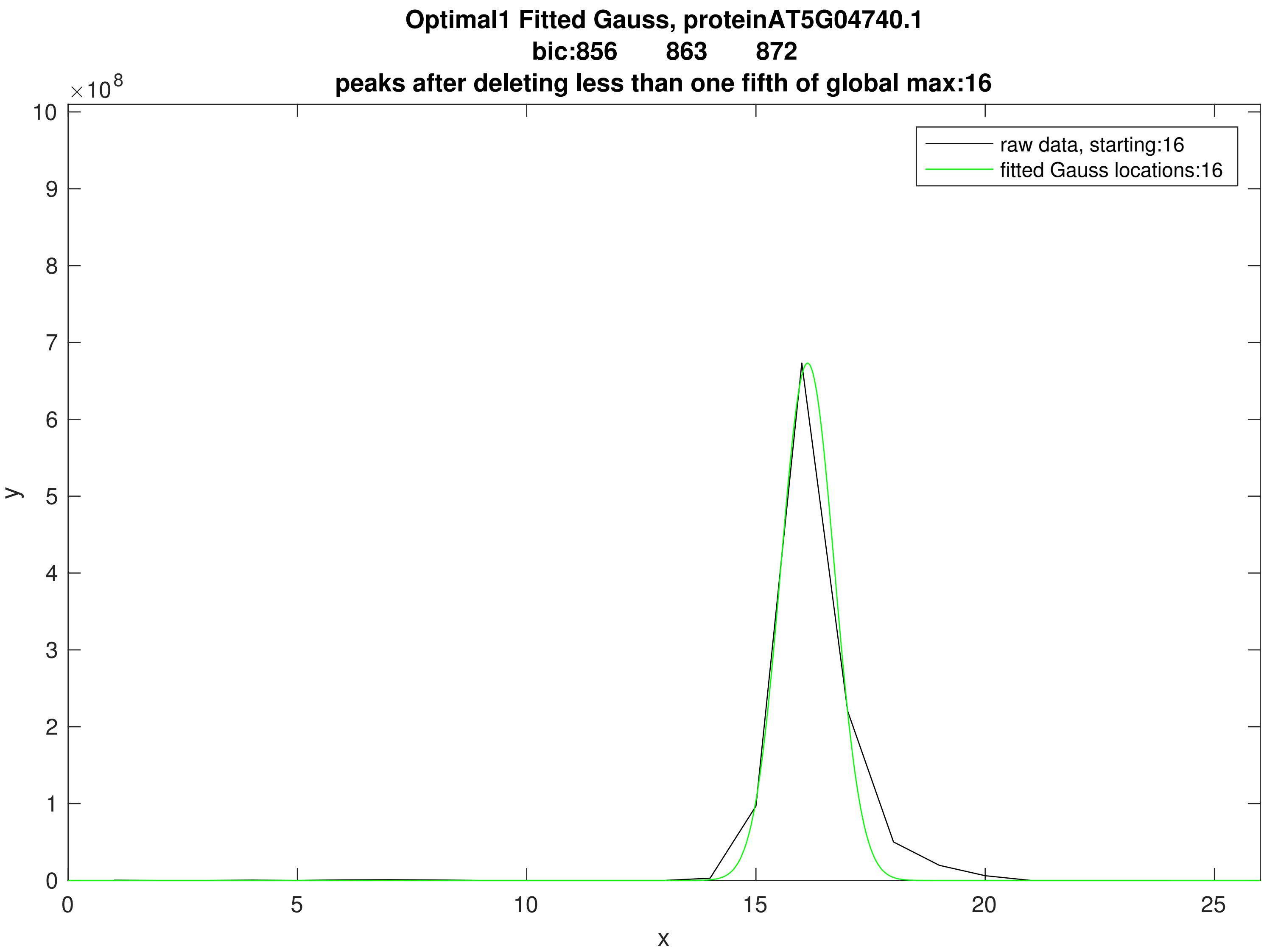
Optimal1 Fitted Gauss, proteinAT5G04740.1
bic:856 863 872
peaks after deleting less than one fifth of global max:16
×10^8
raw data, starting:16
fitted Gauss locations:16
y
x
0
1
2
3
4
5
6
7
8
9
10
0
5
10
15
20
25

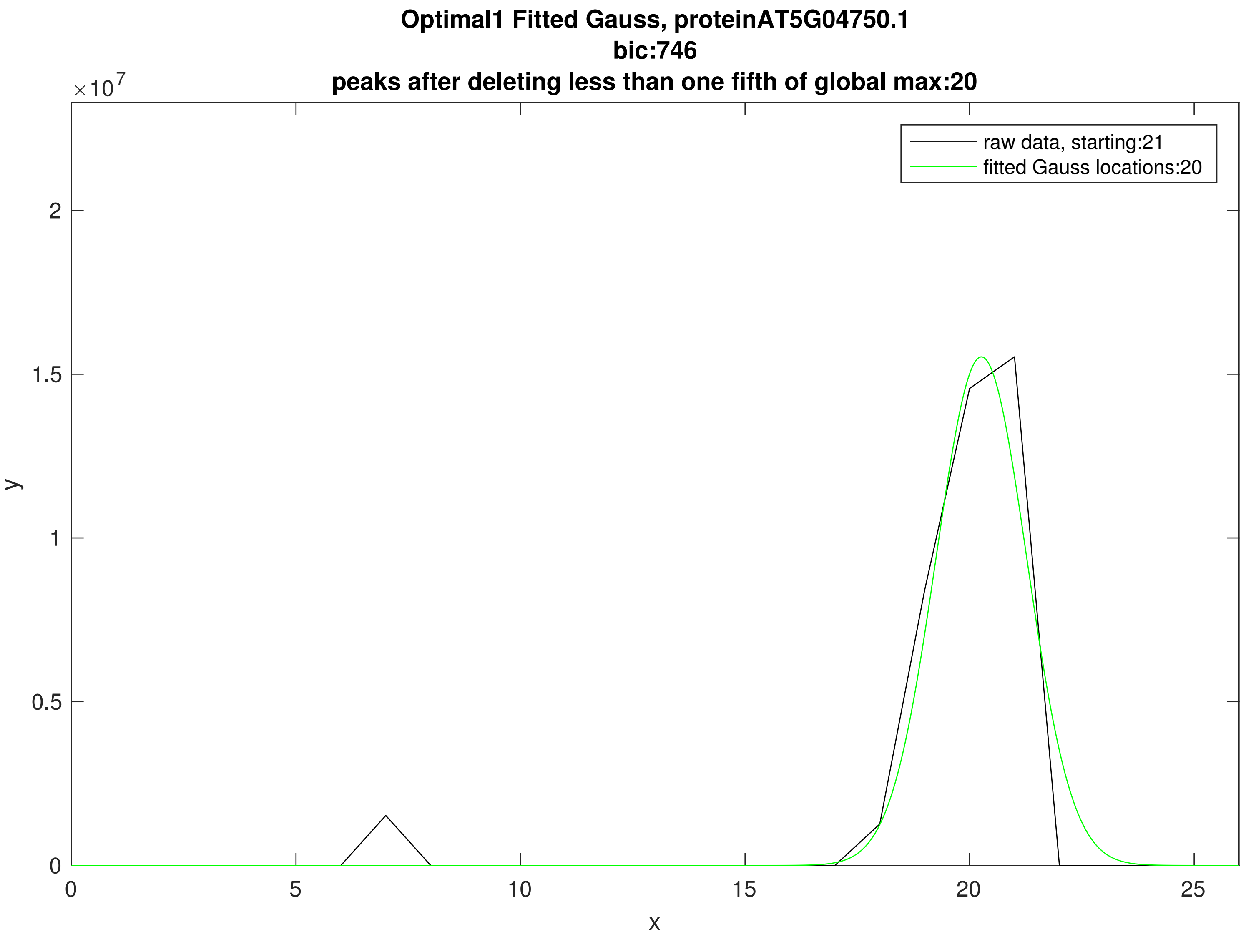

Optimal1 Fitted Gauss, proteinAT5G04750.1
bic:746
peaks after deleting less than one fifth of global max:20
×10^7
raw data, starting:21
fitted Gauss locations:20
y
x
0
0.5
1
1.5
2
0
5
10
15
20
25

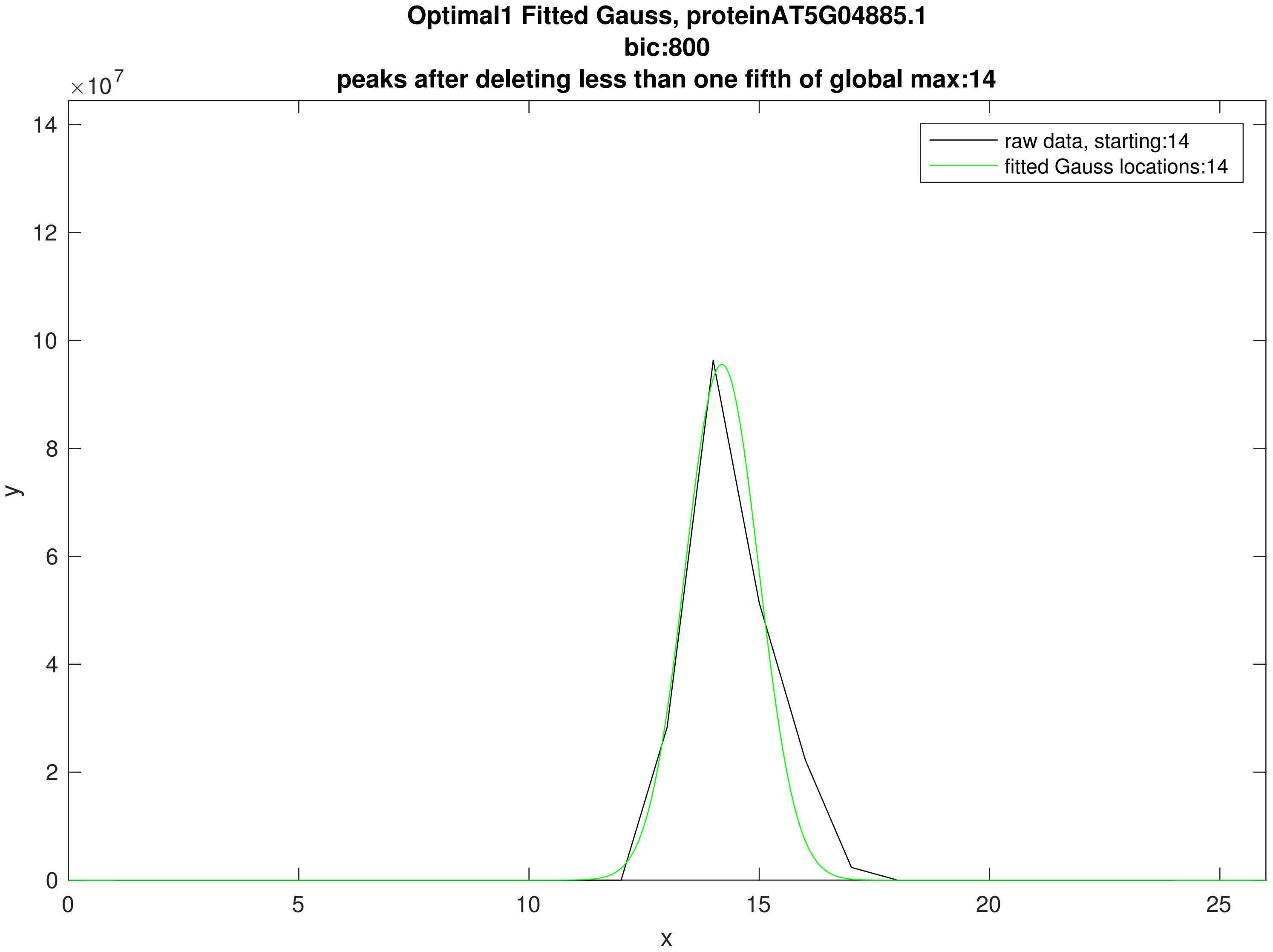

Optimal1 Fitted Gauss, proteinAT5G04885.1
bic:800
peaks after deleting less than one fifth of global max:14
×10^7
14
12
10
8
6
4
2
0
y
0
5
10
15
20
25
x
raw data, starting:14
fitted Gauss locations:14

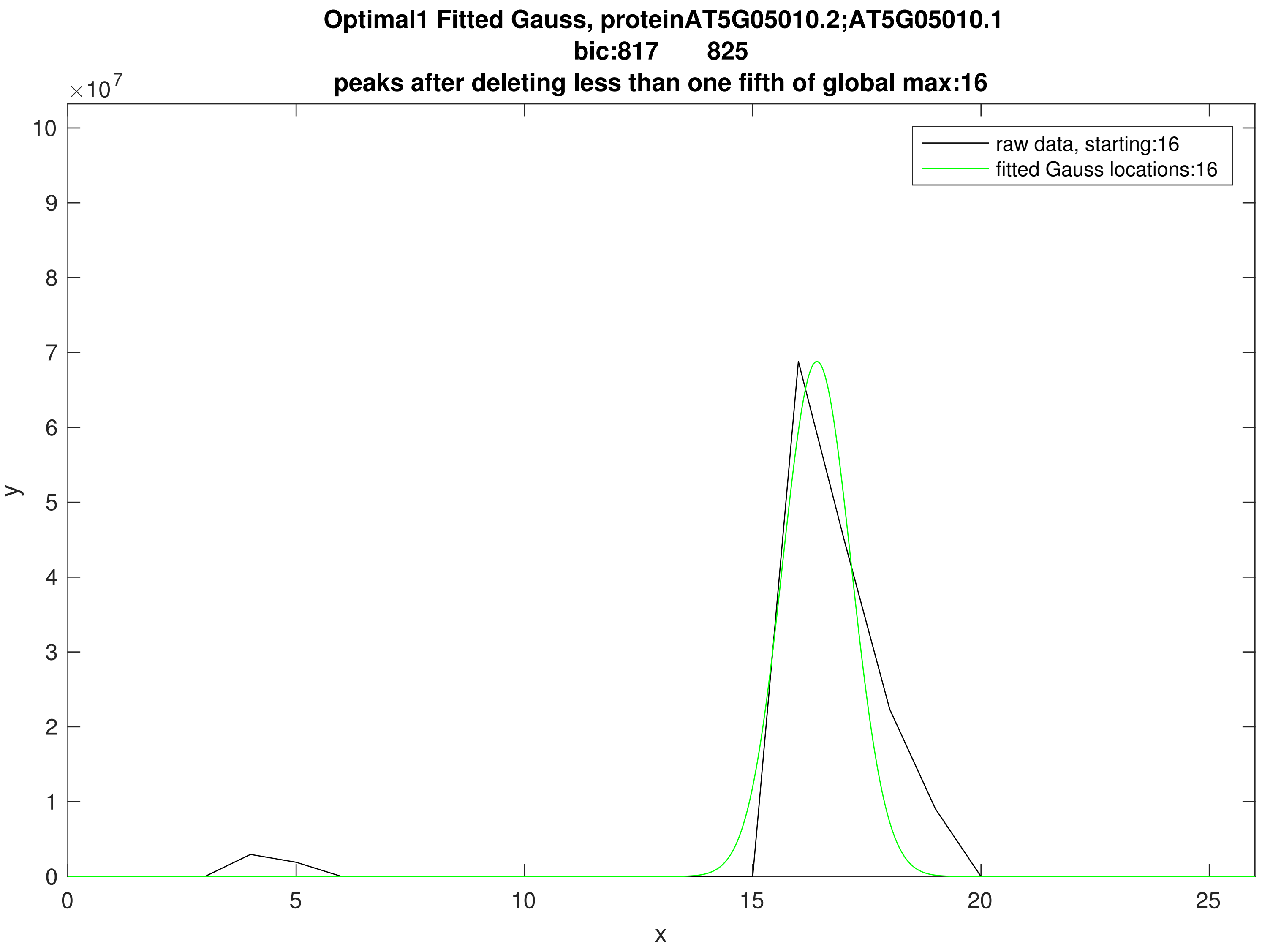
Optimal1 Fitted Gauss, proteinAT5G05010.2;AT5G05010.1
bic:817 825
peaks after deleting less than one fifth of global max:16
×10^7
raw data, starting:16
fitted Gauss locations:16
y
x
0
1
2
3
4
5
6
7
8
9
10
0
5
10
15
20
25

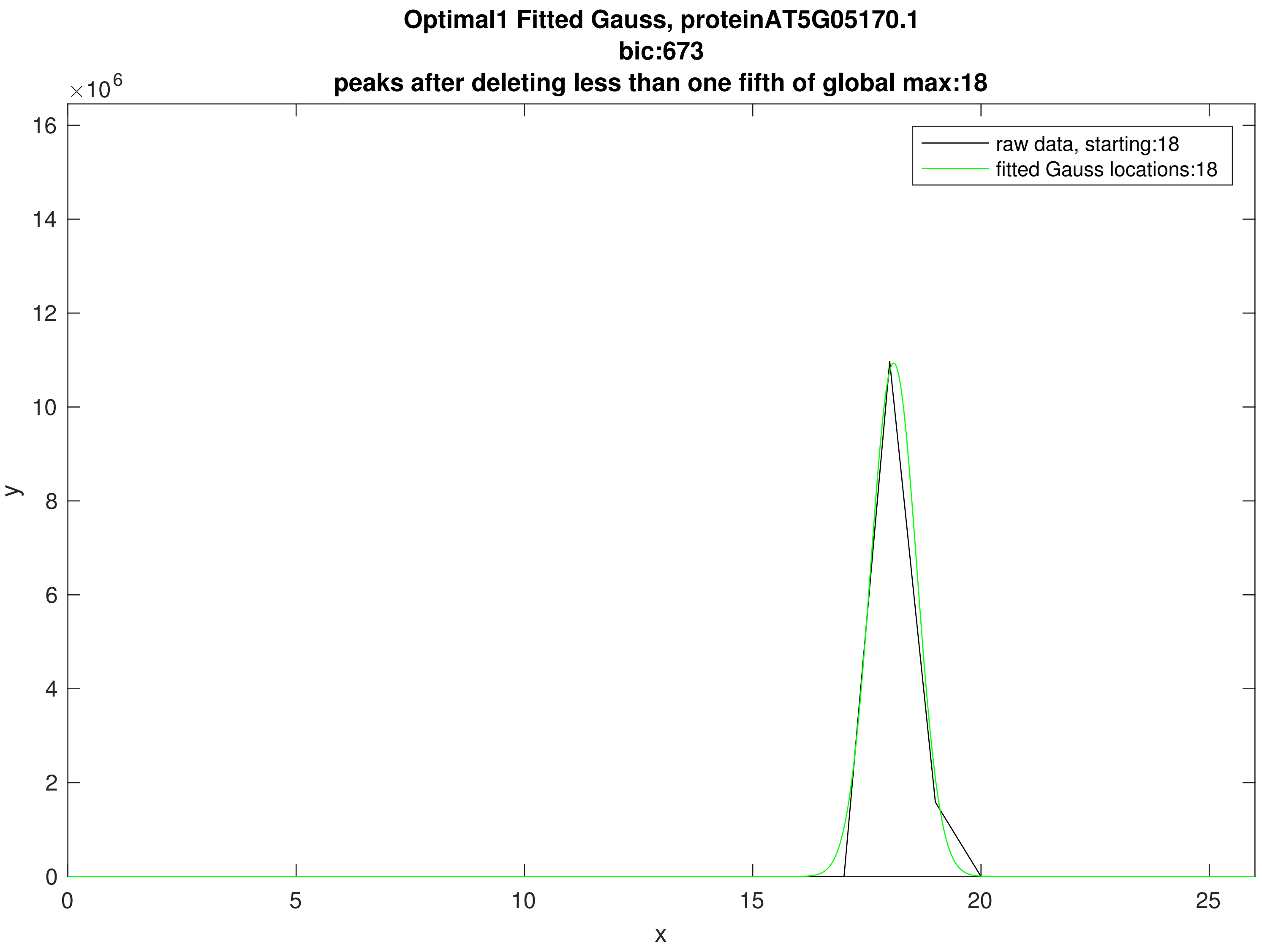
Optimal1 Fitted Gauss, proteinAT5G05170.1
bic:673
peaks after deleting less than one fifth of global max:18
×10^6
16
14
12
10
8
6
4
2
0
y
0
5
10
15
20
25
x
raw data, starting:18
fitted Gauss locations:18

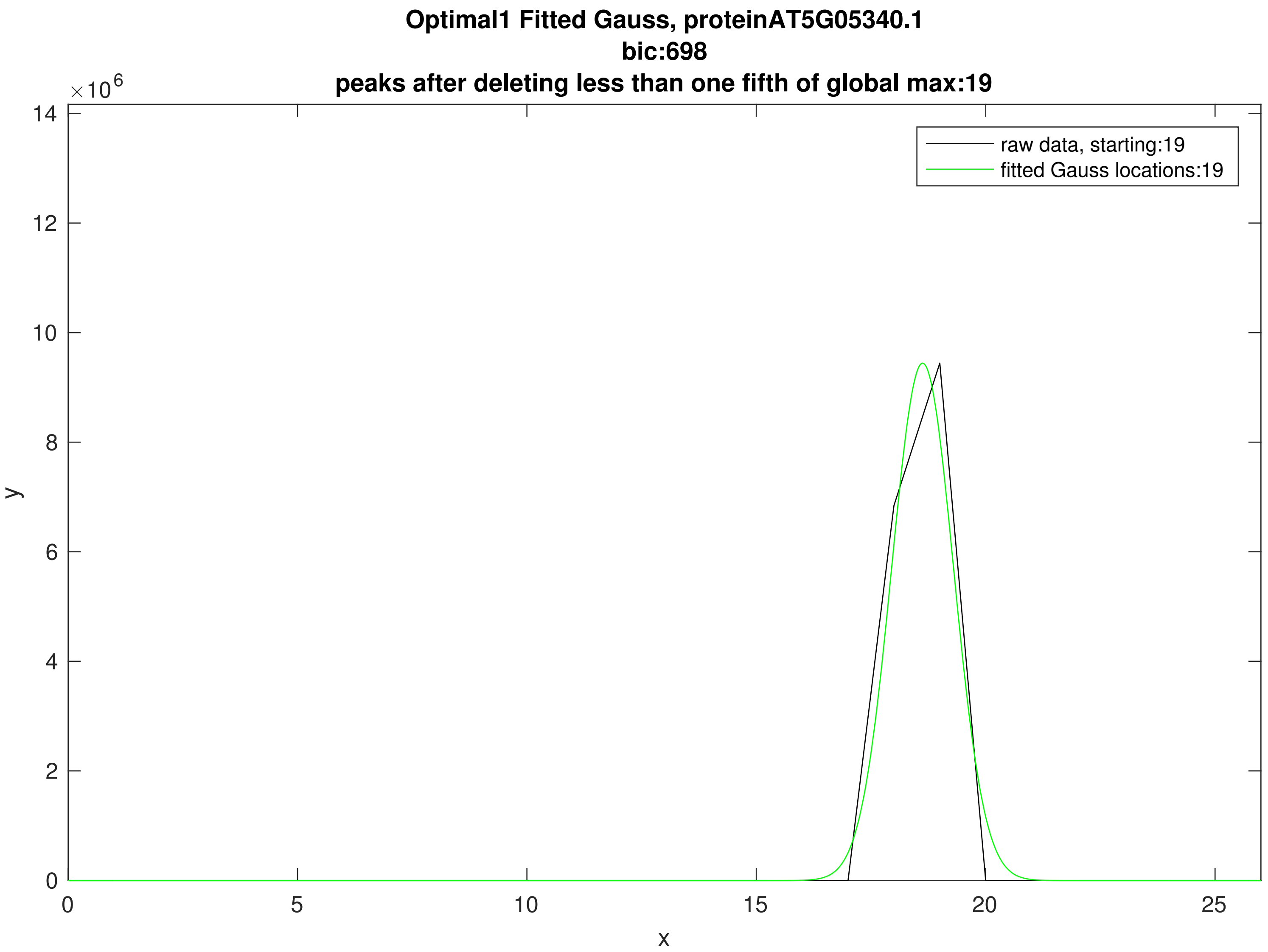

Optimal1 Fitted Gauss, proteinAT5G05340.1
bic:698
peaks after deleting less than one fifth of global max:19
×10^6
raw data, starting:19
fitted Gauss locations:19
y
x
0
2
4
6
8
10
12
14
5
15
20
25

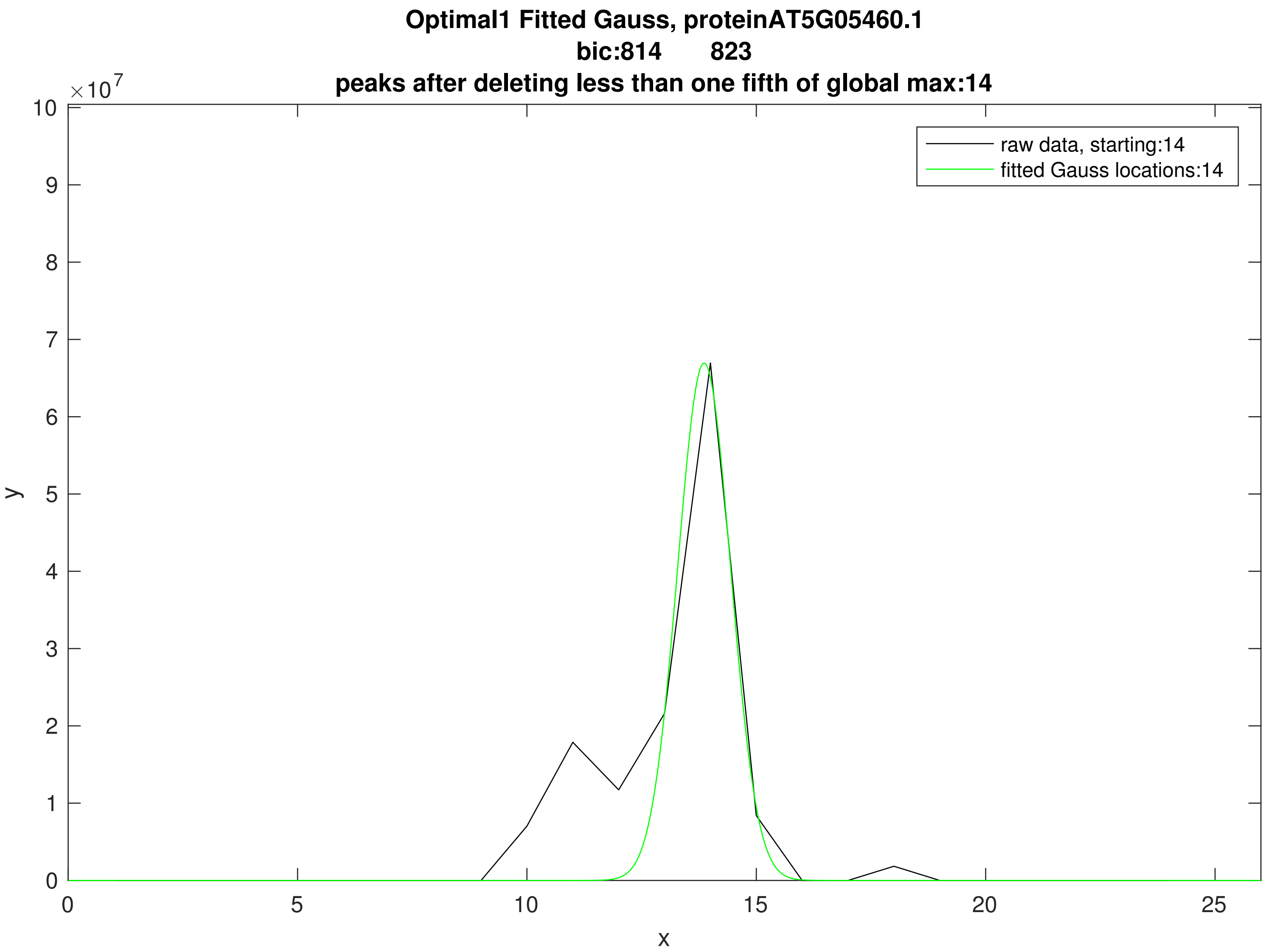
Optimal1 Fitted Gauss, proteinAT5G05460.1
bic:814 823
peaks after deleting less than one fifth of global max:14
×10^7
10
9
8
7
6
5
4
3
2
1
0
y
0
5
10
15
20
25
x
raw data, starting:14
fitted Gauss locations:14

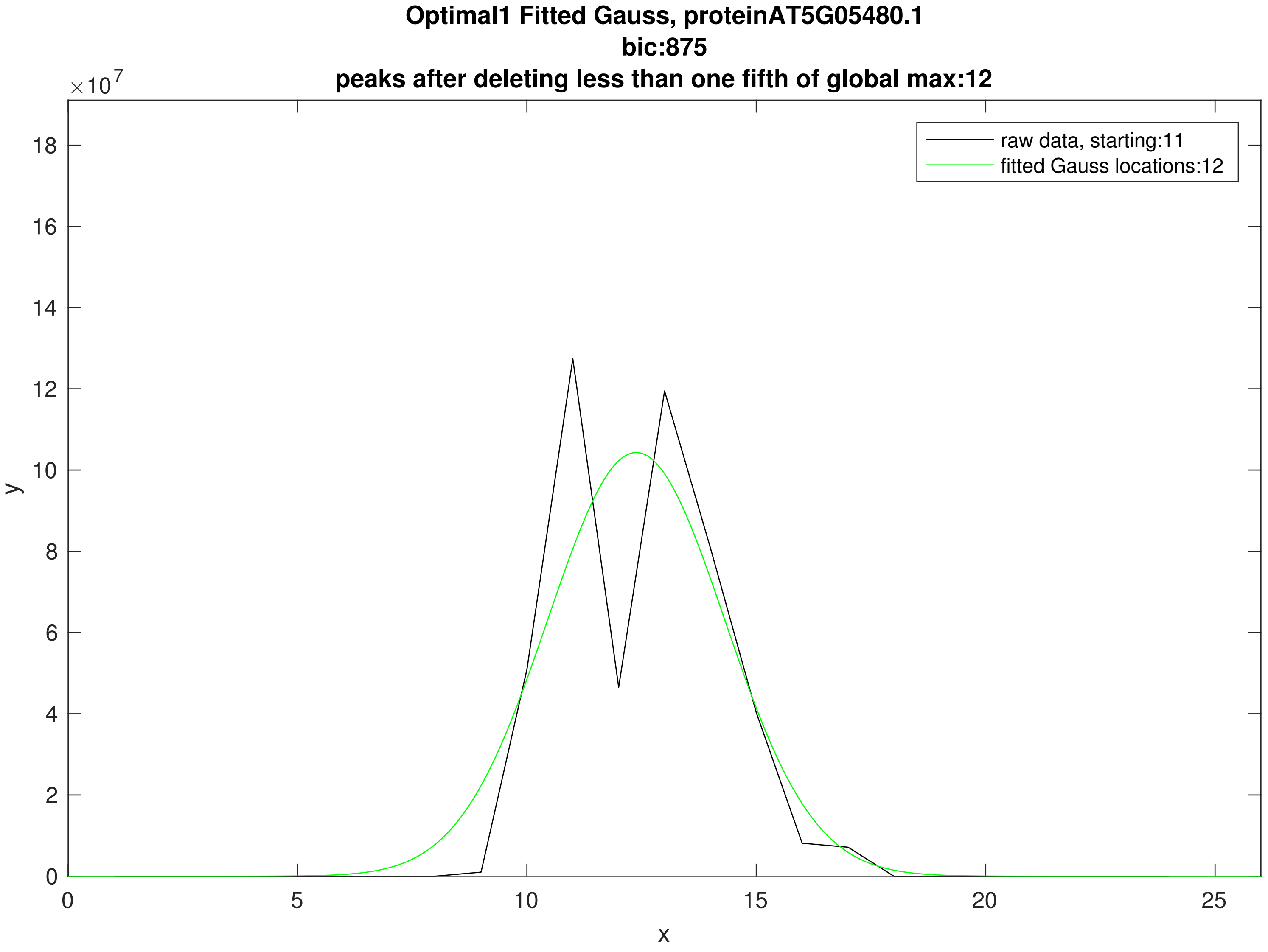
Optimal1 Fitted Gauss, proteinAT5G05480.1
bic:875
peaks after deleting less than one fifth of global max:12
×10^7
raw data, starting:11
fitted Gauss locations:12
y
x
0
2
4
6
8
10
12
14
16
18
0
5
10
15
20
25

**Optimal2 Fitted Gauss, proteinAT5G05730.1;AT5G05730.2**
**bic:878 816**
**peaks after deleting less than one fifth of global max:6.1 13**

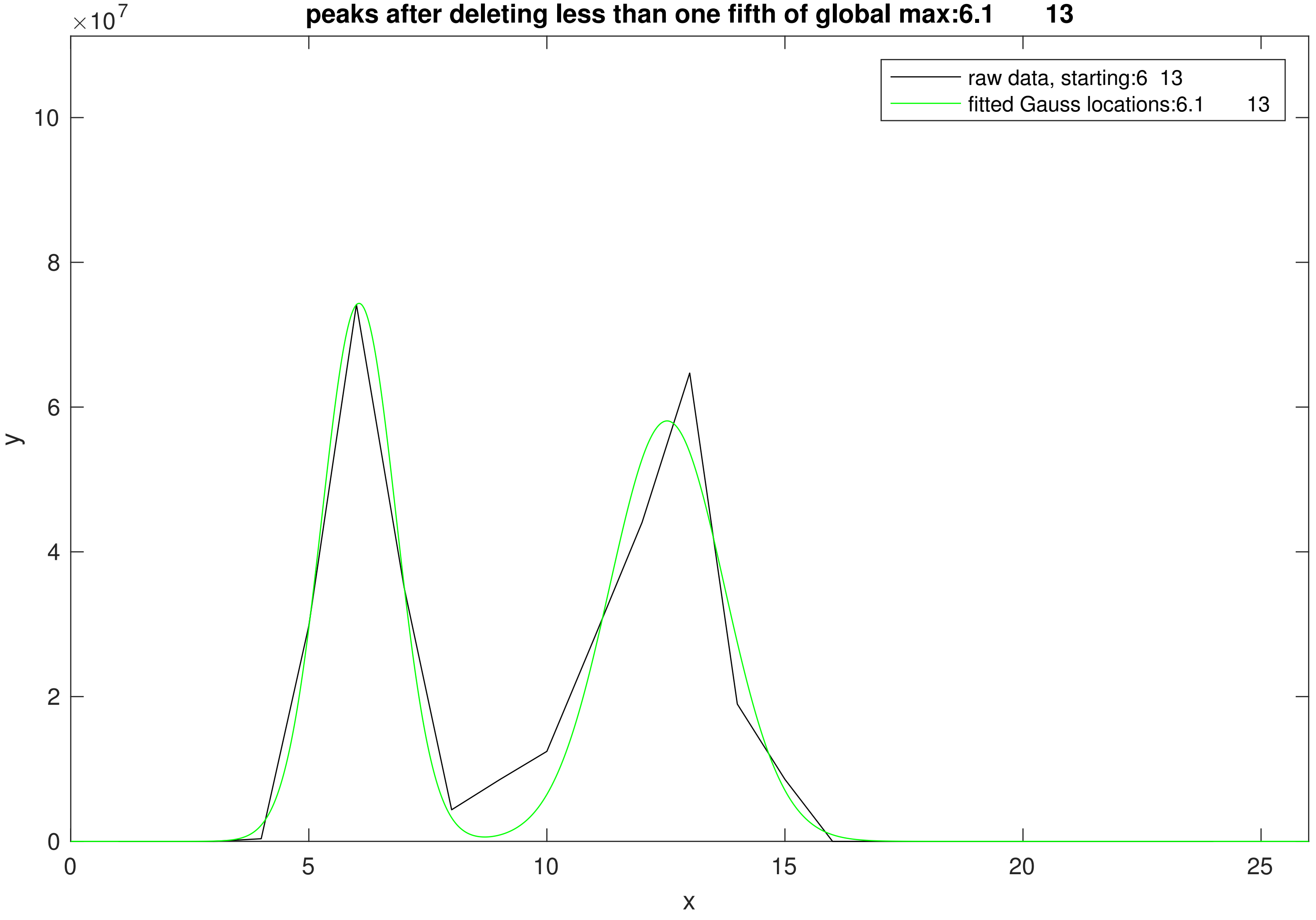

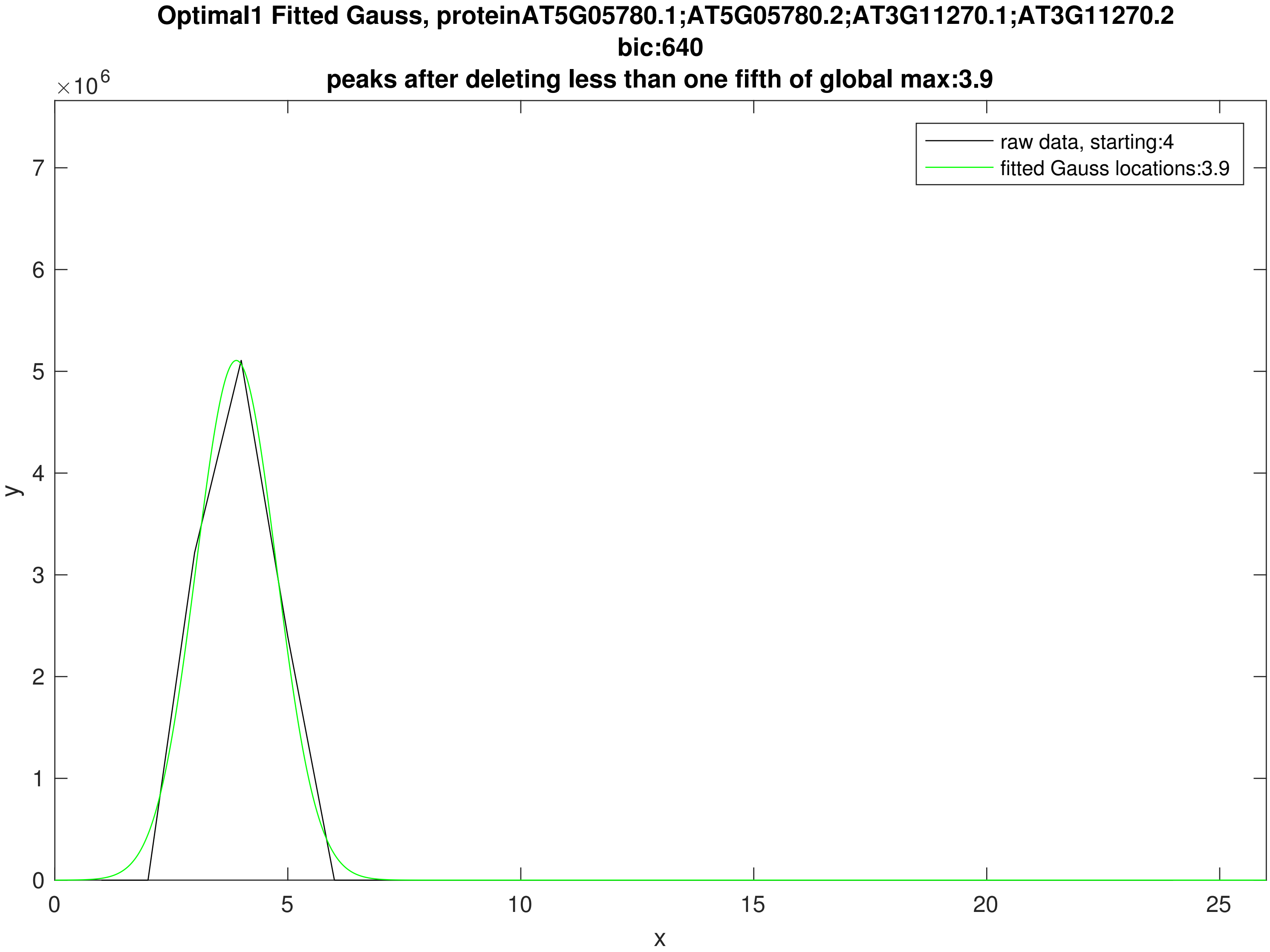
Optimal1 Fitted Gauss, proteinAT5G05780.1;AT5G05780.2;AT3G11270.1;AT3G11270.2
bic:640
peaks after deleting less than one fifth of global max:3.9
×10^6
raw data, starting:4
fitted Gauss locations:3.9
y
x
0
1
2
3
4
5
6
7
0
5
10
15
20
25

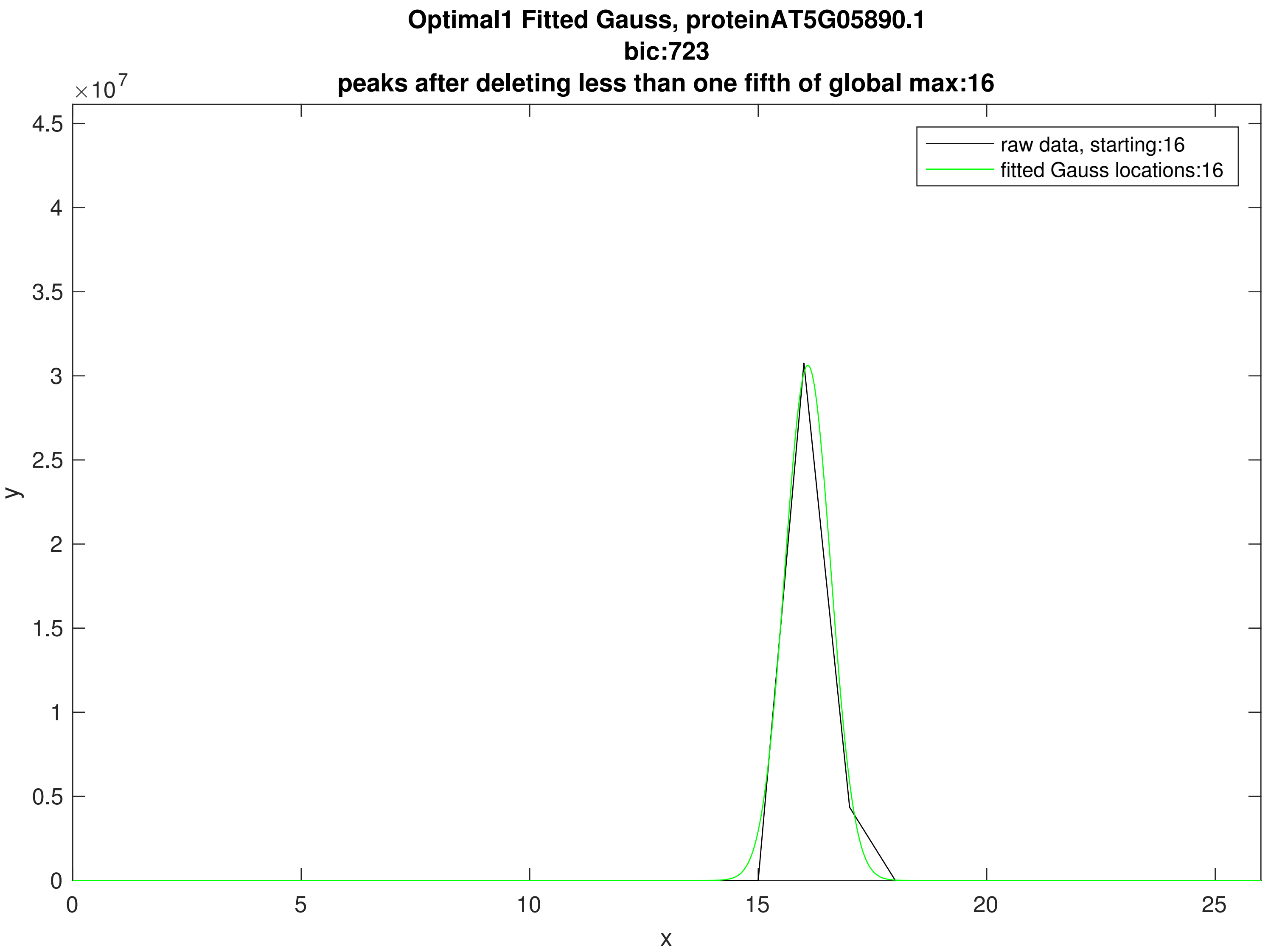

Optimal1 Fitted Gauss, proteinAT5G05890.1
bic:723
peaks after deleting less than one fifth of global max:16
×10^7
raw data, starting:16
fitted Gauss locations:16
y
x

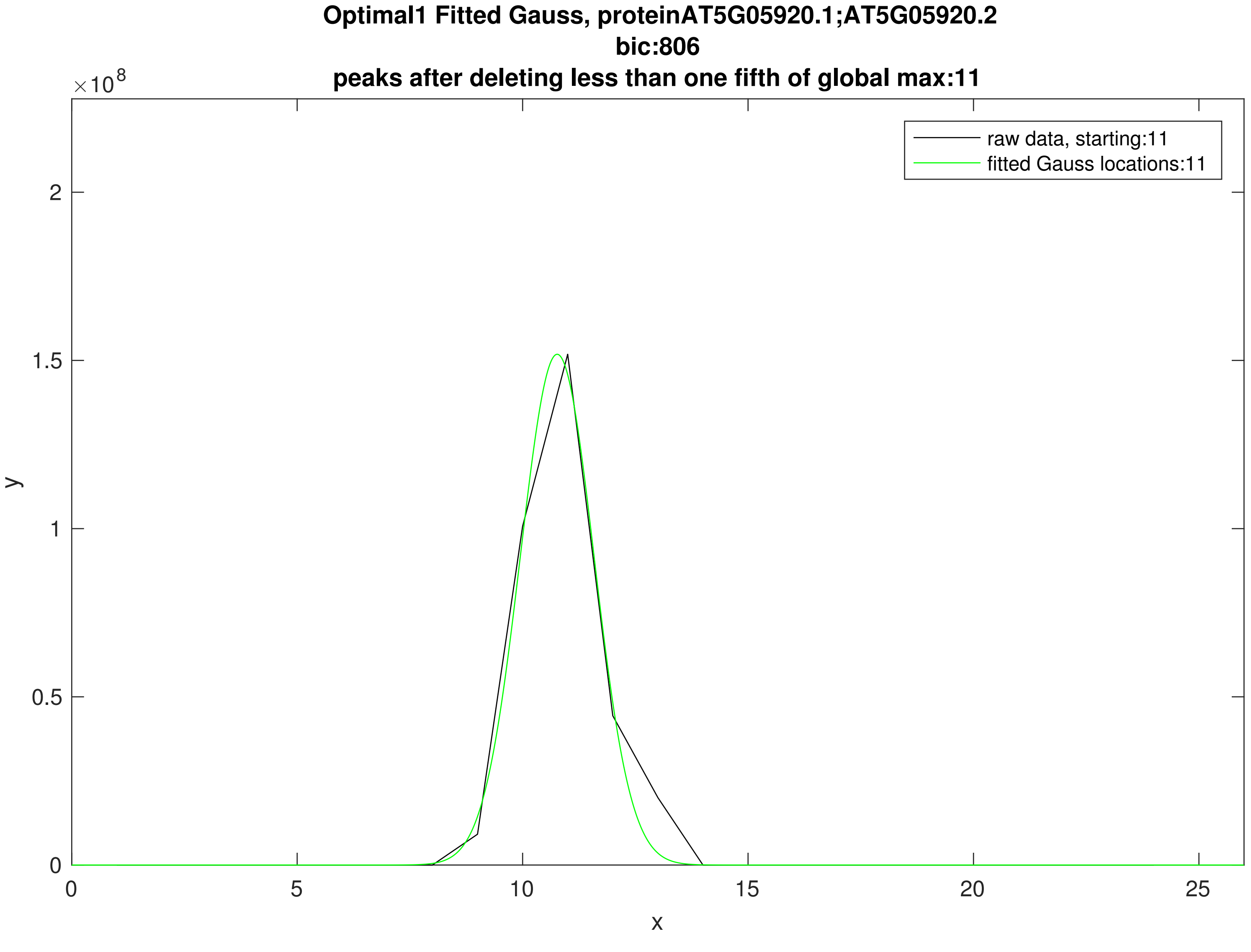

Optimal1 Fitted Gauss, proteinAT5G05920.1;AT5G05920.2
bic:806
peaks after deleting less than one fifth of global max:11
×10^8
raw data, starting:11
fitted Gauss locations:11
y
x
0
0.5
1
1.5
2
0
5
10
15
20
25

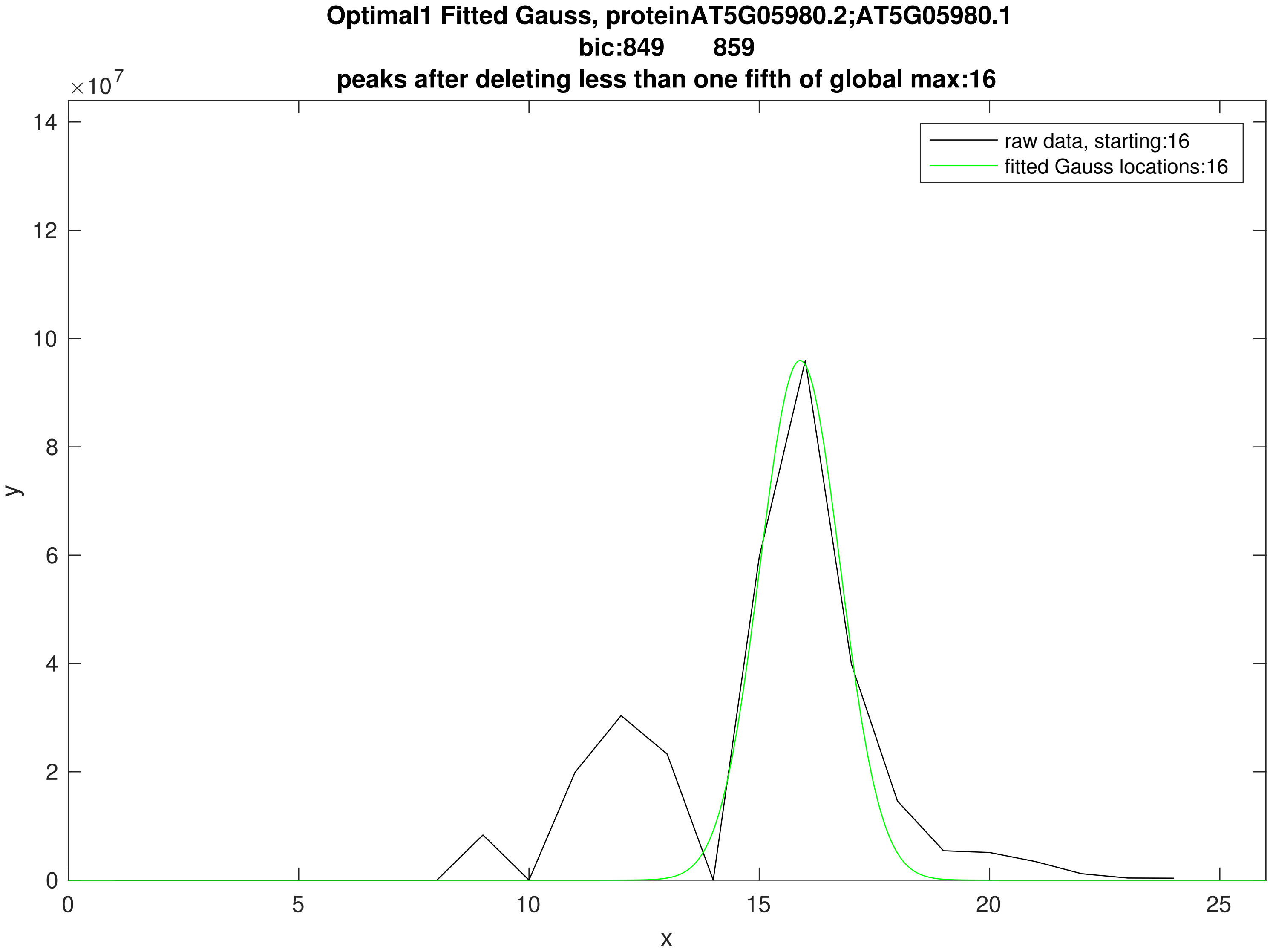
Optimal1 Fitted Gauss, proteinAT5G05980.2;AT5G05980.1
bic:849 859
peaks after deleting less than one fifth of global max:16
×10^7
raw data, starting:16
fitted Gauss locations:16
y
x
0
2
4
6
8
10
12
14
0
5
10
15
20
25

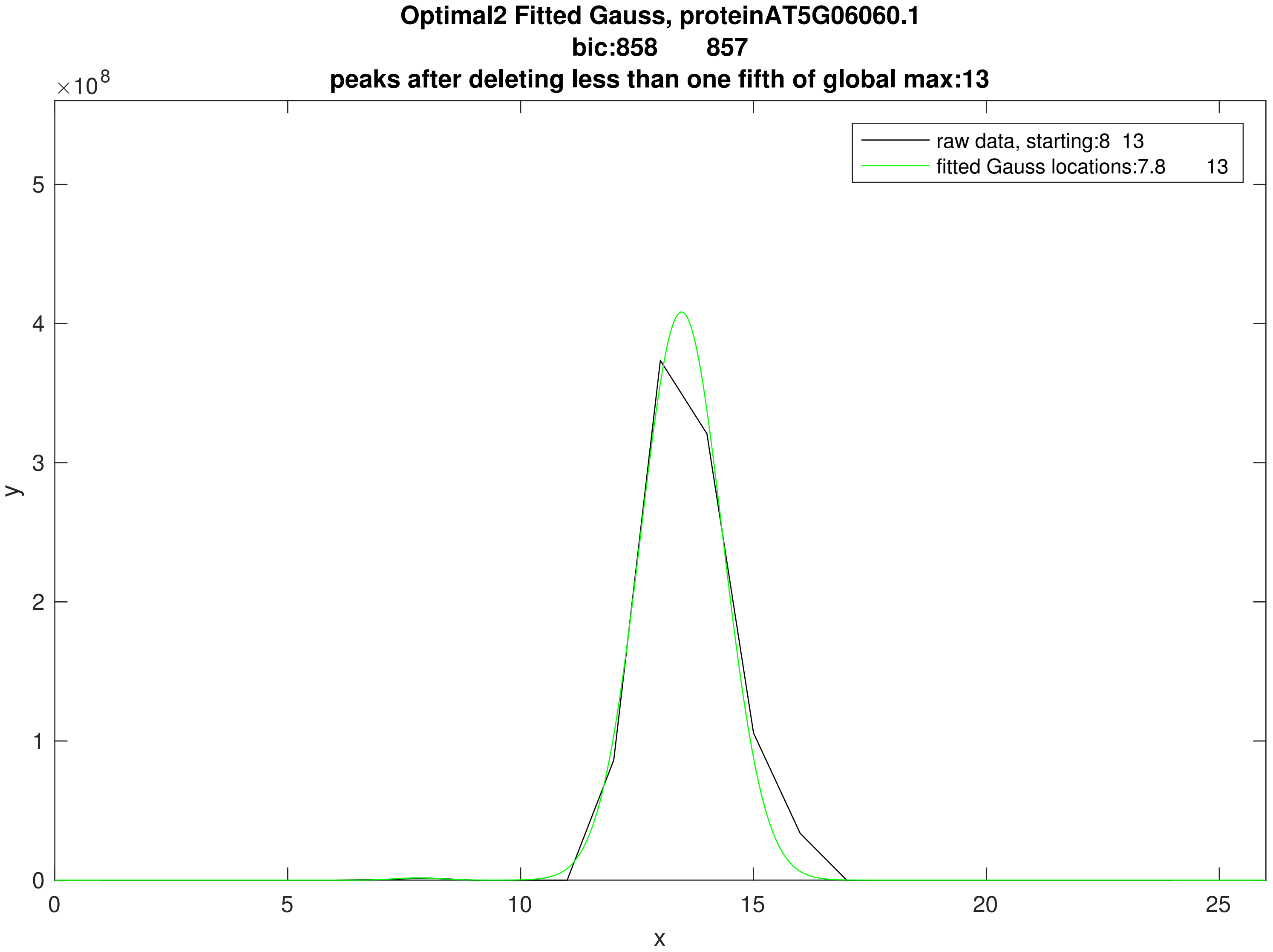
Optimal2 Fitted Gauss, proteinAT5G06060.1
bic:858 857
peaks after deleting less than one fifth of global max:13
×10^8
raw data, starting:8 13
fitted Gauss locations:7.8 13
y
x
0
1
2
3
4
5
0
5
10
15
20
25

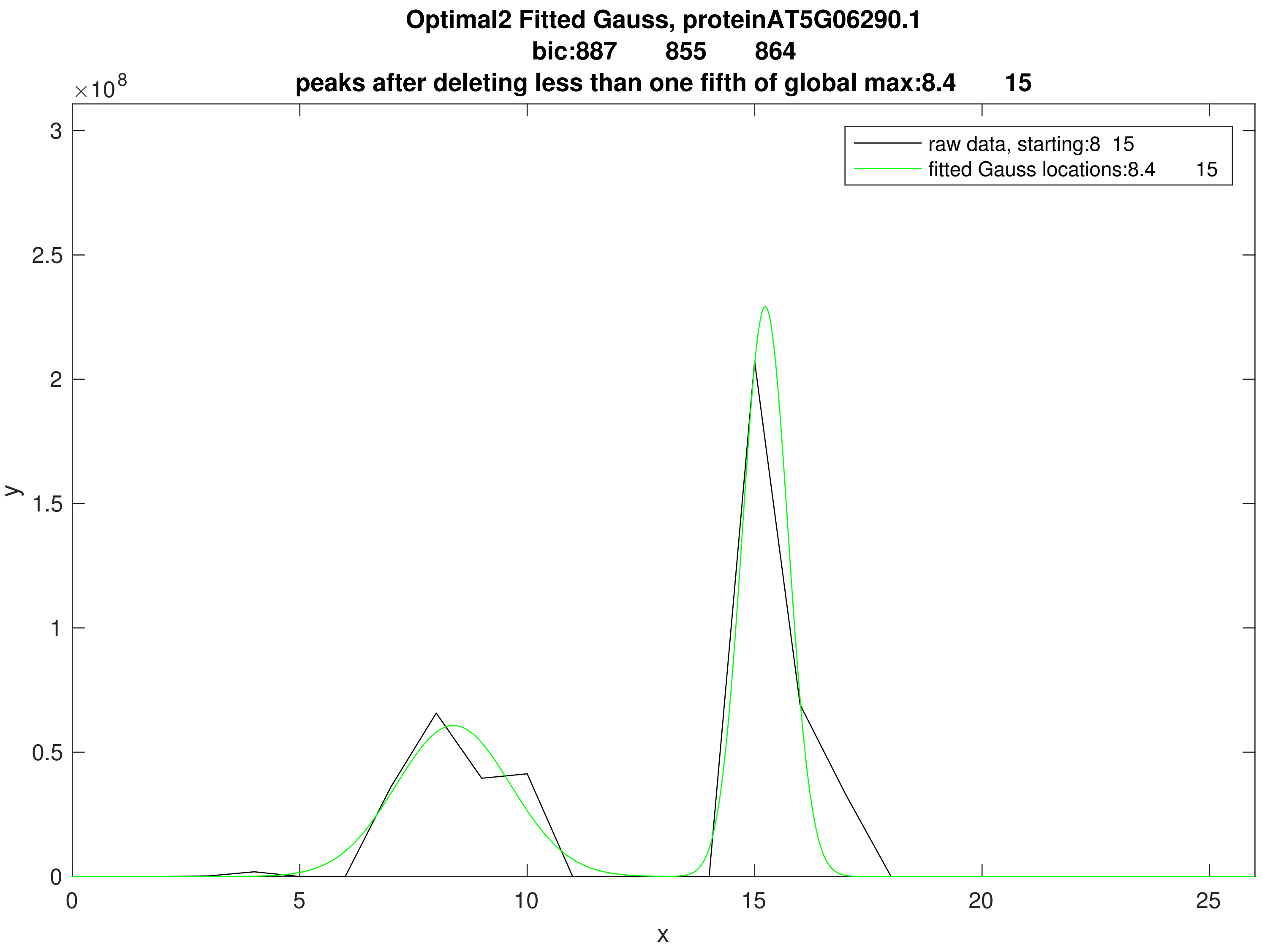

Optimal2 Fitted Gauss, proteinAT5G06290.1
bic:887 855 864
peaks after deleting less than one fifth of global max:8.4 15
×10^8
3
2.5
2
1.5
1
0.5
0
y
0
5
10
15
20
25
x
raw data, starting:8 15
fitted Gauss locations:8.4 15

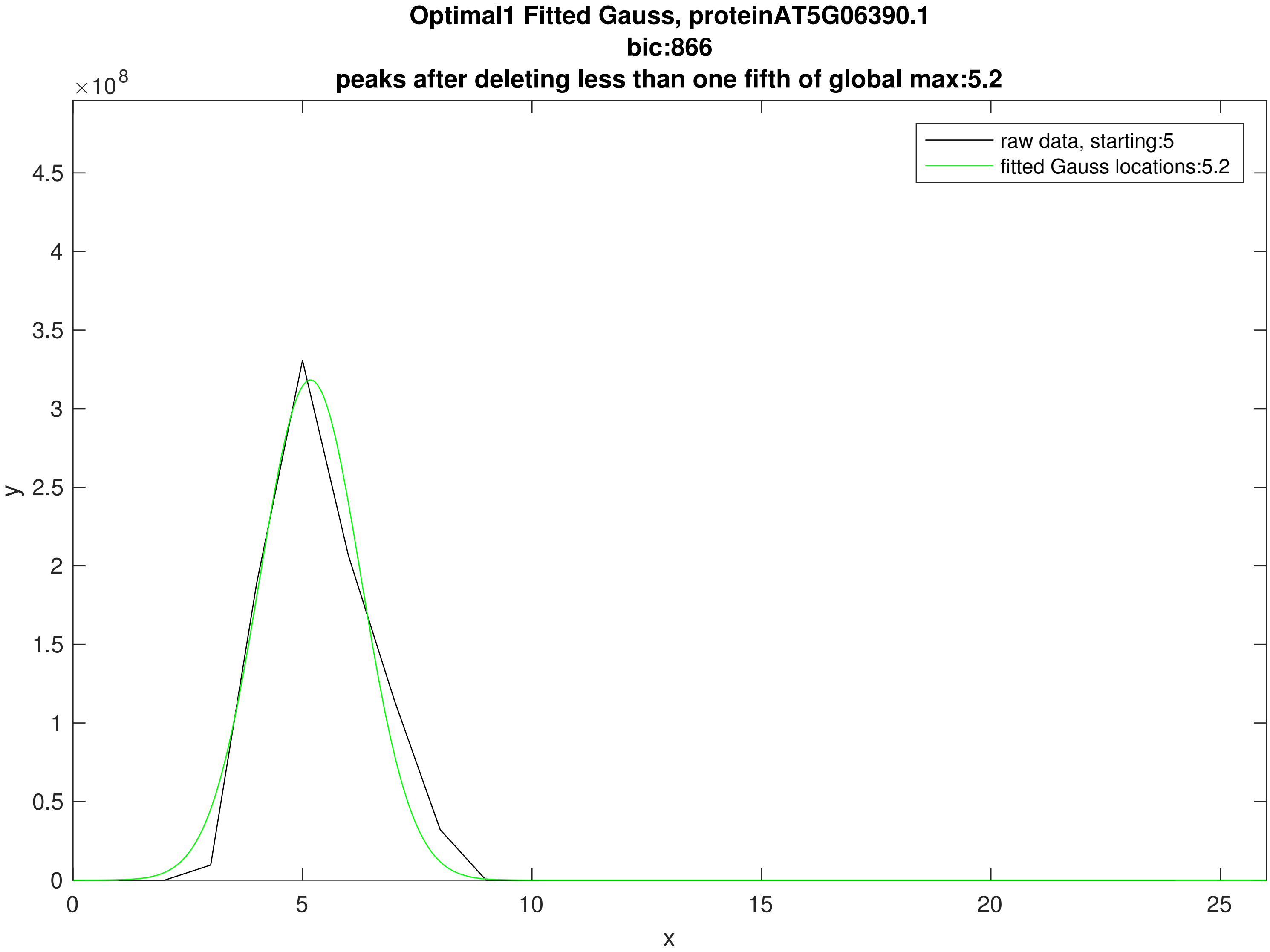

Optimal1 Fitted Gauss, proteinAT5G06390.1
bic:866
peaks after deleting less than one fifth of global max:5.2
×10^8
raw data, starting:5
fitted Gauss locations:5.2
4.5
4
3.5
3
2.5
2
1.5
1
0.5
0
y
0
5
10
15
20
25
x

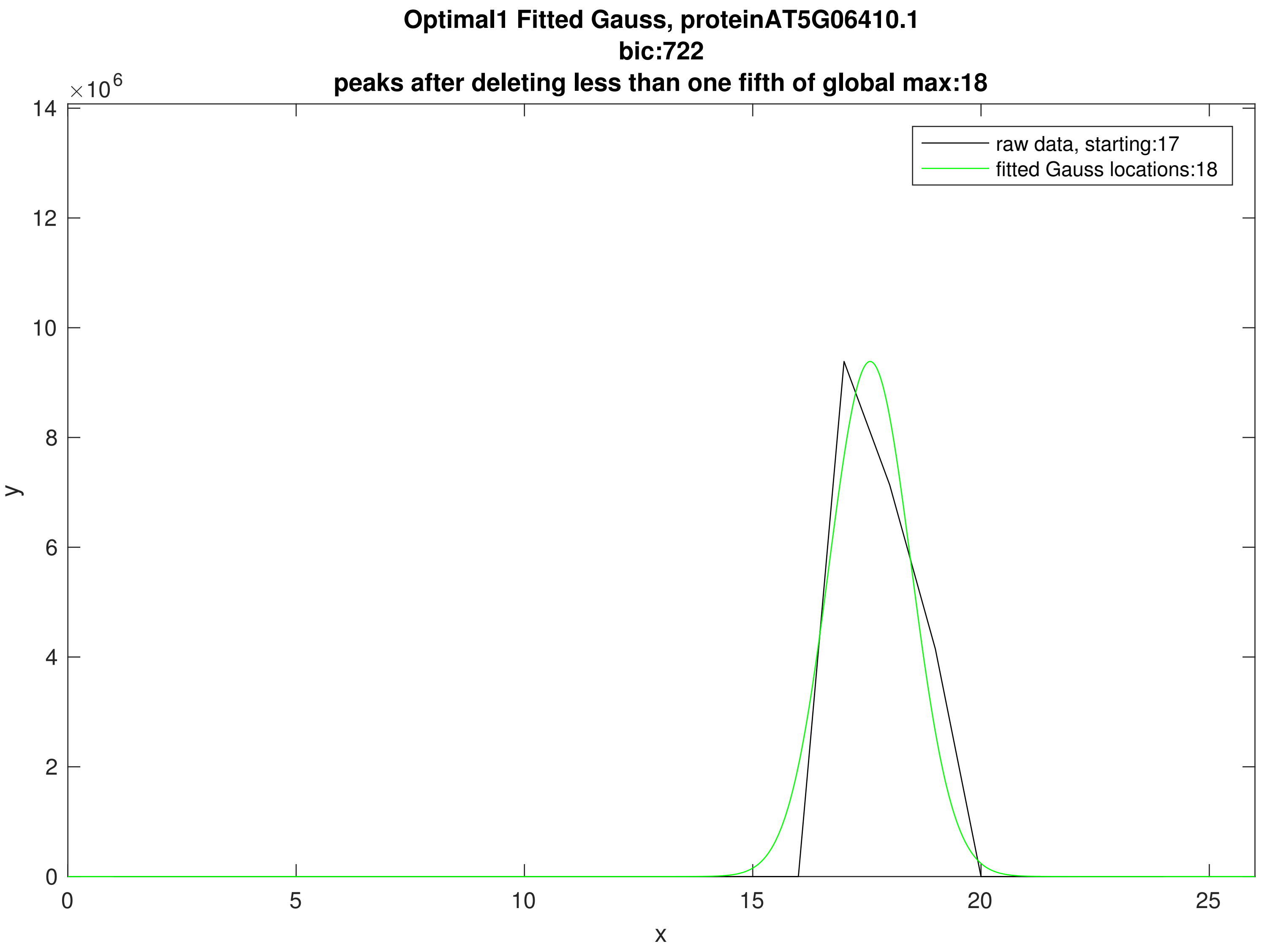
Optimal1 Fitted Gauss, proteinAT5G06410.1
bic:722
peaks after deleting less than one fifth of global max:18
×10^6
14
12
10
8
6
4
2
0
y
0
5
10
15
20
25
x
raw data, starting:17
fitted Gauss locations:18

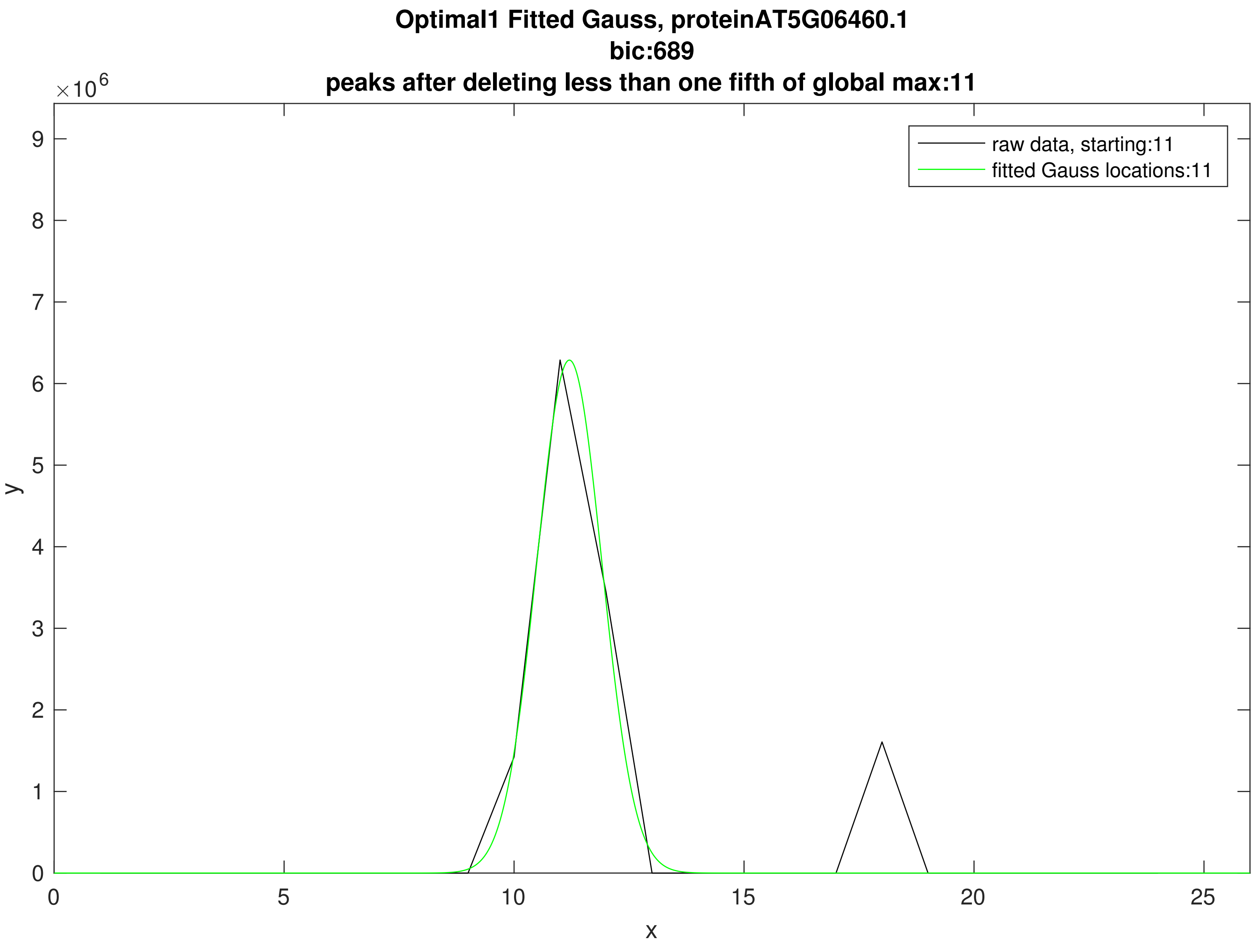
Optimal1 Fitted Gauss, proteinAT5G06460.1
bic:689
peaks after deleting less than one fifth of global max:11
×10^6
raw data, starting:11
fitted Gauss locations:11
y
x
0
1
2
3
4
5
6
7
8
9
0
5
10
15
20
25

**Optimal1 Fitted Gauss, proteinAT5G06580.1**
**bic:801**
**peaks after deleting less than one fifth of global max:12**

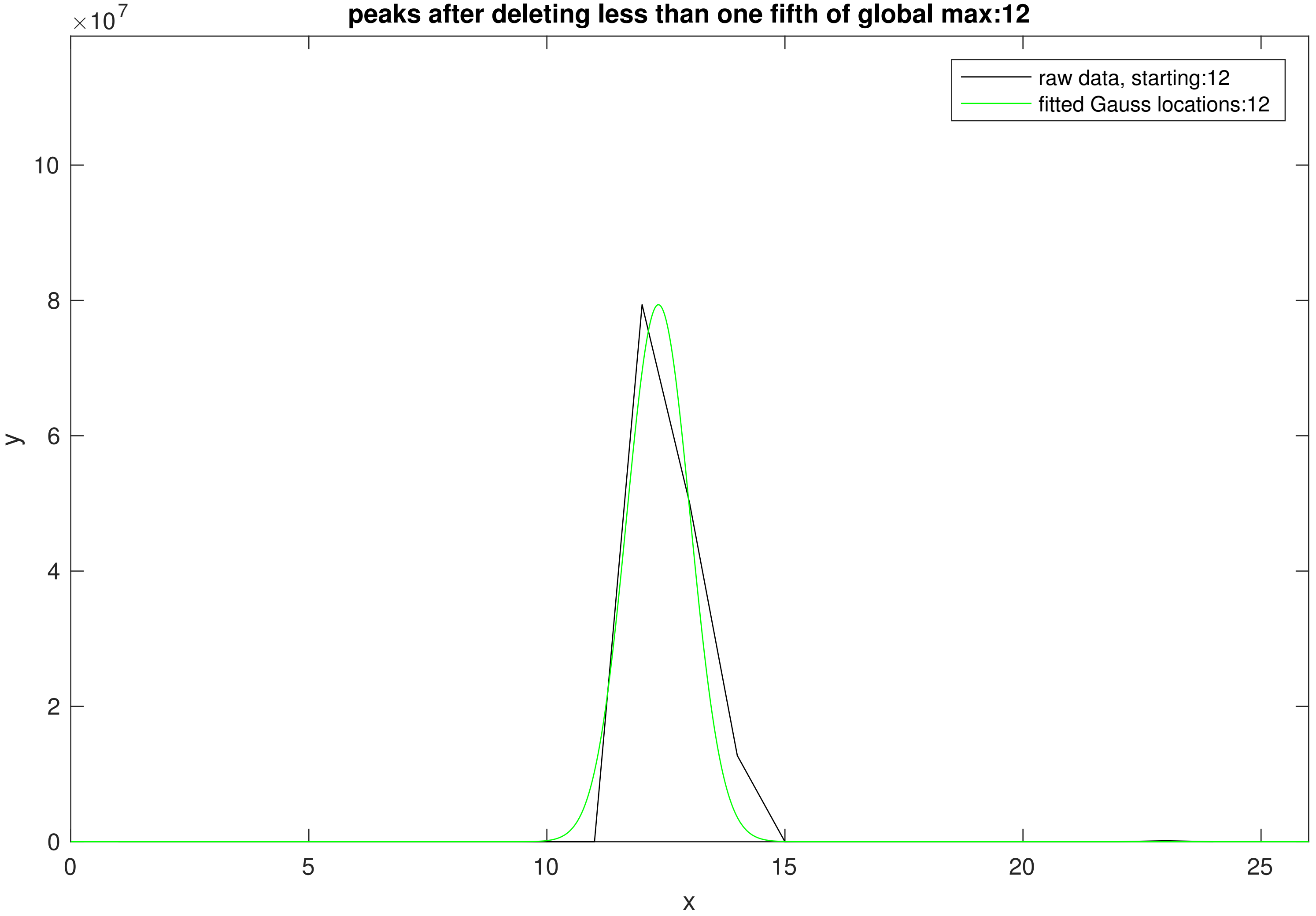

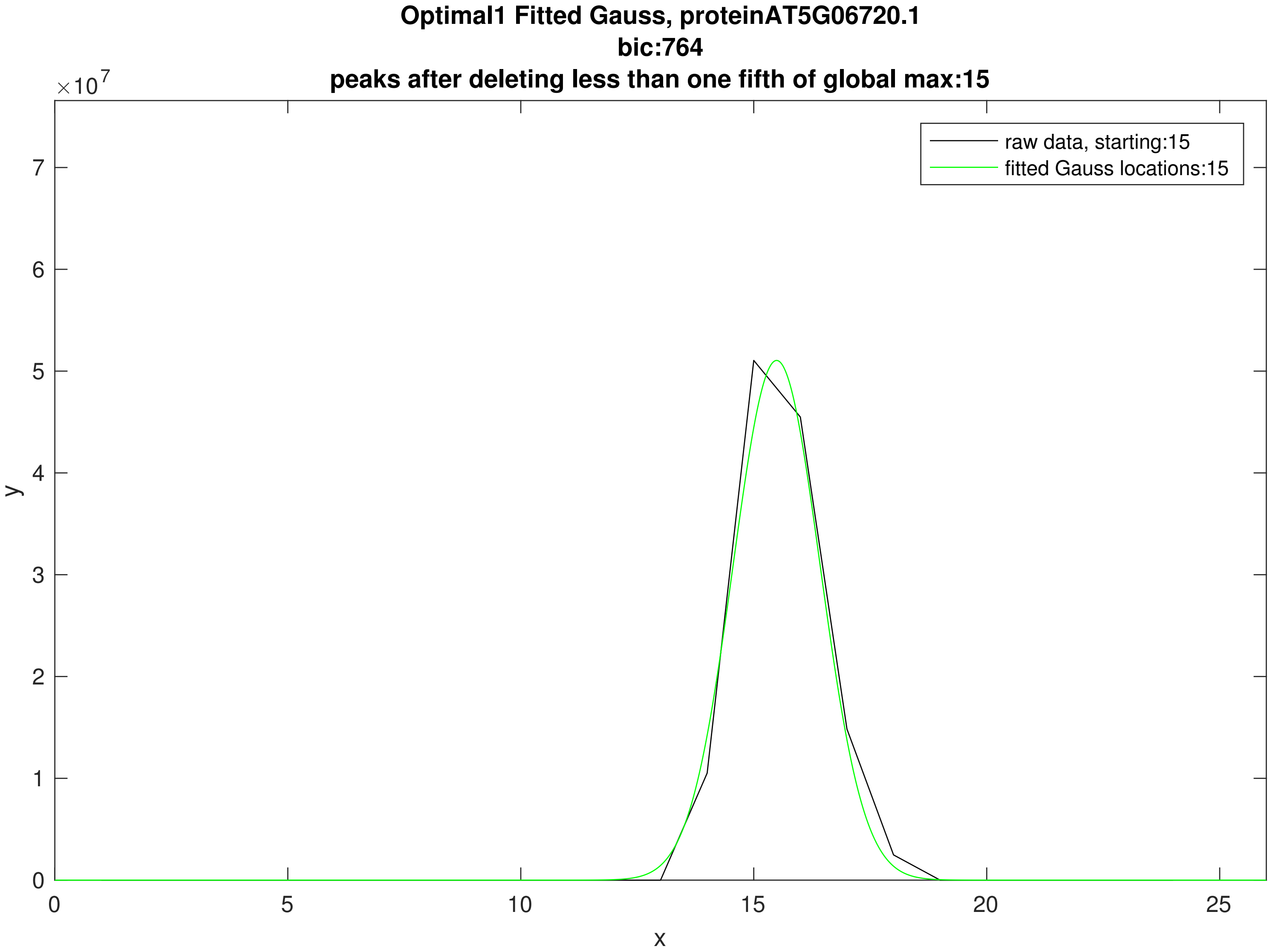

Optimal1 Fitted Gauss, proteinAT5G06720.1
bic:764
peaks after deleting less than one fifth of global max:15
×10^7
raw data, starting:15
fitted Gauss locations:15
0
1
2
3
4
5
6
7
y
0
5
10
15
20
25
x

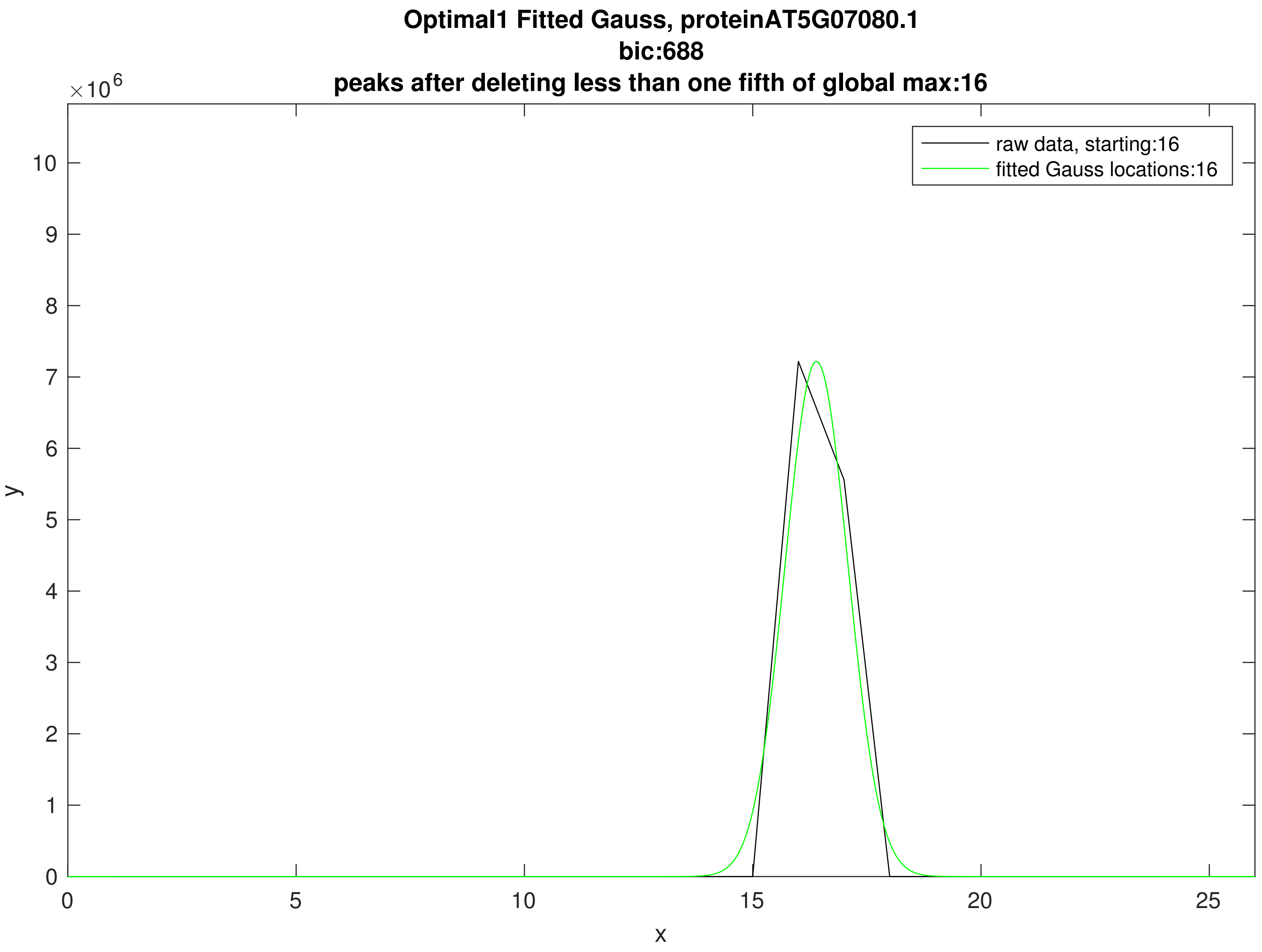

Optimal1 Fitted Gauss, proteinAT5G07080.1
bic:688
peaks after deleting less than one fifth of global max:16
×10^6
raw data, starting:16
fitted Gauss locations:16
y
x
0
1
2
3
4
5
6
7
8
9
10
0
5
10
15
20
25

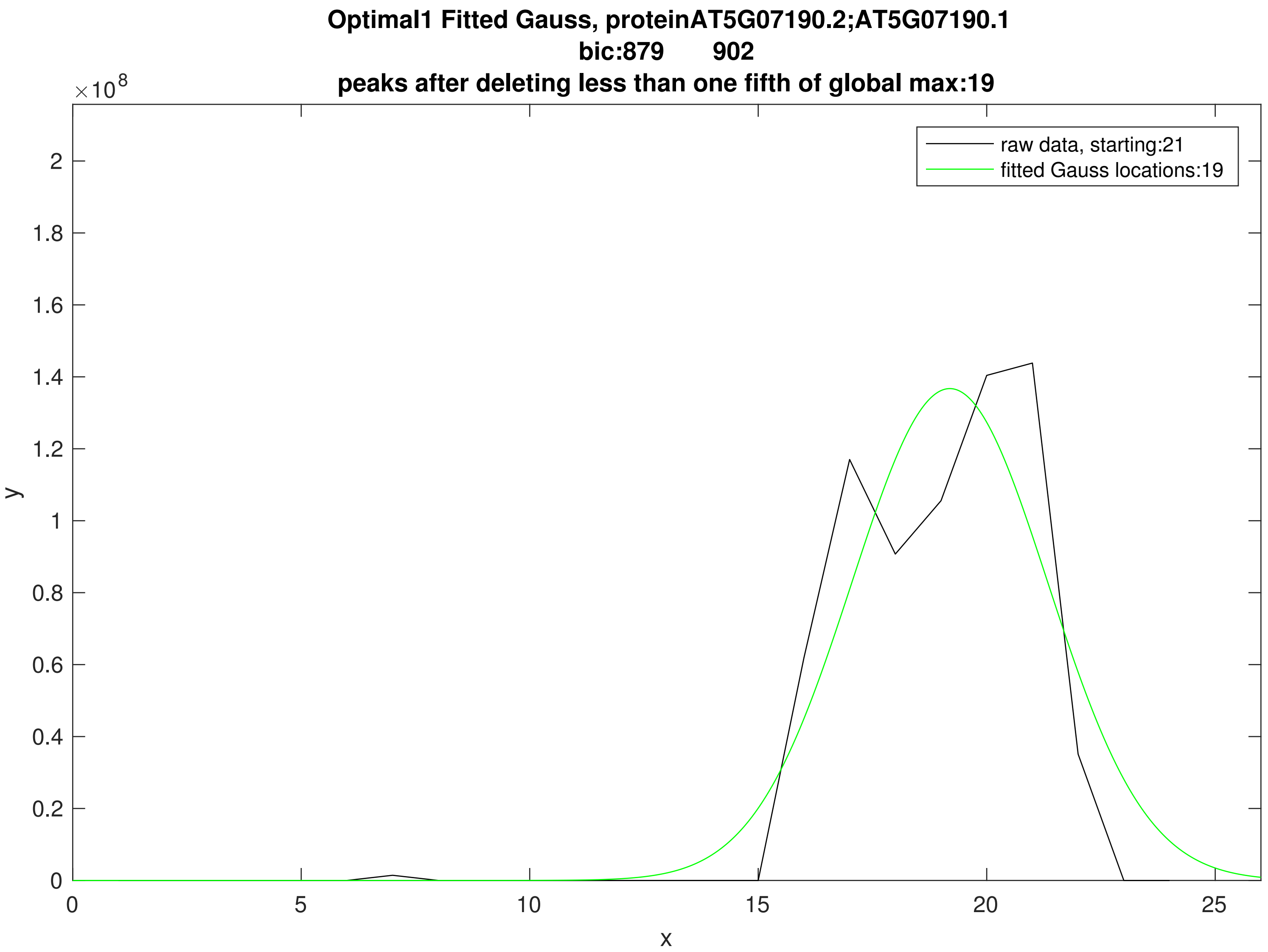
Optimal1 Fitted Gauss, proteinAT5G07190.2;AT5G07190.1
bic:879 902
peaks after deleting less than one fifth of global max:19
×10^8
raw data, starting:21
fitted Gauss locations:19
y
x
0
0.2
0.4
0.6
0.8
1
1.2
1.4
1.6
1.8
2
5
10
15
20
25

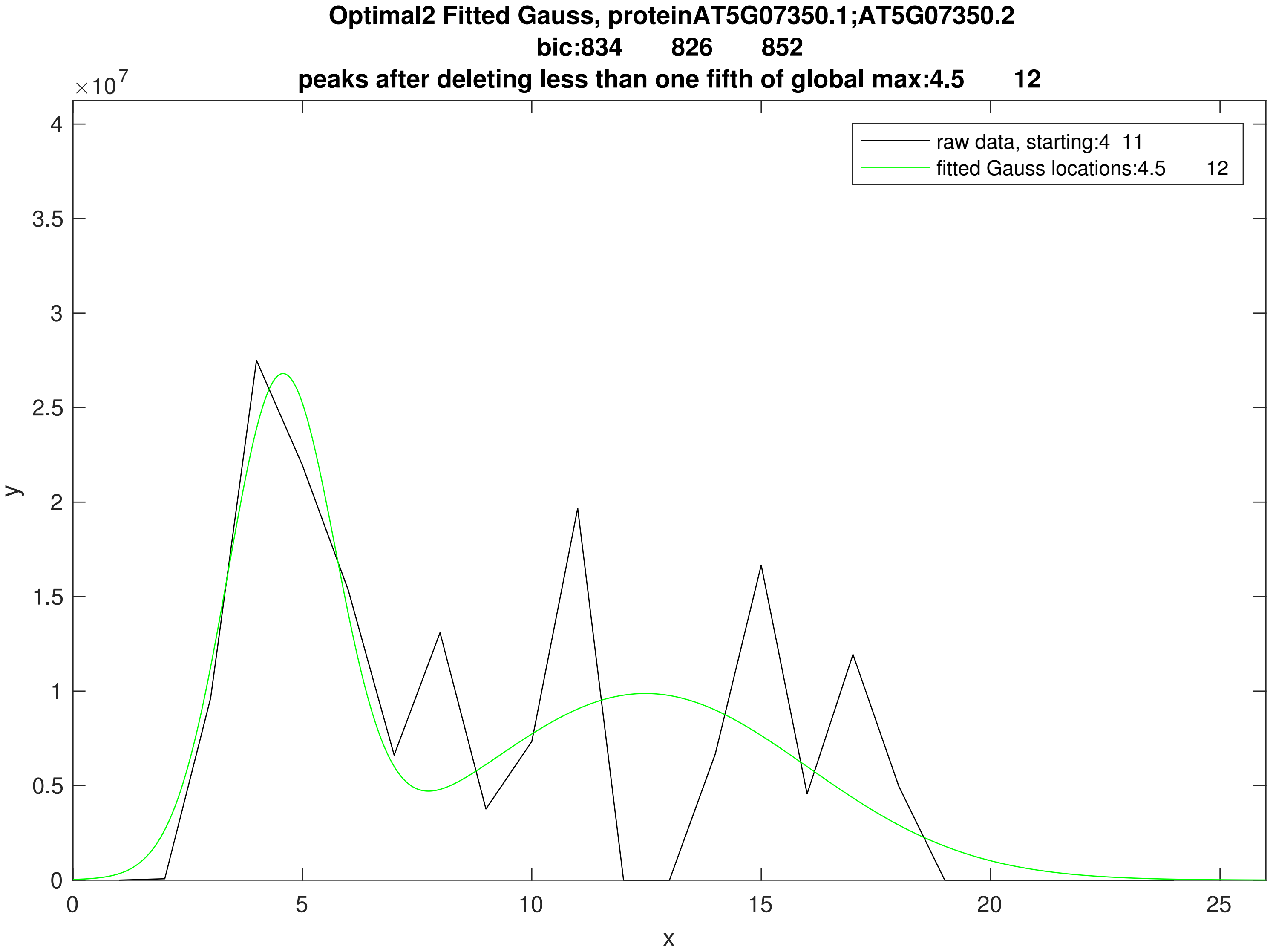

Optimal2 Fitted Gauss, proteinAT5G07350.1;AT5G07350.2
bic:834 826 852
peaks after deleting less than one fifth of global max:4.5 12
×10^7
raw data, starting:4 11
fitted Gauss locations:4.5 12
y
x
0
0.5
1
1.5
2
2.5
3
3.5
4
5
10
15
20
25

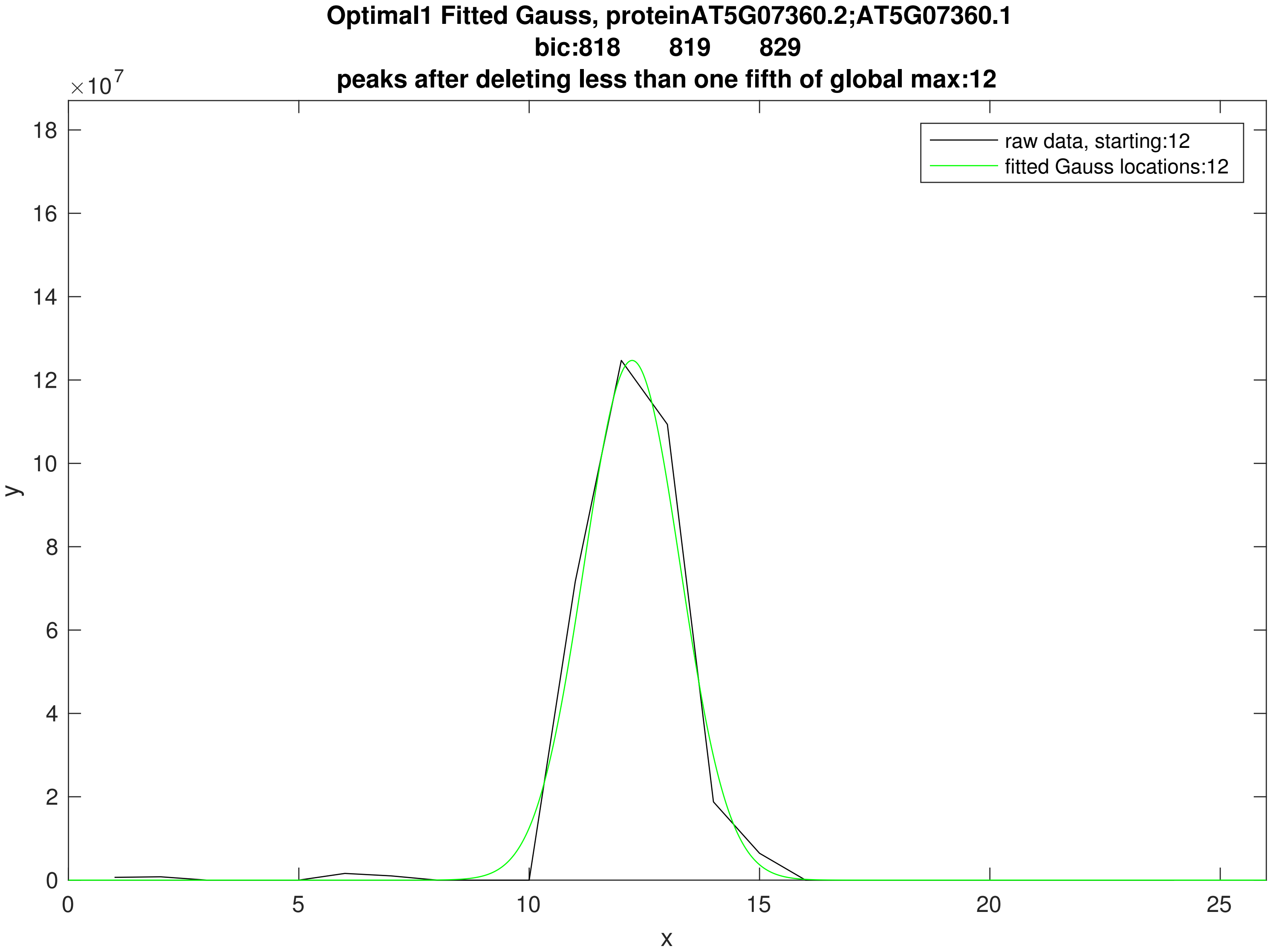
Optimal1 Fitted Gauss, proteinAT5G07360.2;AT5G07360.1
bic:818 819 829
peaks after deleting less than one fifth of global max:12
×10^7
raw data, starting:12
fitted Gauss locations:12
y
x
0
2
4
6
8
10
12
14
16
18
0
5
10
15
20
25

**Optimal1 Fitted Gauss, proteinAT5G07440.1;AT5G07440.2;AT5G07440.3**
**bic:1e+03  1.01e+03  1.02e+03  1.03e+03**
**peaks after deleting less than one fifth of global max:9.1**

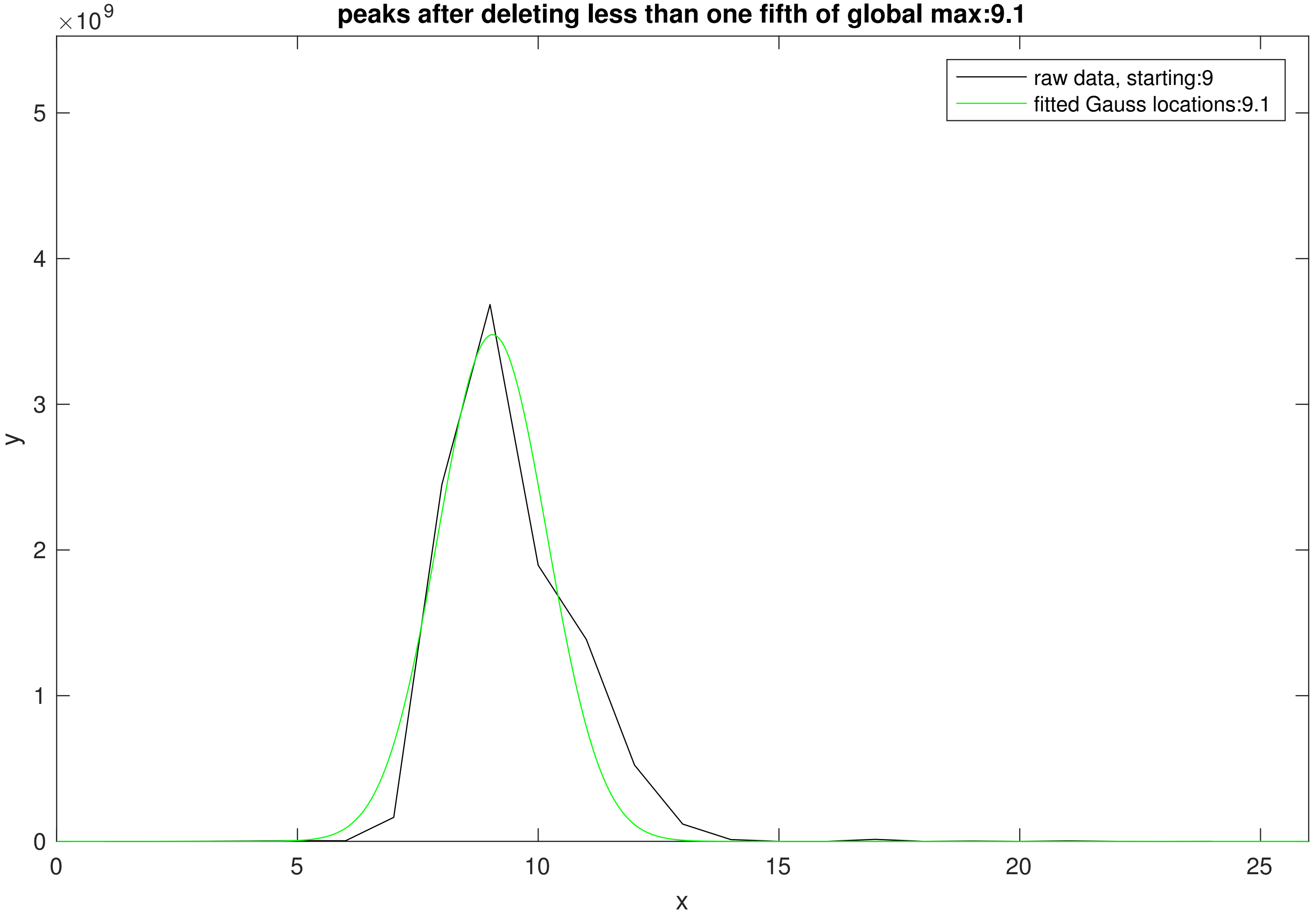

**Optimal1 Fitted Gauss, proteinAT5G07460.1**
**bic:928 934**
**peaks after deleting less than one fifth of global max:16**

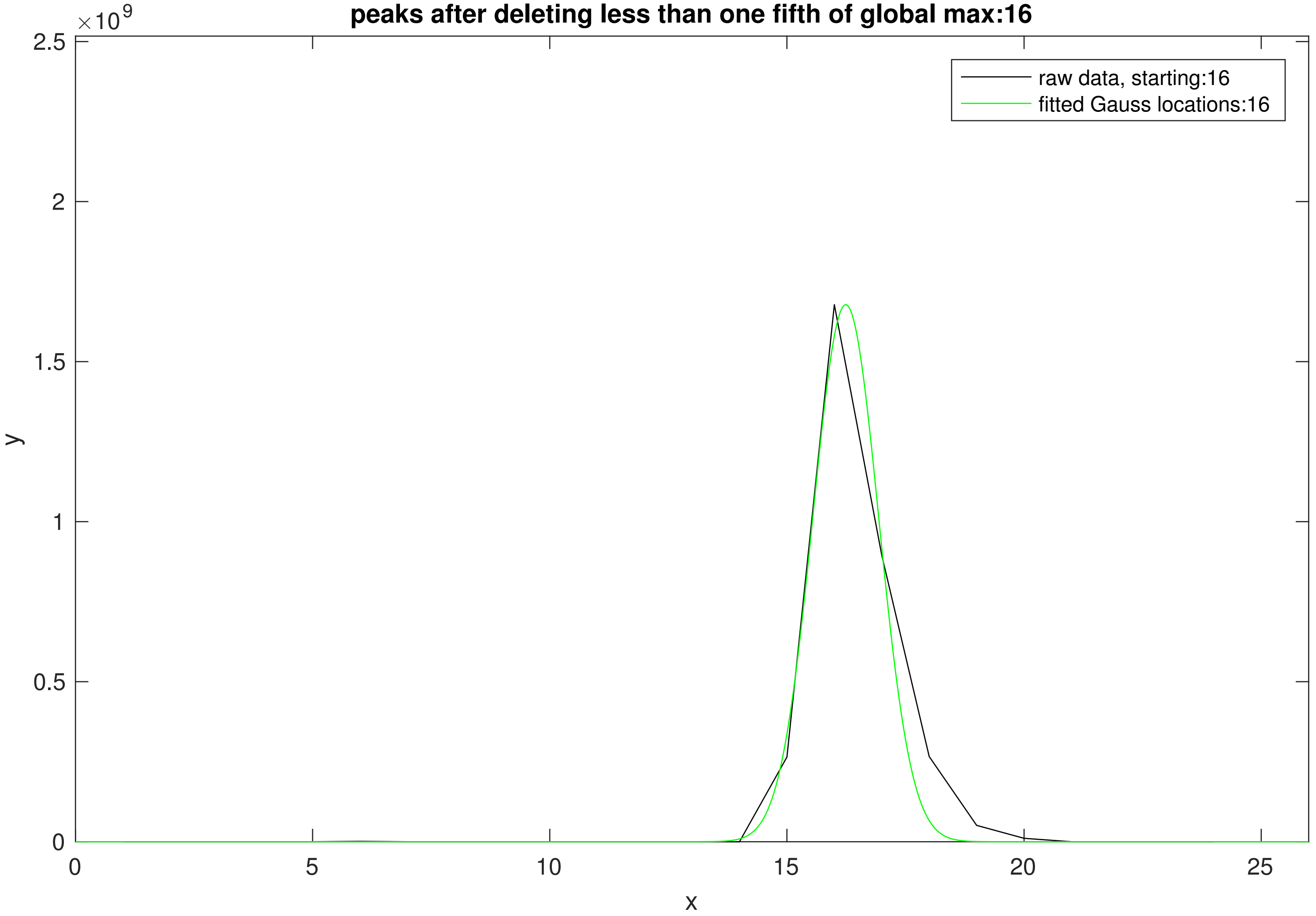

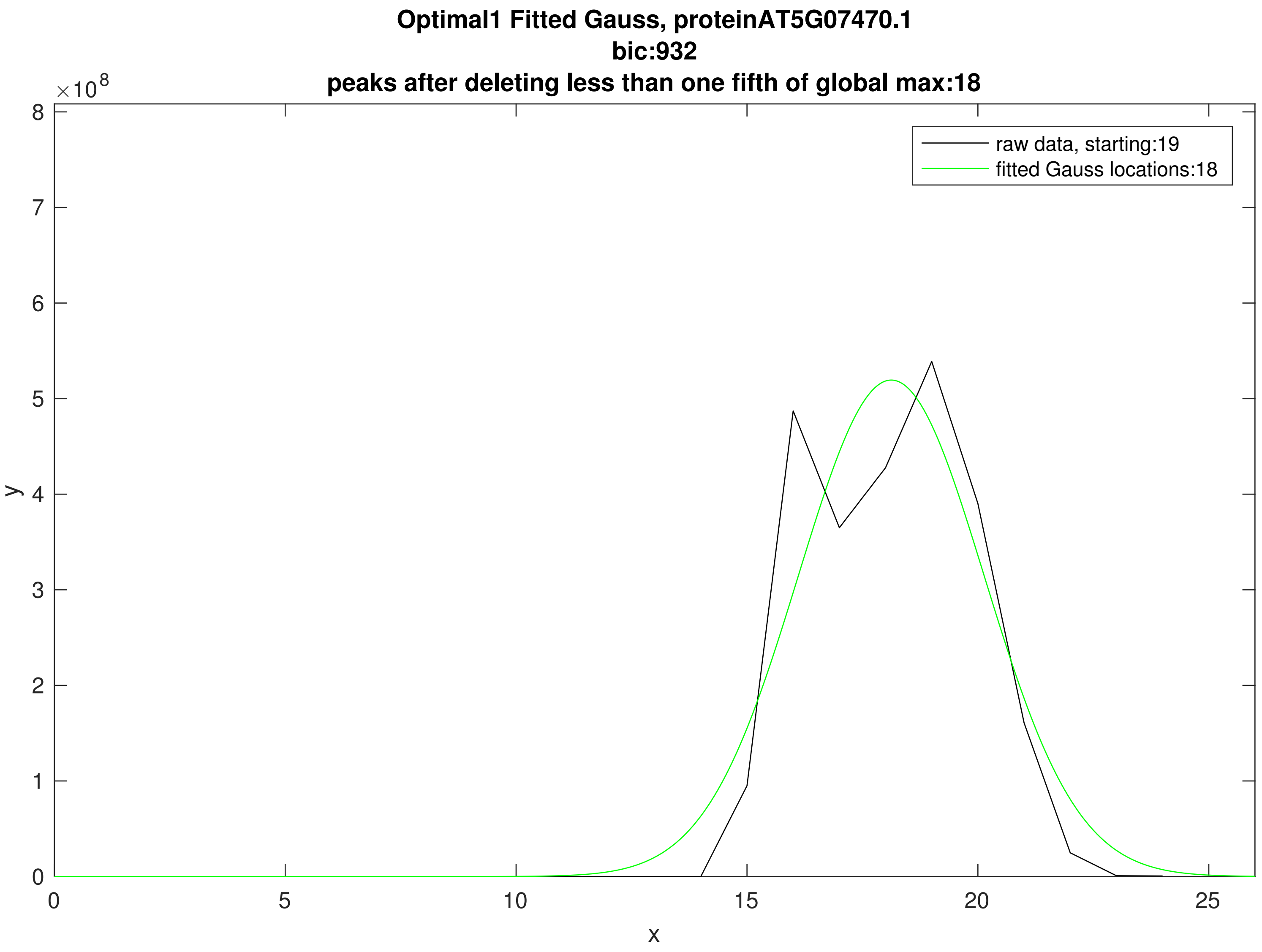

Optimal1 Fitted Gauss, proteinAT5G07470.1
bic:932
peaks after deleting less than one fifth of global max:18
×10^8
raw data, starting:19
fitted Gauss locations:18
y
x
0
1
2
3
4
5
6
7
8
0
5
10
15
20
25

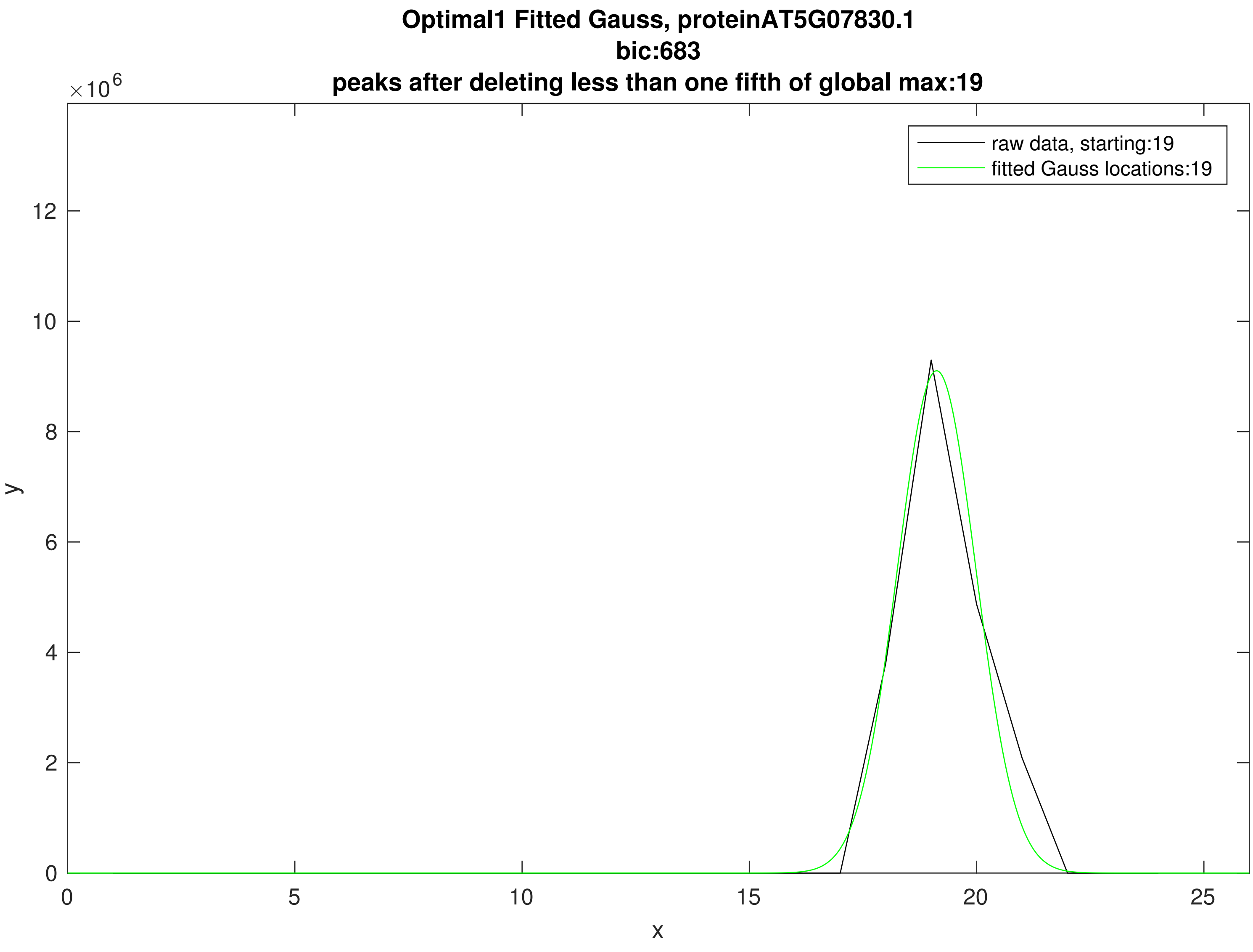
Optimal1 Fitted Gauss, proteinAT5G07830.1
bic:683
peaks after deleting less than one fifth of global max:19
×10^6
raw data, starting:19
fitted Gauss locations:19
y
x
0
2
4
6
8
10
12
0
5
10
15
20
25

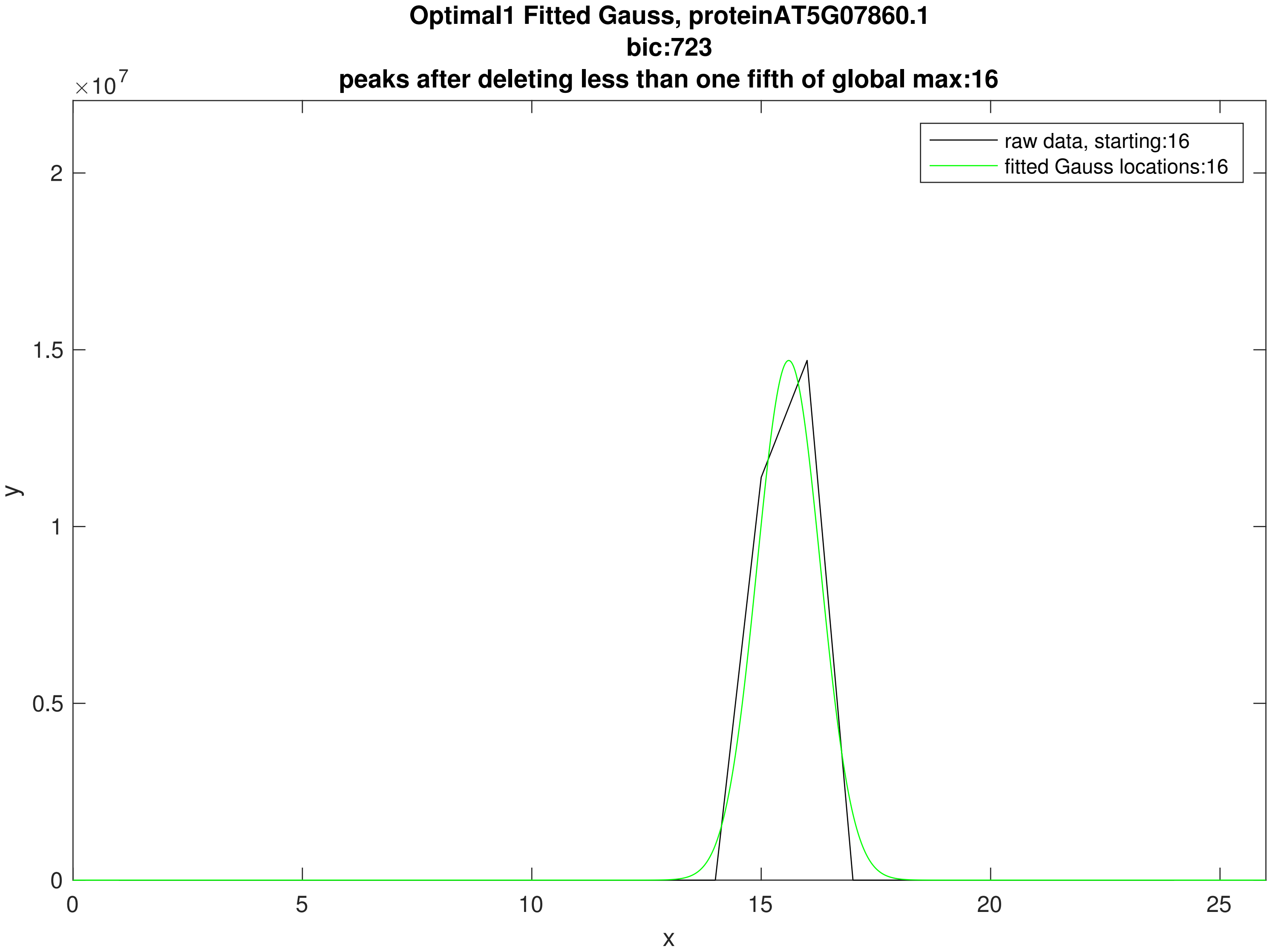

Optimal1 Fitted Gauss, proteinAT5G07860.1
bic:723
peaks after deleting less than one fifth of global max:16
×10^7
raw data, starting:16
fitted Gauss locations:16
y
x
0
0.5
1
1.5
2
5
10
15
20
25

Optimal1 Fitted Gauss, proteinAT5G07870.1
bic:789 791
peaks after deleting less than one fifth of global max:16

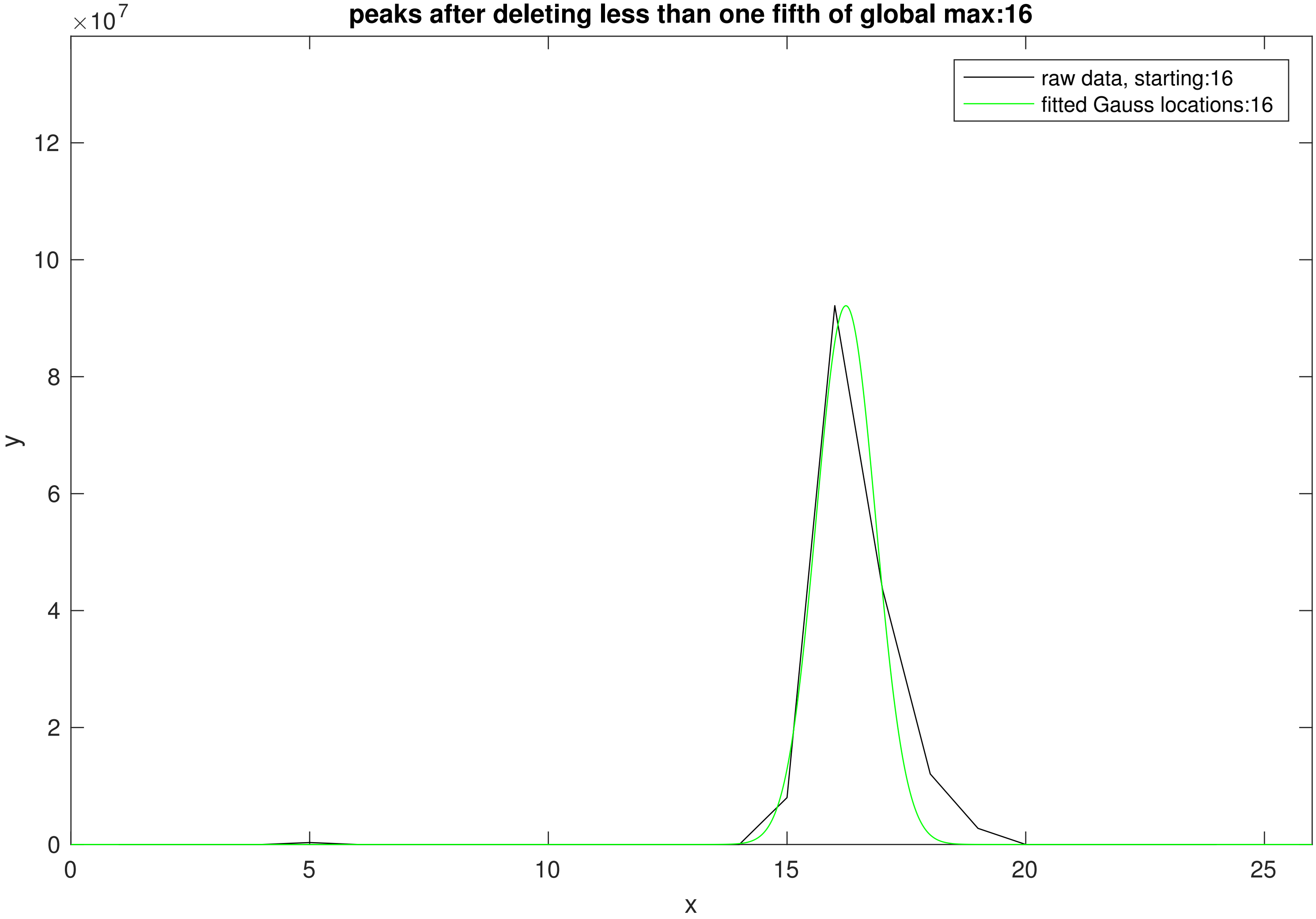

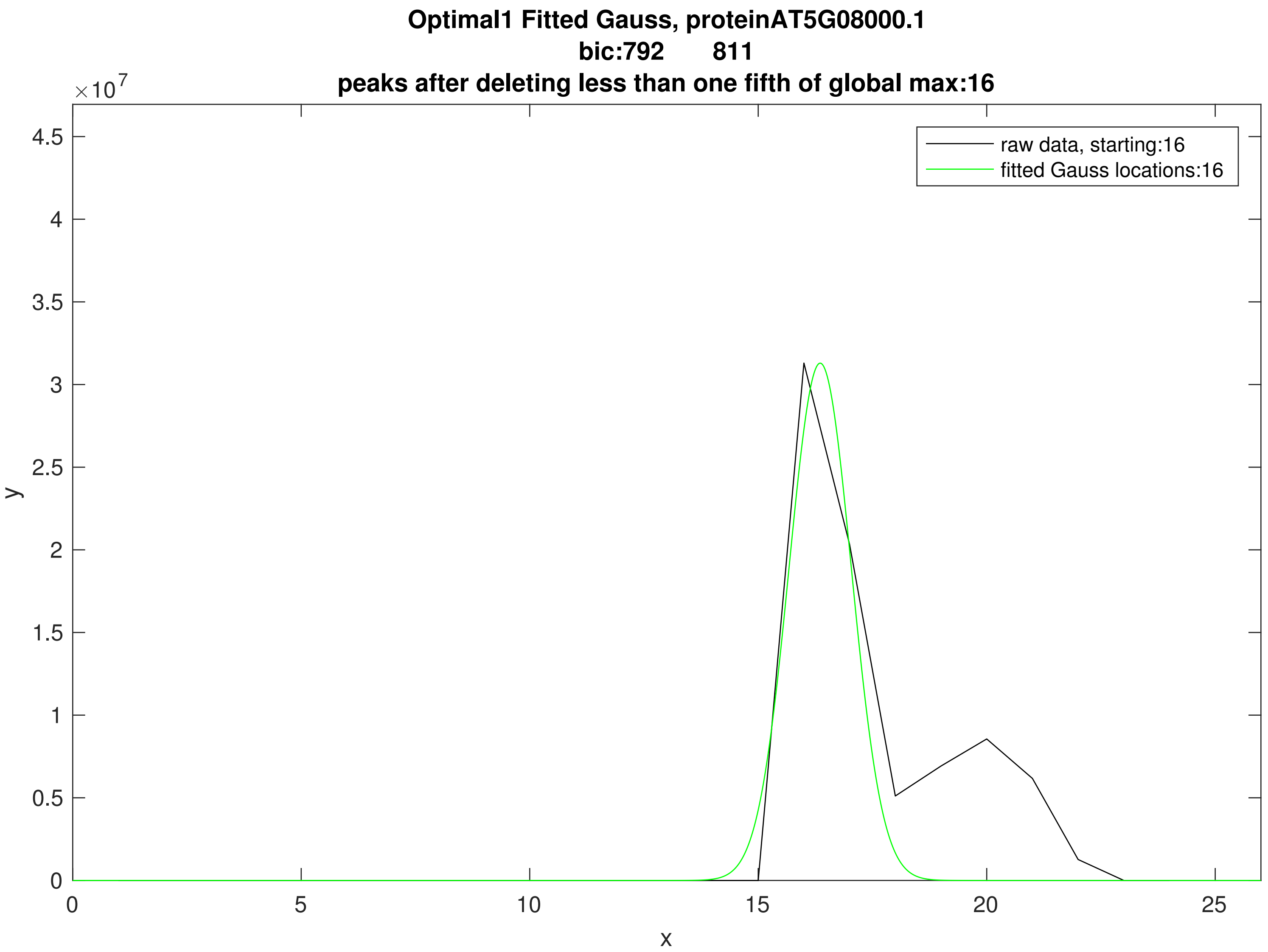

Optimal1 Fitted Gauss, proteinAT5G08000.1
bic:792 811
peaks after deleting less than one fifth of global max:16
×10^7
4.5
4
3.5
3
2.5
2
1.5
1
0.5
0
y
0
5
10
15
20
25
x
raw data, starting:16
fitted Gauss locations:16

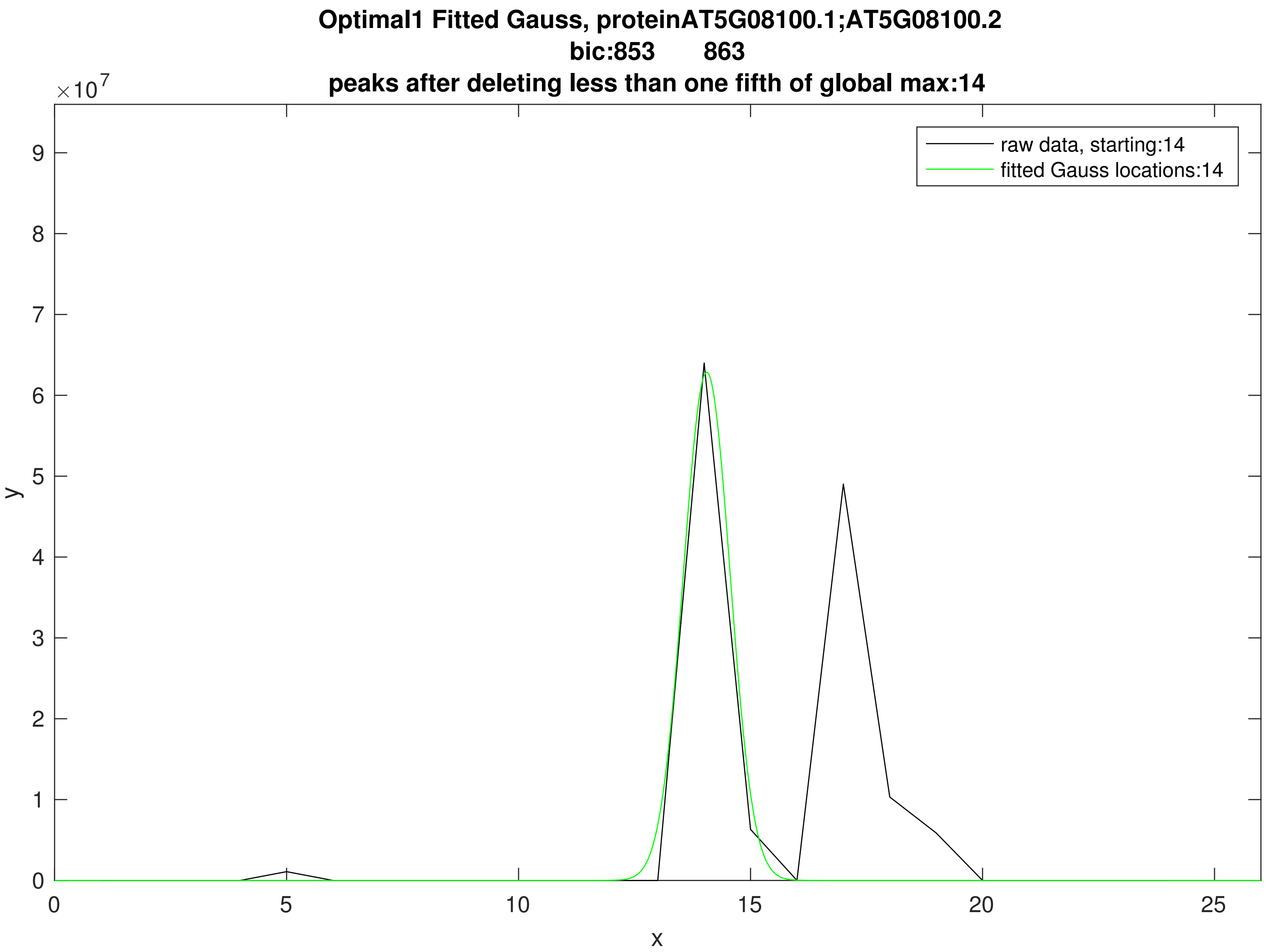
Optimal1 Fitted Gauss, proteinAT5G08100.1;AT5G08100.2
bic:853 863
peaks after deleting less than one fifth of global max:14
×10^7
raw data, starting:14
fitted Gauss locations:14
y
x
0
1
2
3
4
5
6
7
8
9
0
5
10
15
20
25

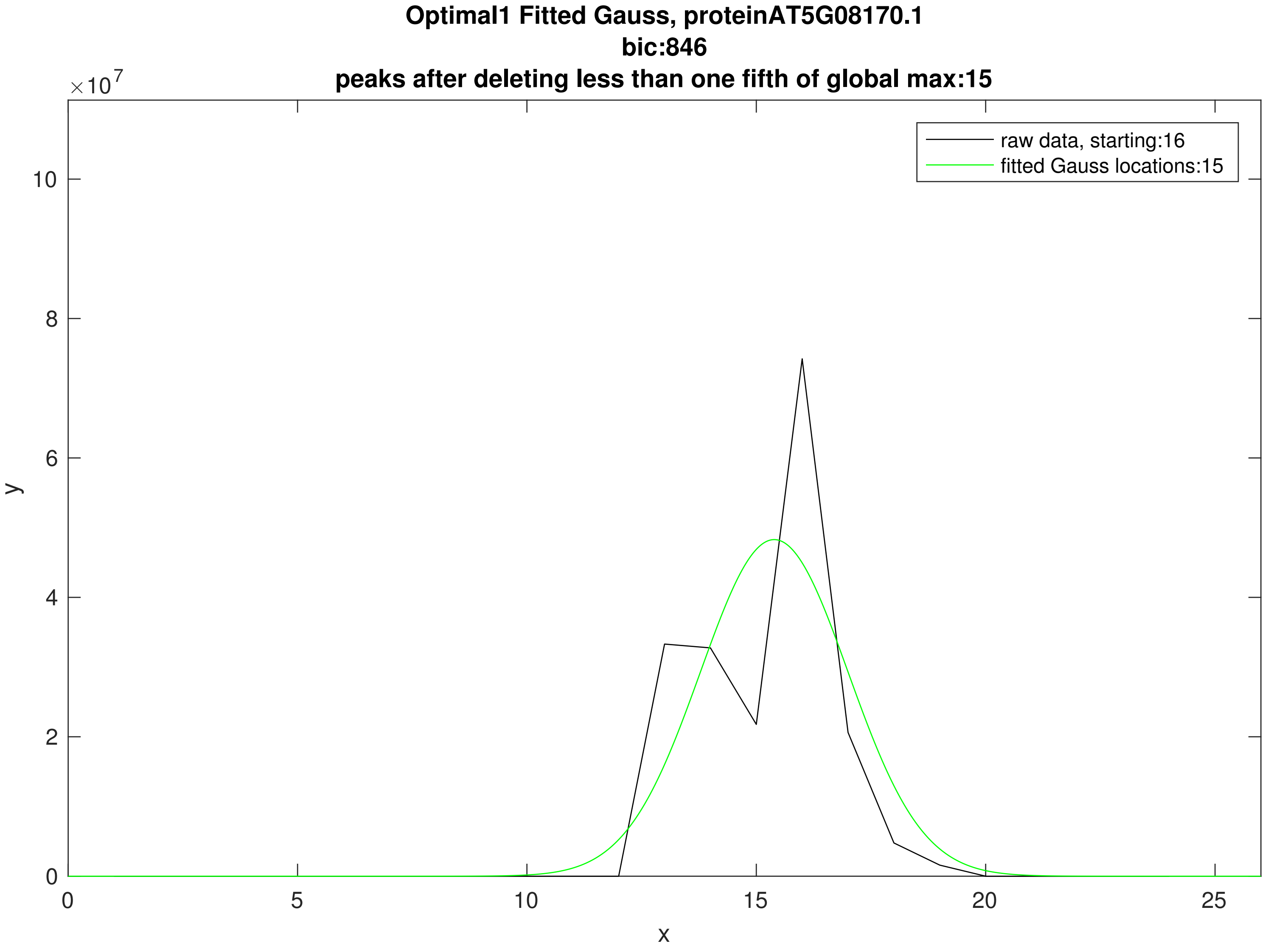

Optimal1 Fitted Gauss, proteinAT5G08170.1
bic:846
peaks after deleting less than one fifth of global max:15
×10^7
raw data, starting:16
fitted Gauss locations:15
y
x
0
2
4
6
8
10
0
5
10
15
20
25

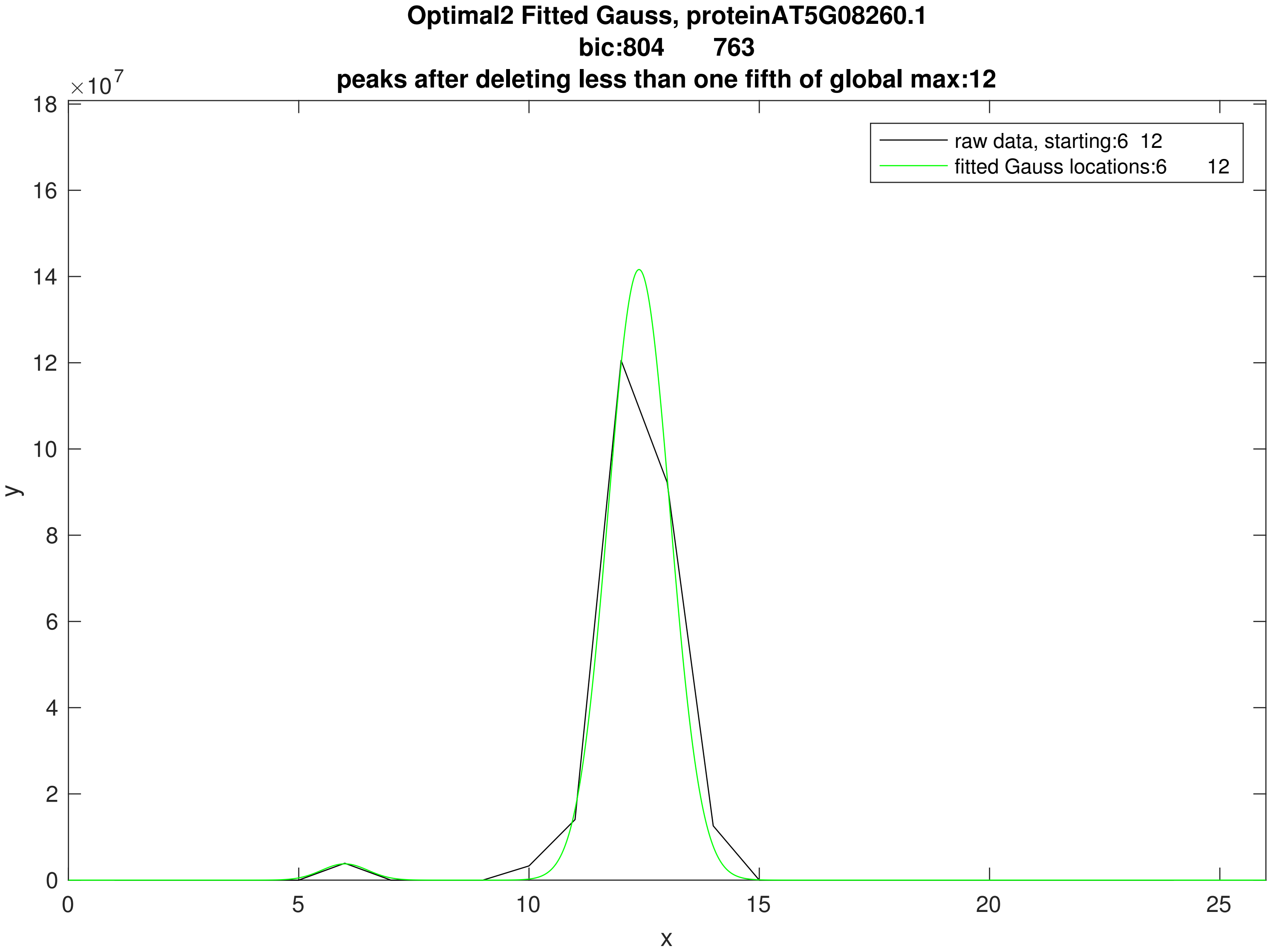

Optimal2 Fitted Gauss, proteinAT5G08260.1
bic:804 763
peaks after deleting less than one fifth of global max:12
×10^7
raw data, starting:6 12
fitted Gauss locations:6 12
y
x
0
2
4
6
8
10
12
14
16
18
0
5
10
15
20
25

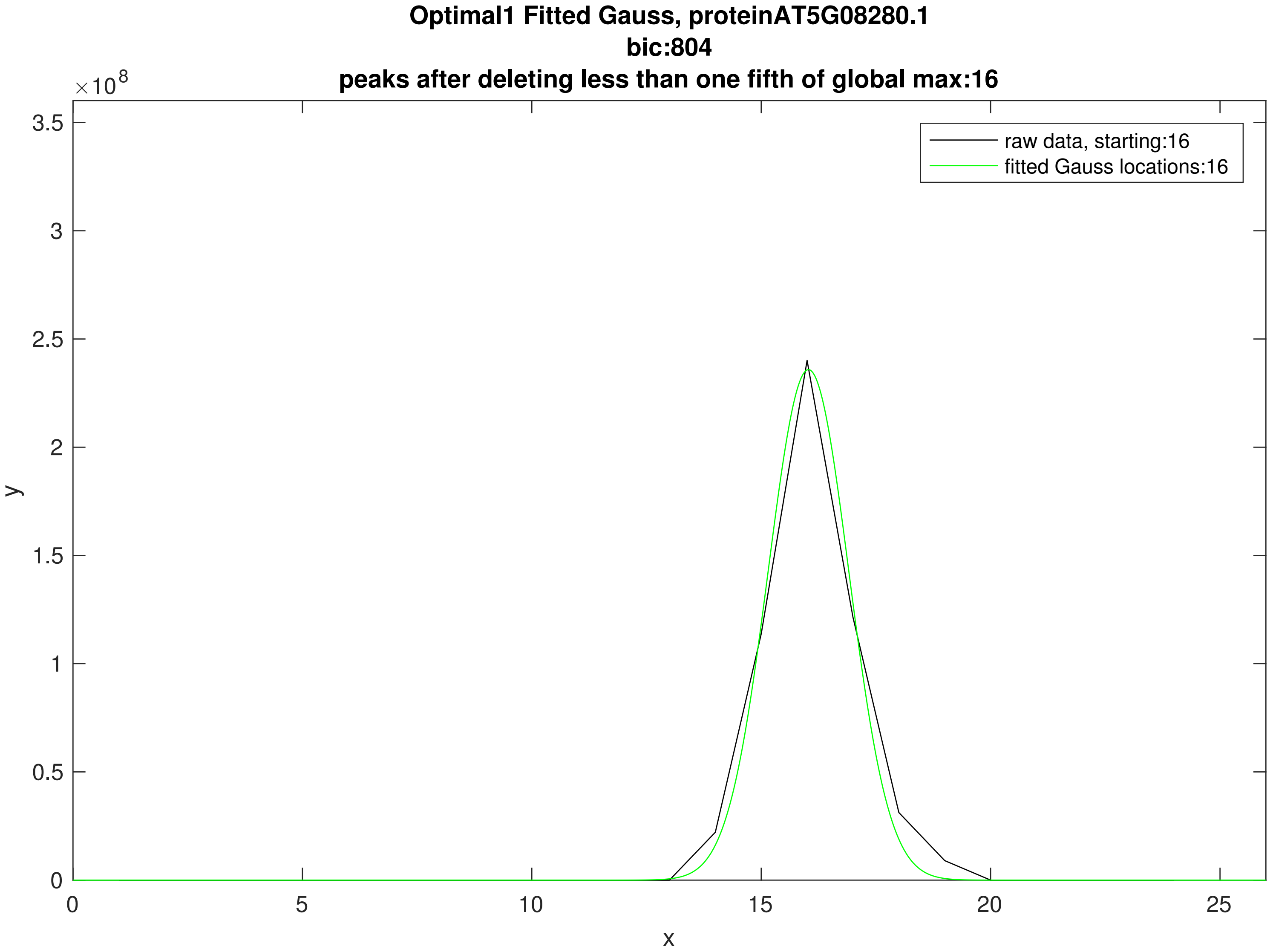

Optimal1 Fitted Gauss, proteinAT5G08280.1
bic:804
peaks after deleting less than one fifth of global max:16
×10^8
raw data, starting:16
fitted Gauss locations:16
y
x
0
0.5
1
1.5
2
2.5
3
3.5
0
5
10
15
20
25

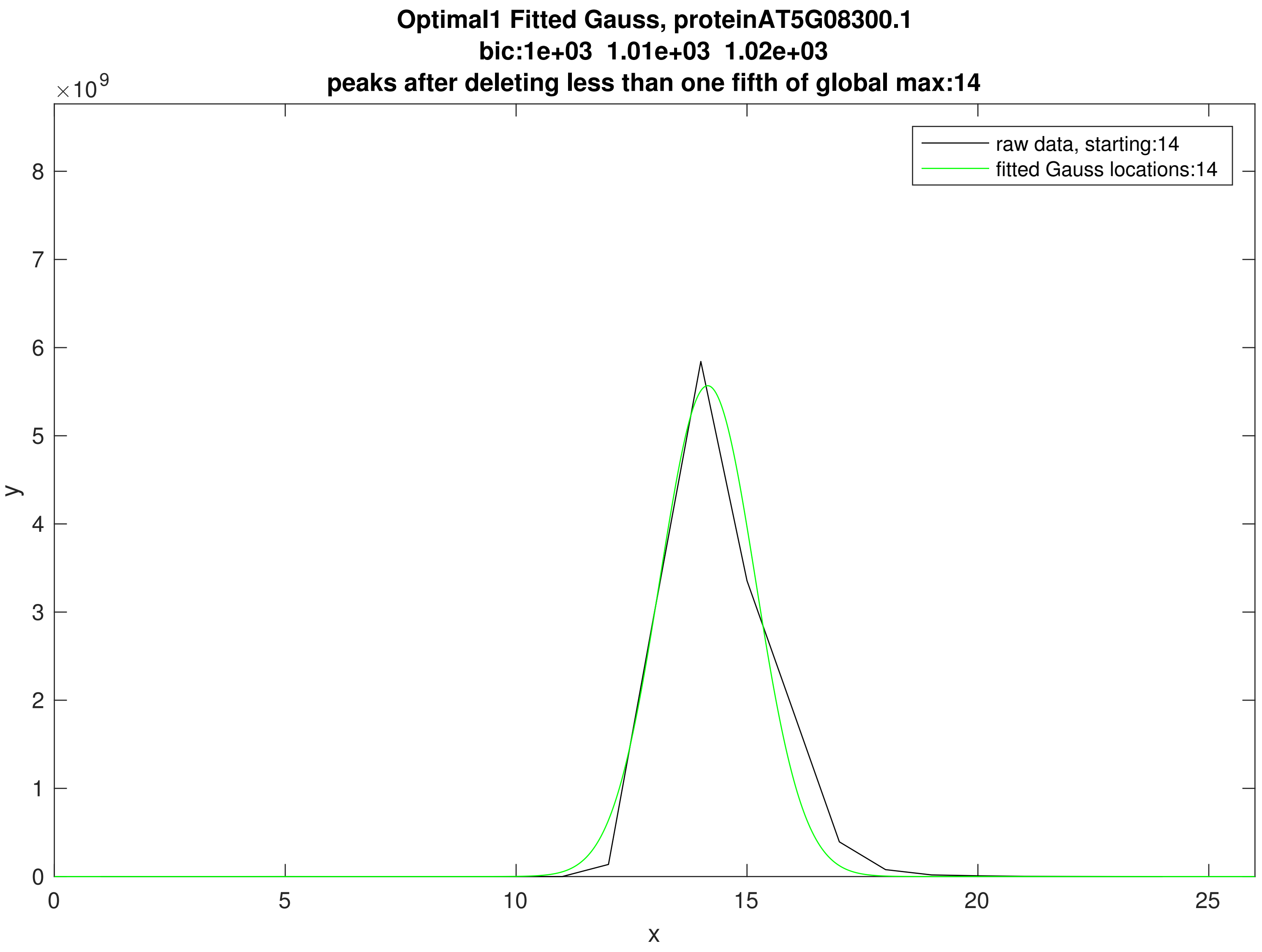
Optimal1 Fitted Gauss, proteinAT5G08300.1
bic:1e+03 1.01e+03 1.02e+03
peaks after deleting less than one fifth of global max:14
×10^9
raw data, starting:14
fitted Gauss locations:14
y
x
0
1
2
3
4
5
6
7
8
0
5
10
15
20
25

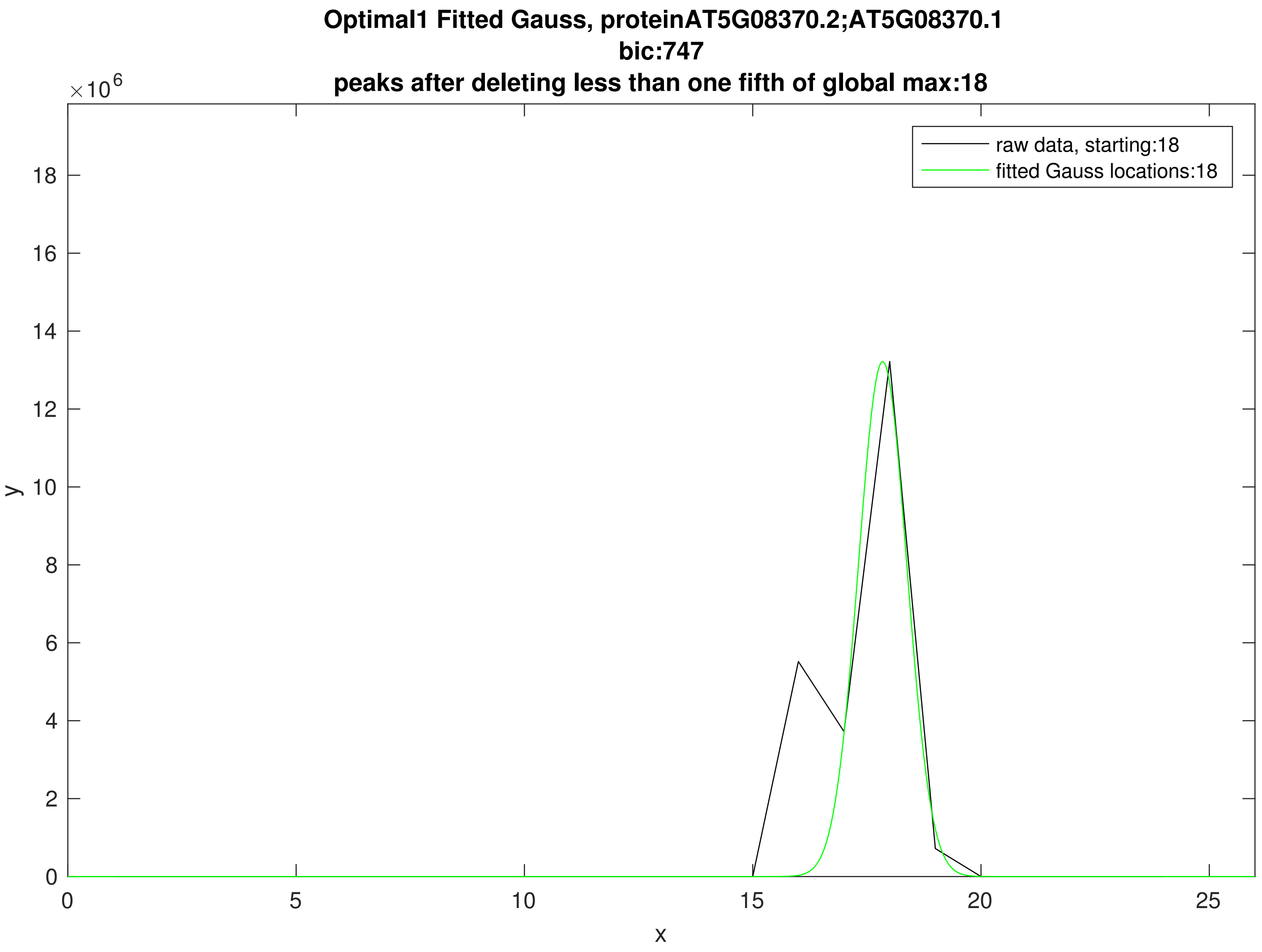

Optimal1 Fitted Gauss, proteinAT5G08370.2;AT5G08370.1
bic:747
peaks after deleting less than one fifth of global max:18
×10^6
raw data, starting:18
fitted Gauss locations:18
y
x
0
2
4
6
8
10
12
14
16
18
0
5
10
15
20
25

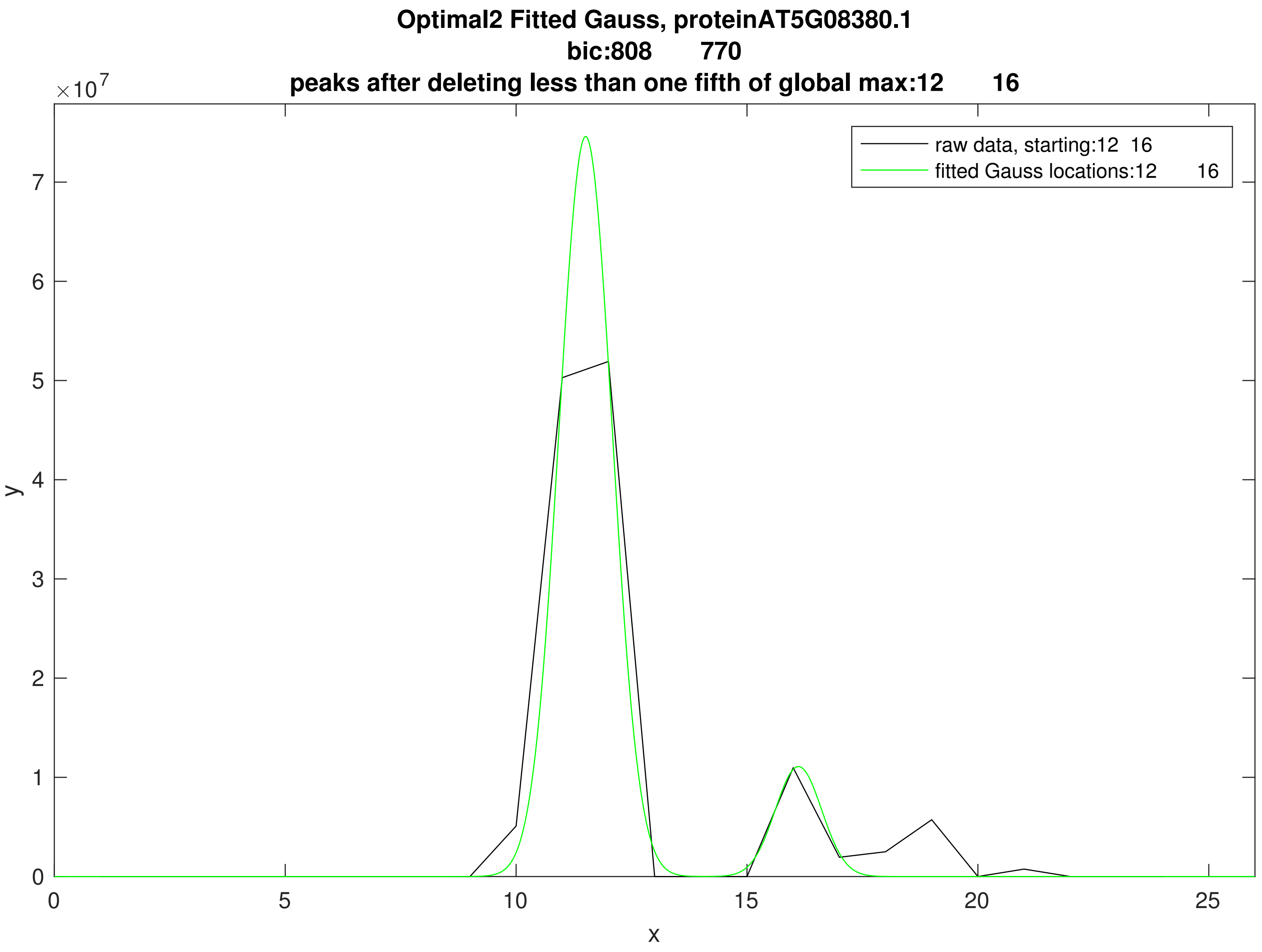

Optimal2 Fitted Gauss, proteinAT5G08380.1
bic:808 770
peaks after deleting less than one fifth of global max:12 16
×10^7
raw data, starting:12 16
fitted Gauss locations:12 16
y
x
0
1
2
3
4
5
6
7
0
5
10
15
20
25

#### Optimal1 Fitted Gauss, proteinAT5G08530.1
#### bic:800
#### peaks after deleting less than one fifth of global max:14

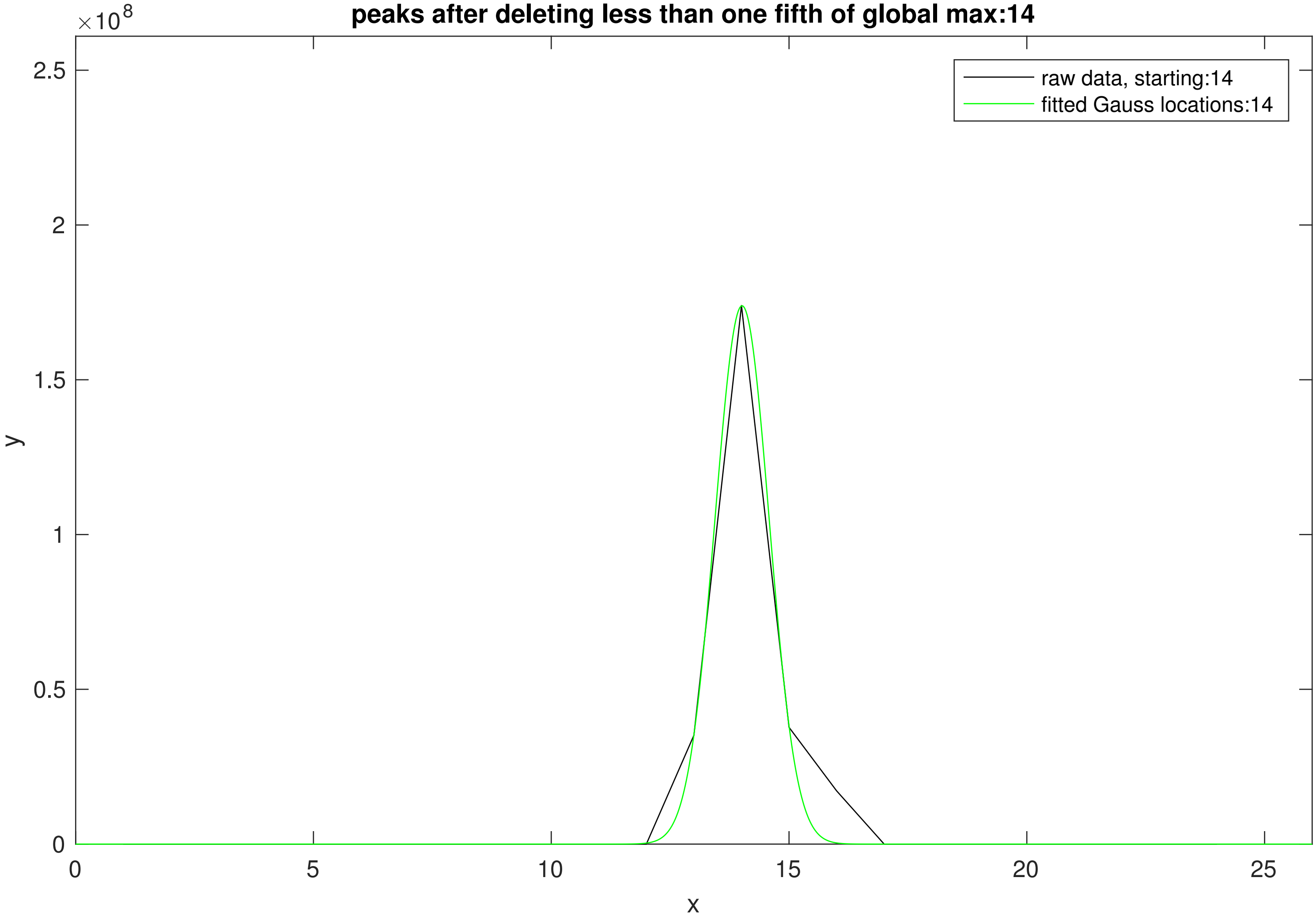

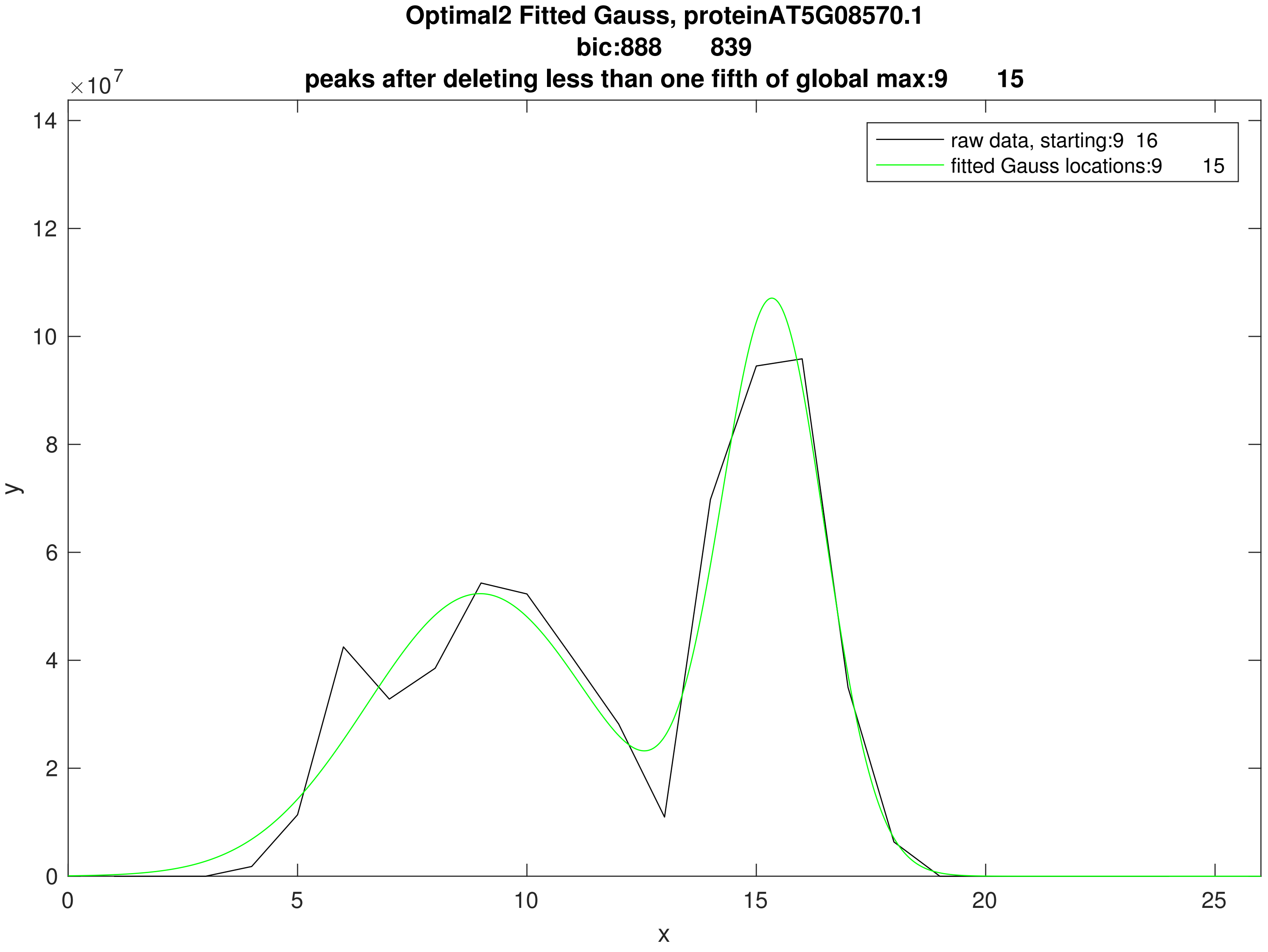

Optimal2 Fitted Gauss, proteinAT5G08570.1
bic:888 839
peaks after deleting less than one fifth of global max:9 15
×10^7
raw data, starting:9 16
fitted Gauss locations:9 15
y
x
0
2
4
6
8
10
12
14
0
5
10
15
20
25

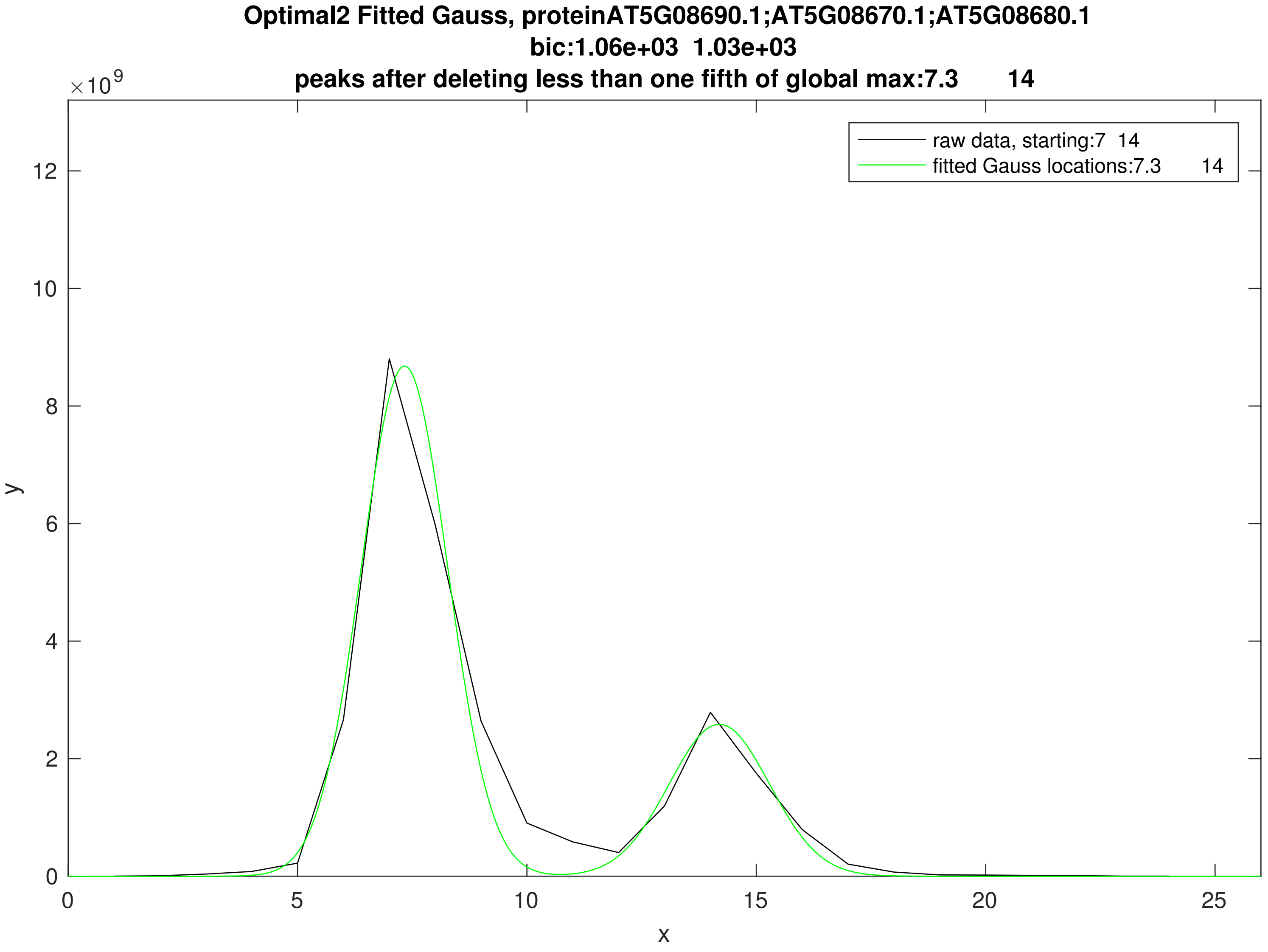

Optimal2 Fitted Gauss, proteinAT5G08690.1;AT5G08670.1;AT5G08680.1
bic:1.06e+03 1.03e+03
peaks after deleting less than one fifth of global max:7.3 14
×10^9
raw data, starting:7 14
fitted Gauss locations:7.3 14
y
x
0
2
4
6
8
10
12
0
5
10
15
20
25

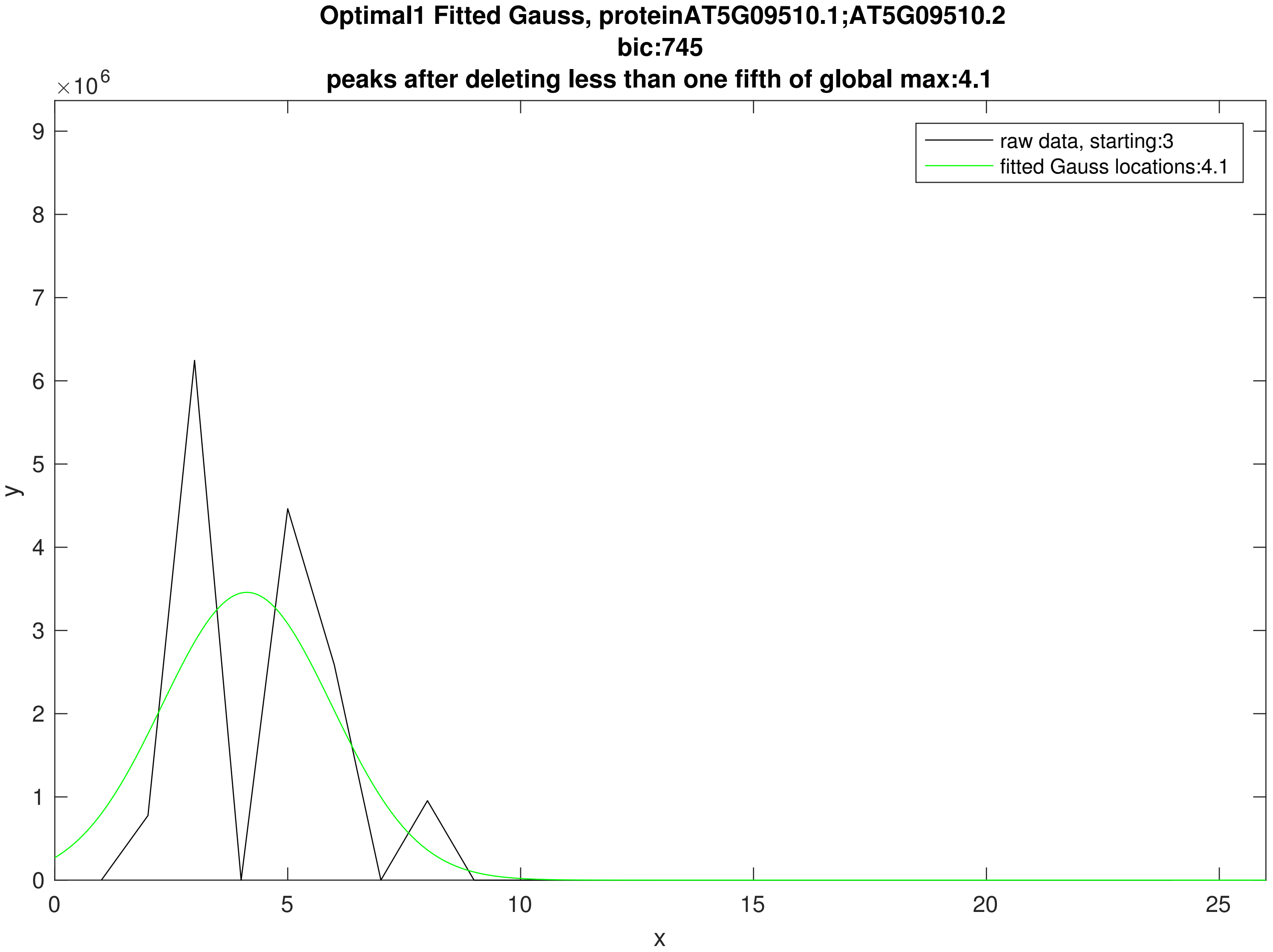
Optimal1 Fitted Gauss, proteinAT5G09510.1;AT5G09510.2
bic:745
peaks after deleting less than one fifth of global max:4.1
×10^6
raw data, starting:3
fitted Gauss locations:4.1
y
x
0
1
2
3
4
5
6
7
8
9
10
15
20
25

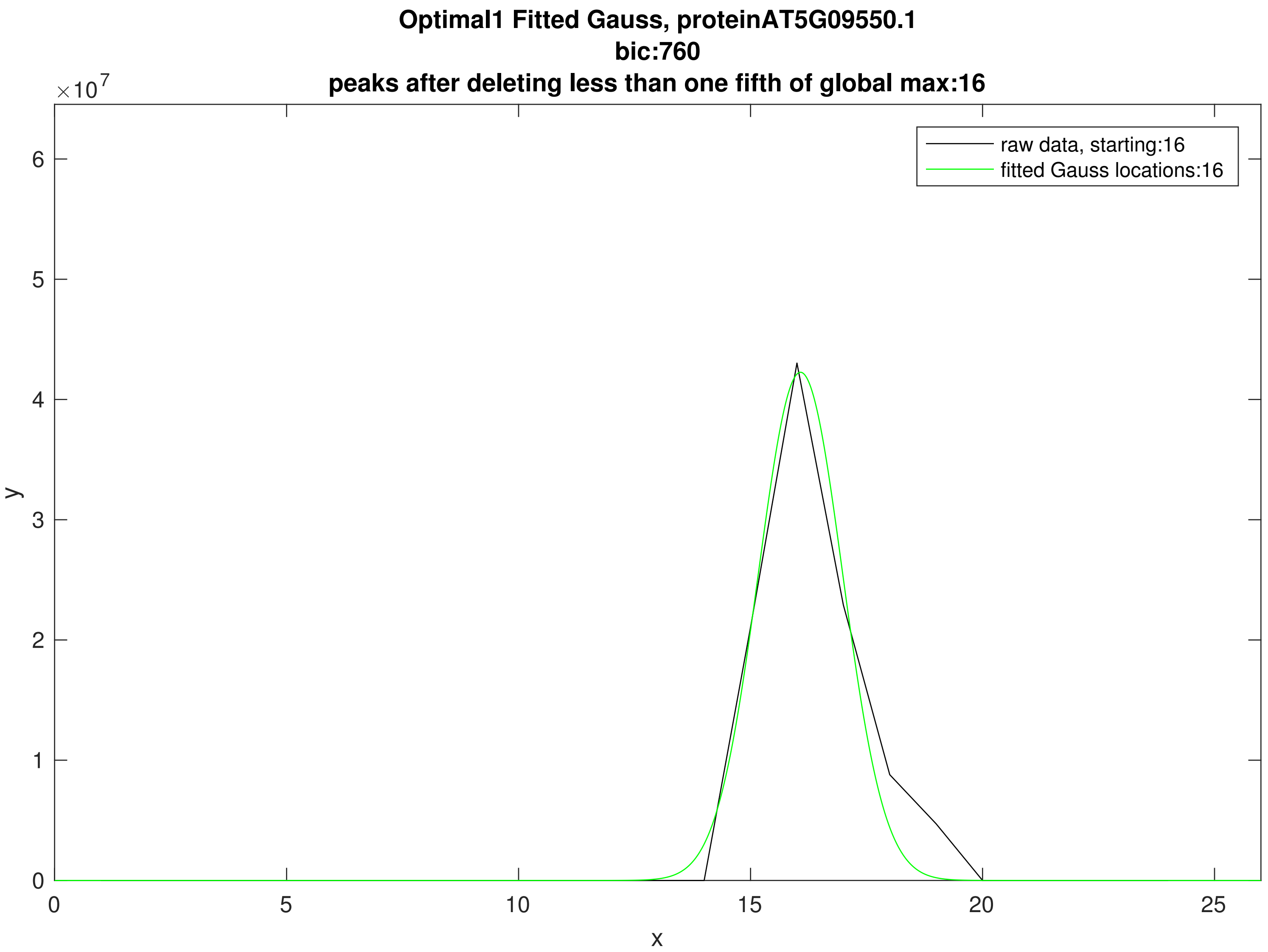

Optimal1 Fitted Gauss, proteinAT5G09550.1
bic:760
peaks after deleting less than one fifth of global max:16
×10^7
raw data, starting:16
fitted Gauss locations:16
y
x
0
1
2
3
4
5
6
0
5
10
15
20
25

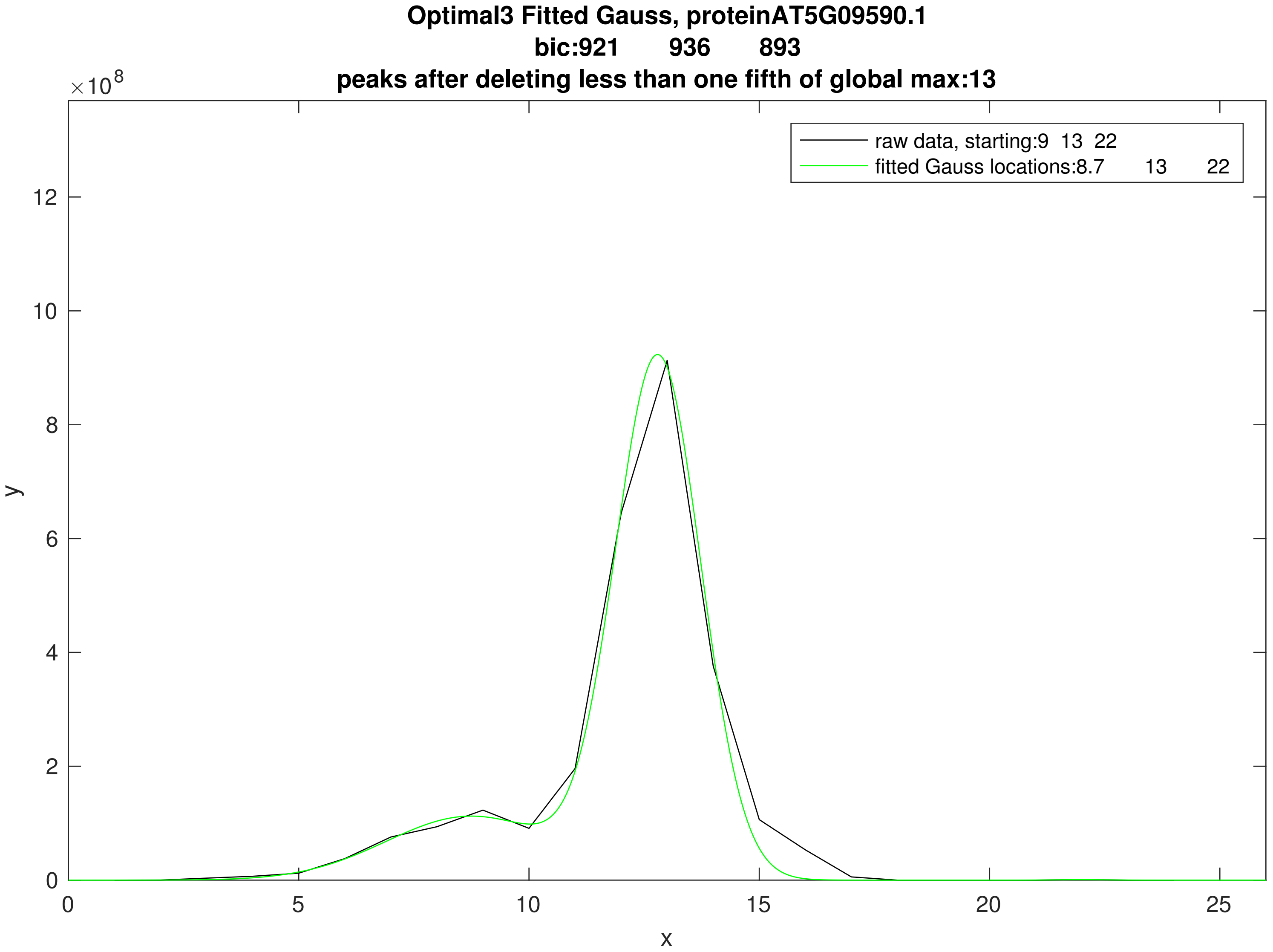

Optimal3 Fitted Gauss, proteinAT5G09590.1
bic:921 936 893
peaks after deleting less than one fifth of global max:13
×10^8
raw data, starting:9 13 22
fitted Gauss locations:8.7 13 22
y
x
0
2
4
6
8
10
12
0
5
10
15
20
25

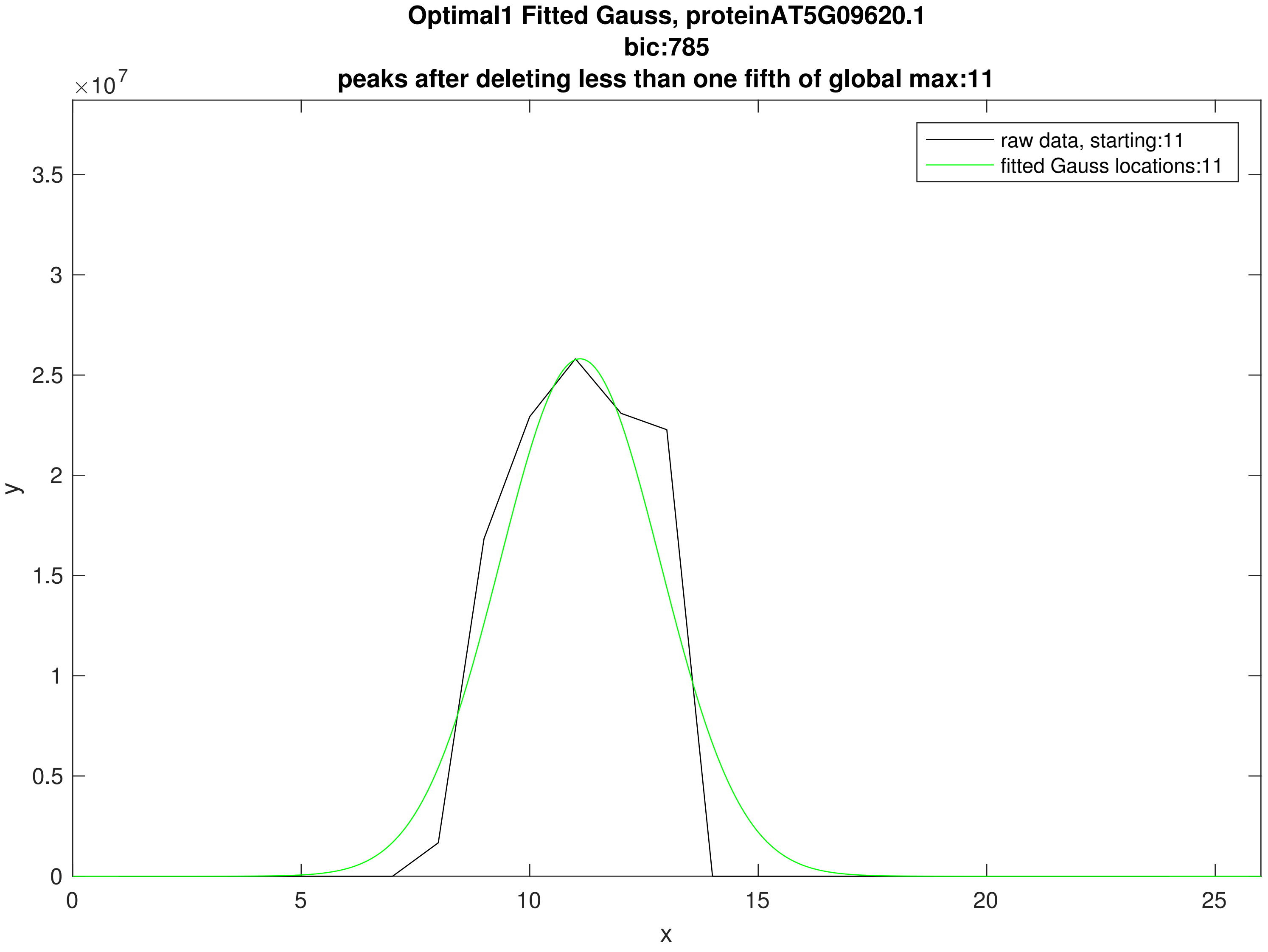
Optimal1 Fitted Gauss, proteinAT5G09620.1
bic:785
peaks after deleting less than one fifth of global max:11
×10^7
3.5
3
2.5
2
1.5
1
0.5
0
y
0
5
10
15
20
25
x
raw data, starting:11
fitted Gauss locations:11

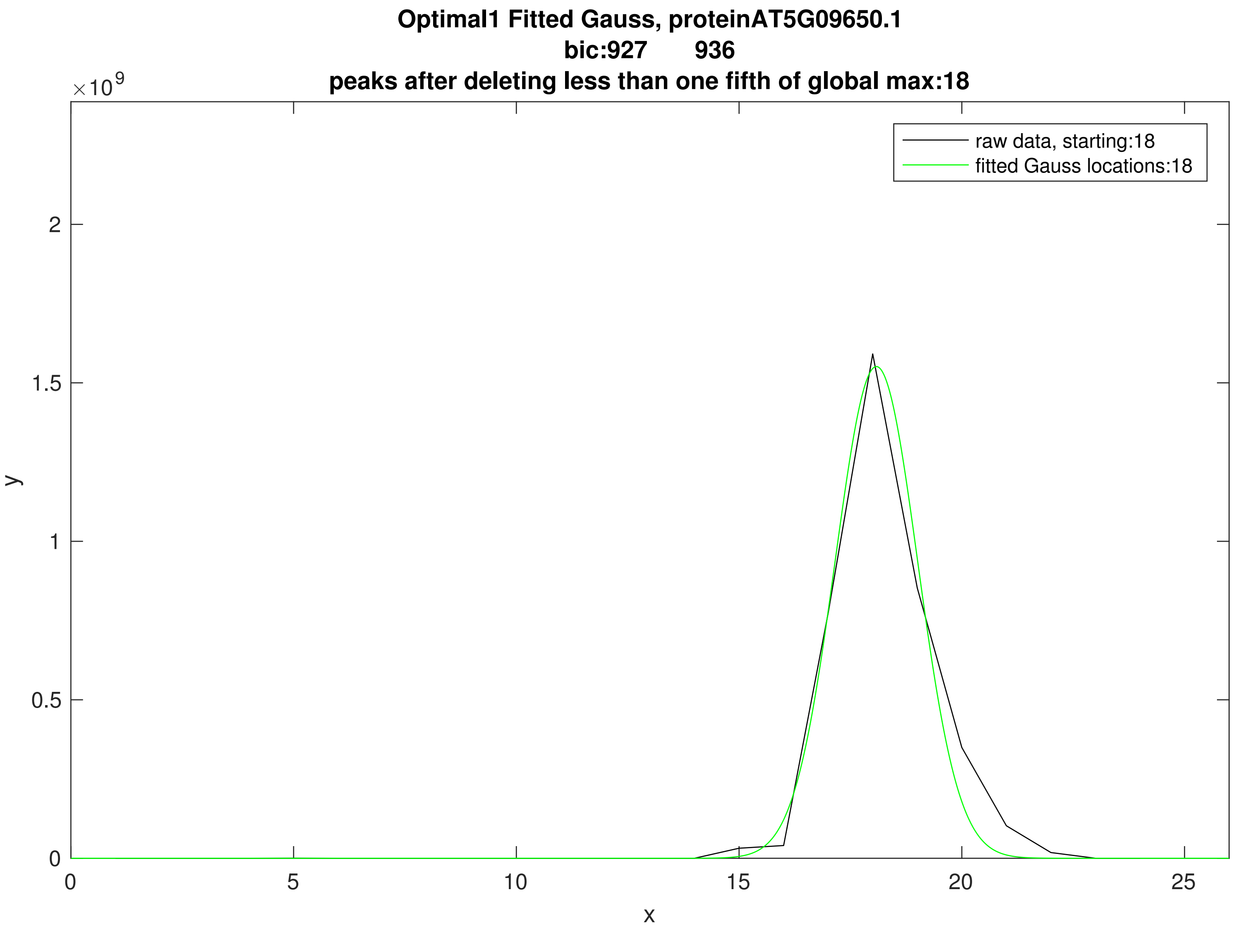

Optimal1 Fitted Gauss, proteinAT5G09650.1
bic:927 936
peaks after deleting less than one fifth of global max:18
×10^9
raw data, starting:18
fitted Gauss locations:18
2
1.5
1
0.5
0
y
0
5
10
15
20
25
x

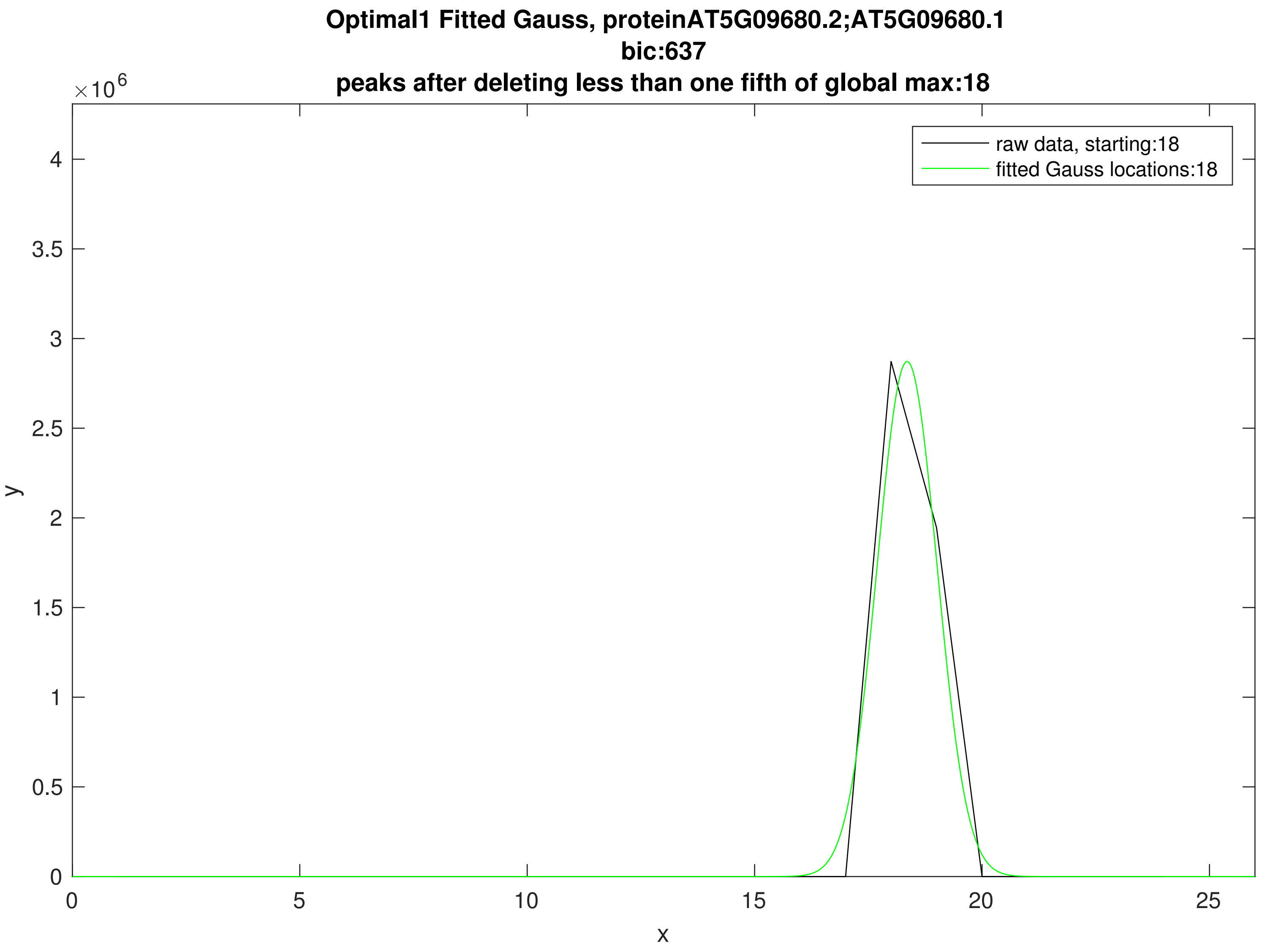

Optimal1 Fitted Gauss, proteinAT5G09680.2;AT5G09680.1
bic:637
peaks after deleting less than one fifth of global max:18
×10^6
raw data, starting:18
fitted Gauss locations:18
y
x
0
0.5
1
1.5
2
2.5
3
3.5
4
0
5
10
15
20
25

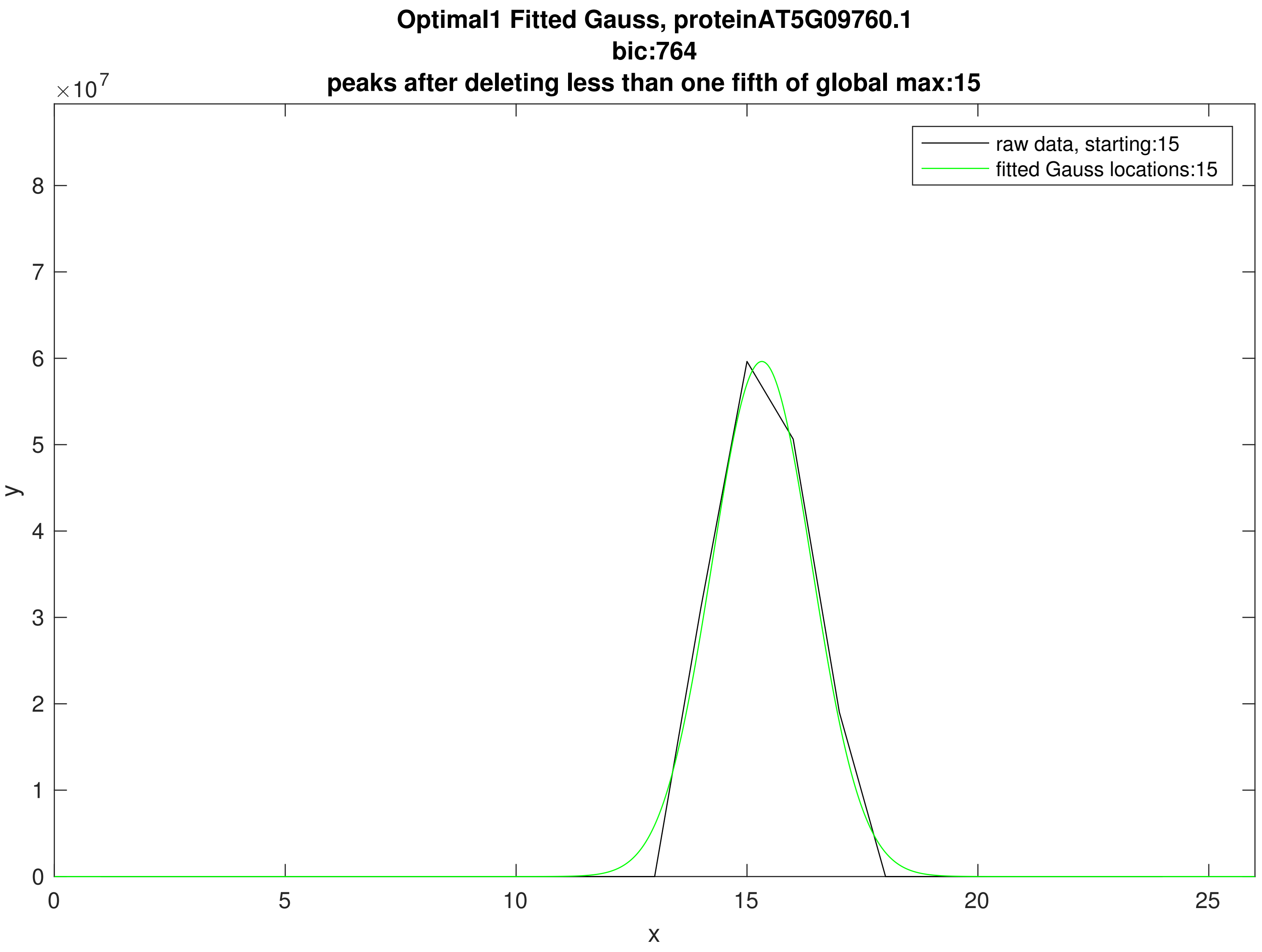

Optimal1 Fitted Gauss, proteinAT5G09760.1
bic:764
peaks after deleting less than one fifth of global max:15
×10^7
raw data, starting:15
fitted Gauss locations:15
y
x
0
1
2
3
4
5
6
7
8
0
5
10
15
20
25

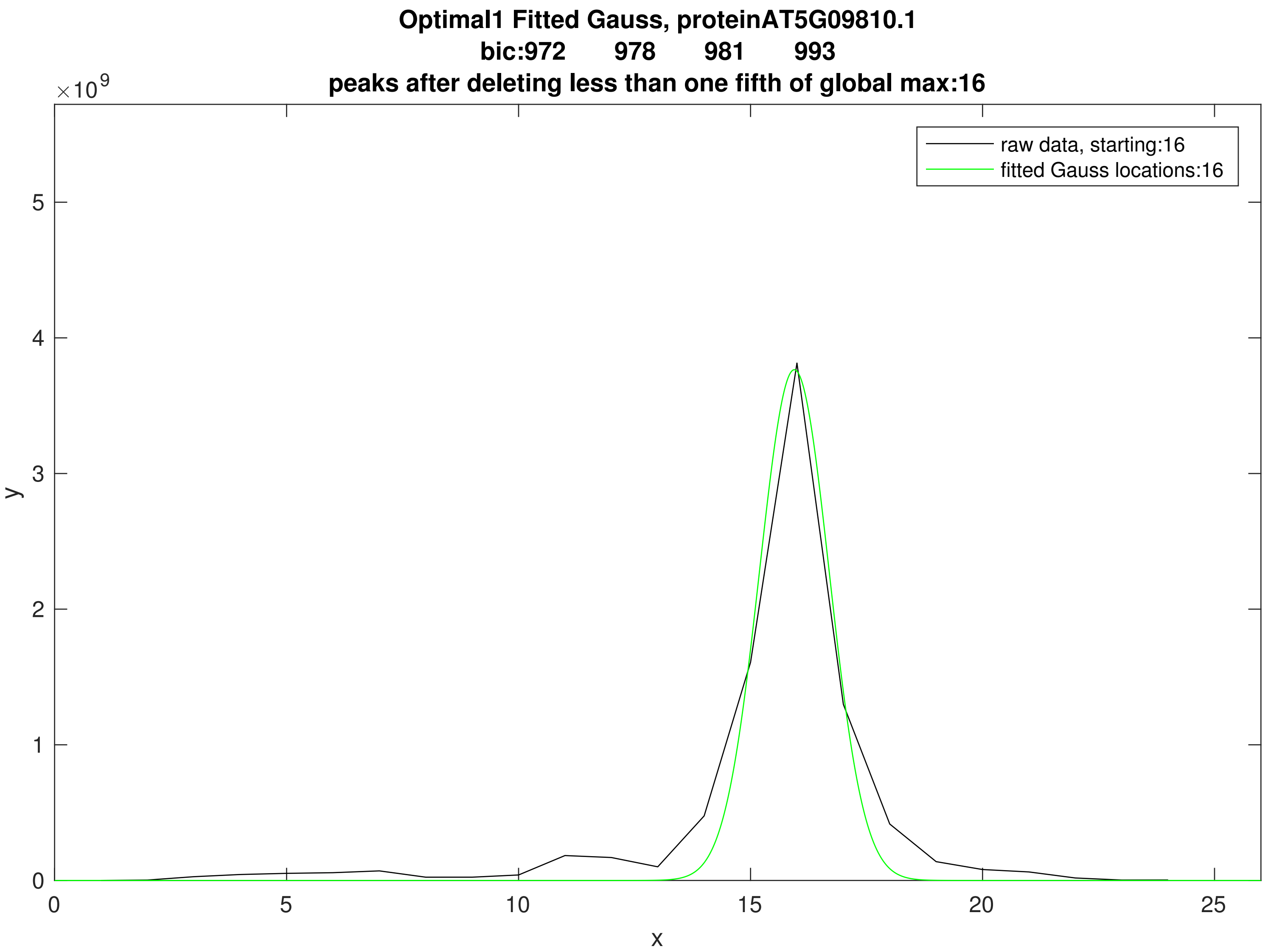
Optimal1 Fitted Gauss, proteinAT5G09810.1
bic:972 978 981 993
peaks after deleting less than one fifth of global max:16
×10^9
raw data, starting:16
fitted Gauss locations:16
y
x
0
1
2
3
4
5
0
5
10
15
20
25

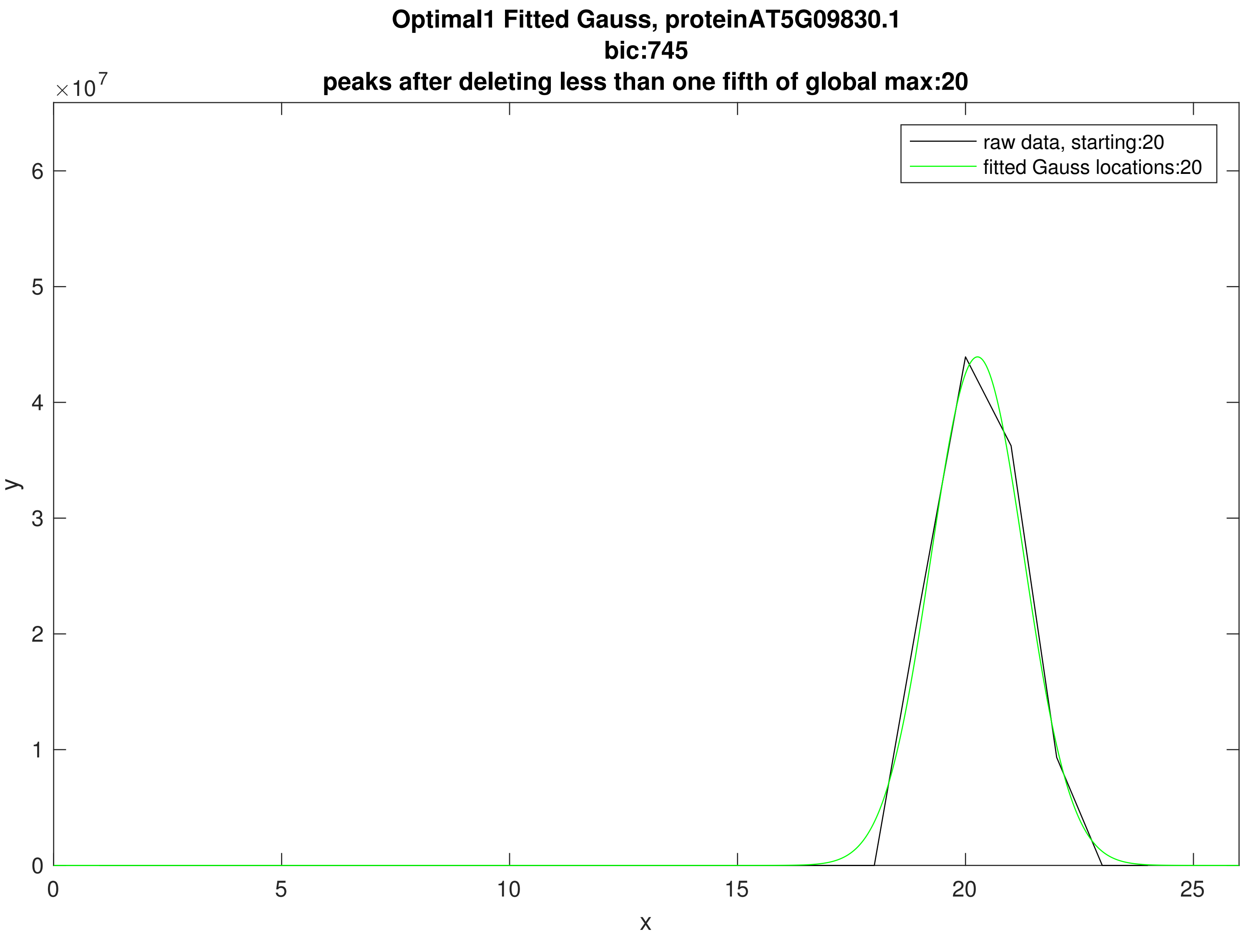
Optimal1 Fitted Gauss, proteinAT5G09830.1
bic:745
peaks after deleting less than one fifth of global max:20
×10^7
raw data, starting:20
fitted Gauss locations:20
6
5
4
3
2
1
0
y
0
5
10
15
20
25
x

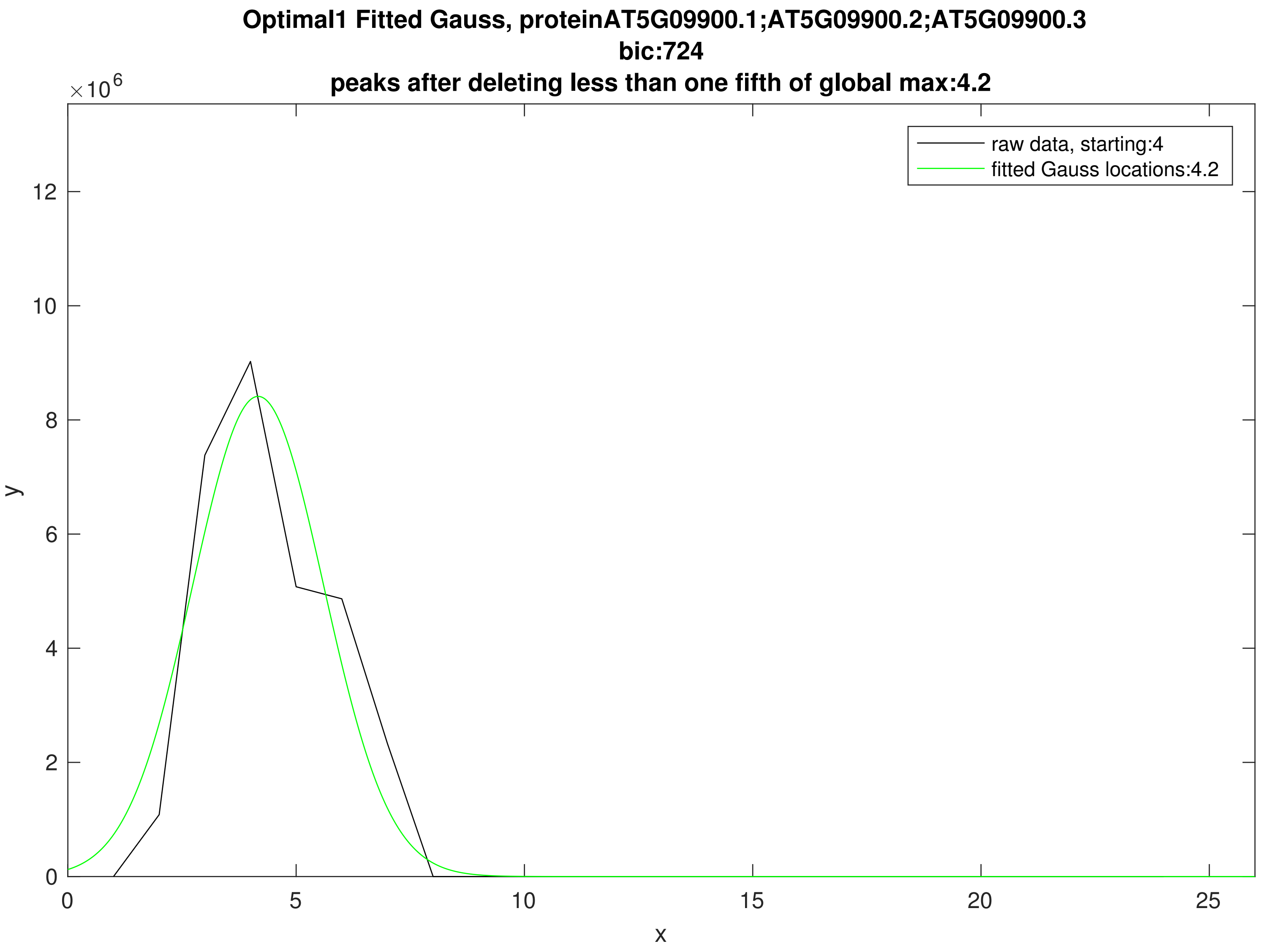

Optimal1 Fitted Gauss, proteinAT5G09900.1;AT5G09900.2;AT5G09900.3
bic:724
peaks after deleting less than one fifth of global max:4.2
×10^6
raw data, starting:4
fitted Gauss locations:4.2
y
x
0
2
4
6
8
10
12
0
5
10
15
20
25

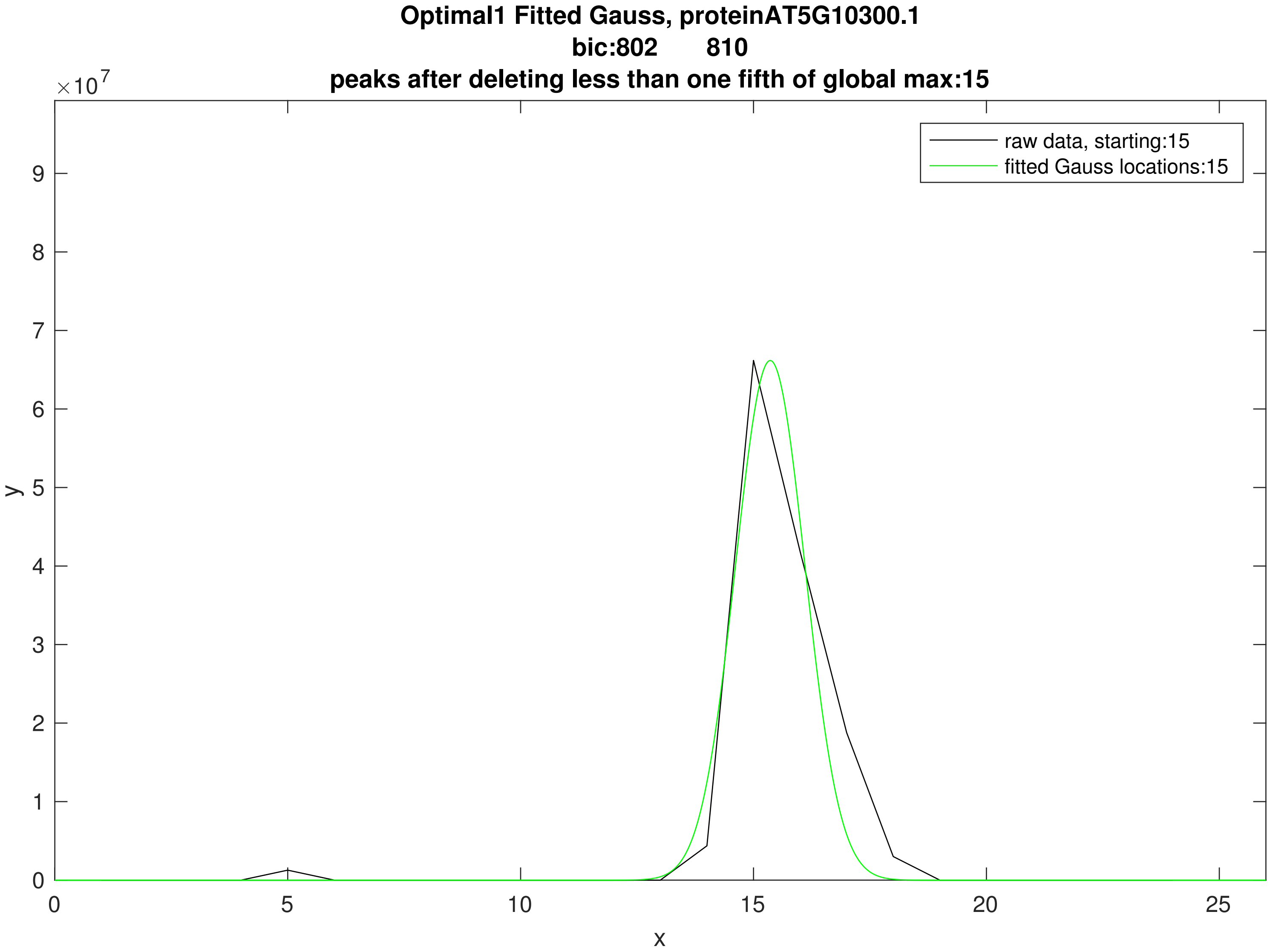

Optimal1 Fitted Gauss, proteinAT5G10300.1
bic:802 810
peaks after deleting less than one fifth of global max:15
×10^7
raw data, starting:15
fitted Gauss locations:15
y
x
0
1
2
3
4
5
6
7
8
9
10
15
20
25

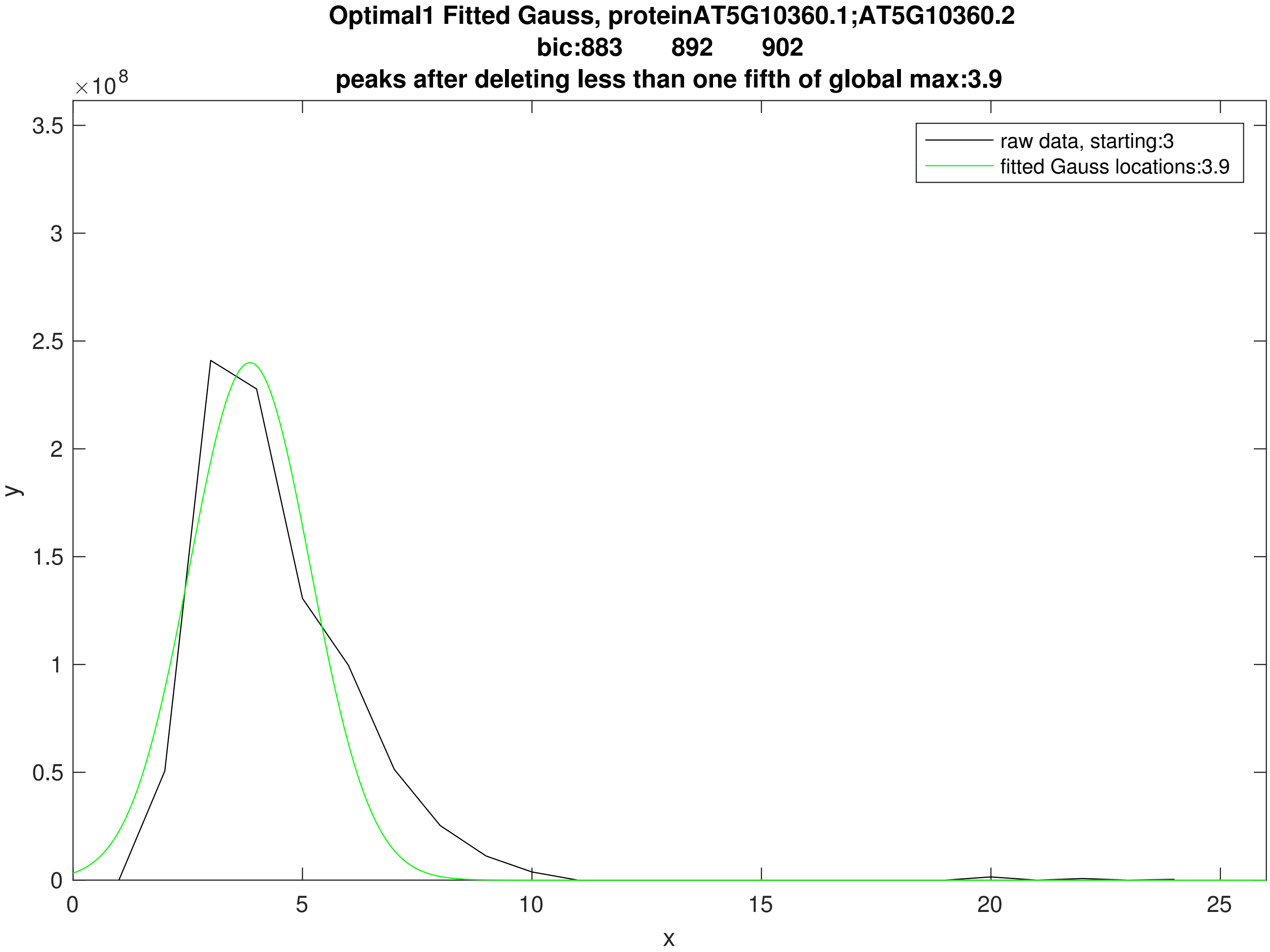

Optimal1 Fitted Gauss, proteinAT5G10360.1;AT5G10360.2
bic:883 892 902
peaks after deleting less than one fifth of global max:3.9
×10^8
3.5
3
2.5
2
1.5
1
0.5
0
y
0
5
10
15
20
25
x
raw data, starting:3
fitted Gauss locations:3.9

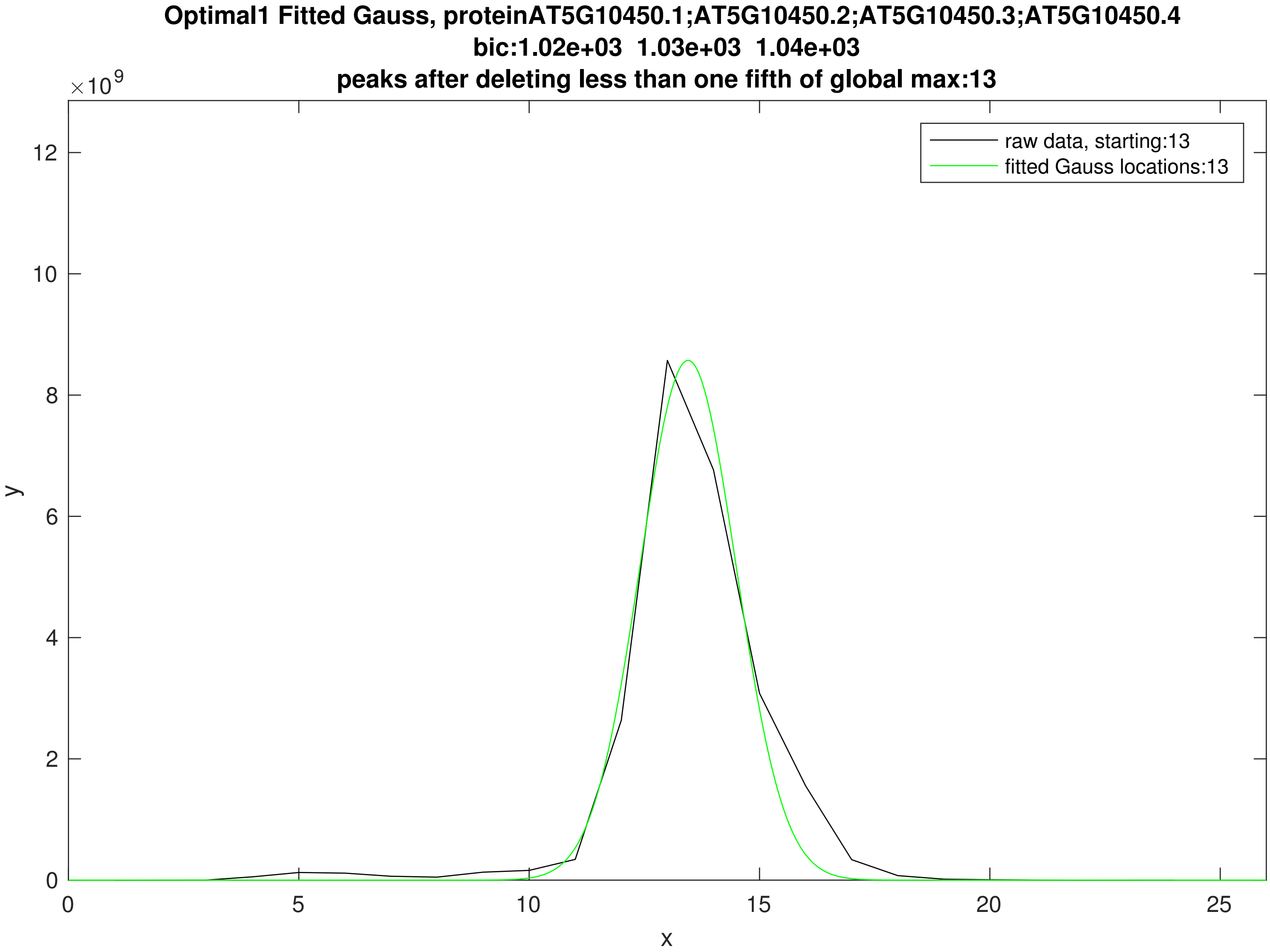
Optimal1 Fitted Gauss, proteinAT5G10450.1;AT5G10450.2;AT5G10450.3;AT5G10450.4
bic:1.02e+03  1.03e+03  1.04e+03
peaks after deleting less than one fifth of global max:13
×10^9
raw data, starting:13
fitted Gauss locations:13
y
x
0
2
4
6
8
10
12
5
15
20
25

#### Optimal1 Fitted Gauss, proteinAT5G10540.1
#### bic:993 1e+03
#### peaks after deleting less than one fifth of global max:14

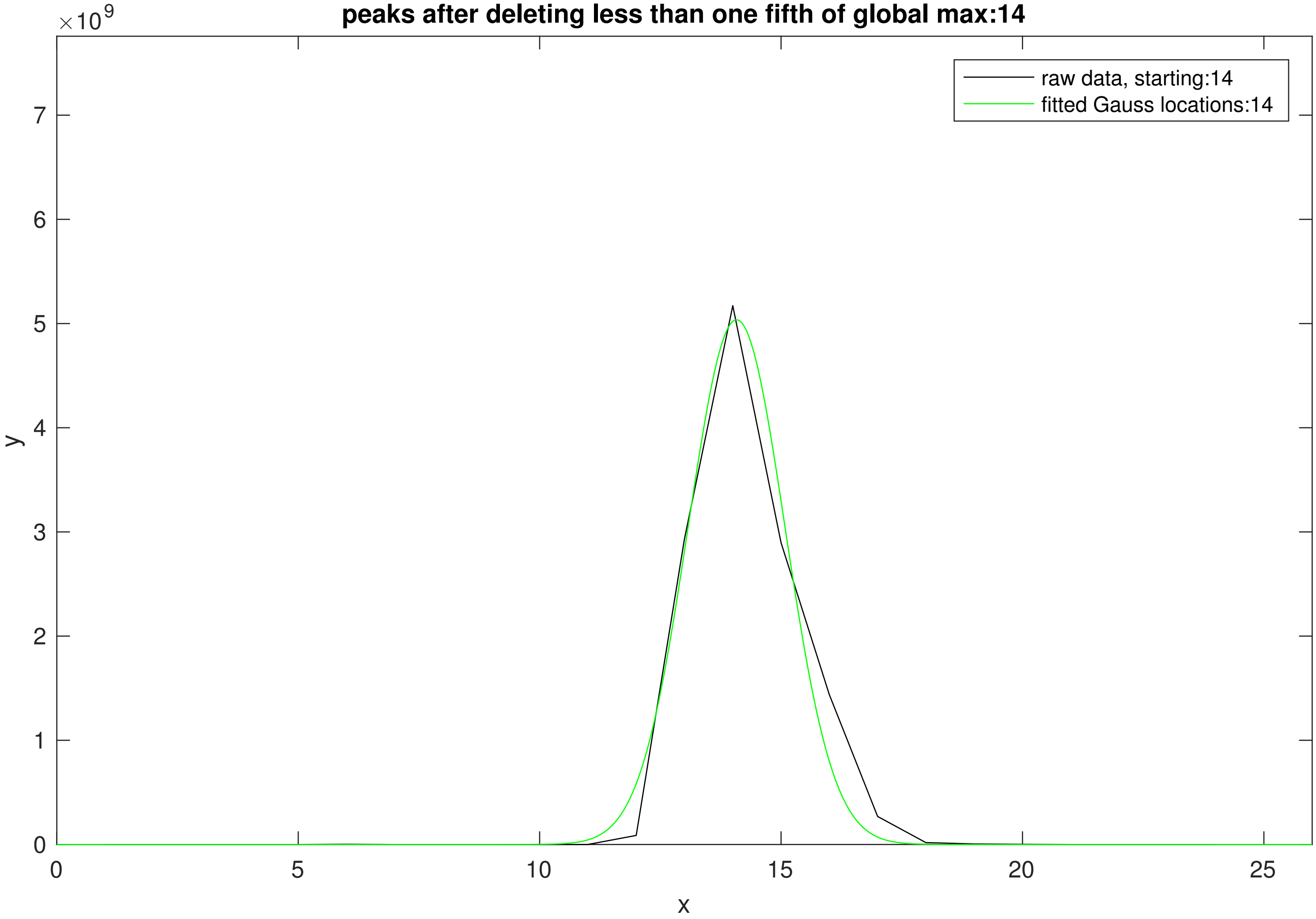

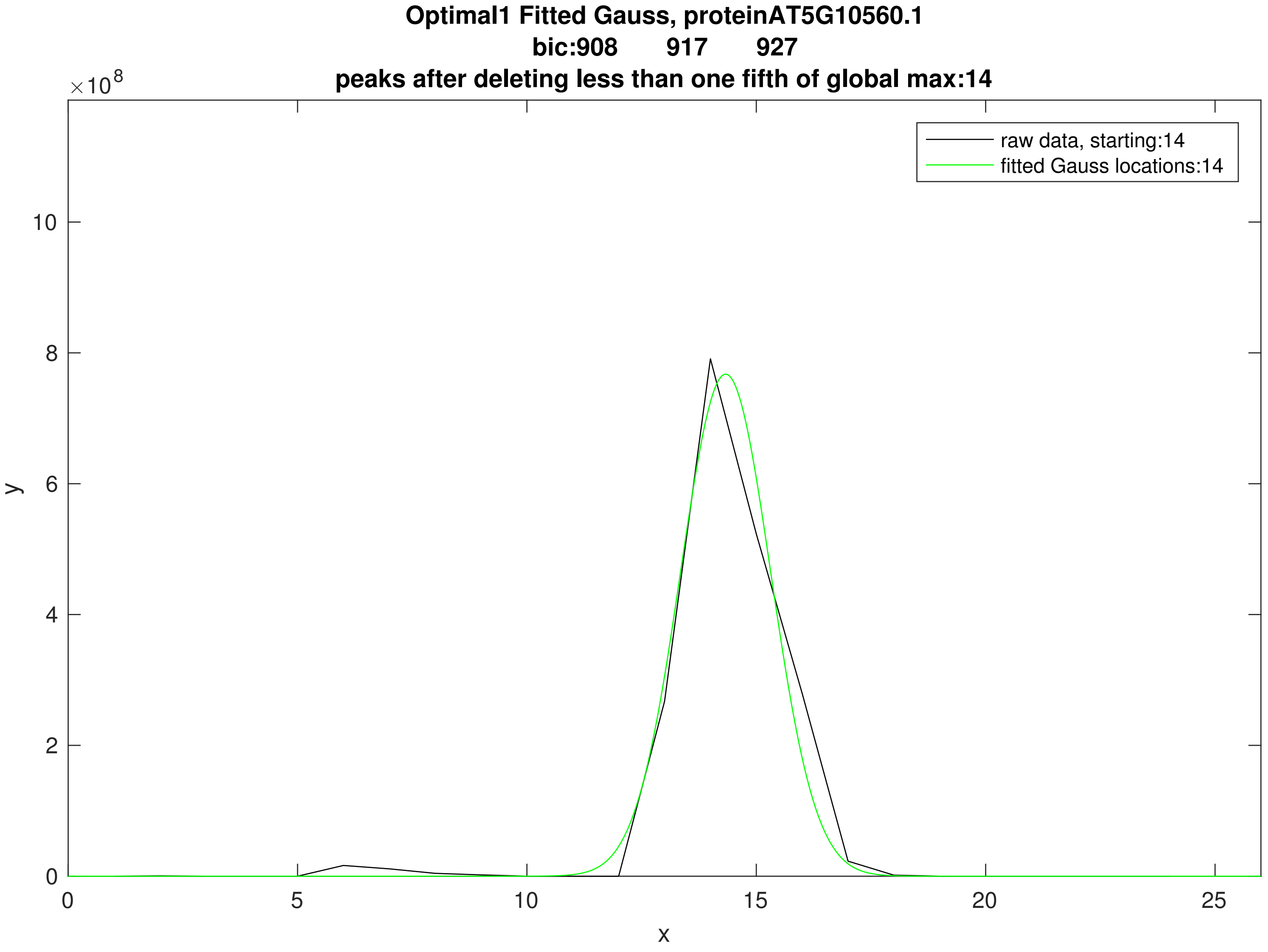

Optimal1 Fitted Gauss, proteinAT5G10560.1
bic:908 917 927
peaks after deleting less than one fifth of global max:14
×10^8
raw data, starting:14
fitted Gauss locations:14
y
x
0
2
4
6
8
10
0
5
10
15
20
25

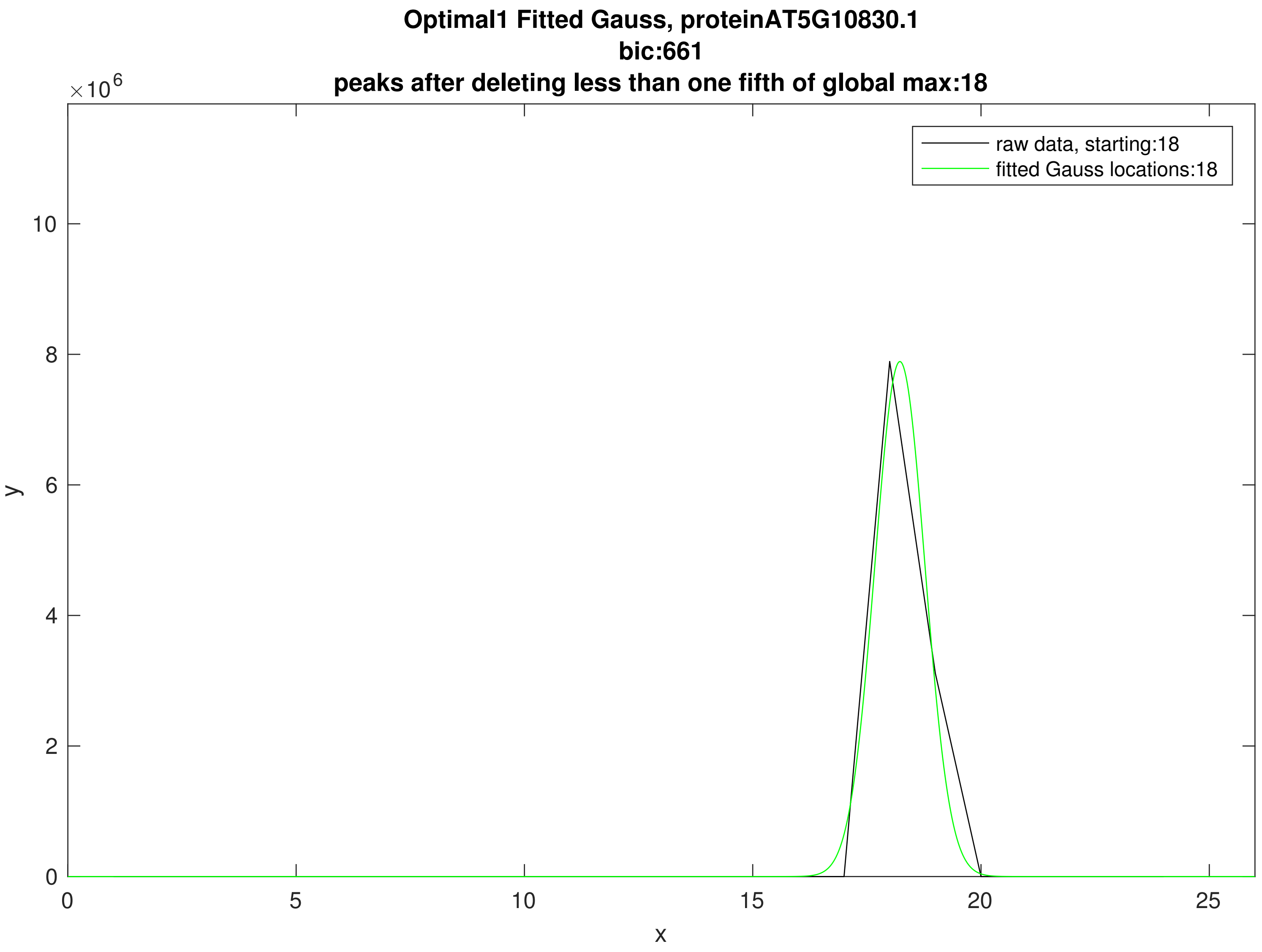

Optimal1 Fitted Gauss, proteinAT5G10830.1
bic:661
peaks after deleting less than one fifth of global max:18
×10^6
raw data, starting:18
fitted Gauss locations:18
y
x
0
2
4
6
8
10
0
5
10
15
20
25

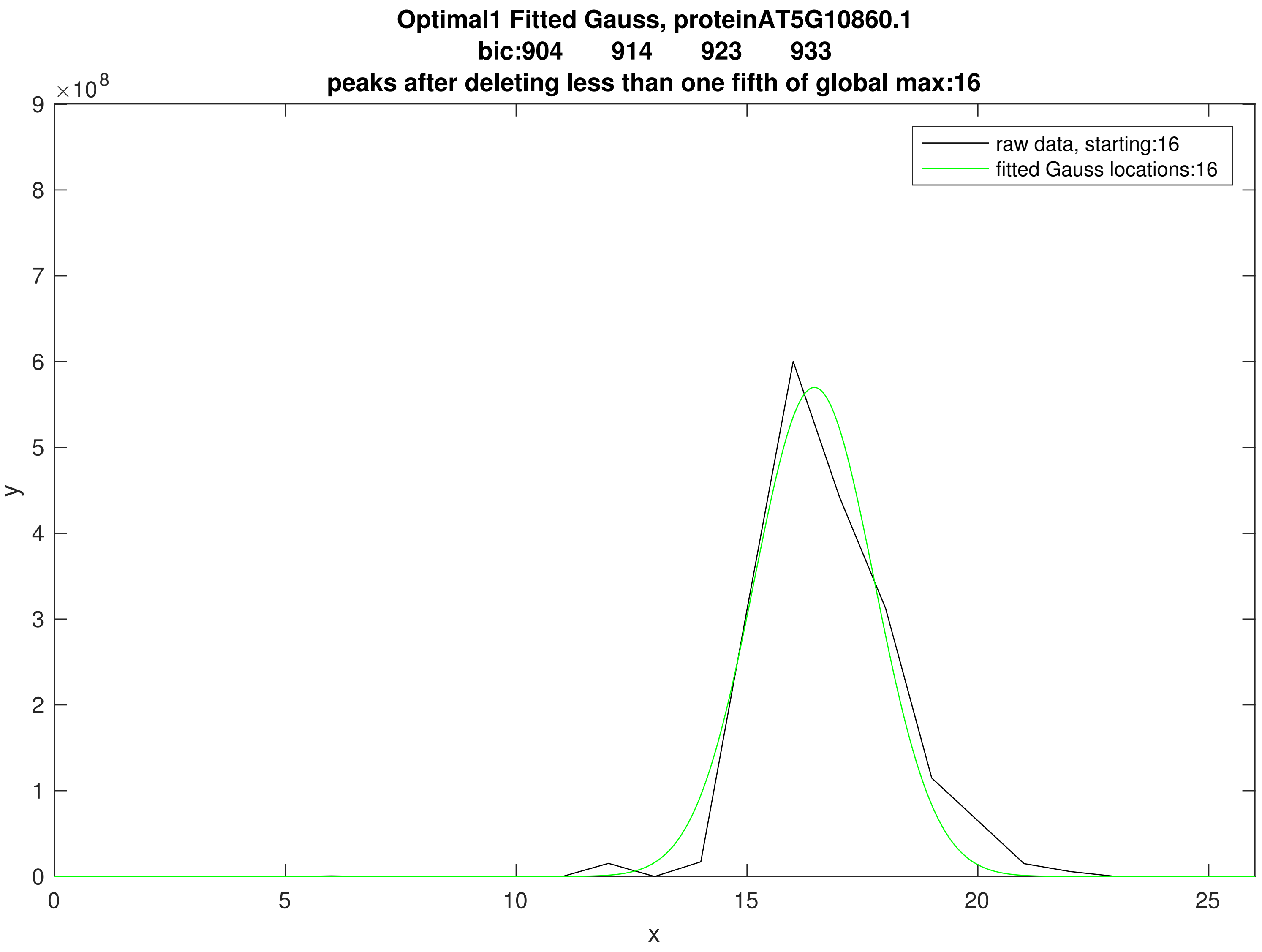
Optimal1 Fitted Gauss, proteinAT5G10860.1
bic:904 914 923 933
peaks after deleting less than one fifth of global max:16
×10^8
raw data, starting:16
fitted Gauss locations:16
y
x
0
1
2
3
4
5
6
7
8
9
0
5
10
15
20
25

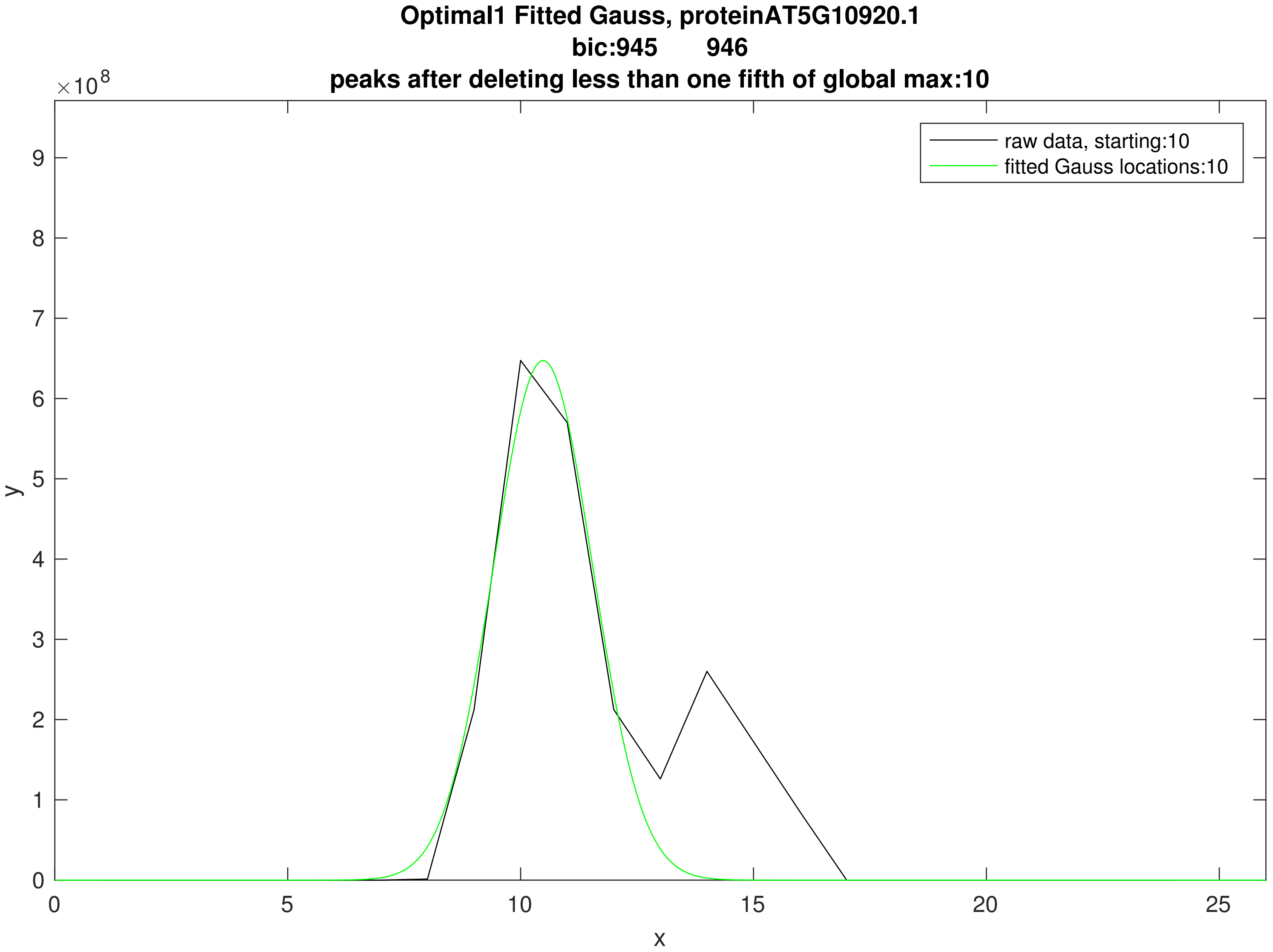
Optimal1 Fitted Gauss, proteinAT5G10920.1
bic:945 946
peaks after deleting less than one fifth of global max:10
×10^8
raw data, starting:10
fitted Gauss locations:10
y
x
0
1
2
3
4
5
6
7
8
9
0
5
10
15
20
25

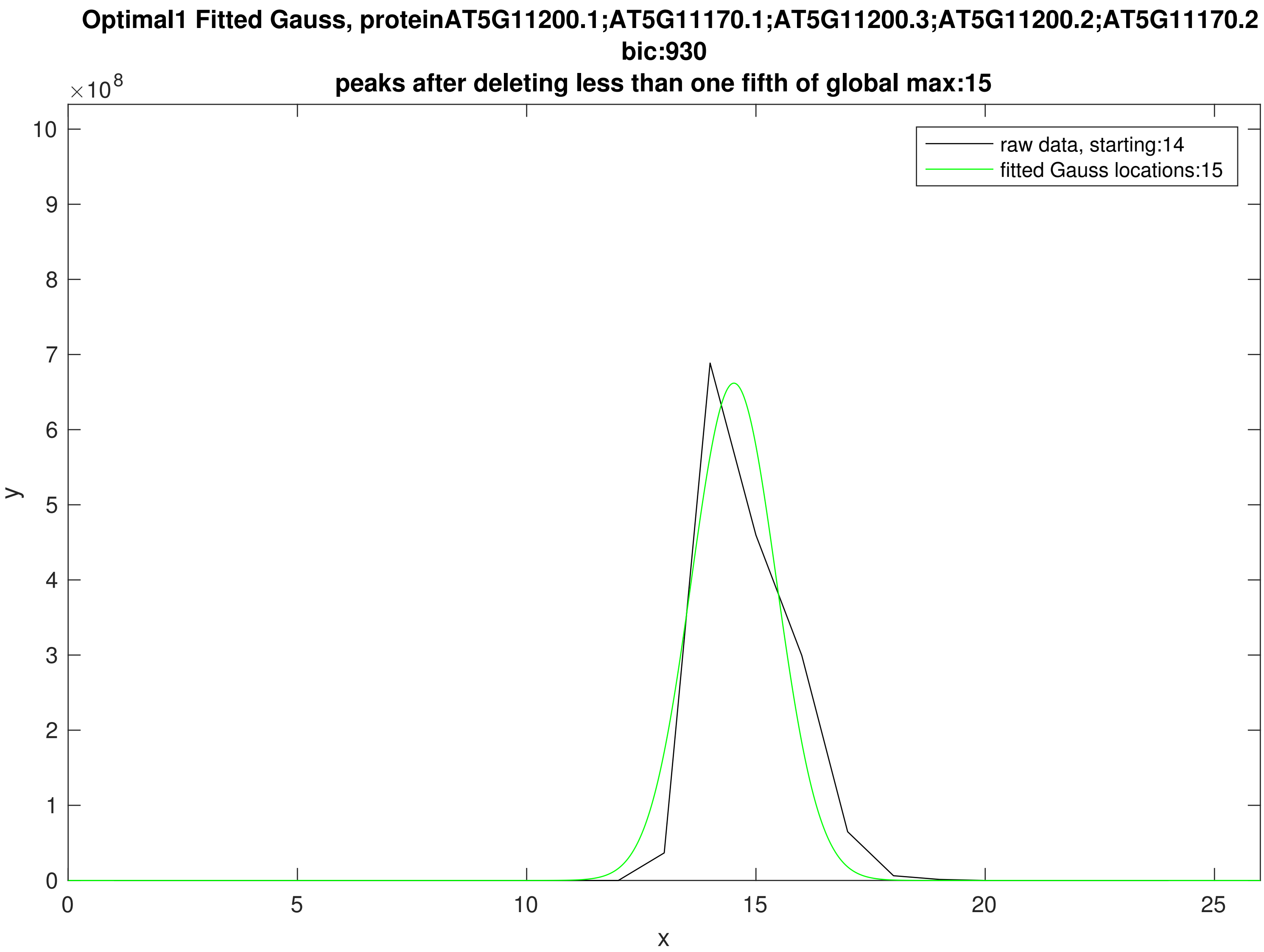

Optimal1 Fitted Gauss, proteinAT5G11200.1;AT5G11170.1;AT5G11200.3;AT5G11200.2;AT5G11170.2
bic:930
peaks after deleting less than one fifth of global max:15
×10^8
raw data, starting:14
fitted Gauss locations:15
y
x
0
1
2
3
4
5
6
7
8
9
10
0
5
10
15
20
25

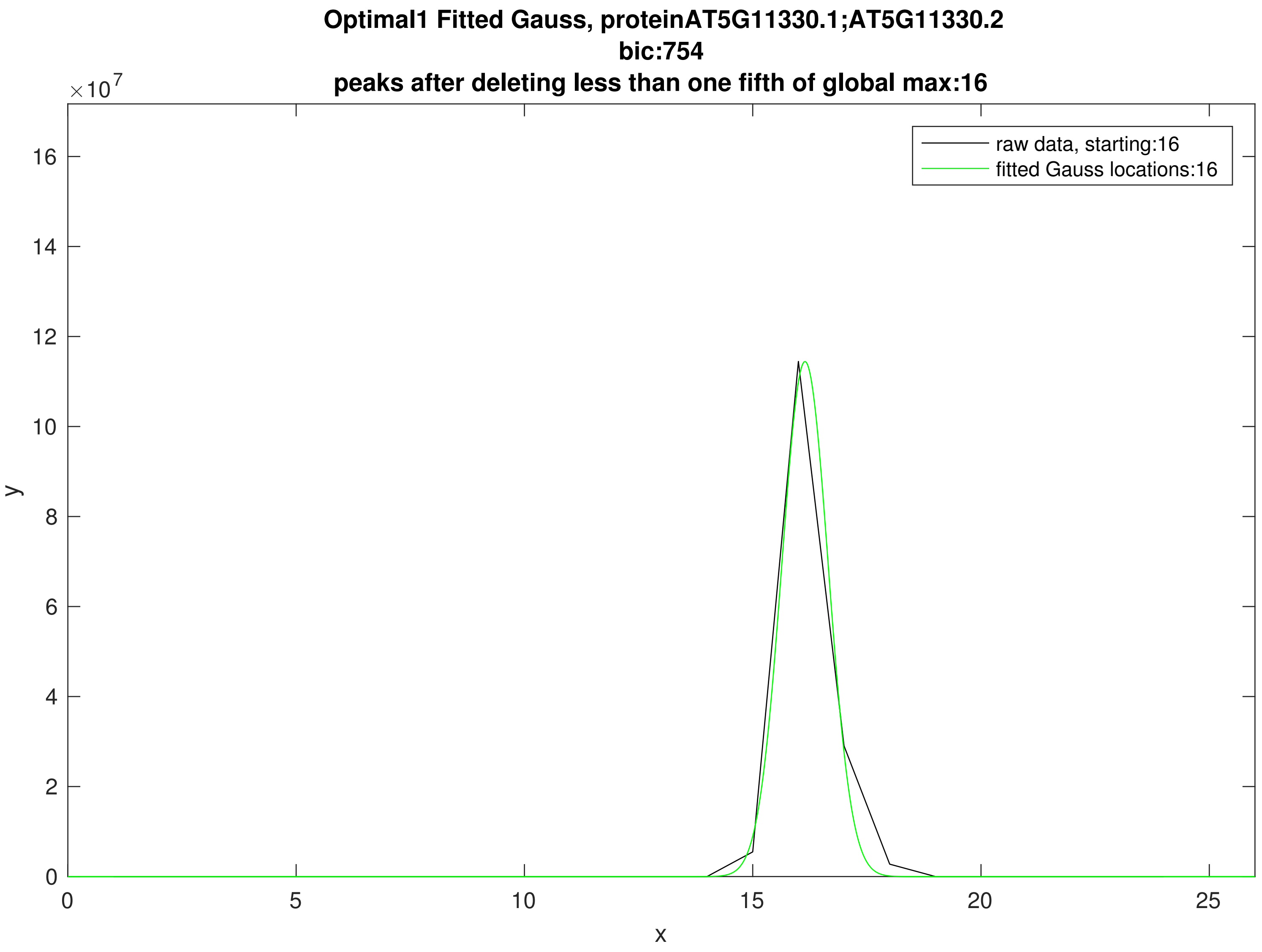
Optimal1 Fitted Gauss, proteinAT5G11330.1;AT5G11330.2
bic:754
peaks after deleting less than one fifth of global max:16
×10^7
16
14
12
10
8
6
4
2
0
y
0
5
10
15
20
25
x
raw data, starting:16
fitted Gauss locations:16

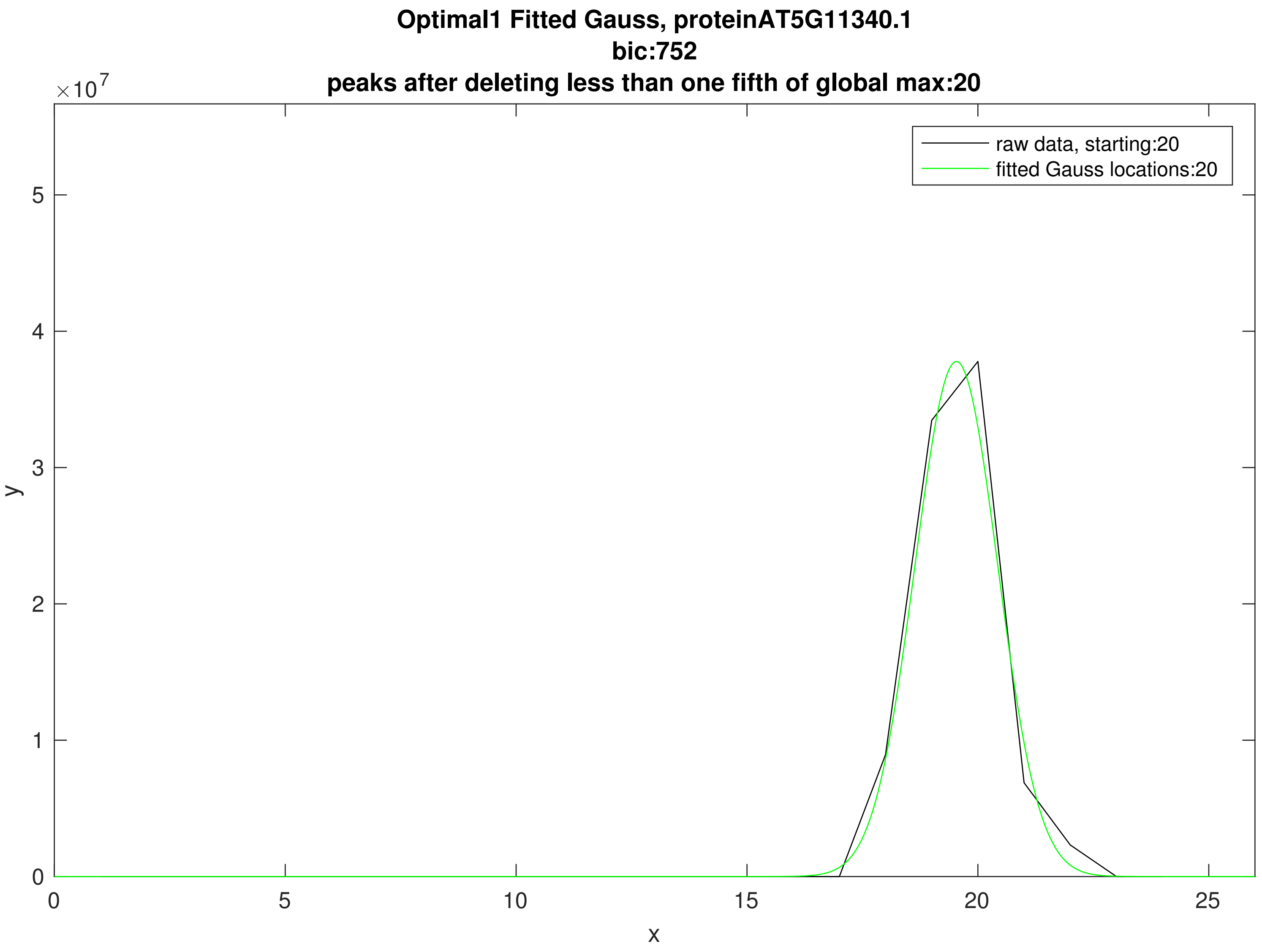

Optimal1 Fitted Gauss, proteinAT5G11340.1
bic:752
peaks after deleting less than one fifth of global max:20
×10^7
raw data, starting:20
fitted Gauss locations:20
y
x
0
1
2
3
4
5
0
5
10
15
20
25

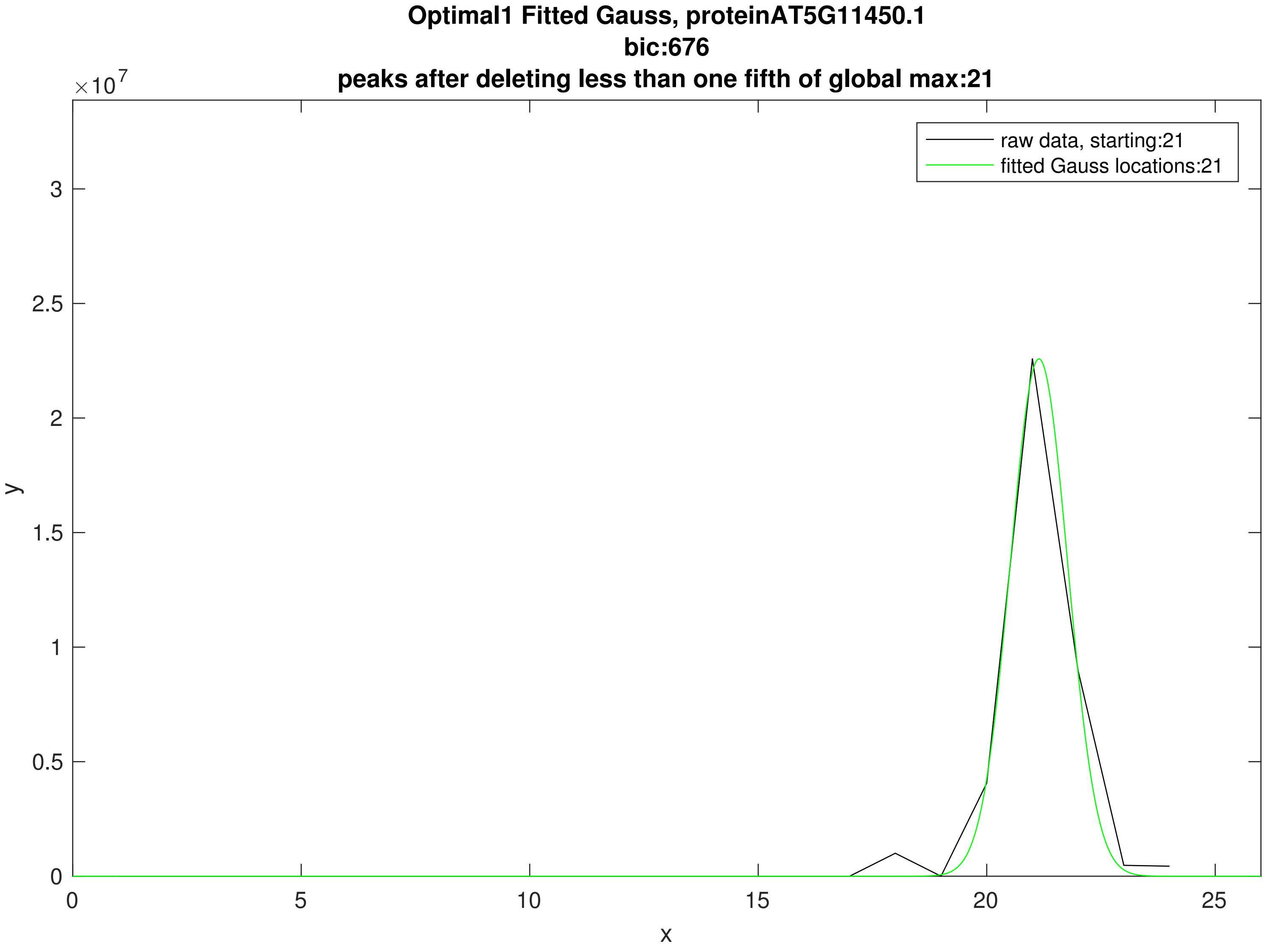
Optimal1 Fitted Gauss, proteinAT5G11450.1
bic:676
peaks after deleting less than one fifth of global max:21
×10^7
raw data, starting:21
fitted Gauss locations:21
y
x
0
0.5
1
1.5
2
2.5
3
5
10
15
20
25

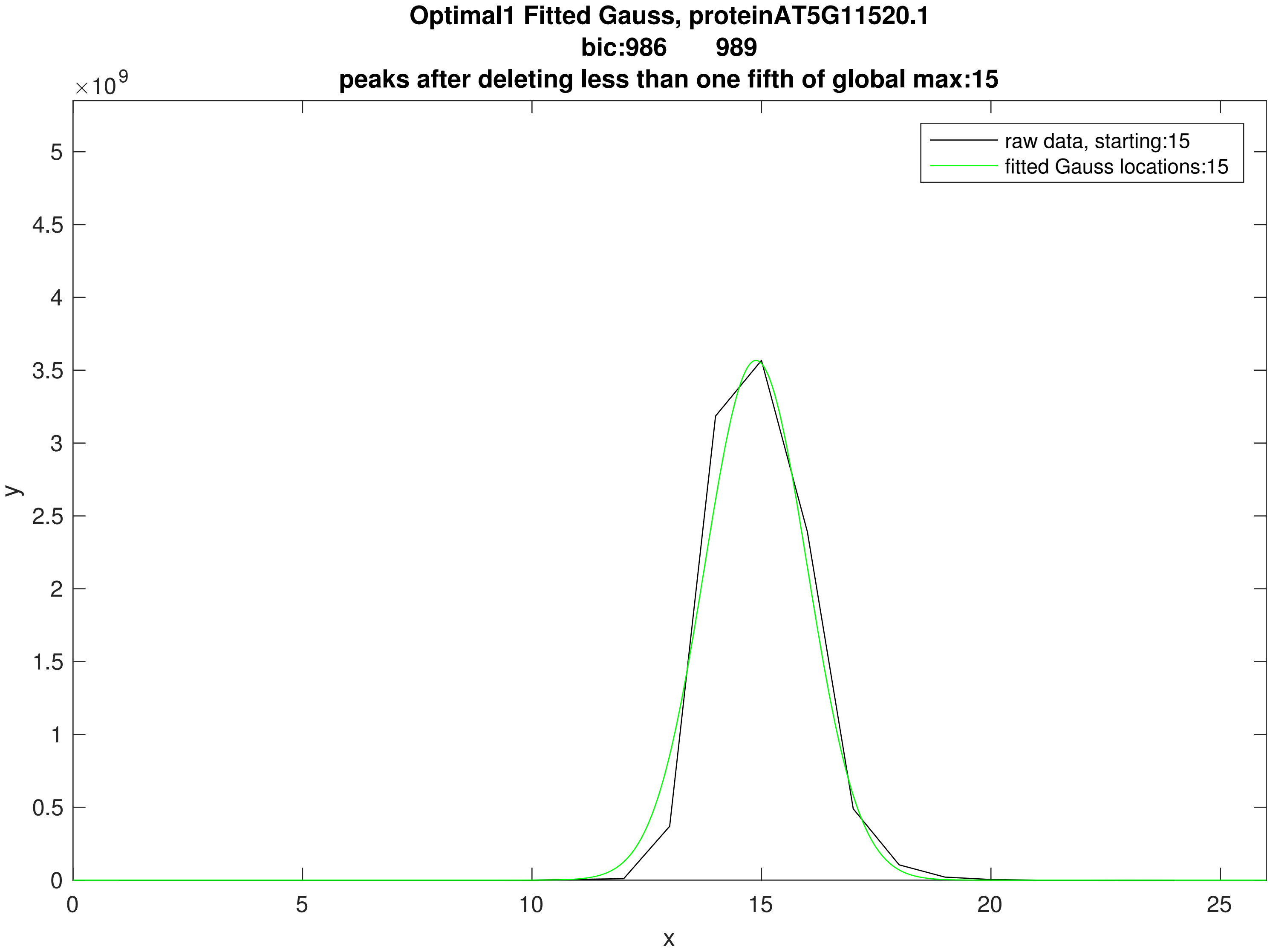
Optimal1 Fitted Gauss, proteinAT5G11520.1
bic:986 989
peaks after deleting less than one fifth of global max:15
×10^9
raw data, starting:15
fitted Gauss locations:15
y
x
0
0.5
1
1.5
2
2.5
3
3.5
4
4.5
5
0
5
10
15
20
25

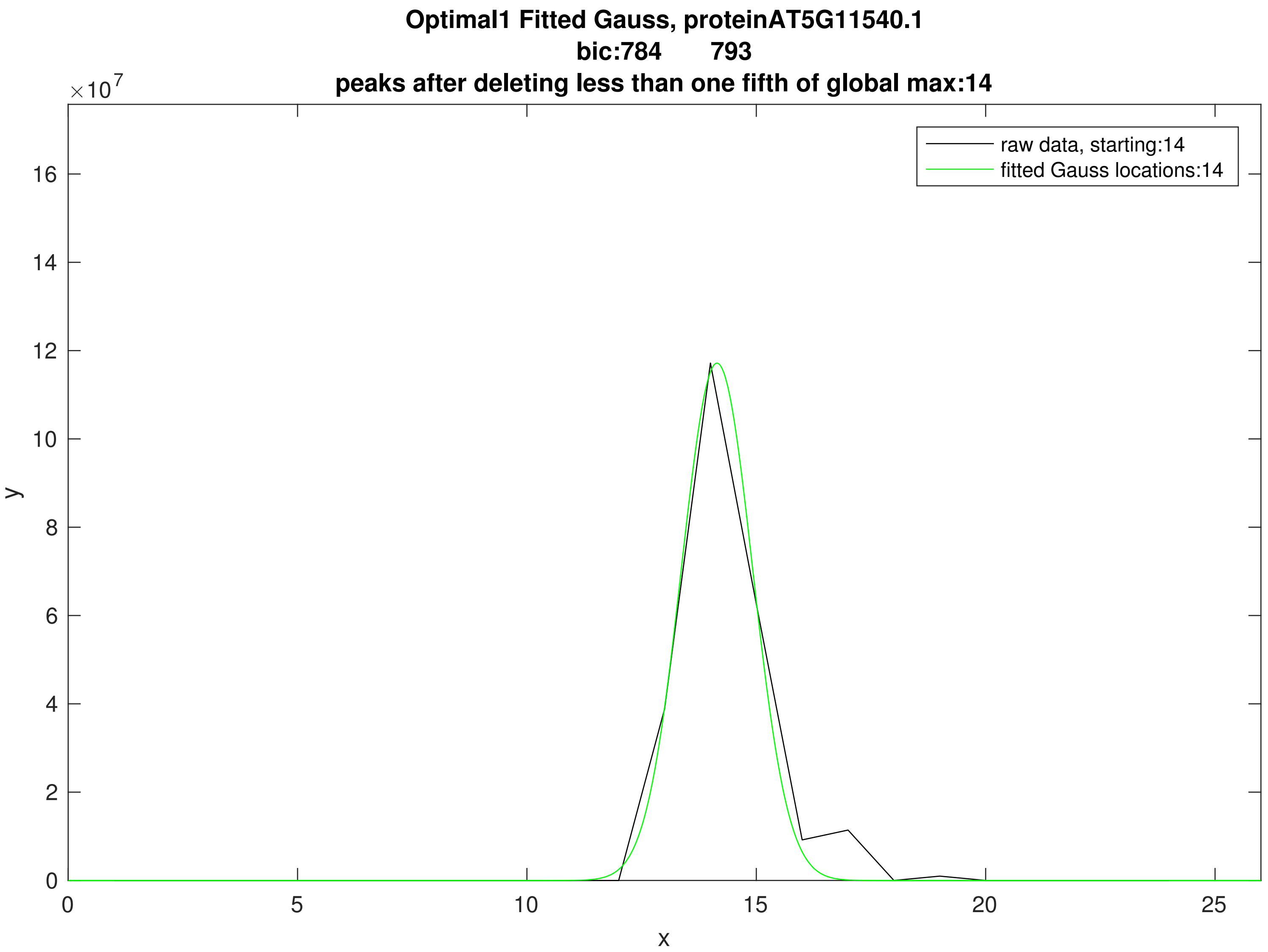
Optimal1 Fitted Gauss, proteinAT5G11540.1
bic:784 793
peaks after deleting less than one fifth of global max:14
×10^7
raw data, starting:14
fitted Gauss locations:14
16
14
12
10
8
6
4
2
0
y
0
5
10
15
20
25
x

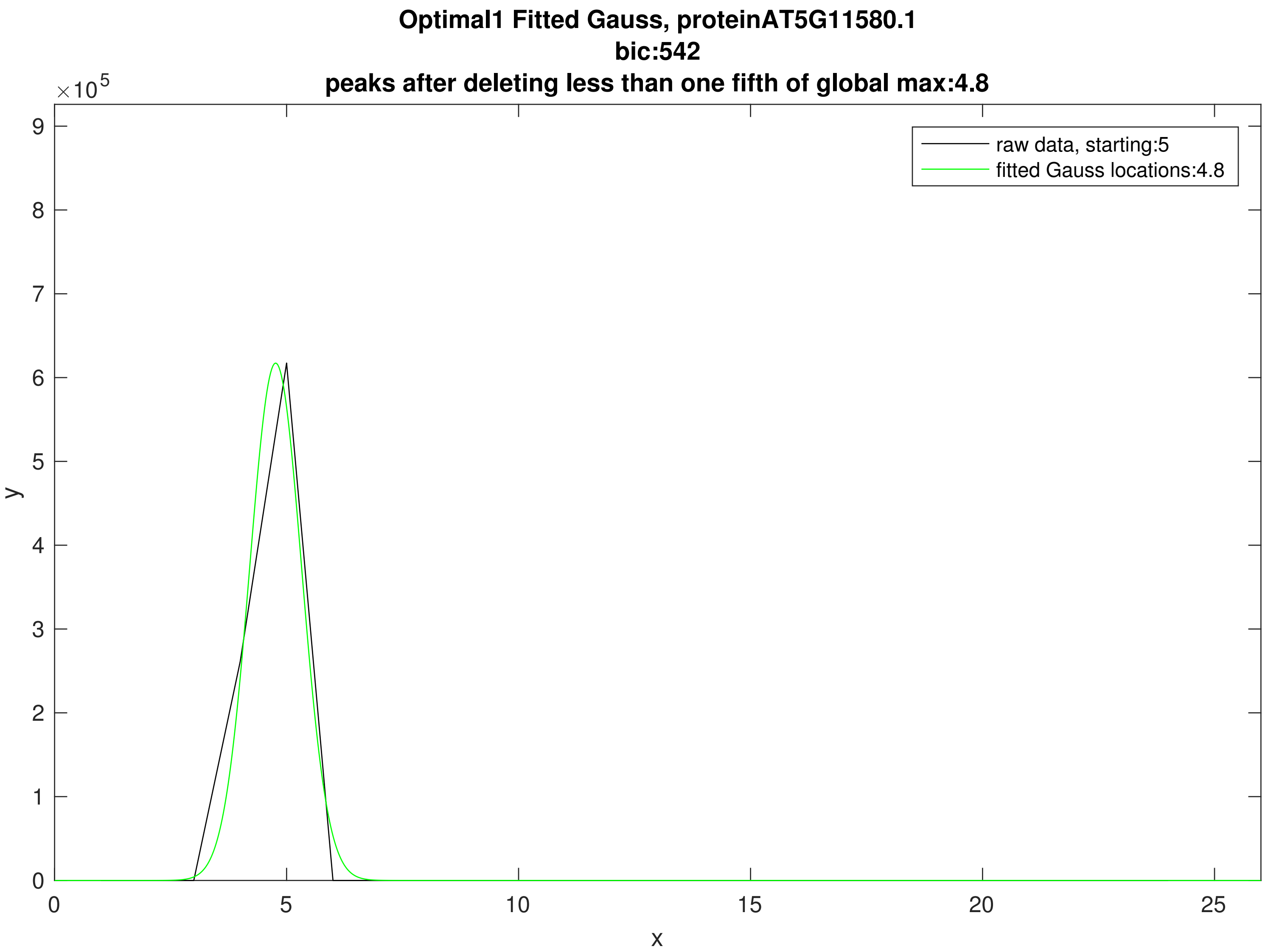

Optimal1 Fitted Gauss, proteinAT5G11580.1
bic:542
peaks after deleting less than one fifth of global max:4.8
×10^5
raw data, starting:5
fitted Gauss locations:4.8
y
x
0
1
2
3
4
5
6
7
8
9
10
15
20
25

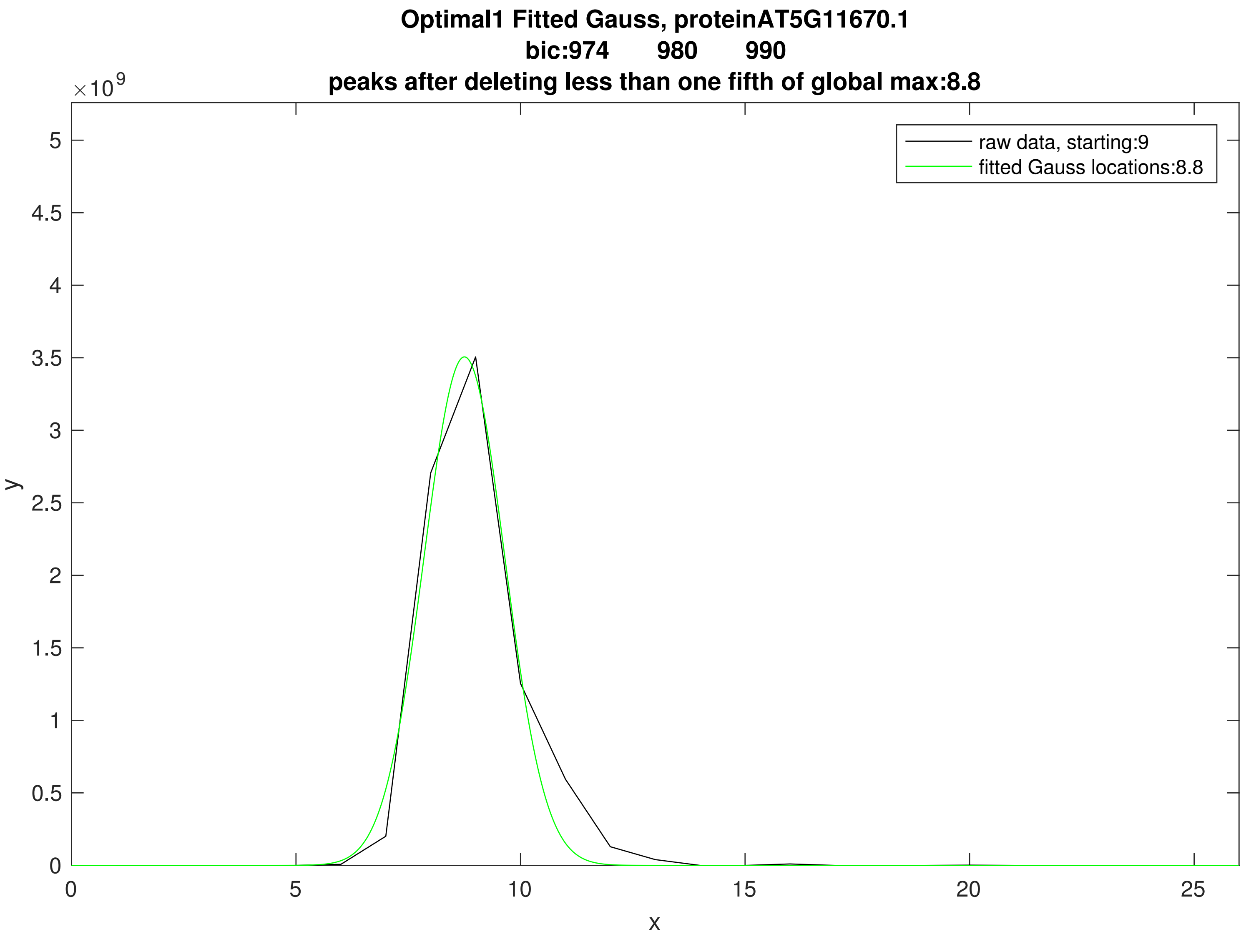

Optimal1 Fitted Gauss, proteinAT5G11670.1
bic:974 980 990
peaks after deleting less than one fifth of global max:8.8
×10^9
raw data, starting:9
fitted Gauss locations:8.8
y
x
0
0.5
1
1.5
2
2.5
3
3.5
4
4.5
5
0
5
10
15
20
25

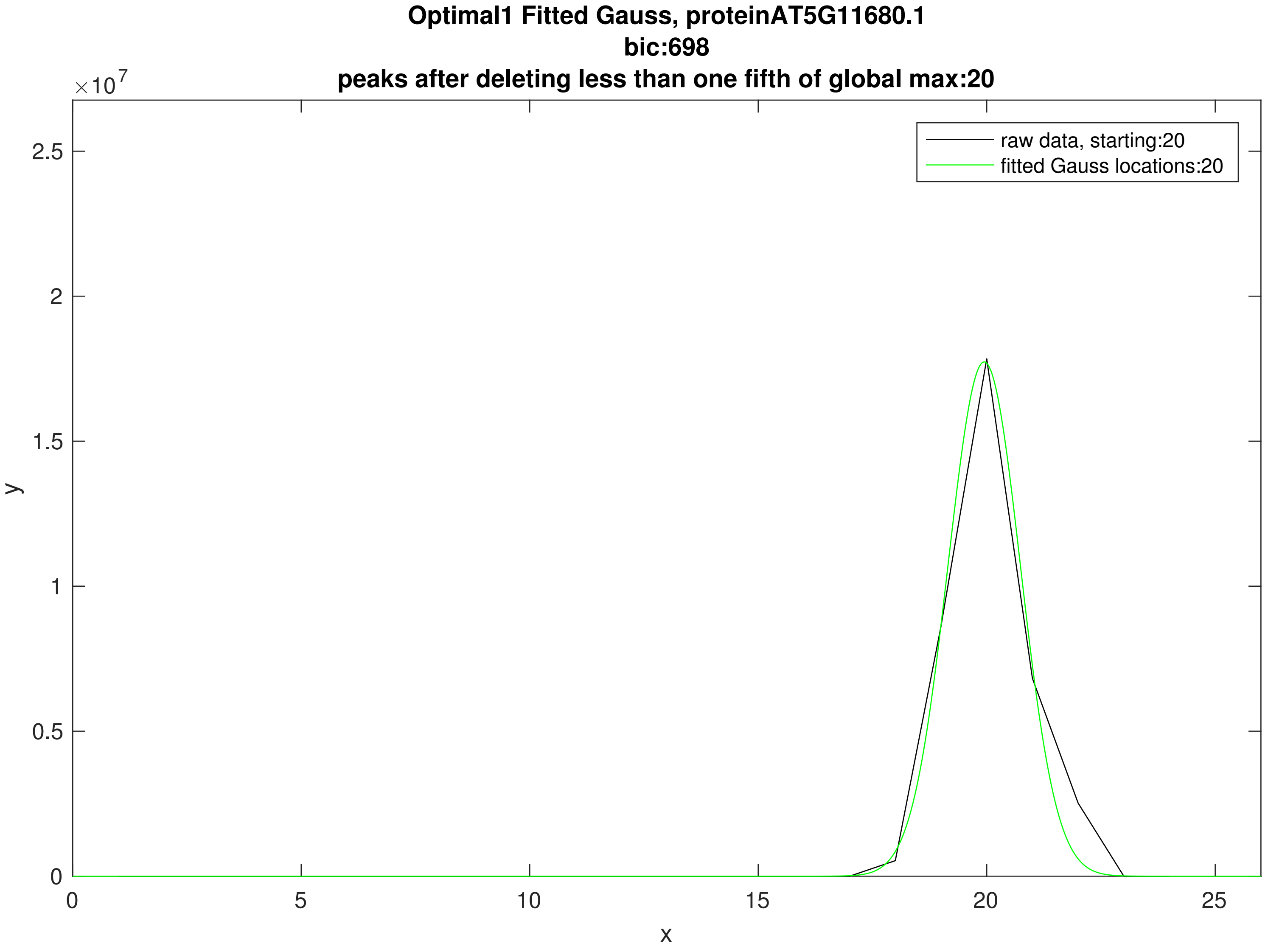

Optimal1 Fitted Gauss, proteinAT5G11680.1
bic:698
peaks after deleting less than one fifth of global max:20
×10^7
2.5
2
1.5
1
0.5
0
y
0
5
10
15
20
25
x
raw data, starting:20
fitted Gauss locations:20

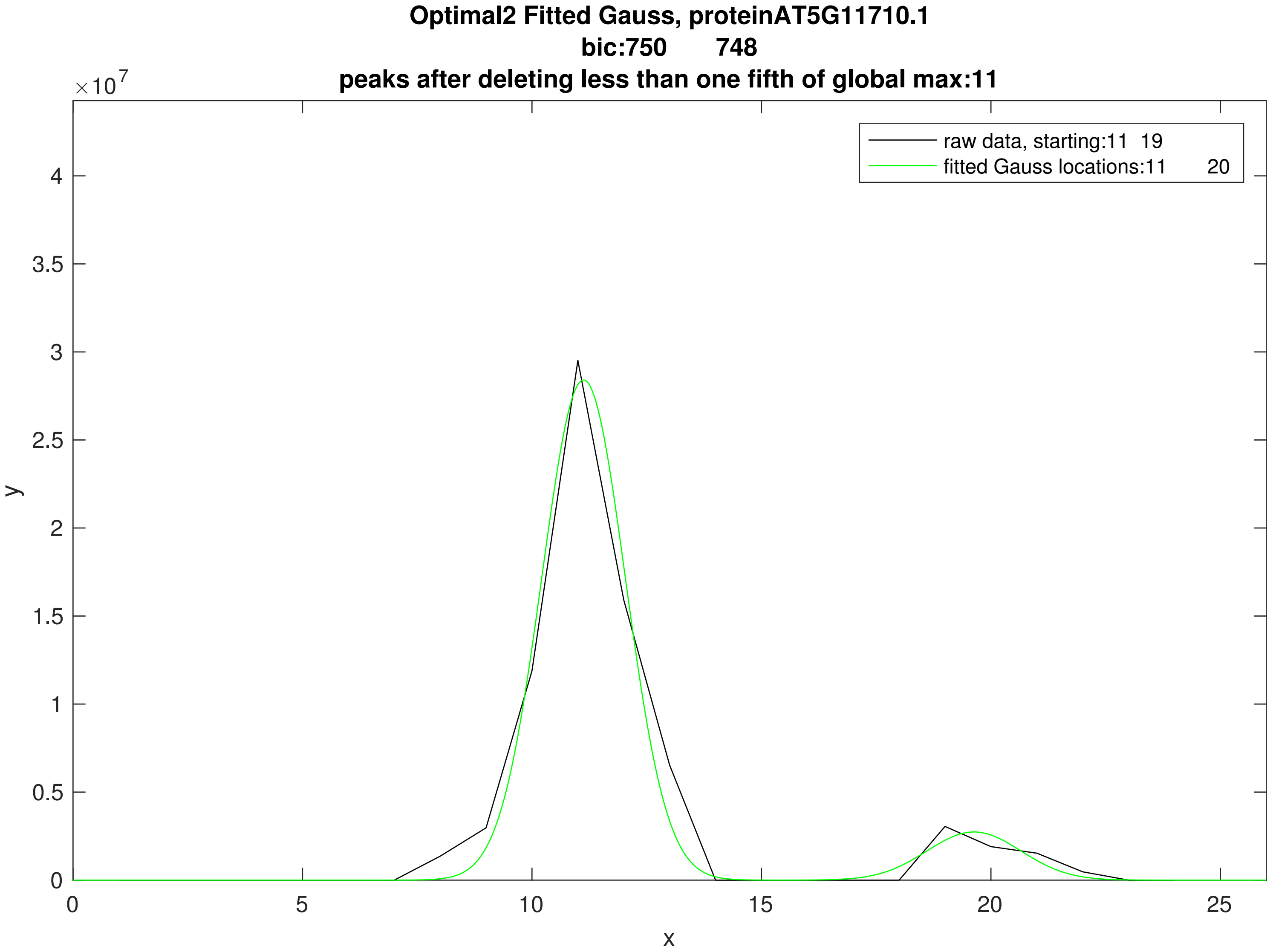

Optimal2 Fitted Gauss, proteinAT5G11710.1
bic:750 748
peaks after deleting less than one fifth of global max:11
×10^7
raw data, starting:11 19
fitted Gauss locations:11 20
4
3.5
3
2.5
2
1.5
1
0.5
0
y
0
5
10
15
20
25
x

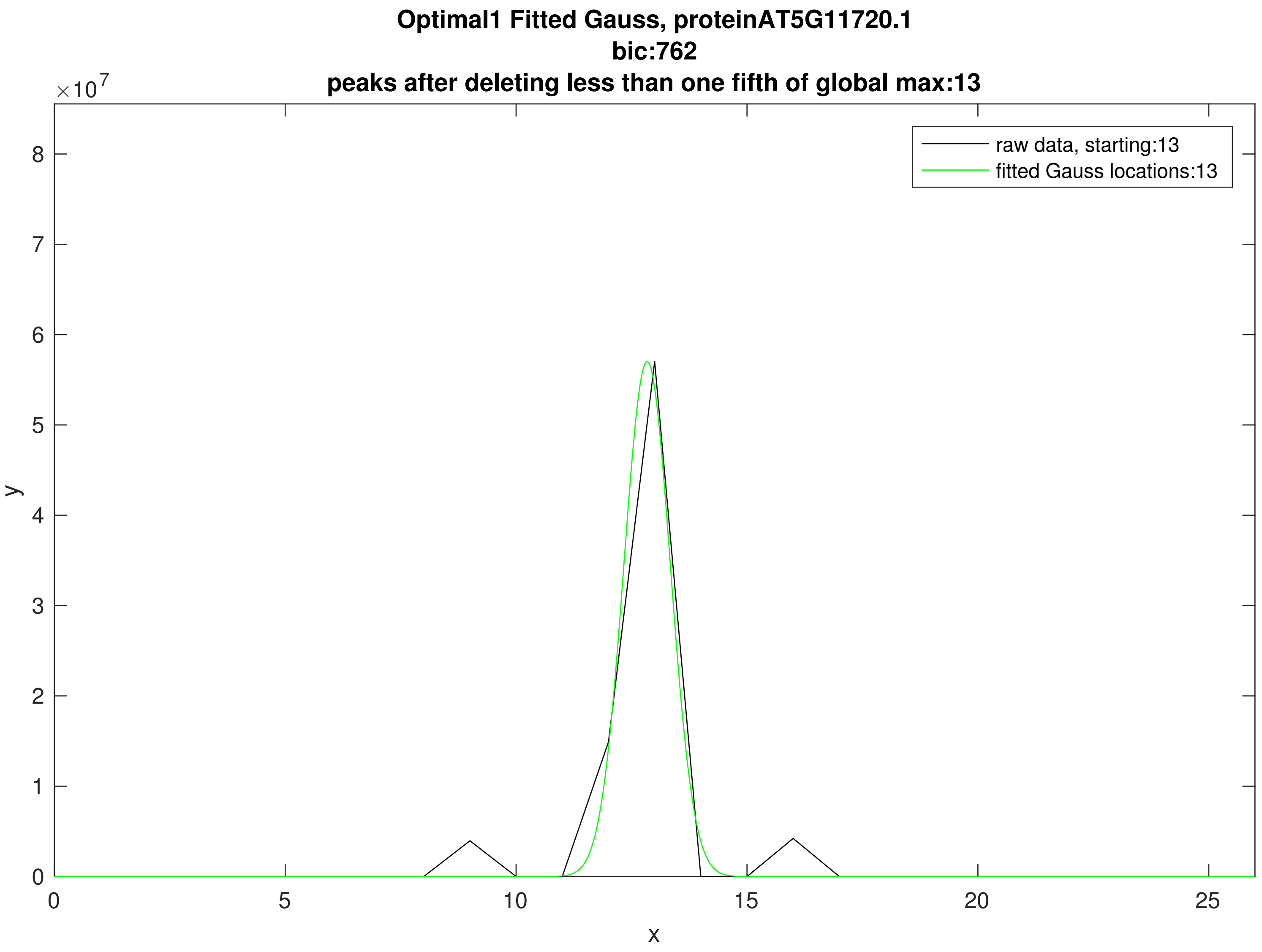

Optimal1 Fitted Gauss, proteinAT5G11720.1
bic:762
peaks after deleting less than one fifth of global max:13
×10^7
raw data, starting:13
fitted Gauss locations:13
y
x
0
1
2
3
4
5
6
7
8
0
5
10
15
20
25

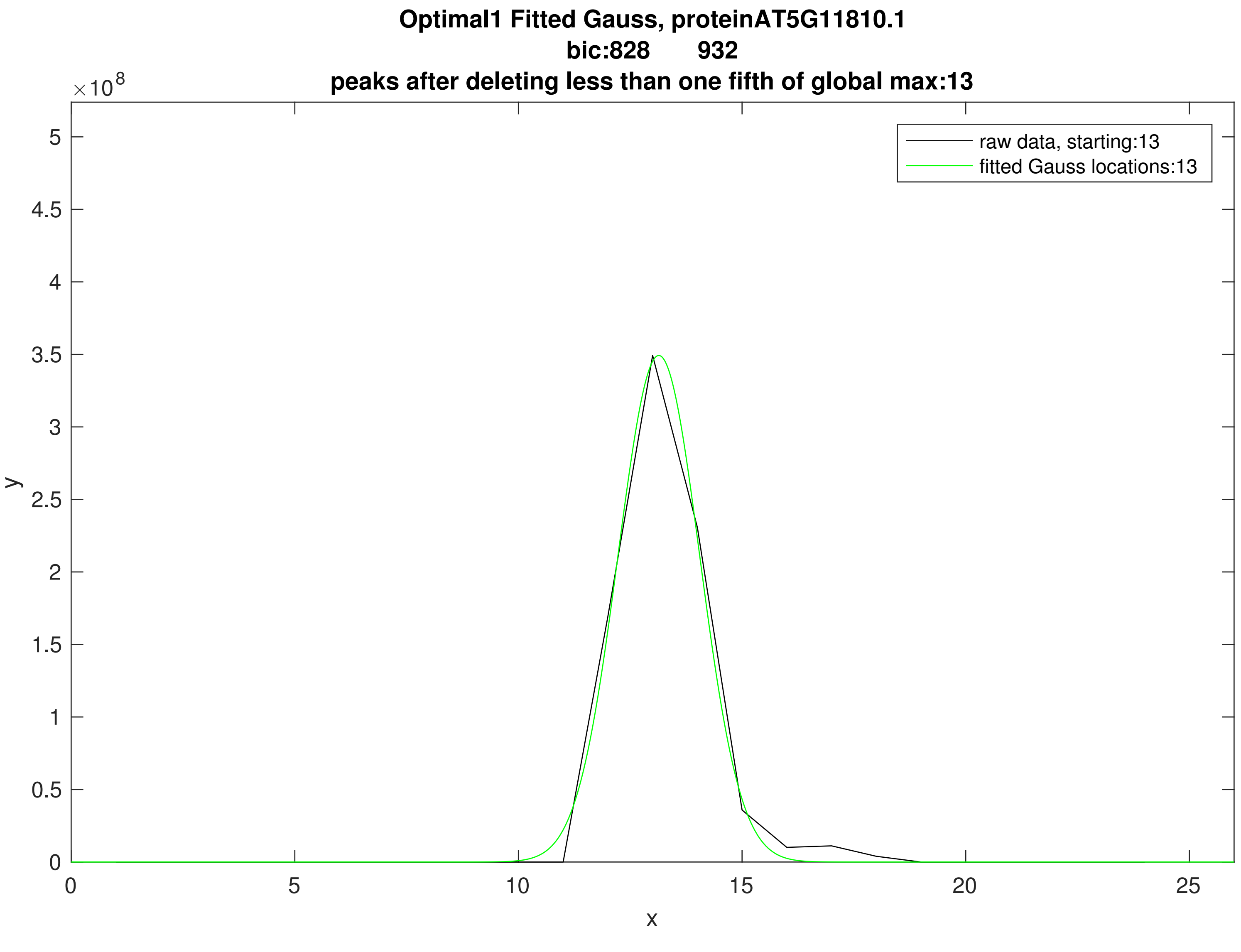
Optimal1 Fitted Gauss, proteinAT5G11810.1
bic:828 932
peaks after deleting less than one fifth of global max:13
$\times 10^8$
raw data, starting:13
fitted Gauss locations:13
y
x
0
0.5
1
1.5
2
2.5
3
3.5
4
4.5
5
0
5
10
15
20
25

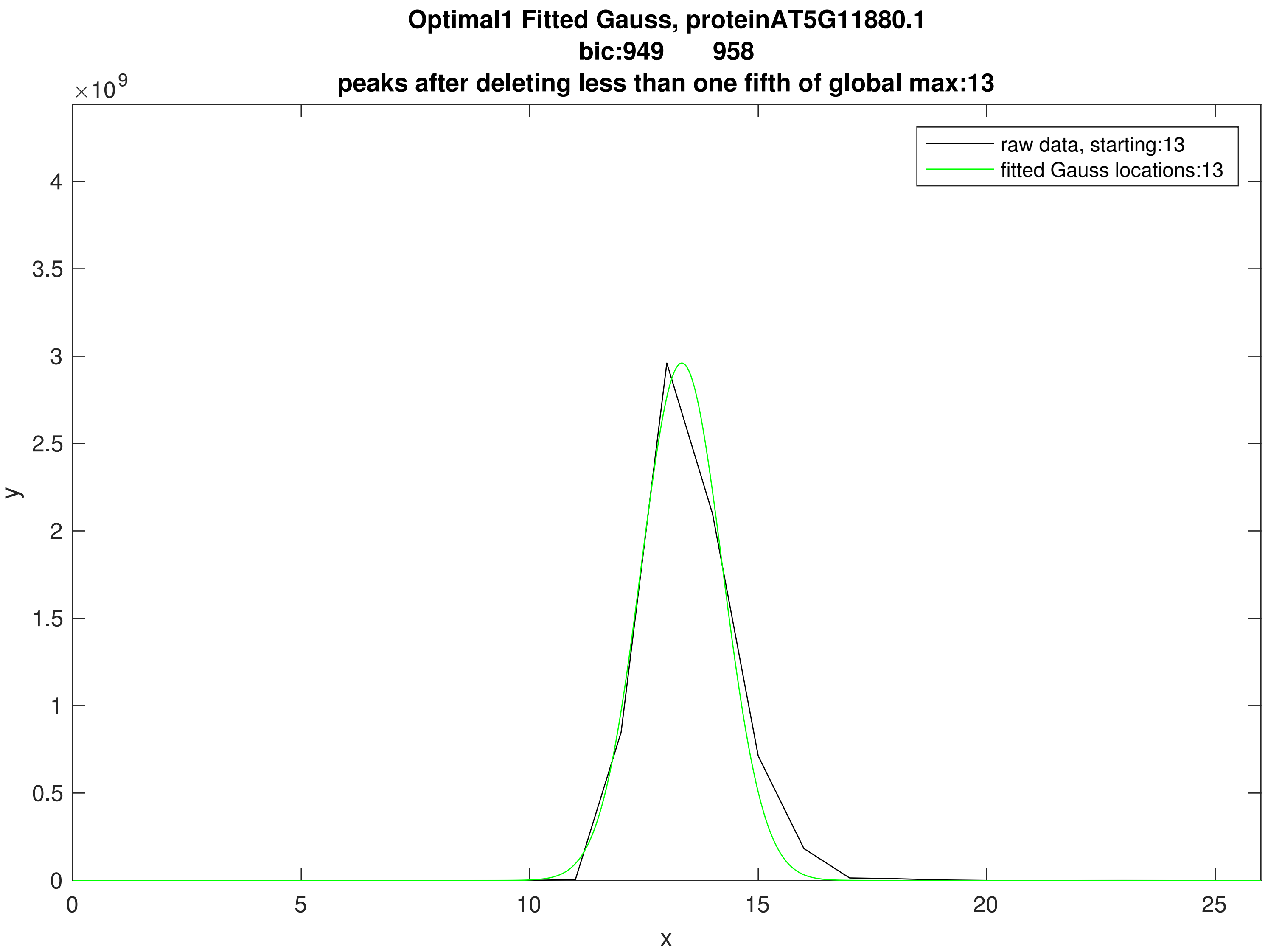
Optimal1 Fitted Gauss, proteinAT5G11880.1
bic:949 958
peaks after deleting less than one fifth of global max:13
×10^9
raw data, starting:13
fitted Gauss locations:13
y
x
0
0.5
1
1.5
2
2.5
3
3.5
4
0
5
10
15
20
25

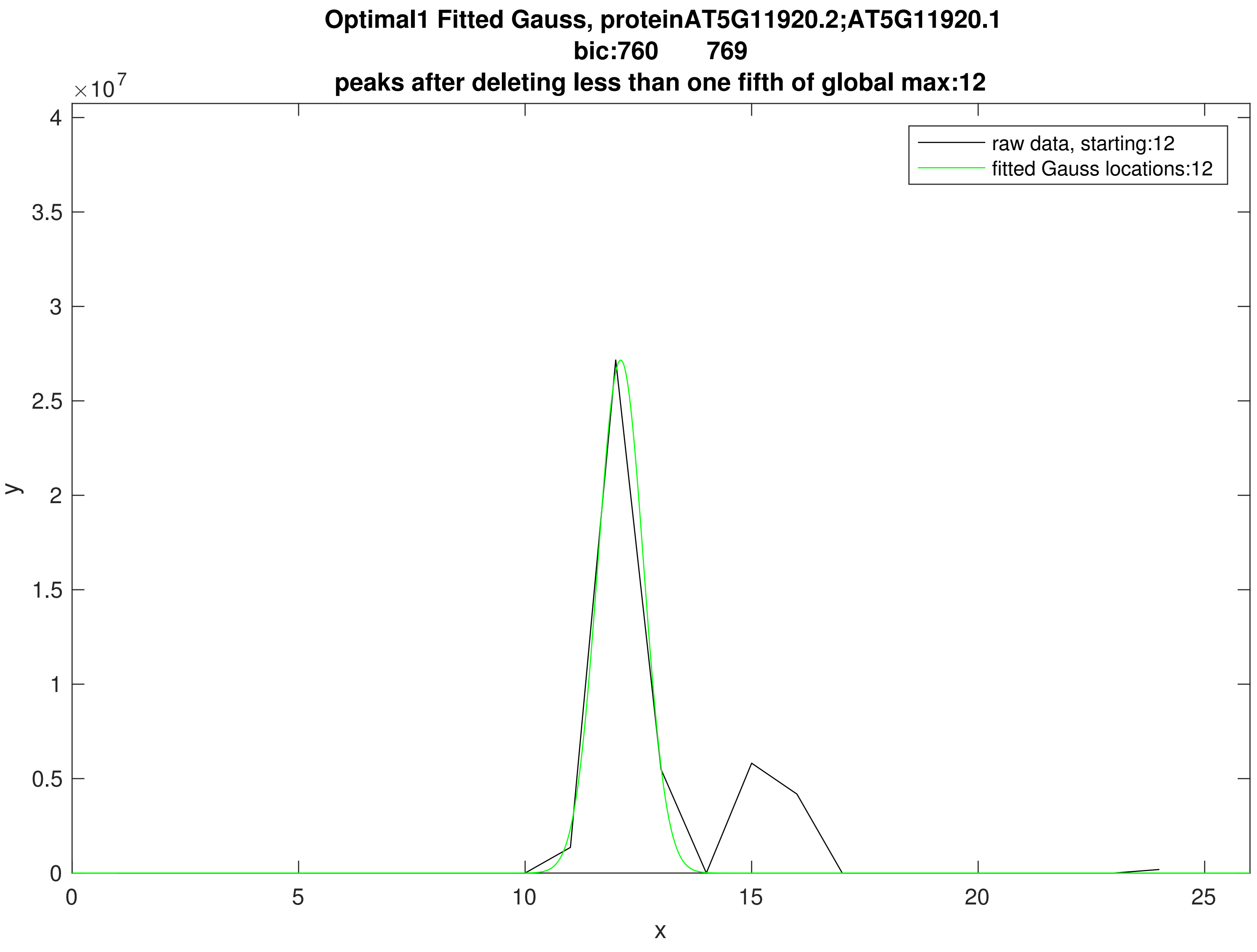
Optimal1 Fitted Gauss, proteinAT5G11920.2;AT5G11920.1
bic:760 769
peaks after deleting less than one fifth of global max:12
×10^7
4
3.5
3
2.5
2
1.5
1
0.5
0
y
0
5
10
15
20
25
x
raw data, starting:12
fitted Gauss locations:12

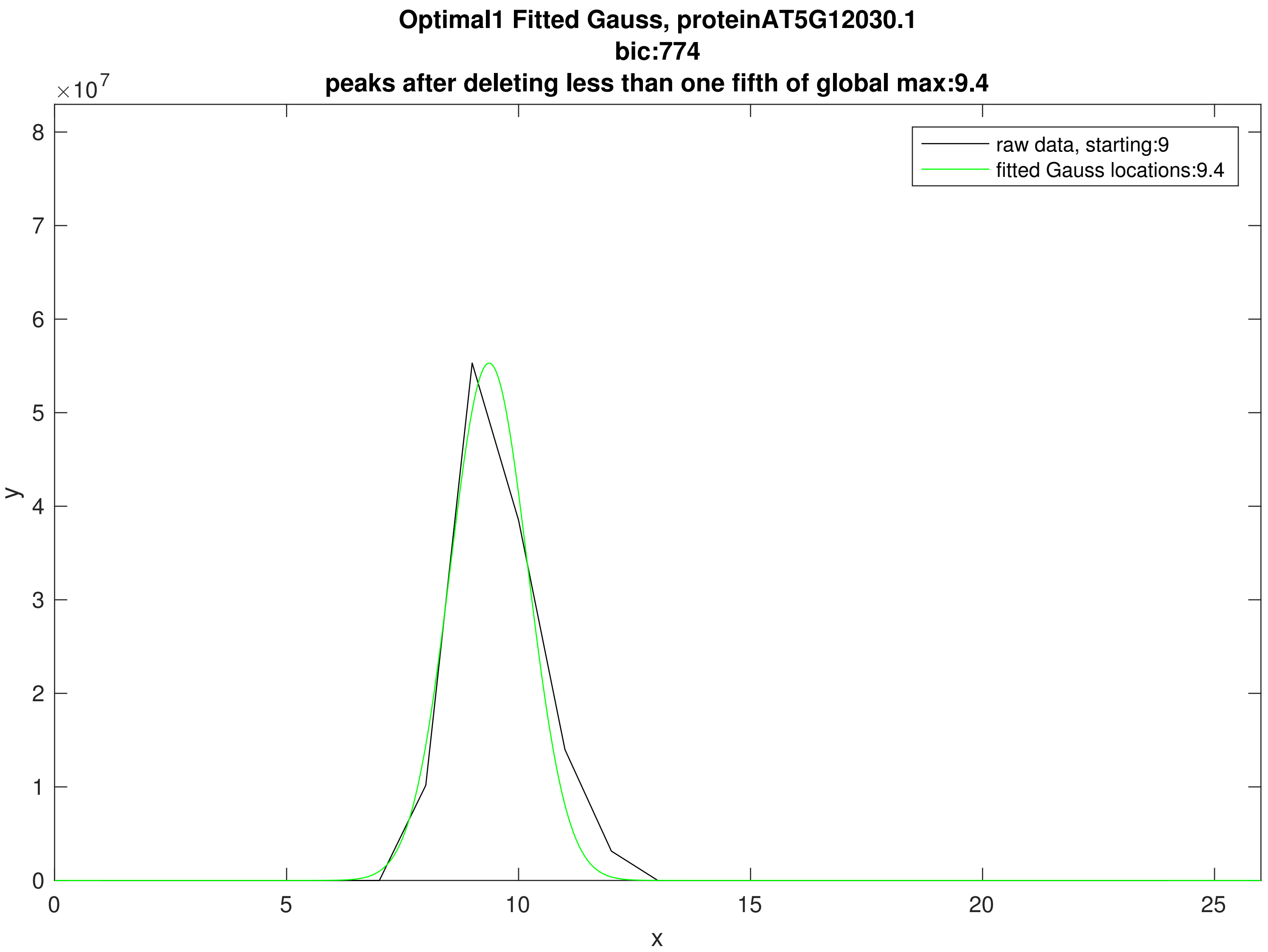
Optimal1 Fitted Gauss, proteinAT5G12030.1
bic:774
peaks after deleting less than one fifth of global max:9.4
×10^7
8
7
6
5
4
3
2
1
0
y
0
5
10
15
20
25
x
raw data, starting:9
fitted Gauss locations:9.4

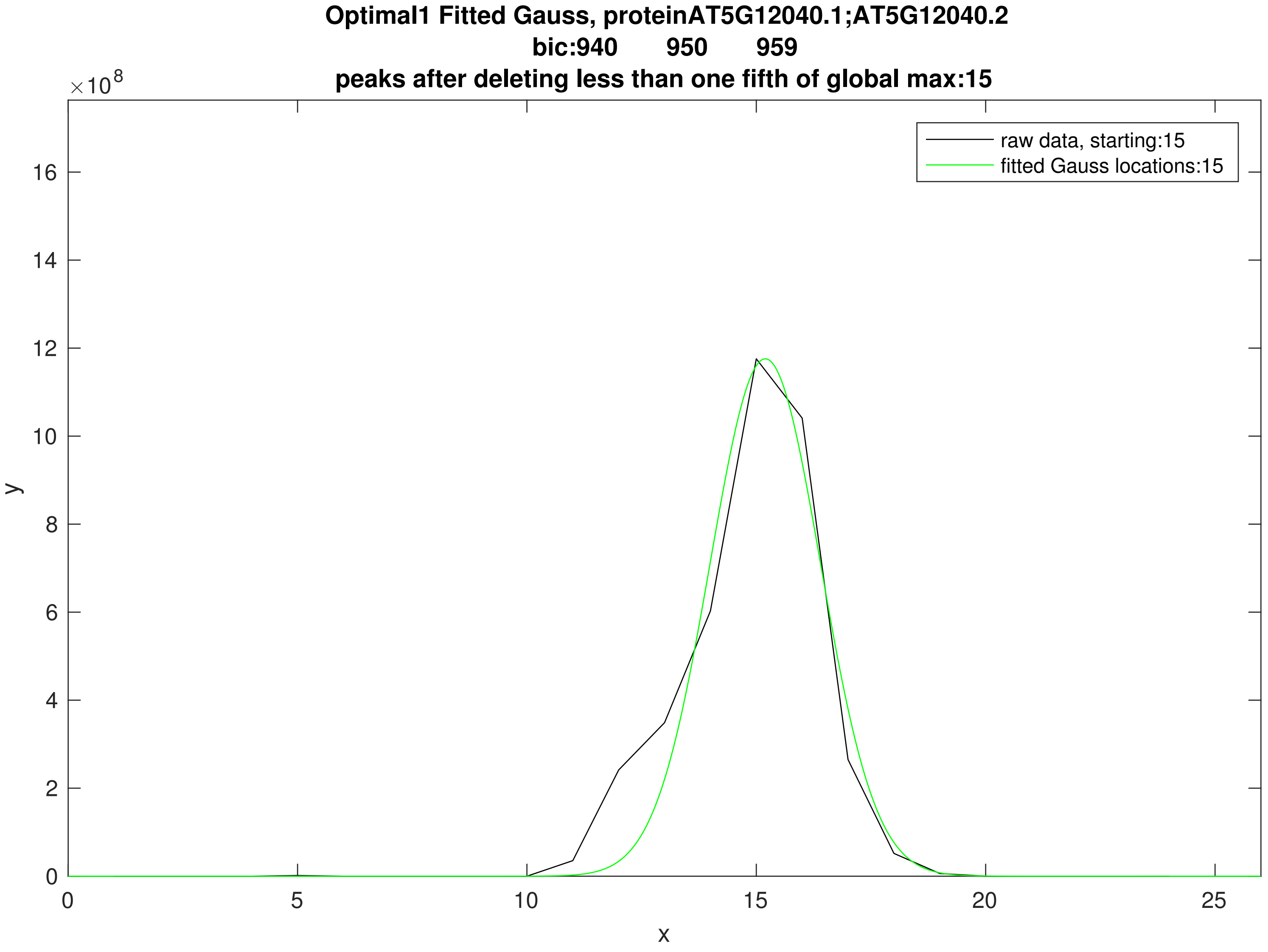

Optimal1 Fitted Gauss, proteinAT5G12040.1;AT5G12040.2
bic:940 950 959
peaks after deleting less than one fifth of global max:15
×10^8
raw data, starting:15
fitted Gauss locations:15
y
x
0
2
4
6
8
10
12
14
16
0
5
10
15
20
25

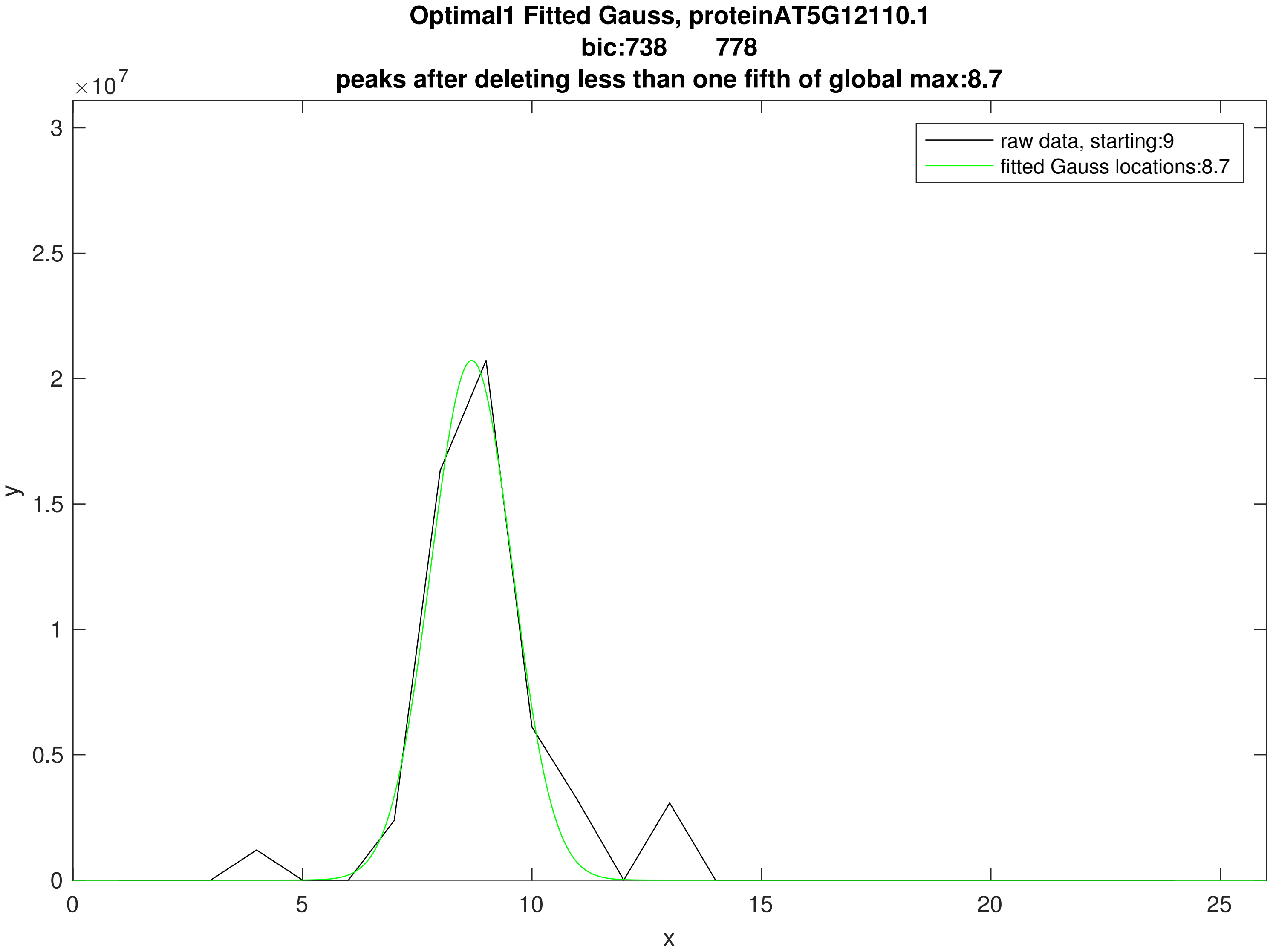

Optimal1 Fitted Gauss, proteinAT5G12110.1
bic:738 778
peaks after deleting less than one fifth of global max:8.7
×10^7
raw data, starting:9
fitted Gauss locations:8.7
3
2.5
2
1.5
1
0.5
0
y
0
5
10
15
20
25
x

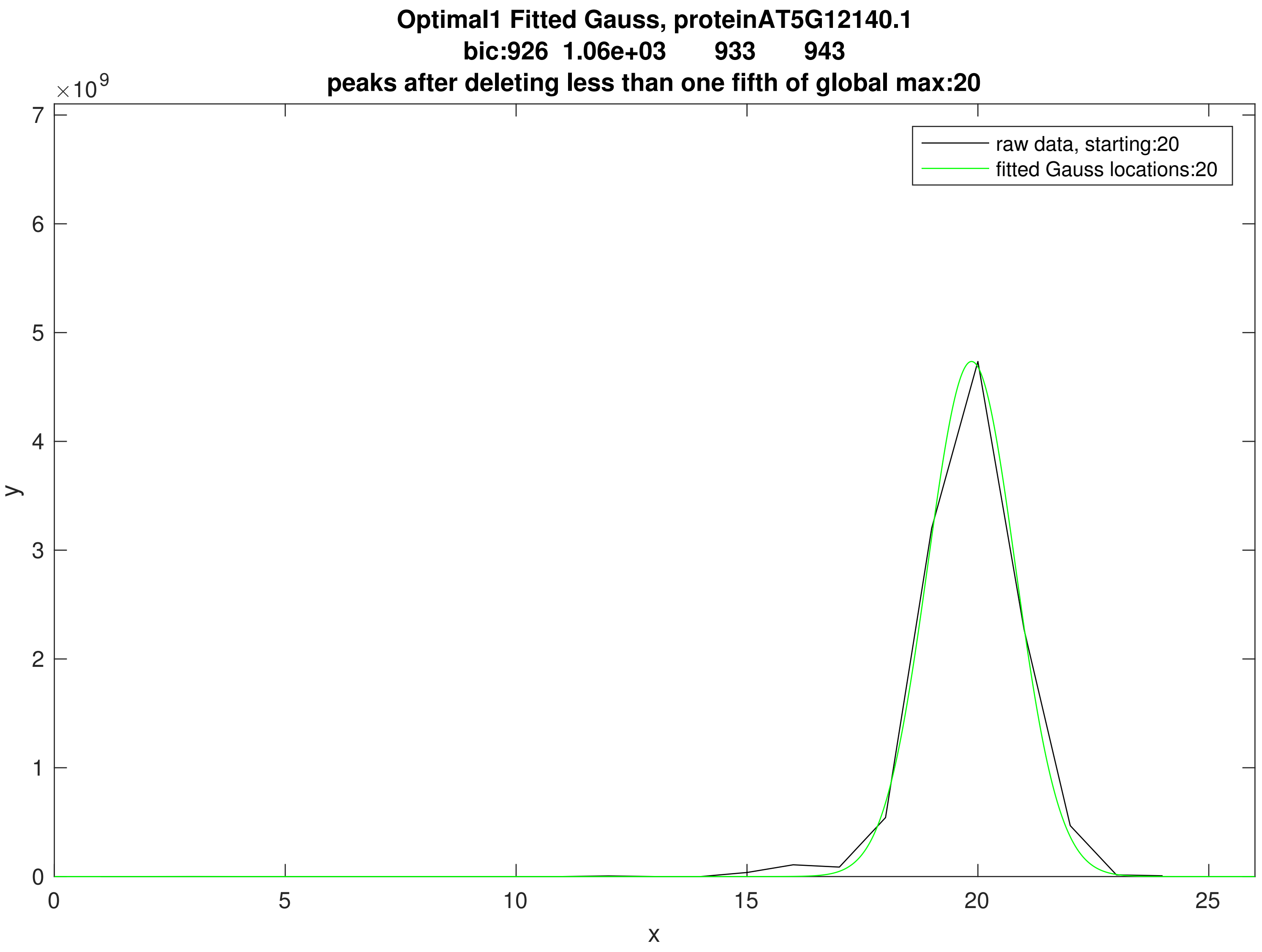
Optimal1 Fitted Gauss, proteinAT5G12140.1
bic:926 1.06e+03 933 943
peaks after deleting less than one fifth of global max:20
×10^9
raw data, starting:20
fitted Gauss locations:20
y
x

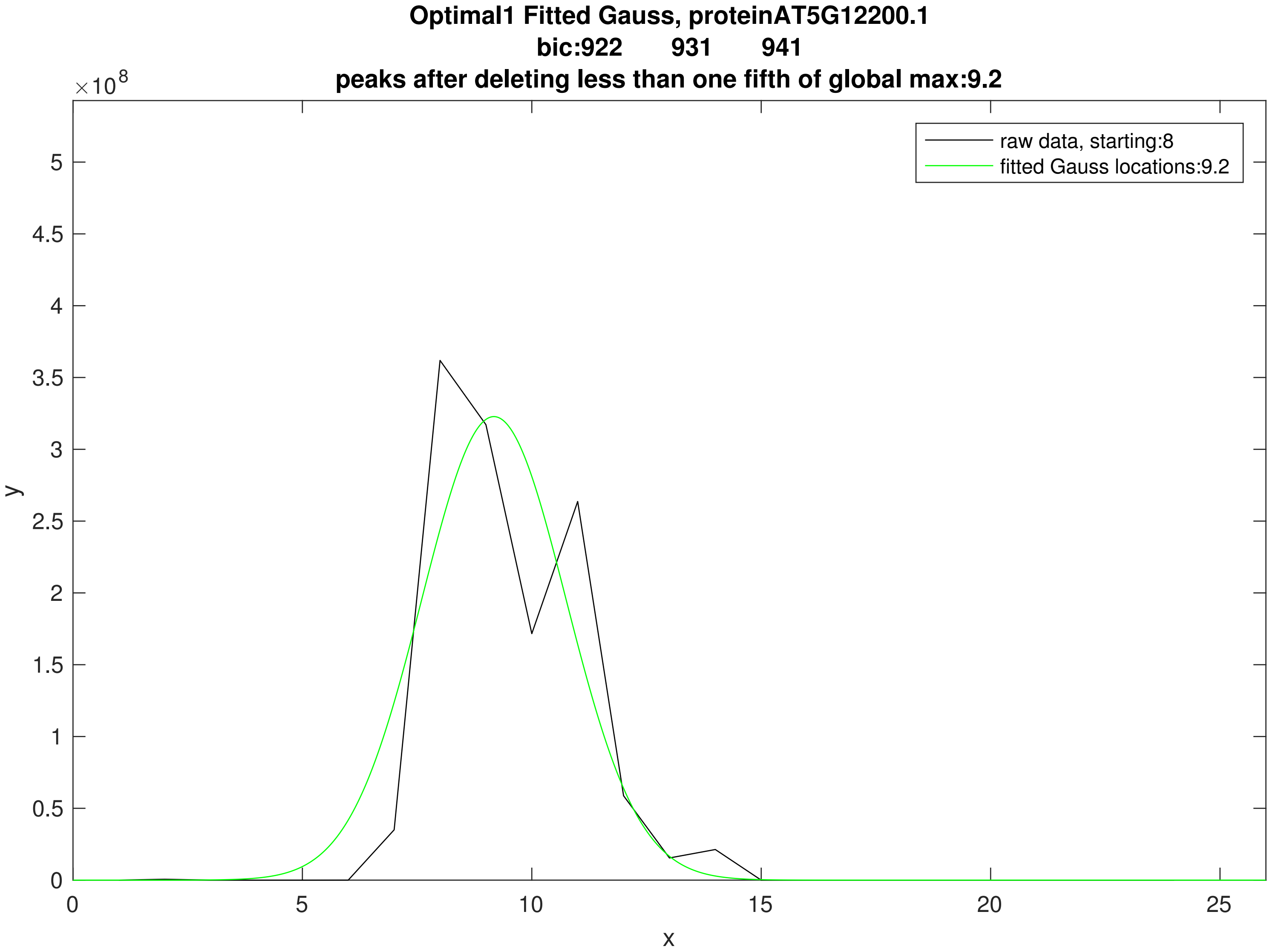
Optimal1 Fitted Gauss, proteinAT5G12200.1
bic:922 931 941
peaks after deleting less than one fifth of global max:9.2
×10^8
raw data, starting:8
fitted Gauss locations:9.2
y
x
0
0.5
1
1.5
2
2.5
3
3.5
4
4.5
5
0
5
10
15
20
25

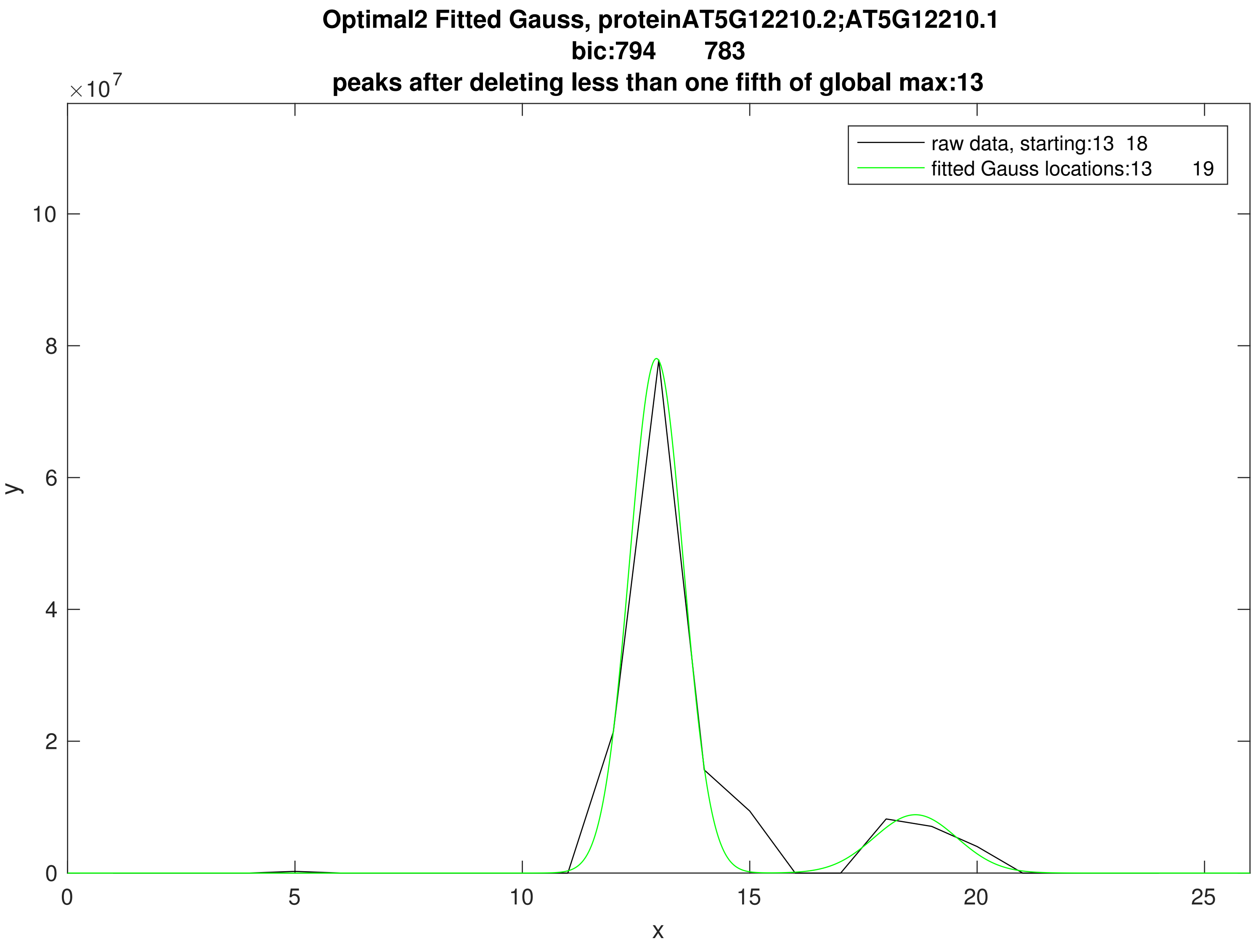
Optimal2 Fitted Gauss, proteinAT5G12210.2;AT5G12210.1
bic:794 783
peaks after deleting less than one fifth of global max:13
×10^7
raw data, starting:13 18
fitted Gauss locations:13 19
10
8
6
4
2
0
y
0
5
10
15
20
25
x

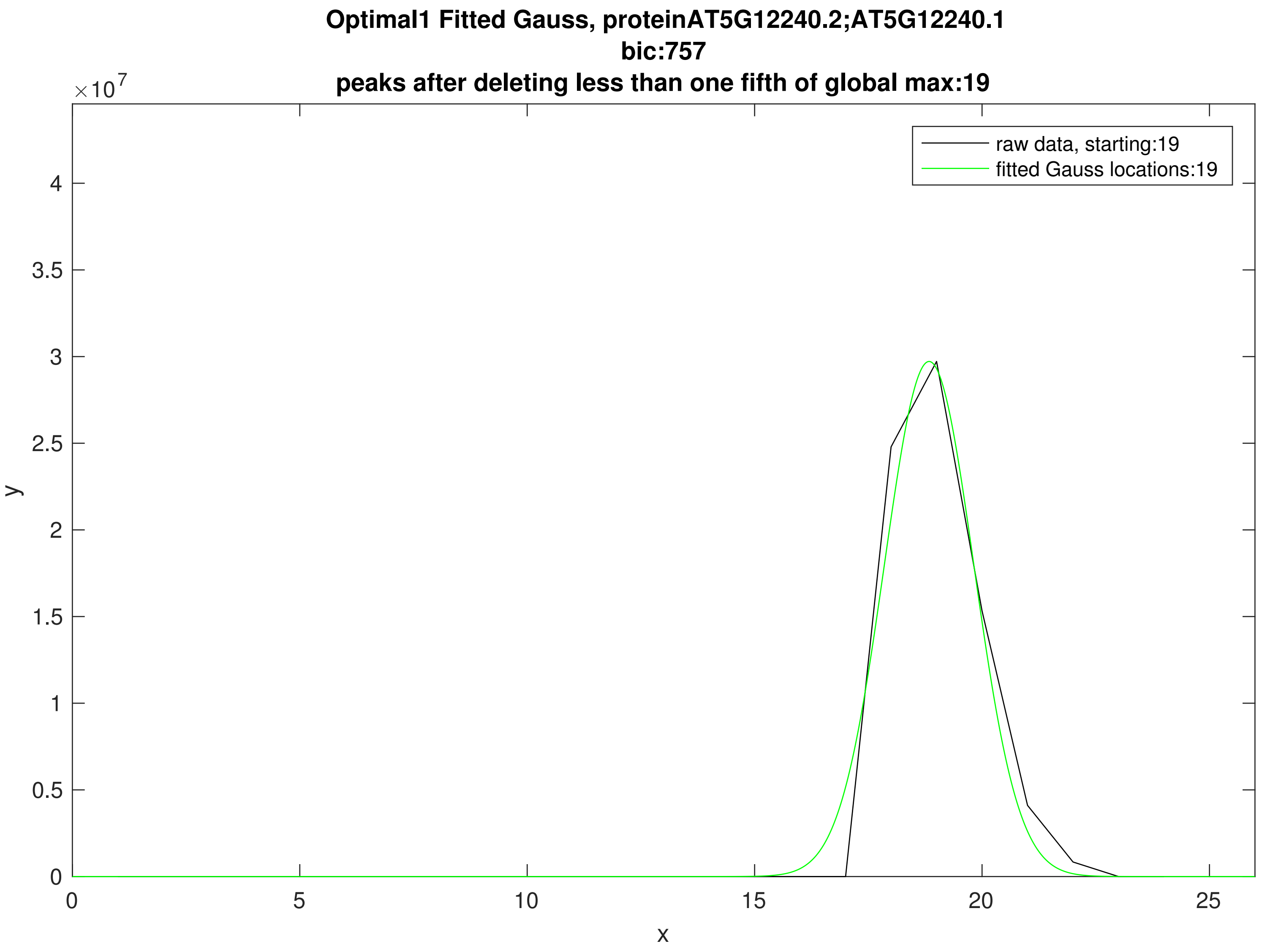
Optimal1 Fitted Gauss, proteinAT5G12240.2;AT5G12240.1
bic:757
peaks after deleting less than one fifth of global max:19
×10^7
raw data, starting:19
fitted Gauss locations:19
y
x
0
0.5
1
1.5
2
2.5
3
3.5
4
0
5
10
15
20
25

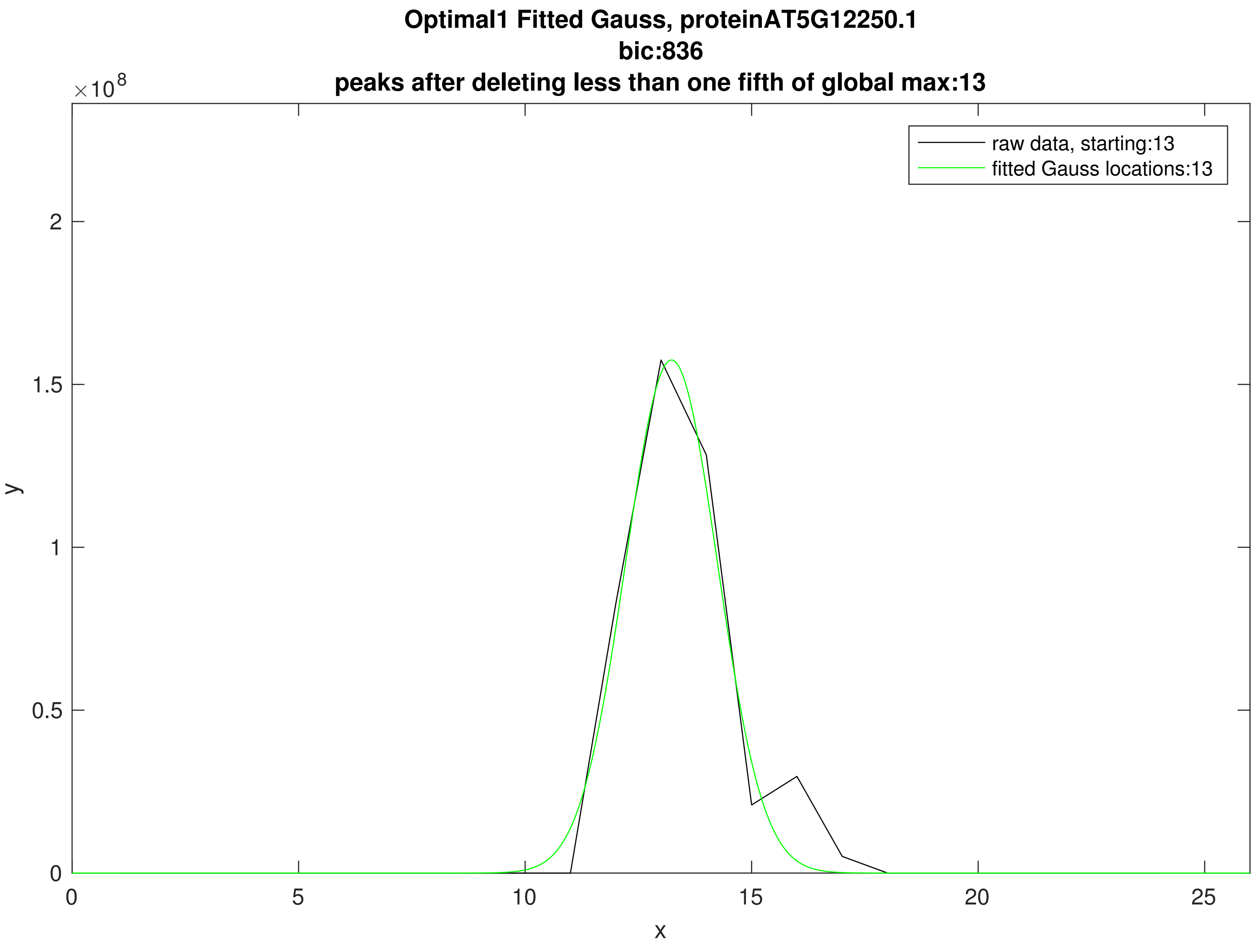
Optimal1 Fitted Gauss, proteinAT5G12250.1
bic:836
peaks after deleting less than one fifth of global max:13
×10^8
raw data, starting:13
fitted Gauss locations:13
y
x
0
0.5
1
1.5
2
0
5
10
15
20
25

**Optimal1 Fitted Gauss, proteinAT5G12320.1**
**bic:775**
**peaks after deleting less than one fifth of global max:19**

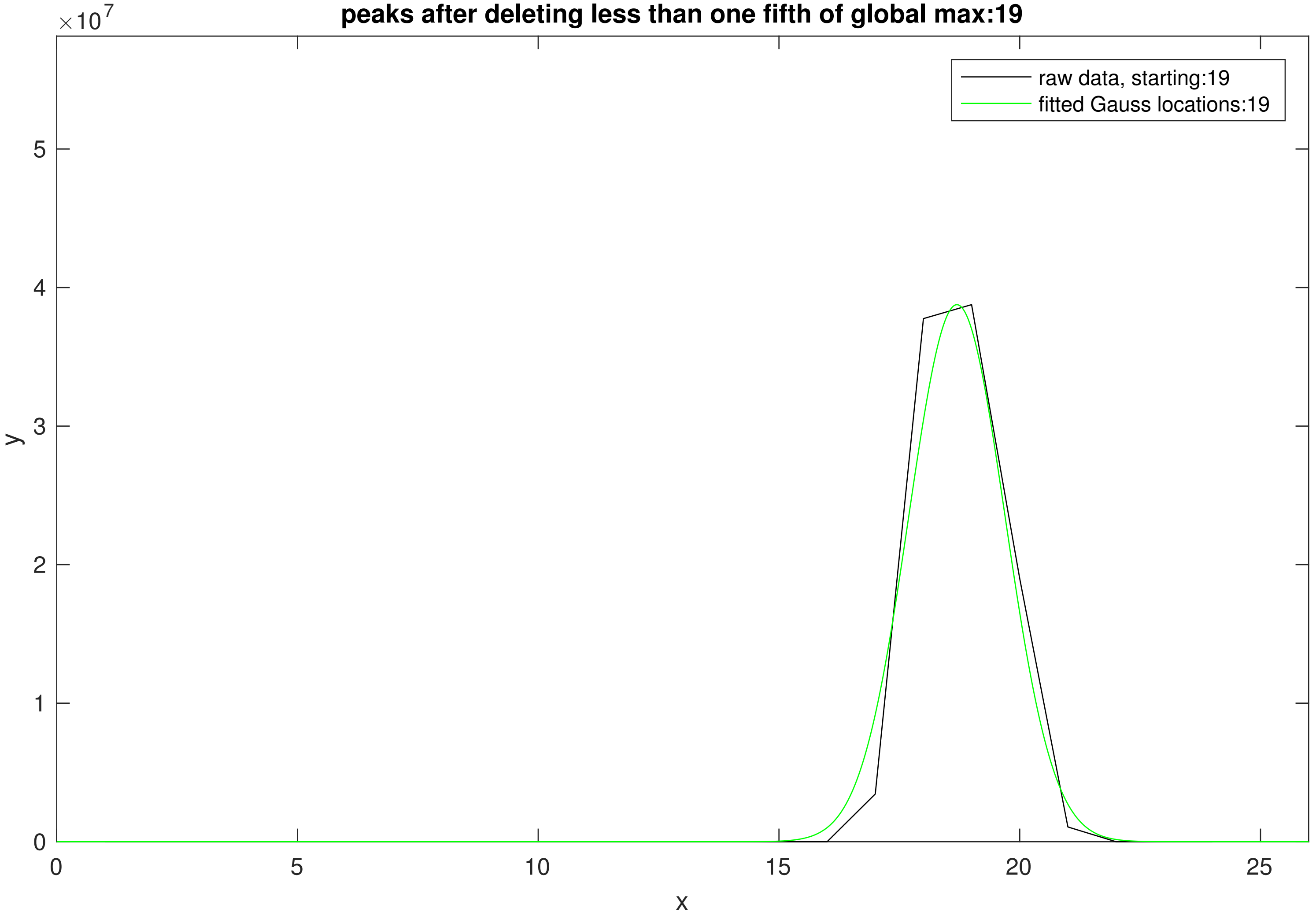

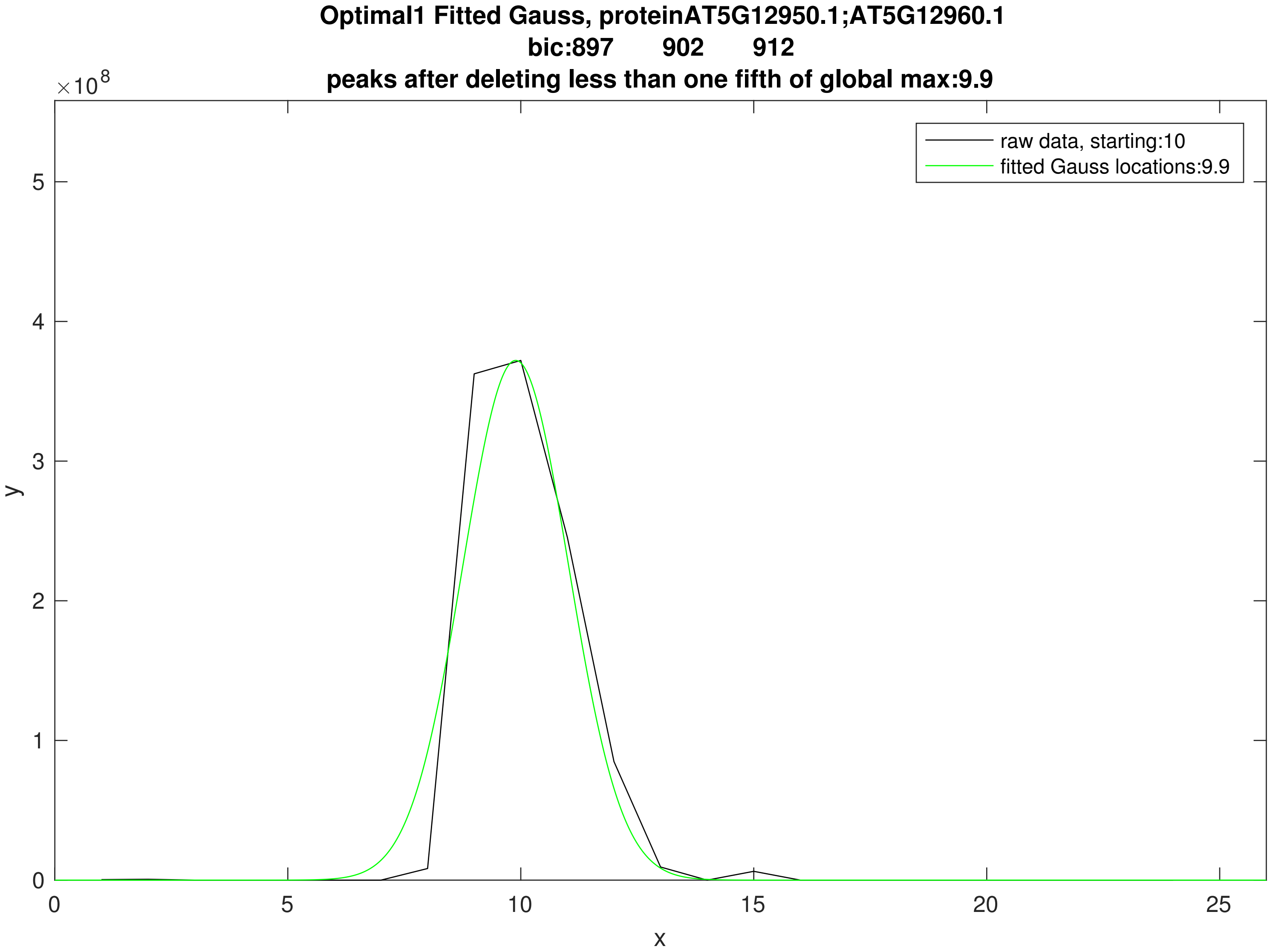

Optimal1 Fitted Gauss, proteinAT5G12950.1;AT5G12960.1
bic:897 902 912
peaks after deleting less than one fifth of global max:9.9
×10^8
raw data, starting:10
fitted Gauss locations:9.9
y
x
0
1
2
3
4
5
10
15
20
25

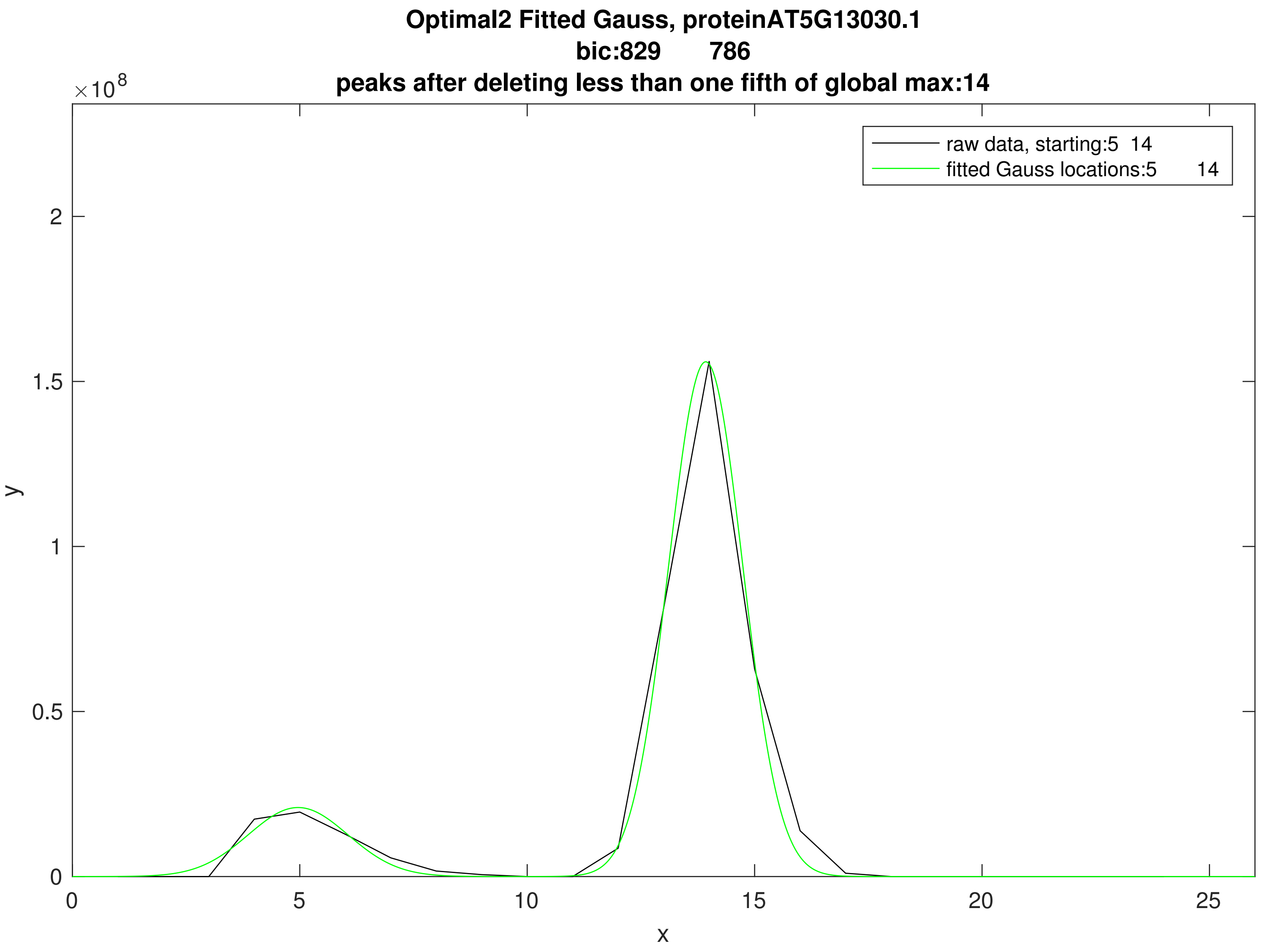
Optimal2 Fitted Gauss, proteinAT5G13030.1
bic:829 786
peaks after deleting less than one fifth of global max:14
×10^8
raw data, starting:5 14
fitted Gauss locations:5 14
y
x
0
0.5
1
1.5
2
5
10
15
20
25

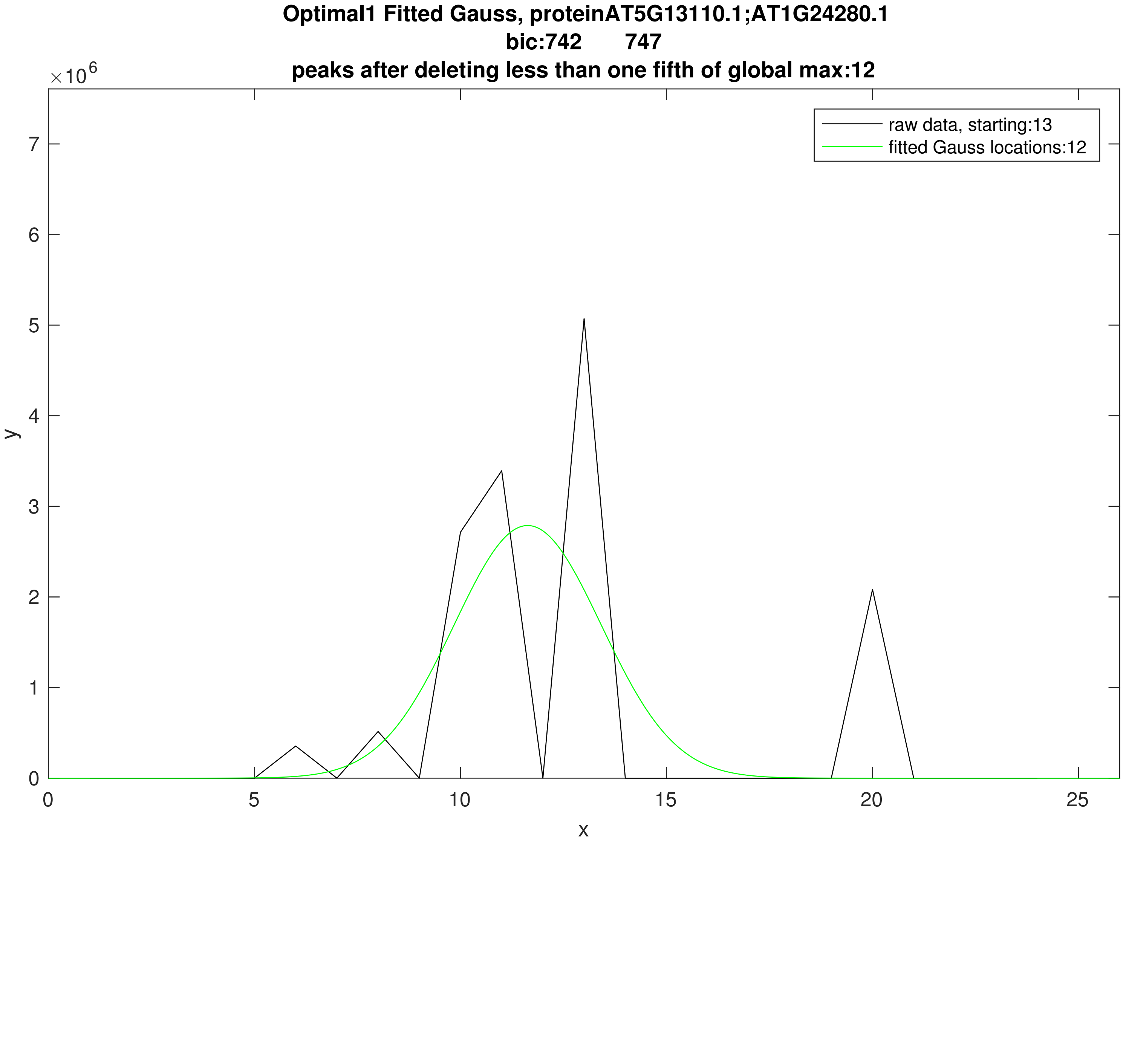
Optimal1 Fitted Gauss, proteinAT5G13110.1;AT1G24280.1
bic:742 747
peaks after deleting less than one fifth of global max:12
×10^6
raw data, starting:13
fitted Gauss locations:12
y
x
0
1
2
3
4
5
6
7
0
5
10
15
20
25

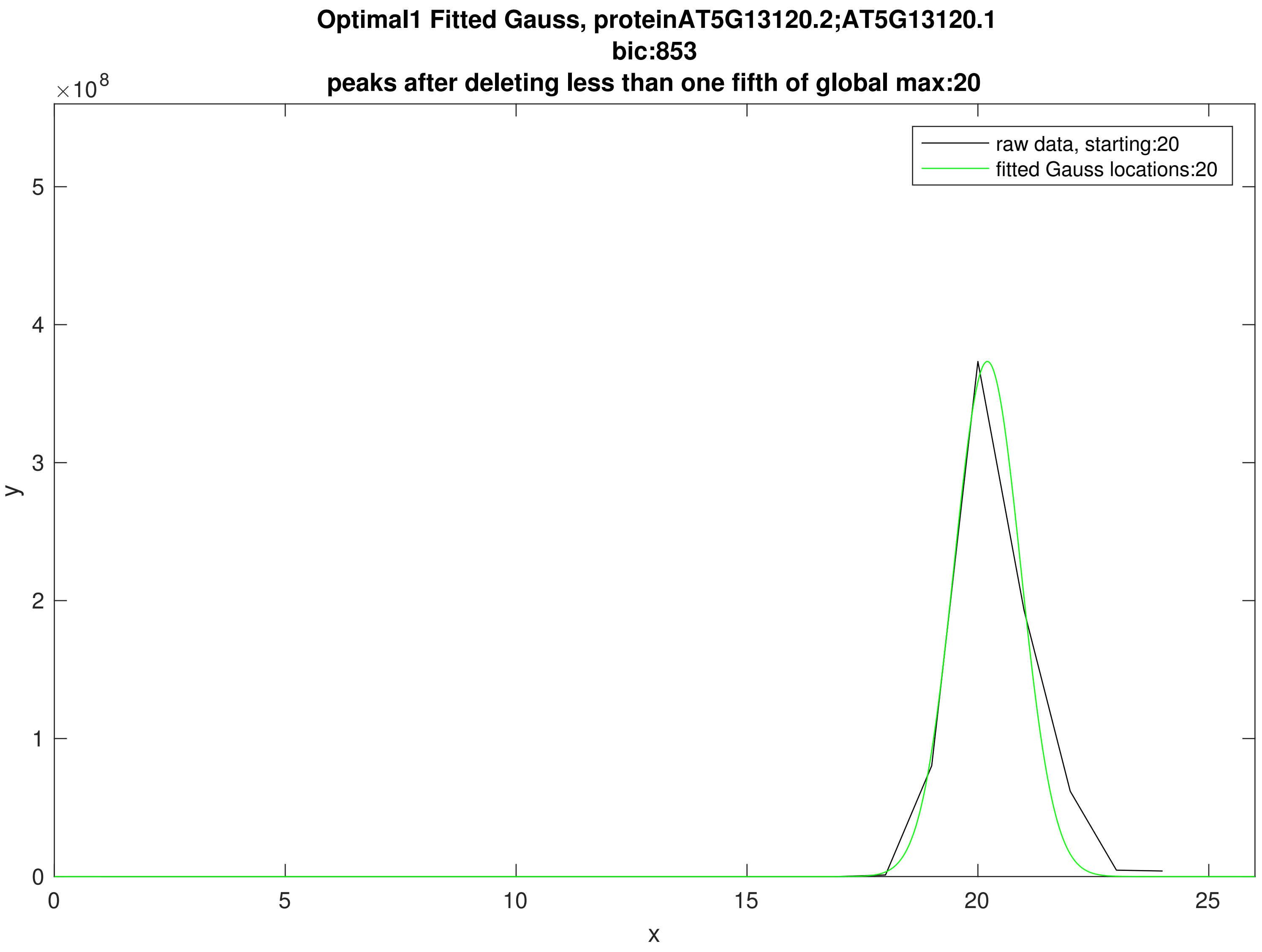

Optimal1 Fitted Gauss, proteinAT5G13120.2;AT5G13120.1
bic:853
peaks after deleting less than one fifth of global max:20
×10^8
raw data, starting:20
fitted Gauss locations:20
y
x
0
1
2
3
4
5
0
5
10
15
20
25

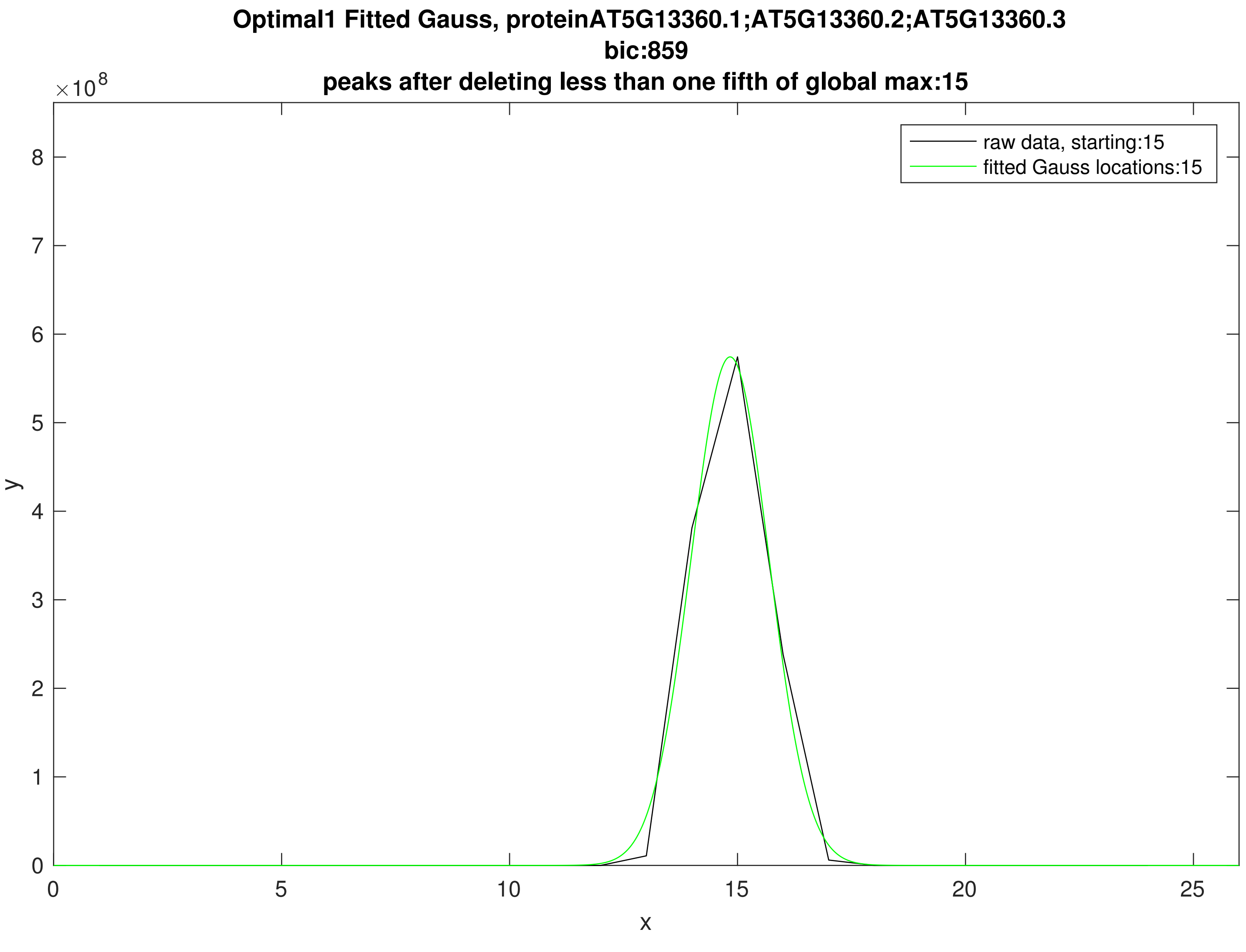
Optimal1 Fitted Gauss, proteinAT5G13360.1;AT5G13360.2;AT5G13360.3
bic:859
peaks after deleting less than one fifth of global max:15
×10^8
raw data, starting:15
fitted Gauss locations:15
y
x
0
1
2
3
4
5
6
7
8
0
5
10
15
20
25

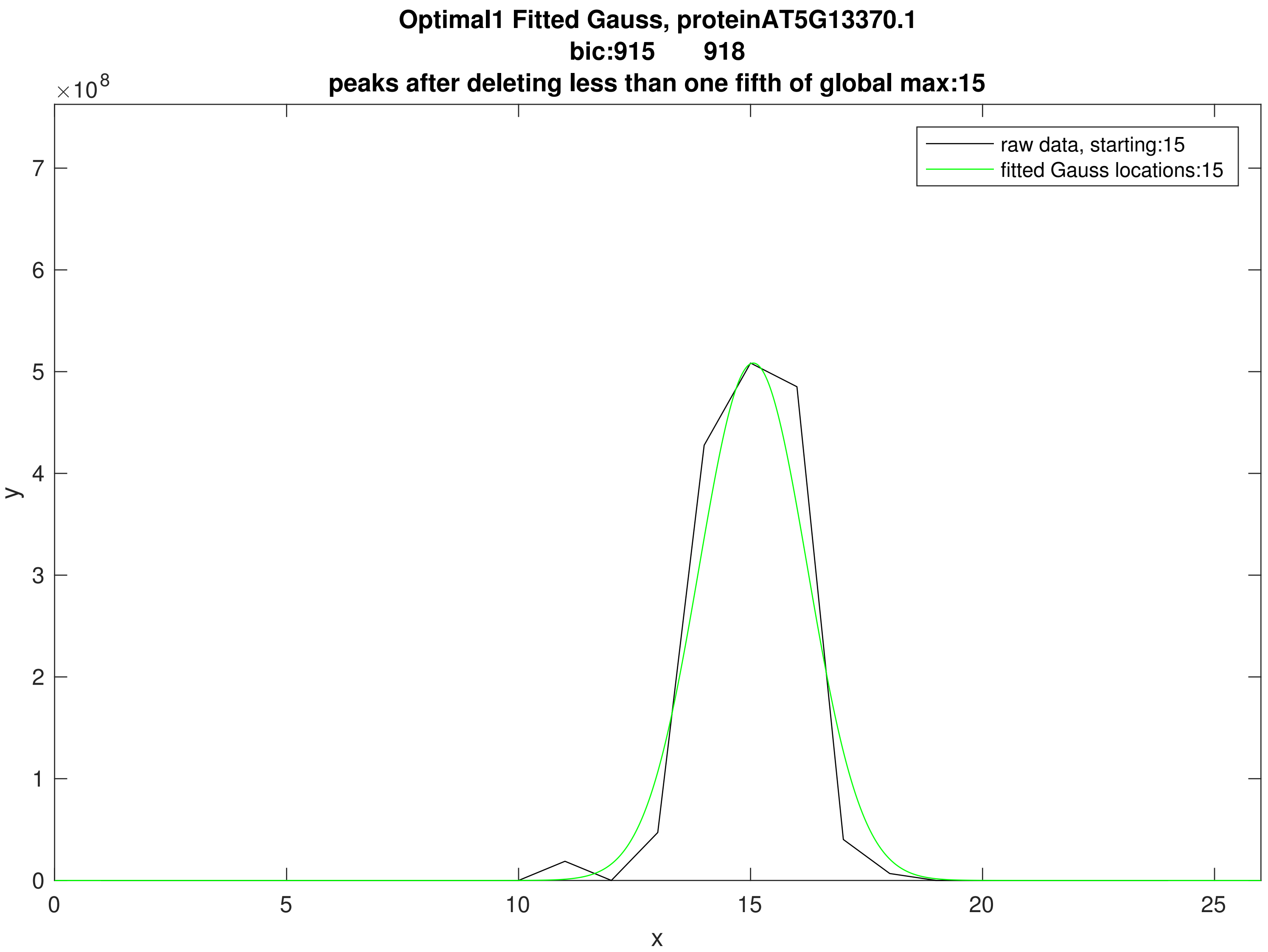

Optimal1 Fitted Gauss, proteinAT5G13370.1
bic:915 918
peaks after deleting less than one fifth of global max:15
×10^8
raw data, starting:15
fitted Gauss locations:15
y
x
0
1
2
3
4
5
6
7
0
5
10
15
20
25

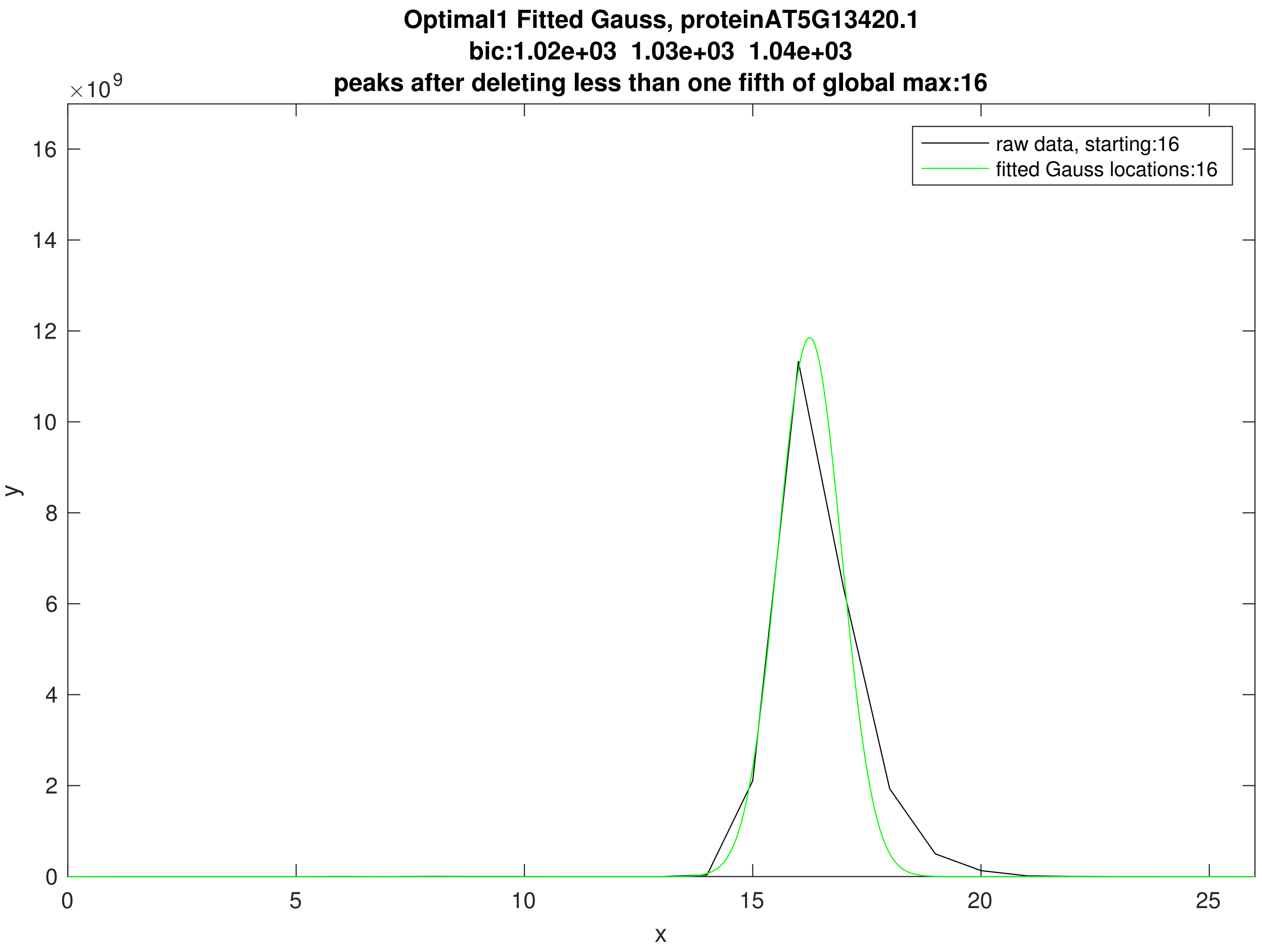
Optimal1 Fitted Gauss, proteinAT5G13420.1
bic:1.02e+03  1.03e+03  1.04e+03
peaks after deleting less than one fifth of global max:16
×10^9
raw data, starting:16
fitted Gauss locations:16
y
x
0
2
4
6
8
10
12
14
16
0
5
10
15
20
25

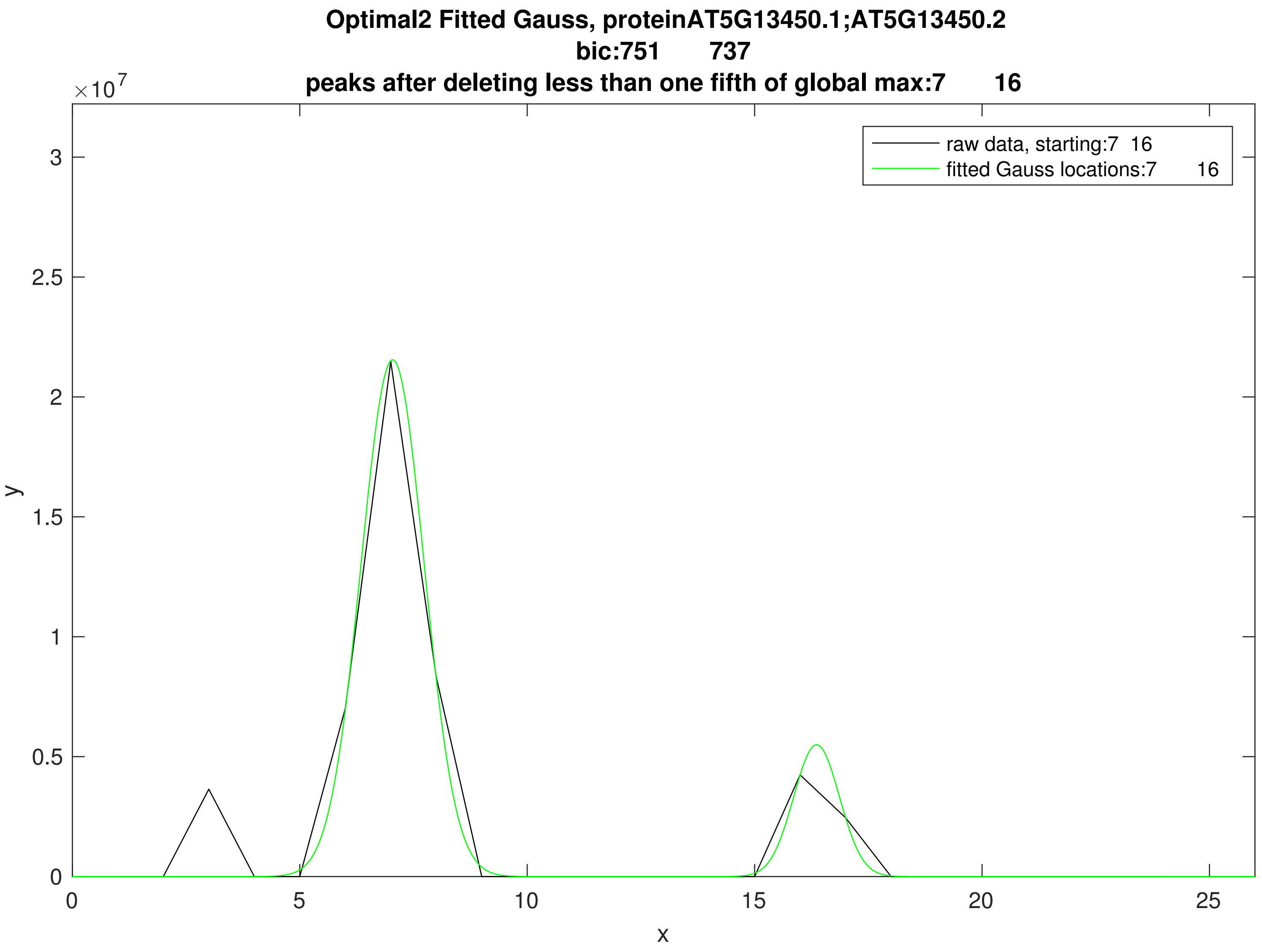
Optimal2 Fitted Gauss, proteinAT5G13450.1;AT5G13450.2
bic:751 737
peaks after deleting less than one fifth of global max:7 16
×10^7
raw data, starting:7 16
fitted Gauss locations:7 16
y
x
0
0.5
1
1.5
2
2.5
3
5
10
15
20
25

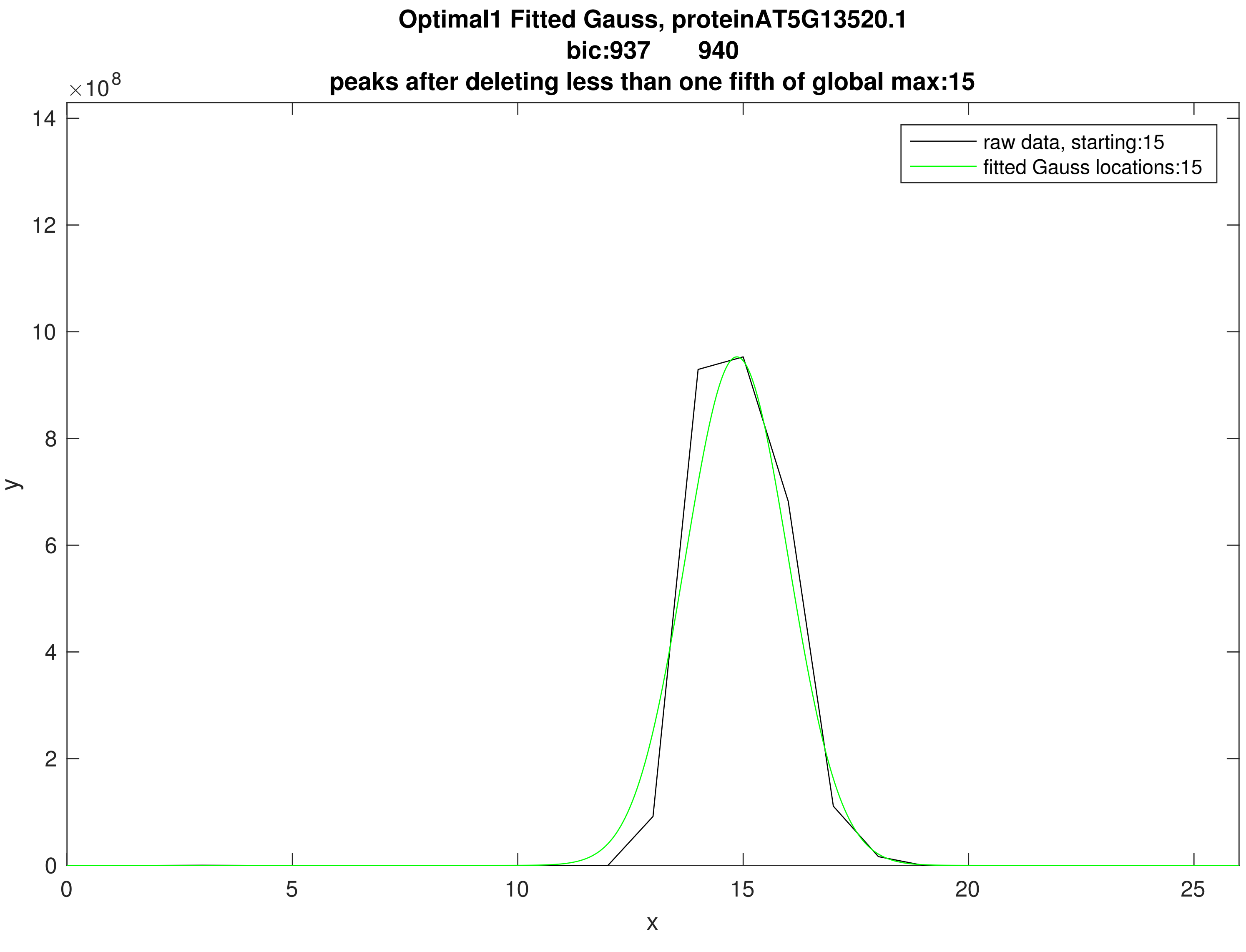

Optimal1 Fitted Gauss, proteinAT5G13520.1
bic:937 940
peaks after deleting less than one fifth of global max:15
×10^8
14
12
10
8
6
4
2
0
y
0
5
10
15
20
25
x
raw data, starting:15
fitted Gauss locations:15

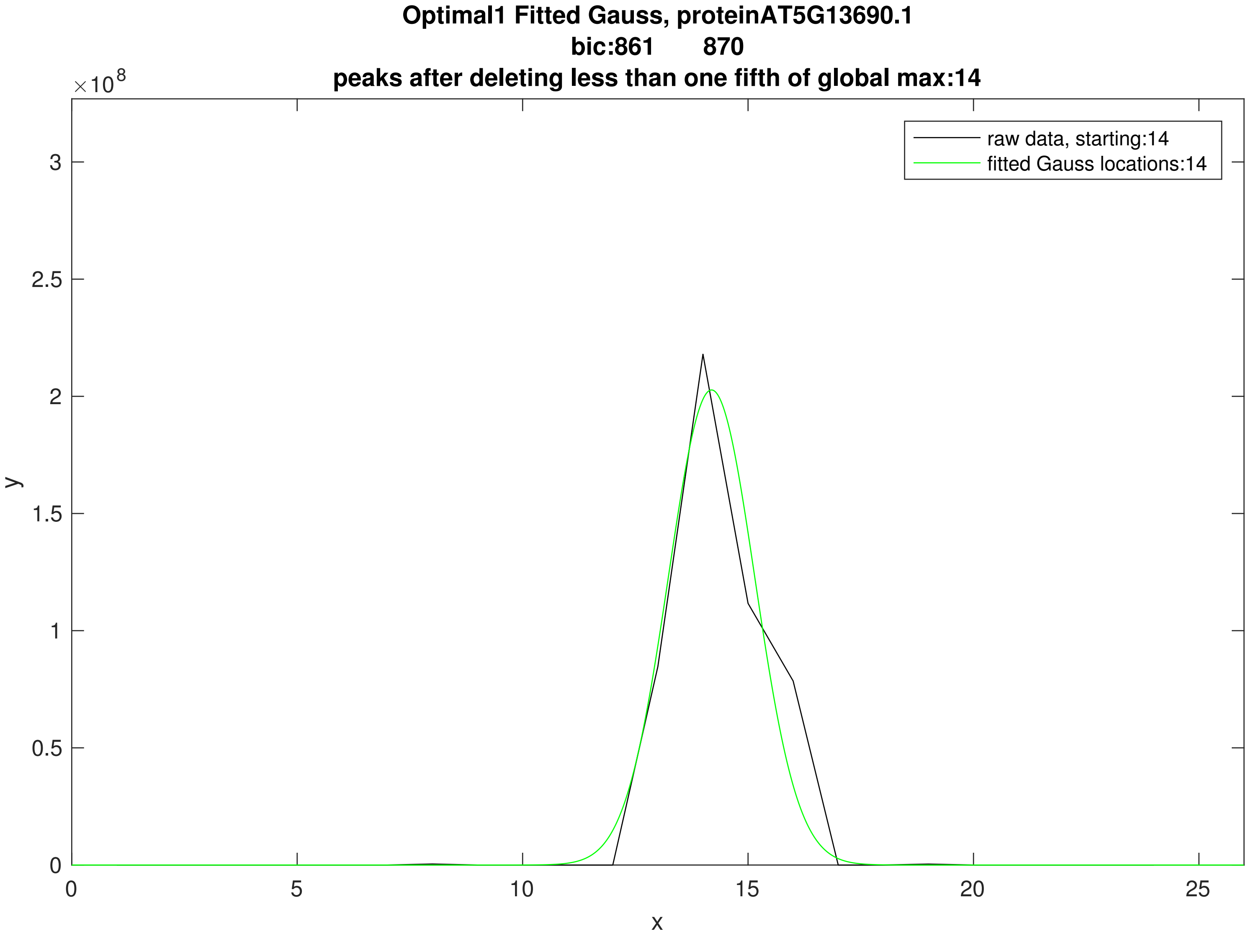

Optimal1 Fitted Gauss, proteinAT5G13690.1
bic:861 870
peaks after deleting less than one fifth of global max:14
×10^8
3
2.5
2
1.5
1
0.5
0
y
0
5
10
15
20
25
x
raw data, starting:14
fitted Gauss locations:14

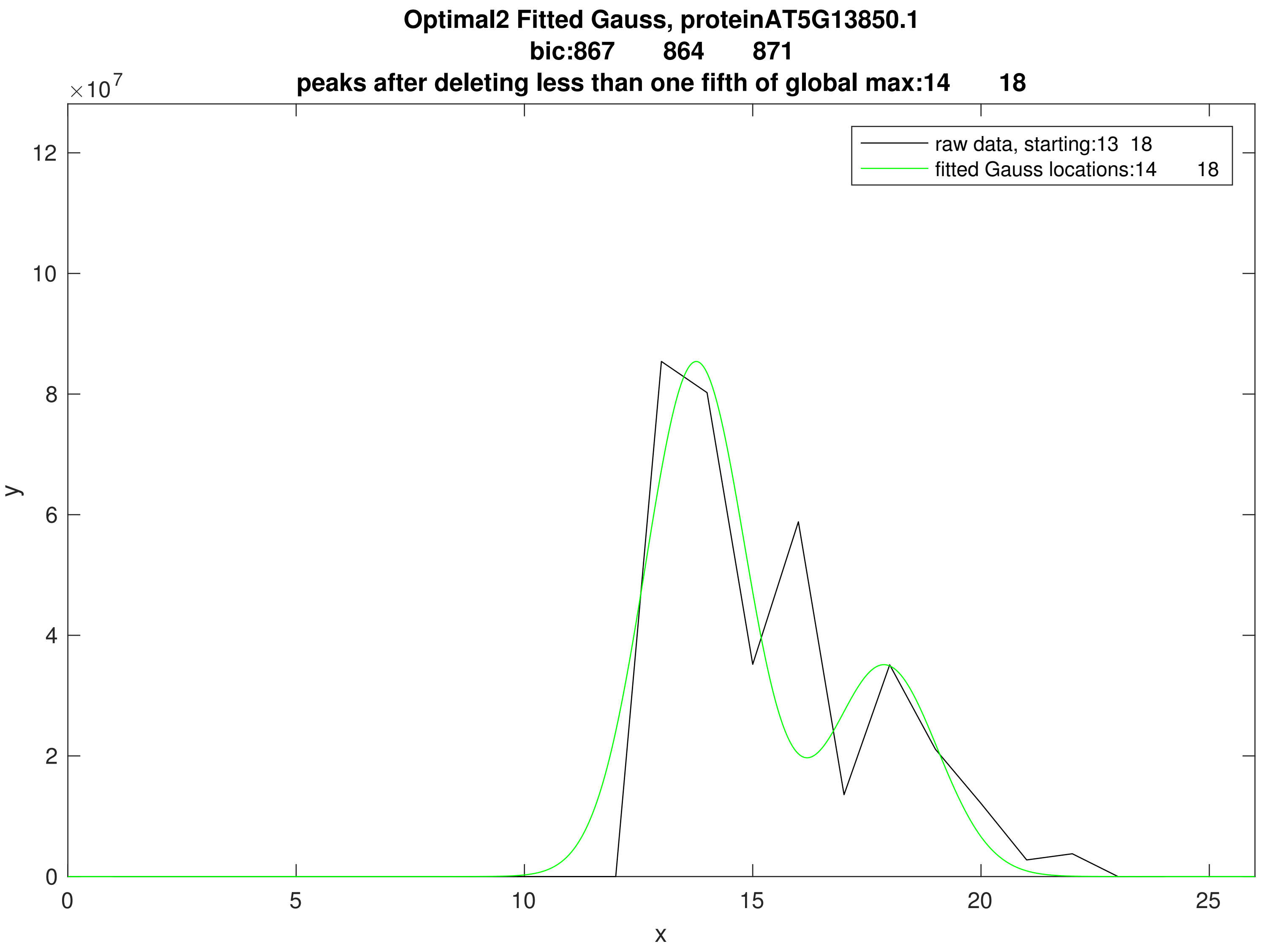
Optimal2 Fitted Gauss, proteinAT5G13850.1
bic:867 864 871
peaks after deleting less than one fifth of global max:14 18
×10^7
raw data, starting:13 18
fitted Gauss locations:14 18
y
x
0
2
4
6
8
10
12
0
5
10
15
20
25

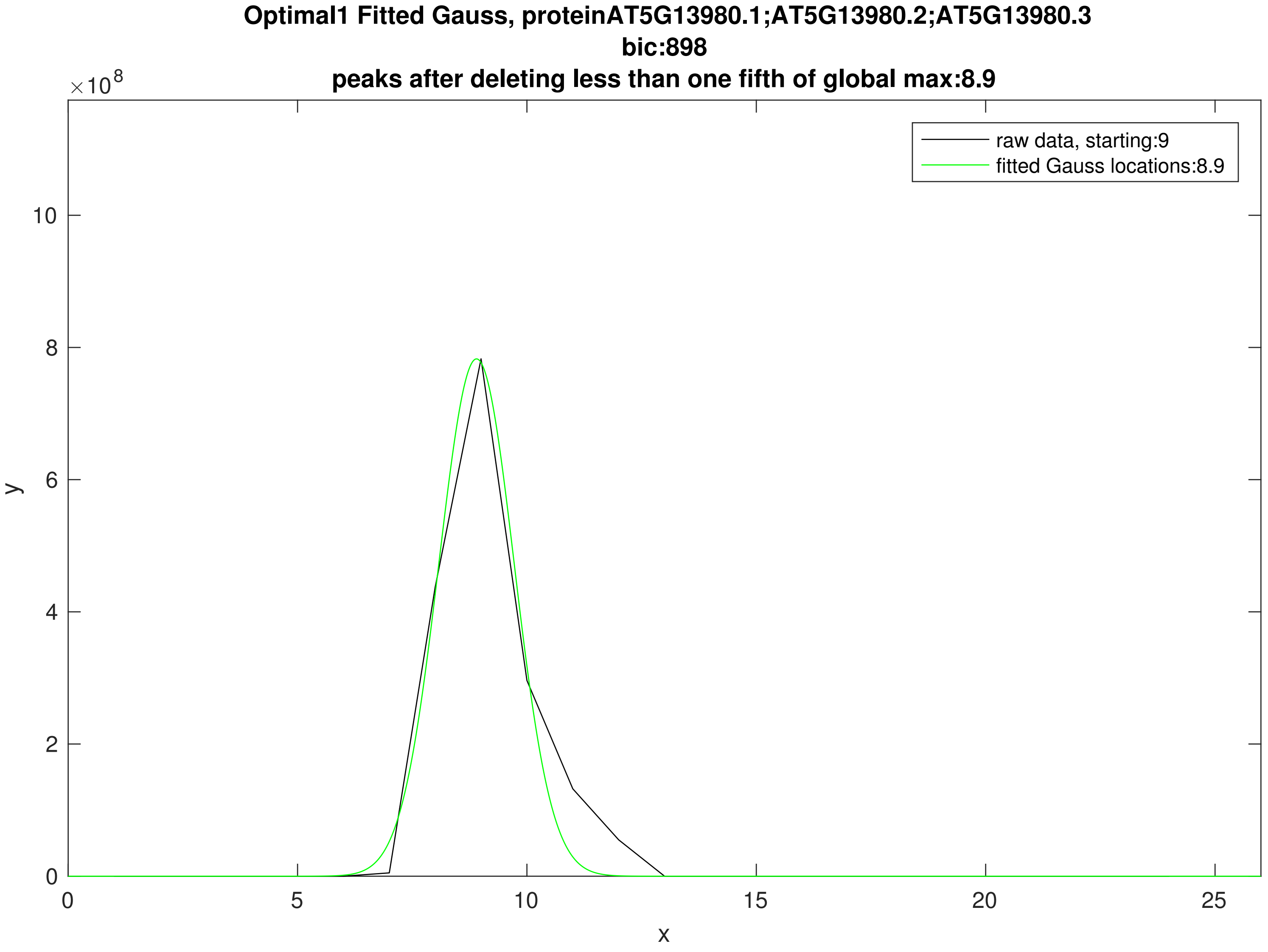

Optimal1 Fitted Gauss, proteinAT5G13980.1;AT5G13980.2;AT5G13980.3
bic:898
peaks after deleting less than one fifth of global max:8.9
×10^8
raw data, starting:9
fitted Gauss locations:8.9
y
x
0
2
4
6
8
10
0
5
10
15
20
25

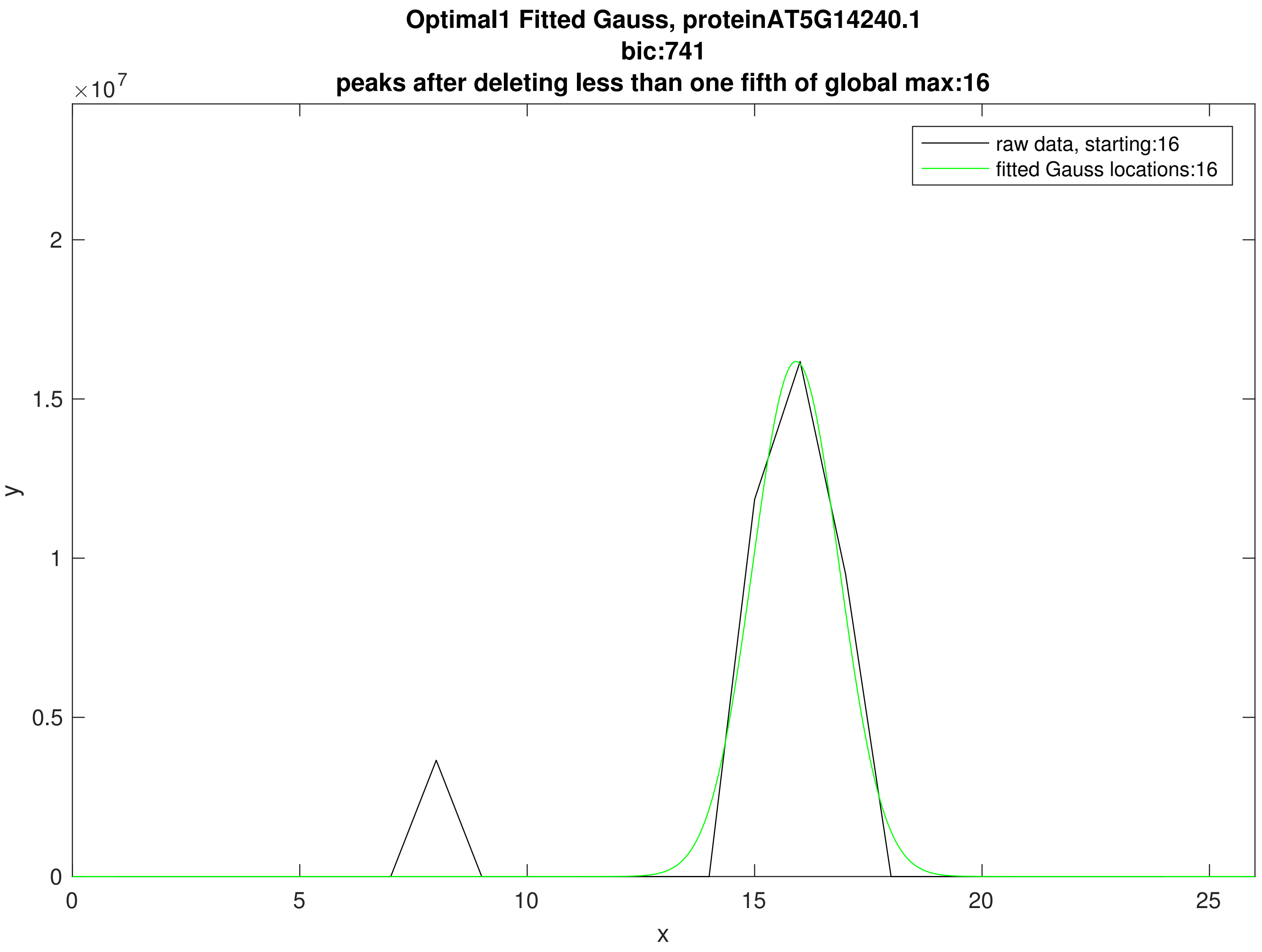

Optimal1 Fitted Gauss, proteinAT5G14240.1
bic:741
peaks after deleting less than one fifth of global max:16
×10^7
raw data, starting:16
fitted Gauss locations:16
y
x
0
0.5
1
1.5
2
0
5
10
15
20
25

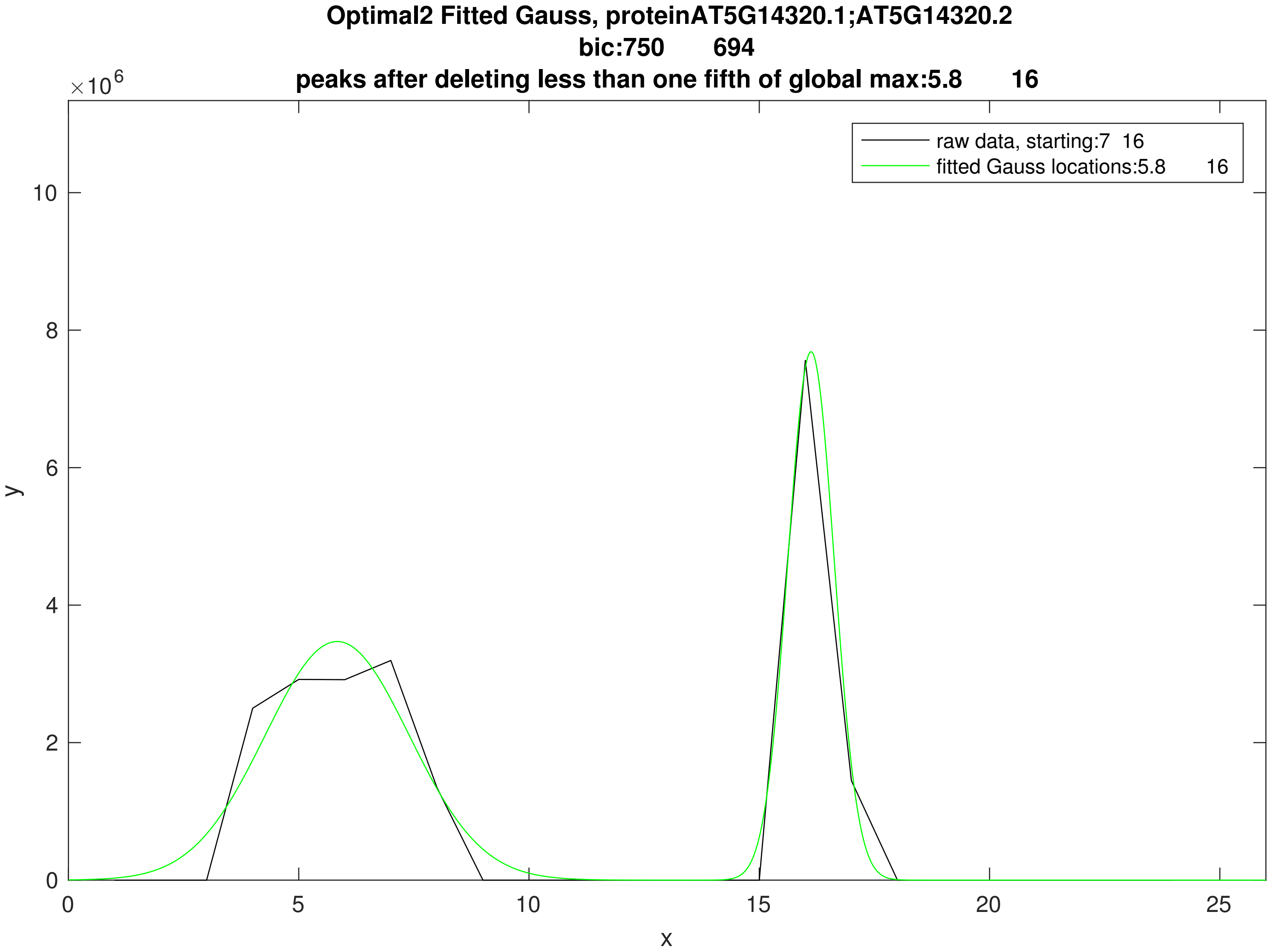
Optimal2 Fitted Gauss, proteinAT5G14320.1;AT5G14320.2
bic:750 694
peaks after deleting less than one fifth of global max:5.8 16
×10^6
raw data, starting:7 16
fitted Gauss locations:5.8 16
10
8
6
4
2
0
y
0
5
10
15
20
25
x

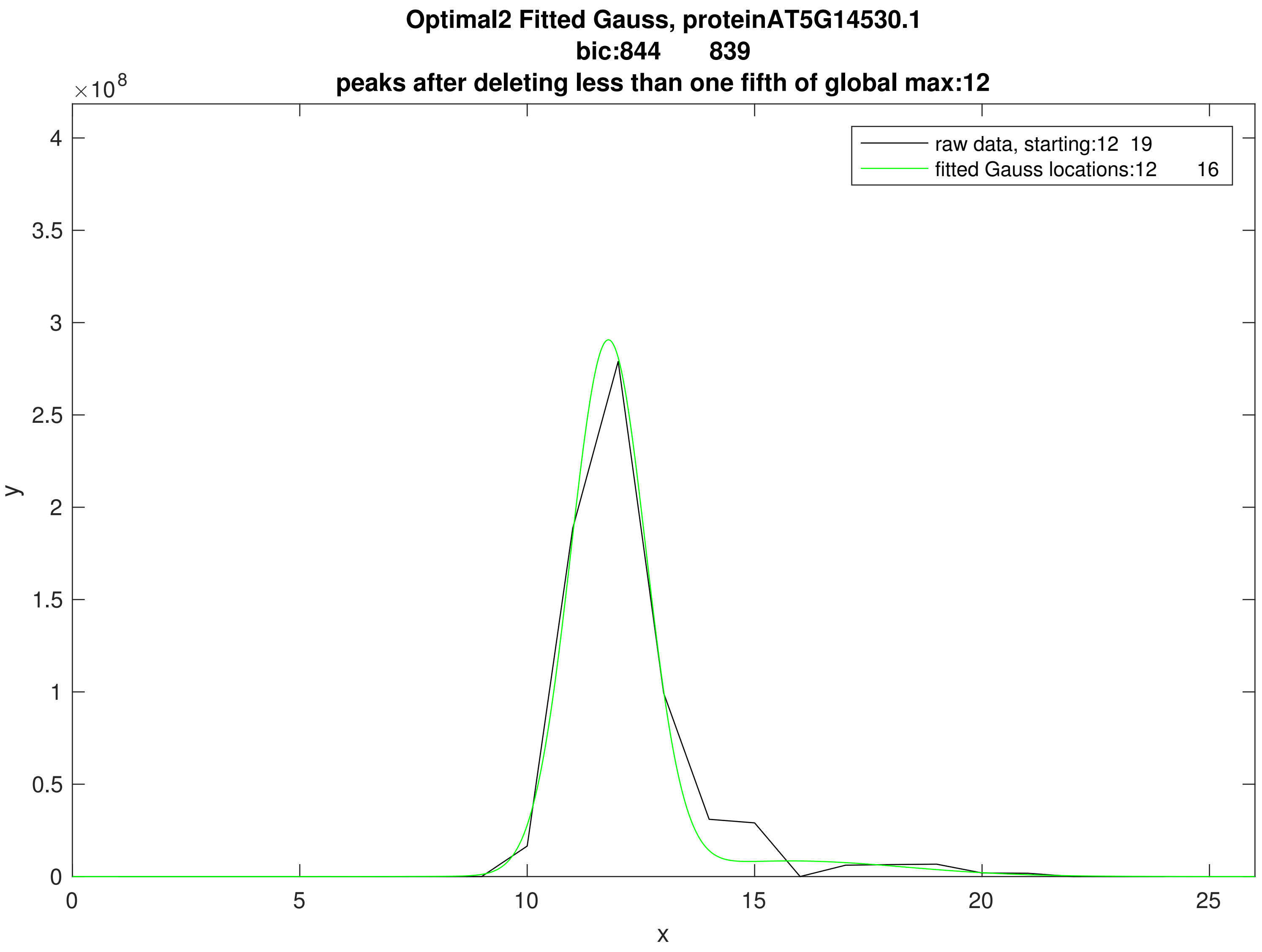
Optimal2 Fitted Gauss, proteinAT5G14530.1
bic:844 839
peaks after deleting less than one fifth of global max:12
×10^8
raw data, starting:12 19
fitted Gauss locations:12 16
y
x
0
0.5
1
1.5
2
2.5
3
3.5
4
0
5
10
15
20
25

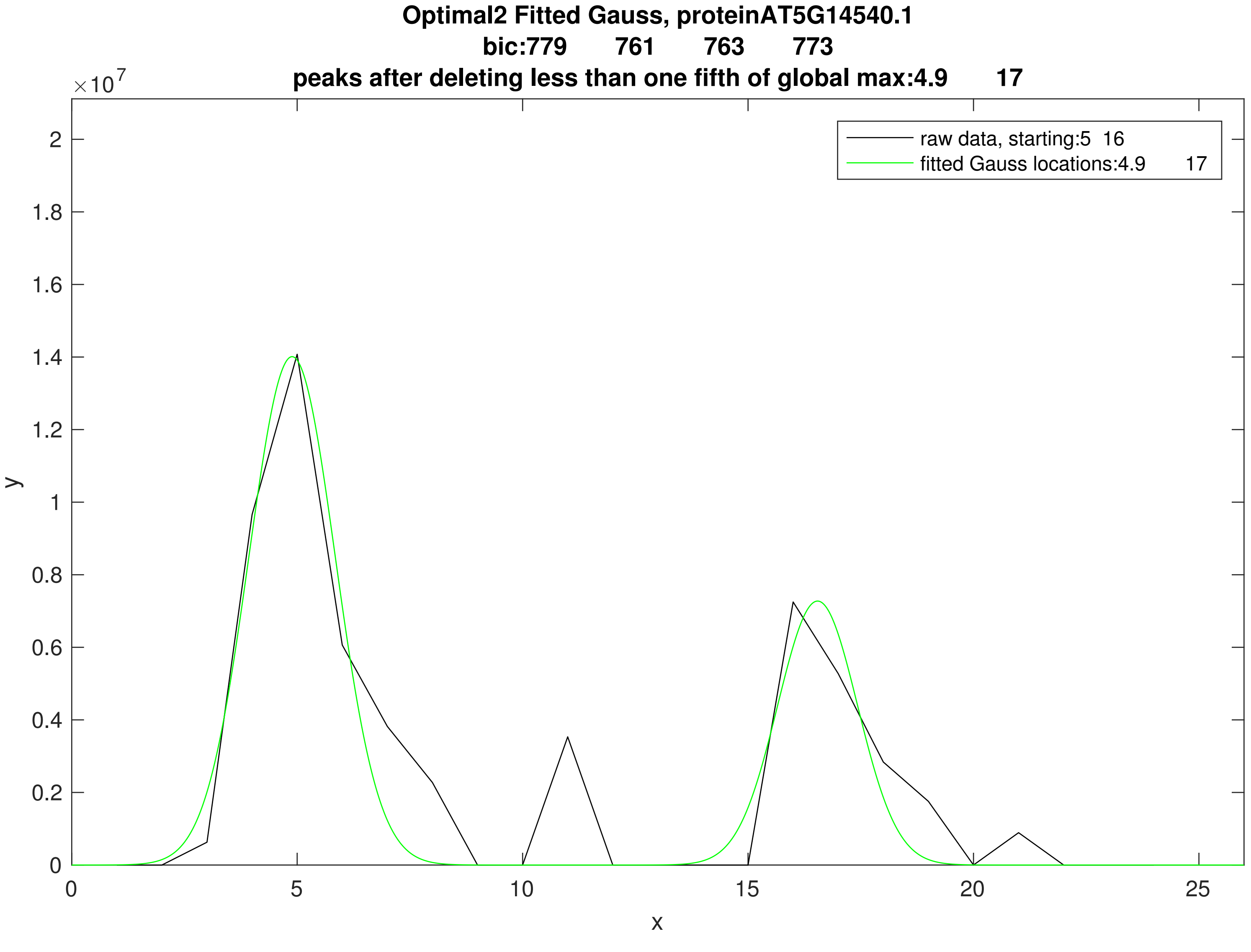
Optimal2 Fitted Gauss, proteinAT5G14540.1
bic:779 761 763 773
peaks after deleting less than one fifth of global max:4.9 17
×10^7
raw data, starting:5 16
fitted Gauss locations:4.9 17
y
x
0
0.2
0.4
0.6
0.8
1
1.2
1.4
1.6
1.8
2
0
5
10
15
20
25

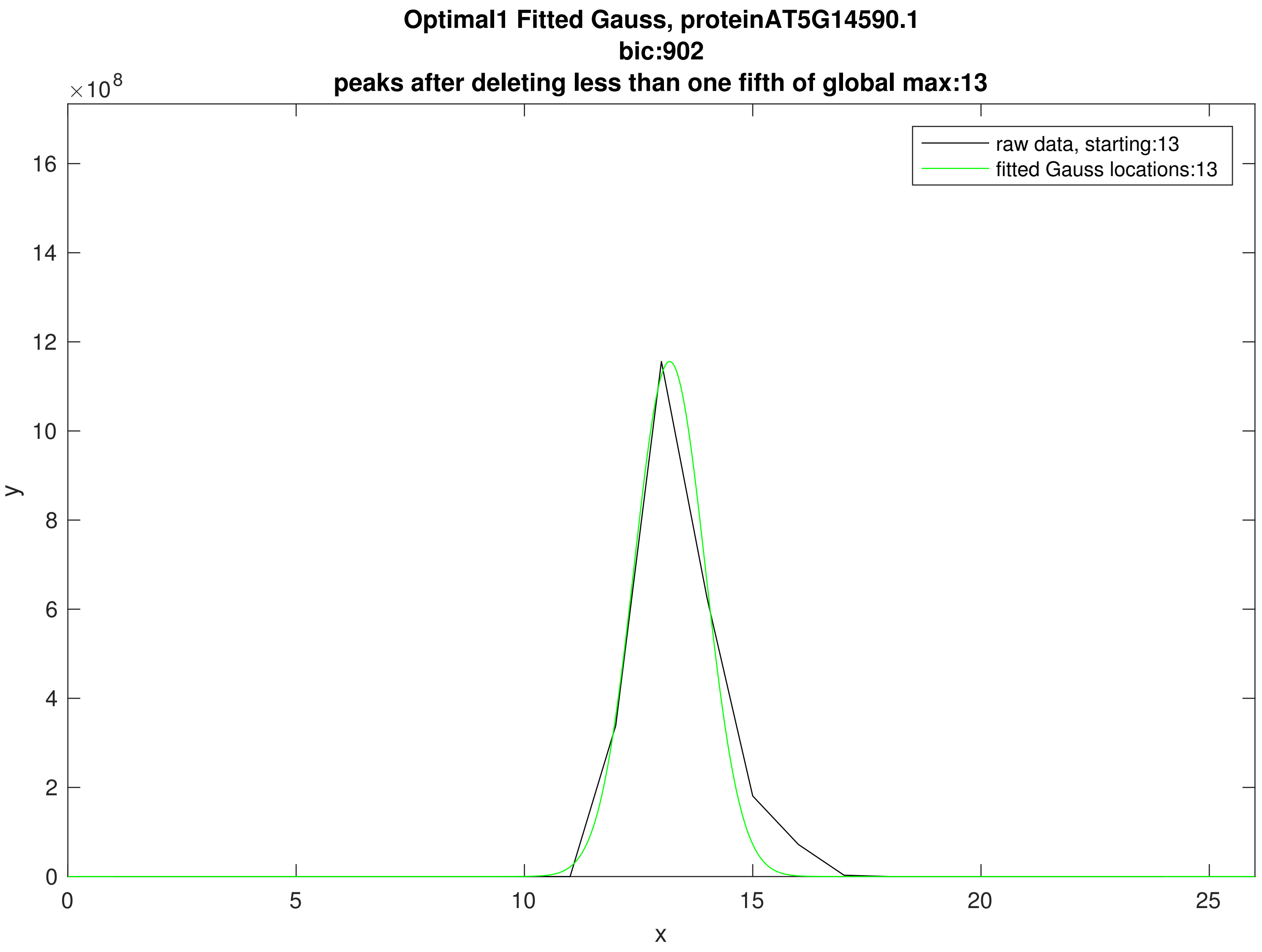

Optimal1 Fitted Gauss, proteinAT5G14590.1
bic:902
peaks after deleting less than one fifth of global max:13
×10^8
raw data, starting:13
fitted Gauss locations:13
y
x
0
2
4
6
8
10
12
14
16
0
5
10
15
20
25

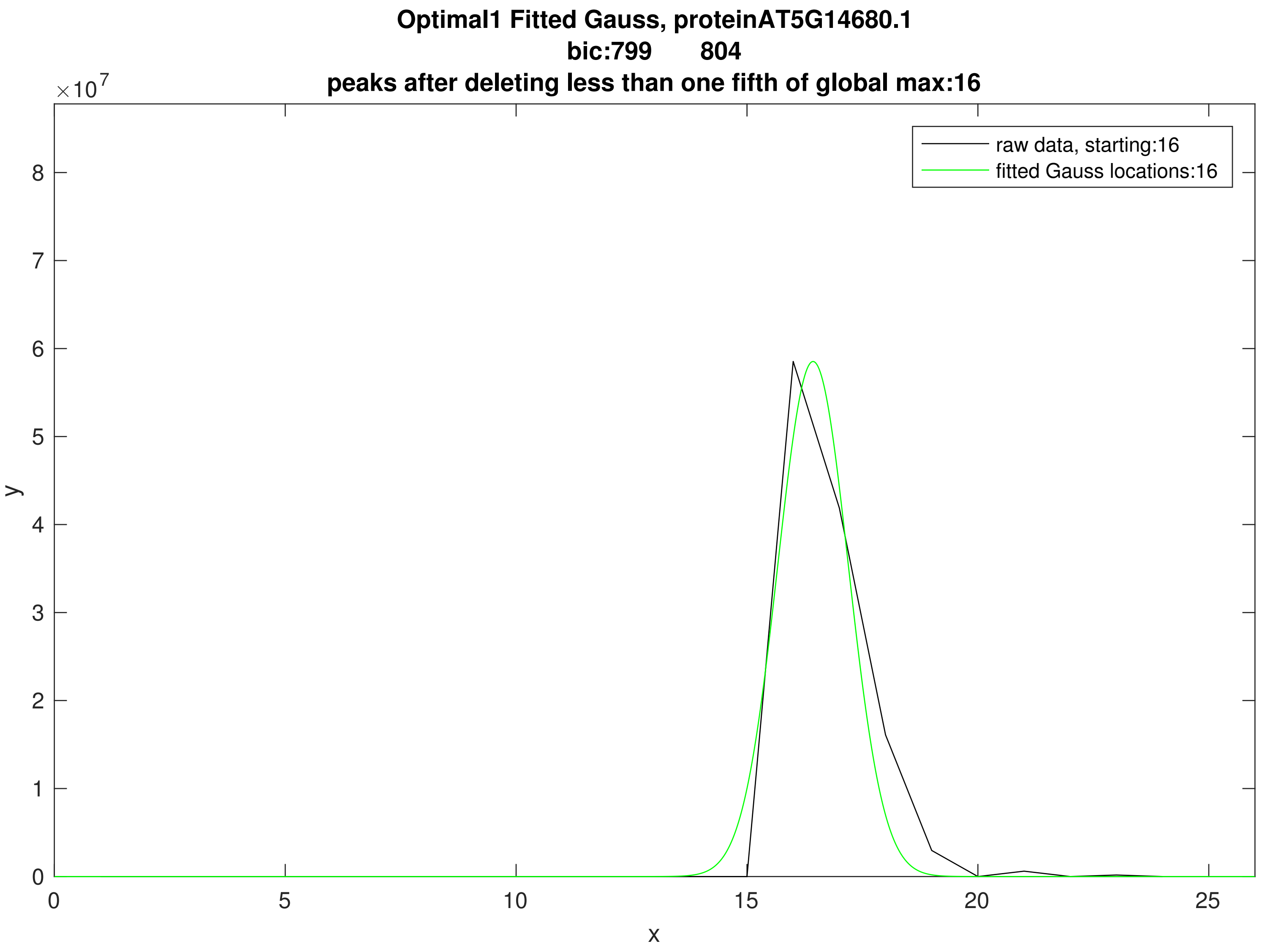
Optimal1 Fitted Gauss, proteinAT5G14680.1
bic:799 804
peaks after deleting less than one fifth of global max:16
×10^7
raw data, starting:16
fitted Gauss locations:16
y
x
0
1
2
3
4
5
6
7
8
0
5
10
15
20
25

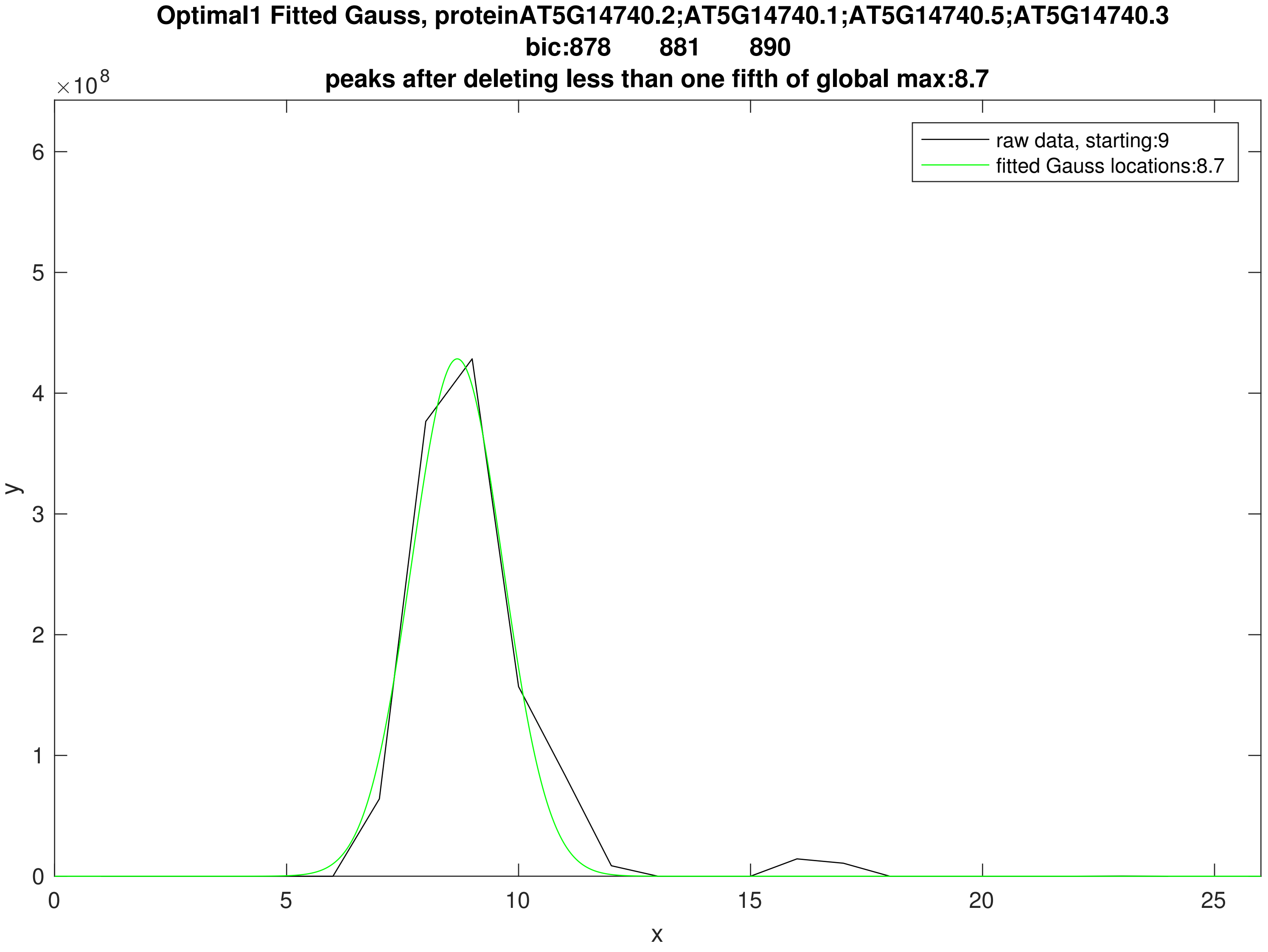
Optimal1 Fitted Gauss, proteinAT5G14740.2;AT5G14740.1;AT5G14740.5;AT5G14740.3
bic:878 881 890
peaks after deleting less than one fifth of global max:8.7
×10^8
raw data, starting:9
fitted Gauss locations:8.7
y
x
0
1
2
3
4
5
6
0
5
10
15
20
25

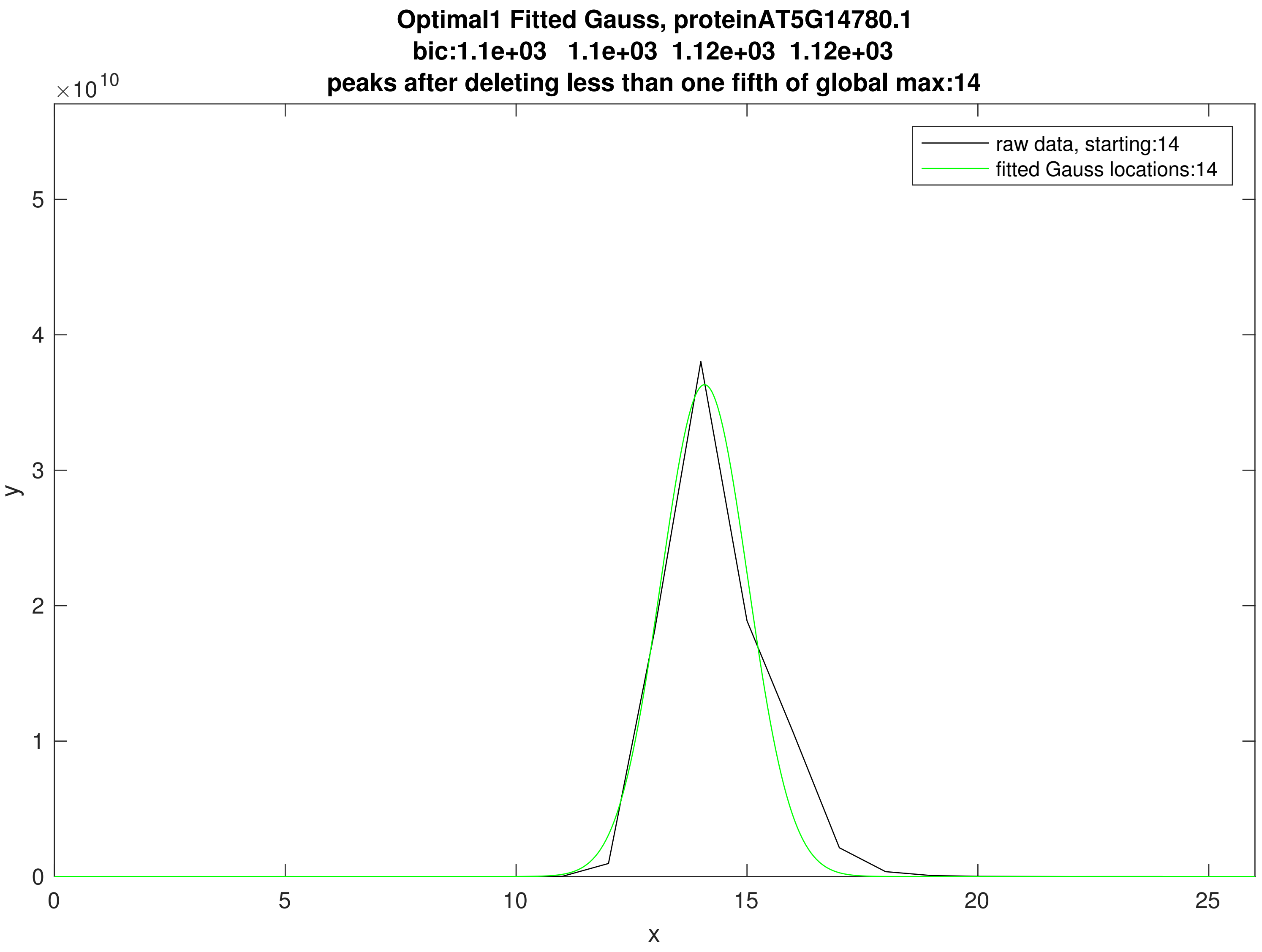
Optimal1 Fitted Gauss, proteinAT5G14780.1
bic:1.1e+03 1.1e+03 1.12e+03 1.12e+03
peaks after deleting less than one fifth of global max:14
×10^10
raw data, starting:14
fitted Gauss locations:14
y
x
0
1
2
3
4
5
0
5
10
15
20
25

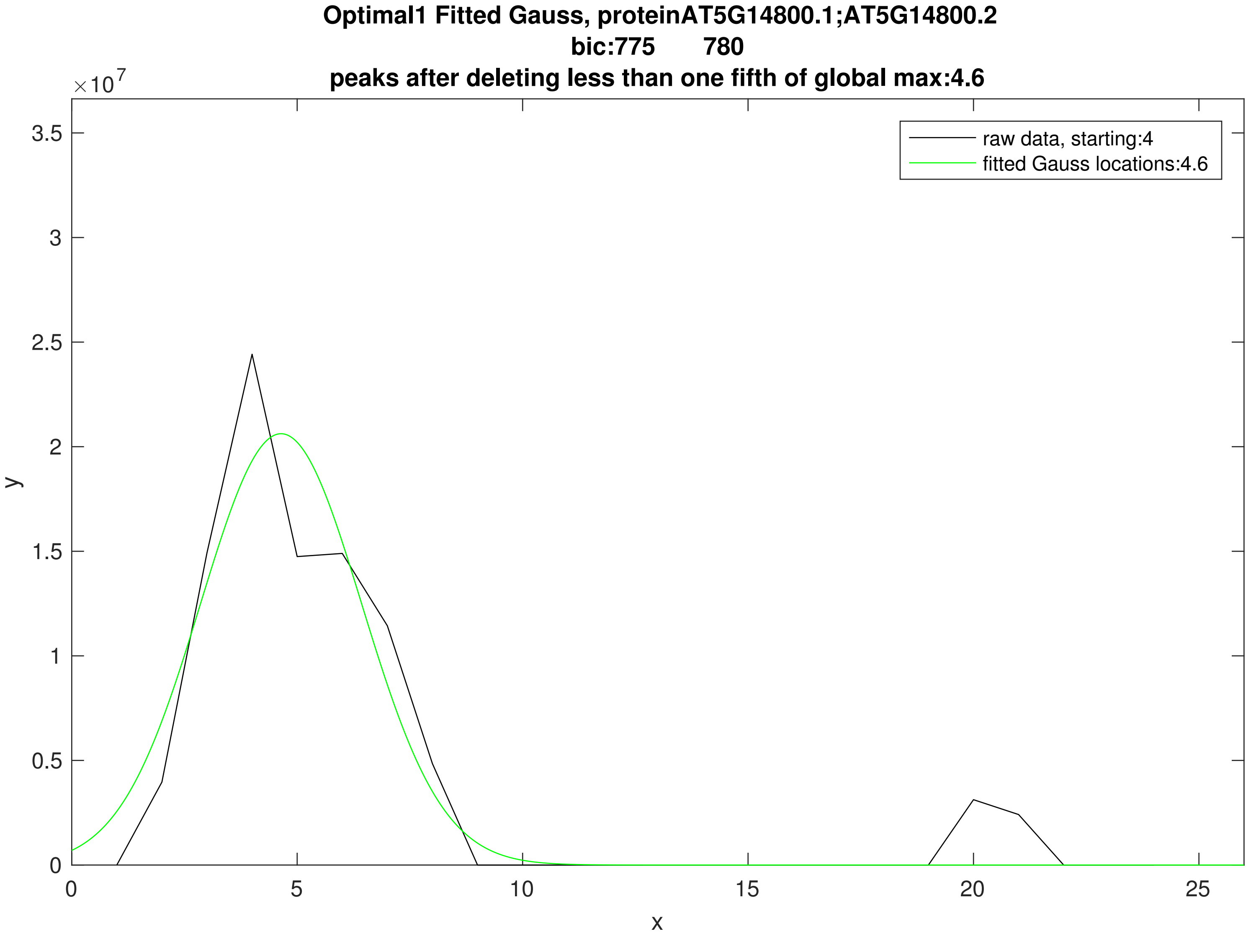
Optimal1 Fitted Gauss, proteinAT5G14800.1;AT5G14800.2
bic:775 780
peaks after deleting less than one fifth of global max:4.6
×10^7
3.5
3
2.5
2
1.5
1
0.5
0
y
0
5
10
15
20
25
x
raw data, starting:4
fitted Gauss locations:4.6

**Optimal1 Fitted Gauss, proteinAT5G14910.1**
**bic:796**
**peaks after deleting less than one fifth of global max:16**

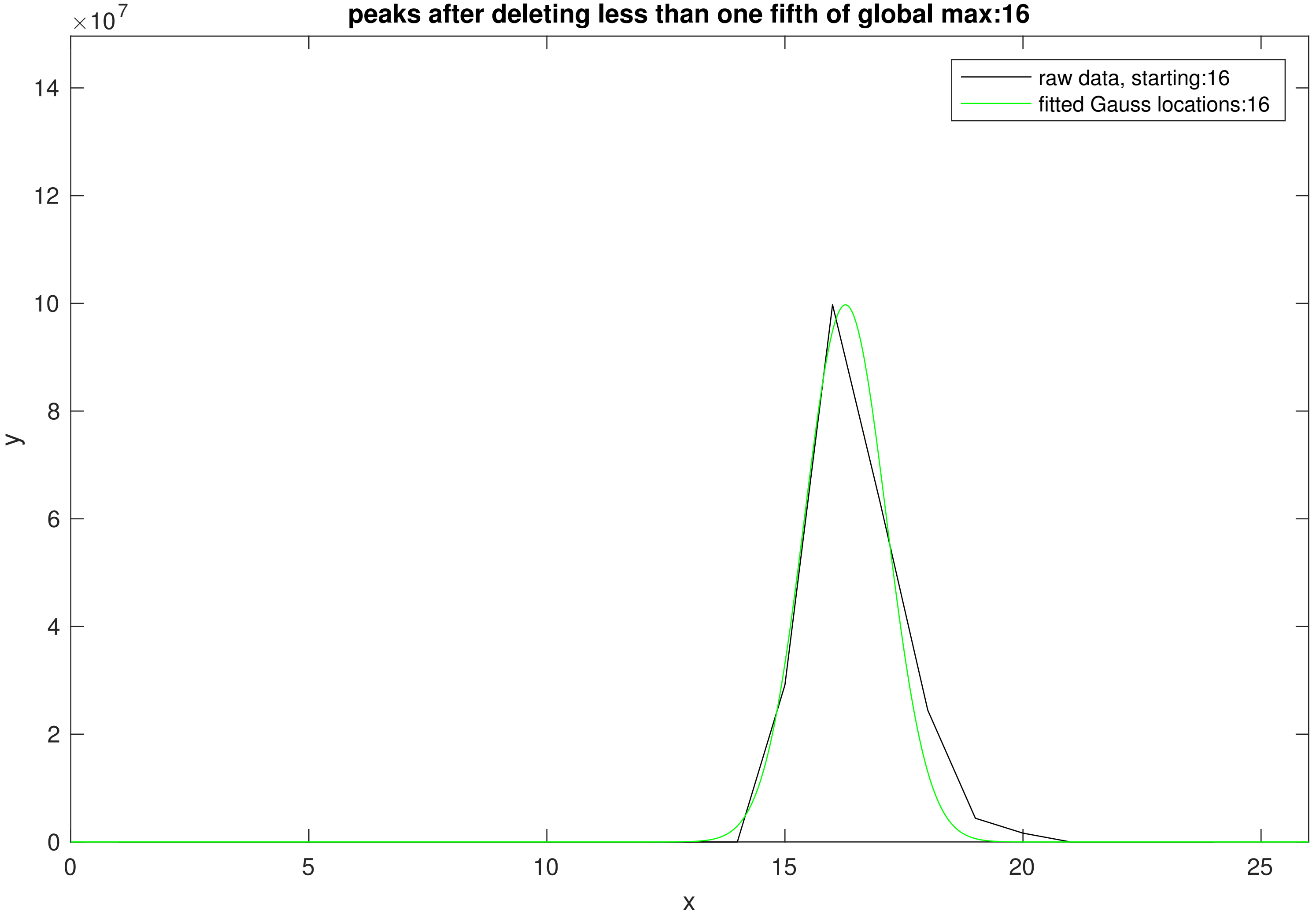

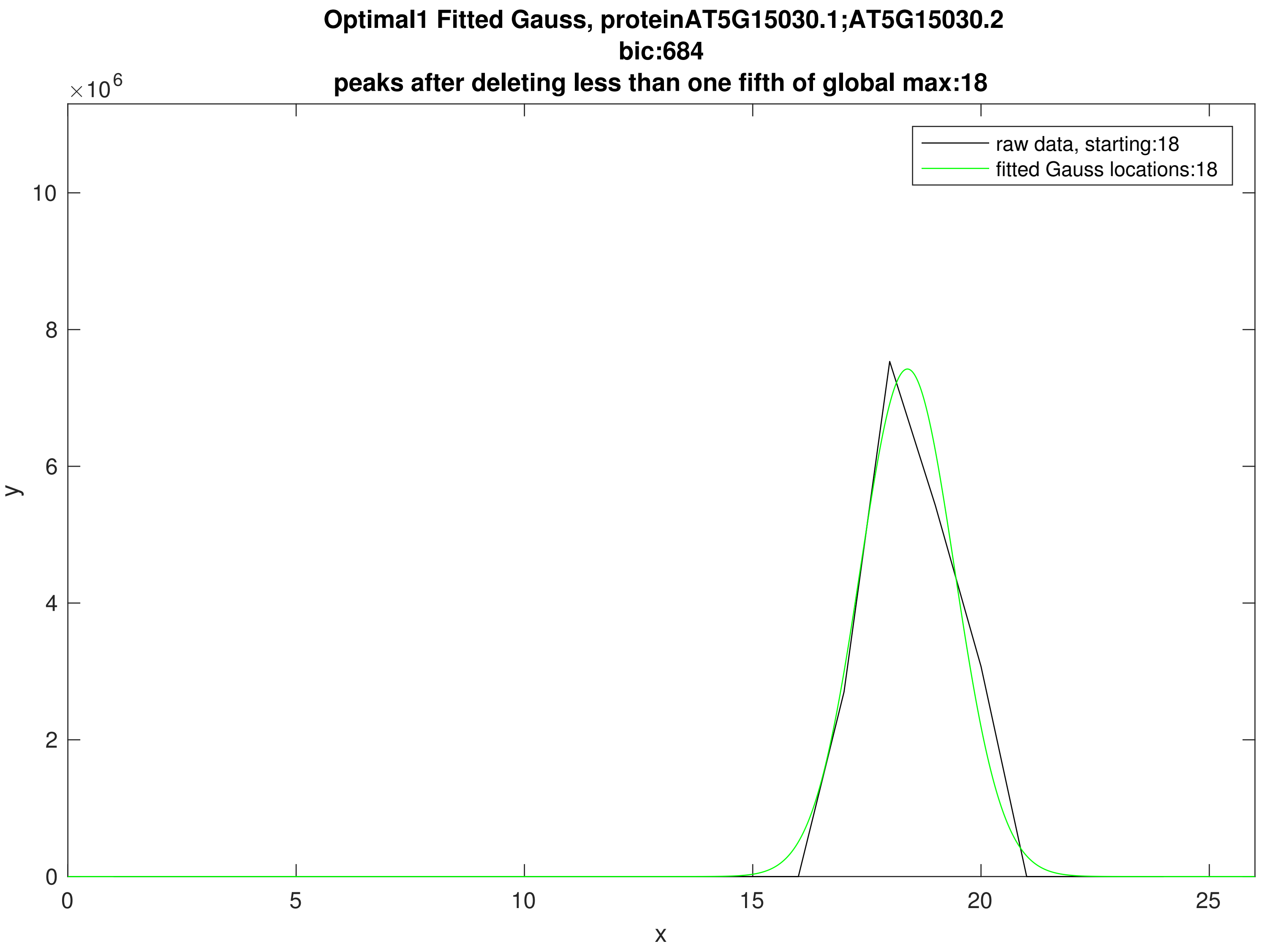

Optimal1 Fitted Gauss, proteinAT5G15030.1;AT5G15030.2
bic:684
peaks after deleting less than one fifth of global max:18
×10^6
raw data, starting:18
fitted Gauss locations:18
10
8
6
4
2
0
y
0
5
10
15
20
25
x

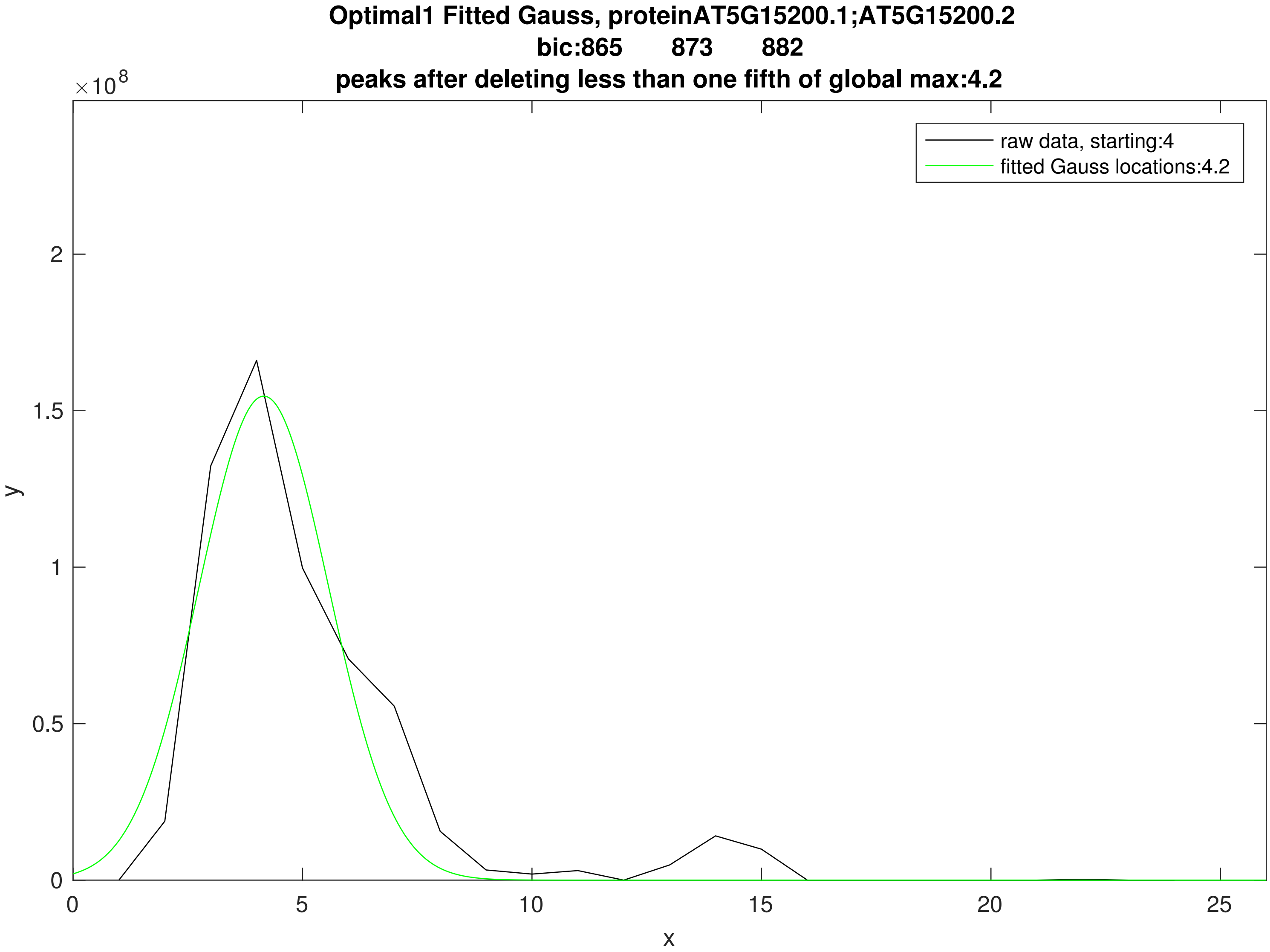
Optimal1 Fitted Gauss, proteinAT5G15200.1;AT5G15200.2
bic:865 873 882
peaks after deleting less than one fifth of global max:4.2
×10^8
raw data, starting:4
fitted Gauss locations:4.2
2
1.5
1
0.5
0
y
0
5
10
15
20
25
x

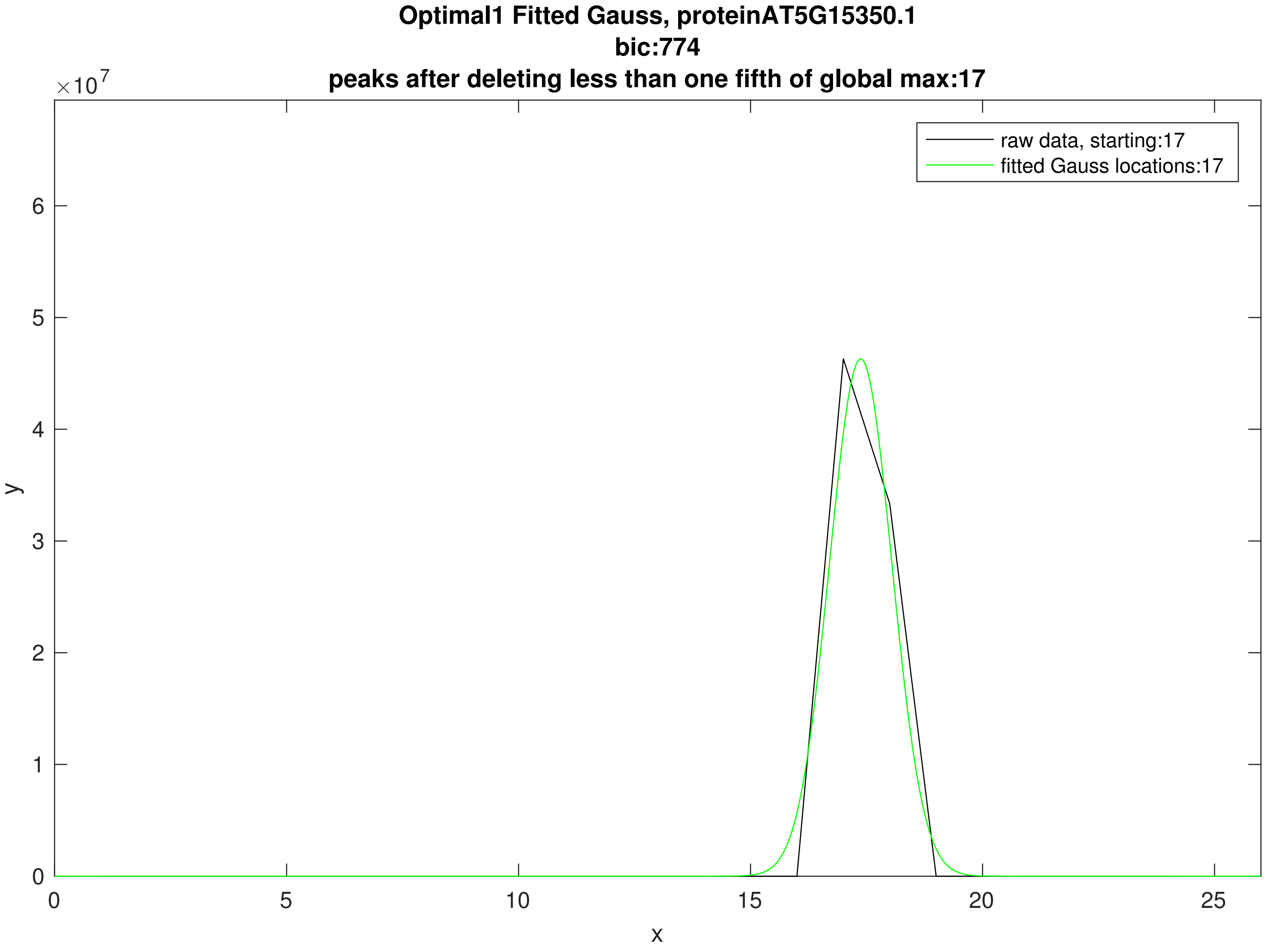
Optimal1 Fitted Gauss, proteinAT5G15350.1
bic:774
peaks after deleting less than one fifth of global max:17
×10^7
raw data, starting:17
fitted Gauss locations:17
y
x
0
1
2
3
4
5
6
0
5
10
15
20
25

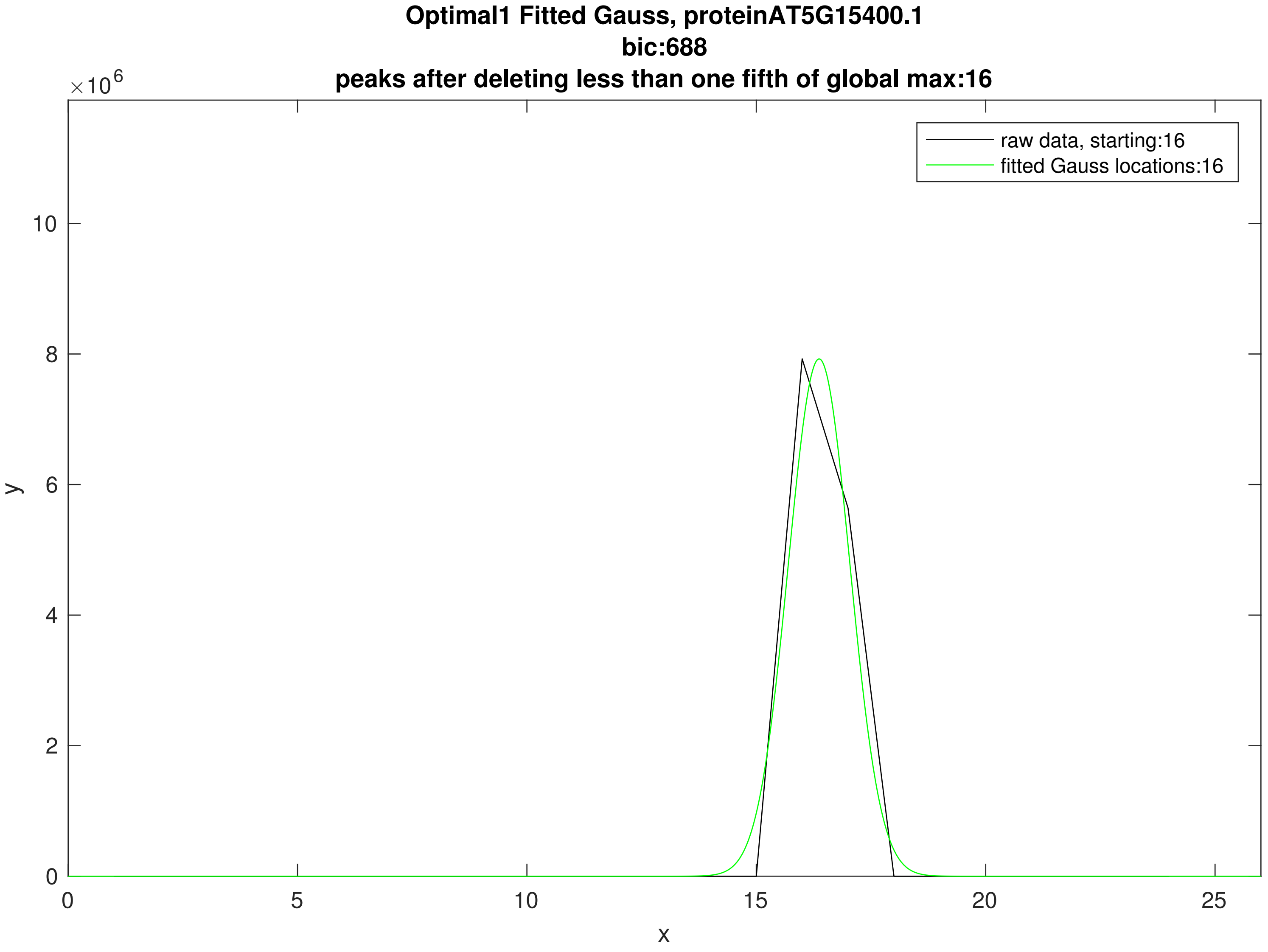
Optimal1 Fitted Gauss, proteinAT5G15400.1
bic:688
peaks after deleting less than one fifth of global max:16
×10^6
raw data, starting:16
fitted Gauss locations:16
10
8
6
4
2
0
y
0
5
10
15
20
25
x

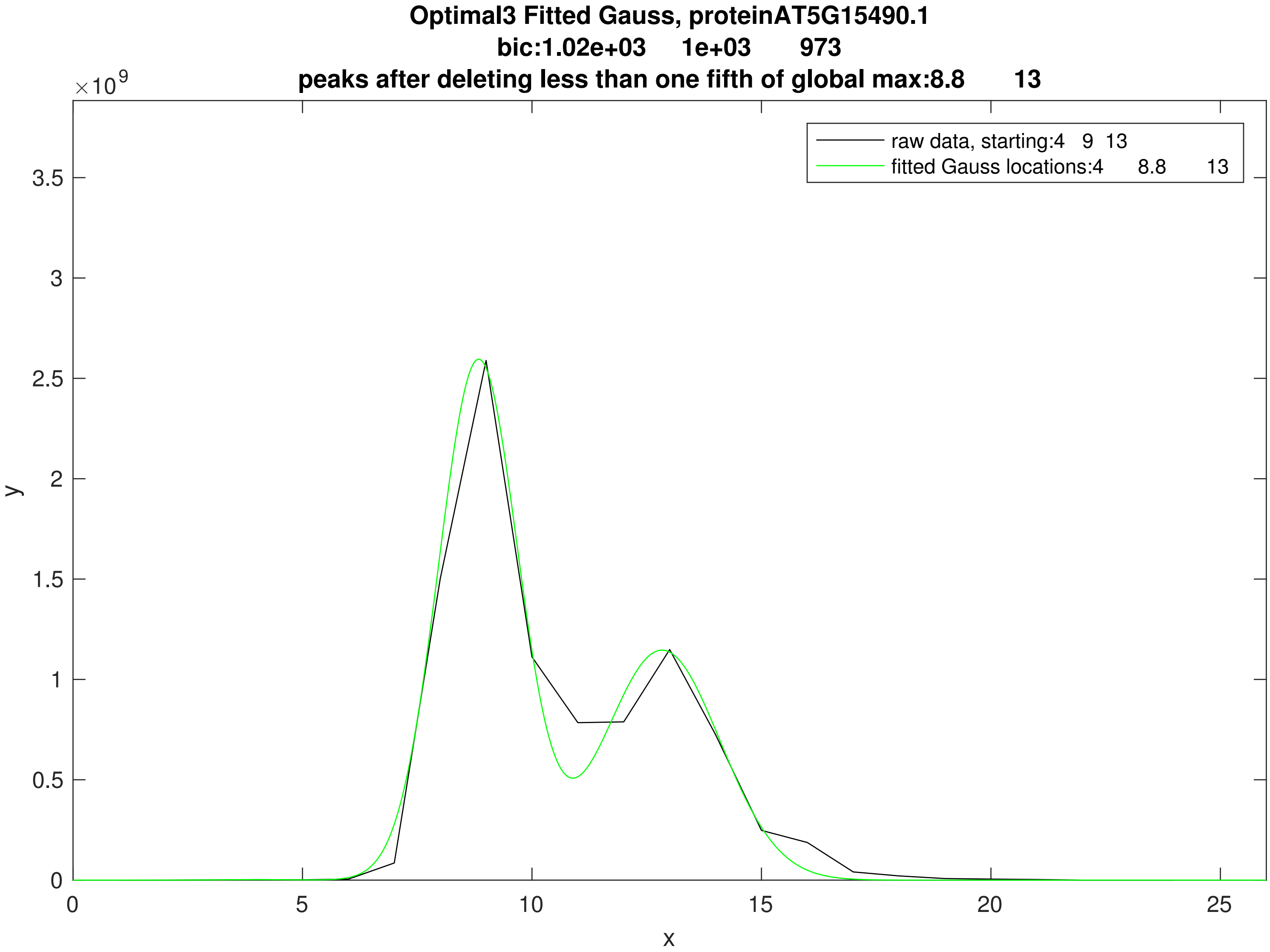
Optimal3 Fitted Gauss, proteinAT5G15490.1
bic:1.02e+03 1e+03 973
peaks after deleting less than one fifth of global max:8.8 13
×10^9
raw data, starting:4 9 13
fitted Gauss locations:4 8.8 13
3.5
3
2.5
2
1.5
1
0.5
0
y
0
5
10
15
20
25
x

**Optimal1 Fitted Gauss, proteinAT5G15650.1**
**bic:956 961 1.01e+03 1.02e+03**
**peaks after deleting less than one fifth of global max:5.9**

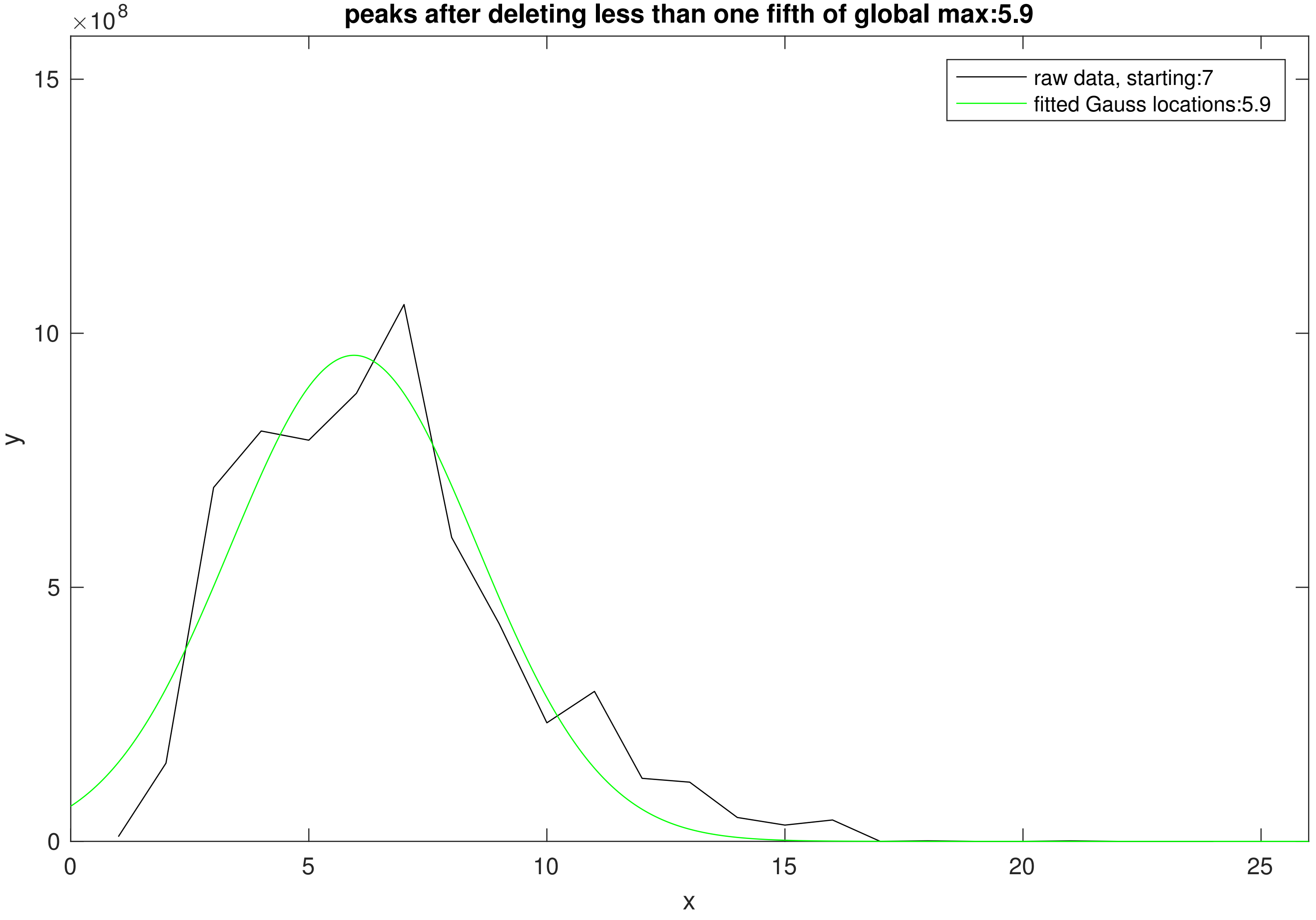

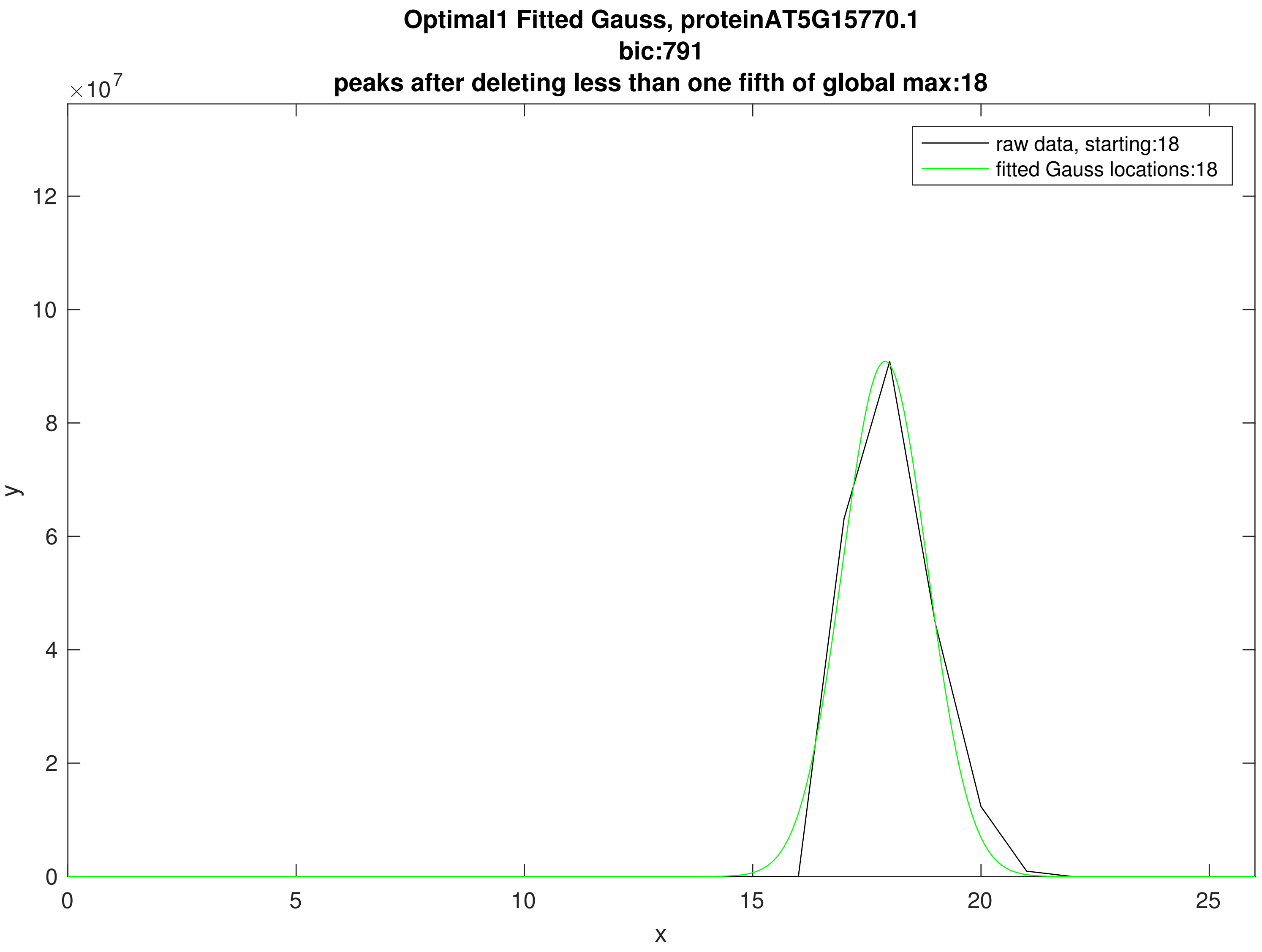
Optimal1 Fitted Gauss, proteinAT5G15770.1
bic:791
peaks after deleting less than one fifth of global max:18
×10^7
raw data, starting:18
fitted Gauss locations:18
12
10
8
6
4
2
0
y
0
5
10
15
20
25
x

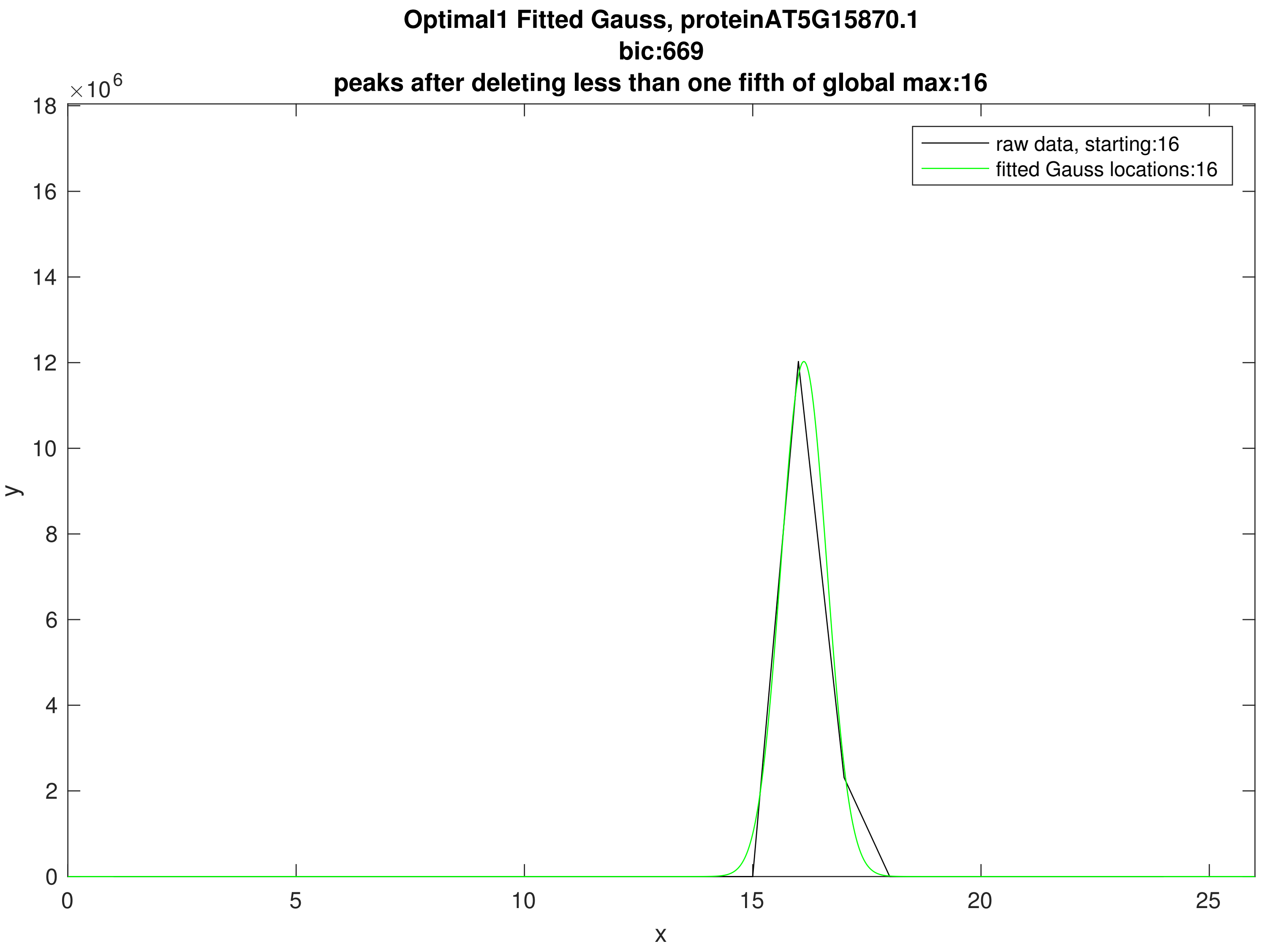

Optimal1 Fitted Gauss, proteinAT5G15870.1
bic:669
peaks after deleting less than one fifth of global max:16
×10^6
18
16
14
12
10
8
6
4
2
0
y
0
5
10
15
20
25
x
raw data, starting:16
fitted Gauss locations:16

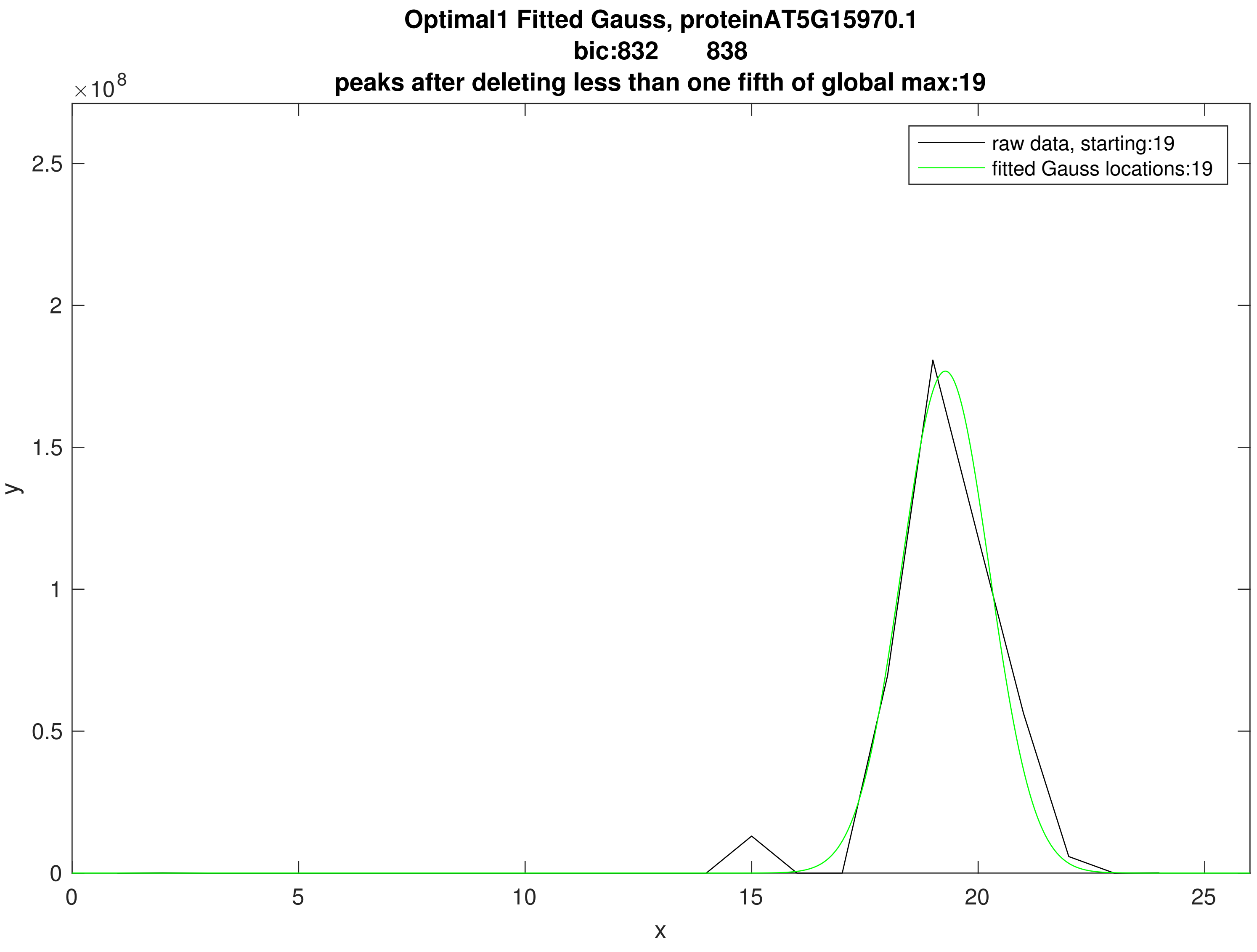
Optimal1 Fitted Gauss, proteinAT5G15970.1
bic:832 838
peaks after deleting less than one fifth of global max:19
×10^8
raw data, starting:19
fitted Gauss locations:19
2.5
2
1.5
1
0.5
0
y
0
5
10
15
20
25
x

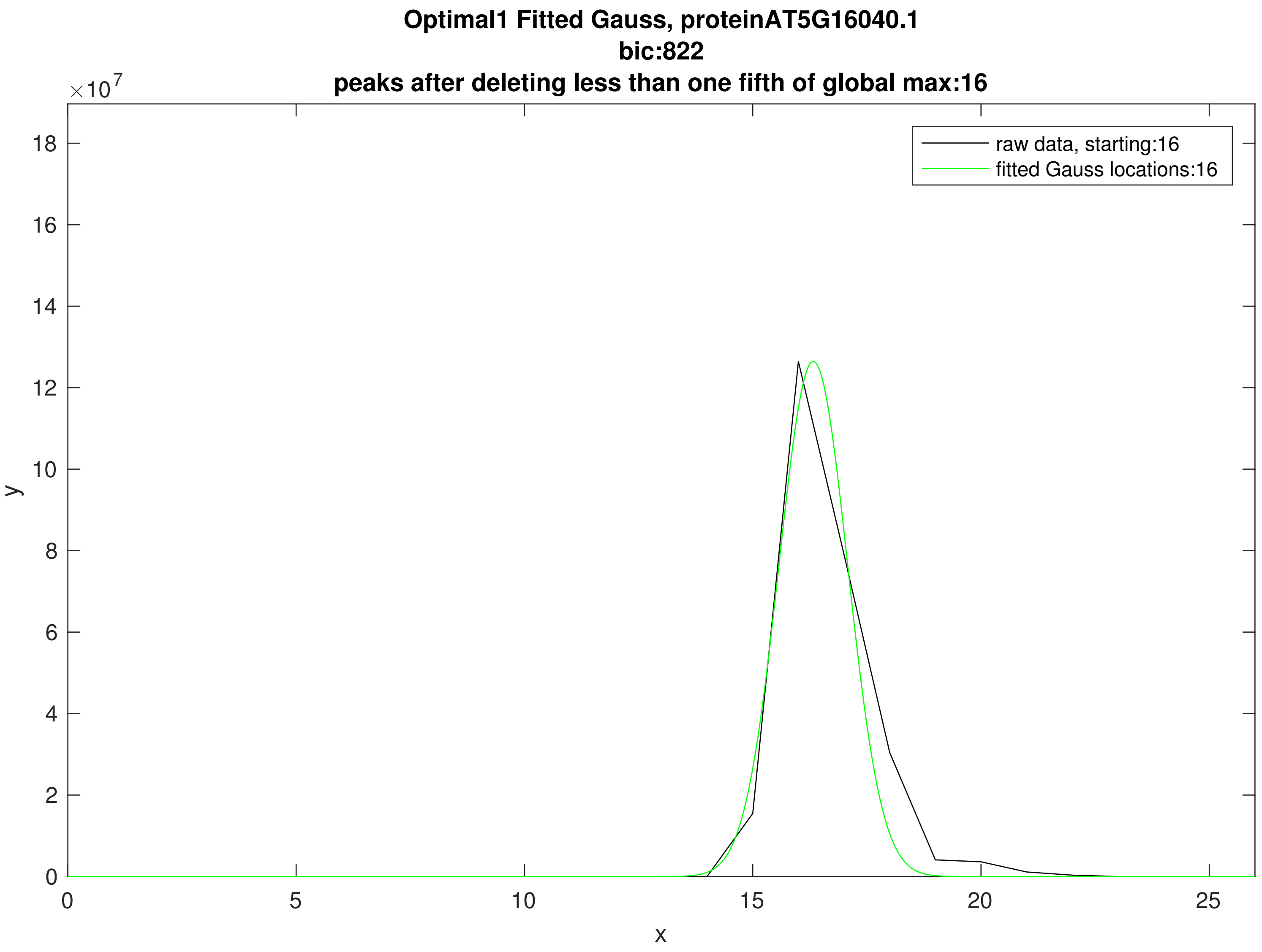
Optimal1 Fitted Gauss, proteinAT5G16040.1
bic:822
peaks after deleting less than one fifth of global max:16
×10^7
raw data, starting:16
fitted Gauss locations:16
y
x
0
2
4
6
8
10
12
14
16
18
0
5
10
15
20
25

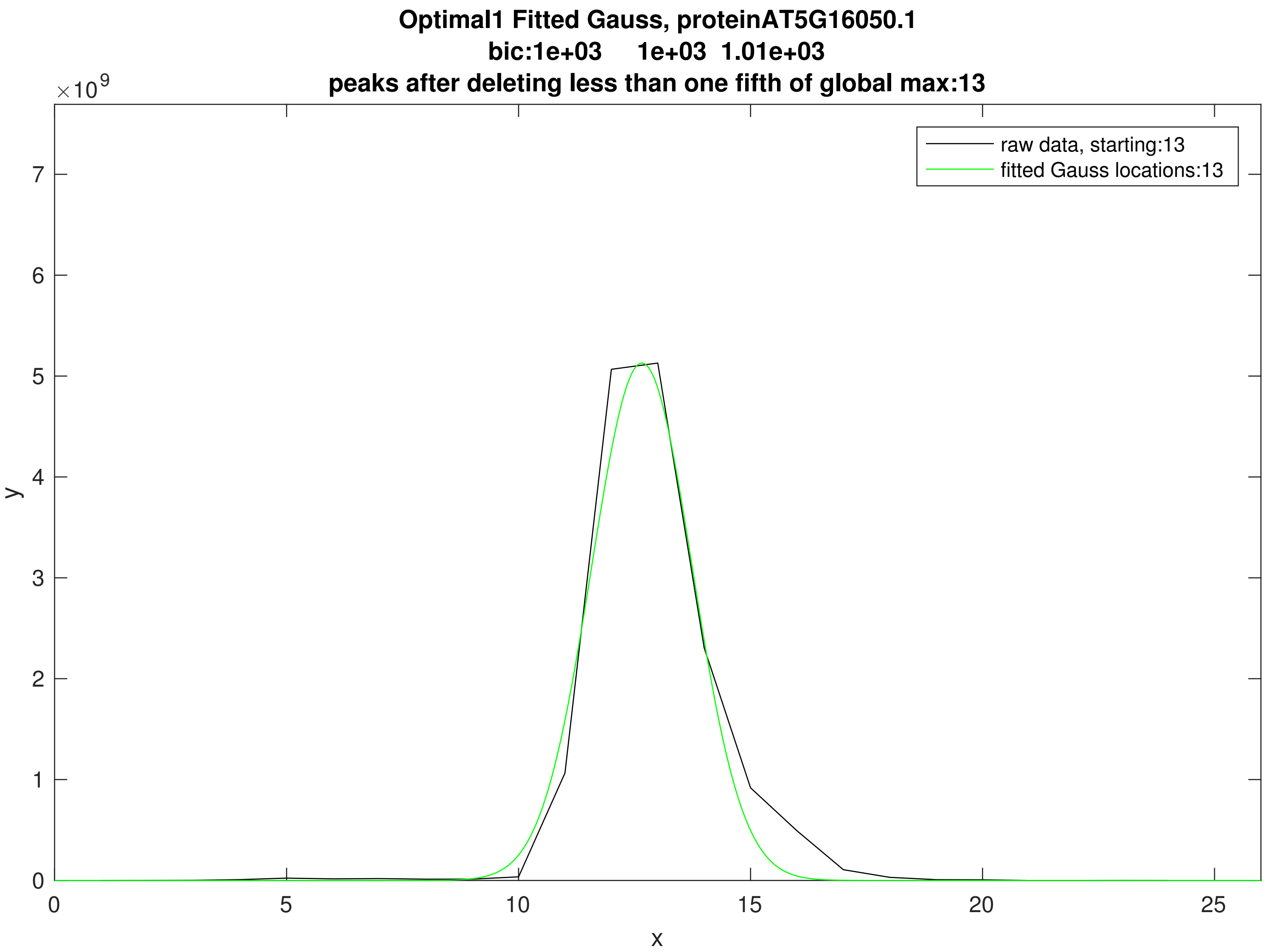
Optimal1 Fitted Gauss, proteinAT5G16050.1
bic:1e+03 1e+03 1.01e+03
peaks after deleting less than one fifth of global max:13
×10^9
raw data, starting:13
fitted Gauss locations:13
y
x
0
1
2
3
4
5
6
7
0
5
10
15
20
25

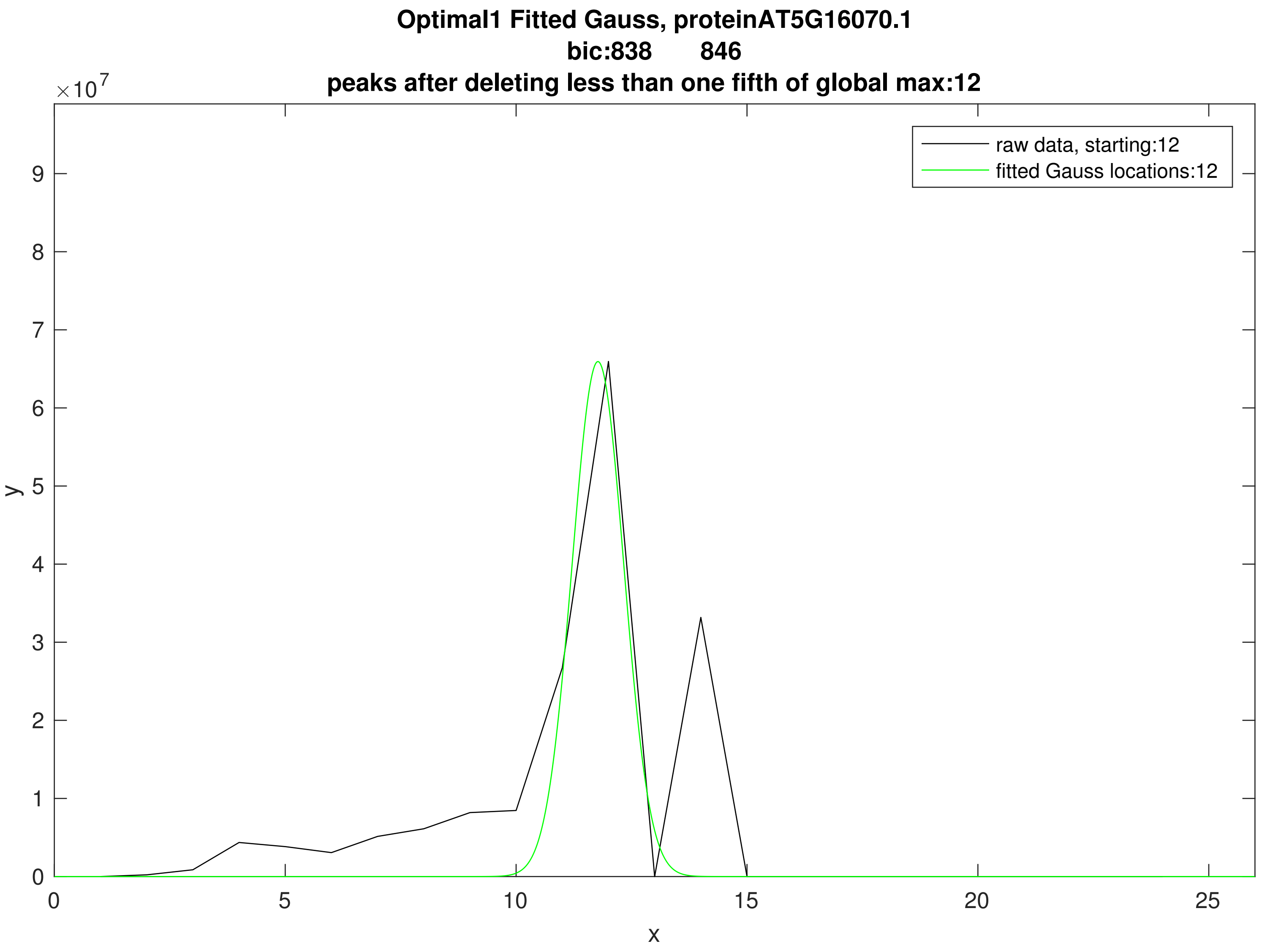

Optimal1 Fitted Gauss, proteinAT5G16070.1
bic:838 846
peaks after deleting less than one fifth of global max:12
×10^7
raw data, starting:12
fitted Gauss locations:12
y
x
0
1
2
3
4
5
6
7
8
9
0
5
10
15
20
25

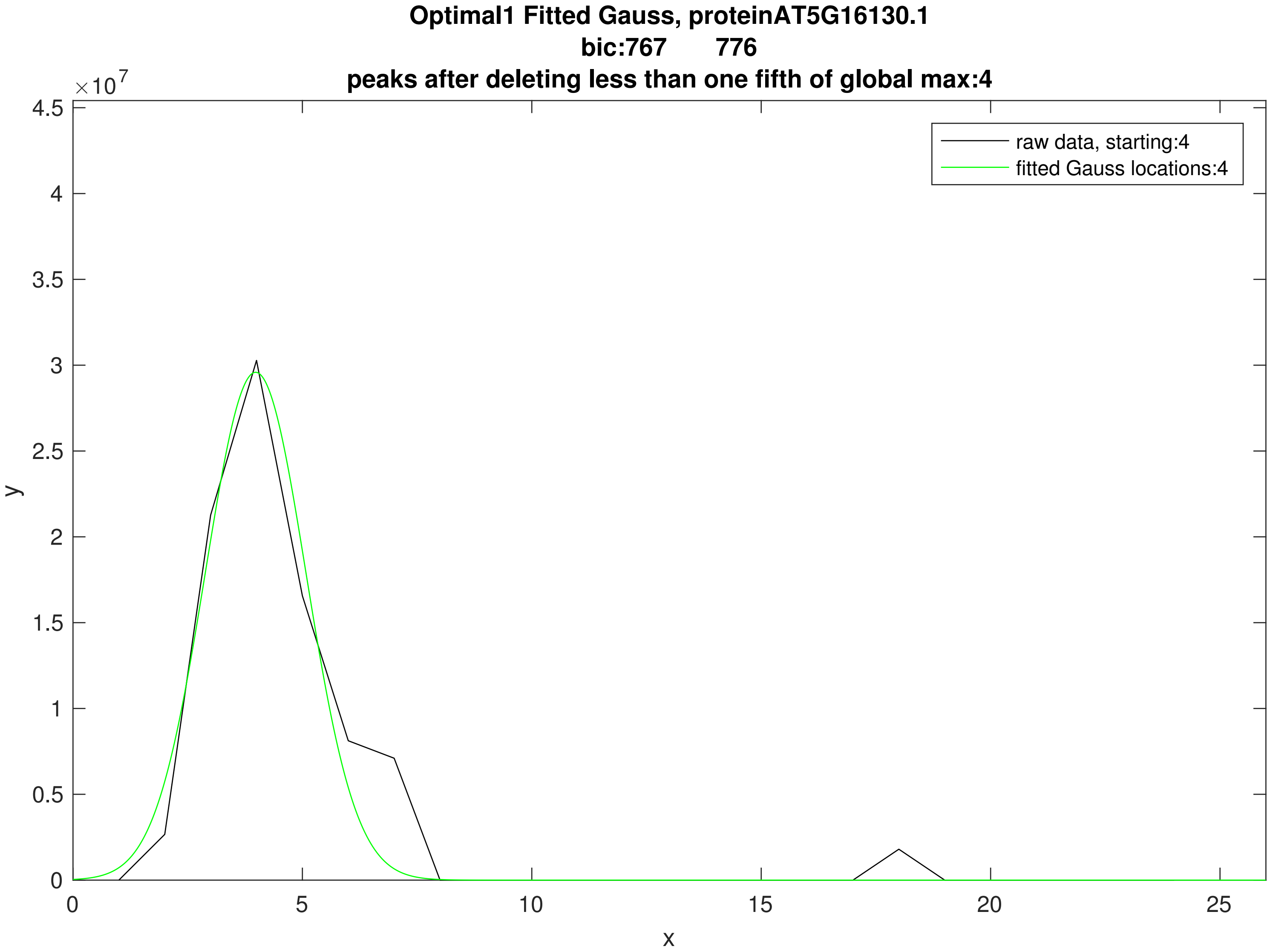

Optimal1 Fitted Gauss, proteinAT5G16130.1
bic:767 776
peaks after deleting less than one fifth of global max:4
×10^7
4.5
4
3.5
3
2.5
2
1.5
1
0.5
0
y
0
5
10
15
20
25
x
raw data, starting:4
fitted Gauss locations:4

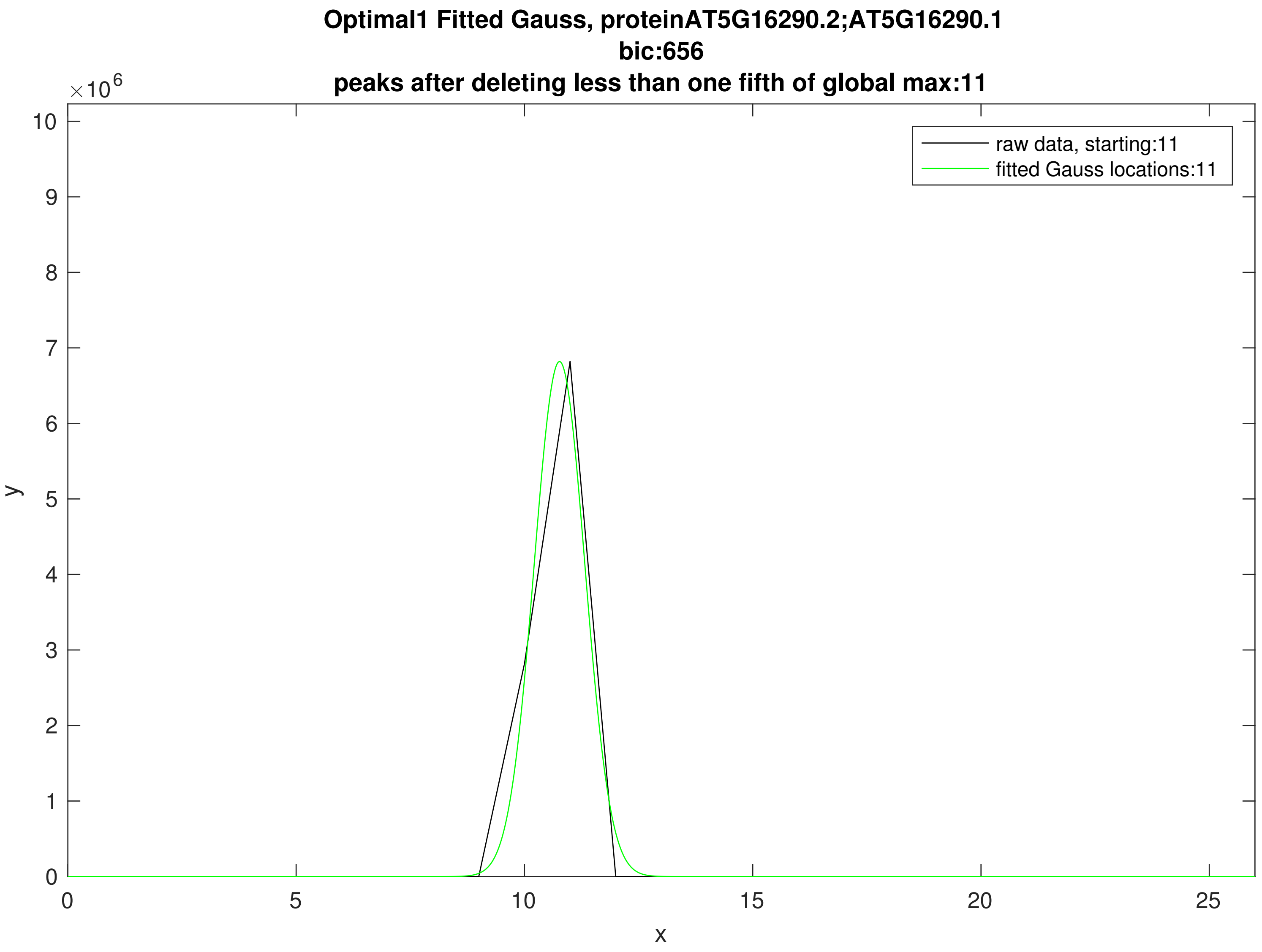

Optimal1 Fitted Gauss, proteinAT5G16290.2;AT5G16290.1
bic:656
peaks after deleting less than one fifth of global max:11
×10^6
10
9
8
7
6
5
4
3
2
1
0
y
0
5
10
15
20
25
x
raw data, starting:11
fitted Gauss locations:11

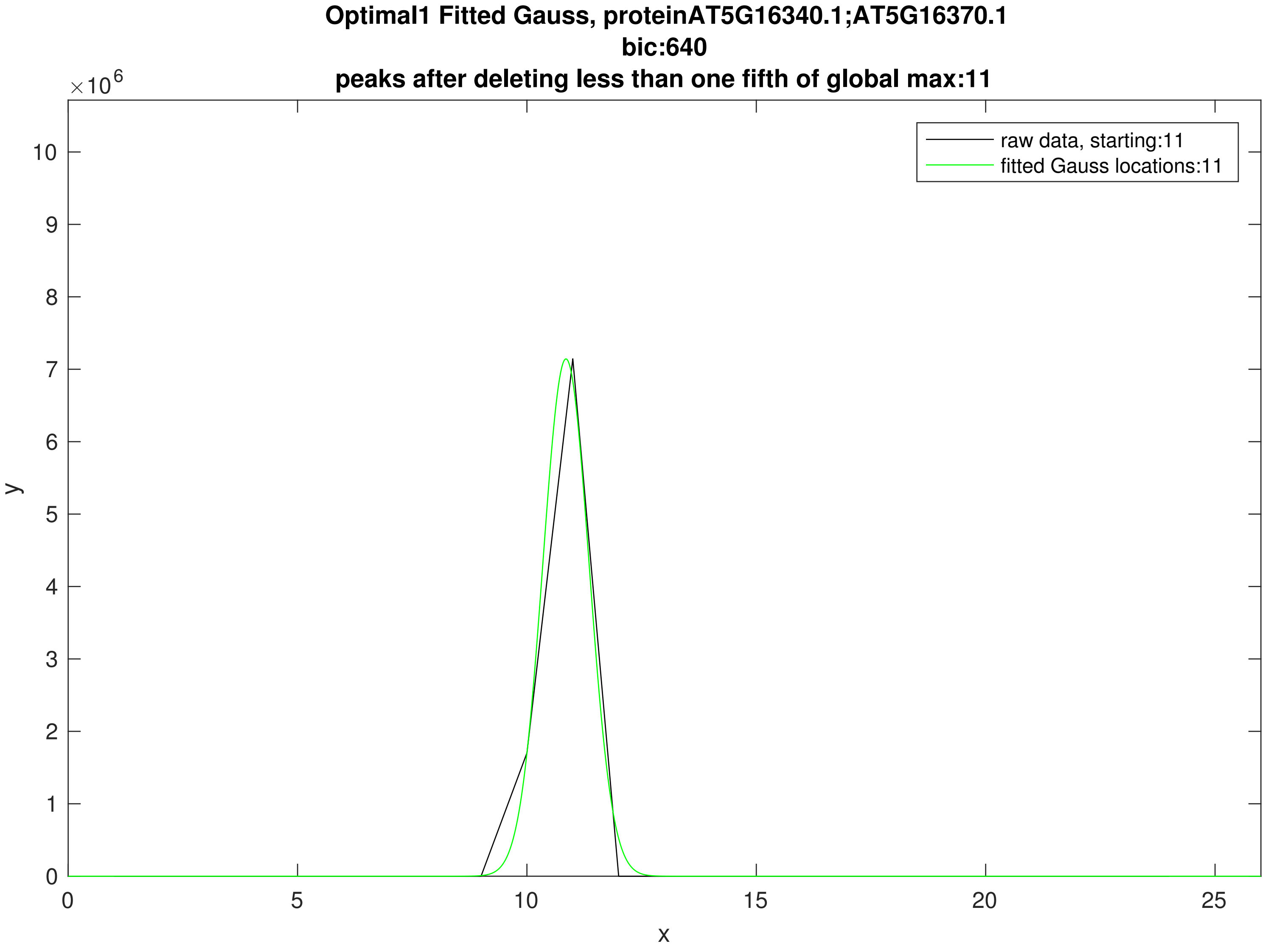
Optimal1 Fitted Gauss, proteinAT5G16340.1;AT5G16370.1
bic:640
peaks after deleting less than one fifth of global max:11
×10^6
raw data, starting:11
fitted Gauss locations:11
y
x
0
1
2
3
4
5
6
7
8
9
10
0
5
10
15
20
25

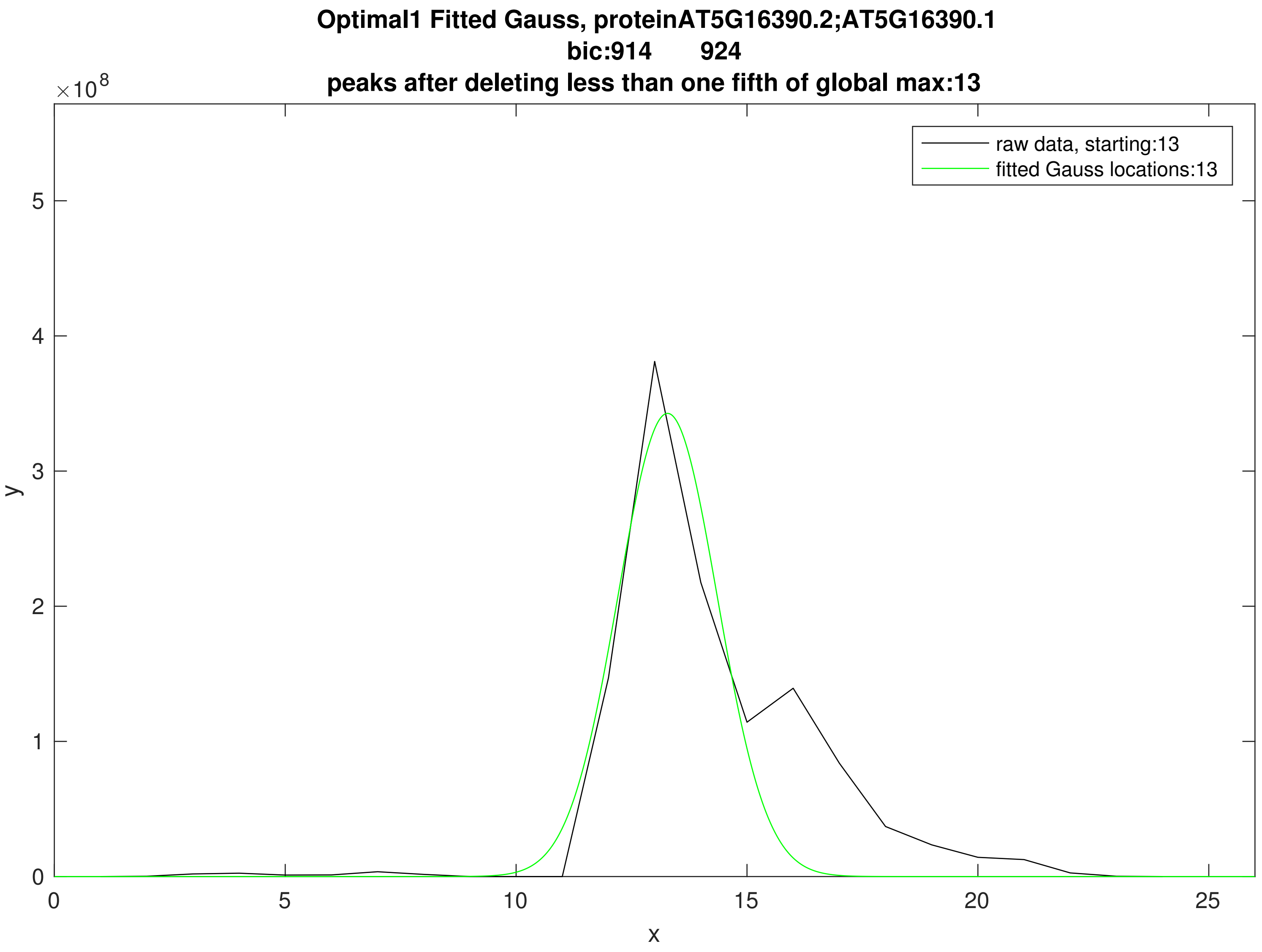
Optimal1 Fitted Gauss, proteinAT5G16390.2;AT5G16390.1
bic:914 924
peaks after deleting less than one fifth of global max:13
×10^8
raw data, starting:13
fitted Gauss locations:13
y
x
0
1
2
3
4
5
0
5
10
15
20
25

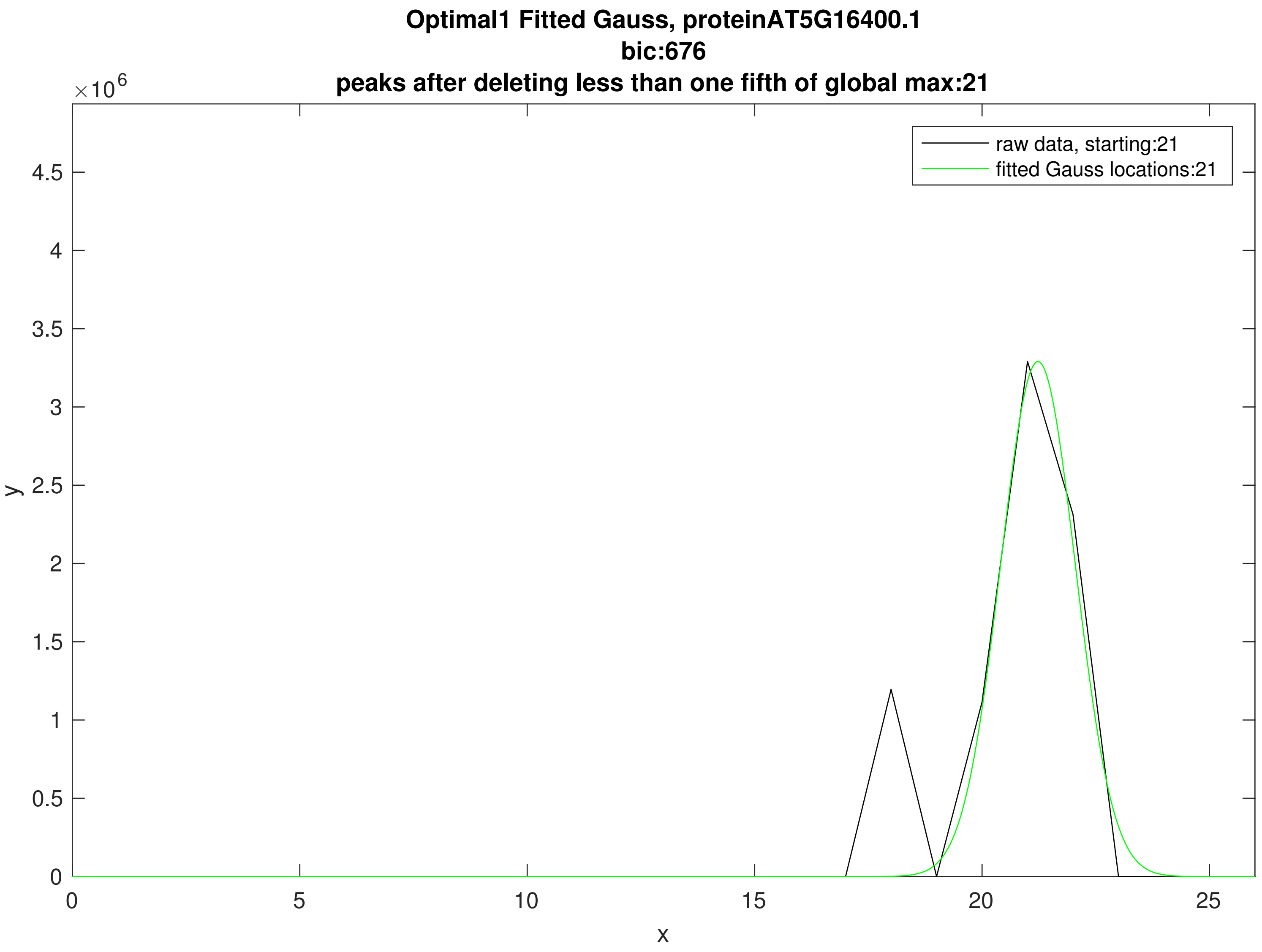
Optimal1 Fitted Gauss, proteinAT5G16400.1
bic:676
peaks after deleting less than one fifth of global max:21
×10^6
raw data, starting:21
fitted Gauss locations:21
y
x
0
0.5
1
1.5
2
2.5
3
3.5
4
4.5
0
5
10
15
20
25

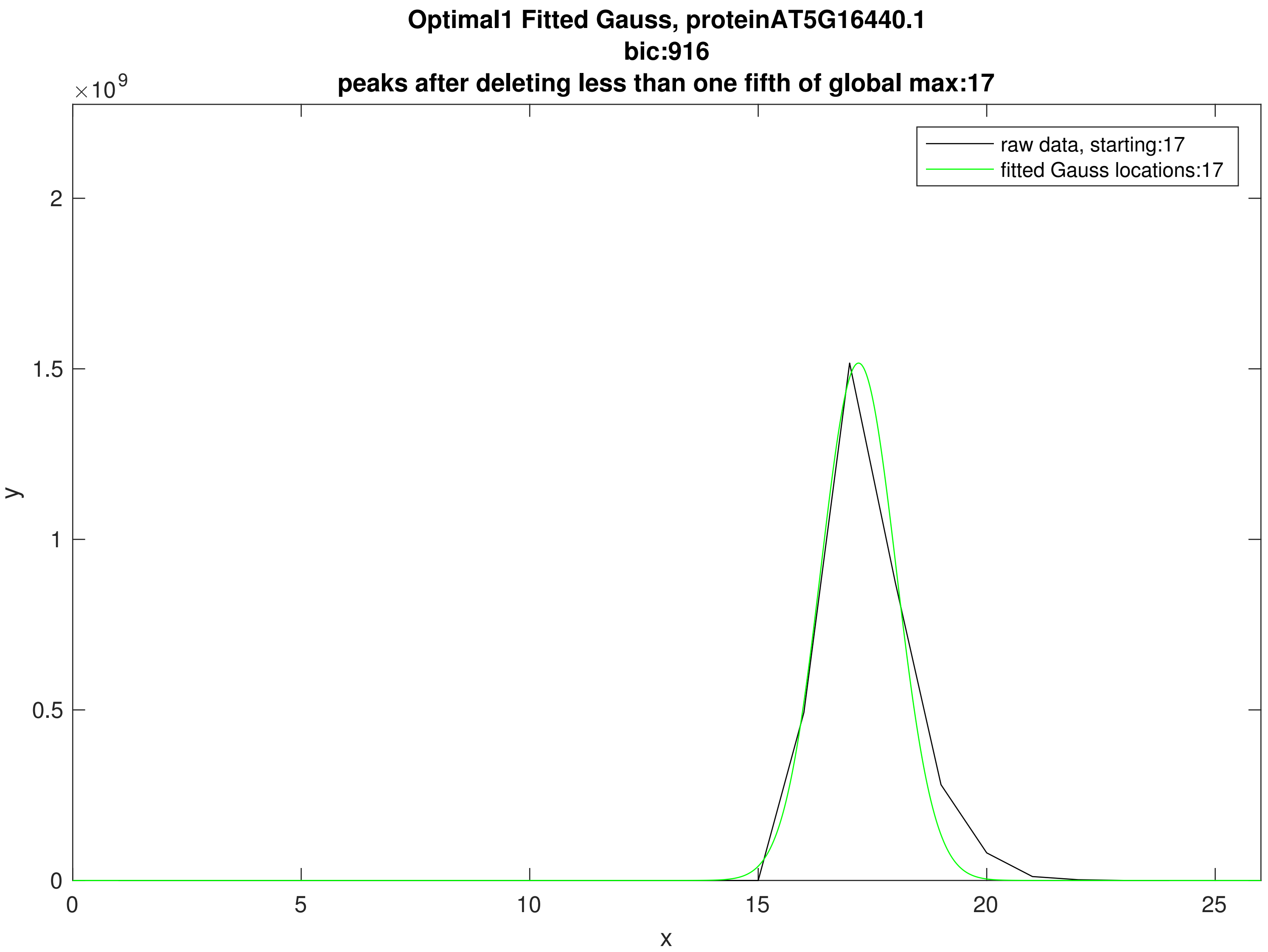
Optimal1 Fitted Gauss, proteinAT5G16440.1
bic:916
peaks after deleting less than one fifth of global max:17
×10^9
raw data, starting:17
fitted Gauss locations:17
y
x
0
0.5
1
1.5
2
0
5
10
15
20
25

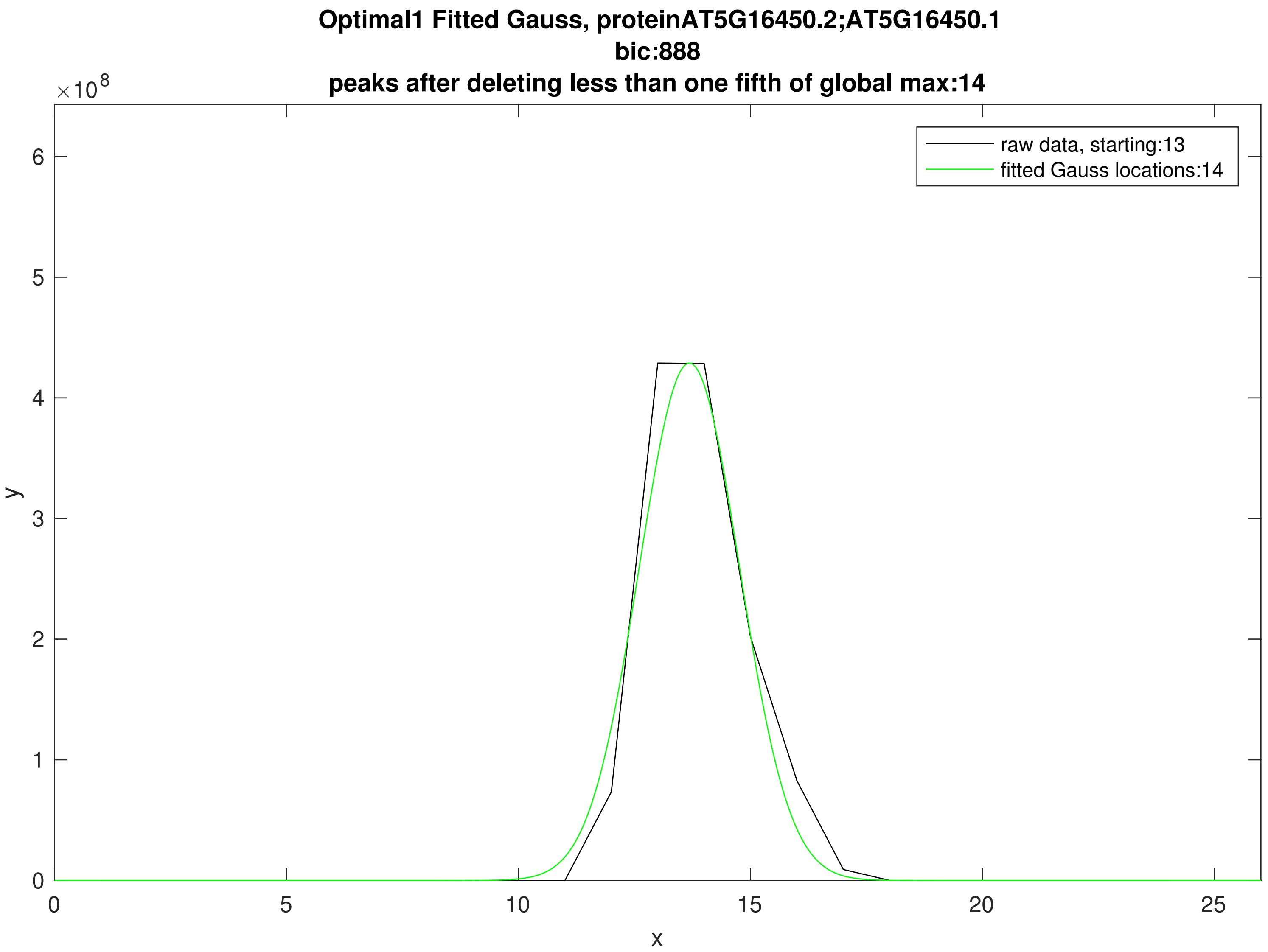
Optimal1 Fitted Gauss, proteinAT5G16450.2;AT5G16450.1
bic:888
peaks after deleting less than one fifth of global max:14
×10^8
raw data, starting:13
fitted Gauss locations:14
y
x
0
1
2
3
4
5
6
0
5
10
15
20
25

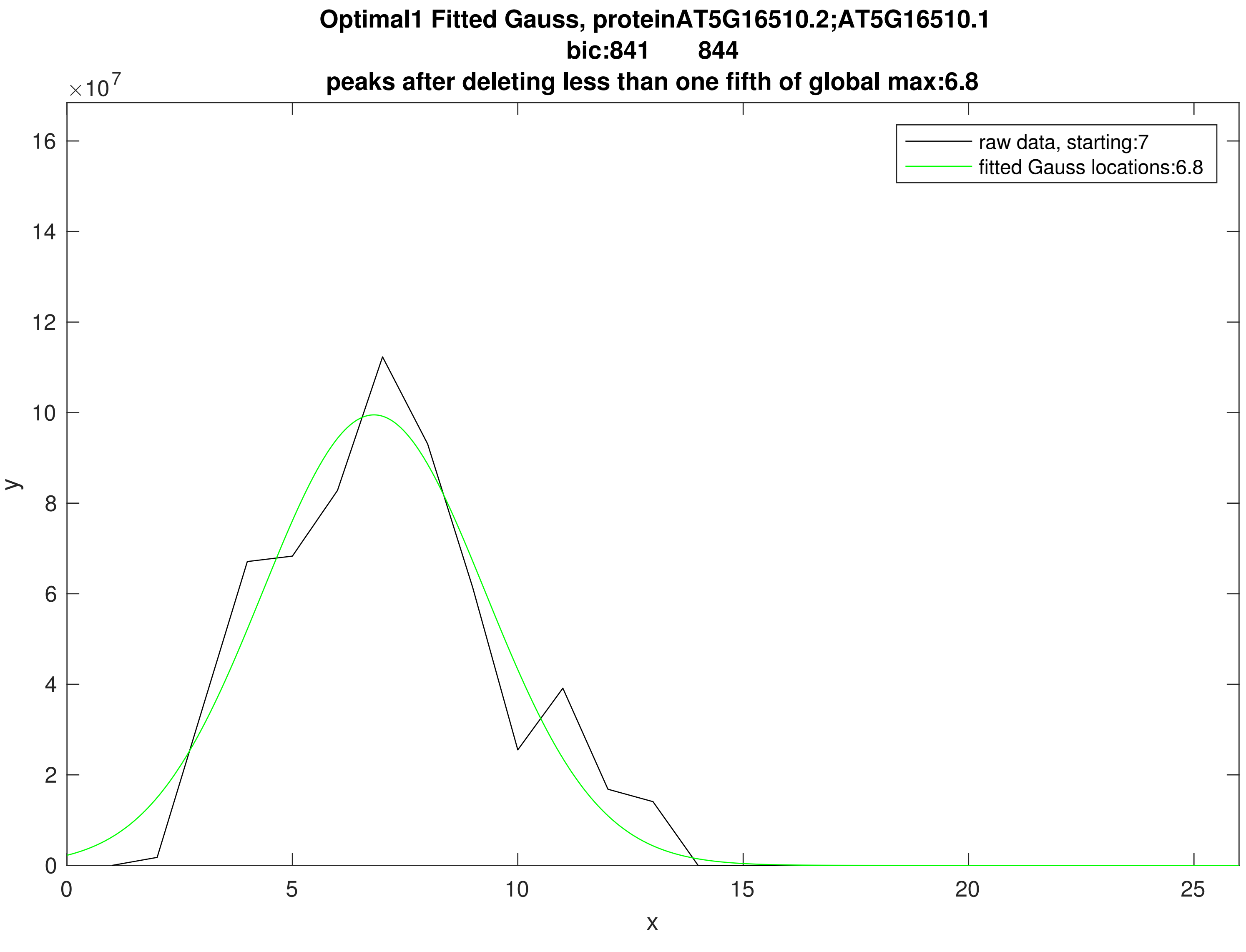

Optimal1 Fitted Gauss, proteinAT5G16510.2;AT5G16510.1
bic:841 844
peaks after deleting less than one fifth of global max:6.8
×10^7
raw data, starting:7
fitted Gauss locations:6.8
y
x
0
2
4
6
8
10
12
14
16
5
15
20
25

Optimal1 Fitted Gauss, proteinAT5G16590.1
bic:842 846 855
peaks after deleting less than one fifth of global max:16

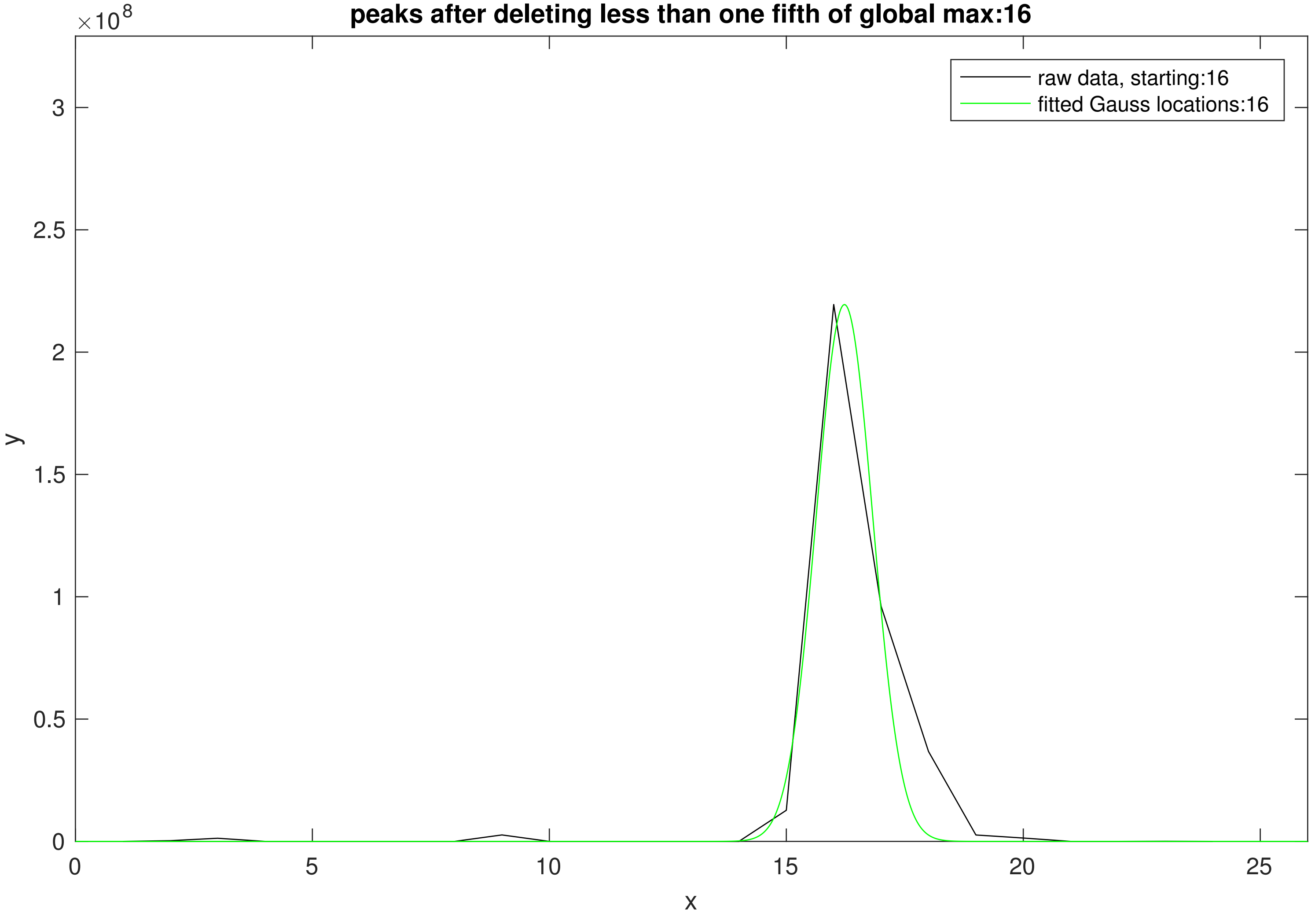

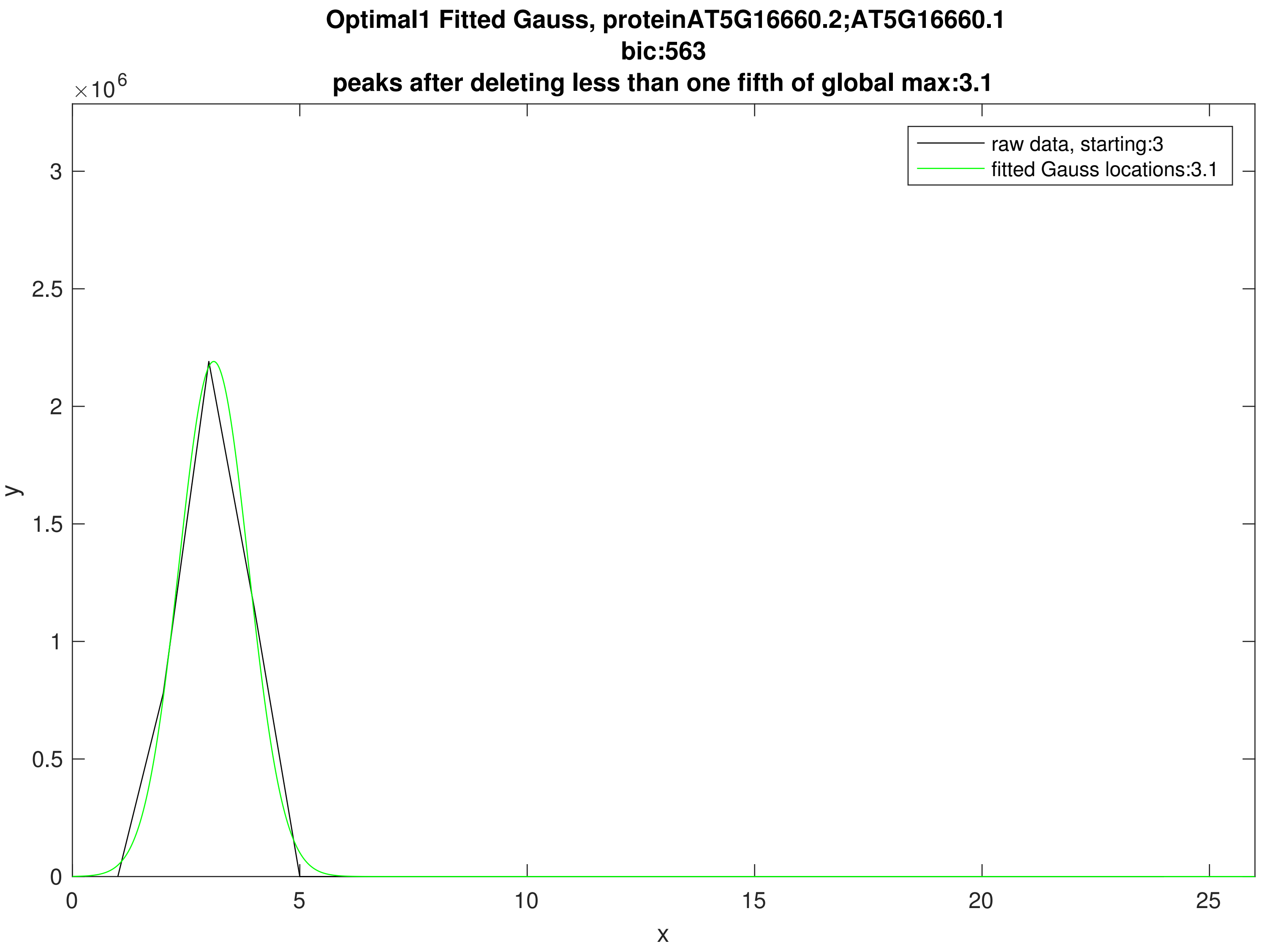

Optimal1 Fitted Gauss, proteinAT5G16660.2;AT5G16660.1
bic:563
peaks after deleting less than one fifth of global max:3.1
×10^6
raw data, starting:3
fitted Gauss locations:3.1
y
x
0
0.5
1
1.5
2
2.5
3
0
5
10
15
20
25

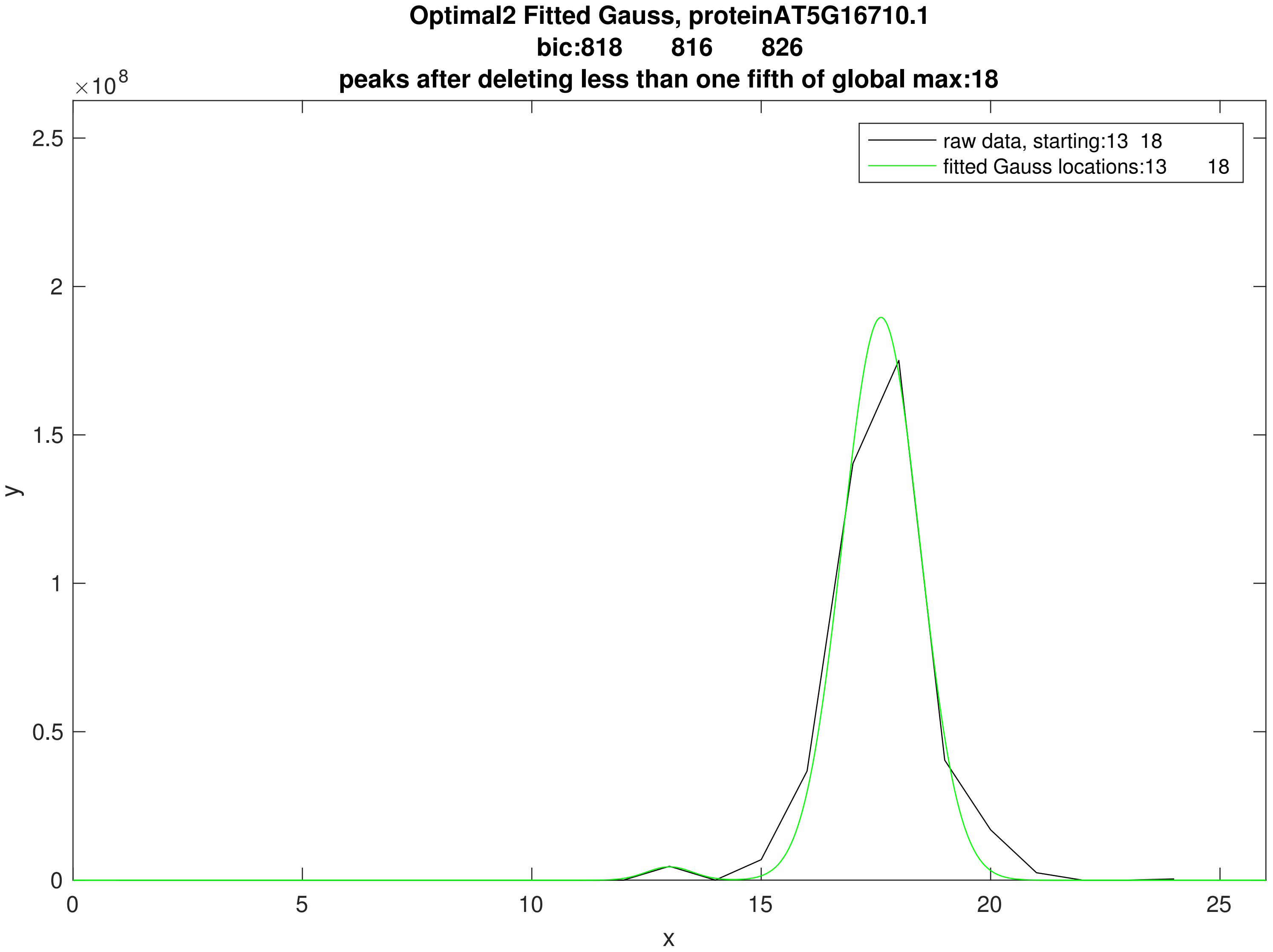
Optimal2 Fitted Gauss, proteinAT5G16710.1
bic:818 816 826
peaks after deleting less than one fifth of global max:18
×10^8
2.5
2
1.5
1
0.5
0
y
0
5
10
15
20
25
x
raw data, starting:13 18
fitted Gauss locations:13 18

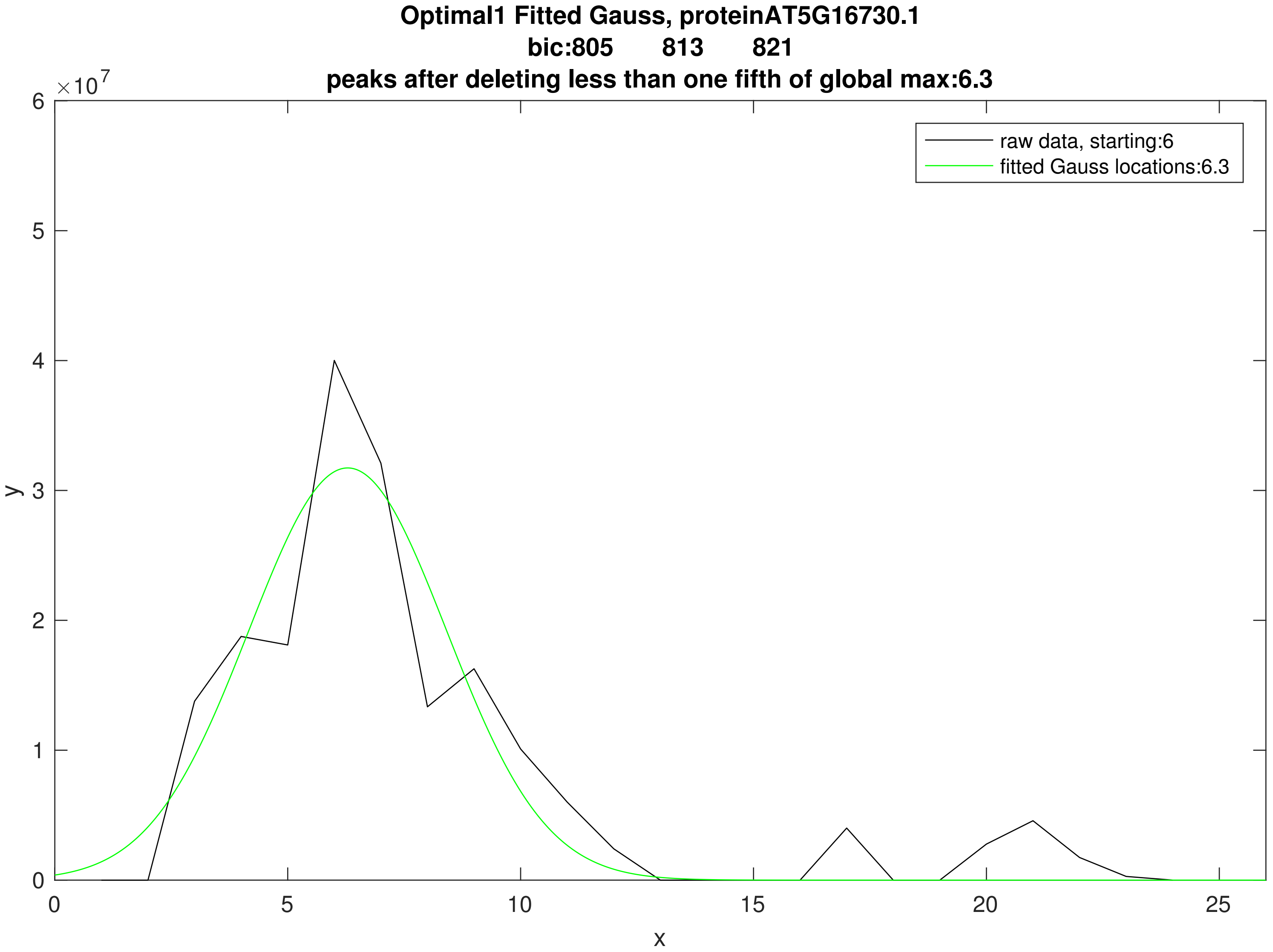
Optimal1 Fitted Gauss, proteinAT5G16730.1
bic:805 813 821
peaks after deleting less than one fifth of global max:6.3
×10^7
raw data, starting:6
fitted Gauss locations:6.3
y
x
0
1
2
3
4
5
6
0
5
10
15
20
25

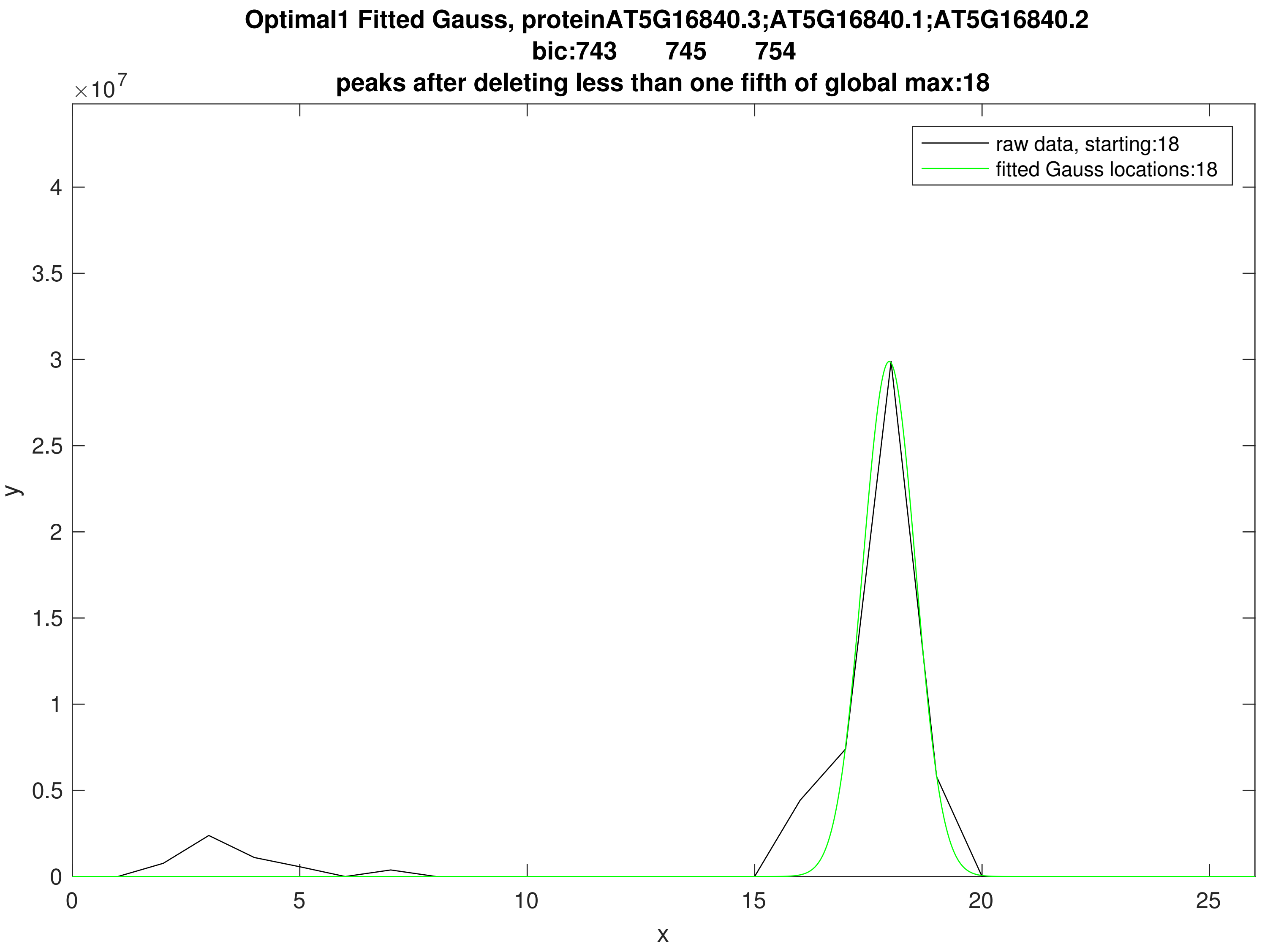

Optimal1 Fitted Gauss, proteinAT5G16840.3;AT5G16840.1;AT5G16840.2
bic:743 745 754
peaks after deleting less than one fifth of global max:18
×10^7
raw data, starting:18
fitted Gauss locations:18
y
x
0
0.5
1
1.5
2
2.5
3
3.5
4
5
10
15
20
25

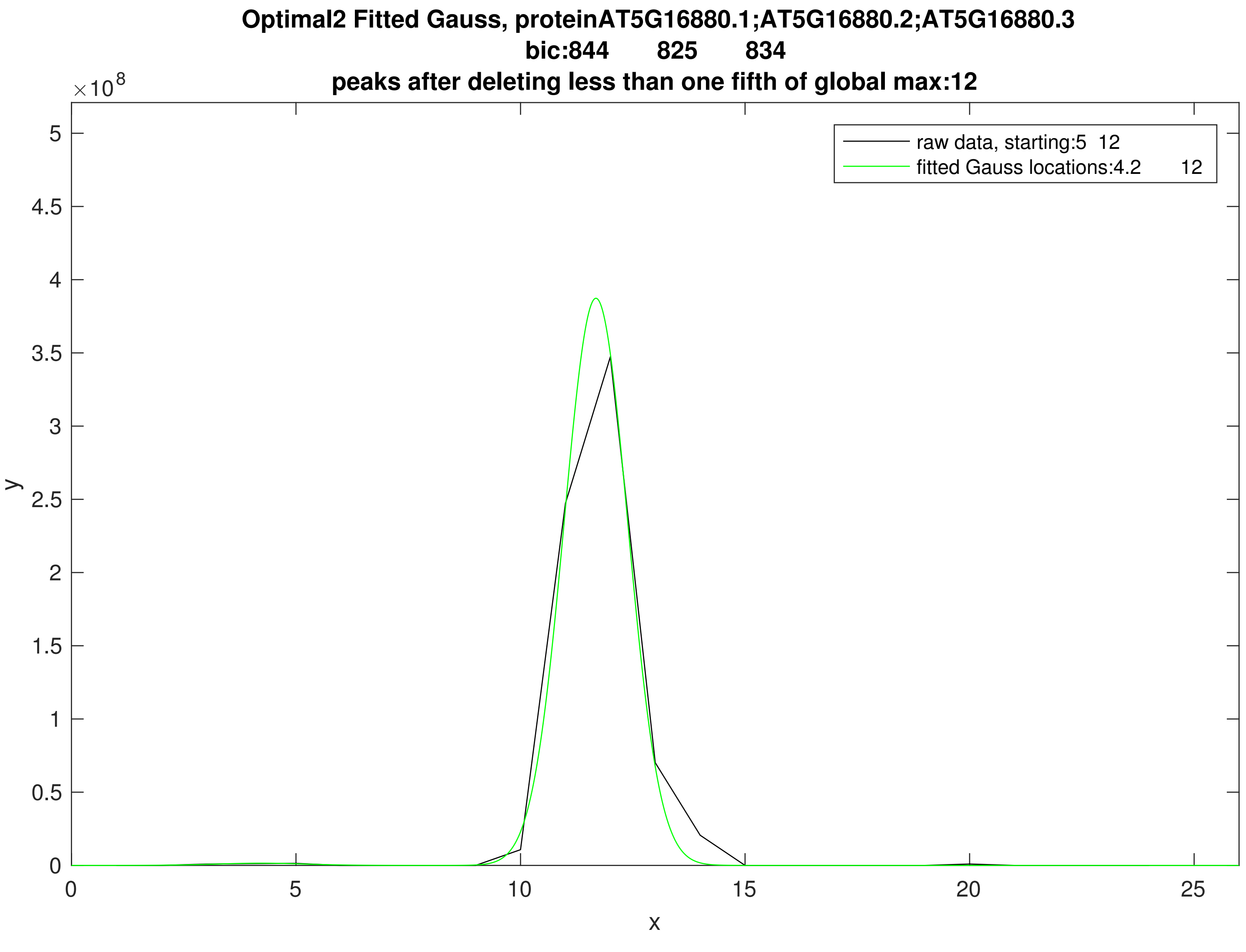
Optimal2 Fitted Gauss, proteinAT5G16880.1;AT5G16880.2;AT5G16880.3
bic:844 825 834
peaks after deleting less than one fifth of global max:12
×10^8
raw data, starting:5 12
fitted Gauss locations:4.2 12
y
x
0
0.5
1
1.5
2
2.5
3
3.5
4
4.5
5
0
5
10
15
20
25

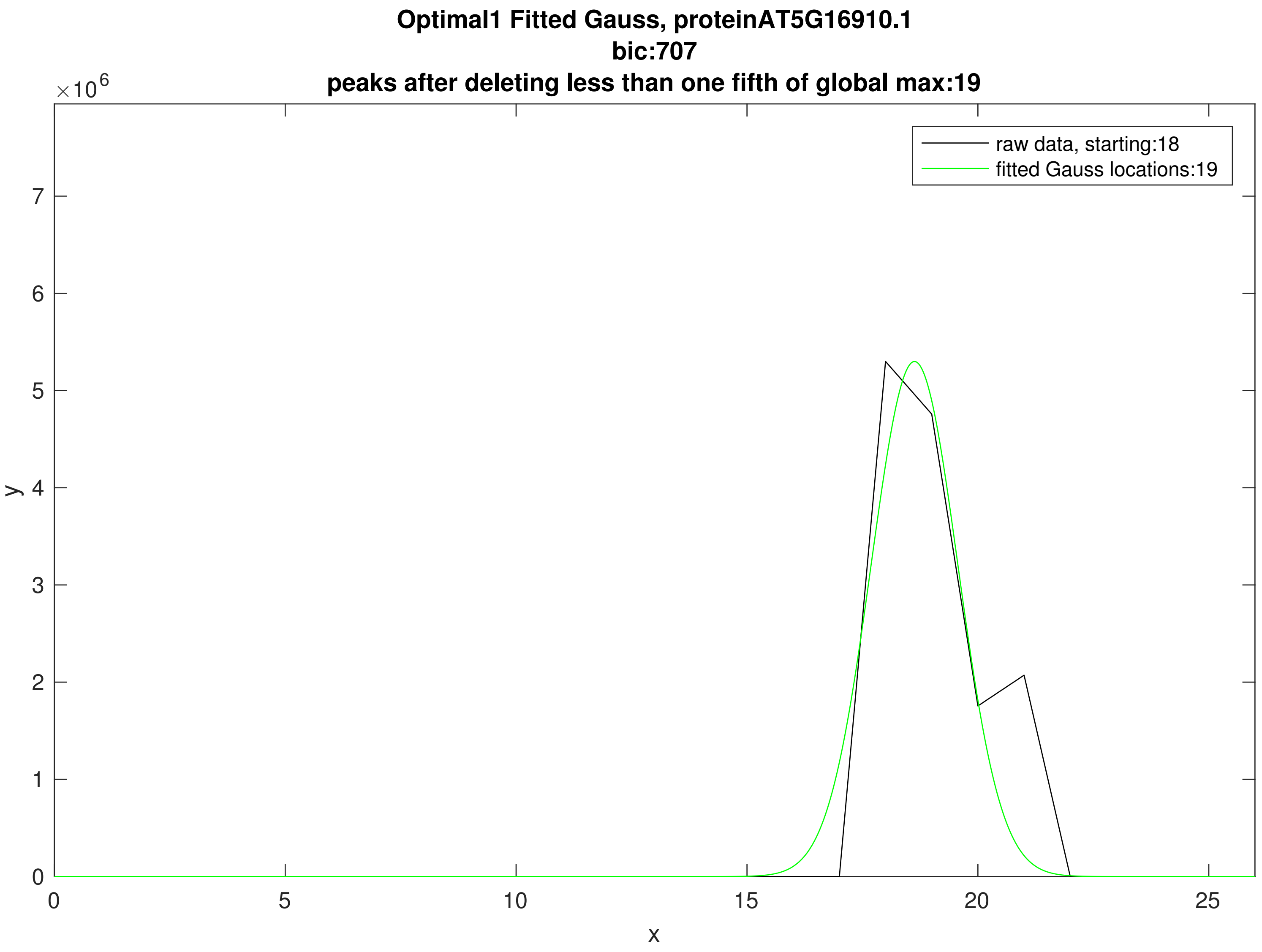
Optimal1 Fitted Gauss, proteinAT5G16910.1
bic:707
peaks after deleting less than one fifth of global max:19
×10^6
raw data, starting:18
fitted Gauss locations:19
y
x
0
1
2
3
4
5
6
7
0
5
10
15
20
25

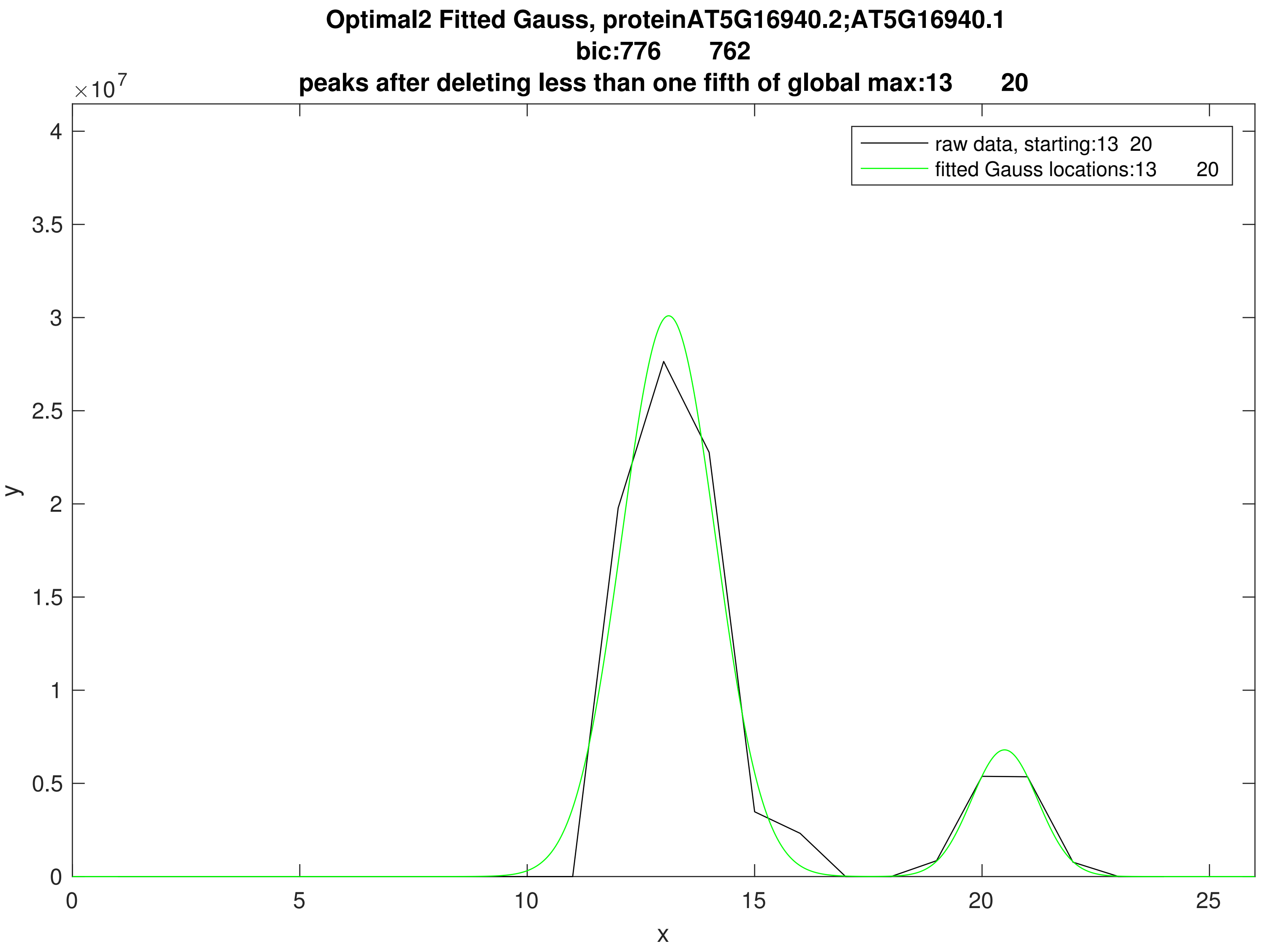

Optimal2 Fitted Gauss, proteinAT5G16940.2;AT5G16940.1
bic:776 762
peaks after deleting less than one fifth of global max:13 20
×10^7
raw data, starting:13 20
fitted Gauss locations:13 20
y
x
0
0.5
1
1.5
2
2.5
3
3.5
4
5
10
15
20
25

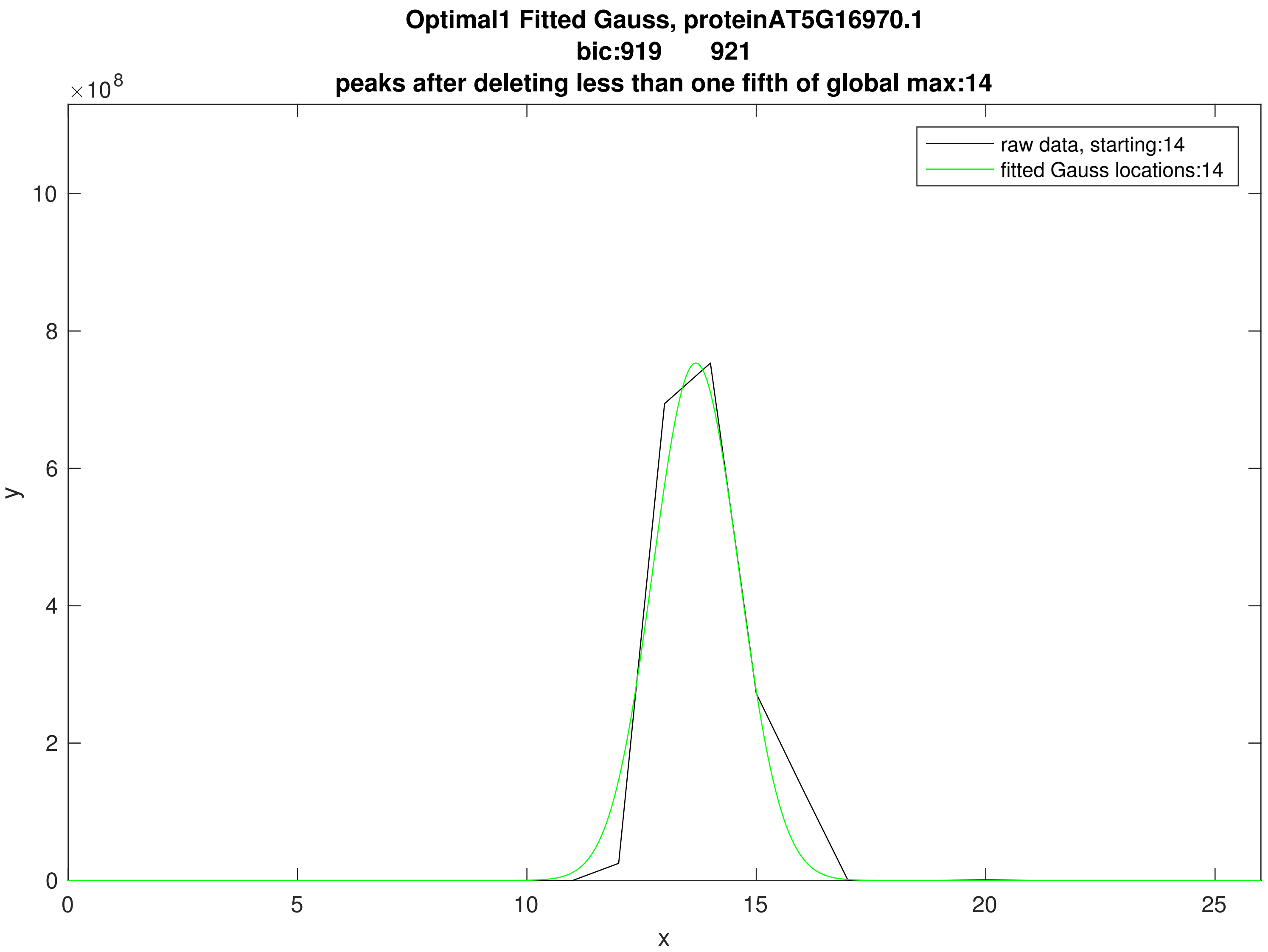

Optimal1 Fitted Gauss, proteinAT5G16970.1
bic:919 921
peaks after deleting less than one fifth of global max:14
×10^8
raw data, starting:14
fitted Gauss locations:14
y
x
0
2
4
6
8
10
0
5
10
15
20
25

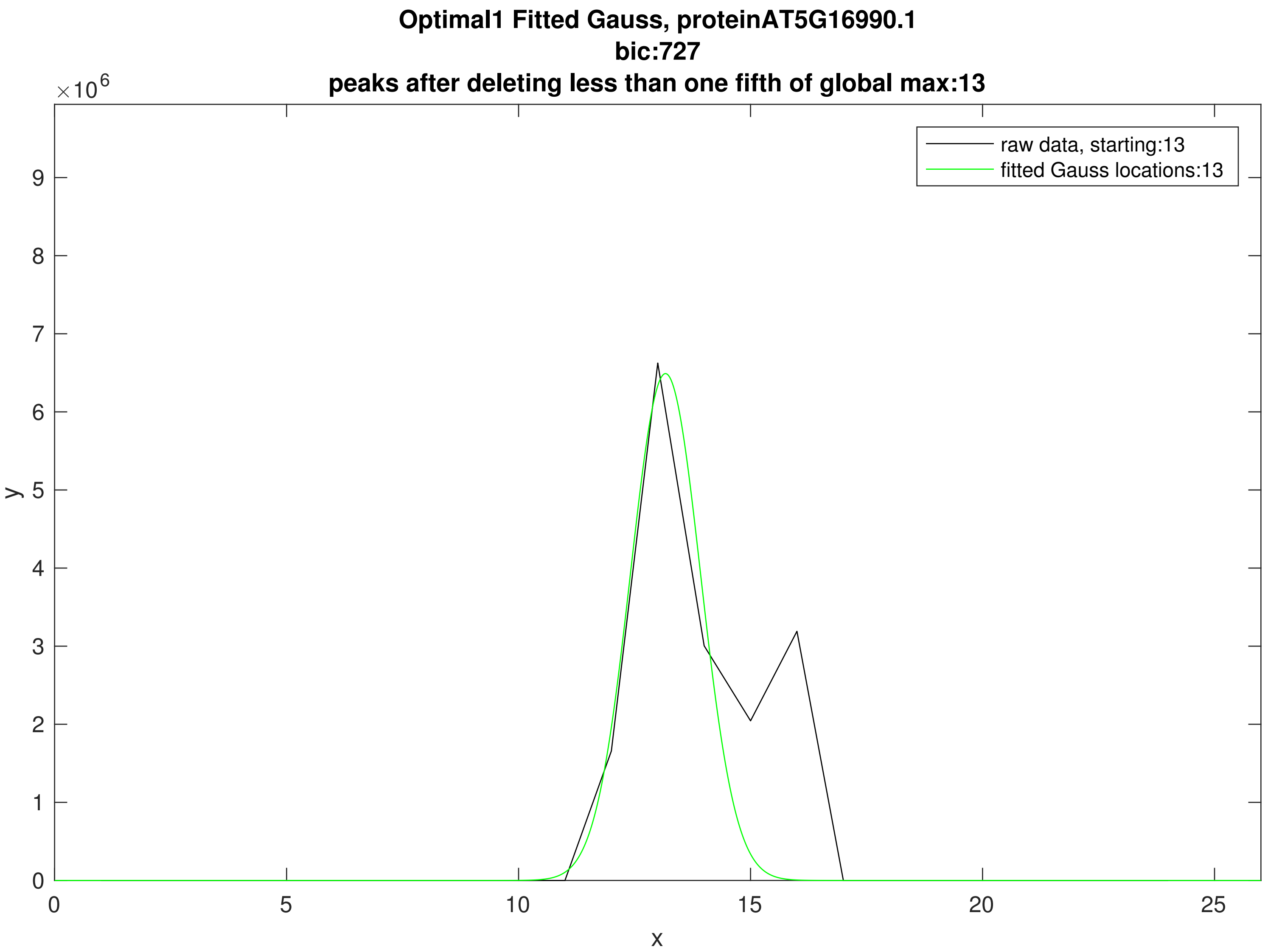
Optimal1 Fitted Gauss, proteinAT5G16990.1
bic:727
peaks after deleting less than one fifth of global max:13
×10^6
raw data, starting:13
fitted Gauss locations:13
y
x
0
1
2
3
4
5
6
7
8
9
0
5
10
15
20
25

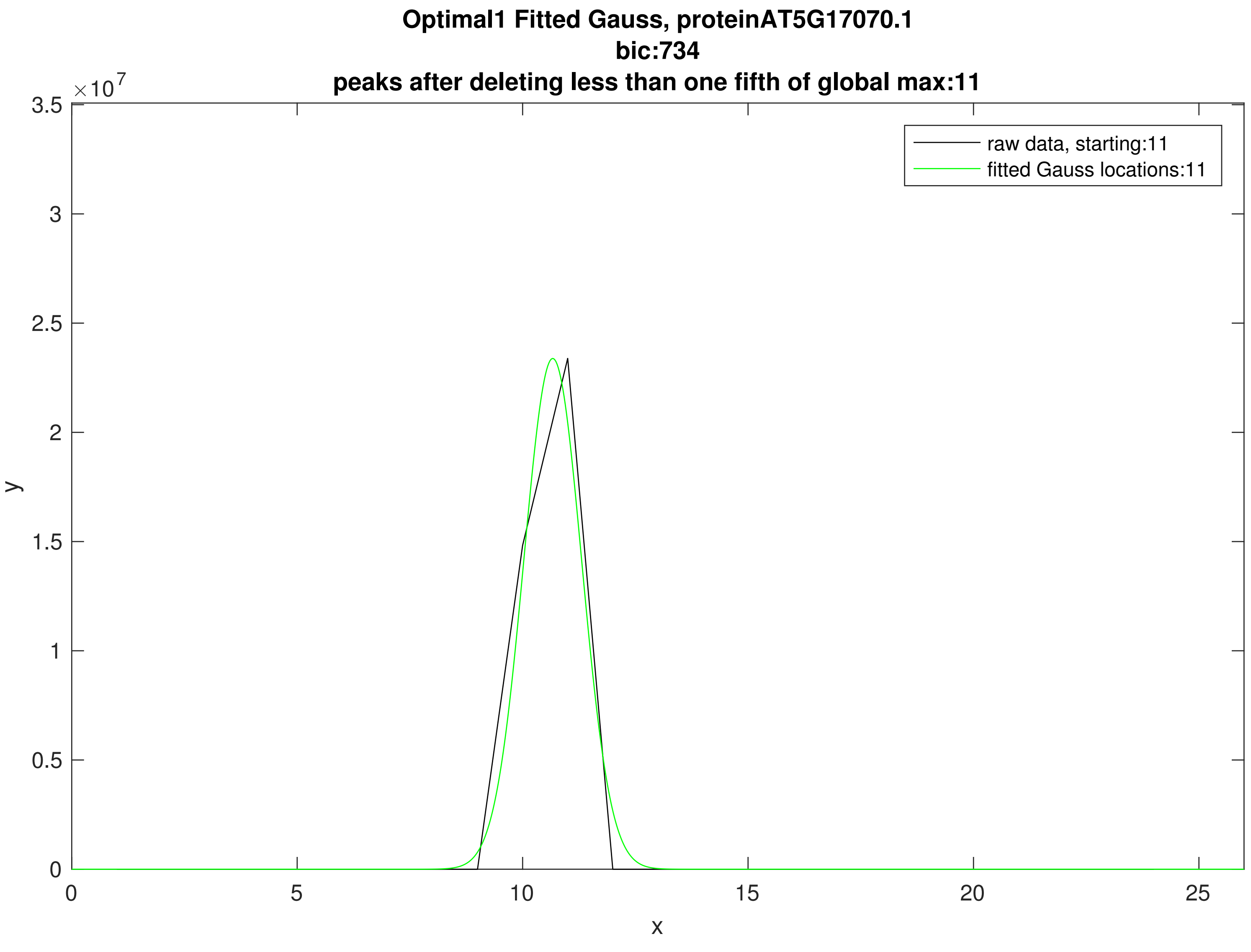
Optimal1 Fitted Gauss, proteinAT5G17070.1
bic:734
peaks after deleting less than one fifth of global max:11
raw data, starting:11
fitted Gauss locations:11
×10^7
3.5
3
2.5
2
1.5
1
0.5
0
y
0
5
10
15
20
25
x

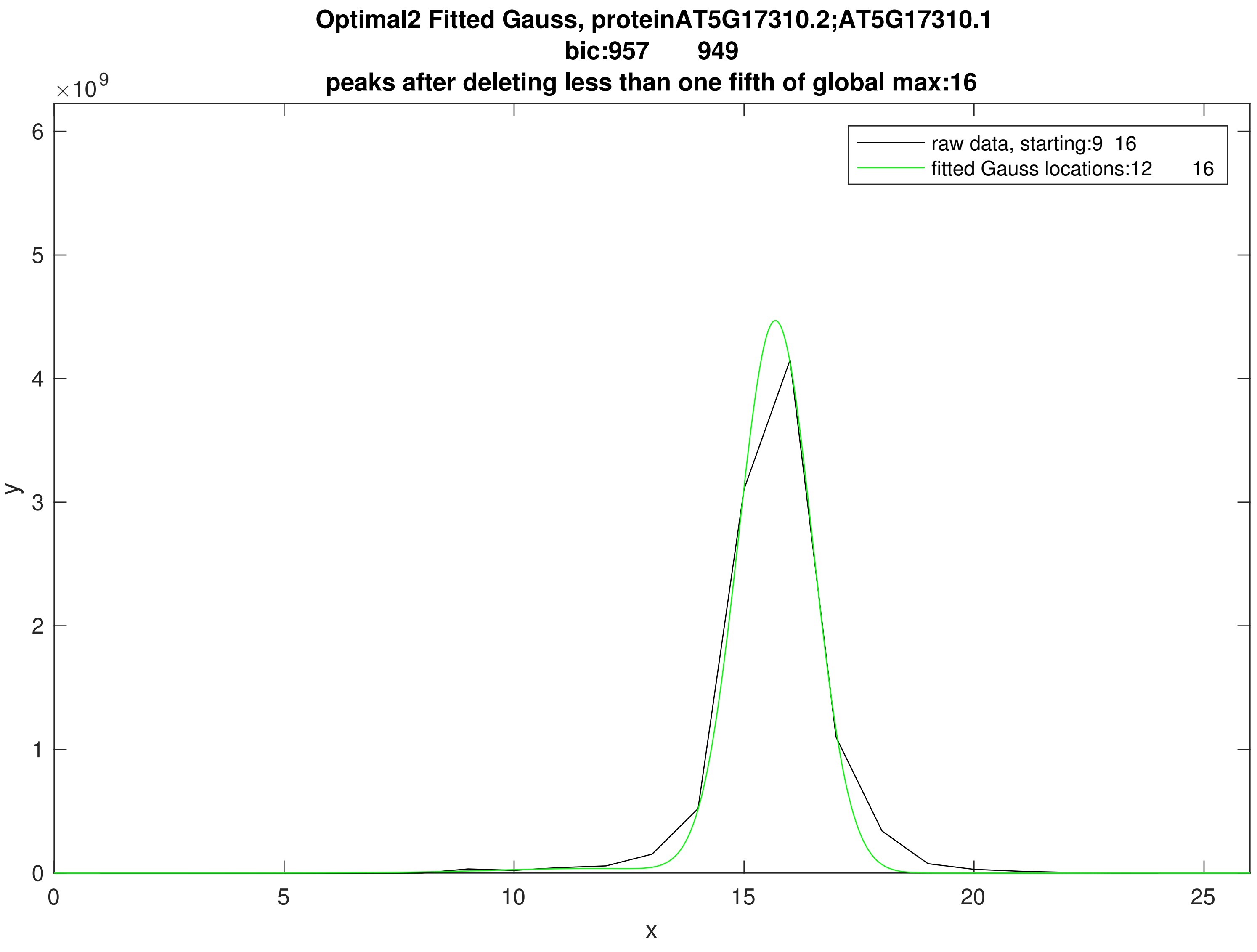
Optimal2 Fitted Gauss, proteinAT5G17310.2;AT5G17310.1
bic:957 949
peaks after deleting less than one fifth of global max:16
×10^9
raw data, starting:9 16
fitted Gauss locations:12 16
y
x
0
1
2
3
4
5
6
0
5
10
15
20
25

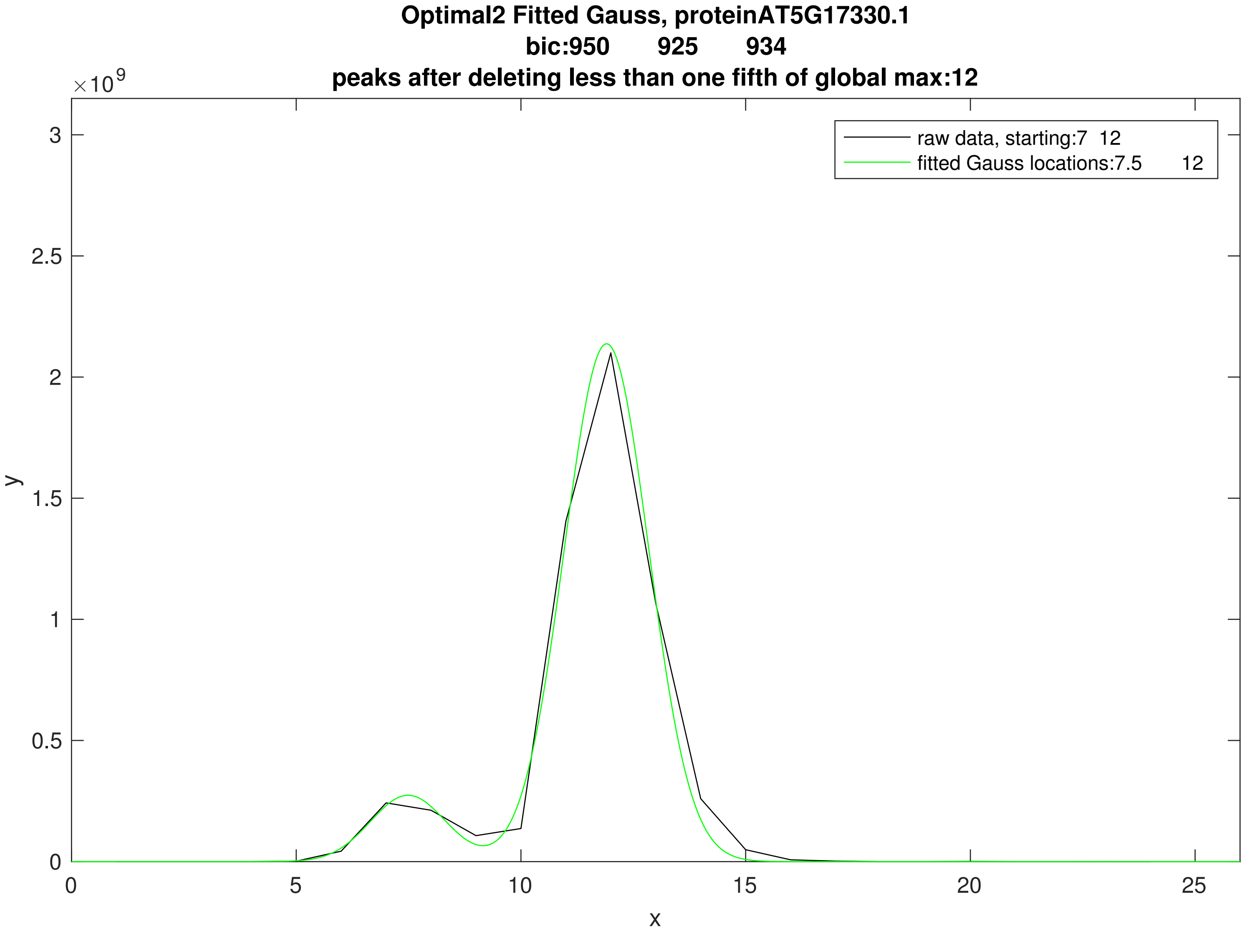

Optimal2 Fitted Gauss, proteinAT5G17330.1
bic:950 925 934
peaks after deleting less than one fifth of global max:12
×10^9
3
2.5
2
1.5
1
0.5
0
y
0
5
10
15
20
25
x
raw data, starting:7 12
fitted Gauss locations:7.5 12

**Optimal1 Fitted Gauss, proteinAT5G17380.1**
**bic:978 987**
**peaks after deleting less than one fifth of global max:12**

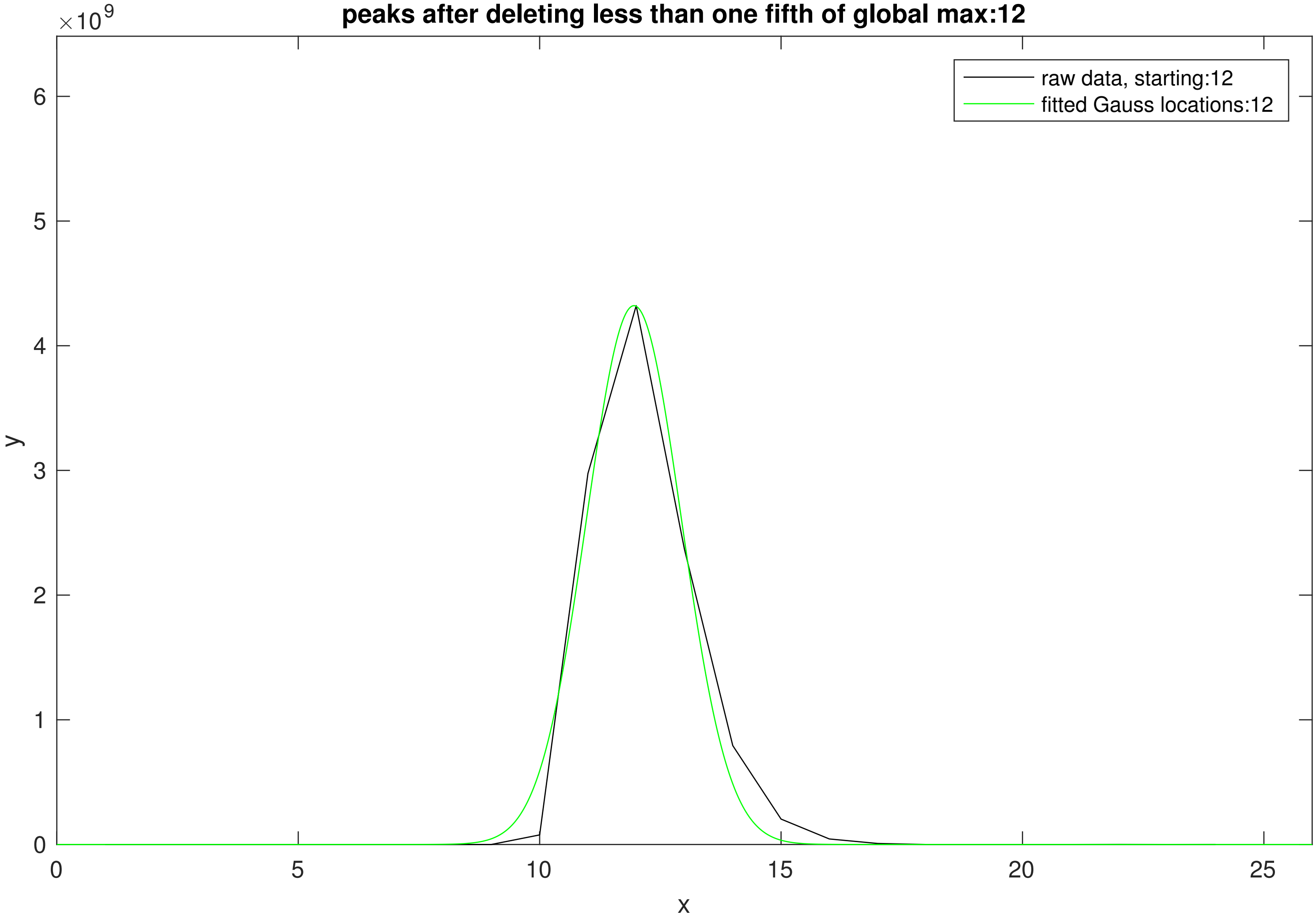

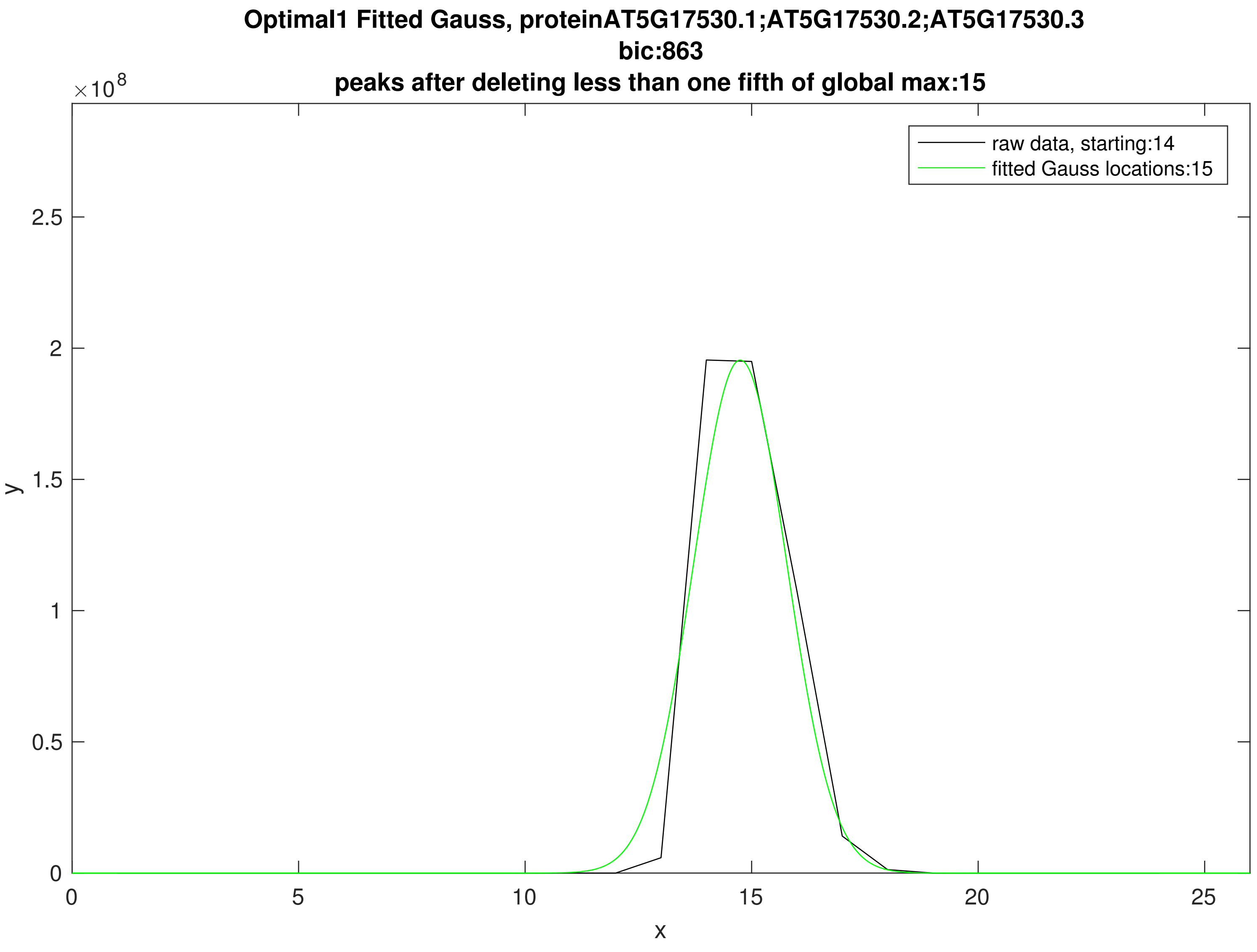
Optimal1 Fitted Gauss, proteinAT5G17530.1;AT5G17530.2;AT5G17530.3
bic:863
peaks after deleting less than one fifth of global max:15
×10^8
raw data, starting:14
fitted Gauss locations:15
2.5
2
1.5
1
0.5
0
y
0
5
10
15
20
25
x

#### Optimal1 Fitted Gauss, proteinAT5G17550.1
#### bic:764
#### peaks after deleting less than one fifth of global max:14

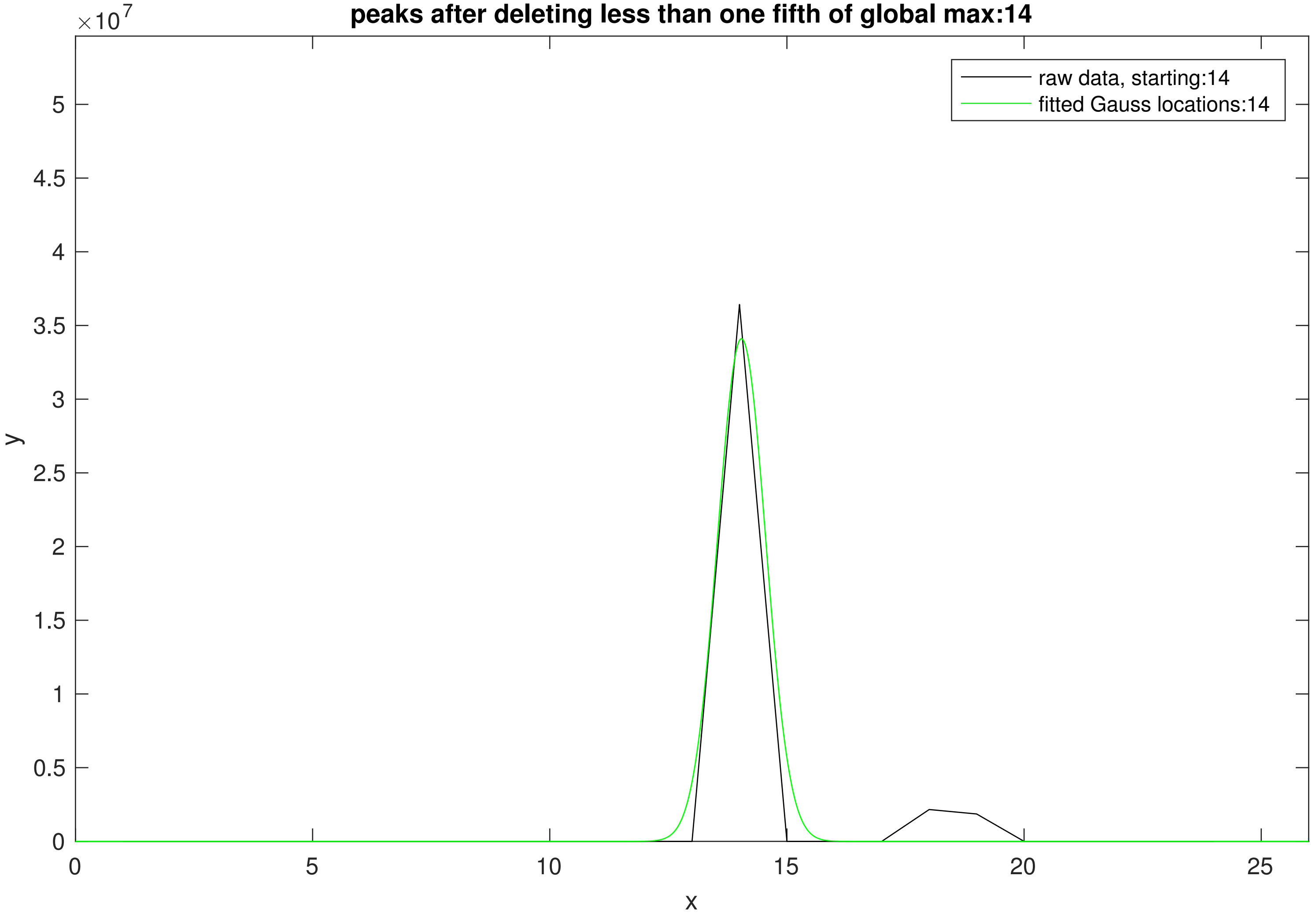

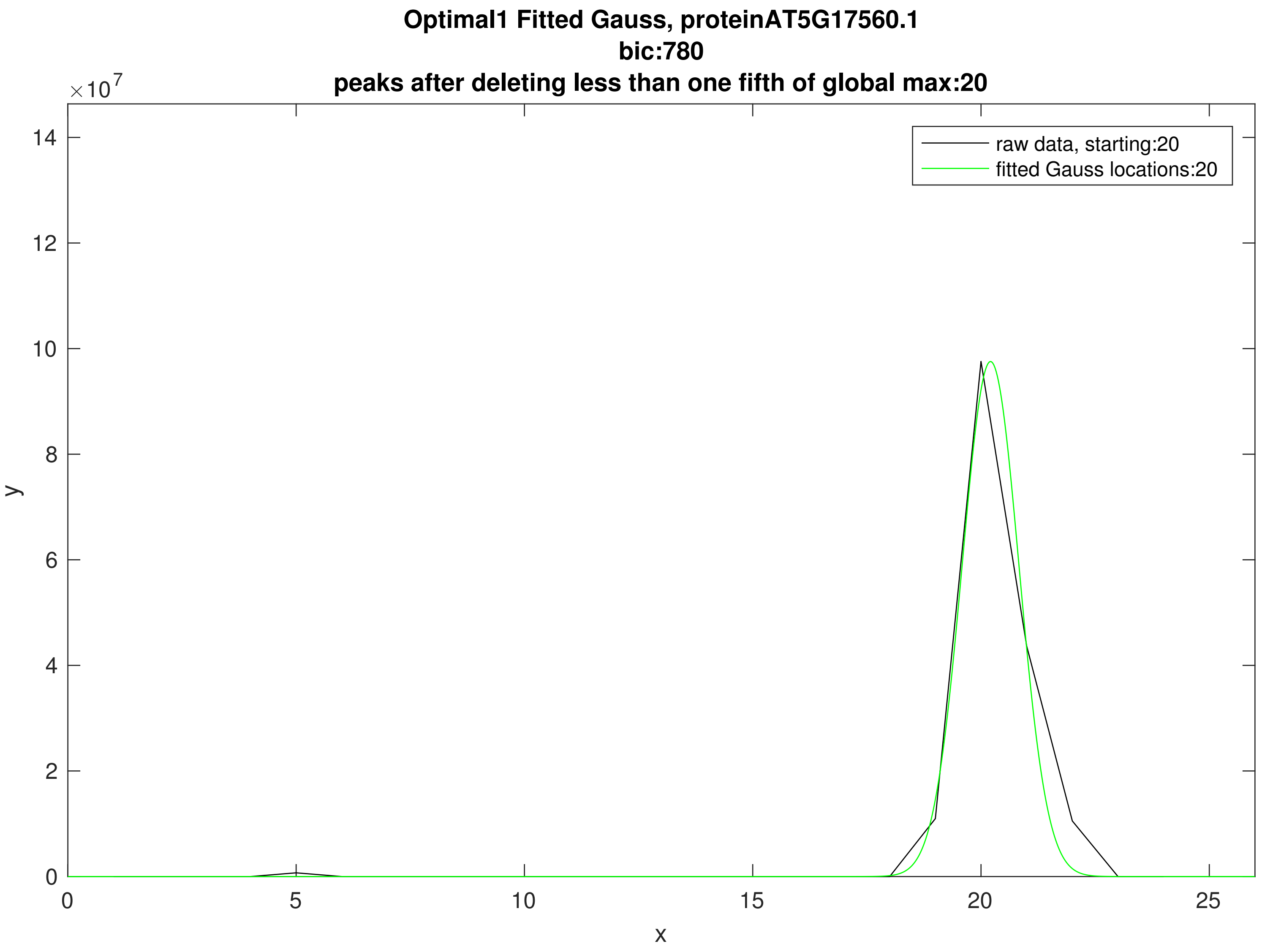
Optimal1 Fitted Gauss, proteinAT5G17560.1
bic:780
peaks after deleting less than one fifth of global max:20
×10^7
14
12
10
8
6
4
2
0
y
0
5
10
15
20
25
x
raw data, starting:20
fitted Gauss locations:20

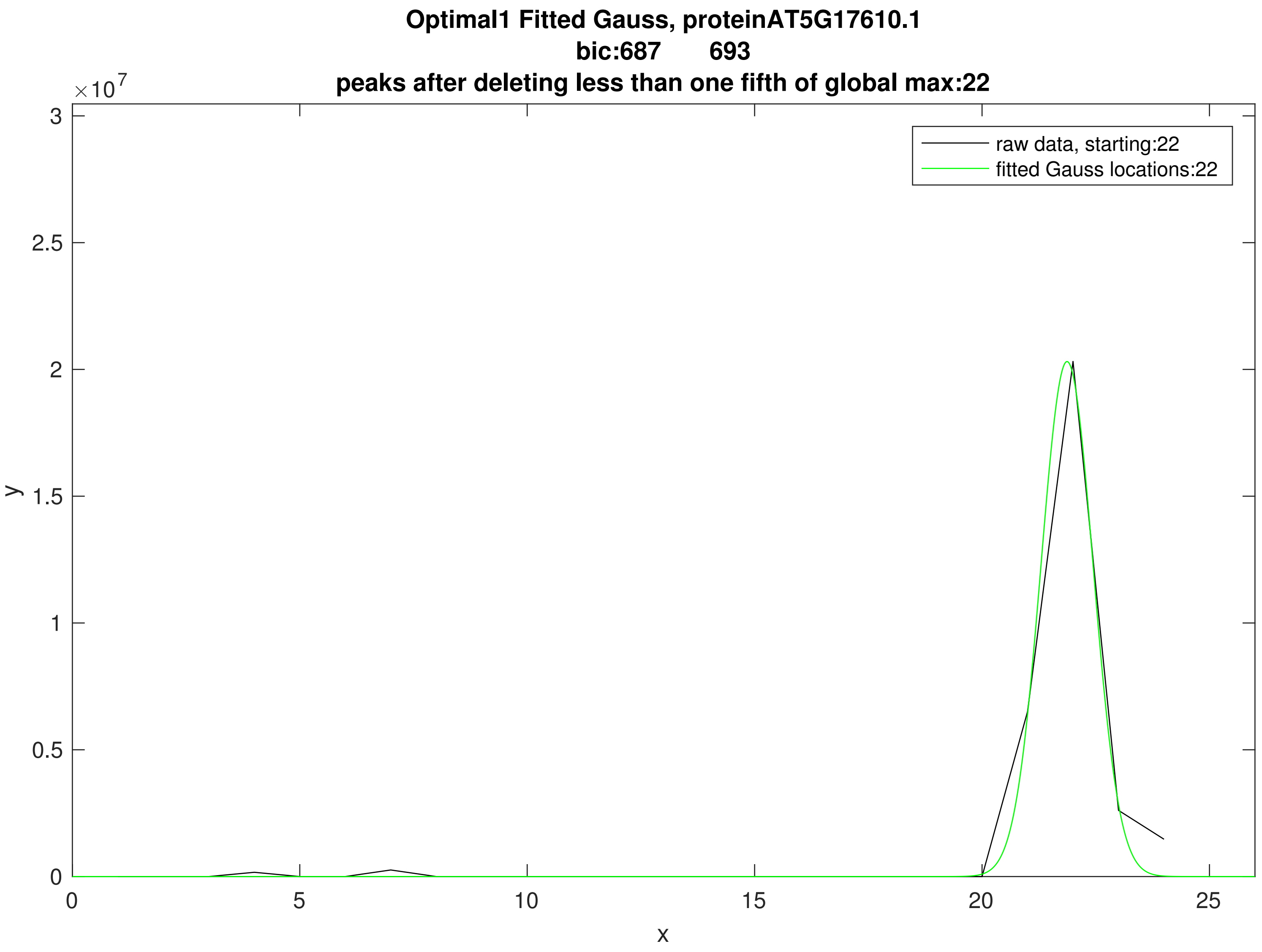

Optimal1 Fitted Gauss, proteinAT5G17610.1
bic:687 693
peaks after deleting less than one fifth of global max:22
×10^7
raw data, starting:22
fitted Gauss locations:22
y
x
0
0.5
1
1.5
2
2.5
3
0
5
10
15
20
25

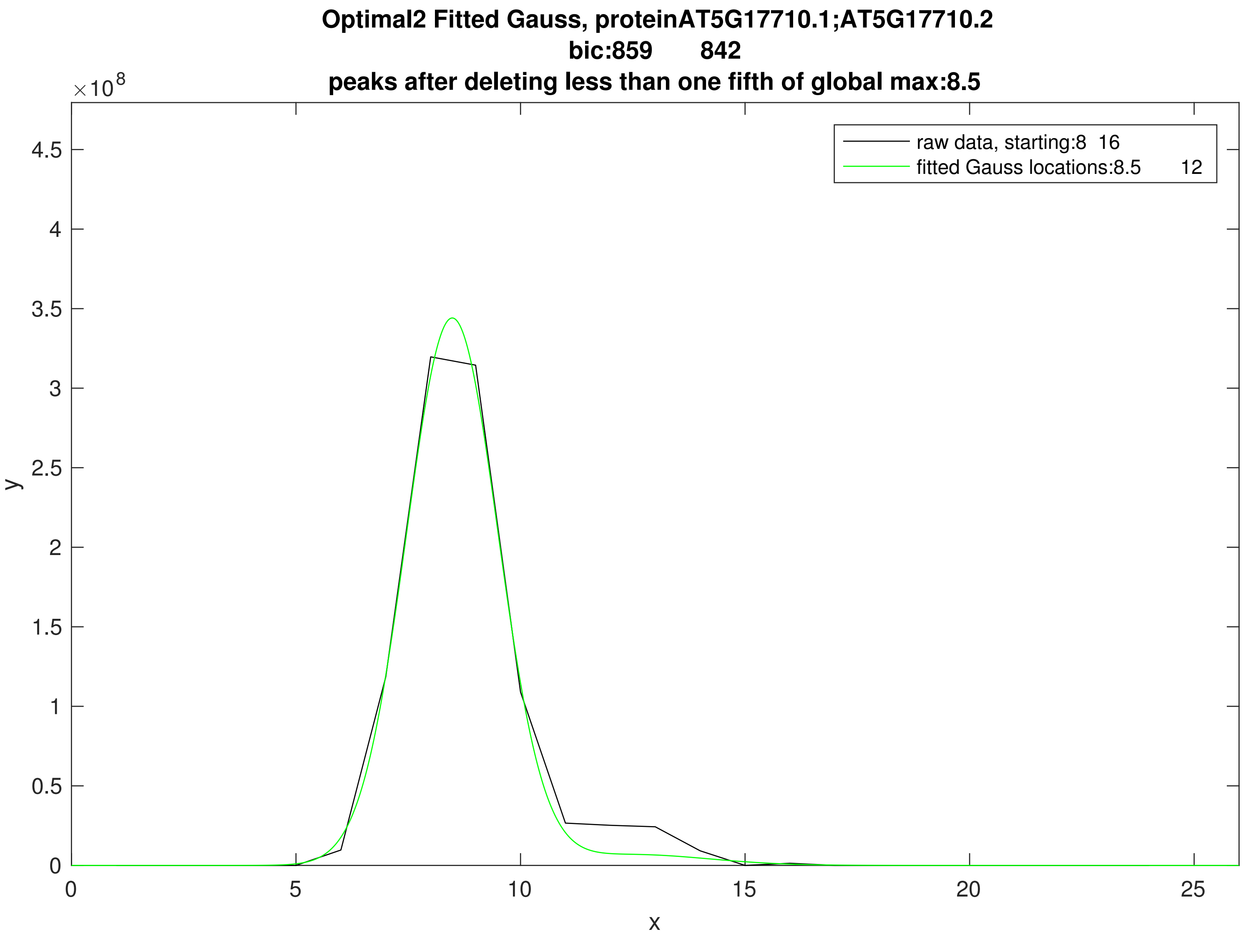

Optimal2 Fitted Gauss, proteinAT5G17710.1;AT5G17710.2
bic:859 842
peaks after deleting less than one fifth of global max:8.5
×10^8
4.5
4
3.5
3
2.5
2
1.5
1
0.5
0
y
0
5
10
15
20
25
x
raw data, starting:8 16
fitted Gauss locations:8.5 12

#### Optimal1 Fitted Gauss, proteinAT5G17910.2;AT5G17910.1
#### bic:788
#### peaks after deleting less than one fifth of global max:5.4

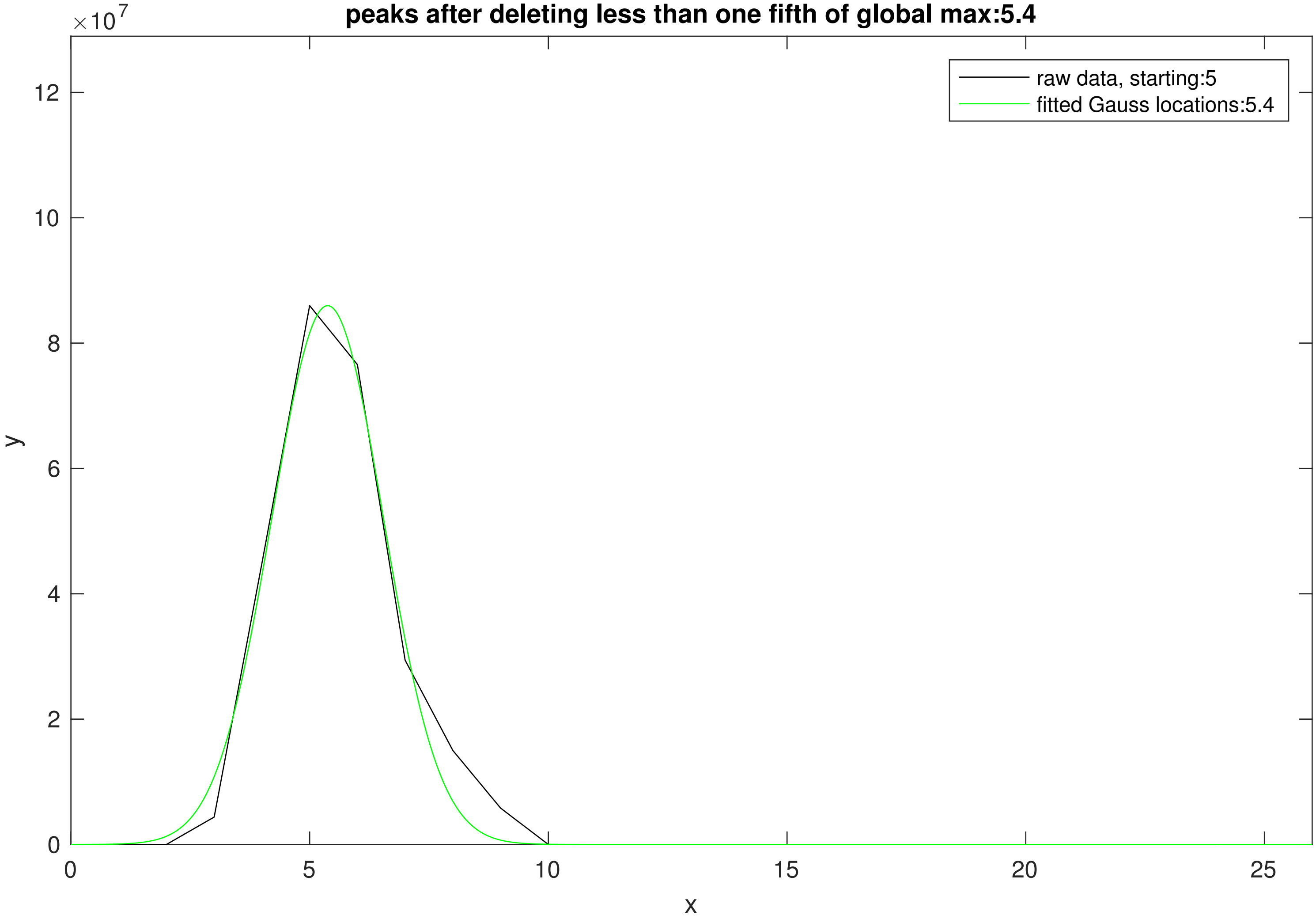

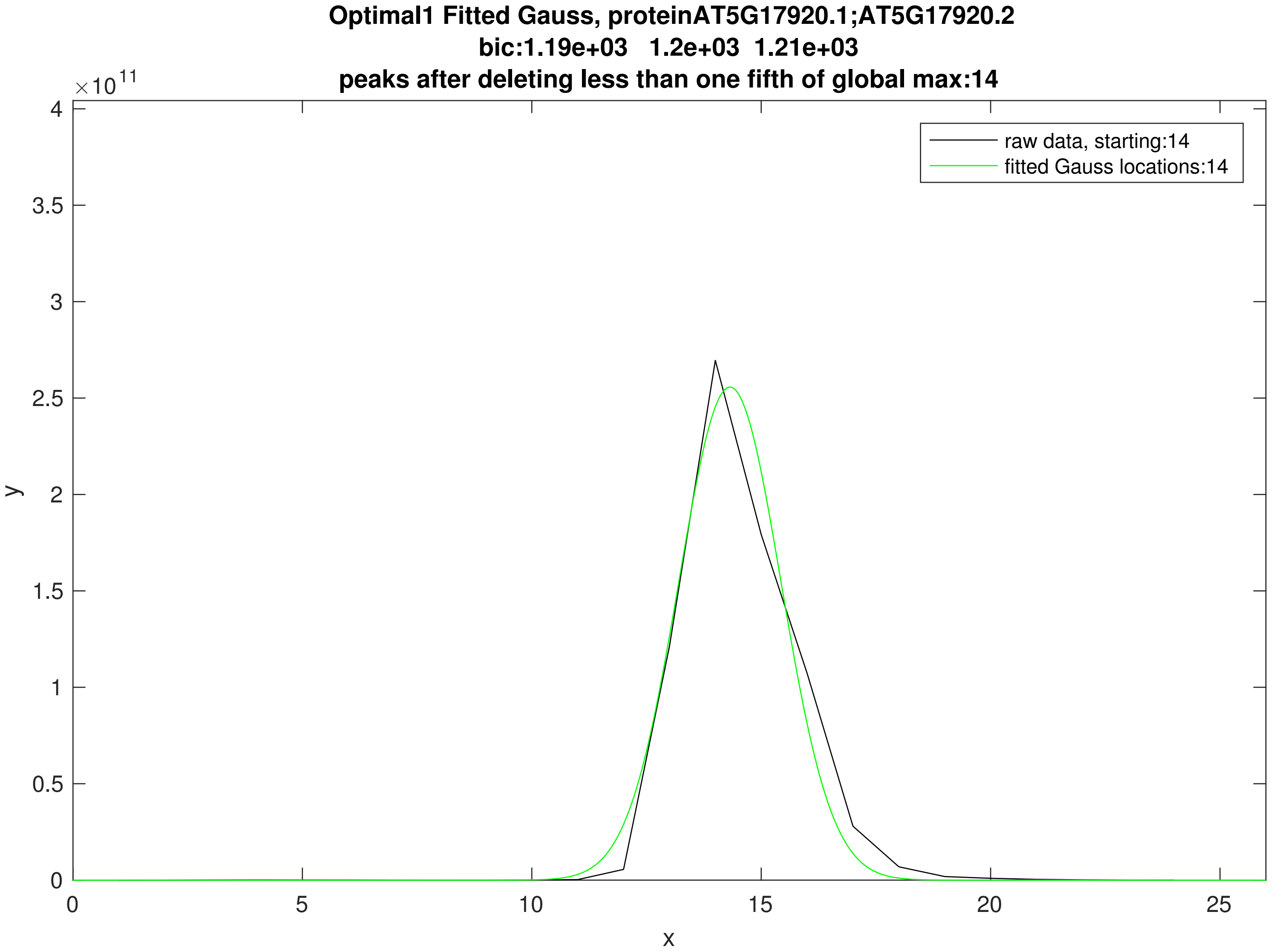

Optimal1 Fitted Gauss, proteinAT5G17920.1;AT5G17920.2
bic:1.19e+03 1.2e+03 1.21e+03
peaks after deleting less than one fifth of global max:14
×10^11
4
3.5
3
2.5
2
1.5
1
0.5
0
y
0
5
10
15
20
25
x
raw data, starting:14
fitted Gauss locations:14

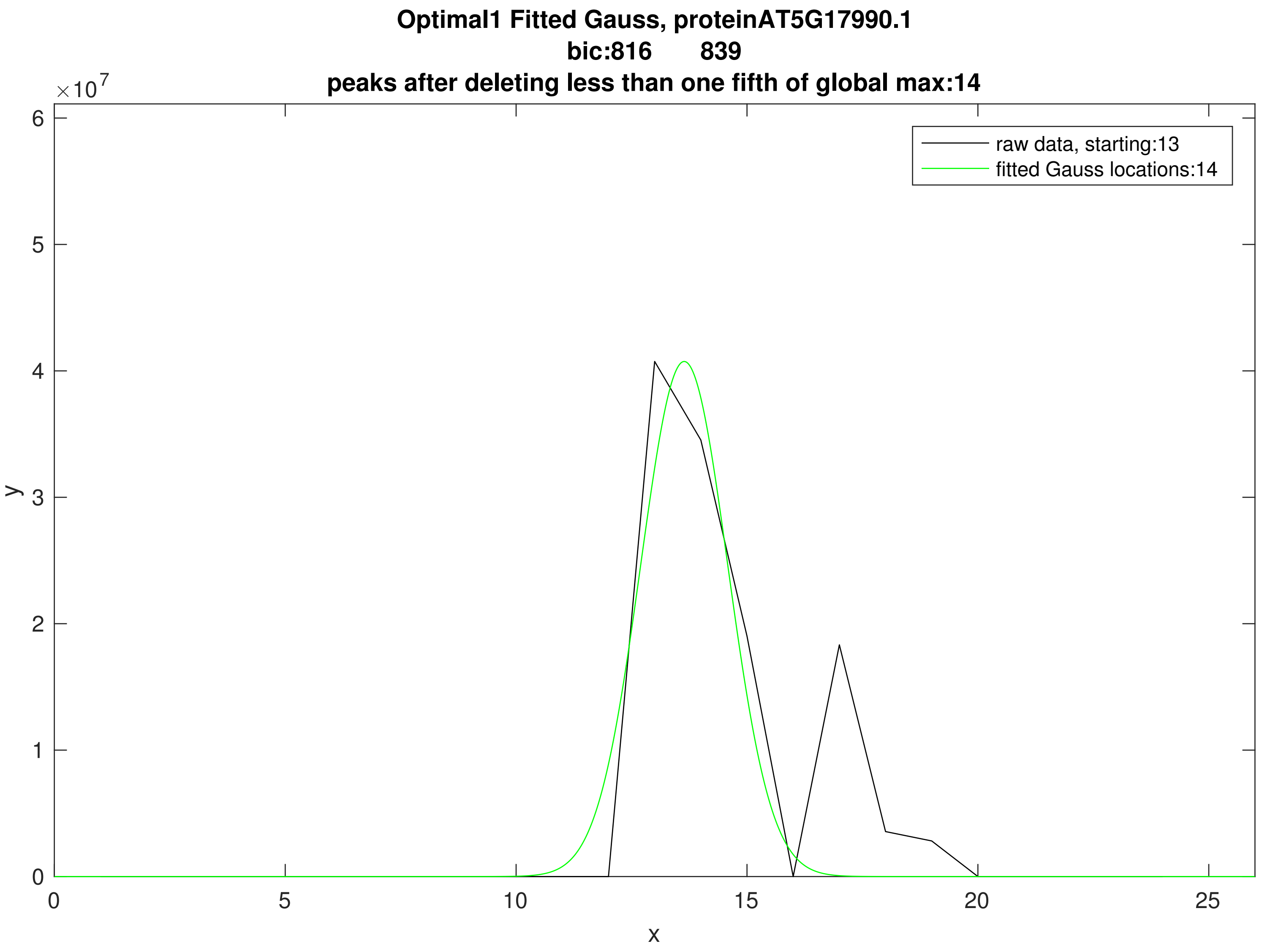
Optimal1 Fitted Gauss, proteinAT5G17990.1
bic:816 839
peaks after deleting less than one fifth of global max:14
×10^7
6
5
4
3
2
1
0
y
0
5
10
15
20
25
x
raw data, starting:13
fitted Gauss locations:14

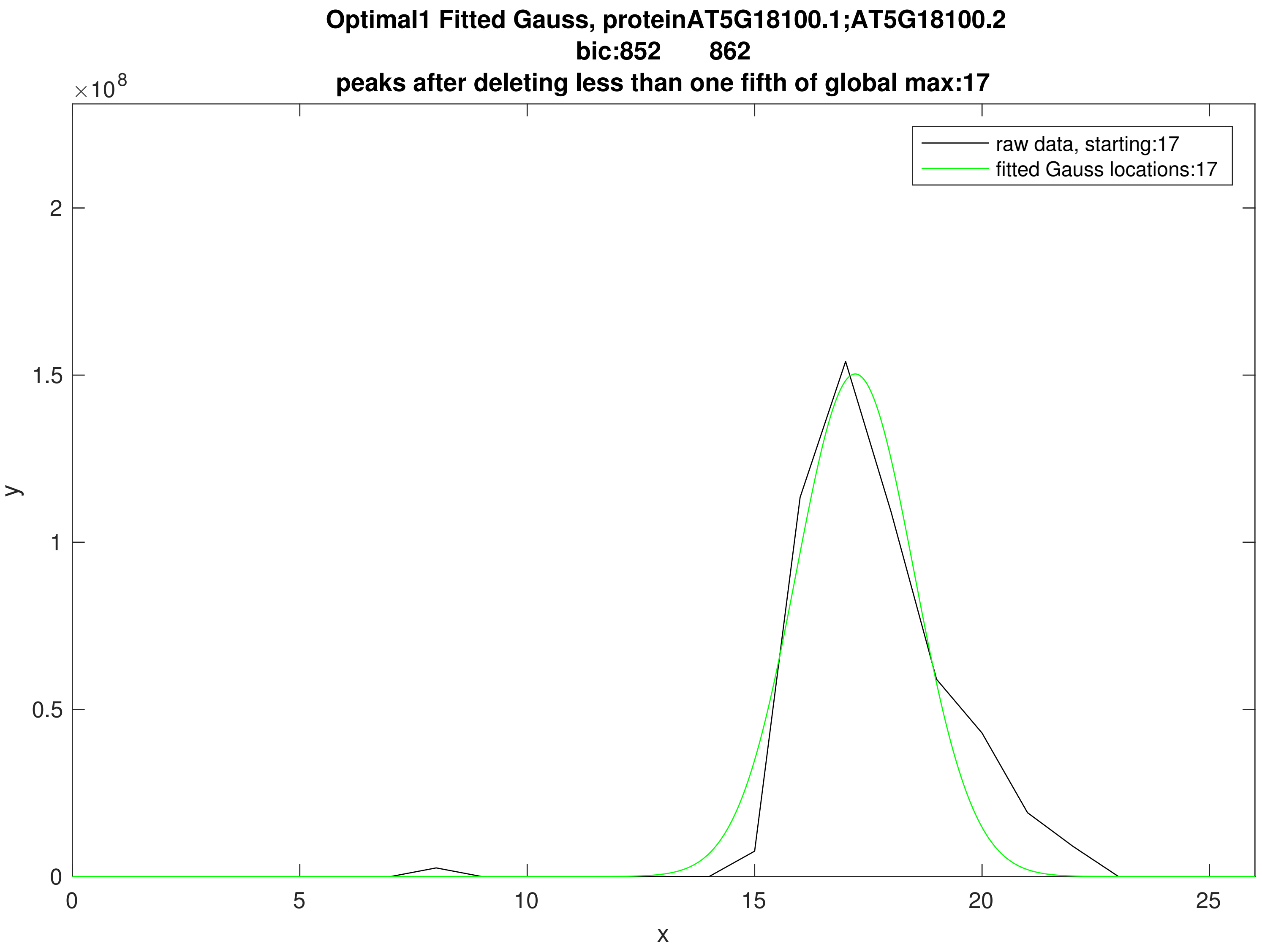

Optimal1 Fitted Gauss, proteinAT5G18100.1;AT5G18100.2
bic:852 862
peaks after deleting less than one fifth of global max:17
×10^8
raw data, starting:17
fitted Gauss locations:17
y
x
0
0.5
1
1.5
2
5
10
15
20
25

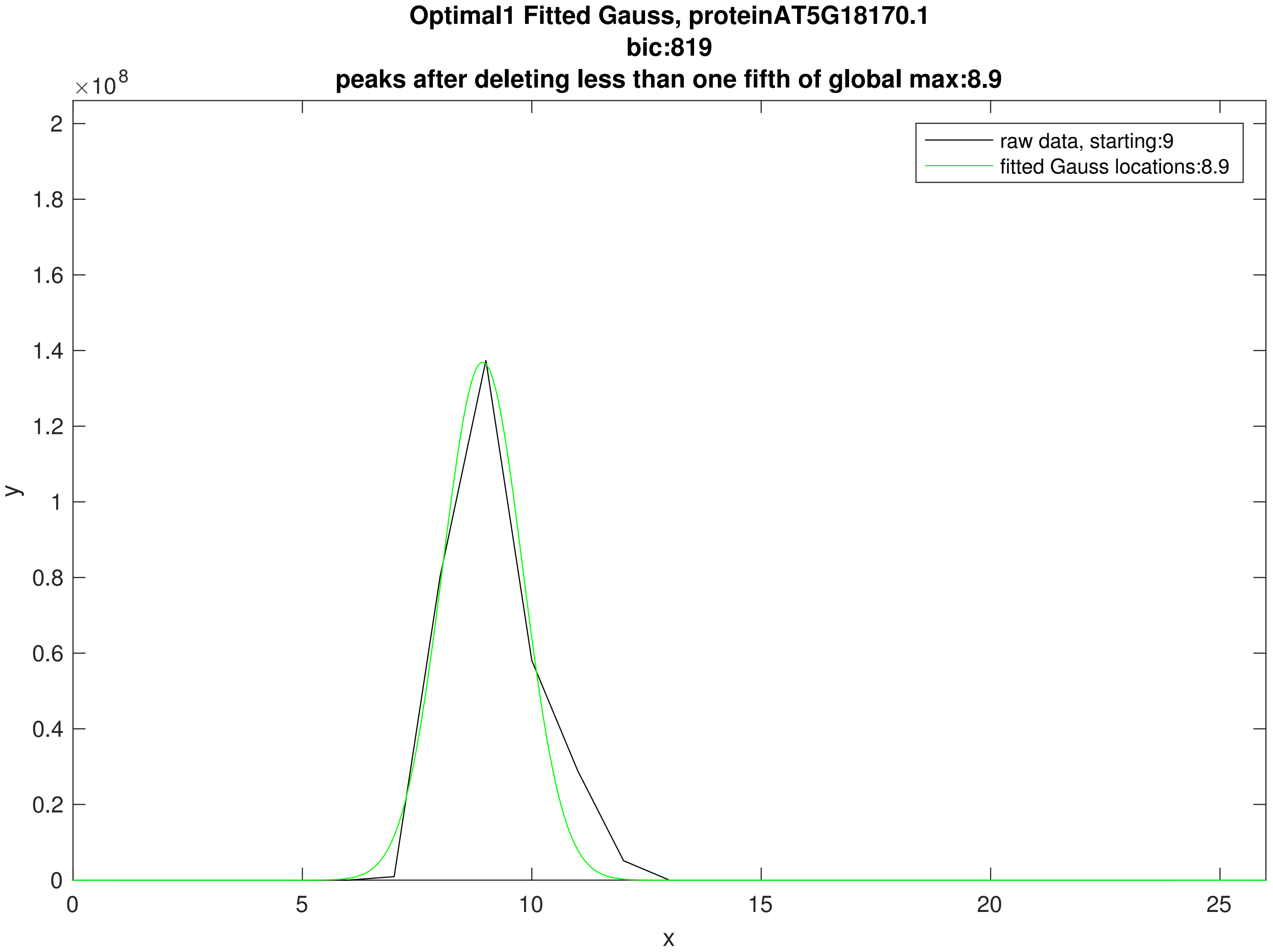

Optimal1 Fitted Gauss, proteinAT5G18170.1
bic:819
peaks after deleting less than one fifth of global max:8.9
×10^8
raw data, starting:9
fitted Gauss locations:8.9
y
x
0
0.2
0.4
0.6
0.8
1
1.2
1.4
1.6
1.8
2
0
5
10
15
20
25

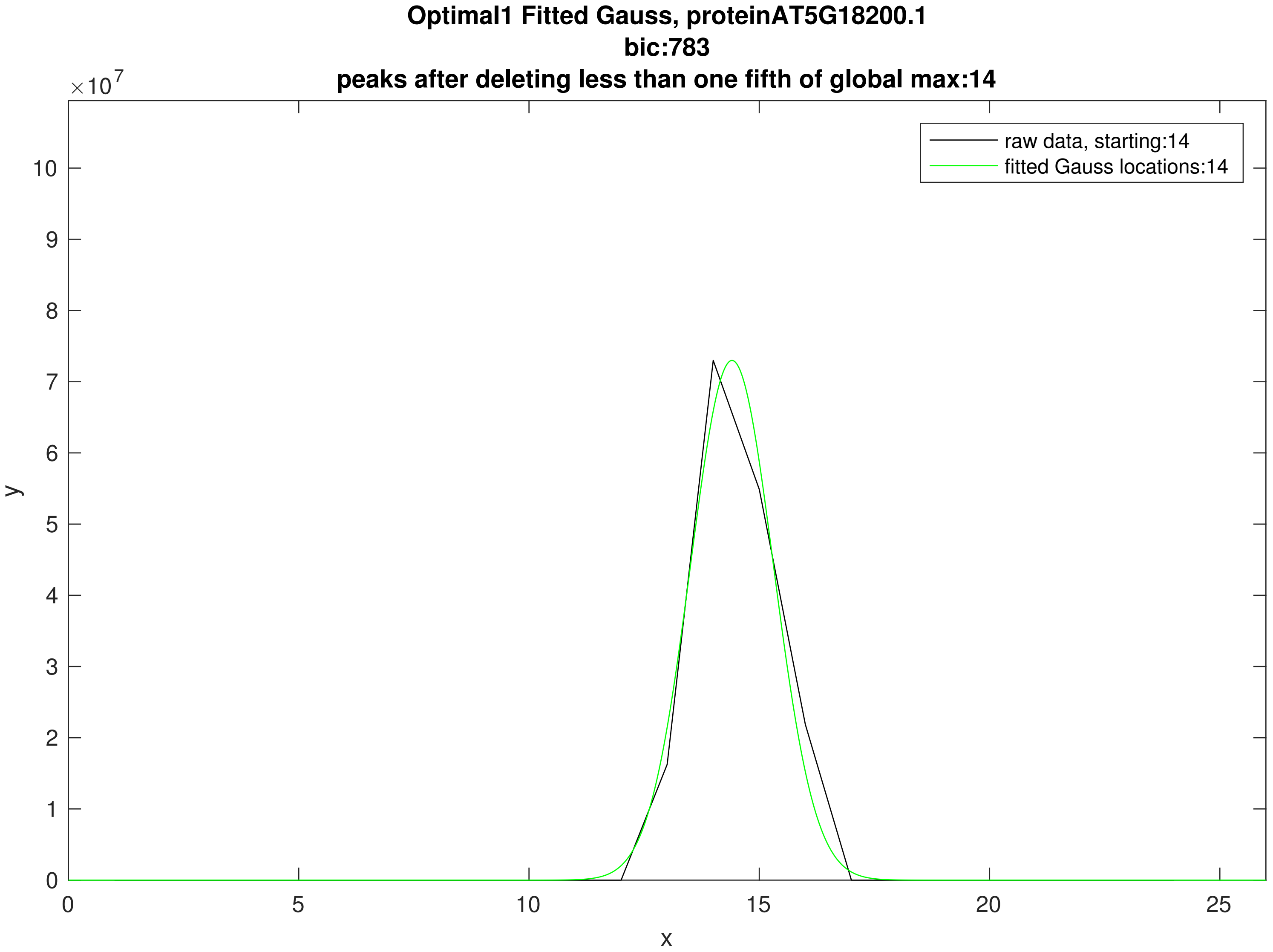
Optimal1 Fitted Gauss, proteinAT5G18200.1
bic:783
peaks after deleting less than one fifth of global max:14
×10^7
raw data, starting:14
fitted Gauss locations:14
y
x
0
1
2
3
4
5
6
7
8
9
10
0
5
10
15
20
25

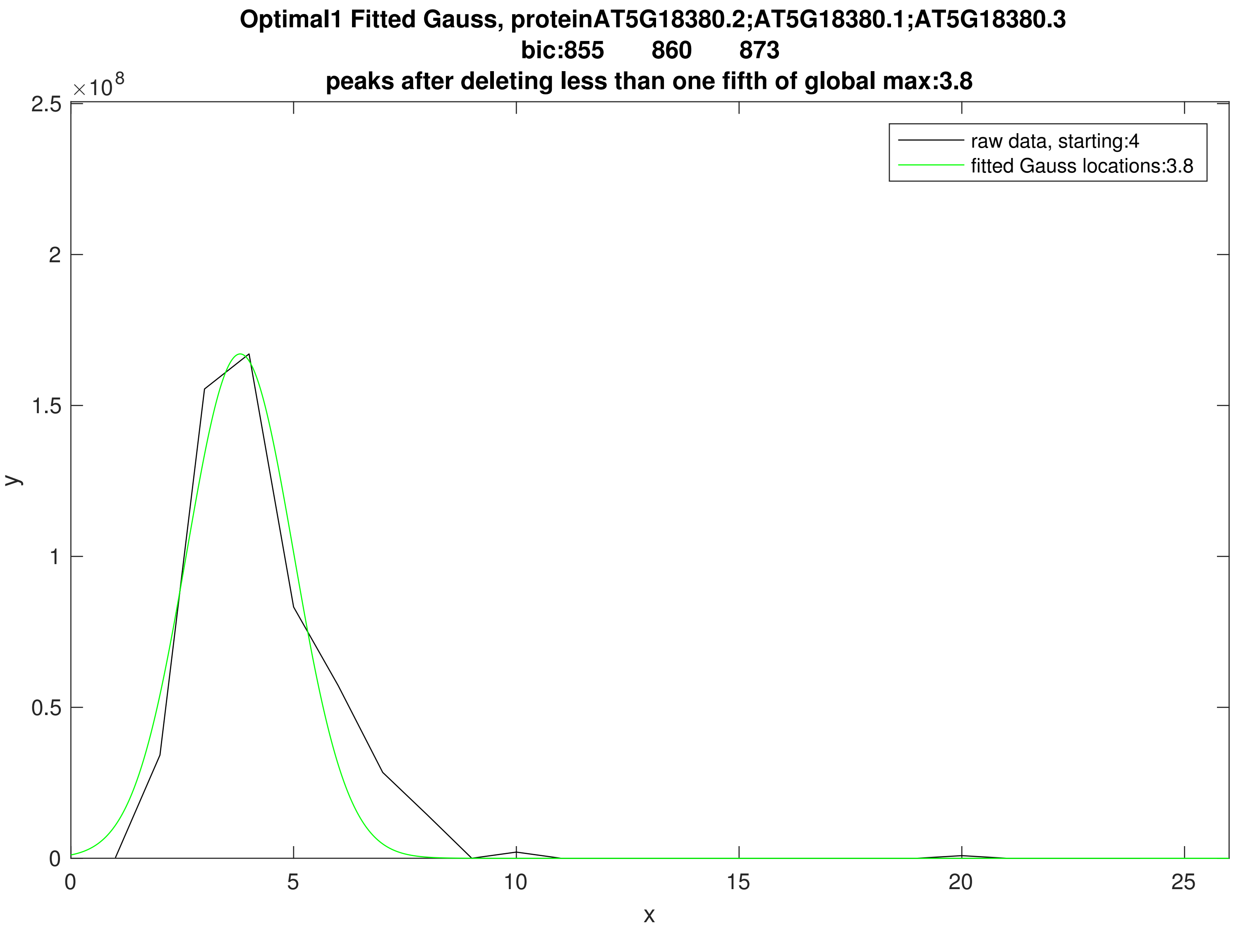
Optimal1 Fitted Gauss, proteinAT5G18380.2;AT5G18380.1;AT5G18380.3
bic:855 860 873
peaks after deleting less than one fifth of global max:3.8
×10^8
2.5
2
1.5
1
0.5
0
y
0
5
10
15
20
25
x
raw data, starting:4
fitted Gauss locations:3.8

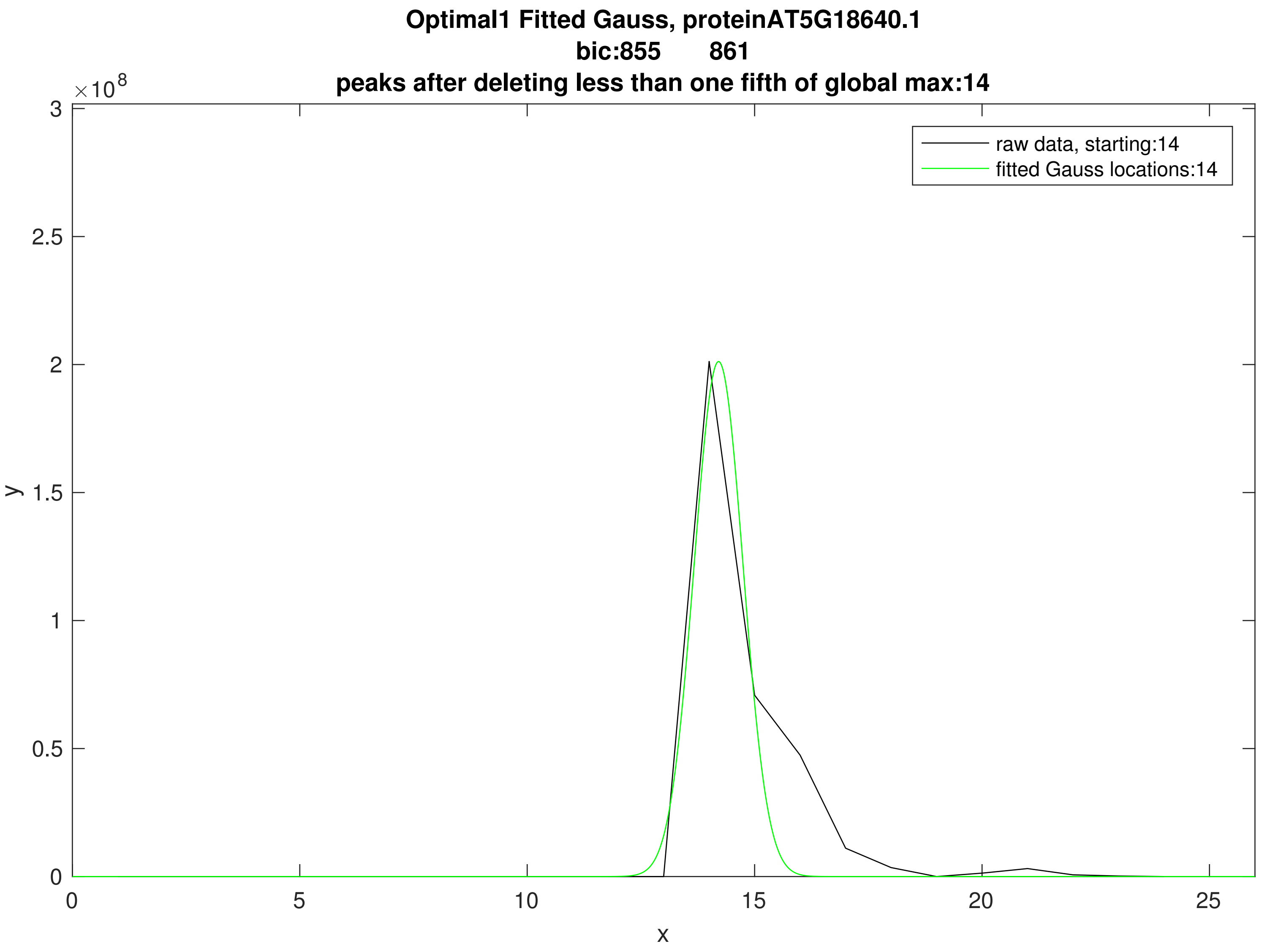

Optimal1 Fitted Gauss, proteinAT5G18640.1
bic:855 861
peaks after deleting less than one fifth of global max:14
×10^8
3
2.5
2
1.5
1
0.5
0
y
0
5
10
15
20
25
x
raw data, starting:14
fitted Gauss locations:14

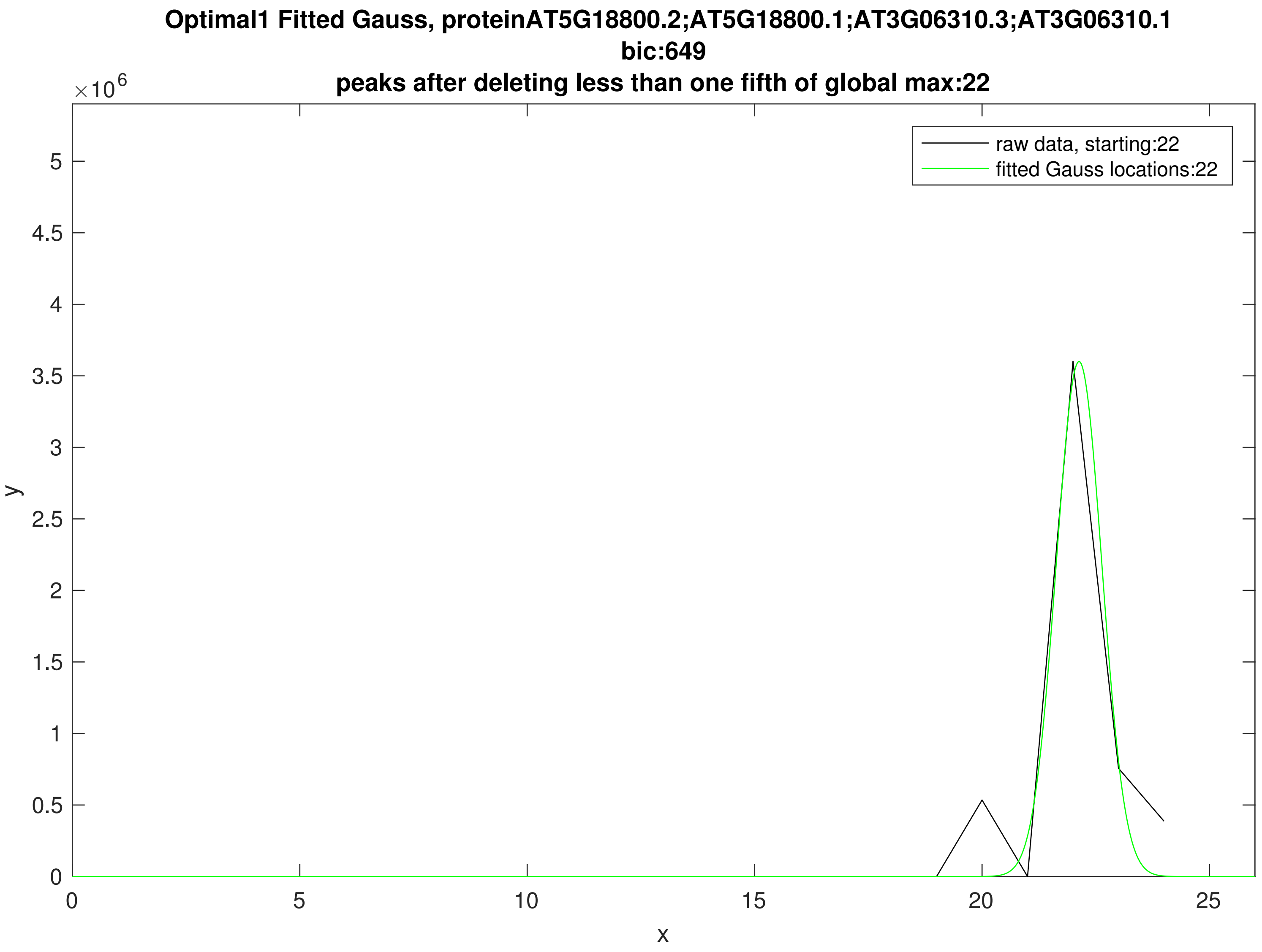
Optimal1 Fitted Gauss, proteinAT5G18800.2;AT5G18800.1;AT3G06310.3;AT3G06310.1
bic:649
peaks after deleting less than one fifth of global max:22
×10^6
raw data, starting:22
fitted Gauss locations:22
y
x
0
5
10
15
20
25
0.5
1
1.5
2
2.5
3
3.5
4
4.5

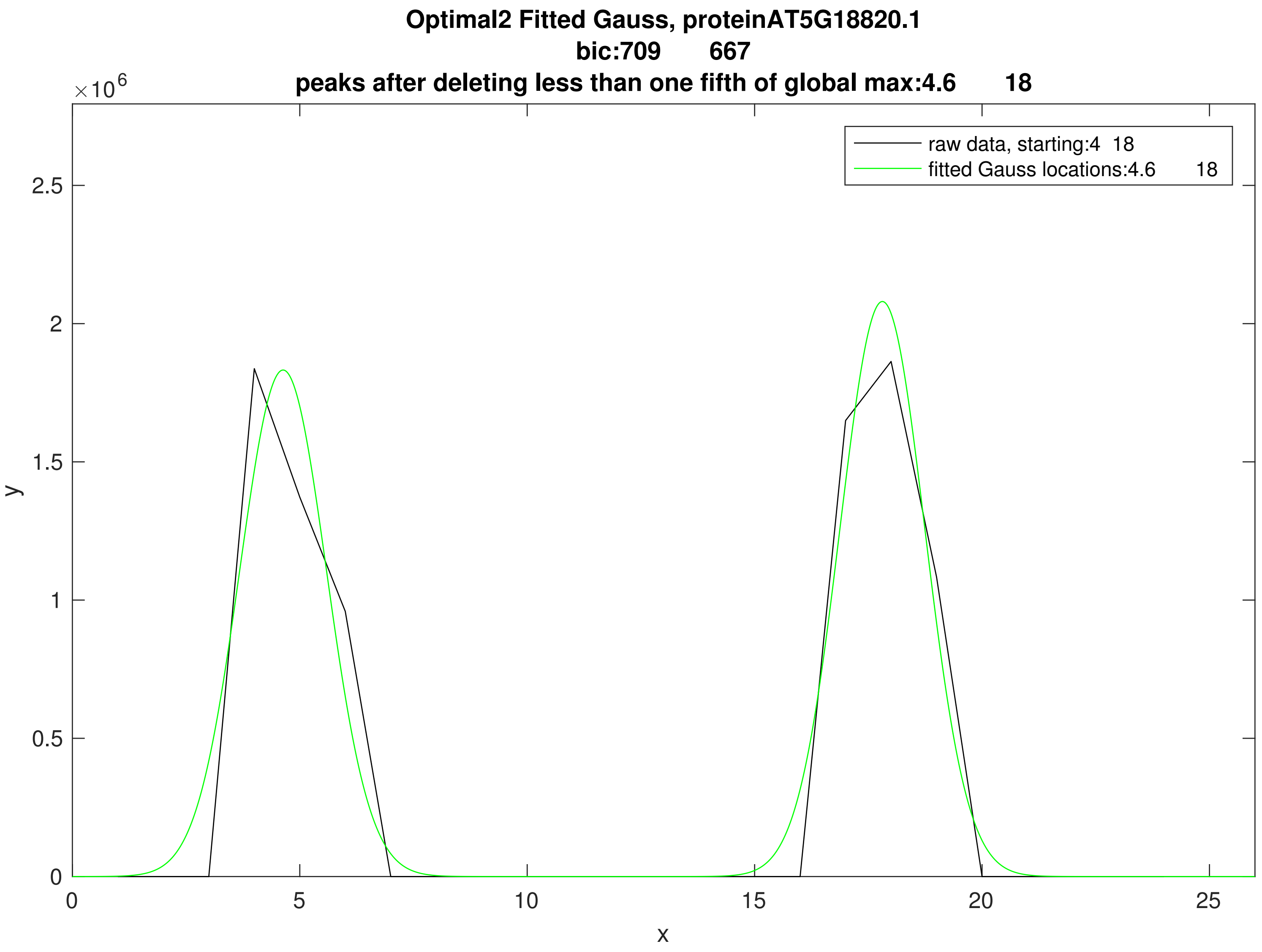
Optimal2 Fitted Gauss, proteinAT5G18820.1
bic:709 667
peaks after deleting less than one fifth of global max:4.6 18
×10^6
raw data, starting:4 18
fitted Gauss locations:4.6 18
y
x
0
0.5
1
1.5
2
2.5
0
5
10
15
20
25

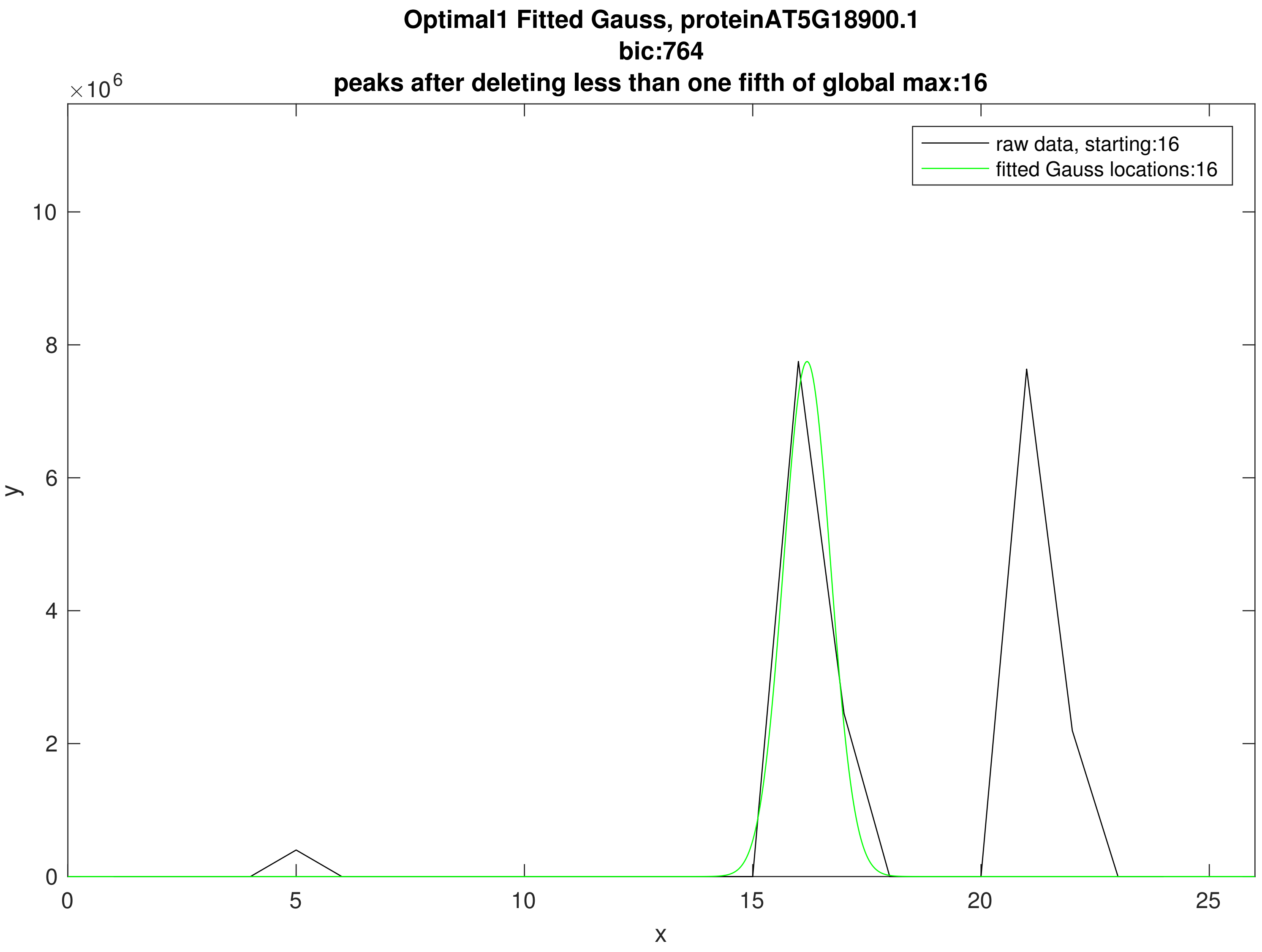
Optimal1 Fitted Gauss, proteinAT5G18900.1
bic:764
peaks after deleting less than one fifth of global max:16
×10^6
raw data, starting:16
fitted Gauss locations:16
y
x
0
2
4
6
8
10
0
5
10
15
20
25

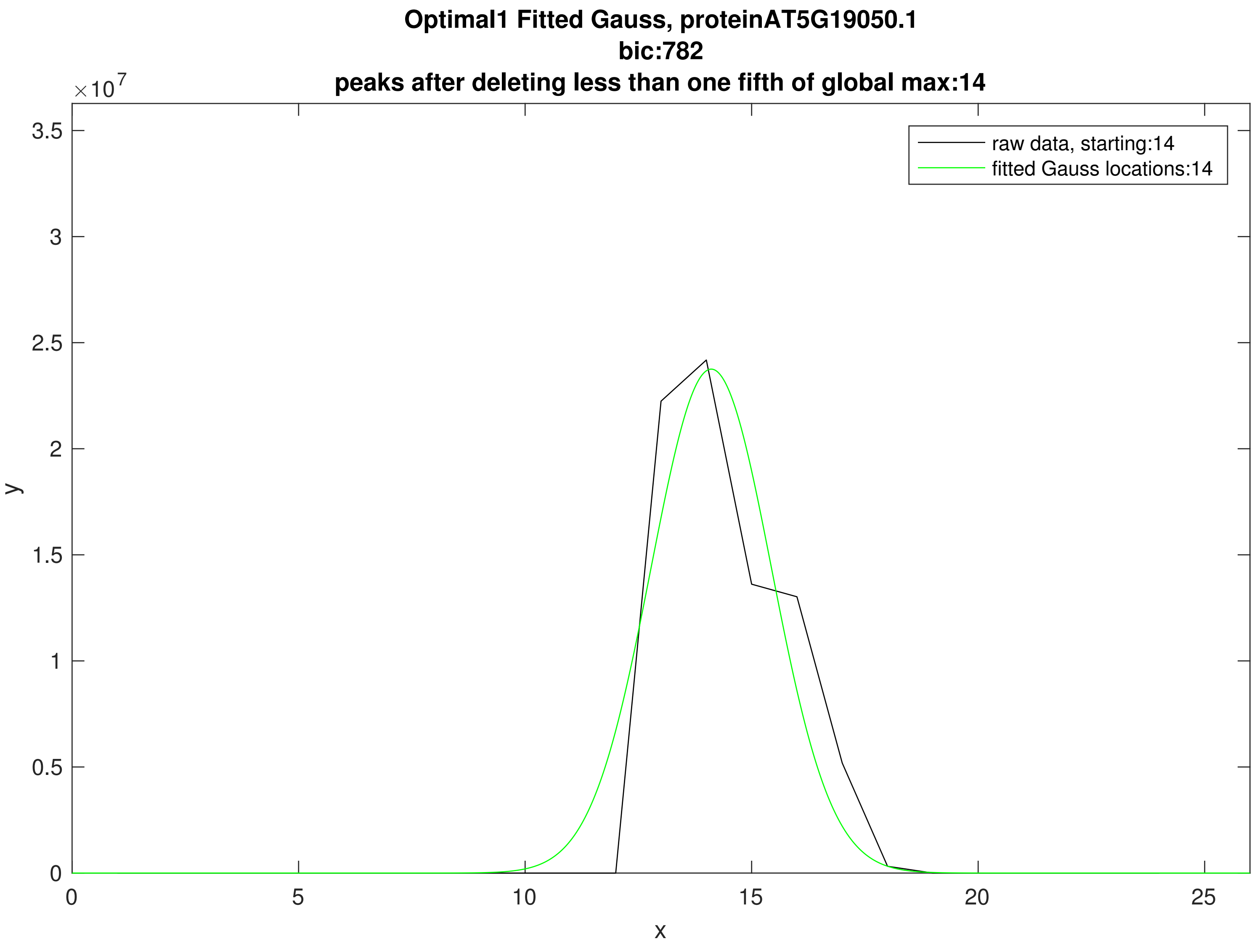

Optimal1 Fitted Gauss, proteinAT5G19050.1
bic:782
peaks after deleting less than one fifth of global max:14
×10^7
raw data, starting:14
fitted Gauss locations:14
3.5
3
2.5
2
1.5
1
0.5
0
y
0
5
10
15
20
25
x

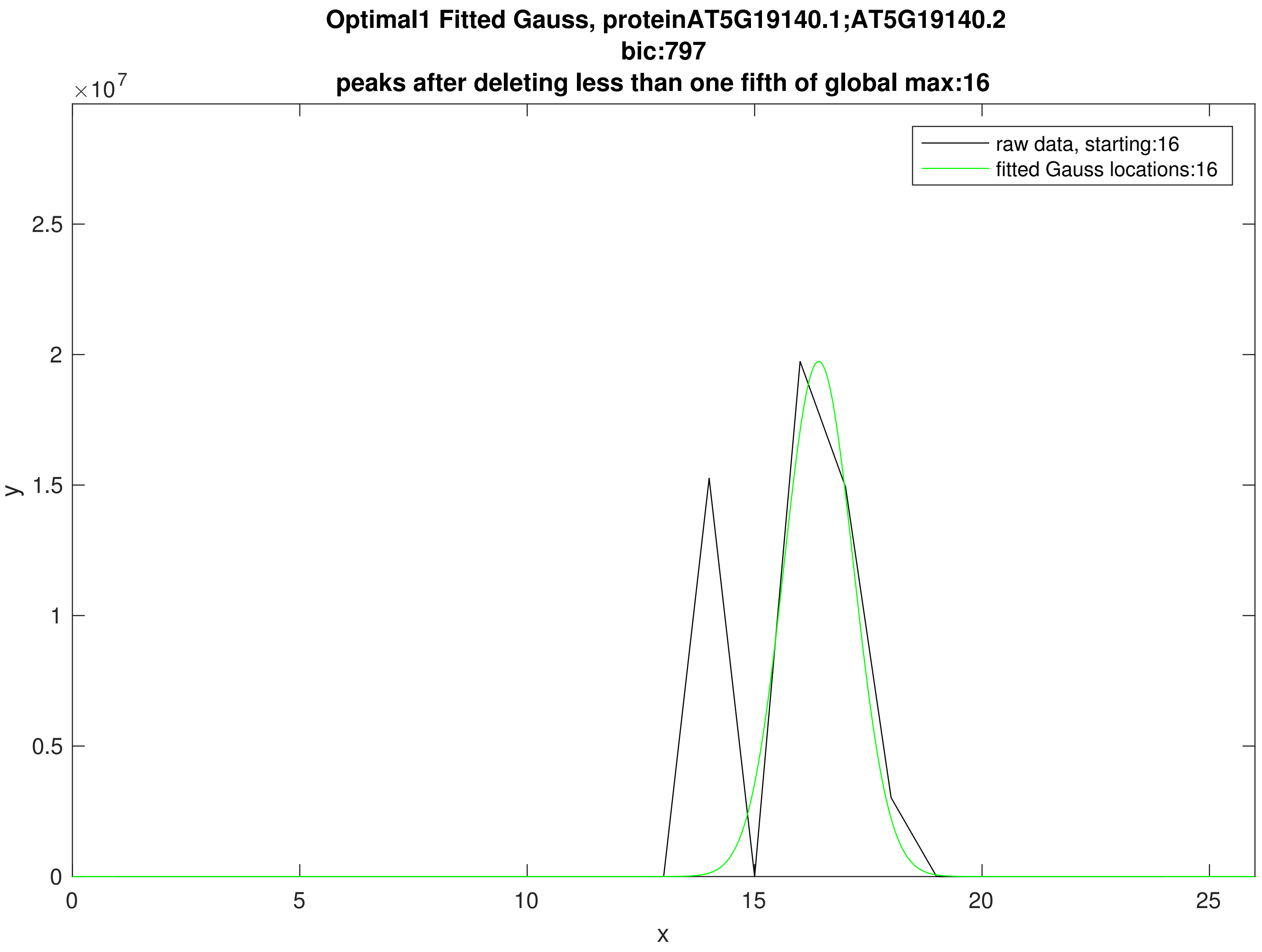
Optimal1 Fitted Gauss, proteinAT5G19140.1;AT5G19140.2
bic:797
peaks after deleting less than one fifth of global max:16
×10^7
raw data, starting:16
fitted Gauss locations:16
y
x
0
0.5
1
1.5
2
2.5
0
5
10
15
20
25

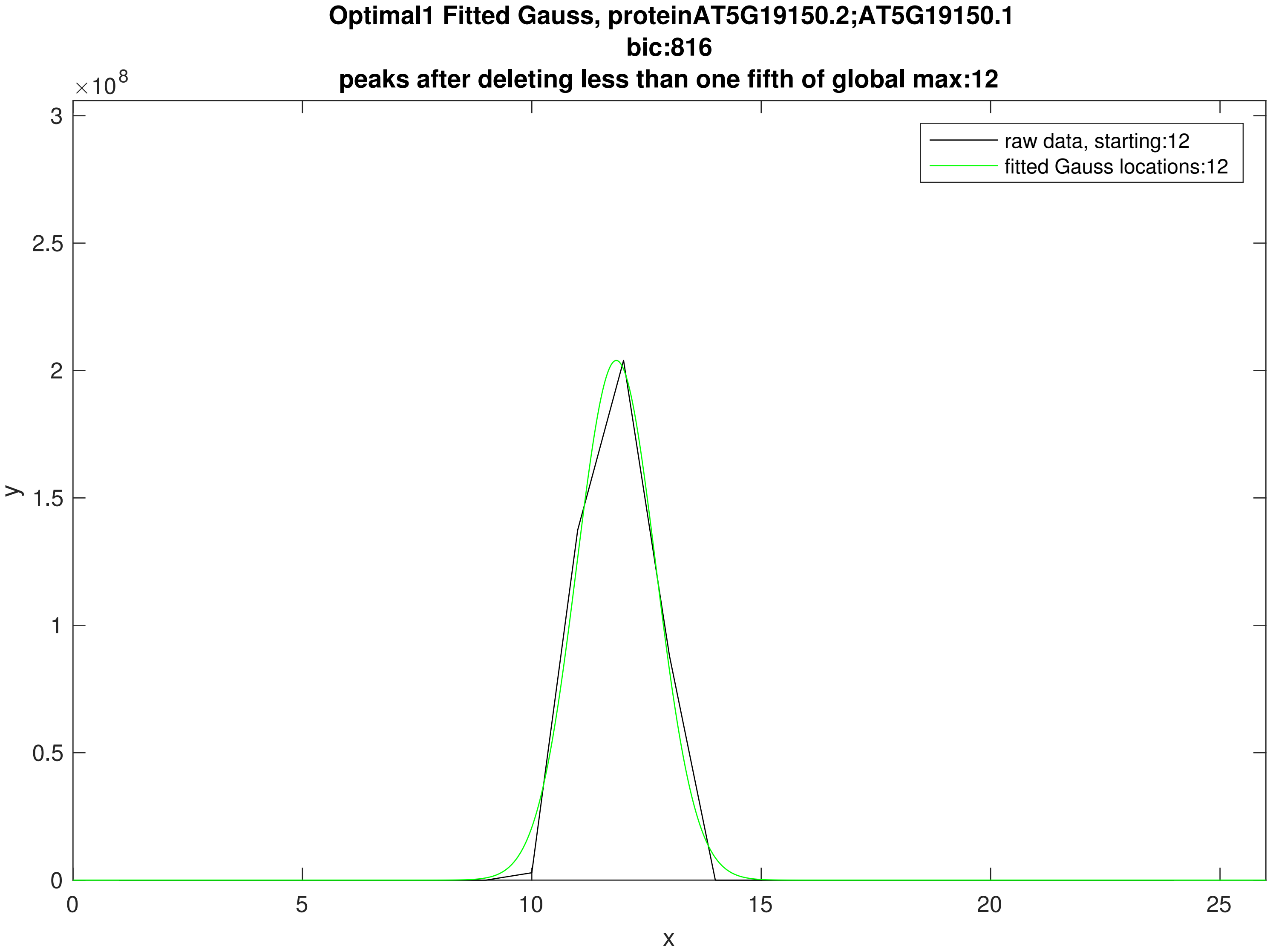

Optimal1 Fitted Gauss, proteinAT5G19150.2;AT5G19150.1
bic:816
peaks after deleting less than one fifth of global max:12
×10^8
3
2.5
2
1.5
1
0.5
0
y
0
5
10
15
20
25
x
raw data, starting:12
fitted Gauss locations:12

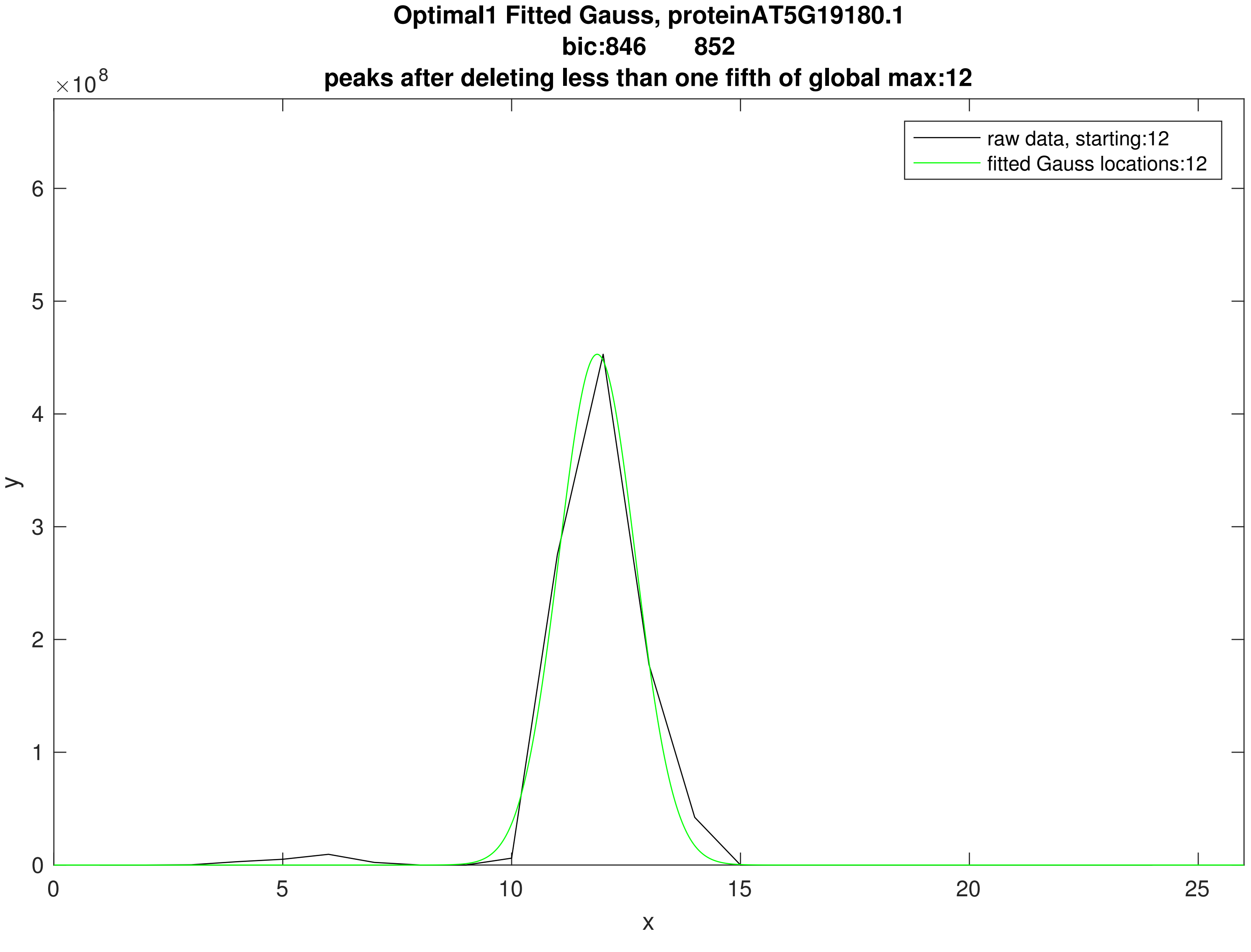

Optimal1 Fitted Gauss, proteinAT5G19180.1
bic:846 852
peaks after deleting less than one fifth of global max:12
×10^8
raw data, starting:12
fitted Gauss locations:12
y
x
0
1
2
3
4
5
6
0
5
10
15
20
25

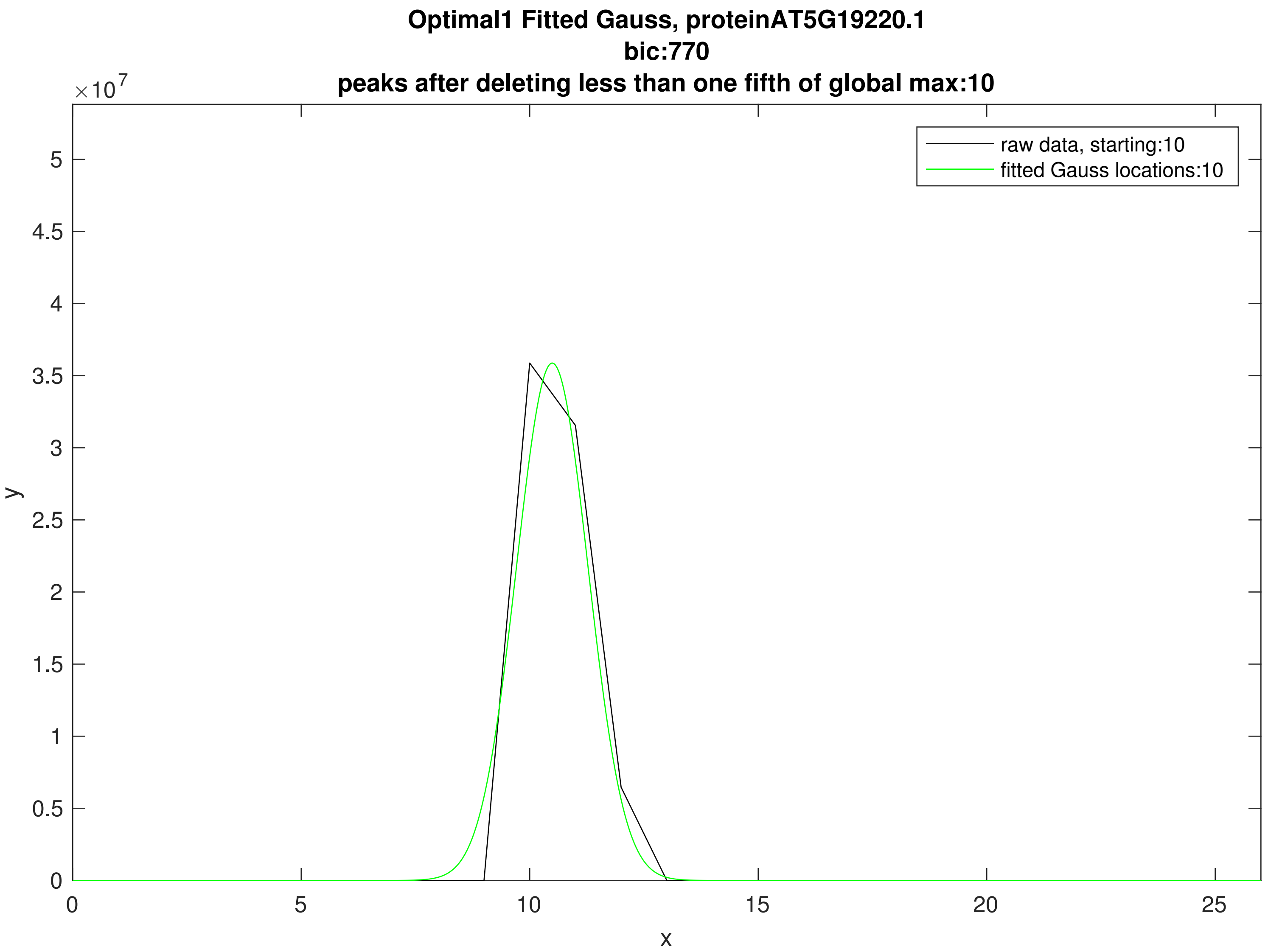
Optimal1 Fitted Gauss, proteinAT5G19220.1
bic:770
peaks after deleting less than one fifth of global max:10
×10^7
raw data, starting:10
fitted Gauss locations:10
y
x
0
0.5
1
1.5
2
2.5
3
3.5
4
4.5
5
0
5
10
15
20
25

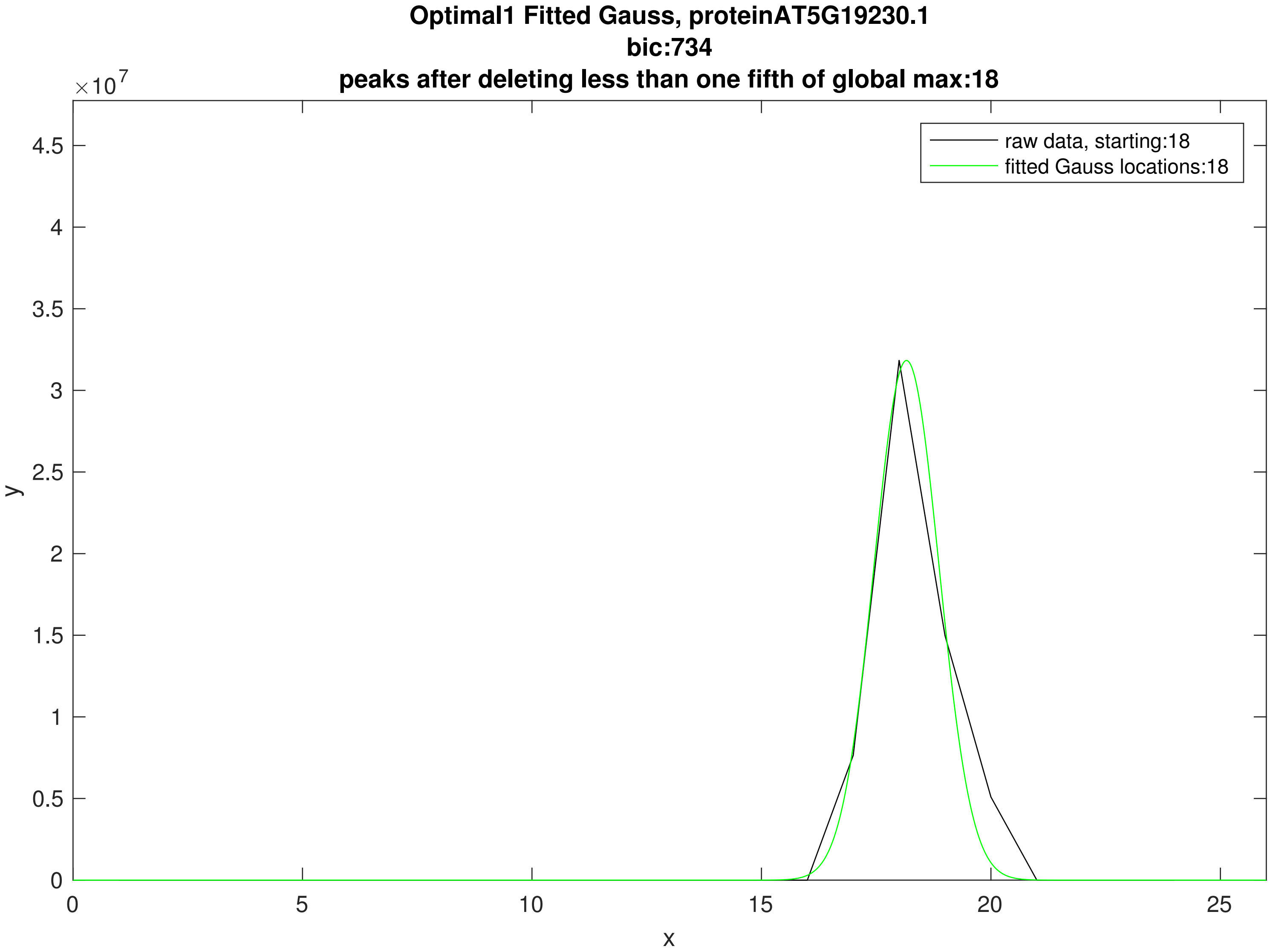

Optimal1 Fitted Gauss, proteinAT5G19230.1
bic:734
peaks after deleting less than one fifth of global max:18
×10^7
4.5
4
3.5
3
2.5
2
1.5
1
0.5
0
y
0
5
10
15
20
25
x
raw data, starting:18
fitted Gauss locations:18

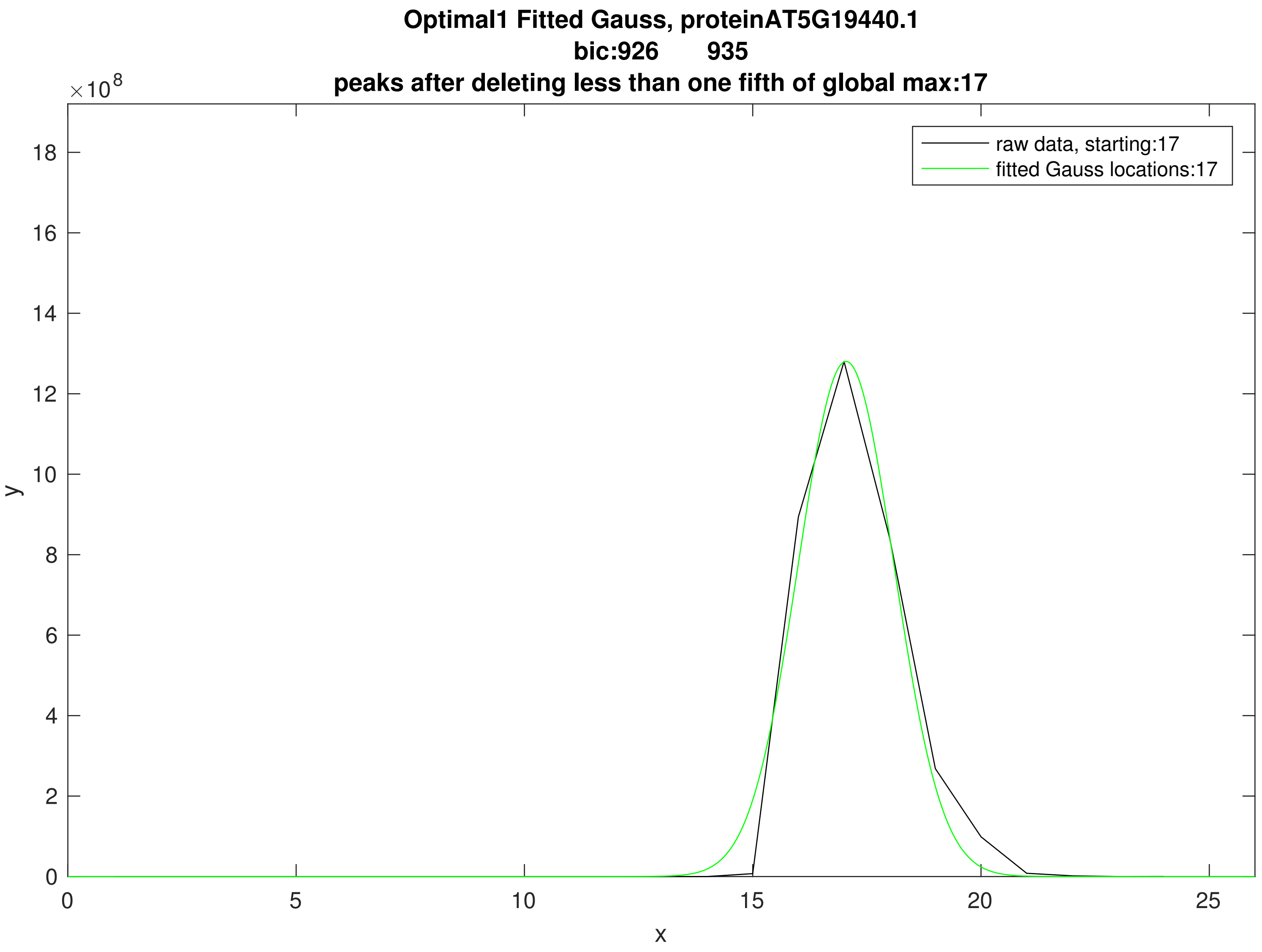
Optimal1 Fitted Gauss, proteinAT5G19440.1
bic:926 935
peaks after deleting less than one fifth of global max:17
×10^8
raw data, starting:17
fitted Gauss locations:17
y
x
0
2
4
6
8
10
12
14
16
18
0
5
10
15
20
25

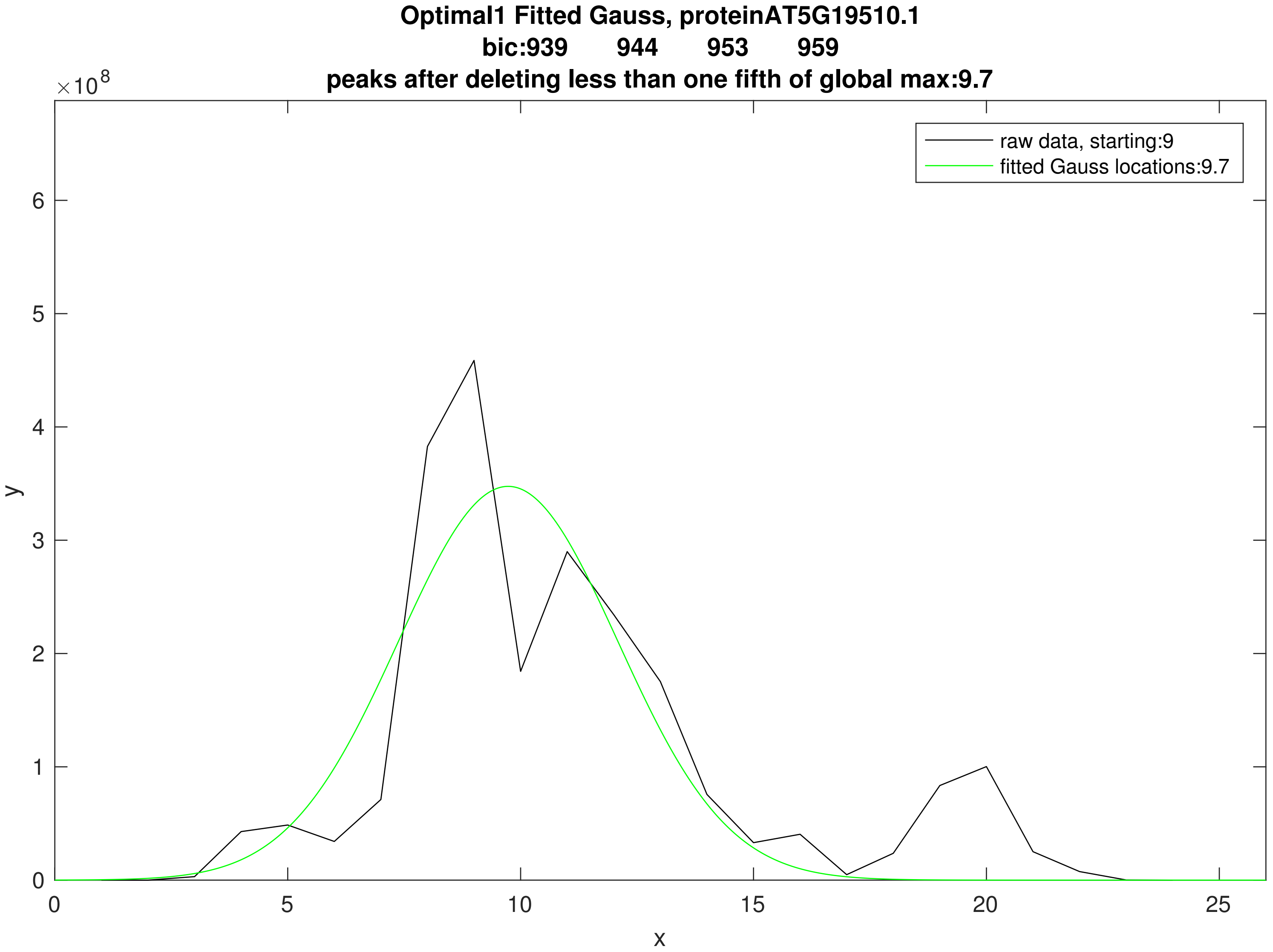
Optimal1 Fitted Gauss, proteinAT5G19510.1
bic:939 944 953 959
peaks after deleting less than one fifth of global max:9.7
×10^8
raw data, starting:9
fitted Gauss locations:9.7
y
x
0
1
2
3
4
5
6
0
5
10
15
20
25

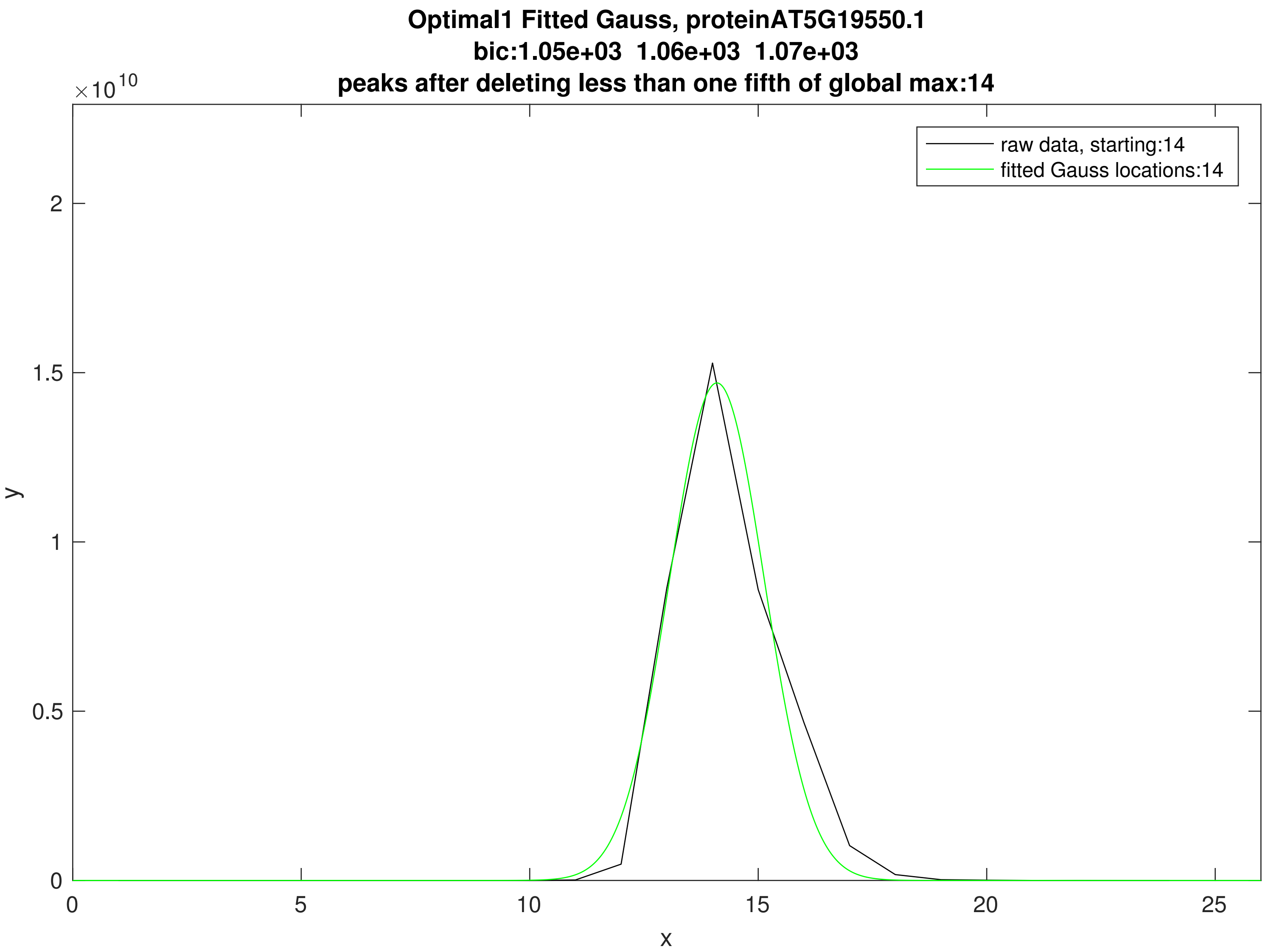
Optimal1 Fitted Gauss, proteinAT5G19550.1
bic:1.05e+03 1.06e+03 1.07e+03
peaks after deleting less than one fifth of global max:14
×10^10
raw data, starting:14
fitted Gauss locations:14
y
x
0
0.5
1
1.5
2
5
10
15
20
25

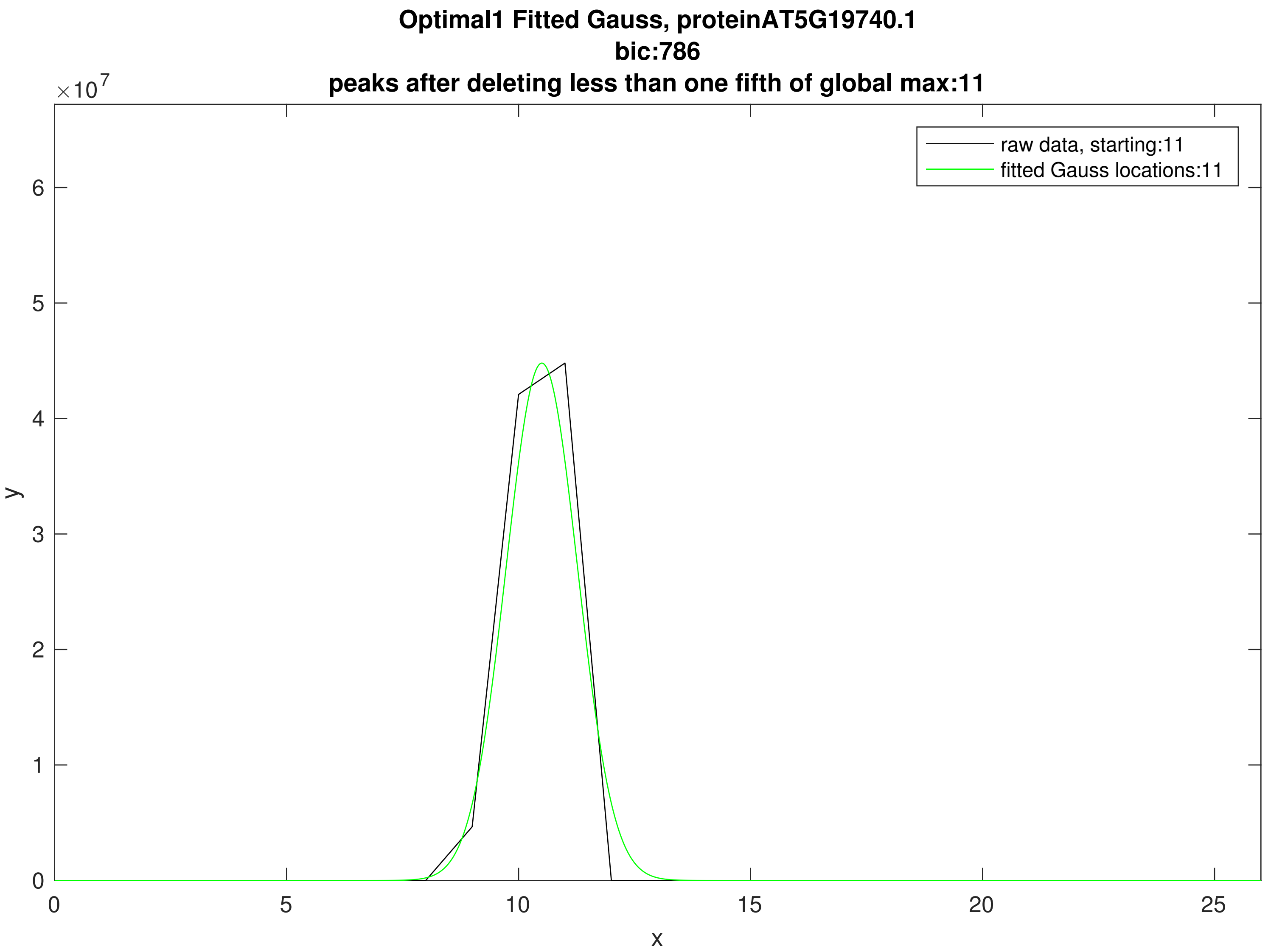

Optimal1 Fitted Gauss, proteinAT5G19740.1
bic:786
peaks after deleting less than one fifth of global max:11
×10^7
raw data, starting:11
fitted Gauss locations:11
y
x
0
1
2
3
4
5
6
0
5
10
15
20
25

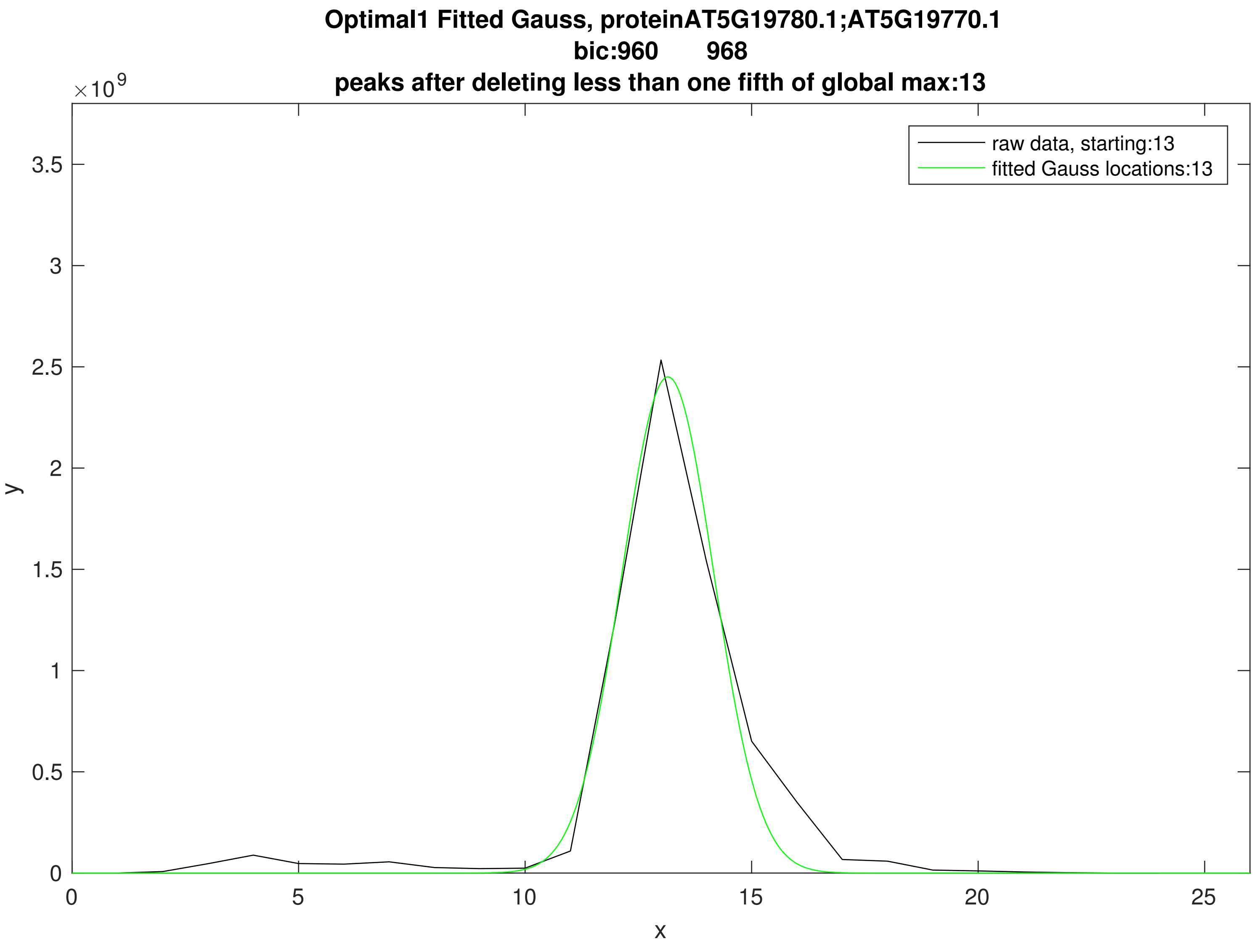
Optimal1 Fitted Gauss, proteinAT5G19780.1;AT5G19770.1
bic:960 968
peaks after deleting less than one fifth of global max:13
×10^9
3.5
3
2.5
2
1.5
1
0.5
0
y
0
5
10
15
20
25
x
raw data, starting:13
fitted Gauss locations:13

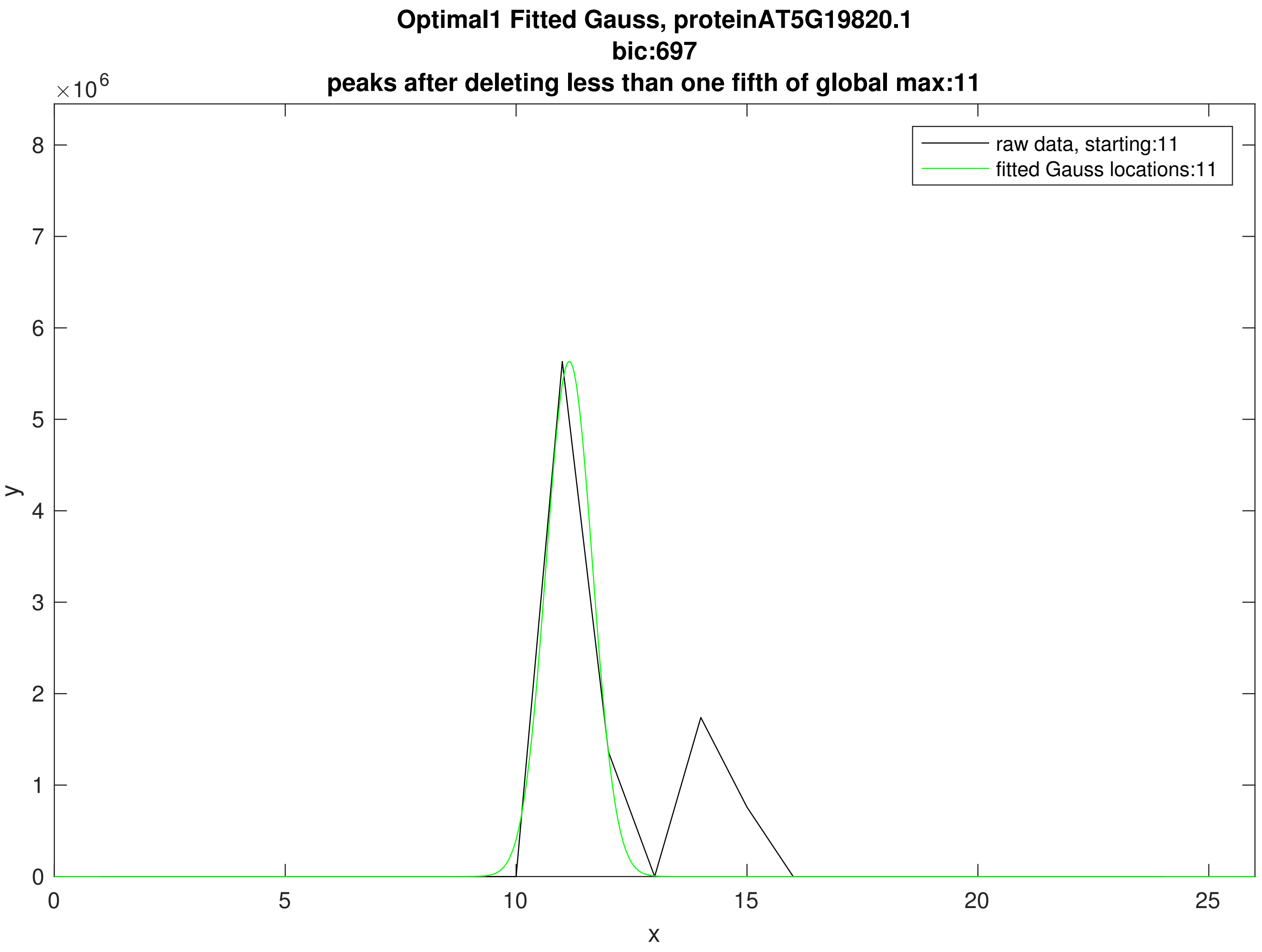
Optimal1 Fitted Gauss, proteinAT5G19820.1
bic:697
peaks after deleting less than one fifth of global max:11
×10^6
raw data, starting:11
fitted Gauss locations:11
y
x
0
1
2
3
4
5
6
7
8
0
5
10
15
20
25

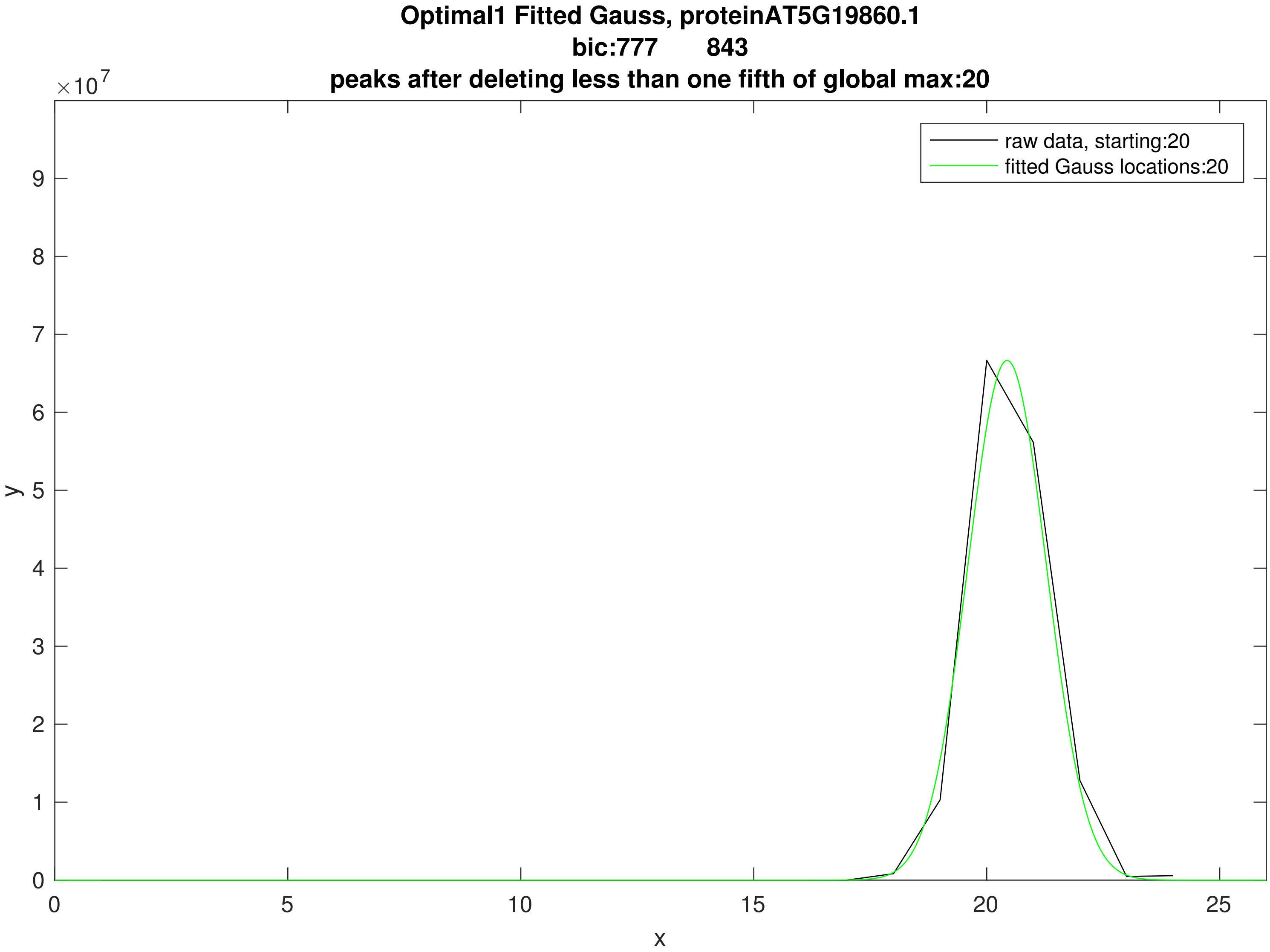
Optimal1 Fitted Gauss, proteinAT5G19860.1
bic:777 843
peaks after deleting less than one fifth of global max:20
×10^7
raw data, starting:20
fitted Gauss locations:20
y
x
0
1
2
3
4
5
6
7
8
9
0
5
10
15
20
25

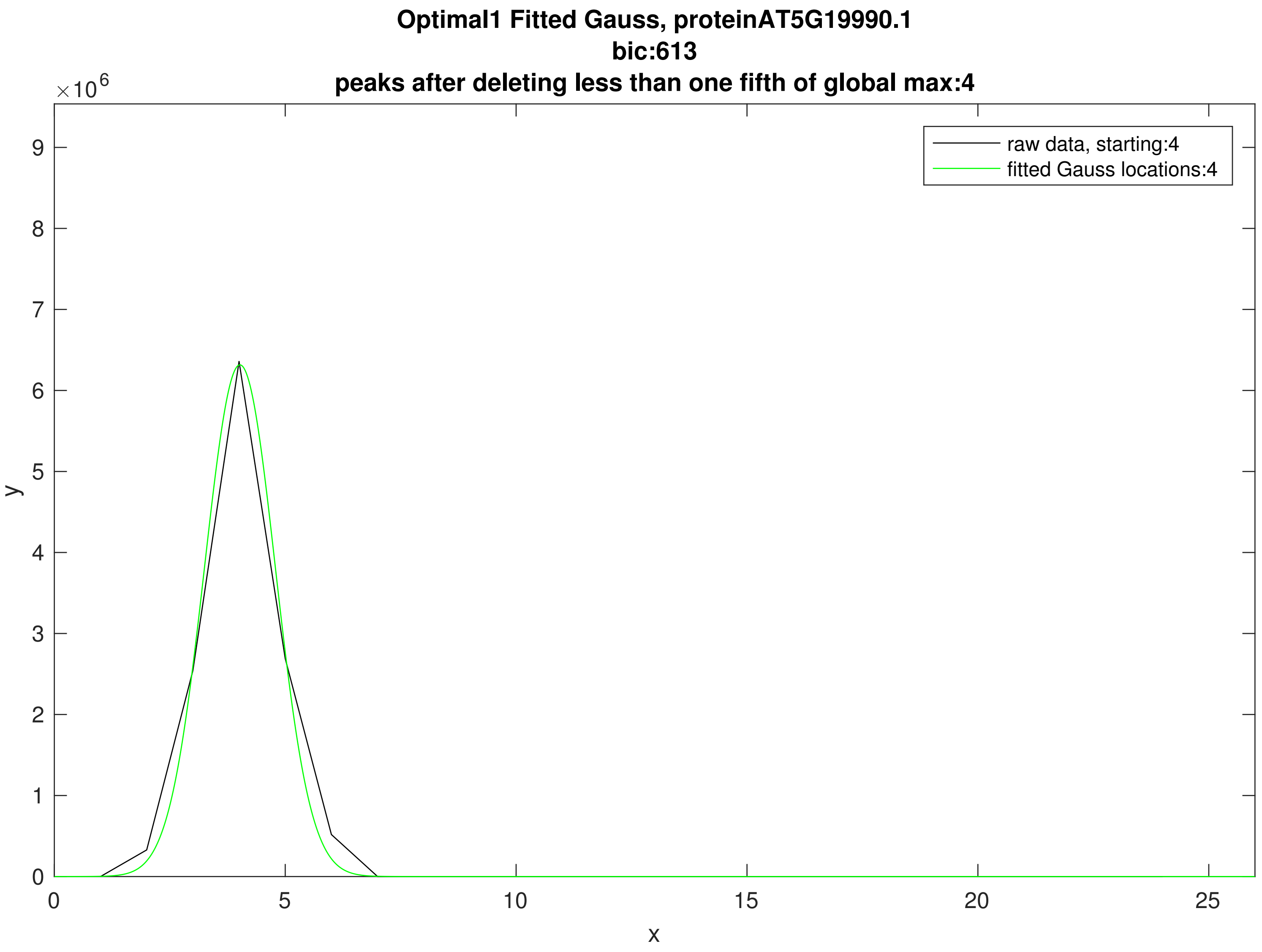

Optimal1 Fitted Gauss, proteinAT5G19990.1
bic:613
peaks after deleting less than one fifth of global max:4
×10^6
raw data, starting:4
fitted Gauss locations:4
y
x
0
1
2
3
4
5
6
7
8
9
0
5
10
15
20
25

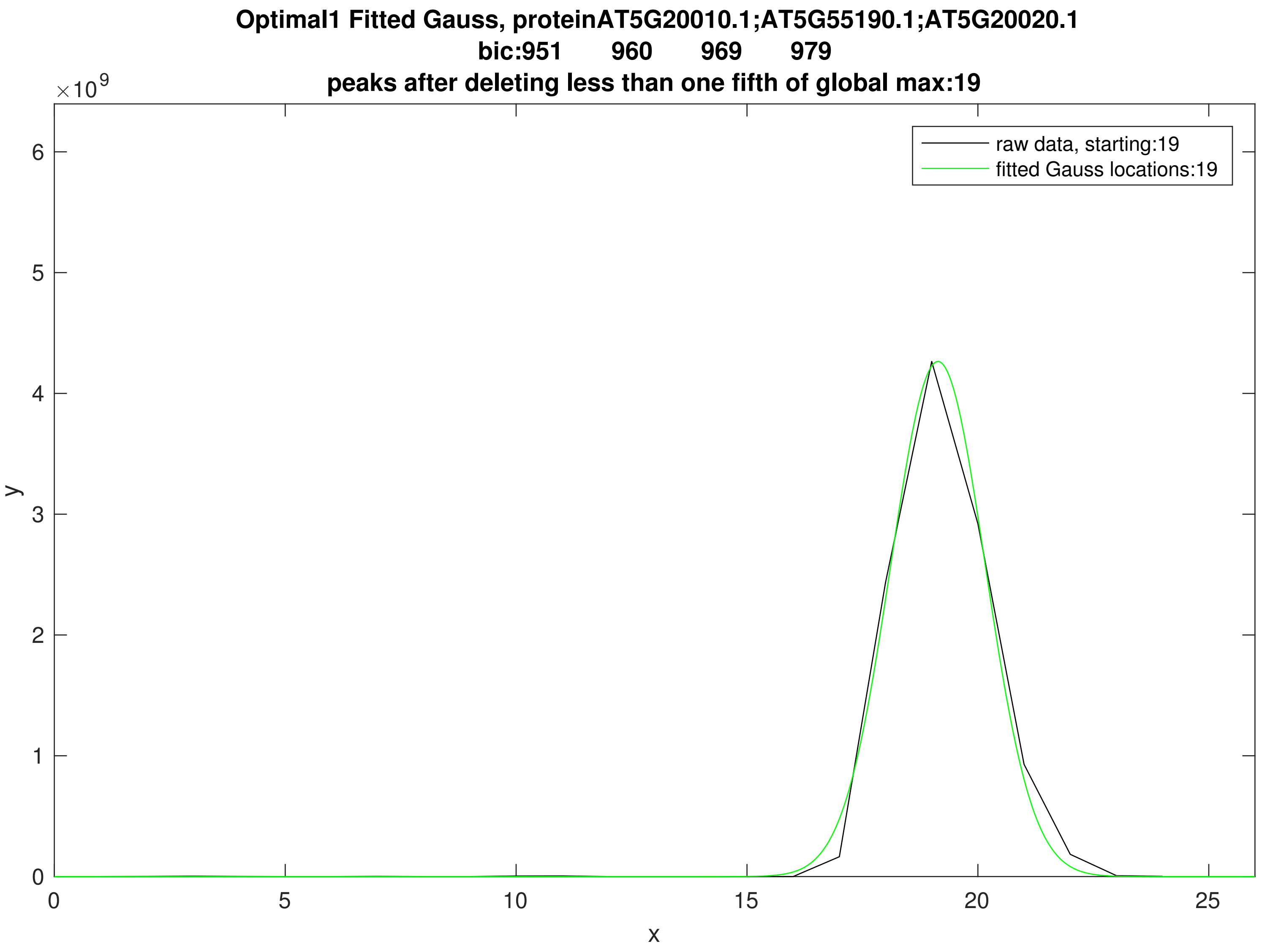
Optimal1 Fitted Gauss, proteinAT5G20010.1;AT5G55190.1;AT5G20020.1
bic:951 960 969 979
peaks after deleting less than one fifth of global max:19
×10^9
raw data, starting:19
fitted Gauss locations:19
y
x
0
1
2
3
4
5
6
0
5
10
15
20
25

**Optimal1 Fitted Gauss, proteinAT5G20060.3;AT5G20060.2;AT5G20060.1**
**bic:798 804**
**peaks after deleting less than one fifth of global max:19**

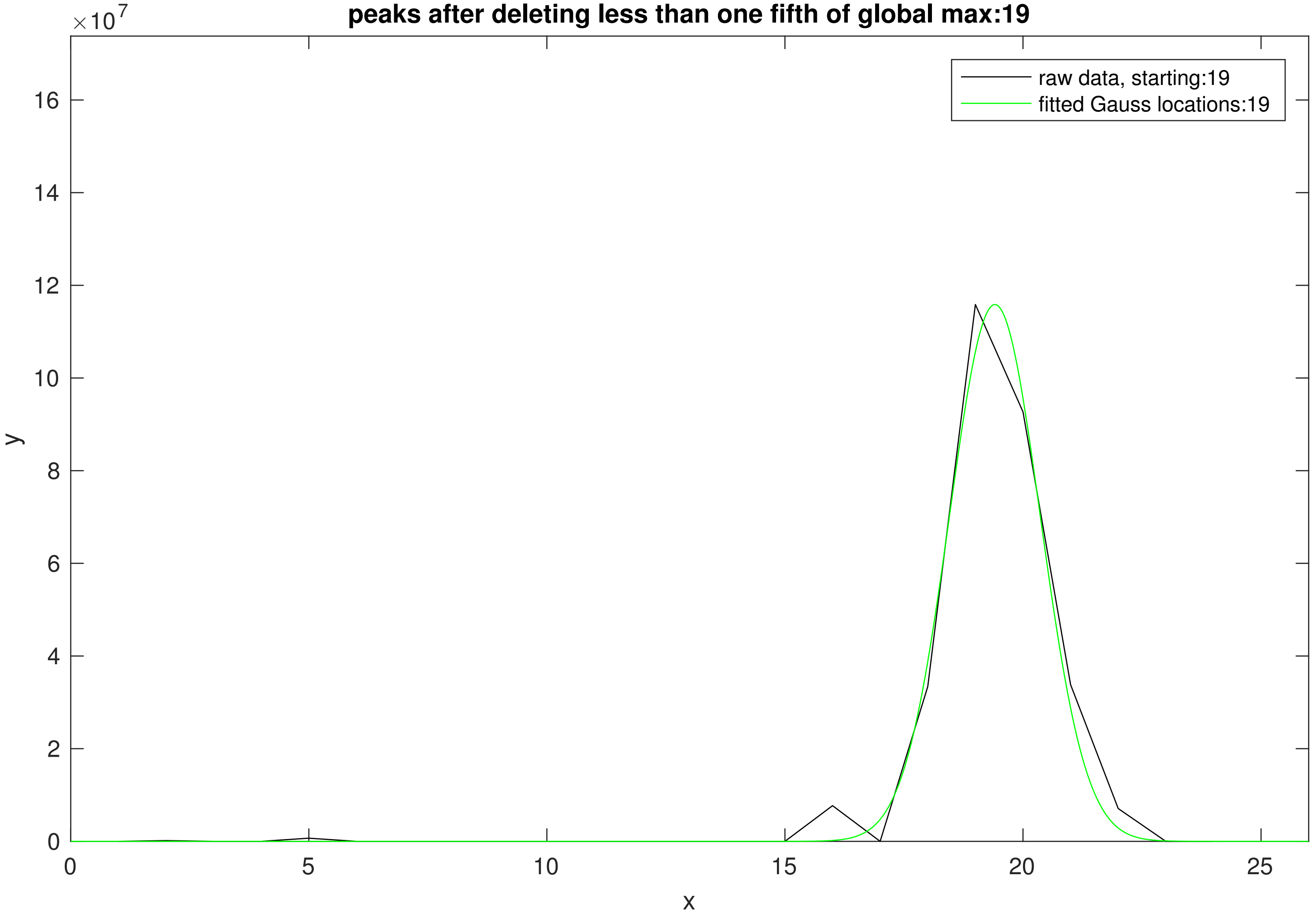

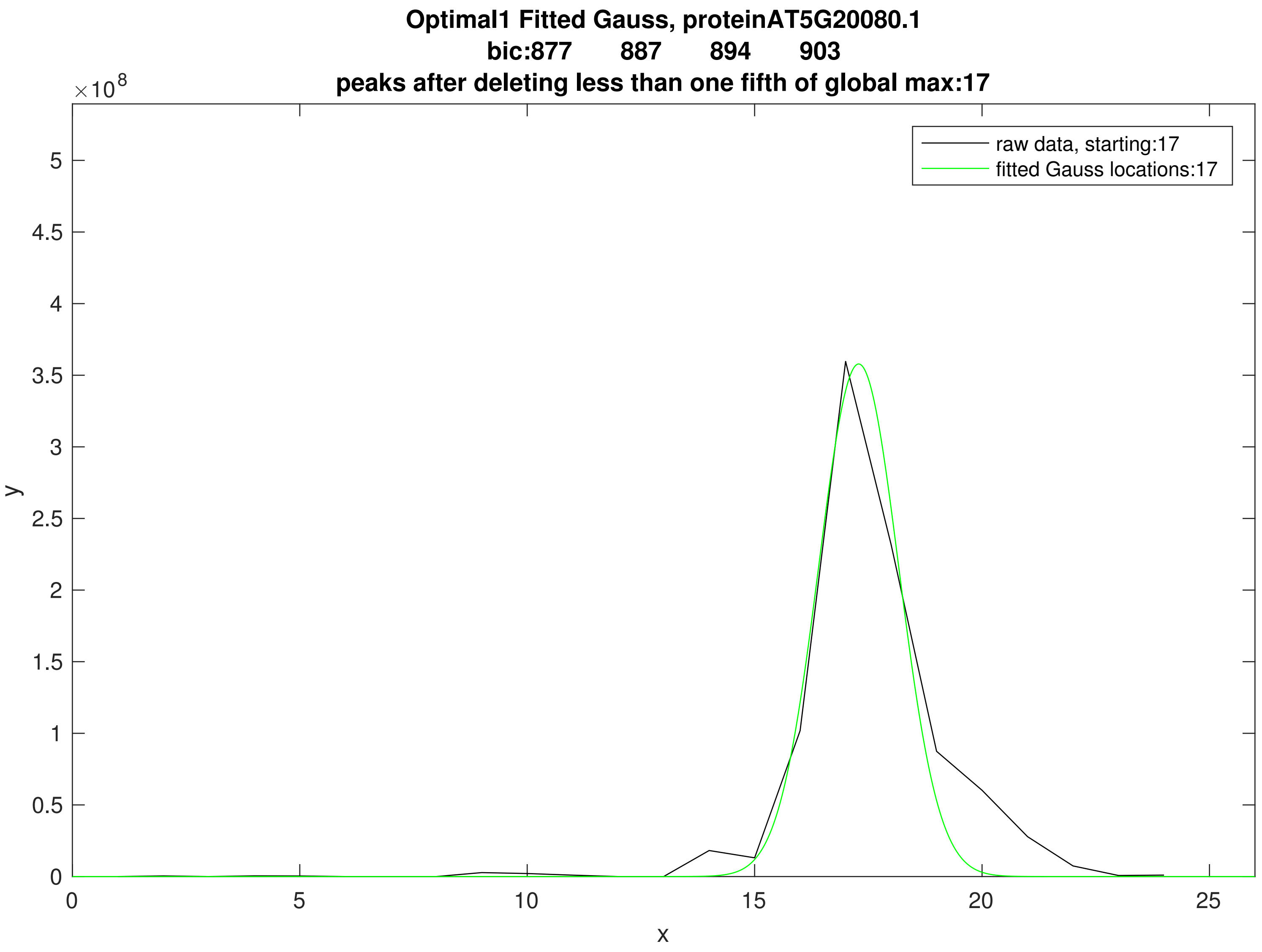
Optimal1 Fitted Gauss, proteinAT5G20080.1
bic:877 887 894 903
peaks after deleting less than one fifth of global max:17
×10^8
raw data, starting:17
fitted Gauss locations:17
5
4.5
4
3.5
3
2.5
2
1.5
1
0.5
0
y
0
5
10
15
20
25
x

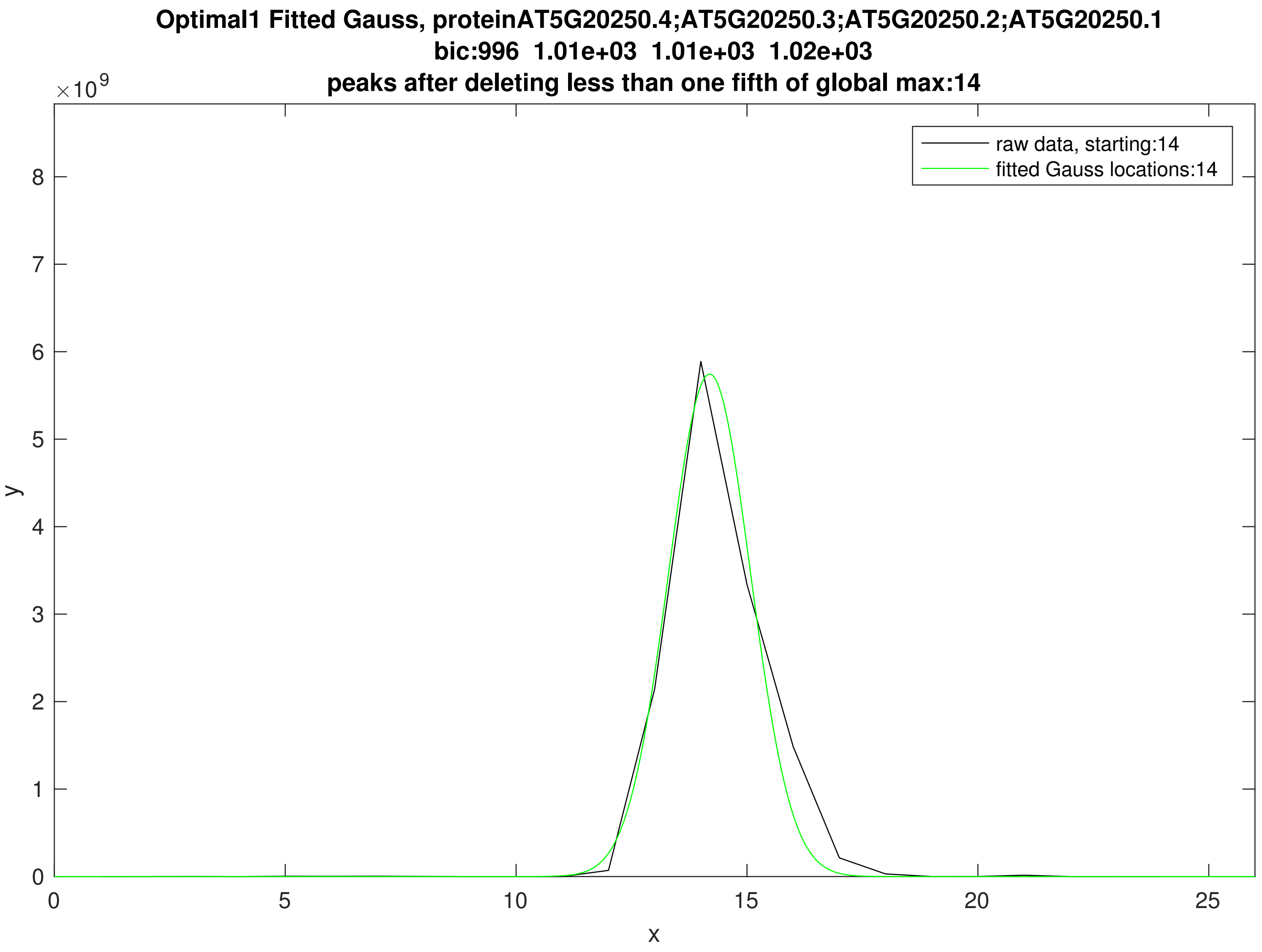

Optimal1 Fitted Gauss, proteinAT5G20250.4;AT5G20250.3;AT5G20250.2;AT5G20250.1
bic:996 1.01e+03 1.01e+03 1.02e+03
peaks after deleting less than one fifth of global max:14
×10^9
raw data, starting:14
fitted Gauss locations:14
y
x
0
1
2
3
4
5
6
7
8
0
5
10
15
20
25

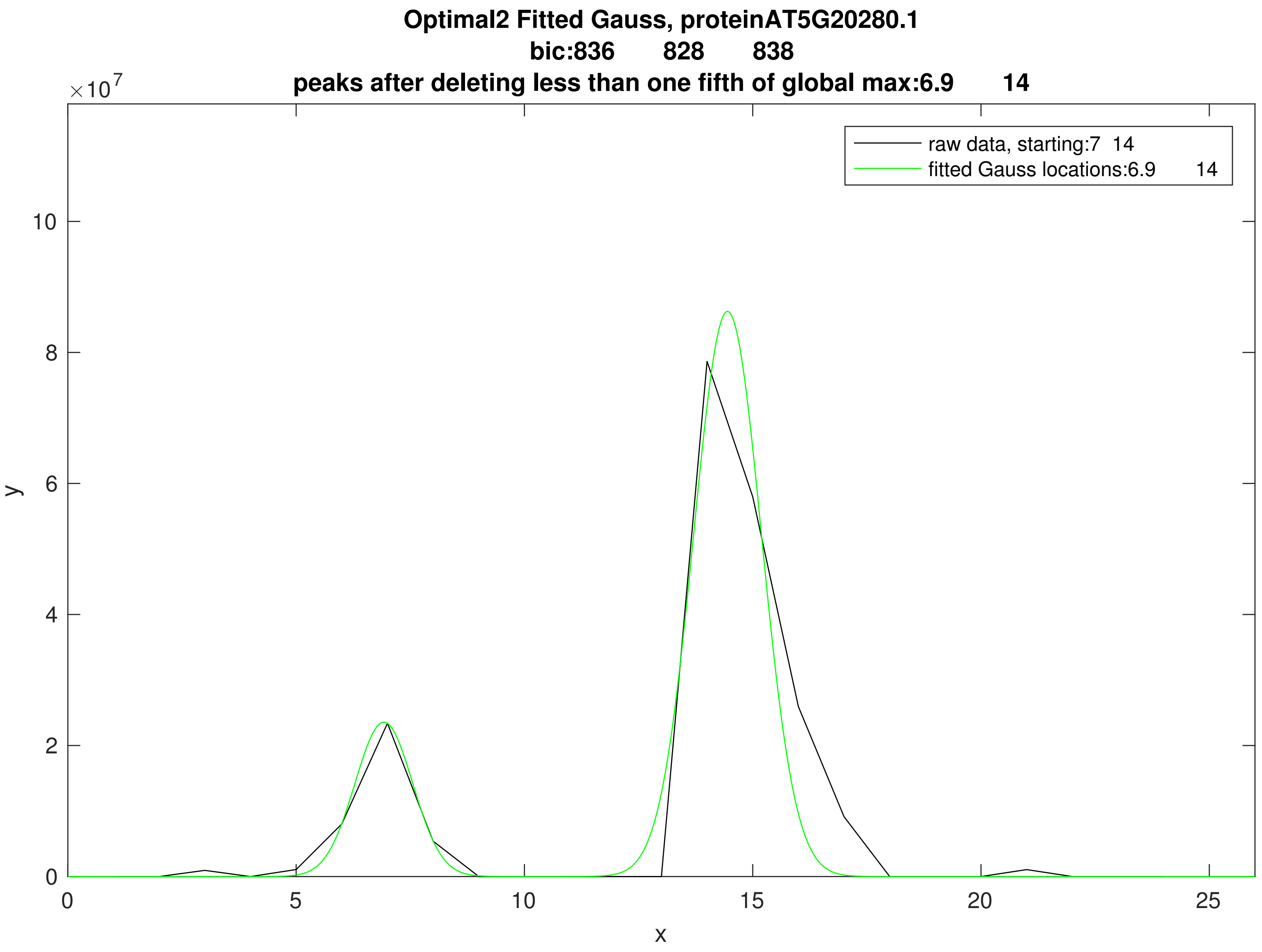
Optimal2 Fitted Gauss, proteinAT5G20280.1
bic:836 828 838
peaks after deleting less than one fifth of global max:6.9 14
×10^7
raw data, starting:7 14
fitted Gauss locations:6.9 14
10
8
6
4
2
0
y
0
5
10
15
20
25
x

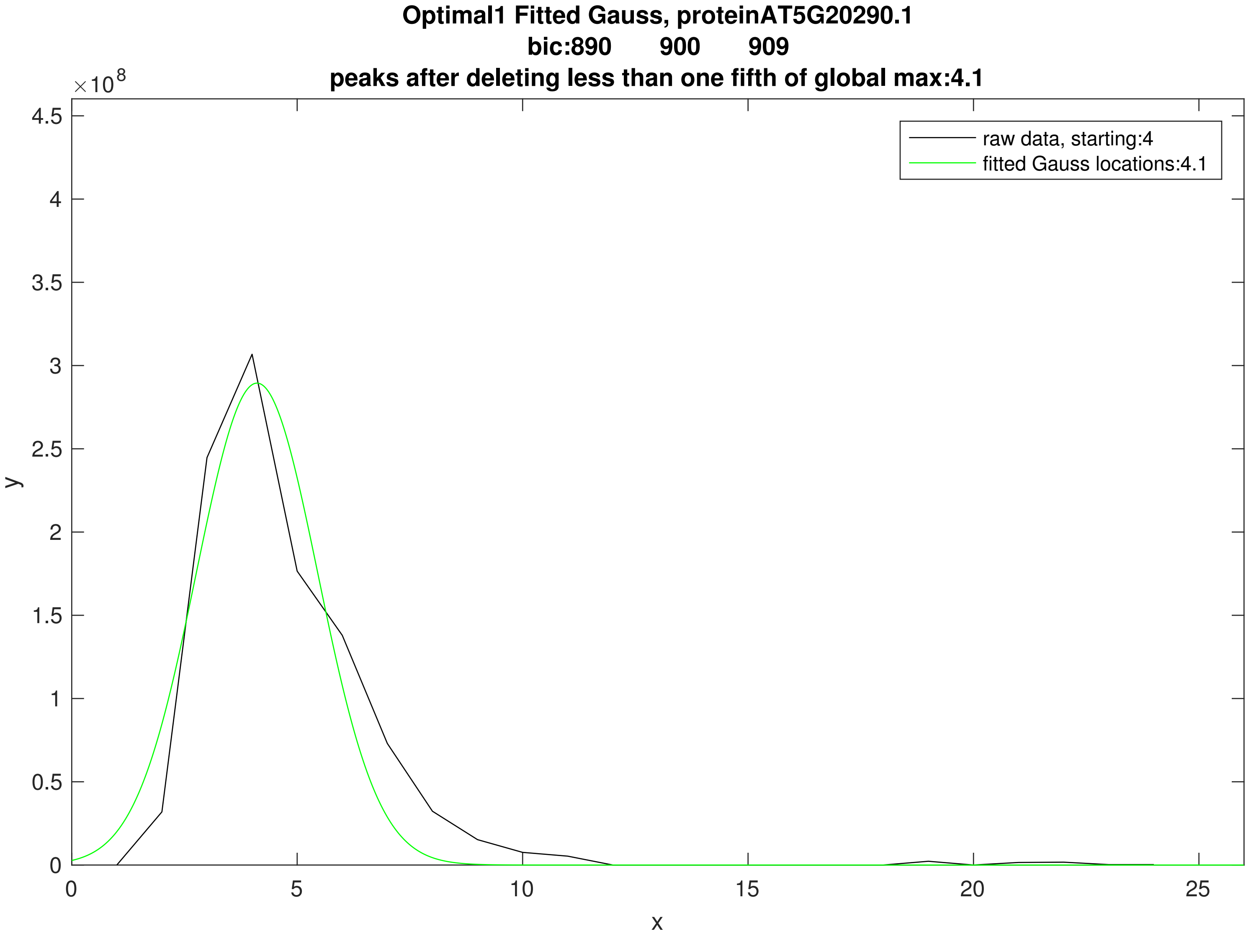

Optimal1 Fitted Gauss, proteinAT5G20290.1
bic:890 900 909
peaks after deleting less than one fifth of global max:4.1
×10^8
4.5
4
3.5
3
2.5
2
1.5
1
0.5
0
y
0
5
10
15
20
25
x
raw data, starting:4
fitted Gauss locations:4.1

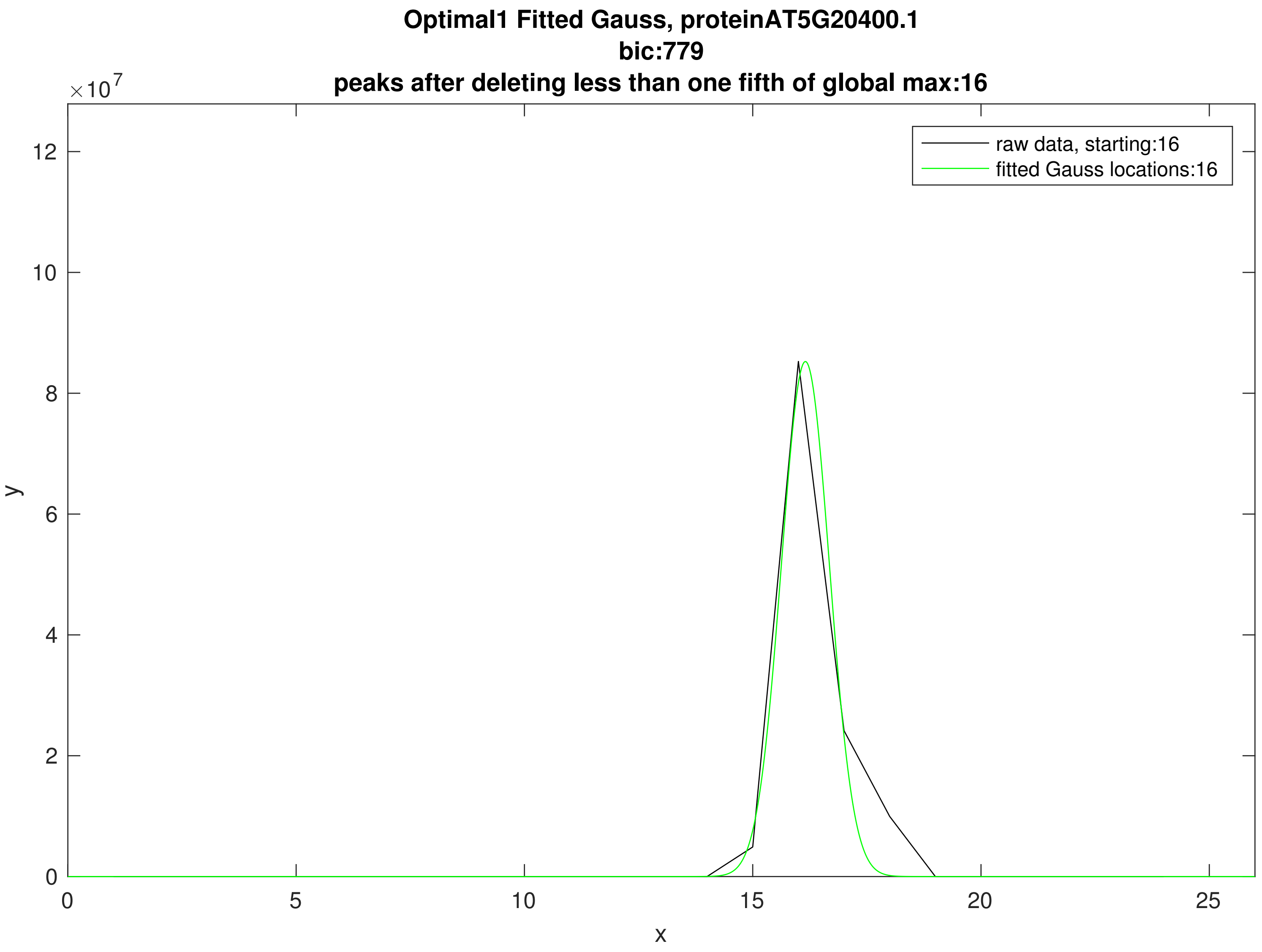

Optimal1 Fitted Gauss, proteinAT5G20400.1
bic:779
peaks after deleting less than one fifth of global max:16
×10^7
raw data, starting:16
fitted Gauss locations:16
y
x
0
2
4
6
8
10
12
0
5
10
15
20
25

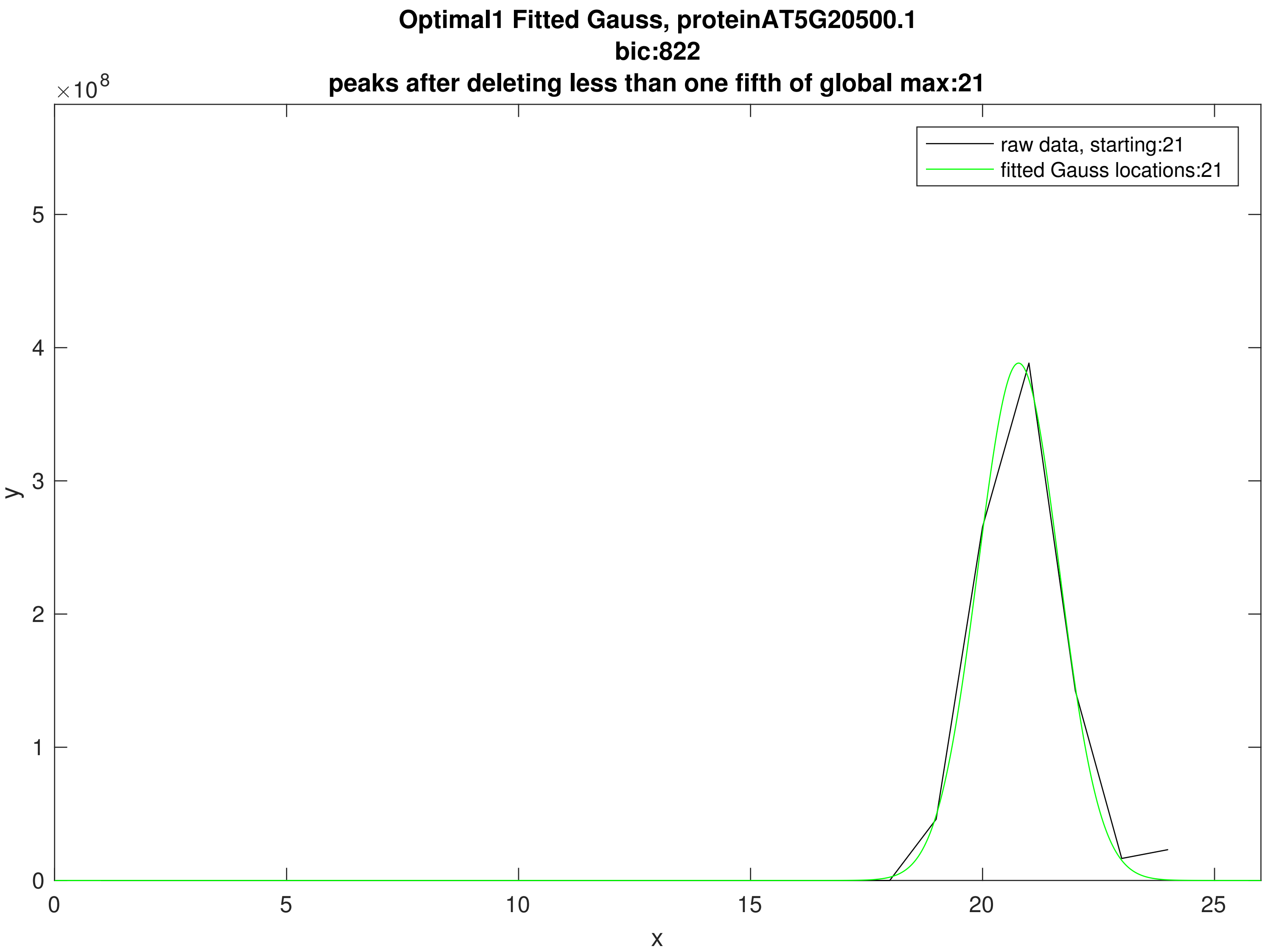
Optimal1 Fitted Gauss, proteinAT5G20500.1
bic:822
peaks after deleting less than one fifth of global max:21
×10^8
raw data, starting:21
fitted Gauss locations:21
5
4
3
2
1
0
y
0
5
10
15
20
25
x

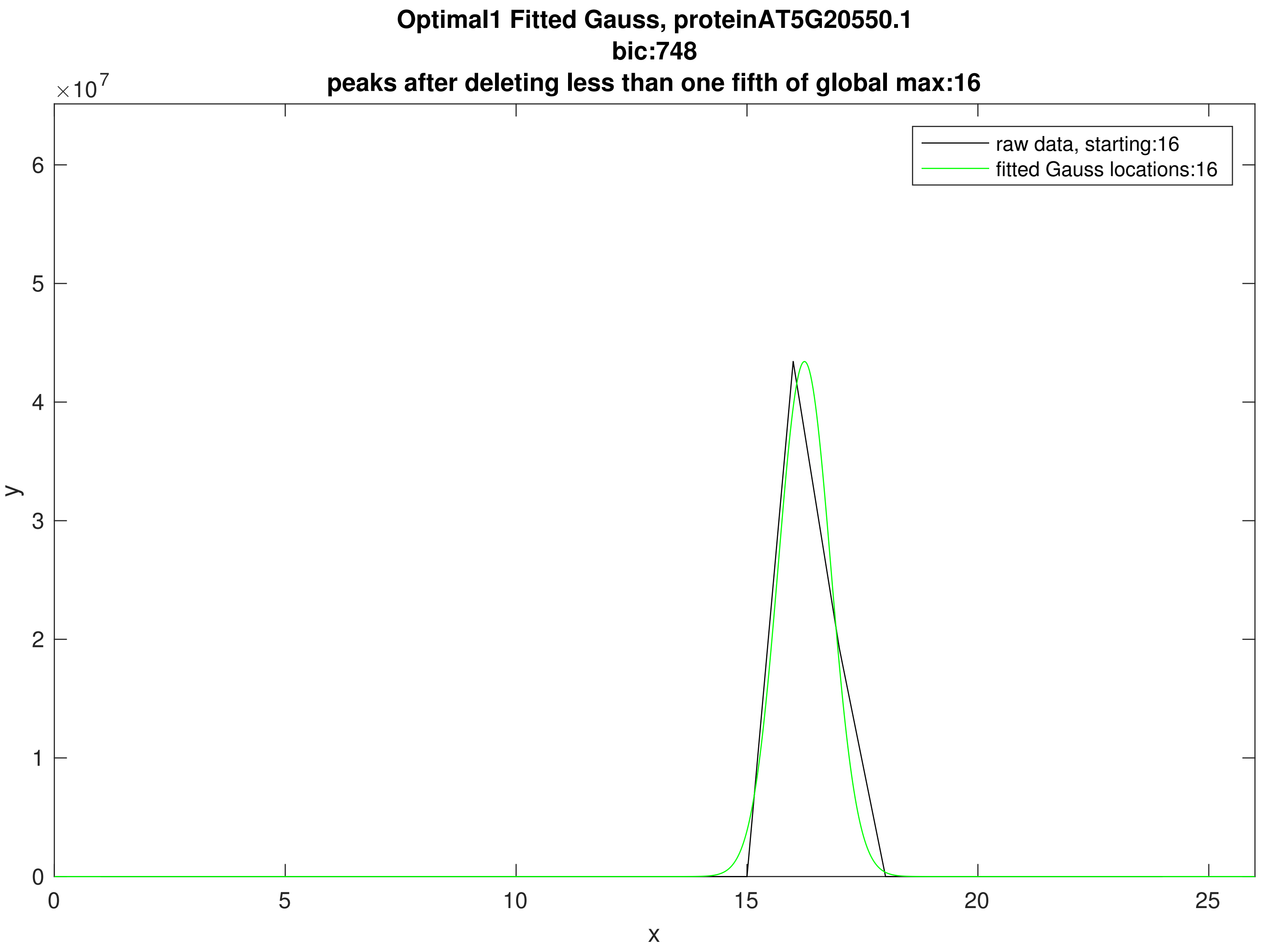

Optimal1 Fitted Gauss, proteinAT5G20550.1
bic:748
peaks after deleting less than one fifth of global max:16
×10^7
raw data, starting:16
fitted Gauss locations:16
y
x
0
1
2
3
4
5
6
0
5
10
15
20
25

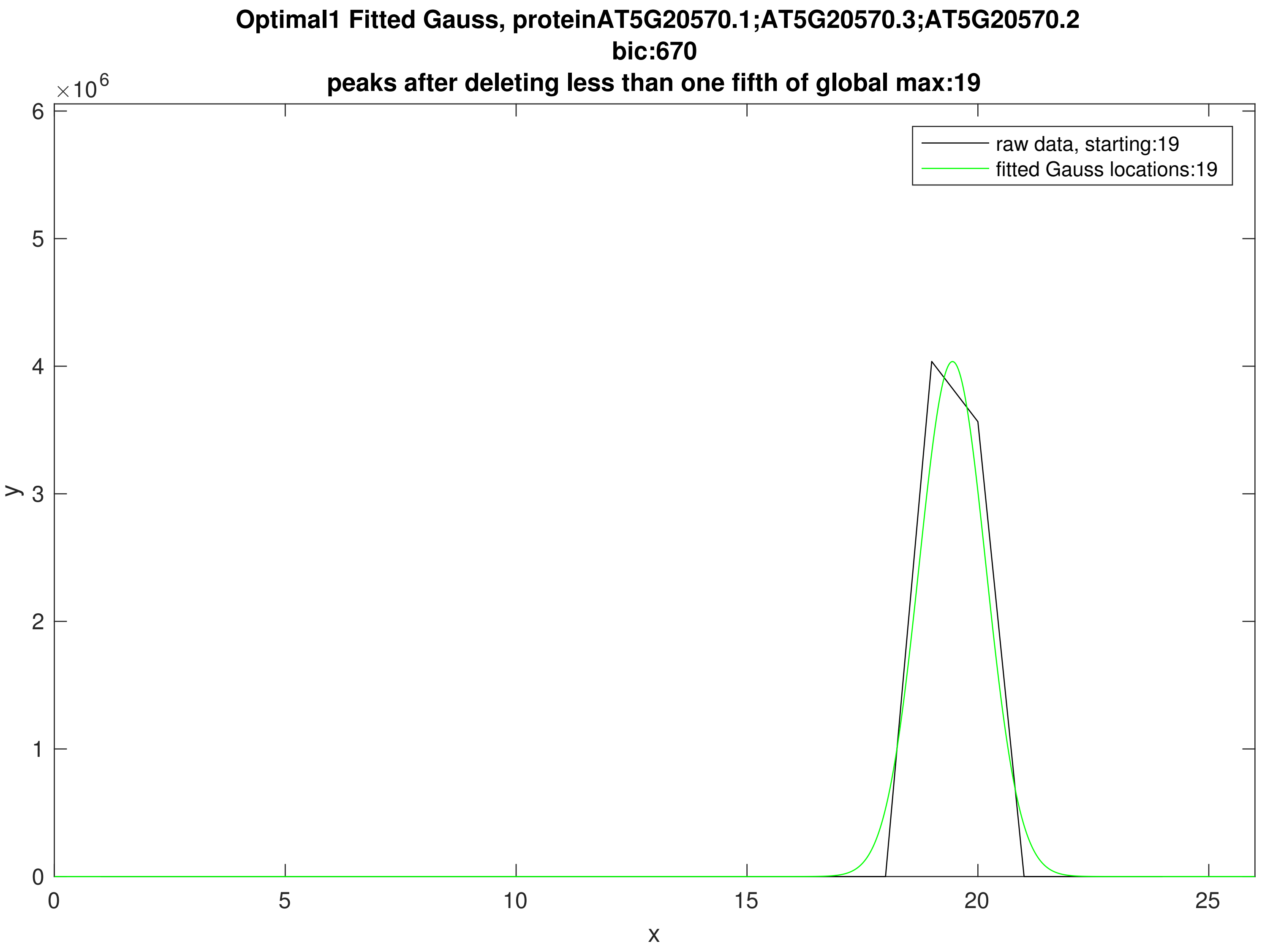
Optimal1 Fitted Gauss, proteinAT5G20570.1;AT5G20570.3;AT5G20570.2
bic:670
peaks after deleting less than one fifth of global max:19
×10^6
raw data, starting:19
fitted Gauss locations:19
y
x
0
1
2
3
4
5
6
0
5
10
15
20
25

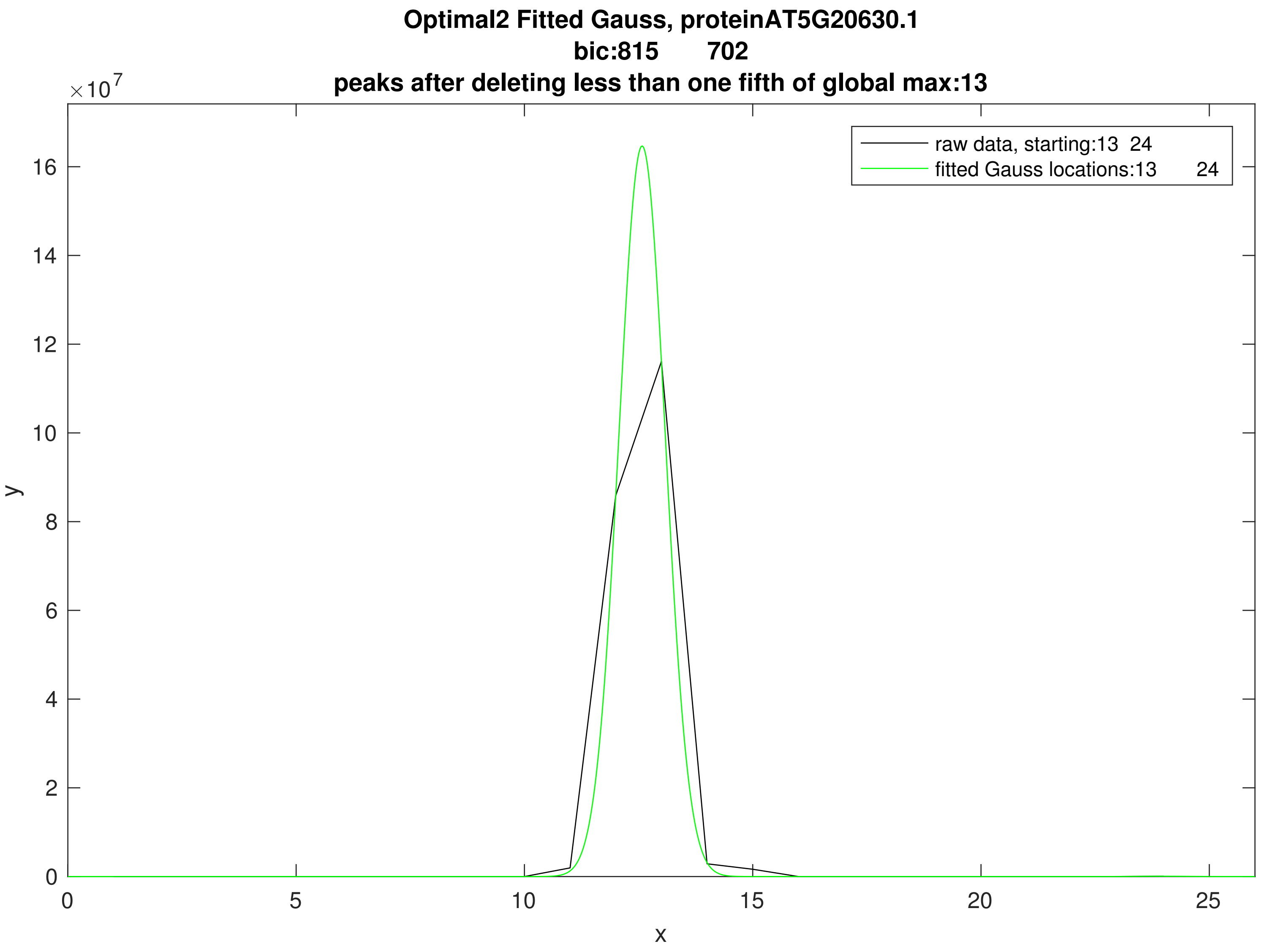

Optimal2 Fitted Gauss, proteinAT5G20630.1
bic:815 702
peaks after deleting less than one fifth of global max:13
×10^7
raw data, starting:13 24
fitted Gauss locations:13 24
16
14
12
10
8
6
4
2
0
y
0
5
10
15
20
25
x

**Optimal1 Fitted Gauss, proteinAT5G20720.3;AT5G20720.2;AT5G20720.1**
**bic:1e+03 1.01e+03 1.02e+03**
**peaks after deleting less than one fifth of global max:13**

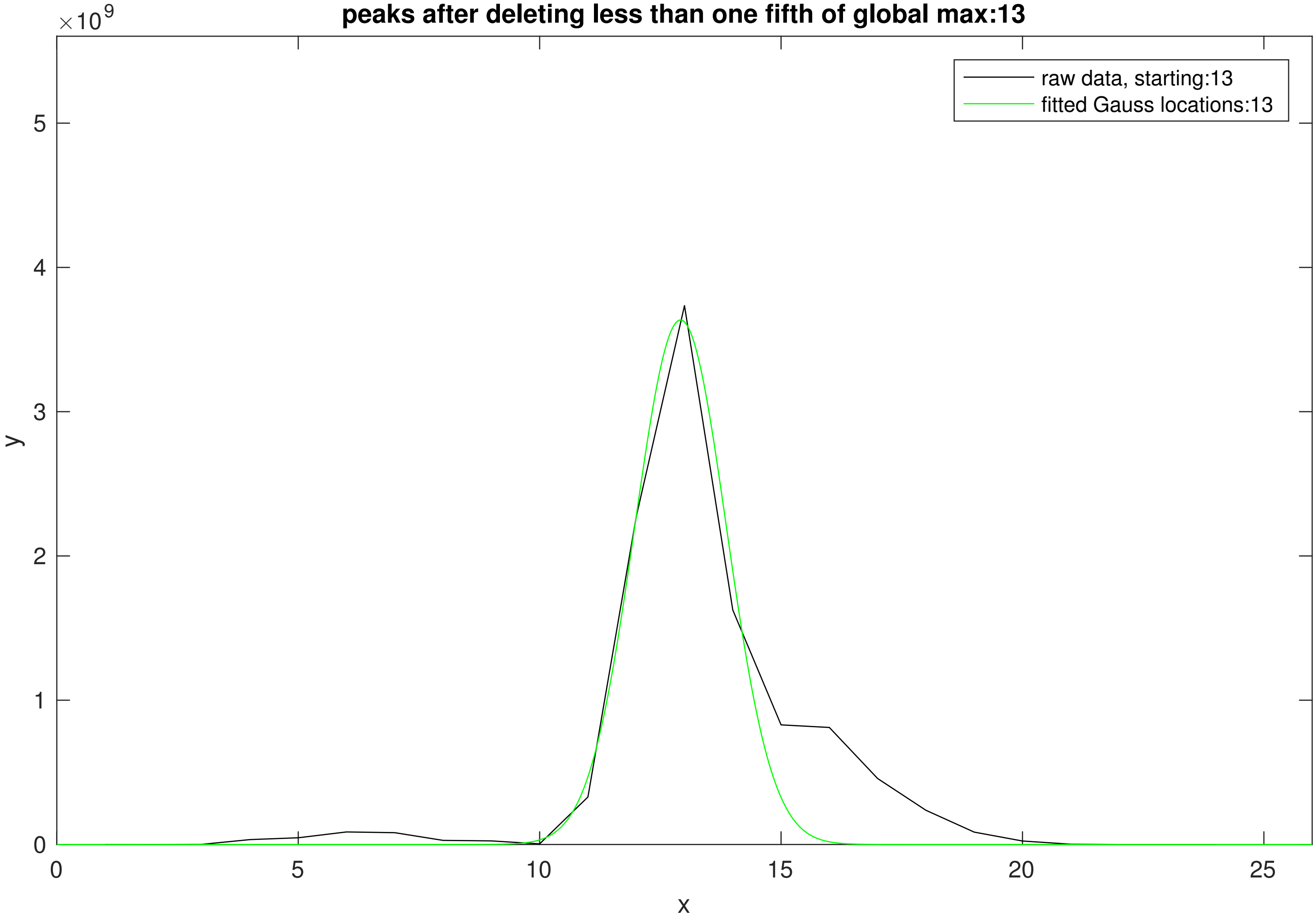

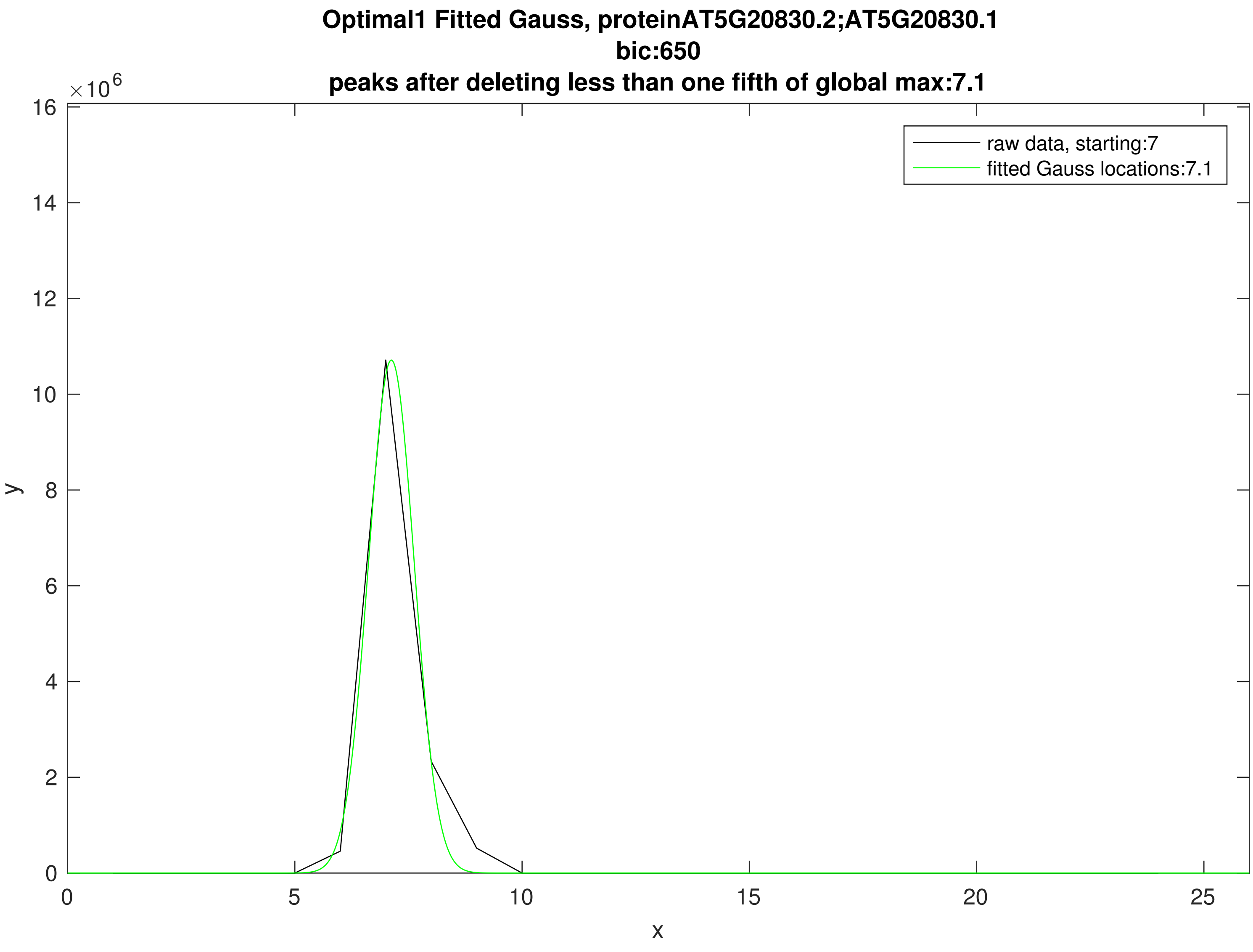
Optimal1 Fitted Gauss, proteinAT5G20830.2;AT5G20830.1
bic:650
peaks after deleting less than one fifth of global max:7.1
×10^6
16
14
12
10
8
6
4
2
0
y
0
5
10
15
20
25
x
raw data, starting:7
fitted Gauss locations:7.1

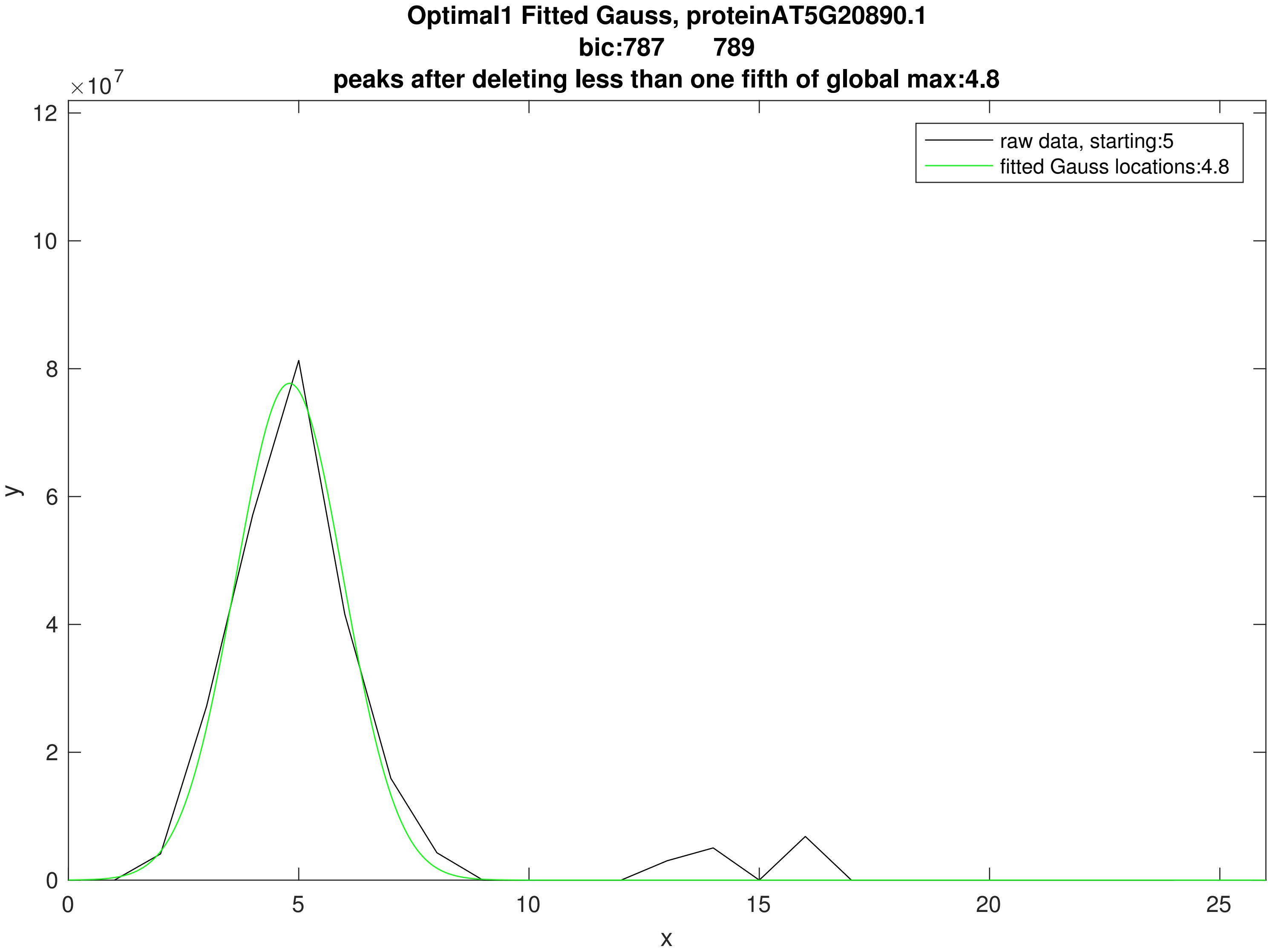

Optimal1 Fitted Gauss, proteinAT5G20890.1
bic:787 789
peaks after deleting less than one fifth of global max:4.8
×10^7
raw data, starting:5
fitted Gauss locations:4.8
y
x
0
2
4
6
8
10
12
5
15
20
25

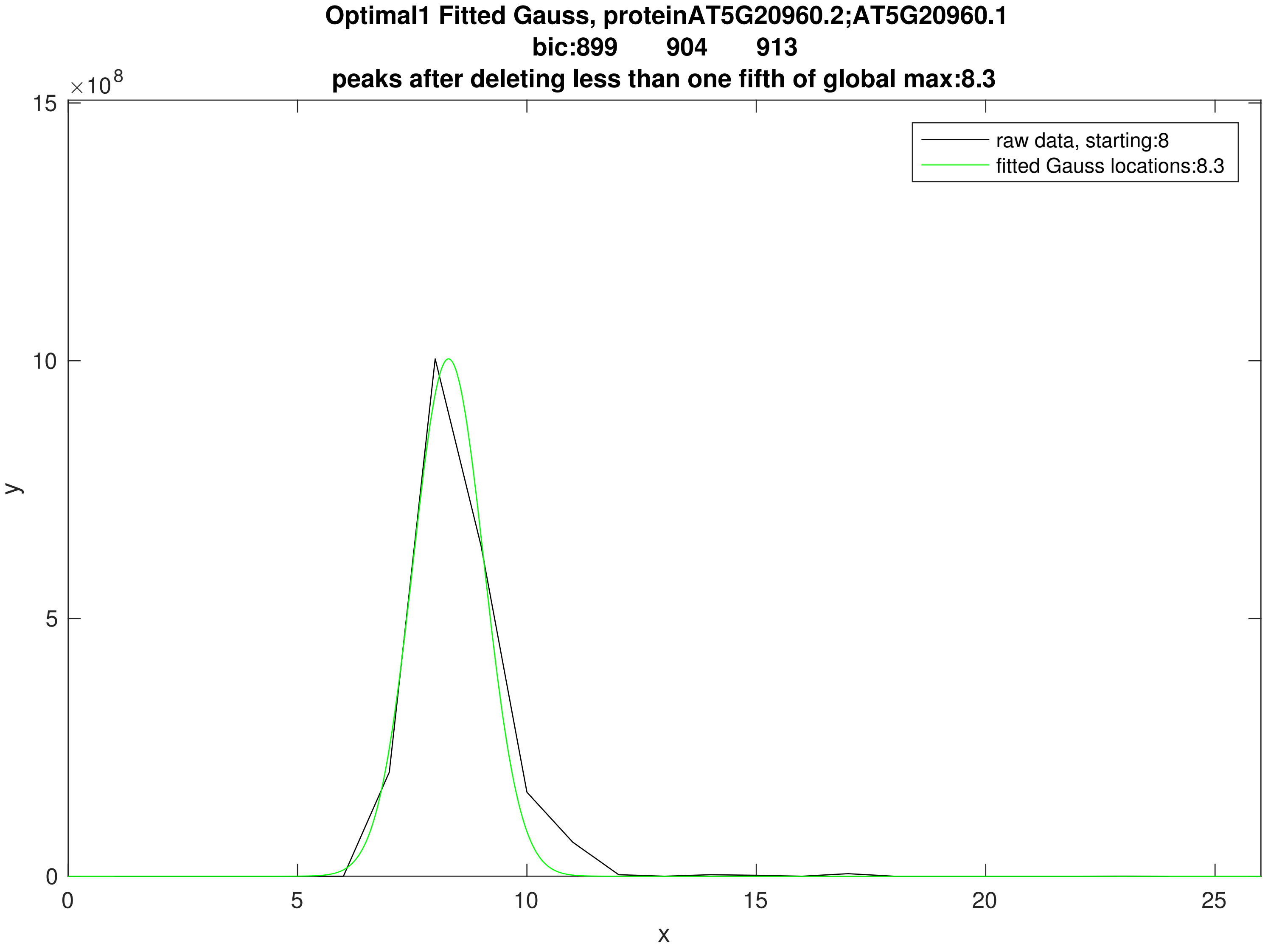
Optimal1 Fitted Gauss, proteinAT5G20960.2;AT5G20960.1
bic:899 904 913
peaks after deleting less than one fifth of global max:8.3
×10^8
15
10
5
0
y
0
5
10
15
20
25
x
raw data, starting:8
fitted Gauss locations:8.3

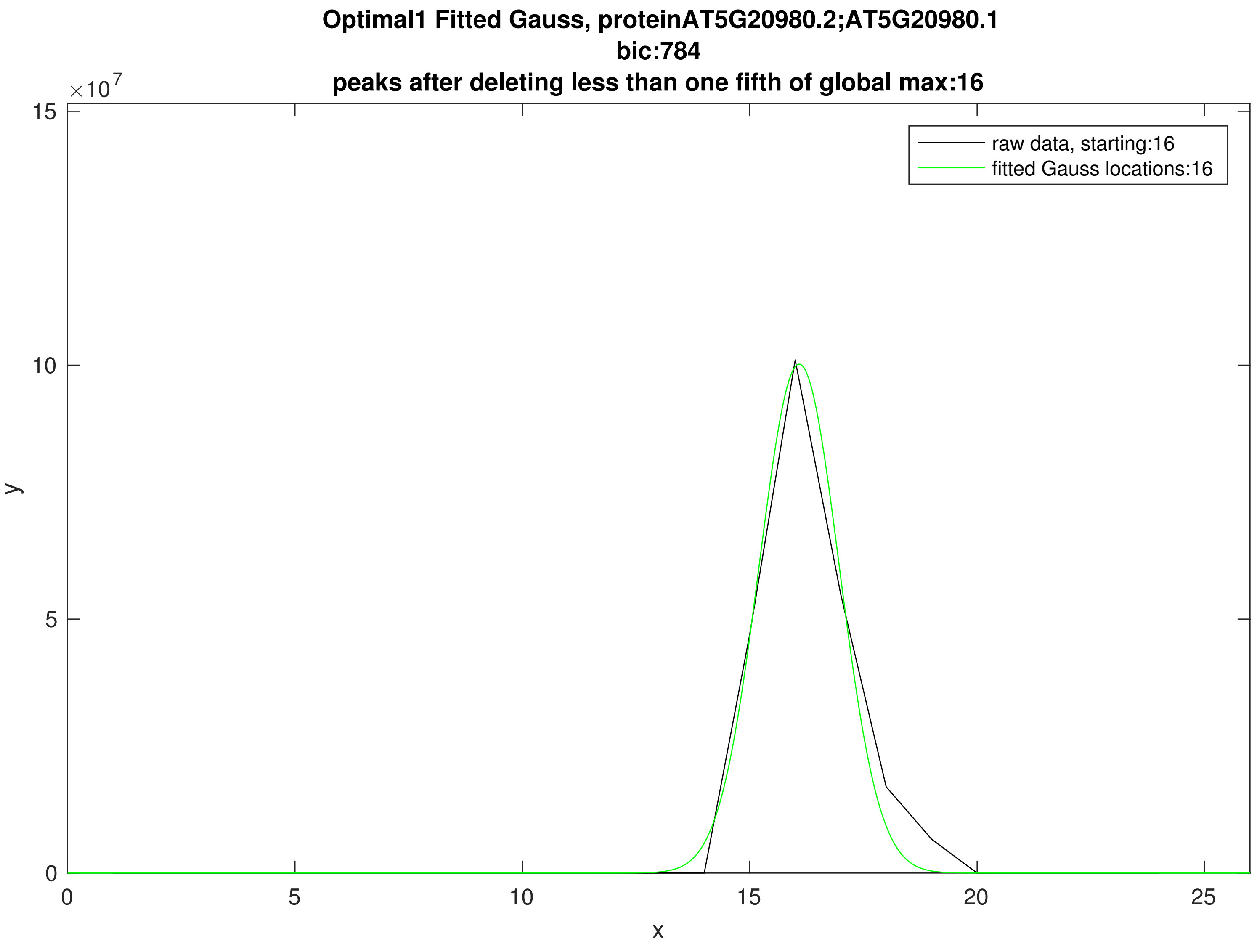

Optimal1 Fitted Gauss, proteinAT5G20980.2;AT5G20980.1
bic:784
peaks after deleting less than one fifth of global max:16
×10^7
15
10
5
0
y
0
5
10
15
20
25
x
raw data, starting:16
fitted Gauss locations:16

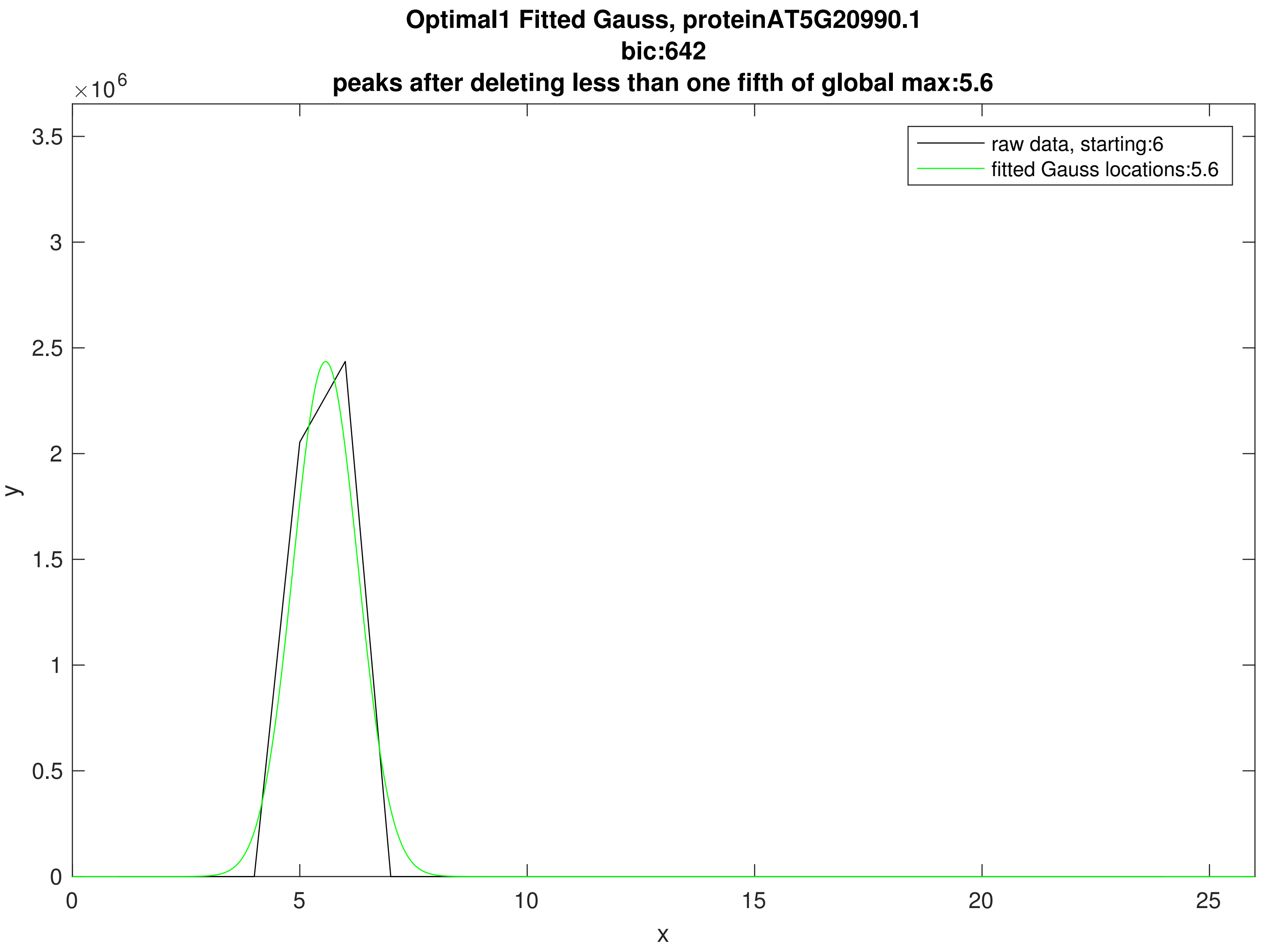

Optimal1 Fitted Gauss, proteinAT5G20990.1
bic:642
peaks after deleting less than one fifth of global max:5.6
×10^6
3.5
3
2.5
2
1.5
1
0.5
0
y
0
5
10
15
20
25
x
raw data, starting:6
fitted Gauss locations:5.6

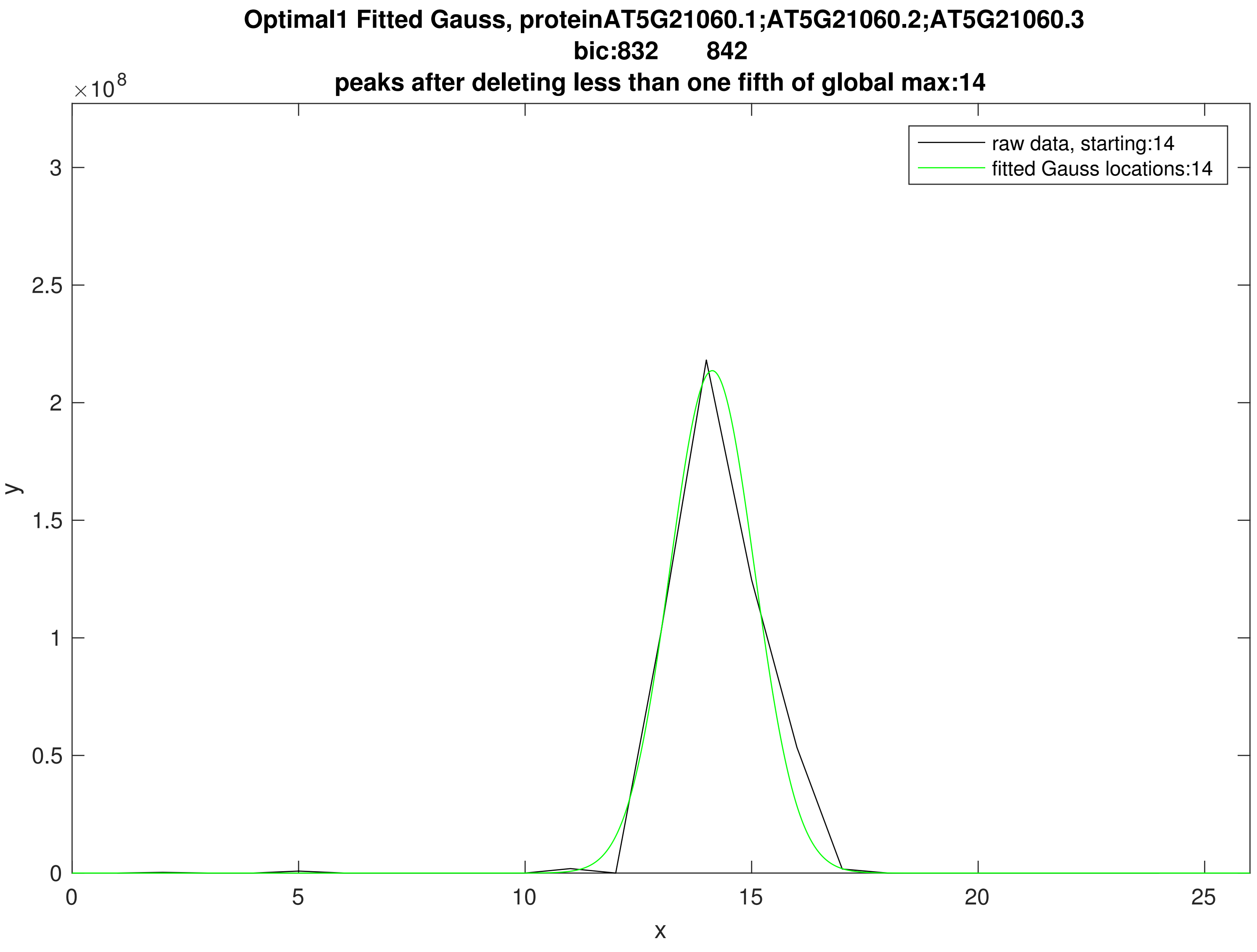

Optimal1 Fitted Gauss, proteinAT5G21060.1;AT5G21060.2;AT5G21060.3
bic:832 842
peaks after deleting less than one fifth of global max:14
×10^8
raw data, starting:14
fitted Gauss locations:14
3
2.5
2
1.5
1
0.5
0
y
0
5
10
15
20
25
x

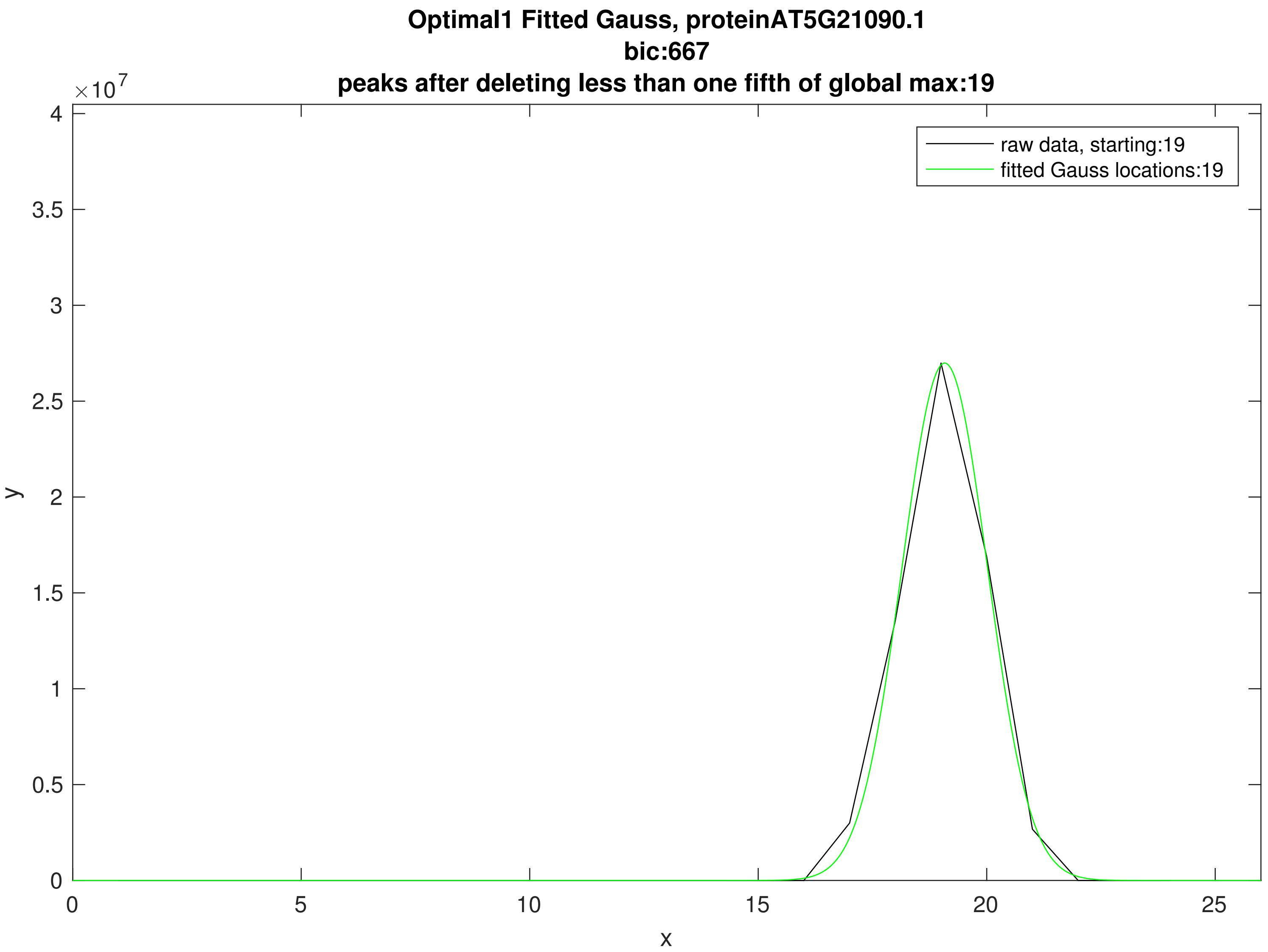

Optimal1 Fitted Gauss, proteinAT5G21090.1
bic:667
peaks after deleting less than one fifth of global max:19
×10^7
4
3.5
3
2.5
2
1.5
1
0.5
0
y
0
5
10
15
20
25
x
raw data, starting:19
fitted Gauss locations:19

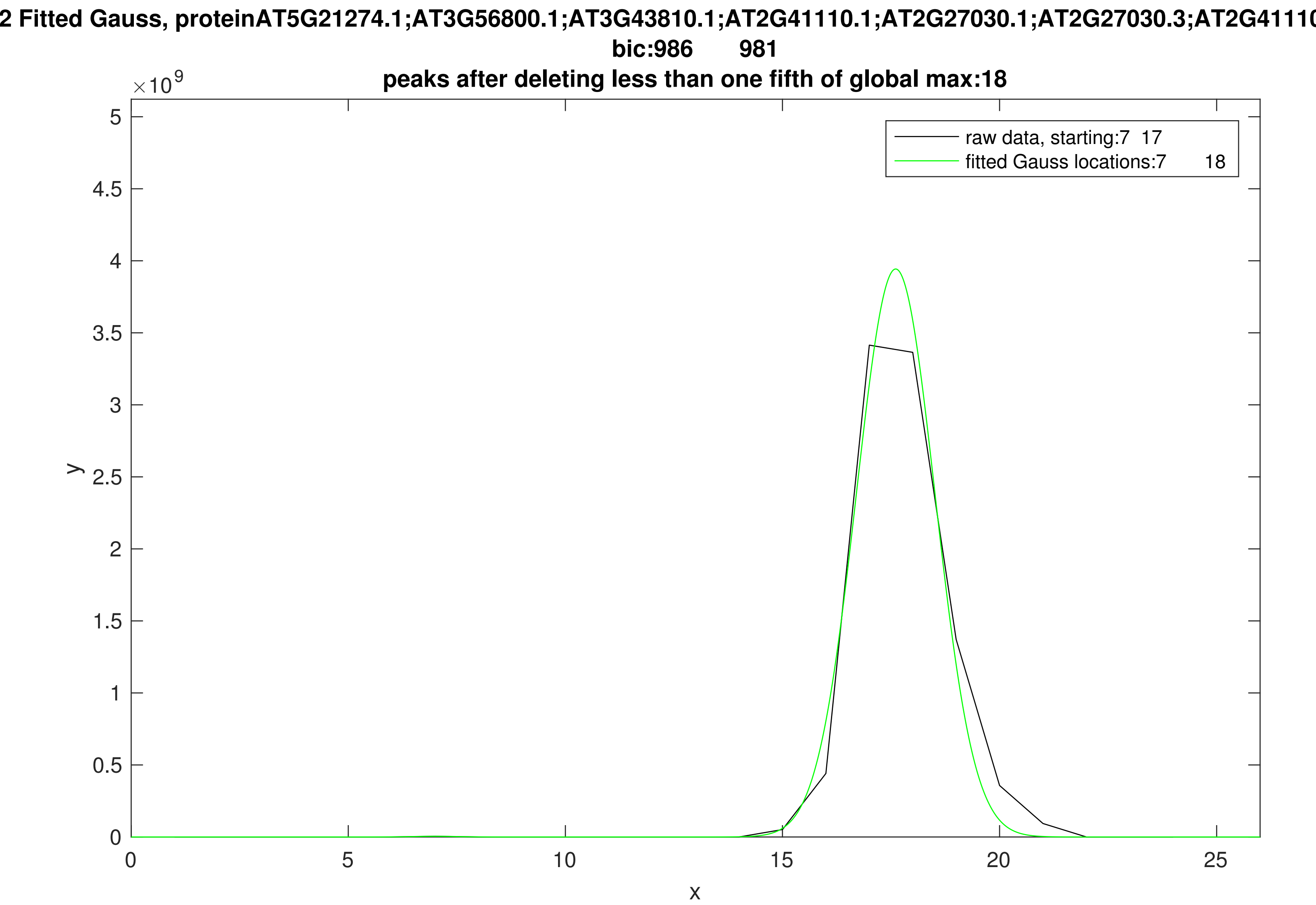

2 Fitted Gauss, proteinAT5G21274.1;AT3G56800.1;AT3G43810.1;AT2G41110.1;AT2G27030.1;AT2G27030.3;AT2G4111
bic:986 981
peaks after deleting less than one fifth of global max:18
×10^9
raw data, starting:7 17
fitted Gauss locations:7 18
y
x

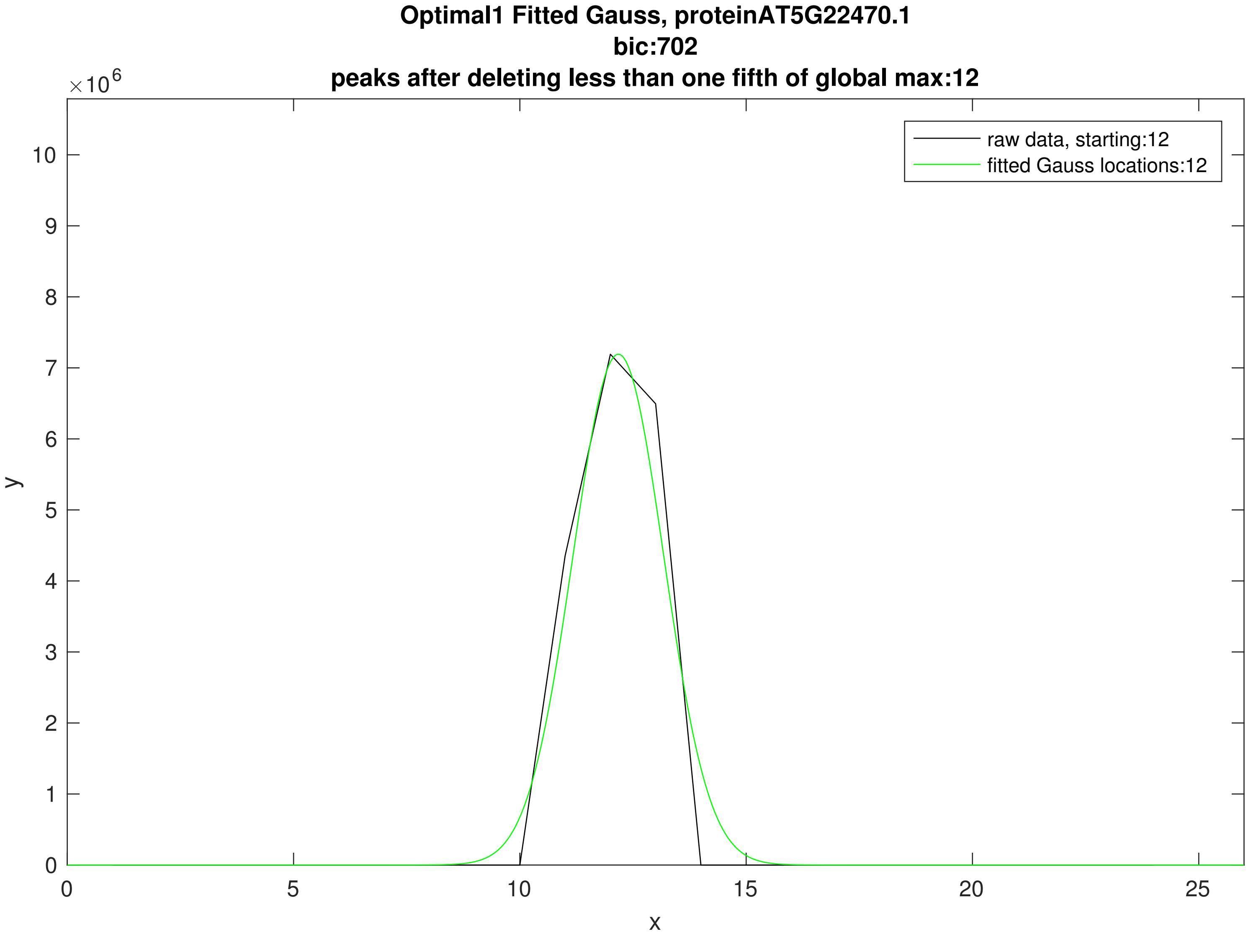

Optimal1 Fitted Gauss, proteinAT5G22470.1
bic:702
peaks after deleting less than one fifth of global max:12
×10^6
raw data, starting:12
fitted Gauss locations:12
y
x
0
1
2
3
4
5
6
7
8
9
10
0
5
10
15
20
25

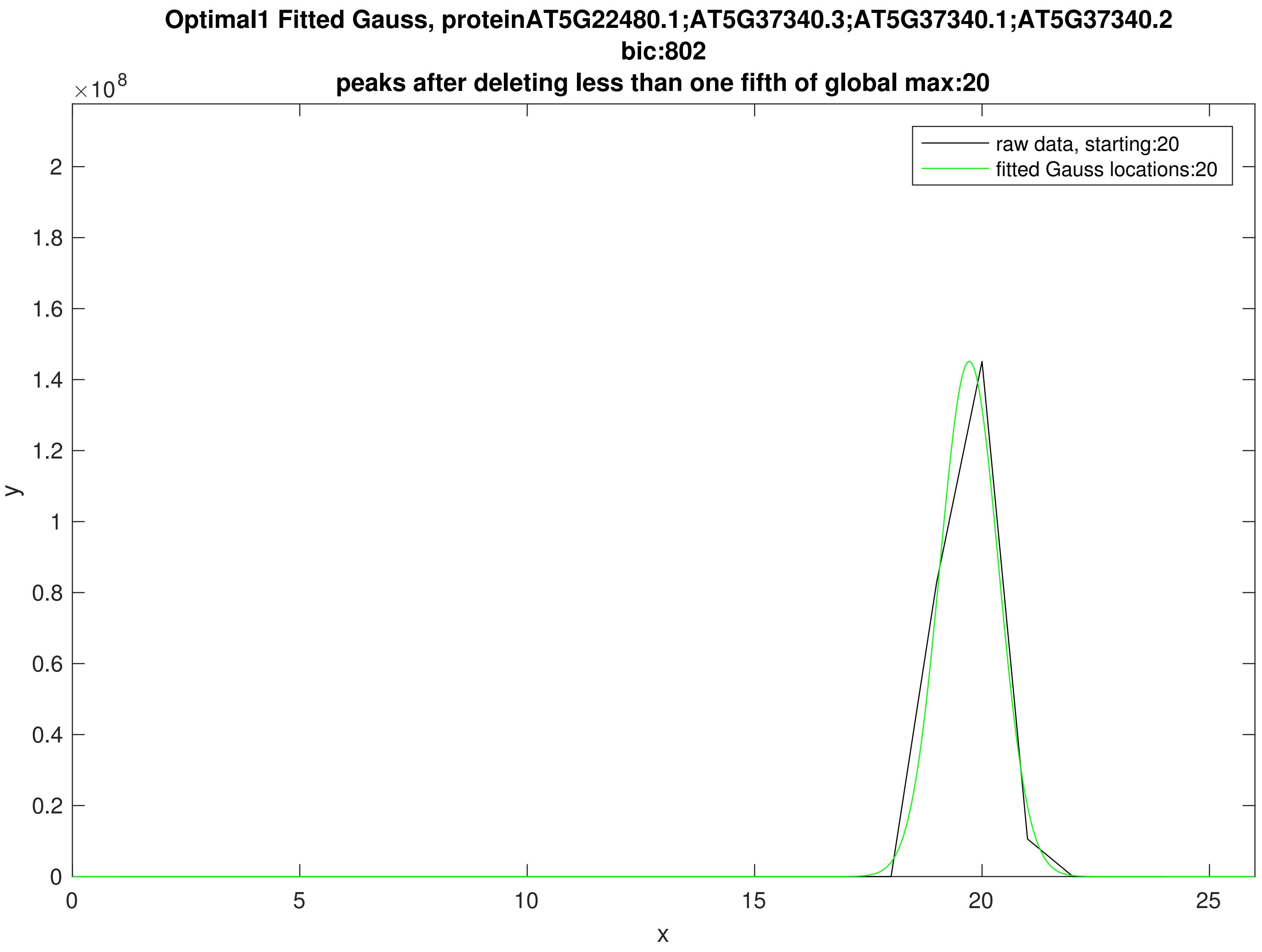
Optimal1 Fitted Gauss, proteinAT5G22480.1;AT5G37340.3;AT5G37340.1;AT5G37340.2
bic:802
peaks after deleting less than one fifth of global max:20
×10^8
raw data, starting:20
fitted Gauss locations:20
y
x
0
0.2
0.4
0.6
0.8
1
1.2
1.4
1.6
1.8
2
0
5
10
15
20
25

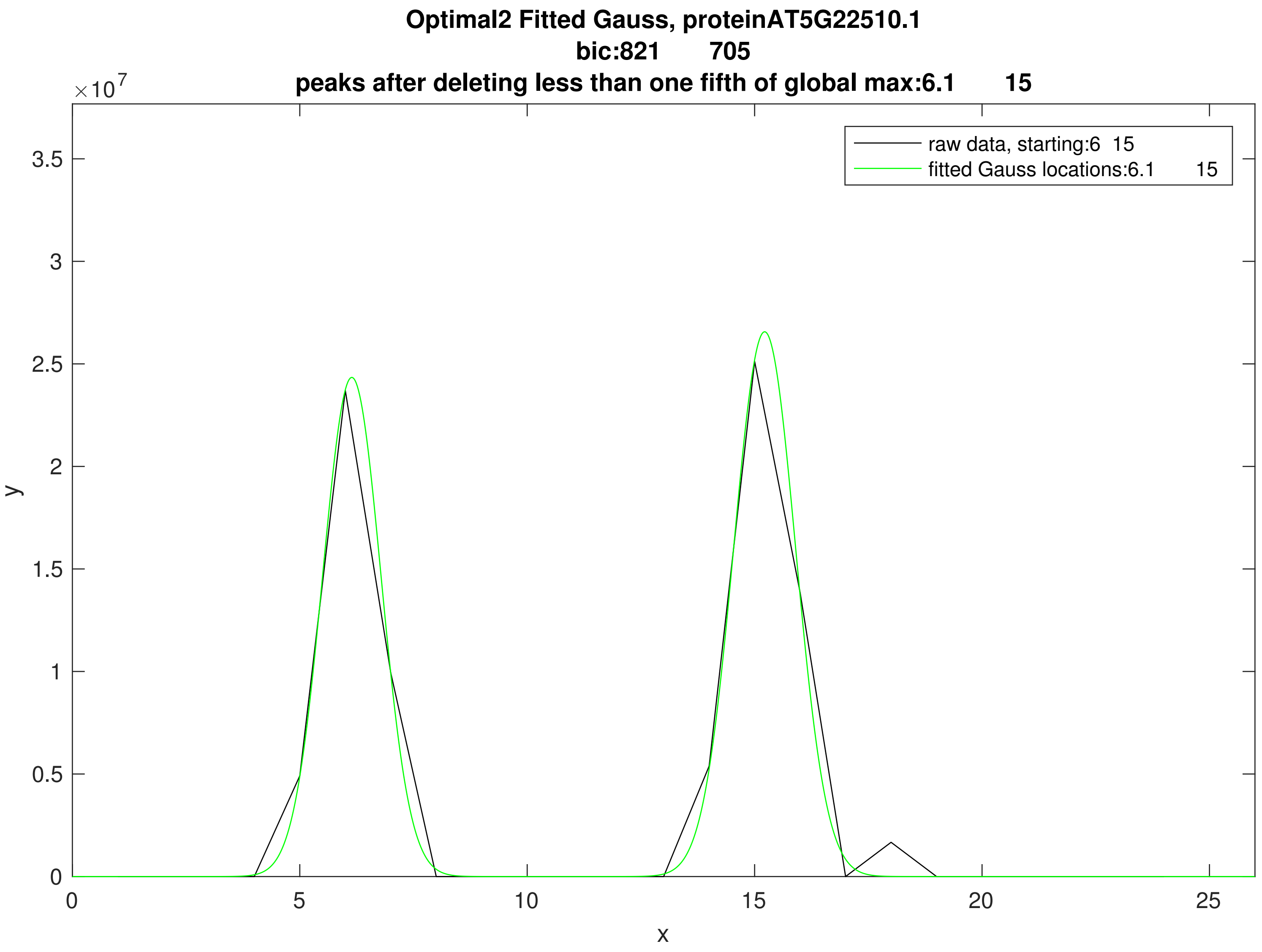

Optimal2 Fitted Gauss, proteinAT5G22510.1
bic:821 705
peaks after deleting less than one fifth of global max:6.1 15
×10^7
raw data, starting:6 15
fitted Gauss locations:6.1 15
3.5
3
2.5
2
1.5
1
0.5
0
y
0
5
10
15
20
25
x

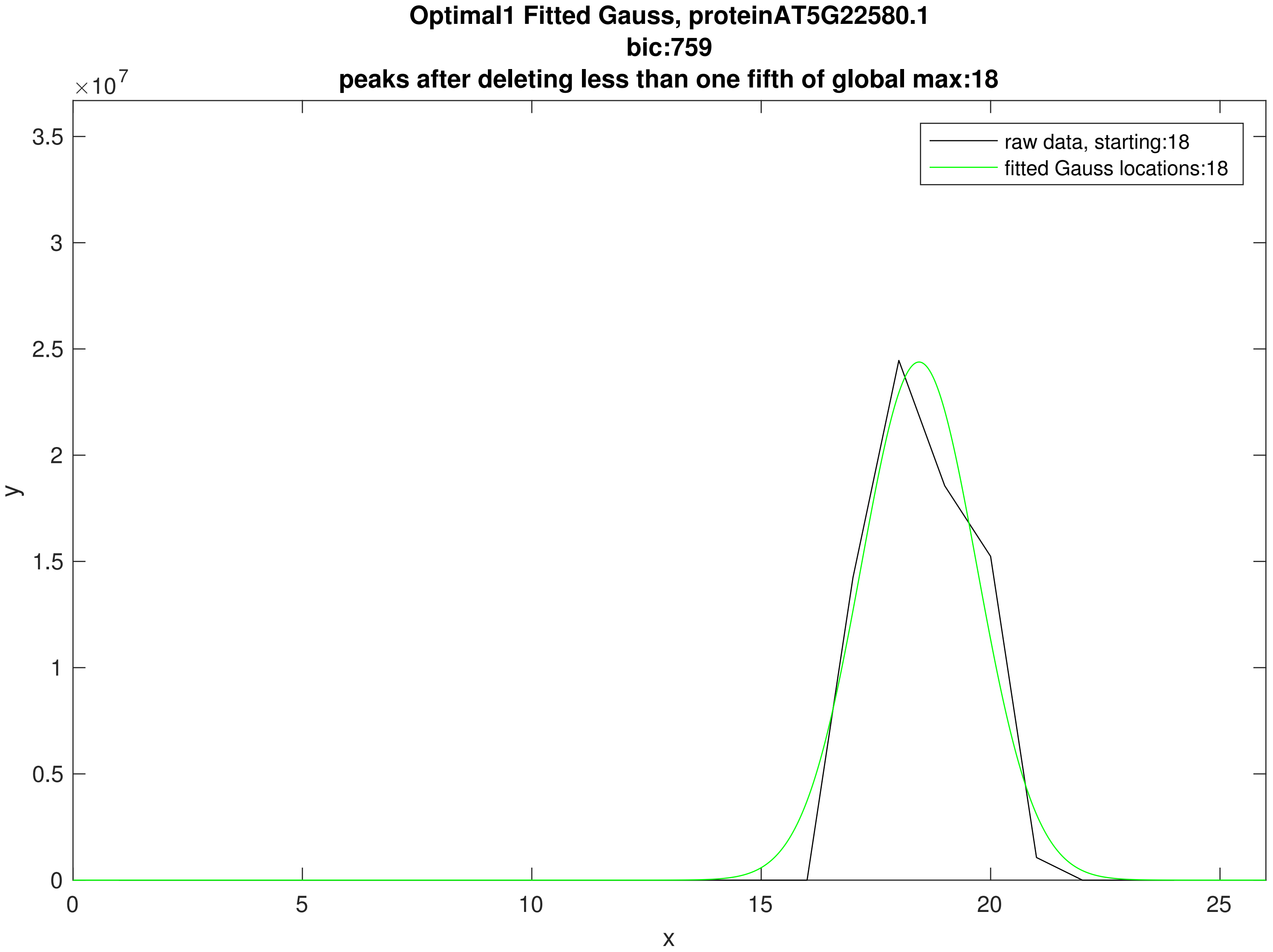
Optimal1 Fitted Gauss, proteinAT5G22580.1
bic:759
peaks after deleting less than one fifth of global max:18
×10^7
3.5
3
2.5
2
1.5
1
0.5
0
y
0
5
10
15
20
25
x
raw data, starting:18
fitted Gauss locations:18

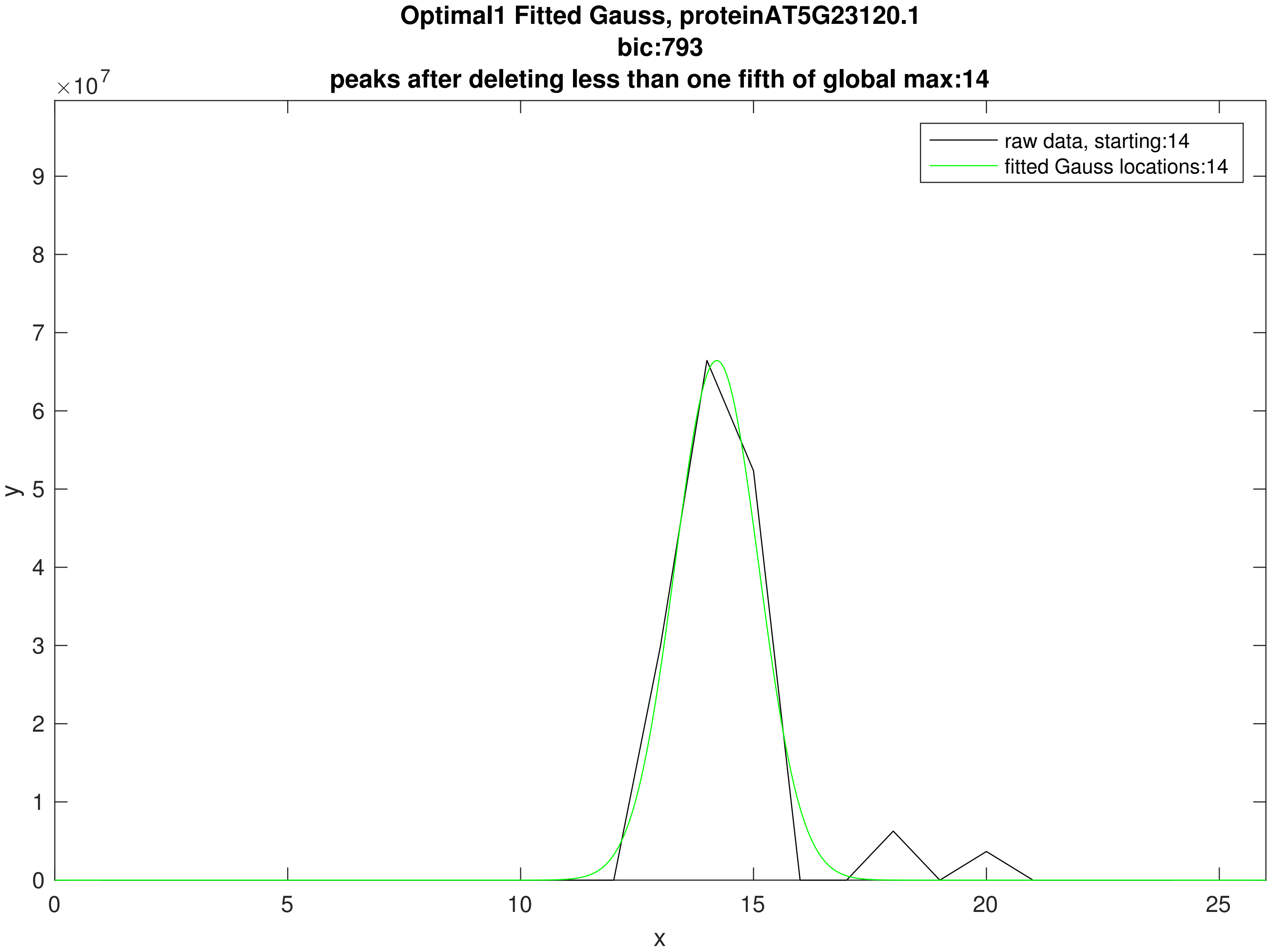
Optimal1 Fitted Gauss, proteinAT5G23120.1
bic:793
peaks after deleting less than one fifth of global max:14
×10^7
raw data, starting:14
fitted Gauss locations:14
y
x
0
1
2
3
4
5
6
7
8
9
0
5
10
15
20
25

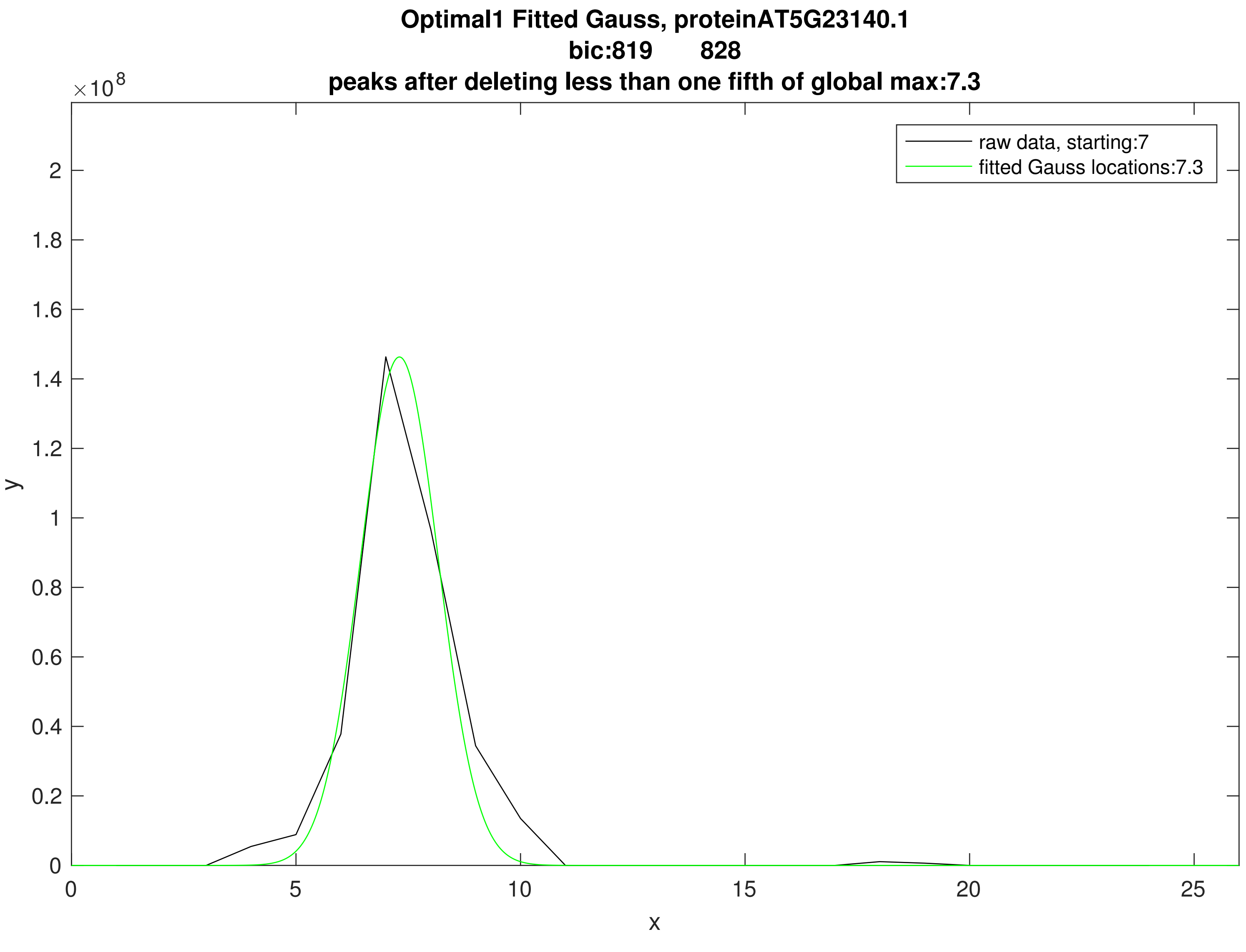

Optimal1 Fitted Gauss, proteinAT5G23140.1
bic:819 828
peaks after deleting less than one fifth of global max:7.3
×10^8
raw data, starting:7
fitted Gauss locations:7.3
y
x
0
0.2
0.4
0.6
0.8
1
1.2
1.4
1.6
1.8
2
5
10
15
20
25

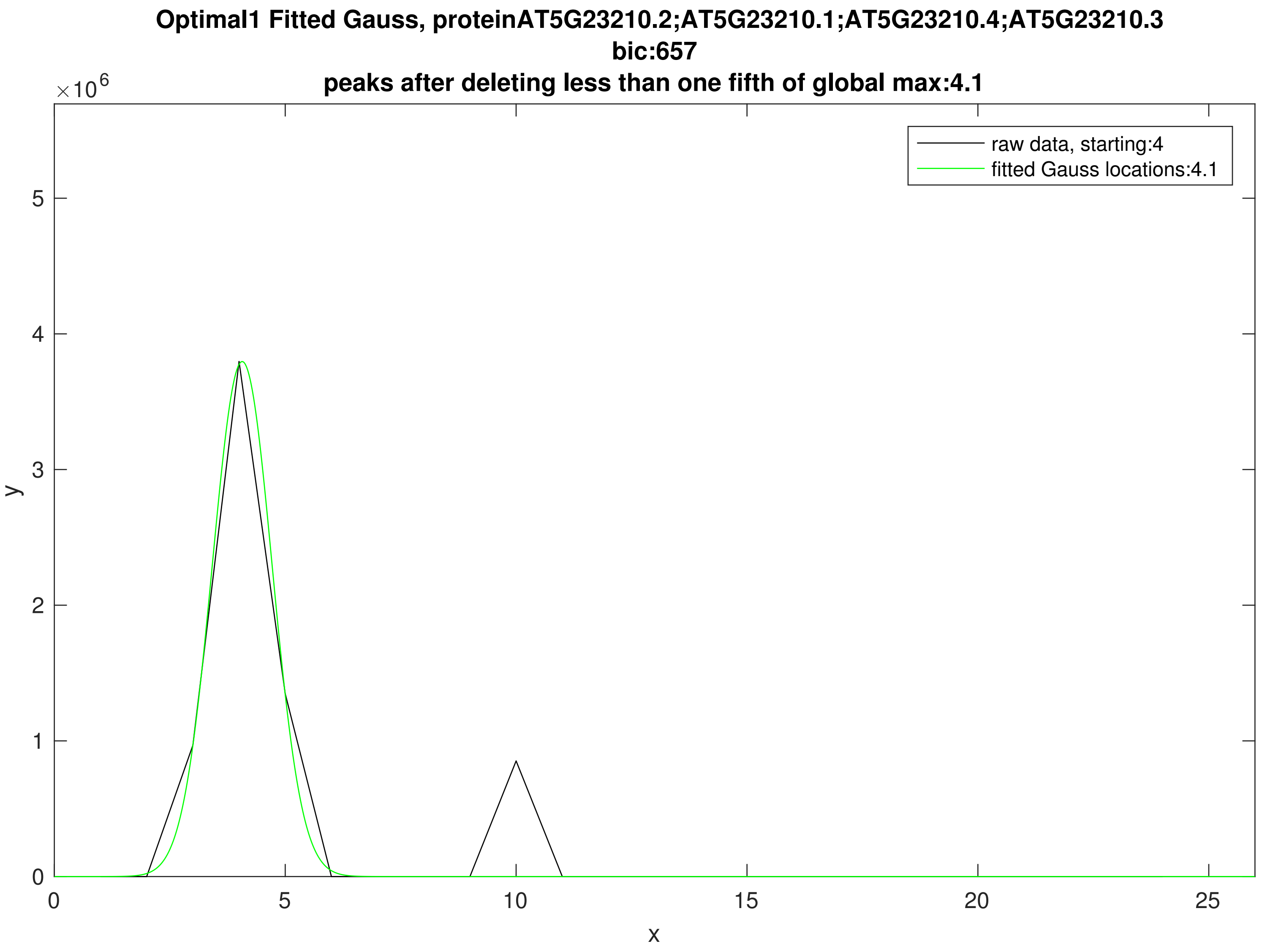
Optimal1 Fitted Gauss, proteinAT5G23210.2;AT5G23210.1;AT5G23210.4;AT5G23210.3
bic:657
peaks after deleting less than one fifth of global max:4.1
×10^6
raw data, starting:4
fitted Gauss locations:4.1
y
x
0
1
2
3
4
5
0
5
10
15
20
25

Optimal1 Fitted Gauss, proteinAT5G23220.1
bic:871
peaks after deleting less than one fifth of global max:17

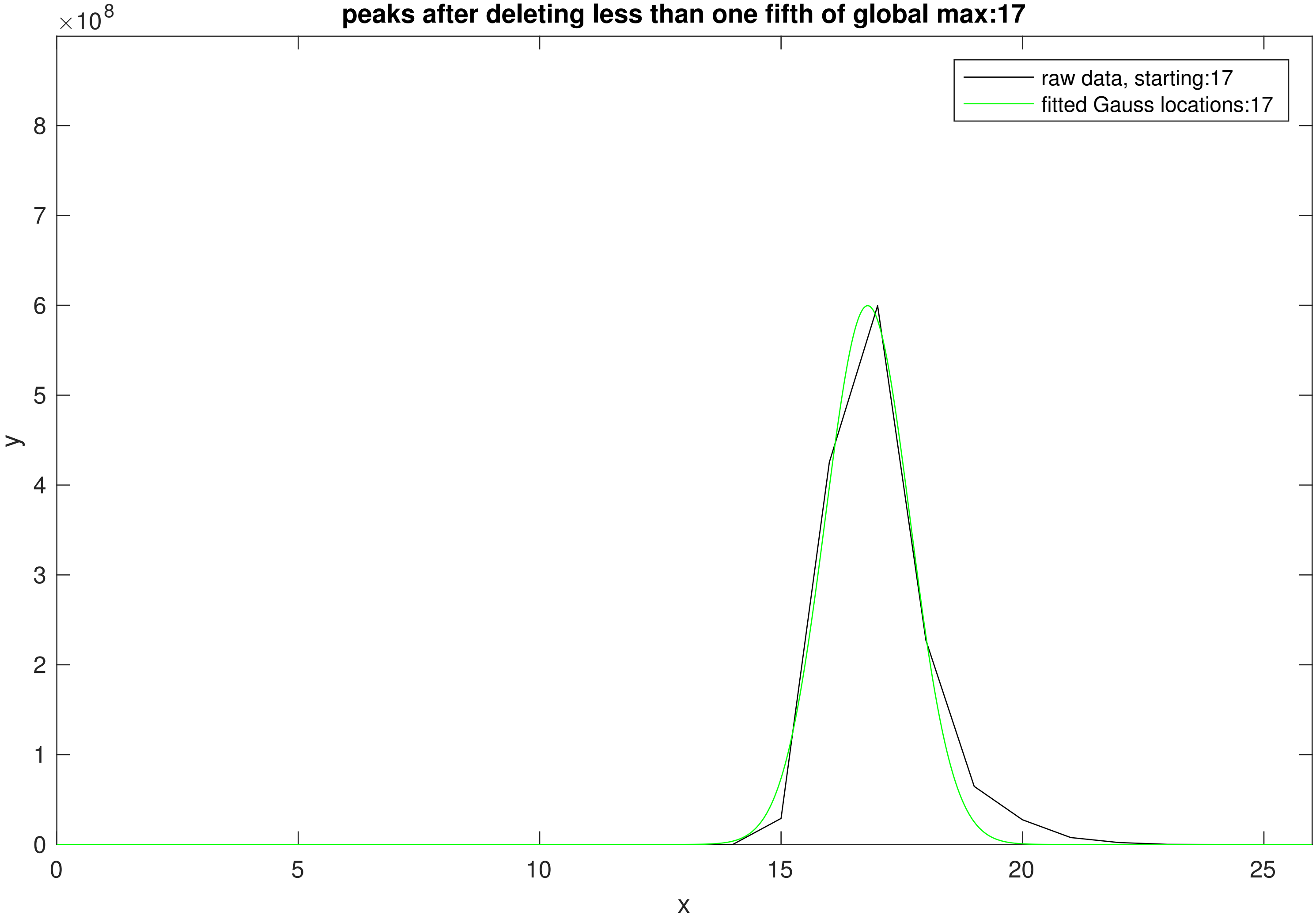

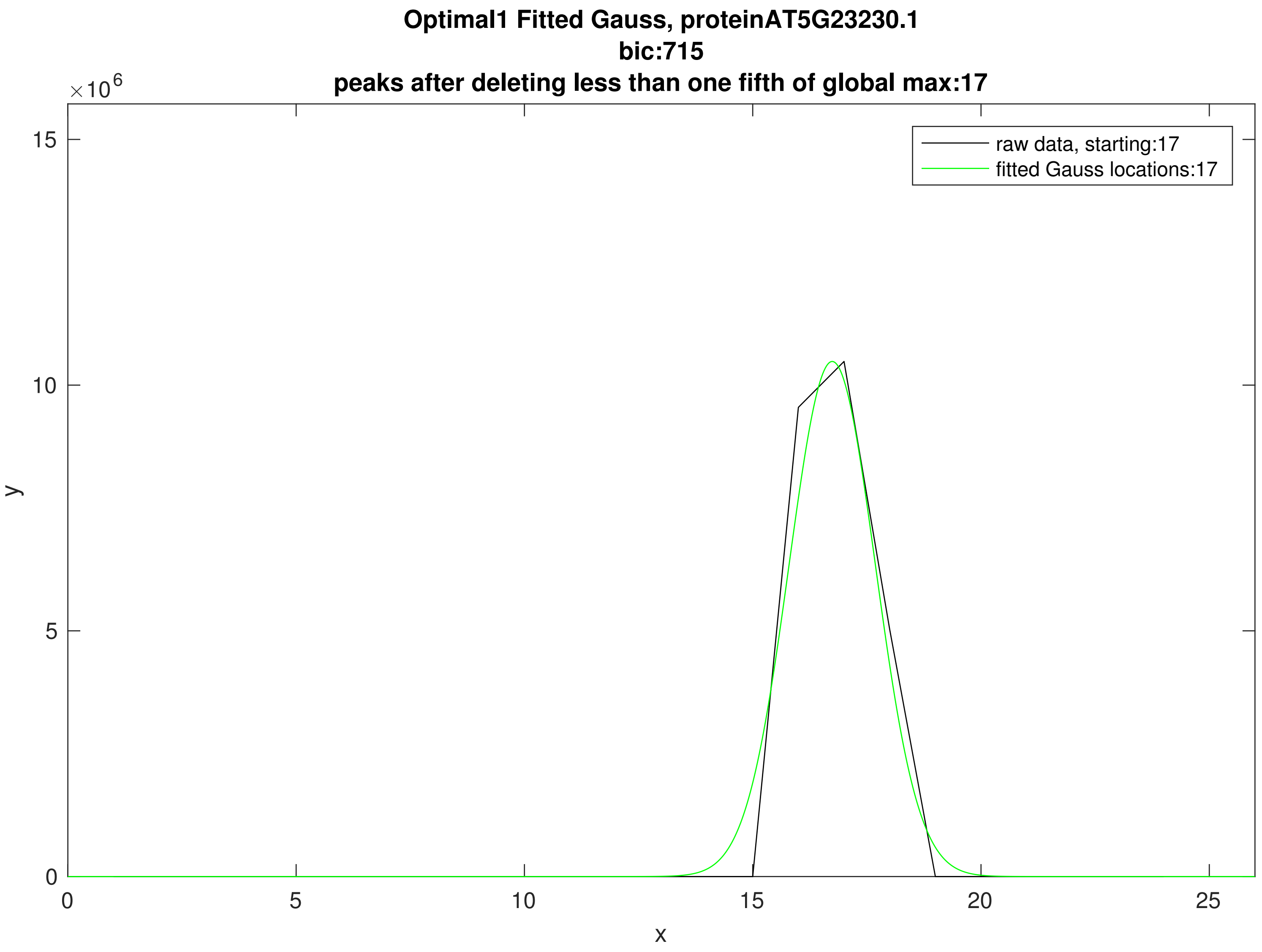

Optimal1 Fitted Gauss, proteinAT5G23230.1
bic:715
peaks after deleting less than one fifth of global max:17
×10^6
15
10
5
0
y
0
5
10
15
20
25
x
raw data, starting:17
fitted Gauss locations:17

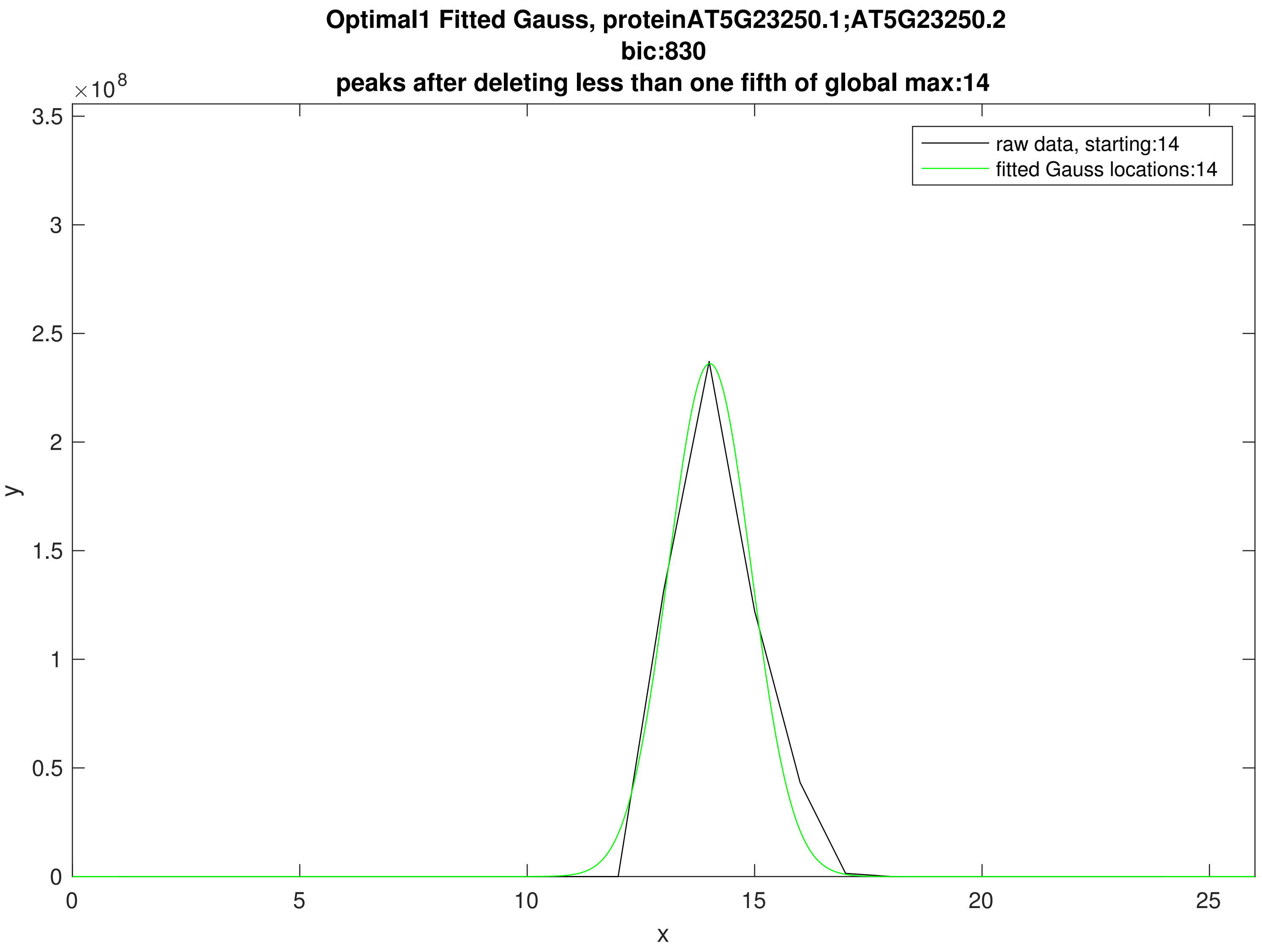
Optimal1 Fitted Gauss, proteinAT5G23250.1;AT5G23250.2
bic:830
peaks after deleting less than one fifth of global max:14
×10^8
3.5
3
2.5
2
1.5
1
0.5
0
y
0
5
10
15
20
25
x
raw data, starting:14
fitted Gauss locations:14

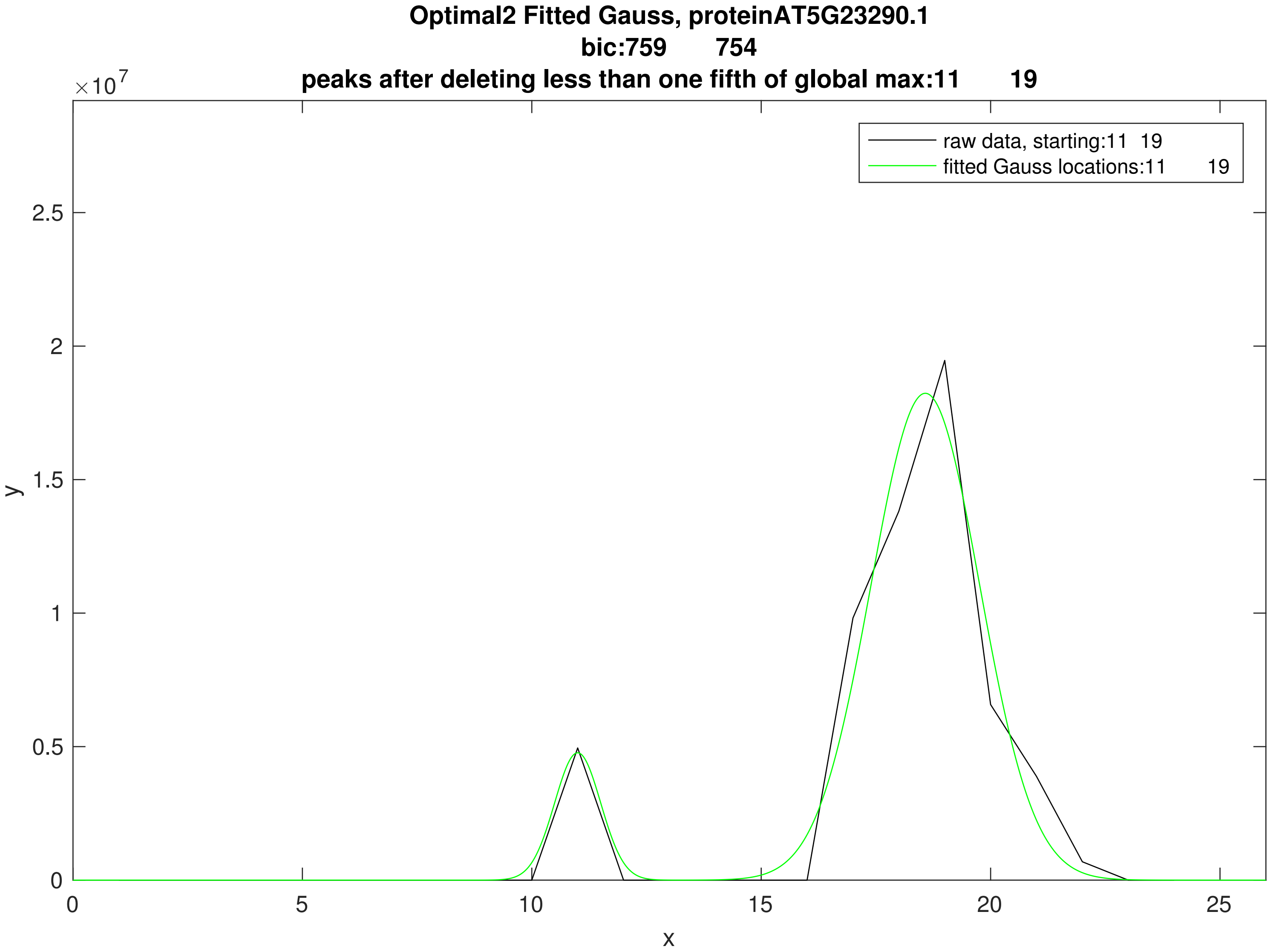
Optimal2 Fitted Gauss, proteinAT5G23290.1
bic:759 754
peaks after deleting less than one fifth of global max:11 19
×10^7
raw data, starting:11 19
fitted Gauss locations:11 19
2.5
2
1.5
1
0.5
0
y
0
5
10
15
20
25
x

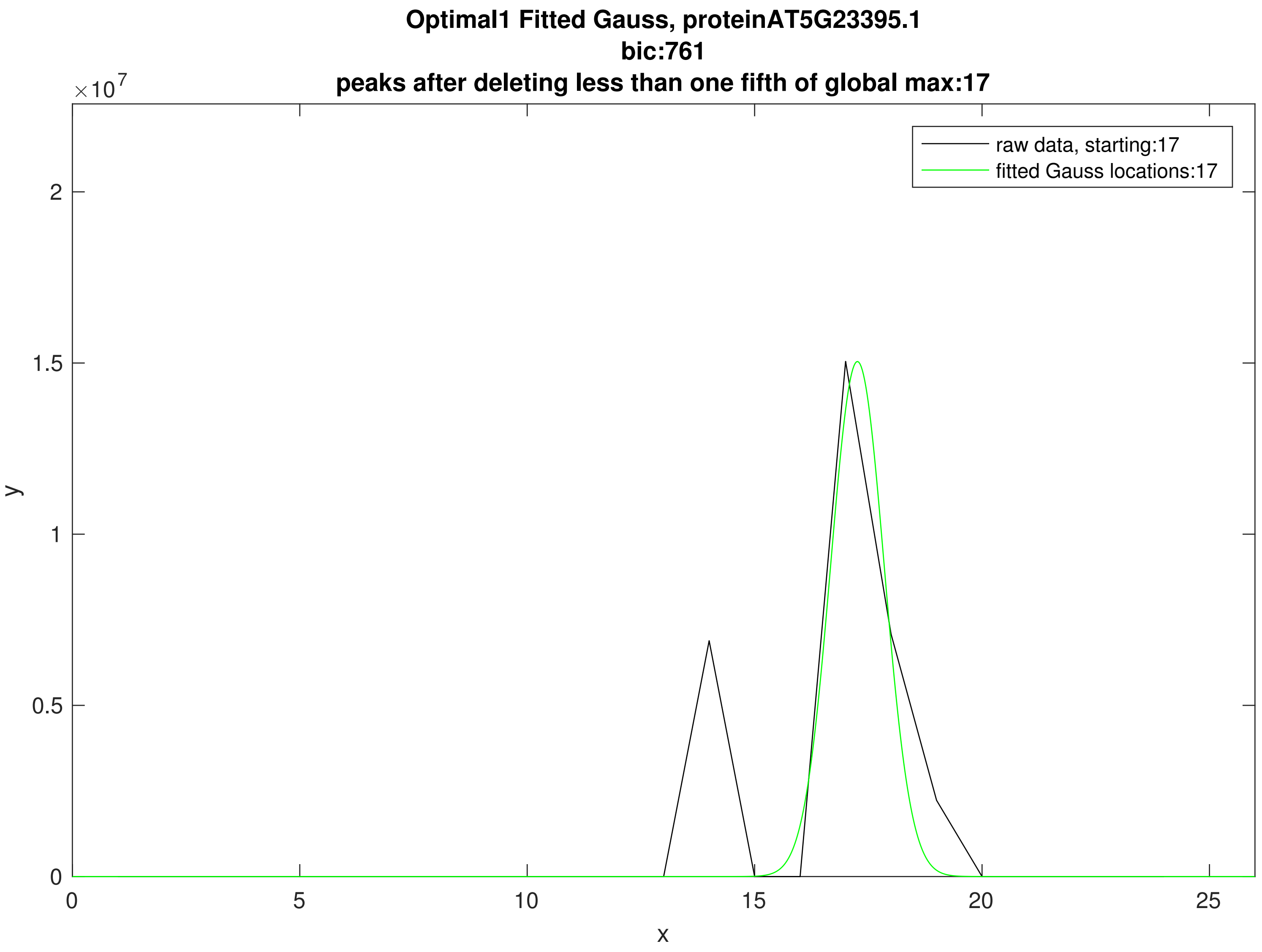

Optimal1 Fitted Gauss, proteinAT5G23395.1
bic:761
peaks after deleting less than one fifth of global max:17
×10^7
raw data, starting:17
fitted Gauss locations:17
y
x
0
0.5
1
1.5
2
0
5
10
15
20
25

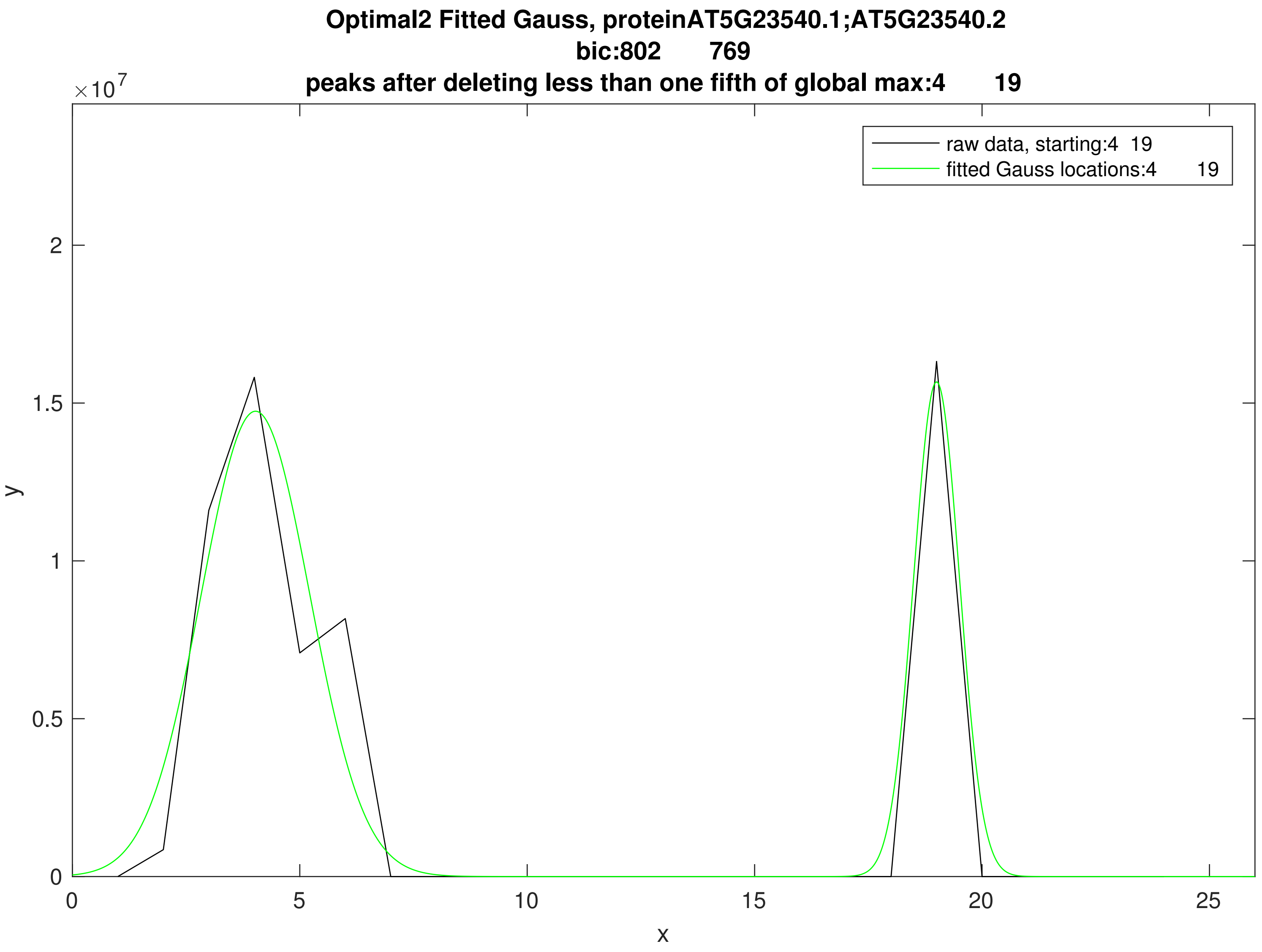
Optimal2 Fitted Gauss, proteinAT5G23540.1;AT5G23540.2
bic:802 769
peaks after deleting less than one fifth of global max:4 19
×10^7
raw data, starting:4 19
fitted Gauss locations:4 19
y
x
0
0.5
1
1.5
2
5
10
15
20
25

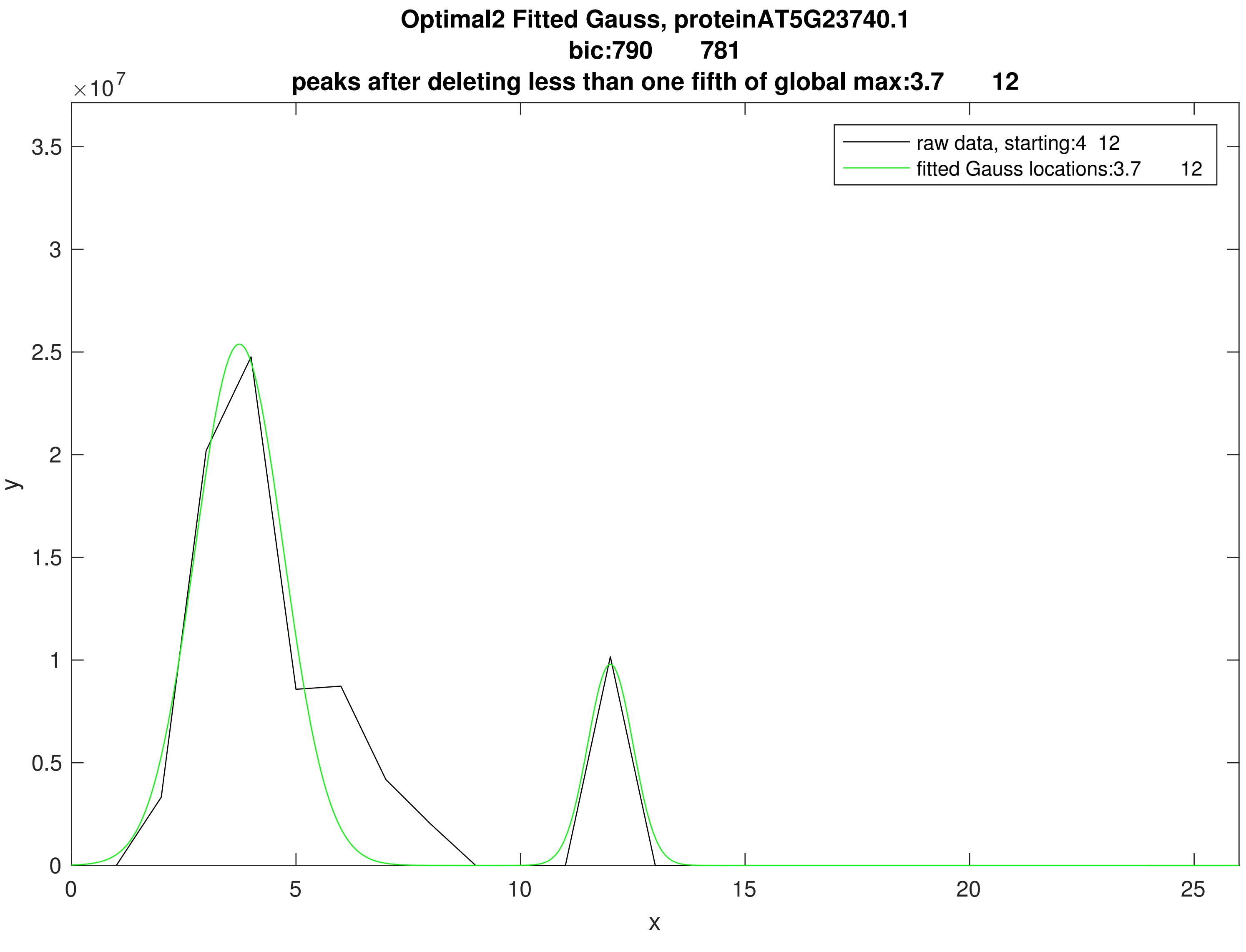
Optimal2 Fitted Gauss, proteinAT5G23740.1
bic:790 781
peaks after deleting less than one fifth of global max:3.7 12
×10^7
3.5
3
2.5
2
1.5
1
0.5
0
y
0
5
10
15
20
25
x
raw data, starting:4 12
fitted Gauss locations:3.7 12

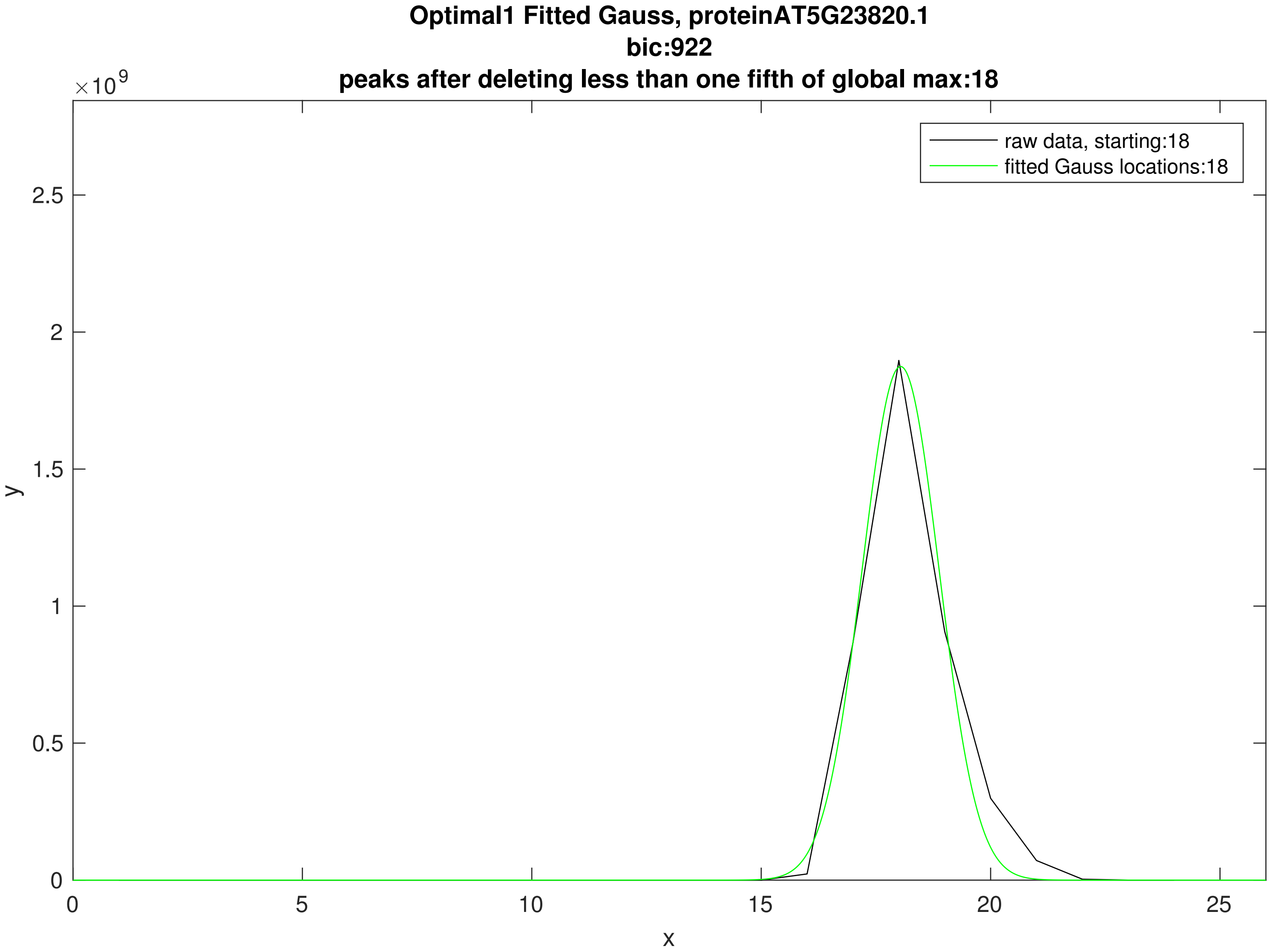
Optimal1 Fitted Gauss, proteinAT5G23820.1
bic:922
peaks after deleting less than one fifth of global max:18
×10^9
raw data, starting:18
fitted Gauss locations:18
y
x
0
0.5
1
1.5
2
2.5
0
5
10
15
20
25

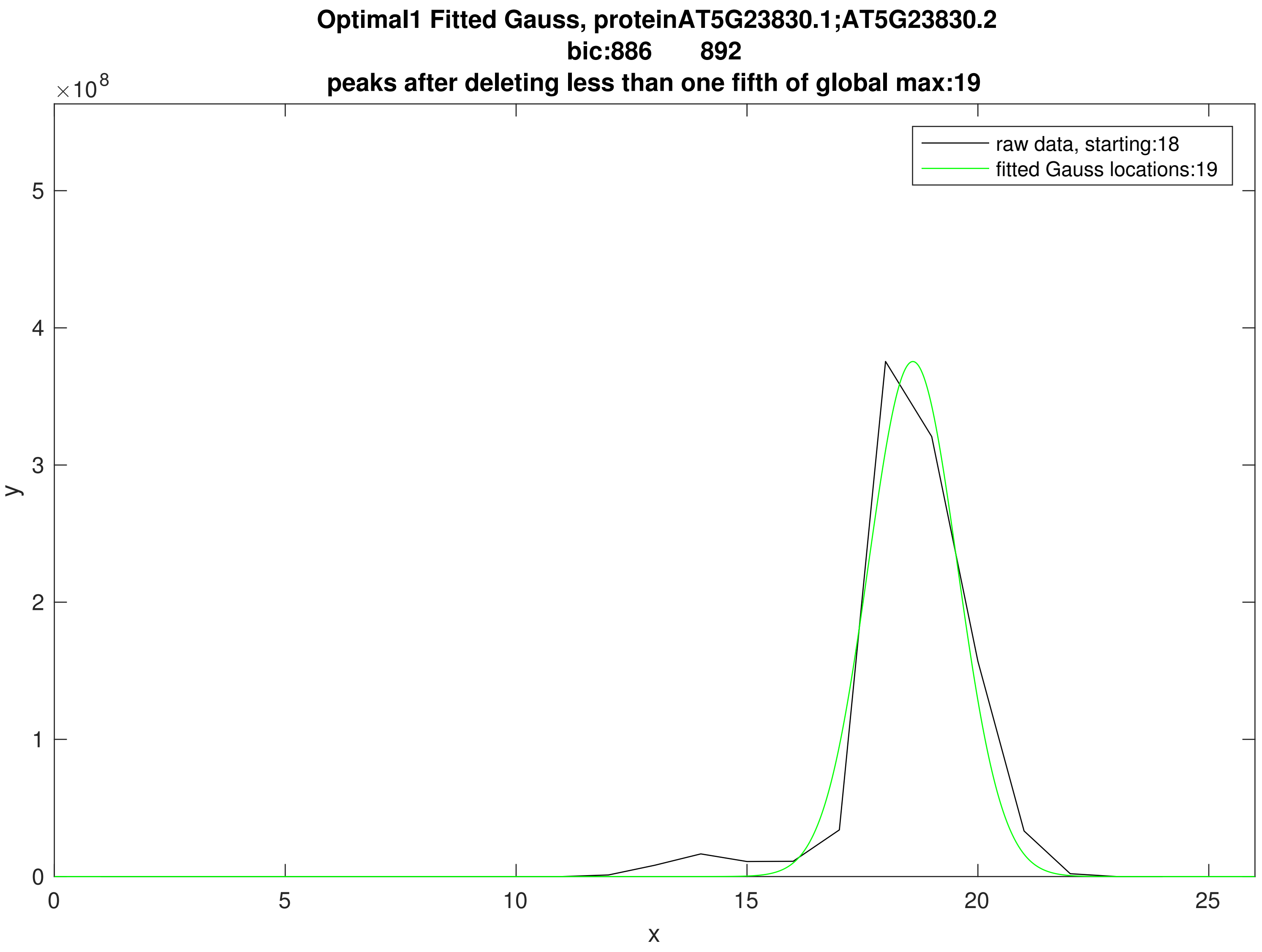

Optimal1 Fitted Gauss, proteinAT5G23830.1;AT5G23830.2
bic:886 892
peaks after deleting less than one fifth of global max:19
×10^8
raw data, starting:18
fitted Gauss locations:19
y
x
0
1
2
3
4
5
0
5
10
15
20
25

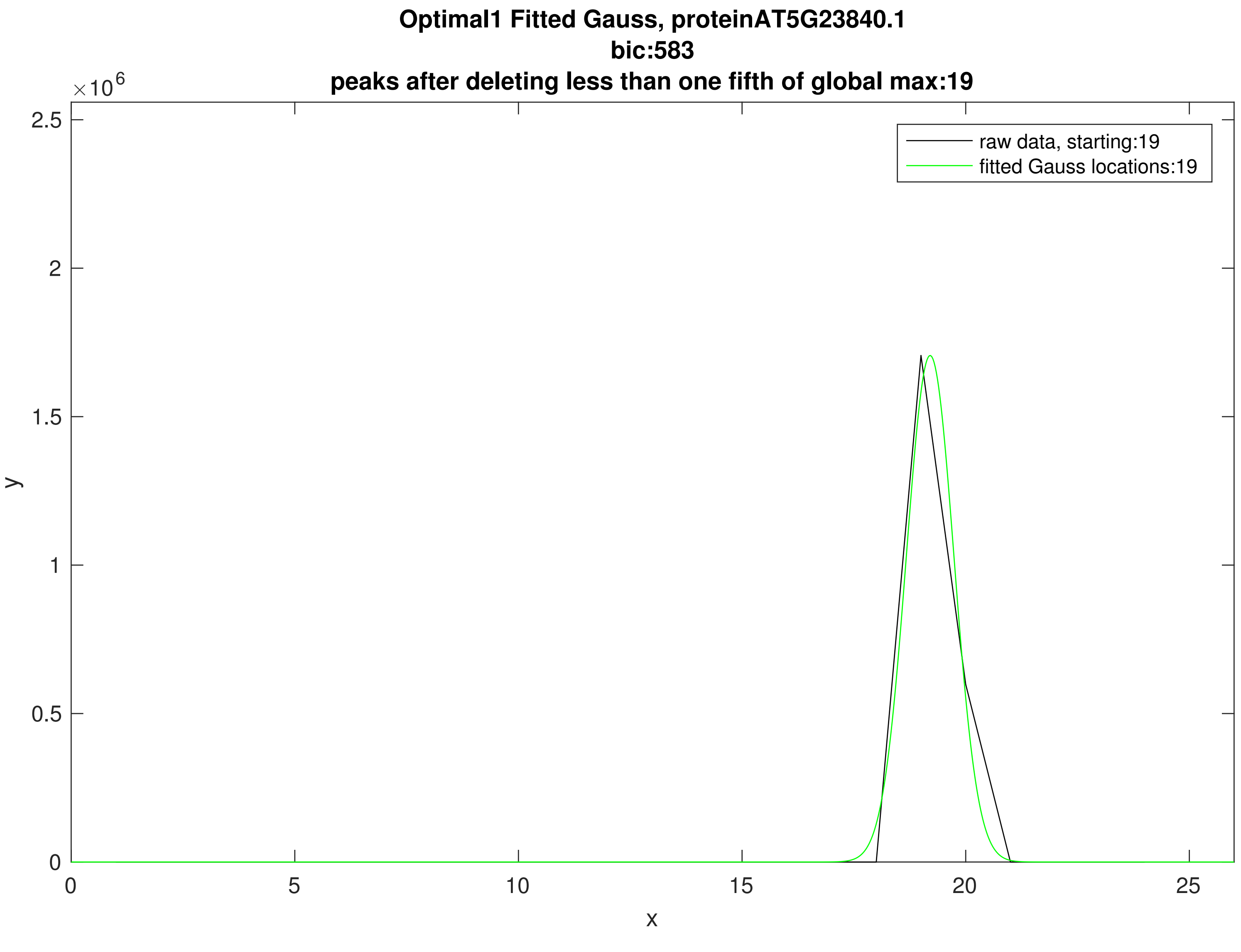

Optimal1 Fitted Gauss, proteinAT5G23840.1
bic:583
peaks after deleting less than one fifth of global max:19
×10^6
raw data, starting:19
fitted Gauss locations:19
y
x
0
0.5
1
1.5
2
2.5
0
5
10
15
20
25

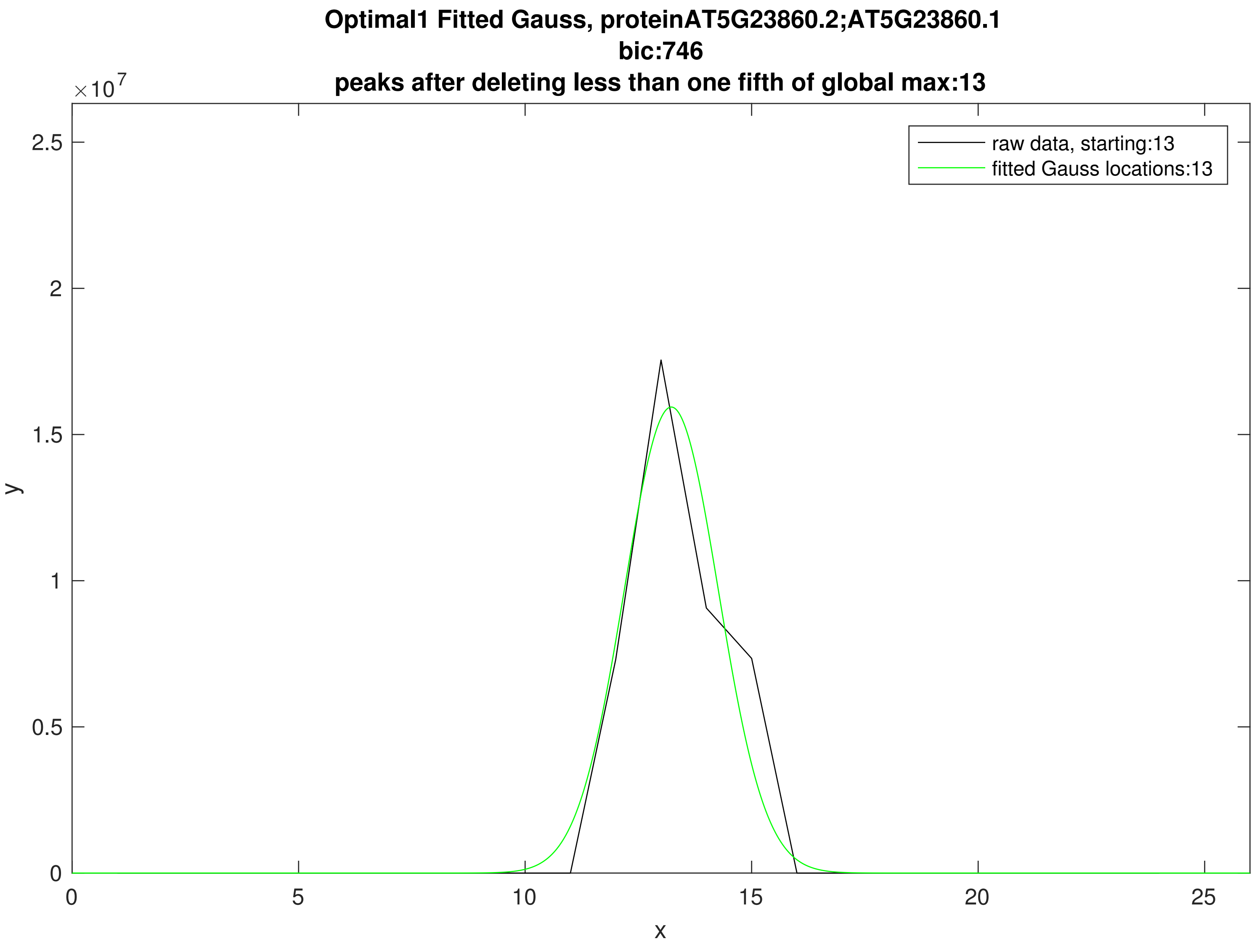
Optimal1 Fitted Gauss, proteinAT5G23860.2;AT5G23860.1
bic:746
peaks after deleting less than one fifth of global max:13
×10^7
raw data, starting:13
fitted Gauss locations:13
y
x
0
0.5
1
1.5
2
2.5
0
5
10
15
20
25

Optimal1 Fitted Gauss, proteinAT5G23900.1
bic:706
peaks after deleting less than one fifth of global max:3.5

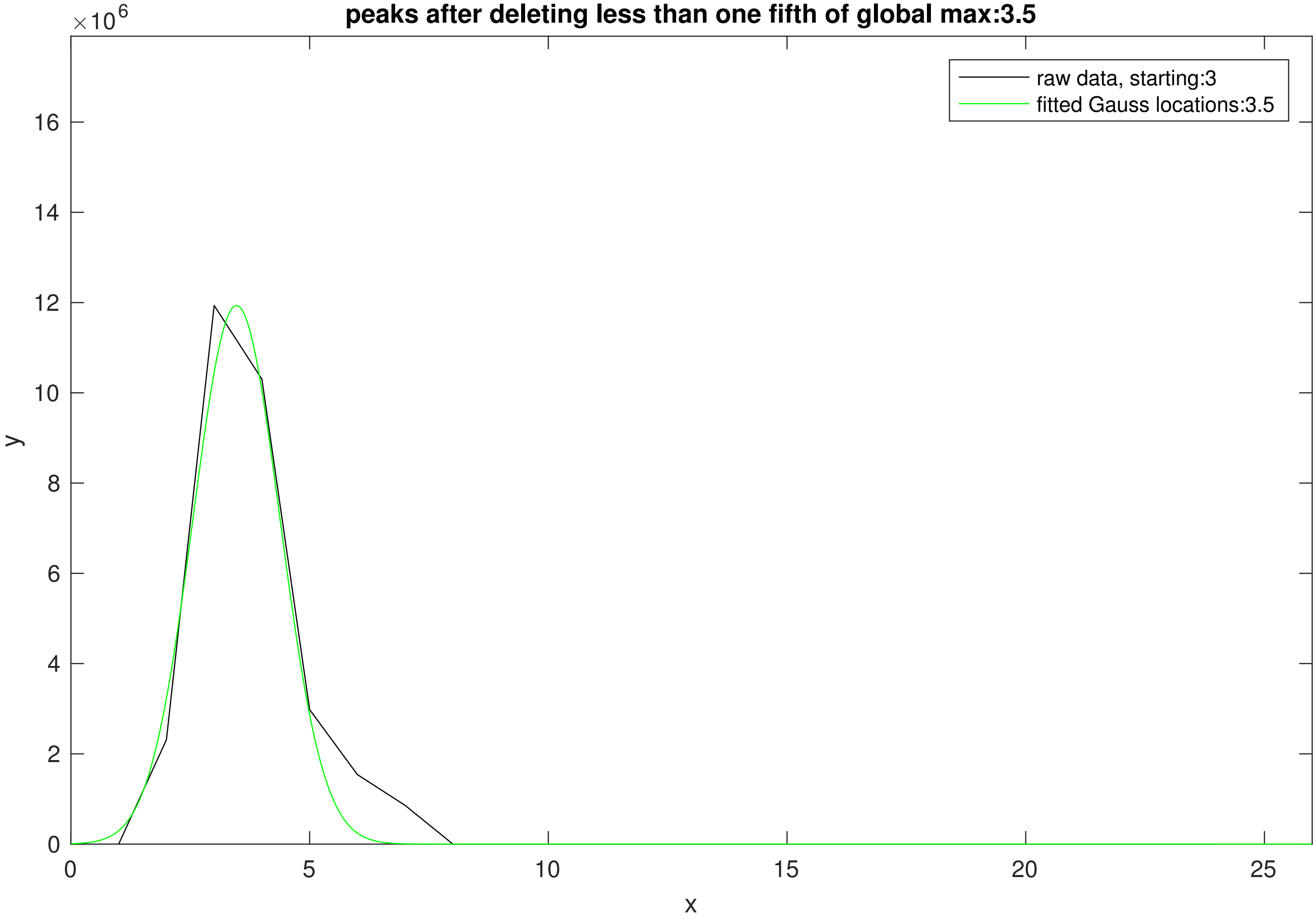

**Optimal1 Fitted Gauss, proteinAT5G23940.1**
**bic:703**
**peaks after deleting less than one fifth of global max:15**

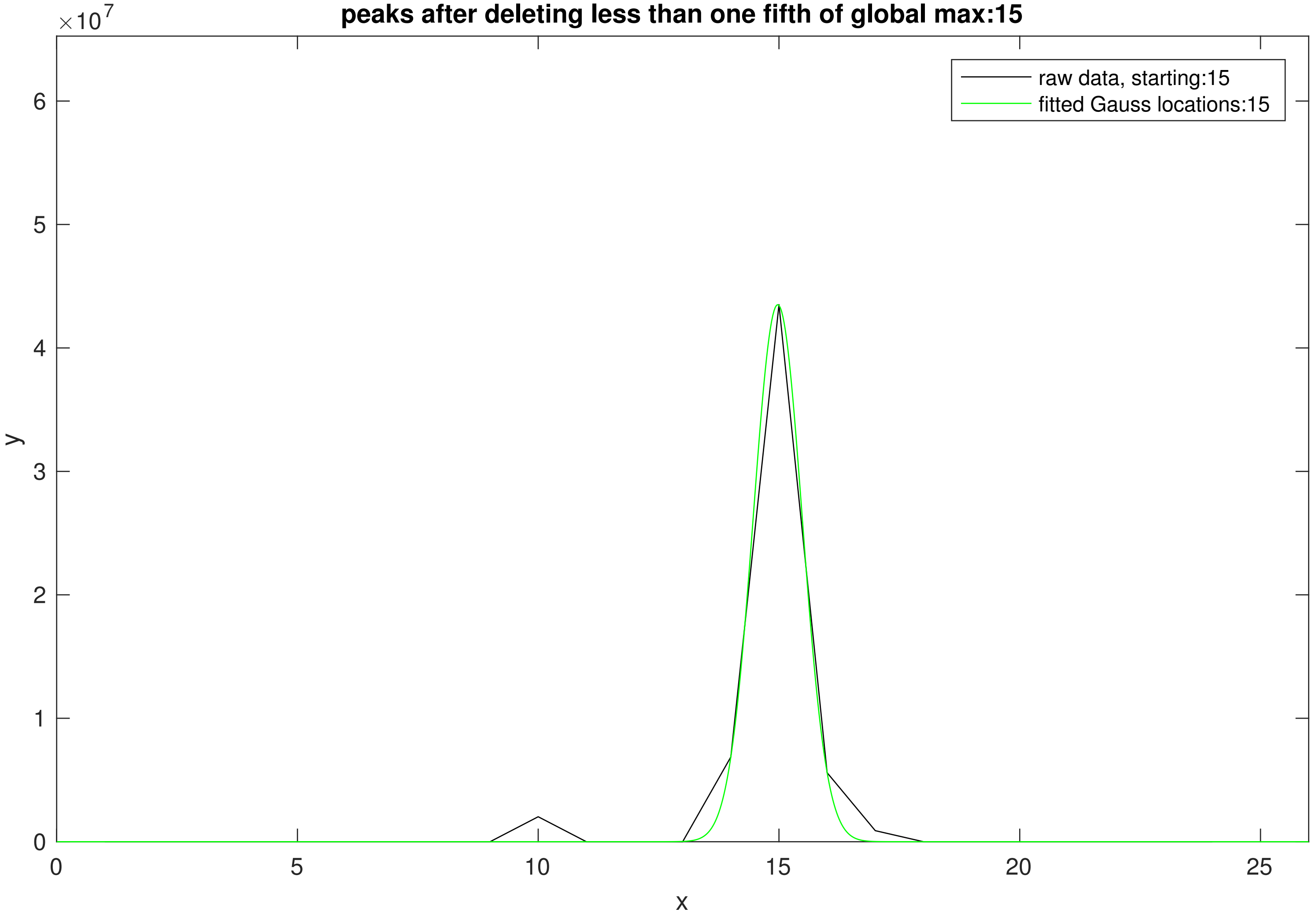

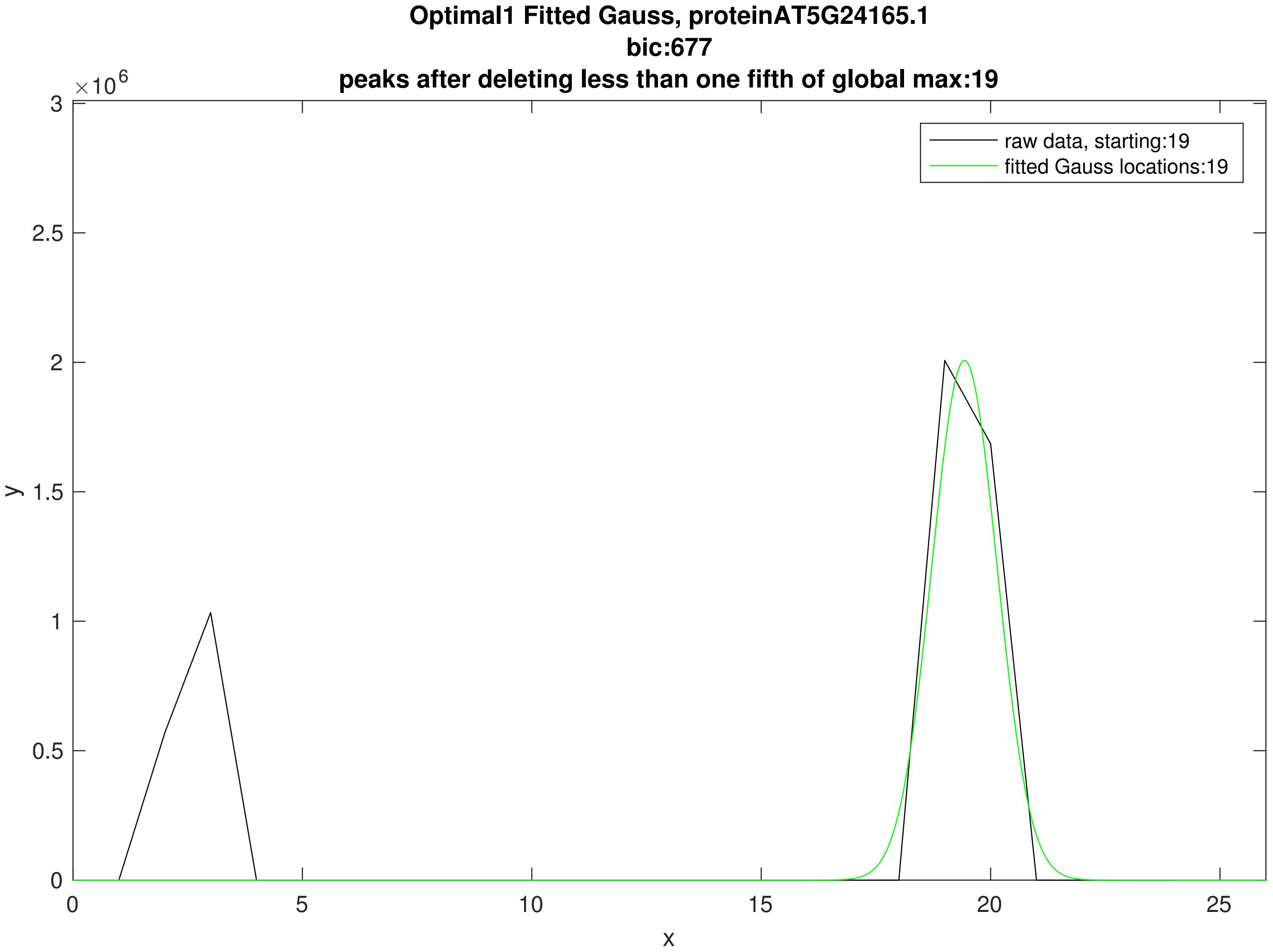

Optimal1 Fitted Gauss, proteinAT5G24165.1
bic:677
peaks after deleting less than one fifth of global max:19
×10^6
raw data, starting:19
fitted Gauss locations:19
y
x
0
0.5
1
1.5
2
2.5
3
0
5
10
15
20
25

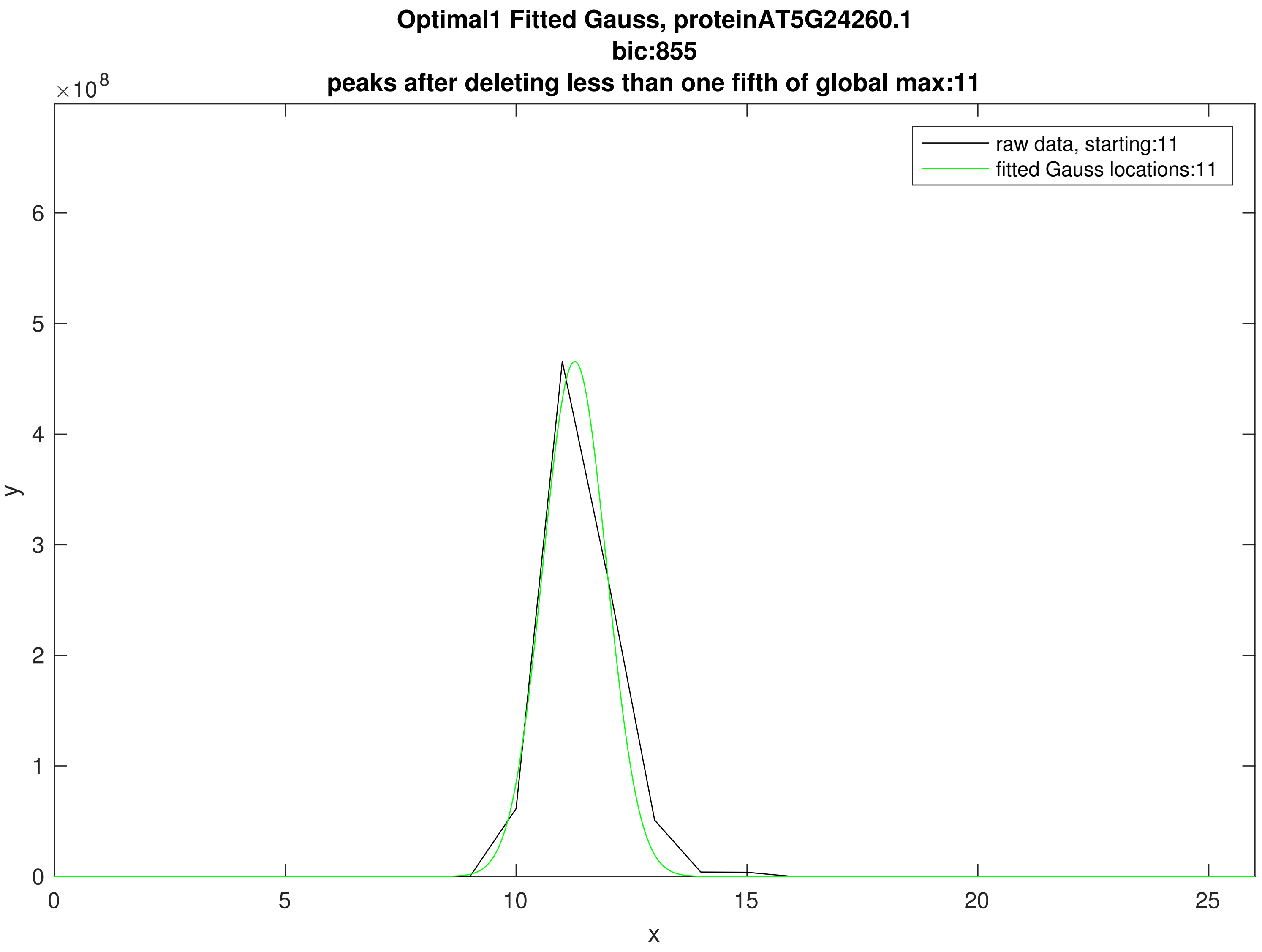

Optimal1 Fitted Gauss, proteinAT5G24260.1
bic:855
peaks after deleting less than one fifth of global max:11
×10^8
raw data, starting:11
fitted Gauss locations:11
y
x
0
1
2
3
4
5
6
0
5
10
15
20
25

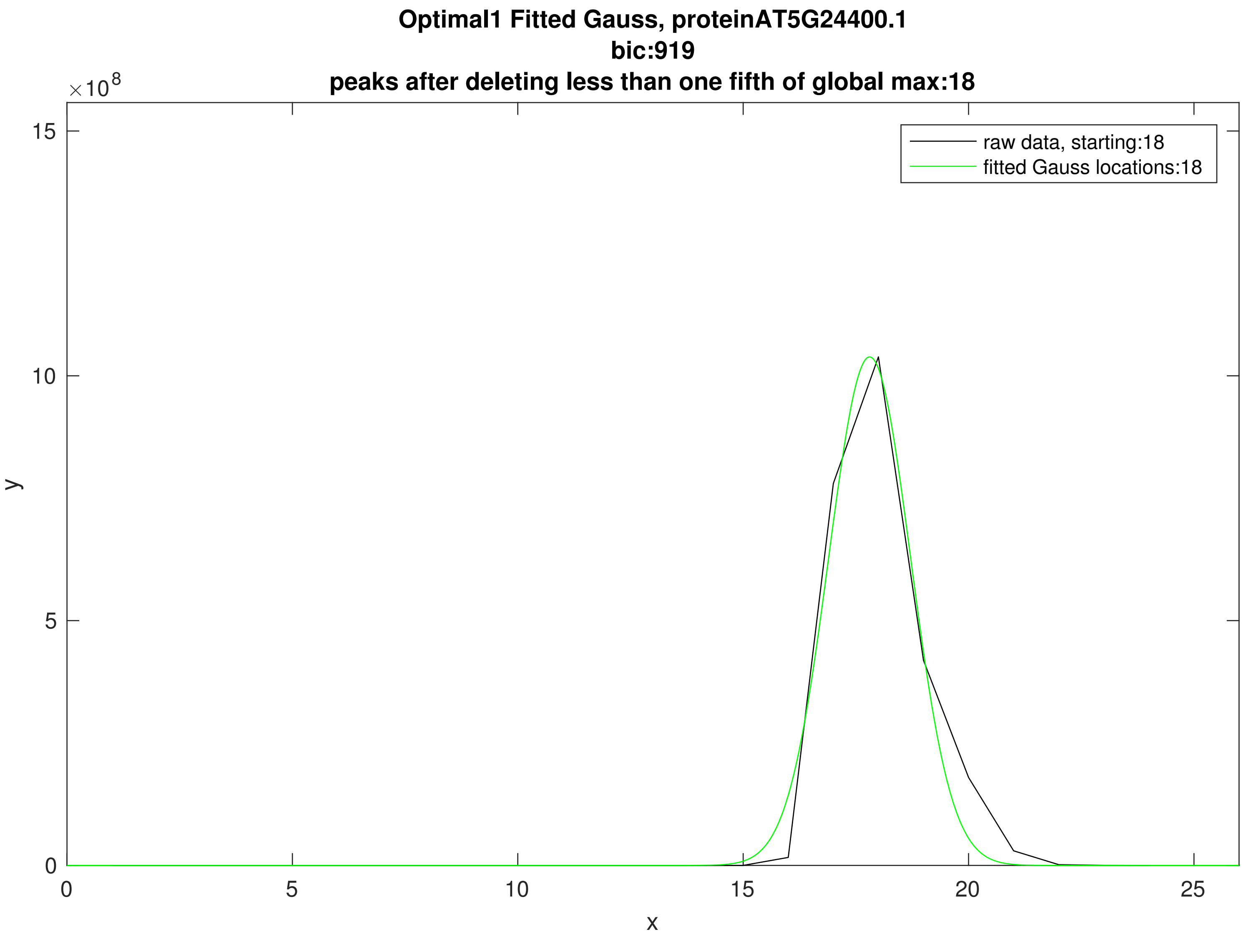

Optimal1 Fitted Gauss, proteinAT5G24400.1
bic:919
peaks after deleting less than one fifth of global max:18
×10^8
15
10
5
0
y
0
5
10
15
20
25
x
raw data, starting:18
fitted Gauss locations:18

Optimal1 Fitted Gauss, proteinAT5G24520.3;AT5G24520.2;AT5G24520.1
bic:774
peaks after deleting less than one fifth of global max:16

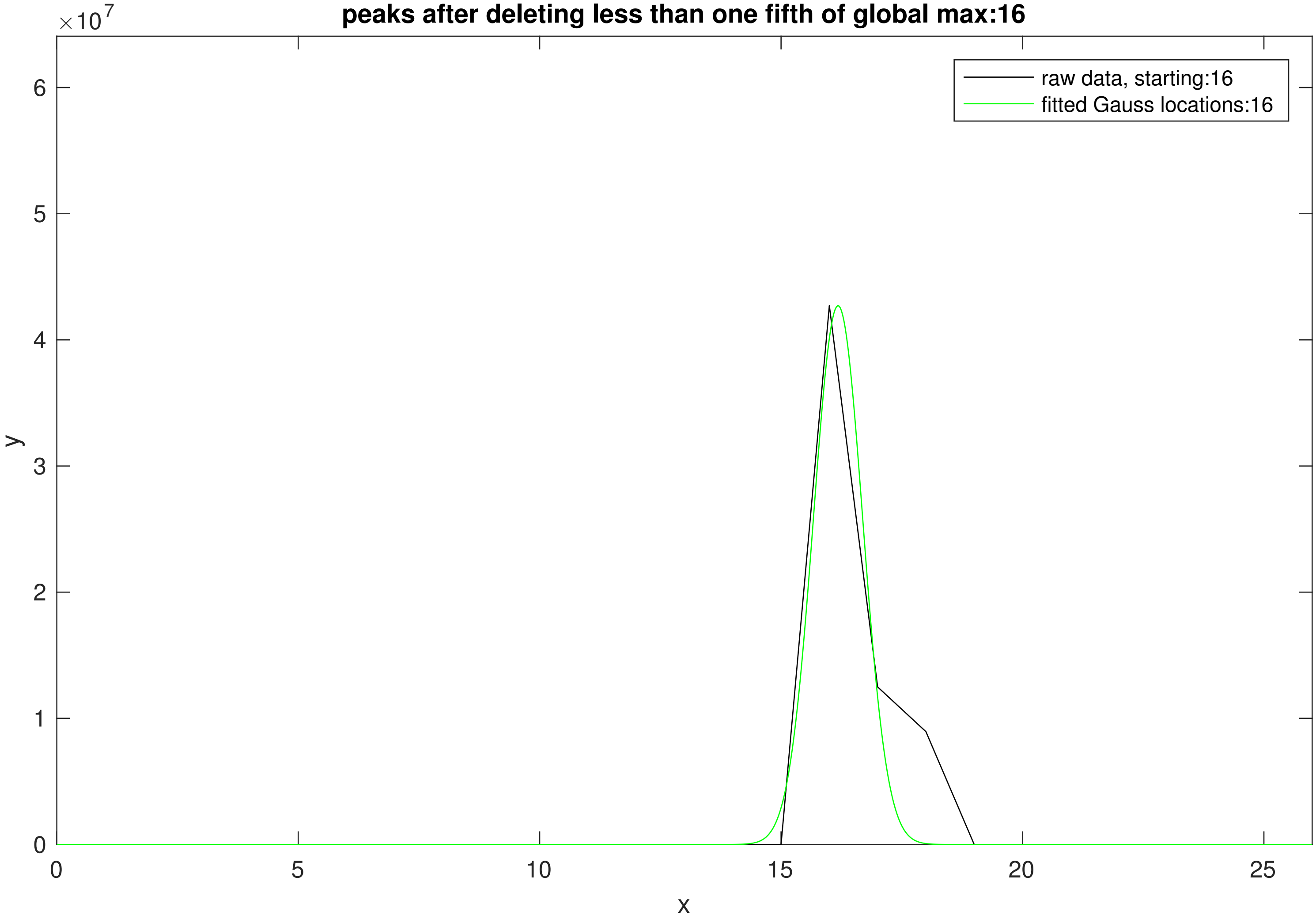

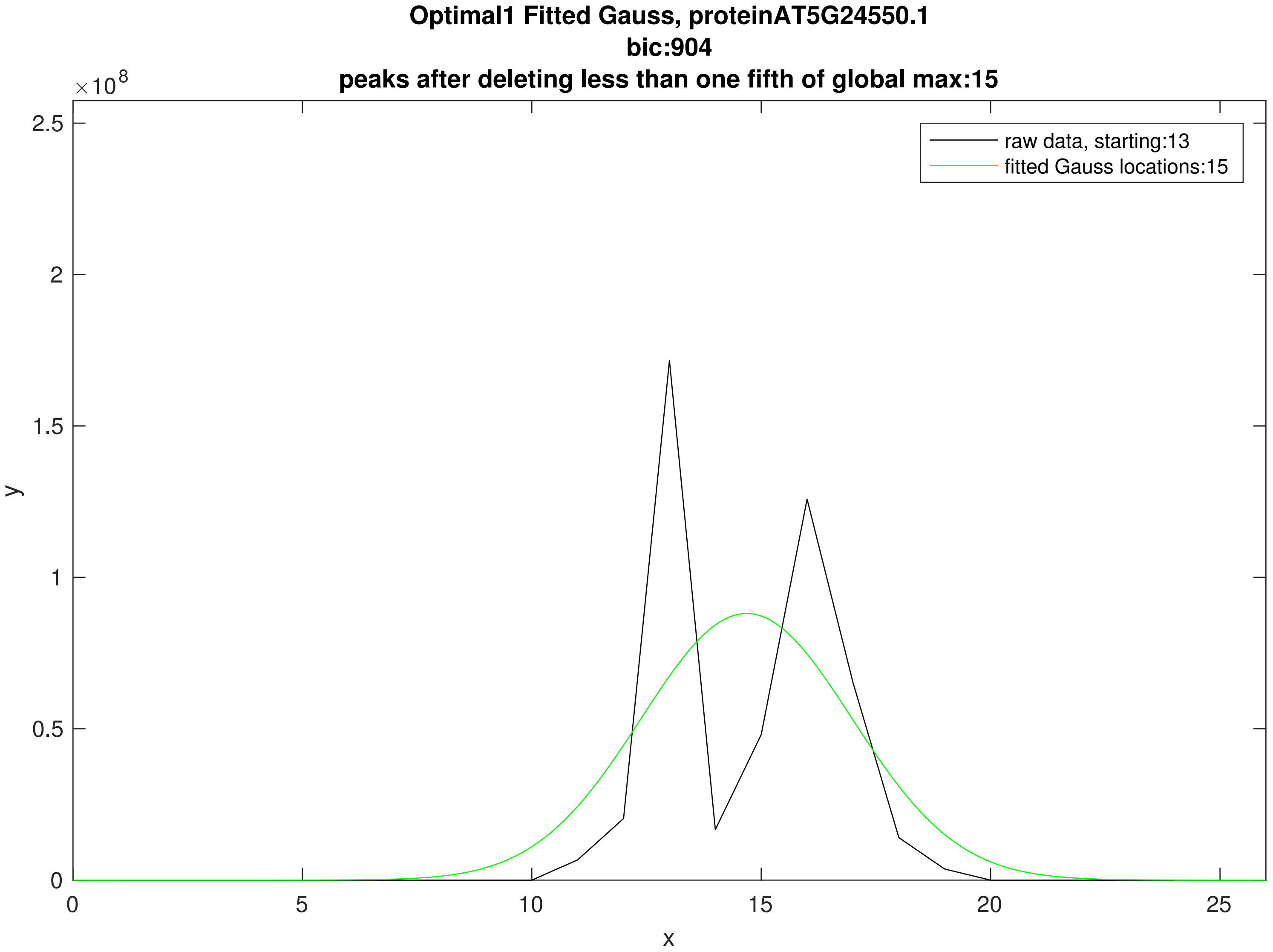

Optimal1 Fitted Gauss, proteinAT5G24550.1
bic:904
peaks after deleting less than one fifth of global max:15
×10^8
2.5
2
1.5
1
0.5
0
y
0
5
10
15
20
25
x
raw data, starting:13
fitted Gauss locations:15

**Optimal1 Fitted Gauss, proteinAT5G24760.1;AT5G24760.3;AT5G24760.2**
**bic:787**
**peaks after deleting less than one fifth of global max:14**

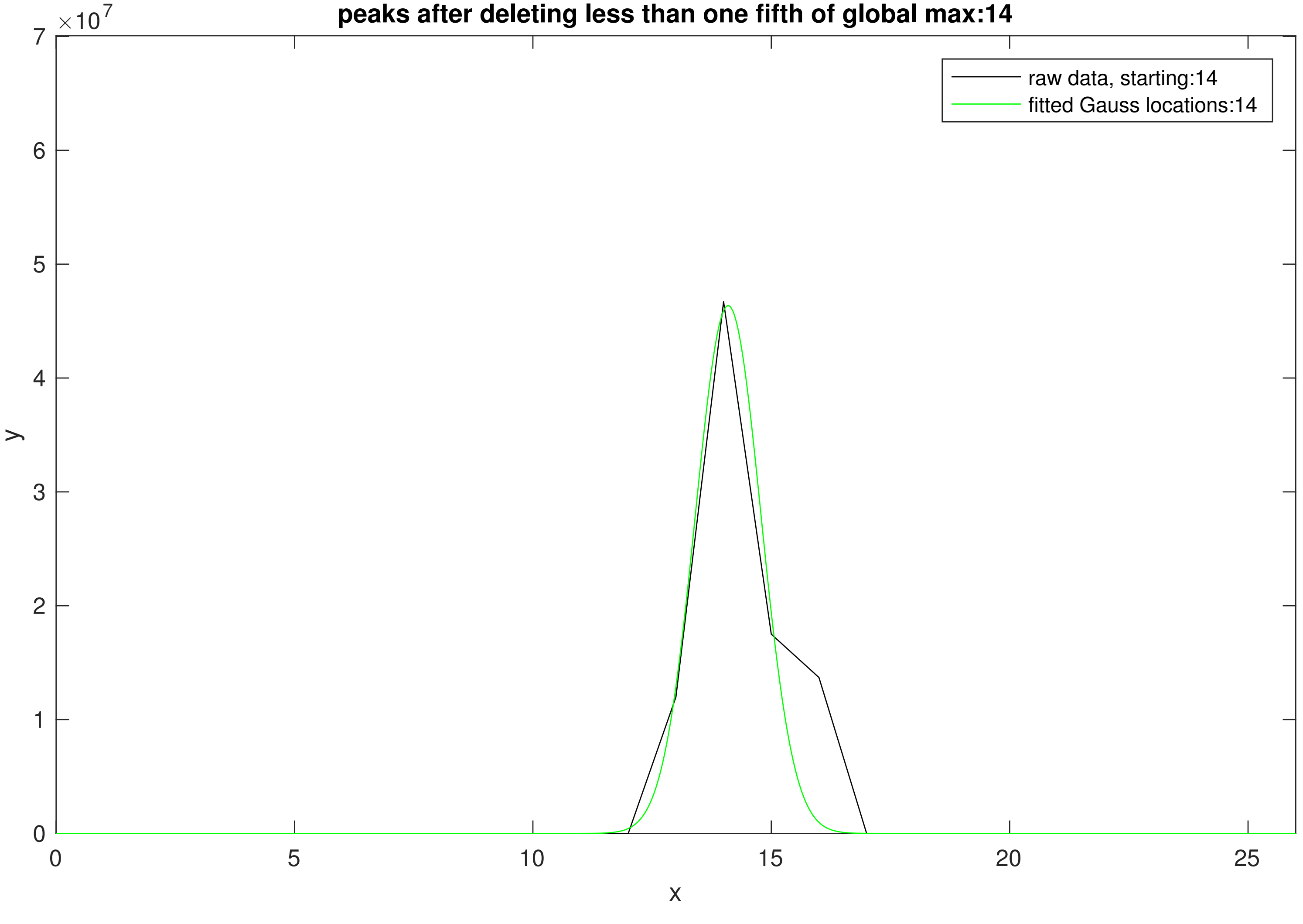

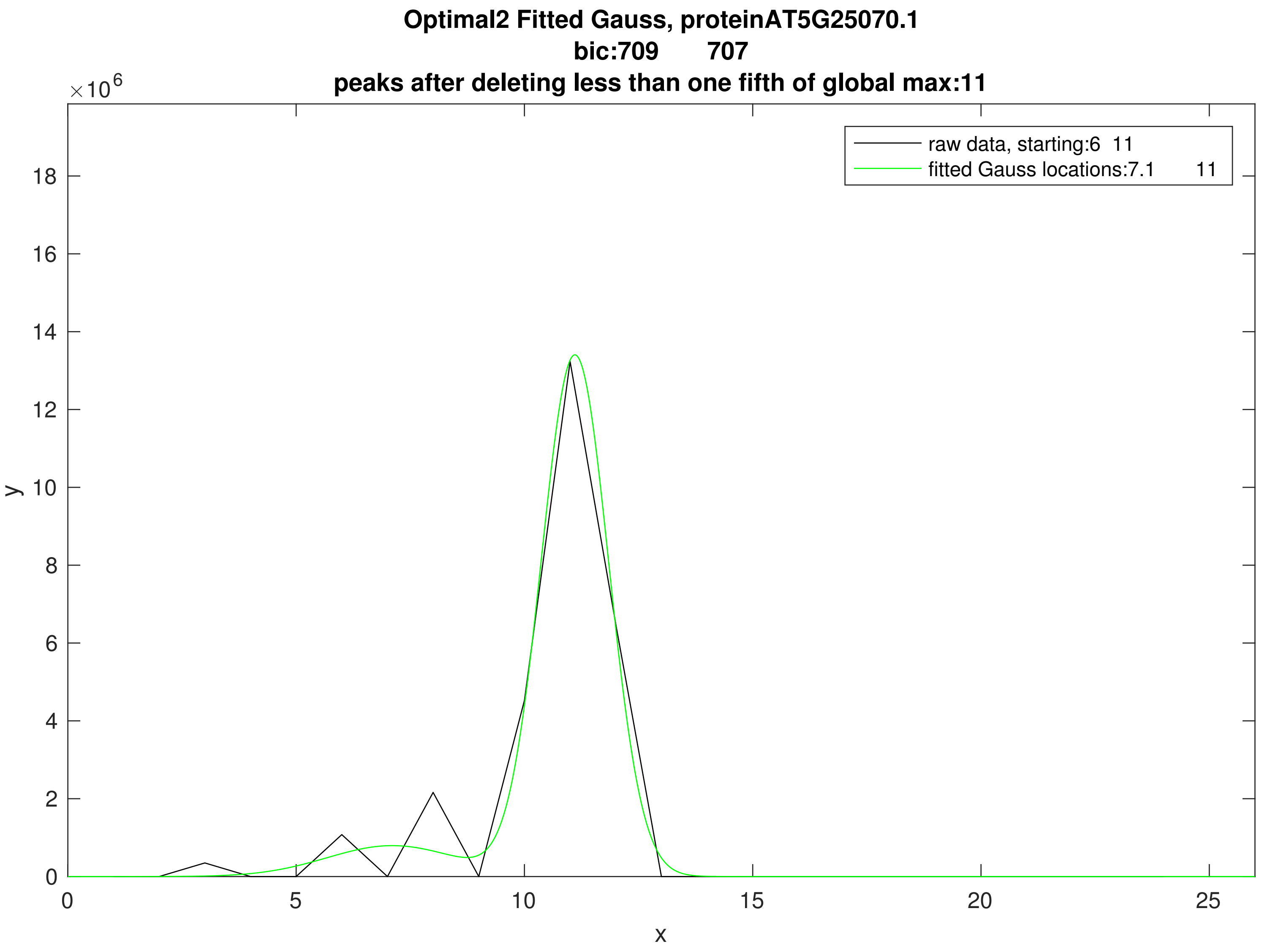
Optimal2 Fitted Gauss, proteinAT5G25070.1
bic:709 707
peaks after deleting less than one fifth of global max:11
×10^6
raw data, starting:6 11
fitted Gauss locations:7.1 11
y
x
0
2
4
6
8
10
12
14
16
18
0
5
10
15
20
25

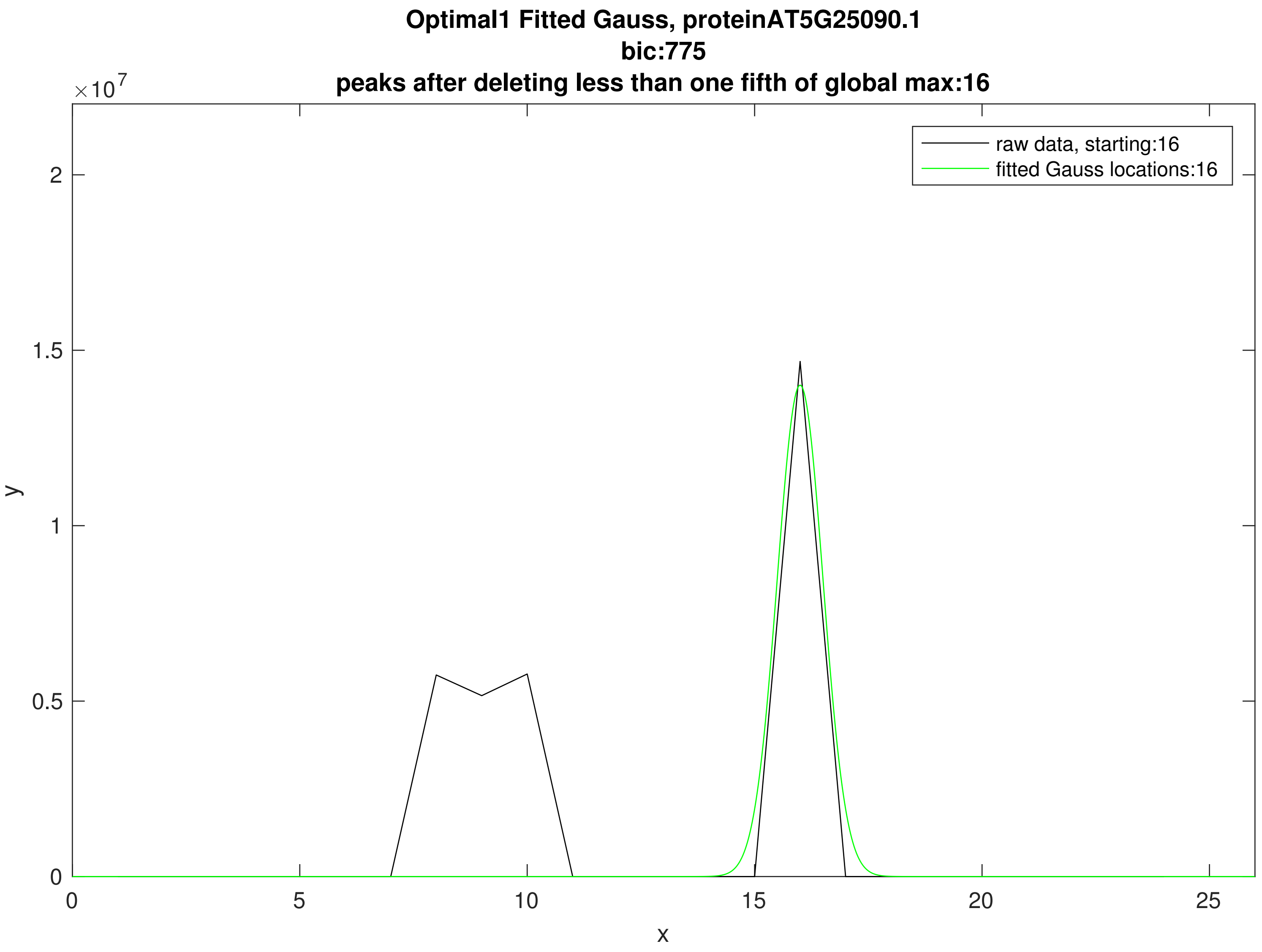
Optimal1 Fitted Gauss, proteinAT5G25090.1
bic:775
peaks after deleting less than one fifth of global max:16
×10^7
raw data, starting:16
fitted Gauss locations:16
y
x
0
0.5
1
1.5
2
0
5
10
15
20
25

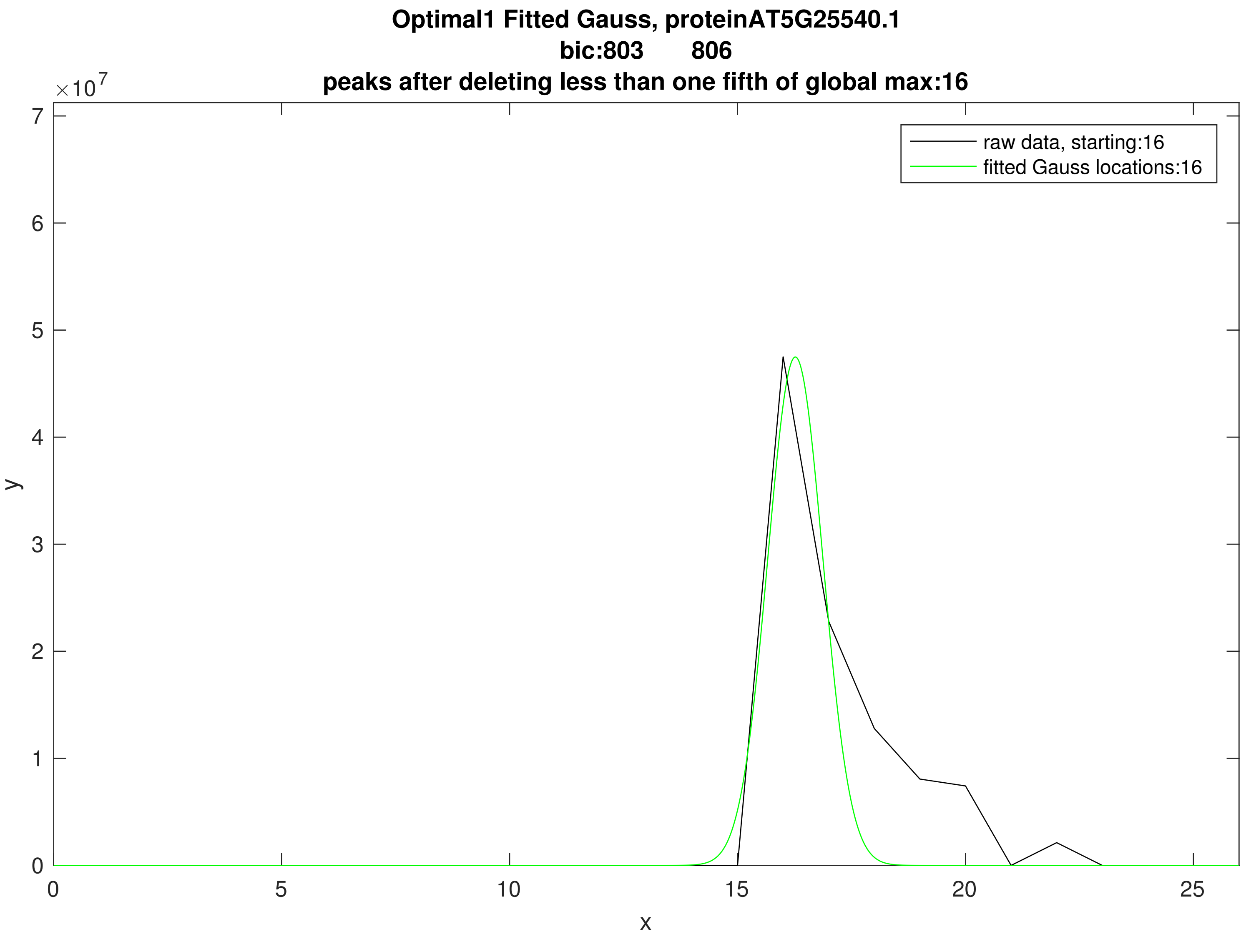

Optimal1 Fitted Gauss, proteinAT5G25540.1
bic:803 806
peaks after deleting less than one fifth of global max:16
×10^7
raw data, starting:16
fitted Gauss locations:16
y
x
0
1
2
3
4
5
6
7
0
5
10
15
20
25

**Optimal2 Fitted Gauss, proteinAT5G25770.1;AT5G25770.2;AT5G25770.3**
**bic:880 878**
**peaks after deleting less than one fifth of global max:17**

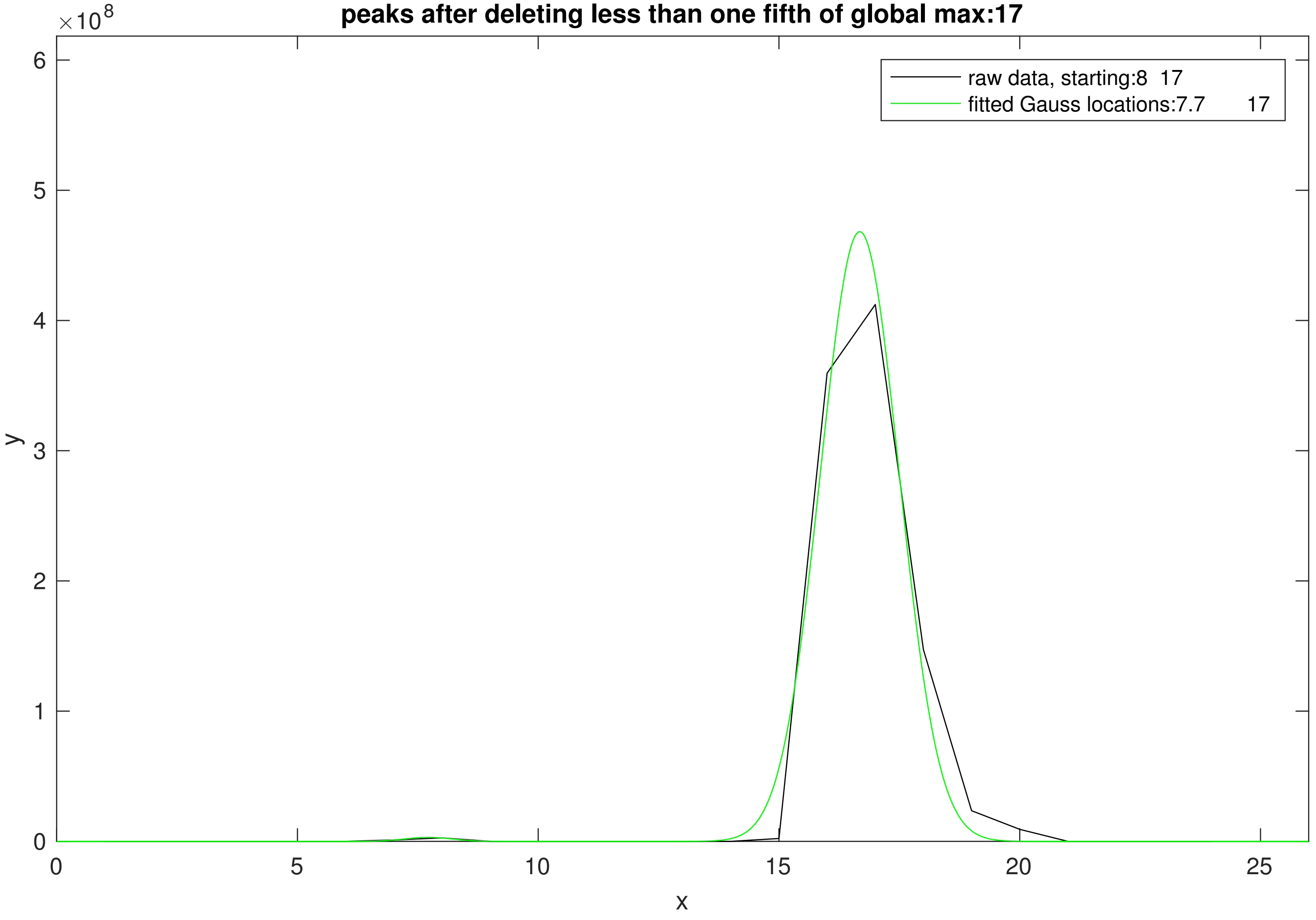

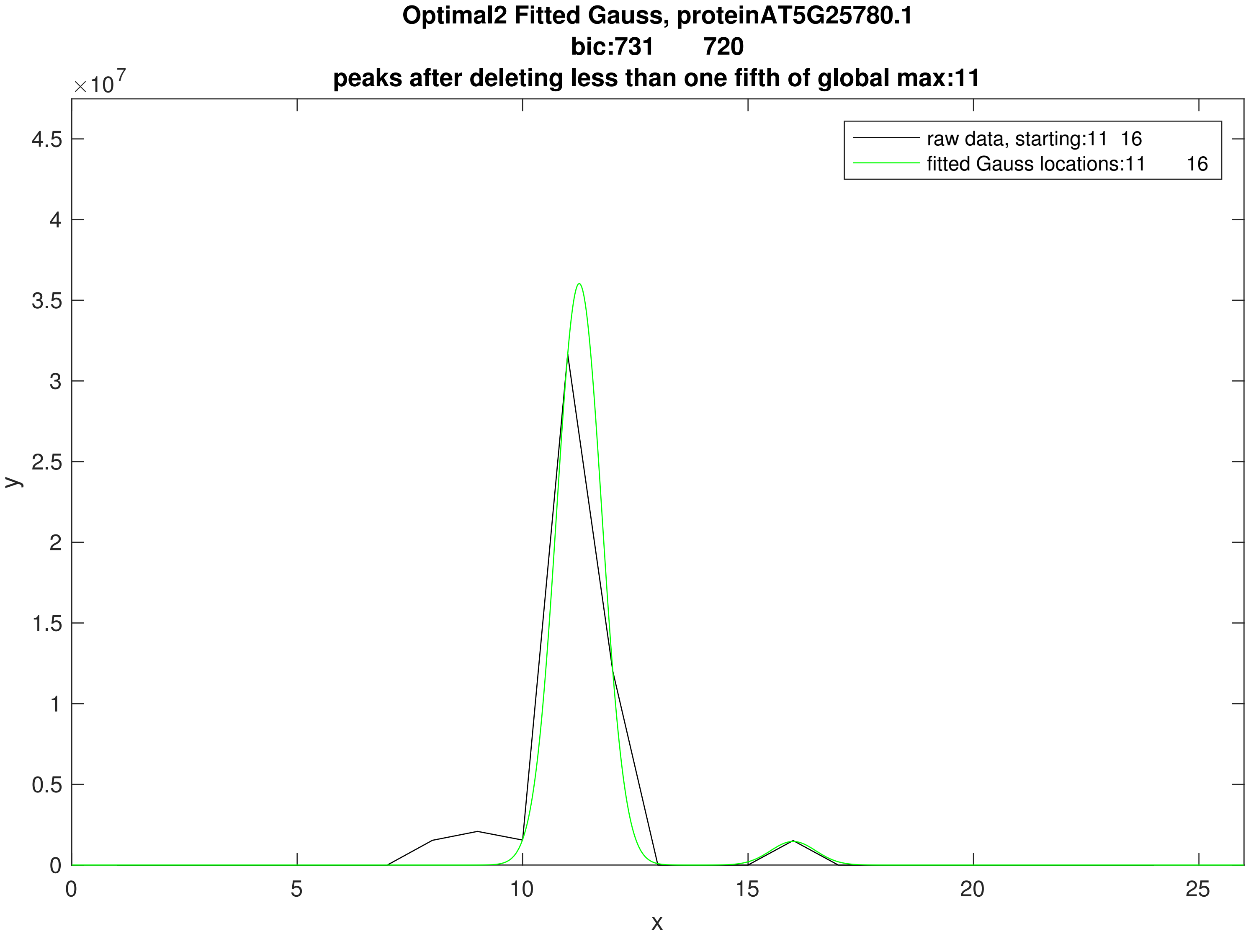
Optimal2 Fitted Gauss, proteinAT5G25780.1
bic:731 720
peaks after deleting less than one fifth of global max:11
×10^7
raw data, starting:11 16
fitted Gauss locations:11 16
y
x
0
0.5
1
1.5
2
2.5
3
3.5
4
4.5
0
5
10
15
20
25

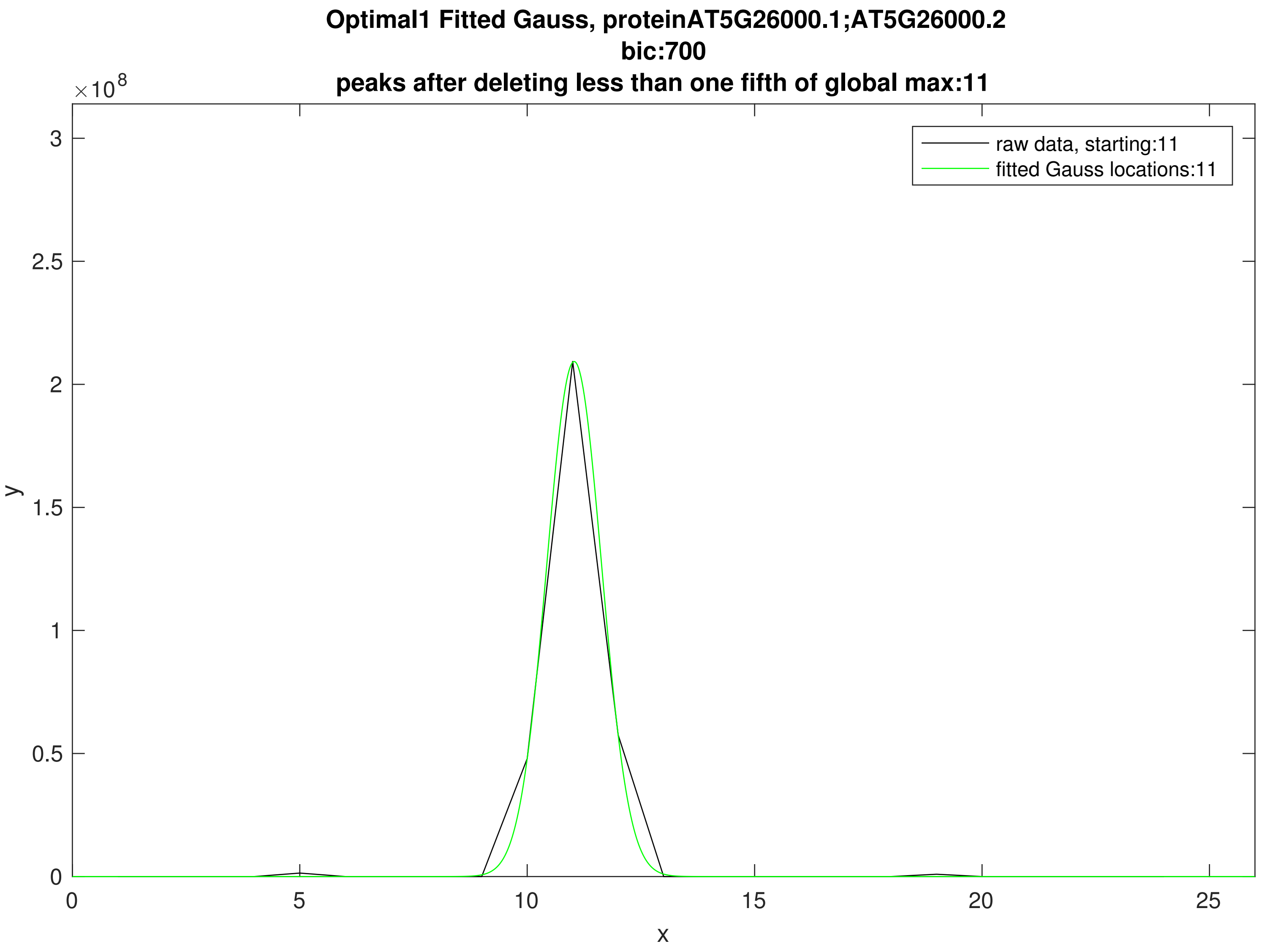
Optimal1 Fitted Gauss, proteinAT5G26000.1;AT5G26000.2
bic:700
peaks after deleting less than one fifth of global max:11
×10^8
3
2.5
2
1.5
1
0.5
0
y
0
5
10
15
20
25
x
raw data, starting:11
fitted Gauss locations:11

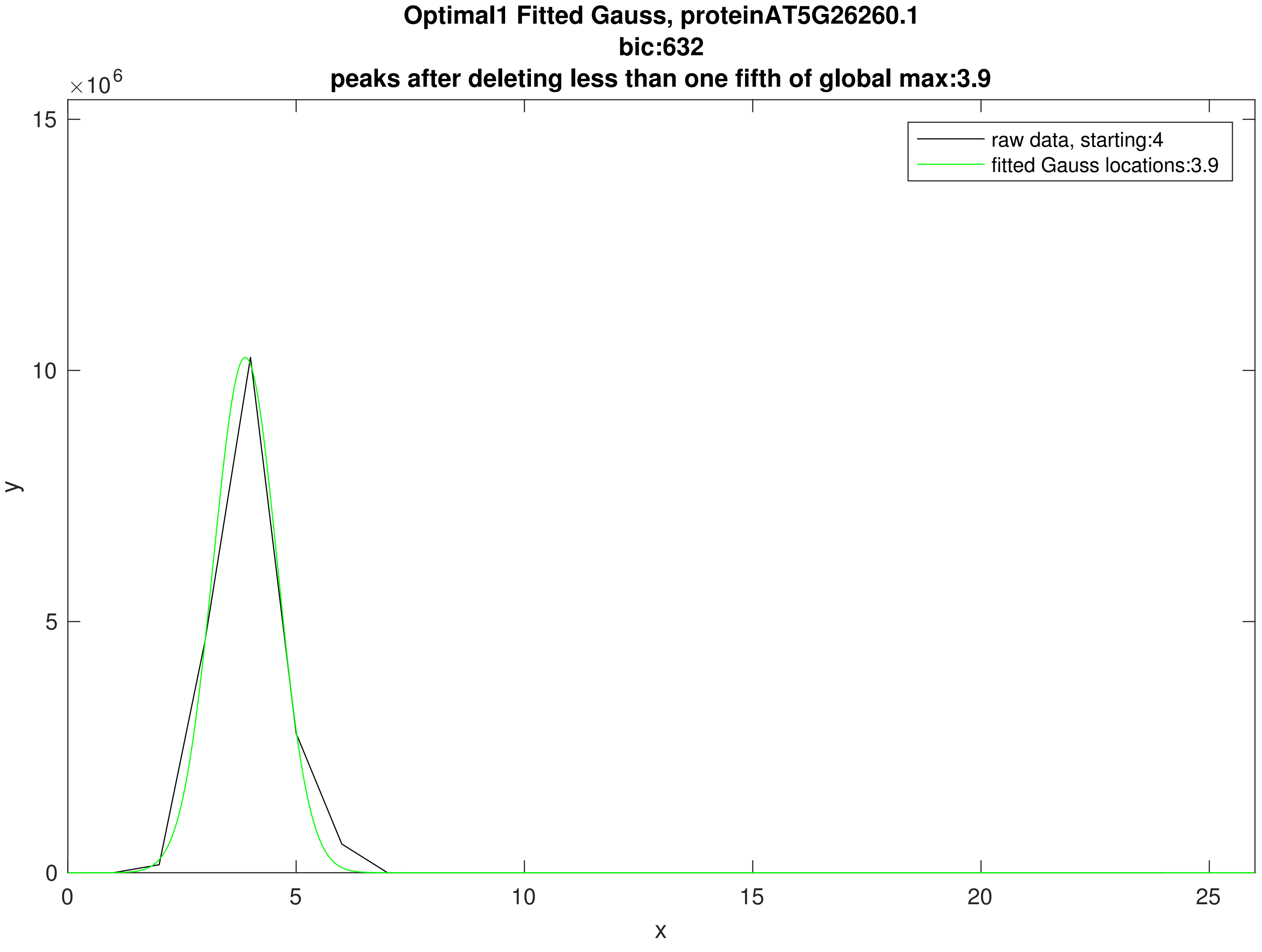
Optimal1 Fitted Gauss, proteinAT5G26260.1
bic:632
peaks after deleting less than one fifth of global max:3.9
×10^6
15
10
5
0
y
0
5
10
15
20
25
x
raw data, starting:4
fitted Gauss locations:3.9

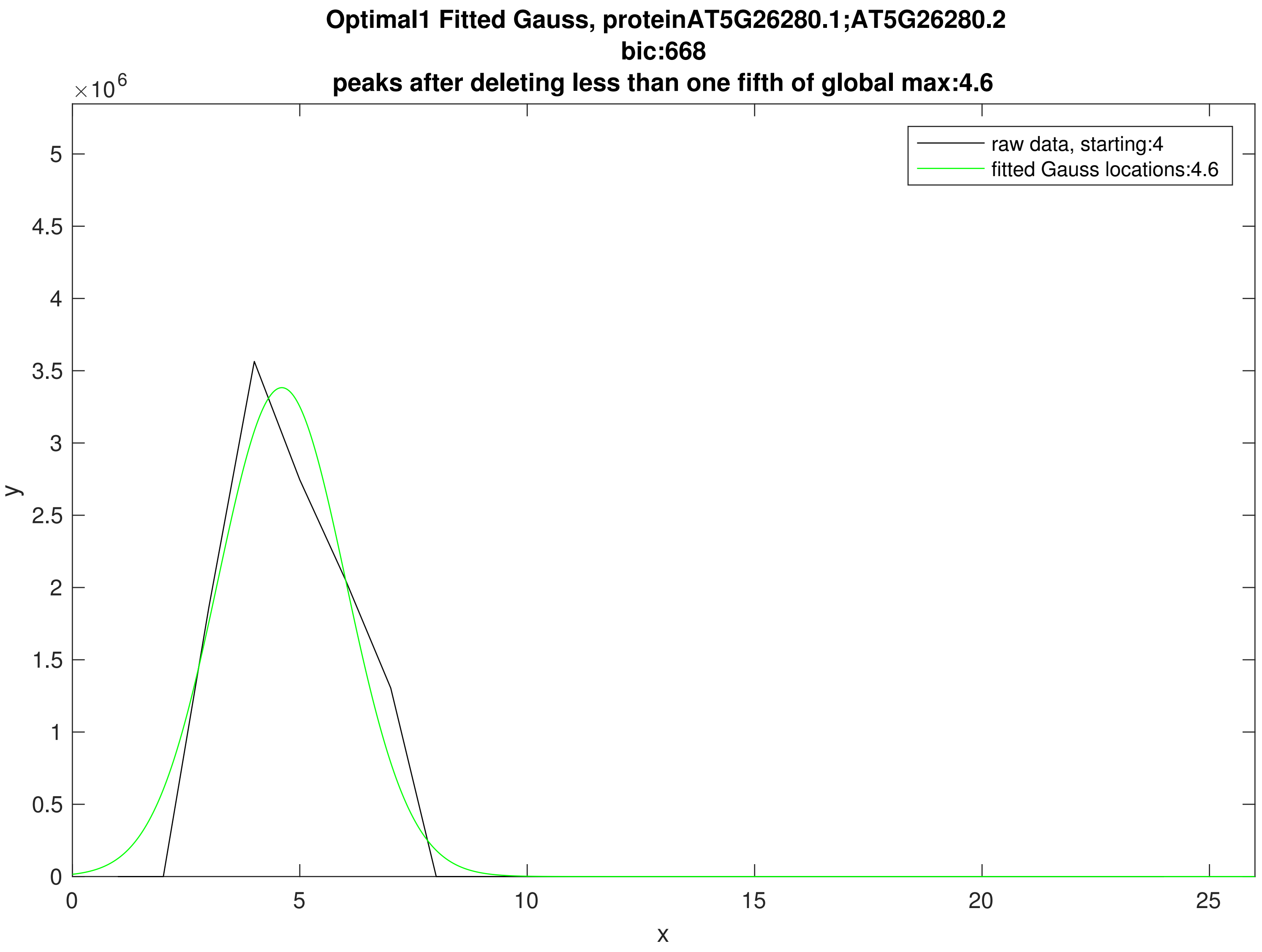

Optimal1 Fitted Gauss, proteinAT5G26280.1;AT5G26280.2
bic:668
peaks after deleting less than one fifth of global max:4.6
×10^6
raw data, starting:4
fitted Gauss locations:4.6
y
x
0
0.5
1
1.5
2
2.5
3
3.5
4
4.5
5
0
5
10
15
20
25

#### Optimal2 Fitted Gauss, proteinAT5G26360.1
#### bic:880 863 872 881
#### peaks after deleting less than one fifth of global max:5.8 18

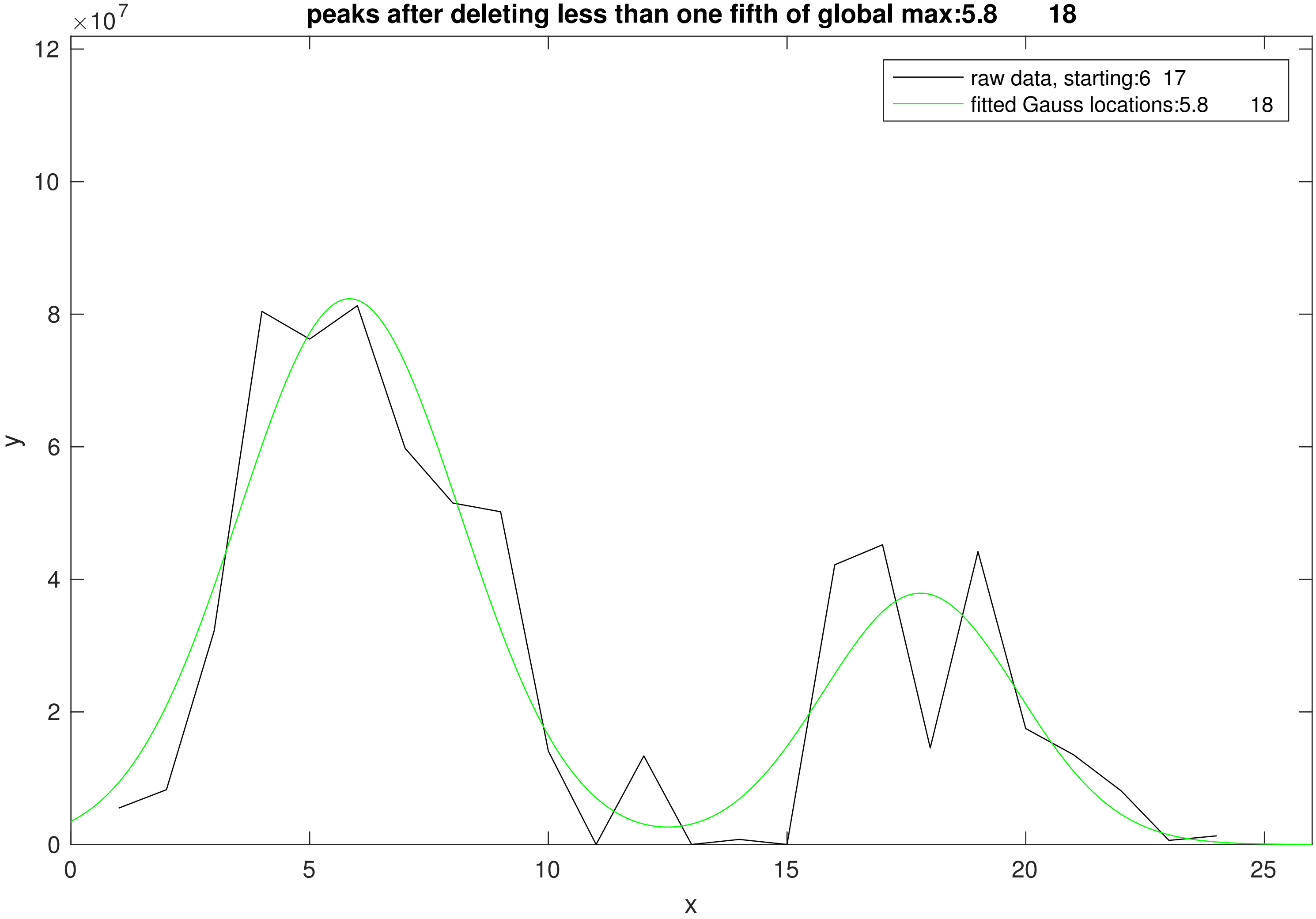

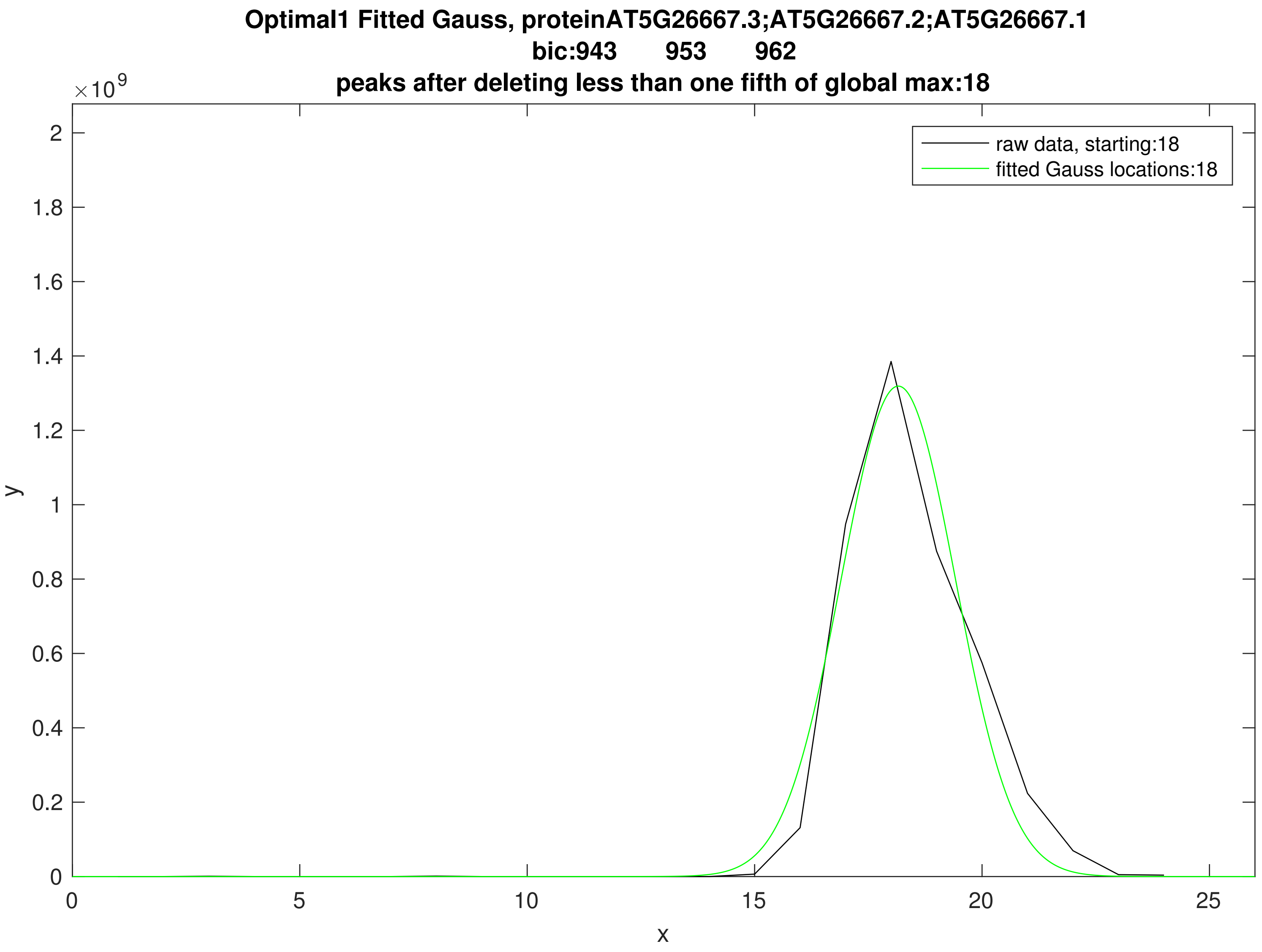

Optimal1 Fitted Gauss, proteinAT5G26667.3;AT5G26667.2;AT5G26667.1
bic:943 953 962
peaks after deleting less than one fifth of global max:18
×10^9
raw data, starting:18
fitted Gauss locations:18
y
x
0
0.2
0.4
0.6
0.8
1
1.2
1.4
1.6
1.8
2
5
10
15
20
25

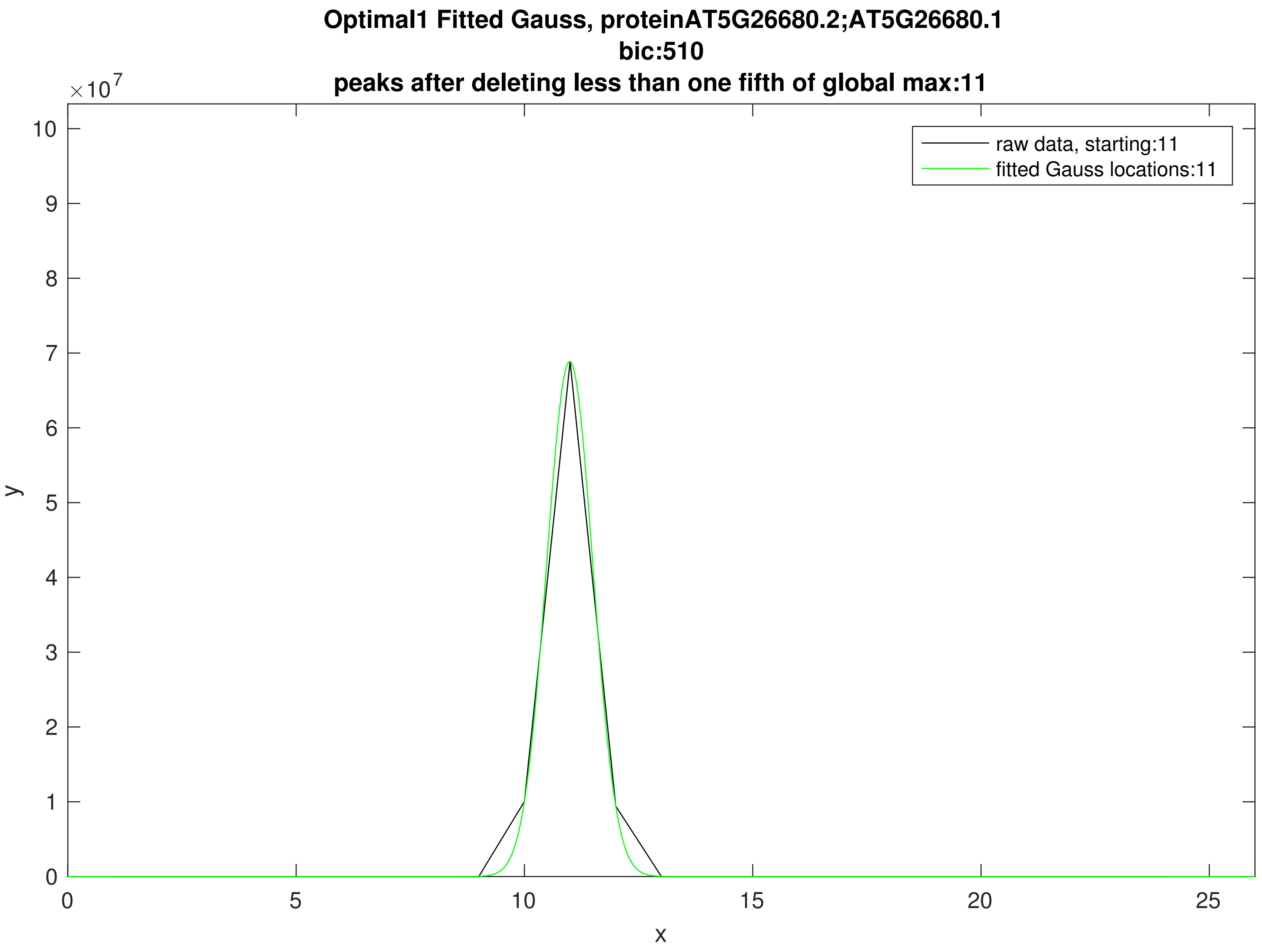
Optimal1 Fitted Gauss, proteinAT5G26680.2;AT5G26680.1
bic:510
peaks after deleting less than one fifth of global max:11
×10^7
raw data, starting:11
fitted Gauss locations:11
y
x
0
1
2
3
4
5
6
7
8
9
10
0
5
10
15
20
25

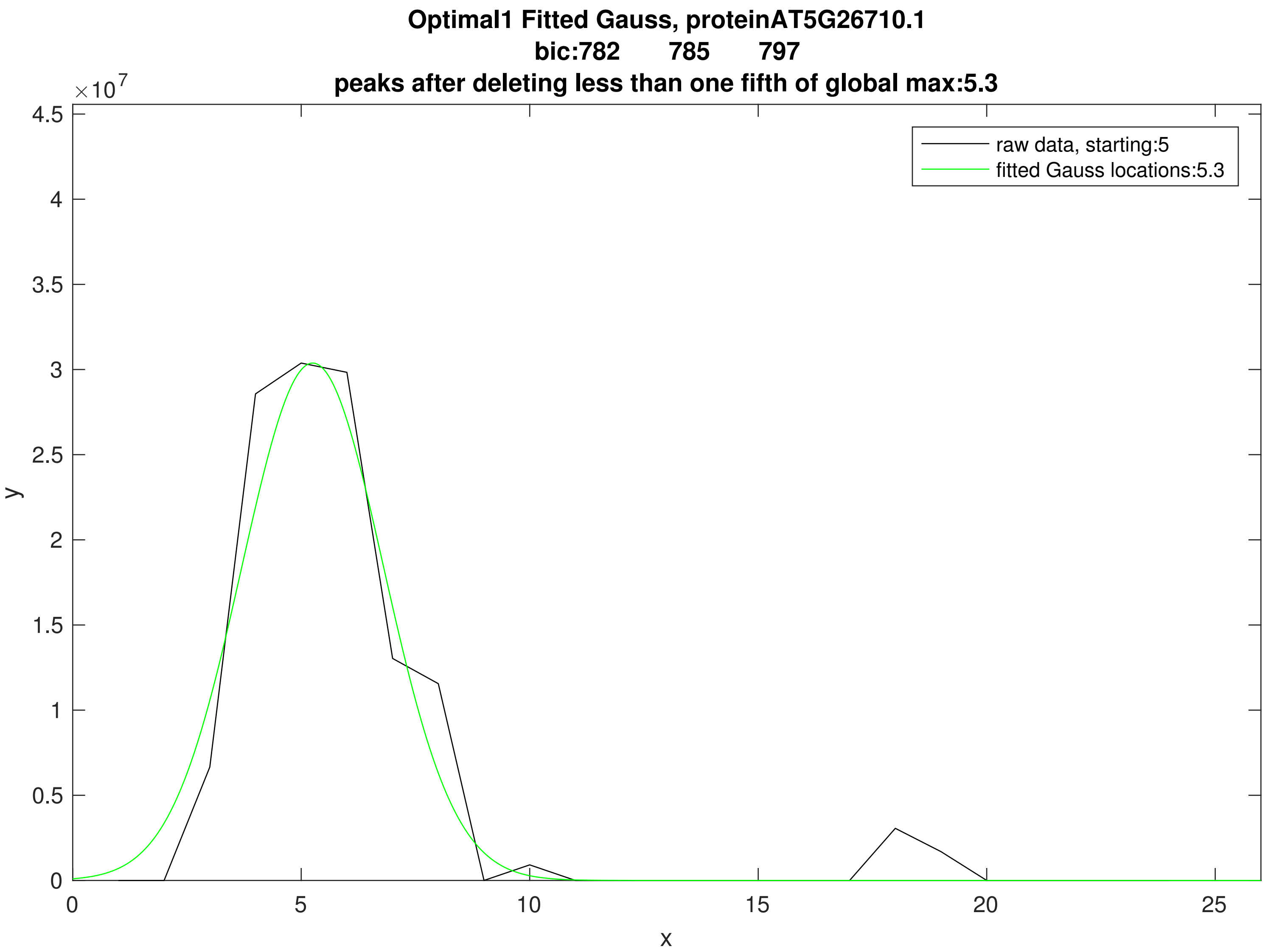
Optimal1 Fitted Gauss, proteinAT5G26710.1
bic:782 785 797
peaks after deleting less than one fifth of global max:5.3
×10^7
4.5
4
3.5
3
2.5
2
1.5
1
0.5
0
y
0
5
10
15
20
25
x
raw data, starting:5
fitted Gauss locations:5.3

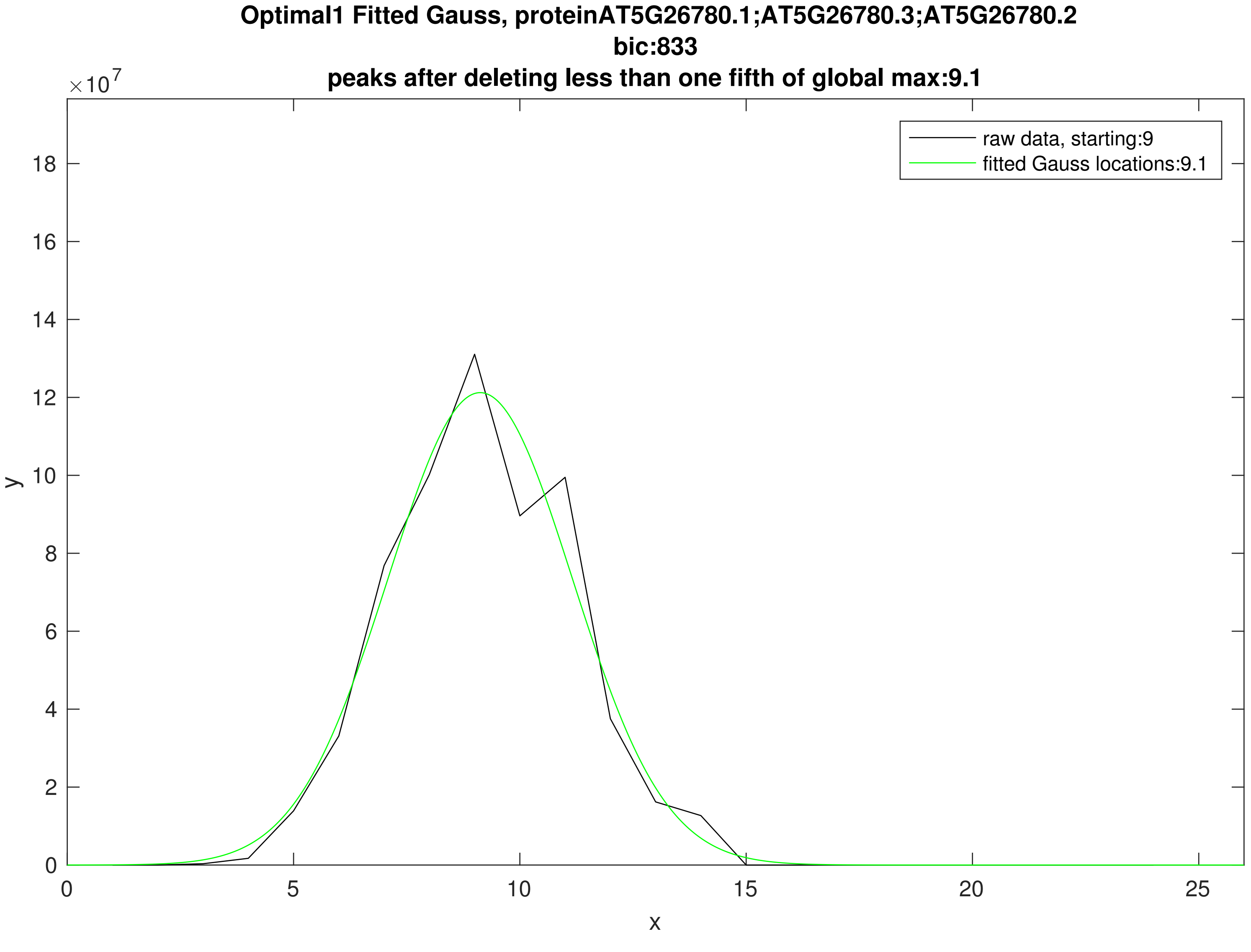

Optimal1 Fitted Gauss, proteinAT5G26780.1;AT5G26780.3;AT5G26780.2
bic:833
peaks after deleting less than one fifth of global max:9.1
×10^7
raw data, starting:9
fitted Gauss locations:9.1
y
x
0
2
4
6
8
10
12
14
16
18
0
5
10
15
20
25

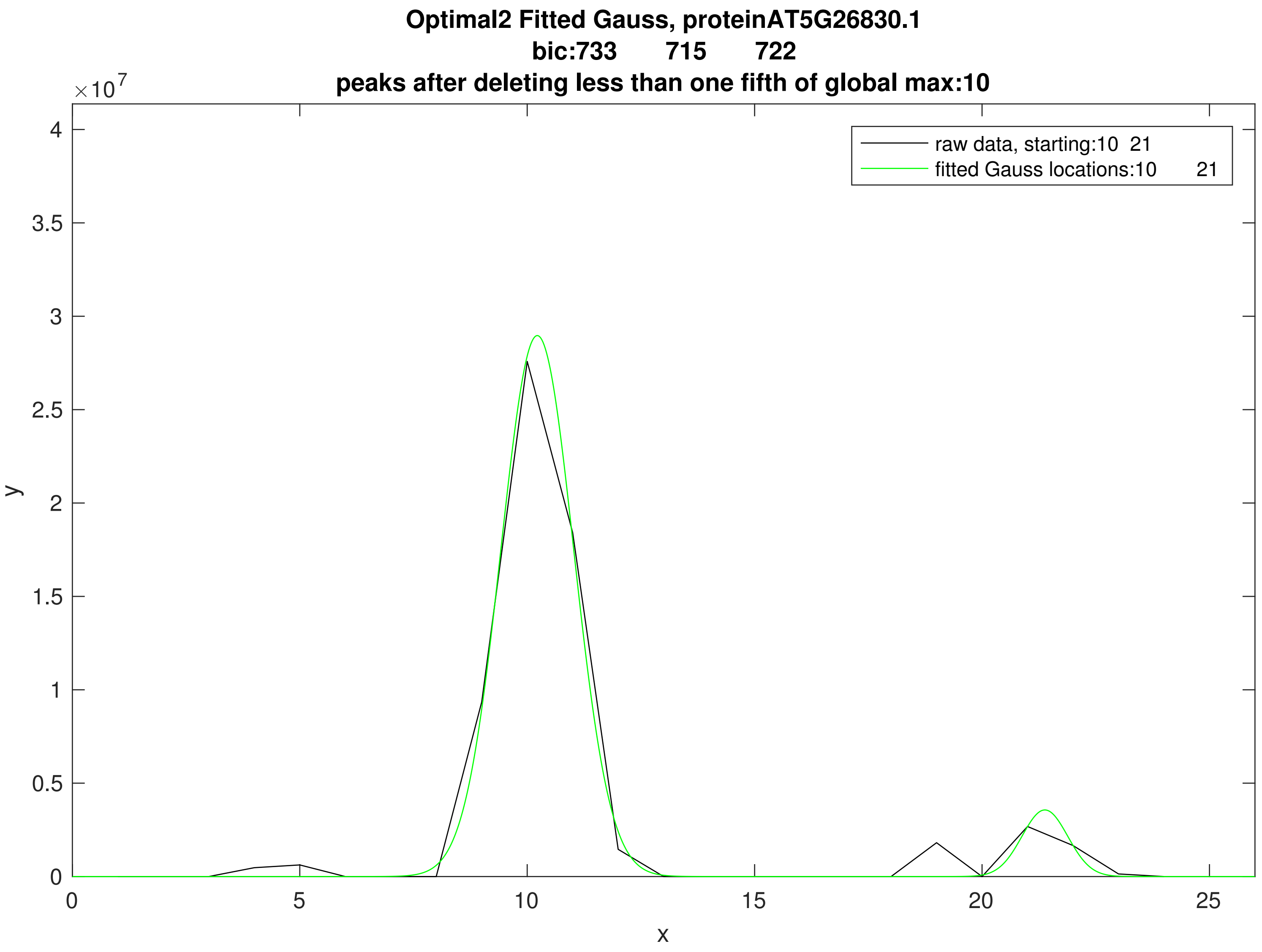
Optimal2 Fitted Gauss, proteinAT5G26830.1
bic:733 715 722
peaks after deleting less than one fifth of global max:10
raw data, starting:10 21
fitted Gauss locations:10 21
×10^7
0
0.5
1
1.5
2
2.5
3
3.5
4
y
0
5
10
15
20
25
x

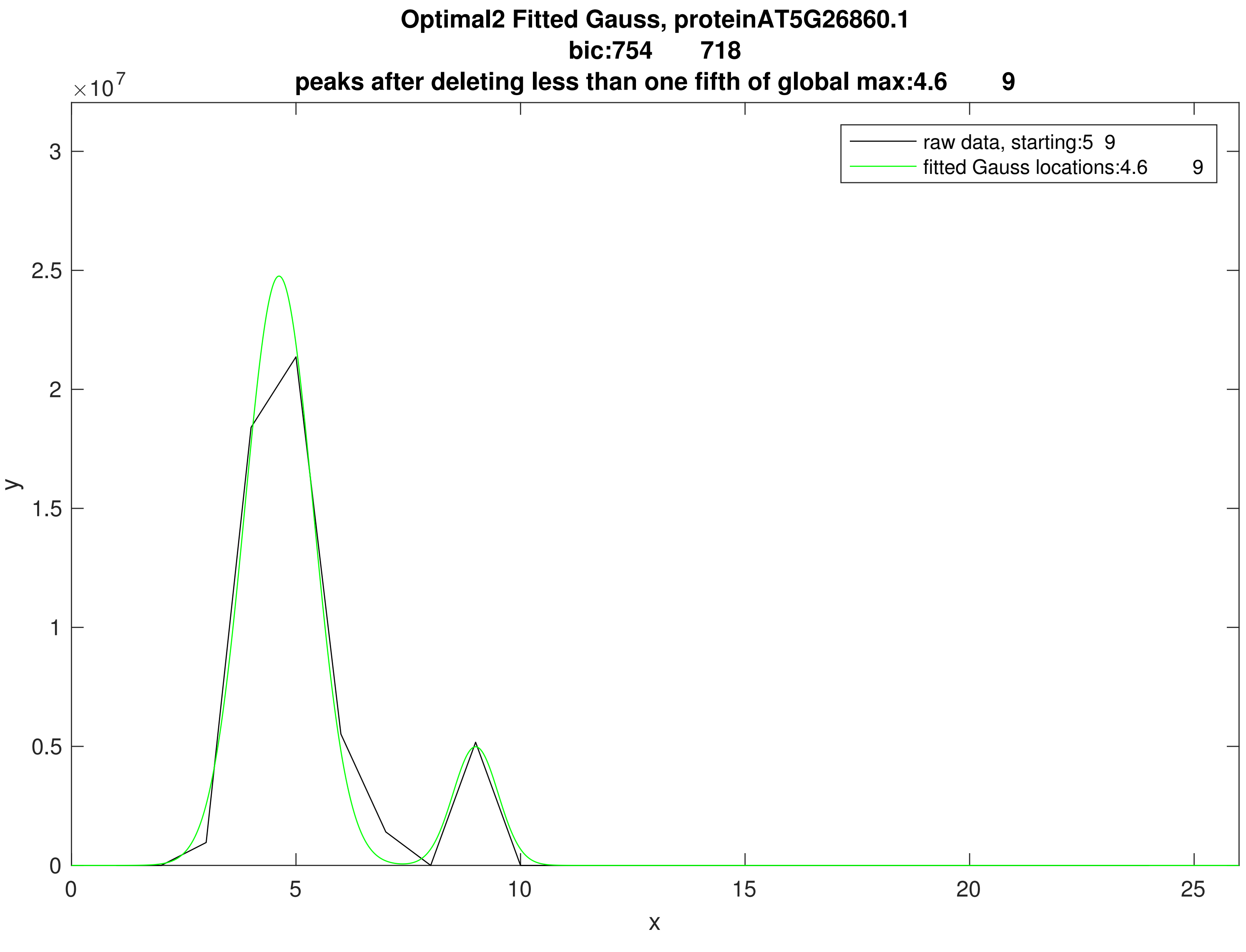

Optimal2 Fitted Gauss, proteinAT5G26860.1
bic:754 718
peaks after deleting less than one fifth of global max:4.6 9
×10^7
3
2.5
2
1.5
1
0.5
0
y
0
5
10
15
20
25
x
raw data, starting:5 9
fitted Gauss locations:4.6 9

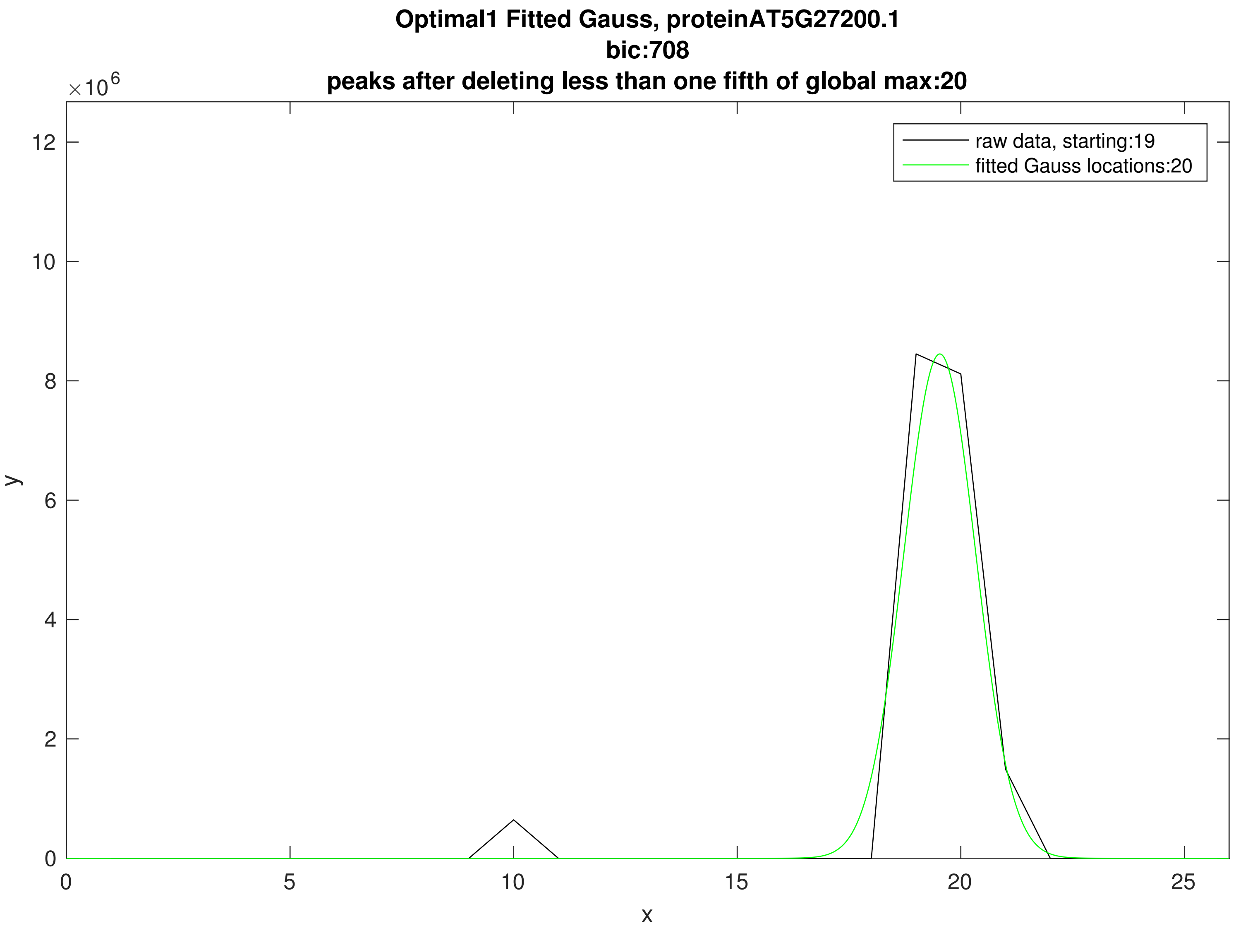
Optimal1 Fitted Gauss, proteinAT5G27200.1
bic:708
peaks after deleting less than one fifth of global max:20
$\times 10^6$
12
10
8
6
4
2
0
y
0
5
10
15
20
25
x
raw data, starting:19
fitted Gauss locations:20

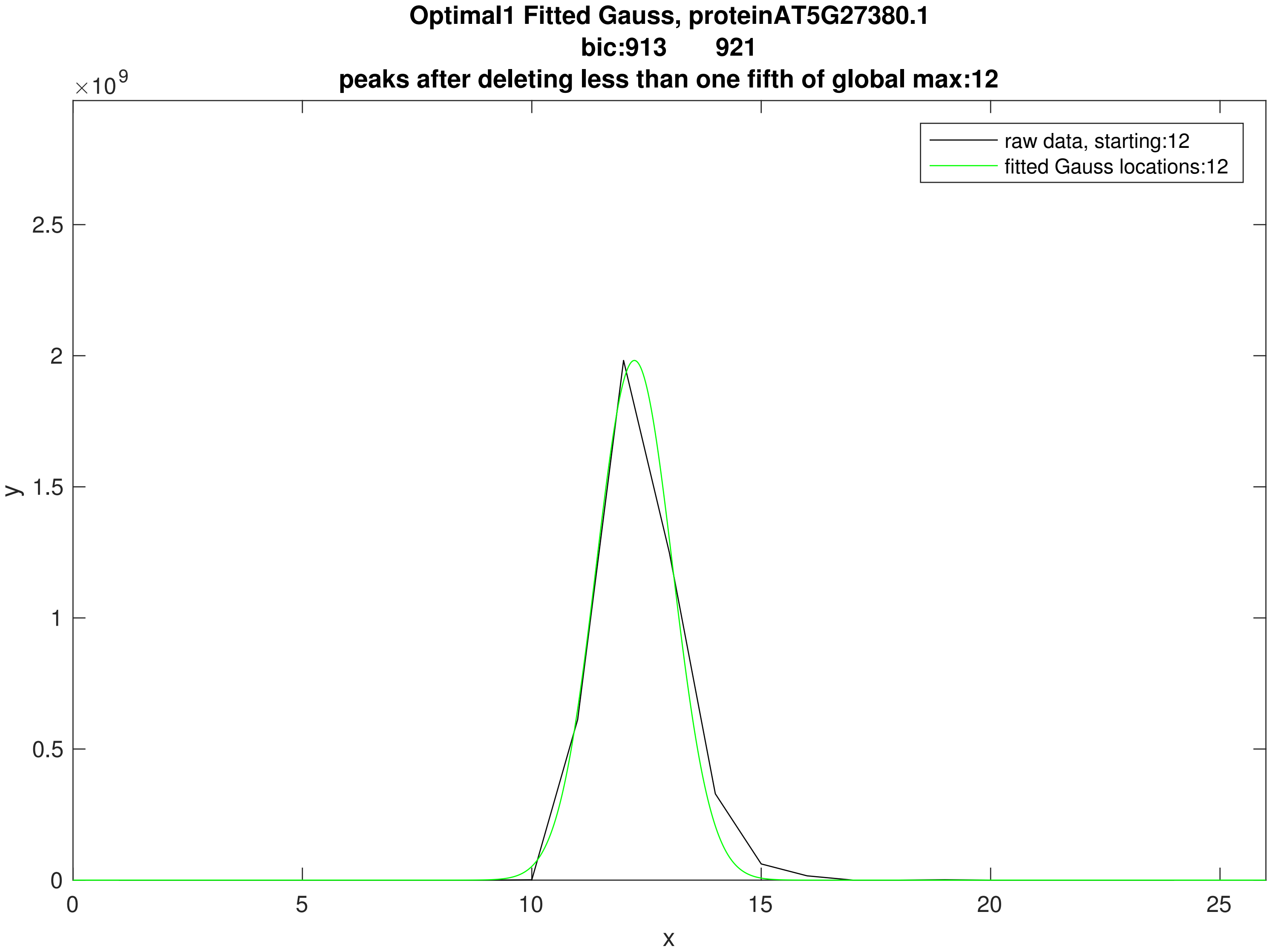
Optimal1 Fitted Gauss, proteinAT5G27380.1
bic:913 921
peaks after deleting less than one fifth of global max:12
×10^9
raw data, starting:12
fitted Gauss locations:12
y
x
0
0.5
1
1.5
2
2.5
0
5
10
15
20
25

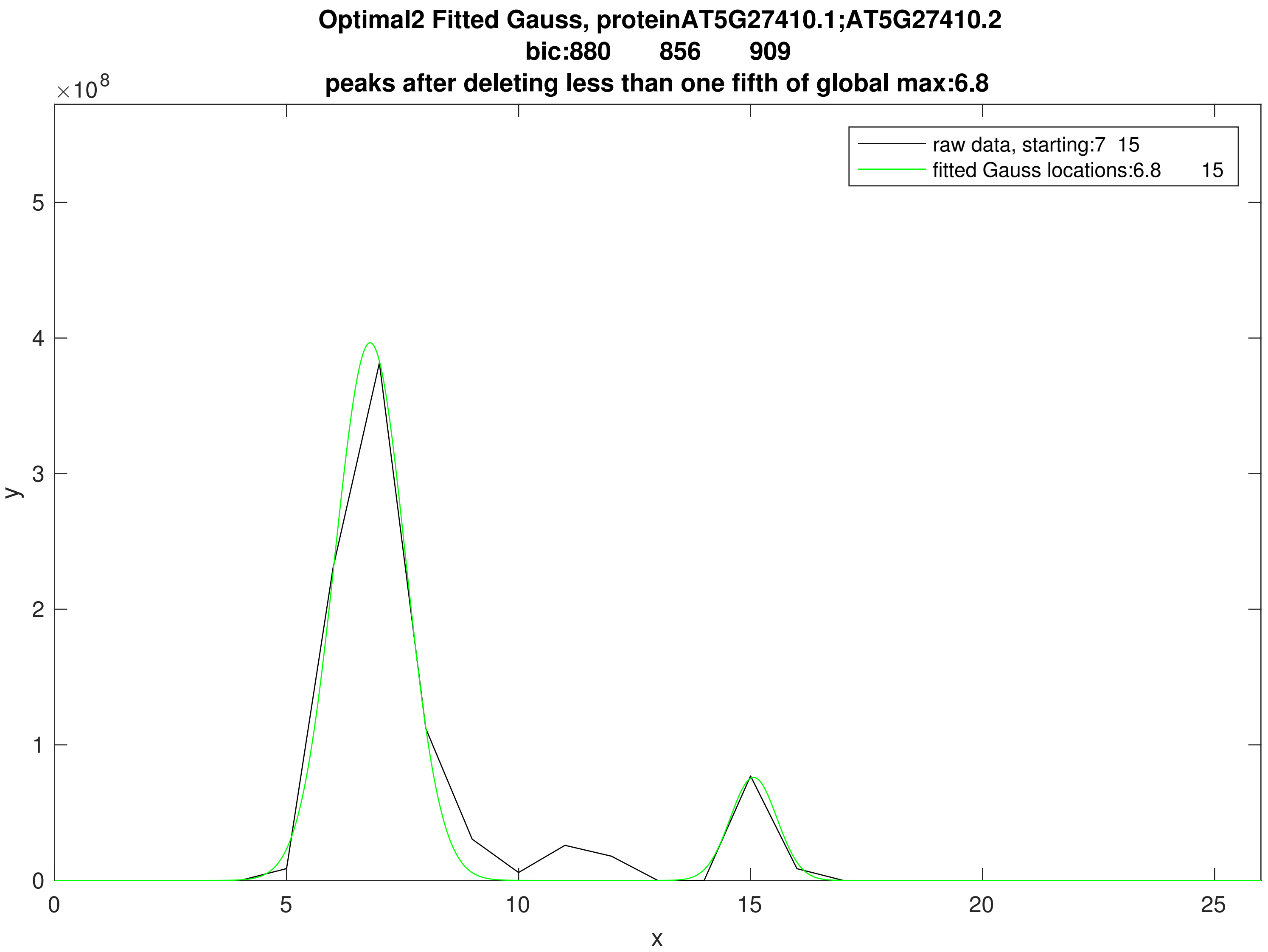

Optimal2 Fitted Gauss, proteinAT5G27410.1;AT5G27410.2
bic:880 856 909
peaks after deleting less than one fifth of global max:6.8
×10^8
raw data, starting:7 15
fitted Gauss locations:6.8 15
y
x
0
1
2
3
4
5
0
5
10
15
20
25

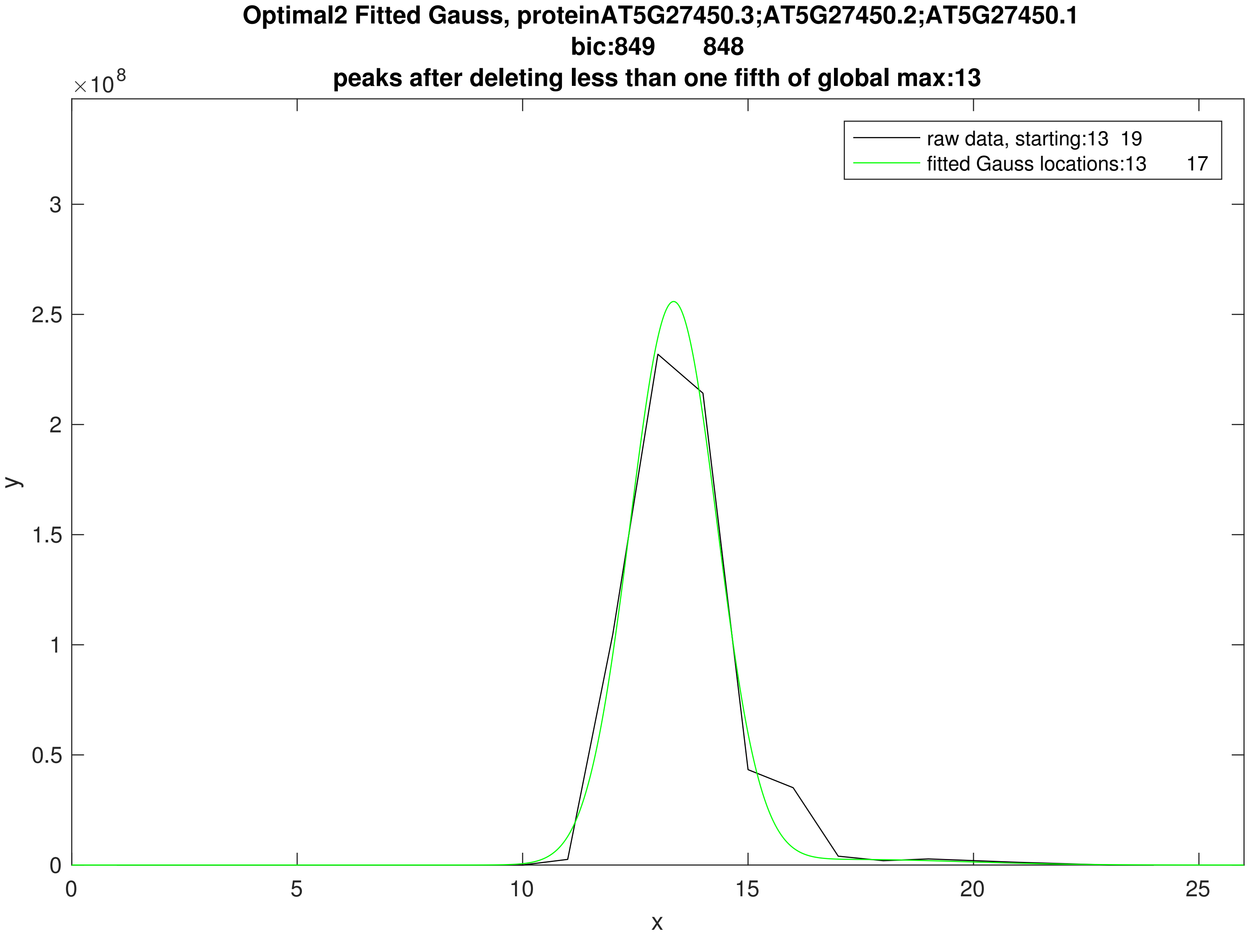

Optimal2 Fitted Gauss, proteinAT5G27450.3;AT5G27450.2;AT5G27450.1
bic:849 848
peaks after deleting less than one fifth of global max:13
×10^8
raw data, starting:13 19
fitted Gauss locations:13 17
y
x
0
0.5
1
1.5
2
2.5
3
5
10
15
20
25

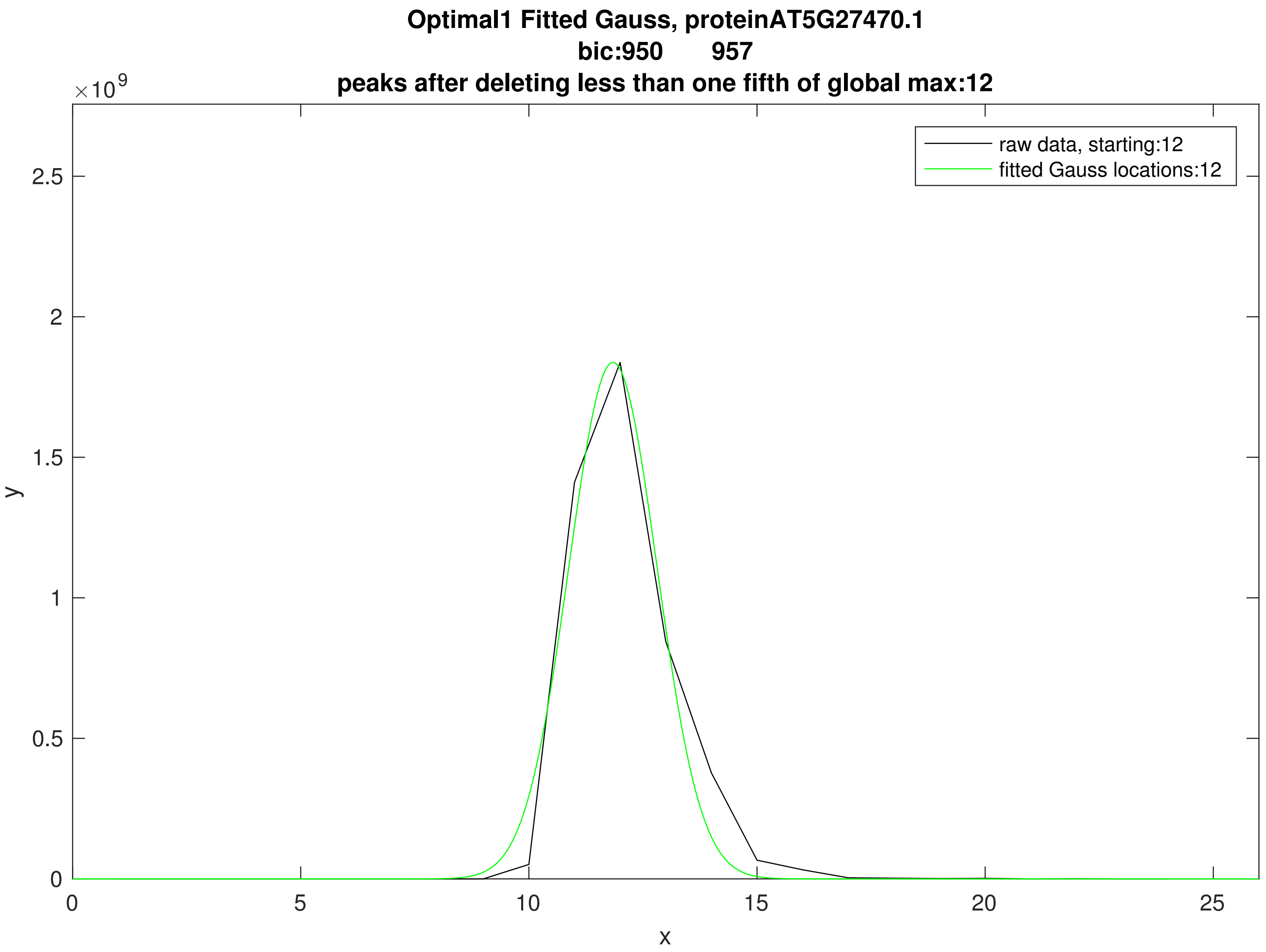
Optimal1 Fitted Gauss, proteinAT5G27470.1
bic:950 957
peaks after deleting less than one fifth of global max:12
×10^9
2.5
2
1.5
1
0.5
0
y
0
5
10
15
20
25
x
raw data, starting:12
fitted Gauss locations:12

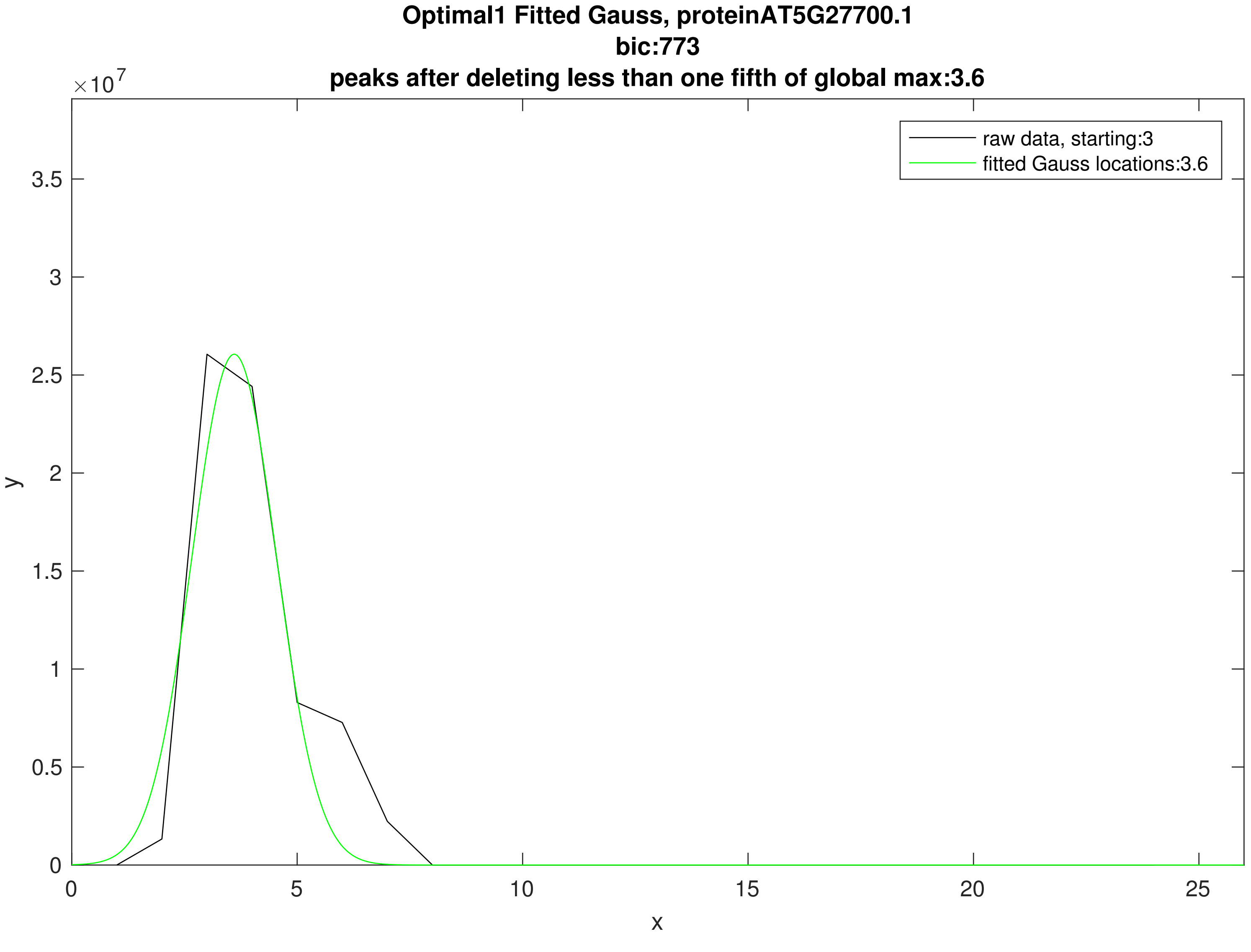
Optimal1 Fitted Gauss, proteinAT5G27700.1
bic:773
peaks after deleting less than one fifth of global max:3.6
×10^7
raw data, starting:3
fitted Gauss locations:3.6
y
x
0
0.5
1
1.5
2
2.5
3
3.5
0
5
10
15
20
25

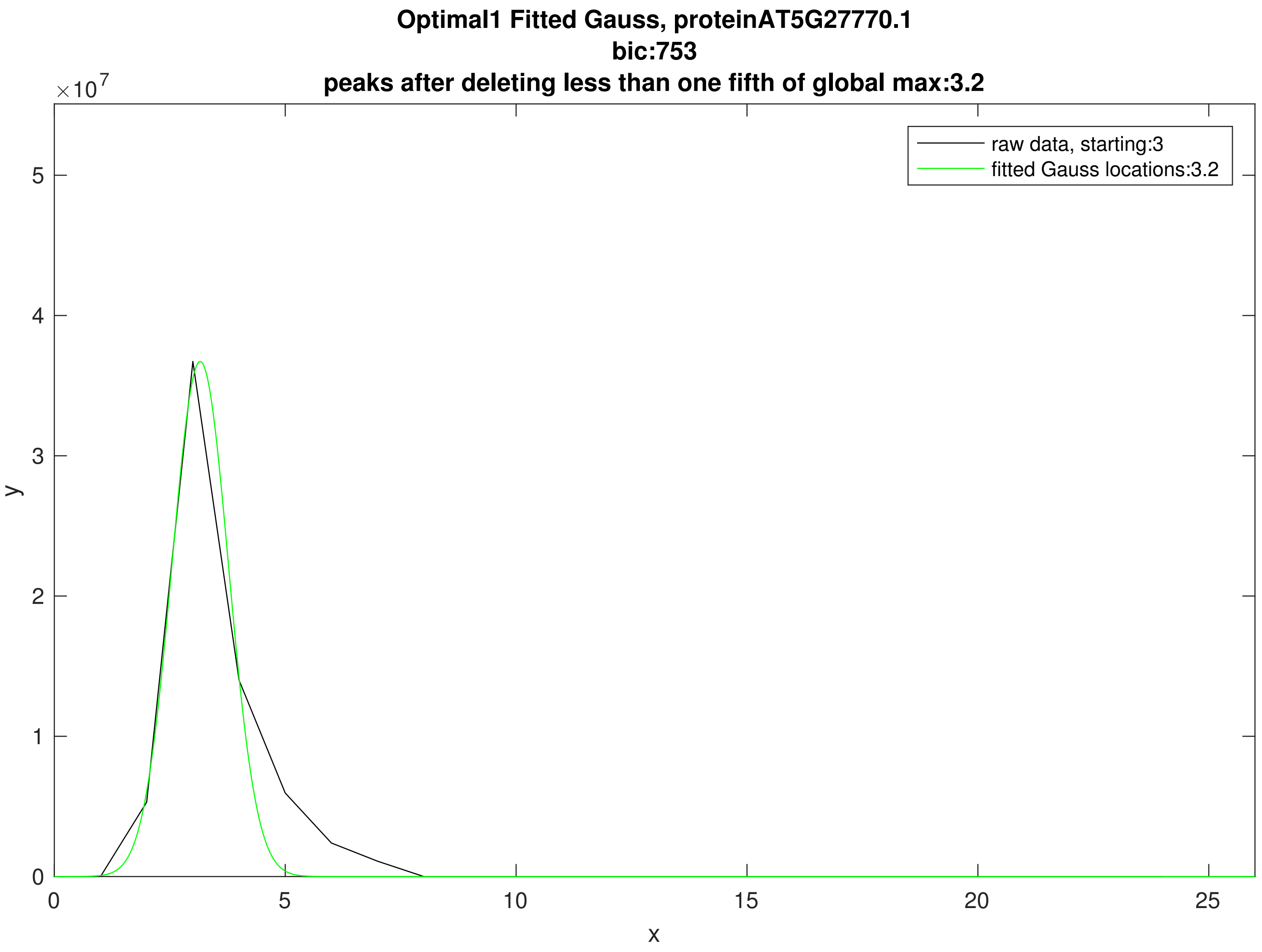
Optimal1 Fitted Gauss, proteinAT5G27770.1
bic:753
peaks after deleting less than one fifth of global max:3.2
×10^7
5
4
3
2
1
0
y
0
5
10
15
20
25
x
raw data, starting:3
fitted Gauss locations:3.2

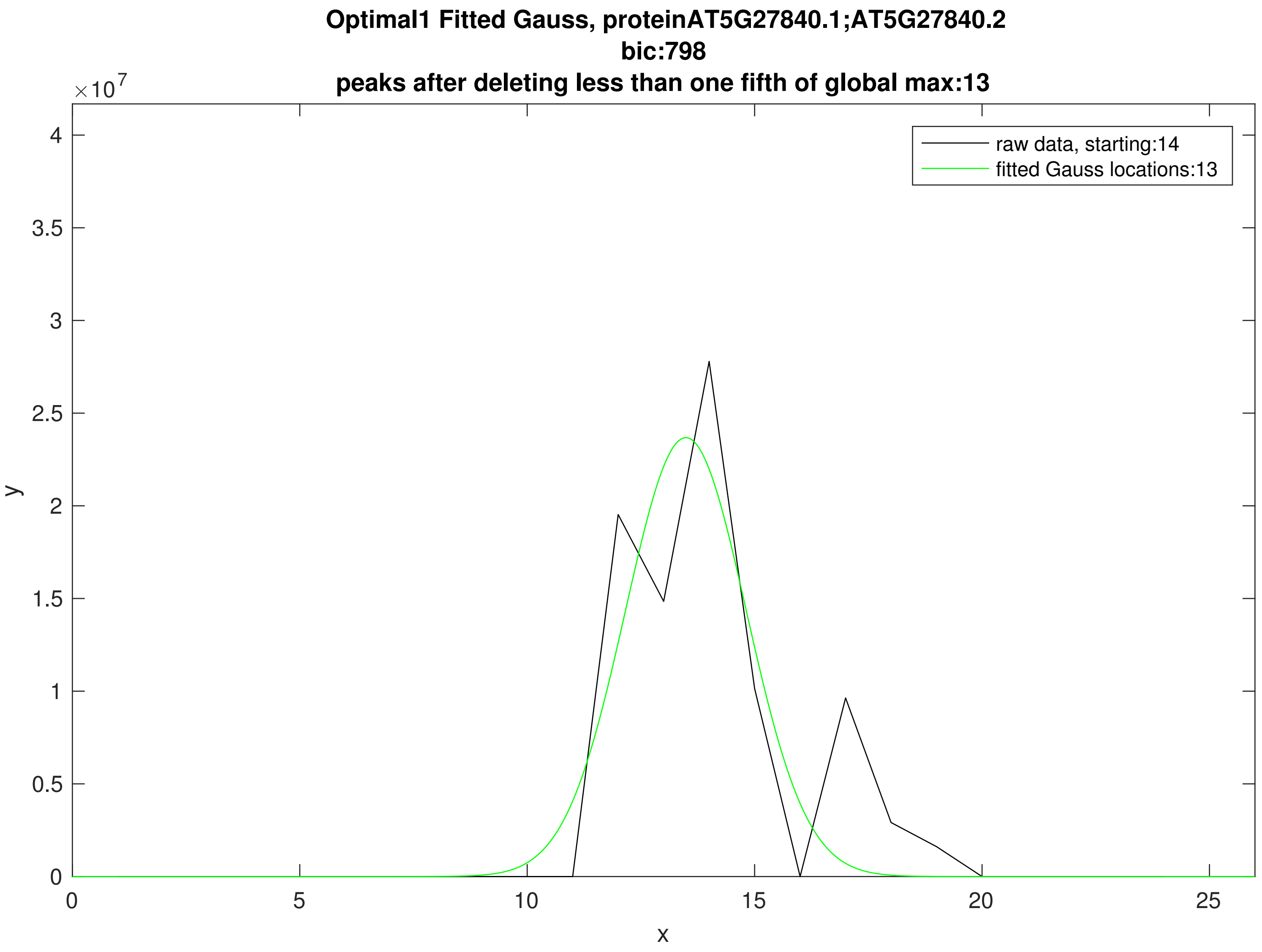
Optimal1 Fitted Gauss, proteinAT5G27840.1;AT5G27840.2
bic:798
peaks after deleting less than one fifth of global max:13
raw data, starting:14
fitted Gauss locations:13
×10^7
y
x
0
5
10
15
20
25
0.5
1
1.5
2
2.5
3
3.5
4

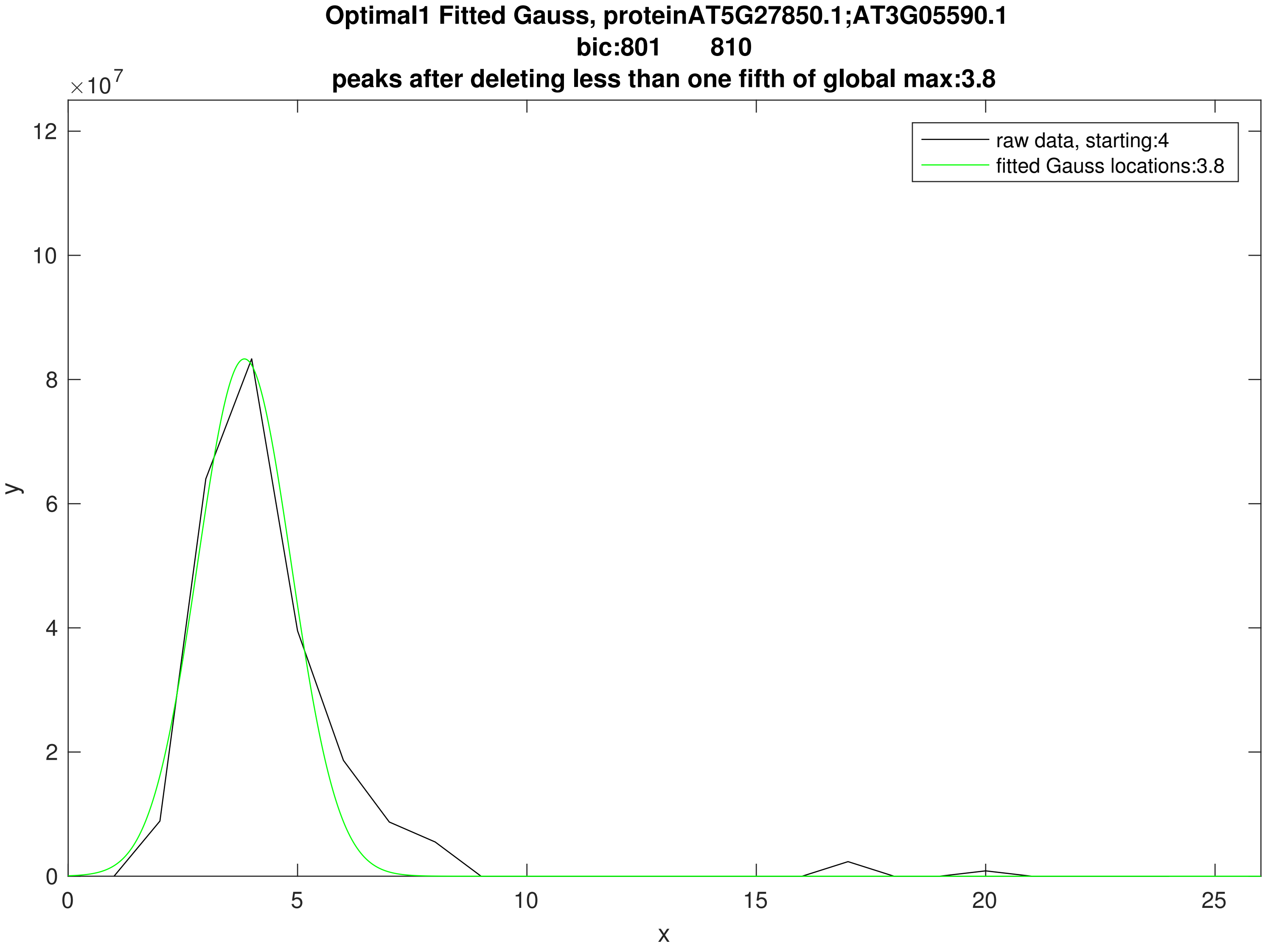

Optimal1 Fitted Gauss, proteinAT5G27850.1;AT3G05590.1
bic:801 810
peaks after deleting less than one fifth of global max:3.8
×10^7
raw data, starting:4
fitted Gauss locations:3.8
y
x
0
2
4
6
8
10
12
5
15
20
25

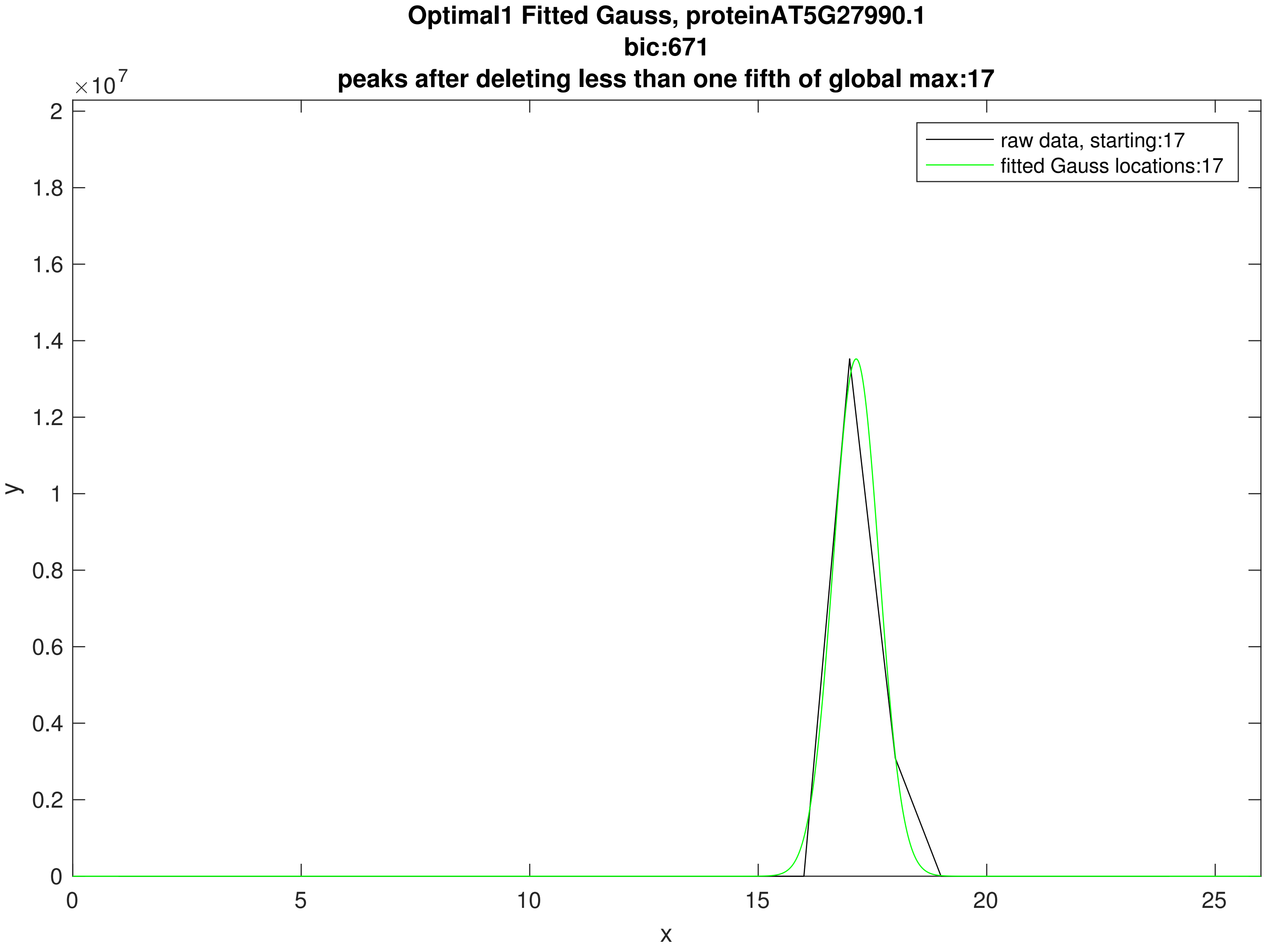
Optimal1 Fitted Gauss, proteinAT5G27990.1
bic:671
peaks after deleting less than one fifth of global max:17
×10^7
2
1.8
1.6
1.4
1.2
1
0.8
0.6
0.4
0.2
0
y
0
5
10
15
20
25
x
raw data, starting:17
fitted Gauss locations:17

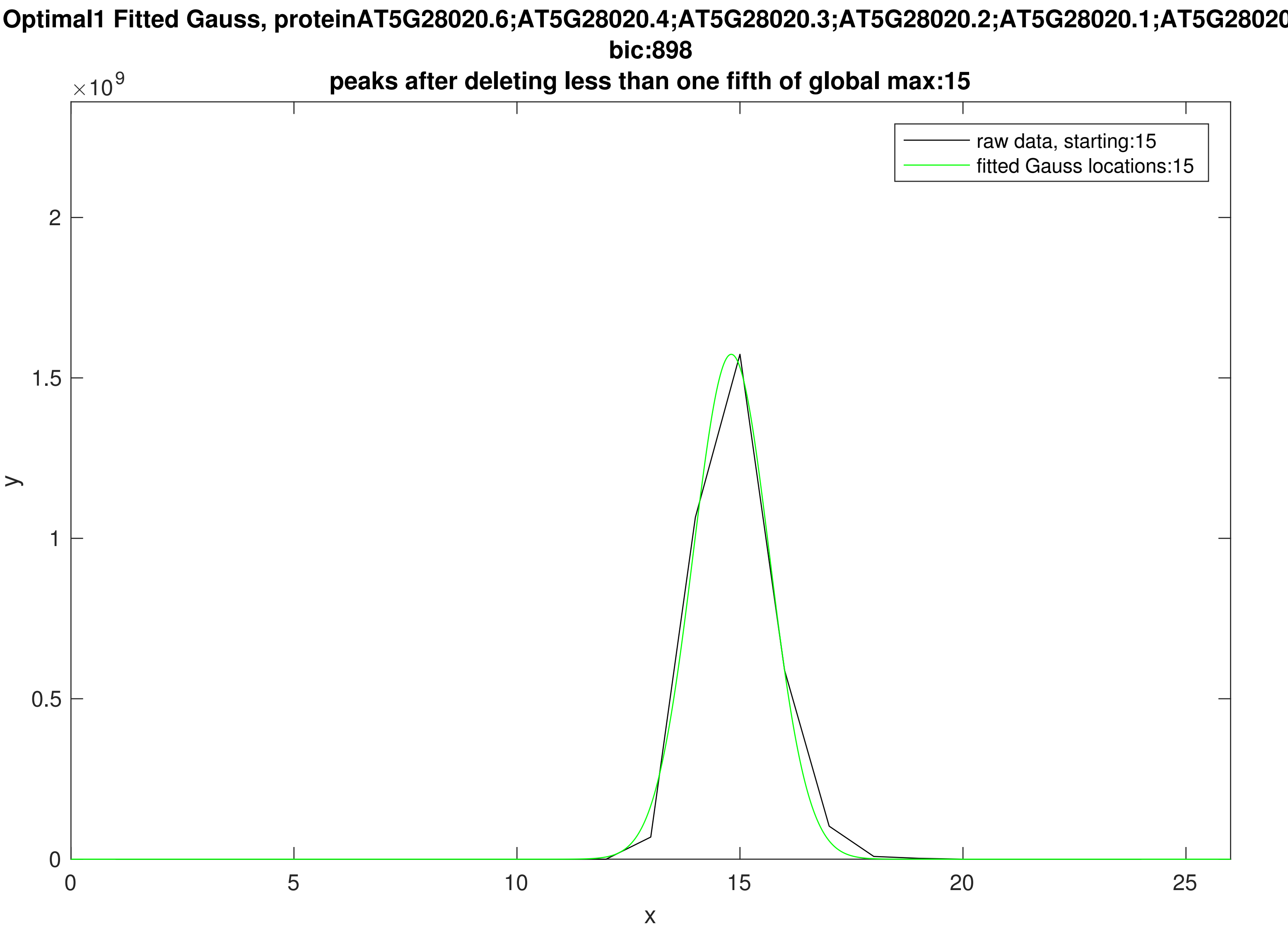
Optimal1 Fitted Gauss, proteinAT5G28020.6;AT5G28020.4;AT5G28020.3;AT5G28020.2;AT5G28020.1;AT5G28020
bic:898
peaks after deleting less than one fifth of global max:15
×10^9
raw data, starting:15
fitted Gauss locations:15
y
x
0
0.5
1
1.5
2
0
5
10
15
20
25

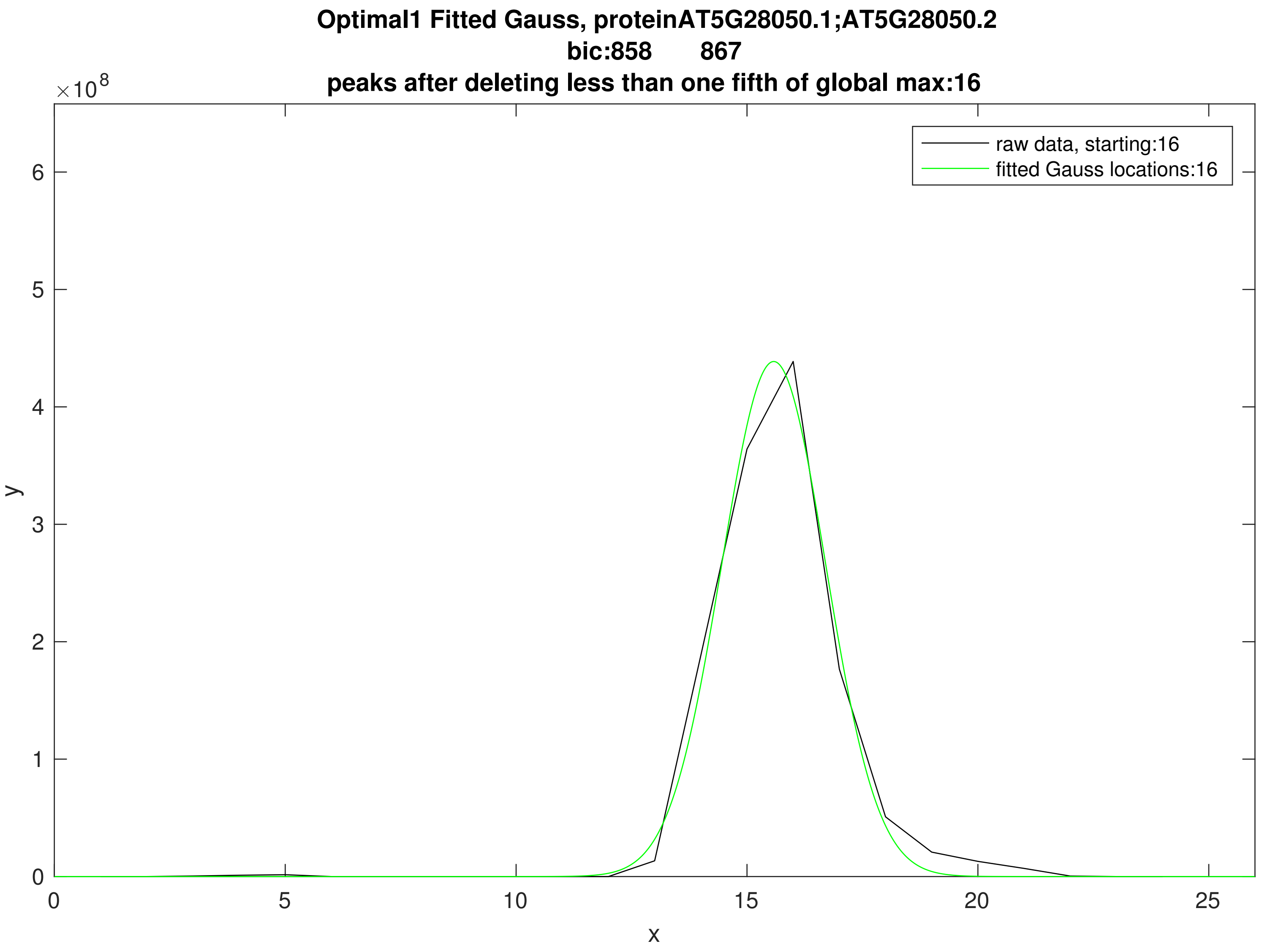

Optimal1 Fitted Gauss, proteinAT5G28050.1;AT5G28050.2
bic:858 867
peaks after deleting less than one fifth of global max:16
×10^8
raw data, starting:16
fitted Gauss locations:16
y
x
0
1
2
3
4
5
6
0
5
10
15
20
25

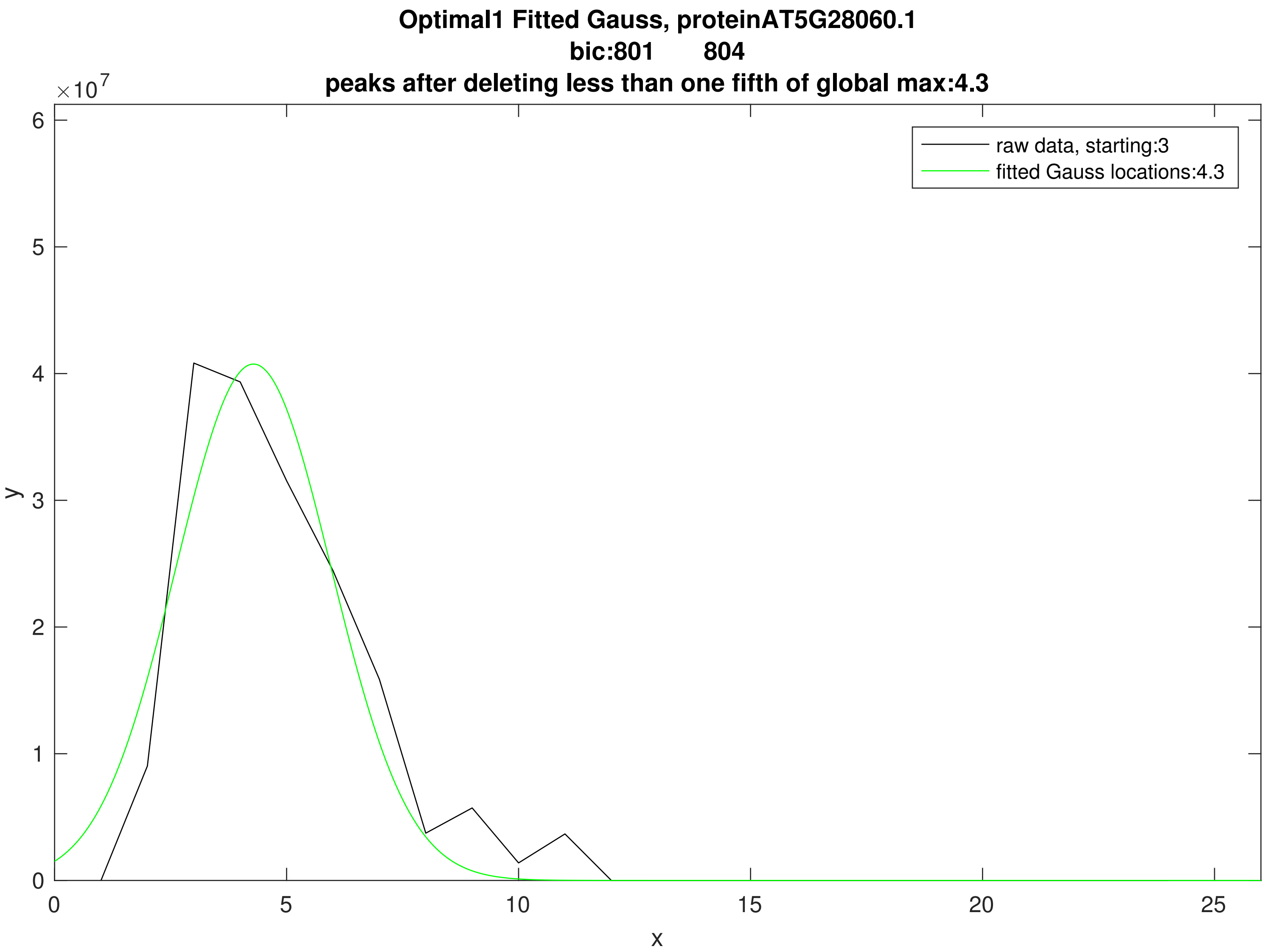

Optimal1 Fitted Gauss, proteinAT5G28060.1
bic:801 804
peaks after deleting less than one fifth of global max:4.3
×10^7
raw data, starting:3
fitted Gauss locations:4.3
y
x

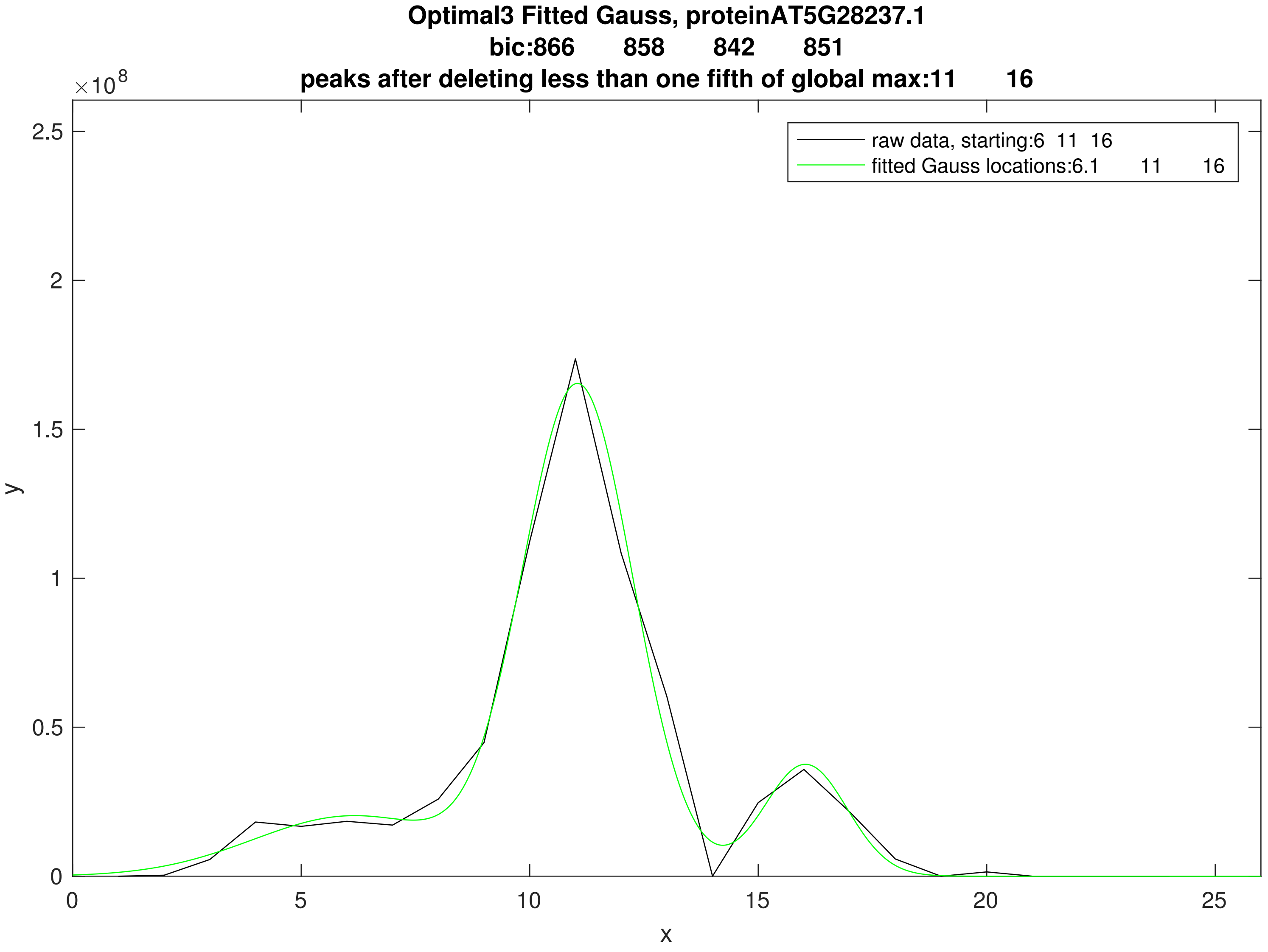
Optimal3 Fitted Gauss, proteinAT5G28237.1
bic:866 858 842 851
peaks after deleting less than one fifth of global max:11 16
×10^8
2.5
2
1.5
1
0.5
0
y
0
5
10
15
20
25
x
raw data, starting:6 11 16
fitted Gauss locations:6.1 11 16

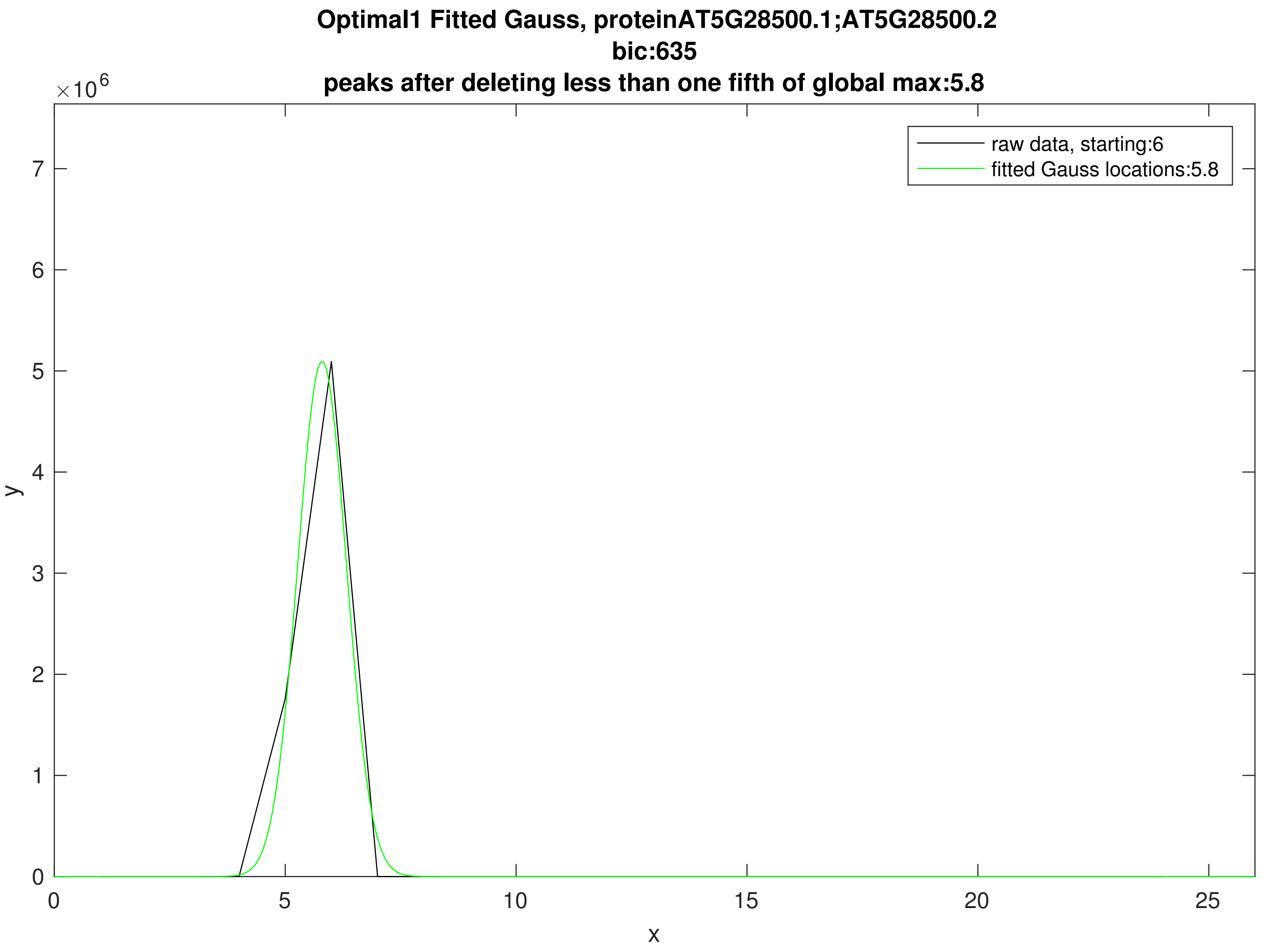

Optimal1 Fitted Gauss, proteinAT5G28500.1;AT5G28500.2
bic:635
peaks after deleting less than one fifth of global max:5.8
×10^6
raw data, starting:6
fitted Gauss locations:5.8
y
x
0
1
2
3
4
5
6
7
0
5
10
15
20
25

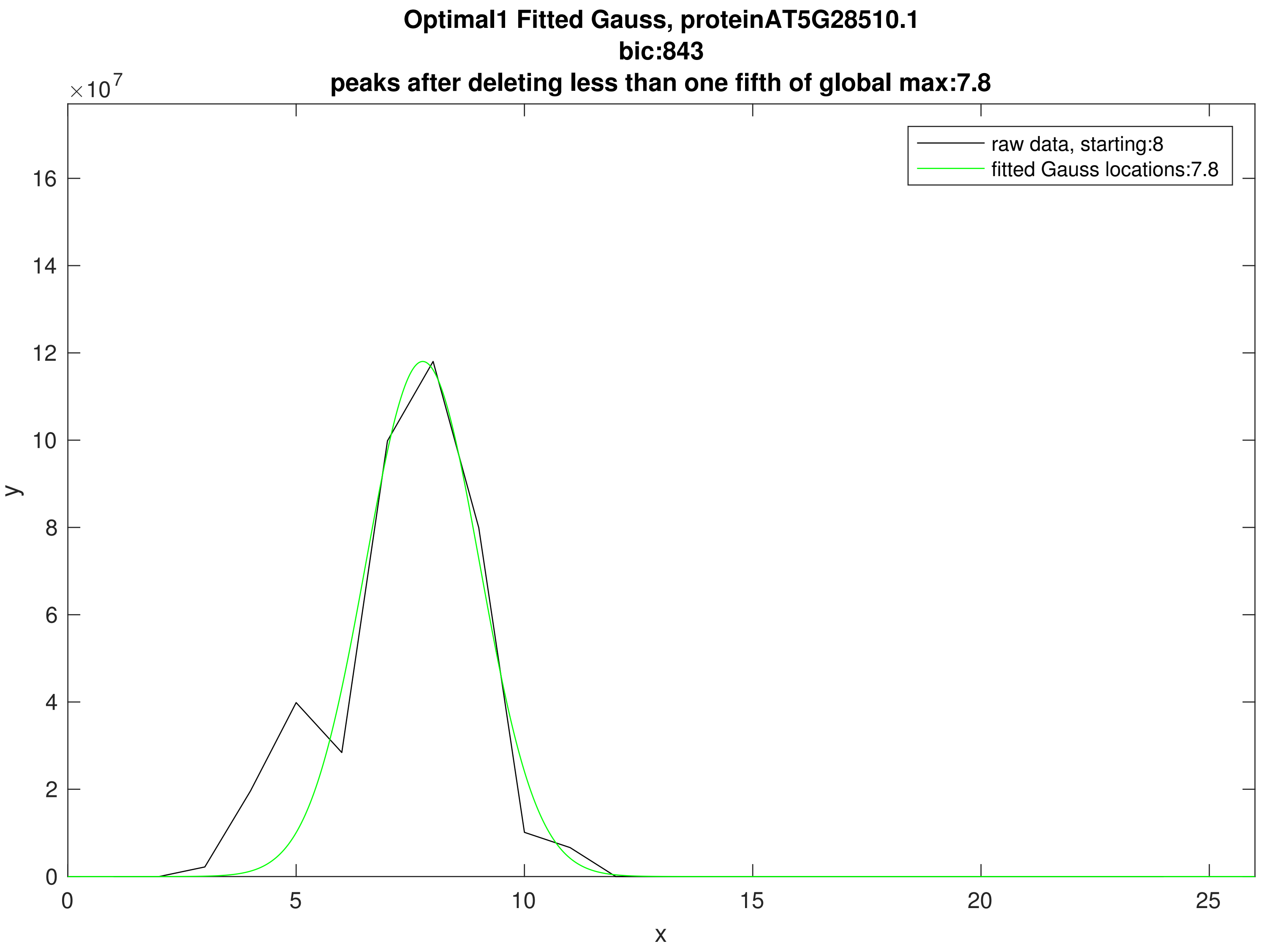
Optimal1 Fitted Gauss, proteinAT5G28510.1
bic:843
peaks after deleting less than one fifth of global max:7.8
×10^7
raw data, starting:8
fitted Gauss locations:7.8
y
x
0
2
4
6
8
10
12
14
16
0
5
10
15
20
25

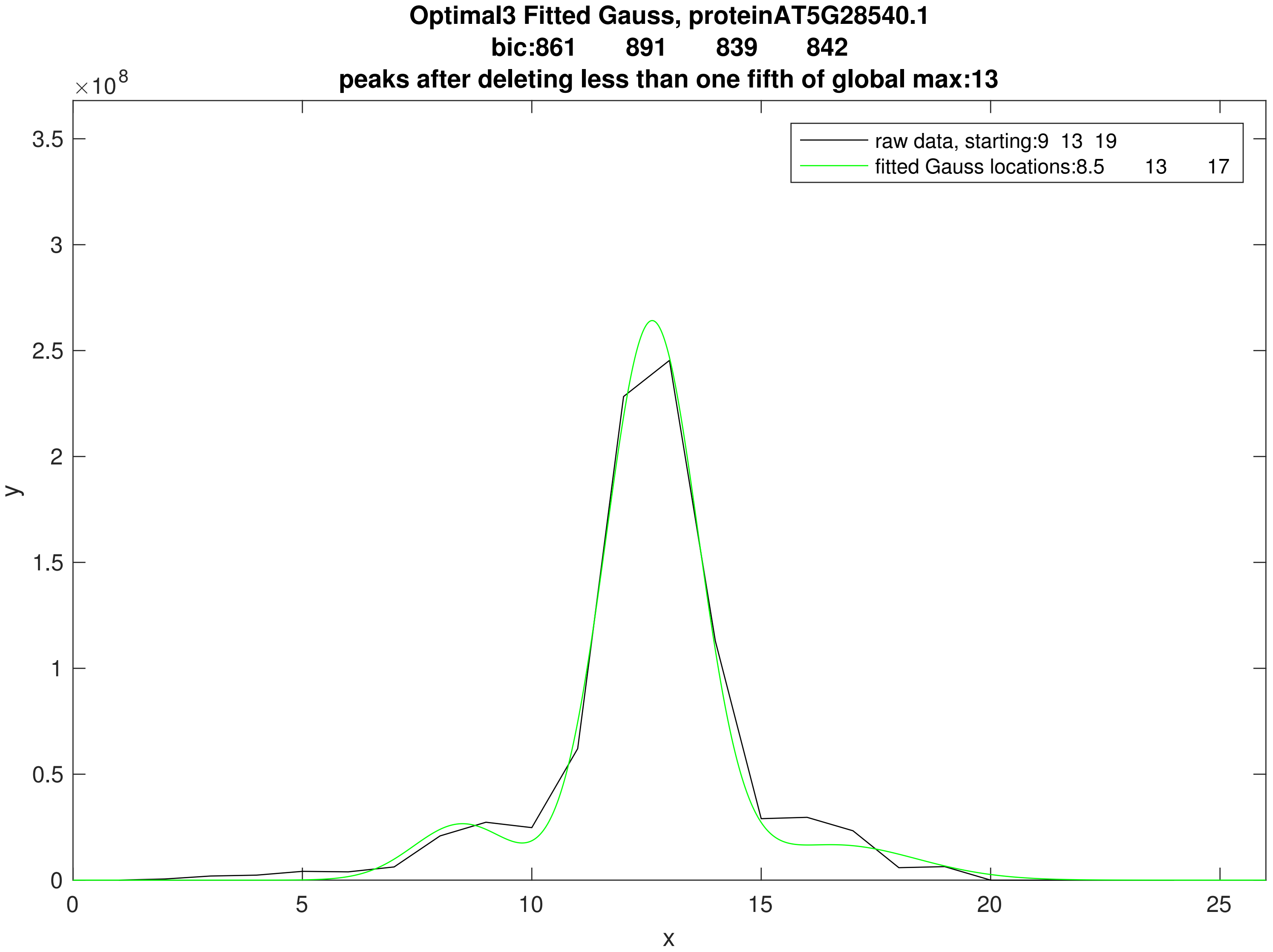

Optimal3 Fitted Gauss, proteinAT5G28540.1
bic:861 891 839 842
peaks after deleting less than one fifth of global max:13
×10^8
raw data, starting:9 13 19
fitted Gauss locations:8.5 13 17
y
x
0
0.5
1
1.5
2
2.5
3
3.5
5
10
15
20
25

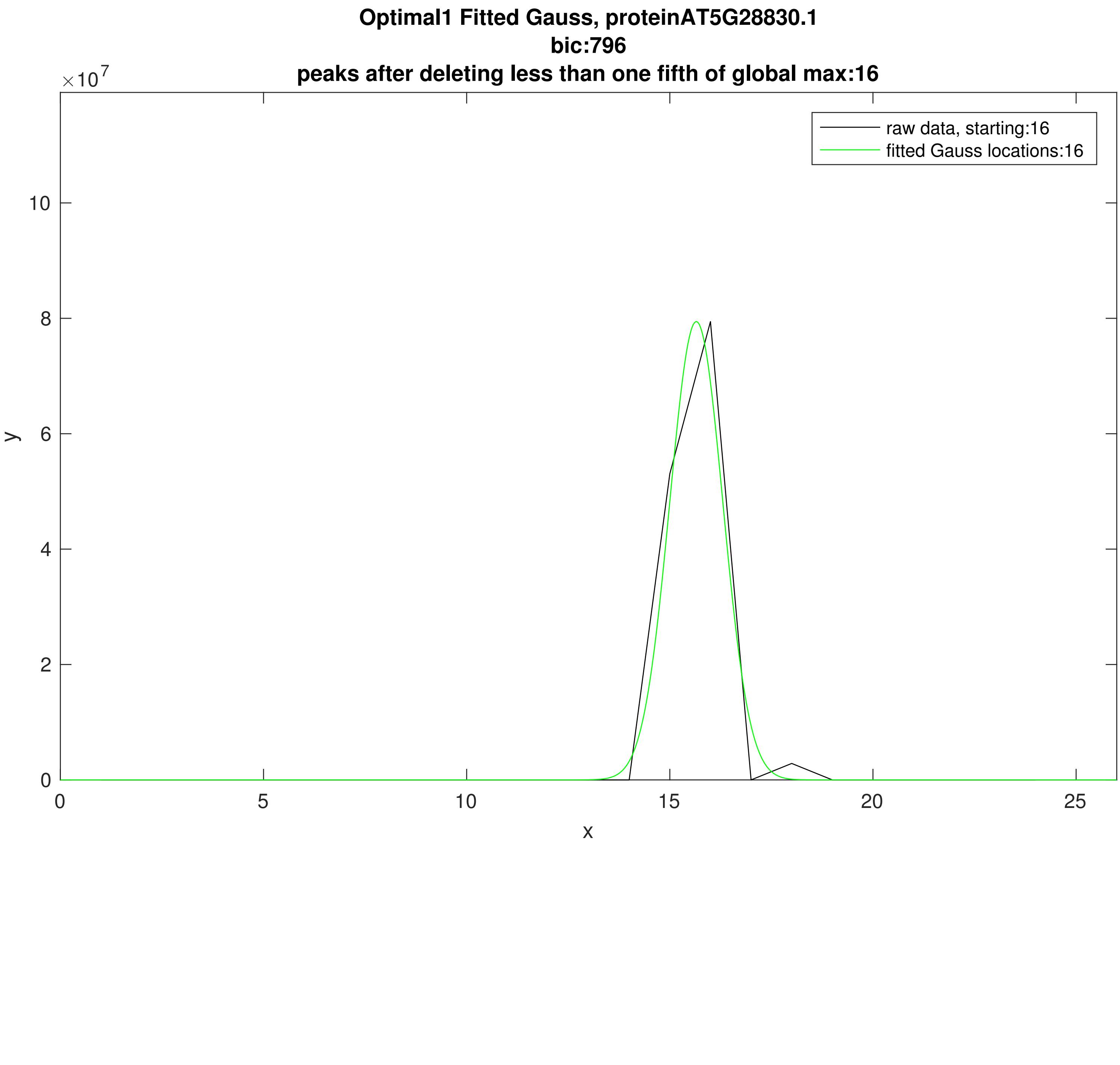
Optimal1 Fitted Gauss, proteinAT5G28830.1
bic:796
peaks after deleting less than one fifth of global max:16
×10^7
raw data, starting:16
fitted Gauss locations:16
10
8
6
4
2
0
y
0
5
10
15
20
25
x

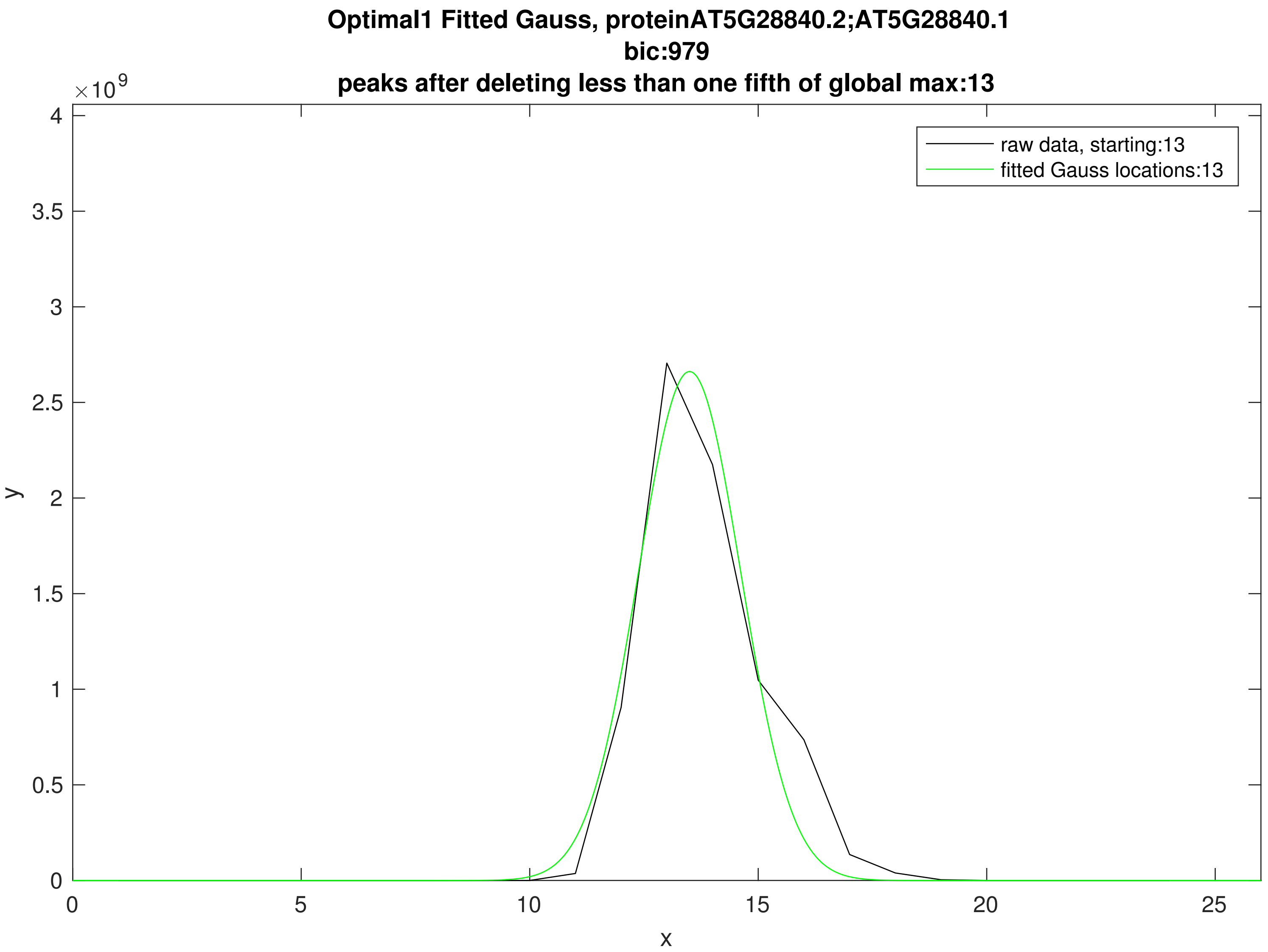
Optimal1 Fitted Gauss, proteinAT5G28840.2;AT5G28840.1
bic:979
peaks after deleting less than one fifth of global max:13
×10^9
raw data, starting:13
fitted Gauss locations:13
y
x
0
0.5
1
1.5
2
2.5
3
3.5
4
0
5
10
15
20
25

#### Optimal1 Fitted Gauss, proteinAT5G30510.1
#### bic:685
#### peaks after deleting less than one fifth of global max:15

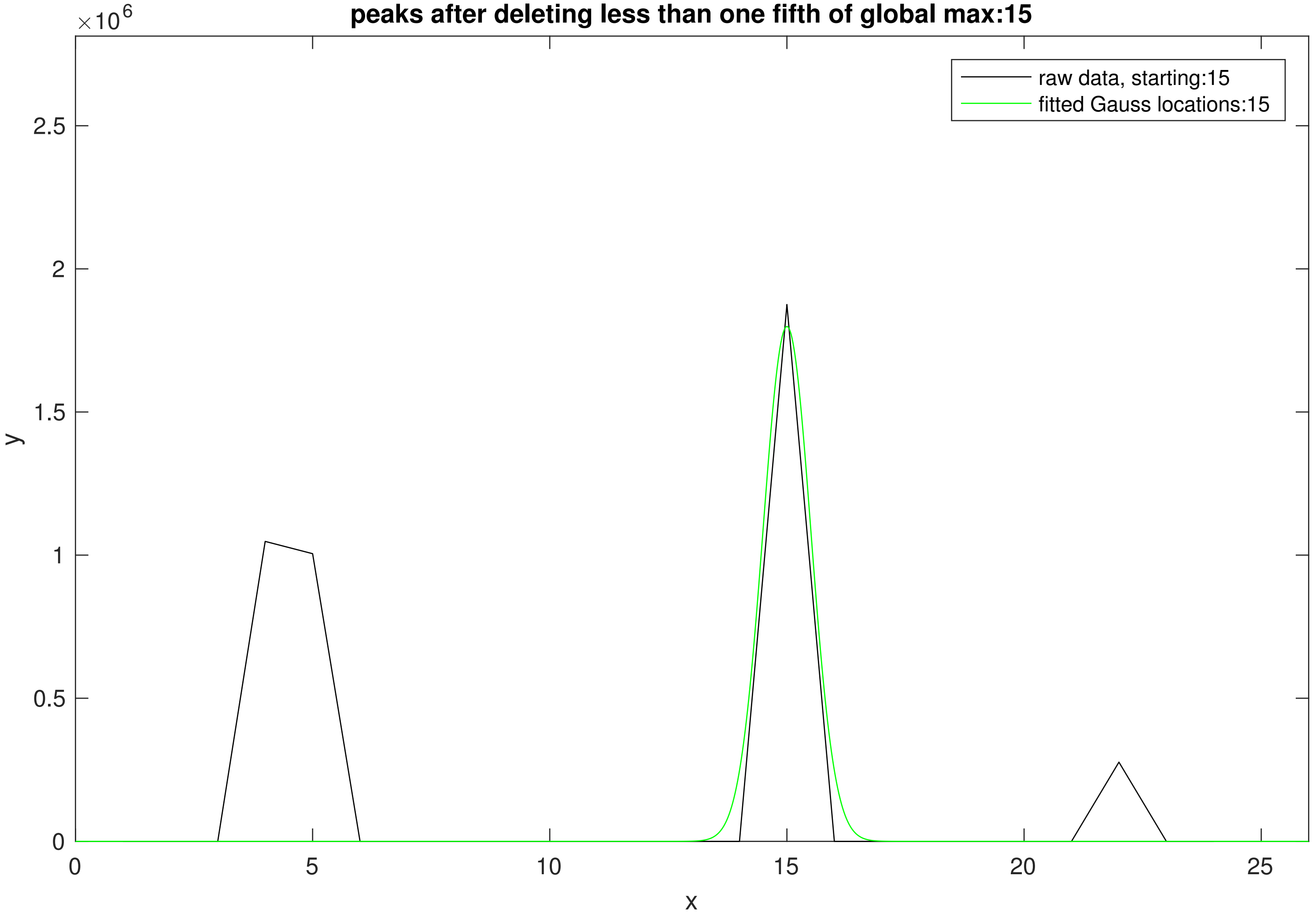

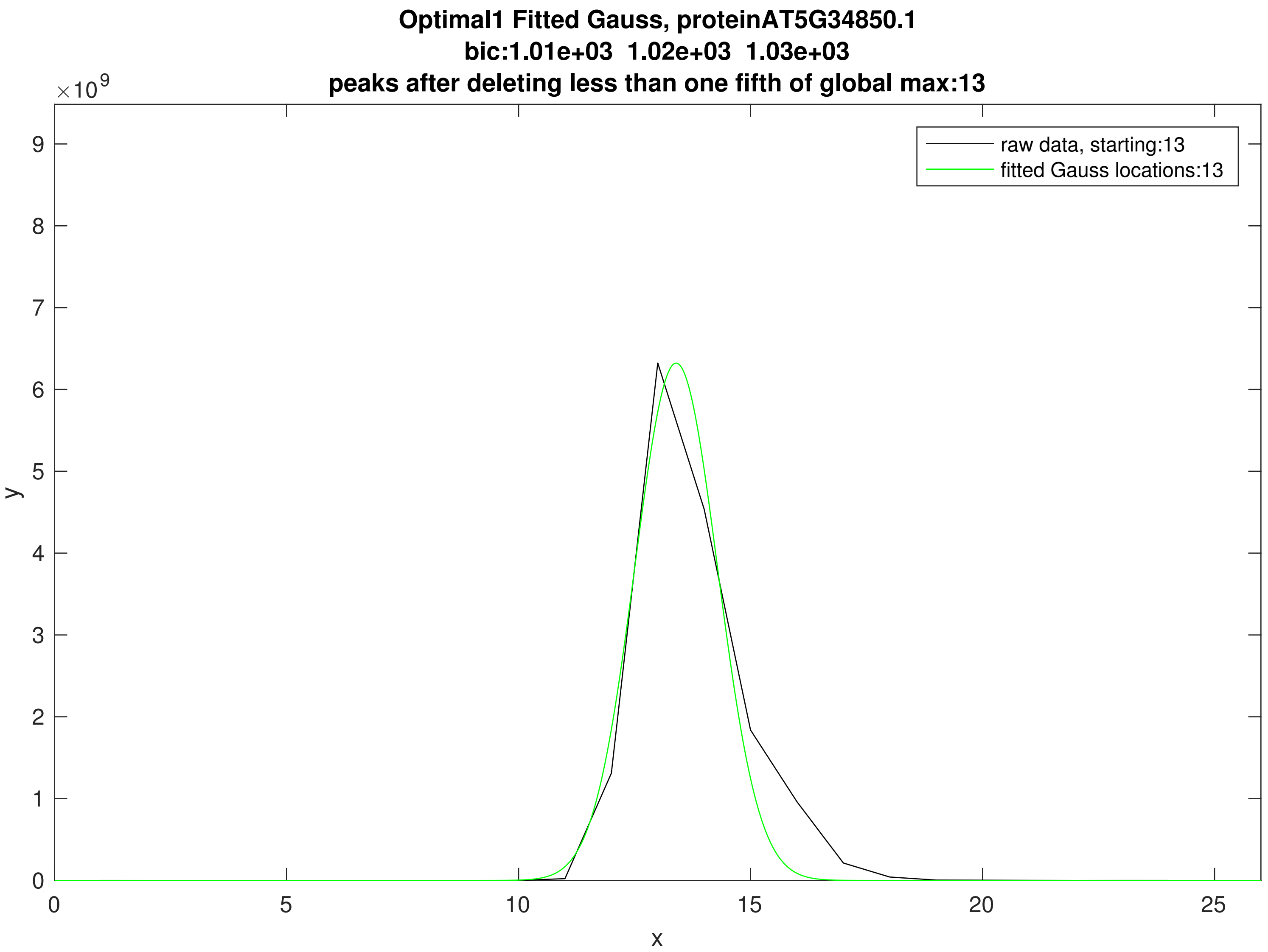

Optimal1 Fitted Gauss, proteinAT5G34850.1
bic:1.01e+03 1.02e+03 1.03e+03
peaks after deleting less than one fifth of global max:13
×10^9
raw data, starting:13
fitted Gauss locations:13
y
x
0
1
2
3
4
5
6
7
8
9
0
5
10
15
20
25

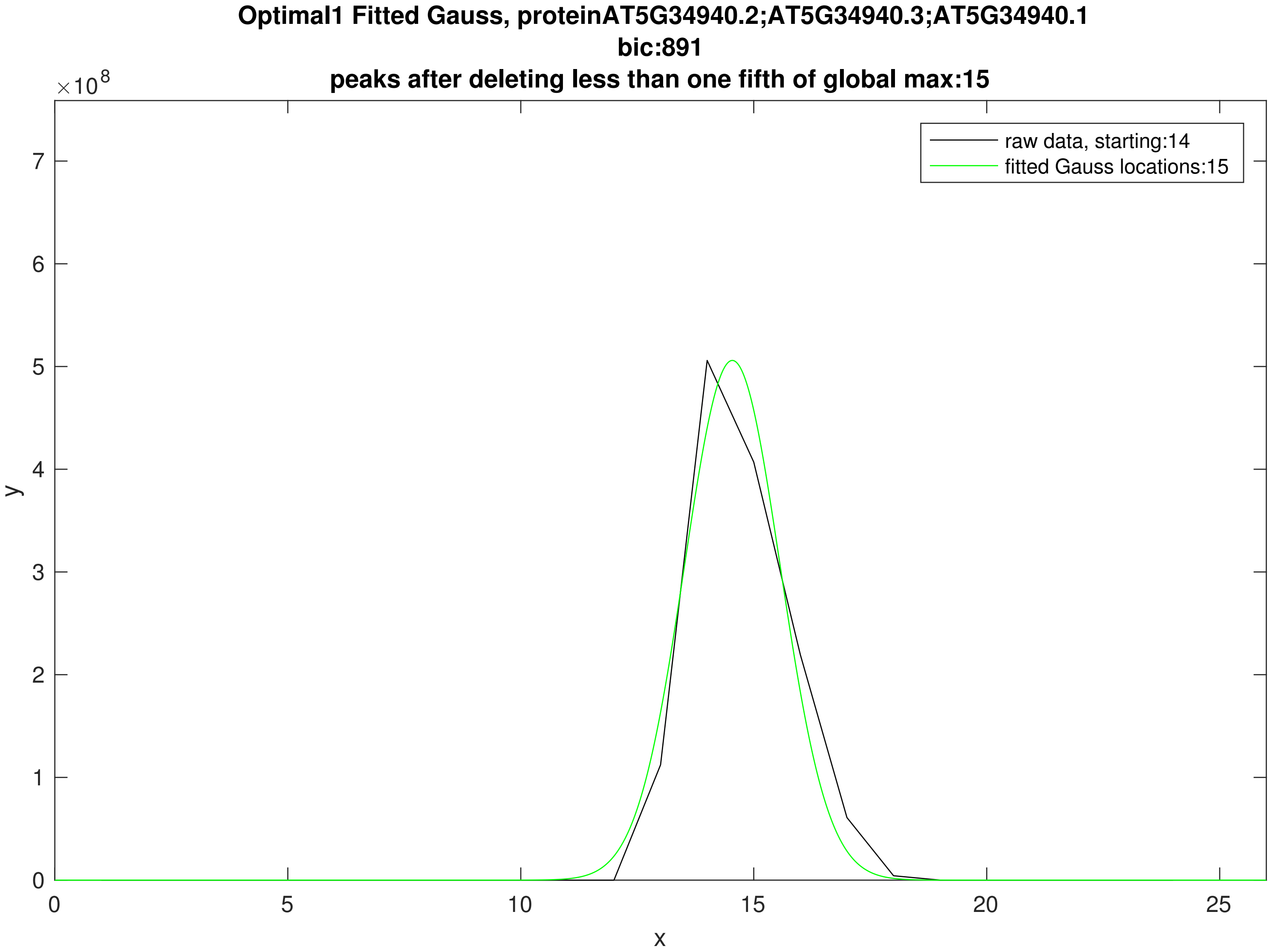

Optimal1 Fitted Gauss, proteinAT5G34940.2;AT5G34940.3;AT5G34940.1
bic:891
peaks after deleting less than one fifth of global max:15
×10^8
raw data, starting:14
fitted Gauss locations:15
y
x
0
1
2
3
4
5
6
7
0
5
10
15
20
25

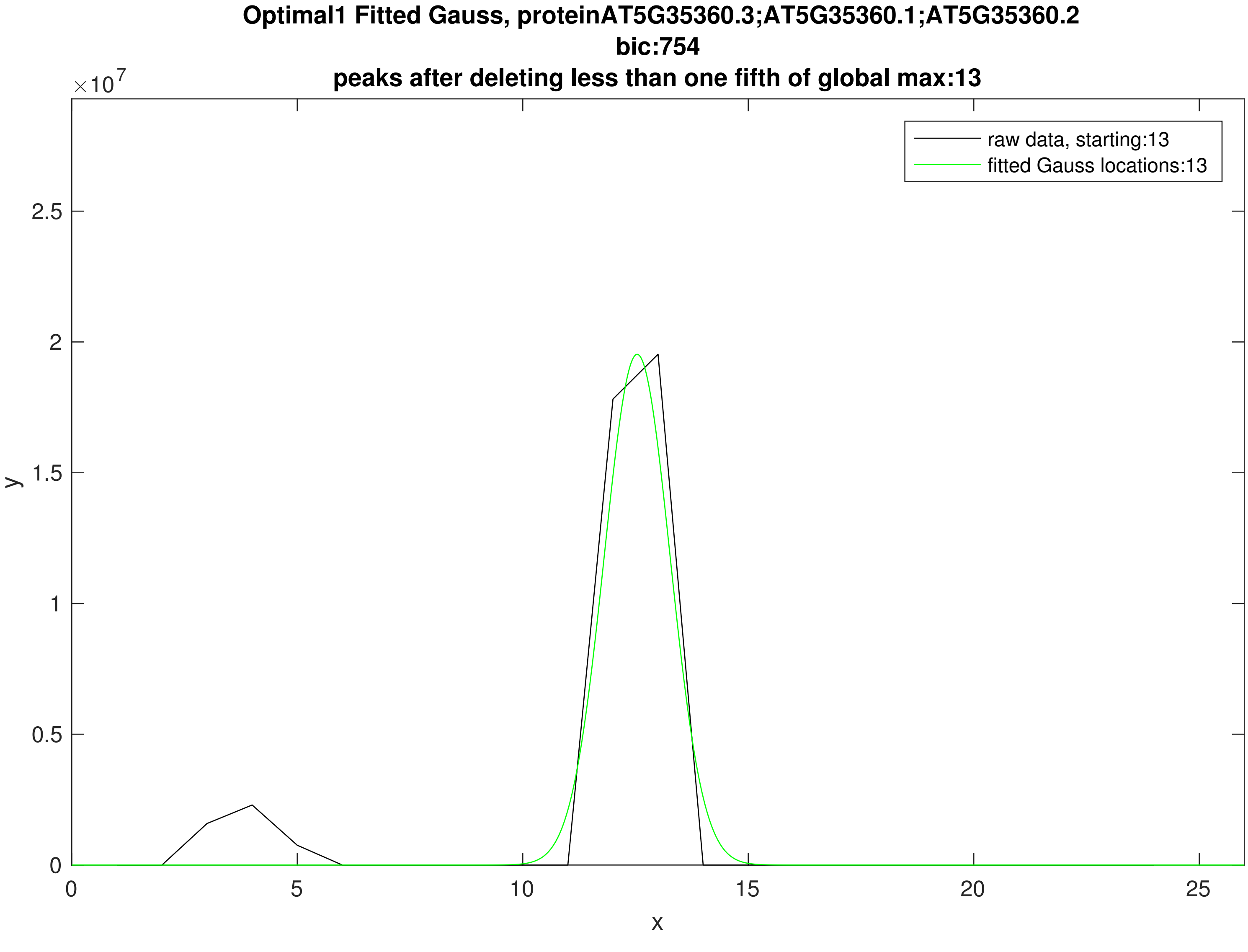
Optimal1 Fitted Gauss, proteinAT5G35360.3;AT5G35360.1;AT5G35360.2
bic:754
peaks after deleting less than one fifth of global max:13
×10^7
raw data, starting:13
fitted Gauss locations:13
y
2.5
2
1.5
1
0.5
0
0
5
10
15
20
25
x

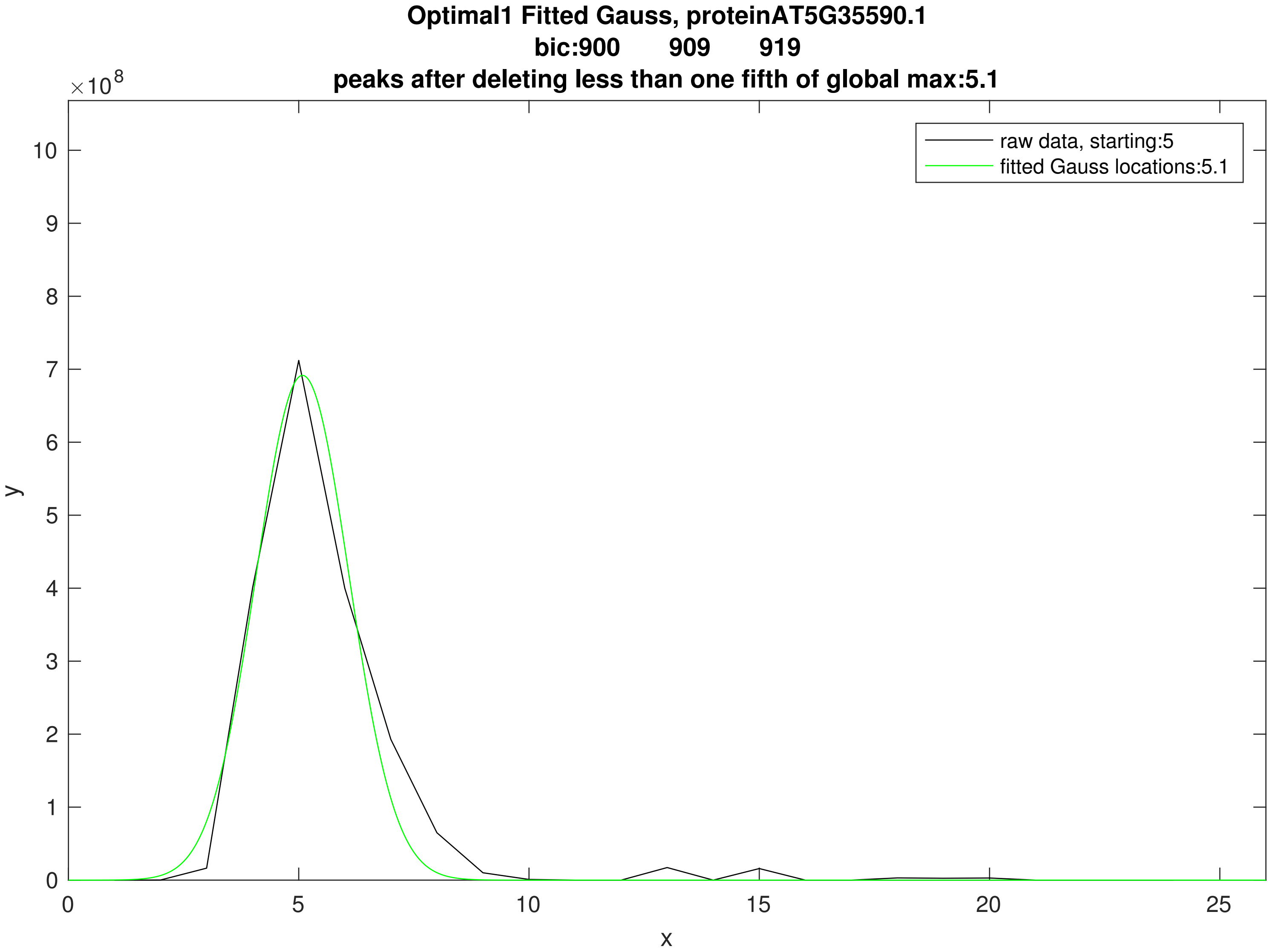

Optimal1 Fitted Gauss, proteinAT5G35590.1
bic:900 909 919
peaks after deleting less than one fifth of global max:5.1
×10^8
raw data, starting:5
fitted Gauss locations:5.1
y
x
0
1
2
3
4
5
6
7
8
9
10
0
5
10
15
20
25

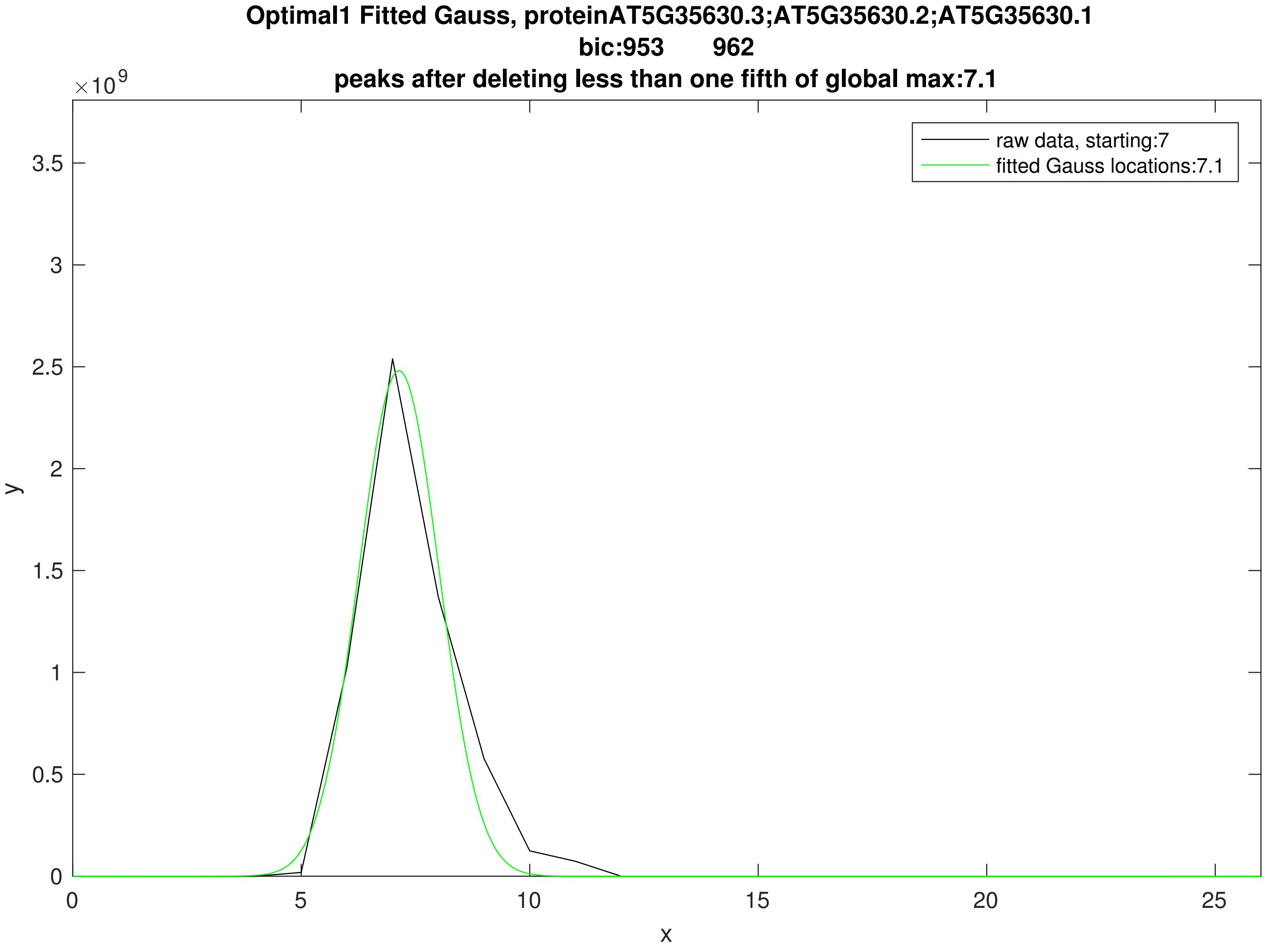

Optimal1 Fitted Gauss, proteinAT5G35630.3;AT5G35630.2;AT5G35630.1
bic:953 962
peaks after deleting less than one fifth of global max:7.1
×10^9
raw data, starting:7
fitted Gauss locations:7.1
3.5
3
2.5
2
1.5
1
0.5
0
y
0
5
10
15
20
25
x

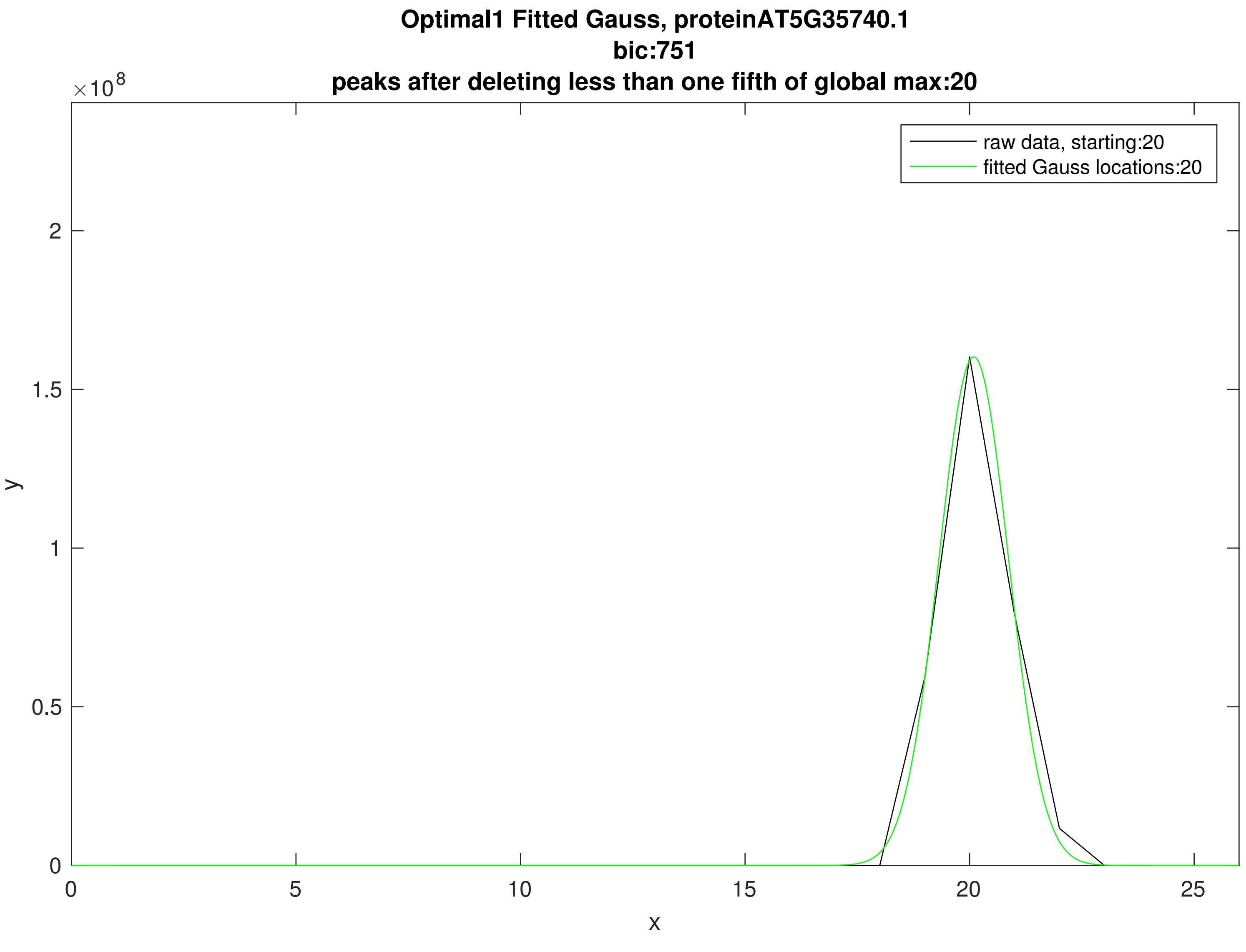
Optimal1 Fitted Gauss, proteinAT5G35740.1
bic:751
peaks after deleting less than one fifth of global max:20
×10^8
raw data, starting:20
fitted Gauss locations:20
y
x
0
0.5
1
1.5
2
5
10
15
20
25

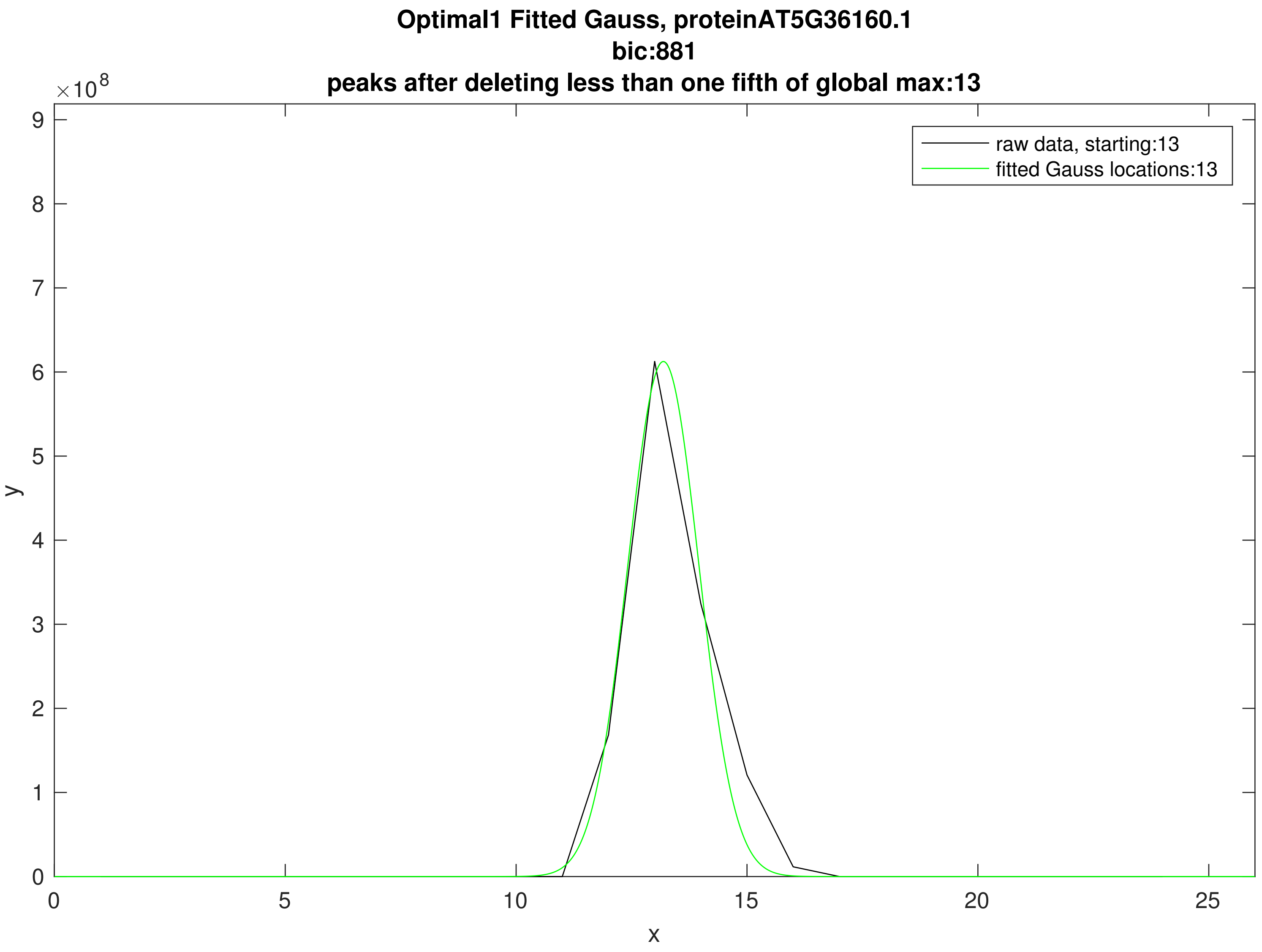
Optimal1 Fitted Gauss, proteinAT5G36160.1
bic:881
peaks after deleting less than one fifth of global max:13
×10^8
raw data, starting:13
fitted Gauss locations:13
y
x
0
1
2
3
4
5
6
7
8
9
0
5
10
15
20
25

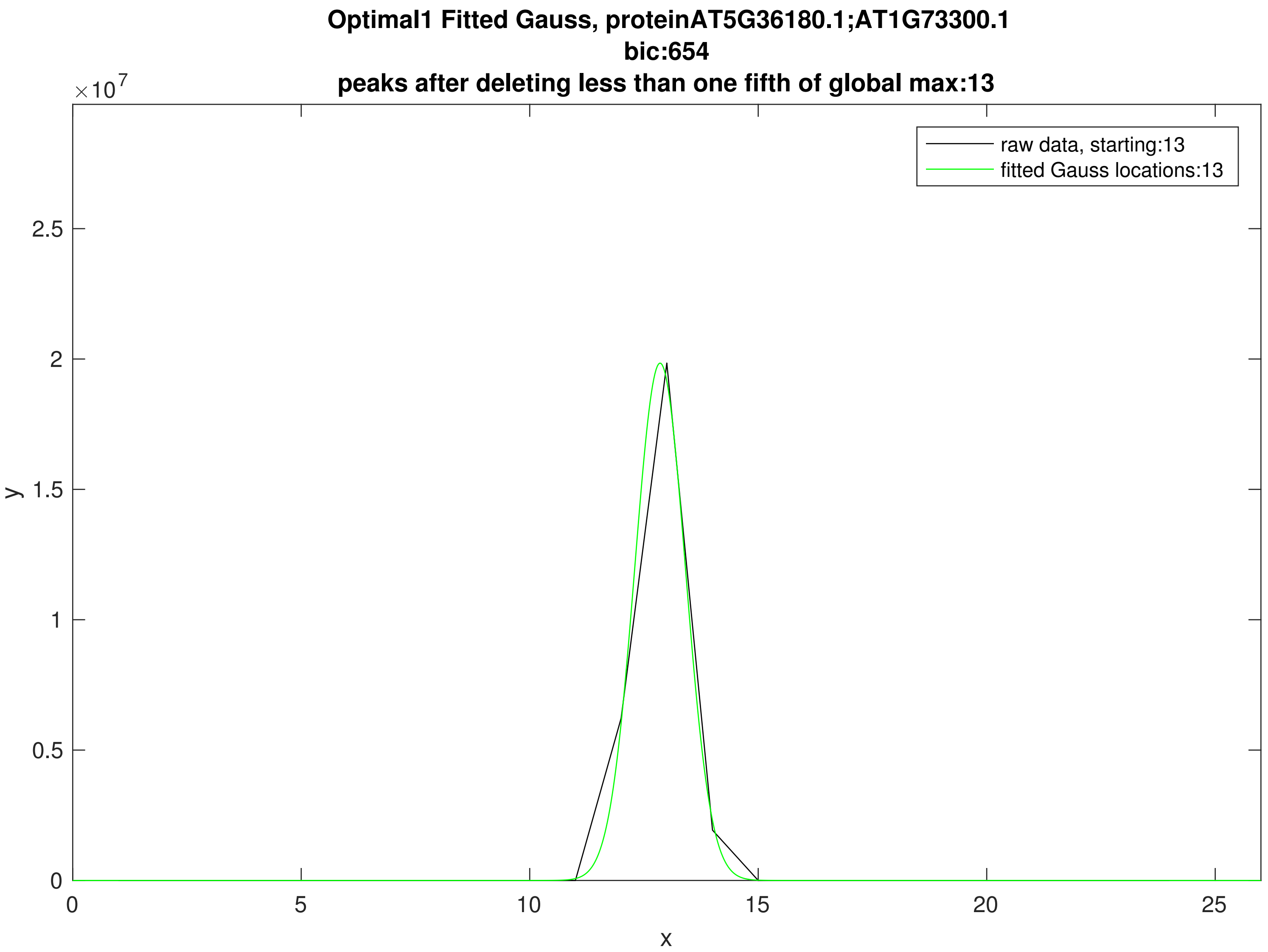

Optimal1 Fitted Gauss, proteinAT5G36180.1;AT1G73300.1
bic:654
peaks after deleting less than one fifth of global max:13
×10^7
raw data, starting:13
fitted Gauss locations:13
2.5
2
1.5
1
0.5
0
y
0
5
10
15
20
25
x

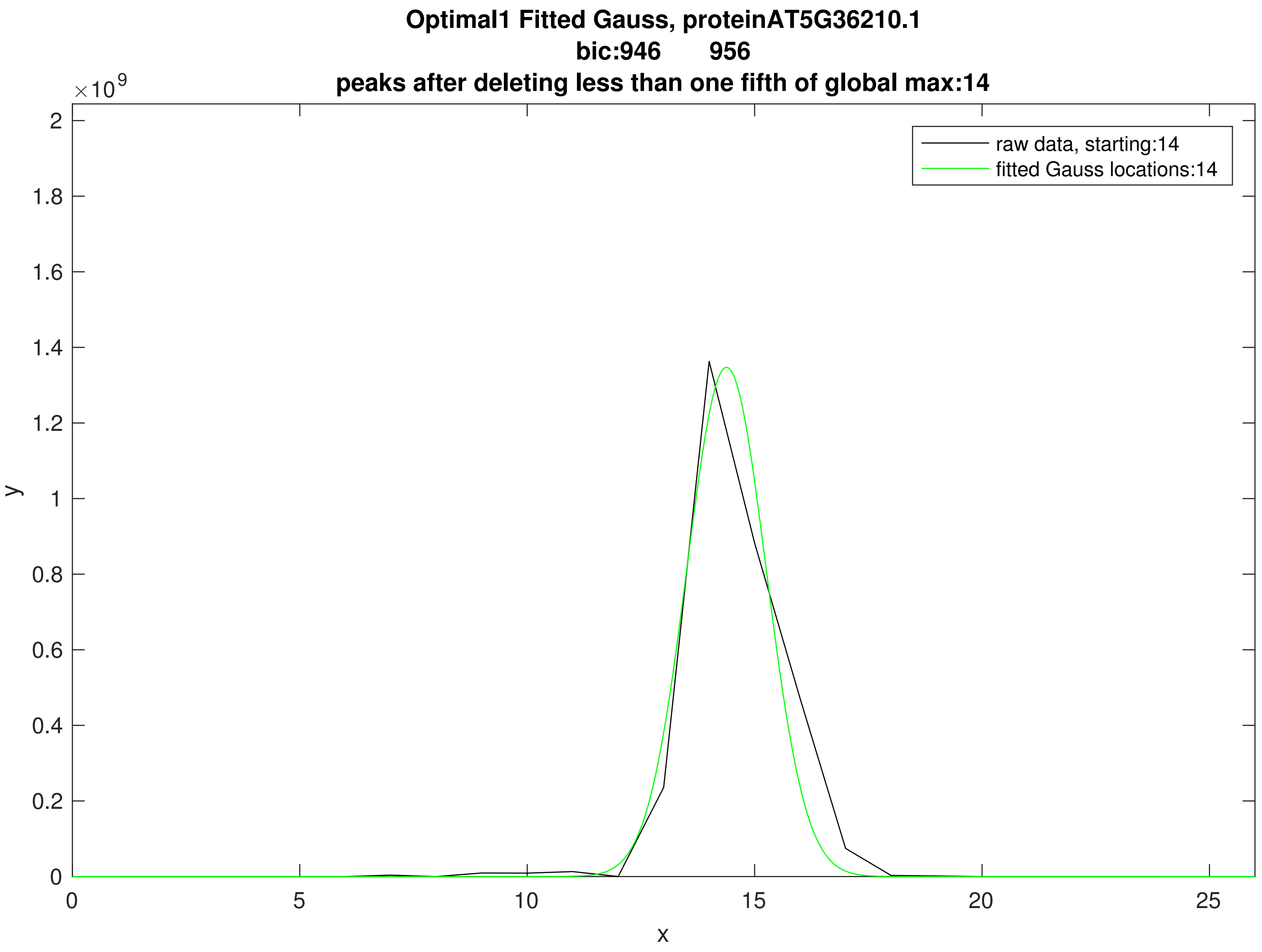

Optimal1 Fitted Gauss, proteinAT5G36210.1
bic:946 956
peaks after deleting less than one fifth of global max:14
×10^9
2
1.8
1.6
1.4
1.2
1
0.8
0.6
0.4
0.2
0
y
0
5
10
15
20
25
x
raw data, starting:14
fitted Gauss locations:14

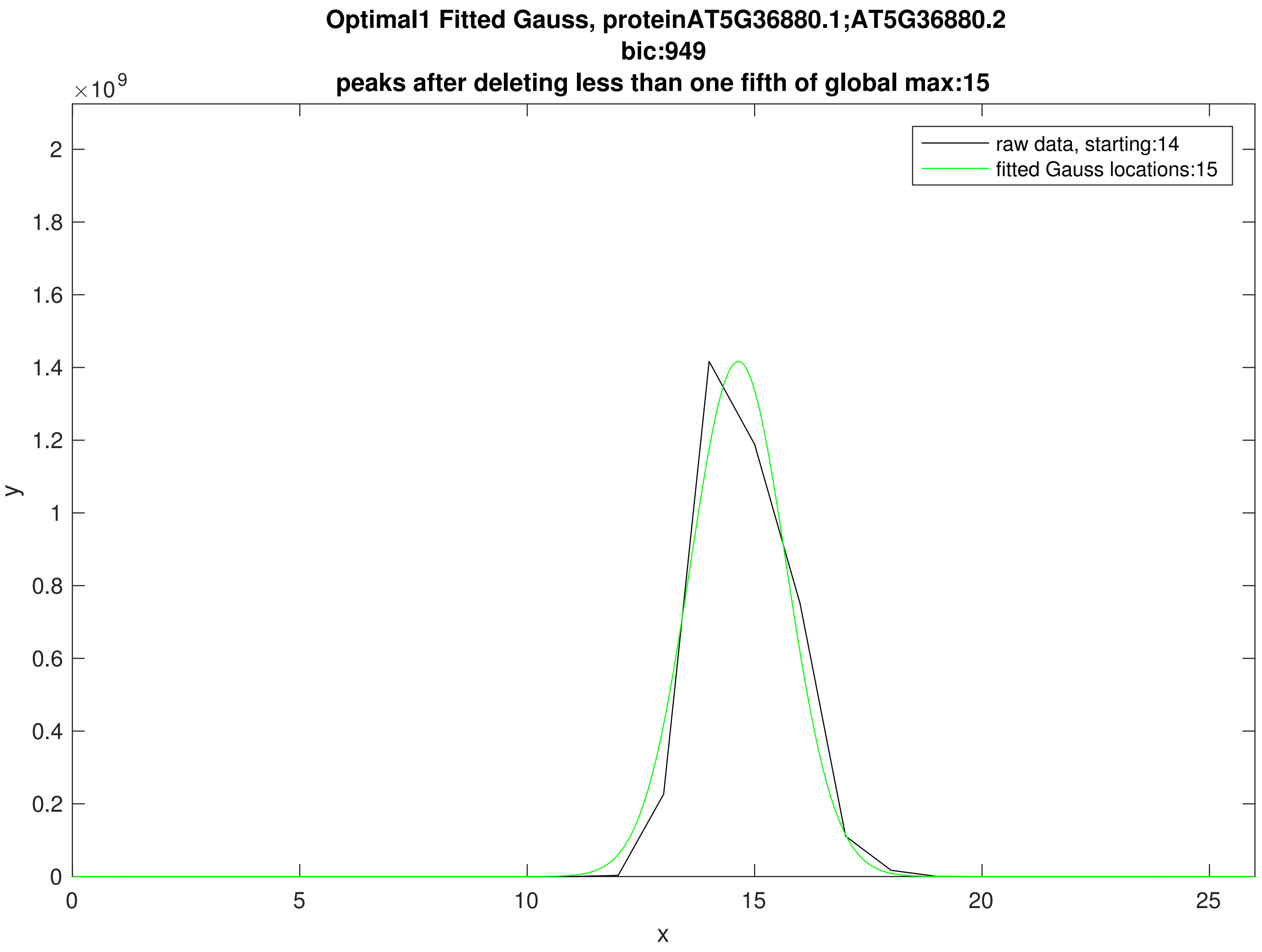
Optimal1 Fitted Gauss, proteinAT5G36880.1;AT5G36880.2
bic:949
peaks after deleting less than one fifth of global max:15
×10^9
raw data, starting:14
fitted Gauss locations:15
y
x
0
0.2
0.4
0.6
0.8
1
1.2
1.4
1.6
1.8
2
0
5
10
15
20
25

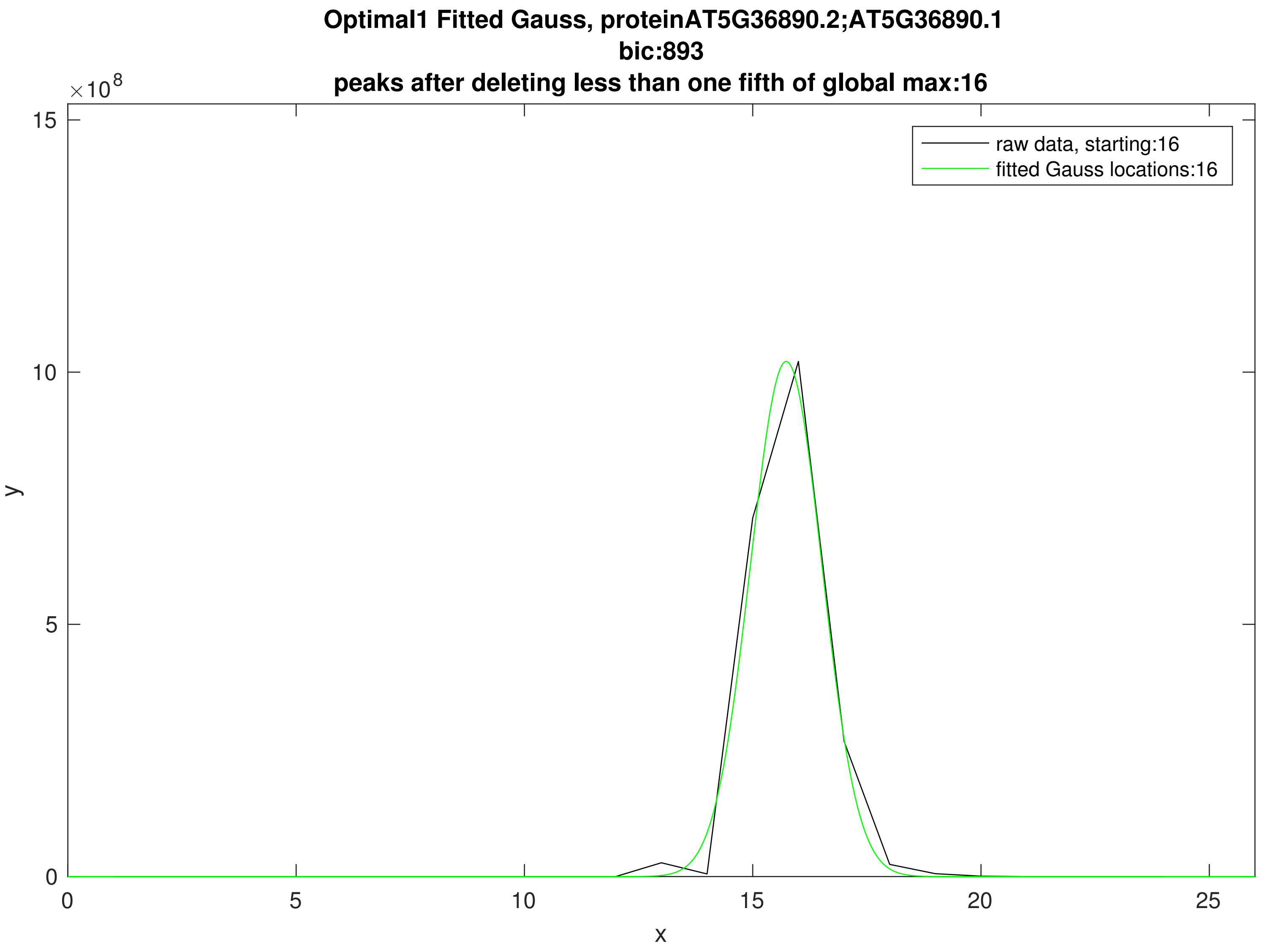
Optimal1 Fitted Gauss, proteinAT5G36890.2;AT5G36890.1
bic:893
peaks after deleting less than one fifth of global max:16
×10^8
15
10
5
0
y
0
5
10
15
20
25
x
raw data, starting:16
fitted Gauss locations:16

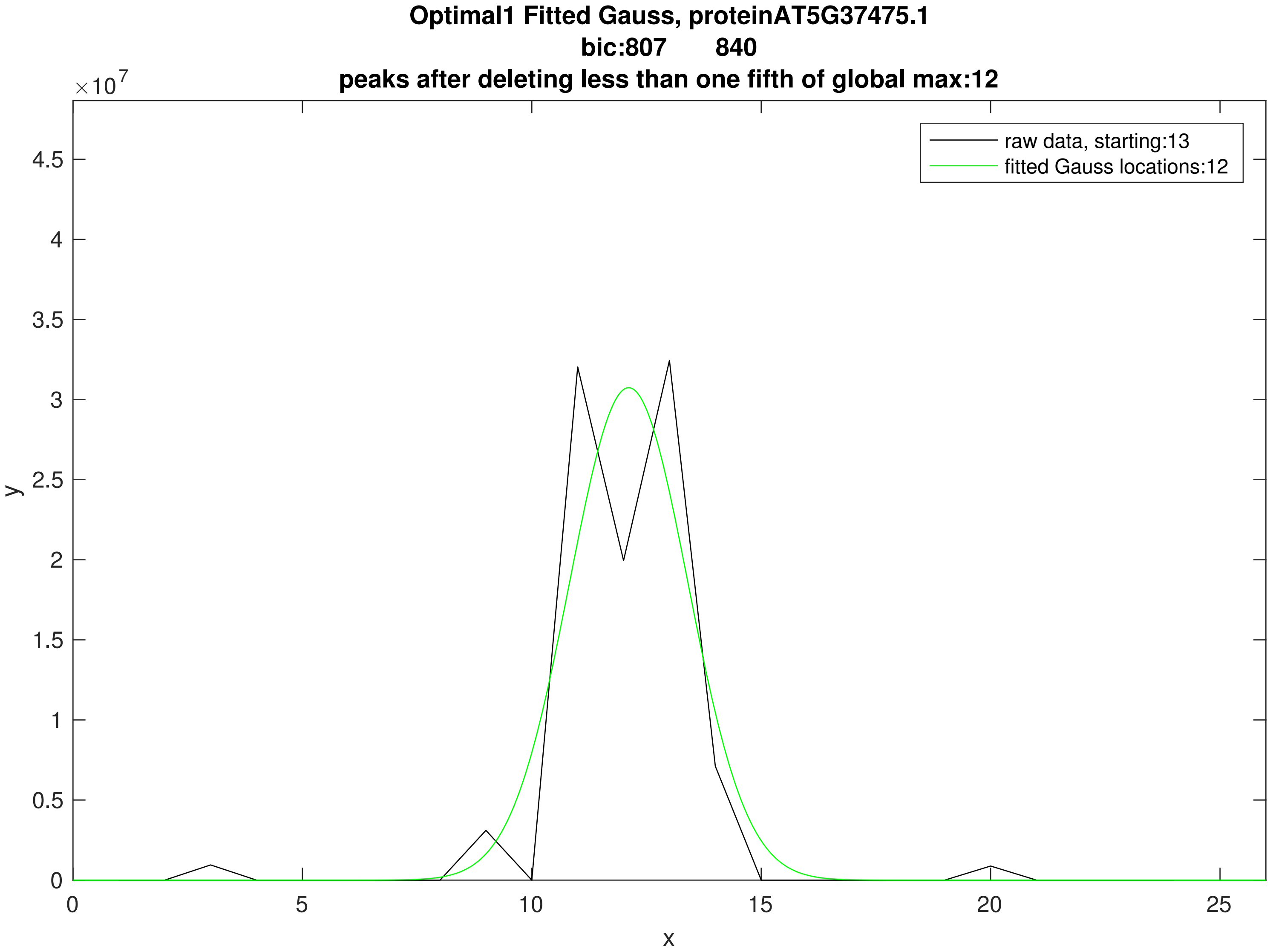
Optimal1 Fitted Gauss, proteinAT5G37475.1
bic:807 840
peaks after deleting less than one fifth of global max:12
×10^7
raw data, starting:13
fitted Gauss locations:12
y
x
0
0.5
1
1.5
2
2.5
3
3.5
4
4.5
5
10
15
20
25

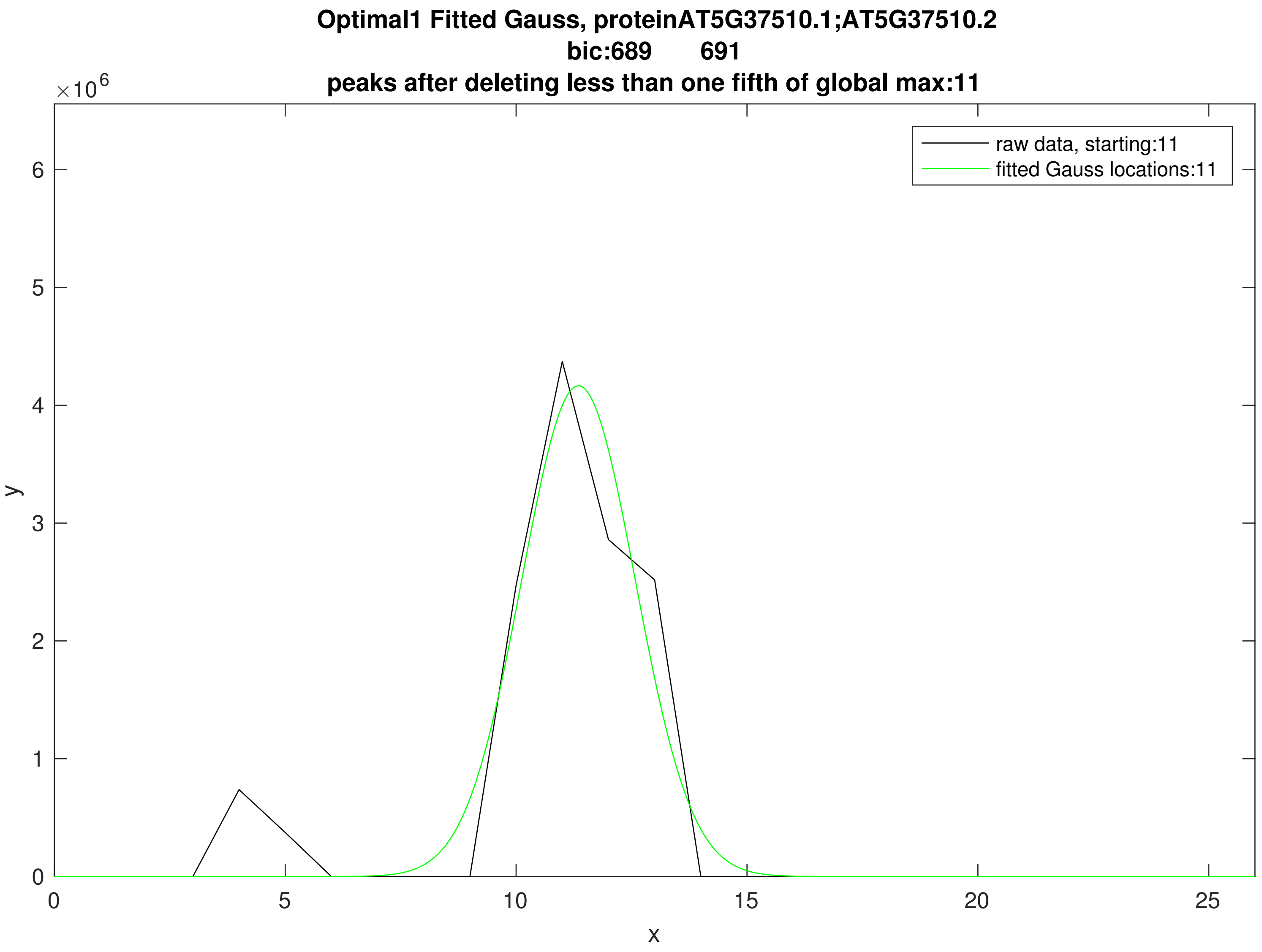
Optimal1 Fitted Gauss, proteinAT5G37510.1;AT5G37510.2
bic:689 691
peaks after deleting less than one fifth of global max:11
×10^6
raw data, starting:11
fitted Gauss locations:11
y
x
0
1
2
3
4
5
6
0
5
10
15
20
25

**Optimal1 Fitted Gauss, proteinAT5G37600.1**
**bic:996 1e+03 1.01e+03**
**peaks after deleting less than one fifth of global max:7.5**

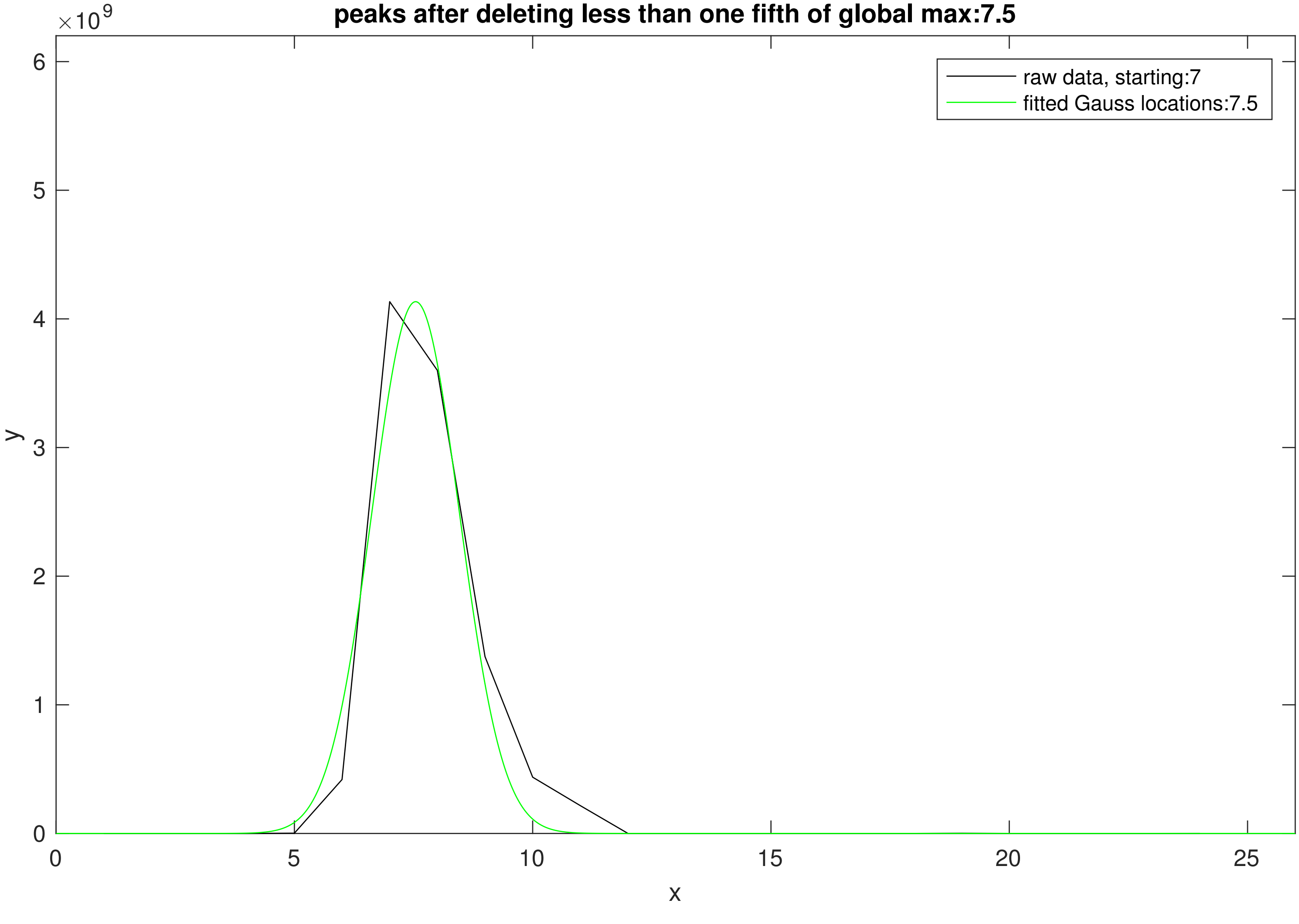

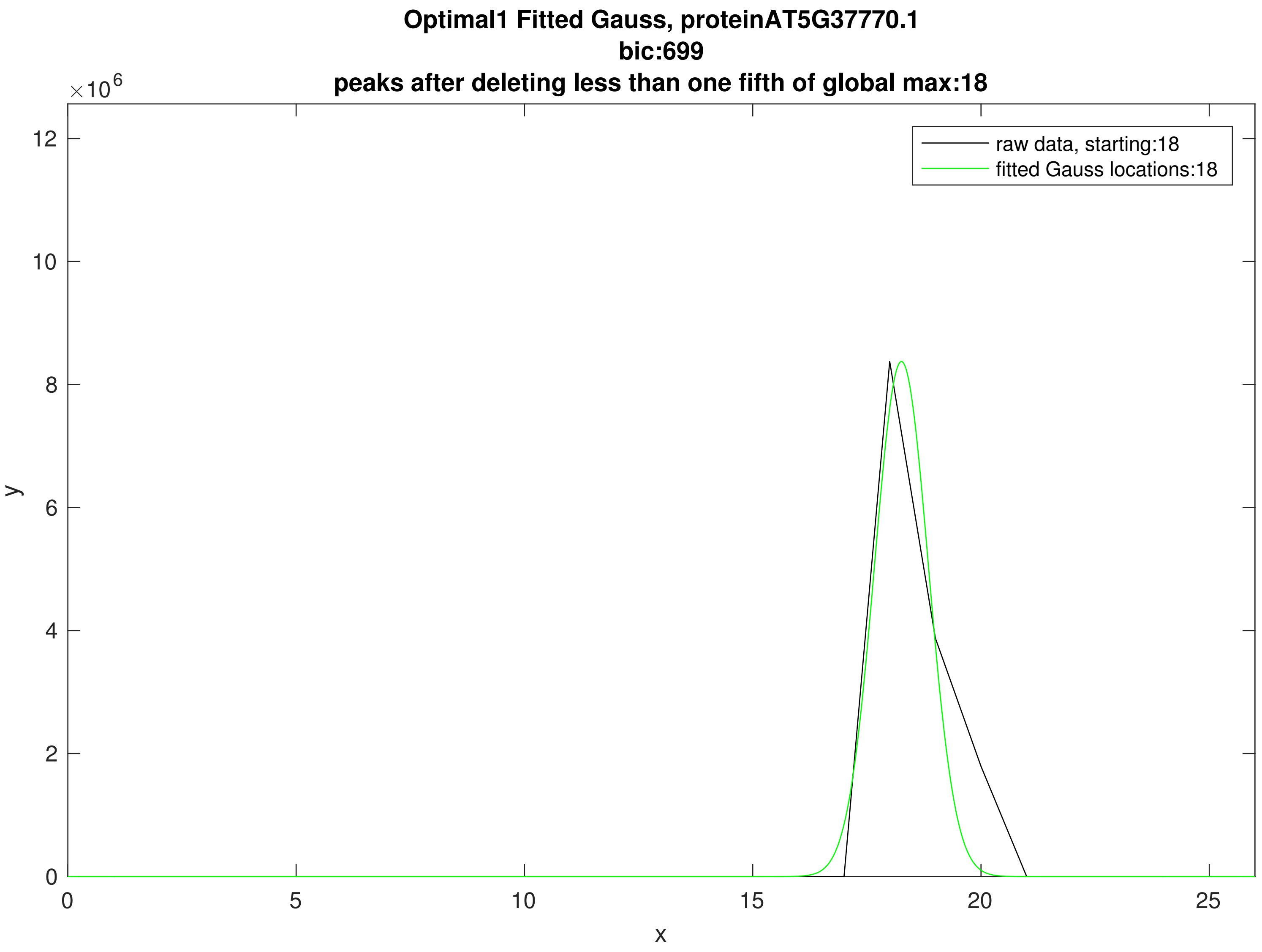

Optimal1 Fitted Gauss, proteinAT5G37770.1
bic:699
peaks after deleting less than one fifth of global max:18
×10^6
raw data, starting:18
fitted Gauss locations:18
y
x
0
2
4
6
8
10
12
0
5
10
15
20
25

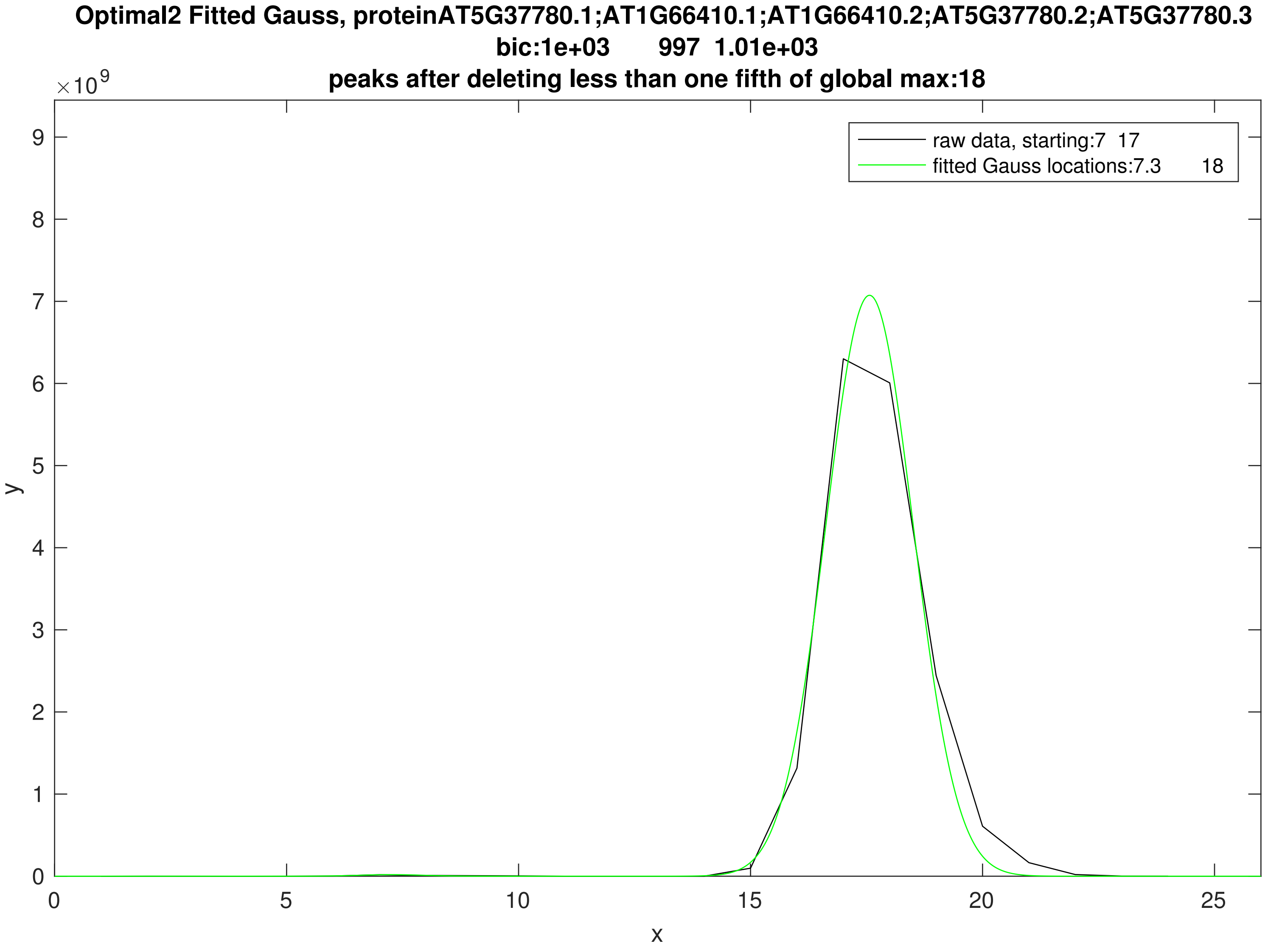

Optimal2 Fitted Gauss, proteinAT5G37780.1;AT1G66410.1;AT1G66410.2;AT5G37780.2;AT5G37780.3
bic:1e+03 997 1.01e+03
peaks after deleting less than one fifth of global max:18
×10^9
raw data, starting:7 17
fitted Gauss locations:7.3 18
y
x
0
1
2
3
4
5
6
7
8
9
0
5
10
15
20
25

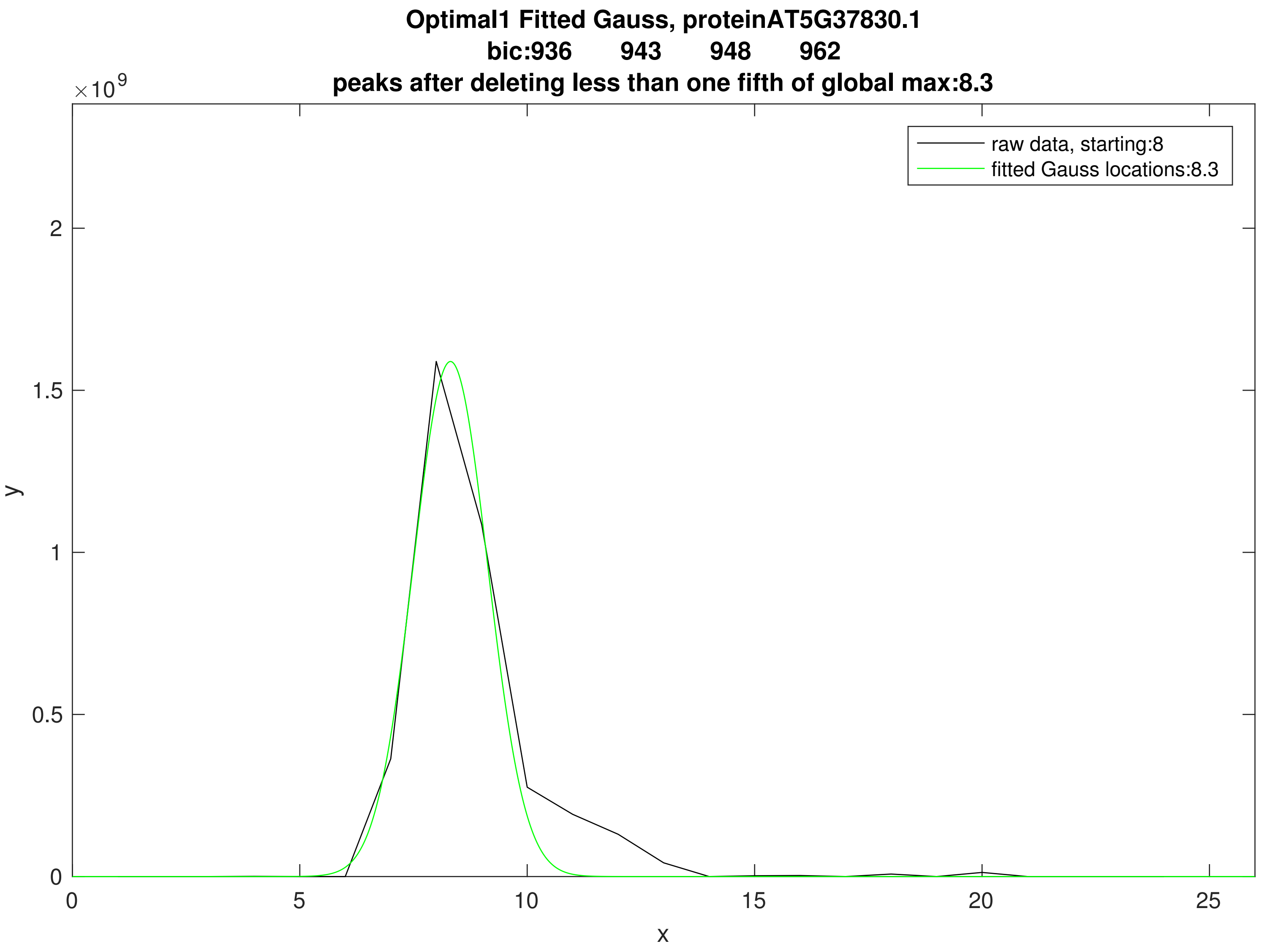
Optimal1 Fitted Gauss, proteinAT5G37830.1
bic:936 943 948 962
peaks after deleting less than one fifth of global max:8.3
×10^9
raw data, starting:8
fitted Gauss locations:8.3
2
1.5
1
0.5
0
y
0
5
10
15
20
25
x

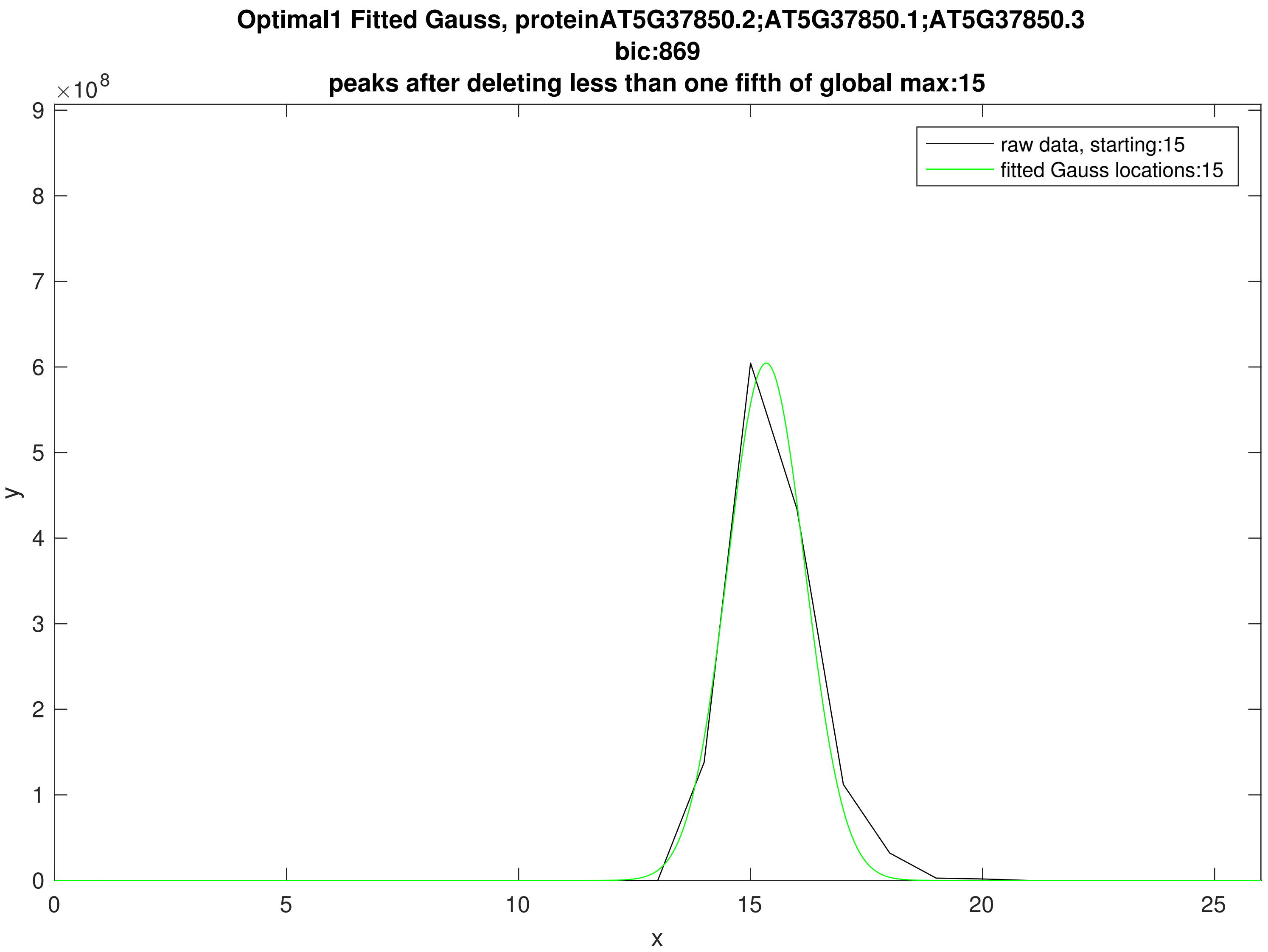
Optimal1 Fitted Gauss, proteinAT5G37850.2;AT5G37850.1;AT5G37850.3
bic:869
peaks after deleting less than one fifth of global max:15
×10^8
raw data, starting:15
fitted Gauss locations:15
y
x
0
1
2
3
4
5
6
7
8
9
0
5
10
15
20
25

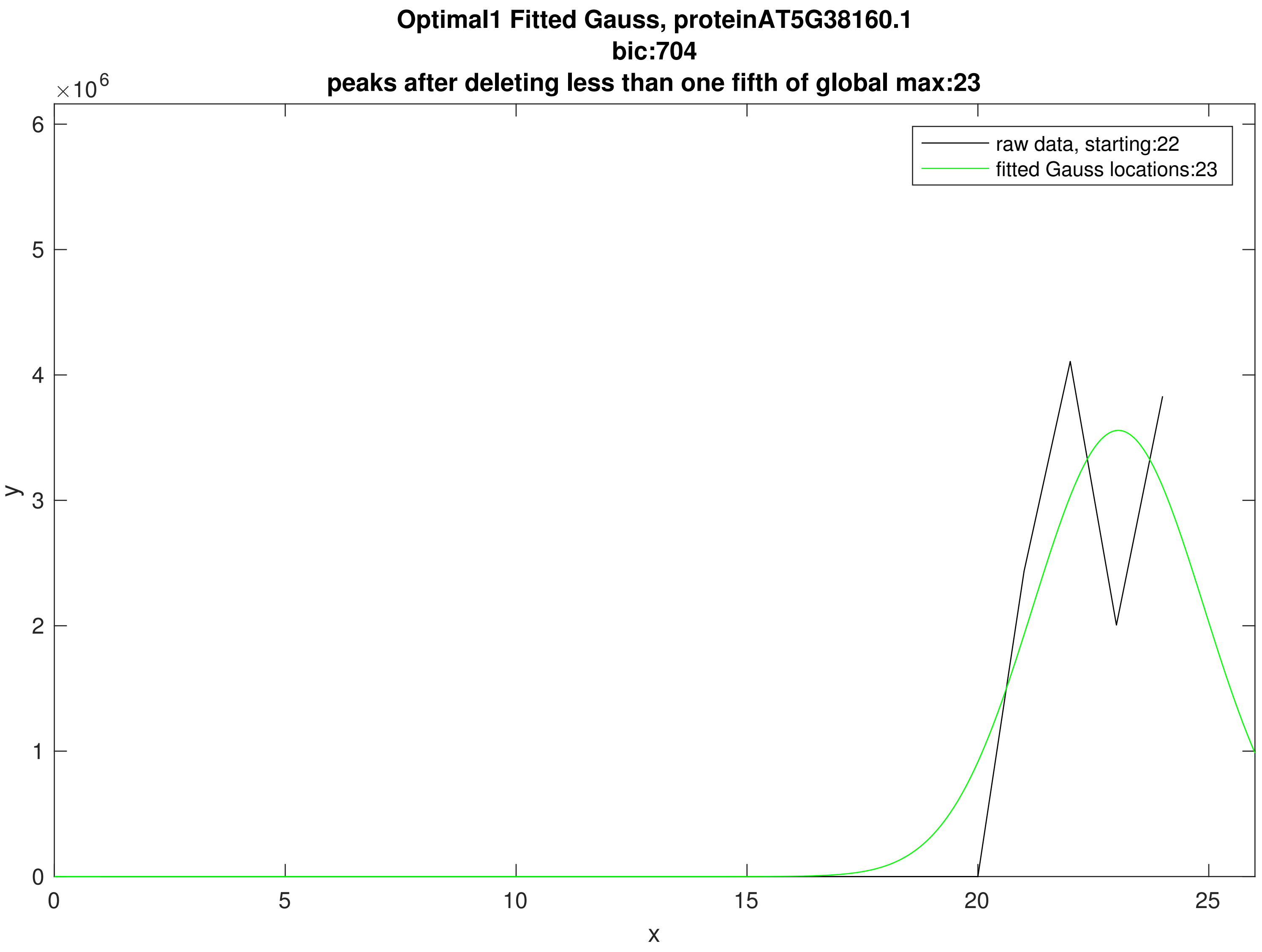
Optimal1 Fitted Gauss, proteinAT5G38160.1
bic:704
peaks after deleting less than one fifth of global max:23
×10^6
raw data, starting:22
fitted Gauss locations:23
y
x
0
1
2
3
4
5
6
0
5
10
15
20
25

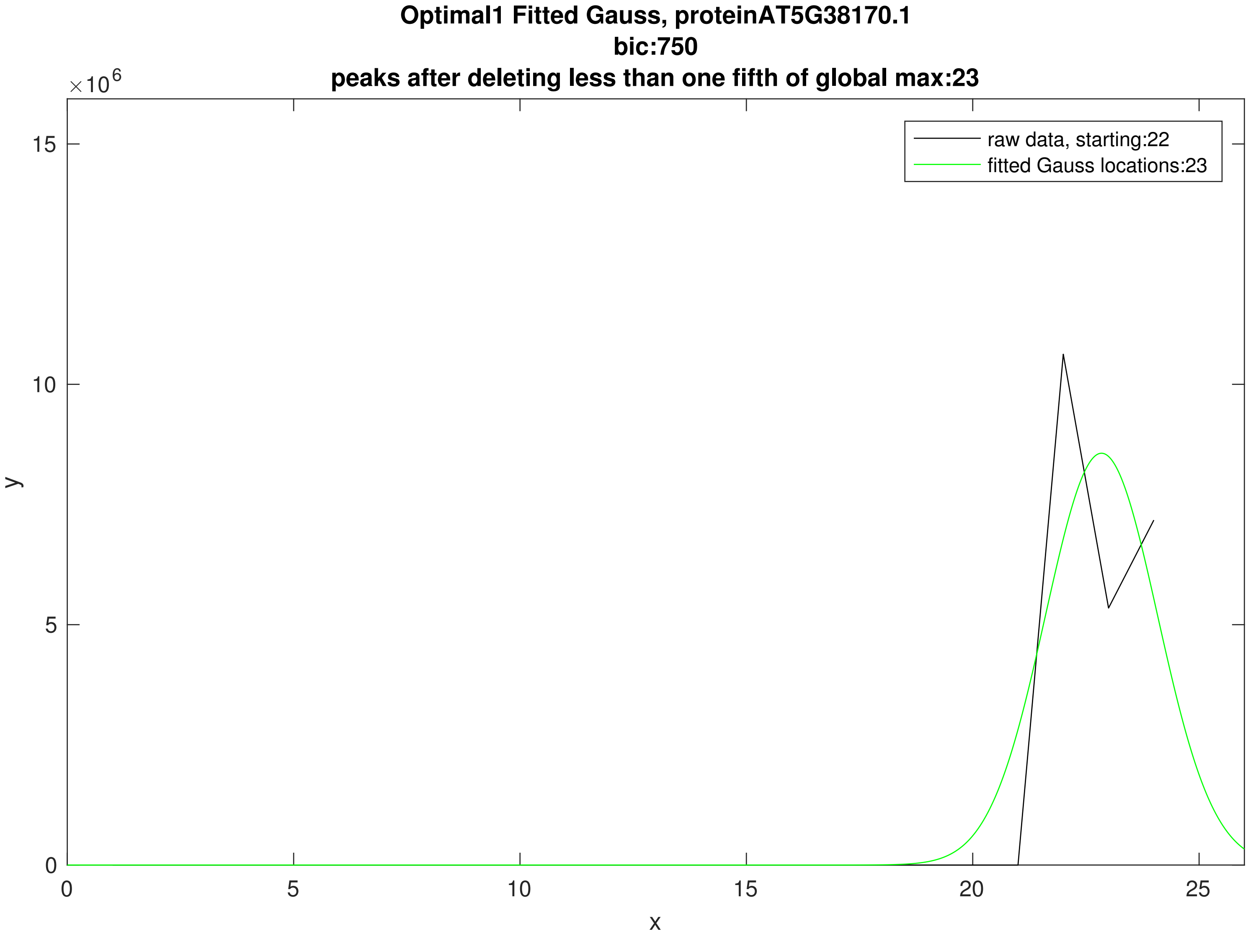
Optimal1 Fitted Gauss, proteinAT5G38170.1
bic:750
peaks after deleting less than one fifth of global max:23
×10^6
15
10
5
0
y
0
5
10
15
20
25
x
raw data, starting:22
fitted Gauss locations:23

**Optimal1 Fitted Gauss, proteinAT5G38420.1;AT5G38410.1;AT5G38410.3;AT5G38410.2**
**bic:994 996 1.01e+03 1.01e+03**
**peaks after deleting less than one fifth of global max:6.7**

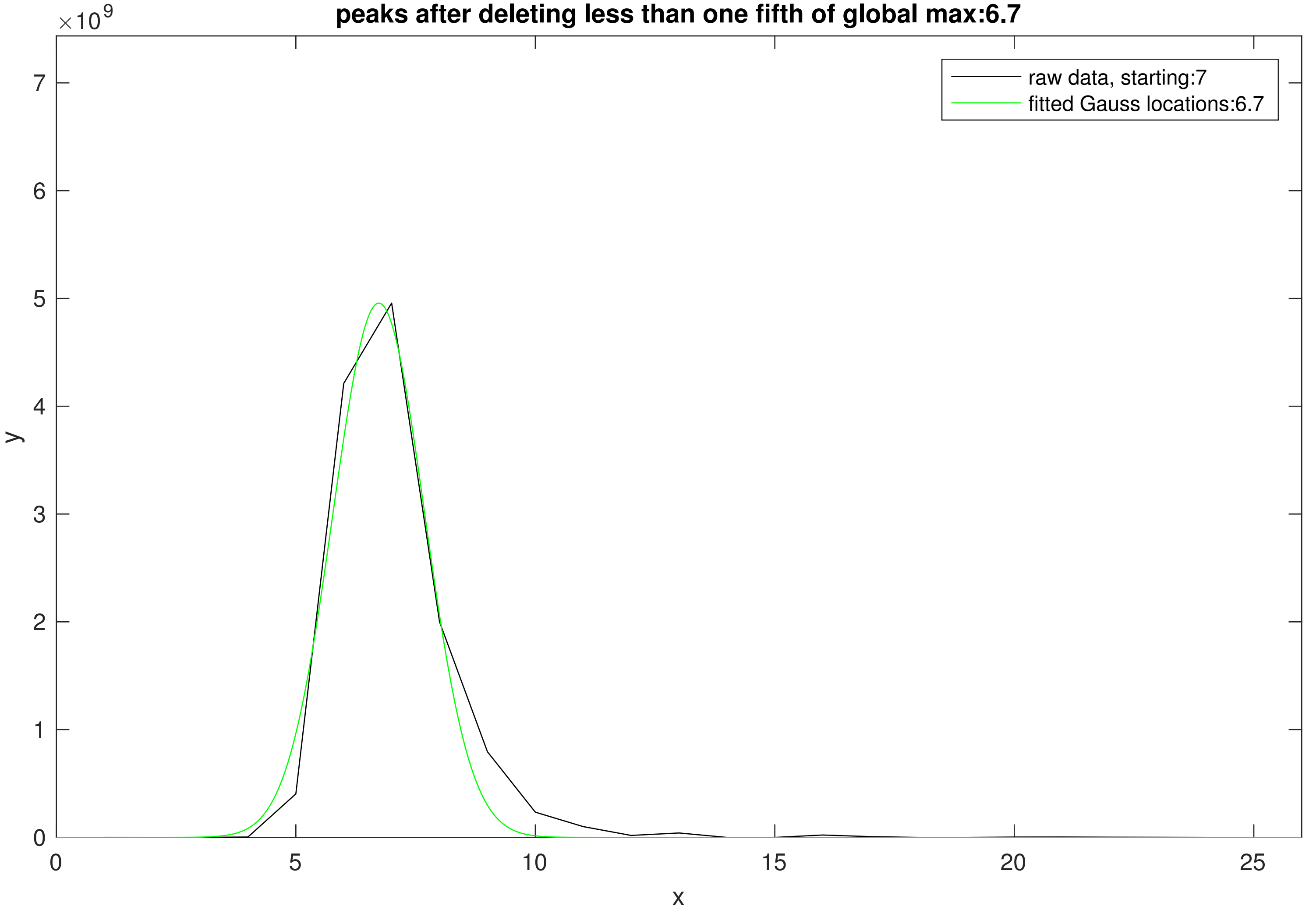

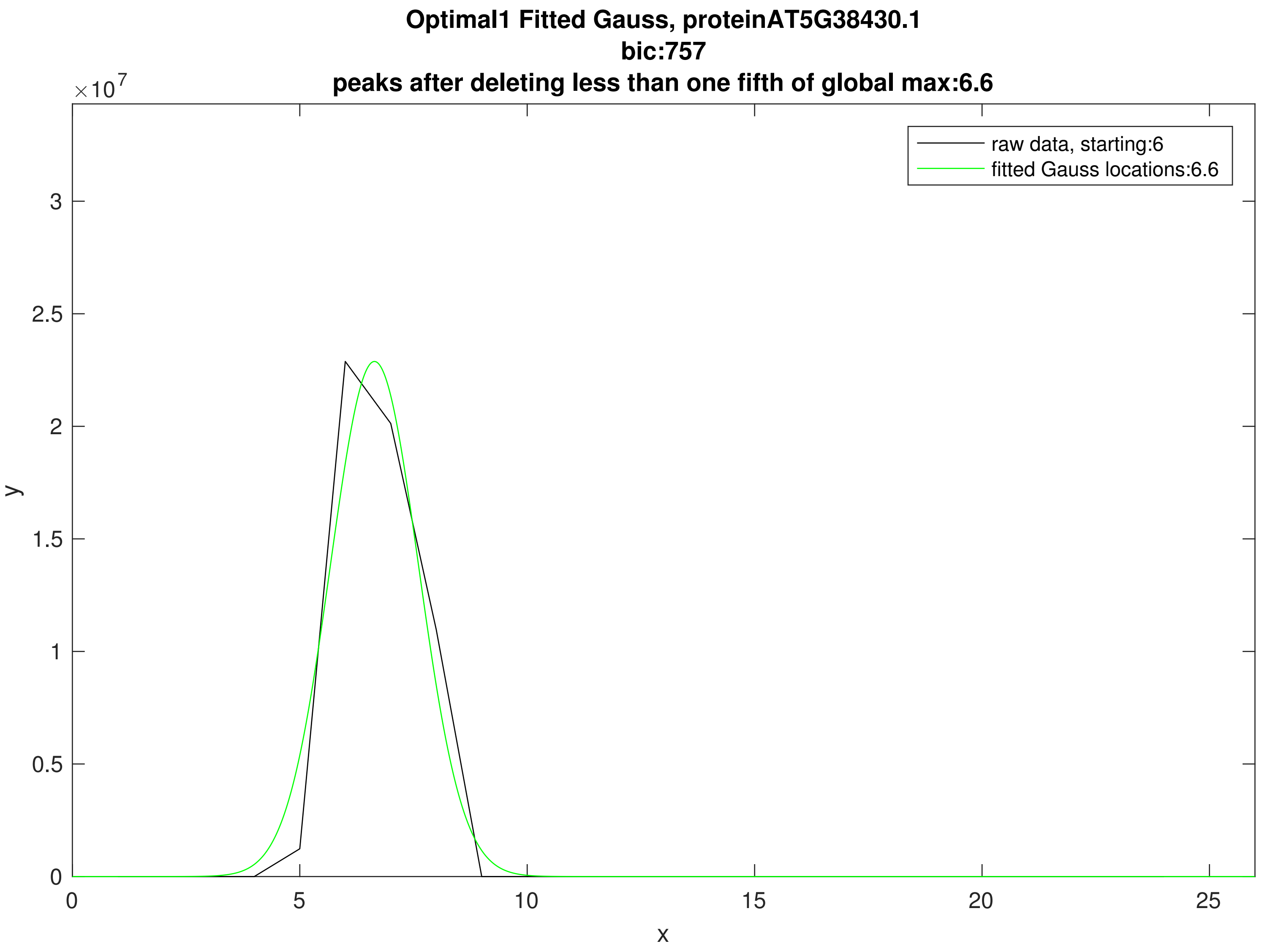
Optimal1 Fitted Gauss, proteinAT5G38430.1
bic:757
peaks after deleting less than one fifth of global max:6.6
×10^7
raw data, starting:6
fitted Gauss locations:6.6
y
x
0
0.5
1
1.5
2
2.5
3
5
10
15
20
25

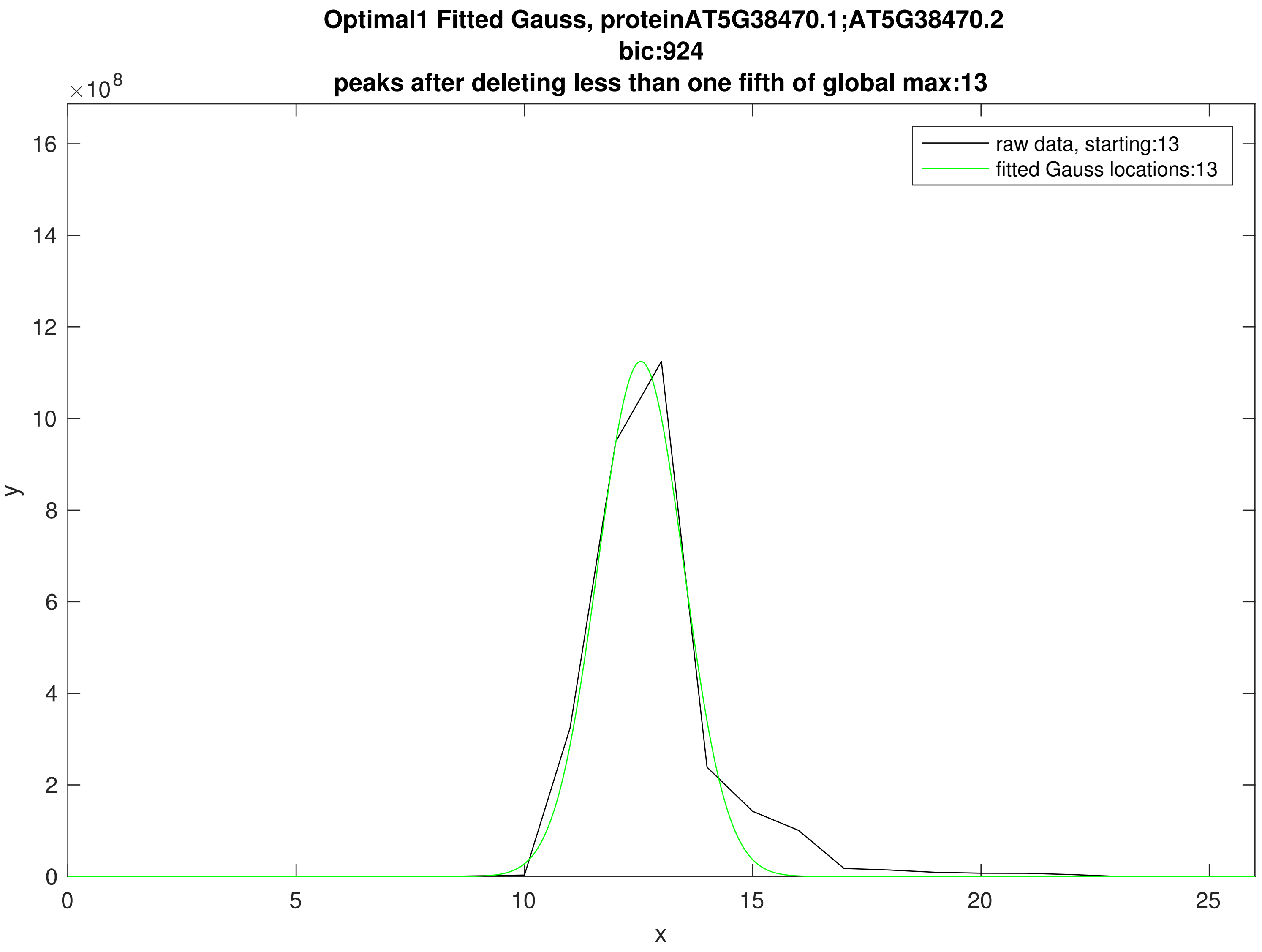
Optimal1 Fitted Gauss, proteinAT5G38470.1;AT5G38470.2
bic:924
peaks after deleting less than one fifth of global max:13
×10^8
raw data, starting:13
fitted Gauss locations:13
y
x
0
2
4
6
8
10
12
14
16
0
5
10
15
20
25

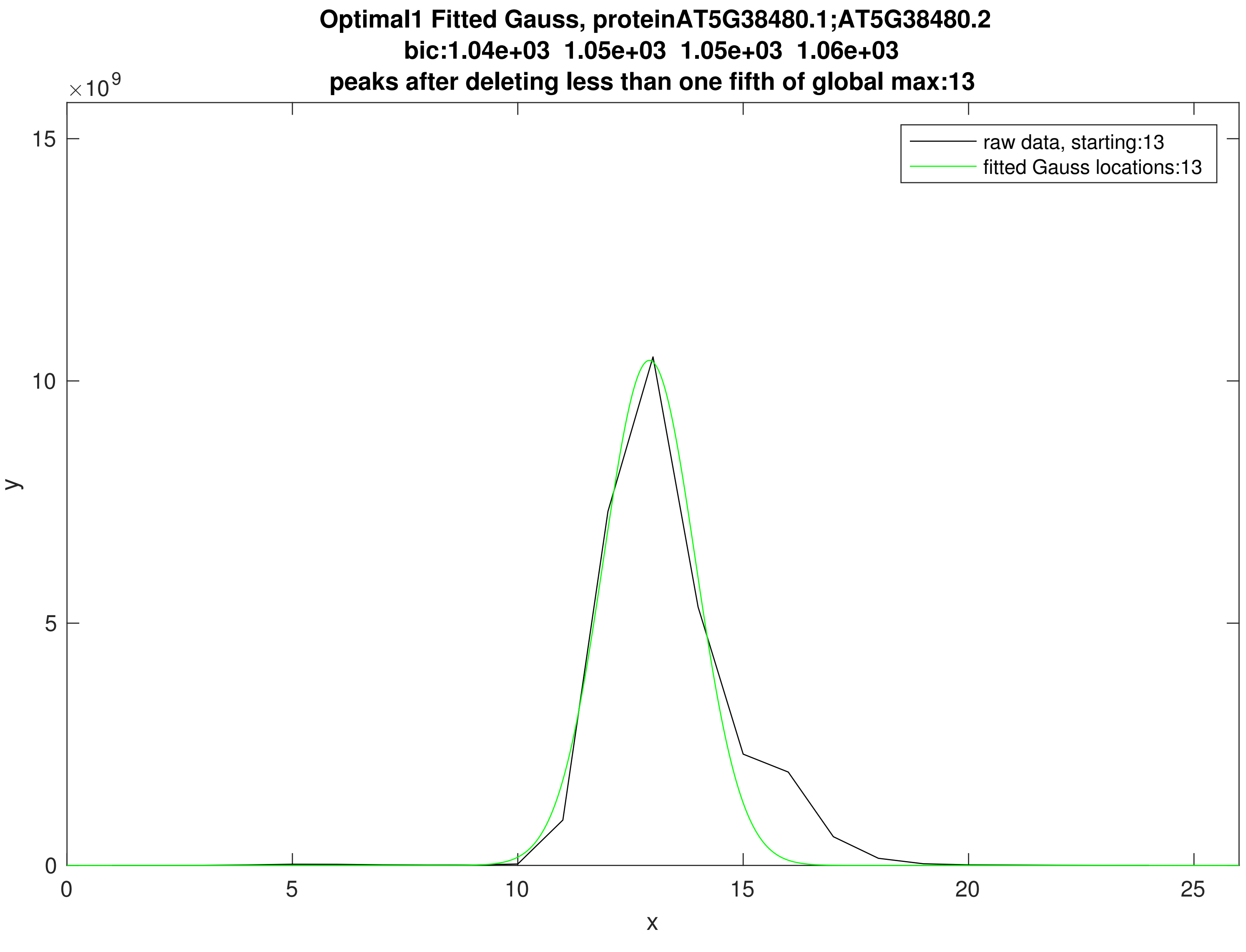

Optimal1 Fitted Gauss, proteinAT5G38480.1;AT5G38480.2
bic:1.04e+03 1.05e+03 1.05e+03 1.06e+03
peaks after deleting less than one fifth of global max:13
×10^9
15
10
5
0
y
0
5
10
15
20
25
x
raw data, starting:13
fitted Gauss locations:13

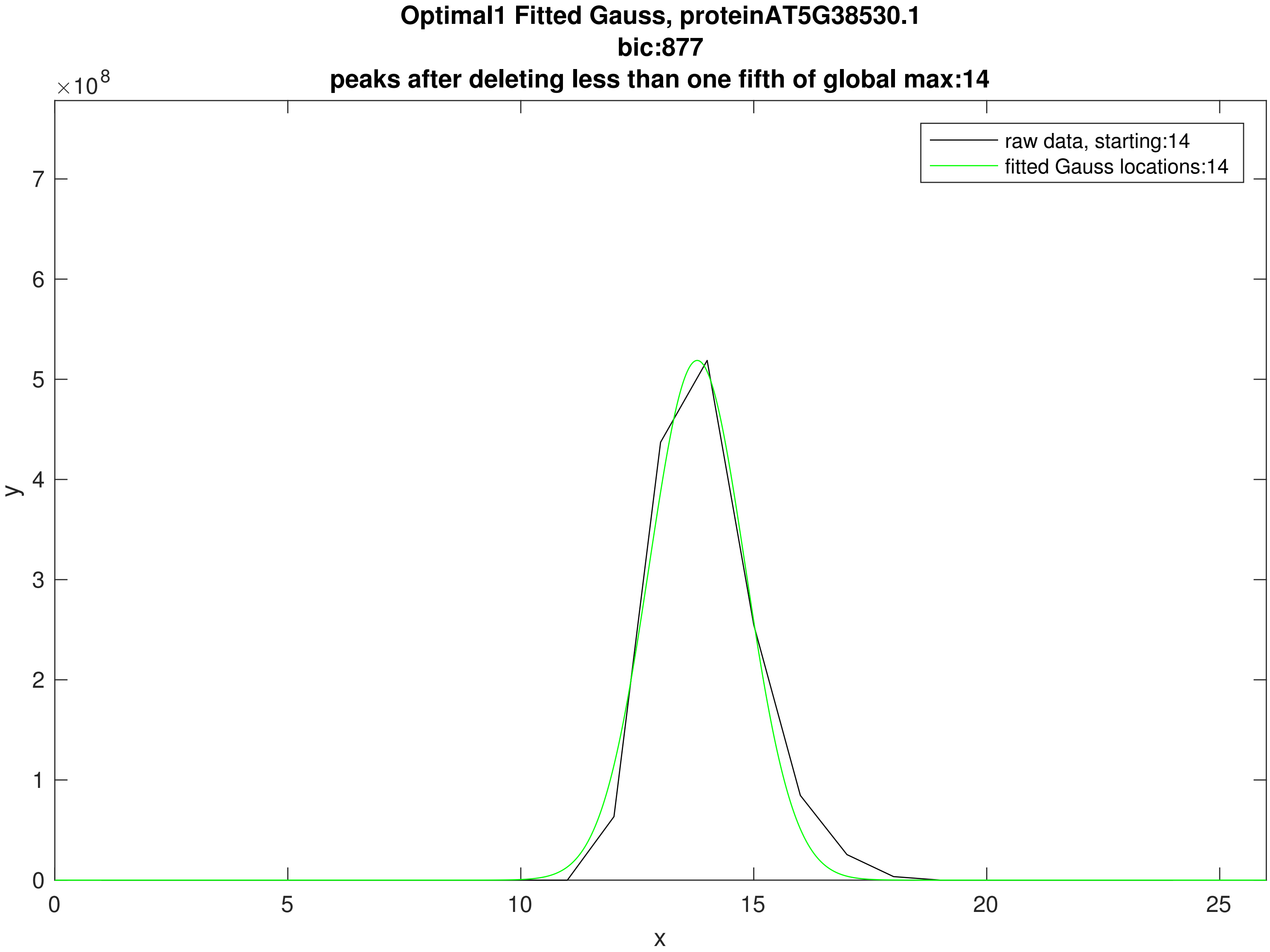
Optimal1 Fitted Gauss, proteinAT5G38530.1
bic:877
peaks after deleting less than one fifth of global max:14
×10^8
raw data, starting:14
fitted Gauss locations:14
y
x
0
1
2
3
4
5
6
7
0
5
10
15
20
25

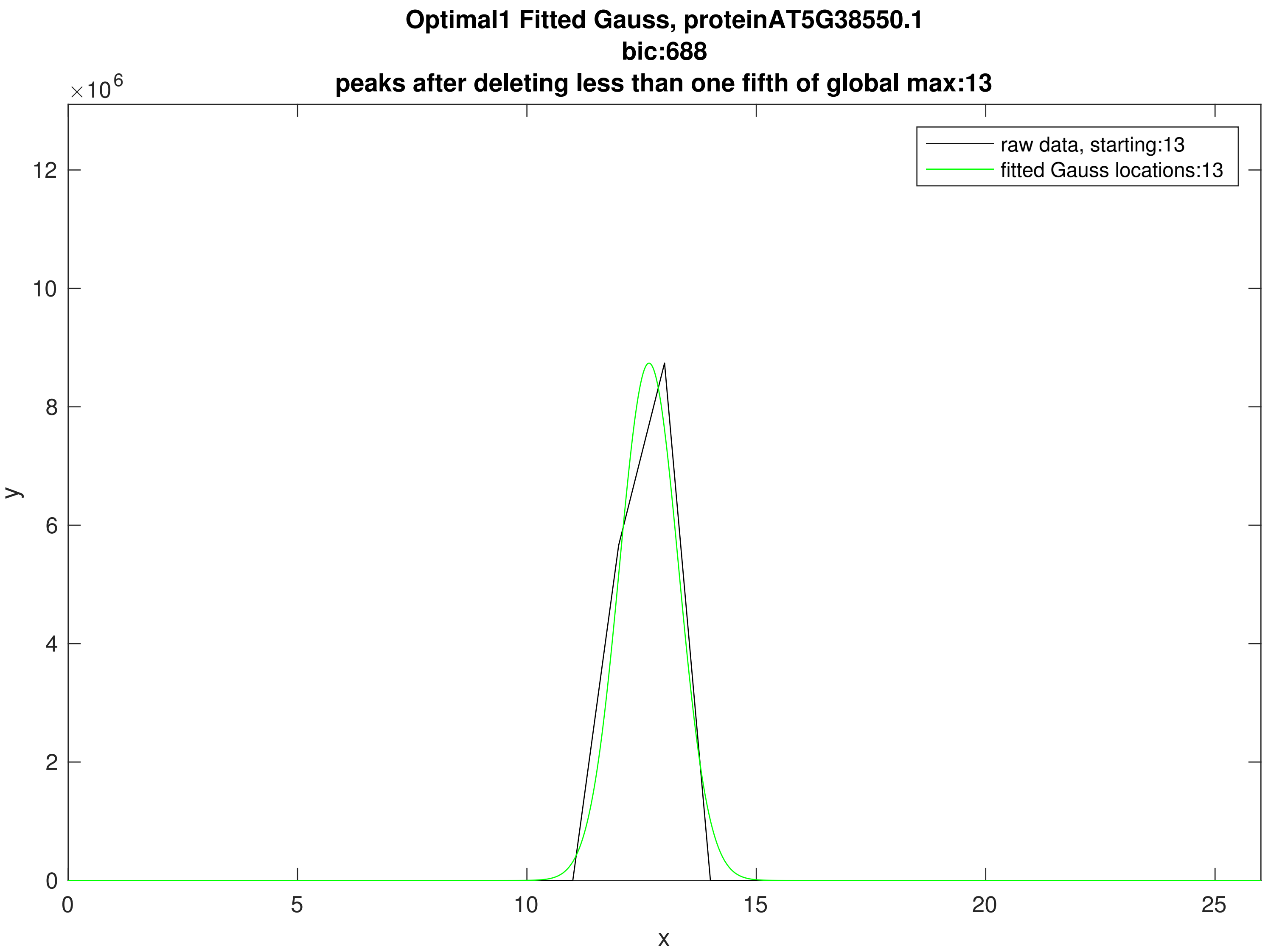
Optimal1 Fitted Gauss, proteinAT5G38550.1
bic:688
peaks after deleting less than one fifth of global max:13
×10^6
raw data, starting:13
fitted Gauss locations:13
y
x
0
2
4
6
8
10
12
0
5
10
15
20
25

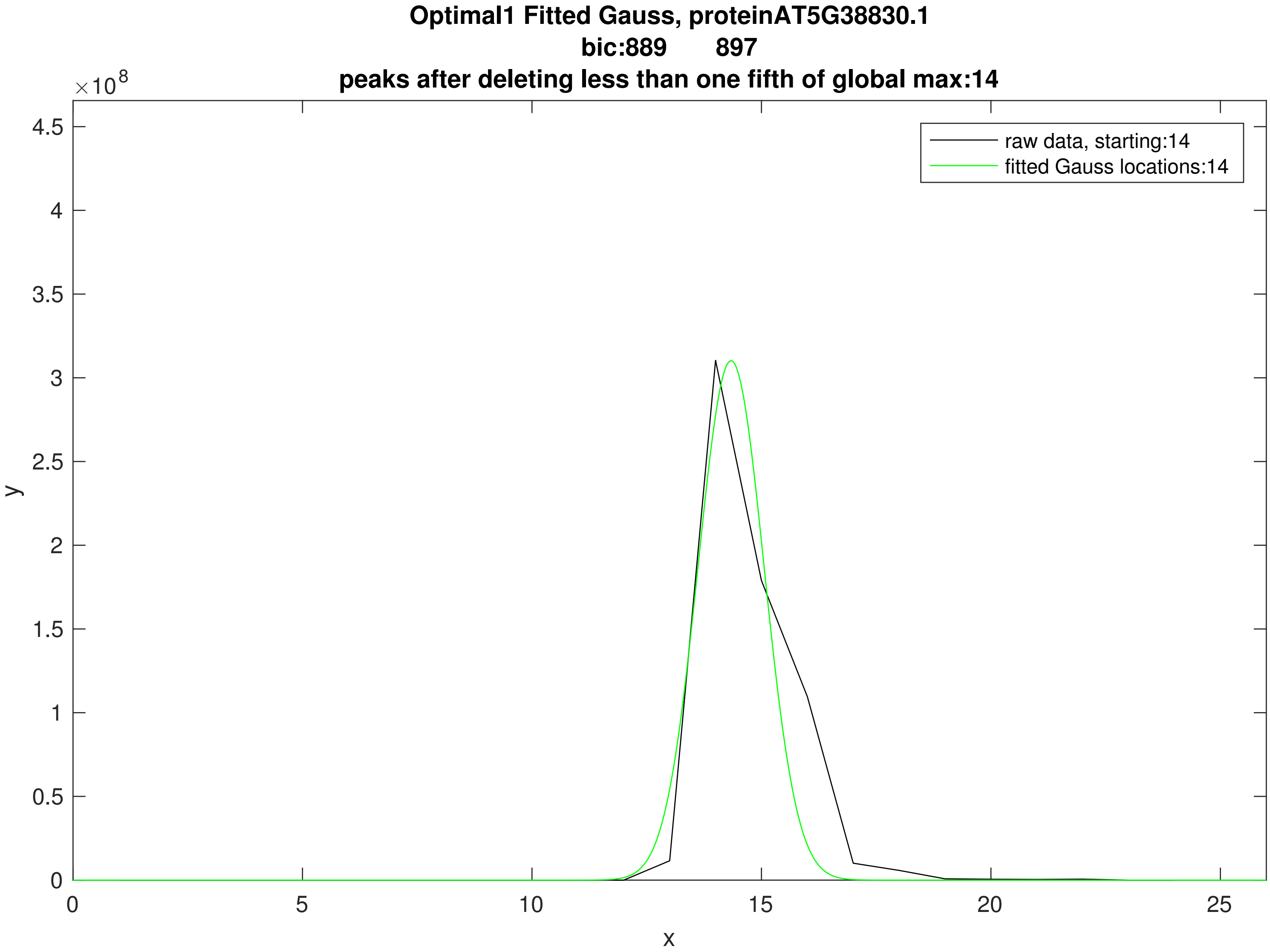
Optimal1 Fitted Gauss, proteinAT5G38830.1
bic:889 897
peaks after deleting less than one fifth of global max:14
×10^8
4.5
4
3.5
3
2.5
2
1.5
1
0.5
0
y
0
5
10
15
20
25
x
raw data, starting:14
fitted Gauss locations:14

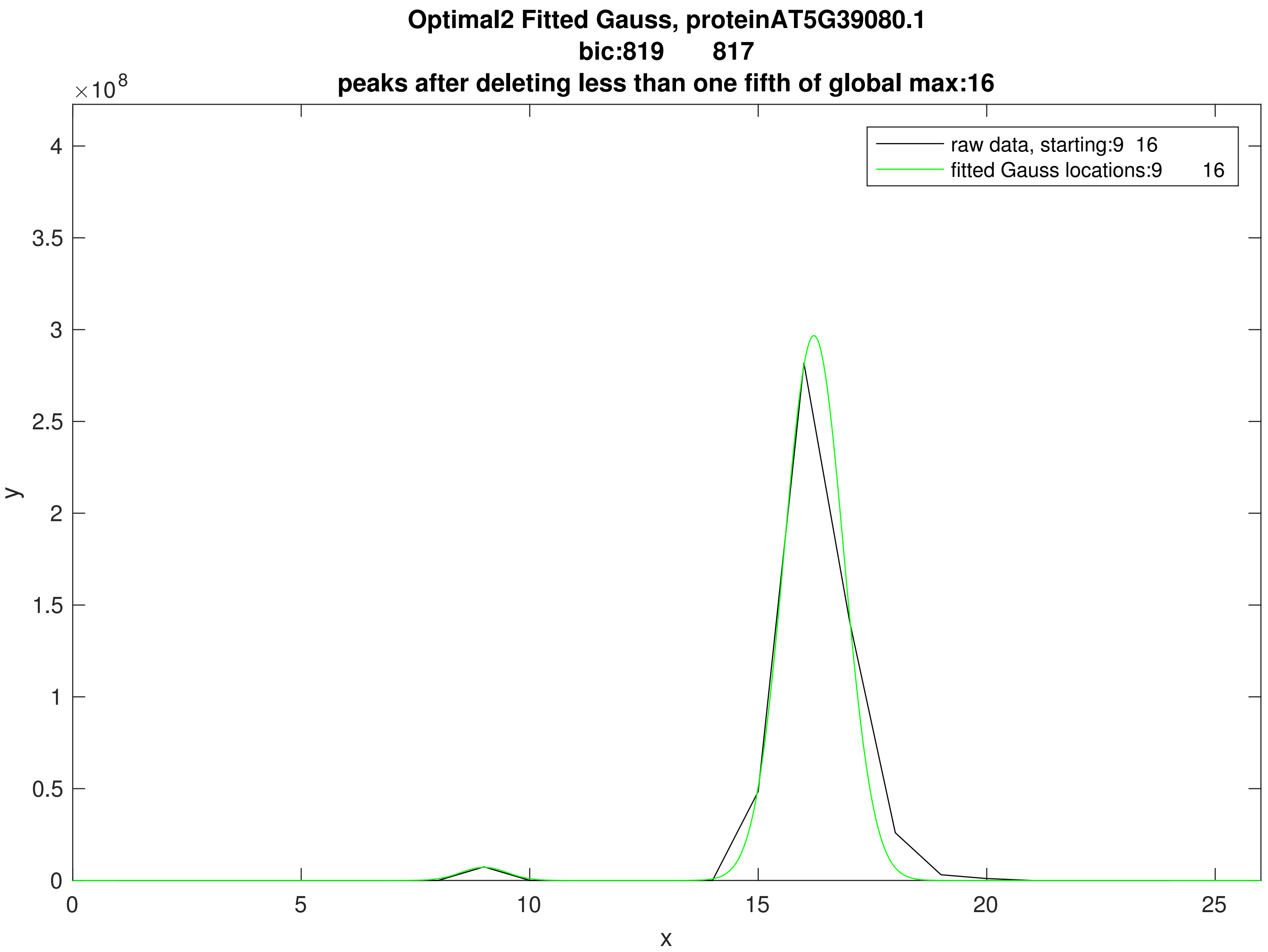
Optimal2 Fitted Gauss, proteinAT5G39080.1
bic:819 817
peaks after deleting less than one fifth of global max:16
×10^8
4
3.5
3
2.5
2
1.5
1
0.5
0
y
0
5
10
15
20
25
x
raw data, starting:9 16
fitted Gauss locations:9 16

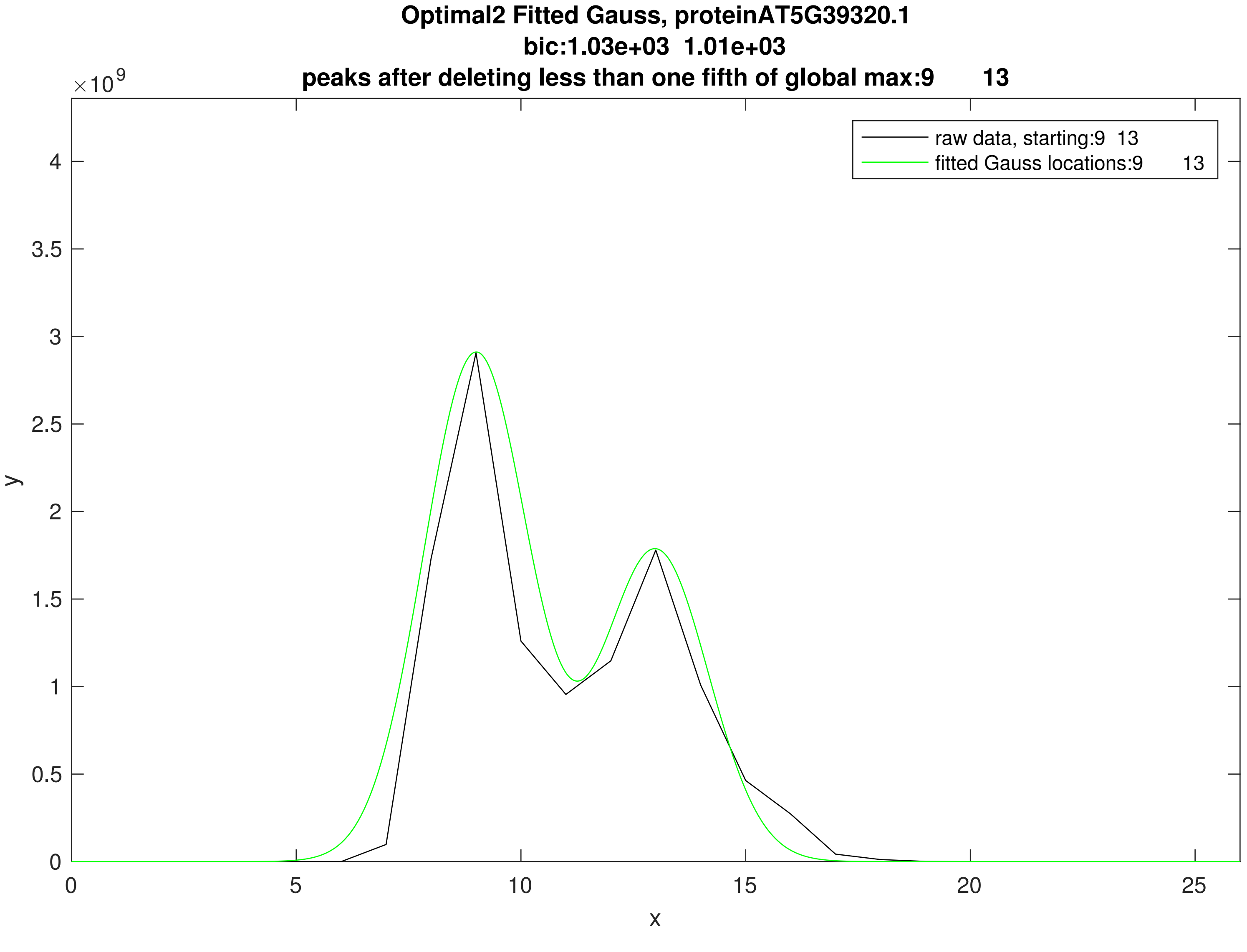
Optimal2 Fitted Gauss, proteinAT5G39320.1
bic:1.03e+03  1.01e+03
peaks after deleting less than one fifth of global max:9       13
×10^9
raw data, starting:9  13
fitted Gauss locations:9       13
y
x
0
0.5
1
1.5
2
2.5
3
3.5
4
5
10
15
20
25

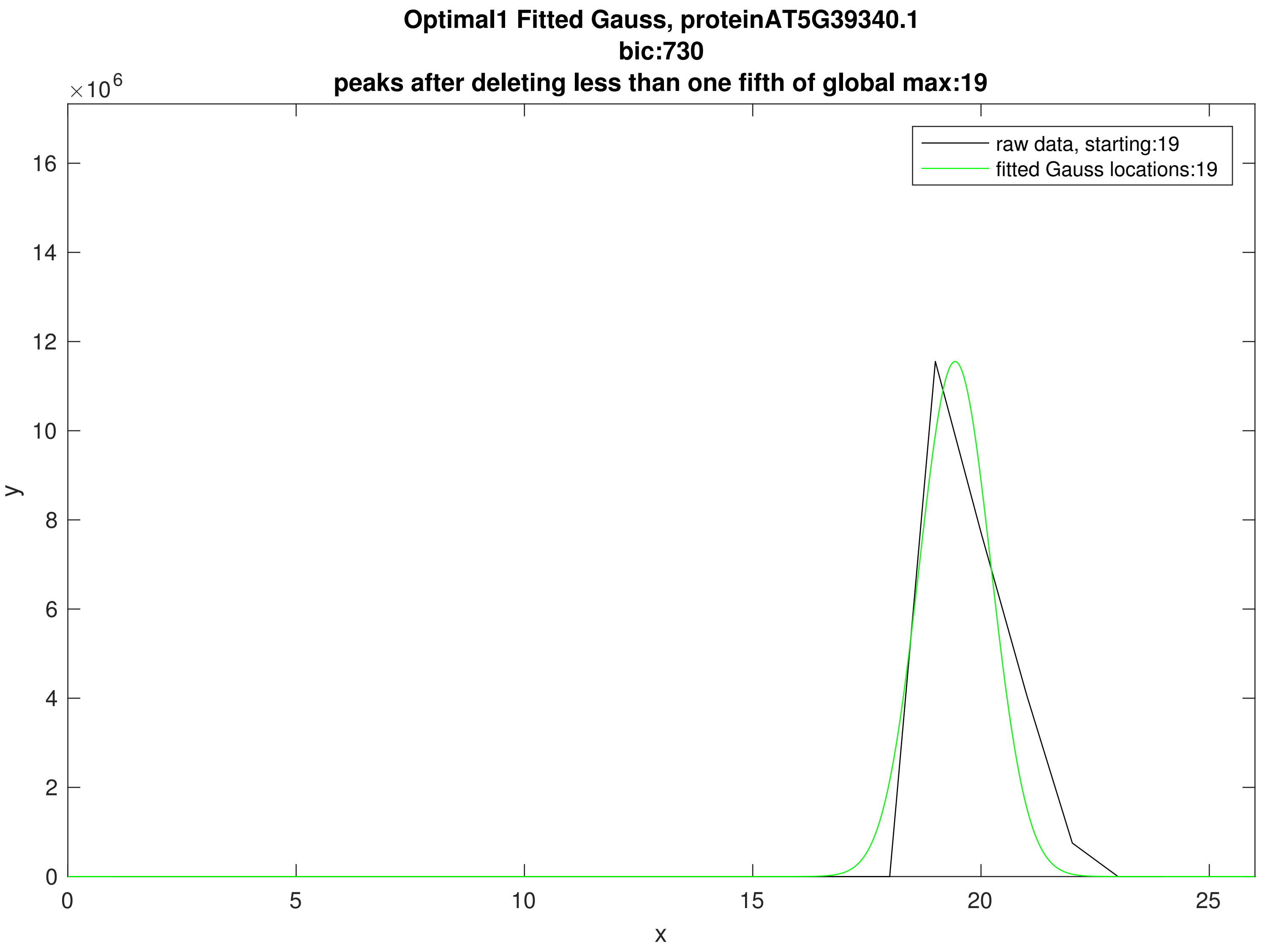

Optimal1 Fitted Gauss, proteinAT5G39340.1
bic:730
peaks after deleting less than one fifth of global max:19
×10^6
raw data, starting:19
fitted Gauss locations:19
y
x
0
2
4
6
8
10
12
14
16
5
15
20
25

**Optimal1 Fitted Gauss, proteinAT5G39510.1**
**bic:729**
**peaks after deleting less than one fifth of global max:19**

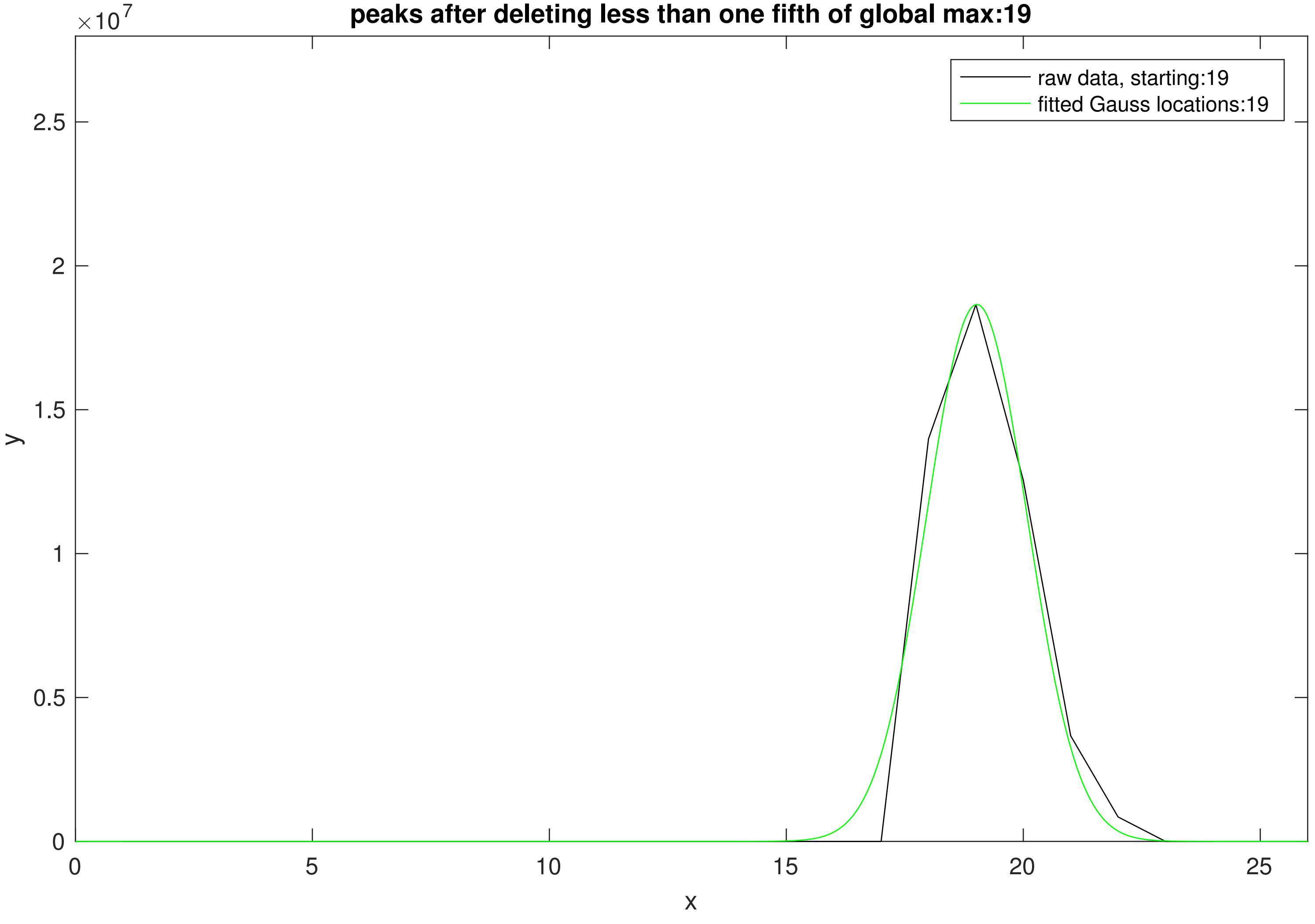

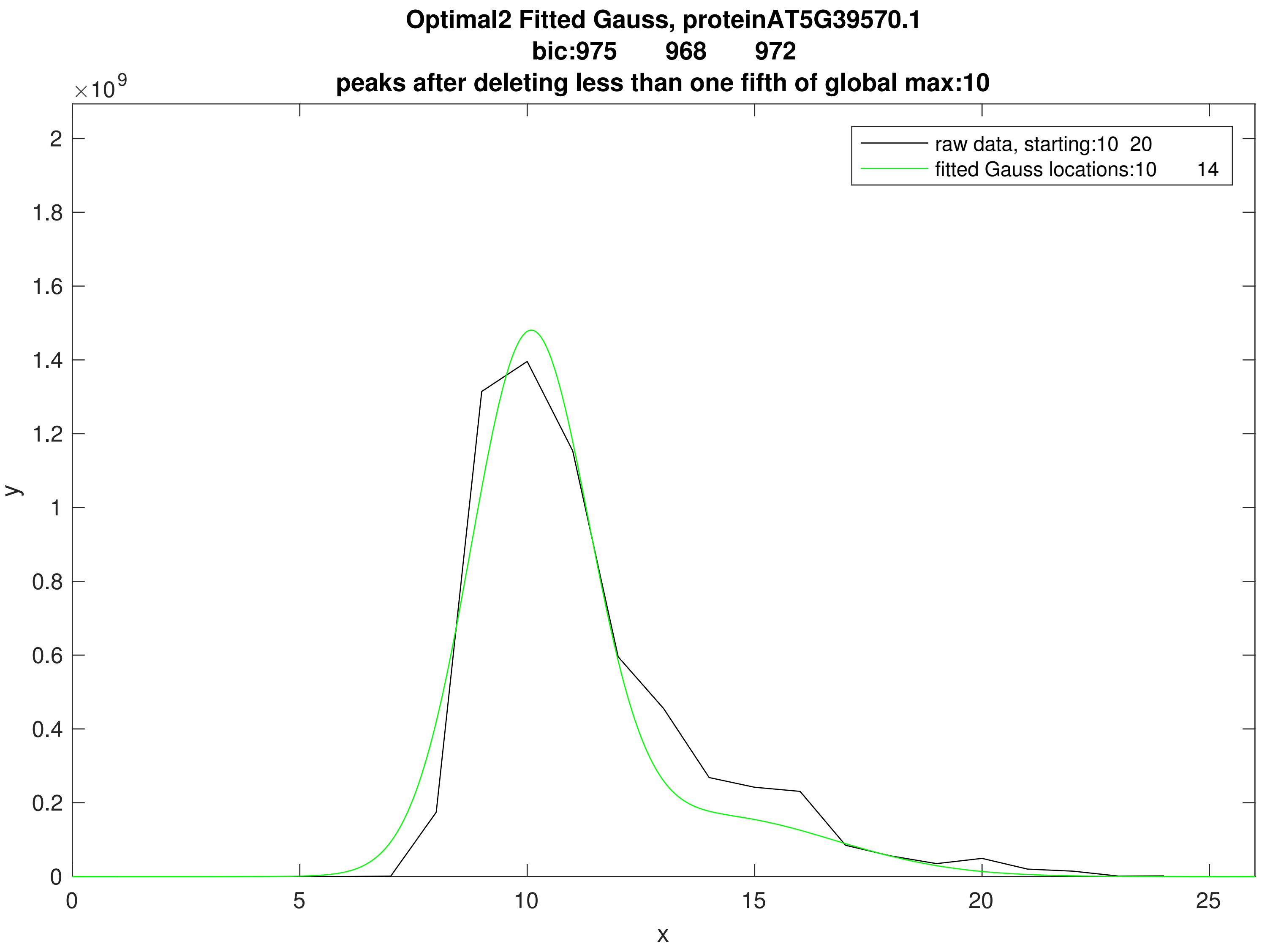

Optimal2 Fitted Gauss, proteinAT5G39570.1
bic:975 968 972
peaks after deleting less than one fifth of global max:10
×10^9
raw data, starting:10 20
fitted Gauss locations:10 14
y
x
0
0.2
0.4
0.6
0.8
1
1.2
1.4
1.6
1.8
2
0
5
10
15
20
25

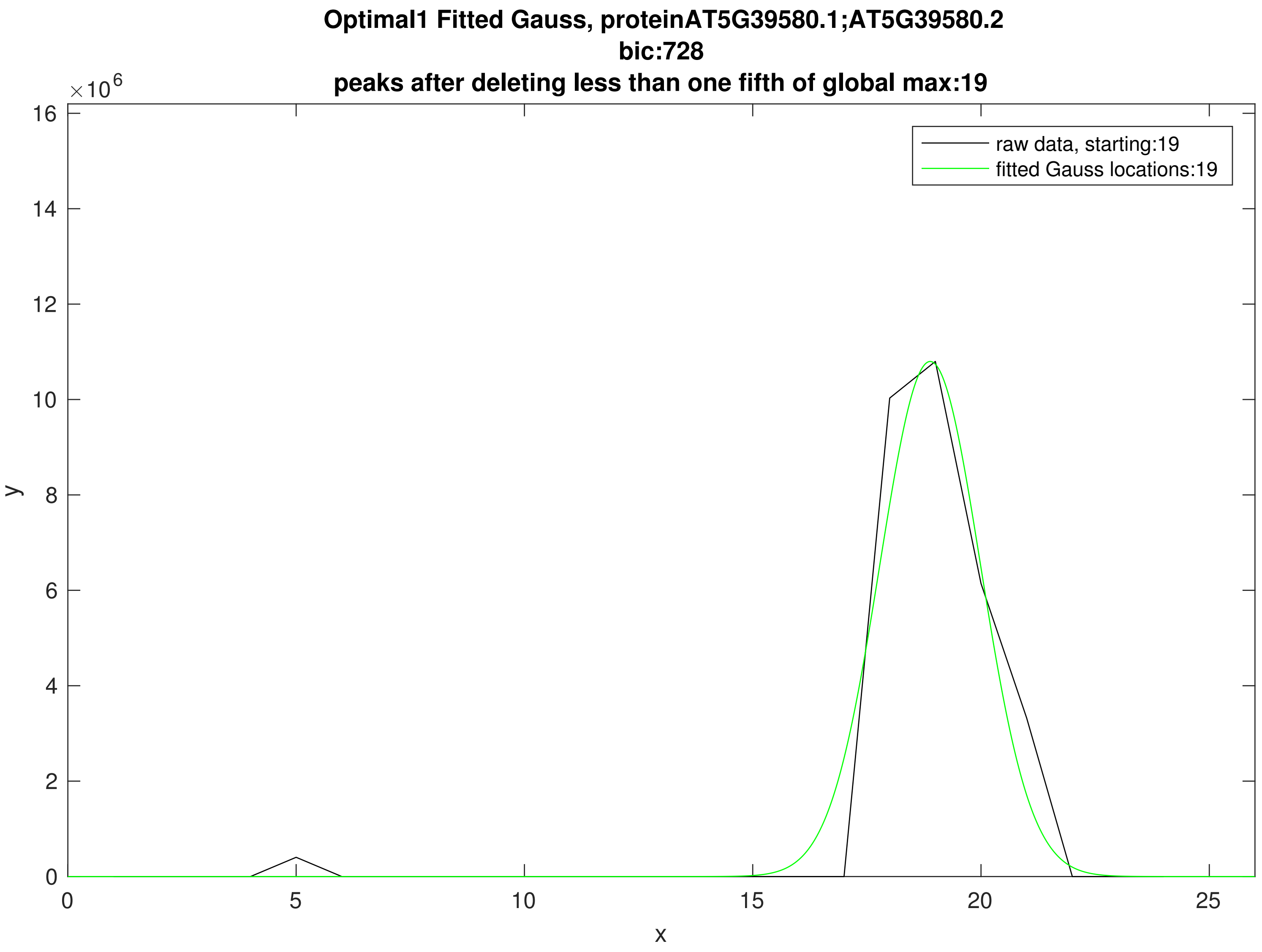

Optimal1 Fitted Gauss, proteinAT5G39580.1;AT5G39580.2
bic:728
peaks after deleting less than one fifth of global max:19
×10^6
16
14
12
10
8
6
4
2
0
y
0
5
10
15
20
25
x
raw data, starting:19
fitted Gauss locations:19

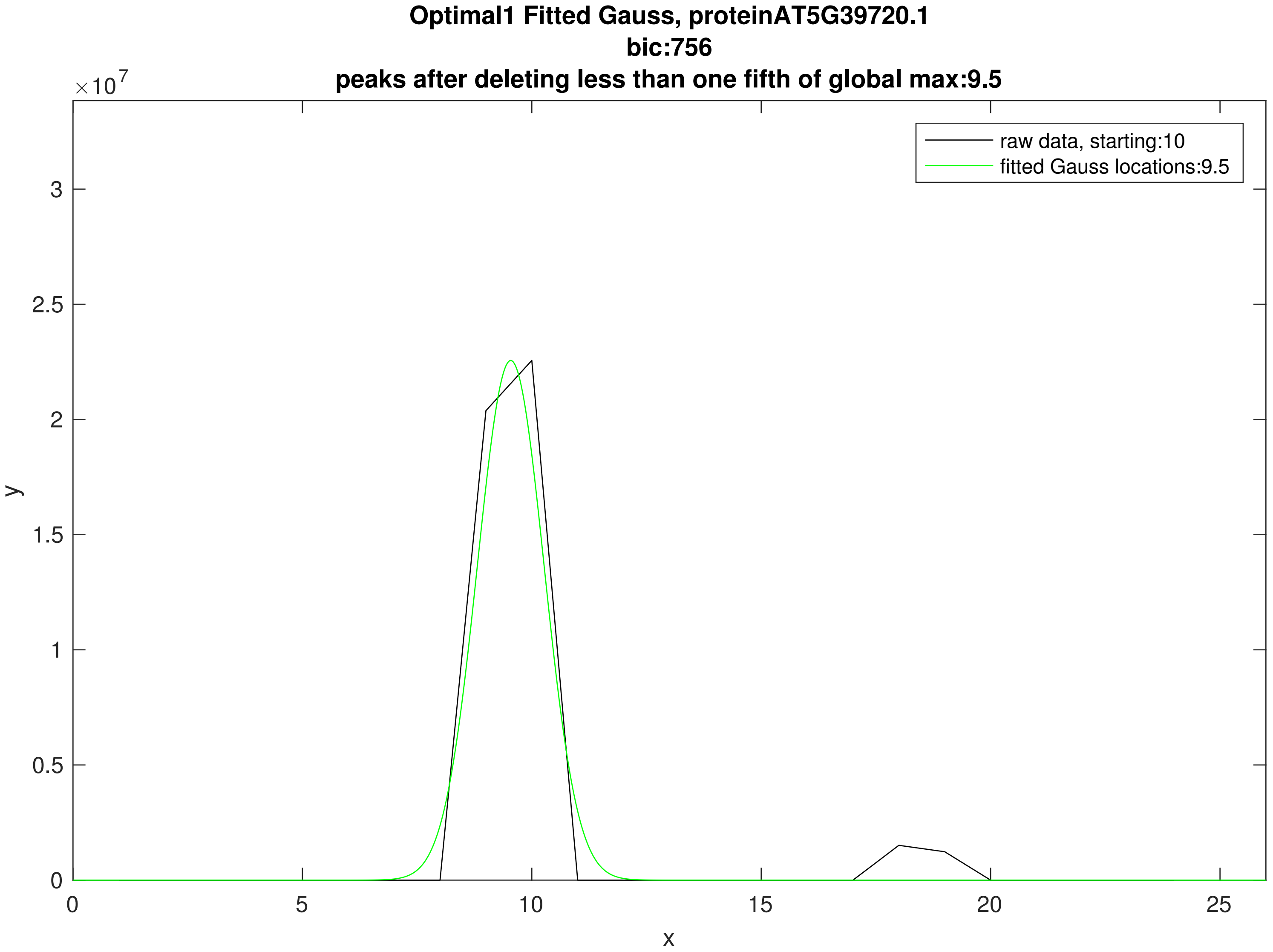
Optimal1 Fitted Gauss, proteinAT5G39720.1
bic:756
peaks after deleting less than one fifth of global max:9.5
×10^7
raw data, starting:10
fitted Gauss locations:9.5
y
x
0
0.5
1
1.5
2
2.5
3
0
5
10
15
20
25

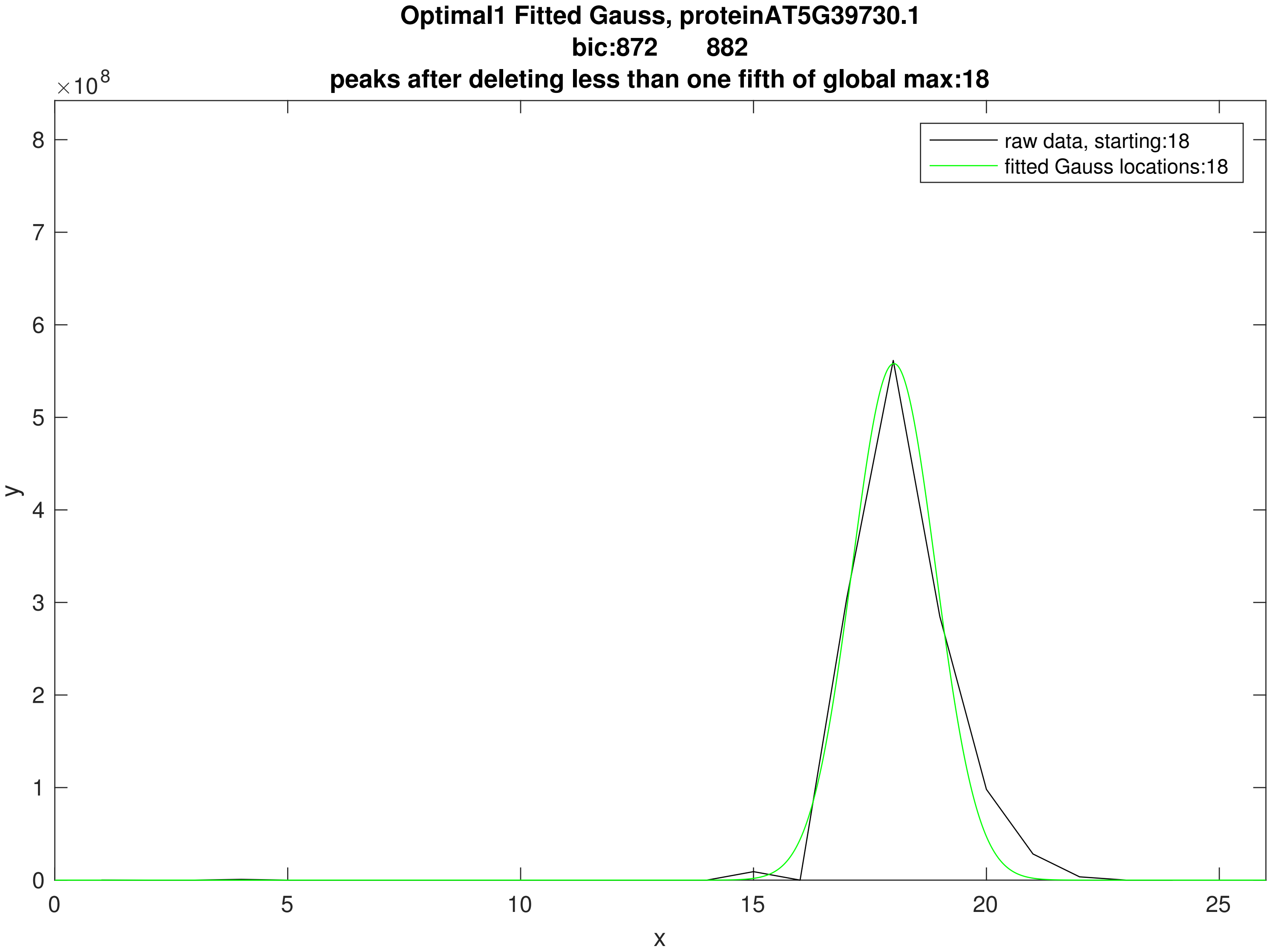
Optimal1 Fitted Gauss, proteinAT5G39730.1
bic:872 882
peaks after deleting less than one fifth of global max:18
×10^8
raw data, starting:18
fitted Gauss locations:18
y
x
0
1
2
3
4
5
6
7
8
0
5
10
15
20
25

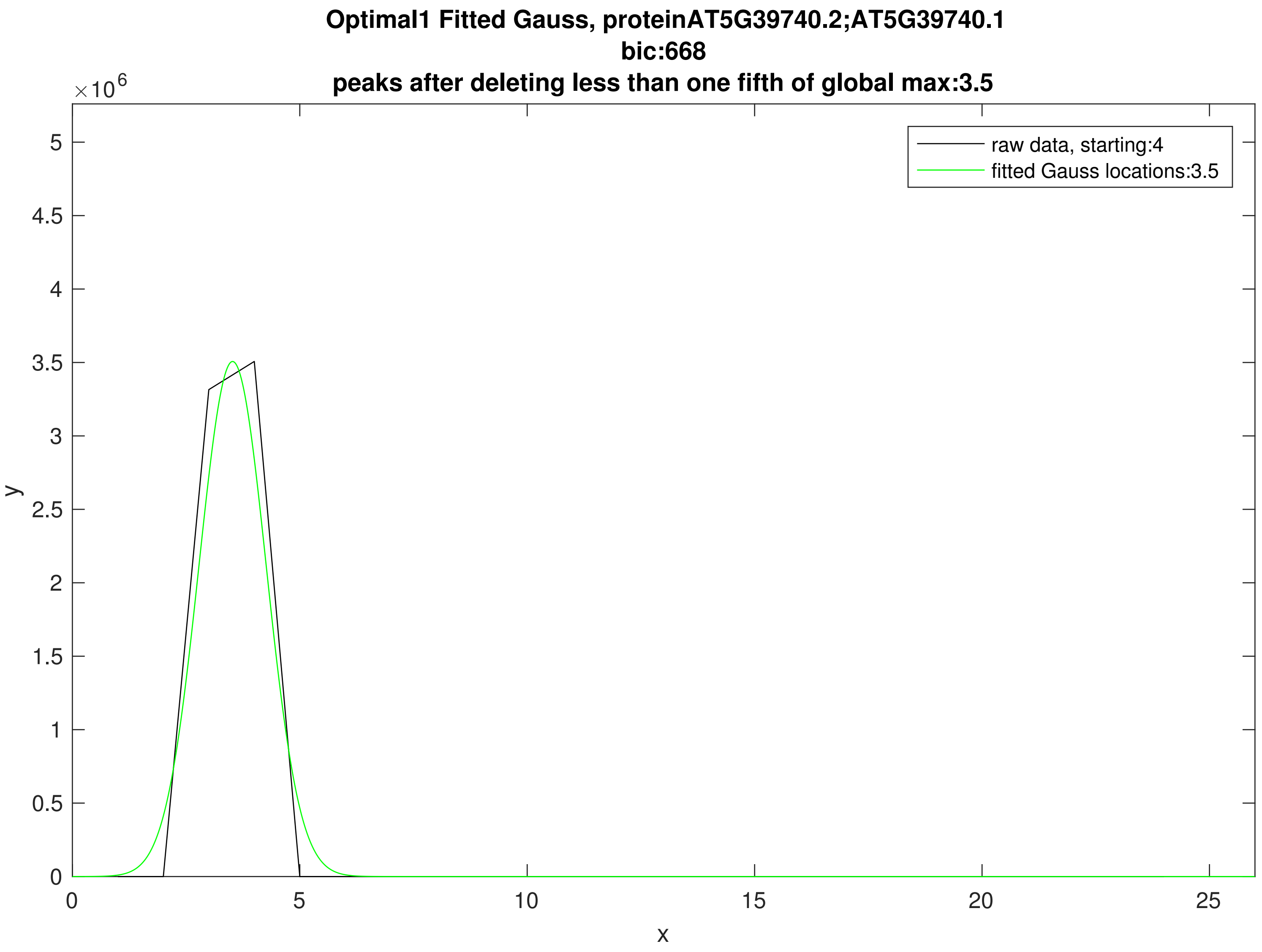
Optimal1 Fitted Gauss, proteinAT5G39740.2;AT5G39740.1
bic:668
peaks after deleting less than one fifth of global max:3.5
×10^6
raw data, starting:4
fitted Gauss locations:3.5
y
x
0
0.5
1
1.5
2
2.5
3
3.5
4
4.5
5
0
5
10
15
20
25

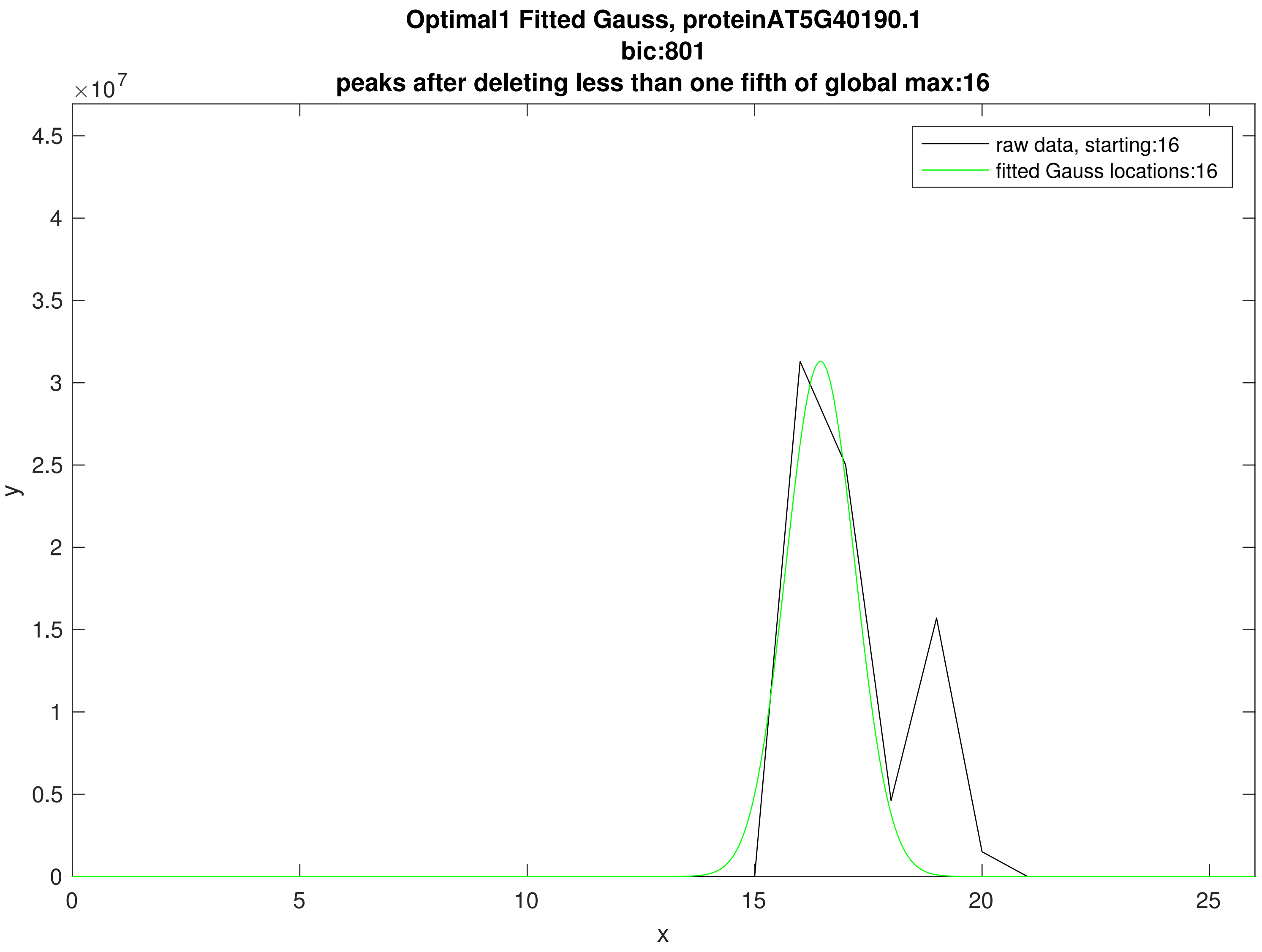
Optimal1 Fitted Gauss, proteinAT5G40190.1
bic:801
peaks after deleting less than one fifth of global max:16
×10^7
4.5
4
3.5
3
2.5
2
1.5
1
0.5
0
y
0
5
10
15
20
25
x
raw data, starting:16
fitted Gauss locations:16

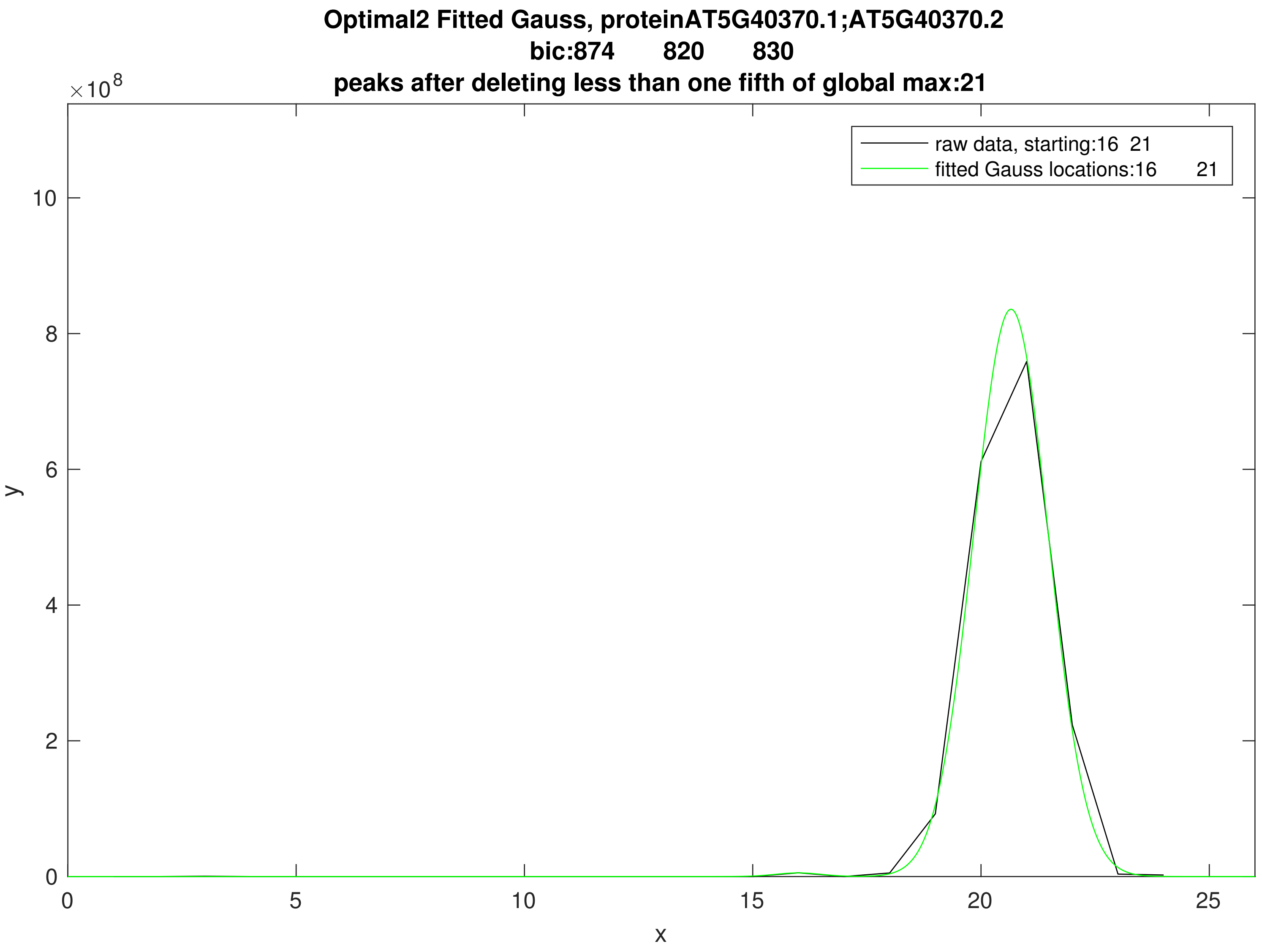
Optimal2 Fitted Gauss, proteinAT5G40370.1;AT5G40370.2
bic:874 820 830
peaks after deleting less than one fifth of global max:21
×10^8
raw data, starting:16 21
fitted Gauss locations:16 21
y
x
0
2
4
6
8
10
0
5
10
15
20
25

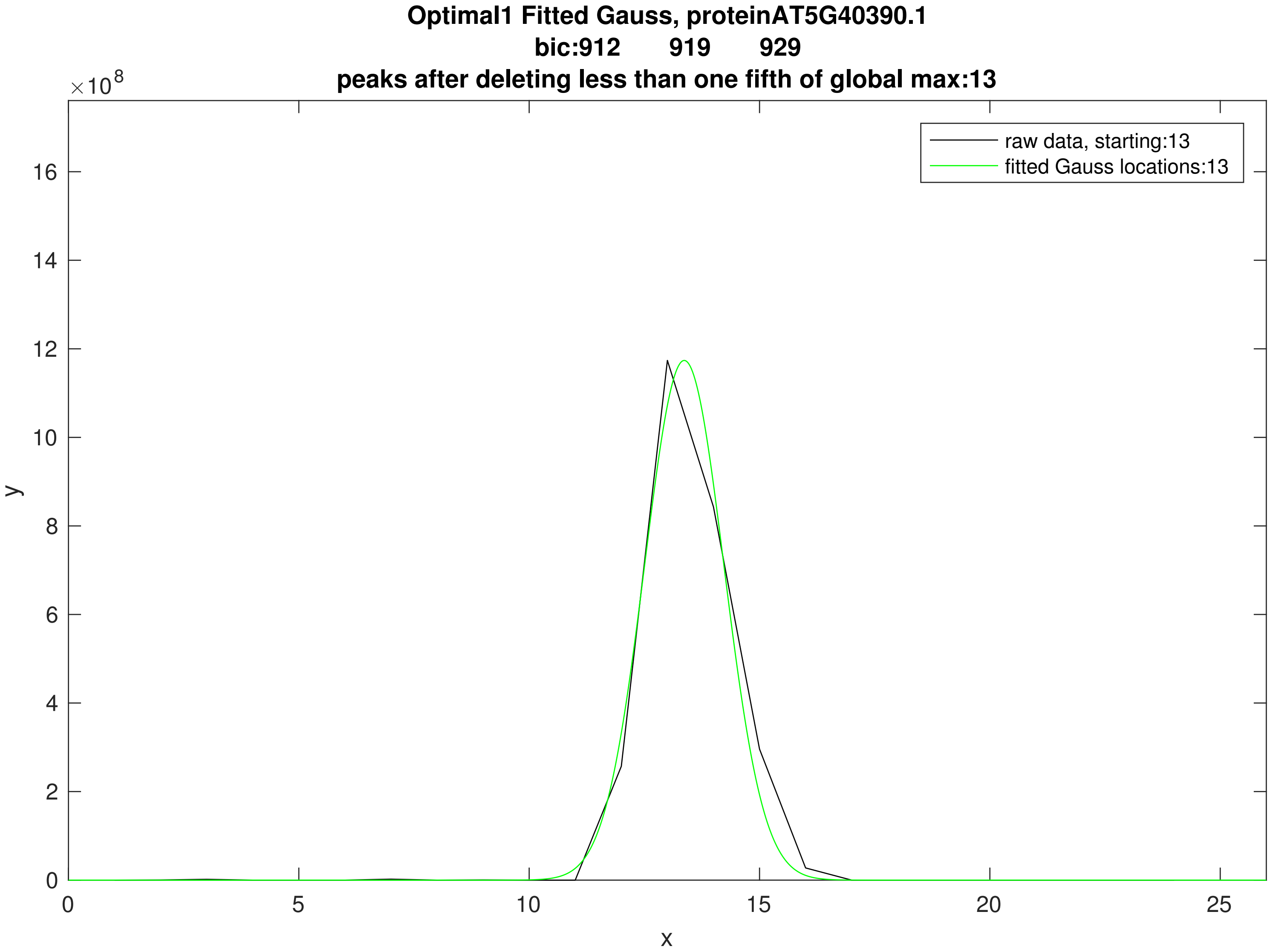
Optimal1 Fitted Gauss, proteinAT5G40390.1
bic:912 919 929
peaks after deleting less than one fifth of global max:13
×10^8
raw data, starting:13
fitted Gauss locations:13
16
14
12
10
8
6
4
2
0
y
0
5
10
15
20
25
x

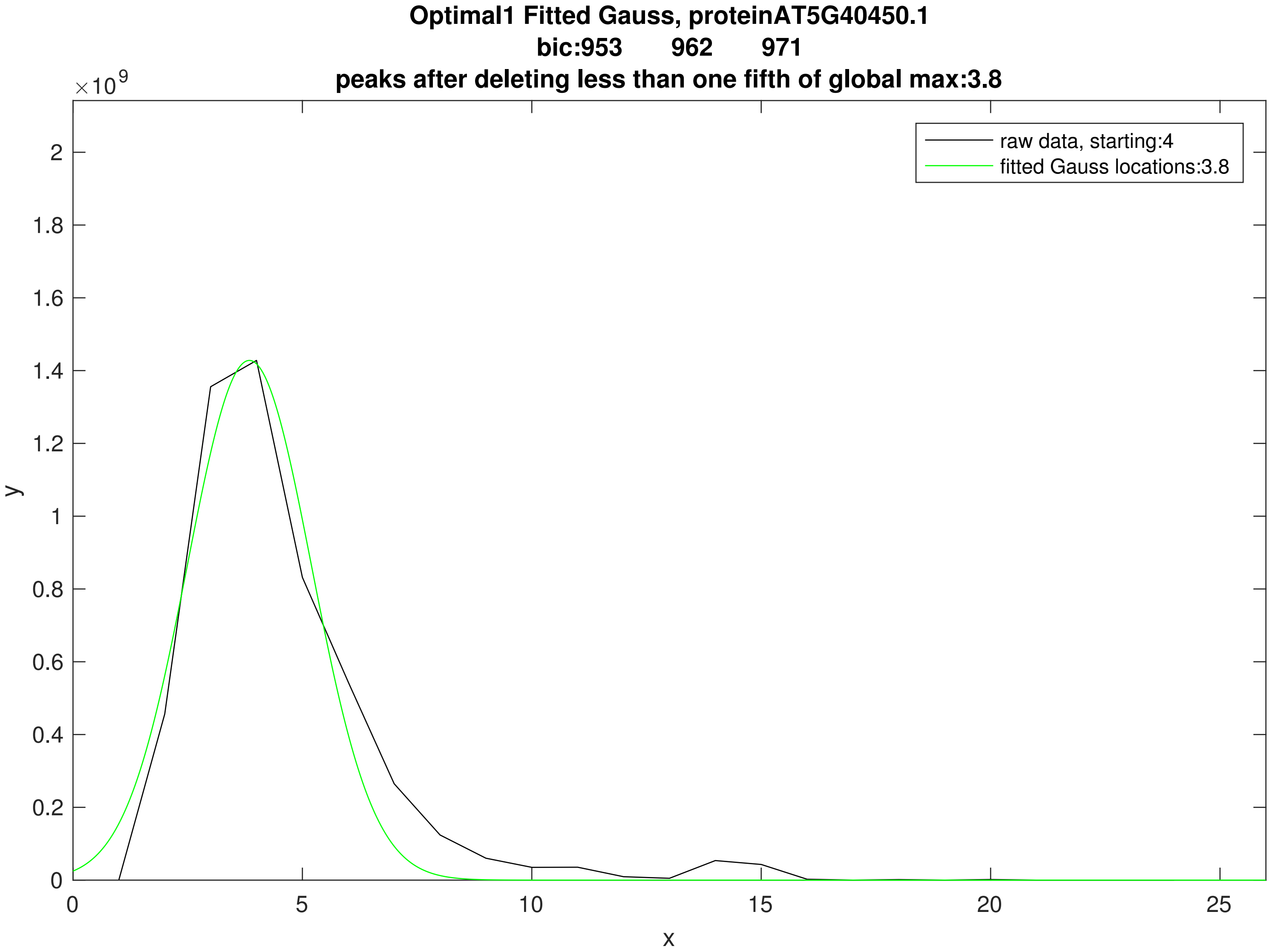
Optimal1 Fitted Gauss, proteinAT5G40450.1
bic:953 962 971
peaks after deleting less than one fifth of global max:3.8
×10^9
raw data, starting:4
fitted Gauss locations:3.8
y
x
0
0.2
0.4
0.6
0.8
1
1.2
1.4
1.6
1.8
2
0
5
10
15
20
25

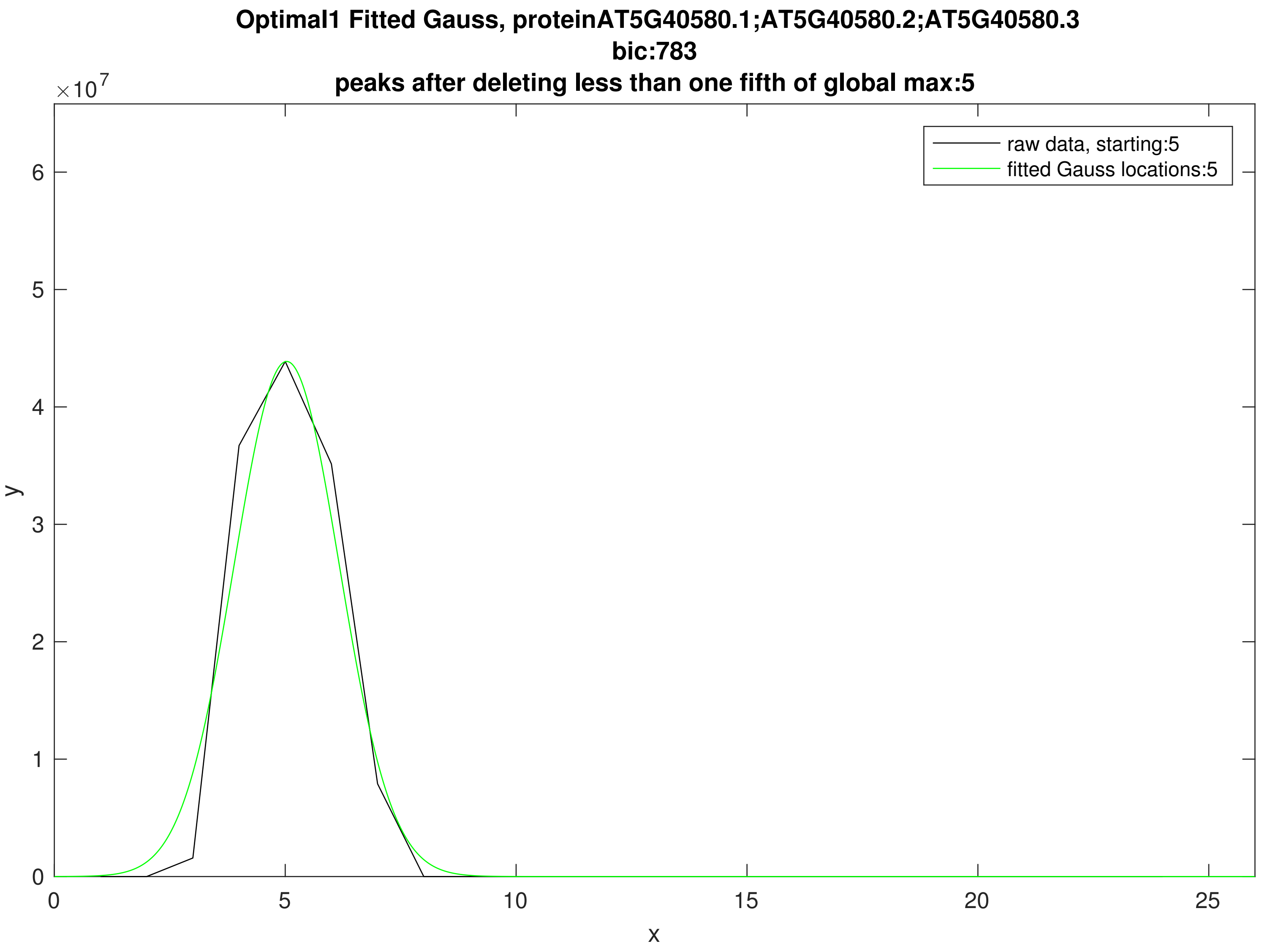
Optimal1 Fitted Gauss, proteinAT5G40580.1;AT5G40580.2;AT5G40580.3
bic:783
peaks after deleting less than one fifth of global max:5
×10^7
raw data, starting:5
fitted Gauss locations:5
y
x

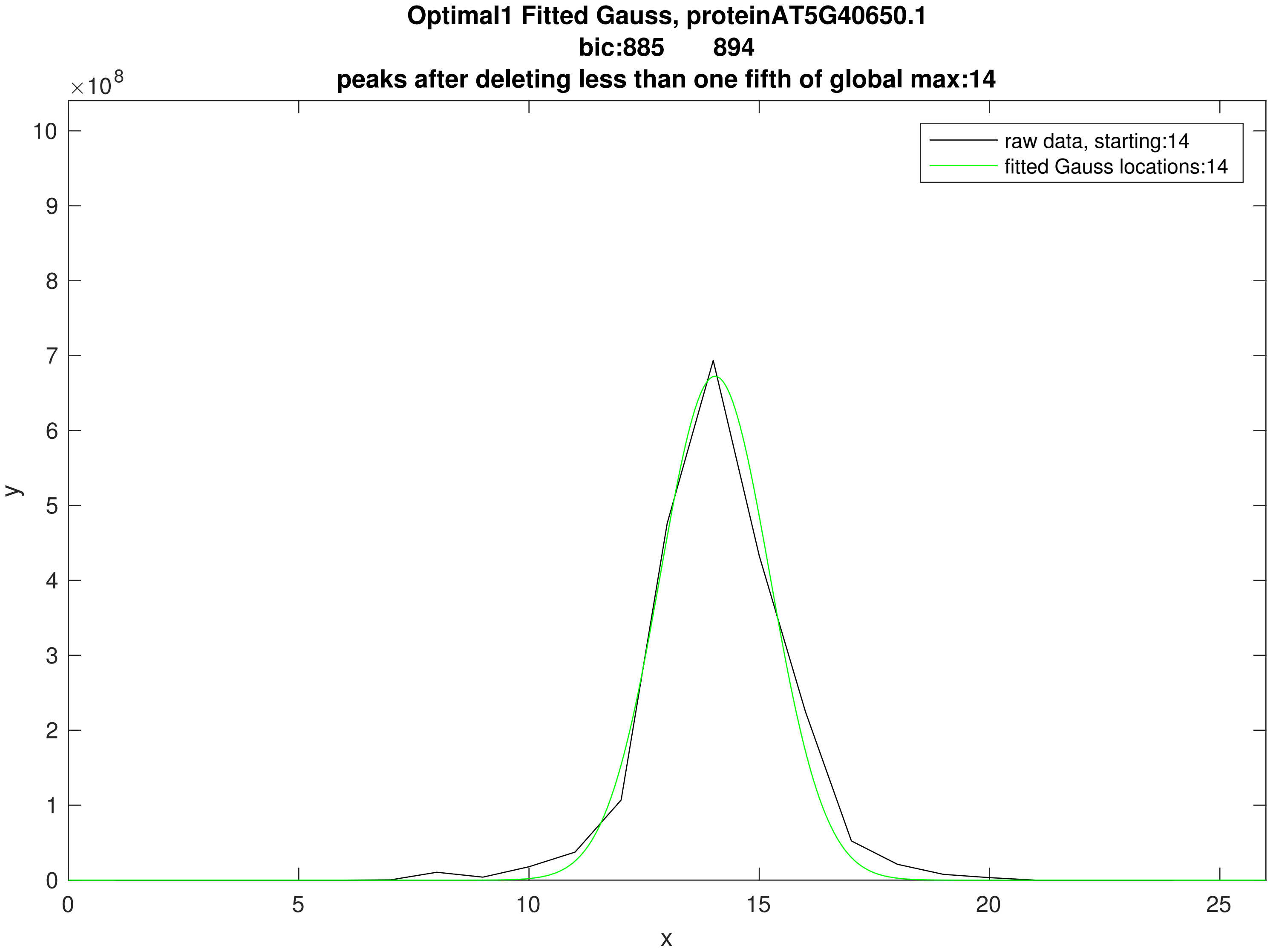

Optimal1 Fitted Gauss, proteinAT5G40650.1
bic:885 894
peaks after deleting less than one fifth of global max:14
×10^8
raw data, starting:14
fitted Gauss locations:14
y
x
0
5
10
15
20
25
1
2
3
4
6
7
8
9

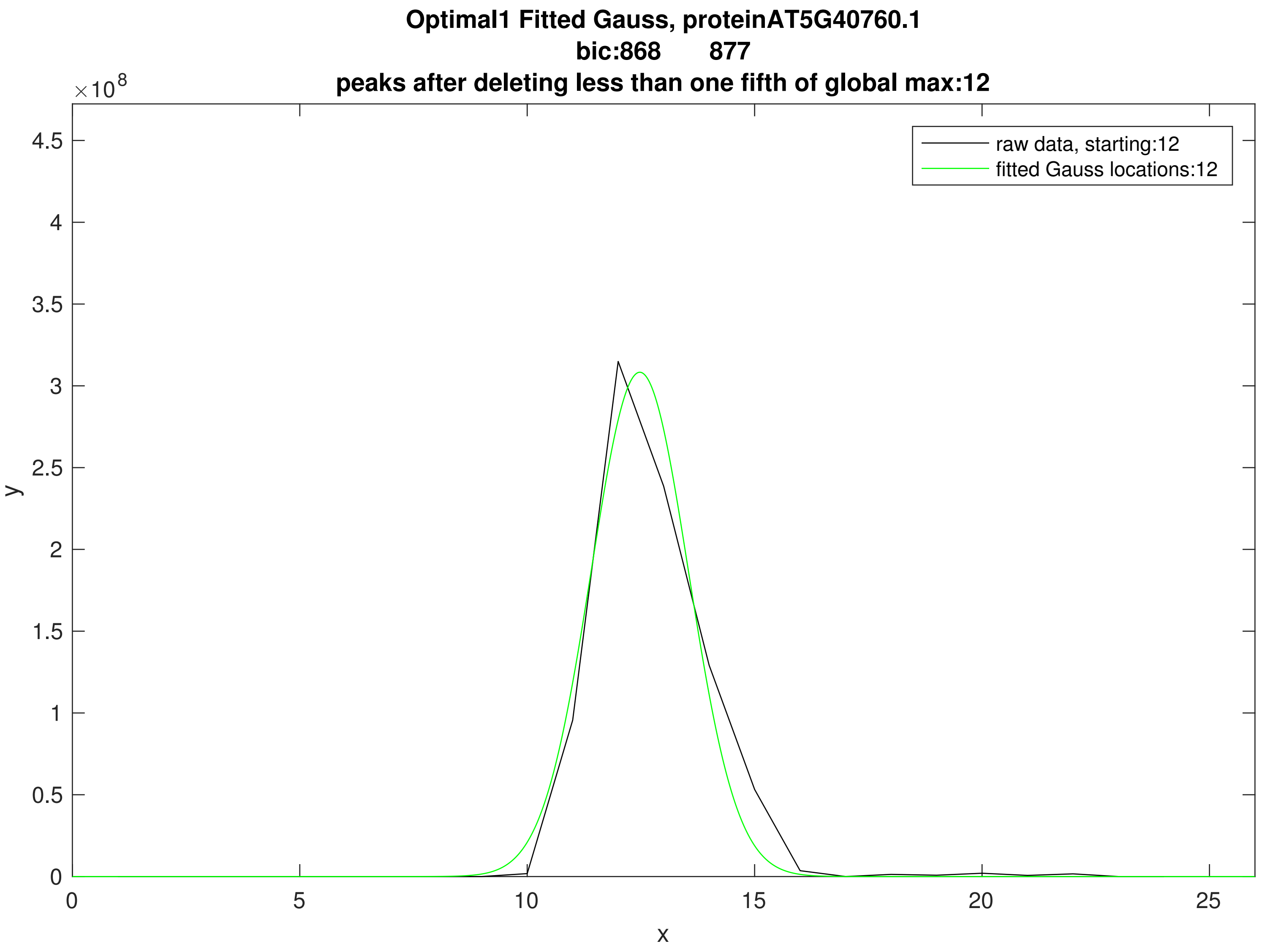
Optimal1 Fitted Gauss, proteinAT5G40760.1
bic:868 877
peaks after deleting less than one fifth of global max:12
×10^8
4.5
4
3.5
3
2.5
2
1.5
1
0.5
0
y
0
5
10
15
20
25
x
raw data, starting:12
fitted Gauss locations:12

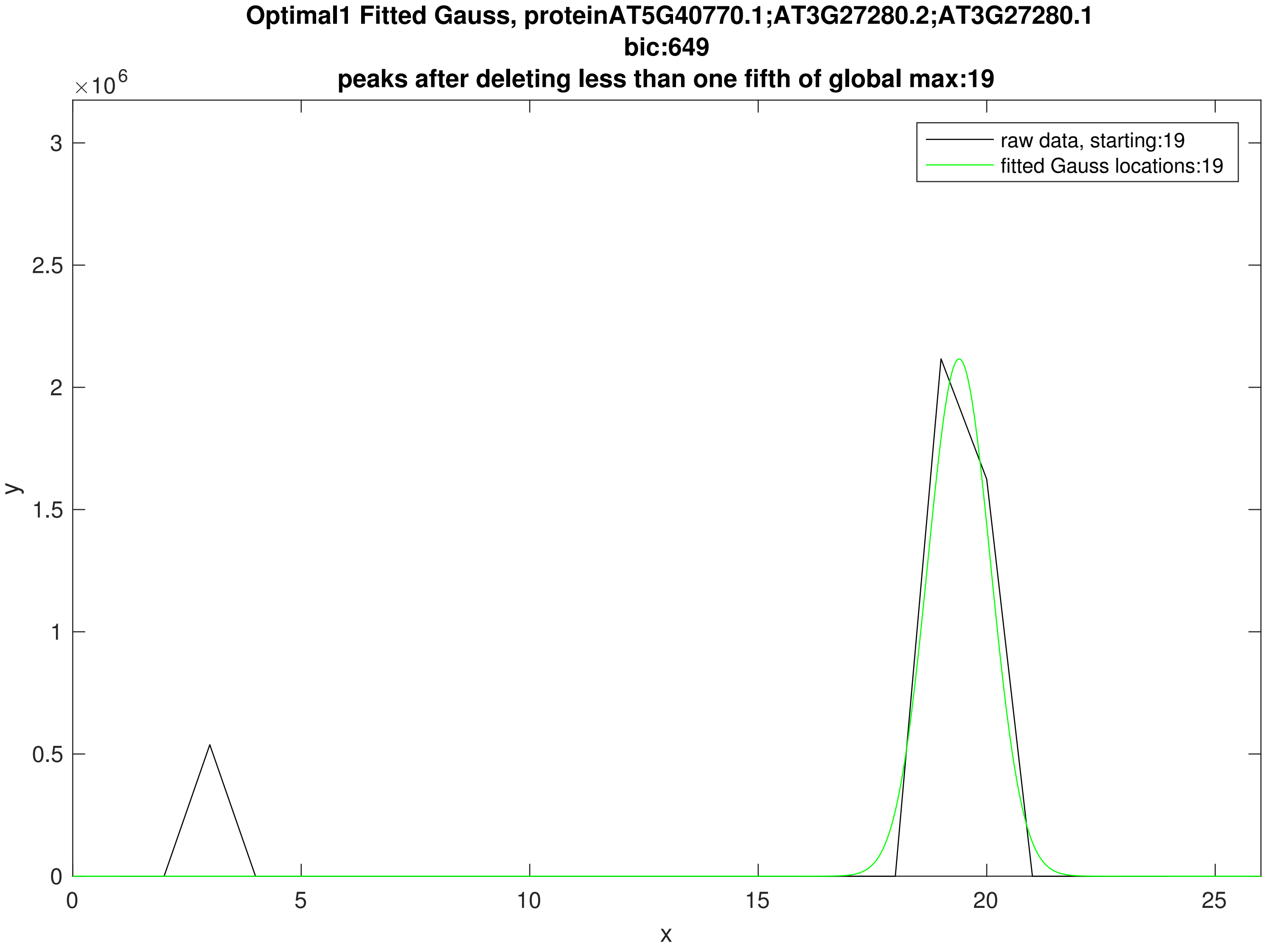
Optimal1 Fitted Gauss, proteinAT5G40770.1;AT3G27280.2;AT3G27280.1
bic:649
peaks after deleting less than one fifth of global max:19
×10^6
raw data, starting:19
fitted Gauss locations:19
y
x
0
0.5
1
1.5
2
2.5
3
0
5
10
15
20
25

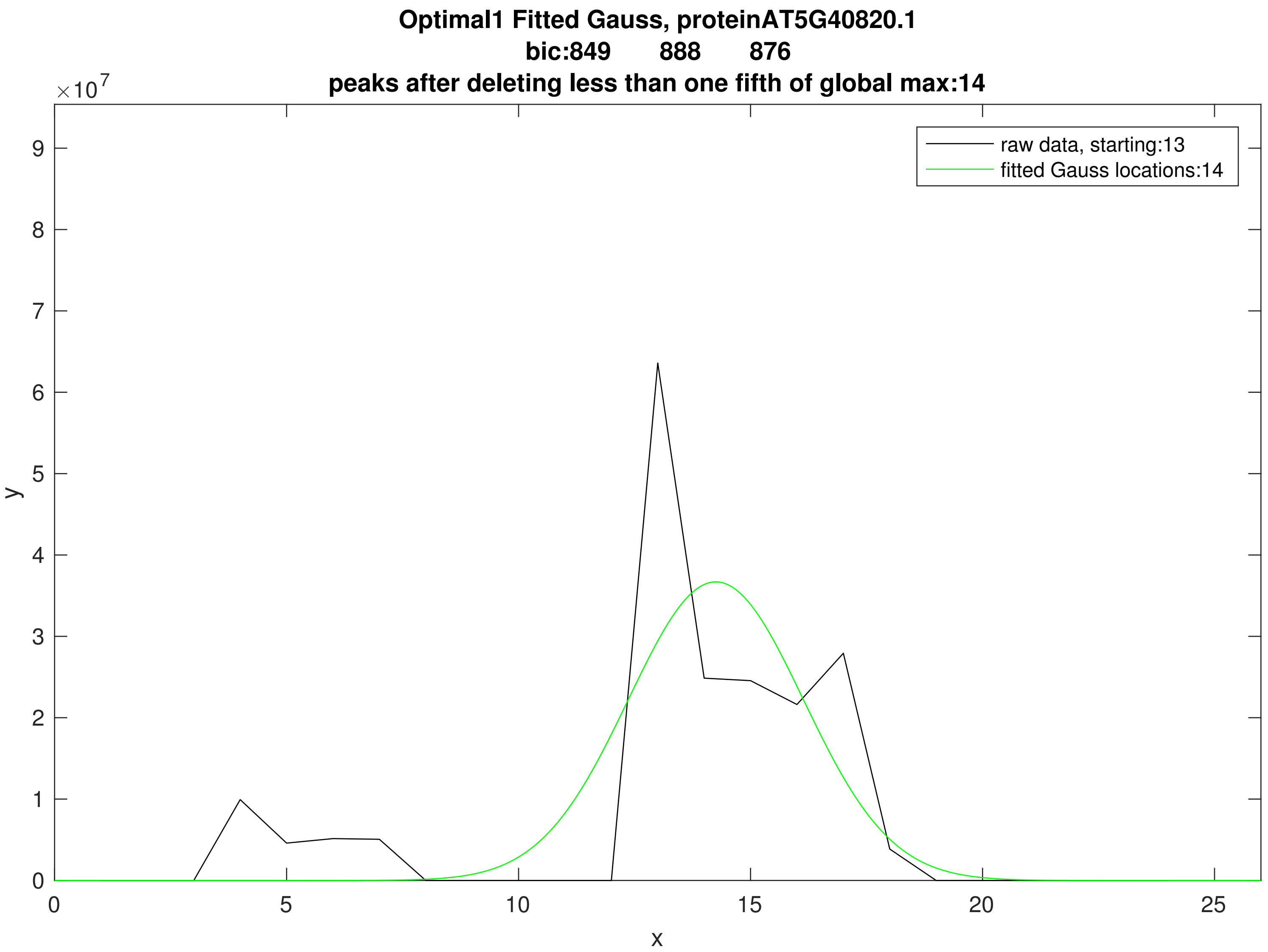
Optimal1 Fitted Gauss, proteinAT5G40820.1
bic:849 888 876
peaks after deleting less than one fifth of global max:14
×10^7
raw data, starting:13
fitted Gauss locations:14
y
x
0
1
2
3
4
5
6
7
8
9
10
15
20
25

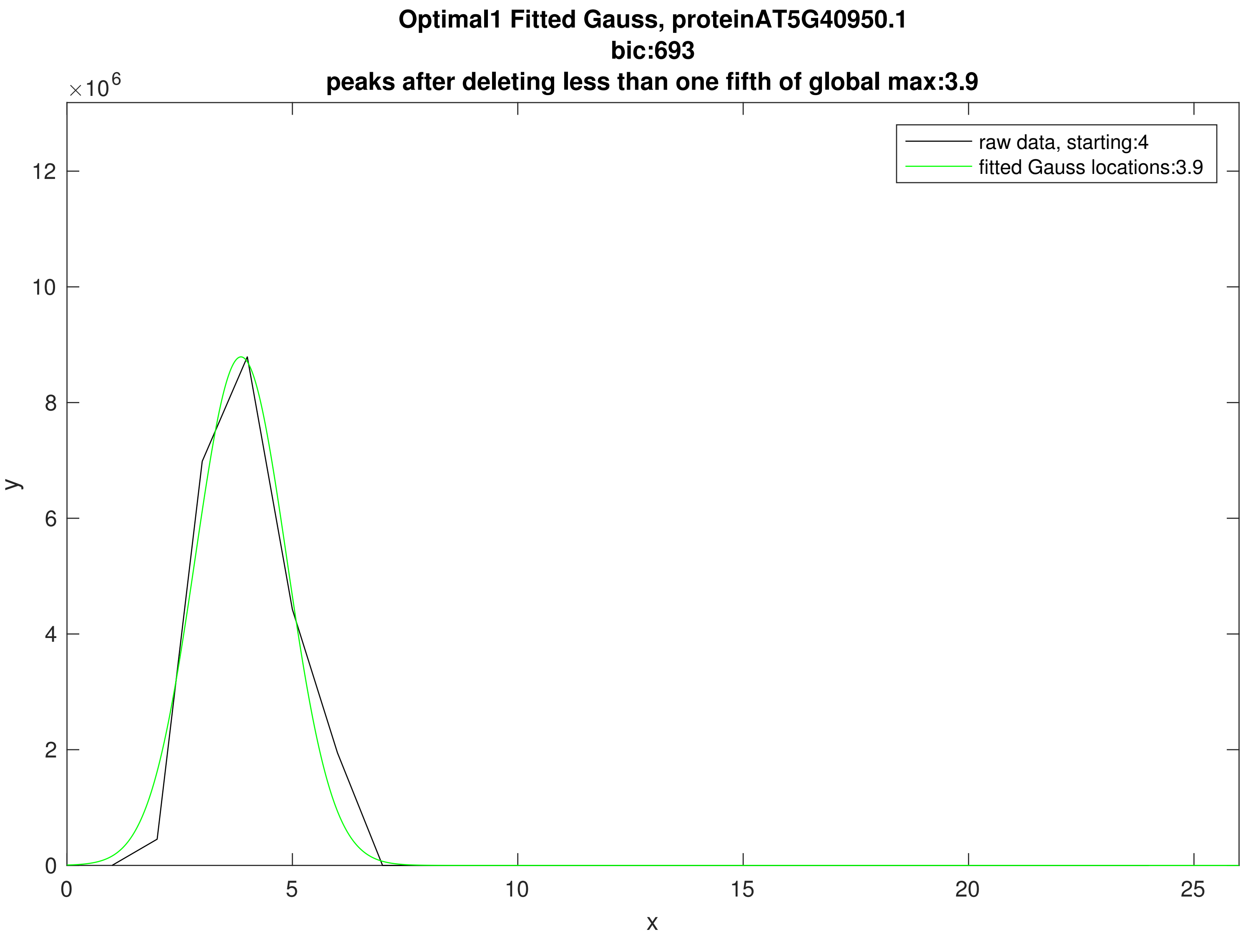

Optimal1 Fitted Gauss, proteinAT5G40950.1
bic:693
peaks after deleting less than one fifth of global max:3.9
×10^6
raw data, starting:4
fitted Gauss locations:3.9
y
x
0
2
4
6
8
10
12
0
5
10
15
20
25

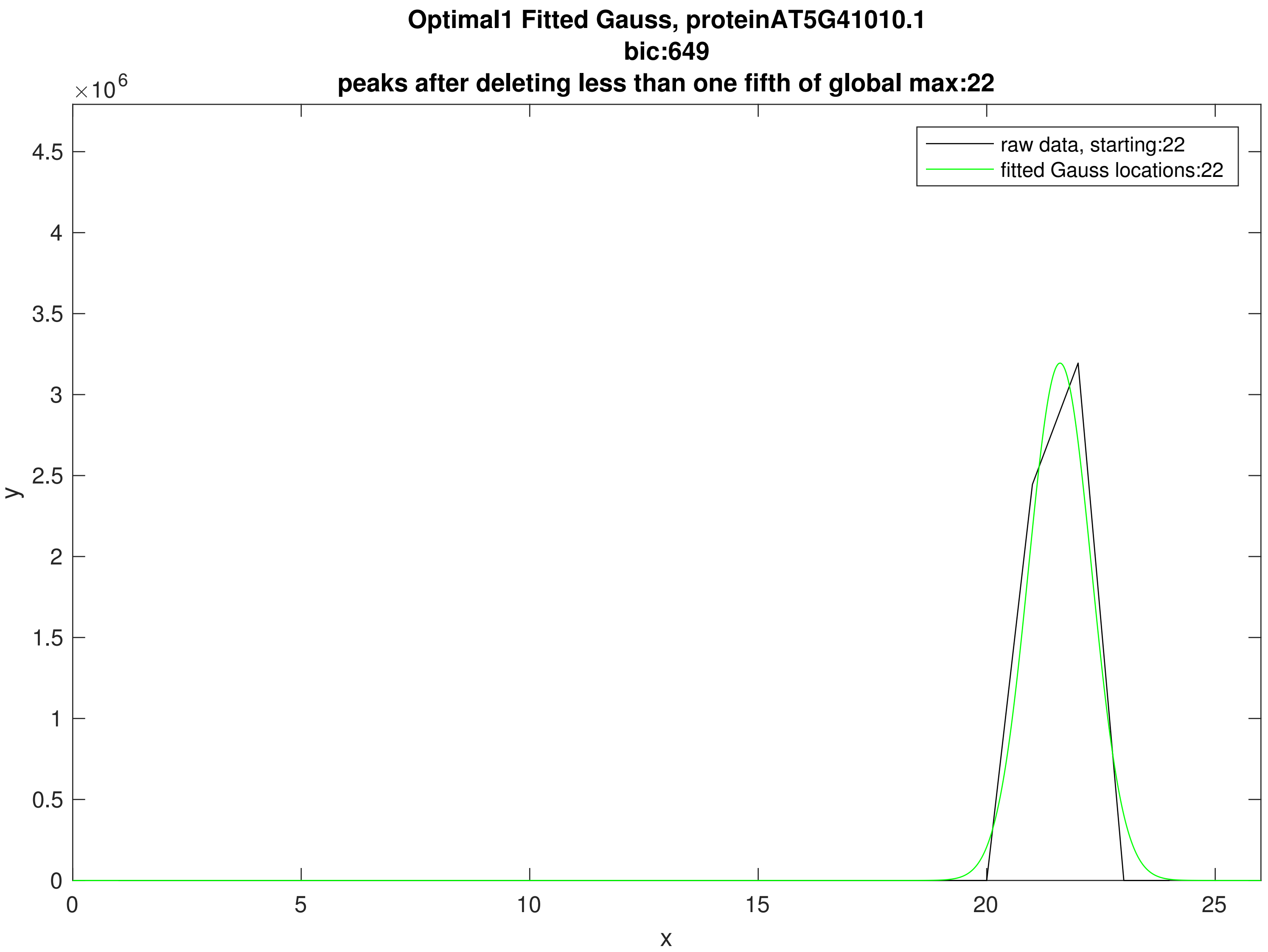
Optimal1 Fitted Gauss, proteinAT5G41010.1
bic:649
peaks after deleting less than one fifth of global max:22
×10^6
raw data, starting:22
fitted Gauss locations:22
y
x
0
0.5
1
1.5
2
2.5
3
3.5
4
4.5
0
5
10
15
20
25

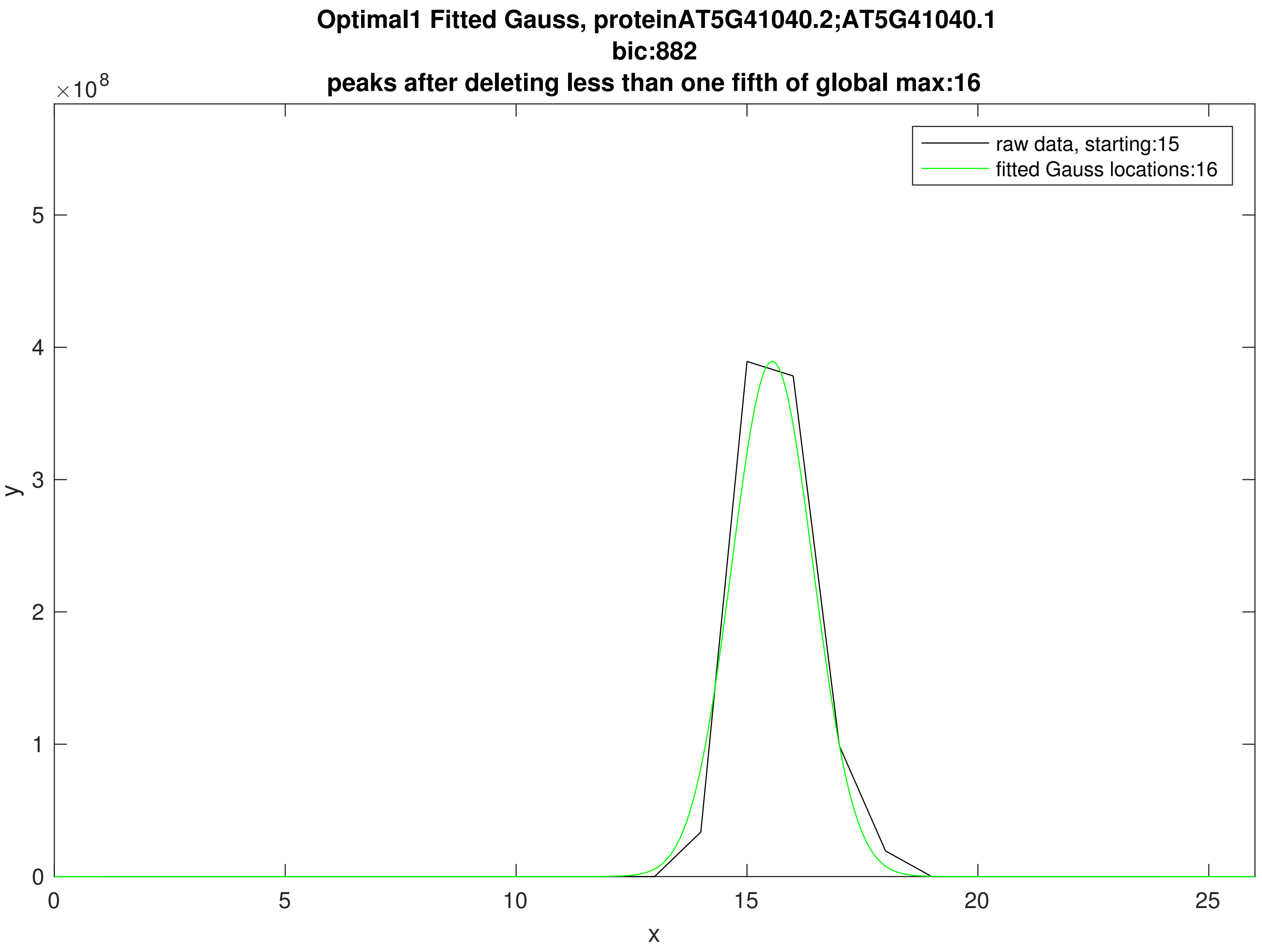

Optimal1 Fitted Gauss, proteinAT5G41040.2;AT5G41040.1
bic:882
peaks after deleting less than one fifth of global max:16
×10^8
raw data, starting:15
fitted Gauss locations:16
y
x
0
1
2
3
4
5
0
5
10
15
20
25

Optimal1 Fitted Gauss, proteinAT5G41520.1;AT5G41520.2
bic:760
peaks after deleting less than one fifth of global max:3.7

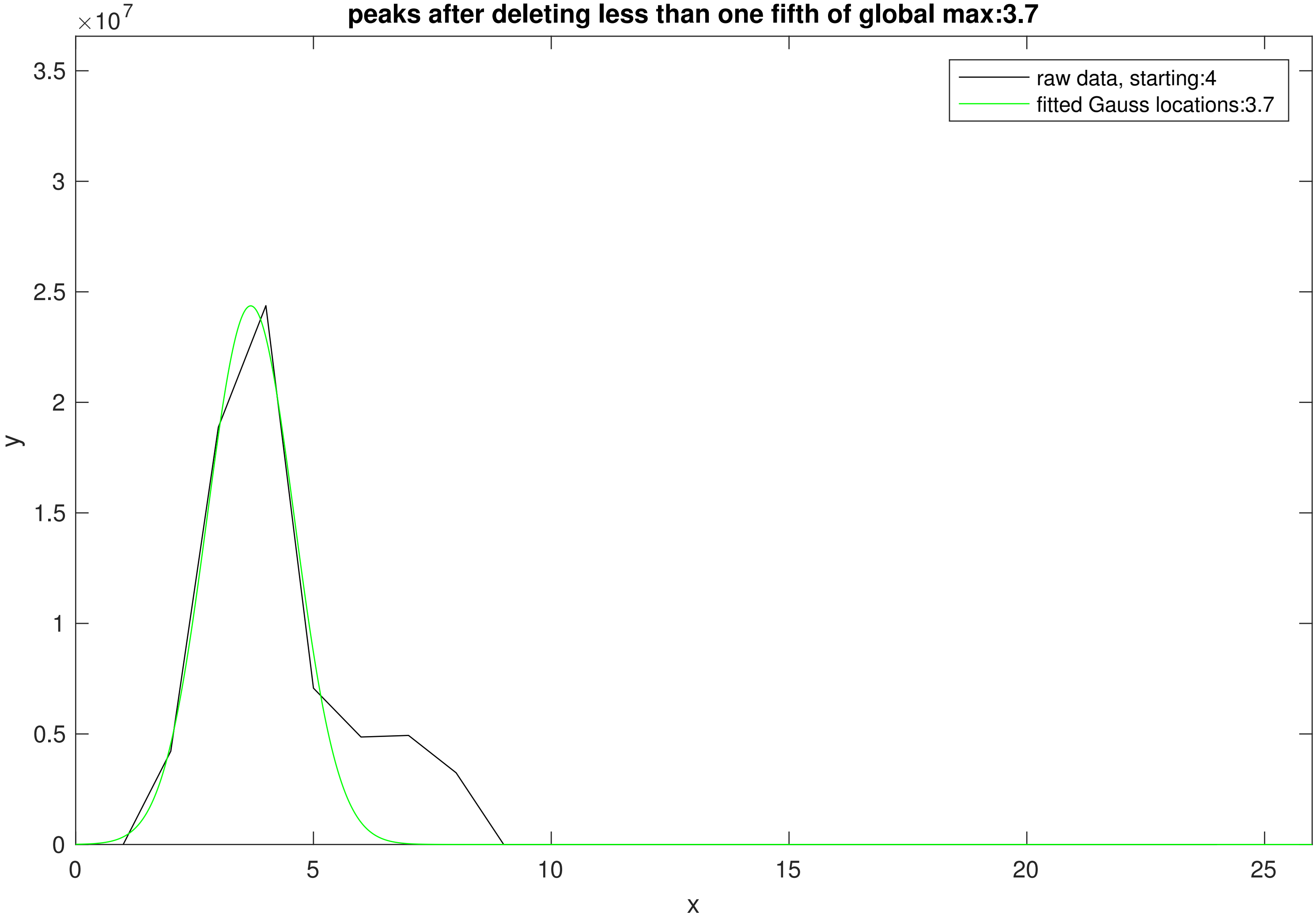

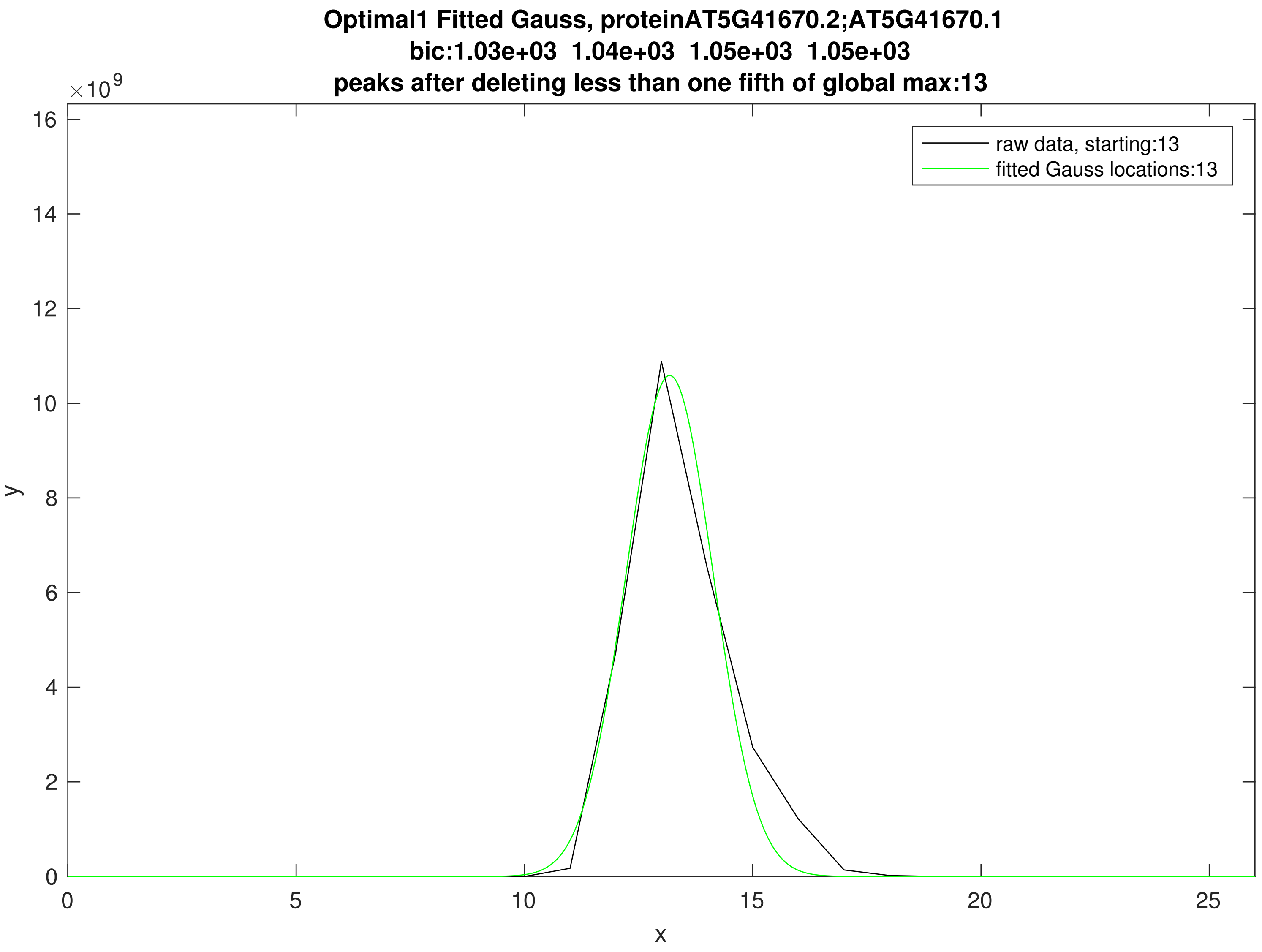
Optimal1 Fitted Gauss, proteinAT5G41670.2;AT5G41670.1
bic:1.03e+03 1.04e+03 1.05e+03 1.05e+03
peaks after deleting less than one fifth of global max:13
×10^9
16
14
12
10
8
6
4
2
0
y
0
5
10
15
20
25
x
raw data, starting:13
fitted Gauss locations:13

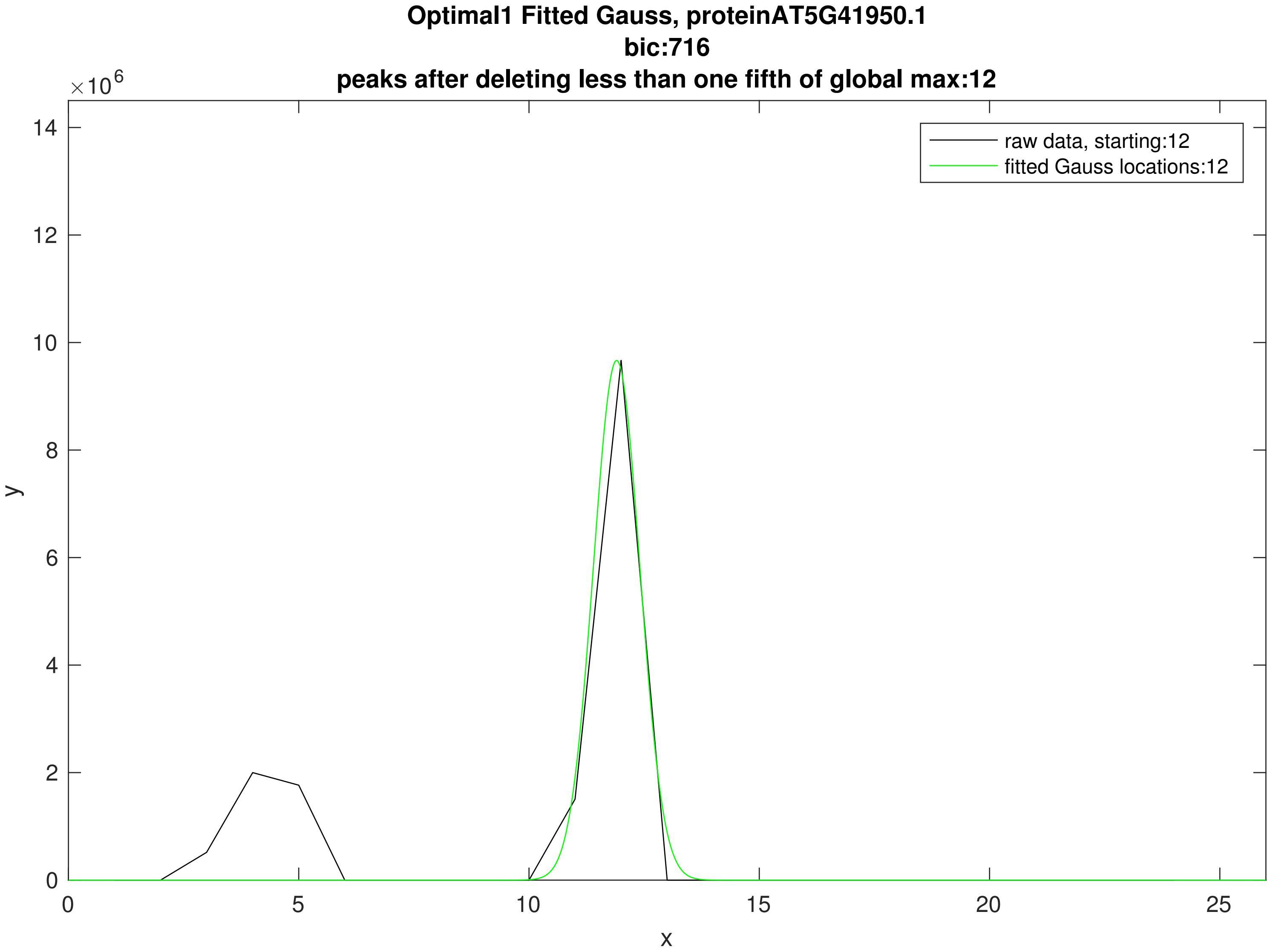
Optimal1 Fitted Gauss, proteinAT5G41950.1
bic:716
peaks after deleting less than one fifth of global max:12
×10^6
raw data, starting:12
fitted Gauss locations:12
y
x
0
2
4
6
8
10
12
14
0
5
10
15
20
25

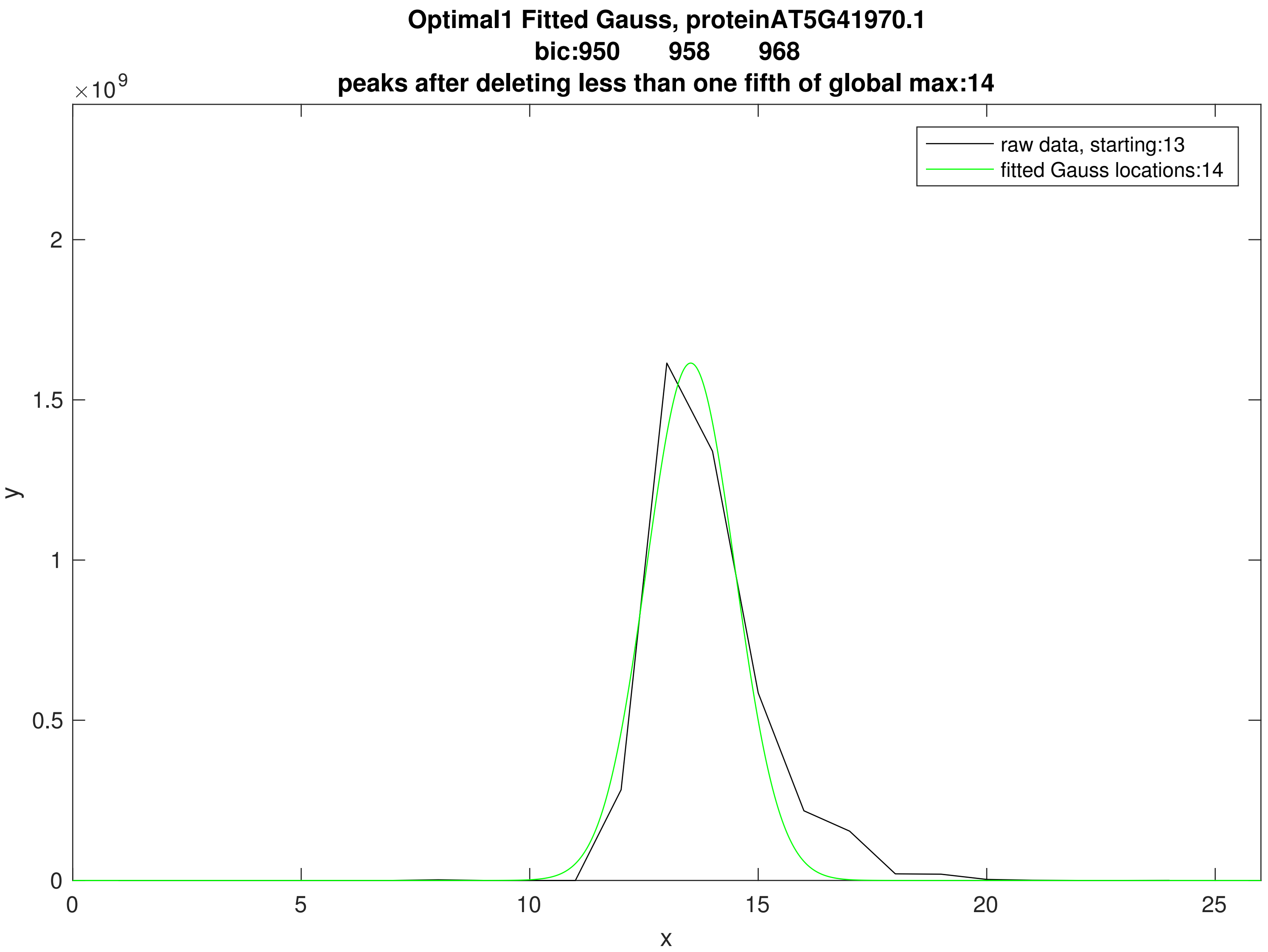

Optimal1 Fitted Gauss, proteinAT5G41970.1
bic:950 958 968
peaks after deleting less than one fifth of global max:14
×10^9
raw data, starting:13
fitted Gauss locations:14
y
x
0
0.5
1
1.5
2
5
10
15
20
25

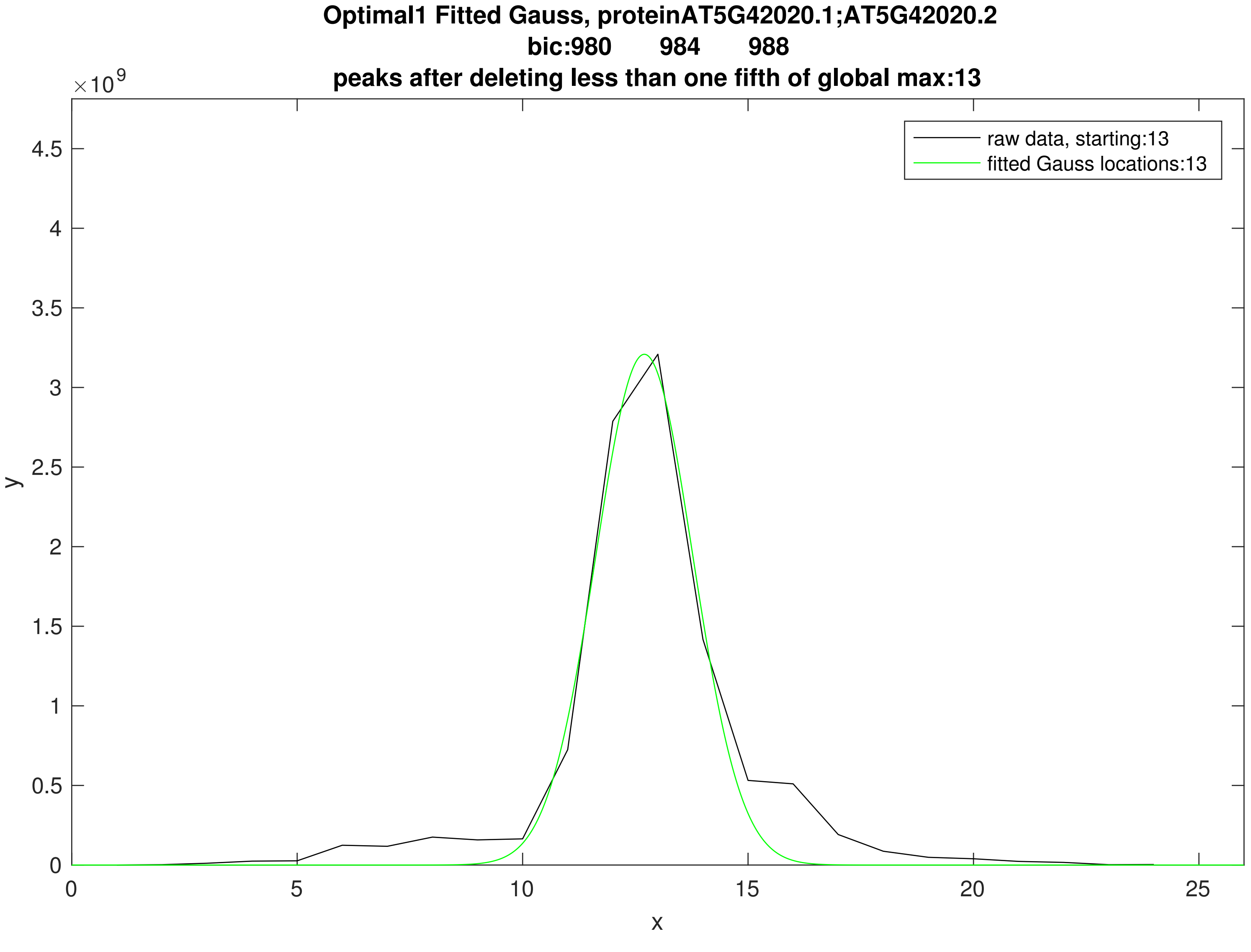
Optimal1 Fitted Gauss, proteinAT5G42020.1;AT5G42020.2
bic:980 984 988
peaks after deleting less than one fifth of global max:13
raw data, starting:13
fitted Gauss locations:13
y
x

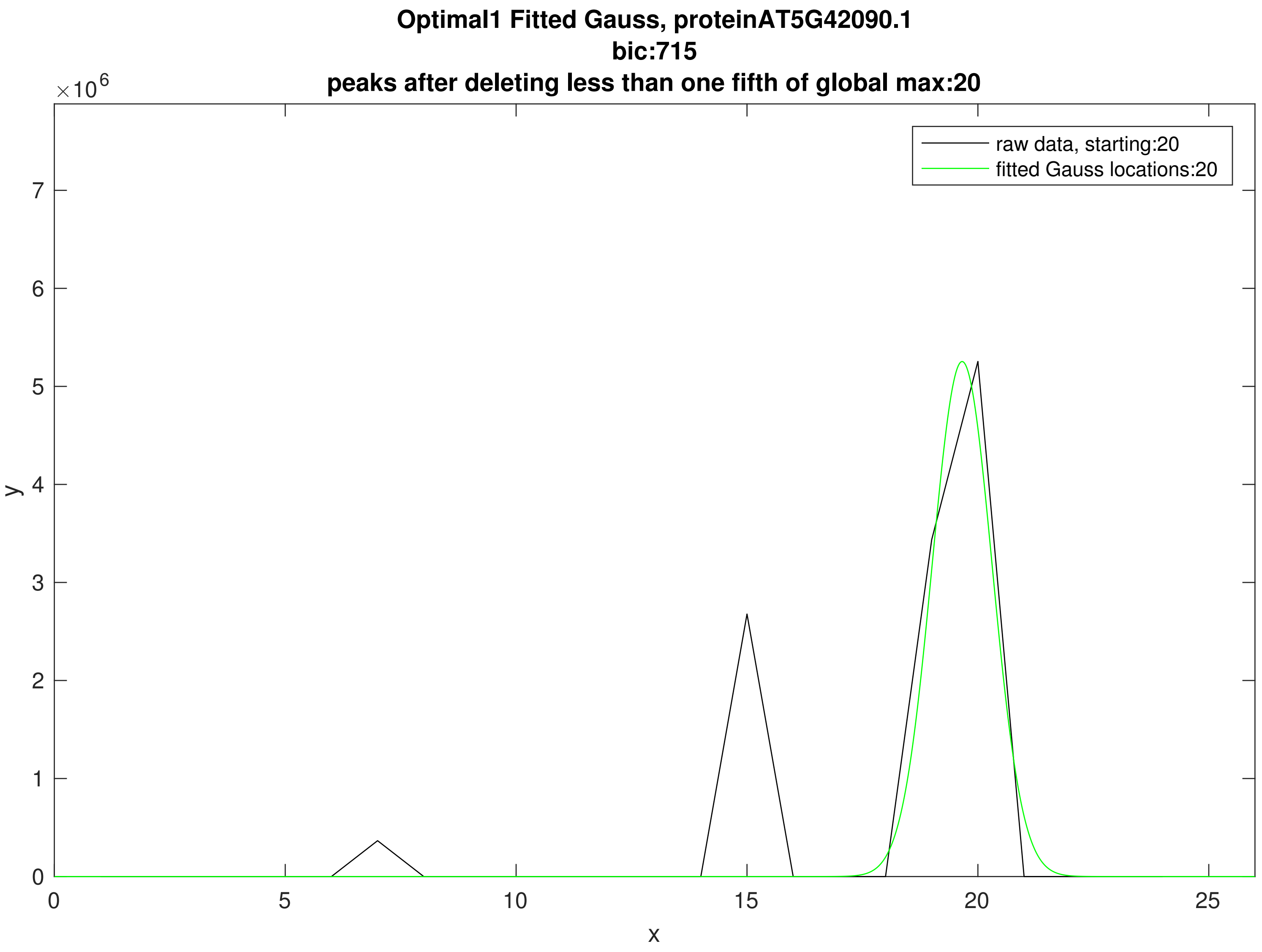
Optimal1 Fitted Gauss, proteinAT5G42090.1
bic:715
peaks after deleting less than one fifth of global max:20
×10^6
raw data, starting:20
fitted Gauss locations:20
y
x
0
1
2
3
4
5
6
7
0
5
10
15
20
25

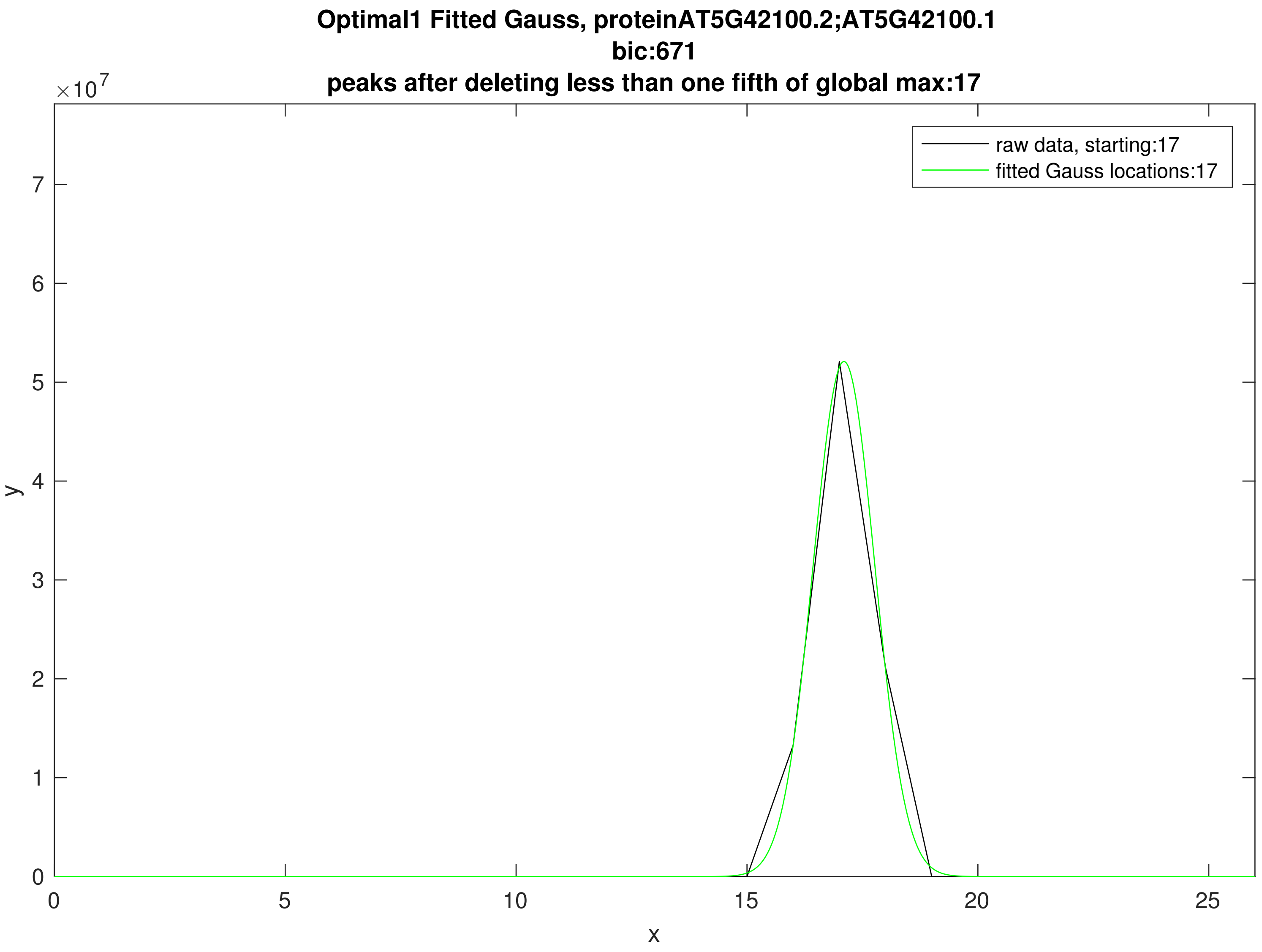

Optimal1 Fitted Gauss, proteinAT5G42100.2;AT5G42100.1
bic:671
peaks after deleting less than one fifth of global max:17
×10^7
raw data, starting:17
fitted Gauss locations:17
y
x
0
1
2
3
4
5
6
7
0
5
10
15
20
25

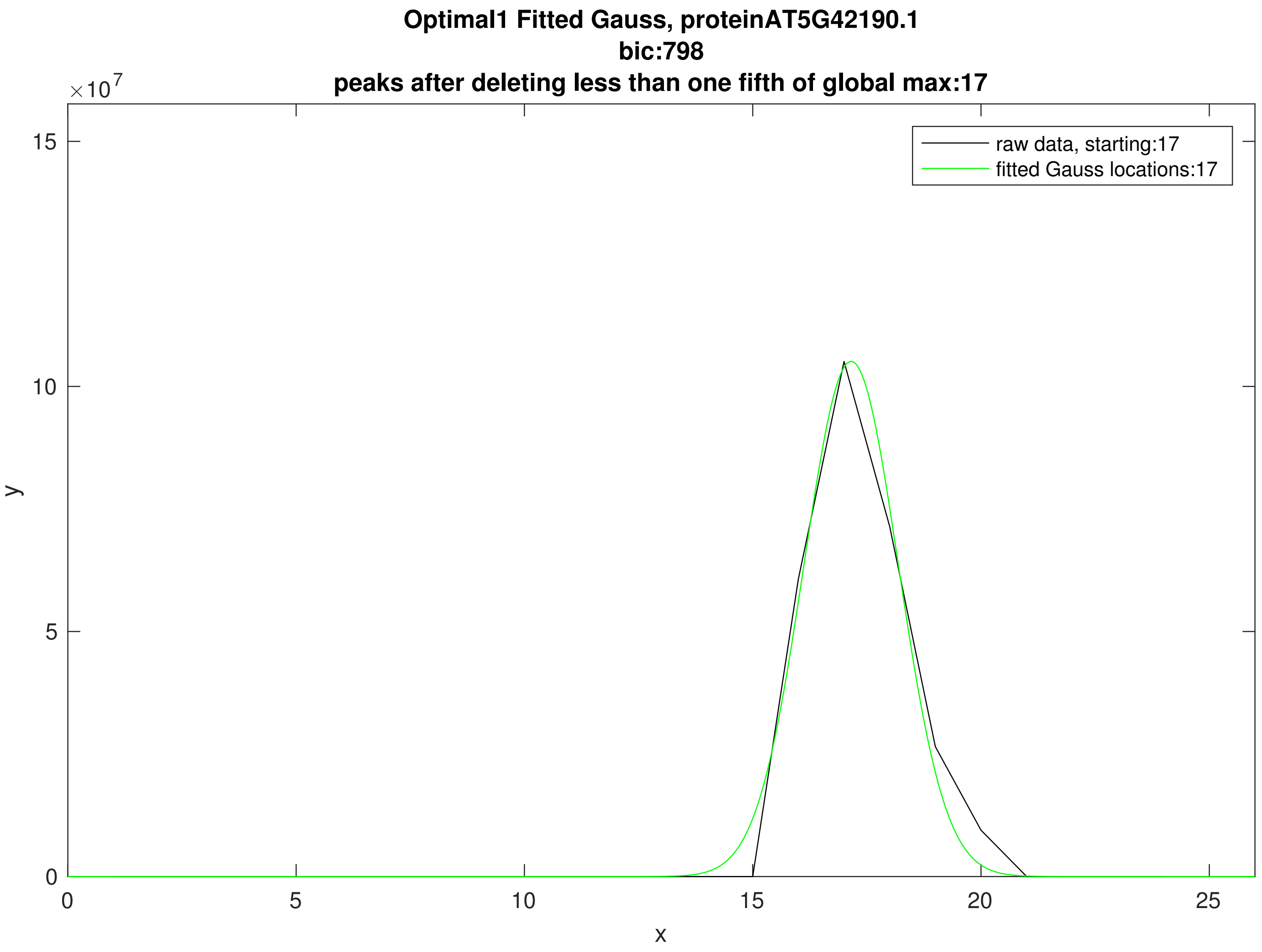

Optimal1 Fitted Gauss, proteinAT5G42190.1
bic:798
peaks after deleting less than one fifth of global max:17
×10^7
raw data, starting:17
fitted Gauss locations:17
y
x
0
5
10
15
20
25

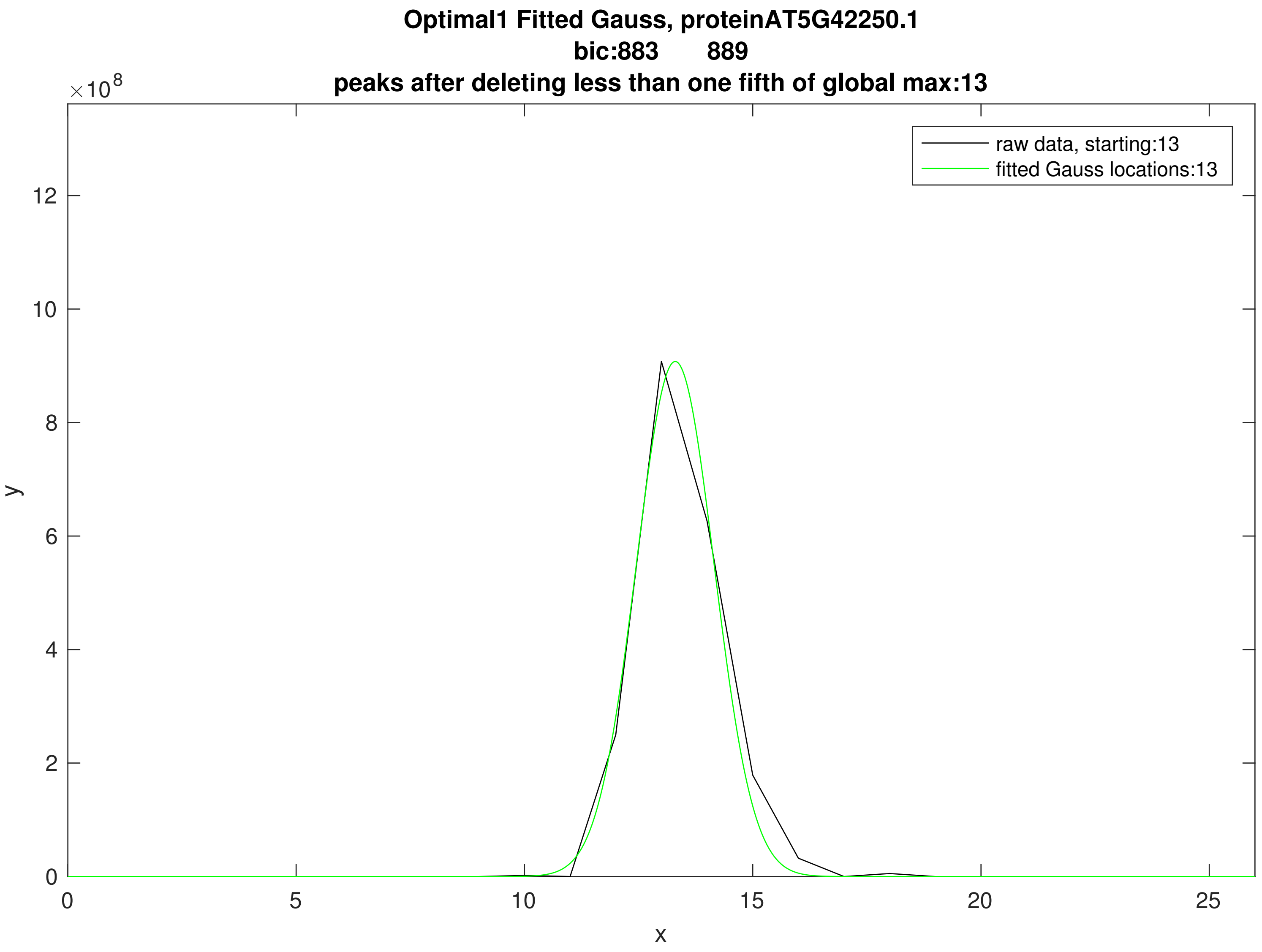

Optimal1 Fitted Gauss, proteinAT5G42250.1
bic:883 889
peaks after deleting less than one fifth of global max:13
×10^8
raw data, starting:13
fitted Gauss locations:13
y
x
0
2
4
6
8
10
12
0
5
10
15
20
25

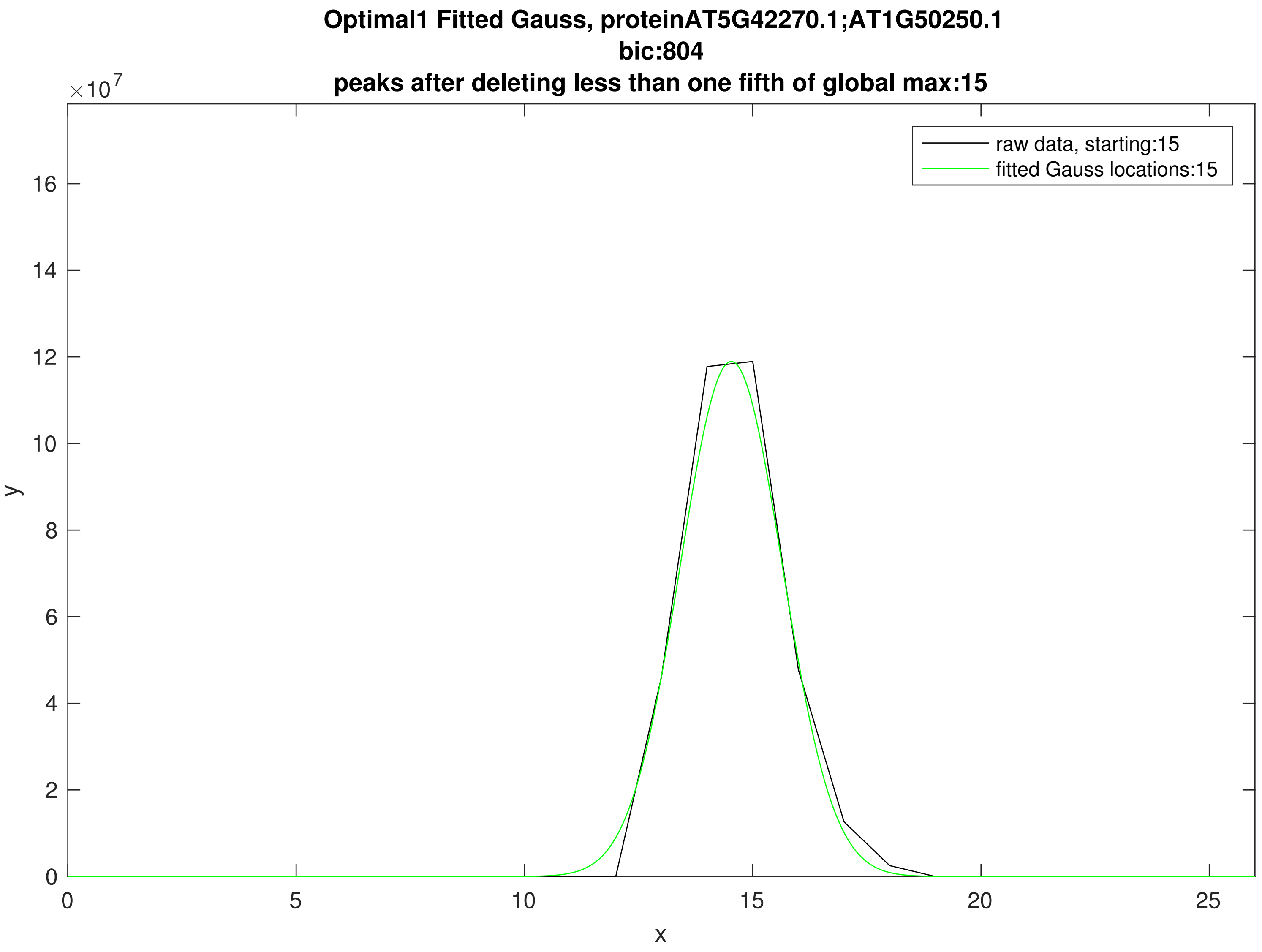
Optimal1 Fitted Gauss, proteinAT5G42270.1;AT1G50250.1
bic:804
peaks after deleting less than one fifth of global max:15
×10^7
raw data, starting:15
fitted Gauss locations:15
y
x
0
2
4
6
8
10
12
14
16
0
5
10
15
20
25

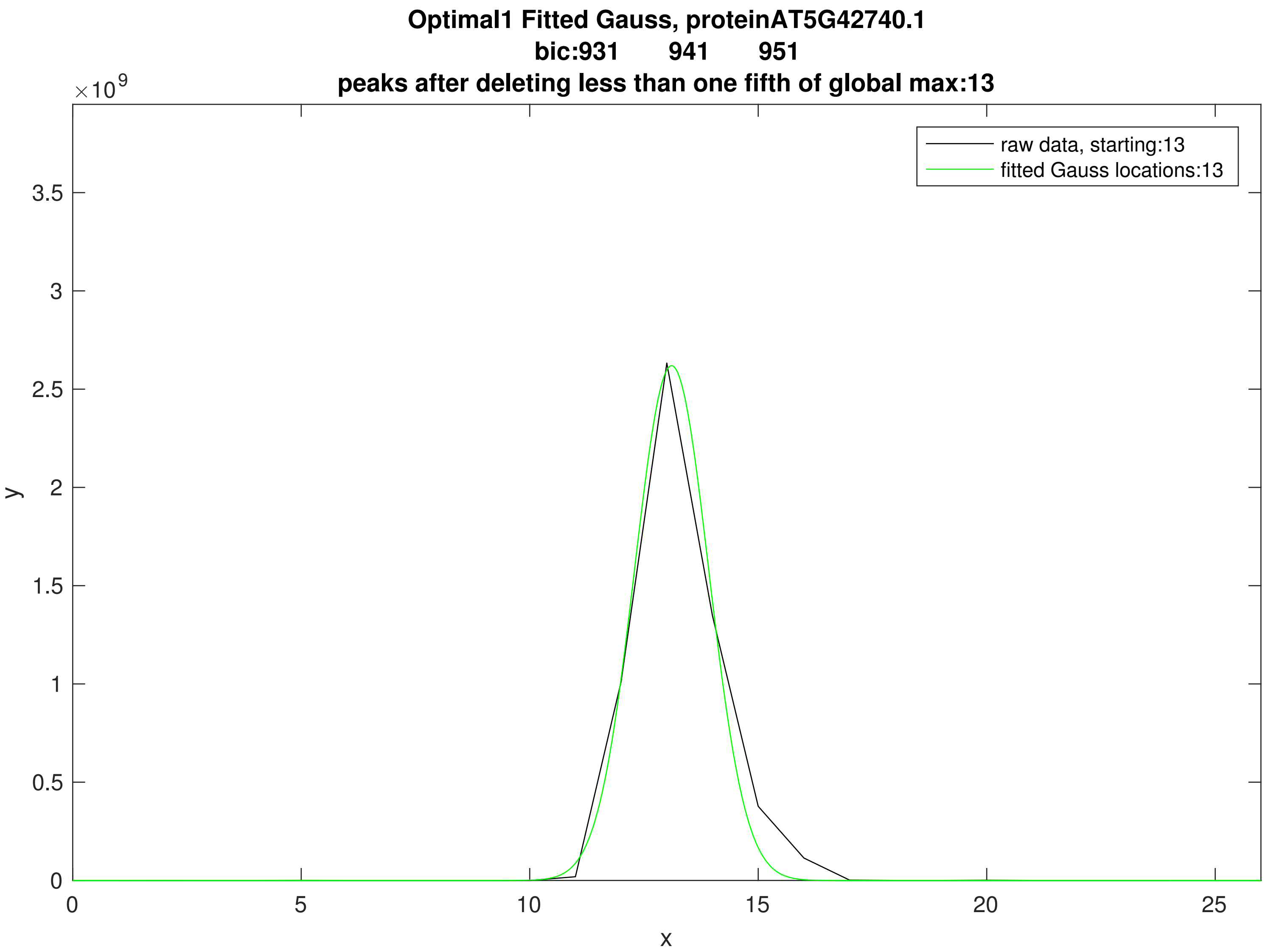

Optimal1 Fitted Gauss, proteinAT5G42740.1
bic:931 941 951
peaks after deleting less than one fifth of global max:13
×10^9
raw data, starting:13
fitted Gauss locations:13
3.5
3
2.5
2
1.5
1
0.5
0
y
0
5
10
15
20
25
x

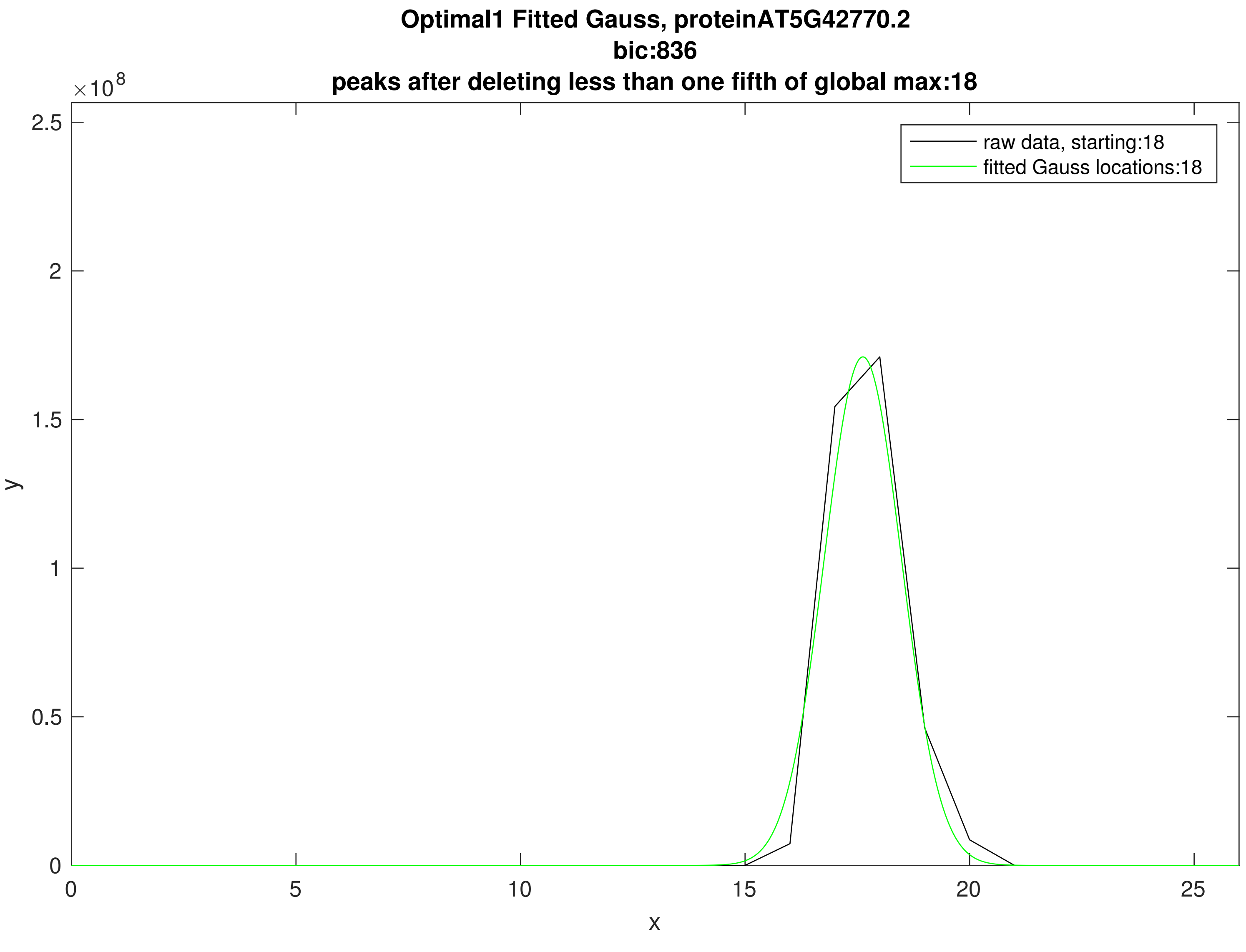
Optimal1 Fitted Gauss, proteinAT5G42770.2
bic:836
peaks after deleting less than one fifth of global max:18
raw data, starting:18
fitted Gauss locations:18
×10^8
2.5
2
1.5
1
0.5
0
5
10
15
20
25
x
y

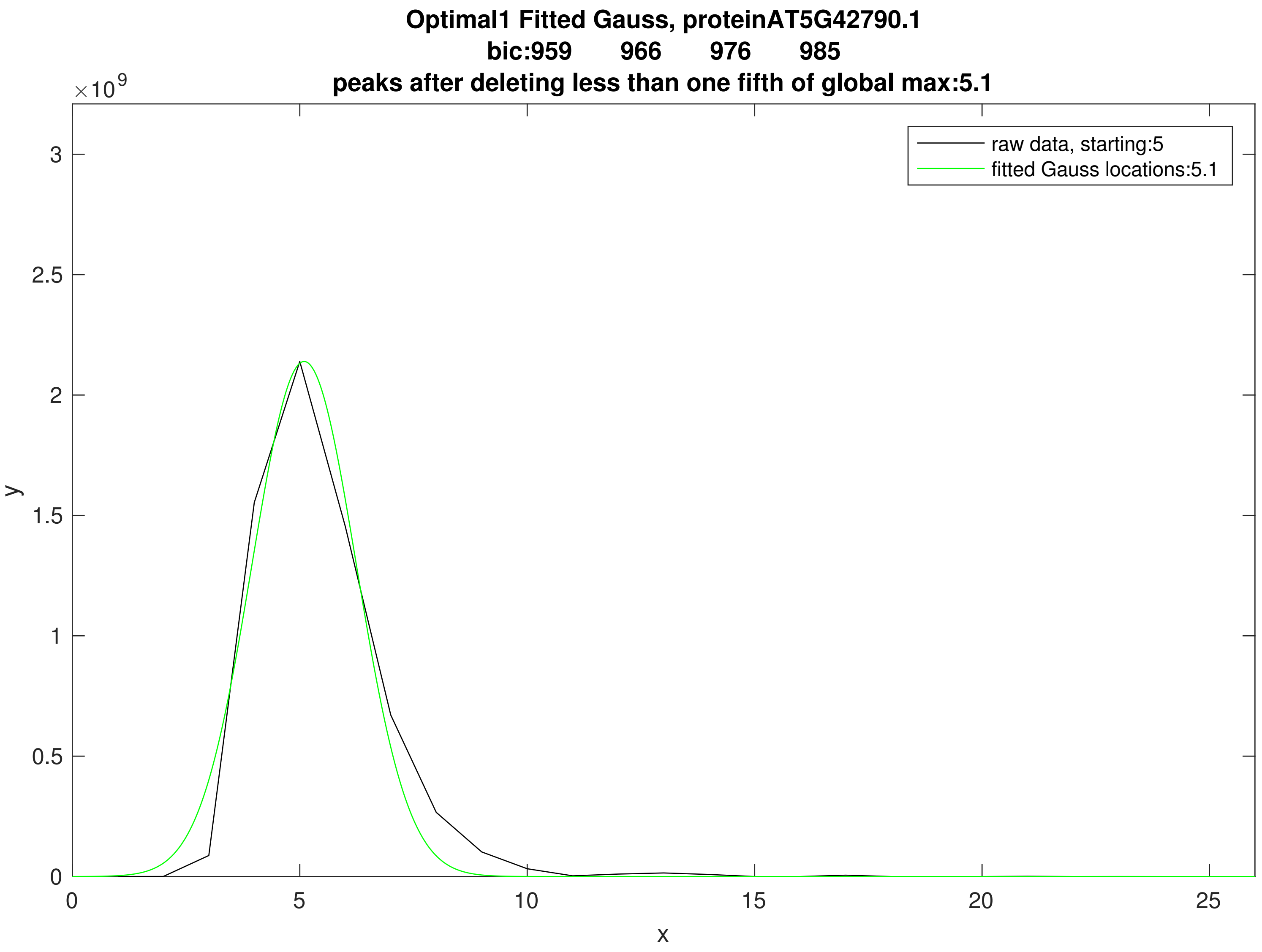

Optimal1 Fitted Gauss, proteinAT5G42790.1
bic:959 966 976 985
peaks after deleting less than one fifth of global max:5.1
×10^9
3
2.5
2
1.5
1
0.5
0
y
0
5
10
15
20
25
x
raw data, starting:5
fitted Gauss locations:5.1

#### Optimal1 Fitted Gauss, proteinAT5G42850.2;AT5G42850.1
#### bic:767
#### peaks after deleting less than one fifth of global max:20

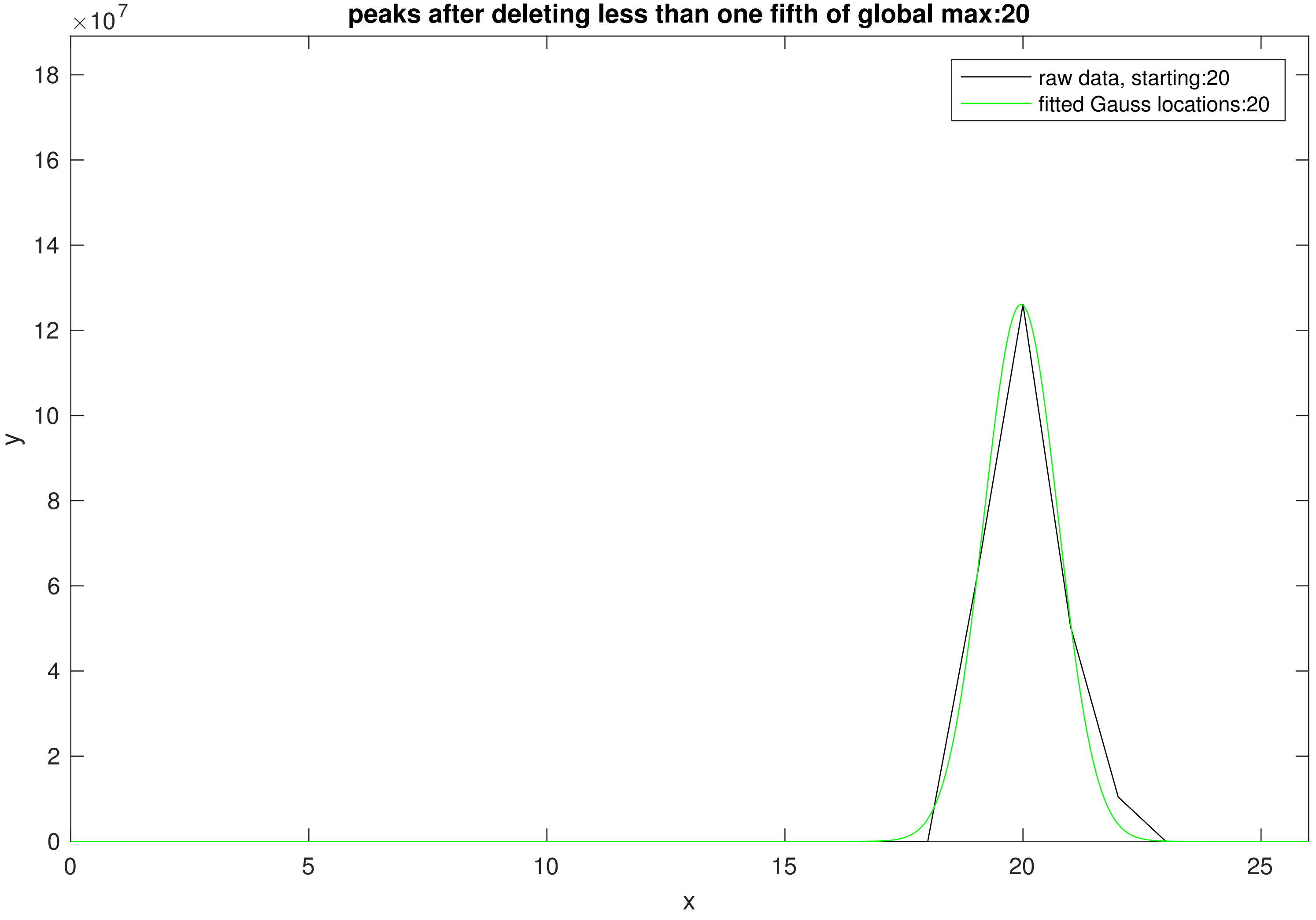

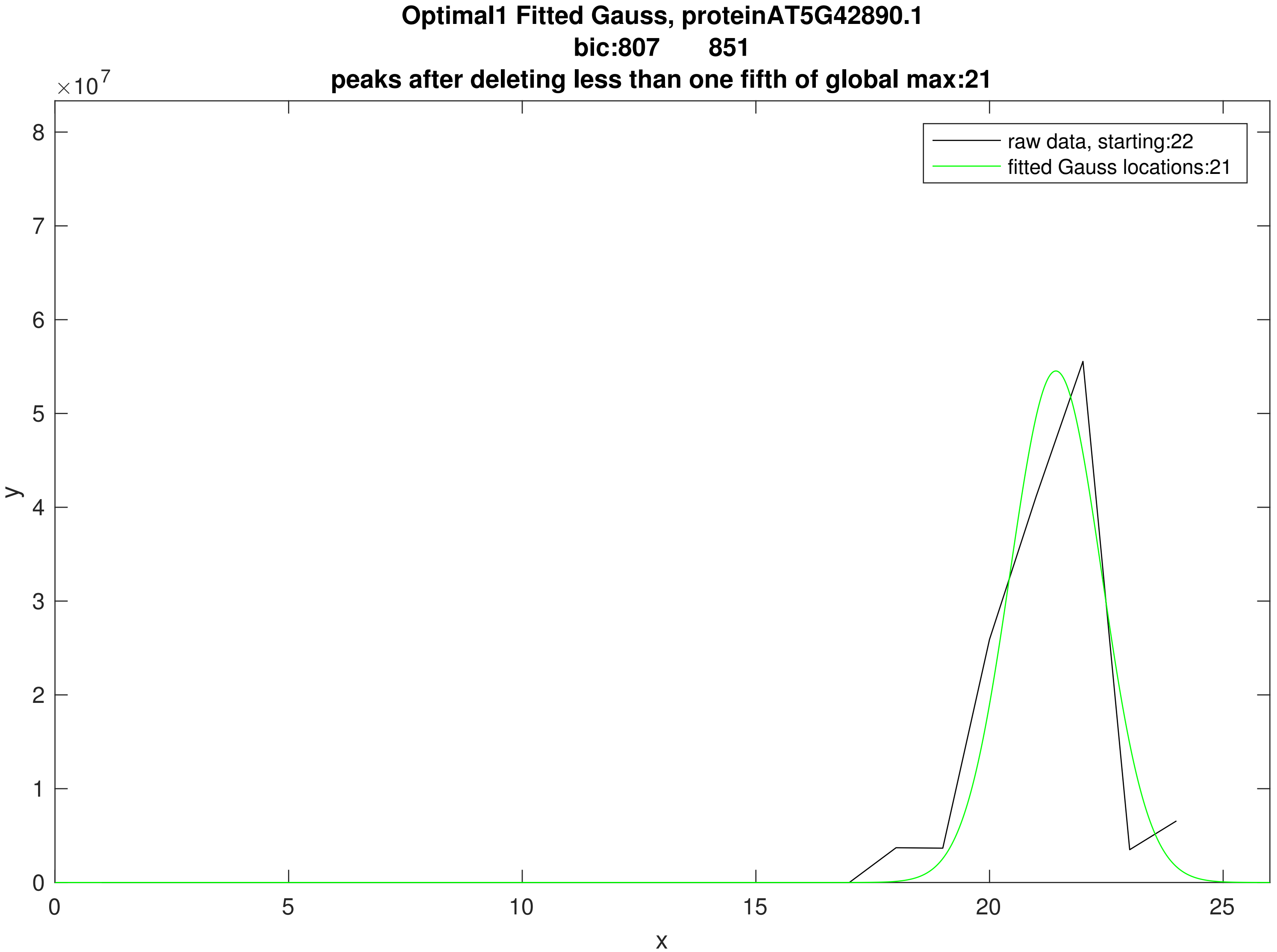
Optimal1 Fitted Gauss, proteinAT5G42890.1
bic:807 851
peaks after deleting less than one fifth of global max:21
×10^7
raw data, starting:22
fitted Gauss locations:21
y
x
0
1
2
3
4
5
6
7
8
0
5
10
15
20
25

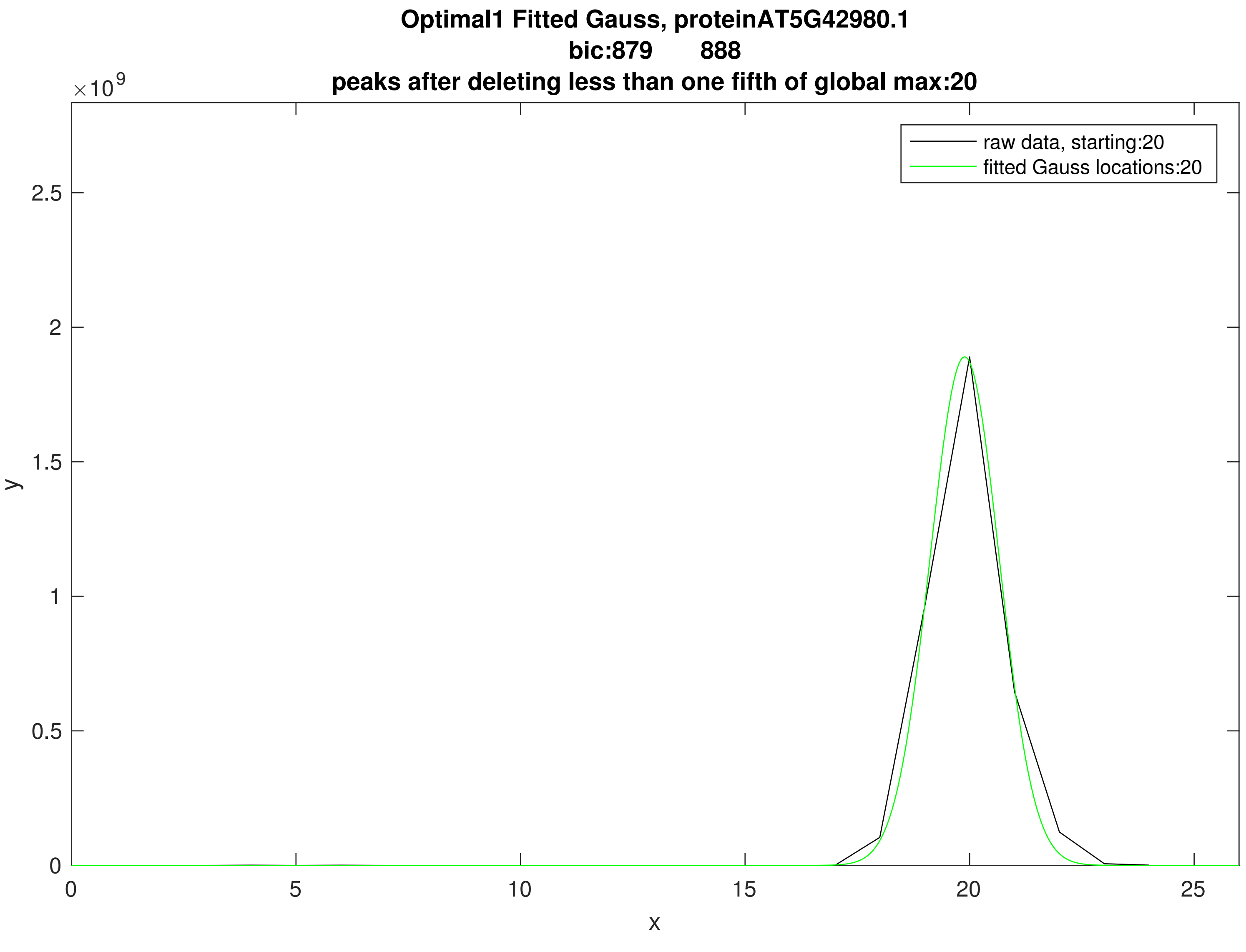

Optimal1 Fitted Gauss, proteinAT5G42980.1
bic:879 888
peaks after deleting less than one fifth of global max:20
×10^9
raw data, starting:20
fitted Gauss locations:20
2.5
2
1.5
1
0.5
0
y
0
5
10
15
20
25
x

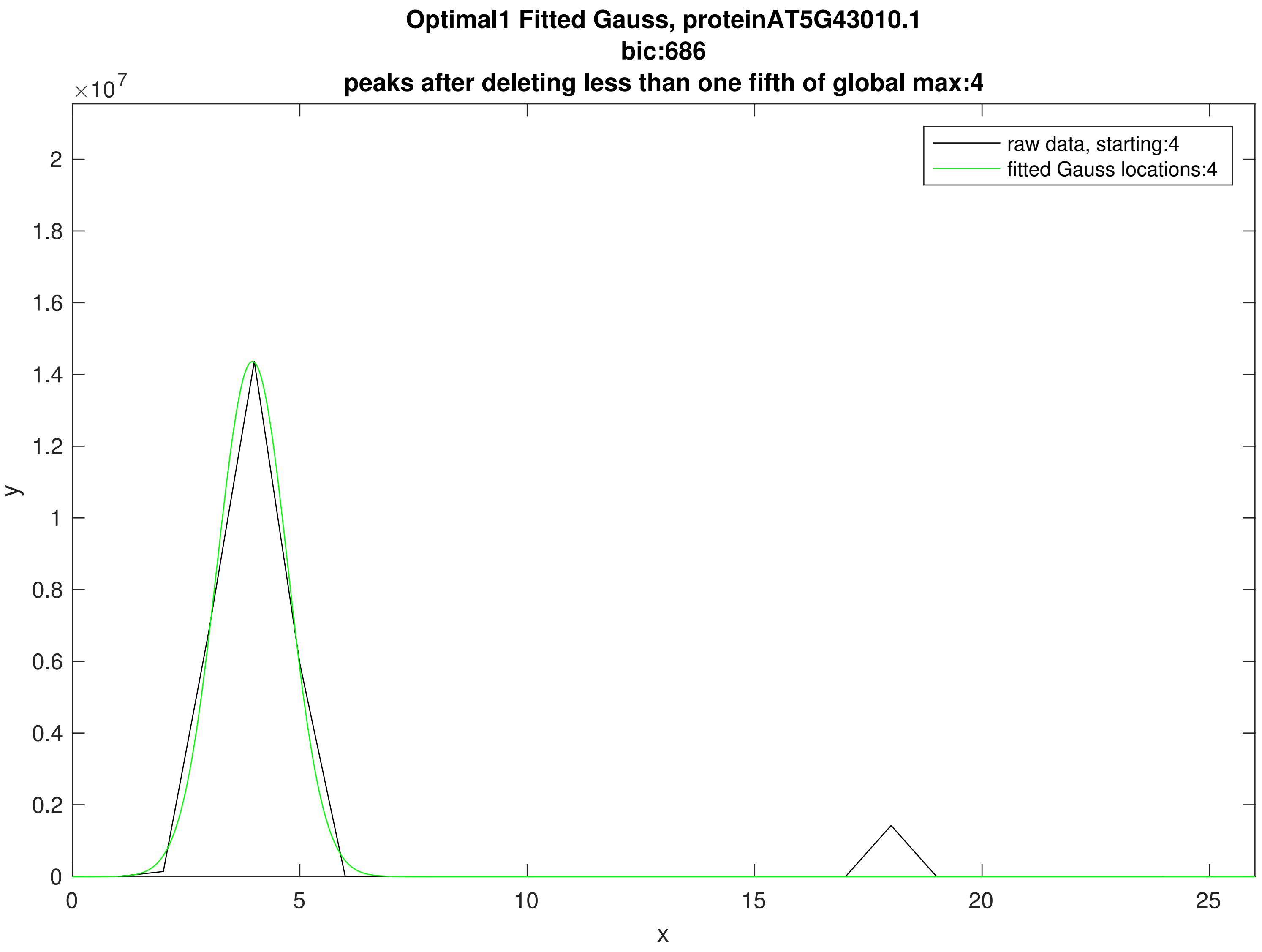

Optimal1 Fitted Gauss, proteinAT5G43010.1
bic:686
peaks after deleting less than one fifth of global max:4
×10^7
raw data, starting:4
fitted Gauss locations:4
y
x

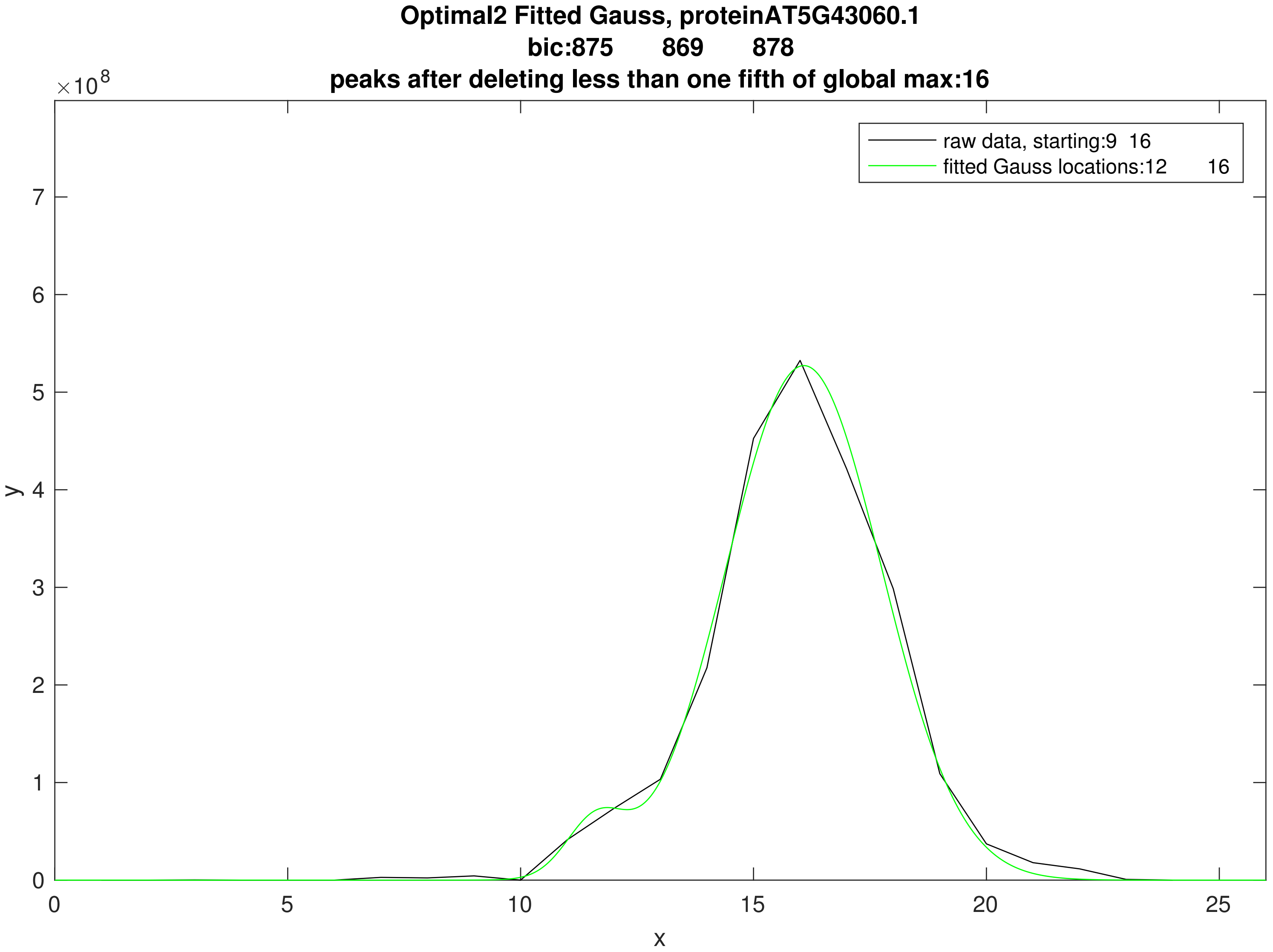

Optimal2 Fitted Gauss, proteinAT5G43060.1
bic:875 869 878
peaks after deleting less than one fifth of global max:16
×10^8
raw data, starting:9 16
fitted Gauss locations:12 16
y
x
0
1
2
3
4
5
6
7
0
5
10
15
20
25

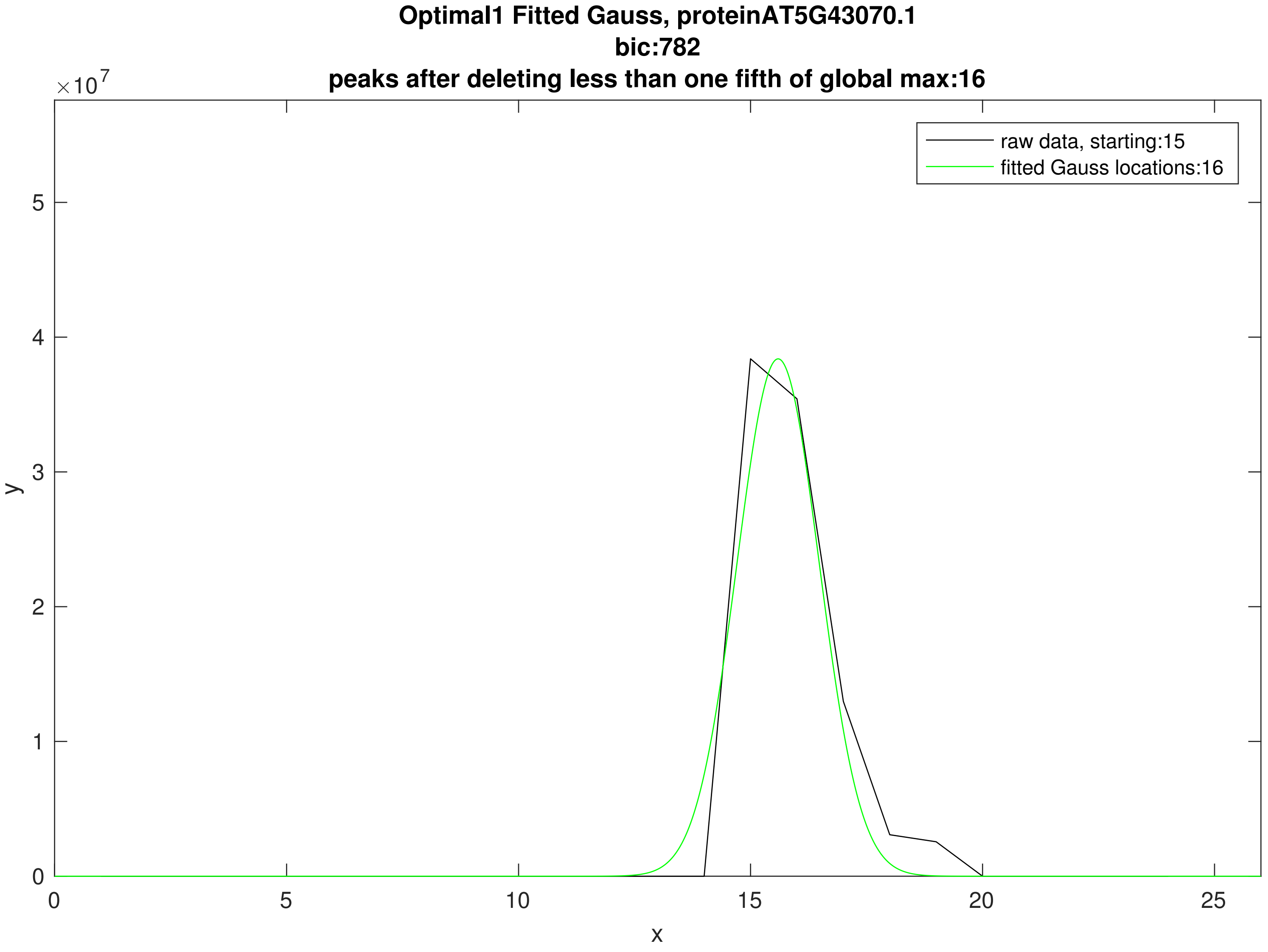
Optimal1 Fitted Gauss, proteinAT5G43070.1
bic:782
peaks after deleting less than one fifth of global max:16
×10^7
raw data, starting:15
fitted Gauss locations:16
0
1
2
3
4
5
y
0
5
10
15
20
25
x

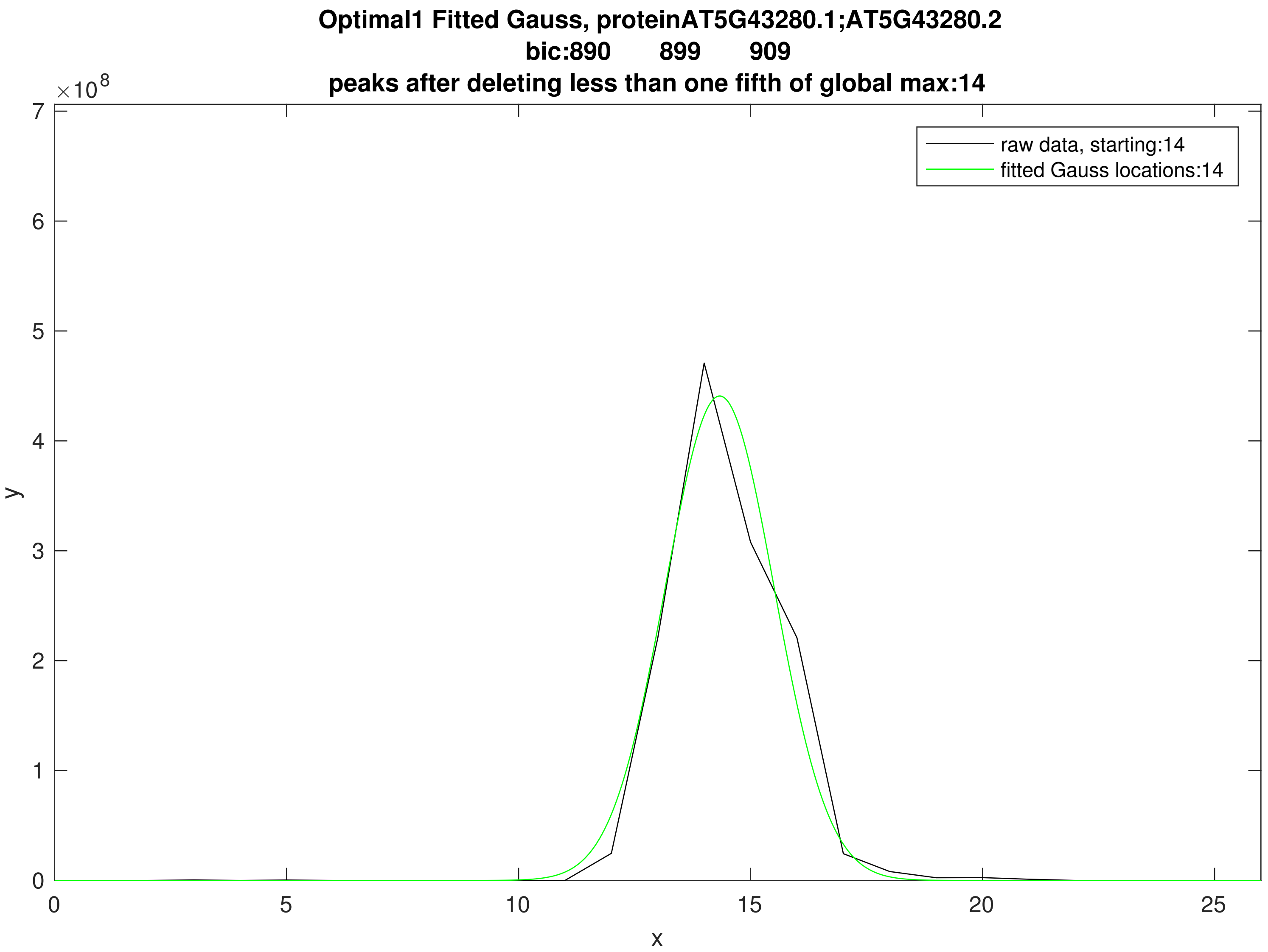

Optimal1 Fitted Gauss, proteinAT5G43280.1;AT5G43280.2
bic:890 899 909
peaks after deleting less than one fifth of global max:14
×10^8
7
6
5
4
3
2
1
0
y
0
5
10
15
20
25
x
raw data, starting:14
fitted Gauss locations:14

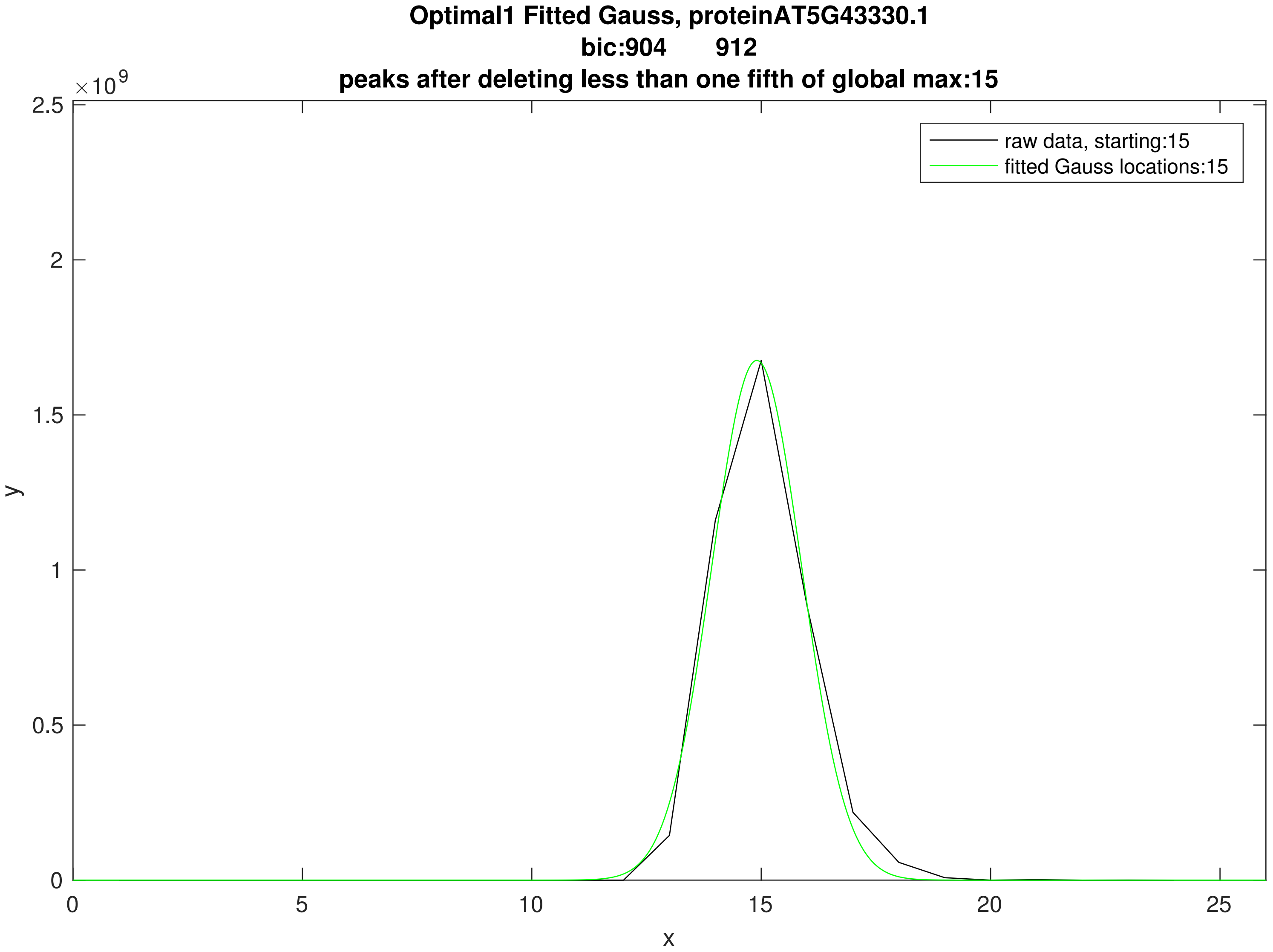

Optimal1 Fitted Gauss, proteinAT5G43330.1
bic:904 912
peaks after deleting less than one fifth of global max:15
×10^9
2.5
2
1.5
1
0.5
0
y
0
5
10
15
20
25
x
raw data, starting:15
fitted Gauss locations:15

**Optimal1 Fitted Gauss, proteinAT5G43400.1**
**bic:765**
**peaks after deleting less than one fifth of global max:13**

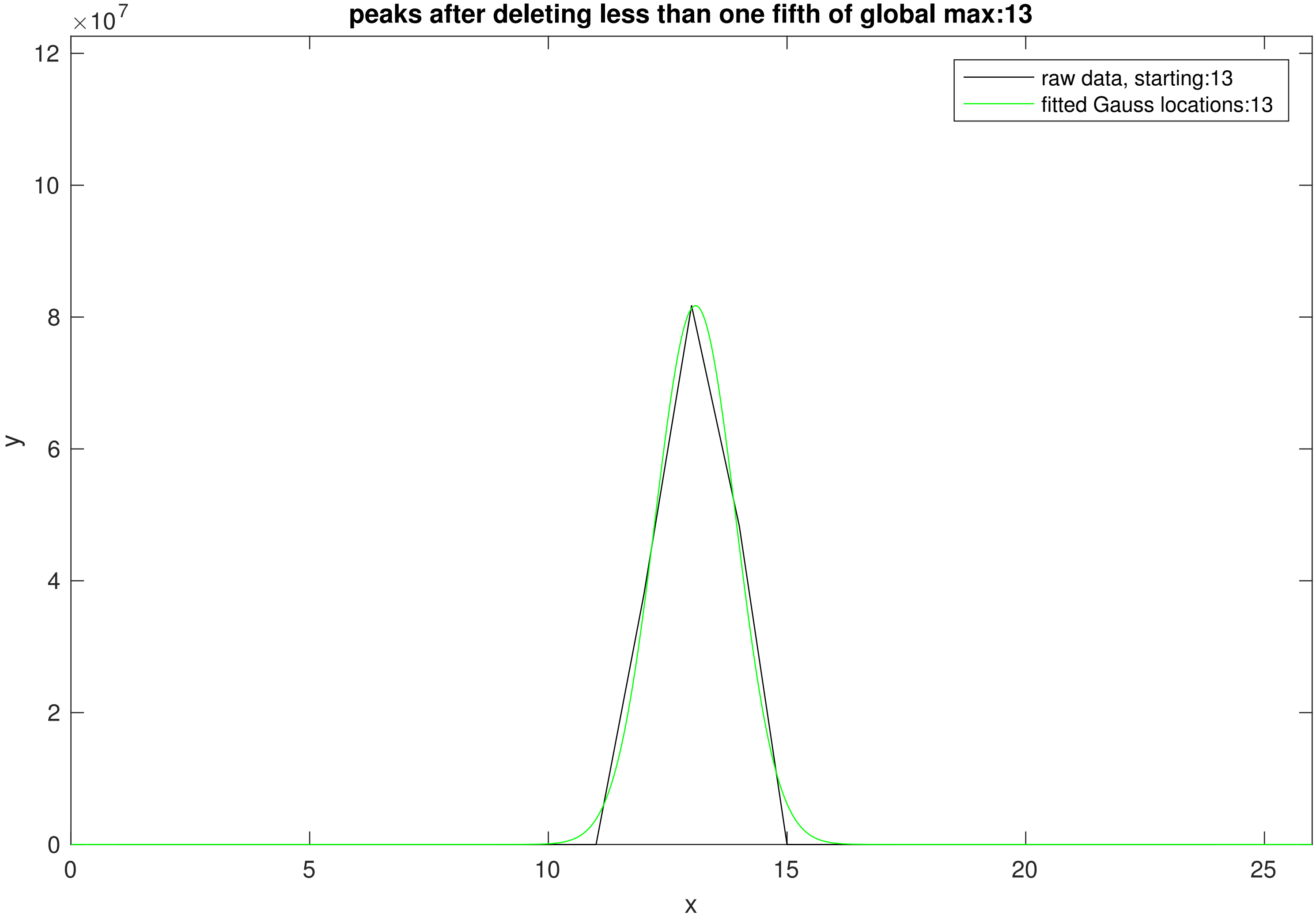

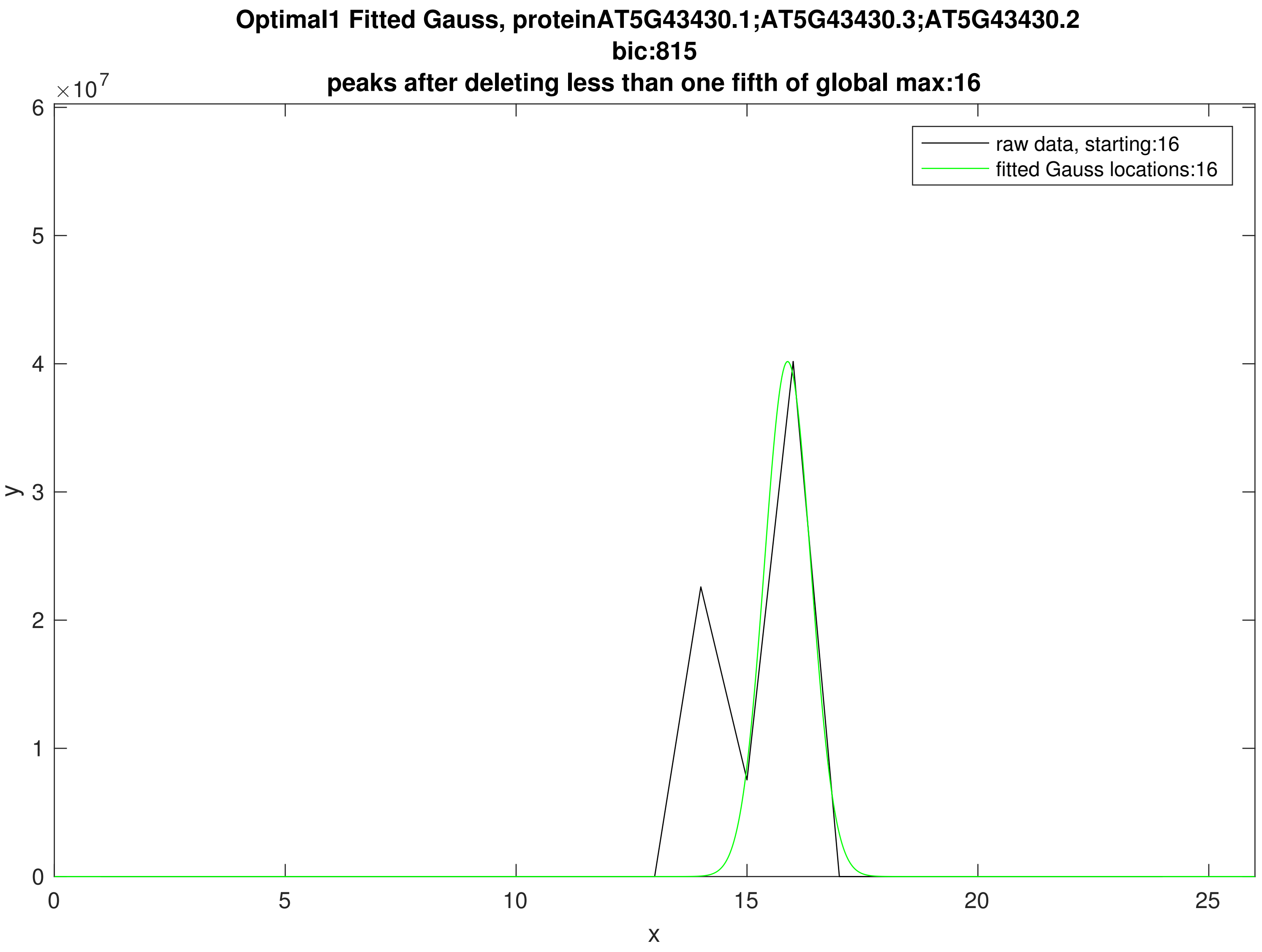

Optimal1 Fitted Gauss, proteinAT5G43430.1;AT5G43430.3;AT5G43430.2
bic:815
peaks after deleting less than one fifth of global max:16
×10^7
6
5
4
3
2
1
0
y
0
5
10
15
20
25
x
raw data, starting:16
fitted Gauss locations:16

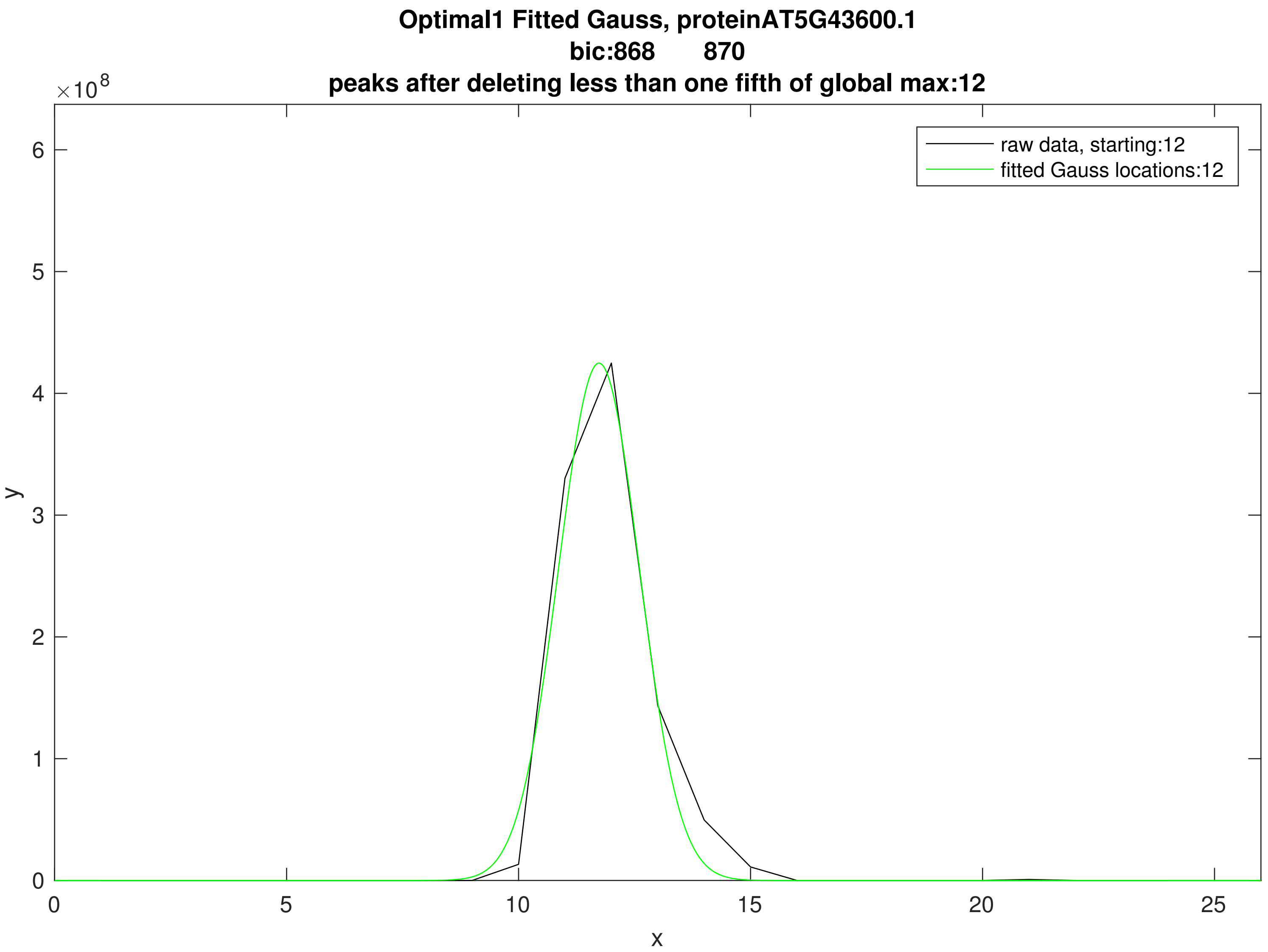
Optimal1 Fitted Gauss, proteinAT5G43600.1
bic:868 870
peaks after deleting less than one fifth of global max:12
×10^8
raw data, starting:12
fitted Gauss locations:12
y
x
0
1
2
3
4
5
6
0
5
10
15
20
25

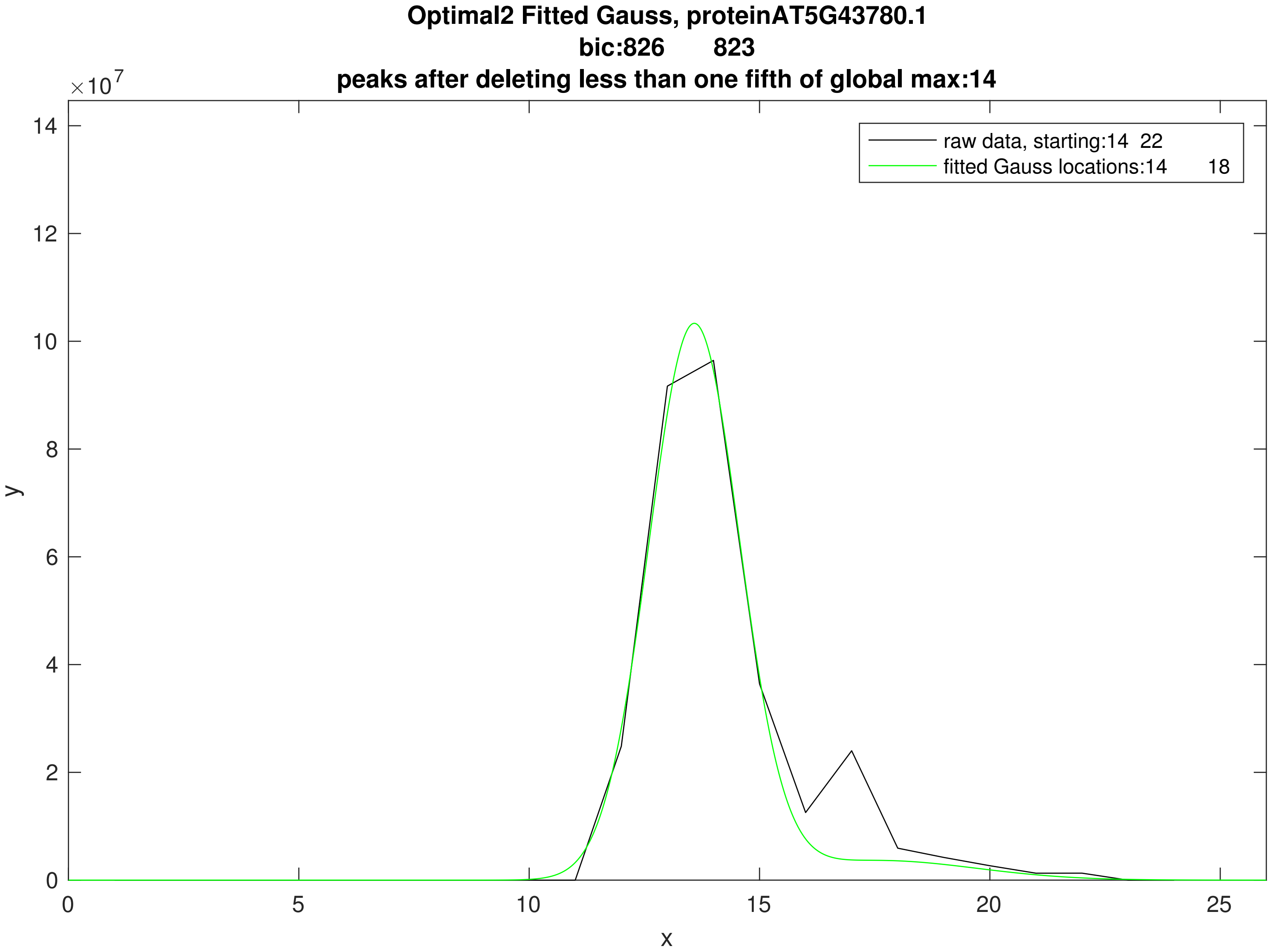
Optimal2 Fitted Gauss, proteinAT5G43780.1
bic:826 823
peaks after deleting less than one fifth of global max:14
×10^7
raw data, starting:14 22
fitted Gauss locations:14 18
y
x
0
2
4
6
8
10
12
14
0
5
10
15
20
25

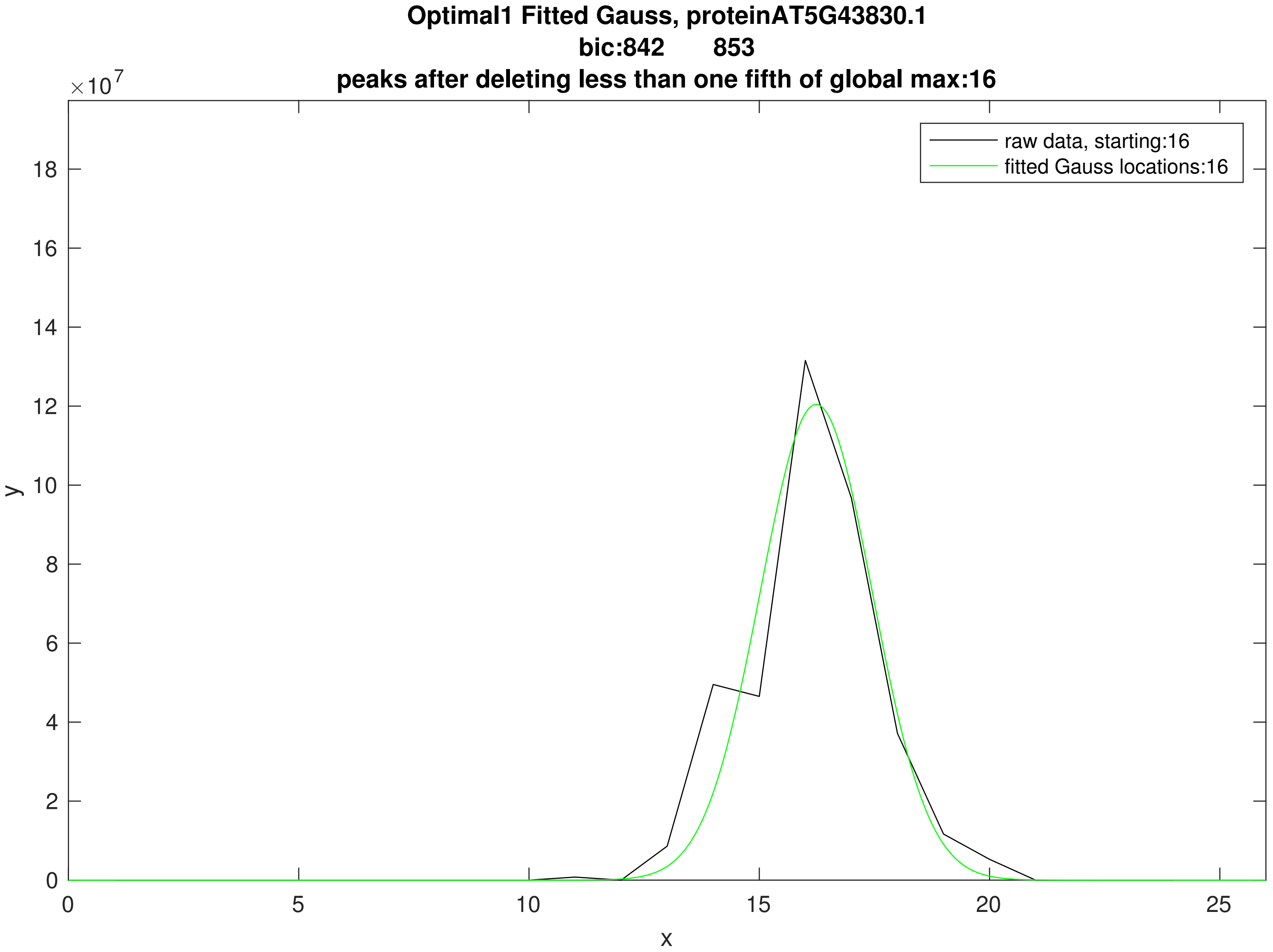
Optimal1 Fitted Gauss, proteinAT5G43830.1
bic:842 853
peaks after deleting less than one fifth of global max:16
×10^7
raw data, starting:16
fitted Gauss locations:16
y
x
0
2
4
6
8
10
12
14
16
18
0
5
10
15
20
25

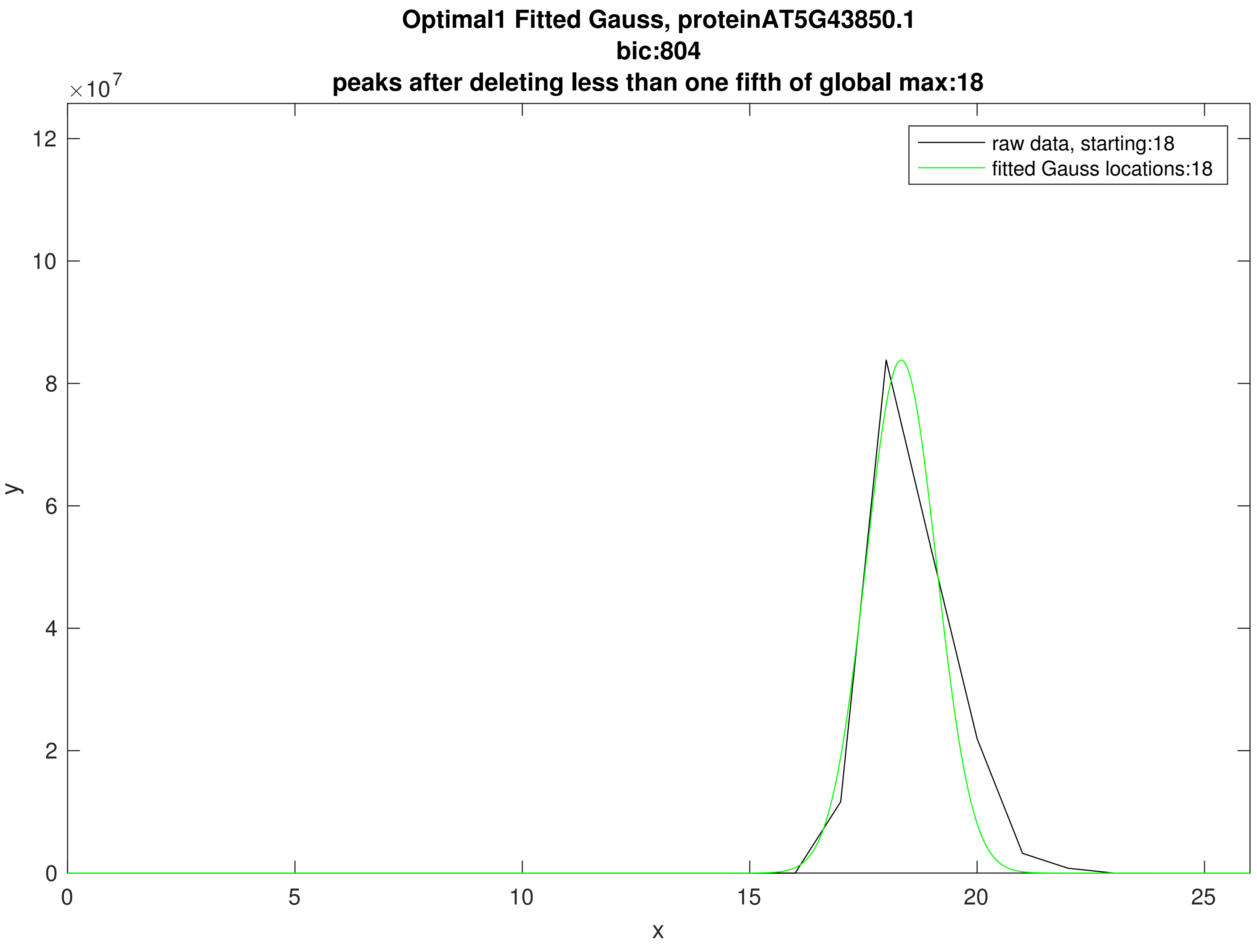
Optimal1 Fitted Gauss, proteinAT5G43850.1
bic:804
peaks after deleting less than one fifth of global max:18
×10^7
raw data, starting:18
fitted Gauss locations:18
y
x
0
2
4
6
8
10
12
0
5
10
15
20
25

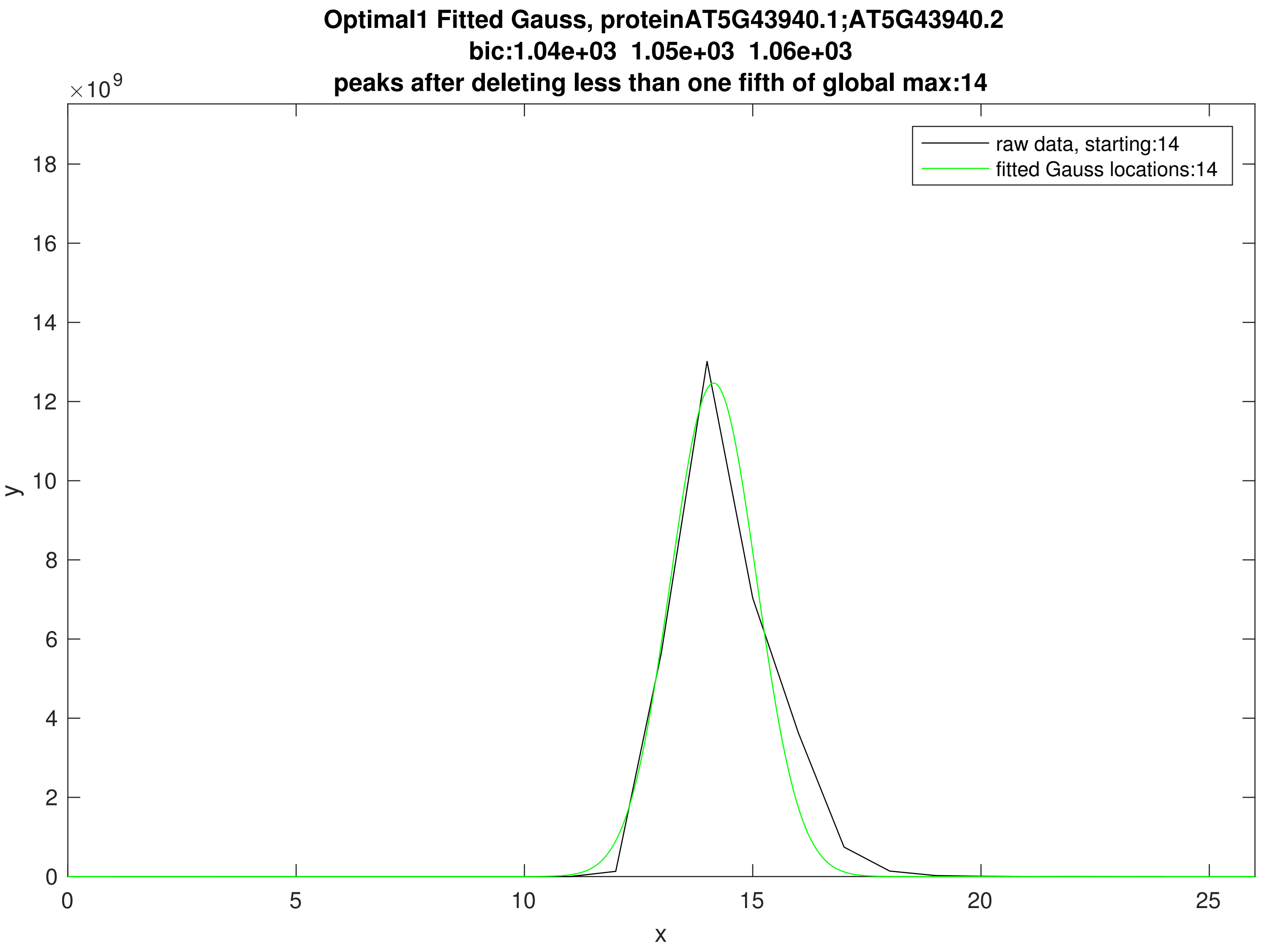
Optimal1 Fitted Gauss, proteinAT5G43940.1;AT5G43940.2
bic:1.04e+03 1.05e+03 1.06e+03
peaks after deleting less than one fifth of global max:14
×10^9
raw data, starting:14
fitted Gauss locations:14
y
x
0
2
4
6
8
10
12
14
16
18
5
15
20
25

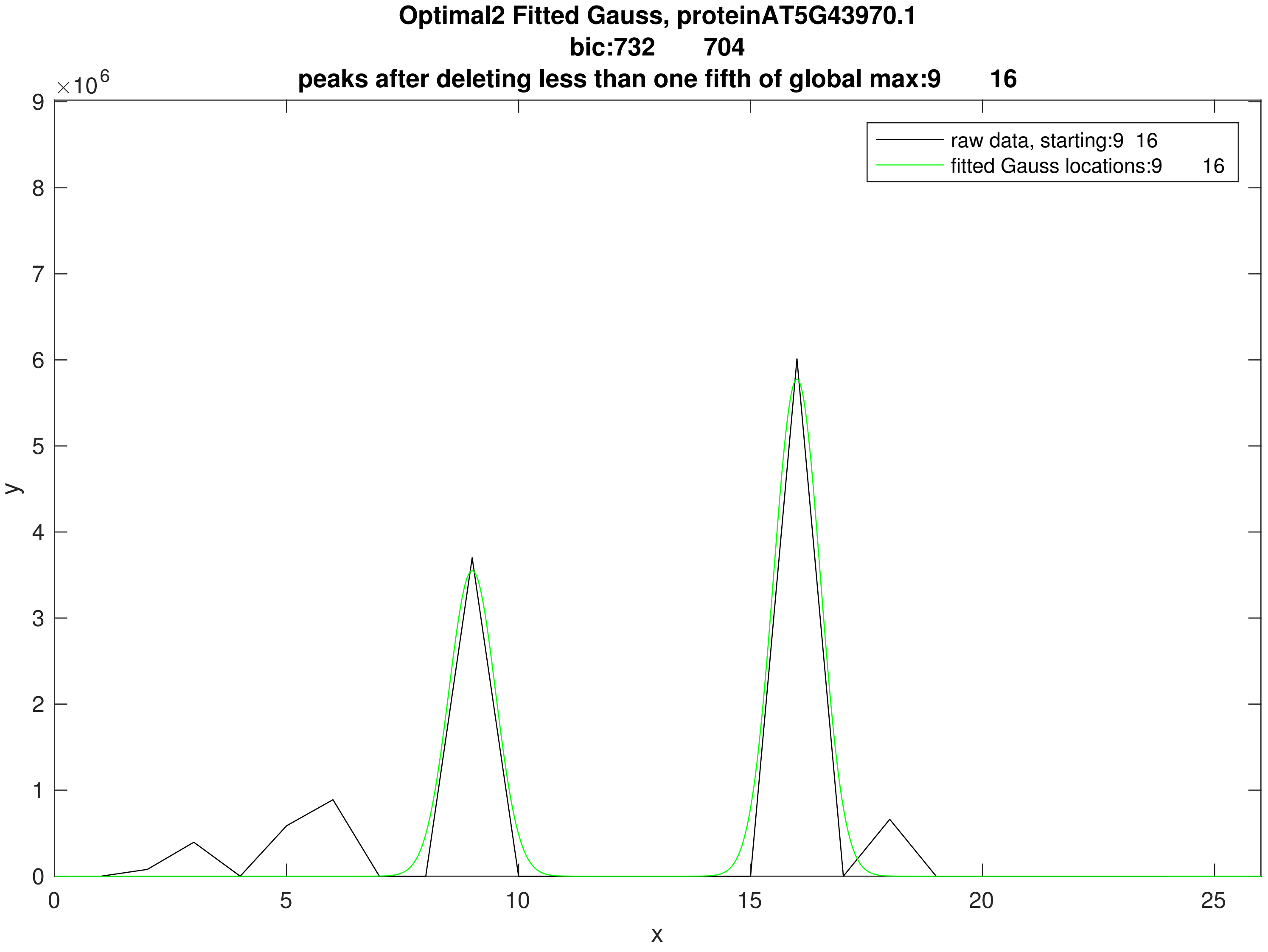
Optimal2 Fitted Gauss, proteinAT5G43970.1
bic:732 704
peaks after deleting less than one fifth of global max:9 16
×10^6
raw data, starting:9 16
fitted Gauss locations:9 16
y
x
0
1
2
3
4
5
6
7
8
9
0
5
10
15
20
25

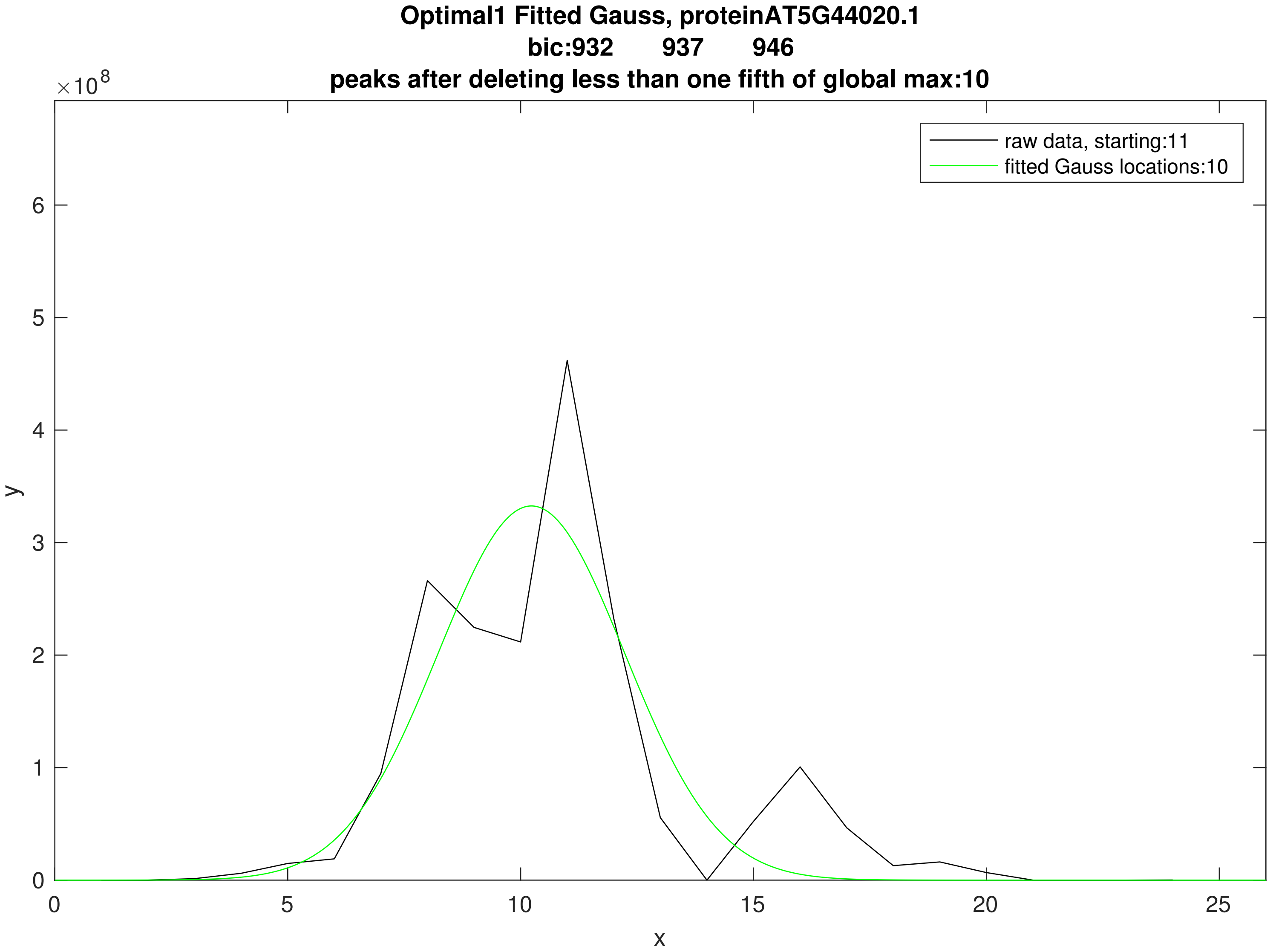
Optimal1 Fitted Gauss, proteinAT5G44020.1
bic:932 937 946
peaks after deleting less than one fifth of global max:10
×10^8
raw data, starting:11
fitted Gauss locations:10
y
x
0
1
2
3
4
5
6
10
15
20
25

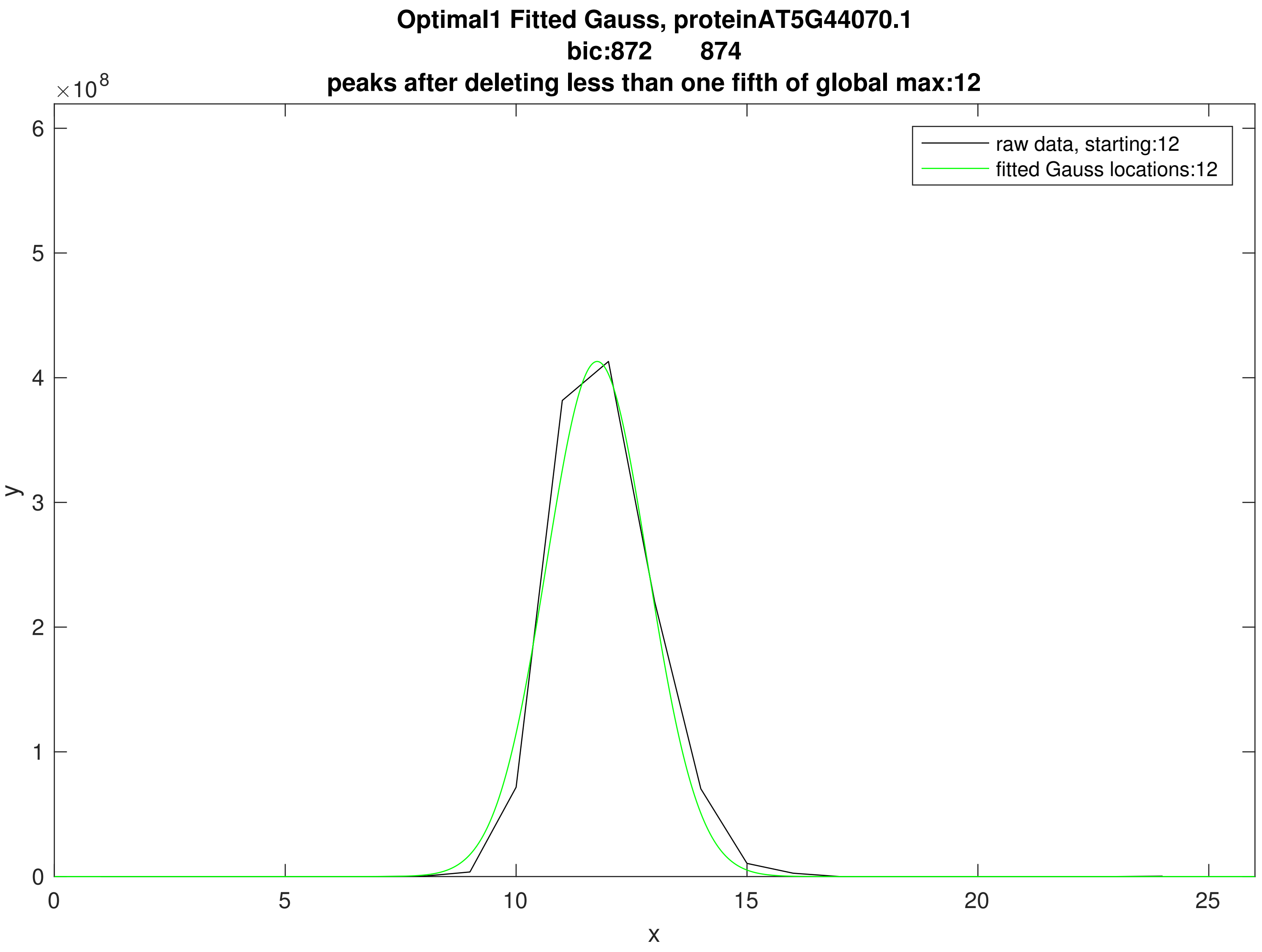

Optimal1 Fitted Gauss, proteinAT5G44070.1
bic:872 874
peaks after deleting less than one fifth of global max:12
×10^8
raw data, starting:12
fitted Gauss locations:12
y
x

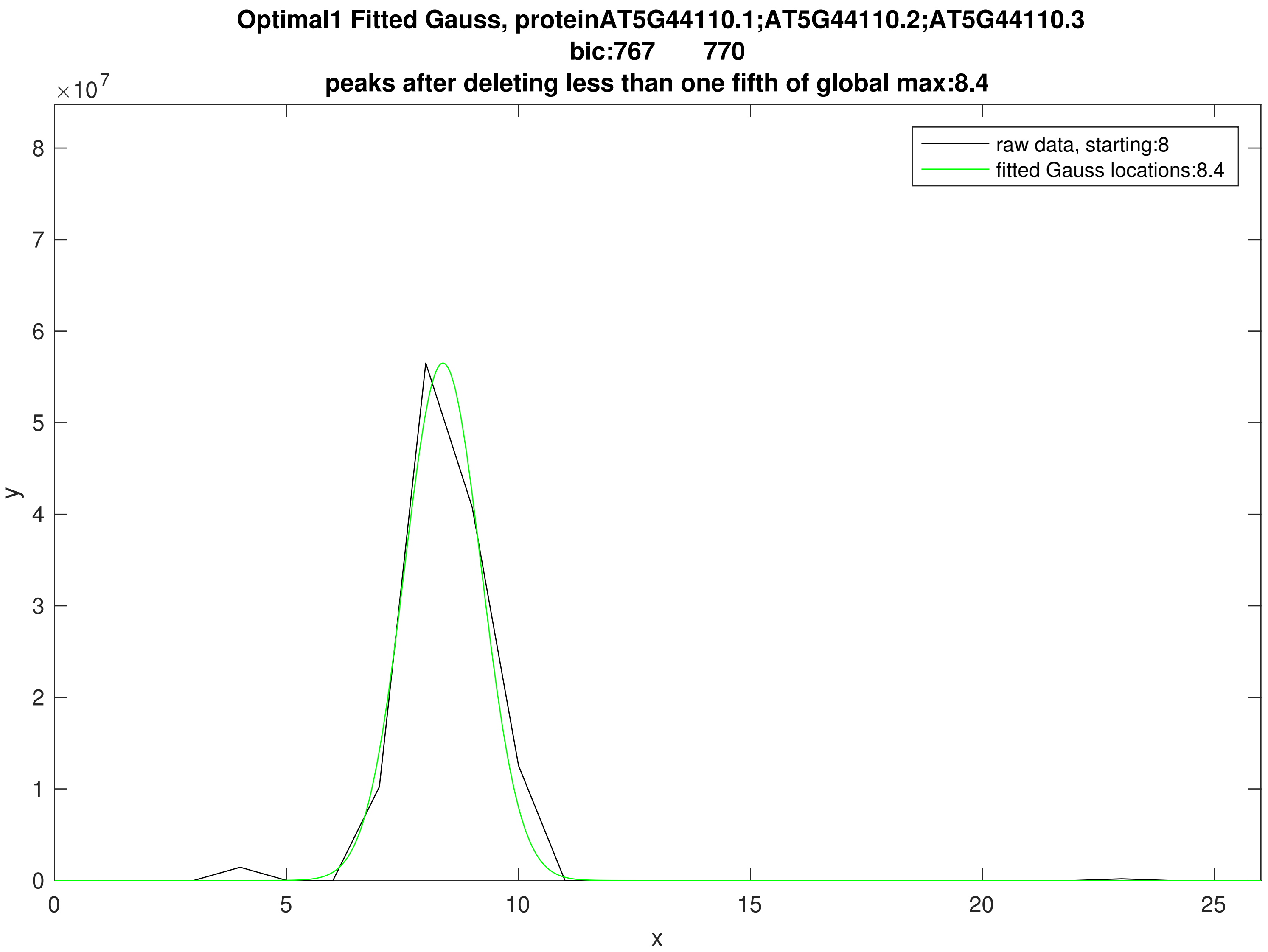
Optimal1 Fitted Gauss, proteinAT5G44110.1;AT5G44110.2;AT5G44110.3
bic:767 770
peaks after deleting less than one fifth of global max:8.4
×10^7
8
7
6
5
4
3
2
1
0
y
0
5
10
15
20
25
x
raw data, starting:8
fitted Gauss locations:8.4

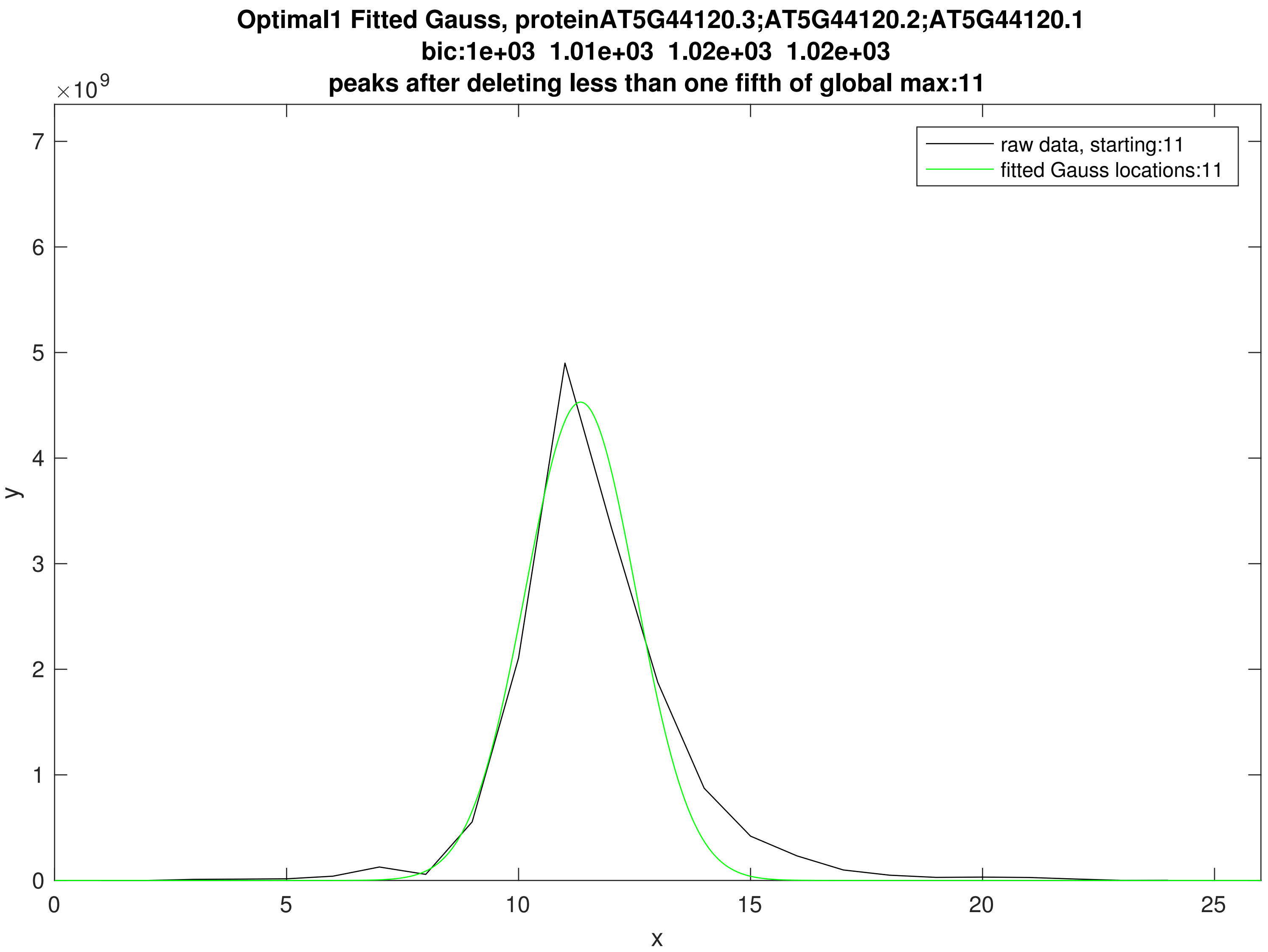

Optimal1 Fitted Gauss, proteinAT5G44120.3;AT5G44120.2;AT5G44120.1
bic:1e+03 1.01e+03 1.02e+03 1.02e+03
peaks after deleting less than one fifth of global max:11
×10^9
raw data, starting:11
fitted Gauss locations:11
y
x
0
1
2
3
4
5
6
7
0
5
10
15
20
25

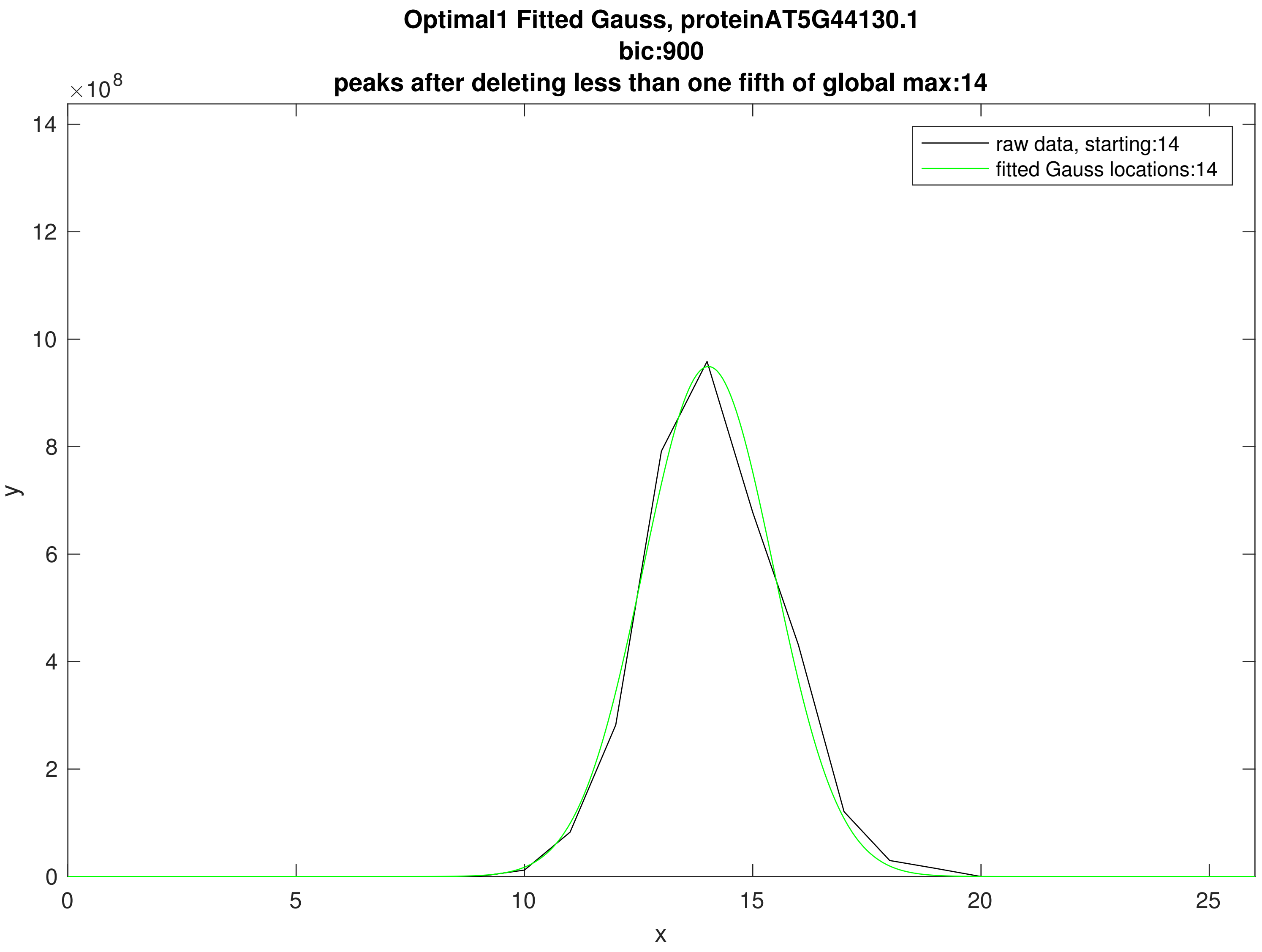

Optimal1 Fitted Gauss, proteinAT5G44130.1
bic:900
peaks after deleting less than one fifth of global max:14
×10^8
raw data, starting:14
fitted Gauss locations:14
y
x
0
2
4
6
8
10
12
14
5
15
20
25

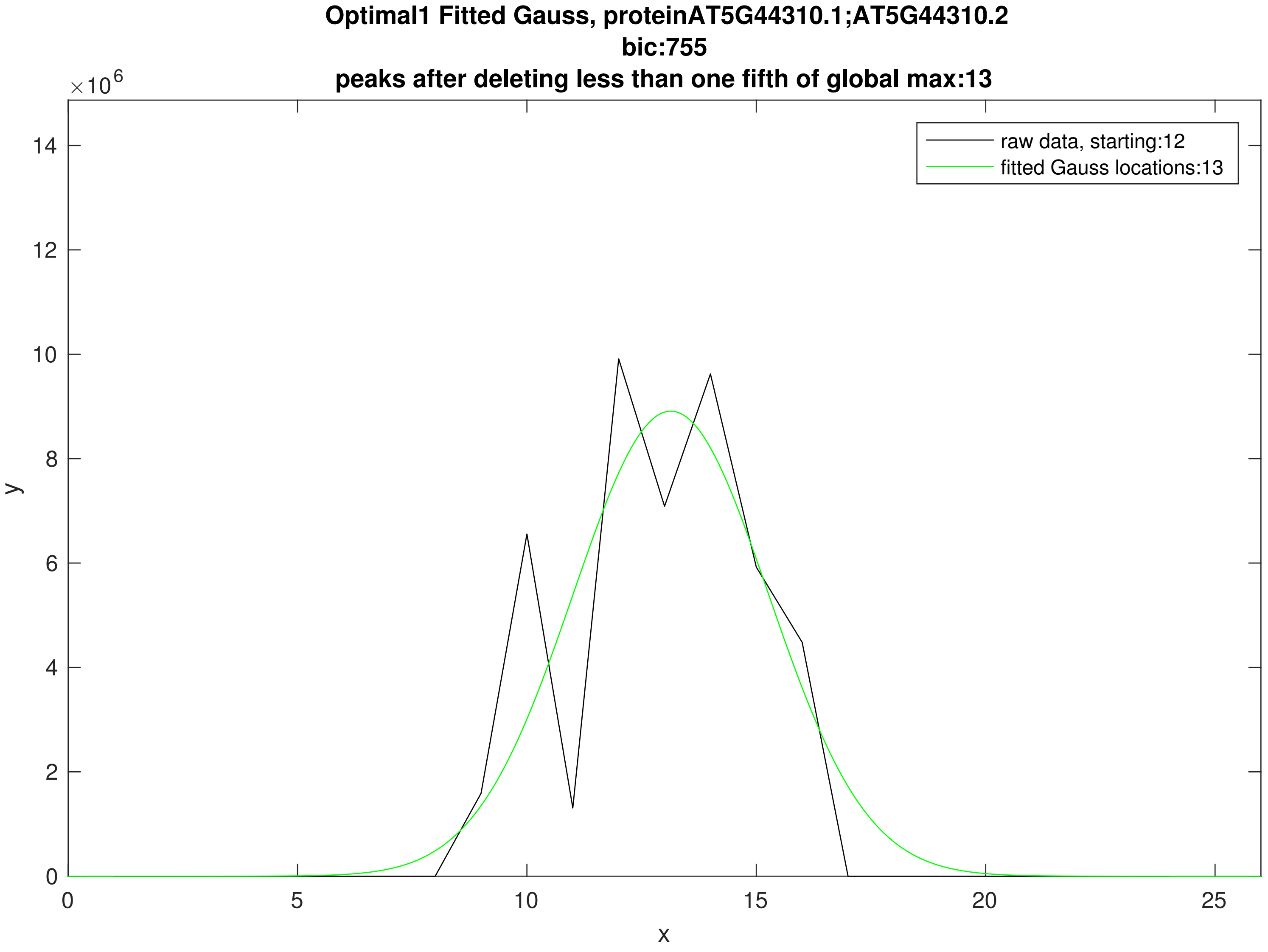
Optimal1 Fitted Gauss, proteinAT5G44310.1;AT5G44310.2
bic:755
peaks after deleting less than one fifth of global max:13
$\times 10^6$
raw data, starting:12
fitted Gauss locations:13
y
x
0
2
4
6
8
10
12
14
0
5
10
15
20
25

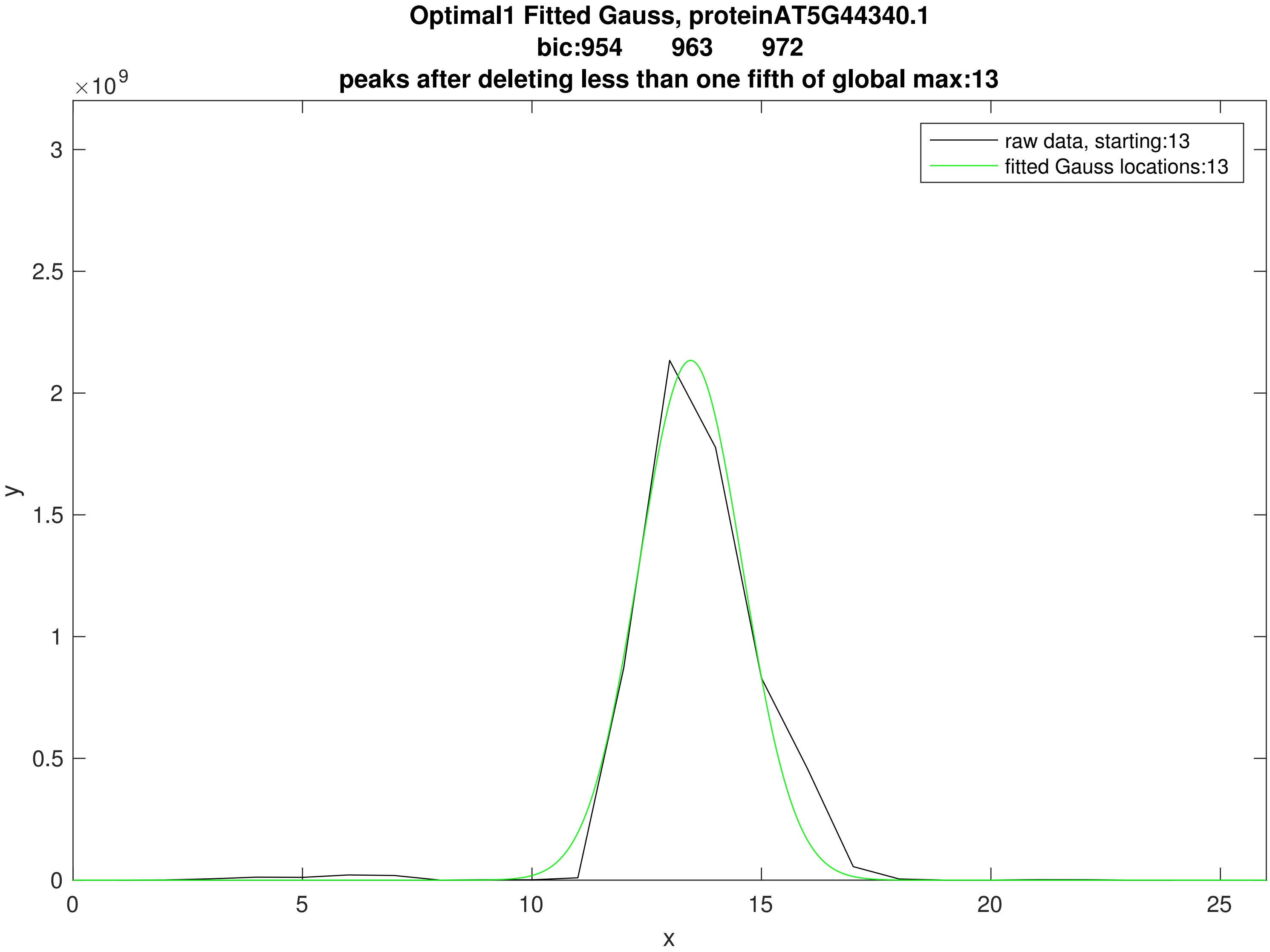
Optimal1 Fitted Gauss, proteinAT5G44340.1
bic:954 963 972
peaks after deleting less than one fifth of global max:13
×10^9
3
2.5
2
1.5
1
0.5
0
y
0
5
10
15
20
25
x
raw data, starting:13
fitted Gauss locations:13

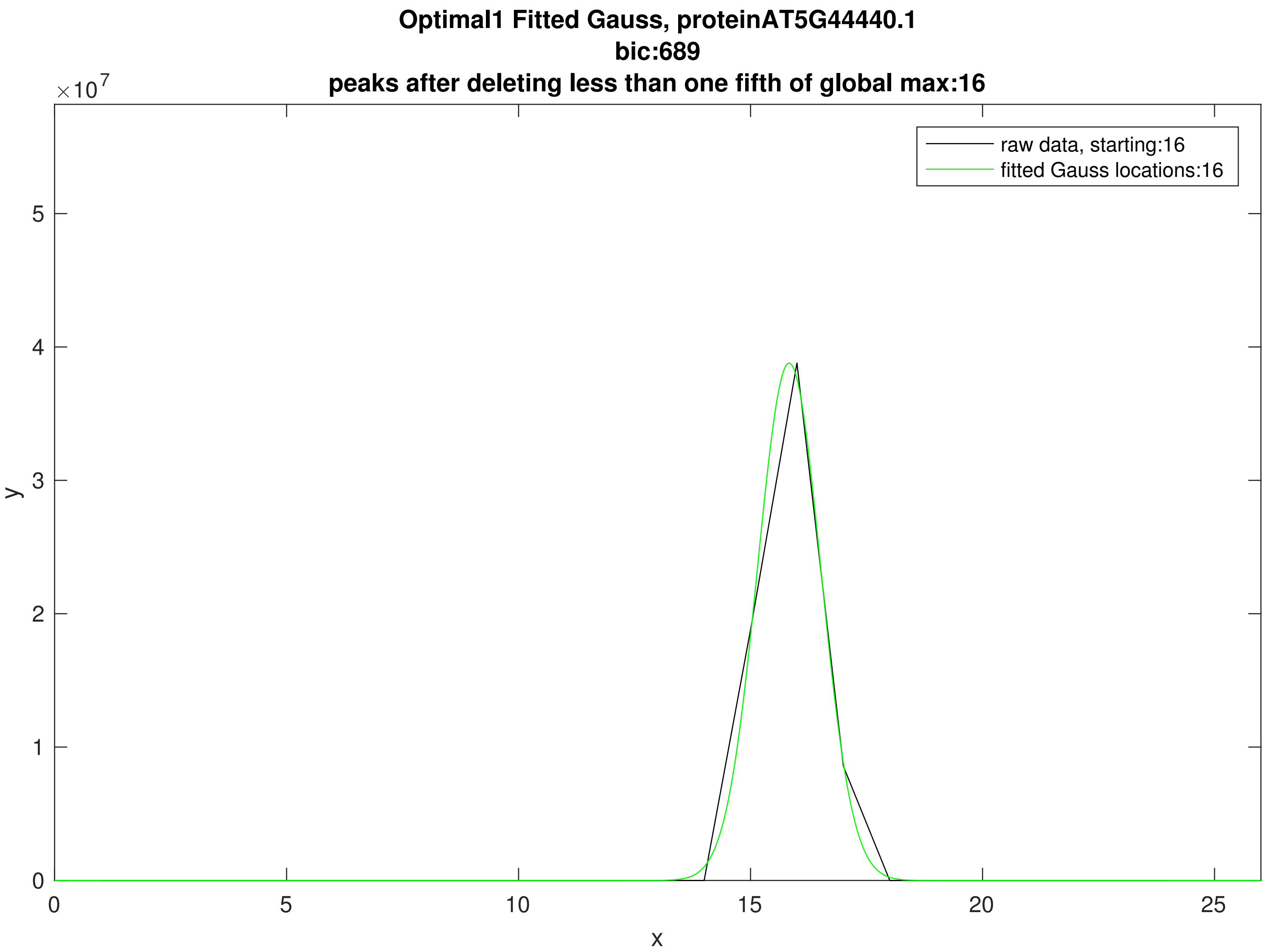

Optimal1 Fitted Gauss, proteinAT5G44440.1
bic:689
peaks after deleting less than one fifth of global max:16
×10^7
raw data, starting:16
fitted Gauss locations:16
y
x
0
1
2
3
4
5
0
5
10
15
20
25

#### Optimal1 Fitted Gauss, proteinAT5G44460.1
#### bic:784
#### peaks after deleting less than one fifth of global max:18

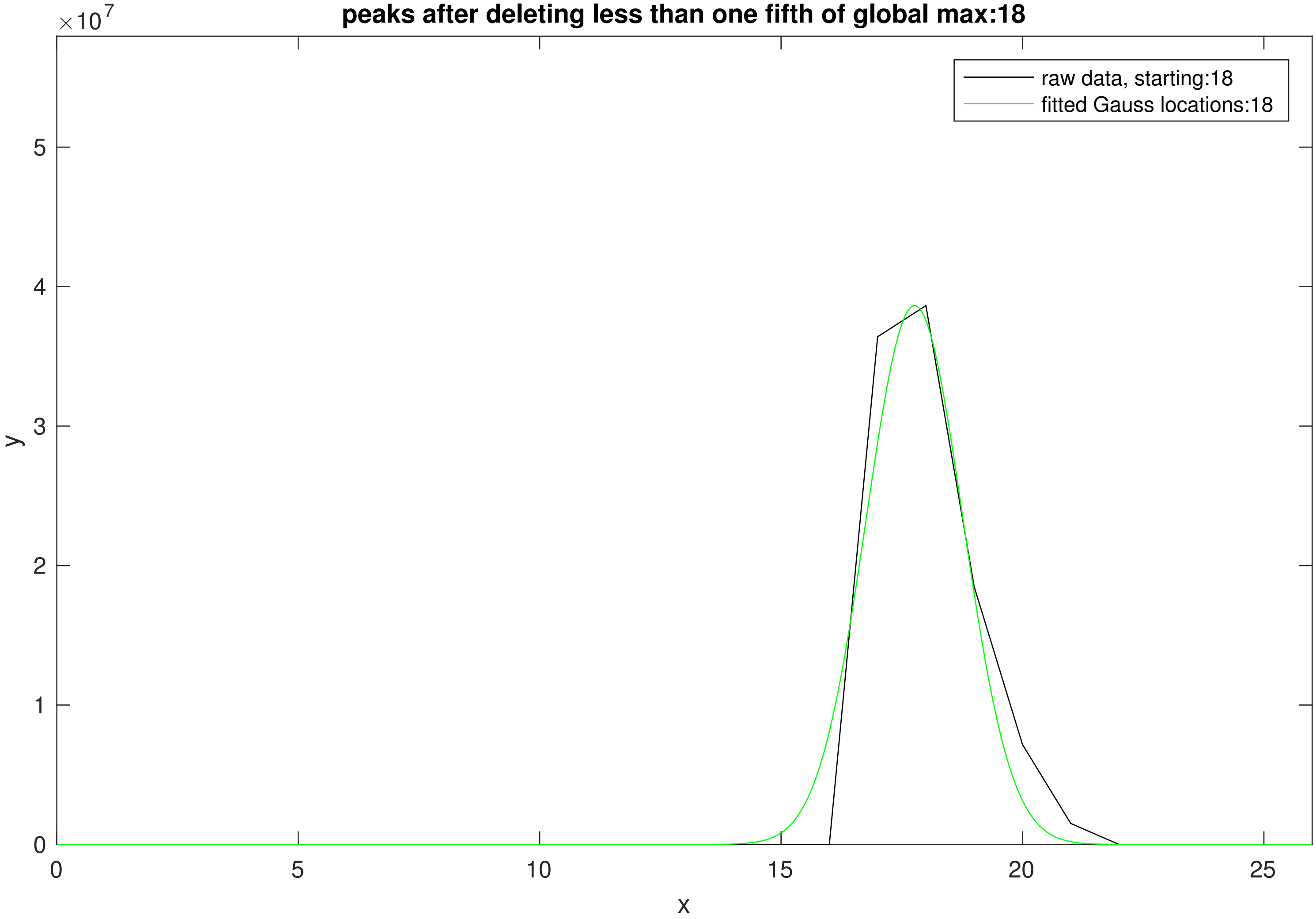

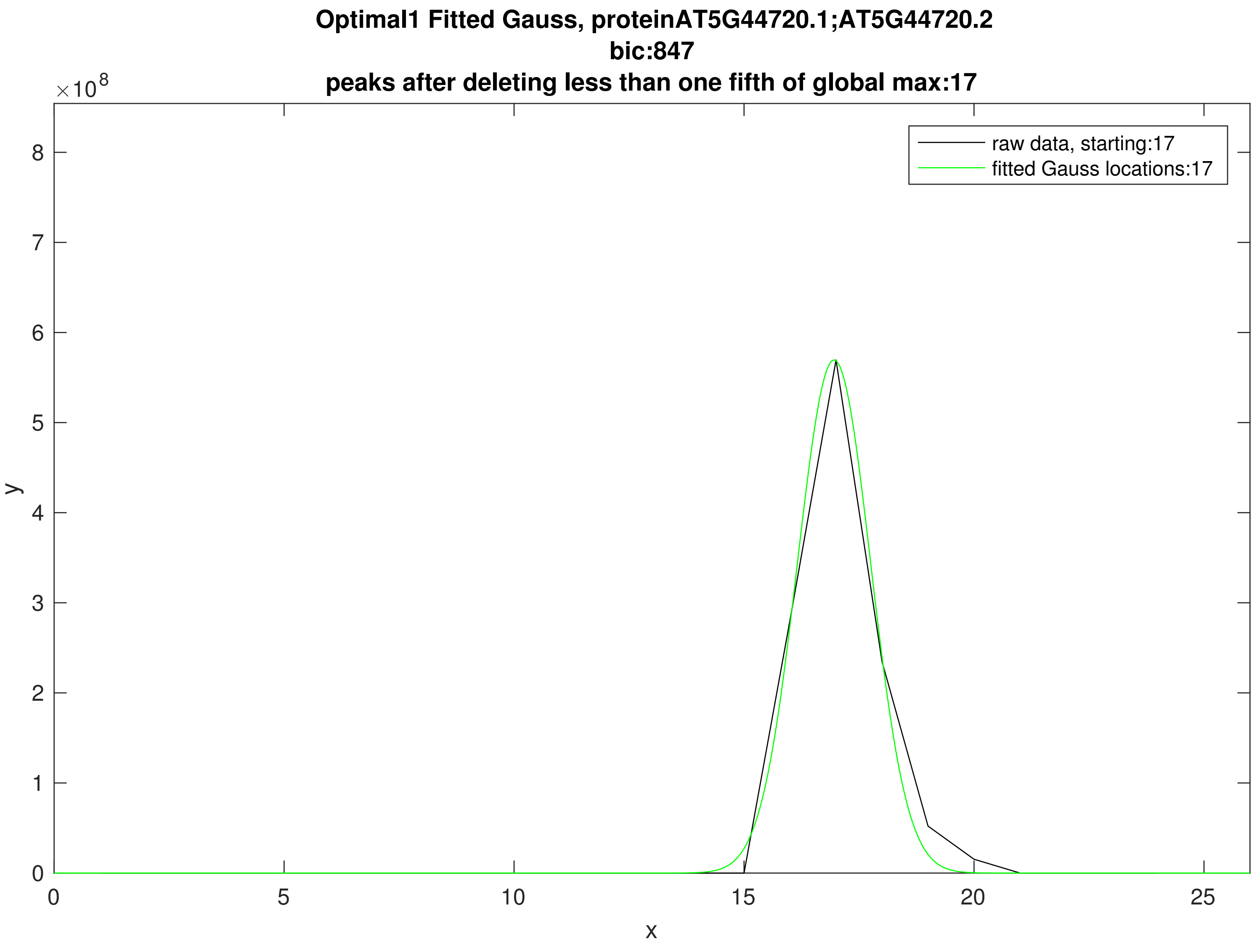

Optimal1 Fitted Gauss, proteinAT5G44720.1;AT5G44720.2
bic:847
peaks after deleting less than one fifth of global max:17
×10^8
raw data, starting:17
fitted Gauss locations:17
y
x
0
1
2
3
4
5
6
7
8
0
5
10
15
20
25

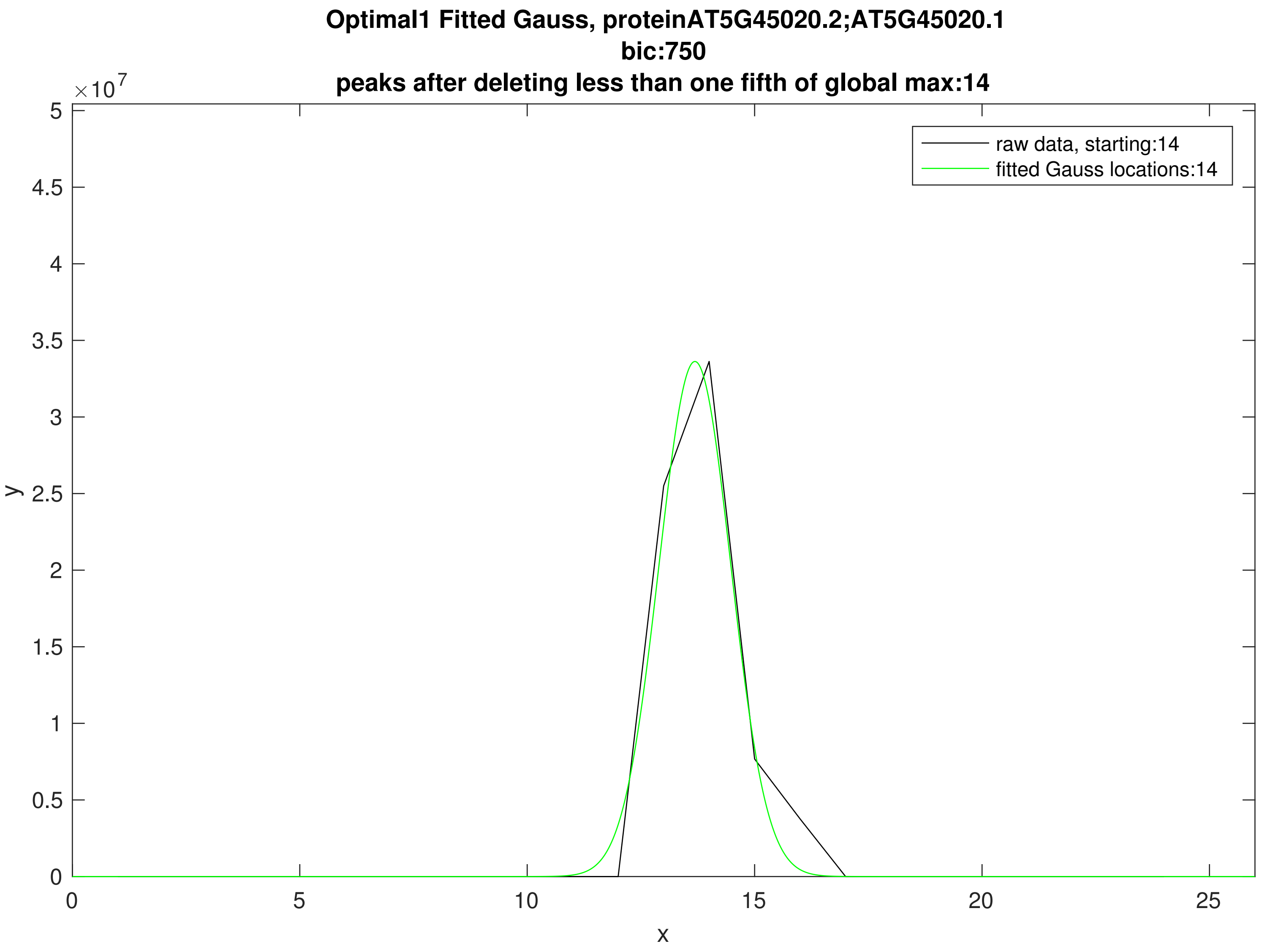
Optimal1 Fitted Gauss, proteinAT5G45020.2;AT5G45020.1
bic:750
peaks after deleting less than one fifth of global max:14
×10^7
5
4.5
4
3.5
3
2.5
2
1.5
1
0.5
0
y
0
5
10
15
20
25
x
raw data, starting:14
fitted Gauss locations:14

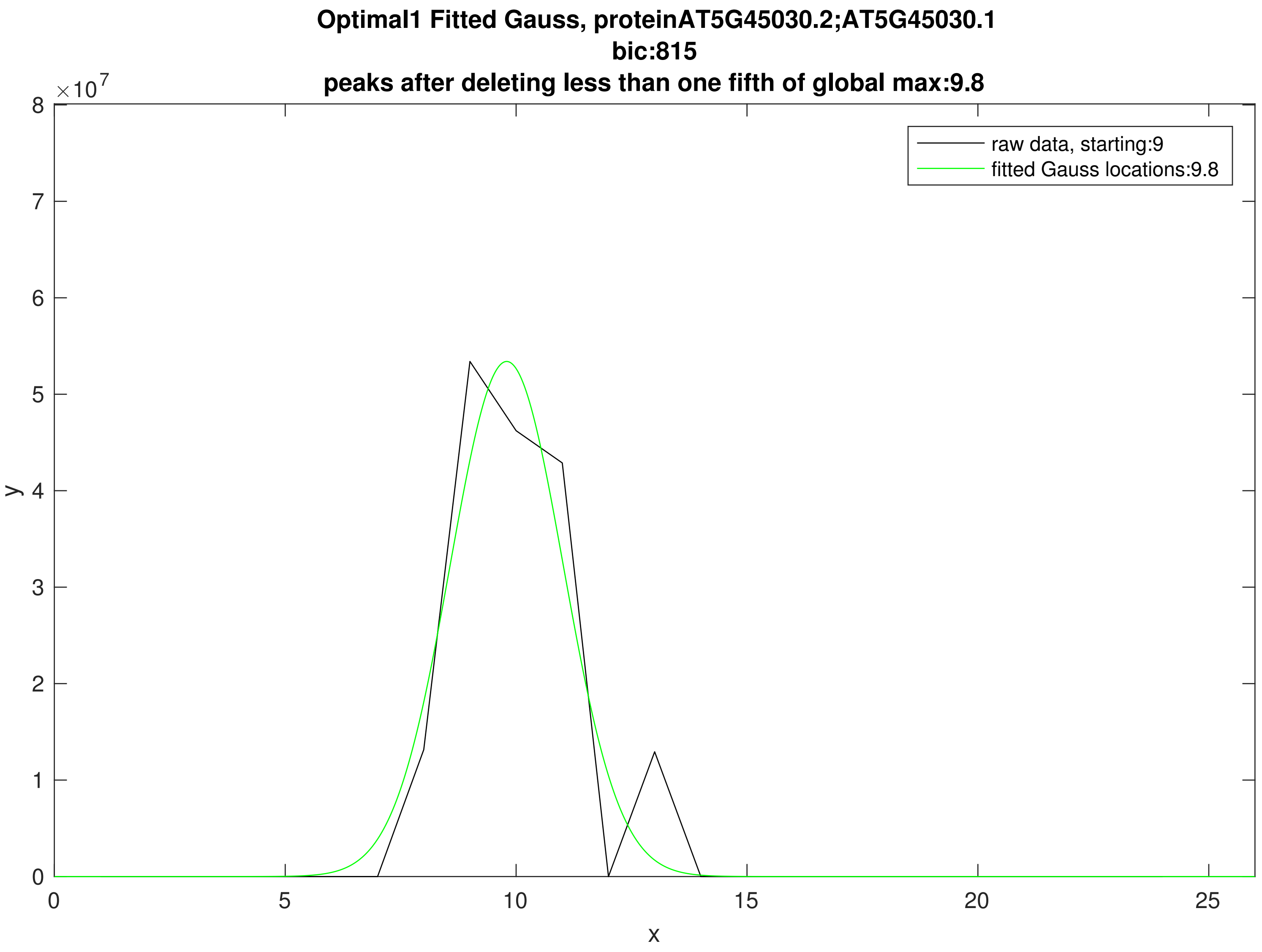
Optimal1 Fitted Gauss, proteinAT5G45030.2;AT5G45030.1
bic:815
peaks after deleting less than one fifth of global max:9.8
×10^7
8
7
6
5
4
3
2
1
0
y
0
5
10
15
20
25
x
raw data, starting:9
fitted Gauss locations:9.8

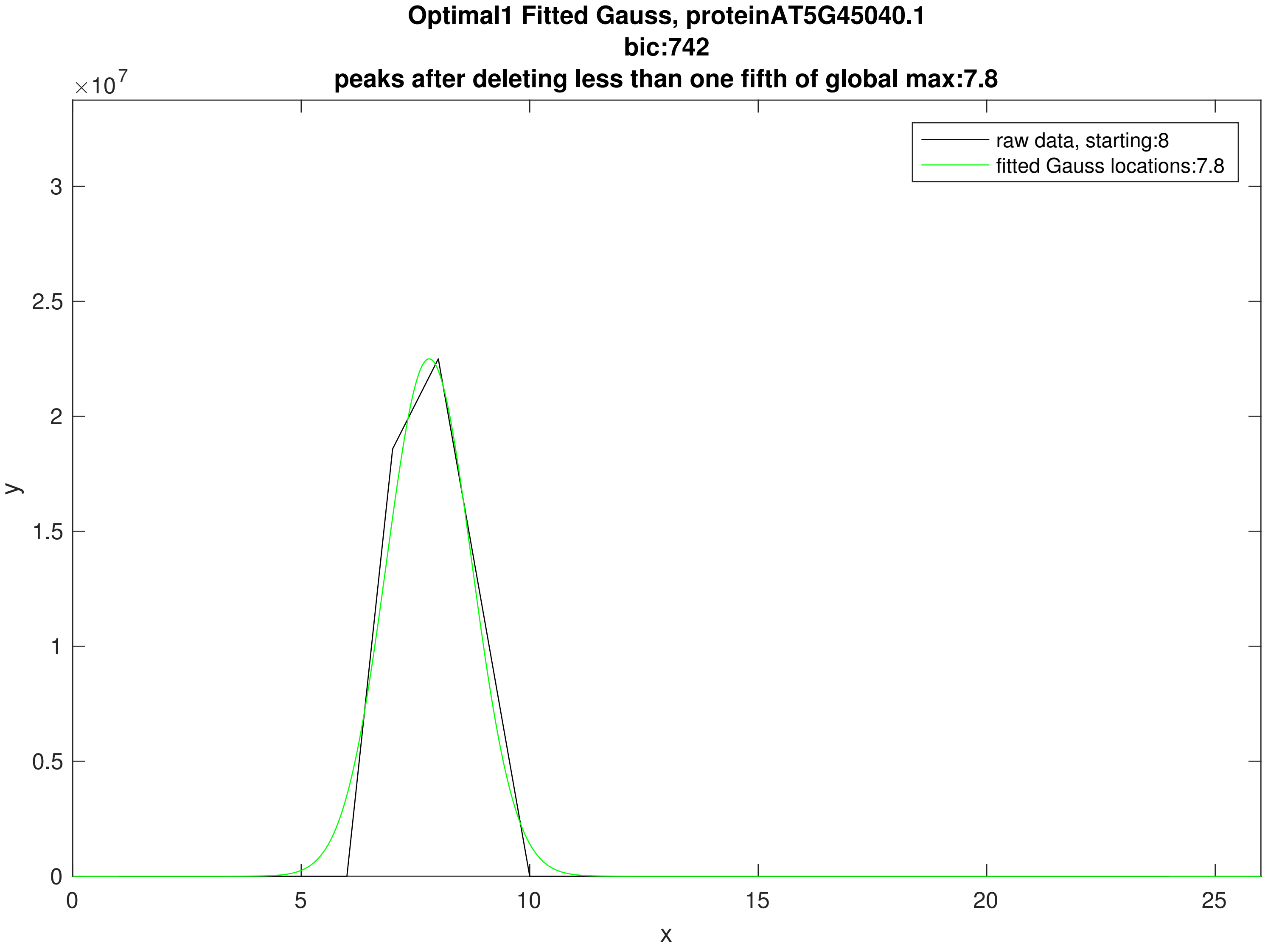
Optimal1 Fitted Gauss, proteinAT5G45040.1
bic:742
peaks after deleting less than one fifth of global max:7.8
×10^7
raw data, starting:8
fitted Gauss locations:7.8
y
x
0
0.5
1
1.5
2
2.5
3
0
5
10
15
20
25

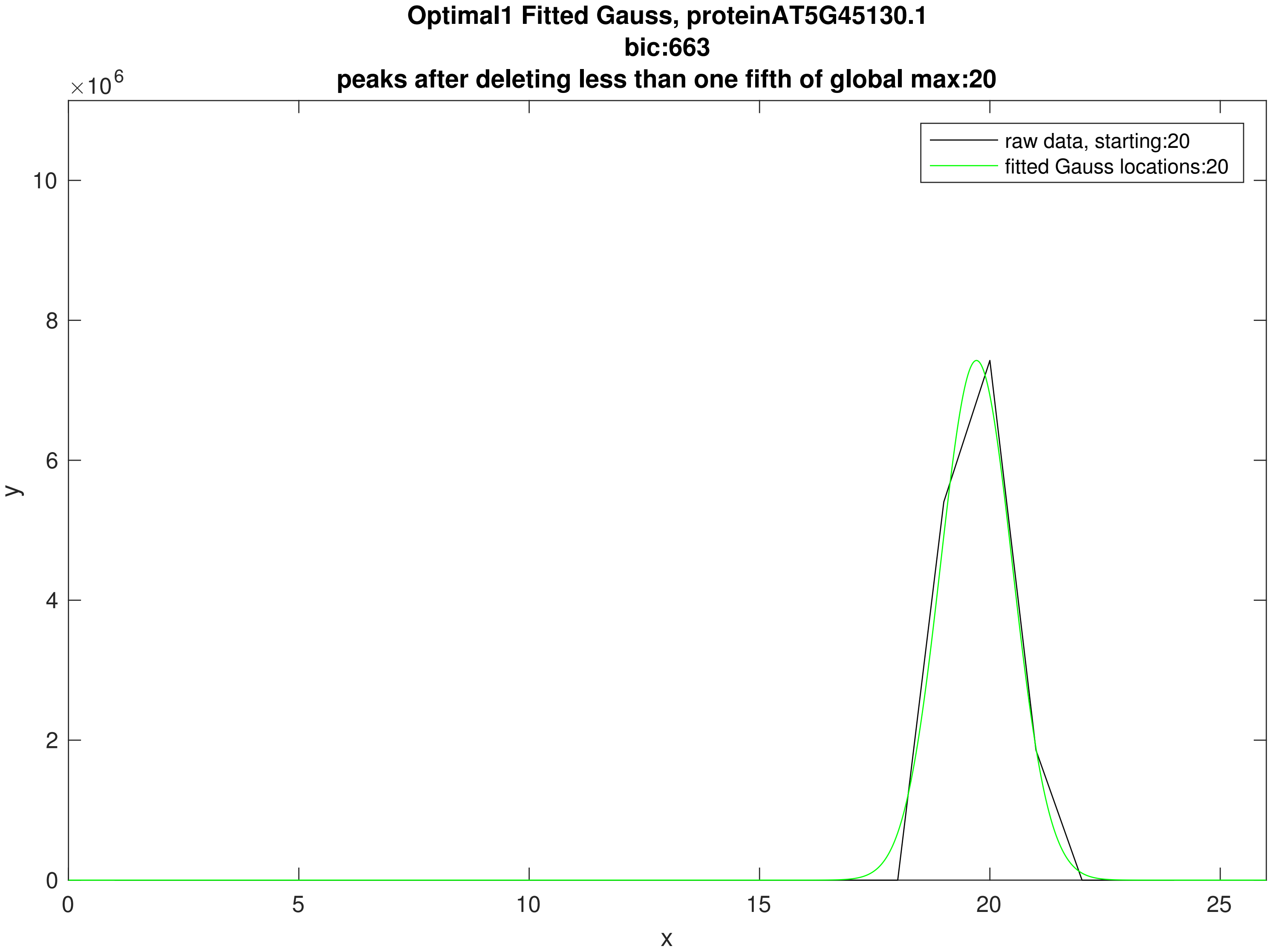

Optimal1 Fitted Gauss, proteinAT5G45130.1
bic:663
peaks after deleting less than one fifth of global max:20
×10^6
raw data, starting:20
fitted Gauss locations:20
10
8
6
4
2
0
y
0
5
10
15
20
25
x

**Optimal1 Fitted Gauss, proteinAT5G45280.2;AT5G45280.1**
**bic:912 921**
**peaks after deleting less than one fifth of global max:18**

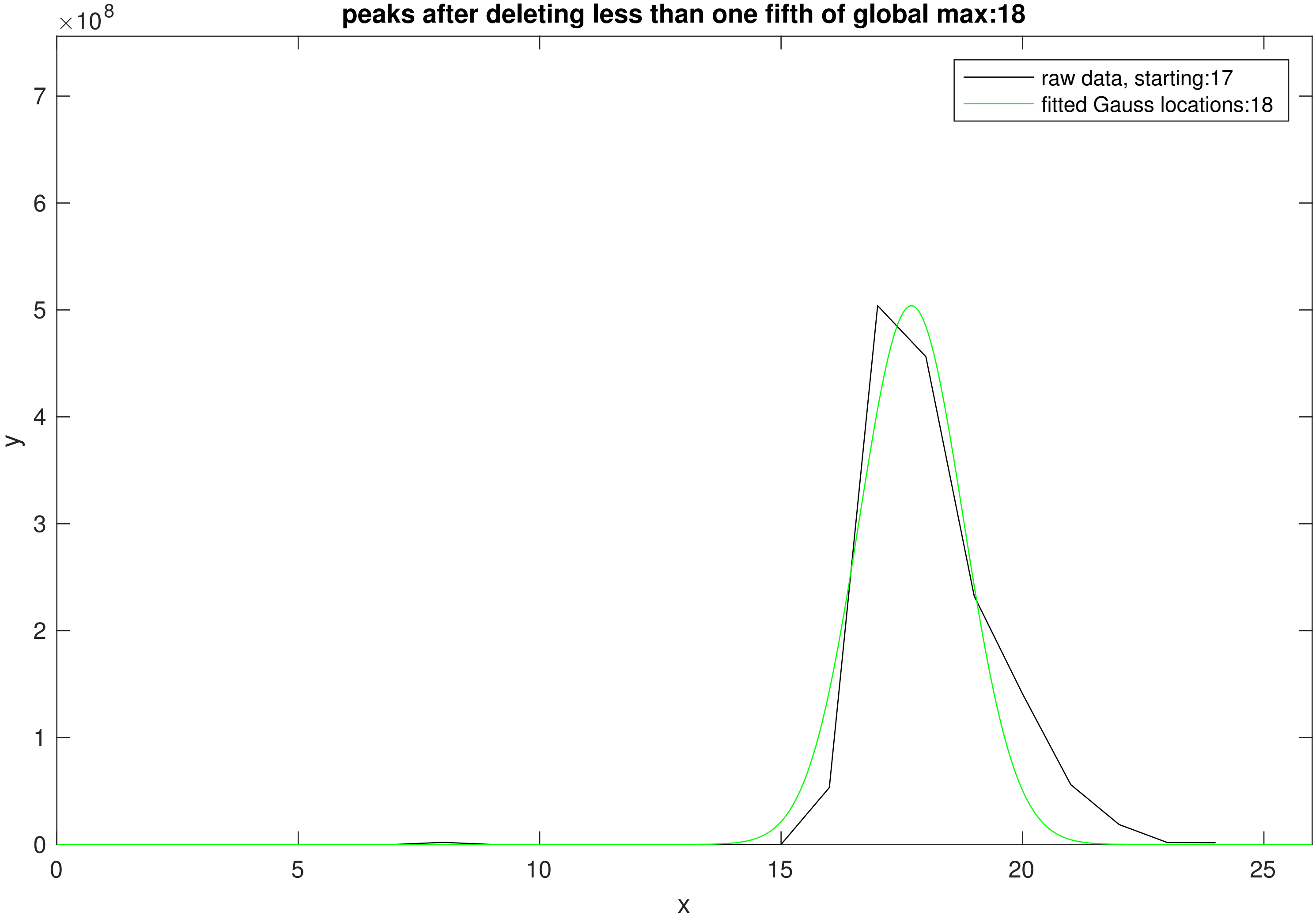

Optimal1 Fitted Gauss, proteinAT5G45390.1
bic:833 842
peaks after deleting less than one fifth of global max:8.5

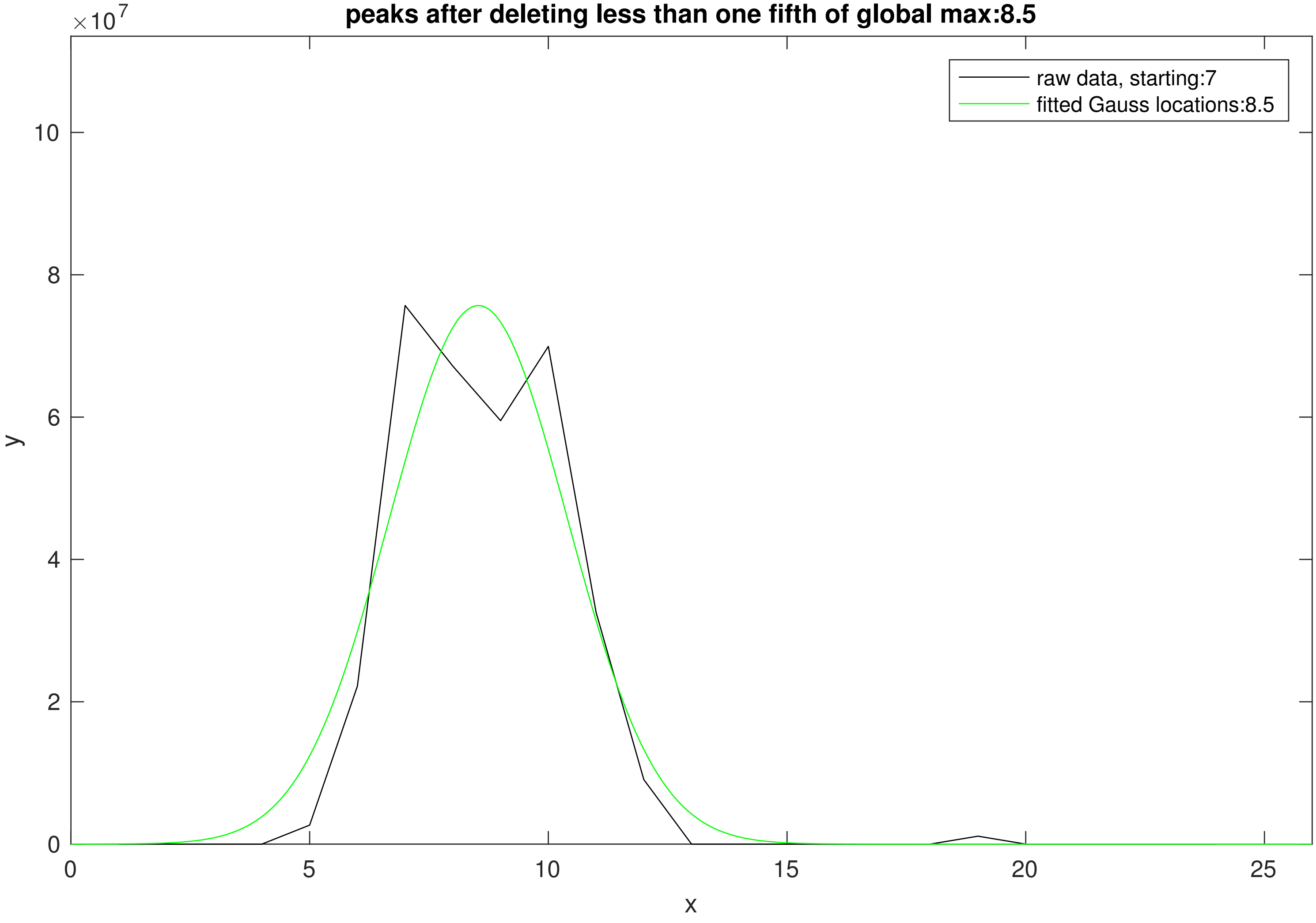

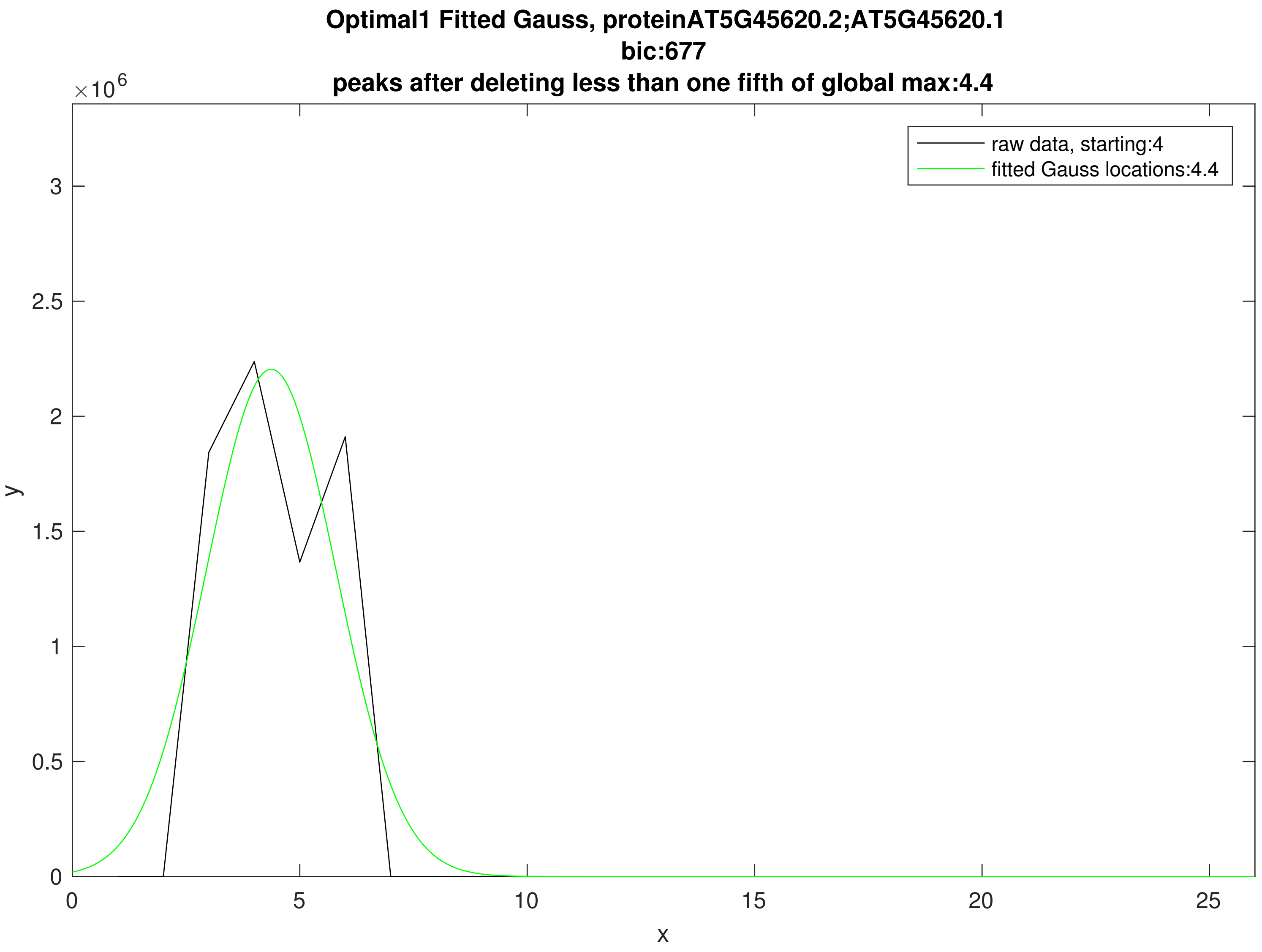
Optimal1 Fitted Gauss, proteinAT5G45620.2;AT5G45620.1
bic:677
peaks after deleting less than one fifth of global max:4.4
×10^6
raw data, starting:4
fitted Gauss locations:4.4
y
x
0
0.5
1
1.5
2
2.5
3
0
5
10
15
20
25

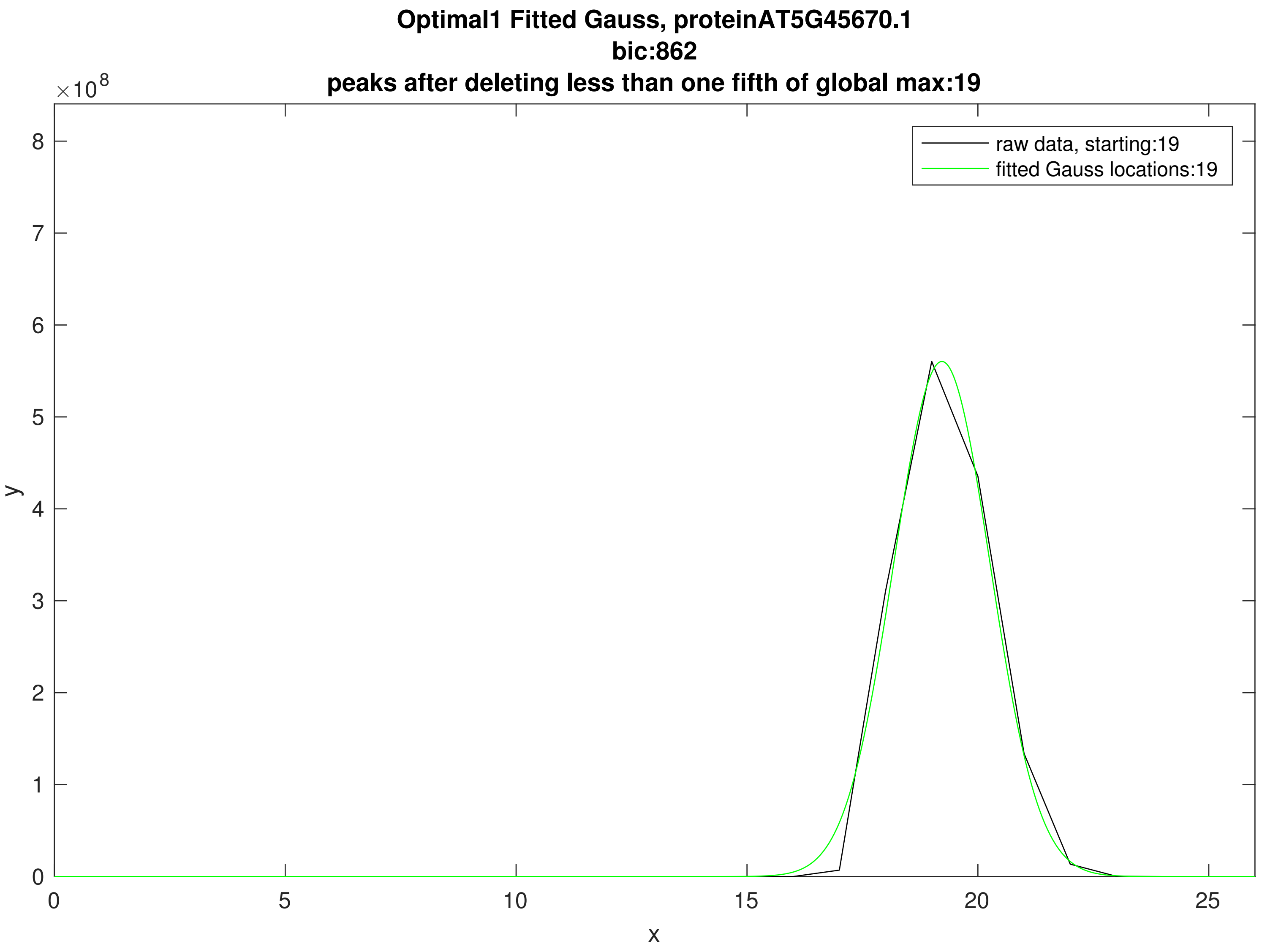

Optimal1 Fitted Gauss, proteinAT5G45670.1
bic:862
peaks after deleting less than one fifth of global max:19
×10^8
raw data, starting:19
fitted Gauss locations:19
y
x
0
1
2
3
4
5
6
7
8
0
5
10
15
20
25

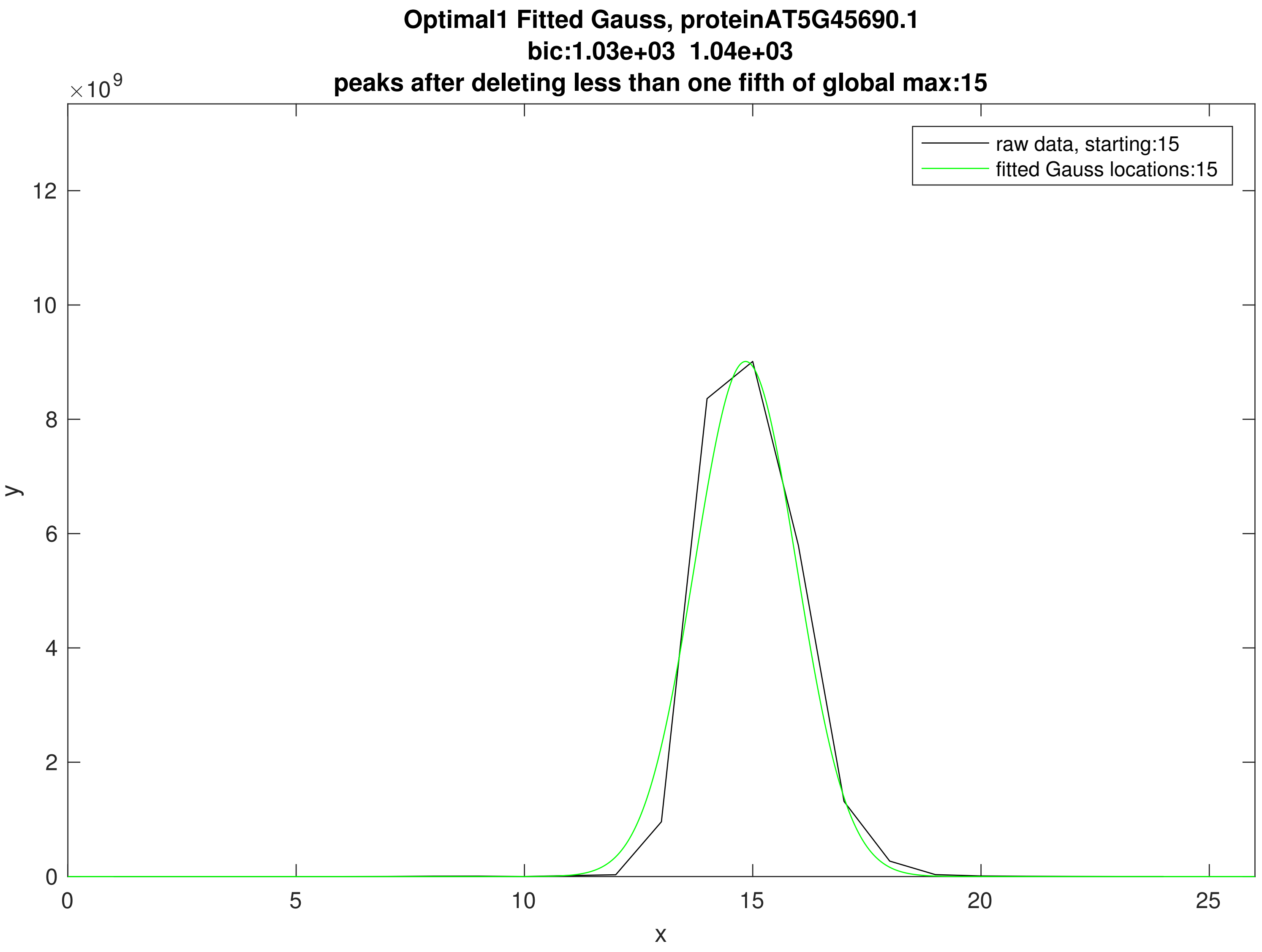

Optimal1 Fitted Gauss, proteinAT5G45690.1
bic:1.03e+03  1.04e+03
peaks after deleting less than one fifth of global max:15
×10^9
raw data, starting:15
fitted Gauss locations:15
12
10
8
6
4
2
0
y
0
5
10
15
20
25
x

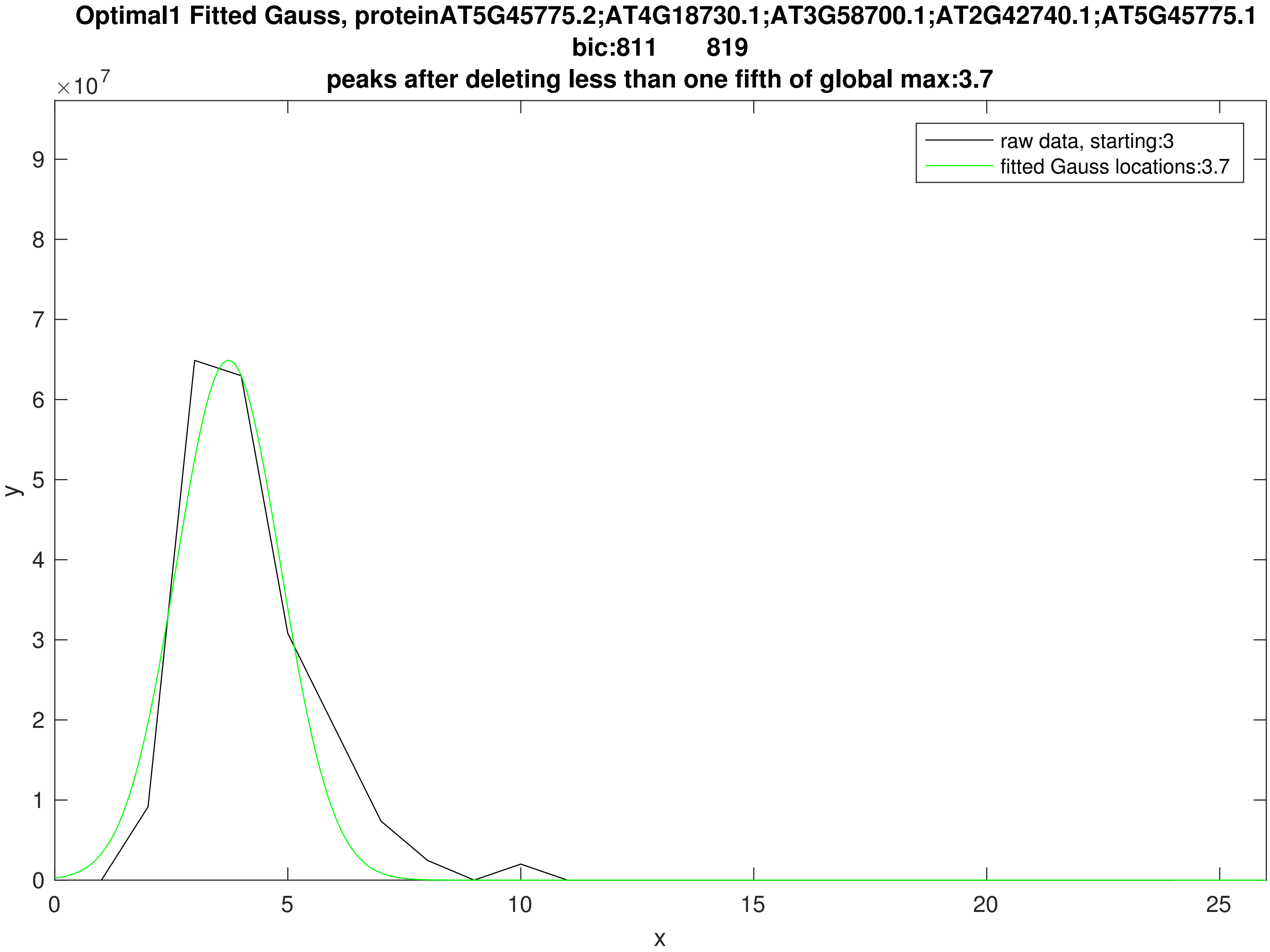
Optimal1 Fitted Gauss, proteinAT5G45775.2;AT4G18730.1;AT3G58700.1;AT2G42740.1;AT5G45775.1
bic:811 819
peaks after deleting less than one fifth of global max:3.7
×10^7
raw data, starting:3
fitted Gauss locations:3.7
y
x
0
1
2
3
4
5
6
7
8
9
0
5
10
15
20
25

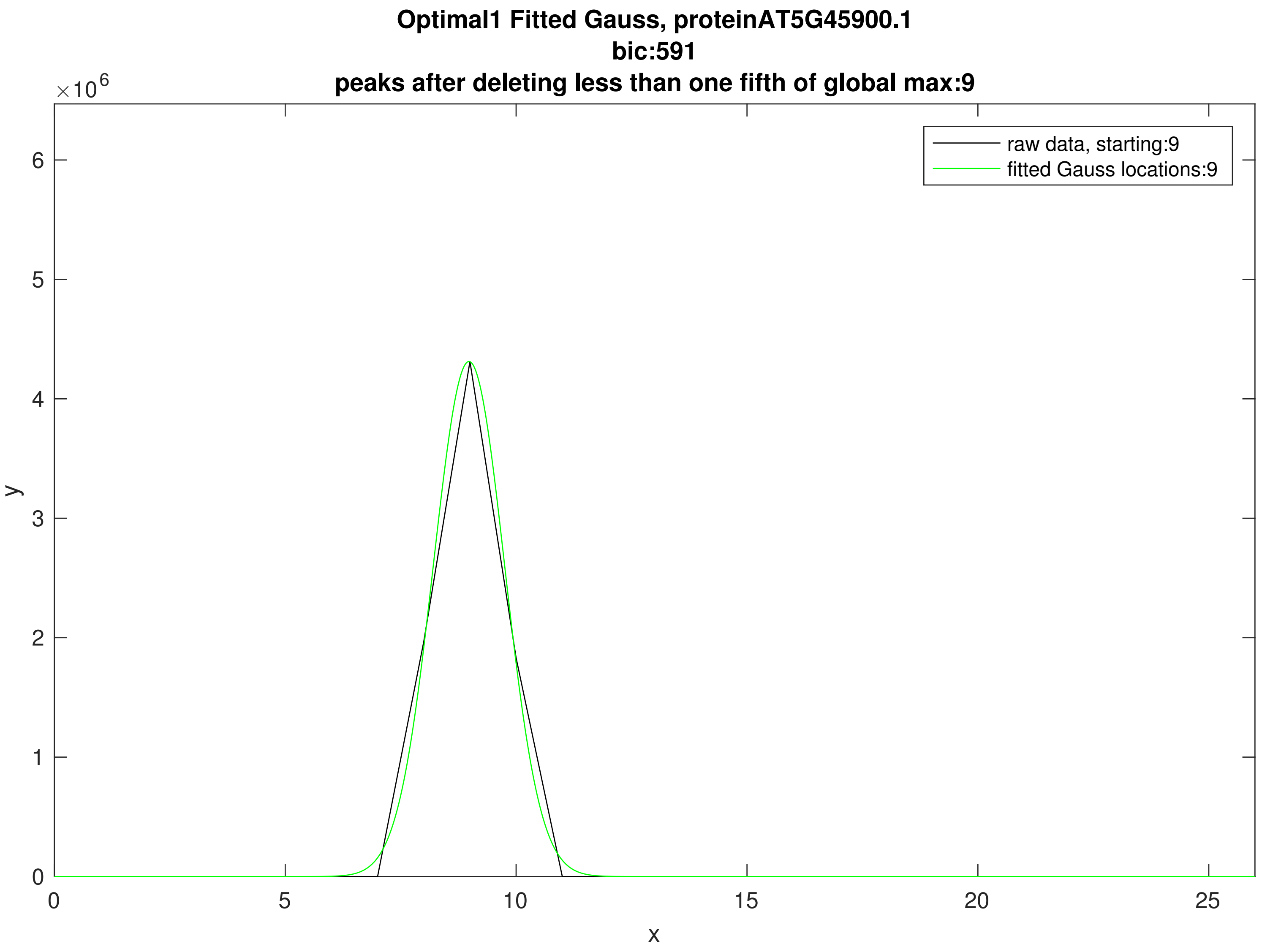
Optimal1 Fitted Gauss, proteinAT5G45900.1
bic:591
peaks after deleting less than one fifth of global max:9
×10^6
raw data, starting:9
fitted Gauss locations:9
y
x
0
1
2
3
4
5
6
0
5
10
15
20
25

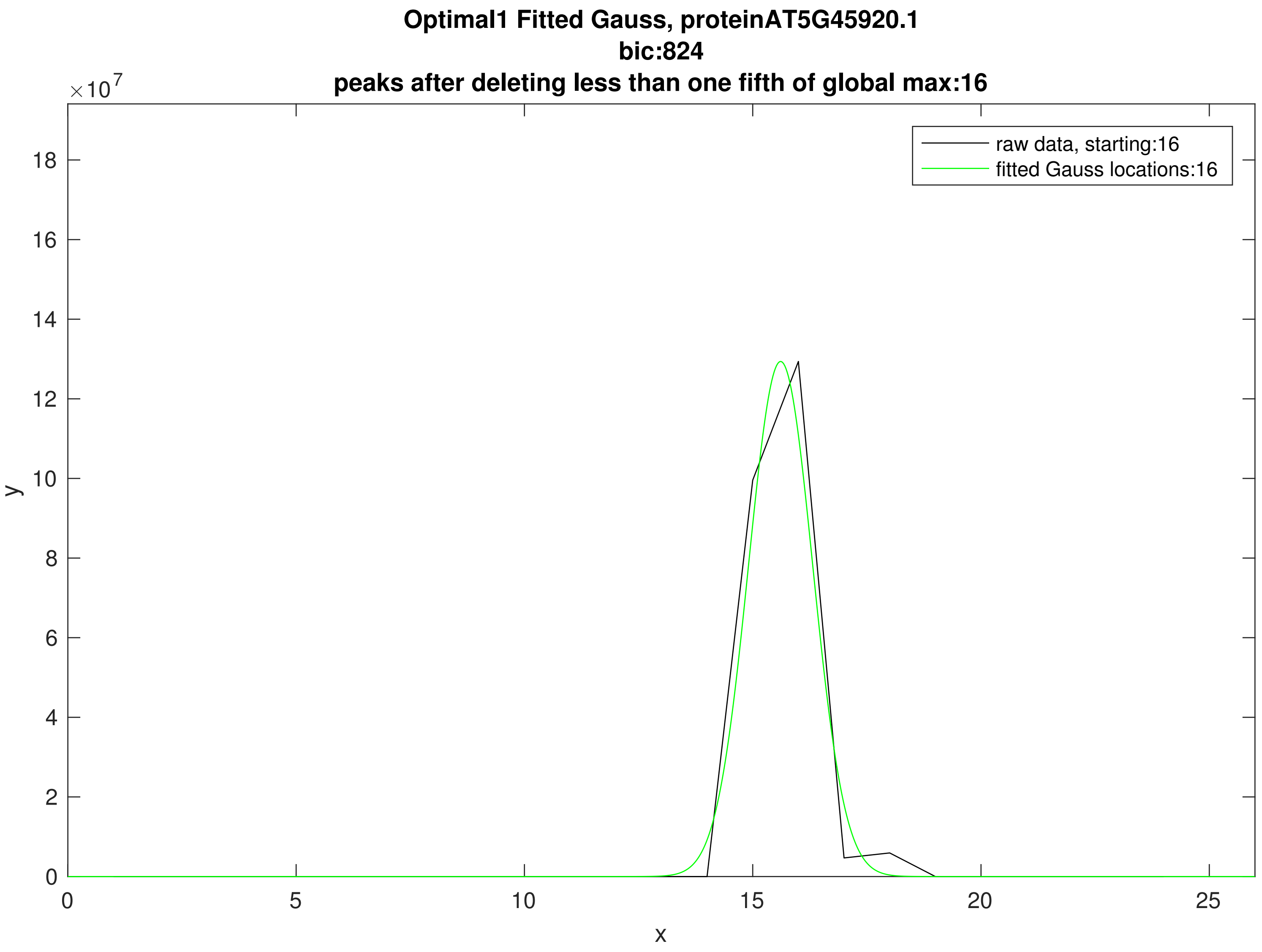

Optimal1 Fitted Gauss, proteinAT5G45920.1
bic:824
peaks after deleting less than one fifth of global max:16
×10^7
raw data, starting:16
fitted Gauss locations:16
y
x
0
2
4
6
8
10
12
14
16
18
0
5
10
15
20
25

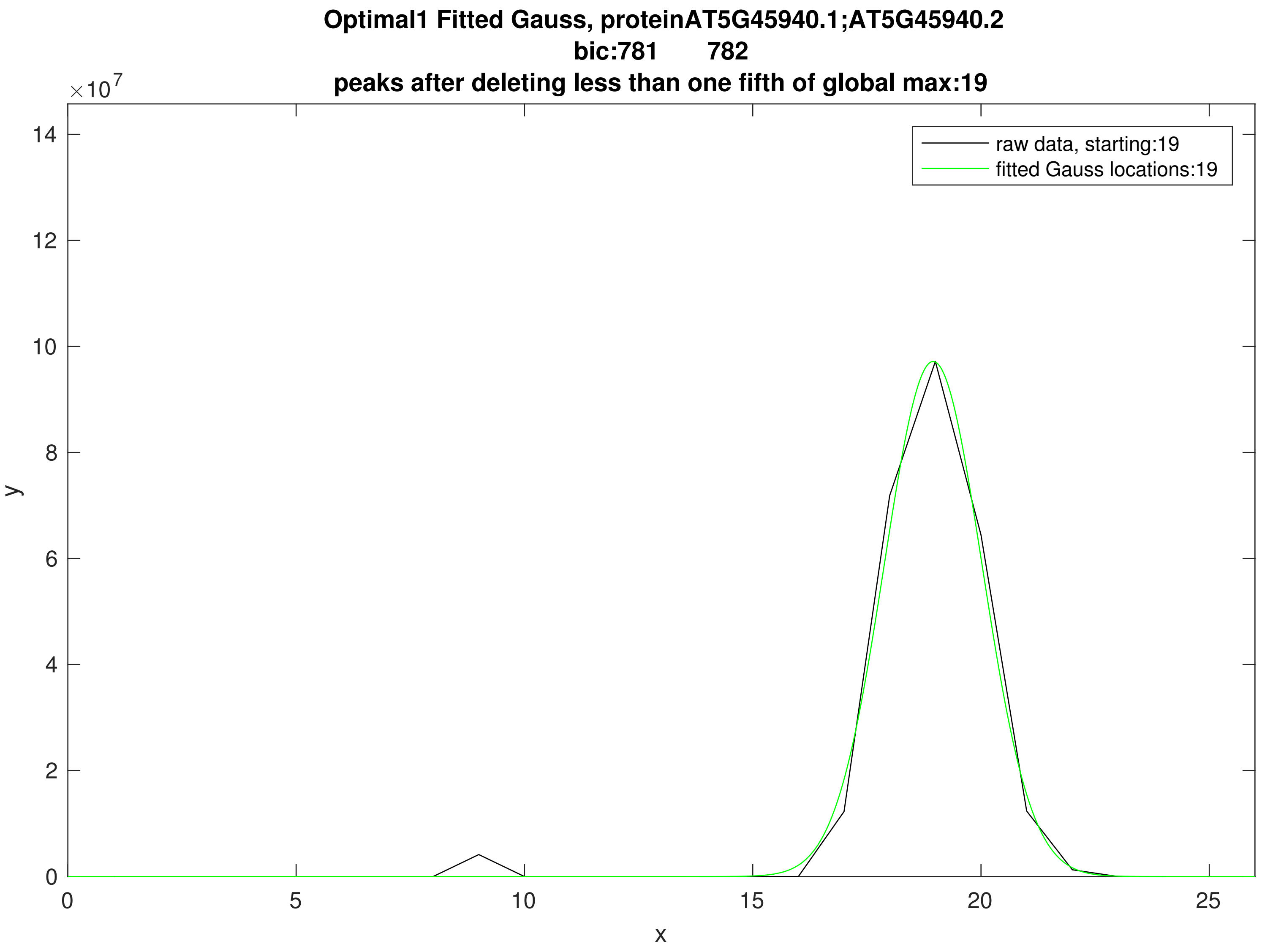

Optimal1 Fitted Gauss, proteinAT5G45940.1;AT5G45940.2
bic:781 782
peaks after deleting less than one fifth of global max:19
×10^7
raw data, starting:19
fitted Gauss locations:19
y
x
0
2
4
6
8
10
12
14
5
15
20
25

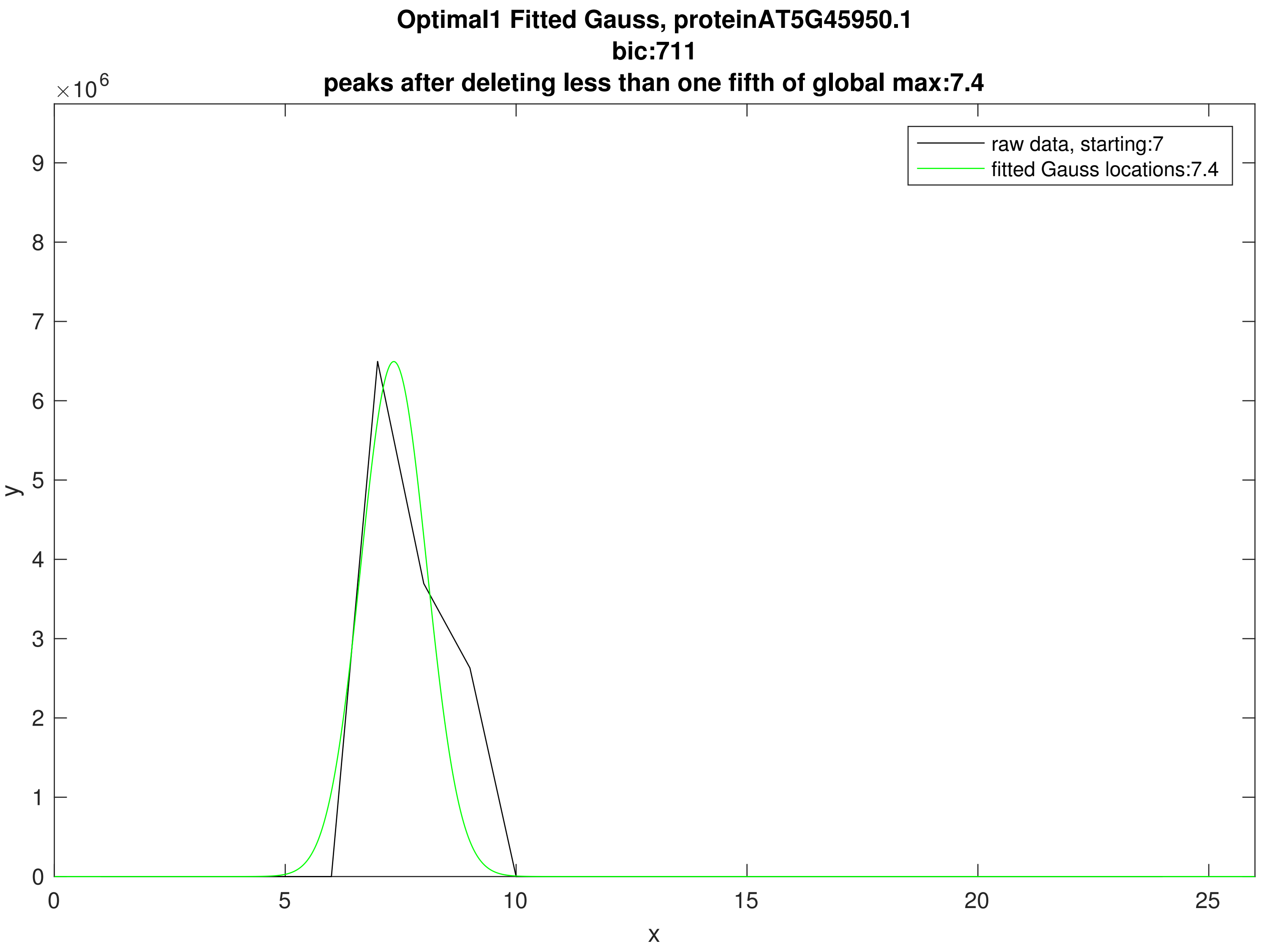

Optimal1 Fitted Gauss, proteinAT5G45950.1
bic:711
peaks after deleting less than one fifth of global max:7.4
×10^6
raw data, starting:7
fitted Gauss locations:7.4
y
x
0
1
2
3
4
5
6
7
8
9
0
5
10
15
20
25

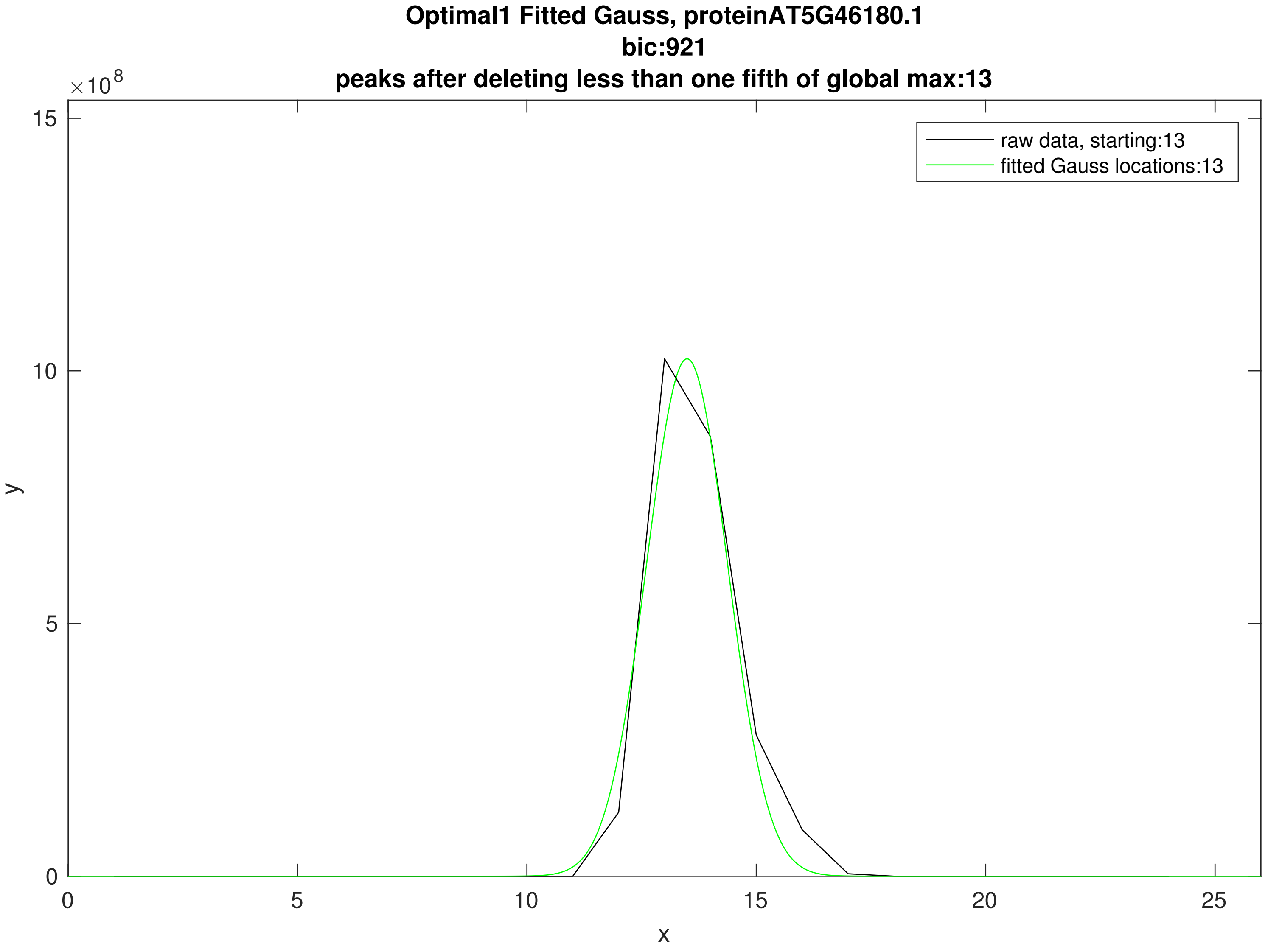
Optimal1 Fitted Gauss, proteinAT5G46180.1
bic:921
peaks after deleting less than one fifth of global max:13
×10^8
15
10
5
0
y
0
5
10
15
20
25
x
raw data, starting:13
fitted Gauss locations:13

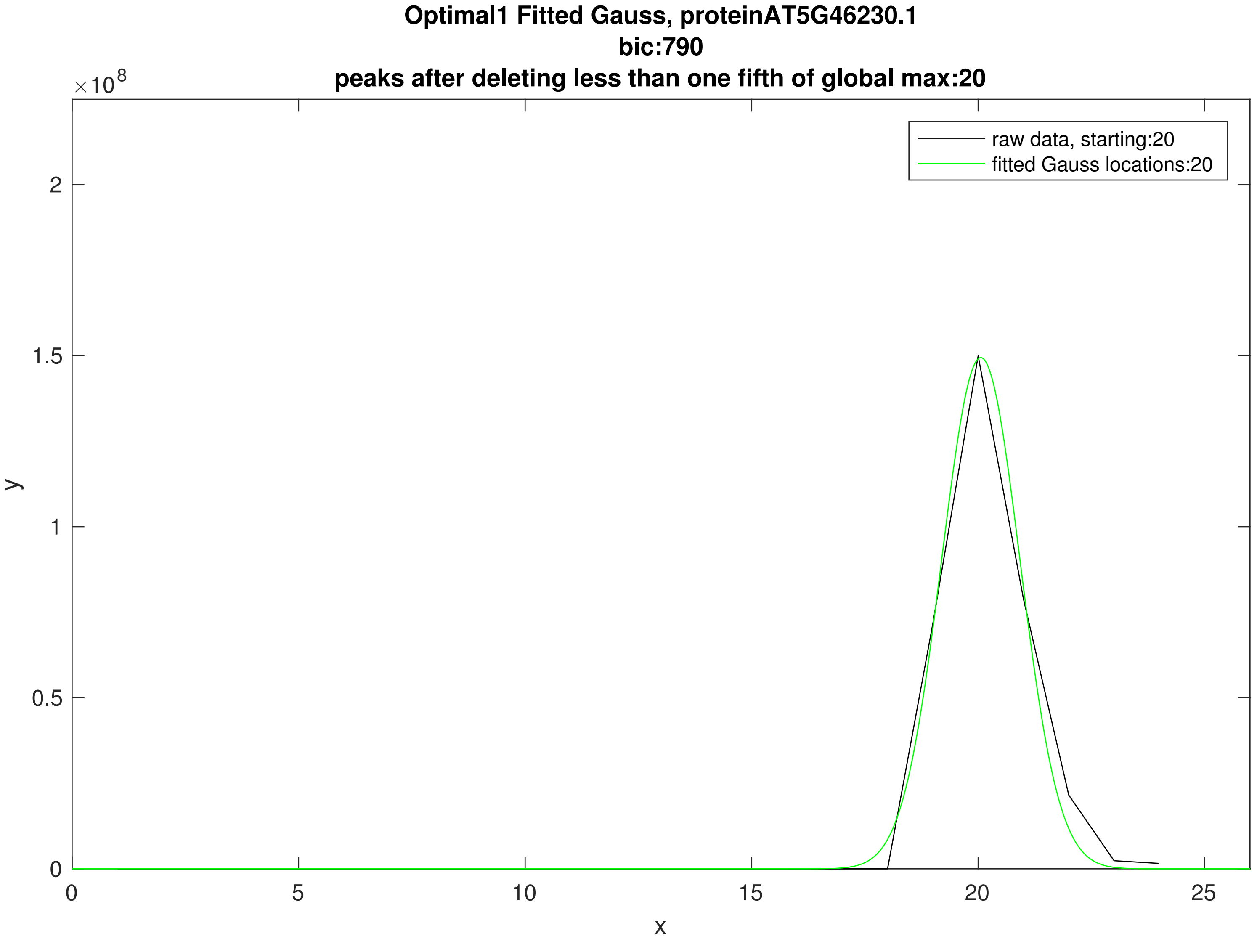
Optimal1 Fitted Gauss, proteinAT5G46230.1
bic:790
peaks after deleting less than one fifth of global max:20
×10^8
raw data, starting:20
fitted Gauss locations:20
2
1.5
1
0.5
0
y
0
5
10
15
20
25
x

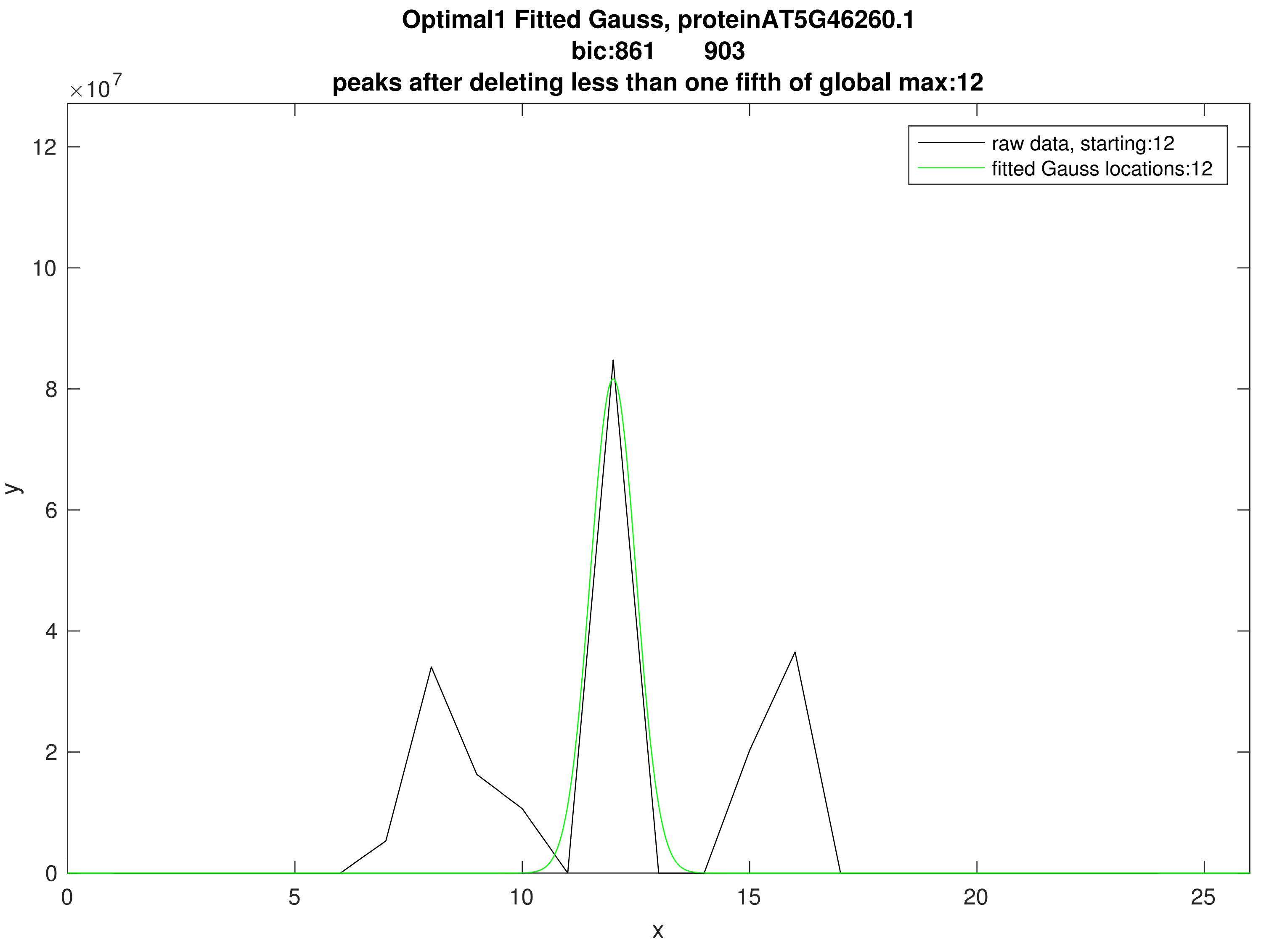
Optimal1 Fitted Gauss, proteinAT5G46260.1
bic:861 903
peaks after deleting less than one fifth of global max:12
×10^7
raw data, starting:12
fitted Gauss locations:12
y
x
0
2
4
6
8
10
12
0
5
10
15
20
25

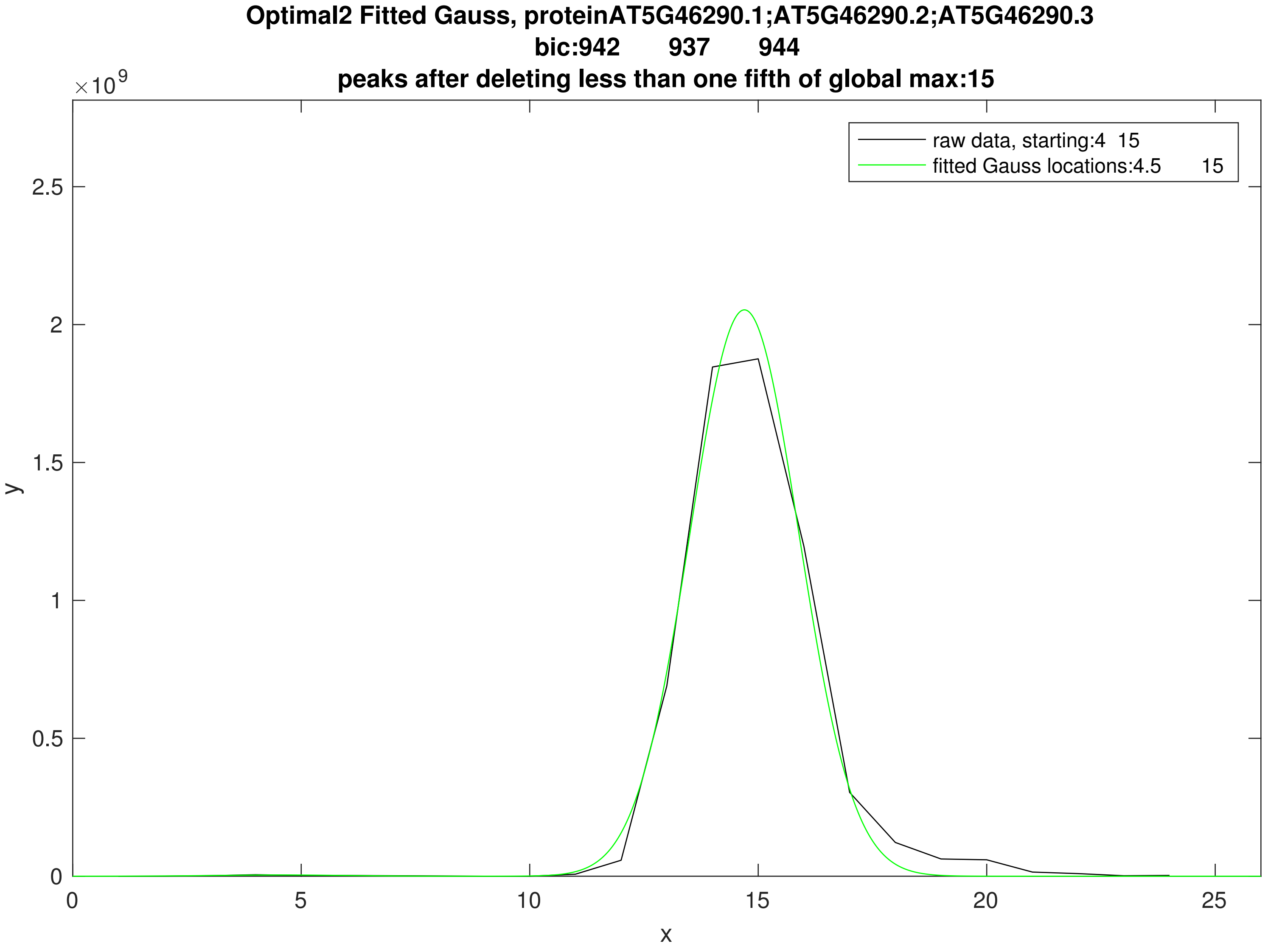
Optimal2 Fitted Gauss, proteinAT5G46290.1;AT5G46290.2;AT5G46290.3
bic:942 937 944
peaks after deleting less than one fifth of global max:15
×10^9
raw data, starting:4 15
fitted Gauss locations:4.5 15
y
x
0
0.5
1
1.5
2
2.5
5
10
15
20
25

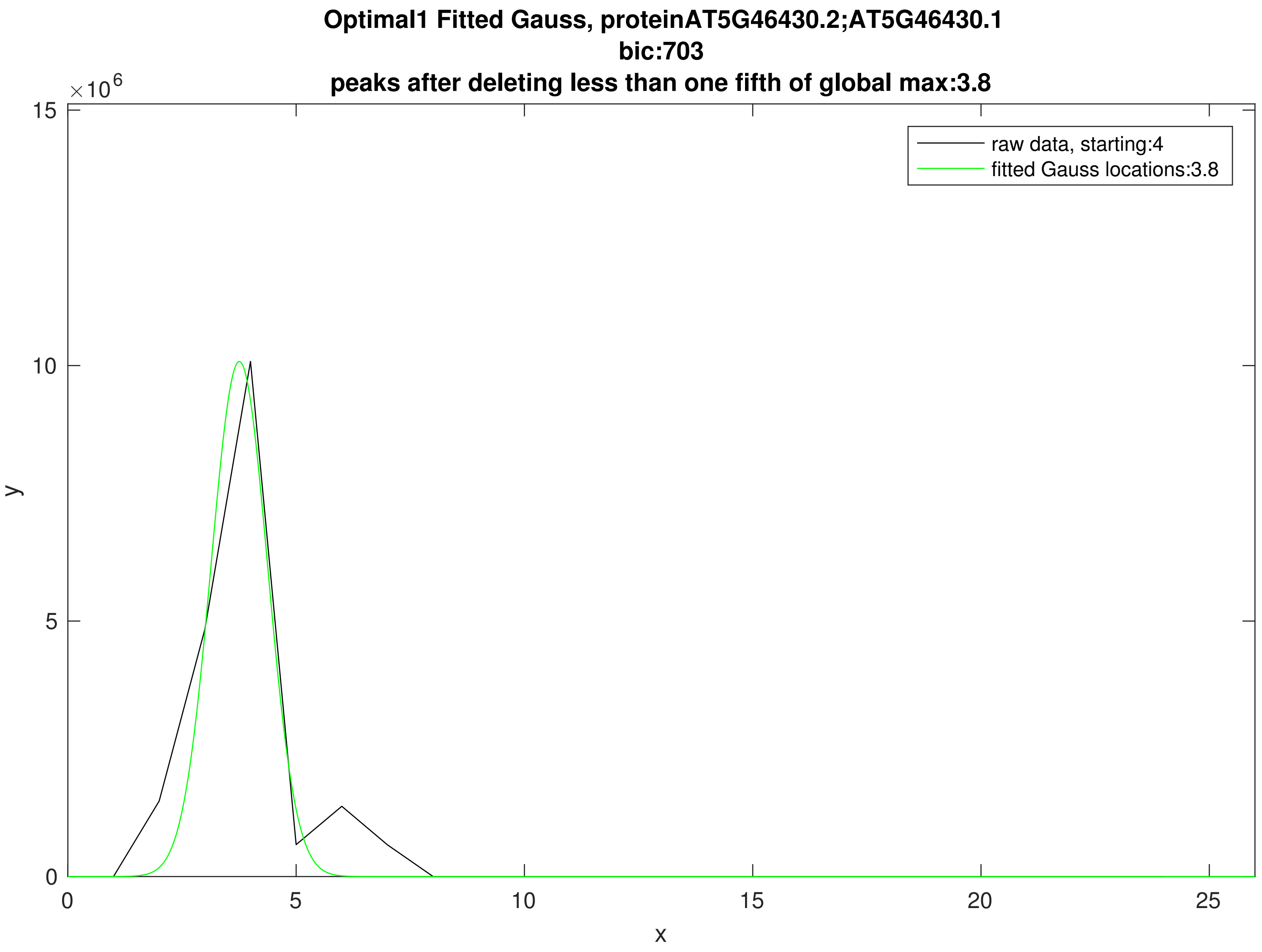

Optimal1 Fitted Gauss, proteinAT5G46430.2;AT5G46430.1
bic:703
peaks after deleting less than one fifth of global max:3.8
×10^6
15
10
5
0
y
0
5
10
15
20
25
x
raw data, starting:4
fitted Gauss locations:3.8

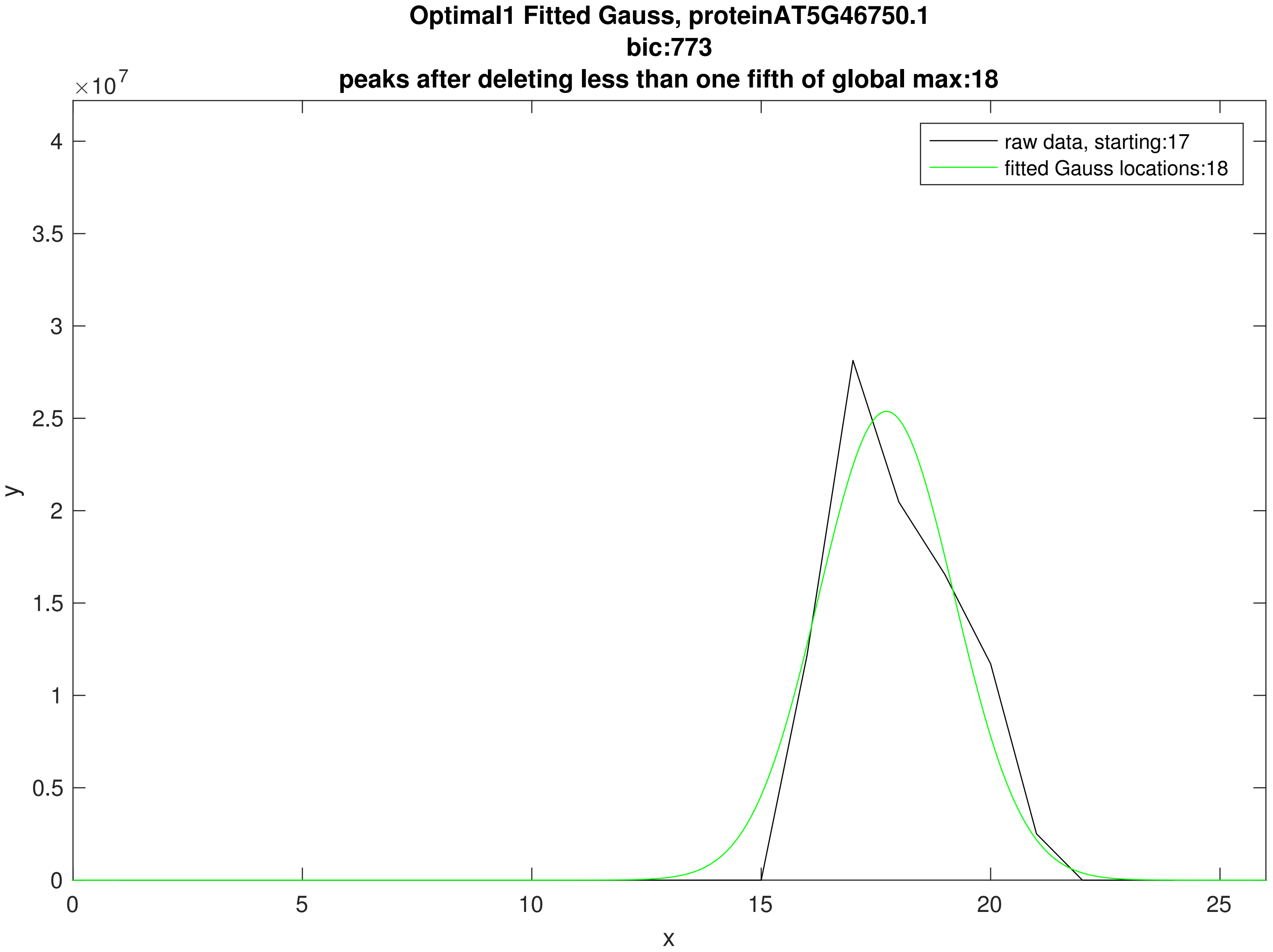
Optimal1 Fitted Gauss, proteinAT5G46750.1
bic:773
peaks after deleting less than one fifth of global max:18
×10^7
raw data, starting:17
fitted Gauss locations:18
y
x
0
0.5
1
1.5
2
2.5
3
3.5
4
0
5
10
15
20
25

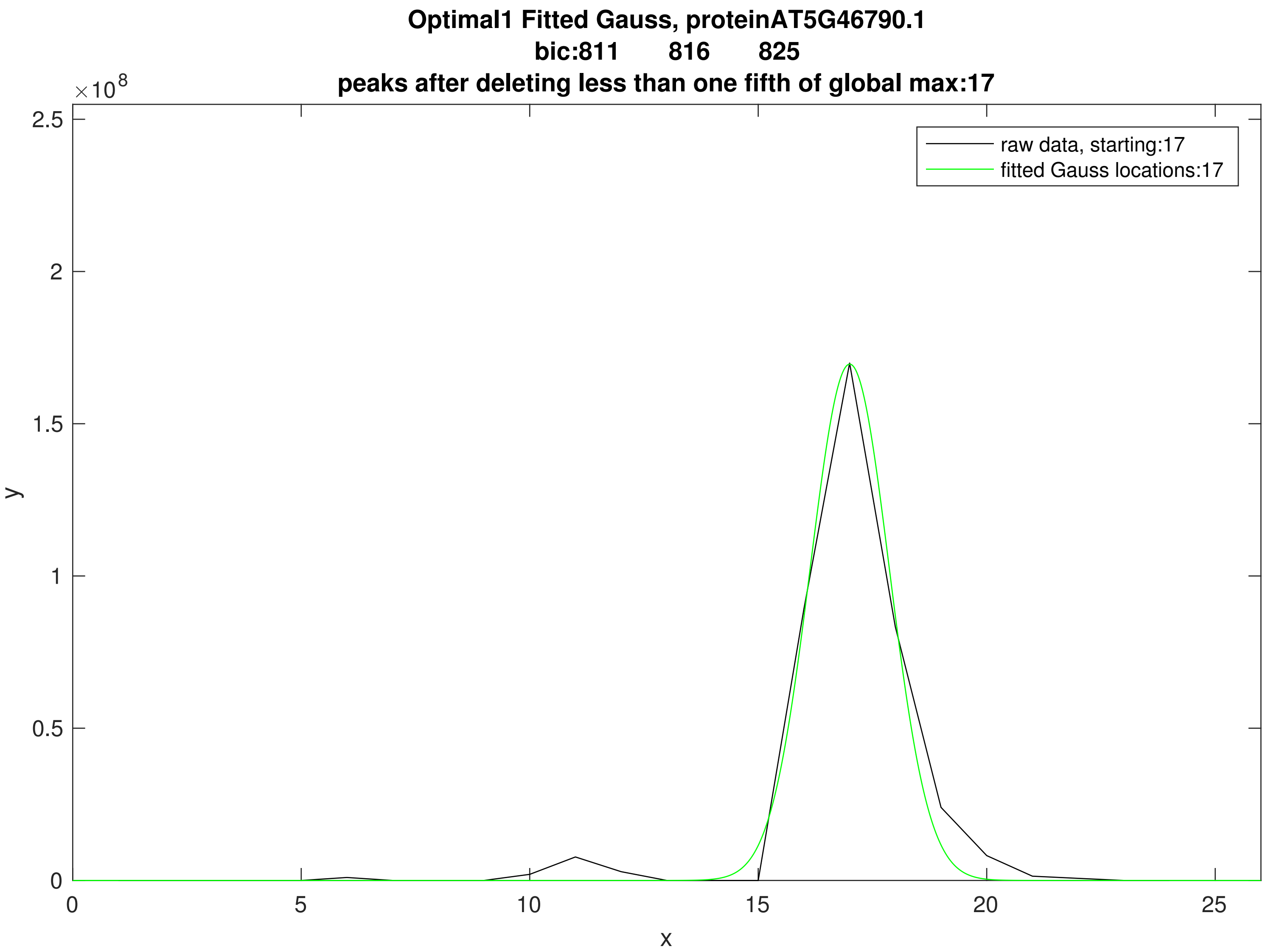

Optimal1 Fitted Gauss, proteinAT5G46790.1
bic:811 816 825
peaks after deleting less than one fifth of global max:17
×10^8
2.5
2
1.5
1
0.5
0
y
0
5
10
15
20
25
x
raw data, starting:17
fitted Gauss locations:17

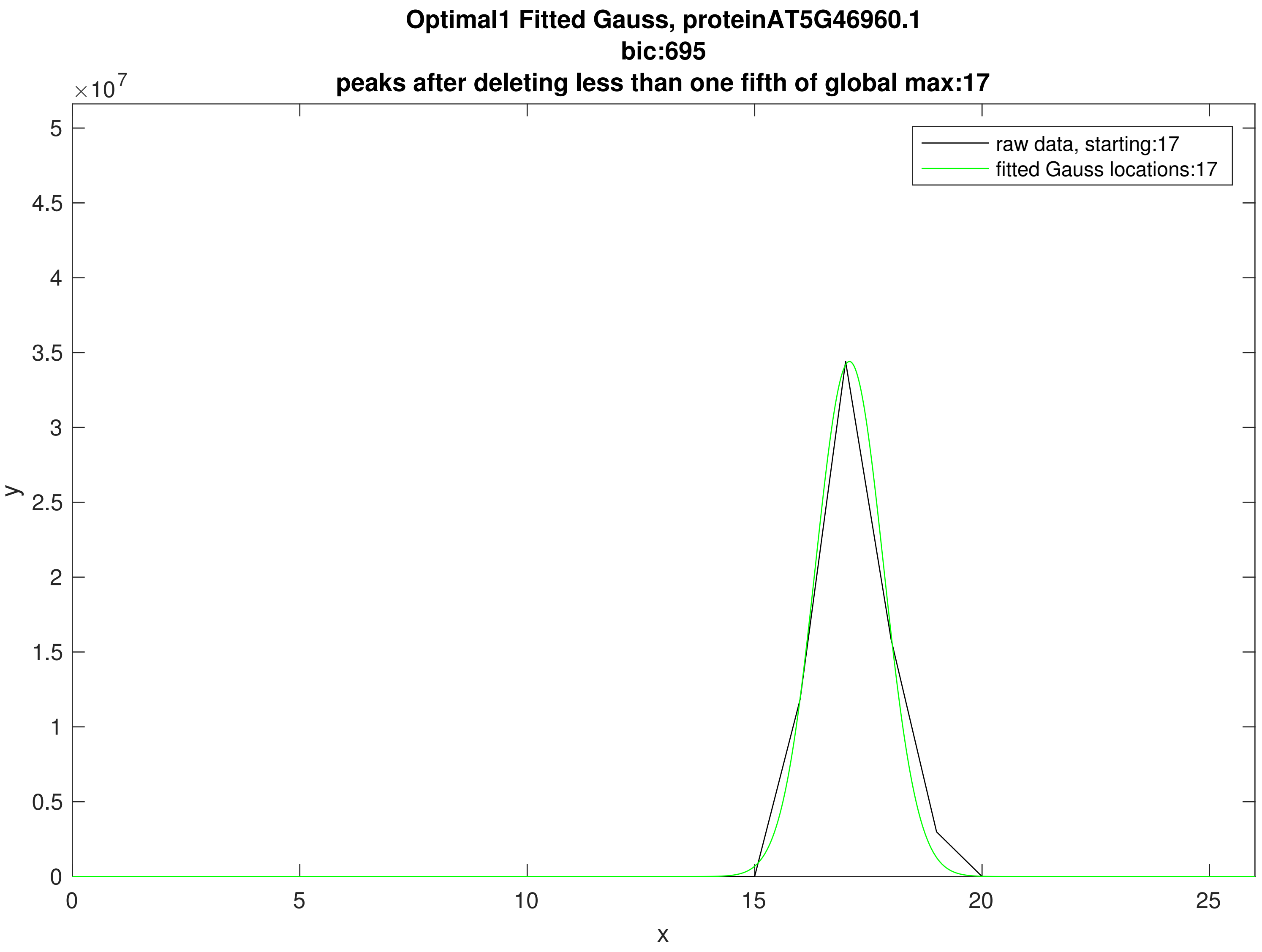

Optimal1 Fitted Gauss, proteinAT5G46960.1
bic:695
peaks after deleting less than one fifth of global max:17
×10^7
raw data, starting:17
fitted Gauss locations:17
5
4.5
4
3.5
3
2.5
2
1.5
1
0.5
0
y
0
5
10
15
20
25
x

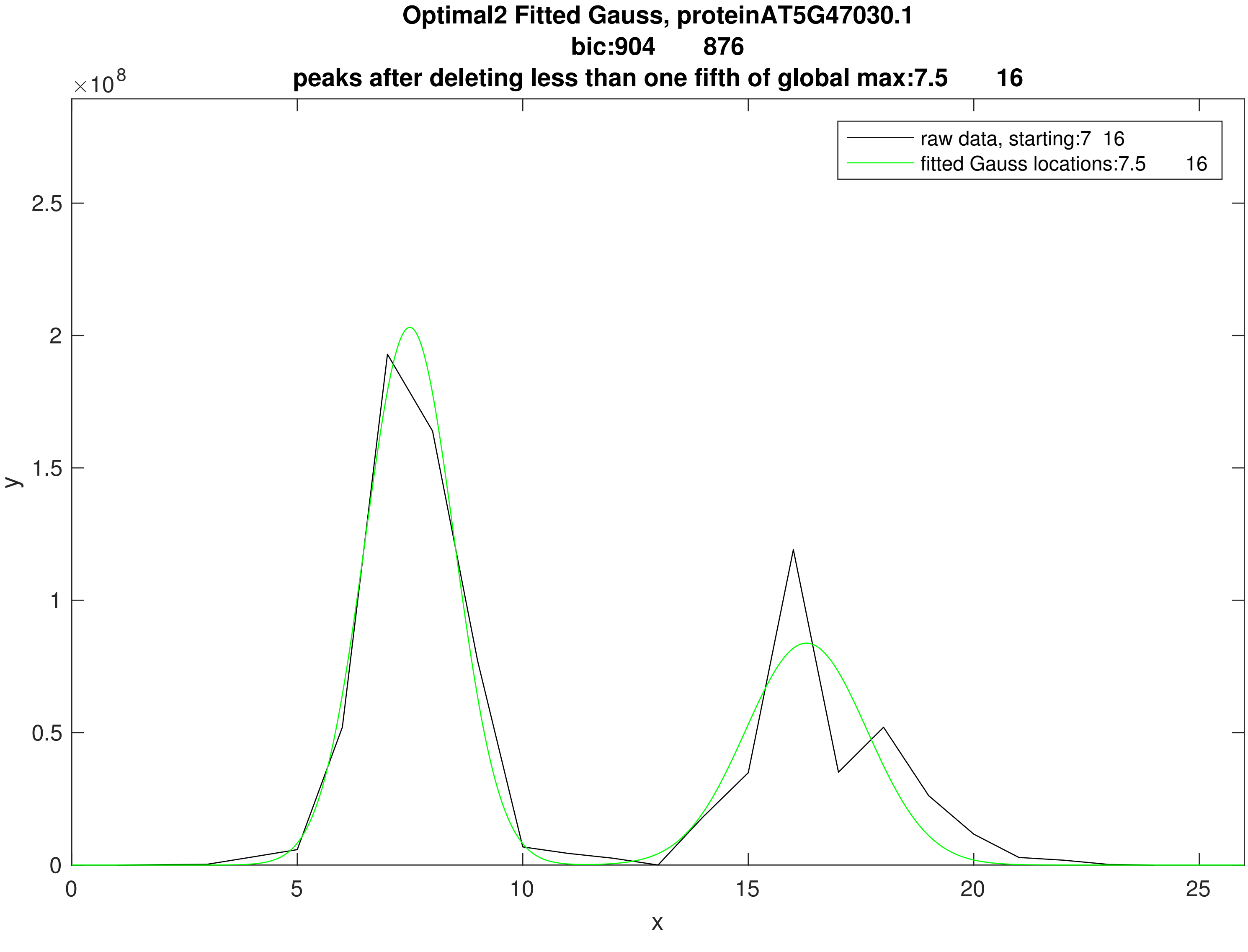
Optimal2 Fitted Gauss, proteinAT5G47030.1
bic:904 876
peaks after deleting less than one fifth of global max:7.5 16
×10^8
raw data, starting:7 16
fitted Gauss locations:7.5 16
y
x
0
0.5
1
1.5
2
2.5
5
10
15
20
25

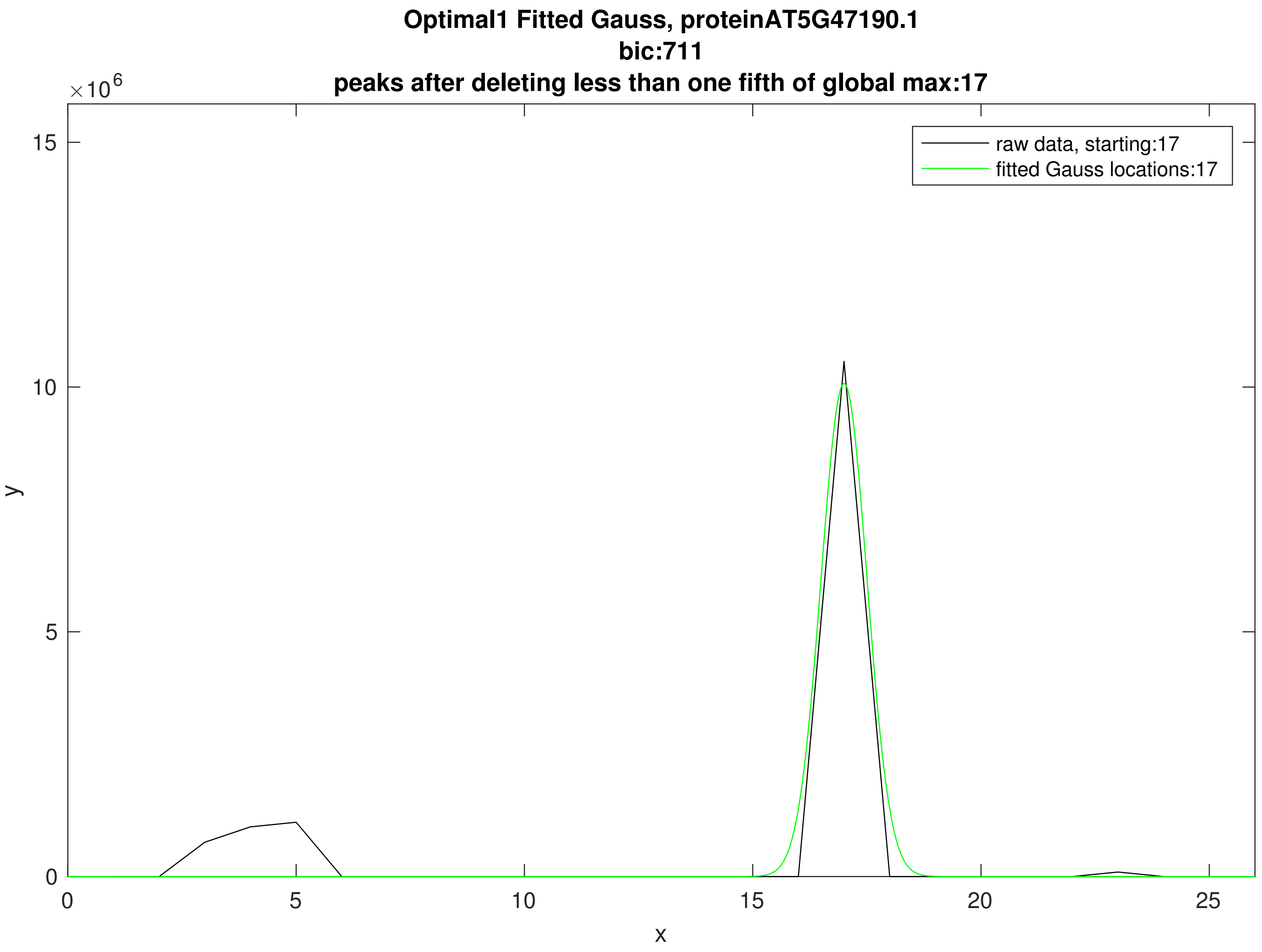
Optimal1 Fitted Gauss, proteinAT5G47190.1
bic:711
peaks after deleting less than one fifth of global max:17
×10^6
raw data, starting:17
fitted Gauss locations:17
y
x
0
5
10
15
20
25

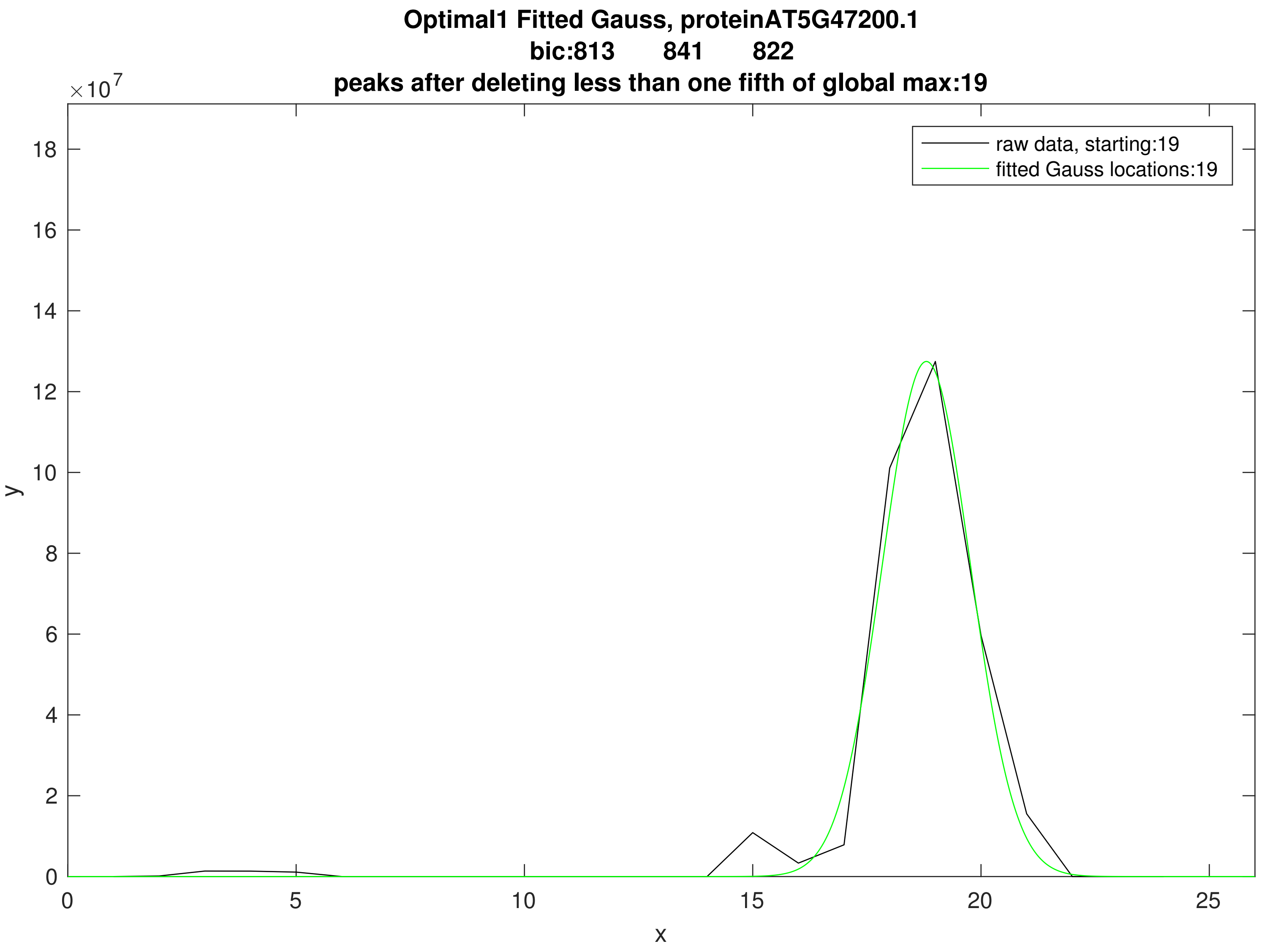

Optimal1 Fitted Gauss, proteinAT5G47200.1
bic:813 841 822
peaks after deleting less than one fifth of global max:19
×10^7
raw data, starting:19
fitted Gauss locations:19
y
x
0
2
4
6
8
10
12
14
16
18
0
5
10
15
20
25

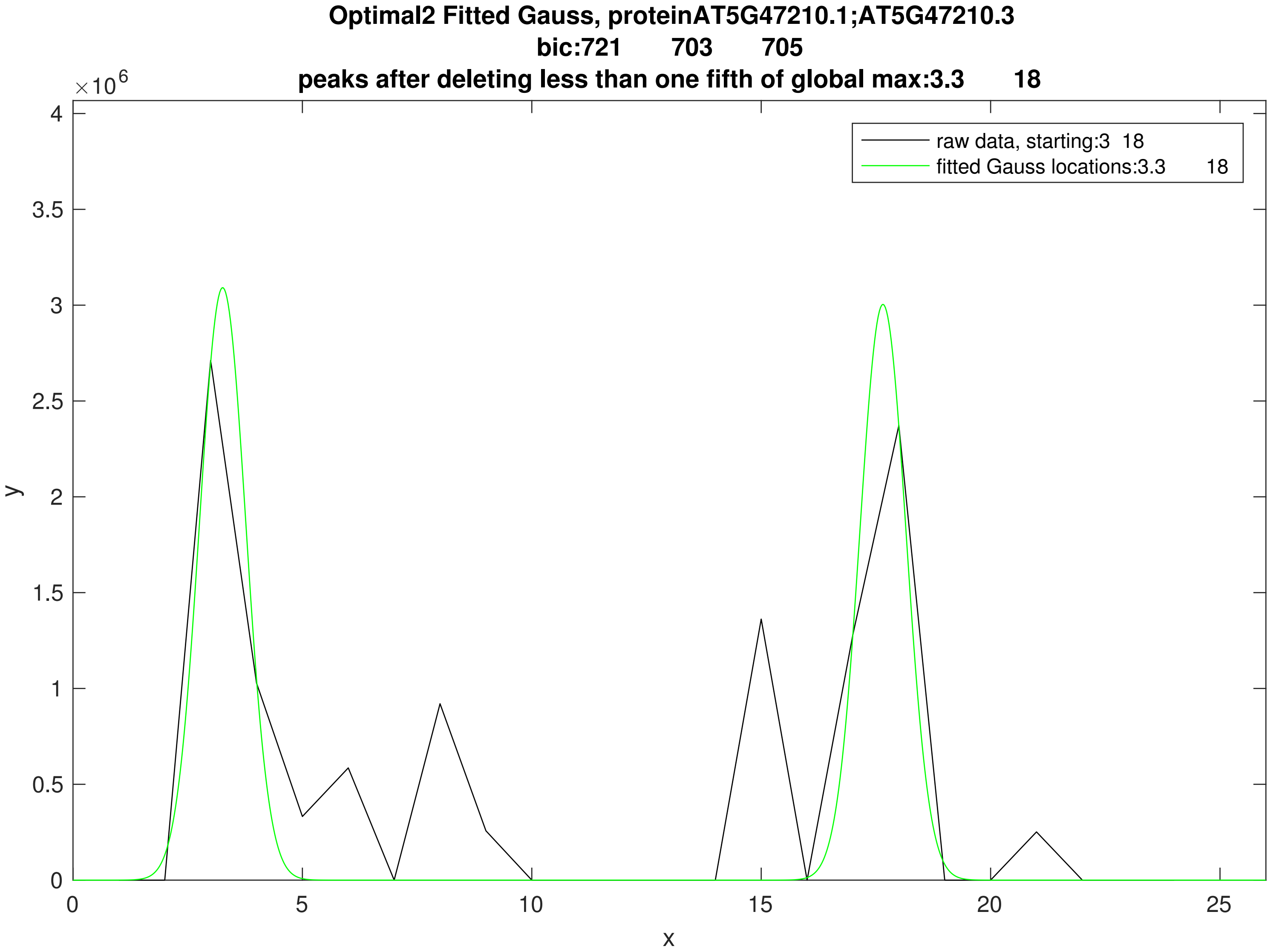
Optimal2 Fitted Gauss, proteinAT5G47210.1;AT5G47210.3
bic:721 703 705
peaks after deleting less than one fifth of global max:3.3 18
×10^6
raw data, starting:3 18
fitted Gauss locations:3.3 18
y
x
0
0.5
1
1.5
2
2.5
3
3.5
4
5
10
15
20
25

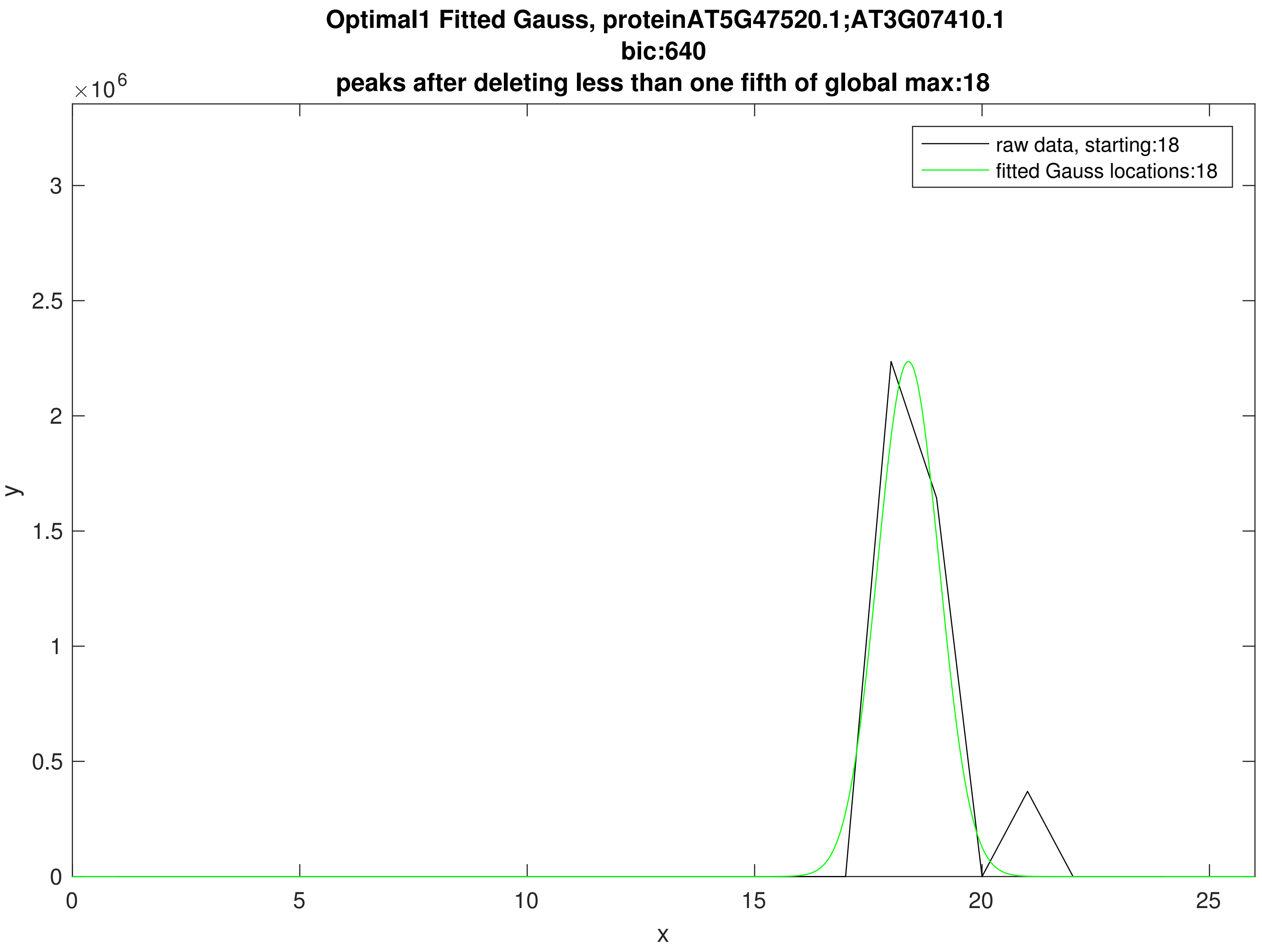

Optimal1 Fitted Gauss, proteinAT5G47520.1;AT3G07410.1
bic:640
peaks after deleting less than one fifth of global max:18
×10^6
raw data, starting:18
fitted Gauss locations:18
y
x
0
0.5
1
1.5
2
2.5
3
0
5
10
15
20
25

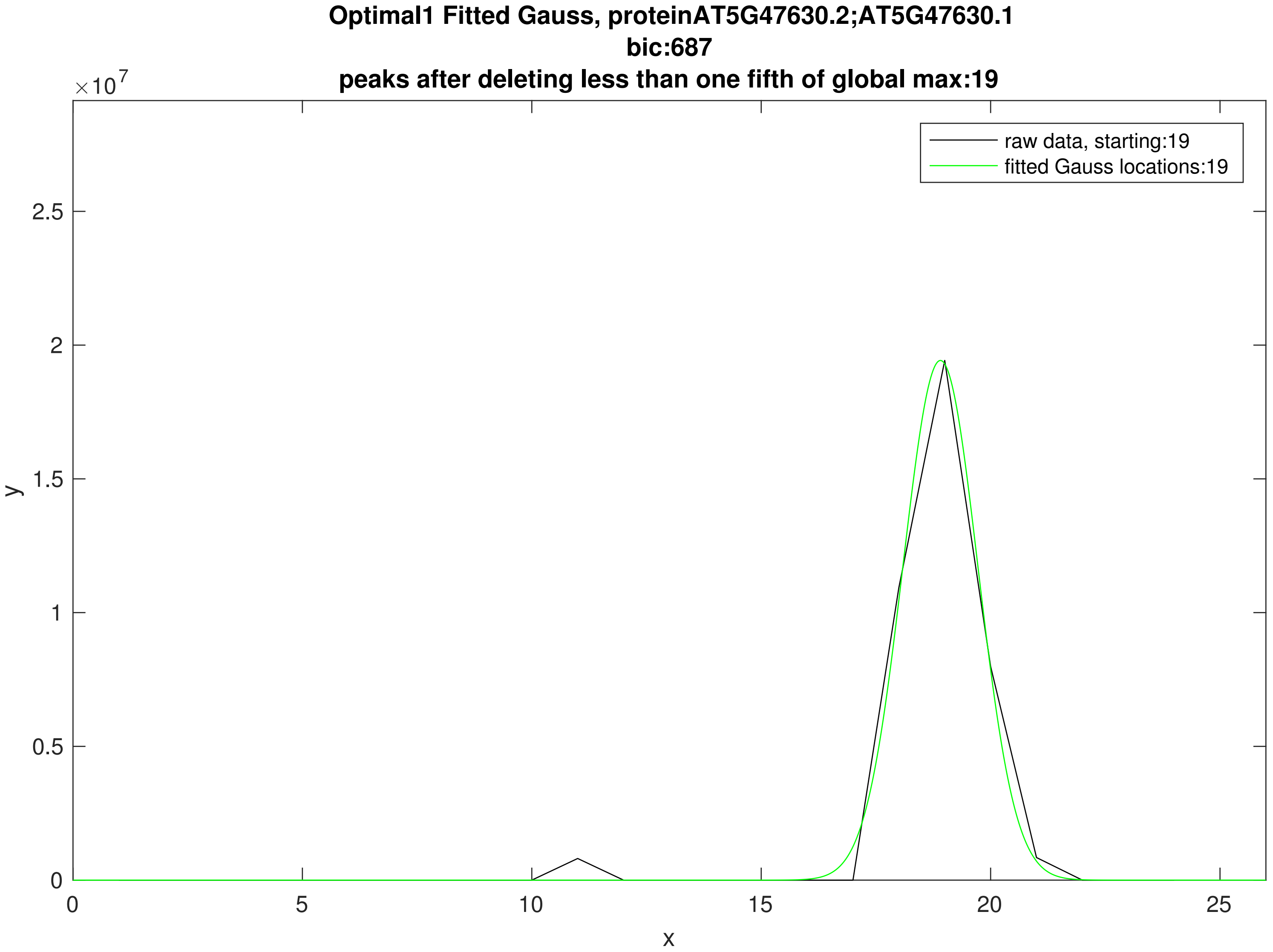

Optimal1 Fitted Gauss, proteinAT5G47630.2;AT5G47630.1
bic:687
peaks after deleting less than one fifth of global max:19
×10^7
raw data, starting:19
fitted Gauss locations:19
y
x
0
0.5
1
1.5
2
2.5
0
5
10
15
20
25

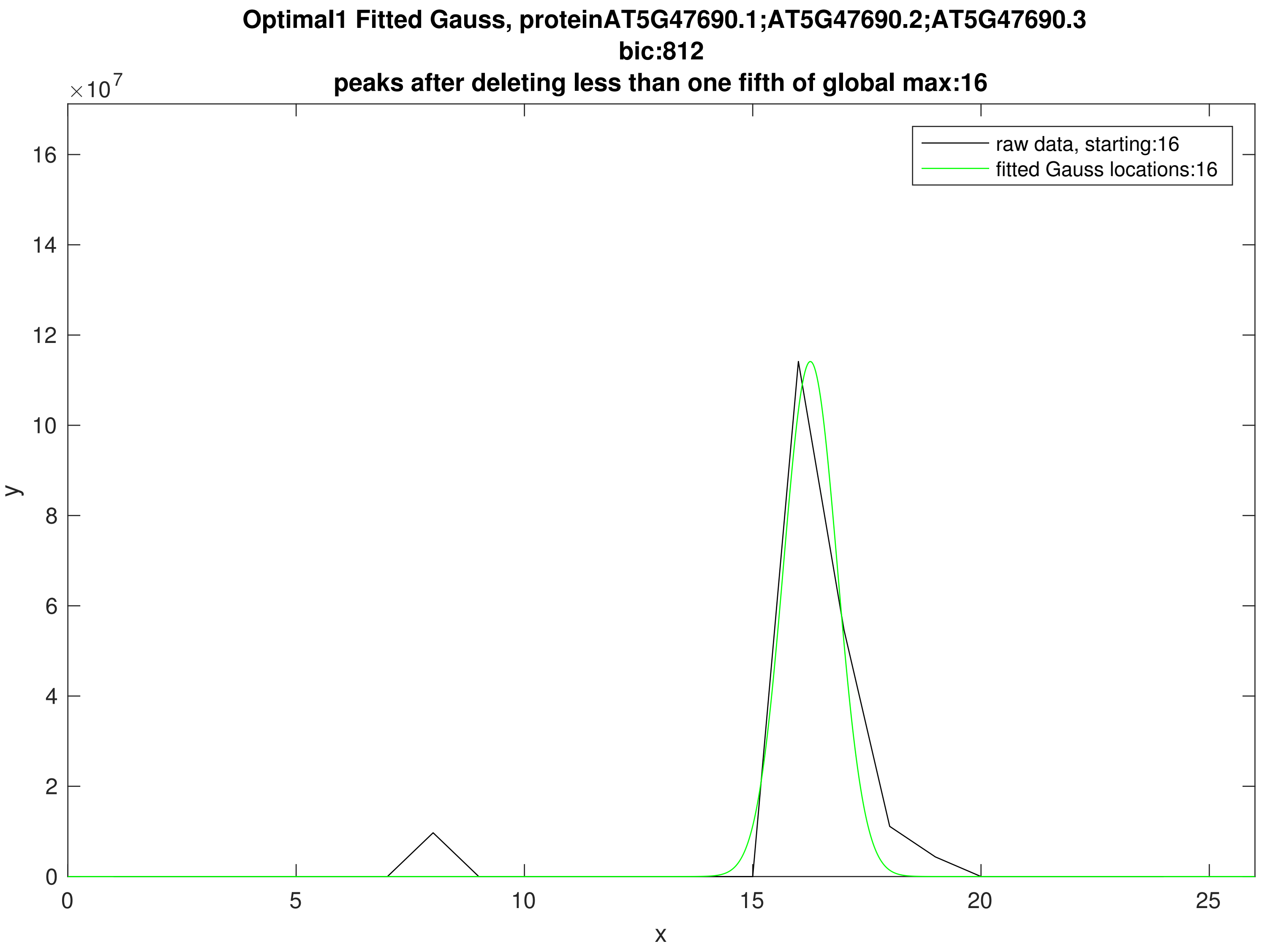

Optimal1 Fitted Gauss, proteinAT5G47690.1;AT5G47690.2;AT5G47690.3
bic:812
peaks after deleting less than one fifth of global max:16
×10^7
raw data, starting:16
fitted Gauss locations:16
y
x
0
2
4
6
8
10
12
14
16
0
5
10
15
20
25

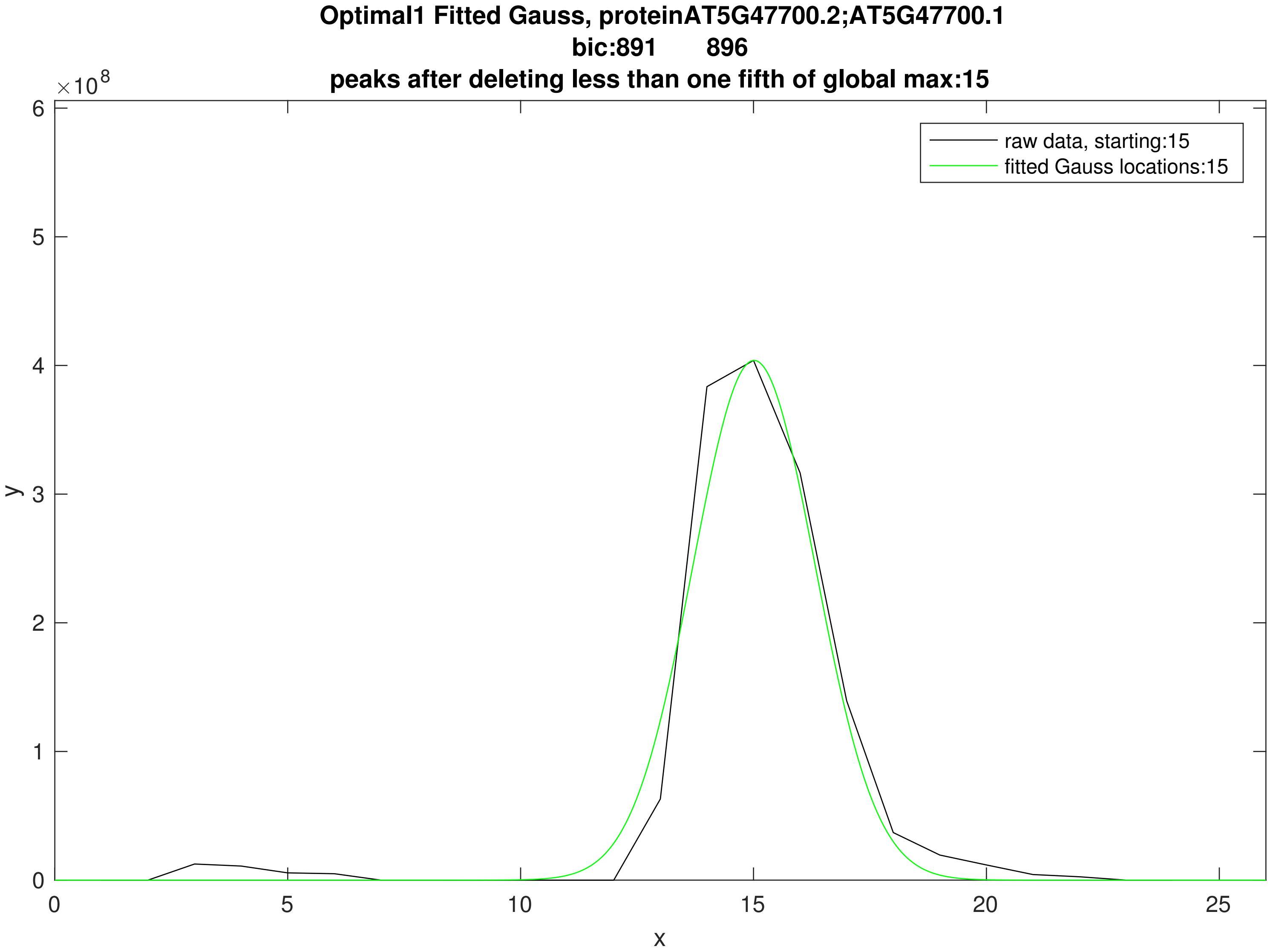

Optimal1 Fitted Gauss, proteinAT5G47700.2;AT5G47700.1
bic:891 896
peaks after deleting less than one fifth of global max:15
×10^8
6
5
4
3
2
1
0
y
0
5
10
15
20
25
x
raw data, starting:15
fitted Gauss locations:15

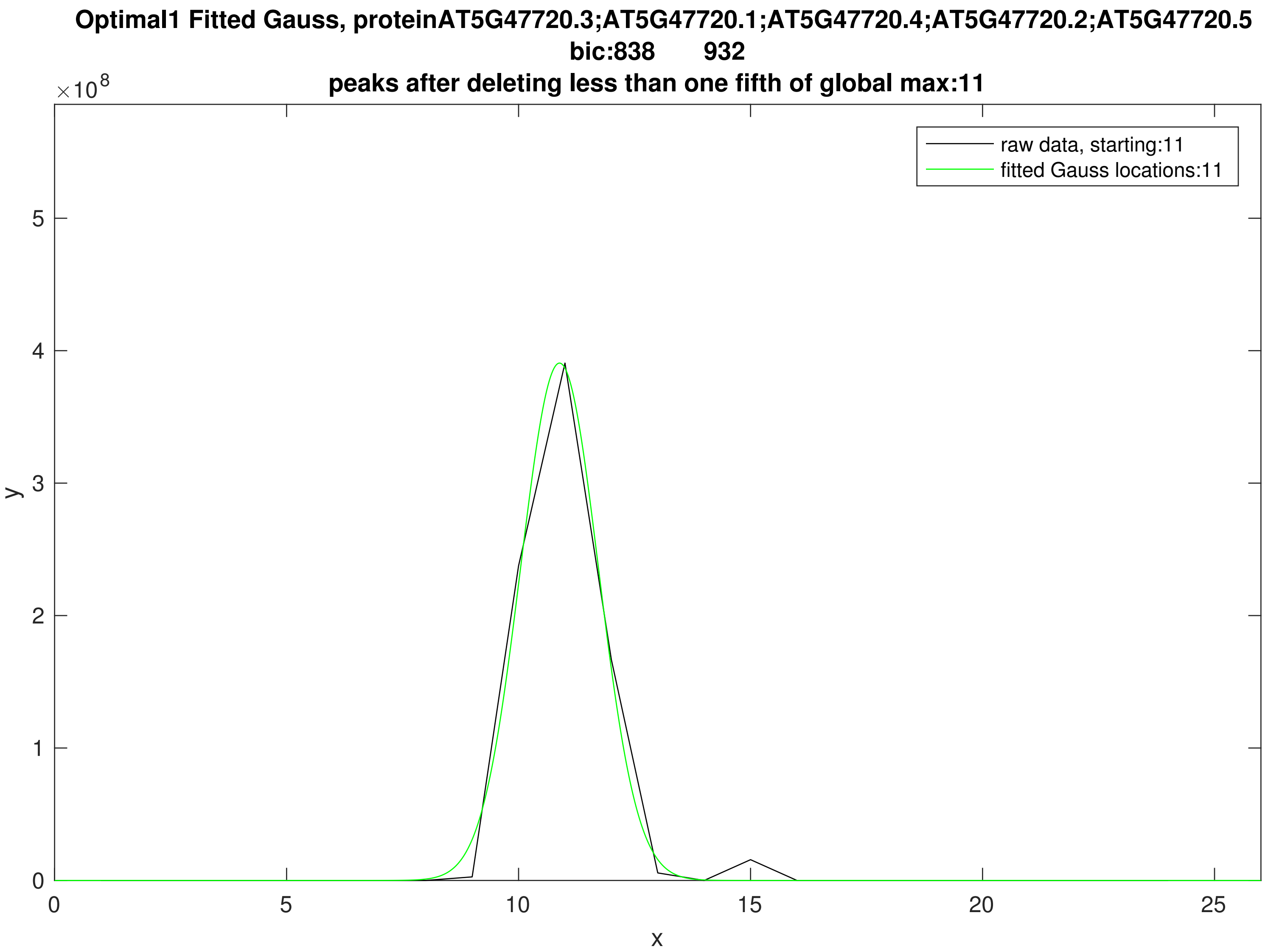

Optimal1 Fitted Gauss, proteinAT5G47720.3;AT5G47720.1;AT5G47720.4;AT5G47720.2;AT5G47720.5
bic:838 932
peaks after deleting less than one fifth of global max:11
×10^8
raw data, starting:11
fitted Gauss locations:11
y
x
0
1
2
3
4
5
0
5
10
15
20
25

**Optimal1 Fitted Gauss, proteinAT5G47760.1**
**bic:833**
**peaks after deleting less than one fifth of global max:15**

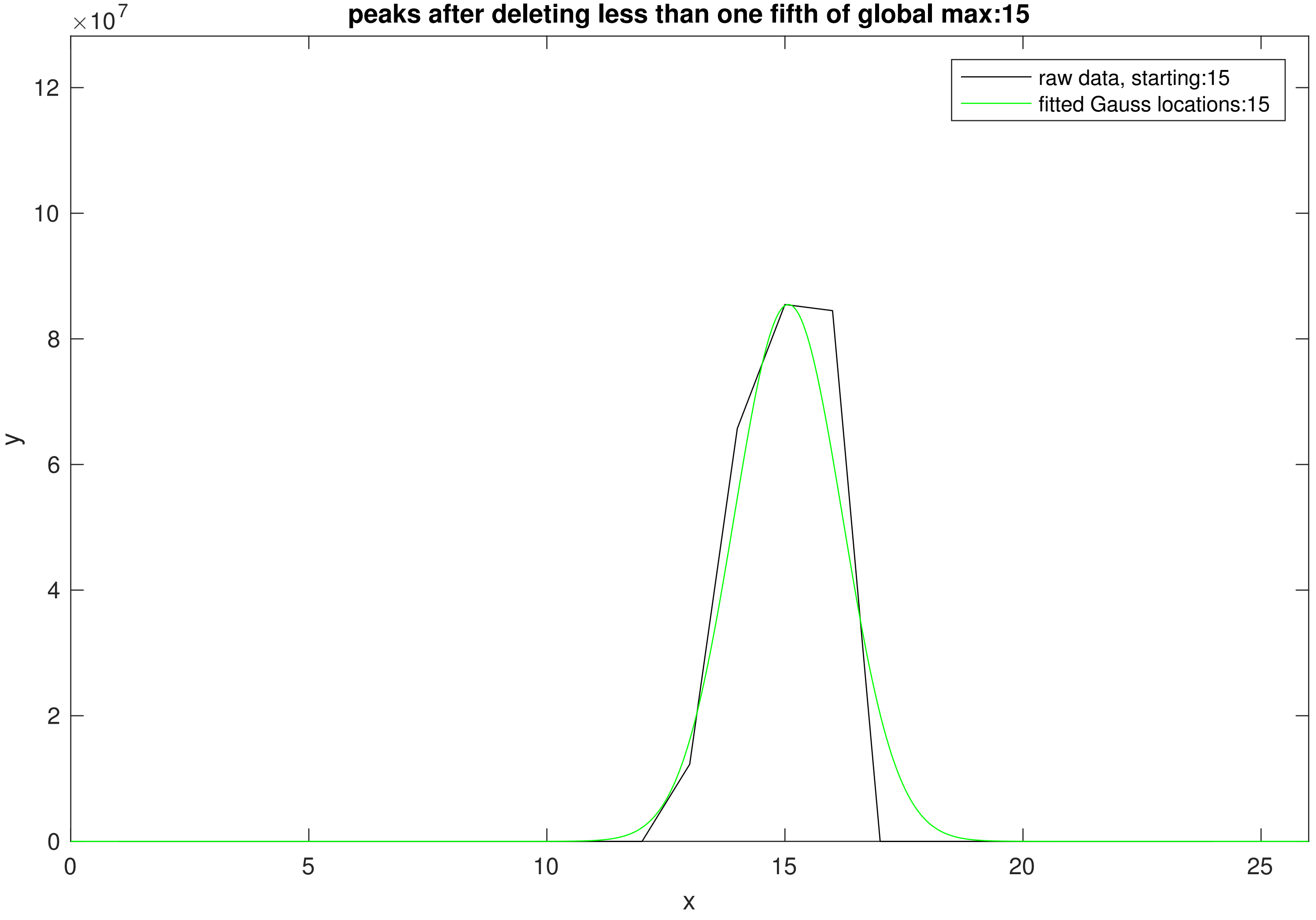

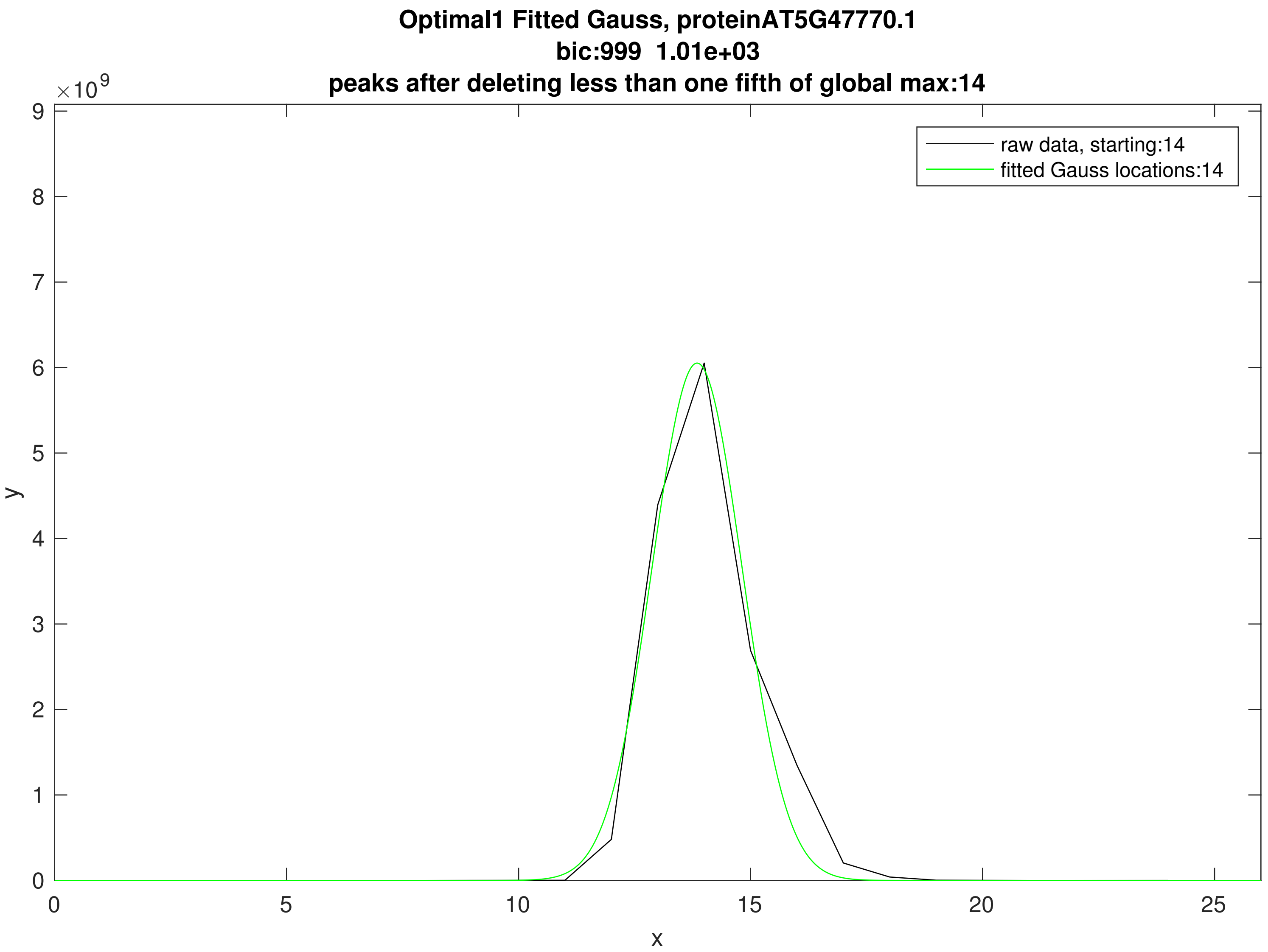

Optimal1 Fitted Gauss, proteinAT5G47770.1
bic:999 1.01e+03
peaks after deleting less than one fifth of global max:14
×10^9
9
8
7
6
5
4
3
2
1
0
y
0
5
10
15
20
25
x
raw data, starting:14
fitted Gauss locations:14

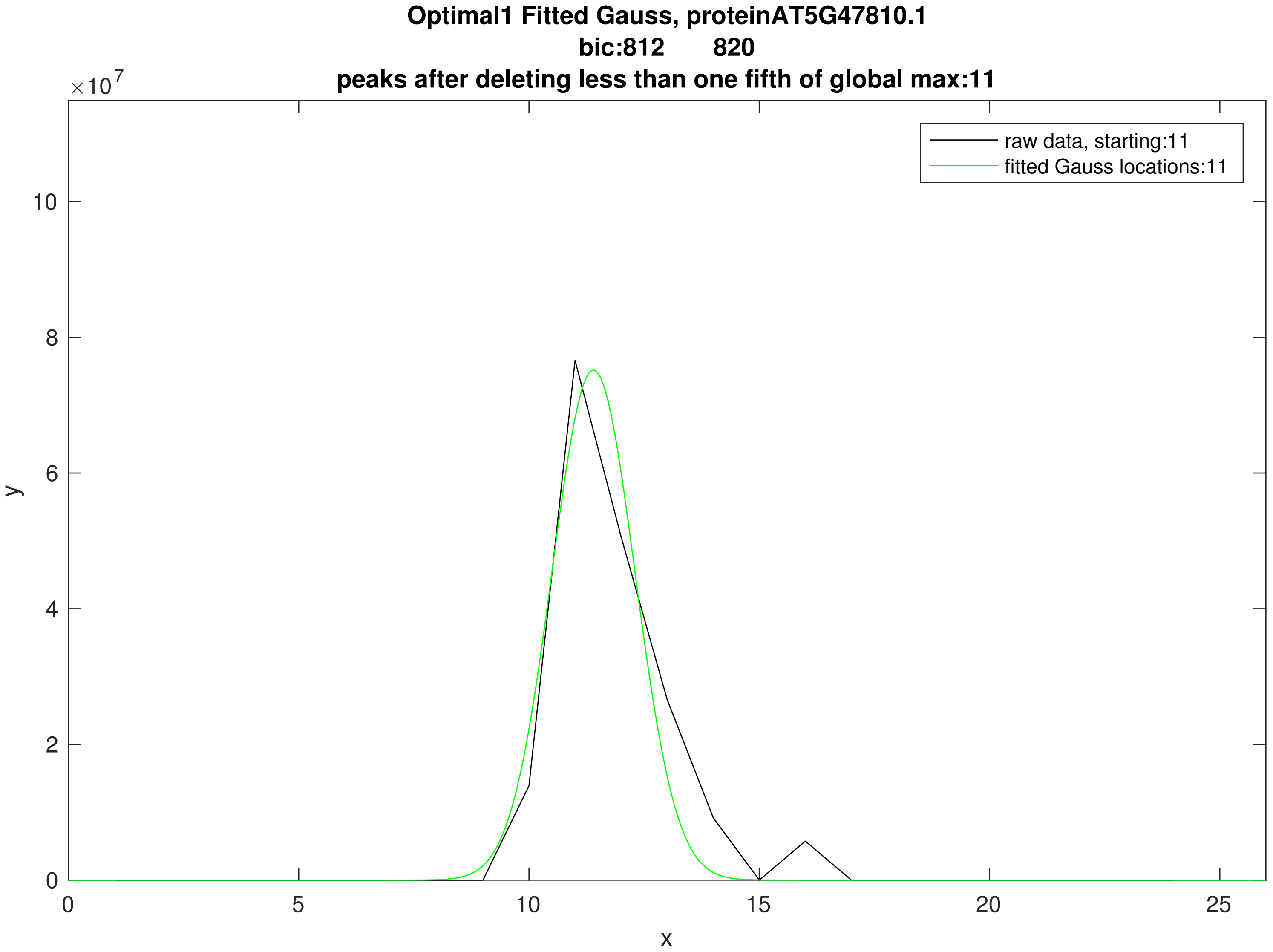
Optimal1 Fitted Gauss, proteinAT5G47810.1
bic:812 820
peaks after deleting less than one fifth of global max:11
×10^7
raw data, starting:11
fitted Gauss locations:11
y
x
0
2
4
6
8
10
0
5
10
15
20
25

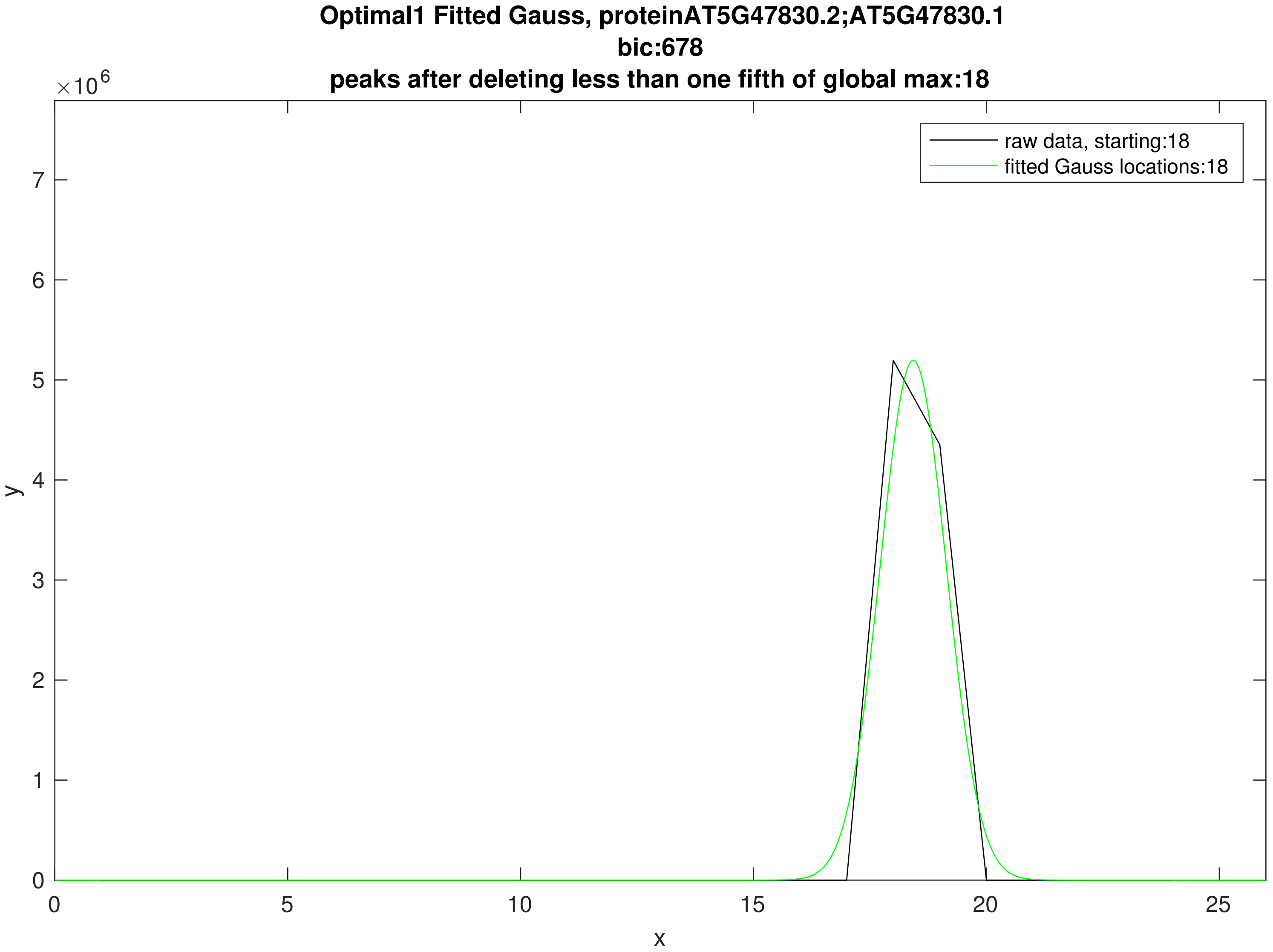
Optimal1 Fitted Gauss, proteinAT5G47830.2;AT5G47830.1
bic:678
peaks after deleting less than one fifth of global max:18
×10^6
raw data, starting:18
fitted Gauss locations:18
y
x
0
1
2
3
4
5
6
7
0
5
10
15
20
25

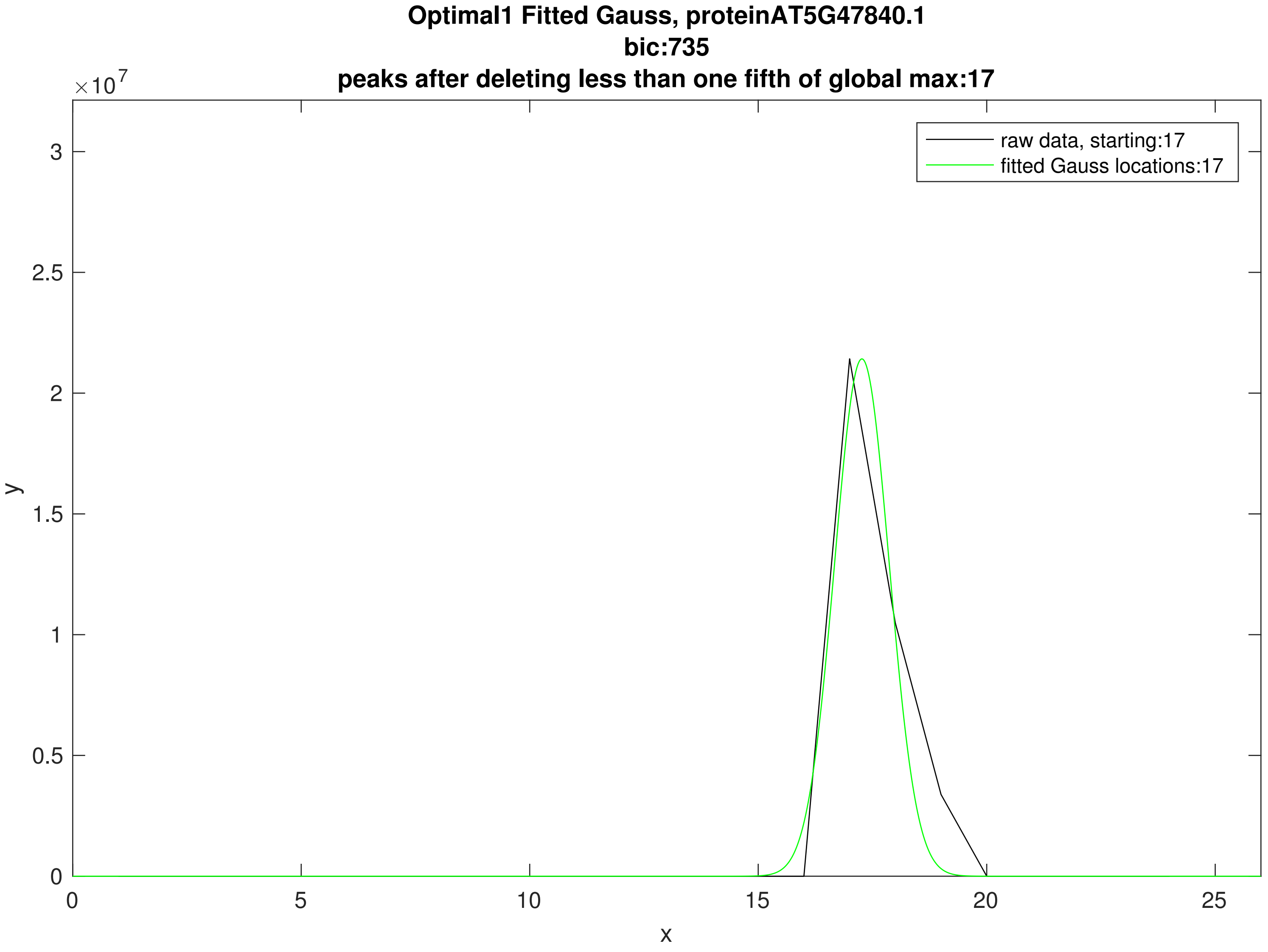
Optimal1 Fitted Gauss, proteinAT5G47840.1
bic:735
peaks after deleting less than one fifth of global max:17
×10^7
raw data, starting:17
fitted Gauss locations:17
y
x
0
0.5
1
1.5
2
2.5
3
0
5
10
15
20
25

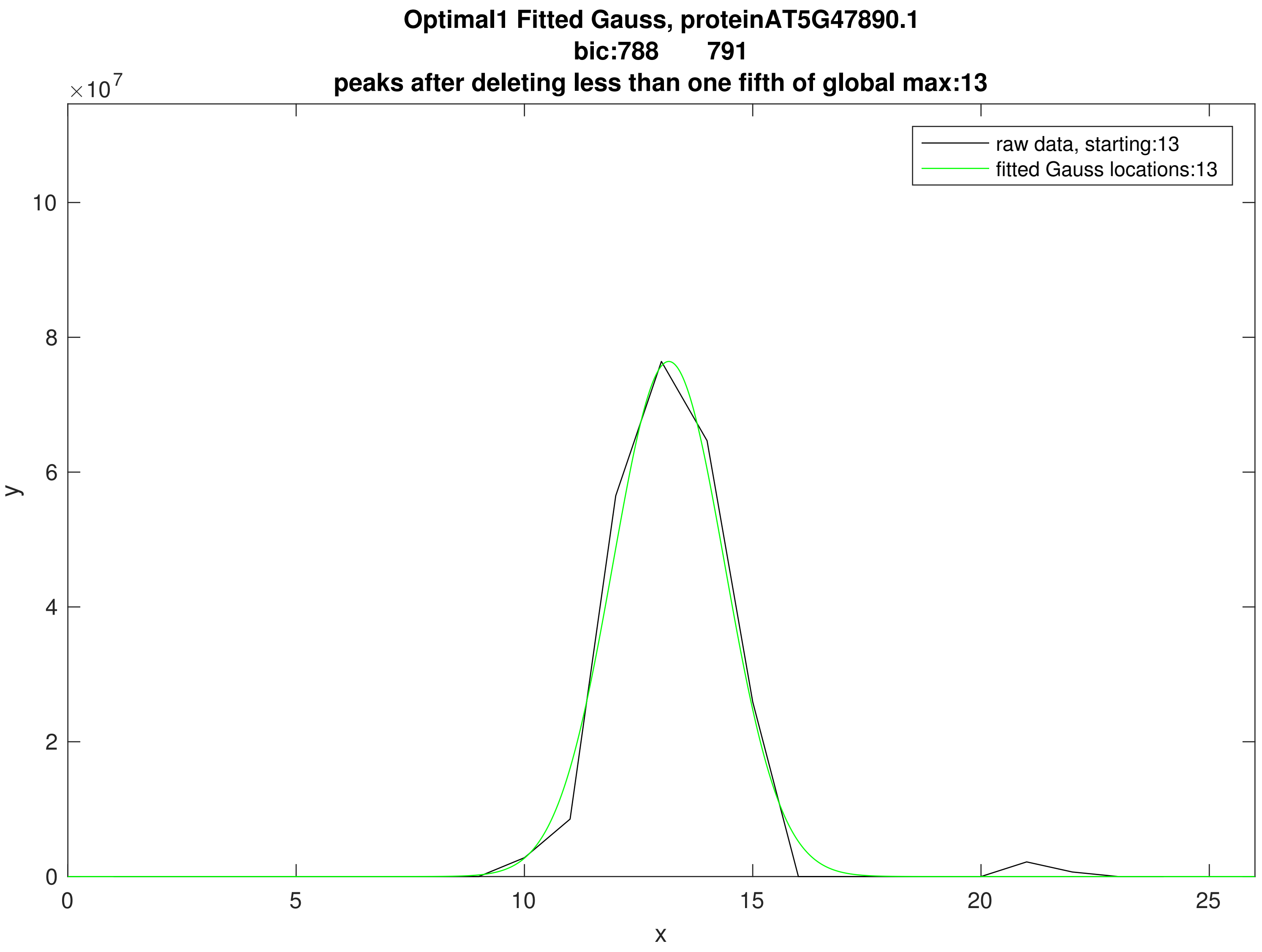

Optimal1 Fitted Gauss, proteinAT5G47890.1
bic:788 791
peaks after deleting less than one fifth of global max:13
×10^7
raw data, starting:13
fitted Gauss locations:13
y
x
0
2
4
6
8
10
0
5
10
15
20
25

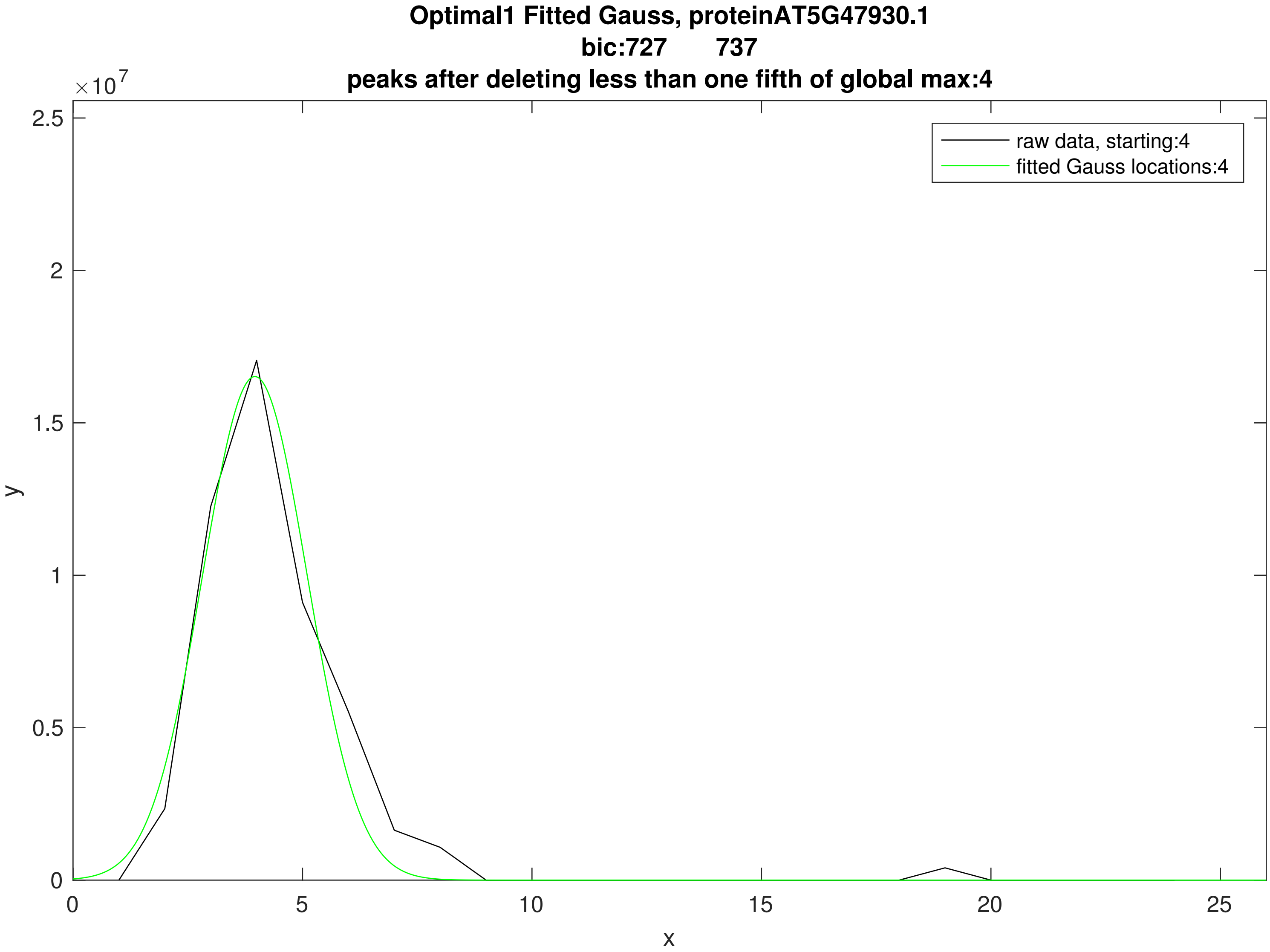

Optimal1 Fitted Gauss, proteinAT5G47930.1
bic:727 737
peaks after deleting less than one fifth of global max:4
×10^7
raw data, starting:4
fitted Gauss locations:4
y
x
0
0.5
1
1.5
2
2.5
5
10
15
20
25

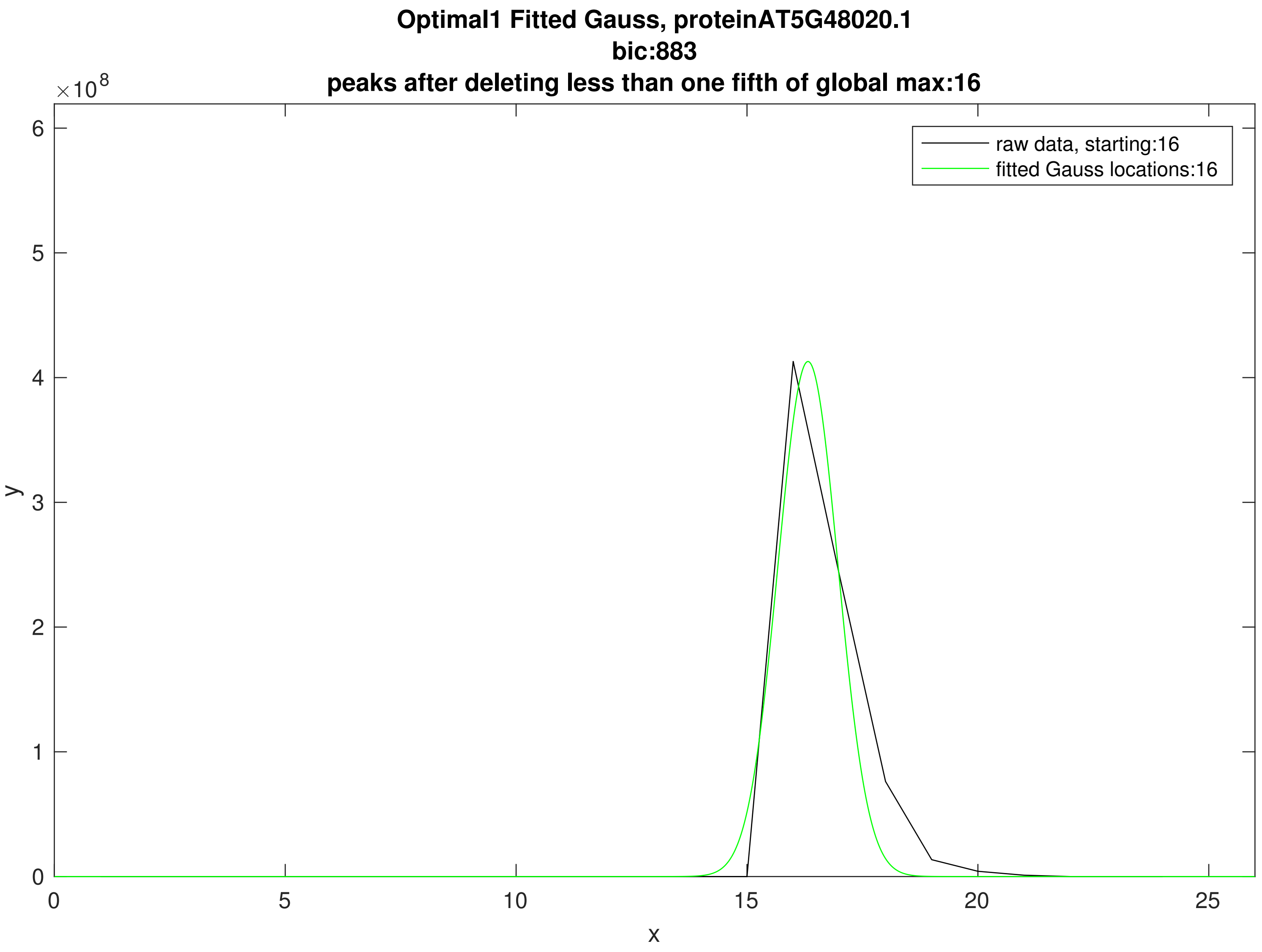
Optimal1 Fitted Gauss, proteinAT5G48020.1
bic:883
peaks after deleting less than one fifth of global max:16
×10^8
6
5
4
3
2
1
0
y
0
5
10
15
20
25
x
raw data, starting:16
fitted Gauss locations:16

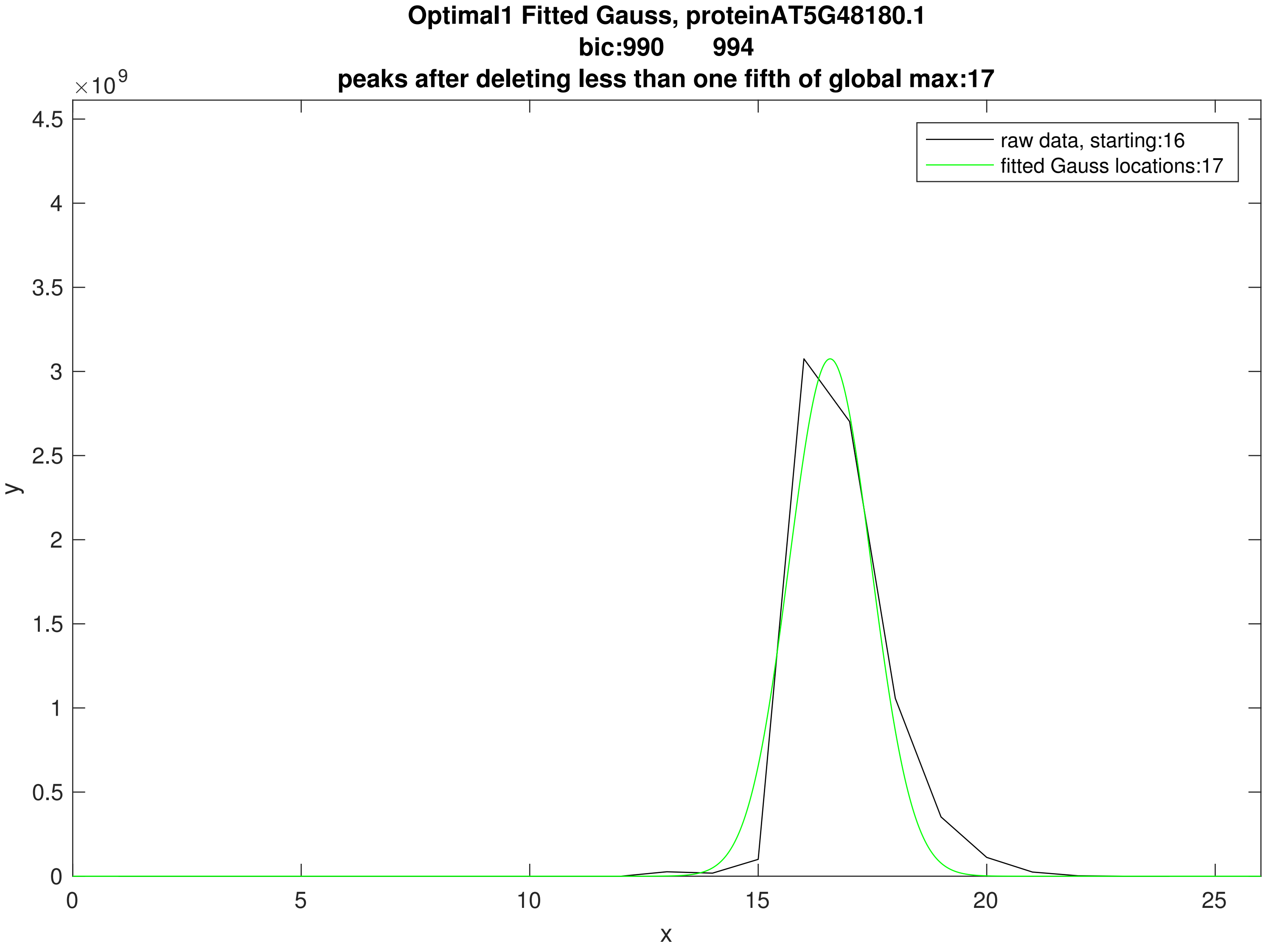

Optimal1 Fitted Gauss, proteinAT5G48180.1
bic:990 994
peaks after deleting less than one fifth of global max:17
×10^9
4.5
4
3.5
3
2.5
2
1.5
1
0.5
0
y
0
5
10
15
20
25
x
raw data, starting:16
fitted Gauss locations:17

**Optimal1 Fitted Gauss, proteinAT5G48220.1;AT5G48220.3;AT5G48220.2**
**bic:673**
**peaks after deleting less than one fifth of global max:17**

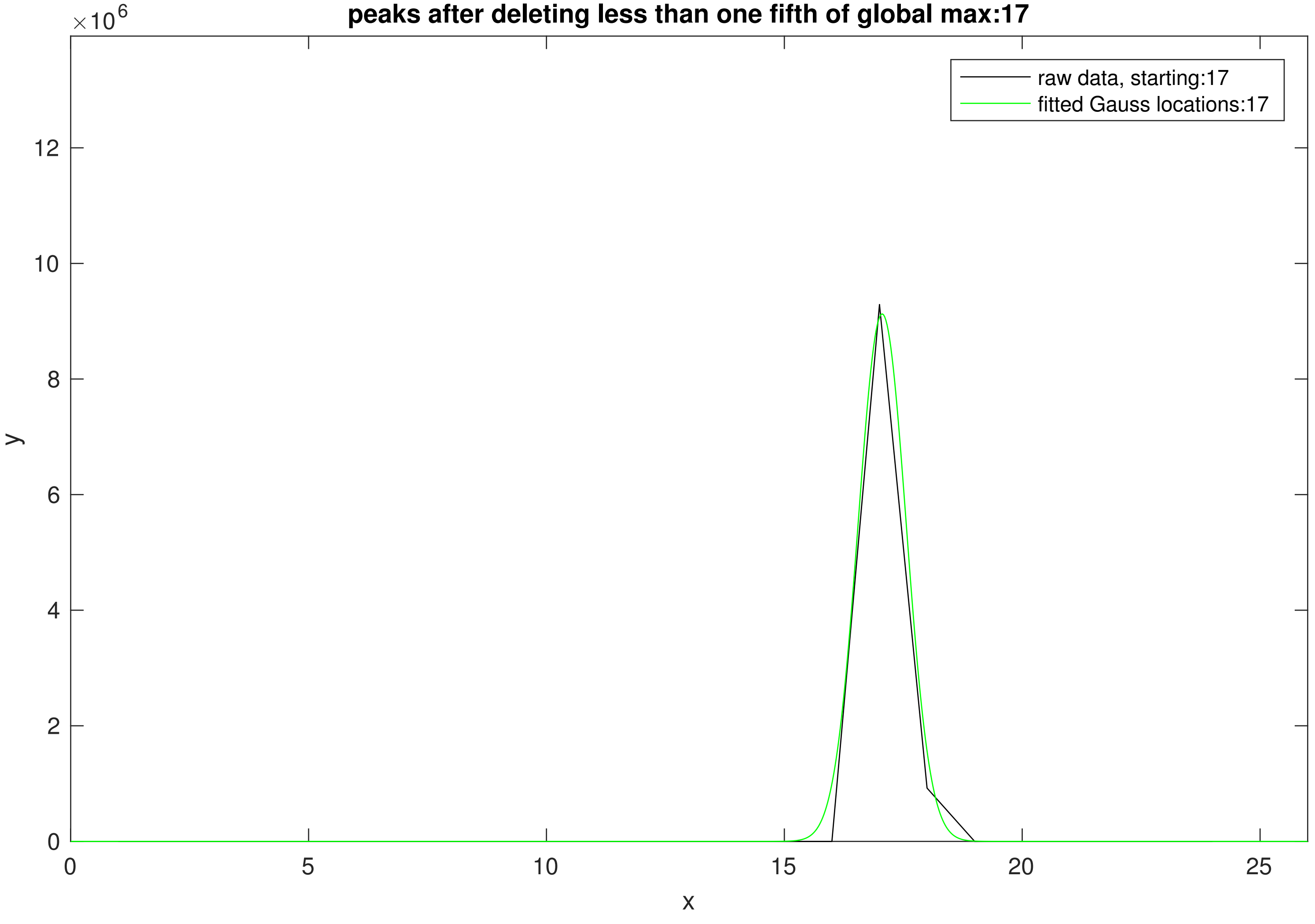

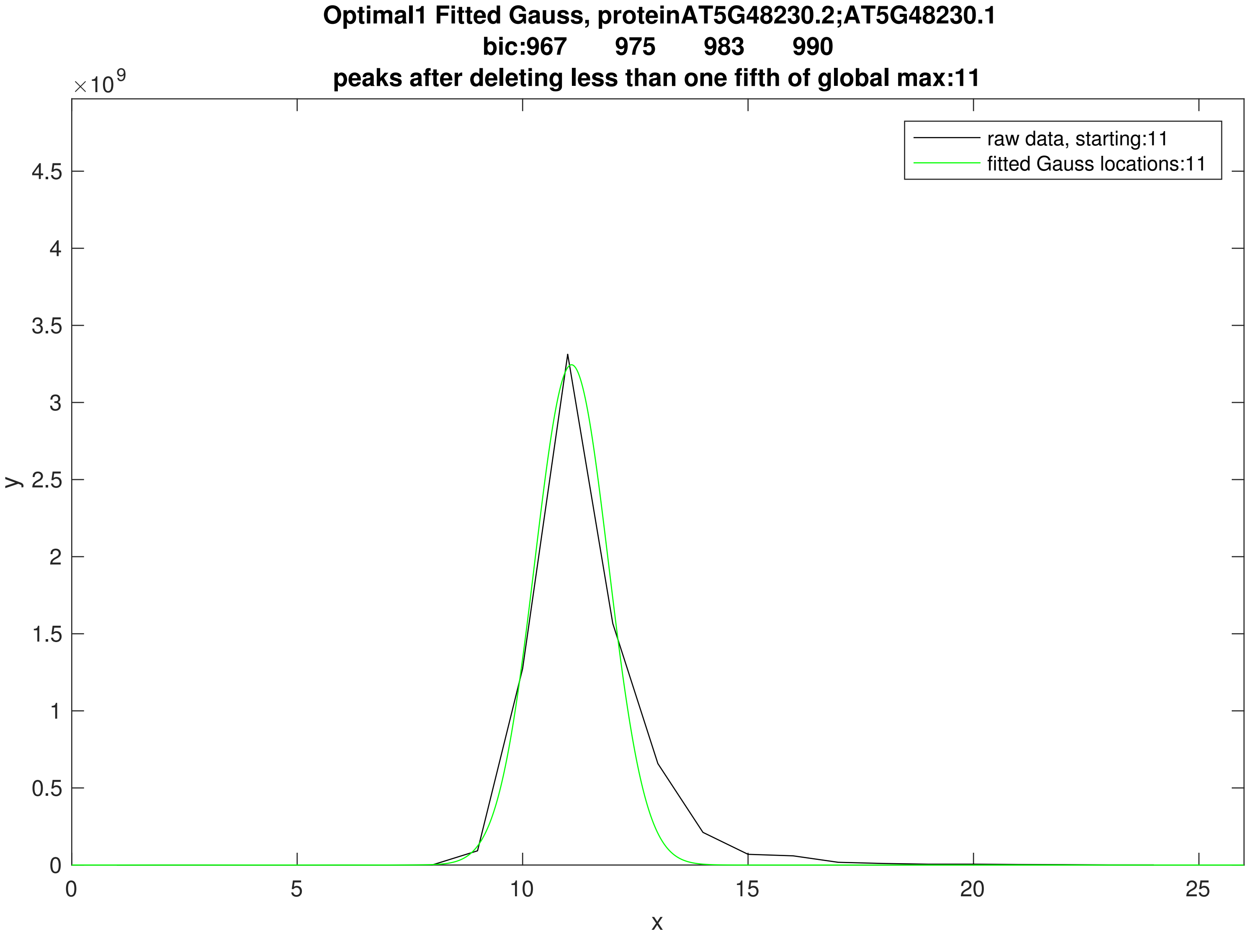
Optimal1 Fitted Gauss, proteinAT5G48230.2;AT5G48230.1
bic:967 975 983 990
peaks after deleting less than one fifth of global max:11
×10^9
4.5
4
3.5
3
2.5
2
1.5
1
0.5
0
y
0
5
10
15
20
25
x
raw data, starting:11
fitted Gauss locations:11

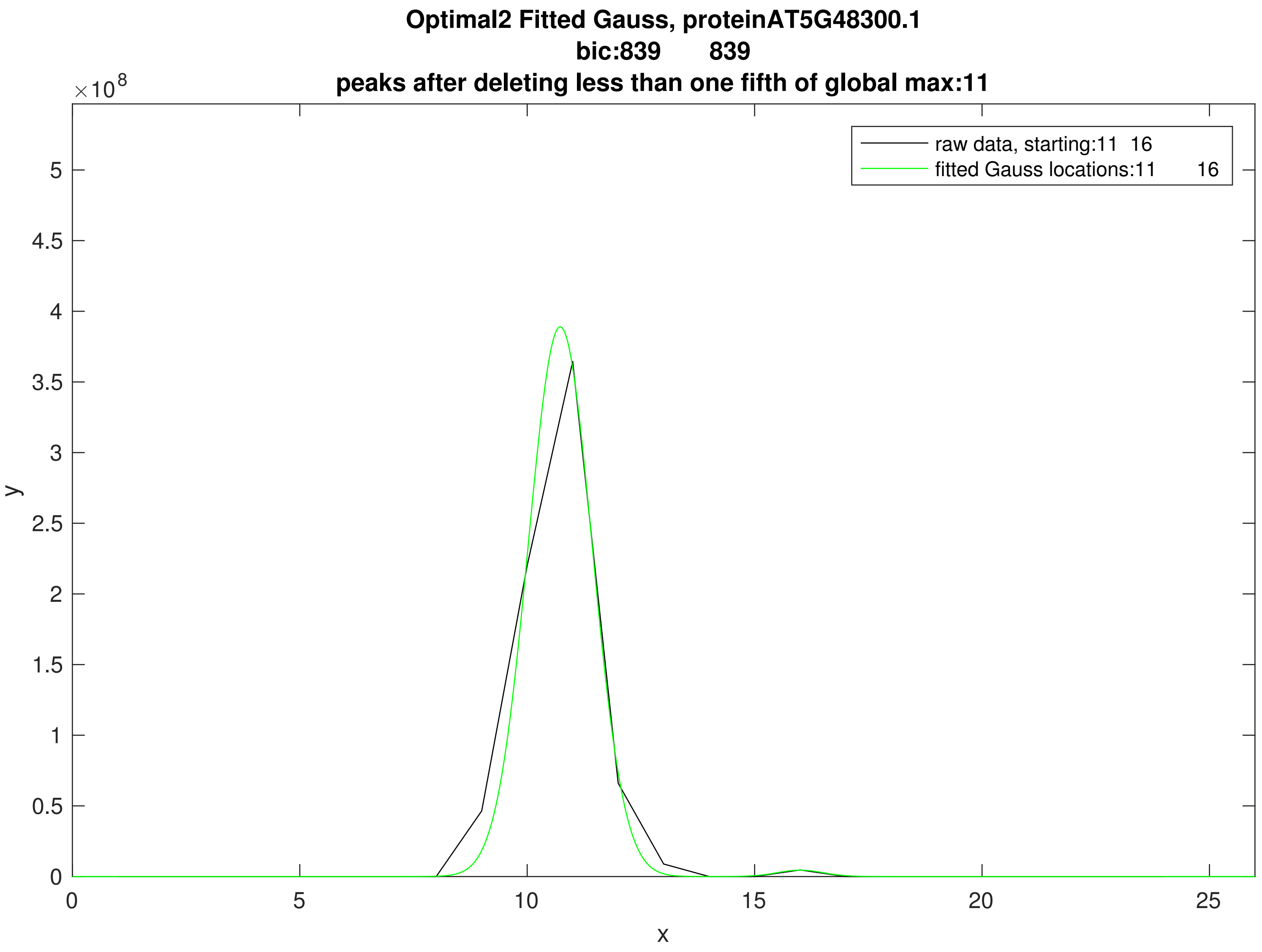

Optimal2 Fitted Gauss, proteinAT5G48300.1
bic:839 839
peaks after deleting less than one fifth of global max:11
×10^8
5
4.5
4
3.5
3
2.5
2
1.5
1
0.5
0
y
0
5
10
15
20
25
x
raw data, starting:11 16
fitted Gauss locations:11 16

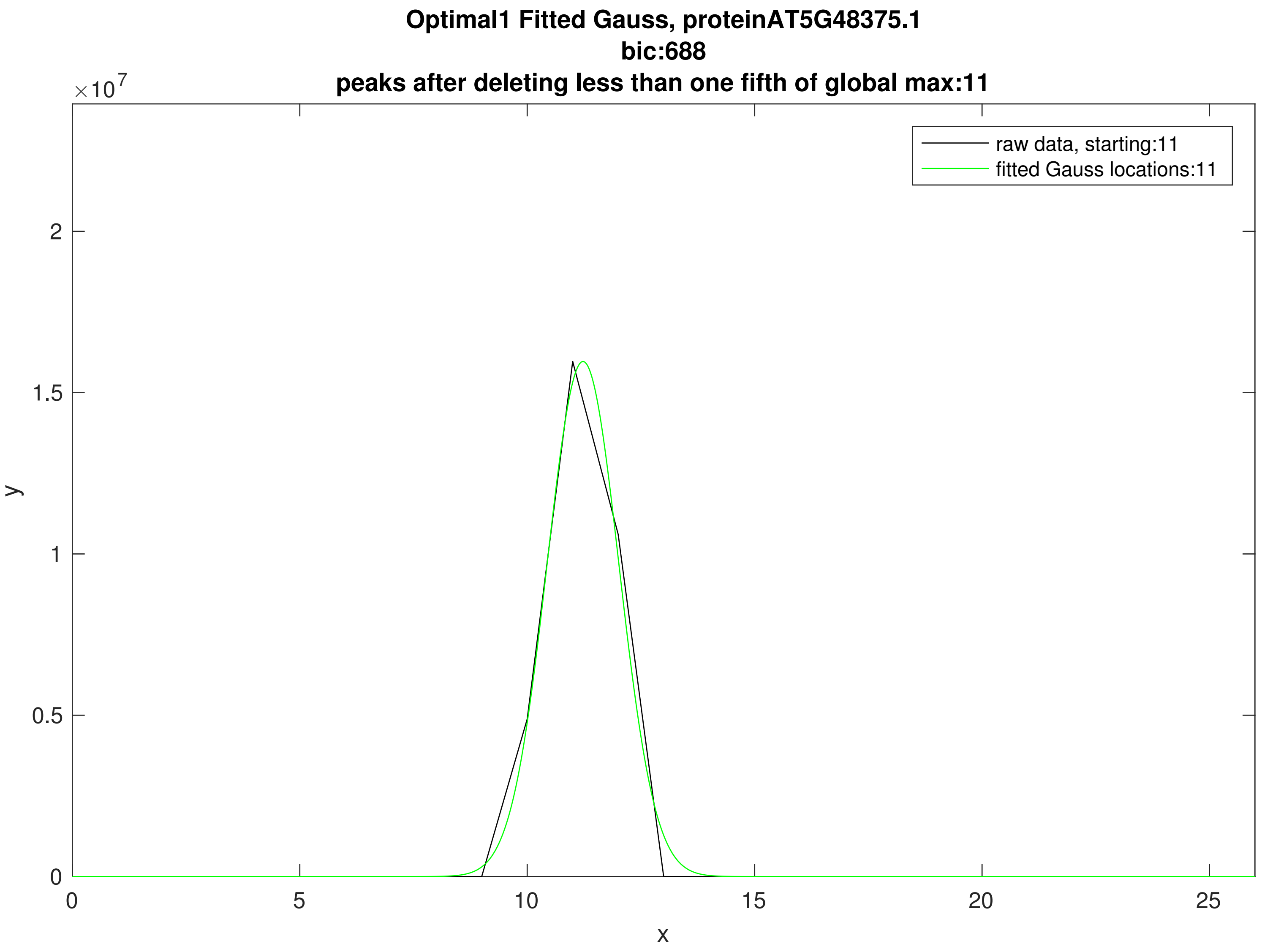
Optimal1 Fitted Gauss, proteinAT5G48375.1
bic:688
peaks after deleting less than one fifth of global max:11
×10^7
raw data, starting:11
fitted Gauss locations:11
y
x
0
0.5
1
1.5
2
0
5
10
15
20
25

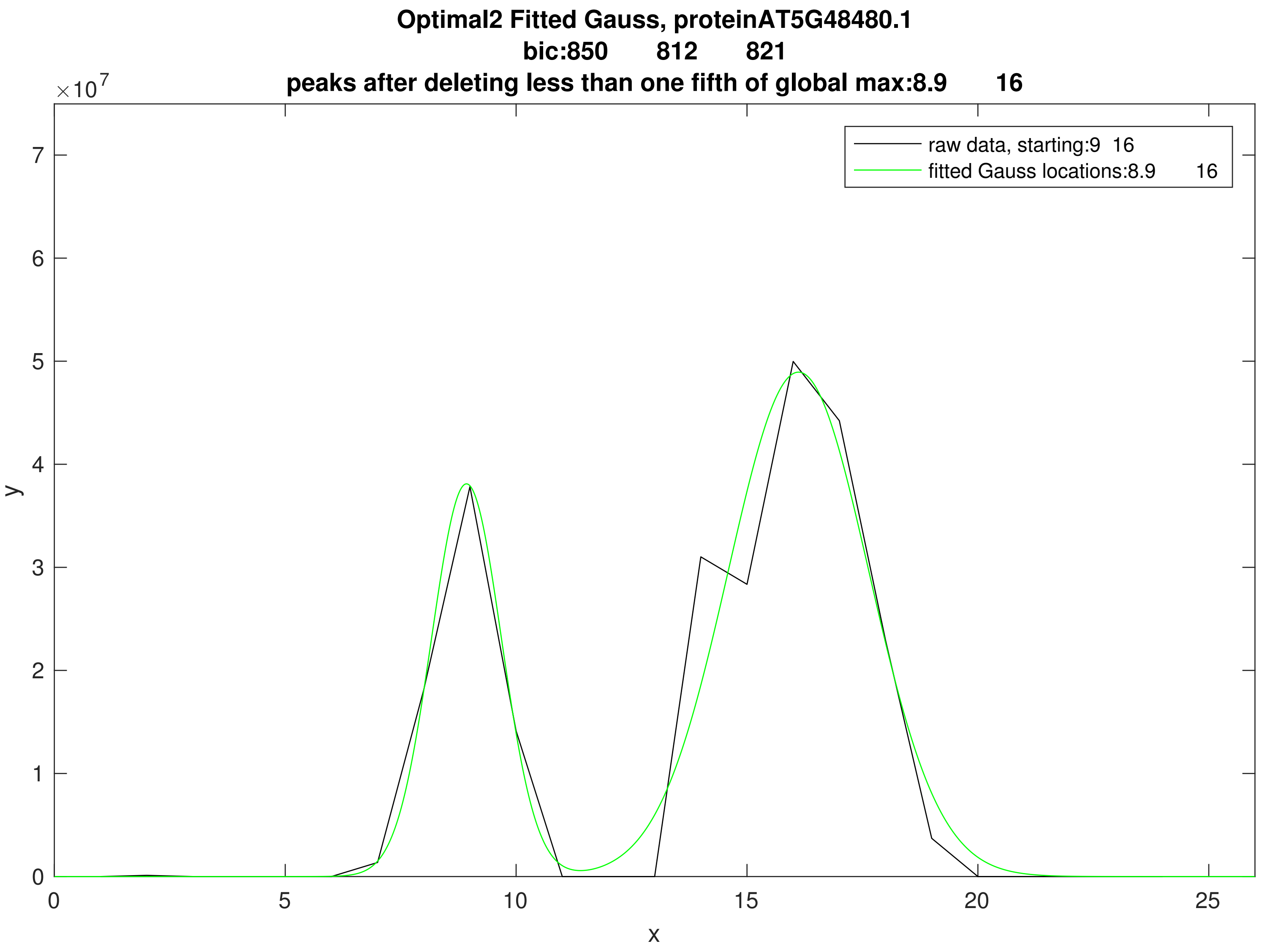
Optimal2 Fitted Gauss, proteinAT5G48480.1
bic:850 812 821
peaks after deleting less than one fifth of global max:8.9 16
×10^7
raw data, starting:9 16
fitted Gauss locations:8.9 16
y
x
0
5
10
15
20
25
1
2
3
4
5
6
7

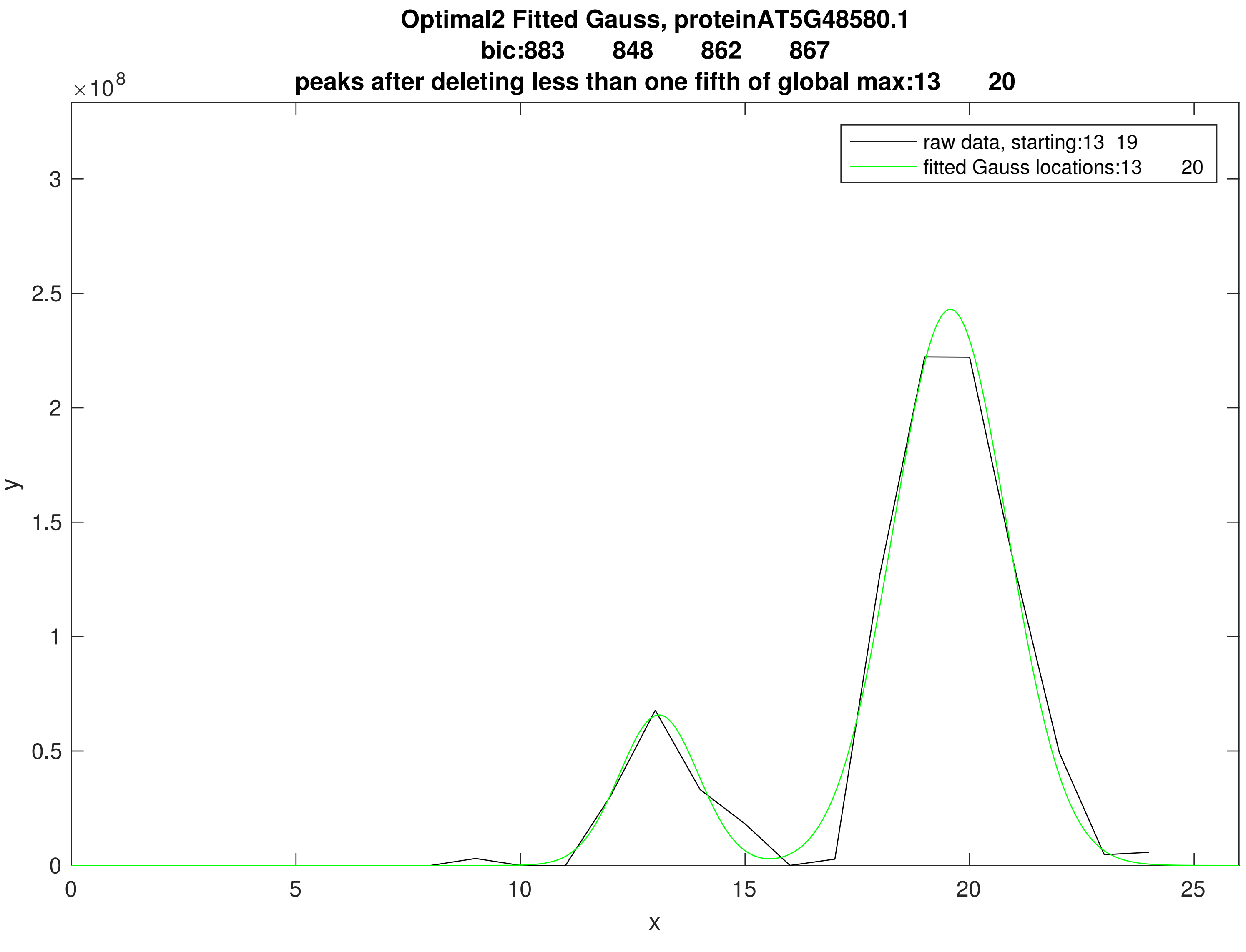
Optimal2 Fitted Gauss, proteinAT5G48580.1
bic:883 848 862 867
peaks after deleting less than one fifth of global max:13 20
×10^8
raw data, starting:13 19
fitted Gauss locations:13 20
y
x
0
0.5
1
1.5
2
2.5
3
0
5
10
15
20
25

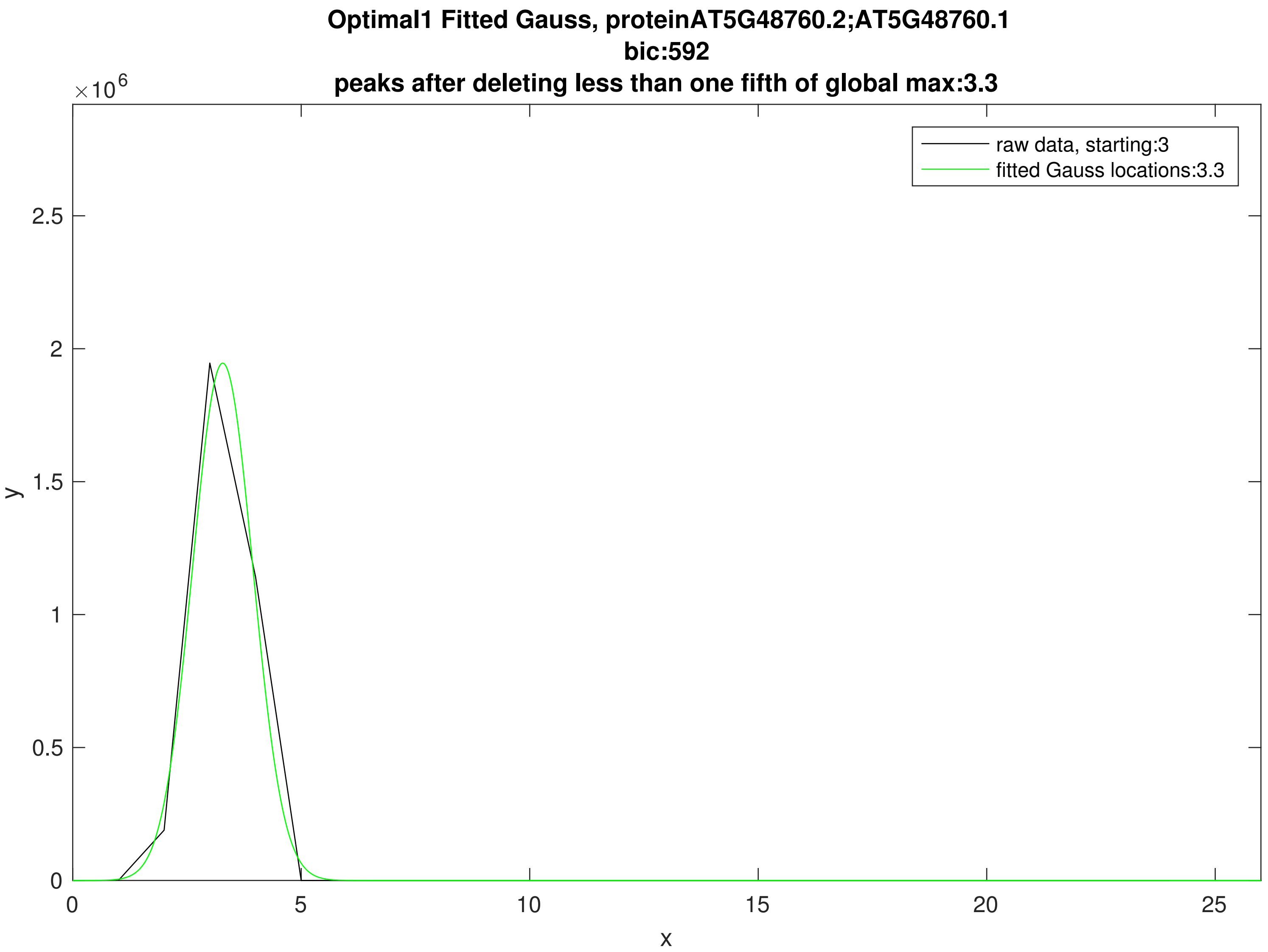
Optimal1 Fitted Gauss, proteinAT5G48760.2;AT5G48760.1
bic:592
peaks after deleting less than one fifth of global max:3.3
×10^6
raw data, starting:3
fitted Gauss locations:3.3
y
x
0
0.5
1
1.5
2
2.5
5
10
15
20
25

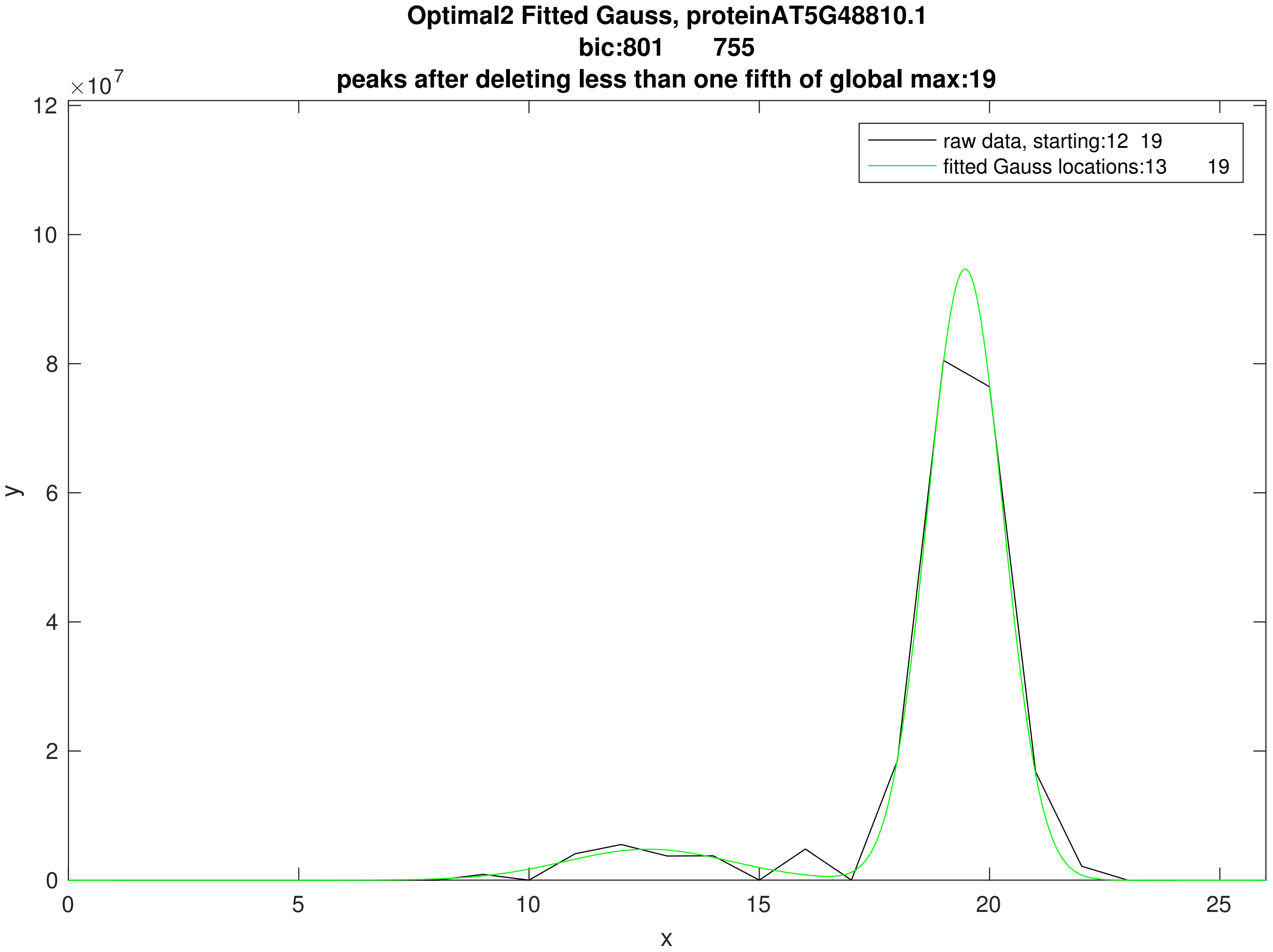
Optimal2 Fitted Gauss, proteinAT5G48810.1
bic:801 755
peaks after deleting less than one fifth of global max:19
×10^7
raw data, starting:12 19
fitted Gauss locations:13 19
y
x
0
2
4
6
8
10
12
0
5
10
15
20
25

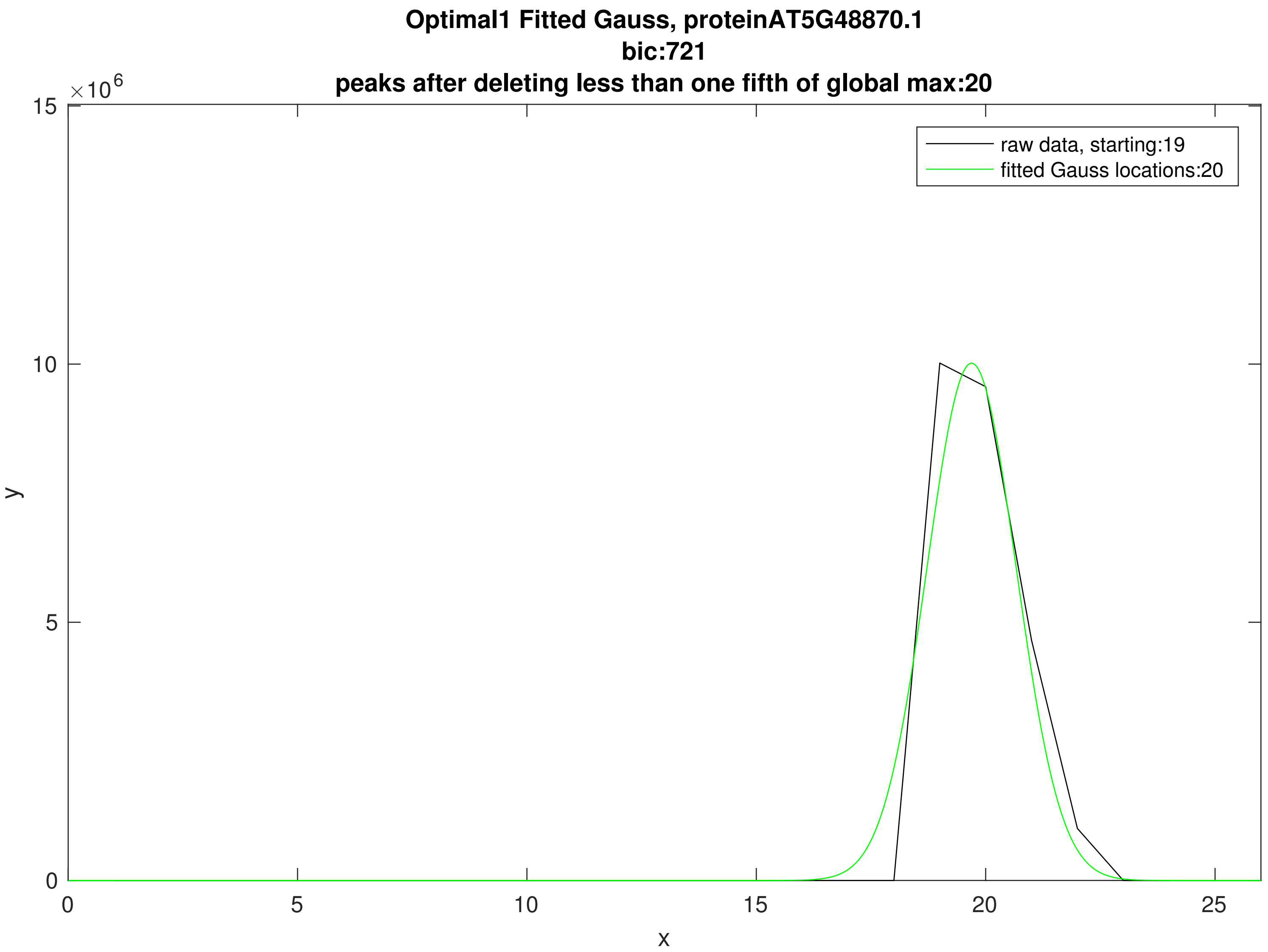
Optimal1 Fitted Gauss, proteinAT5G48870.1
bic:721
peaks after deleting less than one fifth of global max:20
×10^6
15
10
5
0
y
0
5
10
15
20
25
x
raw data, starting:19
fitted Gauss locations:20

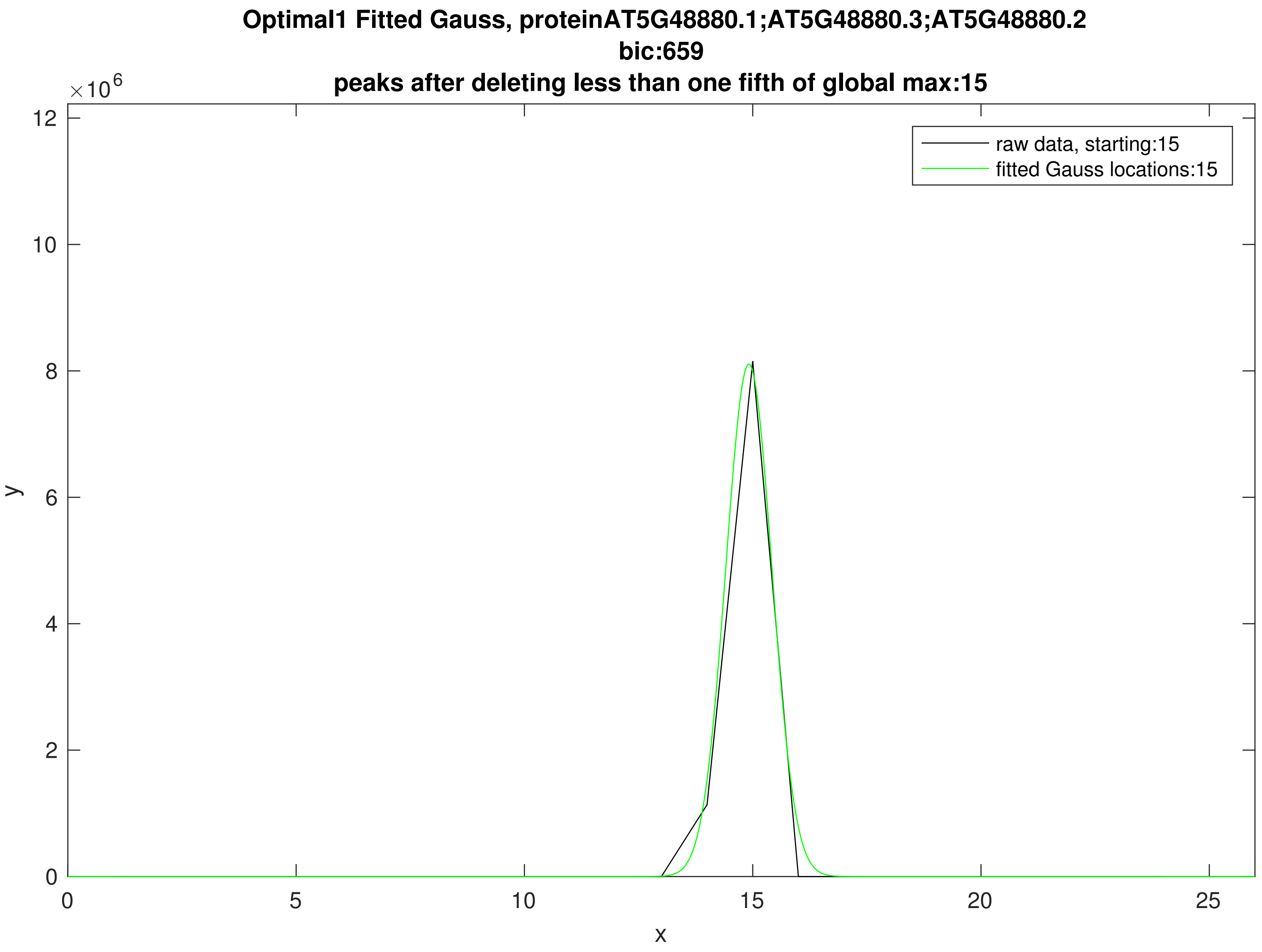

Optimal1 Fitted Gauss, proteinAT5G48880.1;AT5G48880.3;AT5G48880.2
bic:659
peaks after deleting less than one fifth of global max:15
×10^6
raw data, starting:15
fitted Gauss locations:15
y
x
0
2
4
6
8
10
12
0
5
10
15
20
25

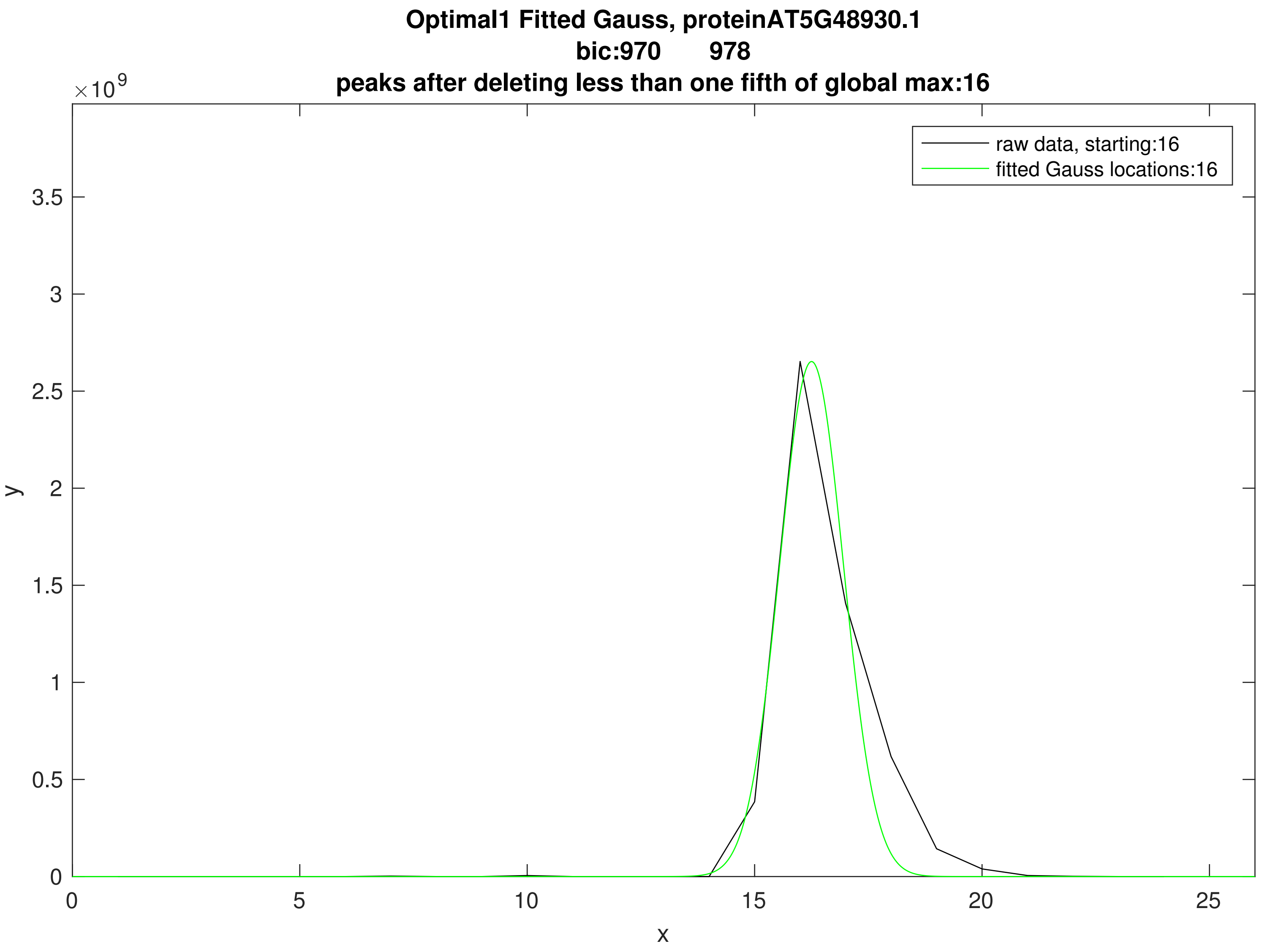

Optimal1 Fitted Gauss, proteinAT5G48930.1
bic:970 978
peaks after deleting less than one fifth of global max:16
×10^9
raw data, starting:16
fitted Gauss locations:16
3.5
3
2.5
2
1.5
1
0.5
0
y
0
5
10
15
20
25
x

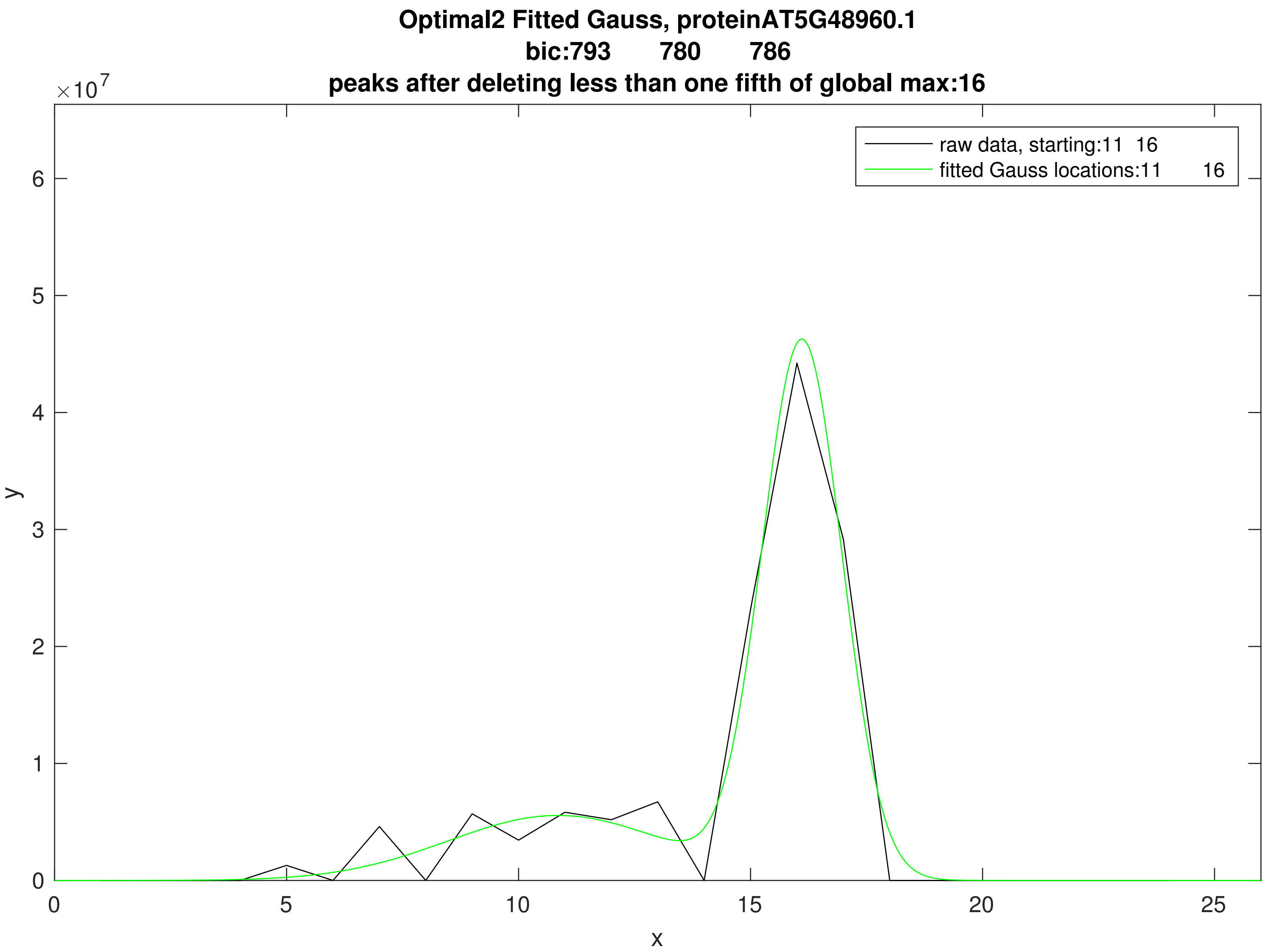
Optimal2 Fitted Gauss, proteinAT5G48960.1
bic:793 780 786
peaks after deleting less than one fifth of global max:16
×10^7
raw data, starting:11 16
fitted Gauss locations:11 16
y
x
0
1
2
3
4
5
6
0
5
10
15
20
25

**Optimal1 Fitted Gauss, proteinAT5G49020.1;AT5G49020.2**
**bic:697**
**peaks after deleting less than one fifth of global max:6.4**

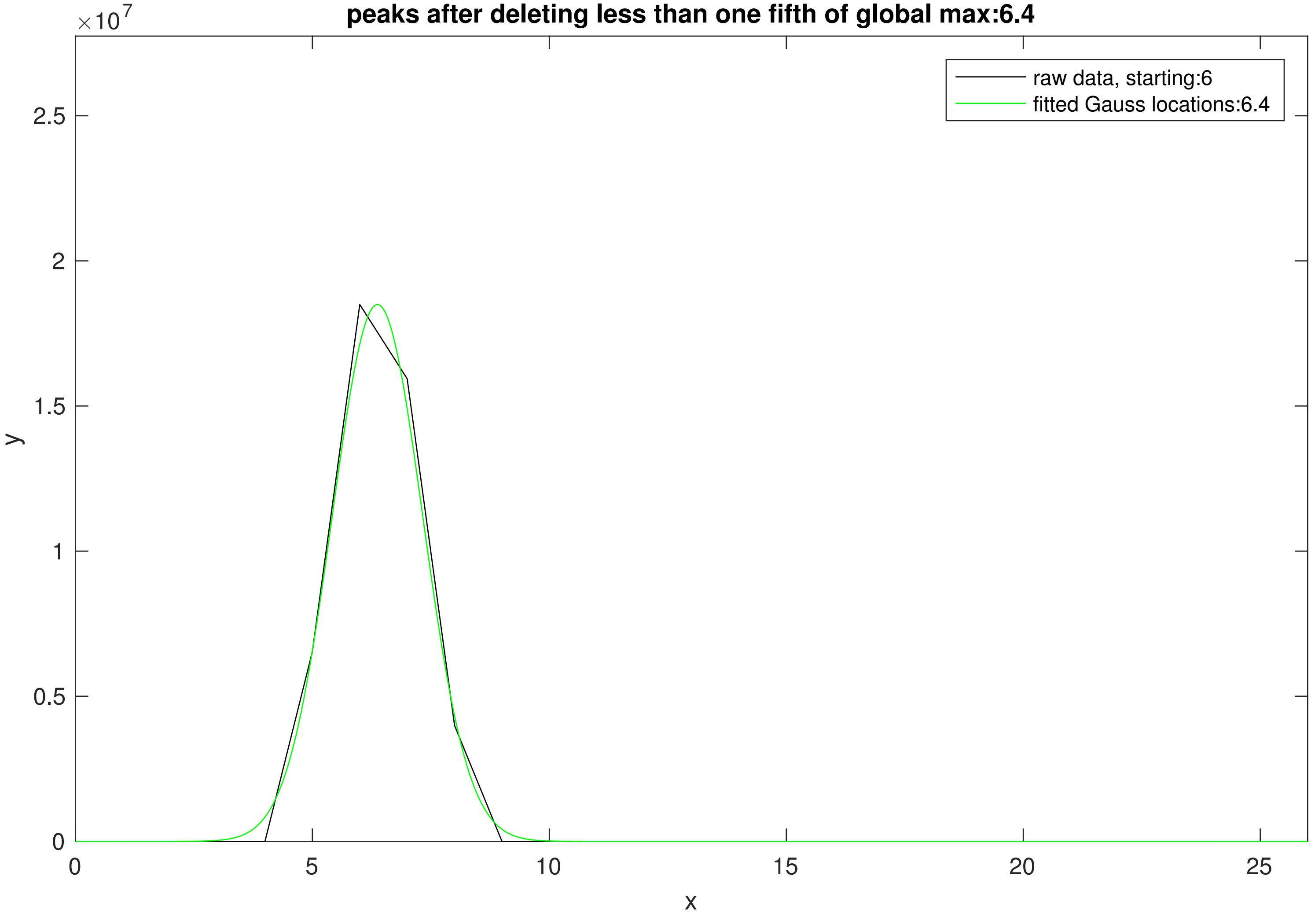

#### Optimal1 Fitted Gauss, proteinAT5G49030.3;AT5G49030.1;AT5G49030.2
#### bic:761
#### peaks after deleting less than one fifth of global max:11

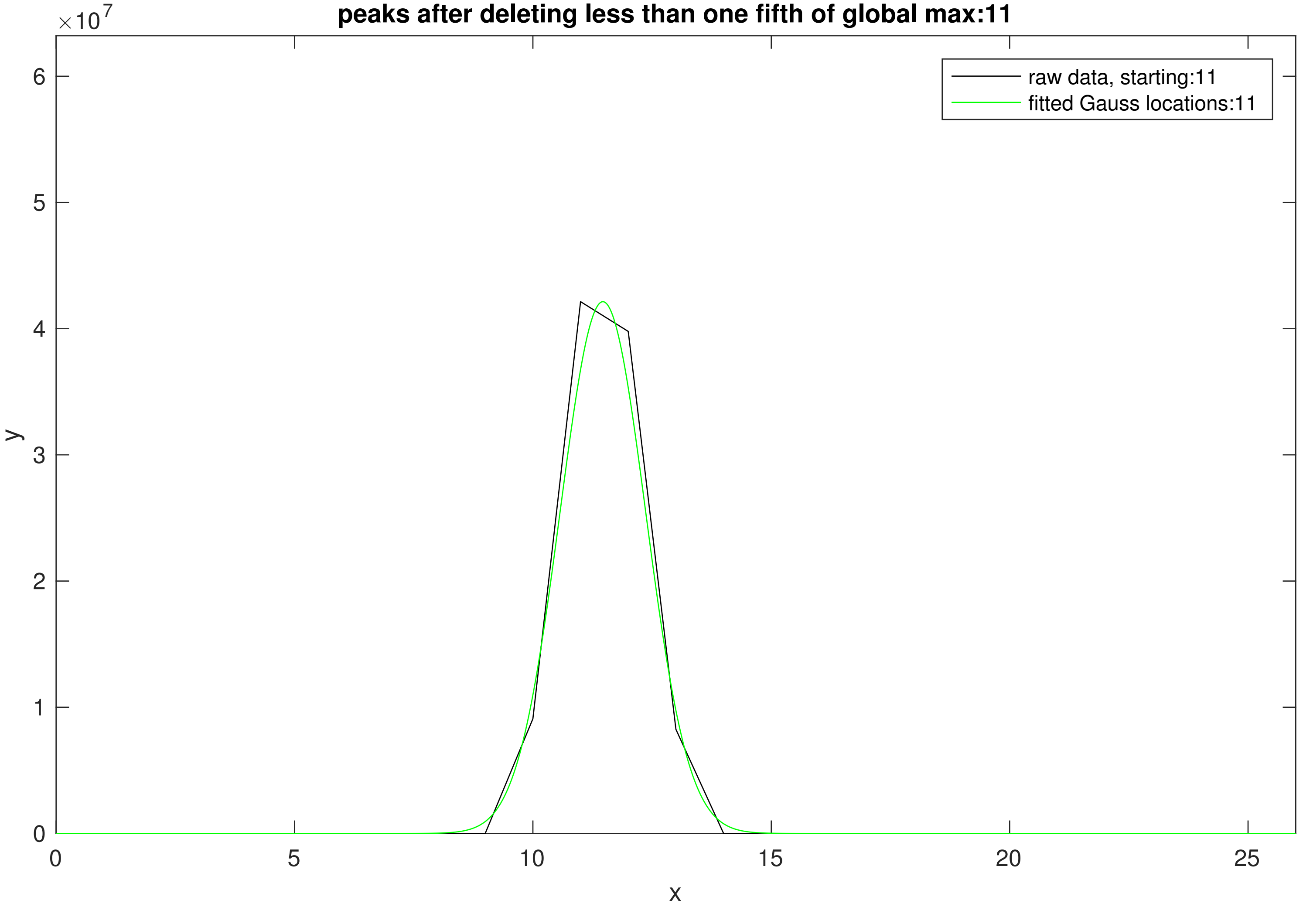

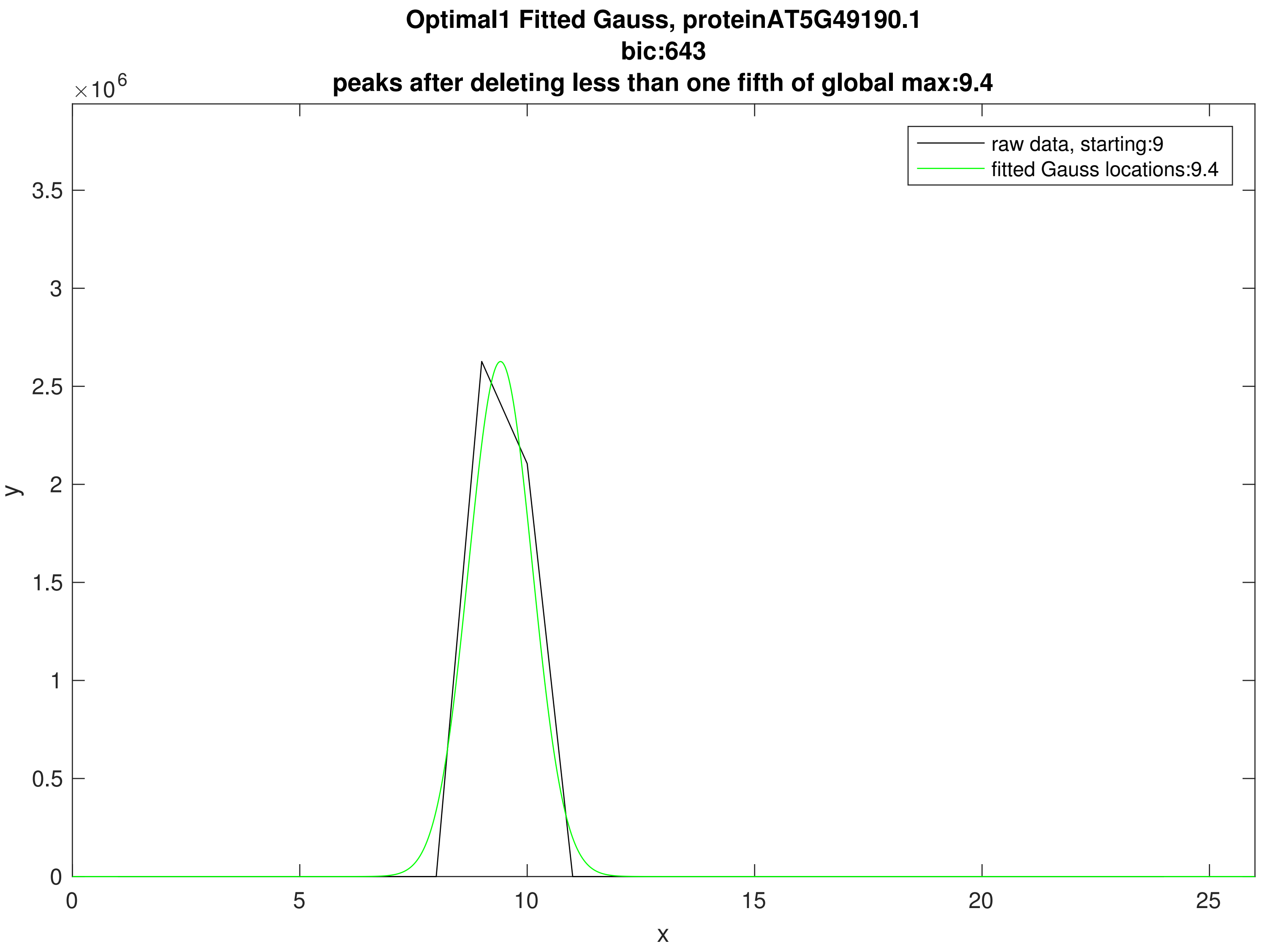

Optimal1 Fitted Gauss, proteinAT5G49190.1
bic:643
peaks after deleting less than one fifth of global max:9.4
×10^6
raw data, starting:9
fitted Gauss locations:9.4
y
x
0
0.5
1
1.5
2
2.5
3
3.5
0
5
10
15
20
25

#### Optimal1 Fitted Gauss, proteinAT5G49215.2;AT5G49215.1
#### bic:798
#### peaks after deleting less than one fifth of global max:15

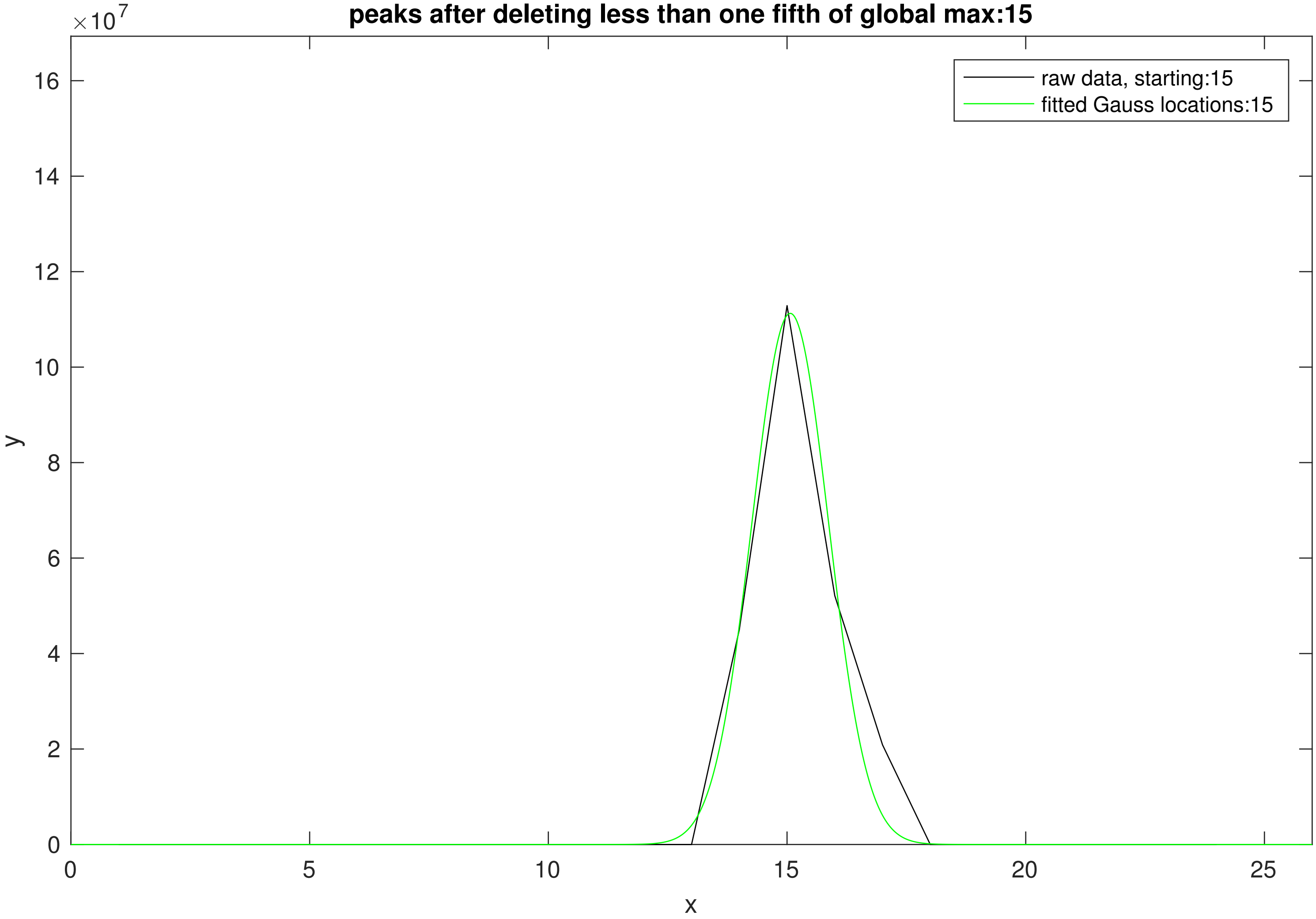

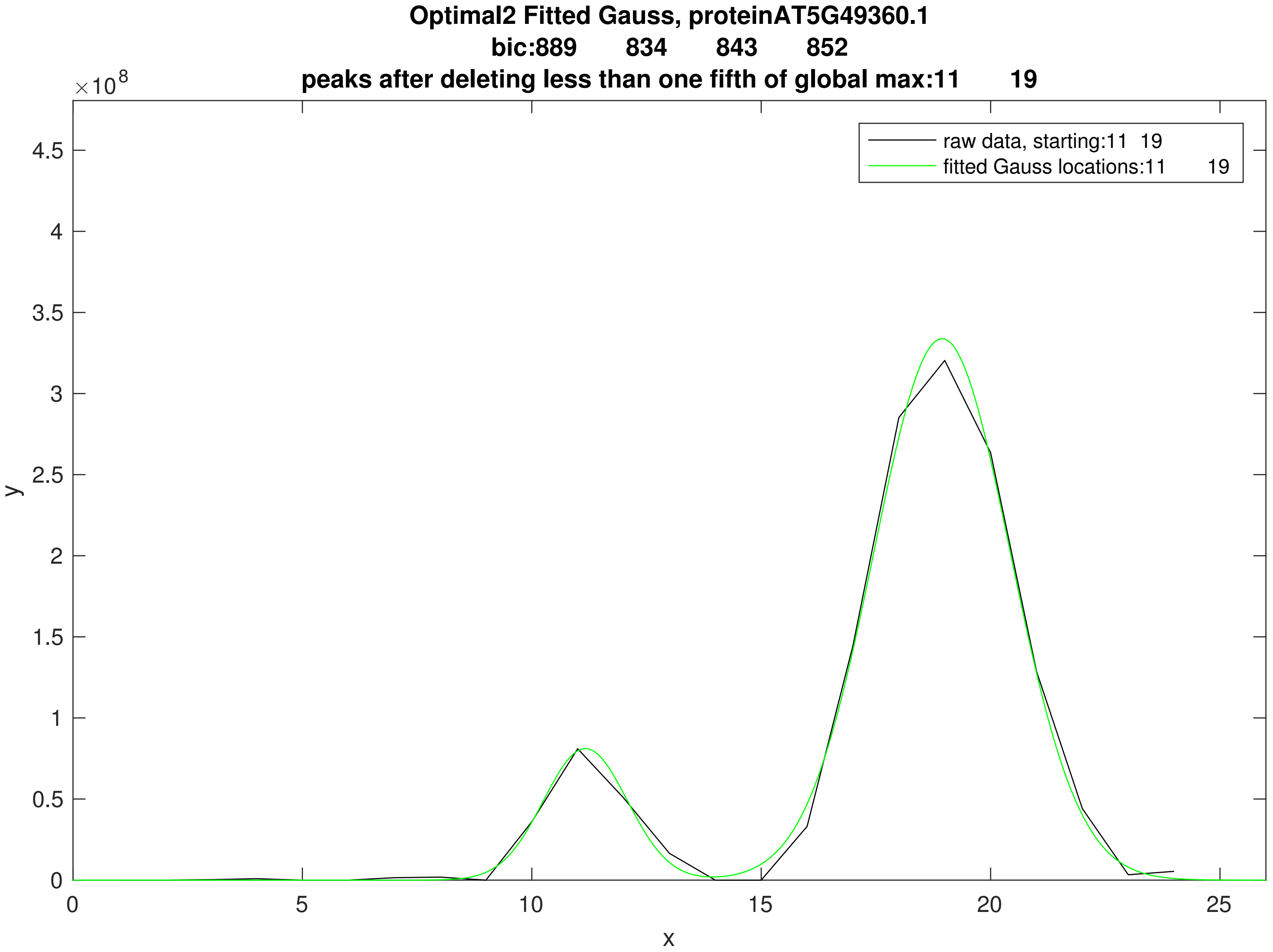
Optimal2 Fitted Gauss, proteinAT5G49360.1
bic:889 834 843 852
peaks after deleting less than one fifth of global max:11 19
×10^8
raw data, starting:11 19
fitted Gauss locations:11 19
y
x
0
0.5
1
1.5
2
2.5
3
3.5
4
4.5
0
5
10
15
20
25

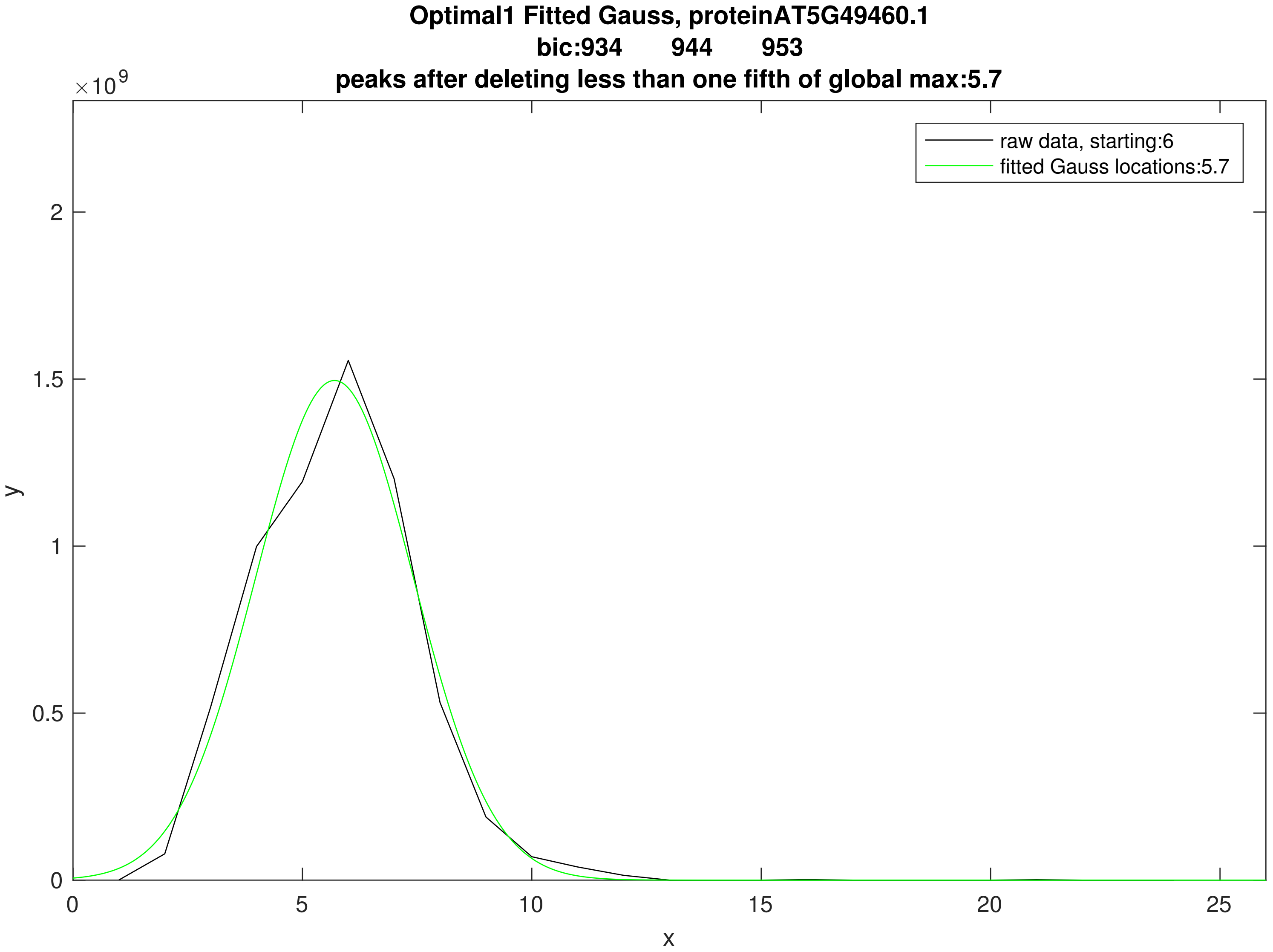
Optimal1 Fitted Gauss, proteinAT5G49460.1
bic:934 944 953
peaks after deleting less than one fifth of global max:5.7
×10^9
raw data, starting:6
fitted Gauss locations:5.7
y
x
0
0.5
1
1.5
2
0
5
10
15
20
25

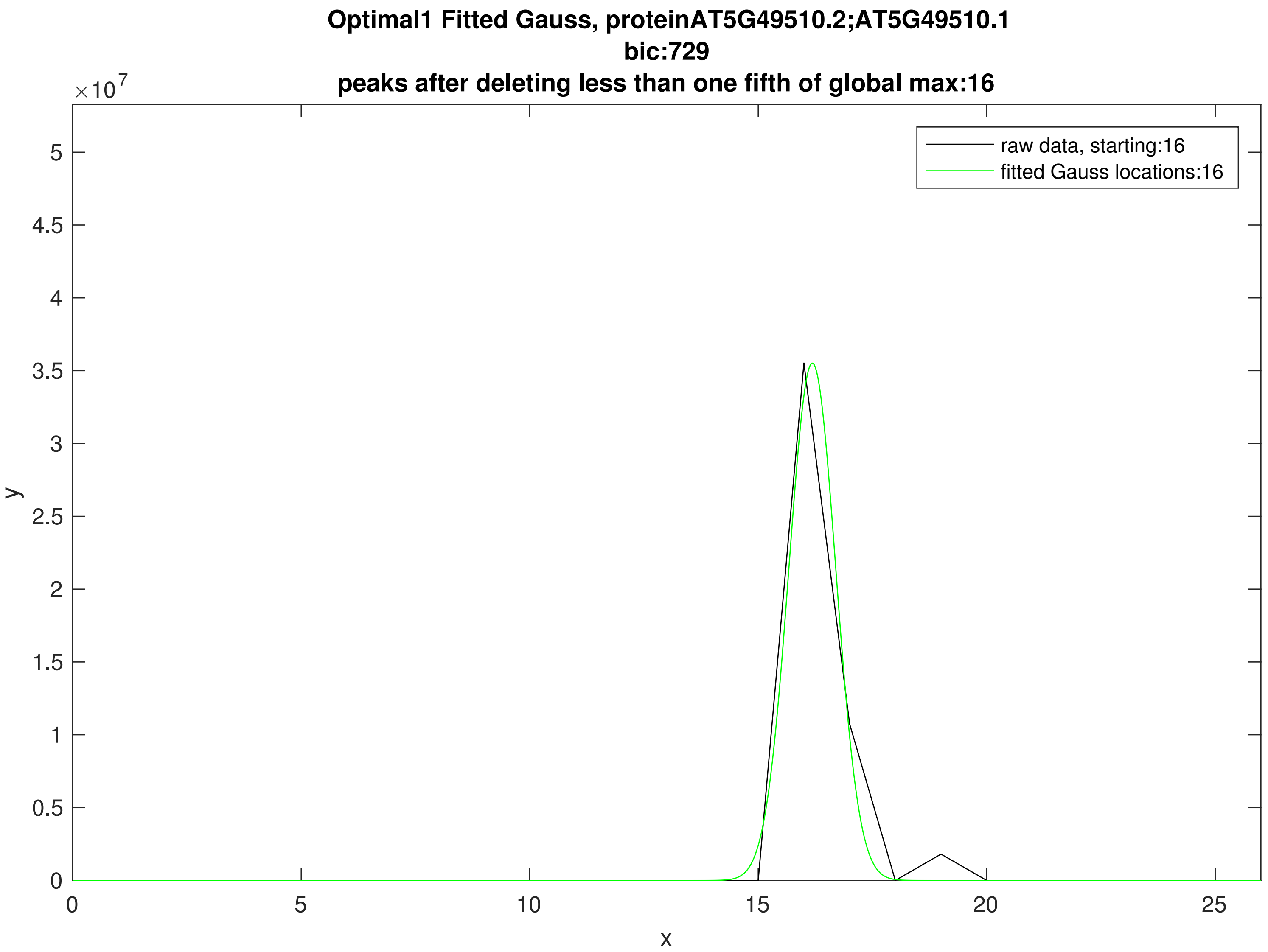

Optimal1 Fitted Gauss, proteinAT5G49510.2;AT5G49510.1
bic:729
peaks after deleting less than one fifth of global max:16
×10^7
raw data, starting:16
fitted Gauss locations:16
y
x
0
0.5
1
1.5
2
2.5
3
3.5
4
4.5
5
0
5
10
15
20
25

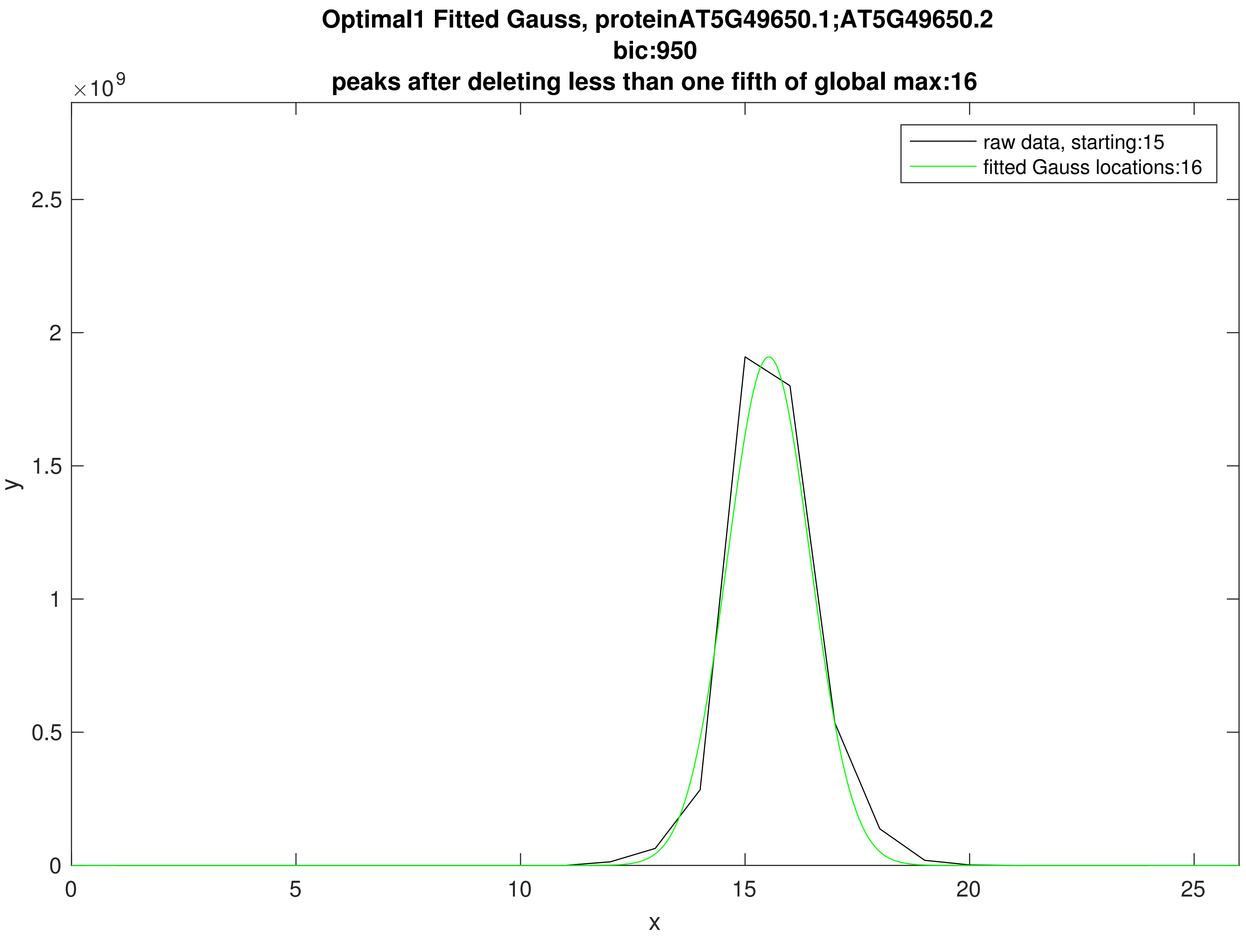

Optimal1 Fitted Gauss, proteinAT5G49650.1;AT5G49650.2
bic:950
peaks after deleting less than one fifth of global max:16
×10^9
raw data, starting:15
fitted Gauss locations:16
2.5
2
1.5
1
0.5
0
y
0
5
10
15
20
25
x

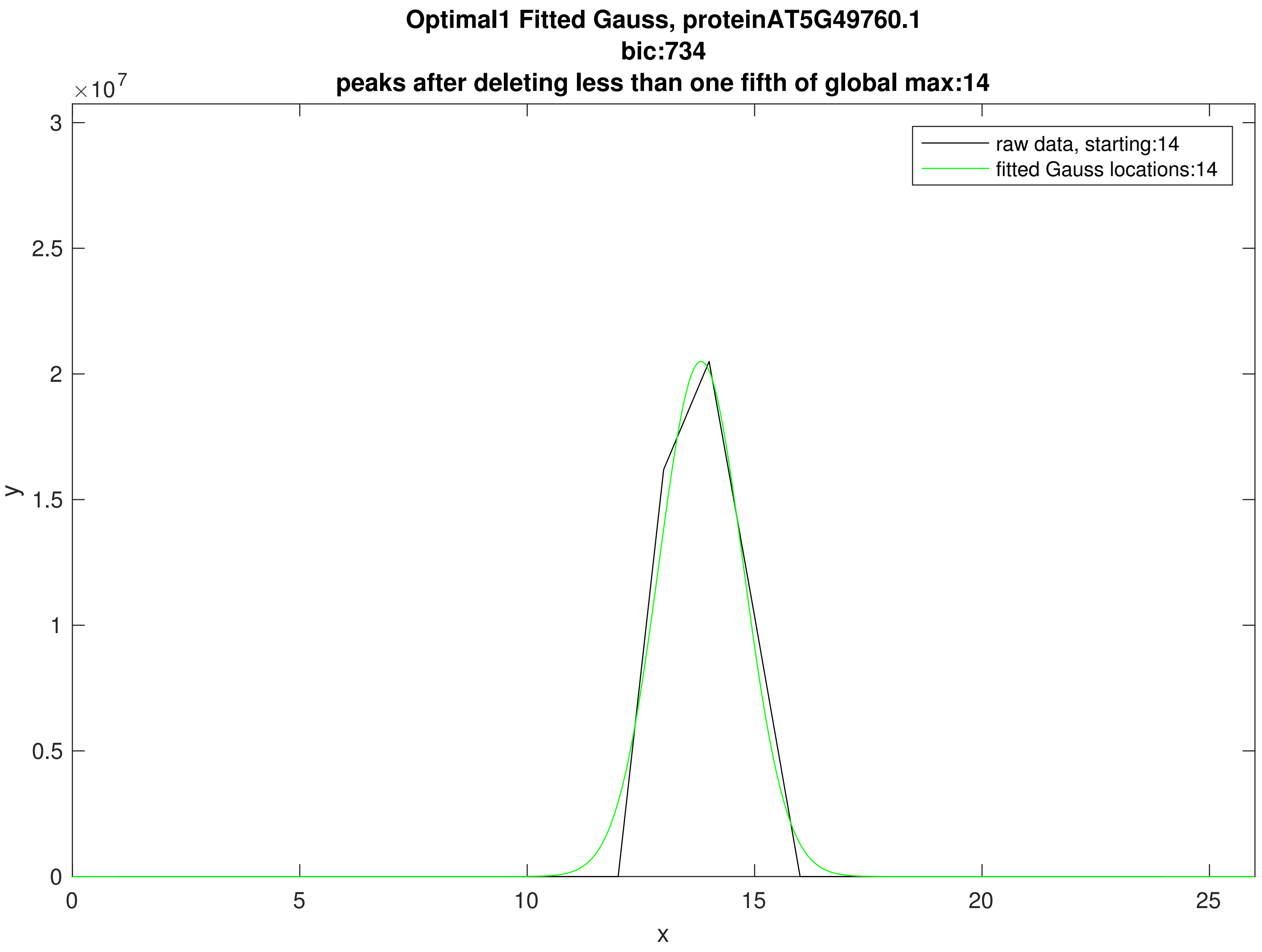
Optimal1 Fitted Gauss, proteinAT5G49760.1
bic:734
peaks after deleting less than one fifth of global max:14
×10^7
3
2.5
2
1.5
1
0.5
0
y
0
5
10
15
20
25
x
raw data, starting:14
fitted Gauss locations:14

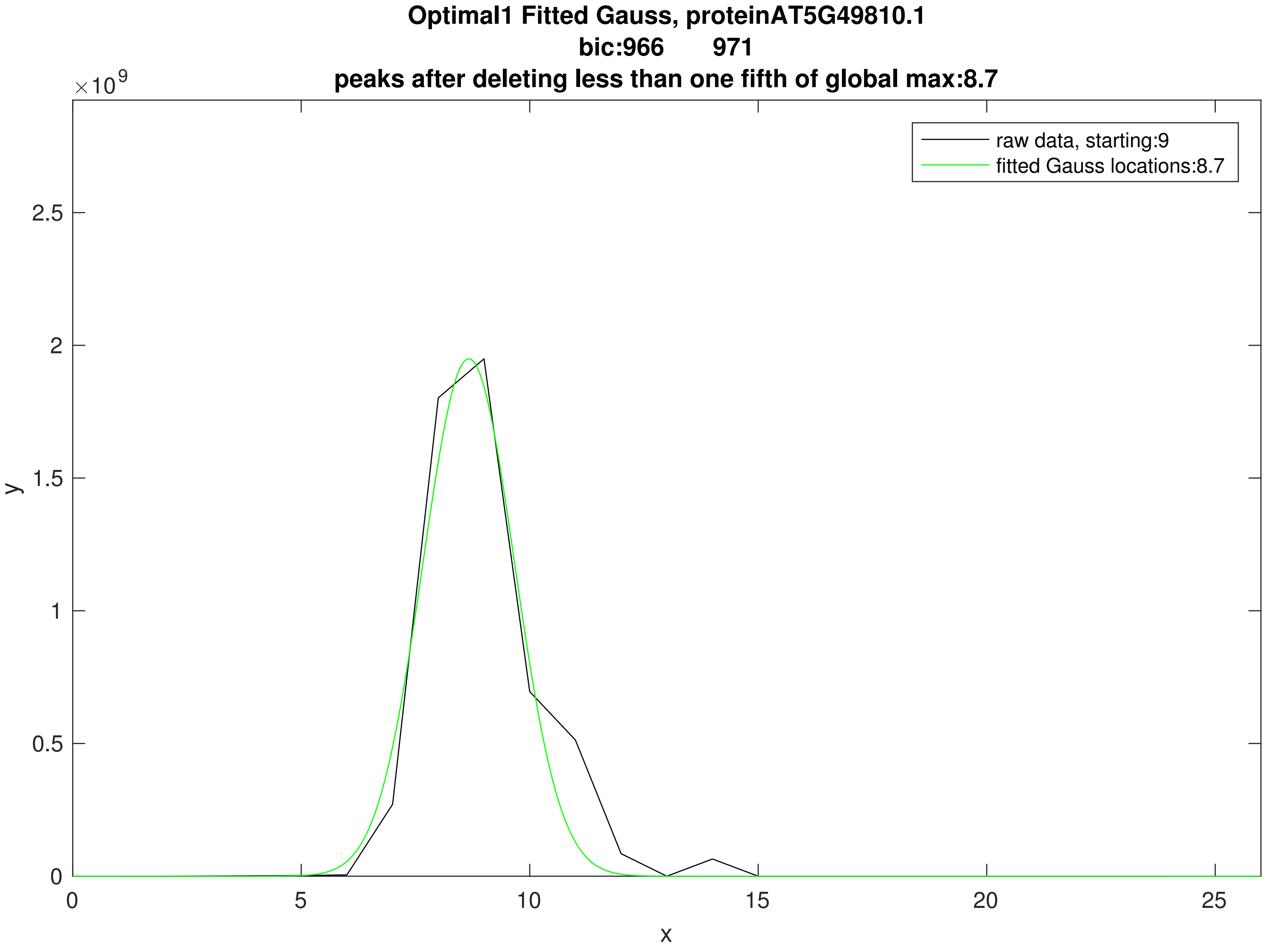

Optimal1 Fitted Gauss, proteinAT5G49810.1
bic:966 971
peaks after deleting less than one fifth of global max:8.7
×10^9
raw data, starting:9
fitted Gauss locations:8.7
y
0
0.5
1
1.5
2
2.5
x
0
5
10
15
20
25

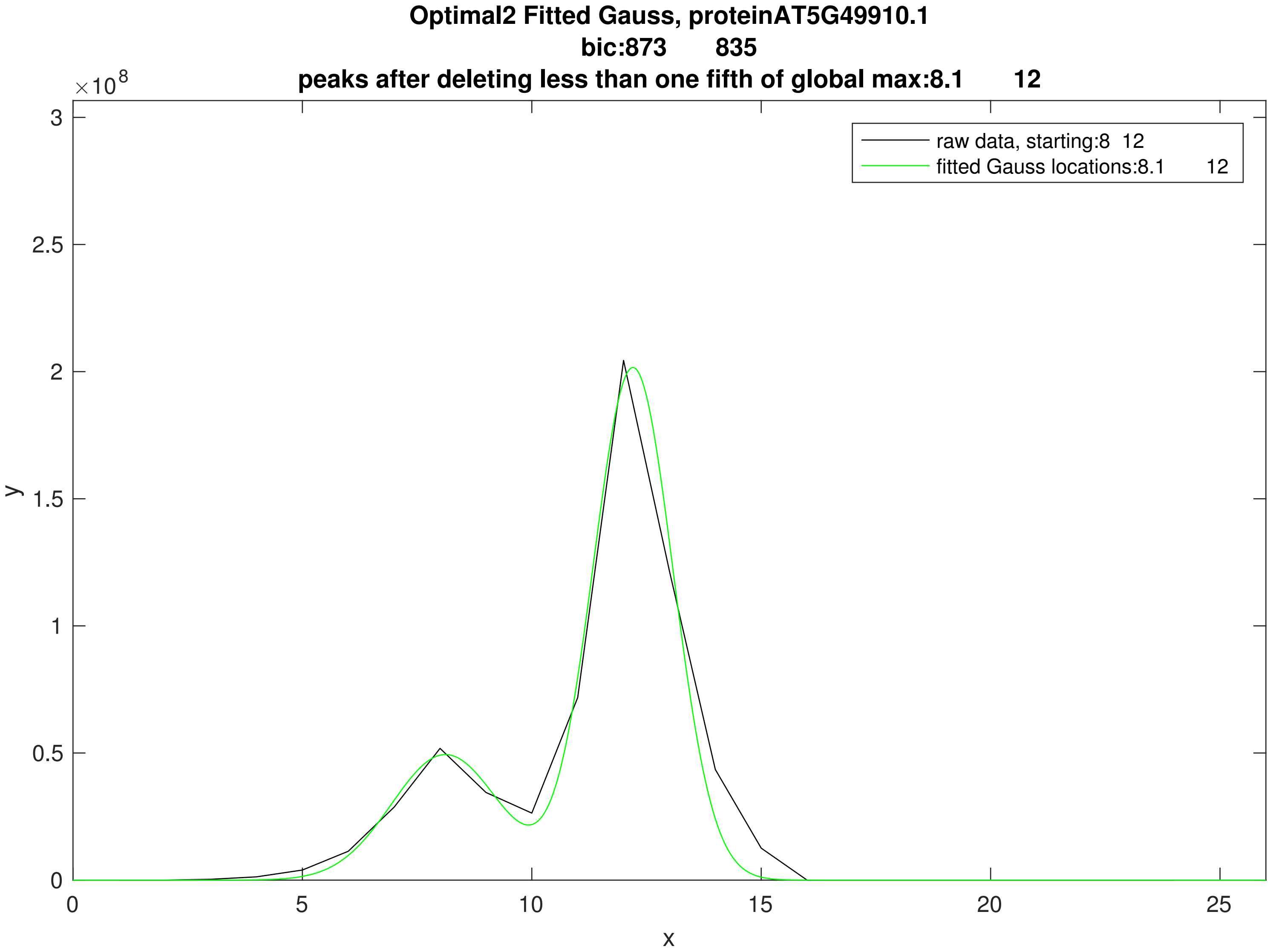

Optimal2 Fitted Gauss, proteinAT5G49910.1
bic:873 835
peaks after deleting less than one fifth of global max:8.1 12
×10^8
3
2.5
2
1.5
1
0.5
0
y
0
5
10
15
20
25
x
raw data, starting:8 12
fitted Gauss locations:8.1 12

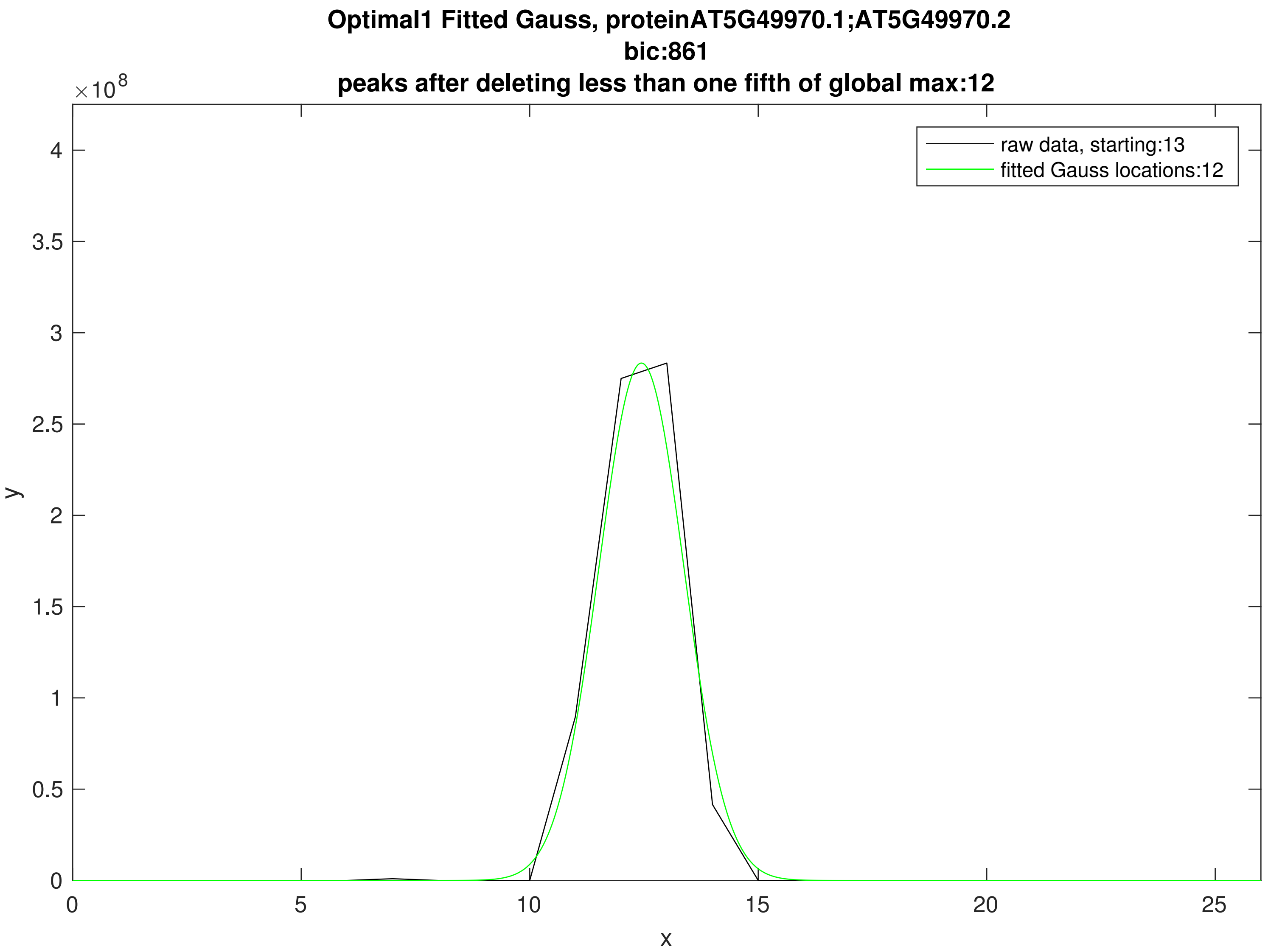

Optimal1 Fitted Gauss, proteinAT5G49970.1;AT5G49970.2
bic:861
peaks after deleting less than one fifth of global max:12
×10^8
4
3.5
3
2.5
2
1.5
1
0.5
0
y
0
5
10
15
20
25
x
raw data, starting:13
fitted Gauss locations:12

Optimal1 Fitted Gauss, proteinAT5G49980.1
bic:707
peaks after deleting less than one fifth of global max:8.6

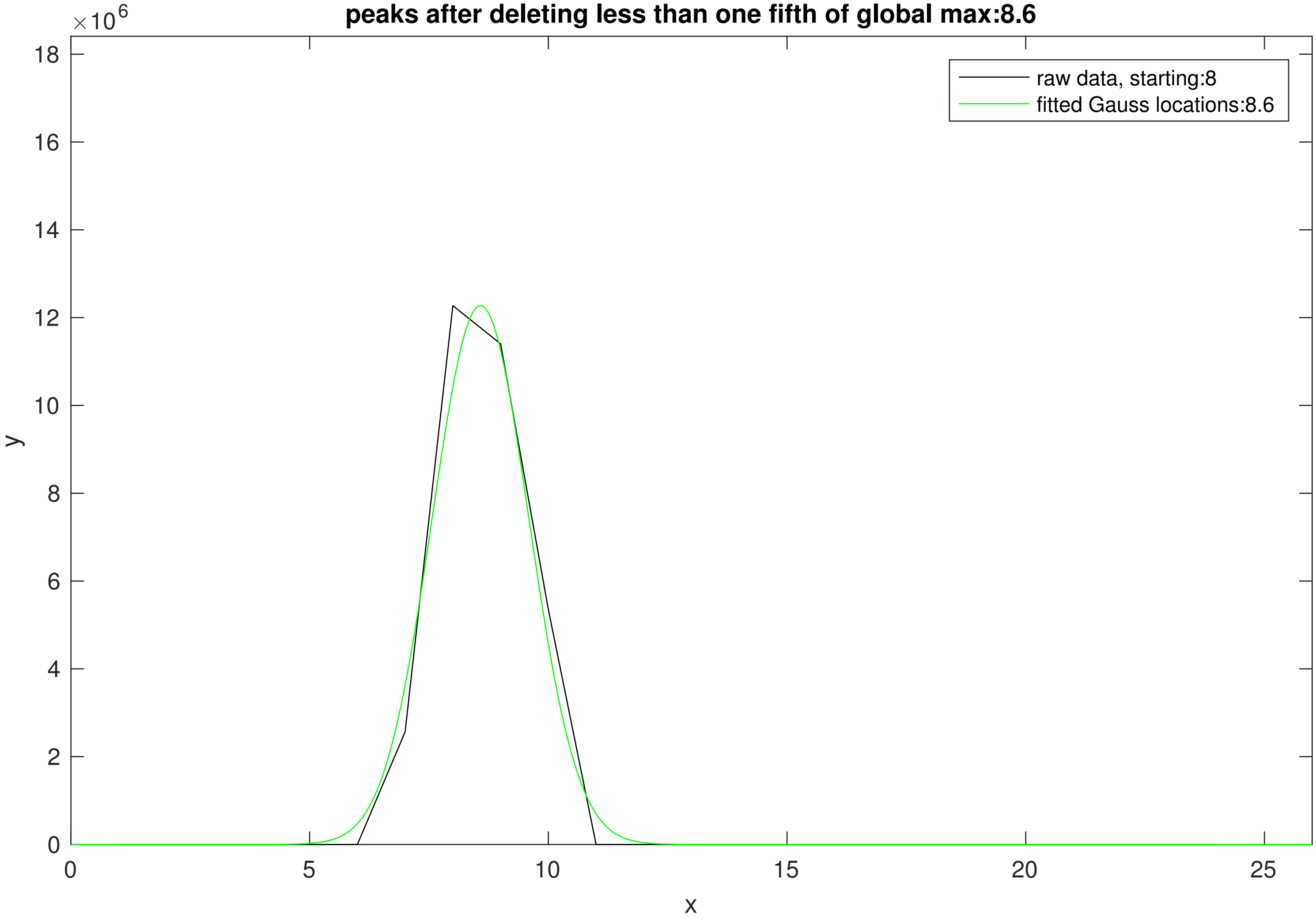

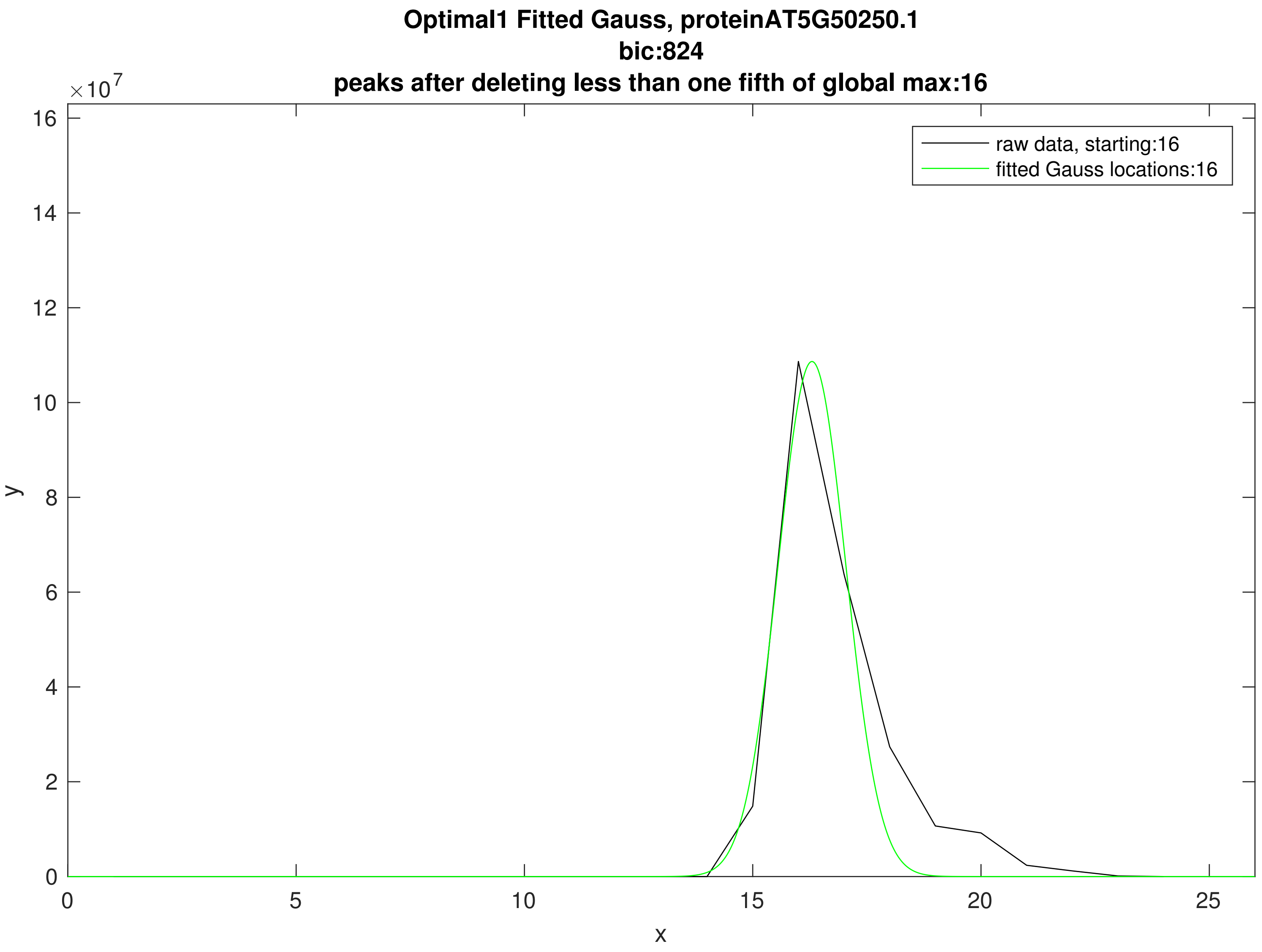

Optimal1 Fitted Gauss, proteinAT5G50250.1
bic:824
peaks after deleting less than one fifth of global max:16
×10^7
16
14
12
10
8
6
4
2
0
y
0
5
10
15
20
25
x
raw data, starting:16
fitted Gauss locations:16

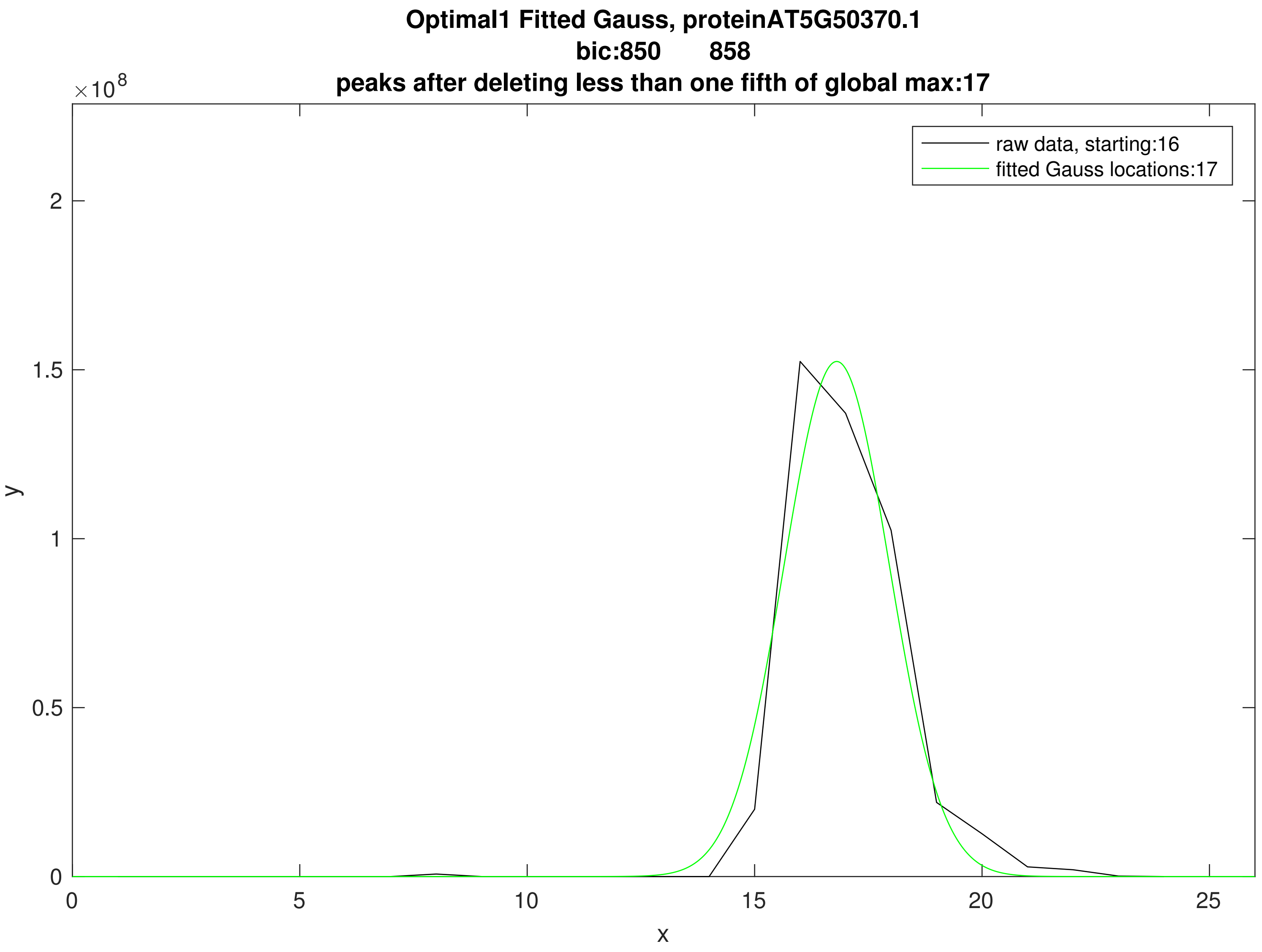
Optimal1 Fitted Gauss, proteinAT5G50370.1
bic:850 858
peaks after deleting less than one fifth of global max:17
×10^8
raw data, starting:16
fitted Gauss locations:17
y
x
0
0.5
1
1.5
2
0
5
10
15
20
25

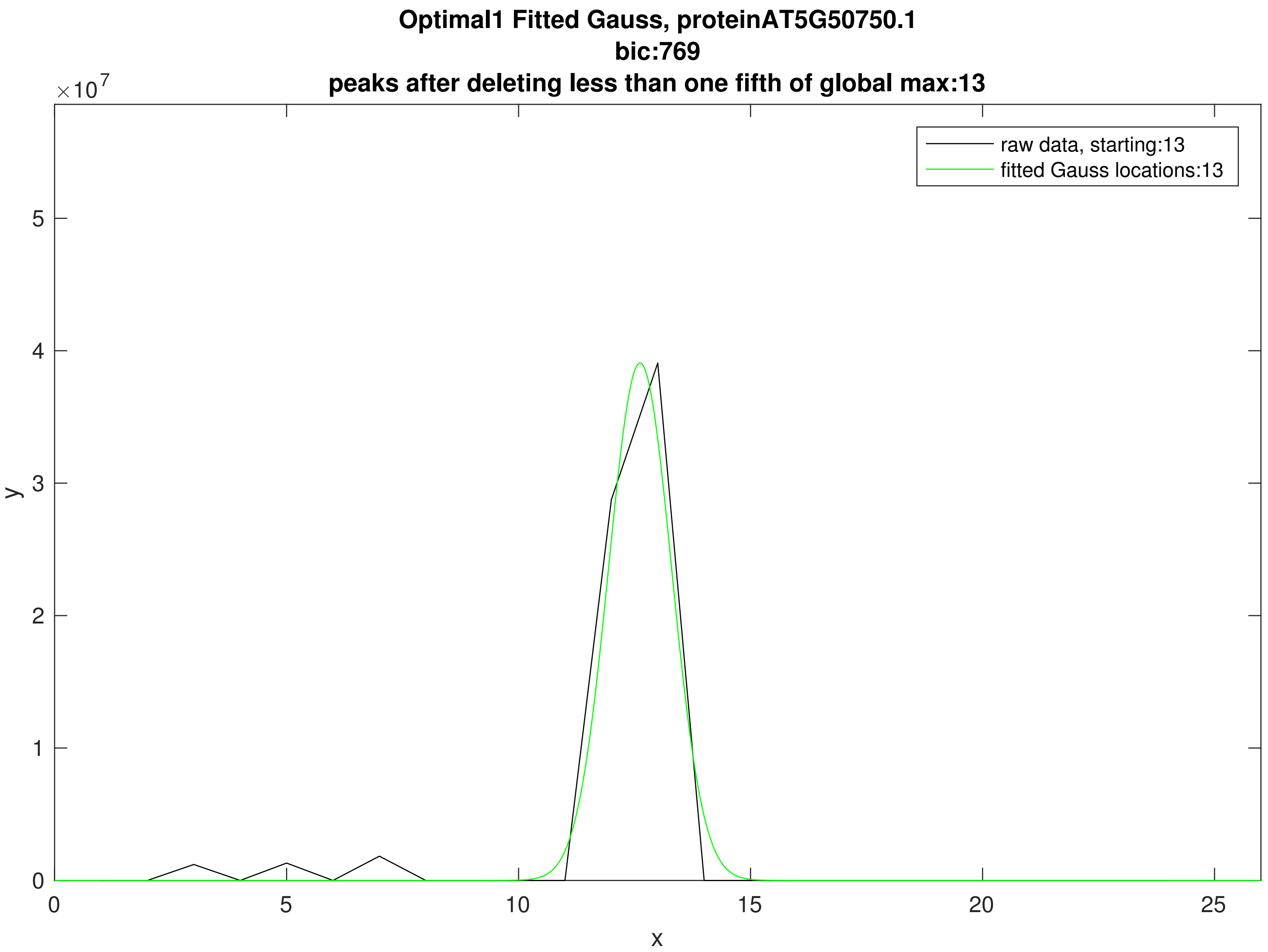
Optimal1 Fitted Gauss, proteinAT5G50750.1
bic:769
peaks after deleting less than one fifth of global max:13
×10^7
raw data, starting:13
fitted Gauss locations:13
y
x
0
1
2
3
4
5
0
5
10
15
20
25

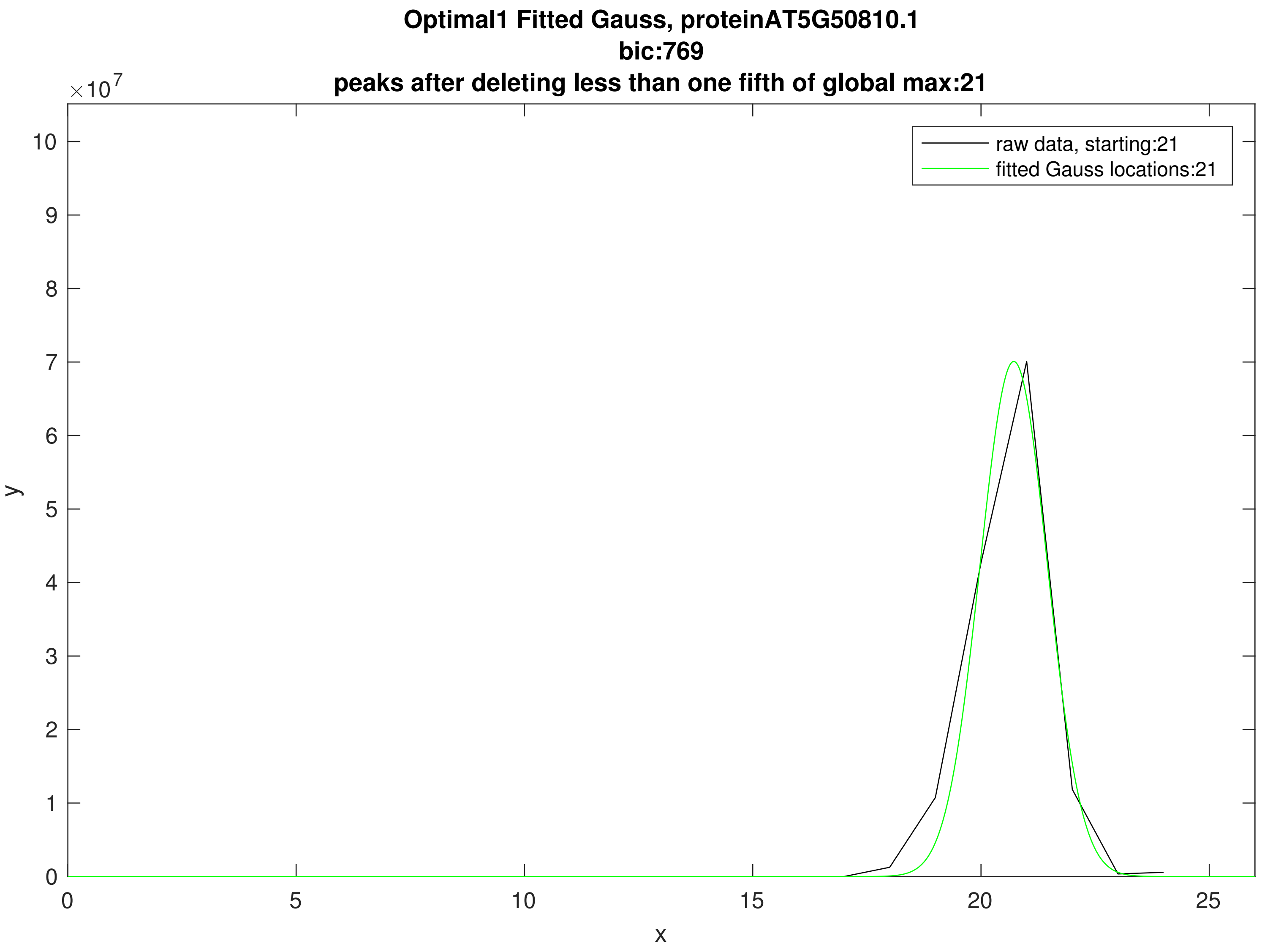
Optimal1 Fitted Gauss, proteinAT5G50810.1
bic:769
peaks after deleting less than one fifth of global max:21
×10^7
raw data, starting:21
fitted Gauss locations:21
y
x
0
1
2
3
4
5
6
7
8
9
10
0
5
10
15
20
25

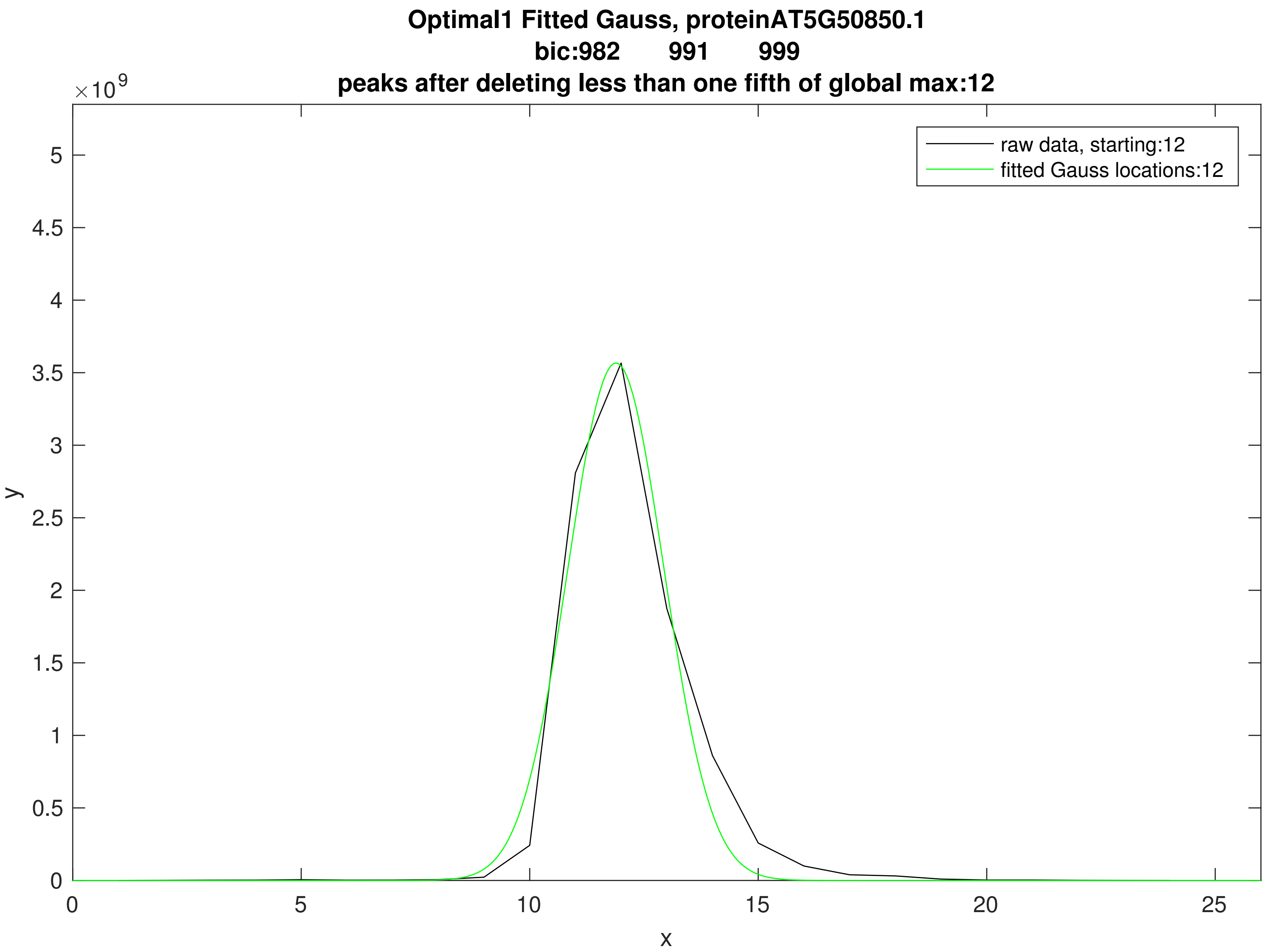
Optimal1 Fitted Gauss, proteinAT5G50850.1
bic:982 991 999
peaks after deleting less than one fifth of global max:12
×10^9
raw data, starting:12
fitted Gauss locations:12
y
x
0
0.5
1
1.5
2
2.5
3
3.5
4
4.5
5
0
5
10
15
20
25

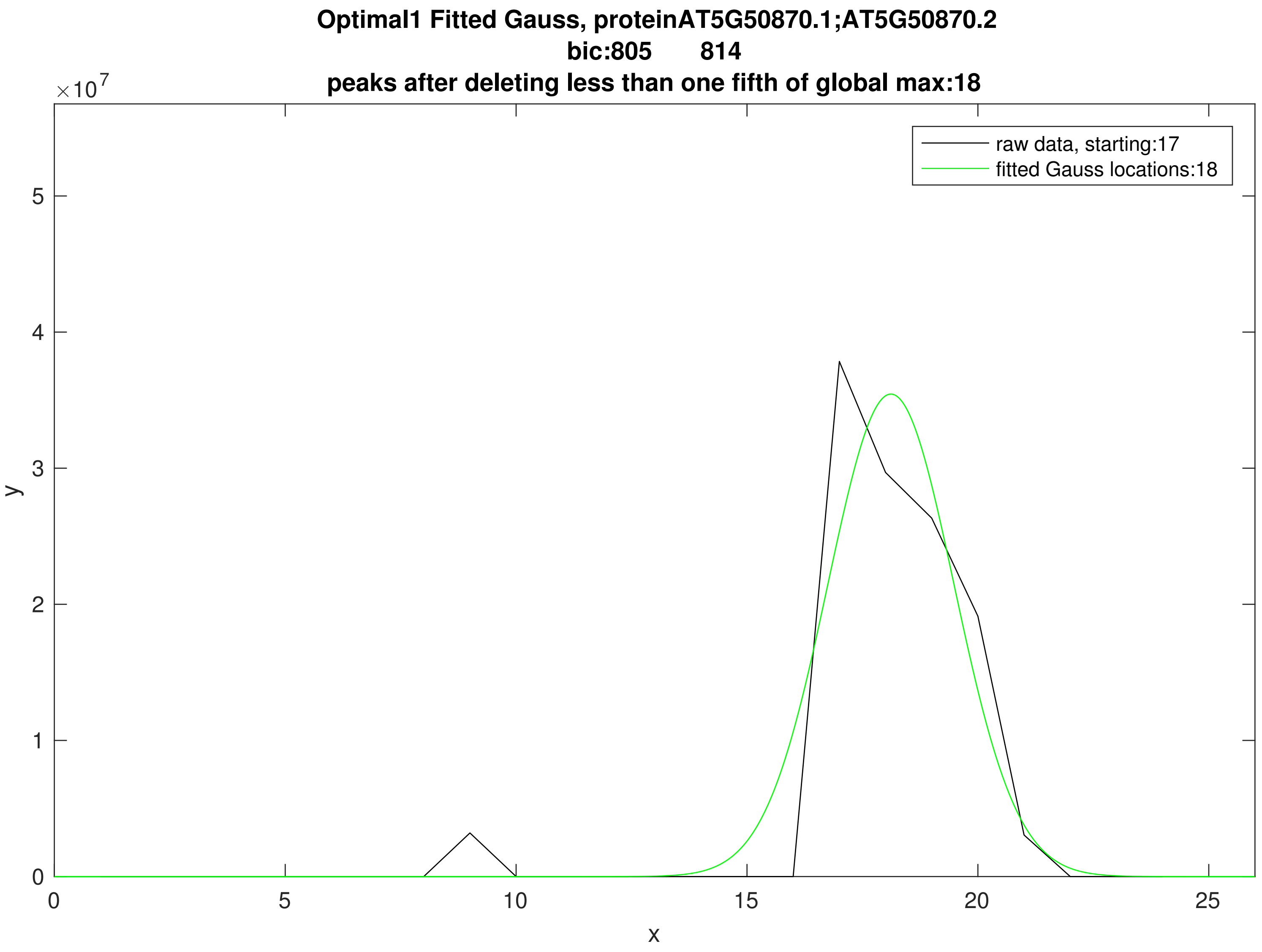
Optimal1 Fitted Gauss, proteinAT5G50870.1;AT5G50870.2
bic:805 814
peaks after deleting less than one fifth of global max:18
×10^7
raw data, starting:17
fitted Gauss locations:18
y
x
0
1
2
3
4
5
0
5
10
15
20
25

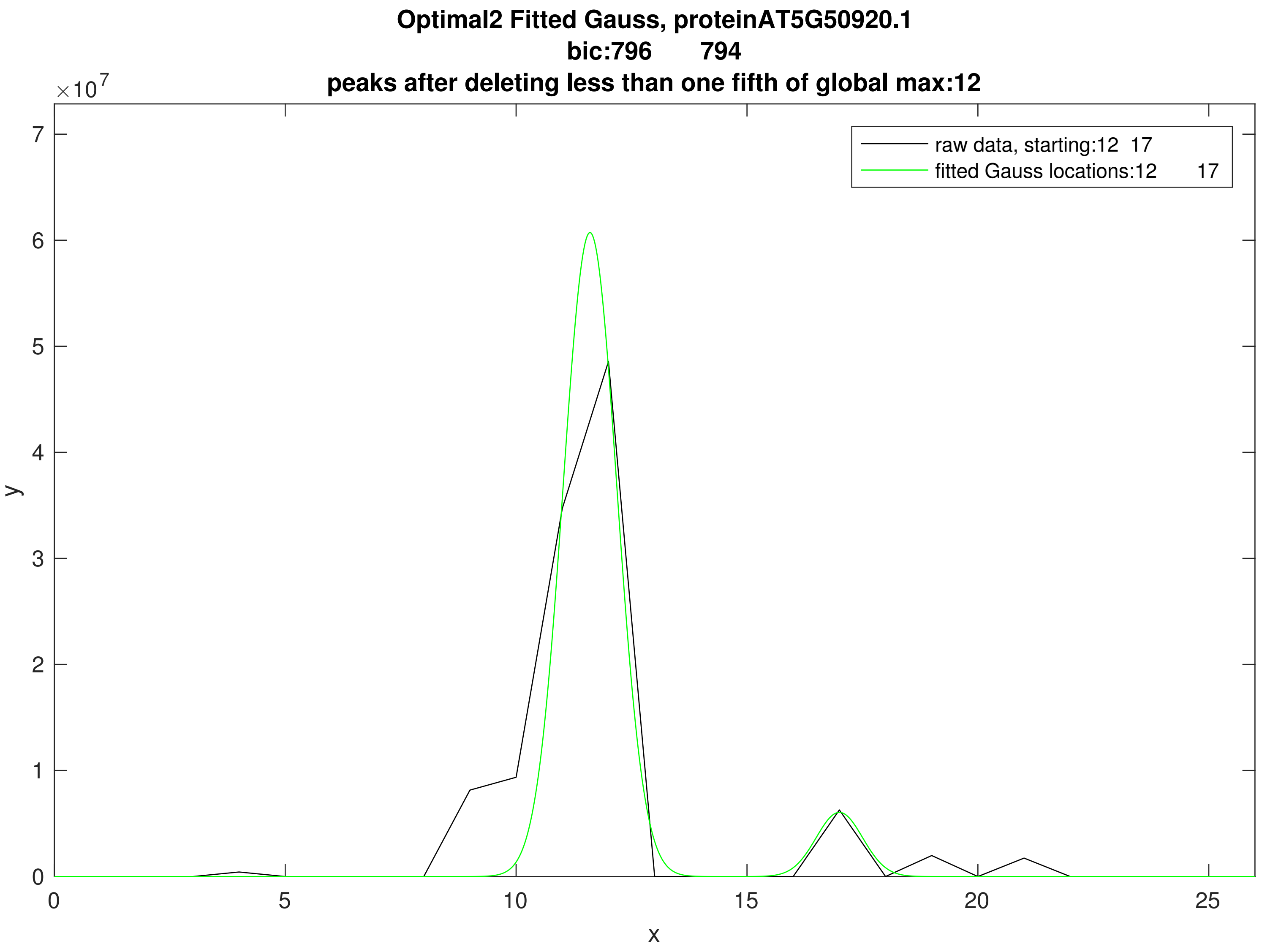
Optimal2 Fitted Gauss, proteinAT5G50920.1
bic:796 794
peaks after deleting less than one fifth of global max:12
×10^7
raw data, starting:12 17
fitted Gauss locations:12 17
y
x
0
1
2
3
4
5
6
7
10
15
20
25

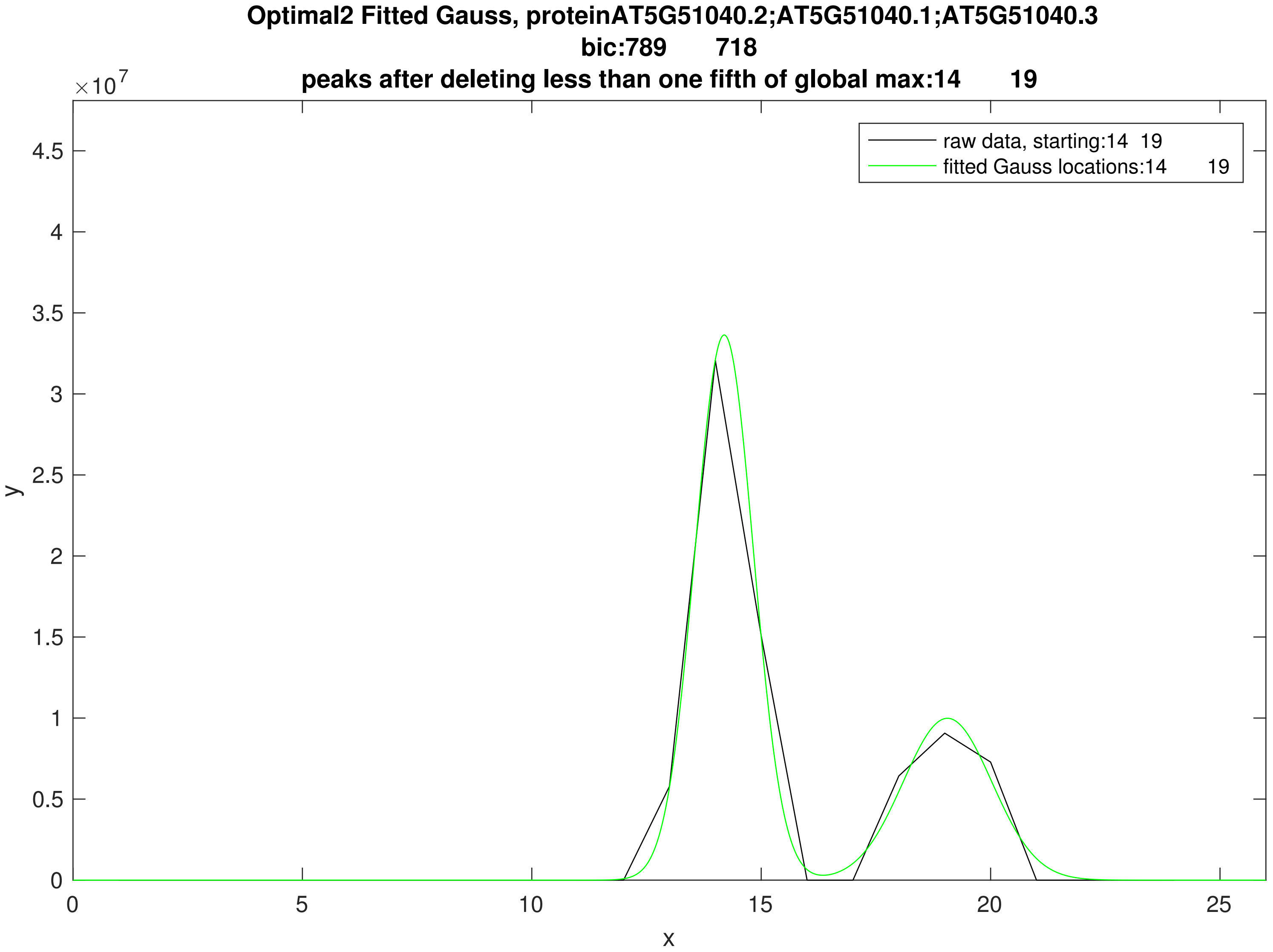
Optimal2 Fitted Gauss, proteinAT5G51040.2;AT5G51040.1;AT5G51040.3
bic:789 718
peaks after deleting less than one fifth of global max:14 19
×10^7
raw data, starting:14 19
fitted Gauss locations:14 19
y
x
0
0.5
1
1.5
2
2.5
3
3.5
4
4.5
5
10
15
20
25

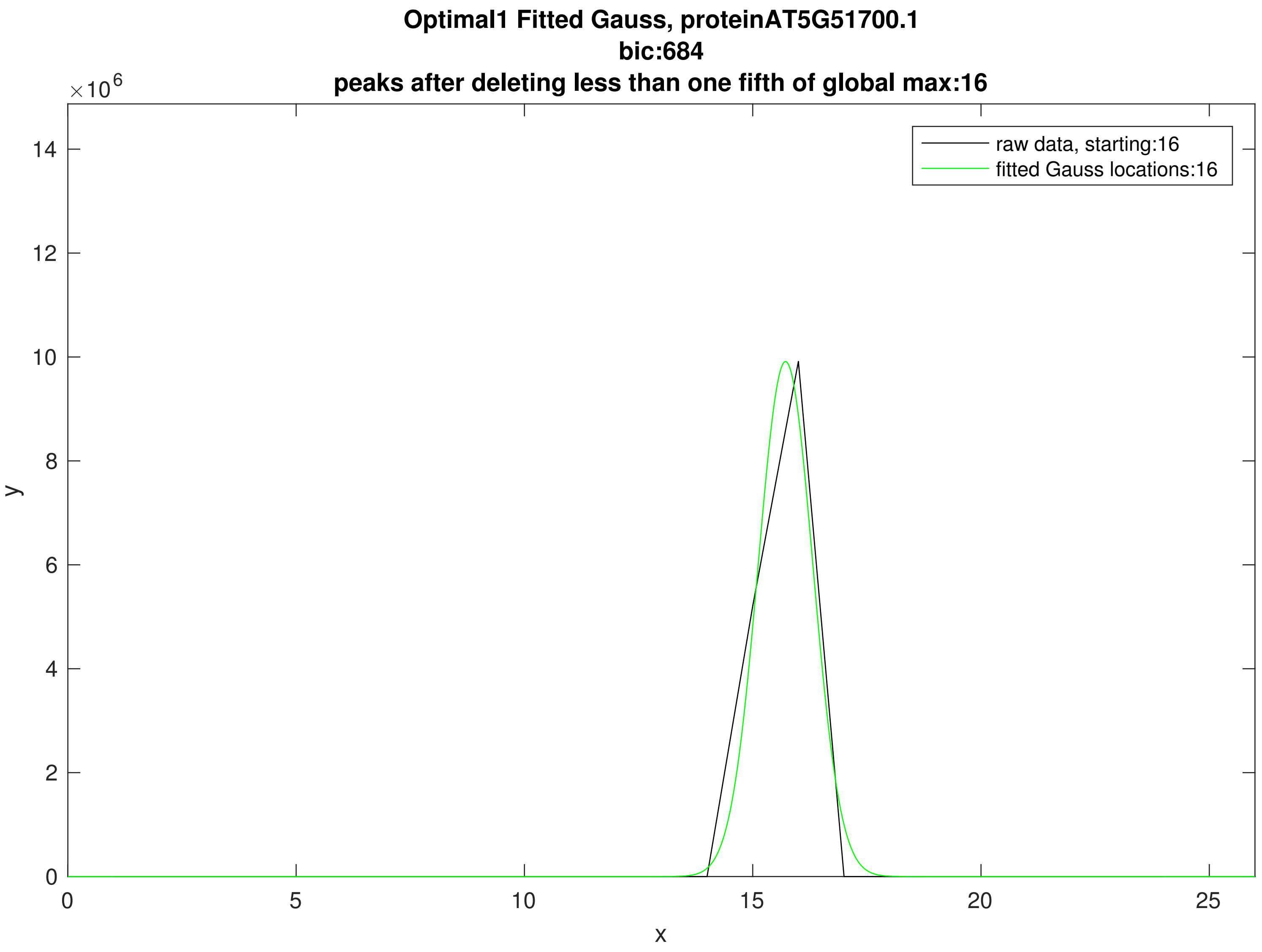

Optimal1 Fitted Gauss, proteinAT5G51700.1
bic:684
peaks after deleting less than one fifth of global max:16
×10^6
raw data, starting:16
fitted Gauss locations:16
y
x
0
2
4
6
8
10
12
14
0
5
10
15
20
25

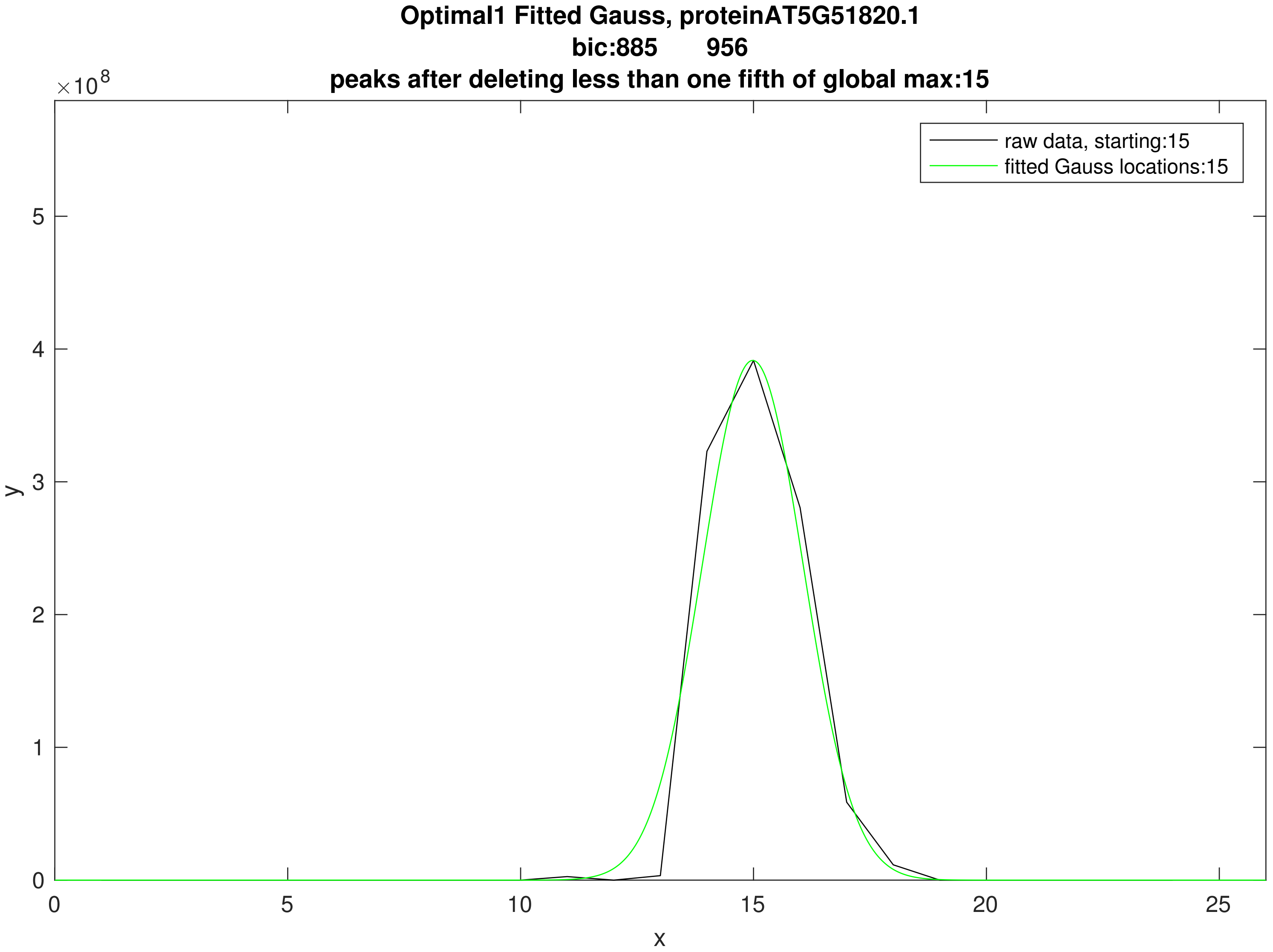
Optimal1 Fitted Gauss, proteinAT5G51820.1
bic:885 956
peaks after deleting less than one fifth of global max:15
×10^8
raw data, starting:15
fitted Gauss locations:15
y
x
0
1
2
3
4
5
0
5
10
15
20
25

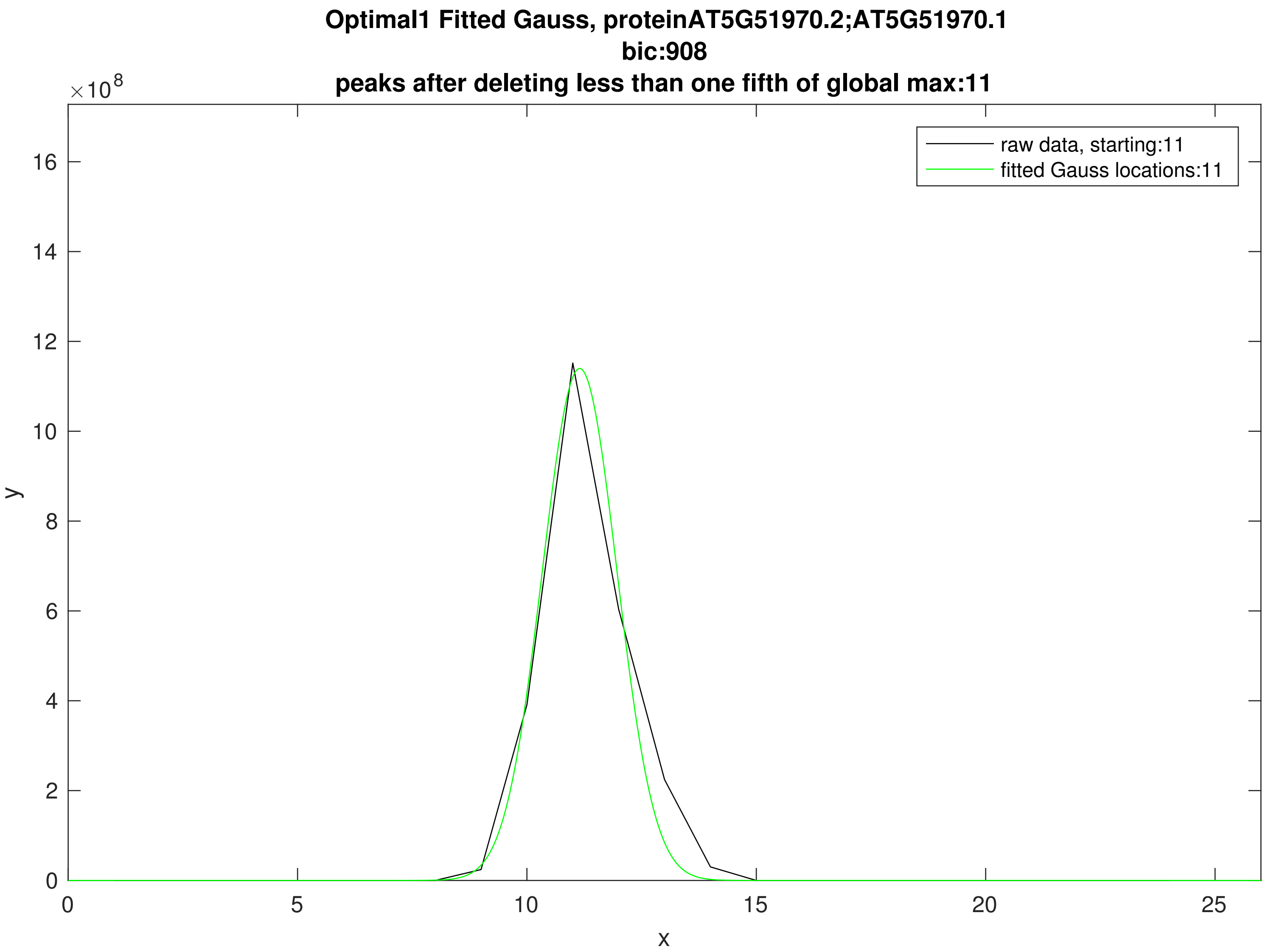

Optimal1 Fitted Gauss, proteinAT5G51970.2;AT5G51970.1
bic:908
peaks after deleting less than one fifth of global max:11
×10^8
16
14
12
10
8
6
4
2
0
y
0
5
10
15
20
25
x
raw data, starting:11
fitted Gauss locations:11

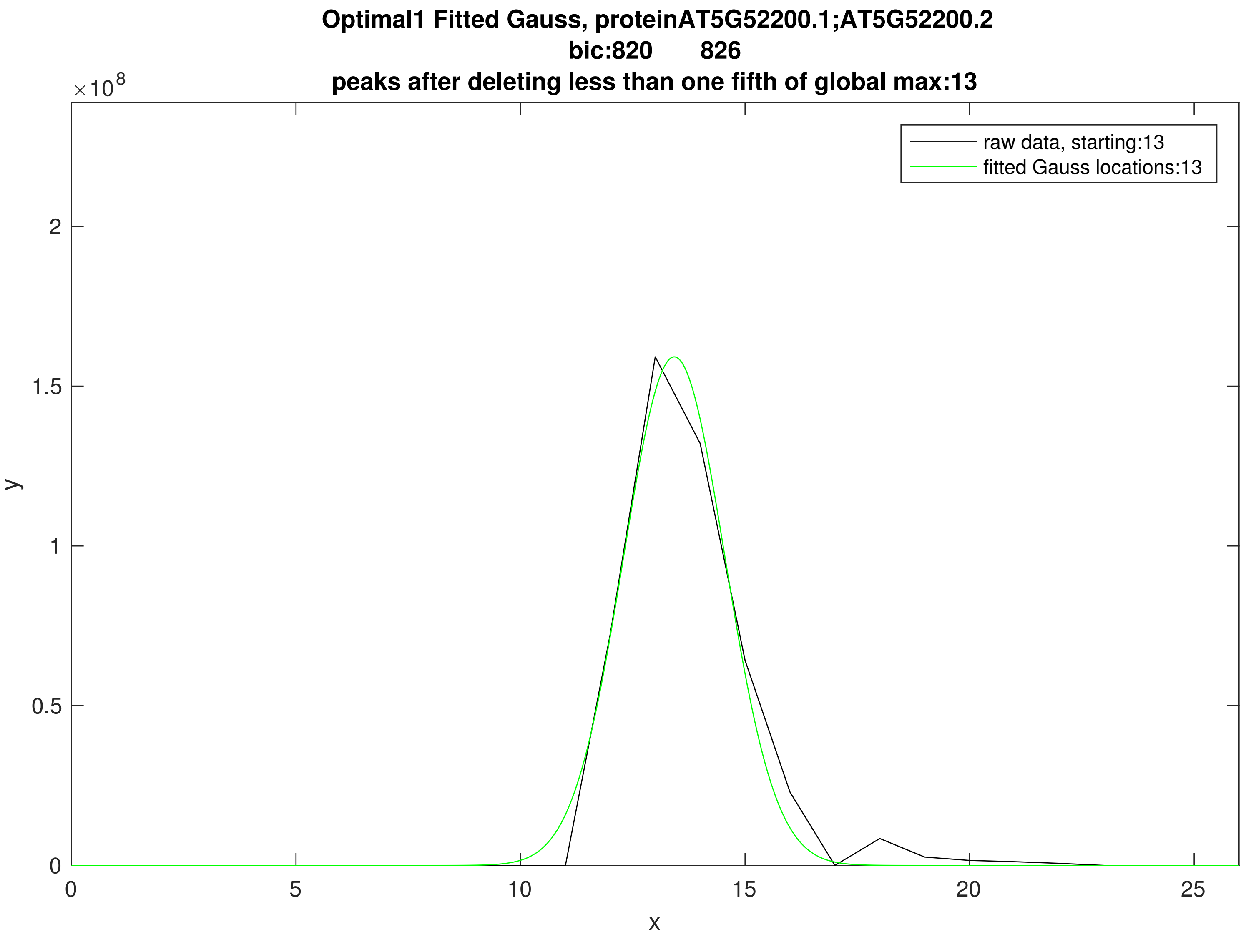
Optimal1 Fitted Gauss, proteinAT5G52200.1;AT5G52200.2
bic:820 826
peaks after deleting less than one fifth of global max:13
×10^8
raw data, starting:13
fitted Gauss locations:13
y
x
0
0.5
1
1.5
2
0
5
10
15
20
25

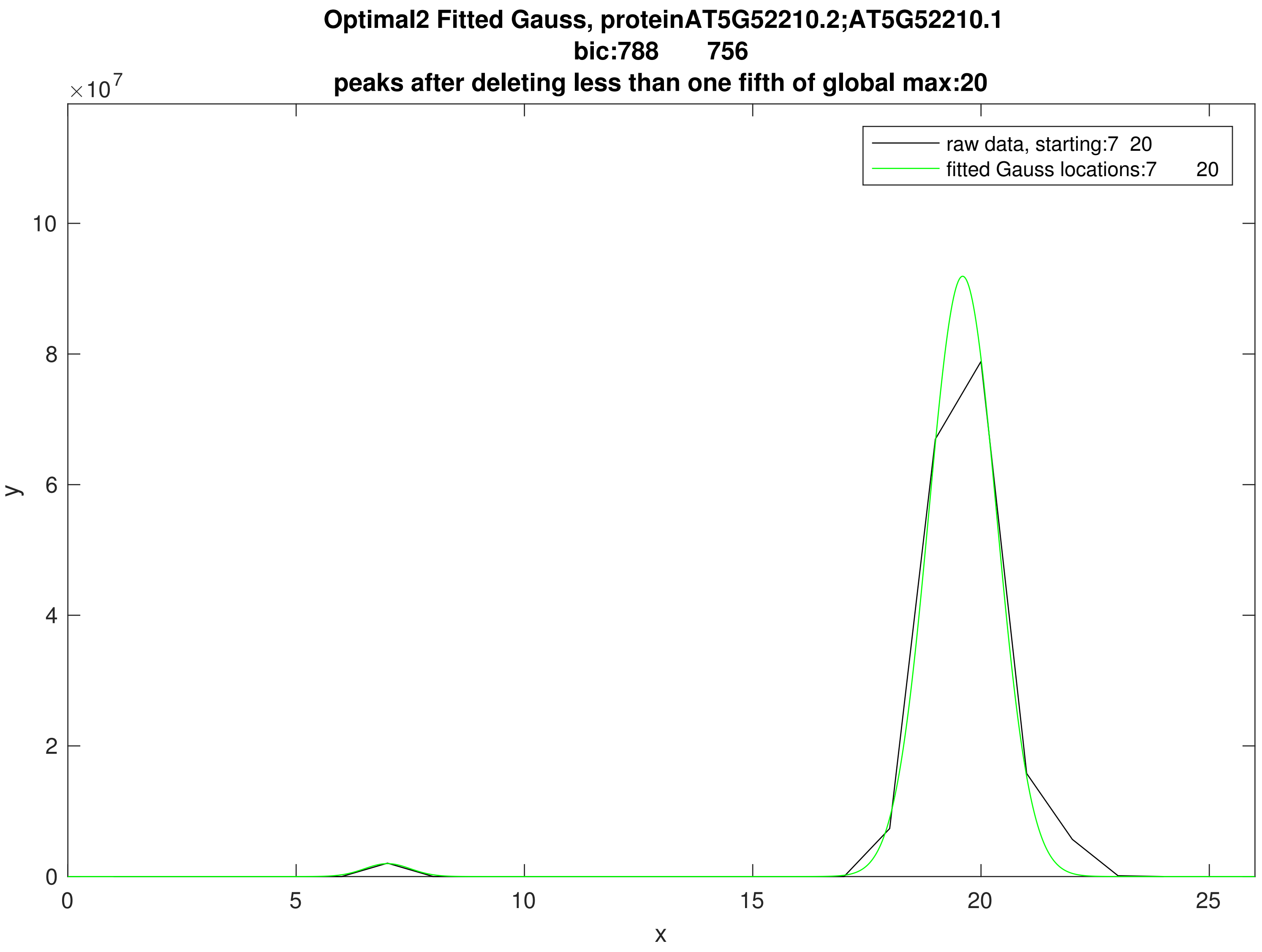
Optimal2 Fitted Gauss, proteinAT5G52210.2;AT5G52210.1
bic:788 756
peaks after deleting less than one fifth of global max:20
×10^7
raw data, starting:7 20
fitted Gauss locations:7 20
10
8
6
4
2
0
y
0
5
10
15
20
25
x

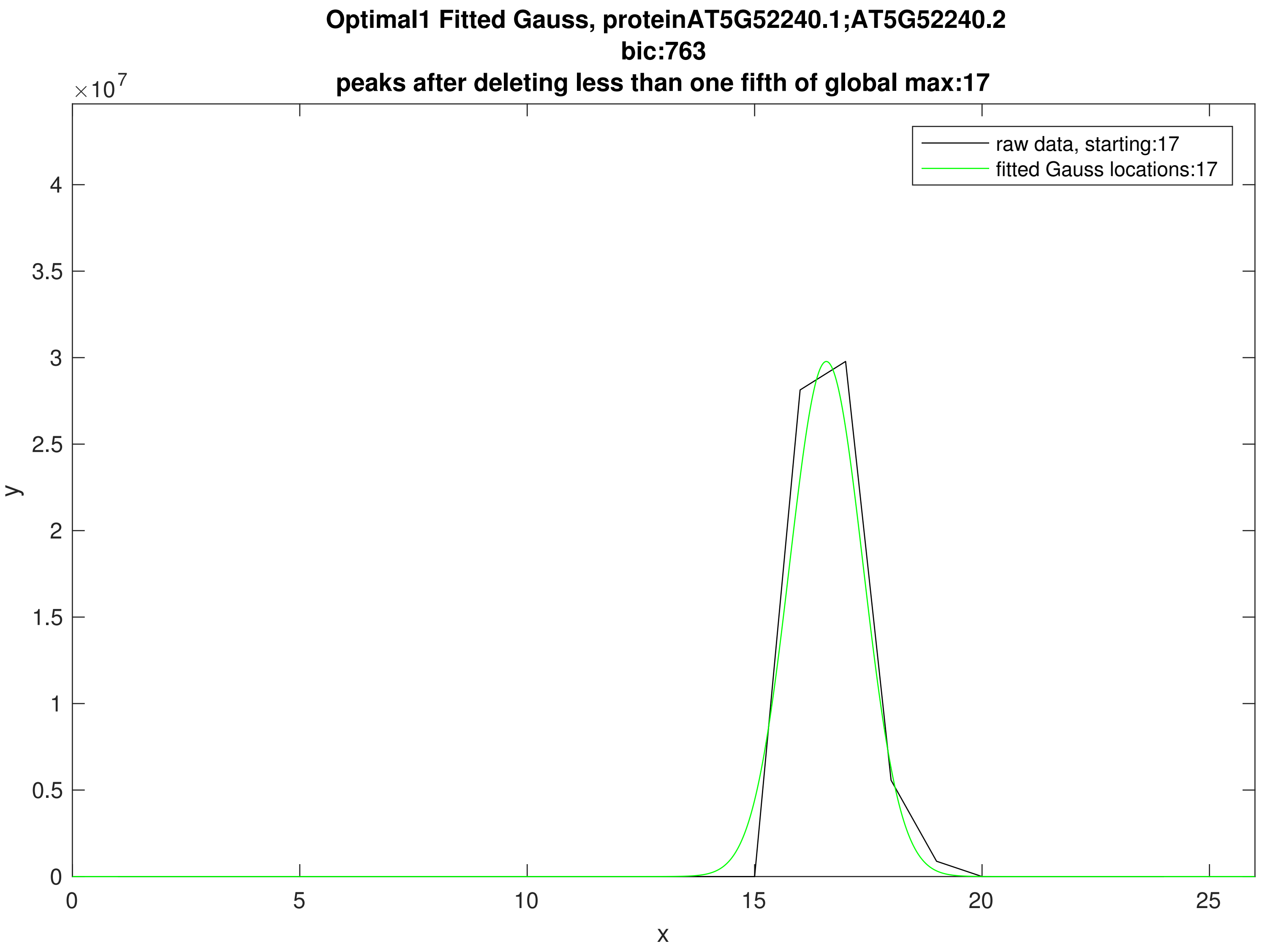
Optimal1 Fitted Gauss, proteinAT5G52240.1;AT5G52240.2
bic:763
peaks after deleting less than one fifth of global max:17
×10^7
raw data, starting:17
fitted Gauss locations:17
4
3.5
3
2.5
2
1.5
1
0.5
0
y
0
5
10
15
20
25
x

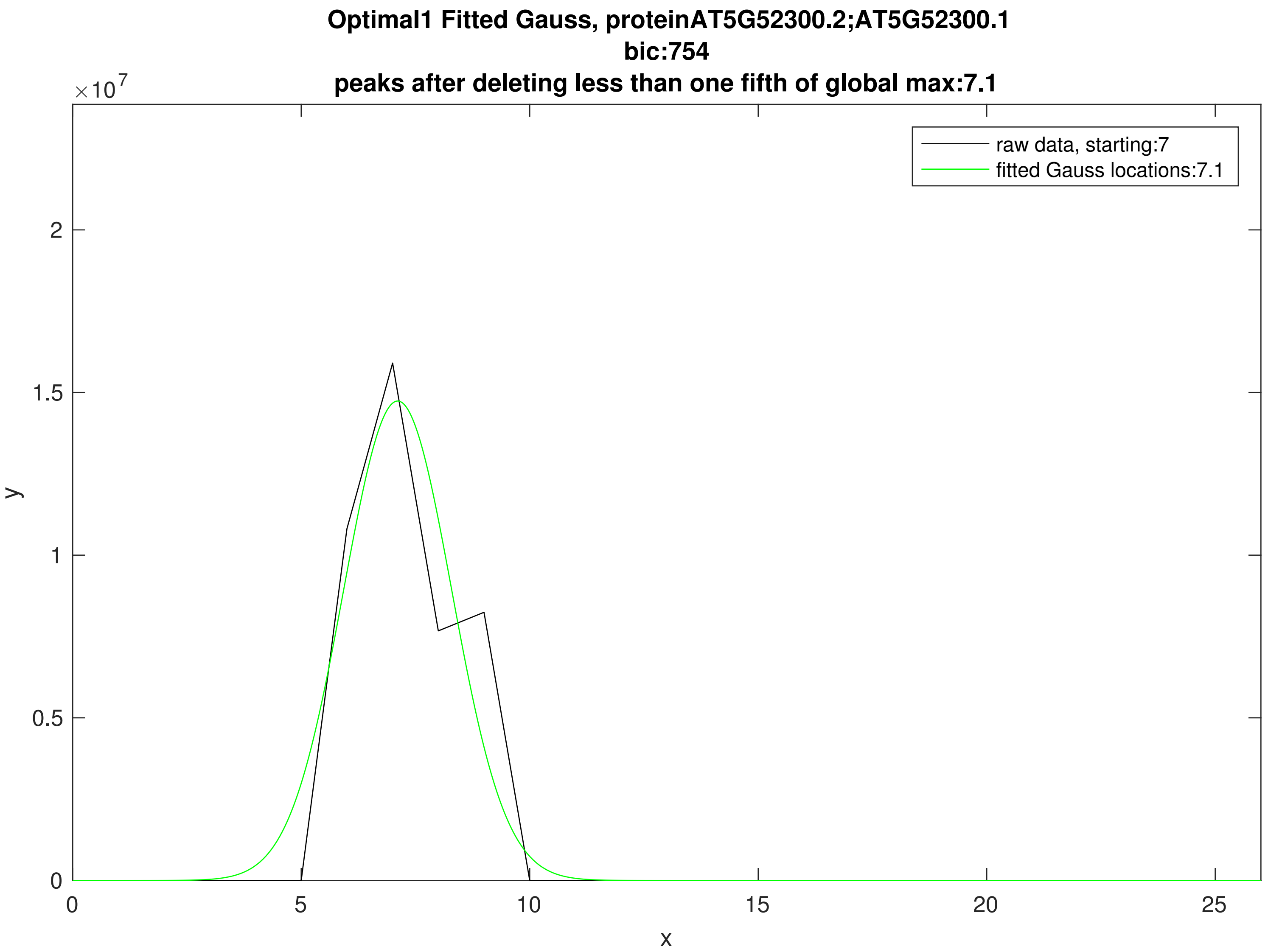

Optimal1 Fitted Gauss, proteinAT5G52300.2;AT5G52300.1
bic:754
peaks after deleting less than one fifth of global max:7.1
×10^7
raw data, starting:7
fitted Gauss locations:7.1
2
1.5
1
0.5
0
y
0
5
10
15
20
25
x

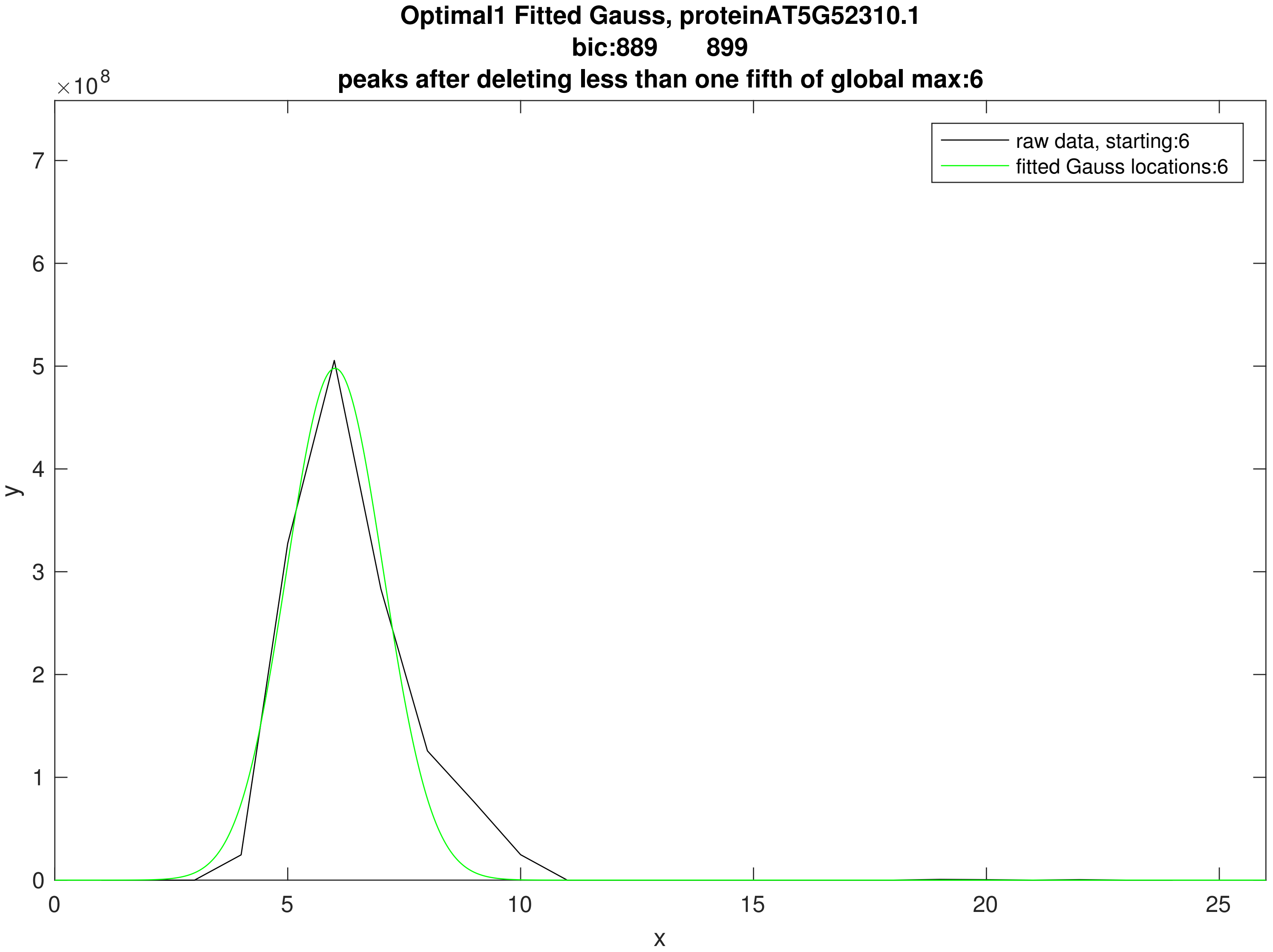

Optimal1 Fitted Gauss, proteinAT5G52310.1
bic:889 899
peaks after deleting less than one fifth of global max:6
×10^8
raw data, starting:6
fitted Gauss locations:6
y
x
0
1
2
3
4
5
6
7
0
5
10
15
20
25

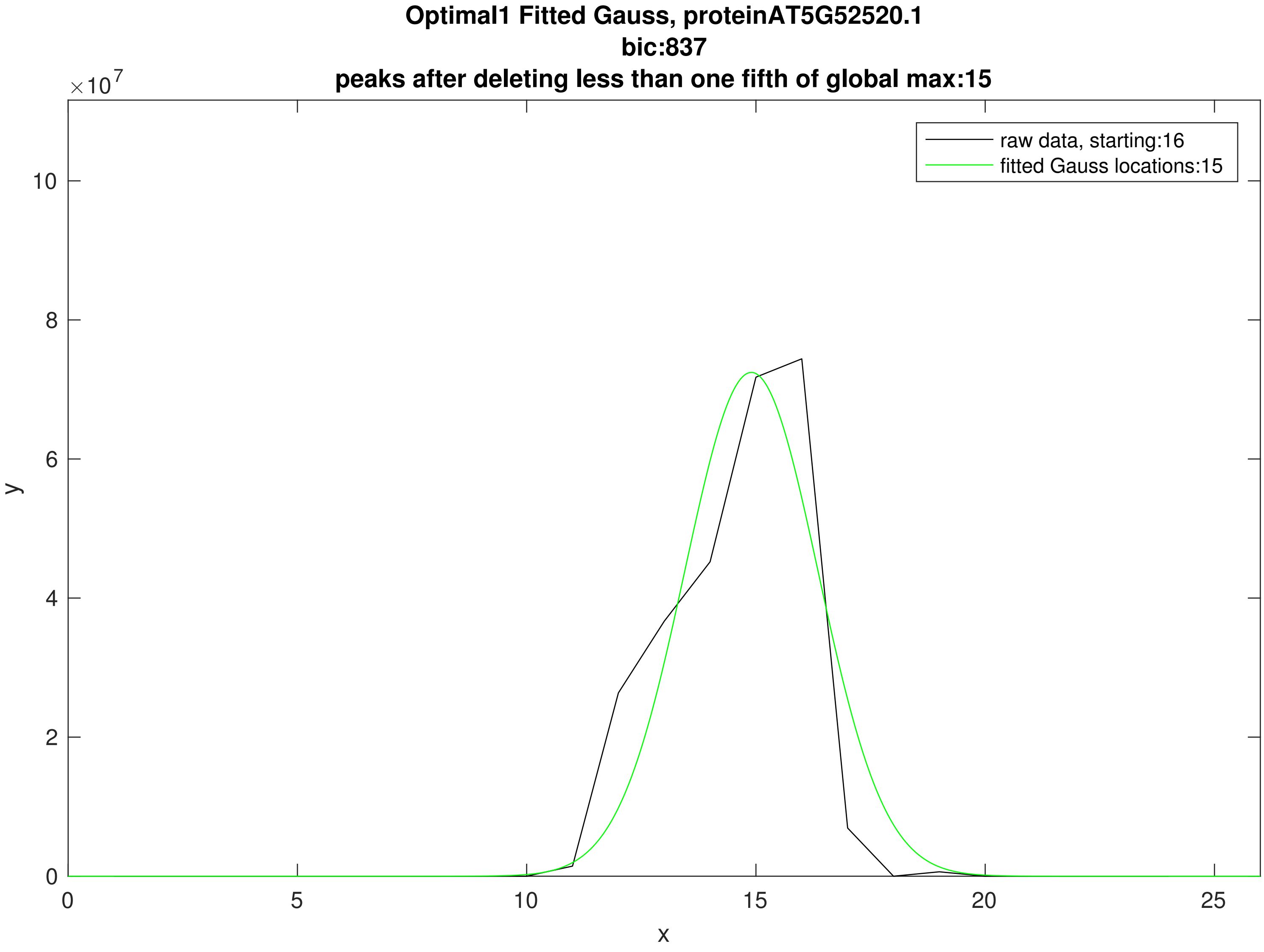
Optimal1 Fitted Gauss, proteinAT5G52520.1
bic:837
peaks after deleting less than one fifth of global max:15
×10^7
raw data, starting:16
fitted Gauss locations:15
10
8
6
4
2
0
y
0
5
10
15
20
25
x

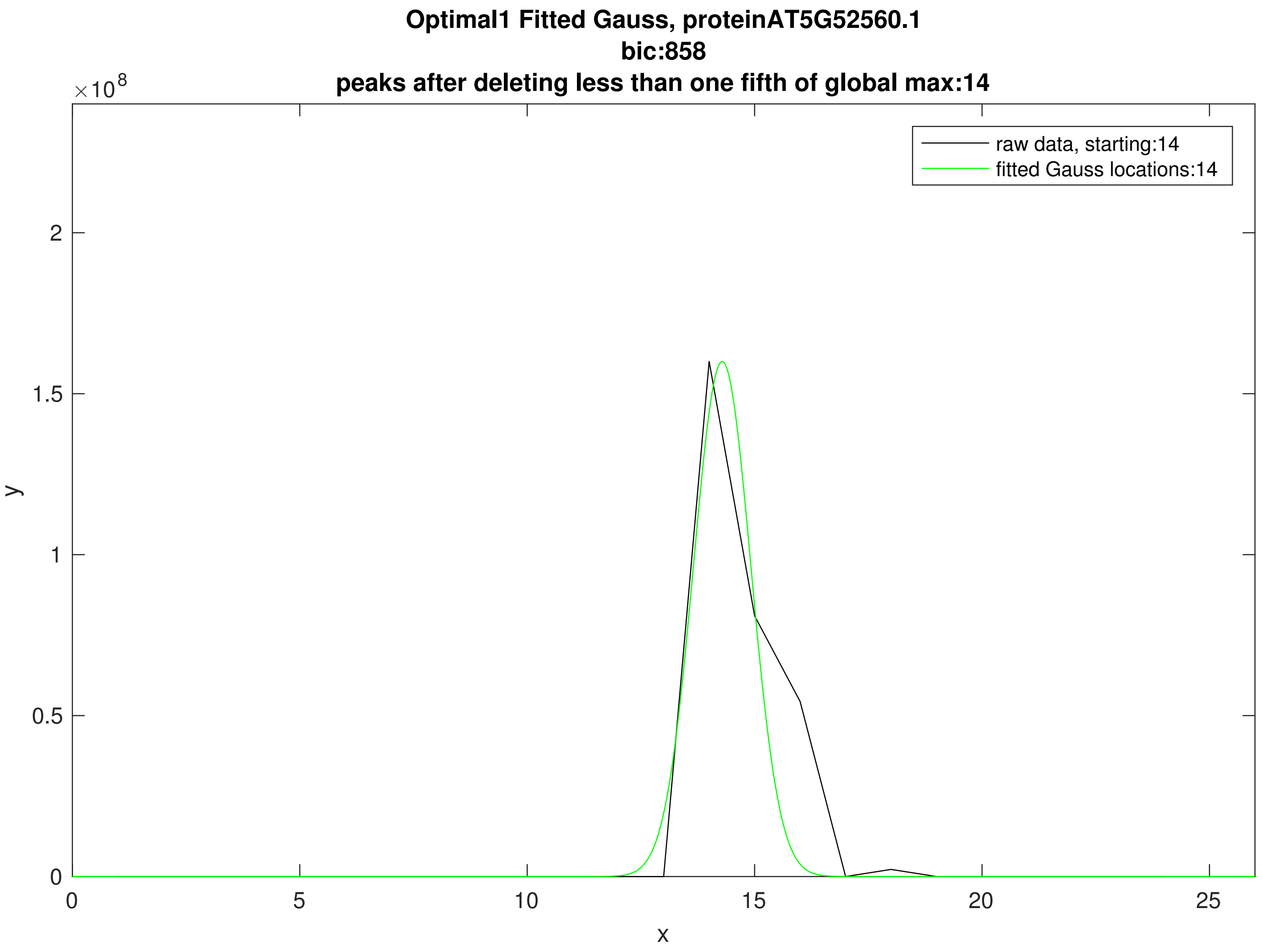
Optimal1 Fitted Gauss, proteinAT5G52560.1
bic:858
peaks after deleting less than one fifth of global max:14
×10^8
raw data, starting:14
fitted Gauss locations:14
y
x
0
0.5
1
1.5
2
0
5
10
15
20
25

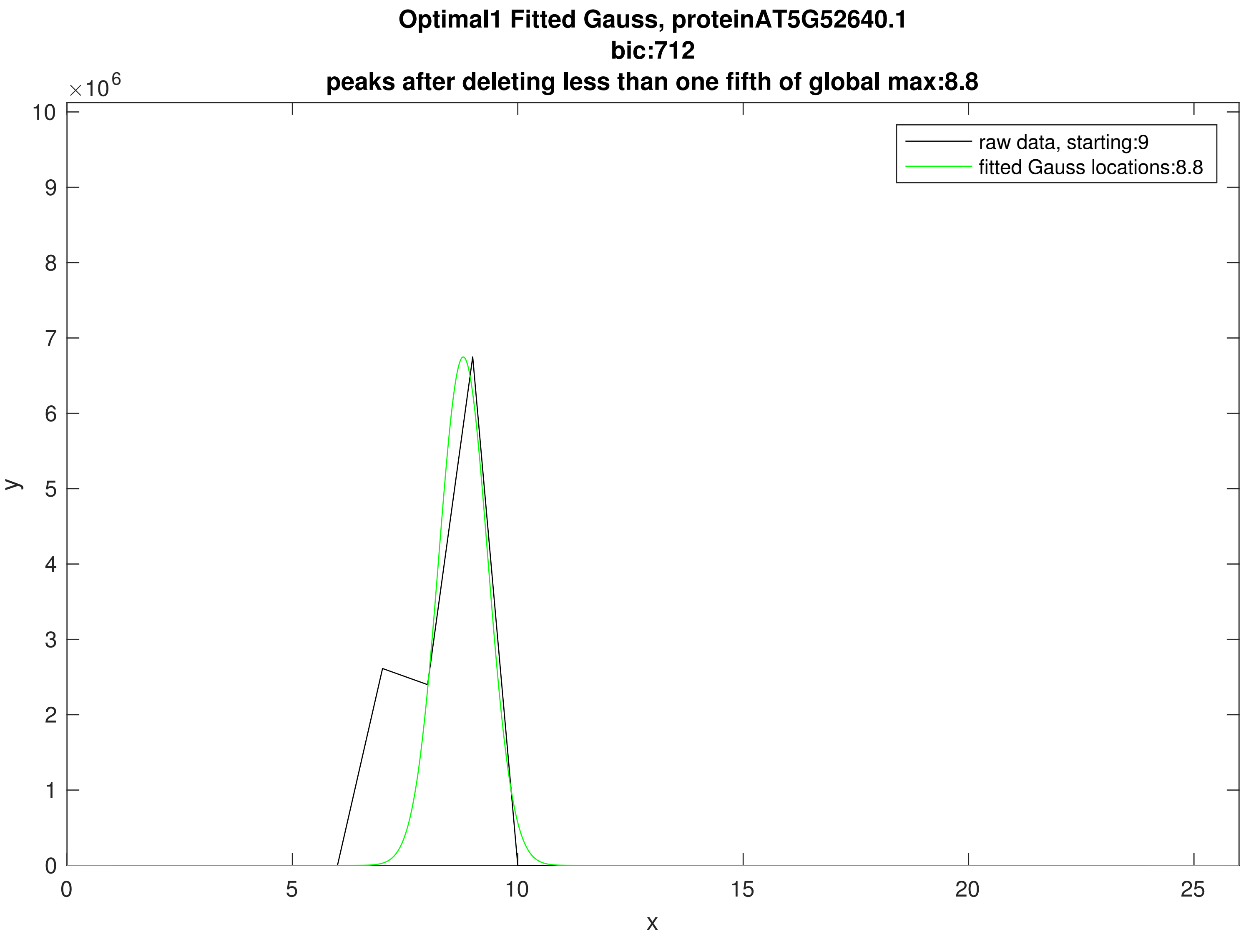

Optimal1 Fitted Gauss, proteinAT5G52640.1
bic:712
peaks after deleting less than one fifth of global max:8.8
×10^6
raw data, starting:9
fitted Gauss locations:8.8
y
x
0
1
2
3
4
5
6
7
8
9
10
0
5
10
15
20
25

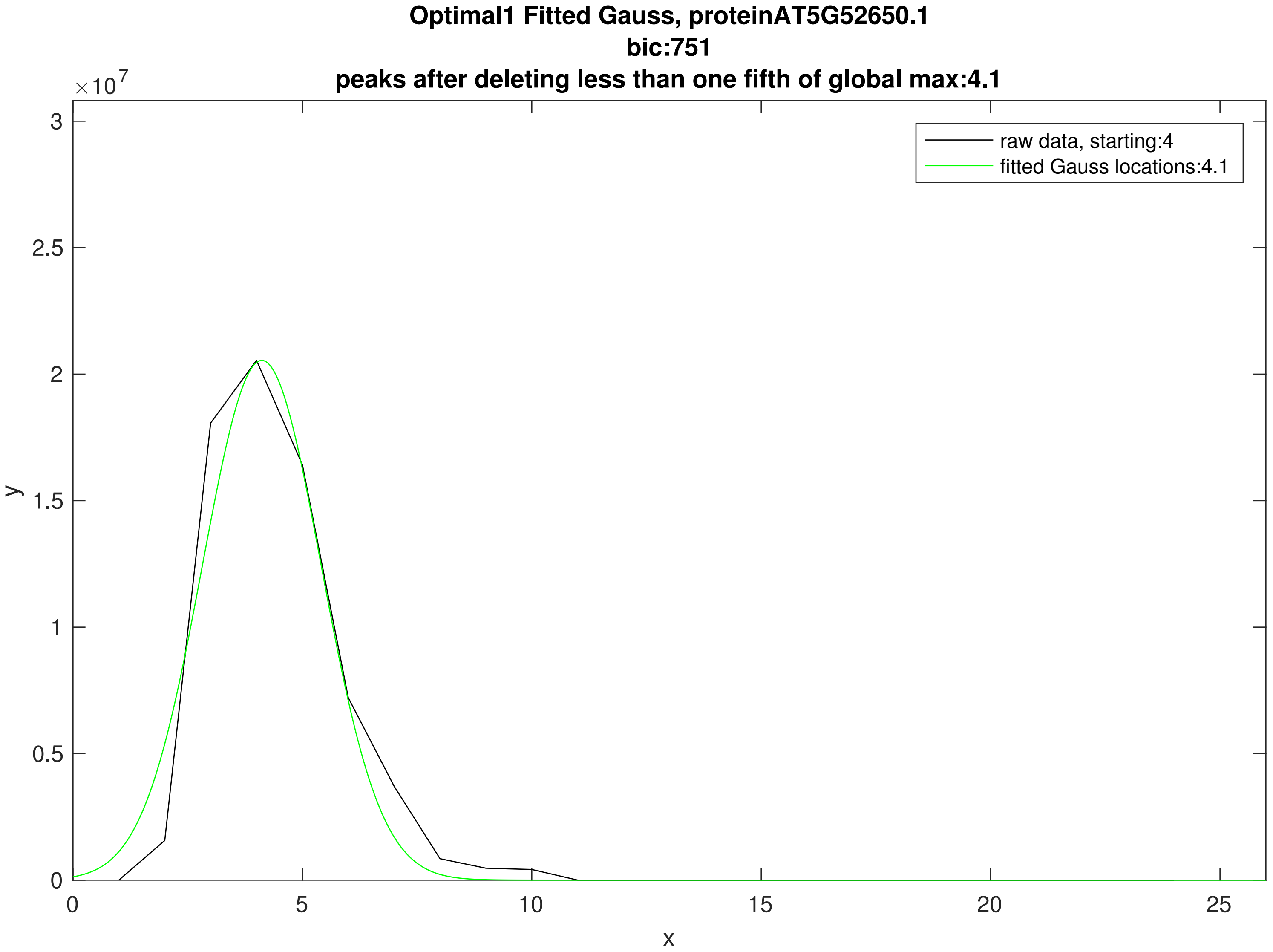

Optimal1 Fitted Gauss, proteinAT5G52650.1
bic:751
peaks after deleting less than one fifth of global max:4.1
×10^7
raw data, starting:4
fitted Gauss locations:4.1
y
x
0
0.5
1
1.5
2
2.5
3
0
5
10
15
20
25

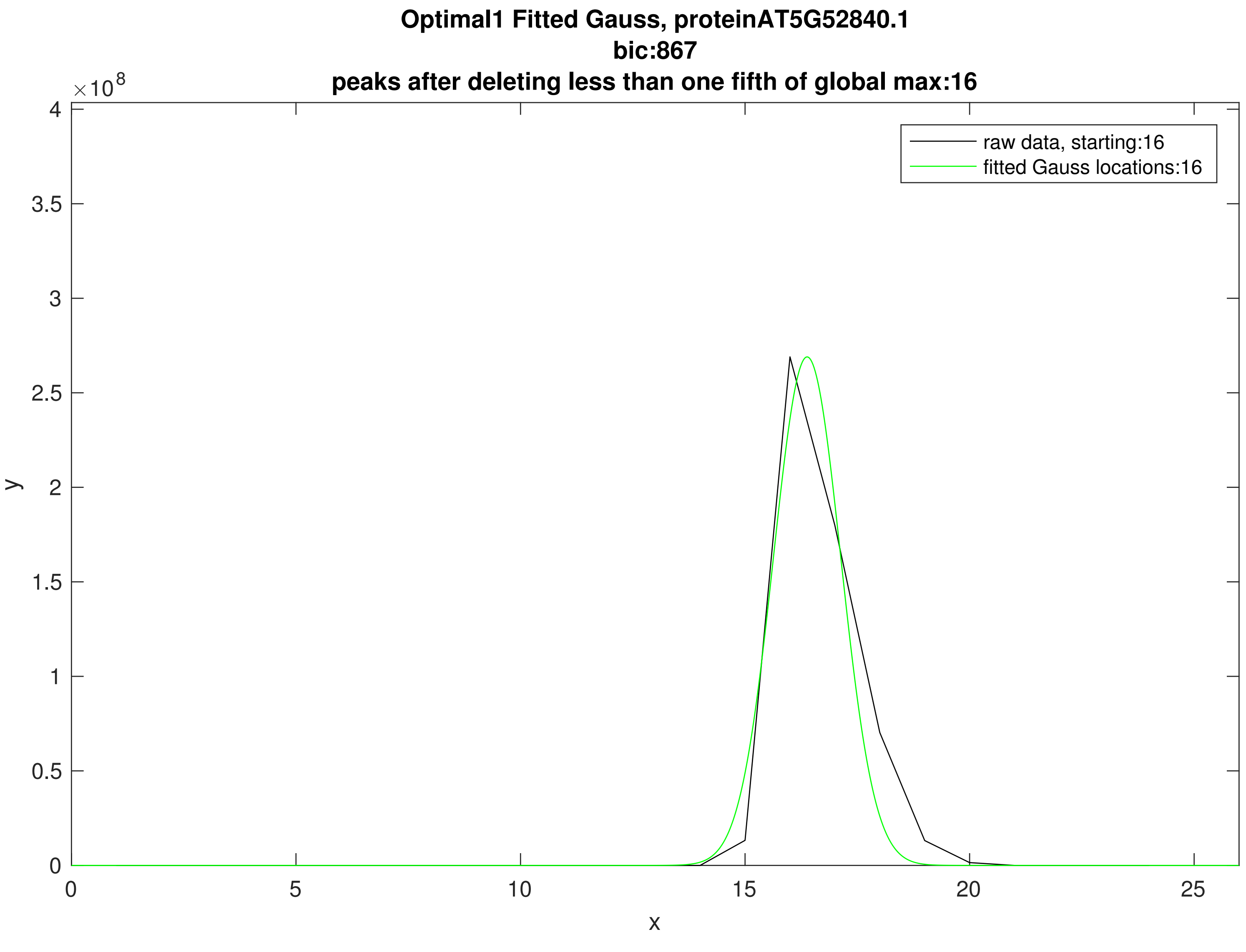

Optimal1 Fitted Gauss, proteinAT5G52840.1
bic:867
peaks after deleting less than one fifth of global max:16
×10^8
y
x
raw data, starting:16
fitted Gauss locations:16

**Optimal2 Fitted Gauss, proteinAT5G52920.1**
**bic:872 838 848**
**peaks after deleting less than one fifth of global max:8.3**

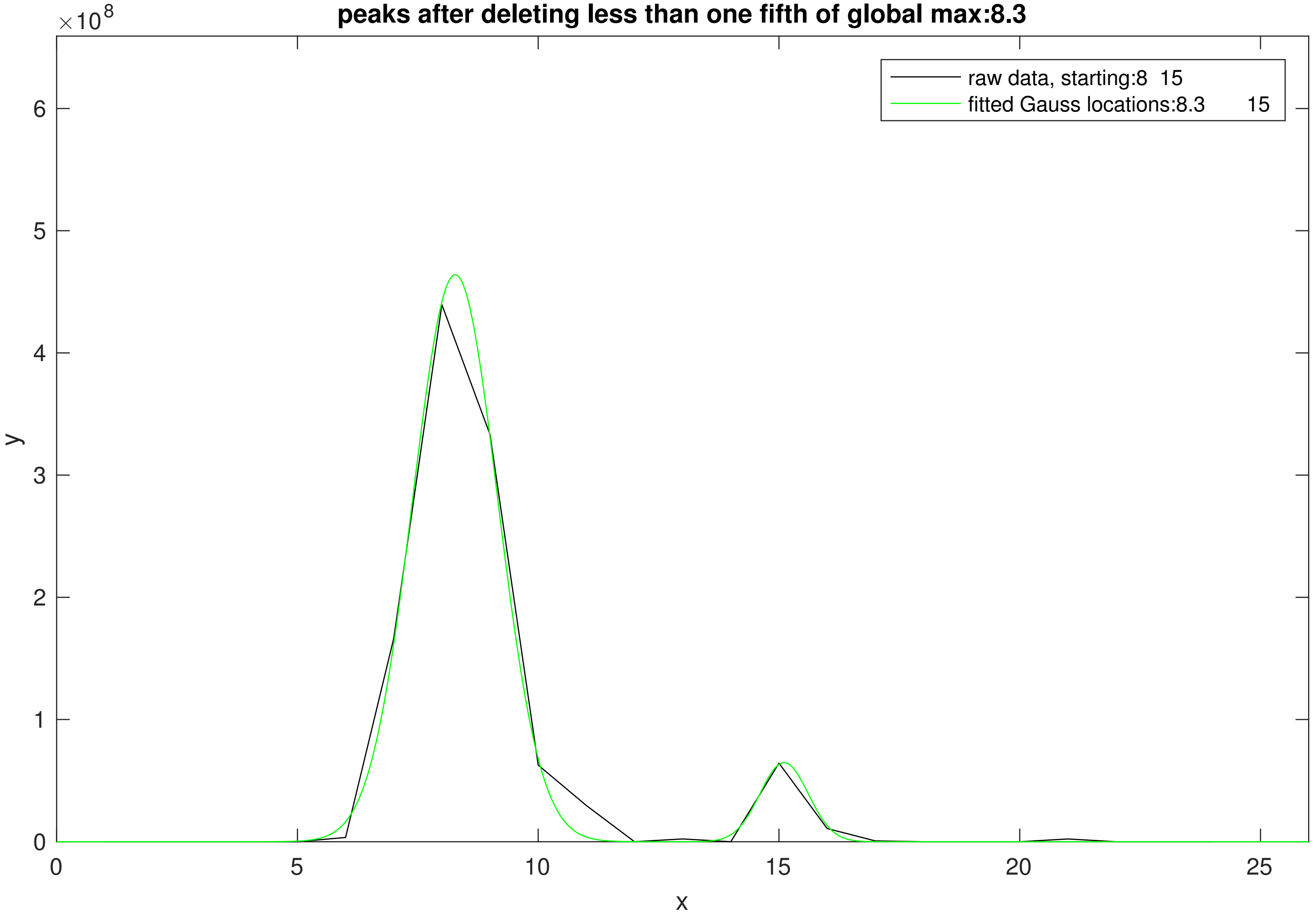

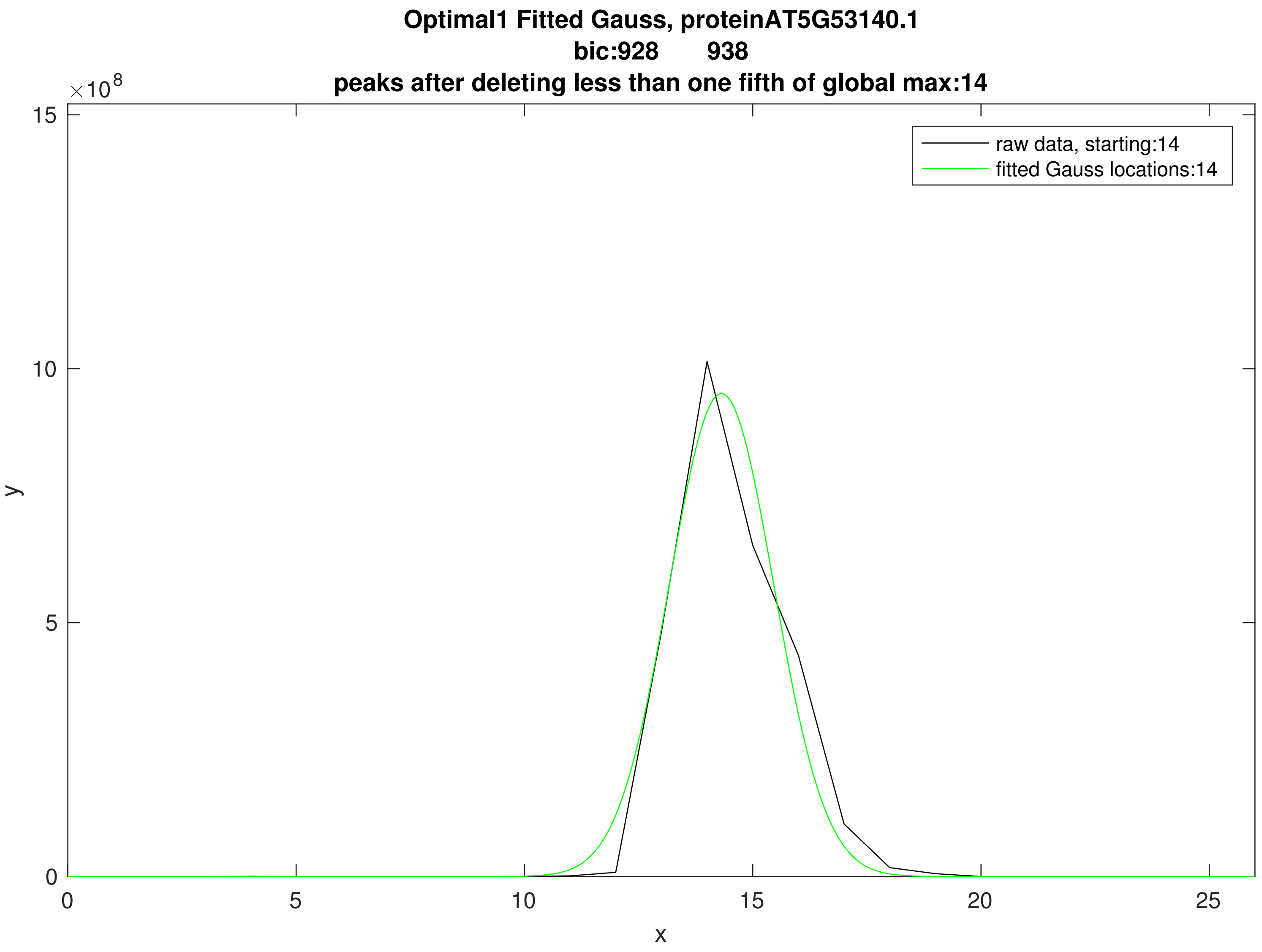

Optimal1 Fitted Gauss, proteinAT5G53140.1
bic:928 938
peaks after deleting less than one fifth of global max:14
×10^8
15
10
5
0
y
0
5
10
15
20
25
x
raw data, starting:14
fitted Gauss locations:14

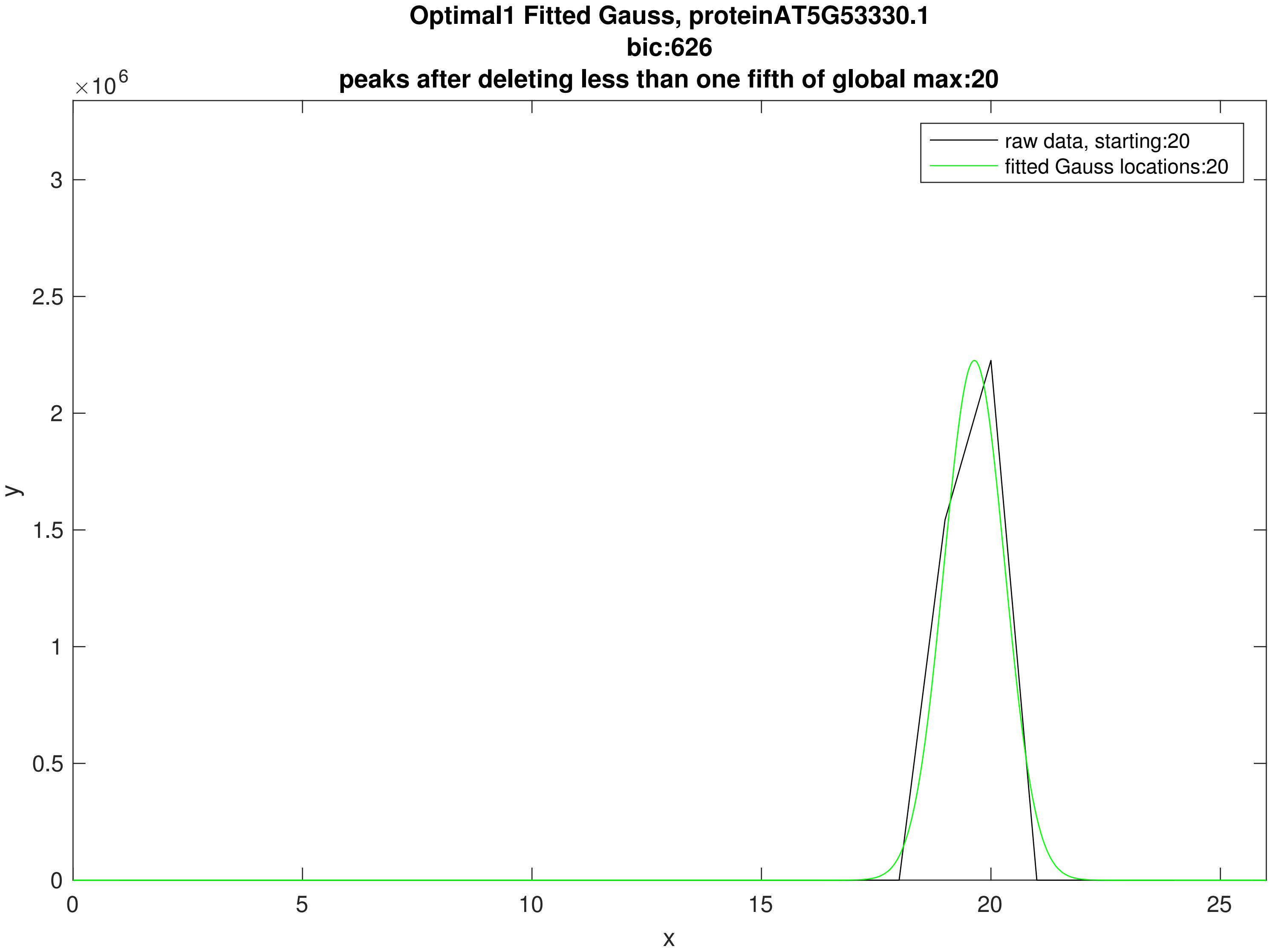
Optimal1 Fitted Gauss, proteinAT5G53330.1
bic:626
peaks after deleting less than one fifth of global max:20
×10^6
raw data, starting:20
fitted Gauss locations:20
3
2.5
2
1.5
1
0.5
0
y
0
5
10
15
20
25
x

**Optimal1 Fitted Gauss, proteinAT5G53460.3;AT5G53460.2;AT5G53460.1**
**bic:1.01e+03  1.02e+03  1.02e+03**
**peaks after deleting less than one fifth of global max:8**

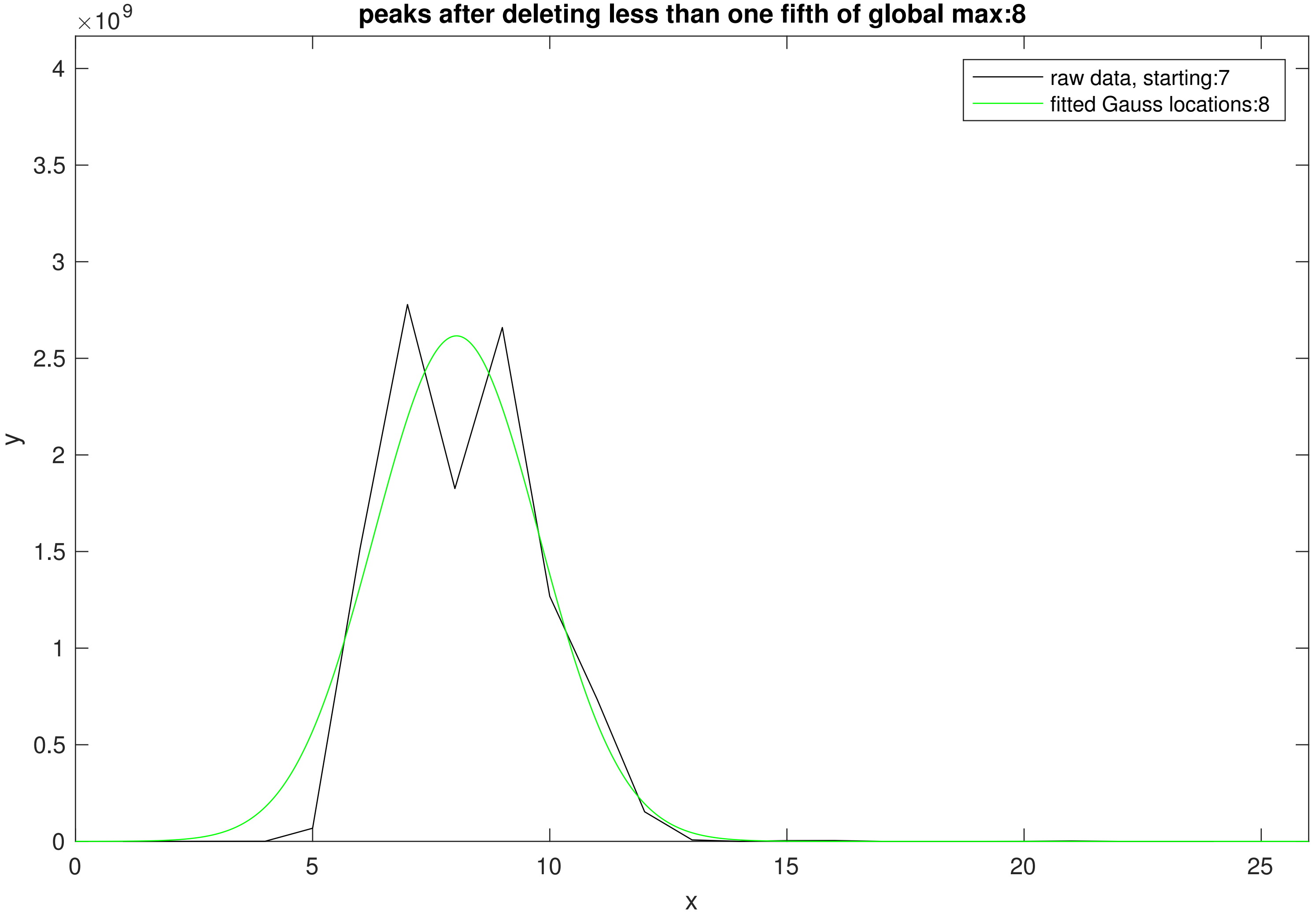

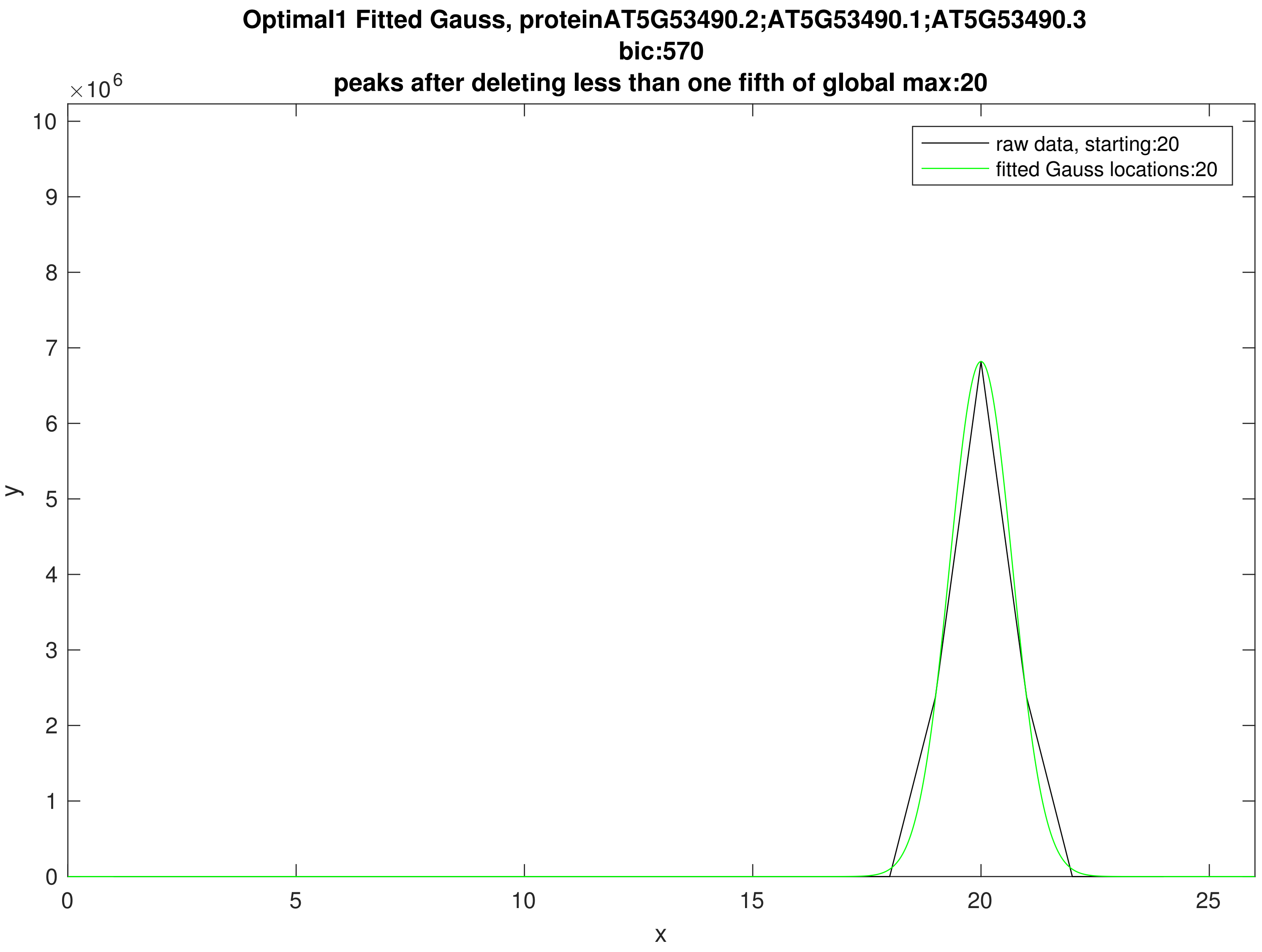
Optimal1 Fitted Gauss, proteinAT5G53490.2;AT5G53490.1;AT5G53490.3
bic:570
peaks after deleting less than one fifth of global max:20
×10^6
raw data, starting:20
fitted Gauss locations:20
y
x
0
5
10
15
20
25
1
2
3
4
6
7
8
9
10

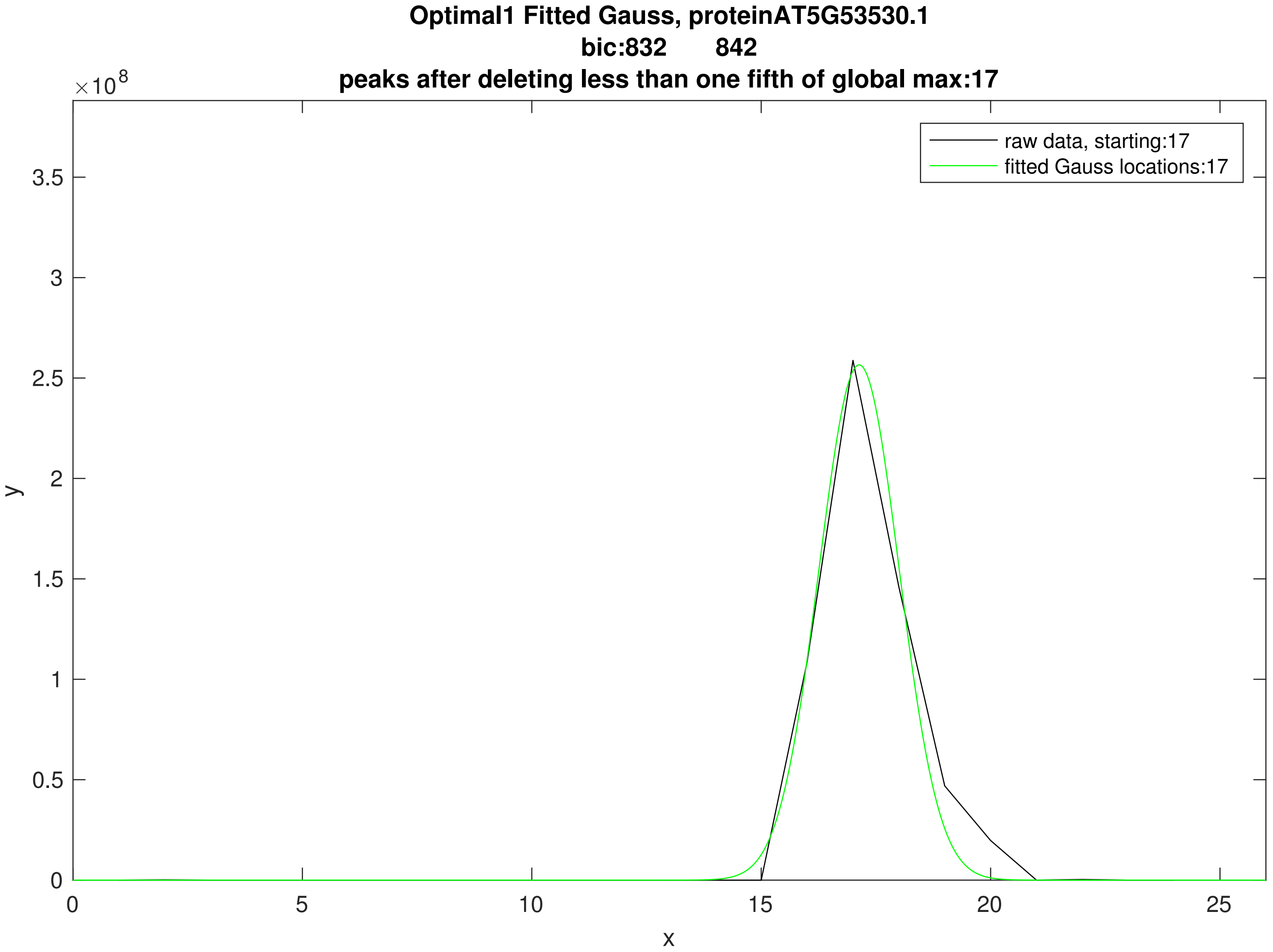
Optimal1 Fitted Gauss, proteinAT5G53530.1
bic:832 842
peaks after deleting less than one fifth of global max:17
×10^8
raw data, starting:17
fitted Gauss locations:17
y
x
0
0.5
1
1.5
2
2.5
3
3.5
5
10
15
20
25

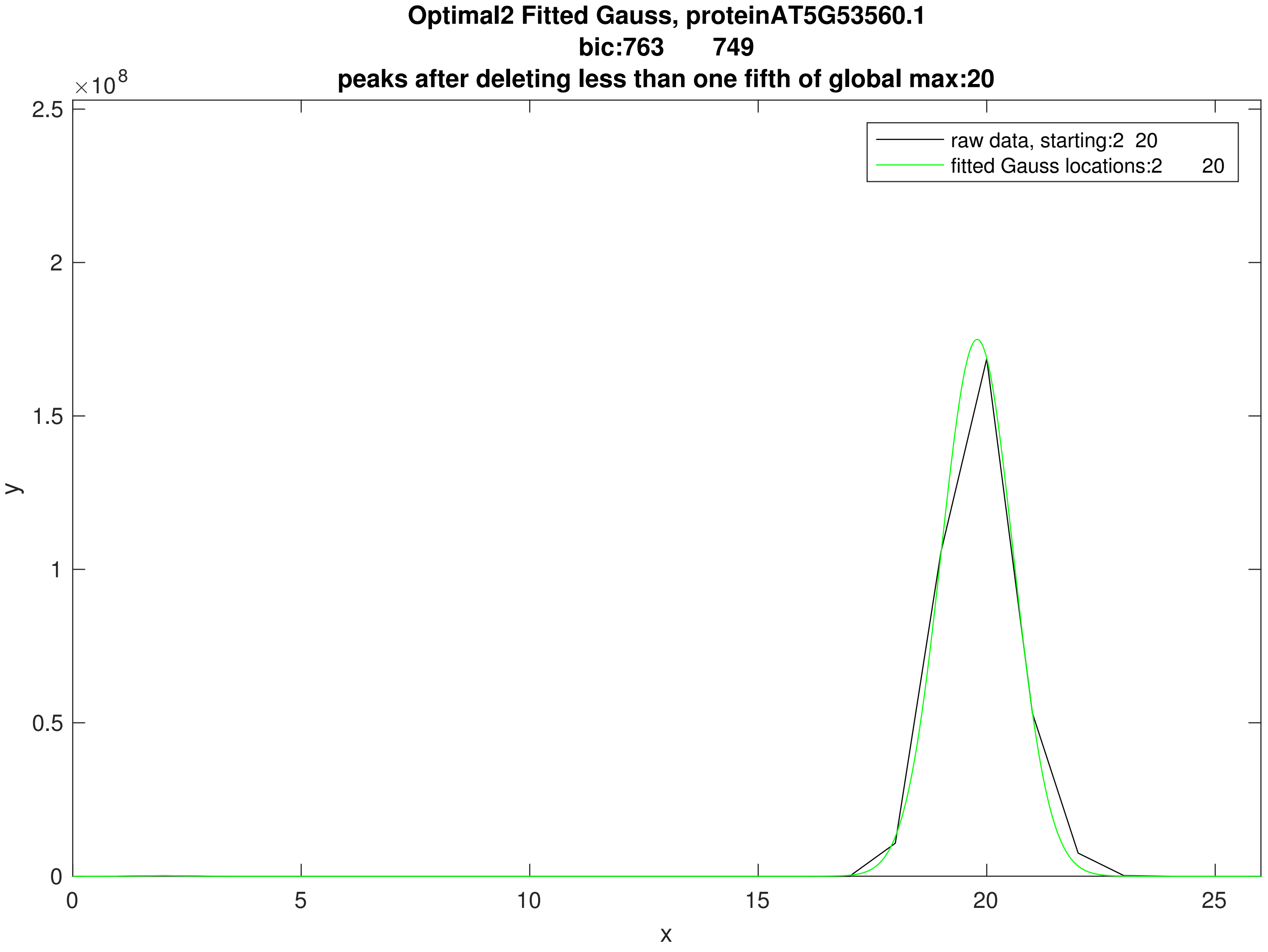

Optimal2 Fitted Gauss, proteinAT5G53560.1
bic:763 749
peaks after deleting less than one fifth of global max:20
×10^8
raw data, starting:2 20
fitted Gauss locations:2 20
y
x

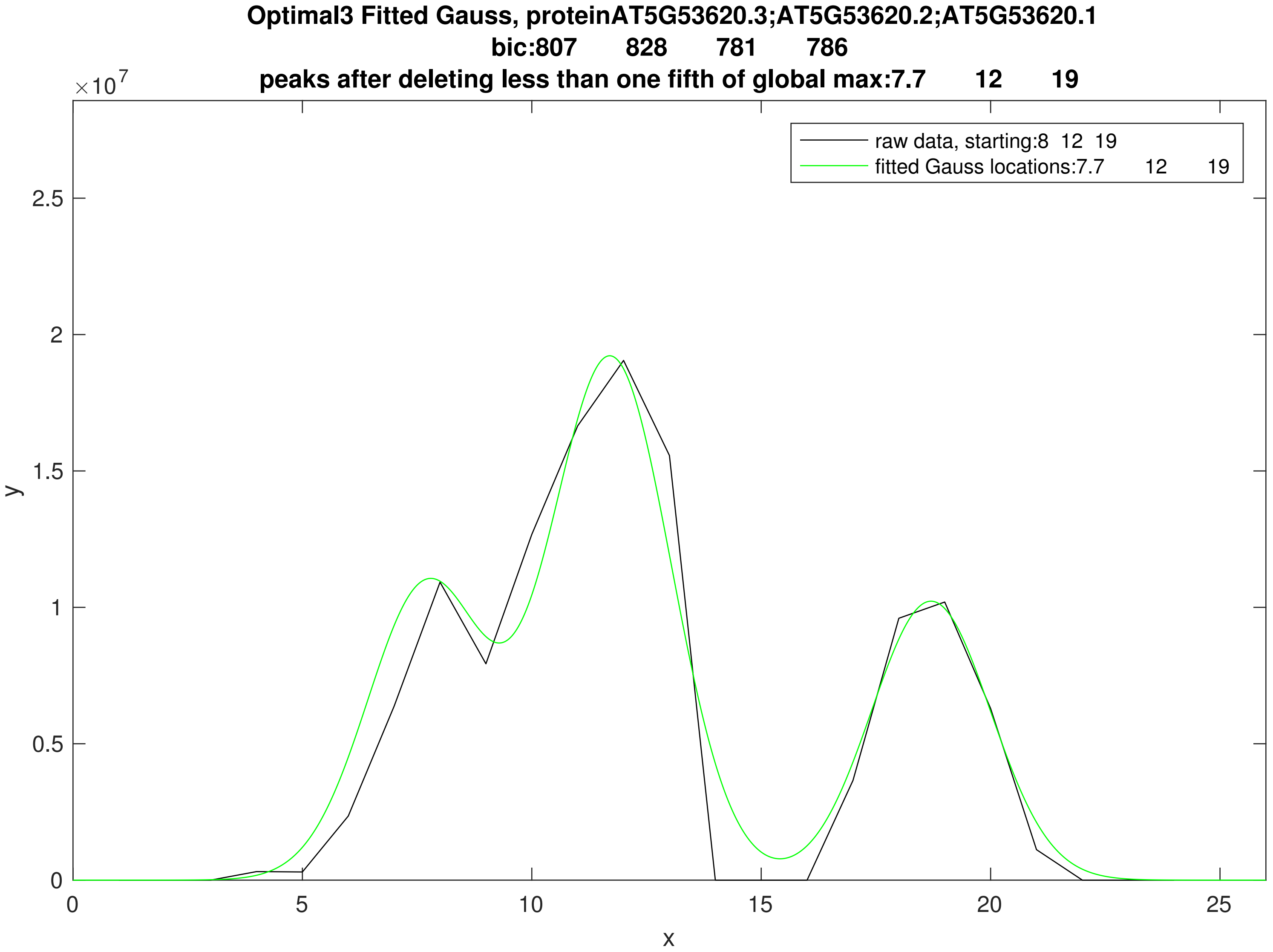

Optimal3 Fitted Gauss, proteinAT5G53620.3;AT5G53620.2;AT5G53620.1
bic:807 828 781 786
peaks after deleting less than one fifth of global max:7.7 12 19
×10^7
raw data, starting:8 12 19
fitted Gauss locations:7.7 12 19
y
x
0
0.5
1
1.5
2
2.5
5
10
15
20
25

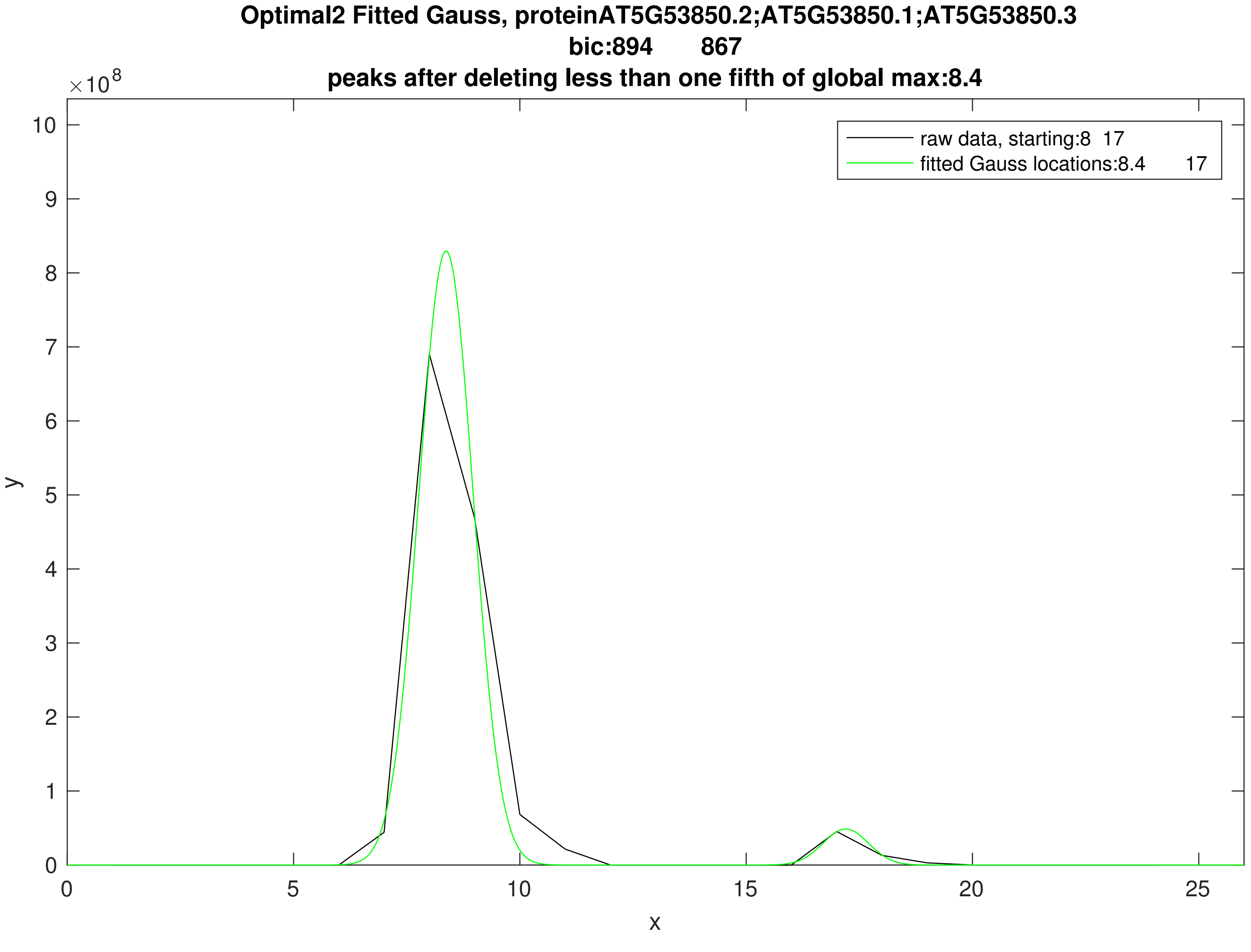

Optimal2 Fitted Gauss, proteinAT5G53850.2;AT5G53850.1;AT5G53850.3
bic:894 867
peaks after deleting less than one fifth of global max:8.4
×10^8
raw data, starting:8 17
fitted Gauss locations:8.4 17
y
x
0
1
2
3
4
5
6
7
8
9
10
0
5
10
15
20
25

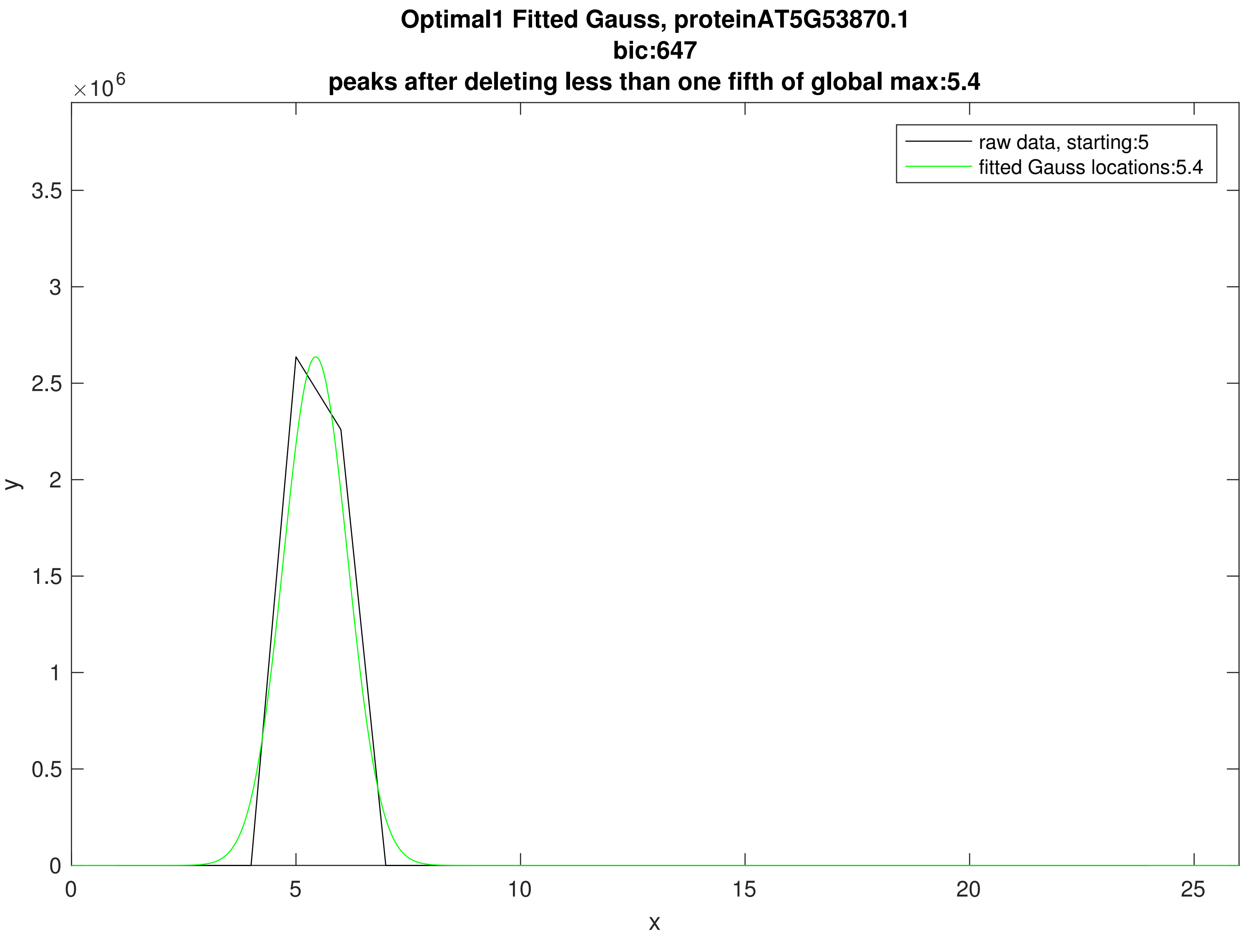
Optimal1 Fitted Gauss, proteinAT5G53870.1
bic:647
peaks after deleting less than one fifth of global max:5.4
×10^6
raw data, starting:5
fitted Gauss locations:5.4
y
x
0
0.5
1
1.5
2
2.5
3
3.5
0
5
10
15
20
25

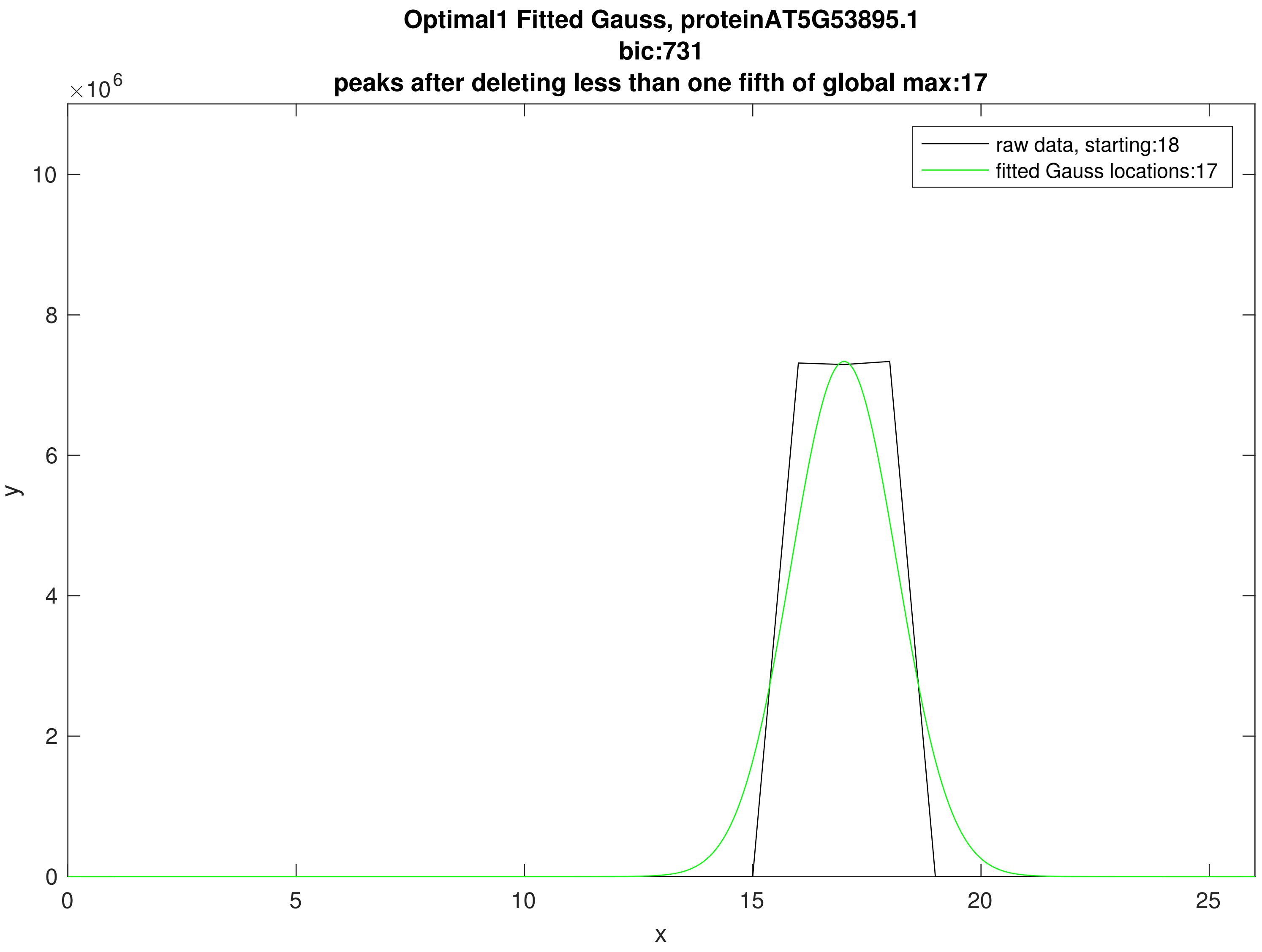
Optimal1 Fitted Gauss, proteinAT5G53895.1
bic:731
peaks after deleting less than one fifth of global max:17
×10^6
raw data, starting:18
fitted Gauss locations:17
10
8
6
4
2
0
y
0
5
10
15
20
25
x

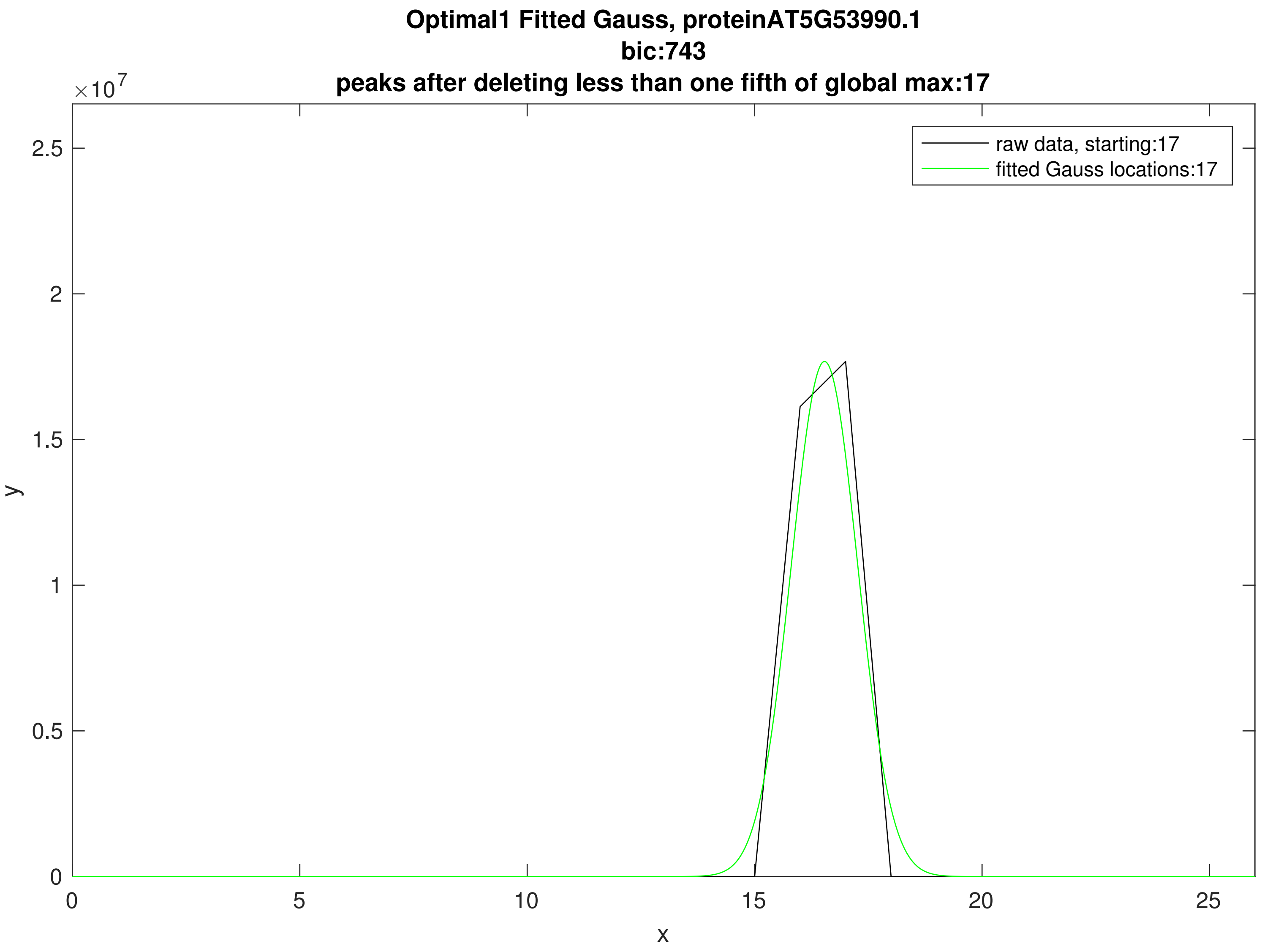
Optimal1 Fitted Gauss, proteinAT5G53990.1
bic:743
peaks after deleting less than one fifth of global max:17
×10^7
2.5
2
1.5
1
0.5
0
y
0
5
10
15
20
25
x
raw data, starting:17
fitted Gauss locations:17

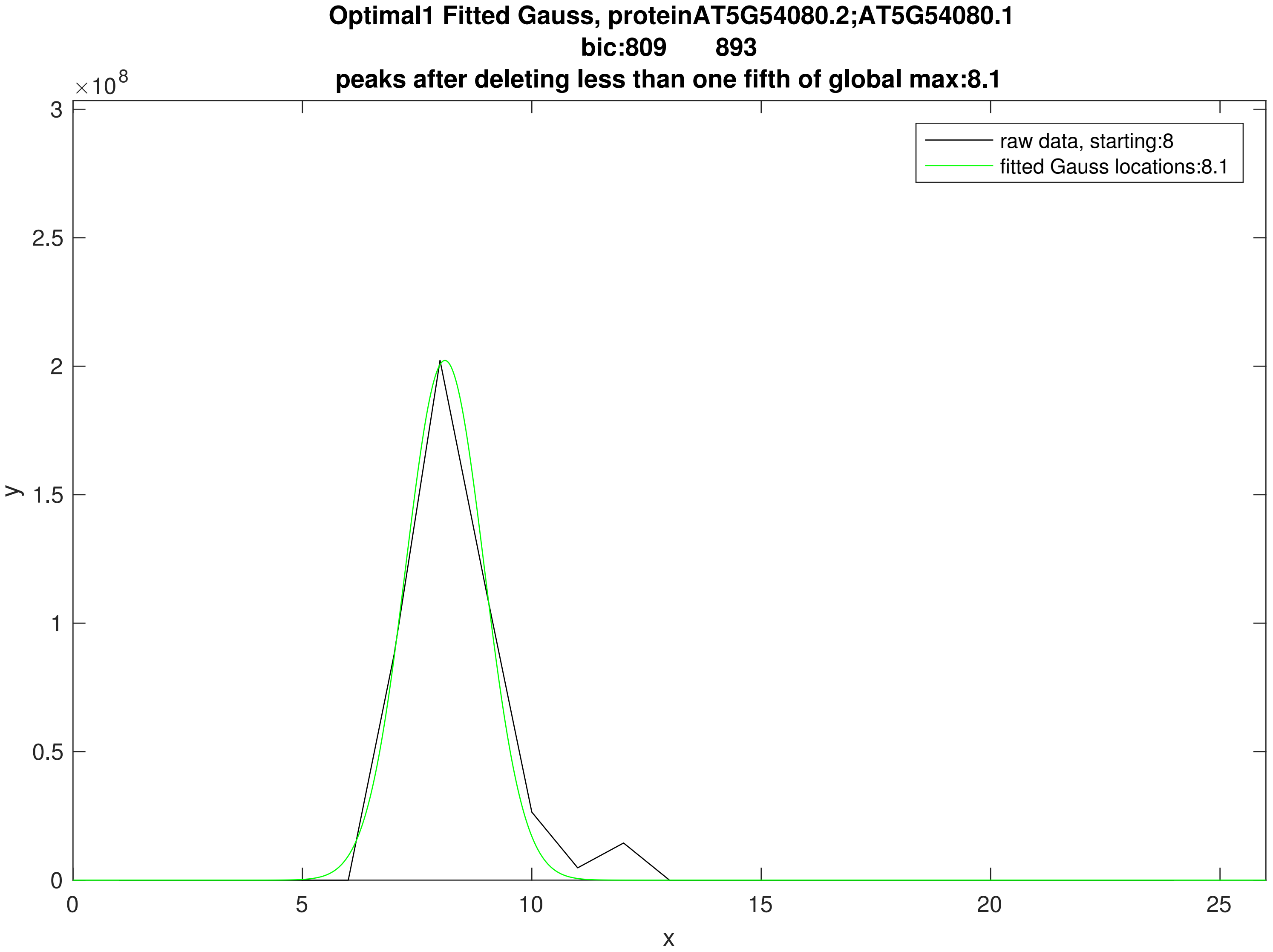

Optimal1 Fitted Gauss, proteinAT5G54080.2;AT5G54080.1
bic:809 893
peaks after deleting less than one fifth of global max:8.1
×10^8
3
2.5
2
1.5
1
0.5
0
y
0
5
10
15
20
25
x
raw data, starting:8
fitted Gauss locations:8.1

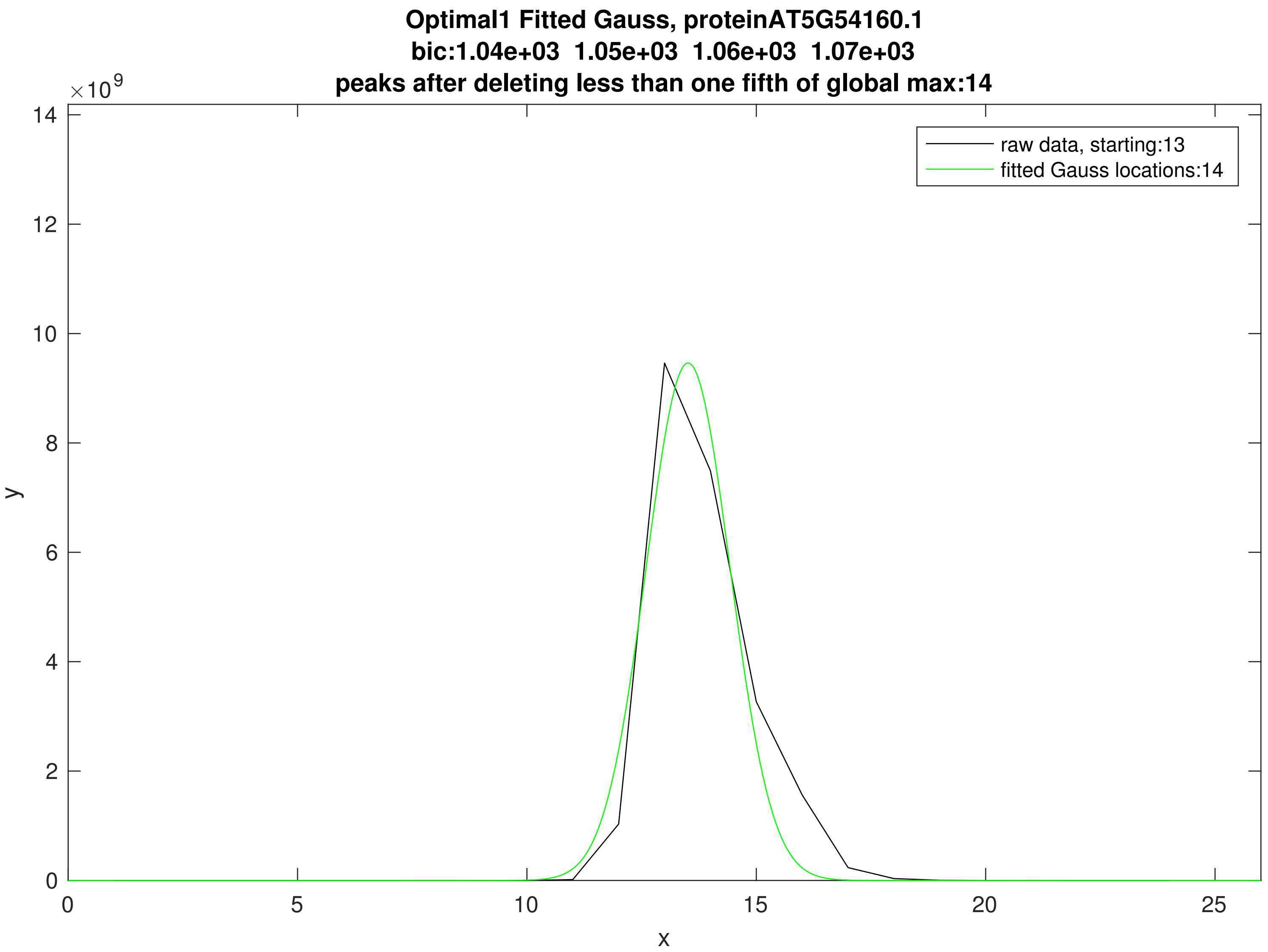

Optimal1 Fitted Gauss, proteinAT5G54160.1
bic:1.04e+03  1.05e+03  1.06e+03  1.07e+03
peaks after deleting less than one fifth of global max:14
×10^9
raw data, starting:13
fitted Gauss locations:14
y
x
0
2
4
6
8
10
12
14
0
5
10
15
20
25

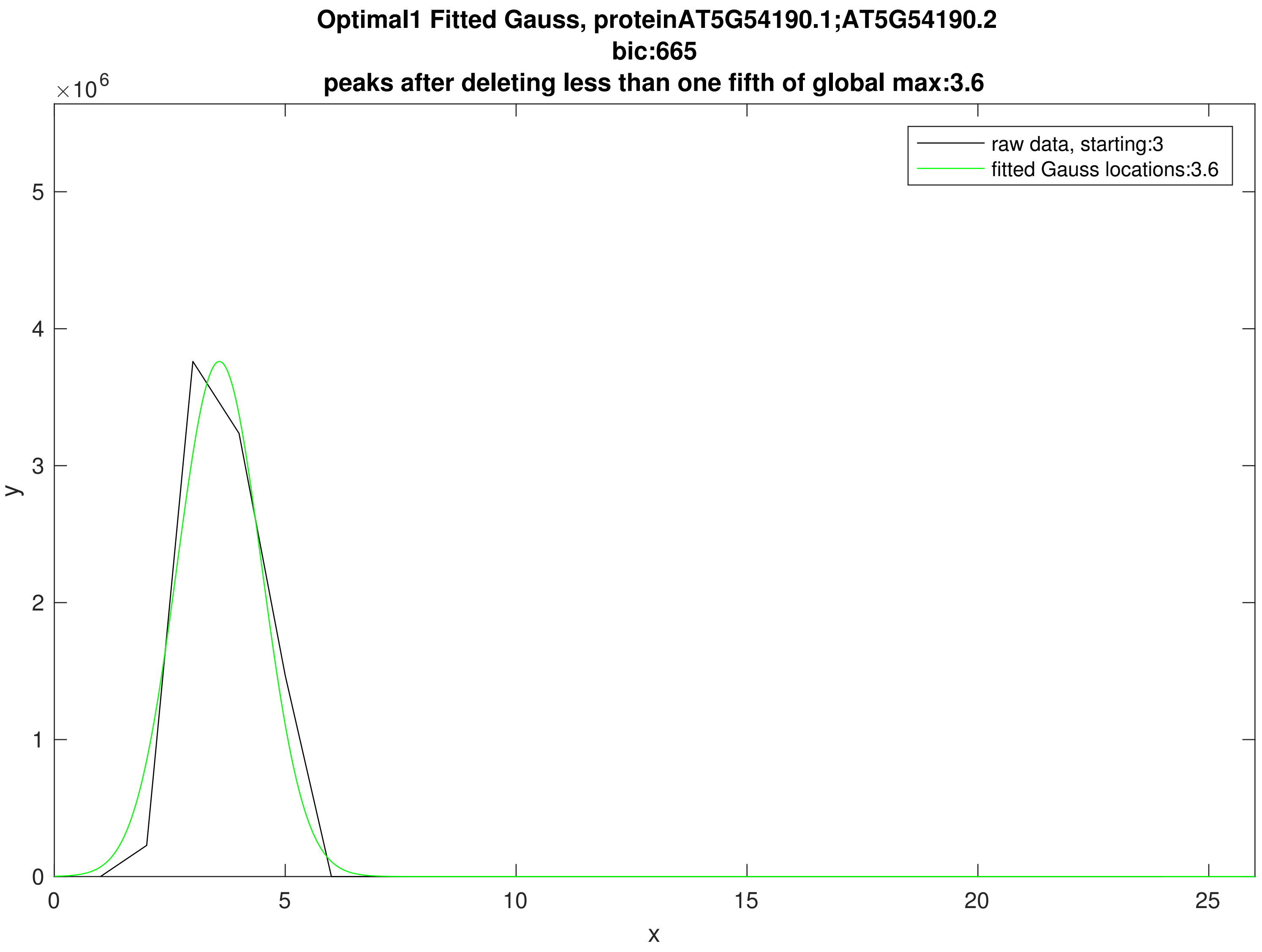
Optimal1 Fitted Gauss, proteinAT5G54190.1;AT5G54190.2
bic:665
peaks after deleting less than one fifth of global max:3.6
×10^6
raw data, starting:3
fitted Gauss locations:3.6
y
x
0
1
2
3
4
5
0
5
10
15
20
25

**Optimal1 Fitted Gauss, proteinAT5G54310.1**
**bic:775**
**peaks after deleting less than one fifth of global max:12**

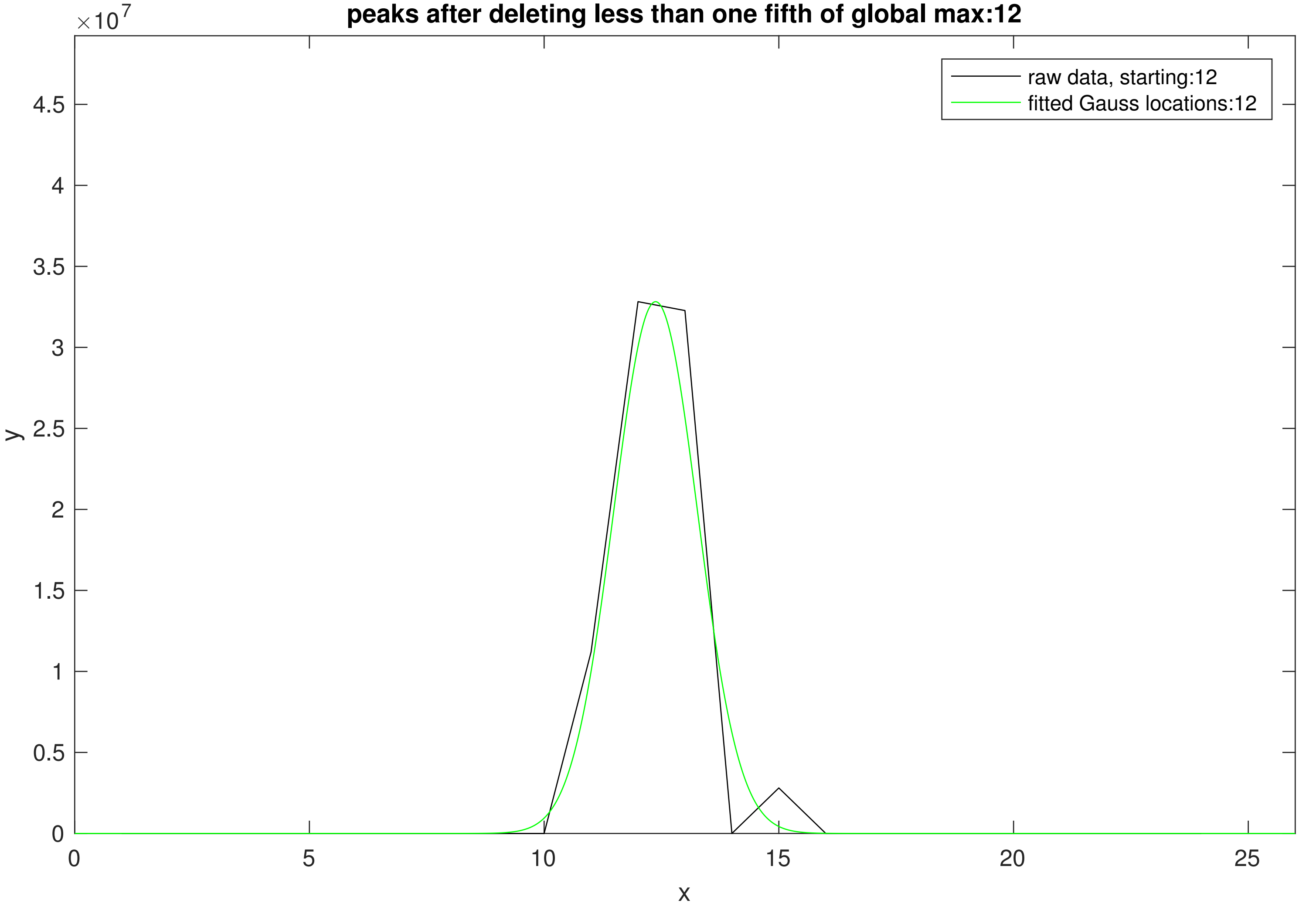

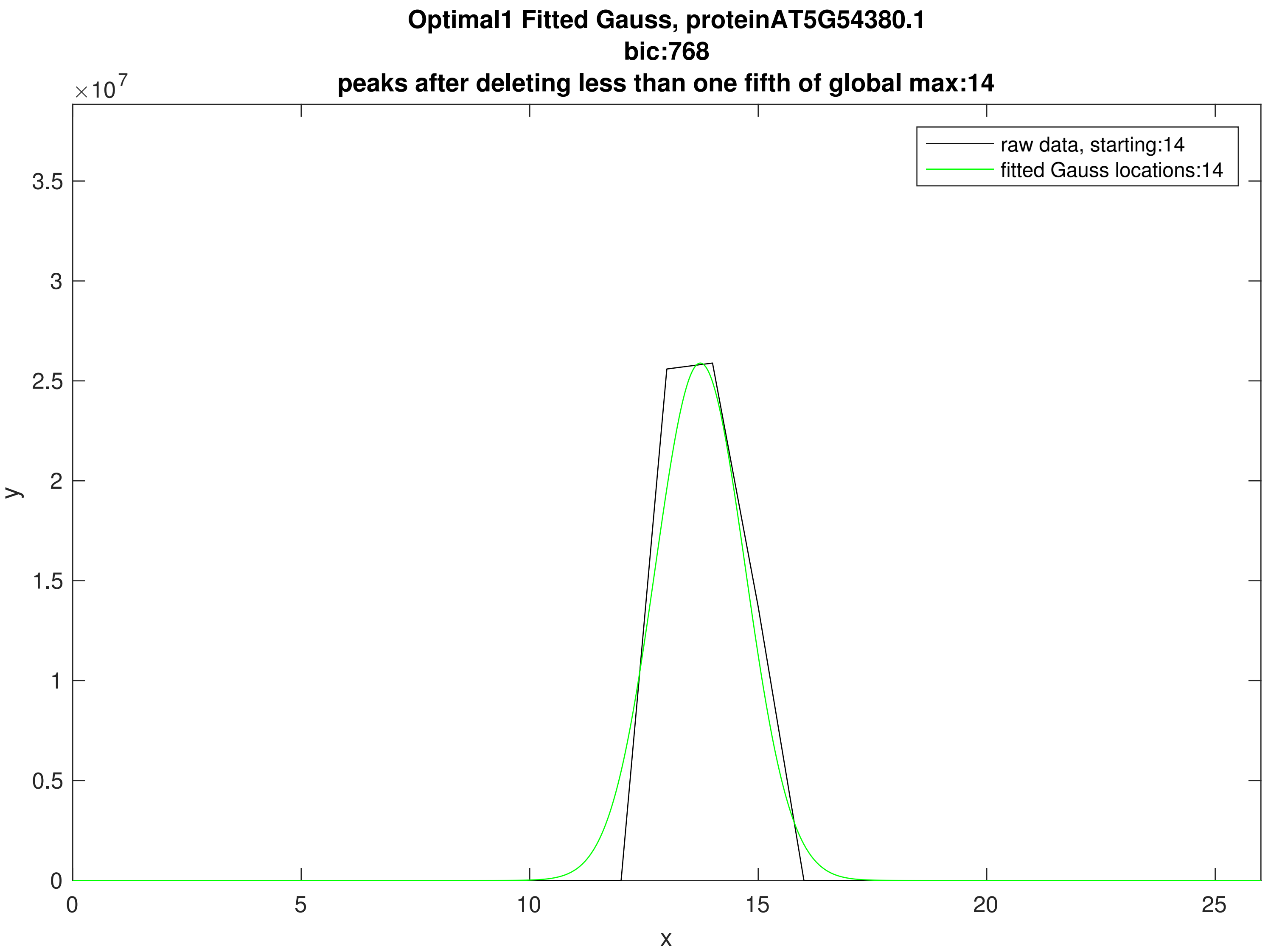
Optimal1 Fitted Gauss, proteinAT5G54380.1
bic:768
peaks after deleting less than one fifth of global max:14
×10^7
raw data, starting:14
fitted Gauss locations:14
3.5
3
2.5
2
1.5
1
0.5
0
y
0
5
10
15
20
25
x

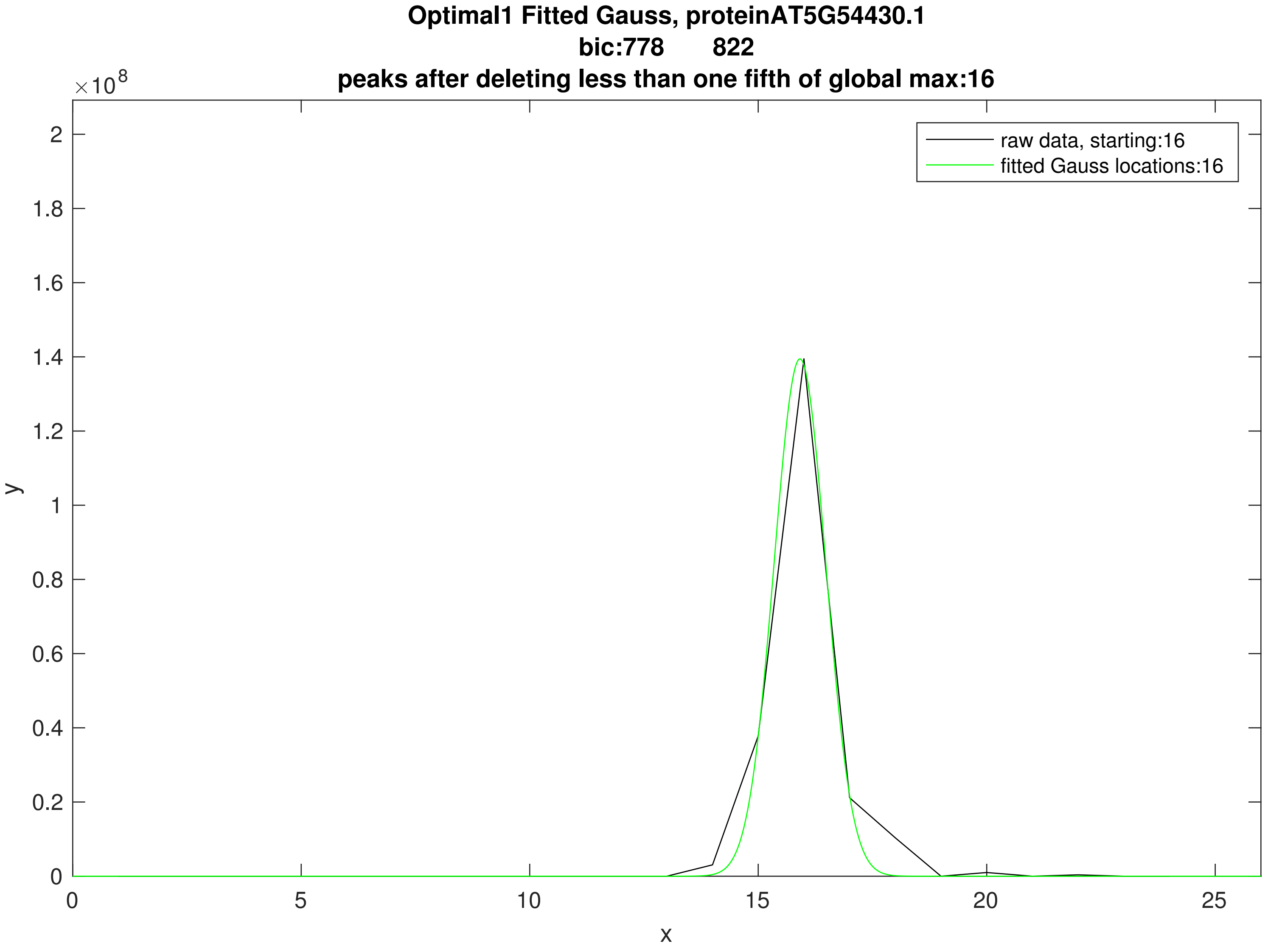
Optimal1 Fitted Gauss, proteinAT5G54430.1
bic:778 822
peaks after deleting less than one fifth of global max:16
×10^8
raw data, starting:16
fitted Gauss locations:16
y
x
0
0.2
0.4
0.6
0.8
1
1.2
1.4
1.6
1.8
2
0
5
10
15
20
25

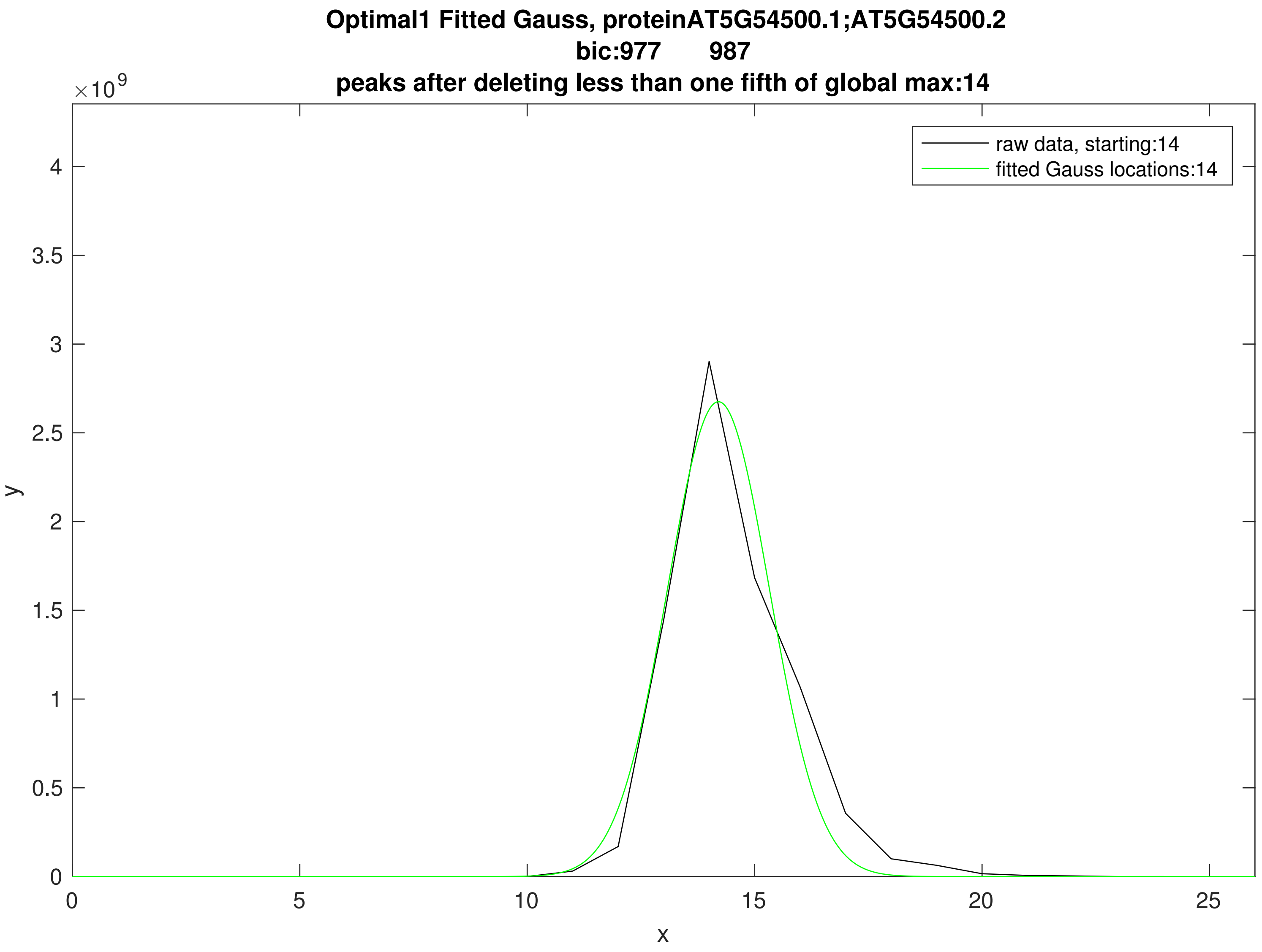

Optimal1 Fitted Gauss, proteinAT5G54500.1;AT5G54500.2
bic:977 987
peaks after deleting less than one fifth of global max:14
×10^9
raw data, starting:14
fitted Gauss locations:14
y
x
0
0.5
1
1.5
2
2.5
3
3.5
4
0
5
10
15
20
25

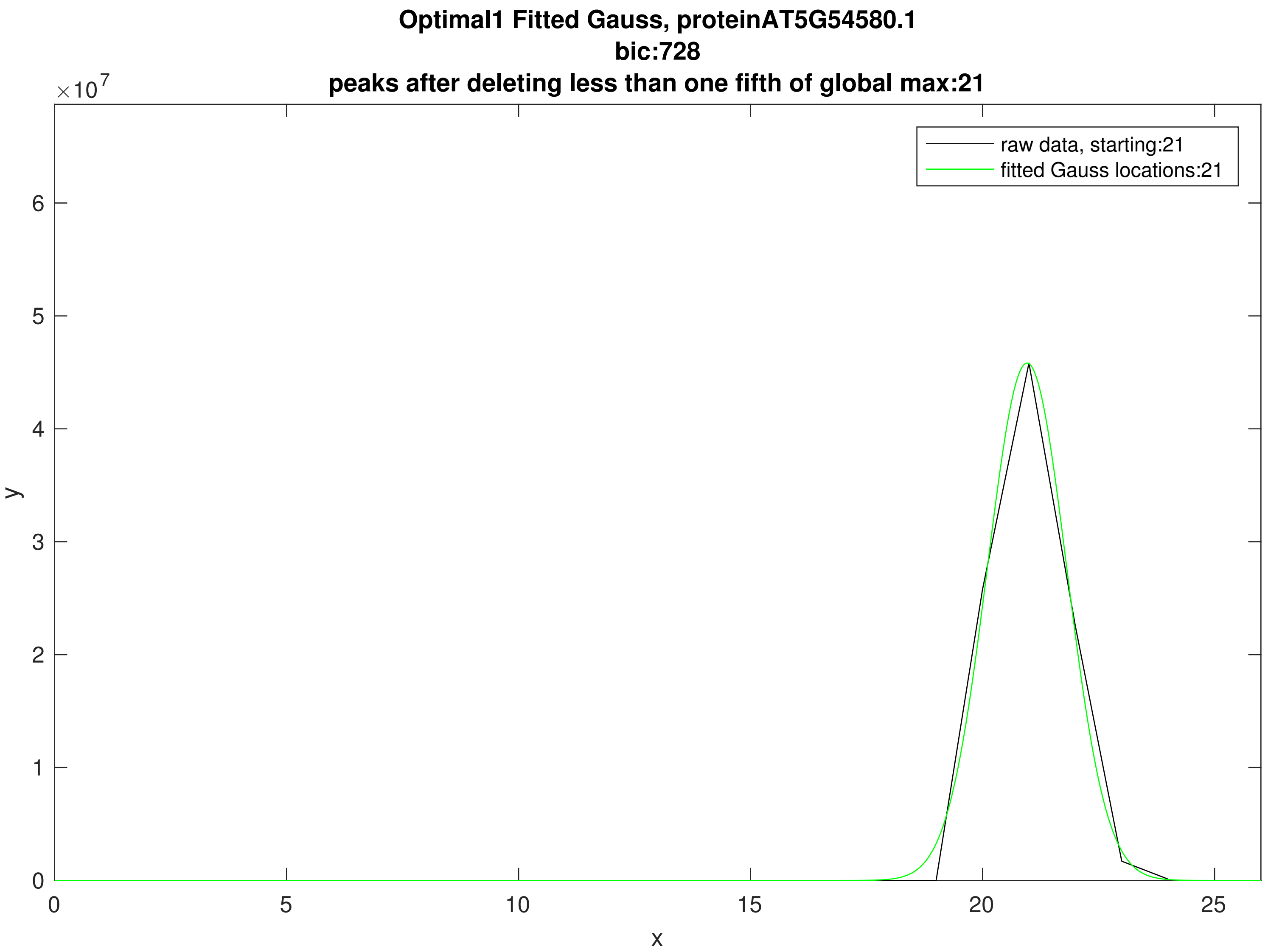
Optimal1 Fitted Gauss, proteinAT5G54580.1
bic:728
peaks after deleting less than one fifth of global max:21
×10^7
raw data, starting:21
fitted Gauss locations:21
y
x
0
1
2
3
4
5
6
0
5
10
15
20
25

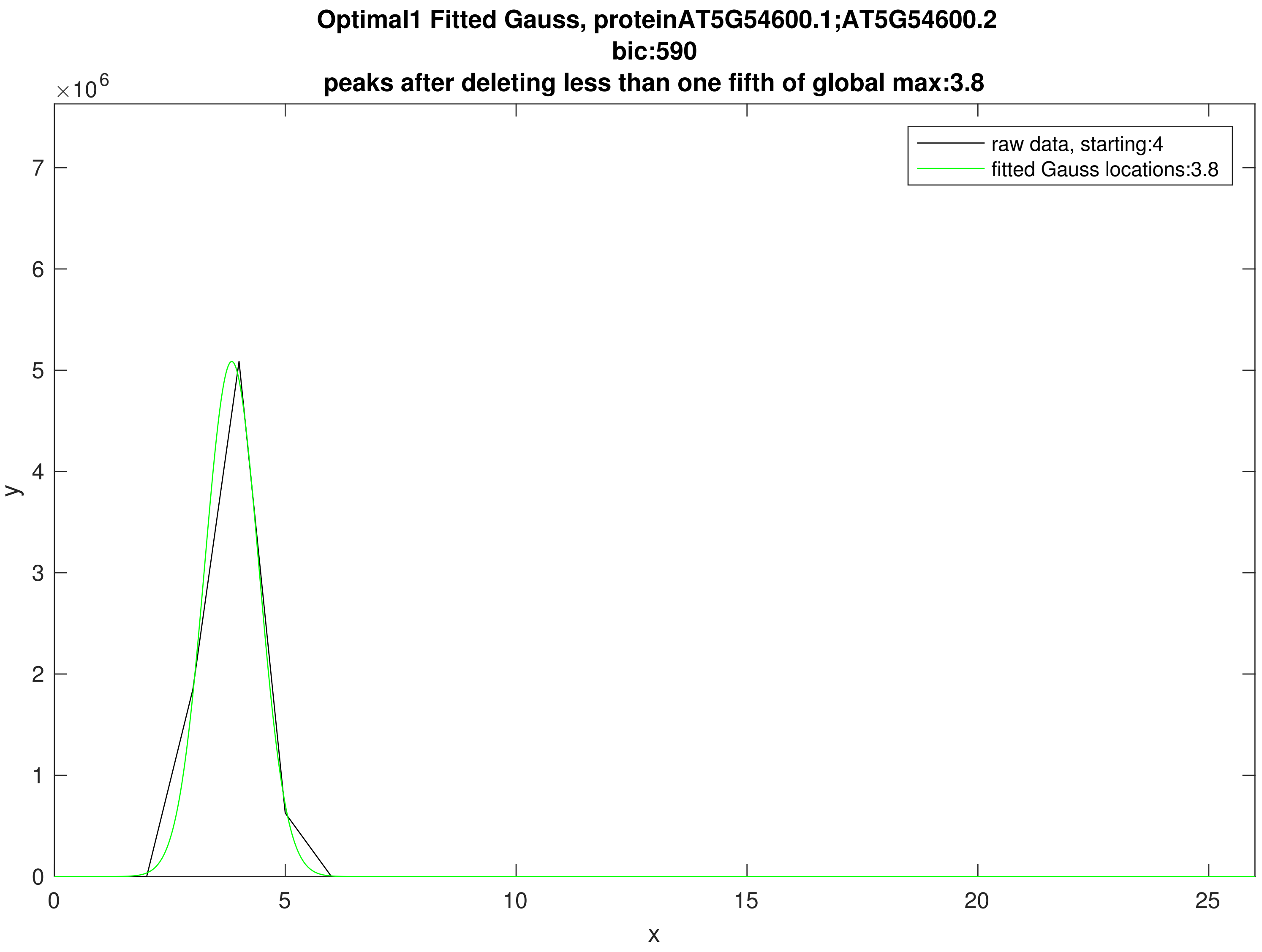
Optimal1 Fitted Gauss, proteinAT5G54600.1;AT5G54600.2
bic:590
peaks after deleting less than one fifth of global max:3.8
×10^6
raw data, starting:4
fitted Gauss locations:3.8
y
x
0
1
2
3
4
5
6
7
0
5
10
15
20
25

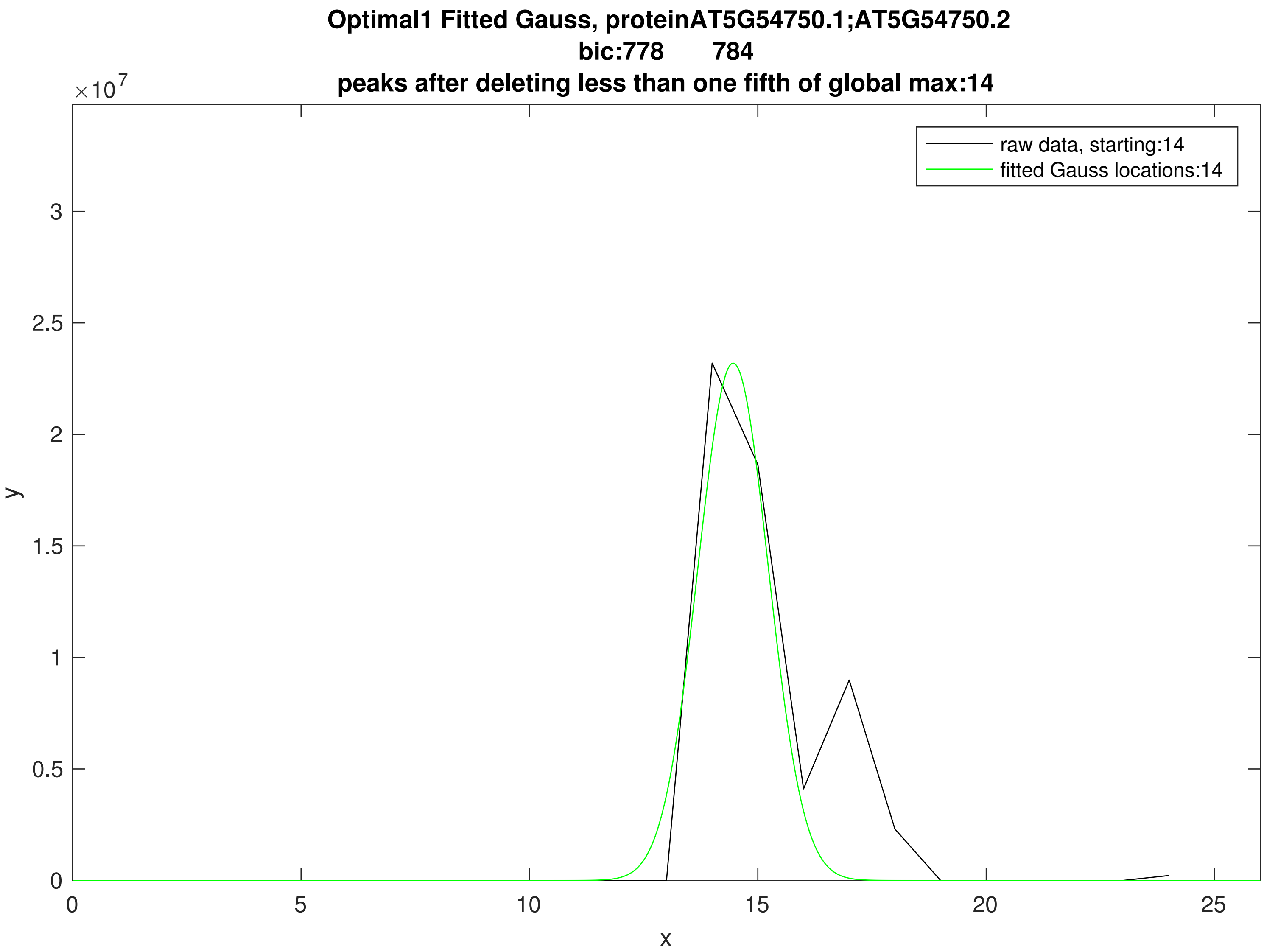
Optimal1 Fitted Gauss, proteinAT5G54750.1;AT5G54750.2
bic:778 784
peaks after deleting less than one fifth of global max:14
×10^7
raw data, starting:14
fitted Gauss locations:14
y
x
0
0.5
1
1.5
2
2.5
3
5
10
15
20
25

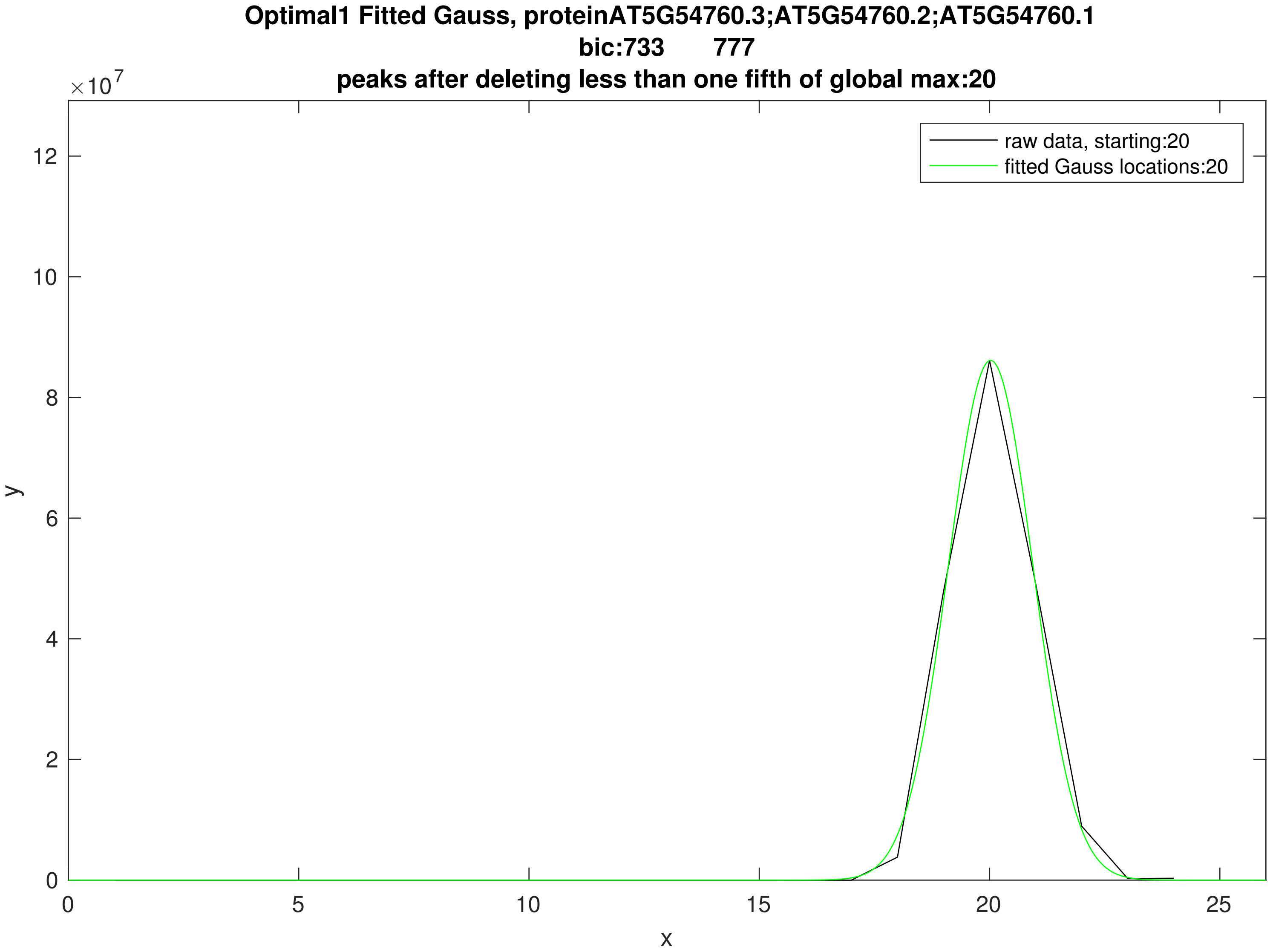
Optimal1 Fitted Gauss, proteinAT5G54760.3;AT5G54760.2;AT5G54760.1
bic:733 777
peaks after deleting less than one fifth of global max:20
×10^7
raw data, starting:20
fitted Gauss locations:20
y
x

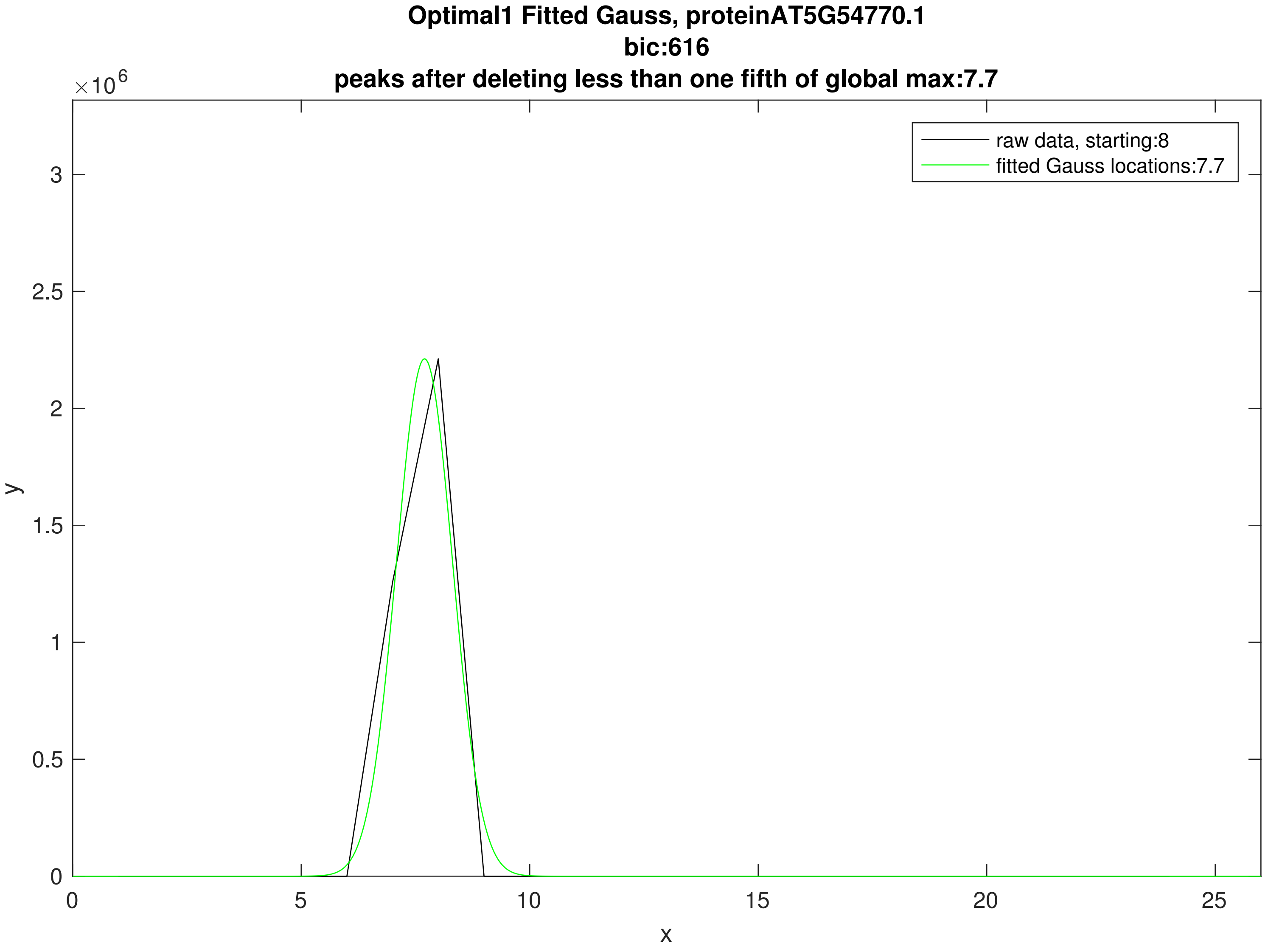

Optimal1 Fitted Gauss, proteinAT5G54770.1
bic:616
peaks after deleting less than one fifth of global max:7.7
×10^6
3
2.5
2
1.5
1
0.5
0
y
0
5
10
15
20
25
x
raw data, starting:8
fitted Gauss locations:7.7

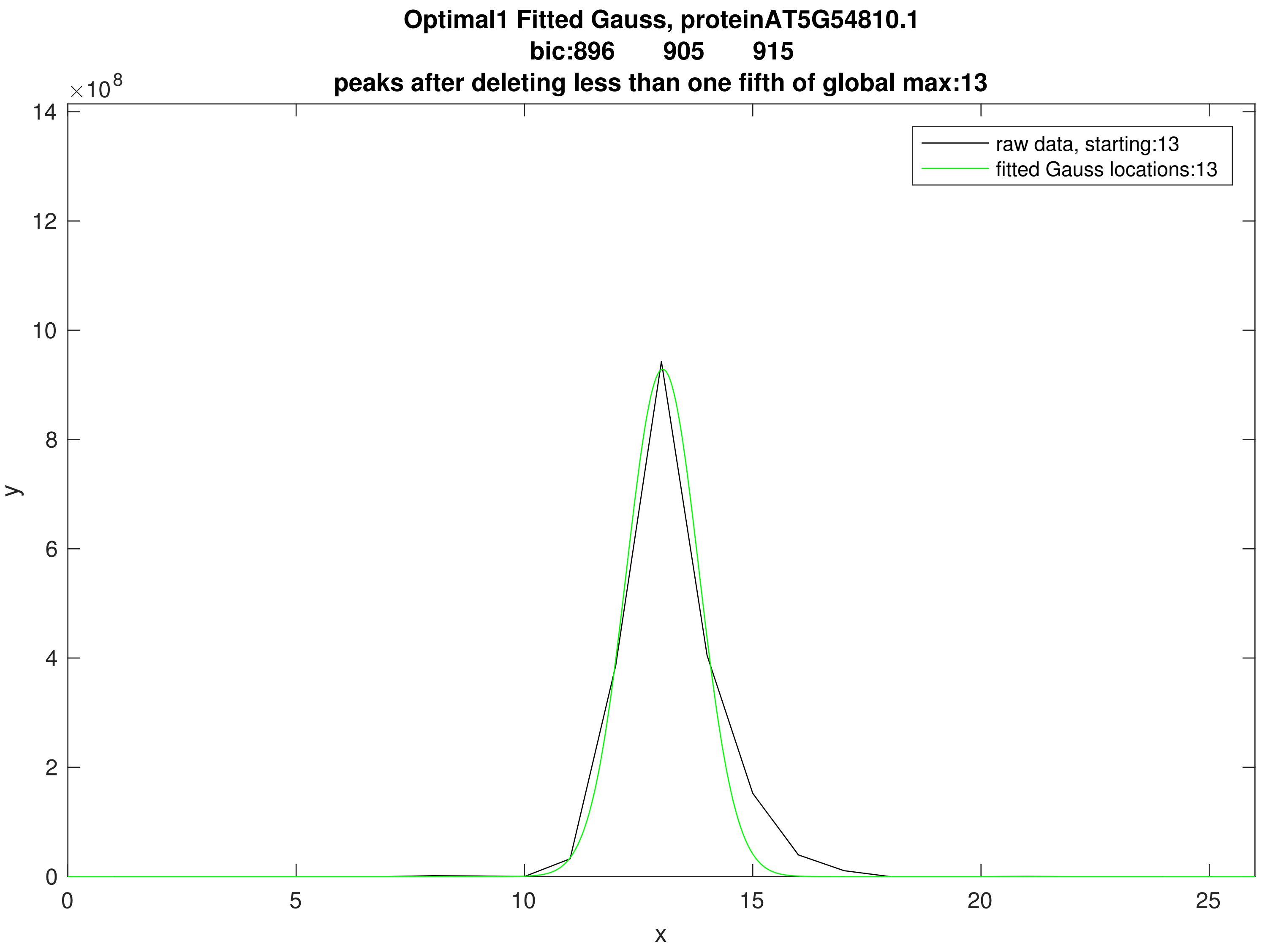
Optimal1 Fitted Gauss, proteinAT5G54810.1
bic:896 905 915
peaks after deleting less than one fifth of global max:13
×10^8
14
12
10
8
6
4
2
0
y
0
5
10
15
20
25
x
raw data, starting:13
fitted Gauss locations:13

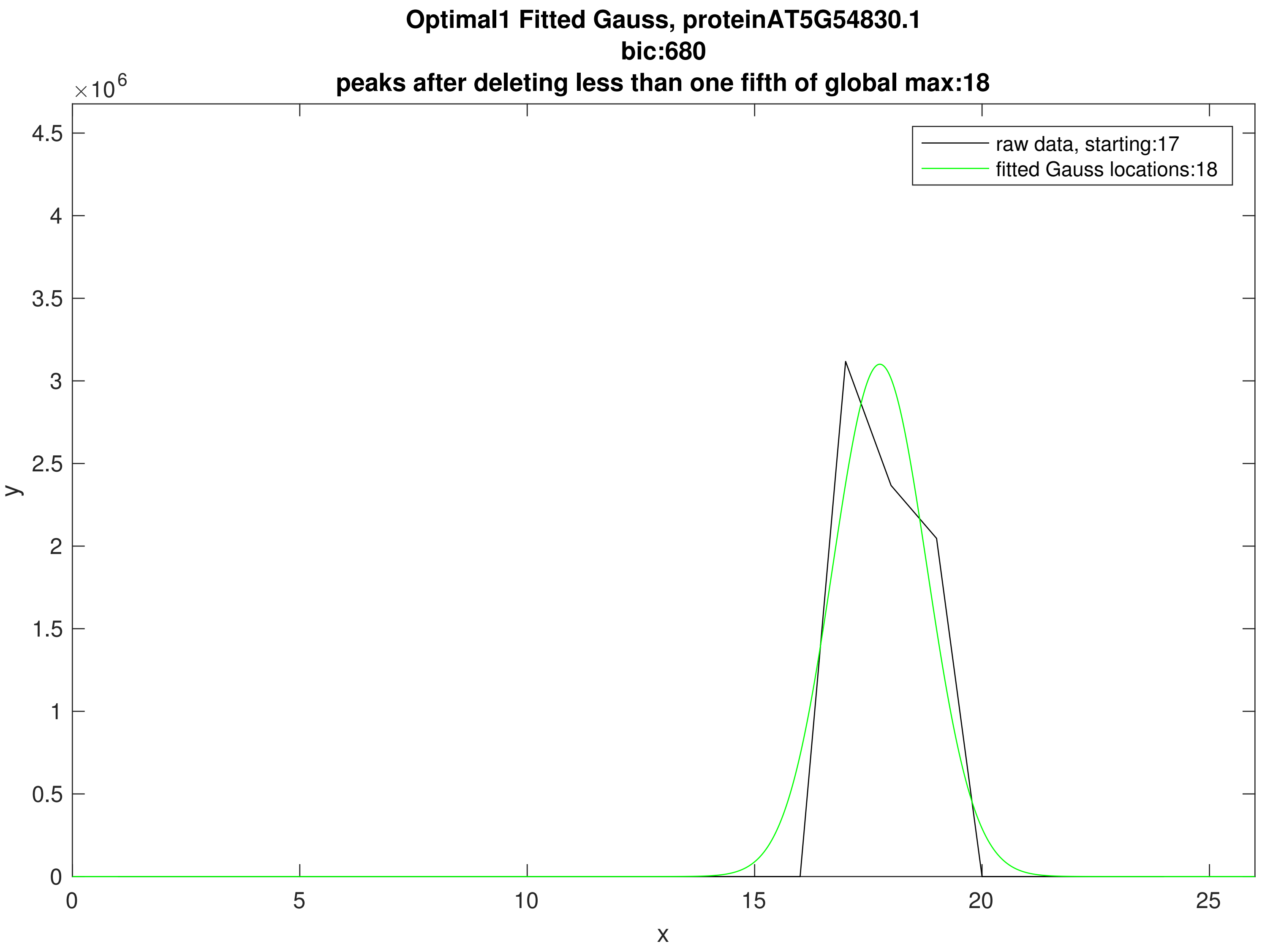

Optimal1 Fitted Gauss, proteinAT5G54830.1
bic:680
peaks after deleting less than one fifth of global max:18
×10^6
4.5
4
3.5
3
2.5
2
1.5
1
0.5
0
y
0
5
10
15
20
25
x
raw data, starting:17
fitted Gauss locations:18

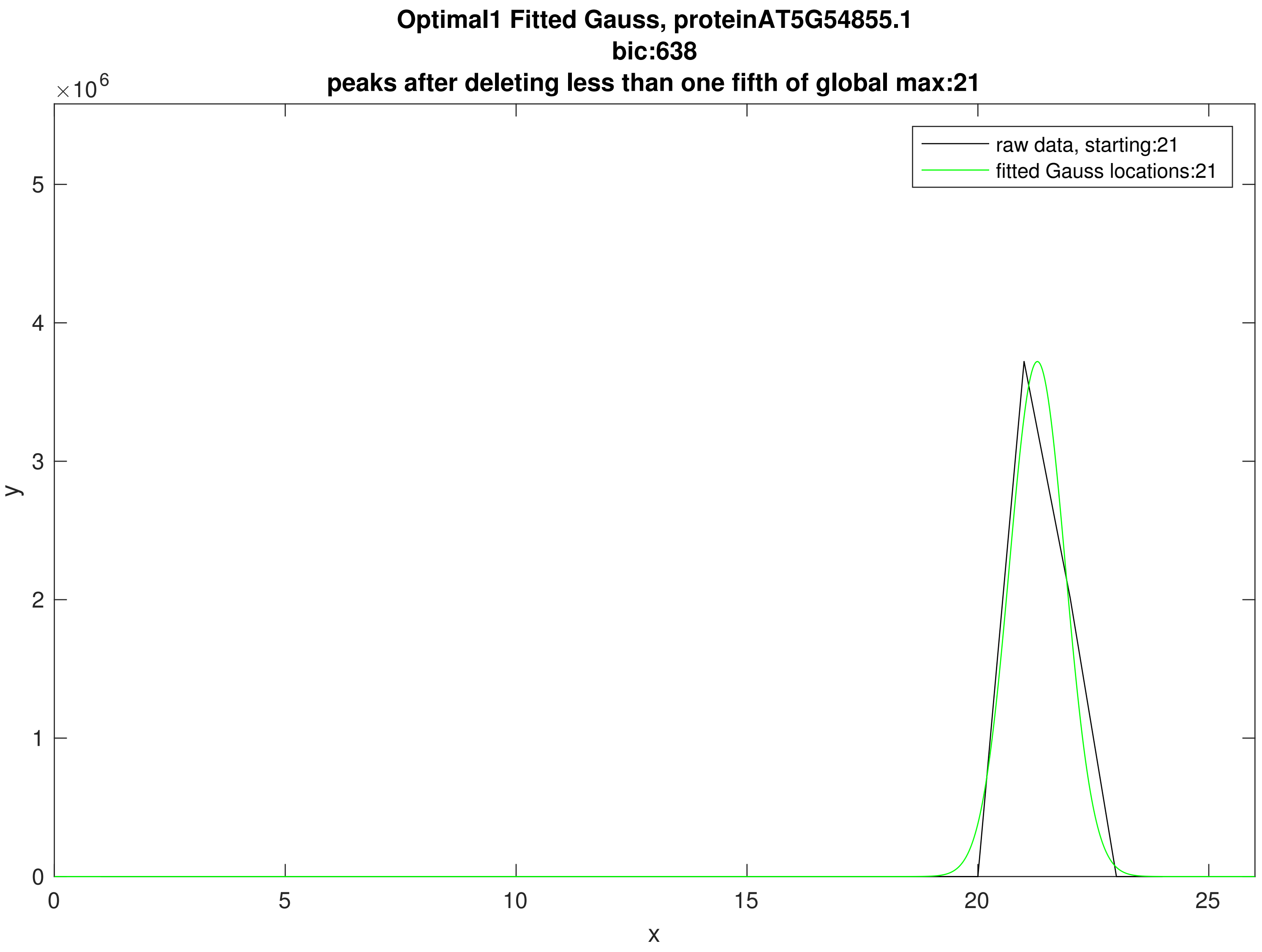
Optimal1 Fitted Gauss, proteinAT5G54855.1
bic:638
peaks after deleting less than one fifth of global max:21
×10^6
raw data, starting:21
fitted Gauss locations:21
y
x
0
1
2
3
4
5
0
5
10
15
20
25

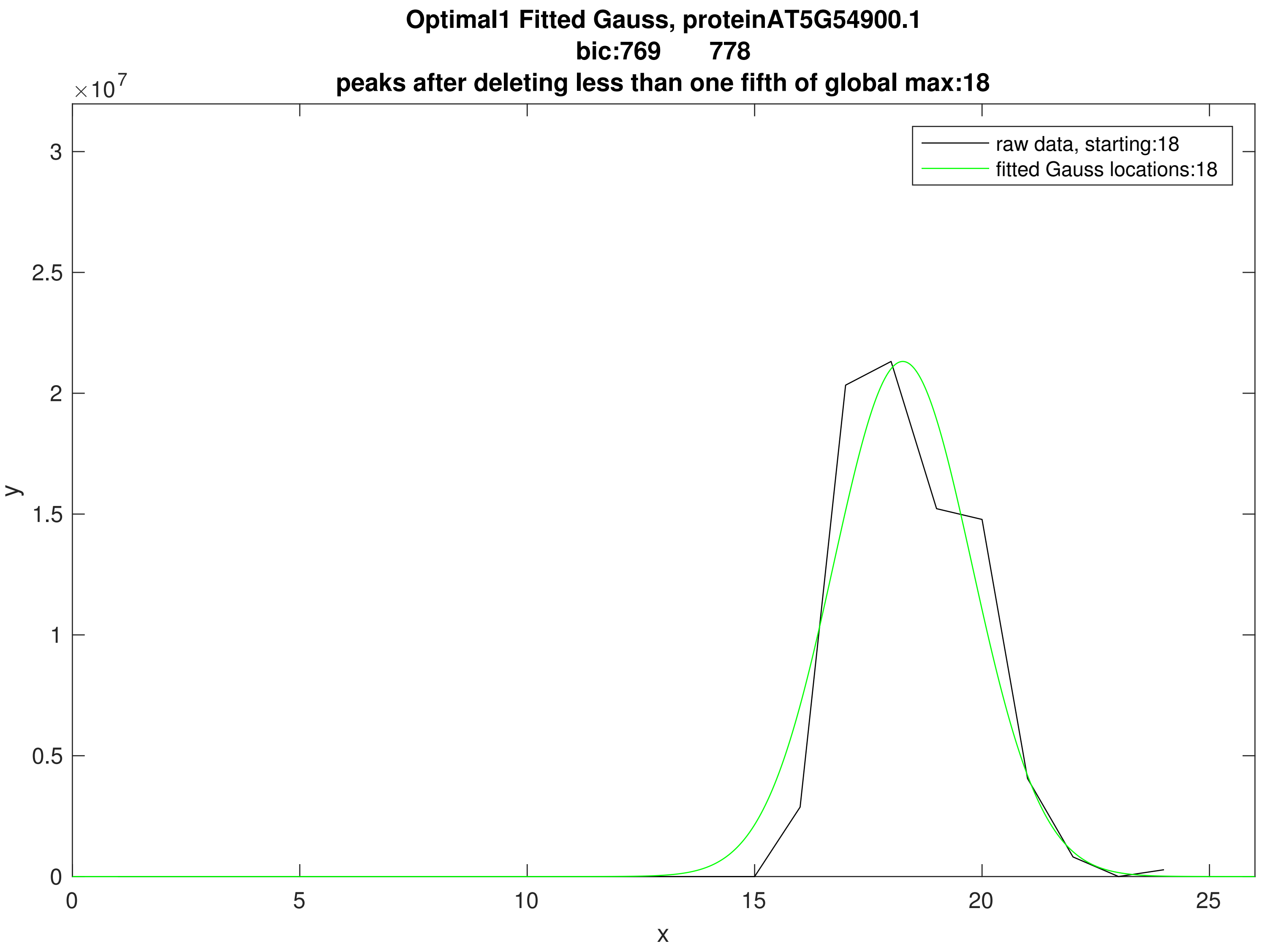

Optimal1 Fitted Gauss, proteinAT5G54900.1
bic:769 778
peaks after deleting less than one fifth of global max:18
×10^7
3
2.5
2
1.5
1
0.5
0
y
0
5
10
15
20
25
x
raw data, starting:18
fitted Gauss locations:18

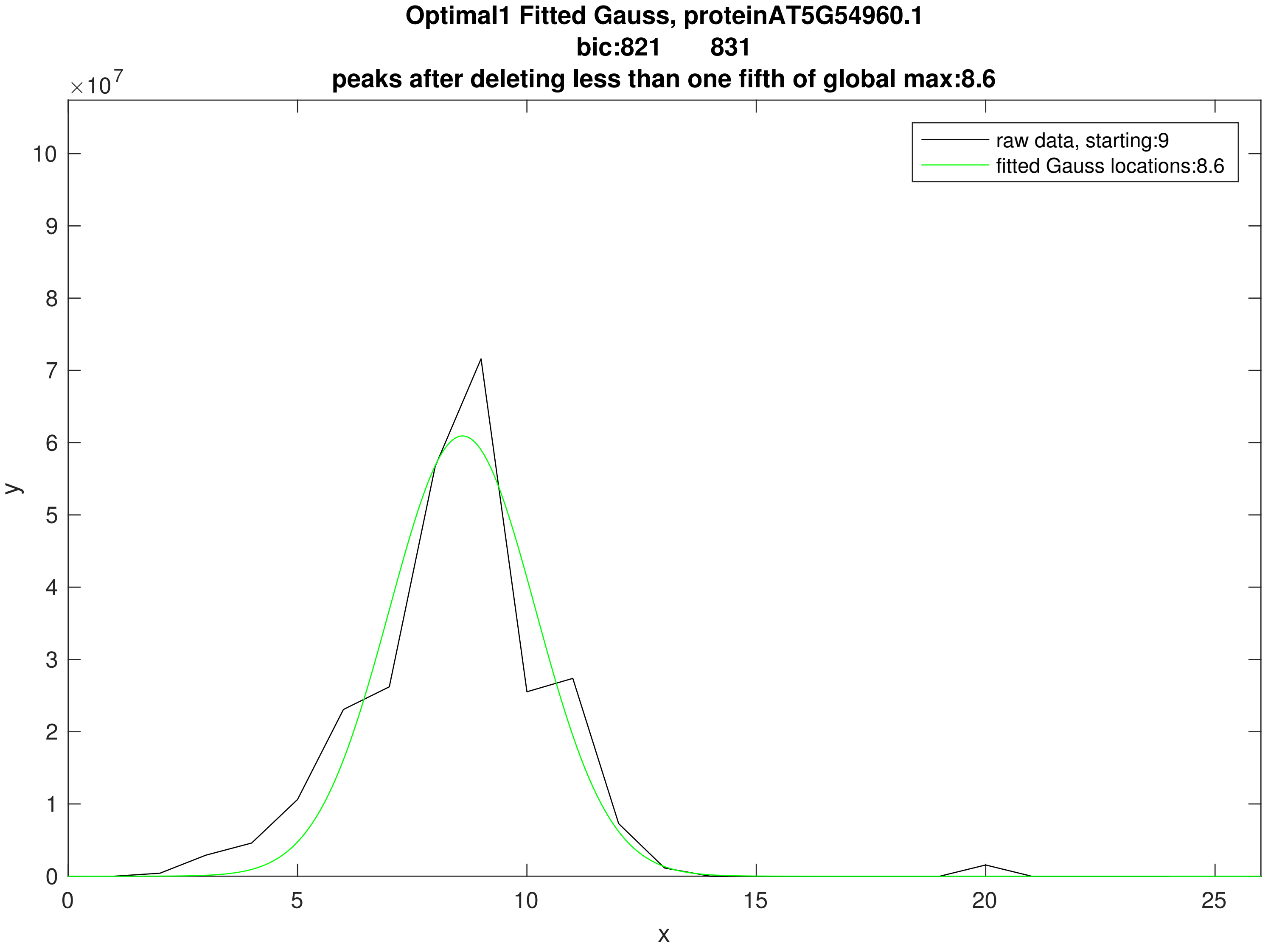

Optimal1 Fitted Gauss, proteinAT5G54960.1
bic:821 831
peaks after deleting less than one fifth of global max:8.6
×10^7
raw data, starting:9
fitted Gauss locations:8.6
y
x
0
1
2
3
4
5
6
7
8
9
10
0
5
10
15
20
25

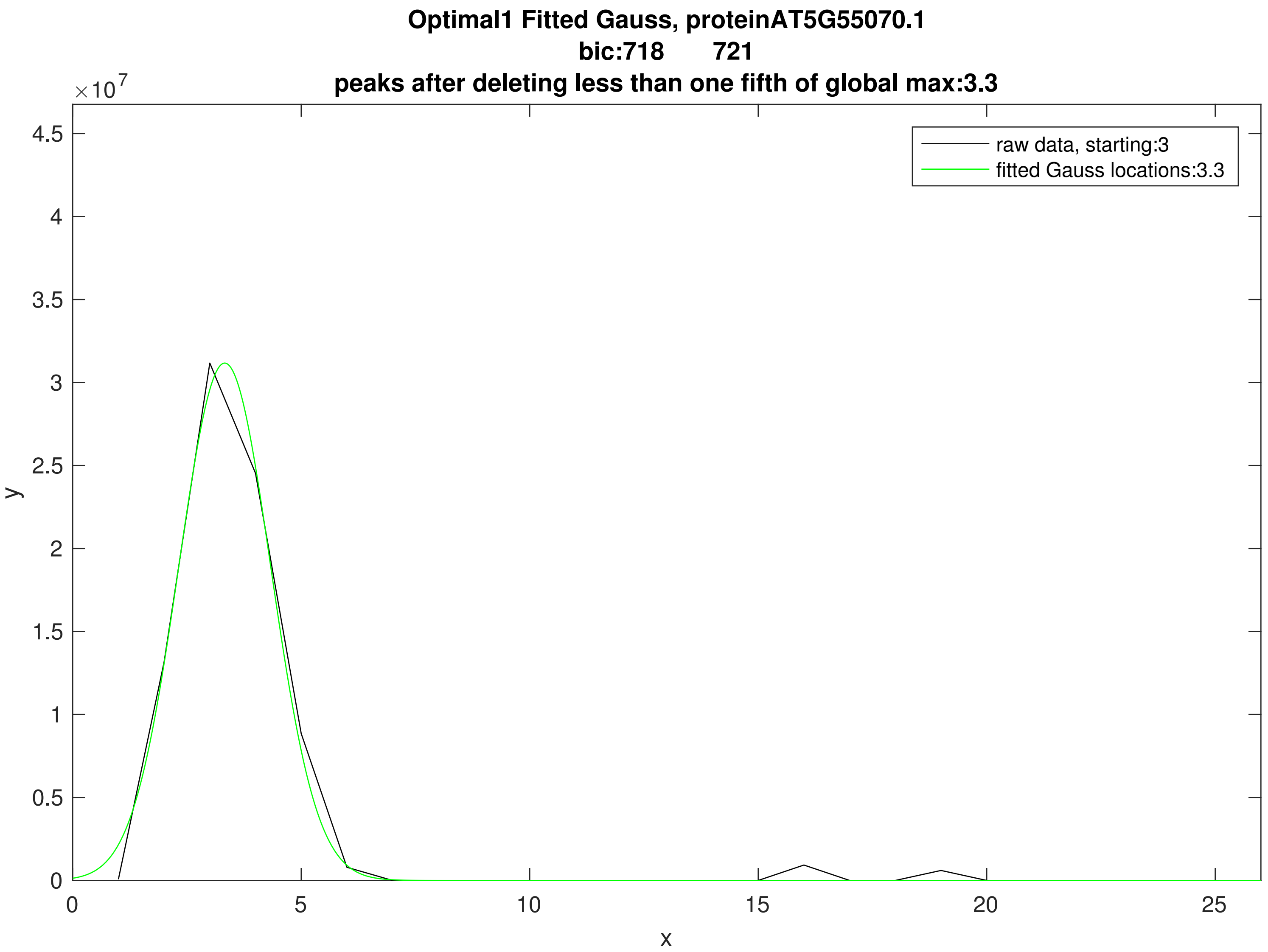
Optimal1 Fitted Gauss, proteinAT5G55070.1
bic:718 721
peaks after deleting less than one fifth of global max:3.3
×10^7
raw data, starting:3
fitted Gauss locations:3.3
y
x
0
0.5
1
1.5
2
2.5
3
3.5
4
4.5
5
10
15
20
25

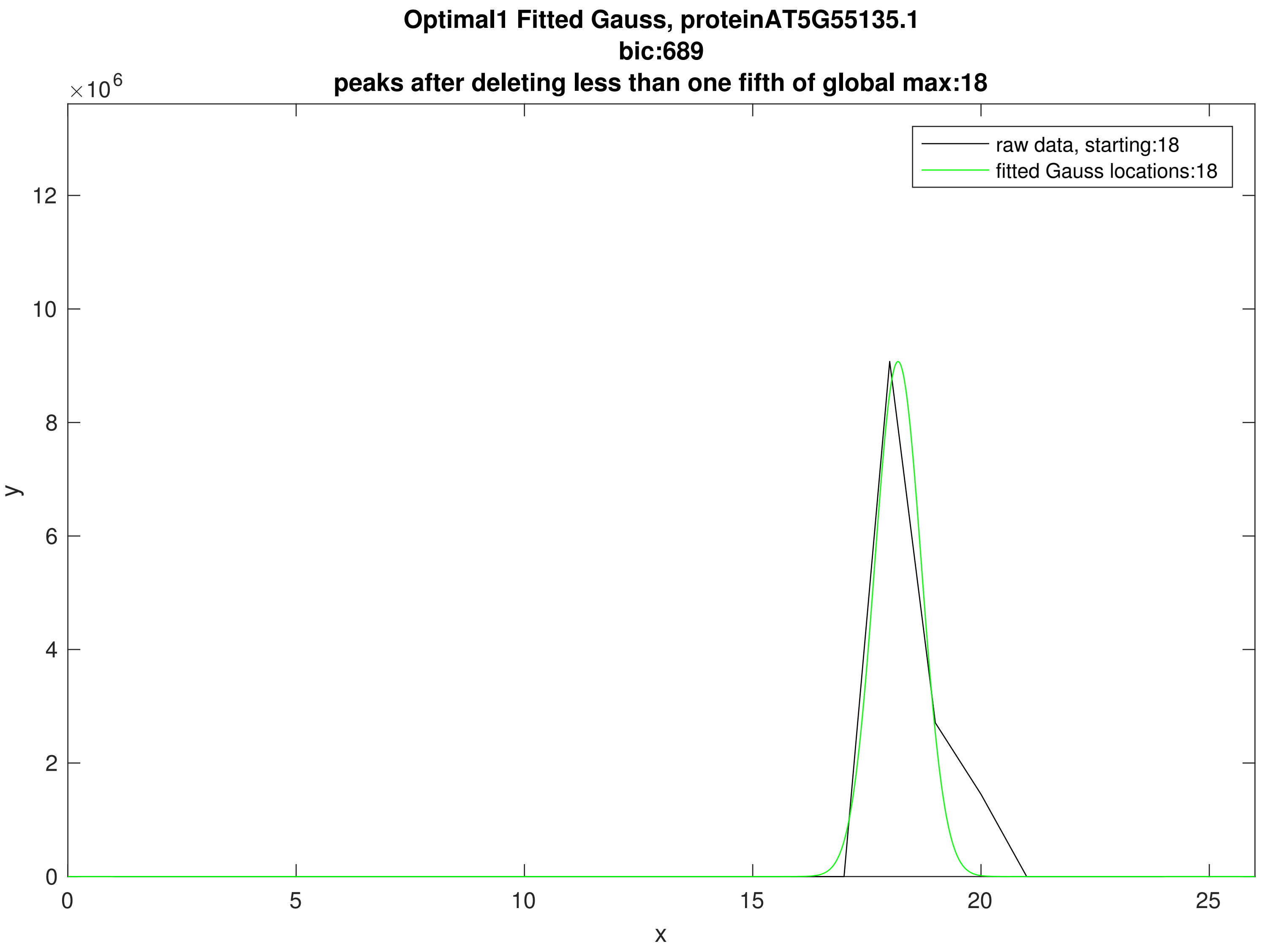
Optimal1 Fitted Gauss, proteinAT5G55135.1
bic:689
peaks after deleting less than one fifth of global max:18
×10^6
raw data, starting:18
fitted Gauss locations:18
y
x
0
2
4
6
8
10
12
0
5
10
15
20
25

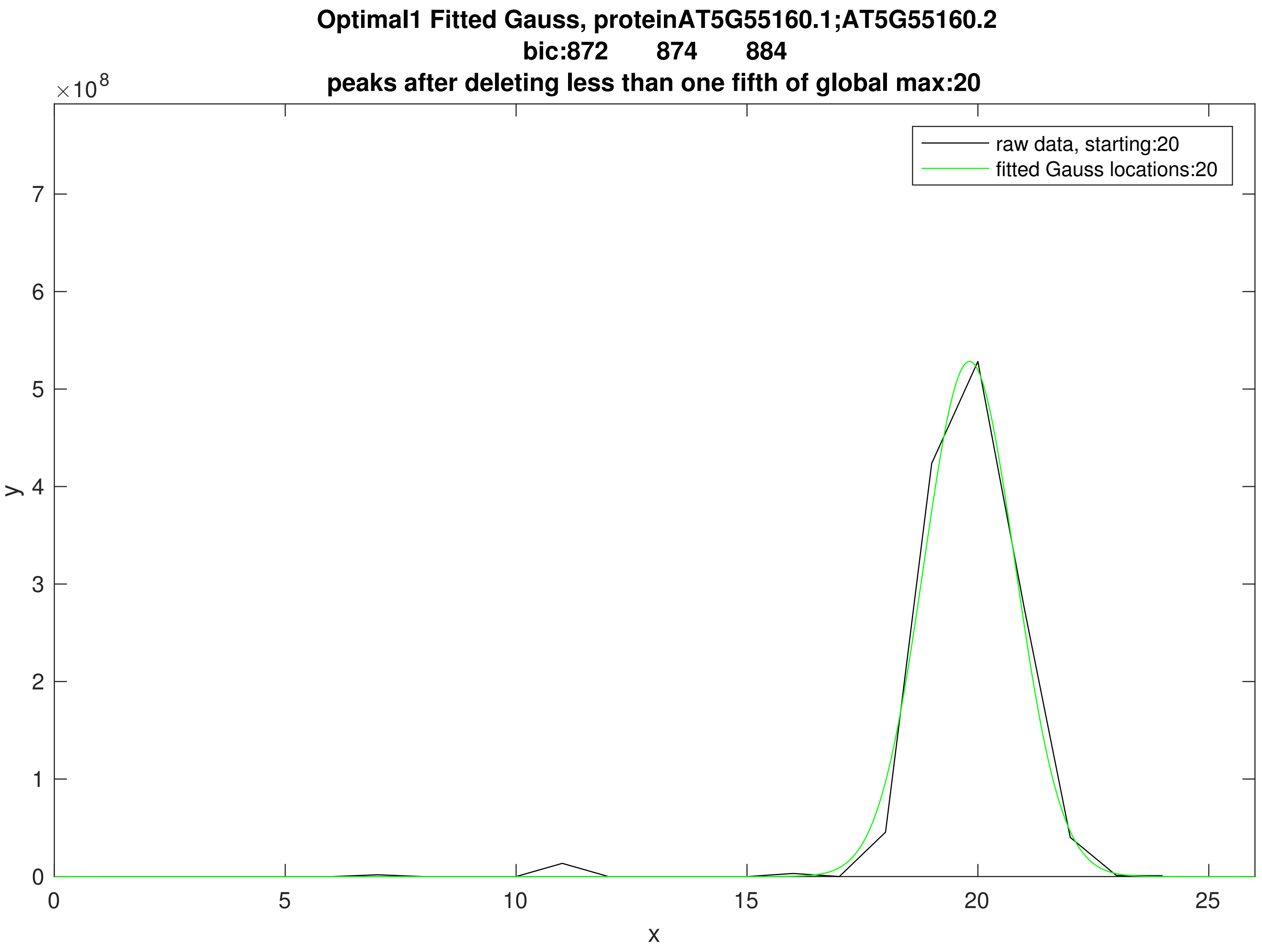
Optimal1 Fitted Gauss, proteinAT5G55160.1;AT5G55160.2
bic:872 874 884
peaks after deleting less than one fifth of global max:20
×10^8
raw data, starting:20
fitted Gauss locations:20
y
x
0
1
2
3
4
5
6
7
0
5
10
15
20
25

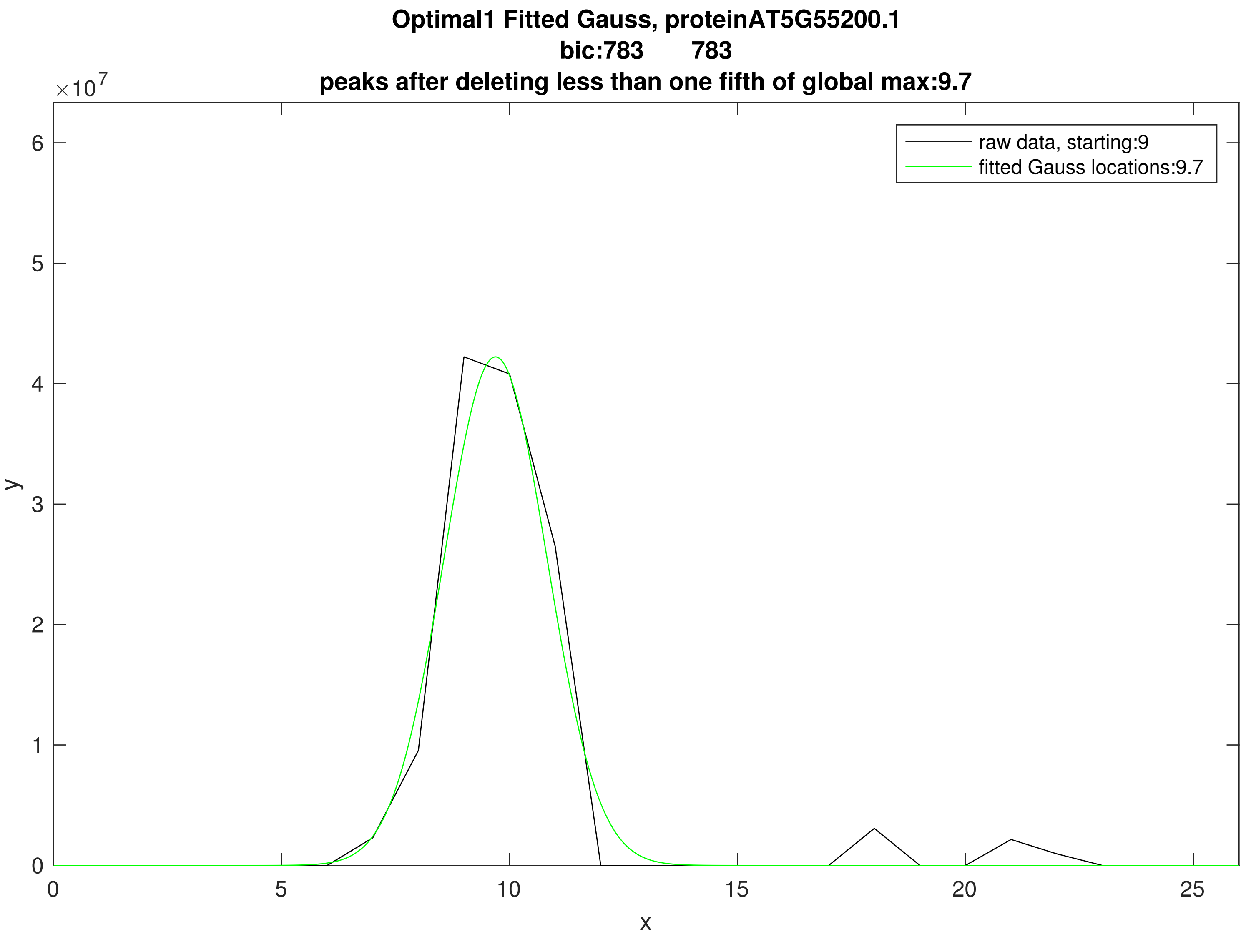
Optimal1 Fitted Gauss, proteinAT5G55200.1
bic:783 783
peaks after deleting less than one fifth of global max:9.7
×10^7
raw data, starting:9
fitted Gauss locations:9.7
y
x
0
1
2
3
4
5
6
0
5
10
15
20
25

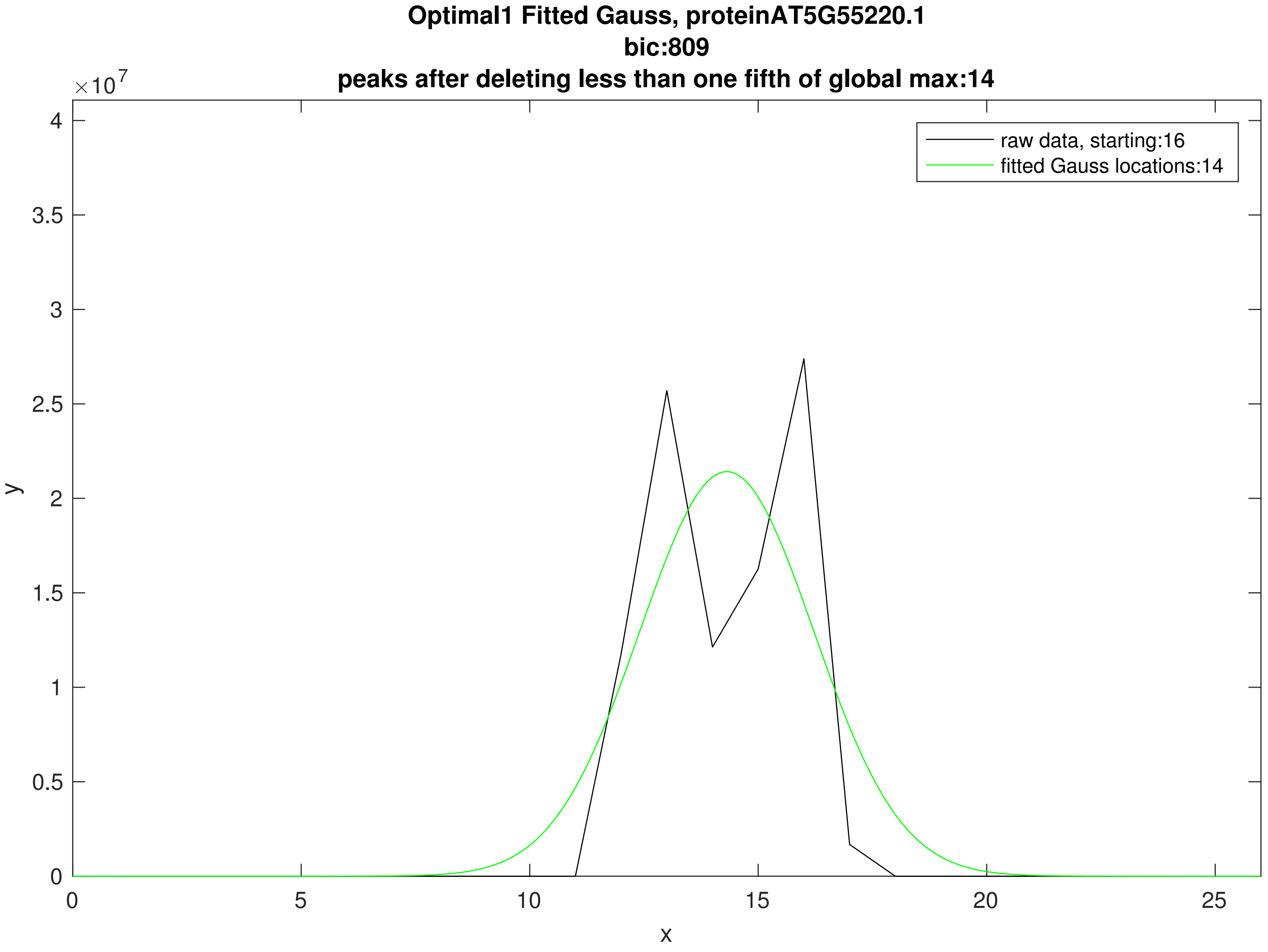

Optimal1 Fitted Gauss, proteinAT5G55220.1
bic:809
peaks after deleting less than one fifth of global max:14
×10^7
raw data, starting:16
fitted Gauss locations:14
y
x
0
0.5
1
1.5
2
2.5
3
3.5
4
0
5
10
15
20
25

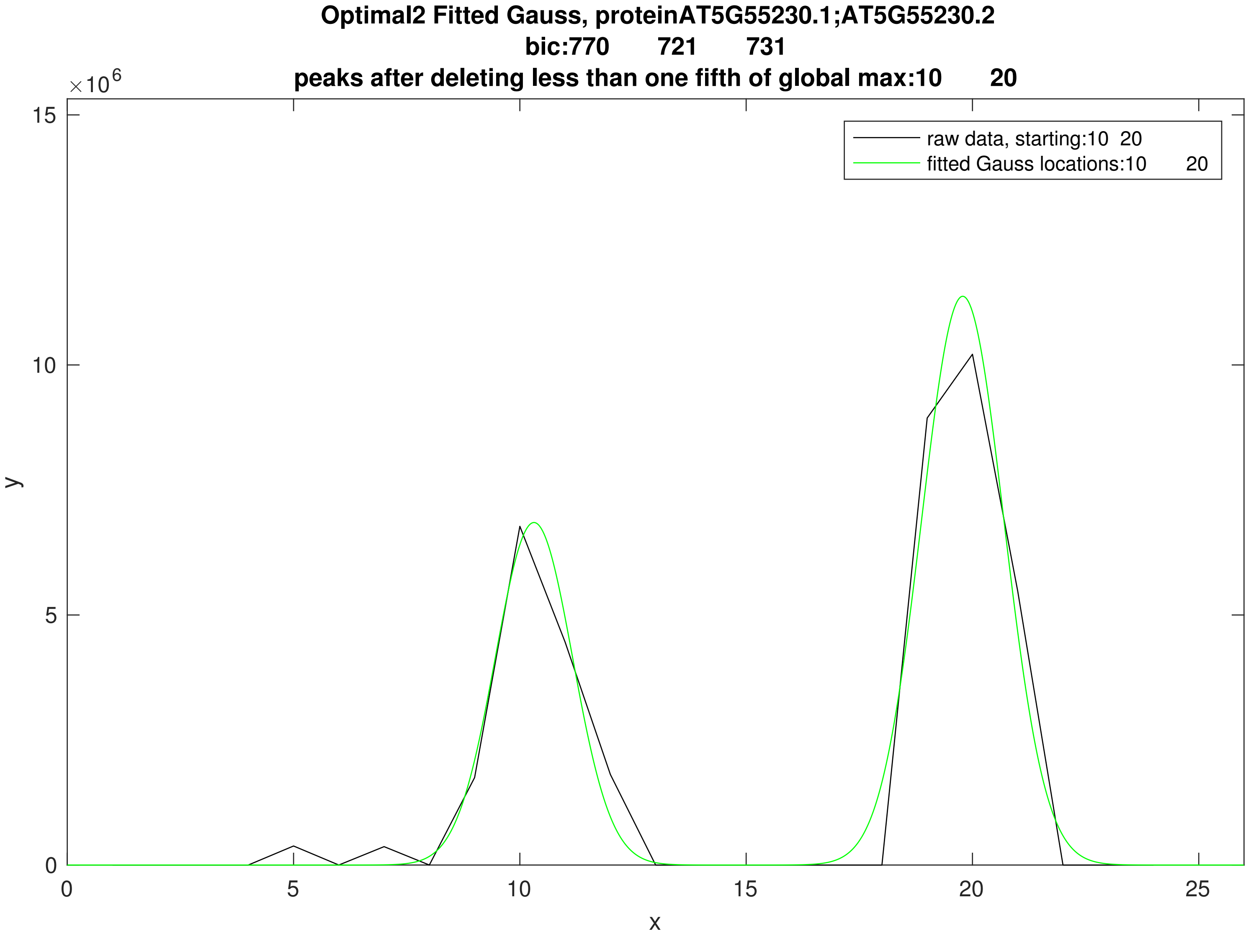
Optimal2 Fitted Gauss, proteinAT5G55230.1;AT5G55230.2
bic:770 721 731
peaks after deleting less than one fifth of global max:10 20
×10^6
raw data, starting:10 20
fitted Gauss locations:10 20
y
x
0
5
10
15
20
25

**Optimal1 Fitted Gauss, proteinAT5G55260.1**
**bic:705**
**peaks after deleting less than one fifth of global max:18**

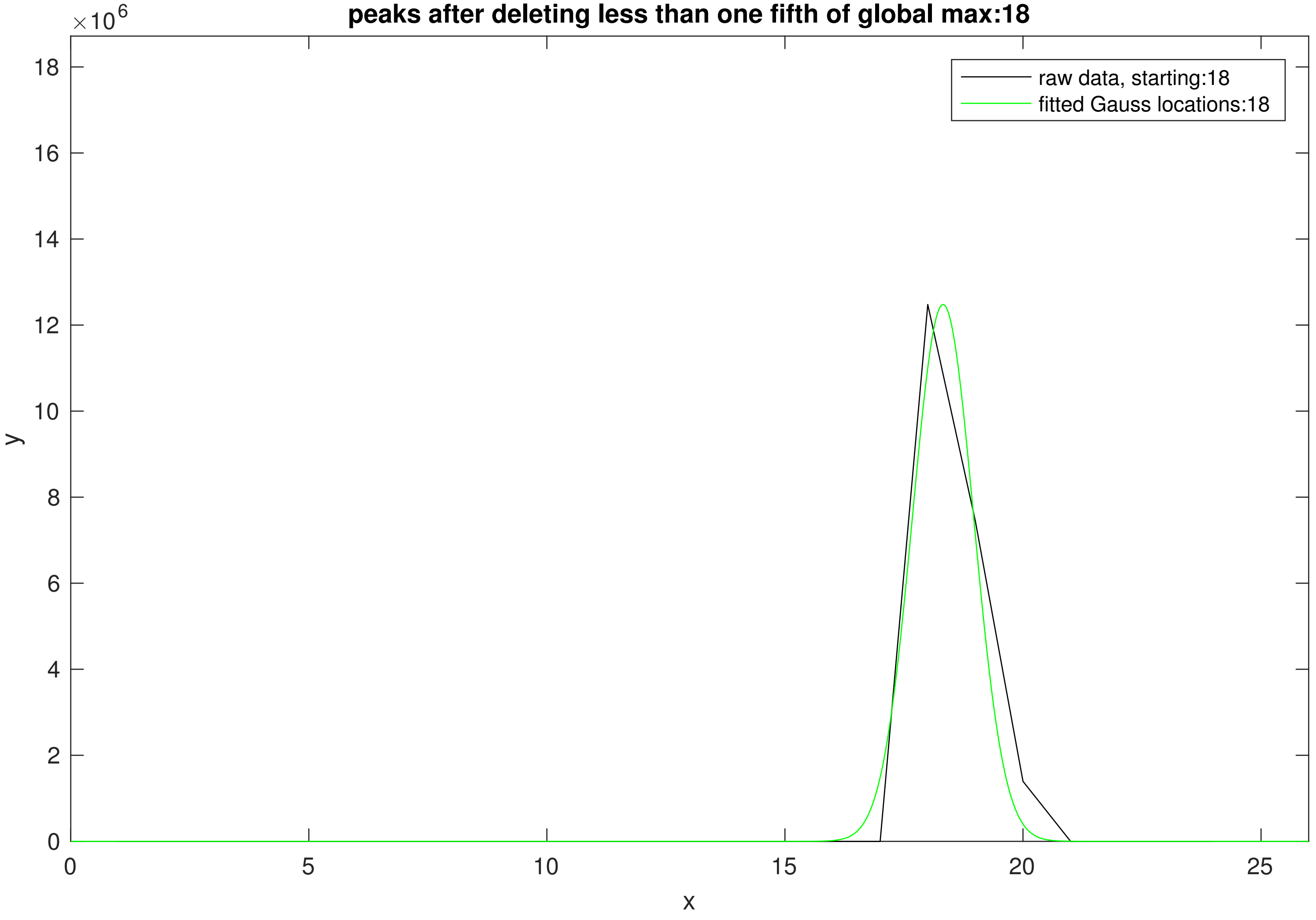

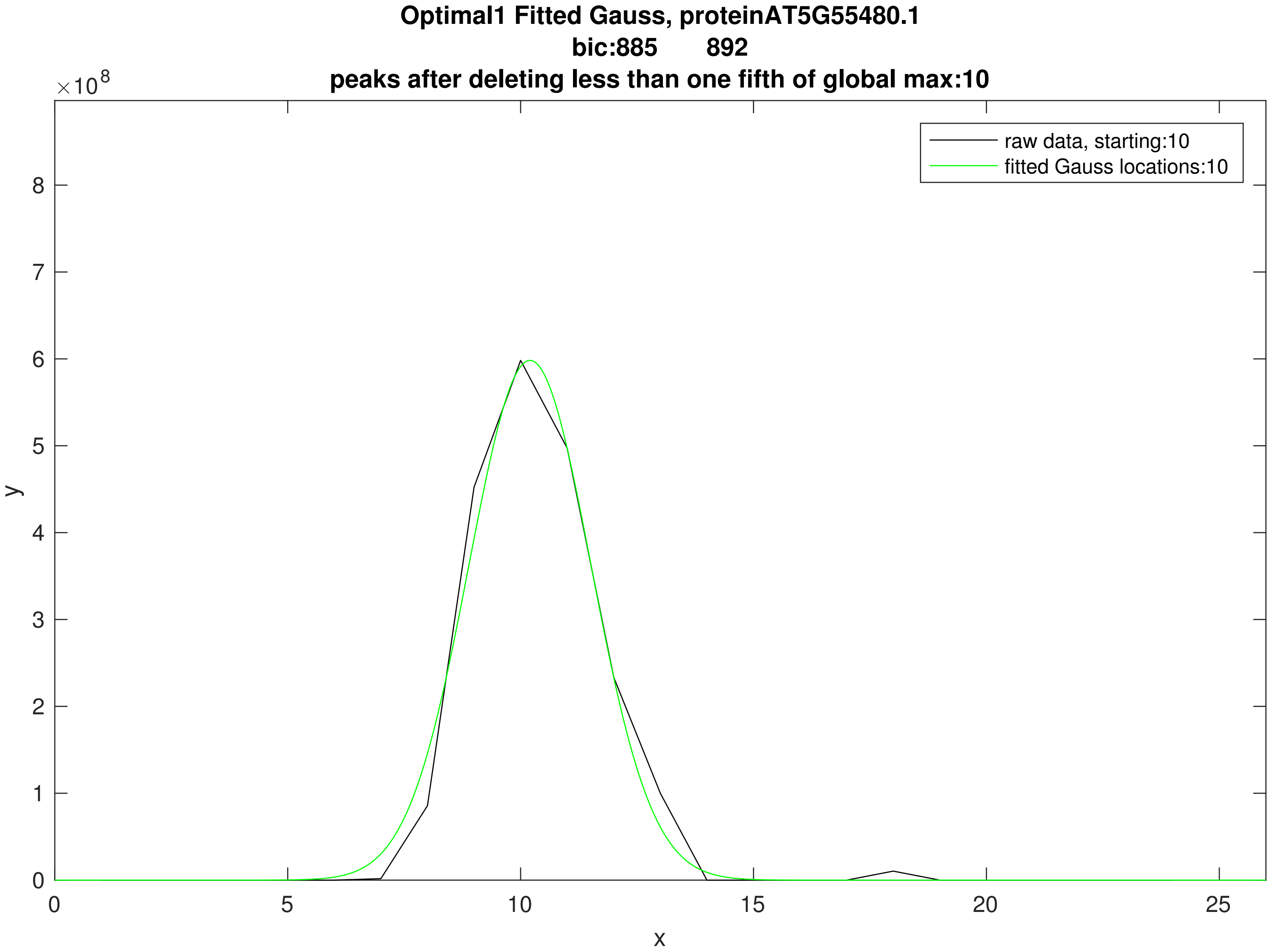
Optimal1 Fitted Gauss, proteinAT5G55480.1
bic:885 892
peaks after deleting less than one fifth of global max:10
×10^8
raw data, starting:10
fitted Gauss locations:10
y
x
0
1
2
3
4
5
6
7
8
0
5
10
15
20
25

#### Optimal2 Fitted Gauss, proteinAT5G55730.2;AT5G55730.1
#### bic:978 937 947 956
#### peaks after deleting less than one fifth of global max:10 17

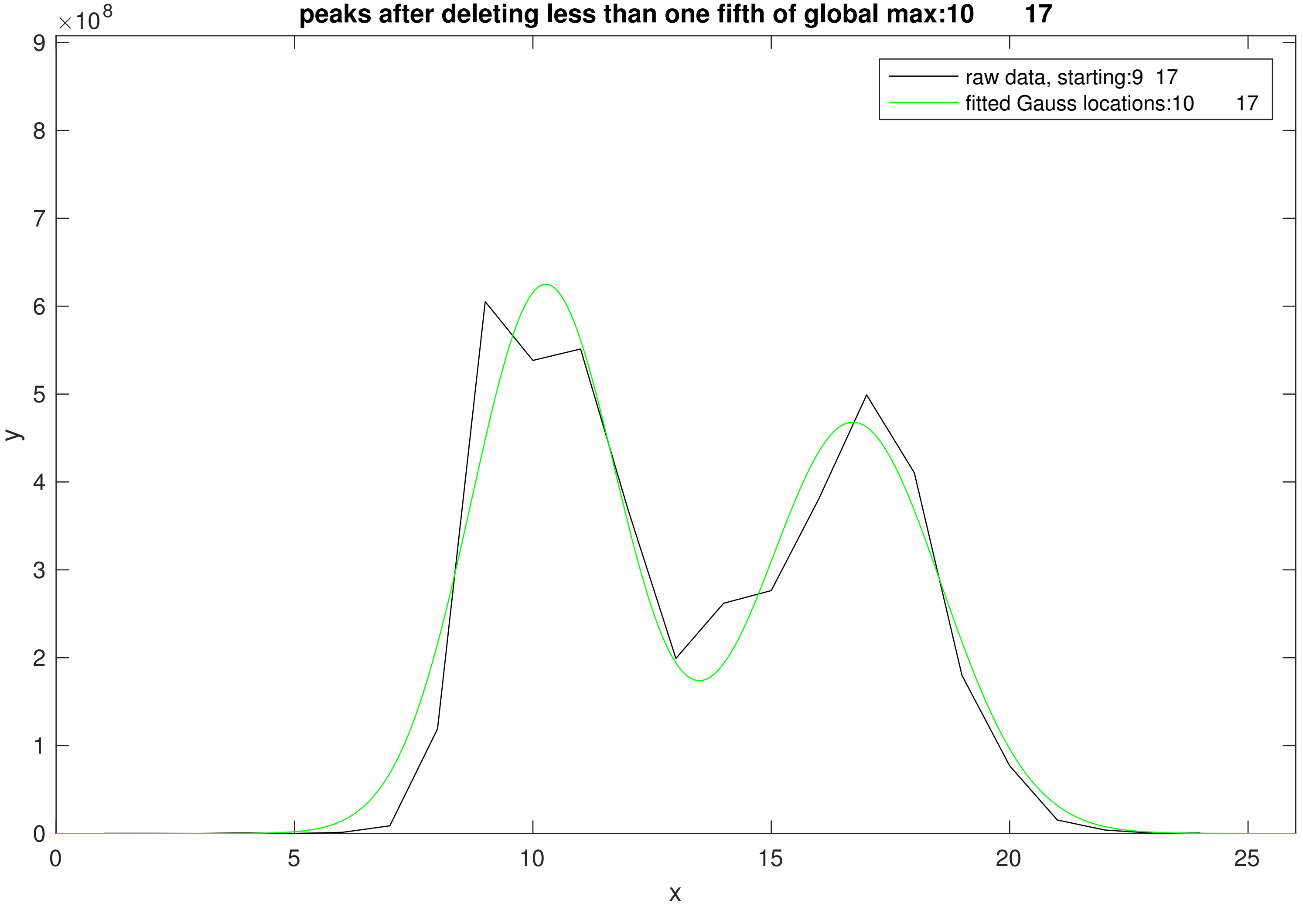

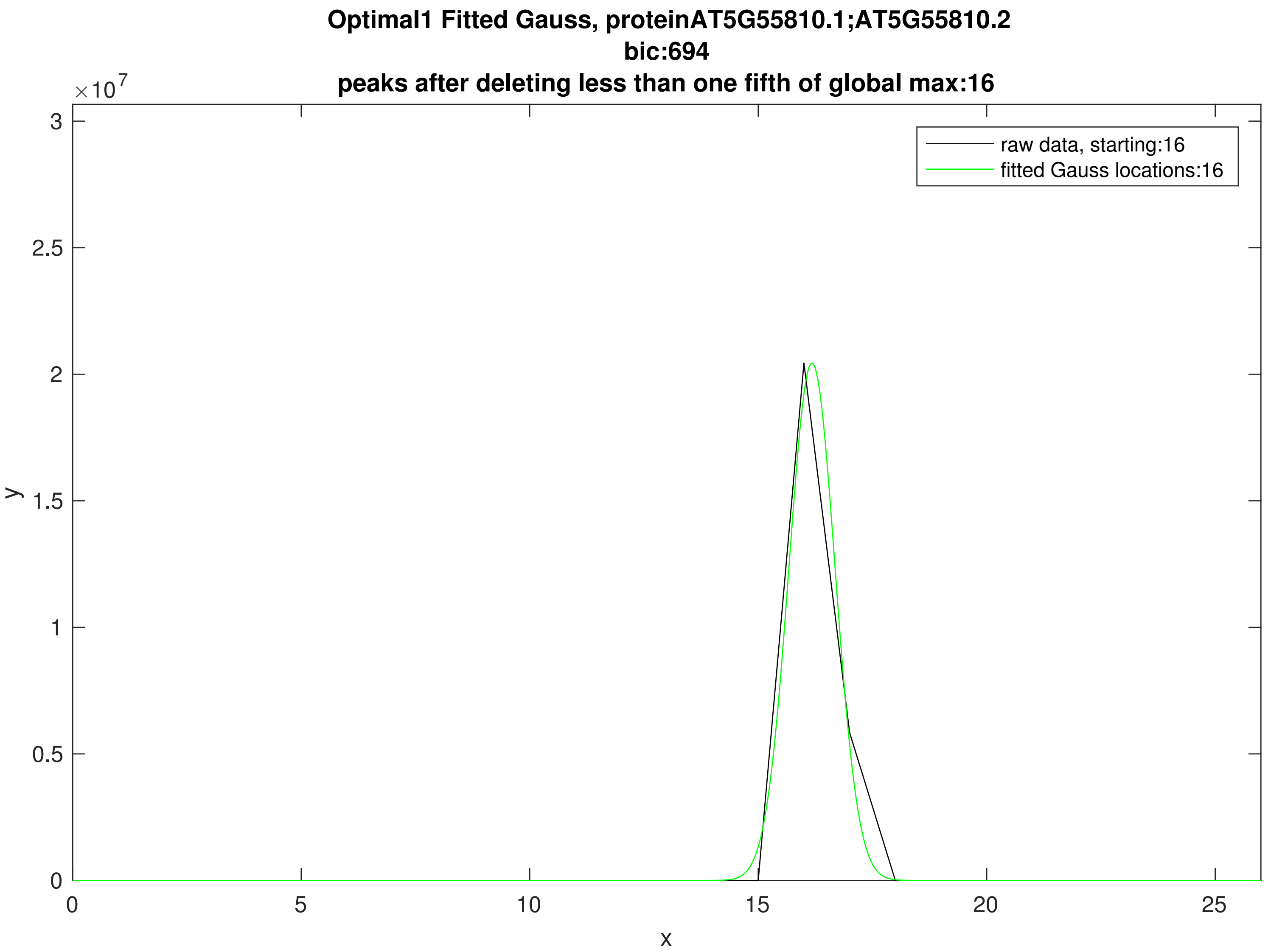

Optimal1 Fitted Gauss, proteinAT5G55810.1;AT5G55810.2
bic:694
peaks after deleting less than one fifth of global max:16
×10^7
raw data, starting:16
fitted Gauss locations:16
y
x
0
0.5
1
1.5
2
2.5
3
0
5
10
15
20
25

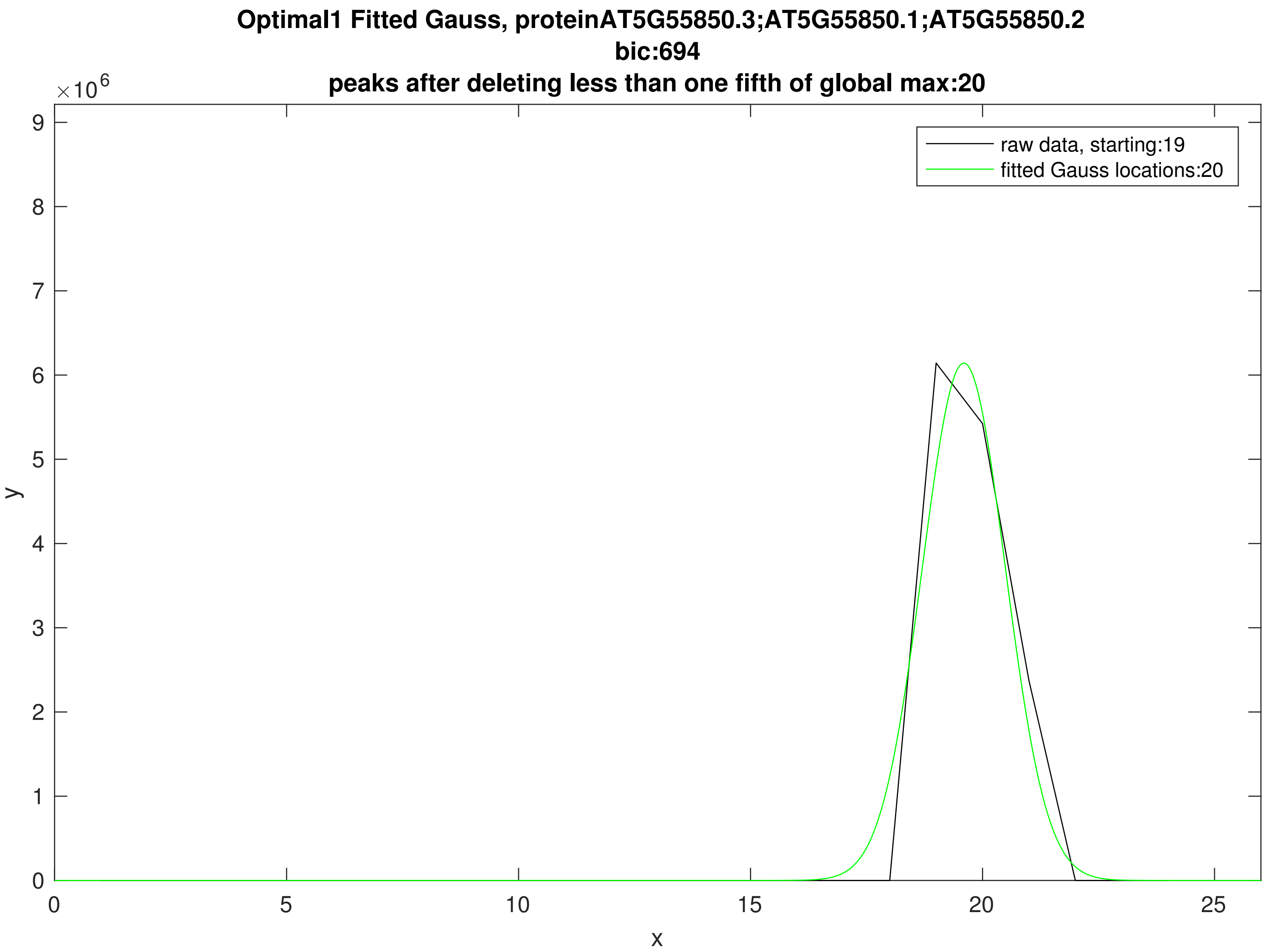
Optimal1 Fitted Gauss, proteinAT5G55850.3;AT5G55850.1;AT5G55850.2
bic:694
peaks after deleting less than one fifth of global max:20
×10^6
raw data, starting:19
fitted Gauss locations:20
y
x
0
1
2
3
4
5
6
7
8
9
0
5
10
15
20
25

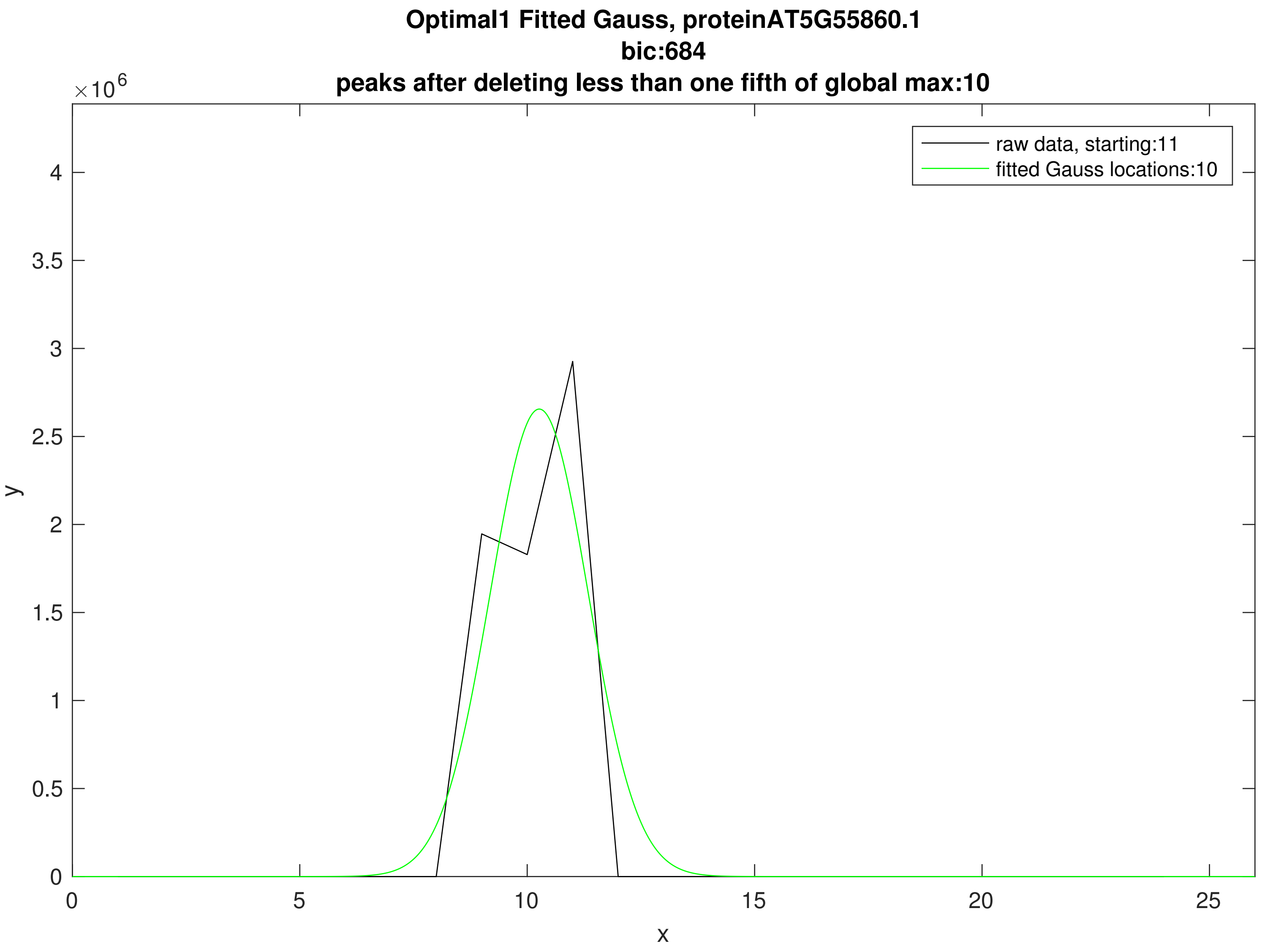
Optimal1 Fitted Gauss, proteinAT5G55860.1
bic:684
peaks after deleting less than one fifth of global max:10
×10^6
raw data, starting:11
fitted Gauss locations:10
y
x
0
0.5
1
1.5
2
2.5
3
3.5
4
0
5
10
15
20
25

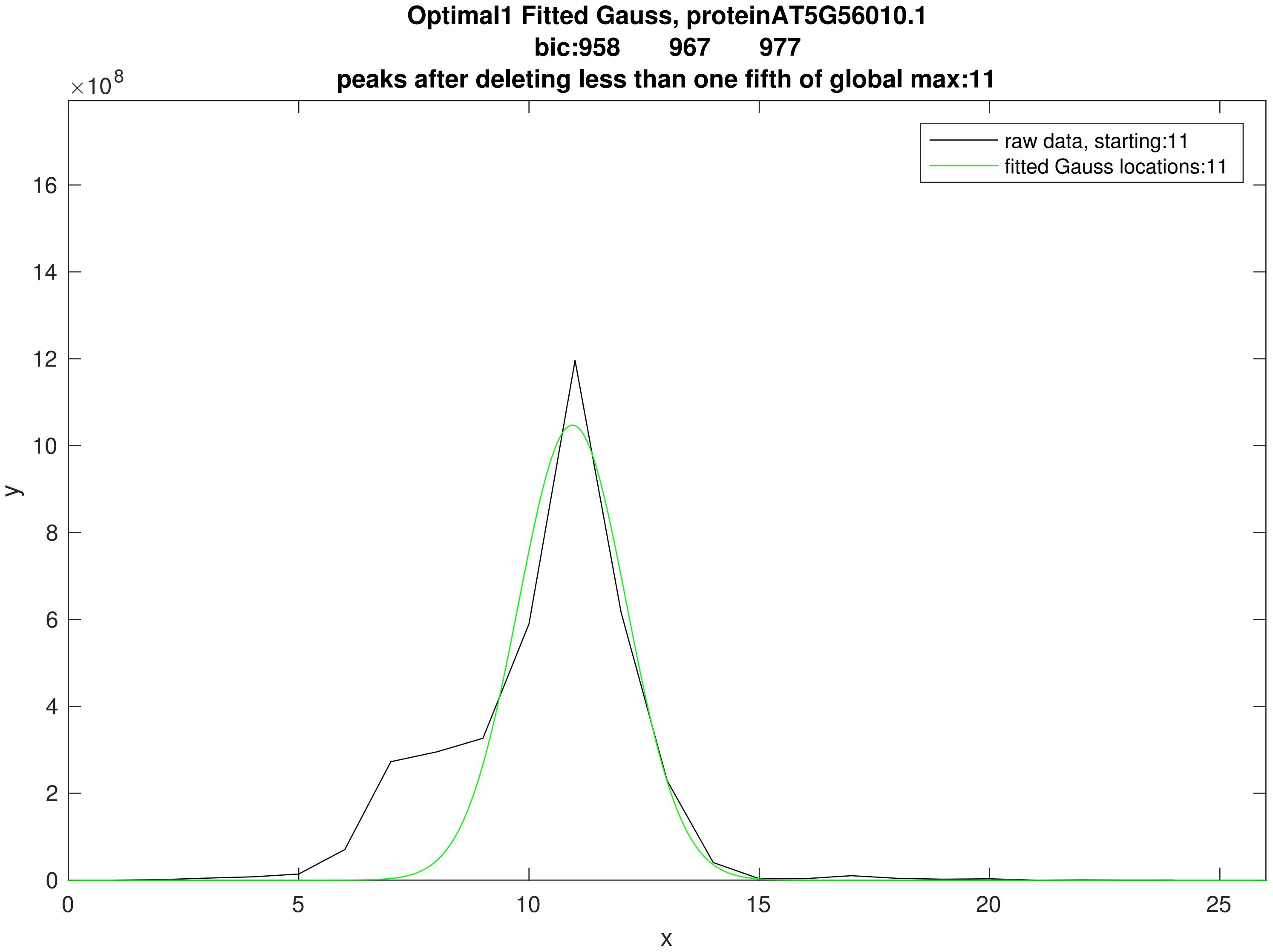
Optimal1 Fitted Gauss, proteinAT5G56010.1
bic:958 967 977
peaks after deleting less than one fifth of global max:11
×10^8
raw data, starting:11
fitted Gauss locations:11
y
x
0
2
4
6
8
10
12
14
16
0
5
10
15
20
25

**Optimal1 Fitted Gauss, proteinAT5G56030.1;AT5G56030.2**
**bic:779**
**peaks after deleting less than one fifth of global max:9**

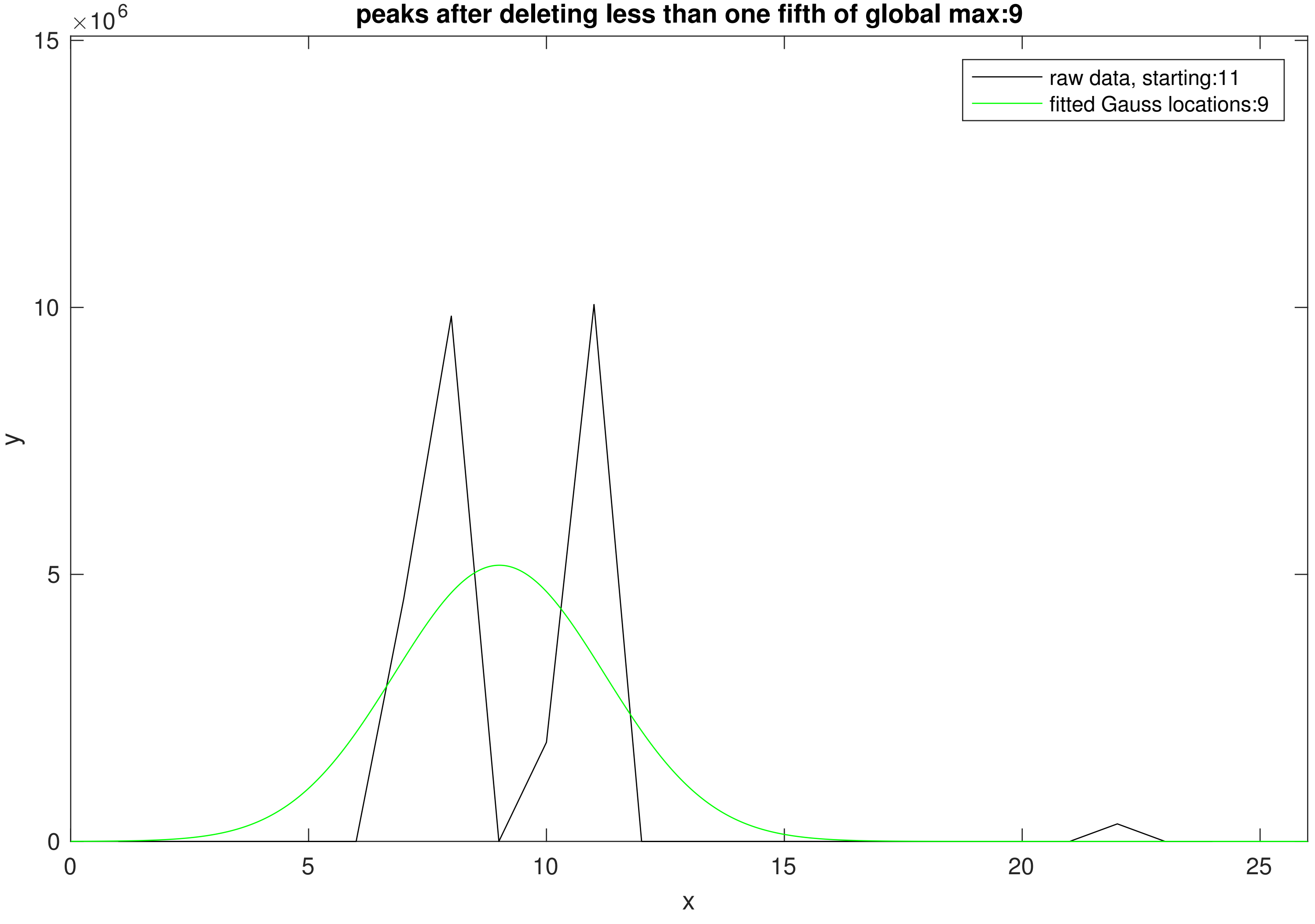

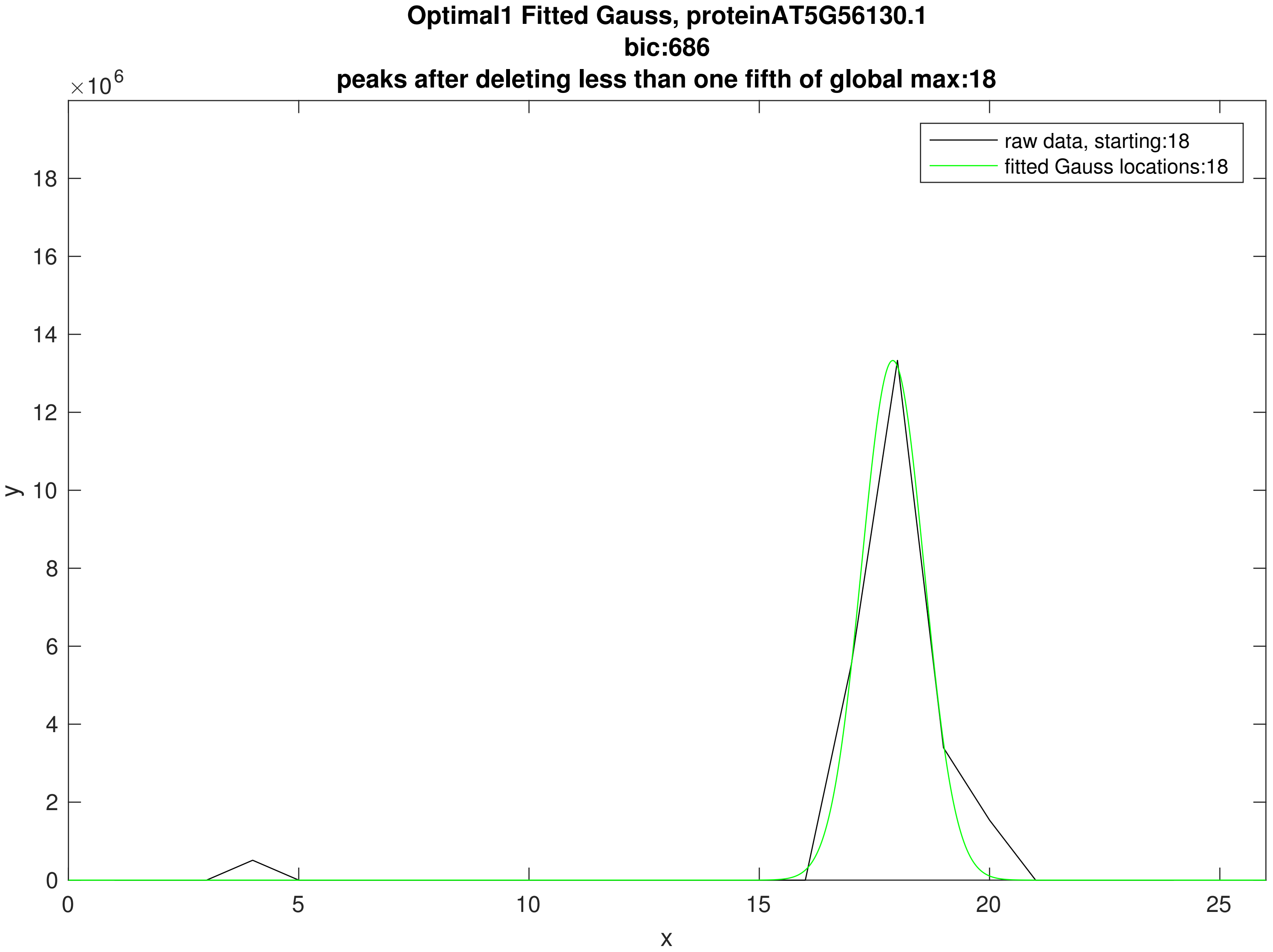
Optimal1 Fitted Gauss, proteinAT5G56130.1
bic:686
peaks after deleting less than one fifth of global max:18
×10^6
raw data, starting:18
fitted Gauss locations:18
y
x
0
2
4
6
8
10
12
14
16
18
0
5
10
15
20
25

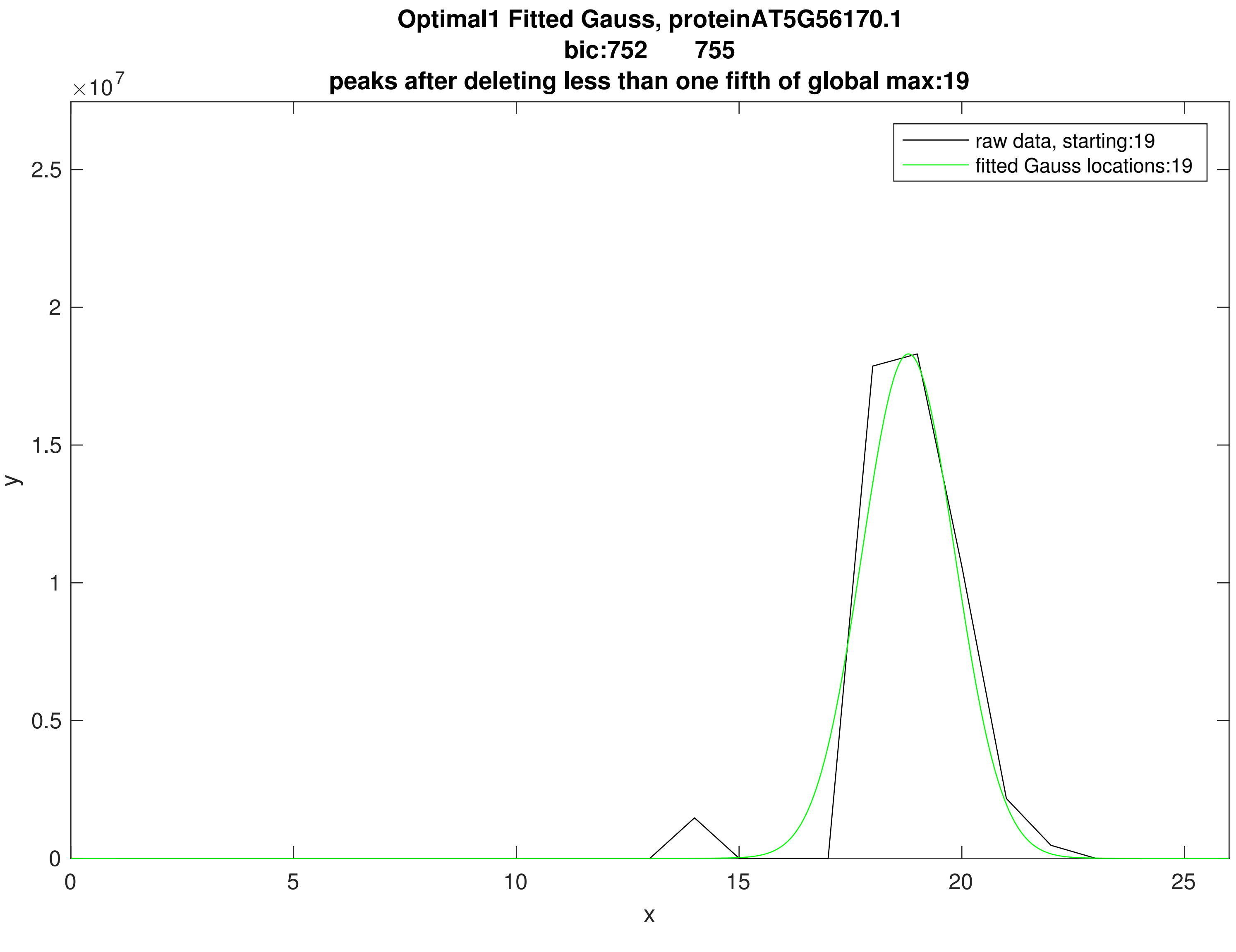

Optimal1 Fitted Gauss, proteinAT5G56170.1
bic:752 755
peaks after deleting less than one fifth of global max:19
×10^7
2.5
2
1.5
1
0.5
0
y
0
5
10
15
20
25
x
raw data, starting:19
fitted Gauss locations:19

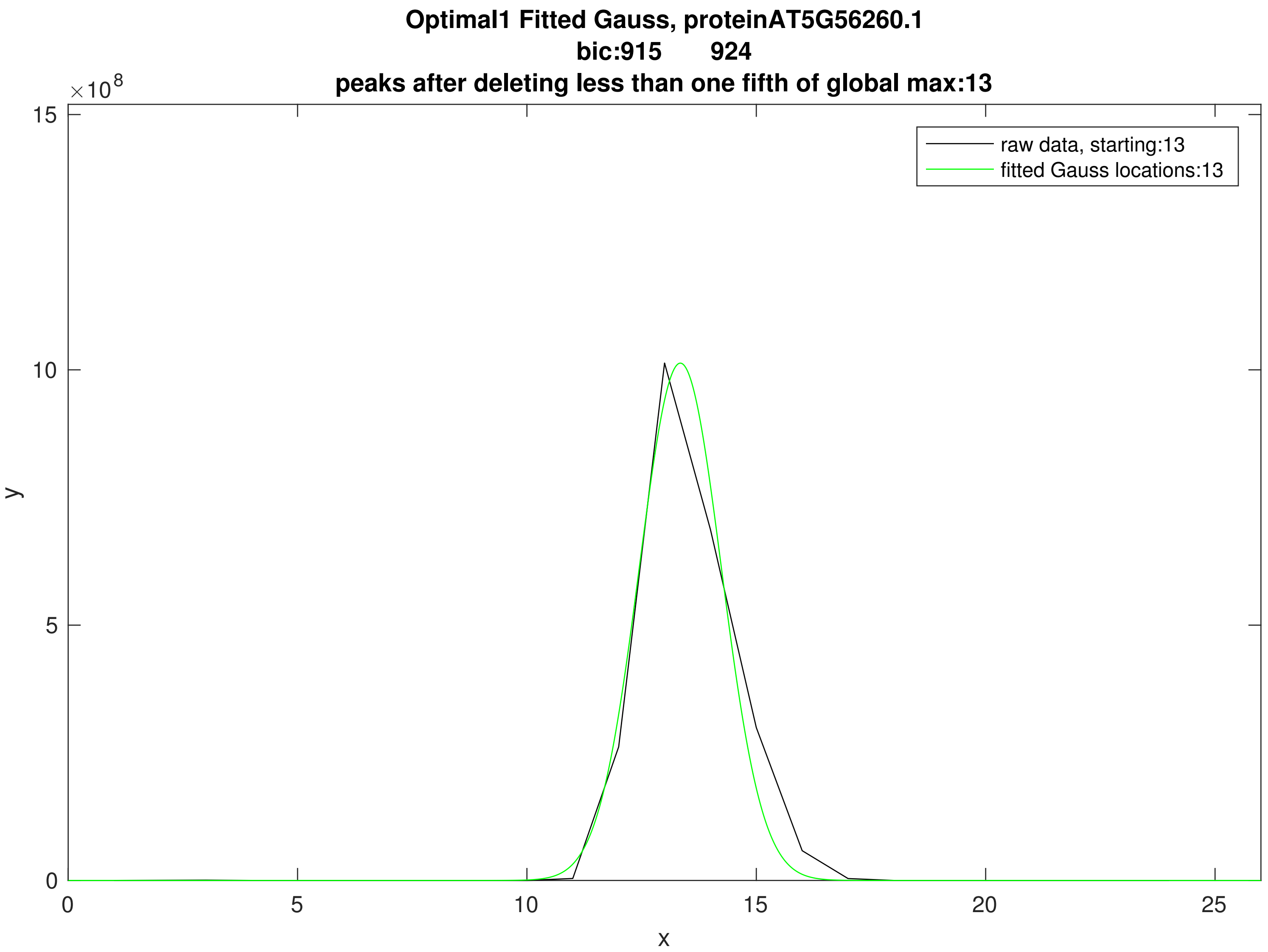

Optimal1 Fitted Gauss, proteinAT5G56260.1
bic:915 924
peaks after deleting less than one fifth of global max:13
×10^8
15
10
5
0
0
5
10
15
20
25
x
y
raw data, starting:13
fitted Gauss locations:13

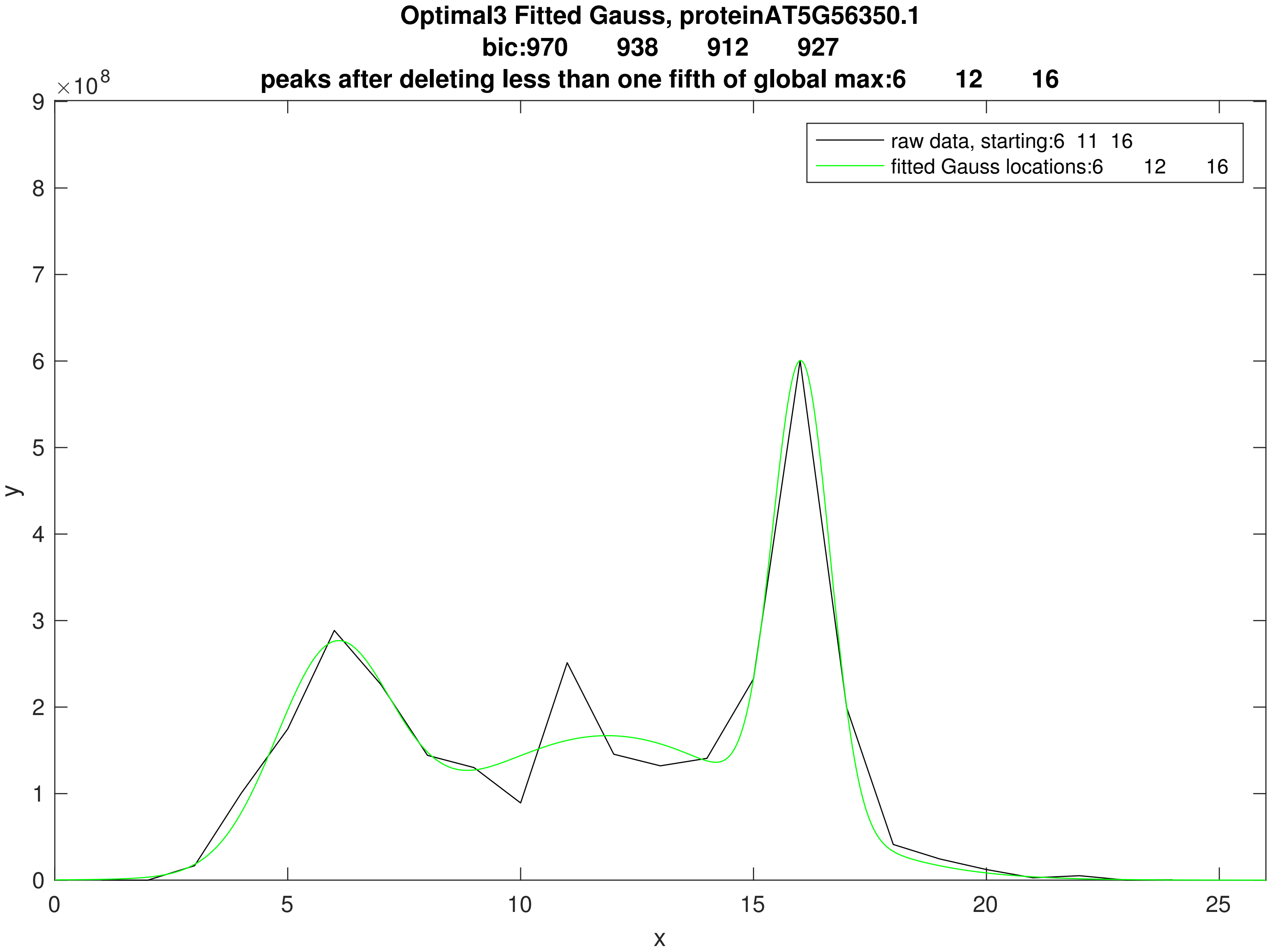
Optimal3 Fitted Gauss, proteinAT5G56350.1
bic:970 938 912 927
peaks after deleting less than one fifth of global max:6 12 16
raw data, starting:6 11 16
fitted Gauss locations:6 12 16
×10^8
y
x
0
1
2
3
4
5
6
7
8
9
0
5
10
15
20
25

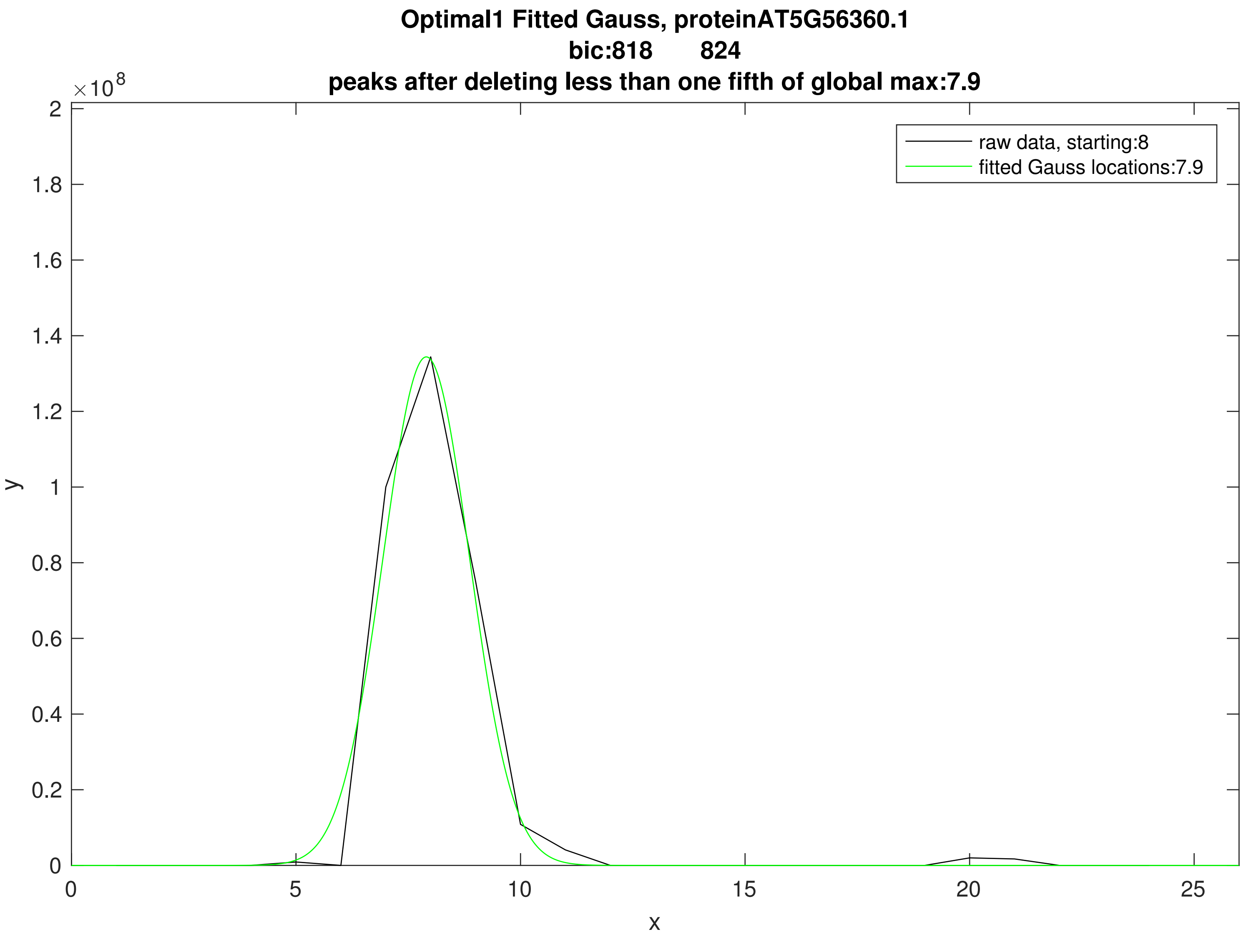
Optimal1 Fitted Gauss, proteinAT5G56360.1
bic:818 824
peaks after deleting less than one fifth of global max:7.9
×10^8
raw data, starting:8
fitted Gauss locations:7.9
y
x
0
0.2
0.4
0.6
0.8
1
1.2
1.4
1.6
1.8
2
0
5
10
15
20
25

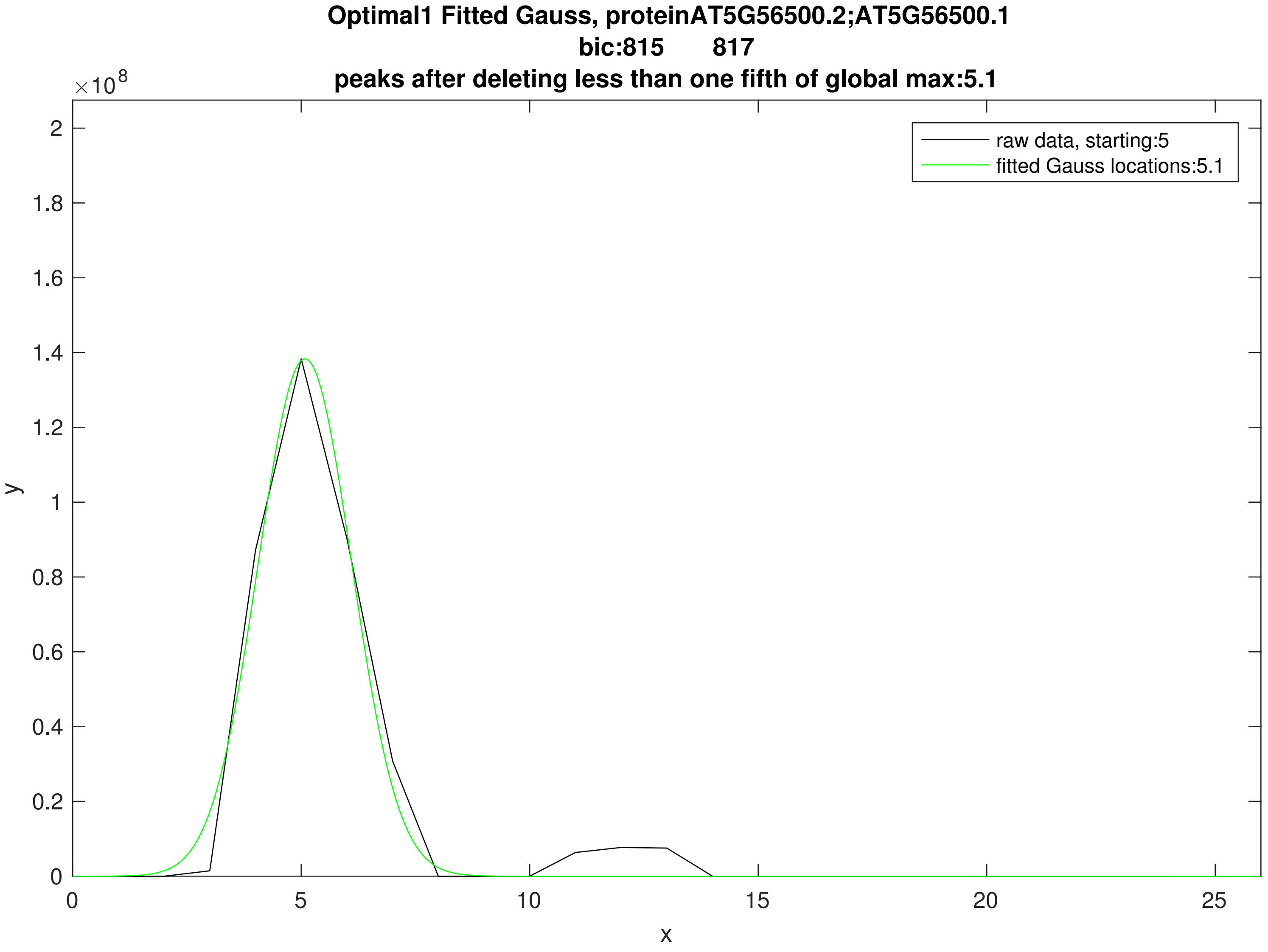
Optimal1 Fitted Gauss, proteinAT5G56500.2;AT5G56500.1
bic:815 817
peaks after deleting less than one fifth of global max:5.1
×10^8
2
1.8
1.6
1.4
1.2
1
0.8
0.6
0.4
0.2
0
y
0
5
10
15
20
25
x
raw data, starting:5
fitted Gauss locations:5.1

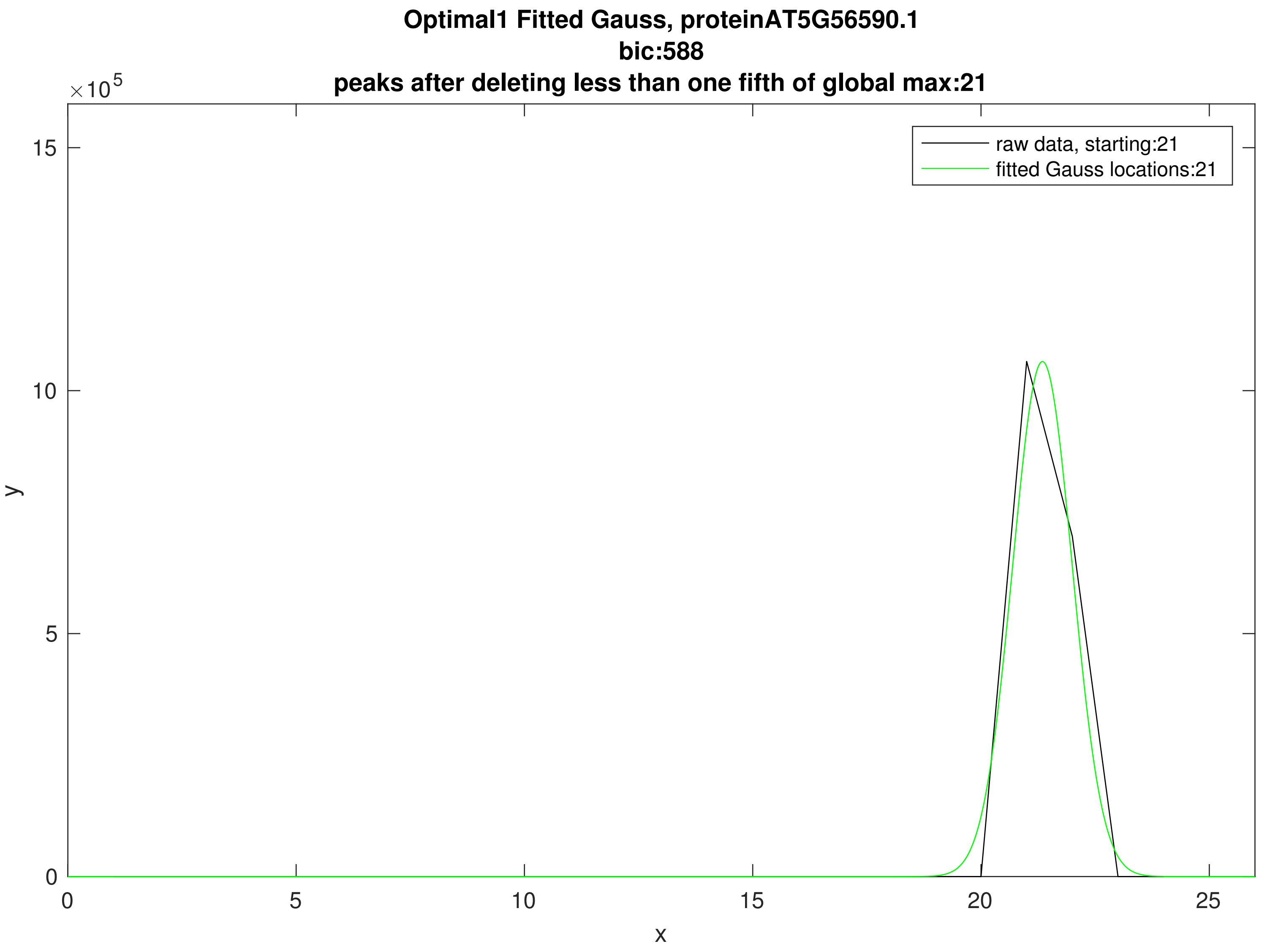
Optimal1 Fitted Gauss, proteinAT5G56590.1
bic:588
peaks after deleting less than one fifth of global max:21
×10^5
raw data, starting:21
fitted Gauss locations:21
15
10
5
0
0
5
10
15
20
25
y
x

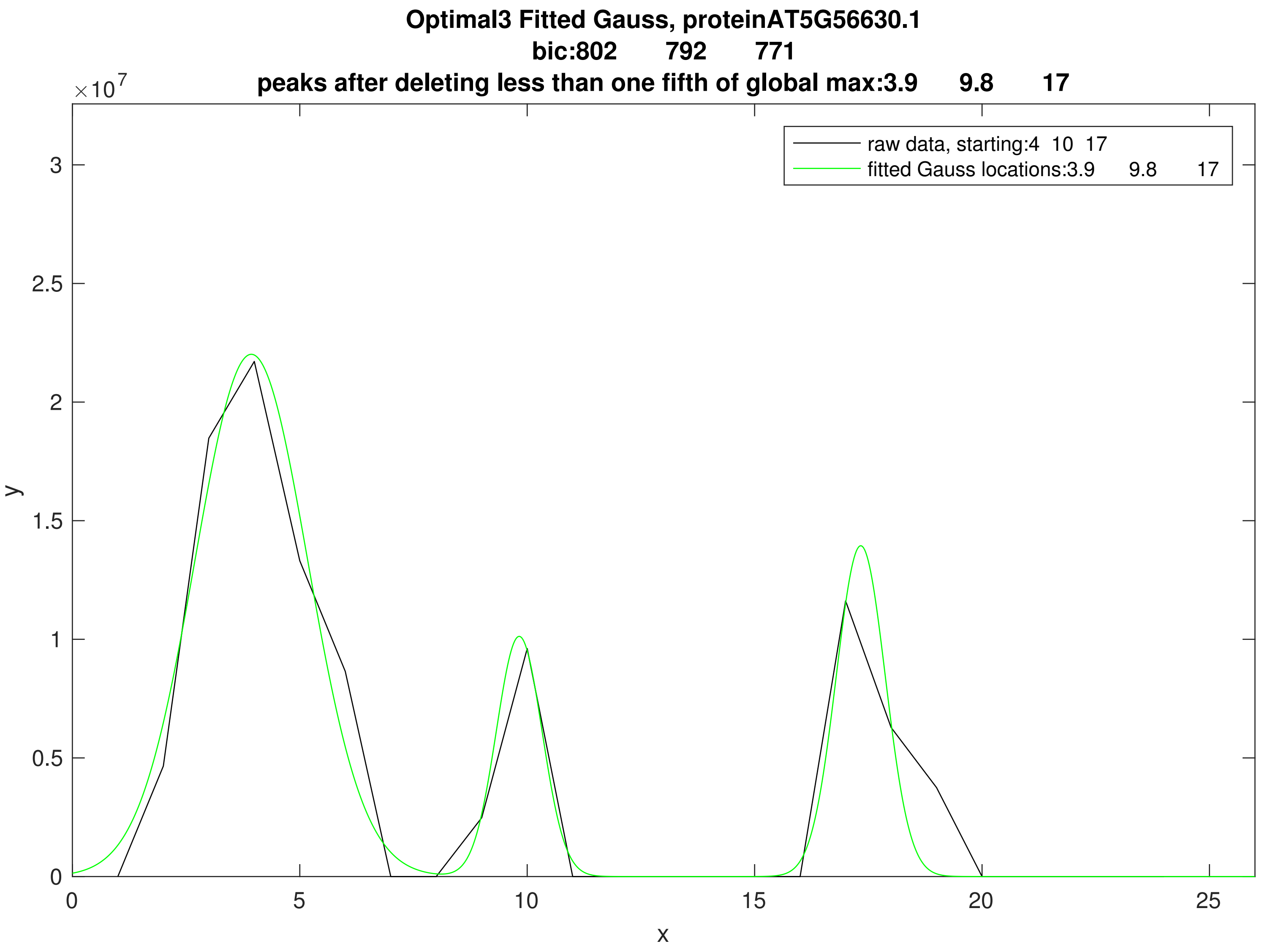

Optimal3 Fitted Gauss, proteinAT5G56630.1
bic:802 792 771
peaks after deleting less than one fifth of global max:3.9 9.8 17
×10^7
raw data, starting:4 10 17
fitted Gauss locations:3.9 9.8 17
y
x
0
0.5
1
1.5
2
2.5
3
5
10
15
20
25

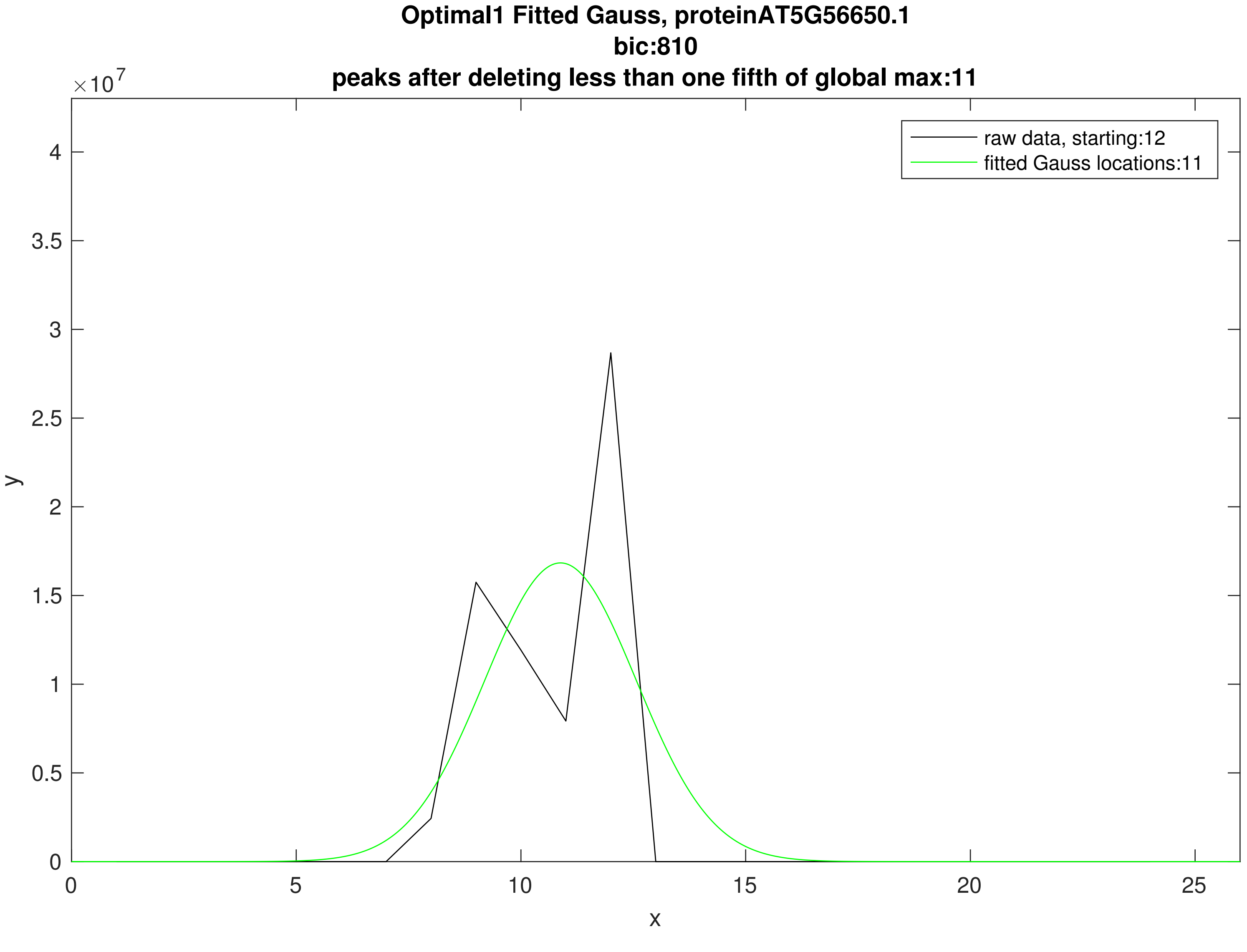

Optimal1 Fitted Gauss, proteinAT5G56650.1
bic:810
peaks after deleting less than one fifth of global max:11
×10^7
raw data, starting:12
fitted Gauss locations:11
y
x
0
0.5
1
1.5
2
2.5
3
3.5
4
0
5
10
15
20
25

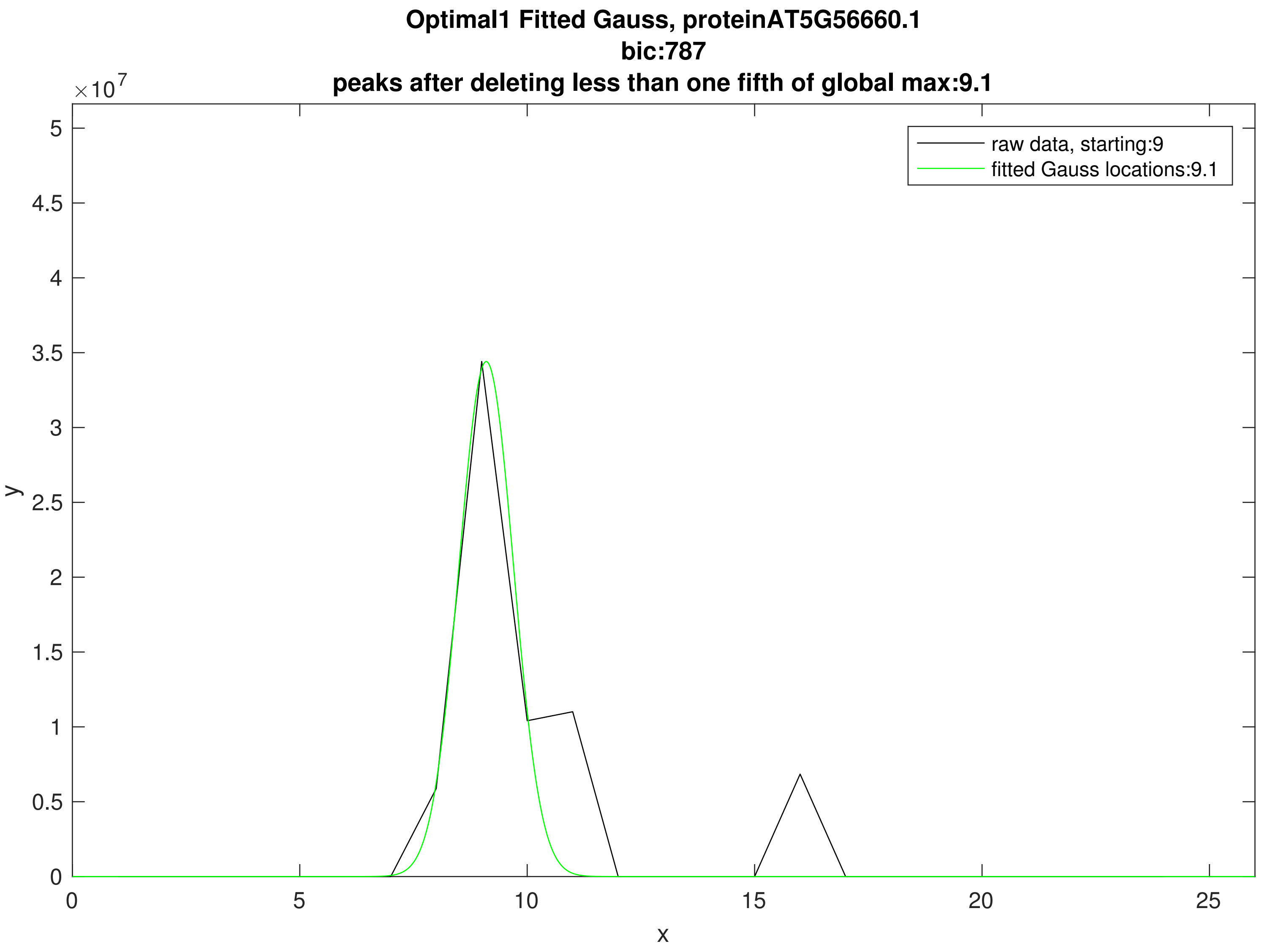
Optimal1 Fitted Gauss, proteinAT5G56660.1
bic:787
peaks after deleting less than one fifth of global max:9.1
×10^7
raw data, starting:9
fitted Gauss locations:9.1
y
x
0
0.5
1
1.5
2
2.5
3
3.5
4
4.5
5
0
5
10
15
20
25

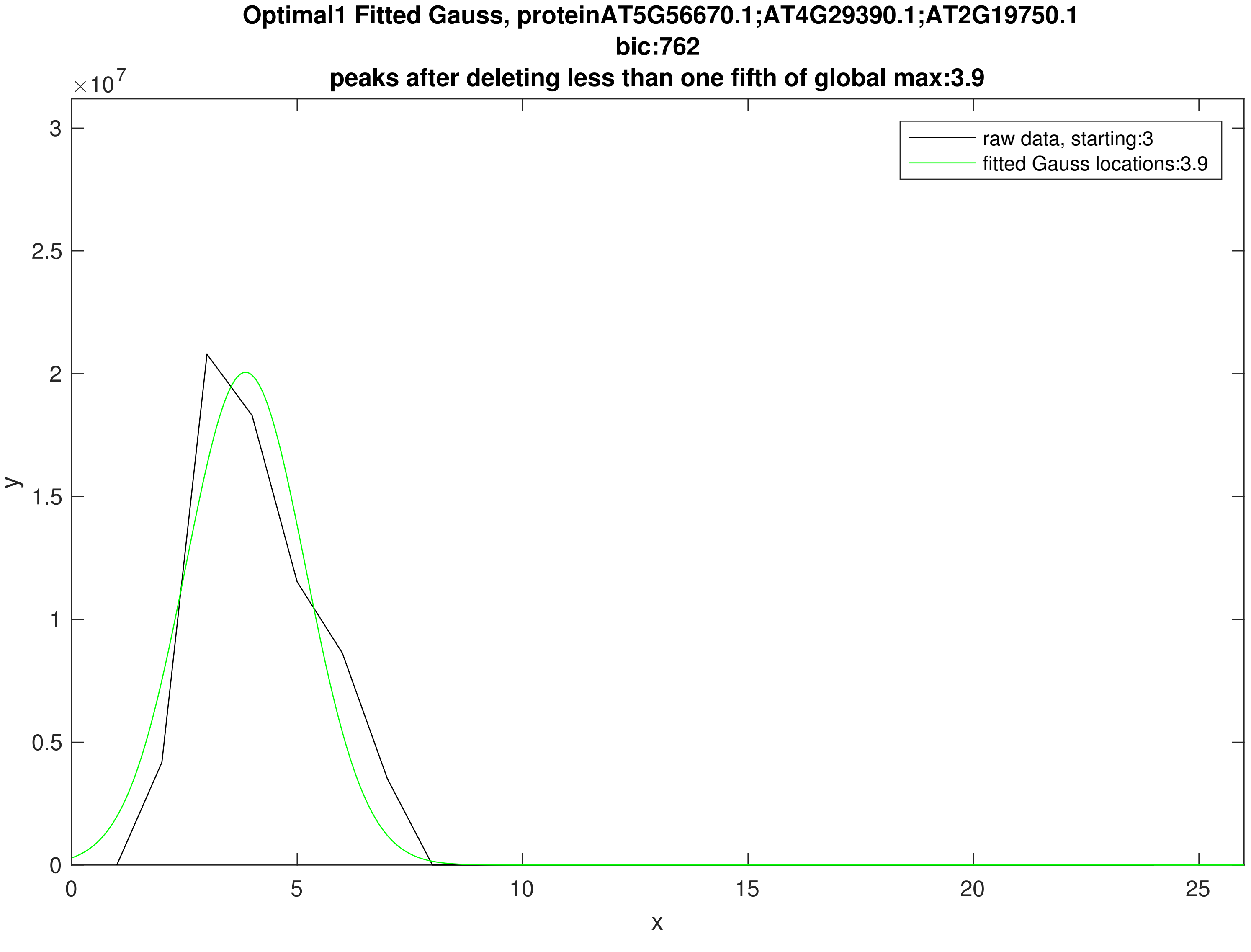

Optimal1 Fitted Gauss, proteinAT5G56670.1;AT4G29390.1;AT2G19750.1
bic:762
peaks after deleting less than one fifth of global max:3.9
×10^7
raw data, starting:3
fitted Gauss locations:3.9
y
x
0
0.5
1
1.5
2
2.5
3
5
10
15
20
25

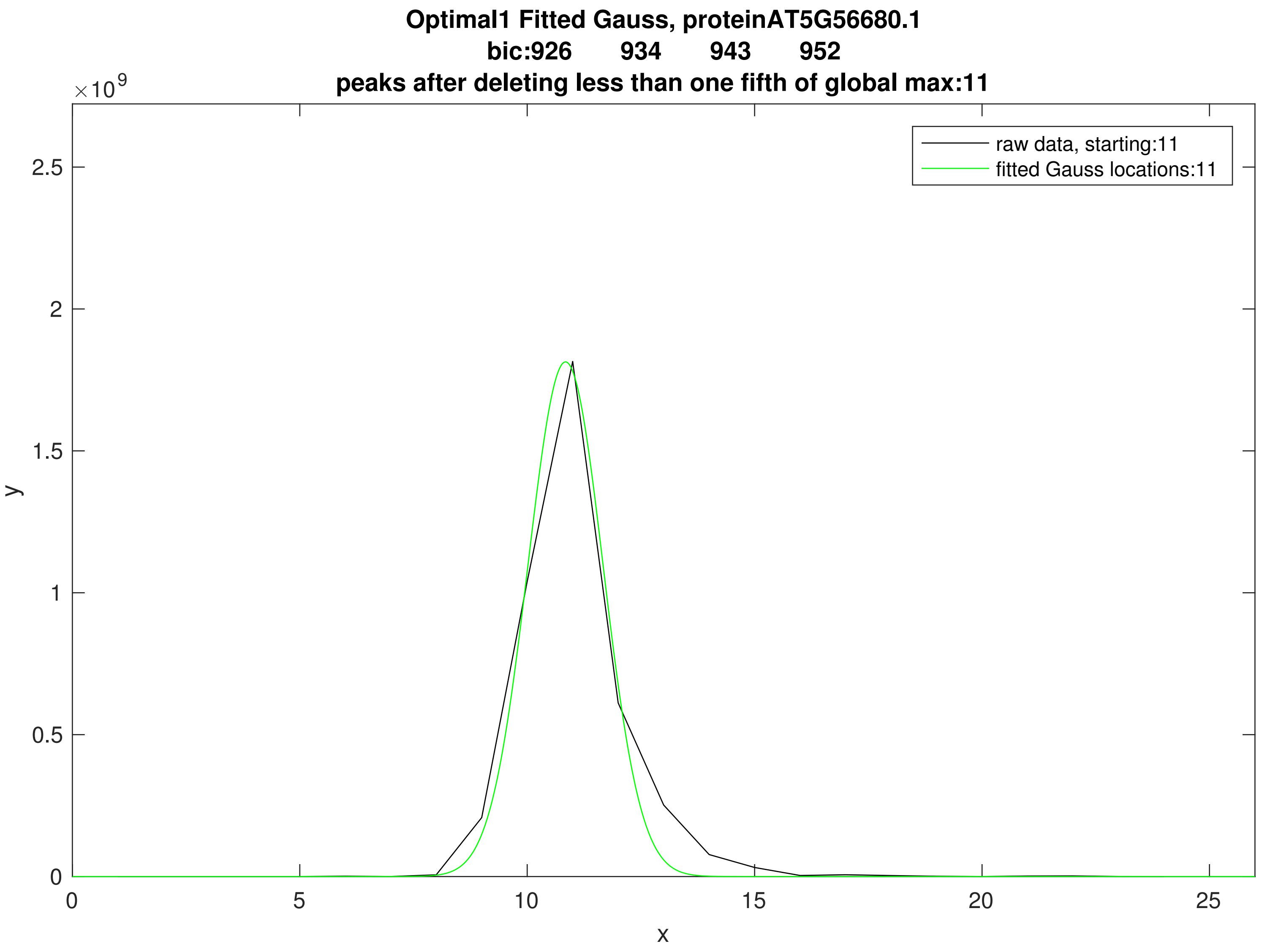
Optimal1 Fitted Gauss, proteinAT5G56680.1
bic:926 934 943 952
peaks after deleting less than one fifth of global max:11
×10^9
raw data, starting:11
fitted Gauss locations:11
y
x
0
0.5
1
1.5
2
2.5
0
5
10
15
20
25

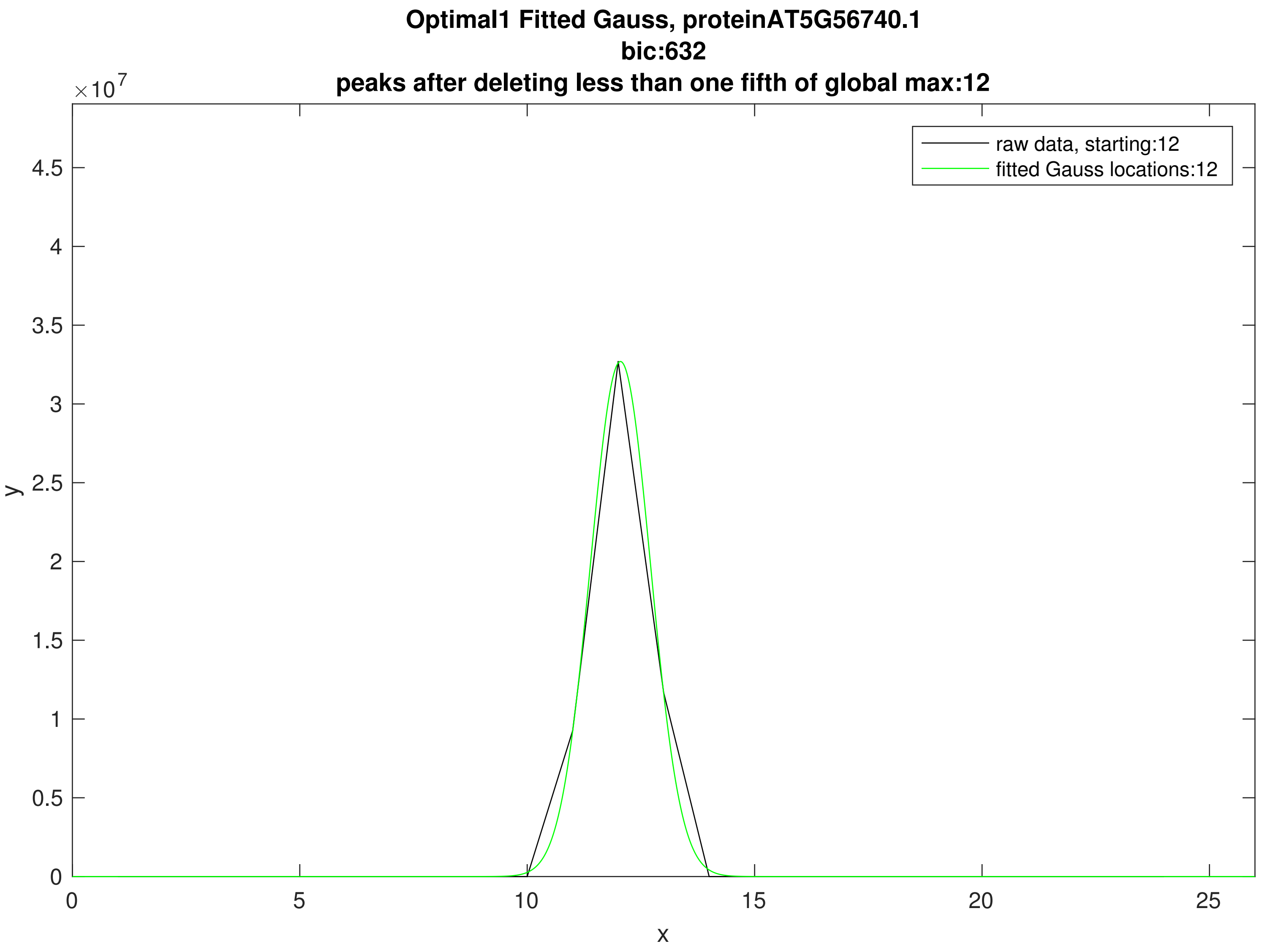
Optimal1 Fitted Gauss, proteinAT5G56740.1
bic:632
peaks after deleting less than one fifth of global max:12
×10^7
y
x
raw data, starting:12
fitted Gauss locations:12
0
0.5
1
1.5
2
2.5
3
3.5
4
4.5
0
5
10
15
20
25

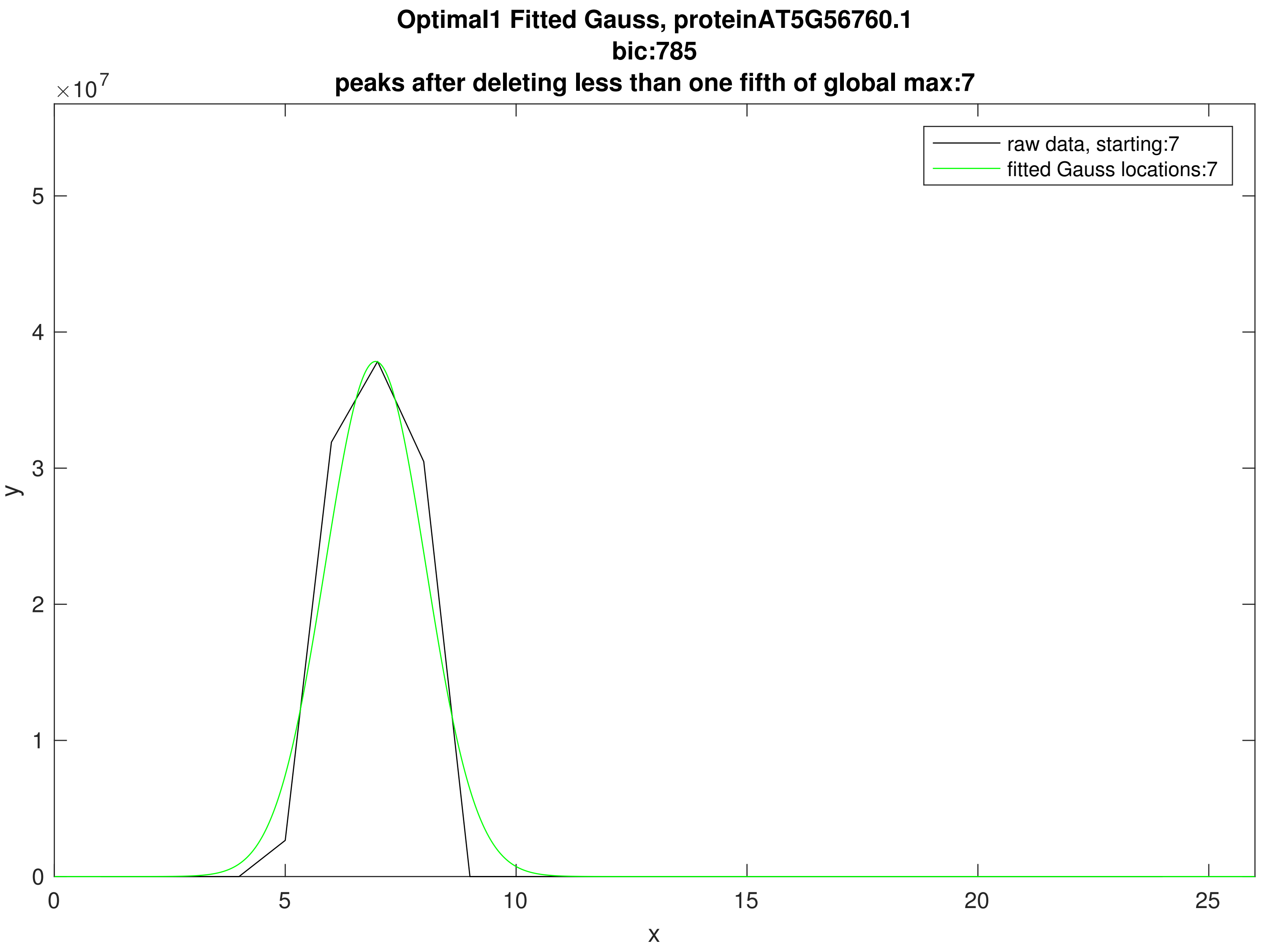
Optimal1 Fitted Gauss, proteinAT5G56760.1
bic:785
peaks after deleting less than one fifth of global max:7
×10^7
raw data, starting:7
fitted Gauss locations:7
y
x
0
1
2
3
4
5
0
5
10
15
20
25

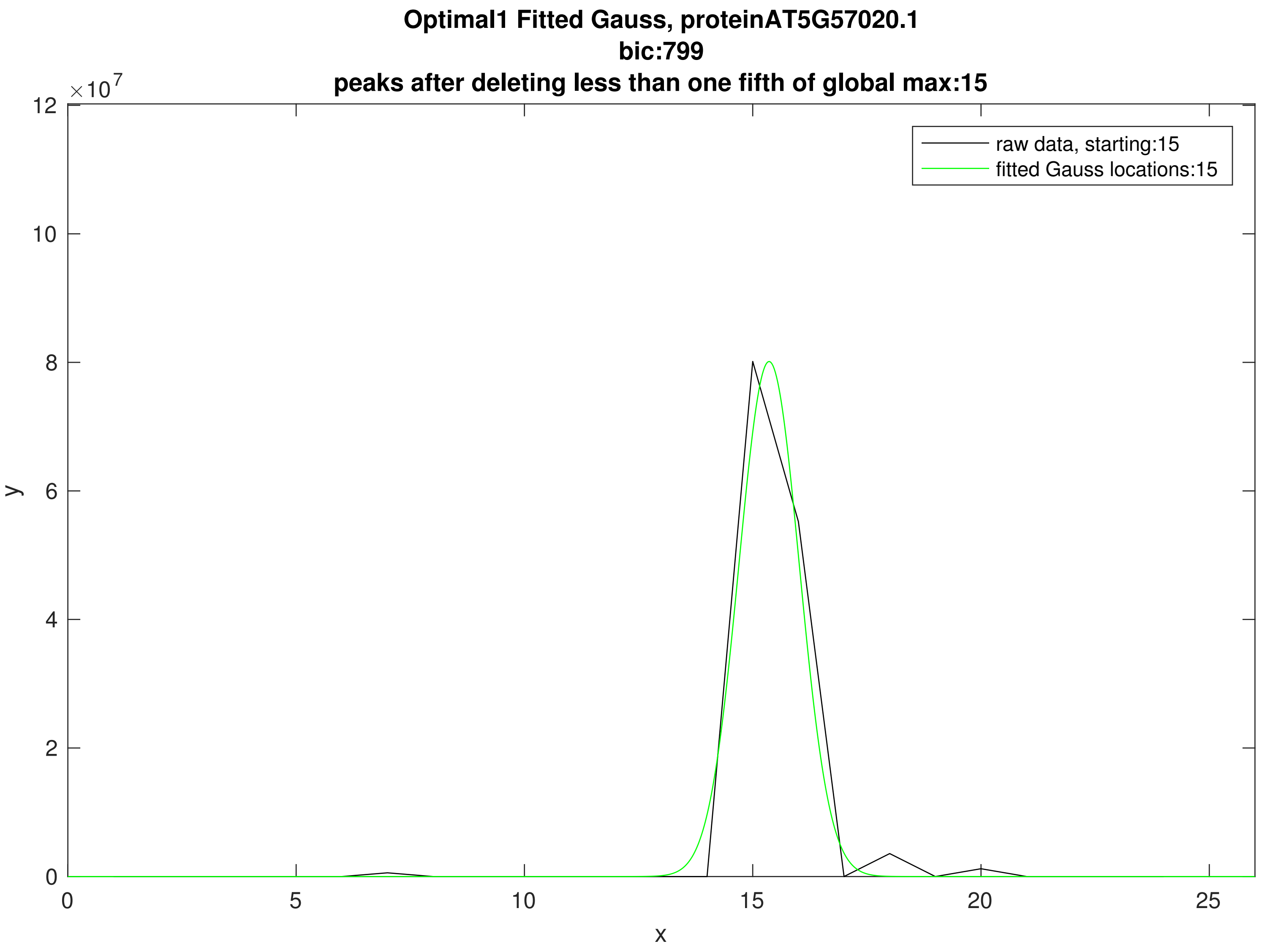
Optimal1 Fitted Gauss, proteinAT5G57020.1
bic:799
peaks after deleting less than one fifth of global max:15
×10^7
raw data, starting:15
fitted Gauss locations:15
y
x
0
2
4
6
8
10
12
0
5
10
15
20
25

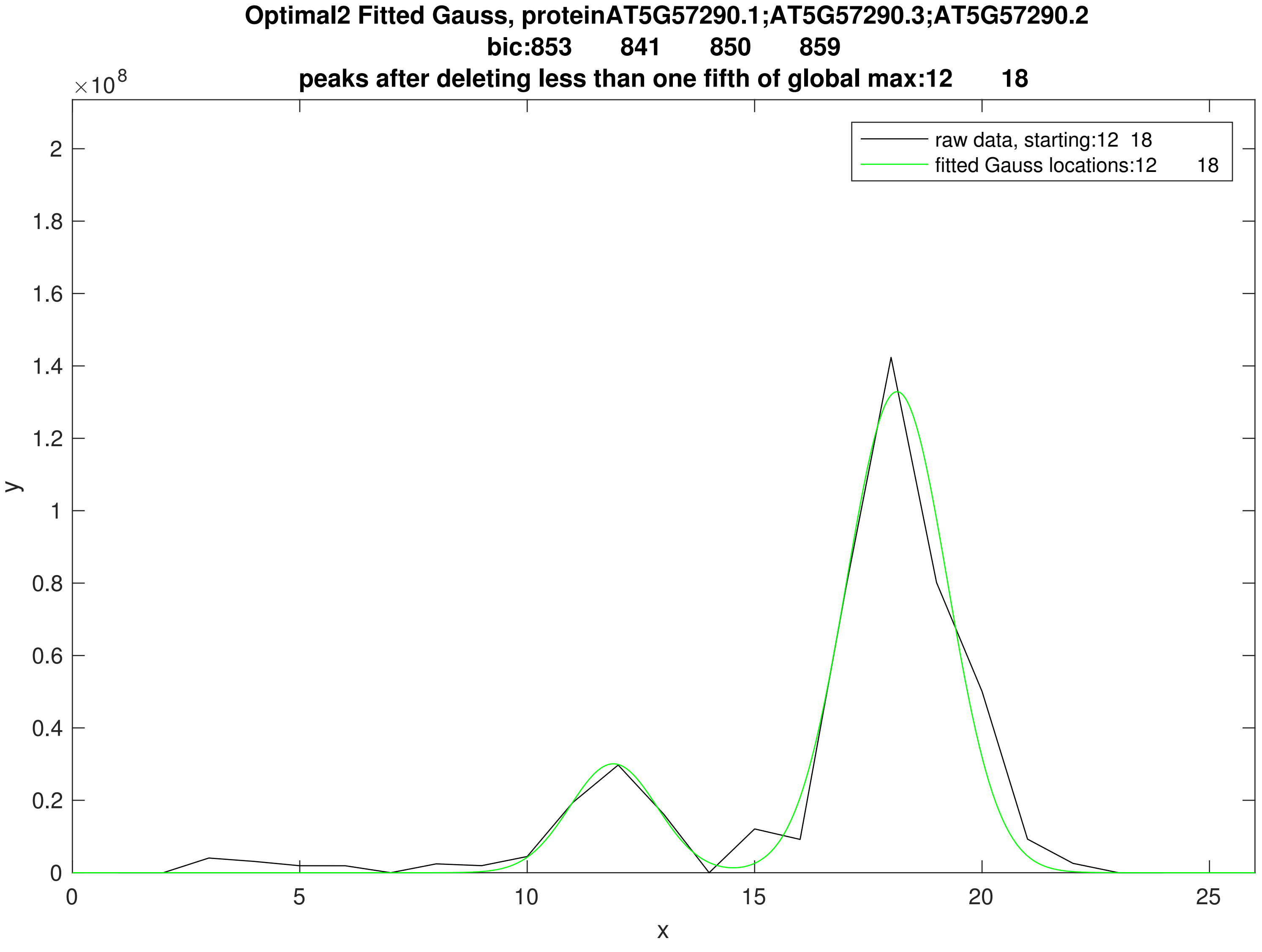

Optimal2 Fitted Gauss, proteinAT5G57290.1;AT5G57290.3;AT5G57290.2
bic:853 841 850 859
peaks after deleting less than one fifth of global max:12 18
×10^8
raw data, starting:12 18
fitted Gauss locations:12 18
y
x
0
0.2
0.4
0.6
0.8
1
1.2
1.4
1.6
1.8
2
0
5
10
15
20
25

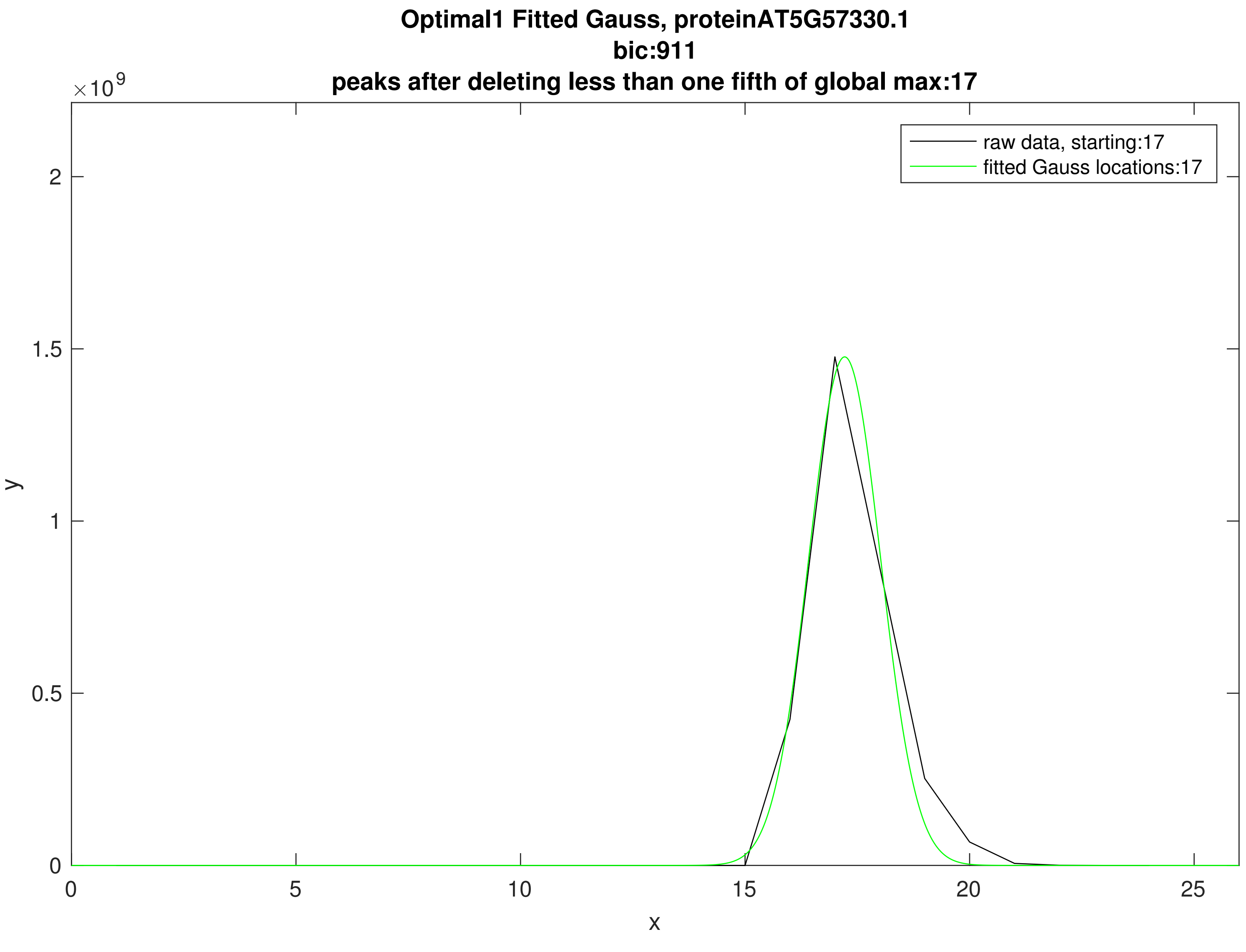
Optimal1 Fitted Gauss, proteinAT5G57330.1
bic:911
peaks after deleting less than one fifth of global max:17
×10^9
raw data, starting:17
fitted Gauss locations:17
2
1.5
1
0.5
0
y
0
5
10
15
20
25
x

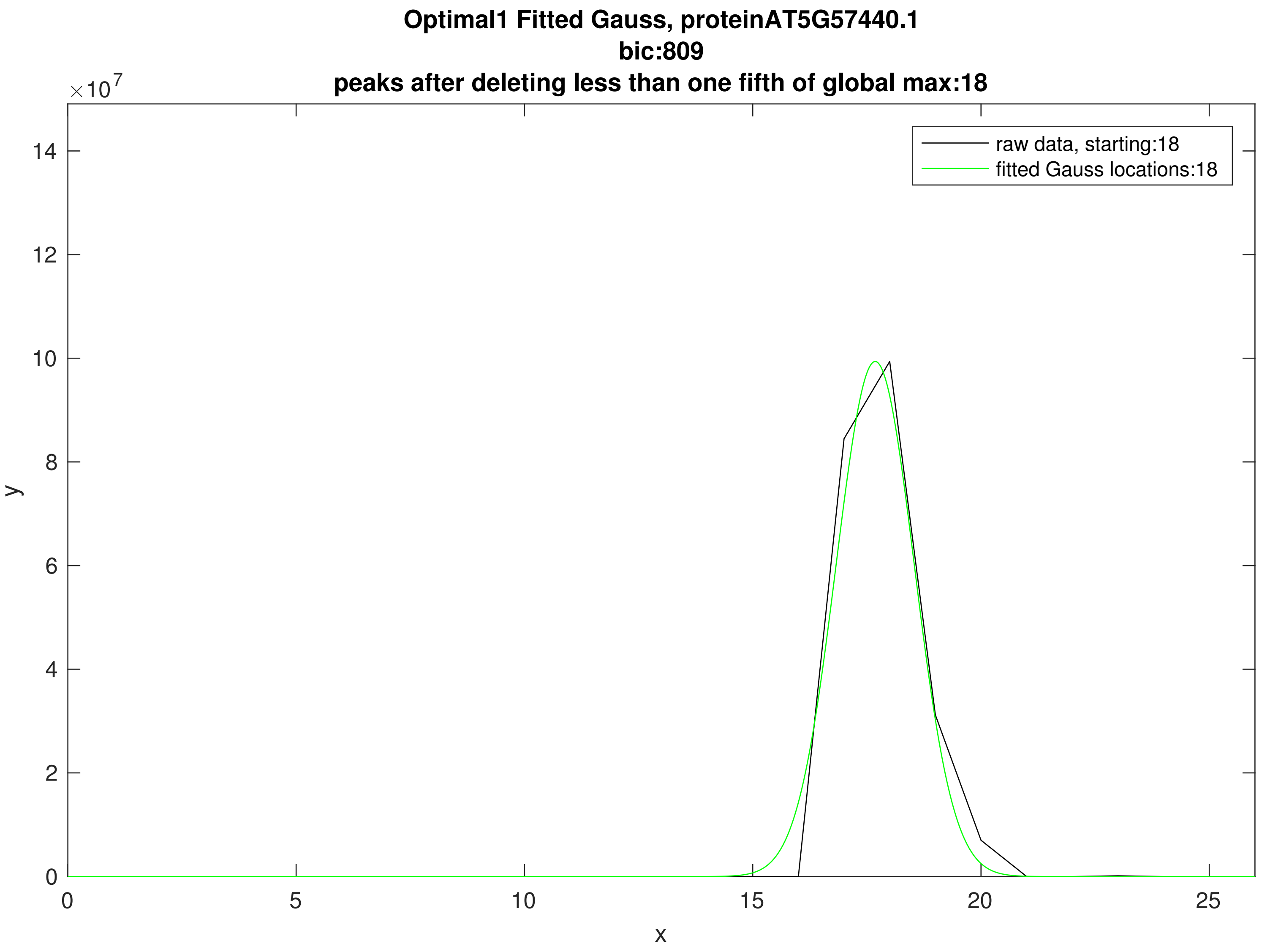
Optimal1 Fitted Gauss, proteinAT5G57440.1
bic:809
peaks after deleting less than one fifth of global max:18
×10^7
raw data, starting:18
fitted Gauss locations:18
y
x
0
2
4
6
8
10
12
14
0
5
10
15
20
25

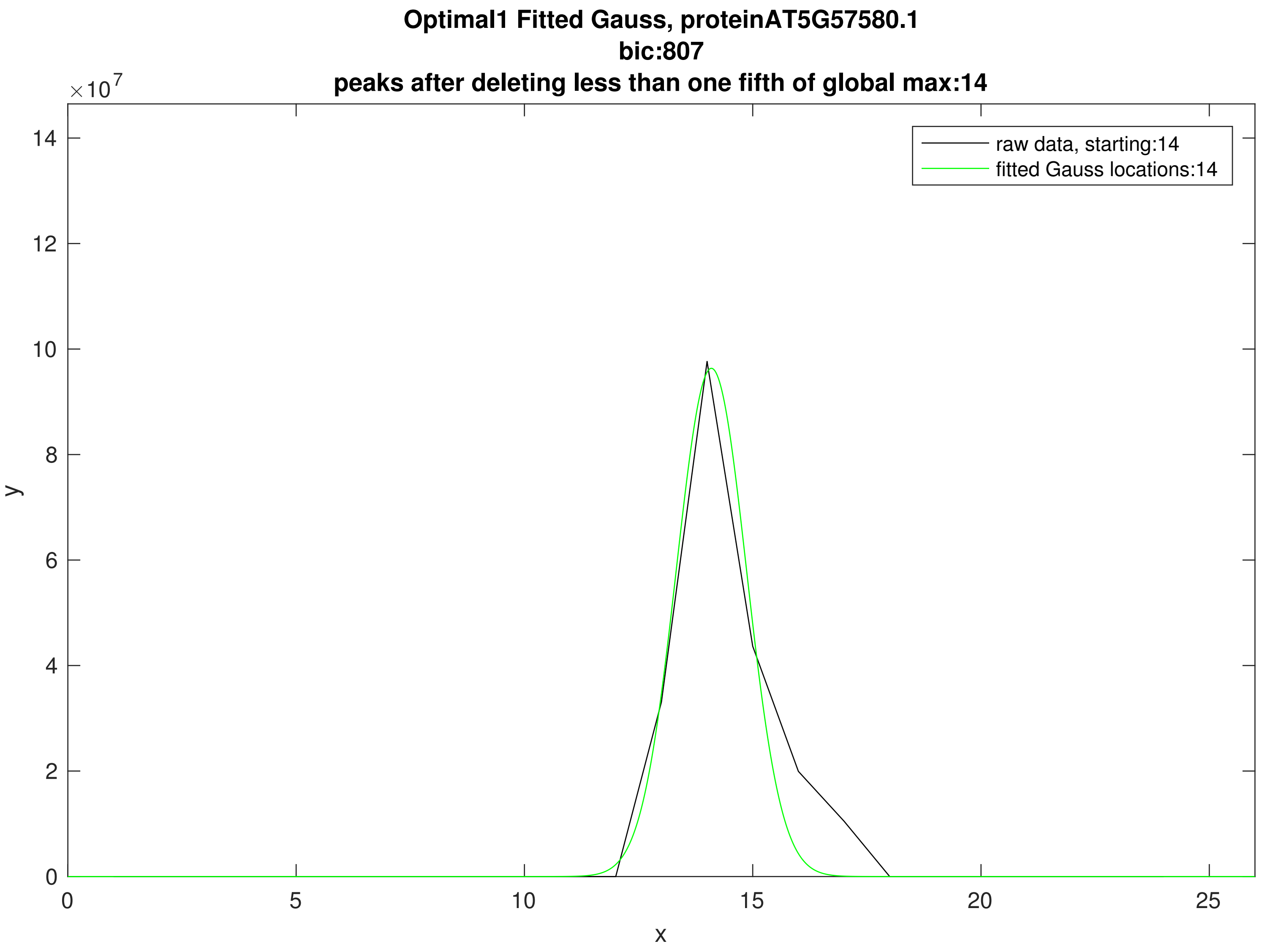
Optimal1 Fitted Gauss, proteinAT5G57580.1
bic:807
peaks after deleting less than one fifth of global max:14
×10^7
raw data, starting:14
fitted Gauss locations:14
y
x
0
2
4
6
8
10
12
14
0
5
10
15
20
25

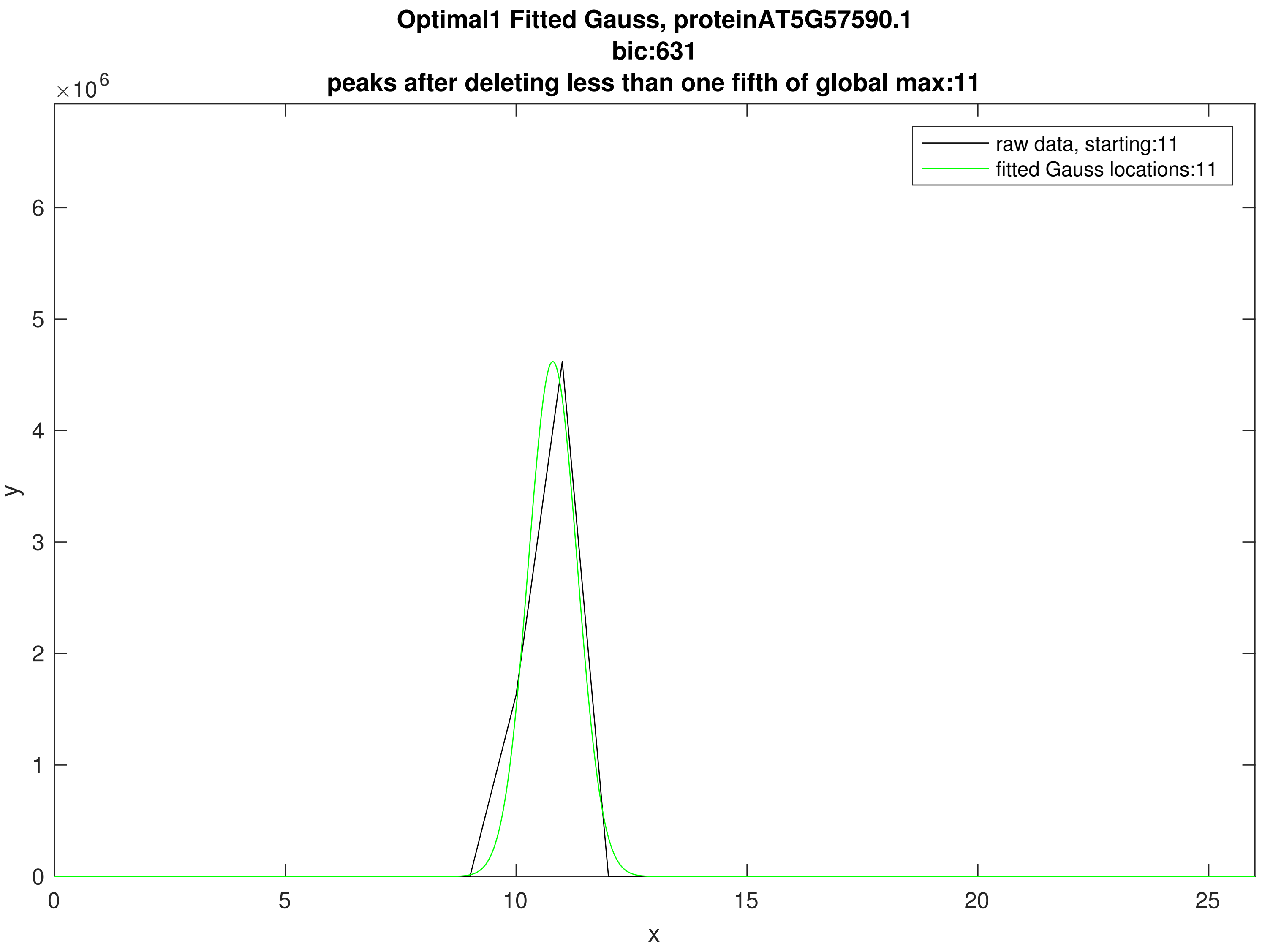

Optimal1 Fitted Gauss, proteinAT5G57590.1
bic:631
peaks after deleting less than one fifth of global max:11
×10^6
raw data, starting:11
fitted Gauss locations:11
y
x
0
1
2
3
4
5
6
0
5
10
15
20
25

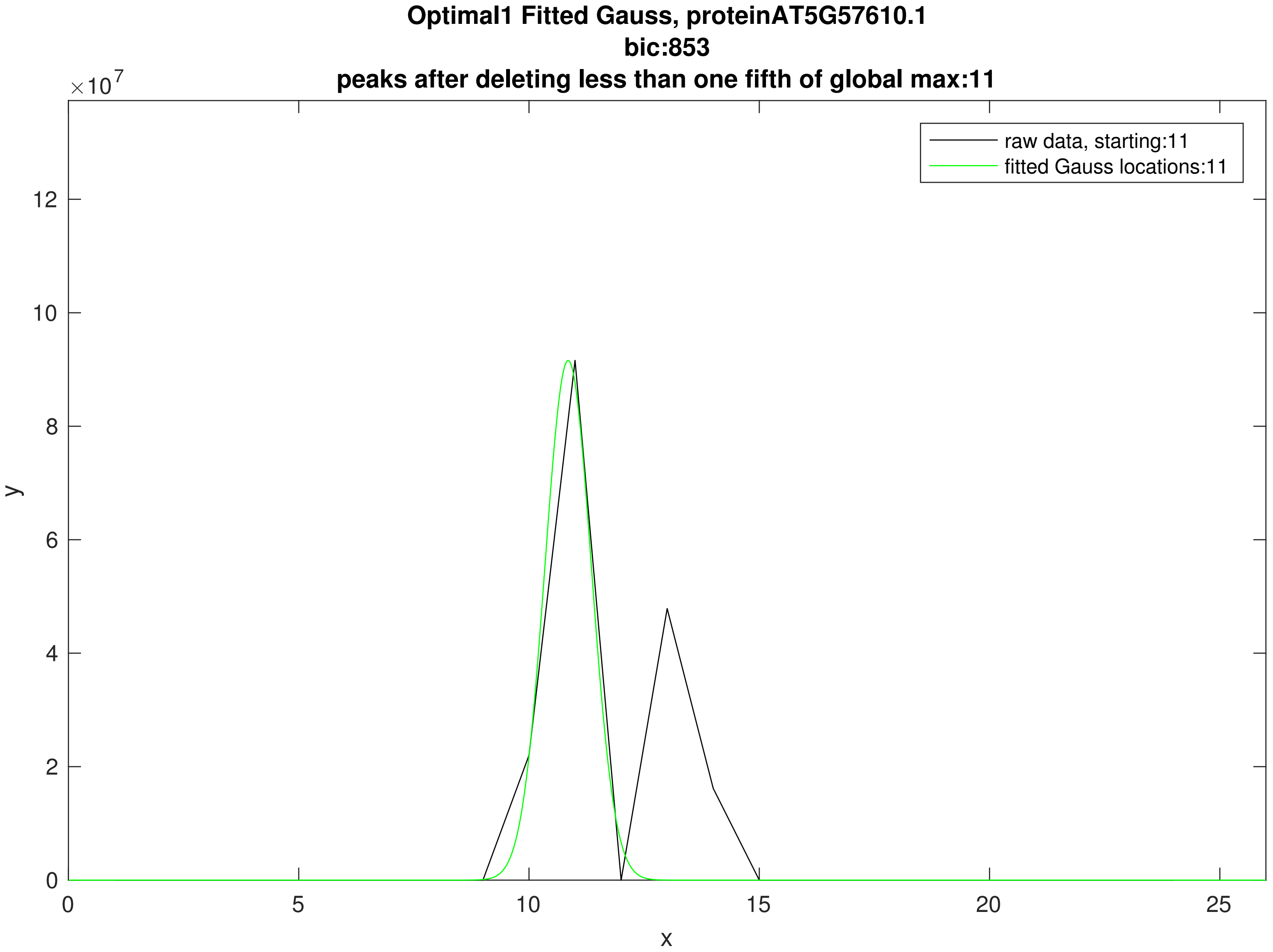

Optimal1 Fitted Gauss, proteinAT5G57610.1
bic:853
peaks after deleting less than one fifth of global max:11
×10^7
raw data, starting:11
fitted Gauss locations:11
y
x
0
2
4
6
8
10
12
0
5
10
15
20
25

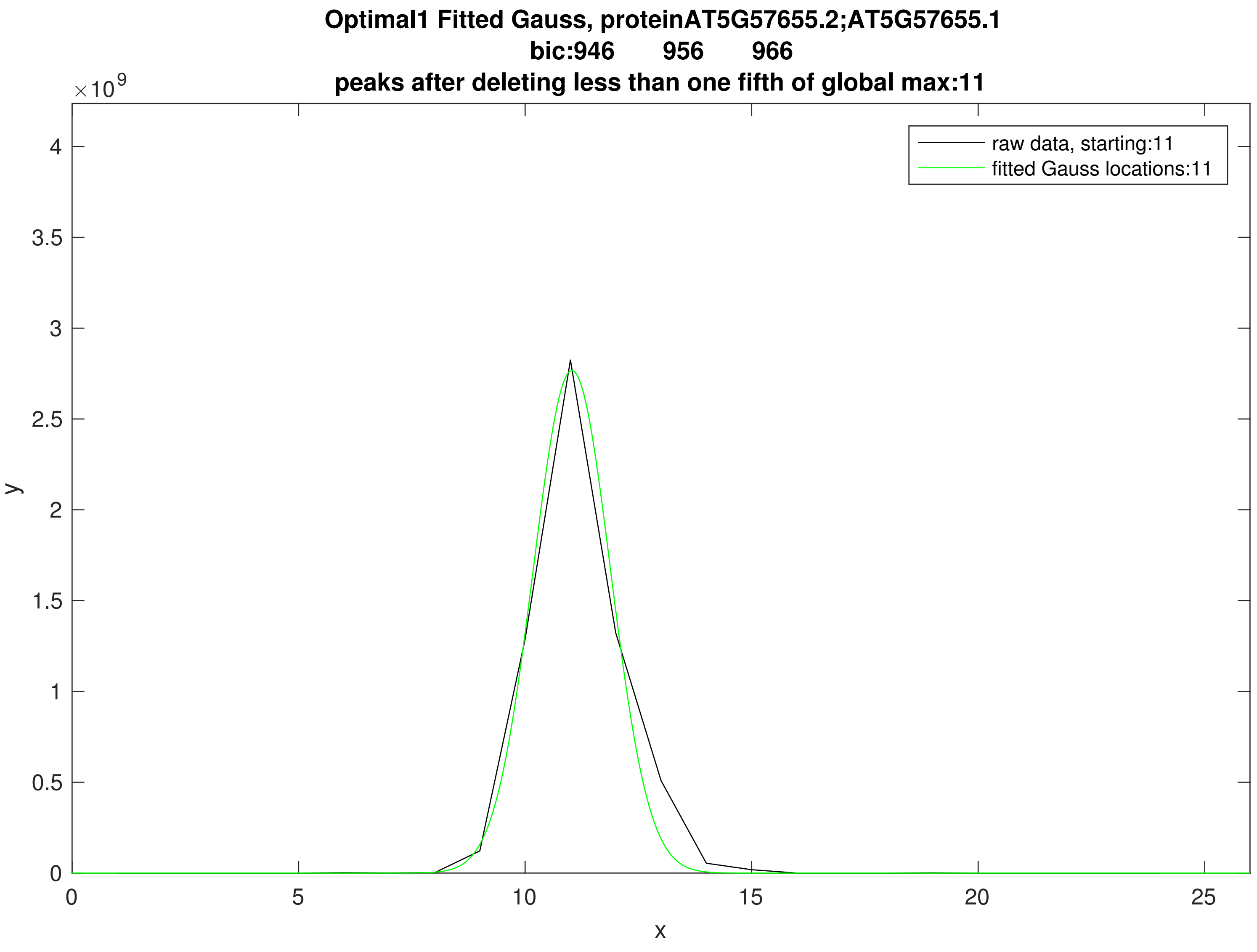
Optimal1 Fitted Gauss, proteinAT5G57655.2;AT5G57655.1
bic:946 956 966
peaks after deleting less than one fifth of global max:11
×10^9
raw data, starting:11
fitted Gauss locations:11
y
x
0
0.5
1
1.5
2
2.5
3
3.5
4
0
5
10
15
20
25

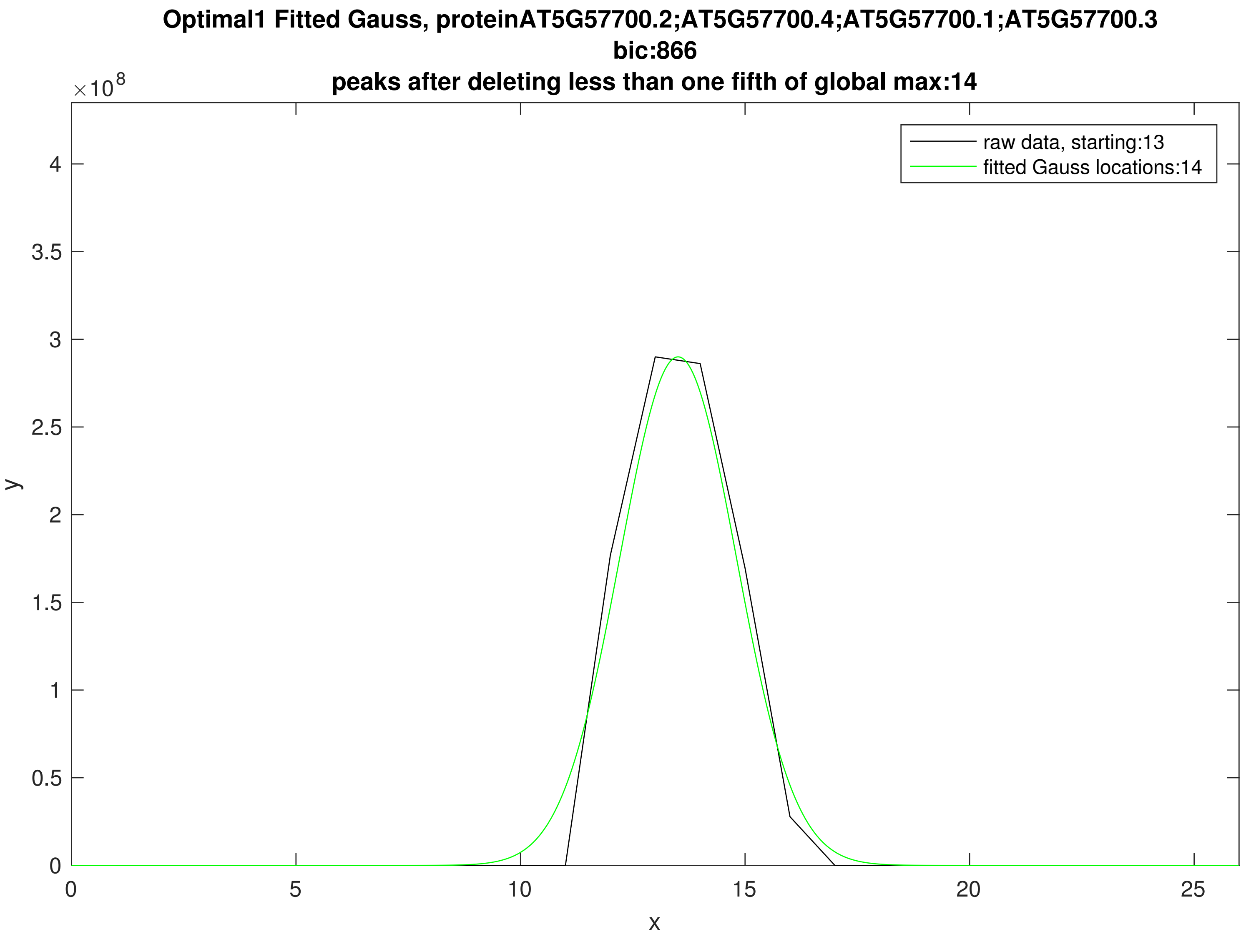

Optimal1 Fitted Gauss, proteinAT5G57700.2;AT5G57700.4;AT5G57700.1;AT5G57700.3
bic:866
peaks after deleting less than one fifth of global max:14
×10^8
raw data, starting:13
fitted Gauss locations:14
4
3.5
3
2.5
2
1.5
1
0.5
0
y
0
5
10
15
20
25
x

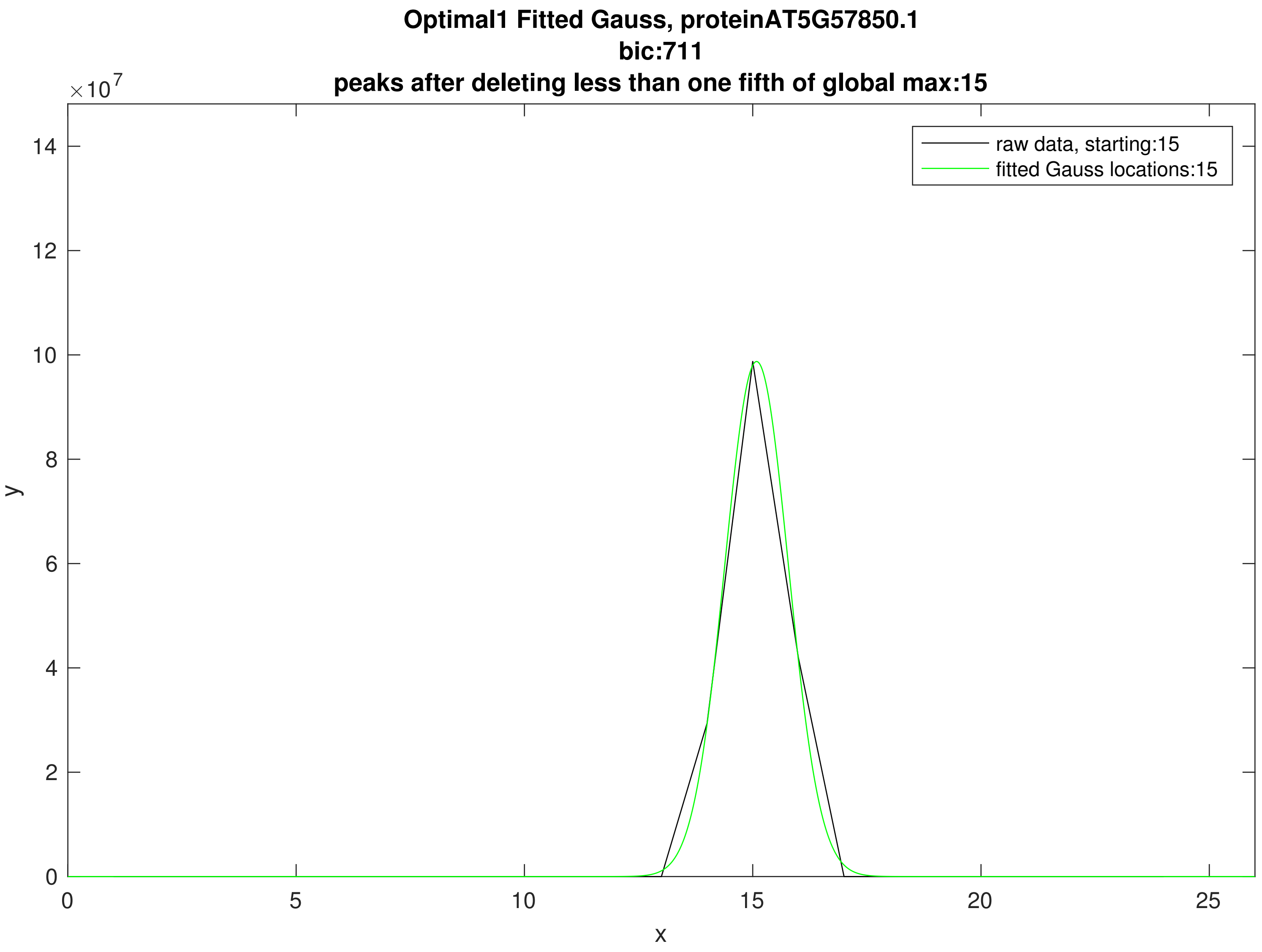
Optimal1 Fitted Gauss, proteinAT5G57850.1
bic:711
peaks after deleting less than one fifth of global max:15
×10^7
14
12
10
8
6
4
2
0
y
0
5
10
15
20
25
x
raw data, starting:15
fitted Gauss locations:15

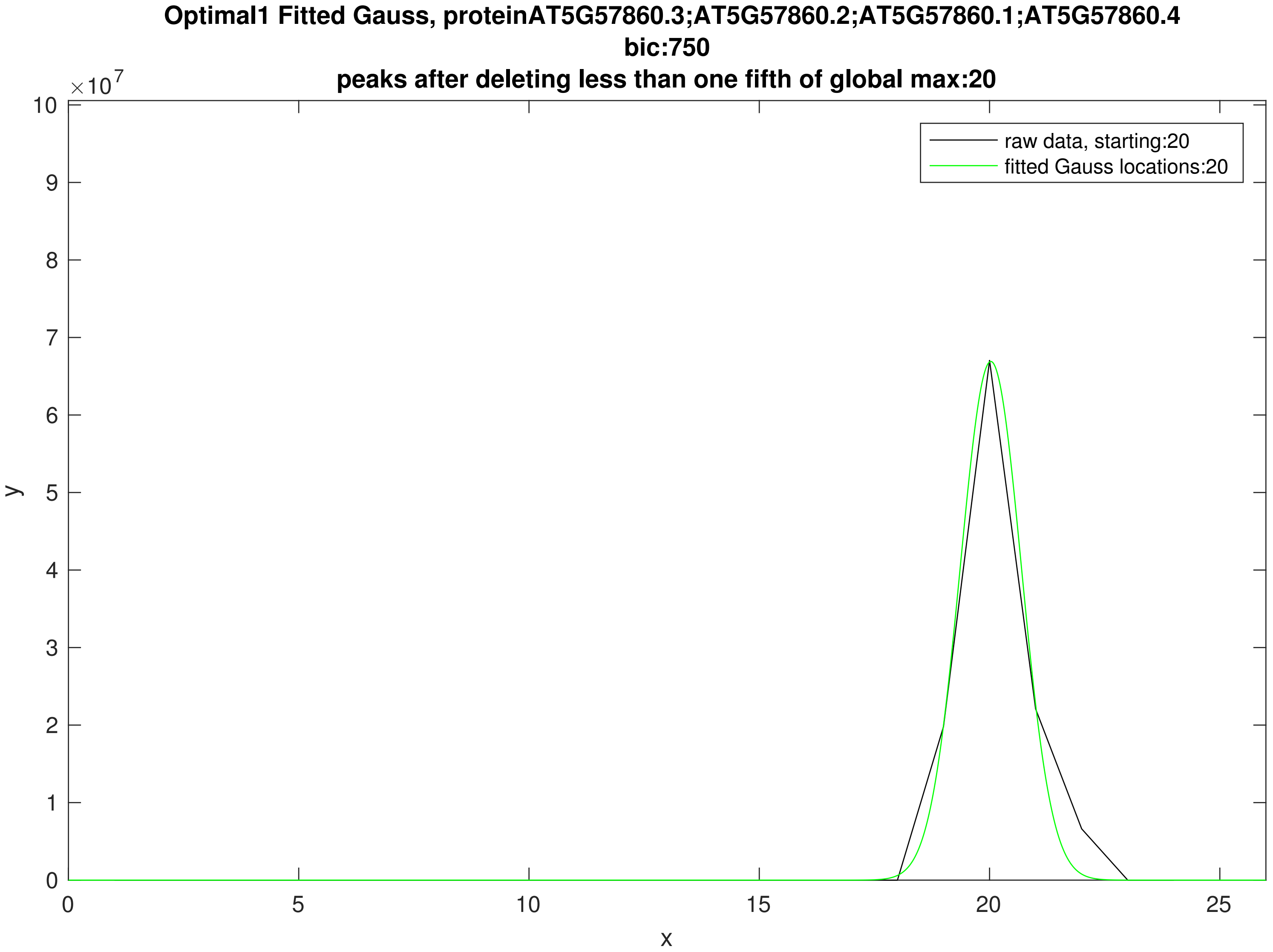
Optimal1 Fitted Gauss, proteinAT5G57860.3;AT5G57860.2;AT5G57860.1;AT5G57860.4
bic:750
peaks after deleting less than one fifth of global max:20
×10^7
raw data, starting:20
fitted Gauss locations:20
y
x
0
5
10
15
20
25
1
2
3
4
6
7
8
9
10

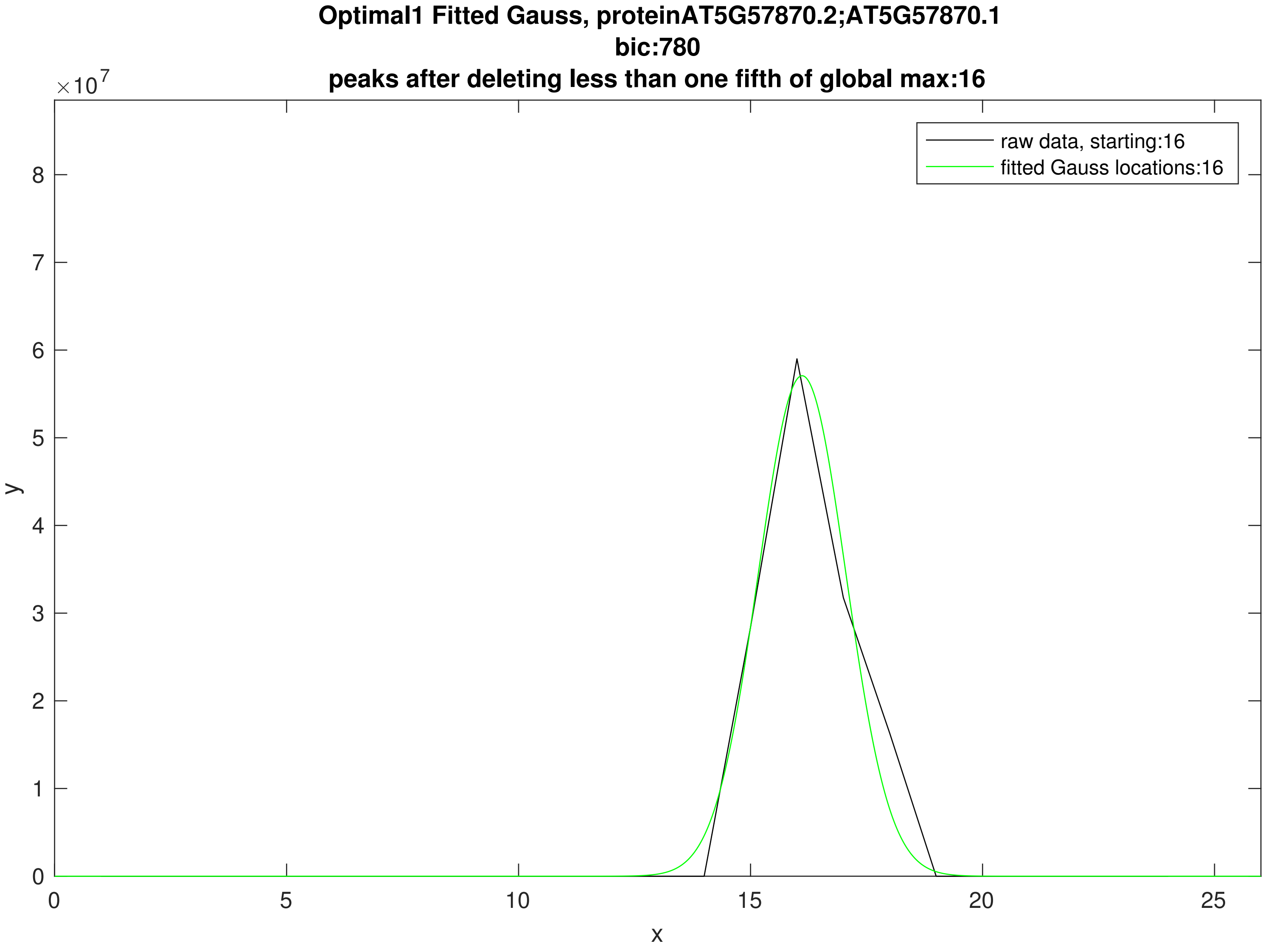
Optimal1 Fitted Gauss, proteinAT5G57870.2;AT5G57870.1
bic:780
peaks after deleting less than one fifth of global max:16
×10^7
raw data, starting:16
fitted Gauss locations:16
y
x
0
1
2
3
4
5
6
7
8
0
5
10
15
20
25

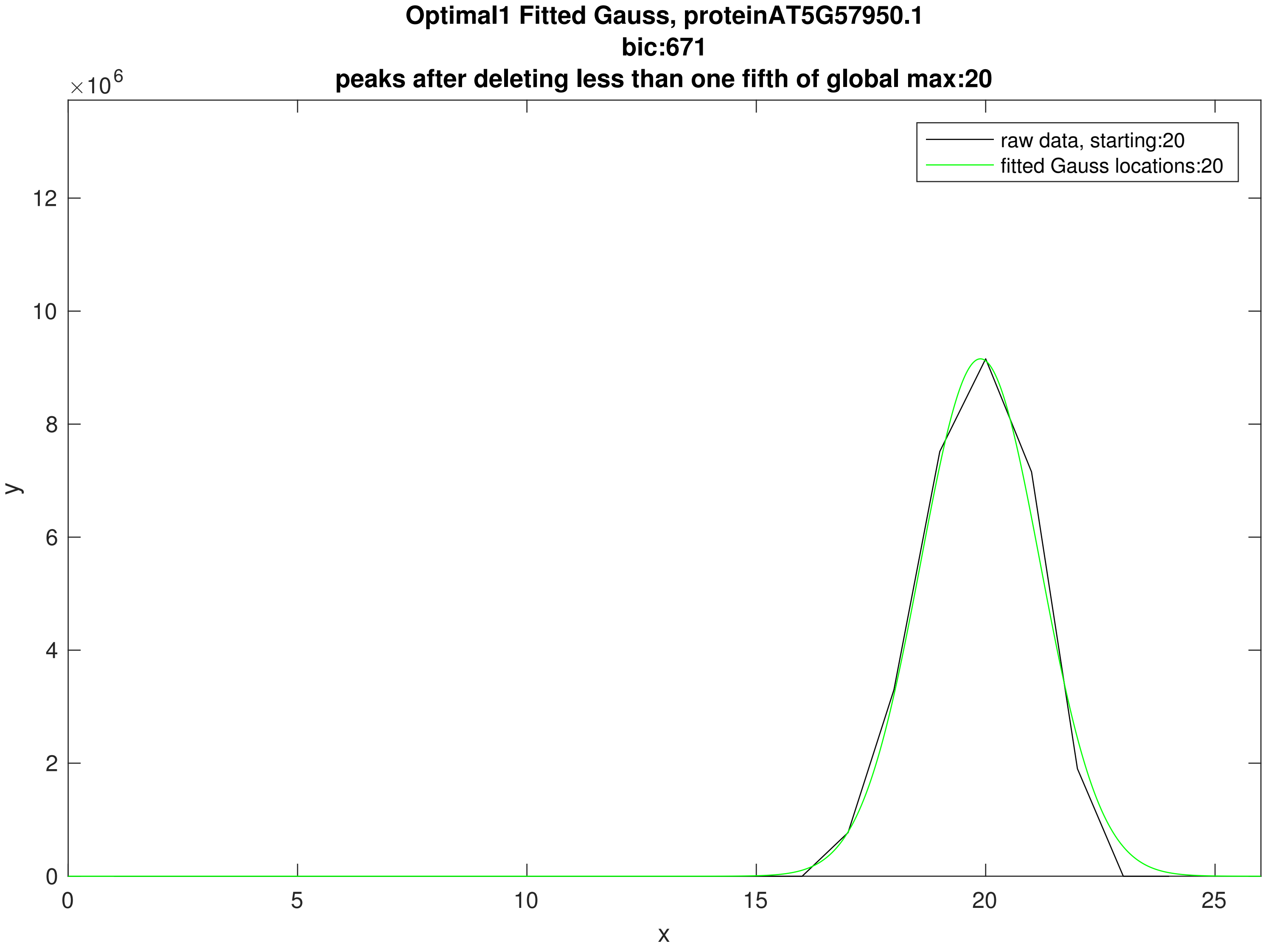
Optimal1 Fitted Gauss, proteinAT5G57950.1
bic:671
peaks after deleting less than one fifth of global max:20
×10^6
raw data, starting:20
fitted Gauss locations:20
y
x
0
2
4
6
8
10
12
0
5
10
15
20
25

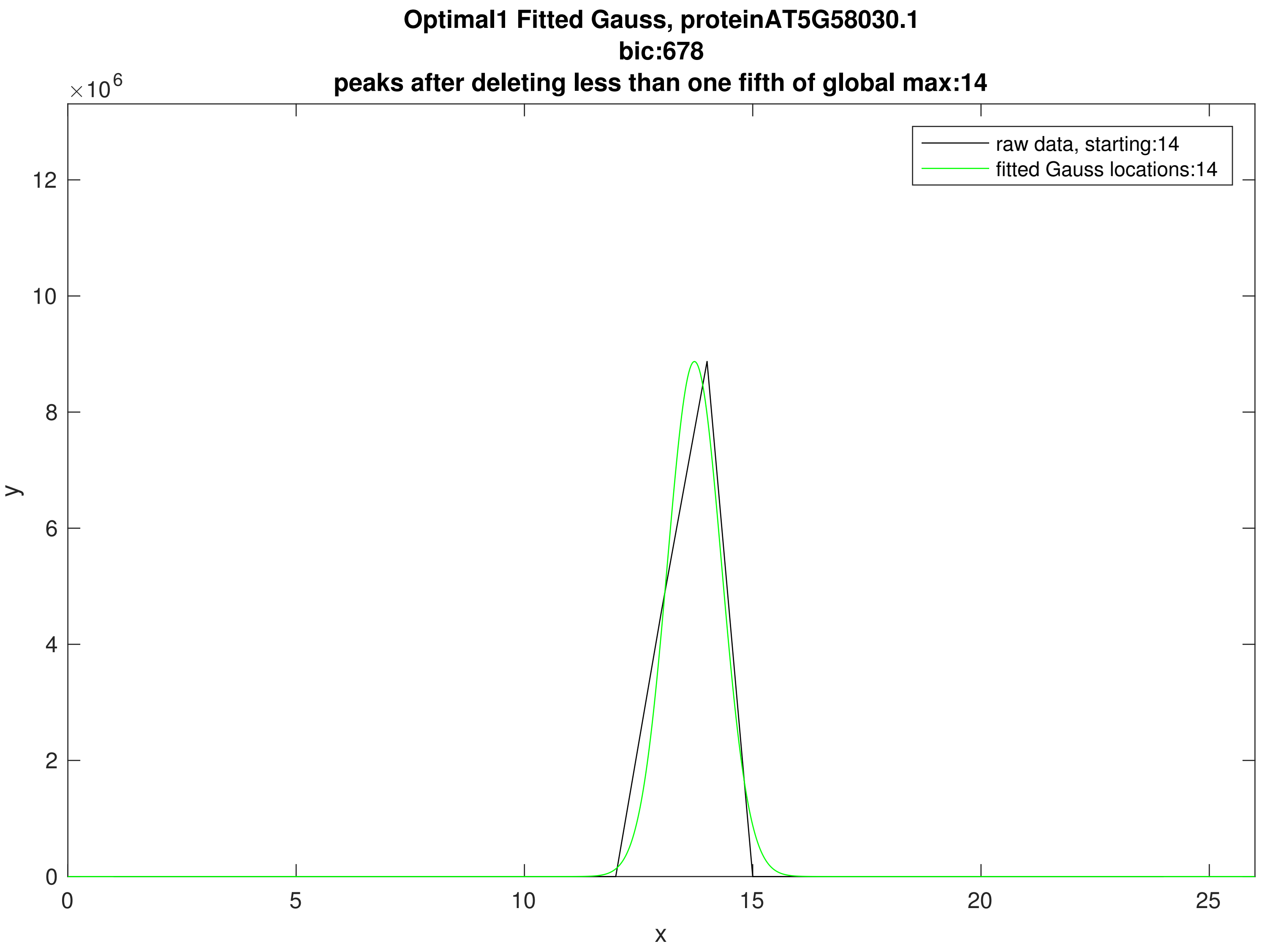
Optimal1 Fitted Gauss, proteinAT5G58030.1
bic:678
peaks after deleting less than one fifth of global max:14
×10^6
raw data, starting:14
fitted Gauss locations:14
y
x
0
2
4
6
8
10
12
0
5
10
15
20
25

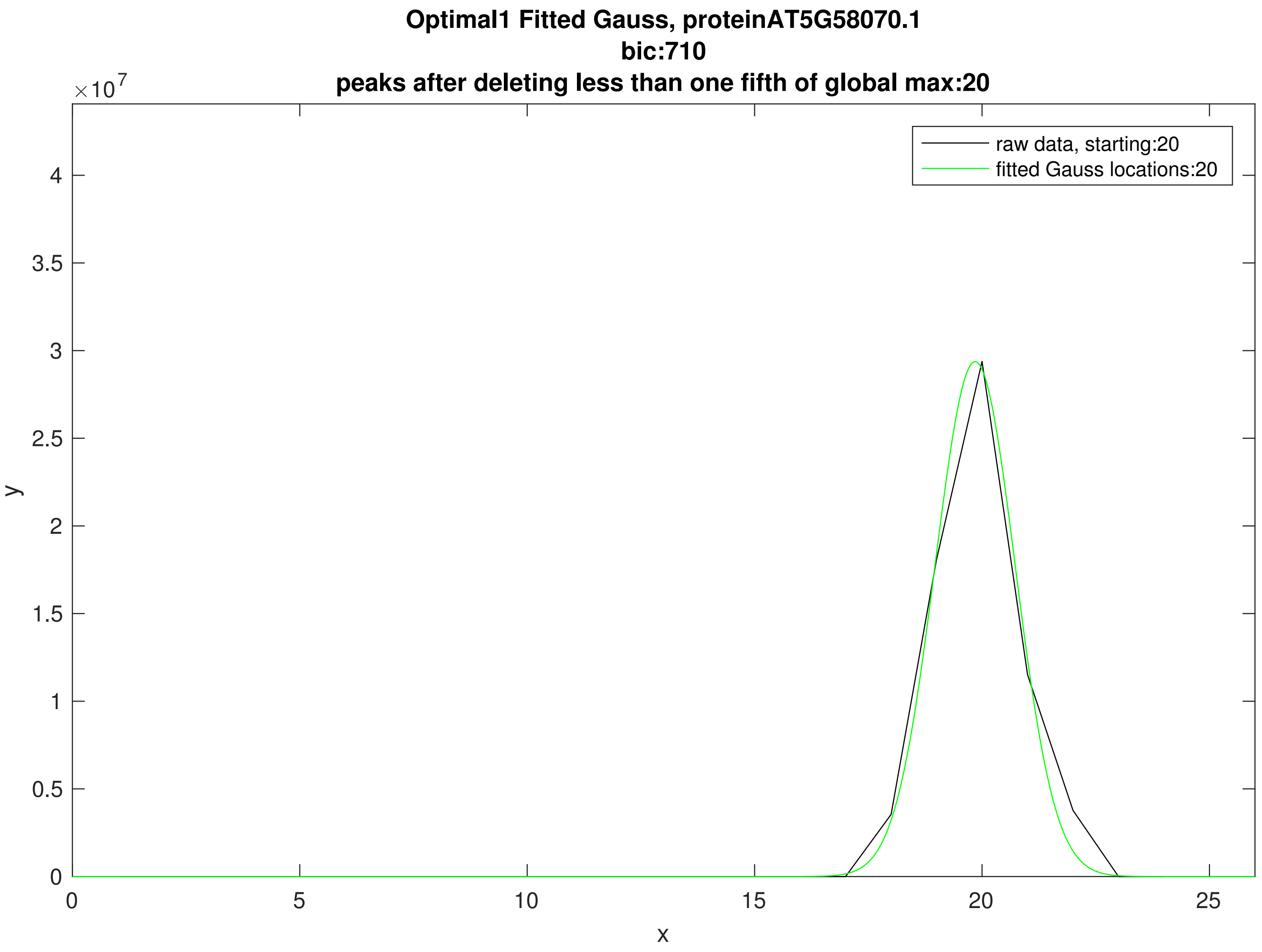
Optimal1 Fitted Gauss, proteinAT5G58070.1
bic:710
peaks after deleting less than one fifth of global max:20
×10^7
raw data, starting:20
fitted Gauss locations:20
y
x
0
0.5
1
1.5
2
2.5
3
3.5
4
0
5
10
15
20
25

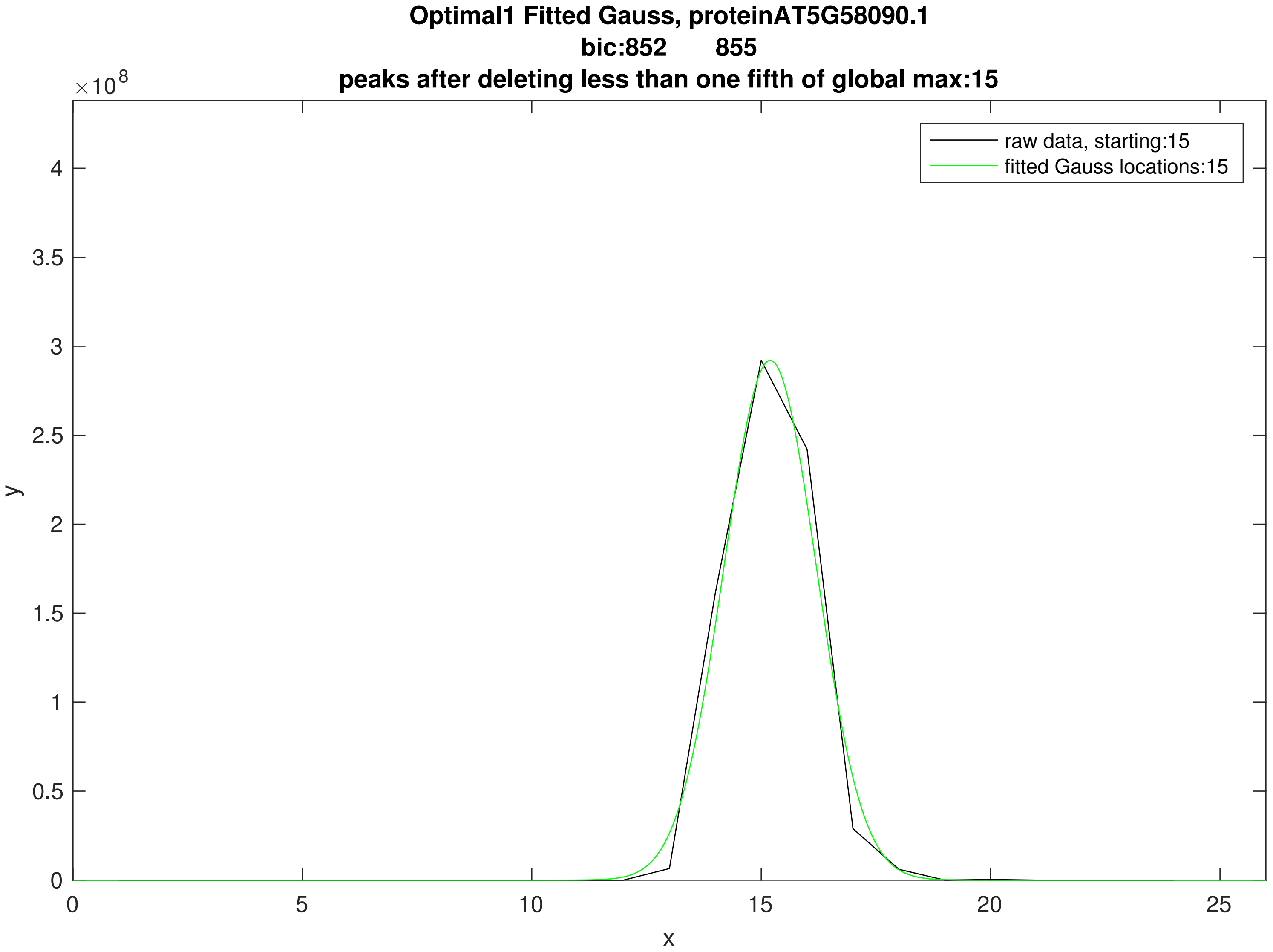

Optimal1 Fitted Gauss, proteinAT5G58090.1
bic:852 855
peaks after deleting less than one fifth of global max:15
×10^8
raw data, starting:15
fitted Gauss locations:15
y
x
0
0.5
1
1.5
2
2.5
3
3.5
4
0
5
10
15
20
25

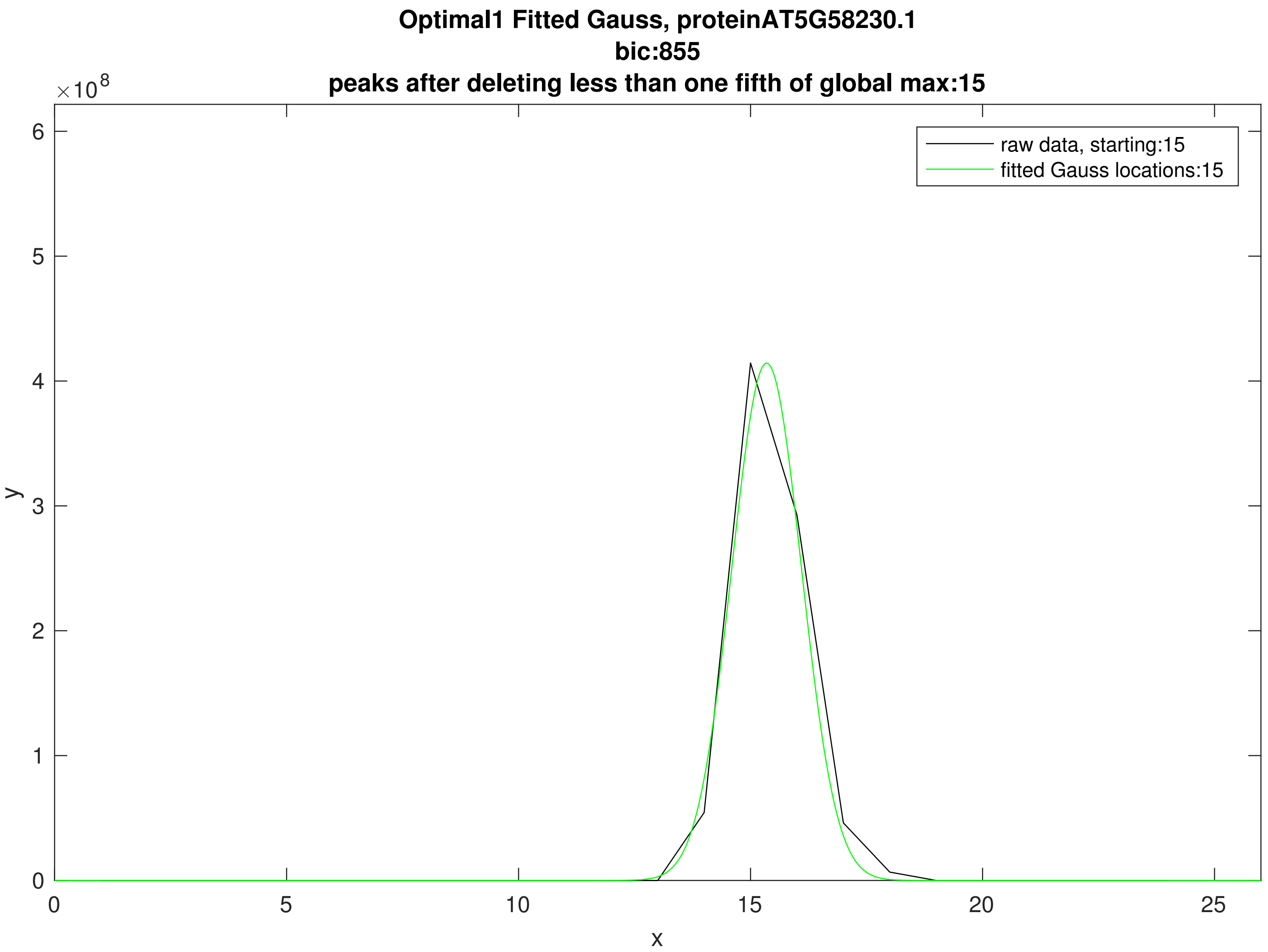

Optimal1 Fitted Gauss, proteinAT5G58230.1
bic:855
peaks after deleting less than one fifth of global max:15
×10^8
raw data, starting:15
fitted Gauss locations:15
y
x
0
1
2
3
4
5
6
0
5
10
15
20
25

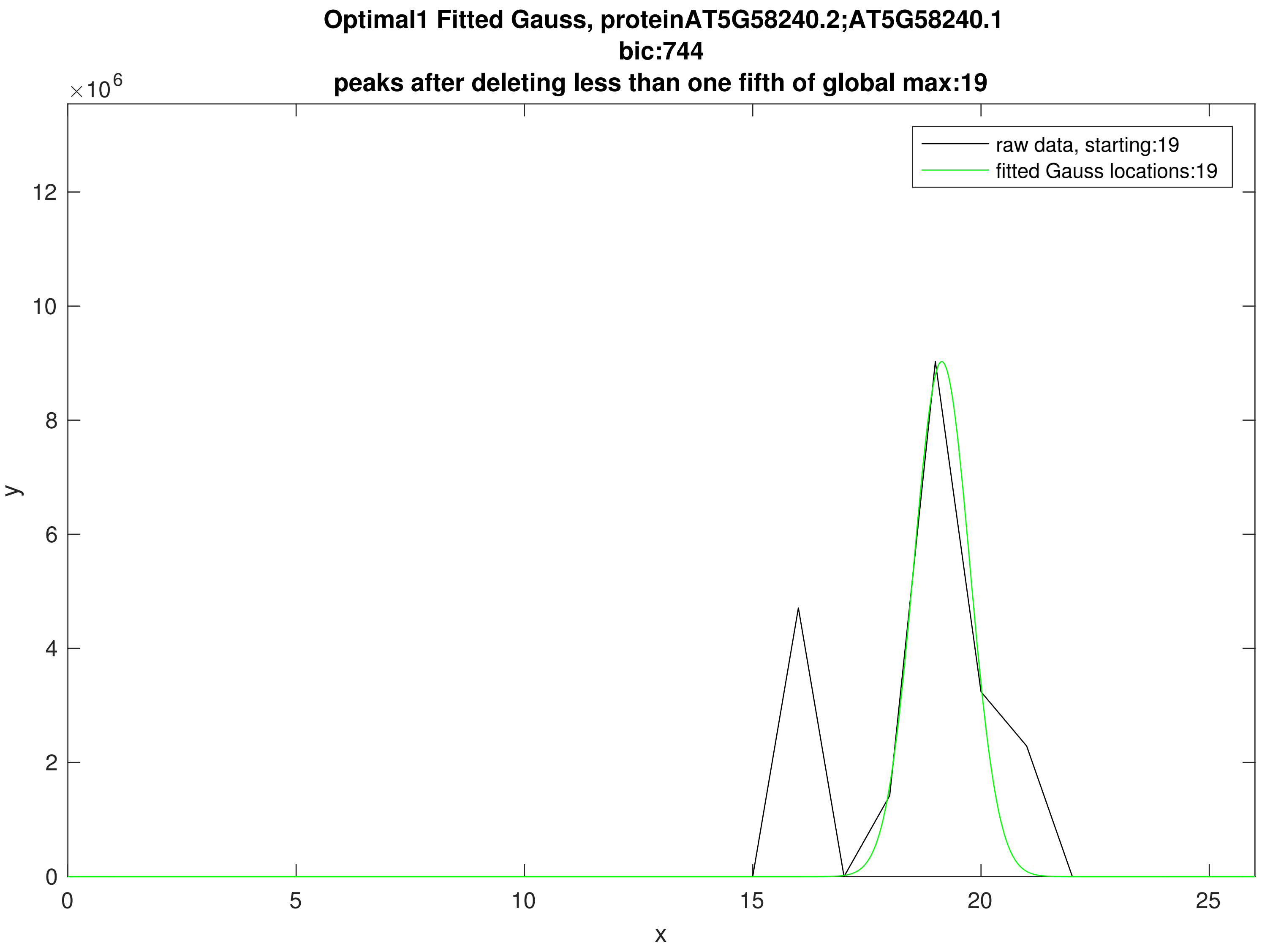

Optimal1 Fitted Gauss, proteinAT5G58240.2;AT5G58240.1
bic:744
peaks after deleting less than one fifth of global max:19
×10^6
raw data, starting:19
fitted Gauss locations:19
y
x
0
2
4
6
8
10
12
0
5
10
15
20
25

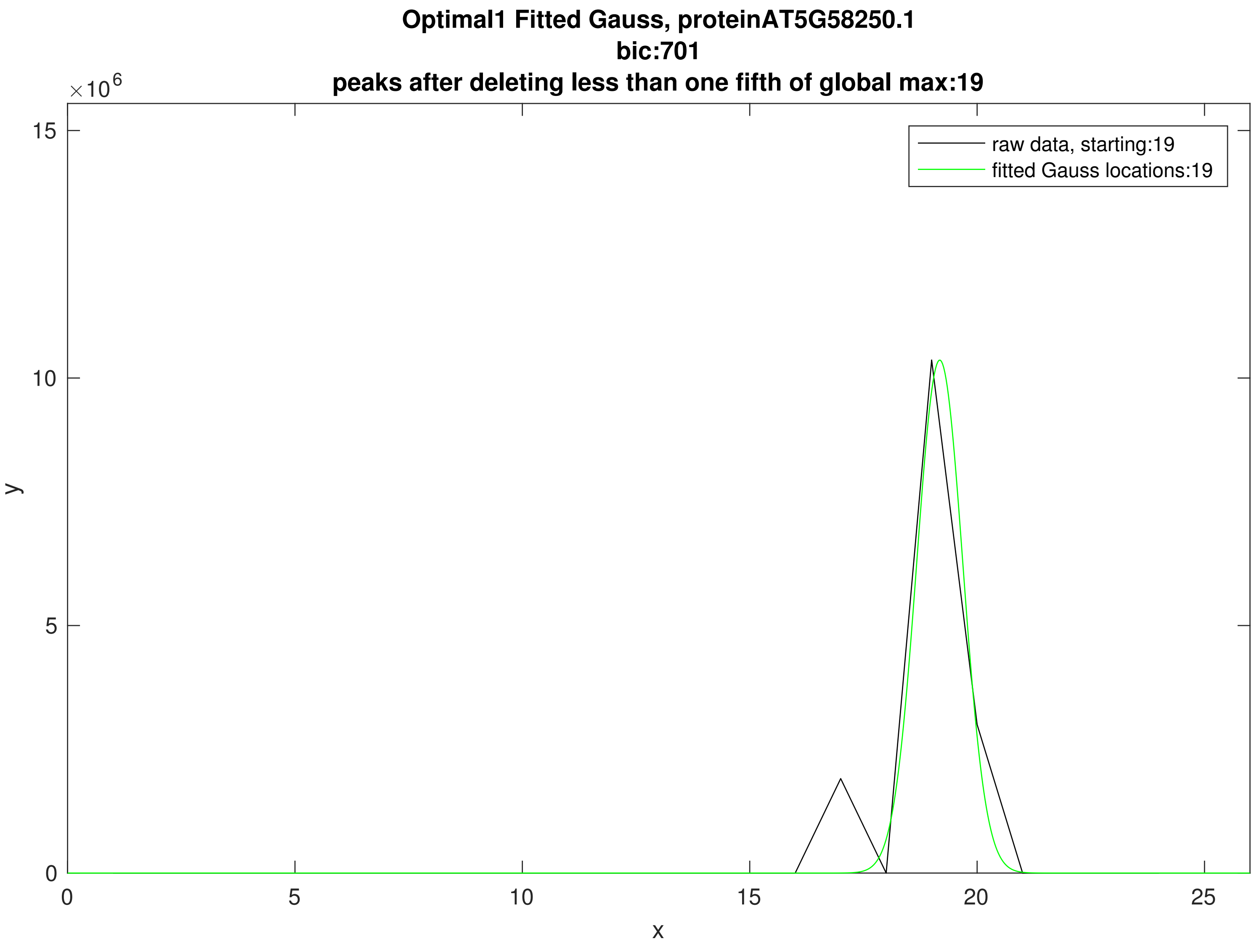

Optimal1 Fitted Gauss, proteinAT5G58250.1
bic:701
peaks after deleting less than one fifth of global max:19
×10^6
15
10
5
0
y
0
5
10
15
20
25
x
raw data, starting:19
fitted Gauss locations:19

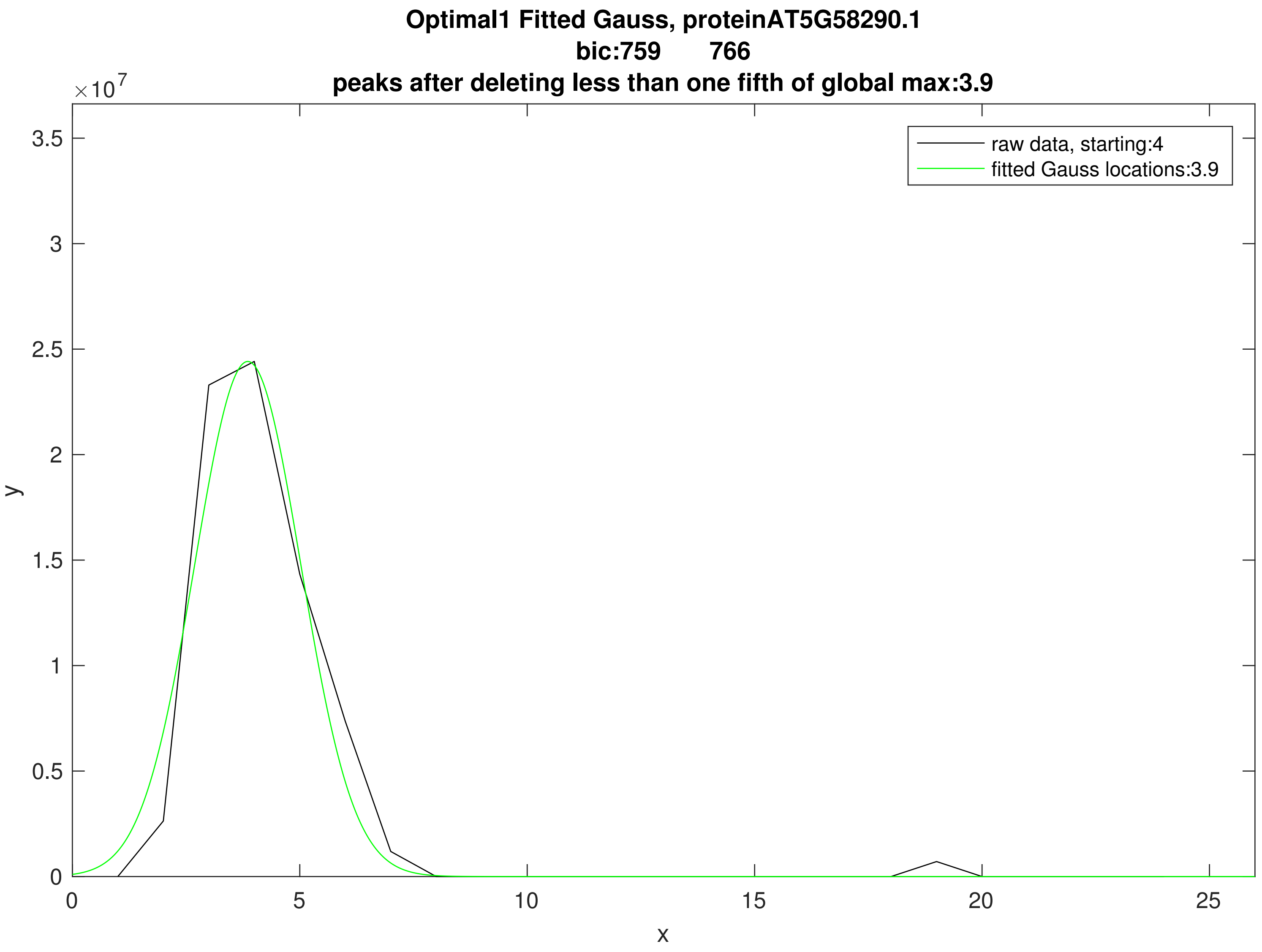

Optimal1 Fitted Gauss, proteinAT5G58290.1
bic:759 766
peaks after deleting less than one fifth of global max:3.9
×10^7
raw data, starting:4
fitted Gauss locations:3.9
y
x
0
0.5
1
1.5
2
2.5
3
3.5
5
10
15
20
25

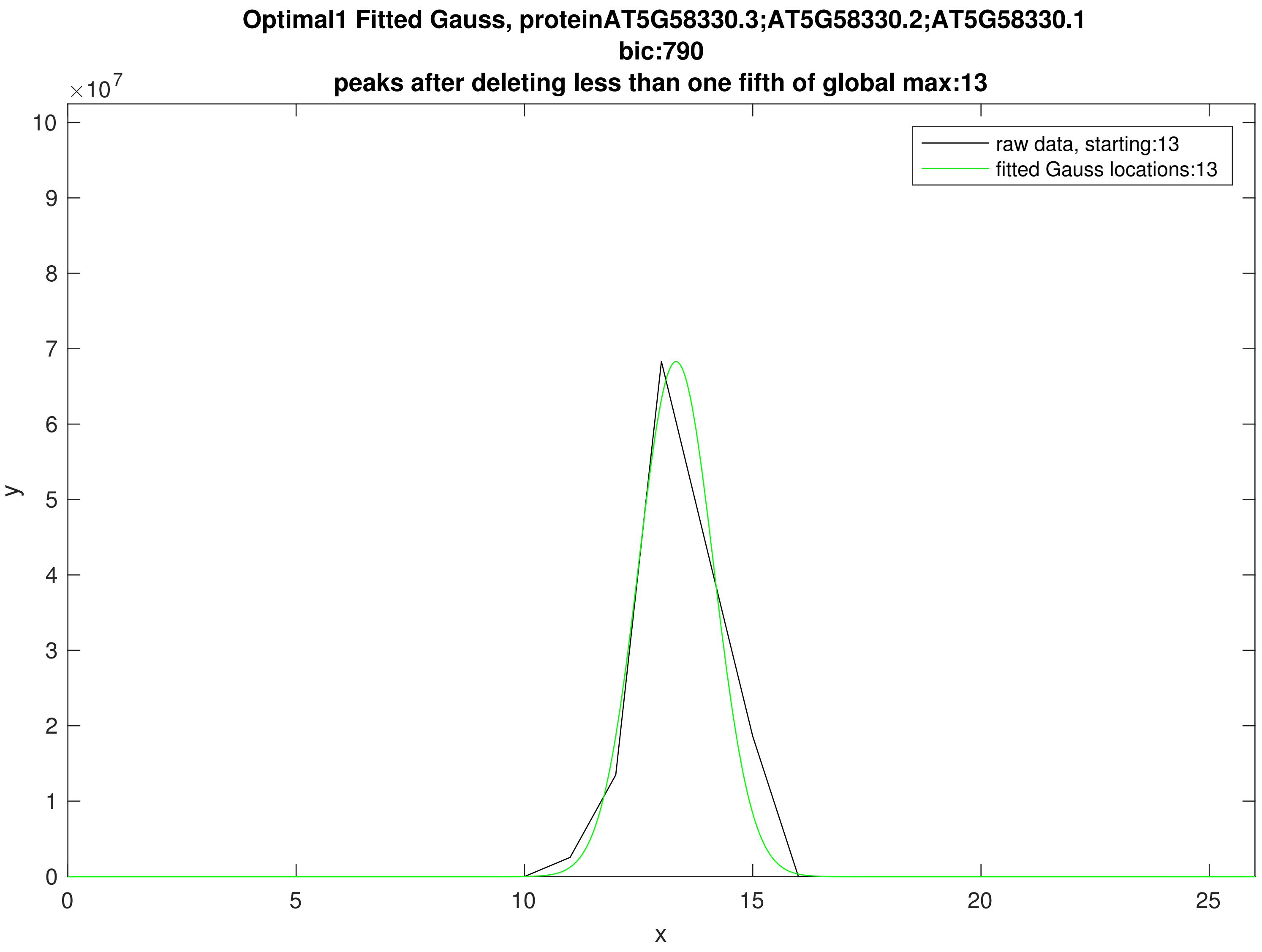
Optimal1 Fitted Gauss, proteinAT5G58330.3;AT5G58330.2;AT5G58330.1
bic:790
peaks after deleting less than one fifth of global max:13
×10^7
raw data, starting:13
fitted Gauss locations:13
y
x
0
1
2
3
4
5
6
7
8
9
10
0
5
10
15
20
25

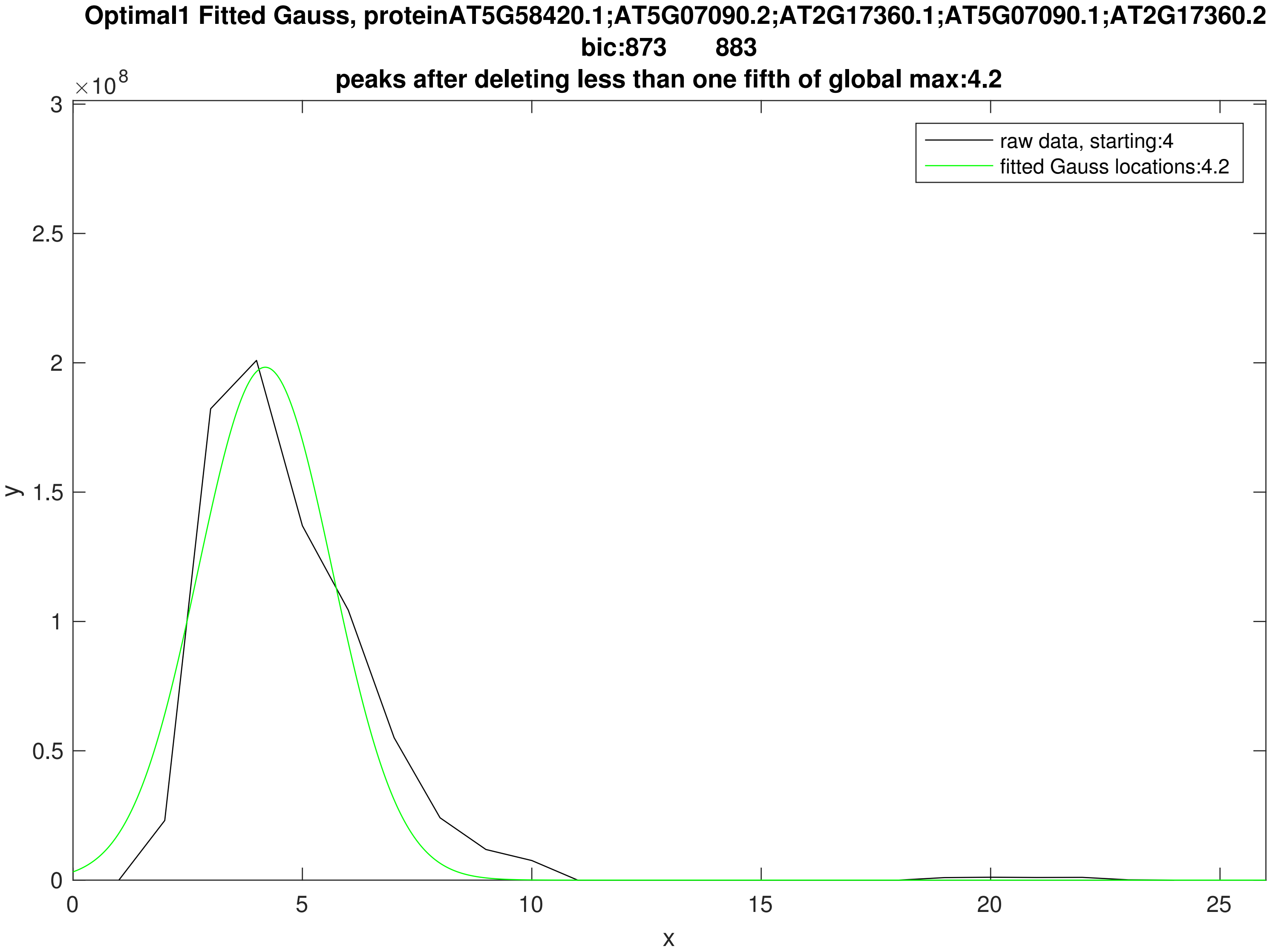

Optimal1 Fitted Gauss, proteinAT5G58420.1;AT5G07090.2;AT2G17360.1;AT5G07090.1;AT2G17360.2
bic:873 883
peaks after deleting less than one fifth of global max:4.2
×10^8
3
2.5
2
1.5
1
0.5
0
y
0
5
10
15
20
25
x
raw data, starting:4
fitted Gauss locations:4.2

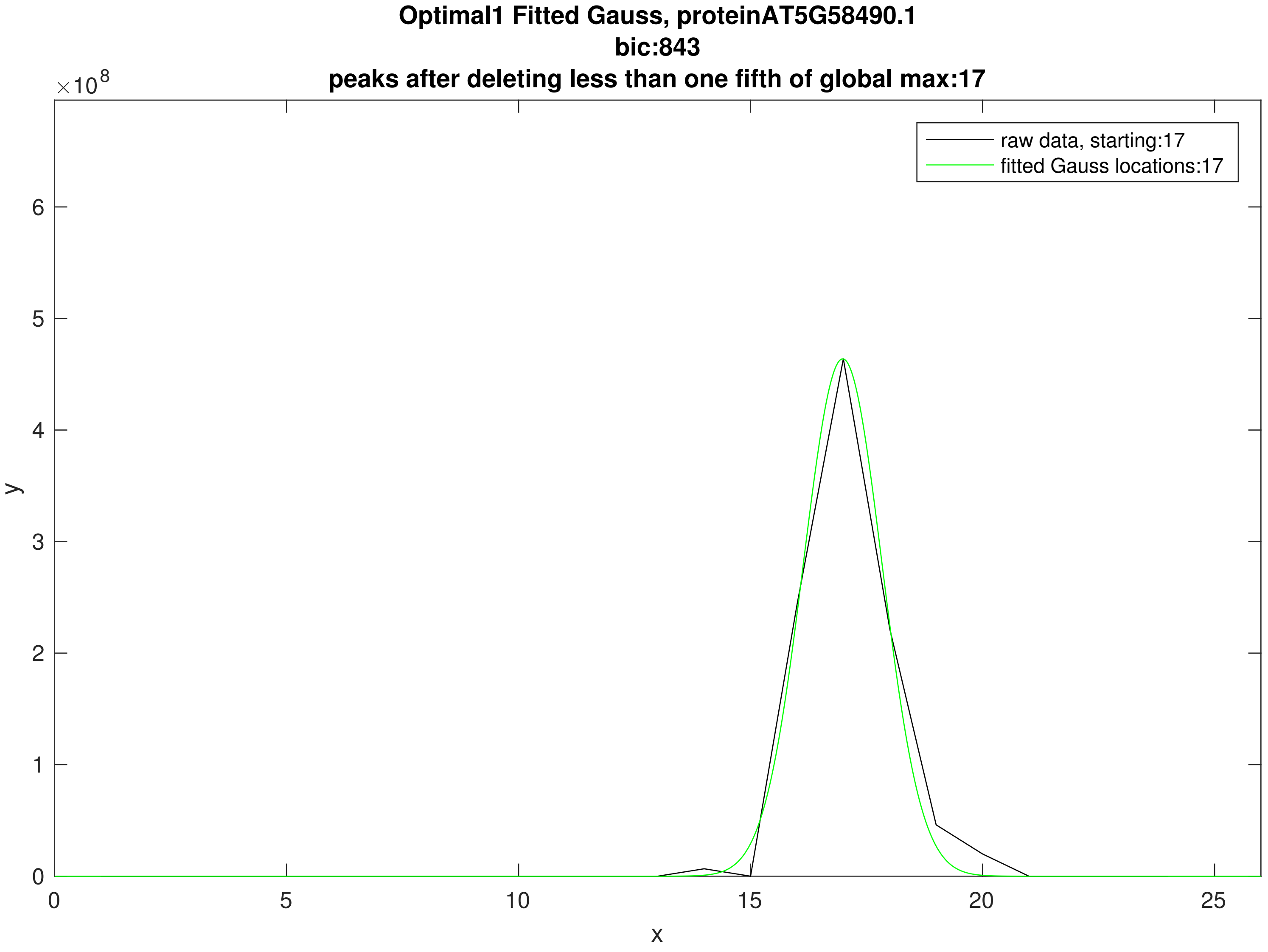

Optimal1 Fitted Gauss, proteinAT5G58490.1
bic:843
peaks after deleting less than one fifth of global max:17
×10^8
raw data, starting:17
fitted Gauss locations:17
y
x
0
1
2
3
4
5
6
0
5
10
15
20
25

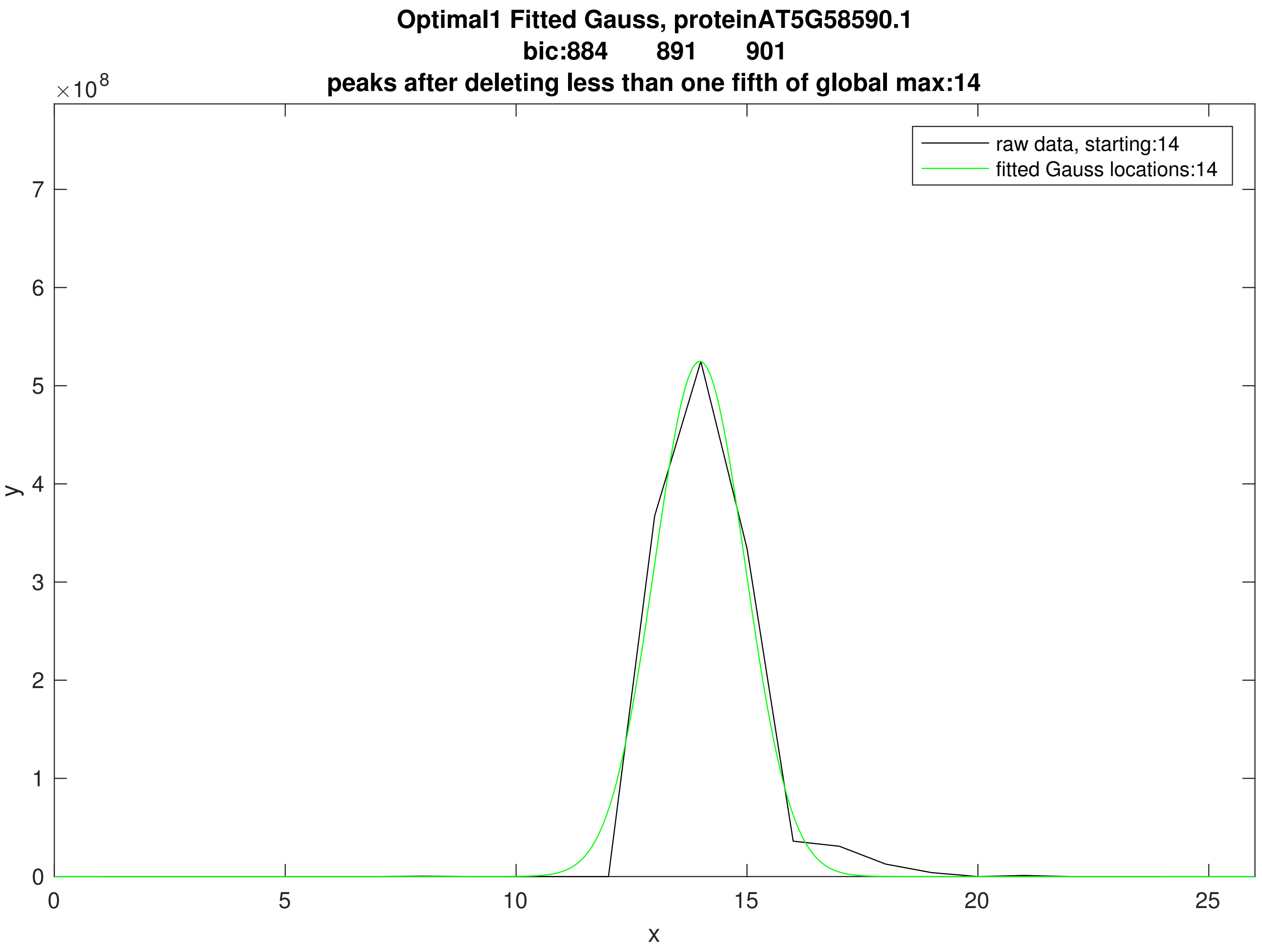

Optimal1 Fitted Gauss, proteinAT5G58590.1
bic:884 891 901
peaks after deleting less than one fifth of global max:14
×10^8
raw data, starting:14
fitted Gauss locations:14
y
x
0
1
2
3
4
5
6
7
0
5
10
15
20
25

**Optimal1 Fitted Gauss, proteinAT5G58600.1;AT5G58600.2**
**bic:640**
**peaks after deleting less than one fifth of global max:18**

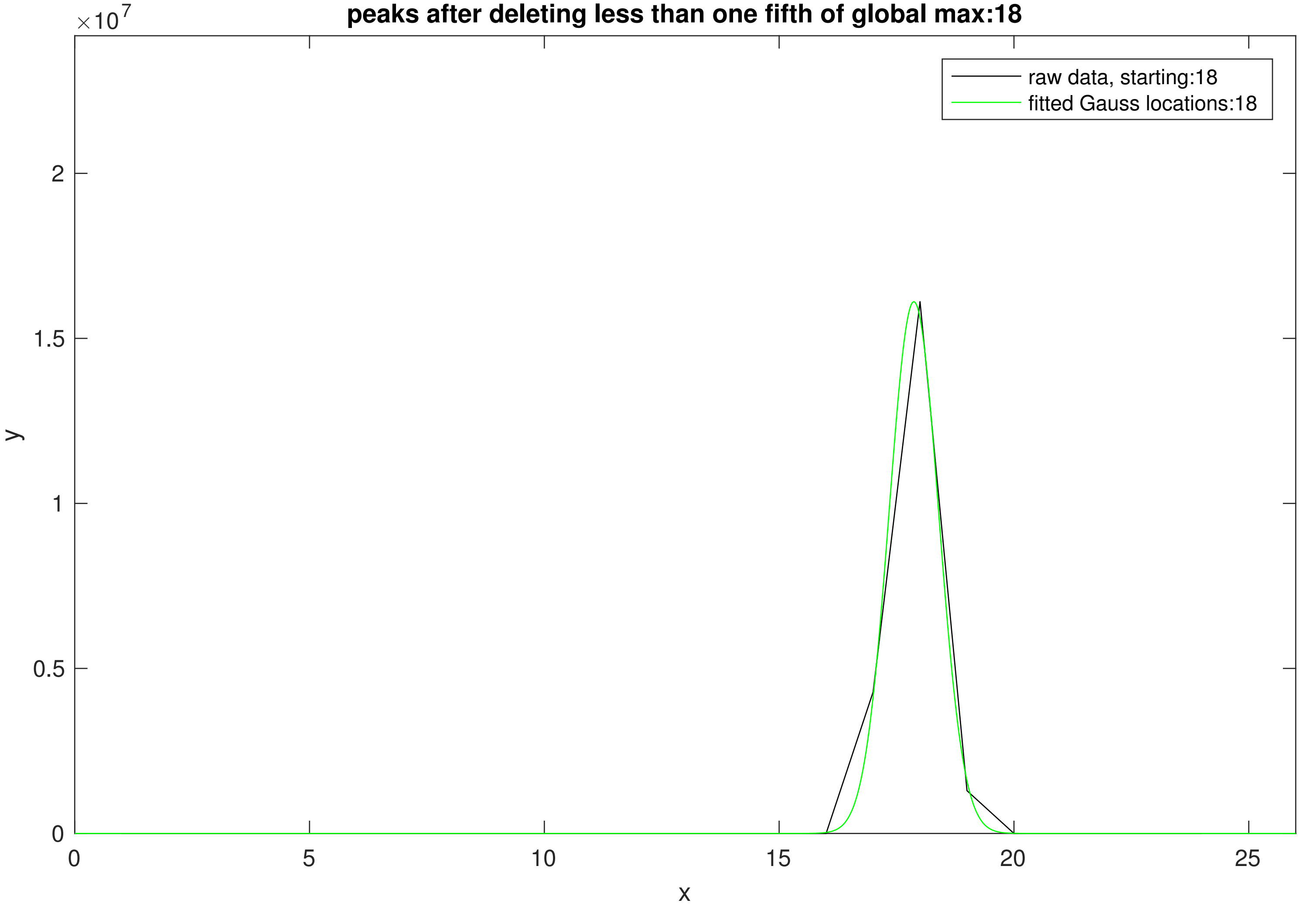

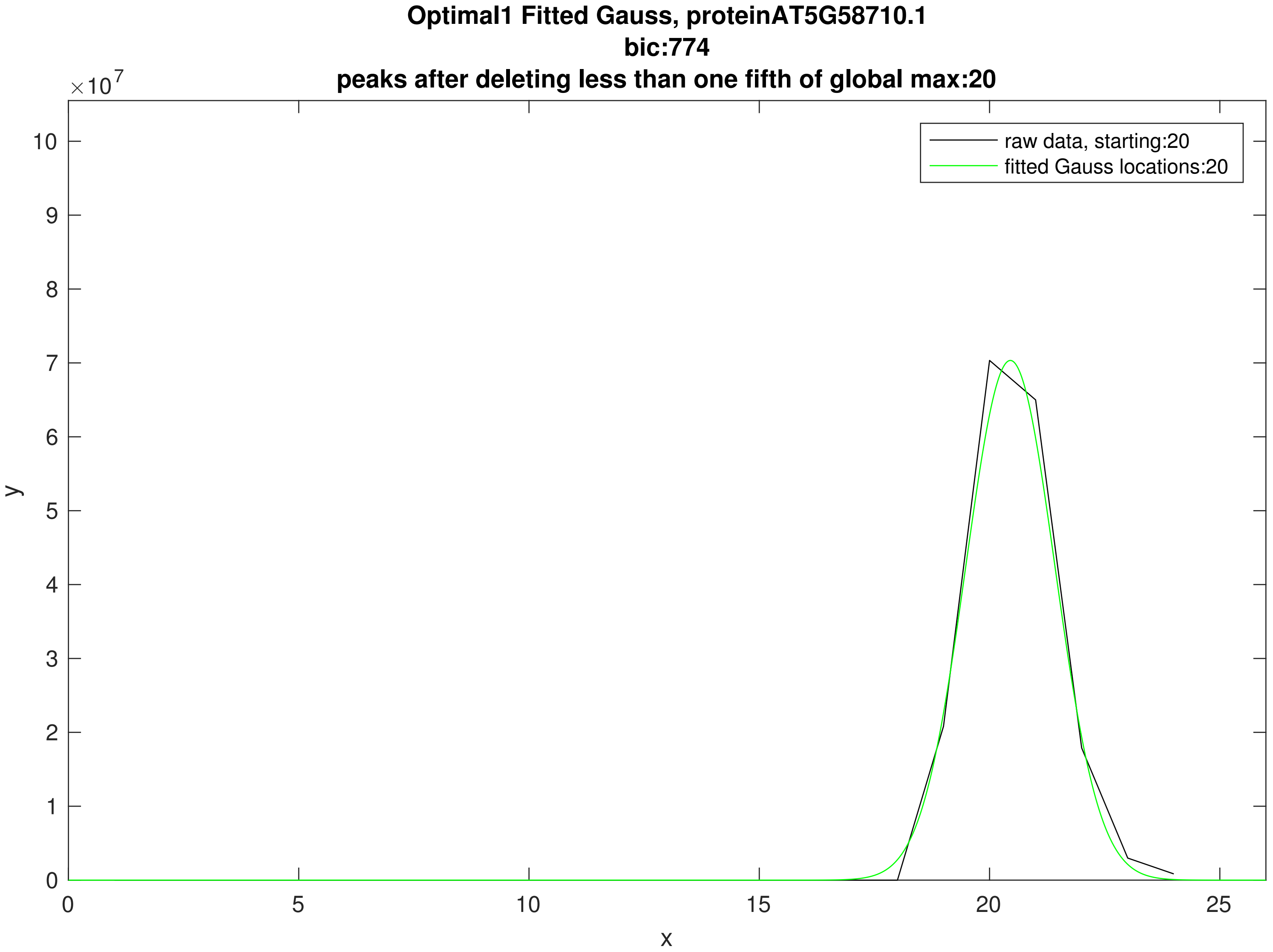
Optimal1 Fitted Gauss, proteinAT5G58710.1
bic:774
peaks after deleting less than one fifth of global max:20
×10^7
raw data, starting:20
fitted Gauss locations:20
y
x
0
1
2
3
4
5
6
7
8
9
10
0
5
10
15
20
25

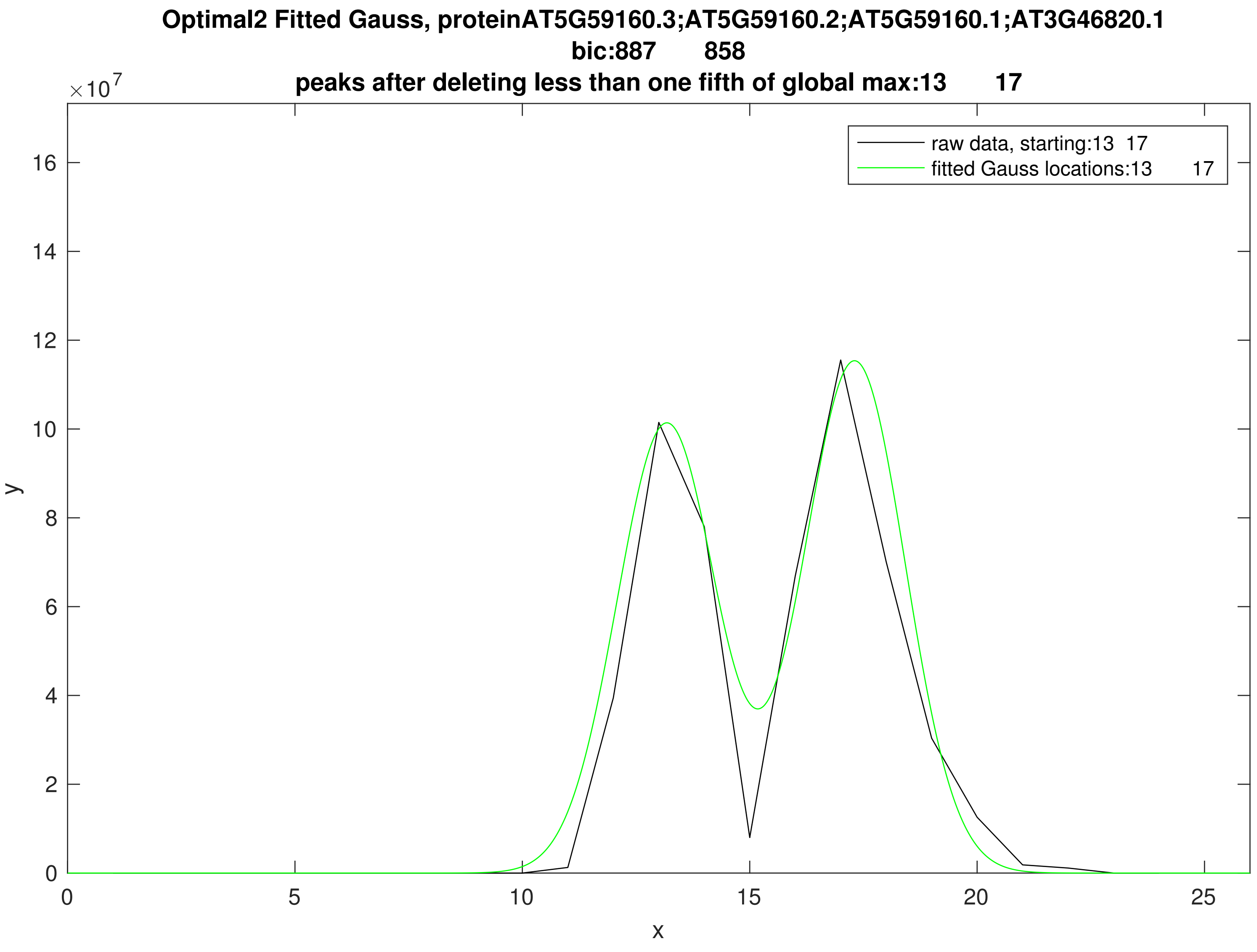
Optimal2 Fitted Gauss, proteinAT5G59160.3;AT5G59160.2;AT5G59160.1;AT3G46820.1
bic:887 858
peaks after deleting less than one fifth of global max:13 17
×10^7
raw data, starting:13 17
fitted Gauss locations:13 17
0
2
4
6
8
10
12
14
16
y
0
5
10
15
20
25
x

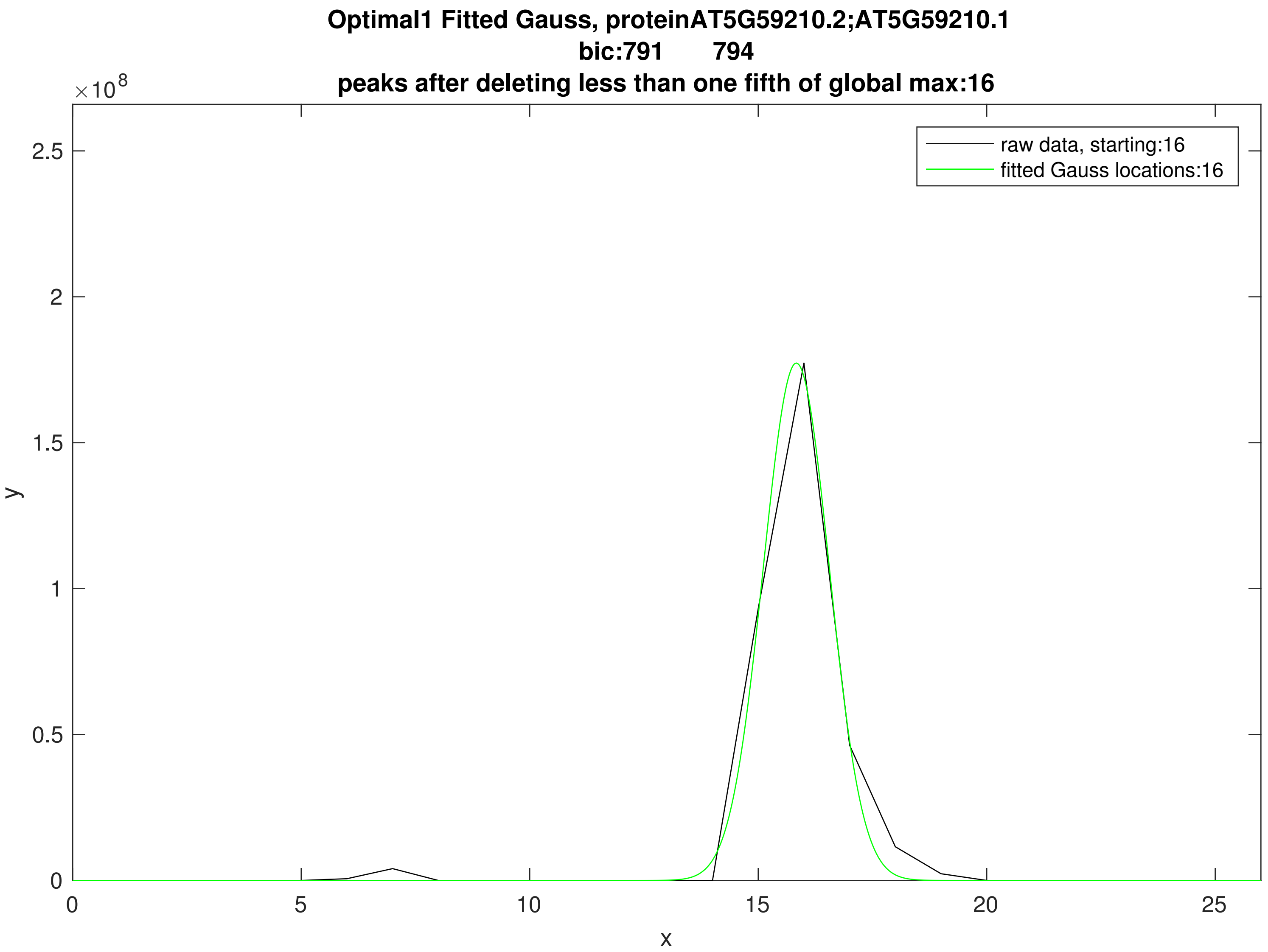

Optimal1 Fitted Gauss, proteinAT5G59210.2;AT5G59210.1
bic:791 794
peaks after deleting less than one fifth of global max:16
×10^8
2.5
2
1.5
1
0.5
0
0
5
10
15
20
25
x
y
raw data, starting:16
fitted Gauss locations:16

**Optimal1 Fitted Gauss, proteinAT5G59290.1**
**bic:1.04e+03  1.04e+03  1.05e+03**
**peaks after deleting less than one fifth of global max:9.7**

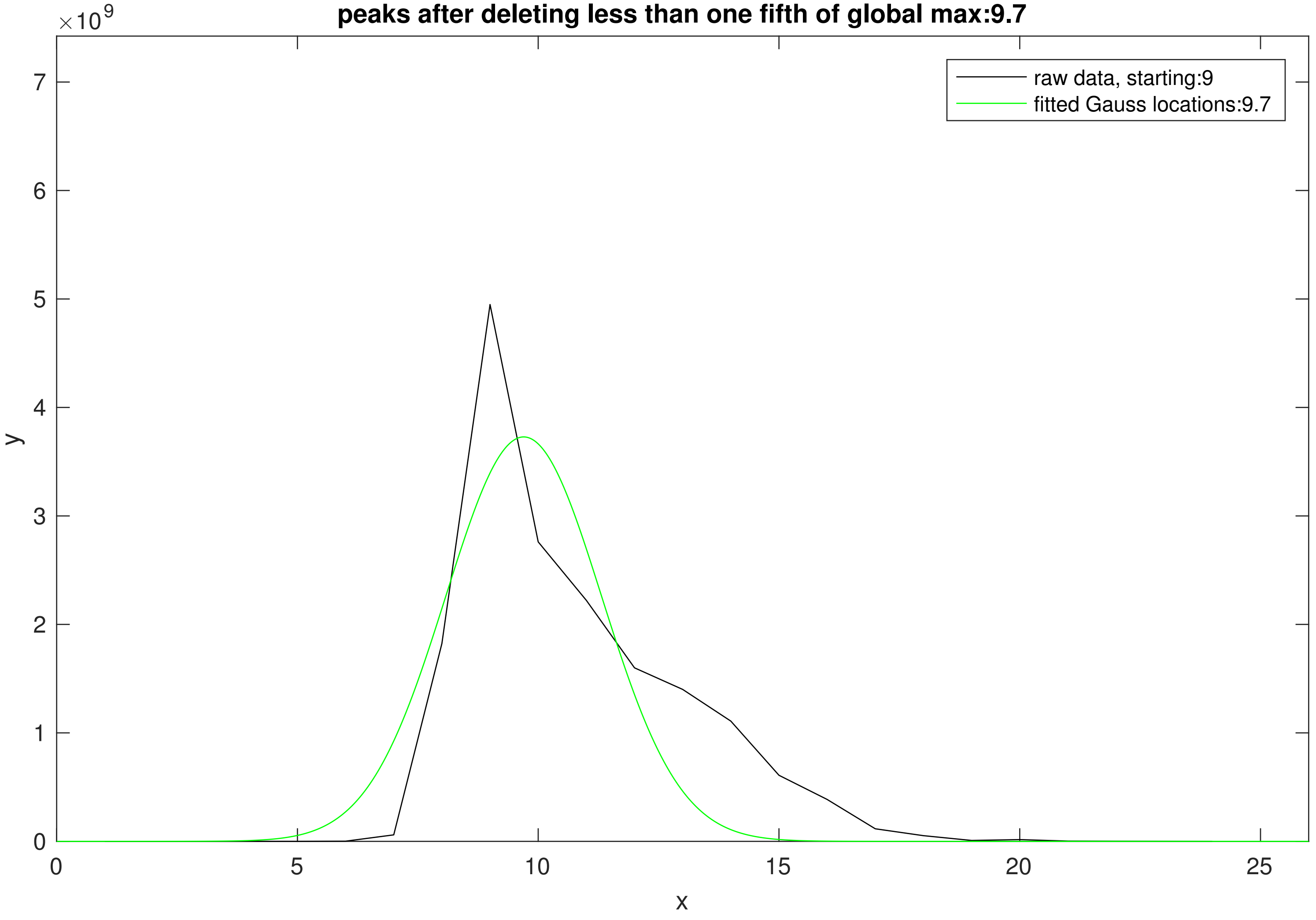

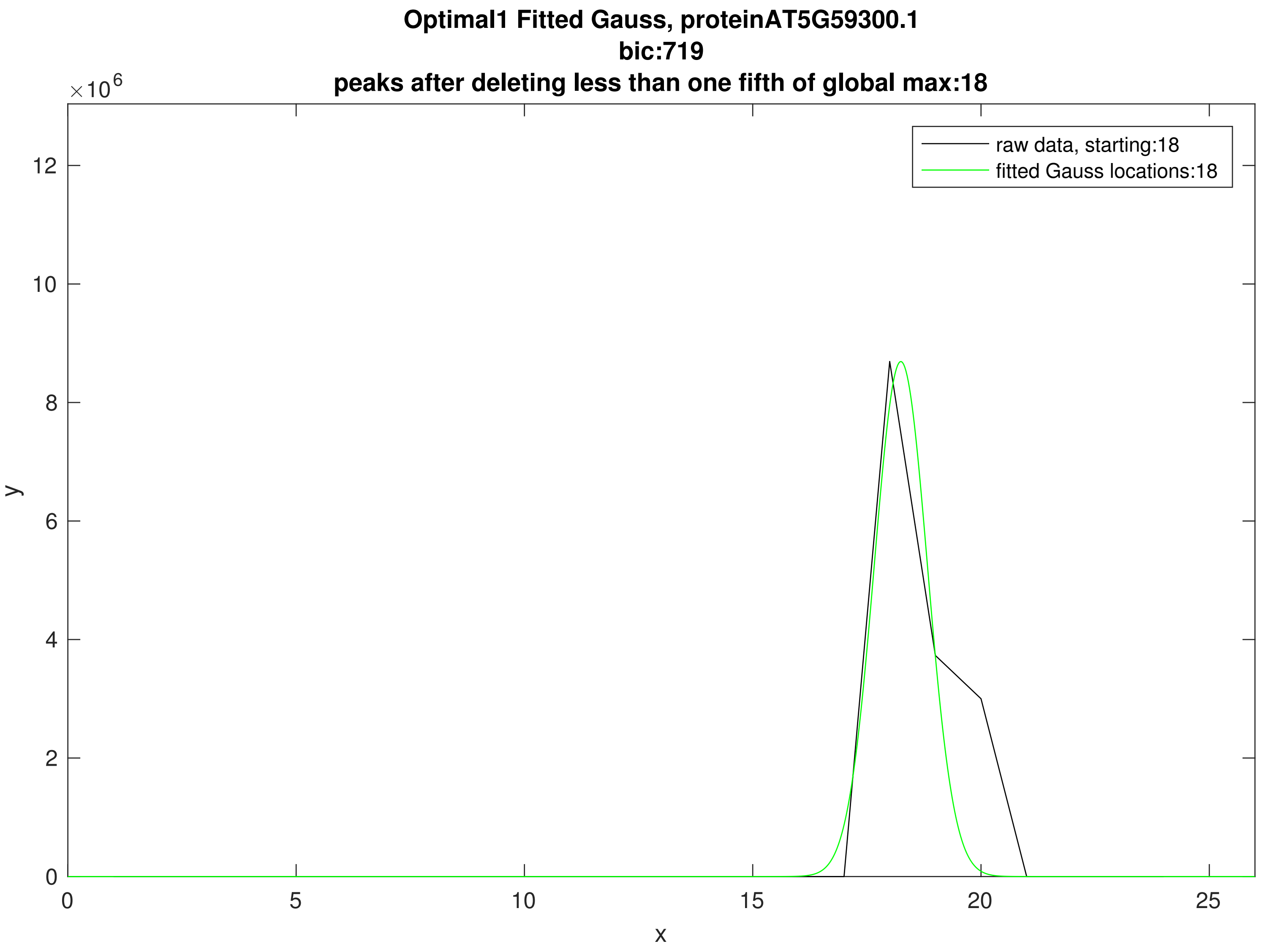

Optimal1 Fitted Gauss, proteinAT5G59300.1
bic:719
peaks after deleting less than one fifth of global max:18
×10^6
raw data, starting:18
fitted Gauss locations:18
12
10
8
6
4
2
0
y
0
5
10
15
20
25
x

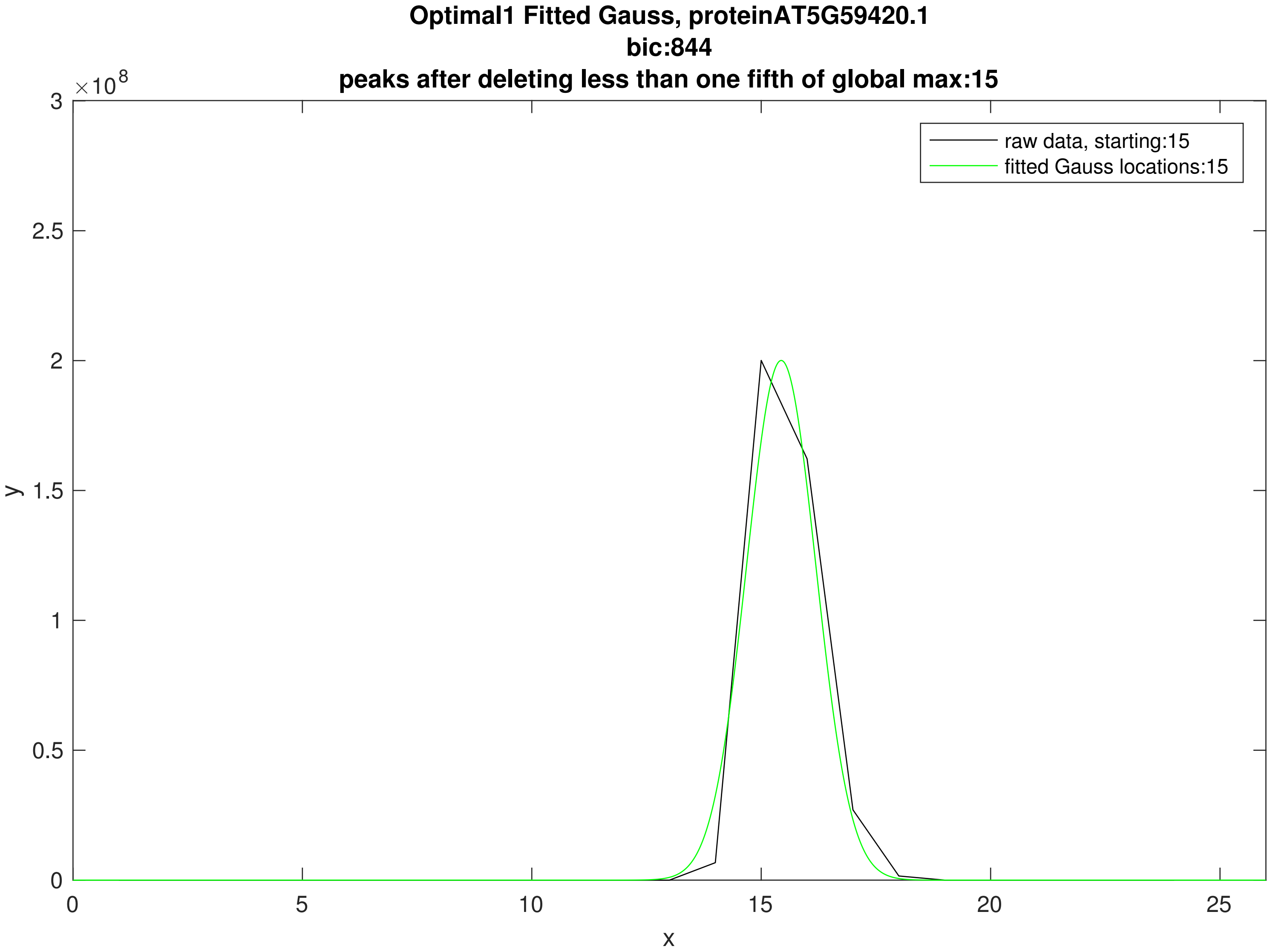
Optimal1 Fitted Gauss, proteinAT5G59420.1
bic:844
peaks after deleting less than one fifth of global max:15
$\times 10^8$
3
2.5
2
1.5
1
0.5
0
y
0
5
10
15
20
25
x
raw data, starting:15
fitted Gauss locations:15

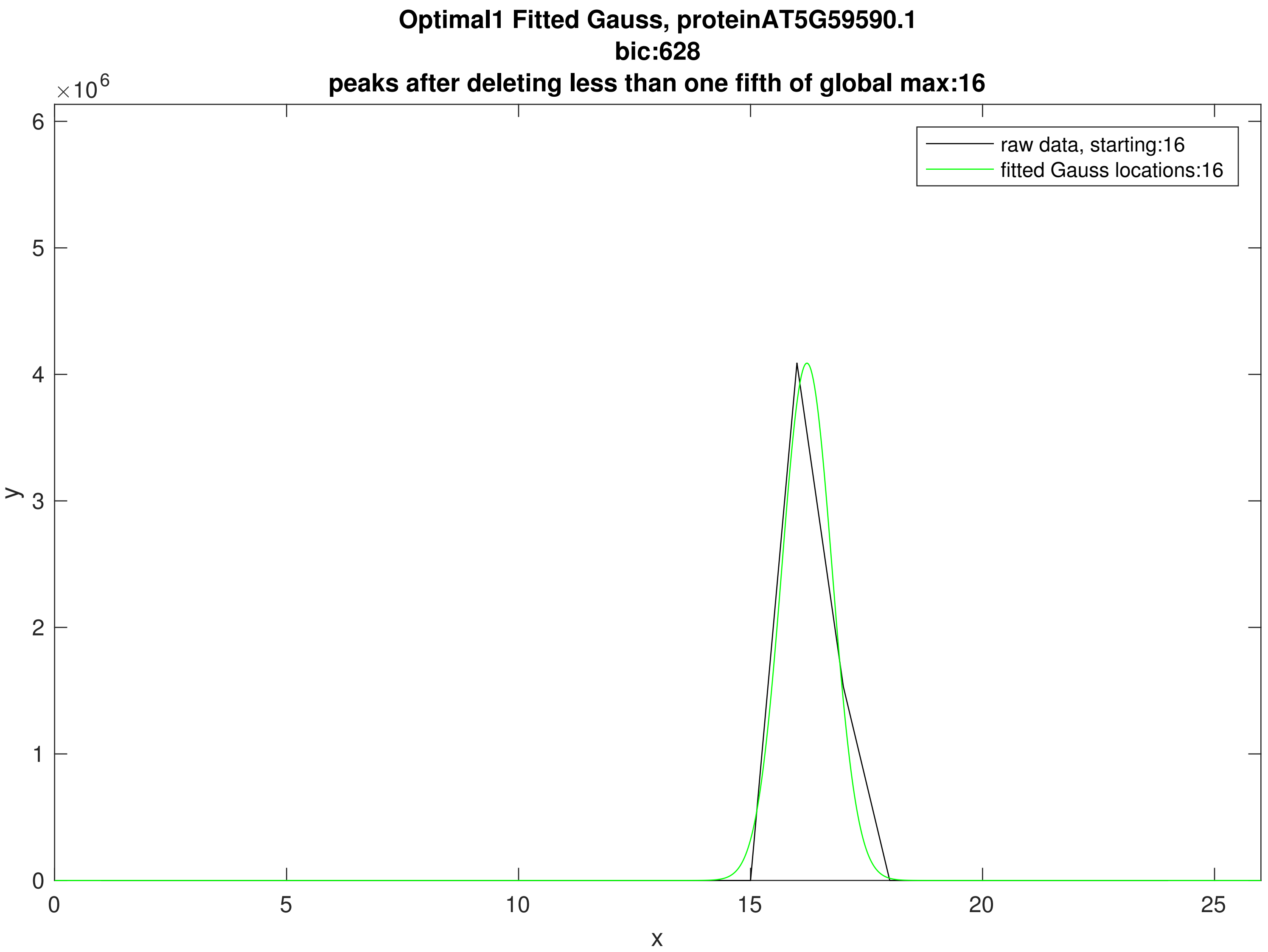
Optimal1 Fitted Gauss, proteinAT5G59590.1
bic:628
peaks after deleting less than one fifth of global max:16
×10^6
raw data, starting:16
fitted Gauss locations:16
y
x
0
1
2
3
4
5
6
0
5
10
15
20
25

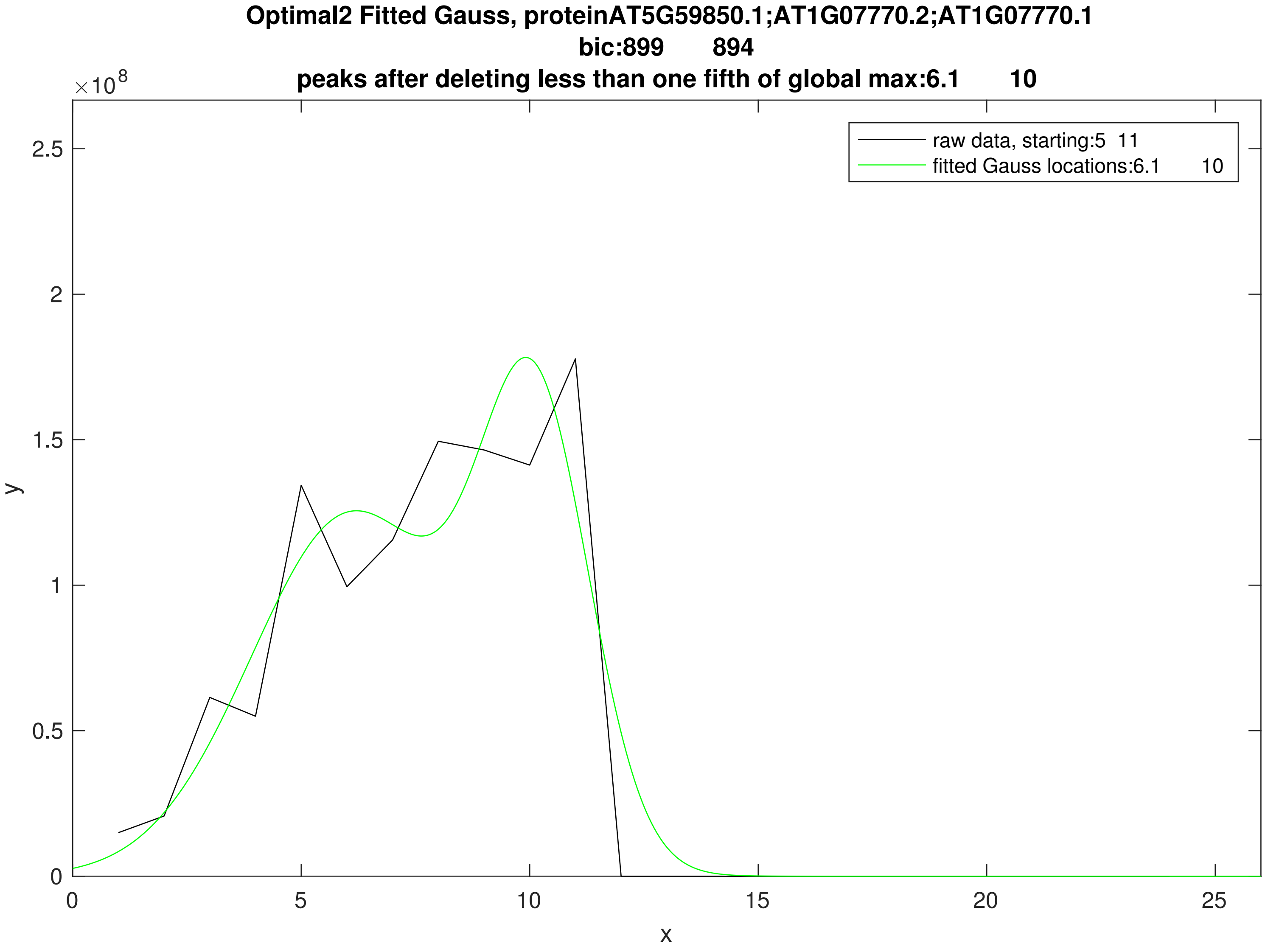

Optimal2 Fitted Gauss, proteinAT5G59850.1;AT1G07770.2;AT1G07770.1
bic:899 894
peaks after deleting less than one fifth of global max:6.1 10
raw data, starting:5 11
fitted Gauss locations:6.1 10
$\times 10^8$
2.5
2
1.5
1
0.5
0
y
0
5
10
15
20
25
x

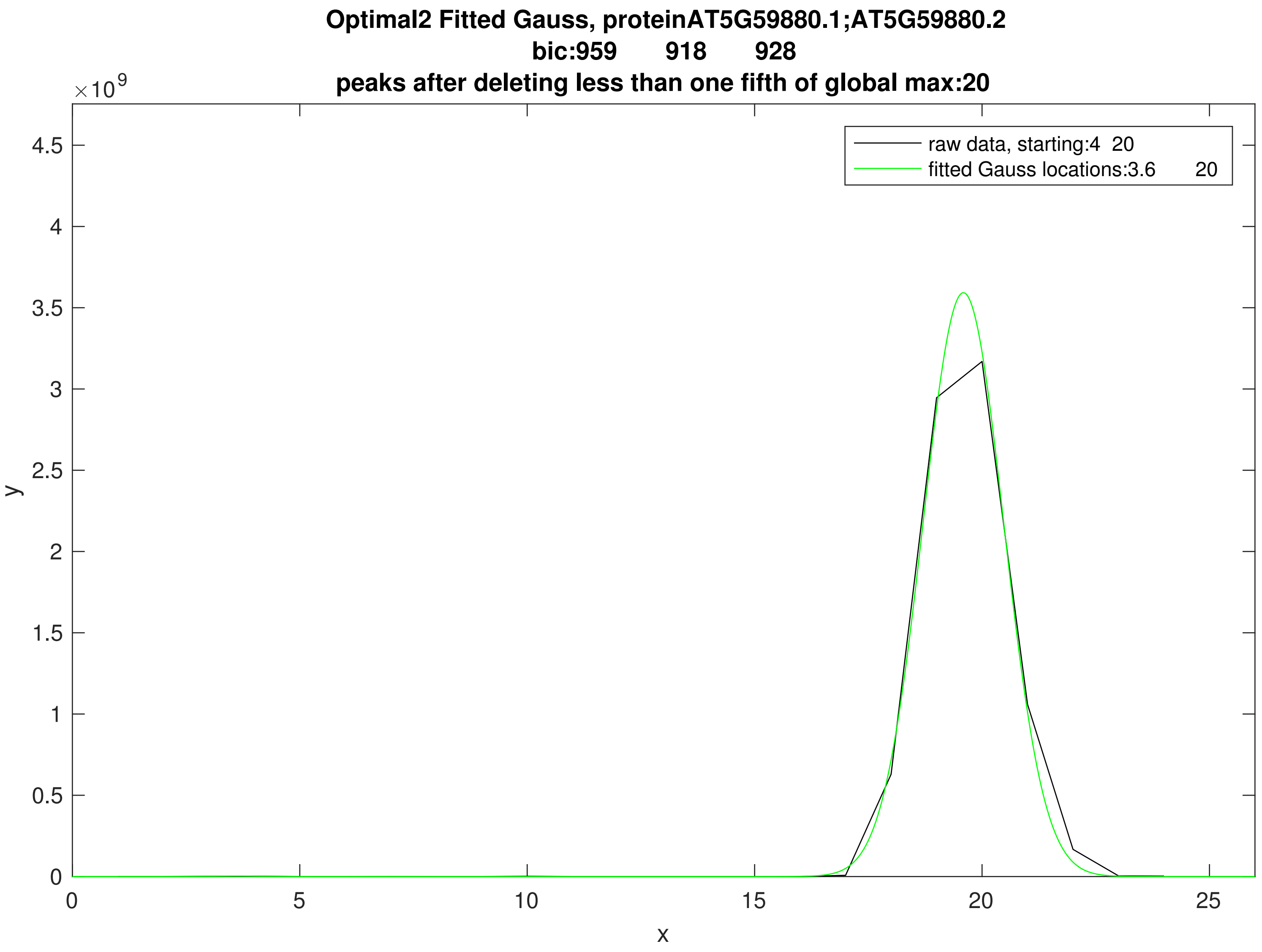
Optimal2 Fitted Gauss, proteinAT5G59880.1;AT5G59880.2
bic:959 918 928
peaks after deleting less than one fifth of global max:20
×10^9
raw data, starting:4 20
fitted Gauss locations:3.6 20
y
x
0
0.5
1
1.5
2
2.5
3
3.5
4
4.5
5
10
15
20
25

#### Optimal1 Fitted Gauss, proteinAT5G59890.1;AT5G59890.2
#### bic:848
#### peaks after deleting less than one fifth of global max:20

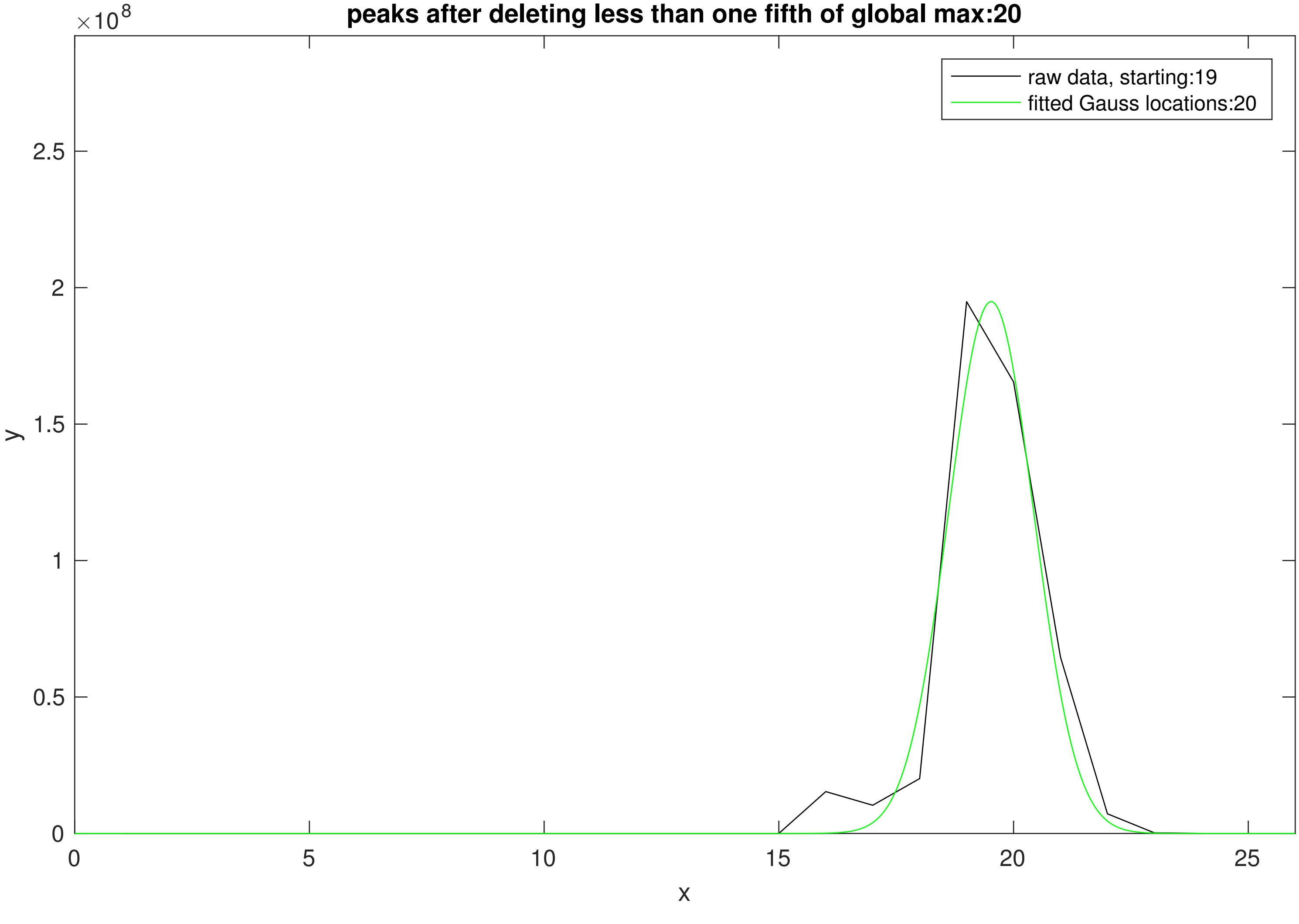

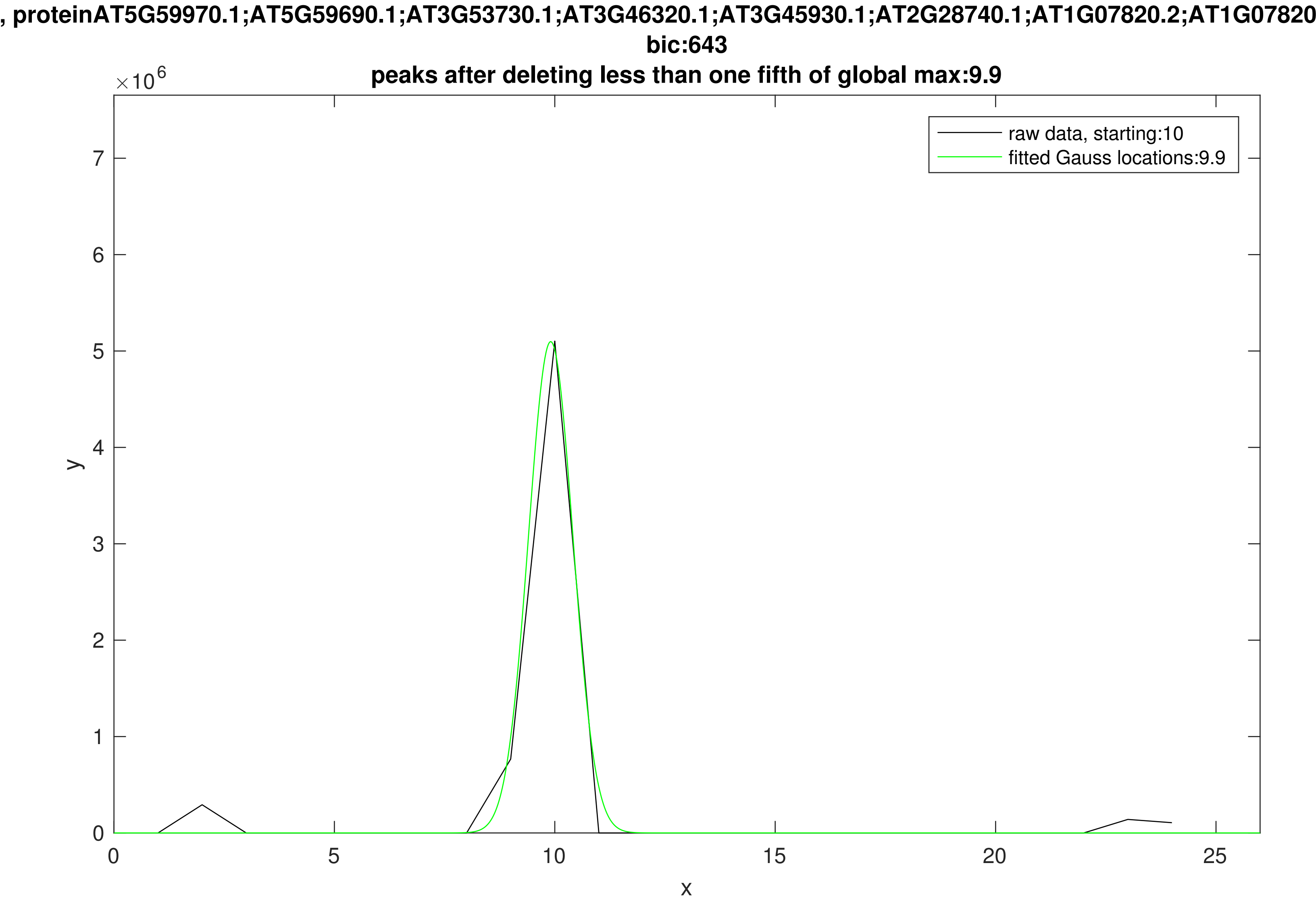

, proteinAT5G59970.1;AT5G59690.1;AT3G53730.1;AT3G46320.1;AT3G45930.1;AT2G28740.1;AT1G07820.2;AT1G07820
bic:643
peaks after deleting less than one fifth of global max:9.9
×10^6
raw data, starting:10
fitted Gauss locations:9.9
y
x
0
1
2
3
4
5
6
7
0
5
10
15
20
25

#### Optimal1 Fitted Gauss, proteinAT5G60160.1
#### bic:894
#### peaks after deleting less than one fifth of global max:6.7

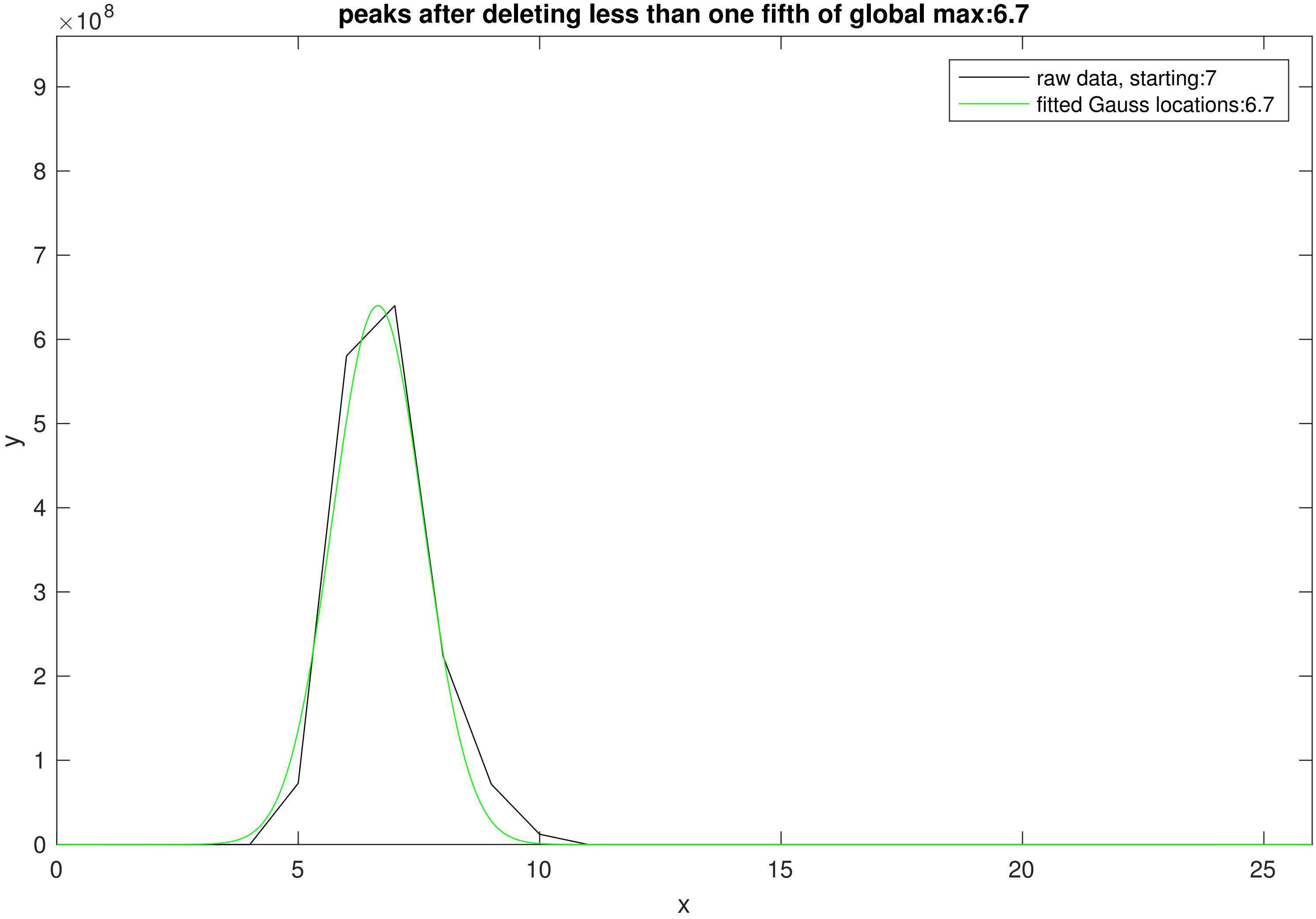

#### Optimal1 Fitted Gauss, proteinAT5G60340.1
#### bic:706
#### peaks after deleting less than one fifth of global max:19

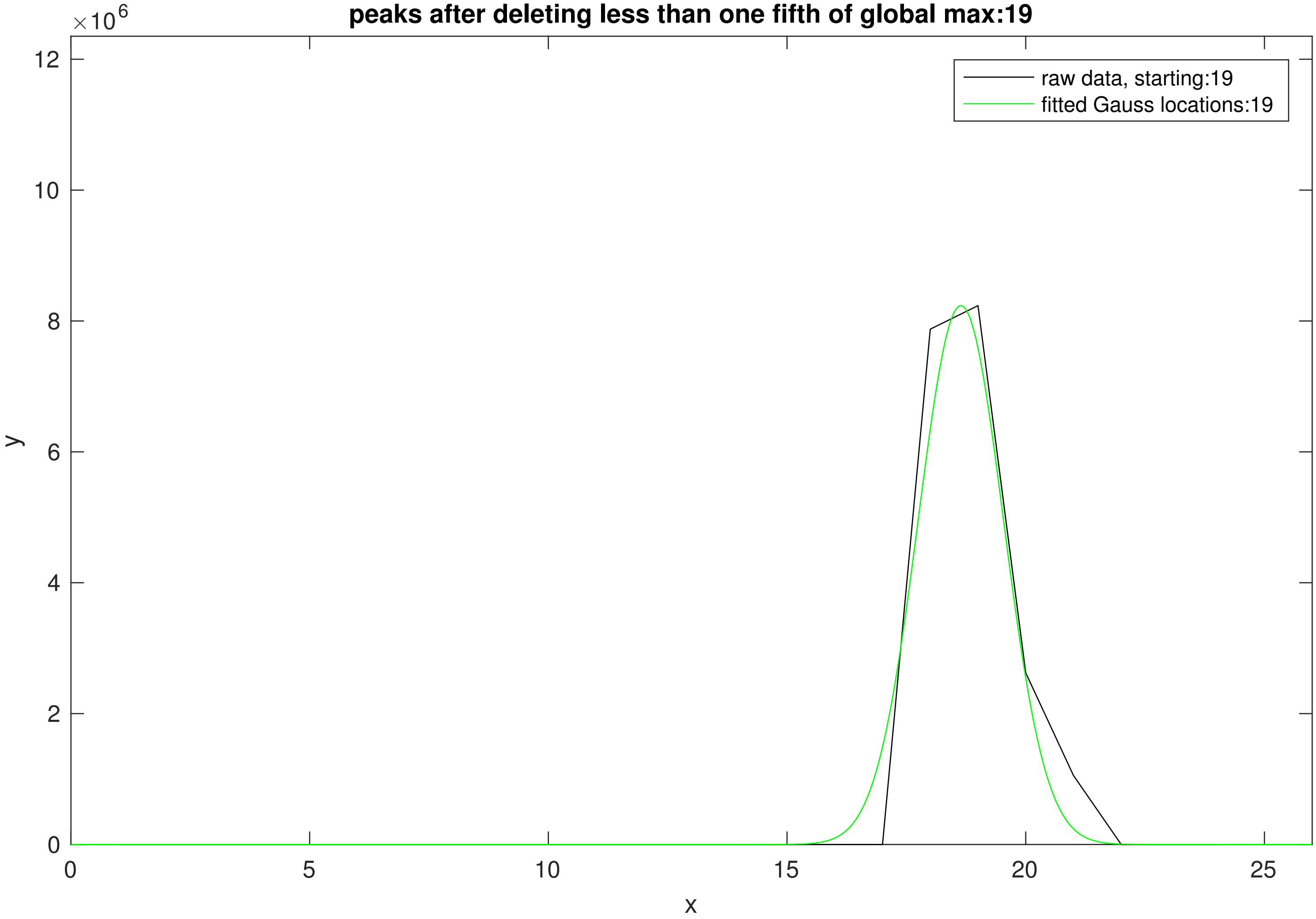

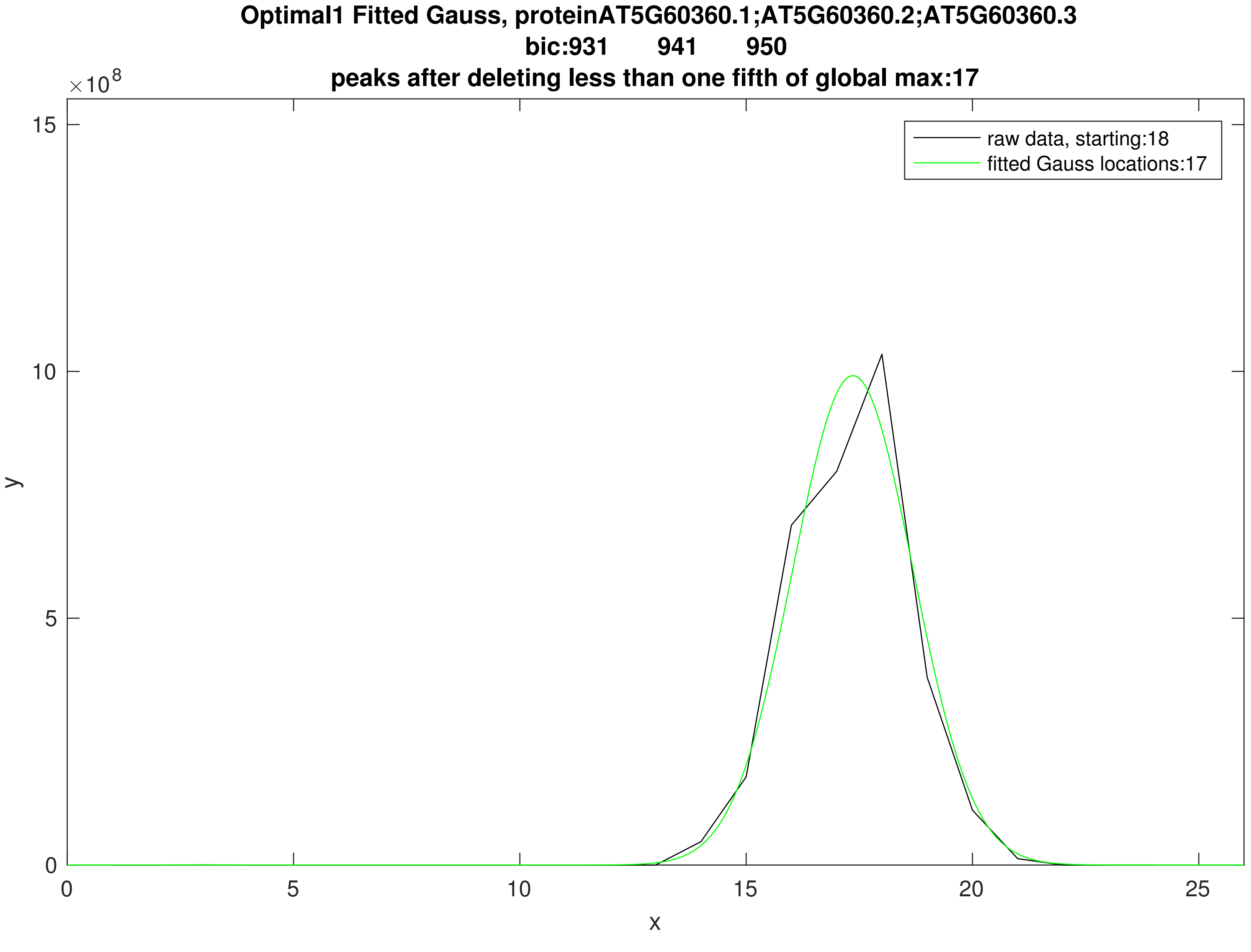

Optimal1 Fitted Gauss, proteinAT5G60360.1;AT5G60360.2;AT5G60360.3
bic:931 941 950
peaks after deleting less than one fifth of global max:17
×10^8
raw data, starting:18
fitted Gauss locations:17
y
x
0
5
10
15
20
25

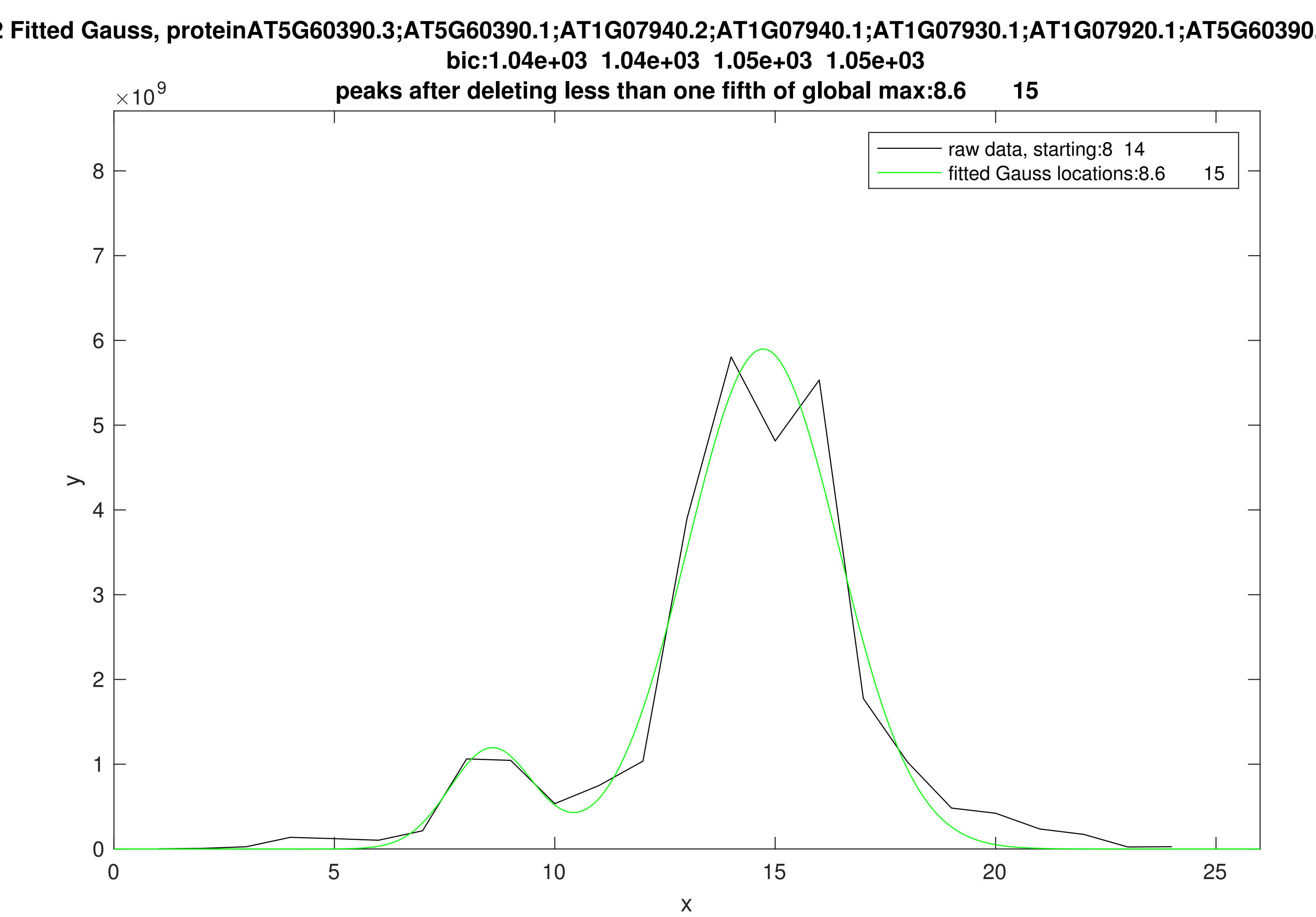
2 Fitted Gauss, proteinAT5G60390.3;AT5G60390.1;AT1G07940.2;AT1G07940.1;AT1G07930.1;AT1G07920.1;AT5G60390.
bic:1.04e+03 1.04e+03 1.05e+03 1.05e+03
peaks after deleting less than one fifth of global max:8.6 15
×10^9
raw data, starting:8 14
fitted Gauss locations:8.6 15
y
x
0
1
2
3
4
5
6
7
8
0
5
10
15
20
25

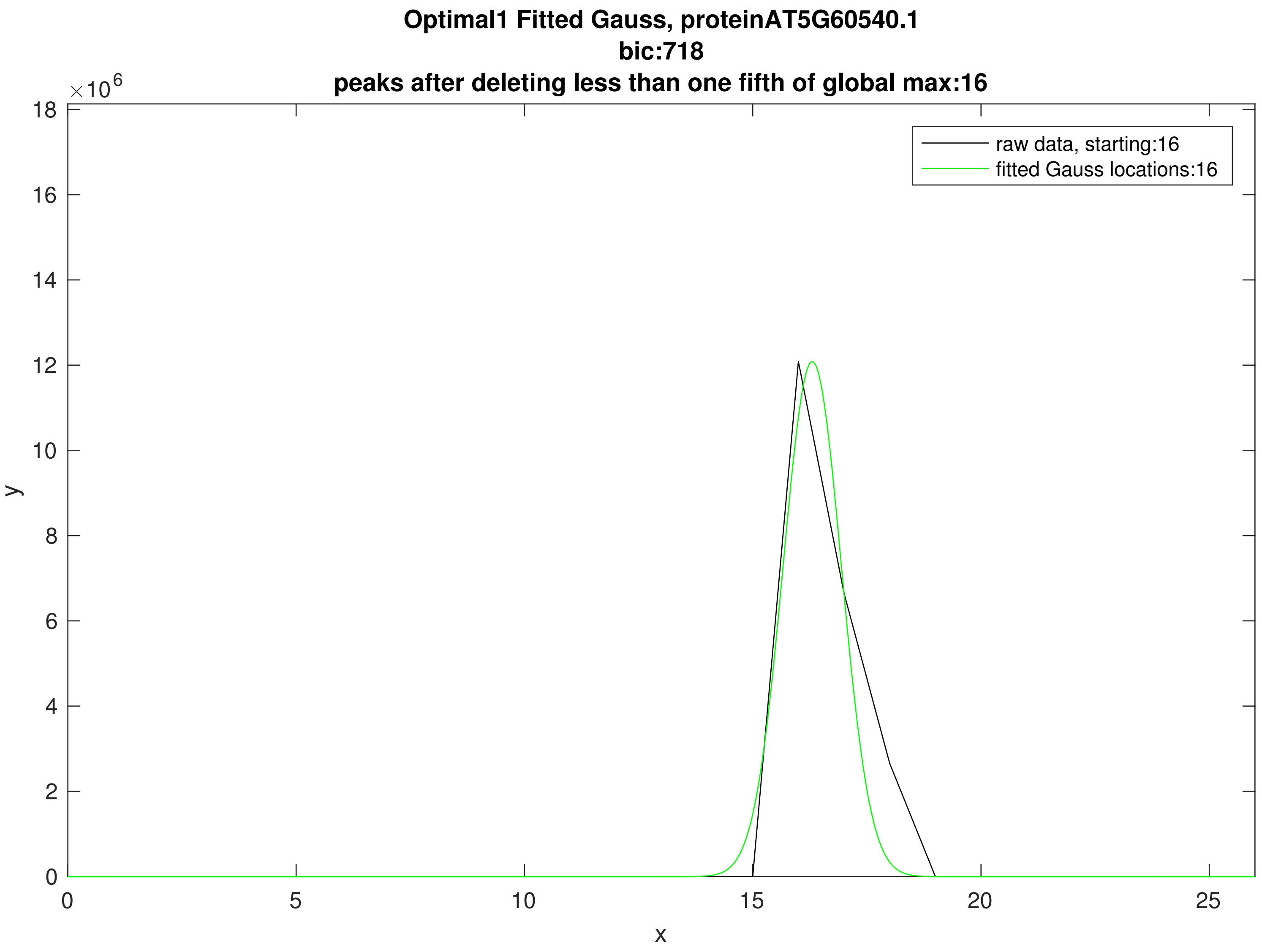

Optimal1 Fitted Gauss, proteinAT5G60540.1
bic:718
peaks after deleting less than one fifth of global max:16
×10^6
18
16
14
12
10
8
6
4
2
0
y
0
5
10
15
20
25
x
raw data, starting:16
fitted Gauss locations:16

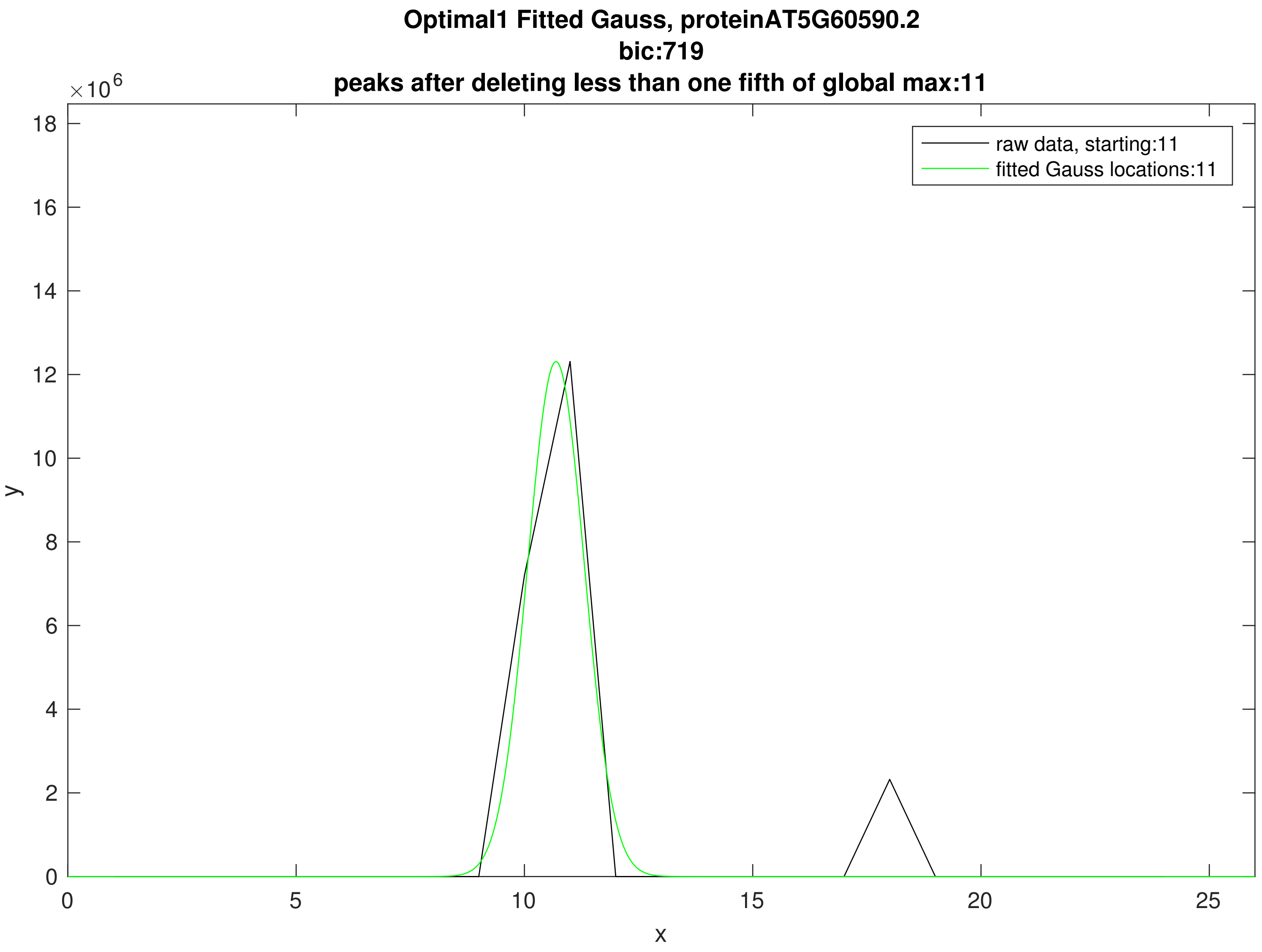

Optimal1 Fitted Gauss, proteinAT5G60590.2
bic:719
peaks after deleting less than one fifth of global max:11
×10^6
18
16
14
12
10
8
6
4
2
0
y
0
5
10
15
20
25
x
raw data, starting:11
fitted Gauss locations:11

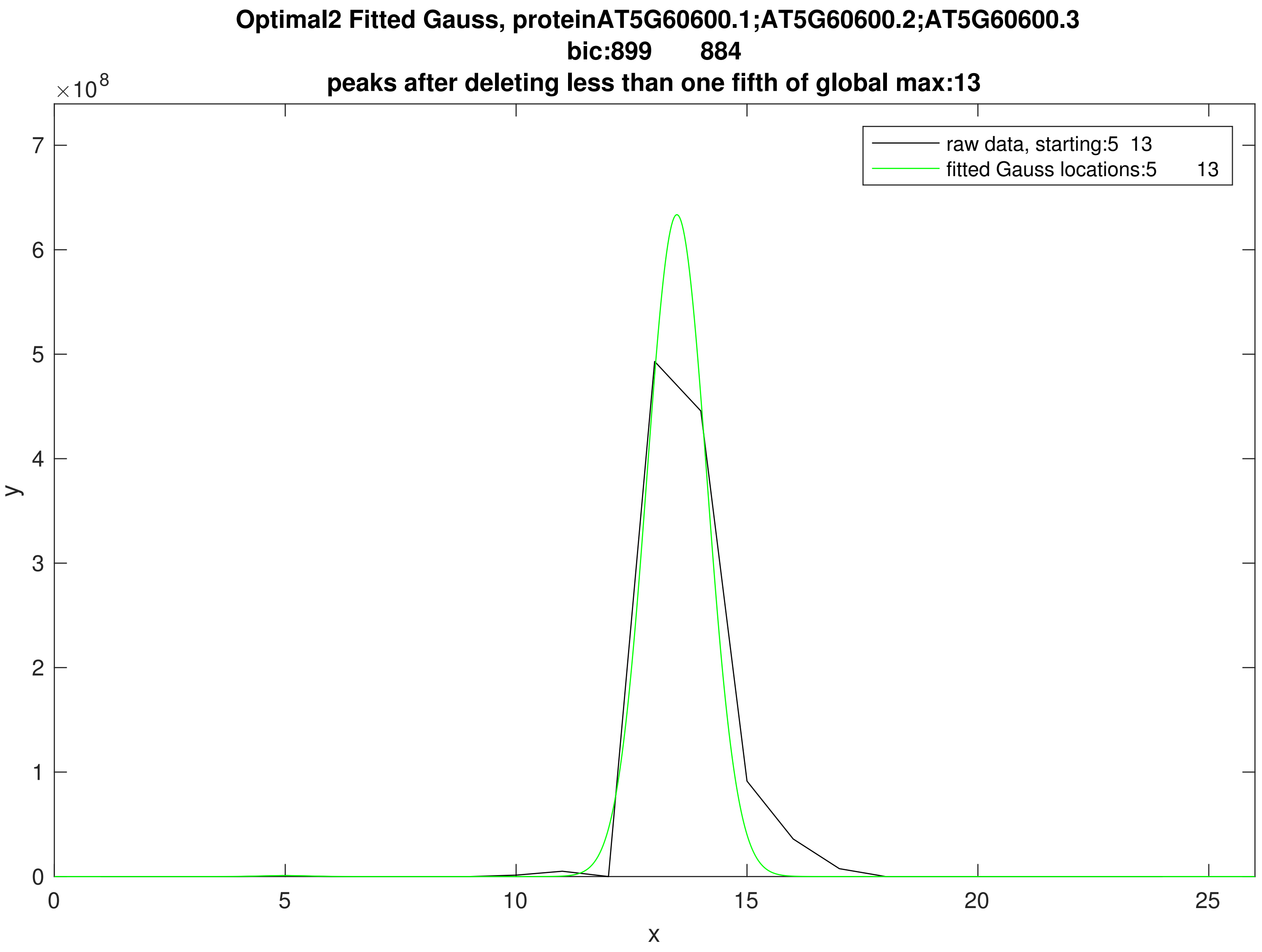
Optimal2 Fitted Gauss, proteinAT5G60600.1;AT5G60600.2;AT5G60600.3
bic:899 884
peaks after deleting less than one fifth of global max:13
×10^8
raw data, starting:5 13
fitted Gauss locations:5 13
y
x
0
1
2
3
4
5
6
7
0
5
10
15
20
25

**Optimal1 Fitted Gauss, proteinAT5G60640.1;AT5G60640.3;AT5G60640.2**
**bic:917 921 930**
**peaks after deleting less than one fifth of global max:11**

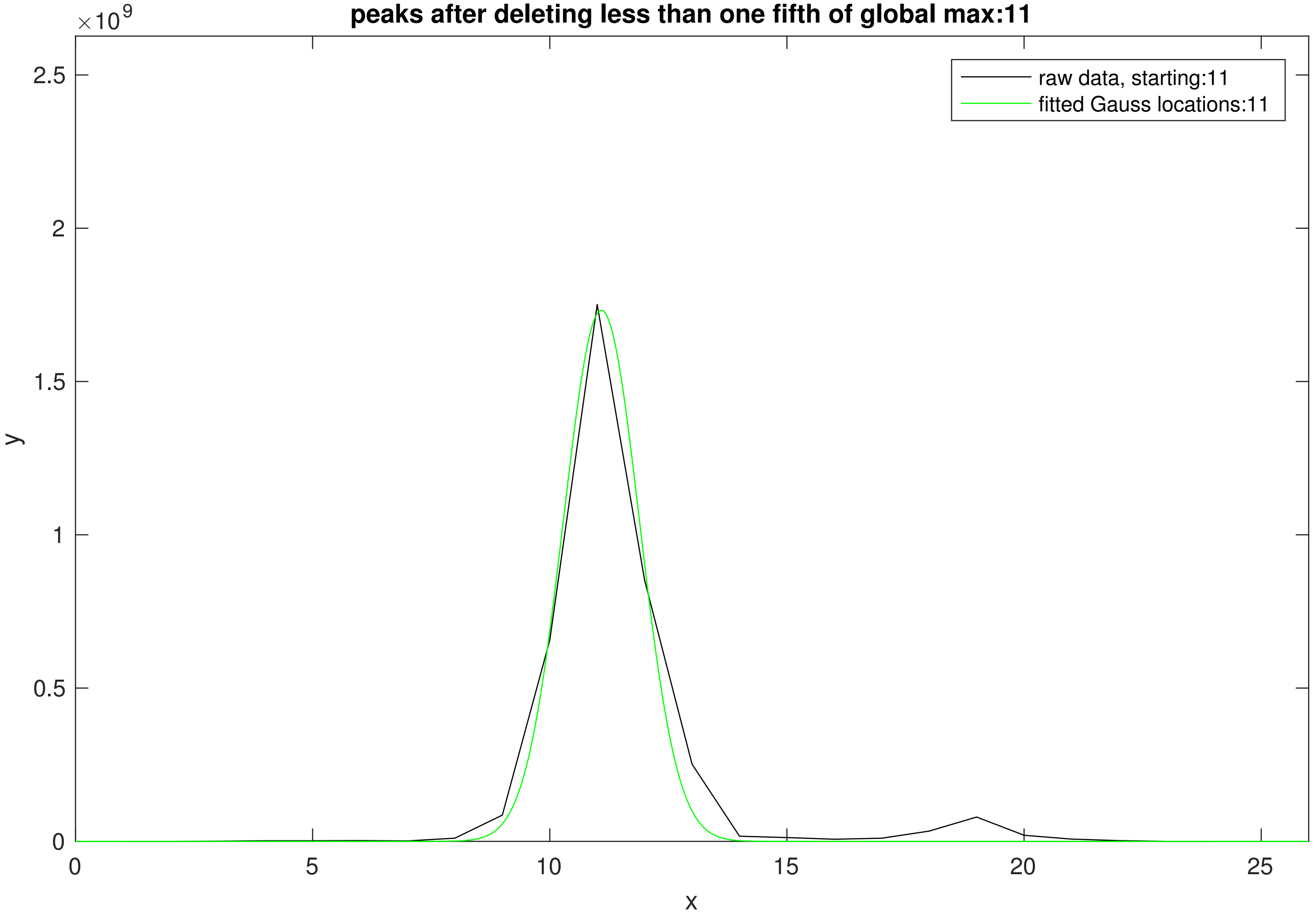

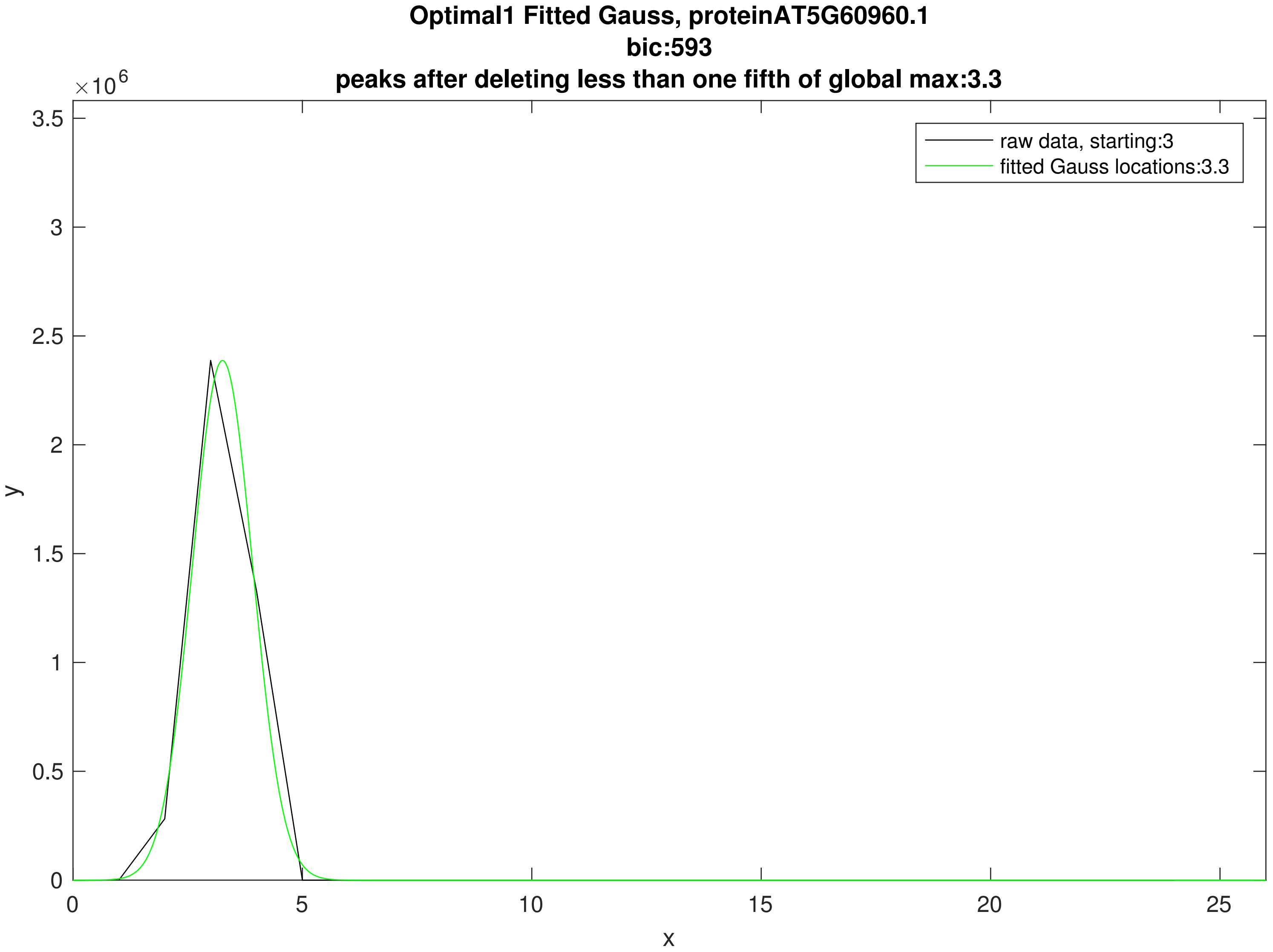
Optimal1 Fitted Gauss, proteinAT5G60960.1
bic:593
peaks after deleting less than one fifth of global max:3.3
×10^6
raw data, starting:3
fitted Gauss locations:3.3
3.5
3
2.5
2
1.5
1
0.5
0
y
0
5
10
15
20
25
x

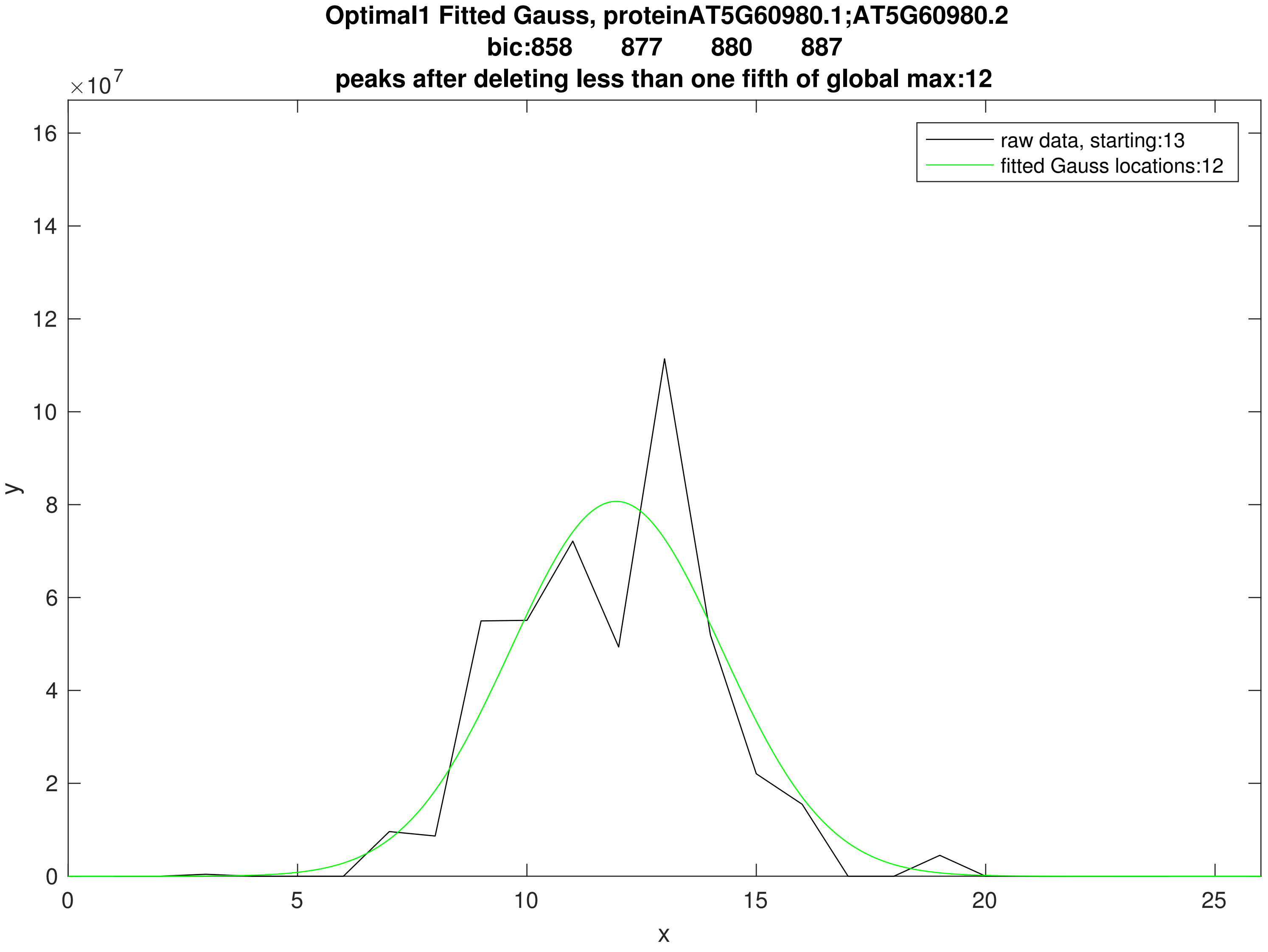

Optimal1 Fitted Gauss, proteinAT5G60980.1;AT5G60980.2
bic:858 877 880 887
peaks after deleting less than one fifth of global max:12
×10^7
raw data, starting:13
fitted Gauss locations:12
0
2
4
6
8
10
12
14
16
y
0
5
10
15
20
25
x

**Optimal2 Fitted Gauss, proteinAT5G61030.1**
**bic:786 780**
**peaks after deleting less than one fifth of global max:17**

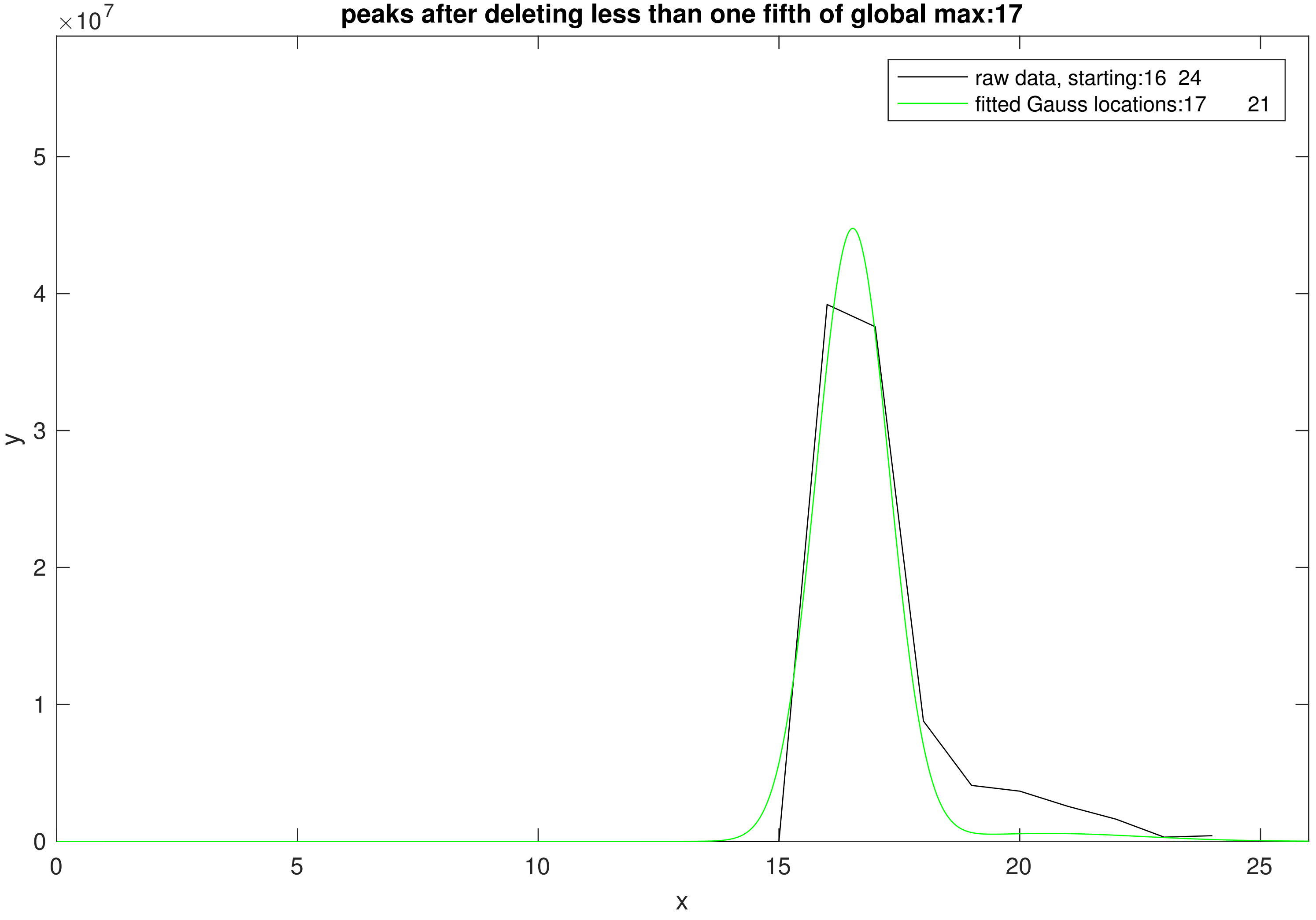

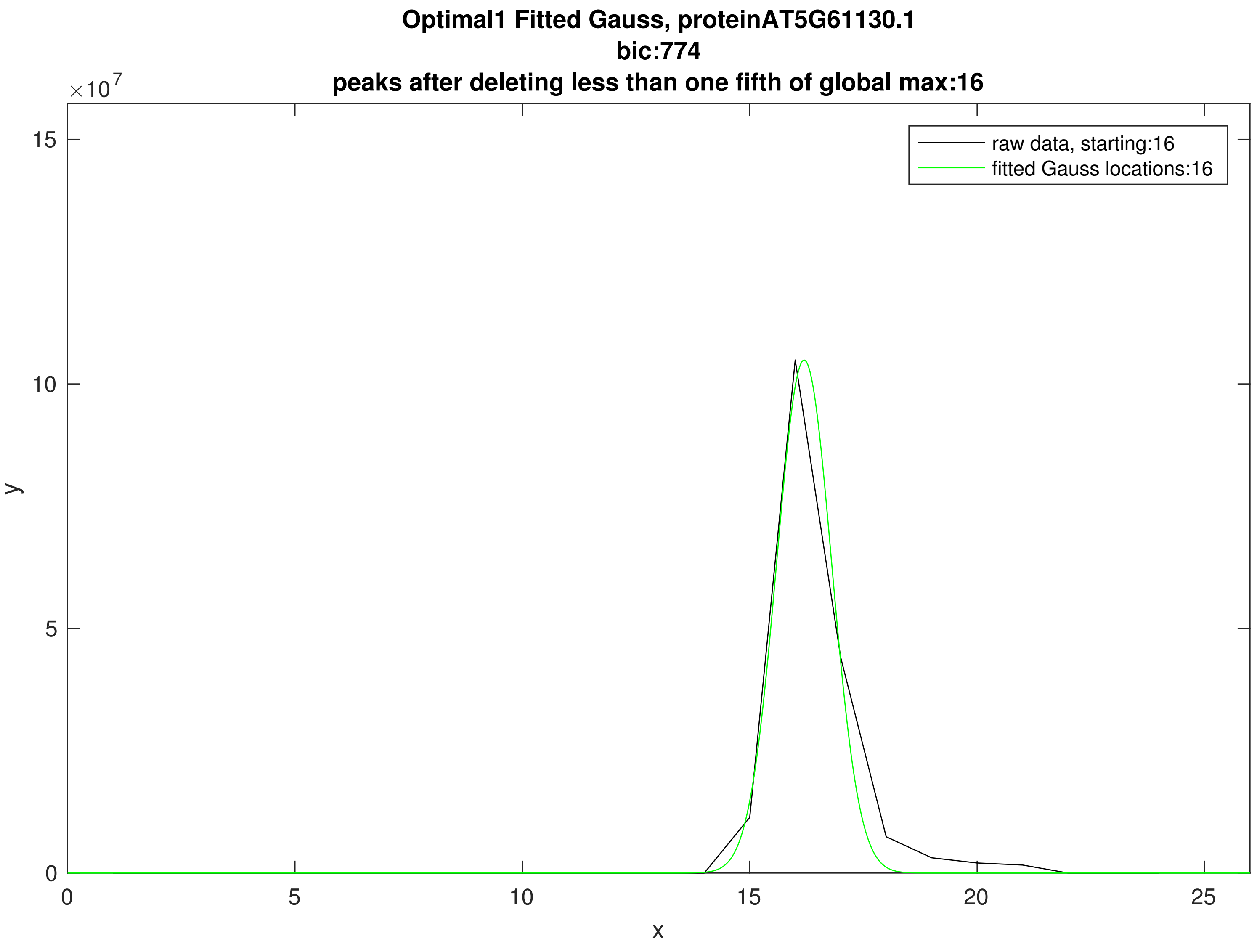
Optimal1 Fitted Gauss, proteinAT5G61130.1
bic:774
peaks after deleting less than one fifth of global max:16
×10^7
15
10
5
0
y
0
5
10
15
20
25
x
raw data, starting:16
fitted Gauss locations:16

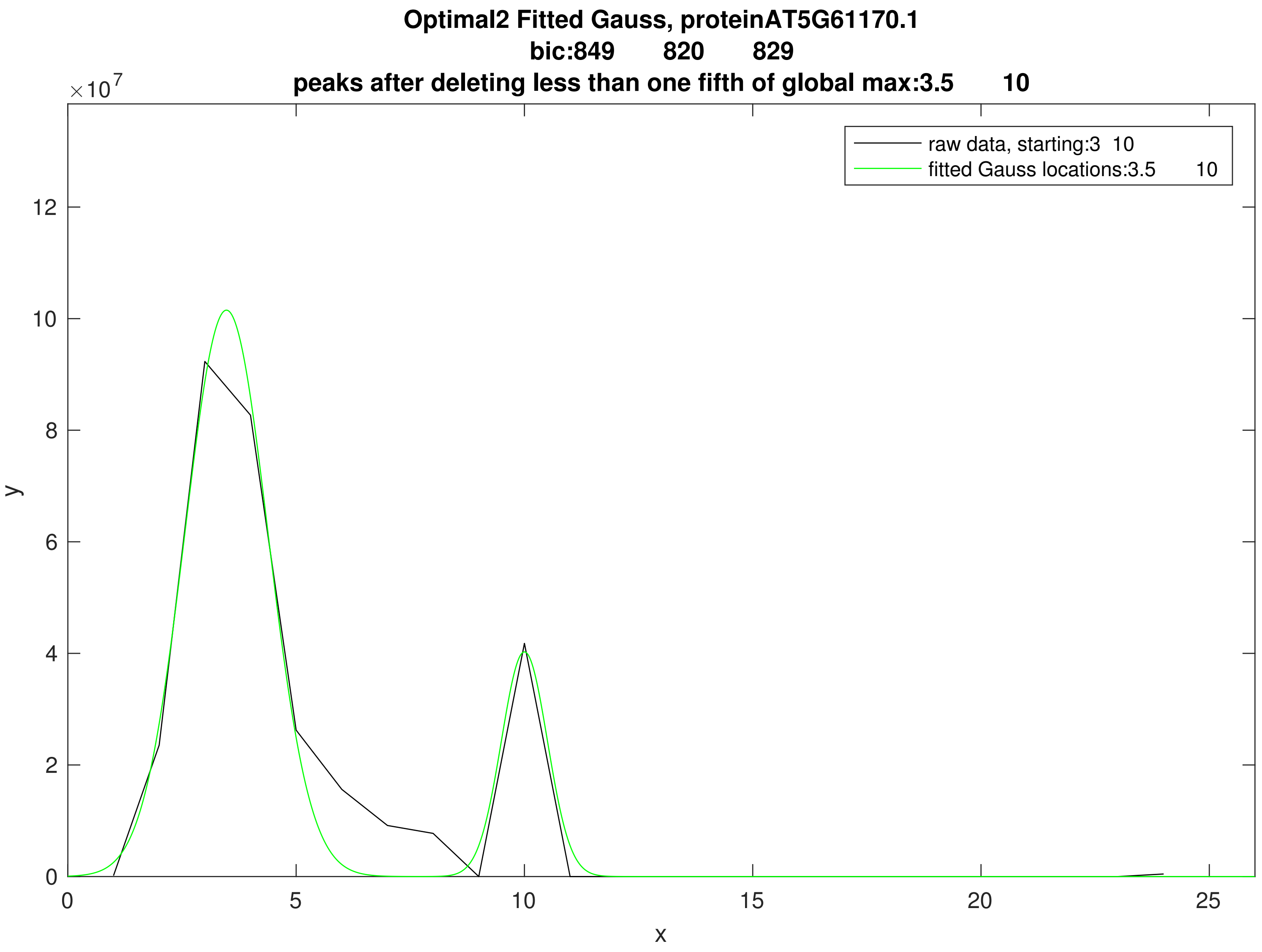

Optimal2 Fitted Gauss, proteinAT5G61170.1
bic:849 820 829
peaks after deleting less than one fifth of global max:3.5 10
×10^7
raw data, starting:3 10
fitted Gauss locations:3.5 10
y
x
0
2
4
6
8
10
12
0
5
10
15
20
25

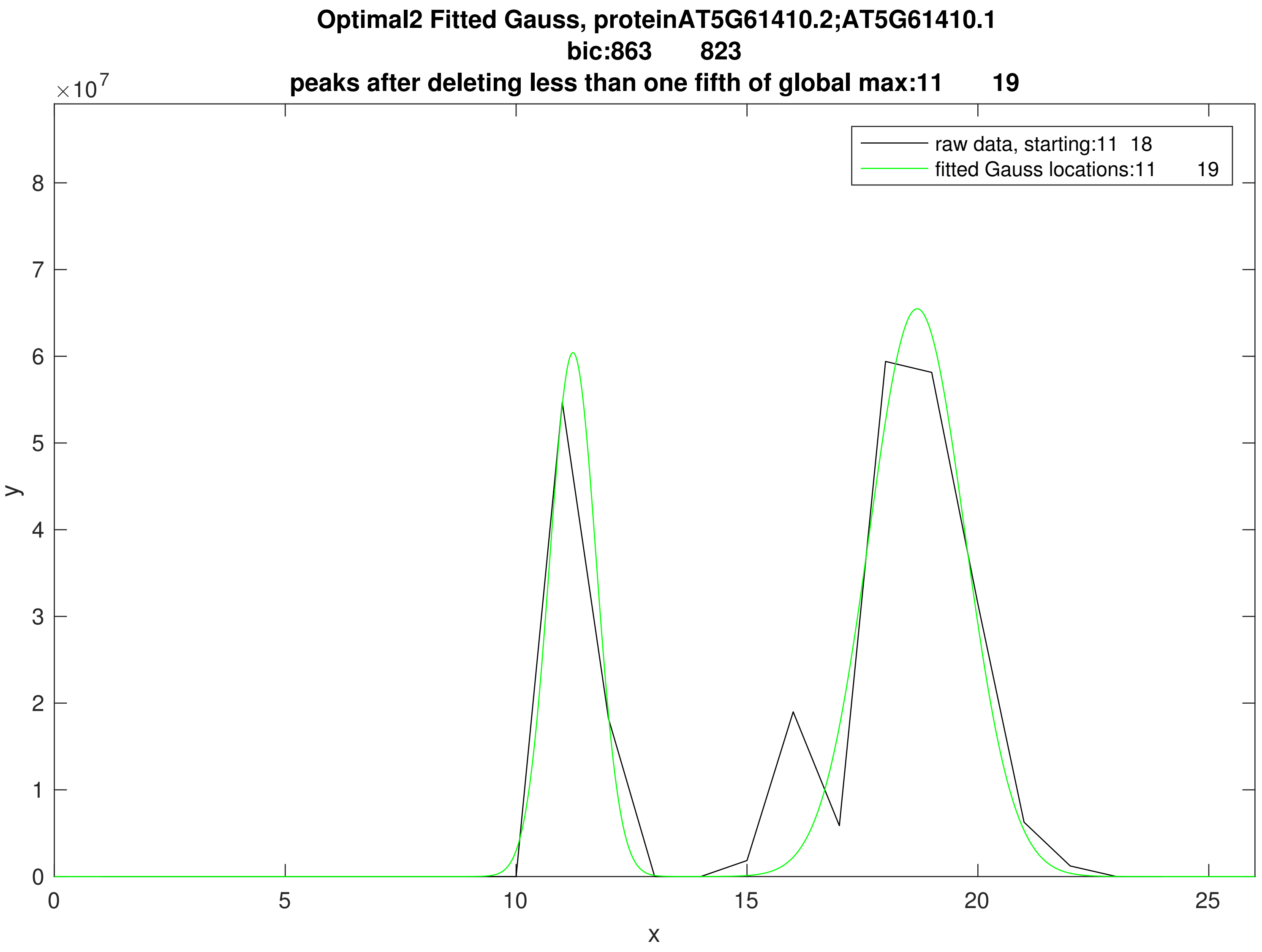

Optimal2 Fitted Gauss, proteinAT5G61410.2;AT5G61410.1
bic:863 823
peaks after deleting less than one fifth of global max:11 19
×10^7
raw data, starting:11 18
fitted Gauss locations:11 19
y
x
0
1
2
3
4
5
6
7
8
0
5
10
15
20
25

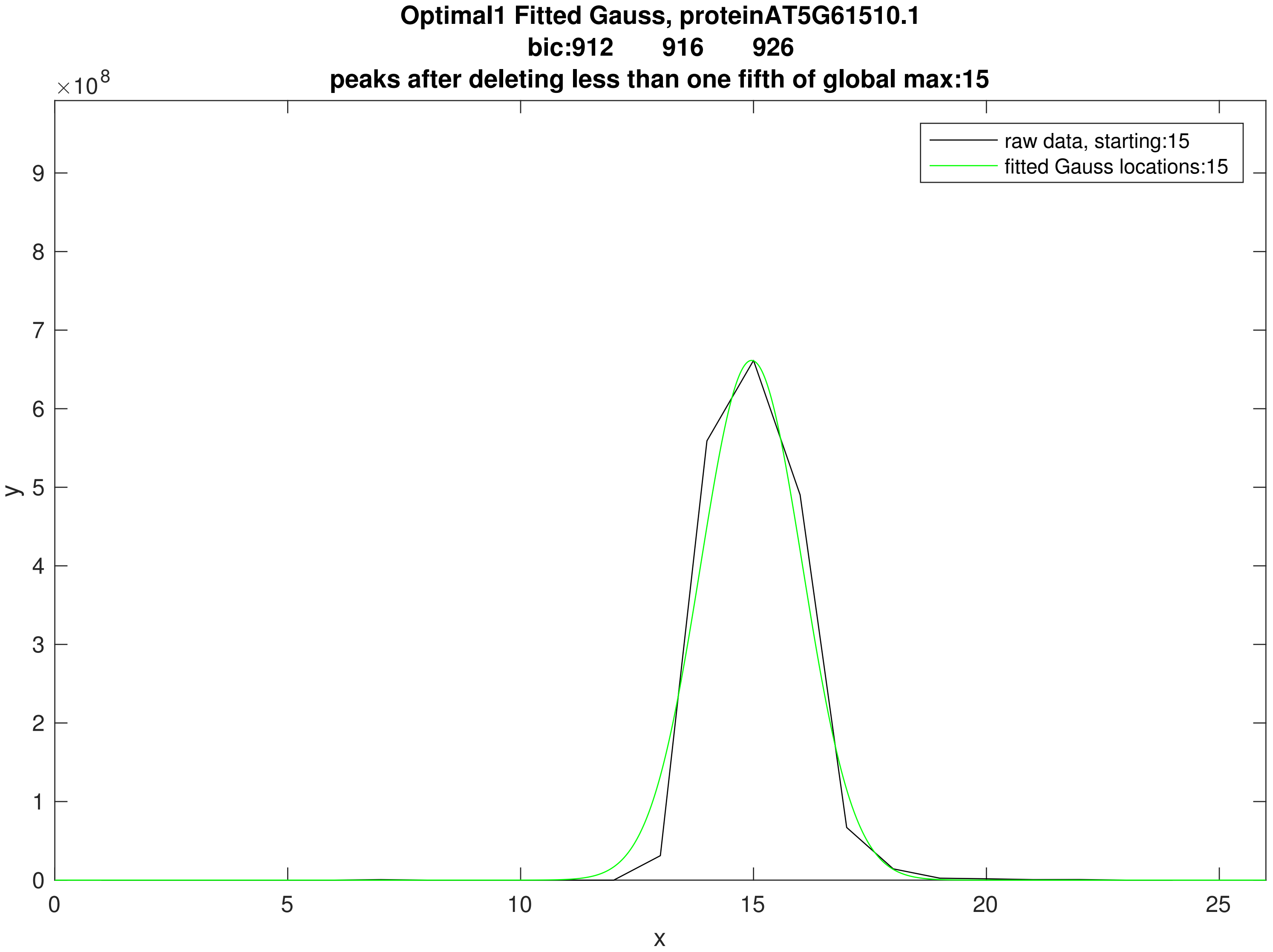

Optimal1 Fitted Gauss, proteinAT5G61510.1
bic:912 916 926
peaks after deleting less than one fifth of global max:15
×10^8
raw data, starting:15
fitted Gauss locations:15
y
x
0
1
2
3
4
5
6
7
8
9
0
5
10
15
20
25

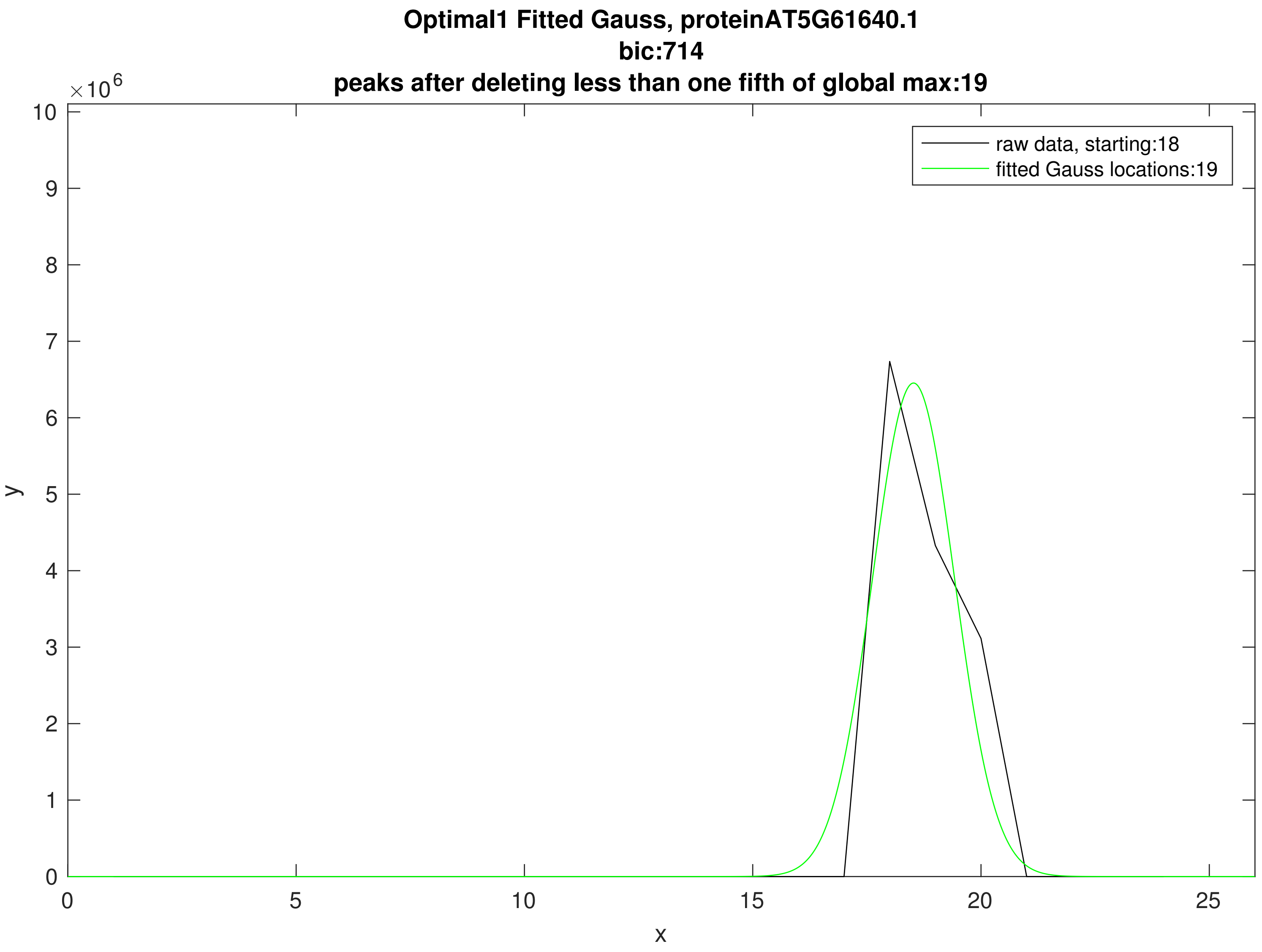
Optimal1 Fitted Gauss, proteinAT5G61640.1
bic:714
peaks after deleting less than one fifth of global max:19
×10^6
10
9
8
7
6
5
4
3
2
1
0
y
0
5
10
15
20
25
x
raw data, starting:18
fitted Gauss locations:19

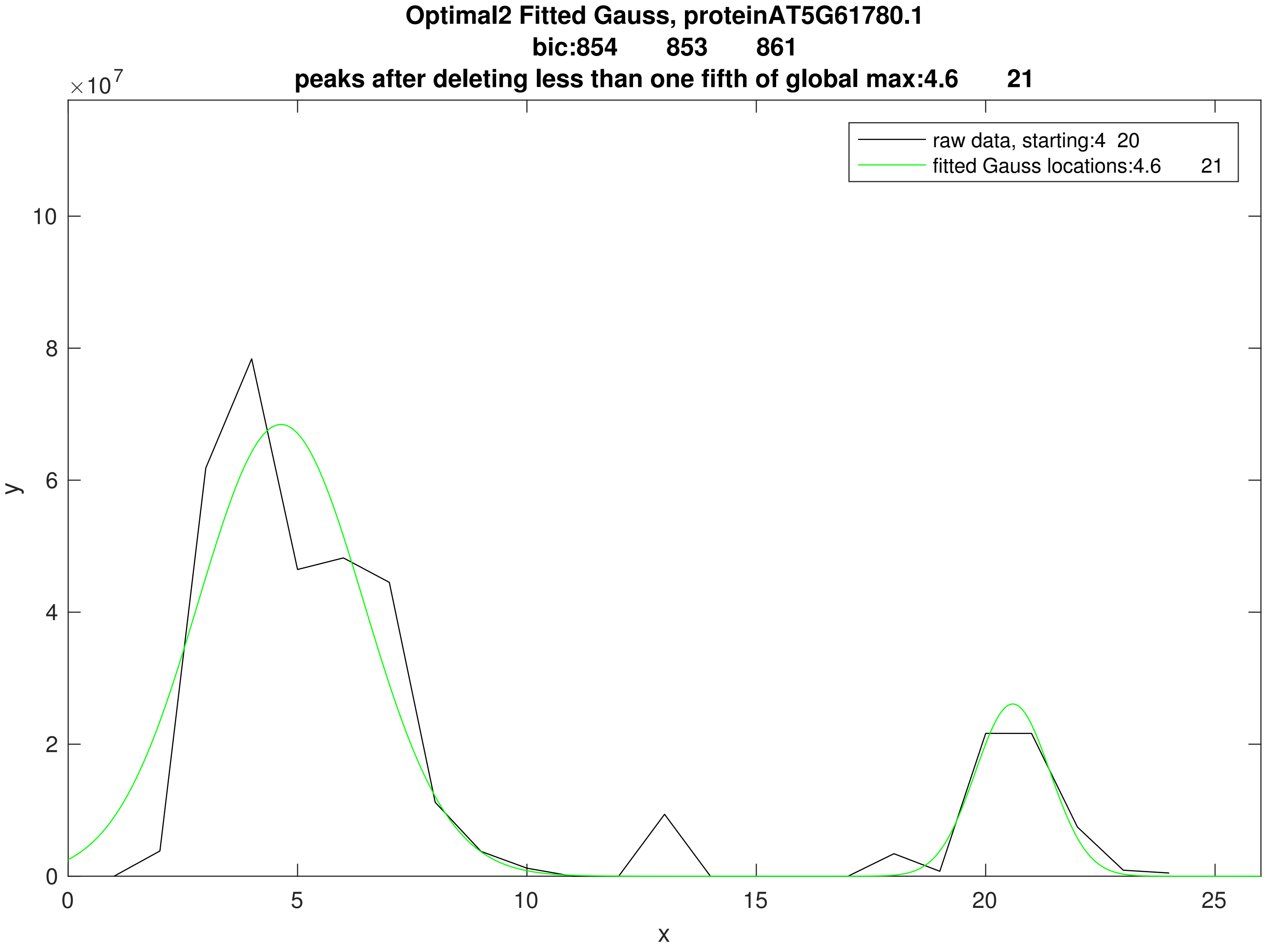

Optimal2 Fitted Gauss, proteinAT5G61780.1
bic:854 853 861
peaks after deleting less than one fifth of global max:4.6 21
×10^7
raw data, starting:4 20
fitted Gauss locations:4.6 21
y
x
0
2
4
6
8
10
0
5
10
15
20
25

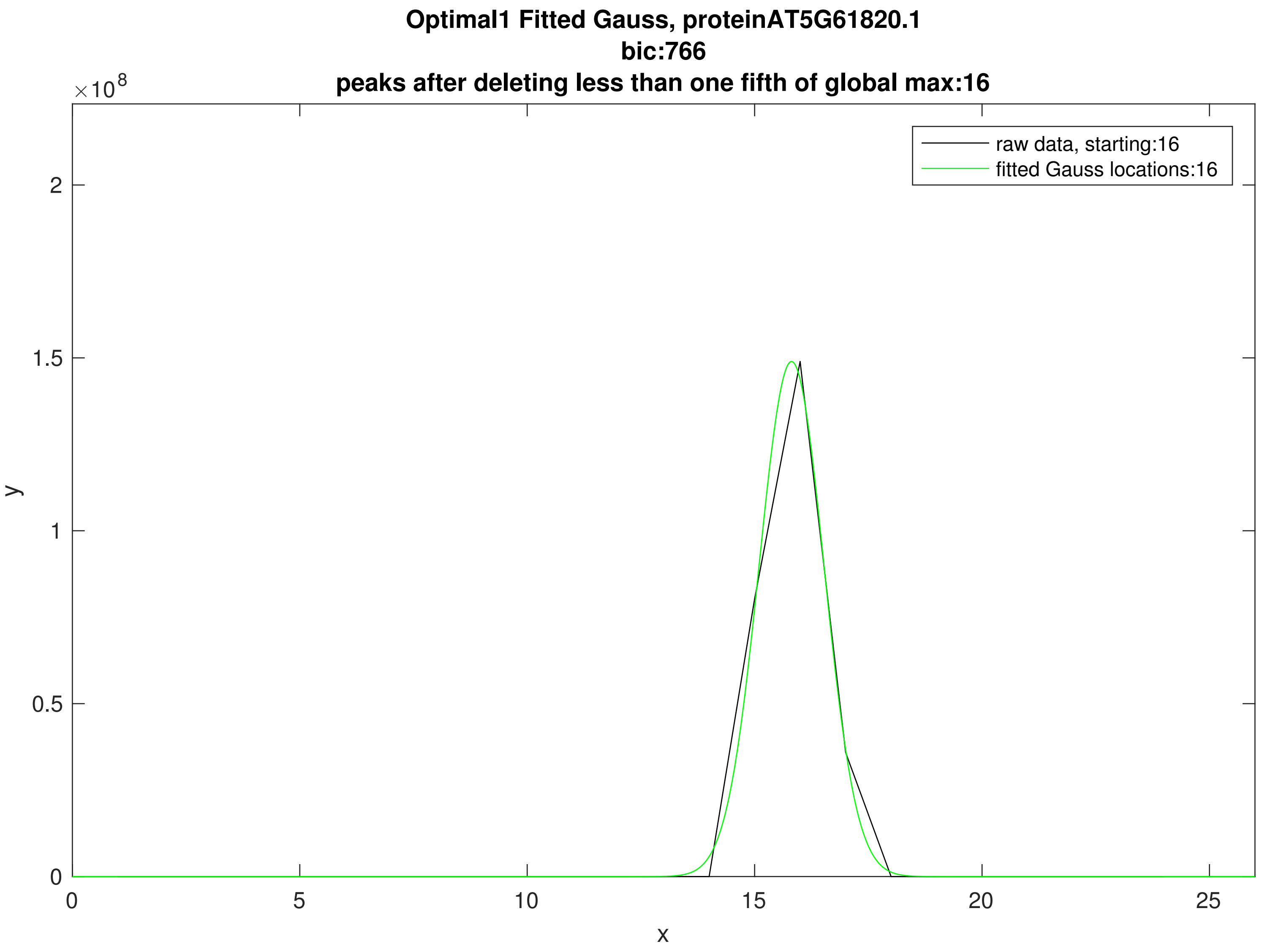

Optimal1 Fitted Gauss, proteinAT5G61820.1
bic:766
peaks after deleting less than one fifth of global max:16
×10^8
raw data, starting:16
fitted Gauss locations:16
2
1.5
1
0.5
0
y
0
5
10
15
20
25
x

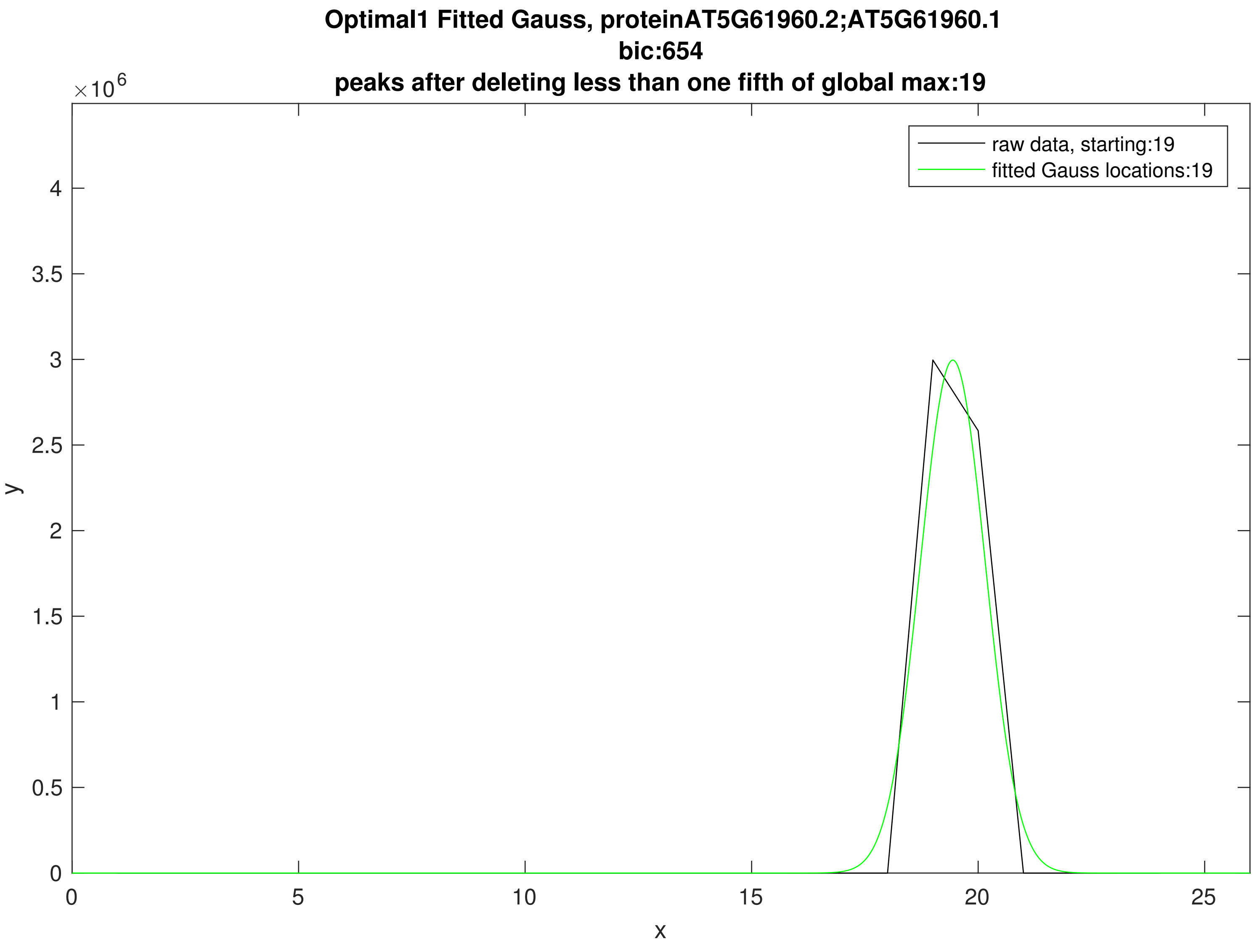
Optimal1 Fitted Gauss, proteinAT5G61960.2;AT5G61960.1
bic:654
peaks after deleting less than one fifth of global max:19
×10^6
raw data, starting:19
fitted Gauss locations:19
y
x
0
0.5
1
1.5
2
2.5
3
3.5
4
0
5
10
15
20
25

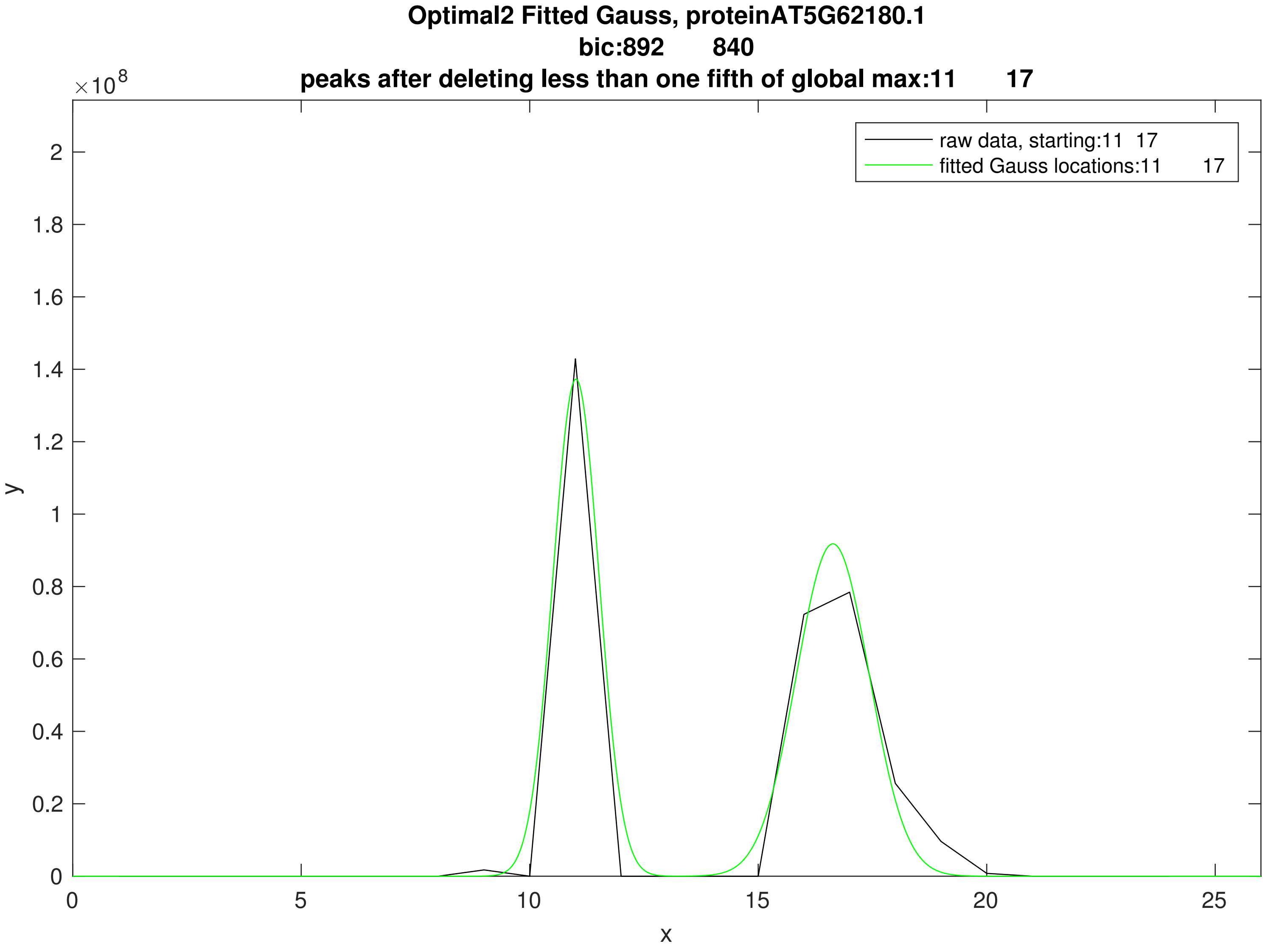

Optimal2 Fitted Gauss, proteinAT5G62180.1
bic:892 840
peaks after deleting less than one fifth of global max:11 17
raw data, starting:11 17
fitted Gauss locations:11 17
×10^8
2
1.8
1.6
1.4
1.2
1
0.8
0.6
0.4
0.2
0
y
0
5
10
15
20
25
x

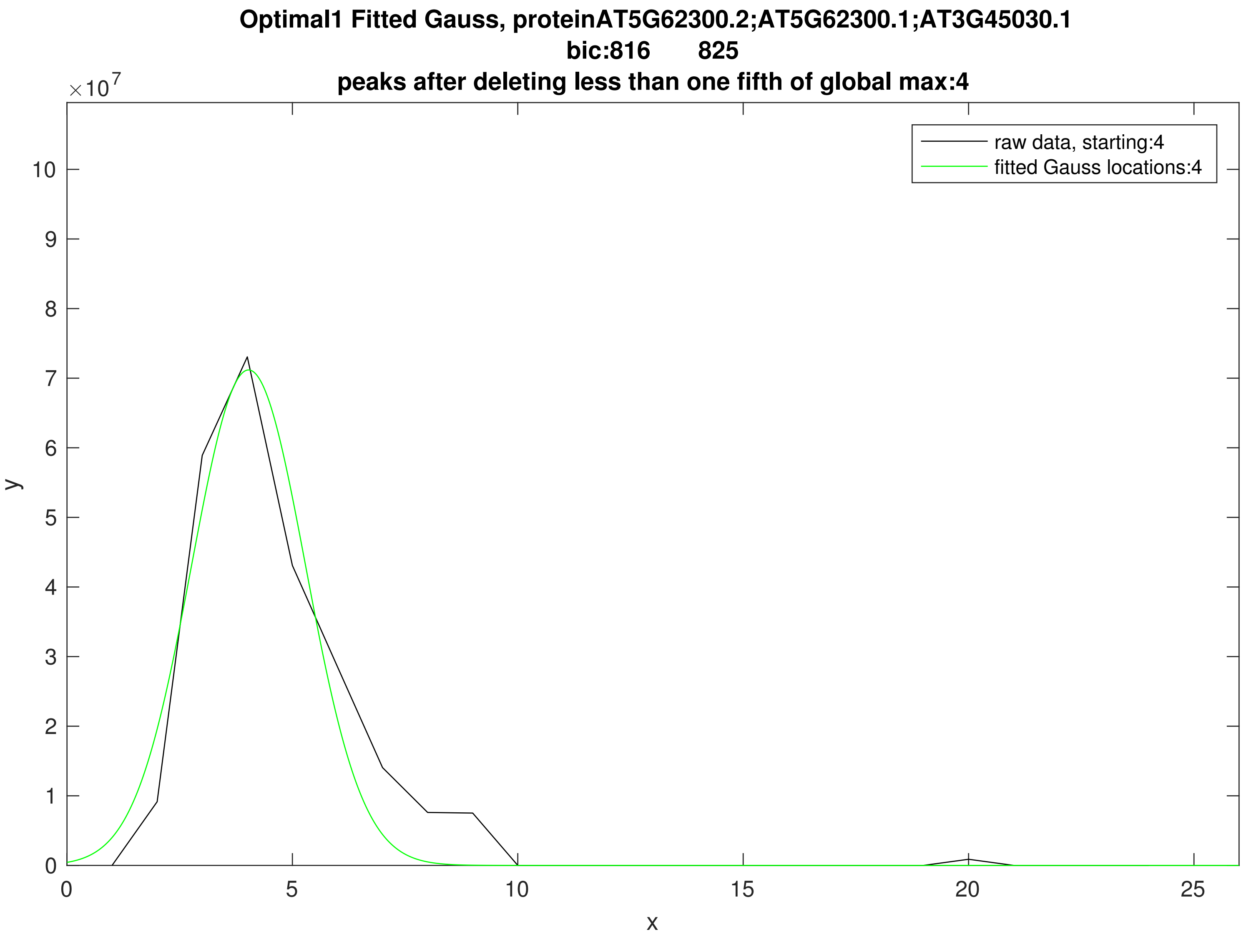
Optimal1 Fitted Gauss, proteinAT5G62300.2;AT5G62300.1;AT3G45030.1
bic:816 825
peaks after deleting less than one fifth of global max:4
×10^7
raw data, starting:4
fitted Gauss locations:4
y
x
0
1
2
3
4
5
6
7
8
9
10
0
5
10
15
20
25

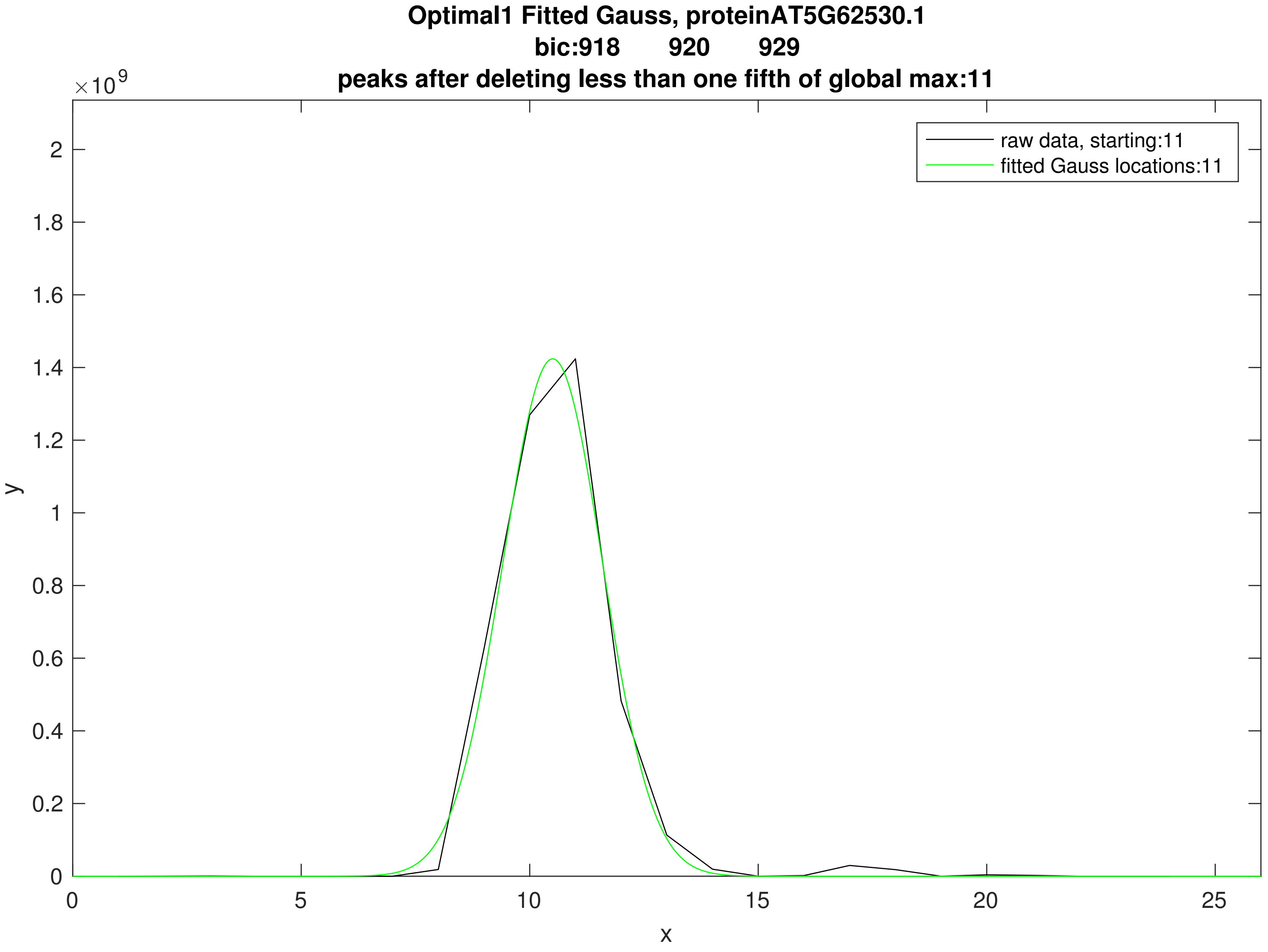
Optimal1 Fitted Gauss, proteinAT5G62530.1
bic:918 920 929
peaks after deleting less than one fifth of global max:11
×10^9
raw data, starting:11
fitted Gauss locations:11
y
x
0
0.2
0.4
0.6
0.8
1
1.2
1.4
1.6
1.8
2
5
10
15
20
25

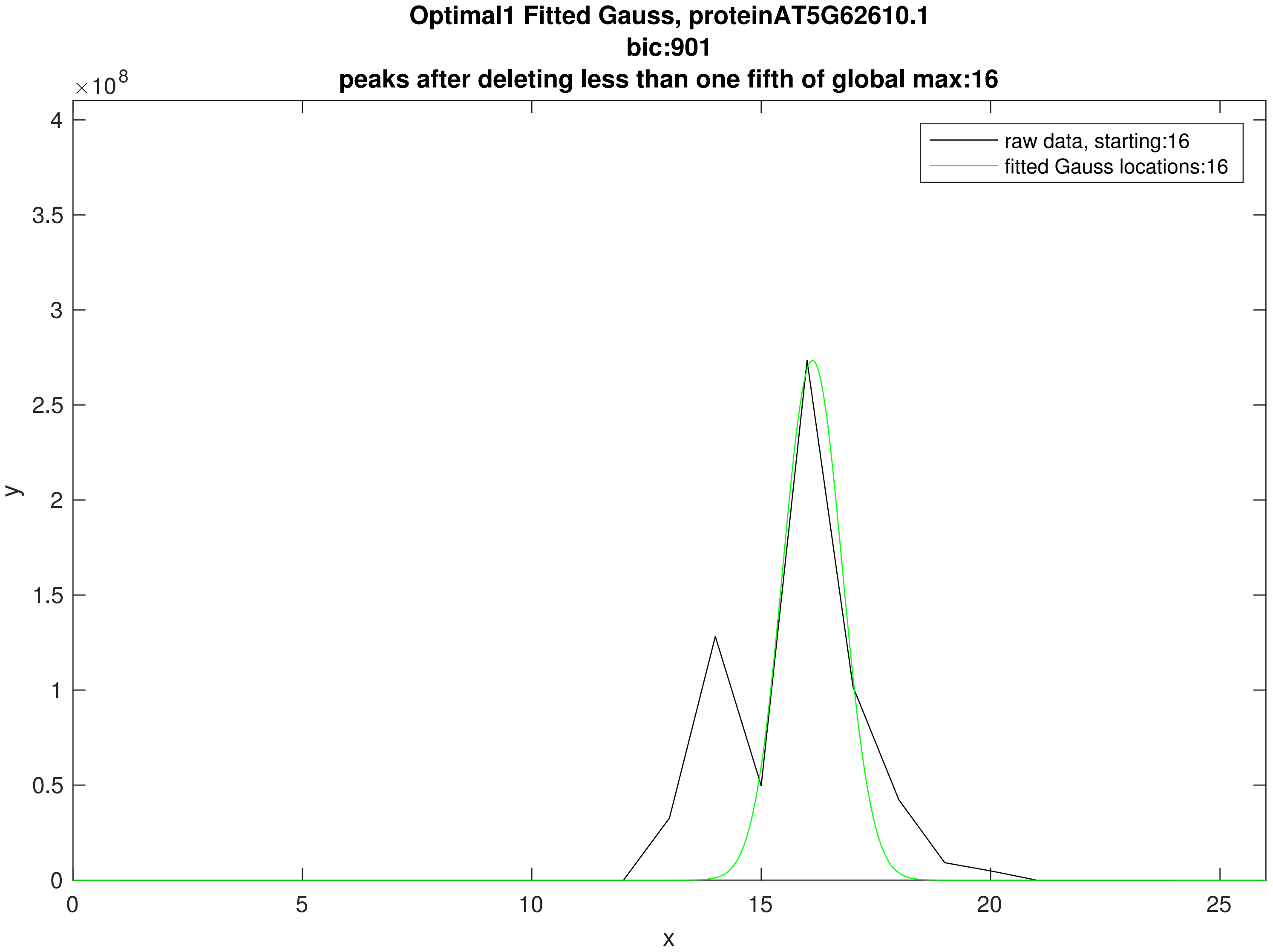

Optimal1 Fitted Gauss, proteinAT5G62610.1
bic:901
peaks after deleting less than one fifth of global max:16
×10^8
raw data, starting:16
fitted Gauss locations:16
y
x
0
0.5
1
1.5
2
2.5
3
3.5
4
0
5
10
15
20
25

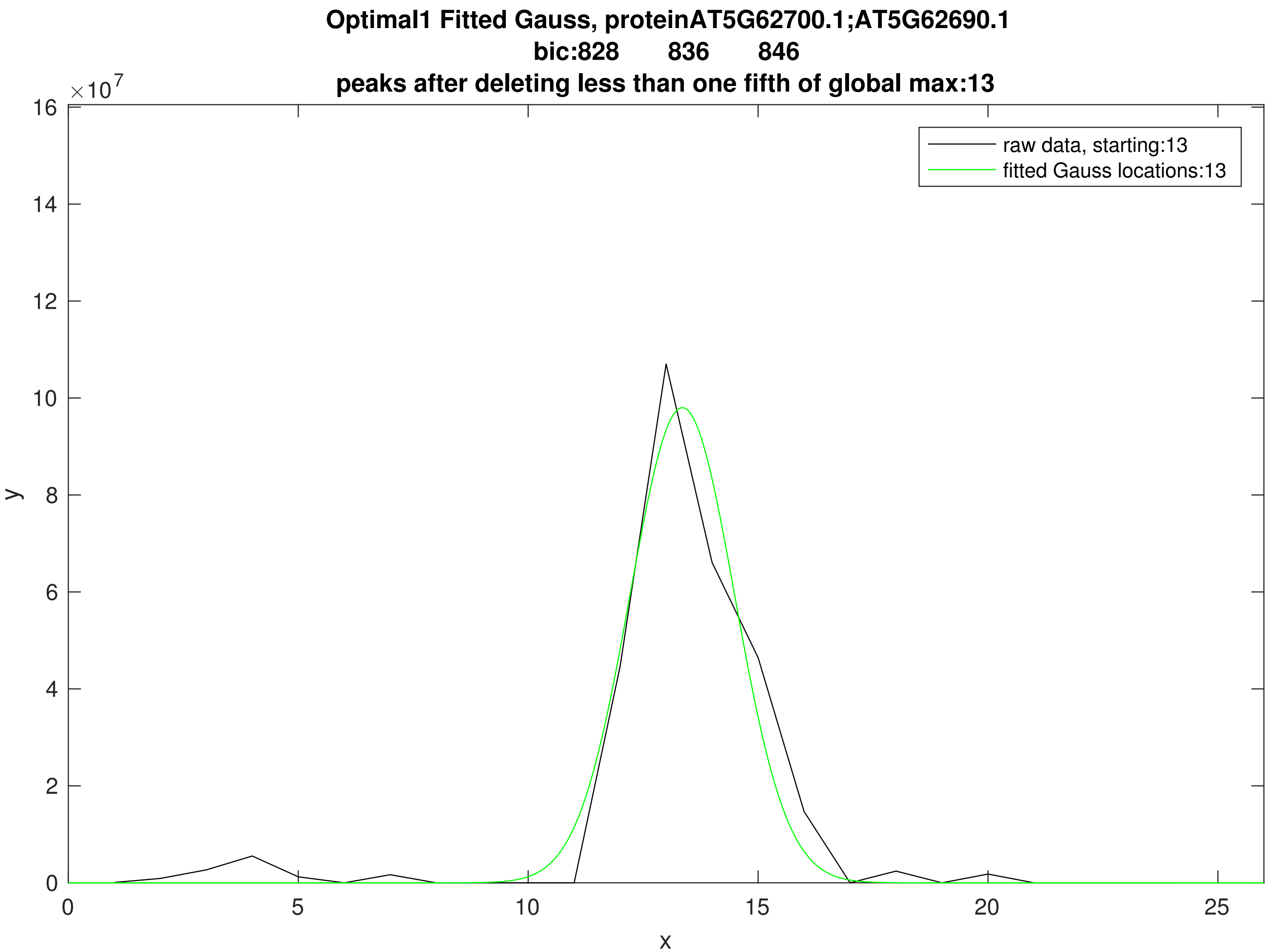
Optimal1 Fitted Gauss, proteinAT5G62700.1;AT5G62690.1
bic:828 836 846
peaks after deleting less than one fifth of global max:13
×10^7
16
14
12
10
8
6
4
2
0
y
0
5
10
15
20
25
x
raw data, starting:13
fitted Gauss locations:13

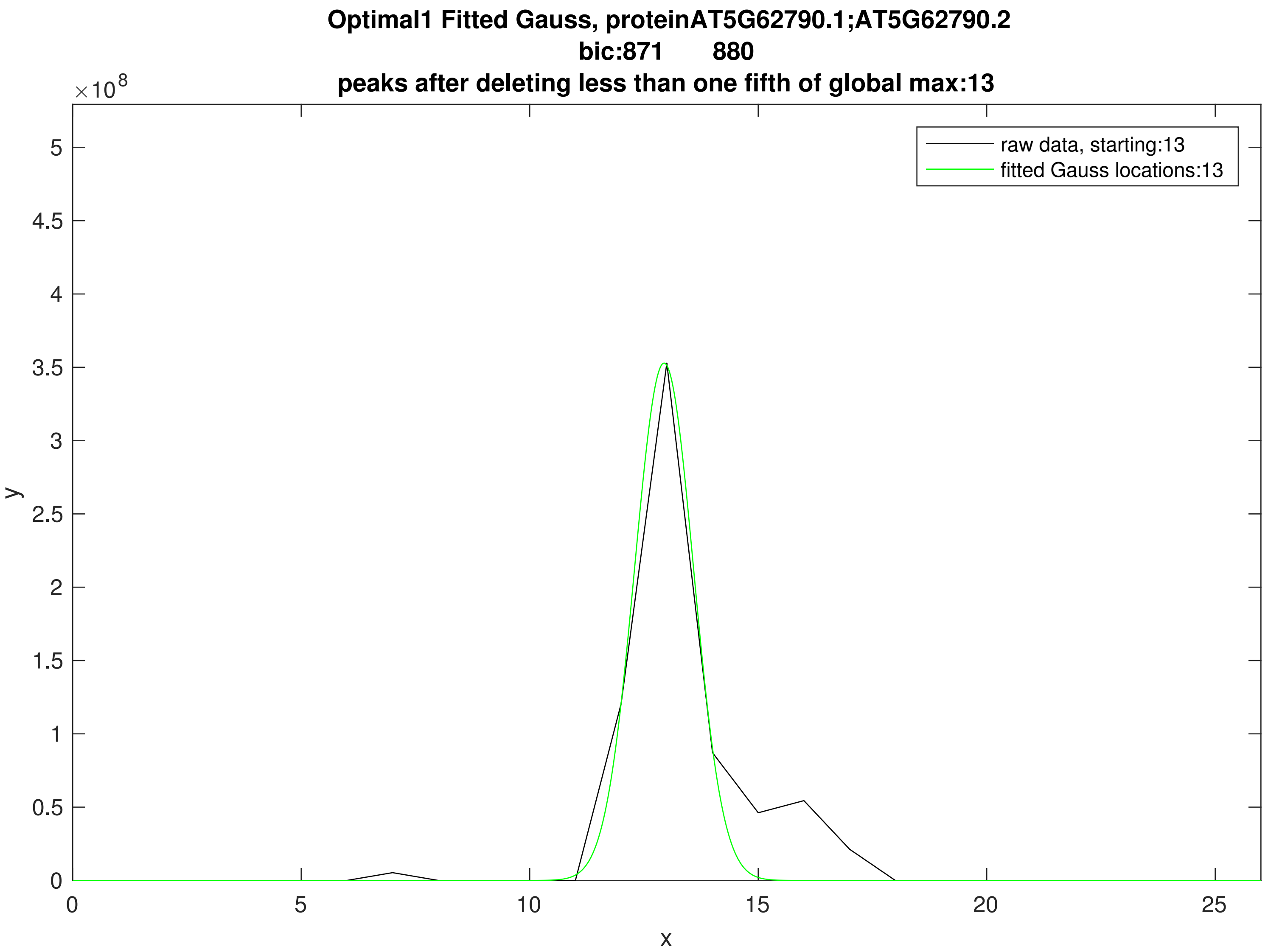
Optimal1 Fitted Gauss, proteinAT5G62790.1;AT5G62790.2
bic:871 880
peaks after deleting less than one fifth of global max:13
×10^8
5
4.5
4
3.5
3
2.5
2
1.5
1
0.5
0
y
0
5
10
15
20
25
x
raw data, starting:13
fitted Gauss locations:13

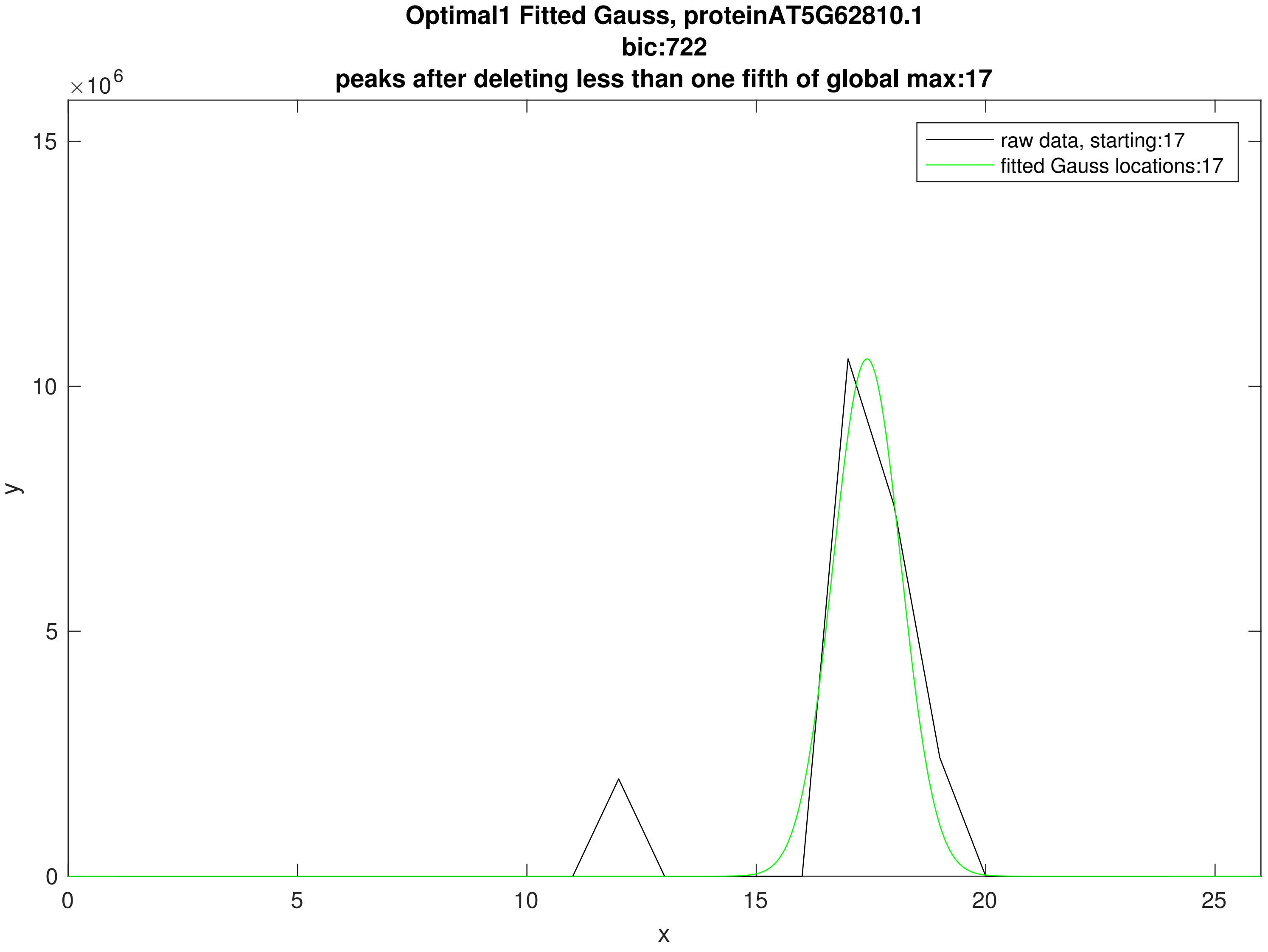

Optimal1 Fitted Gauss, proteinAT5G62810.1
bic:722
peaks after deleting less than one fifth of global max:17
×10^6
15
10
5
0
y
0
5
10
15
20
25
x
raw data, starting:17
fitted Gauss locations:17

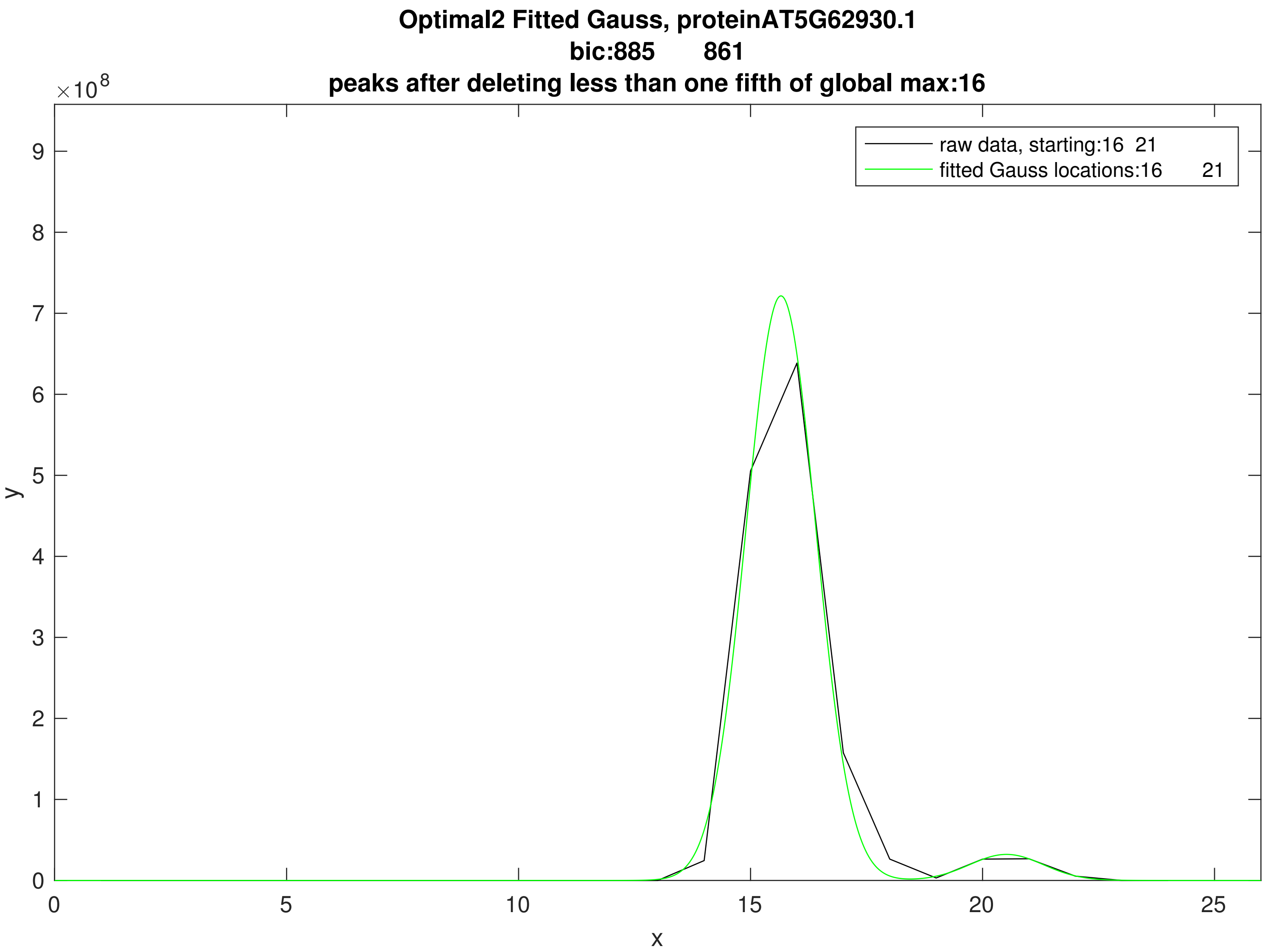
Optimal2 Fitted Gauss, proteinAT5G62930.1
bic:885 861
peaks after deleting less than one fifth of global max:16
×10^8
y
x
raw data, starting:16 21
fitted Gauss locations:16 21
0
1
2
3
4
5
6
7
8
9
0
5
10
15
20
25

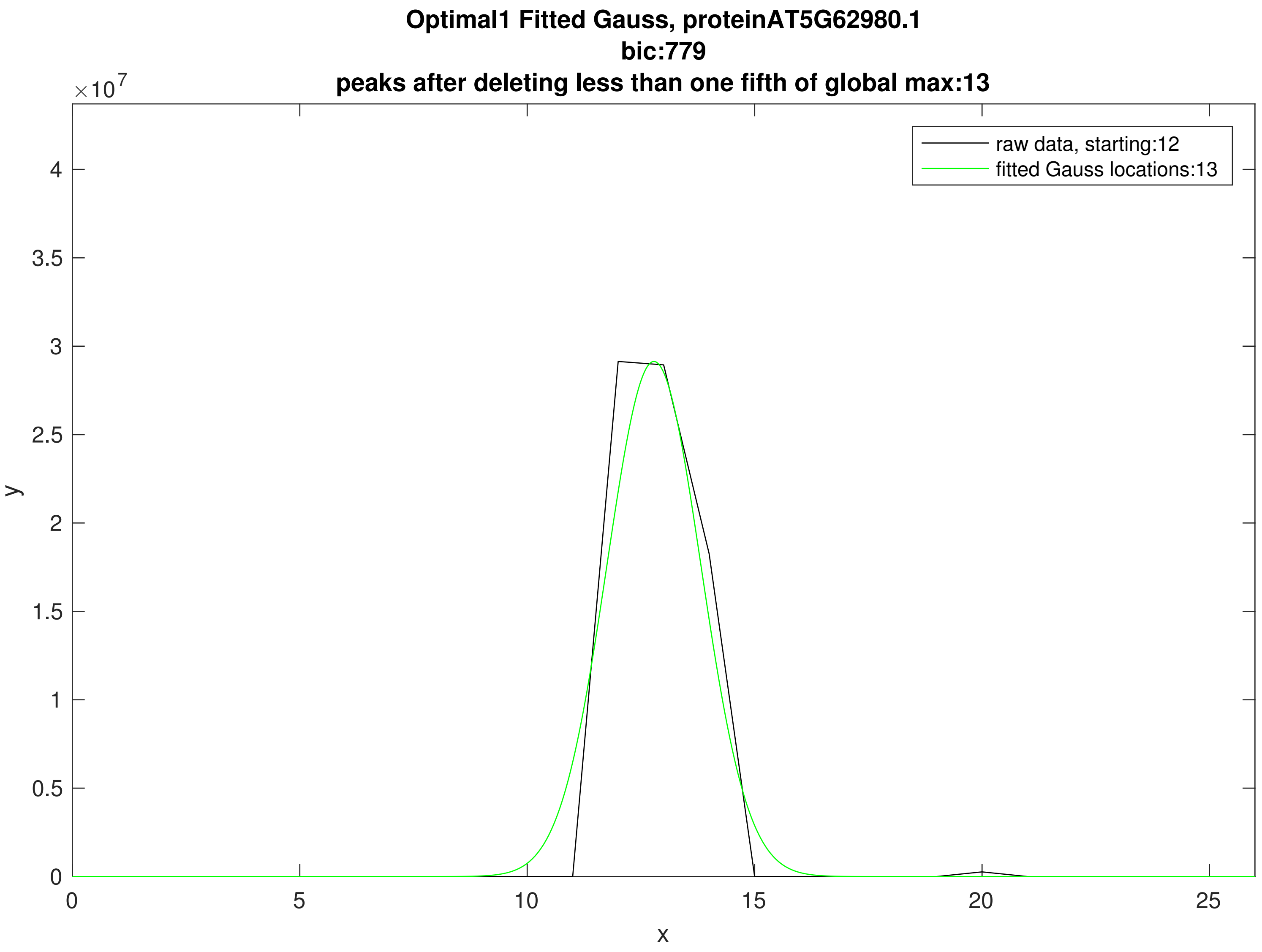
Optimal1 Fitted Gauss, proteinAT5G62980.1
bic:779
peaks after deleting less than one fifth of global max:13
×10^7
raw data, starting:12
fitted Gauss locations:13
4
3.5
3
2.5
2
1.5
1
0.5
0
y
0
5
10
15
20
25
x

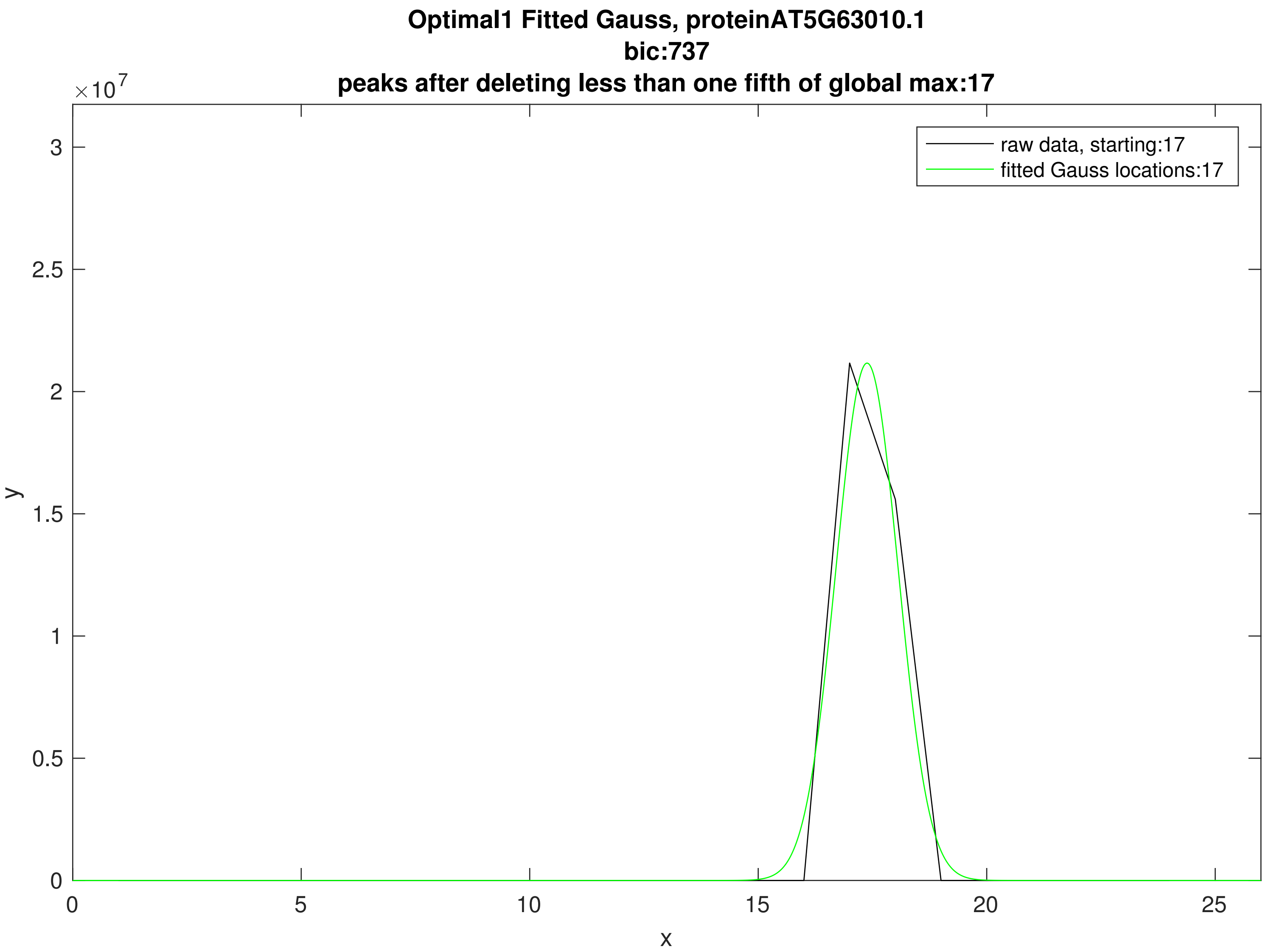
Optimal1 Fitted Gauss, proteinAT5G63010.1
bic:737
peaks after deleting less than one fifth of global max:17
×10^7
raw data, starting:17
fitted Gauss locations:17
y
x
0
0.5
1
1.5
2
2.5
3
0
5
10
15
20
25

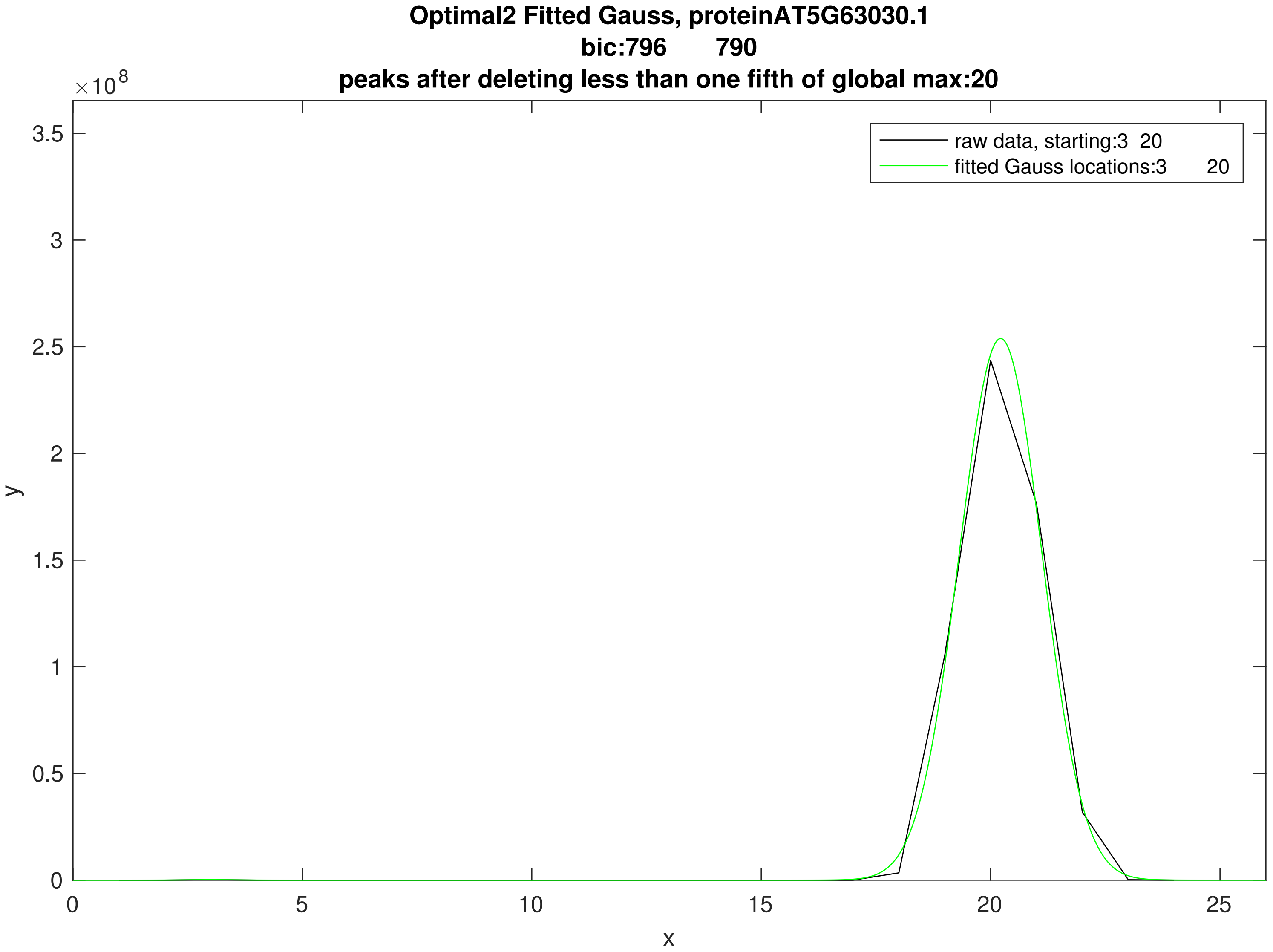
Optimal2 Fitted Gauss, proteinAT5G63030.1
bic:796 790
peaks after deleting less than one fifth of global max:20
×10^8
3.5
3
2.5
2
1.5
1
0.5
0
y
0
5
10
15
20
25
x
raw data, starting:3 20
fitted Gauss locations:3 20

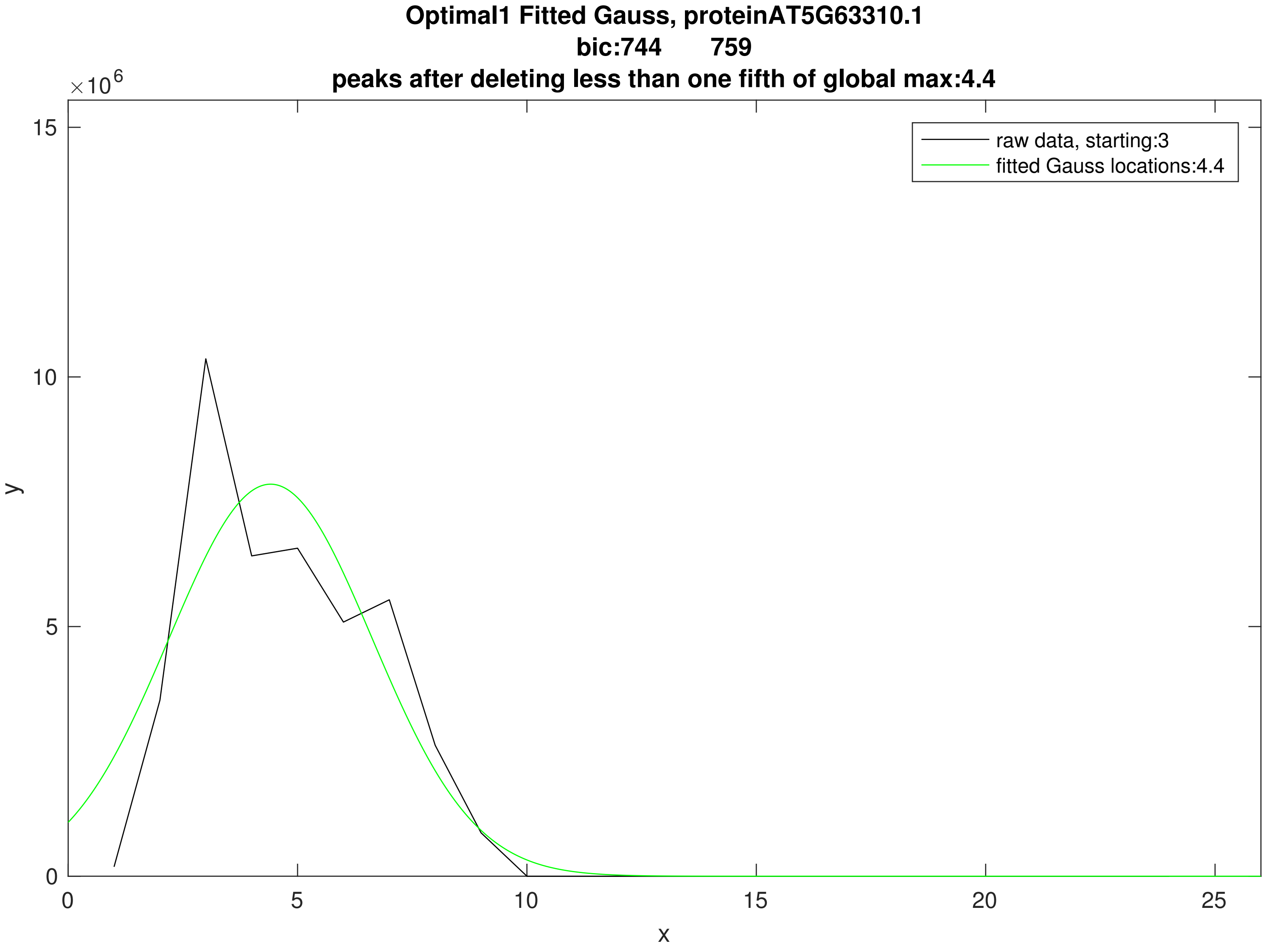
Optimal1 Fitted Gauss, proteinAT5G63310.1
bic:744 759
peaks after deleting less than one fifth of global max:4.4
×10^6
raw data, starting:3
fitted Gauss locations:4.4
y
x
0
5
10
15
0
5
10
15
20
25

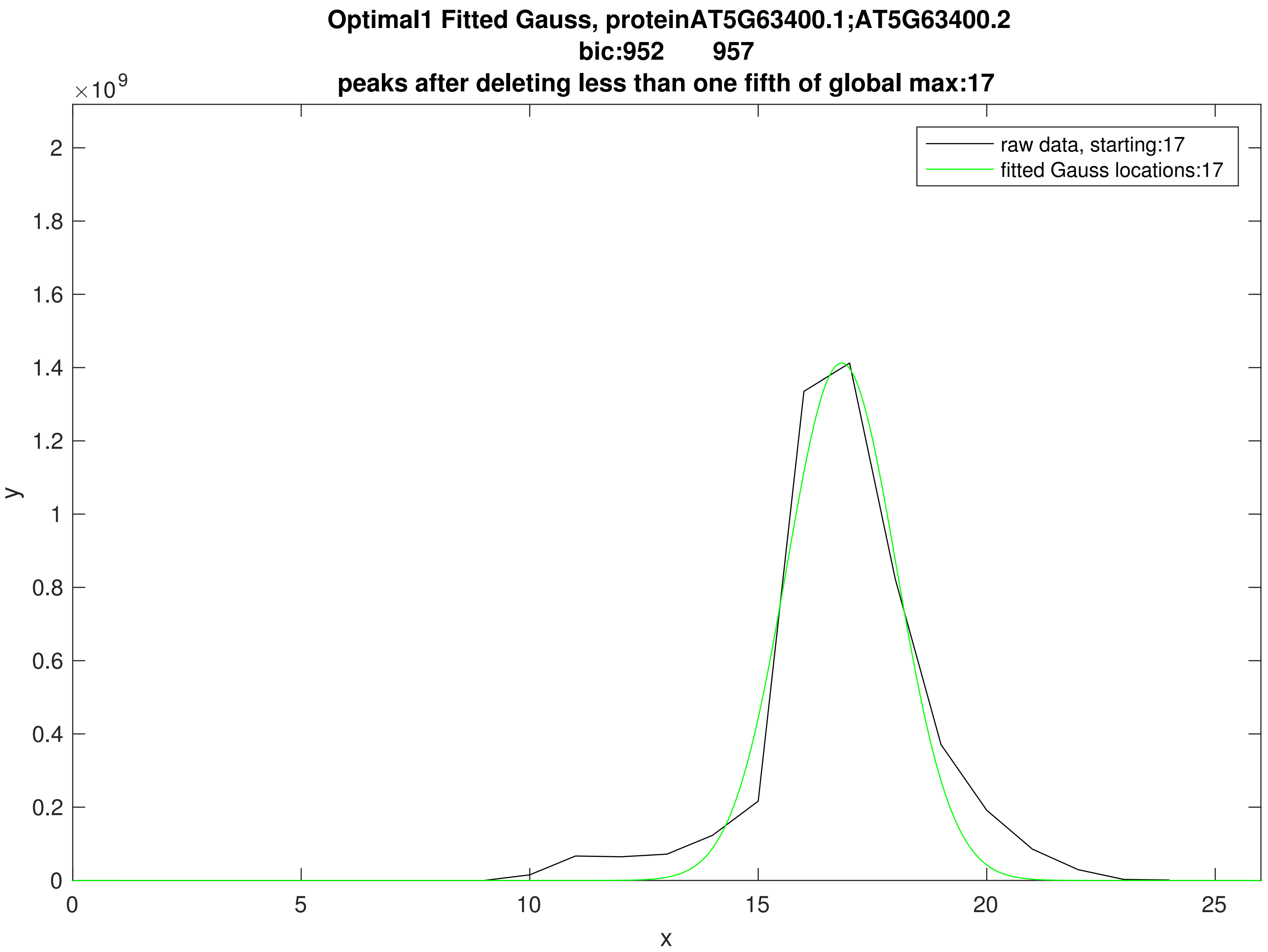

Optimal1 Fitted Gauss, proteinAT5G63400.1;AT5G63400.2
bic:952 957
peaks after deleting less than one fifth of global max:17
×10^9
raw data, starting:17
fitted Gauss locations:17
y
x
0
0.2
0.4
0.6
0.8
1
1.2
1.4
1.6
1.8
2
0
5
10
15
20
25

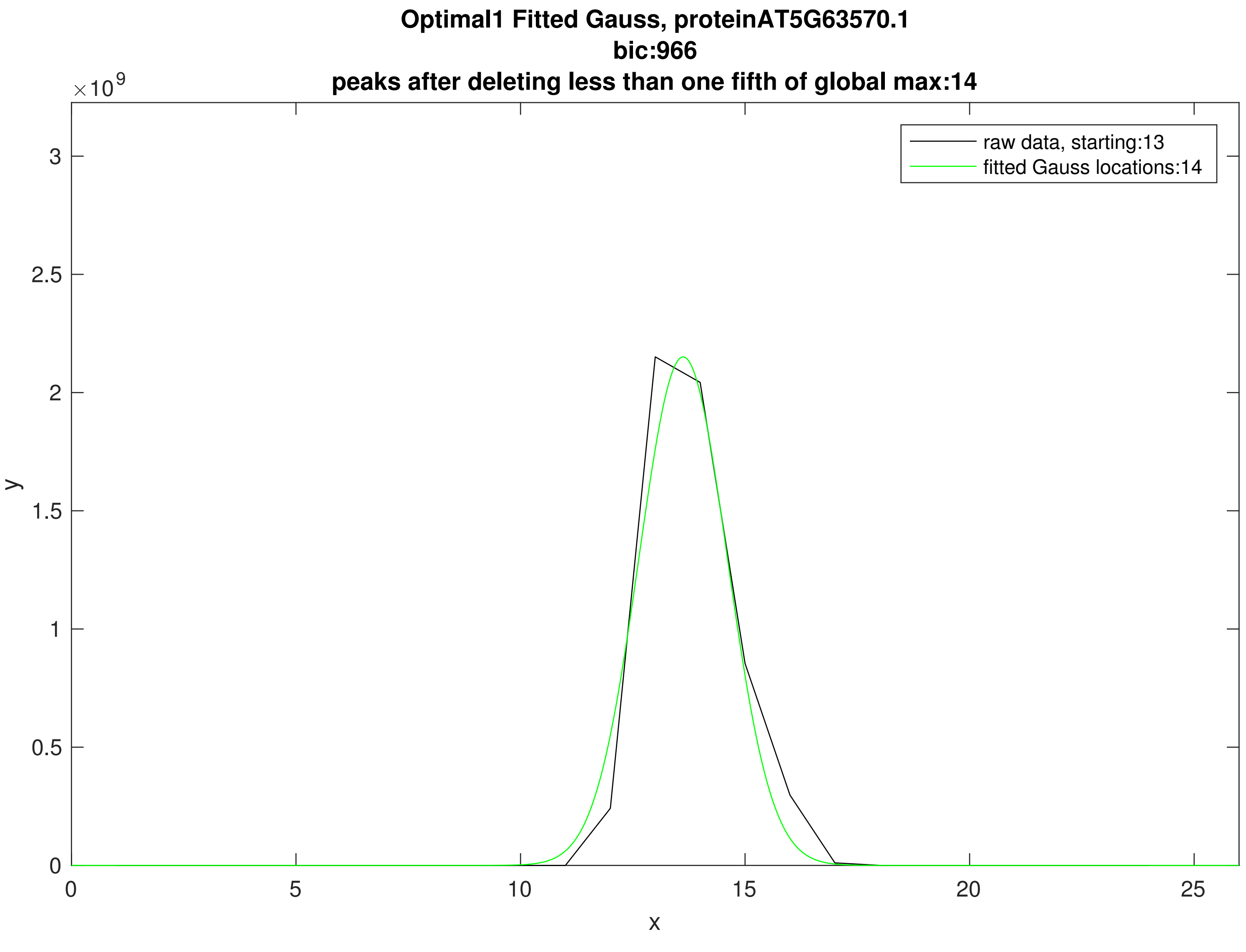
Optimal1 Fitted Gauss, proteinAT5G63570.1
bic:966
peaks after deleting less than one fifth of global max:14
×10^9
raw data, starting:13
fitted Gauss locations:14
y
x
0
0.5
1
1.5
2
2.5
3
0
5
10
15
20
25

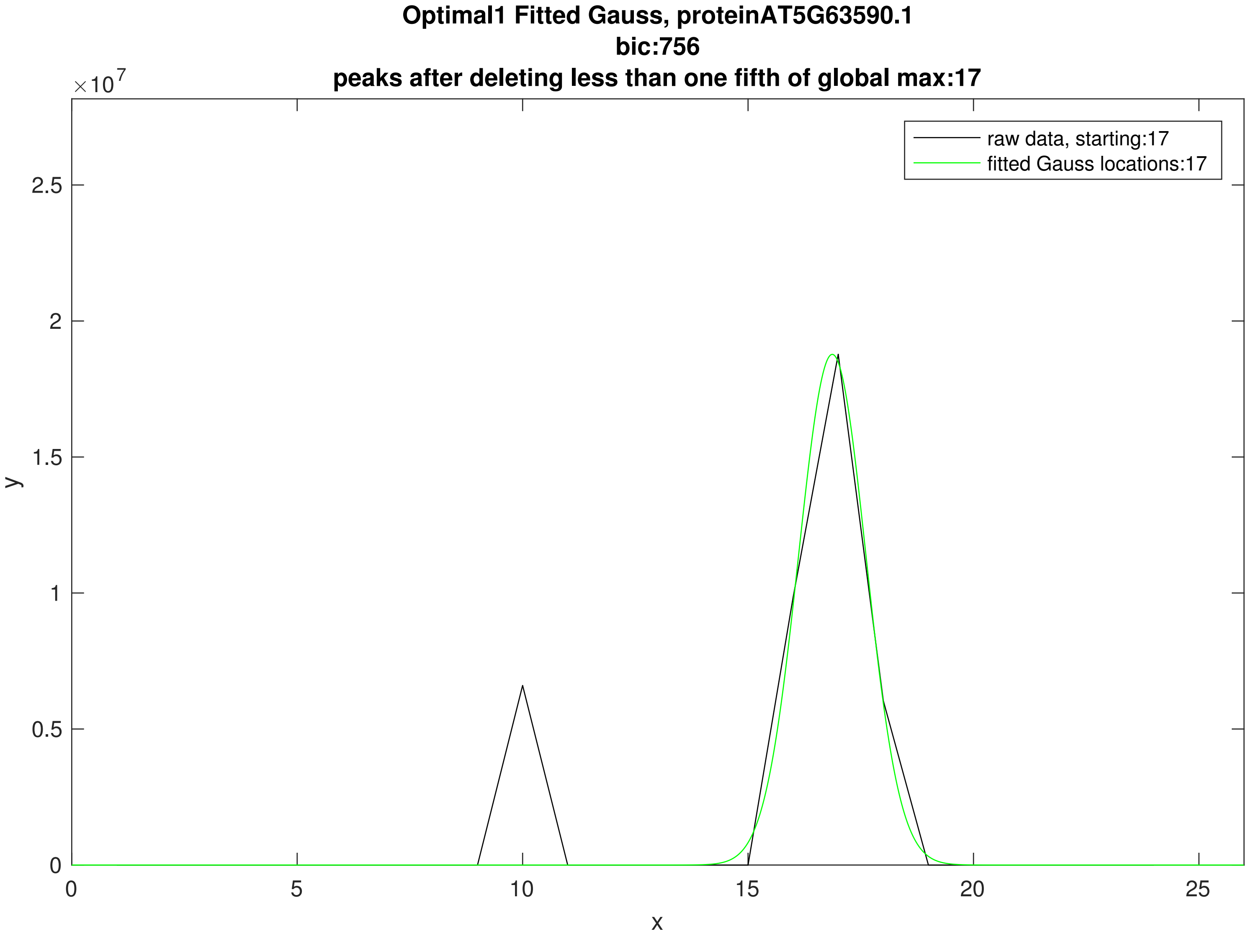
Optimal1 Fitted Gauss, proteinAT5G63590.1
bic:756
peaks after deleting less than one fifth of global max:17
×10^7
raw data, starting:17
fitted Gauss locations:17
y
x
0
0.5
1
1.5
2
2.5
0
5
10
15
20
25

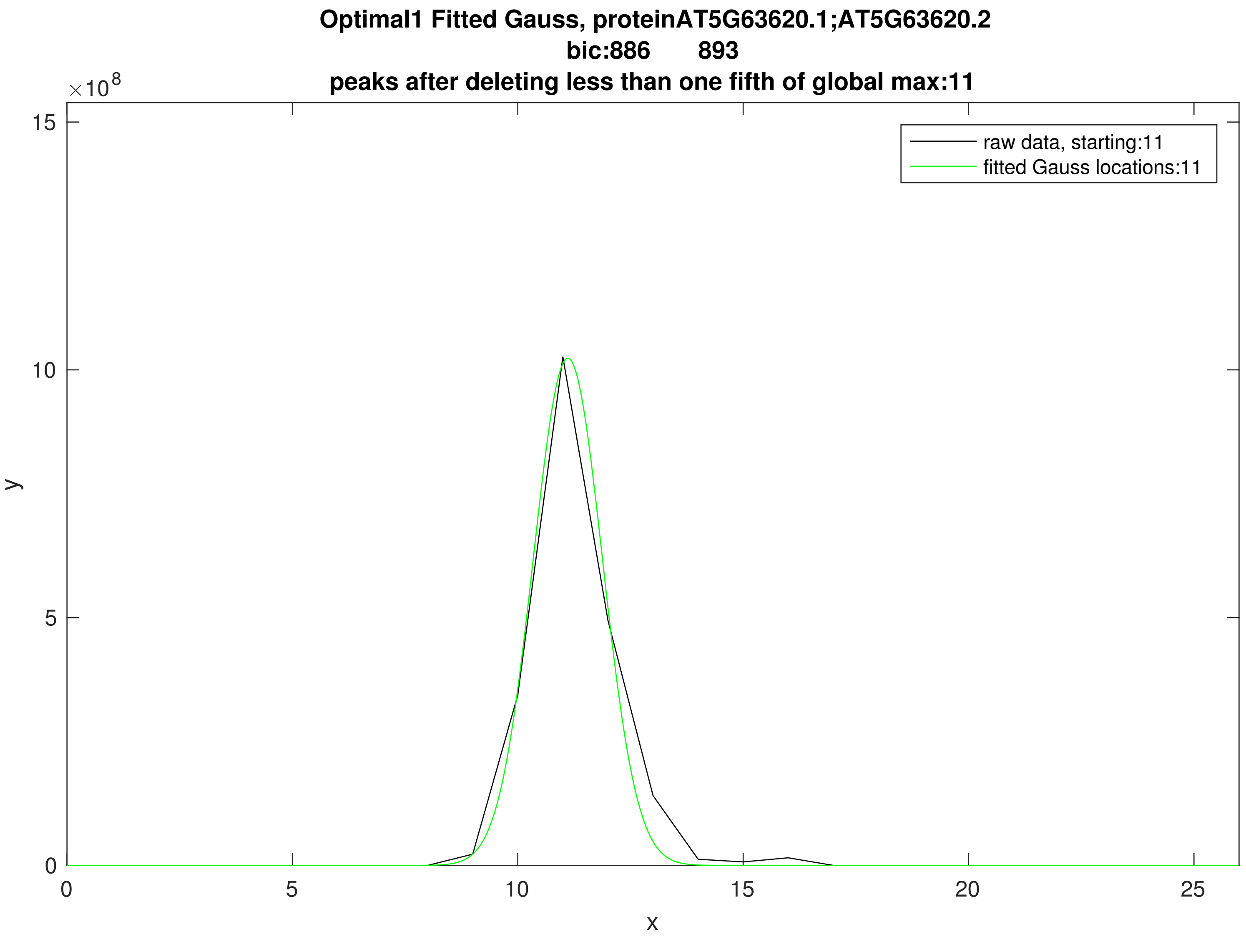

Optimal1 Fitted Gauss, proteinAT5G63620.1;AT5G63620.2
bic:886 893
peaks after deleting less than one fifth of global max:11
×10^8
15
10
5
0
y
0
5
10
15
20
25
x
raw data, starting:11
fitted Gauss locations:11

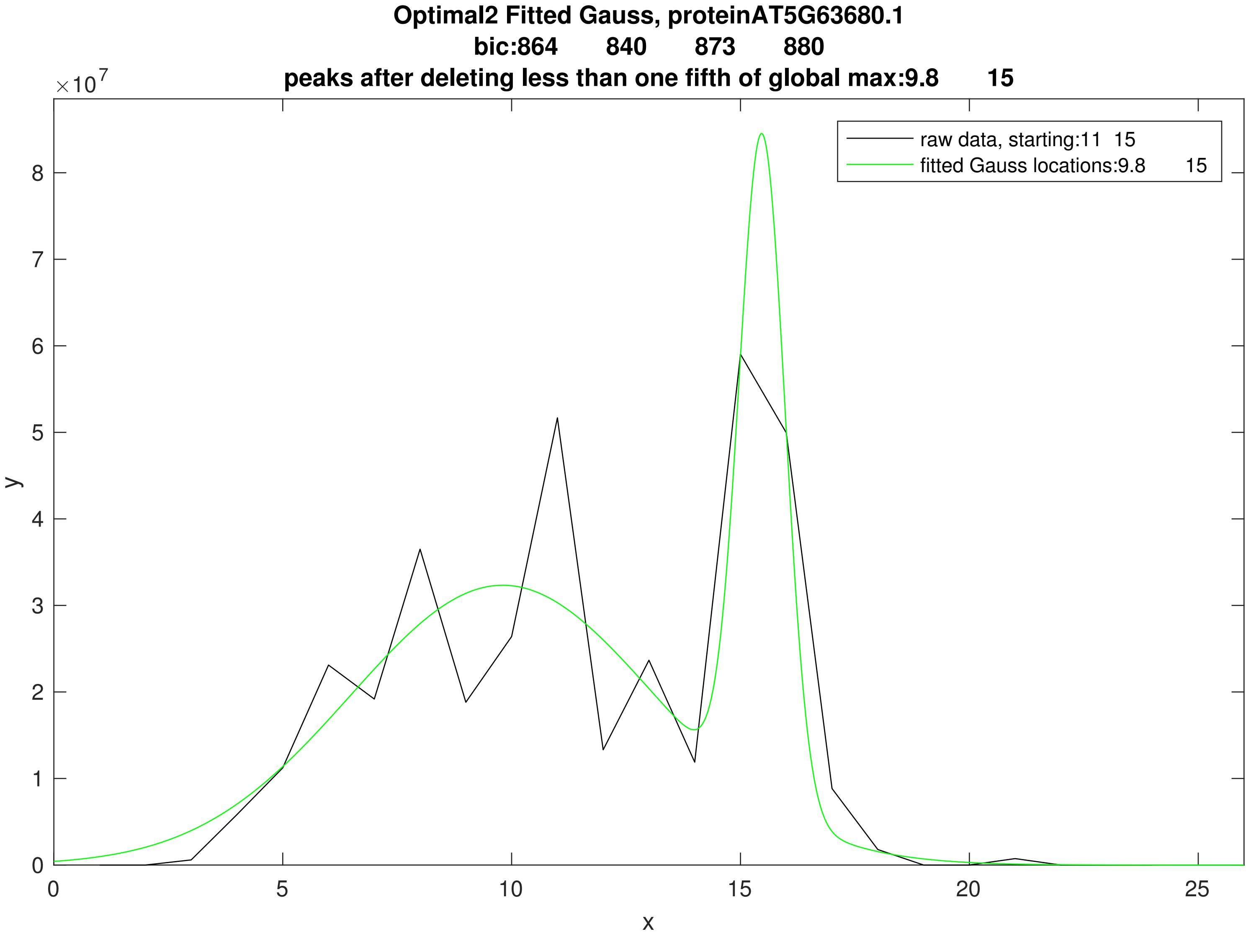
Optimal2 Fitted Gauss, proteinAT5G63680.1
bic:864 840 873 880
peaks after deleting less than one fifth of global max:9.8 15
×10^7
raw data, starting:11 15
fitted Gauss locations:9.8 15
y
x

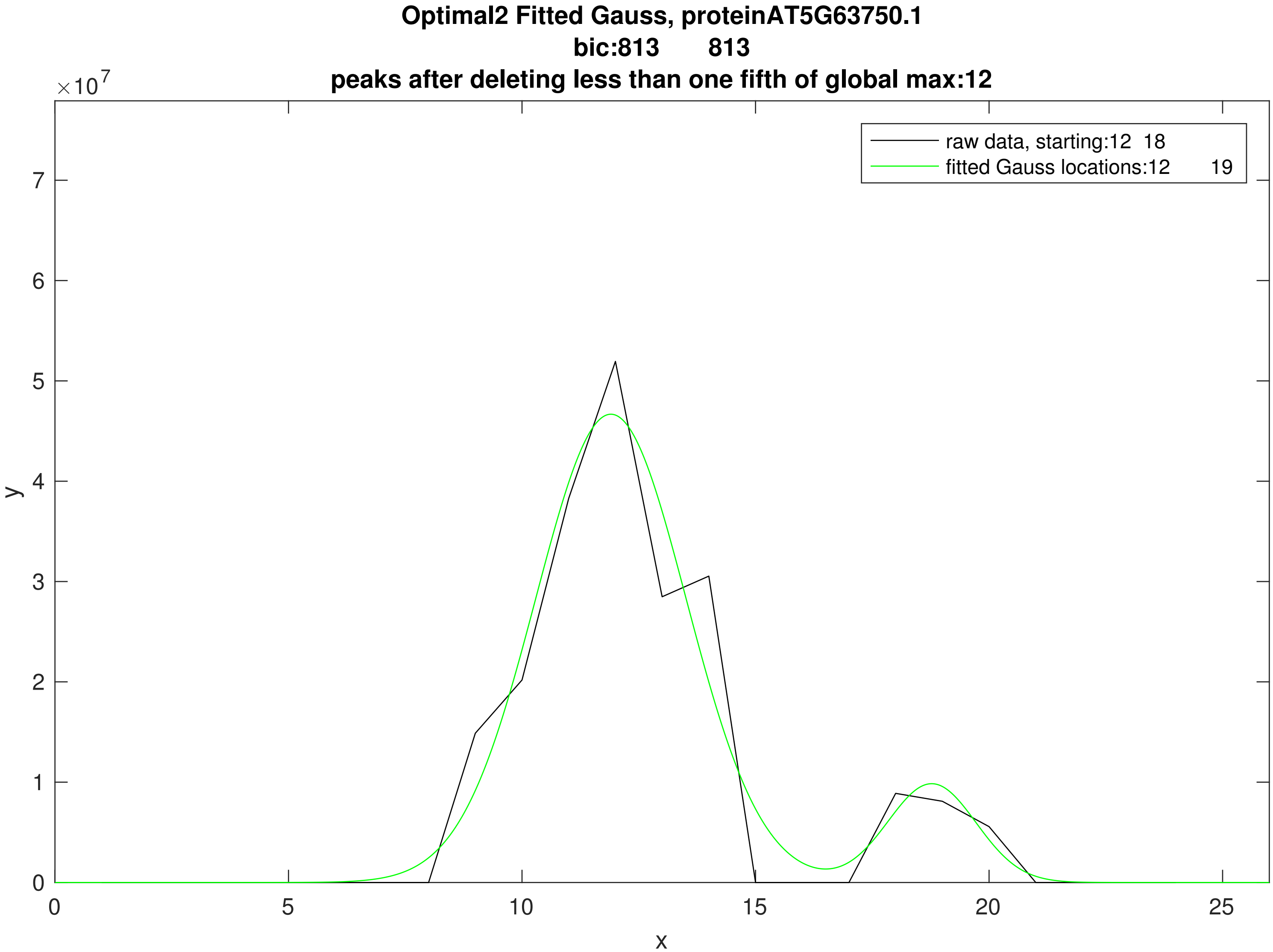
Optimal2 Fitted Gauss, proteinAT5G63750.1
bic:813 813
peaks after deleting less than one fifth of global max:12
×10^7
raw data, starting:12 18
fitted Gauss locations:12 19
y
x
0
1
2
3
4
5
6
7
0
5
10
15
20
25

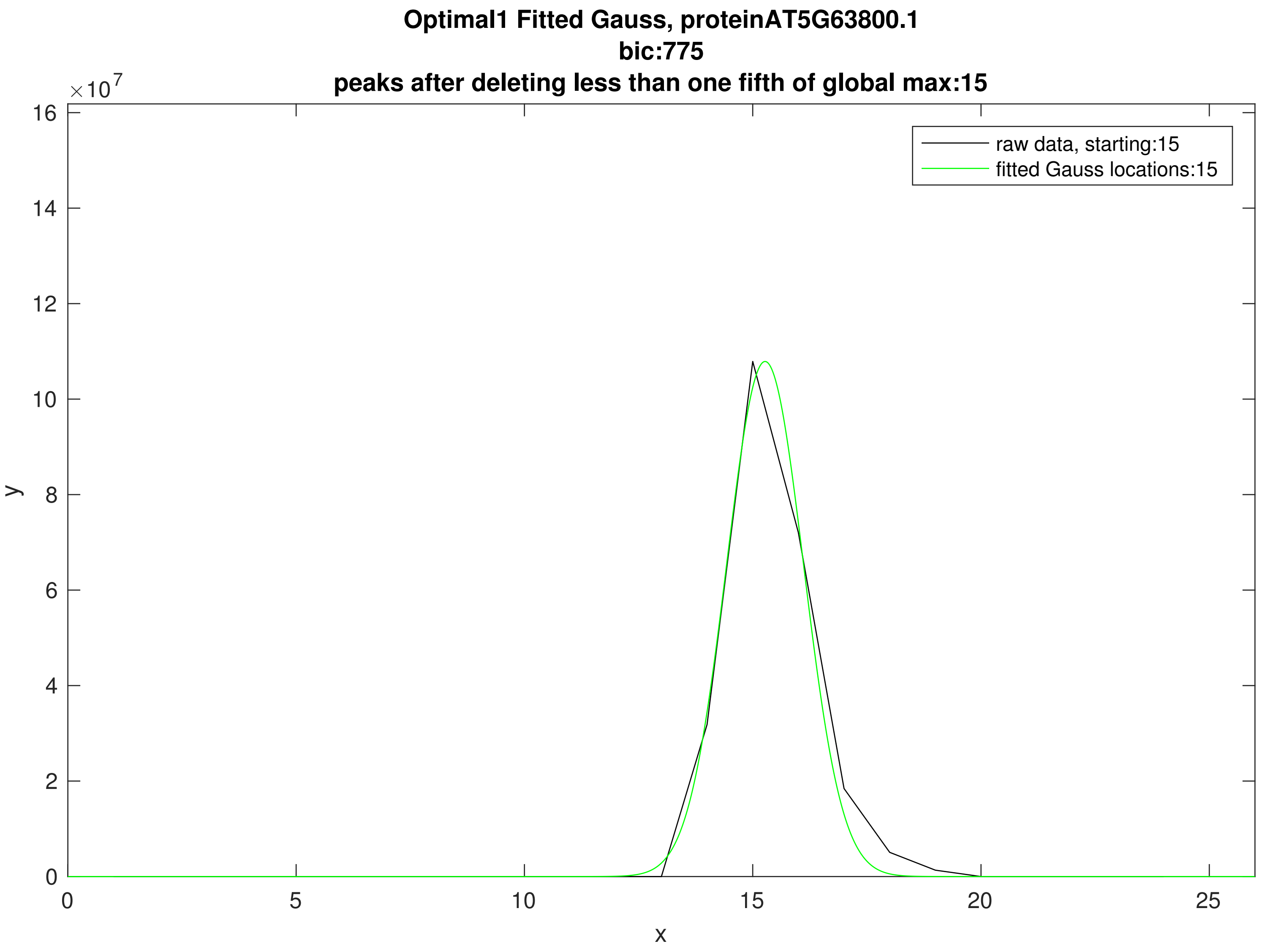
Optimal1 Fitted Gauss, proteinAT5G63800.1
bic:775
peaks after deleting less than one fifth of global max:15
×10^7
raw data, starting:15
fitted Gauss locations:15
y
x
0
2
4
6
8
10
12
14
16
0
5
10
15
20
25

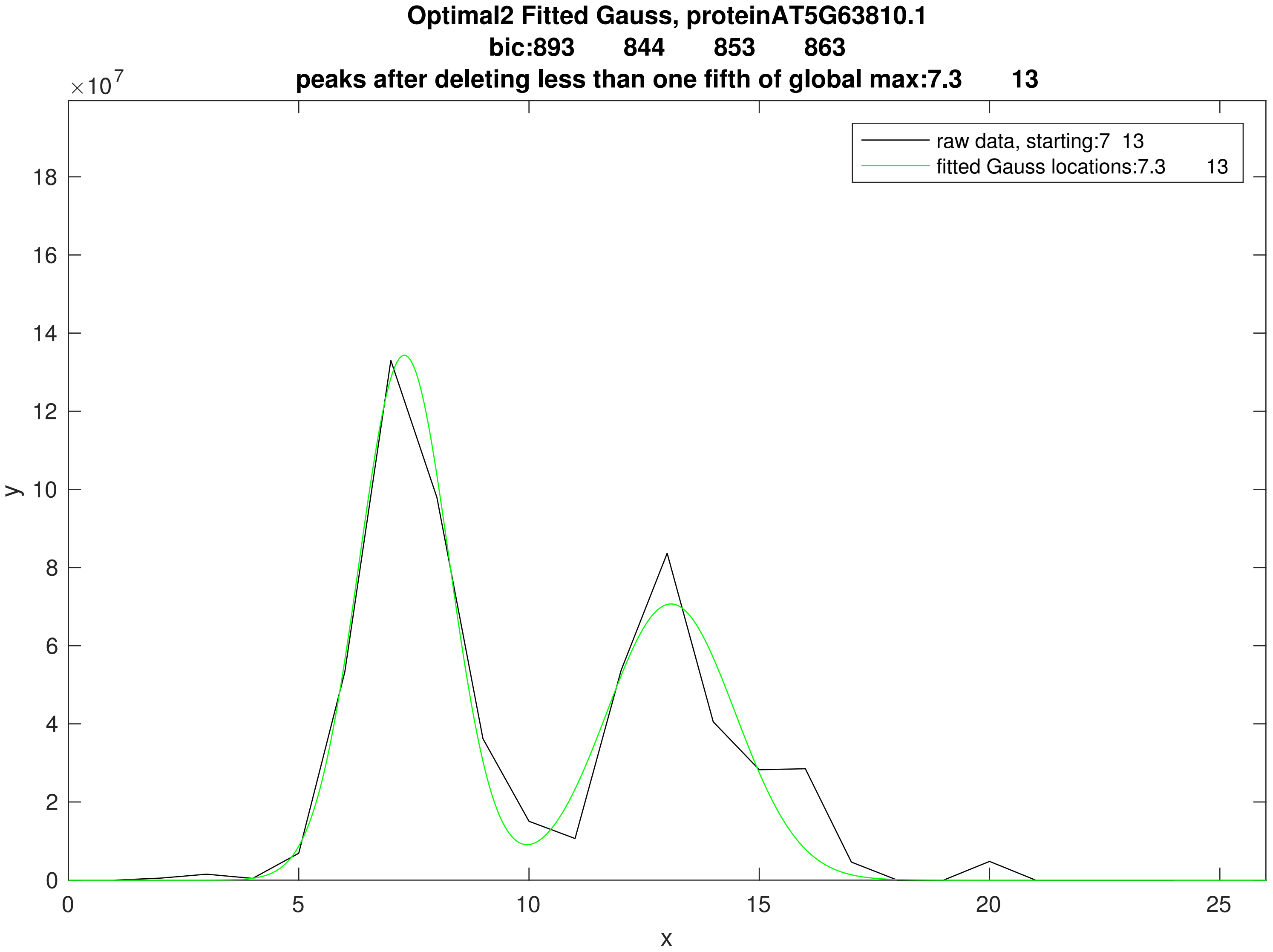
Optimal2 Fitted Gauss, proteinAT5G63810.1
bic:893 844 853 863
peaks after deleting less than one fifth of global max:7.3 13
×10^7
raw data, starting:7 13
fitted Gauss locations:7.3 13
y
x
0
2
4
6
8
10
12
14
16
18
0
5
10
15
20
25

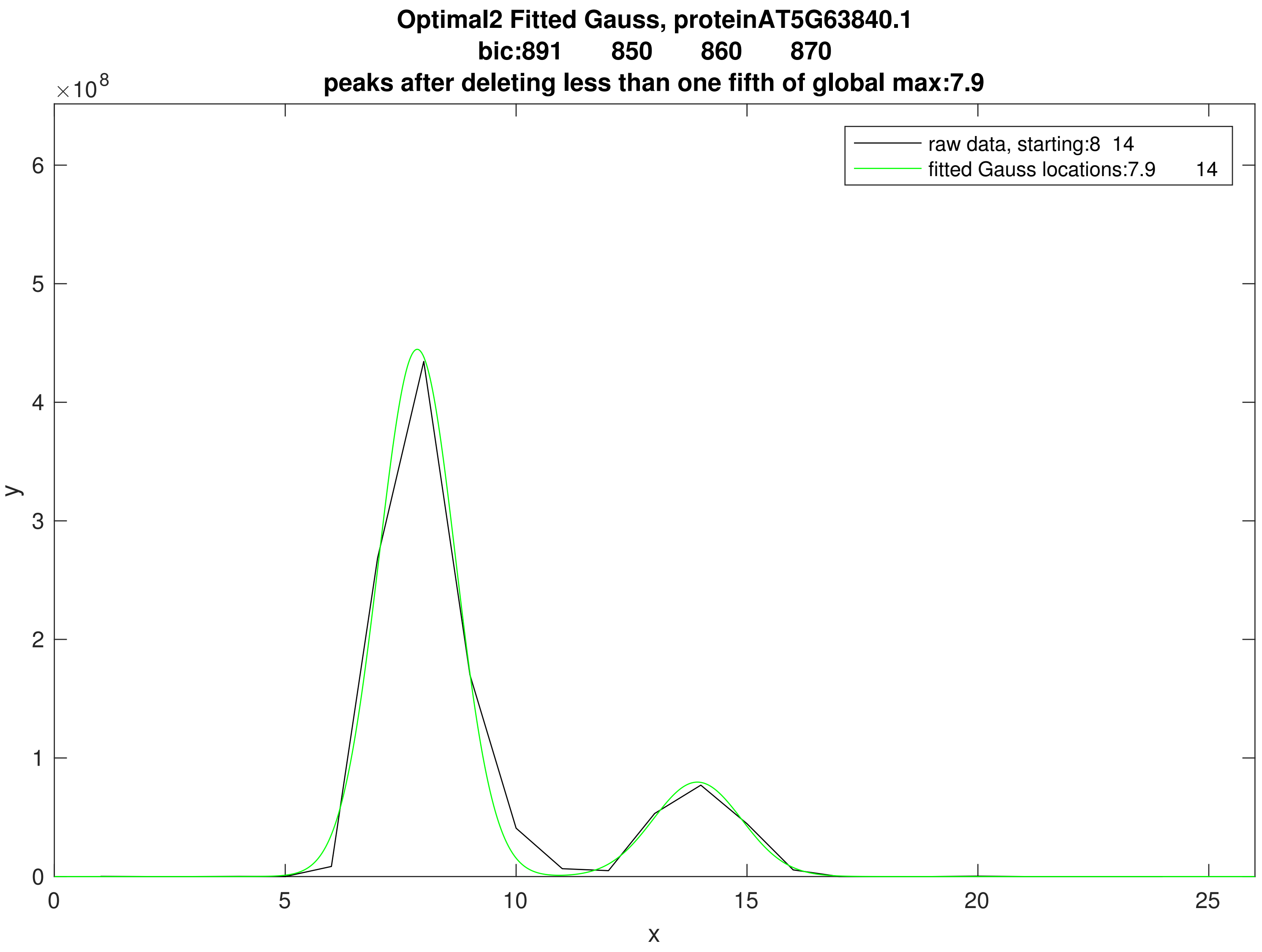
Optimal2 Fitted Gauss, proteinAT5G63840.1
bic:891 850 860 870
peaks after deleting less than one fifth of global max:7.9
×10^8
raw data, starting:8 14
fitted Gauss locations:7.9 14
y
x
0
1
2
3
4
5
6
0
5
10
15
20
25

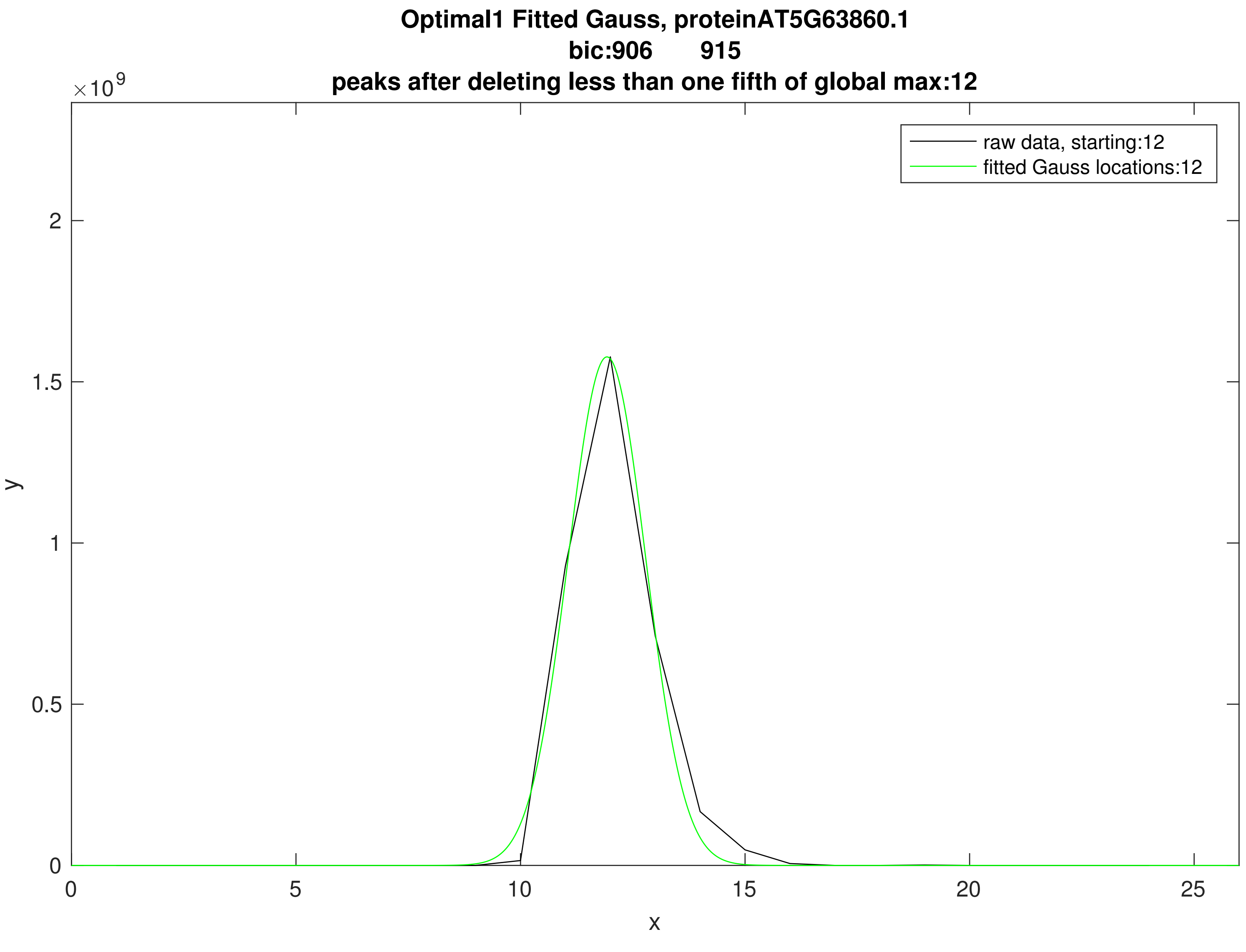
Optimal1 Fitted Gauss, proteinAT5G63860.1
bic:906 915
peaks after deleting less than one fifth of global max:12
×10^9
raw data, starting:12
fitted Gauss locations:12
y
x
0
0.5
1
1.5
2
0
5
10
15
20
25

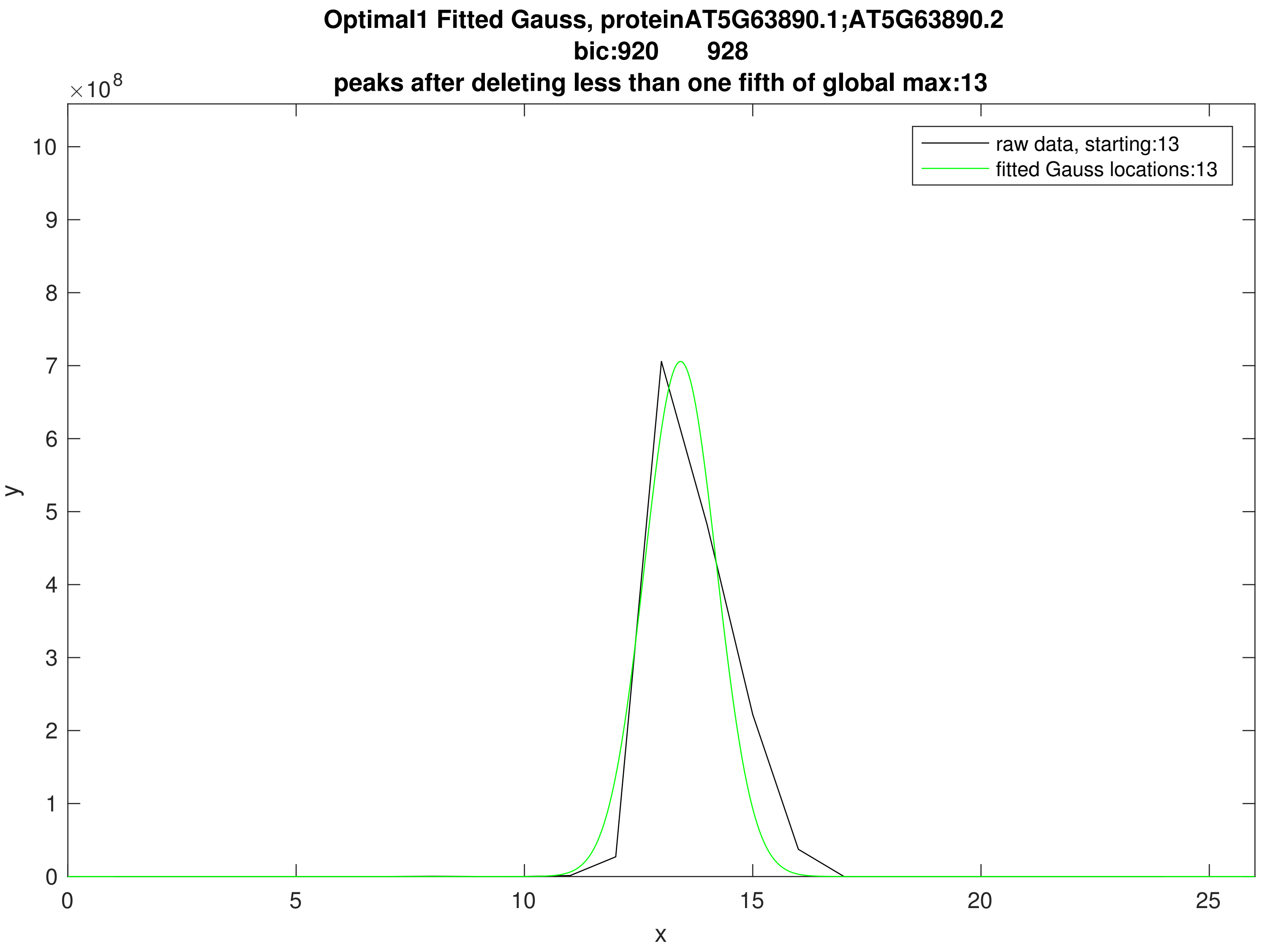

Optimal1 Fitted Gauss, proteinAT5G63890.1;AT5G63890.2
bic:920 928
peaks after deleting less than one fifth of global max:13
×10^8
10
9
8
7
6
5
4
3
2
1
0
y
0
5
10
15
20
25
x
raw data, starting:13
fitted Gauss locations:13

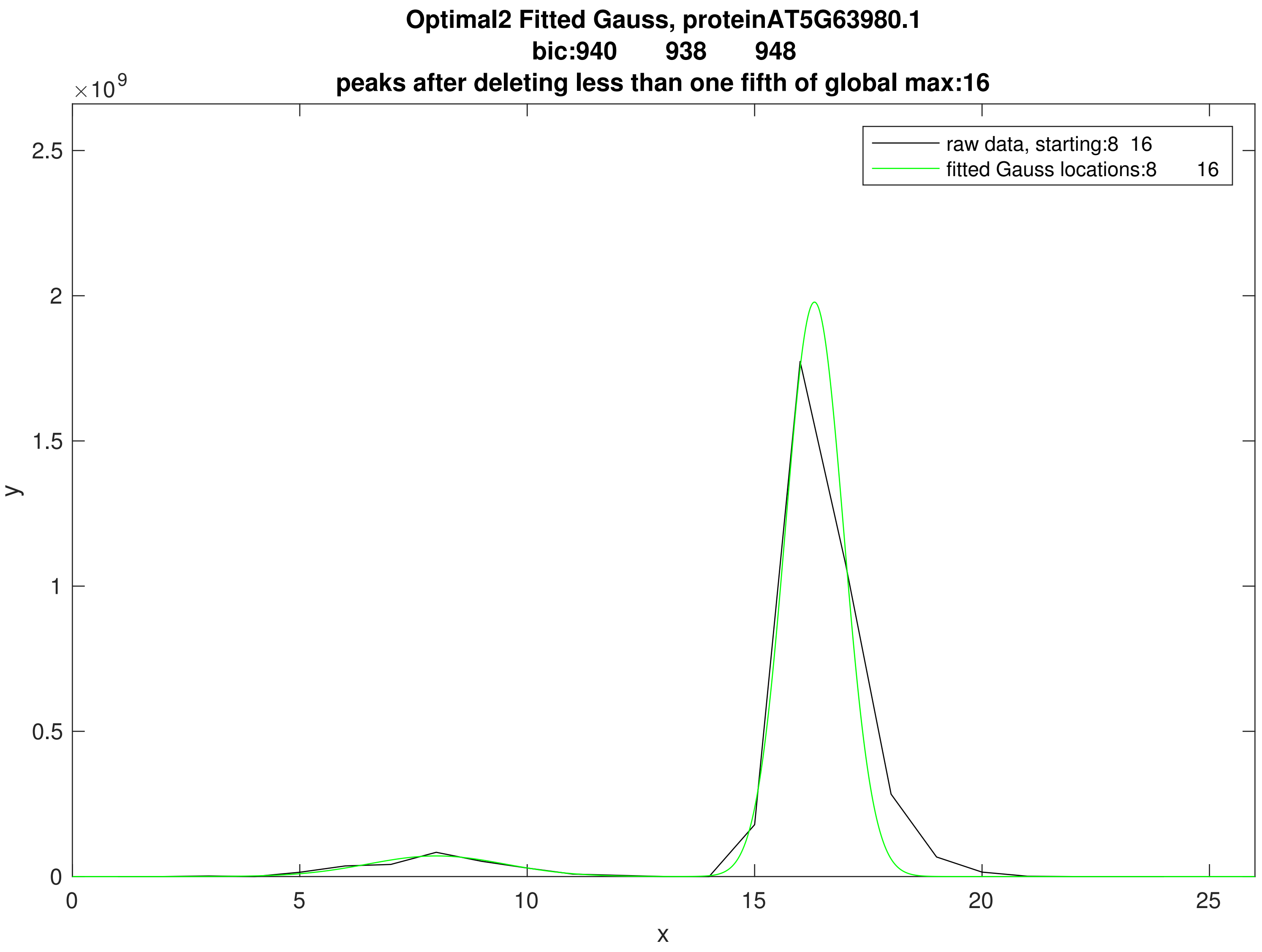
Optimal2 Fitted Gauss, proteinAT5G63980.1
bic:940 938 948
peaks after deleting less than one fifth of global max:16
×10^9
raw data, starting:8 16
fitted Gauss locations:8 16
y
x
0
0.5
1
1.5
2
2.5
5
10
15
20
25

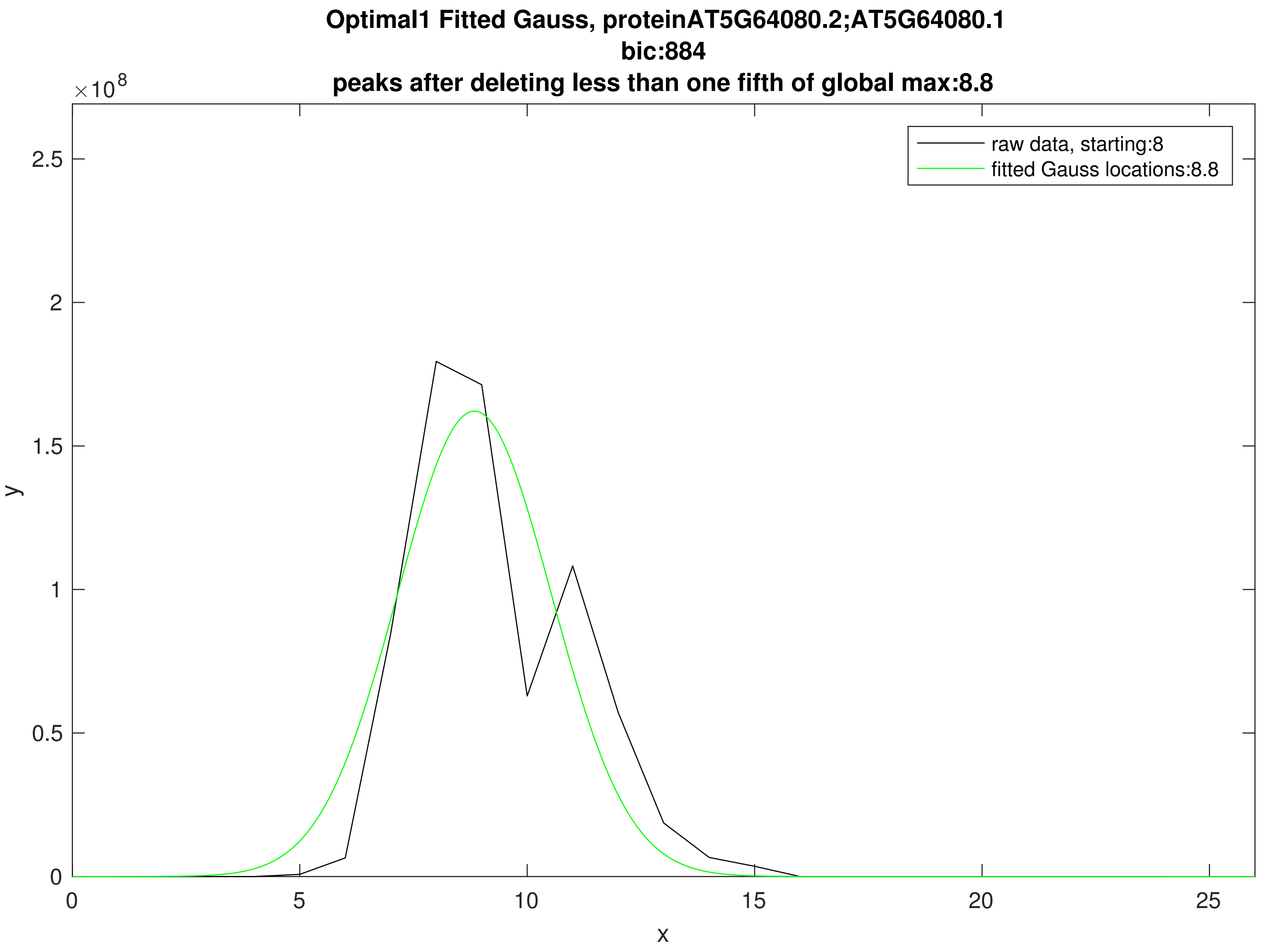

Optimal1 Fitted Gauss, proteinAT5G64080.2;AT5G64080.1
bic:884
peaks after deleting less than one fifth of global max:8.8
×10^8
2.5
2
1.5
1
0.5
0
y
0
5
10
15
20
25
x
raw data, starting:8
fitted Gauss locations:8.8

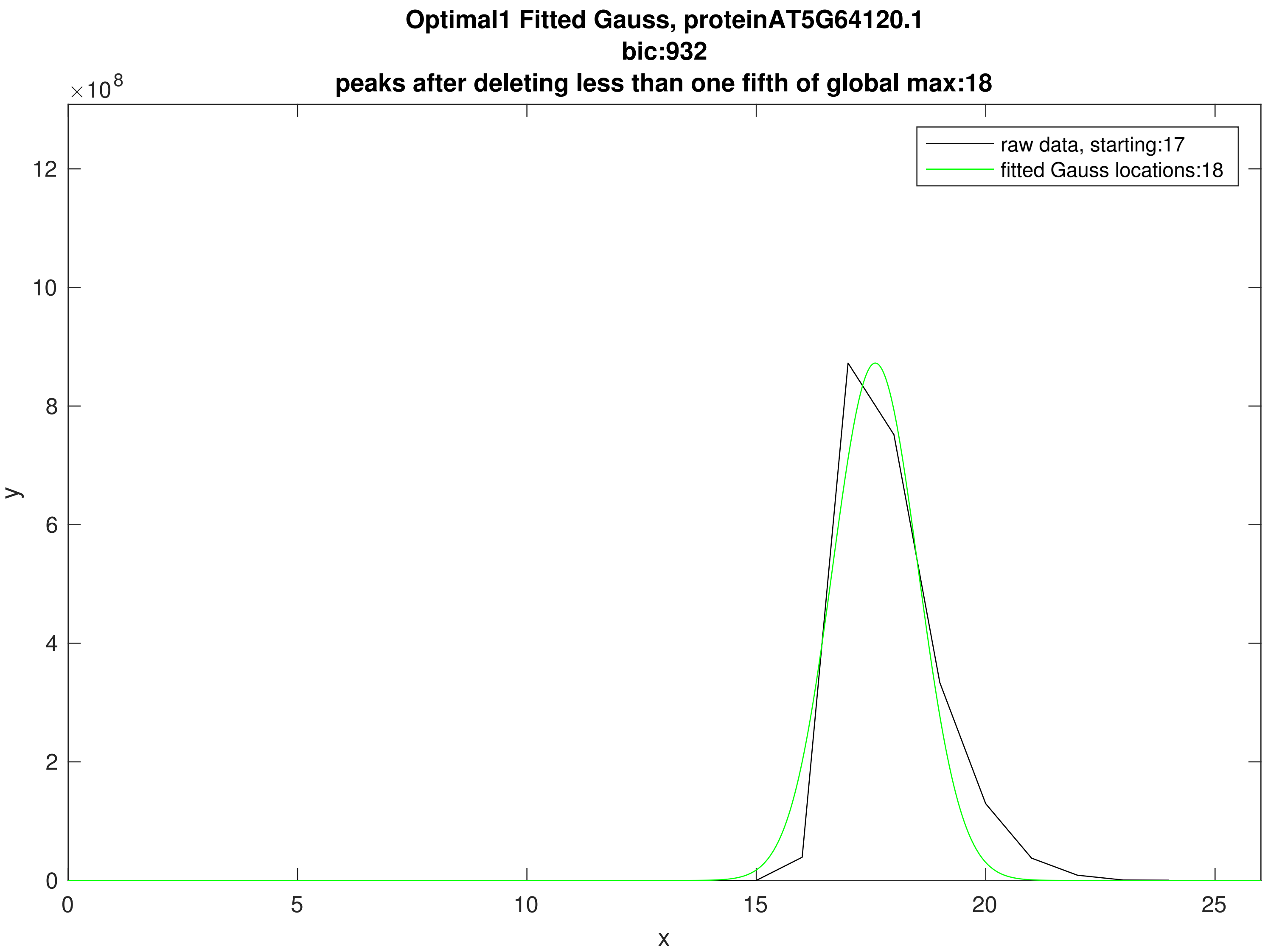
Optimal1 Fitted Gauss, proteinAT5G64120.1
bic:932
peaks after deleting less than one fifth of global max:18
×10^8
raw data, starting:17
fitted Gauss locations:18
12
10
8
6
4
2
0
y
0
5
10
15
20
25
x

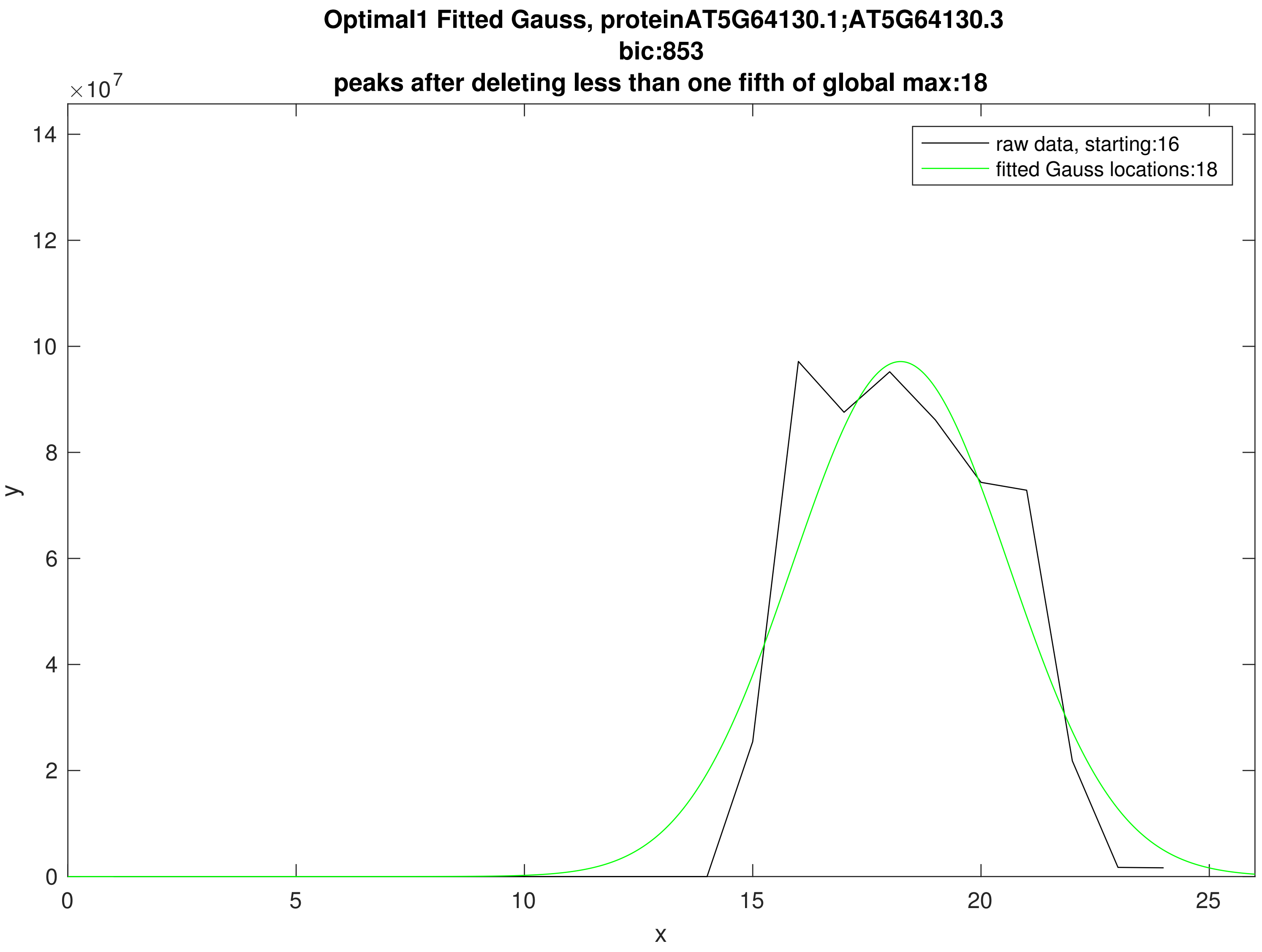
Optimal1 Fitted Gauss, proteinAT5G64130.1;AT5G64130.3
bic:853
peaks after deleting less than one fifth of global max:18
×10^7
14
12
10
8
6
4
2
0
y
0
5
10
15
20
25
x
raw data, starting:16
fitted Gauss locations:18

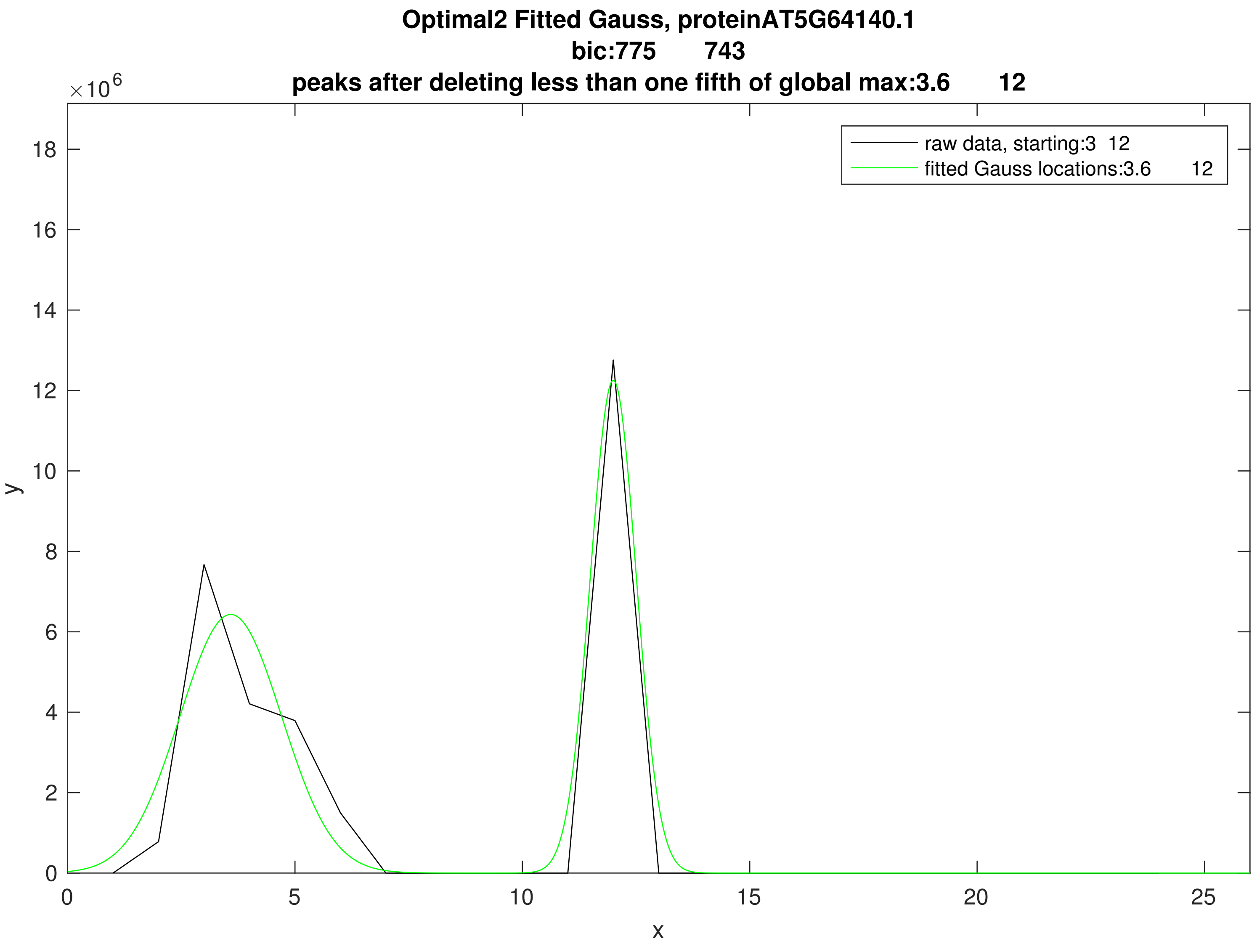

Optimal2 Fitted Gauss, proteinAT5G64140.1
bic:775 743
peaks after deleting less than one fifth of global max:3.6 12
×10^6
raw data, starting:3 12
fitted Gauss locations:3.6 12
y
x
0
2
4
6
8
10
12
14
16
18
0
5
10
15
20
25

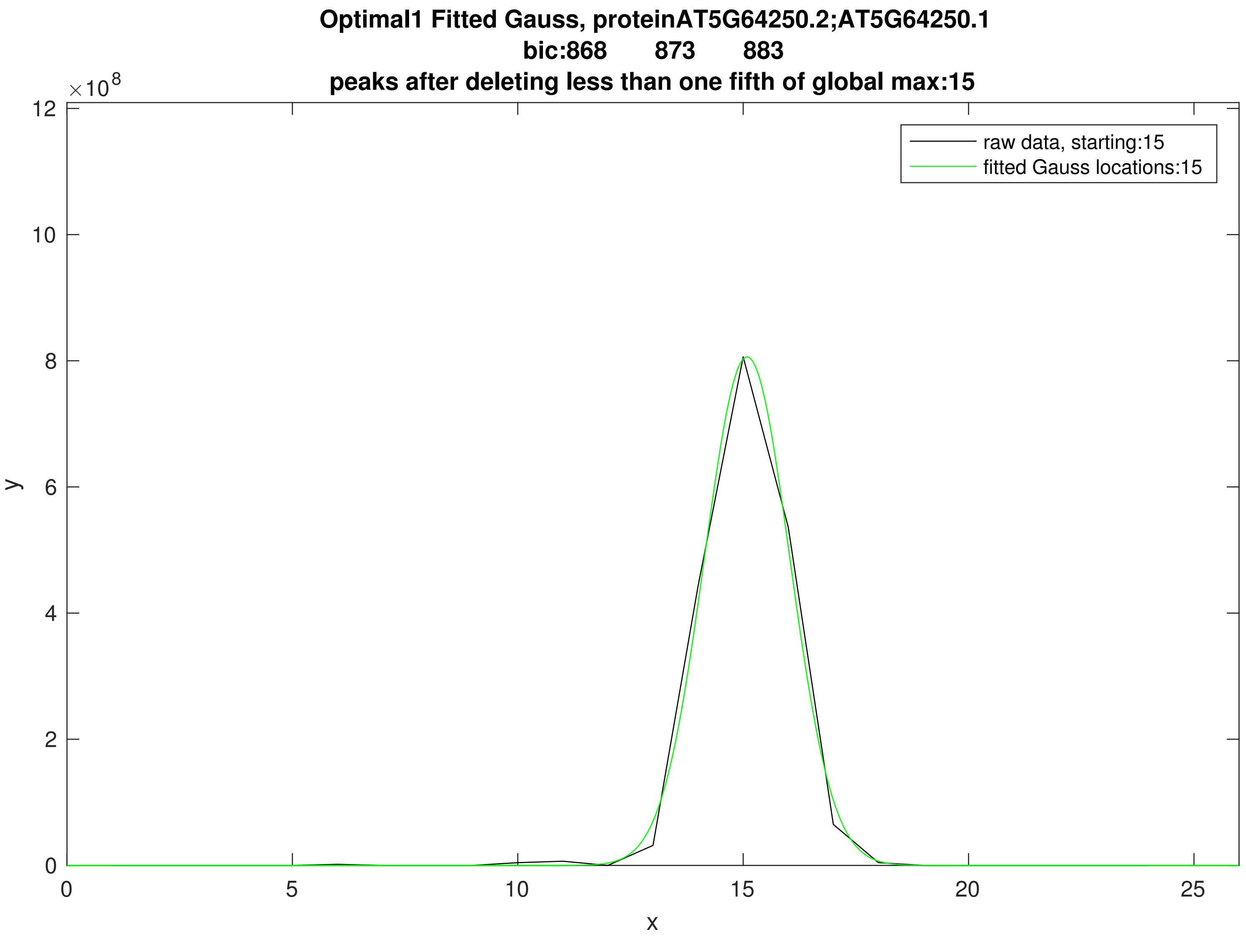
Optimal1 Fitted Gauss, proteinAT5G64250.2;AT5G64250.1
bic:868 873 883
peaks after deleting less than one fifth of global max:15
×10^8
12
10
8
6
4
2
0
y
0
5
10
15
20
25
x
raw data, starting:15
fitted Gauss locations:15

**Optimal1 Fitted Gauss, proteinAT5G64270.1**
**bic:746 755**
**peaks after deleting less than one fifth of global max:18**

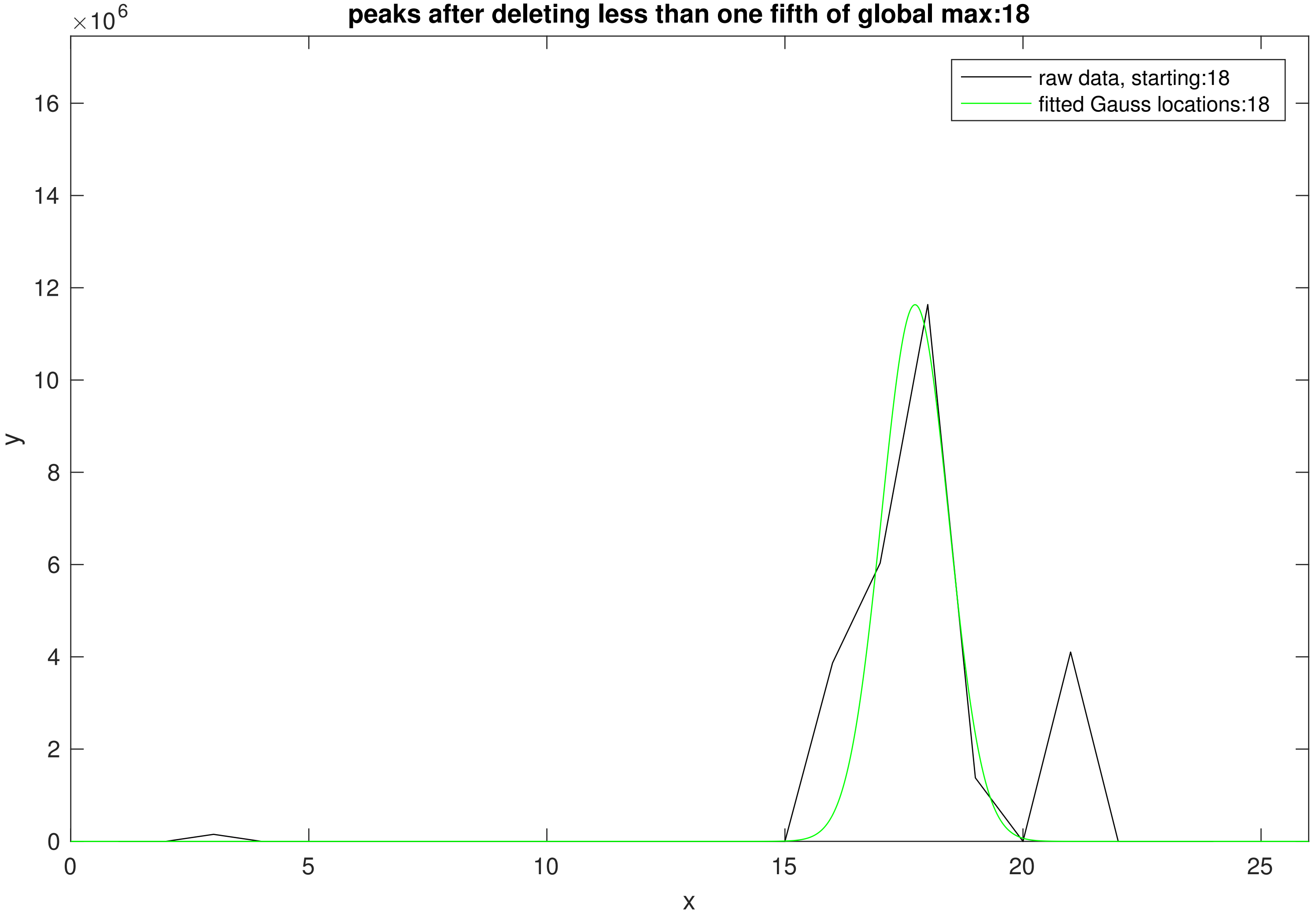

Optimal2 Fitted Gauss, proteinAT5G64300.1
bic:819 790
peaks after deleting less than one fifth of global max:10 14

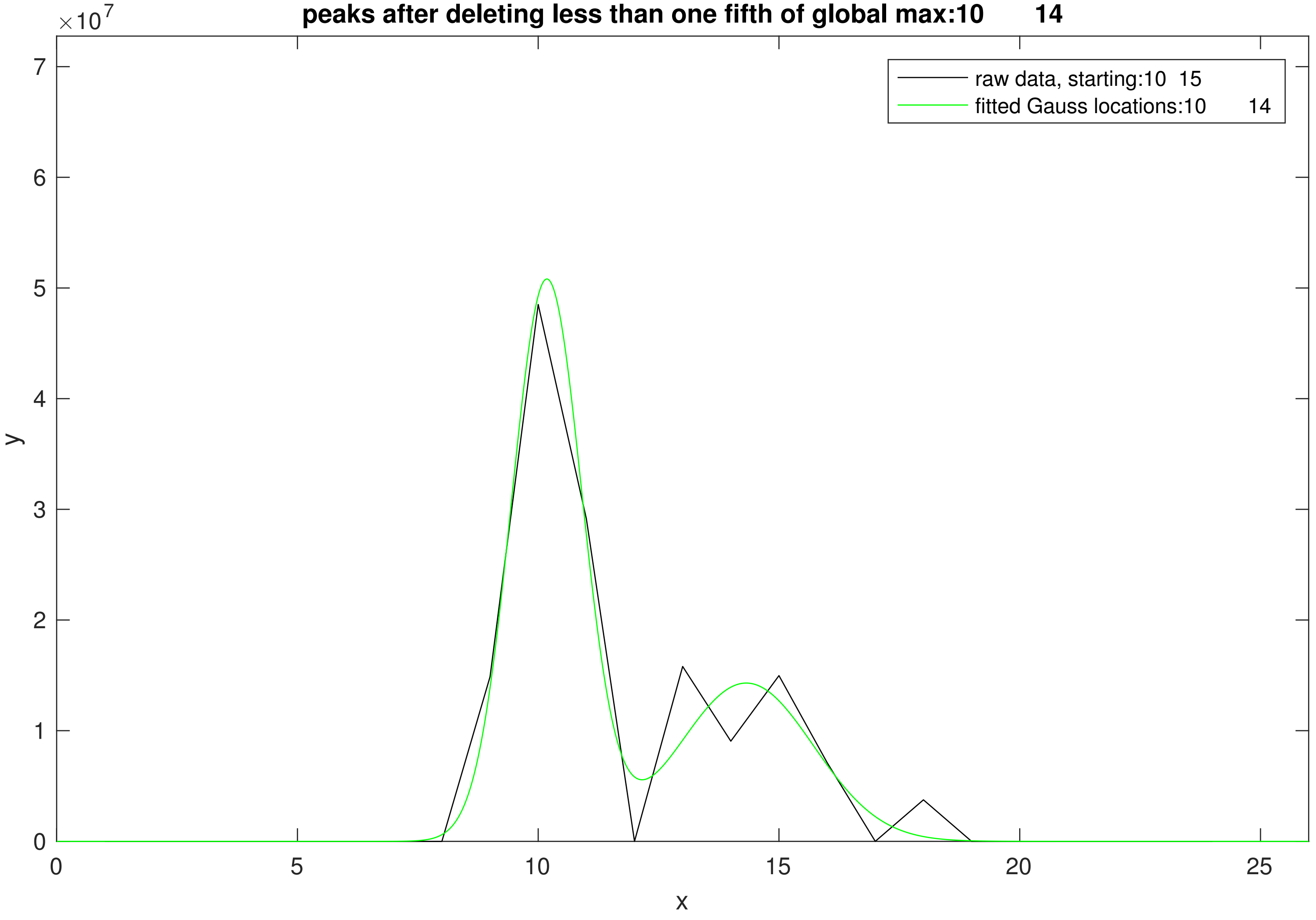

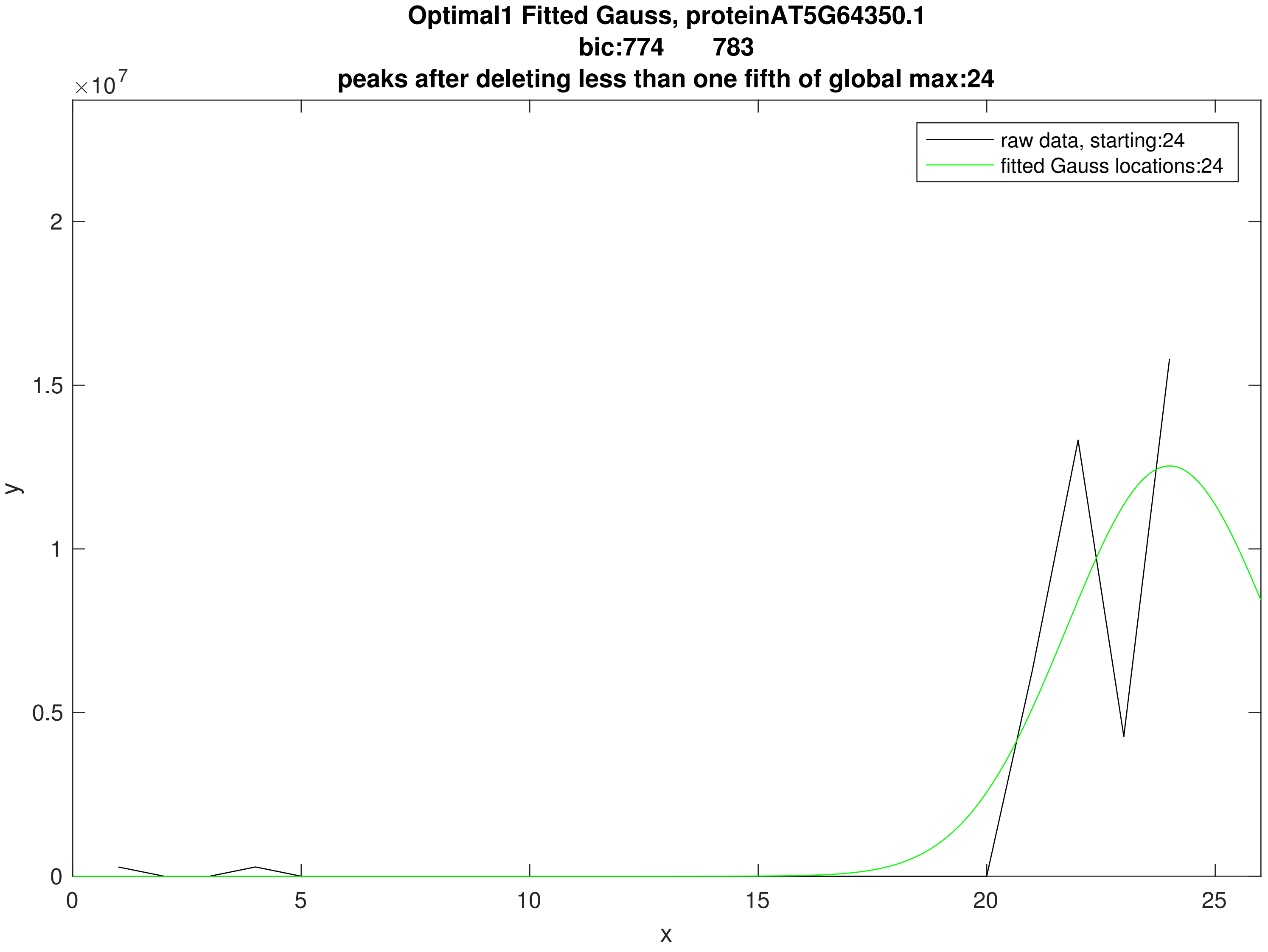

Optimal1 Fitted Gauss, proteinAT5G64350.1
bic:774 783
peaks after deleting less than one fifth of global max:24
×10^7
raw data, starting:24
fitted Gauss locations:24
y
x
0
0.5
1
1.5
2
0
5
10
15
20
25

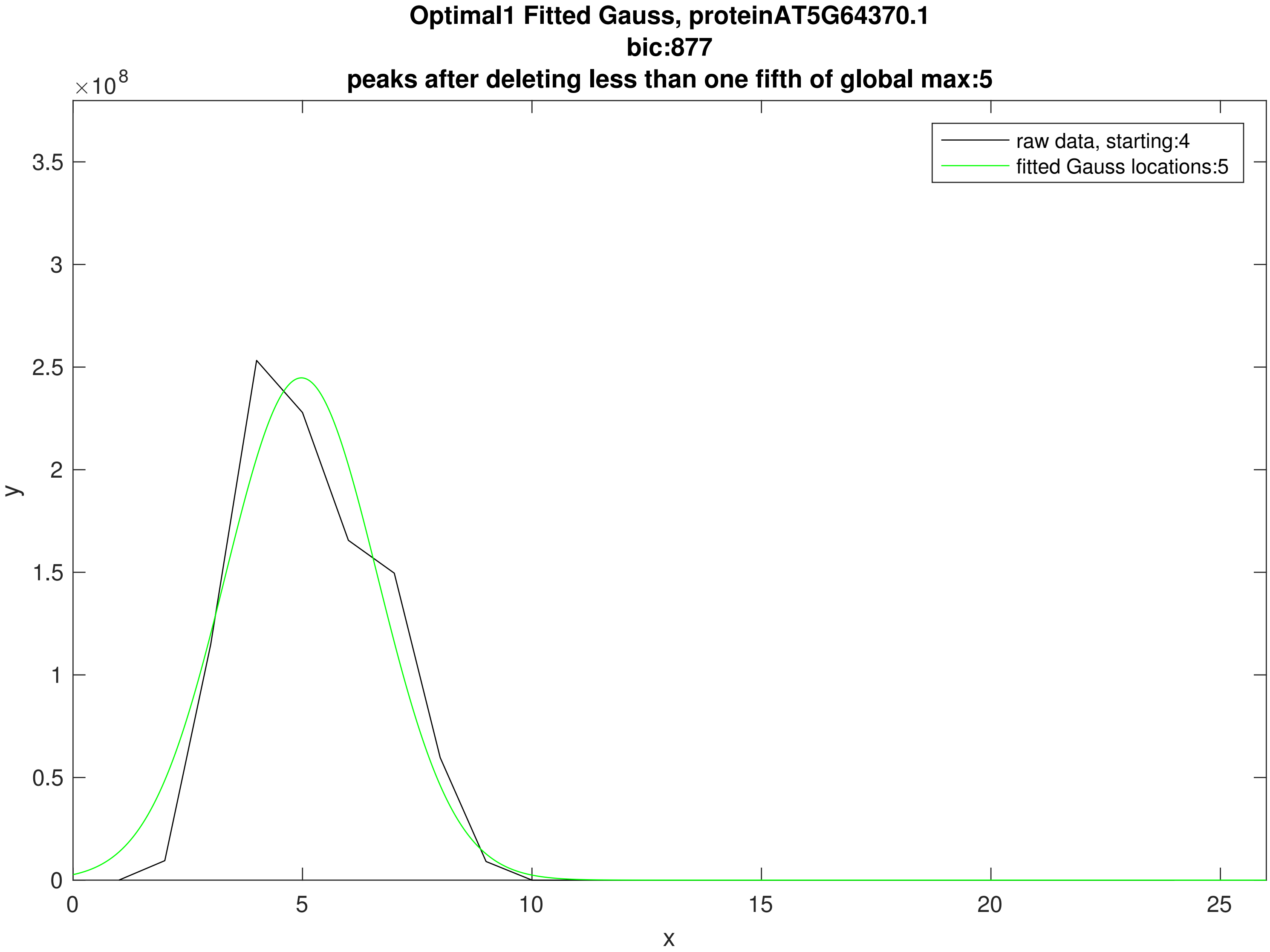

Optimal1 Fitted Gauss, proteinAT5G64370.1
bic:877
peaks after deleting less than one fifth of global max:5
×10^8
raw data, starting:4
fitted Gauss locations:5
y
x
0
0.5
1
1.5
2
2.5
3
3.5
0
5
10
15
20
25

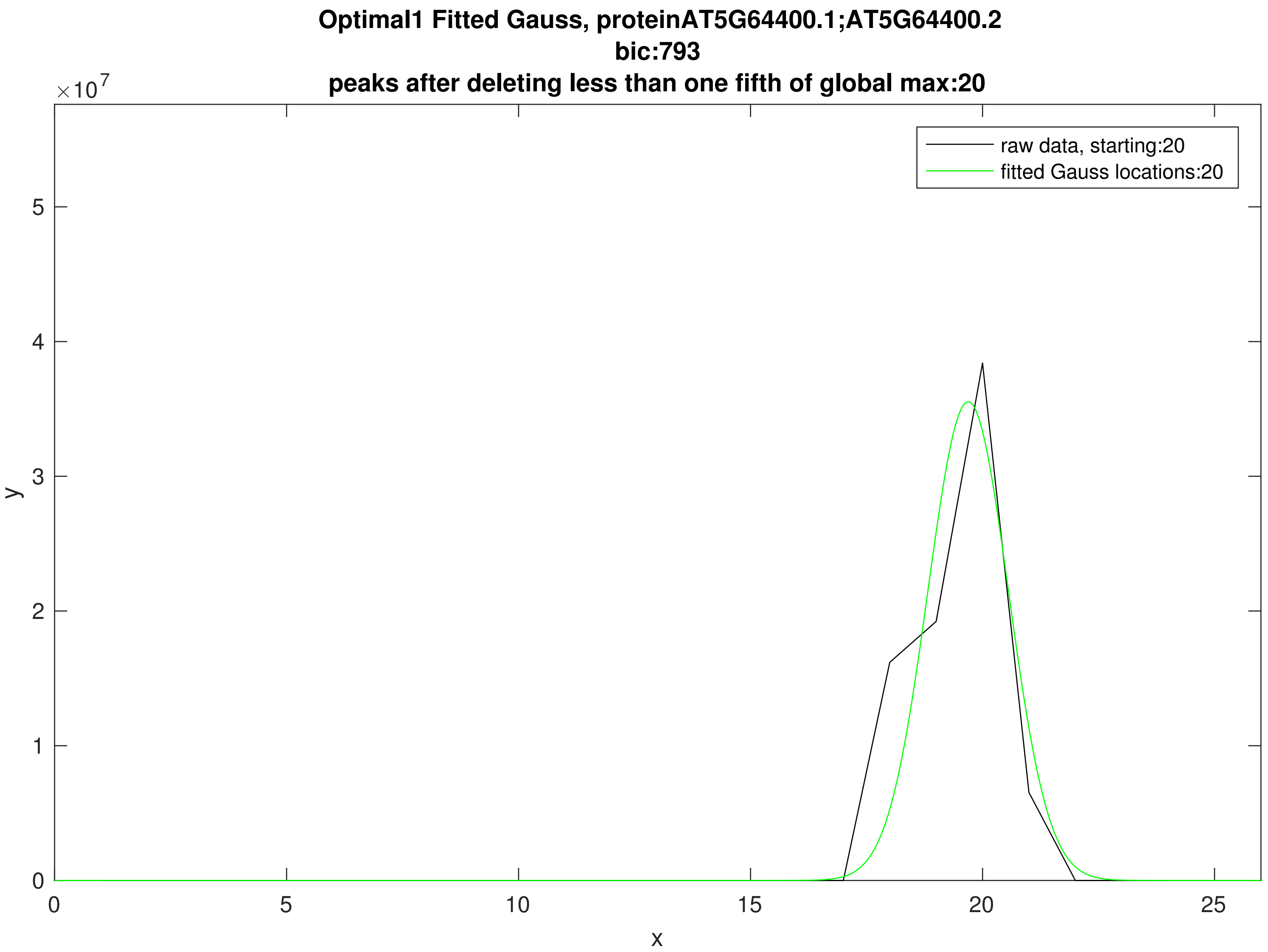

Optimal1 Fitted Gauss, proteinAT5G64400.1;AT5G64400.2
bic:793
peaks after deleting less than one fifth of global max:20
×10^7
raw data, starting:20
fitted Gauss locations:20
y
x
0
1
2
3
4
5
5
10
15
20
25

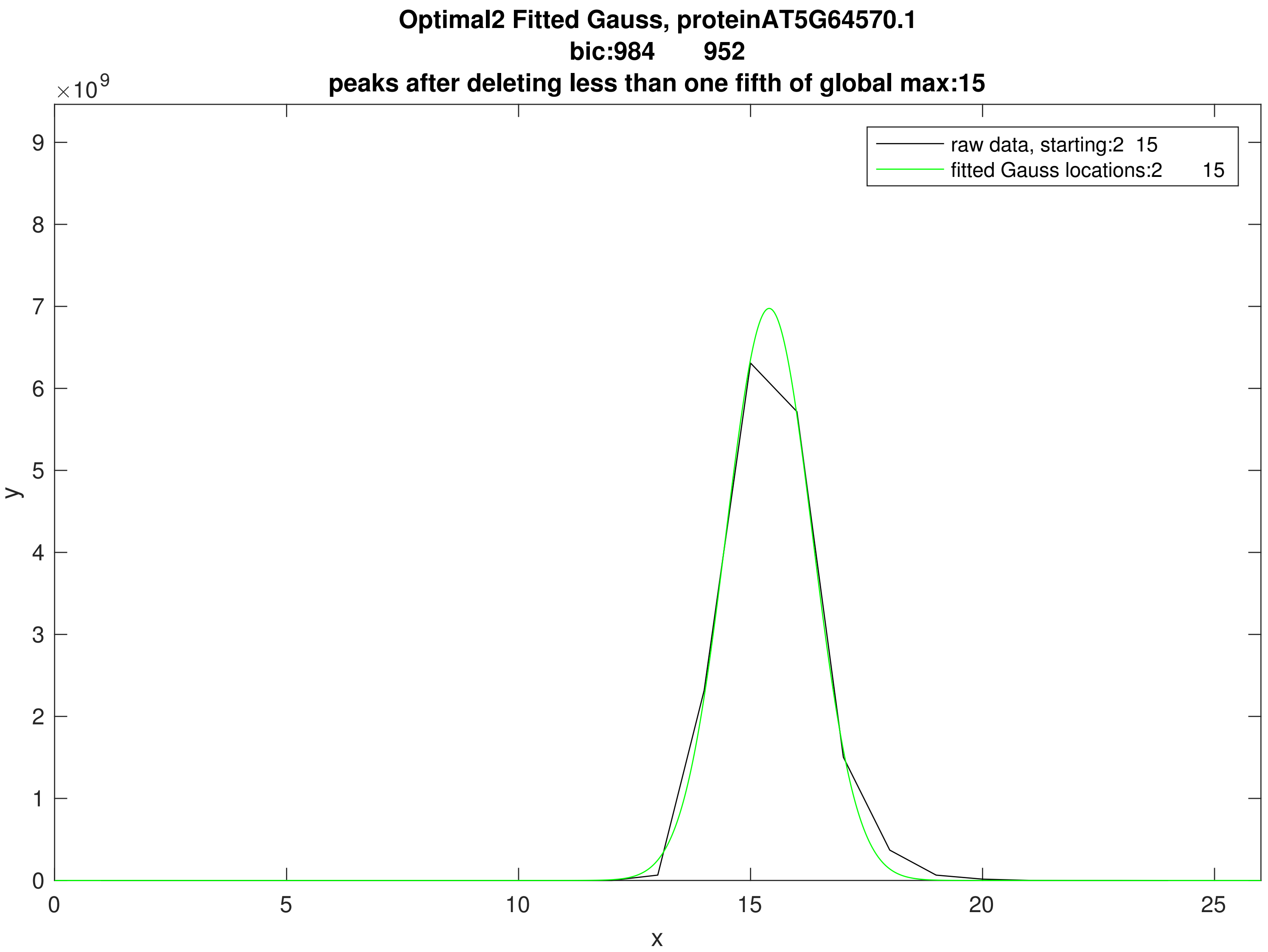

Optimal2 Fitted Gauss, proteinAT5G64570.1
bic:984 952
peaks after deleting less than one fifth of global max:15
×10^9
raw data, starting:2 15
fitted Gauss locations:2 15
y
x
0
1
2
3
4
5
6
7
8
9
0
5
10
15
20
25

Optimal1 Fitted Gauss, proteinAT5G64670.1
bic:711
peaks after deleting less than one fifth of global max:9

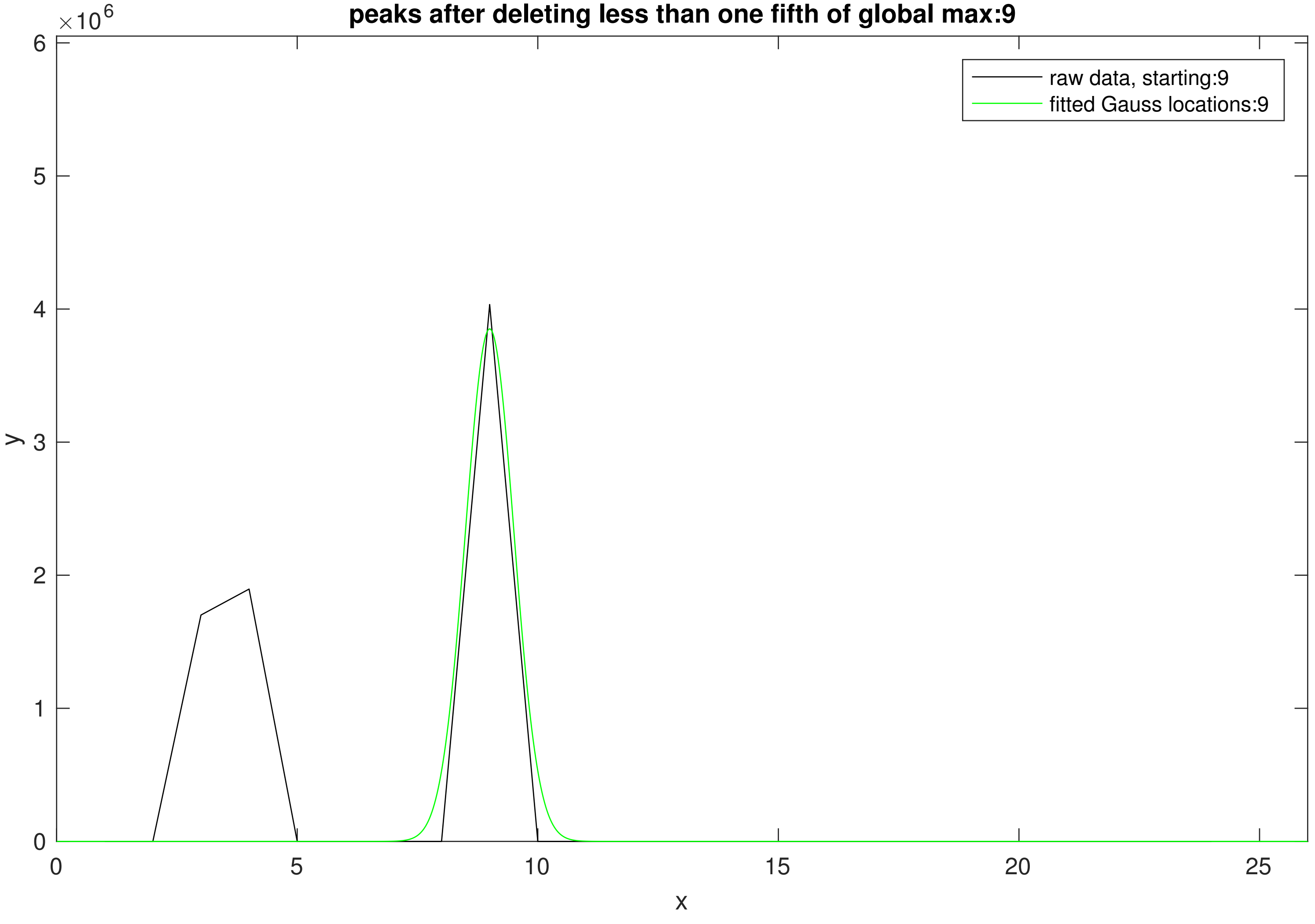

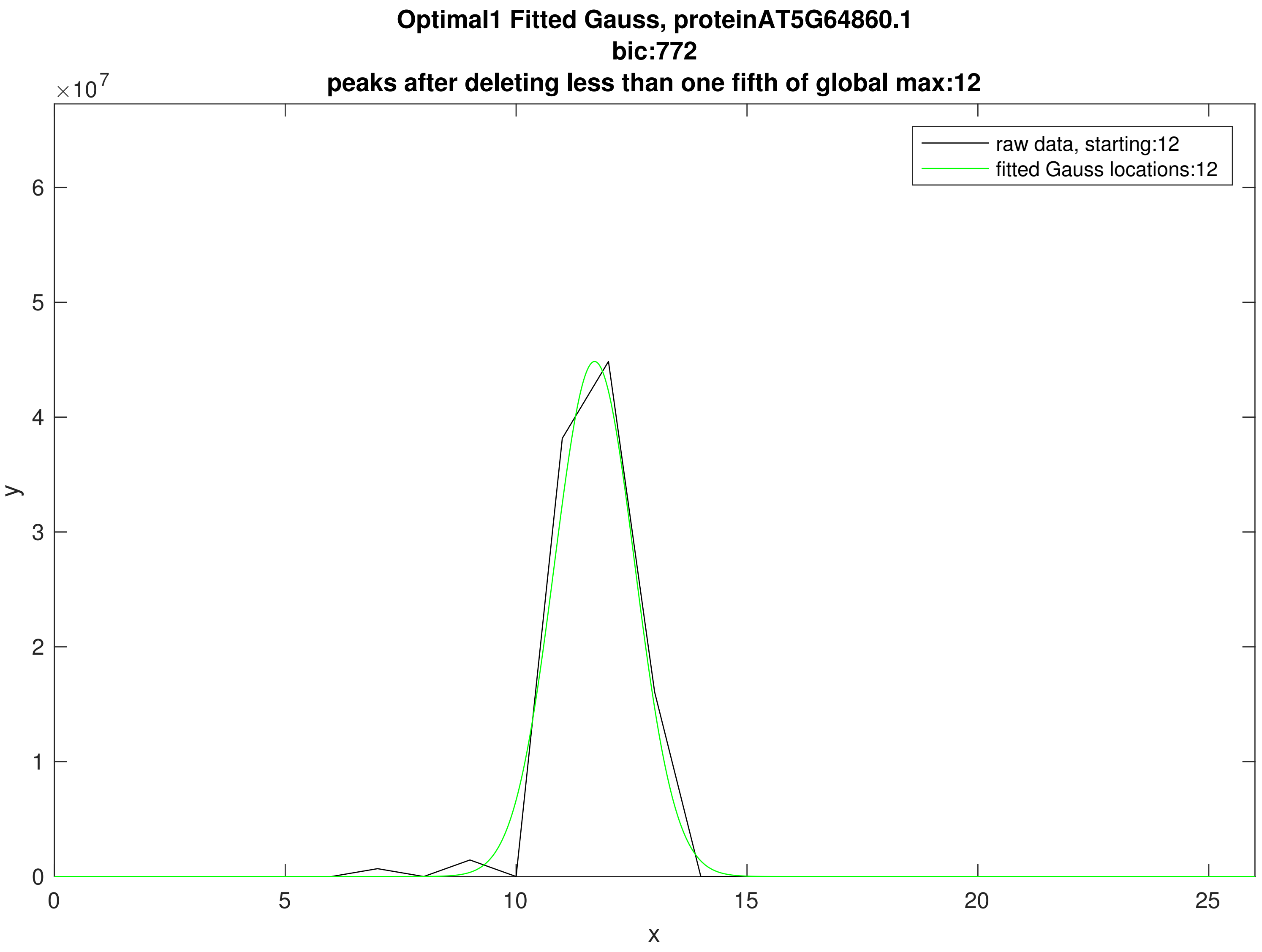

Optimal1 Fitted Gauss, proteinAT5G64860.1
bic:772
peaks after deleting less than one fifth of global max:12
×10^7
raw data, starting:12
fitted Gauss locations:12
y
x
0
1
2
3
4
5
6
0
5
10
15
20
25

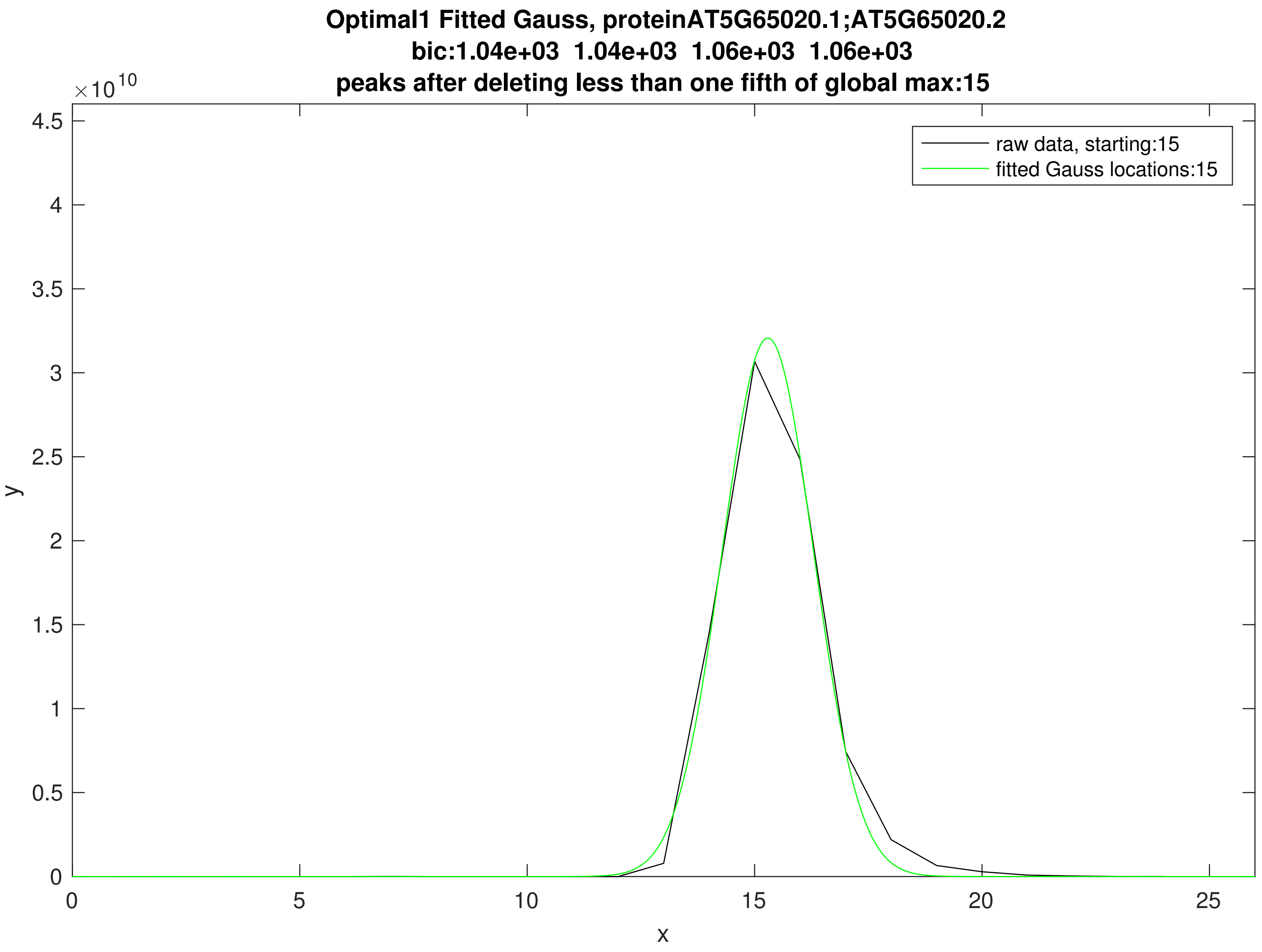

Optimal1 Fitted Gauss, proteinAT5G65020.1;AT5G65020.2
bic:1.04e+03 1.04e+03 1.06e+03 1.06e+03
peaks after deleting less than one fifth of global max:15
×10^10
raw data, starting:15
fitted Gauss locations:15
y
x
0
0.5
1
1.5
2
2.5
3
3.5
4
4.5
5
10
15
20
25

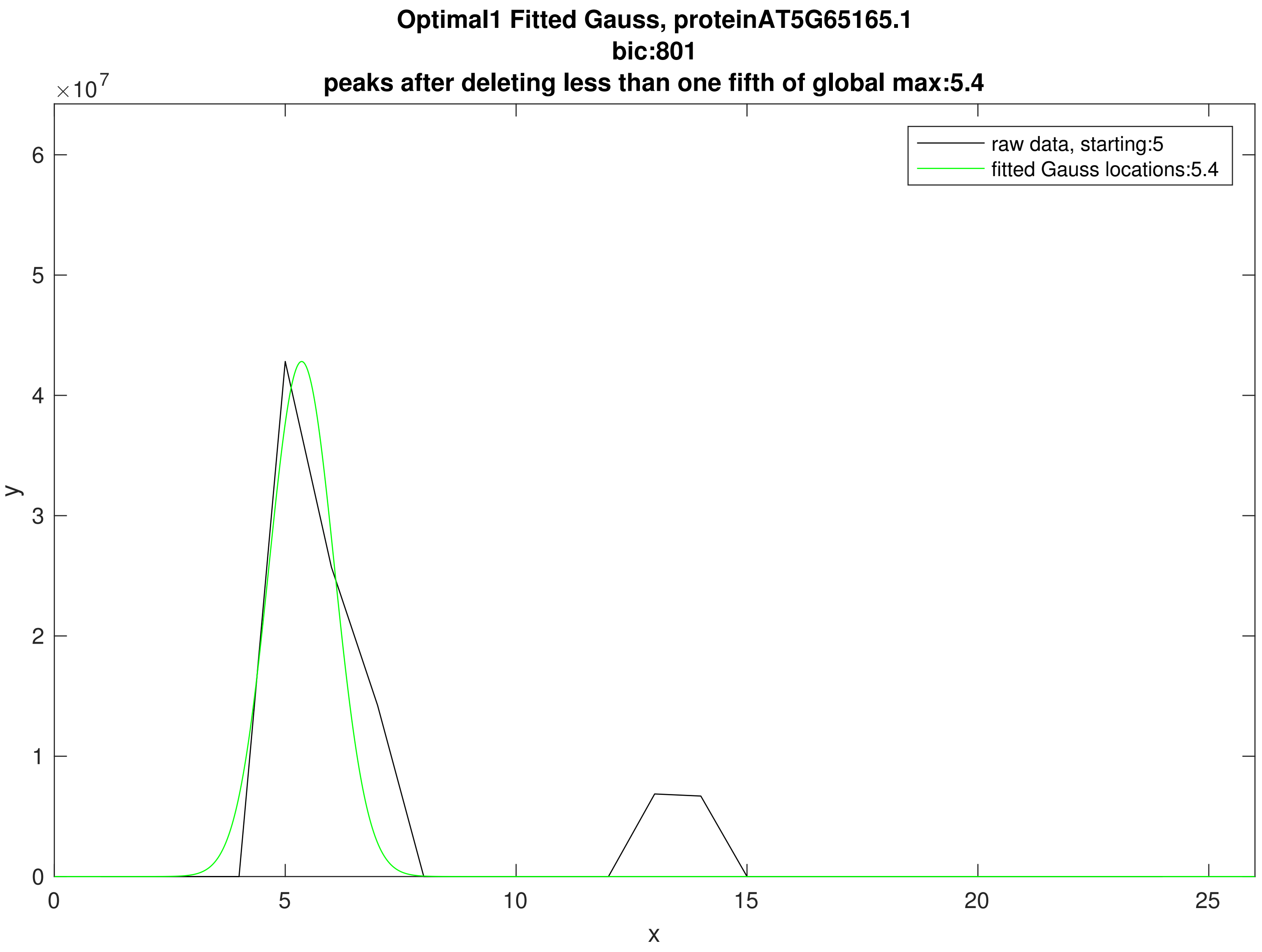

Optimal1 Fitted Gauss, proteinAT5G65165.1
bic:801
peaks after deleting less than one fifth of global max:5.4
×10^7
raw data, starting:5
fitted Gauss locations:5.4
y
x
0
1
2
3
4
5
6
0
5
10
15
20
25

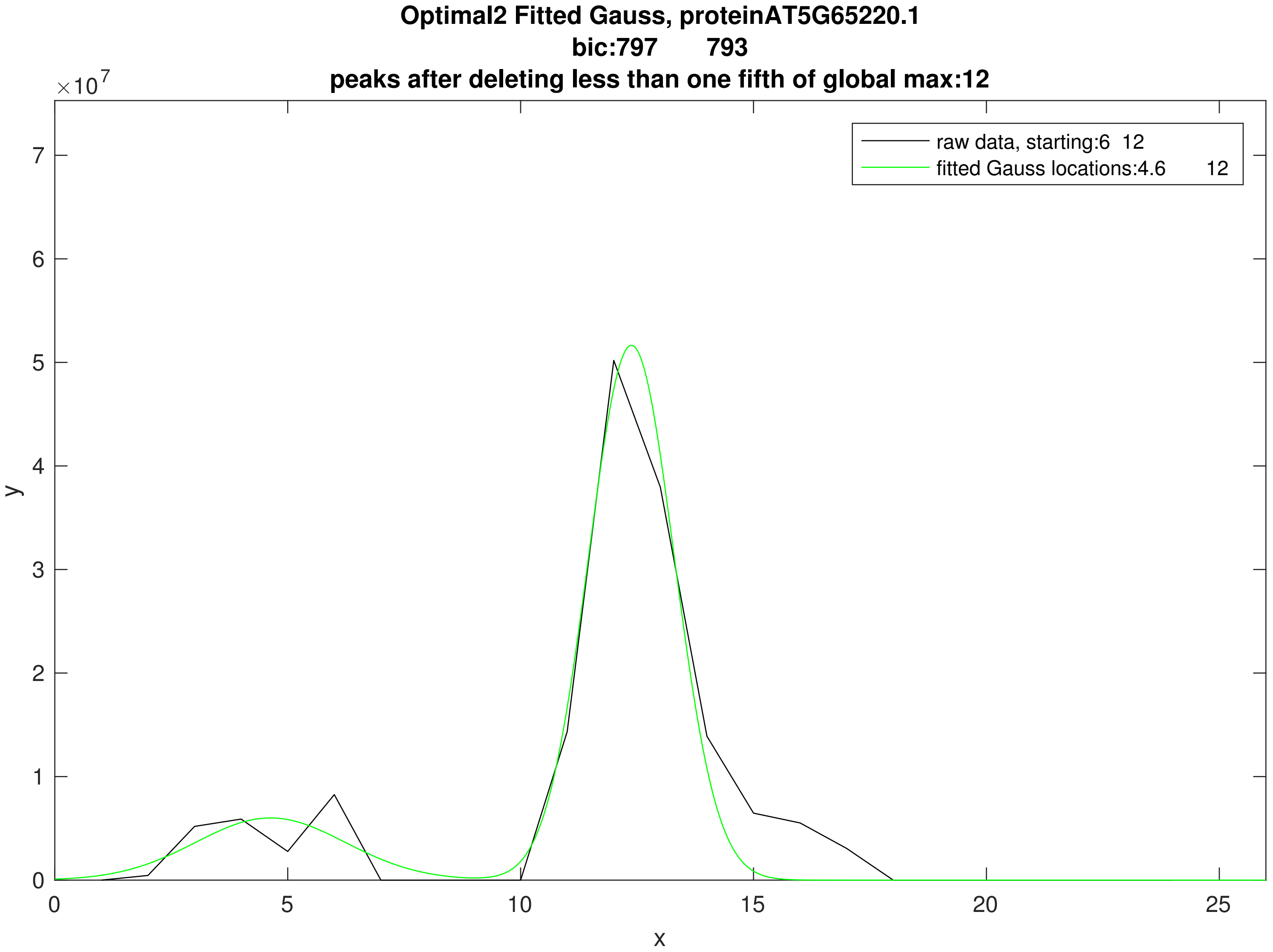

Optimal2 Fitted Gauss, proteinAT5G65220.1
bic:797 793
peaks after deleting less than one fifth of global max:12
×10^7
raw data, starting:6 12
fitted Gauss locations:4.6 12
y
x
0
1
2
3
4
5
6
7
0
5
10
15
20
25

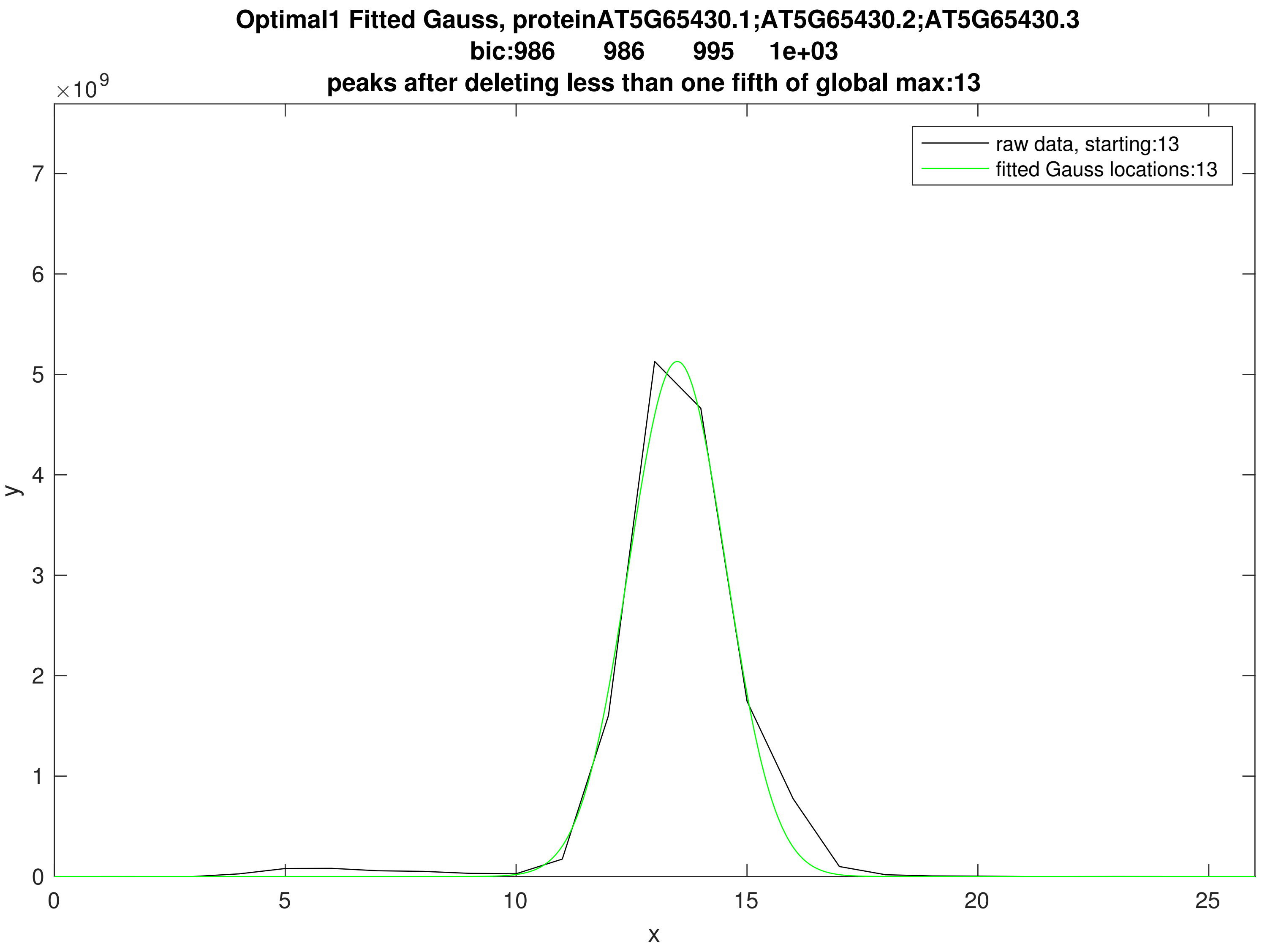

Optimal1 Fitted Gauss, proteinAT5G65430.1;AT5G65430.2;AT5G65430.3
bic:986 986 995 1e+03
peaks after deleting less than one fifth of global max:13
×10^9
raw data, starting:13
fitted Gauss locations:13
y
x
0
1
2
3
4
5
6
7
0
5
10
15
20
25

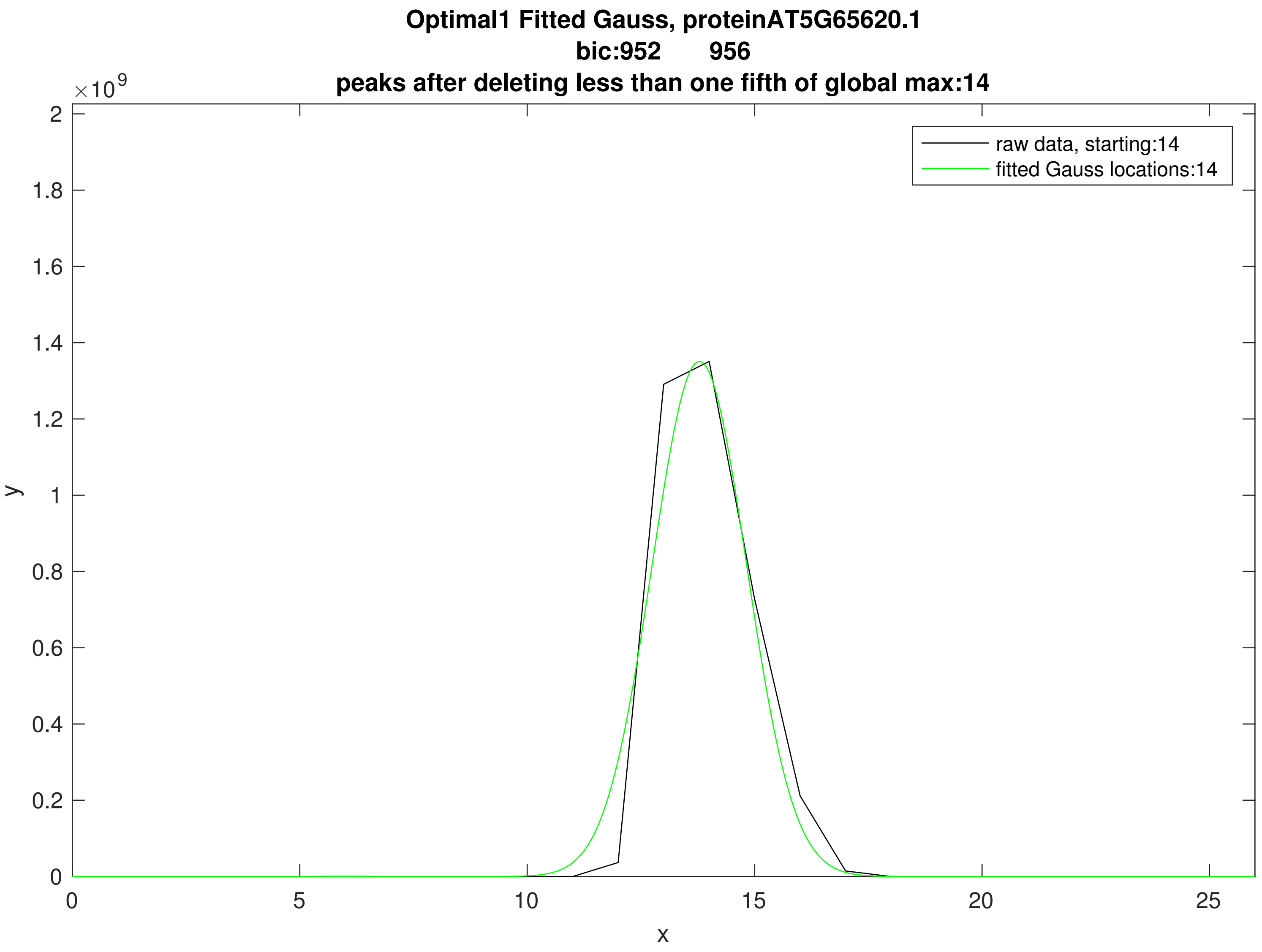
Optimal1 Fitted Gauss, proteinAT5G65620.1
bic:952 956
peaks after deleting less than one fifth of global max:14
×10^9
2
1.8
1.6
1.4
1.2
1
0.8
0.6
0.4
0.2
0
y
0
5
10
15
20
25
x
raw data, starting:14
fitted Gauss locations:14

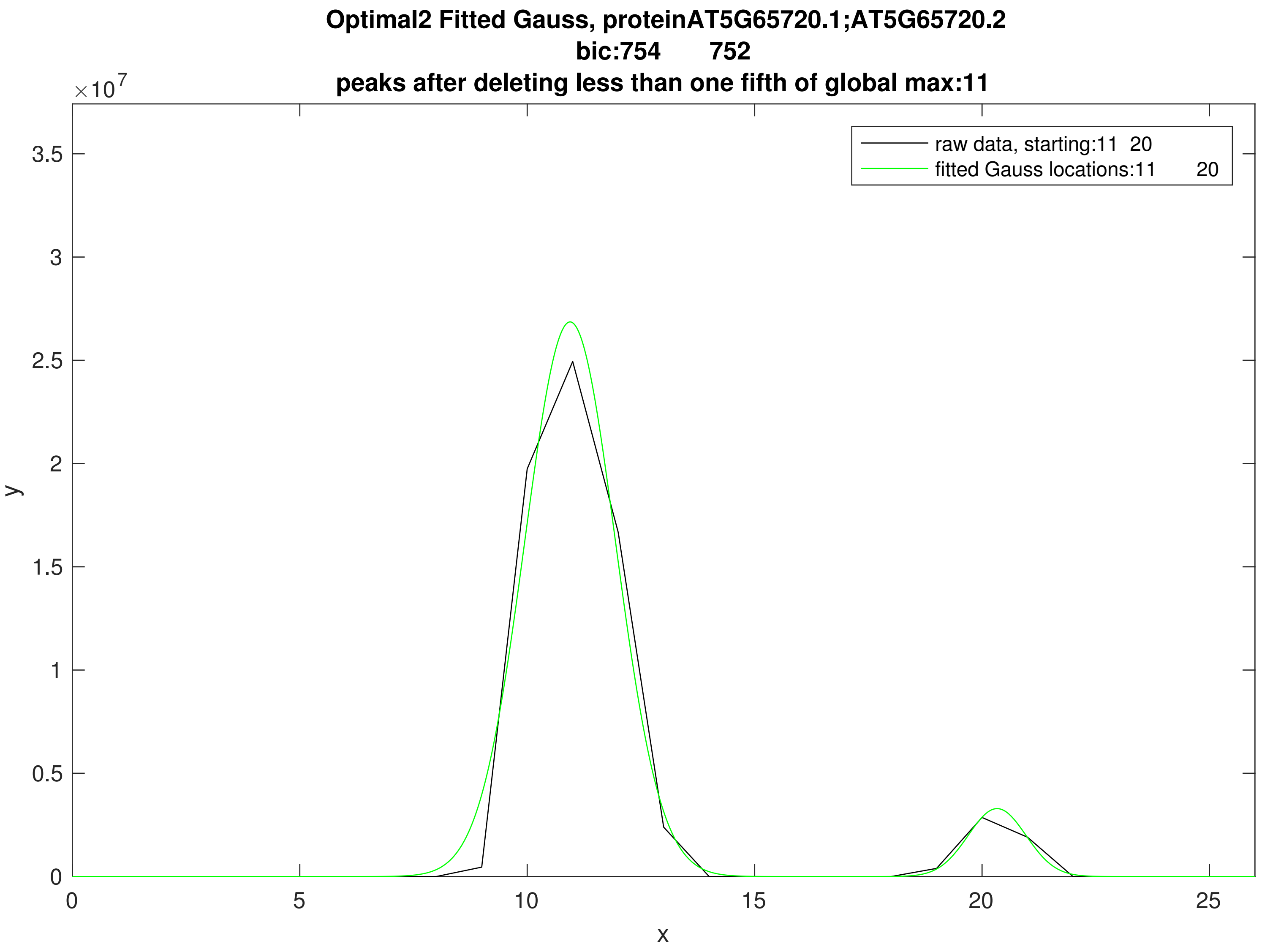

Optimal2 Fitted Gauss, proteinAT5G65720.1;AT5G65720.2
bic:754 752
peaks after deleting less than one fifth of global max:11
×10^7
raw data, starting:11 20
fitted Gauss locations:11 20
y
x
0
0.5
1
1.5
2
2.5
3
3.5
5
10
15
20
25

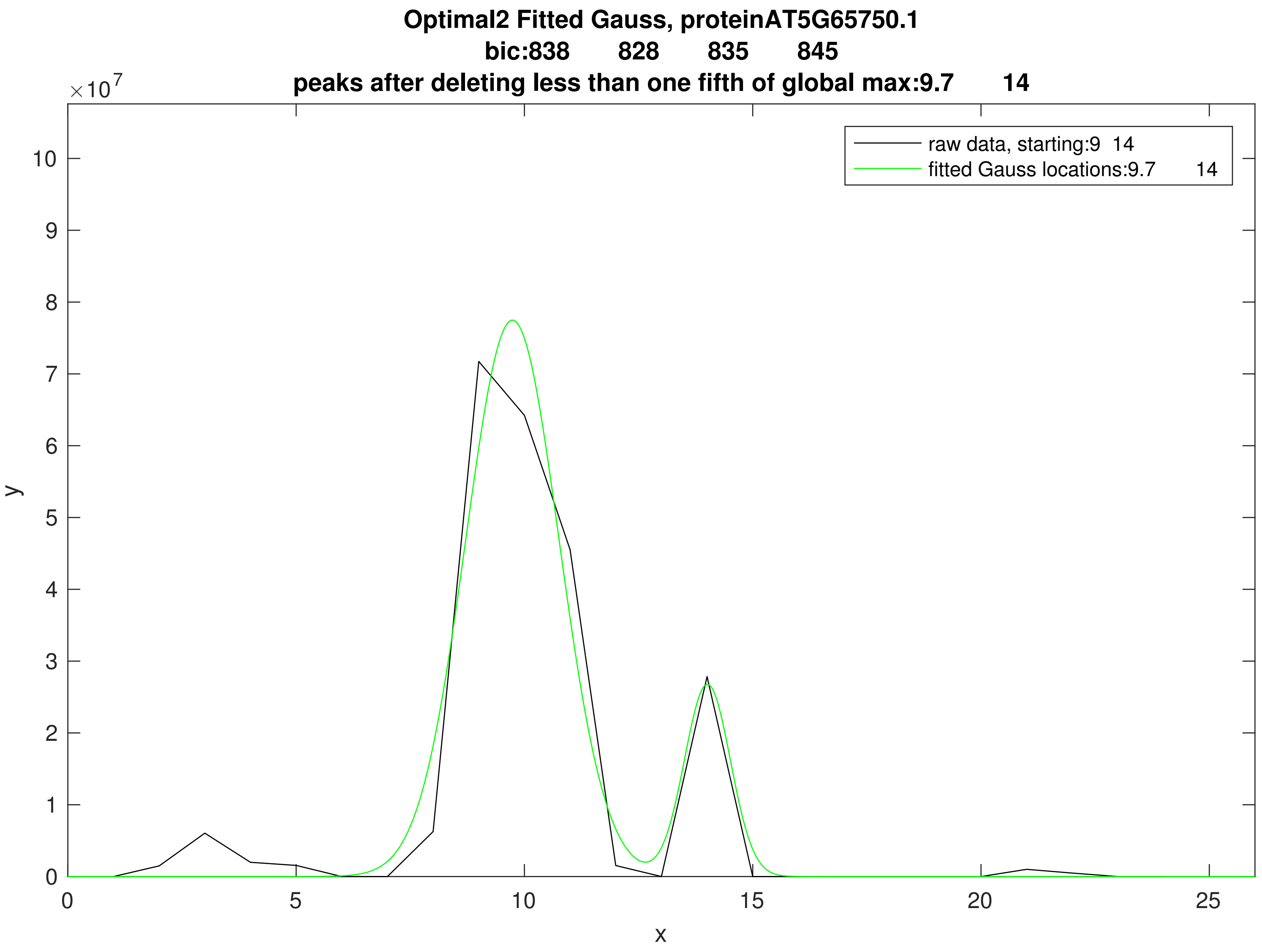

Optimal2 Fitted Gauss, proteinAT5G65750.1
bic:838 828 835 845
peaks after deleting less than one fifth of global max:9.7 14
×10^7
raw data, starting:9 14
fitted Gauss locations:9.7 14
y
x
0
1
2
3
4
5
6
7
8
9
10
0
5
10
15
20
25

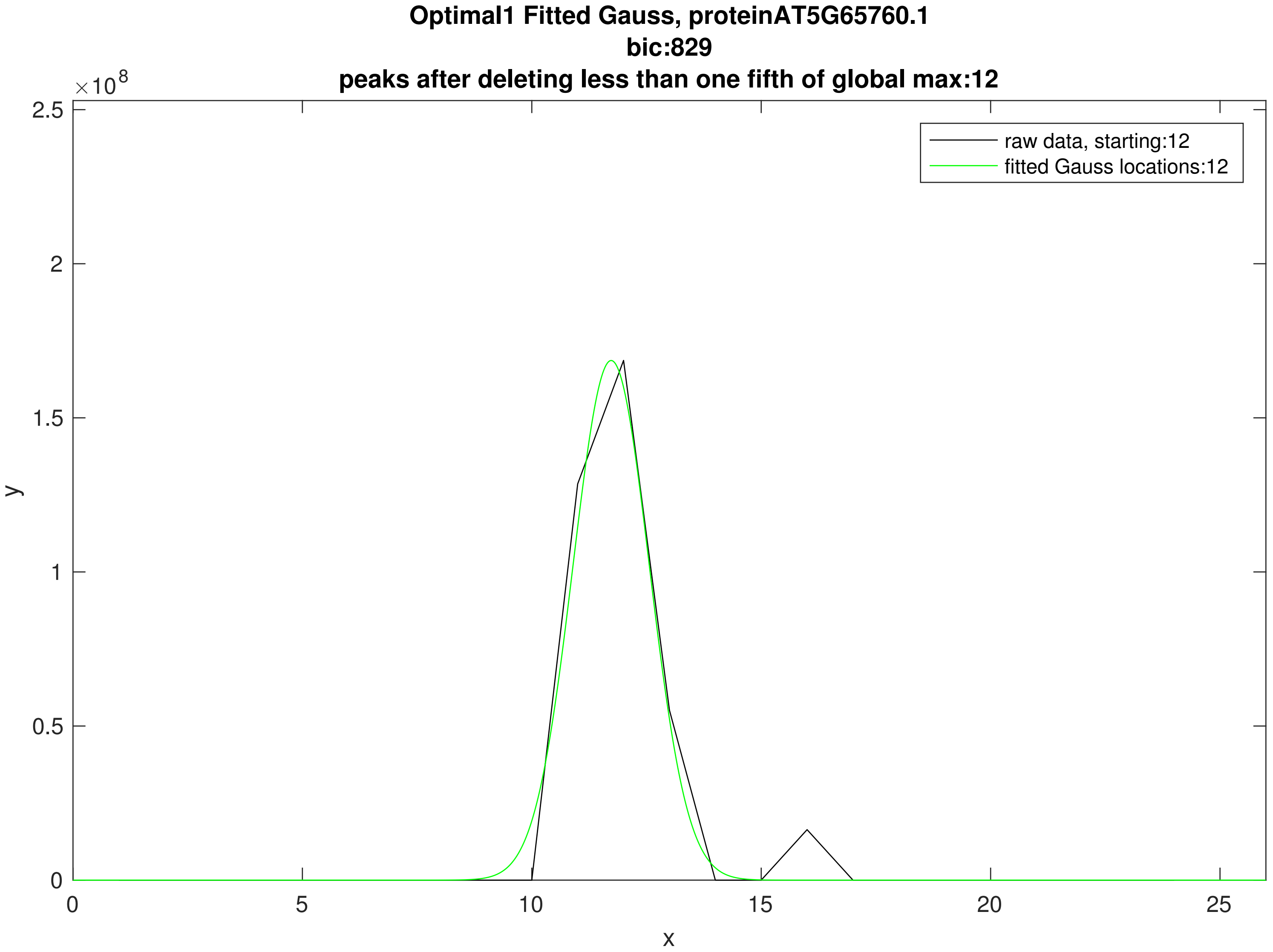
Optimal1 Fitted Gauss, proteinAT5G65760.1
bic:829
peaks after deleting less than one fifth of global max:12
×10^8
2.5
2
1.5
1
0.5
0
y
0
5
10
15
20
25
x
raw data, starting:12
fitted Gauss locations:12

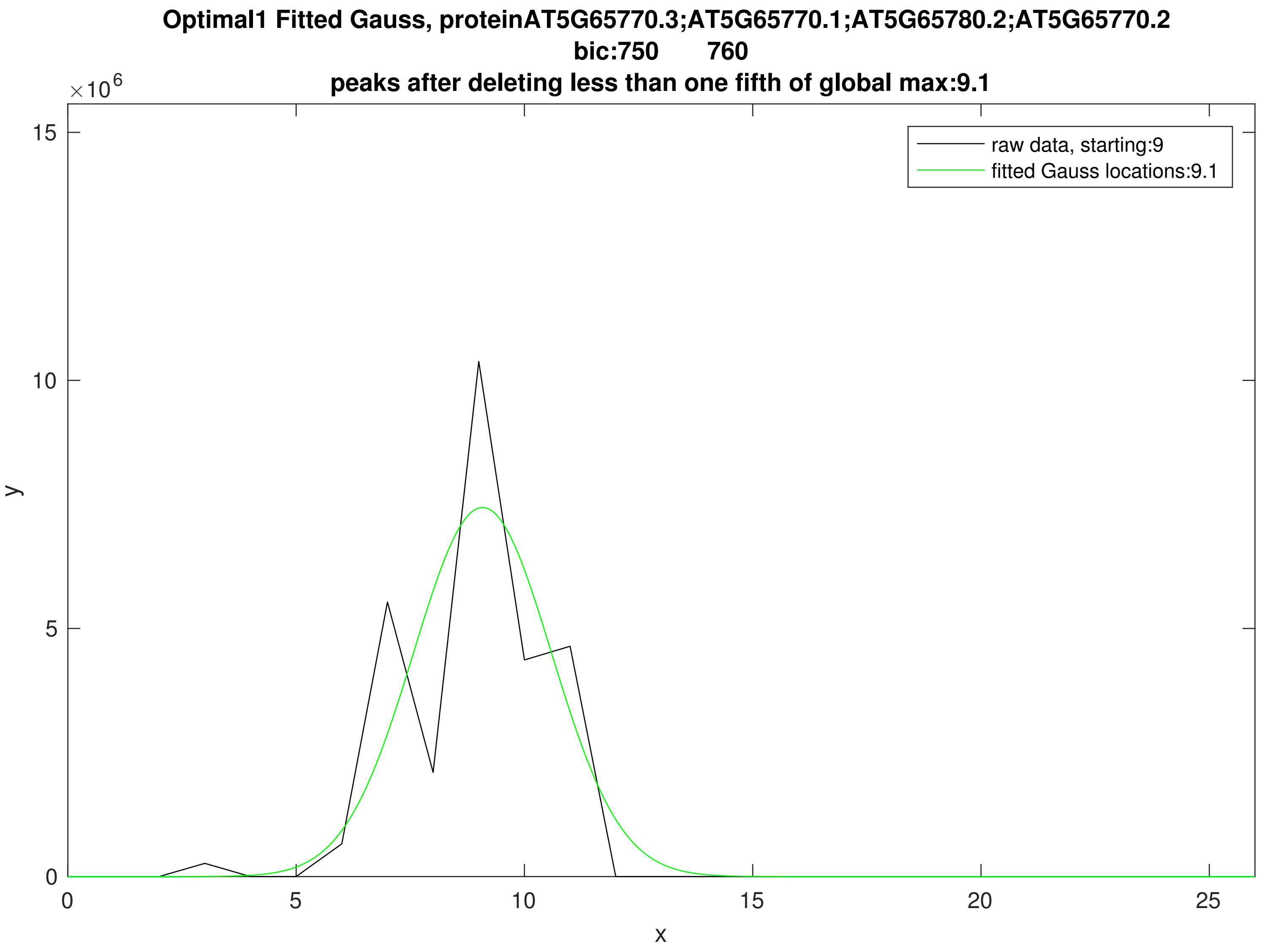

Optimal1 Fitted Gauss, proteinAT5G65770.3;AT5G65770.1;AT5G65780.2;AT5G65770.2
bic:750 760
peaks after deleting less than one fifth of global max:9.1
×10^6
15
10
5
0
0
5
10
15
20
25
x
y
raw data, starting:9
fitted Gauss locations:9.1

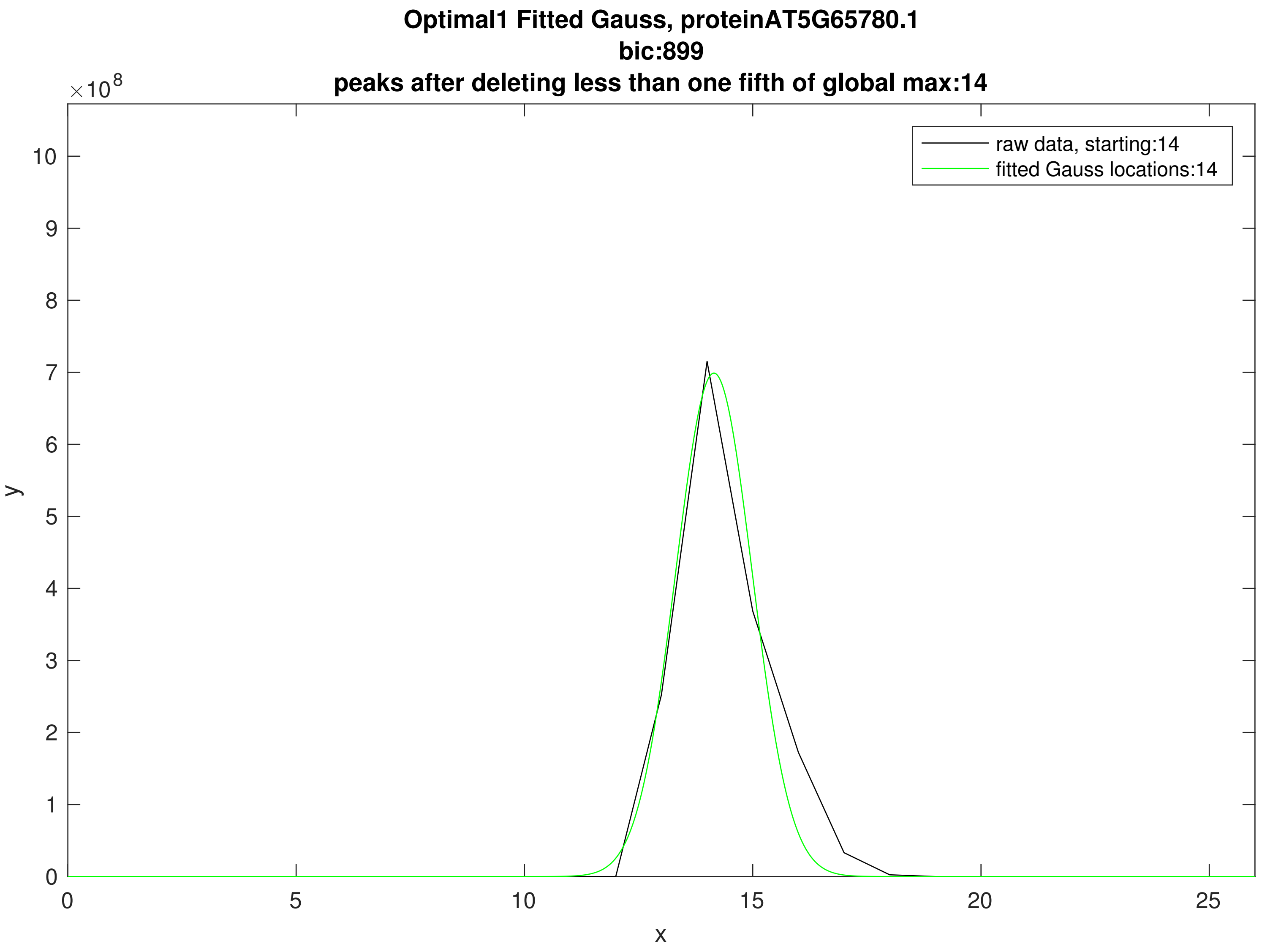

Optimal1 Fitted Gauss, proteinAT5G65780.1
bic:899
peaks after deleting less than one fifth of global max:14
×10^8
raw data, starting:14
fitted Gauss locations:14
y
x
0
1
2
3
4
5
6
7
8
9
10
0
5
10
15
20
25

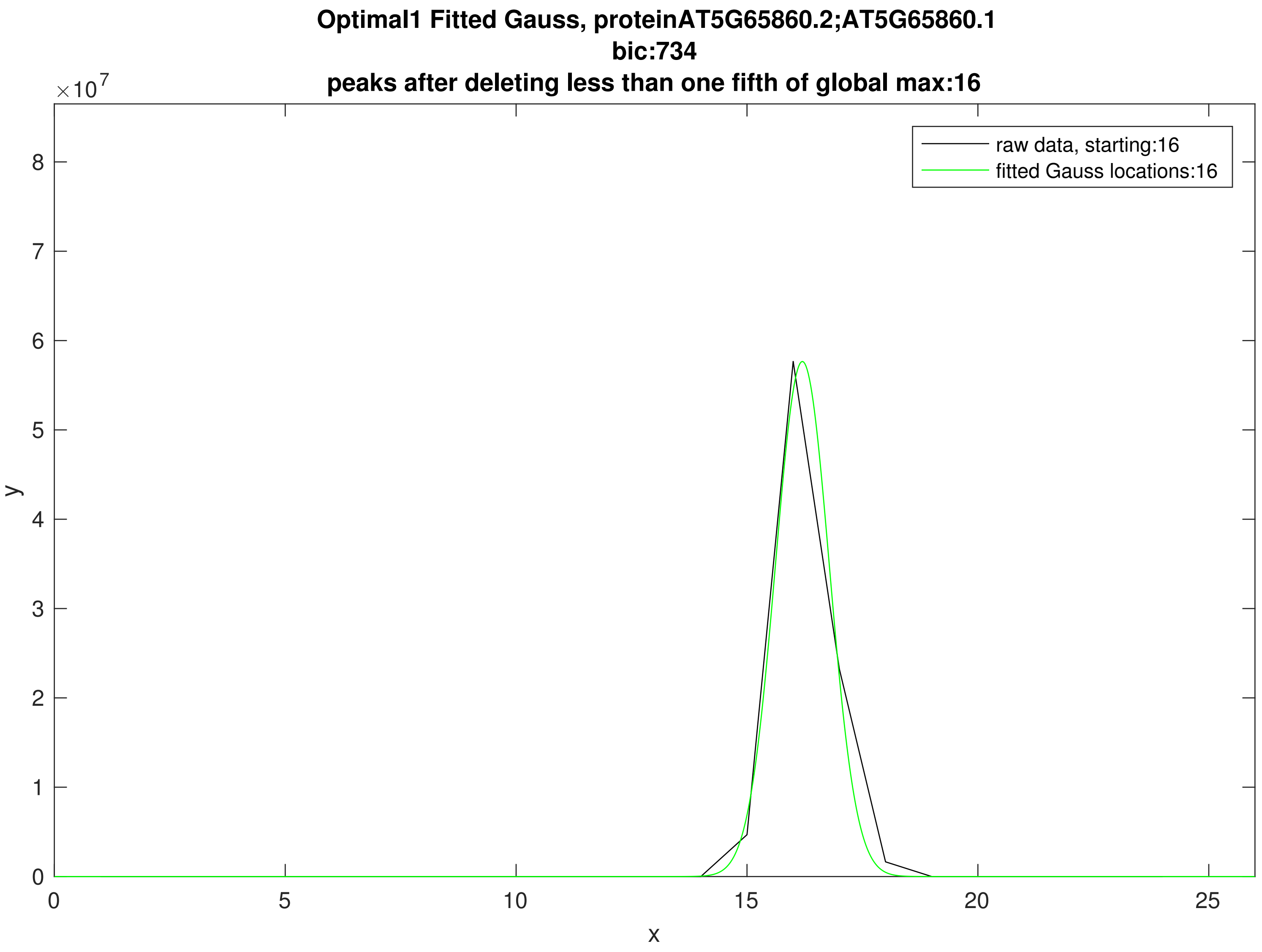
Optimal1 Fitted Gauss, proteinAT5G65860.2;AT5G65860.1
bic:734
peaks after deleting less than one fifth of global max:16
×10^7
raw data, starting:16
fitted Gauss locations:16
y
x
0
1
2
3
4
5
6
7
8
0
5
10
15
20
25

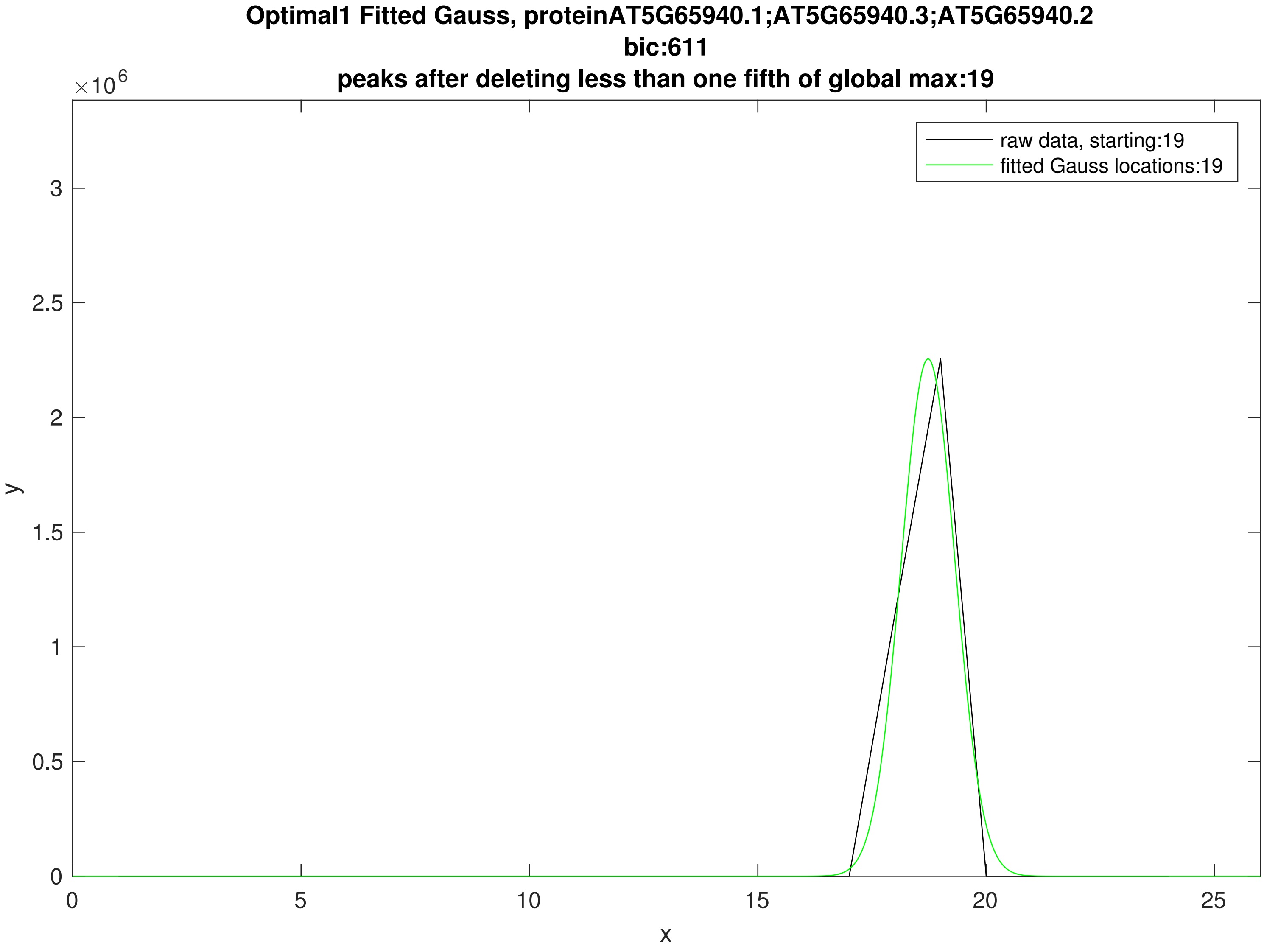
Optimal1 Fitted Gauss, proteinAT5G65940.1;AT5G65940.3;AT5G65940.2
bic:611
peaks after deleting less than one fifth of global max:19
×10^6
raw data, starting:19
fitted Gauss locations:19
y
x
0
0.5
1
1.5
2
2.5
3
0
5
10
15
20
25

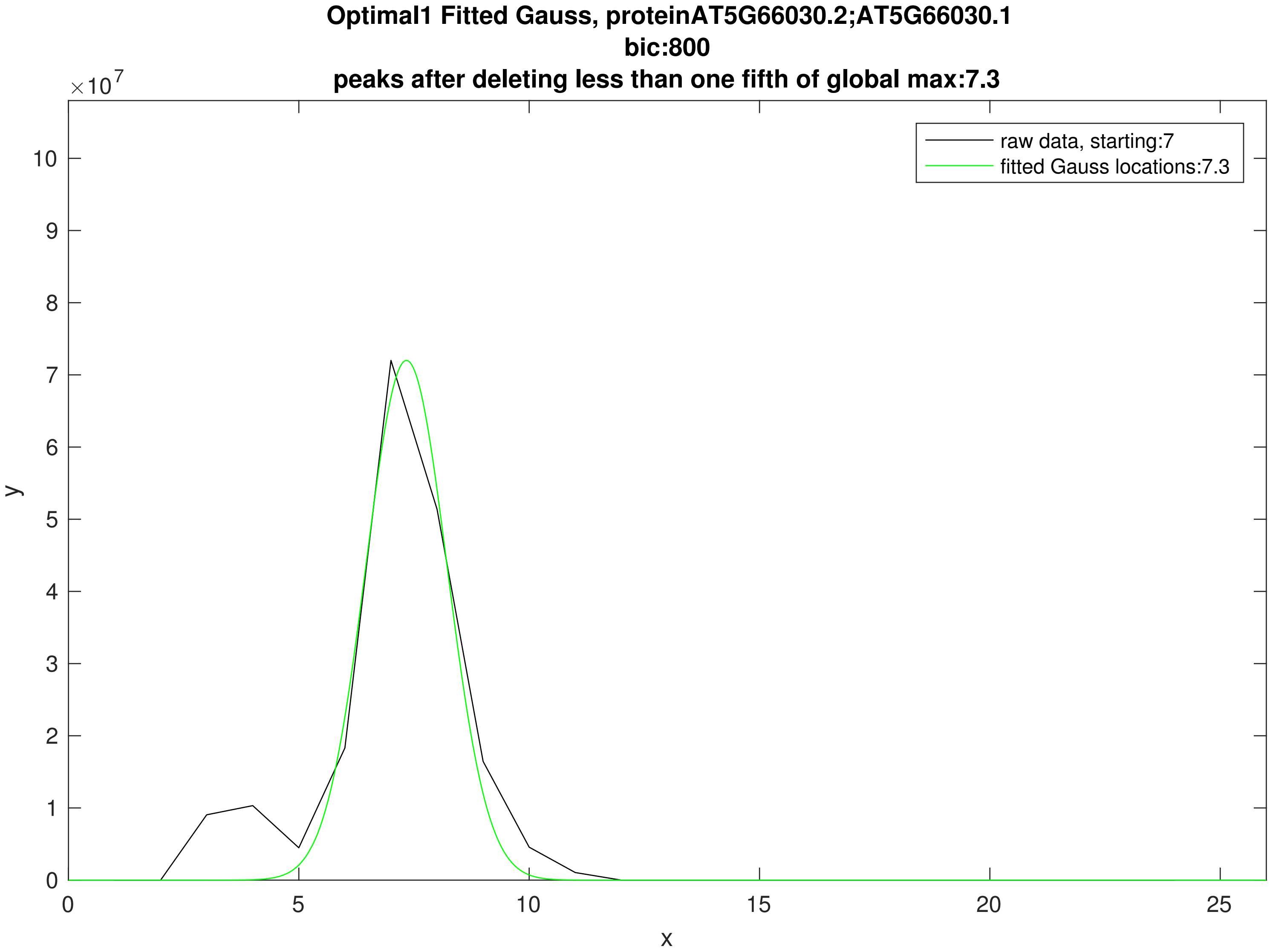

Optimal1 Fitted Gauss, proteinAT5G66030.2;AT5G66030.1
bic:800
peaks after deleting less than one fifth of global max:7.3
×10^7
raw data, starting:7
fitted Gauss locations:7.3
y
x

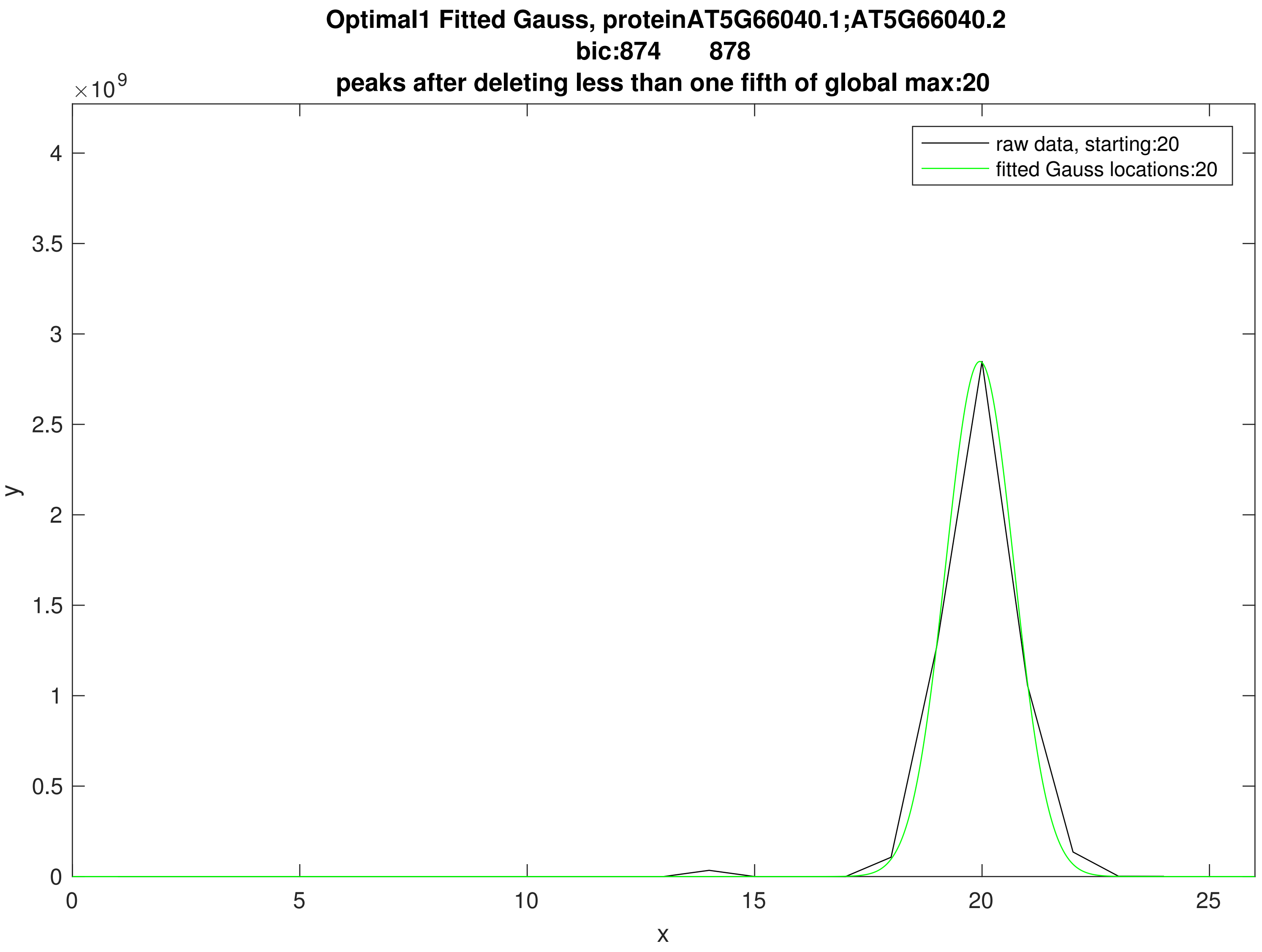

Optimal1 Fitted Gauss, proteinAT5G66040.1;AT5G66040.2
bic:874 878
peaks after deleting less than one fifth of global max:20
×10^9
raw data, starting:20
fitted Gauss locations:20
y
x
0
0.5
1
1.5
2
2.5
3
3.5
4
0
5
10
15
20
25

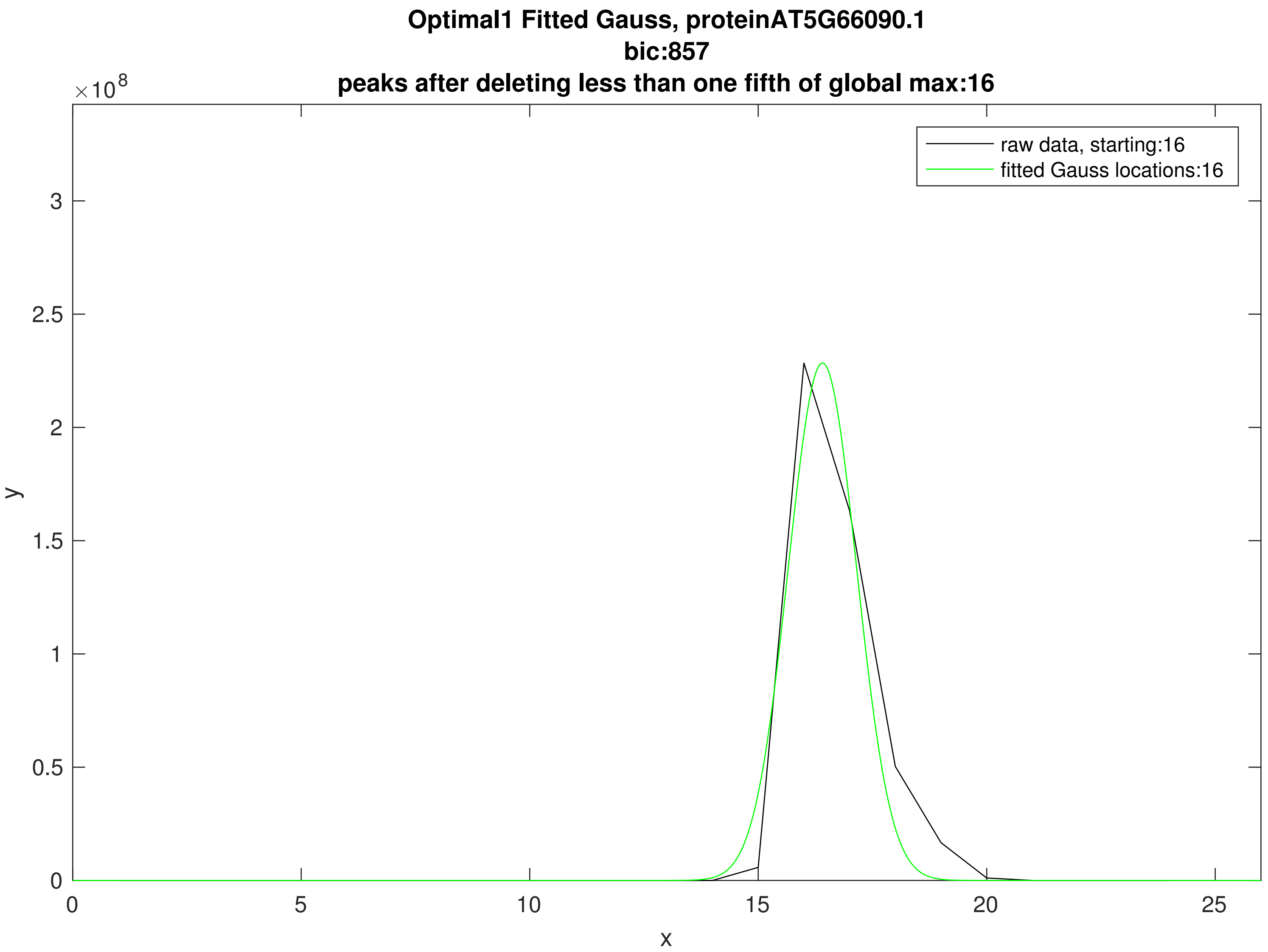
Optimal1 Fitted Gauss, proteinAT5G66090.1
bic:857
peaks after deleting less than one fifth of global max:16
×10^8
raw data, starting:16
fitted Gauss locations:16
y
x
0
0.5
1
1.5
2
2.5
3
0
5
10
15
20
25

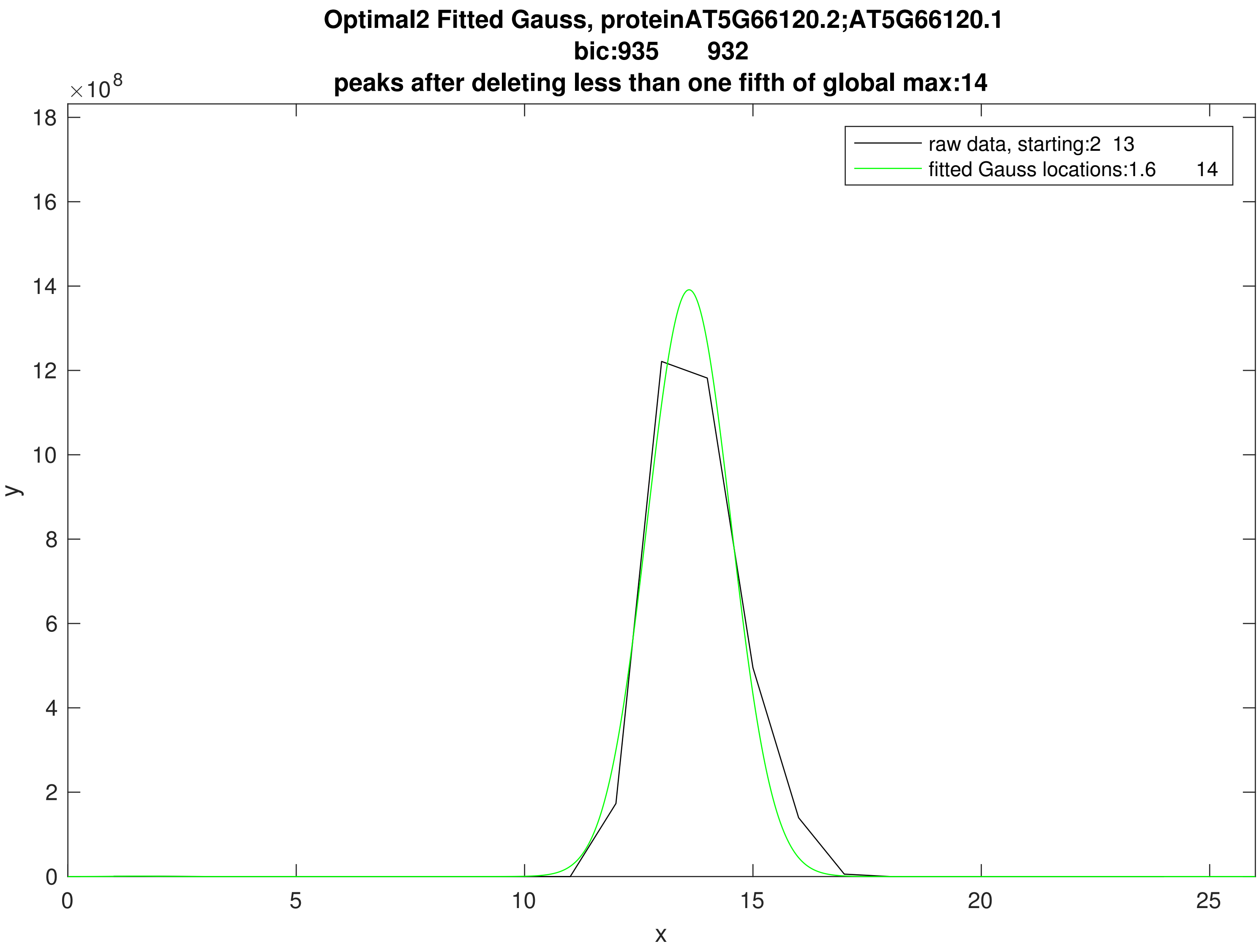

Optimal2 Fitted Gauss, proteinAT5G66120.2;AT5G66120.1
bic:935 932
peaks after deleting less than one fifth of global max:14
×10^8
raw data, starting:2 13
fitted Gauss locations:1.6 14
y
x
0
2
4
6
8
10
12
14
16
18
0
5
10
15
20
25

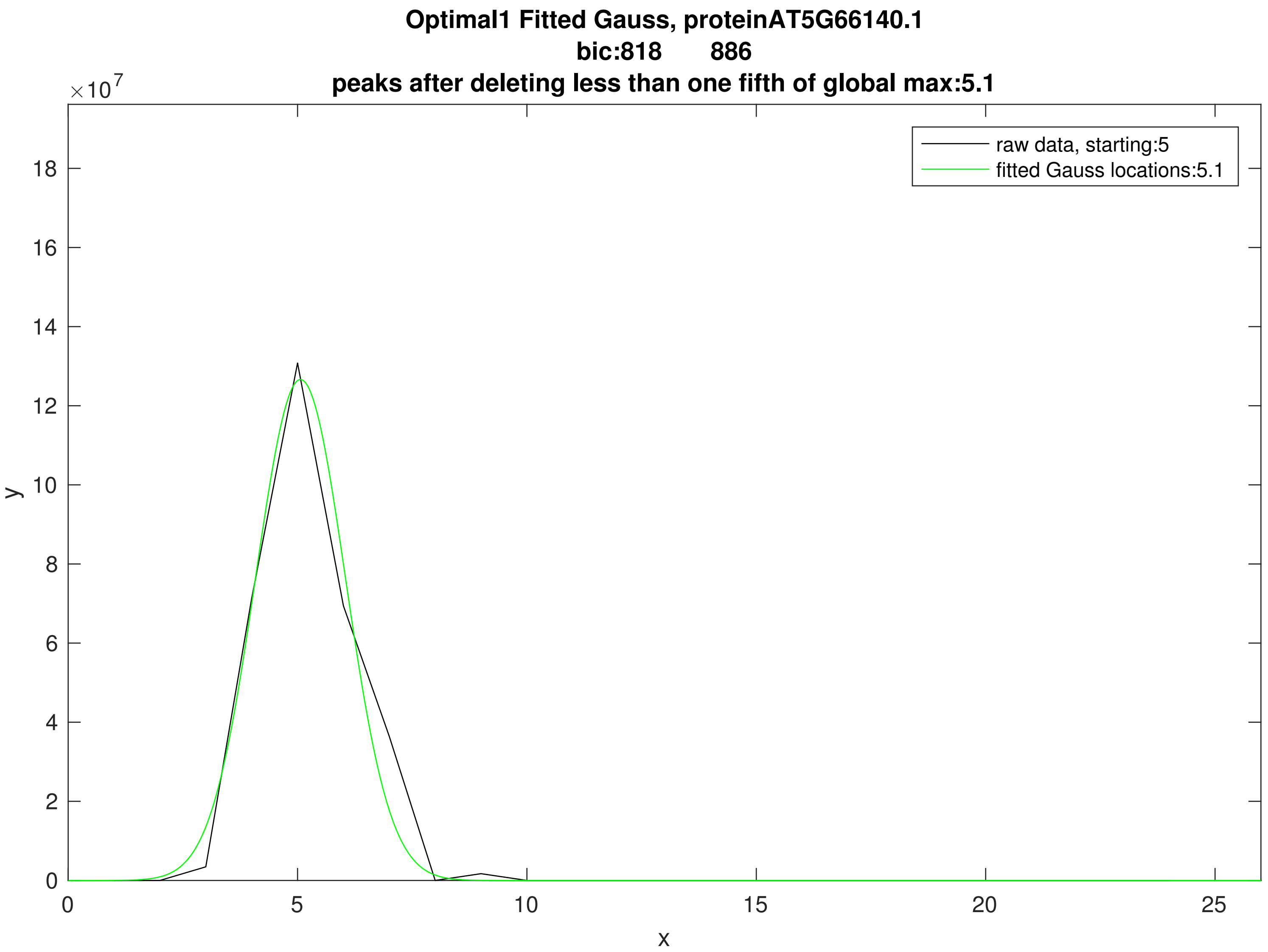
Optimal1 Fitted Gauss, proteinAT5G66140.1
bic:818 886
peaks after deleting less than one fifth of global max:5.1
×10^7
raw data, starting:5
fitted Gauss locations:5.1
y
x
0
2
4
6
8
10
12
14
16
18
0
5
10
15
20
25

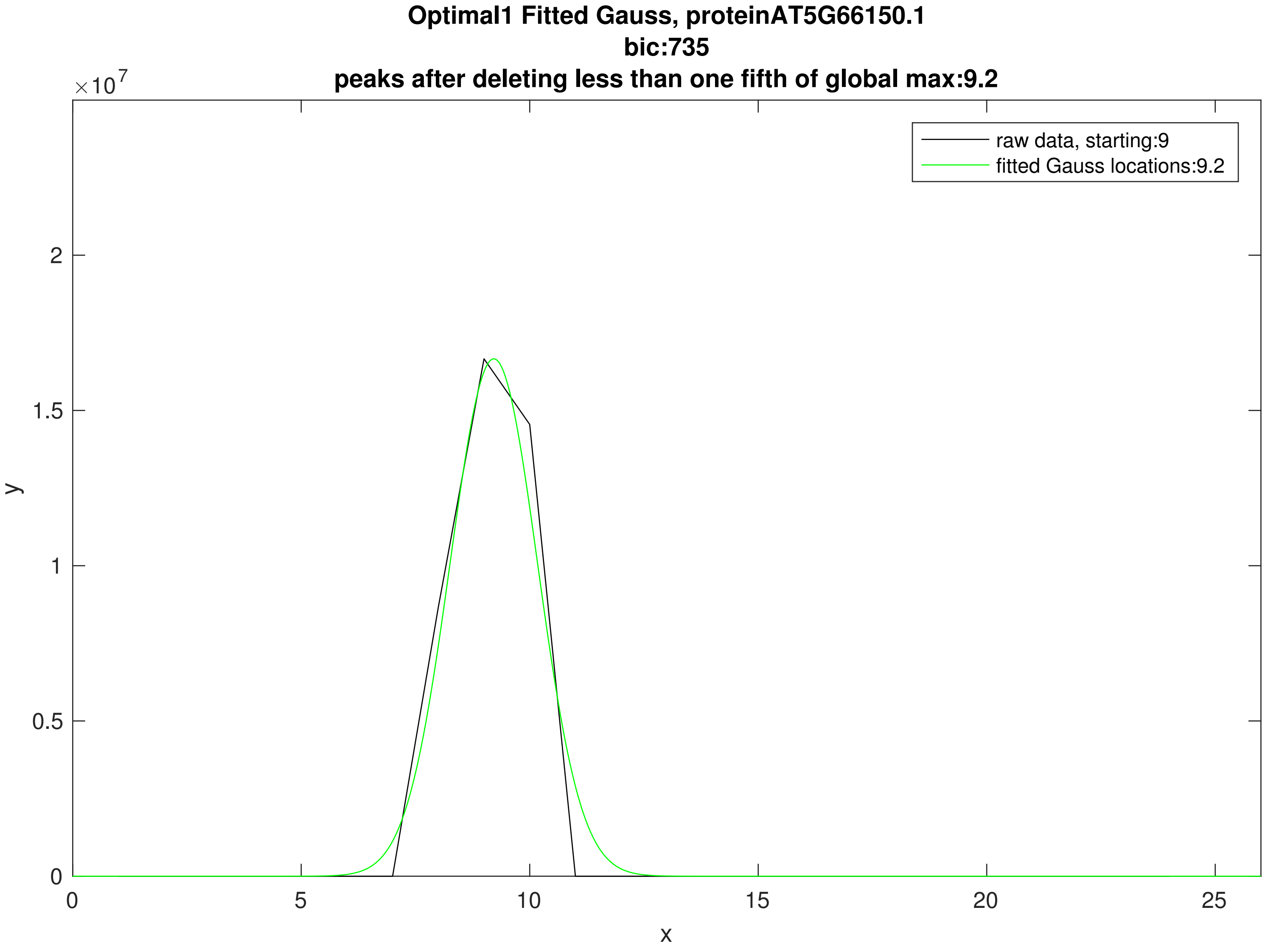
Optimal1 Fitted Gauss, proteinAT5G66150.1
bic:735
peaks after deleting less than one fifth of global max:9.2
×10^7
raw data, starting:9
fitted Gauss locations:9.2
y
x
0
0.5
1
1.5
2
0
5
10
15
20
25

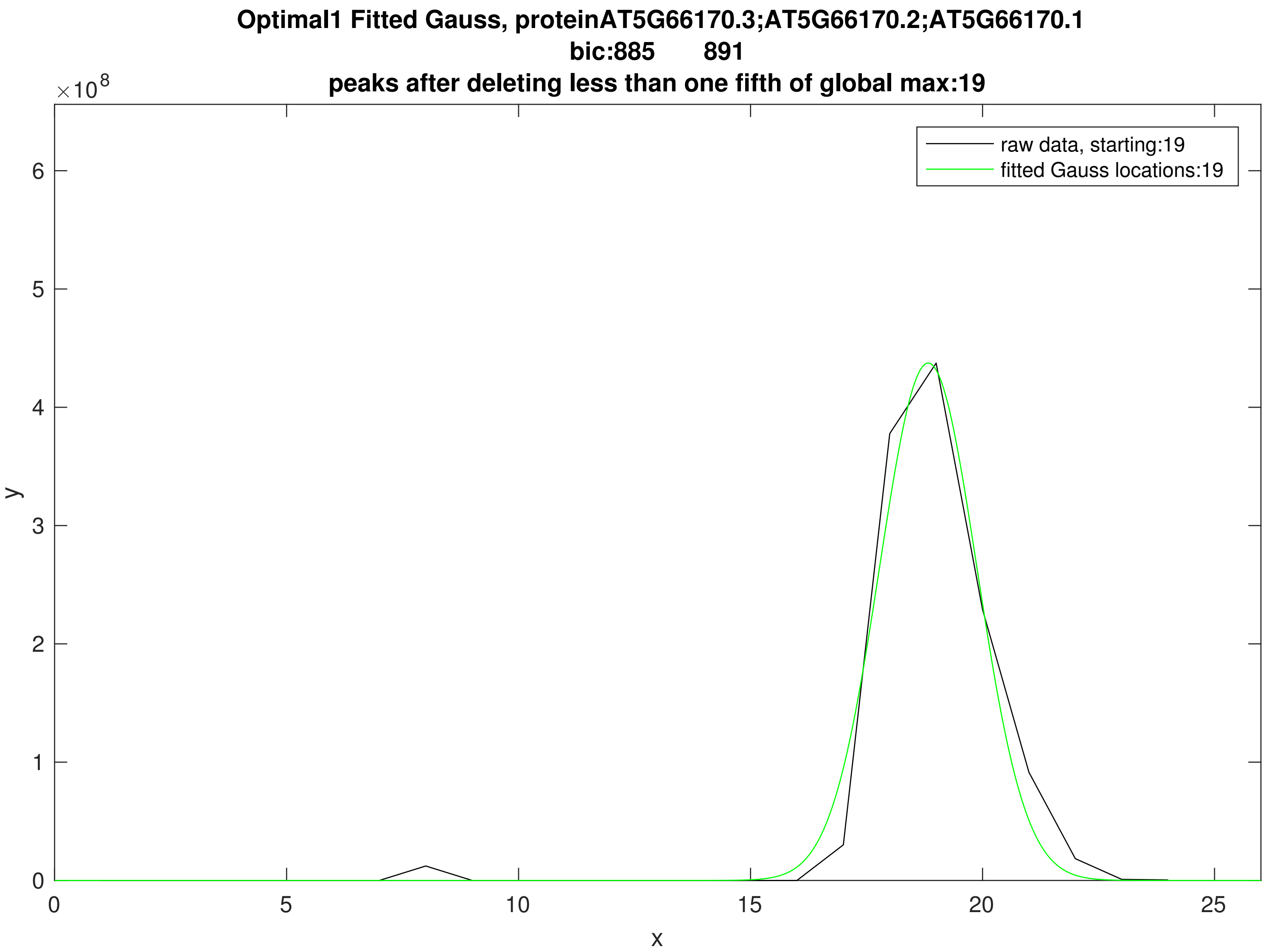

Optimal1 Fitted Gauss, proteinAT5G66170.3;AT5G66170.2;AT5G66170.1
bic:885 891
peaks after deleting less than one fifth of global max:19
×10^8
raw data, starting:19
fitted Gauss locations:19
y
x
0
1
2
3
4
5
6
0
5
10
15
20
25

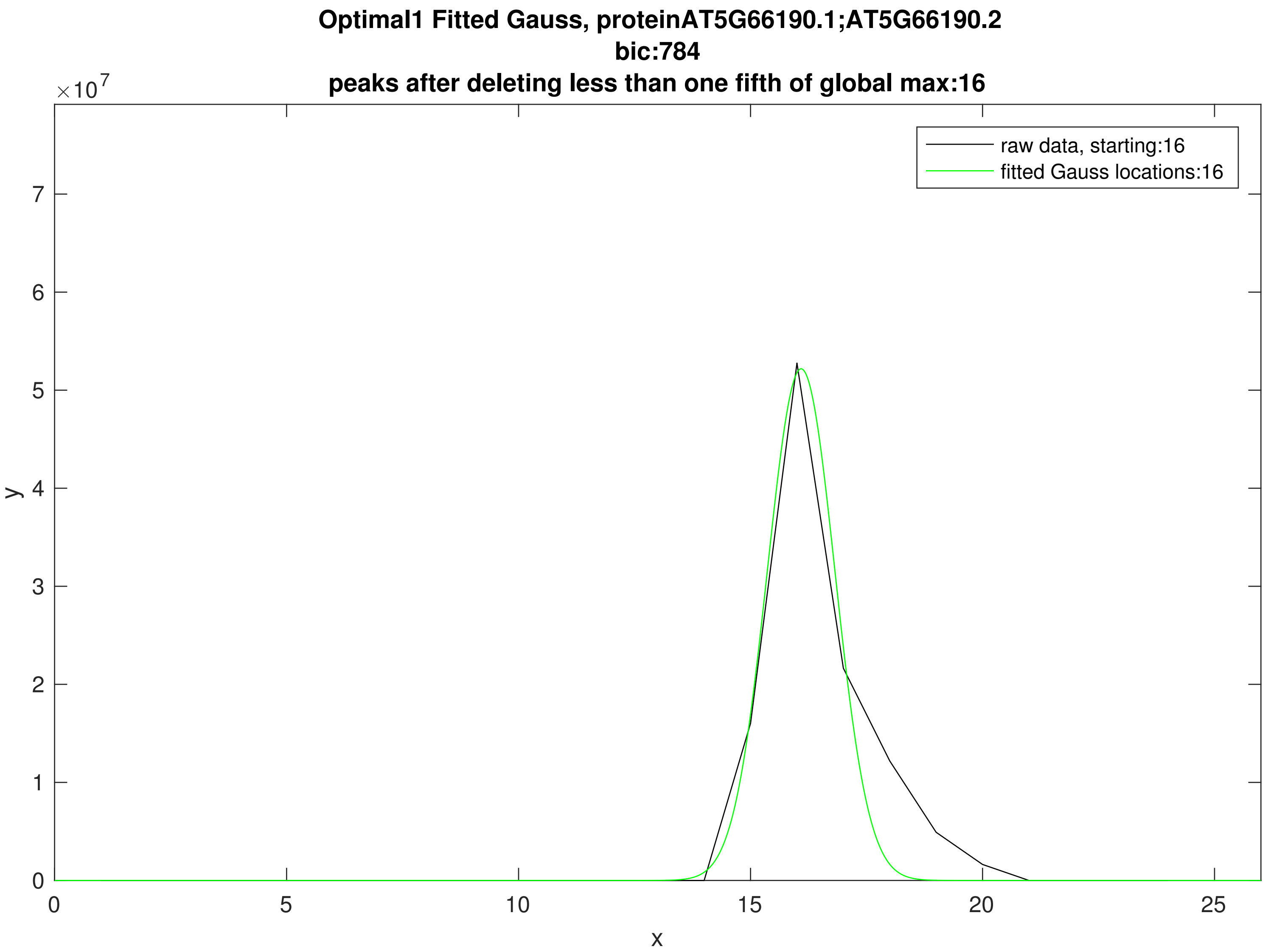

Optimal1 Fitted Gauss, proteinAT5G66190.1;AT5G66190.2
bic:784
peaks after deleting less than one fifth of global max:16
×10^7
raw data, starting:16
fitted Gauss locations:16
y
x
0
1
2
3
4
5
6
7
0
5
10
15
20
25

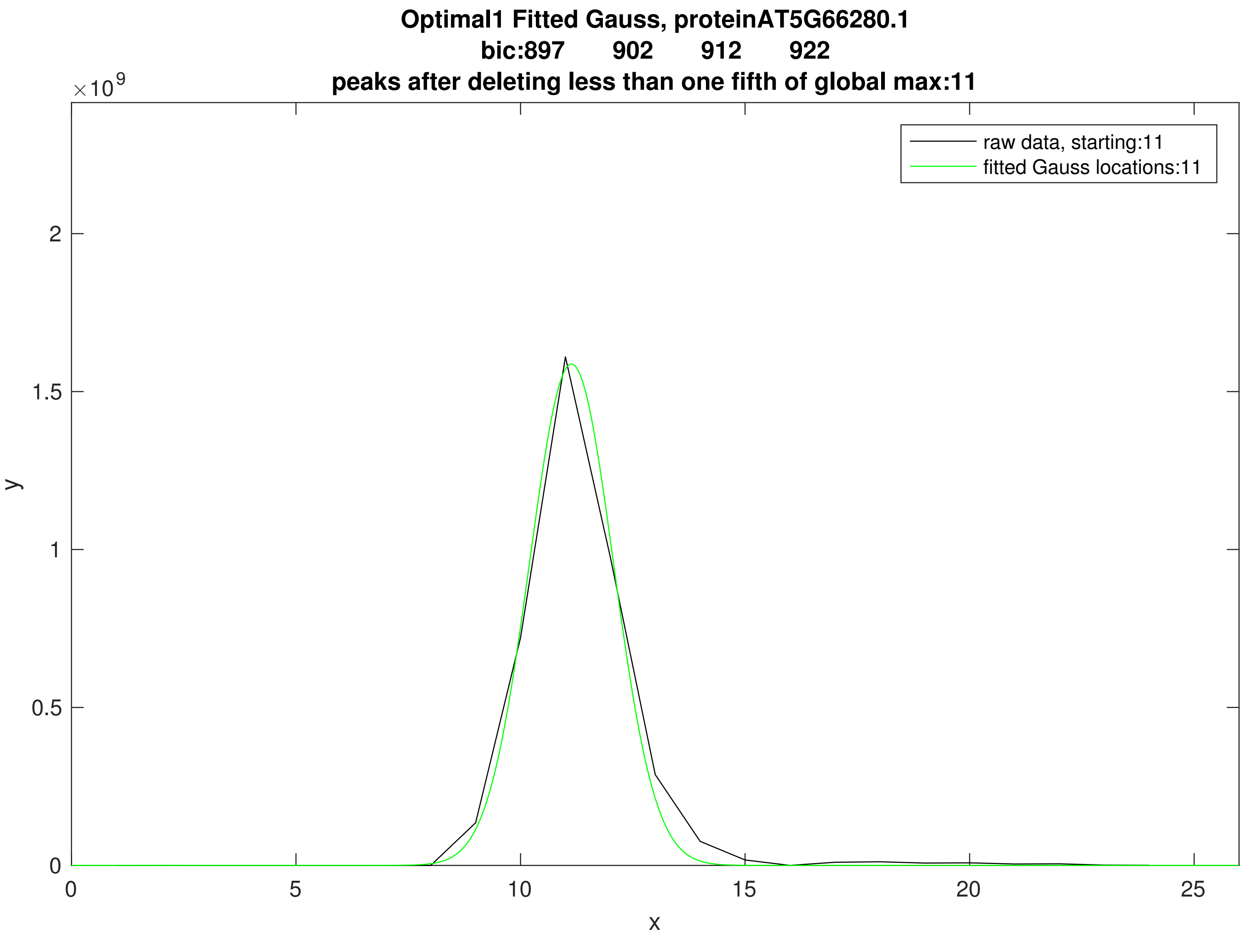
Optimal1 Fitted Gauss, proteinAT5G66280.1
bic:897 902 912 922
peaks after deleting less than one fifth of global max:11
×10^9
raw data, starting:11
fitted Gauss locations:11
y
x
0
0.5
1
1.5
2
0
5
10
15
20
25

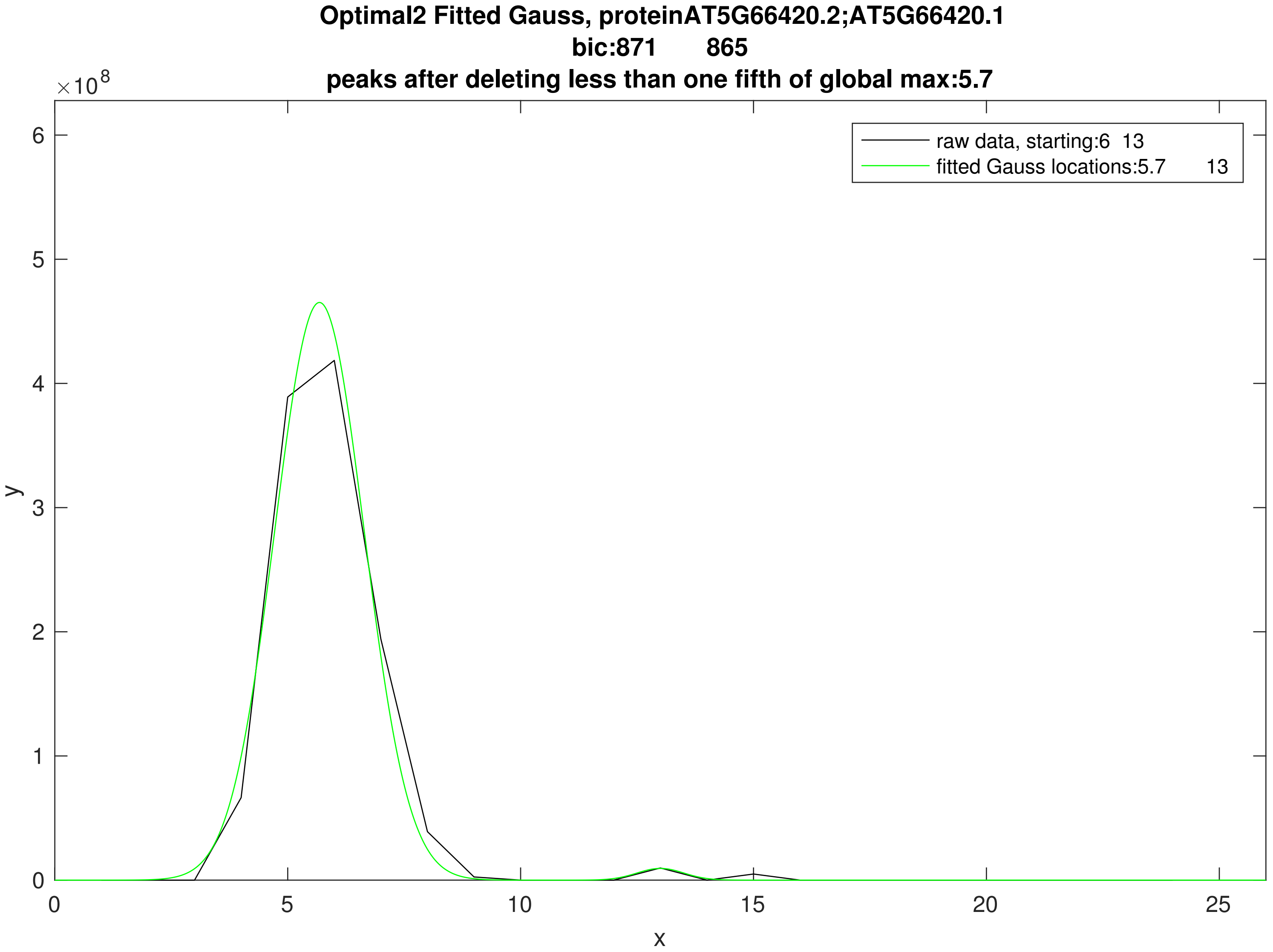
Optimal2 Fitted Gauss, proteinAT5G66420.2;AT5G66420.1
bic:871 865
peaks after deleting less than one fifth of global max:5.7
×10^8
raw data, starting:6 13
fitted Gauss locations:5.7 13
y
x
0
1
2
3
4
5
6
0
5
10
15
20
25

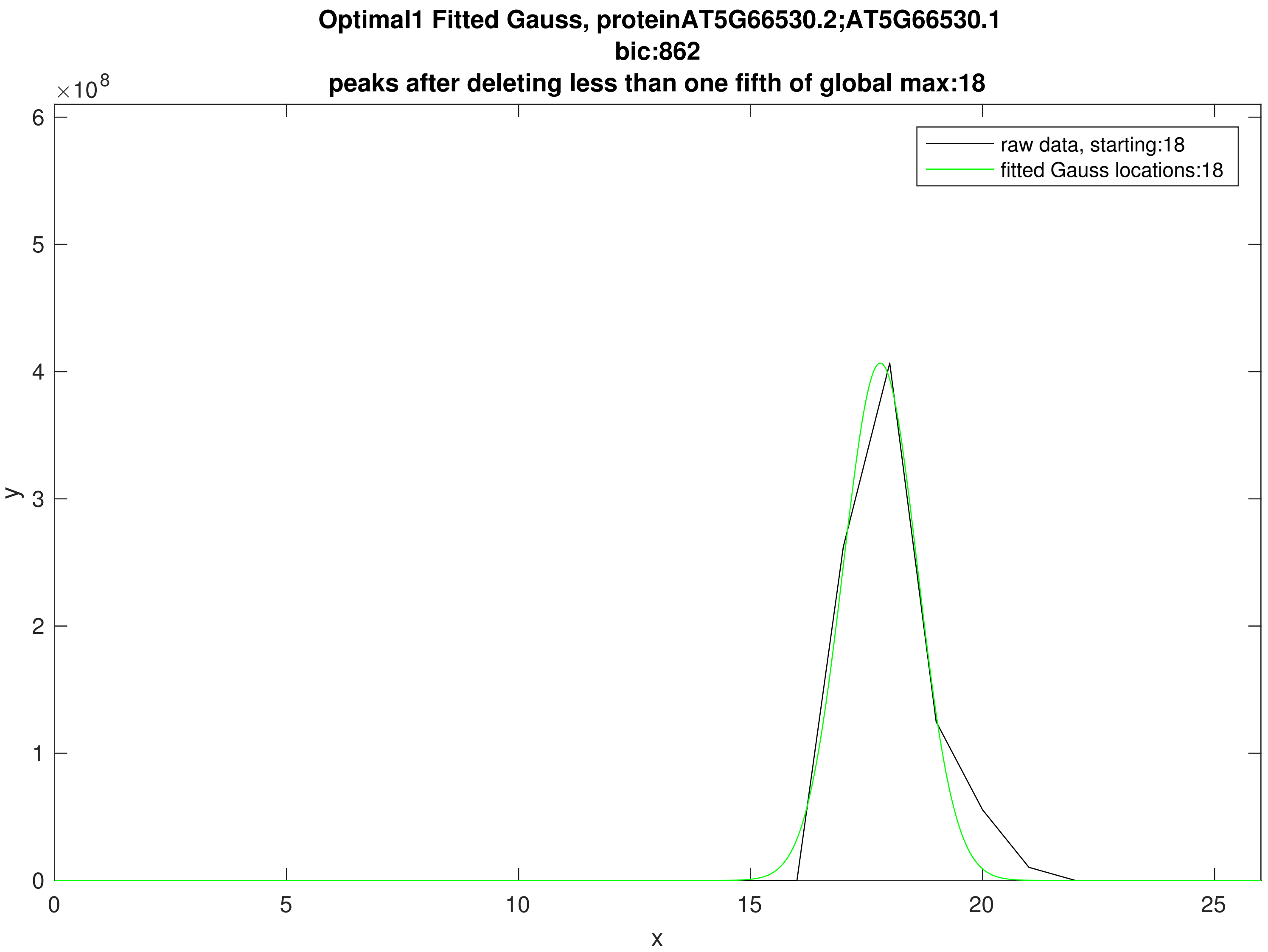
Optimal1 Fitted Gauss, proteinAT5G66530.2;AT5G66530.1
bic:862
peaks after deleting less than one fifth of global max:18
×10^8
6
5
4
3
2
1
0
y
0
5
10
15
20
25
x
raw data, starting:18
fitted Gauss locations:18

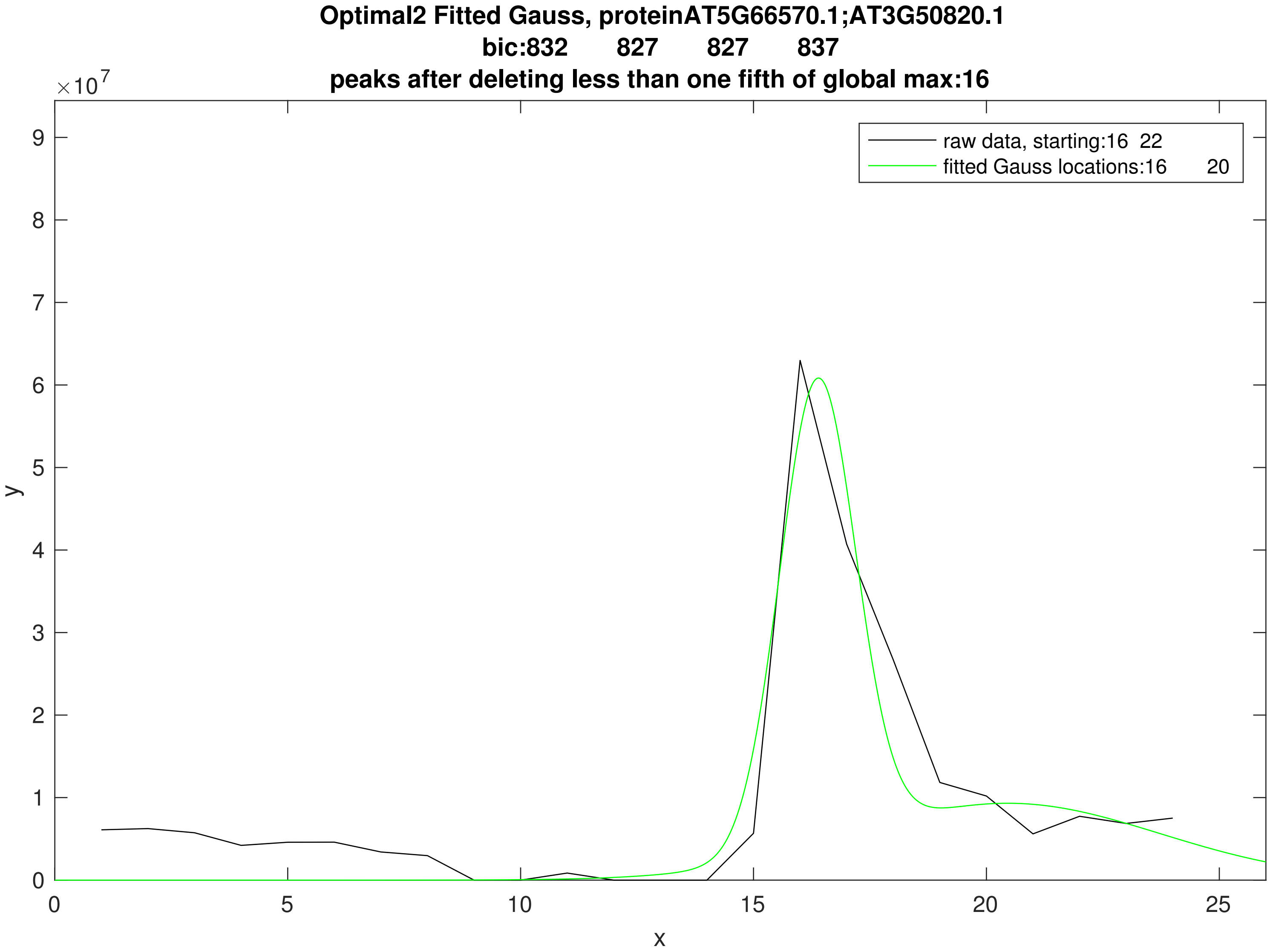

Optimal2 Fitted Gauss, proteinAT5G66570.1;AT3G50820.1
bic:832 827 827 837
peaks after deleting less than one fifth of global max:16
×10^7
raw data, starting:16 22
fitted Gauss locations:16 20
y
x
0
5
10
15
20
25
1
2
3
4
6
7
8
9

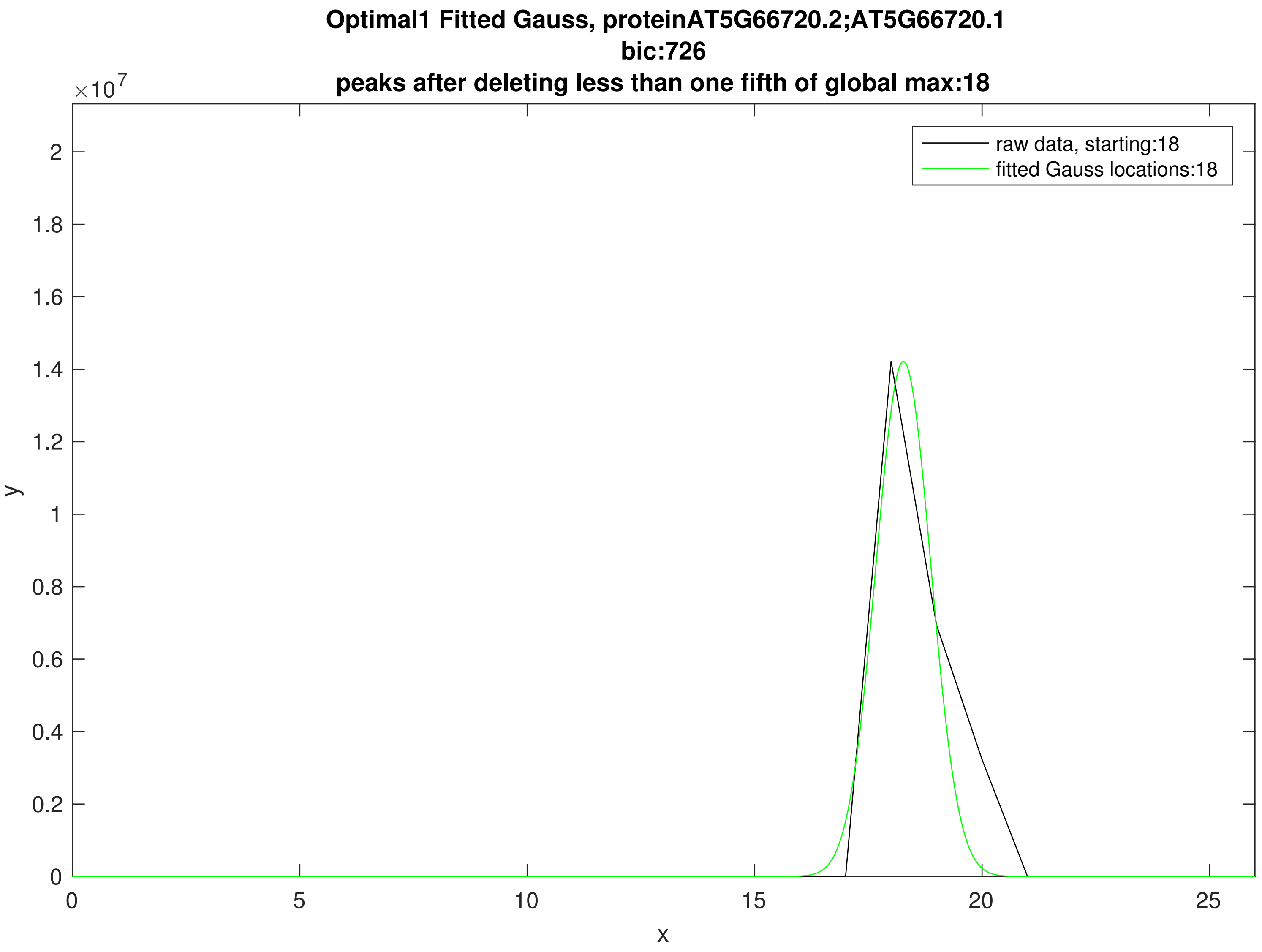
Optimal1 Fitted Gauss, proteinAT5G66720.2;AT5G66720.1
bic:726
peaks after deleting less than one fifth of global max:18
×10^7
raw data, starting:18
fitted Gauss locations:18
y
x
0
0.2
0.4
0.6
0.8
1
1.2
1.4
1.6
1.8
2
5
10
15
20
25

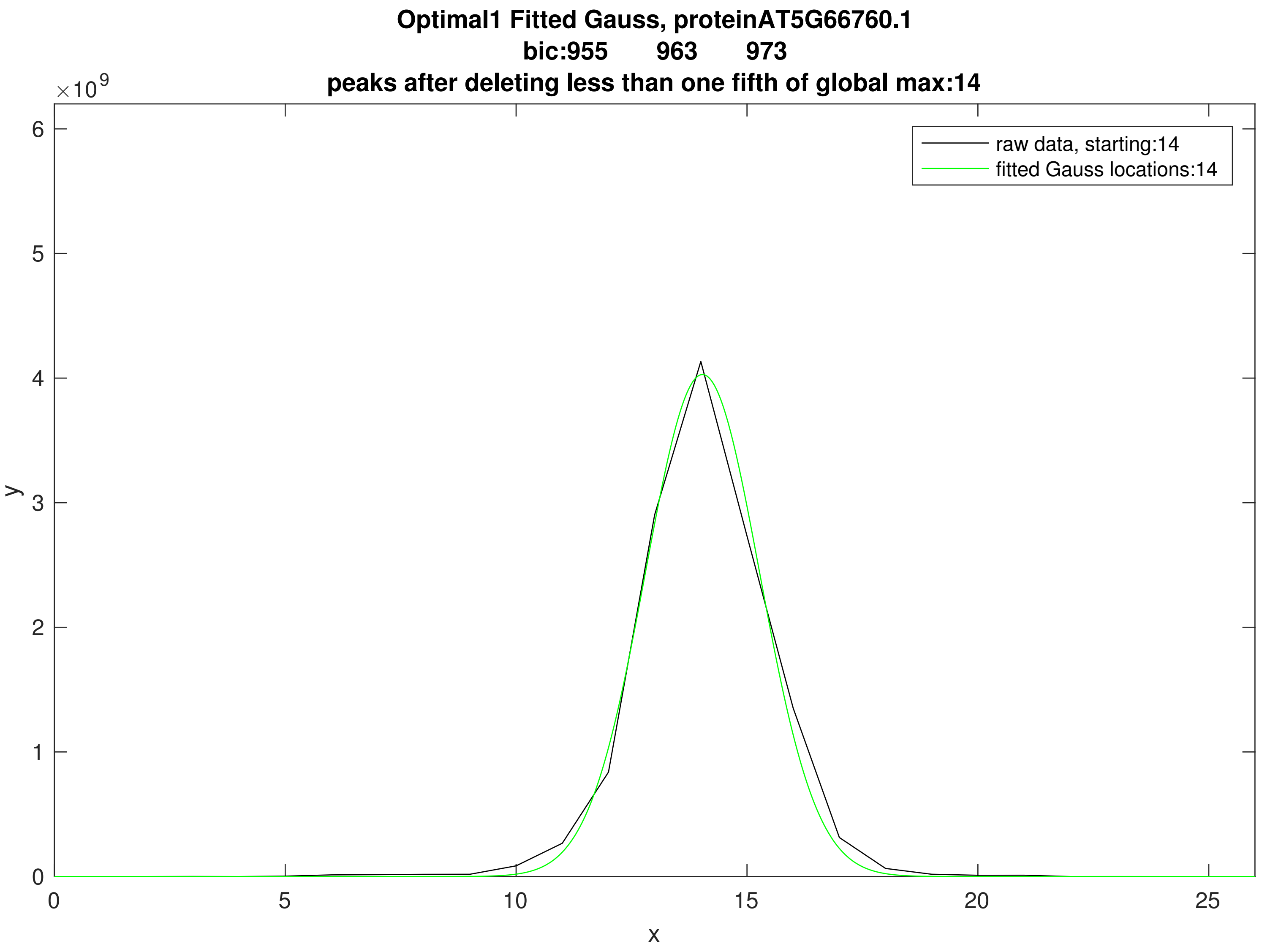

Optimal1 Fitted Gauss, proteinAT5G66760.1
bic:955 963 973
peaks after deleting less than one fifth of global max:14
×10^9
raw data, starting:14
fitted Gauss locations:14
y
x
0
1
2
3
4
5
6
0
5
10
15
20
25

**Optimal1 Fitted Gauss, proteinAT5G66780.1**
**bic:753**
**peaks after deleting less than one fifth of global max:18**

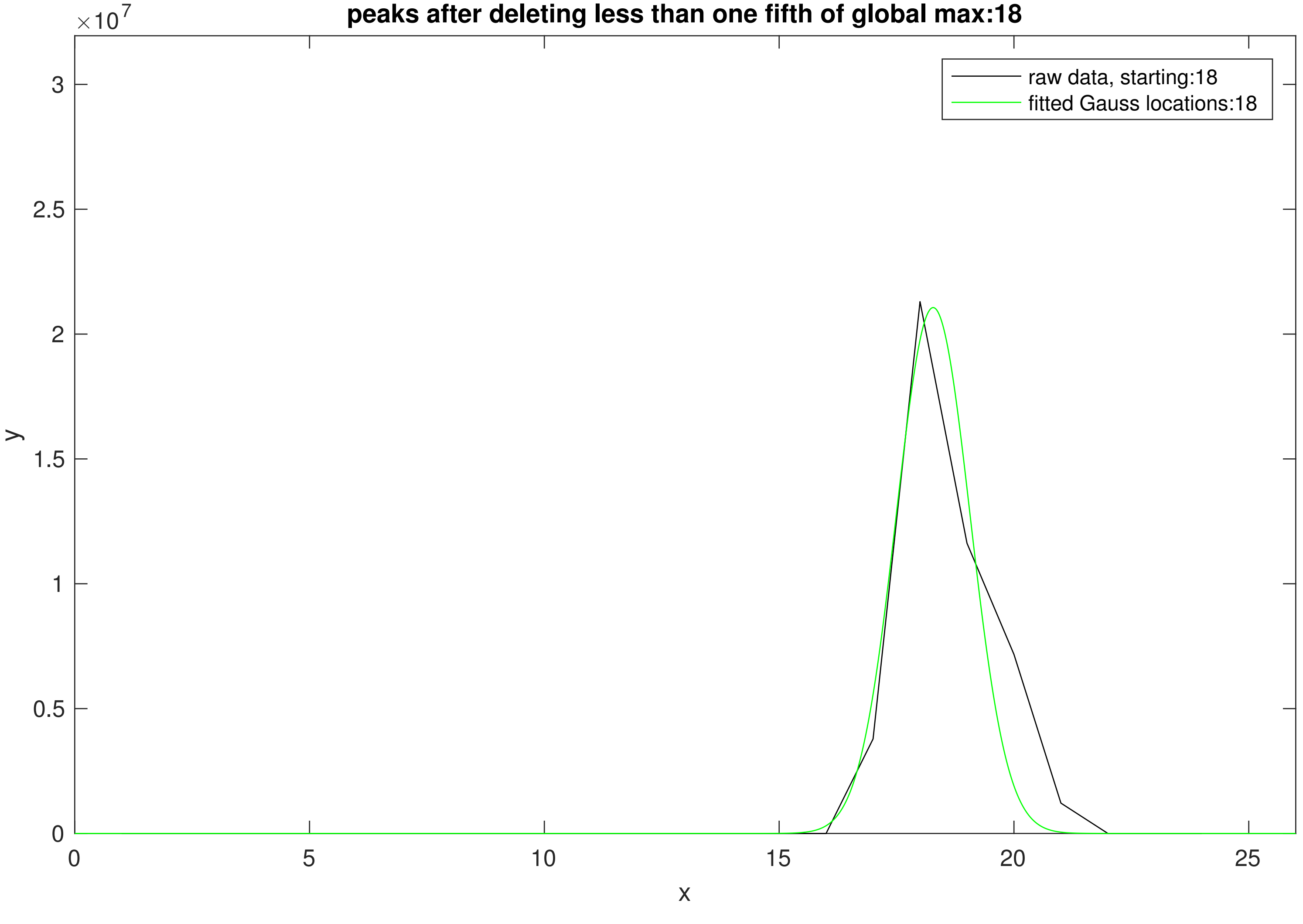

Optimal1 Fitted Gauss, proteinAT5G66860.1
bic:618
peaks after deleting less than one fifth of global max:3.4

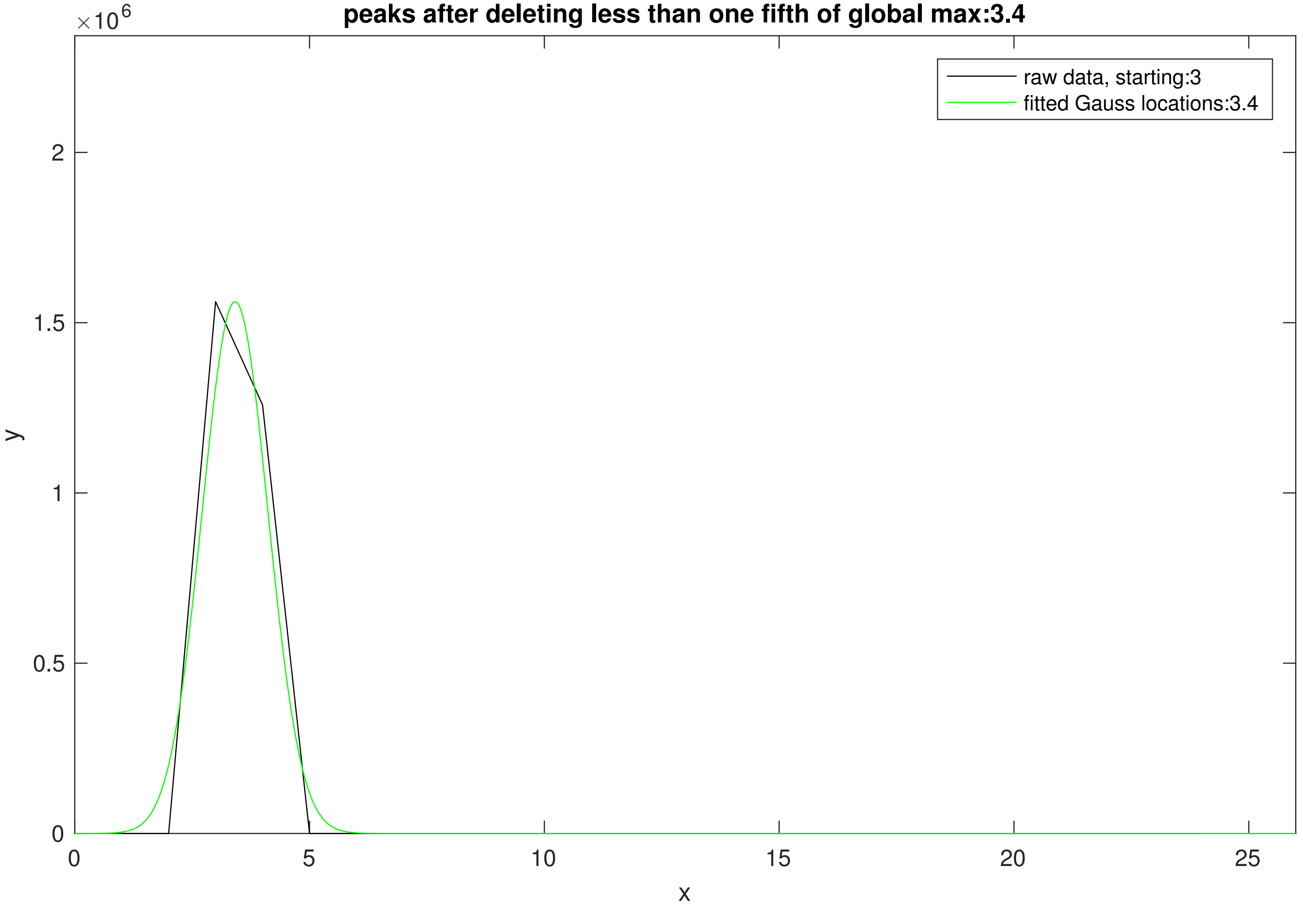

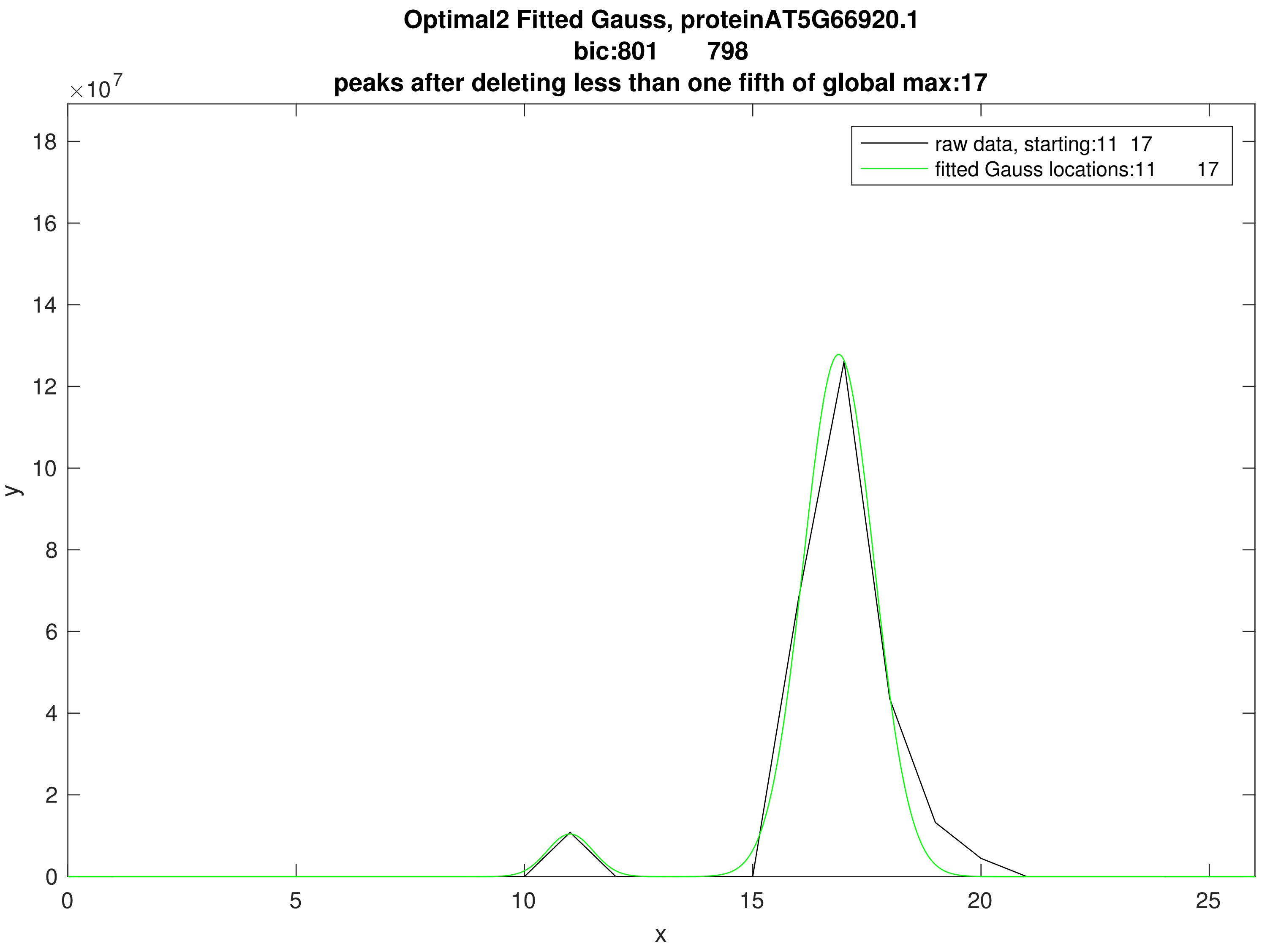
Optimal2 Fitted Gauss, proteinAT5G66920.1
bic:801 798
peaks after deleting less than one fifth of global max:17
×10^7
raw data, starting:11 17
fitted Gauss locations:11 17
y
x
0
2
4
6
8
10
12
14
16
18
5
15
20
25

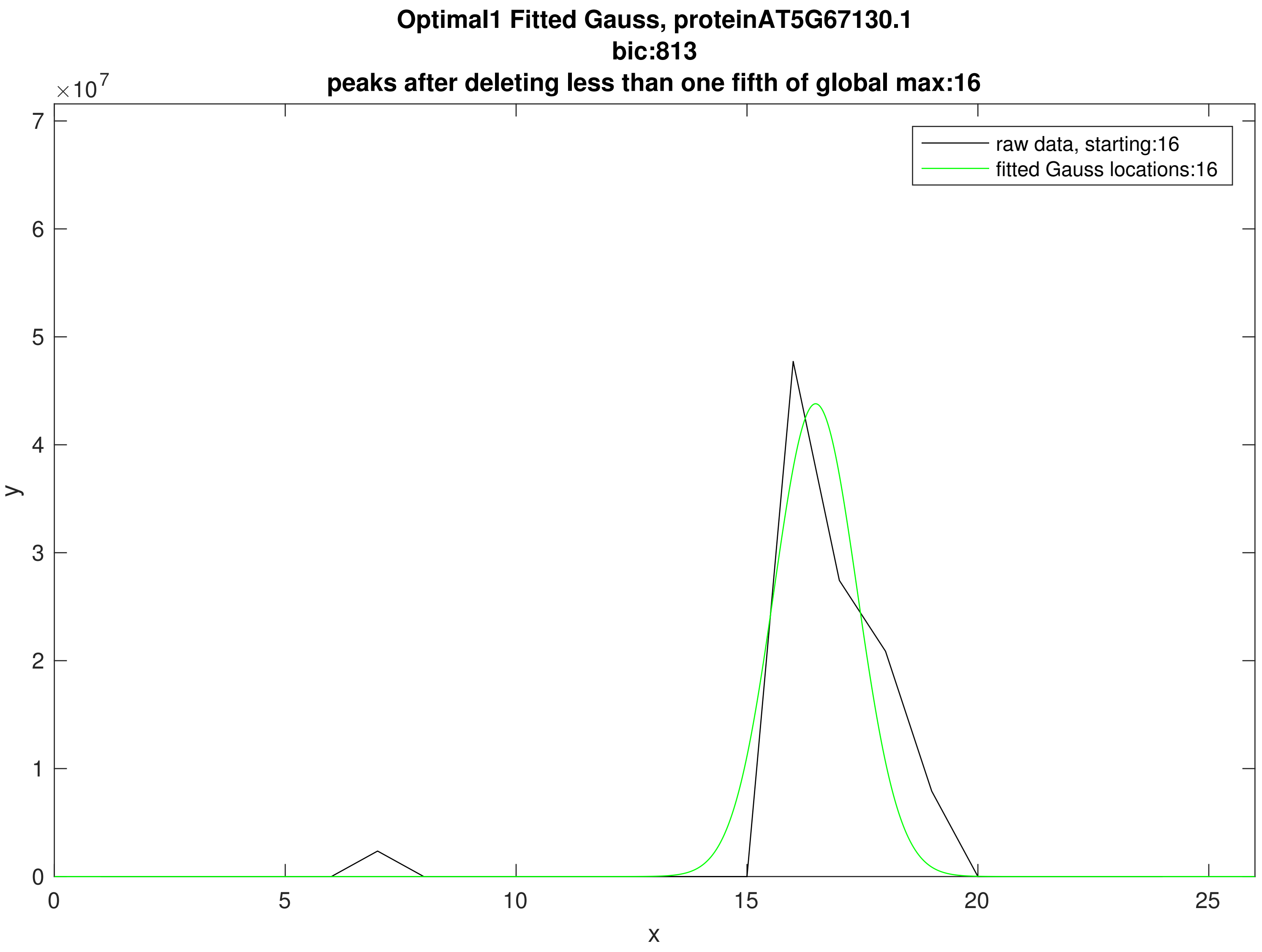
Optimal1 Fitted Gauss, proteinAT5G67130.1
bic:813
peaks after deleting less than one fifth of global max:16
×10^7
raw data, starting:16
fitted Gauss locations:16
y
x

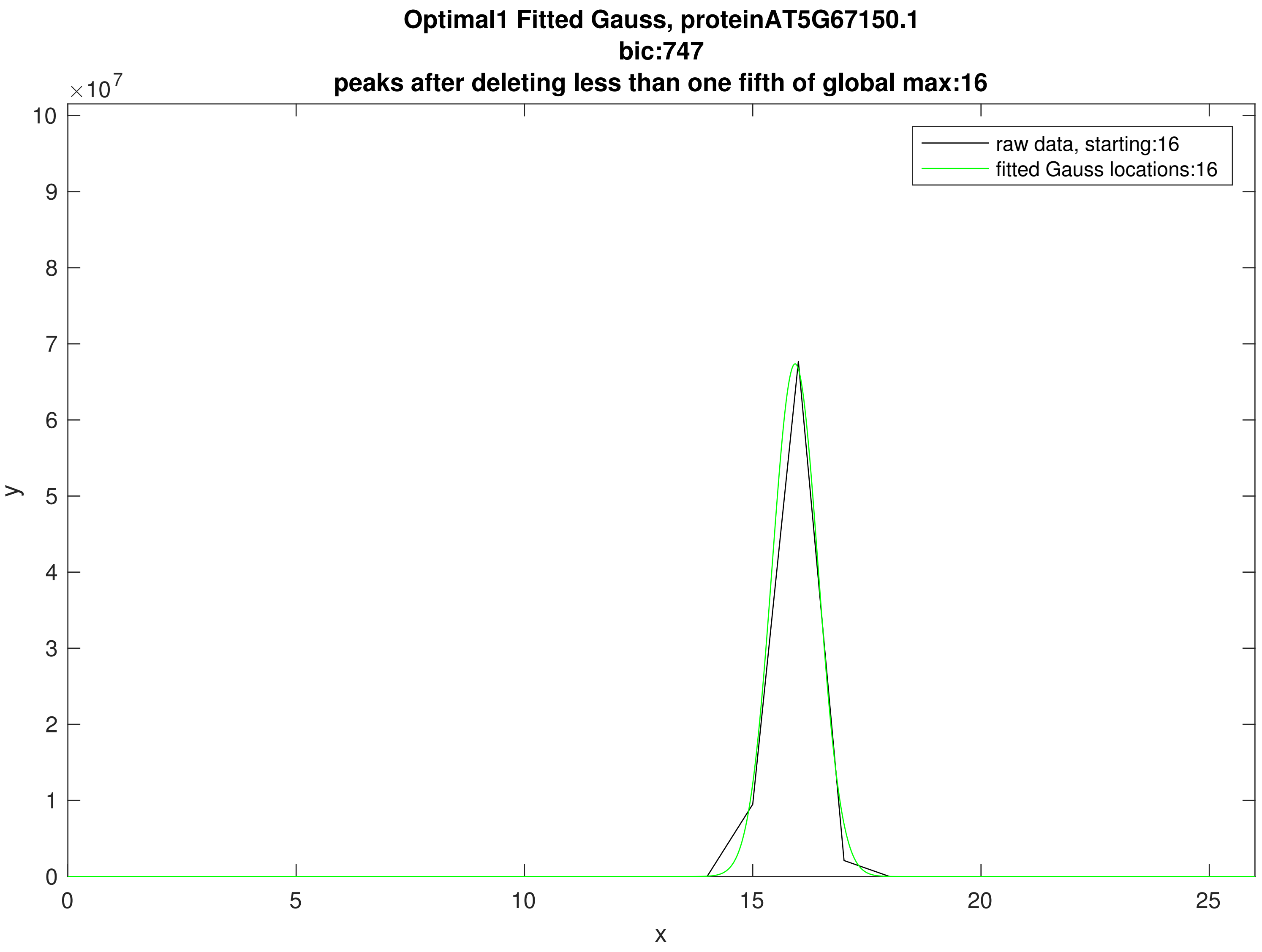
Optimal1 Fitted Gauss, proteinAT5G67150.1
bic:747
peaks after deleting less than one fifth of global max:16
×10^7
raw data, starting:16
fitted Gauss locations:16
y
x
0
1
2
3
4
5
6
7
8
9
10
0
5
10
15
20
25

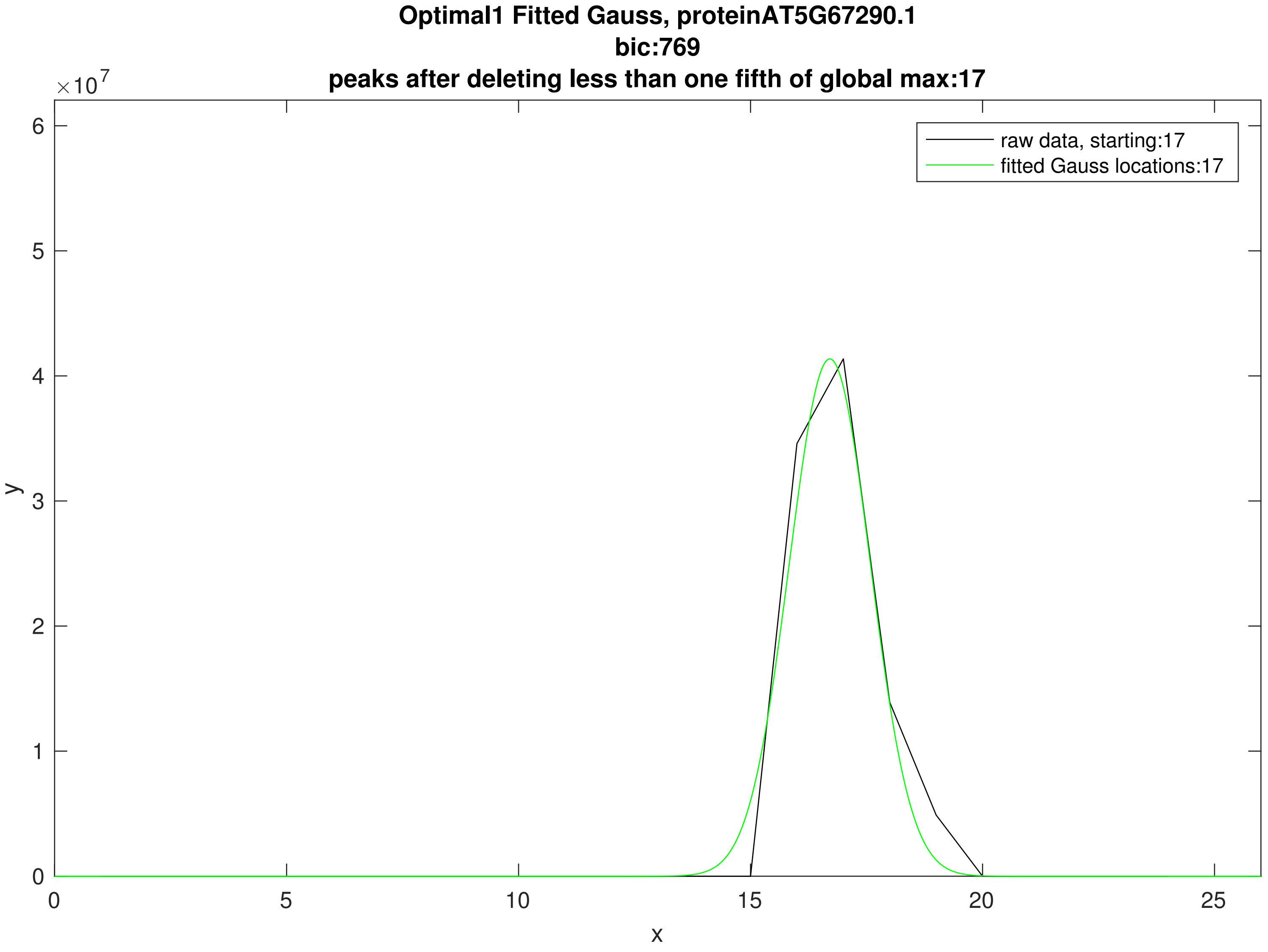
Optimal1 Fitted Gauss, proteinAT5G67290.1
bic:769
peaks after deleting less than one fifth of global max:17
×10^7
raw data, starting:17
fitted Gauss locations:17
y
x
0
1
2
3
4
5
6
0
5
10
15
20
25

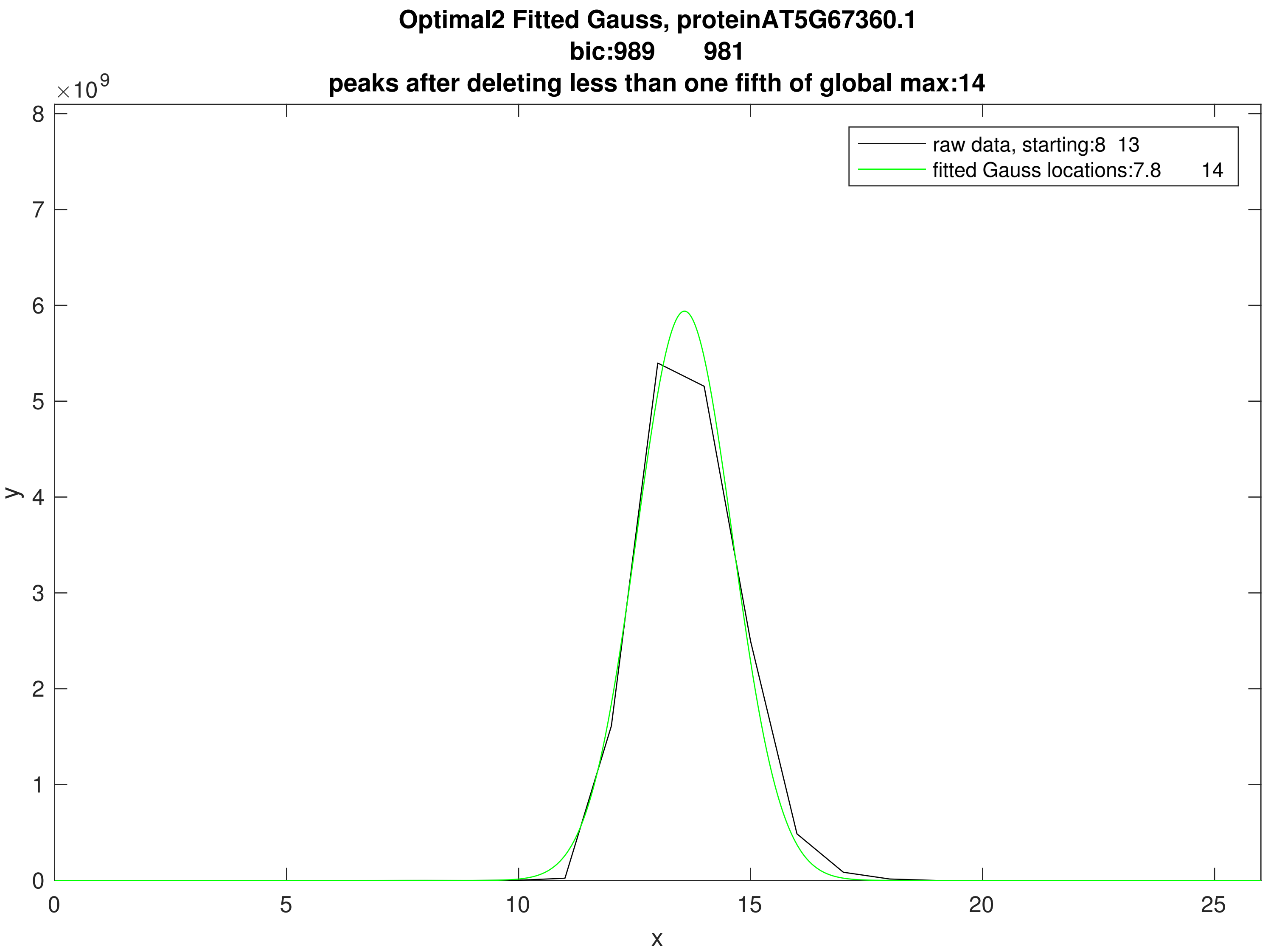
Optimal2 Fitted Gauss, proteinAT5G67360.1
bic:989 981
peaks after deleting less than one fifth of global max:14
×10^9
raw data, starting:8 13
fitted Gauss locations:7.8 14
y
x
0
1
2
3
4
5
6
7
8
0
5
10
15
20
25

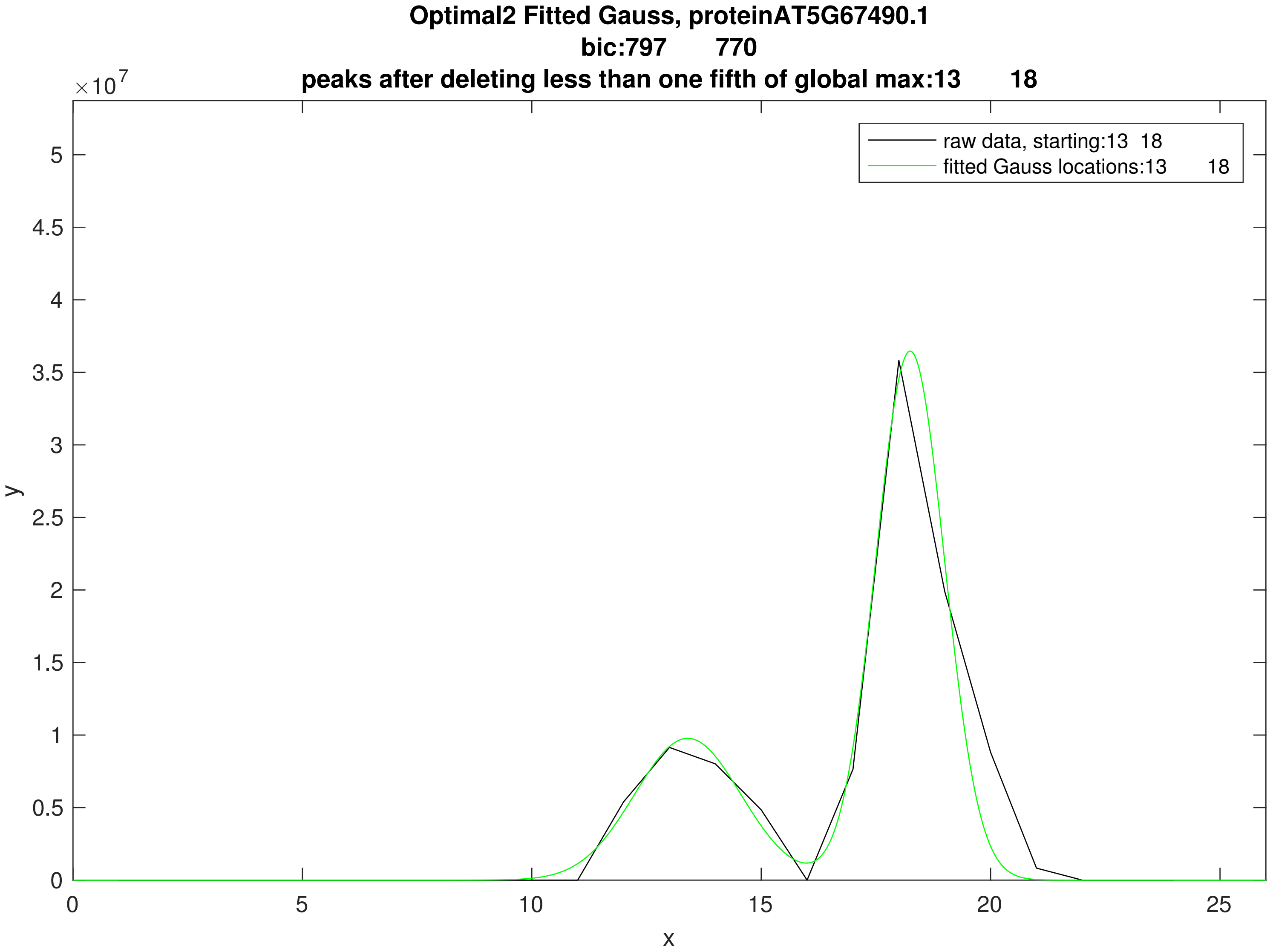

Optimal2 Fitted Gauss, proteinAT5G67490.1
bic:797 770
peaks after deleting less than one fifth of global max:13 18
×10^7
raw data, starting:13 18
fitted Gauss locations:13 18
y
x
0
0.5
1
1.5
2
2.5
3
3.5
4
4.5
5
0
5
10
15
20
25

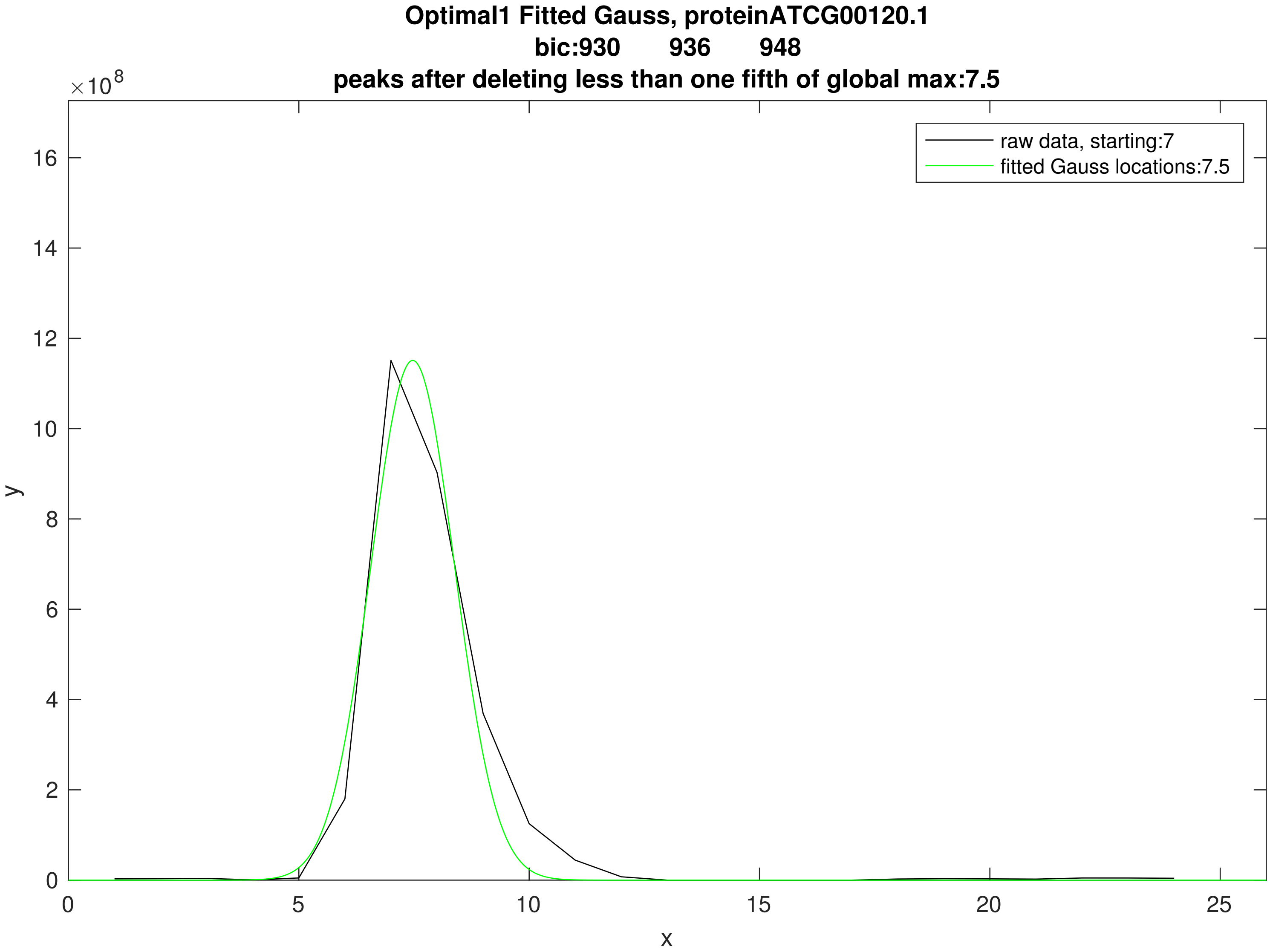

Optimal1 Fitted Gauss, proteinATCG00120.1
bic:930 936 948
peaks after deleting less than one fifth of global max:7.5
×10^8
raw data, starting:7
fitted Gauss locations:7.5
16
14
12
10
8
6
4
2
0
y
0
5
10
15
20
25
x

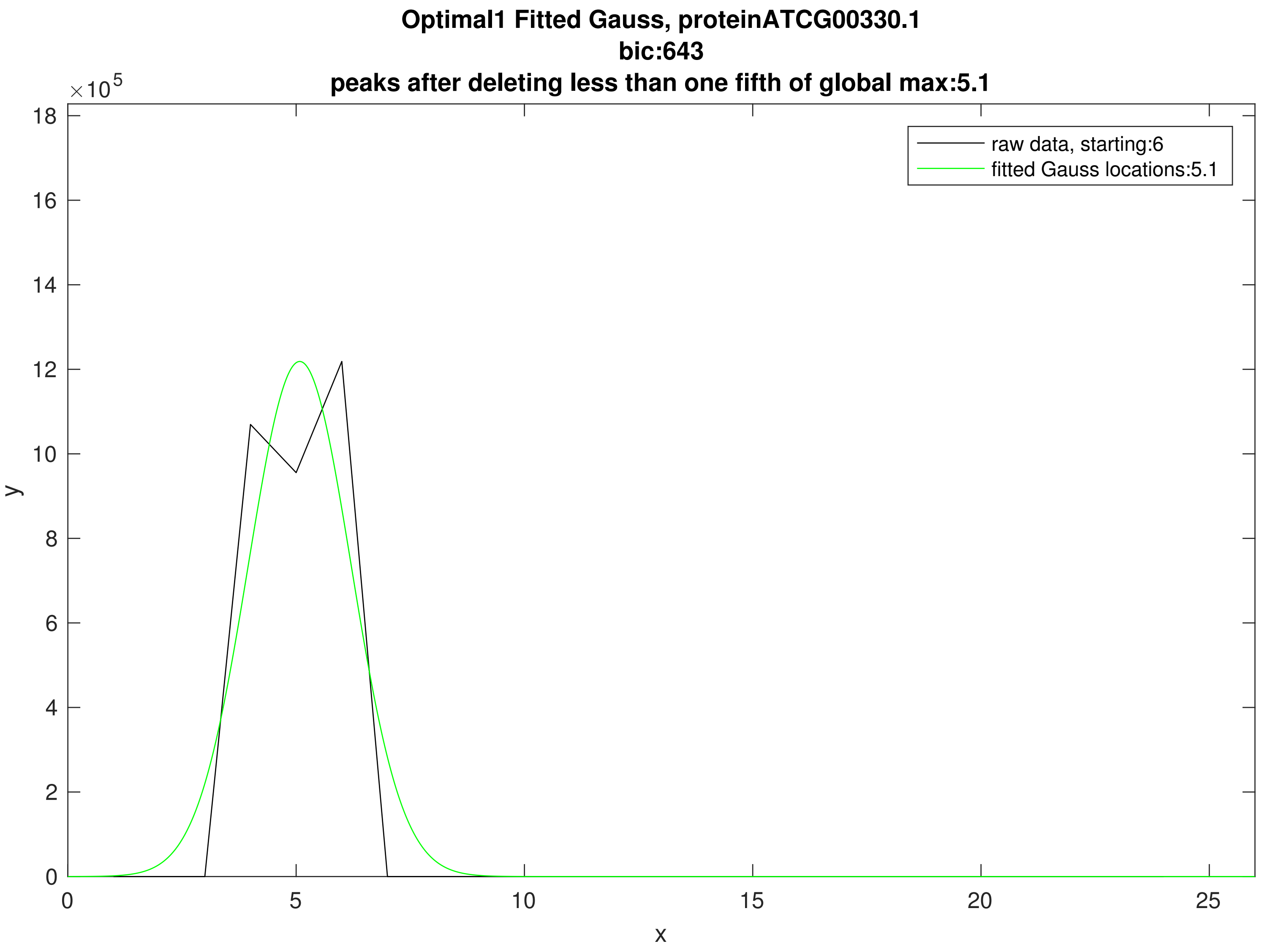
Optimal1 Fitted Gauss, proteinATCG00330.1
bic:643
peaks after deleting less than one fifth of global max:5.1
×10^5
raw data, starting:6
fitted Gauss locations:5.1
18
16
14
12
10
8
6
4
2
0
y
0
5
10
15
20
25
x

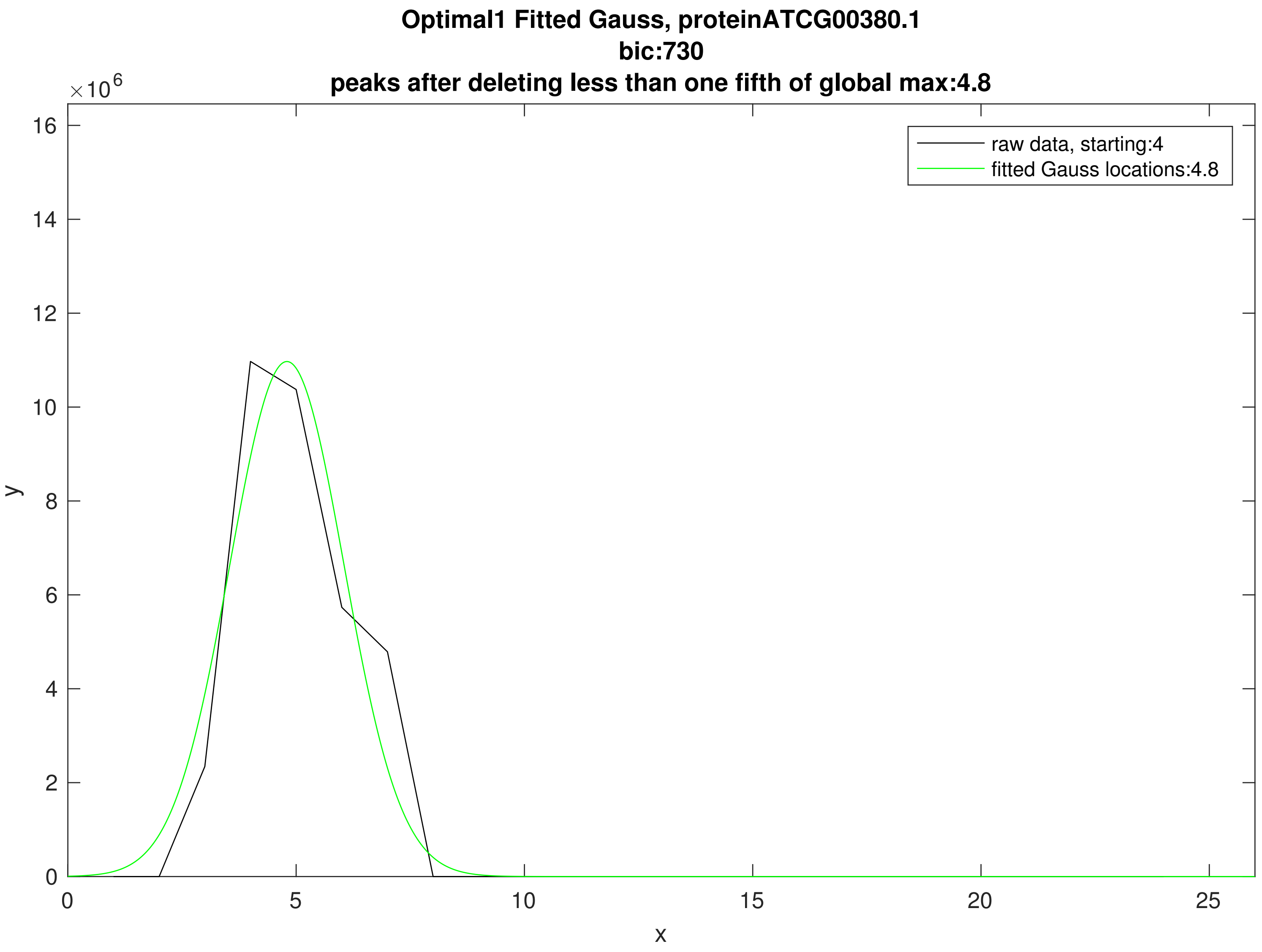
Optimal1 Fitted Gauss, proteinATCG00380.1
bic:730
peaks after deleting less than one fifth of global max:4.8
×10^6
raw data, starting:4
fitted Gauss locations:4.8
y
x
0
2
4
6
8
10
12
14
16
0
5
10
15
20
25

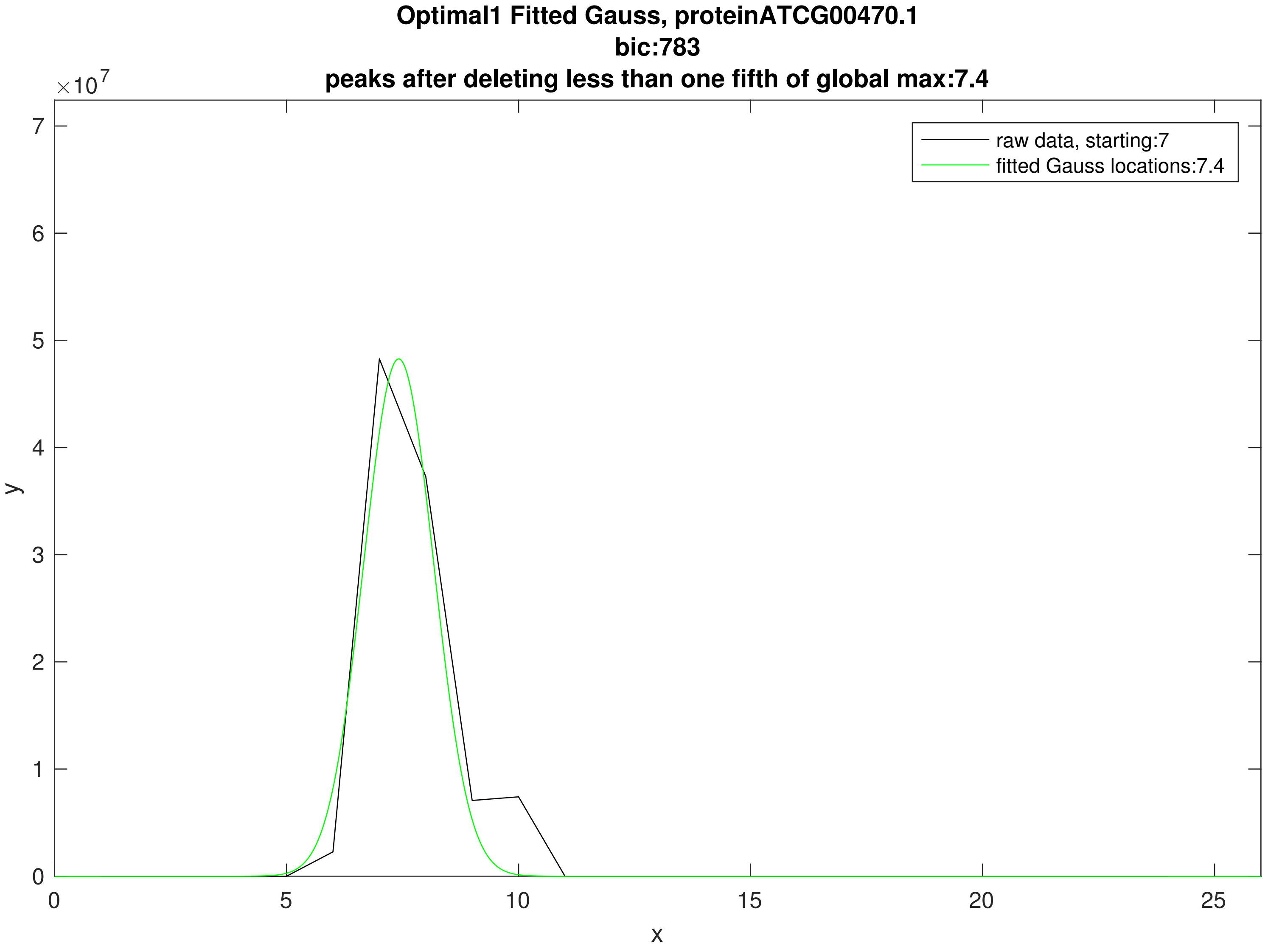

Optimal1 Fitted Gauss, proteinATCG00470.1
bic:783
peaks after deleting less than one fifth of global max:7.4
×10^7
raw data, starting:7
fitted Gauss locations:7.4
y
x
0
1
2
3
4
5
6
7
10
15
20
25

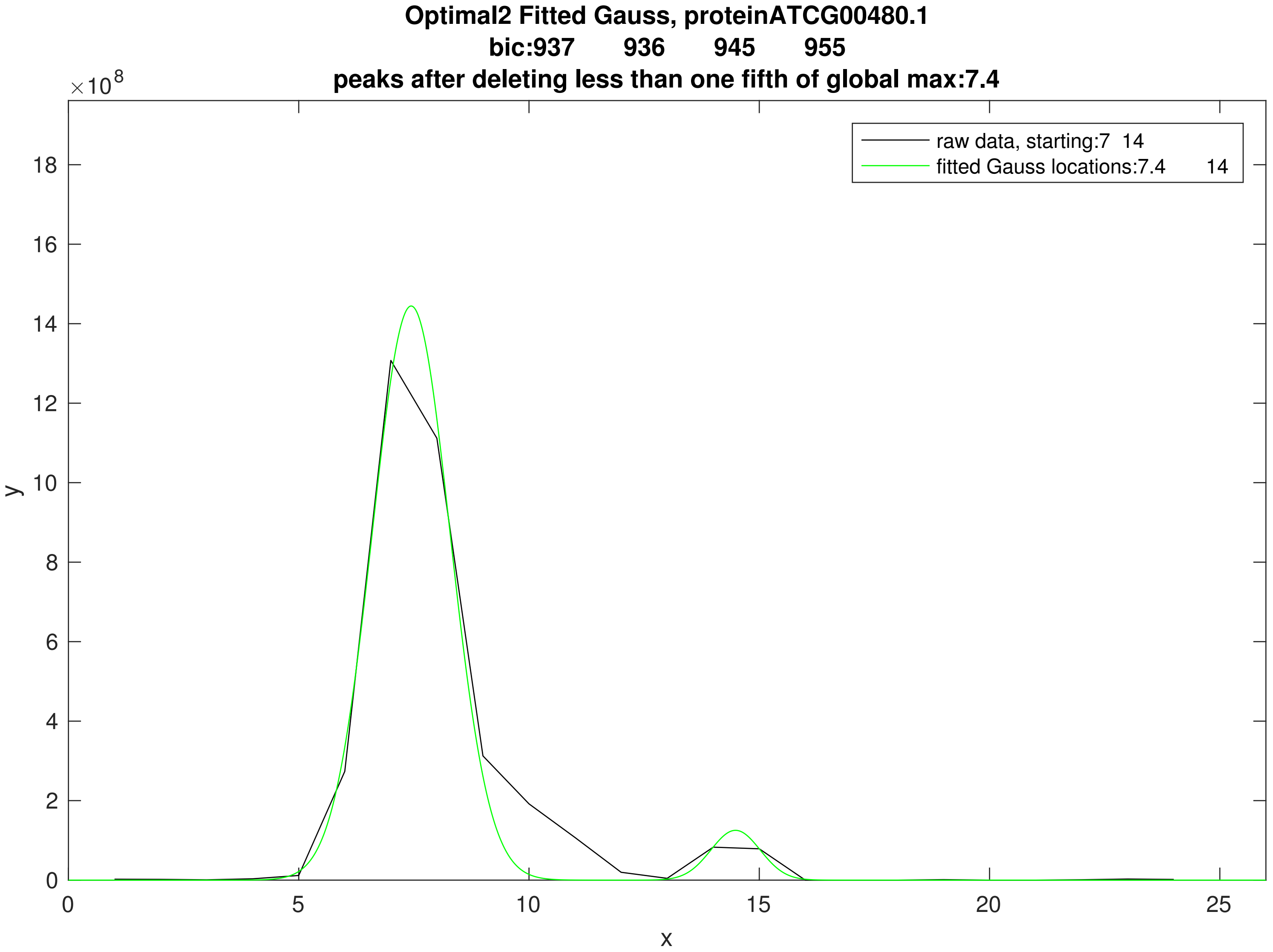
Optimal2 Fitted Gauss, proteinATCG00480.1
bic:937 936 945 955
peaks after deleting less than one fifth of global max:7.4
×10^8
raw data, starting:7 14
fitted Gauss locations:7.4 14
18
16
14
12
10
8
6
4
2
0
y
0
5
10
15
20
25
x

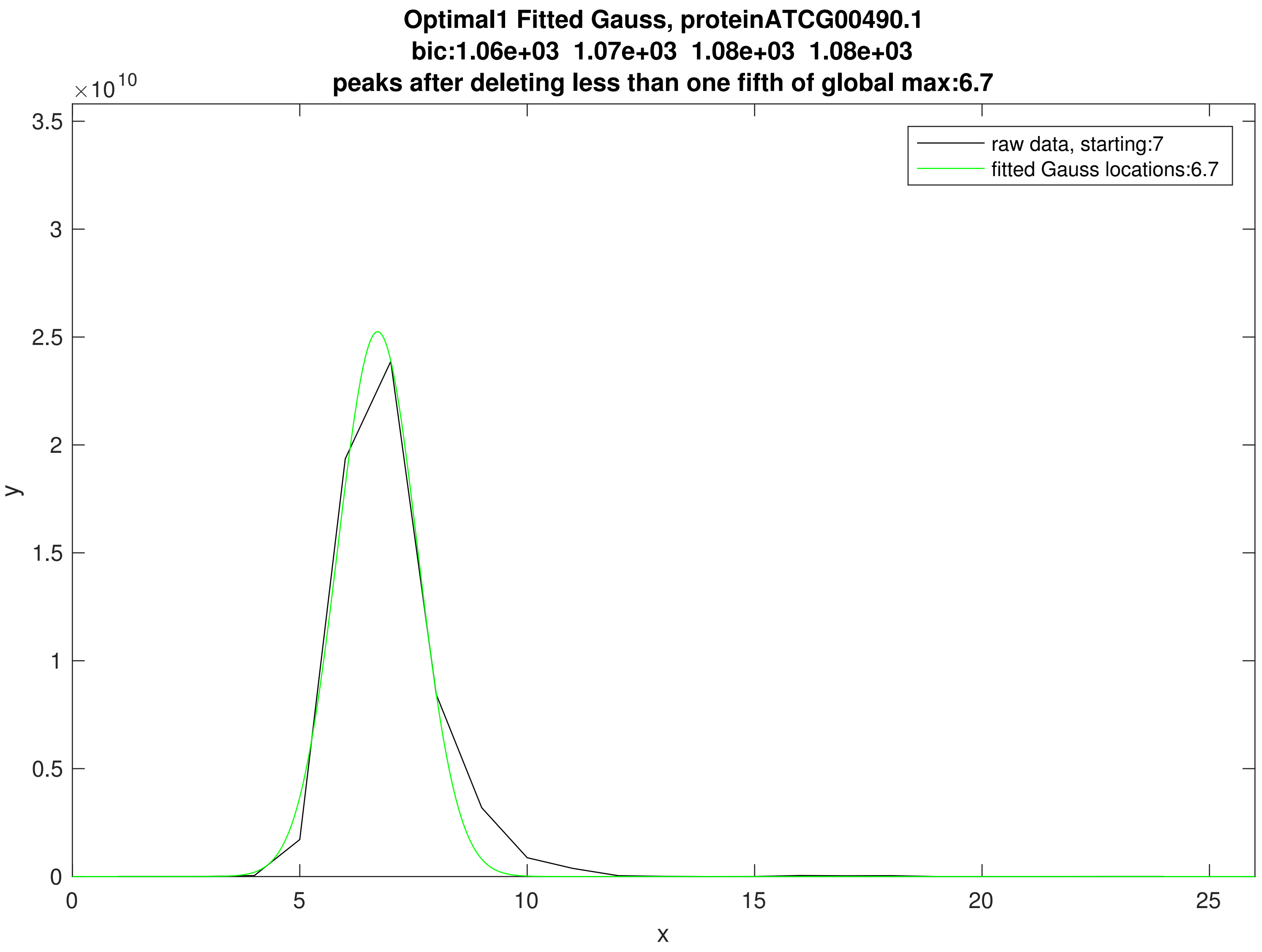

Optimal1 Fitted Gauss, proteinATCG00490.1
bic:1.06e+03 1.07e+03 1.08e+03 1.08e+03
peaks after deleting less than one fifth of global max:6.7
×10^10
3.5
3
2.5
2
1.5
1
0.5
0
y
0
5
10
15
20
25
x
raw data, starting:7
fitted Gauss locations:6.7

**Optimal1 Fitted Gauss, proteinATCG00500.1**
**bic:580**
**peaks after deleting less than one fifth of global max:5.1**

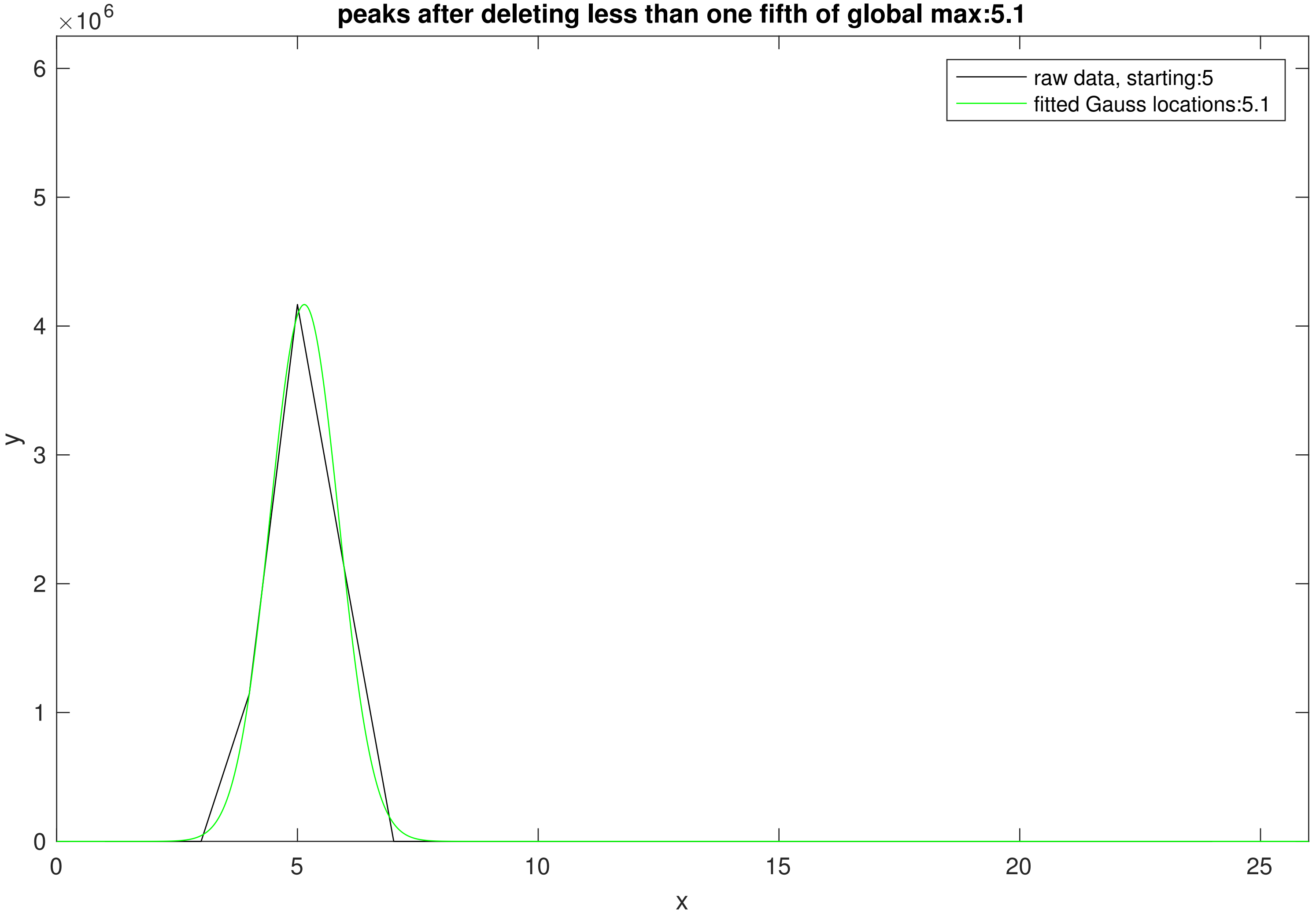

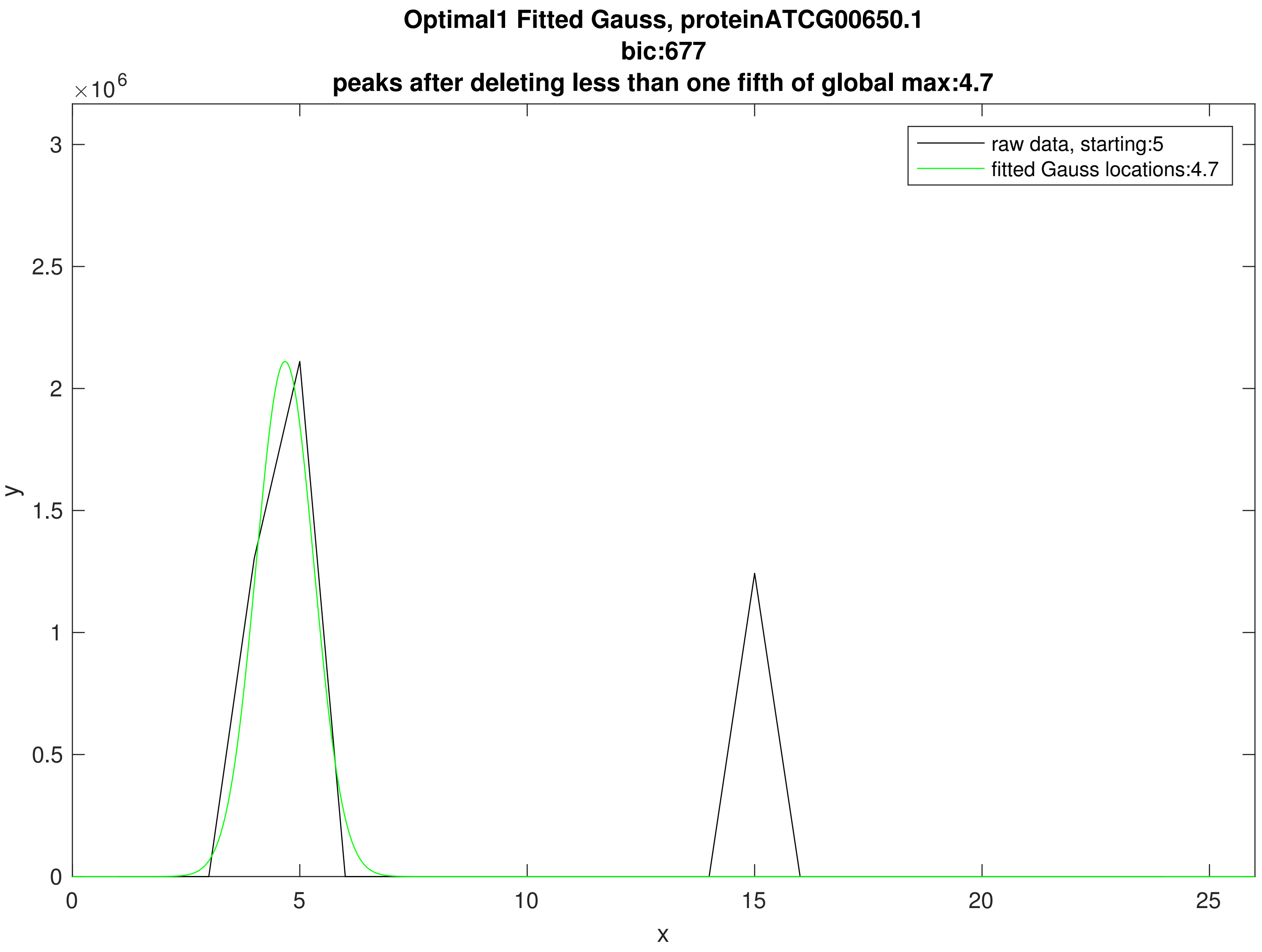

Optimal1 Fitted Gauss, proteinATCG00650.1
bic:677
peaks after deleting less than one fifth of global max:4.7
×10^6
3
2.5
2
1.5
1
0.5
0
y
0
5
10
15
20
25
x
raw data, starting:5
fitted Gauss locations:4.7

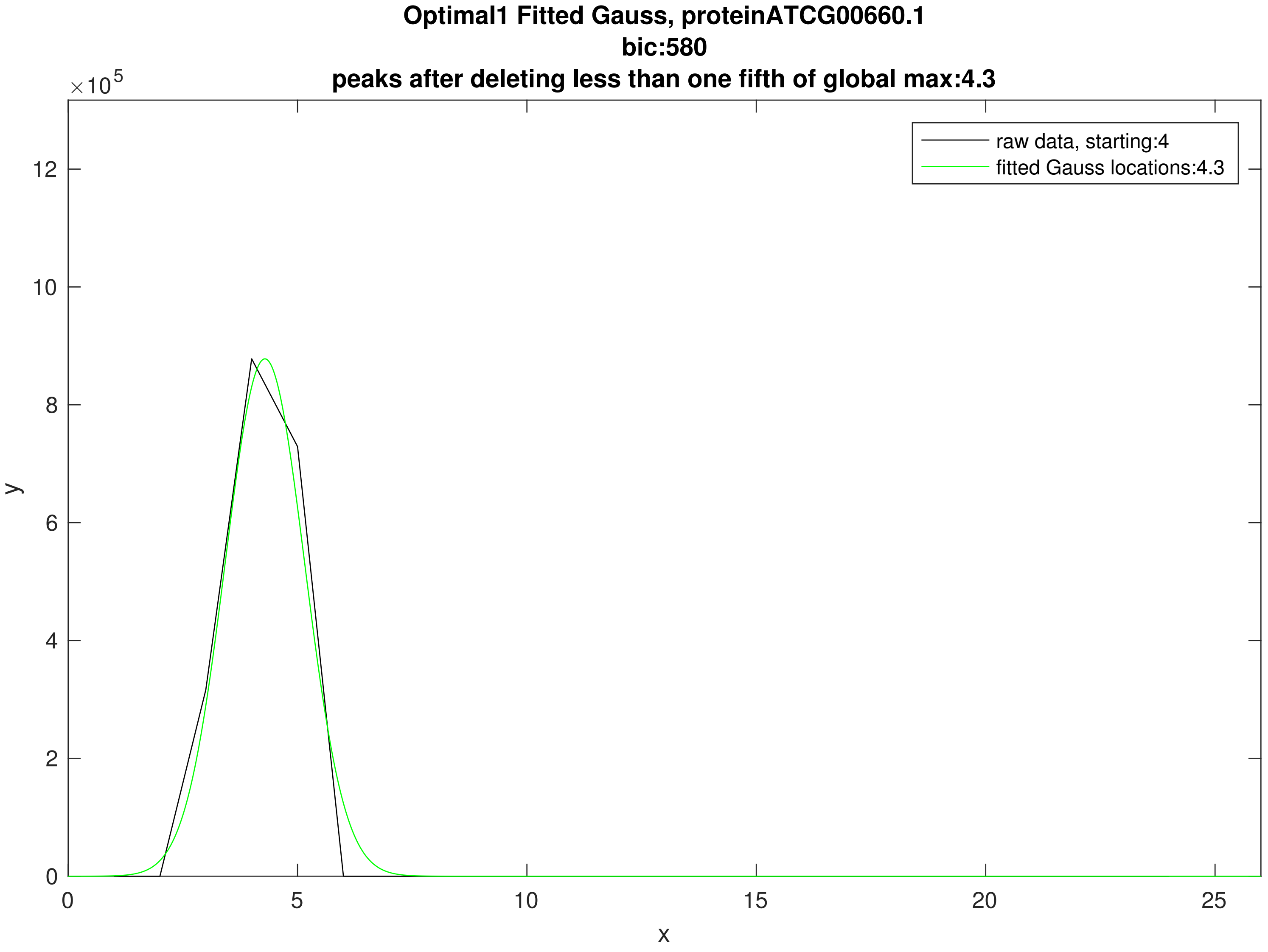

Optimal1 Fitted Gauss, proteinATCG00660.1
bic:580
peaks after deleting less than one fifth of global max:4.3
×10^5
raw data, starting:4
fitted Gauss locations:4.3
12
10
8
6
4
2
0
y
0
5
10
15
20
25
x

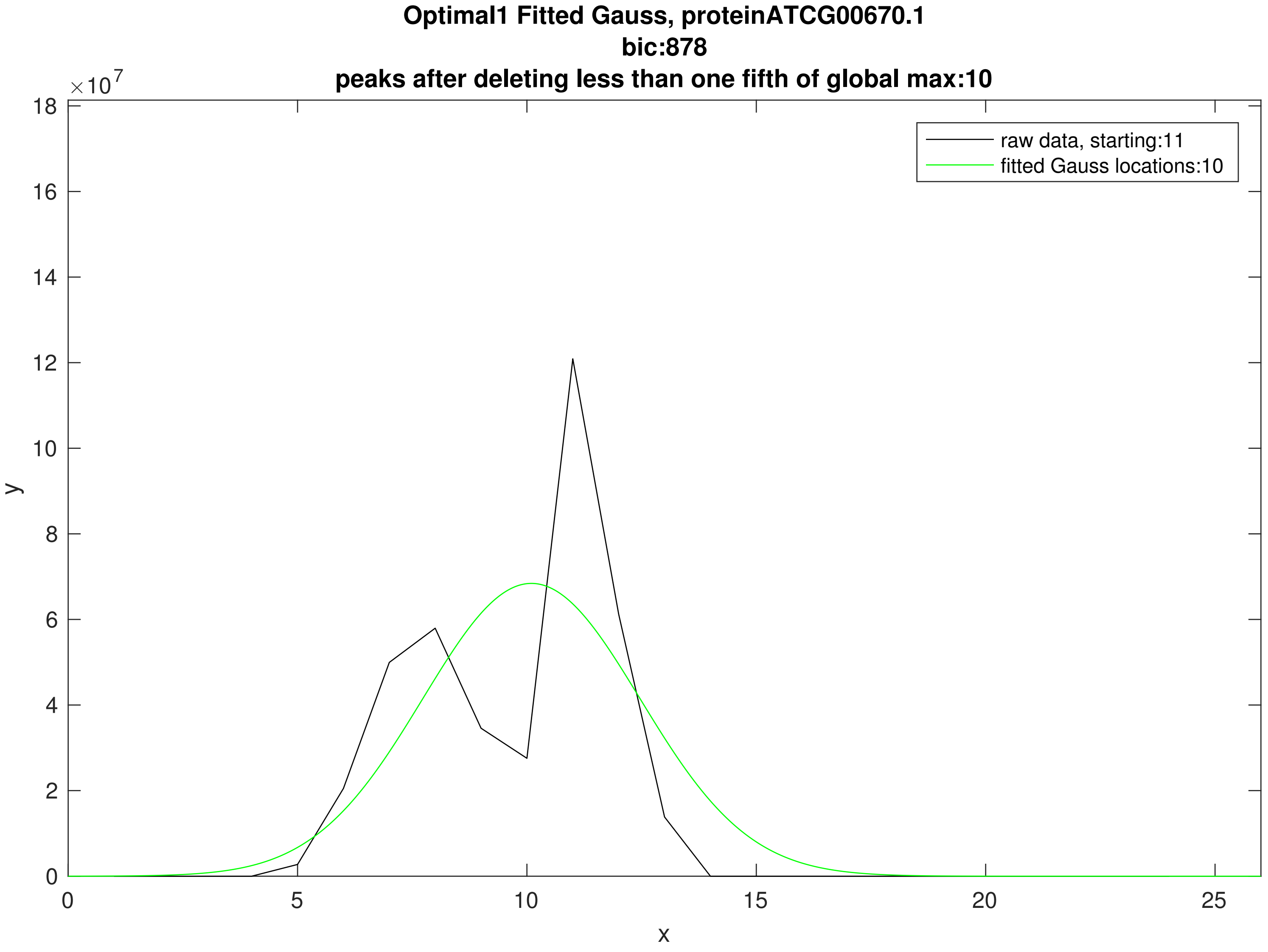
Optimal1 Fitted Gauss, proteinATCG00670.1
bic:878
peaks after deleting less than one fifth of global max:10
×10^7
raw data, starting:11
fitted Gauss locations:10
y
x
0
2
4
6
8
10
12
14
16
18
0
5
10
15
20
25

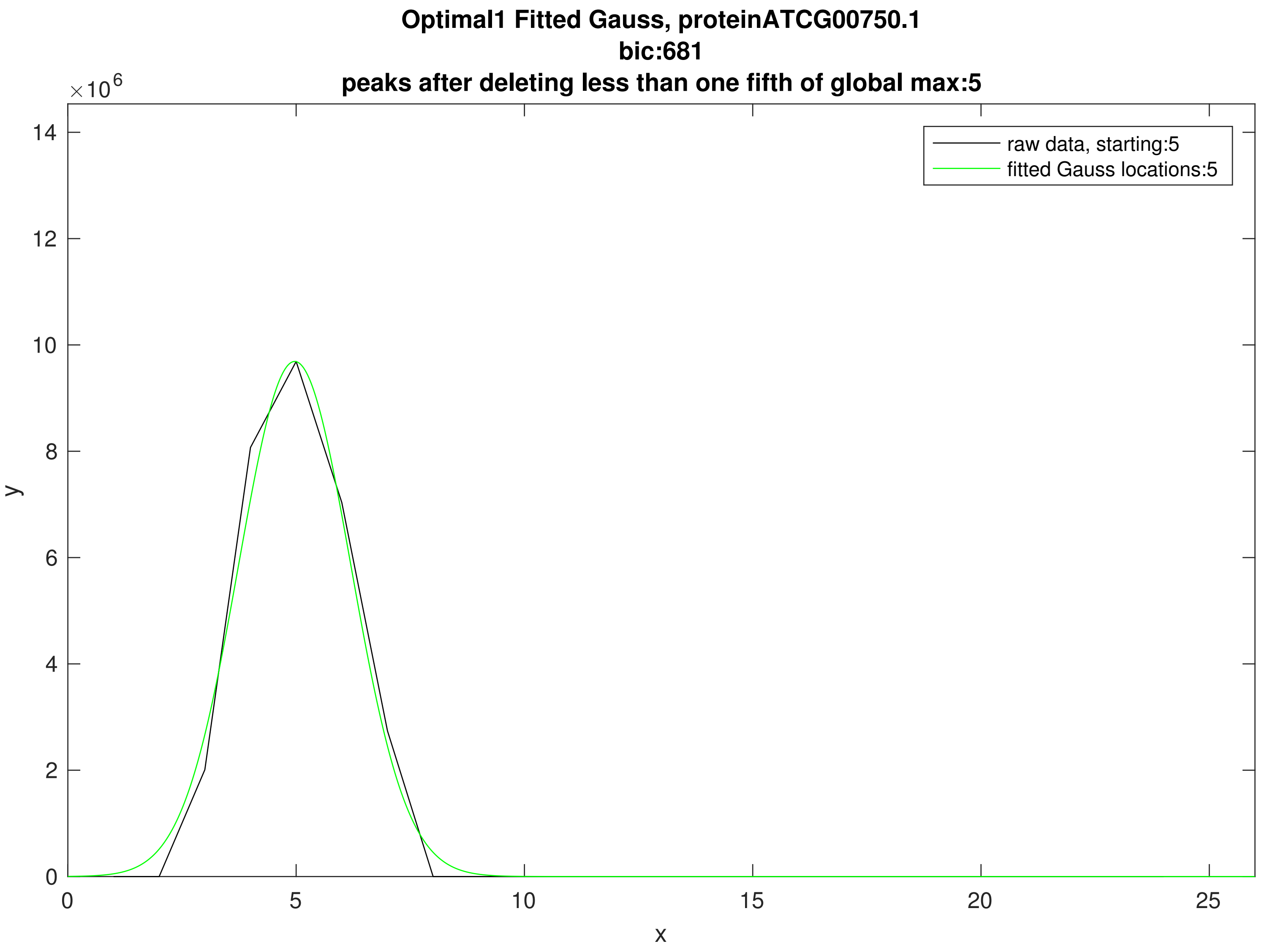

Optimal1 Fitted Gauss, proteinATCG00750.1
bic:681
peaks after deleting less than one fifth of global max:5
×10^6
14
12
10
8
6
4
2
0
y
0
5
10
15
20
25
x
raw data, starting:5
fitted Gauss locations:5

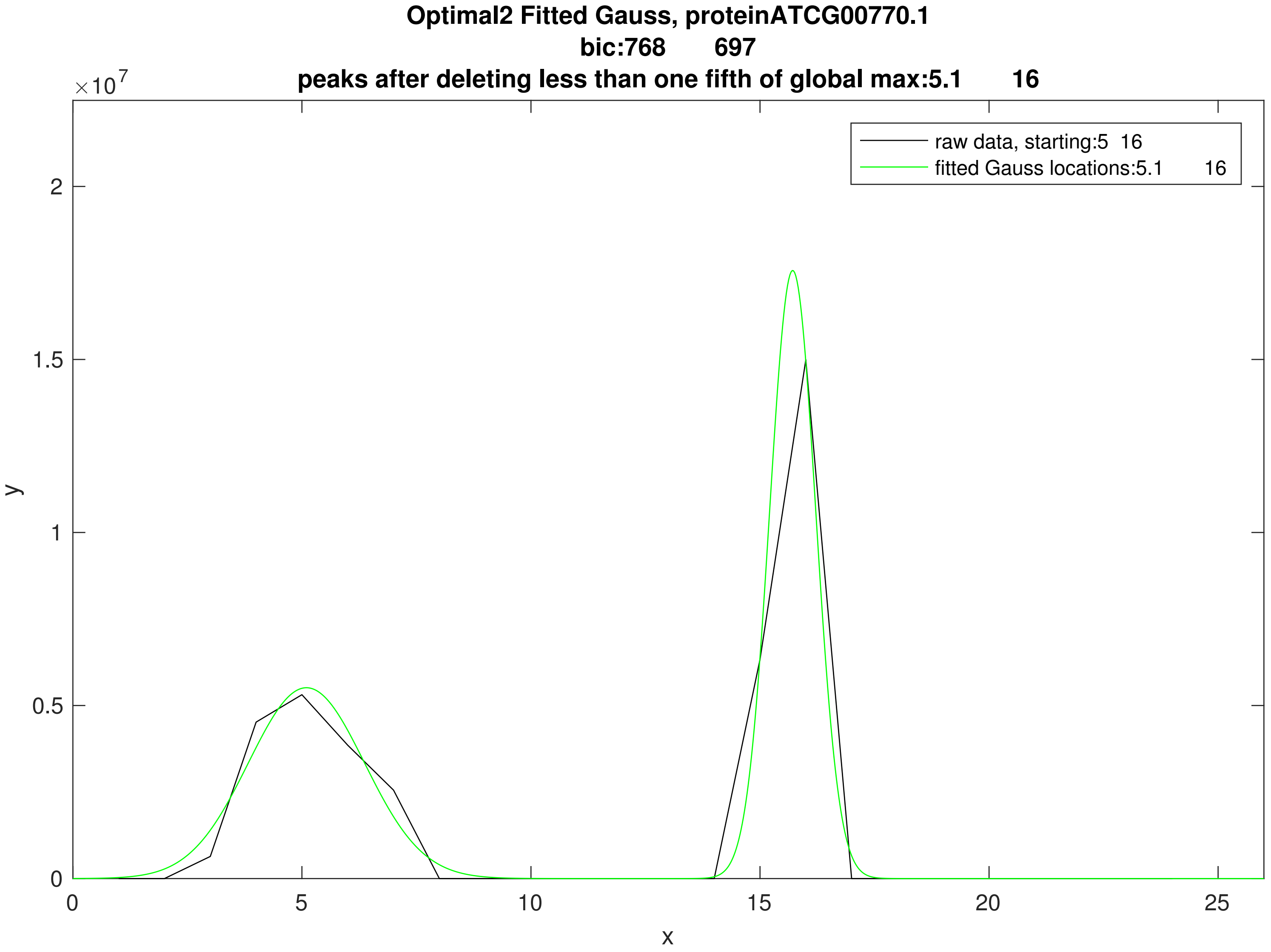
Optimal2 Fitted Gauss, proteinATCG00770.1
bic:768 697
peaks after deleting less than one fifth of global max:5.1 16
×10^7
raw data, starting:5 16
fitted Gauss locations:5.1 16
0
0.5
1
1.5
2
y
0
5
10
15
20
25
x

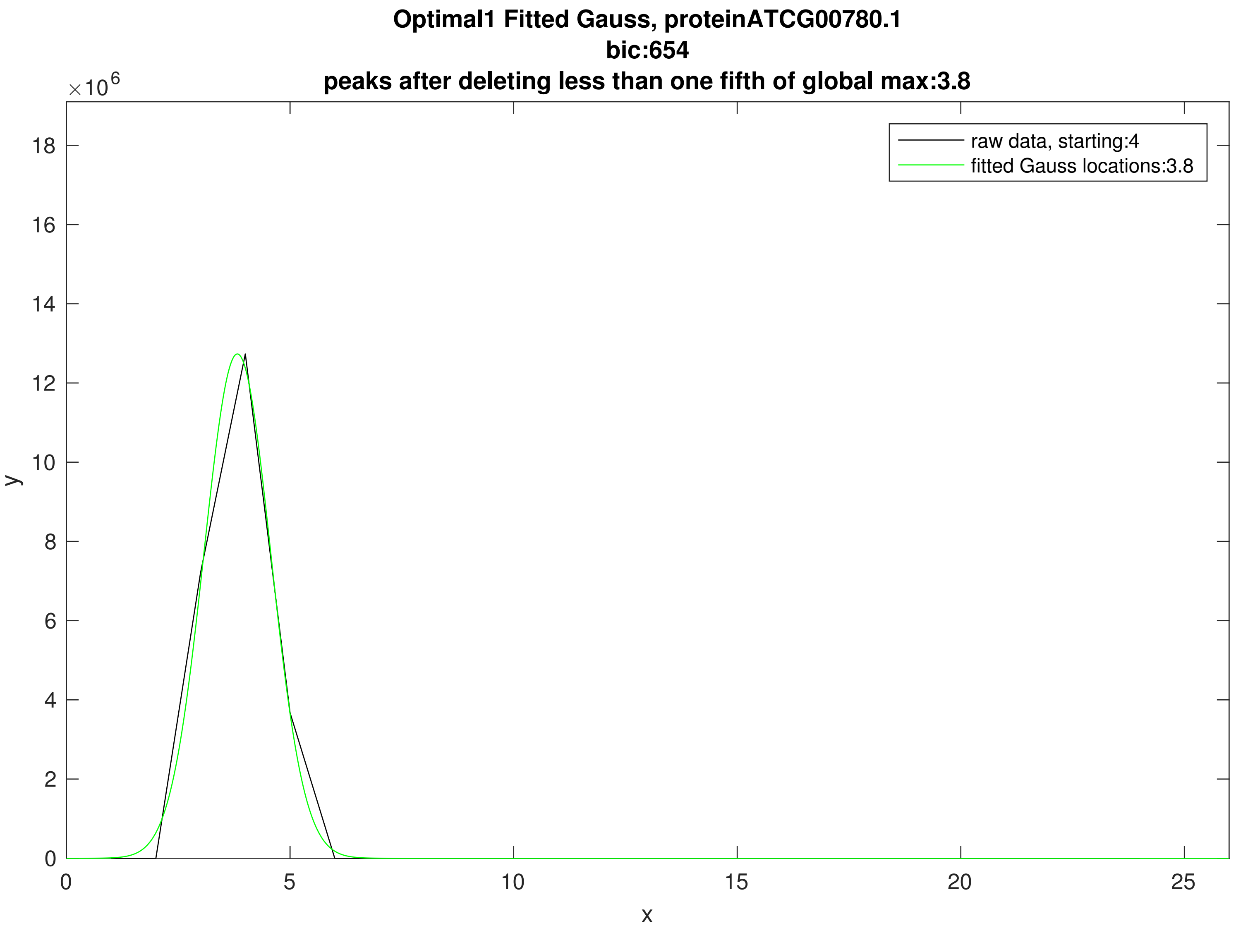

Optimal1 Fitted Gauss, proteinATCG00780.1
bic:654
peaks after deleting less than one fifth of global max:3.8
×10^6
raw data, starting:4
fitted Gauss locations:3.8
y
x
0
2
4
6
8
10
12
14
16
18
0
5
10
15
20
25

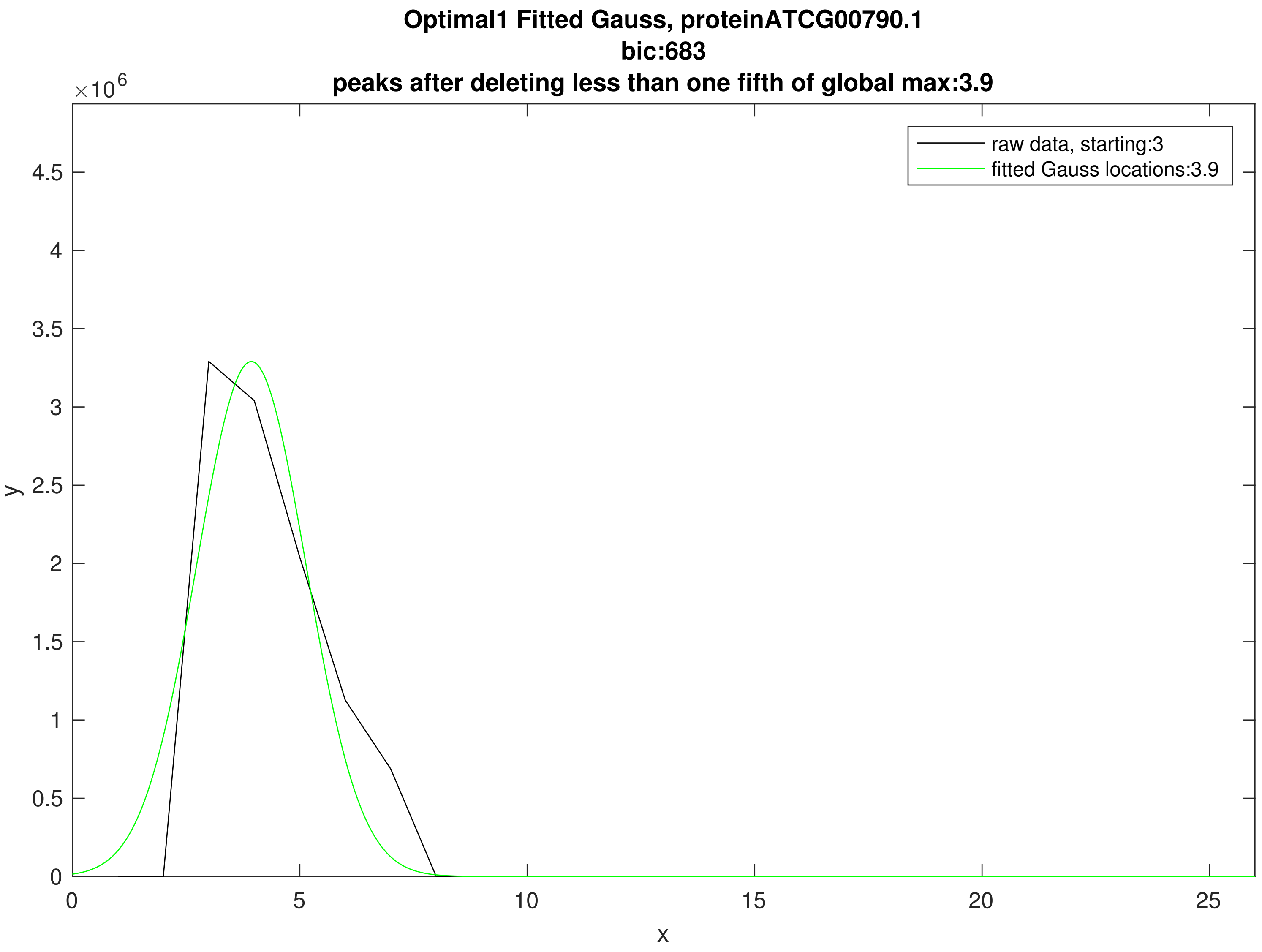
Optimal1 Fitted Gauss, proteinATCG00790.1
bic:683
peaks after deleting less than one fifth of global max:3.9
×10^6
4.5
4
3.5
3
2.5
2
1.5
1
0.5
0
y
0
5
10
15
20
25
x
raw data, starting:3
fitted Gauss locations:3.9

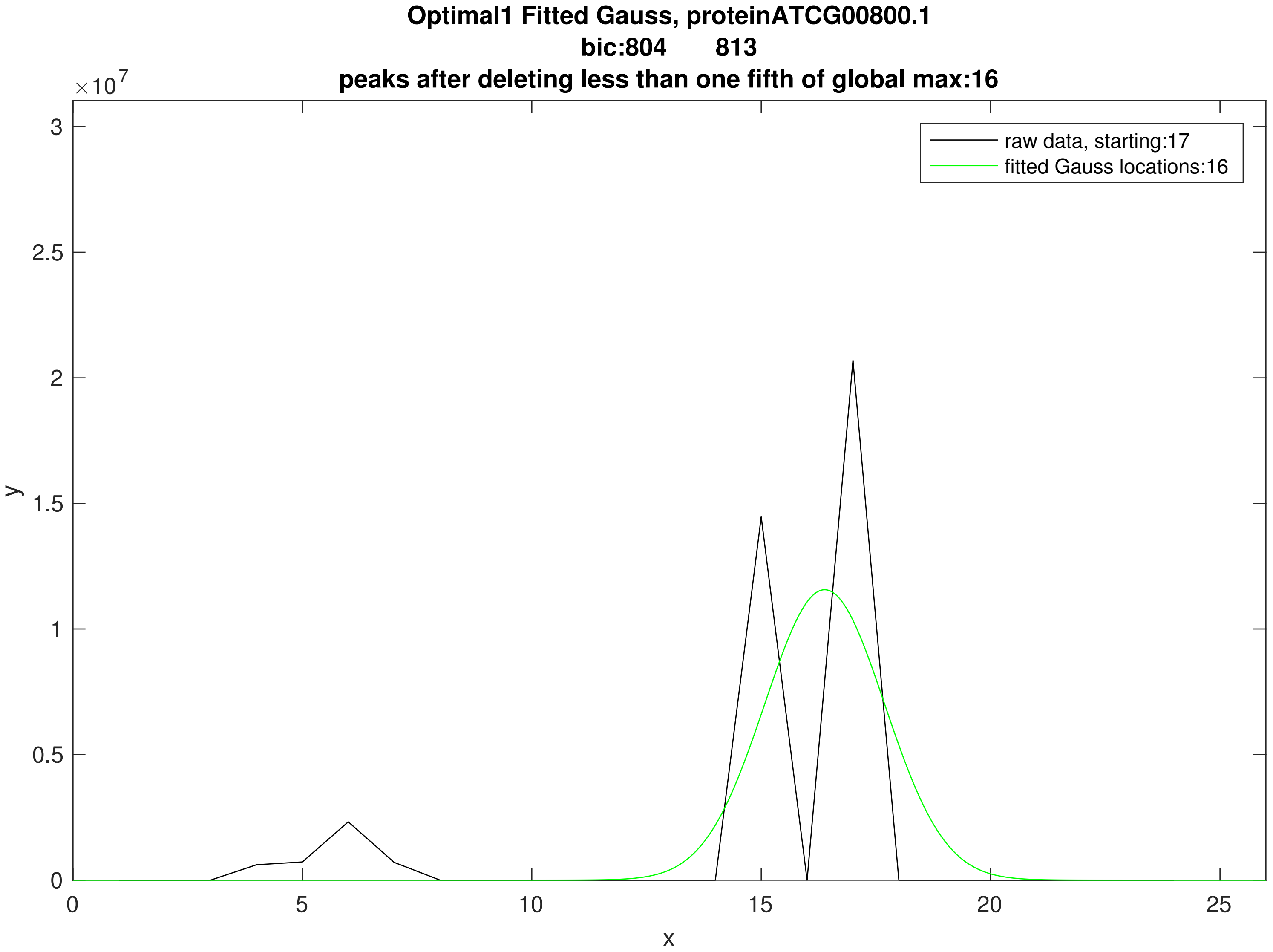

Optimal1 Fitted Gauss, proteinATCG00800.1
bic:804 813
peaks after deleting less than one fifth of global max:16
×10^7
3
2.5
2
1.5
1
0.5
0
y
0
5
10
15
20
25
x
raw data, starting:17
fitted Gauss locations:16

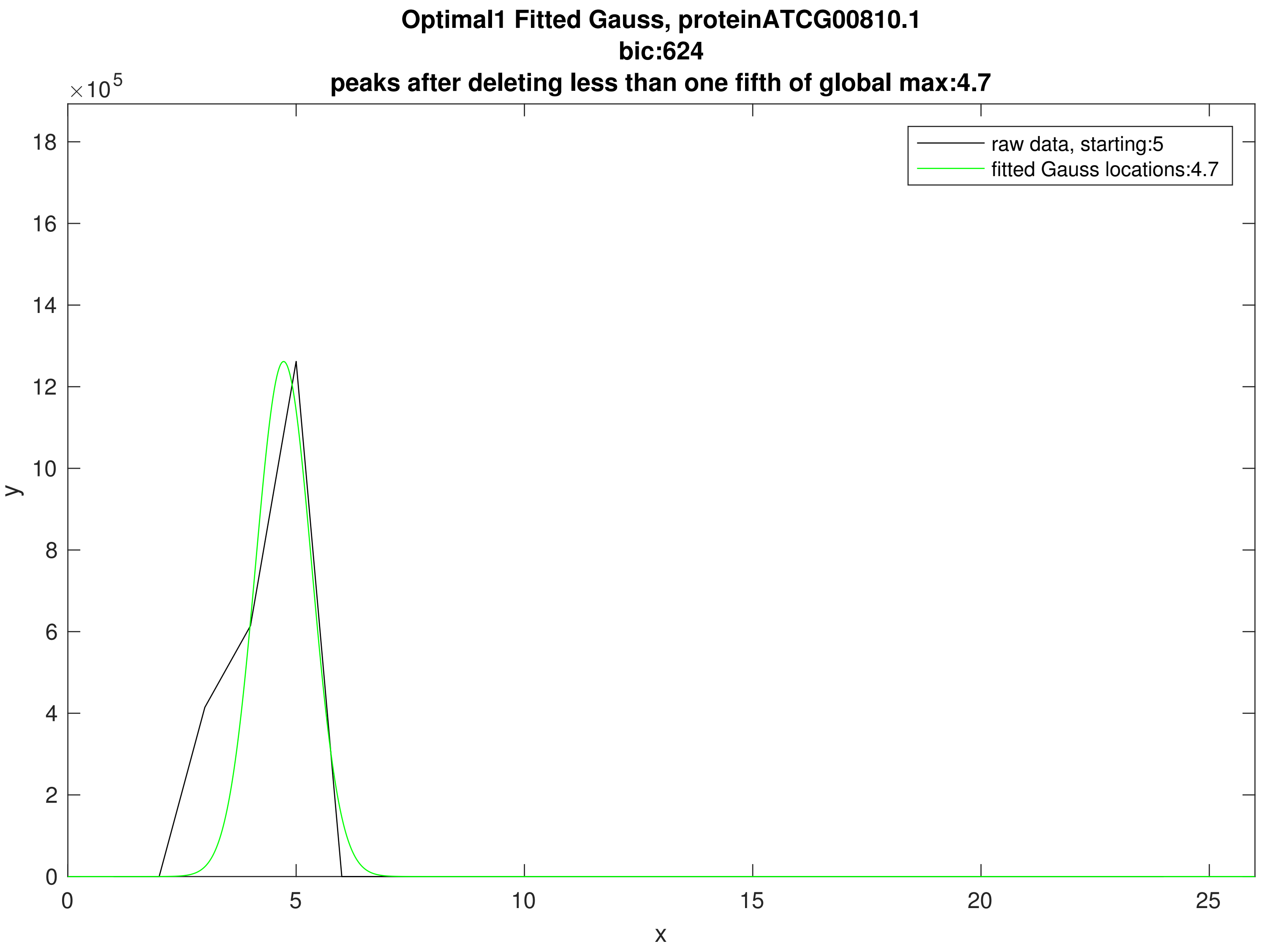
Optimal1 Fitted Gauss, proteinATCG00810.1
bic:624
peaks after deleting less than one fifth of global max:4.7
×10^5
raw data, starting:5
fitted Gauss locations:4.7
y
x
0
2
4
6
8
10
12
14
16
18
0
5
10
15
20
25

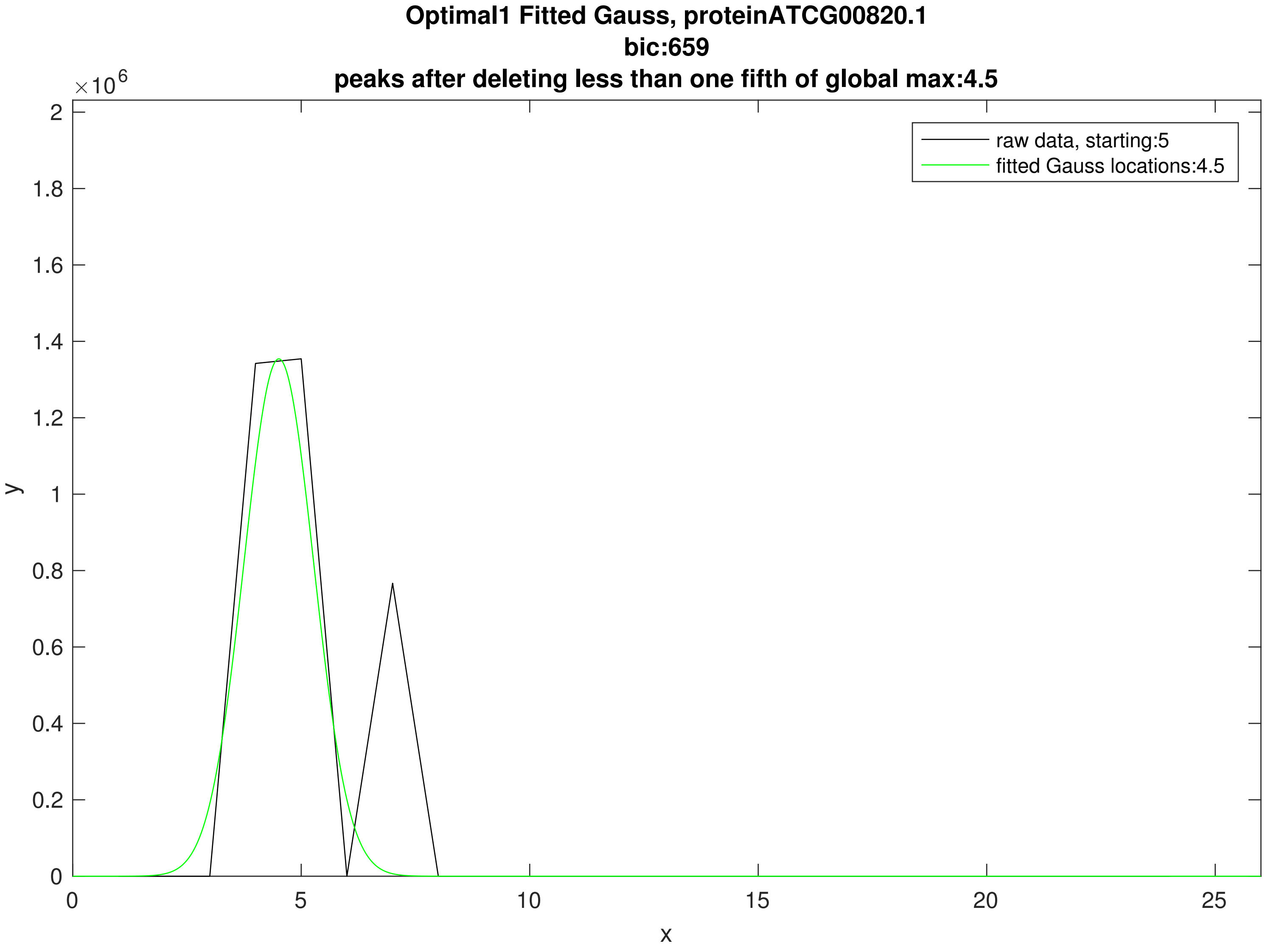

Optimal1 Fitted Gauss, proteinATCG00820.1
bic:659
peaks after deleting less than one fifth of global max:4.5
×10^6
2
1.8
1.6
1.4
1.2
1
0.8
0.6
0.4
0.2
0
y
0
5
10
15
20
25
x
raw data, starting:5
fitted Gauss locations:4.5

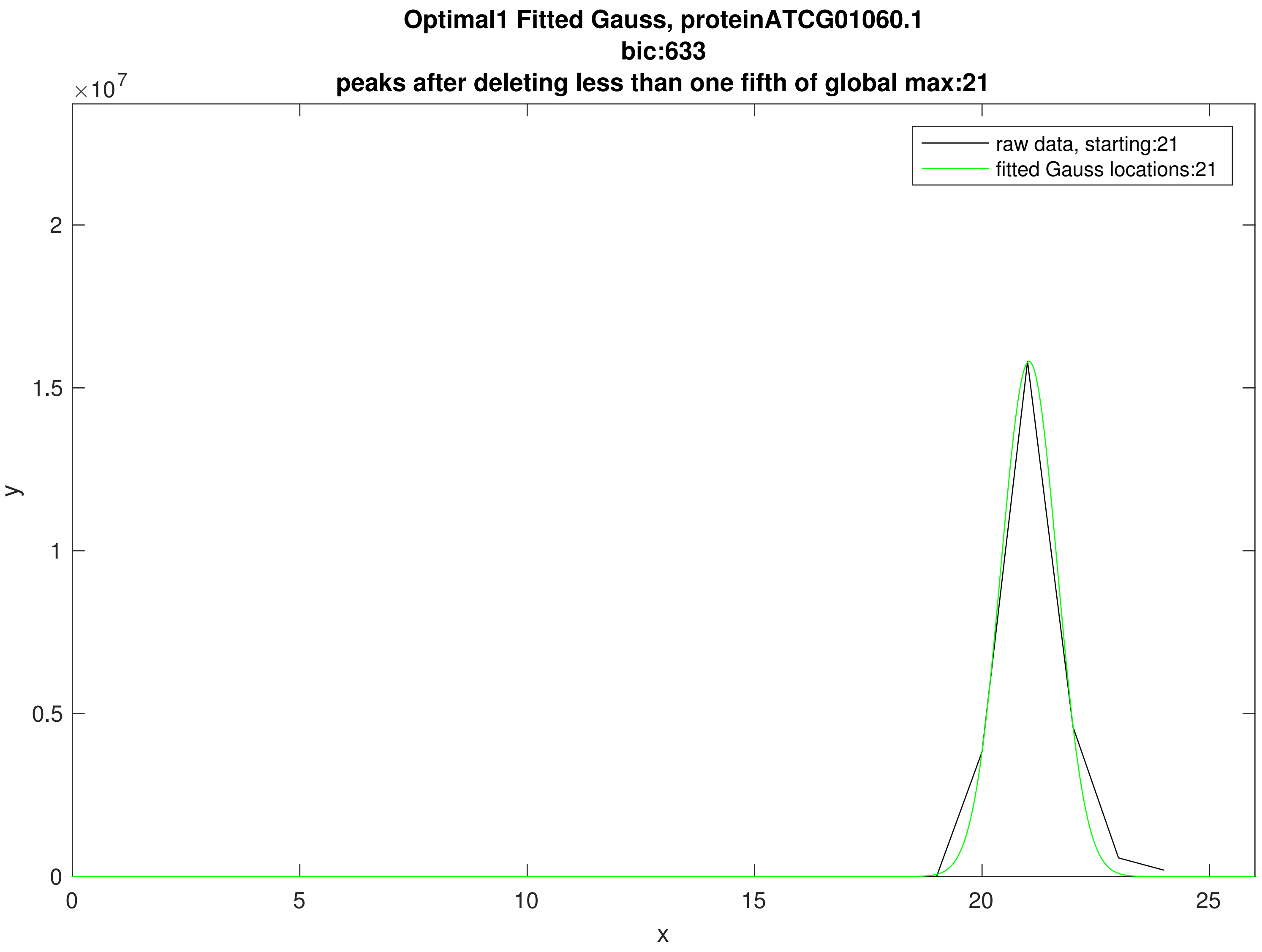
Optimal1 Fitted Gauss, proteinATCG01060.1
bic:633
peaks after deleting less than one fifth of global max:21
×10^7
raw data, starting:21
fitted Gauss locations:21
y
x
0
0.5
1
1.5
2
0
5
10
15
20
25

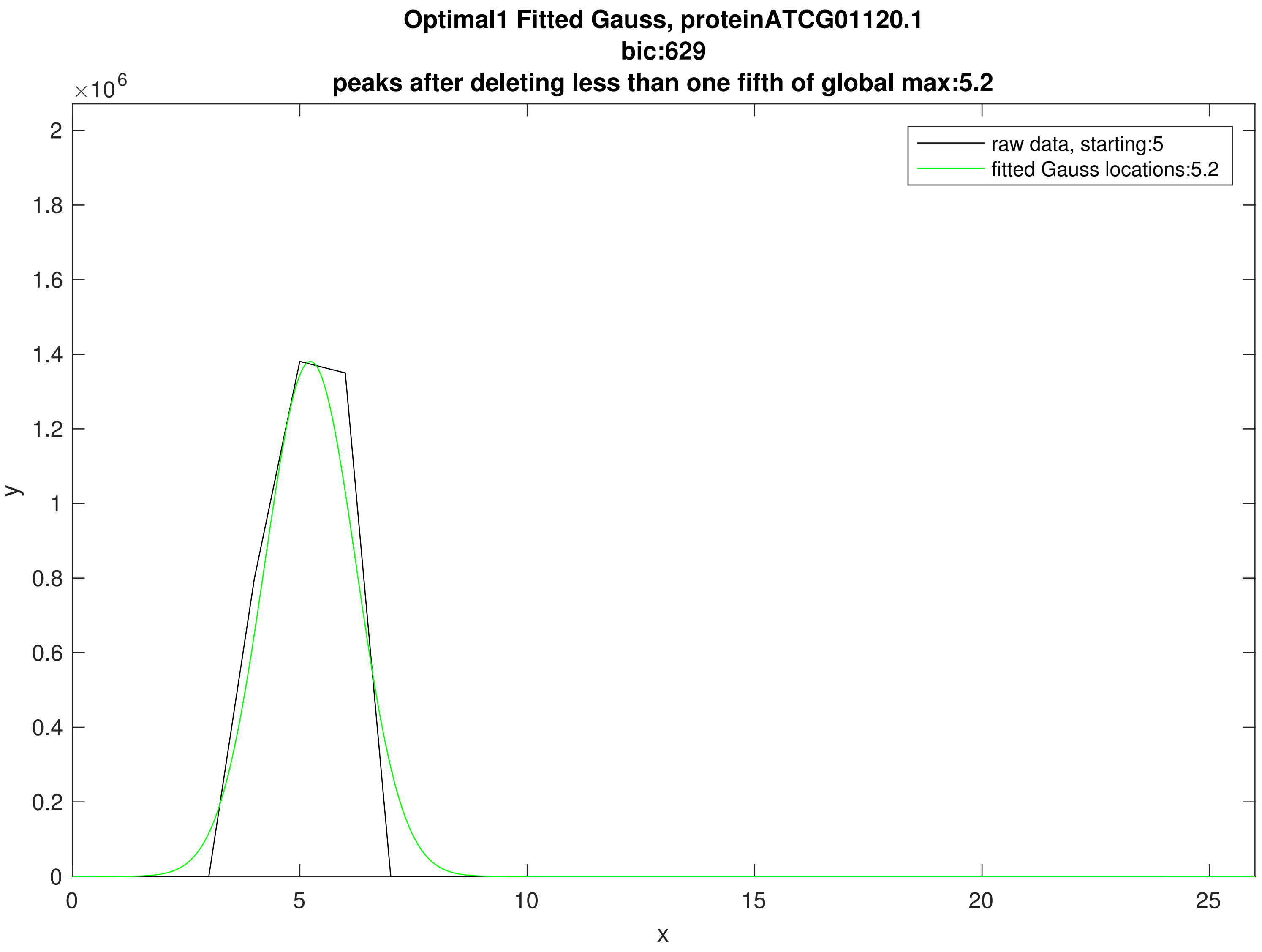

Optimal1 Fitted Gauss, proteinATCG01120.1
bic:629
peaks after deleting less than one fifth of global max:5.2
×10^6
2
1.8
1.6
1.4
1.2
1
0.8
0.6
0.4
0.2
0
y
0
5
10
15
20
25
x
raw data, starting:5
fitted Gauss locations:5.2

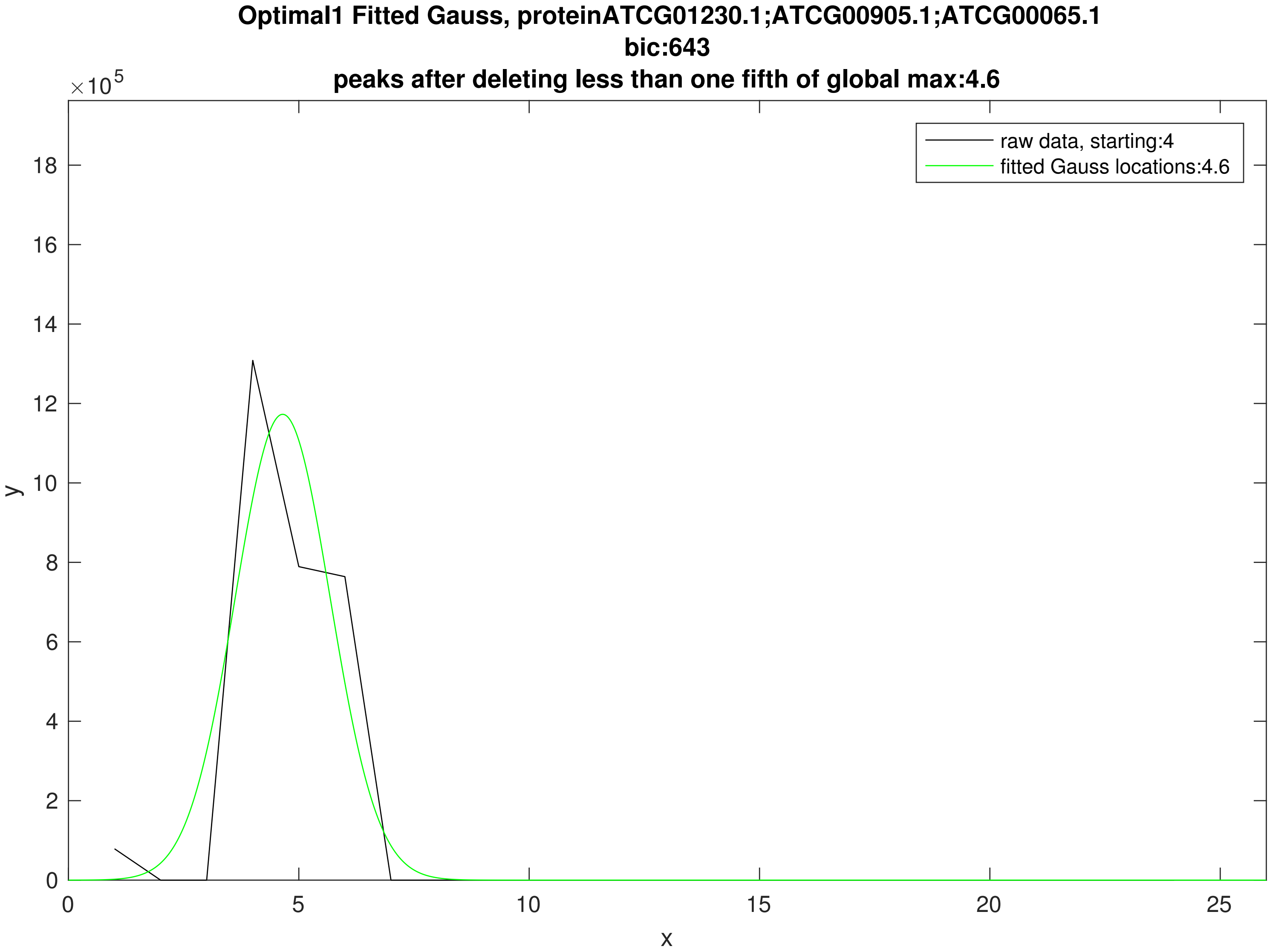

Optimal1 Fitted Gauss, proteinATCG01230.1;ATCG00905.1;ATCG00065.1
bic:643
peaks after deleting less than one fifth of global max:4.6
×10^5
raw data, starting:4
fitted Gauss locations:4.6
y
x
0
2
4
6
8
10
12
14
16
18
0
5
10
15
20
25

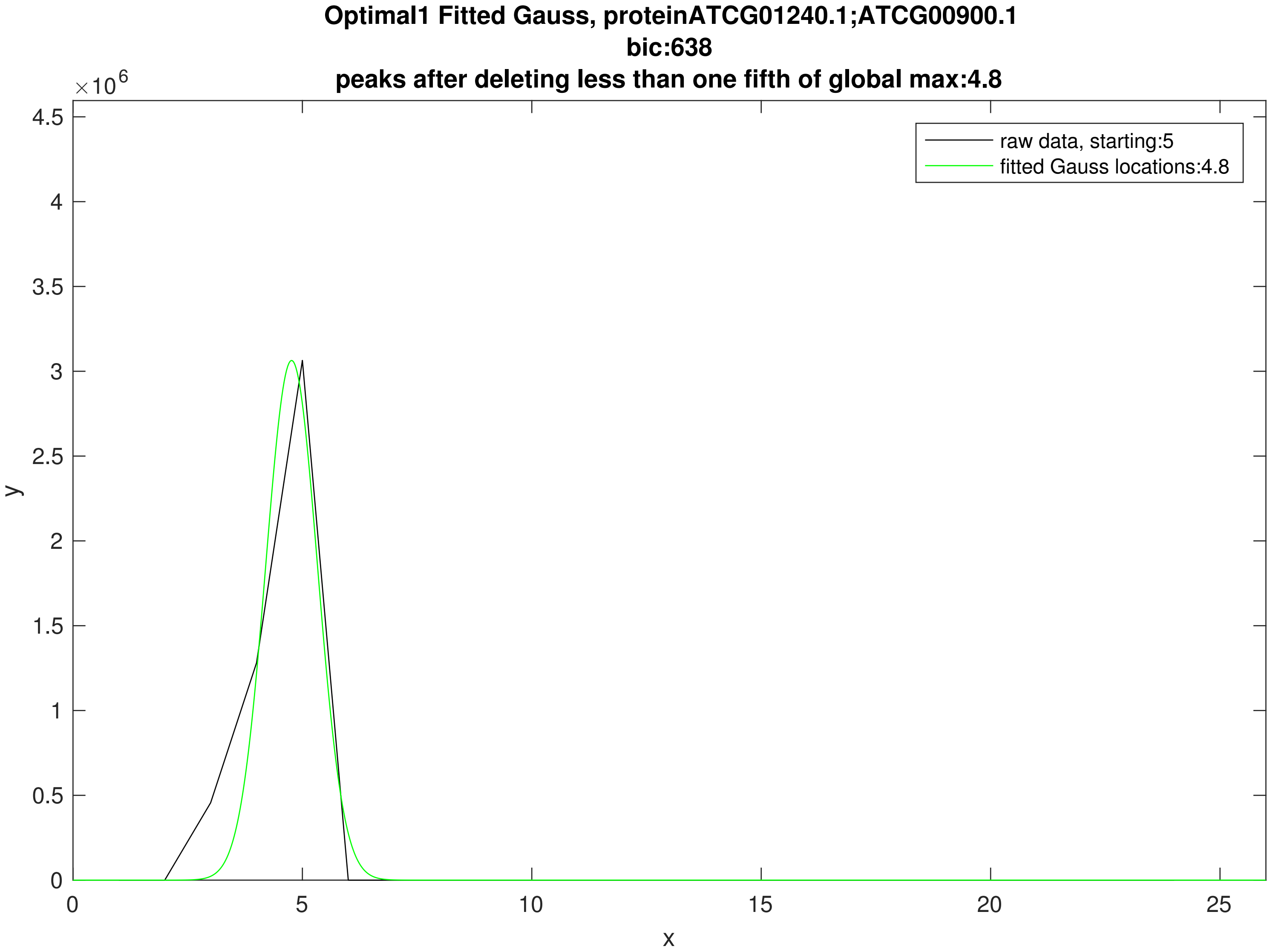
Optimal1 Fitted Gauss, proteinATCG01240.1;ATCG00900.1
bic:638
peaks after deleting less than one fifth of global max:4.8
raw data, starting:5
fitted Gauss locations:4.8
y
x
×10^6
4.5
4
3.5
3
2.5
2
1.5
1
0.5
0
0
5
10
15
20
25

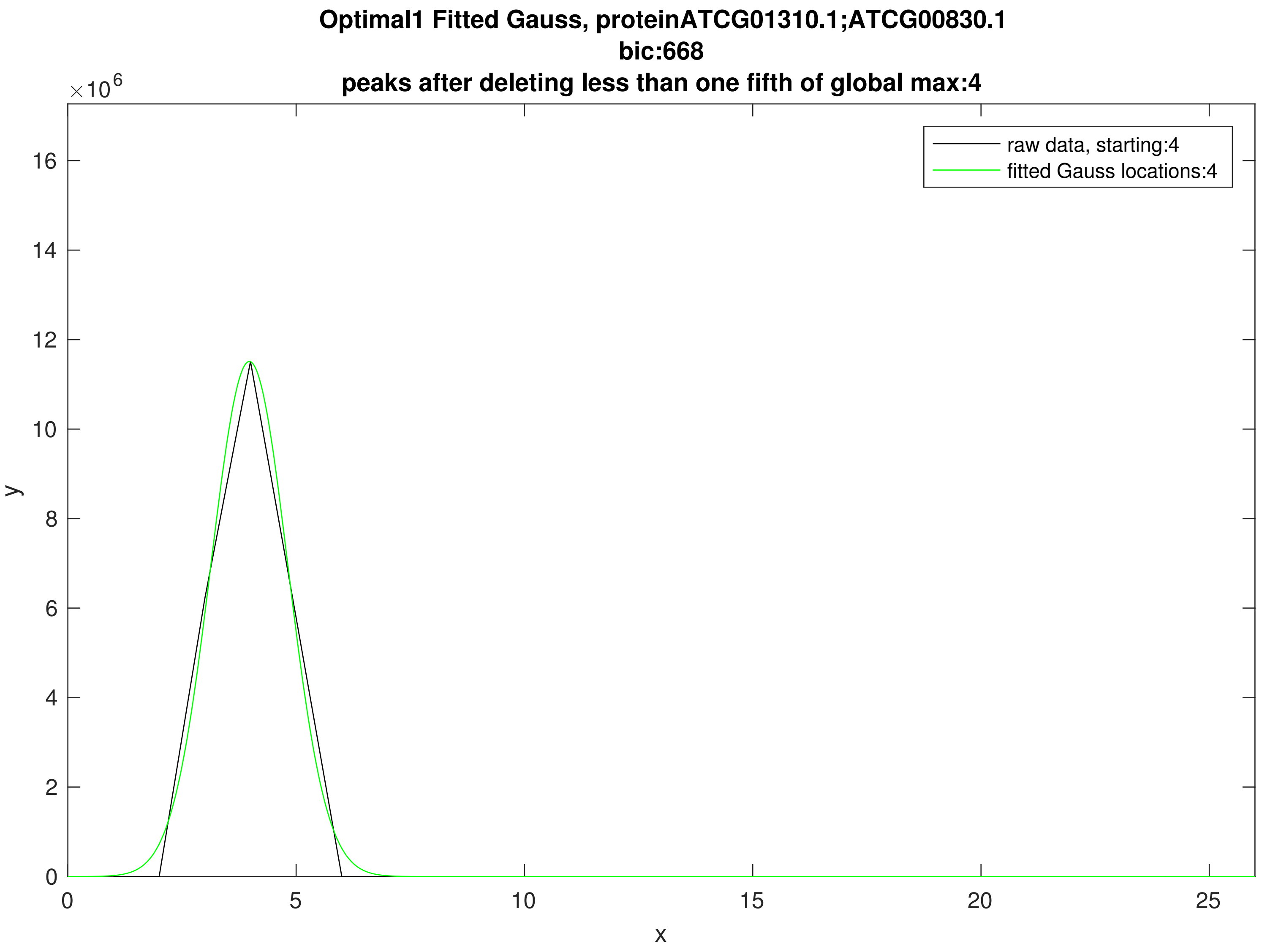
Optimal1 Fitted Gauss, proteinATCG01310.1;ATCG00830.1
bic:668
peaks after deleting less than one fifth of global max:4
×10^6
raw data, starting:4
fitted Gauss locations:4
y
x
0
2
4
6
8
10
12
14
16
5
15
20
25

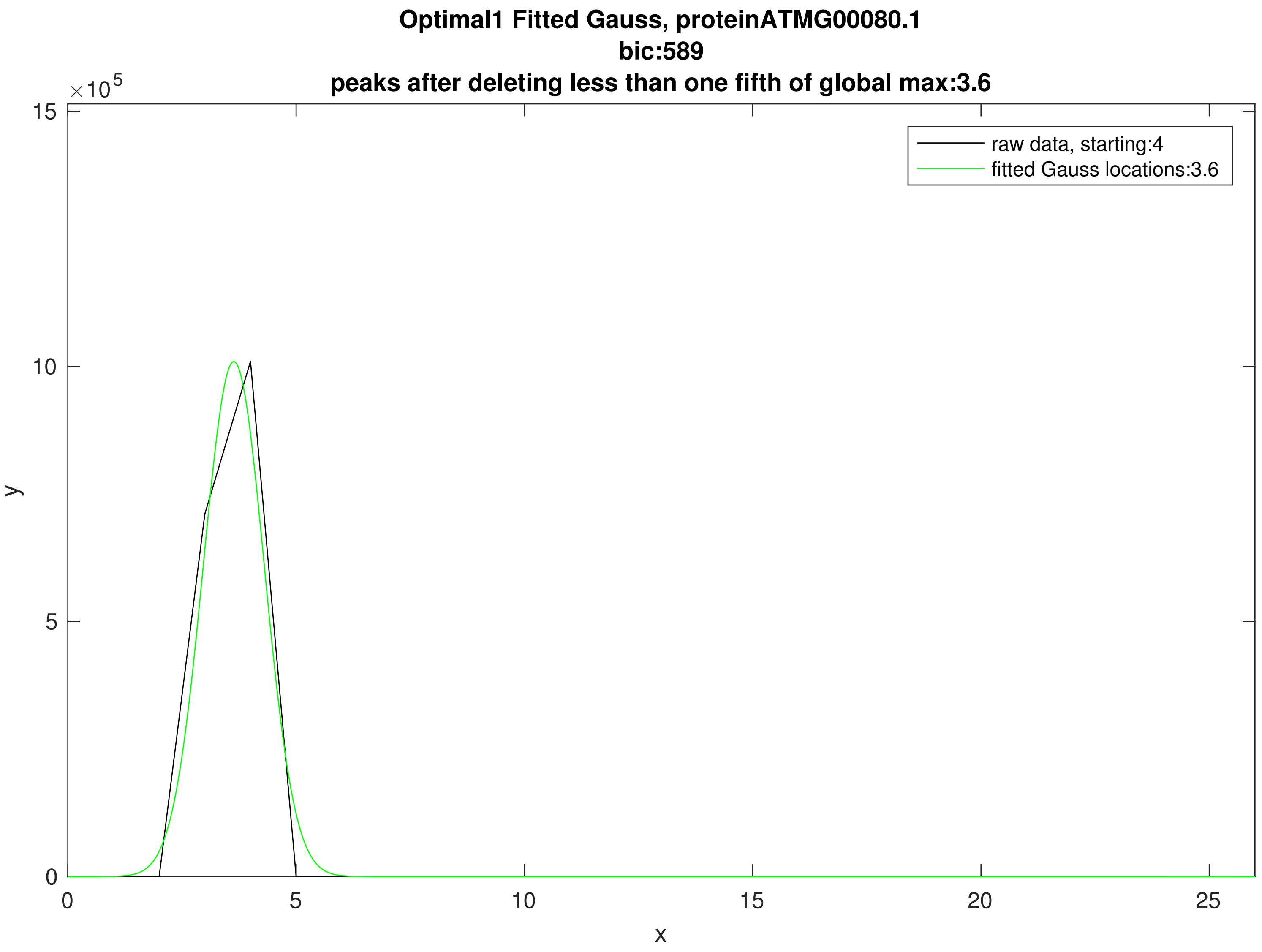
Optimal1 Fitted Gauss, proteinATMG00080.1
bic:589
peaks after deleting less than one fifth of global max:3.6
×10^5
raw data, starting:4
fitted Gauss locations:3.6
y
x
0
5
10
15
20
25

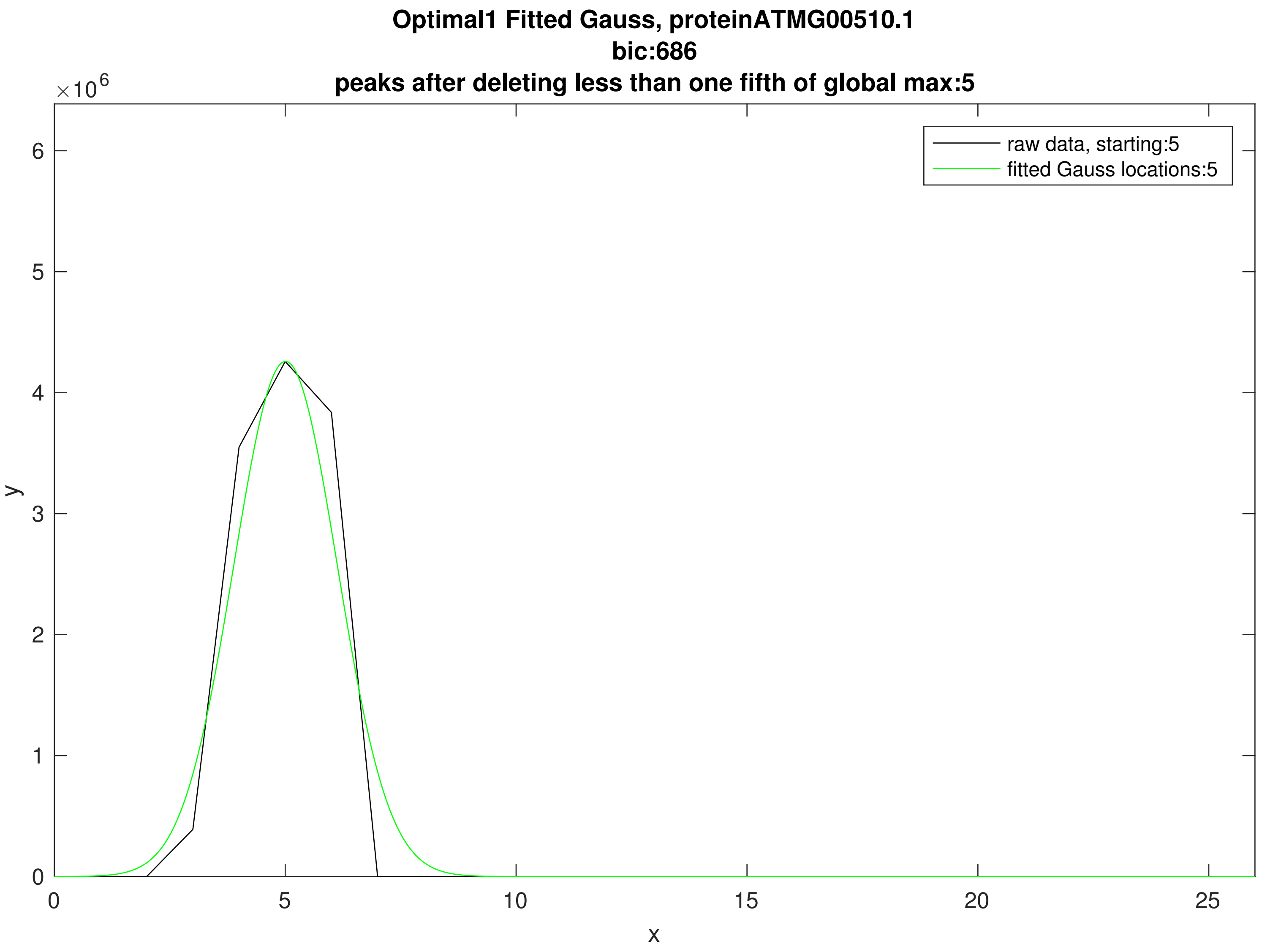
Optimal1 Fitted Gauss, proteinATMG00510.1
bic:686
peaks after deleting less than one fifth of global max:5
×10^6
raw data, starting:5
fitted Gauss locations:5
y
x
0
1
2
3
4
5
6
0
5
10
15
20
25

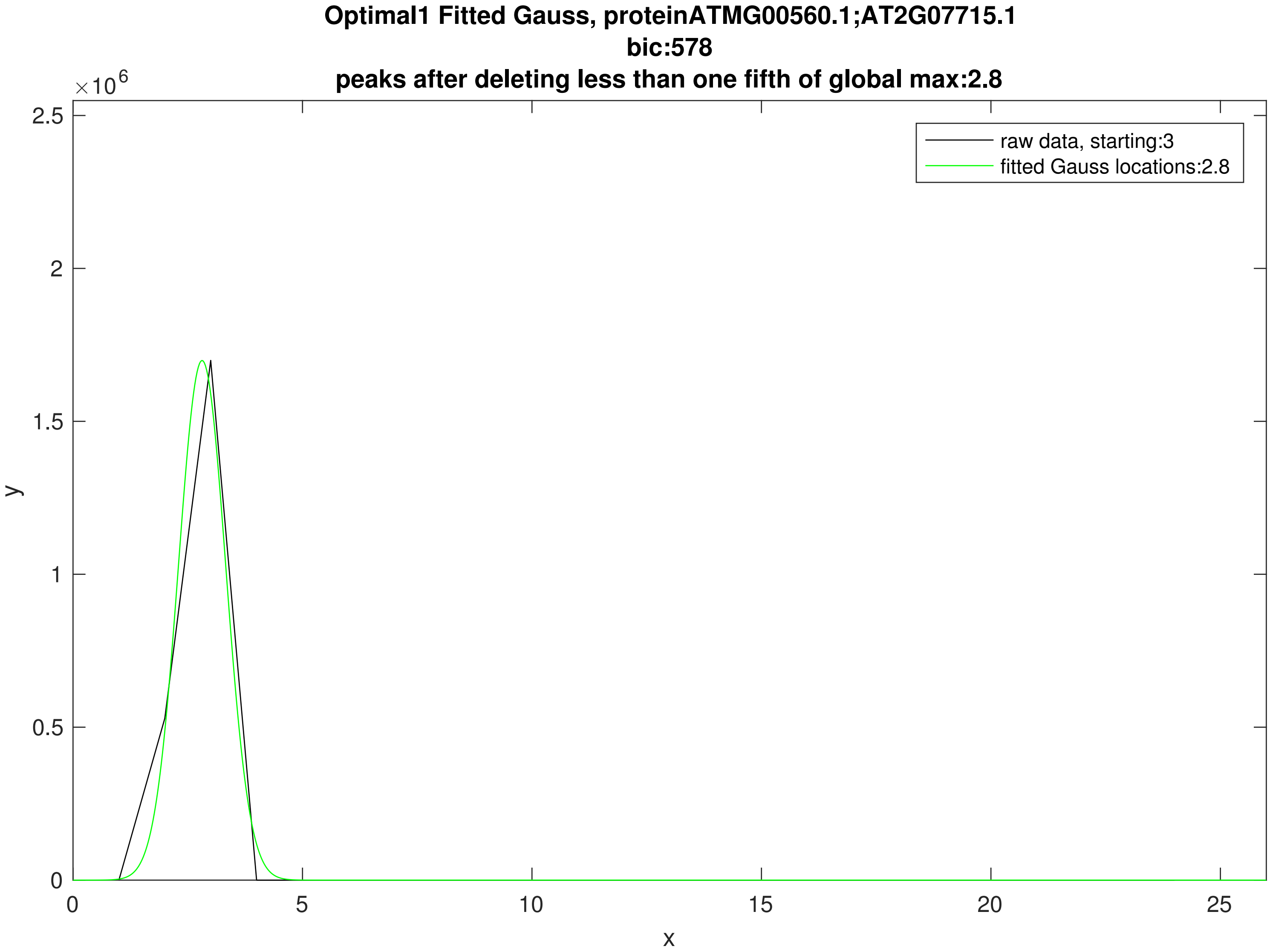
Optimal1 Fitted Gauss, proteinATMG00560.1;AT2G07715.1
bic:578
peaks after deleting less than one fifth of global max:2.8
×10^6
2.5
2
1.5
1
0.5
0
y
0
5
10
15
20
25
x
raw data, starting:3
fitted Gauss locations:2.8

### $SEC_{Ethylene}$ replicate 1 elution profiles

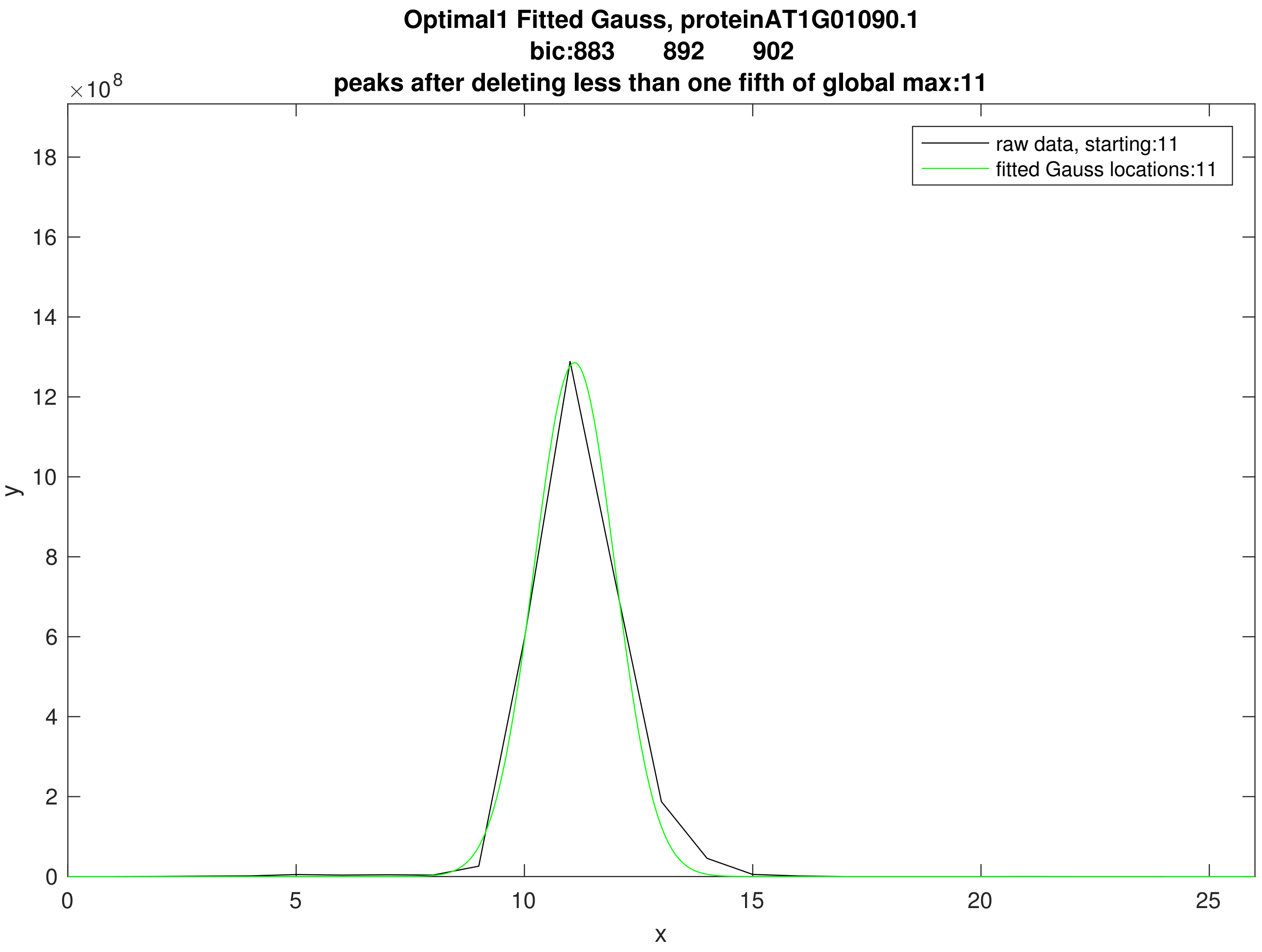
Optimal1 Fitted Gauss, proteinAT1G01090.1
bic:883 892 902
peaks after deleting less than one fifth of global max:11
×10^8
18
16
14
12
10
8
6
4
2
0
y
0
5
10
15
20
25
x
raw data, starting:11
fitted Gauss locations:11

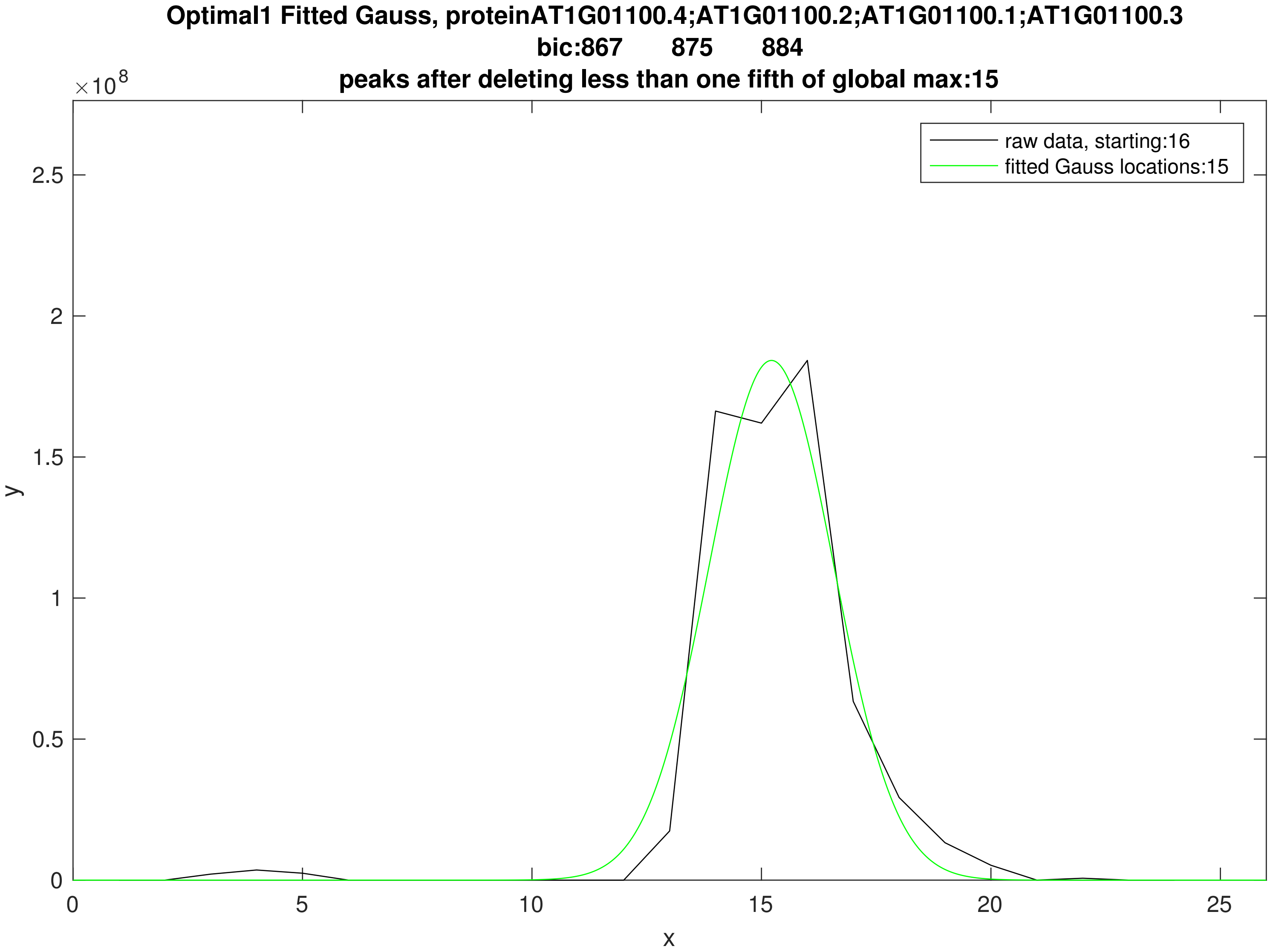
Optimal1 Fitted Gauss, proteinAT1G01100.4;AT1G01100.2;AT1G01100.1;AT1G01100.3
bic:867 875 884
peaks after deleting less than one fifth of global max:15
×10^8
raw data, starting:16
fitted Gauss locations:15
y
x
0
0.5
1
1.5
2
2.5
0
5
10
15
20
25

**Optimal1 Fitted Gauss, proteinAT1G01220.1**
**bic:780**
**peaks after deleting less than one fifth of global max:9.4**

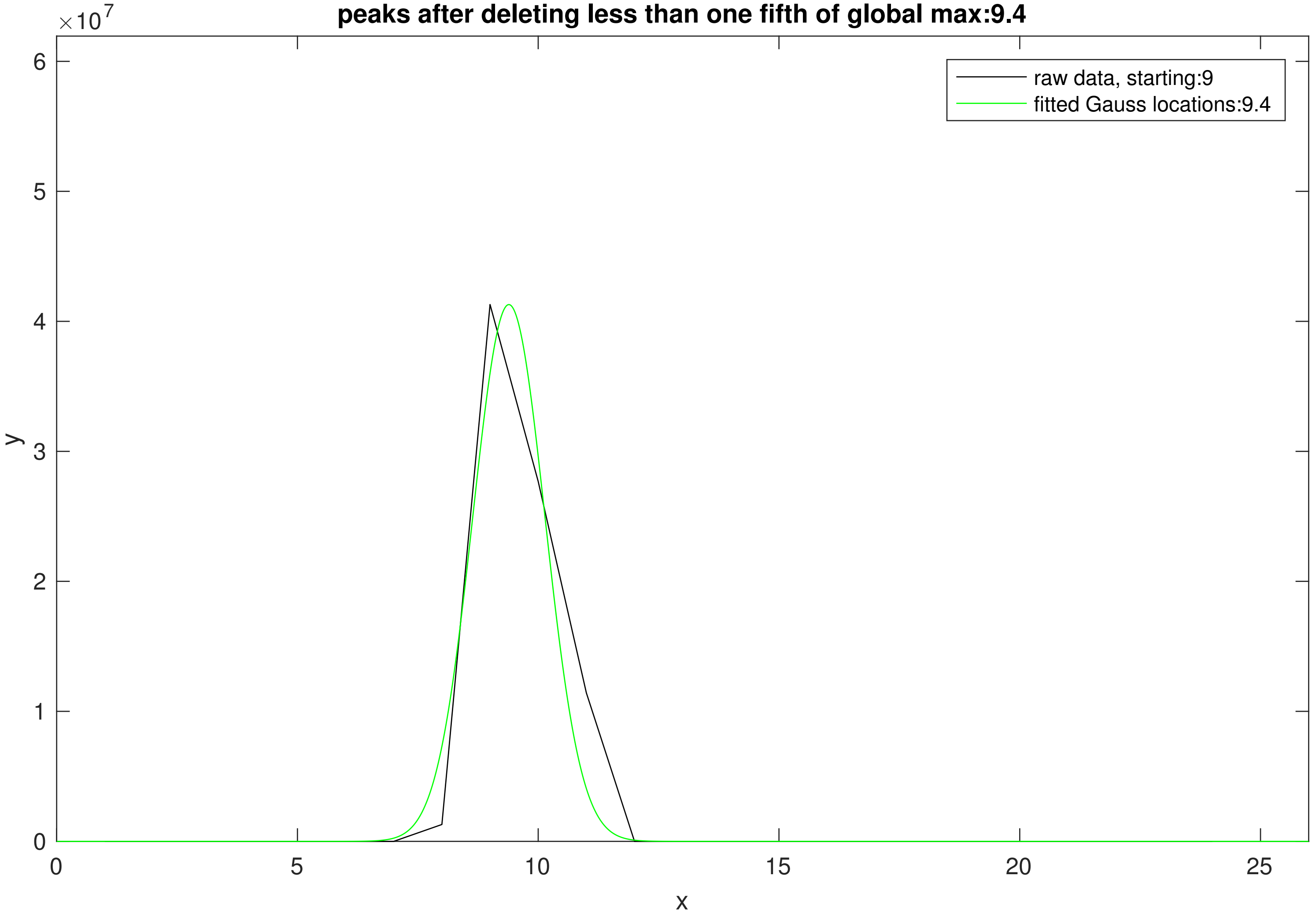

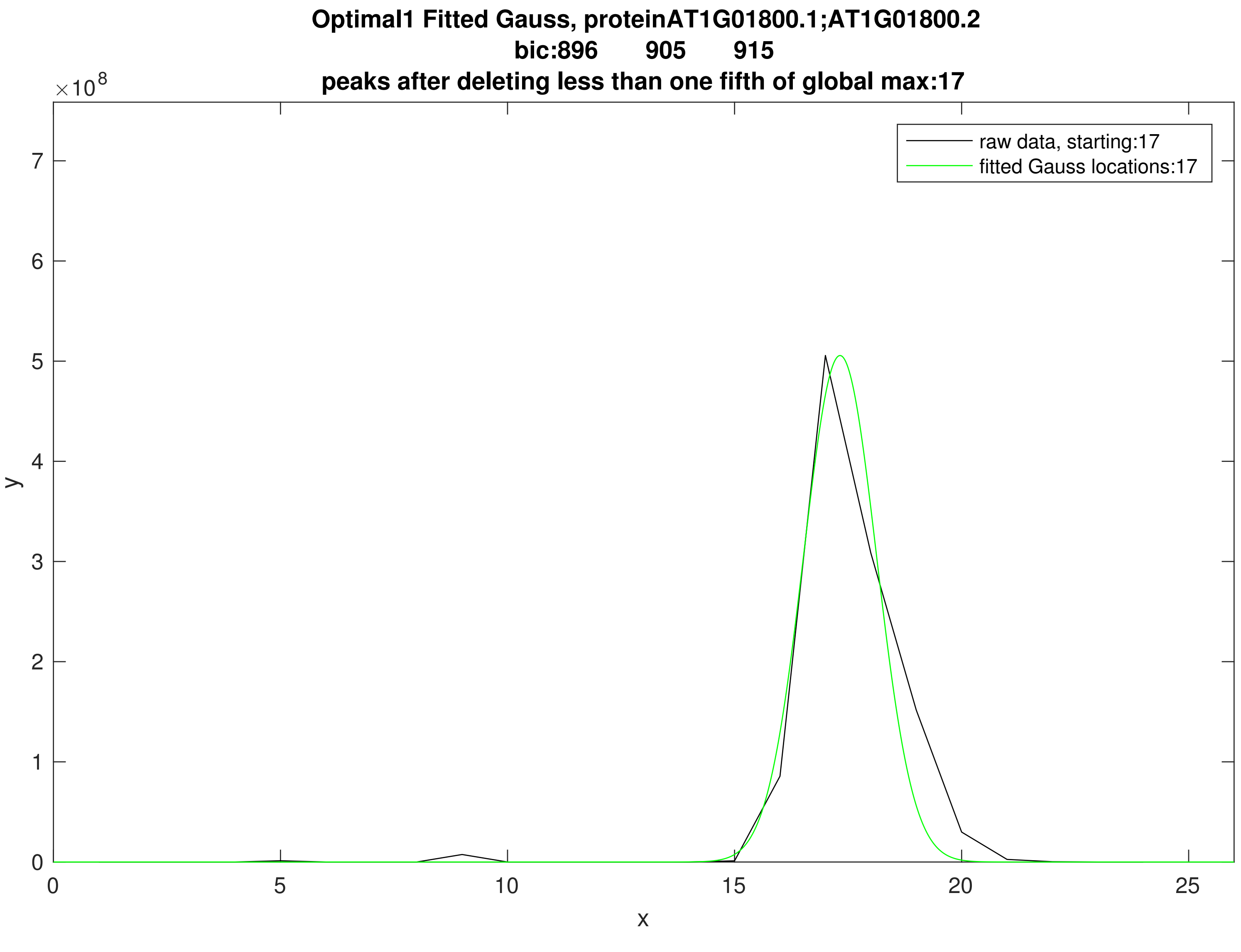
Optimal1 Fitted Gauss, proteinAT1G01800.1;AT1G01800.2
bic:896 905 915
peaks after deleting less than one fifth of global max:17
×10^8
raw data, starting:17
fitted Gauss locations:17
y
x
0
1
2
3
4
5
6
7
0
5
10
15
20
25

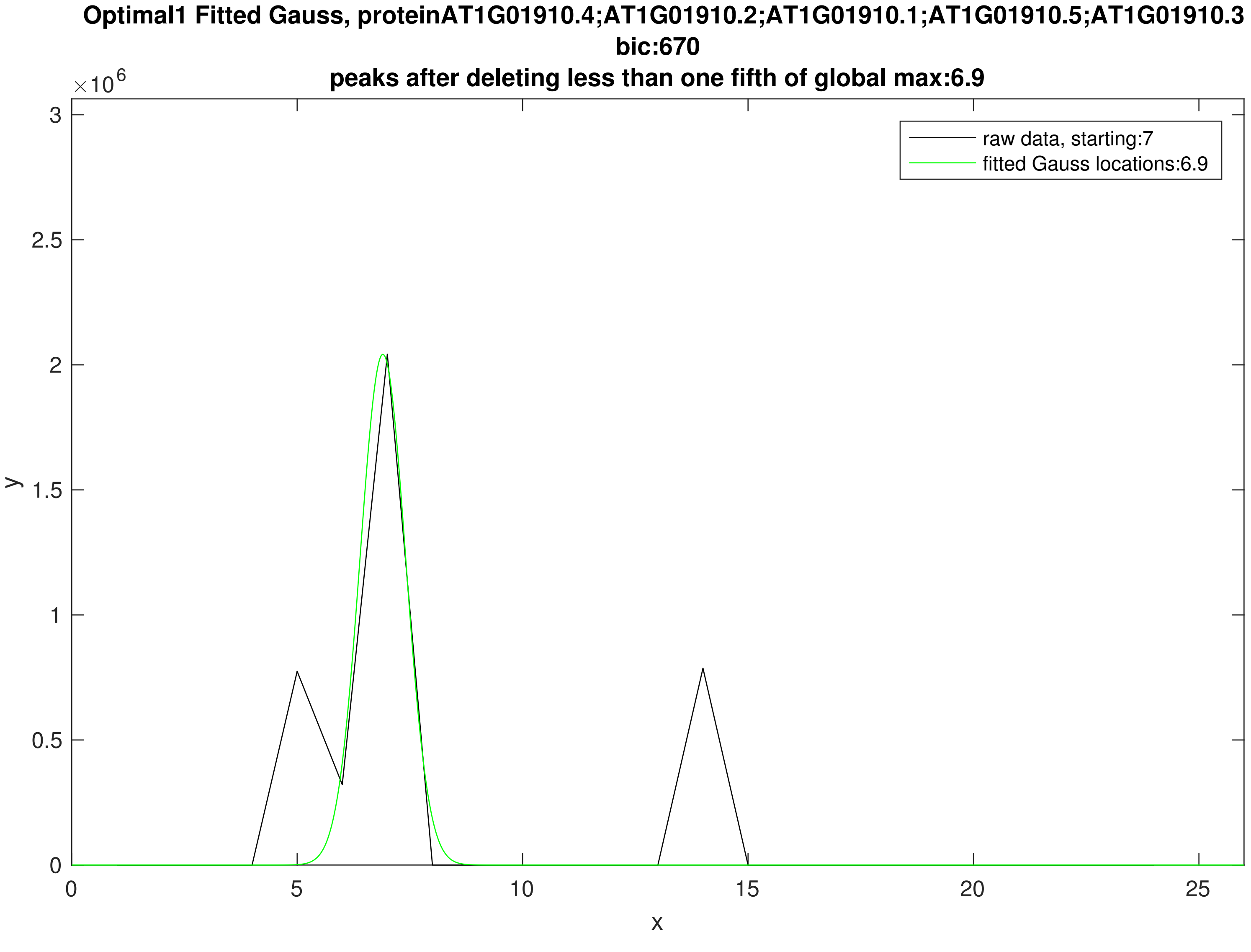

Optimal1 Fitted Gauss, proteinAT1G01910.4;AT1G01910.2;AT1G01910.1;AT1G01910.5;AT1G01910.3
bic:670
peaks after deleting less than one fifth of global max:6.9
×10^6
3
2.5
2
1.5
1
0.5
0
y
0
5
10
15
20
25
x
raw data, starting:7
fitted Gauss locations:6.9

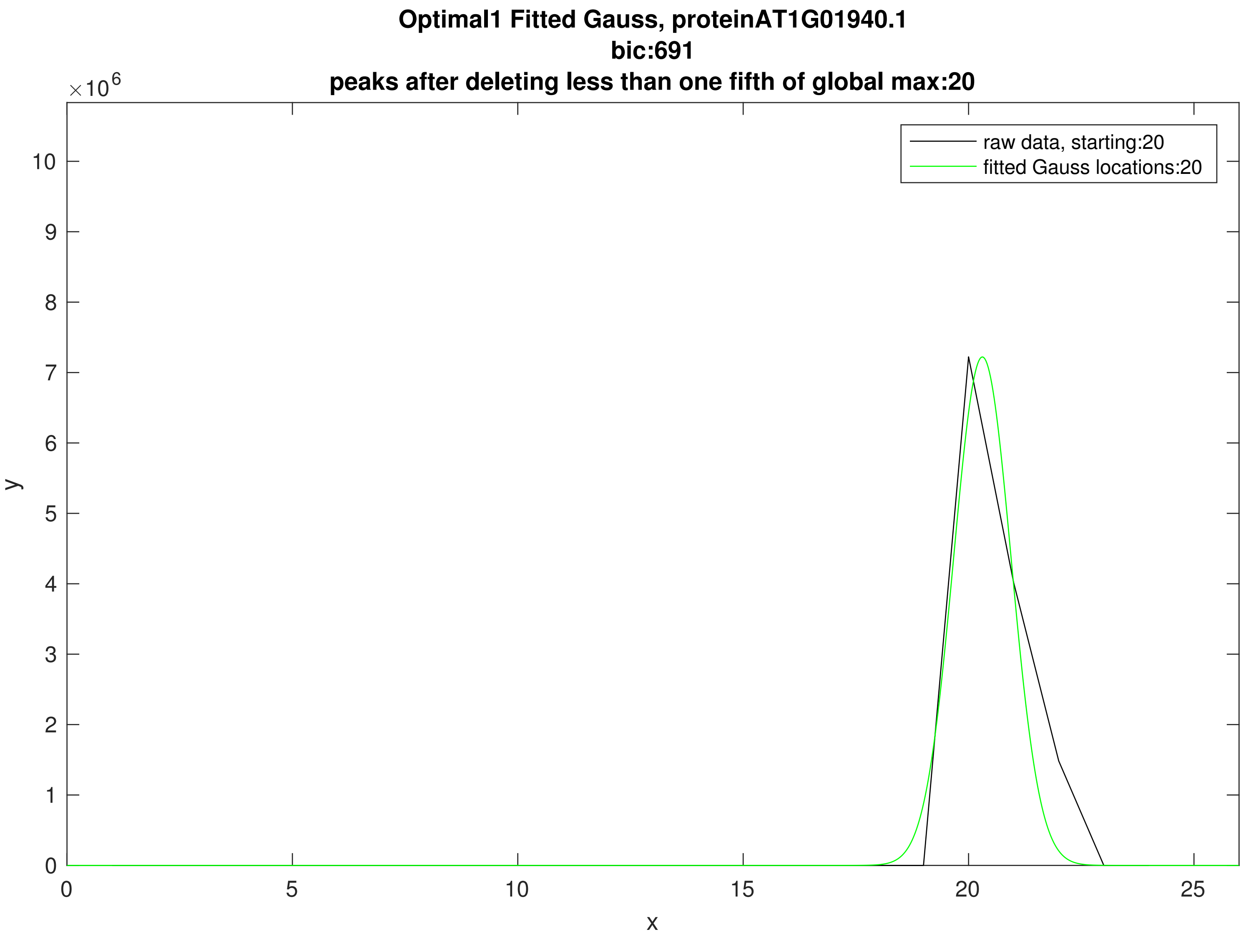
Optimal1 Fitted Gauss, proteinAT1G01940.1
bic:691
peaks after deleting less than one fifth of global max:20
×10^6
raw data, starting:20
fitted Gauss locations:20
y
x
0
1
2
3
4
5
6
7
8
9
10
0
5
10
15
20
25

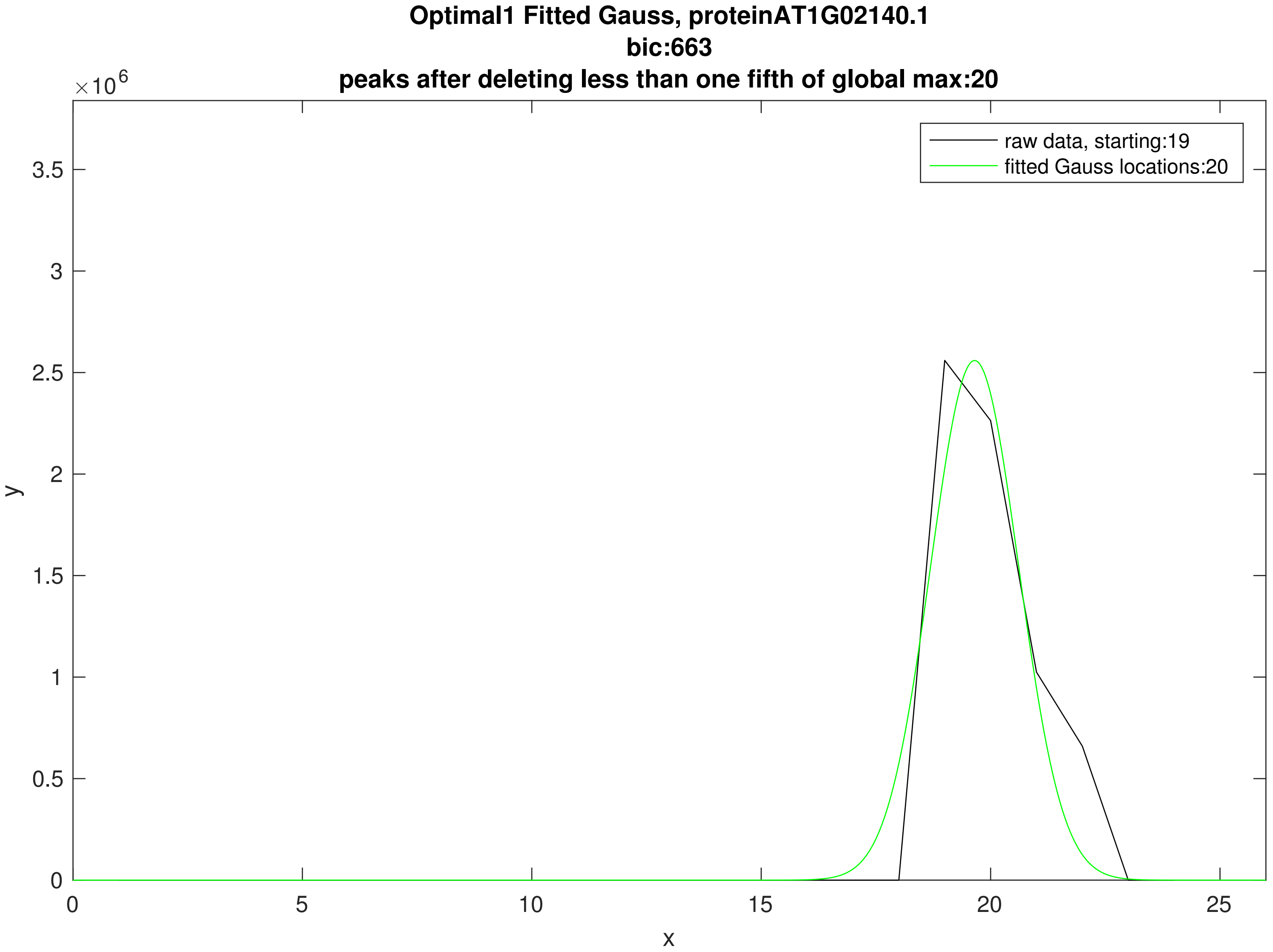
Optimal1 Fitted Gauss, proteinAT1G02140.1
bic:663
peaks after deleting less than one fifth of global max:20
×10^6
raw data, starting:19
fitted Gauss locations:20
3.5
3
2.5
2
1.5
1
0.5
0
y
0
5
10
15
20
25
x

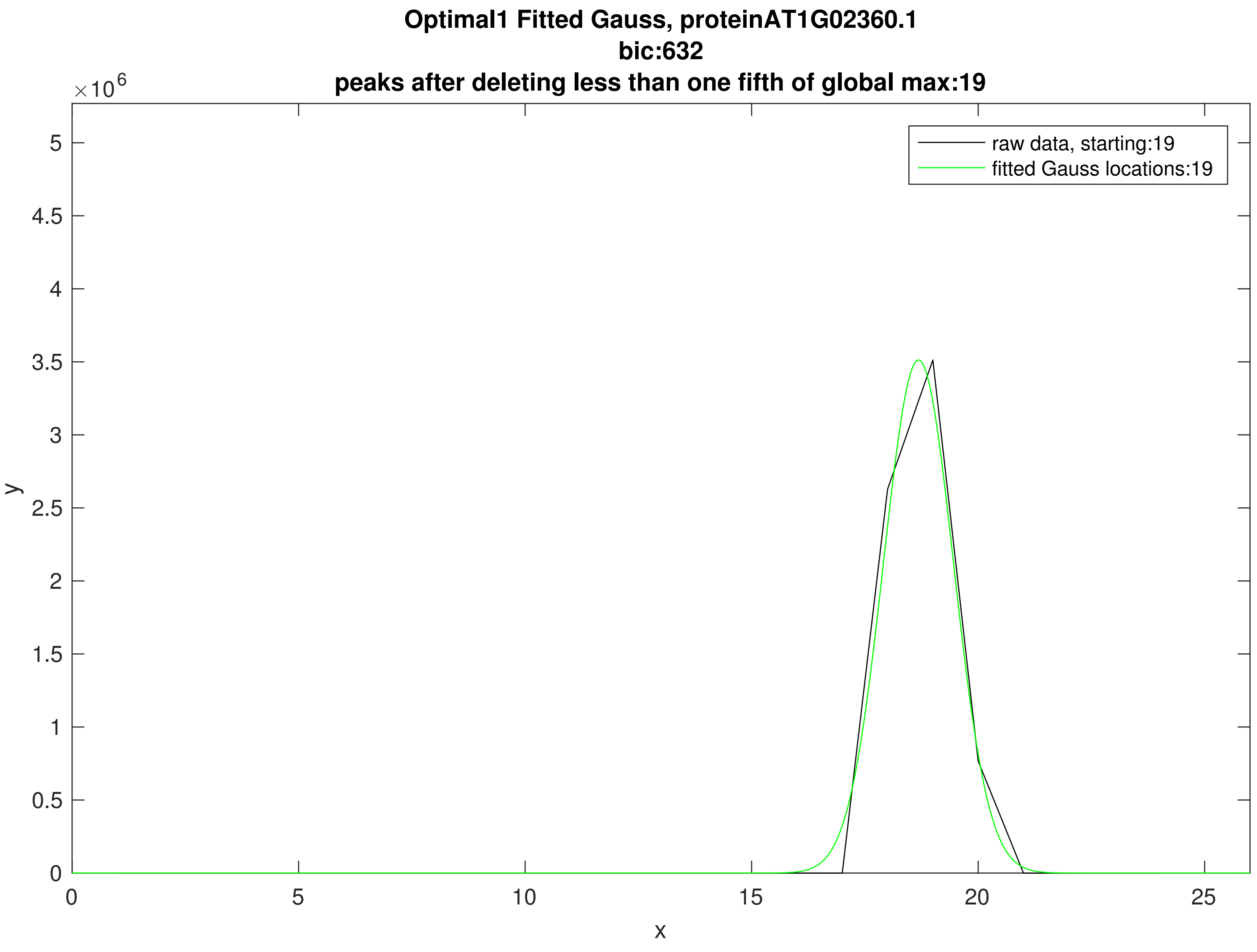
Optimal1 Fitted Gauss, proteinAT1G02360.1
bic:632
peaks after deleting less than one fifth of global max:19
×10^6
raw data, starting:19
fitted Gauss locations:19
y
x
0
0.5
1
1.5
2
2.5
3
3.5
4
4.5
5
0
5
10
15
20
25

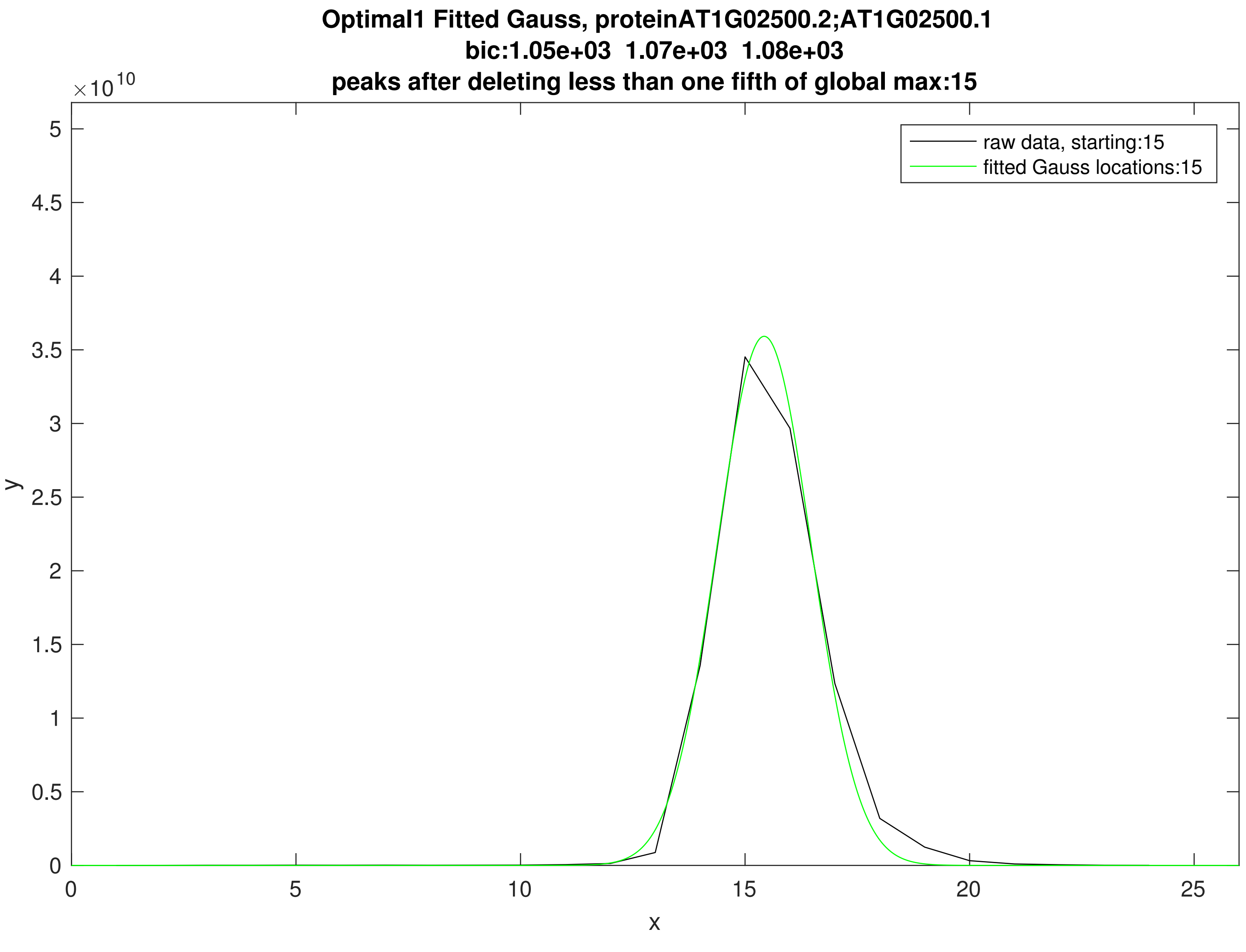

Optimal1 Fitted Gauss, proteinAT1G02500.2;AT1G02500.1
bic:1.05e+03  1.07e+03  1.08e+03
peaks after deleting less than one fifth of global max:15
×10^10
raw data, starting:15
fitted Gauss locations:15
y
x
0
0.5
1
1.5
2
2.5
3
3.5
4
4.5
5
0
5
10
15
20
25

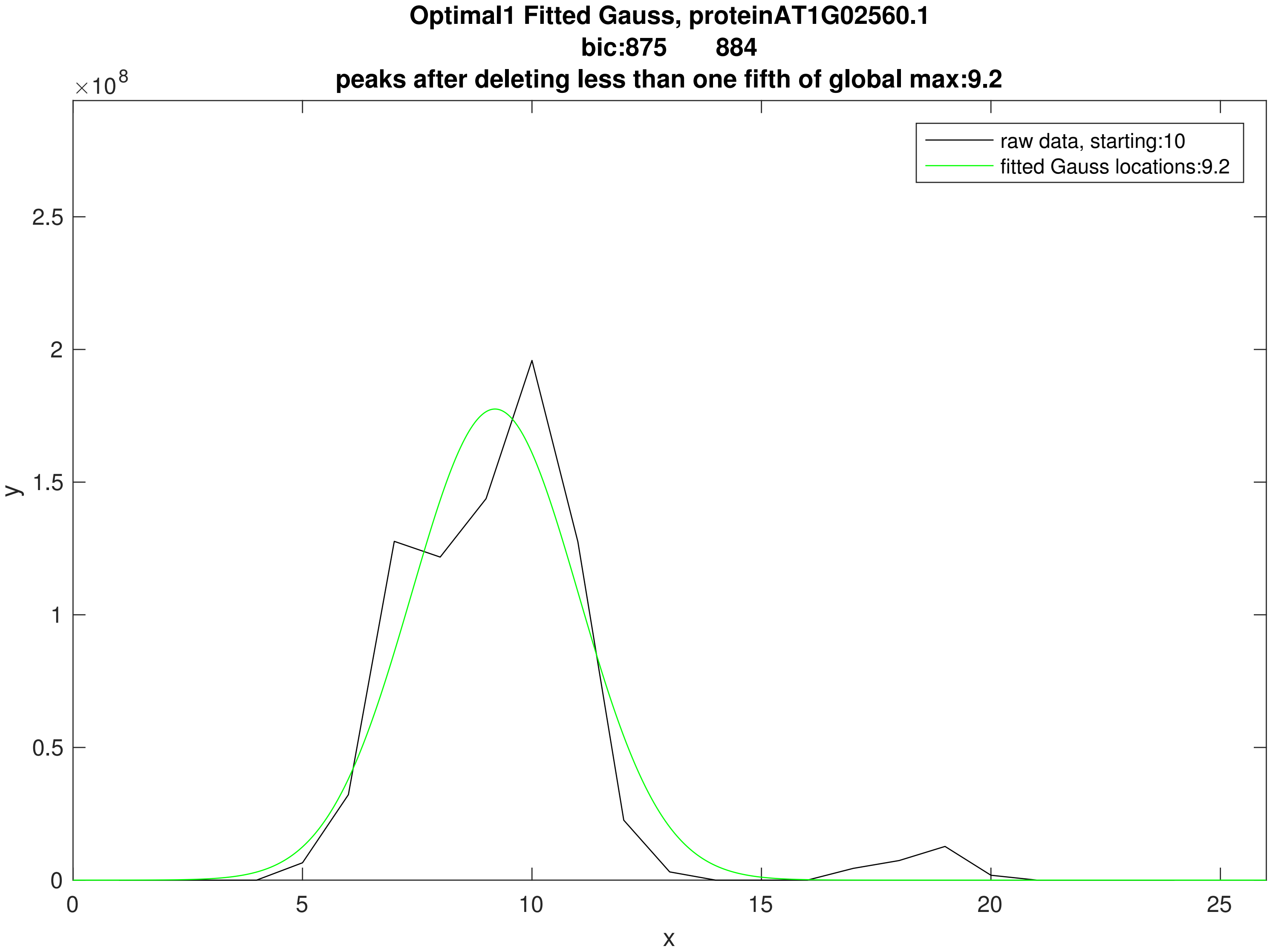

Optimal1 Fitted Gauss, proteinAT1G02560.1
bic:875 884
peaks after deleting less than one fifth of global max:9.2
×10^8
raw data, starting:10
fitted Gauss locations:9.2
y
x
0
0.5
1
1.5
2
2.5
0
5
10
15
20
25

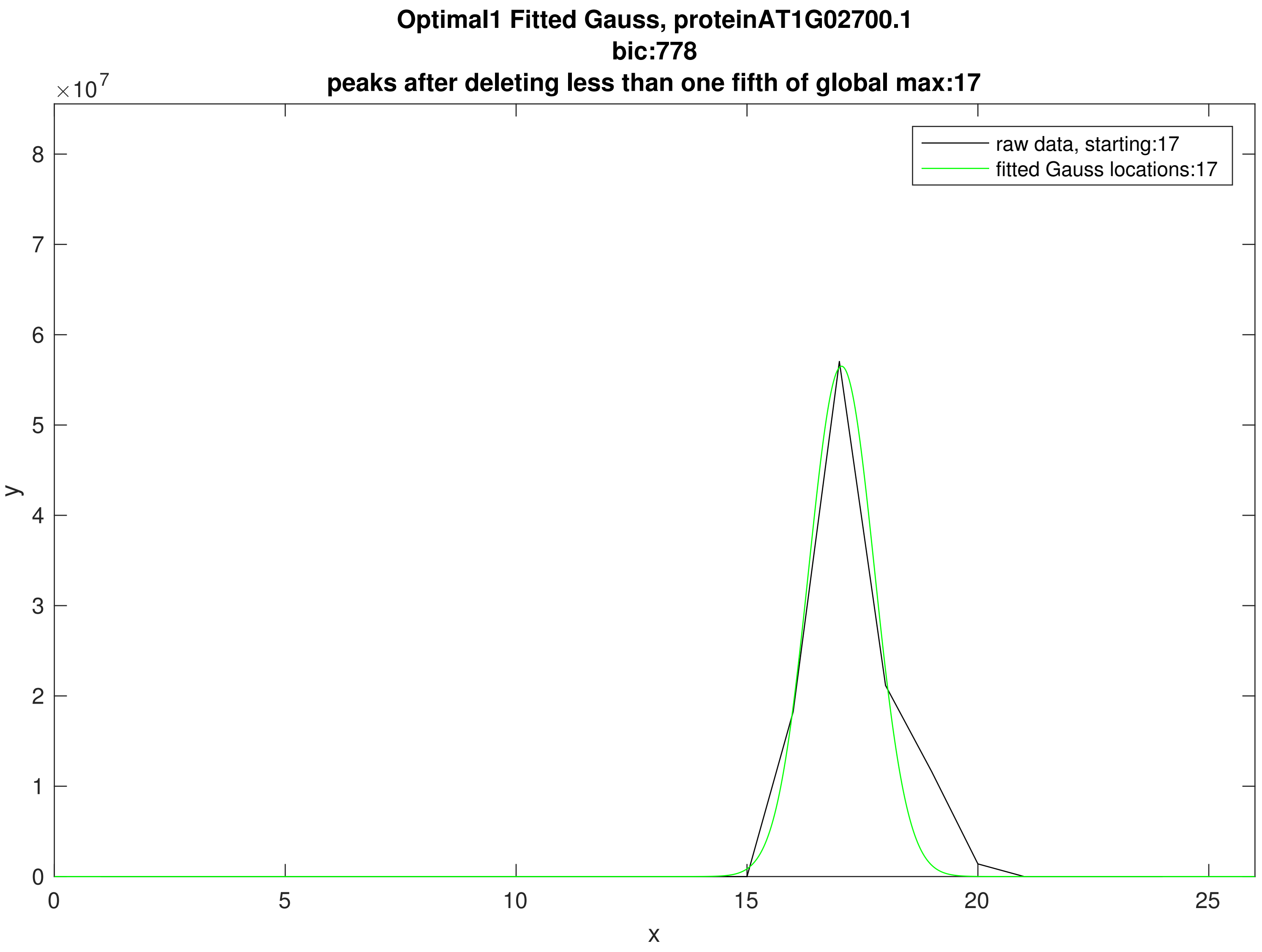
Optimal1 Fitted Gauss, proteinAT1G02700.1
bic:778
peaks after deleting less than one fifth of global max:17
×10^7
raw data, starting:17
fitted Gauss locations:17
y
x
0
1
2
3
4
5
6
7
8
0
5
10
15
20
25

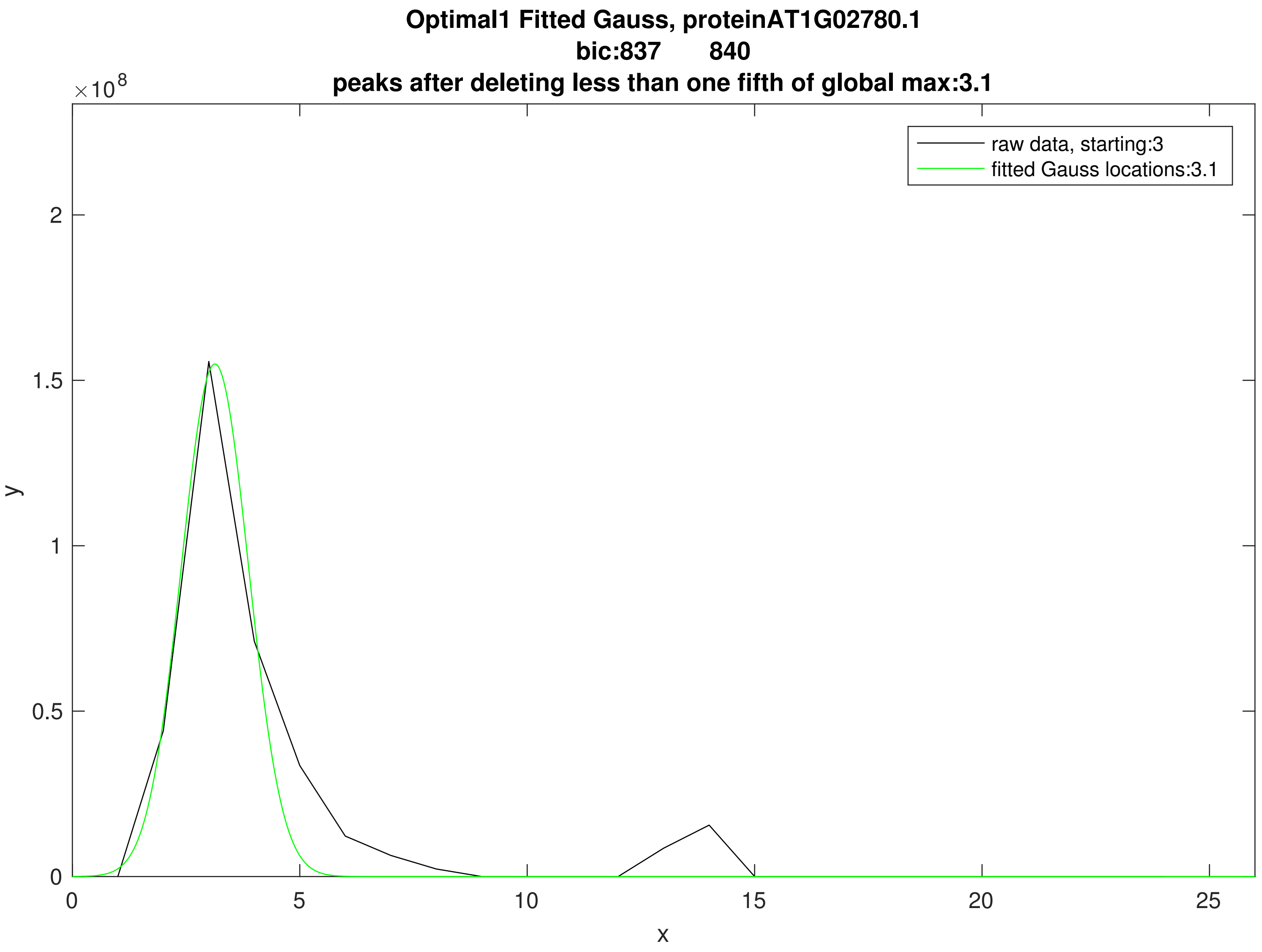
Optimal1 Fitted Gauss, proteinAT1G02780.1
bic:837 840
peaks after deleting less than one fifth of global max:3.1
×10^8
raw data, starting:3
fitted Gauss locations:3.1
y
x
0
0.5
1
1.5
2
0
5
10
15
20
25

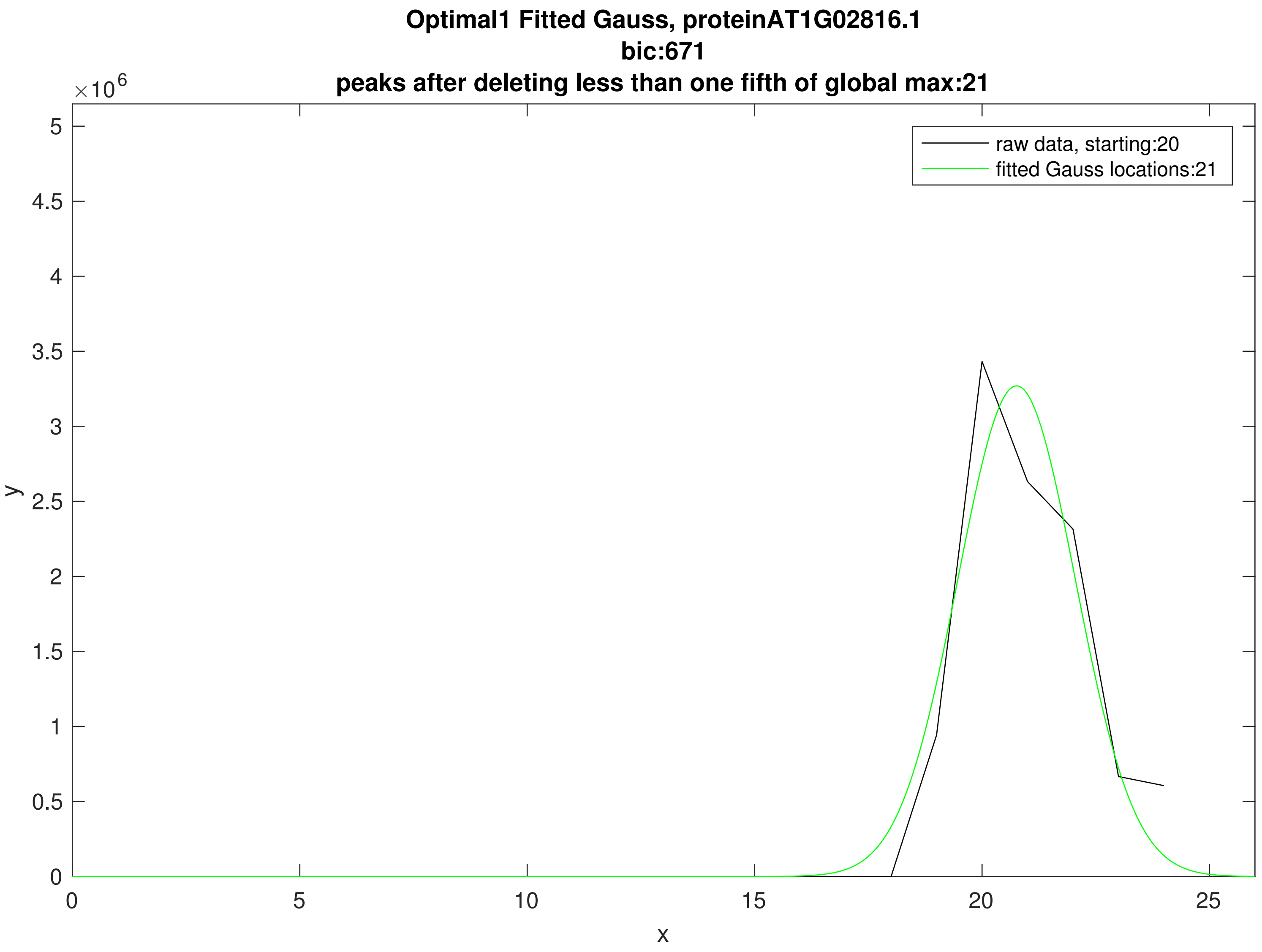

Optimal1 Fitted Gauss, proteinAT1G02816.1
bic:671
peaks after deleting less than one fifth of global max:21
×10^6
raw data, starting:20
fitted Gauss locations:21
y
x
0
0.5
1
1.5
2
2.5
3
3.5
4
4.5
5
0
5
10
15
20
25

Optimal1 Fitted Gauss, proteinAT1G02880.2;AT1G02880.3;AT1G02880.4;AT1G02880.1
bic:782
peaks after deleting less than one fifth of global max:15

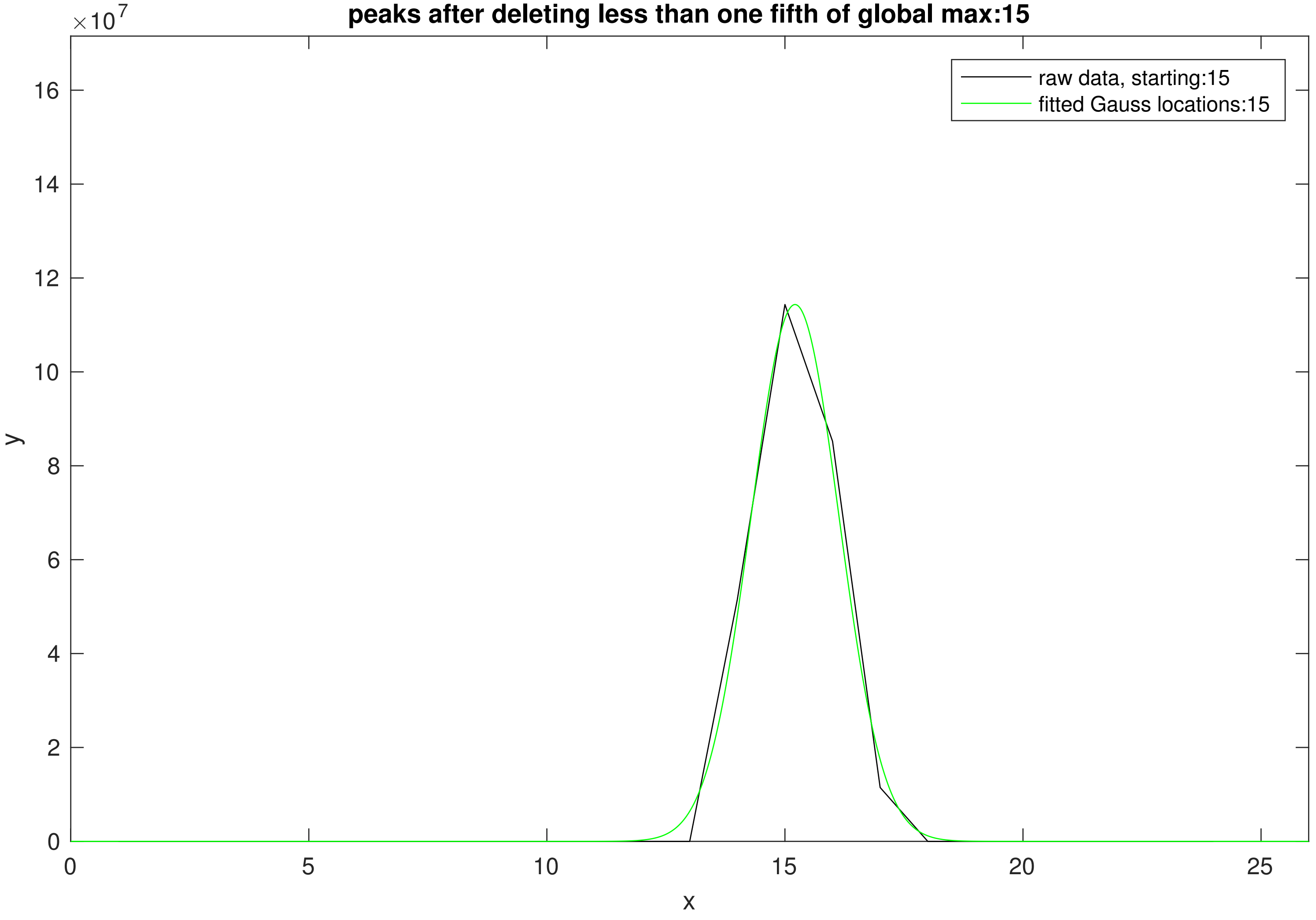

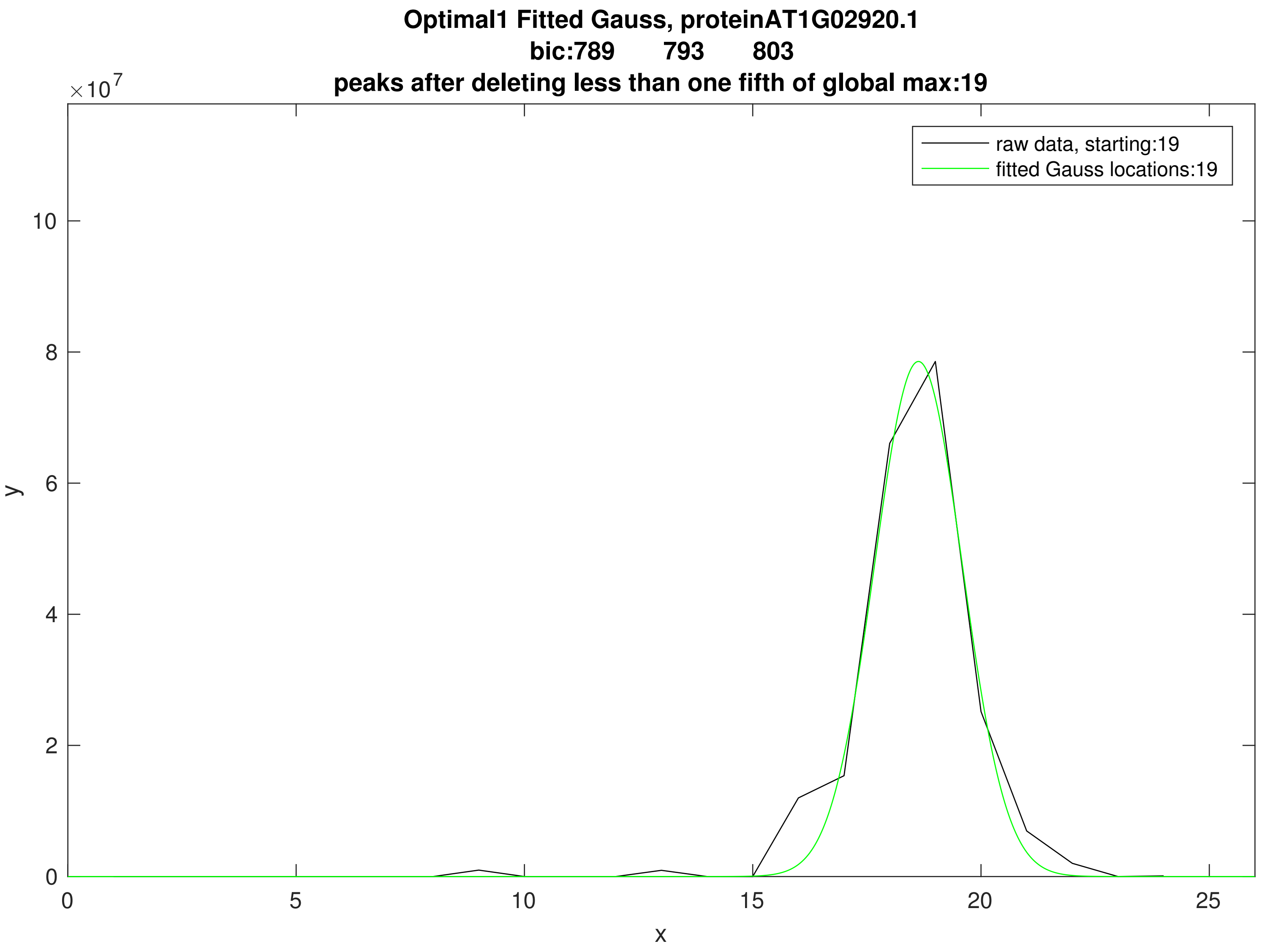
Optimal1 Fitted Gauss, proteinAT1G02920.1
bic:789 793 803
peaks after deleting less than one fifth of global max:19
×10^7
raw data, starting:19
fitted Gauss locations:19
10
8
6
4
2
0
y
0
5
10
15
20
25
x

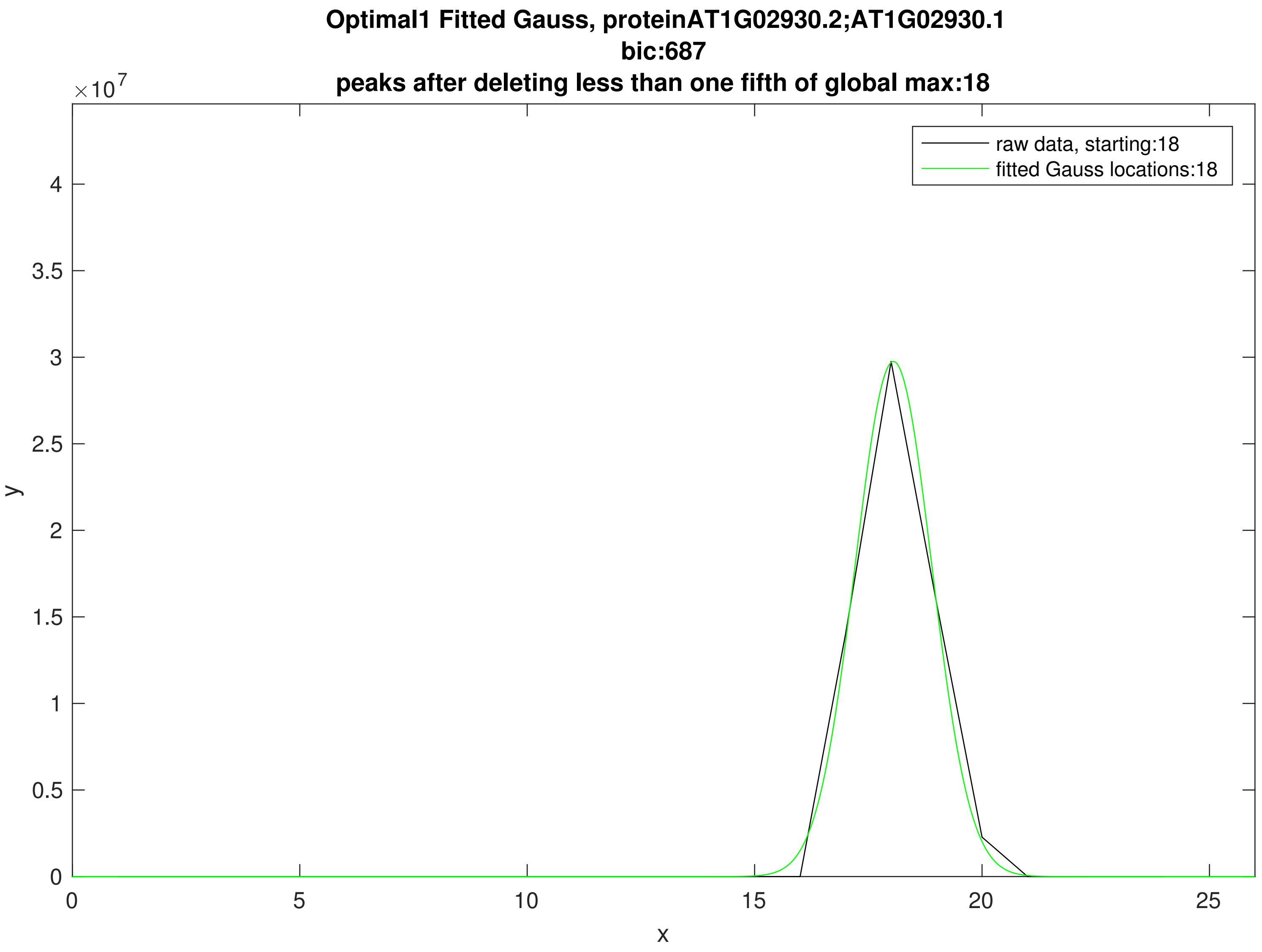

Optimal1 Fitted Gauss, proteinAT1G02930.2;AT1G02930.1
bic:687
peaks after deleting less than one fifth of global max:18
×10^7
raw data, starting:18
fitted Gauss locations:18
y
x
0
0.5
1
1.5
2
2.5
3
3.5
4
0
5
10
15
20
25

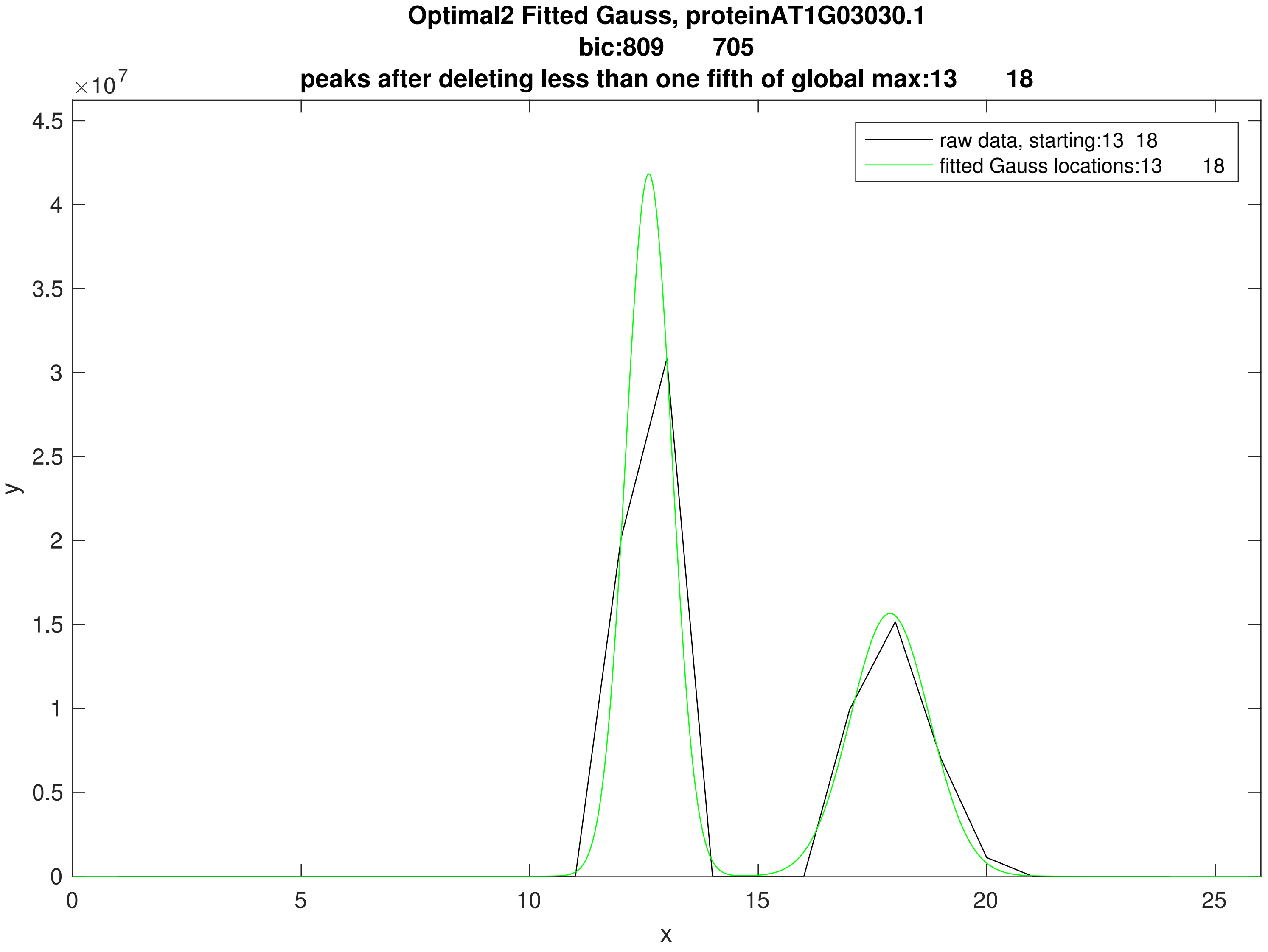

Optimal2 Fitted Gauss, proteinAT1G03030.1
bic:809 705
peaks after deleting less than one fifth of global max:13 18
×10^7
4.5
4
3.5
3
2.5
2
1.5
1
0.5
0
y
0
5
10
15
20
25
x
raw data, starting:13 18
fitted Gauss locations:13 18

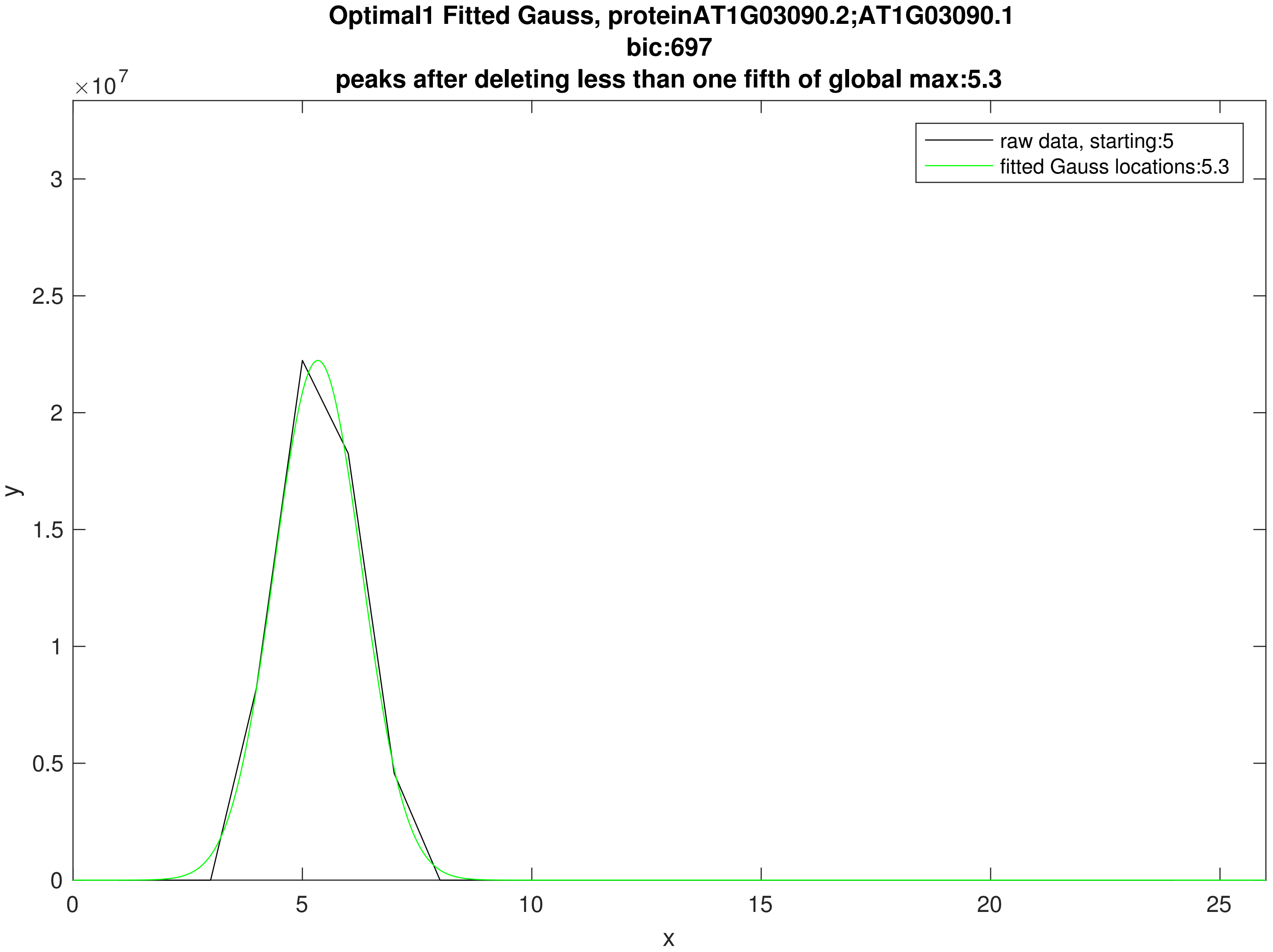

Optimal1 Fitted Gauss, proteinAT1G03090.2;AT1G03090.1
bic:697
peaks after deleting less than one fifth of global max:5.3
×10^7
raw data, starting:5
fitted Gauss locations:5.3
y
x
0
0.5
1
1.5
2
2.5
3
0
5
10
15
20
25

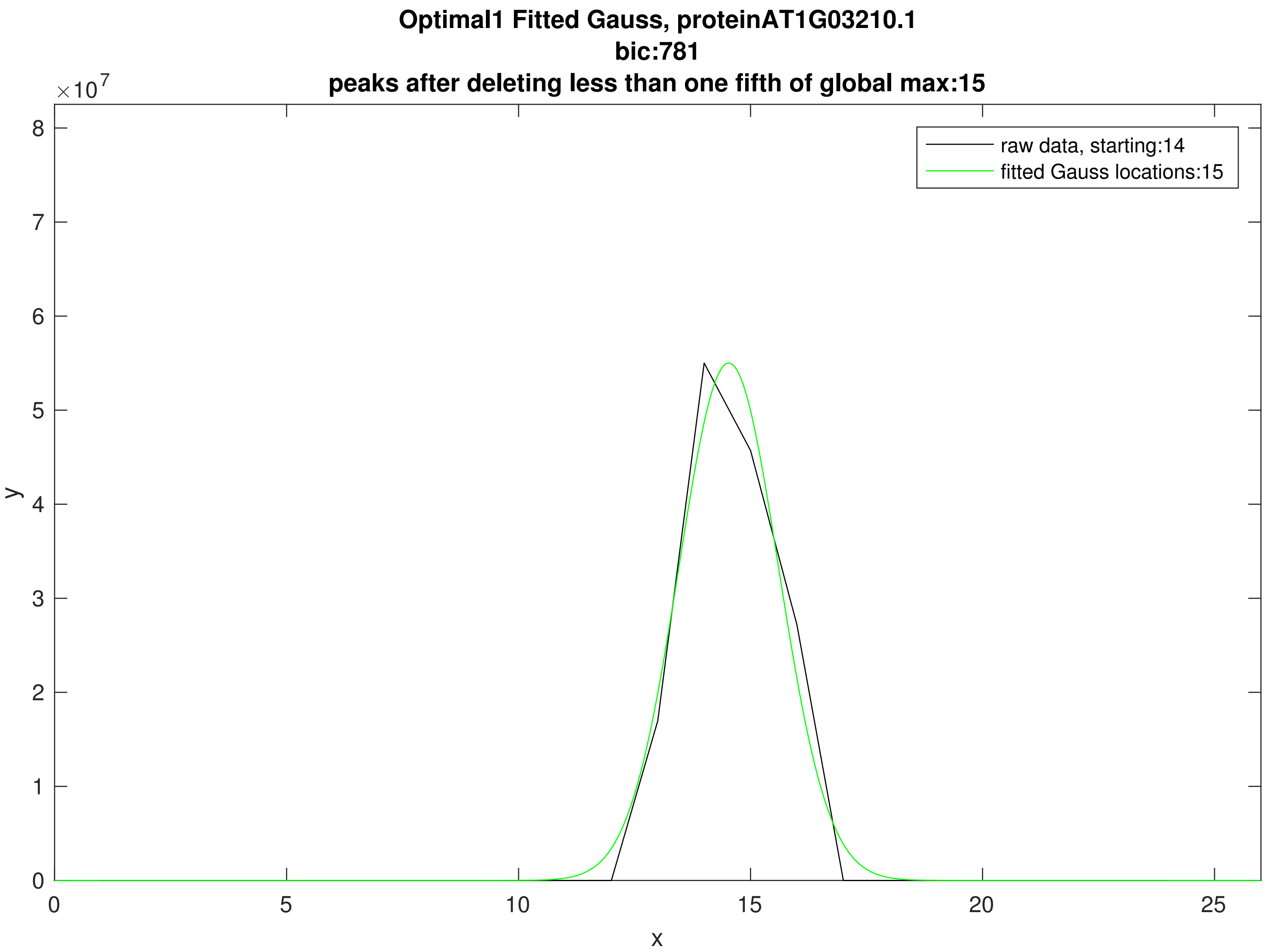
Optimal1 Fitted Gauss, proteinAT1G03210.1
bic:781
peaks after deleting less than one fifth of global max:15
×10^7
raw data, starting:14
fitted Gauss locations:15
y
x
0
1
2
3
4
5
6
7
8
0
5
10
15
20
25

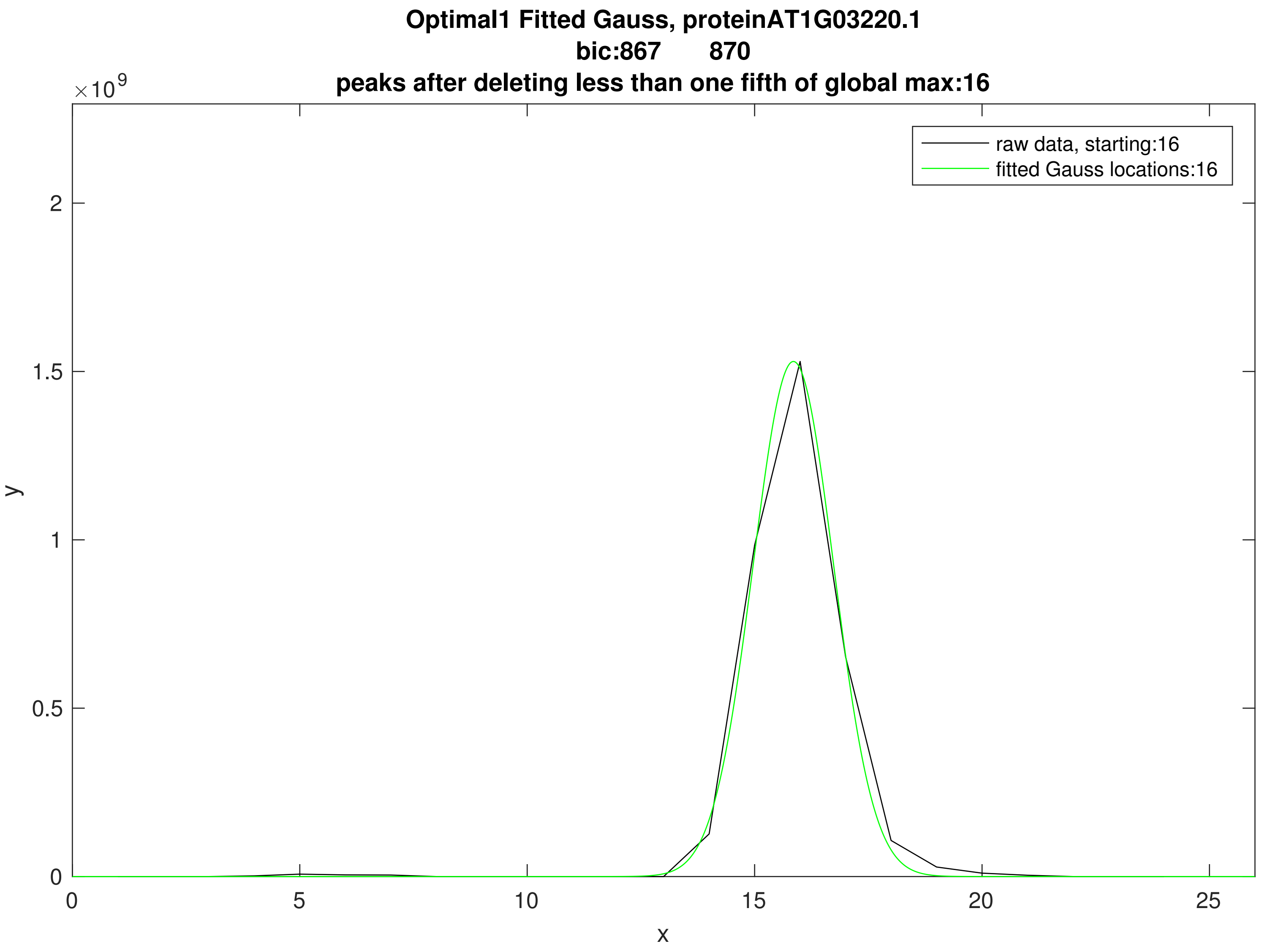

Optimal1 Fitted Gauss, proteinAT1G03220.1
bic:867 870
peaks after deleting less than one fifth of global max:16
×10^9
raw data, starting:16
fitted Gauss locations:16
2
1.5
1
0.5
0
y
0
5
10
15
20
25
x

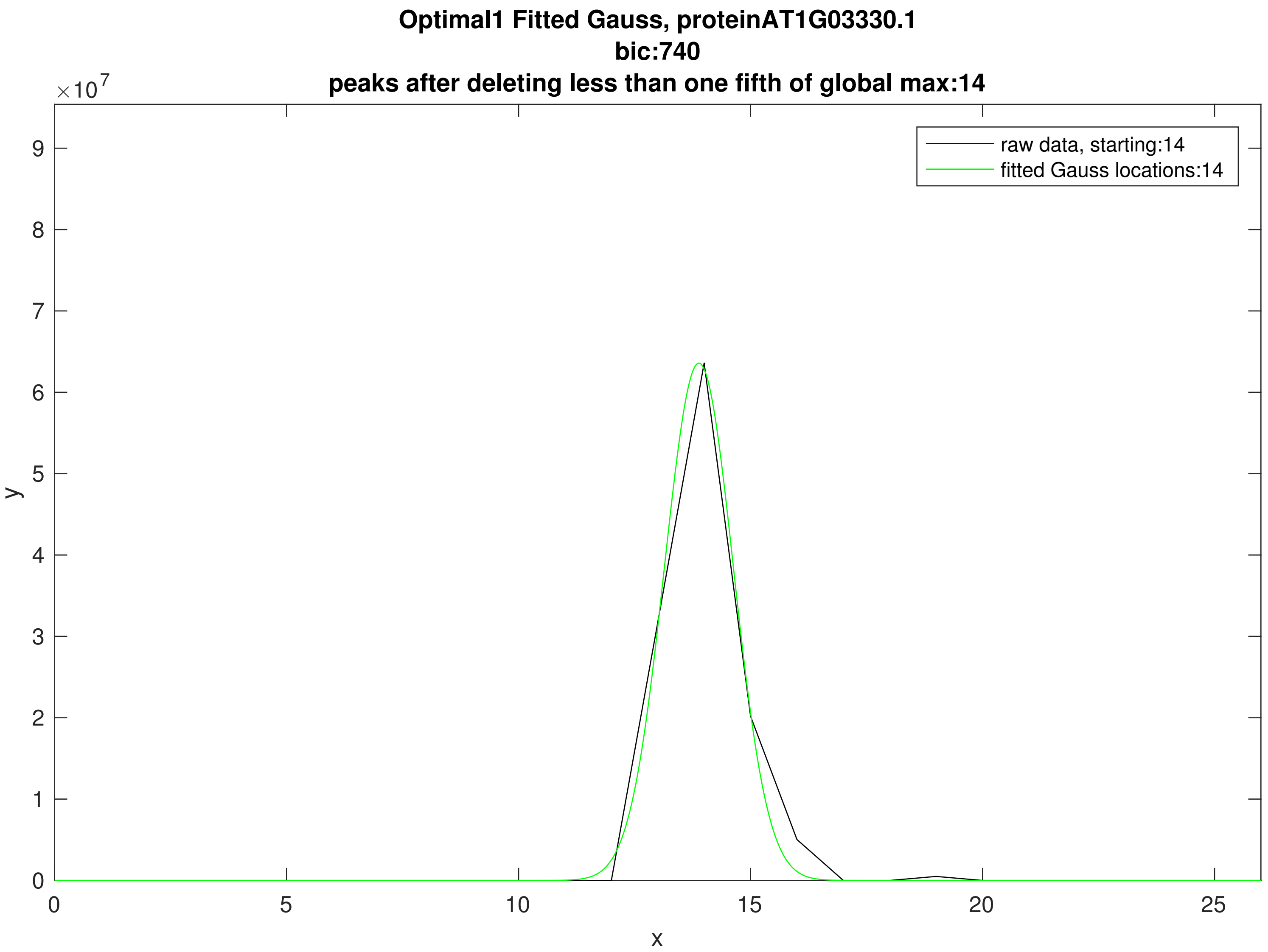

Optimal1 Fitted Gauss, proteinAT1G03330.1
bic:740
peaks after deleting less than one fifth of global max:14
×10^7
raw data, starting:14
fitted Gauss locations:14
y
x
0
1
2
3
4
5
6
7
8
9
0
5
10
15
20
25

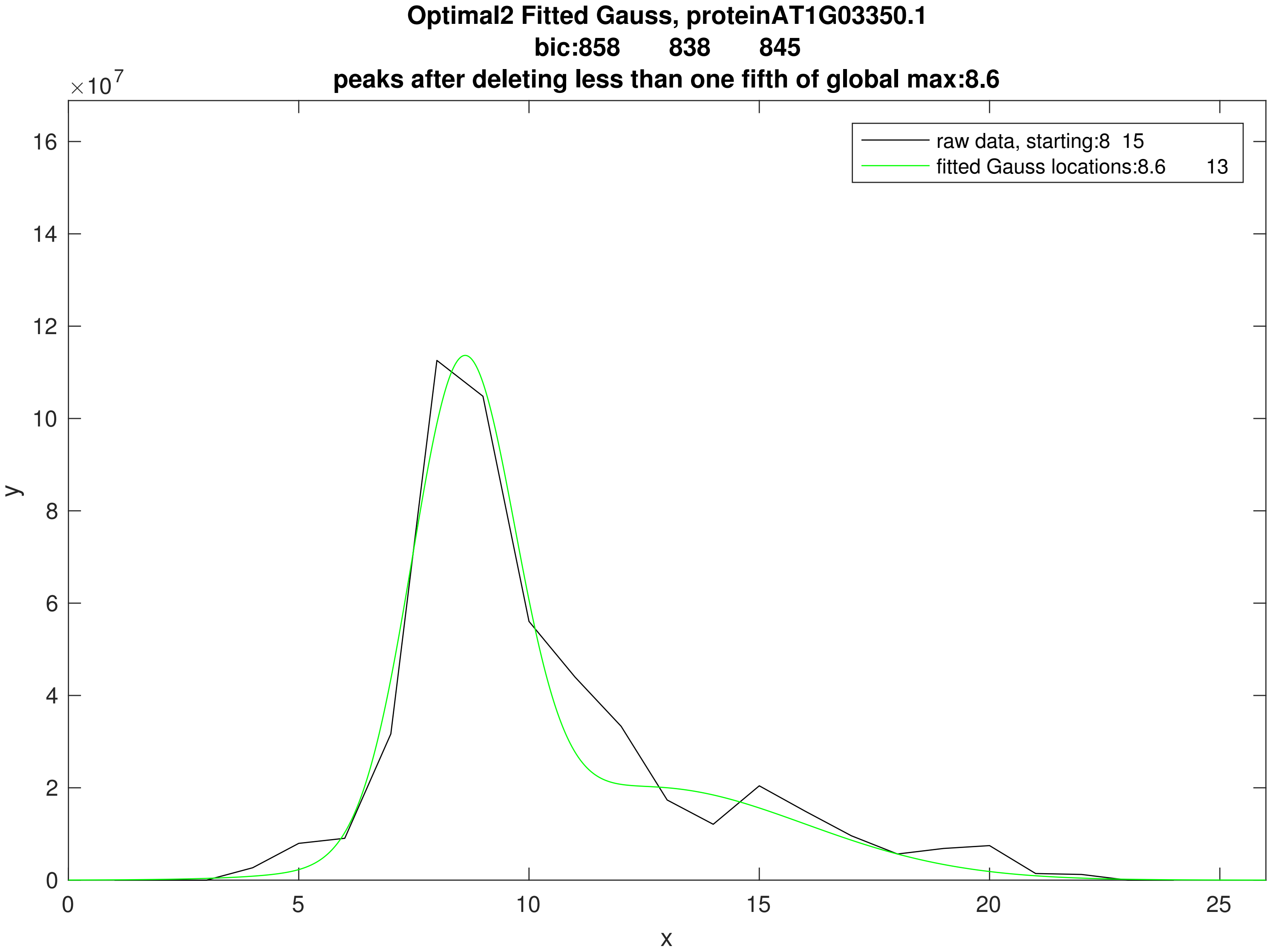

Optimal2 Fitted Gauss, proteinAT1G03350.1
bic:858 838 845
peaks after deleting less than one fifth of global max:8.6
×10^7
raw data, starting:8 15
fitted Gauss locations:8.6 13
y
x
0
2
4
6
8
10
12
14
16
0
5
10
15
20
25

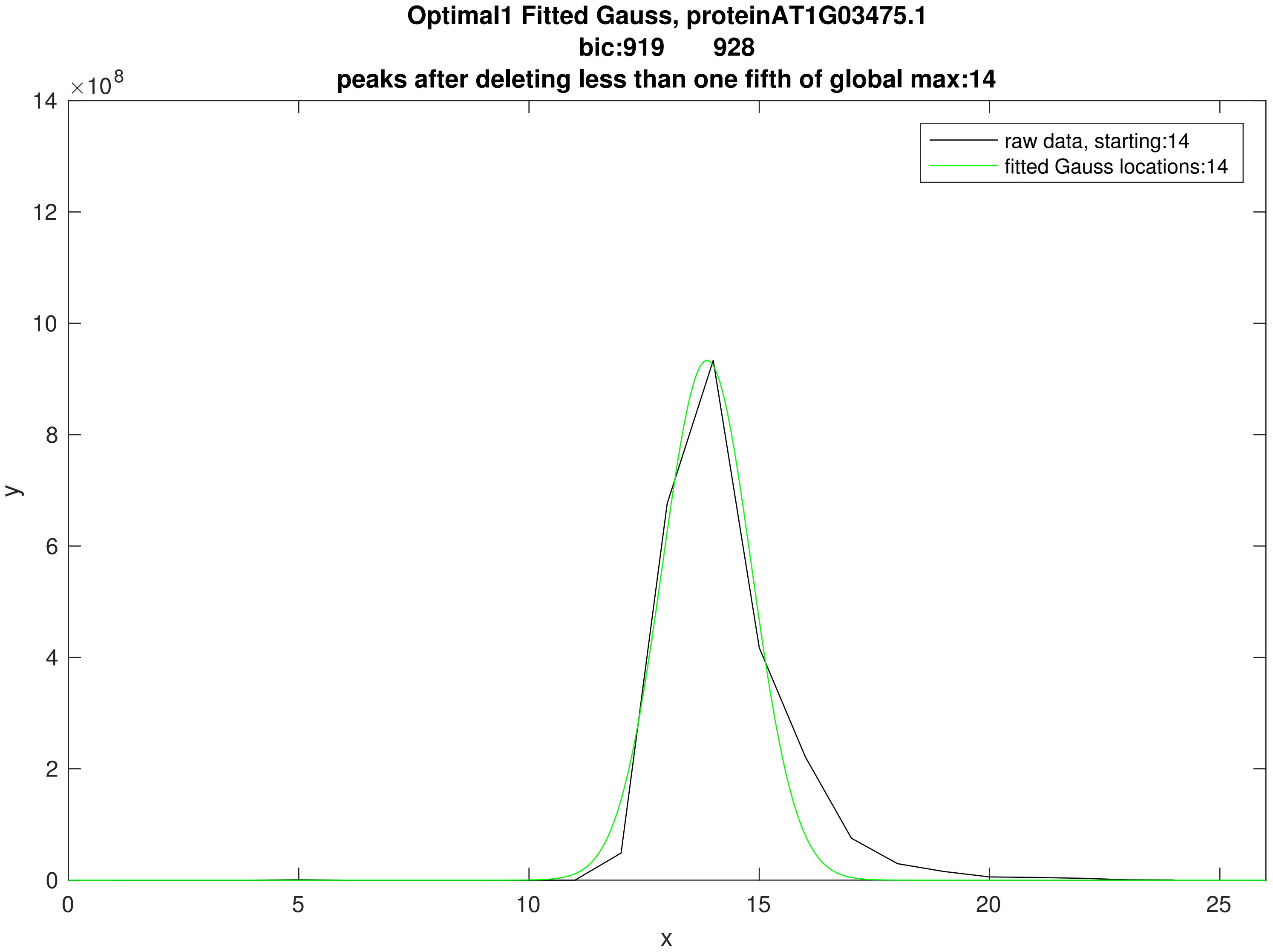
Optimal1 Fitted Gauss, proteinAT1G03475.1
bic:919 928
peaks after deleting less than one fifth of global max:14
×10^8
14
12
10
8
6
4
2
0
y
0
5
10
15
20
25
x
raw data, starting:14
fitted Gauss locations:14

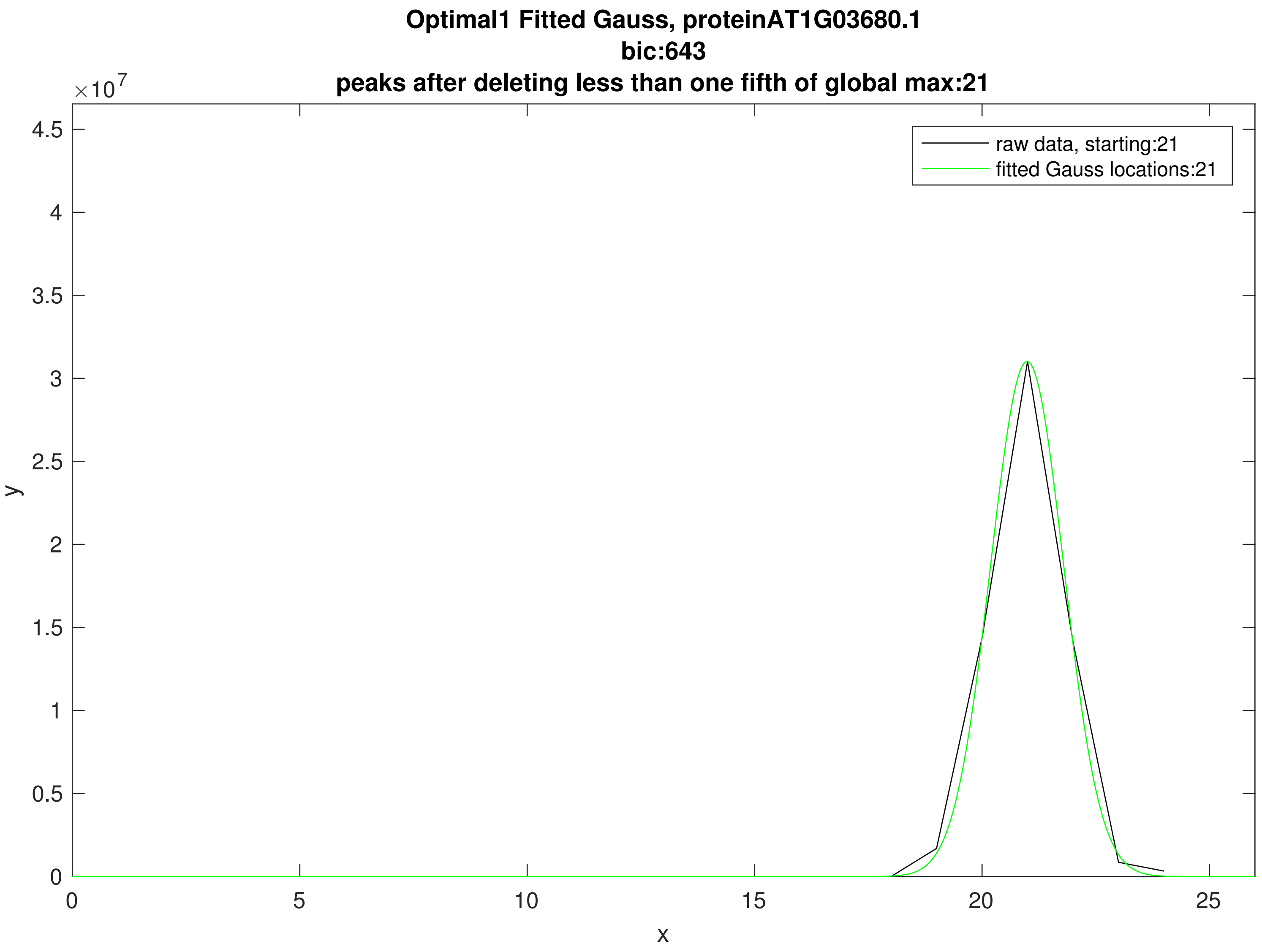

Optimal1 Fitted Gauss, proteinAT1G03680.1
bic:643
peaks after deleting less than one fifth of global max:21
×10^7
4.5
4
3.5
3
2.5
2
1.5
1
0.5
0
y
0
5
10
15
20
25
x
raw data, starting:21
fitted Gauss locations:21

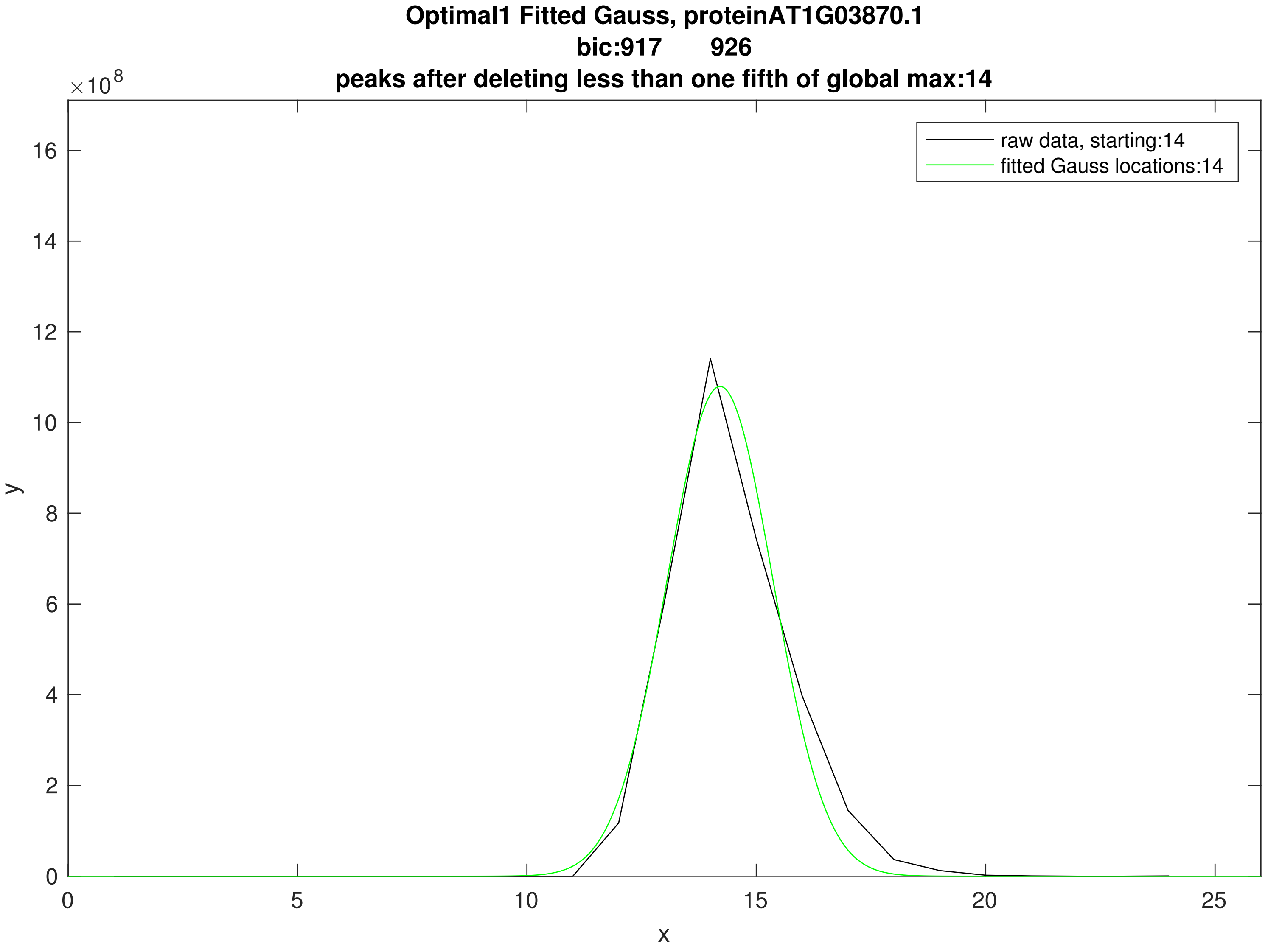
Optimal1 Fitted Gauss, proteinAT1G03870.1
bic:917 926
peaks after deleting less than one fifth of global max:14
raw data, starting:14
fitted Gauss locations:14
y
x
×10^8
0
2
4
6
8
10
12
14
16
0
5
10
15
20
25

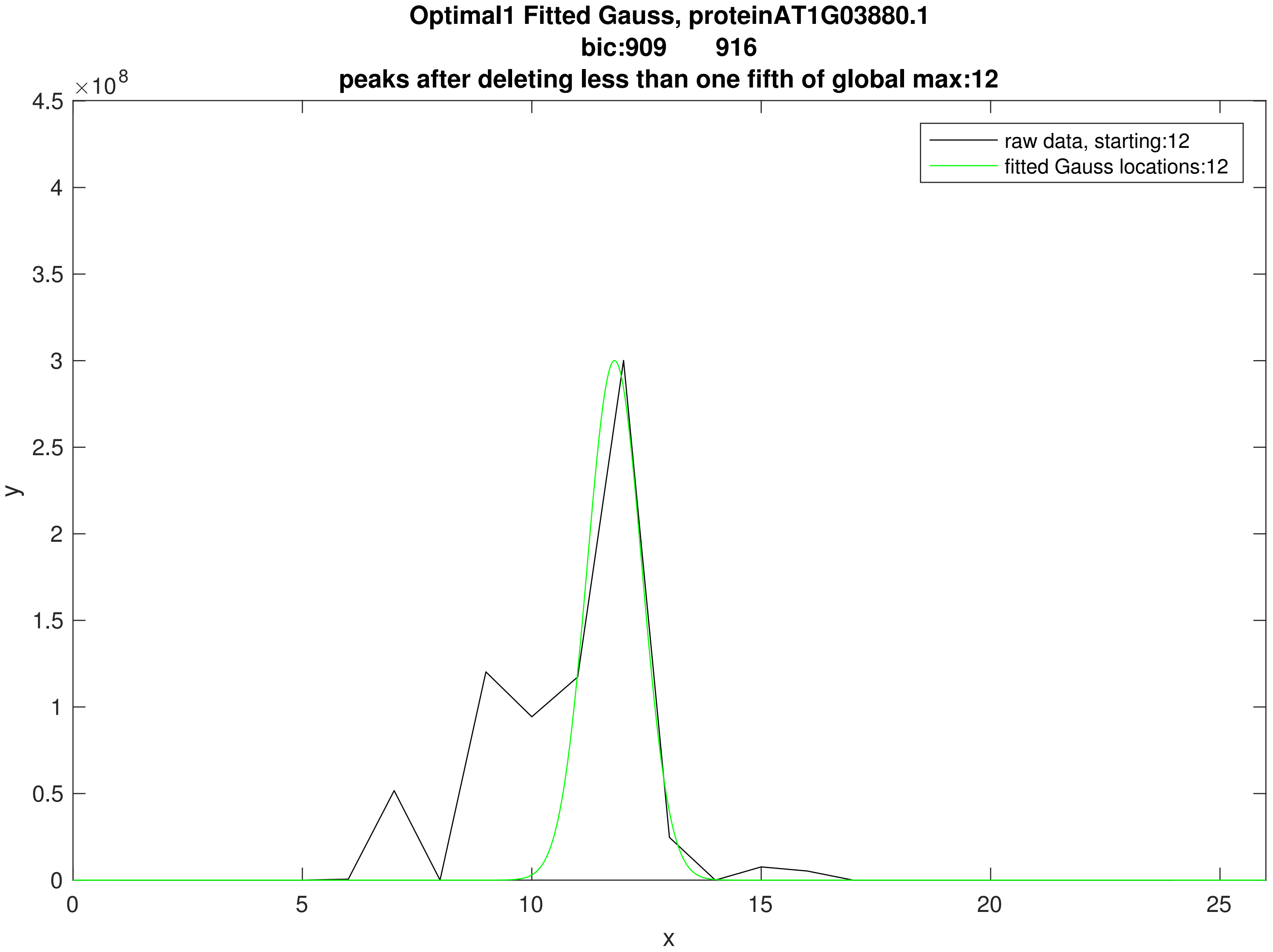

Optimal1 Fitted Gauss, proteinAT1G03880.1
bic:909 916
peaks after deleting less than one fifth of global max:12
×10^8
4.5
4
3.5
3
2.5
2
1.5
1
0.5
0
y
0
5
10
15
20
25
x
raw data, starting:12
fitted Gauss locations:12

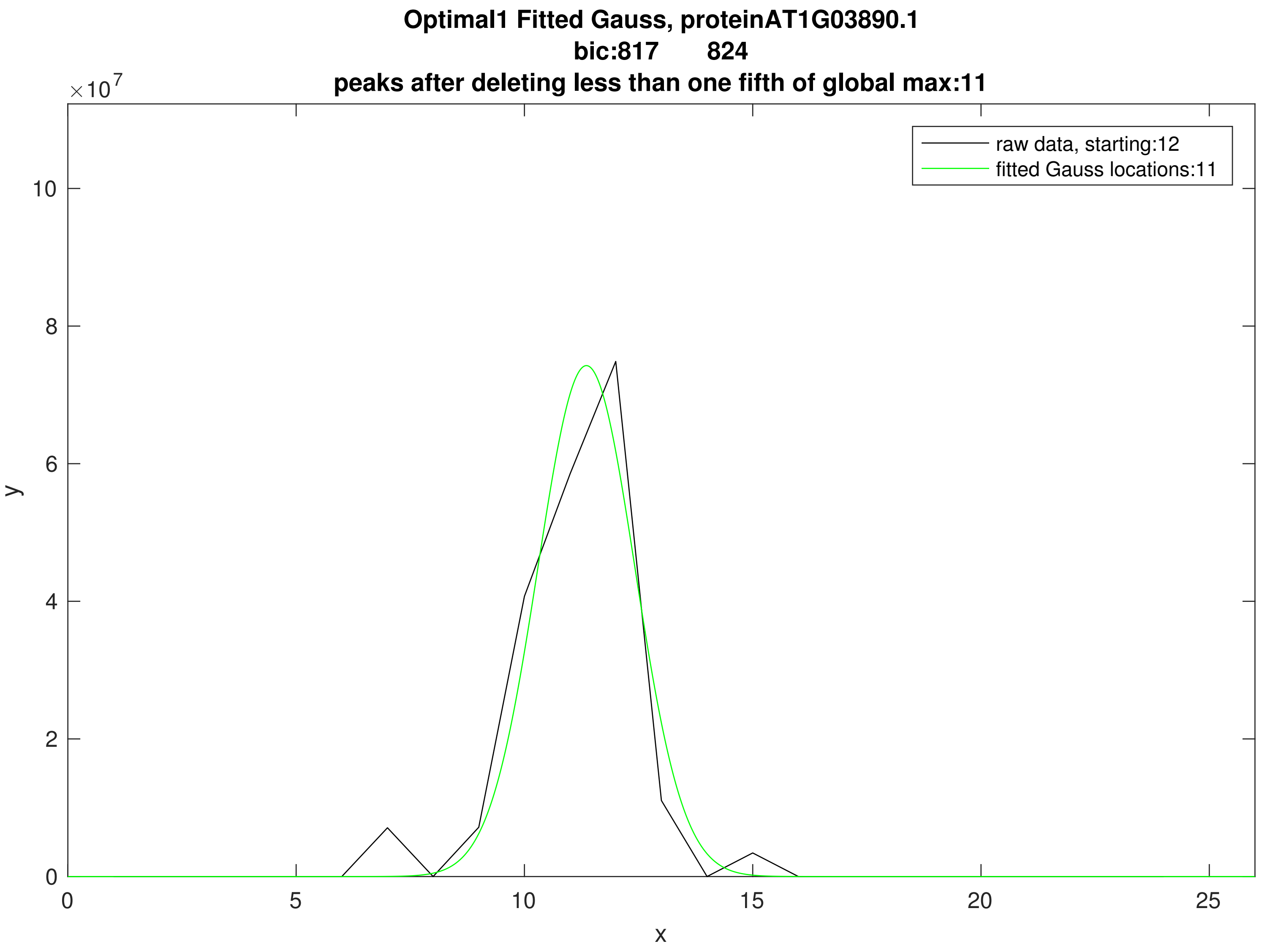
Optimal1 Fitted Gauss, proteinAT1G03890.1
bic:817 824
peaks after deleting less than one fifth of global max:11
×10^7
raw data, starting:12
fitted Gauss locations:11
y
x
0
2
4
6
8
10
0
5
10
15
20
25

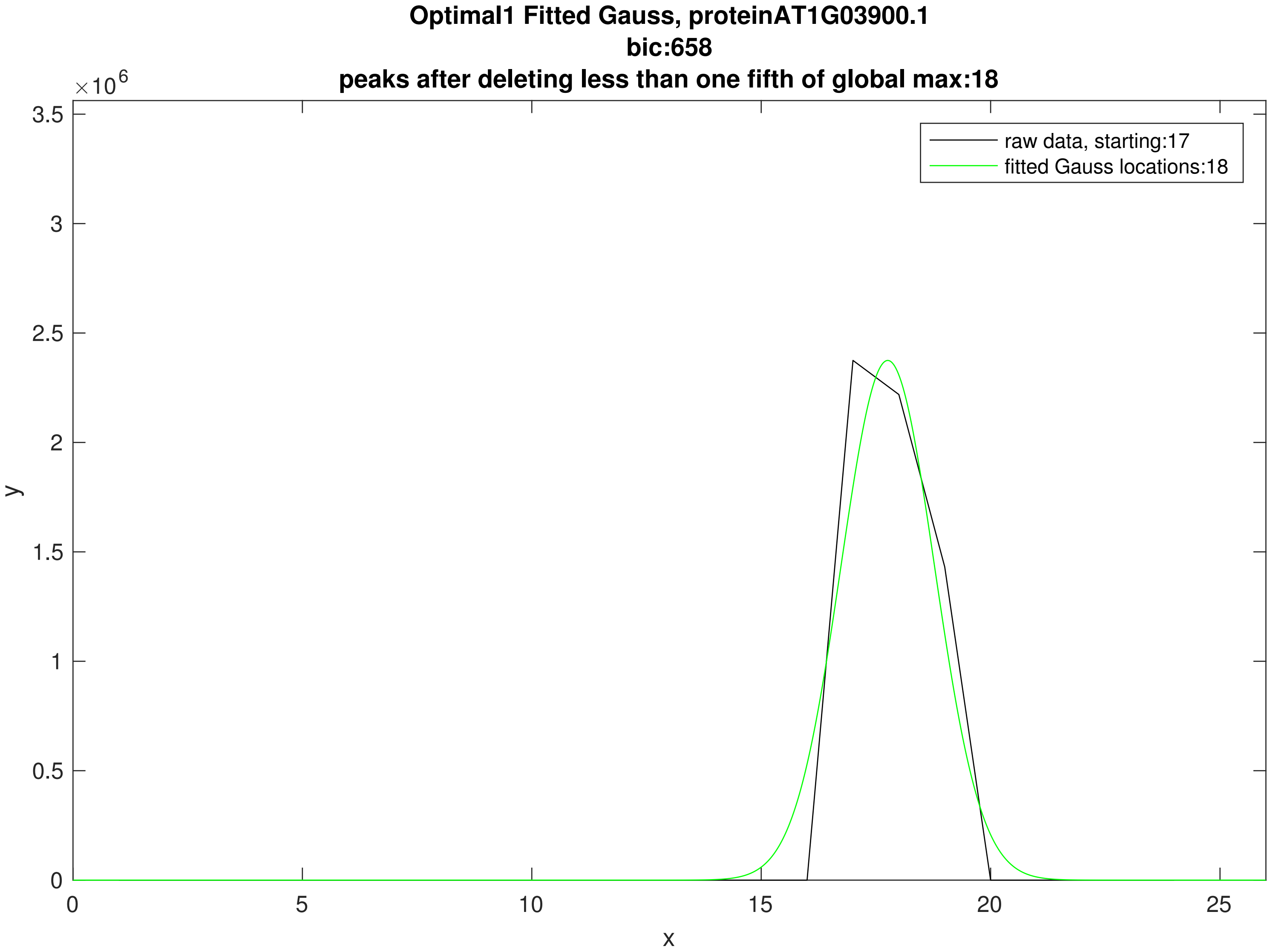

Optimal1 Fitted Gauss, proteinAT1G03900.1
bic:658
peaks after deleting less than one fifth of global max:18
×10^6
3.5
3
2.5
2
1.5
1
0.5
0
y
0
5
10
15
20
25
x
raw data, starting:17
fitted Gauss locations:18

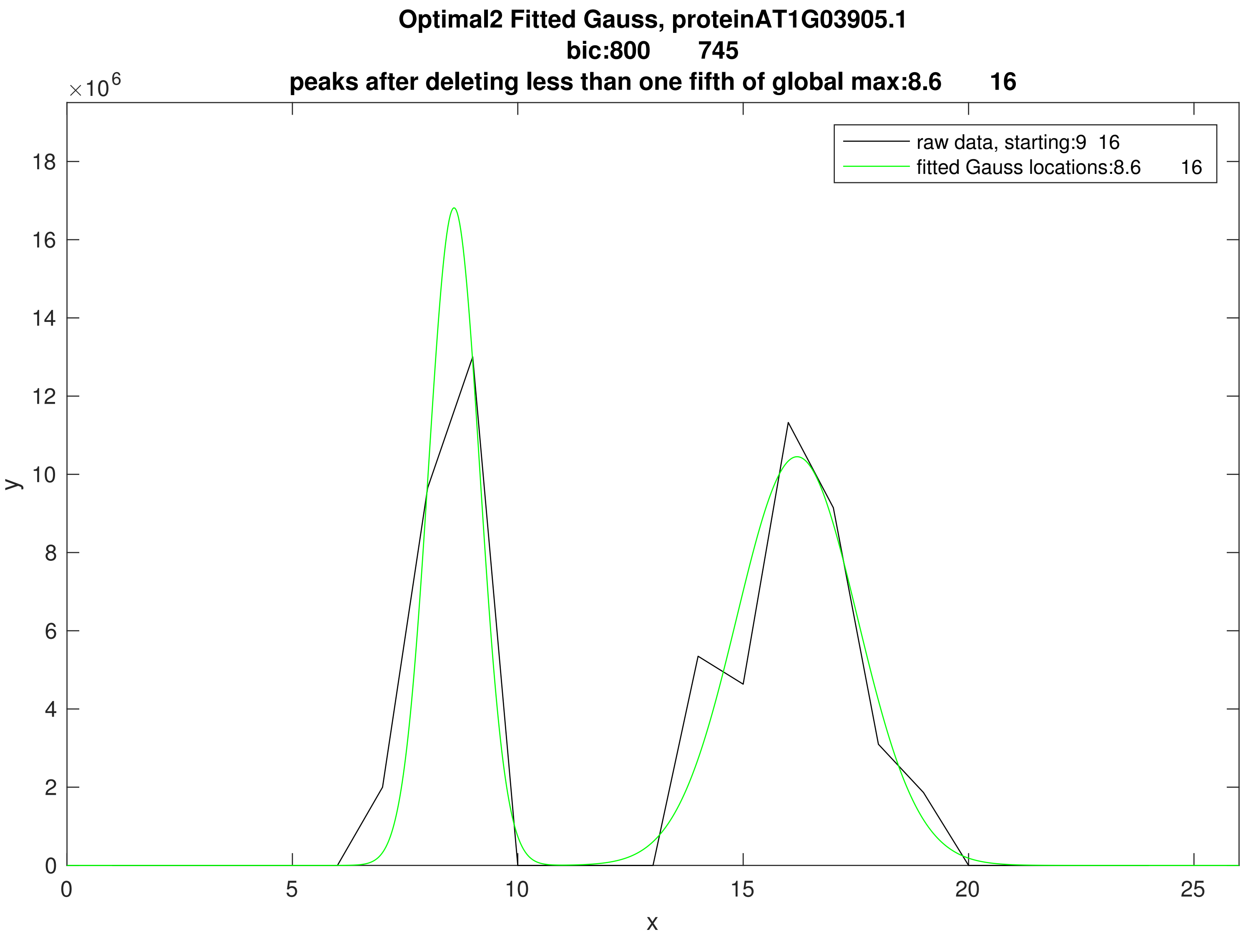

Optimal2 Fitted Gauss, proteinAT1G03905.1
bic:800 745
peaks after deleting less than one fifth of global max:8.6 16
×10^6
raw data, starting:9 16
fitted Gauss locations:8.6 16
y
x
0
2
4
6
8
10
12
14
16
18
0
5
10
15
20
25

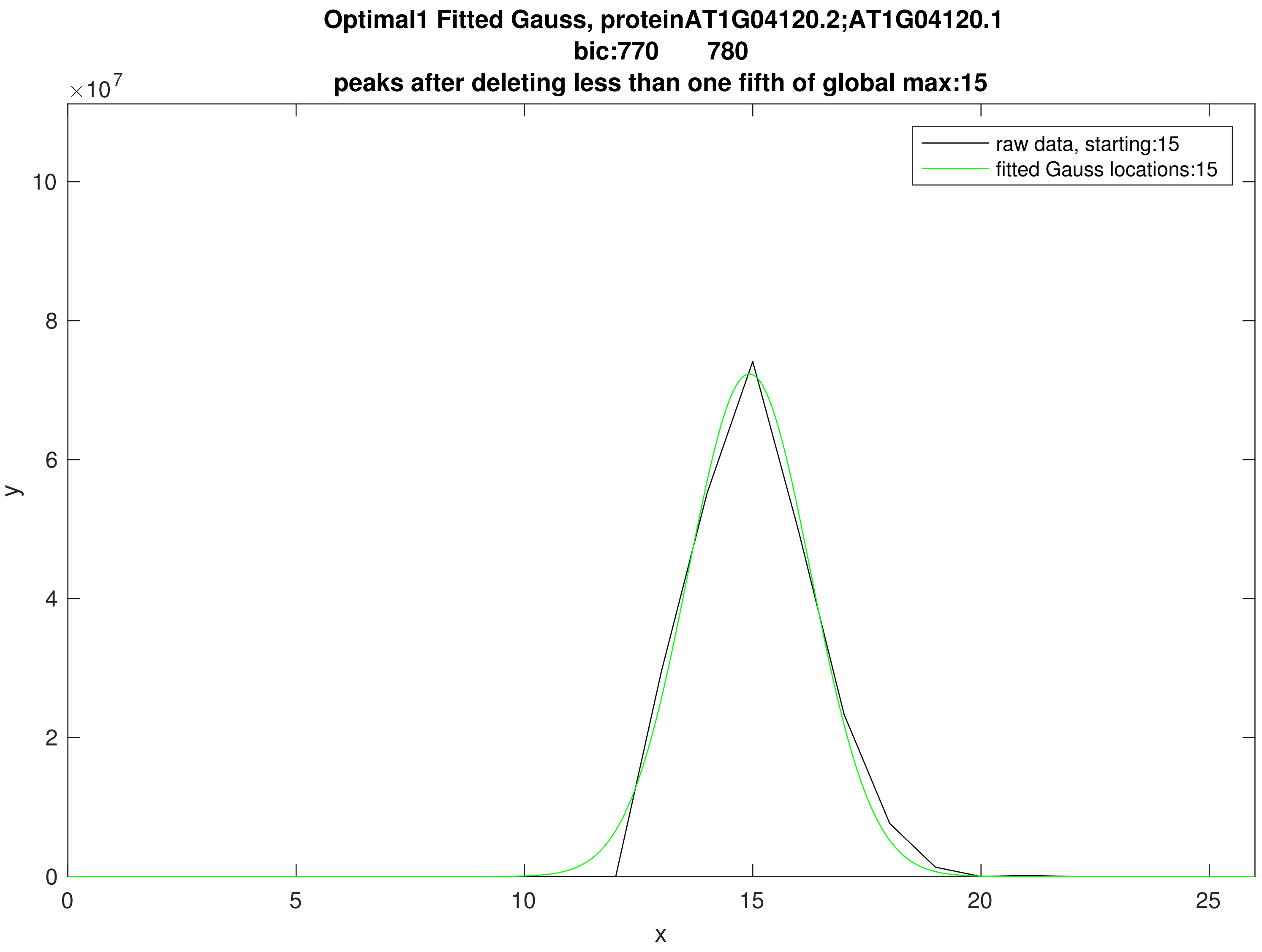
Optimal1 Fitted Gauss, proteinAT1G04120.2;AT1G04120.1
bic:770 780
peaks after deleting less than one fifth of global max:15
×10^7
raw data, starting:15
fitted Gauss locations:15
y
x
0
2
4
6
8
10
0
5
10
15
20
25

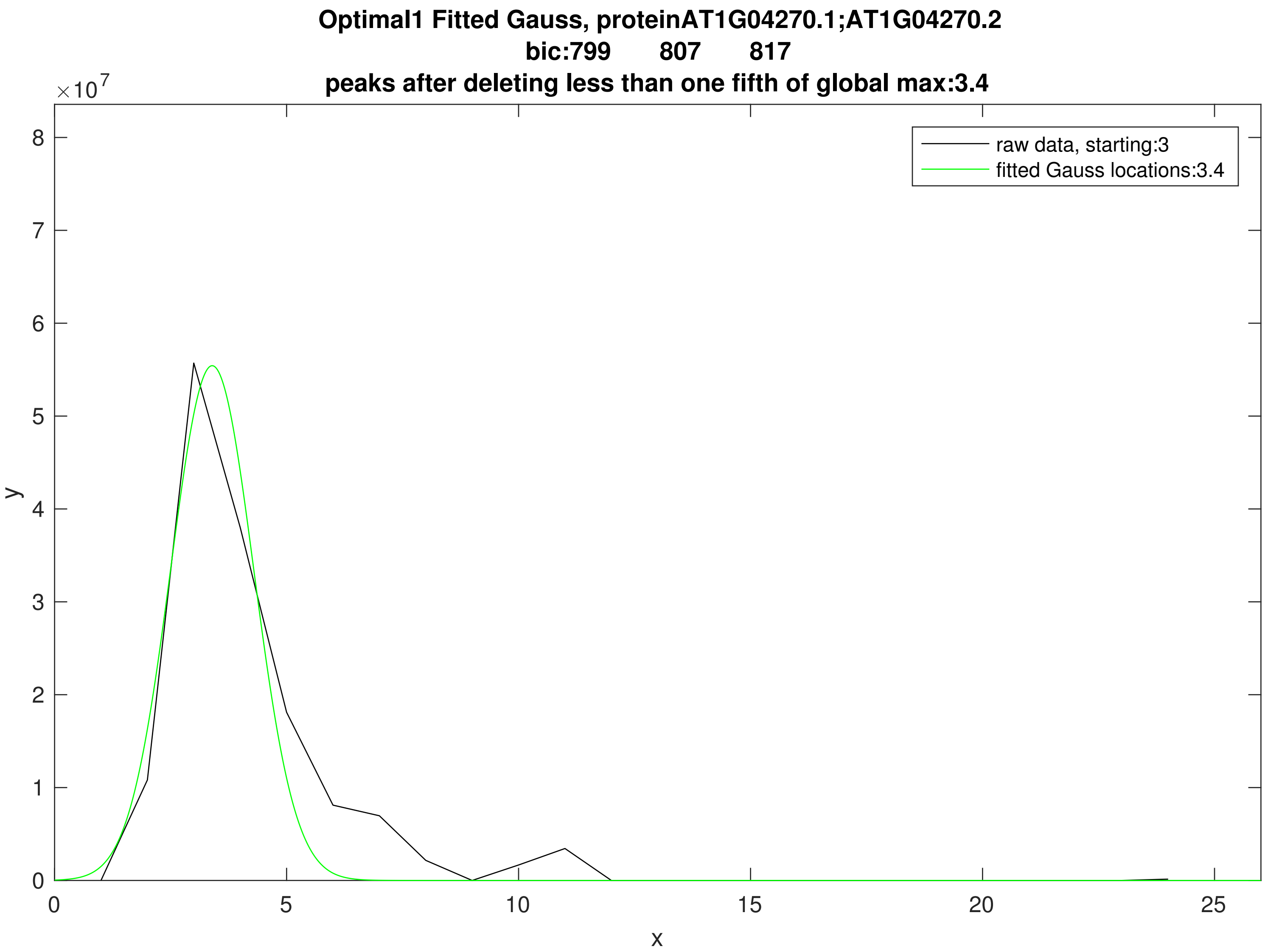

Optimal1 Fitted Gauss, proteinAT1G04270.1;AT1G04270.2
bic:799 807 817
peaks after deleting less than one fifth of global max:3.4
×10^7
raw data, starting:3
fitted Gauss locations:3.4
y
x
0
1
2
3
4
5
6
7
8
0
5
10
15
20
25

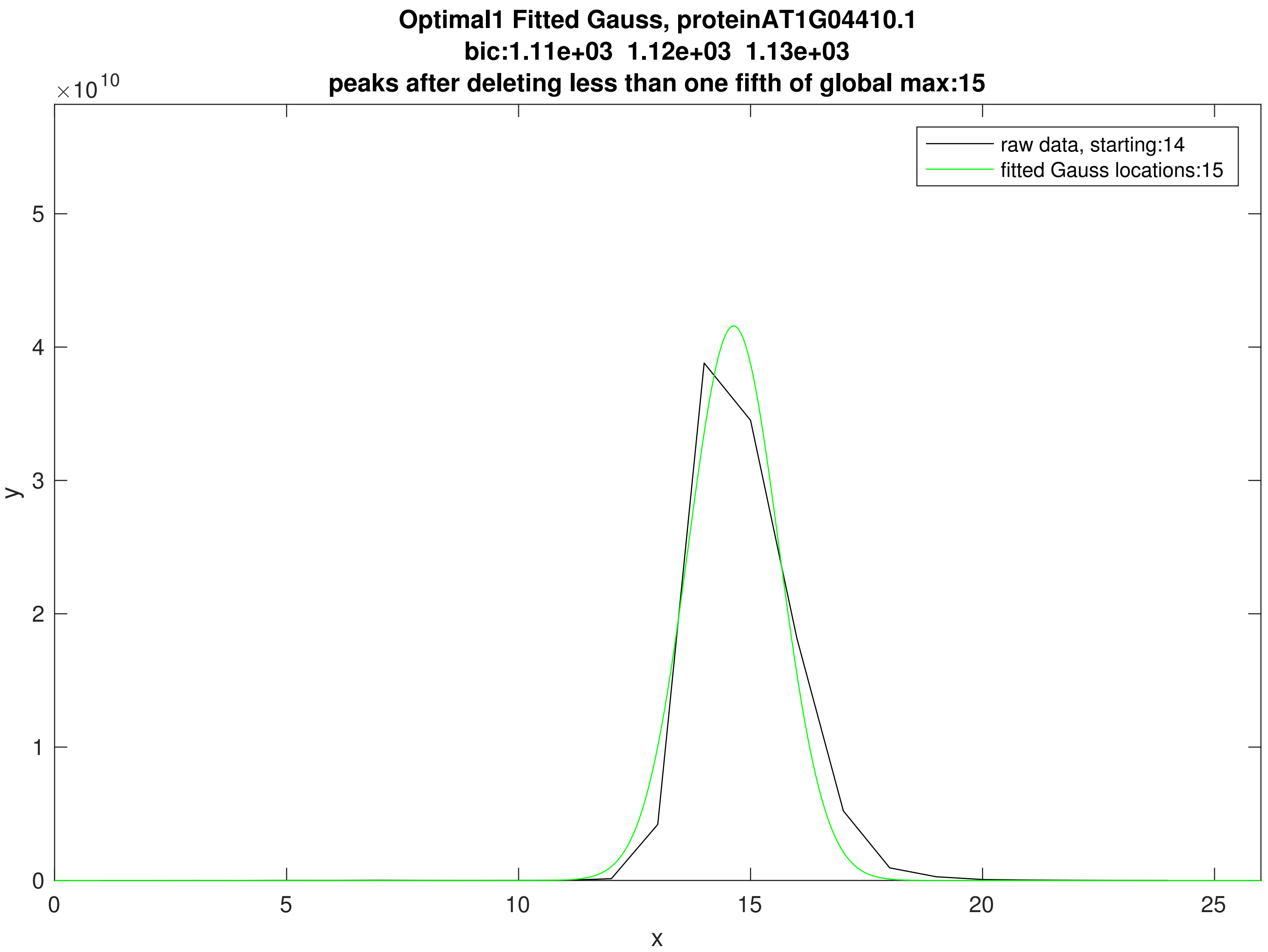

Optimal1 Fitted Gauss, proteinAT1G04410.1
bic:1.11e+03  1.12e+03  1.13e+03
peaks after deleting less than one fifth of global max:15
×10^10
raw data, starting:14
fitted Gauss locations:15
y
x
0
1
2
3
4
5
0
5
10
15
20
25

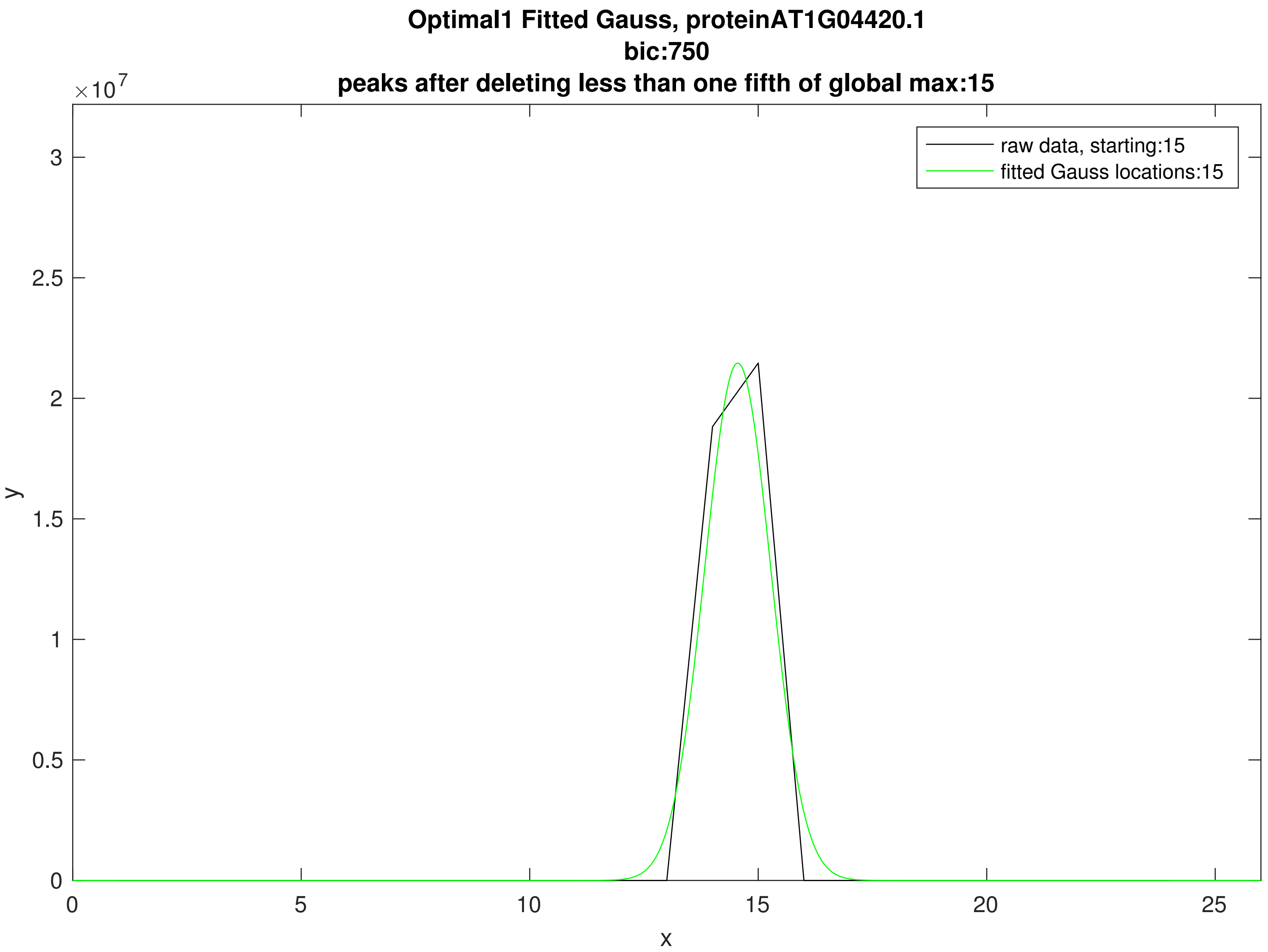

Optimal1 Fitted Gauss, proteinAT1G04420.1
bic:750
peaks after deleting less than one fifth of global max:15
×10^7
raw data, starting:15
fitted Gauss locations:15
y
x
0
0.5
1
1.5
2
2.5
3
0
5
10
15
20
25

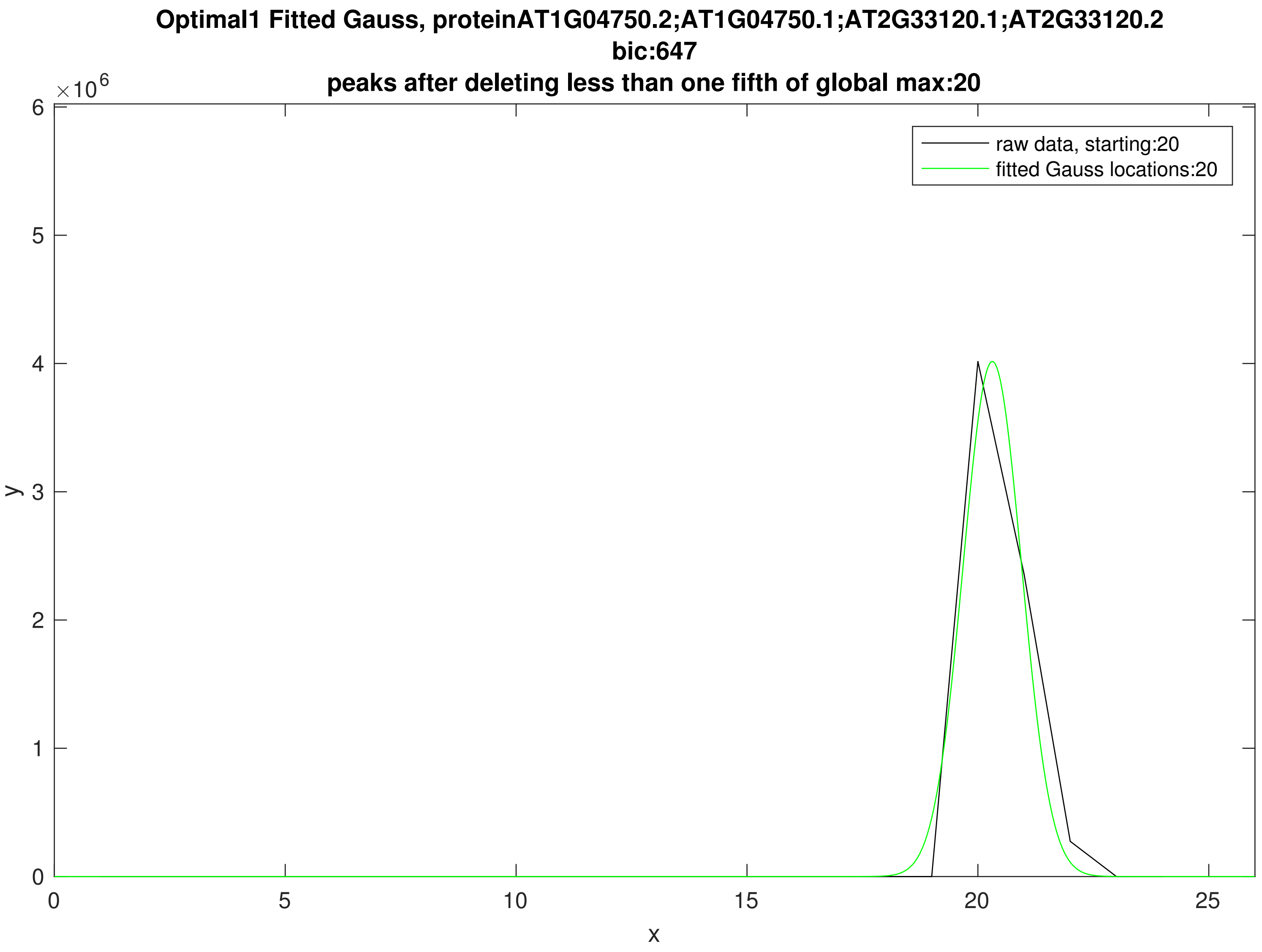

Optimal1 Fitted Gauss, proteinAT1G04750.2;AT1G04750.1;AT2G33120.1;AT2G33120.2
bic:647
peaks after deleting less than one fifth of global max:20
×10^6
6
5
4
3
2
1
0
y
0
5
10
15
20
25
x
raw data, starting:20
fitted Gauss locations:20

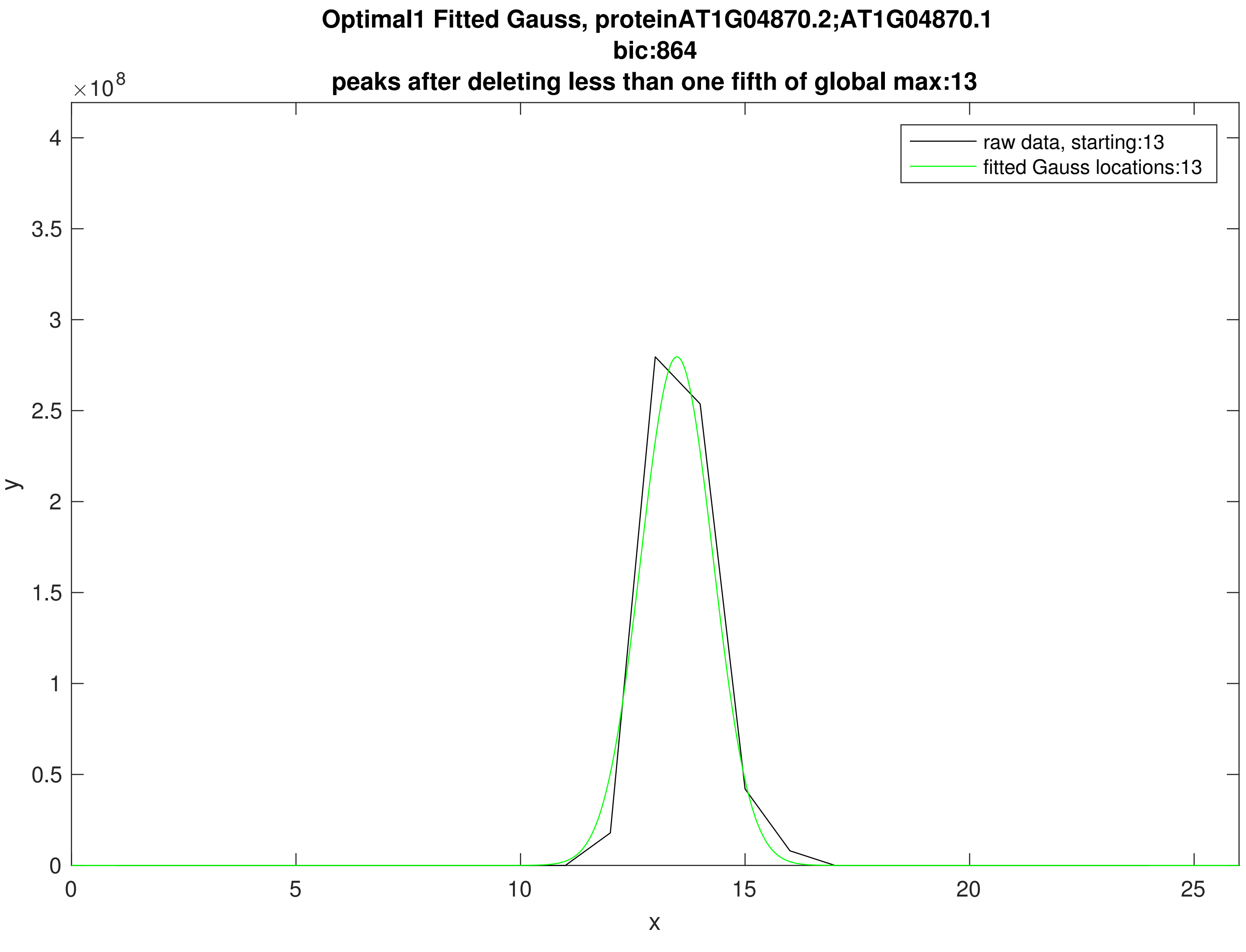

Optimal1 Fitted Gauss, proteinAT1G04870.2;AT1G04870.1
bic:864
peaks after deleting less than one fifth of global max:13
×10^8
raw data, starting:13
fitted Gauss locations:13
y
x
0
0.5
1
1.5
2
2.5
3
3.5
4
0
5
10
15
20
25

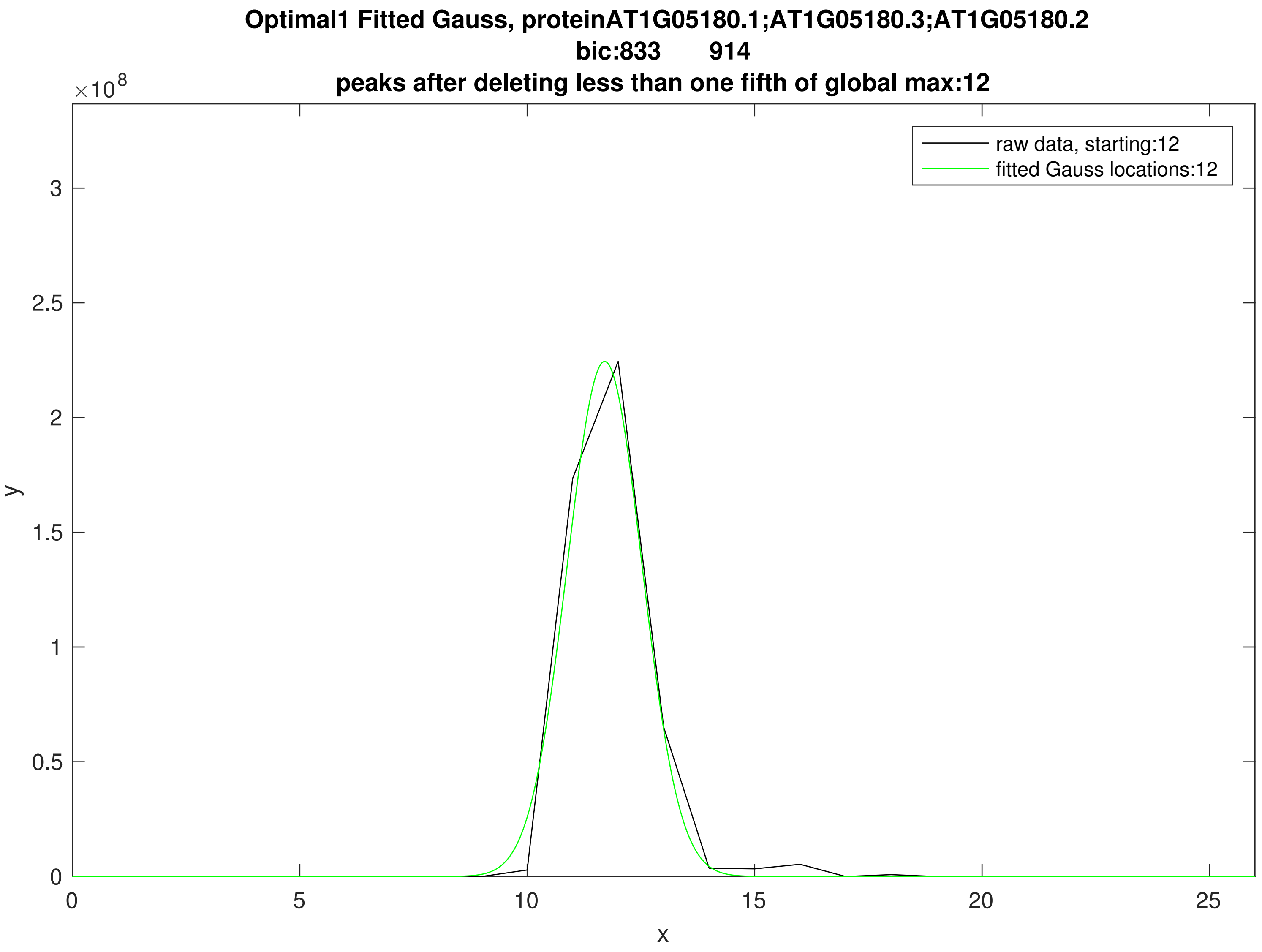

Optimal1 Fitted Gauss, proteinAT1G05180.1;AT1G05180.3;AT1G05180.2
bic:833 914
peaks after deleting less than one fifth of global max:12
×10^8
raw data, starting:12
fitted Gauss locations:12
y
x
0
0.5
1
1.5
2
2.5
3
5
10
15
20
25

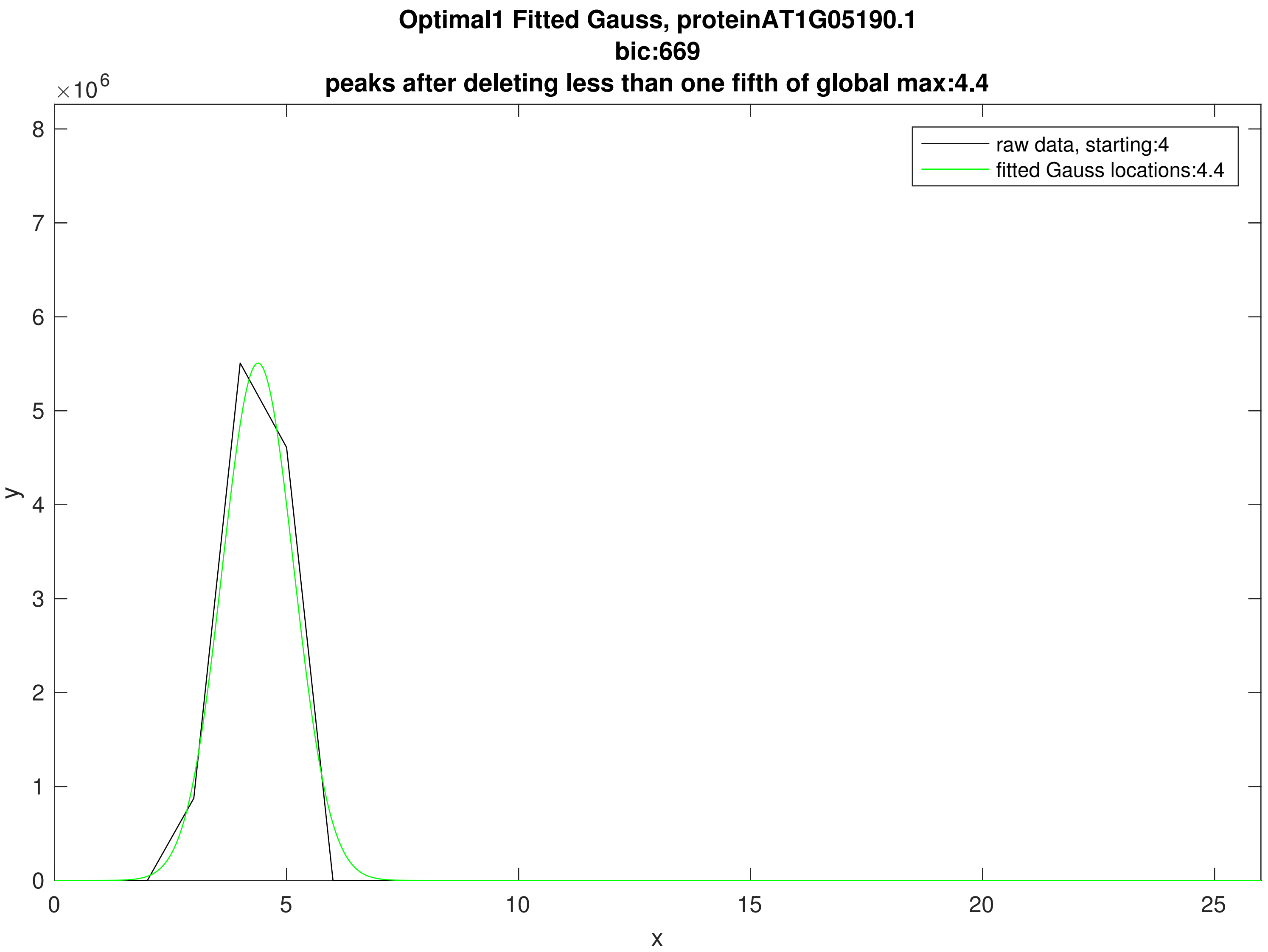

Optimal1 Fitted Gauss, proteinAT1G05190.1
bic:669
peaks after deleting less than one fifth of global max:4.4
×10^6
raw data, starting:4
fitted Gauss locations:4.4
y
x
0
1
2
3
4
5
6
7
8
0
5
10
15
20
25

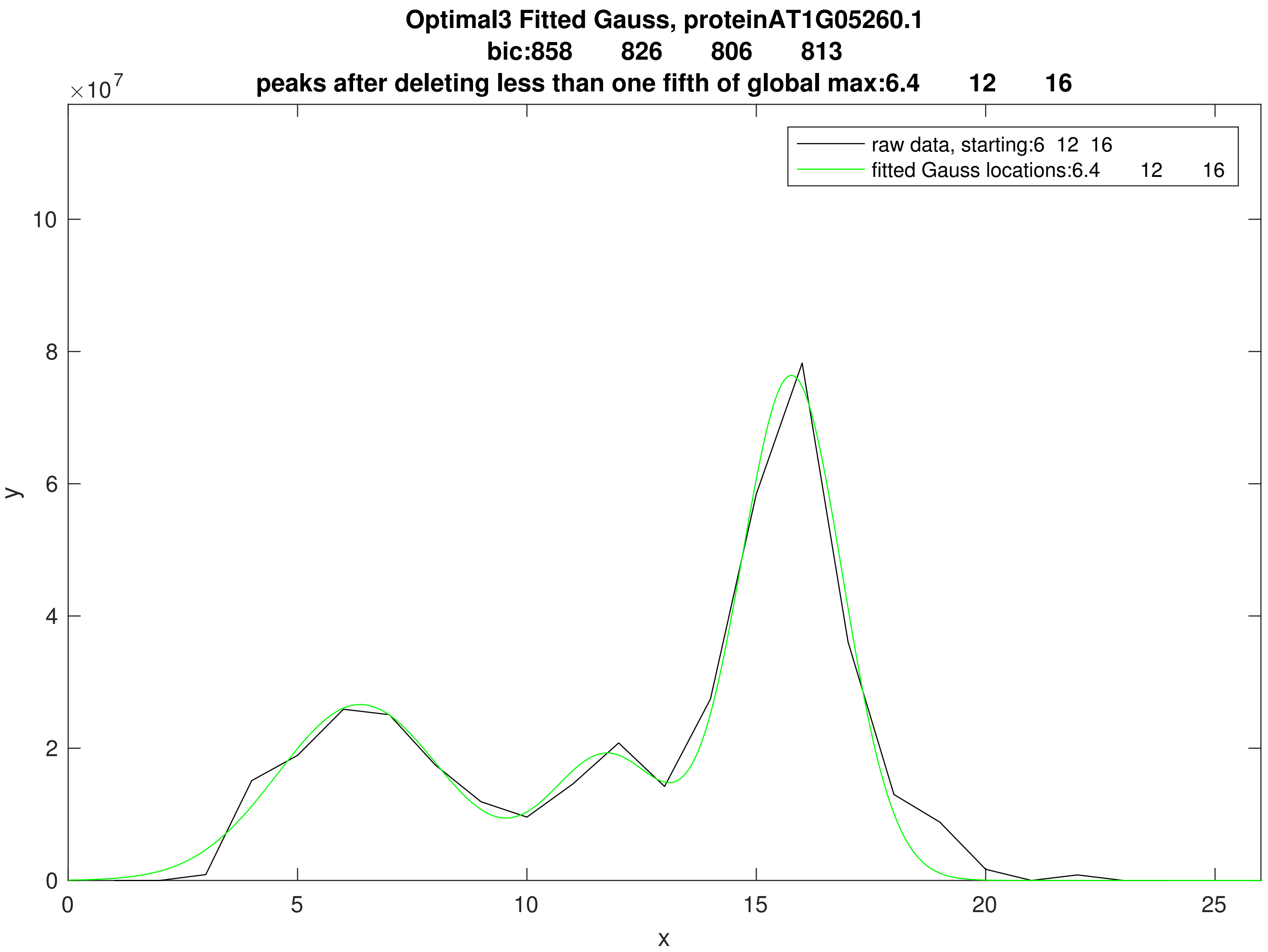
Optimal3 Fitted Gauss, proteinAT1G05260.1
bic:858 826 806 813
peaks after deleting less than one fifth of global max:6.4 12 16
×10^7
raw data, starting:6 12 16
fitted Gauss locations:6.4 12 16
y
x
0
2
4
6
8
10
0
5
10
15
20
25

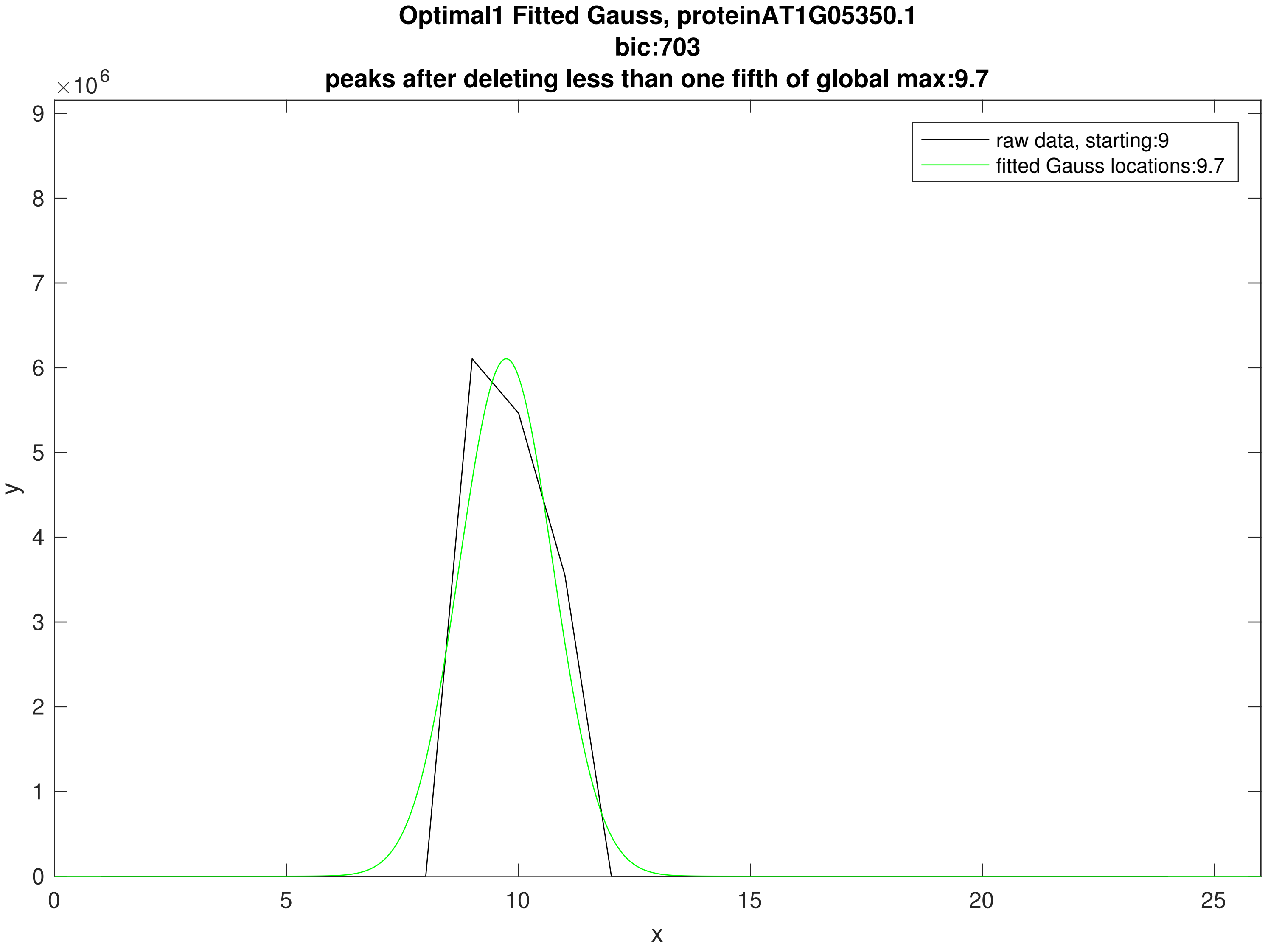

Optimal1 Fitted Gauss, proteinAT1G05350.1
bic:703
peaks after deleting less than one fifth of global max:9.7
×10^6
raw data, starting:9
fitted Gauss locations:9.7
y
x
0
1
2
3
4
5
6
7
8
9
0
5
10
15
20
25

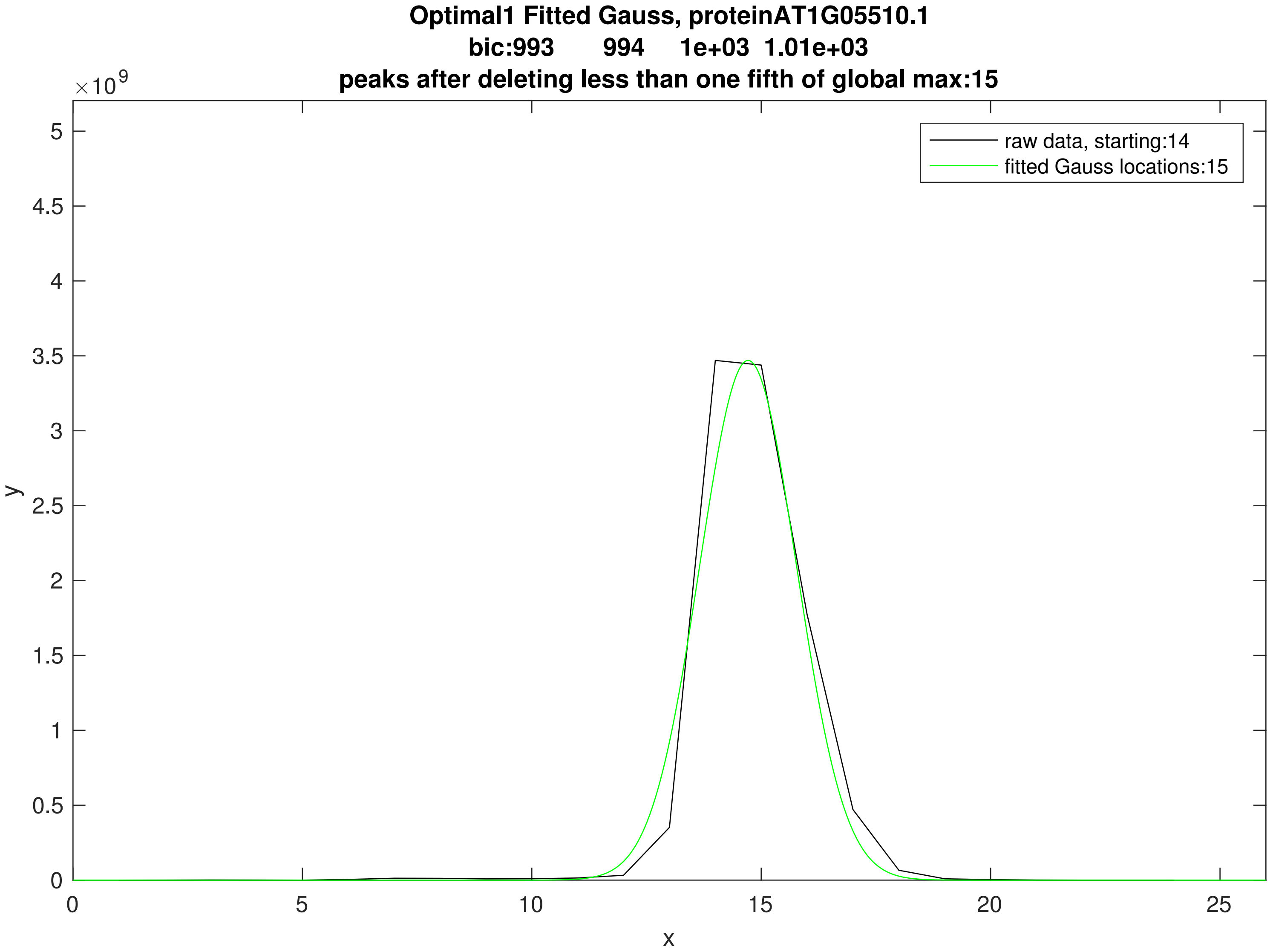

Optimal1 Fitted Gauss, proteinAT1G05510.1
bic:993 994 1e+03 1.01e+03
peaks after deleting less than one fifth of global max:15
×10^9
raw data, starting:14
fitted Gauss locations:15
y
x
0
0.5
1
1.5
2
2.5
3
3.5
4
4.5
5
0
5
10
15
20
25

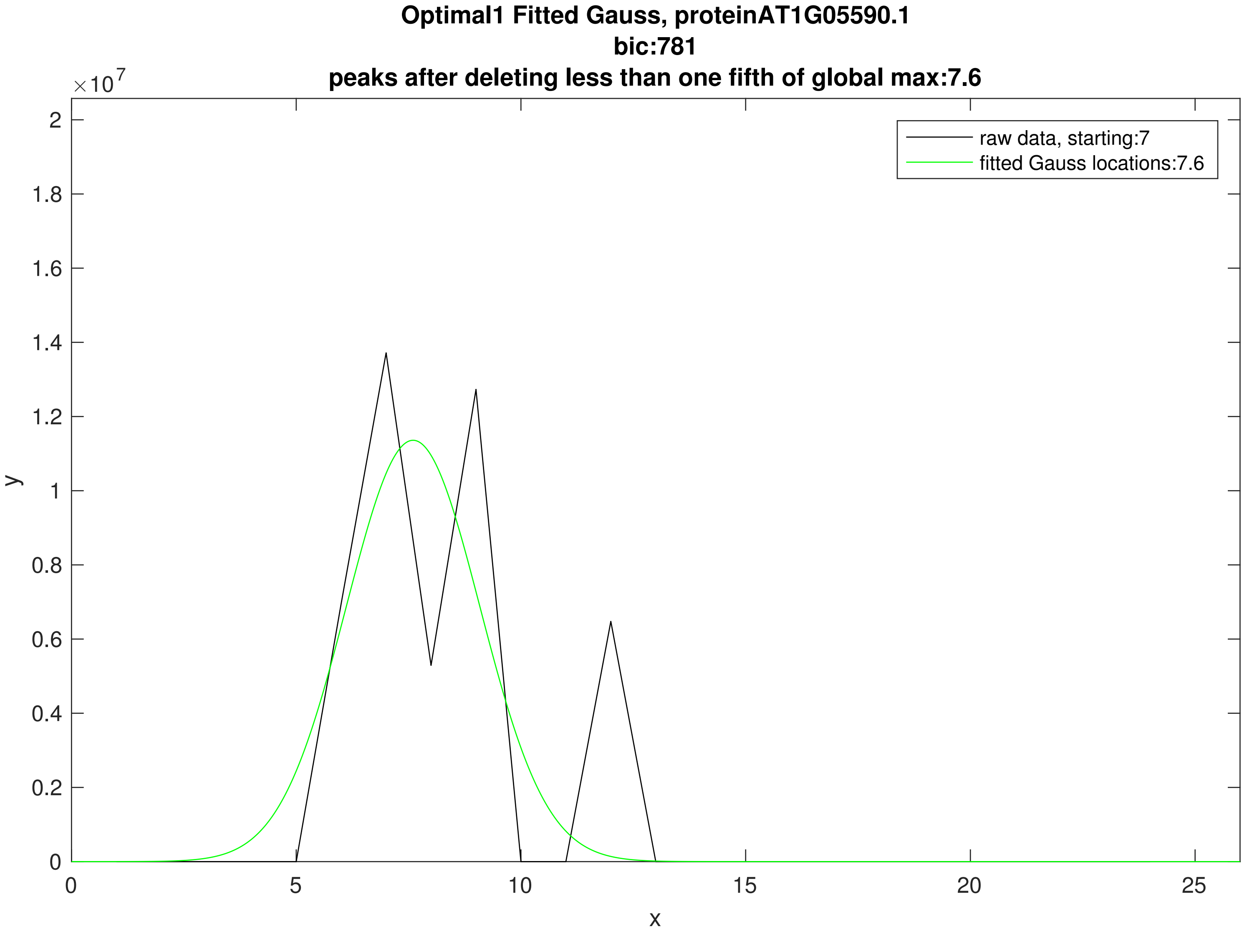

Optimal1 Fitted Gauss, proteinAT1G05590.1
bic:781
peaks after deleting less than one fifth of global max:7.6
×10^7
raw data, starting:7
fitted Gauss locations:7.6
y
x
0
0.2
0.4
0.6
0.8
1
1.2
1.4
1.6
1.8
2
0
5
10
15
20
25

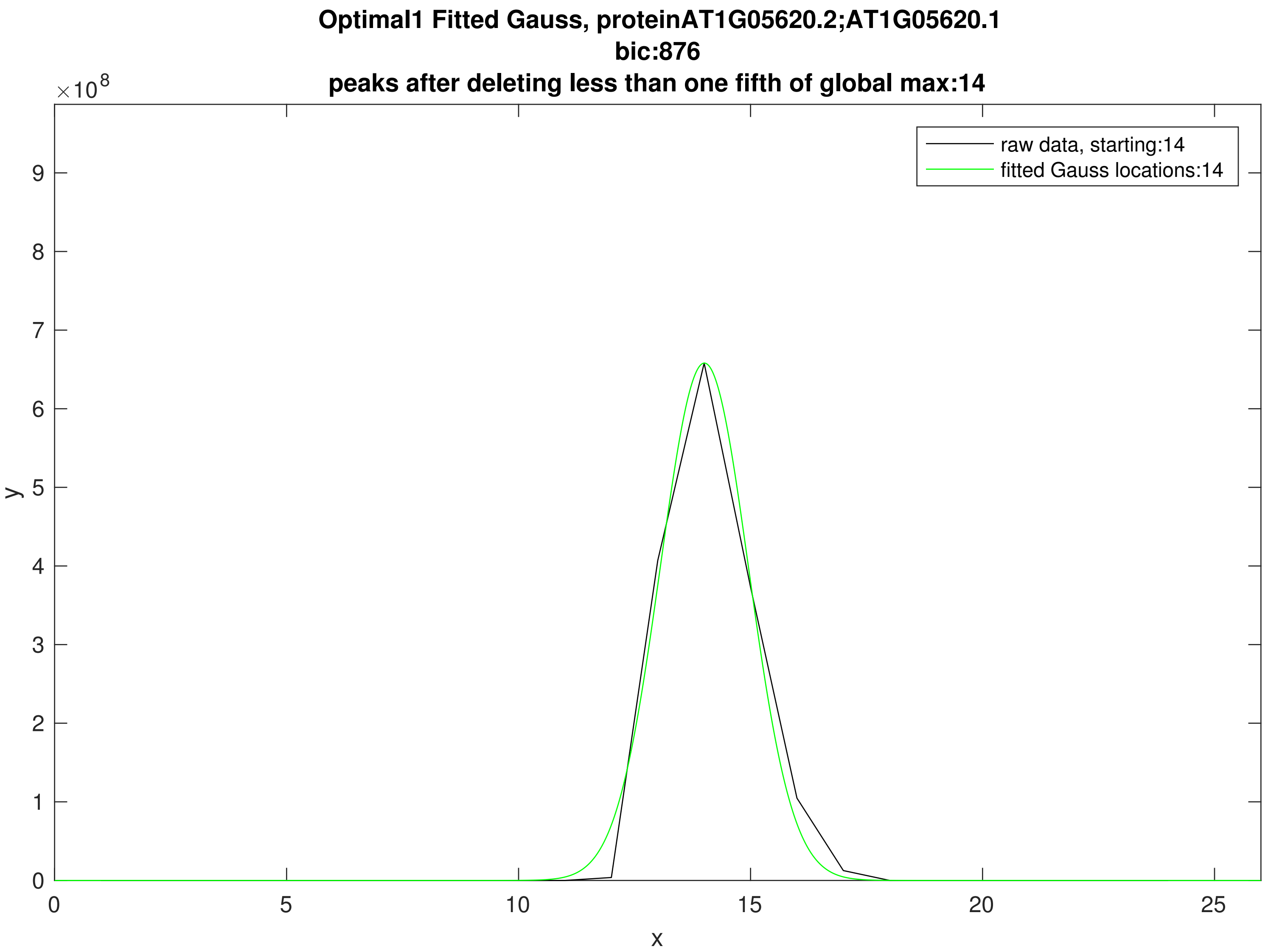
Optimal1 Fitted Gauss, proteinAT1G05620.2;AT1G05620.1
bic:876
peaks after deleting less than one fifth of global max:14
×10^8
raw data, starting:14
fitted Gauss locations:14
y
x
0
1
2
3
4
5
6
7
8
9
0
5
10
15
20
25

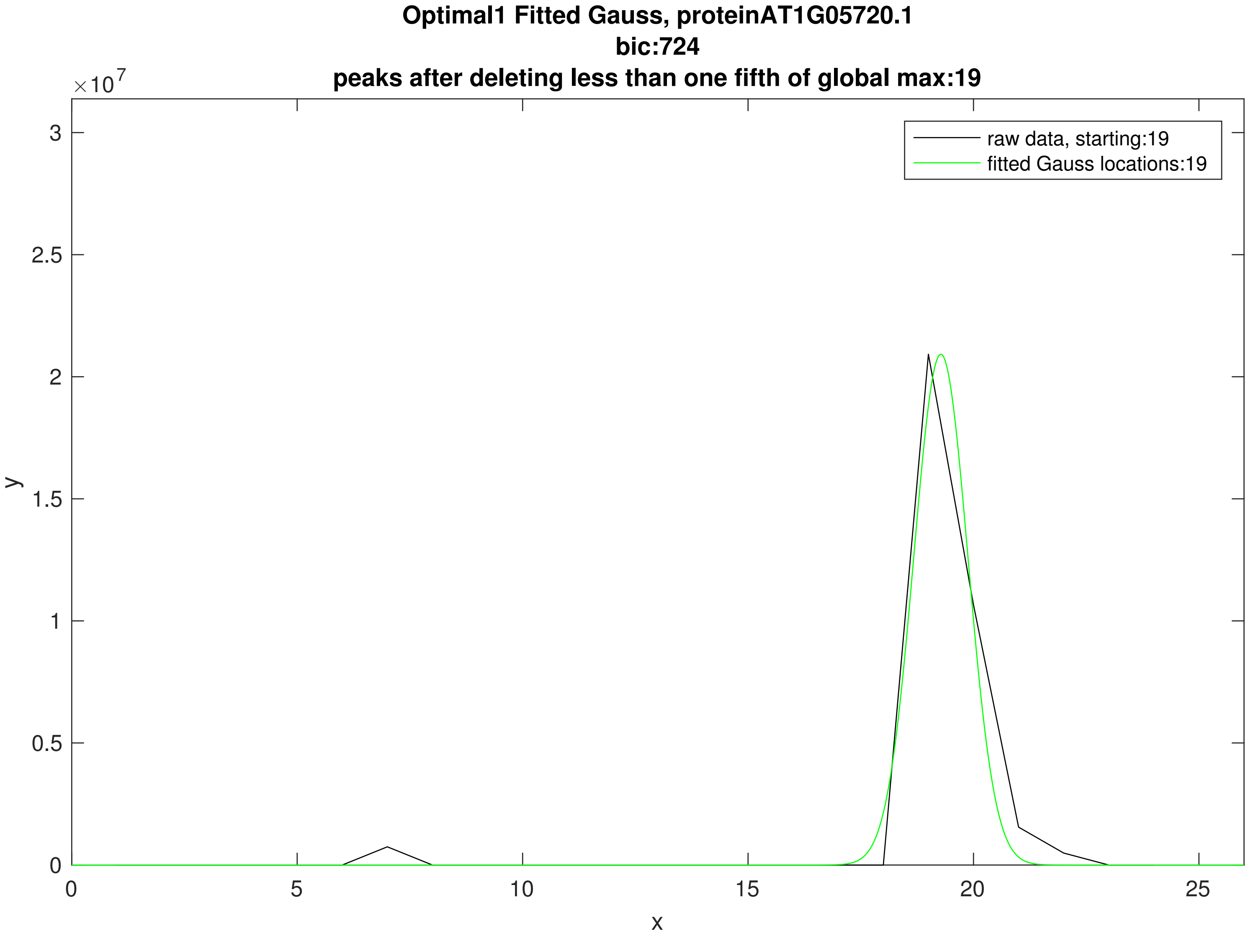

Optimal1 Fitted Gauss, proteinAT1G05720.1
bic:724
peaks after deleting less than one fifth of global max:19
×10^7
3
2.5
2
1.5
1
0.5
0
y
0
5
10
15
20
25
x
raw data, starting:19
fitted Gauss locations:19

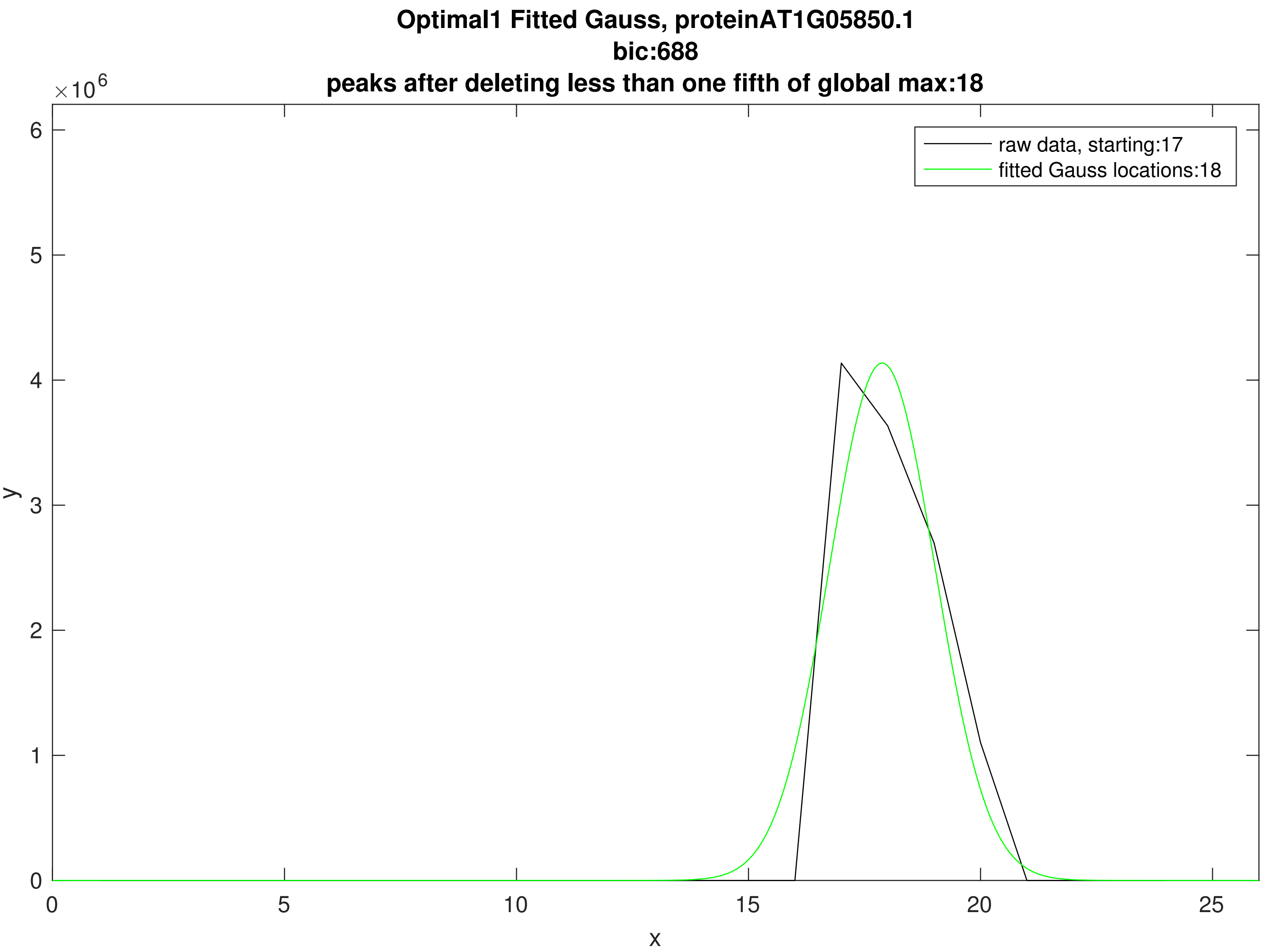
Optimal1 Fitted Gauss, proteinAT1G05850.1
bic:688
peaks after deleting less than one fifth of global max:18
×10^6
raw data, starting:17
fitted Gauss locations:18
y
x
0
1
2
3
4
5
6
0
5
10
15
20
25

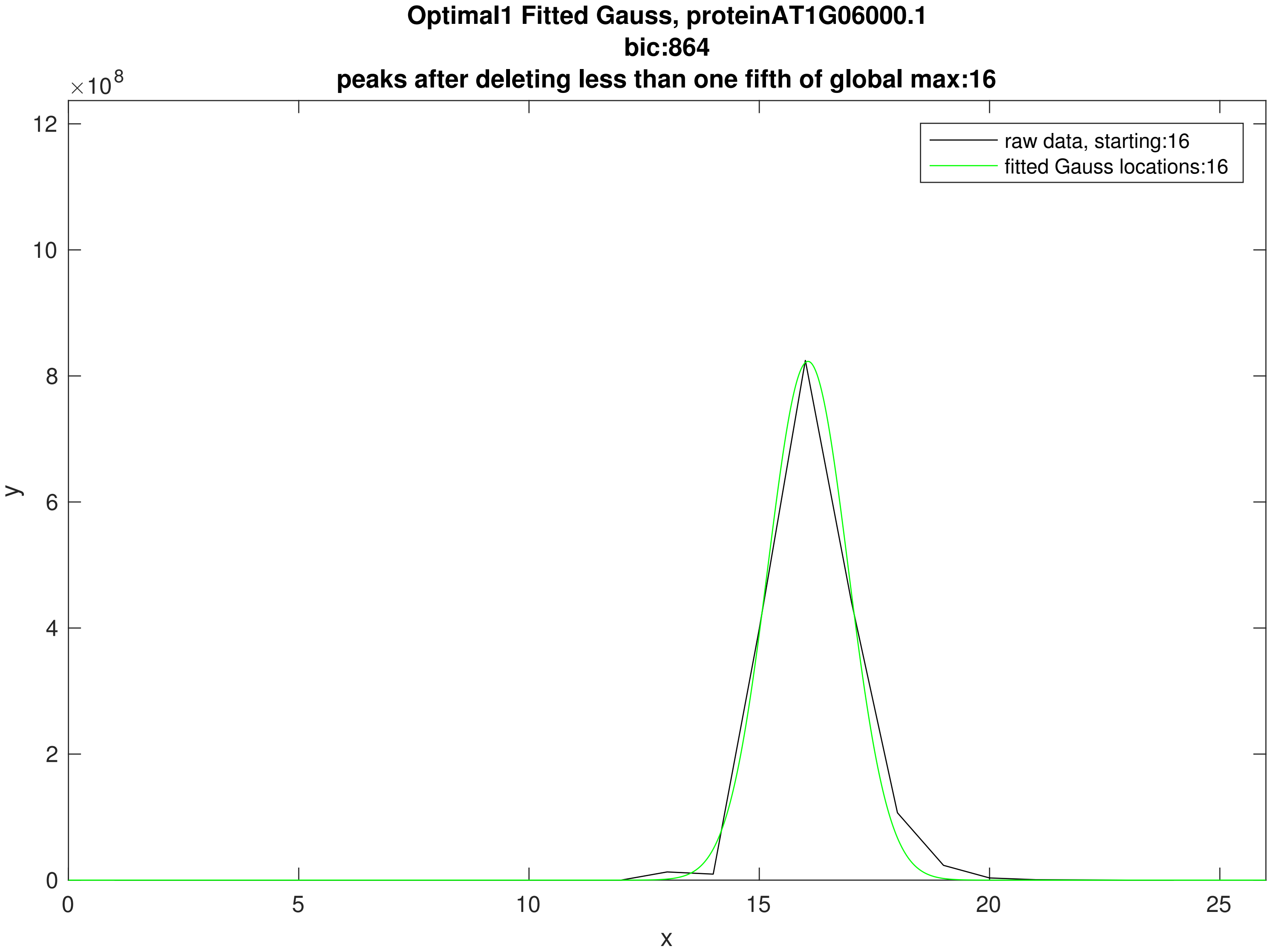
Optimal1 Fitted Gauss, proteinAT1G06000.1
bic:864
peaks after deleting less than one fifth of global max:16
×10^8
raw data, starting:16
fitted Gauss locations:16
12
10
8
6
4
2
0
y
0
5
10
15
20
25
x

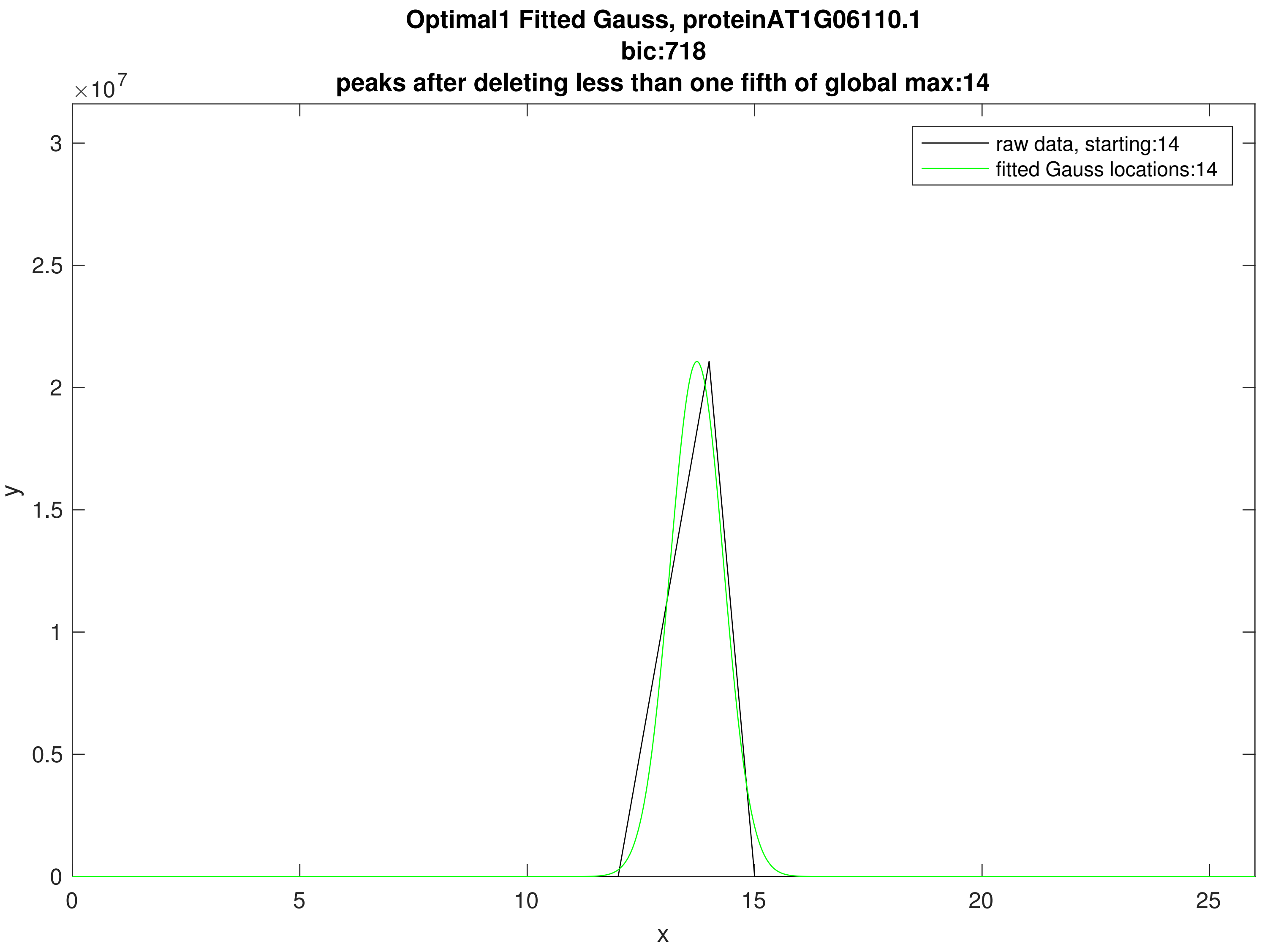

Optimal1 Fitted Gauss, proteinAT1G06110.1
bic:718
peaks after deleting less than one fifth of global max:14
×10^7
3
2.5
2
1.5
1
0.5
0
y
0
5
10
15
20
25
x
raw data, starting:14
fitted Gauss locations:14

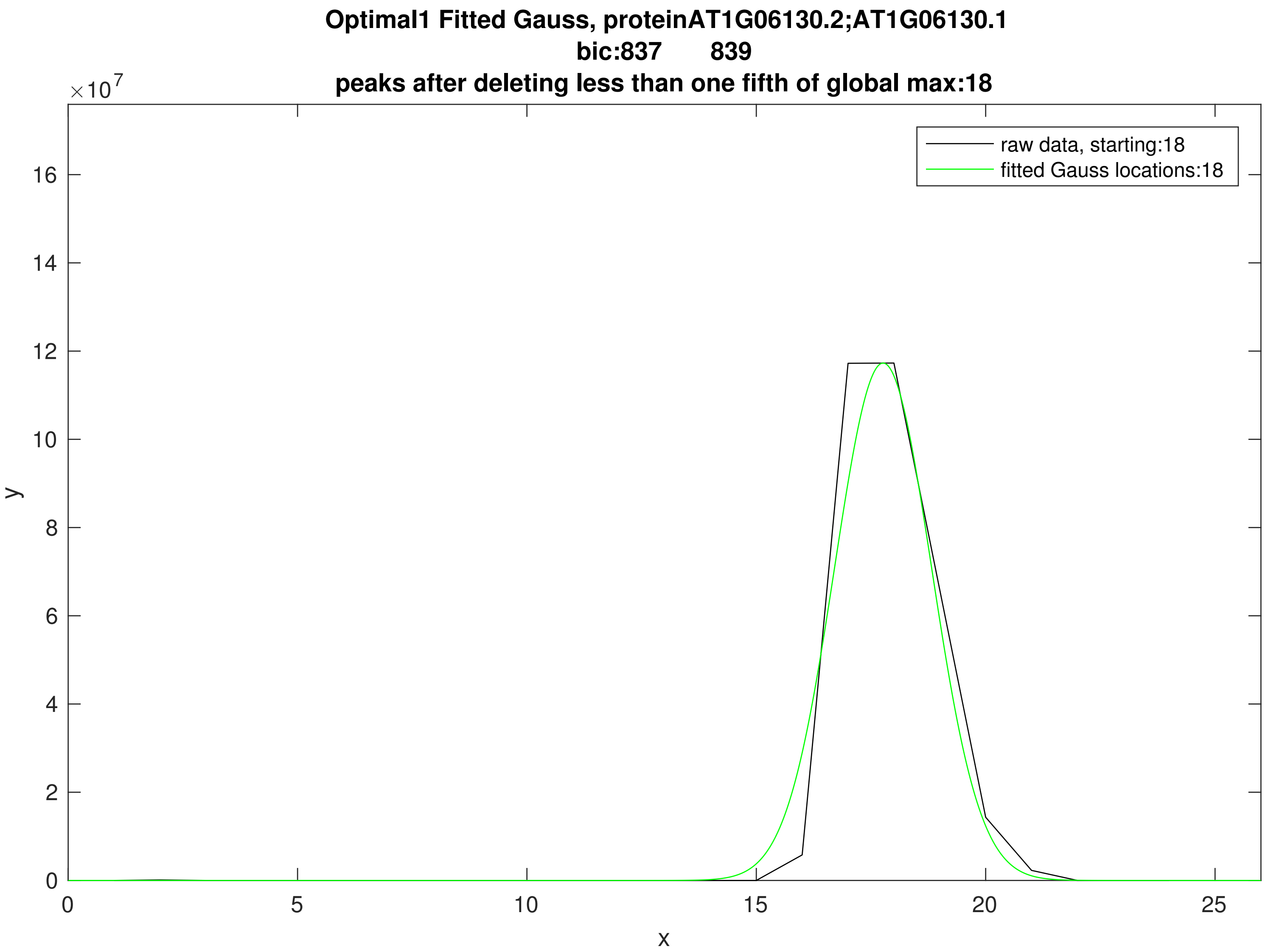
Optimal1 Fitted Gauss, proteinAT1G06130.2;AT1G06130.1
bic:837 839
peaks after deleting less than one fifth of global max:18
×10^7
raw data, starting:18
fitted Gauss locations:18
y
x
0
2
4
6
8
10
12
14
16
0
5
10
15
20
25

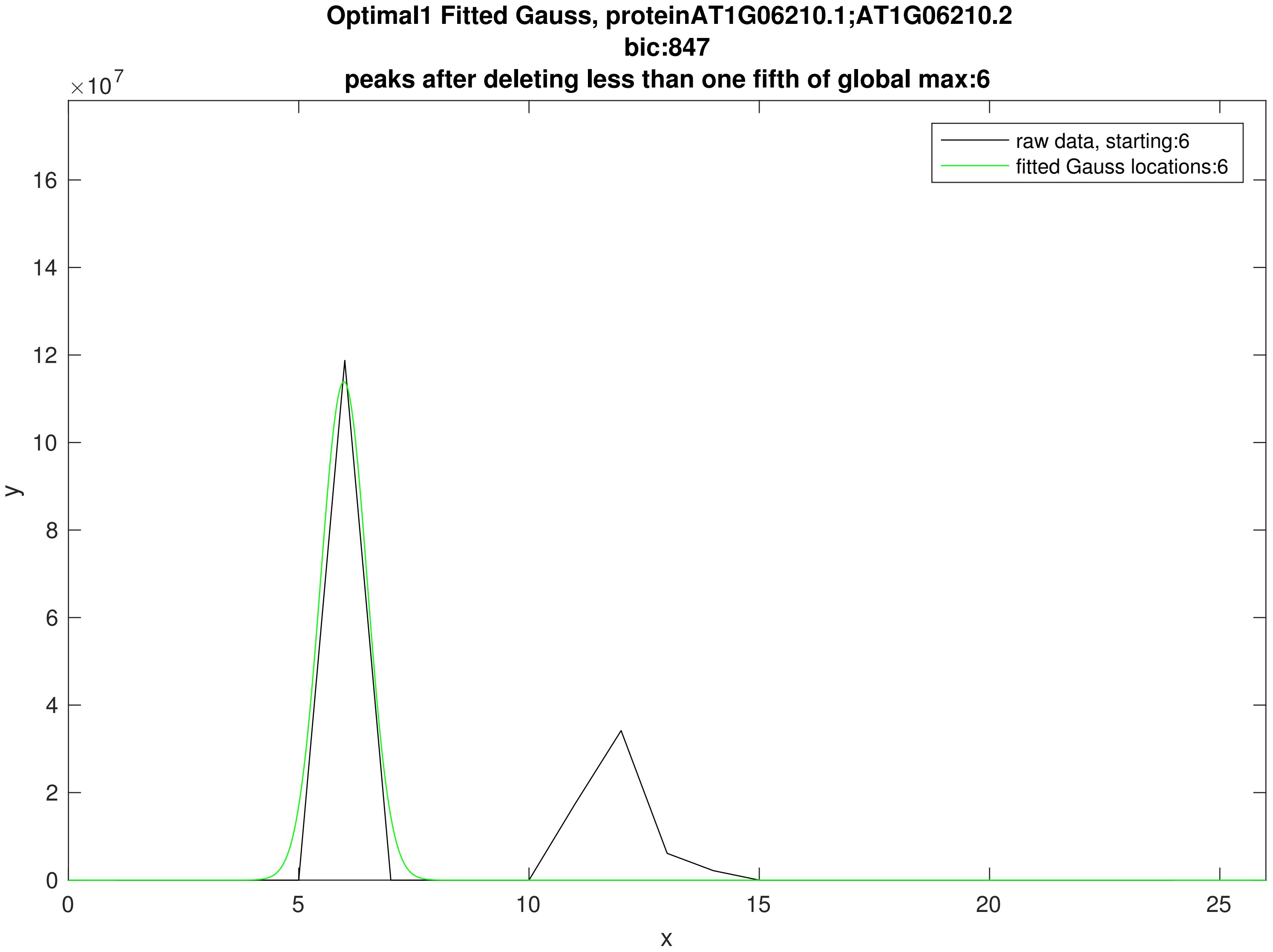
Optimal1 Fitted Gauss, proteinAT1G06210.1;AT1G06210.2
bic:847
peaks after deleting less than one fifth of global max:6
×10^7
16
14
12
10
8
6
4
2
0
y
0
5
10
15
20
25
x
raw data, starting:6
fitted Gauss locations:6

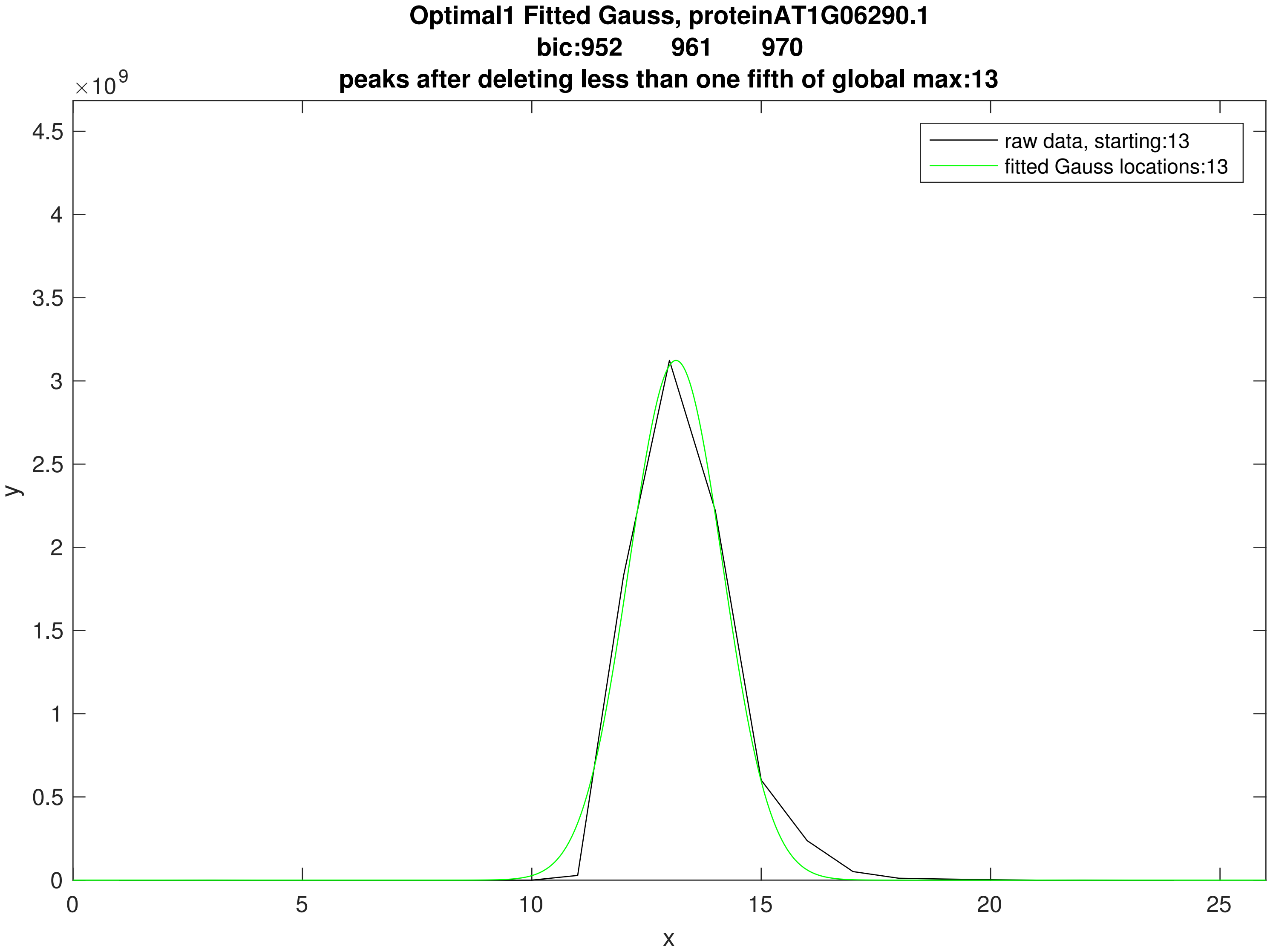
Optimal1 Fitted Gauss, proteinAT1G06290.1
bic:952 961 970
peaks after deleting less than one fifth of global max:13
×10^9
raw data, starting:13
fitted Gauss locations:13
y
x
0
0.5
1
1.5
2
2.5
3
3.5
4
4.5
0
5
10
15
20
25

**Optimal1 Fitted Gauss, proteinAT1G06410.1**
**bic:644**
**peaks after deleting less than one fifth of global max:10**

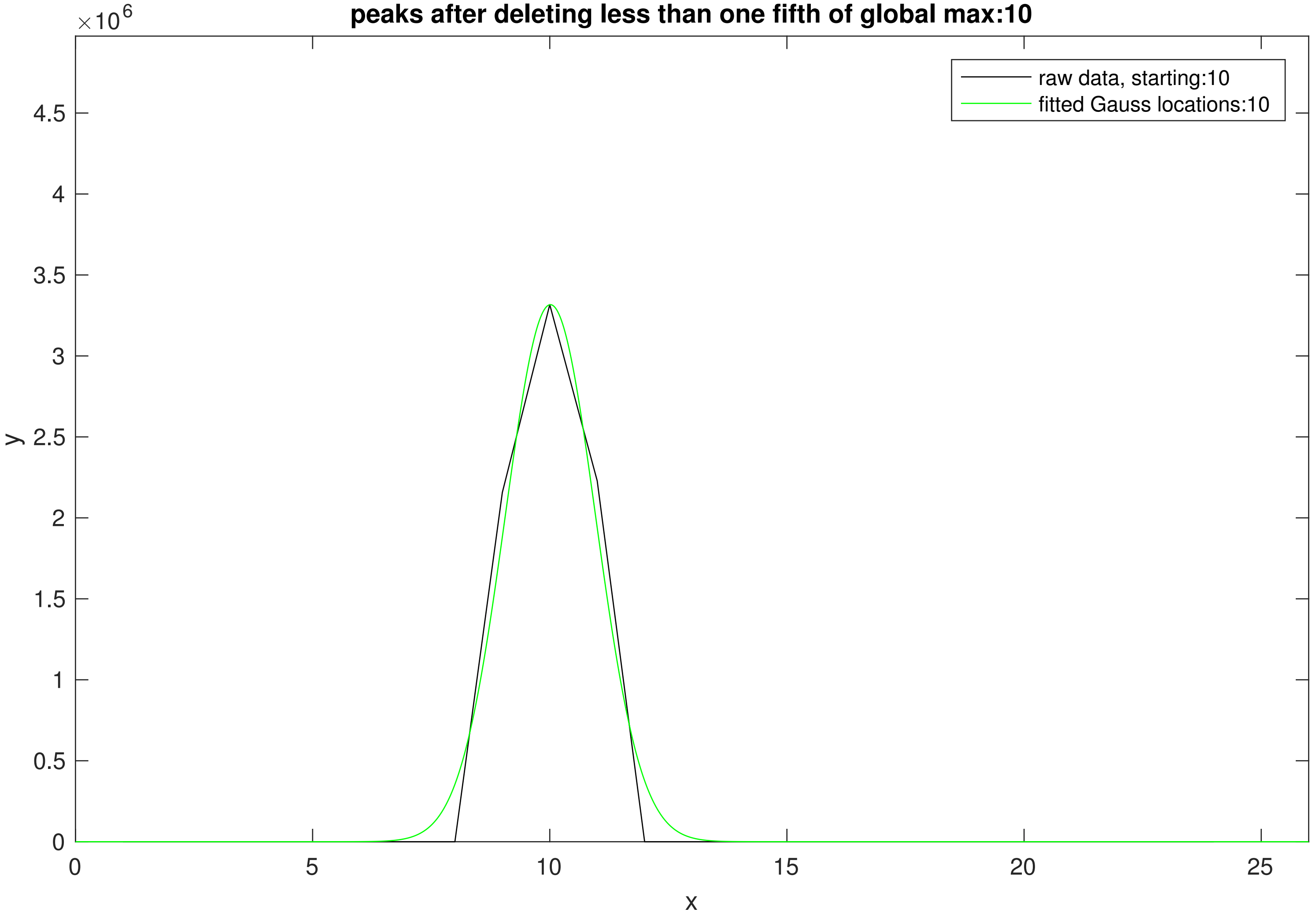

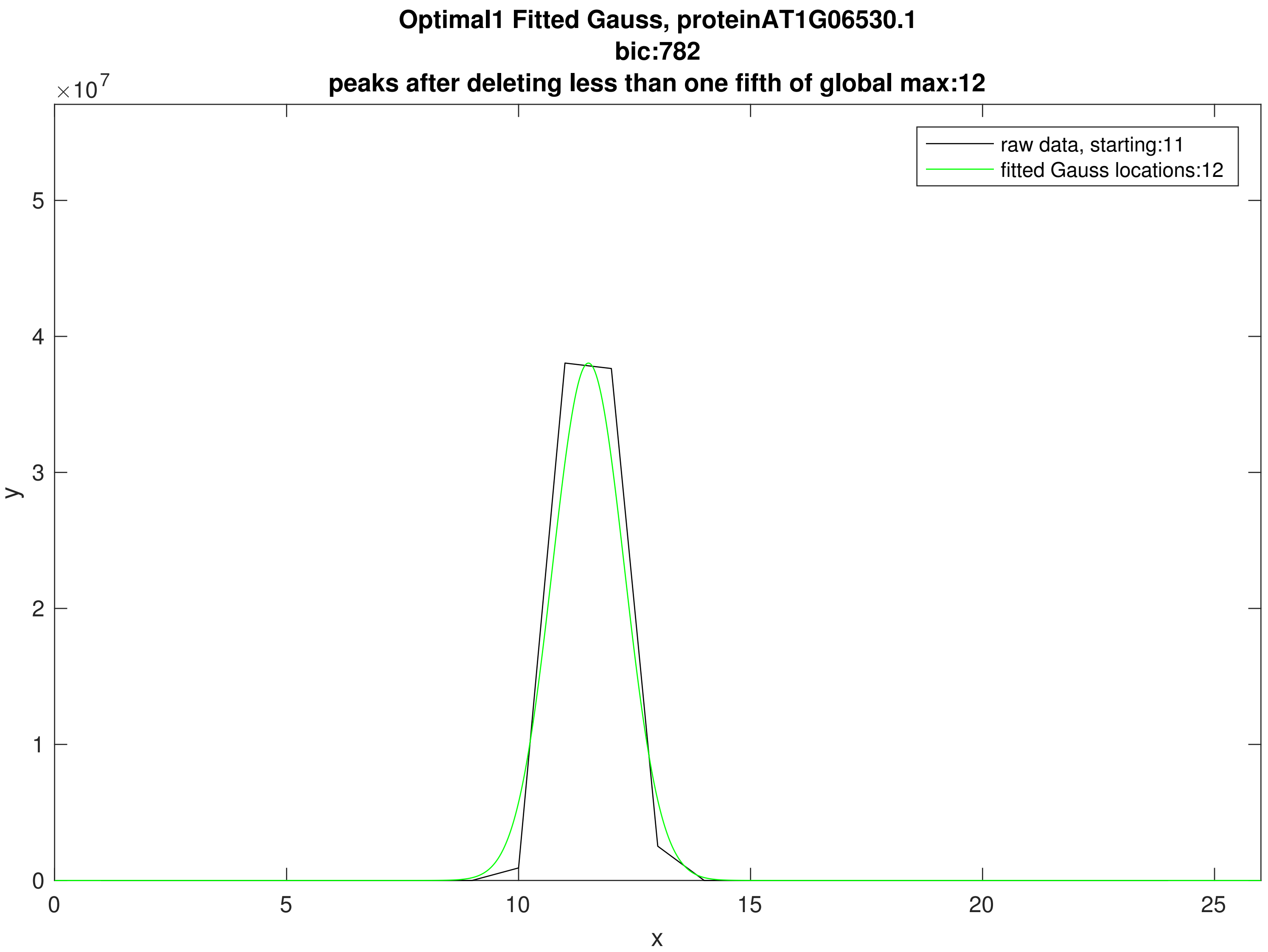

Optimal1 Fitted Gauss, proteinAT1G06530.1
bic:782
peaks after deleting less than one fifth of global max:12
×10^7
raw data, starting:11
fitted Gauss locations:12
y
x
0
1
2
3
4
5
0
5
10
15
20
25

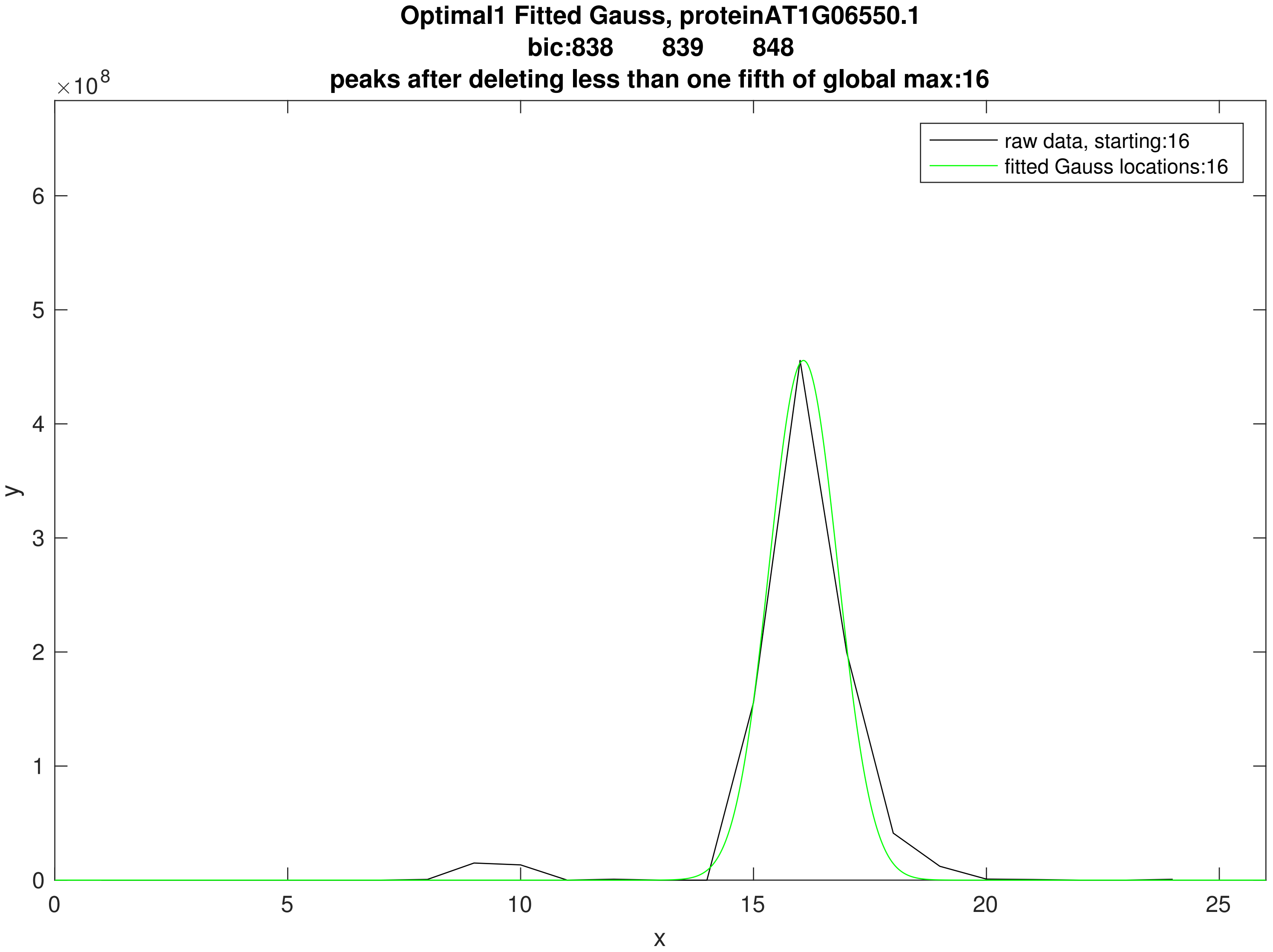

Optimal1 Fitted Gauss, proteinAT1G06550.1
bic:838 839 848
peaks after deleting less than one fifth of global max:16
×10^8
raw data, starting:16
fitted Gauss locations:16
y
x
0
1
2
3
4
5
6
0
5
10
15
20
25

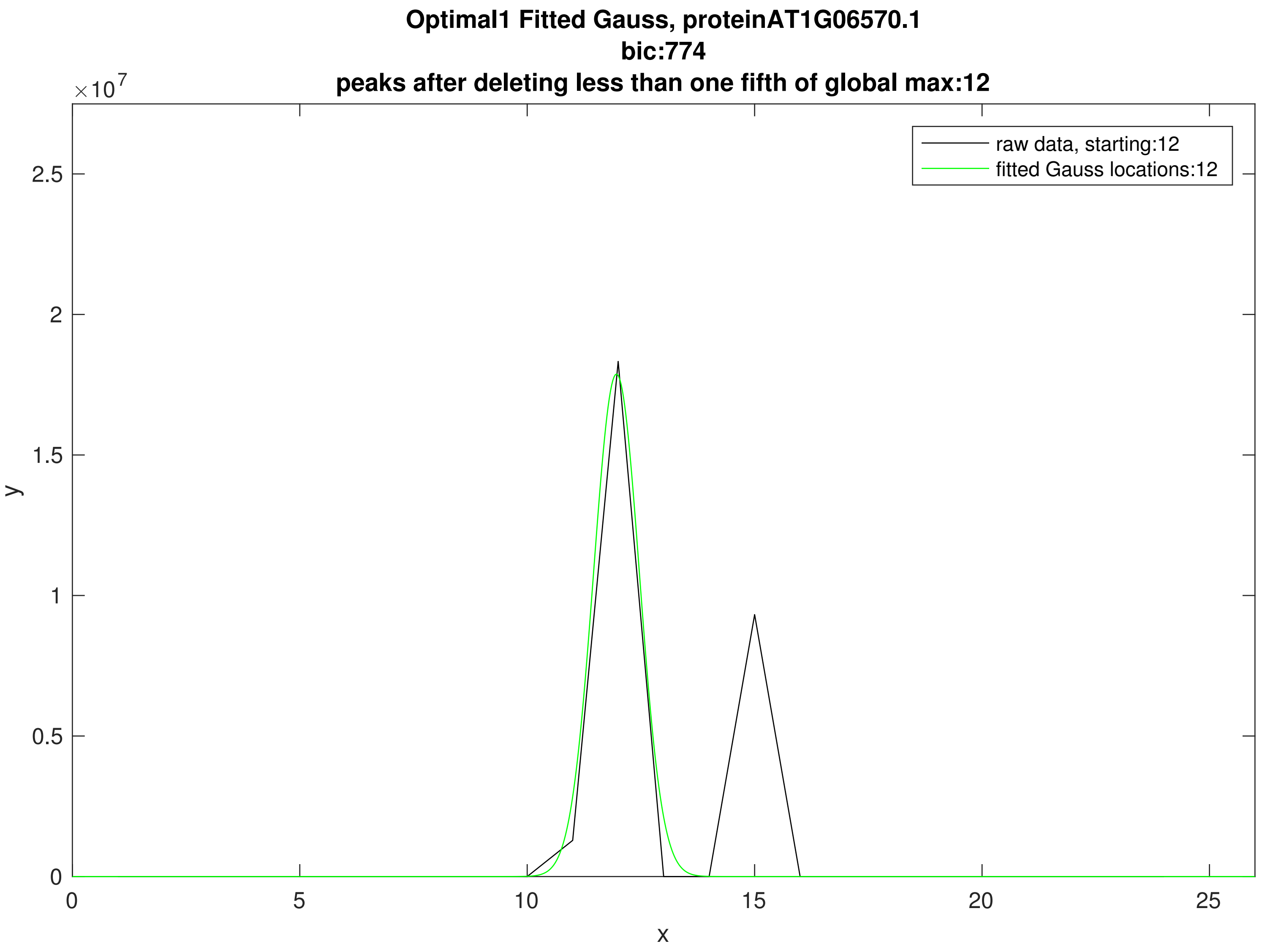
Optimal1 Fitted Gauss, proteinAT1G06570.1
bic:774
peaks after deleting less than one fifth of global max:12
×10^7
raw data, starting:12
fitted Gauss locations:12
y
x
0
0.5
1
1.5
2
2.5
0
5
10
15
20
25

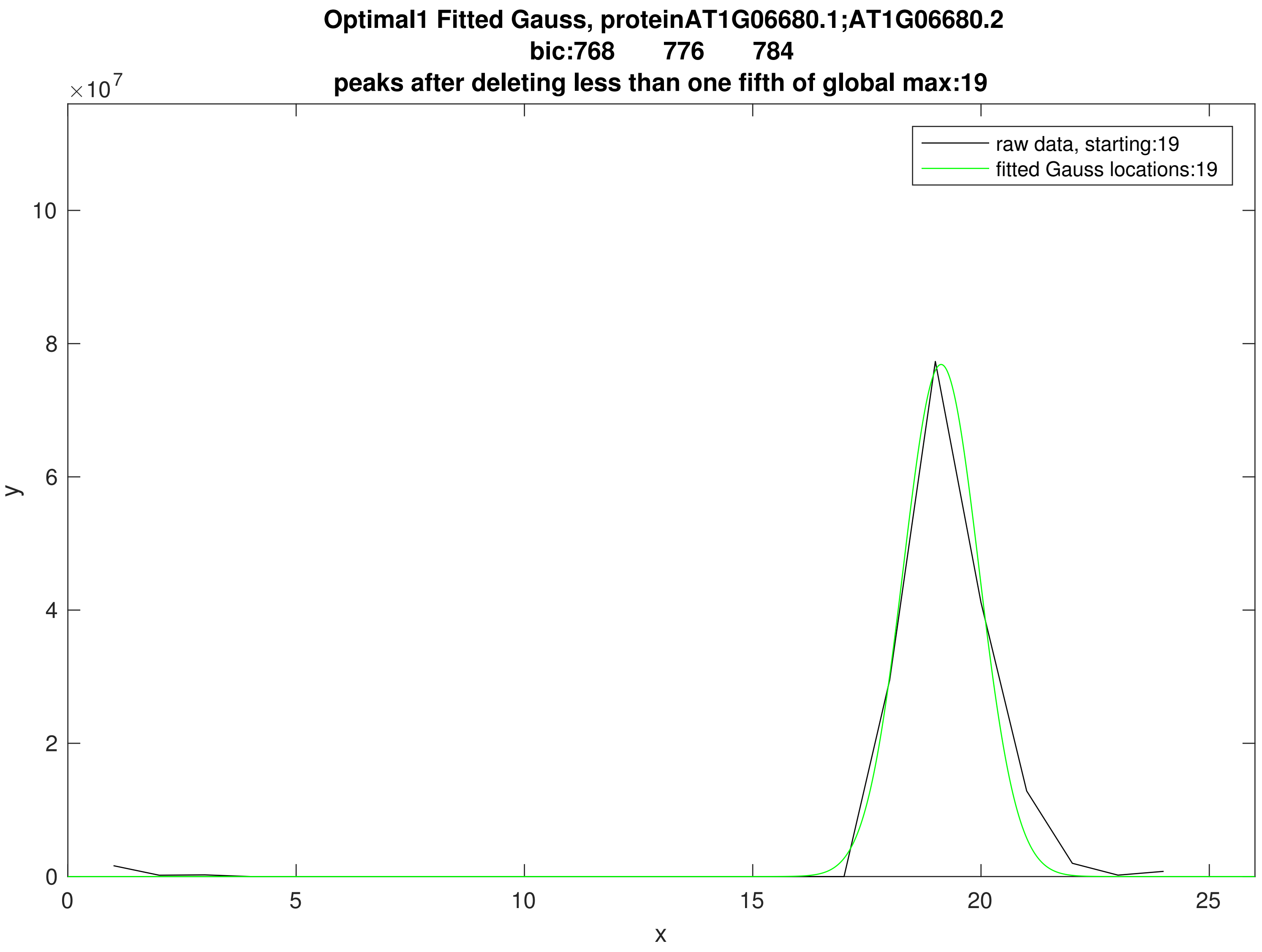

Optimal1 Fitted Gauss, proteinAT1G06680.1;AT1G06680.2
bic:768 776 784
peaks after deleting less than one fifth of global max:19
×10^7
raw data, starting:19
fitted Gauss locations:19
10
8
6
4
2
0
y
0
5
10
15
20
25
x

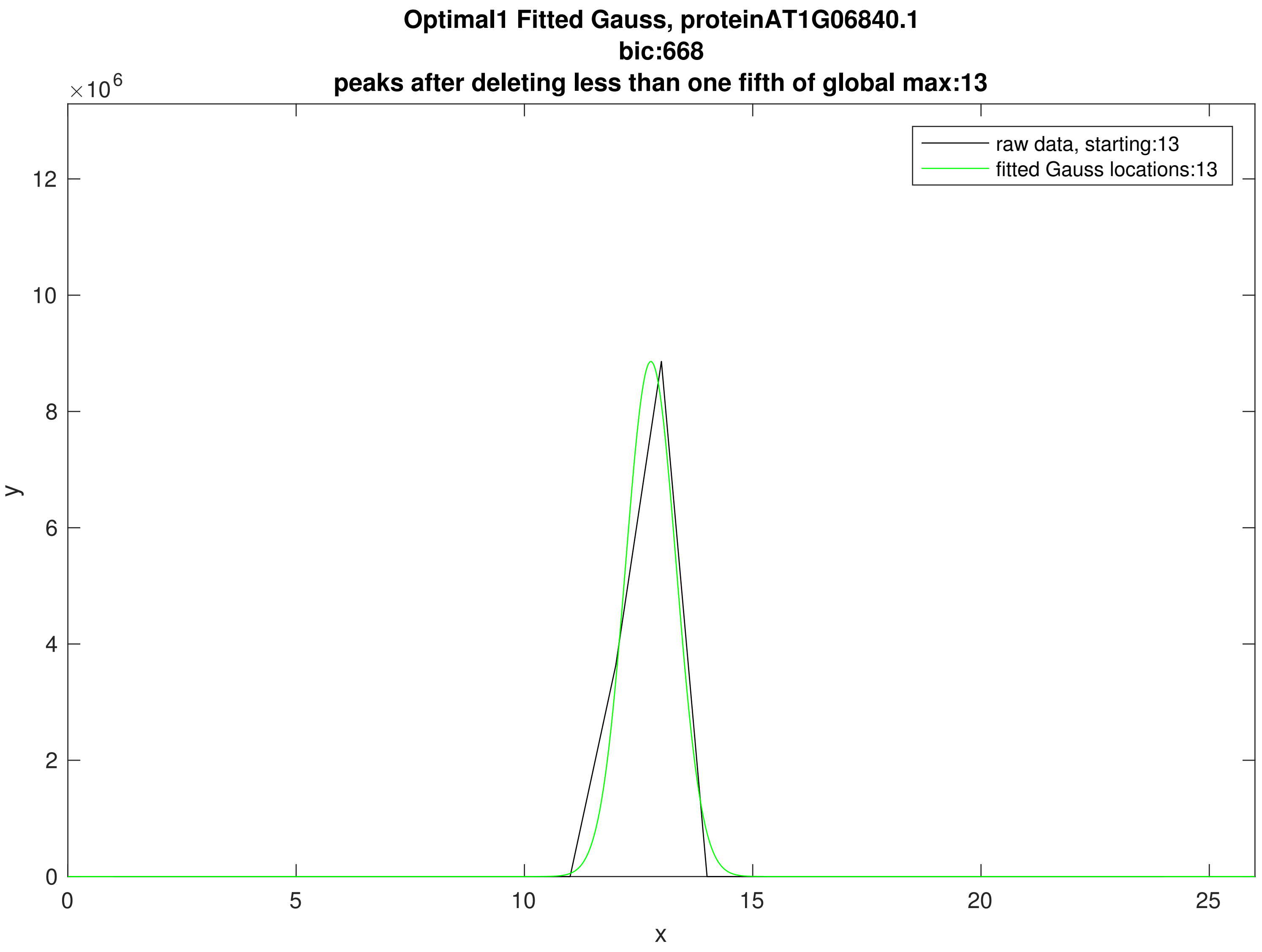
Optimal1 Fitted Gauss, proteinAT1G06840.1
bic:668
peaks after deleting less than one fifth of global max:13
$\times 10^6$
raw data, starting:13
fitted Gauss locations:13
0
2
4
6
8
10
12
y
0
5
10
15
20
25
x

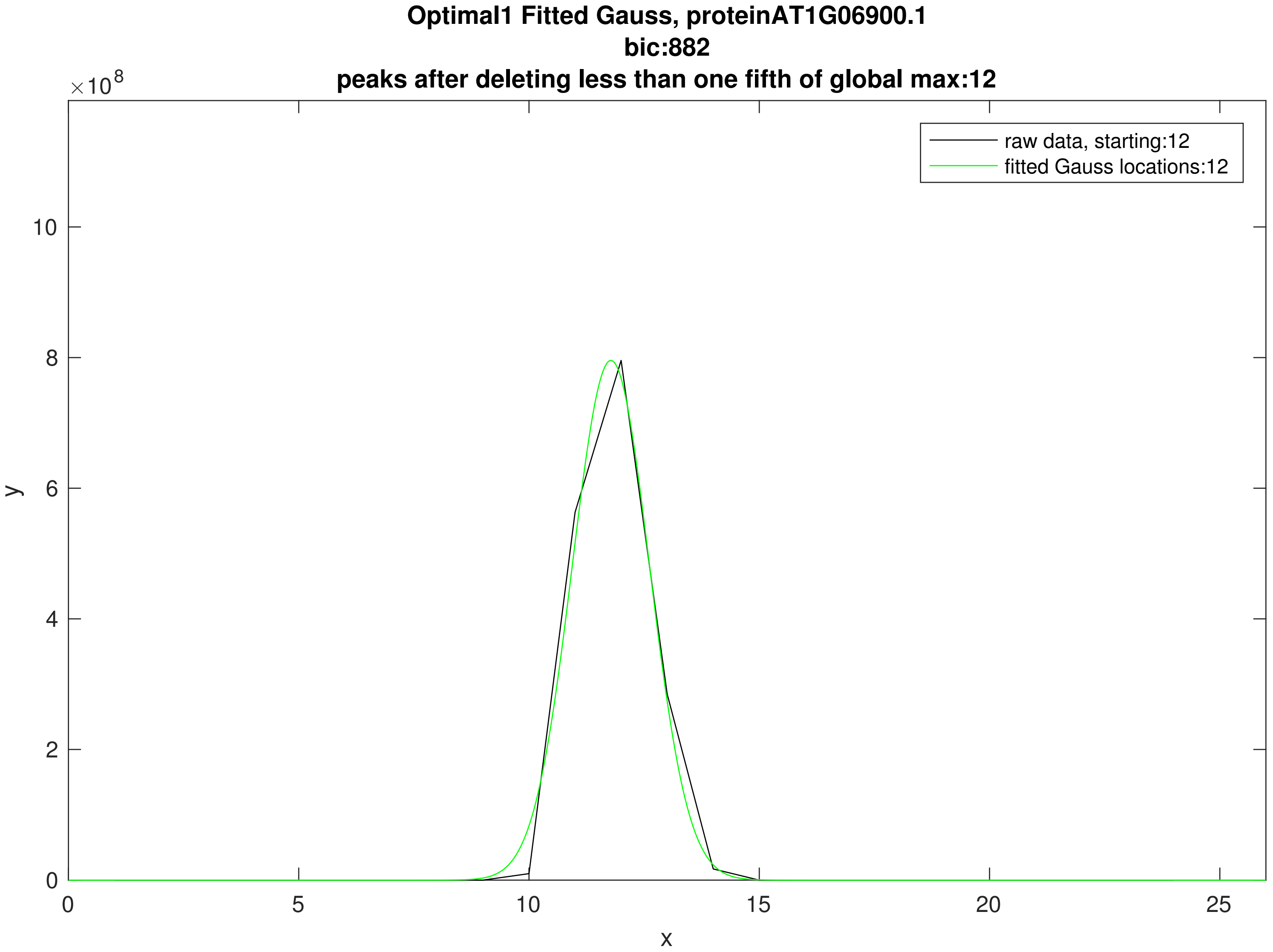
Optimal1 Fitted Gauss, proteinAT1G06900.1
bic:882
peaks after deleting less than one fifth of global max:12
×10^8
raw data, starting:12
fitted Gauss locations:12
y
x
0
2
4
6
8
10
0
5
10
15
20
25

**Optimal1 Fitted Gauss, proteinAT1G07040.1**
**bic:833**
**peaks after deleting less than one fifth of global max:14**

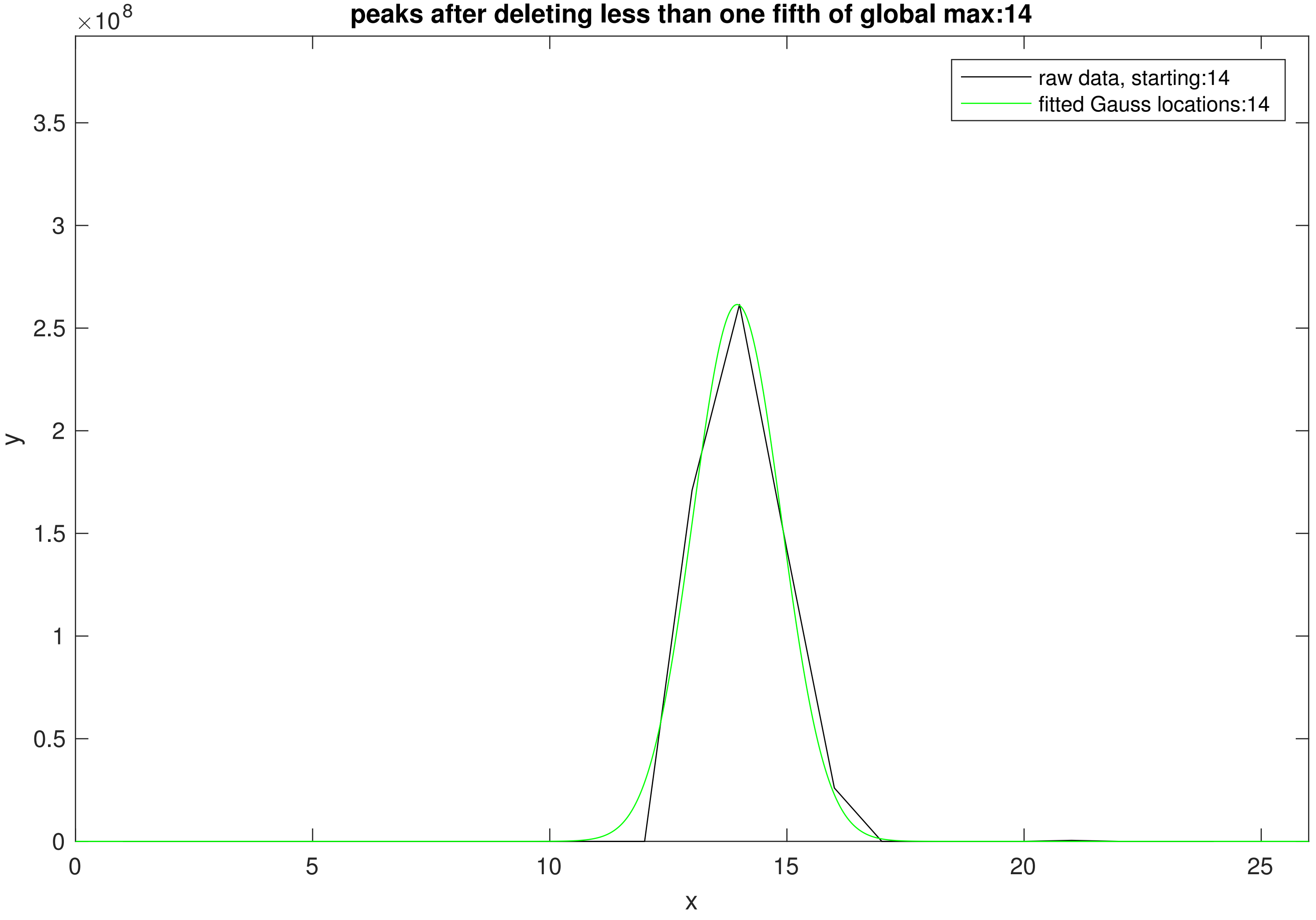

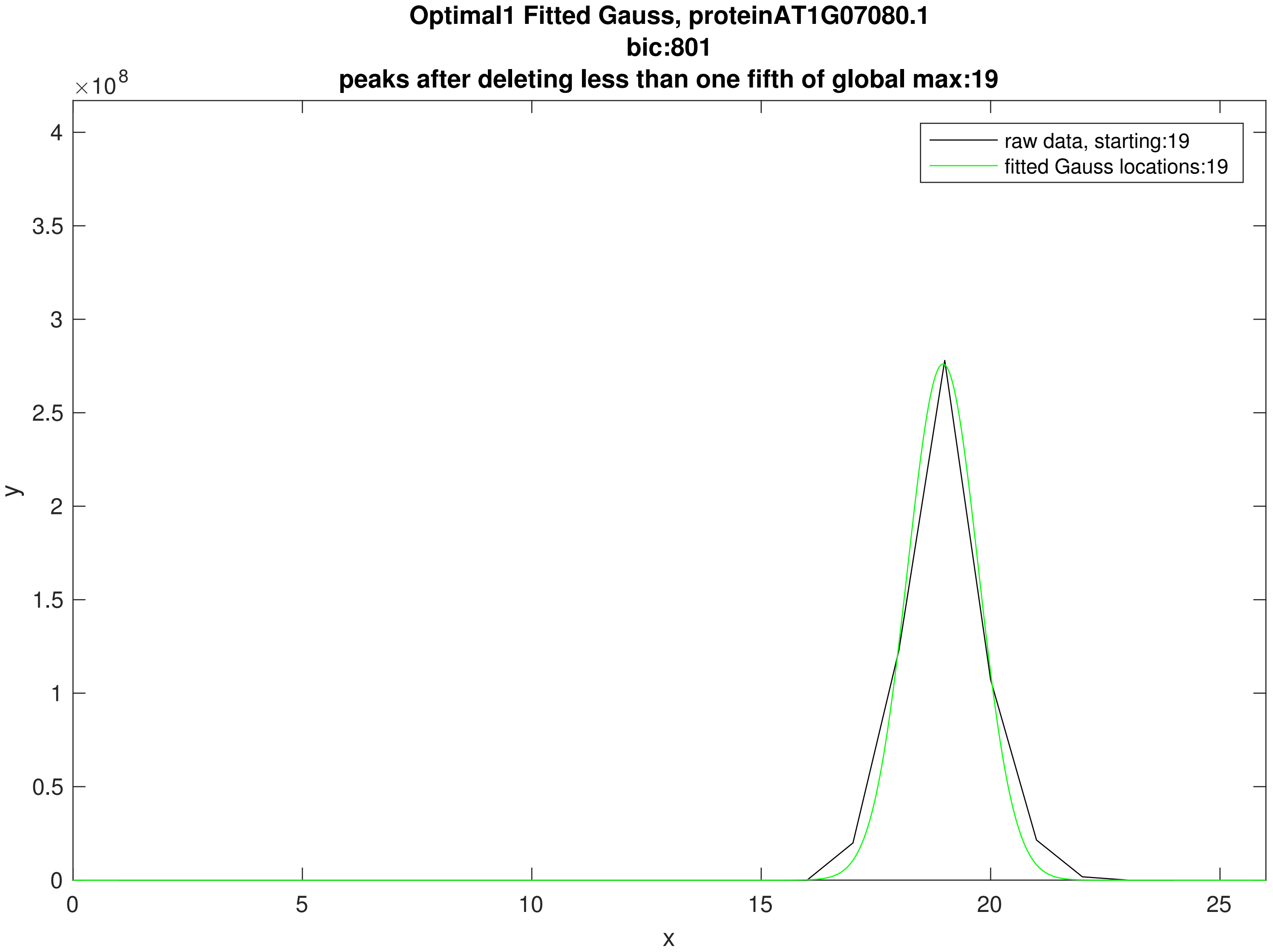
Optimal1 Fitted Gauss, proteinAT1G07080.1
bic:801
peaks after deleting less than one fifth of global max:19
×10^8
4
3.5
3
2.5
2
1.5
1
0.5
0
y
0
5
10
15
20
25
x
raw data, starting:19
fitted Gauss locations:19

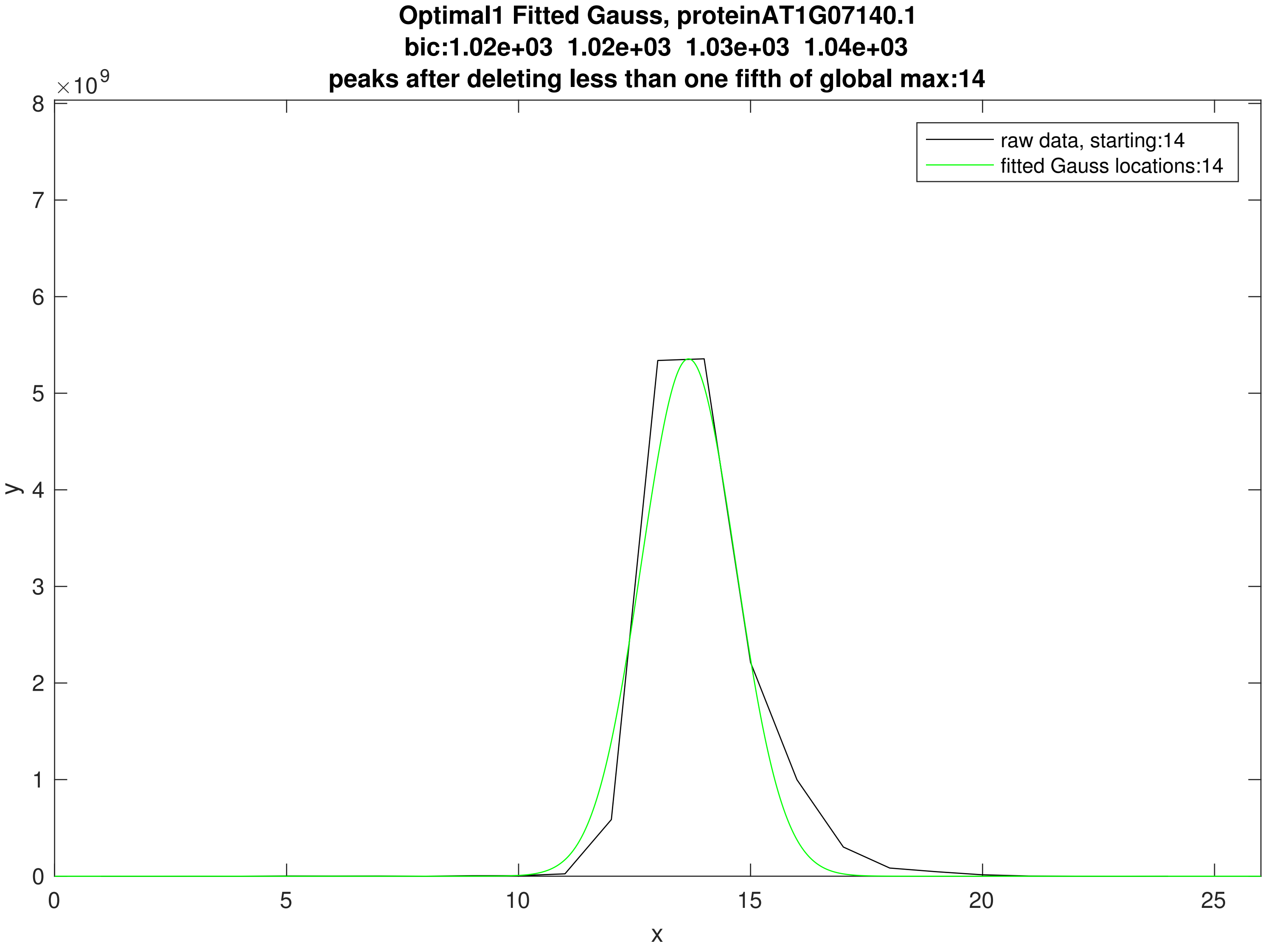
Optimal1 Fitted Gauss, proteinAT1G07140.1
bic:1.02e+03 1.02e+03 1.03e+03 1.04e+03
peaks after deleting less than one fifth of global max:14
×10^9
raw data, starting:14
fitted Gauss locations:14
y
x
0
1
2
3
4
5
6
7
8
0
5
10
15
20
25

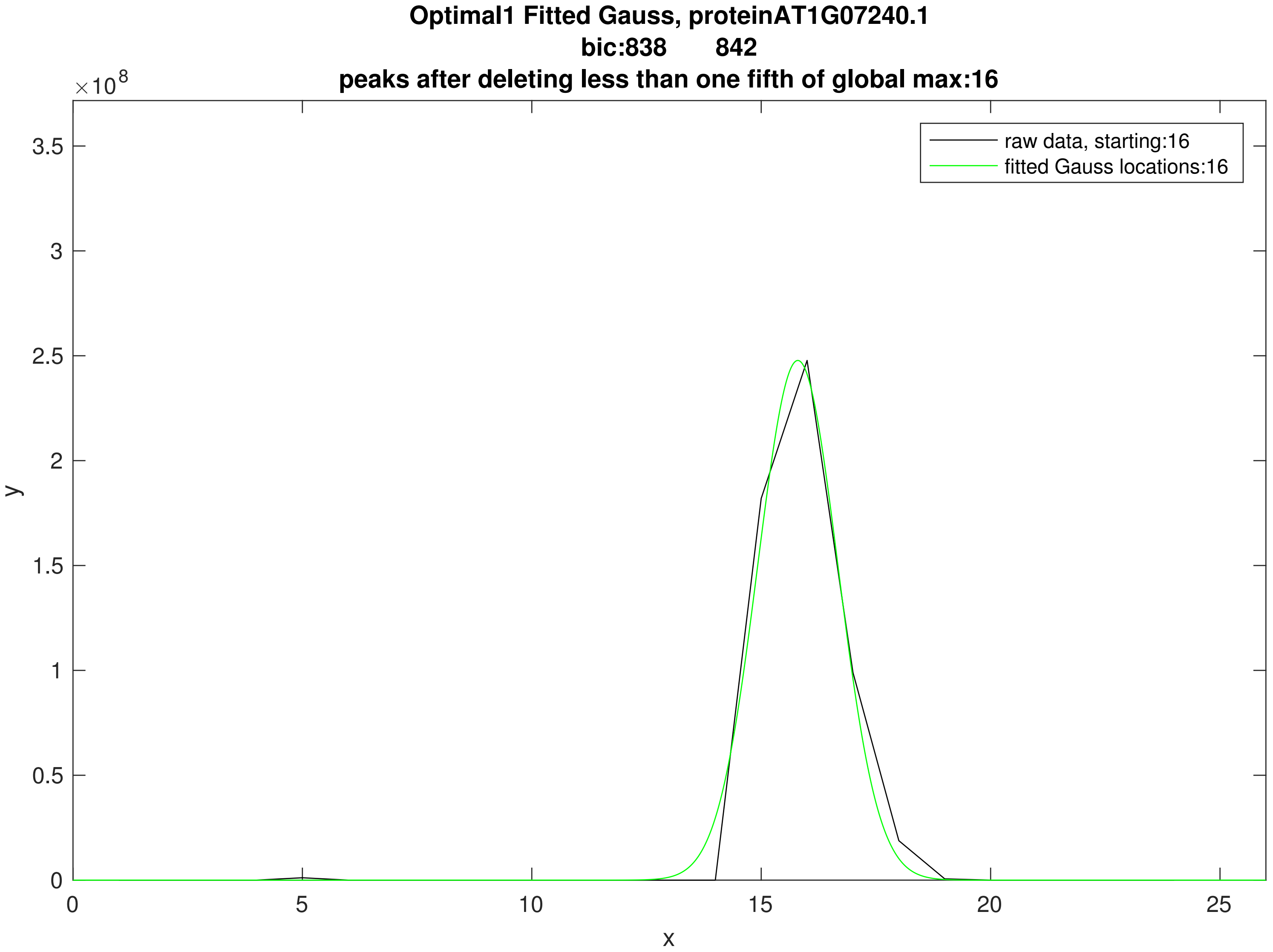
Optimal1 Fitted Gauss, proteinAT1G07240.1
bic:838 842
peaks after deleting less than one fifth of global max:16
raw data, starting:16
fitted Gauss locations:16
×10^8
3.5
3
2.5
2
1.5
1
0.5
0
y
0
5
10
15
20
25
x

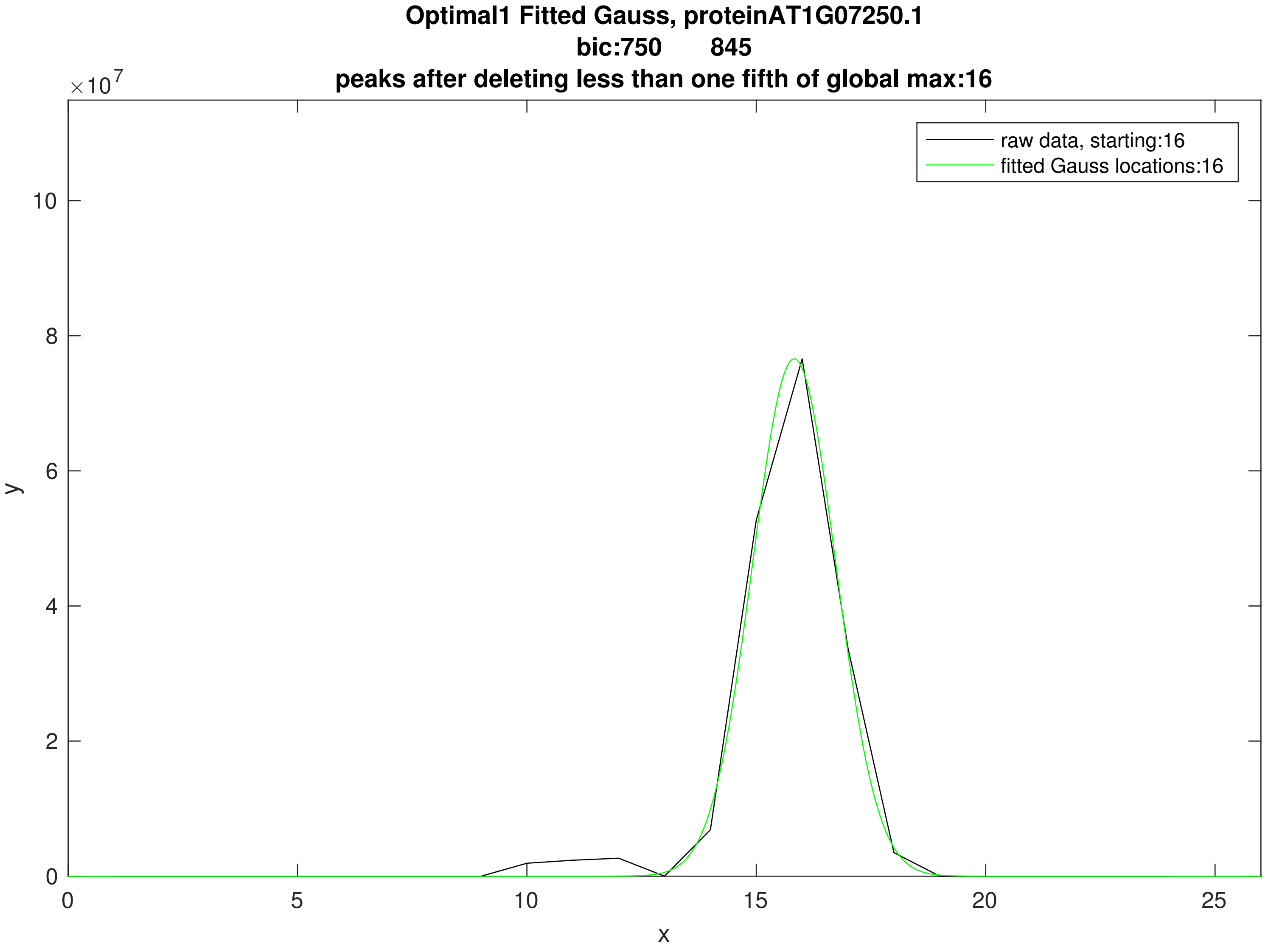
Optimal1 Fitted Gauss, proteinAT1G07250.1
bic:750 845
peaks after deleting less than one fifth of global max:16
×10^7
raw data, starting:16
fitted Gauss locations:16
y
x
0
2
4
6
8
10
0
5
10
15
20
25

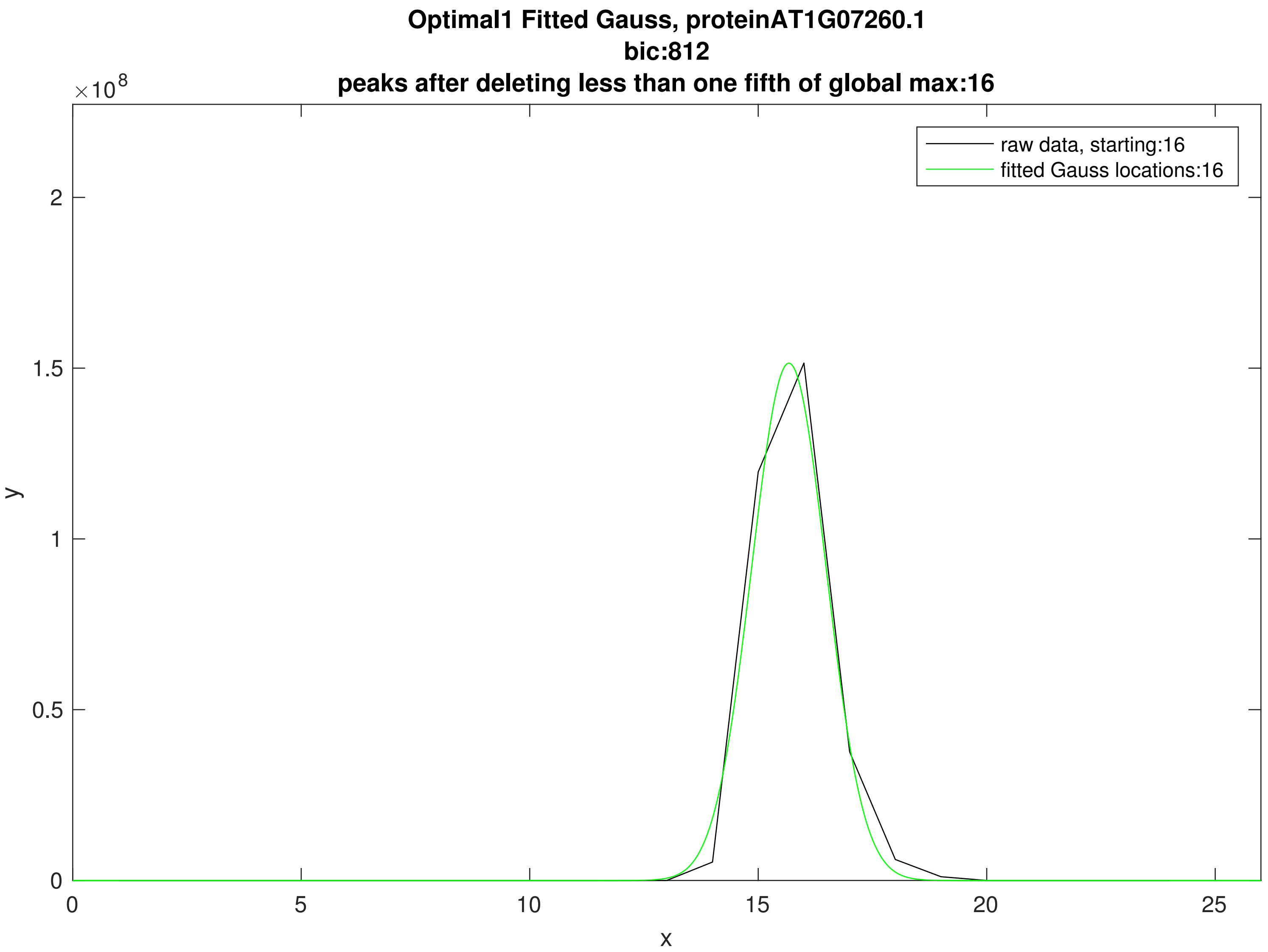
Optimal1 Fitted Gauss, proteinAT1G07260.1
bic:812
peaks after deleting less than one fifth of global max:16
×10^8
2
1.5
1
0.5
0
y
0
5
10
15
20
25
x
raw data, starting:16
fitted Gauss locations:16

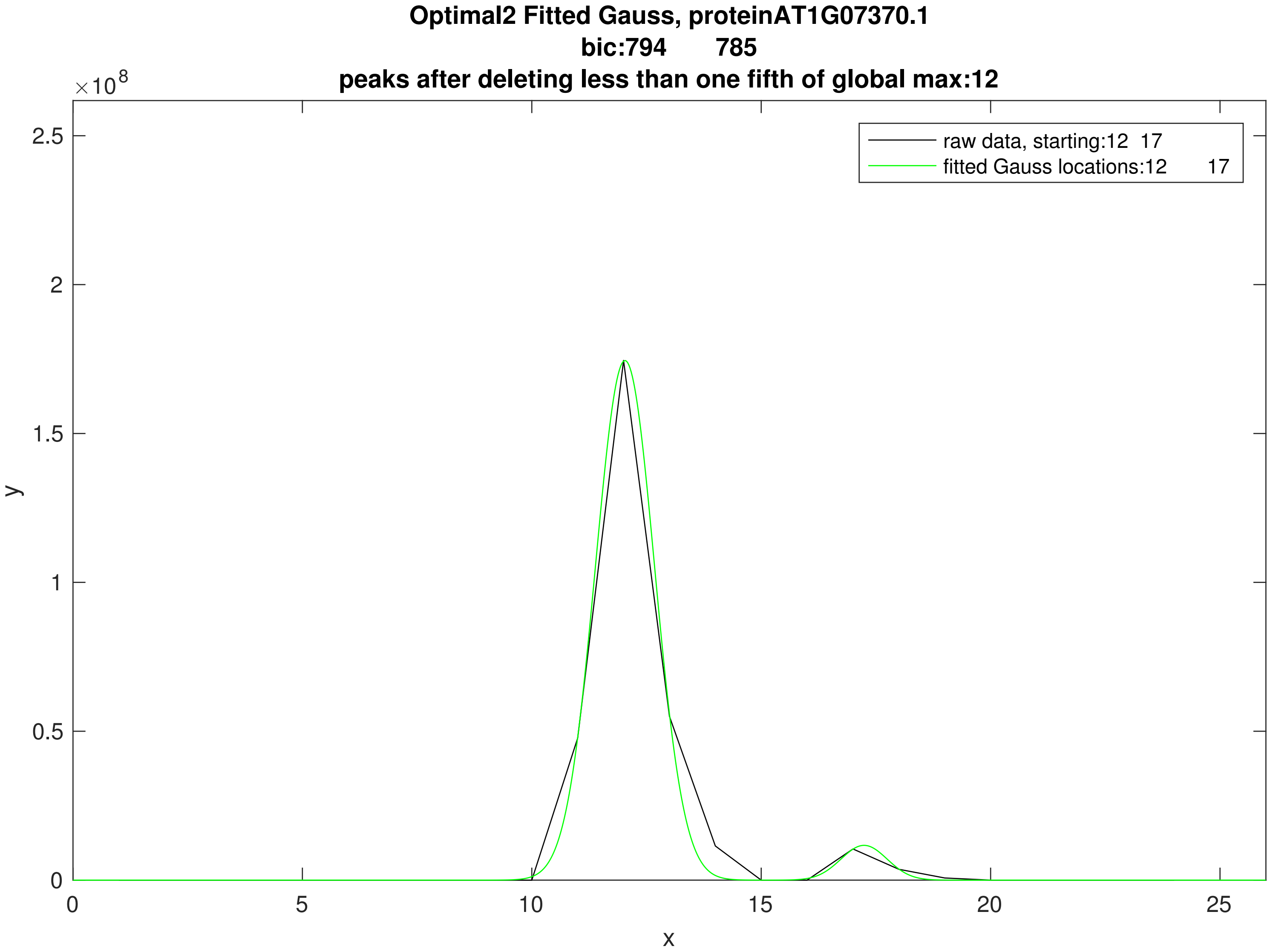

Optimal2 Fitted Gauss, proteinAT1G07370.1
bic:794 785
peaks after deleting less than one fifth of global max:12
×10^8
raw data, starting:12 17
fitted Gauss locations:12 17
y
x

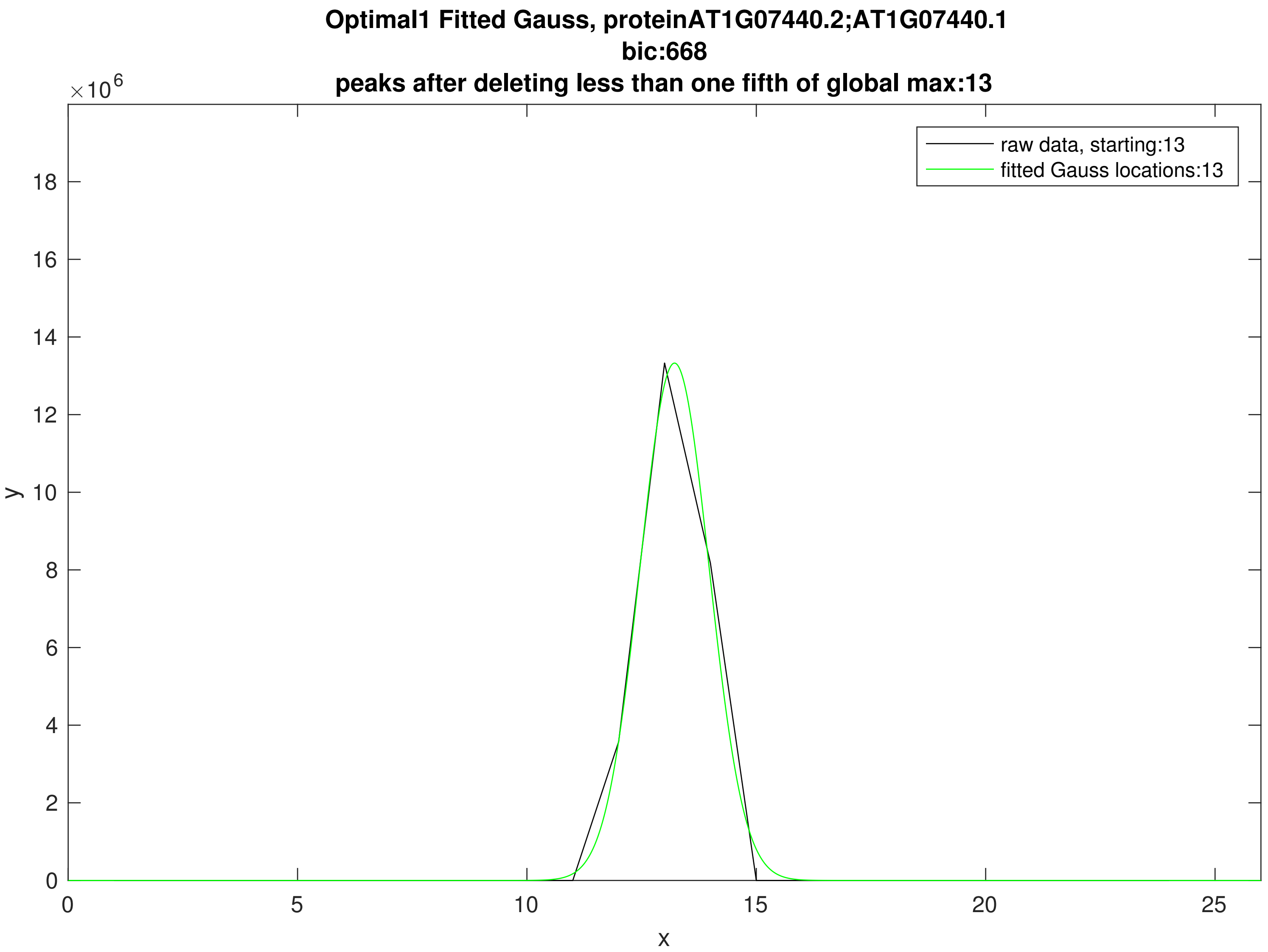
Optimal1 Fitted Gauss, proteinAT1G07440.2;AT1G07440.1
bic:668
peaks after deleting less than one fifth of global max:13
×10^6
raw data, starting:13
fitted Gauss locations:13
y
x
0
2
4
6
8
10
12
14
16
18
0
5
10
15
20
25

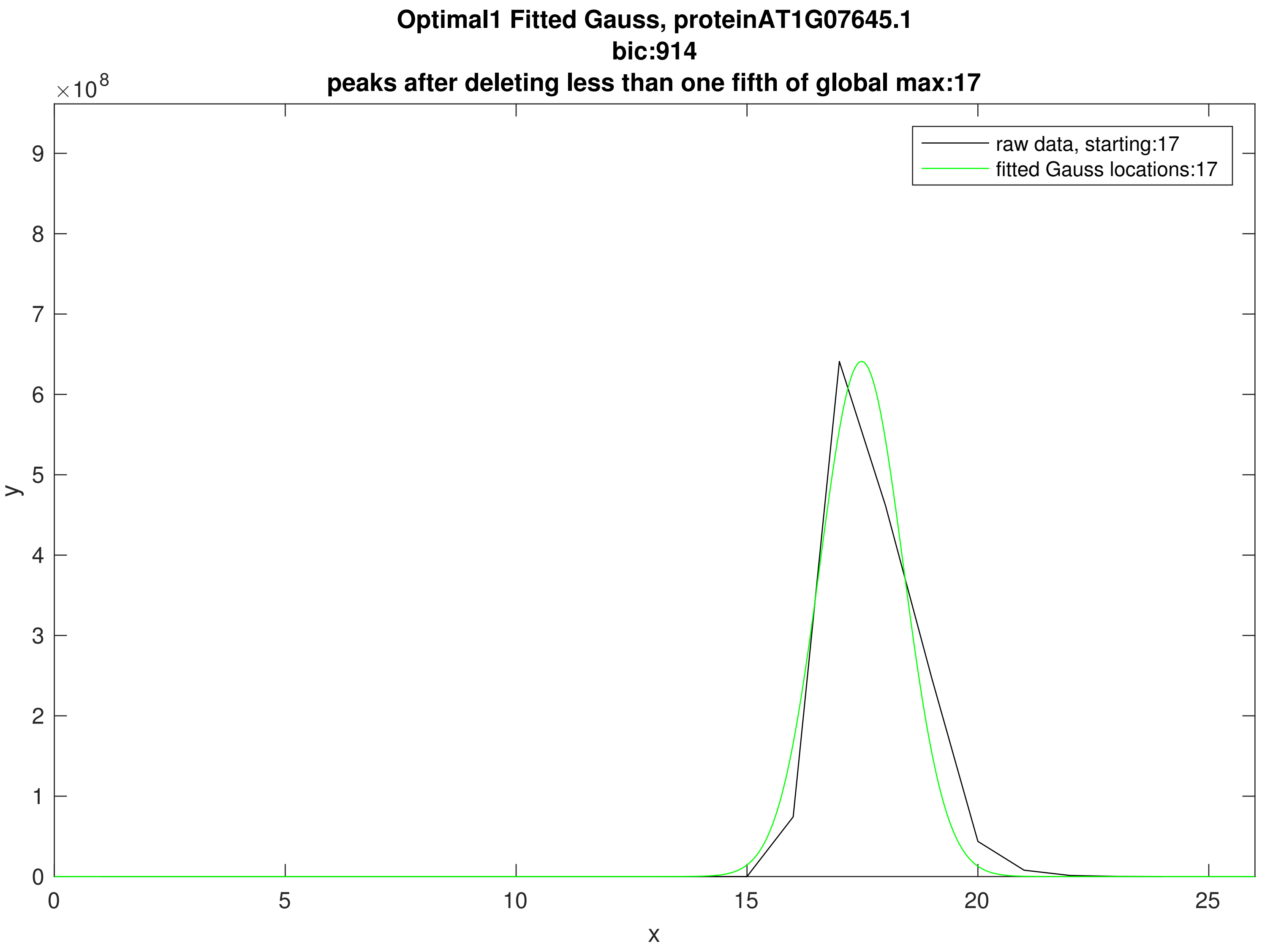
Optimal1 Fitted Gauss, proteinAT1G07645.1
bic:914
peaks after deleting less than one fifth of global max:17
×10^8
raw data, starting:17
fitted Gauss locations:17
y
x
0
1
2
3
4
5
6
7
8
9
0
5
10
15
20
25

, proteinAT1G07660.2;AT5G59970.1;AT5G59690.1;AT3G53730.1;AT3G46320.1;AT3G45930.1;AT2G28740.1;AT1G07820
bic:715
peaks after deleting less than one fifth of global max:19

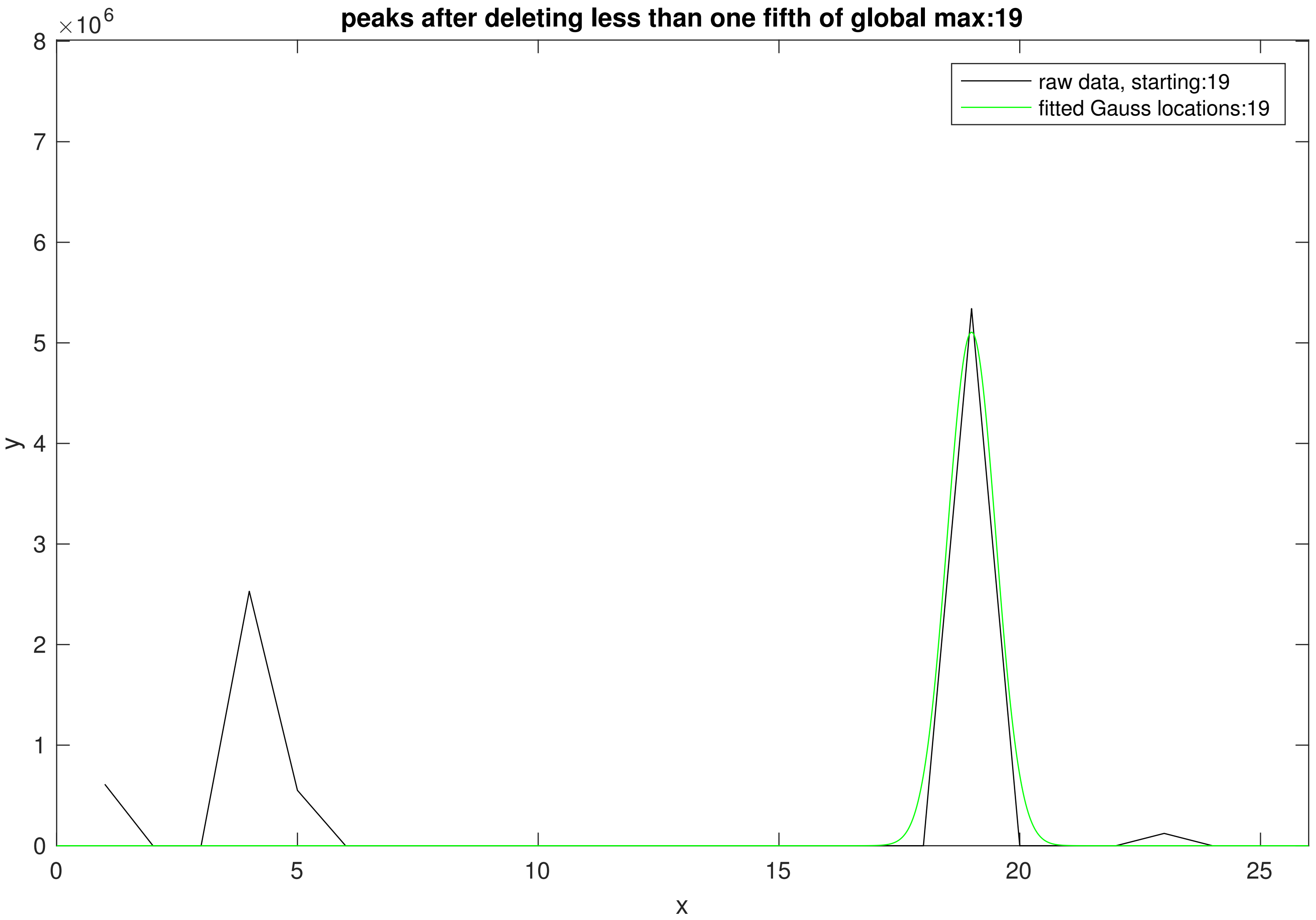

Optimal1 Fitted Gauss, proteinAT1G07750.1
bic:942
peaks after deleting less than one fifth of global max:13

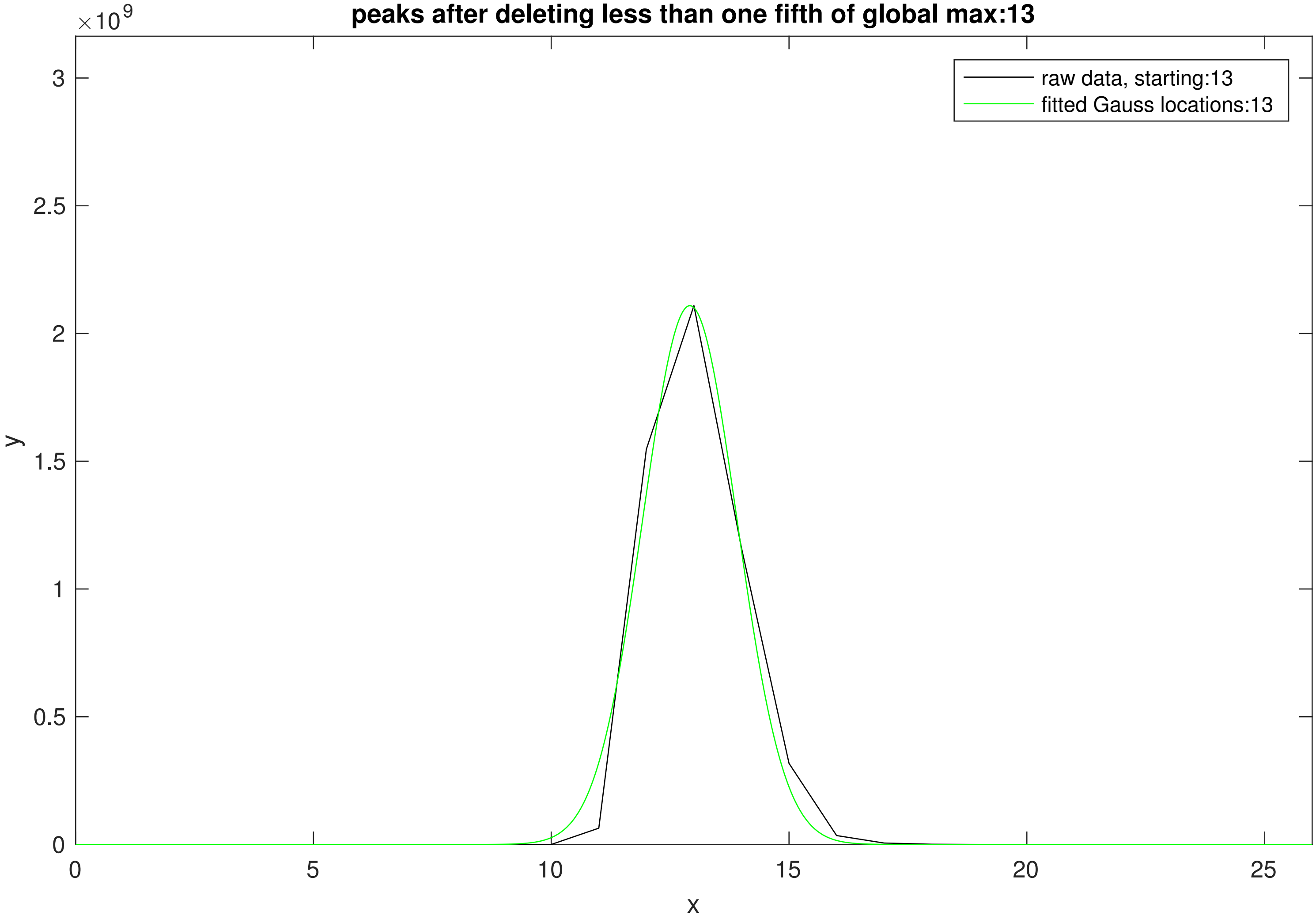

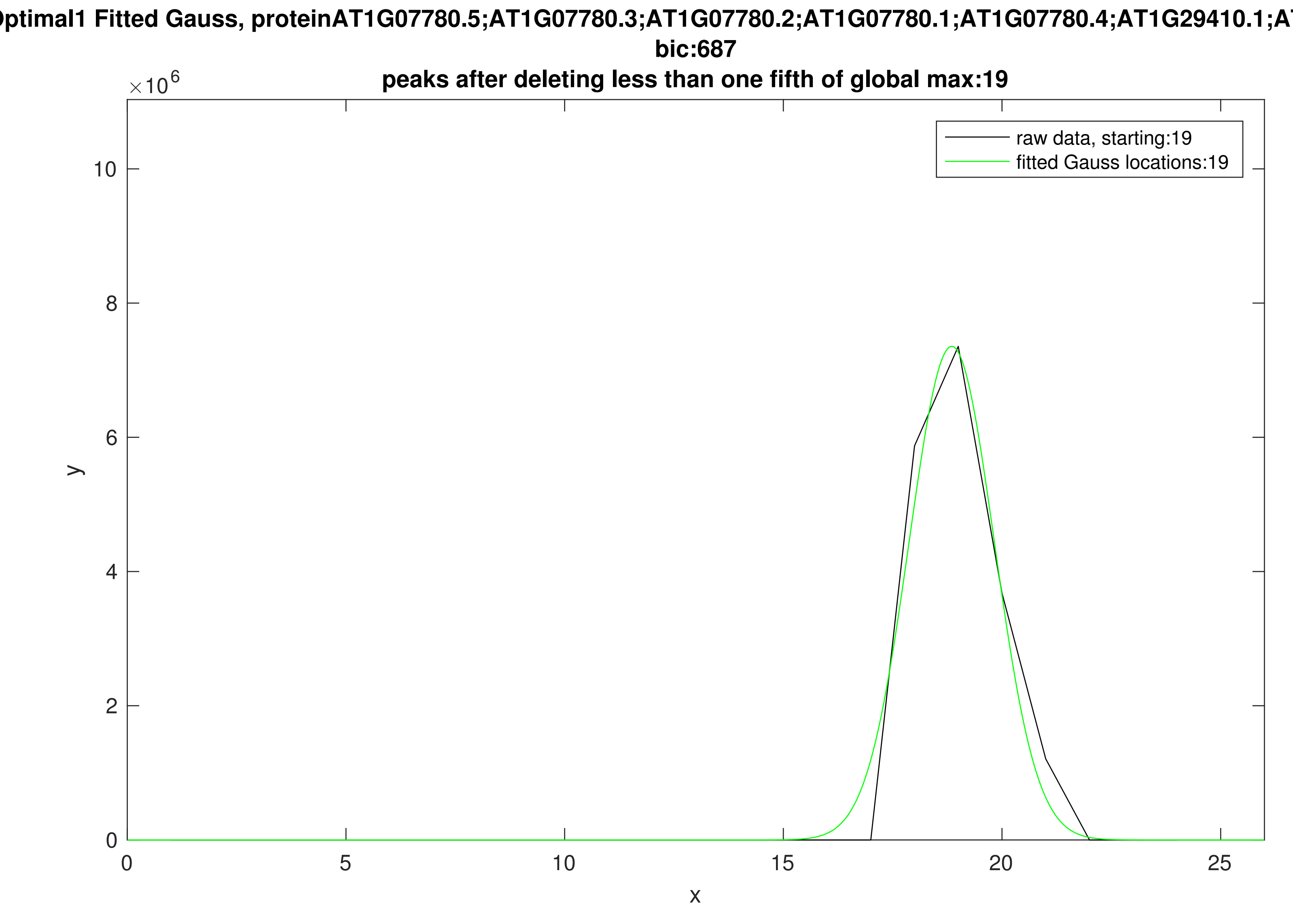

Optimal1 Fitted Gauss, proteinAT1G07780.5;AT1G07780.3;AT1G07780.2;AT1G07780.1;AT1G07780.4;AT1G29410.1;AT1
bic:687
peaks after deleting less than one fifth of global max:19
×10^6
raw data, starting:19
fitted Gauss locations:19
y
x
0
2
4
6
8
10
0
5
10
15
20
25

**1 Fitted Gauss, proteinAT1G07890.8;AT1G07890.7;AT1G07890.5;AT1G07890.4;AT1G07890.3;AT1G07890.2;AT1G0789[illegible]**
**bic:1.05e+03 1.06e+03 1.08e+03**
**peaks after deleting less than one fifth of global max:16**

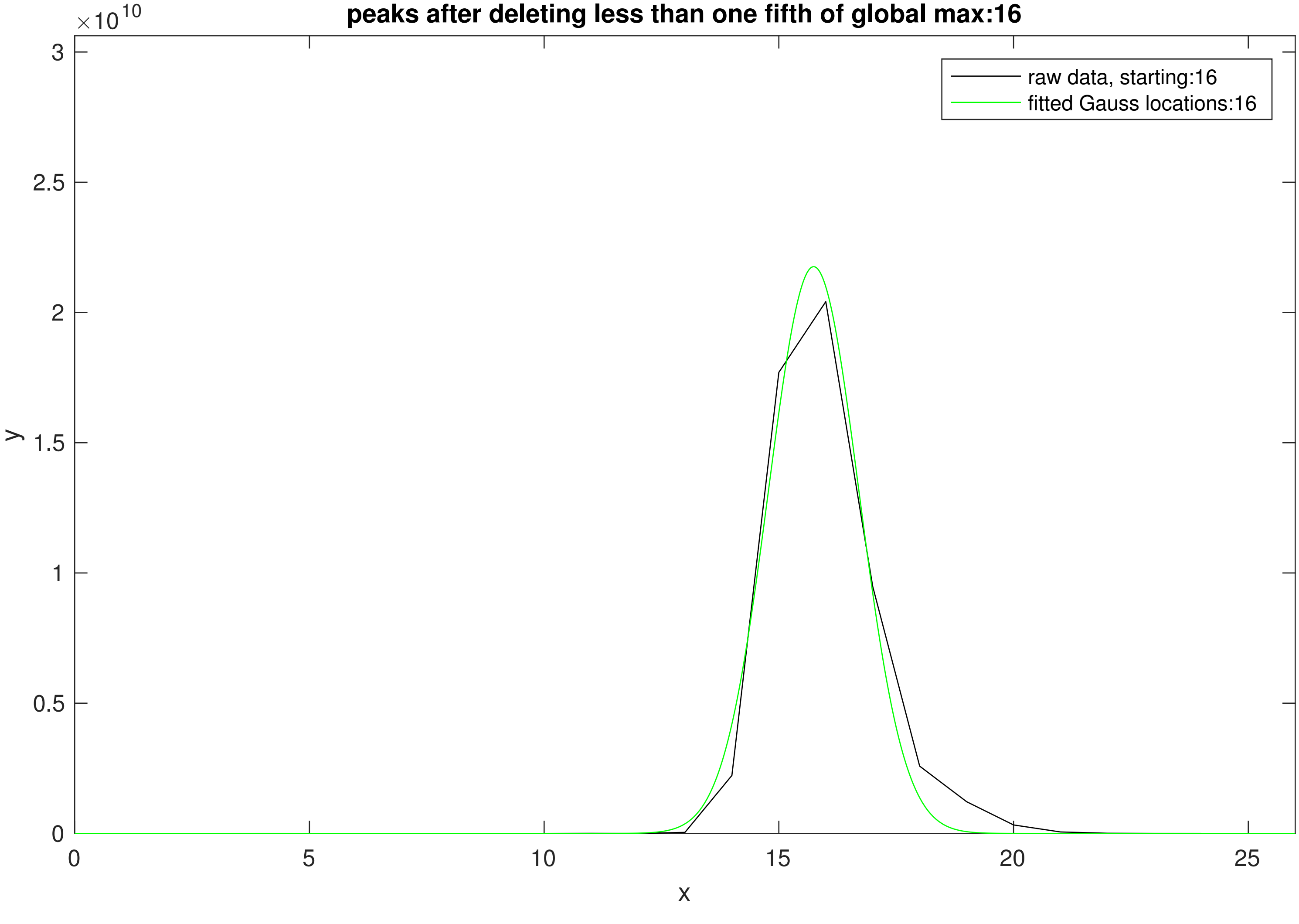

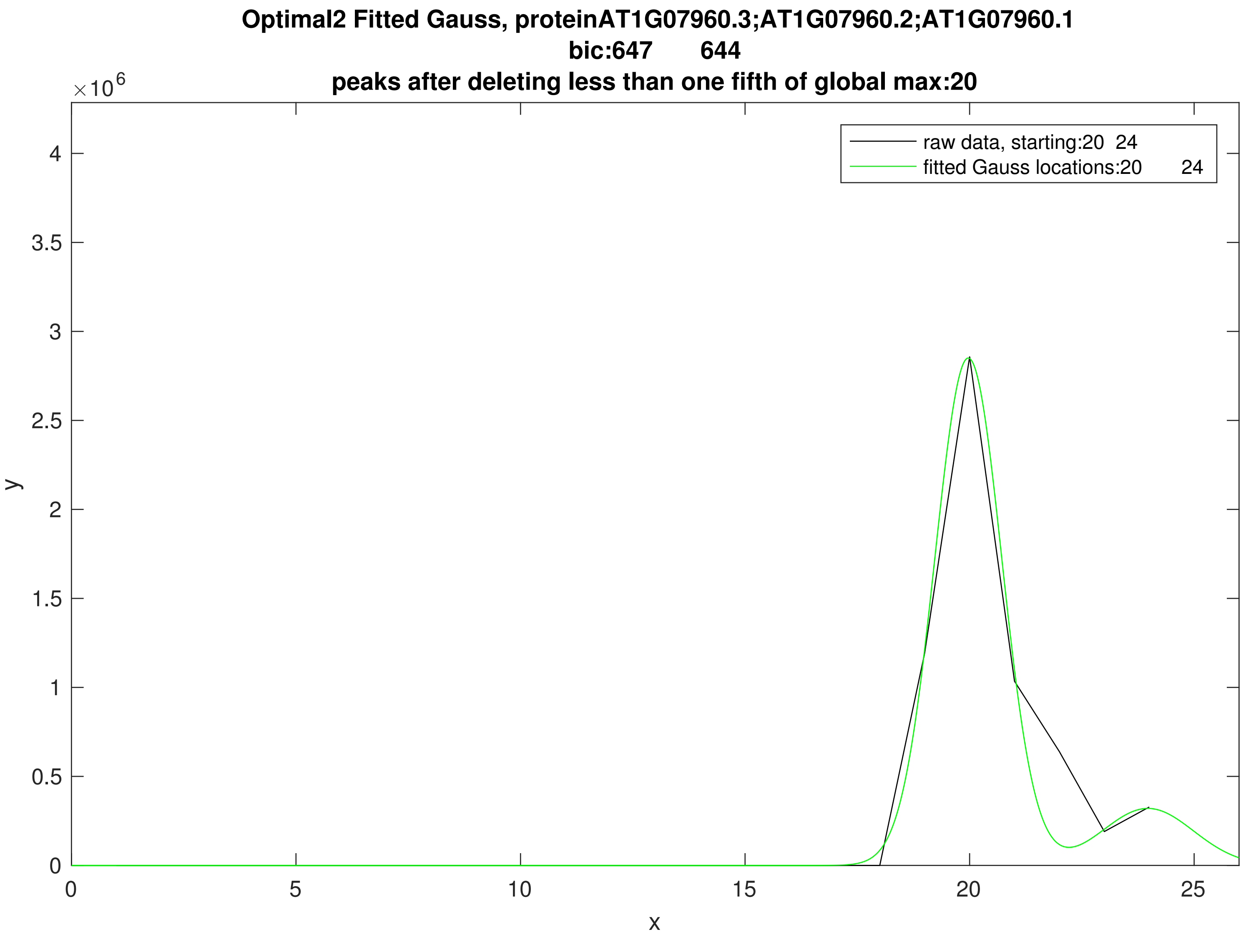

Optimal2 Fitted Gauss, proteinAT1G07960.3;AT1G07960.2;AT1G07960.1
bic:647 644
peaks after deleting less than one fifth of global max:20
×10^6
raw data, starting:20 24
fitted Gauss locations:20 24
4
3.5
3
2.5
2
1.5
1
0.5
0
y
0
5
10
15
20
25
x

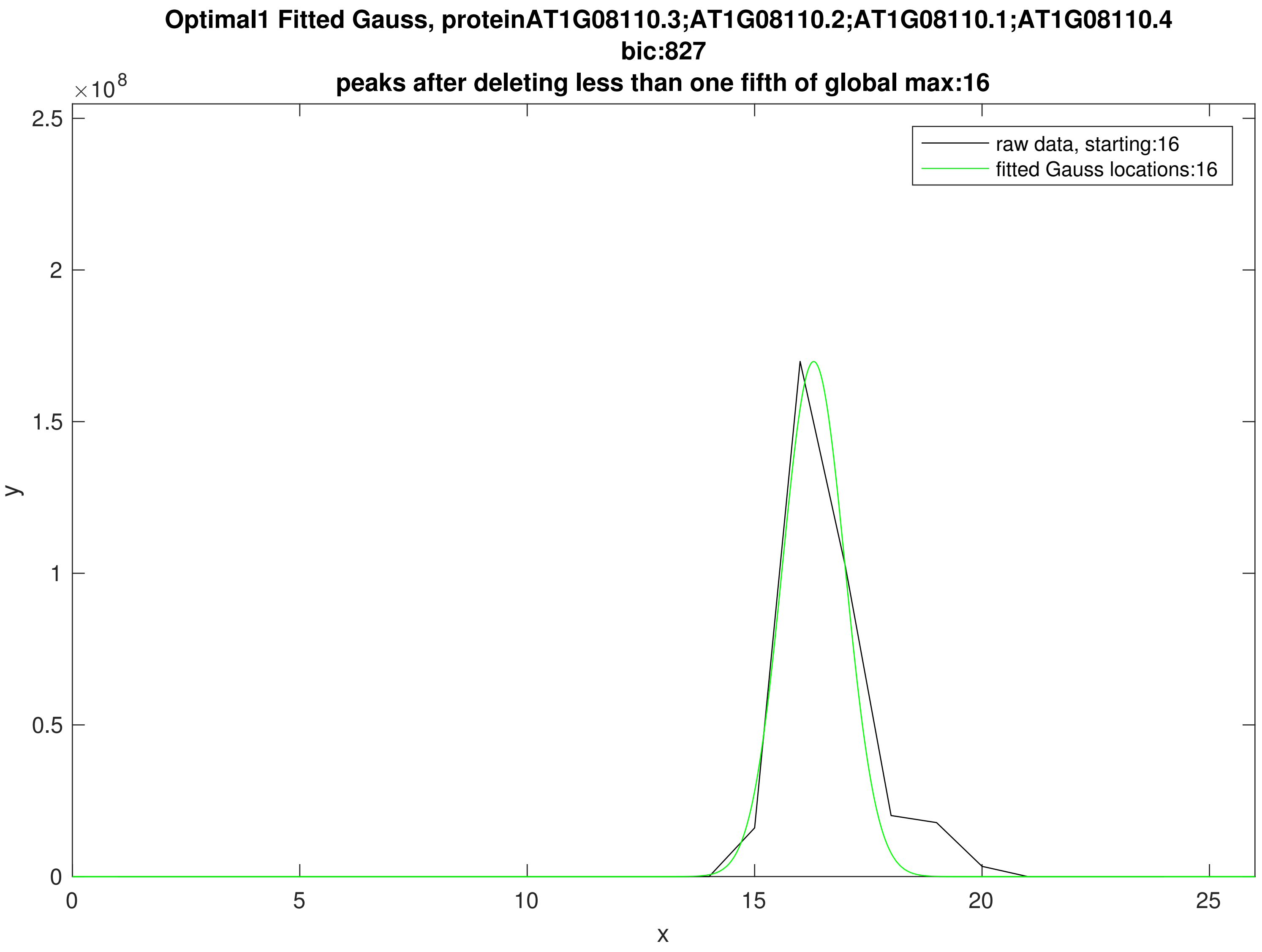
Optimal1 Fitted Gauss, proteinAT1G08110.3;AT1G08110.2;AT1G08110.1;AT1G08110.4
bic:827
peaks after deleting less than one fifth of global max:16
×10^8
2.5
2
1.5
1
0.5
0
y
0
5
10
15
20
25
x
raw data, starting:16
fitted Gauss locations:16

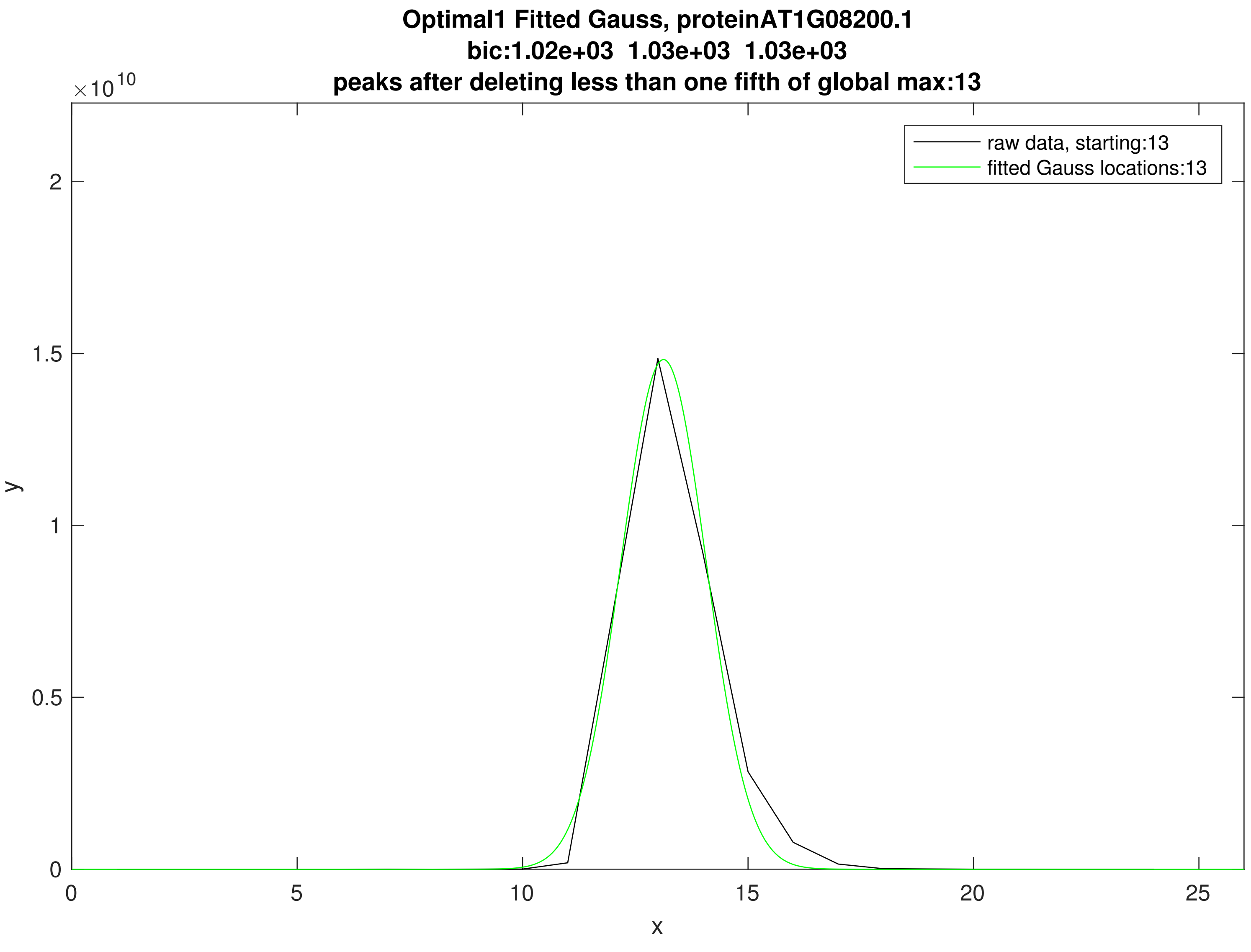
Optimal1 Fitted Gauss, proteinAT1G08200.1
bic:1.02e+03 1.03e+03 1.03e+03
peaks after deleting less than one fifth of global max:13
×10^10
raw data, starting:13
fitted Gauss locations:13
y
x
0
0.5
1
1.5
2
0
5
10
15
20
25

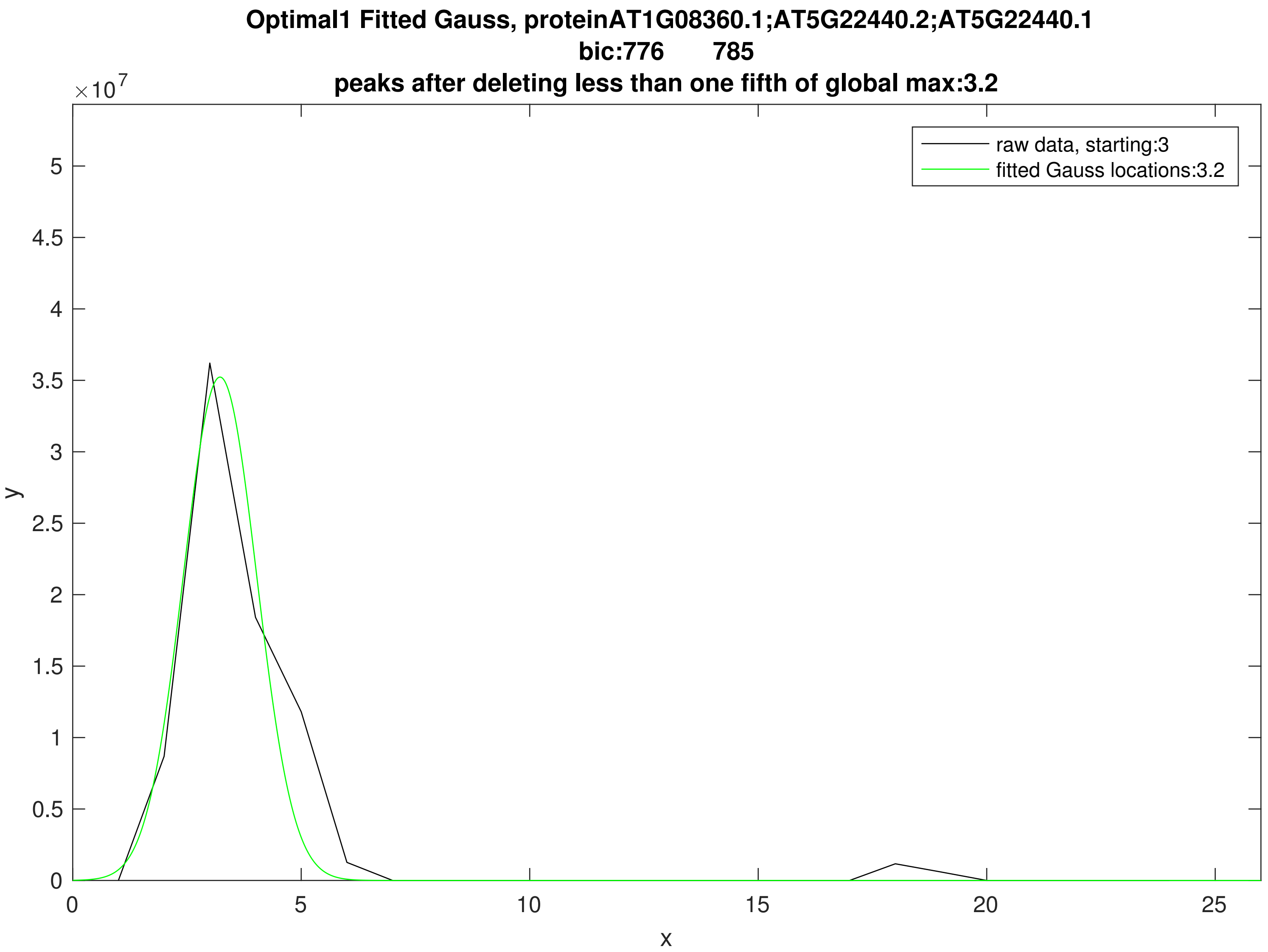

Optimal1 Fitted Gauss, proteinAT1G08360.1;AT5G22440.2;AT5G22440.1
bic:776 785
peaks after deleting less than one fifth of global max:3.2
×10^7
5
4.5
4
3.5
3
2.5
2
1.5
1
0.5
0
y
0
5
10
15
20
25
x
raw data, starting:3
fitted Gauss locations:3.2

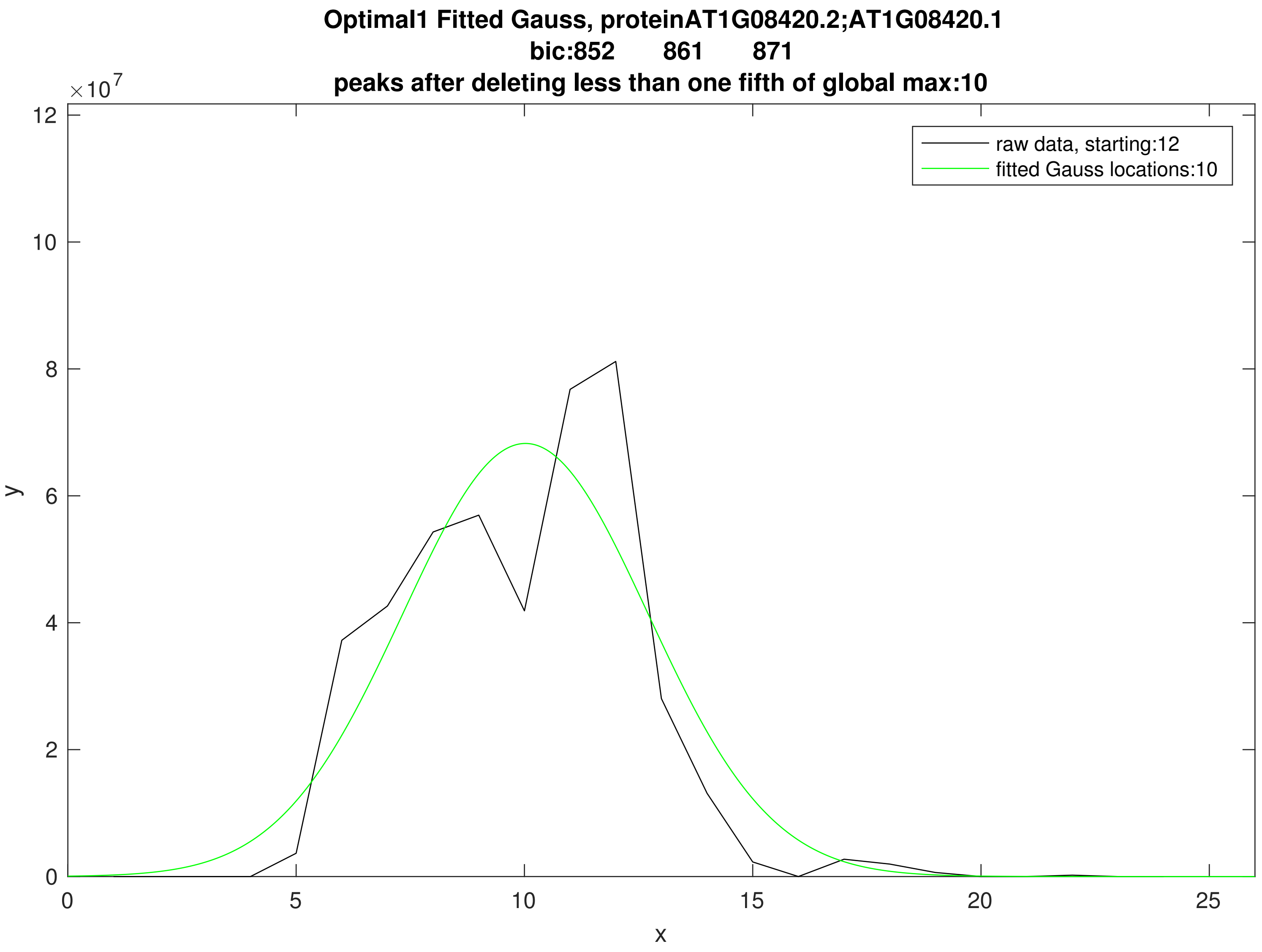
Optimal1 Fitted Gauss, proteinAT1G08420.2;AT1G08420.1
bic:852 861 871
peaks after deleting less than one fifth of global max:10
×10^7
raw data, starting:12
fitted Gauss locations:10
y
x
0
2
4
6
8
10
12
0
5
10
15
20
25

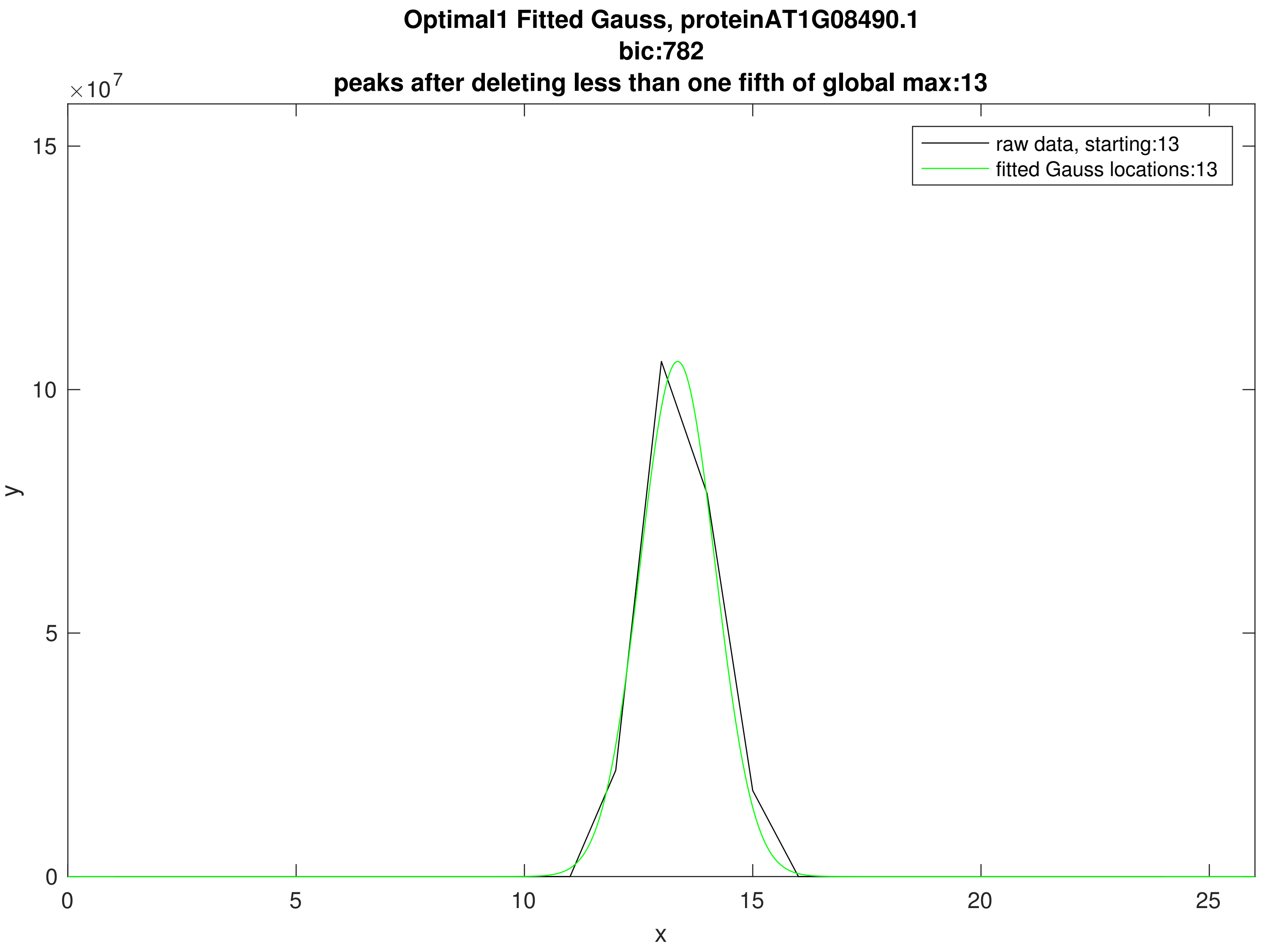
Optimal1 Fitted Gauss, proteinAT1G08490.1
bic:782
peaks after deleting less than one fifth of global max:13
×10^7
15
10
5
0
y
0
5
10
15
20
25
x
raw data, starting:13
fitted Gauss locations:13

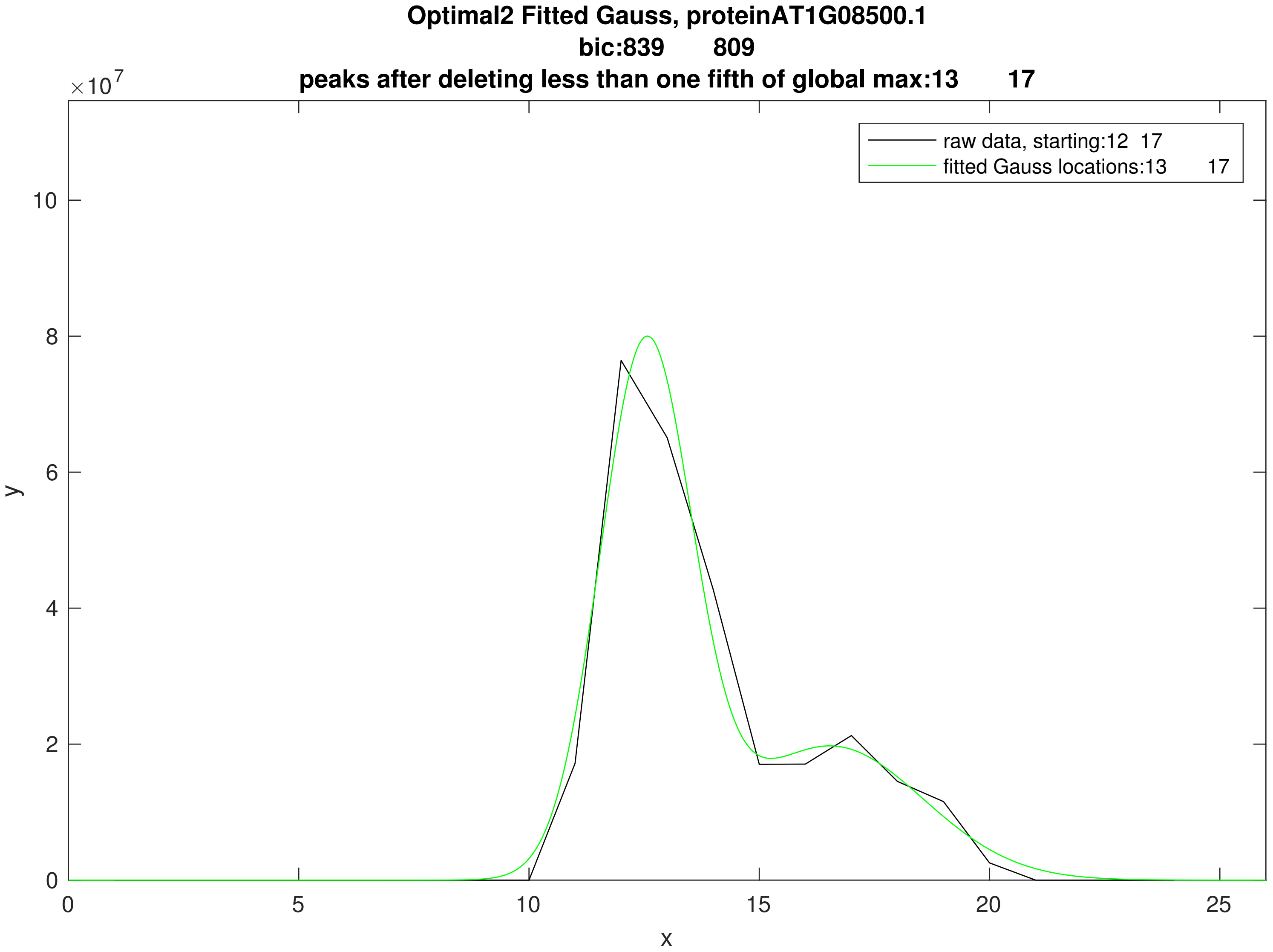
Optimal2 Fitted Gauss, proteinAT1G08500.1
bic:839 809
peaks after deleting less than one fifth of global max:13 17
×10^7
raw data, starting:12 17
fitted Gauss locations:13 17
y
x
0
2
4
6
8
10
0
5
10
15
20
25

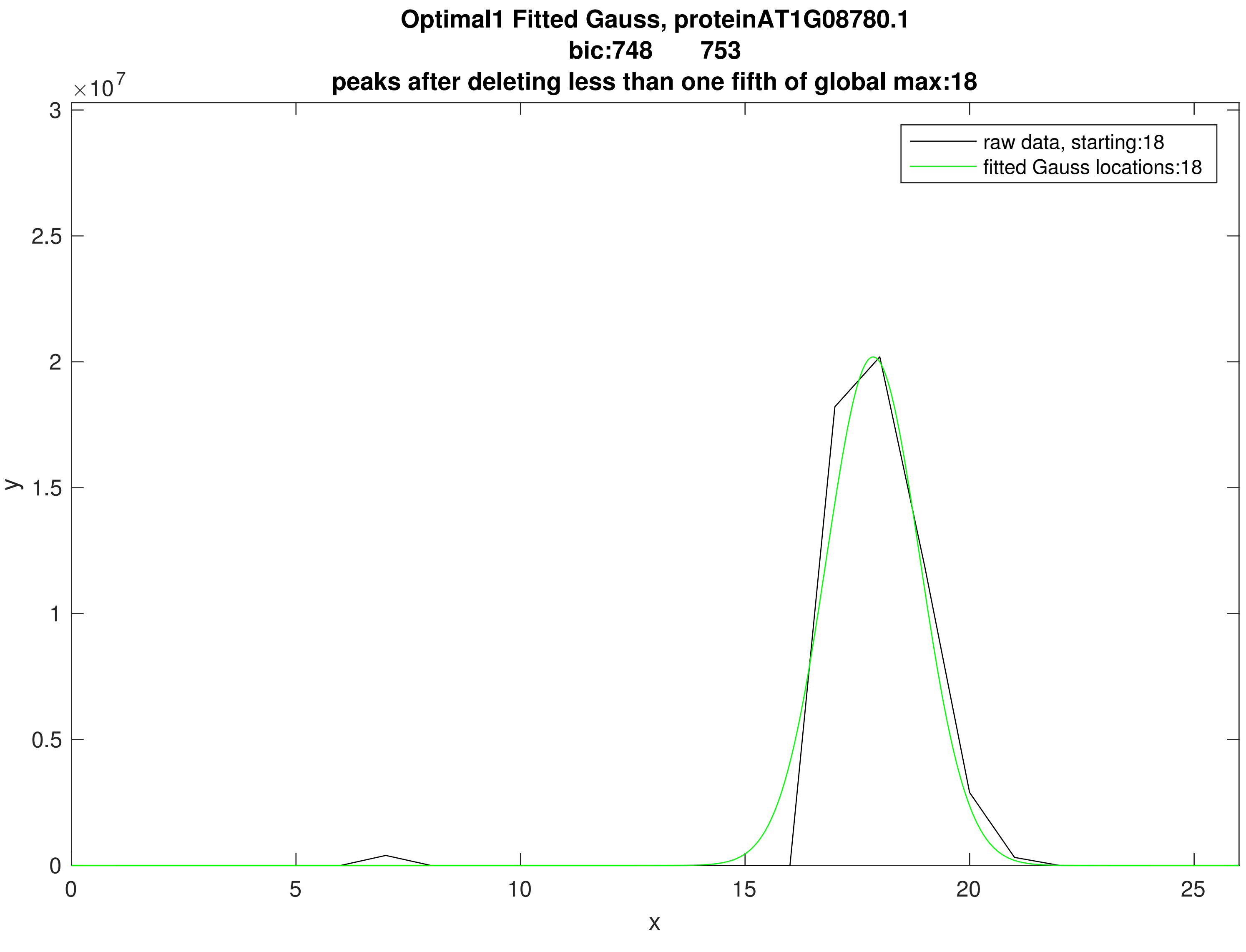
Optimal1 Fitted Gauss, proteinAT1G08780.1
bic:748 753
peaks after deleting less than one fifth of global max:18
×10^7
3
2.5
2
1.5
1
0.5
0
y
0
5
10
15
20
25
x
raw data, starting:18
fitted Gauss locations:18

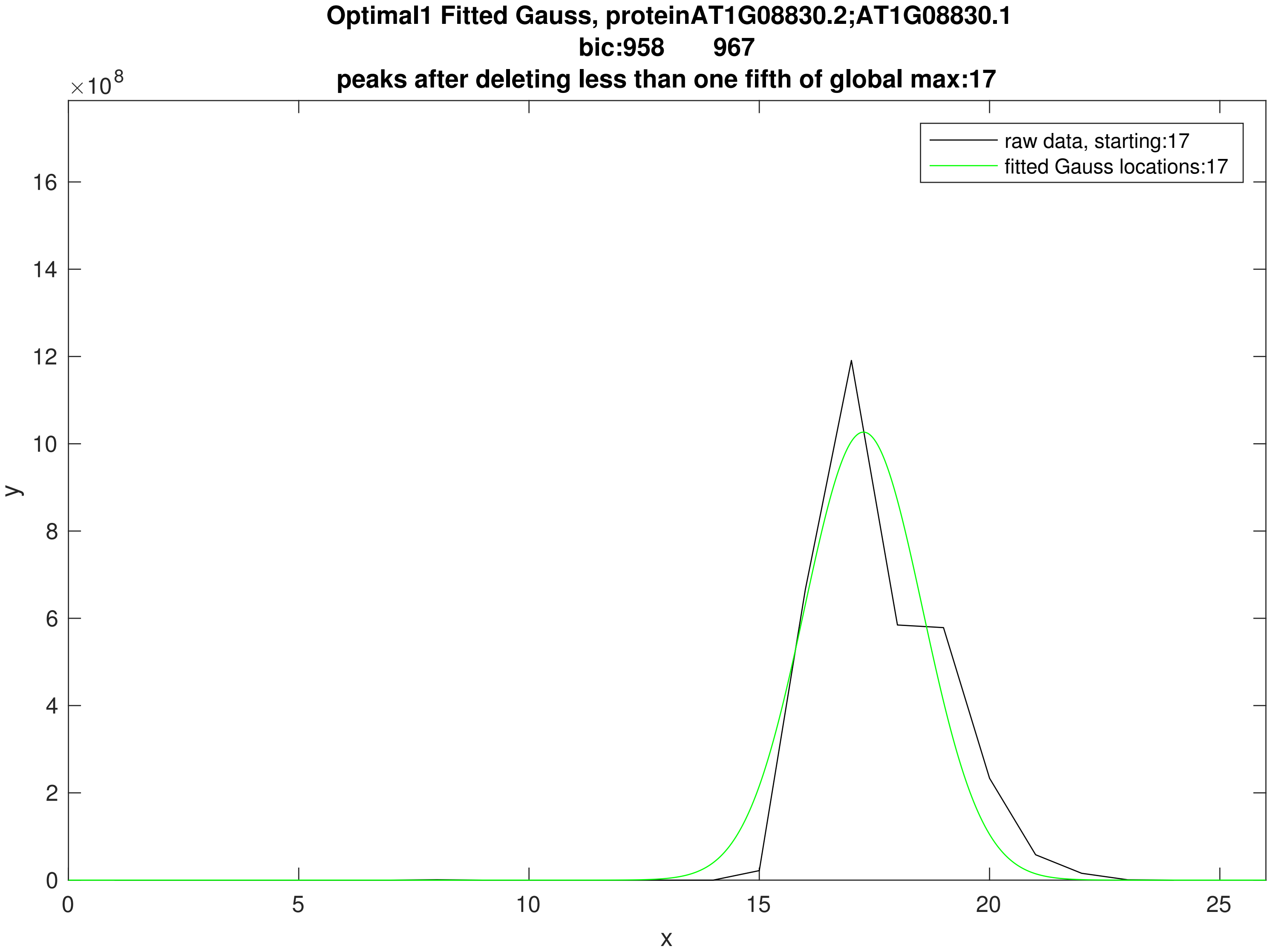
Optimal1 Fitted Gauss, proteinAT1G08830.2;AT1G08830.1
bic:958 967
peaks after deleting less than one fifth of global max:17
×10^8
raw data, starting:17
fitted Gauss locations:17
y
x
0
2
4
6
8
10
12
14
16
0
5
10
15
20
25

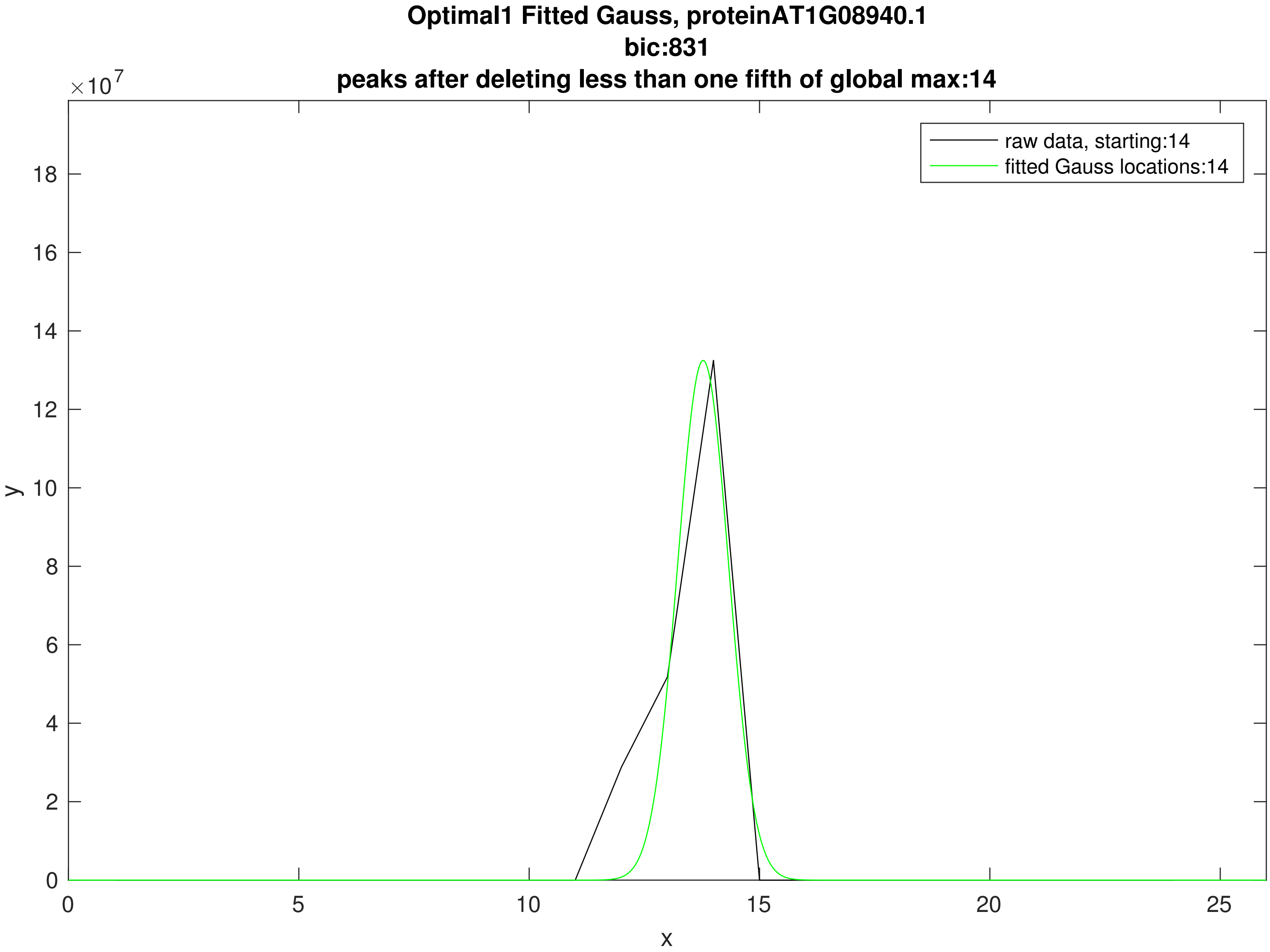

Optimal1 Fitted Gauss, proteinAT1G08940.1
bic:831
peaks after deleting less than one fifth of global max:14
×10^7
raw data, starting:14
fitted Gauss locations:14
18
16
14
12
10
8
6
4
2
0
y
0
5
10
15
20
25
x

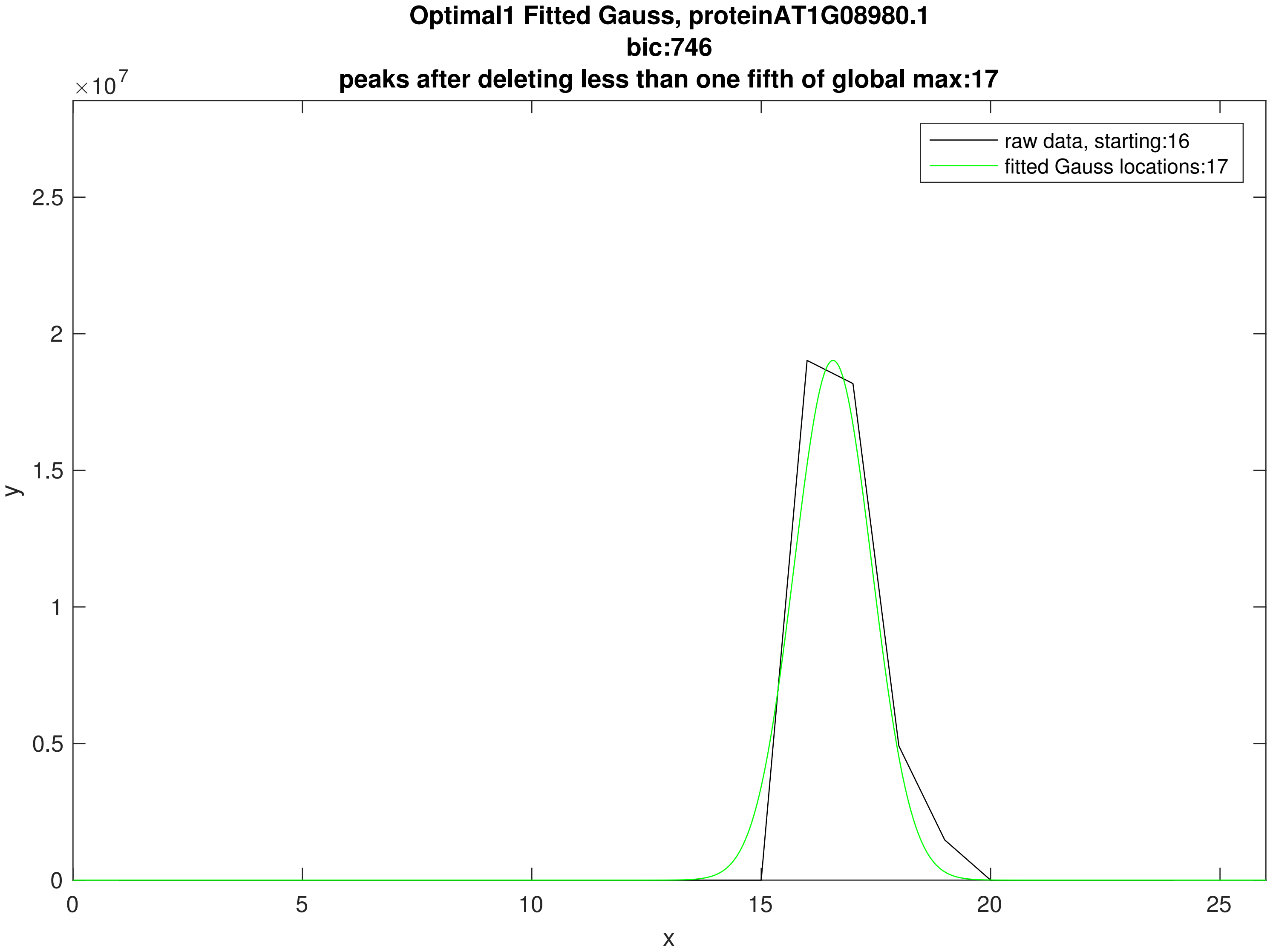
Optimal1 Fitted Gauss, proteinAT1G08980.1
bic:746
peaks after deleting less than one fifth of global max:17
×10^7
raw data, starting:16
fitted Gauss locations:17
y
x
0
0.5
1
1.5
2
2.5
0
5
10
15
20
25

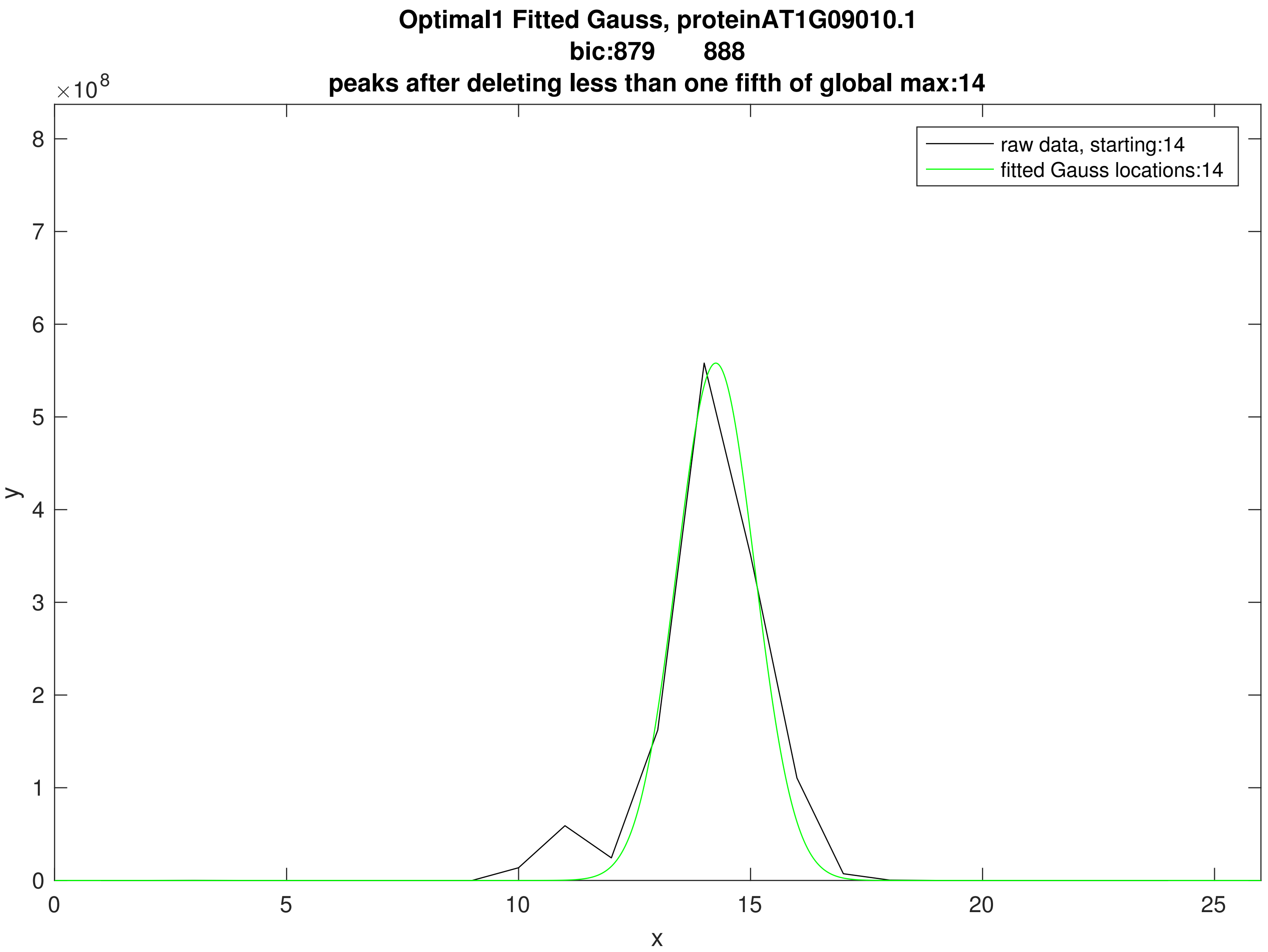
Optimal1 Fitted Gauss, proteinAT1G09010.1
bic:879 888
peaks after deleting less than one fifth of global max:14
×10^8
8
7
6
5
4
3
2
1
0
y
0
5
10
15
20
25
x
raw data, starting:14
fitted Gauss locations:14

**Optimal1 Fitted Gauss, proteinAT1G09130.1;AT1G09130.2;AT1G09130.3**
**bic:856 863**
**peaks after deleting less than one fifth of global max:9.6**

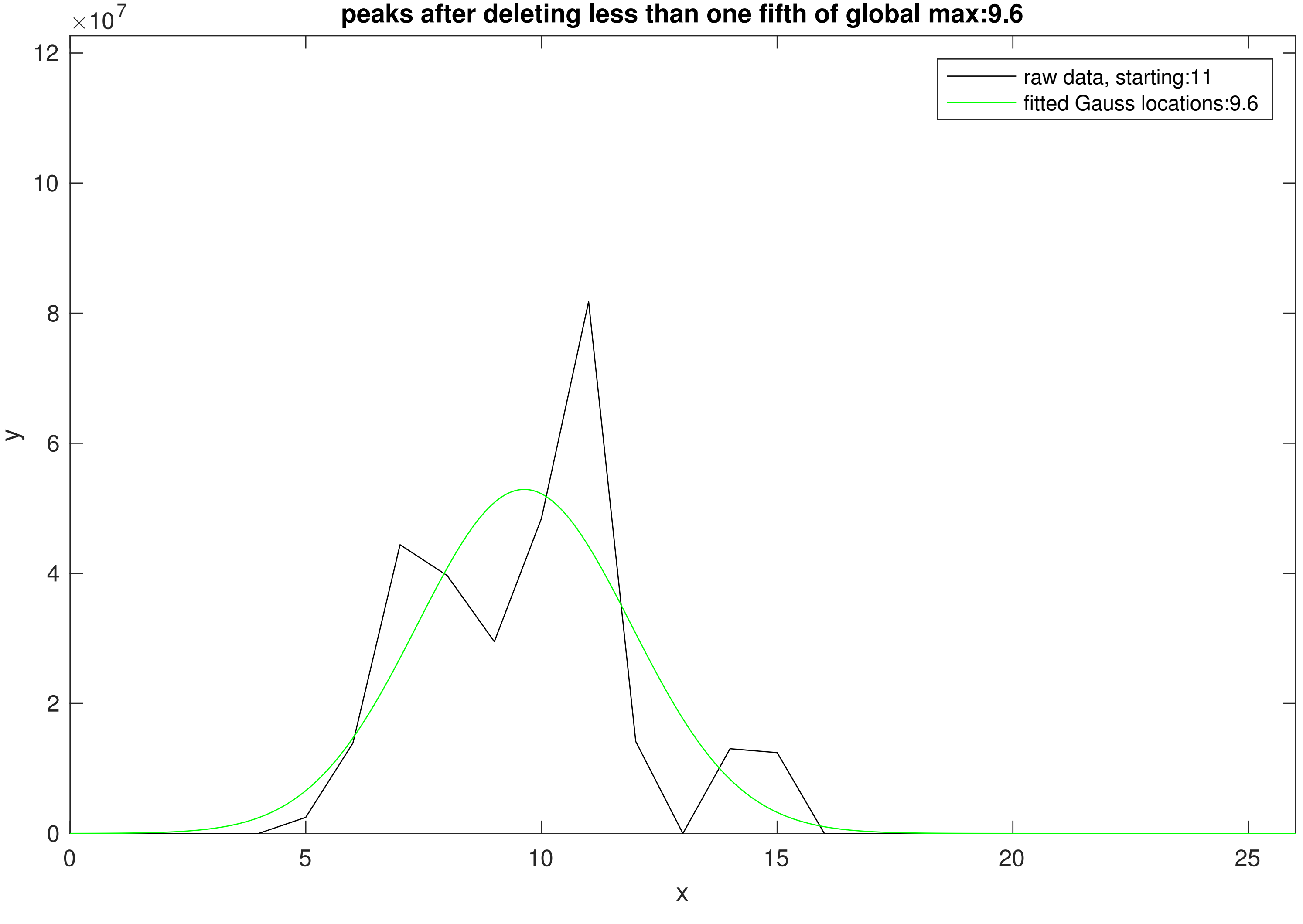

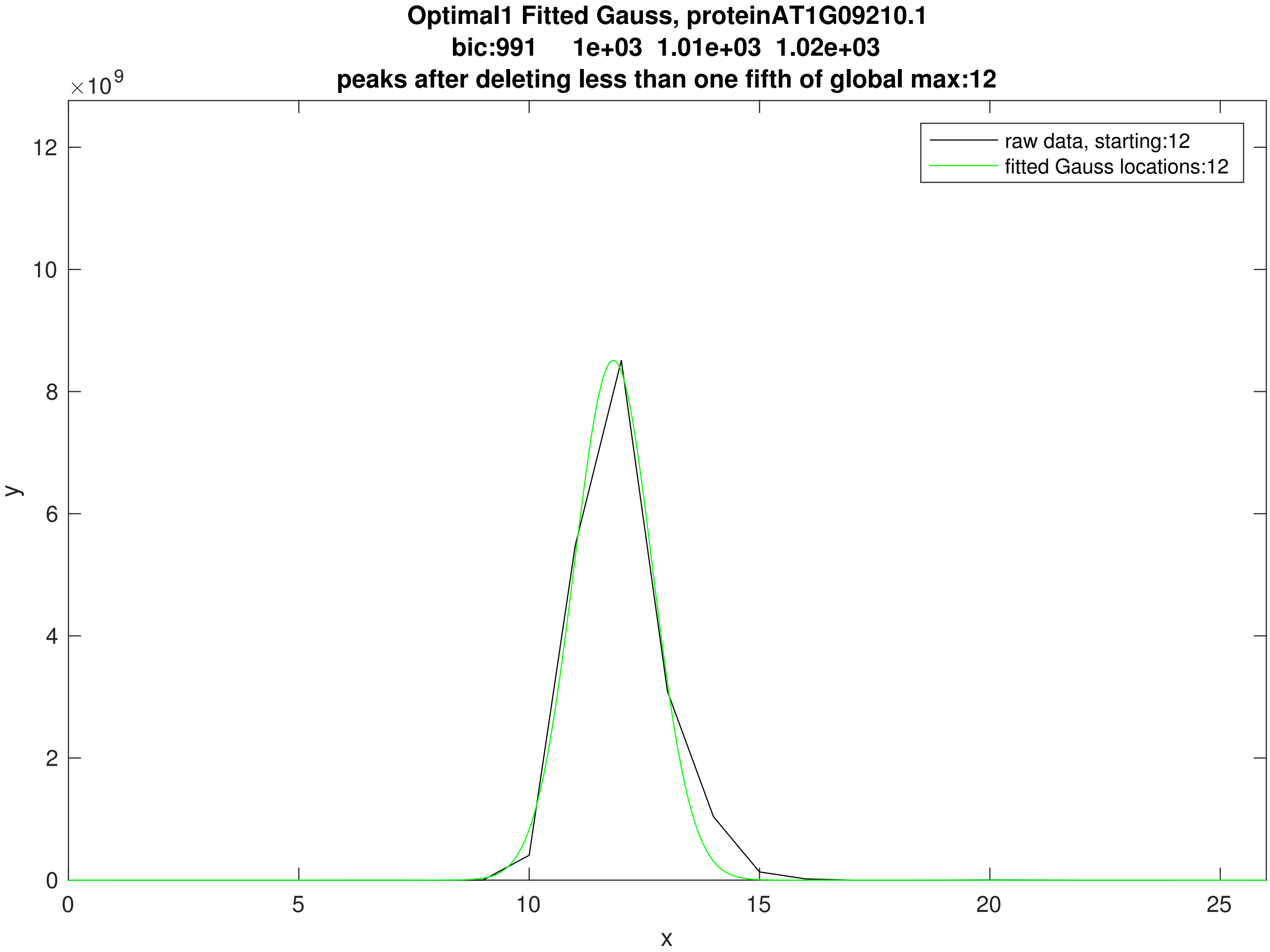
Optimal1 Fitted Gauss, proteinAT1G09210.1
bic:991 1e+03 1.01e+03 1.02e+03
peaks after deleting less than one fifth of global max:12
×10^9
raw data, starting:12
fitted Gauss locations:12
y
x
0
2
4
6
8
10
12
5
15
20
25

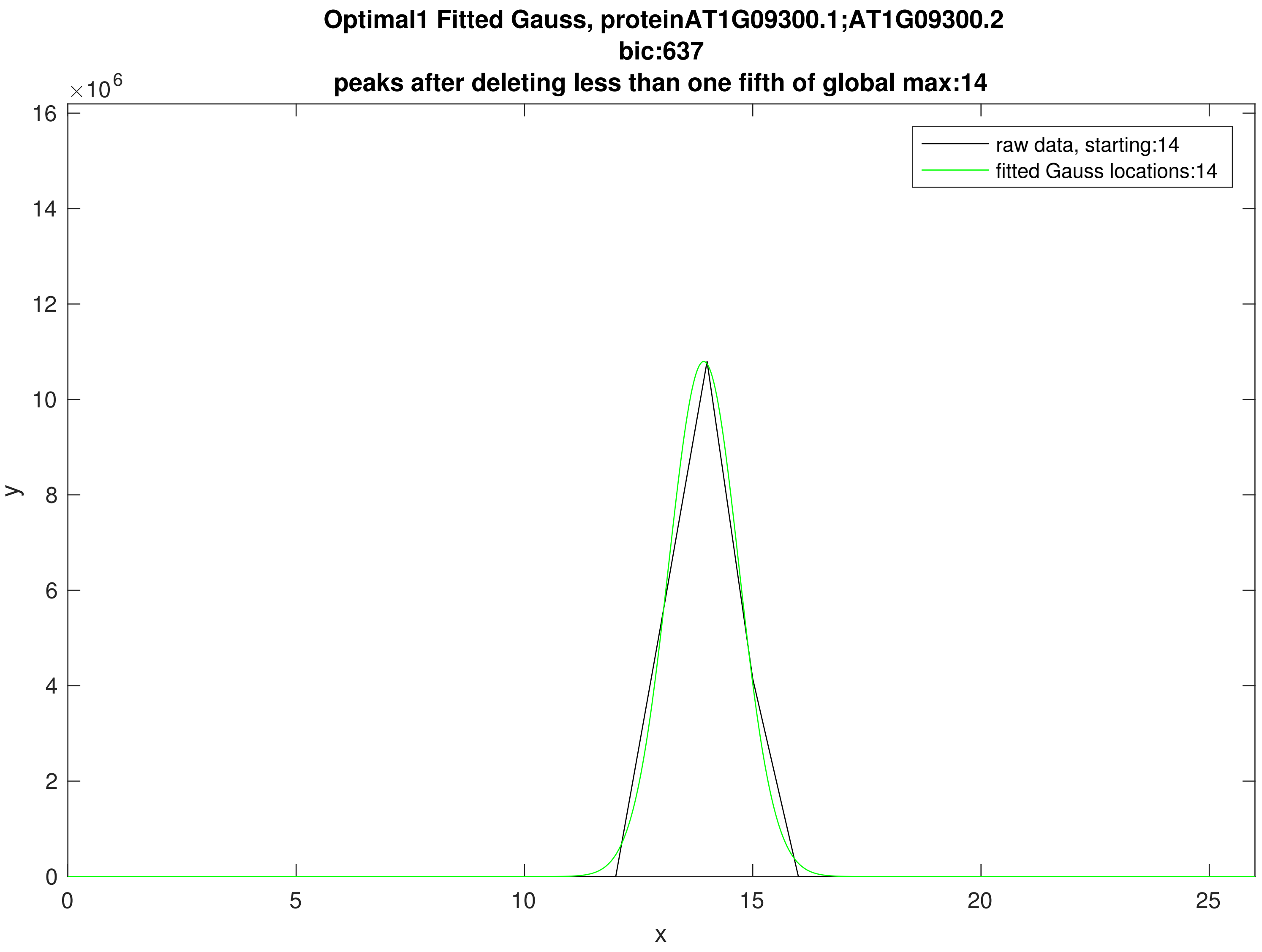
Optimal1 Fitted Gauss, proteinAT1G09300.1;AT1G09300.2
bic:637
peaks after deleting less than one fifth of global max:14
×10^6
16
14
12
10
8
6
4
2
0
y
0
5
10
15
20
25
x
raw data, starting:14
fitted Gauss locations:14

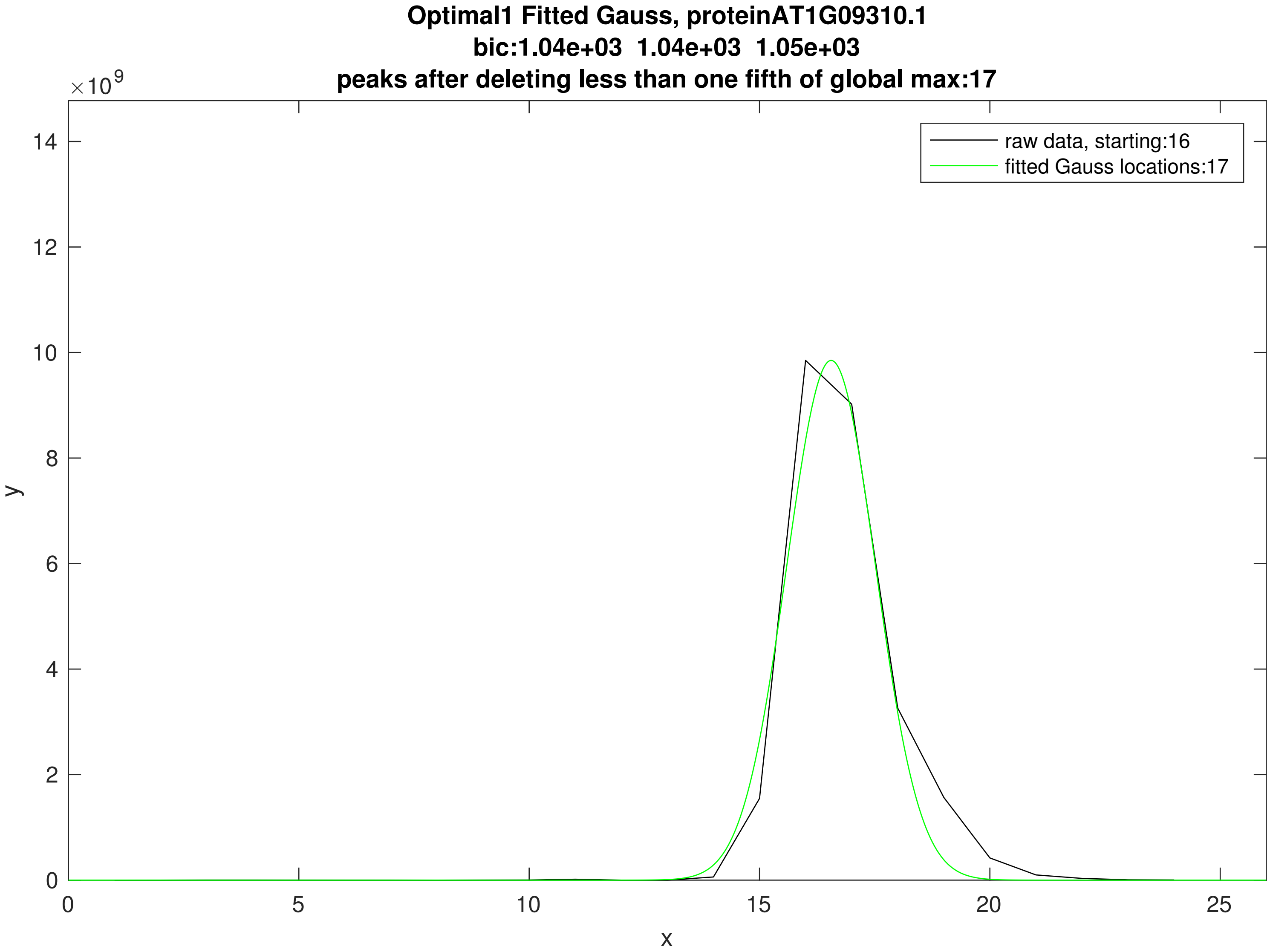

Optimal1 Fitted Gauss, proteinAT1G09310.1
bic:1.04e+03 1.04e+03 1.05e+03
peaks after deleting less than one fifth of global max:17
×10^9
14
12
10
8
6
4
2
0
y
0
5
10
15
20
25
x
raw data, starting:16
fitted Gauss locations:17

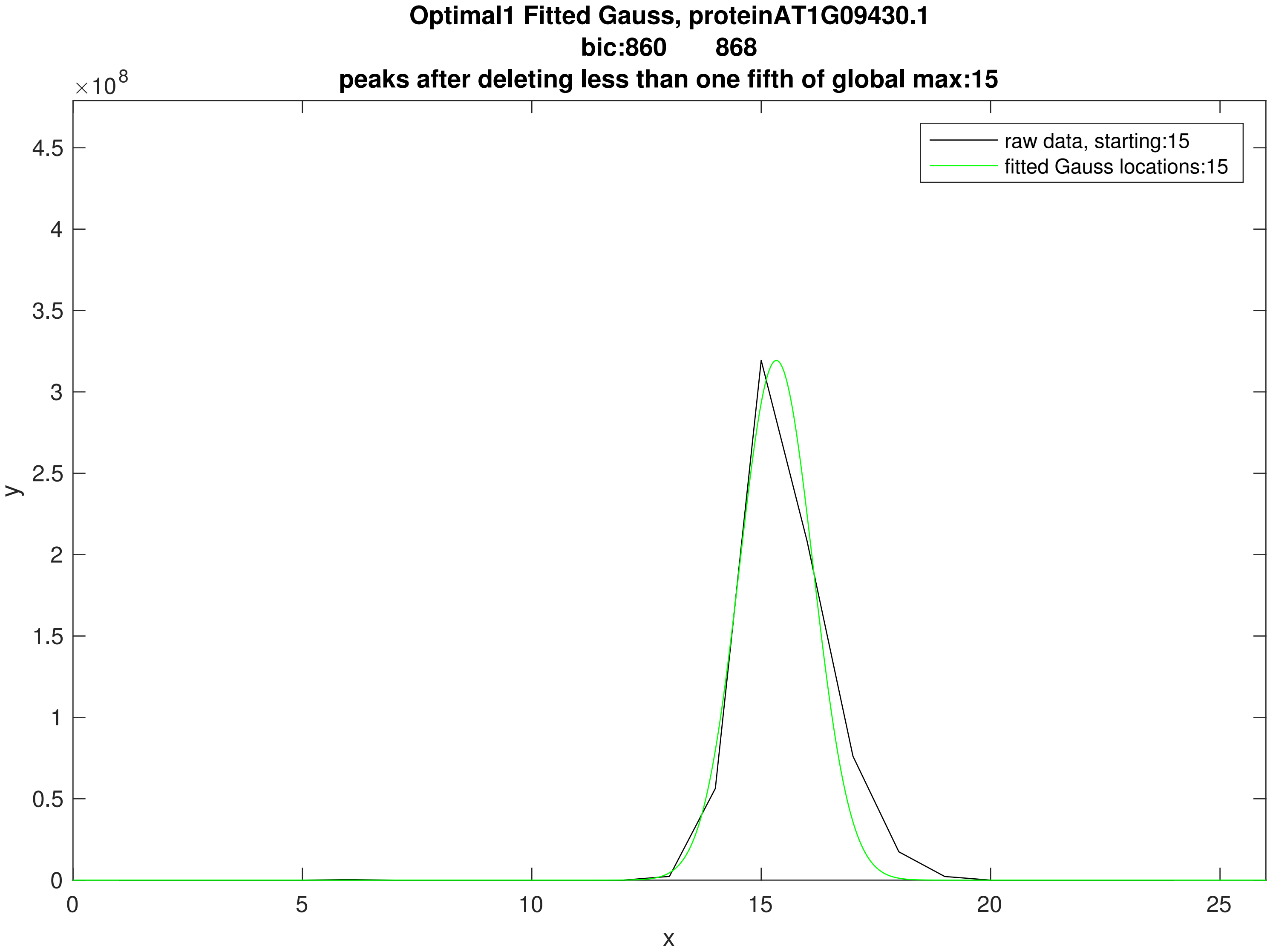
Optimal1 Fitted Gauss, proteinAT1G09430.1
bic:860 868
peaks after deleting less than one fifth of global max:15
×10^8
4.5
4
3.5
3
2.5
2
1.5
1
0.5
0
y
0
5
10
15
20
25
x
raw data, starting:15
fitted Gauss locations:15

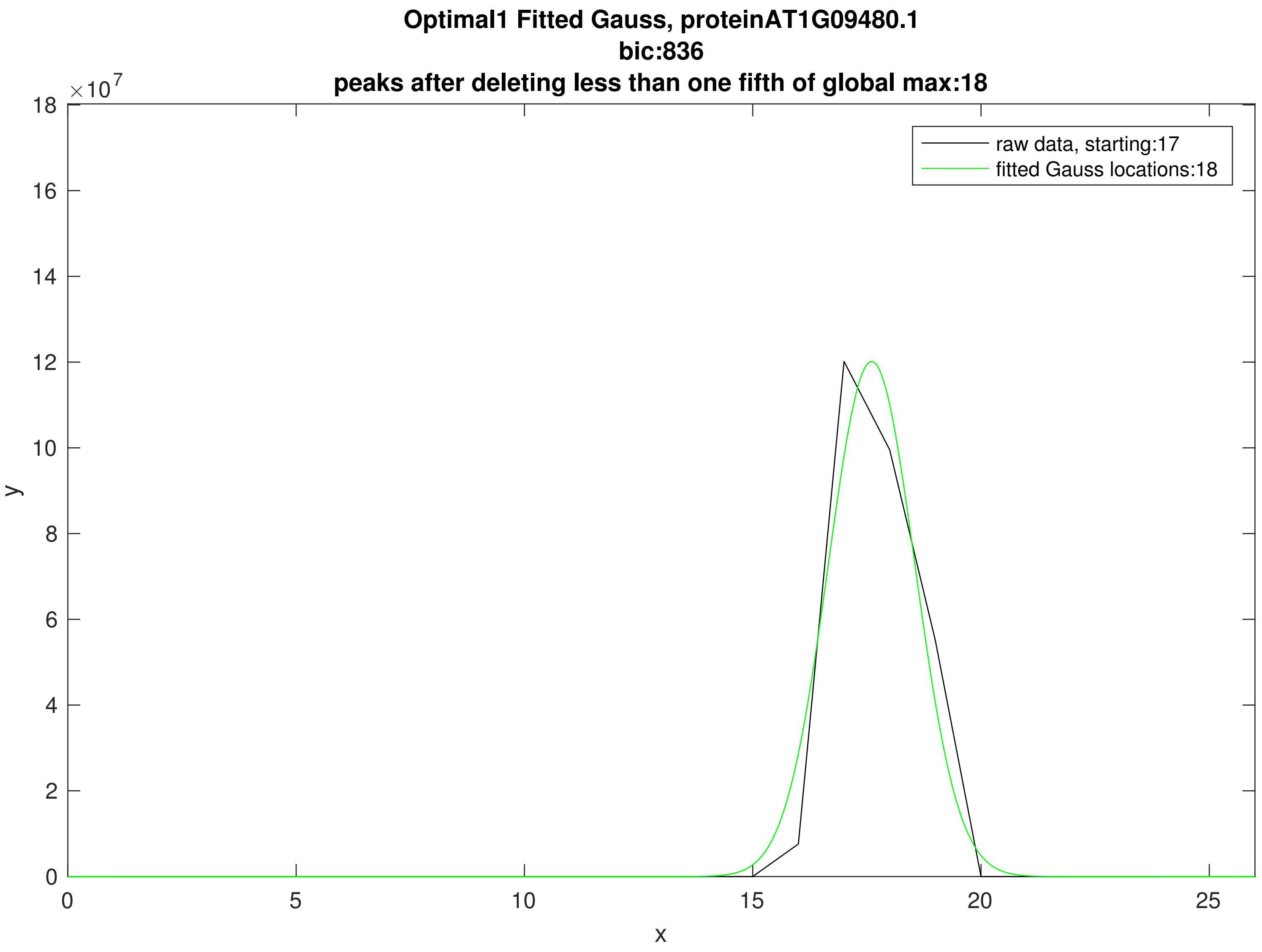
Optimal1 Fitted Gauss, proteinAT1G09480.1
bic:836
peaks after deleting less than one fifth of global max:18
×10^7
raw data, starting:17
fitted Gauss locations:18
y
x
0
2
4
6
8
10
12
14
16
18
0
5
10
15
20
25

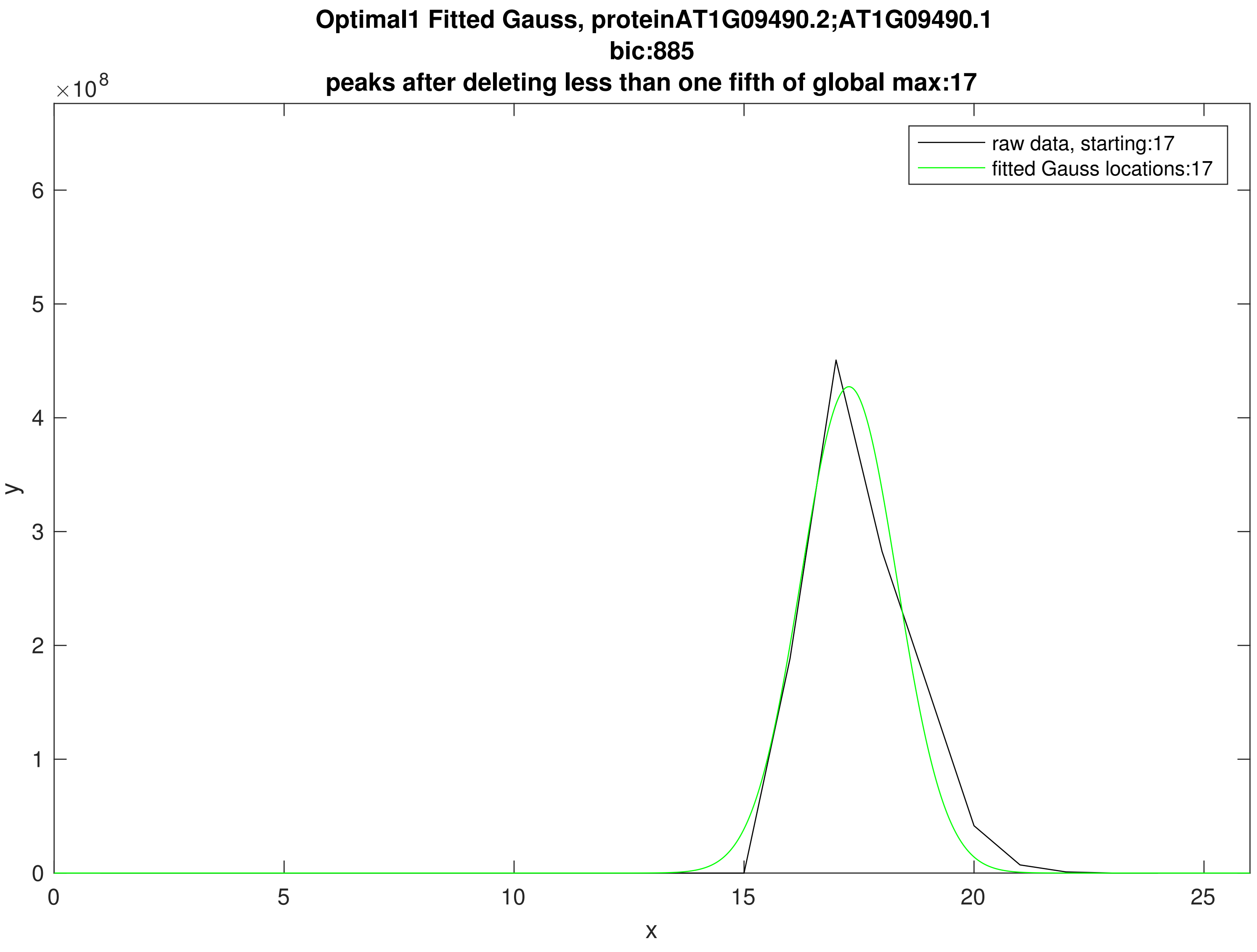
Optimal1 Fitted Gauss, proteinAT1G09490.2;AT1G09490.1
bic:885
peaks after deleting less than one fifth of global max:17
×10^8
raw data, starting:17
fitted Gauss locations:17
y
x
0
1
2
3
4
5
6
0
5
10
15
20
25

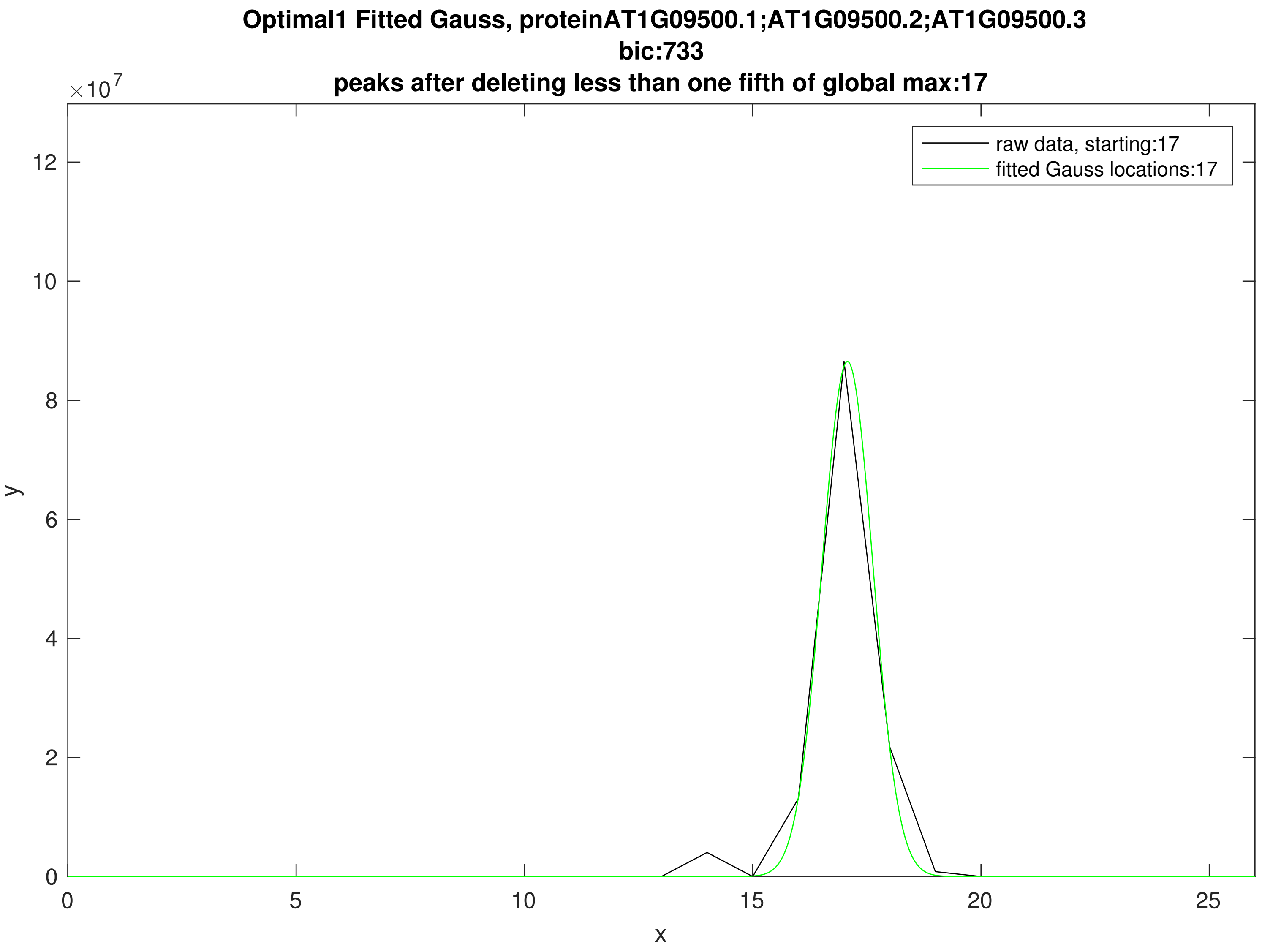
Optimal1 Fitted Gauss, proteinAT1G09500.1;AT1G09500.2;AT1G09500.3
bic:733
peaks after deleting less than one fifth of global max:17
×10^7
raw data, starting:17
fitted Gauss locations:17
y
x
0
2
4
6
8
10
12
0
5
10
15
20
25

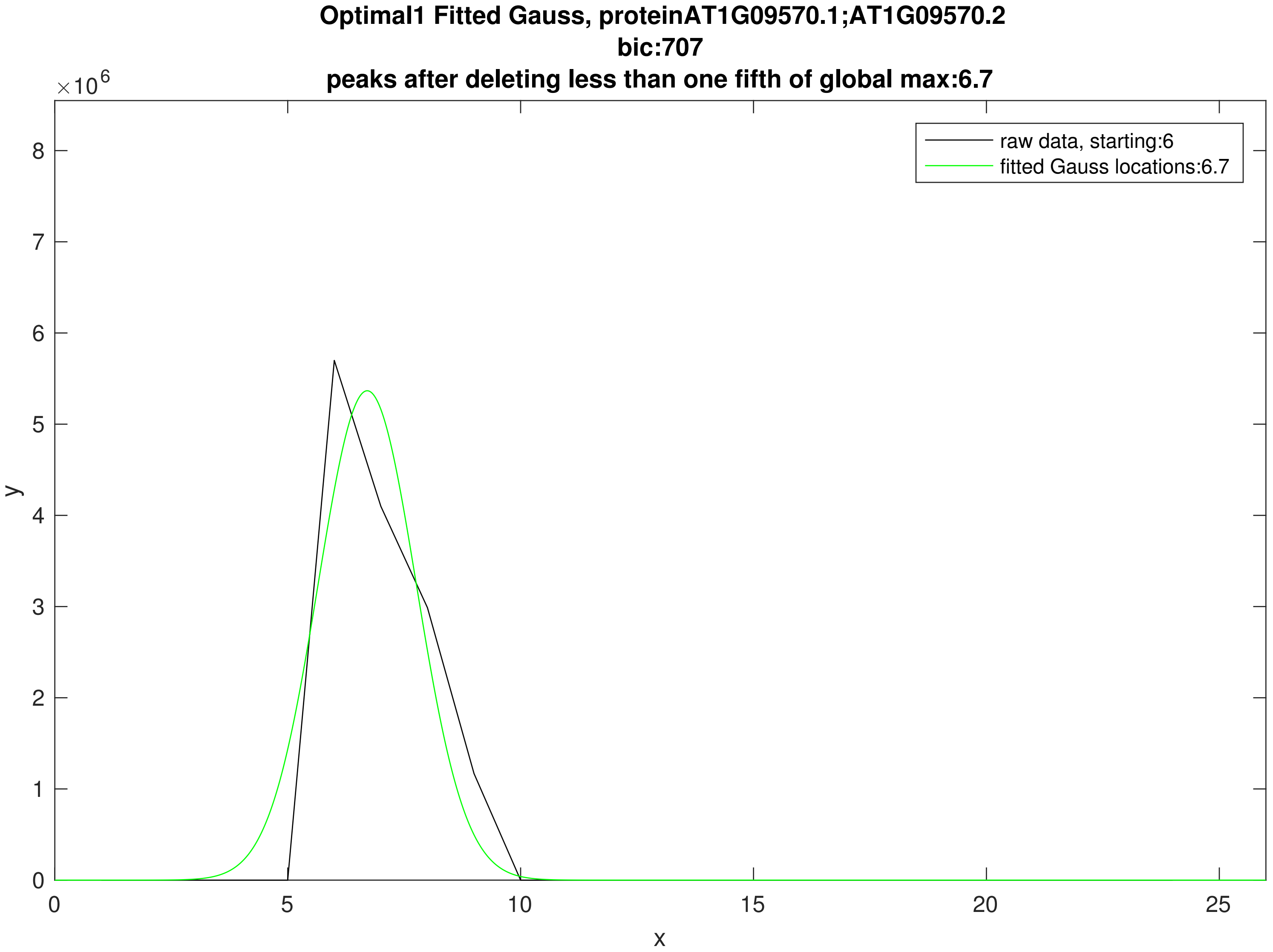
Optimal1 Fitted Gauss, proteinAT1G09570.1;AT1G09570.2
bic:707
peaks after deleting less than one fifth of global max:6.7
×10^6
raw data, starting:6
fitted Gauss locations:6.7
y
x
0
1
2
3
4
5
6
7
8
0
5
10
15
20
25

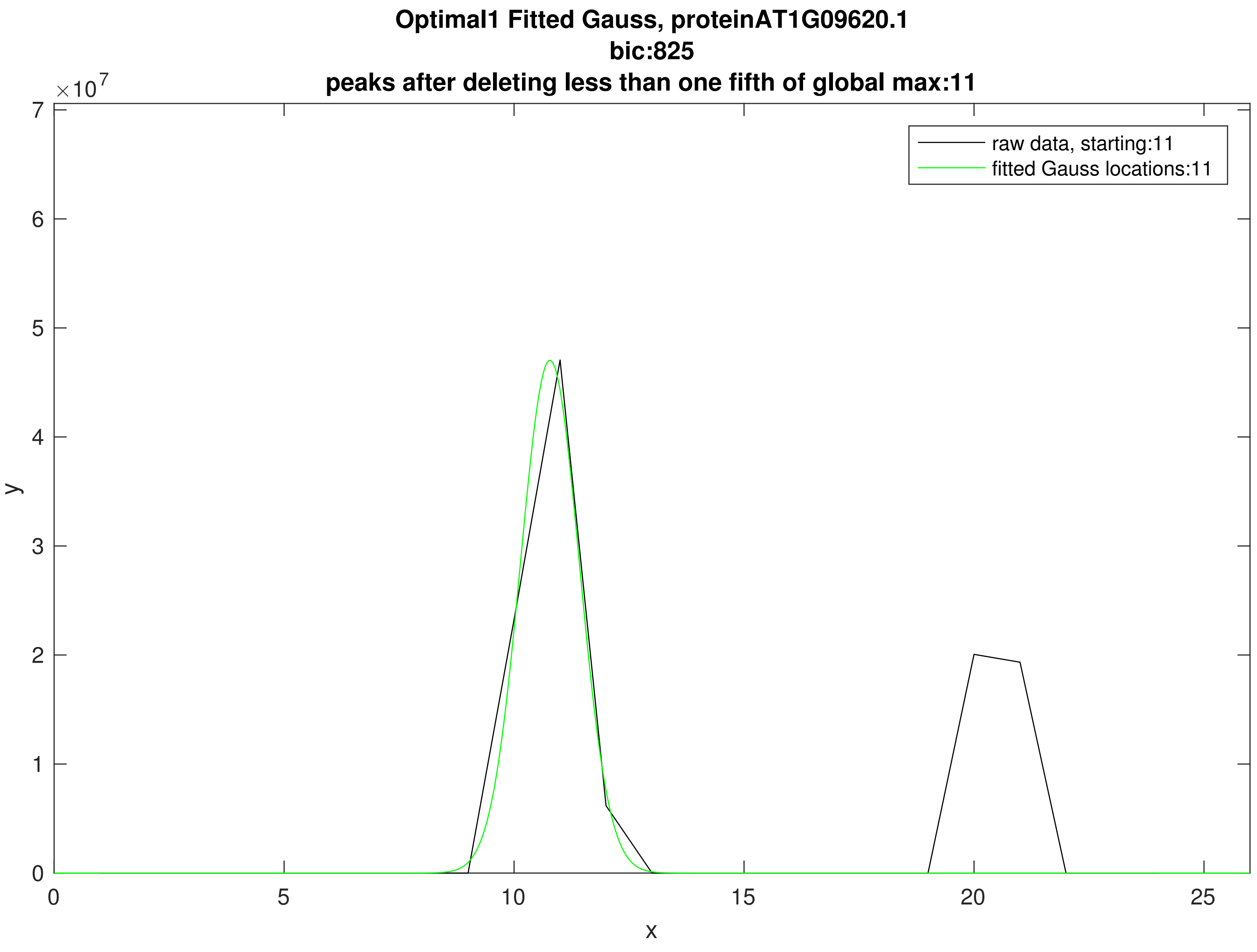
Optimal1 Fitted Gauss, proteinAT1G09620.1
bic:825
peaks after deleting less than one fifth of global max:11
×10^7
raw data, starting:11
fitted Gauss locations:11
y
x
0
1
2
3
4
5
6
7
0
5
10
15
20
25

Optimal1 Fitted Gauss, proteinAT1G09640.1
bic:838 847 857
peaks after deleting less than one fifth of global max:4.9

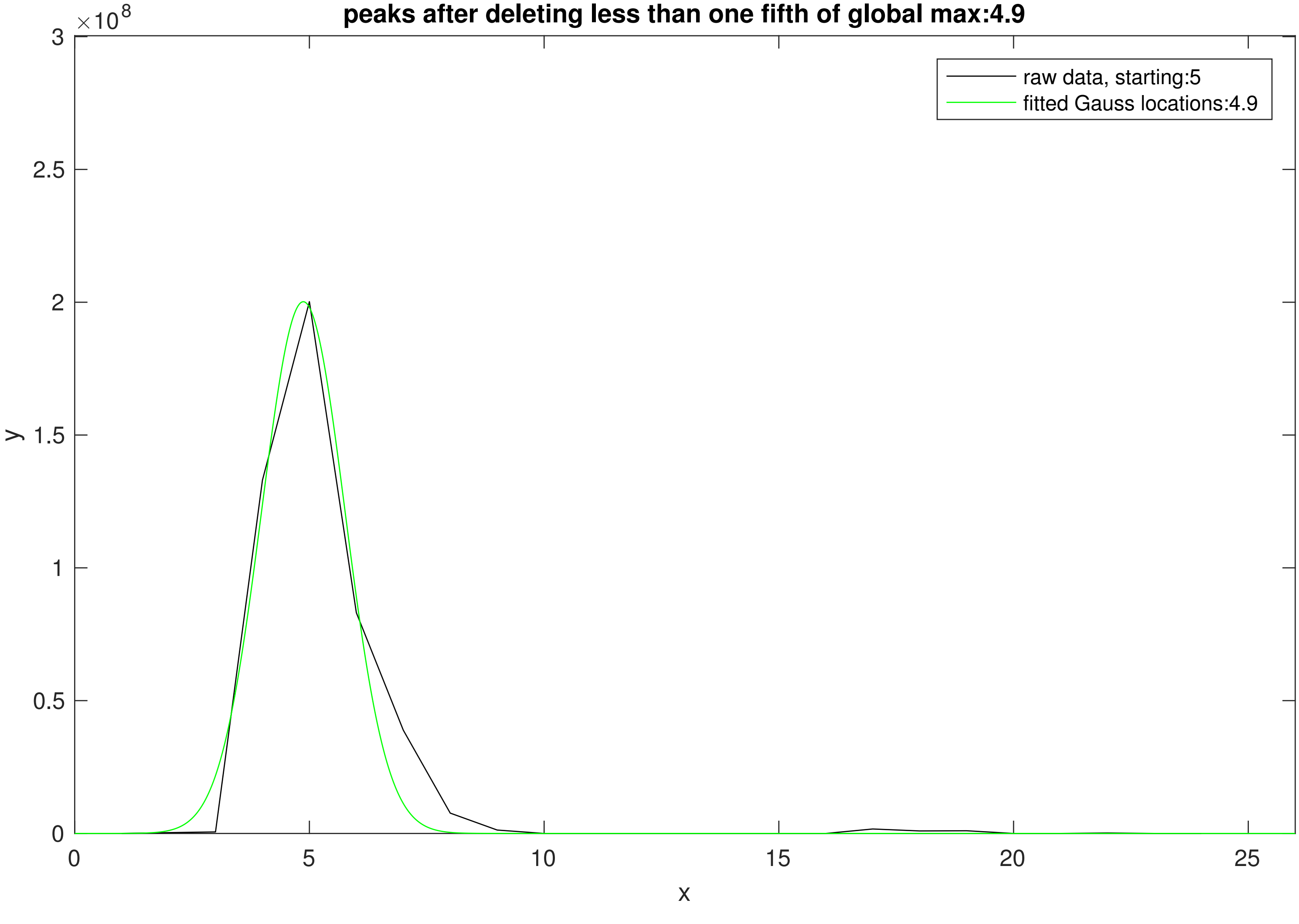

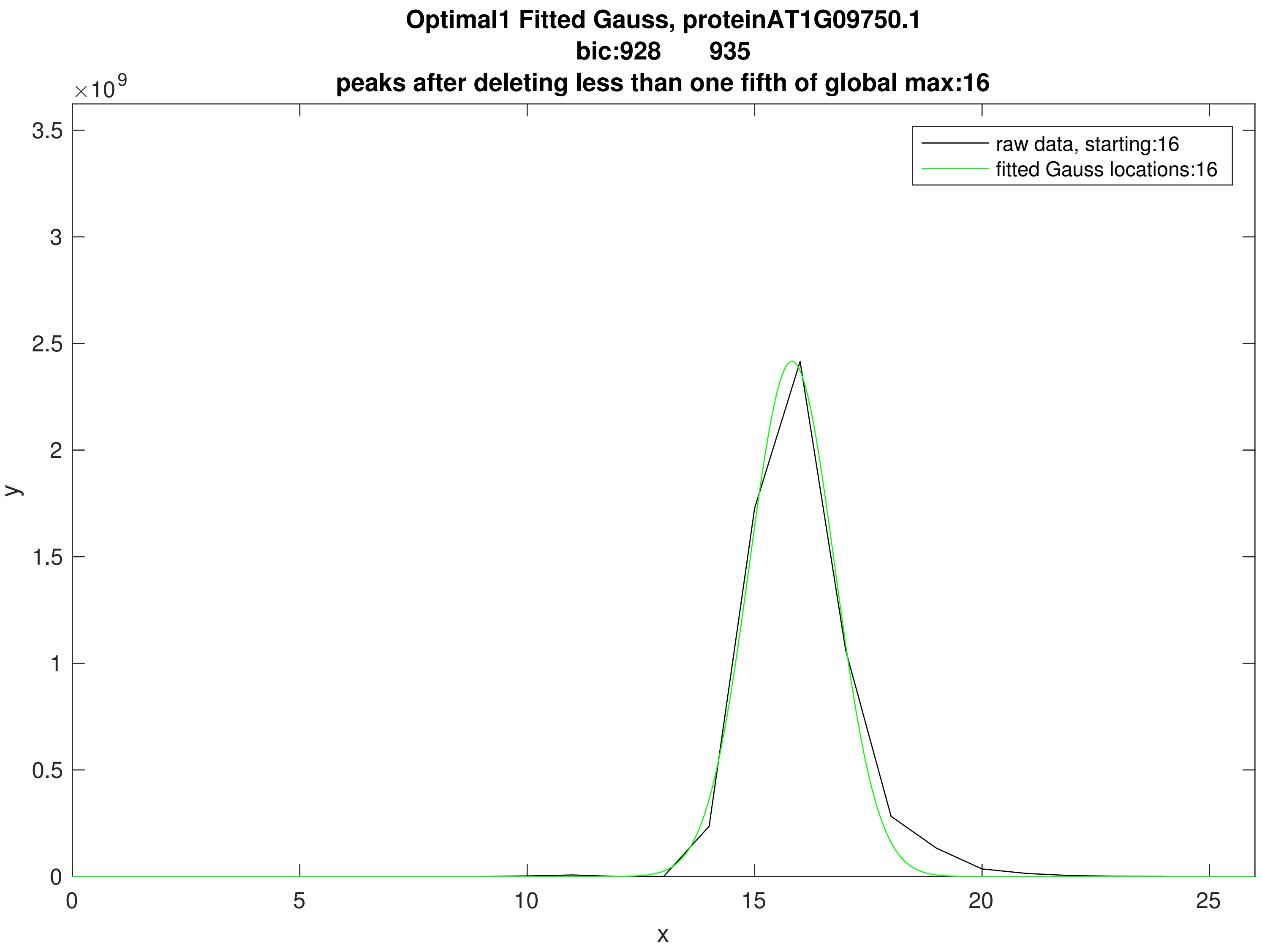
Optimal1 Fitted Gauss, proteinAT1G09750.1
bic:928 935
peaks after deleting less than one fifth of global max:16
×10^9
raw data, starting:16
fitted Gauss locations:16
y
x
0
5
10
15
20
25
0.5
1
1.5
2
2.5
3
3.5

**Optimal1 Fitted Gauss, proteinAT1G09780.1**
**bic:1.04e+03  1.05e+03  1.06e+03**
**peaks after deleting less than one fifth of global max:15**

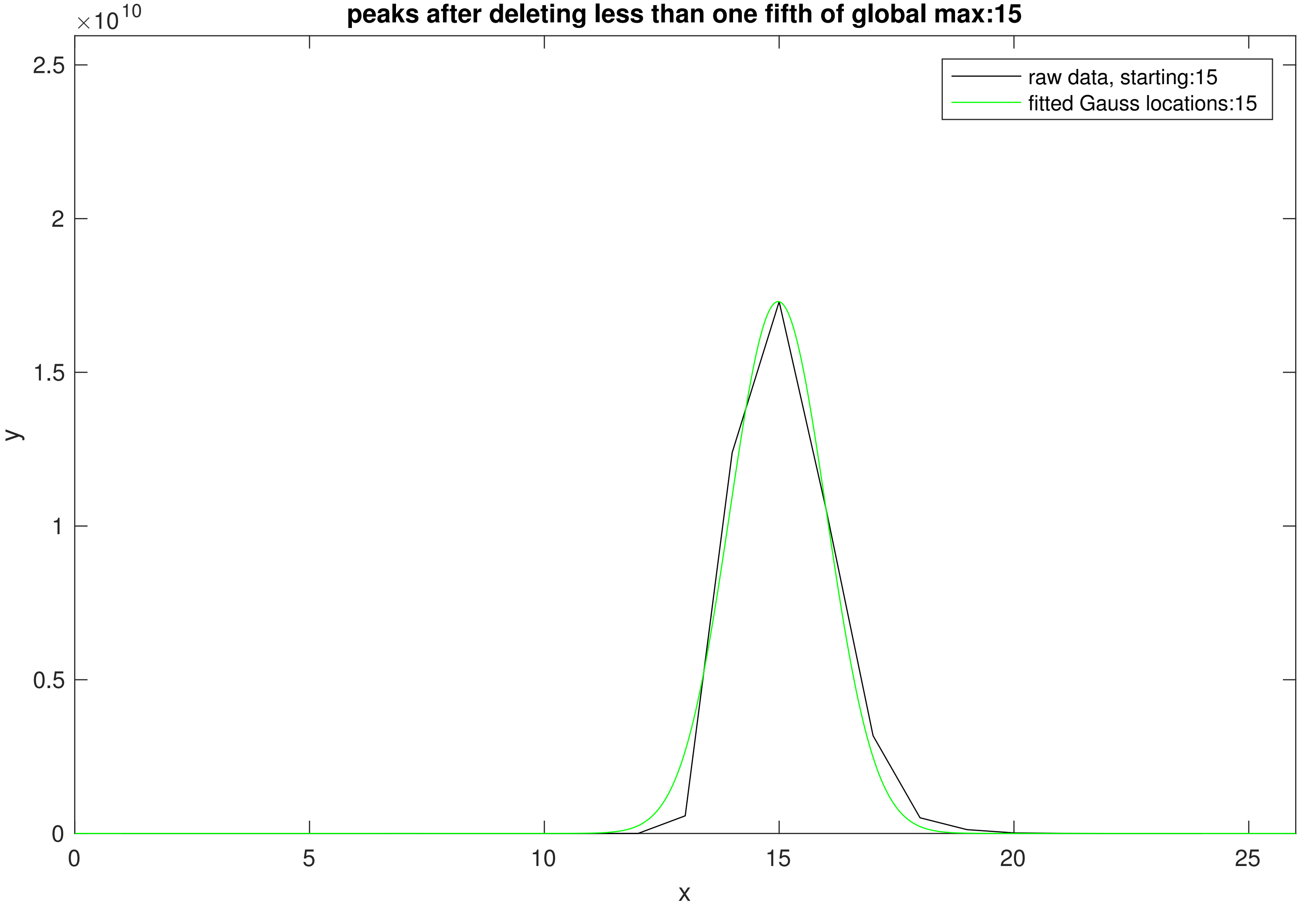

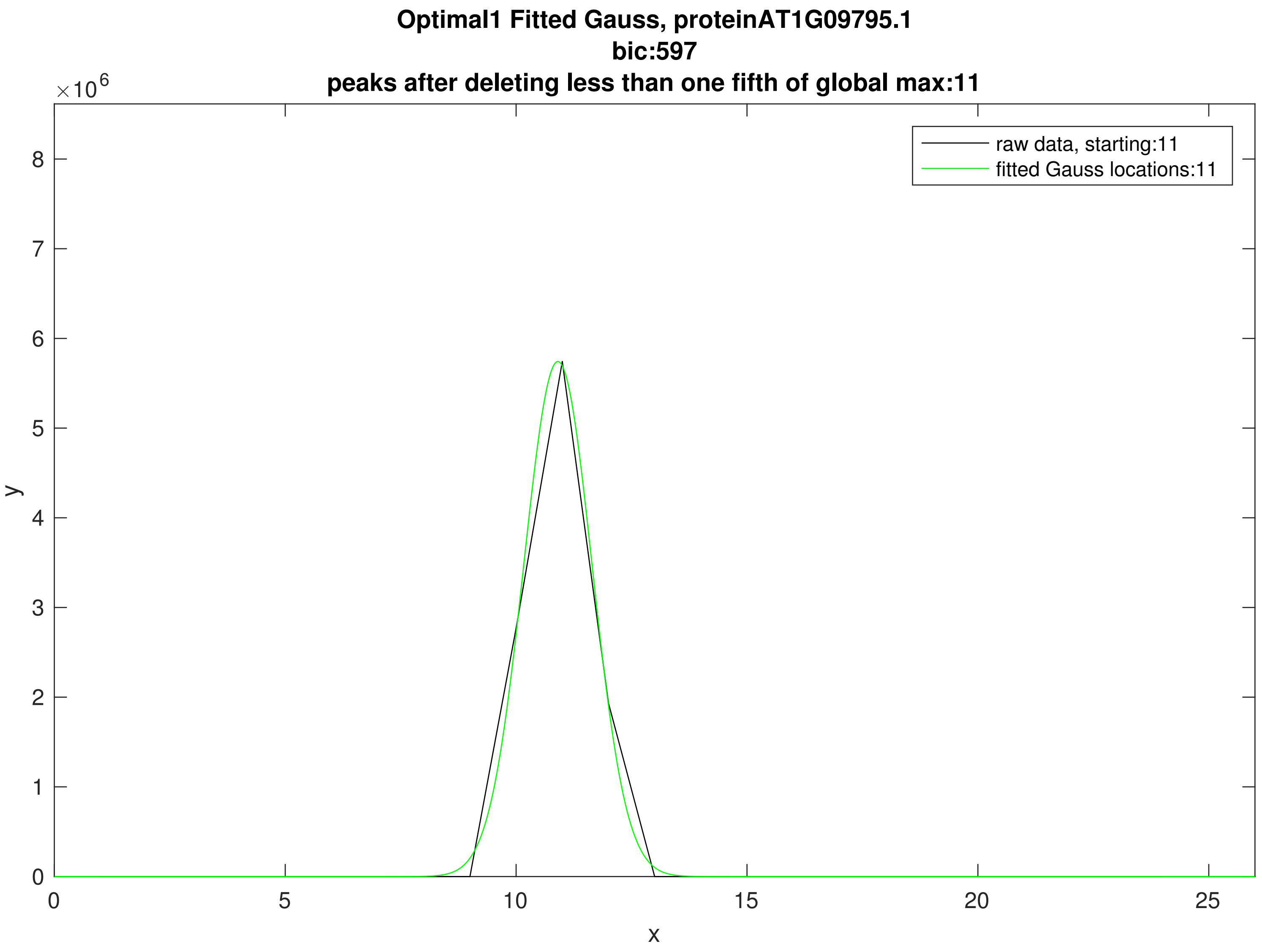
Optimal1 Fitted Gauss, proteinAT1G09795.1
bic:597
peaks after deleting less than one fifth of global max:11
×10^6
raw data, starting:11
fitted Gauss locations:11
y
x
0
1
2
3
4
5
6
7
8
0
5
10
15
20
25

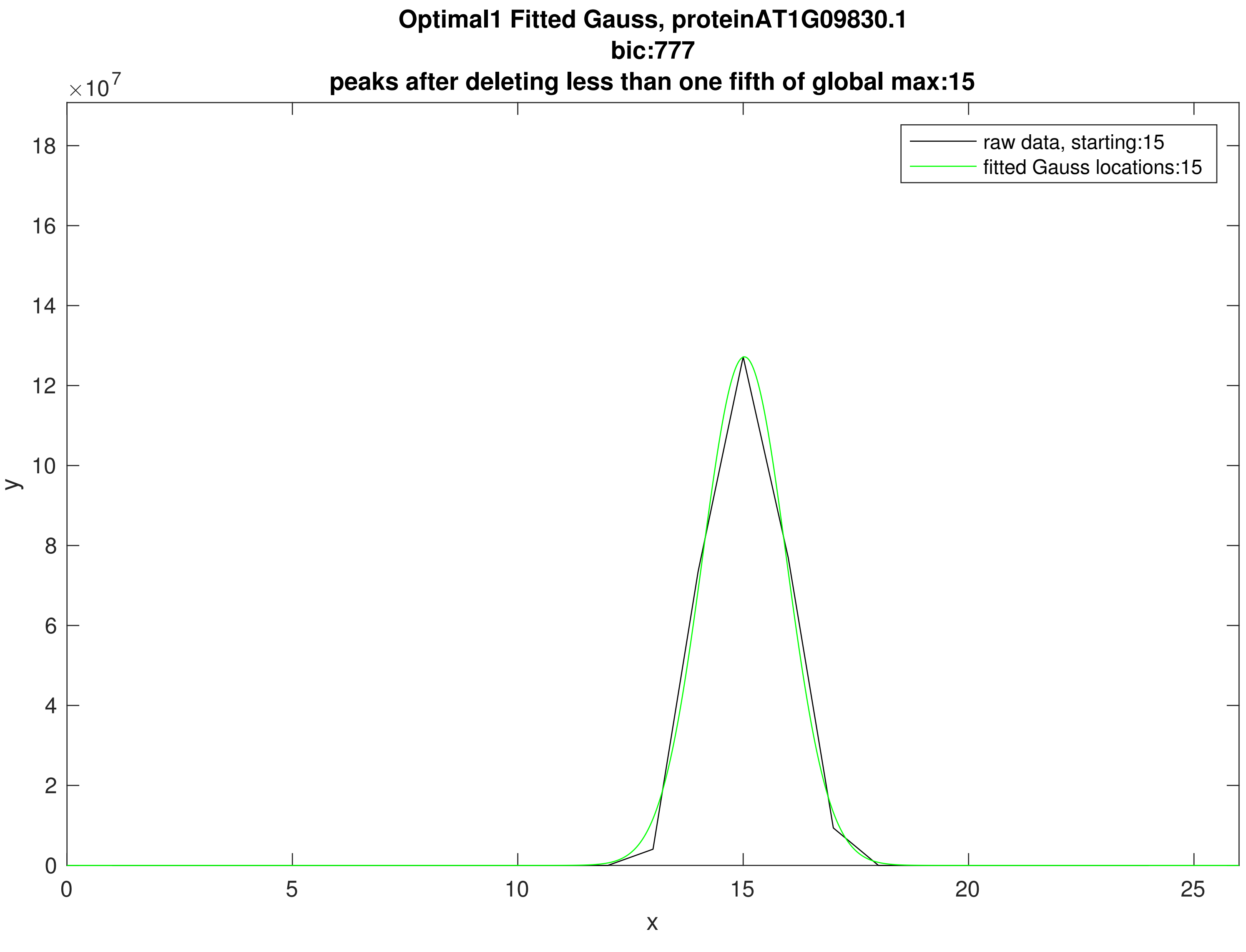

Optimal1 Fitted Gauss, proteinAT1G09830.1
bic:777
peaks after deleting less than one fifth of global max:15
×10^7
raw data, starting:15
fitted Gauss locations:15
y
x
0
2
4
6
8
10
12
14
16
18
0
5
10
15
20
25

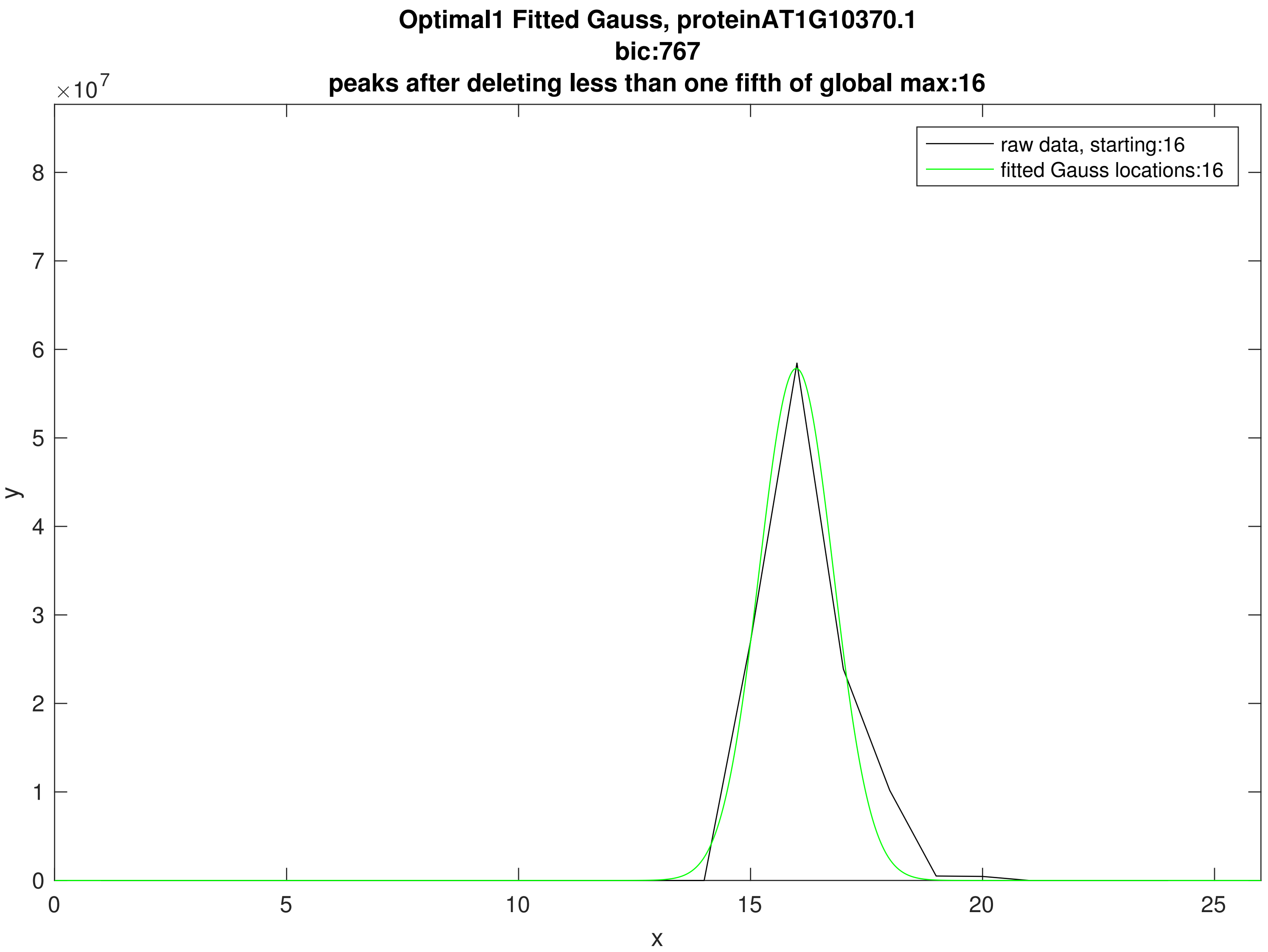

Optimal1 Fitted Gauss, proteinAT1G10370.1
bic:767
peaks after deleting less than one fifth of global max:16
×10^7
8
7
6
5
4
3
2
1
0
y
0
5
10
15
20
25
x
raw data, starting:16
fitted Gauss locations:16

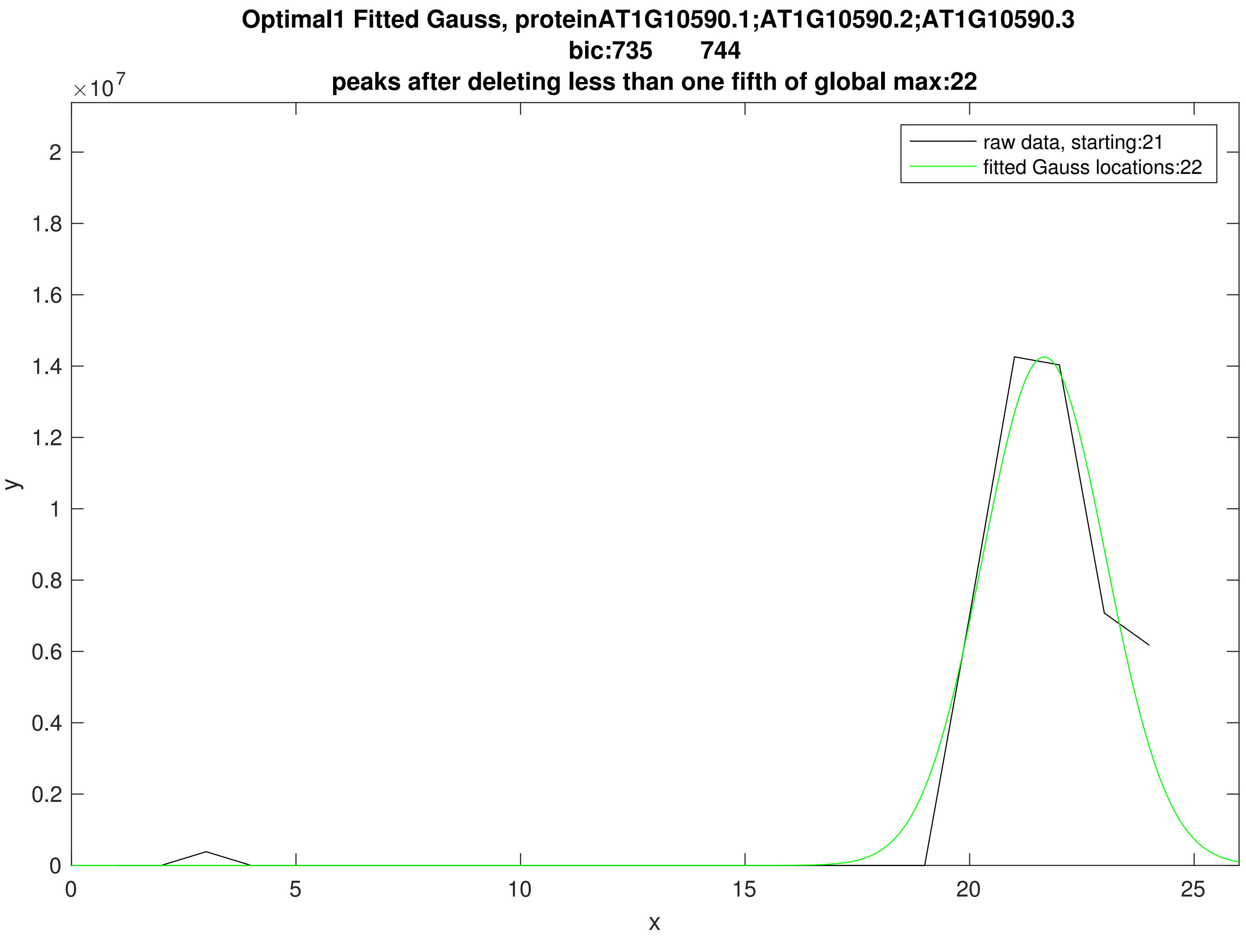

Optimal1 Fitted Gauss, proteinAT1G10590.1;AT1G10590.2;AT1G10590.3
bic:735 744
peaks after deleting less than one fifth of global max:22
×10^7
raw data, starting:21
fitted Gauss locations:22
2
1.8
1.6
1.4
1.2
1
0.8
0.6
0.4
0.2
0
y
0
5
10
15
20
25
x

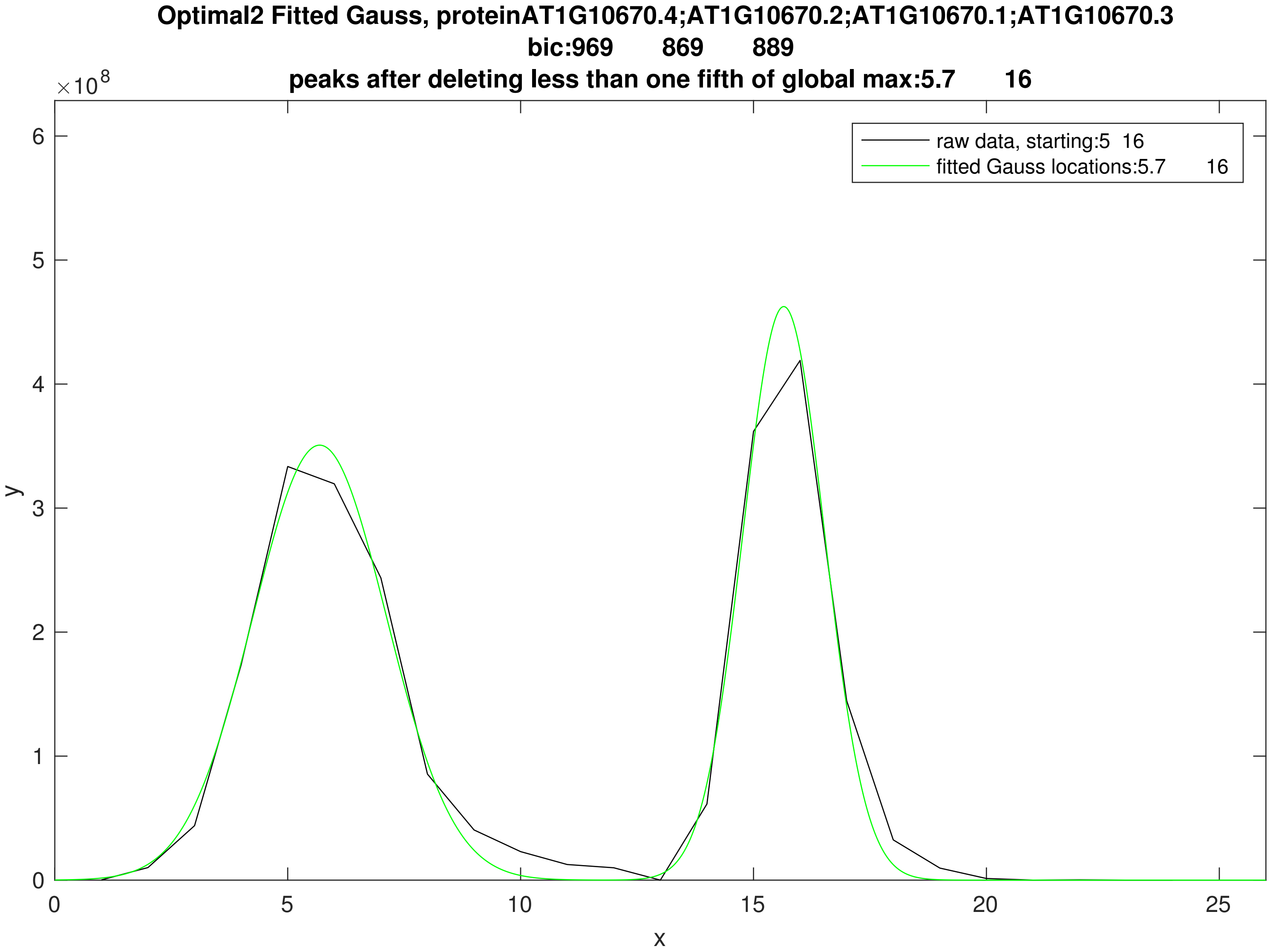

Optimal2 Fitted Gauss, proteinAT1G10670.4;AT1G10670.2;AT1G10670.1;AT1G10670.3
bic:969 869 889
peaks after deleting less than one fifth of global max:5.7 16
×10^8
raw data, starting:5 16
fitted Gauss locations:5.7 16
y
x
0
1
2
3
4
5
6
10
15
20
25

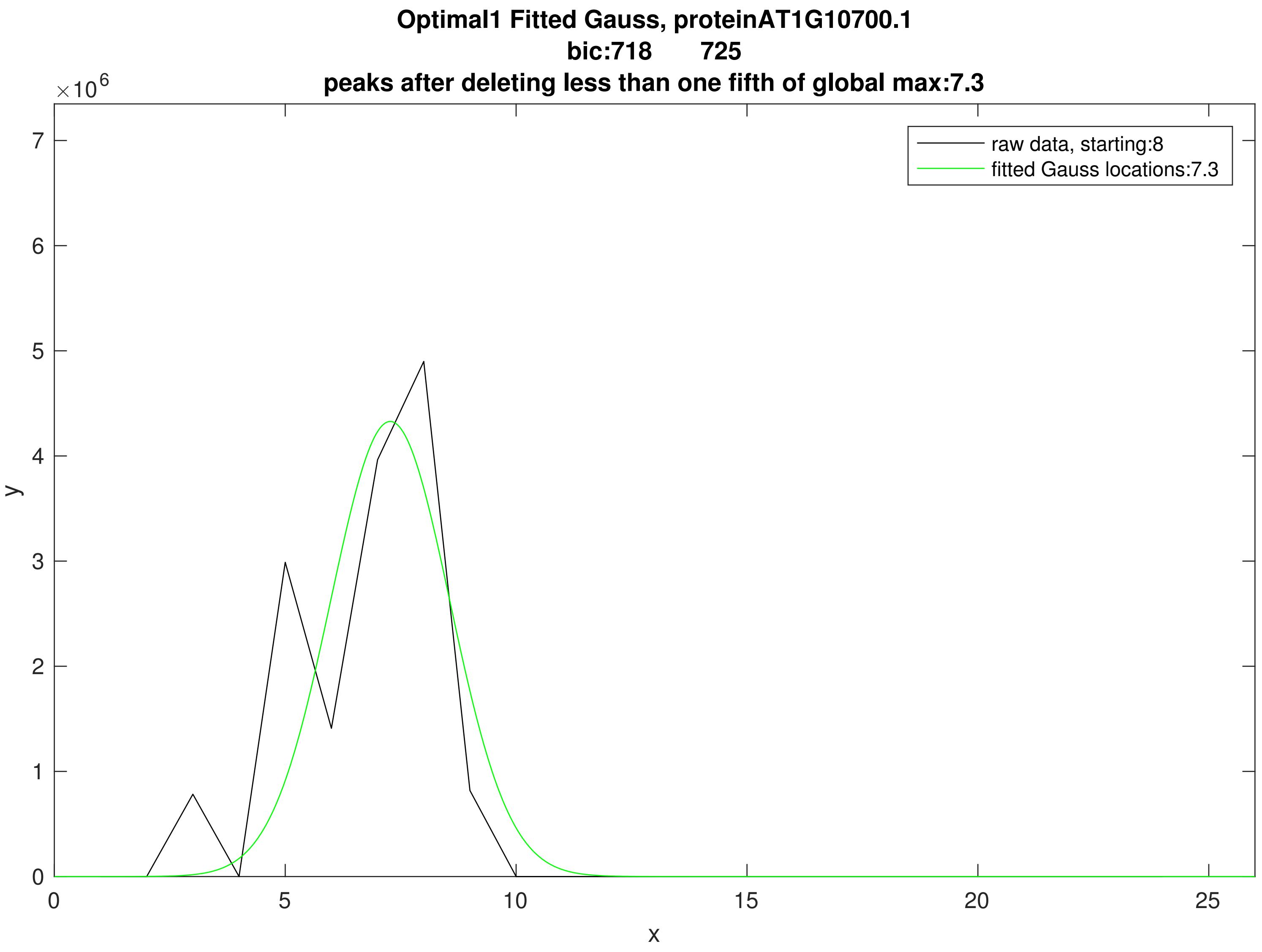
Optimal1 Fitted Gauss, proteinAT1G10700.1
bic:718 725
peaks after deleting less than one fifth of global max:7.3
×10^6
raw data, starting:8
fitted Gauss locations:7.3
y
x
0
1
2
3
4
5
6
7
0
5
10
15
20
25

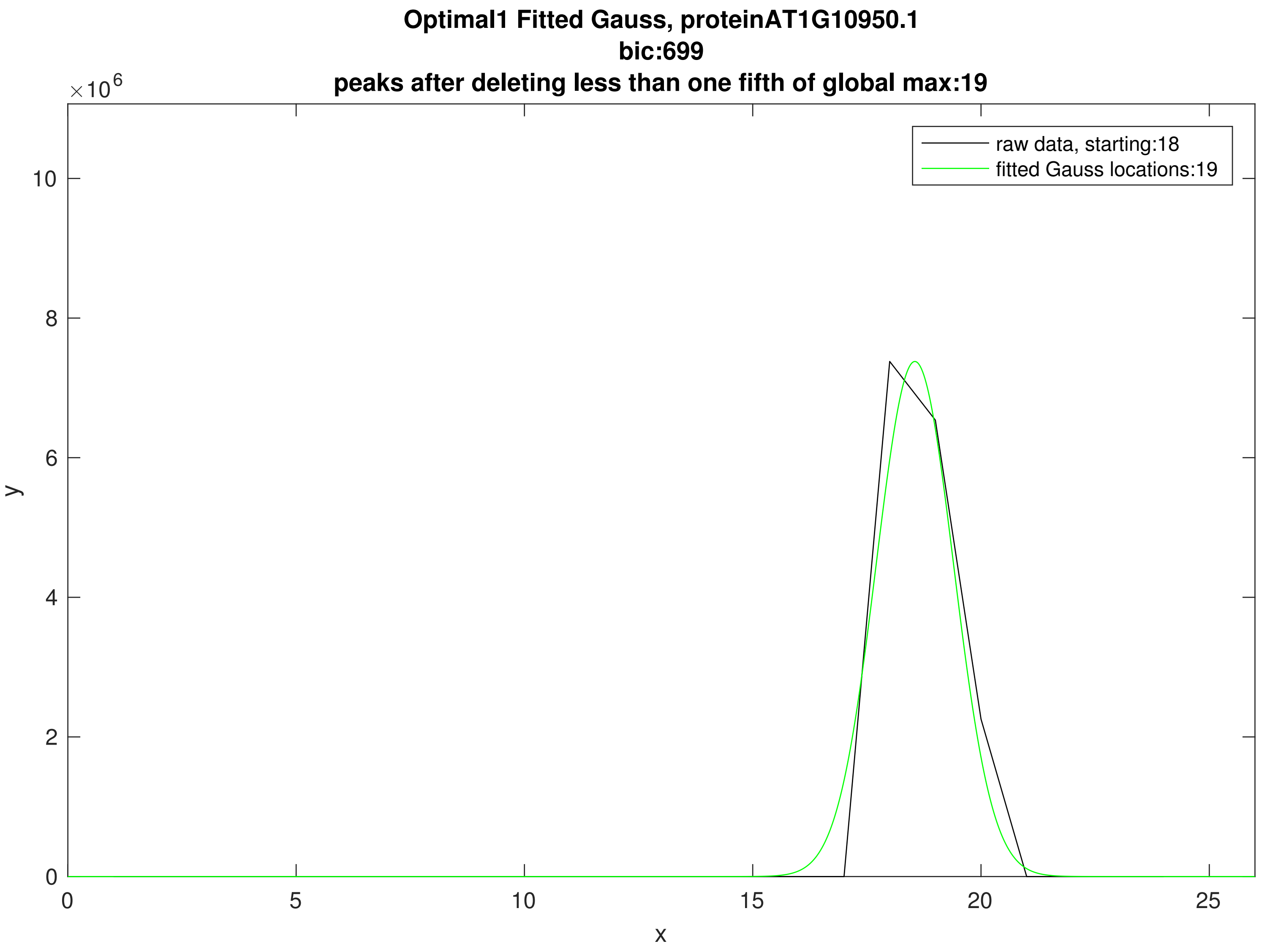
Optimal1 Fitted Gauss, proteinAT1G10950.1
bic:699
peaks after deleting less than one fifth of global max:19
×10^6
raw data, starting:18
fitted Gauss locations:19
y
x
0
2
4
6
8
10
0
5
10
15
20
25

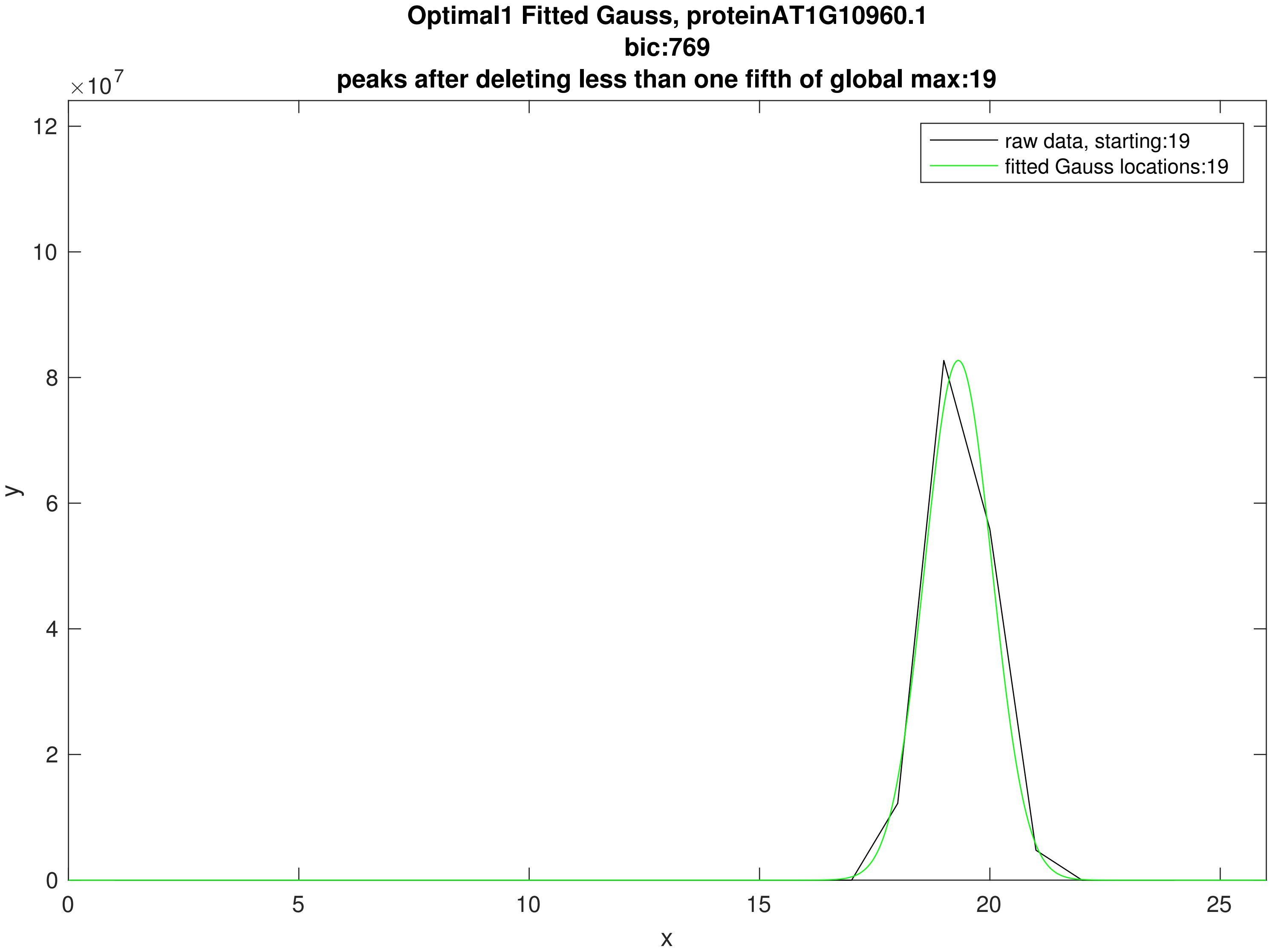

Optimal1 Fitted Gauss, proteinAT1G10960.1
bic:769
peaks after deleting less than one fifth of global max:19
×10^7
raw data, starting:19
fitted Gauss locations:19
y
x
0
2
4
6
8
10
12
0
5
10
15
20
25

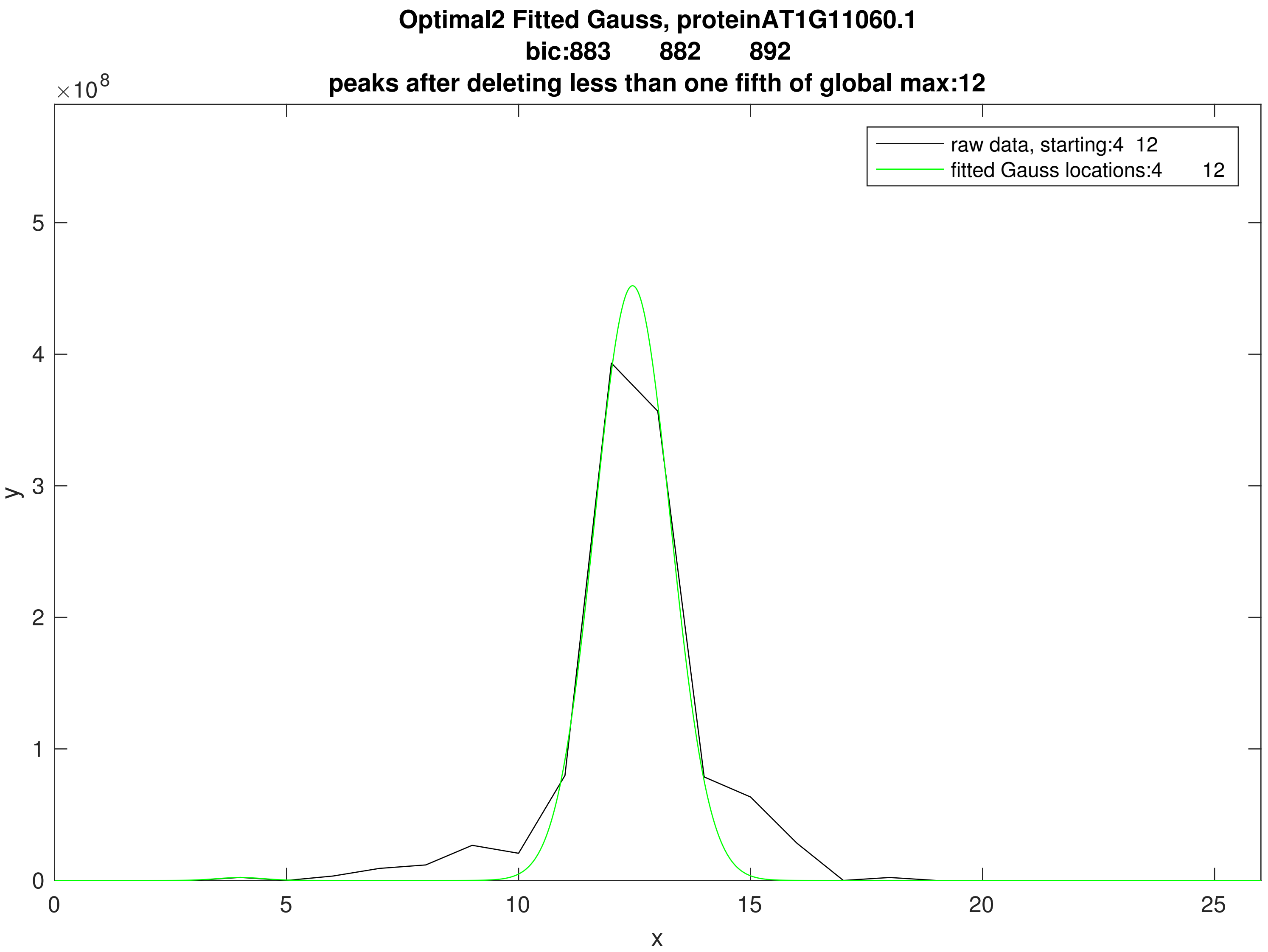

Optimal2 Fitted Gauss, proteinAT1G11060.1
bic:883 882 892
peaks after deleting less than one fifth of global max:12
×10^8
raw data, starting:4 12
fitted Gauss locations:4 12
5
4
3
2
1
0
y
0
5
10
15
20
25
x

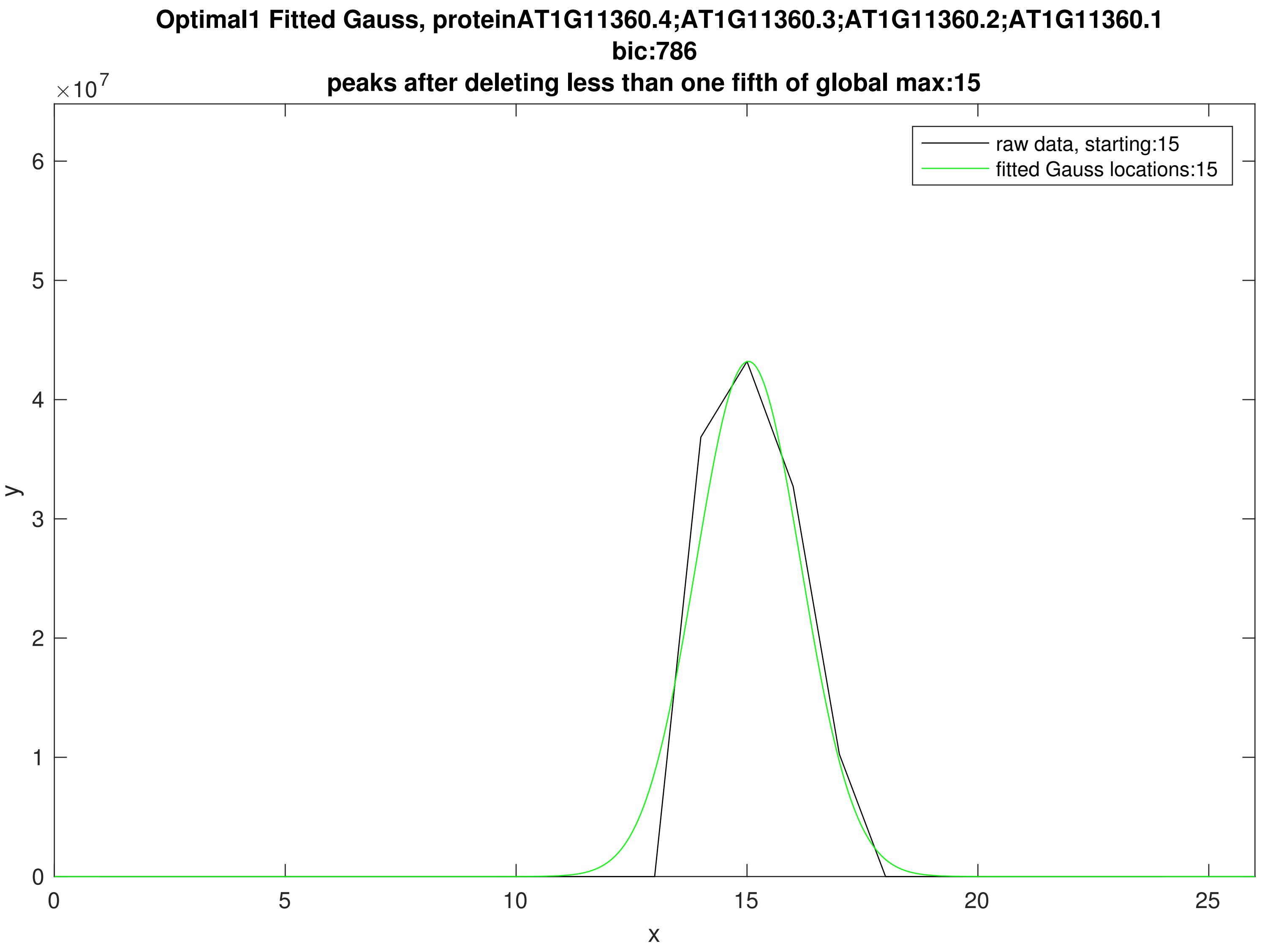

Optimal1 Fitted Gauss, proteinAT1G11360.4;AT1G11360.3;AT1G11360.2;AT1G11360.1
bic:786
peaks after deleting less than one fifth of global max:15
×10^7
raw data, starting:15
fitted Gauss locations:15
y
x
0
1
2
3
4
5
6
0
5
10
15
20
25

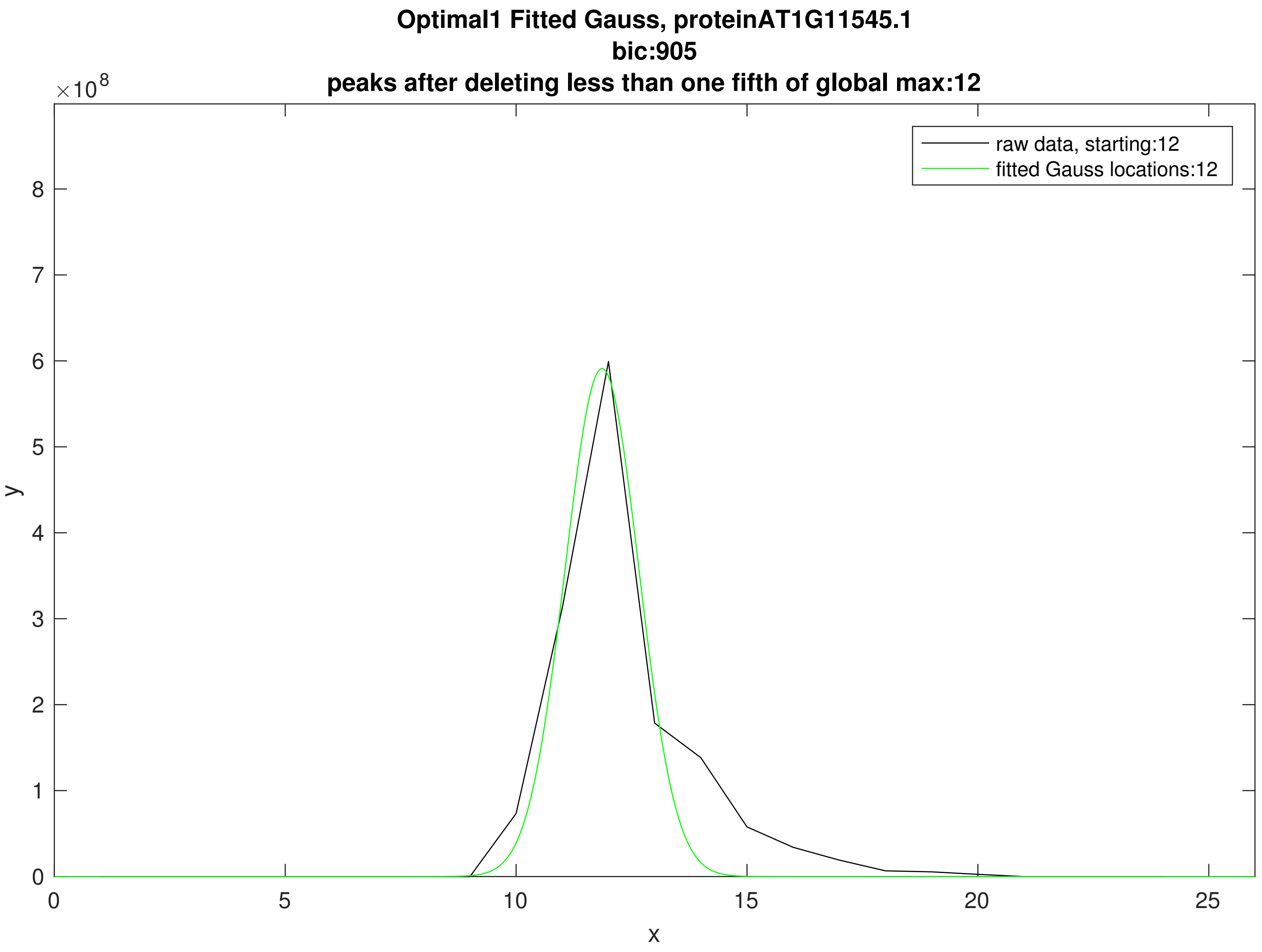

Optimal1 Fitted Gauss, proteinAT1G11545.1
bic:905
peaks after deleting less than one fifth of global max:12
×10^8
raw data, starting:12
fitted Gauss locations:12
y
x
0
1
2
3
4
5
6
7
8
0
5
10
15
20
25

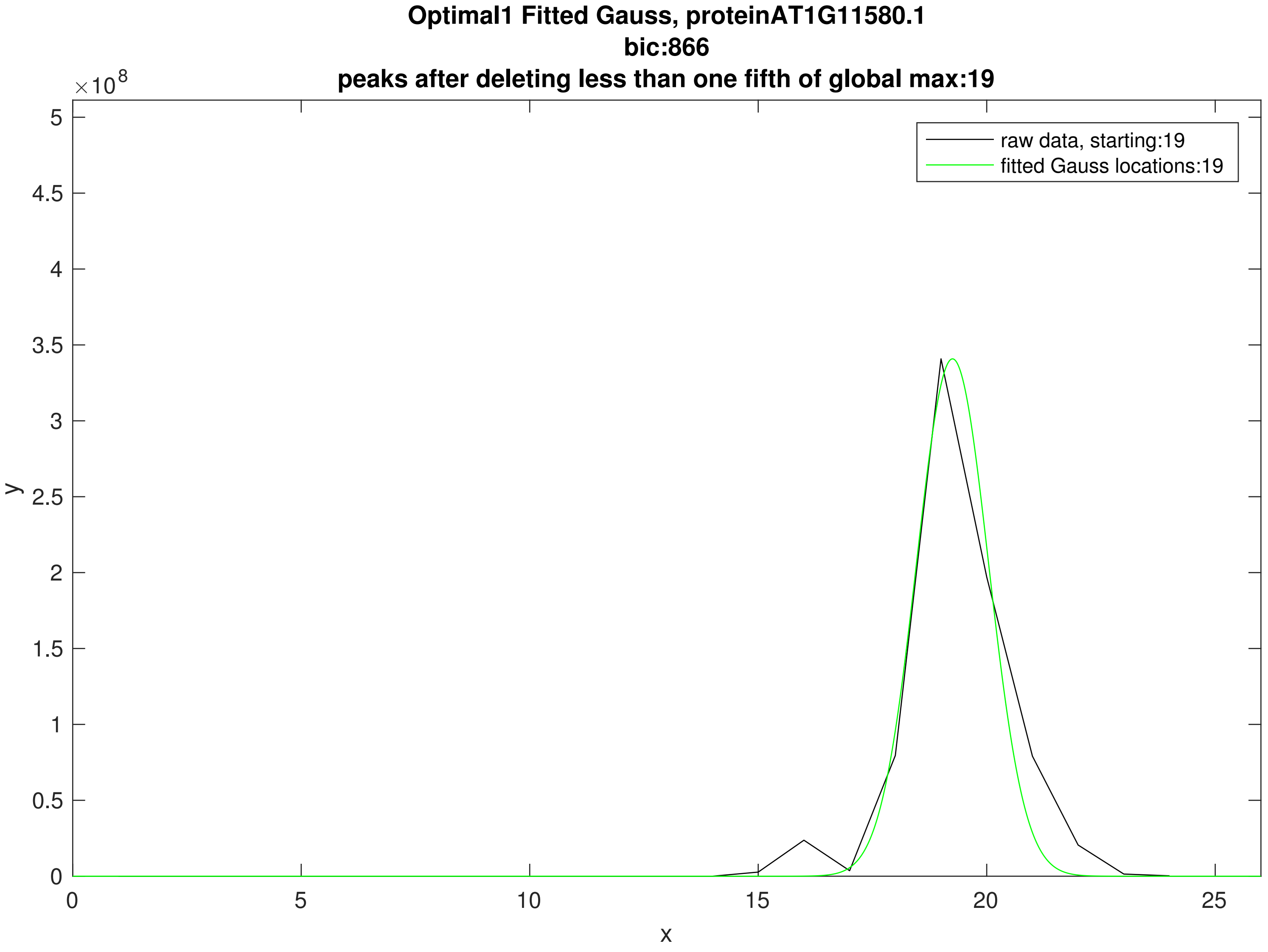

Optimal1 Fitted Gauss, proteinAT1G11580.1
bic:866
peaks after deleting less than one fifth of global max:19
×10^8
raw data, starting:19
fitted Gauss locations:19
y
x
0
0.5
1
1.5
2
2.5
3
3.5
4
4.5
5
0
5
10
15
20
25

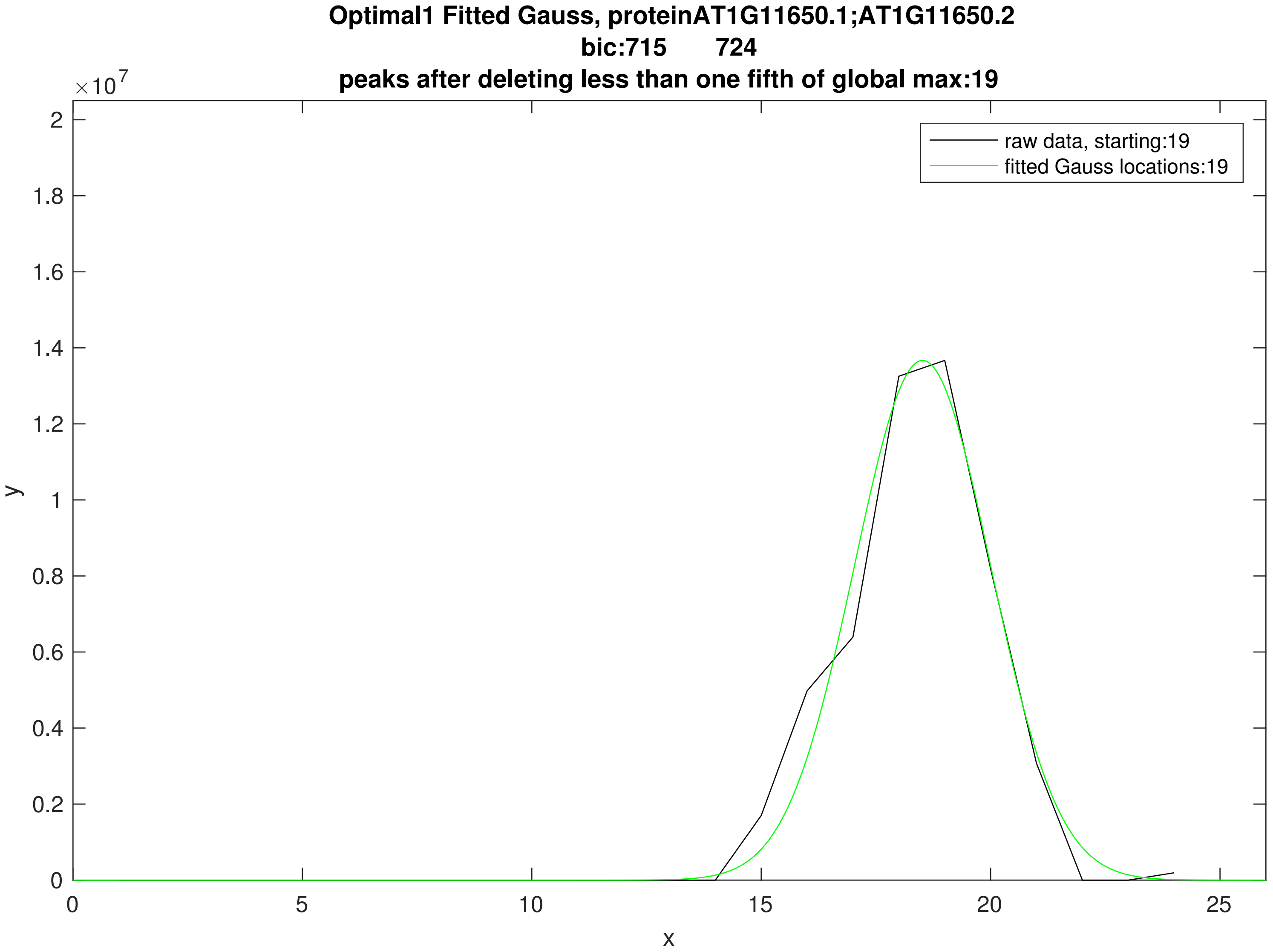
Optimal1 Fitted Gauss, proteinAT1G11650.1;AT1G11650.2
bic:715 724
peaks after deleting less than one fifth of global max:19
×10^7
2
1.8
1.6
1.4
1.2
1
0.8
0.6
0.4
0.2
0
y
0
5
10
15
20
25
x
raw data, starting:19
fitted Gauss locations:19

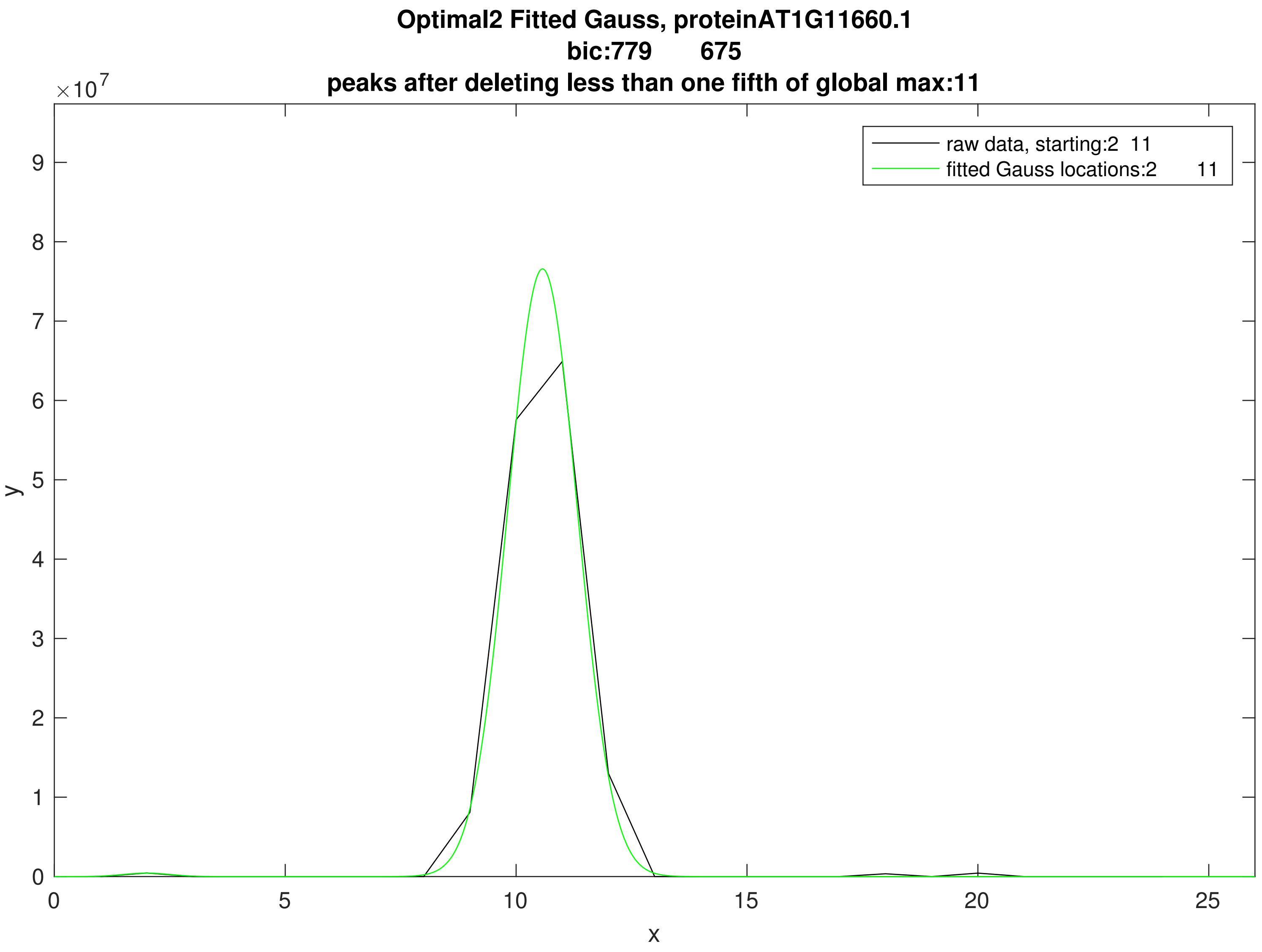
Optimal2 Fitted Gauss, proteinAT1G11660.1
bic:779 675
peaks after deleting less than one fifth of global max:11
raw data, starting:2 11
fitted Gauss locations:2 11
×10^7
y
x
0
1
2
3
4
5
6
7
8
9
0
5
10
15
20
25

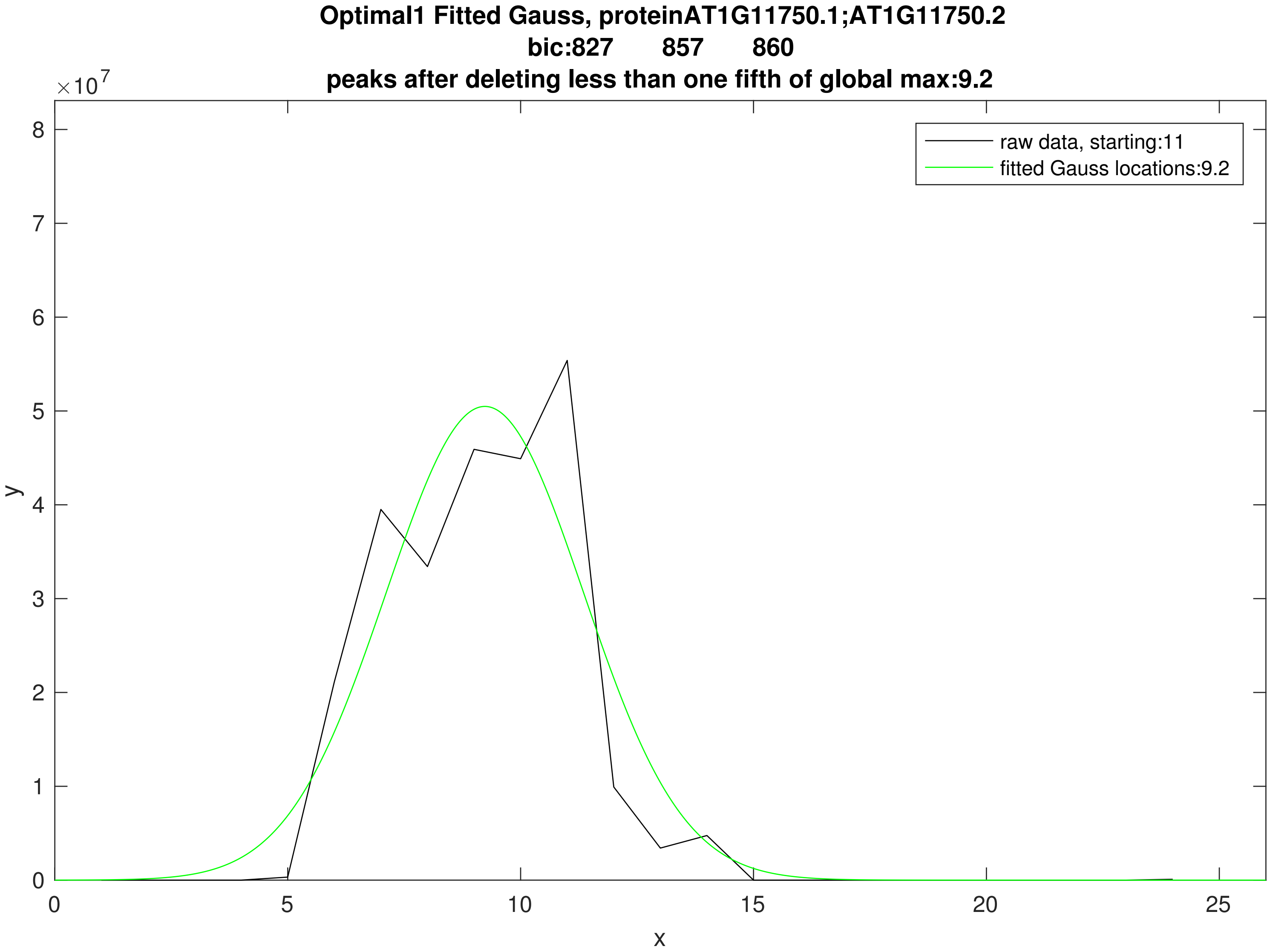
Optimal1 Fitted Gauss, proteinAT1G11750.1;AT1G11750.2
bic:827 857 860
peaks after deleting less than one fifth of global max:9.2
×10^7
raw data, starting:11
fitted Gauss locations:9.2
y
x
0
1
2
3
4
5
6
7
8
0
5
10
15
20
25

Optimal1 Fitted Gauss, proteinAT1G11840.4;AT1G11840.3;AT1G11840.2;AT1G11840.1;AT1G11840.6;AT1G11840.
bic:1e+03 1.01e+03 1.02e+03
peaks after deleting less than one fifth of global max:18

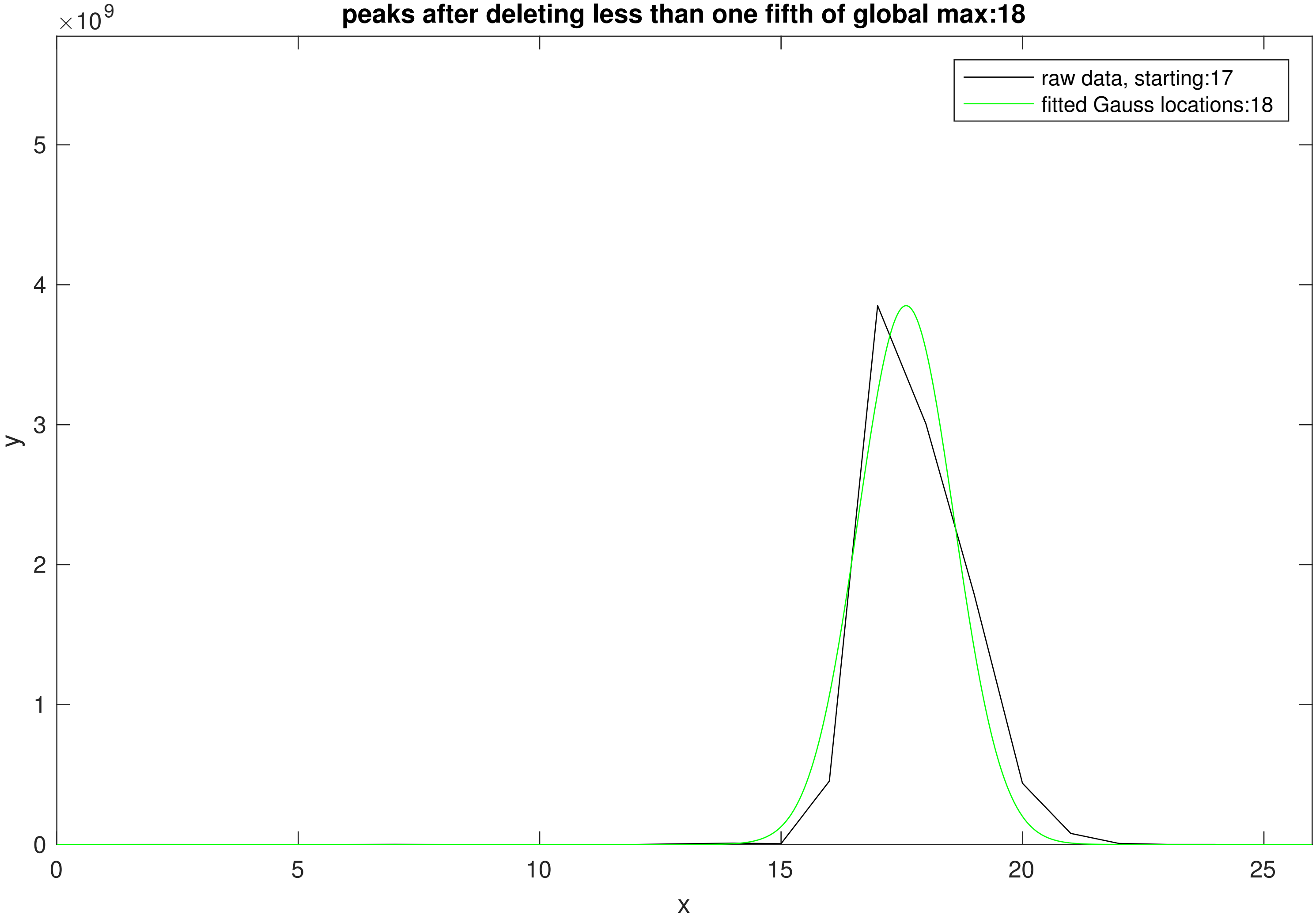

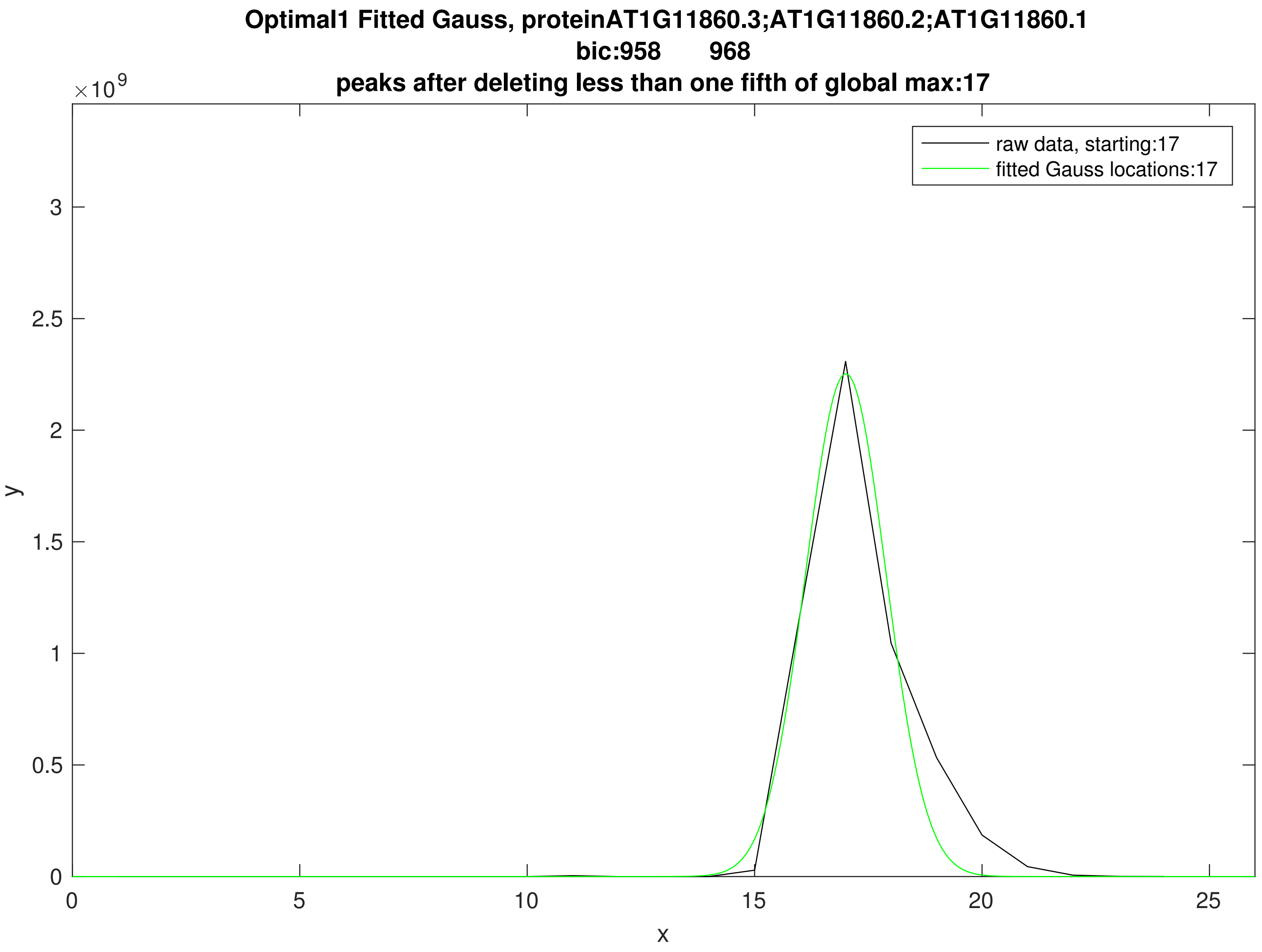

Optimal1 Fitted Gauss, proteinAT1G11860.3;AT1G11860.2;AT1G11860.1
bic:958 968
peaks after deleting less than one fifth of global max:17
×10^9
raw data, starting:17
fitted Gauss locations:17
y
x
0
0.5
1
1.5
2
2.5
3
0
5
10
15
20
25

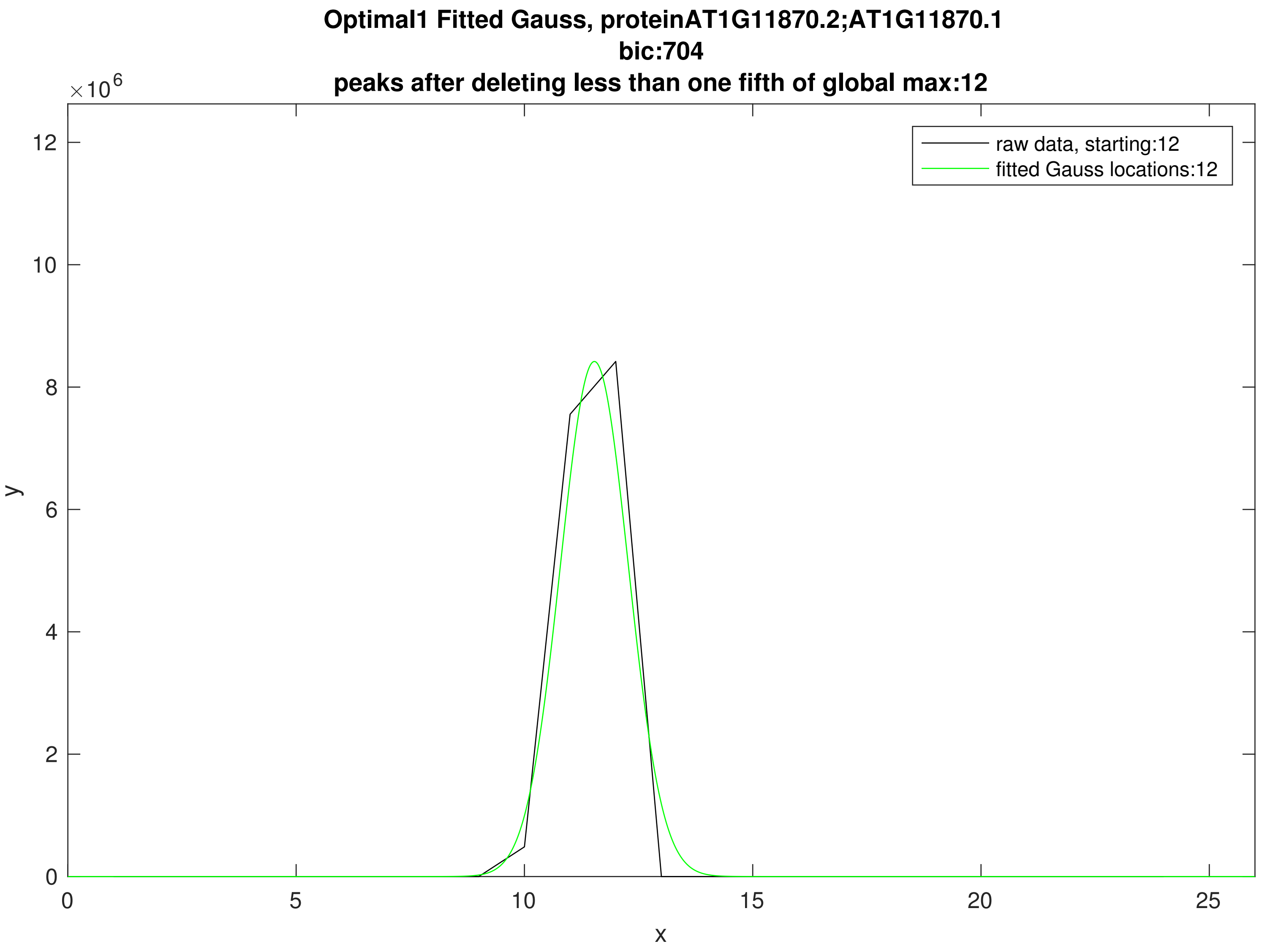

Optimal1 Fitted Gauss, proteinAT1G11870.2;AT1G11870.1
bic:704
peaks after deleting less than one fifth of global max:12
×10^6
raw data, starting:12
fitted Gauss locations:12
y
x
0
2
4
6
8
10
12
0
5
10
15
20
25

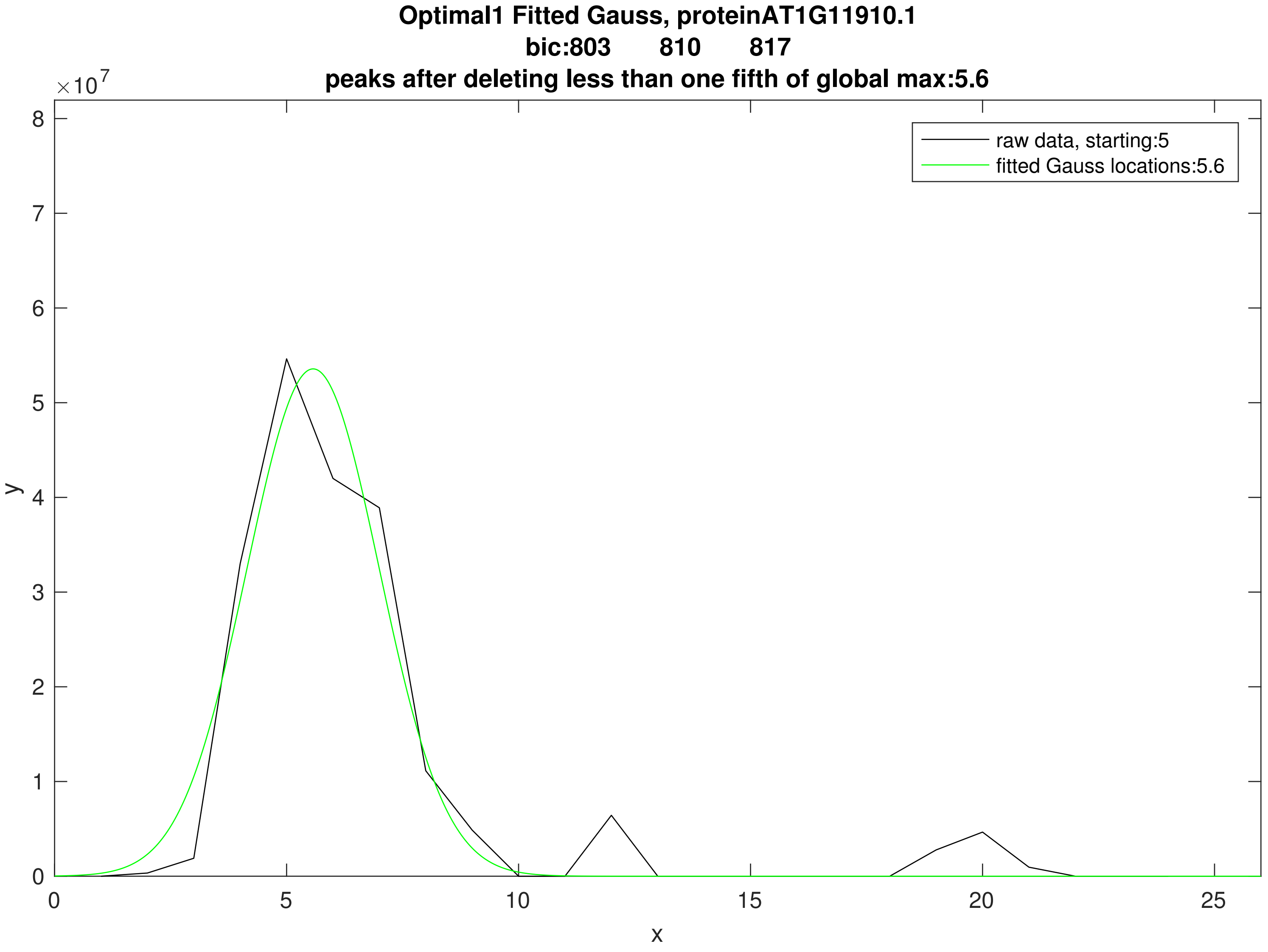

Optimal1 Fitted Gauss, proteinAT1G11910.1
bic:803 810 817
peaks after deleting less than one fifth of global max:5.6
×10^7
raw data, starting:5
fitted Gauss locations:5.6
y
x
0
1
2
3
4
5
6
7
8
10
15
20
25

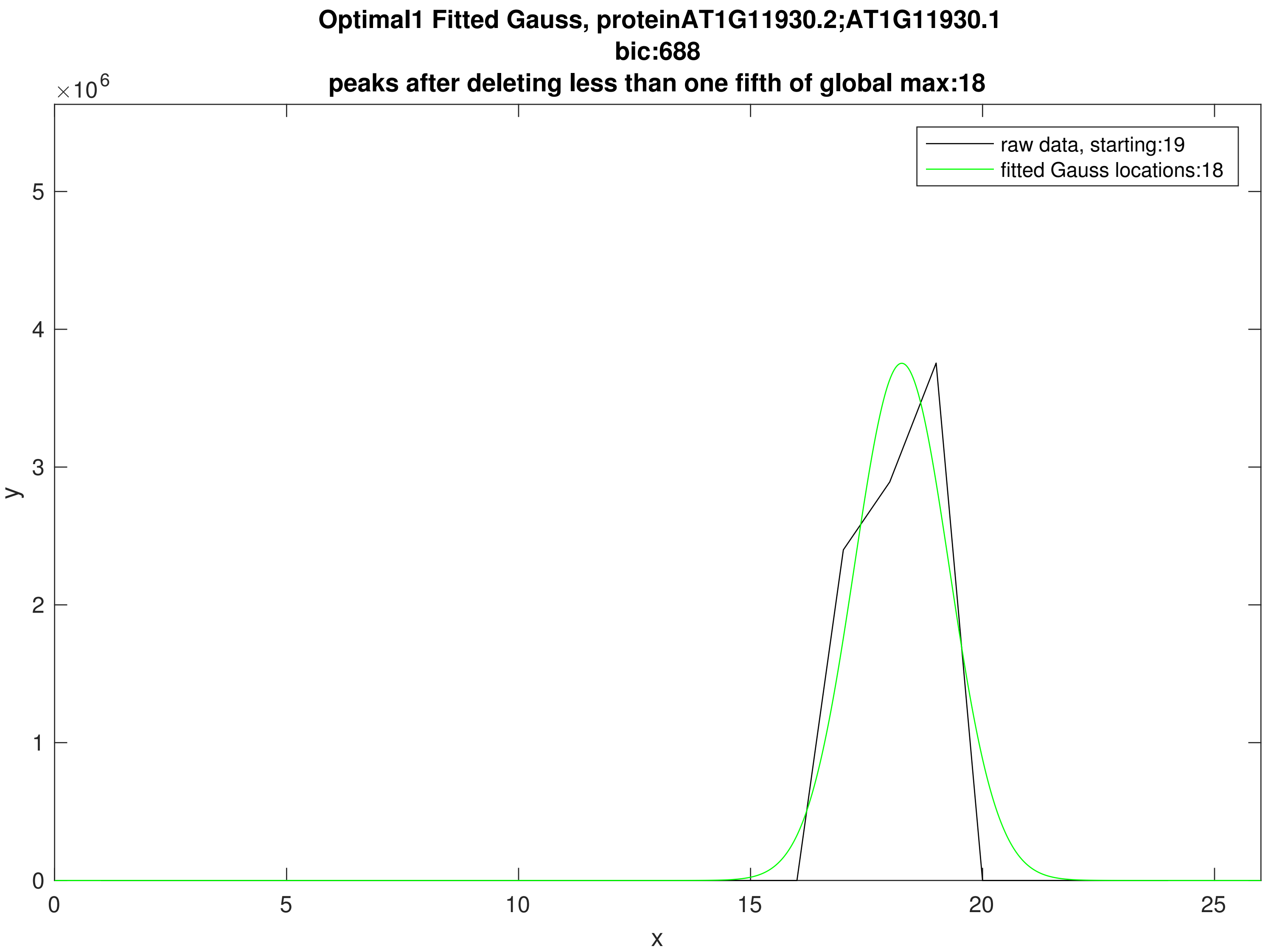

Optimal1 Fitted Gauss, proteinAT1G11930.2;AT1G11930.1
bic:688
peaks after deleting less than one fifth of global max:18
×10^6
raw data, starting:19
fitted Gauss locations:18
y
x
0
1
2
3
4
5
0
5
10
15
20
25

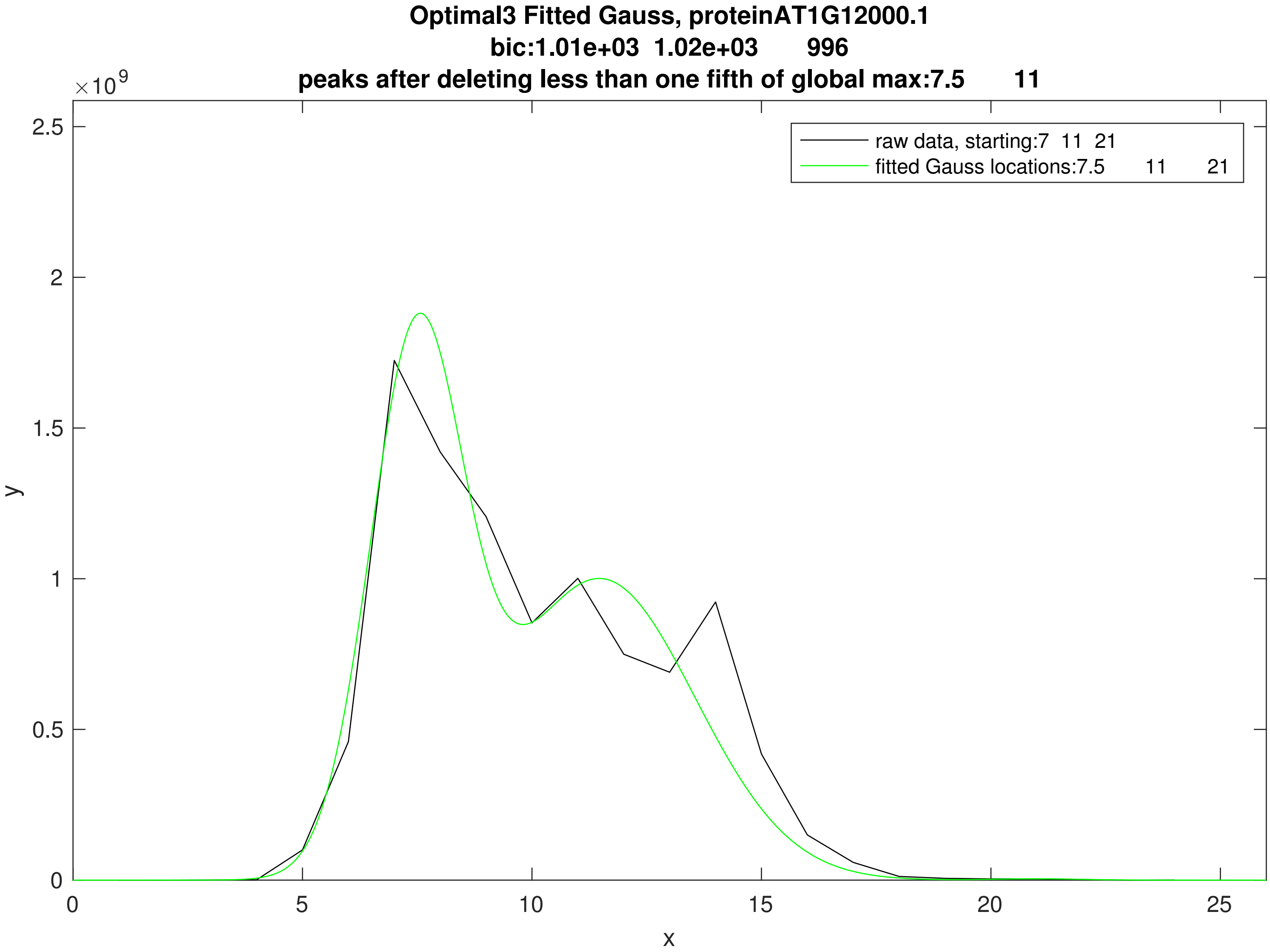

Optimal3 Fitted Gauss, proteinAT1G12000.1
bic:1.01e+03 1.02e+03 996
peaks after deleting less than one fifth of global max:7.5 11
×10^9
raw data, starting:7 11 21
fitted Gauss locations:7.5 11 21
y
x
0
0.5
1
1.5
2
2.5
0
5
10
15
20
25

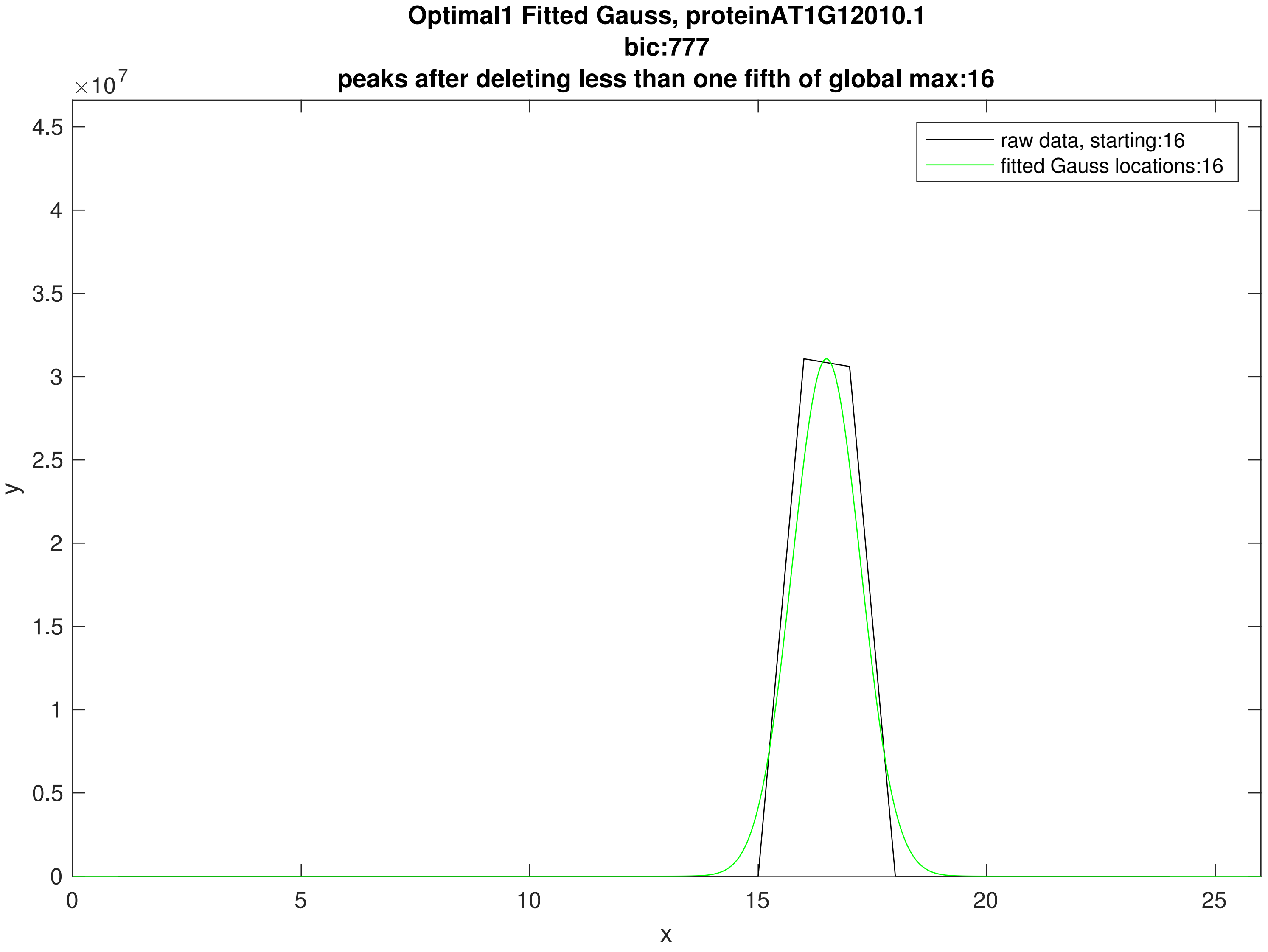
Optimal1 Fitted Gauss, proteinAT1G12010.1
bic:777
peaks after deleting less than one fifth of global max:16
×10^7
4.5
4
3.5
3
2.5
2
1.5
1
0.5
0
y
0
5
10
15
20
25
x
raw data, starting:16
fitted Gauss locations:16

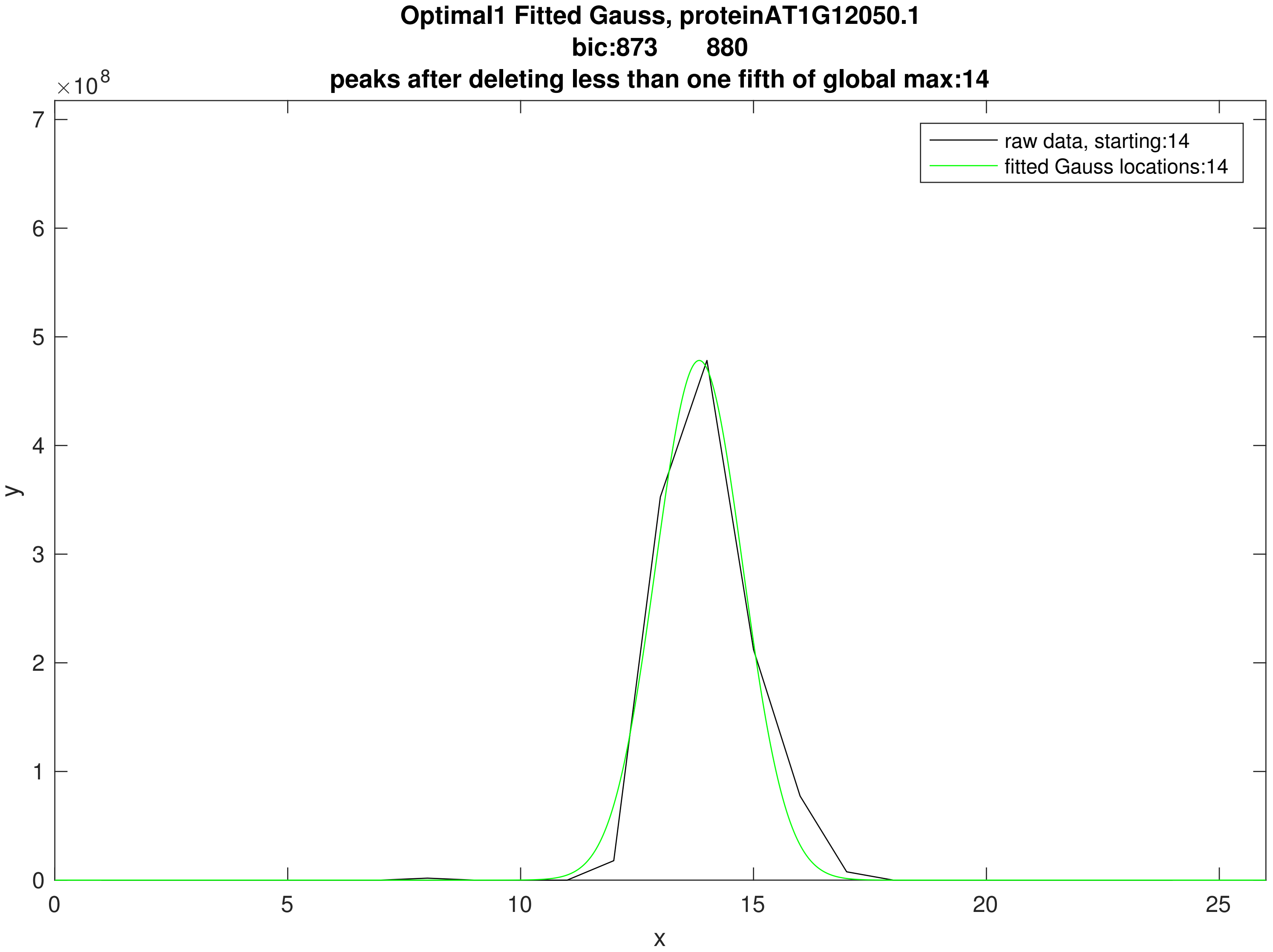
Optimal1 Fitted Gauss, proteinAT1G12050.1
bic:873 880
peaks after deleting less than one fifth of global max:14
×10^8
raw data, starting:14
fitted Gauss locations:14
y
x
0
1
2
3
4
5
6
7
0
5
10
15
20
25

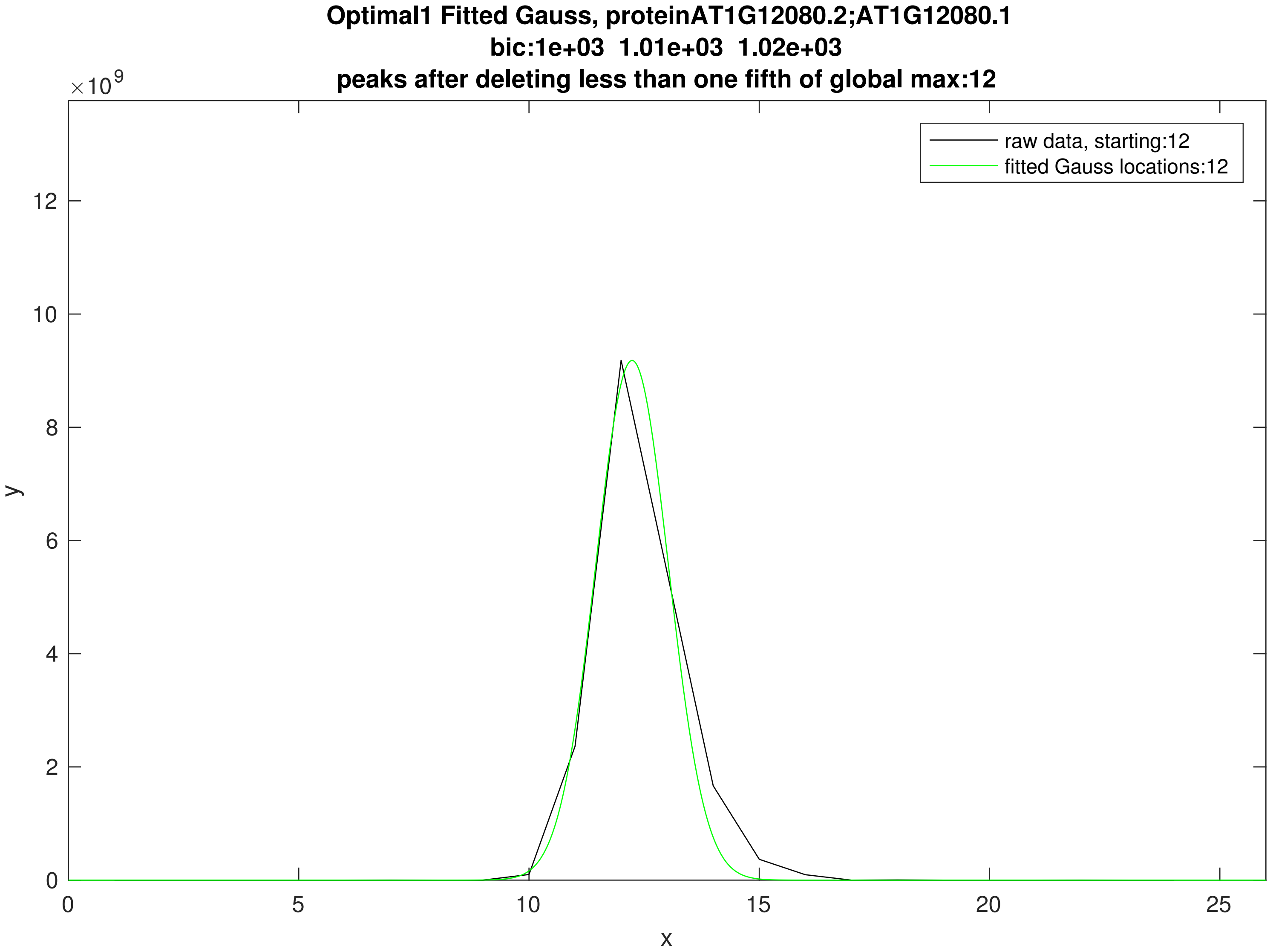
Optimal1 Fitted Gauss, proteinAT1G12080.2;AT1G12080.1
bic:1e+03 1.01e+03 1.02e+03
peaks after deleting less than one fifth of global max:12
×10^9
raw data, starting:12
fitted Gauss locations:12
y
x
0
2
4
6
8
10
12
0
5
10
15
20
25

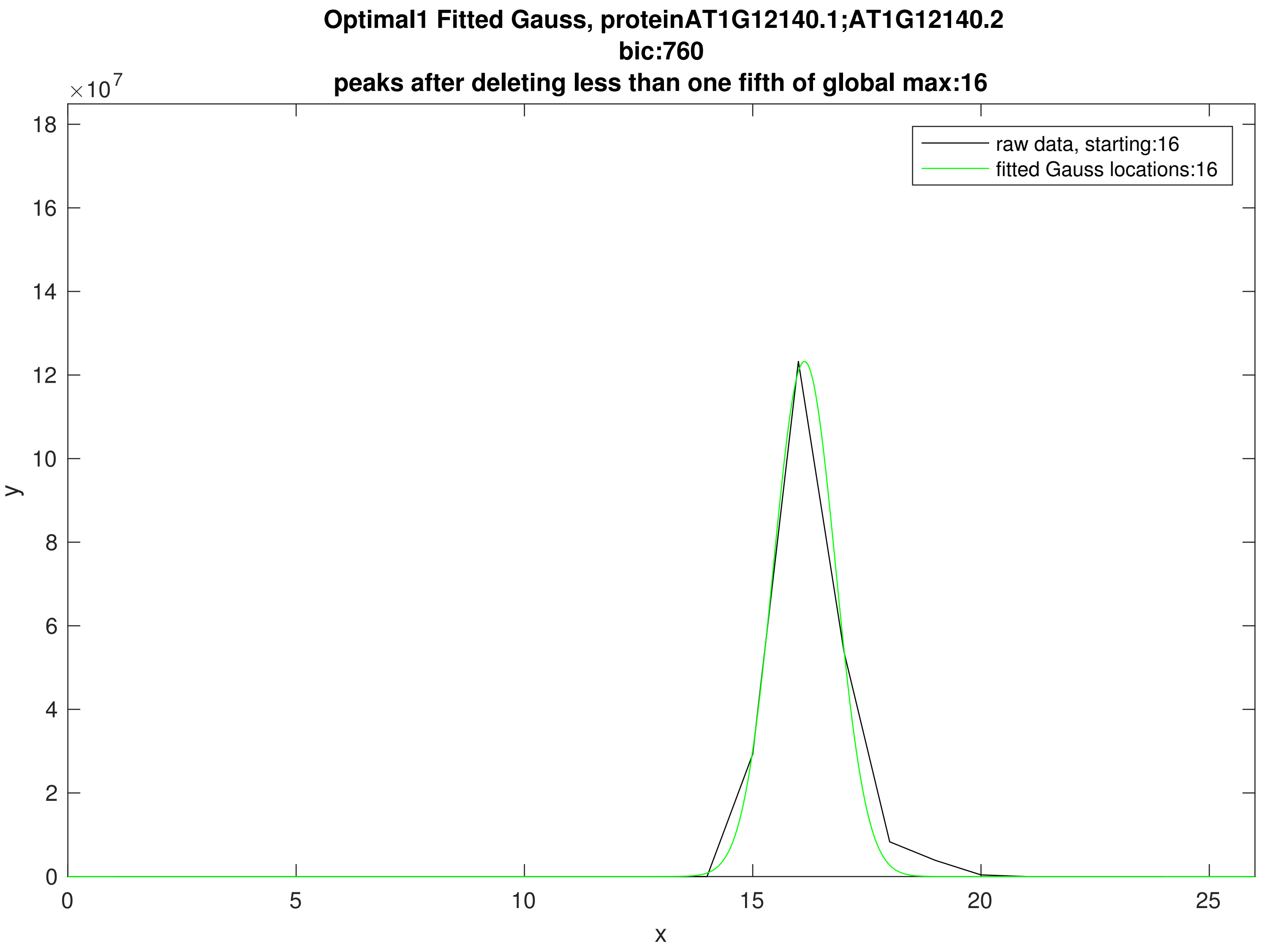
Optimal1 Fitted Gauss, proteinAT1G12140.1;AT1G12140.2
bic:760
peaks after deleting less than one fifth of global max:16
×10^7
raw data, starting:16
fitted Gauss locations:16
y
x
0
2
4
6
8
10
12
14
16
18
0
5
10
15
20
25

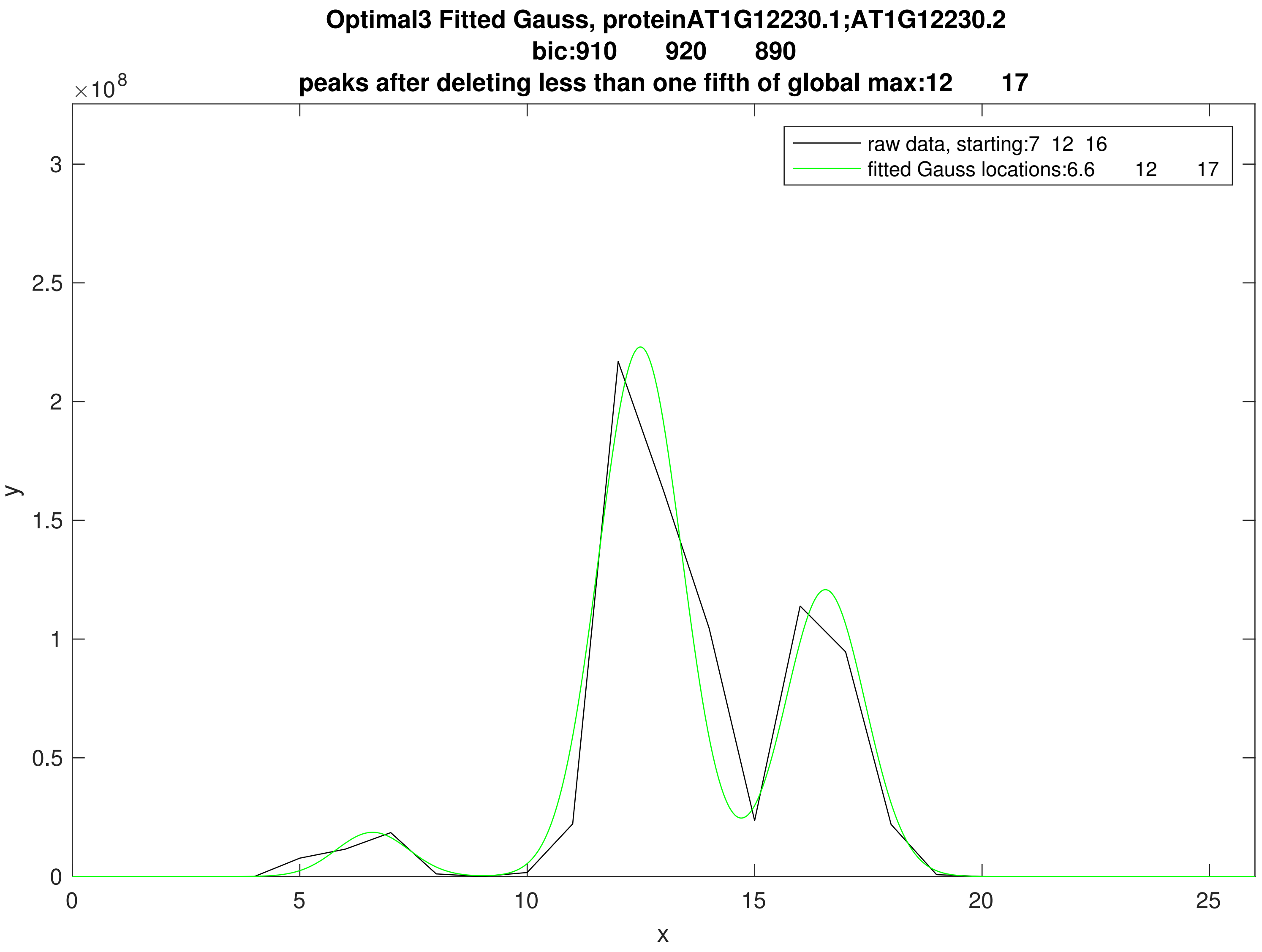
Optimal3 Fitted Gauss, proteinAT1G12230.1;AT1G12230.2
bic:910 920 890
peaks after deleting less than one fifth of global max:12 17
×10^8
raw data, starting:7 12 16
fitted Gauss locations:6.6 12 17
y
x
0
0.5
1
1.5
2
2.5
3
5
10
15
20
25

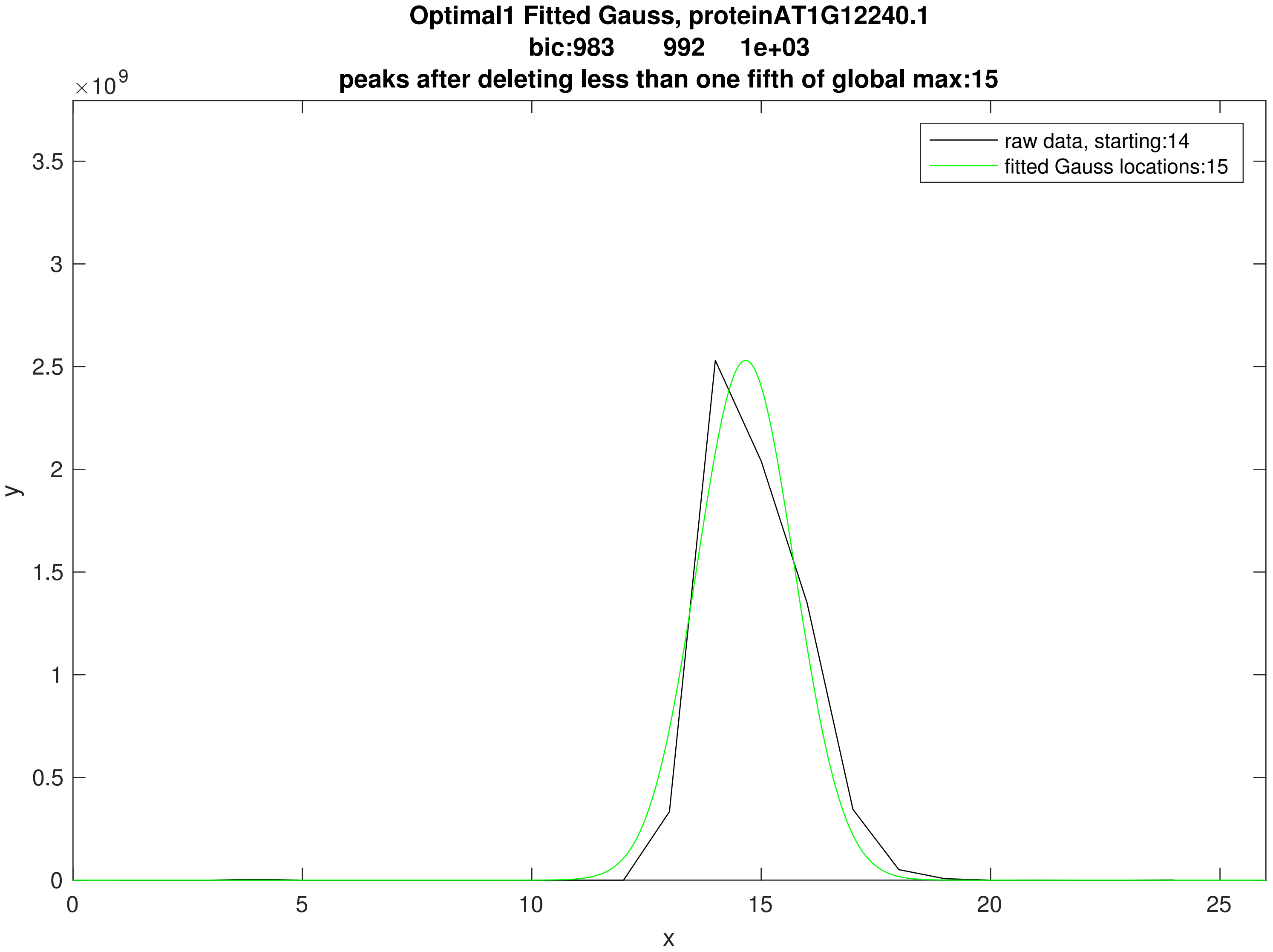
Optimal1 Fitted Gauss, proteinAT1G12240.1
bic:983 992 1e+03
peaks after deleting less than one fifth of global max:15
×10^9
3.5
3
2.5
2
1.5
1
0.5
0
y
0
5
10
15
20
25
x
raw data, starting:14
fitted Gauss locations:15

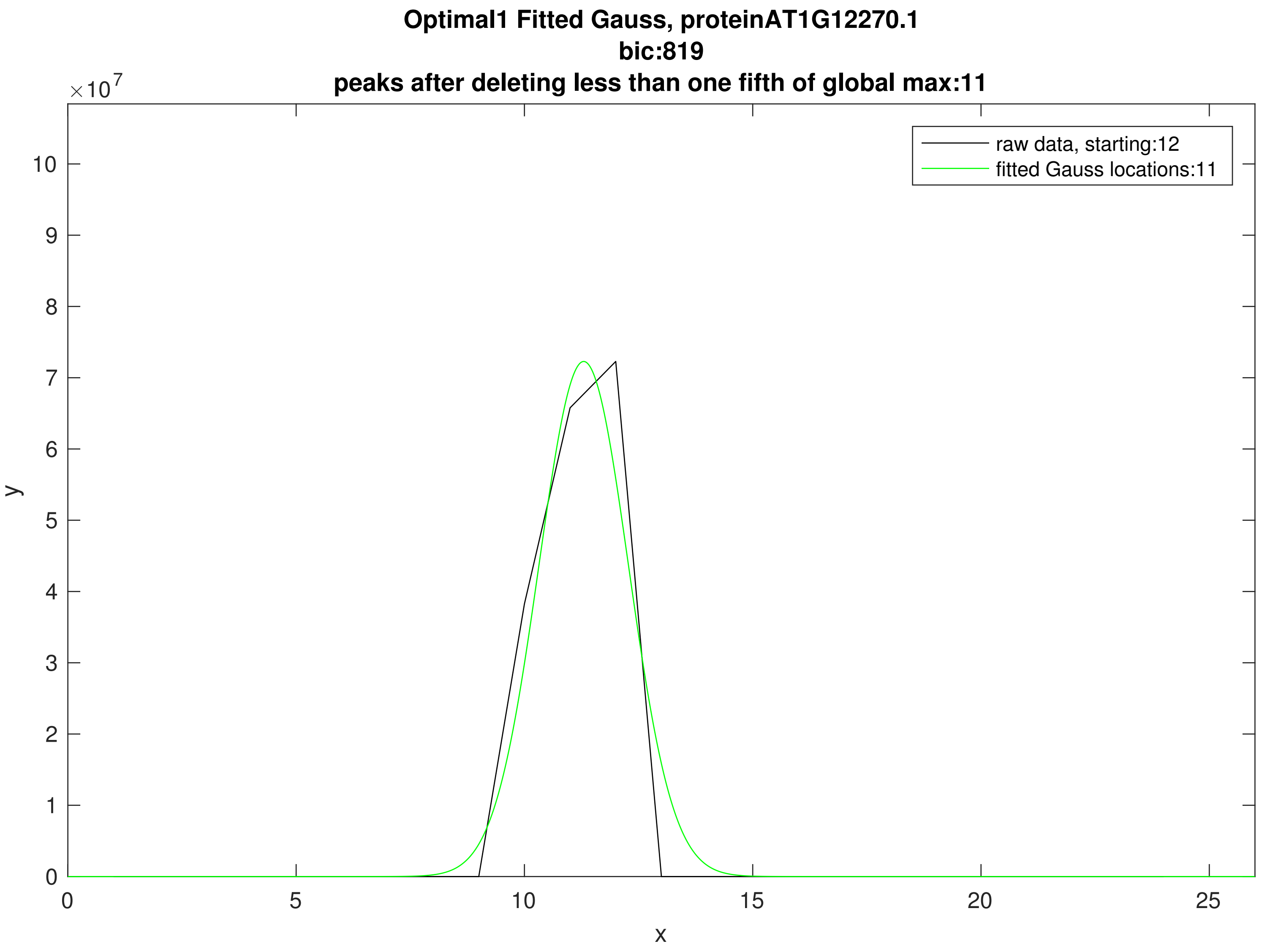

Optimal1 Fitted Gauss, proteinAT1G12270.1
bic:819
peaks after deleting less than one fifth of global max:11
×10^7
raw data, starting:12
fitted Gauss locations:11
y
x
0
1
2
3
4
5
6
7
8
9
10
0
5
10
15
20
25

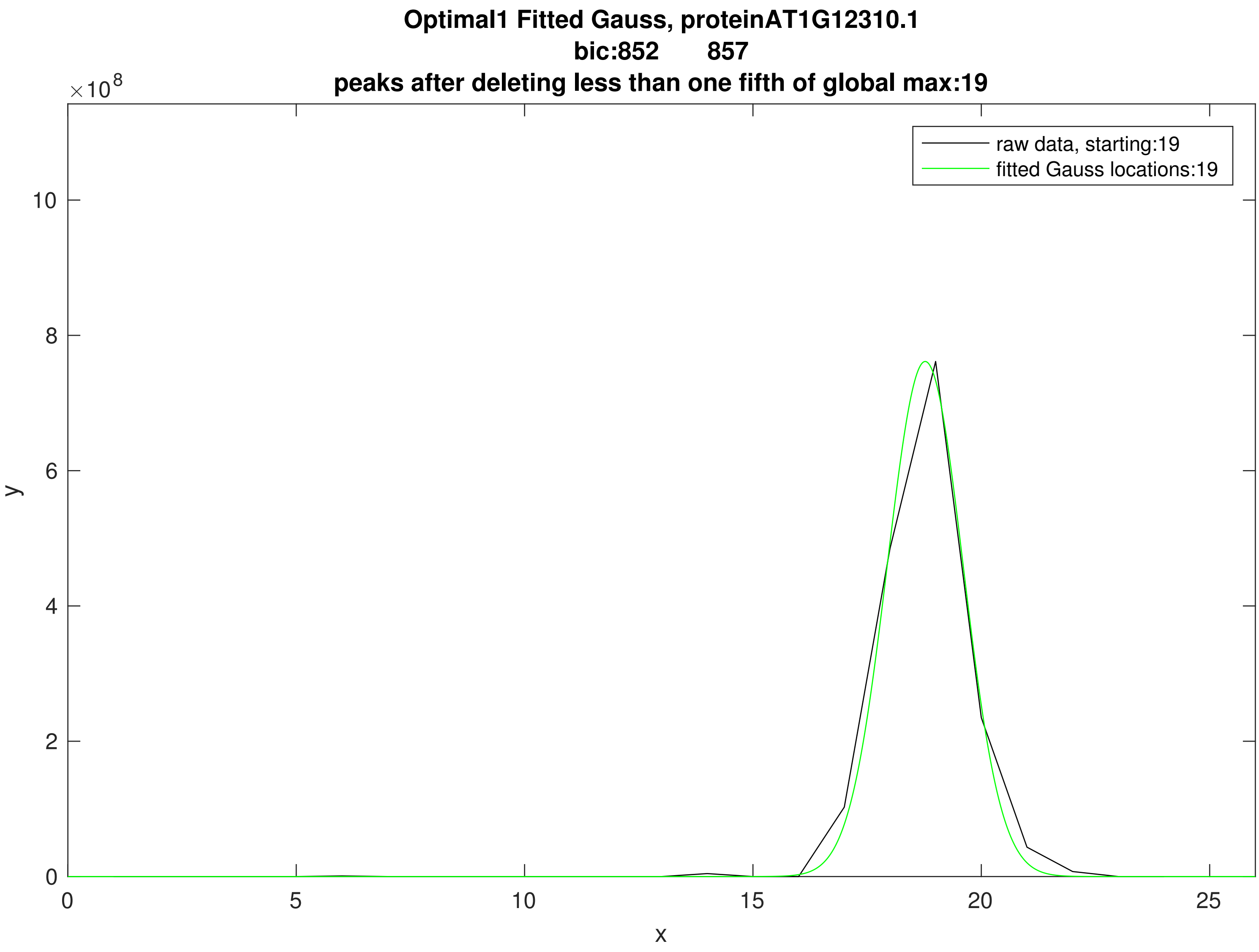
Optimal1 Fitted Gauss, proteinAT1G12310.1
bic:852 857
peaks after deleting less than one fifth of global max:19
×10^8
raw data, starting:19
fitted Gauss locations:19
y
x
0
2
4
6
8
10
0
5
10
15
20
25

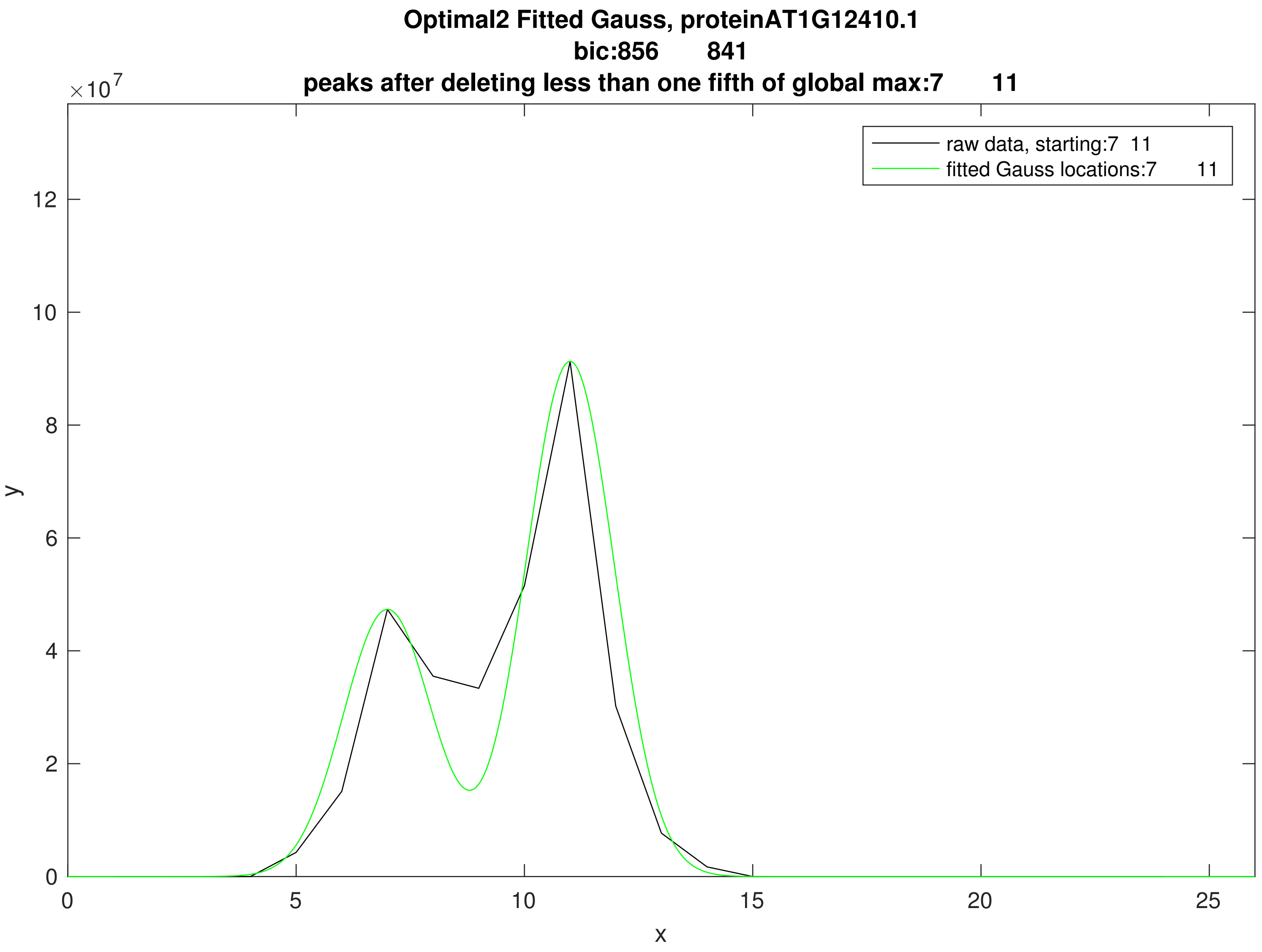

Optimal2 Fitted Gauss, proteinAT1G12410.1
bic:856 841
peaks after deleting less than one fifth of global max:7 11
×10^7
raw data, starting:7 11
fitted Gauss locations:7 11
12
10
8
6
4
2
0
y
0
5
10
15
20
25
x

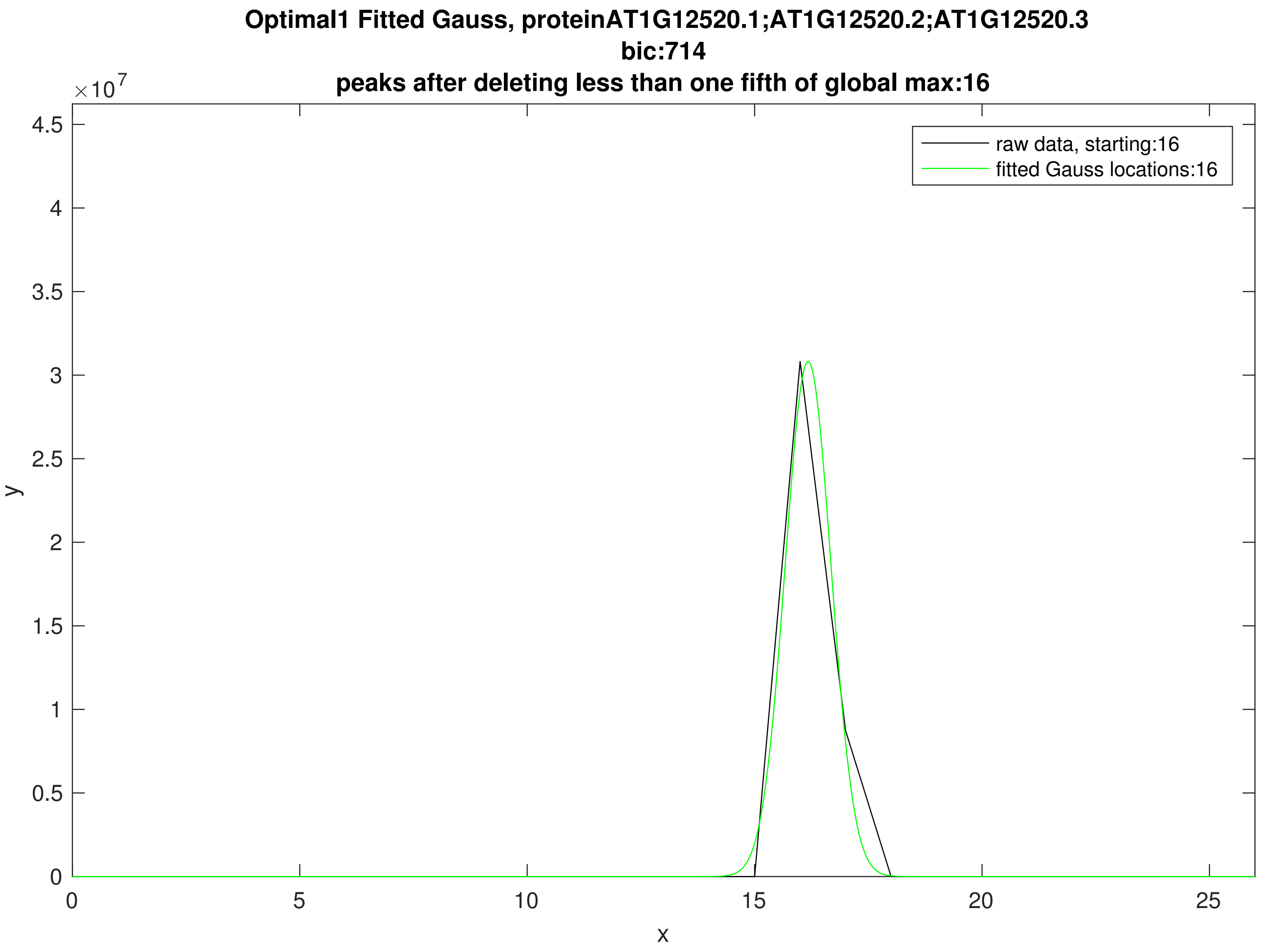
Optimal1 Fitted Gauss, proteinAT1G12520.1;AT1G12520.2;AT1G12520.3
bic:714
peaks after deleting less than one fifth of global max:16
×10^7
4.5
4
3.5
3
2.5
2
1.5
1
0.5
0
y
0
5
10
15
20
25
x
raw data, starting:16
fitted Gauss locations:16

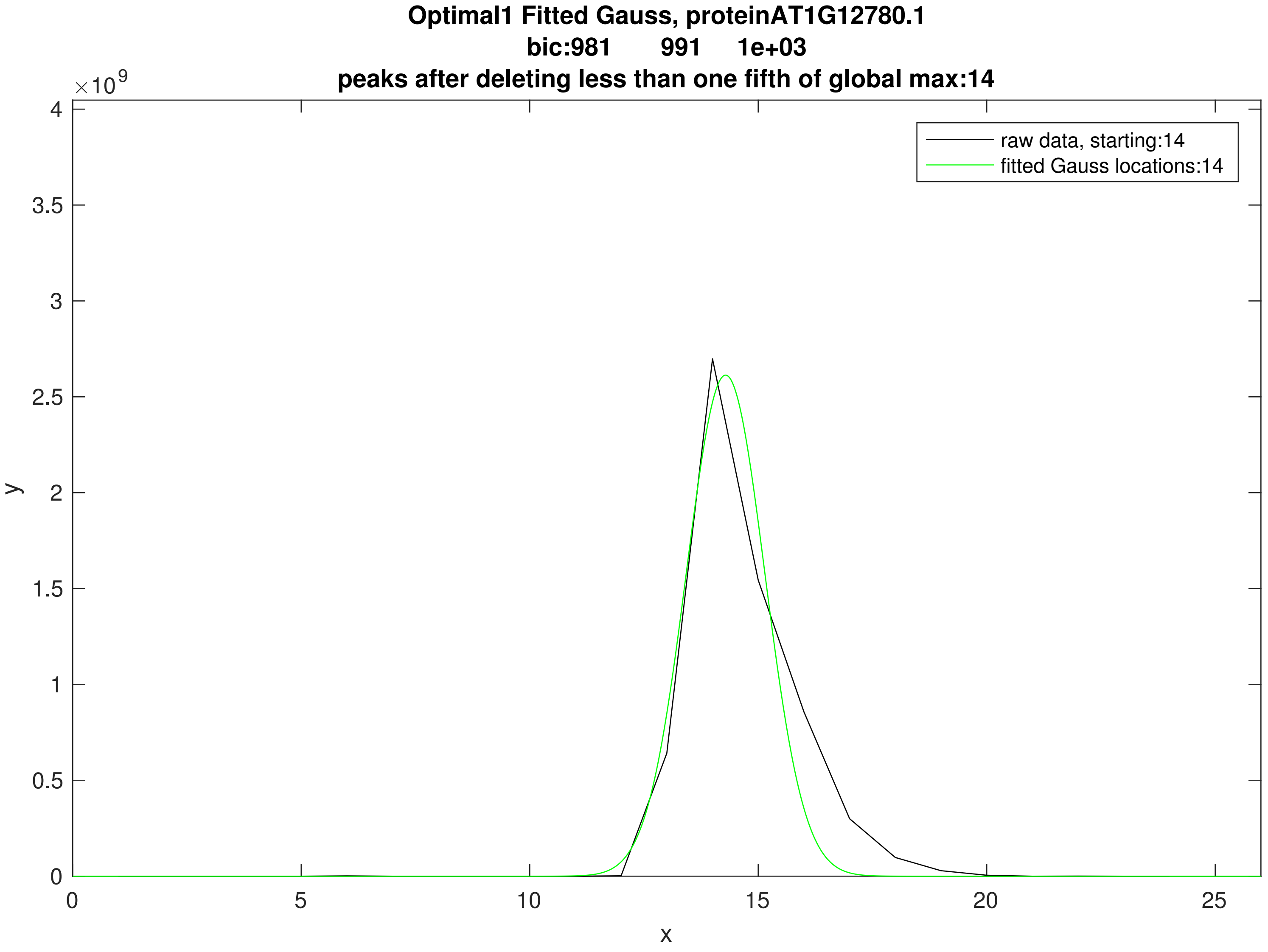

Optimal1 Fitted Gauss, proteinAT1G12780.1
bic:981 991 1e+03
peaks after deleting less than one fifth of global max:14
×10^9
4
3.5
3
2.5
2
1.5
1
0.5
0
y
0
5
10
15
20
25
x
raw data, starting:14
fitted Gauss locations:14

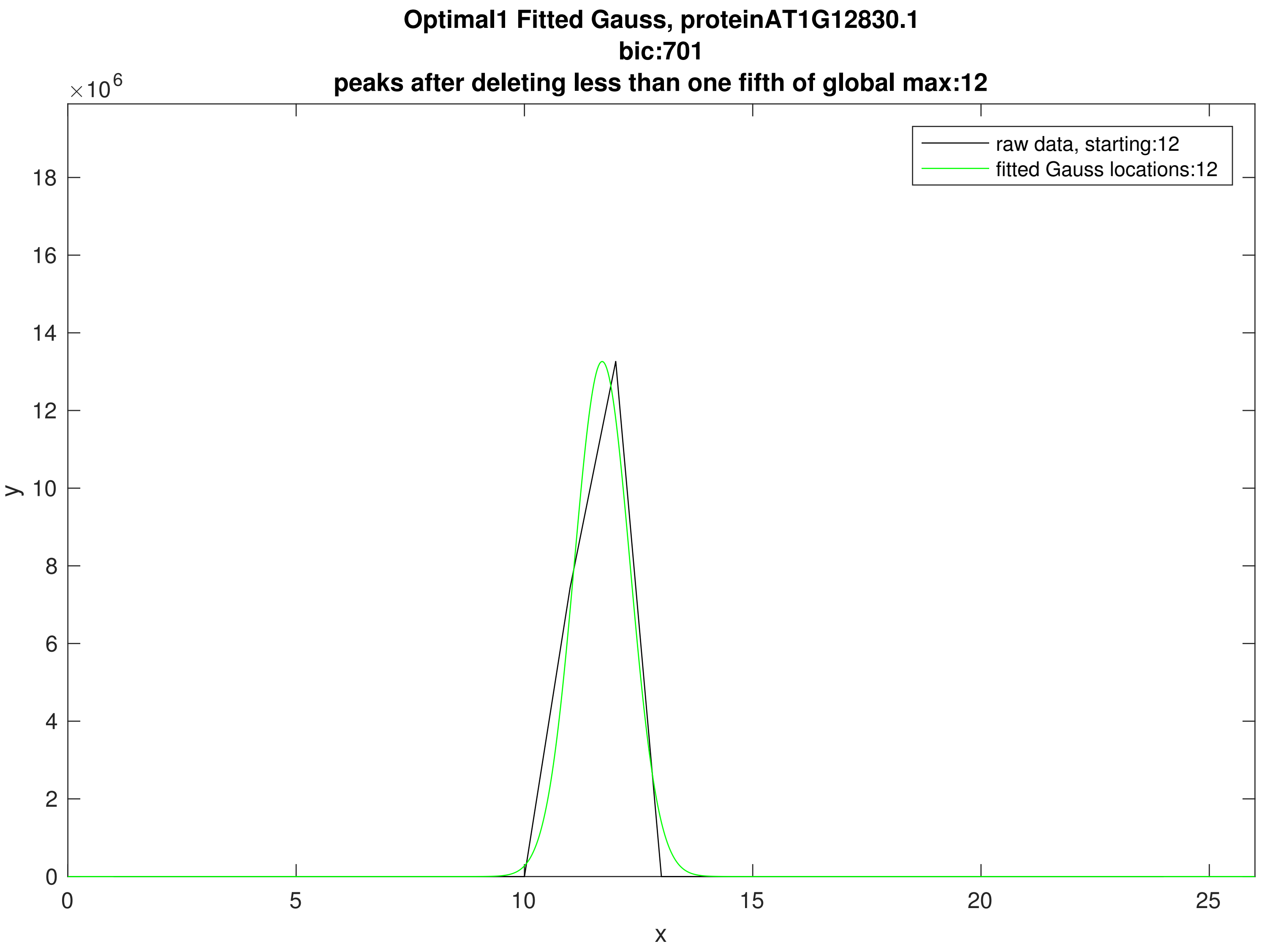
Optimal1 Fitted Gauss, proteinAT1G12830.1
bic:701
peaks after deleting less than one fifth of global max:12
×10^6
raw data, starting:12
fitted Gauss locations:12
y
x
0
2
4
6
8
10
12
14
16
18
0
5
10
15
20
25

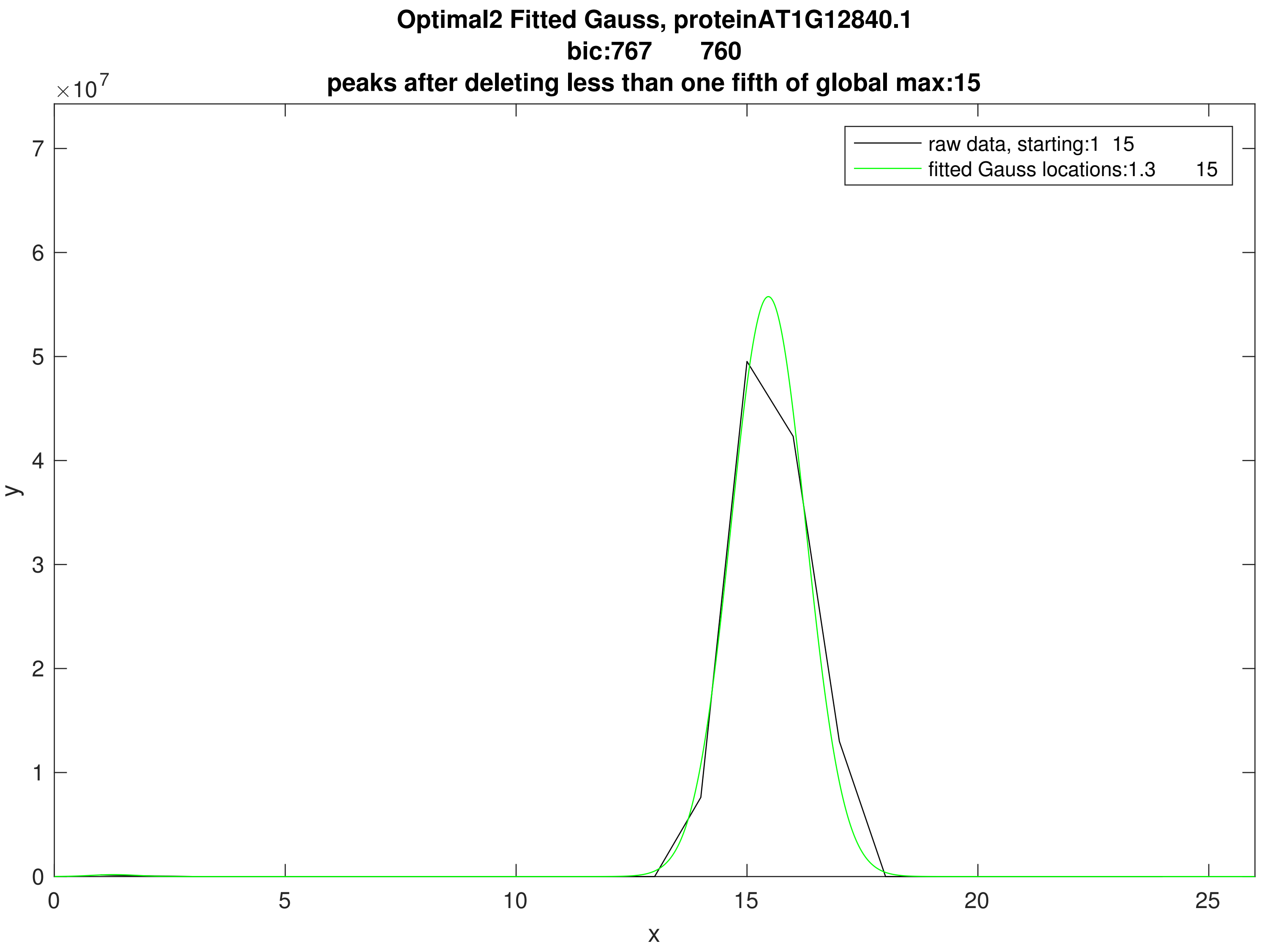
Optimal2 Fitted Gauss, proteinAT1G12840.1
bic:767 760
peaks after deleting less than one fifth of global max:15
×10^7
raw data, starting:1 15
fitted Gauss locations:1.3 15
y
x
0
1
2
3
4
5
6
7
0
5
10
15
20
25

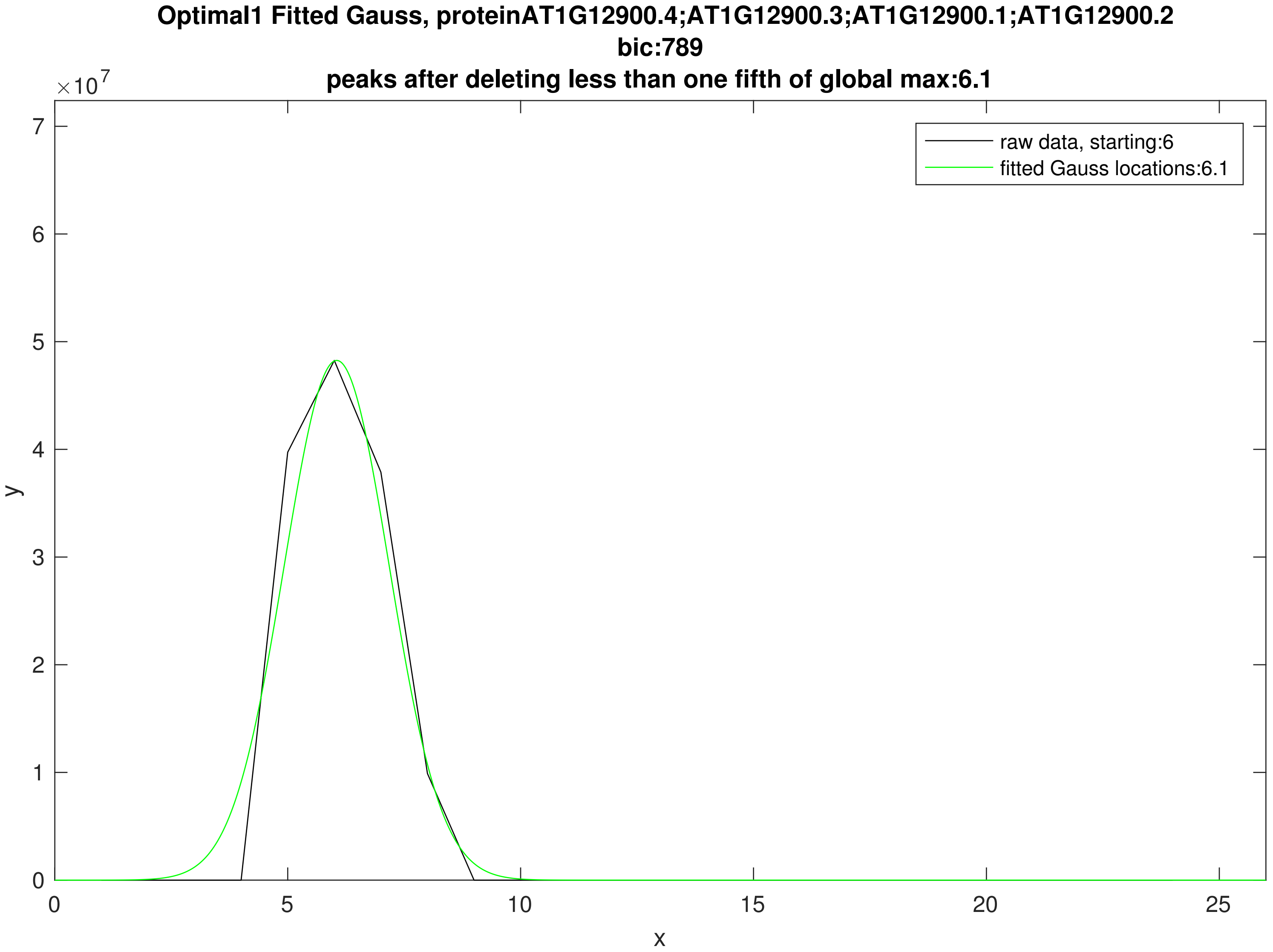

Optimal1 Fitted Gauss, proteinAT1G12900.4;AT1G12900.3;AT1G12900.1;AT1G12900.2
bic:789
peaks after deleting less than one fifth of global max:6.1
×10^7
raw data, starting:6
fitted Gauss locations:6.1
y
x
0
1
2
3
4
5
6
7
0
5
10
15
20
25

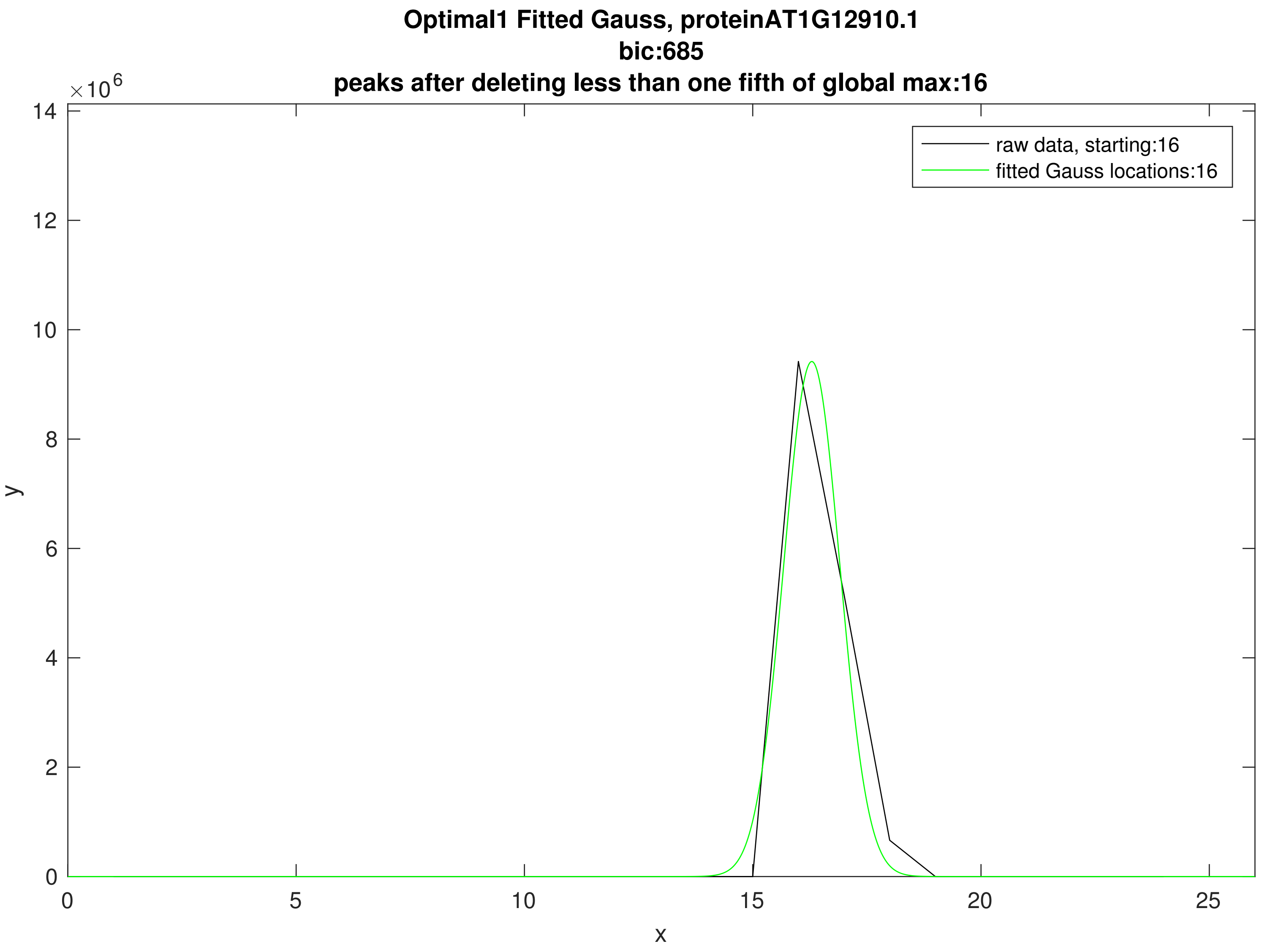
Optimal1 Fitted Gauss, proteinAT1G12910.1
bic:685
peaks after deleting less than one fifth of global max:16
×10^6
14
12
10
8
6
4
2
0
y
0
5
10
15
20
25
x
raw data, starting:16
fitted Gauss locations:16

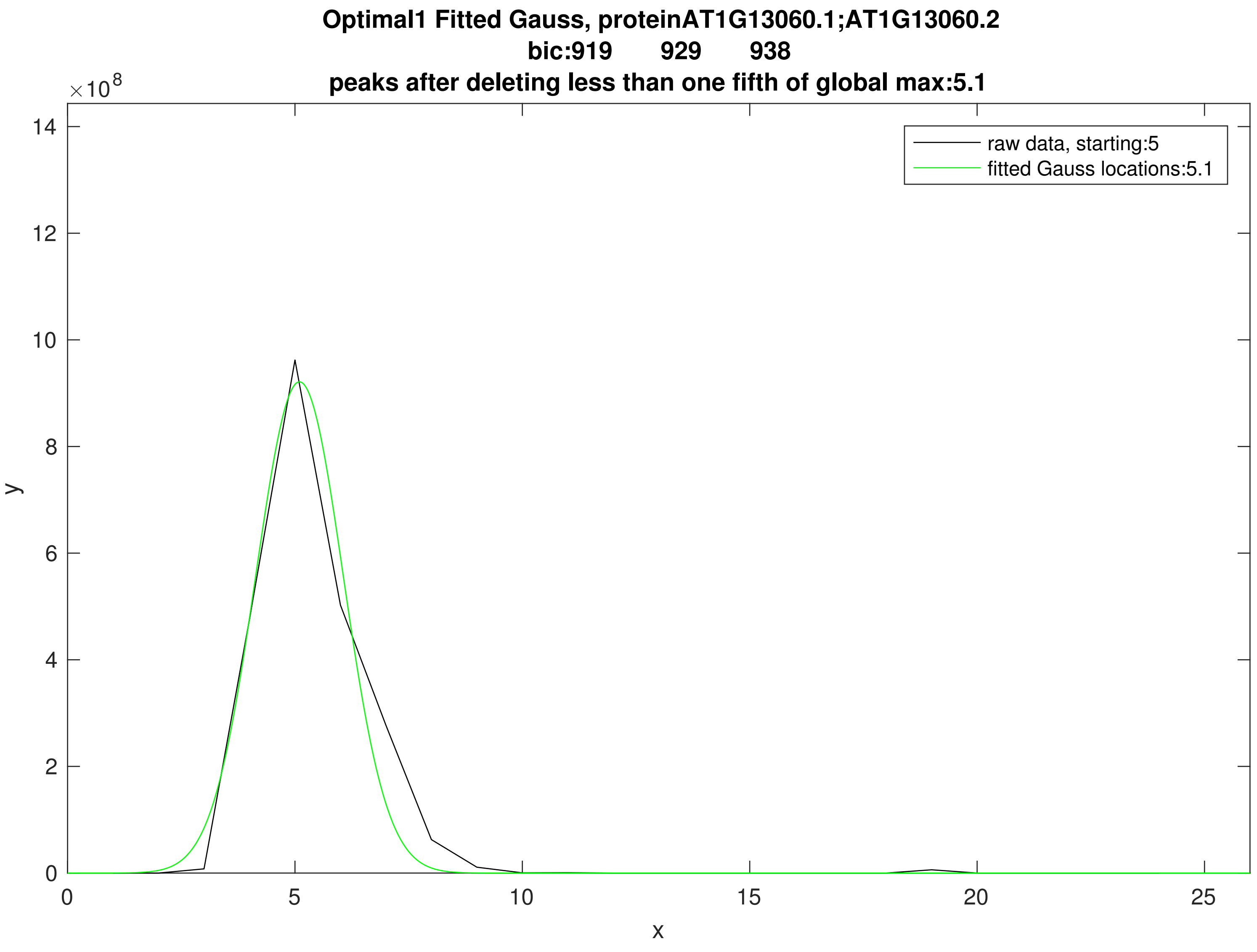
Optimal1 Fitted Gauss, proteinAT1G13060.1;AT1G13060.2
bic:919 929 938
peaks after deleting less than one fifth of global max:5.1
×10^8
14
12
10
8
6
4
2
0
y
0
5
10
15
20
25
x
raw data, starting:5
fitted Gauss locations:5.1

**Optimal1 Fitted Gauss, proteinAT1G13280.1**
**bic:882 888**
**peaks after deleting less than one fifth of global max:15**

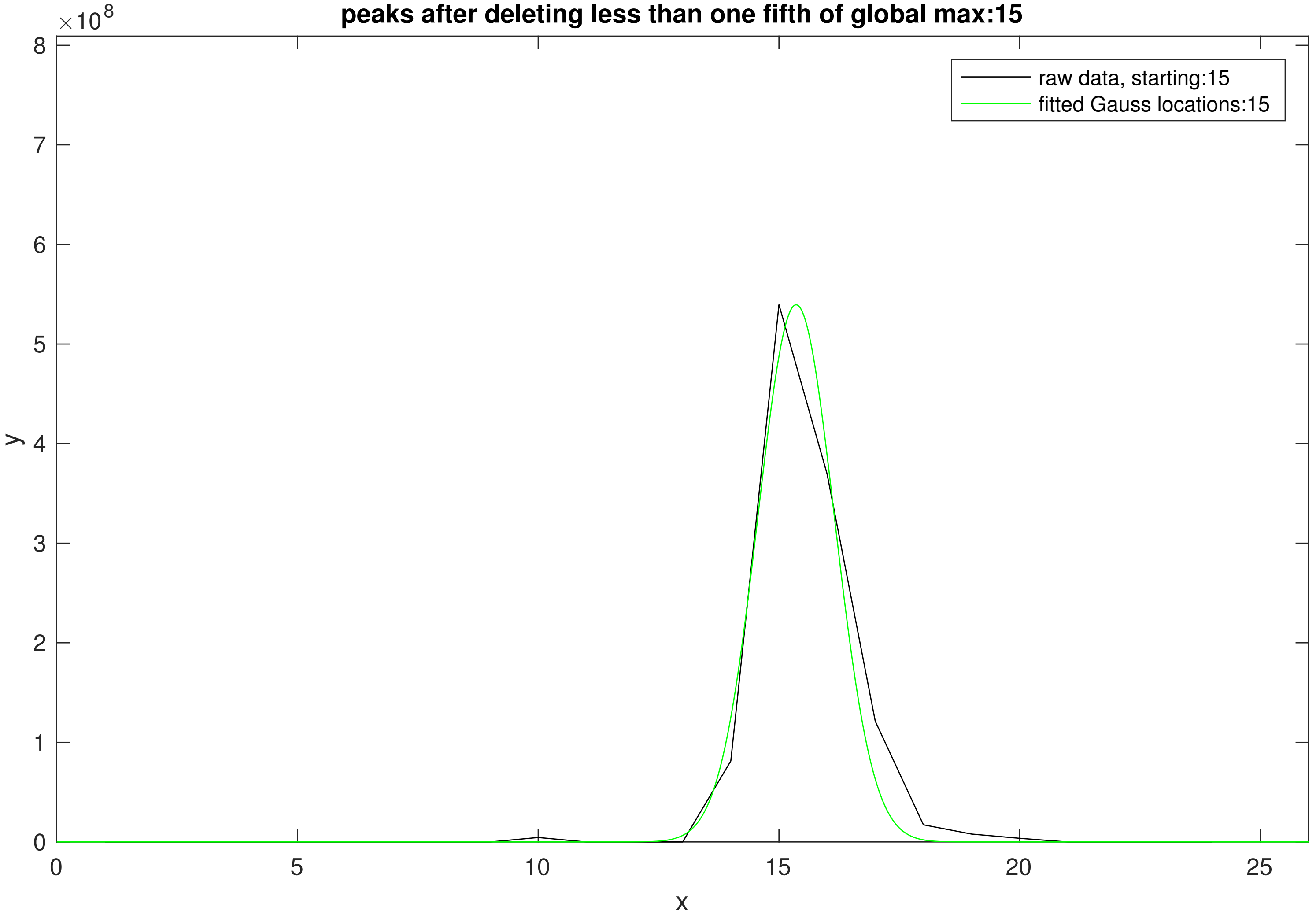

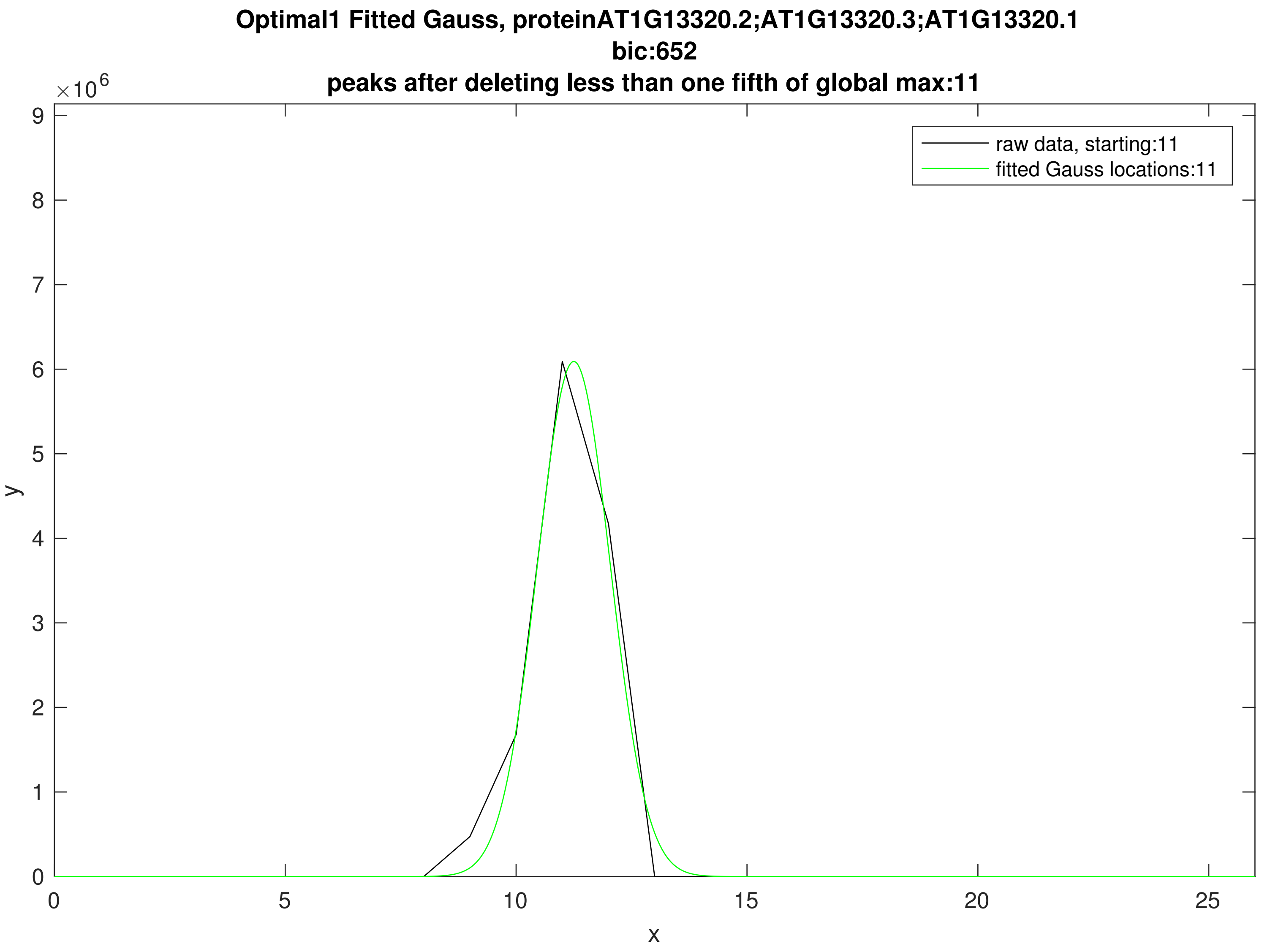
Optimal1 Fitted Gauss, proteinAT1G13320.2;AT1G13320.3;AT1G13320.1
bic:652
peaks after deleting less than one fifth of global max:11
×10^6
raw data, starting:11
fitted Gauss locations:11
y
x
0
5
10
15
20
25
1
2
3
4
6
7
8
9

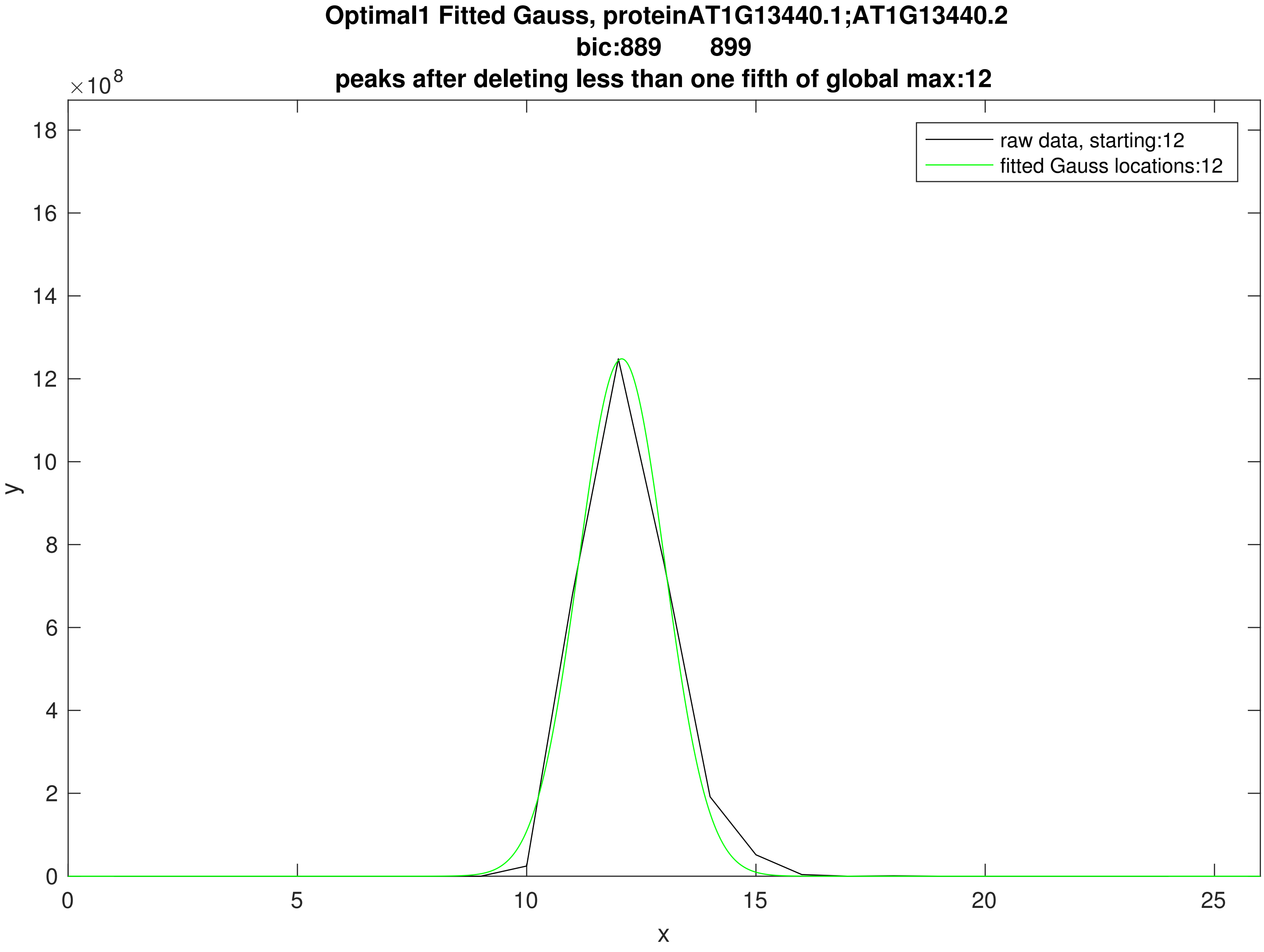
Optimal1 Fitted Gauss, proteinAT1G13440.1;AT1G13440.2
bic:889 899
peaks after deleting less than one fifth of global max:12
×10^8
18
16
14
12
10
8
6
4
2
0
y
0
5
10
15
20
25
x
raw data, starting:12
fitted Gauss locations:12

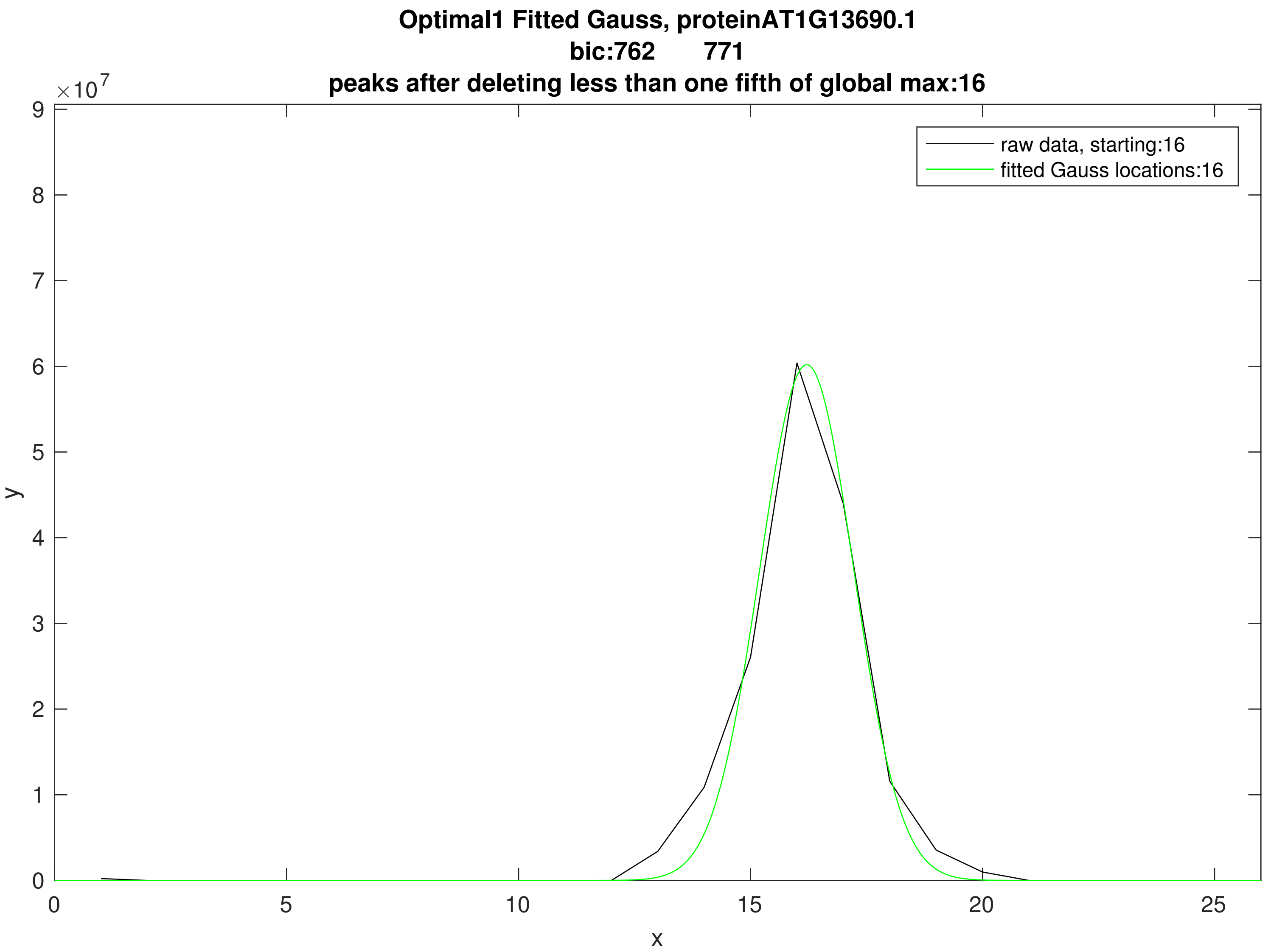

Optimal1 Fitted Gauss, proteinAT1G13690.1
bic:762 771
peaks after deleting less than one fifth of global max:16
×10^7
raw data, starting:16
fitted Gauss locations:16
y
x
0
1
2
3
4
5
6
7
8
9
0
5
10
15
20
25

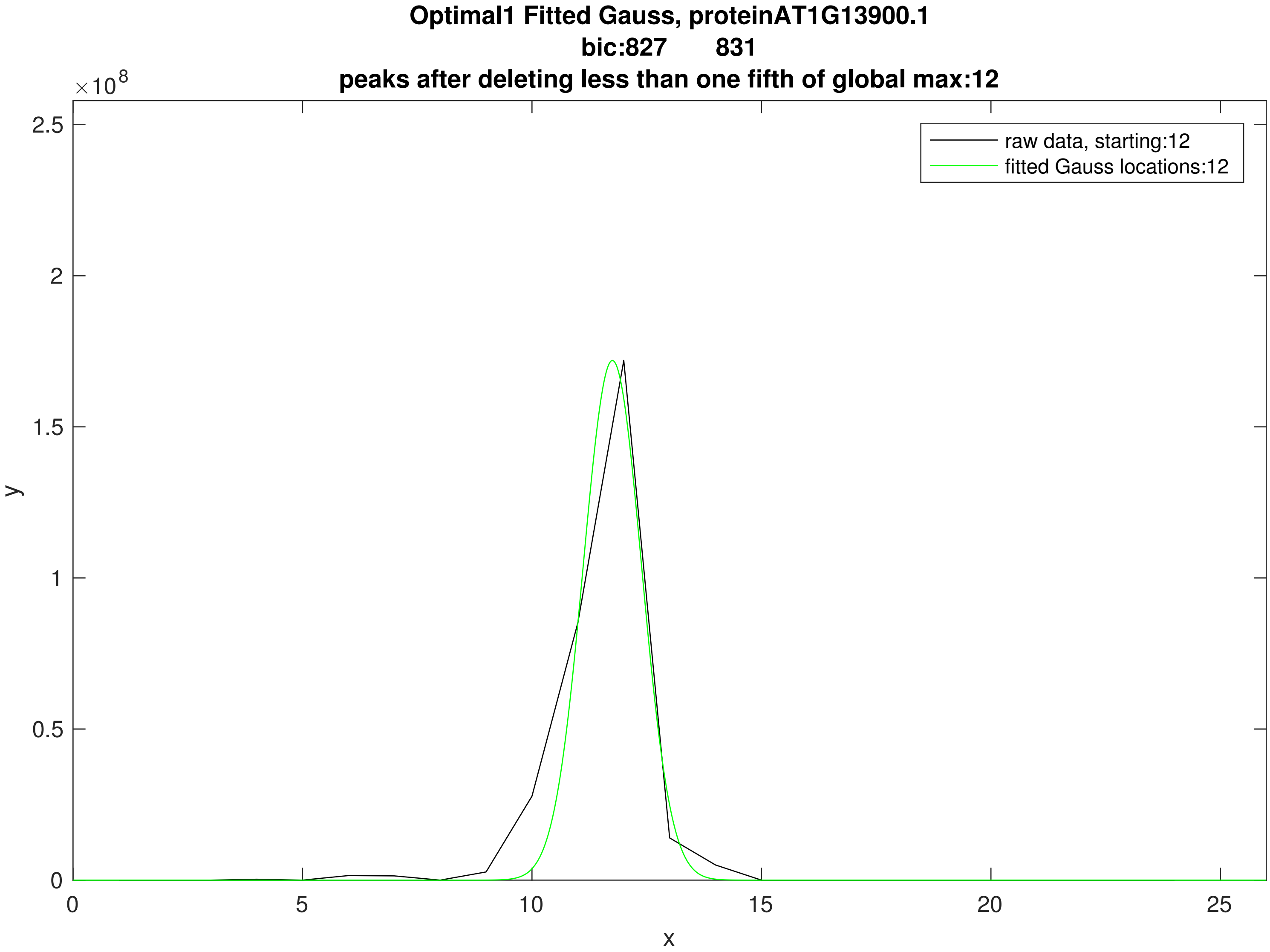
Optimal1 Fitted Gauss, proteinAT1G13900.1
bic:827 831
peaks after deleting less than one fifth of global max:12
×10^8
2.5
2
1.5
1
0.5
0
y
0
5
10
15
20
25
x
raw data, starting:12
fitted Gauss locations:12

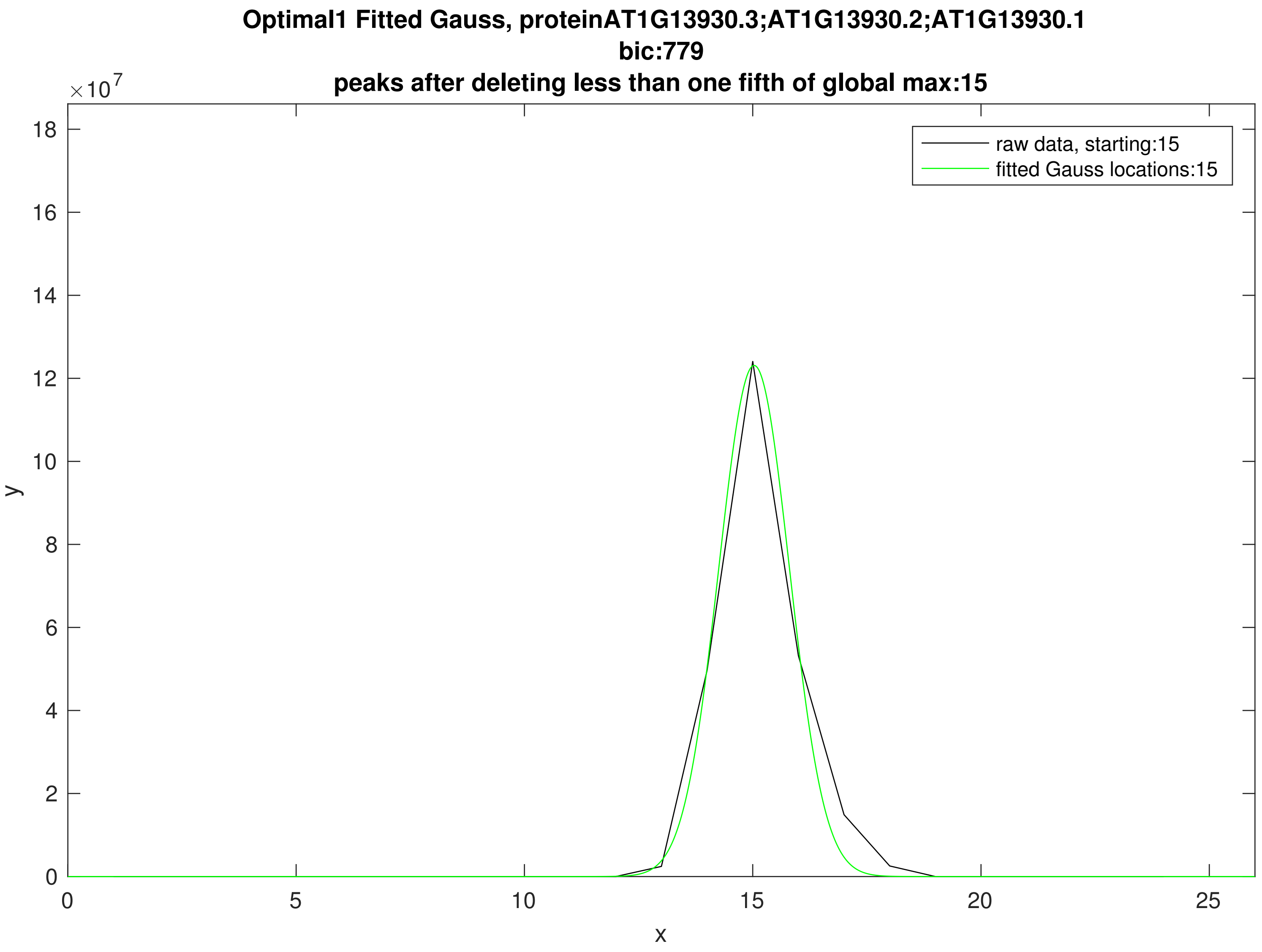
Optimal1 Fitted Gauss, proteinAT1G13930.3;AT1G13930.2;AT1G13930.1
bic:779
peaks after deleting less than one fifth of global max:15
×10^7
18
16
14
12
10
8
6
4
2
0
y
0
5
10
15
20
25
x
raw data, starting:15
fitted Gauss locations:15

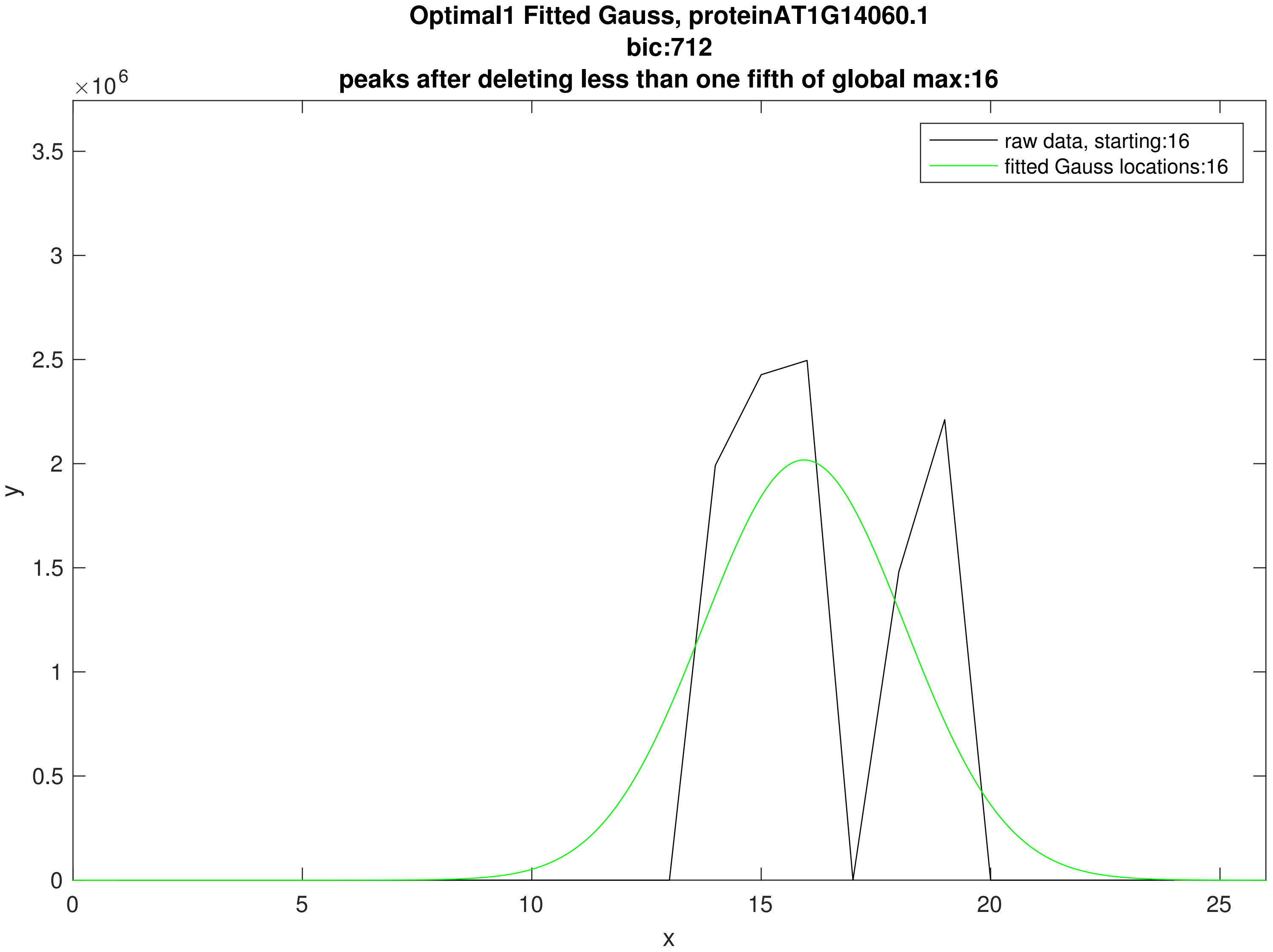
Optimal1 Fitted Gauss, proteinAT1G14060.1
bic:712
peaks after deleting less than one fifth of global max:16
×10^6
3.5
3
2.5
2
1.5
1
0.5
0
y
0
5
10
15
20
25
x
raw data, starting:16
fitted Gauss locations:16

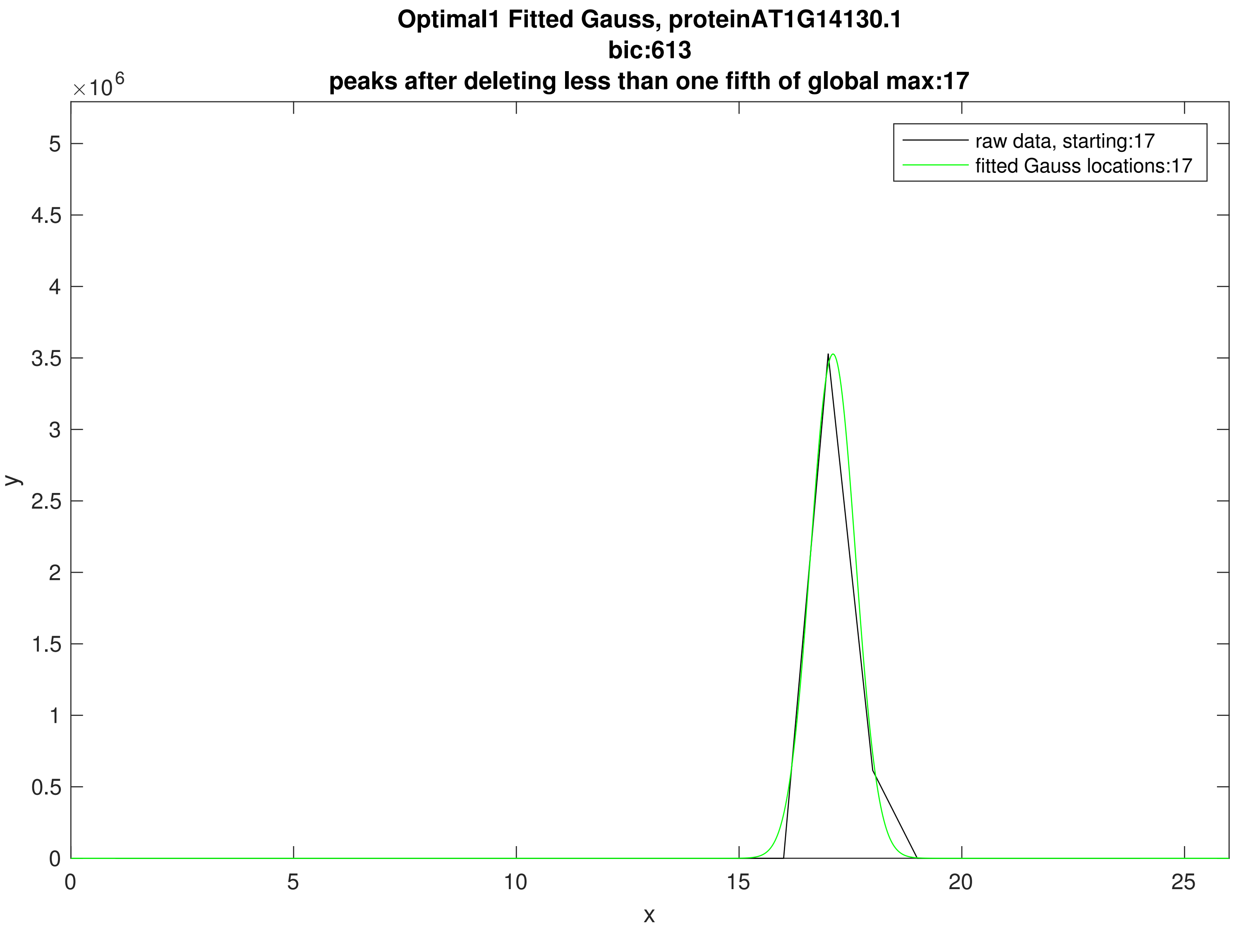

Optimal1 Fitted Gauss, proteinAT1G14130.1
bic:613
peaks after deleting less than one fifth of global max:17
×10^6
raw data, starting:17
fitted Gauss locations:17
y
x
0
0.5
1
1.5
2
2.5
3
3.5
4
4.5
5
0
5
10
15
20
25

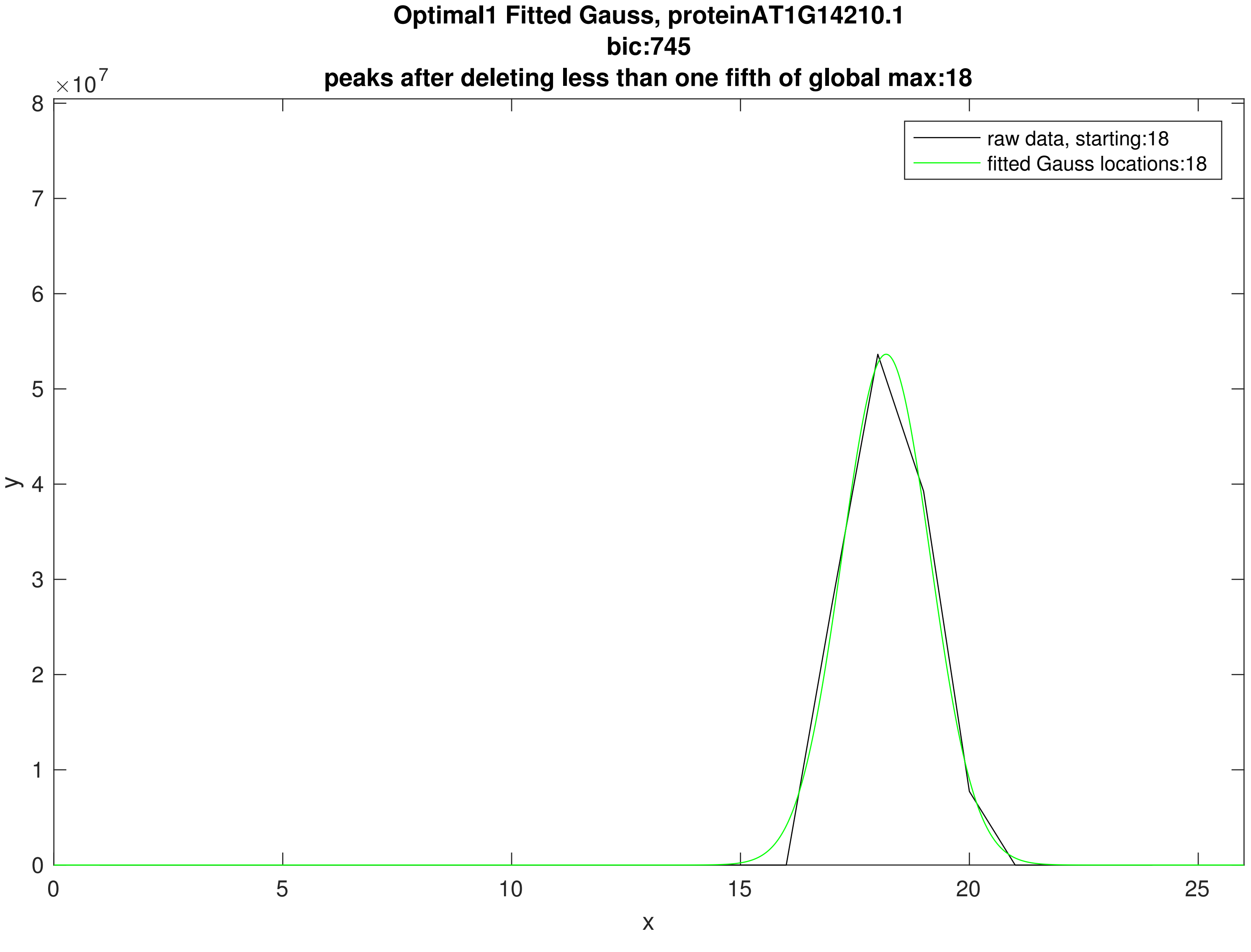

Optimal1 Fitted Gauss, proteinAT1G14210.1
bic:745
peaks after deleting less than one fifth of global max:18
×10^7
raw data, starting:18
fitted Gauss locations:18
y
x
0
1
2
3
4
5
6
7
8
0
5
10
15
20
25

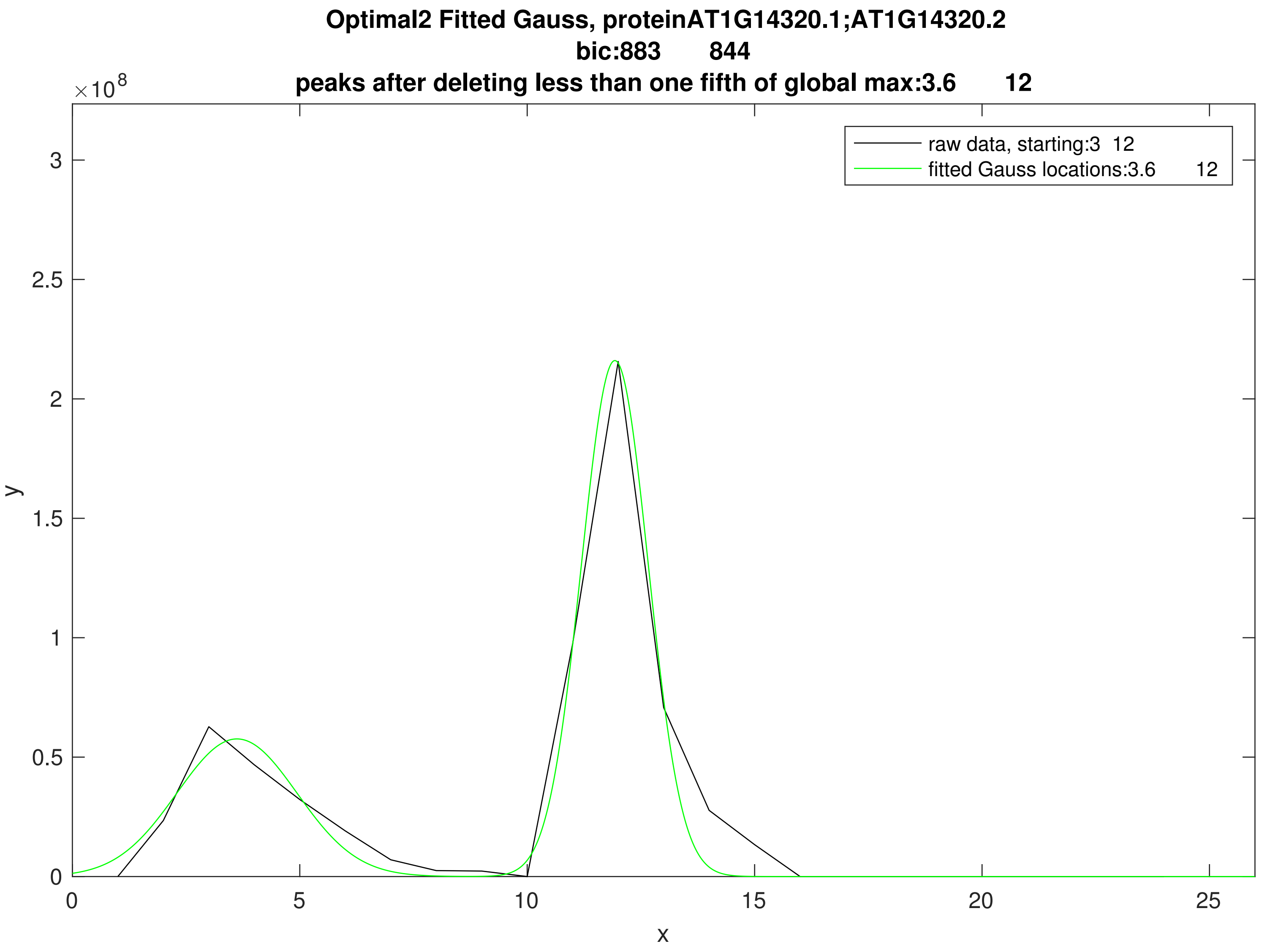
Optimal2 Fitted Gauss, proteinAT1G14320.1;AT1G14320.2
bic:883 844
peaks after deleting less than one fifth of global max:3.6 12
×10^8
raw data, starting:3 12
fitted Gauss locations:3.6 12
y
x
0
0.5
1
1.5
2
2.5
3
0
5
10
15
20
25

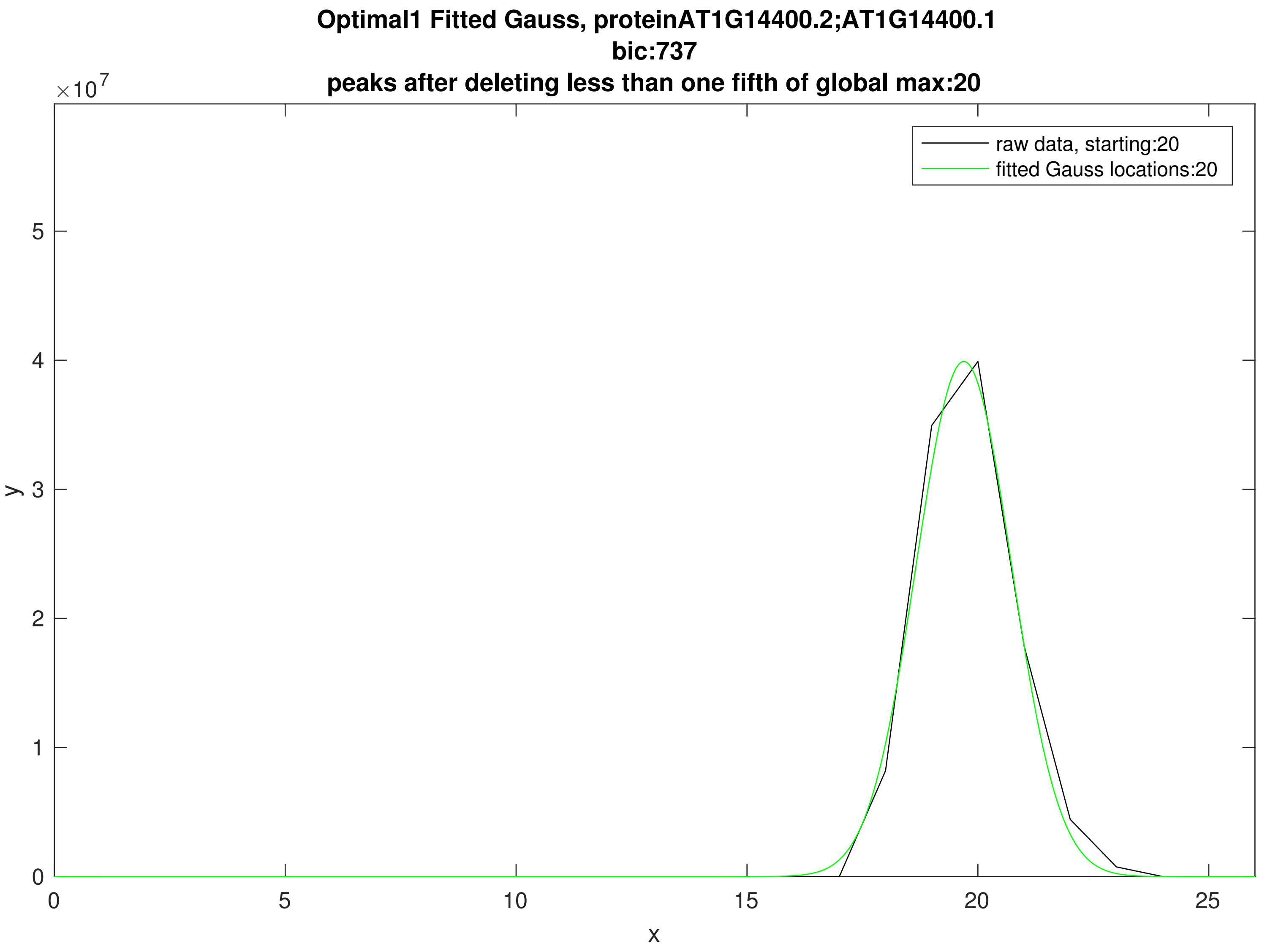

Optimal1 Fitted Gauss, proteinAT1G14400.2;AT1G14400.1
bic:737
peaks after deleting less than one fifth of global max:20
×10^7
raw data, starting:20
fitted Gauss locations:20
y
x
0
1
2
3
4
5
0
5
10
15
20
25

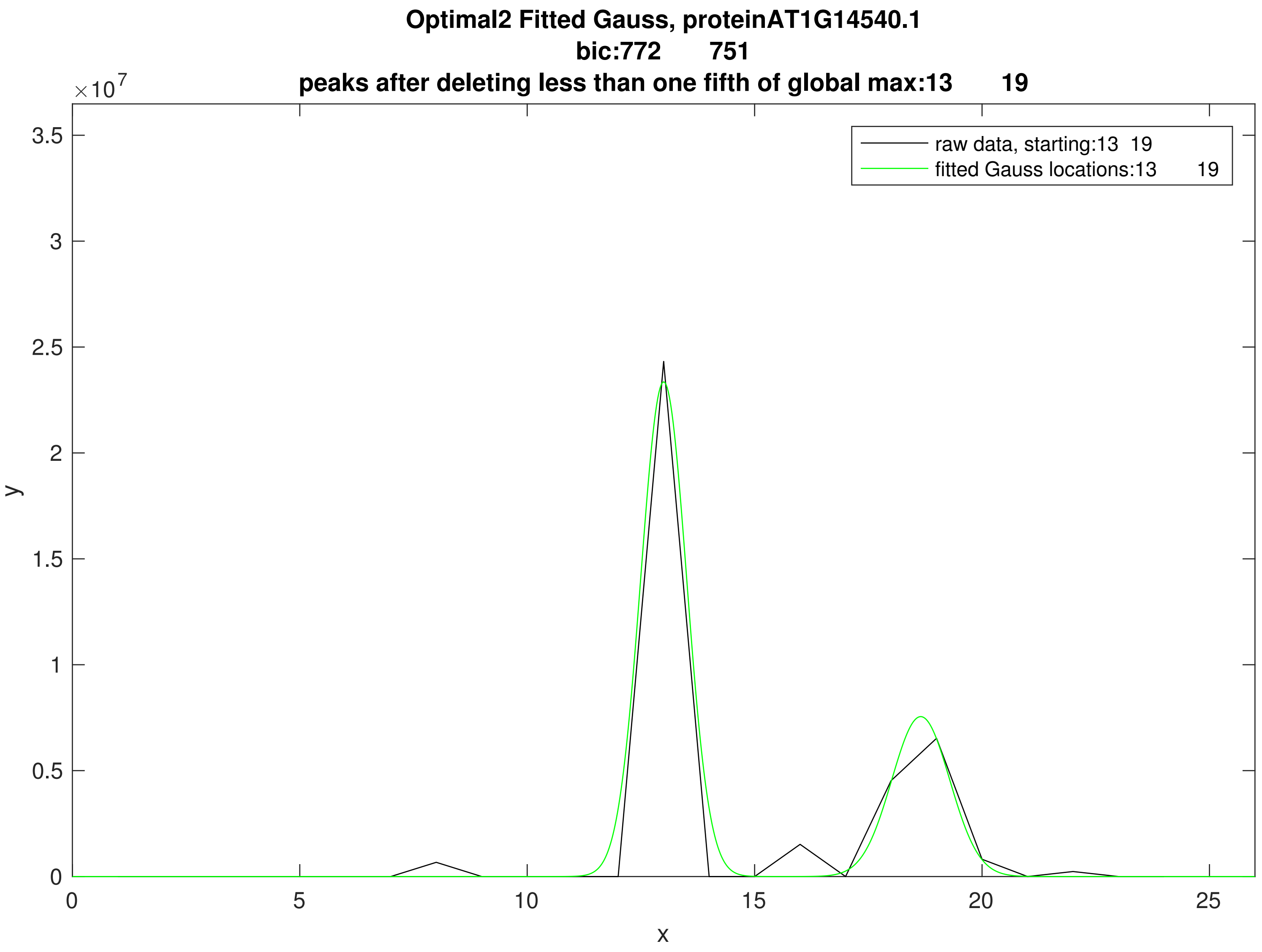

Optimal2 Fitted Gauss, proteinAT1G14540.1
bic:772 751
peaks after deleting less than one fifth of global max:13 19
×10^7
raw data, starting:13 19
fitted Gauss locations:13 19
y
x
0
0.5
1
1.5
2
2.5
3
3.5
0
5
10
15
20
25

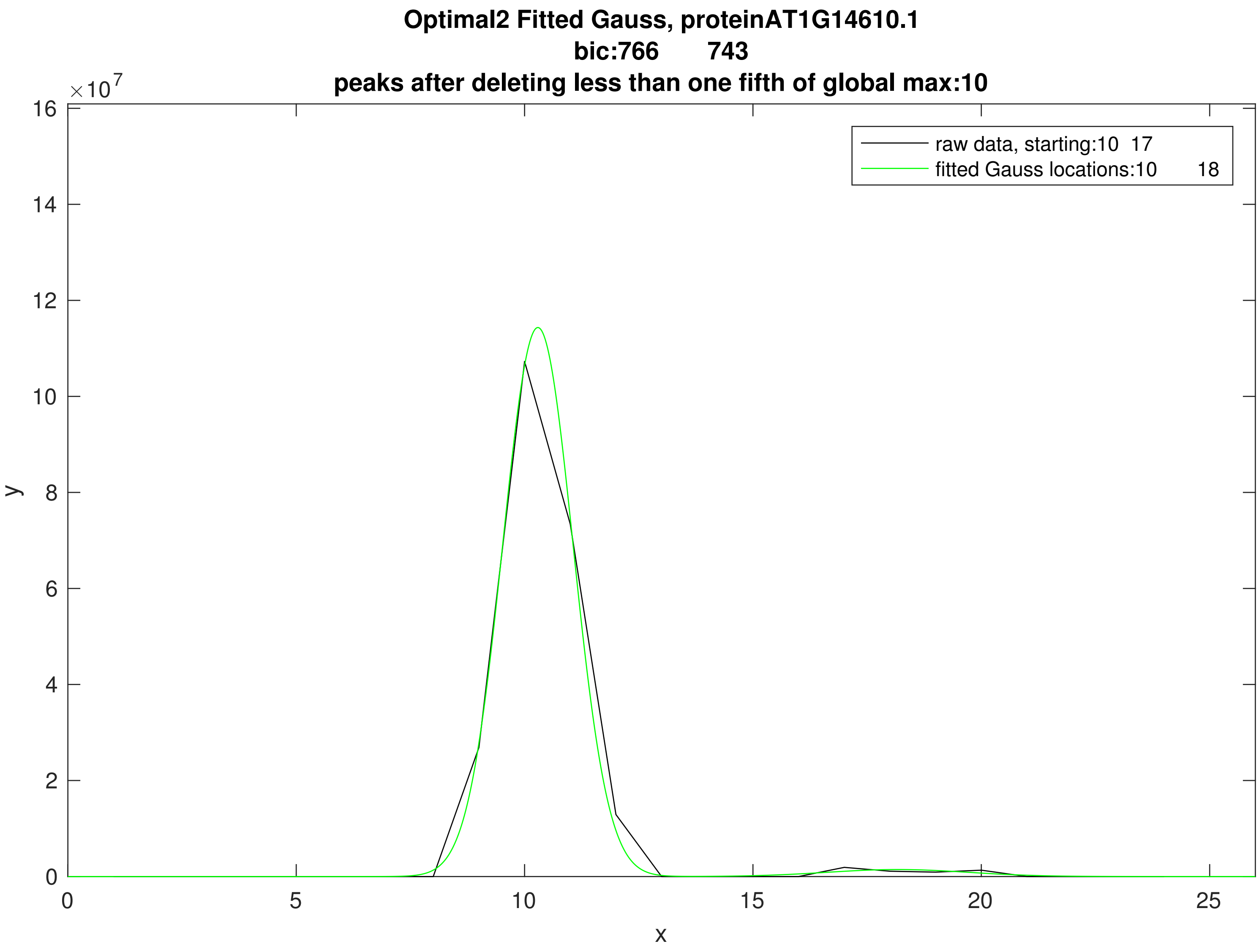
Optimal2 Fitted Gauss, proteinAT1G14610.1
bic:766 743
peaks after deleting less than one fifth of global max:10
×10^7
16
14
12
10
8
6
4
2
0
y
0
5
10
15
20
25
x
raw data, starting:10 17
fitted Gauss locations:10 18

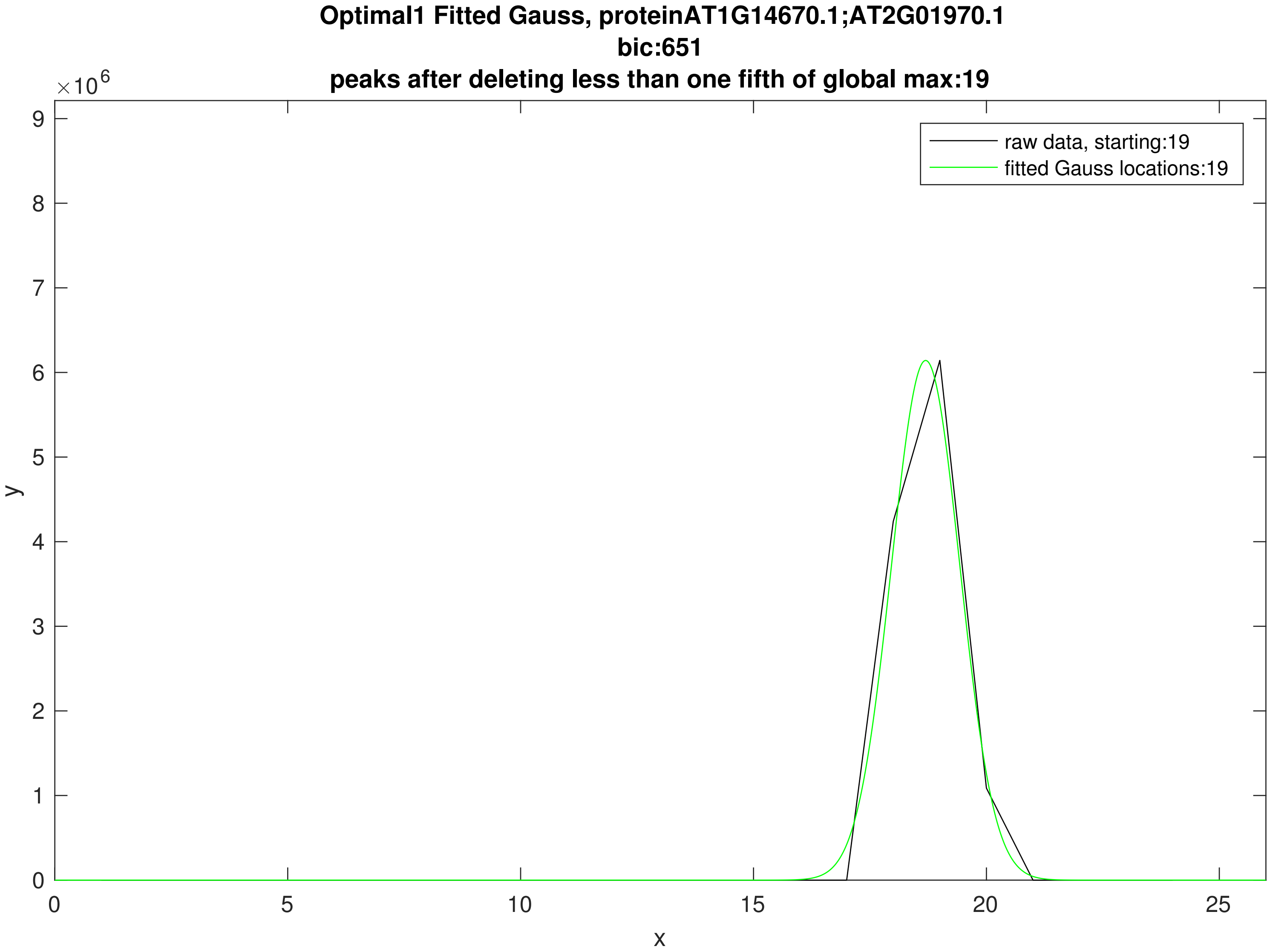

Optimal1 Fitted Gauss, proteinAT1G14670.1;AT2G01970.1
bic:651
peaks after deleting less than one fifth of global max:19
×10^6
0
1
2
3
4
5
6
7
8
9
y
0
5
10
15
20
25
x
raw data, starting:19
fitted Gauss locations:19

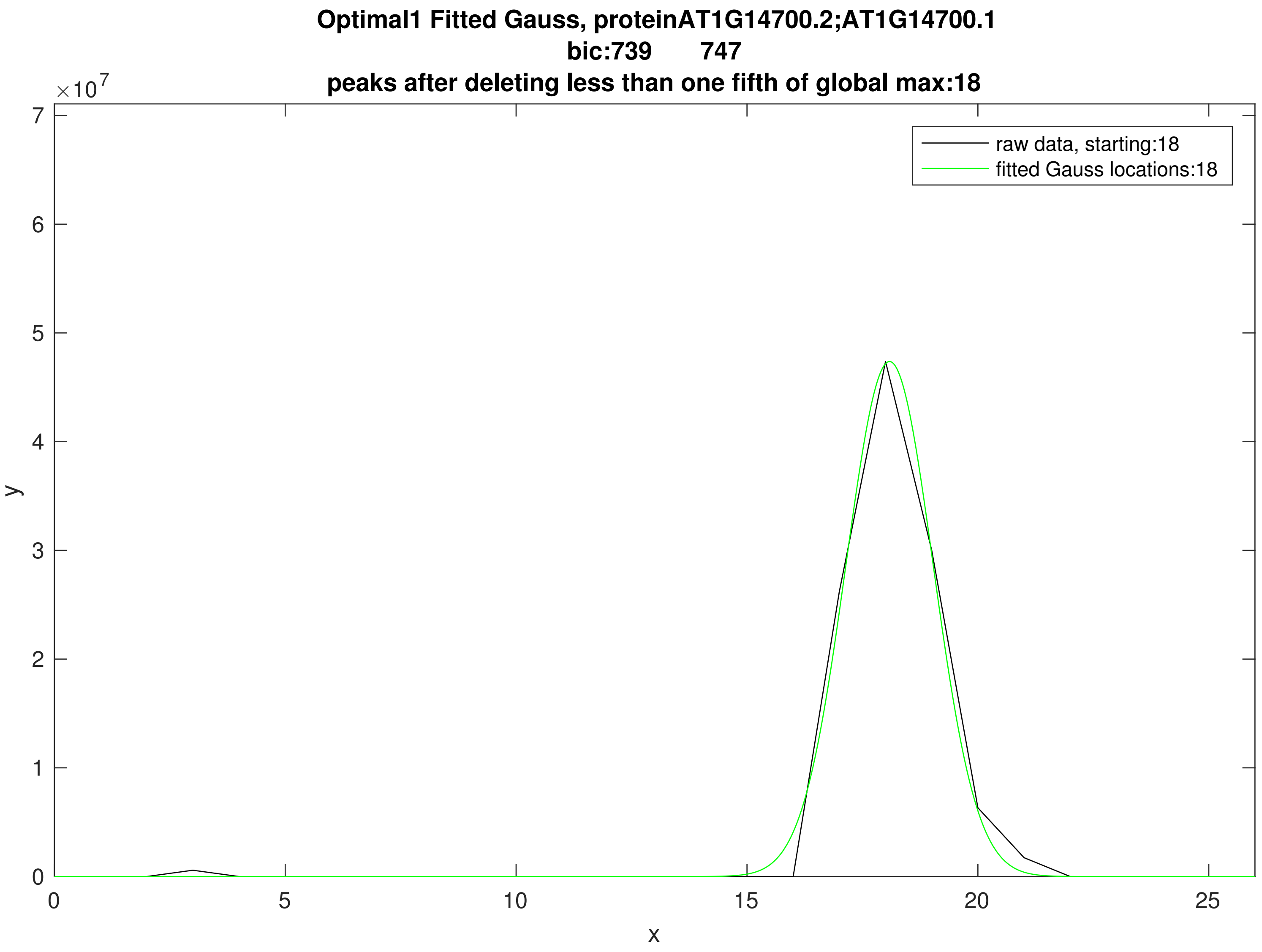
Optimal1 Fitted Gauss, proteinAT1G14700.2;AT1G14700.1
bic:739 747
peaks after deleting less than one fifth of global max:18
×10^7
raw data, starting:18
fitted Gauss locations:18
y
x
0
1
2
3
4
5
6
7
0
5
10
15
20
25

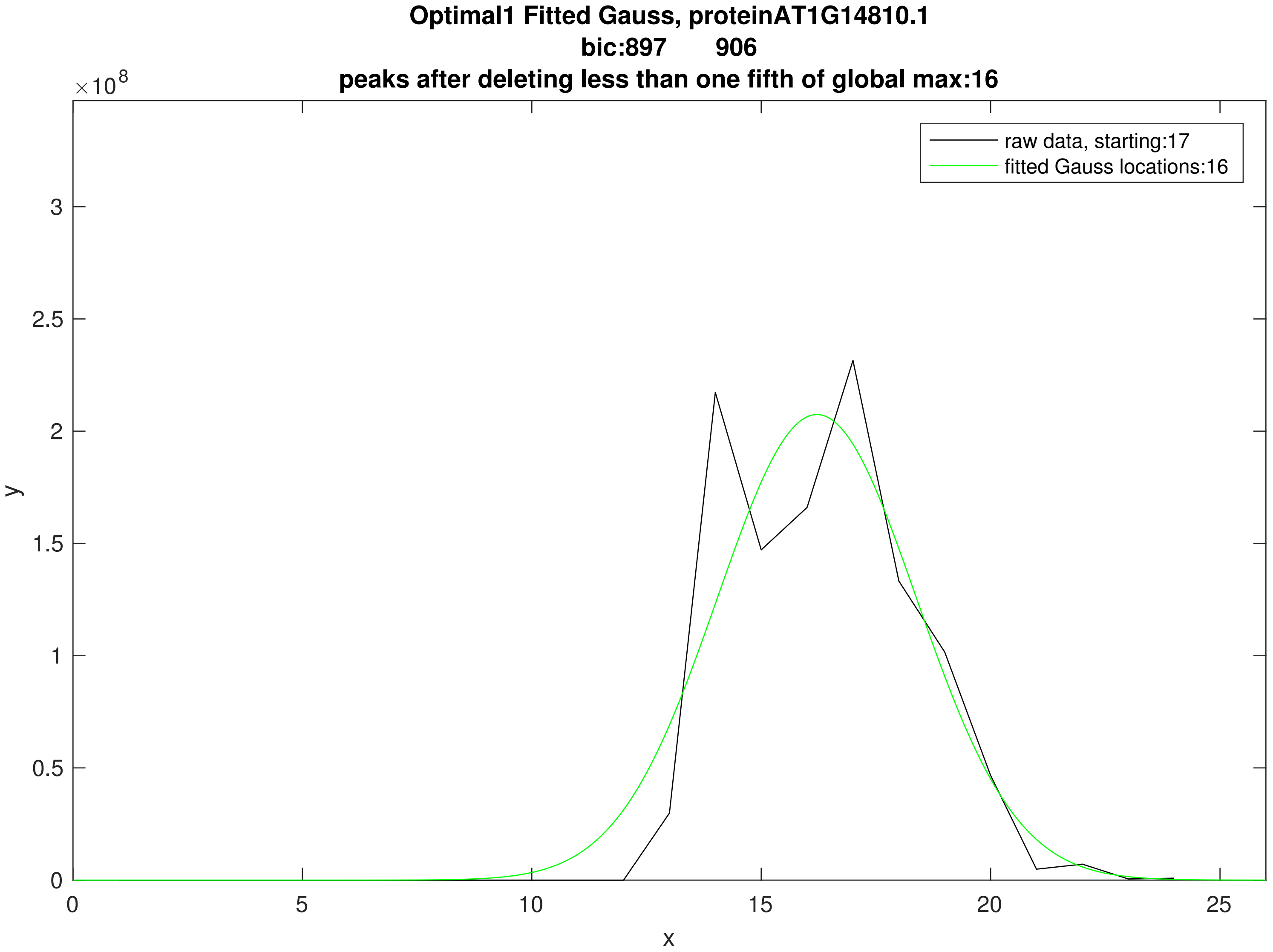

Optimal1 Fitted Gauss, proteinAT1G14810.1
bic:897 906
peaks after deleting less than one fifth of global max:16
×10^8
raw data, starting:17
fitted Gauss locations:16
y
x
0
0.5
1
1.5
2
2.5
3
0
5
10
15
20
25

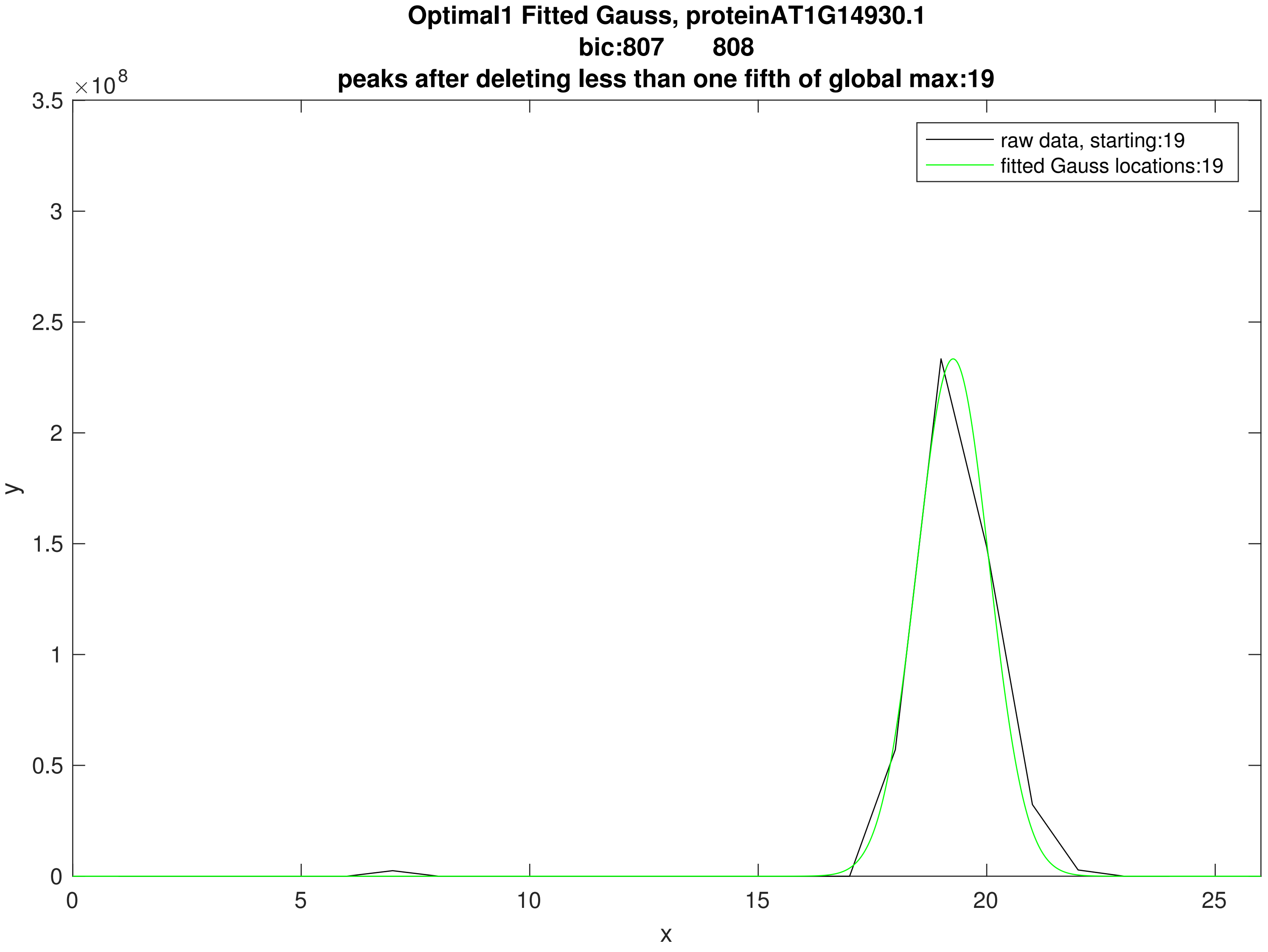
Optimal1 Fitted Gauss, proteinAT1G14930.1
bic:807 808
peaks after deleting less than one fifth of global max:19
×10^8
3.5
3
2.5
2
1.5
1
0.5
0
y
0
5
10
15
20
25
x
raw data, starting:19
fitted Gauss locations:19

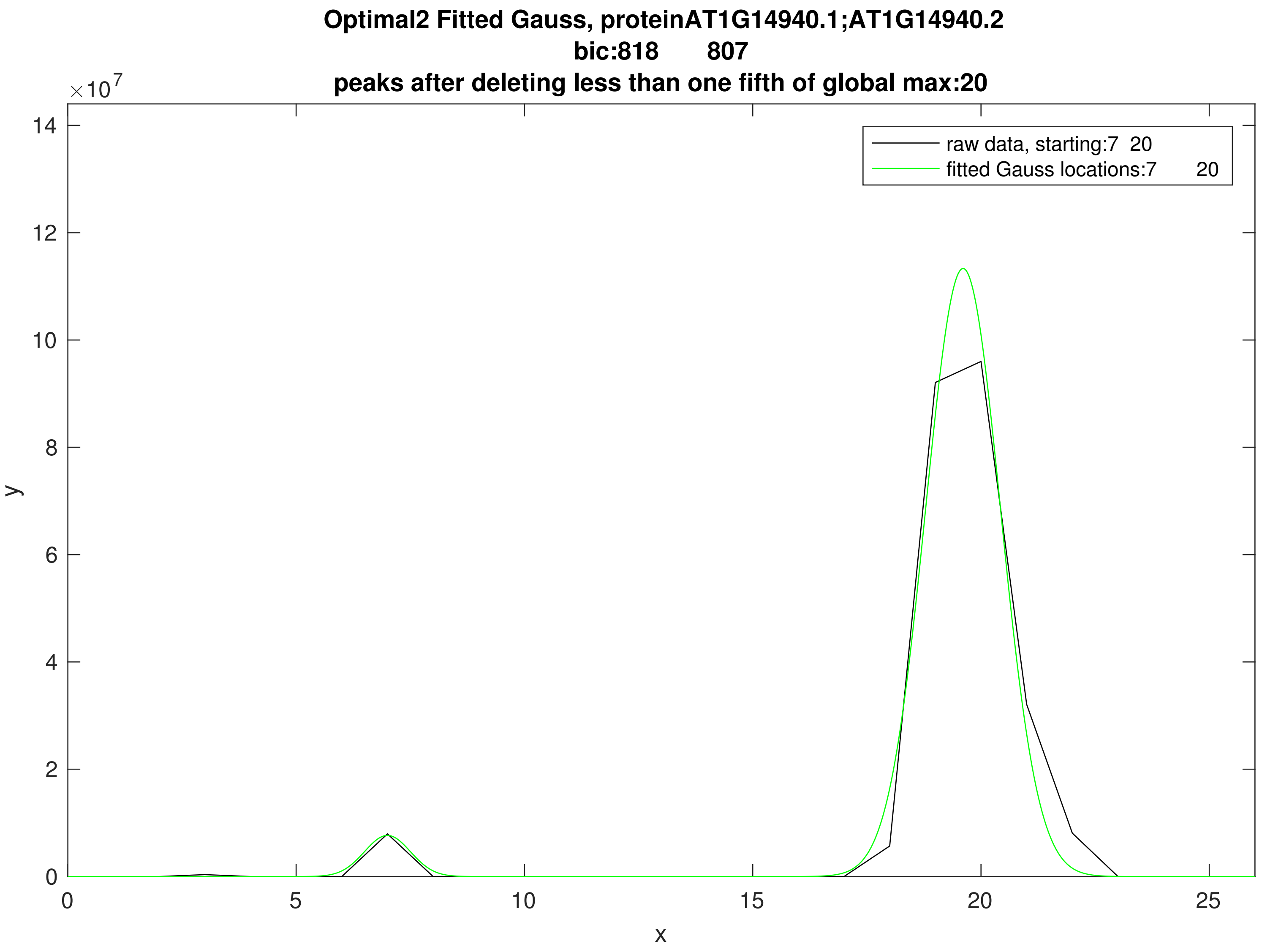

Optimal2 Fitted Gauss, proteinAT1G14940.1;AT1G14940.2
bic:818 807
peaks after deleting less than one fifth of global max:20
×10^7
raw data, starting:7 20
fitted Gauss locations:7 20
y
x
0
2
4
6
8
10
12
14
5
15
20
25

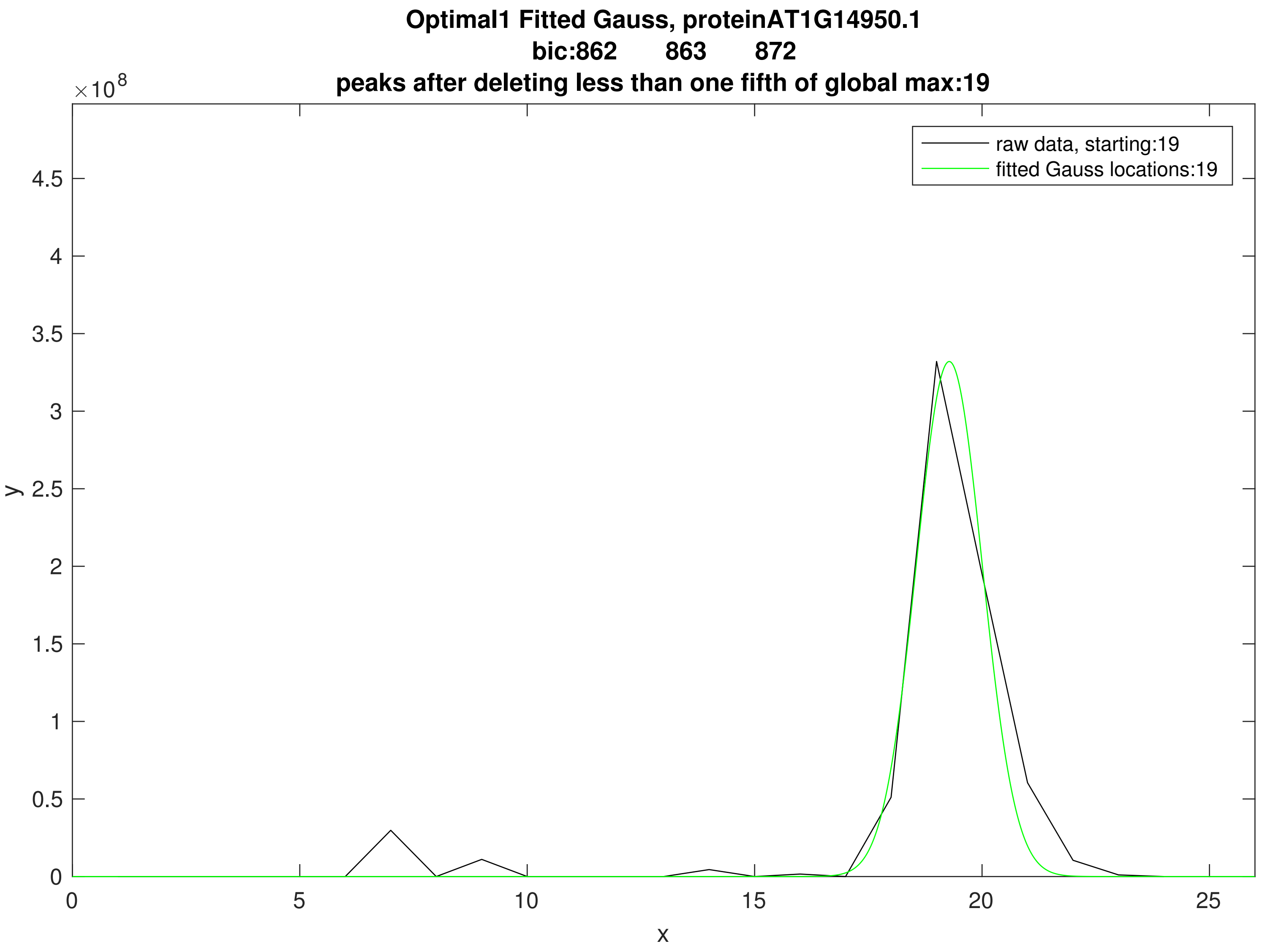
Optimal1 Fitted Gauss, proteinAT1G14950.1
bic:862 863 872
peaks after deleting less than one fifth of global max:19
×10^8
raw data, starting:19
fitted Gauss locations:19
0
0.5
1
1.5
2
2.5
3
3.5
4
4.5
y
0
5
10
15
20
25
x

Optimal1 Fitted Gauss, proteinAT1G14960.1
bic:745
peaks after deleting less than one fifth of global max:18

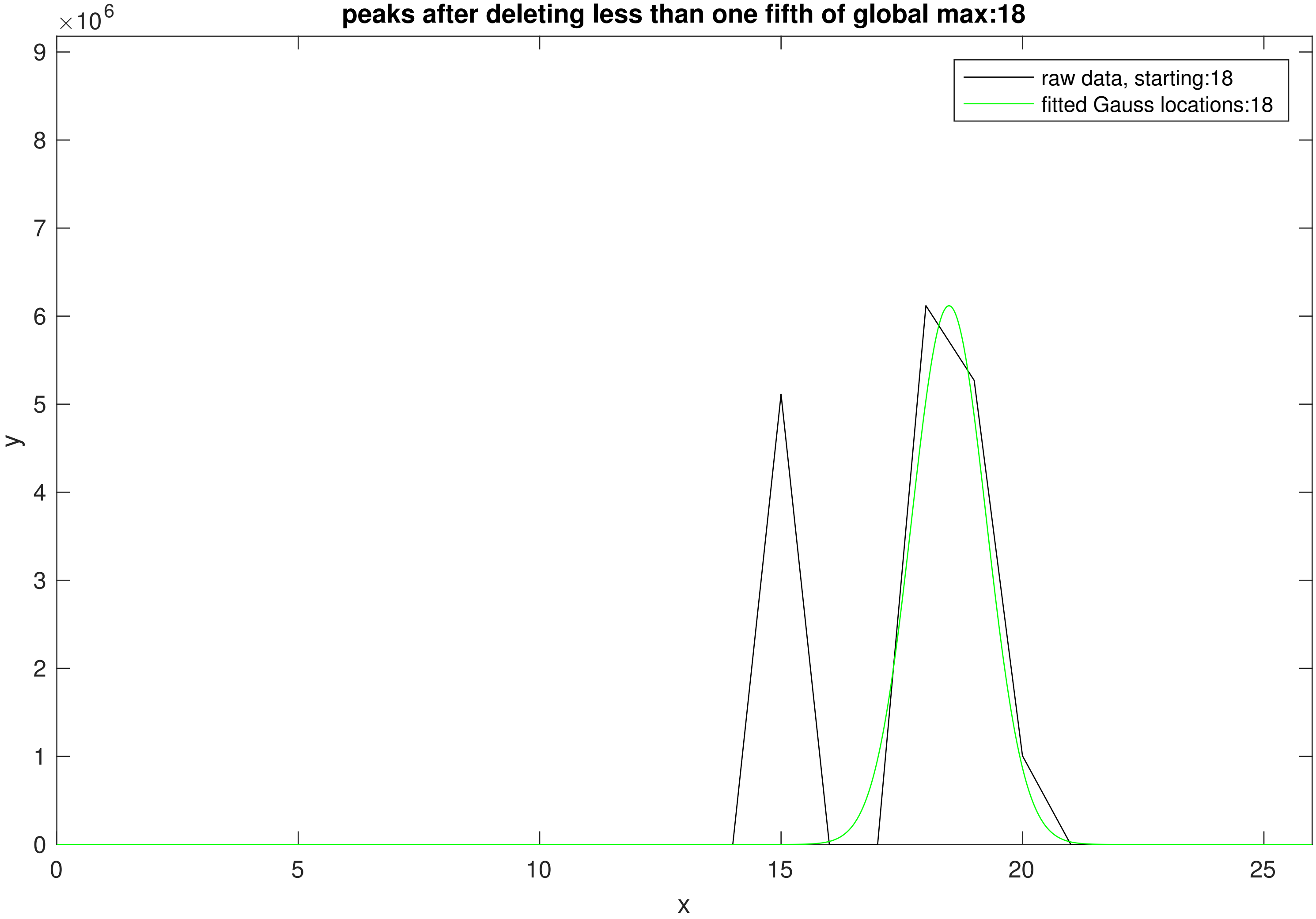

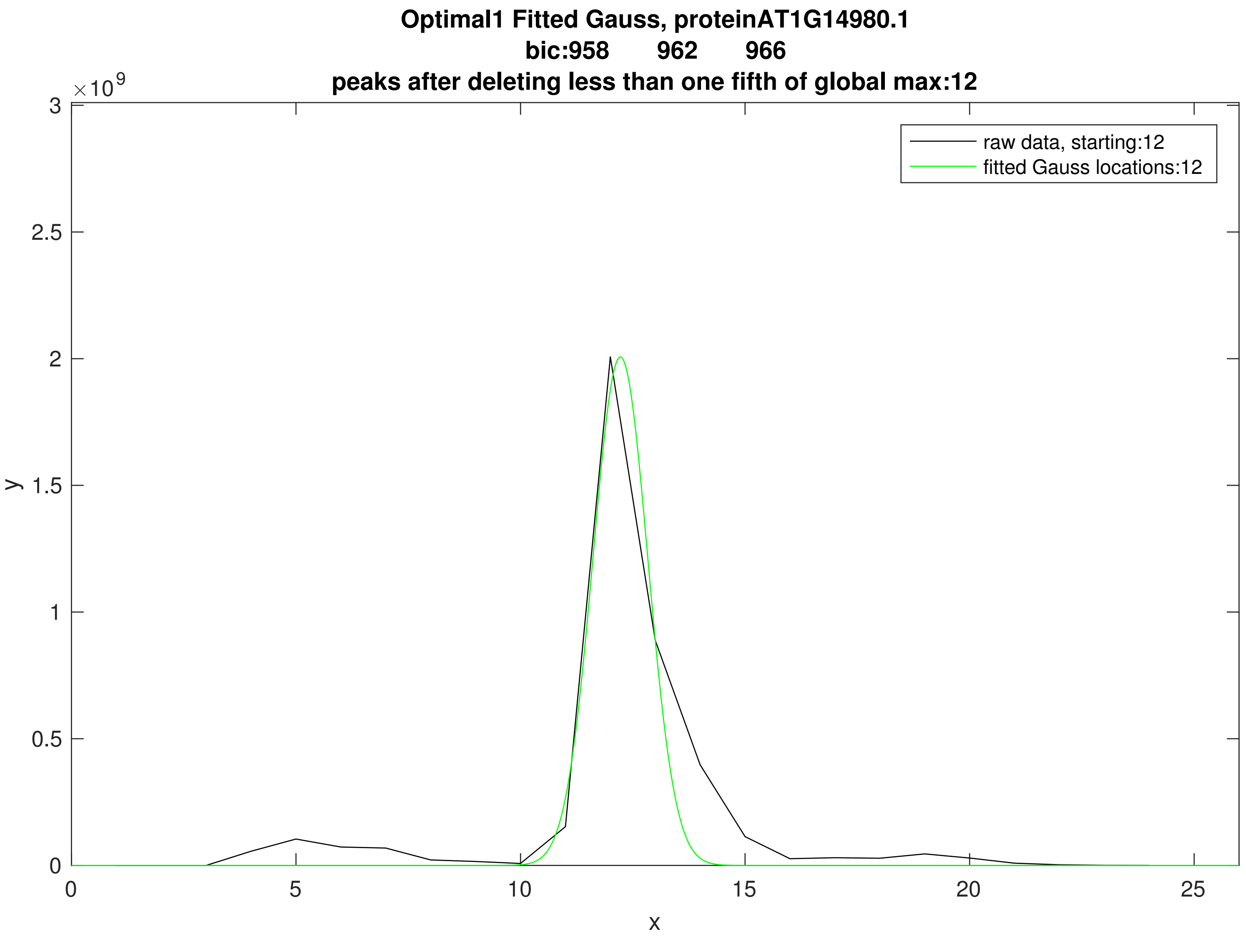

Optimal1 Fitted Gauss, proteinAT1G14980.1
bic:958 962 966
peaks after deleting less than one fifth of global max:12
×10^9
raw data, starting:12
fitted Gauss locations:12
y
x

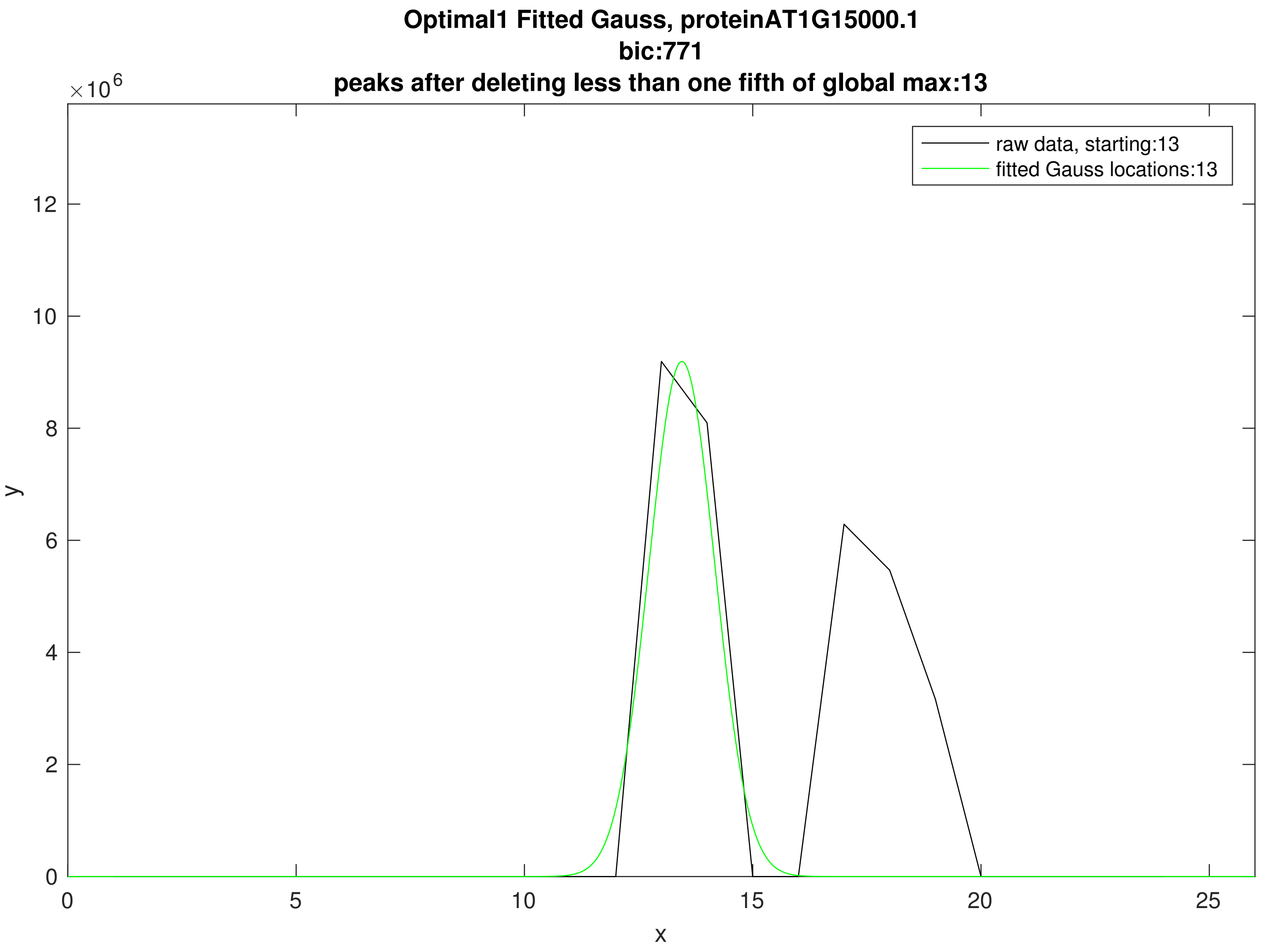

Optimal1 Fitted Gauss, proteinAT1G15000.1
bic:771
peaks after deleting less than one fifth of global max:13
×10^6
raw data, starting:13
fitted Gauss locations:13
12
10
8
6
4
2
0
y
0
5
10
15
20
25
x

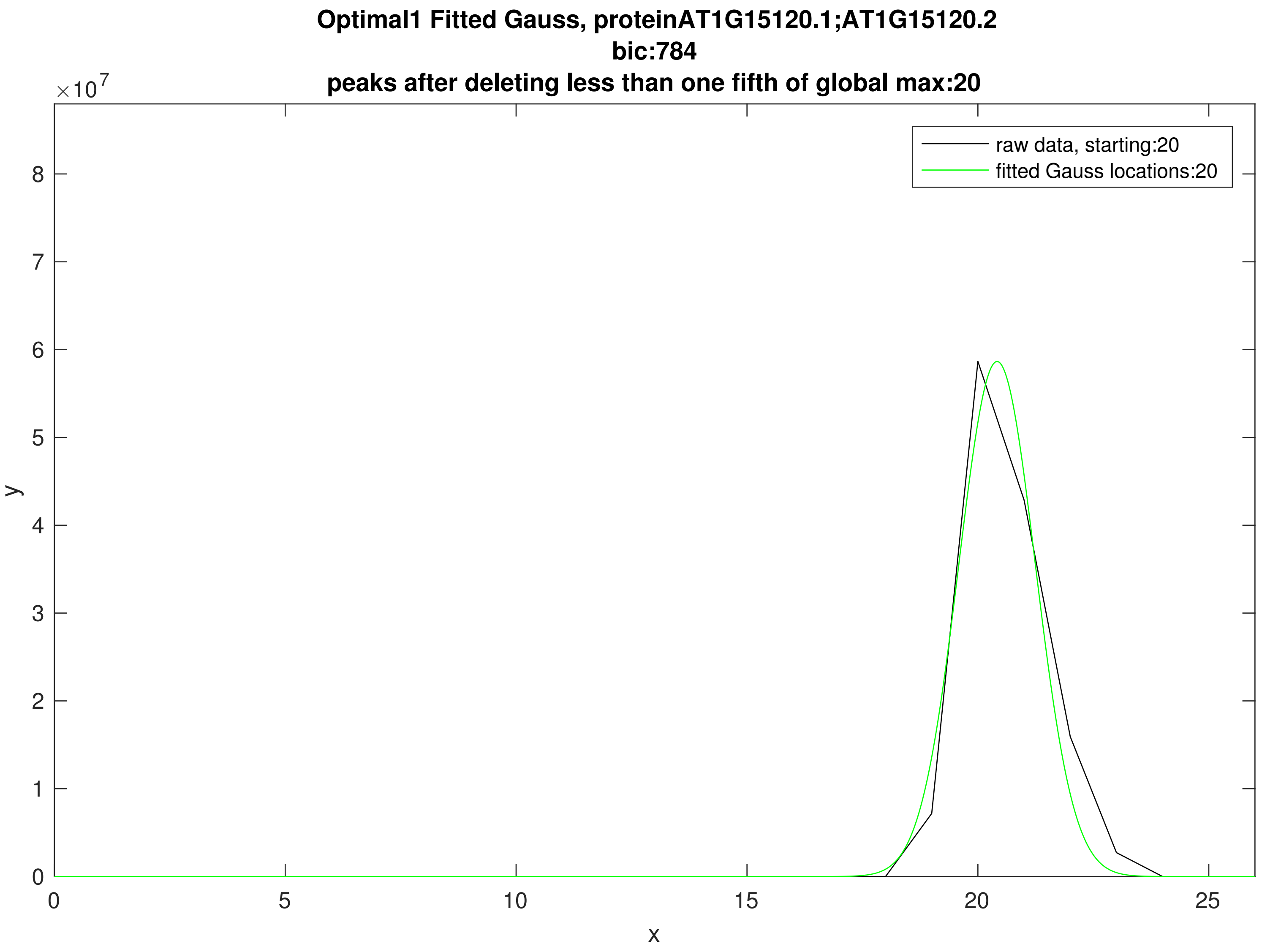

Optimal1 Fitted Gauss, proteinAT1G15120.1;AT1G15120.2
bic:784
peaks after deleting less than one fifth of global max:20
×10^7
raw data, starting:20
fitted Gauss locations:20
y
x
0
1
2
3
4
5
6
7
8
0
5
10
15
20
25

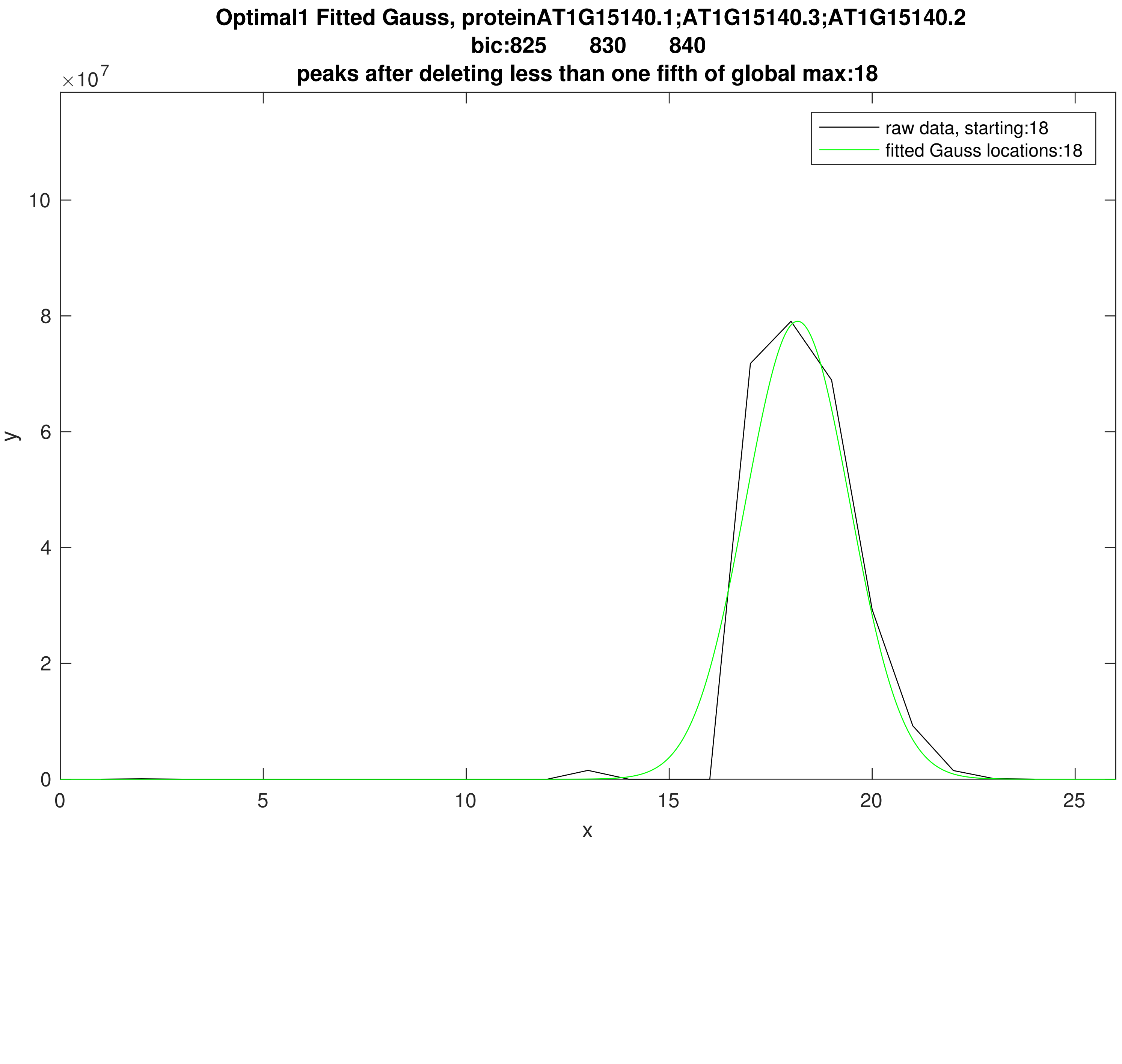
Optimal1 Fitted Gauss, proteinAT1G15140.1;AT1G15140.3;AT1G15140.2
bic:825 830 840
peaks after deleting less than one fifth of global max:18
×10^7
raw data, starting:18
fitted Gauss locations:18
y
x
0
2
4
6
8
10
5
15
20
25

**Optimal1 Fitted Gauss, proteinAT1G15270.1**
**bic:655**
**peaks after deleting less than one fifth of global max:20**

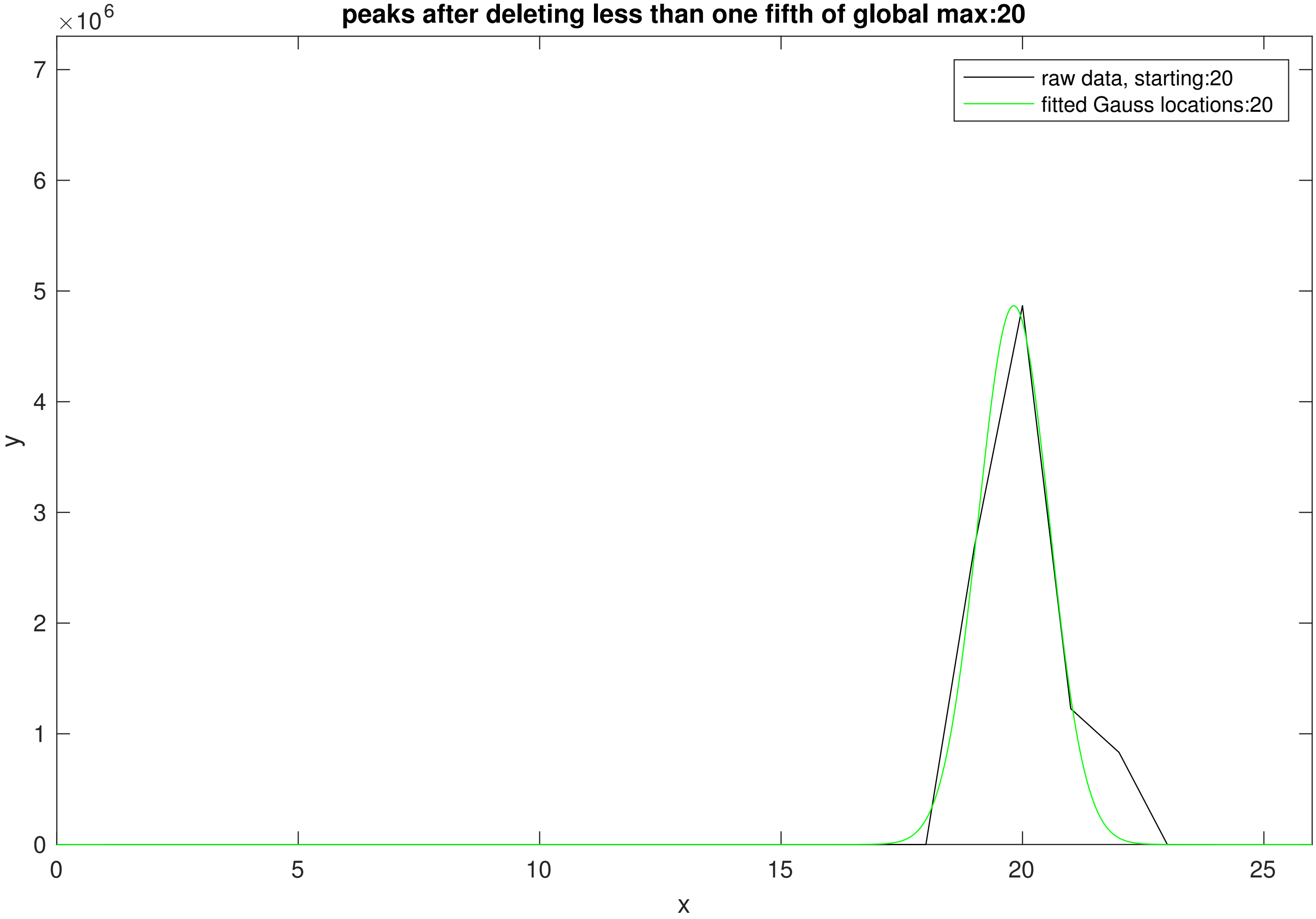

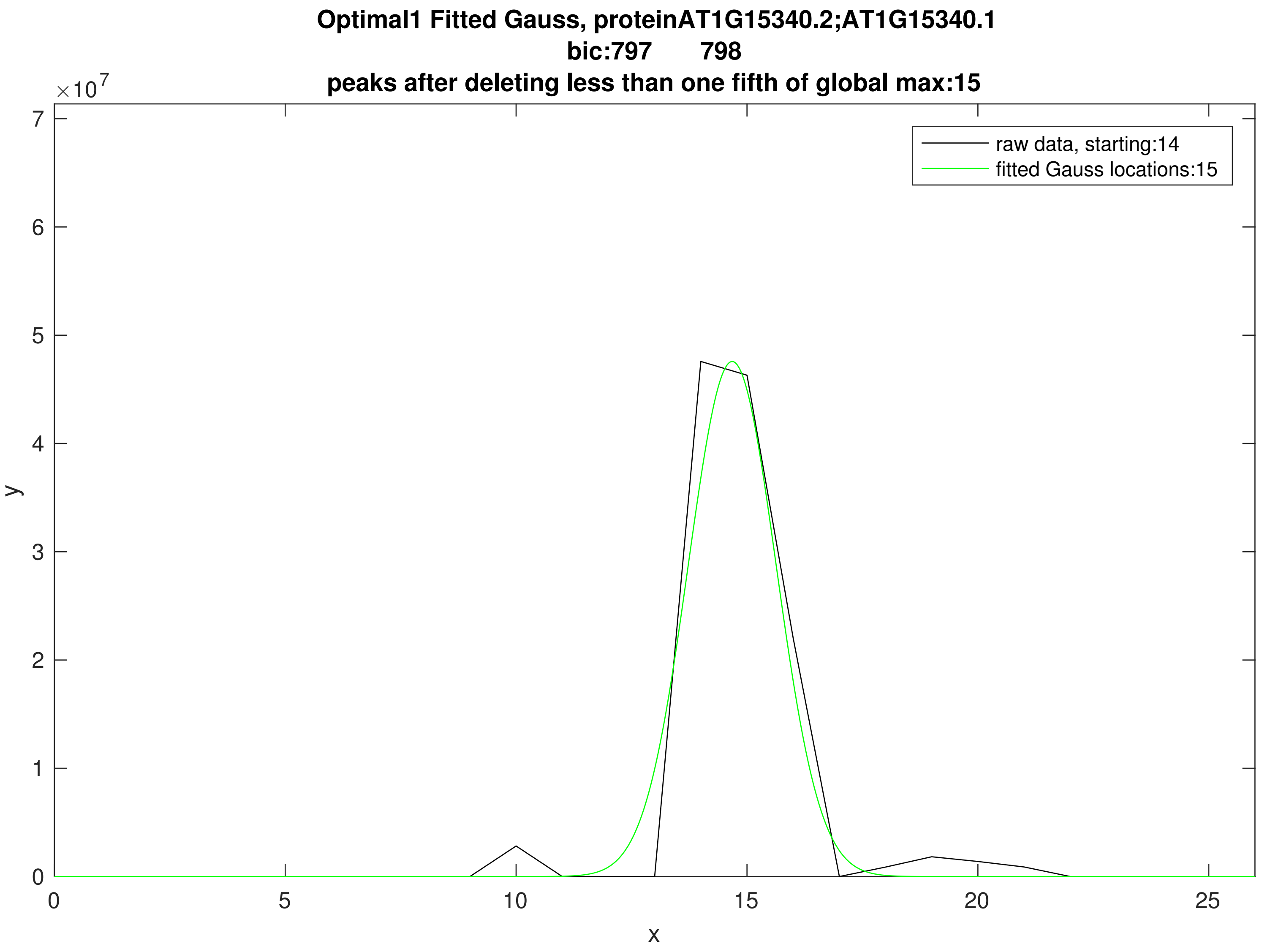
Optimal1 Fitted Gauss, proteinAT1G15340.2;AT1G15340.1
bic:797 798
peaks after deleting less than one fifth of global max:15
×10^7
raw data, starting:14
fitted Gauss locations:15
y
x
0
1
2
3
4
5
6
7
0
5
10
15
20
25

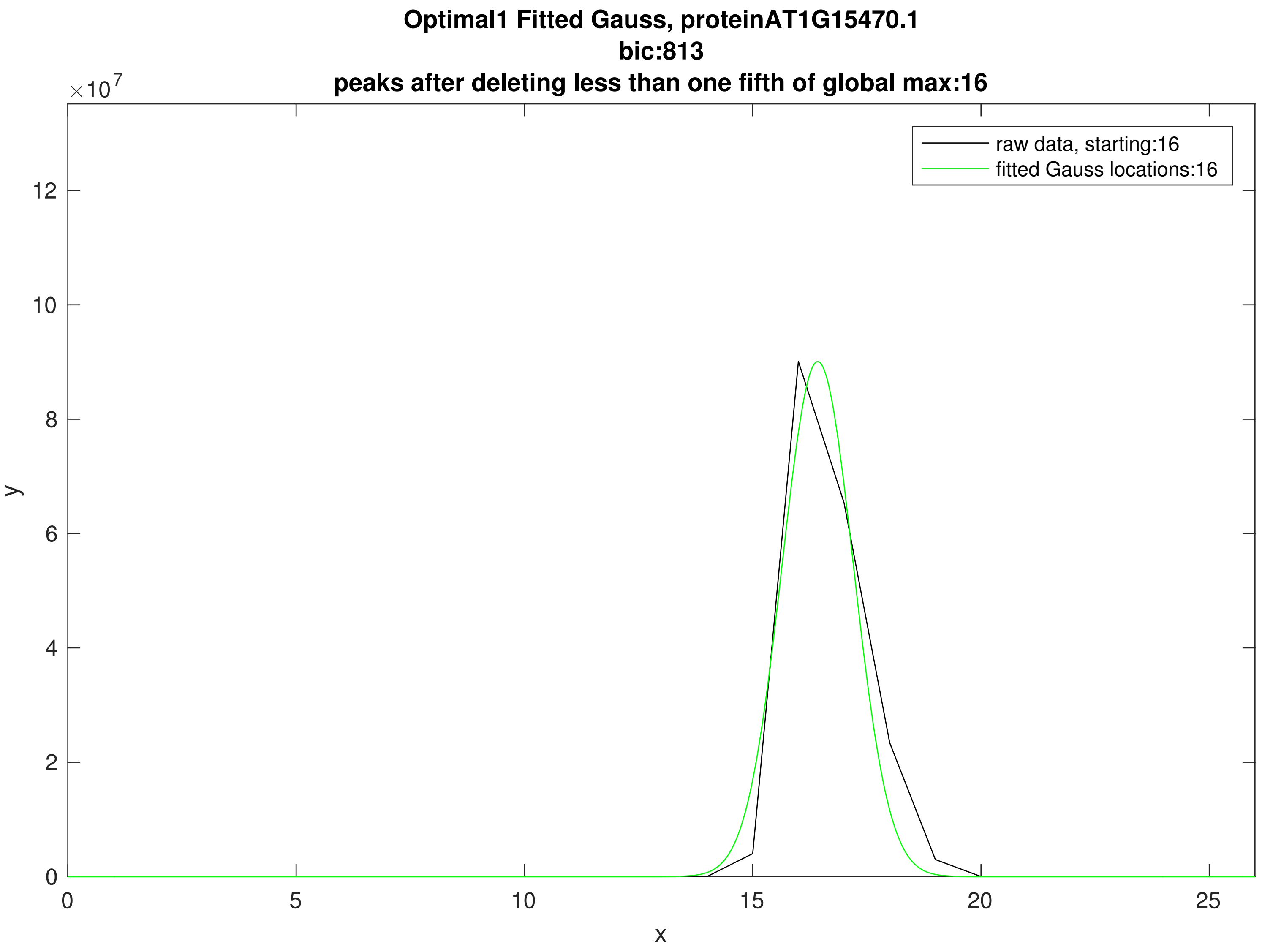
Optimal1 Fitted Gauss, proteinAT1G15470.1
bic:813
peaks after deleting less than one fifth of global max:16
×10^7
raw data, starting:16
fitted Gauss locations:16
12
10
8
6
4
2
0
y
0
5
10
15
20
25
x

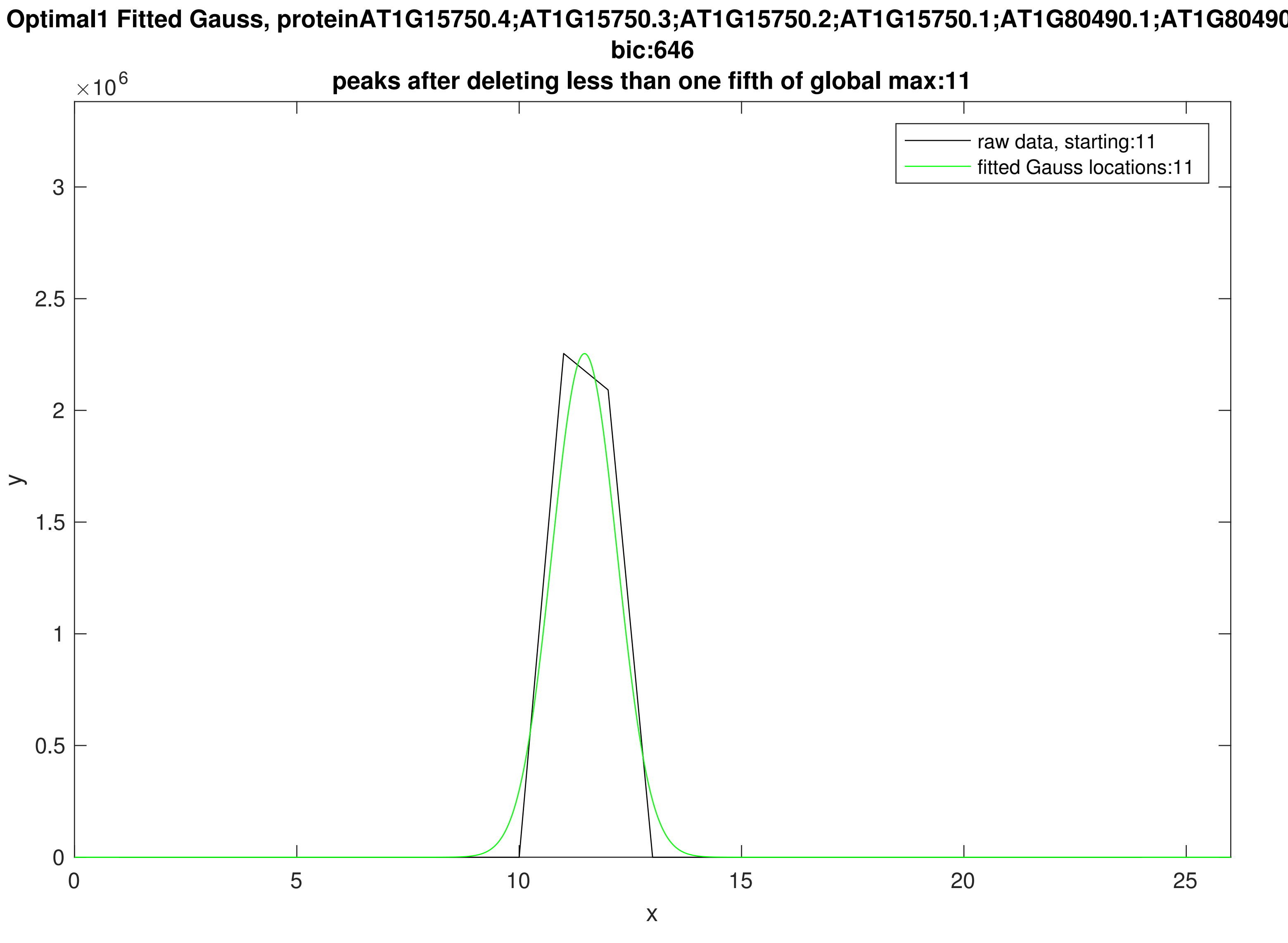

Optimal1 Fitted Gauss, proteinAT1G15750.4;AT1G15750.3;AT1G15750.2;AT1G15750.1;AT1G80490.1;AT1G80490
bic:646
peaks after deleting less than one fifth of global max:11
×10^6
3
2.5
2
1.5
1
0.5
0
y
0
5
10
15
20
25
x
raw data, starting:11
fitted Gauss locations:11

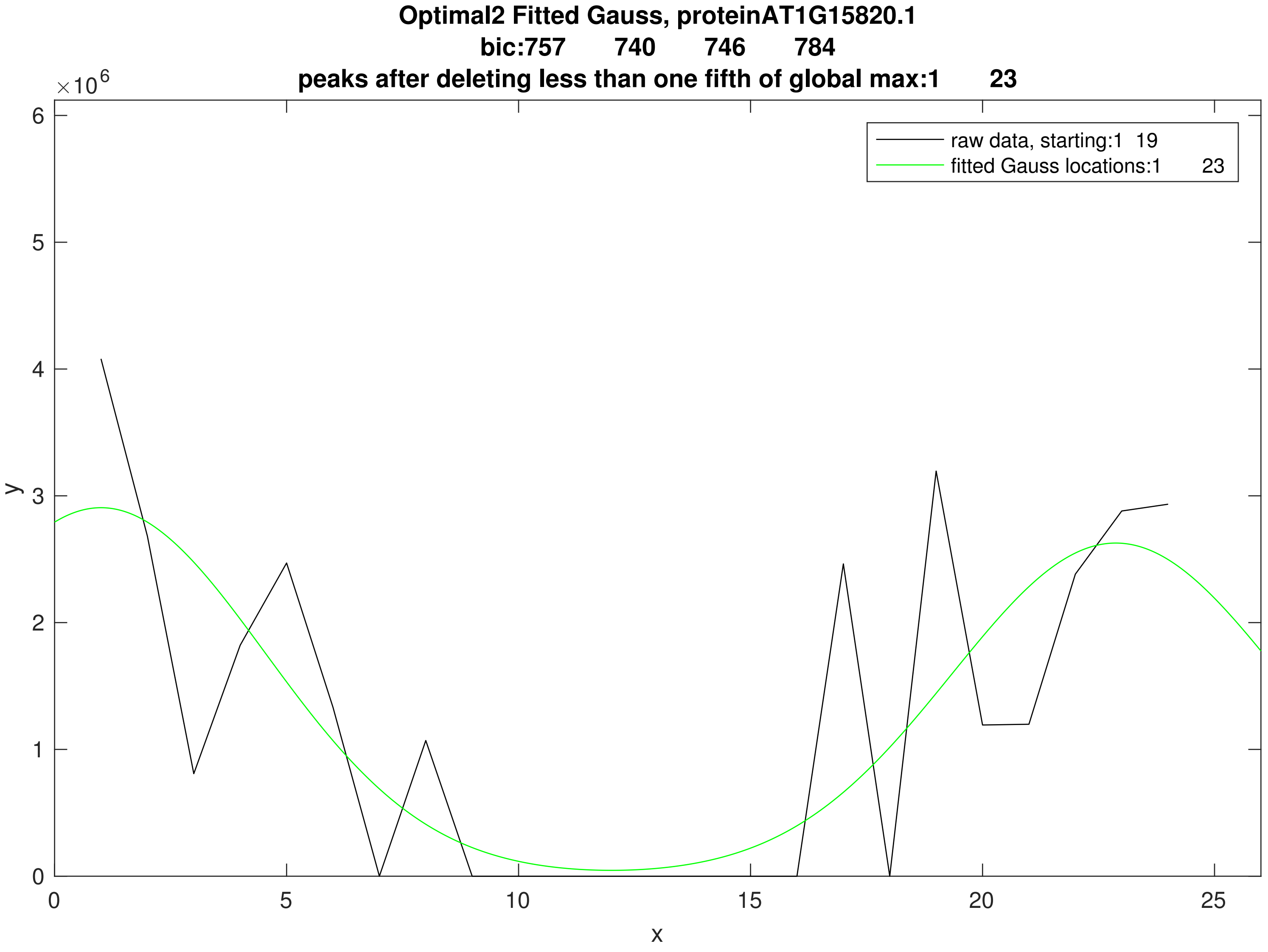
Optimal2 Fitted Gauss, proteinAT1G15820.1
bic:757 740 746 784
peaks after deleting less than one fifth of global max:1 23
×10^6
raw data, starting:1 19
fitted Gauss locations:1 23
y
x
0
1
2
3
4
5
6
10
15
20
25

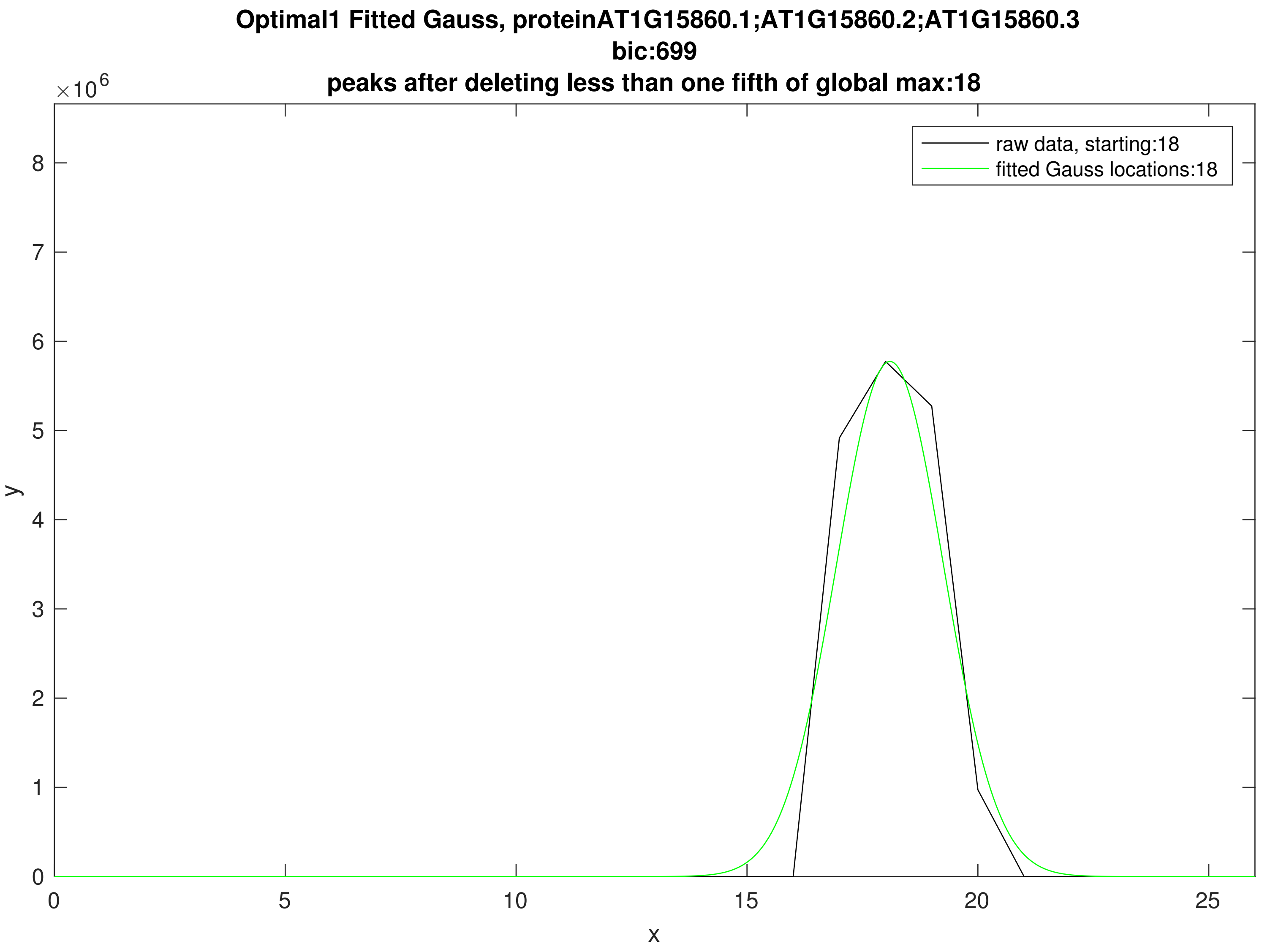
Optimal1 Fitted Gauss, proteinAT1G15860.1;AT1G15860.2;AT1G15860.3
bic:699
peaks after deleting less than one fifth of global max:18
×10^6
raw data, starting:18
fitted Gauss locations:18
y
x
0
1
2
3
4
5
6
7
8
0
5
10
15
20
25

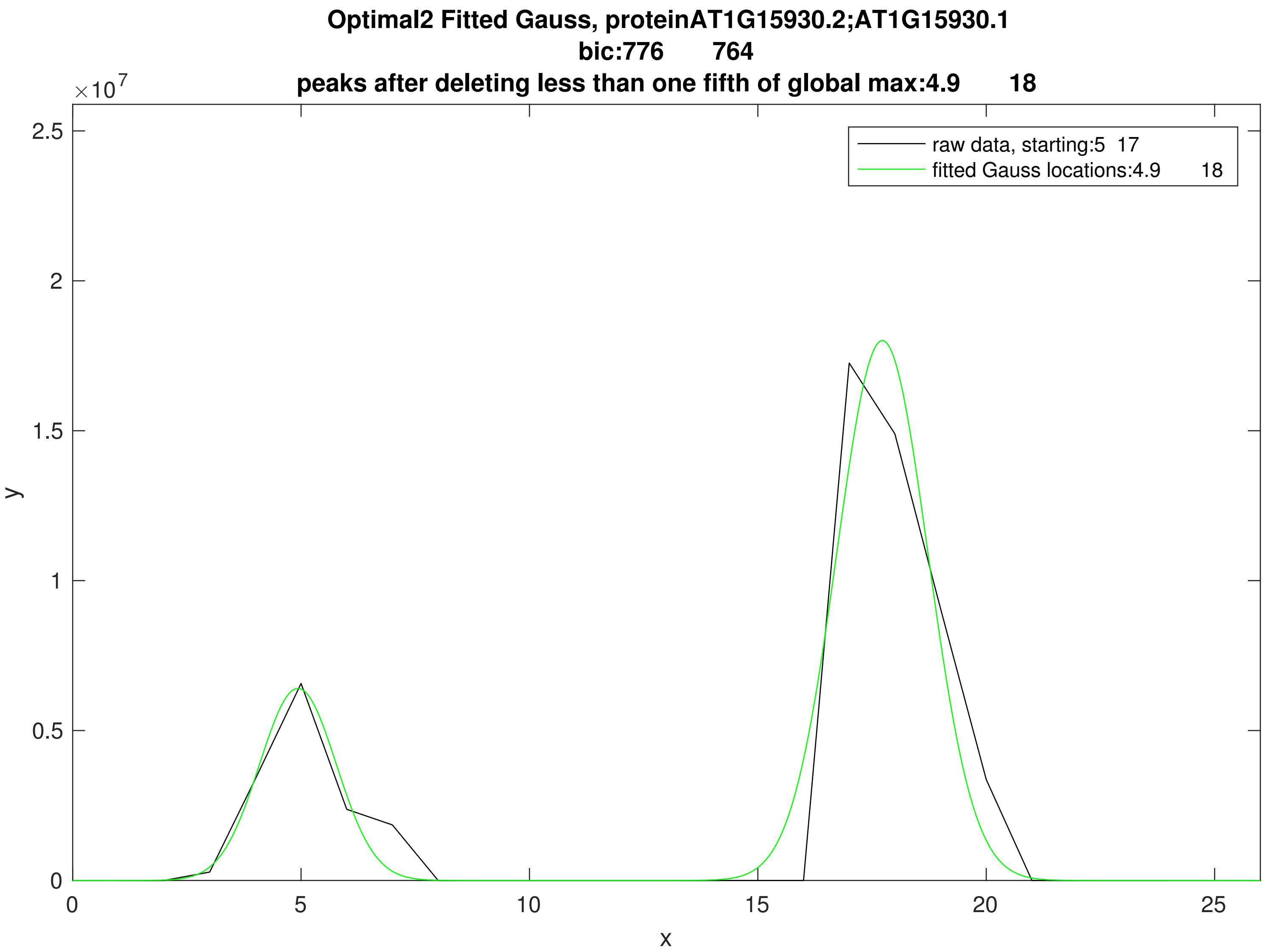
Optimal2 Fitted Gauss, proteinAT1G15930.2;AT1G15930.1
bic:776 764
peaks after deleting less than one fifth of global max:4.9 18
×10^7
raw data, starting:5 17
fitted Gauss locations:4.9 18
2.5
2
1.5
1
0.5
0
y
0
5
10
15
20
25
x

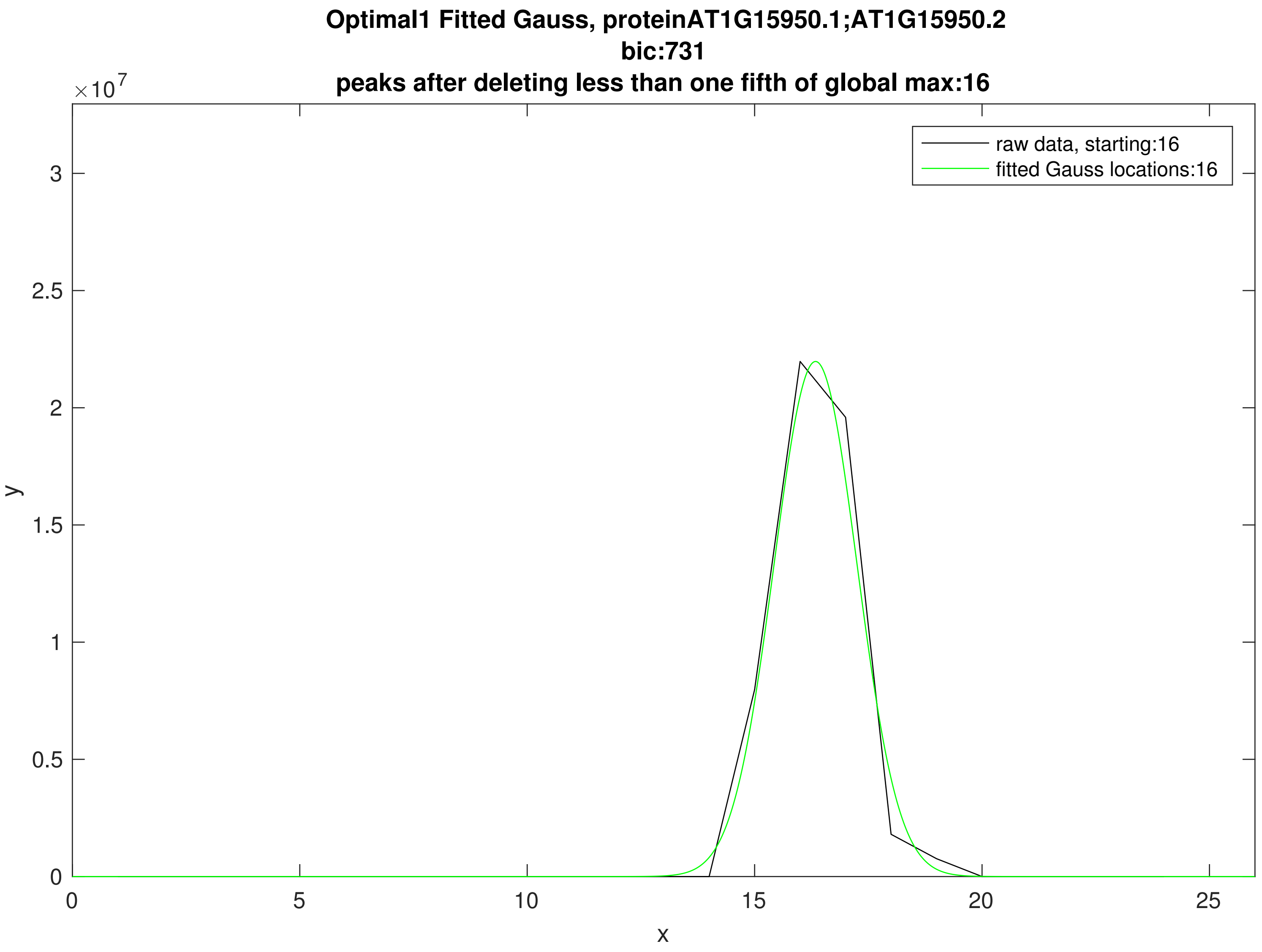

Optimal1 Fitted Gauss, proteinAT1G15950.1;AT1G15950.2
bic:731
peaks after deleting less than one fifth of global max:16
×10^7
raw data, starting:16
fitted Gauss locations:16
y
x
0
0.5
1
1.5
2
2.5
3
0
5
10
15
20
25

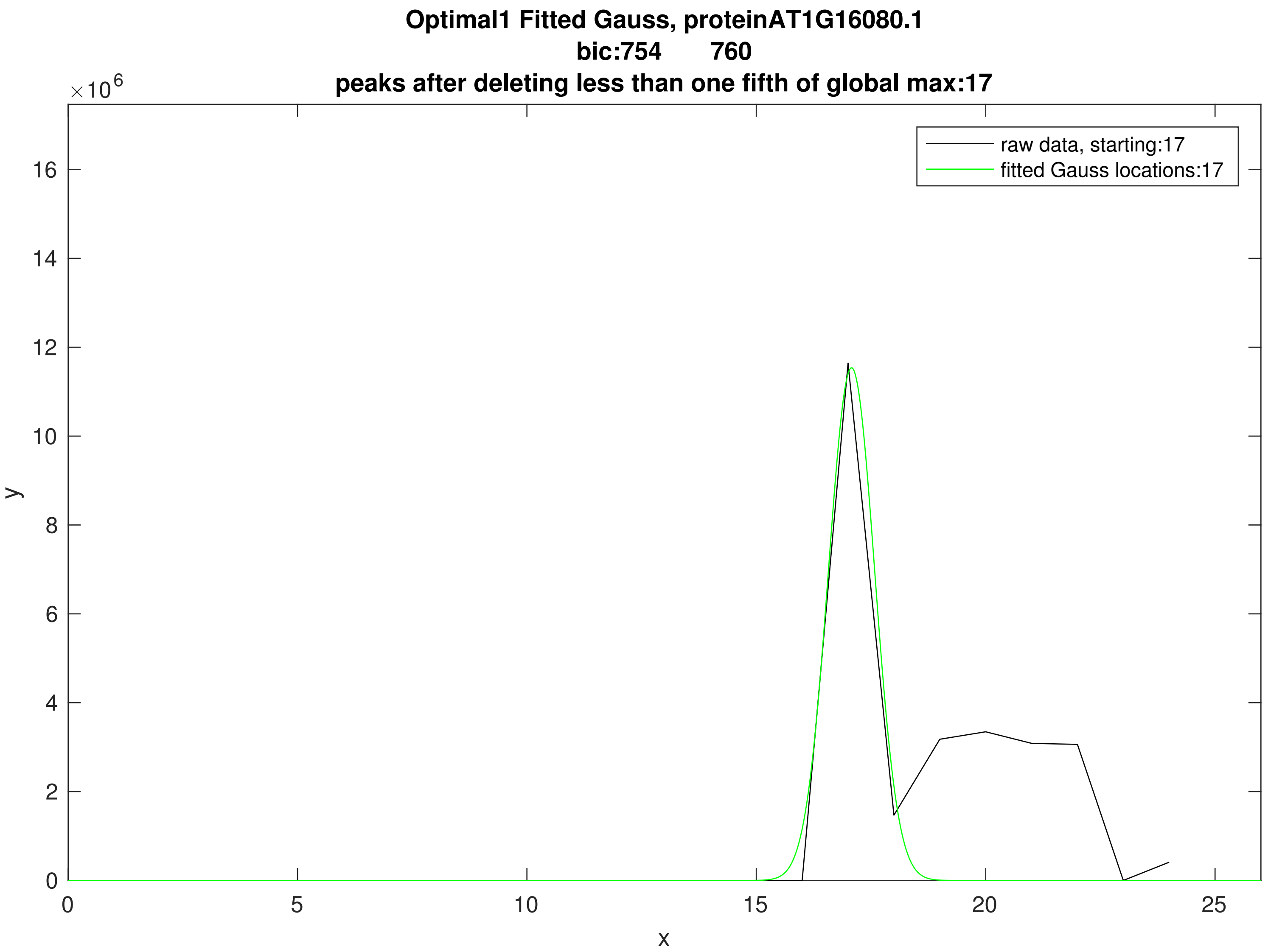

Optimal1 Fitted Gauss, proteinAT1G16080.1
bic:754 760
peaks after deleting less than one fifth of global max:17
×10^6
raw data, starting:17
fitted Gauss locations:17
16
14
12
10
8
6
4
2
0
y
0
5
10
15
20
25
x

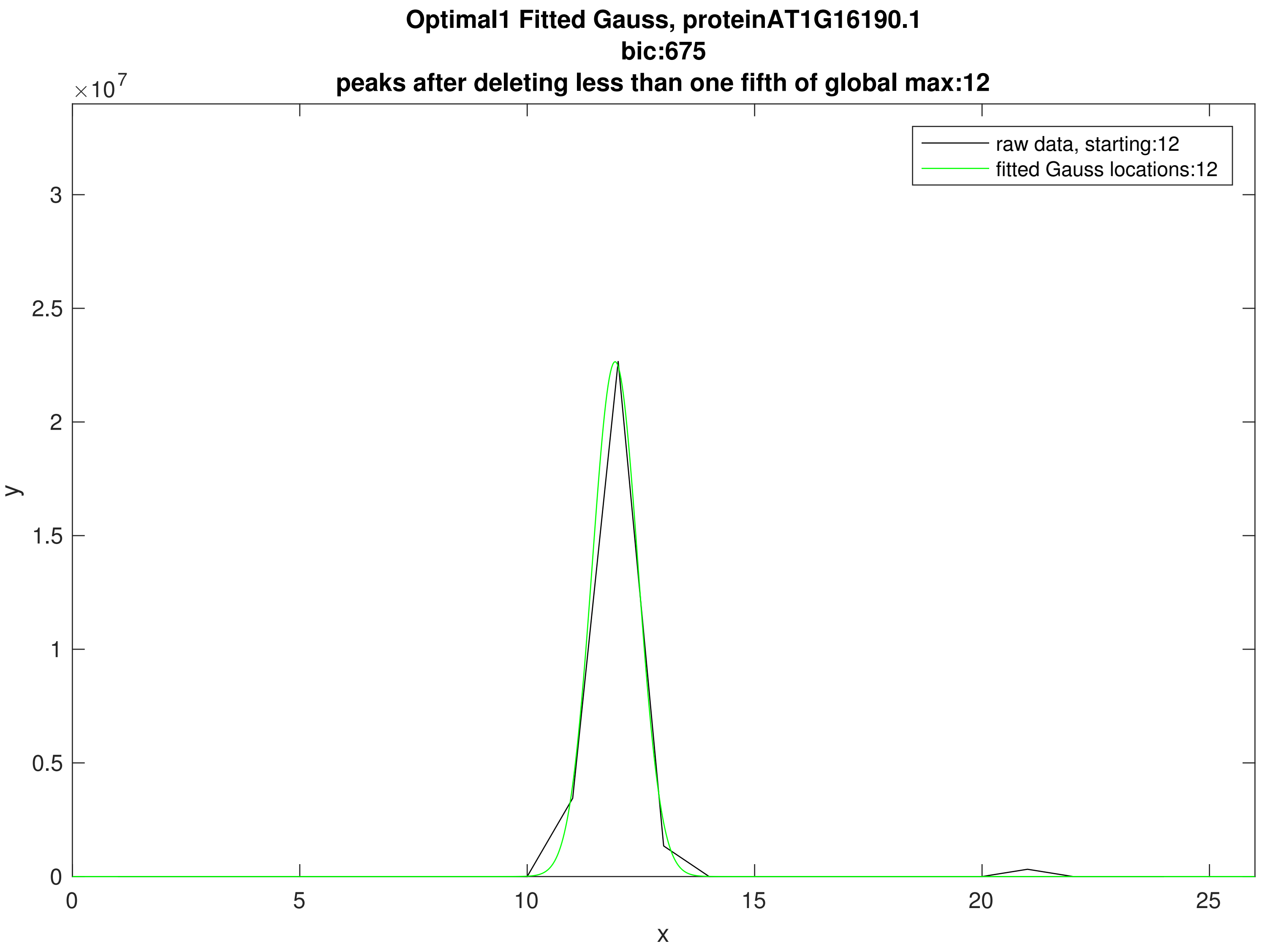
Optimal1 Fitted Gauss, proteinAT1G16190.1
bic:675
peaks after deleting less than one fifth of global max:12
×10^7
raw data, starting:12
fitted Gauss locations:12
y
x
0
0.5
1
1.5
2
2.5
3
0
5
10
15
20
25

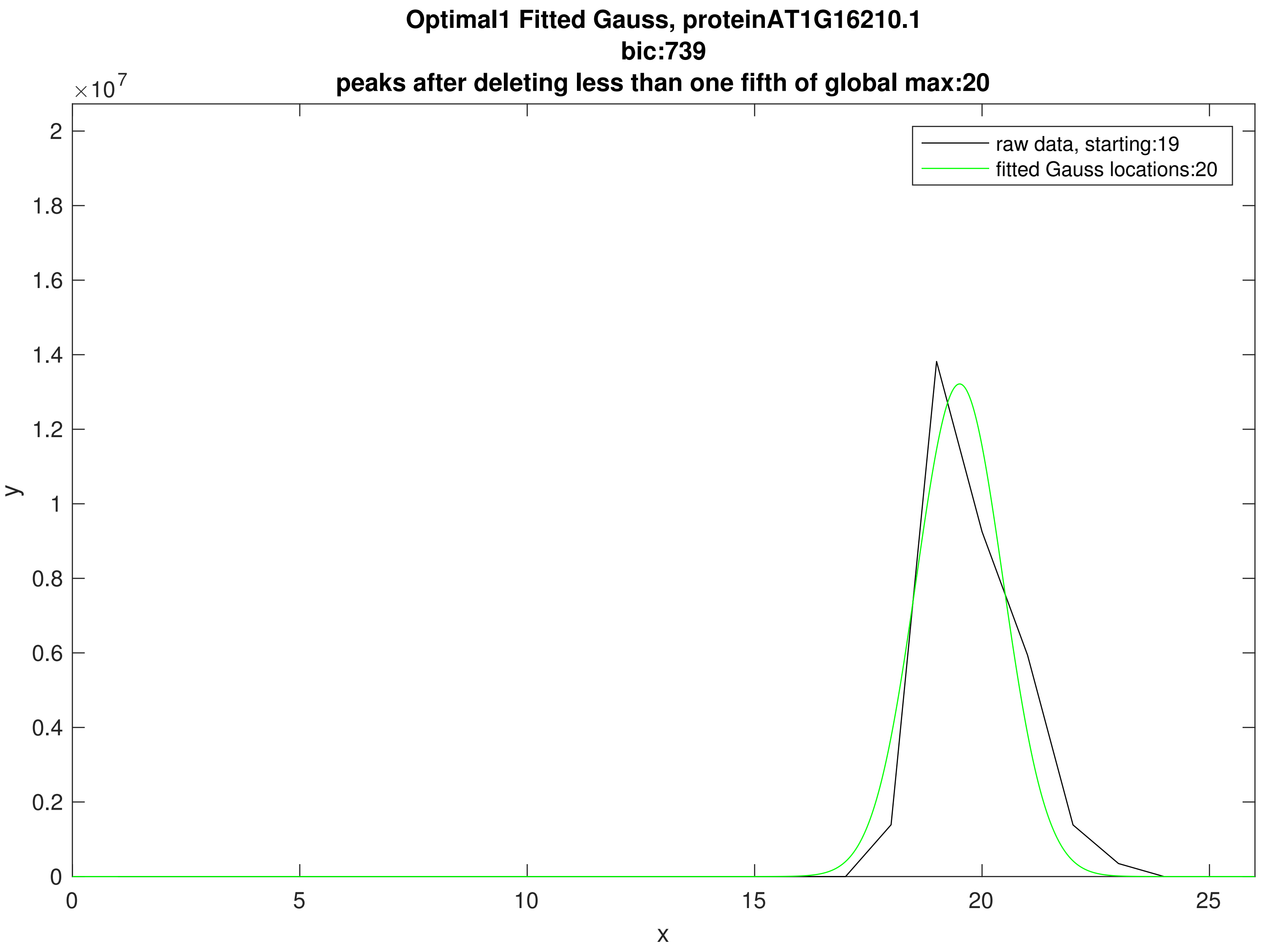
Optimal1 Fitted Gauss, proteinAT1G16210.1
bic:739
peaks after deleting less than one fifth of global max:20
×10^7
y
x
0
0.2
0.4
0.6
0.8
1
1.2
1.4
1.6
1.8
2
0
5
10
15
20
25
raw data, starting:19
fitted Gauss locations:20

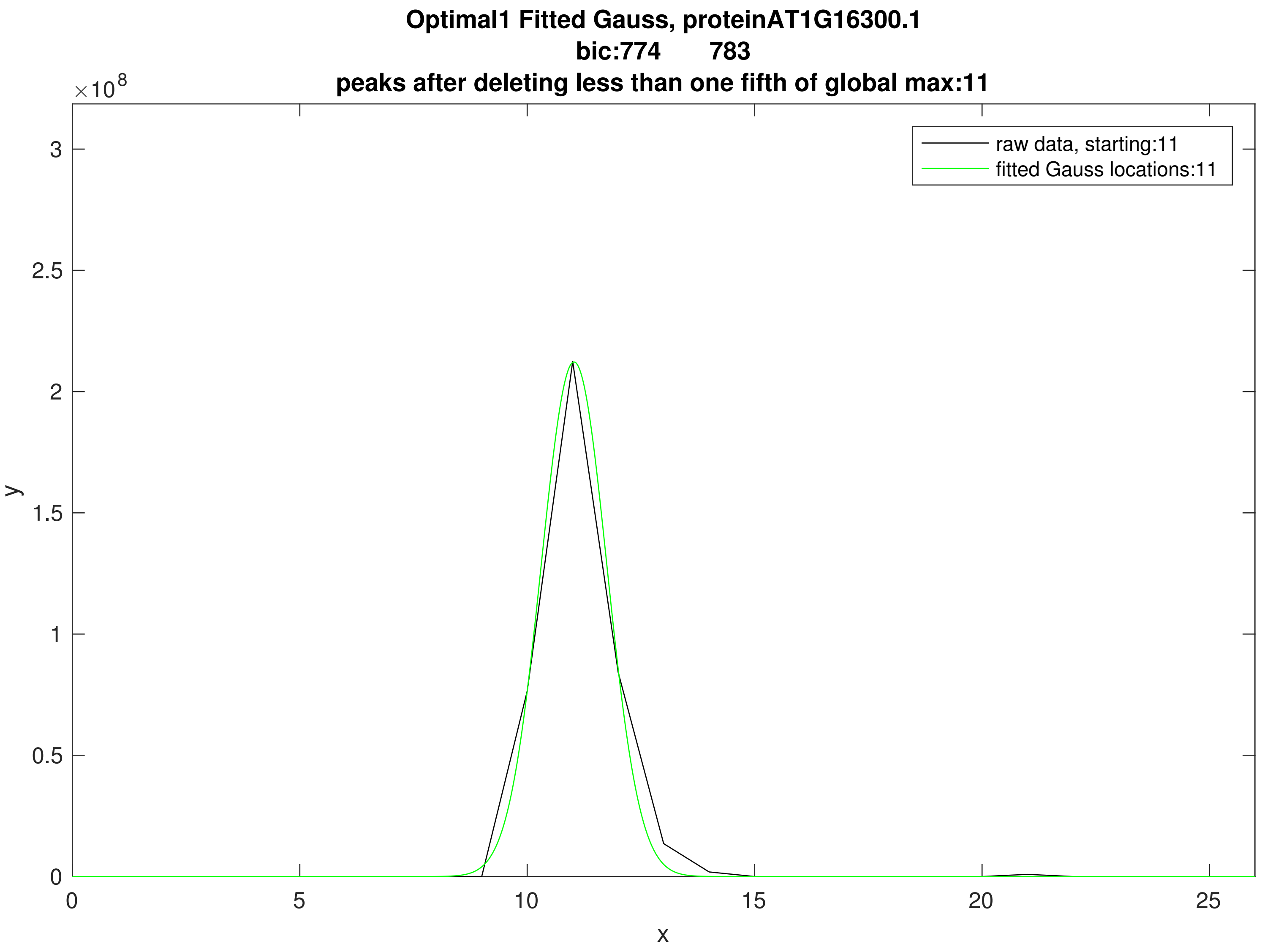

Optimal1 Fitted Gauss, proteinAT1G16300.1
bic:774 783
peaks after deleting less than one fifth of global max:11
×10^8
3
2.5
2
1.5
1
0.5
0
y
0
5
10
15
20
25
x
raw data, starting:11
fitted Gauss locations:11

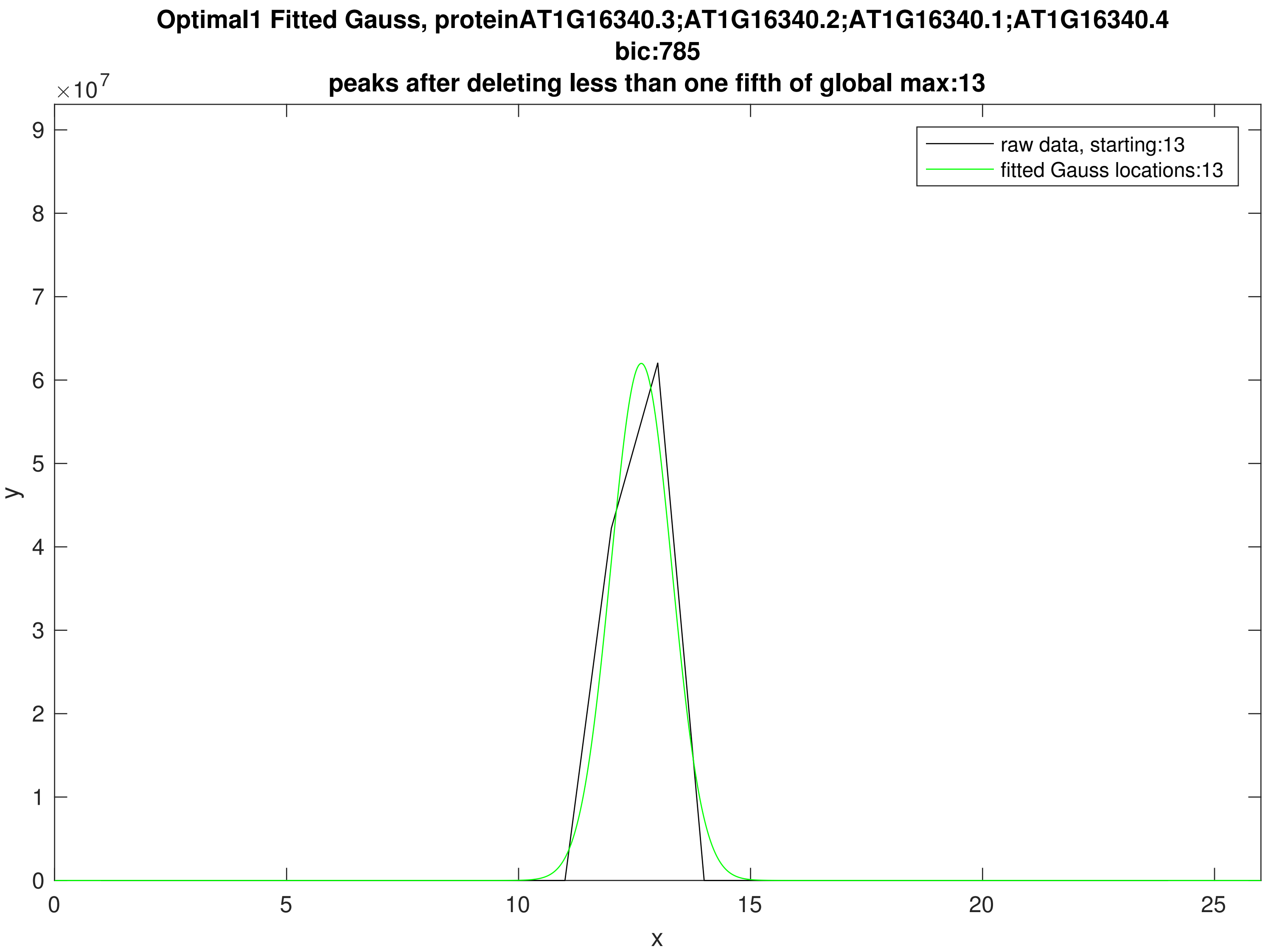

Optimal1 Fitted Gauss, proteinAT1G16340.3;AT1G16340.2;AT1G16340.1;AT1G16340.4
bic:785
peaks after deleting less than one fifth of global max:13
×10^7
raw data, starting:13
fitted Gauss locations:13
y
x
0
1
2
3
4
5
6
7
8
9
0
5
10
15
20
25

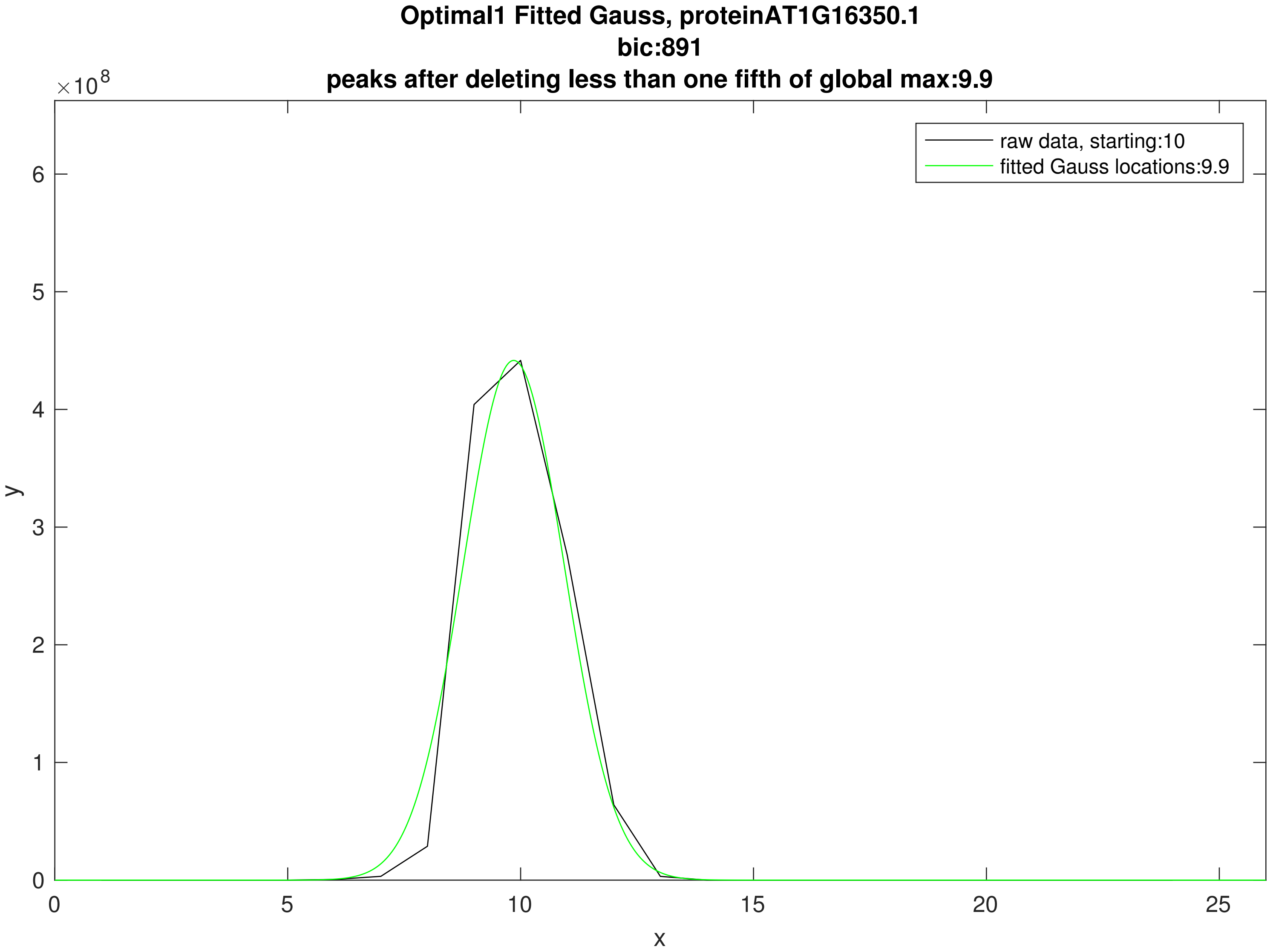

Optimal1 Fitted Gauss, proteinAT1G16350.1
bic:891
peaks after deleting less than one fifth of global max:9.9
×10^8
raw data, starting:10
fitted Gauss locations:9.9
y
x
0
1
2
3
4
5
6
0
5
10
15
20
25

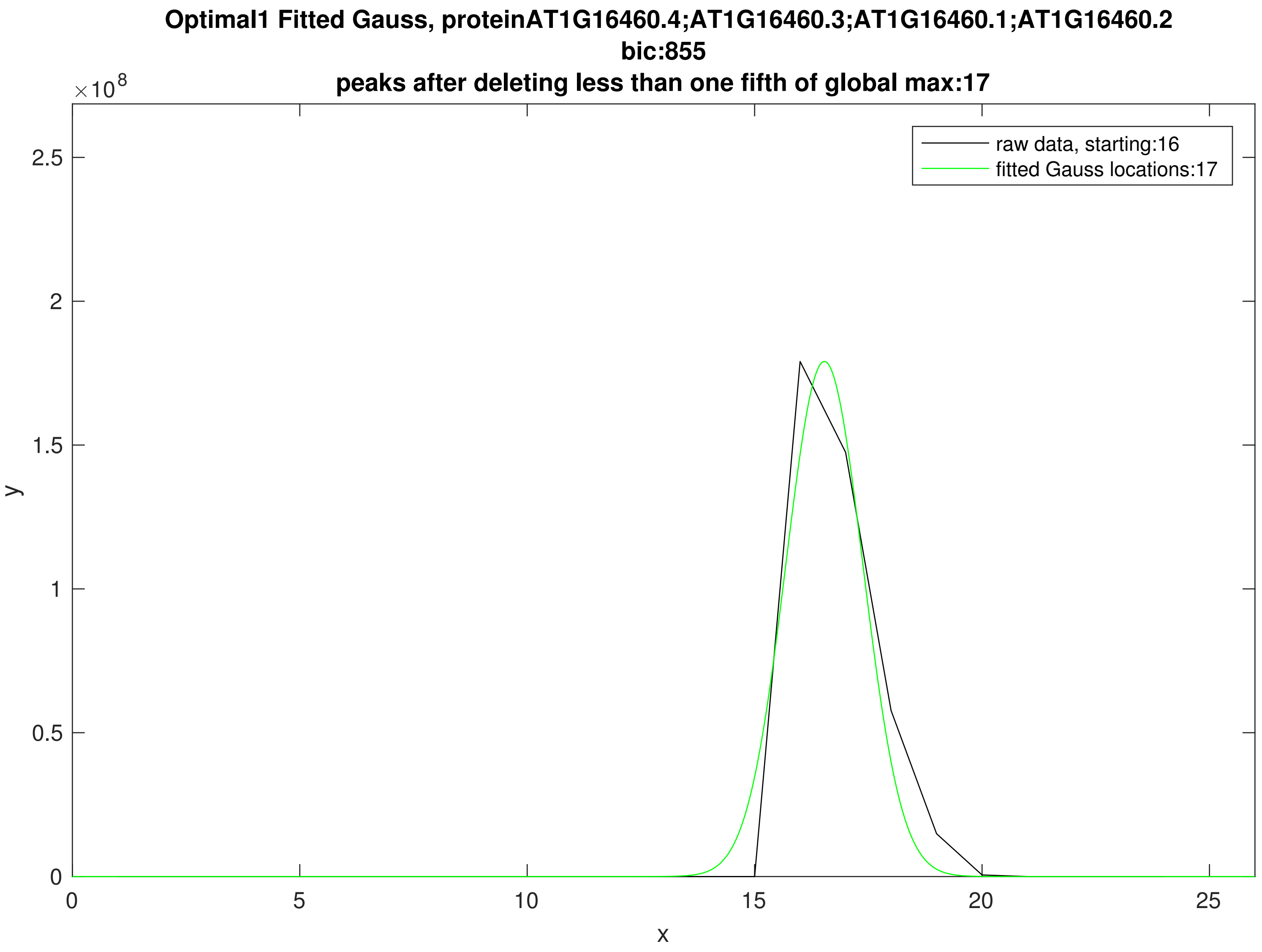
Optimal1 Fitted Gauss, proteinAT1G16460.4;AT1G16460.3;AT1G16460.1;AT1G16460.2
bic:855
peaks after deleting less than one fifth of global max:17
×10^8
2.5
2
1.5
1
0.5
0
y
0
5
10
15
20
25
x
raw data, starting:16
fitted Gauss locations:17

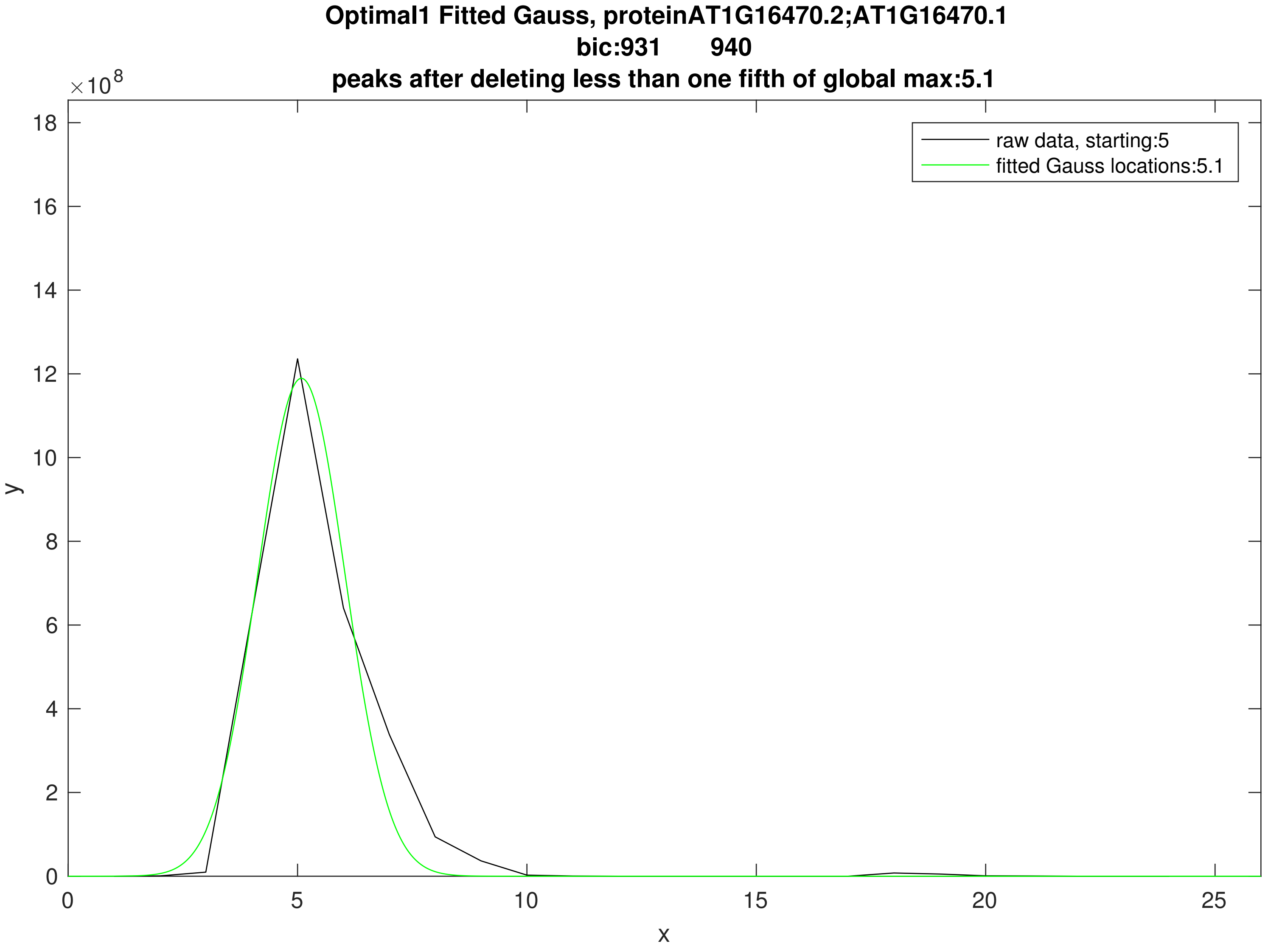
Optimal1 Fitted Gauss, proteinAT1G16470.2;AT1G16470.1
bic:931 940
peaks after deleting less than one fifth of global max:5.1
×10^8
raw data, starting:5
fitted Gauss locations:5.1
y
x
0
2
4
6
8
10
12
14
16
18
0
5
10
15
20
25

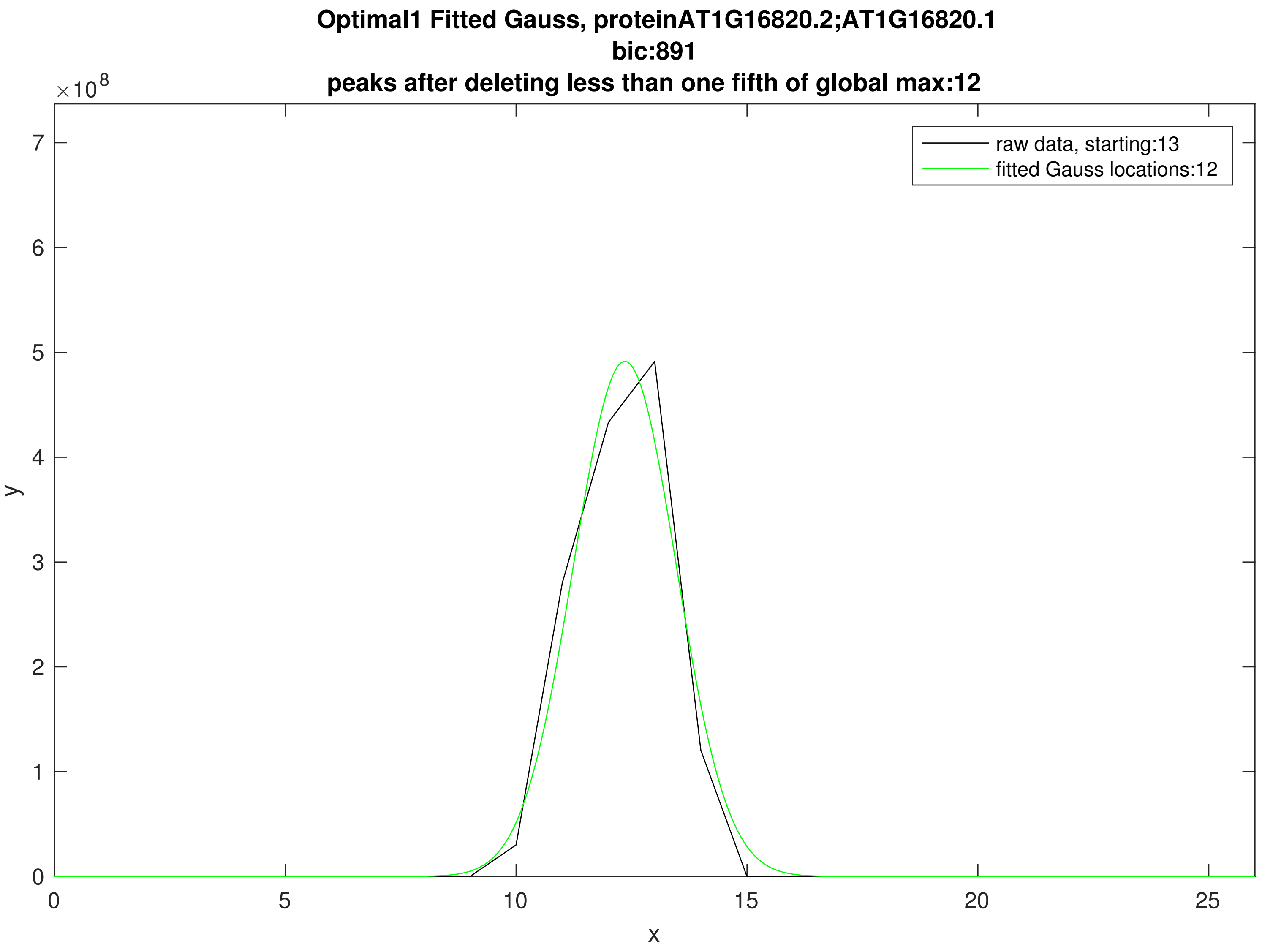

Optimal1 Fitted Gauss, proteinAT1G16820.2;AT1G16820.1
bic:891
peaks after deleting less than one fifth of global max:12
×10^8
raw data, starting:13
fitted Gauss locations:12
y
x
0
1
2
3
4
5
6
7
0
5
10
15
20
25

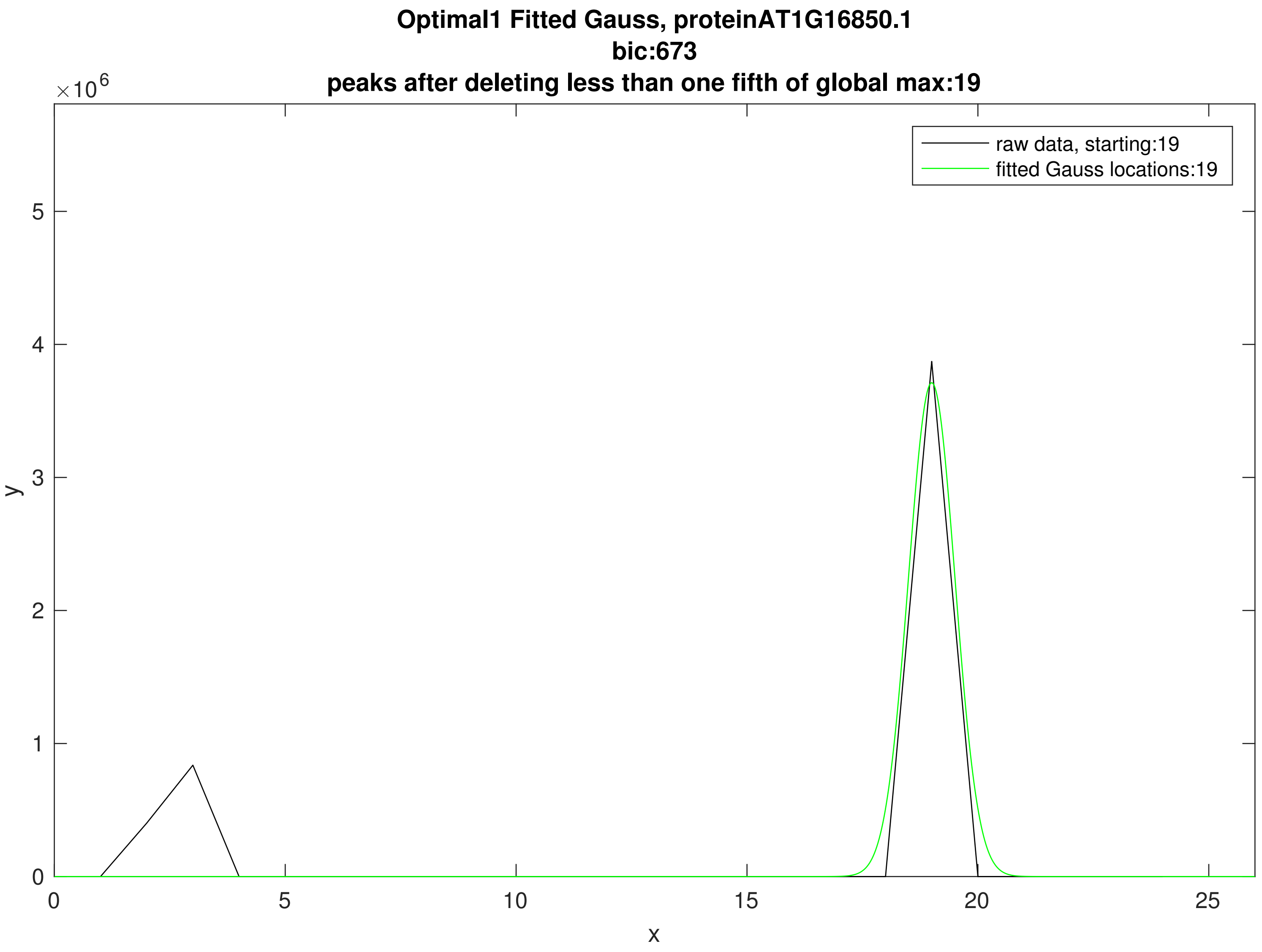

Optimal1 Fitted Gauss, proteinAT1G16850.1
bic:673
peaks after deleting less than one fifth of global max:19
×10^6
raw data, starting:19
fitted Gauss locations:19
y
x
0
1
2
3
4
5
0
5
10
15
20
25

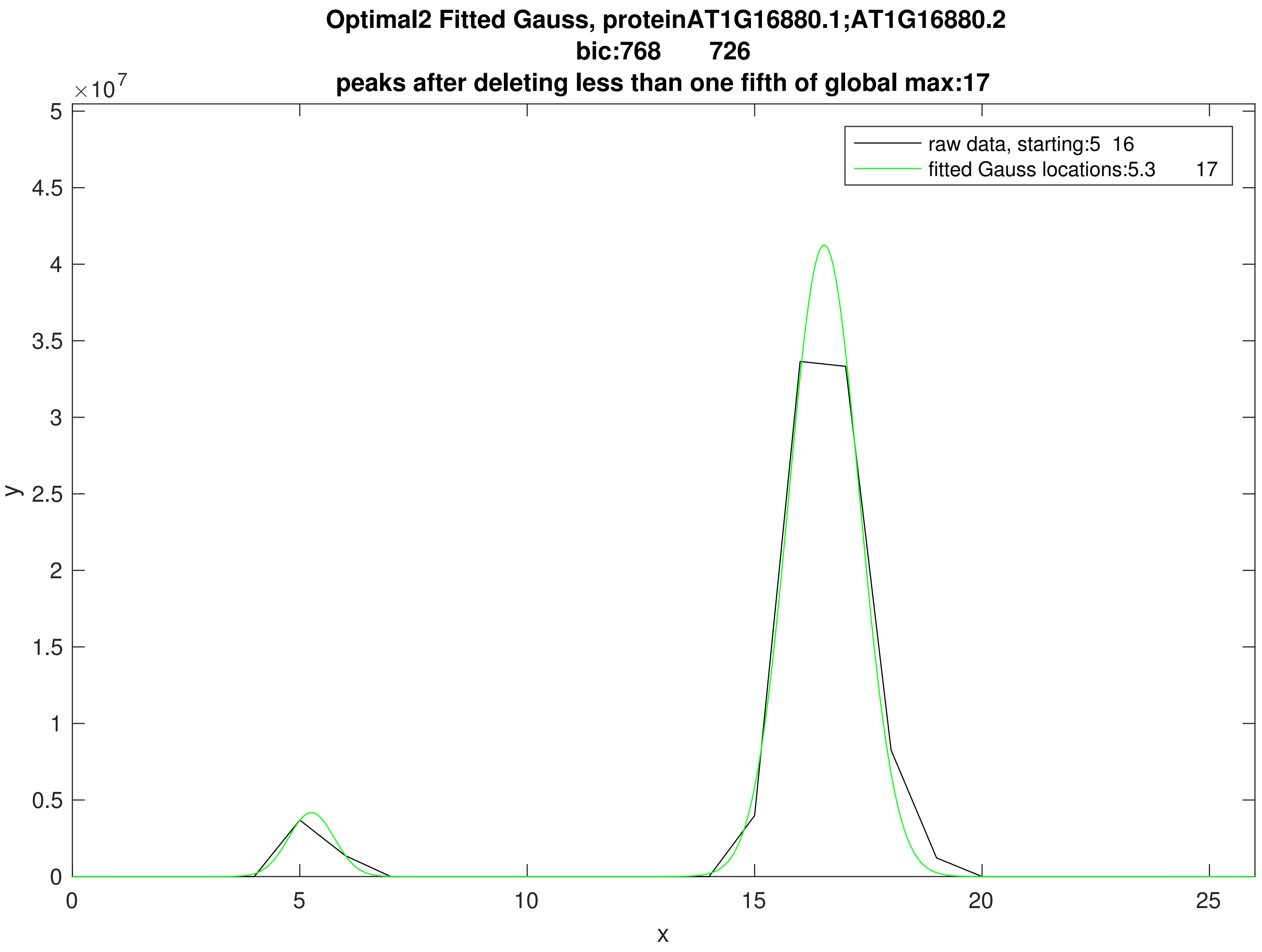

Optimal2 Fitted Gauss, proteinAT1G16880.1;AT1G16880.2
bic:768 726
peaks after deleting less than one fifth of global max:17
×10^7
raw data, starting:5 16
fitted Gauss locations:5.3 17
y
x
0
0.5
1
1.5
2
2.5
3
3.5
4
4.5
5
10
15
20
25

**Optimal1 Fitted Gauss, proteinAT1G16890.2;AT1G16890.3;AT1G16890.1**
**bic:728**
**peaks after deleting less than one fifth of global max:19**

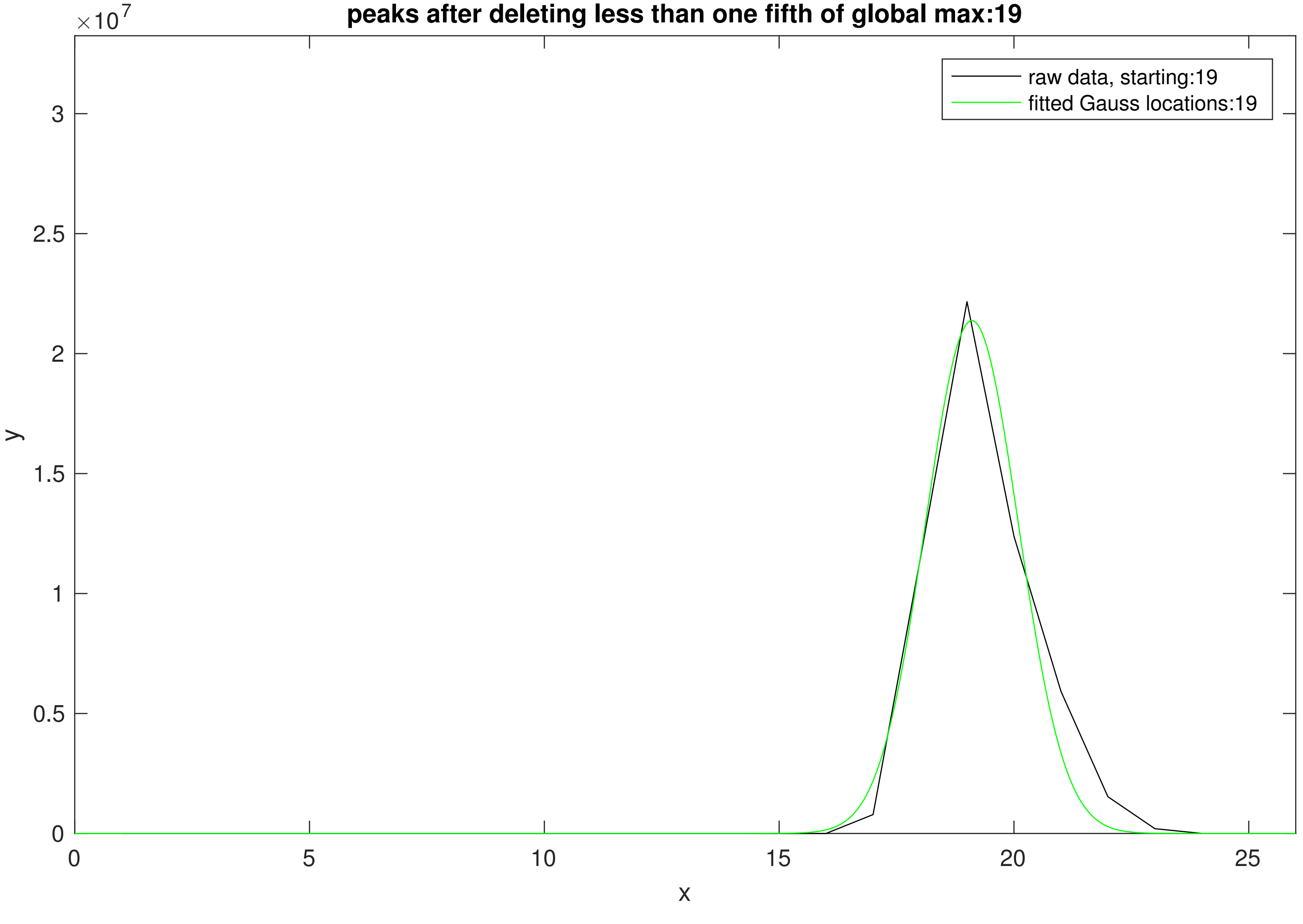

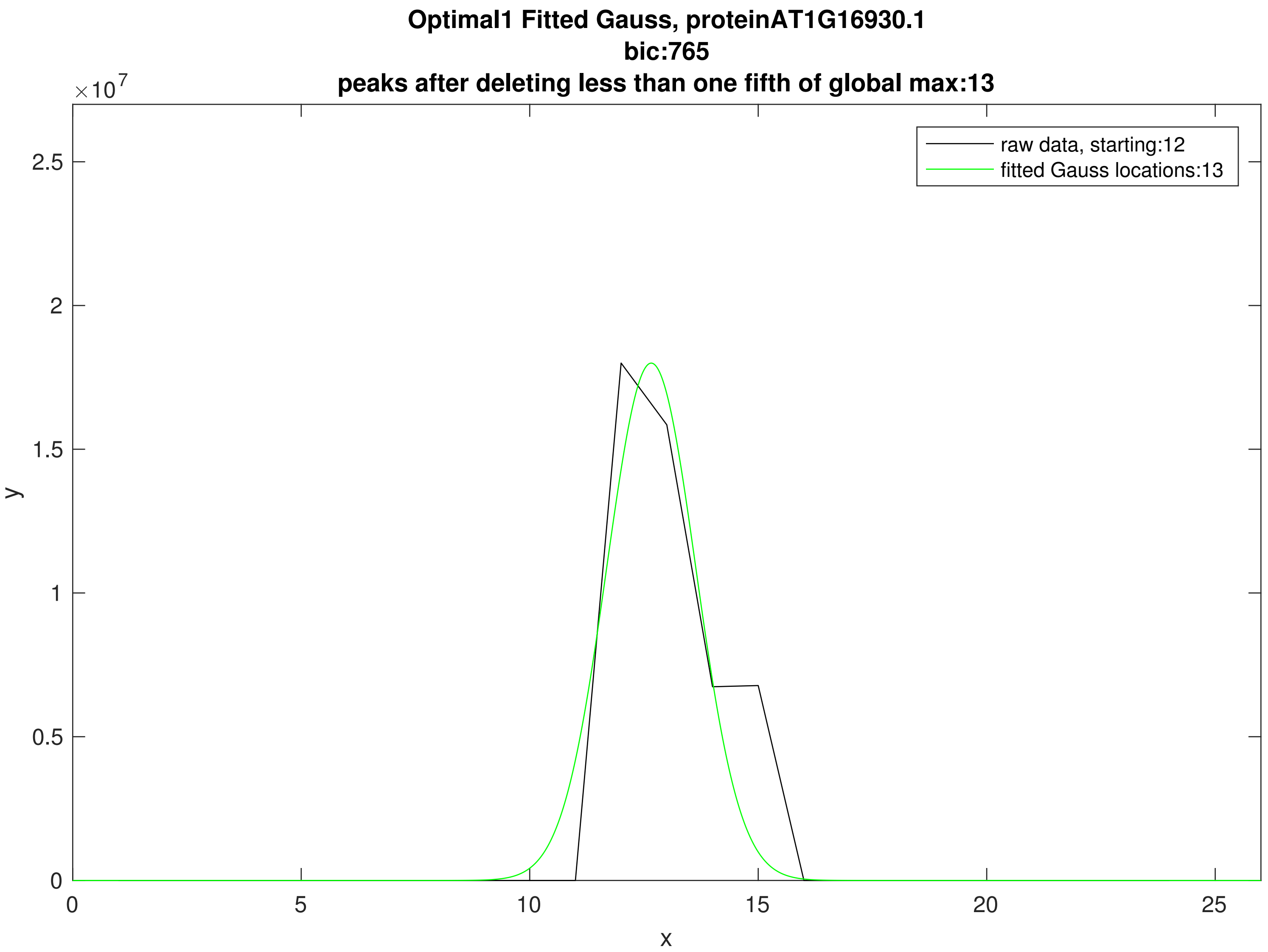
Optimal1 Fitted Gauss, proteinAT1G16930.1
bic:765
peaks after deleting less than one fifth of global max:13
×10^7
raw data, starting:12
fitted Gauss locations:13
y
x
0
0.5
1
1.5
2
2.5
0
5
10
15
20
25

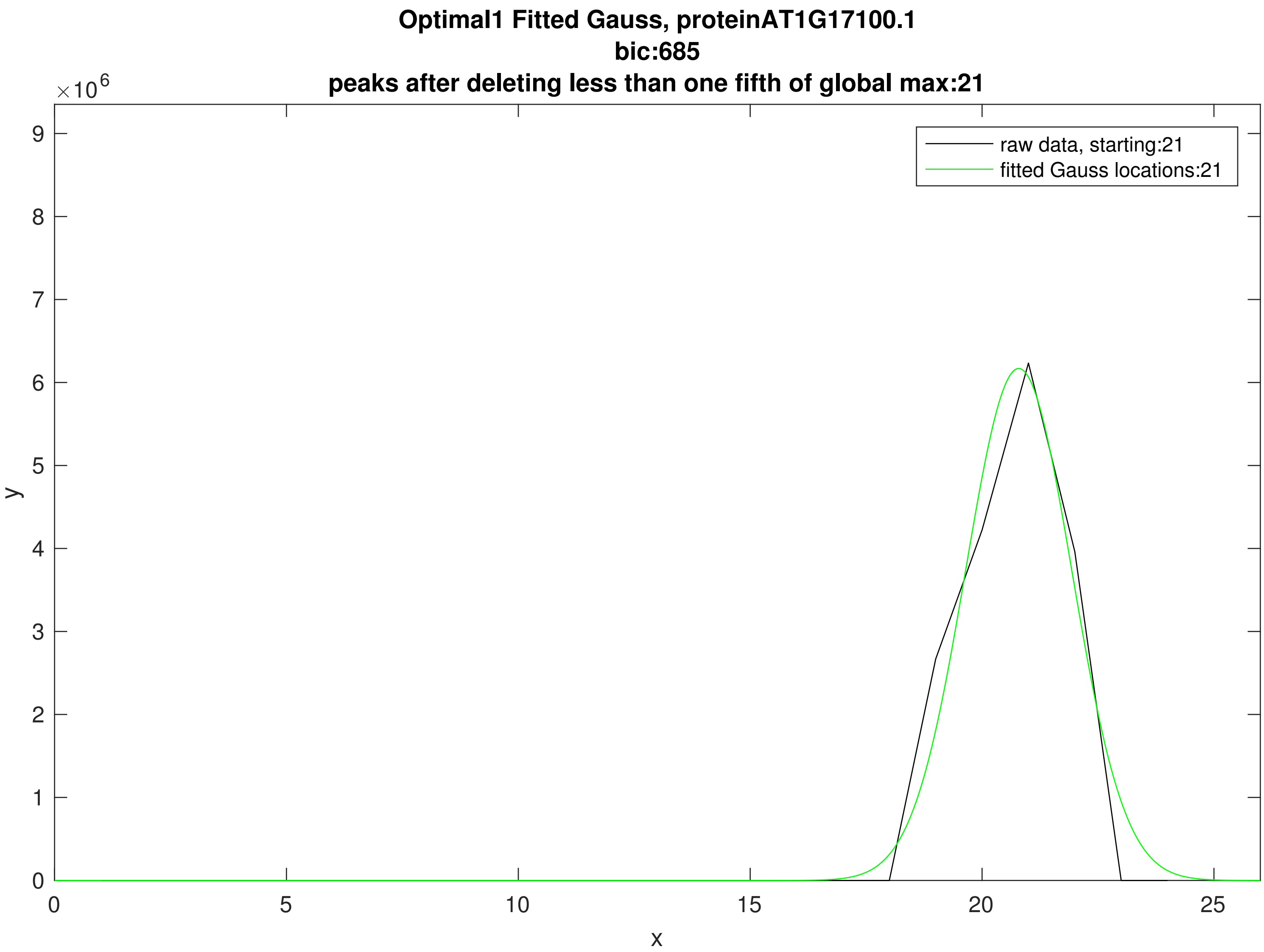
Optimal1 Fitted Gauss, proteinAT1G17100.1
bic:685
peaks after deleting less than one fifth of global max:21
×10^6
9
8
7
6
5
4
3
2
1
0
y
0
5
10
15
20
25
x
raw data, starting:21
fitted Gauss locations:21

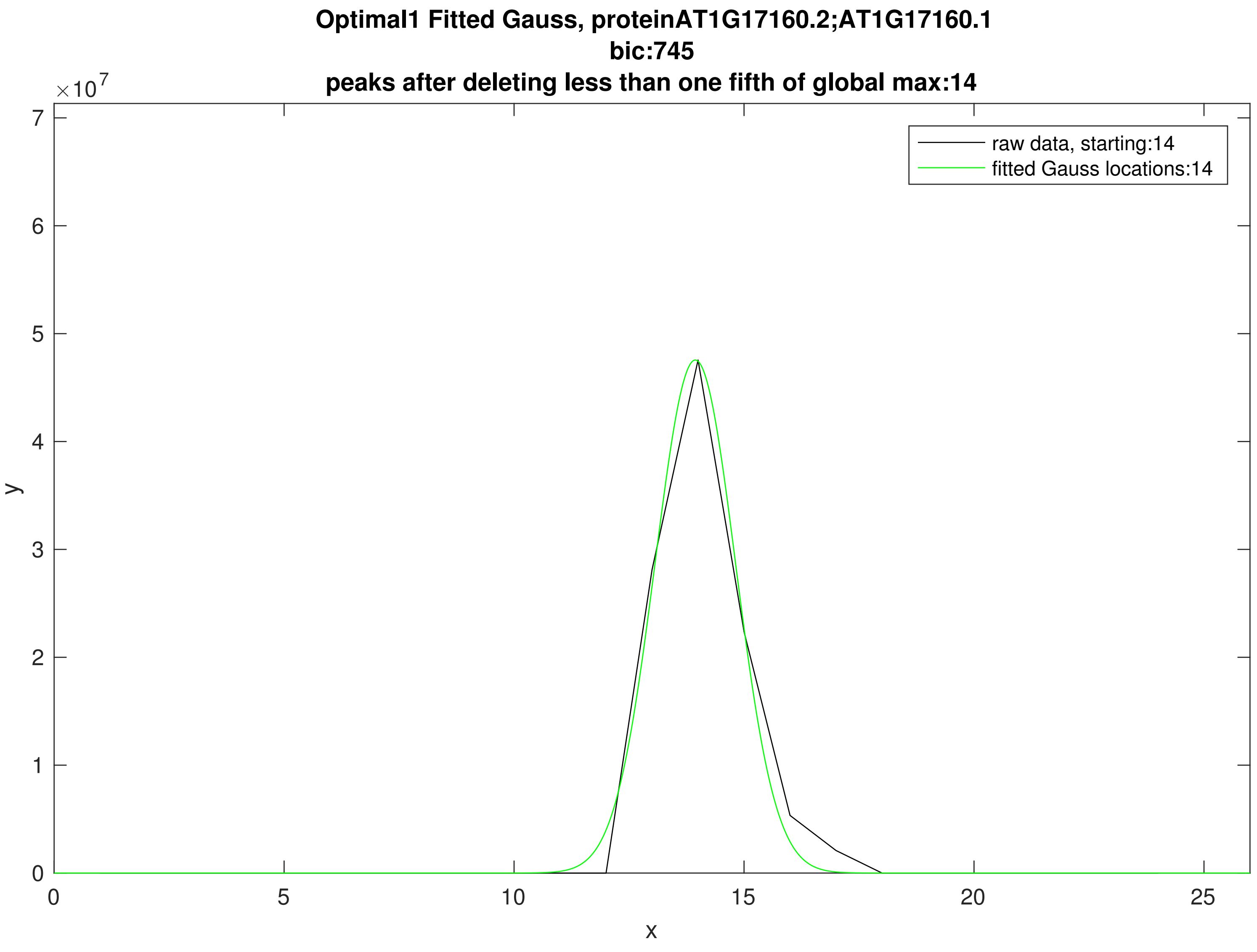
Optimal1 Fitted Gauss, proteinAT1G17160.2;AT1G17160.1
bic:745
peaks after deleting less than one fifth of global max:14
×10^7
raw data, starting:14
fitted Gauss locations:14
y
x
0
1
2
3
4
5
6
7
0
5
10
15
20
25

**Optimal1 Fitted Gauss, proteinAT1G17170.1**
**bic:764**
**peaks after deleting less than one fifth of global max:16**

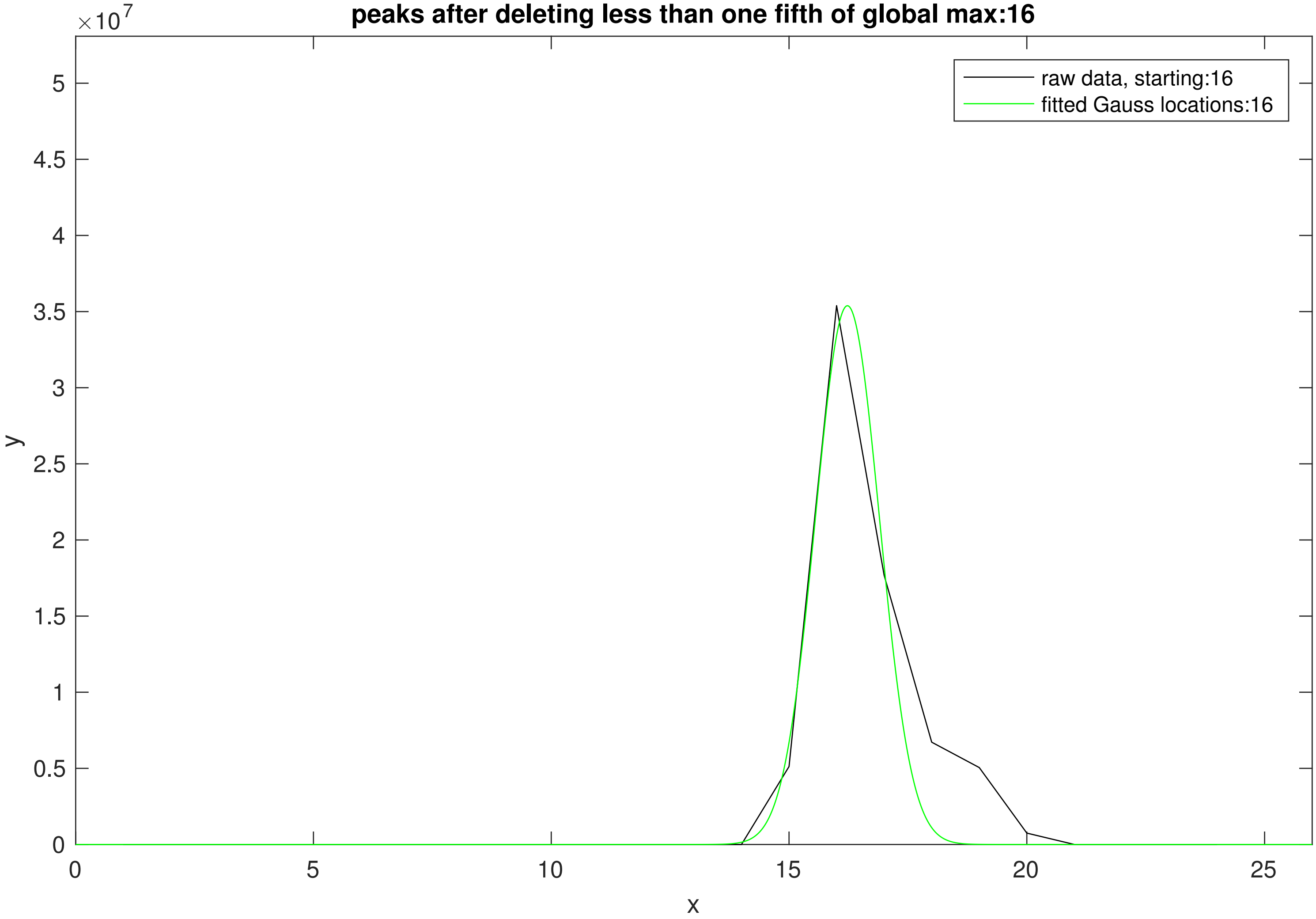

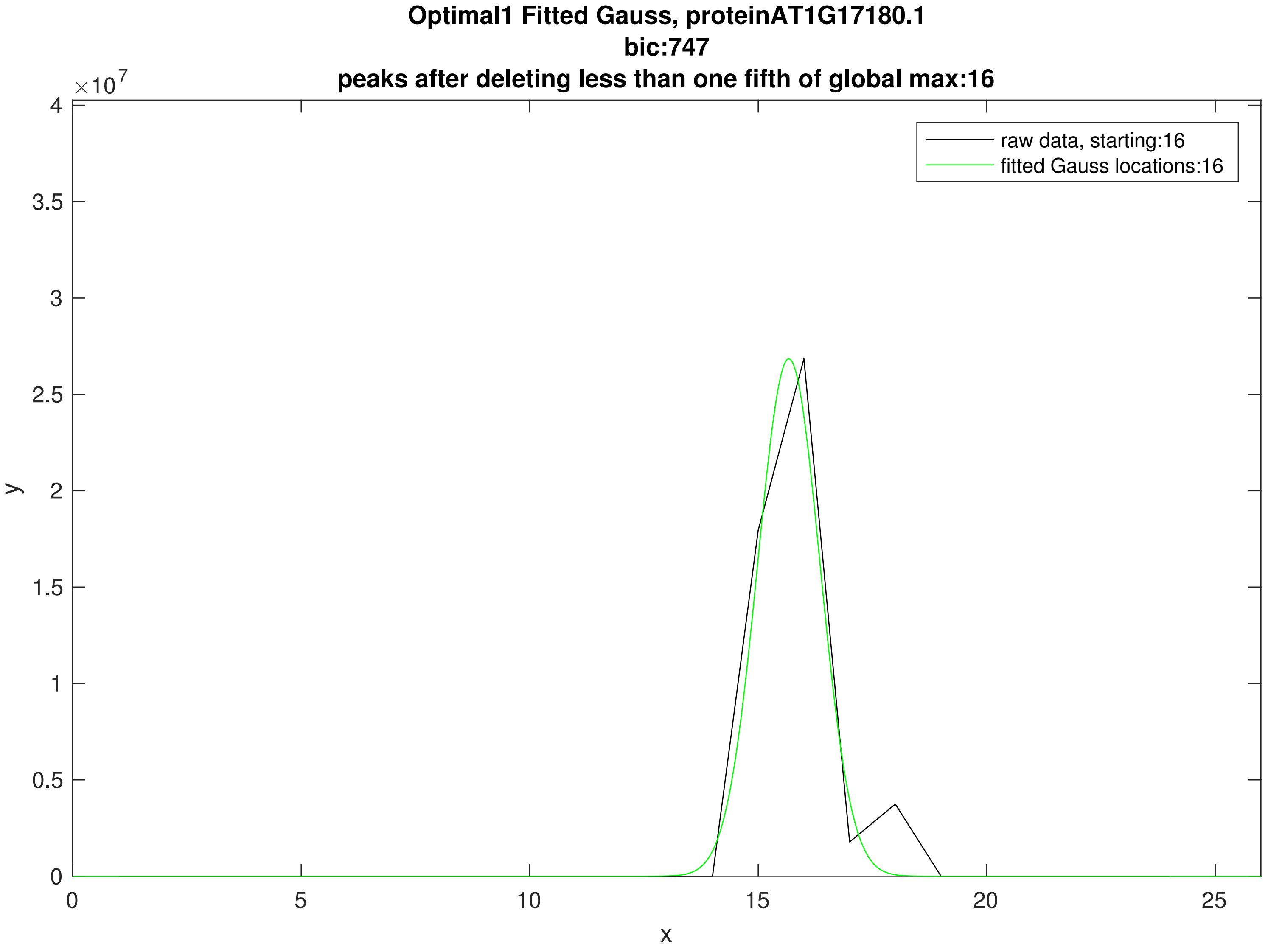
Optimal1 Fitted Gauss, proteinAT1G17180.1
bic:747
peaks after deleting less than one fifth of global max:16
×10^7
4
3.5
3
2.5
2
1.5
1
0.5
0
y
raw data, starting:16
fitted Gauss locations:16
0
5
10
15
20
25
x

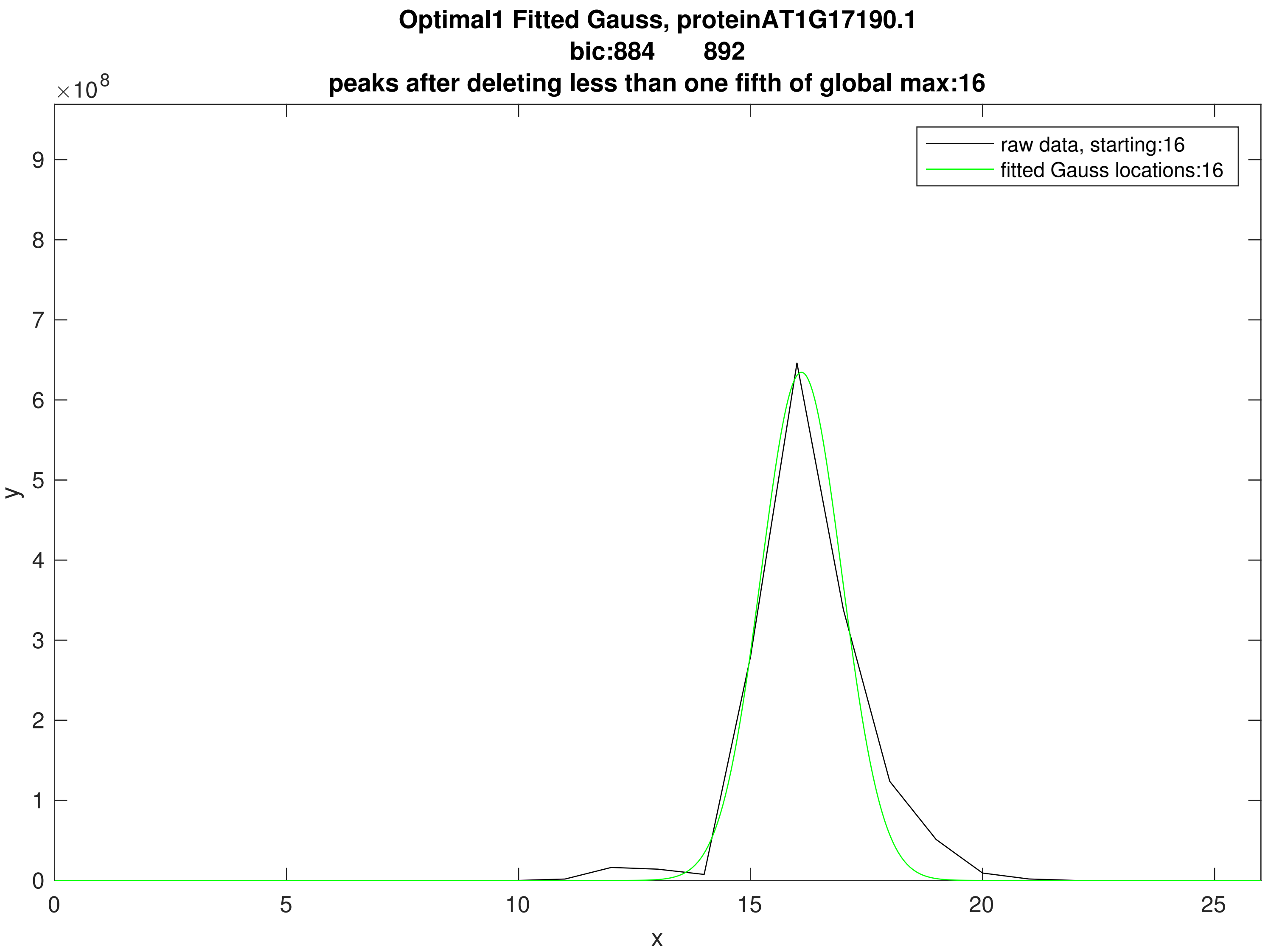
Optimal1 Fitted Gauss, proteinAT1G17190.1
bic:884 892
peaks after deleting less than one fifth of global max:16
×10^8
raw data, starting:16
fitted Gauss locations:16
y
x
0
1
2
3
4
5
6
7
8
9
0
5
10
15
20
25

**Optimal1 Fitted Gauss, proteinAT1G17290.1**
**bic:1e+03  1.01e+03  1.01e+03**
**peaks after deleting less than one fifth of global max:13**

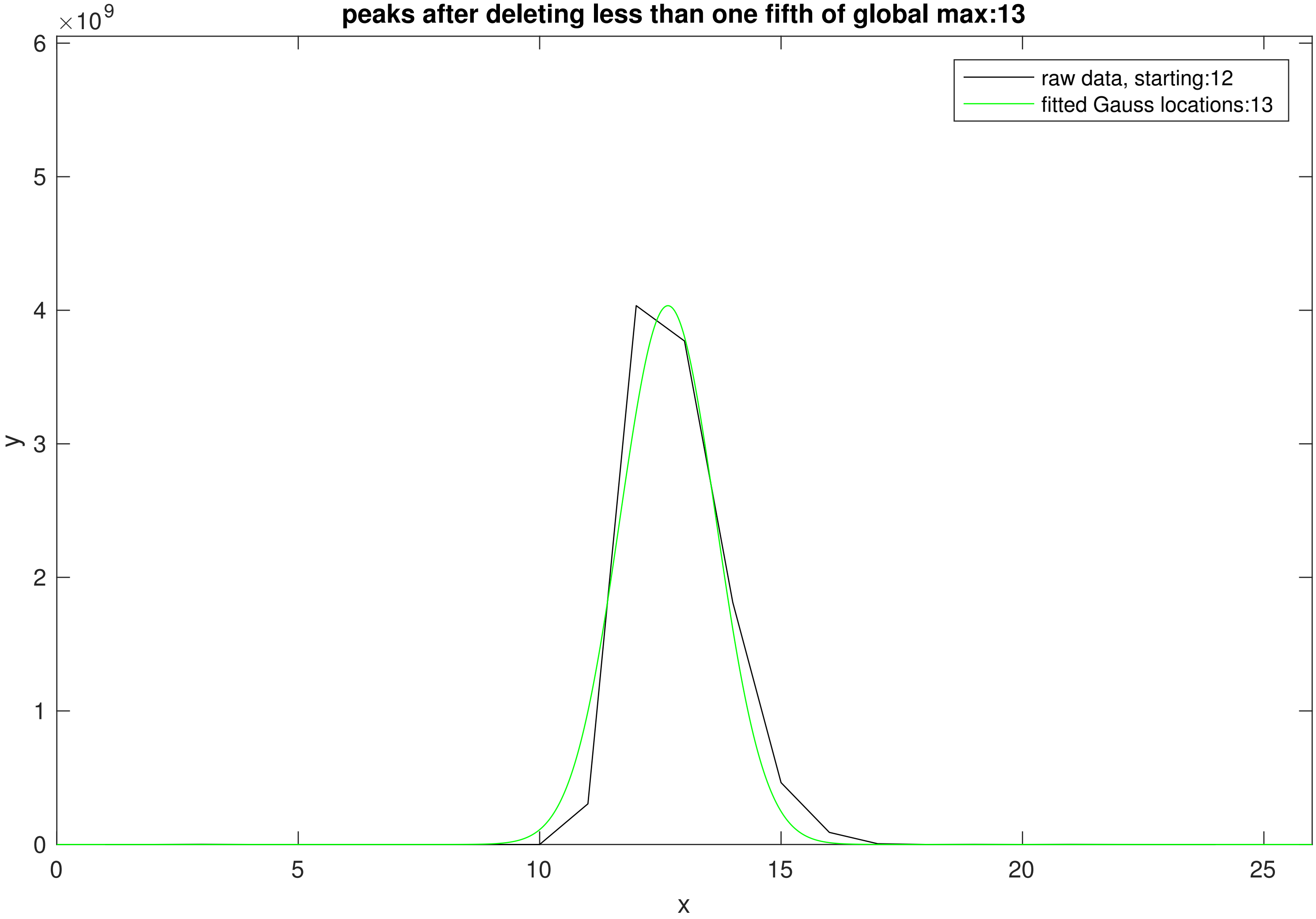

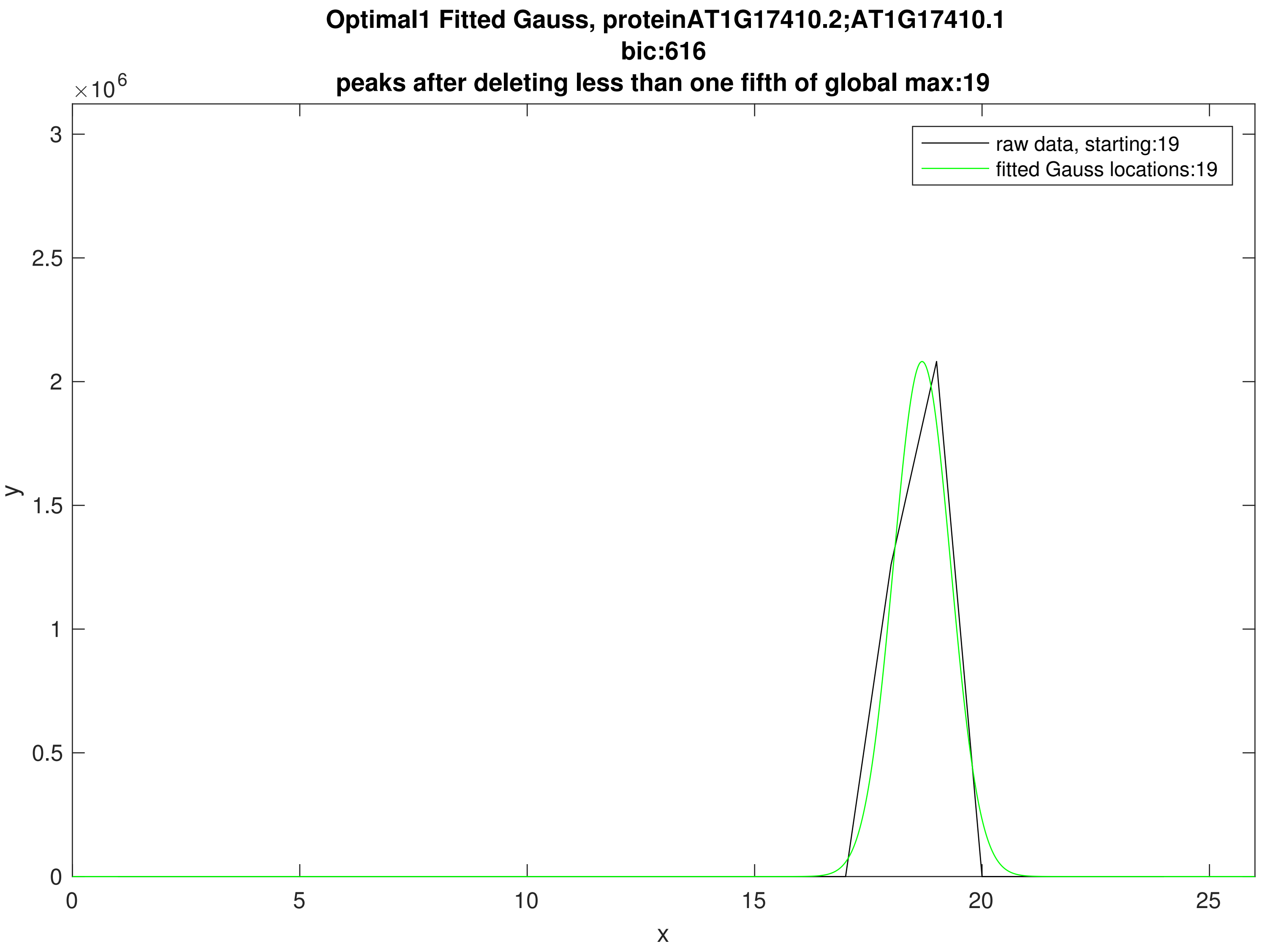
Optimal1 Fitted Gauss, proteinAT1G17410.2;AT1G17410.1
bic:616
peaks after deleting less than one fifth of global max:19
×10^6
3
2.5
2
1.5
1
0.5
0
y
0
5
10
15
20
25
x
raw data, starting:19
fitted Gauss locations:19

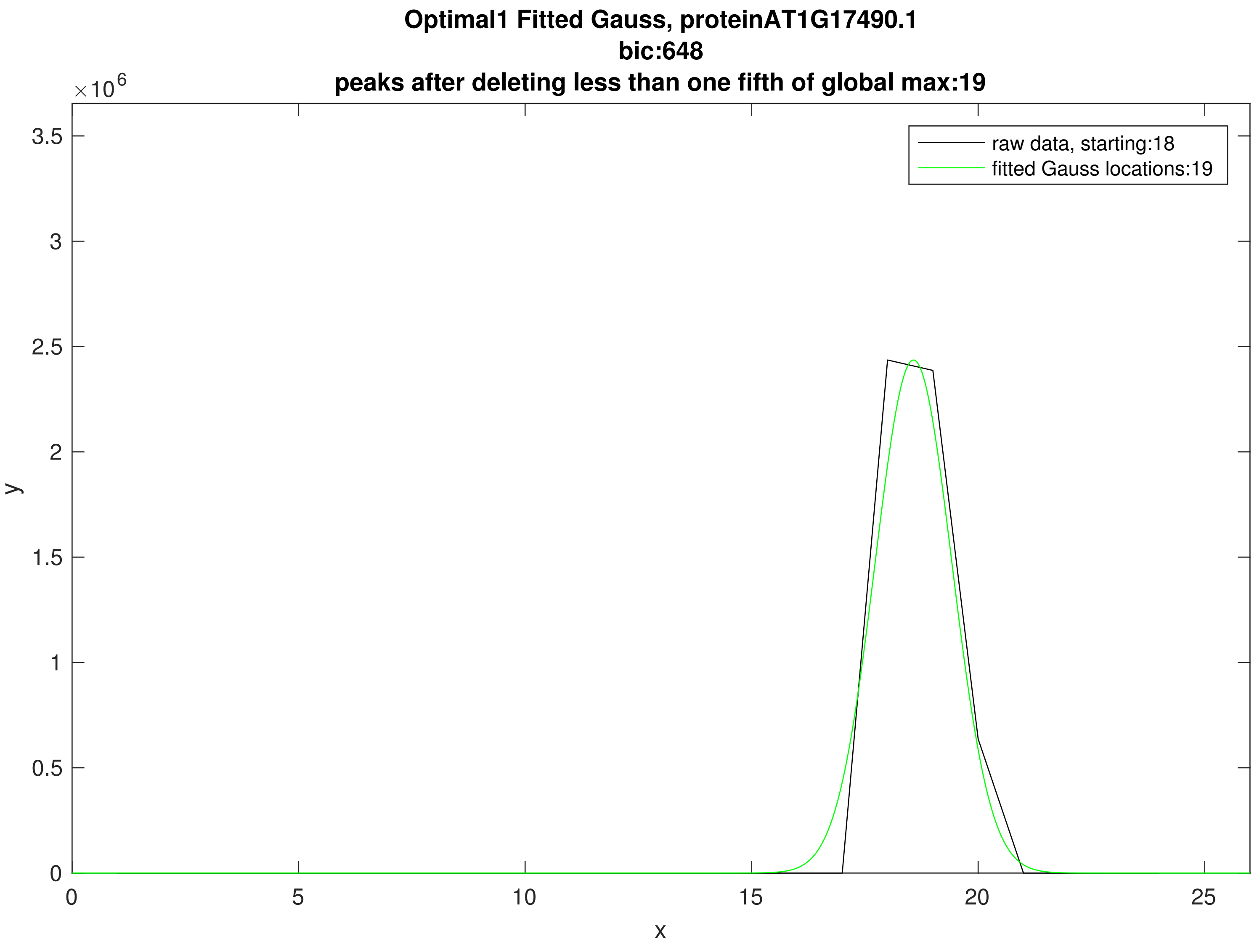
Optimal1 Fitted Gauss, proteinAT1G17490.1
bic:648
peaks after deleting less than one fifth of global max:19
×10^6
3.5
3
2.5
2
1.5
1
0.5
0
y
0
5
10
15
20
25
x
raw data, starting:18
fitted Gauss locations:19

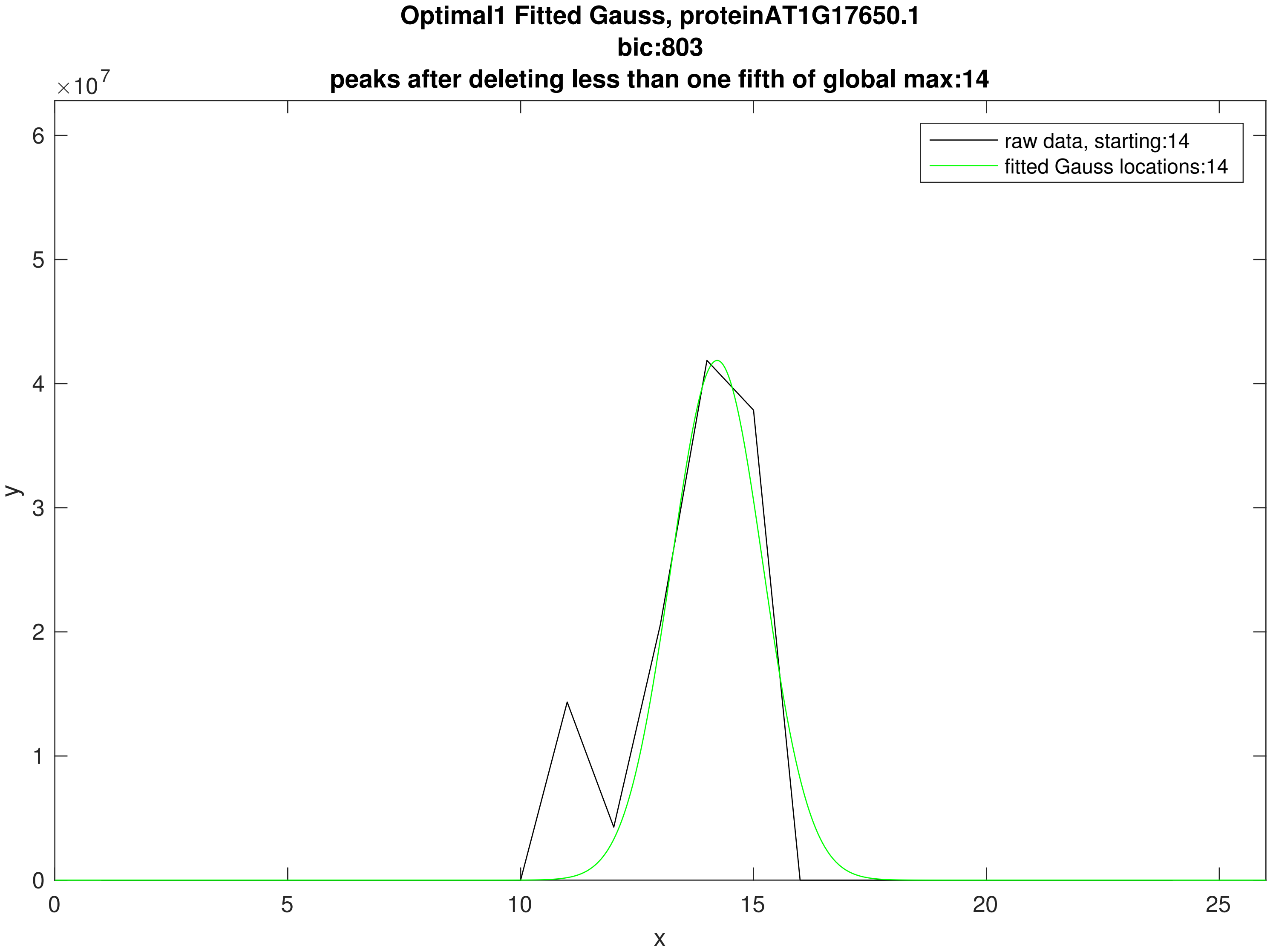
Optimal1 Fitted Gauss, proteinAT1G17650.1
bic:803
peaks after deleting less than one fifth of global max:14
×10^7
raw data, starting:14
fitted Gauss locations:14
y
x
0
1
2
3
4
5
6
0
5
10
15
20
25

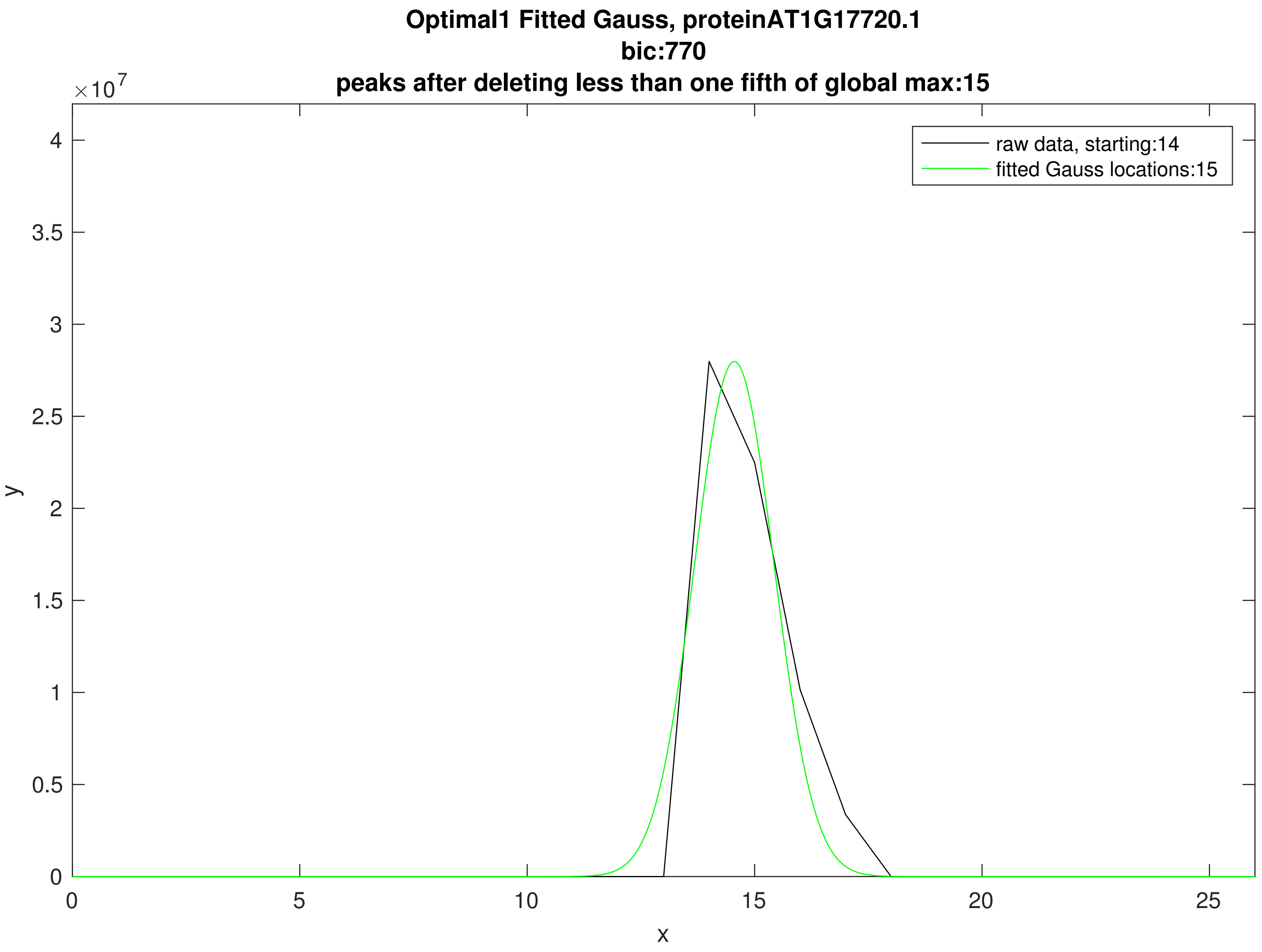
Optimal1 Fitted Gauss, proteinAT1G17720.1
bic:770
peaks after deleting less than one fifth of global max:15
×10^7
raw data, starting:14
fitted Gauss locations:15
y
x
0
0.5
1
1.5
2
2.5
3
3.5
4
5
10
15
20
25

**Optimal2 Fitted Gauss, proteinAT1G17720.2**
**bic:869 859 866**
**peaks after deleting less than one fifth of global max:14**

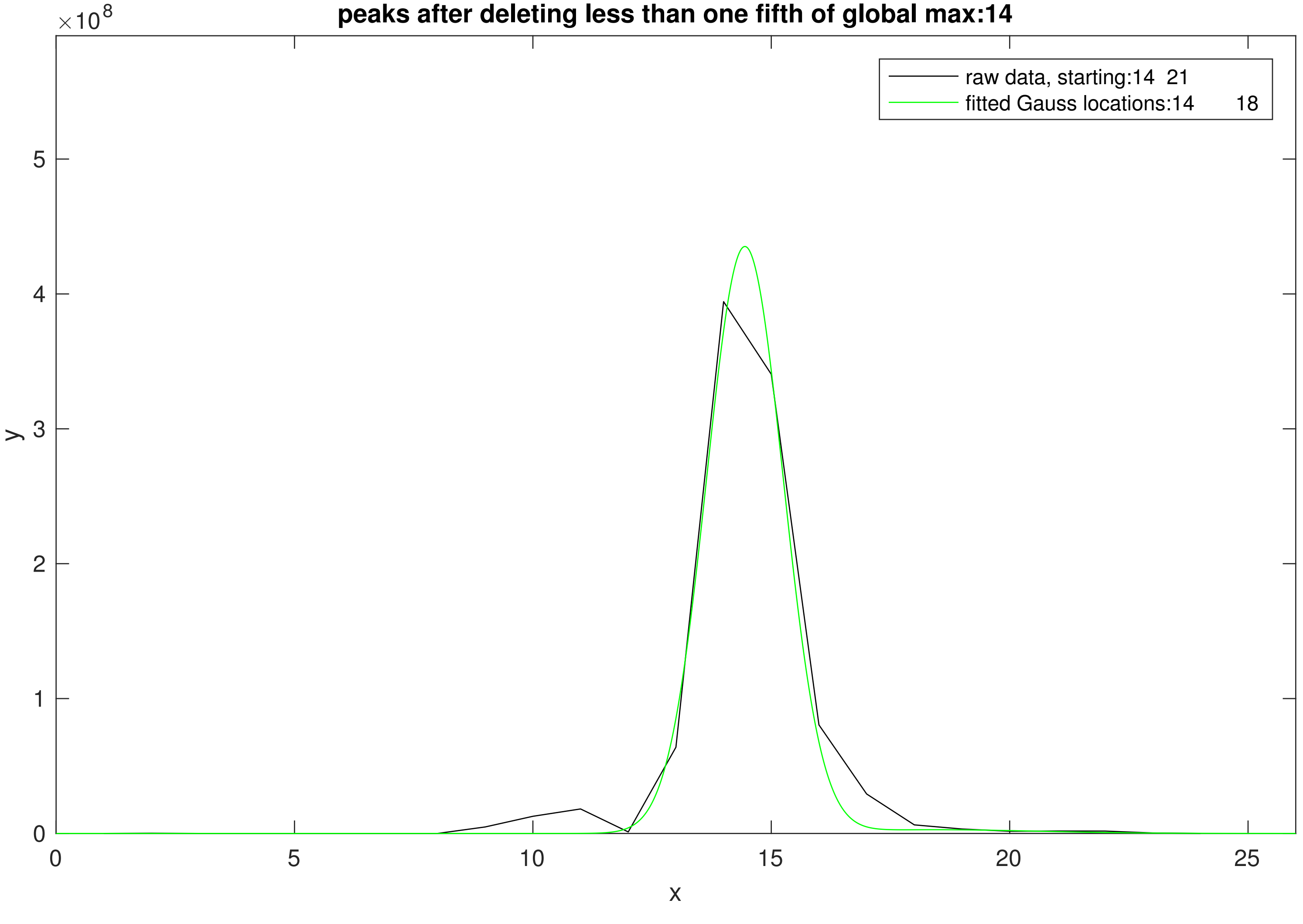

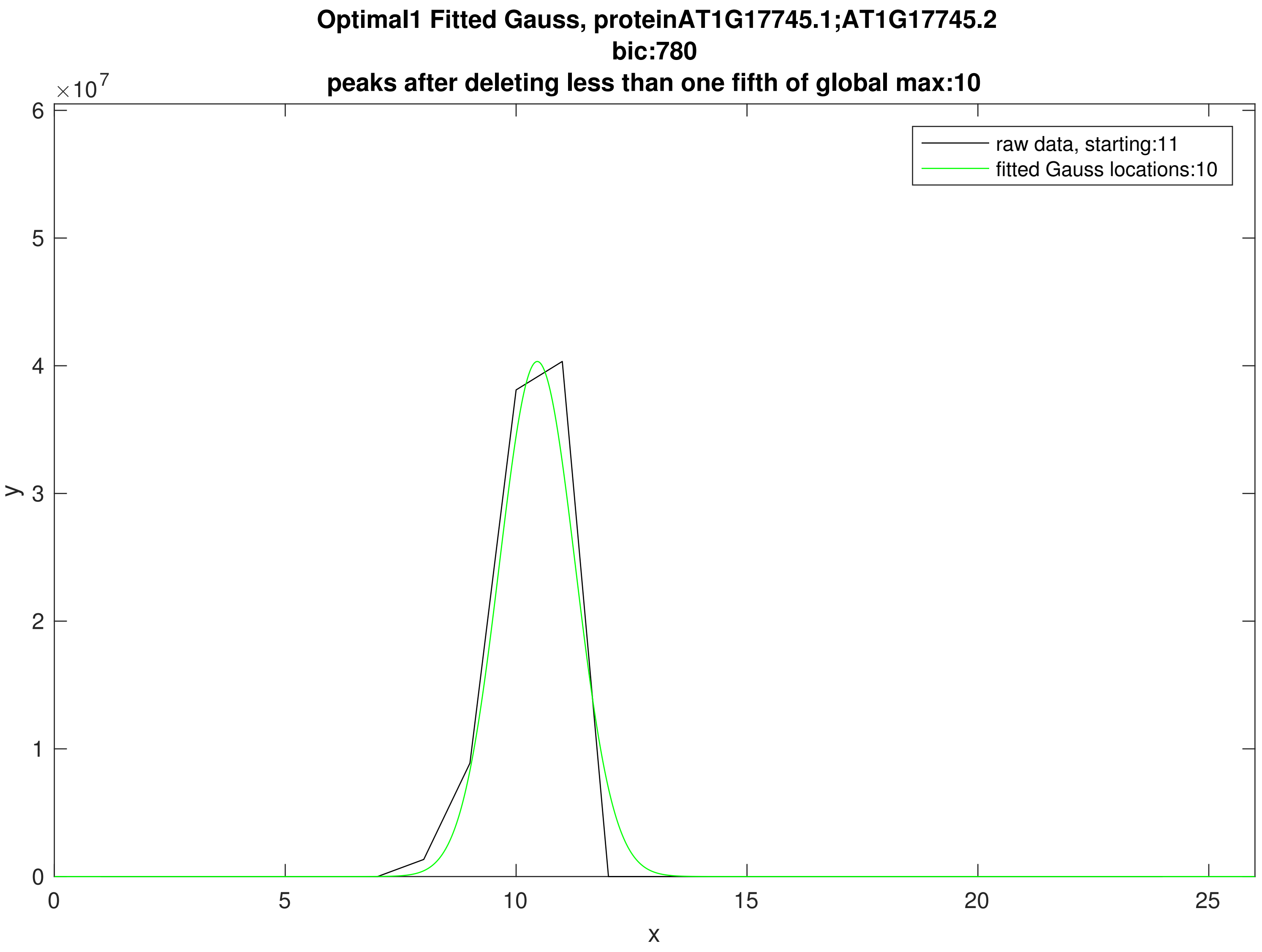
Optimal1 Fitted Gauss, proteinAT1G17745.1;AT1G17745.2
bic:780
peaks after deleting less than one fifth of global max:10
×10^7
6
5
4
3
2
1
0
y
0
5
10
15
20
25
x
raw data, starting:11
fitted Gauss locations:10

**Optimal1 Fitted Gauss, proteinAT1G17860.1**
**bic:685**
**peaks after deleting less than one fifth of global max:21**

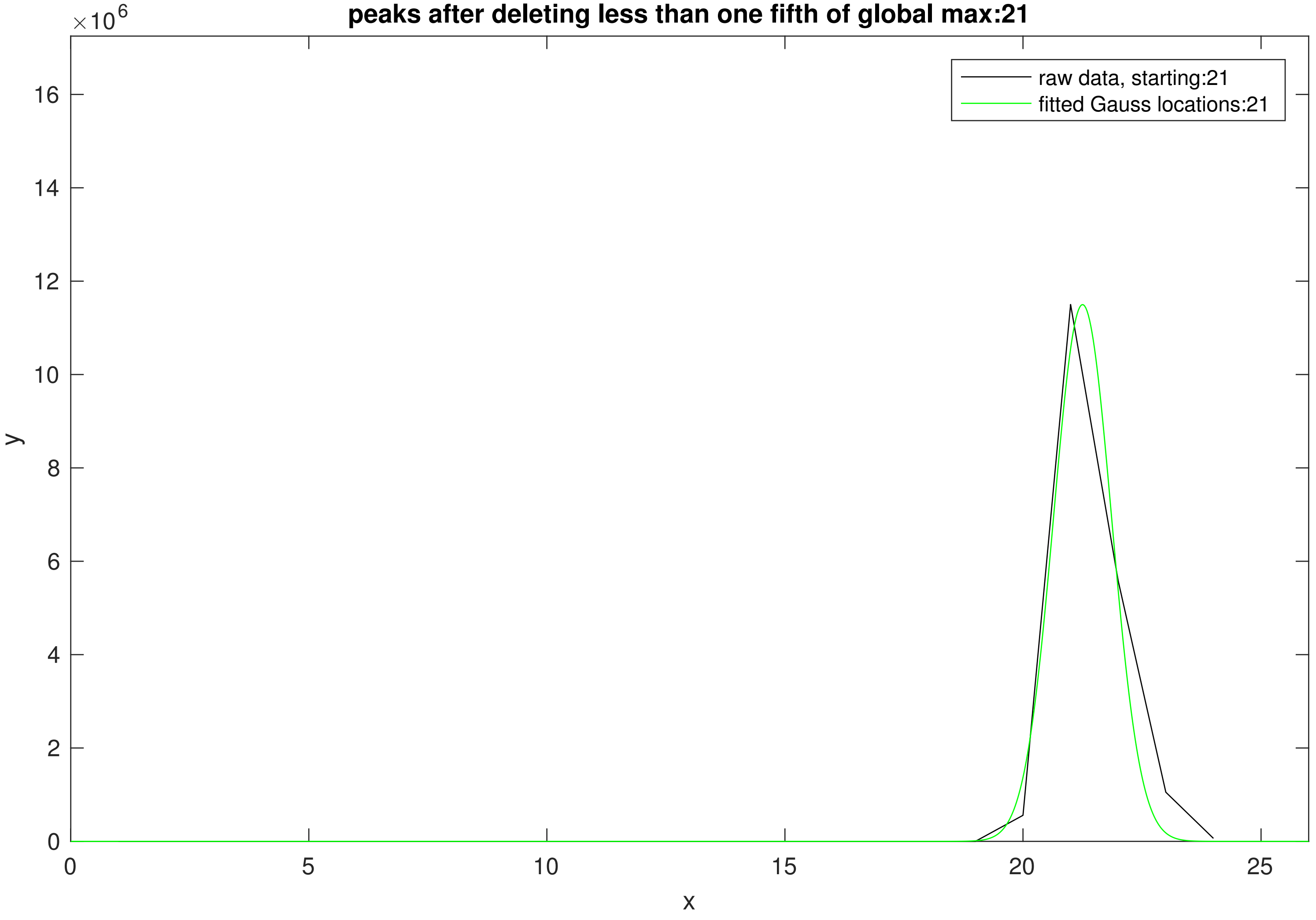

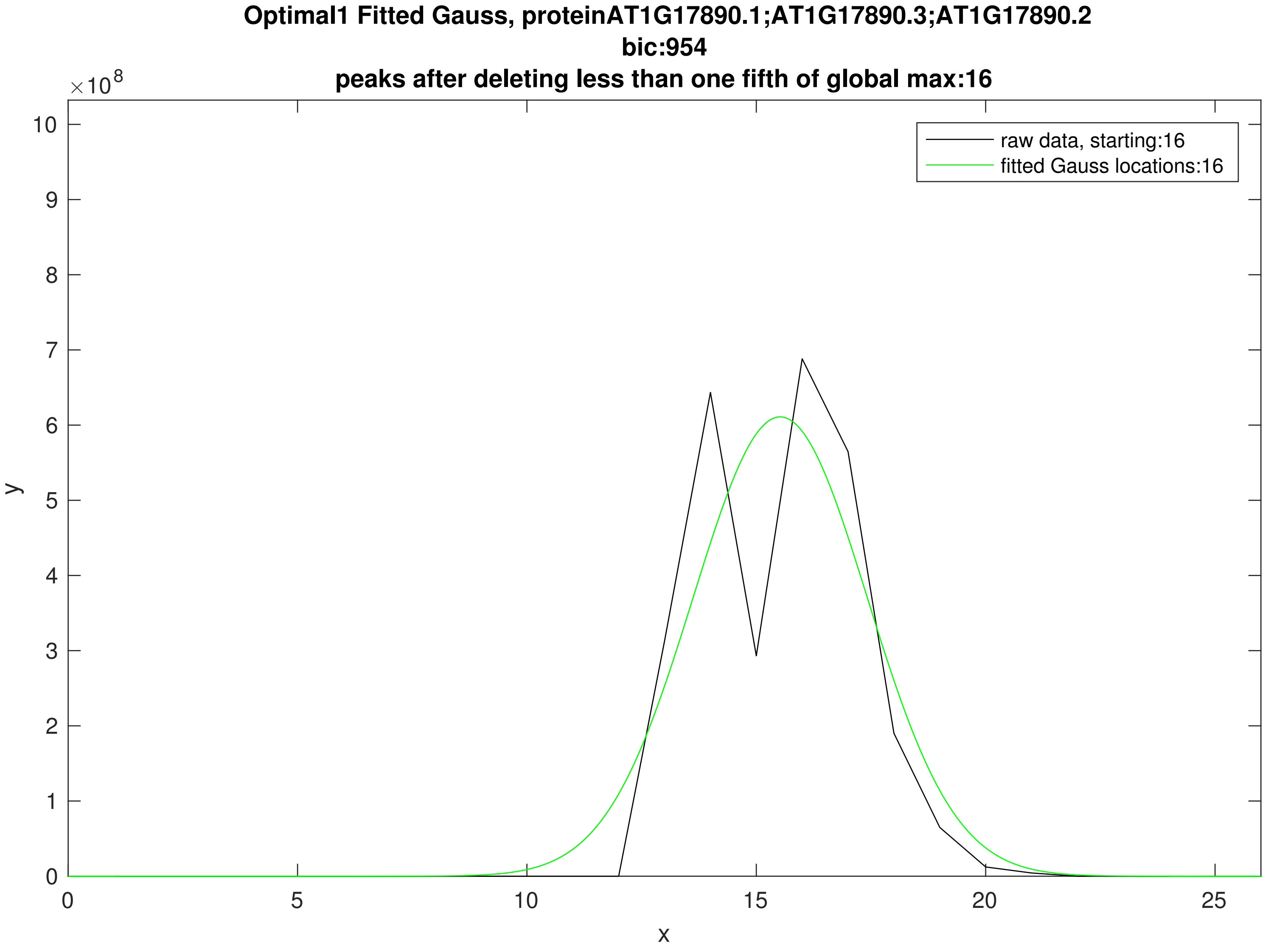
Optimal1 Fitted Gauss, proteinAT1G17890.1;AT1G17890.3;AT1G17890.2
bic:954
peaks after deleting less than one fifth of global max:16
×10^8
raw data, starting:16
fitted Gauss locations:16
y
x
0
1
2
3
4
5
6
7
8
9
10
0
5
10
15
20
25

**Optimal1 Fitted Gauss, proteinAT1G18070.1;AT1G18070.2;AT1G18070.3**
**bic:742 746**
**peaks after deleting less than one fifth of global max:12**

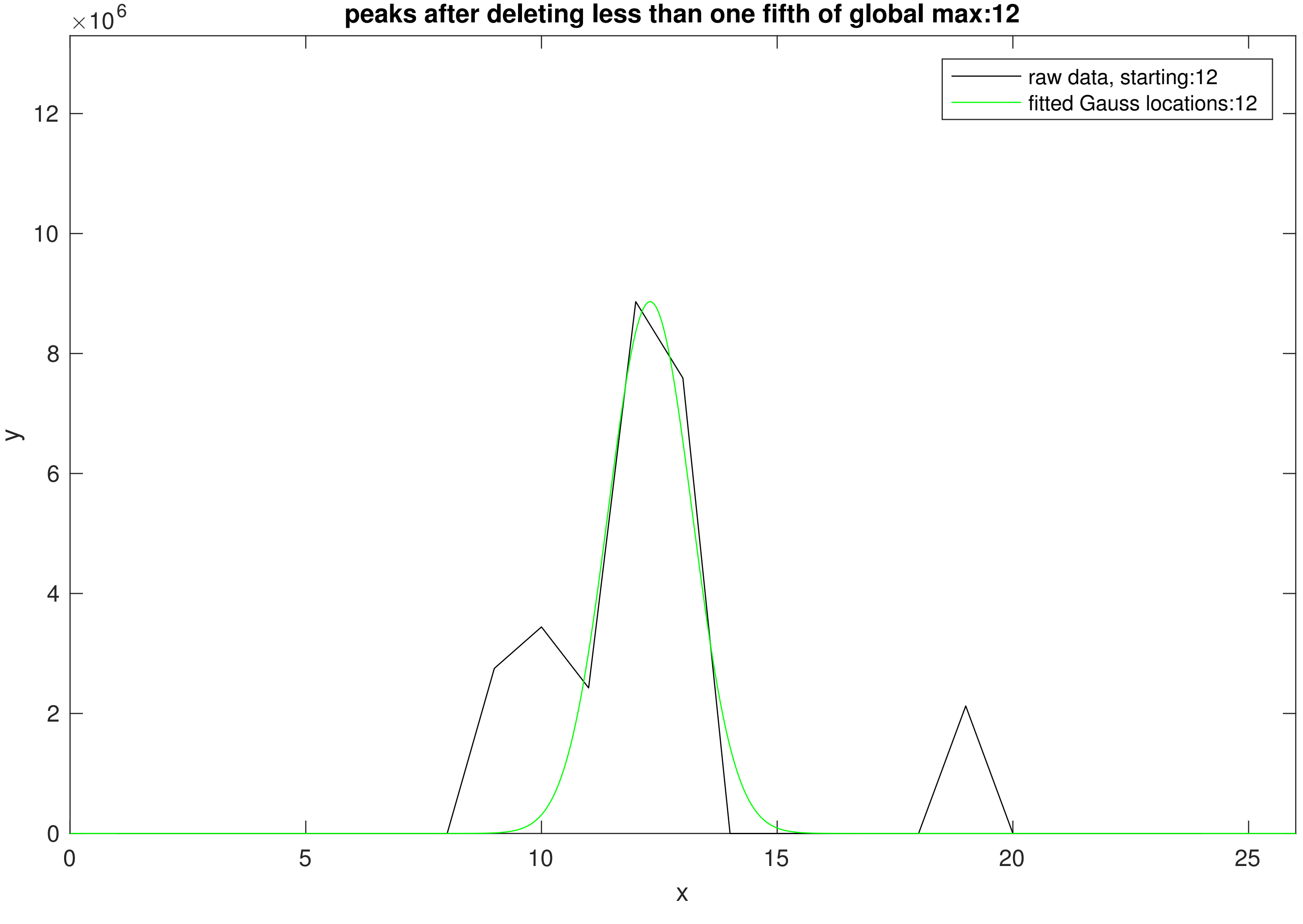

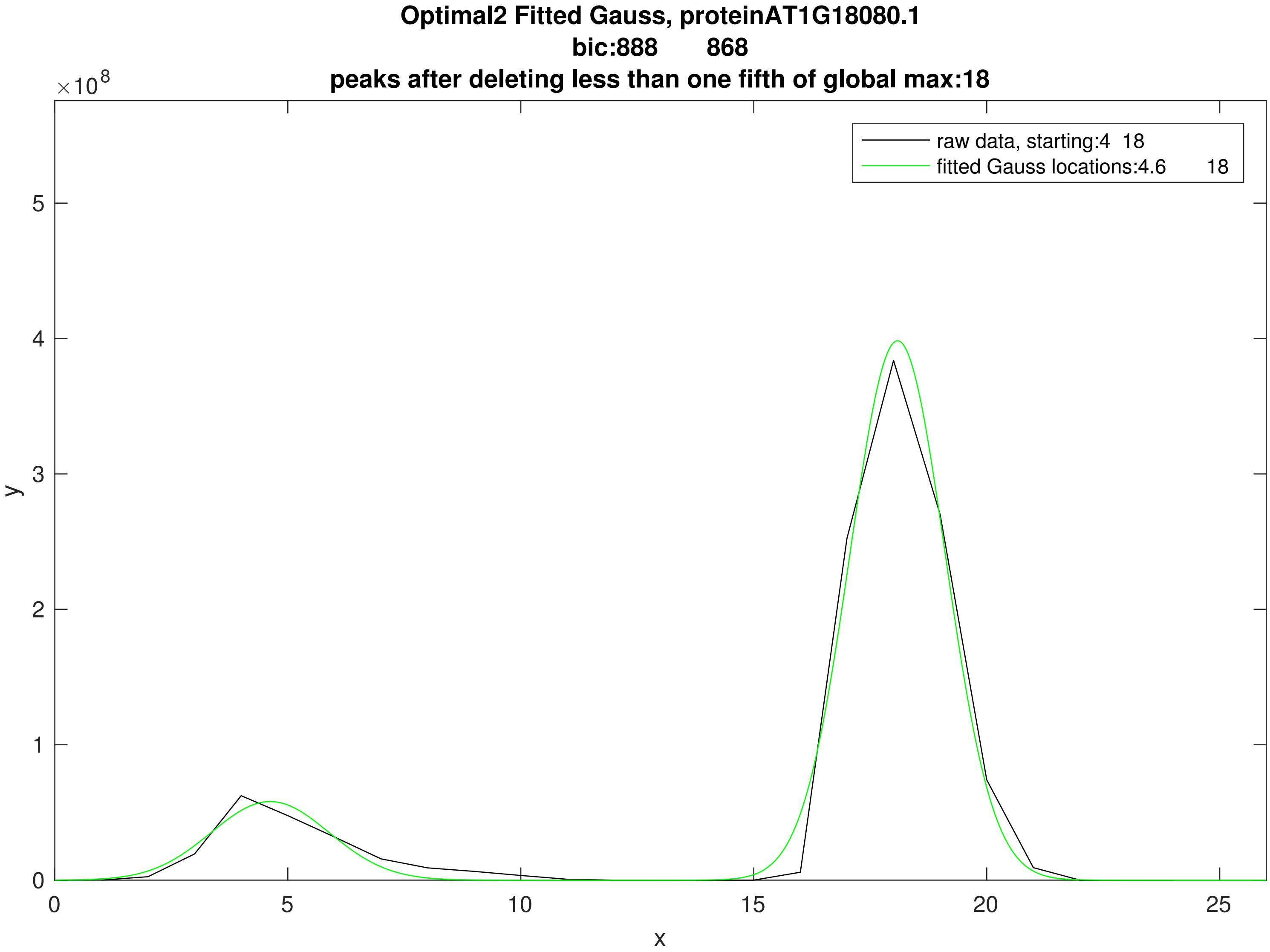
Optimal2 Fitted Gauss, proteinAT1G18080.1
bic:888 868
peaks after deleting less than one fifth of global max:18
×10^8
raw data, starting:4 18
fitted Gauss locations:4.6 18
y
x
0
1
2
3
4
5
0
5
10
15
20
25

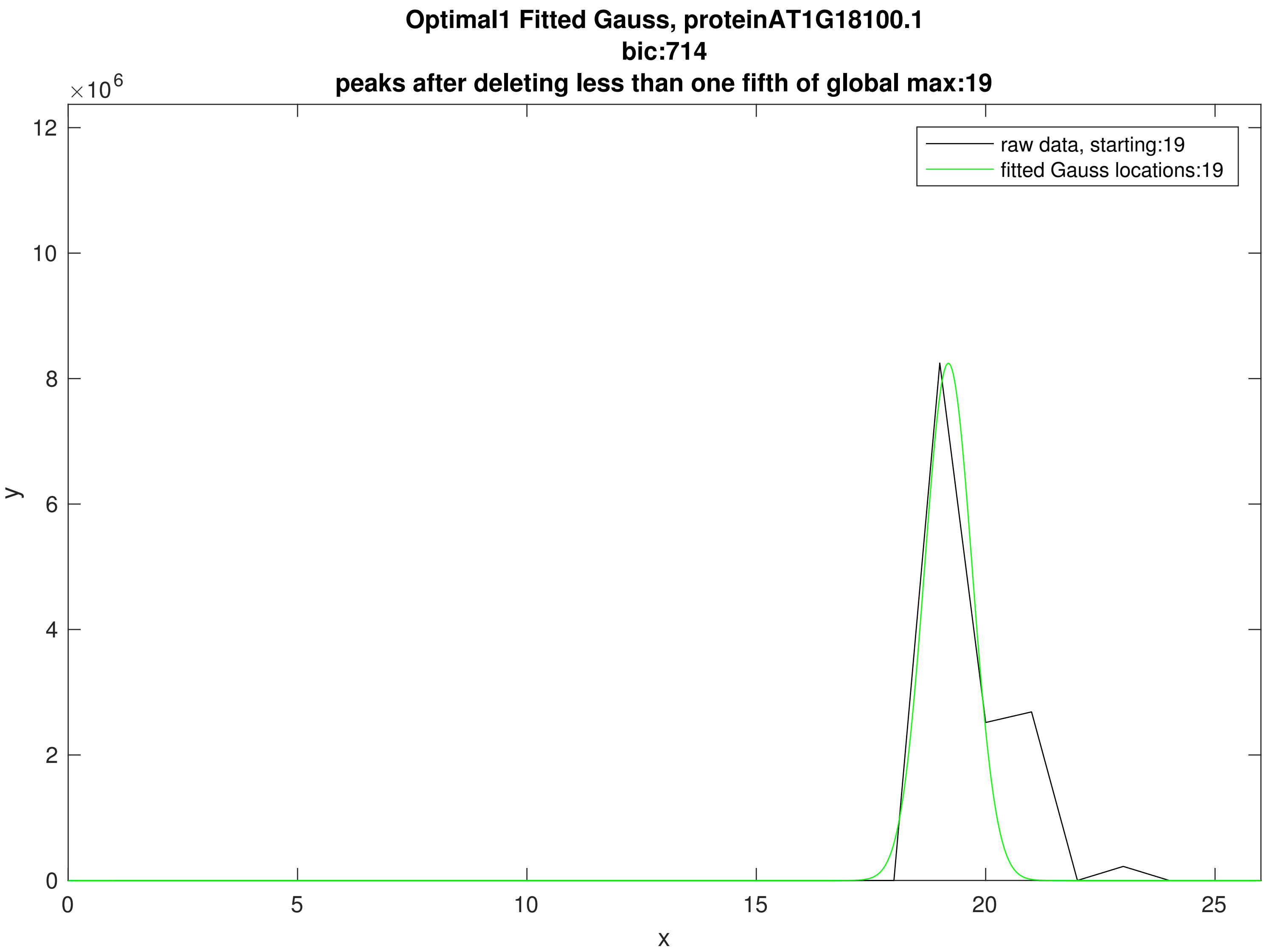

Optimal1 Fitted Gauss, proteinAT1G18100.1
bic:714
peaks after deleting less than one fifth of global max:19
×10^6
raw data, starting:19
fitted Gauss locations:19
y
x
0
2
4
6
8
10
12
0
5
10
15
20
25

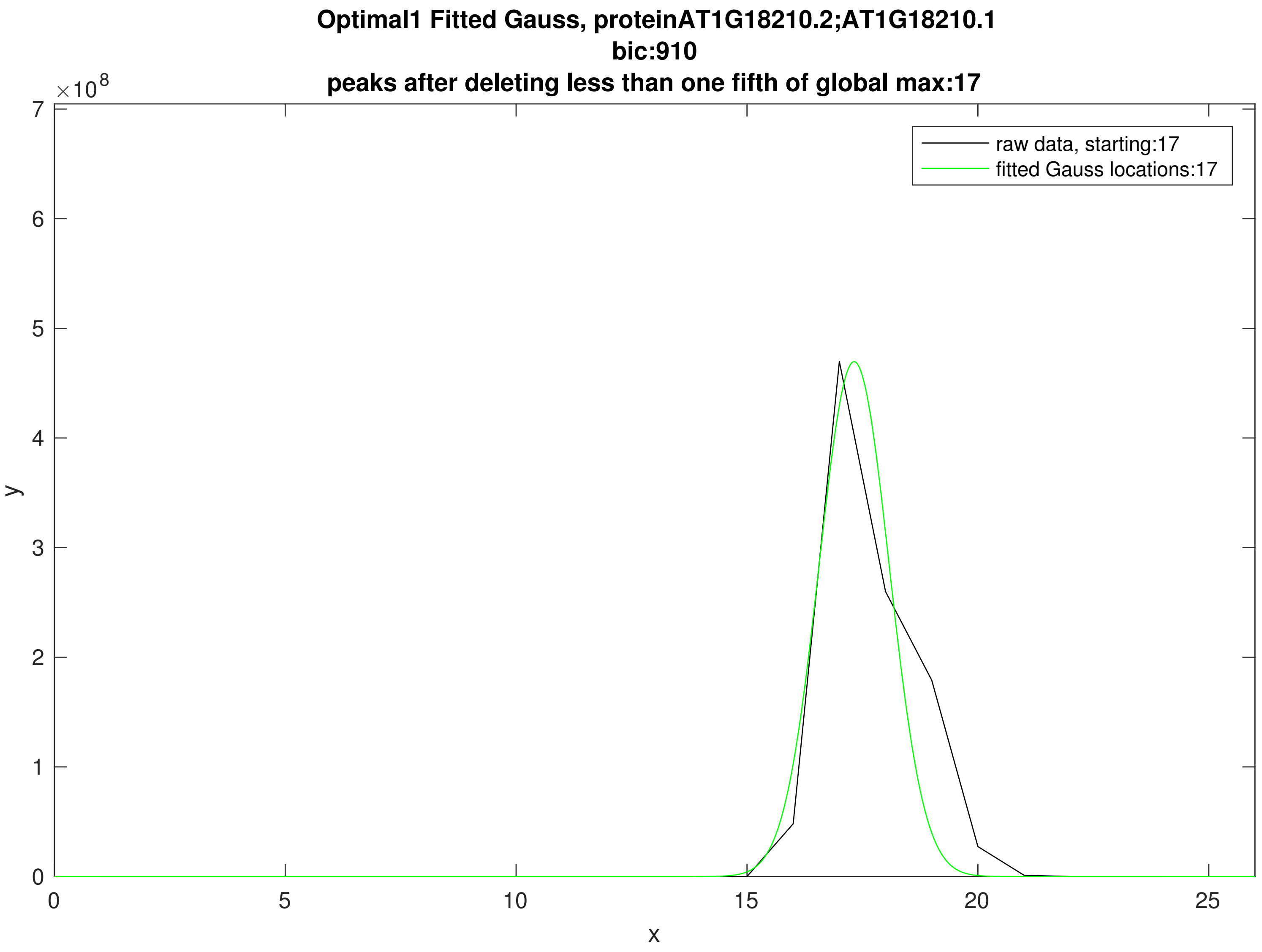

Optimal1 Fitted Gauss, proteinAT1G18210.2;AT1G18210.1
bic:910
peaks after deleting less than one fifth of global max:17
×10^8
raw data, starting:17
fitted Gauss locations:17
y
x

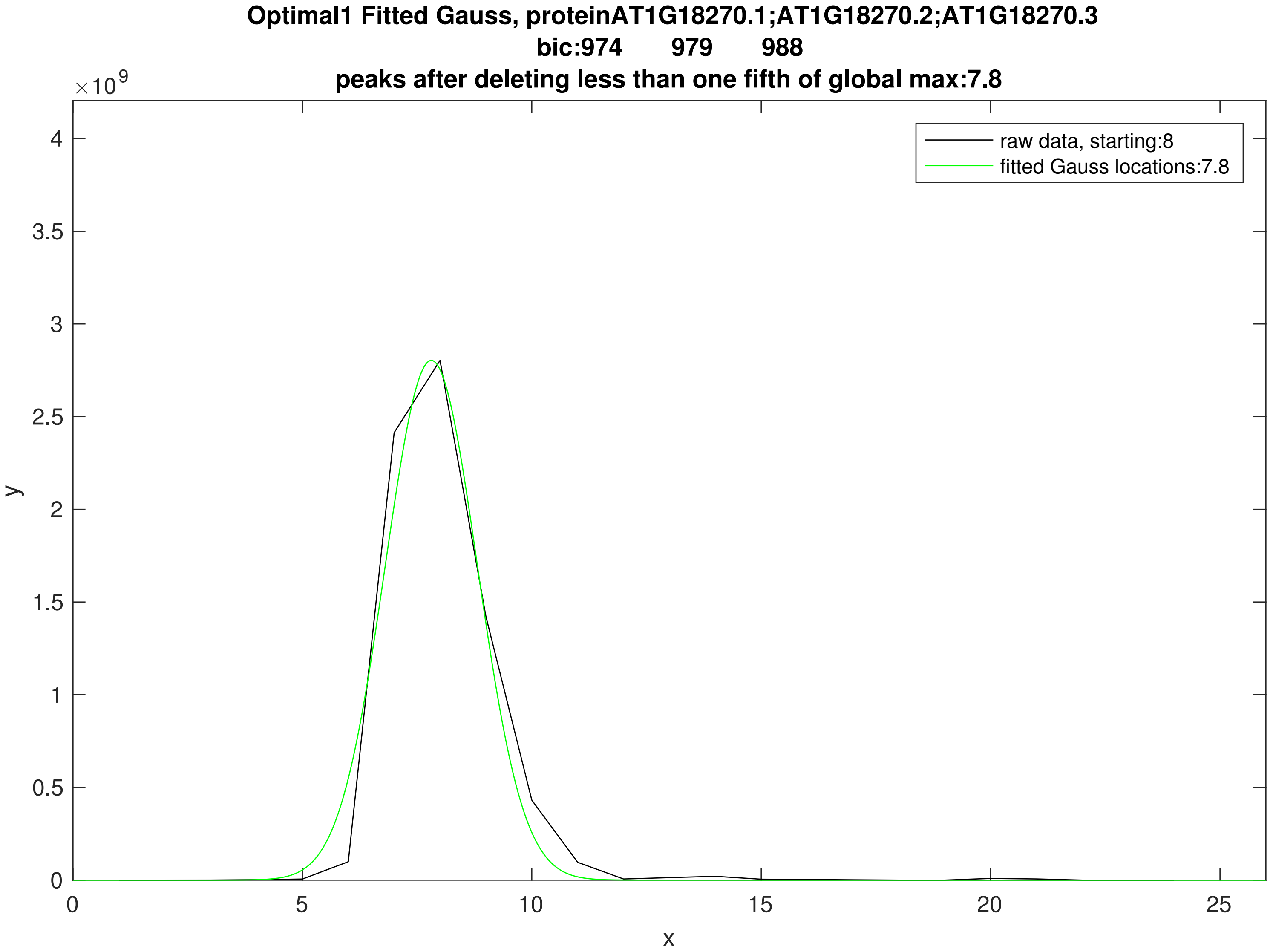

Optimal1 Fitted Gauss, proteinAT1G18270.1;AT1G18270.2;AT1G18270.3
bic:974 979 988
peaks after deleting less than one fifth of global max:7.8
×10^9
raw data, starting:8
fitted Gauss locations:7.8
y
x
0
0.5
1
1.5
2
2.5
3
3.5
4
5
10
15
20
25

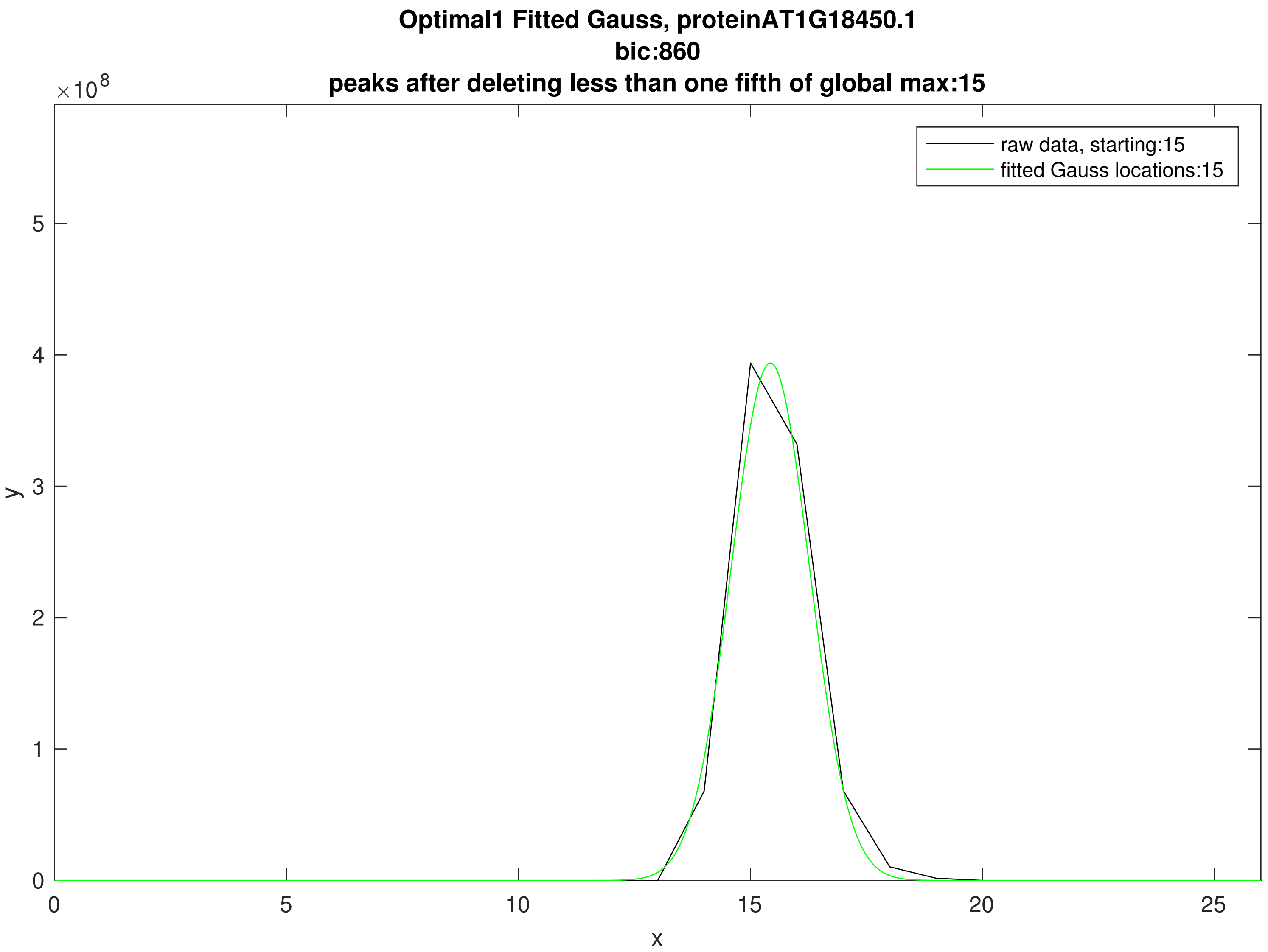
Optimal1 Fitted Gauss, proteinAT1G18450.1
bic:860
peaks after deleting less than one fifth of global max:15
×10^8
raw data, starting:15
fitted Gauss locations:15
y
x
0
1
2
3
4
5
0
5
10
15
20
25

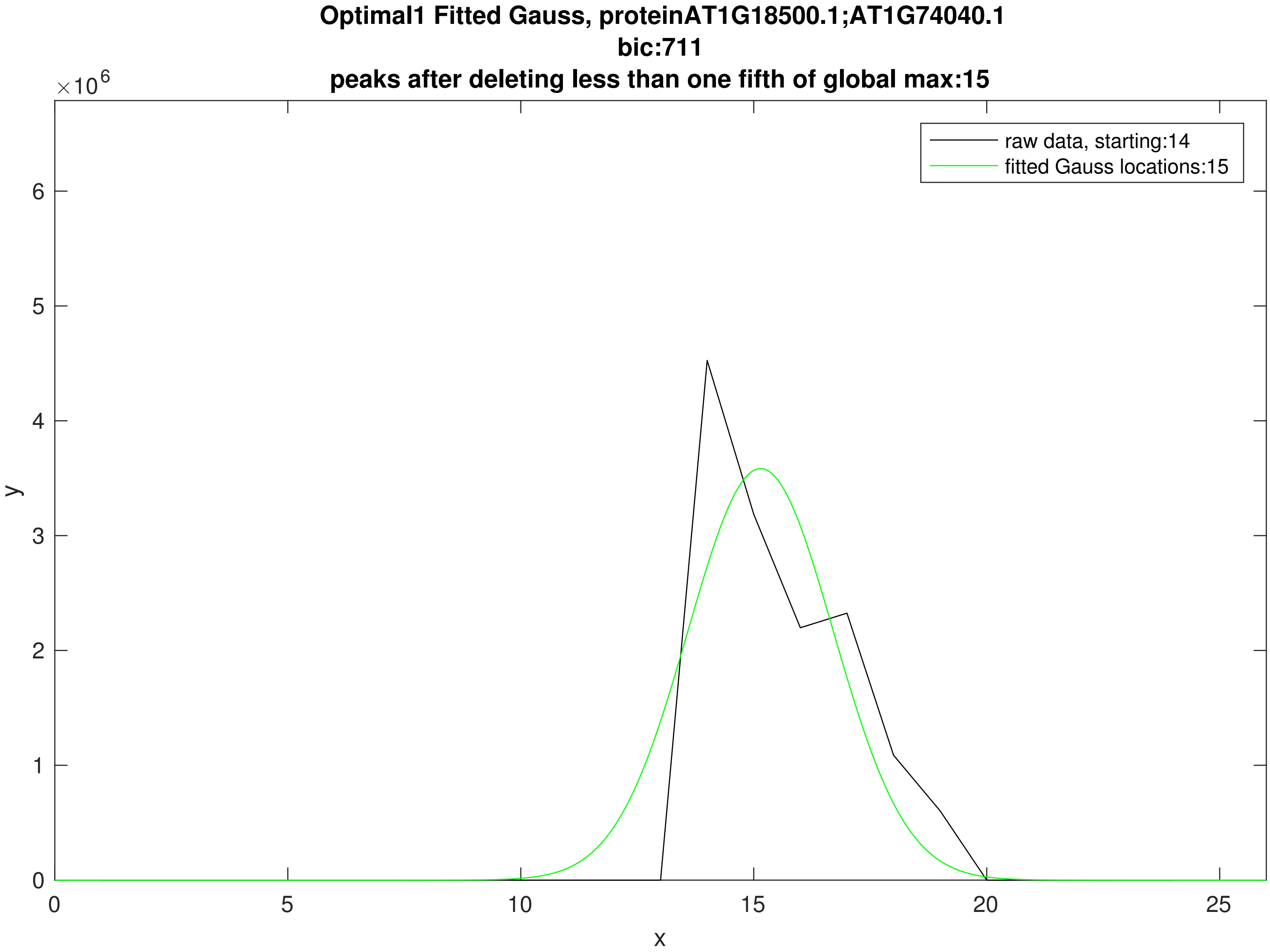

Optimal1 Fitted Gauss, proteinAT1G18500.1;AT1G74040.1
bic:711
peaks after deleting less than one fifth of global max:15
×10^6
raw data, starting:14
fitted Gauss locations:15
y
x
0
1
2
3
4
5
6
0
5
10
15
20
25

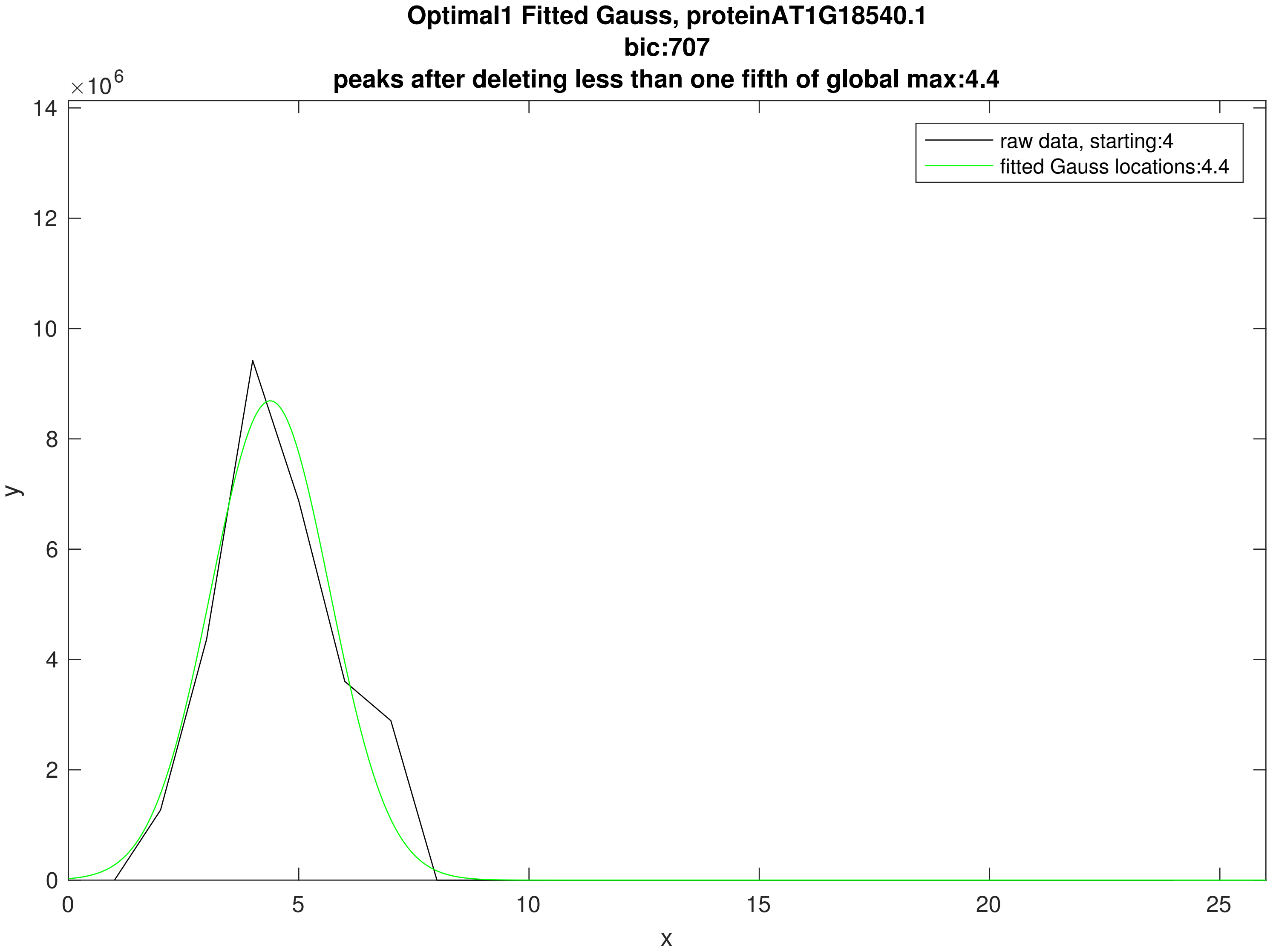
Optimal1 Fitted Gauss, proteinAT1G18540.1
bic:707
peaks after deleting less than one fifth of global max:4.4
×10^6
raw data, starting:4
fitted Gauss locations:4.4
y
x
0
2
4
6
8
10
12
14
0
5
10
15
20
25

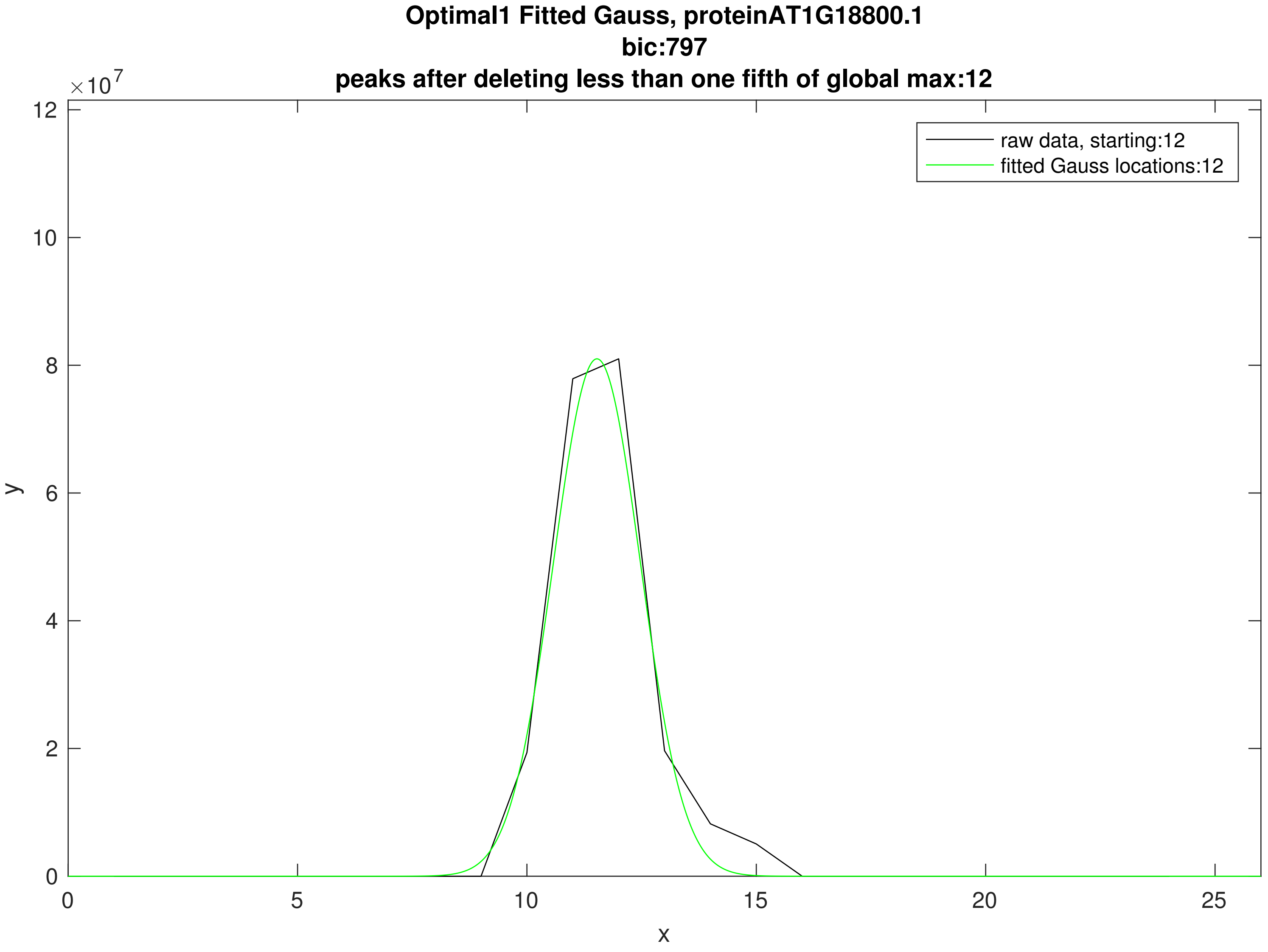

Optimal1 Fitted Gauss, proteinAT1G18800.1
bic:797
peaks after deleting less than one fifth of global max:12
×10^7
12
10
8
6
4
2
0
y
0
5
10
15
20
25
x
raw data, starting:12
fitted Gauss locations:12

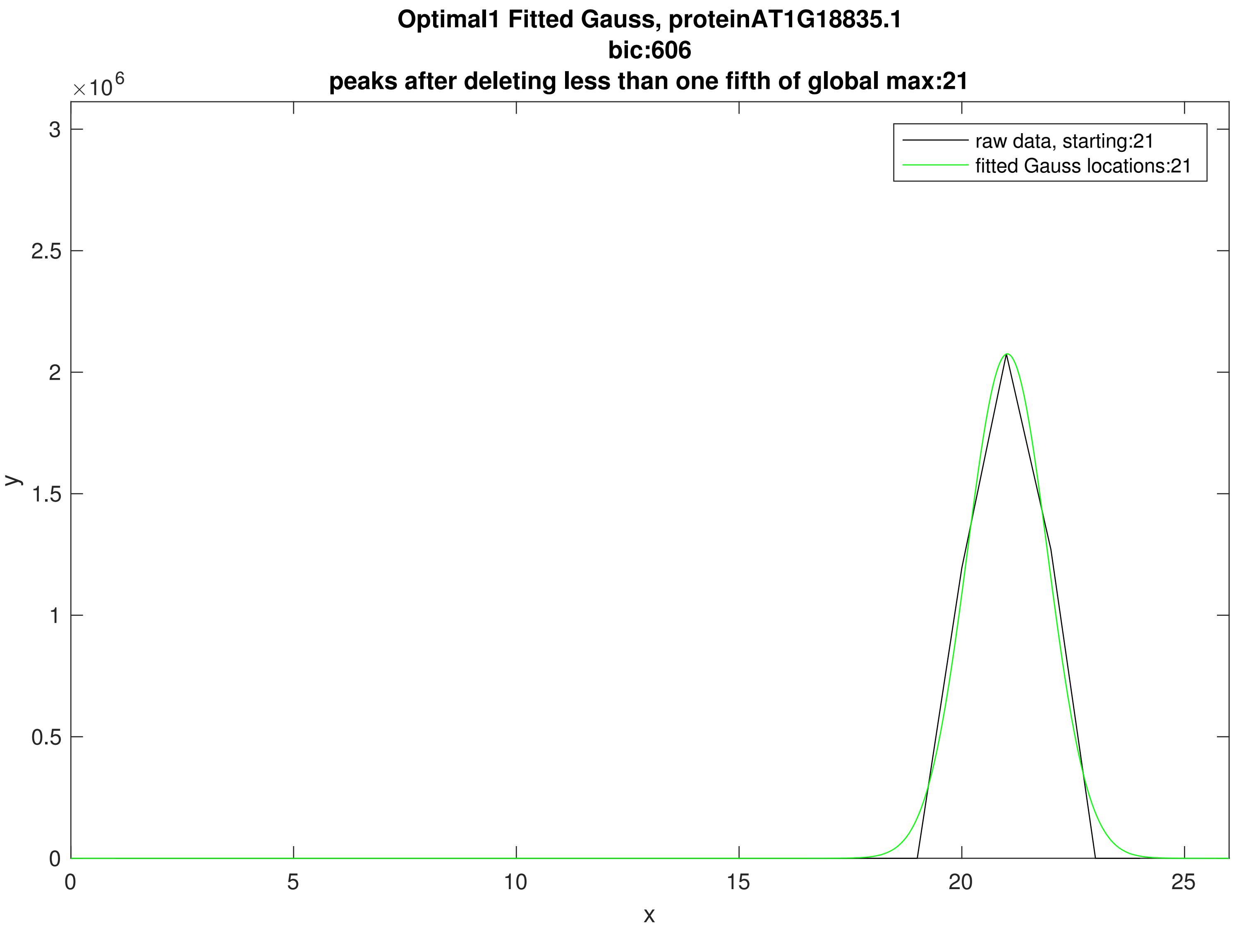
Optimal1 Fitted Gauss, proteinAT1G18835.1
bic:606
peaks after deleting less than one fifth of global max:21
×10^6
raw data, starting:21
fitted Gauss locations:21
y
x
0
0.5
1
1.5
2
2.5
3
0
5
10
15
20
25

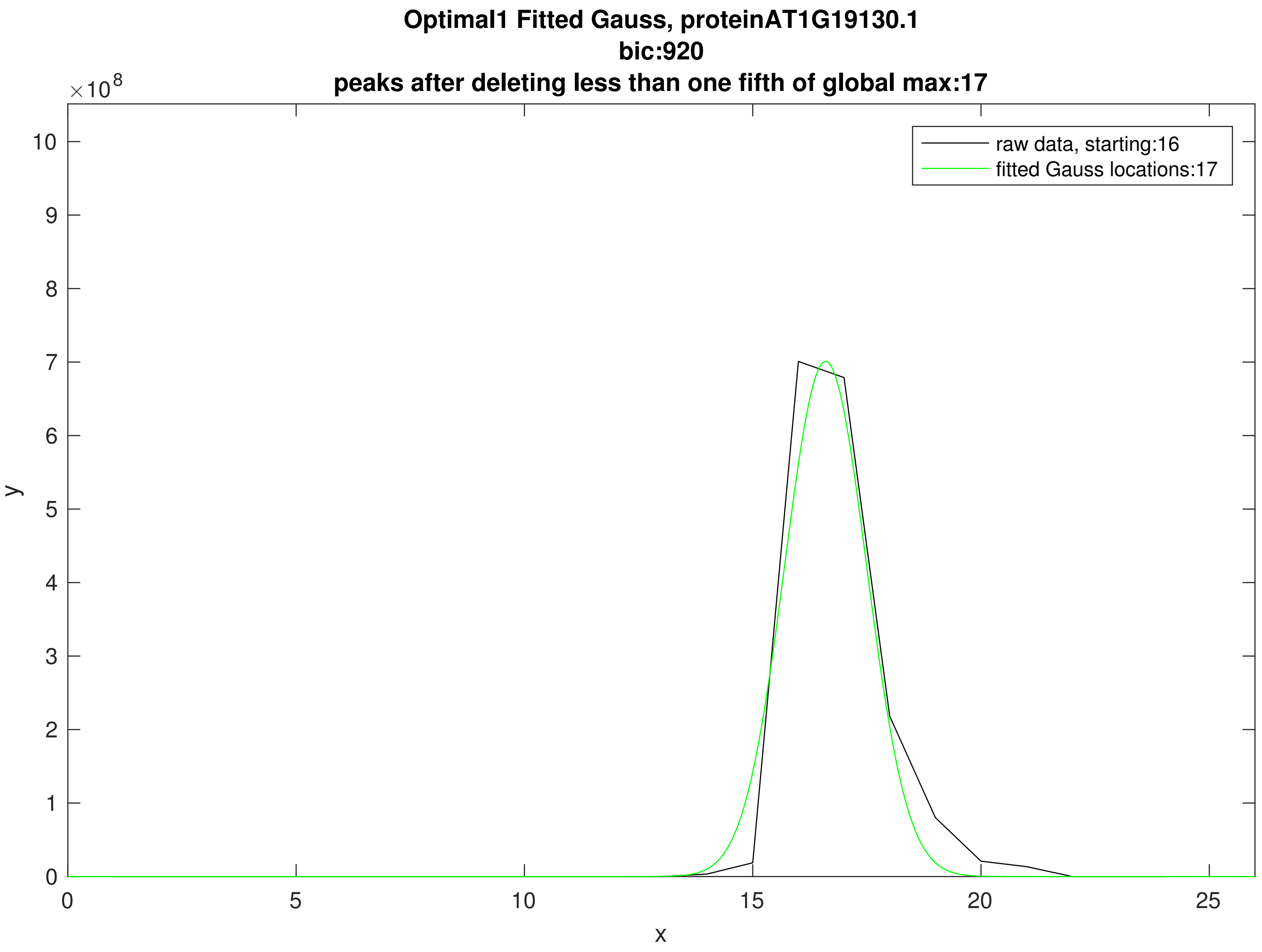

Optimal1 Fitted Gauss, proteinAT1G19130.1
bic:920
peaks after deleting less than one fifth of global max:17
×10^8
raw data, starting:16
fitted Gauss locations:17
y
x
0
1
2
3
4
5
6
7
8
9
10
0
5
10
15
20
25

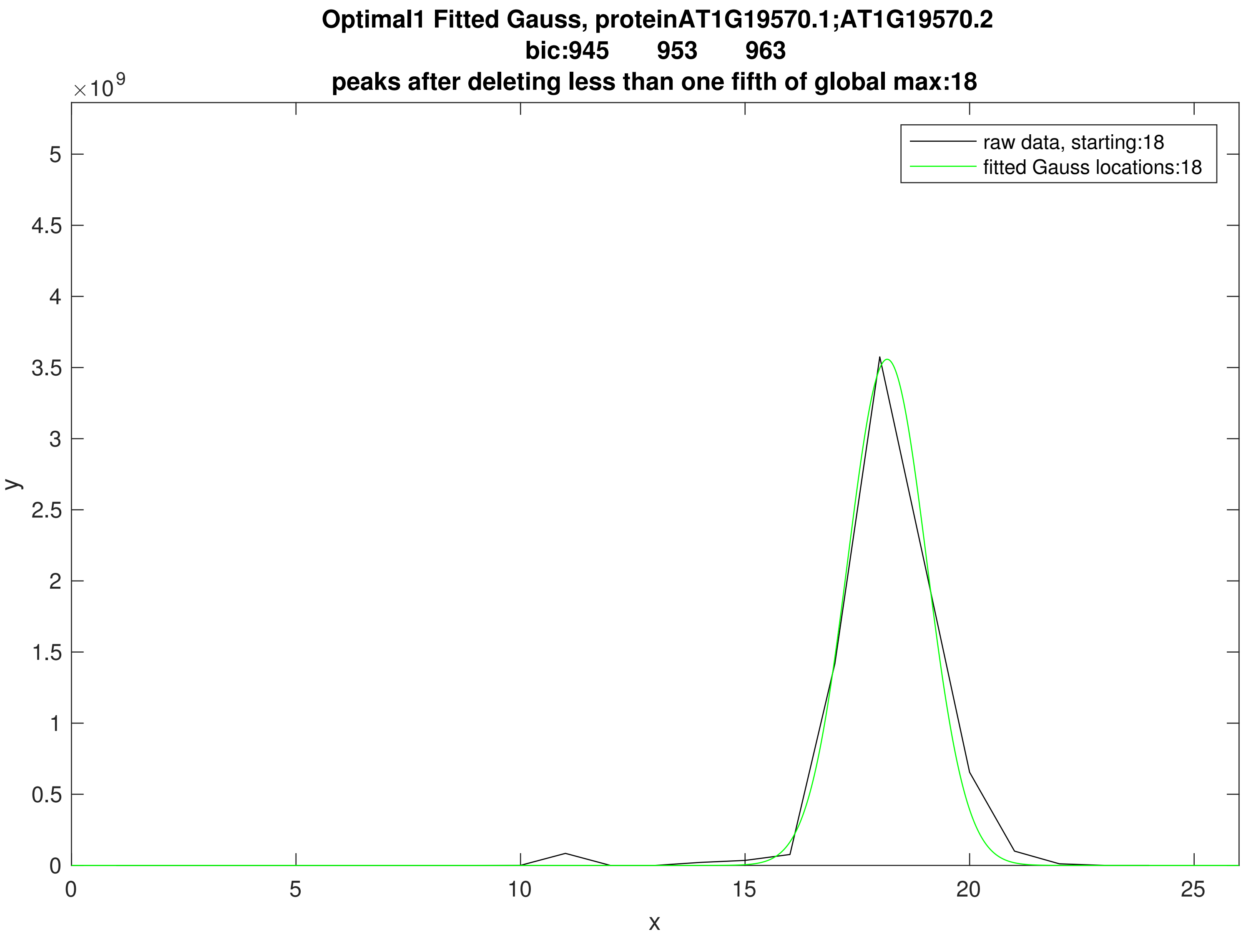
Optimal1 Fitted Gauss, proteinAT1G19570.1;AT1G19570.2
bic:945 953 963
peaks after deleting less than one fifth of global max:18
×10^9
raw data, starting:18
fitted Gauss locations:18
y
x
0
0.5
1
1.5
2
2.5
3
3.5
4
4.5
5
0
5
10
15
20
25

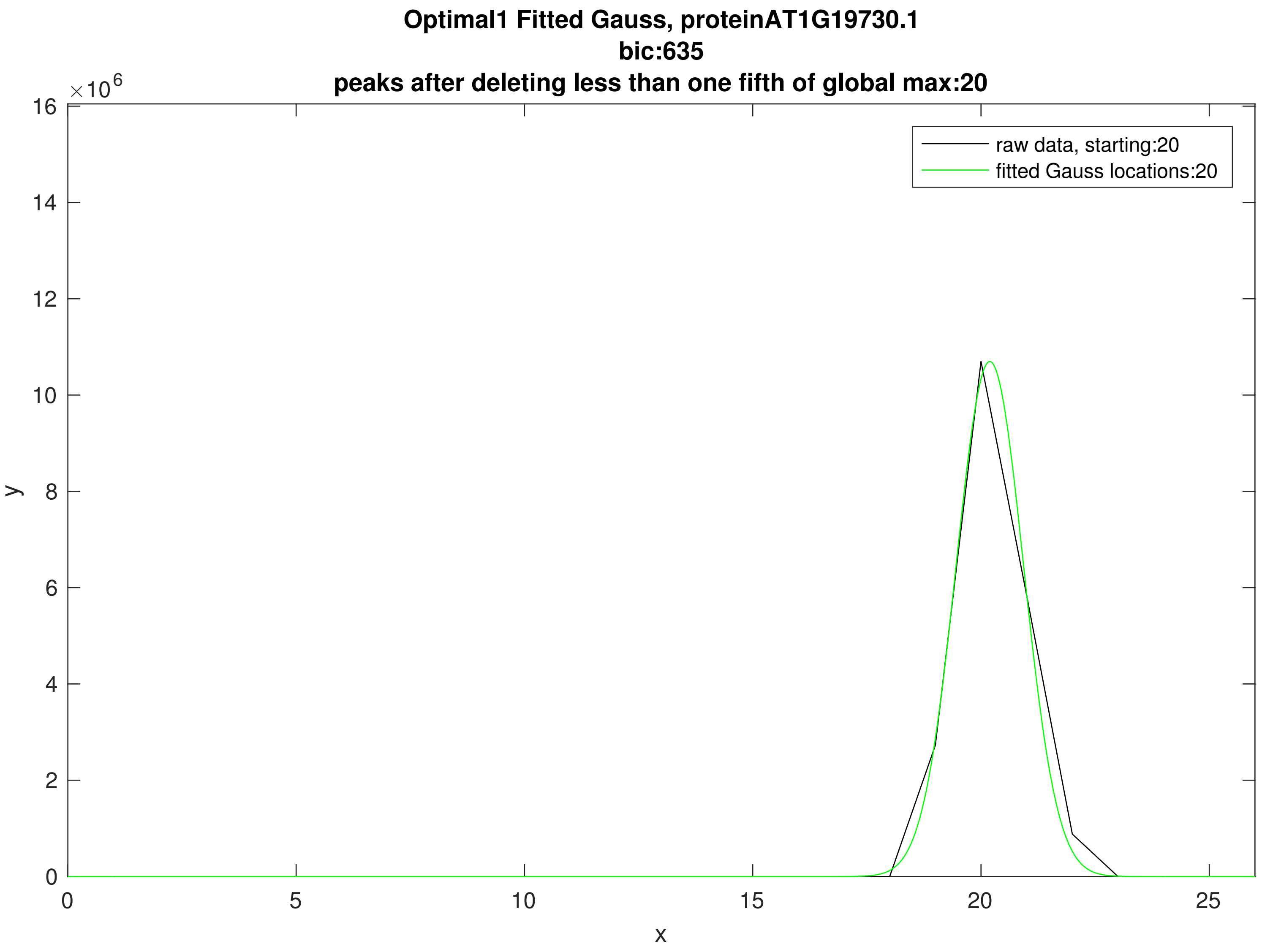
Optimal1 Fitted Gauss, proteinAT1G19730.1
bic:635
peaks after deleting less than one fifth of global max:20
×10^6
16
14
12
10
8
6
4
2
0
y
0
5
10
15
20
25
x
raw data, starting:20
fitted Gauss locations:20

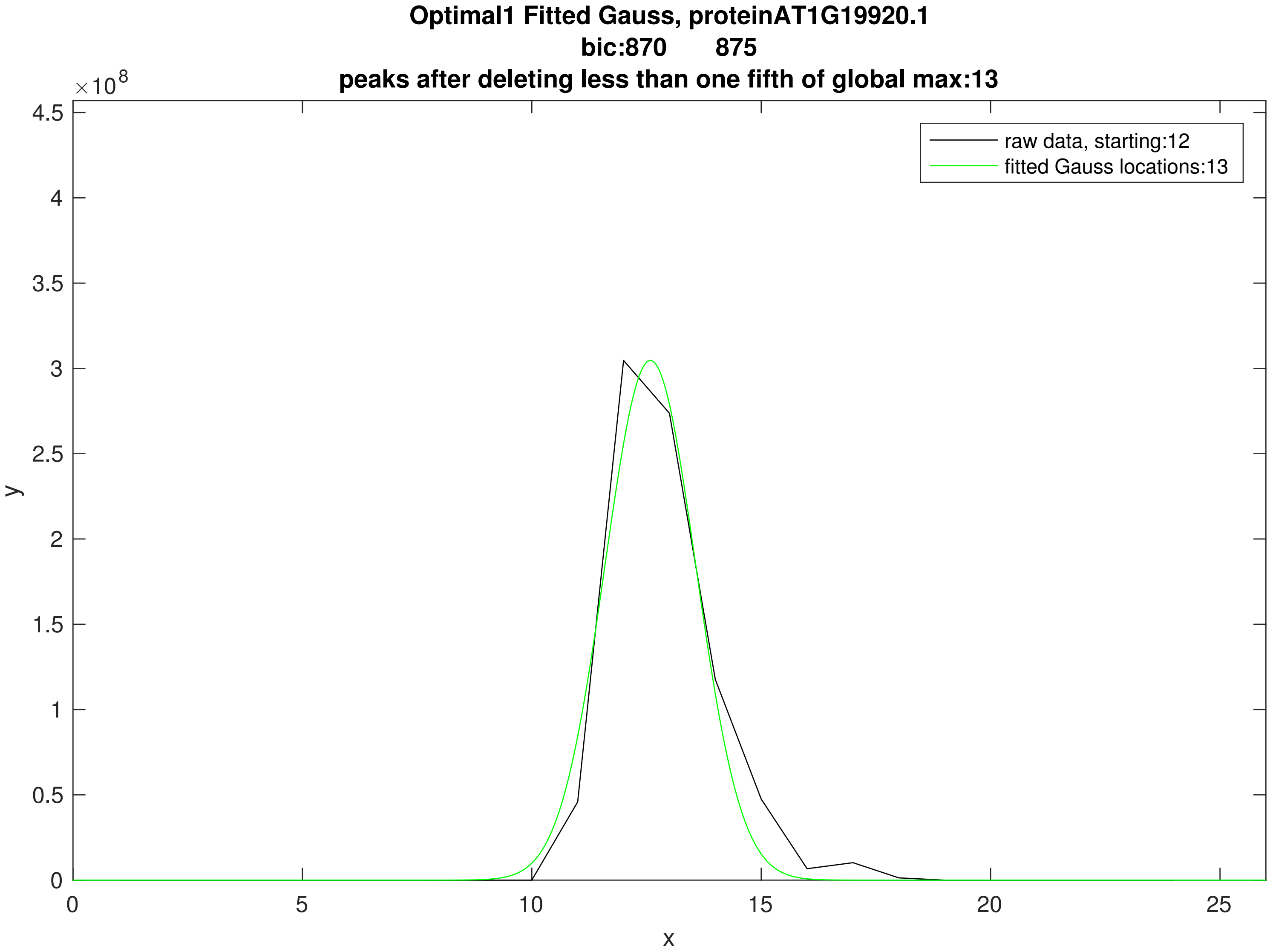

Optimal1 Fitted Gauss, proteinAT1G19920.1
bic:870 875
peaks after deleting less than one fifth of global max:13
×10^8
4.5
4
3.5
3
2.5
2
1.5
1
0.5
0
y
0
5
10
15
20
25
x
raw data, starting:12
fitted Gauss locations:13

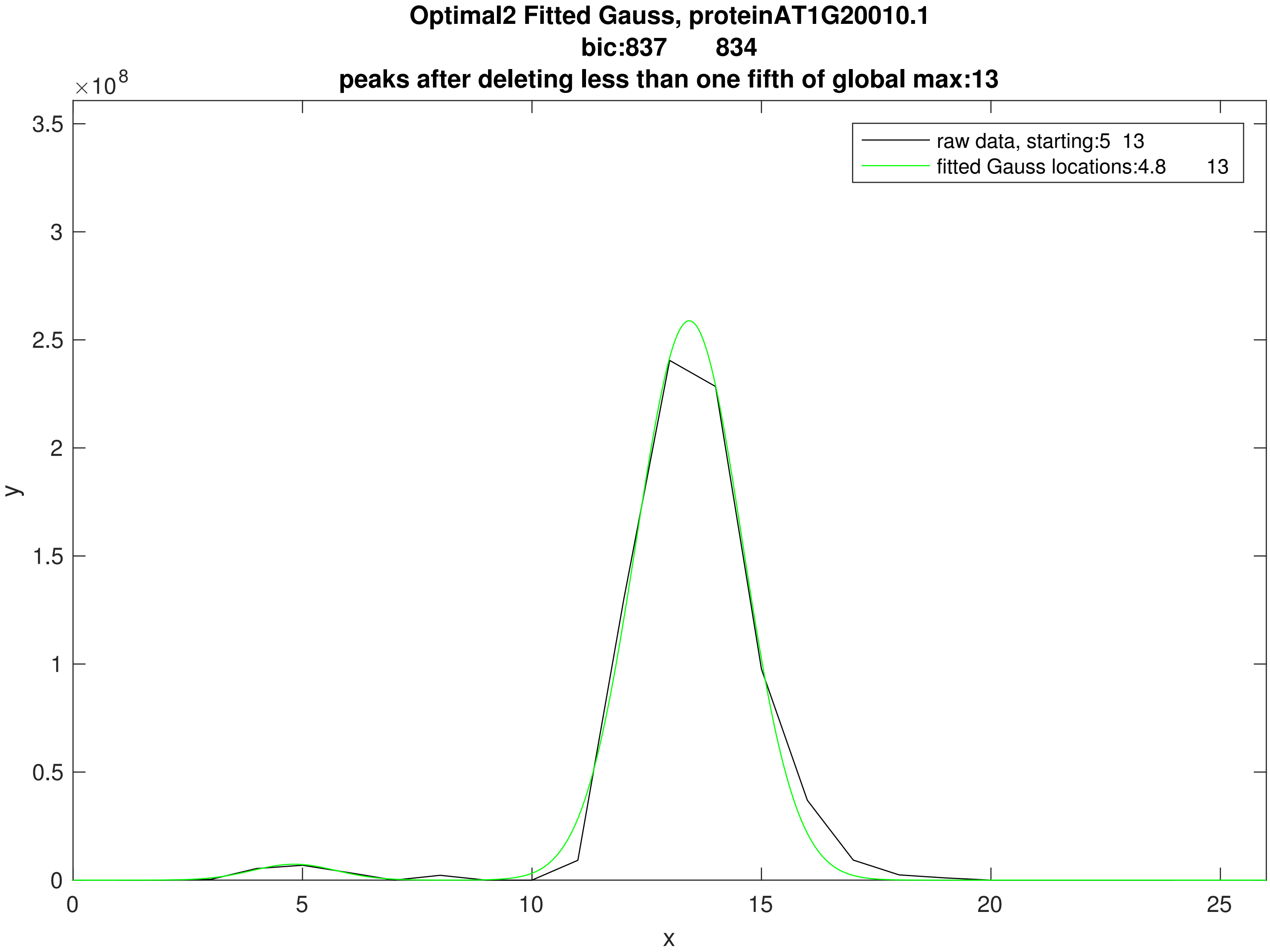
Optimal2 Fitted Gauss, proteinAT1G20010.1
bic:837 834
peaks after deleting less than one fifth of global max:13
×10^8
3.5
3
2.5
2
1.5
1
0.5
0
y
0
5
10
15
20
25
x
raw data, starting:5 13
fitted Gauss locations:4.8 13

#### Optimal2 Fitted Gauss, proteinAT1G20020.2;AT1G20020.1;AT1G20020.3
#### bic:750 734
#### peaks after deleting less than one fifth of global max:16

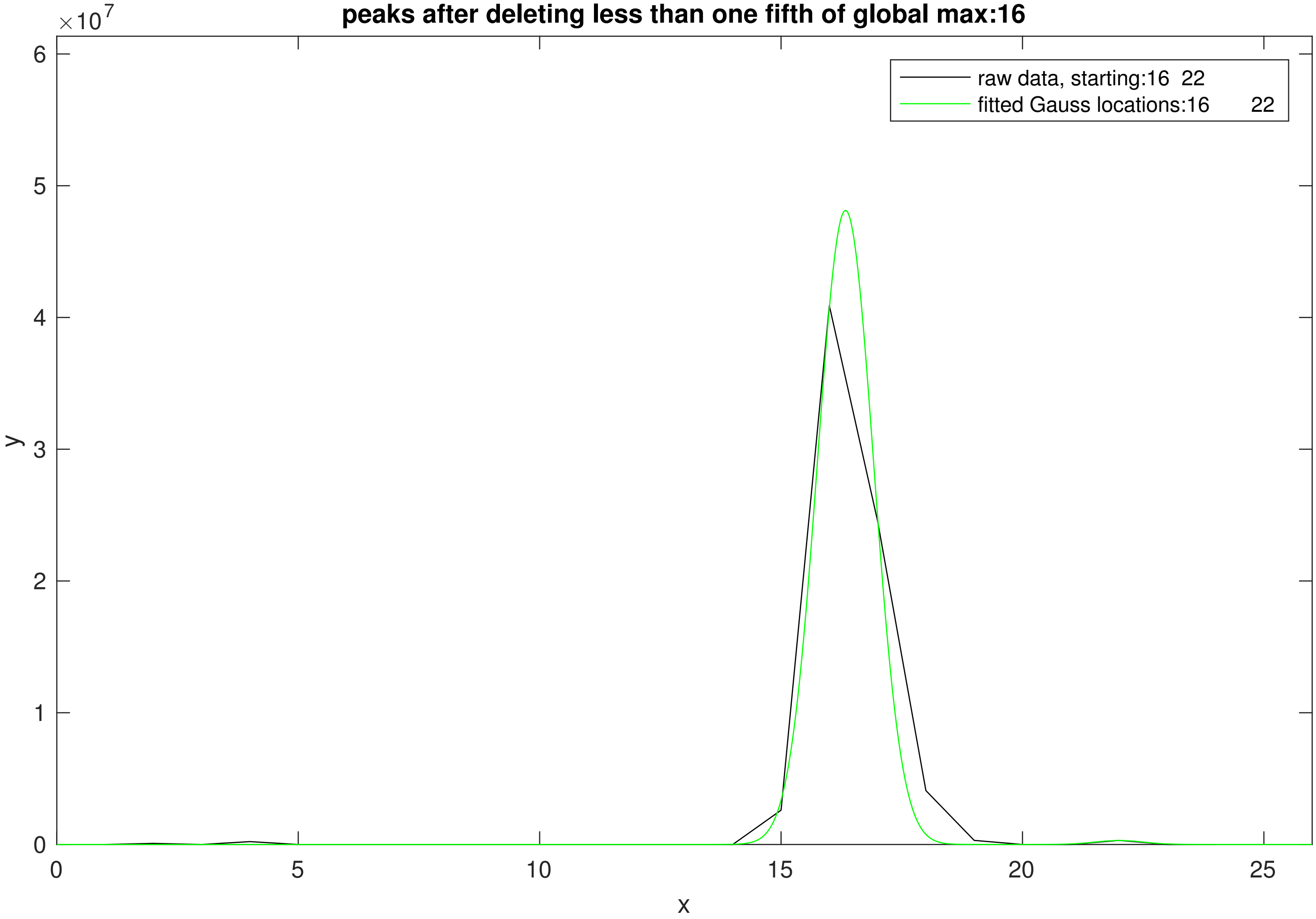

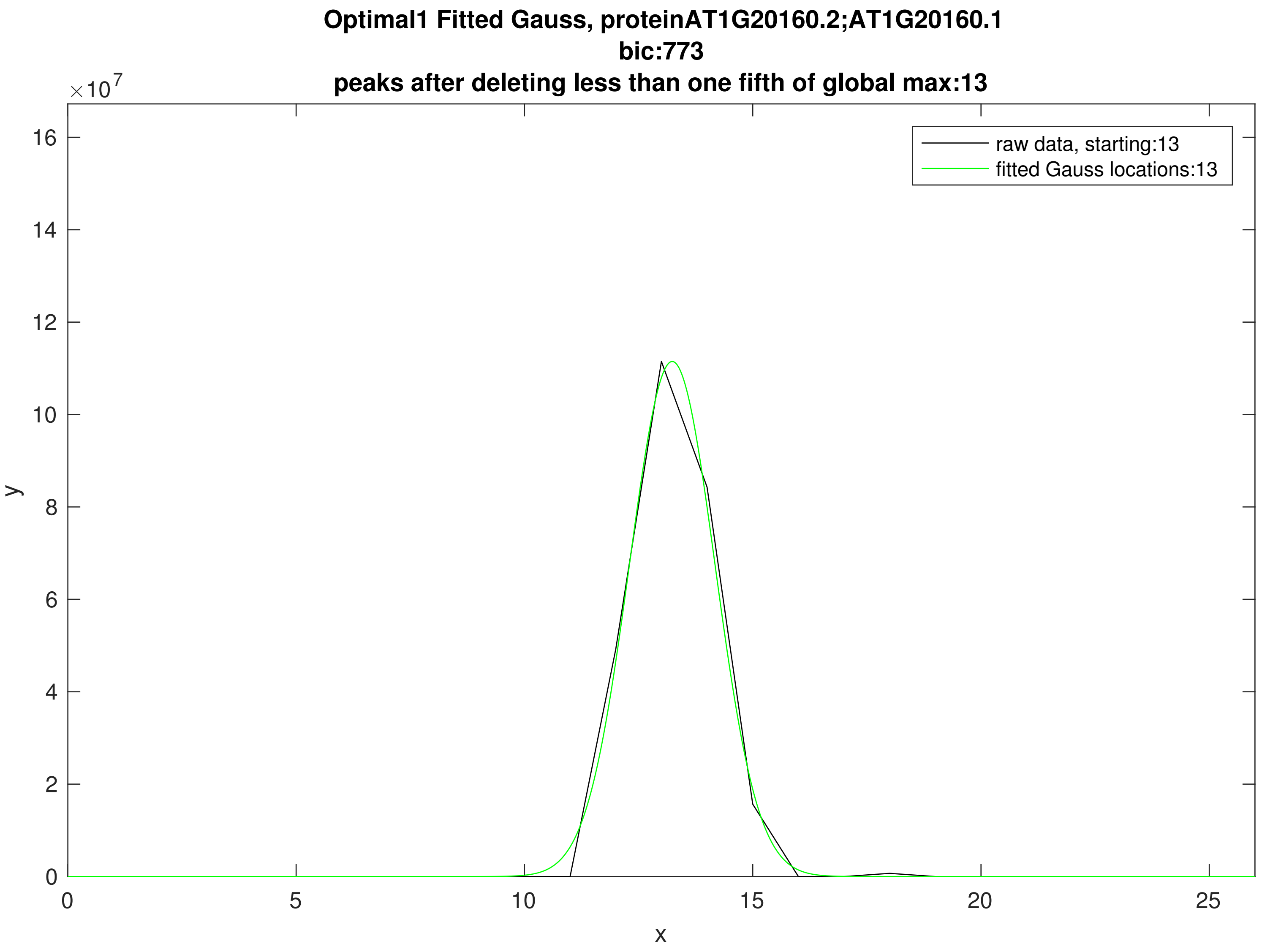
Optimal1 Fitted Gauss, proteinAT1G20160.2;AT1G20160.1
bic:773
peaks after deleting less than one fifth of global max:13
×10^7
raw data, starting:13
fitted Gauss locations:13
y
x
0
2
4
6
8
10
12
14
16
0
5
10
15
20
25

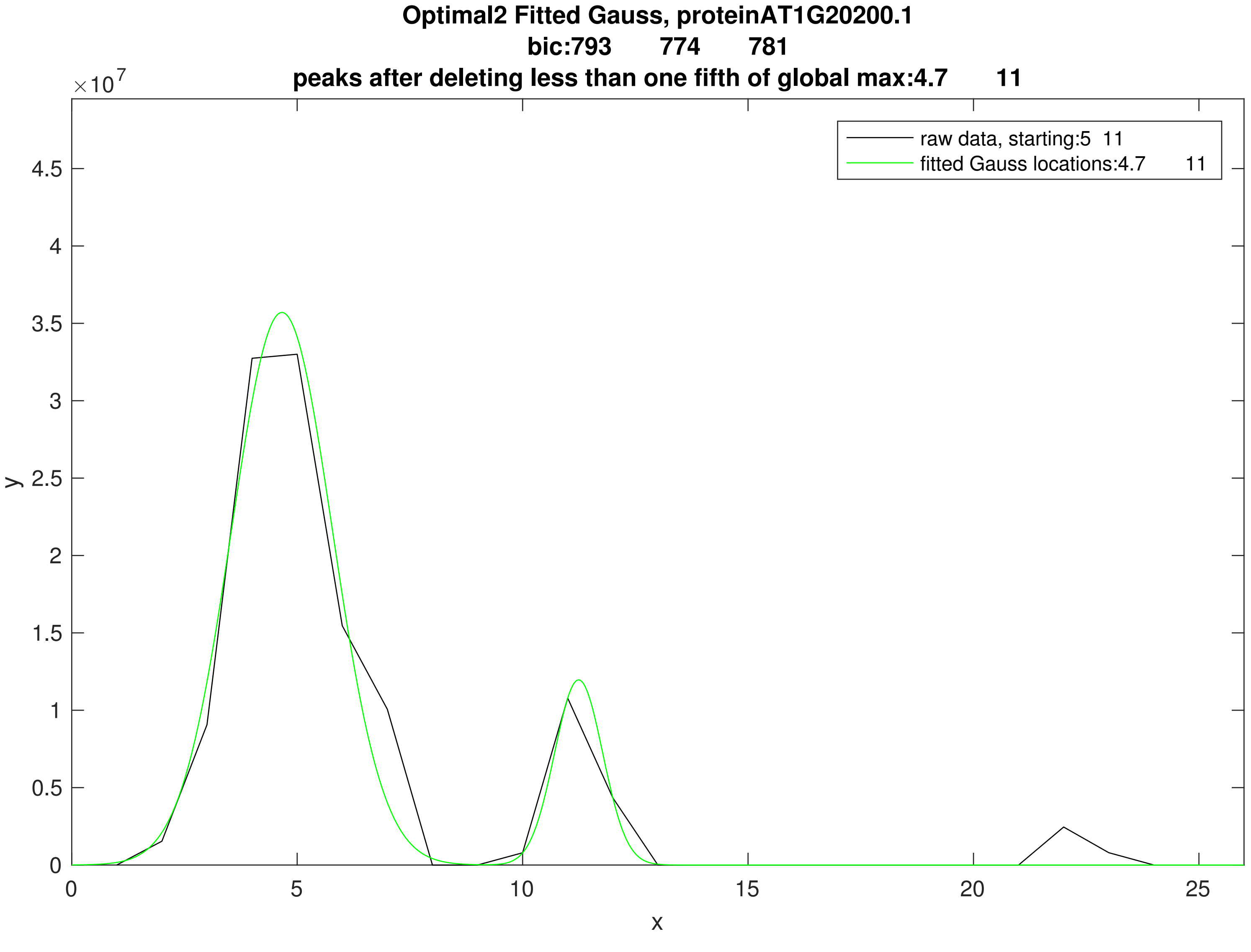
Optimal2 Fitted Gauss, proteinAT1G20200.1
bic:793 774 781
peaks after deleting less than one fifth of global max:4.7 11
×10^7
raw data, starting:5 11
fitted Gauss locations:4.7 11
y
x
0
0.5
1
1.5
2
2.5
3
3.5
4
4.5
5
10
15
20
25

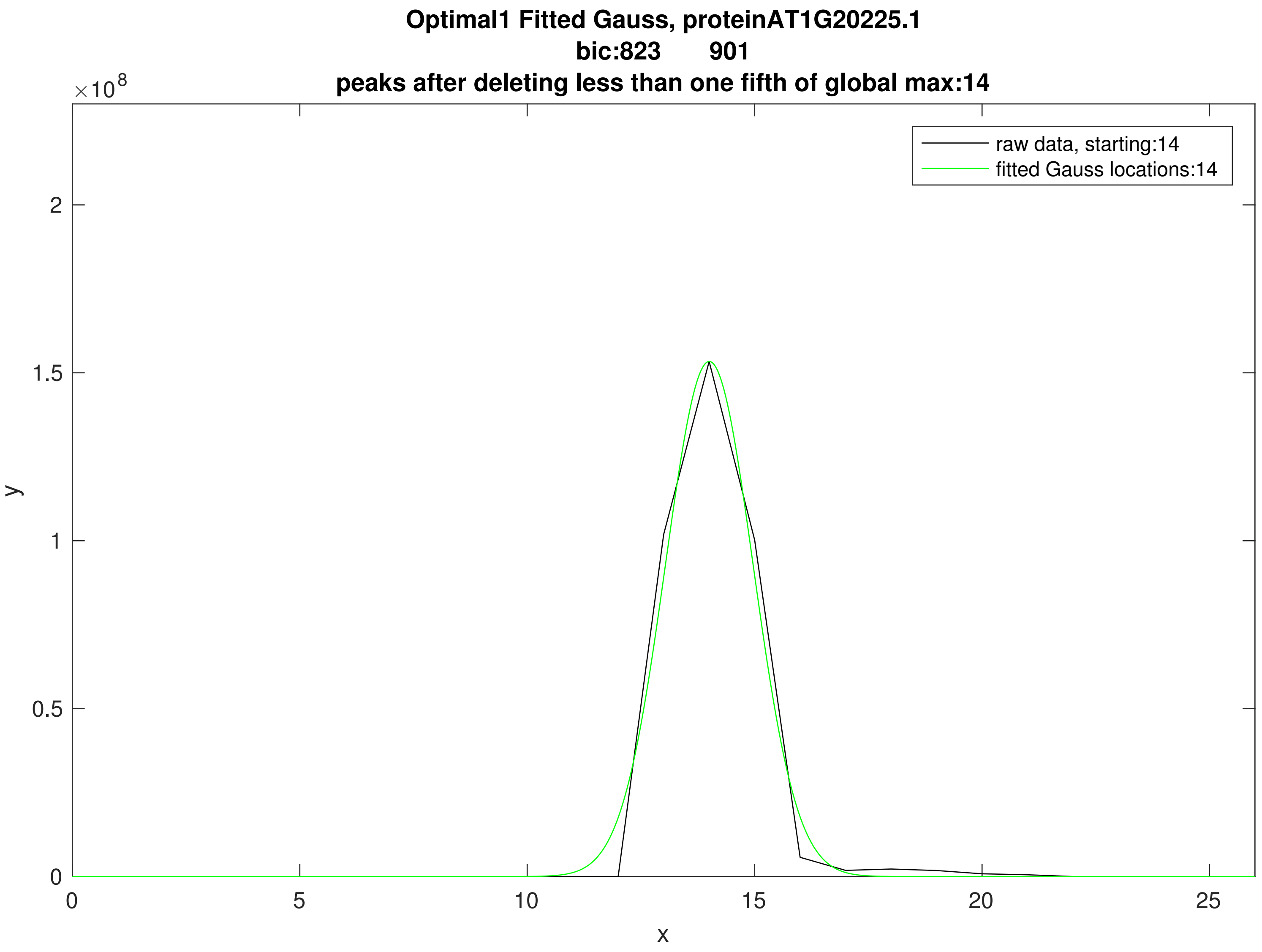
Optimal1 Fitted Gauss, proteinAT1G20225.1
bic:823 901
peaks after deleting less than one fifth of global max:14
×10^8
raw data, starting:14
fitted Gauss locations:14
2
1.5
1
0.5
0
y
0
5
10
15
20
25
x

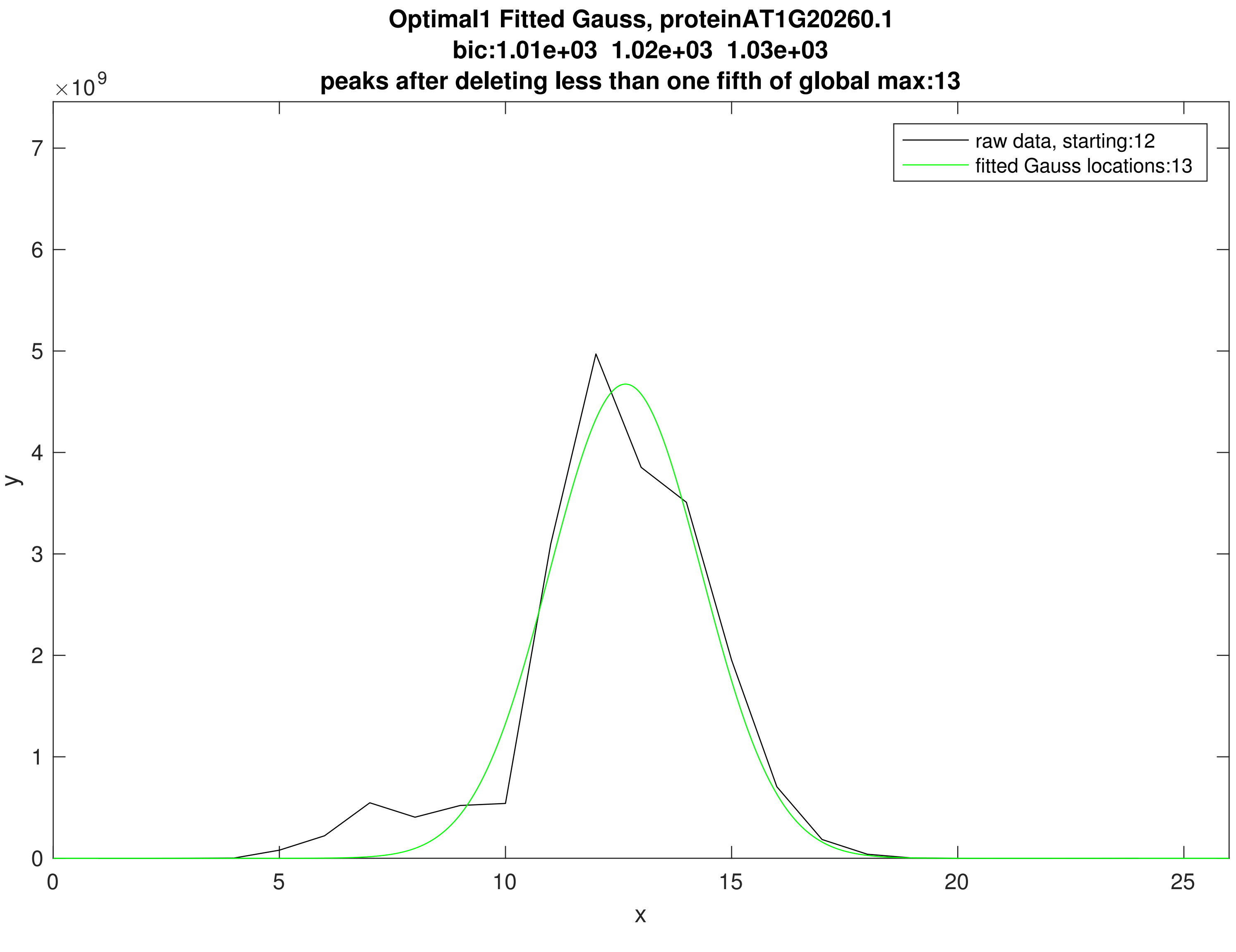

Optimal1 Fitted Gauss, proteinAT1G20260.1
bic:1.01e+03 1.02e+03 1.03e+03
peaks after deleting less than one fifth of global max:13
×10^9
raw data, starting:12
fitted Gauss locations:13
y
x
0
1
2
3
4
5
6
7
0
5
10
15
20
25

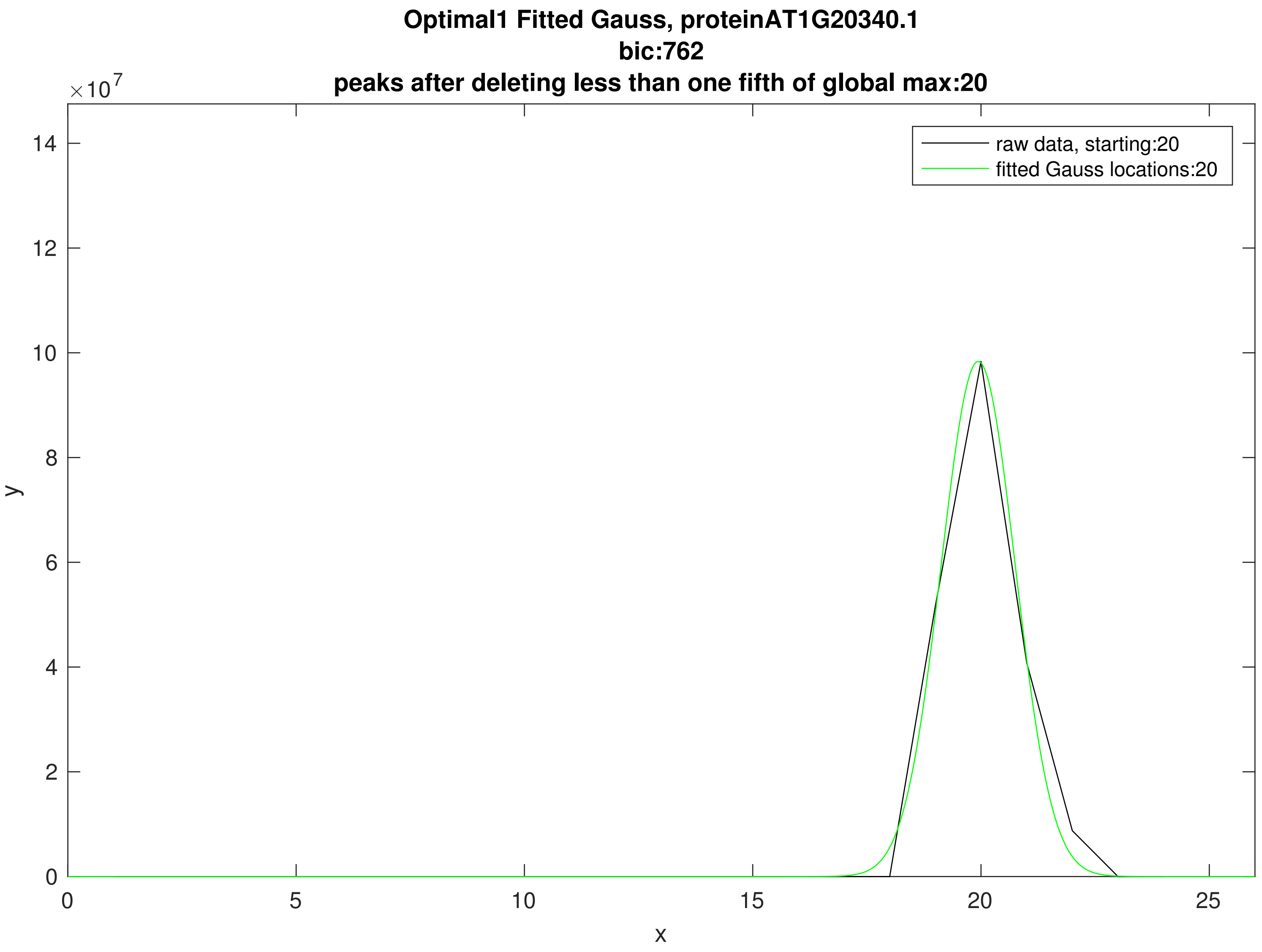
Optimal1 Fitted Gauss, proteinAT1G20340.1
bic:762
peaks after deleting less than one fifth of global max:20
×10^7
raw data, starting:20
fitted Gauss locations:20
y
x
0
2
4
6
8
10
12
14
0
5
10
15
20
25

**Optimal1 Fitted Gauss, proteinAT1G20440.1**
**bic:1.05e+03  1.06e+03  1.06e+03  1.07e+03**
**peaks after deleting less than one fifth of global max:11**

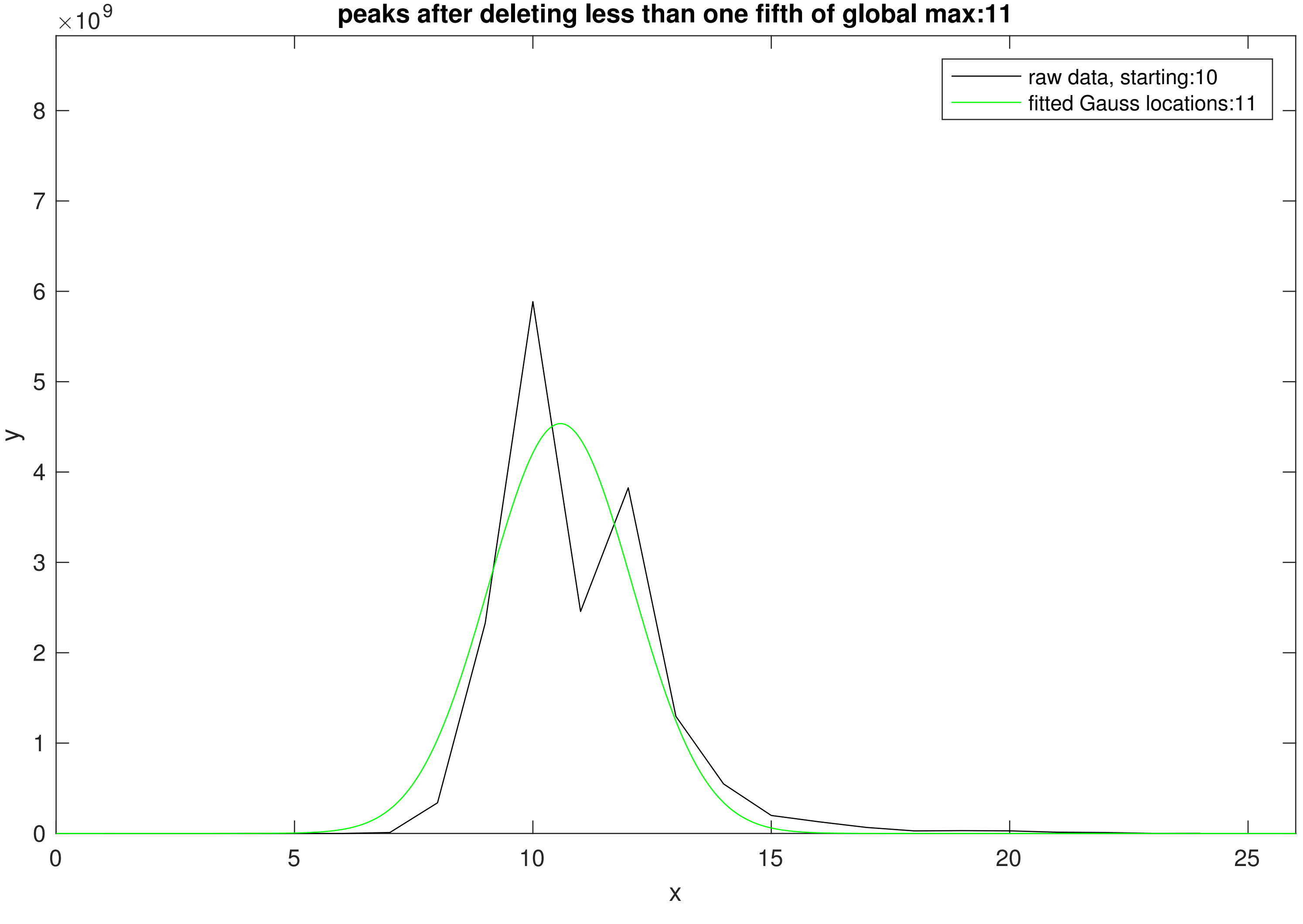

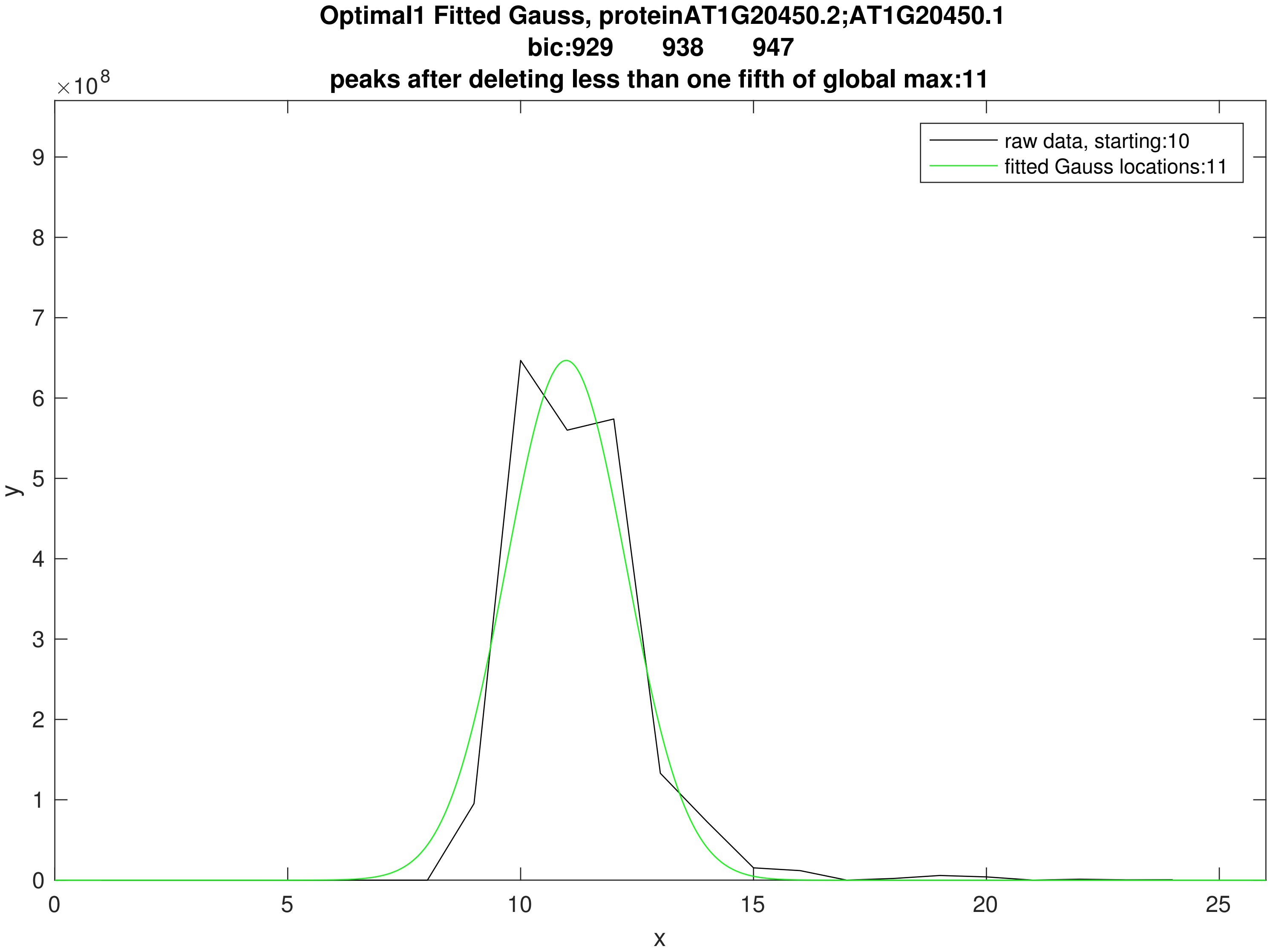

Optimal1 Fitted Gauss, proteinAT1G20450.2;AT1G20450.1
bic:929 938 947
peaks after deleting less than one fifth of global max:11
×10^8
raw data, starting:10
fitted Gauss locations:11
y
x

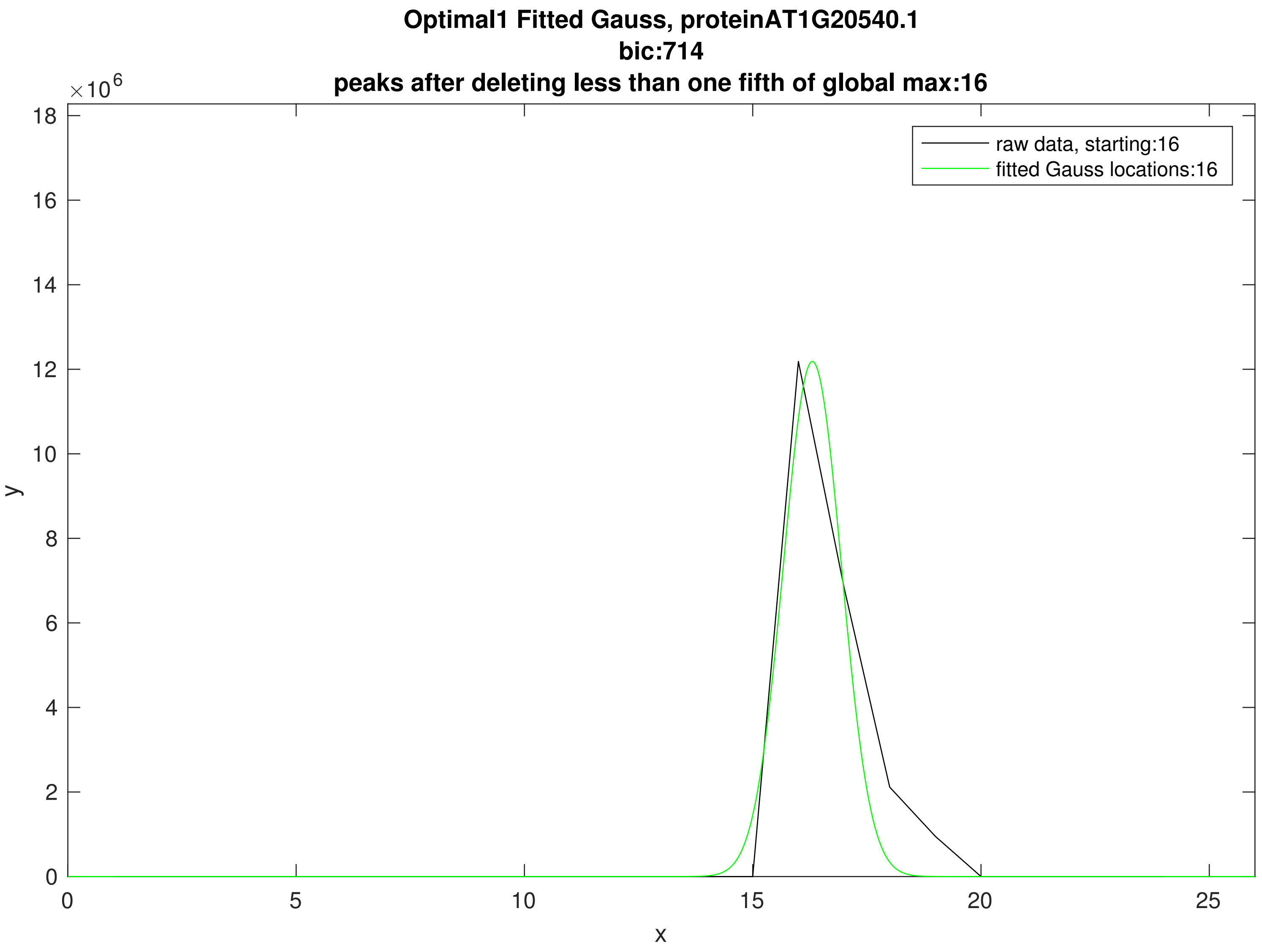

Optimal1 Fitted Gauss, proteinAT1G20540.1
bic:714
peaks after deleting less than one fifth of global max:16
×10^6
18
16
14
12
10
8
6
4
2
0
y
0
5
10
15
20
25
x
raw data, starting:16
fitted Gauss locations:16

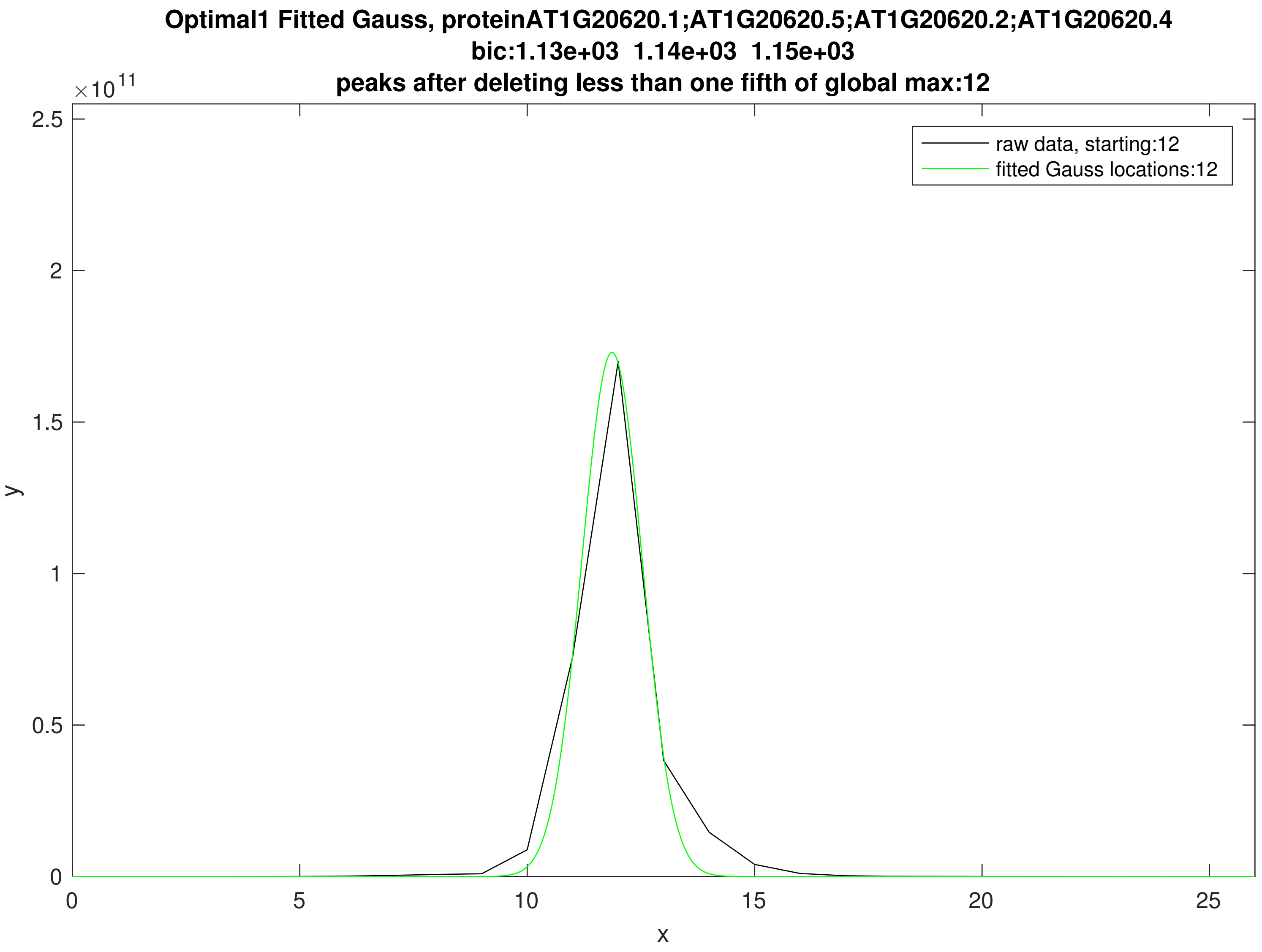

Optimal1 Fitted Gauss, proteinAT1G20620.1;AT1G20620.5;AT1G20620.2;AT1G20620.4
bic:1.13e+03 1.14e+03 1.15e+03
peaks after deleting less than one fifth of global max:12
×10^11
2.5
2
1.5
1
0.5
0
y
0
5
10
15
20
25
x
raw data, starting:12
fitted Gauss locations:12

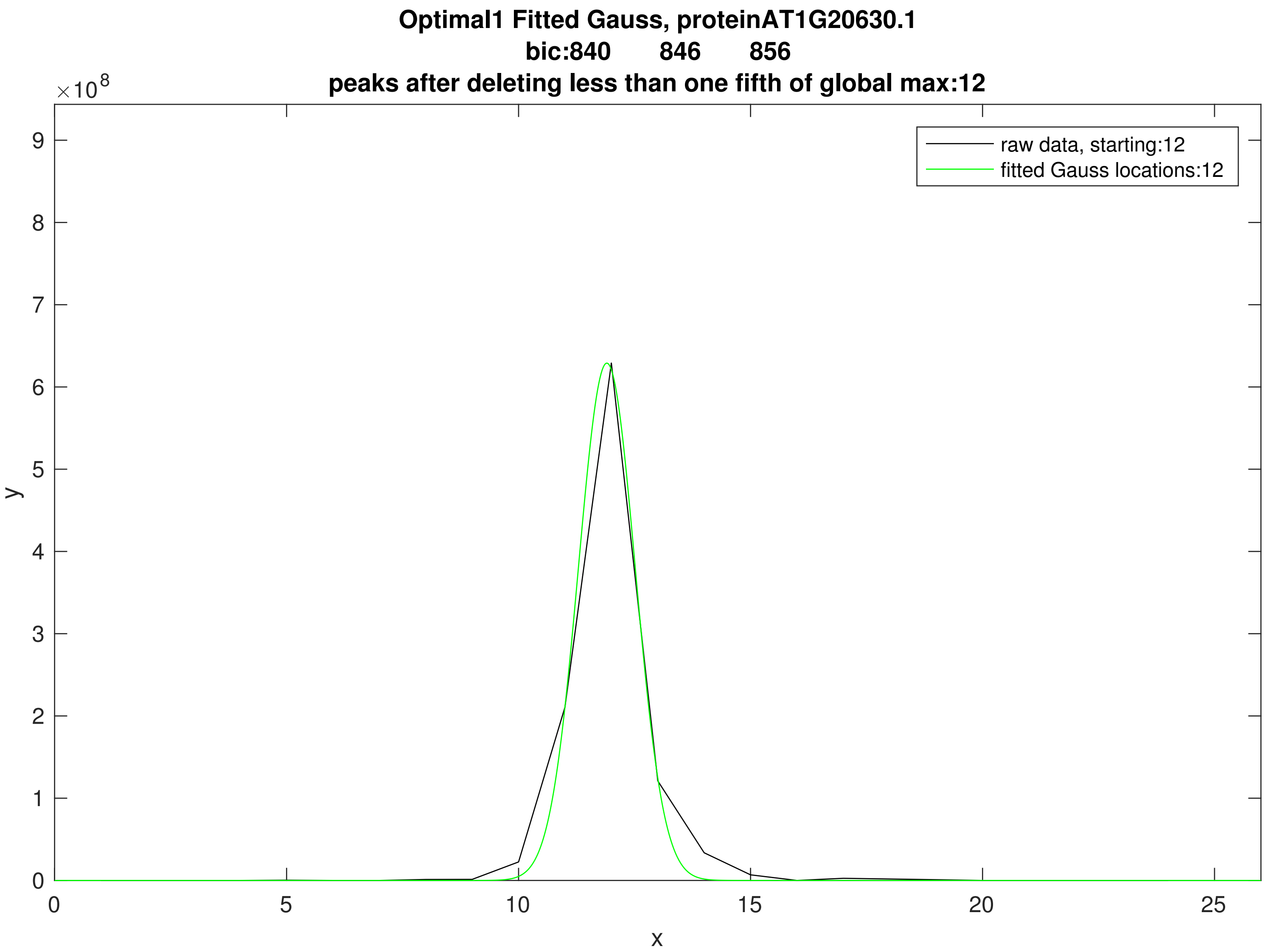

Optimal1 Fitted Gauss, proteinAT1G20630.1
bic:840 846 856
peaks after deleting less than one fifth of global max:12
×10^8
raw data, starting:12
fitted Gauss locations:12
y
x
0
1
2
3
4
5
6
7
8
9
0
5
10
15
20
25

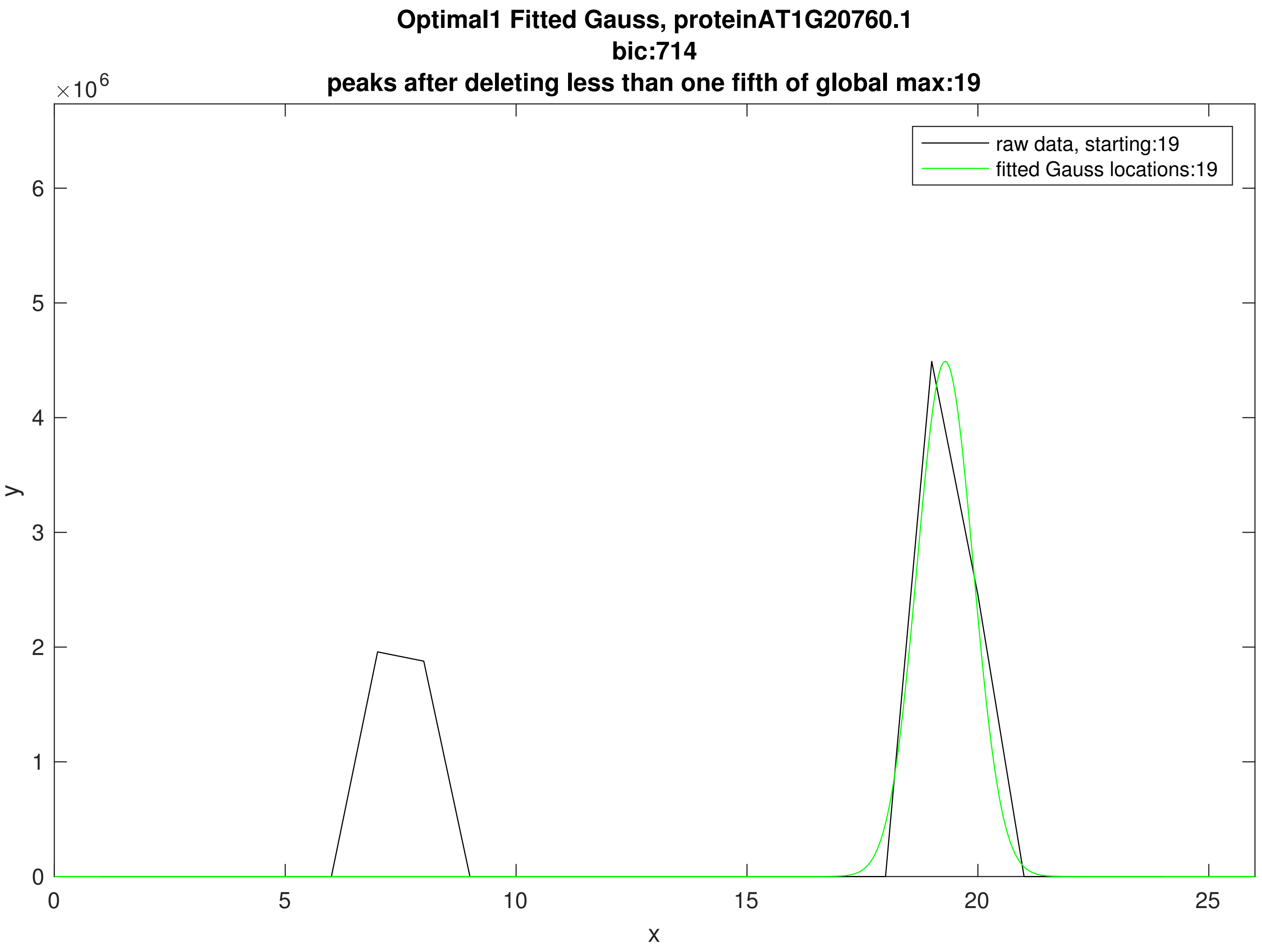
Optimal1 Fitted Gauss, proteinAT1G20760.1
bic:714
peaks after deleting less than one fifth of global max:19
×10^6
raw data, starting:19
fitted Gauss locations:19
y
x
0
1
2
3
4
5
6
0
5
10
15
20
25

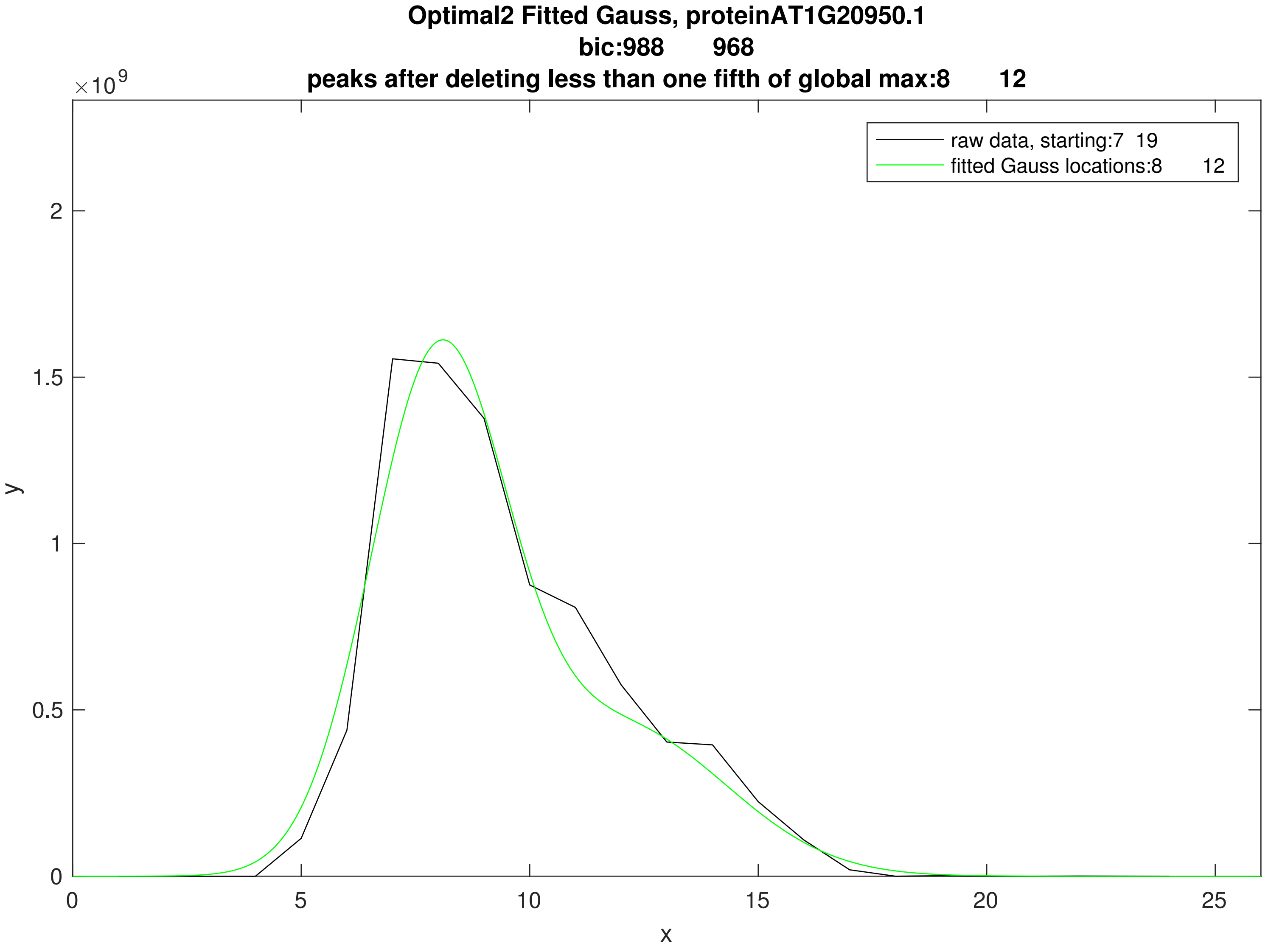
Optimal2 Fitted Gauss, proteinAT1G20950.1
bic:988 968
peaks after deleting less than one fifth of global max:8 12
×10^9
raw data, starting:7 19
fitted Gauss locations:8 12
y
x
0
0.5
1
1.5
2
0
5
10
15
20
25

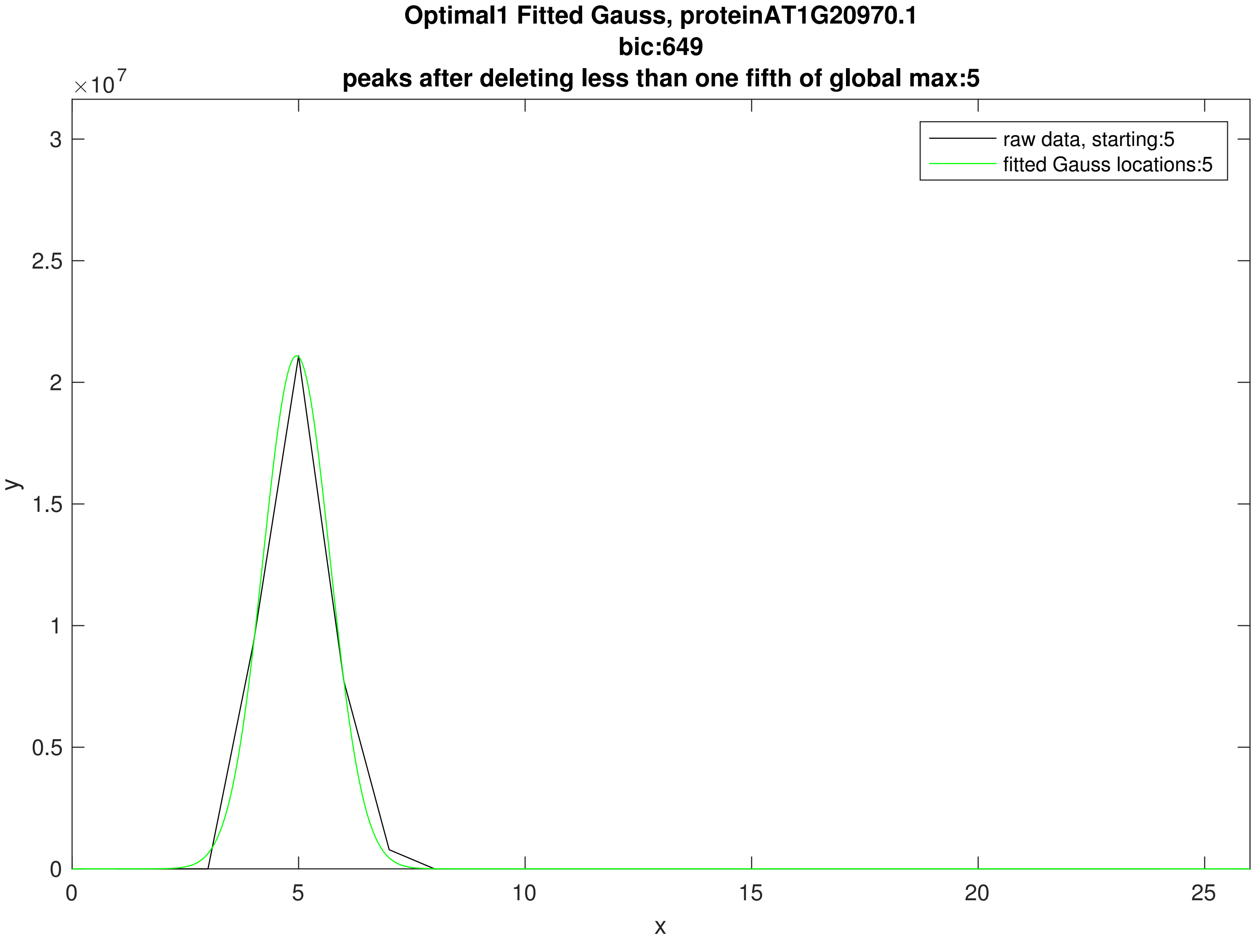
Optimal1 Fitted Gauss, proteinAT1G20970.1
bic:649
peaks after deleting less than one fifth of global max:5
×10^7
raw data, starting:5
fitted Gauss locations:5
3
2.5
2
1.5
1
0.5
0
y
0
5
10
15
20
25
x

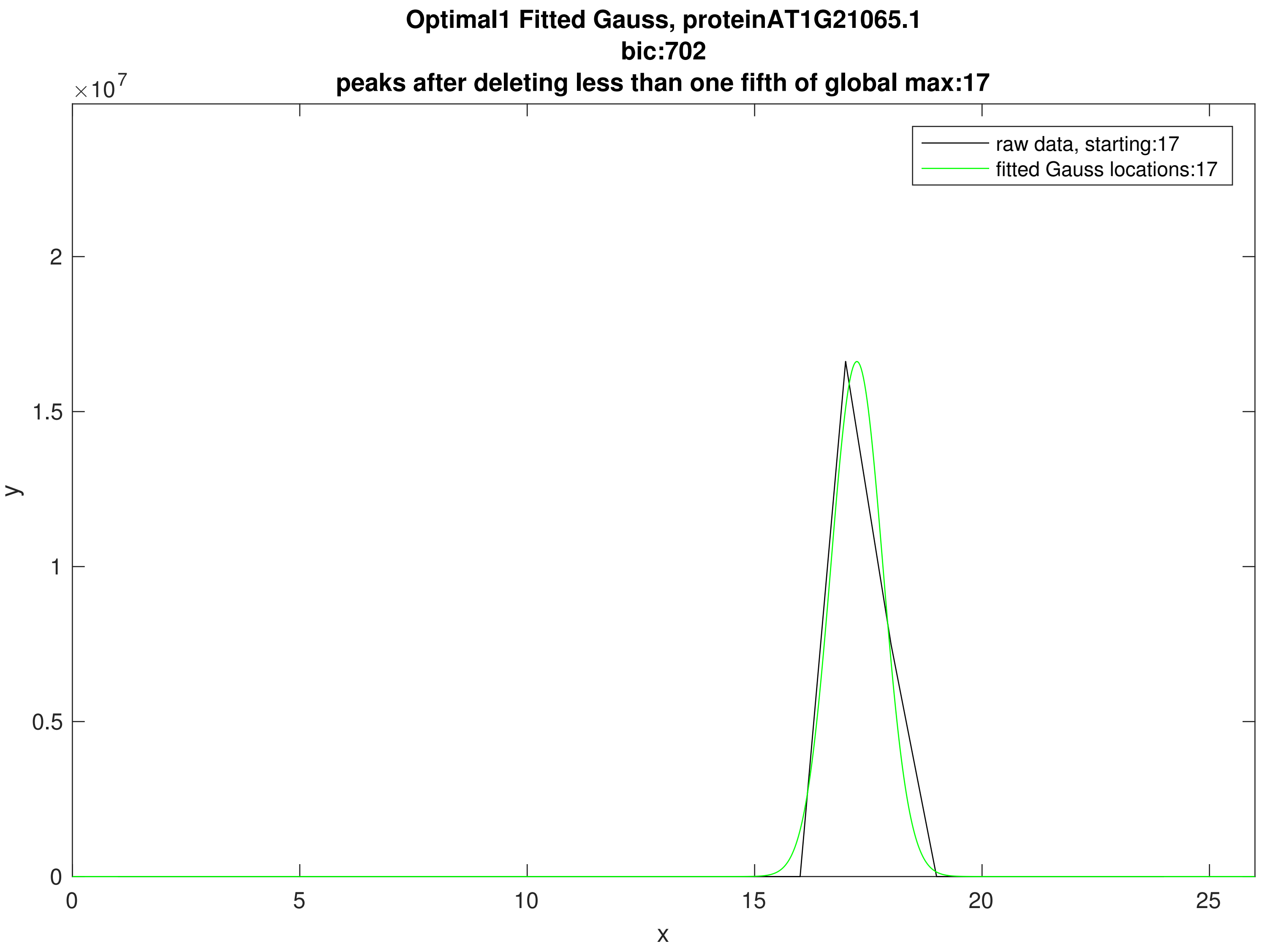
Optimal1 Fitted Gauss, proteinAT1G21065.1
bic:702
peaks after deleting less than one fifth of global max:17
×10^7
raw data, starting:17
fitted Gauss locations:17
y
x
0
0.5
1
1.5
2
0
5
10
15
20
25

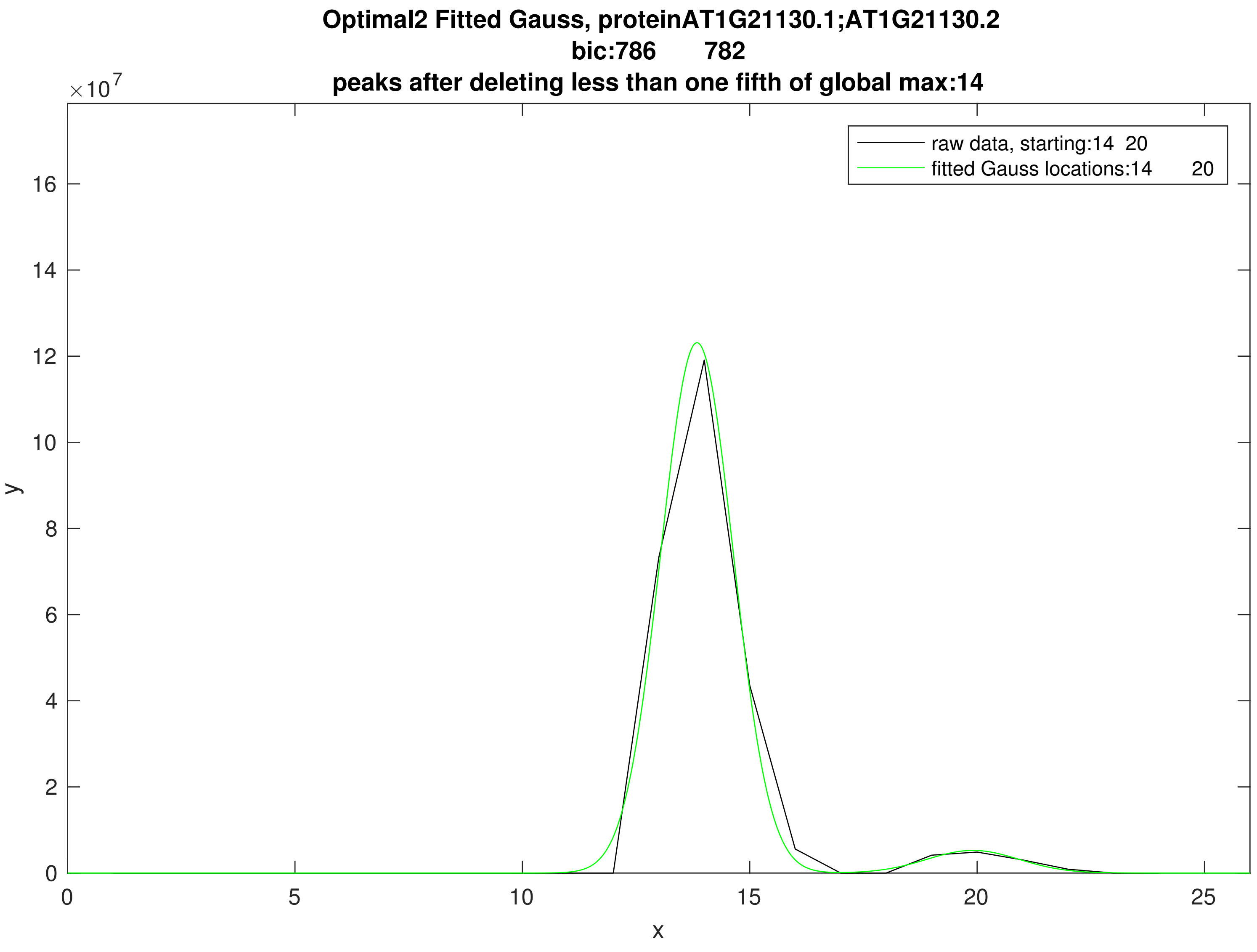
Optimal2 Fitted Gauss, proteinAT1G21130.1;AT1G21130.2
bic:786 782
peaks after deleting less than one fifth of global max:14
×10^7
raw data, starting:14 20
fitted Gauss locations:14 20
y
x
0
2
4
6
8
10
12
14
16
0
5
10
15
20
25

Optimal1 Fitted Gauss, proteinAT1G21350.1;AT1G21350.2;AT1G21350.3
bic:584
peaks after deleting less than one fifth of global max:19

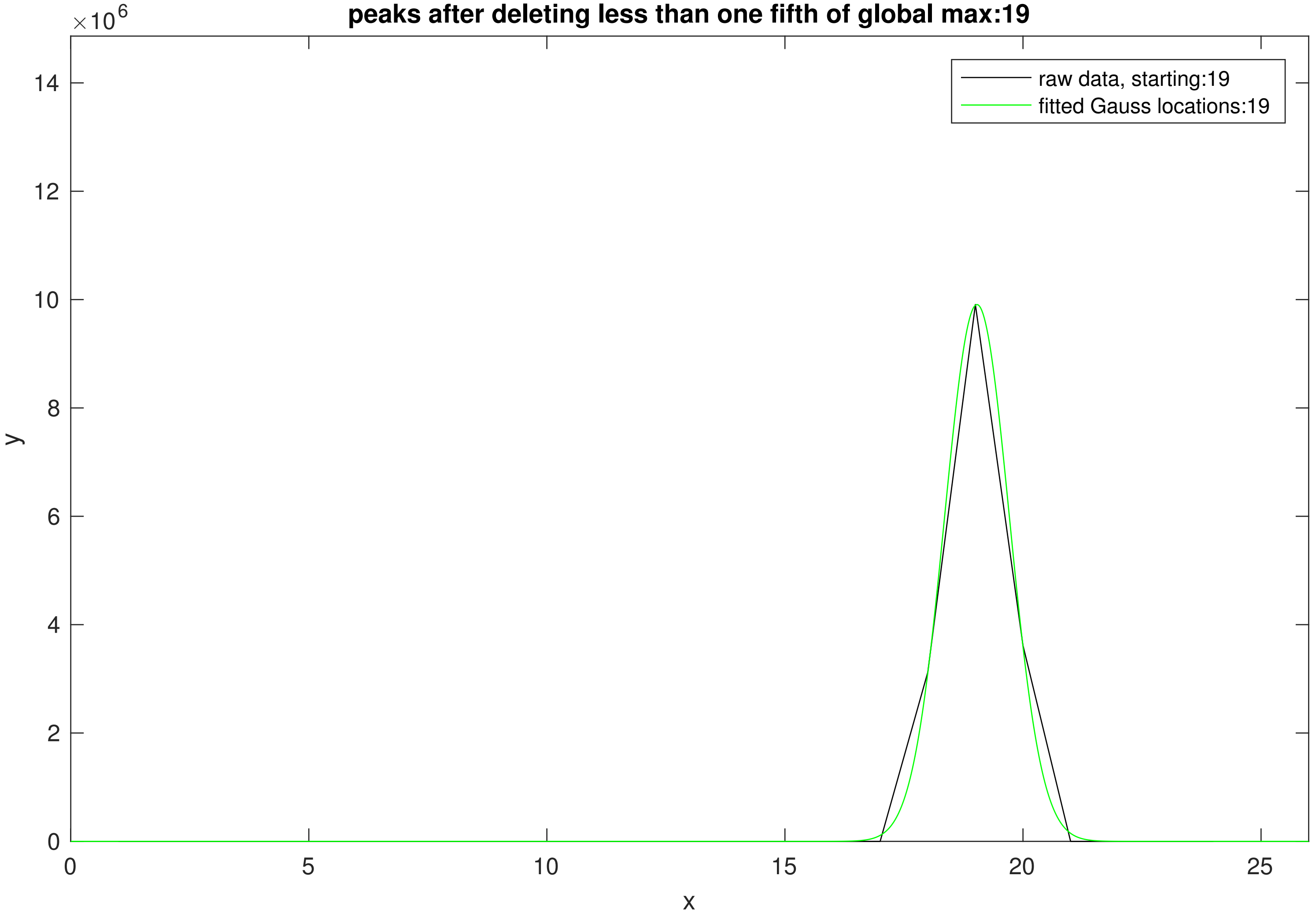

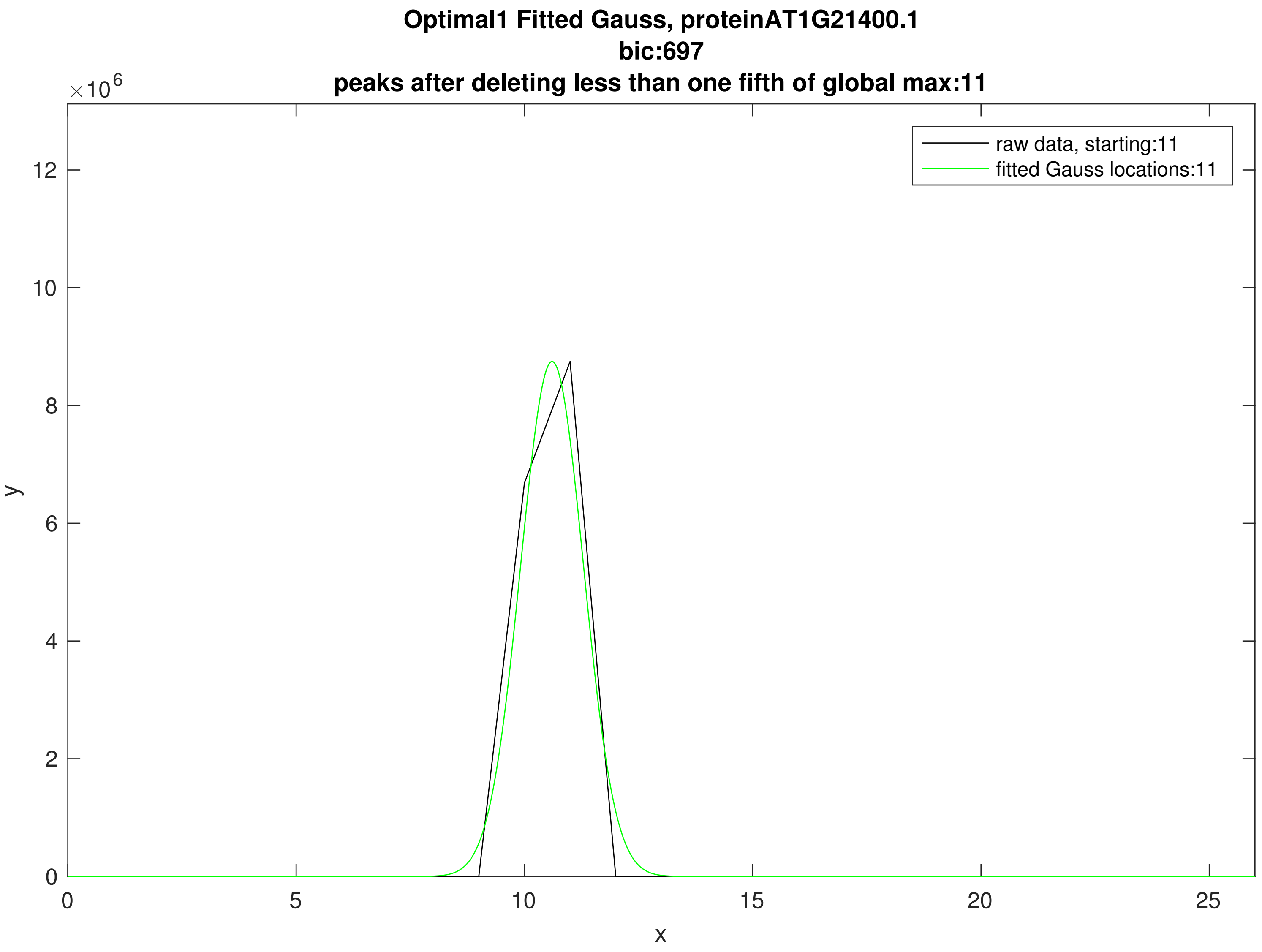
Optimal1 Fitted Gauss, proteinAT1G21400.1
bic:697
peaks after deleting less than one fifth of global max:11
×10^6
12
10
8
6
4
2
0
y
0
5
10
15
20
25
x
raw data, starting:11
fitted Gauss locations:11

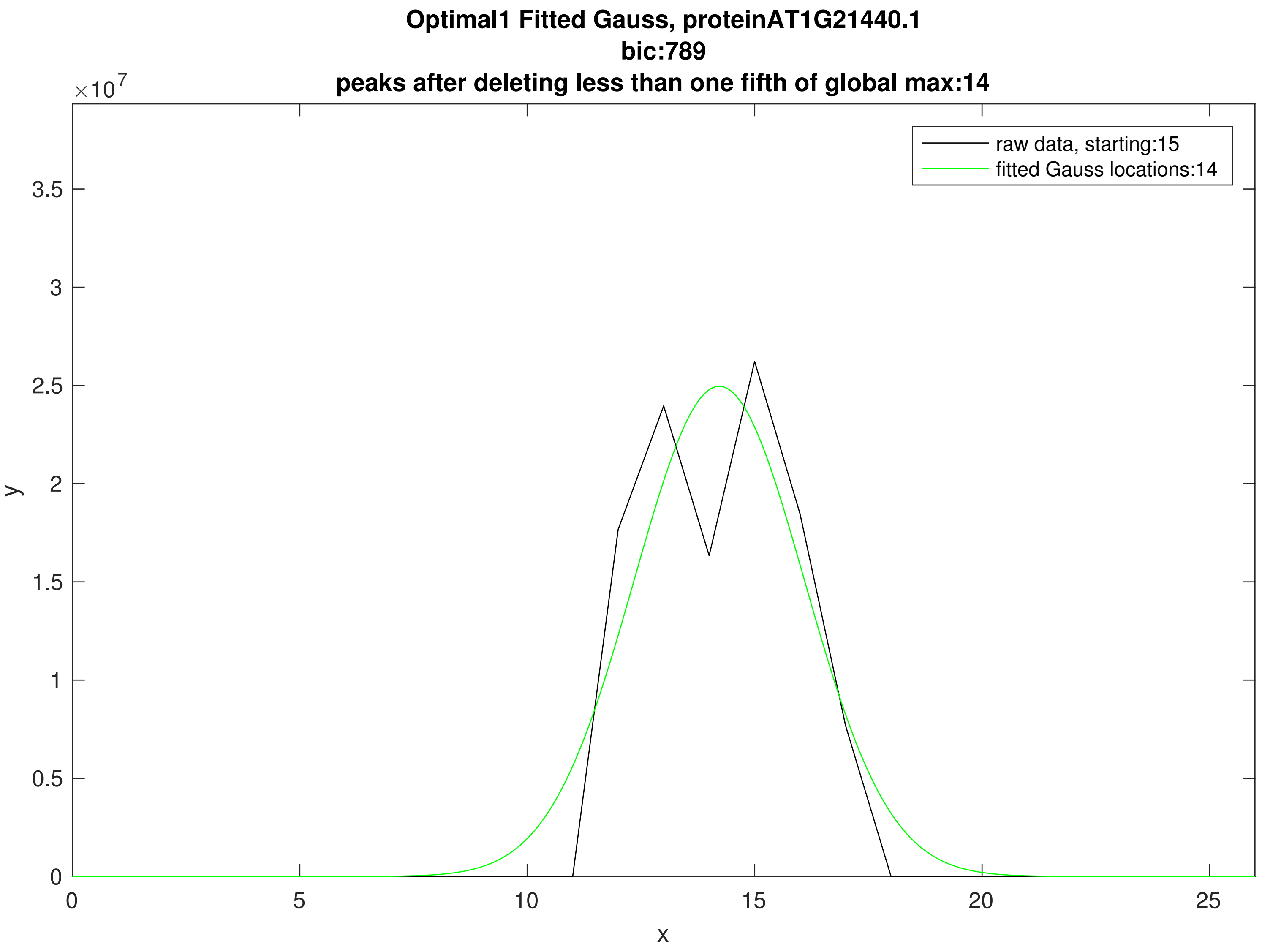
Optimal1 Fitted Gauss, proteinAT1G21440.1
bic:789
peaks after deleting less than one fifth of global max:14
×10^7
3.5
3
2.5
2
1.5
1
0.5
0
y
0
5
10
15
20
25
x
raw data, starting:15
fitted Gauss locations:14

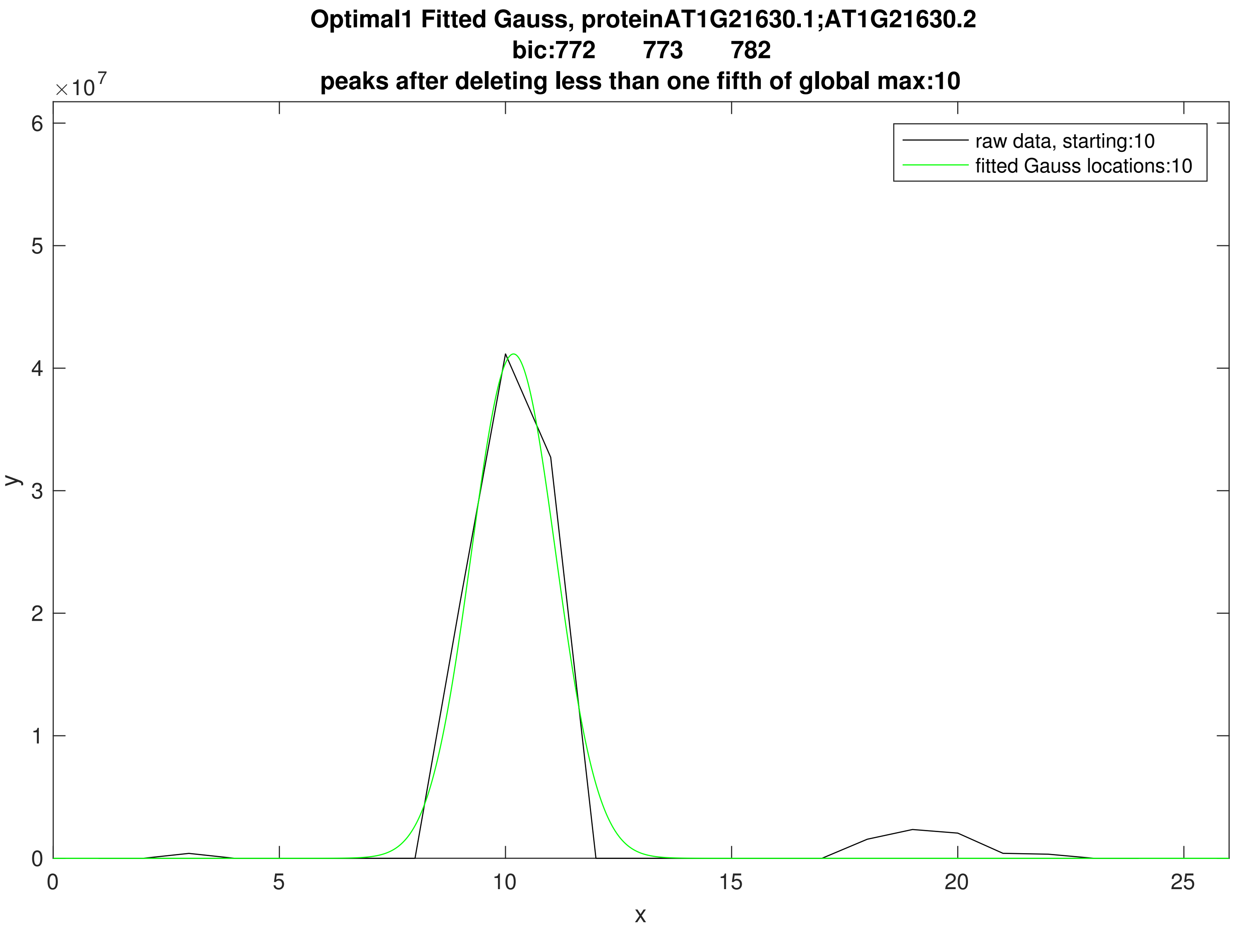
Optimal1 Fitted Gauss, proteinAT1G21630.1;AT1G21630.2
bic:772 773 782
peaks after deleting less than one fifth of global max:10
raw data, starting:10
fitted Gauss locations:10
y
x
×10^7
0
1
2
3
4
5
6
0
5
10
15
20
25

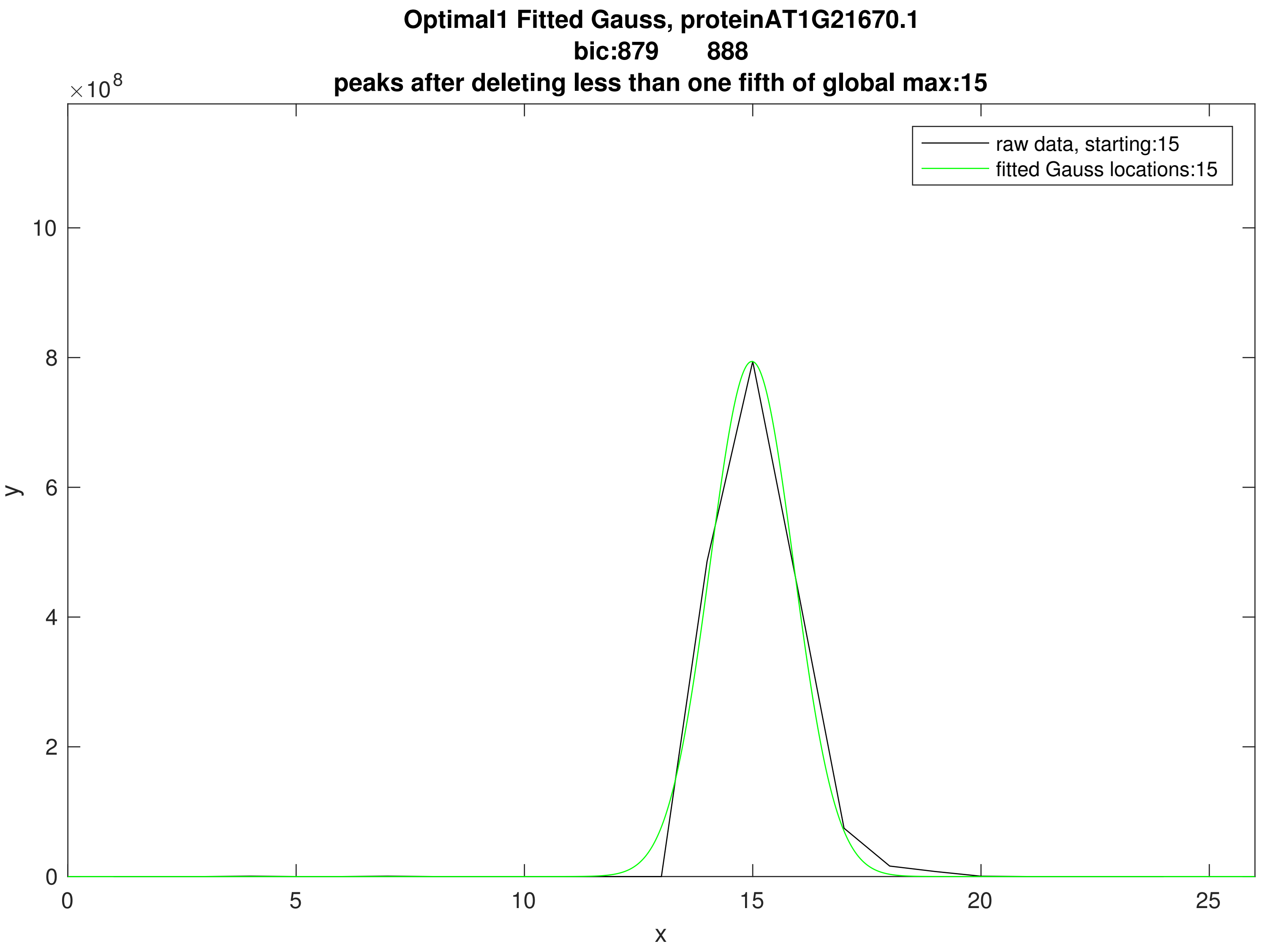

Optimal1 Fitted Gauss, proteinAT1G21670.1
bic:879 888
peaks after deleting less than one fifth of global max:15
×10^8
raw data, starting:15
fitted Gauss locations:15
10
8
6
4
2
0
y
0
5
10
15
20
25
x

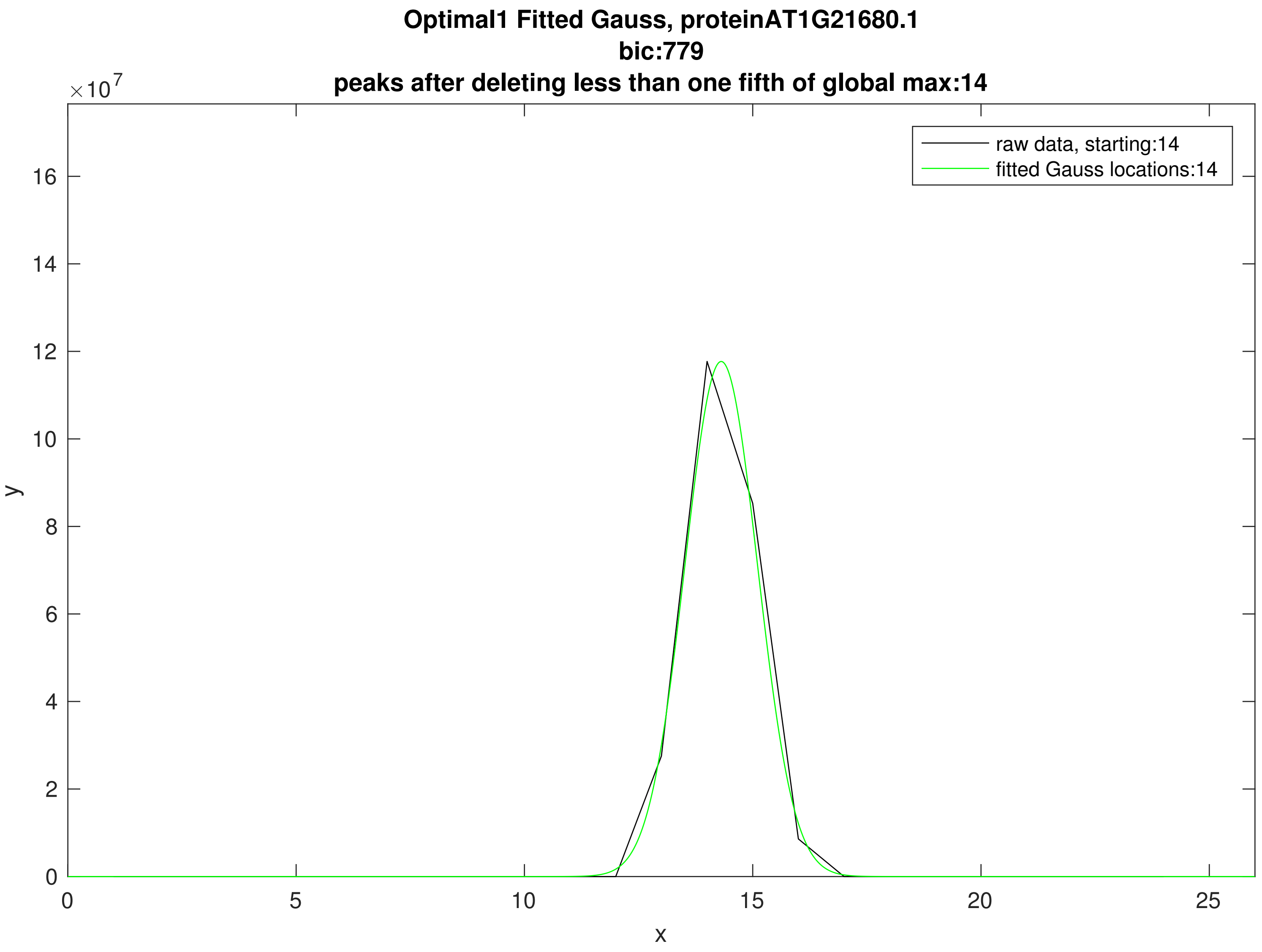

Optimal1 Fitted Gauss, proteinAT1G21680.1
bic:779
peaks after deleting less than one fifth of global max:14
×10^7
16
14
12
10
8
6
4
2
0
y
0
5
10
15
20
25
x
raw data, starting:14
fitted Gauss locations:14

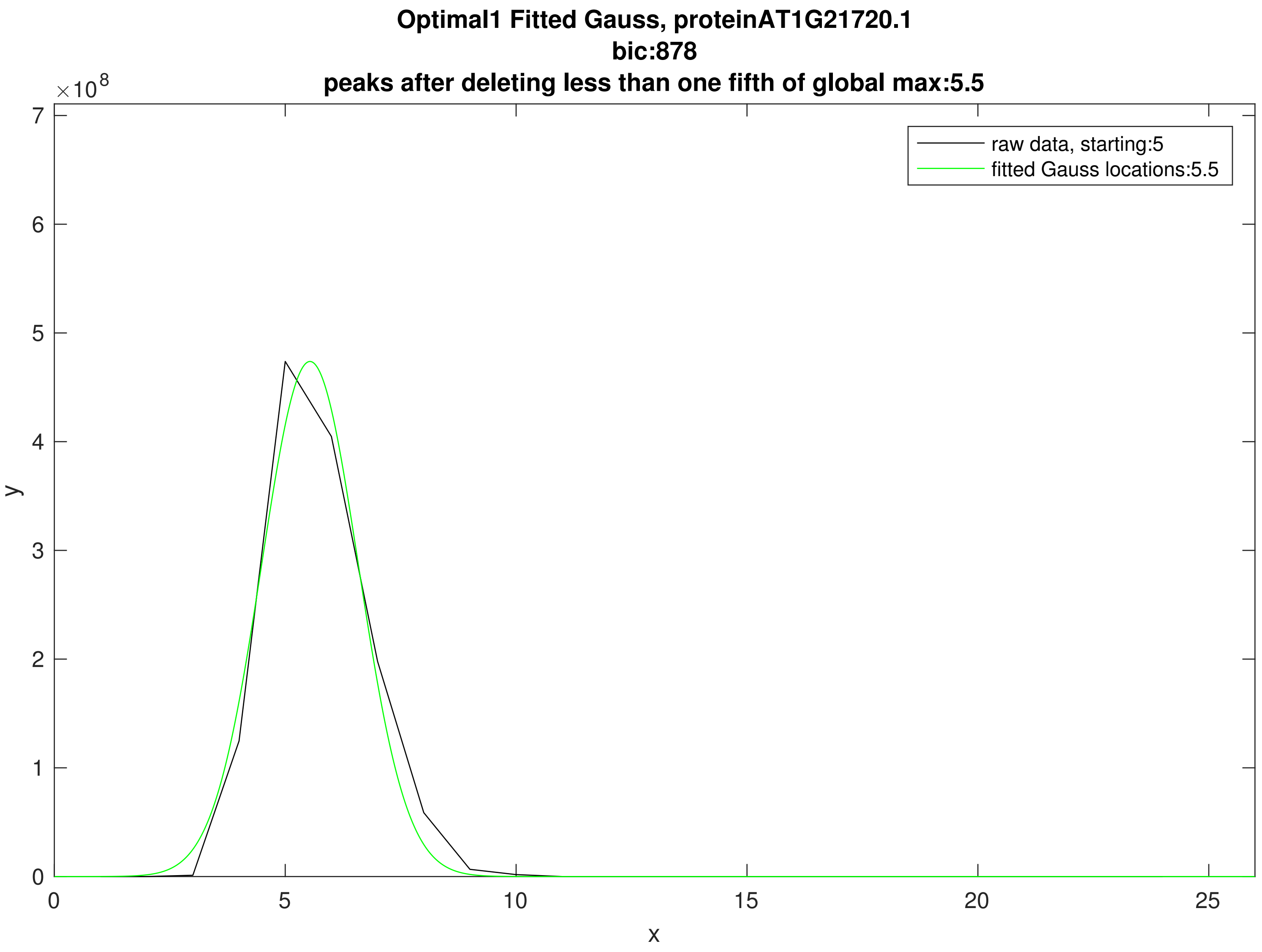
Optimal1 Fitted Gauss, proteinAT1G21720.1
bic:878
peaks after deleting less than one fifth of global max:5.5
×10^8
raw data, starting:5
fitted Gauss locations:5.5
y
x
0
1
2
3
4
5
6
7
0
5
10
15
20
25

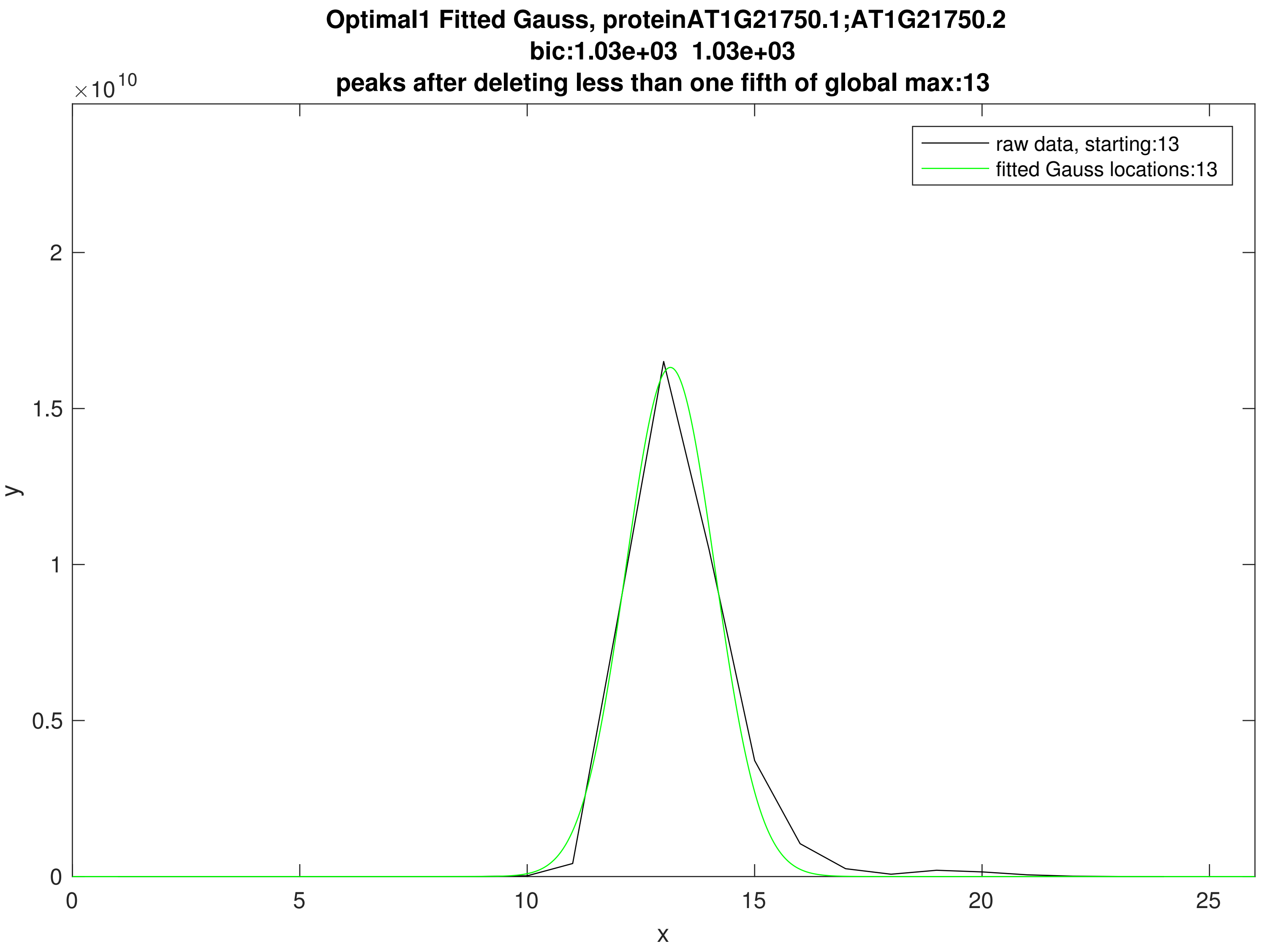

Optimal1 Fitted Gauss, proteinAT1G21750.1;AT1G21750.2
bic:1.03e+03  1.03e+03
peaks after deleting less than one fifth of global max:13
×10^10
raw data, starting:13
fitted Gauss locations:13
y
x
0
0.5
1
1.5
2
5
10
15
20
25

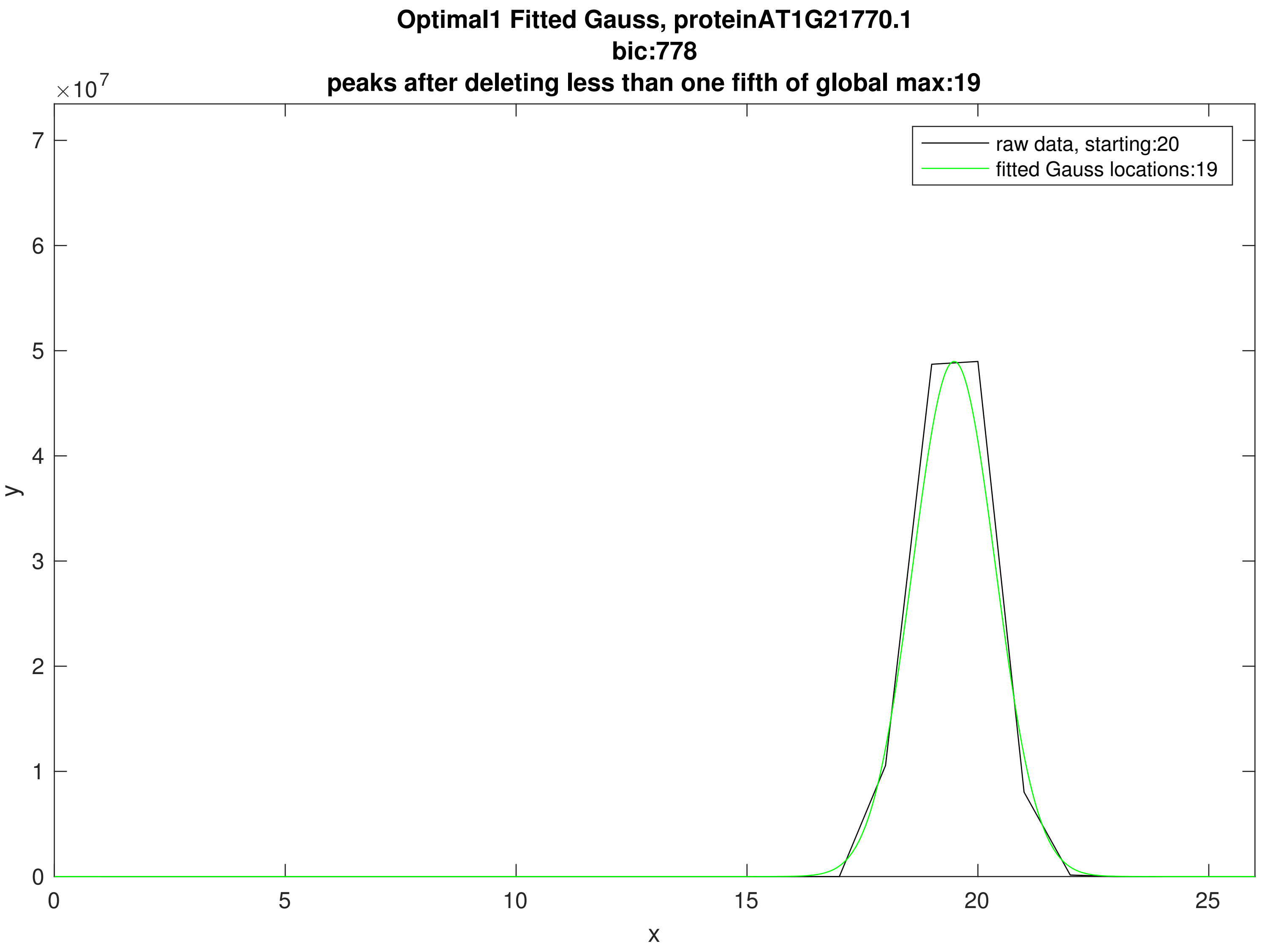
Optimal1 Fitted Gauss, proteinAT1G21770.1
bic:778
peaks after deleting less than one fifth of global max:19
×10^7
7
6
5
4
3
2
1
0
y
0
5
10
15
20
25
x
raw data, starting:20
fitted Gauss locations:19

Optimal1 Fitted Gauss, proteinAT1G21880.2;AT1G21880.1
bic:802
peaks after deleting less than one fifth of global max:7.5

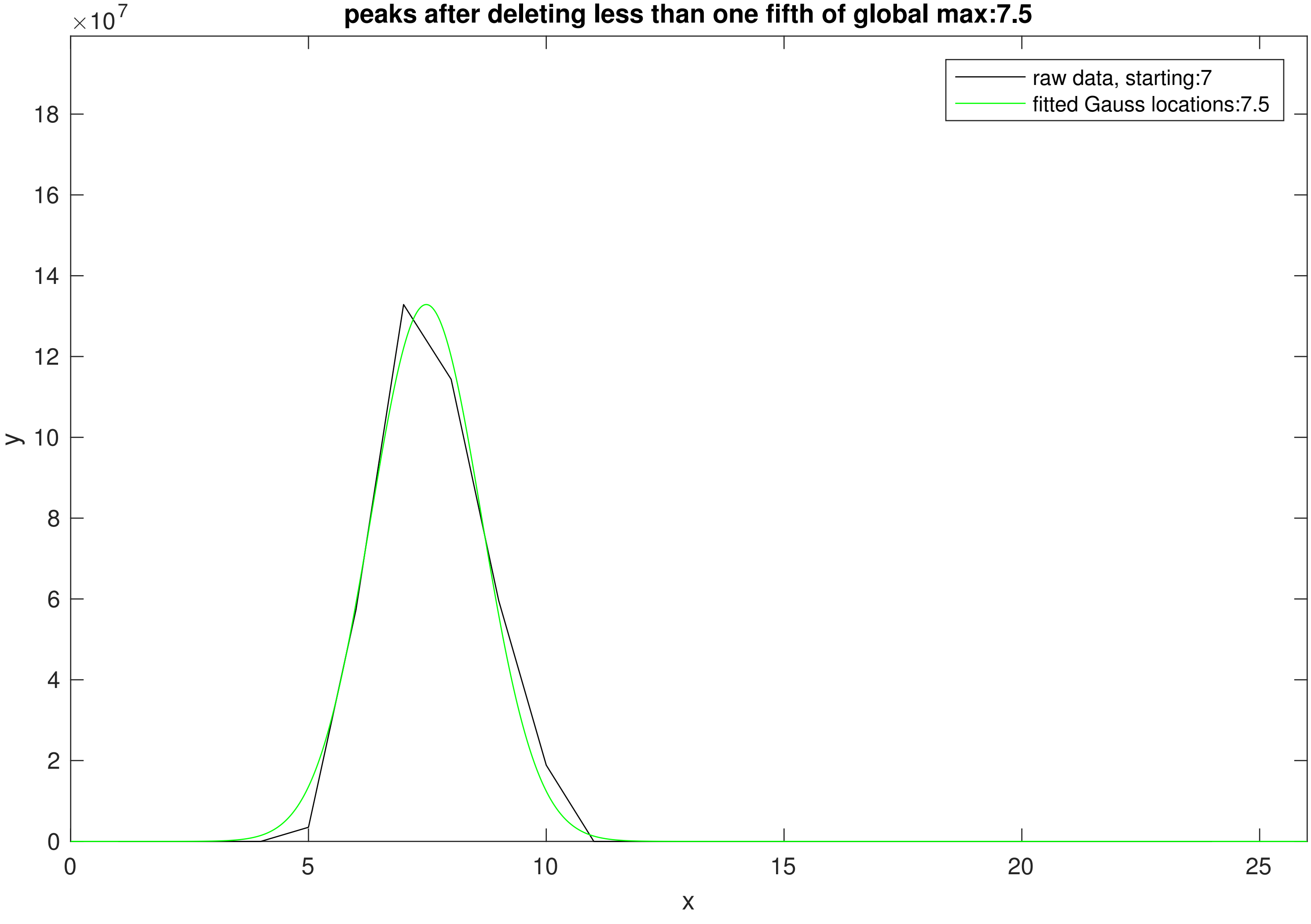

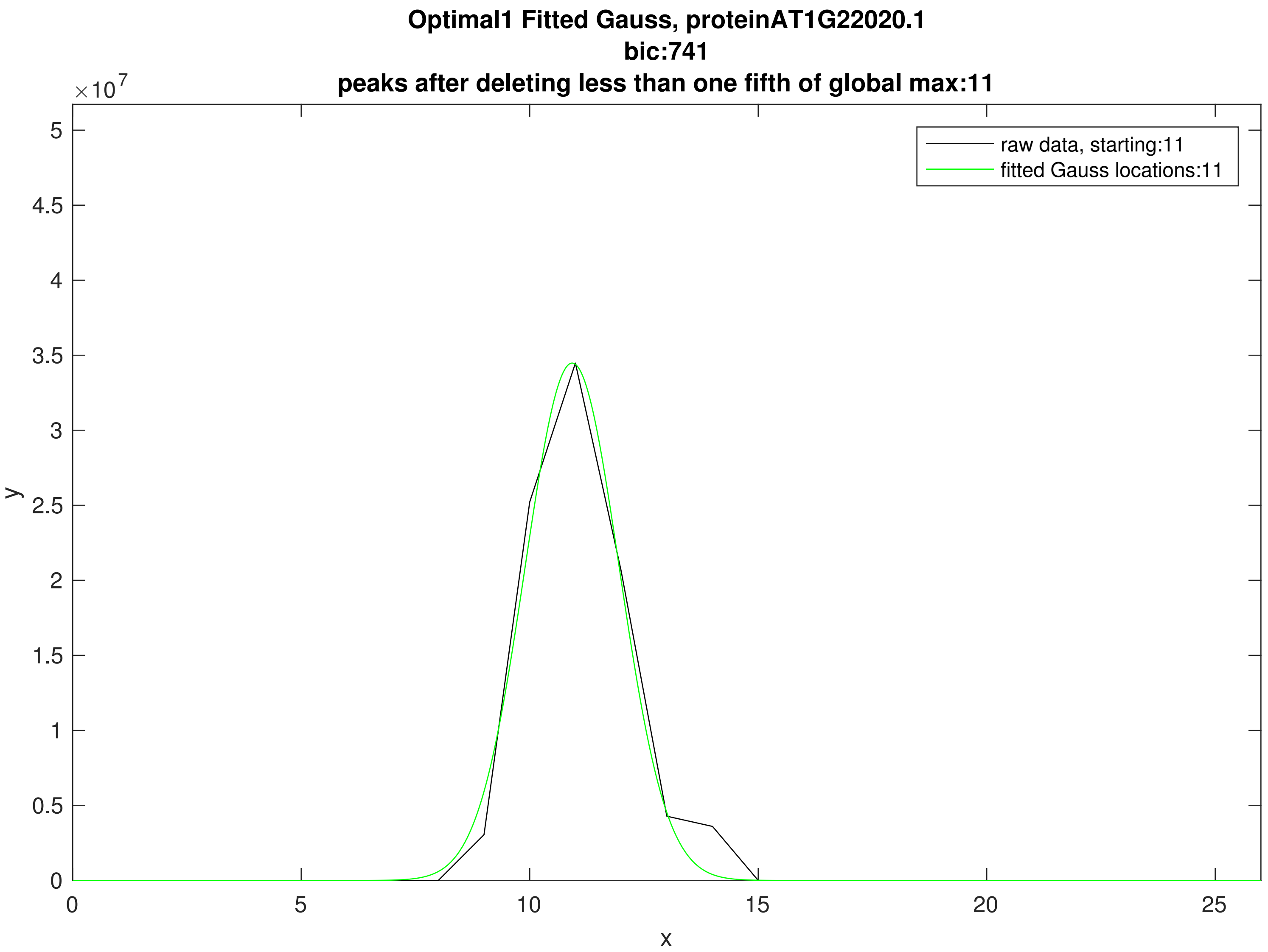

Optimal1 Fitted Gauss, proteinAT1G22020.1
bic:741
peaks after deleting less than one fifth of global max:11
×10^7
raw data, starting:11
fitted Gauss locations:11
y
x
0
0.5
1
1.5
2
2.5
3
3.5
4
4.5
5
0
5
10
15
20
25

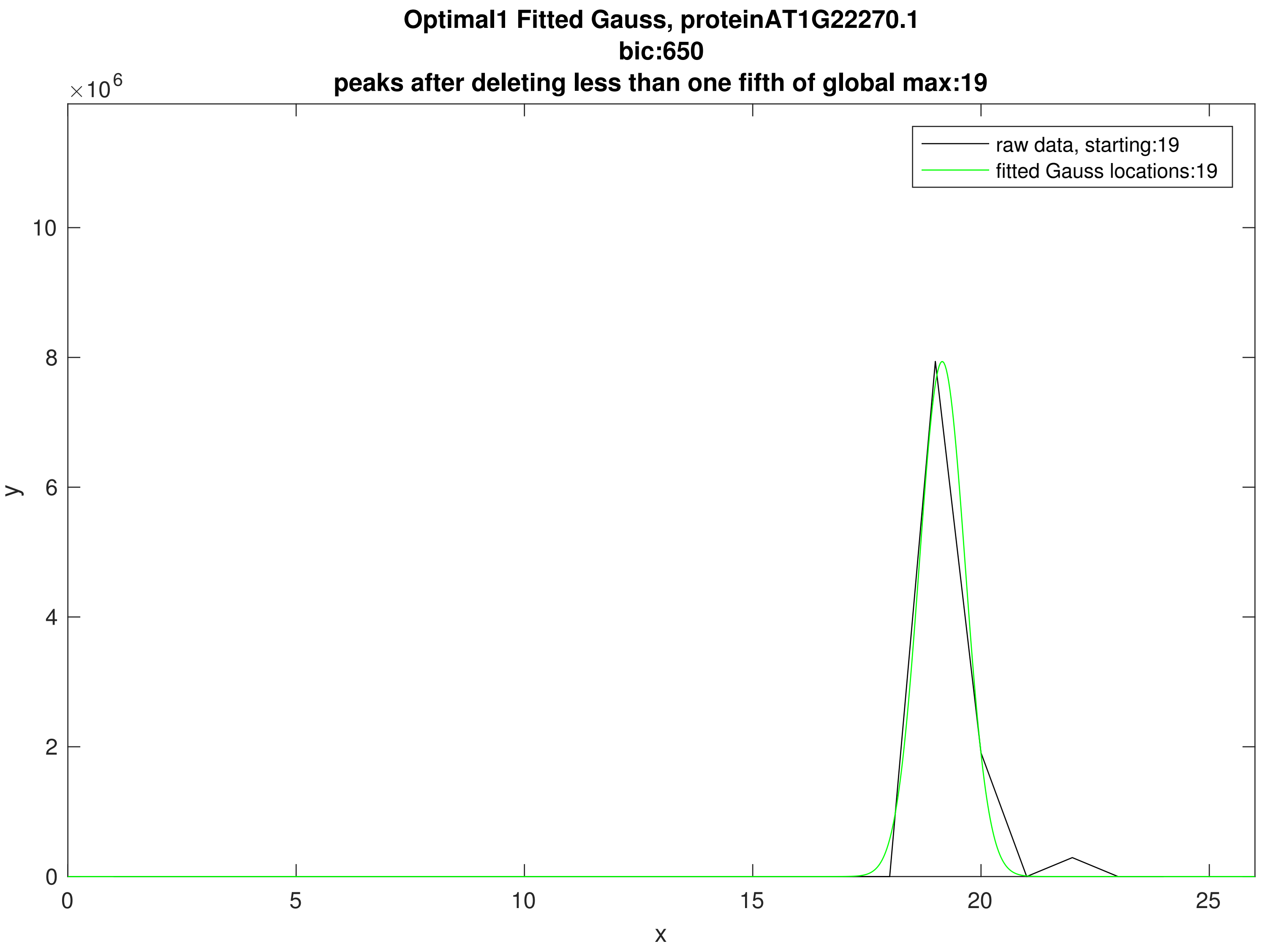

Optimal1 Fitted Gauss, proteinAT1G22270.1
bic:650
peaks after deleting less than one fifth of global max:19
×10^6
raw data, starting:19
fitted Gauss locations:19
10
8
6
4
2
0
y
0
5
10
15
20
25
x

**Optimal1 Fitted Gauss, proteinAT1G22280.1;AT1G22280.3**
**bic:665**
**peaks after deleting less than one fifth of global max:18**

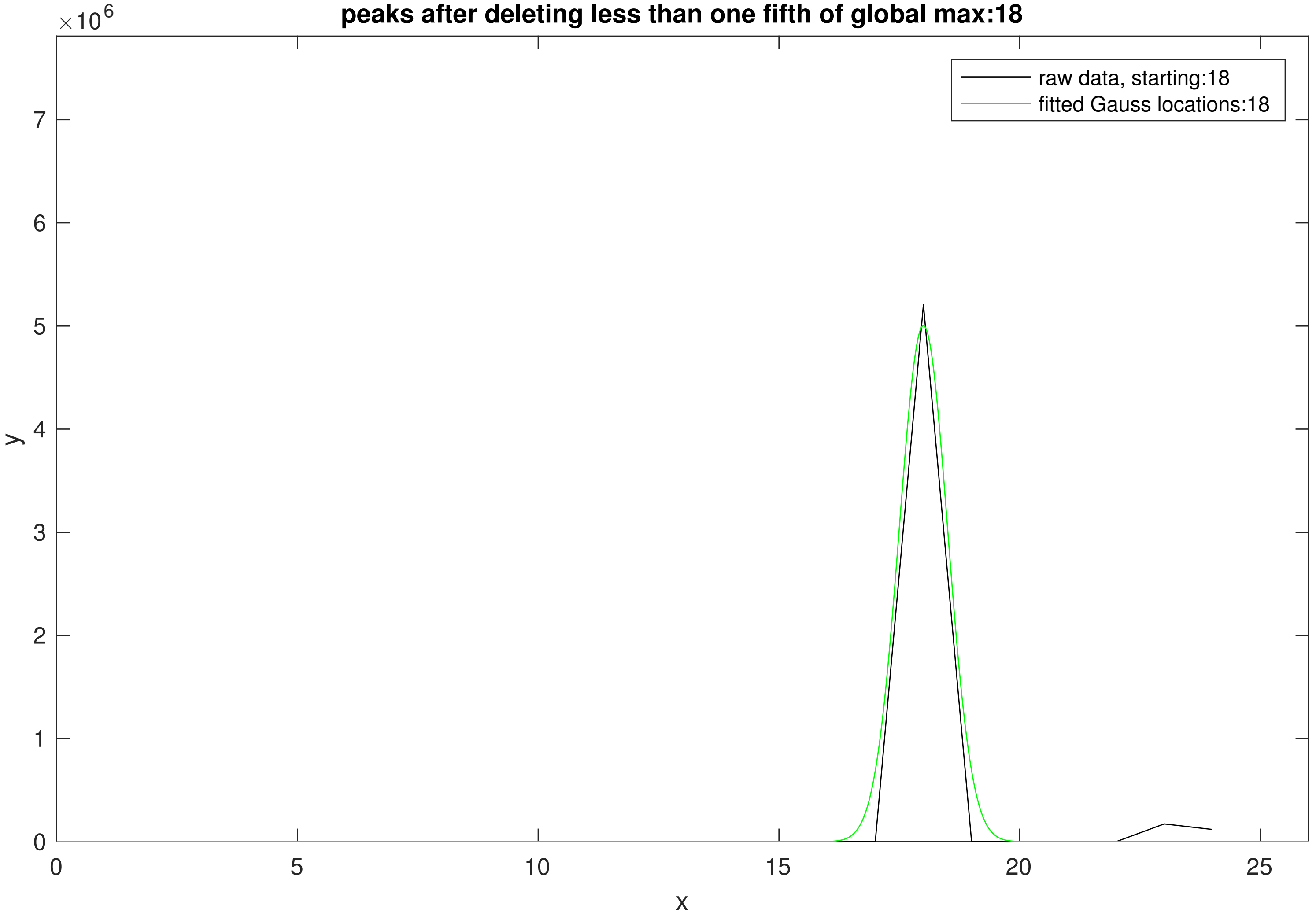

**Optimal1 Fitted Gauss, proteinAT1G22300.1;AT1G22300.2;AT1G22300.3**
**bic:1.03e+03  1.04e+03  1.05e+03**
**peaks after deleting less than one fifth of global max:13**

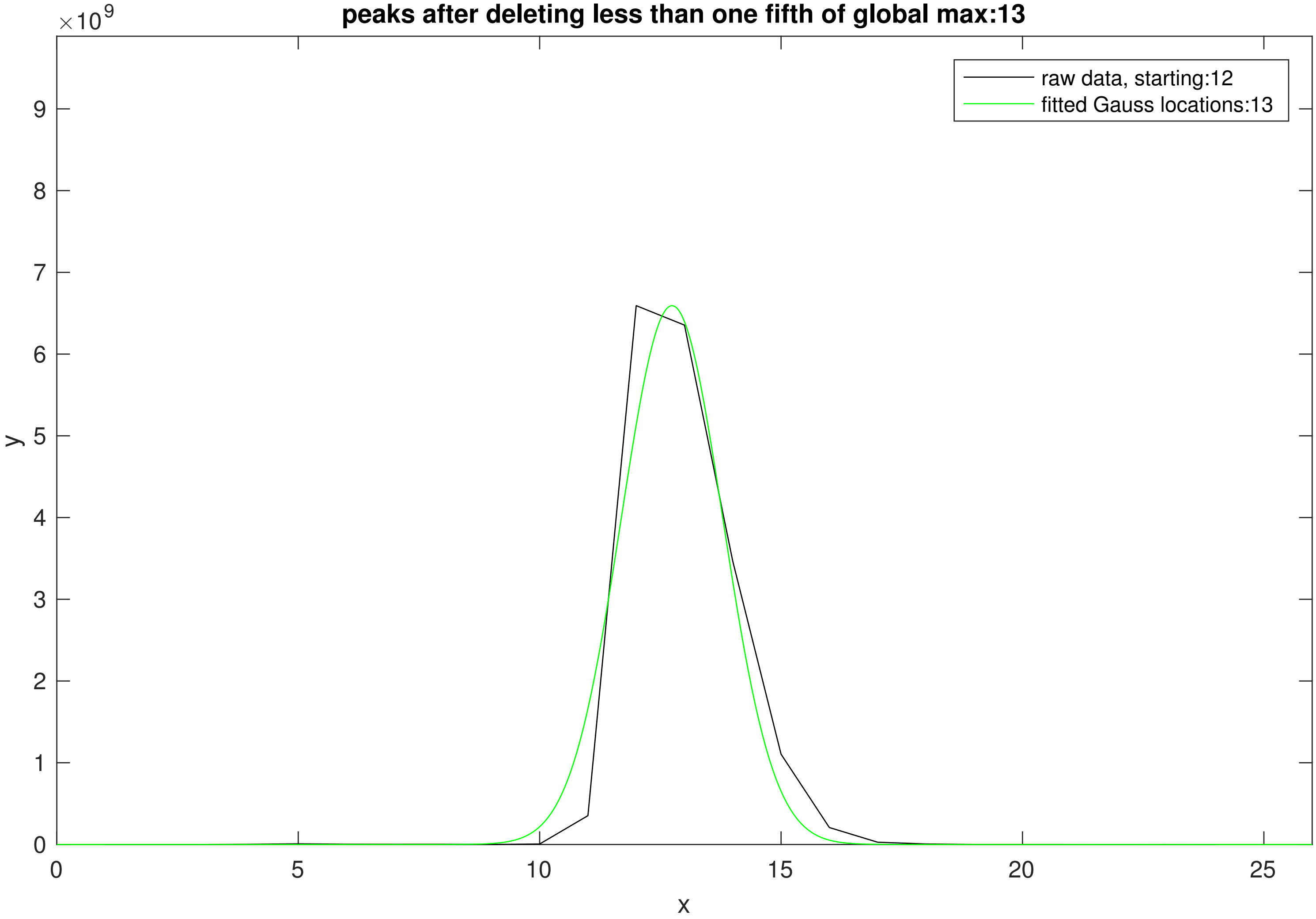

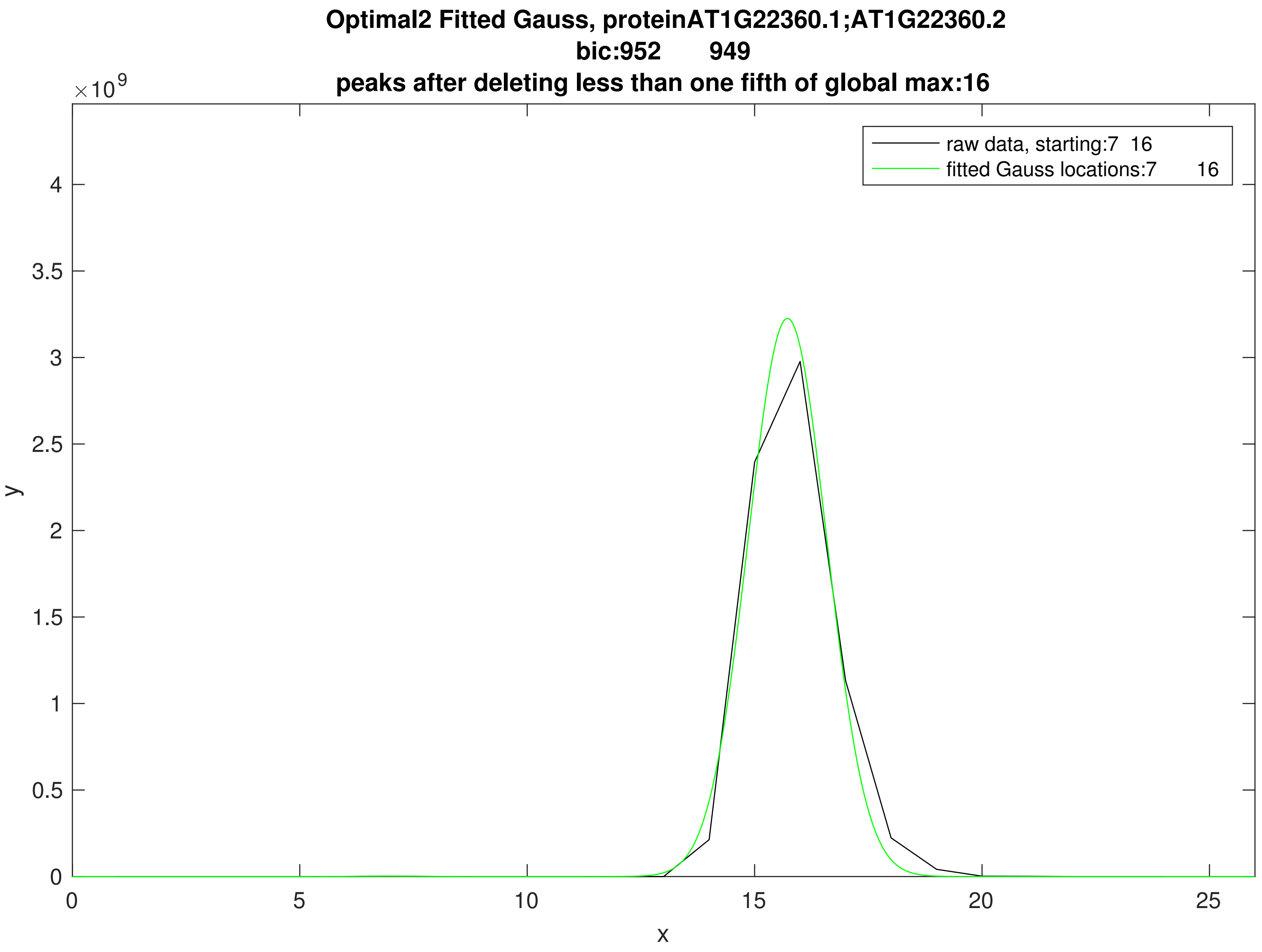

Optimal2 Fitted Gauss, proteinAT1G22360.1;AT1G22360.2
bic:952 949
peaks after deleting less than one fifth of global max:16
×10^9
raw data, starting:7 16
fitted Gauss locations:7 16
y
x
0
0.5
1
1.5
2
2.5
3
3.5
4
5
10
15
20
25

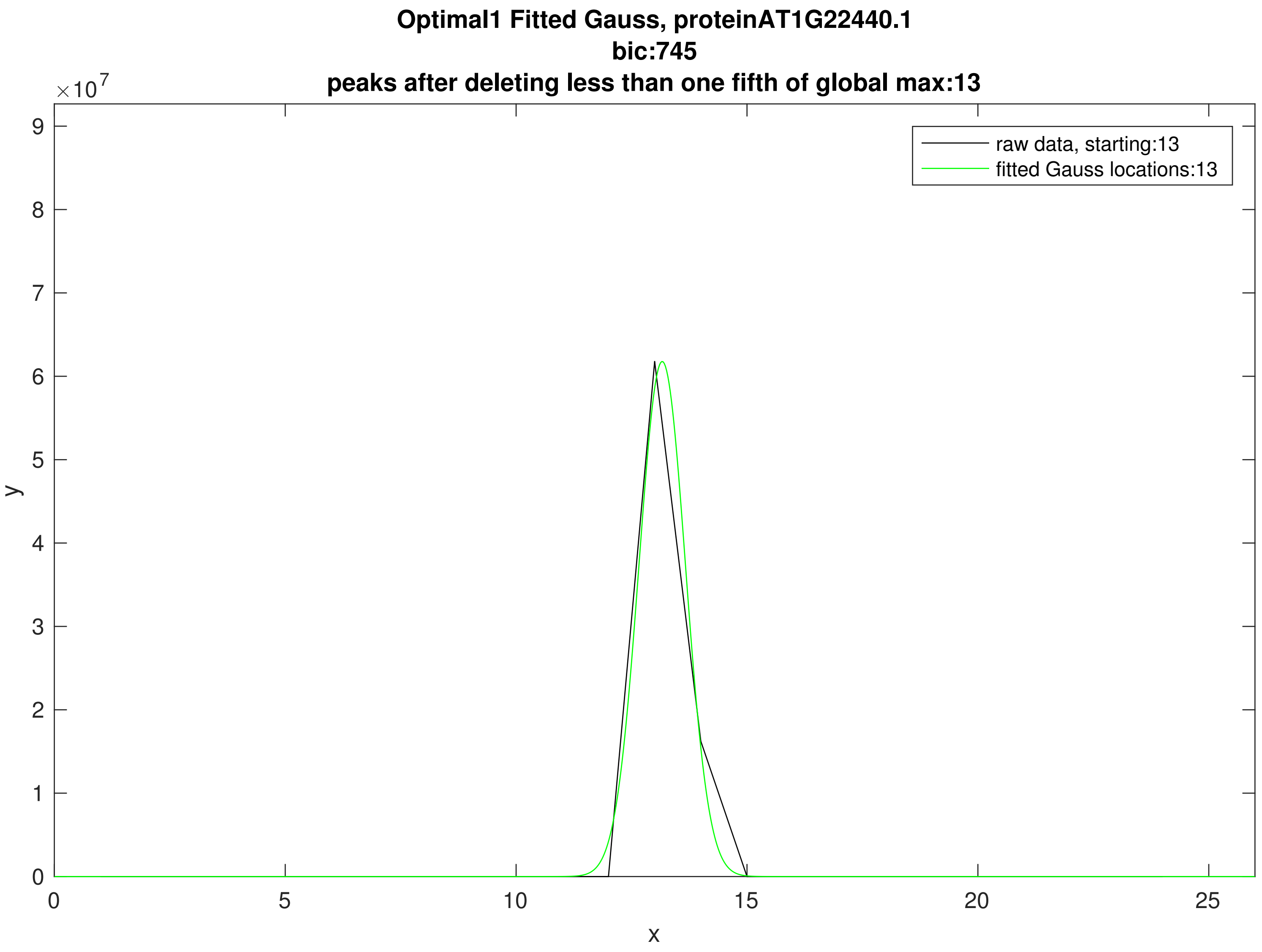
Optimal1 Fitted Gauss, proteinAT1G22440.1
bic:745
peaks after deleting less than one fifth of global max:13
×10^7
raw data, starting:13
fitted Gauss locations:13
y
x
0
1
2
3
4
5
6
7
8
9
0
5
10
15
20
25

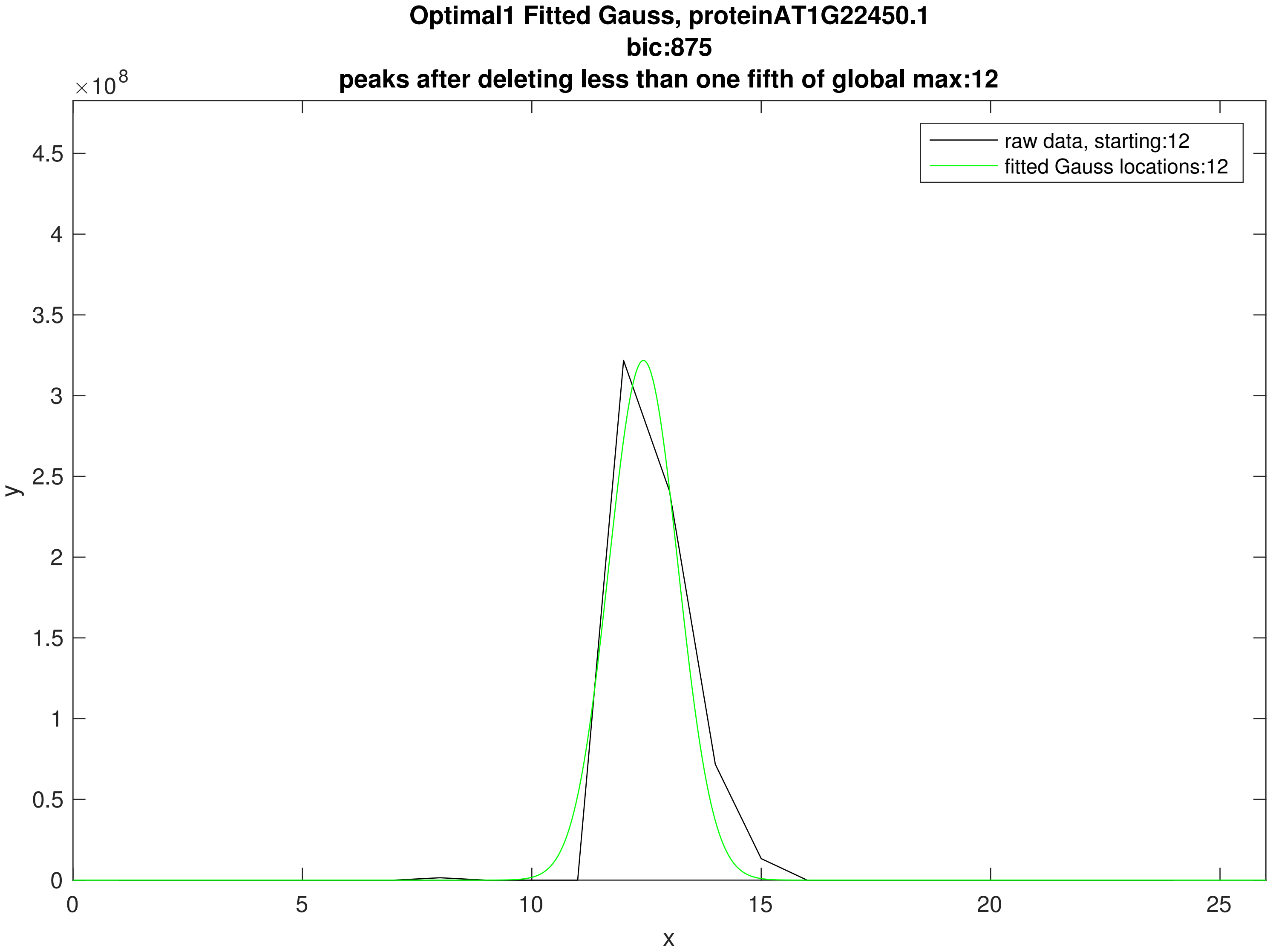

Optimal1 Fitted Gauss, proteinAT1G22450.1
bic:875
peaks after deleting less than one fifth of global max:12
×10^8
4.5
4
3.5
3
2.5
2
1.5
1
0.5
0
y
0
5
10
15
20
25
x
raw data, starting:12
fitted Gauss locations:12

Optimal2 Fitted Gauss, proteinAT1G22530.1
bic:788 778 796
peaks after deleting less than one fifth of global max:8.1 19

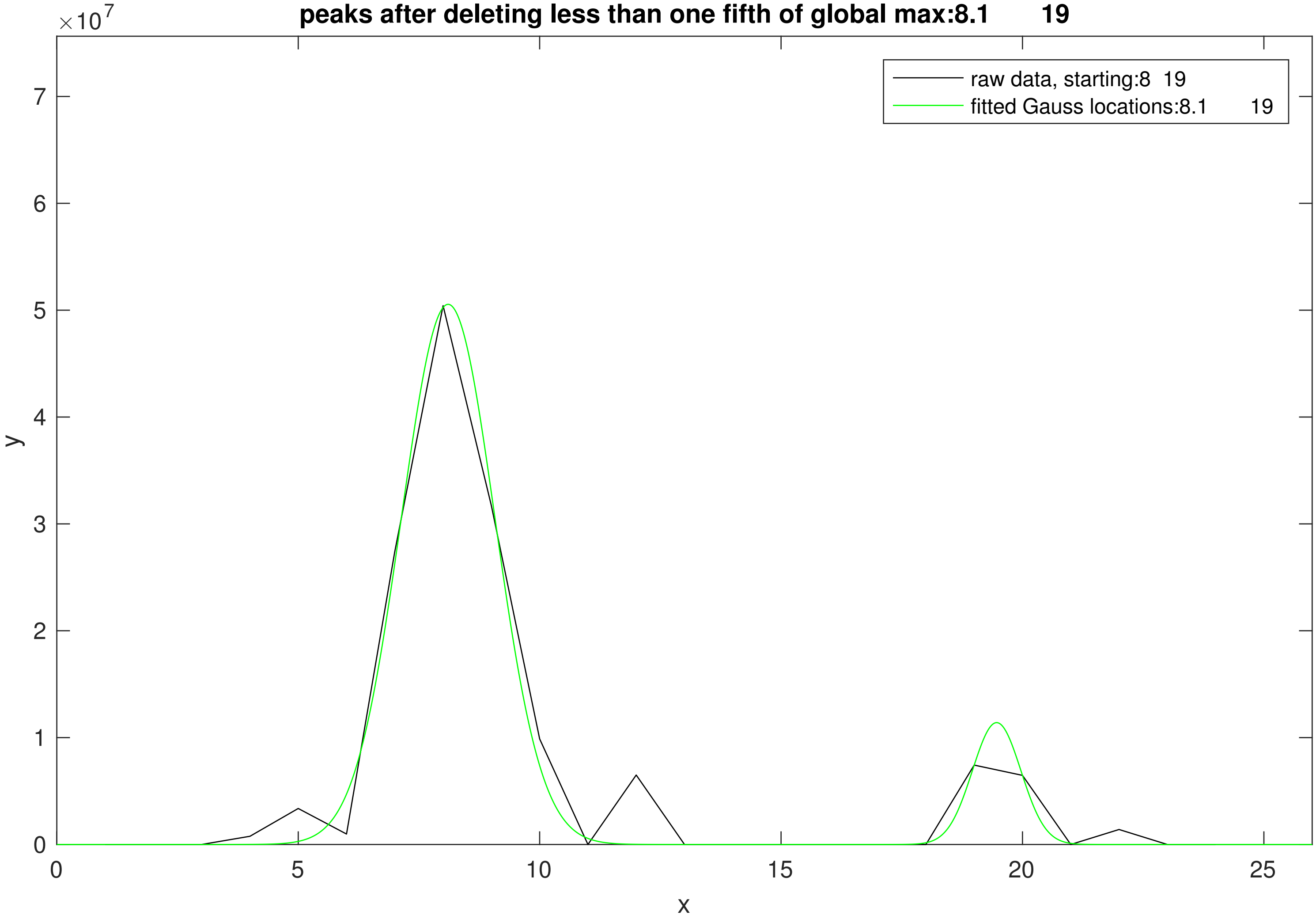

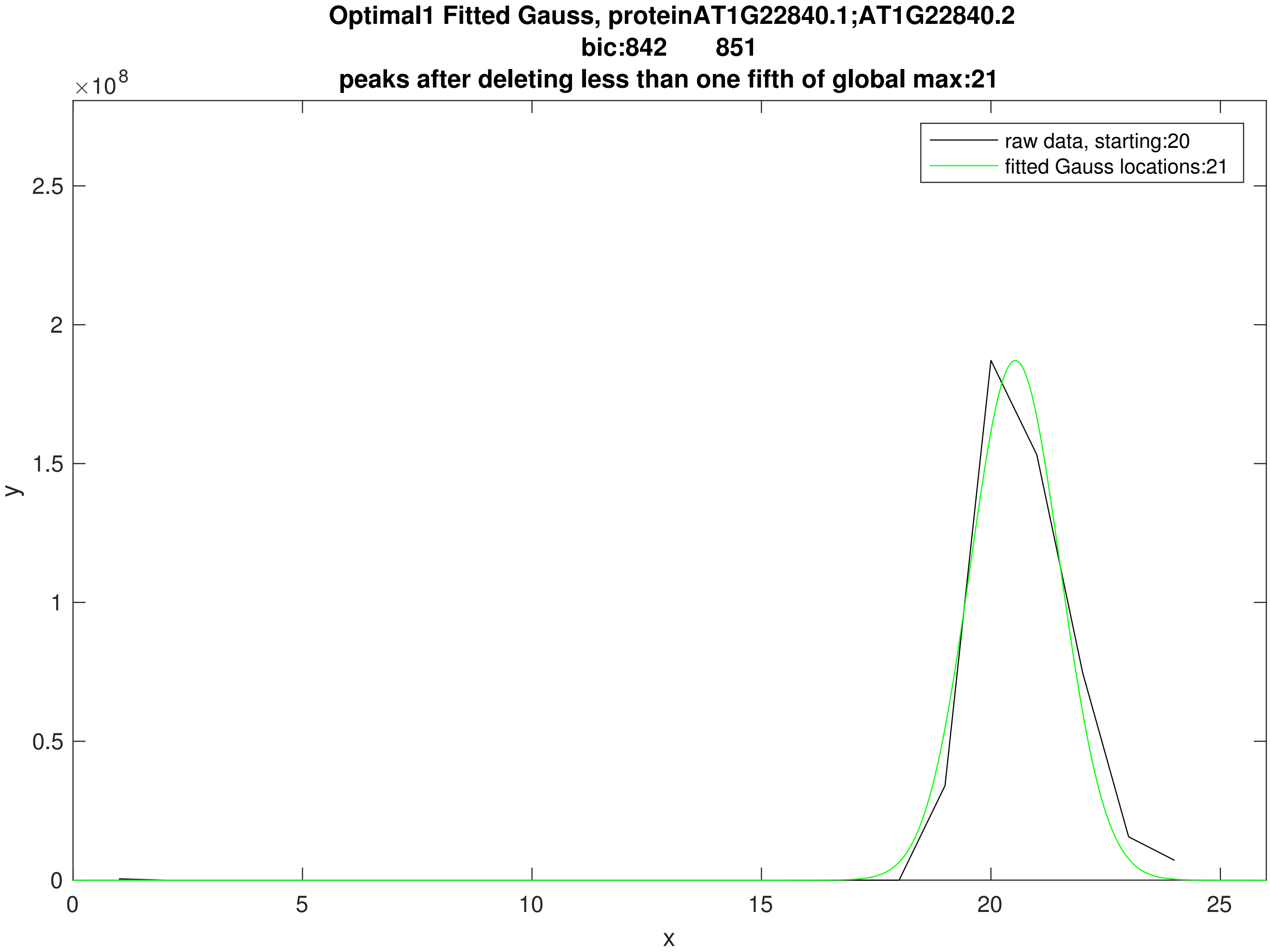
Optimal1 Fitted Gauss, proteinAT1G22840.1;AT1G22840.2
bic:842 851
peaks after deleting less than one fifth of global max:21
×10^8
raw data, starting:20
fitted Gauss locations:21
2.5
2
1.5
1
0.5
0
y
0
5
10
15
20
25
x

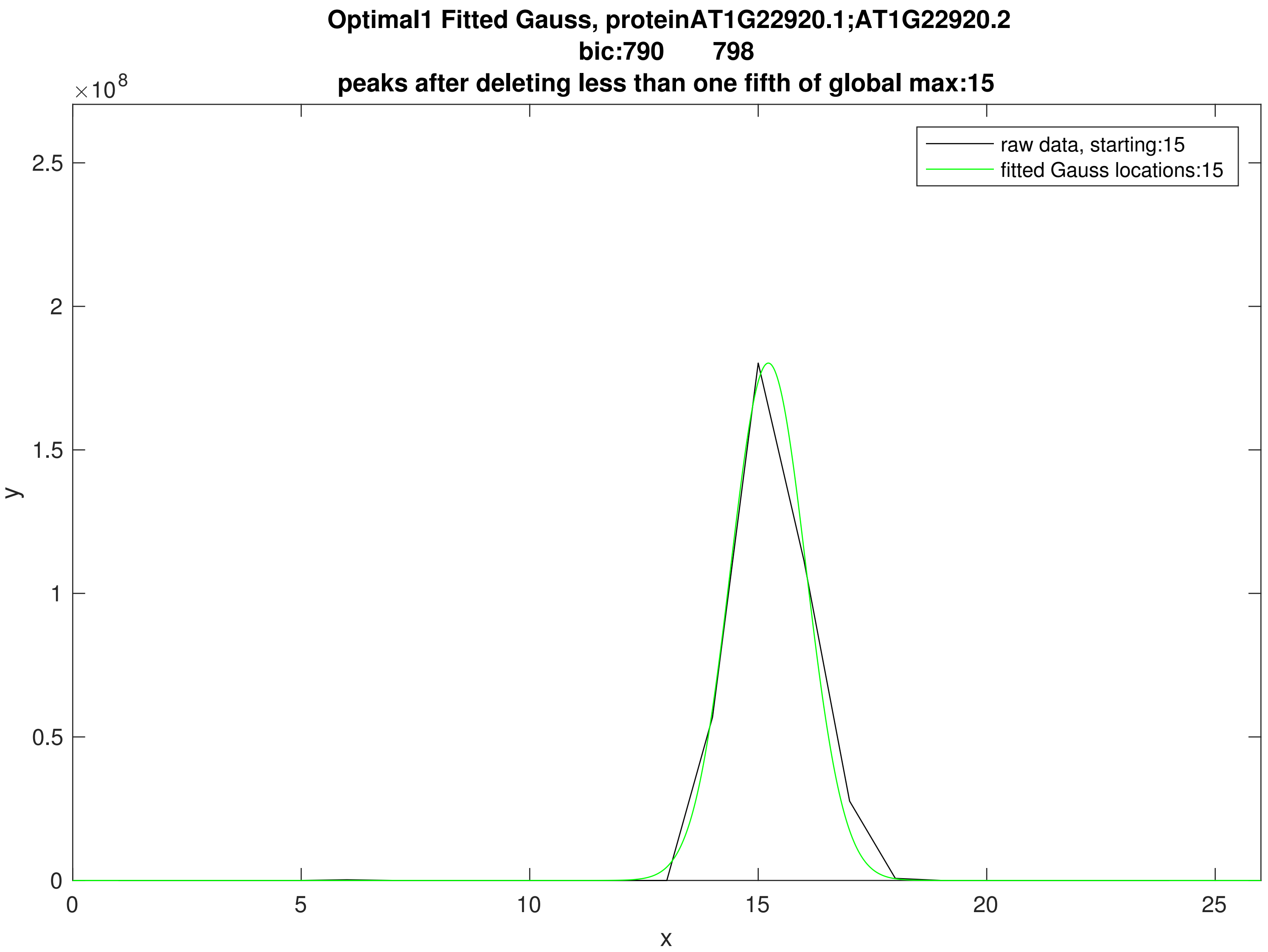
Optimal1 Fitted Gauss, proteinAT1G22920.1;AT1G22920.2
bic:790 798
peaks after deleting less than one fifth of global max:15
×10^8
2.5
2
1.5
1
0.5
0
y
0
5
10
15
20
25
x
raw data, starting:15
fitted Gauss locations:15

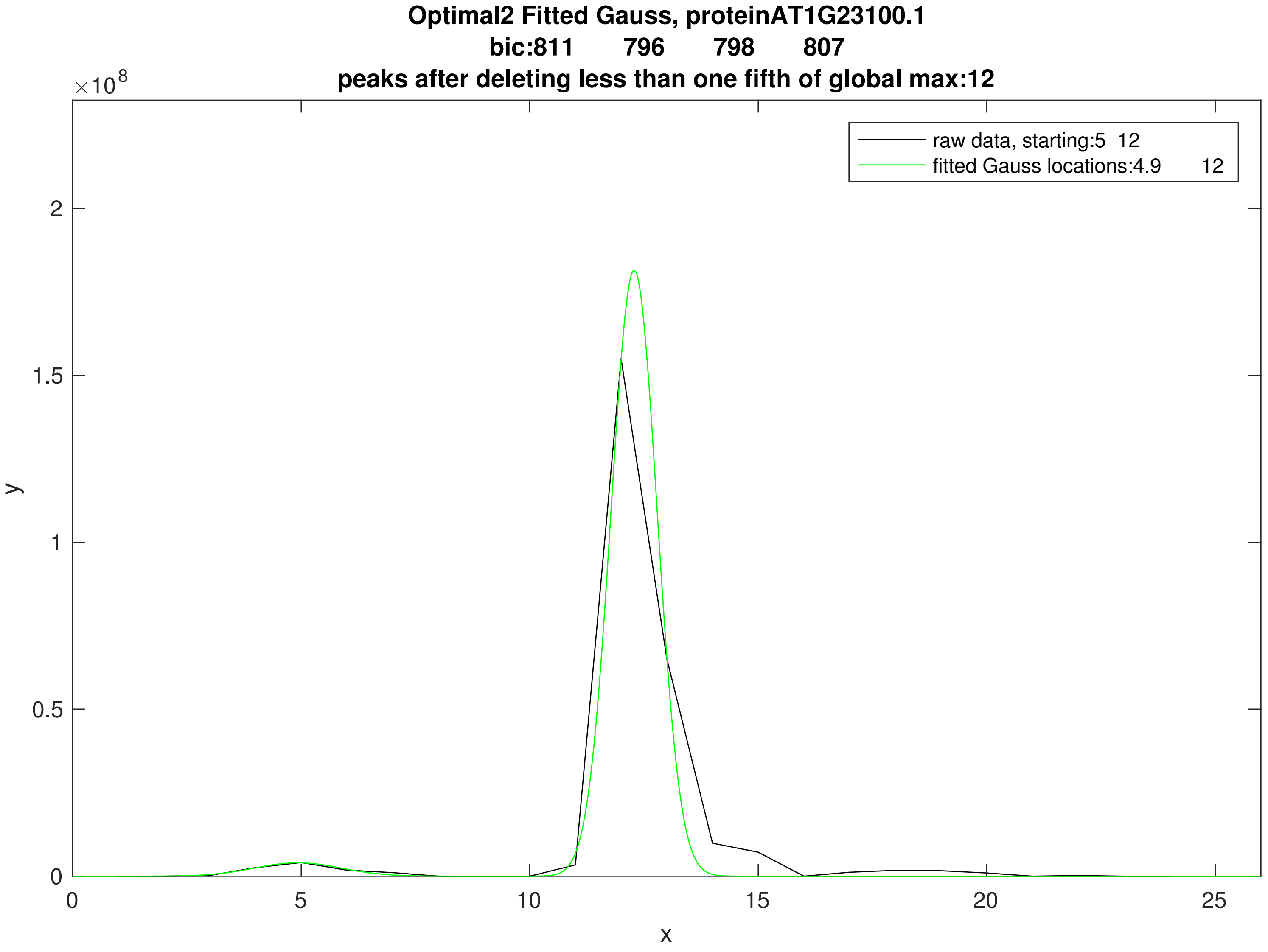

Optimal2 Fitted Gauss, proteinAT1G23100.1
bic:811 796 798 807
peaks after deleting less than one fifth of global max:12
×10^8
raw data, starting:5 12
fitted Gauss locations:4.9 12
2
1.5
1
0.5
0
y
0
5
10
15
20
25
x

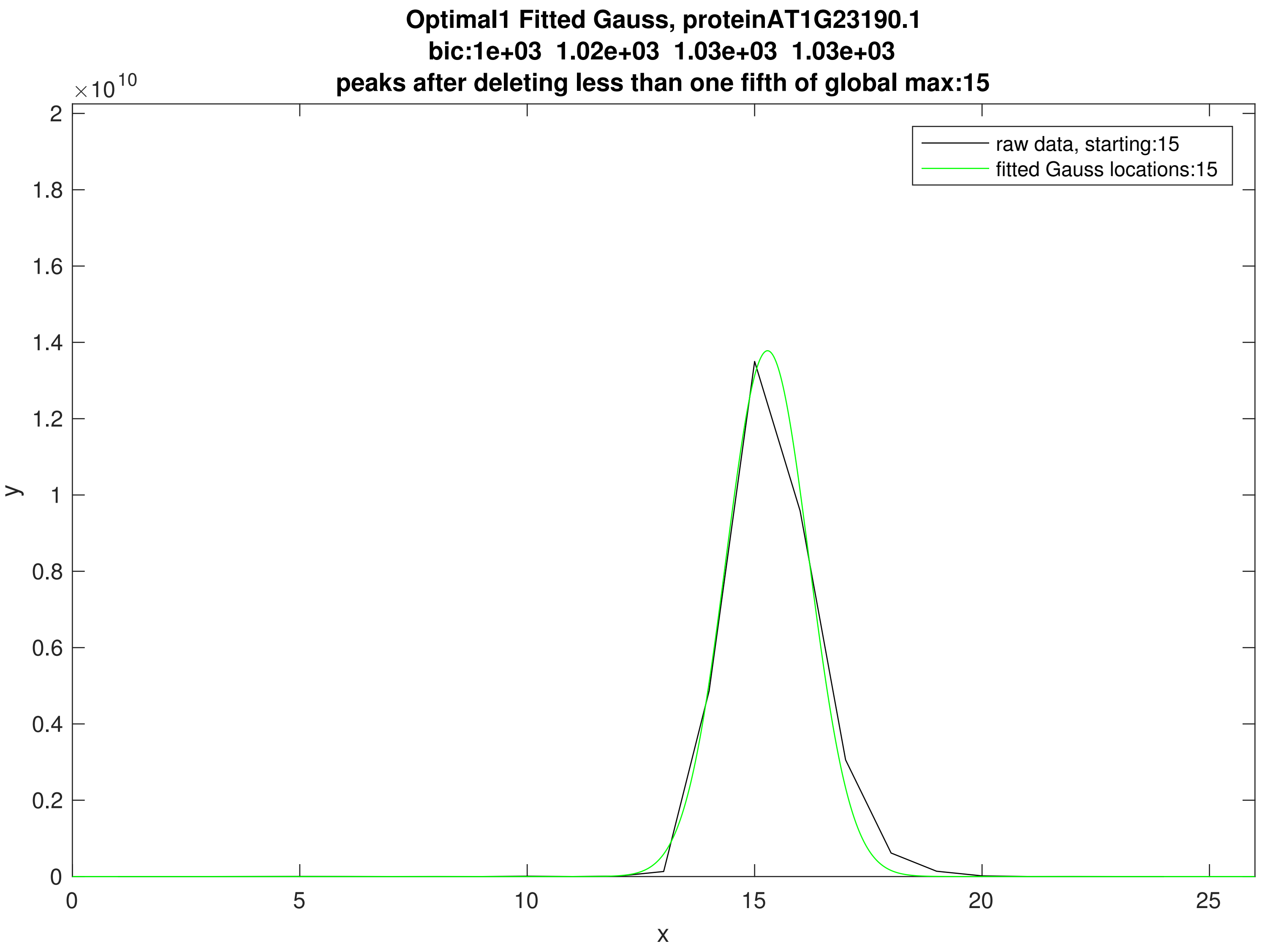

Optimal1 Fitted Gauss, proteinAT1G23190.1
bic:1e+03 1.02e+03 1.03e+03 1.03e+03
peaks after deleting less than one fifth of global max:15
×10^10
2
1.8
1.6
1.4
1.2
1
0.8
0.6
0.4
0.2
0
y
0
5
10
15
20
25
x
raw data, starting:15
fitted Gauss locations:15

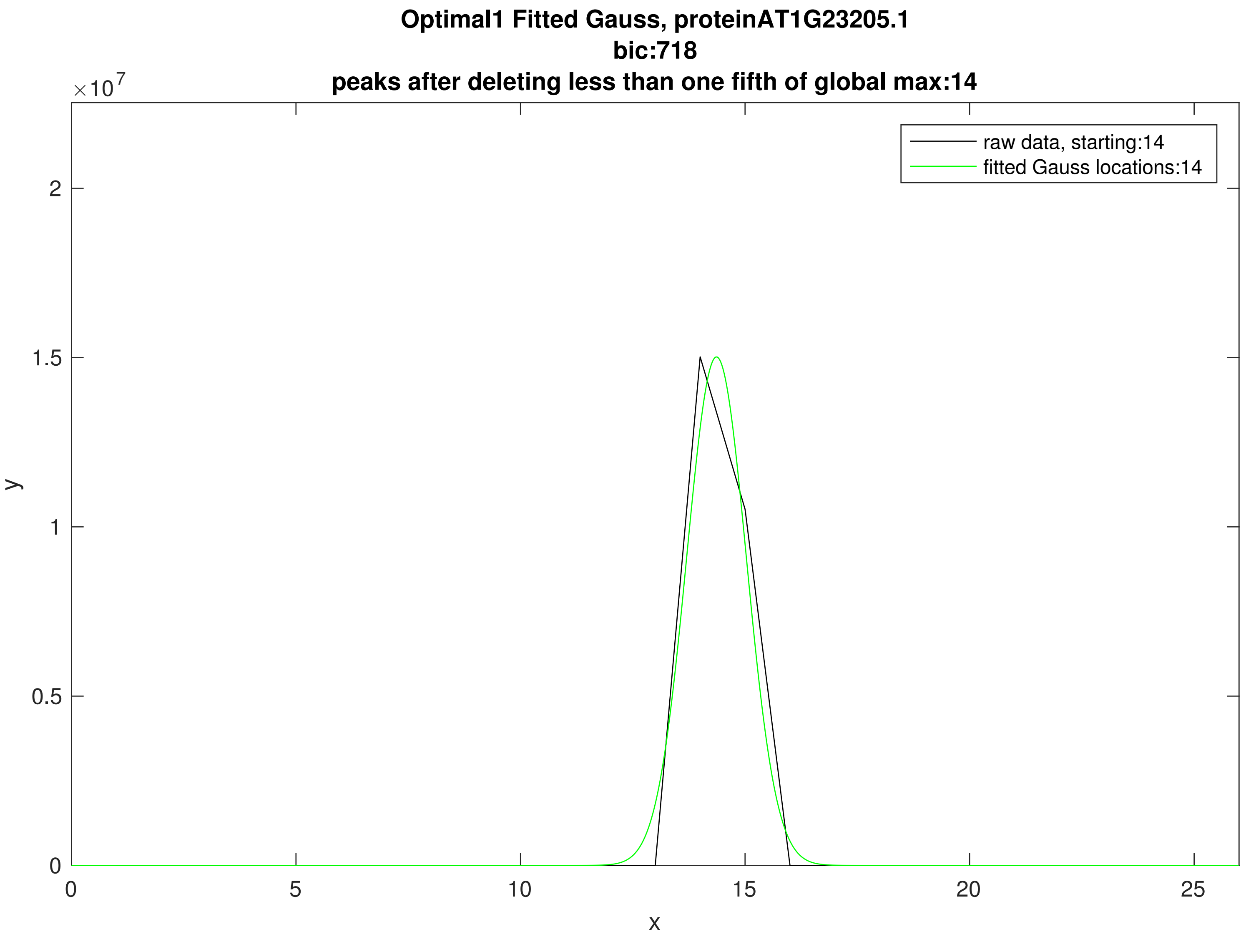
Optimal1 Fitted Gauss, proteinAT1G23205.1
bic:718
peaks after deleting less than one fifth of global max:14
×10^7
raw data, starting:14
fitted Gauss locations:14
2
1.5
1
0.5
0
y
0
5
10
15
20
25
x

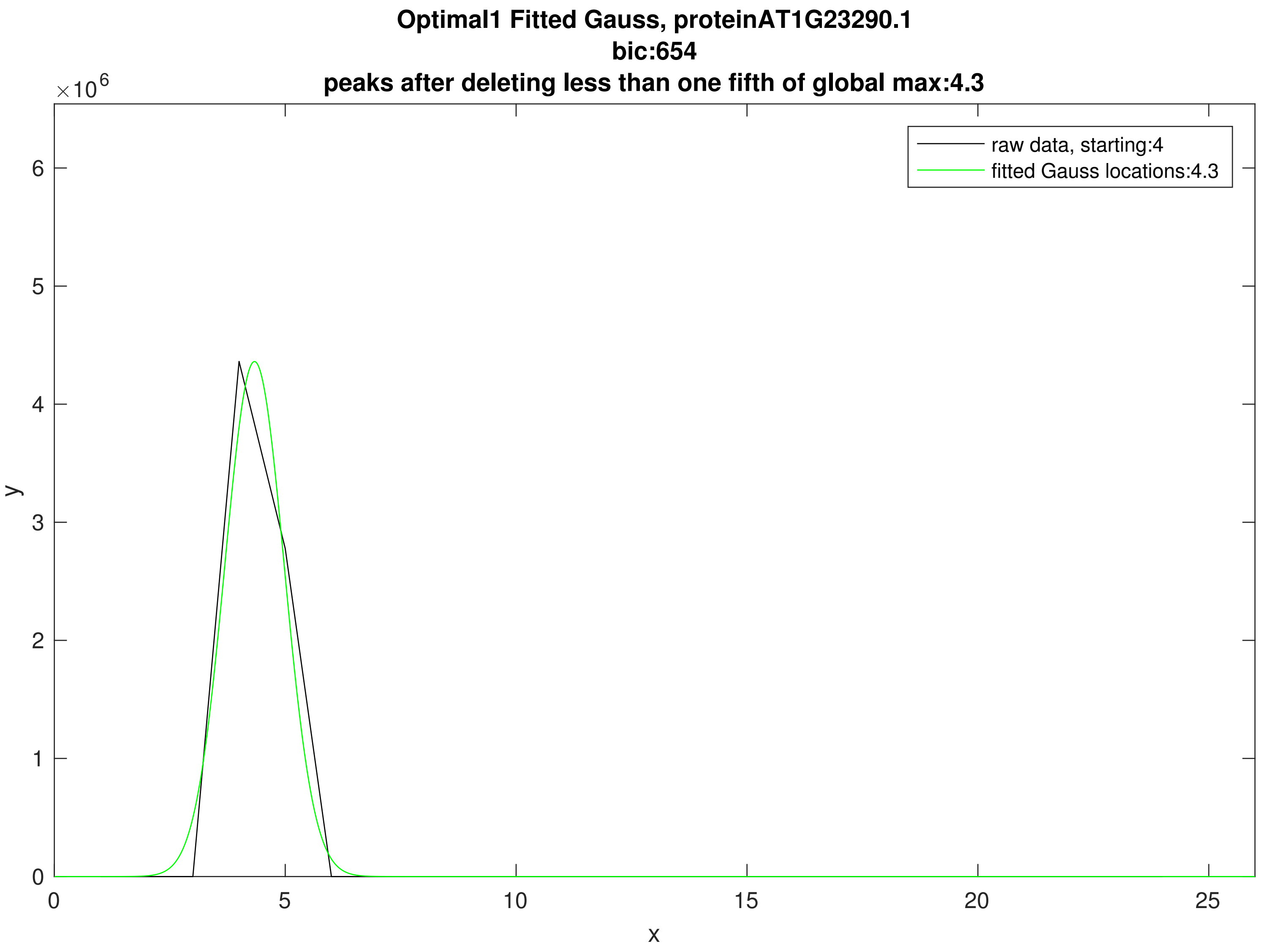

Optimal1 Fitted Gauss, proteinAT1G23290.1
bic:654
peaks after deleting less than one fifth of global max:4.3
×10^6
raw data, starting:4
fitted Gauss locations:4.3
y
x
0
1
2
3
4
5
6
0
5
10
15
20
25

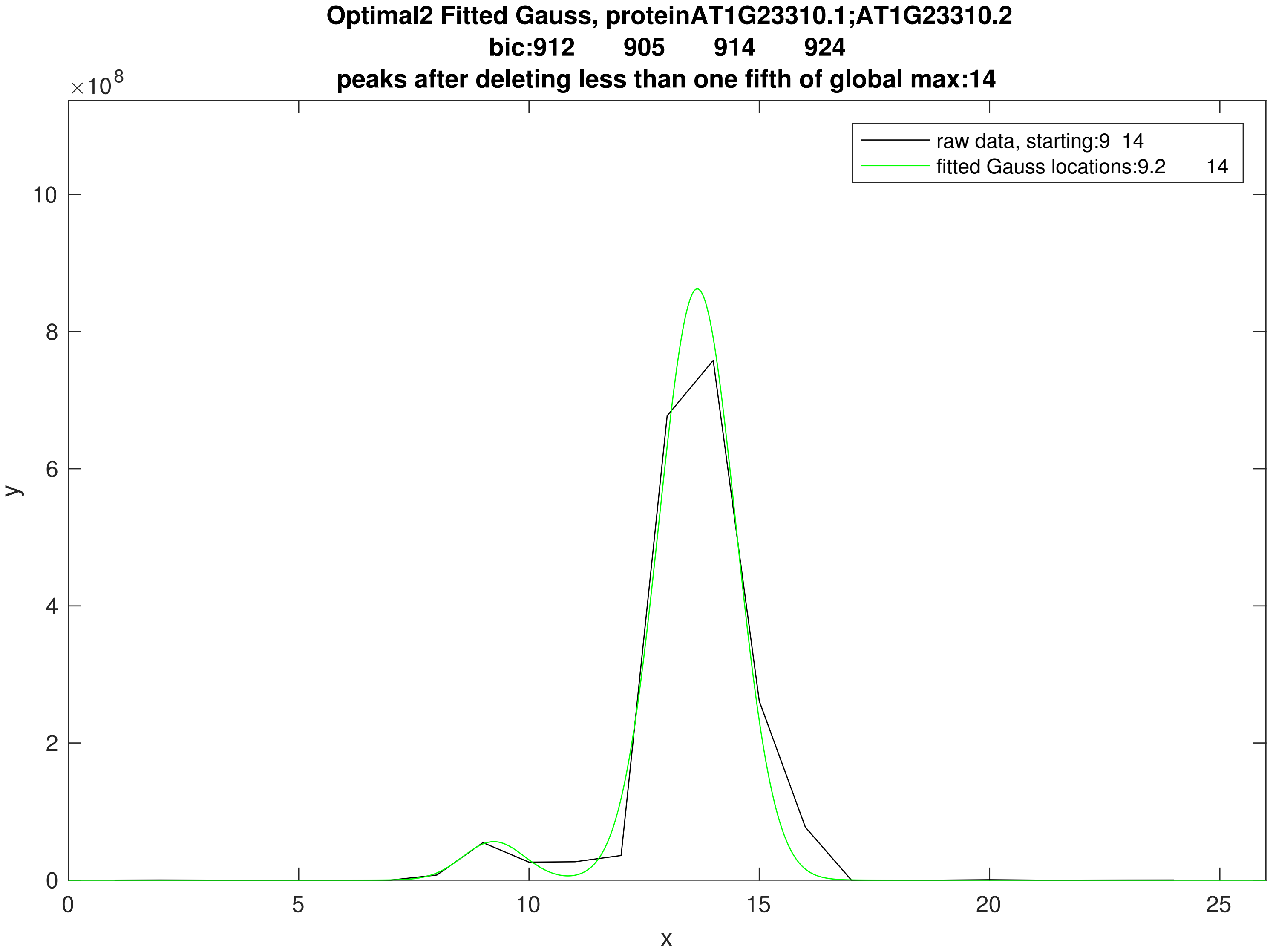

Optimal2 Fitted Gauss, proteinAT1G23310.1;AT1G23310.2
bic:912 905 914 924
peaks after deleting less than one fifth of global max:14
×10^8
raw data, starting:9 14
fitted Gauss locations:9.2 14
y
x
0
2
4
6
8
10
5
15
20
25

#### Optimal1 Fitted Gauss, proteinAT1G23440.1
#### bic:826 834 844
#### peaks after deleting less than one fifth of global max:19

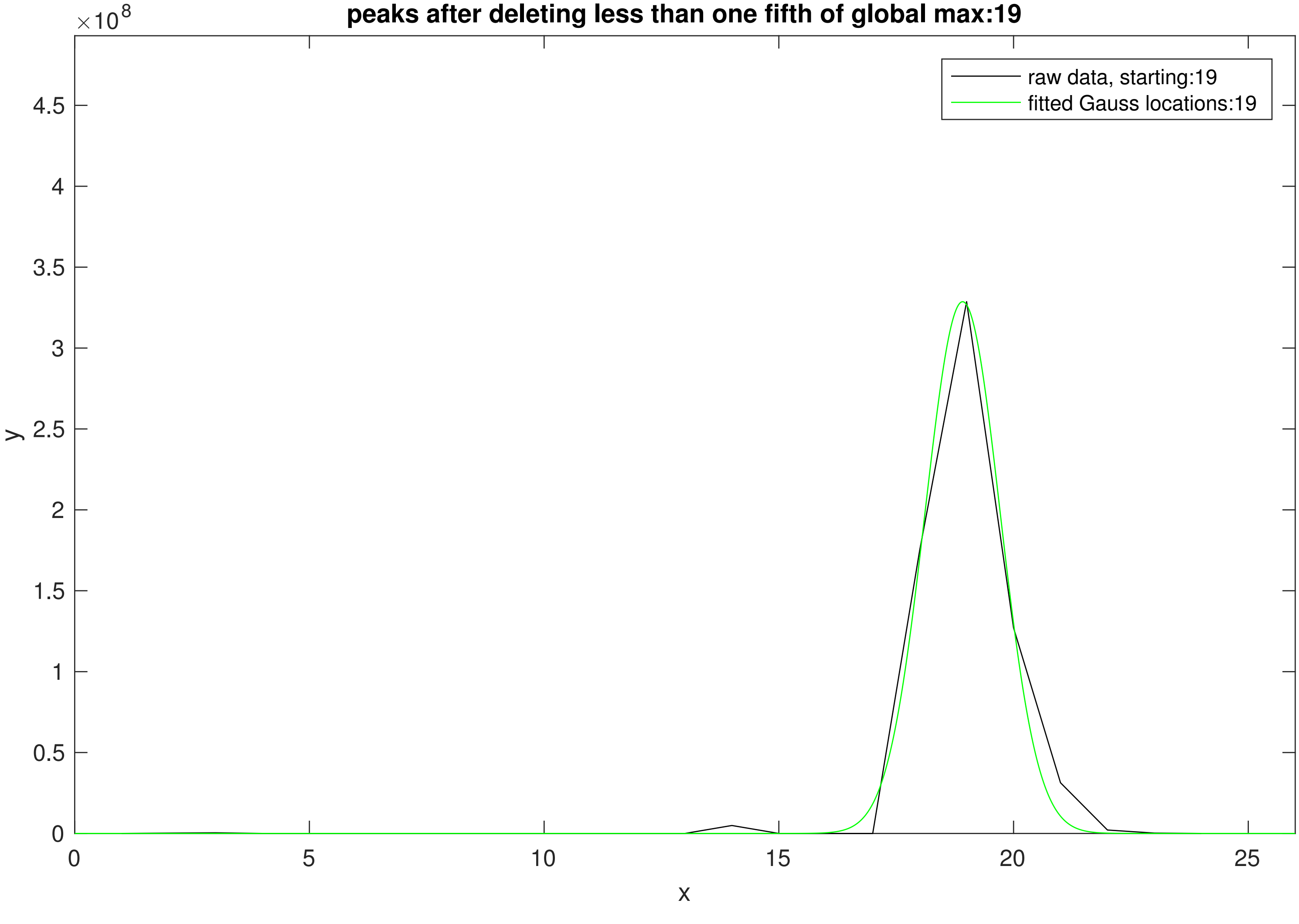

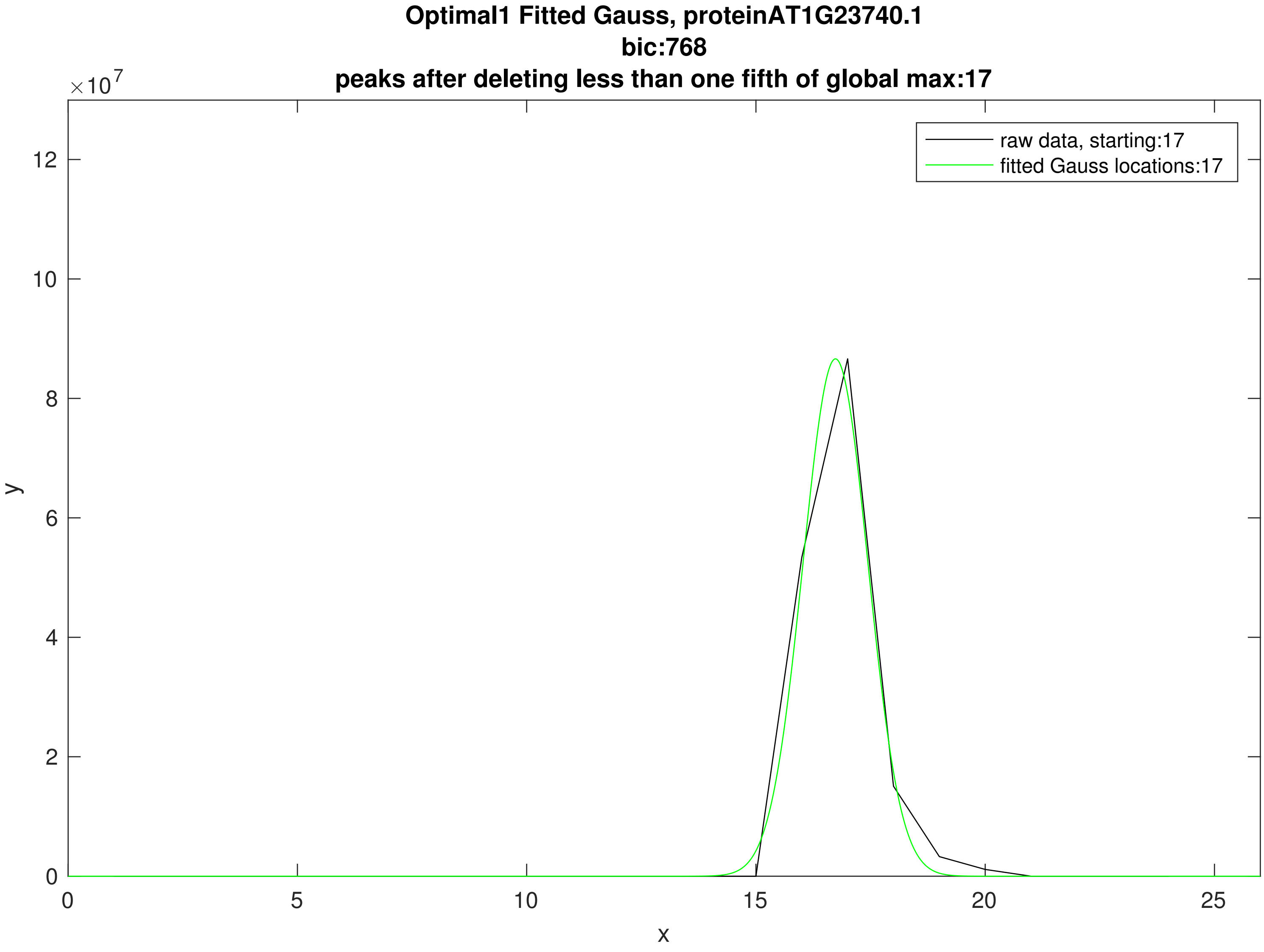

Optimal1 Fitted Gauss, proteinAT1G23740.1
bic:768
peaks after deleting less than one fifth of global max:17
×10^7
raw data, starting:17
fitted Gauss locations:17
y
x
0
2
4
6
8
10
12
0
5
10
15
20
25

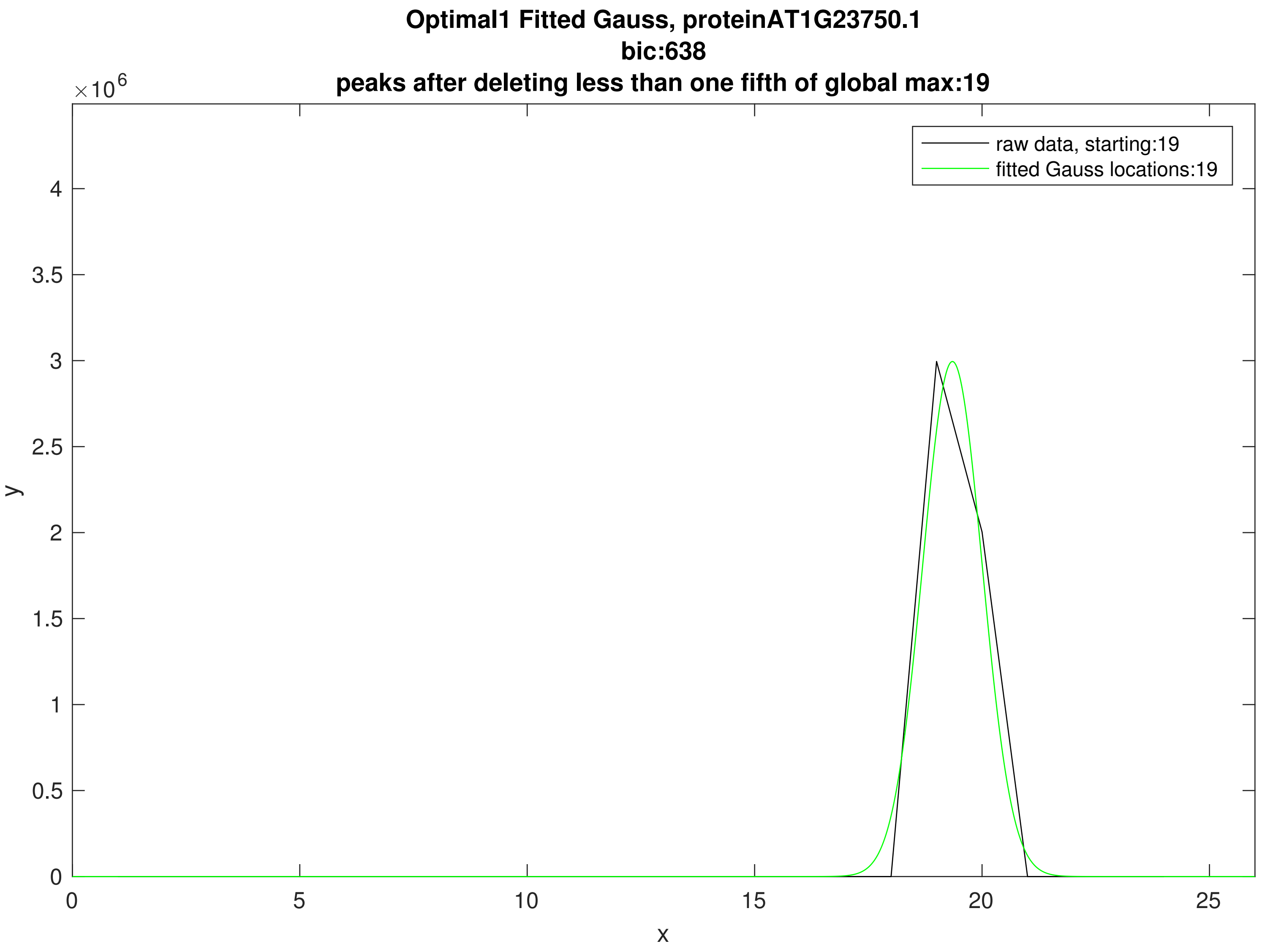
Optimal1 Fitted Gauss, proteinAT1G23750.1
bic:638
peaks after deleting less than one fifth of global max:19
×10^6
raw data, starting:19
fitted Gauss locations:19
y
x
0
0.5
1
1.5
2
2.5
3
3.5
4
0
5
10
15
20
25

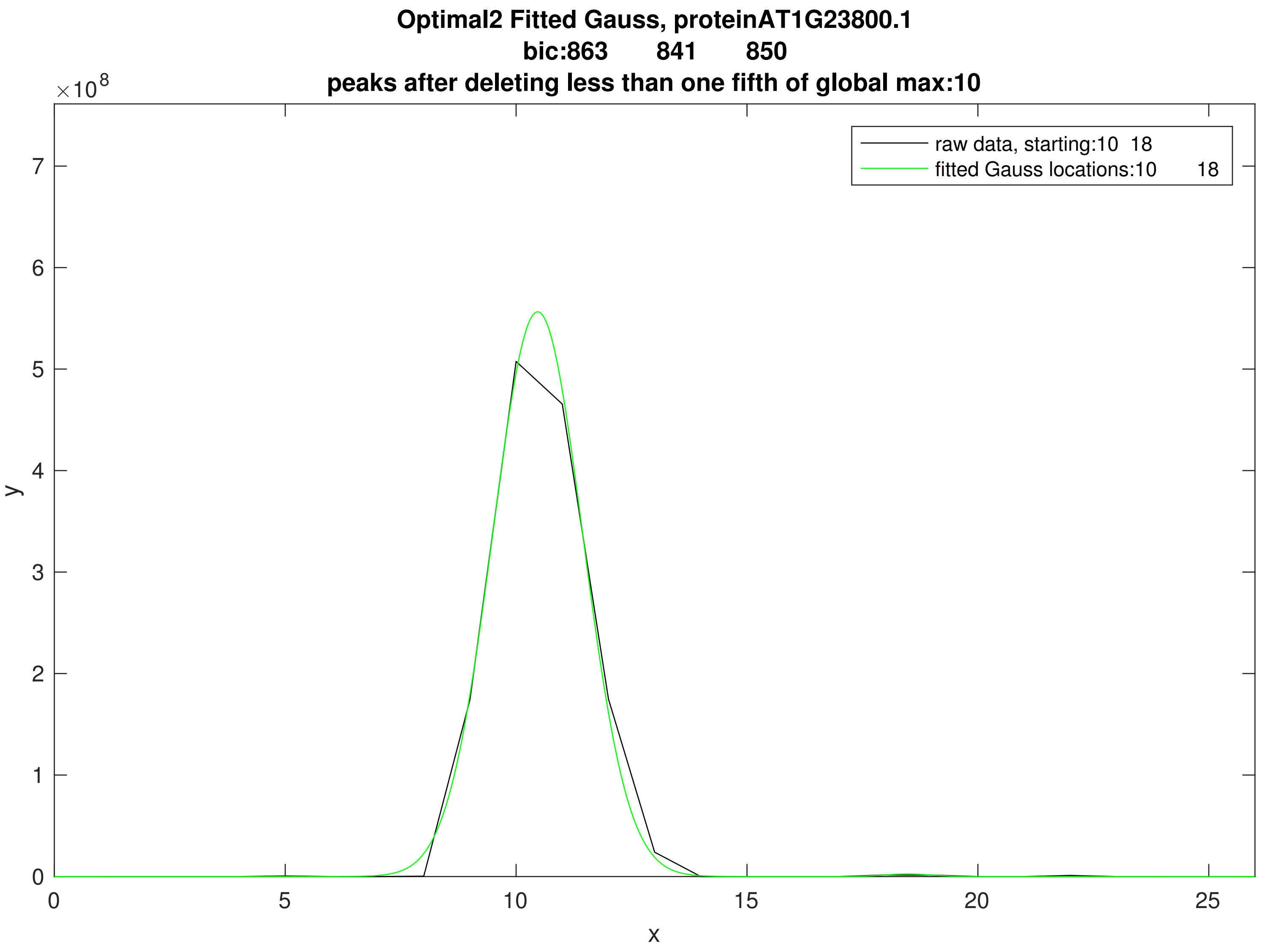

Optimal2 Fitted Gauss, proteinAT1G23800.1
bic:863 841 850
peaks after deleting less than one fifth of global max:10
×10^8
raw data, starting:10 18
fitted Gauss locations:10 18
y
x
0
1
2
3
4
5
6
7
0
5
10
15
20
25

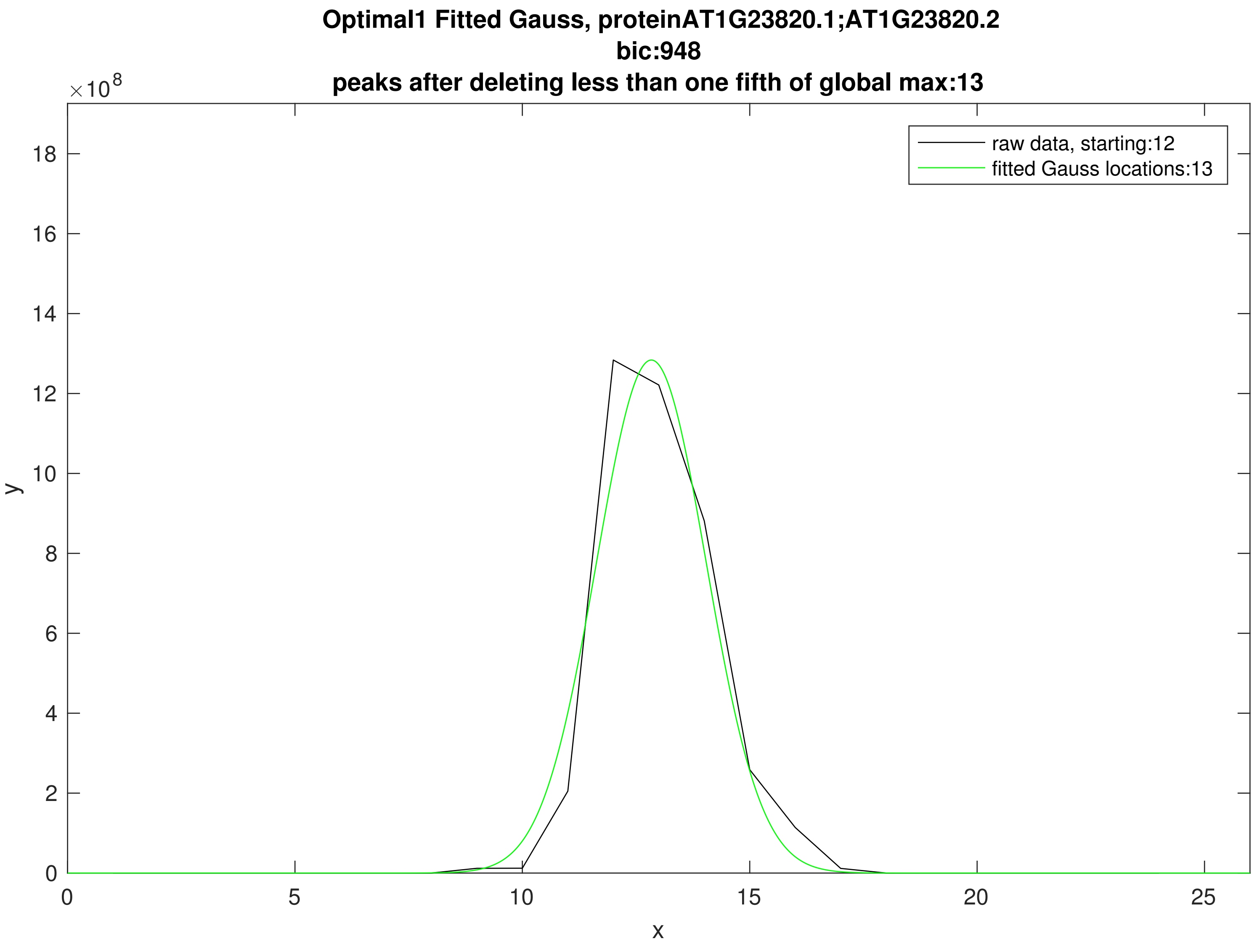

Optimal1 Fitted Gauss, proteinAT1G23820.1;AT1G23820.2
bic:948
peaks after deleting less than one fifth of global max:13
×10^8
raw data, starting:12
fitted Gauss locations:13
y
x
0
2
4
6
8
10
12
14
16
18
0
5
10
15
20
25

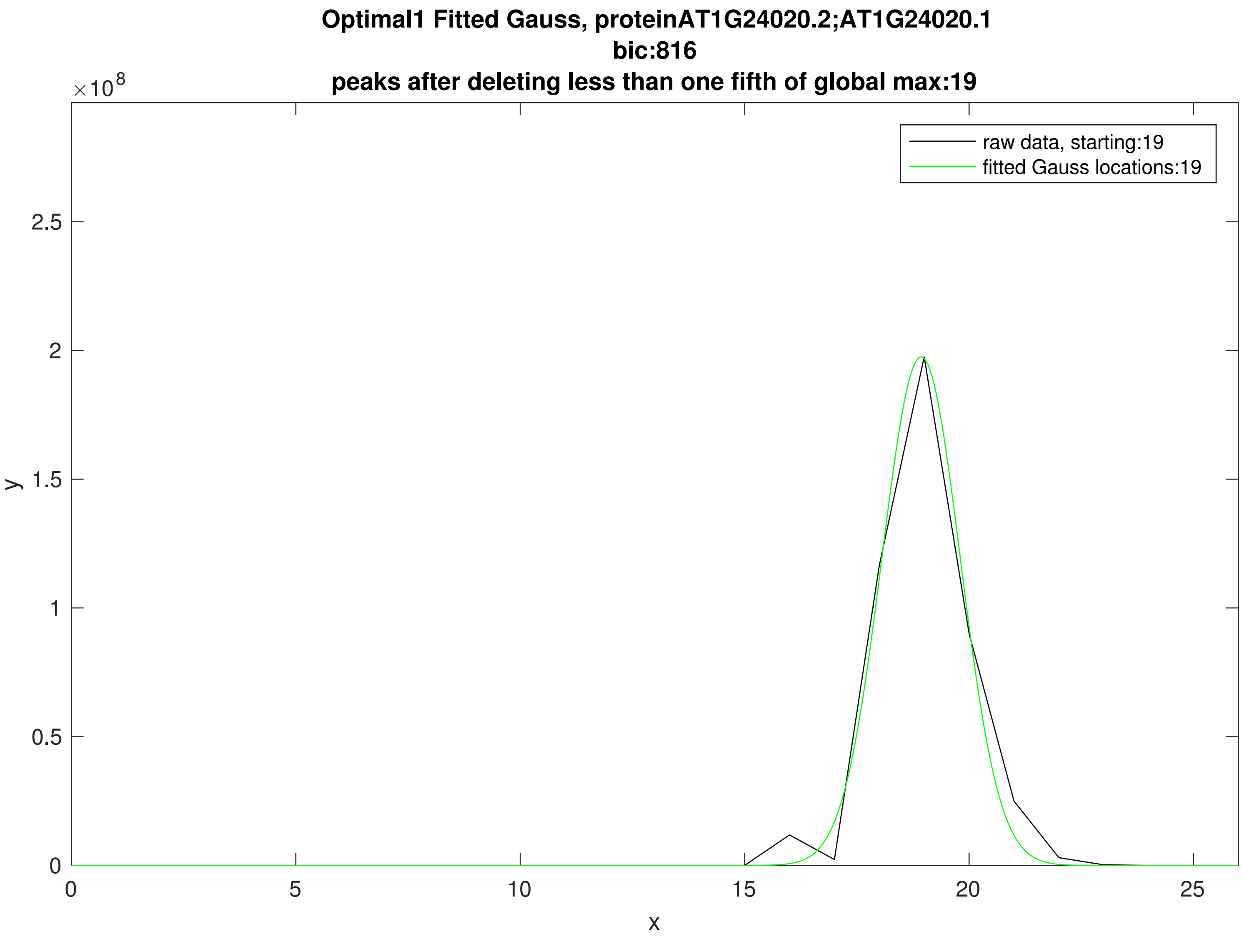

Optimal1 Fitted Gauss, proteinAT1G24020.2;AT1G24020.1
bic:816
peaks after deleting less than one fifth of global max:19
×10^8
raw data, starting:19
fitted Gauss locations:19
2.5
2
1.5
1
0.5
0
y
0
5
10
15
20
25
x

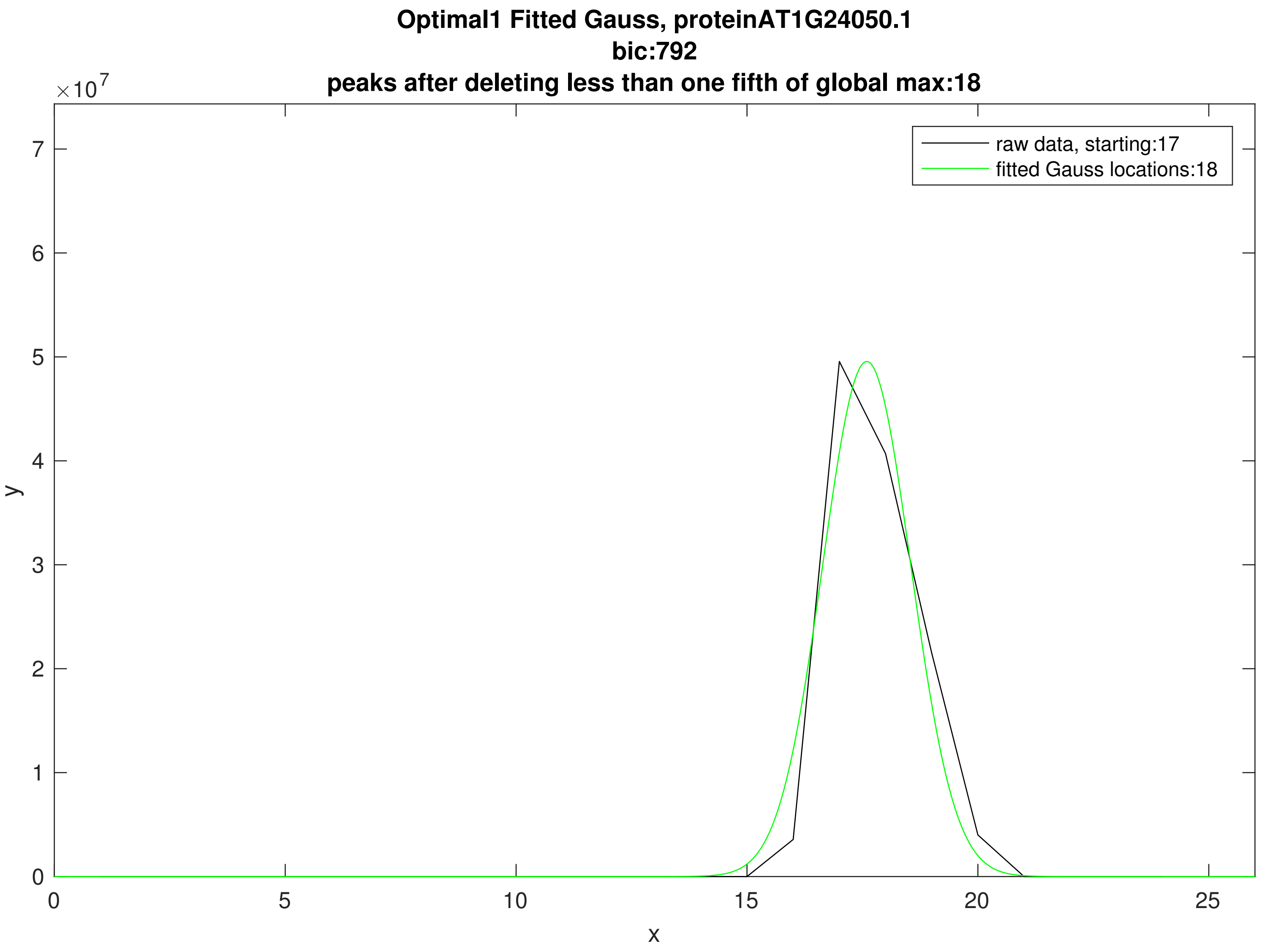
Optimal1 Fitted Gauss, proteinAT1G24050.1
bic:792
peaks after deleting less than one fifth of global max:18
×10^7
raw data, starting:17
fitted Gauss locations:18
y
x
0
1
2
3
4
5
6
7
0
5
10
15
20
25

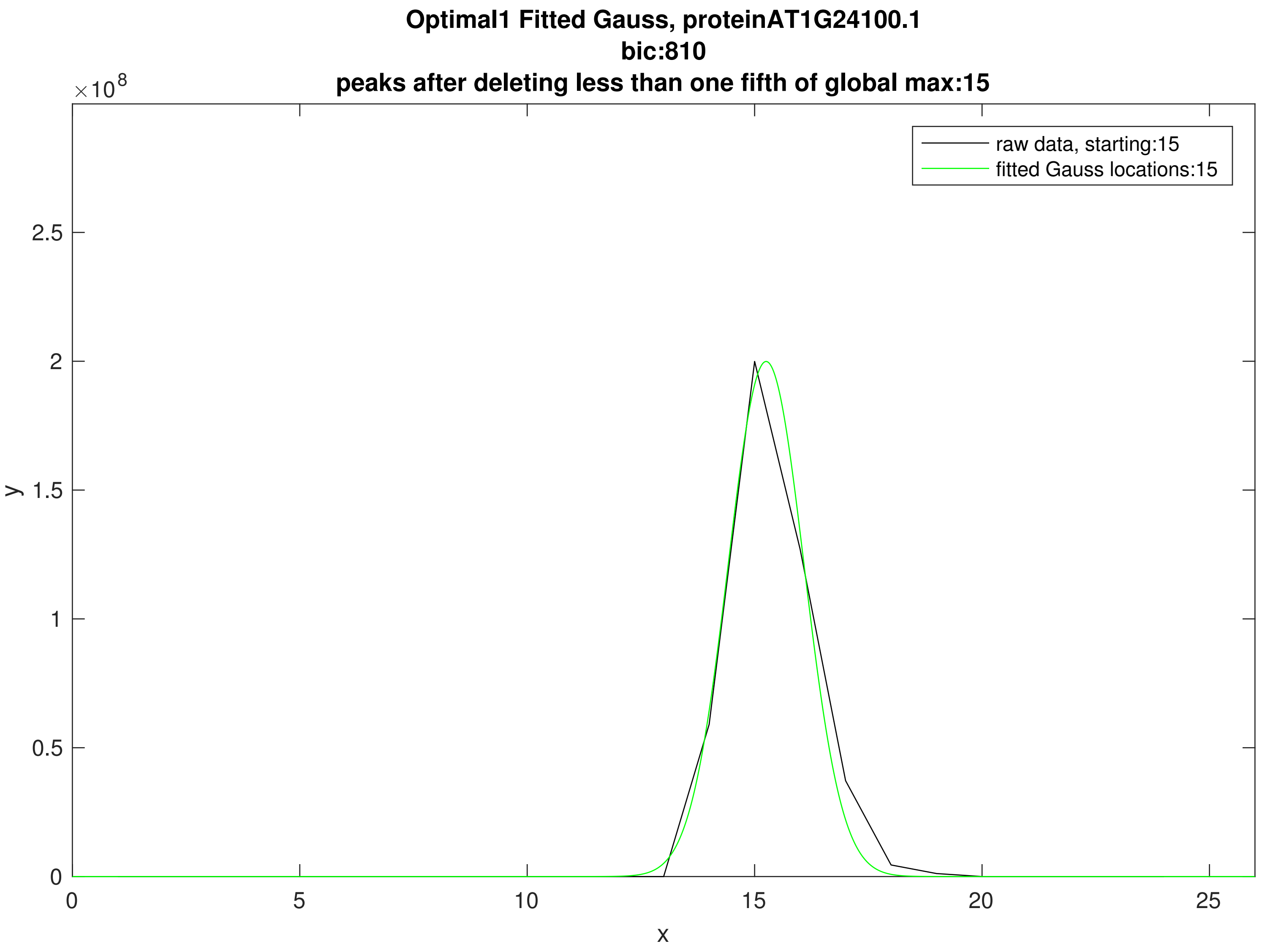
Optimal1 Fitted Gauss, proteinAT1G24100.1
bic:810
peaks after deleting less than one fifth of global max:15
×10^8
raw data, starting:15
fitted Gauss locations:15
2.5
2
y 1.5
1
0.5
0
0
5
10
15
20
25
x

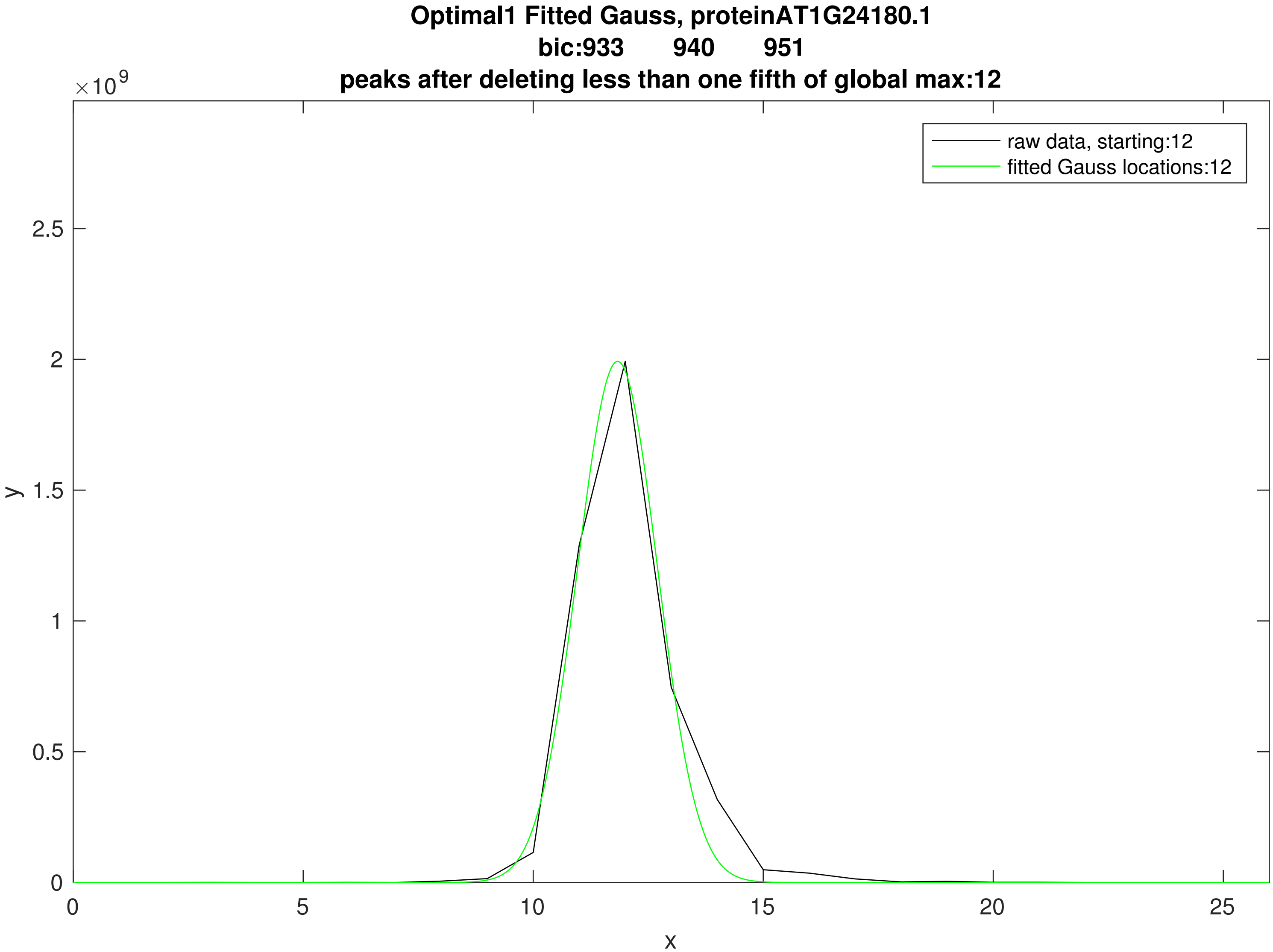
Optimal1 Fitted Gauss, proteinAT1G24180.1
bic:933 940 951
peaks after deleting less than one fifth of global max:12
×10^9
raw data, starting:12
fitted Gauss locations:12
y
x
0
0.5
1
1.5
2
2.5
0
5
10
15
20
25

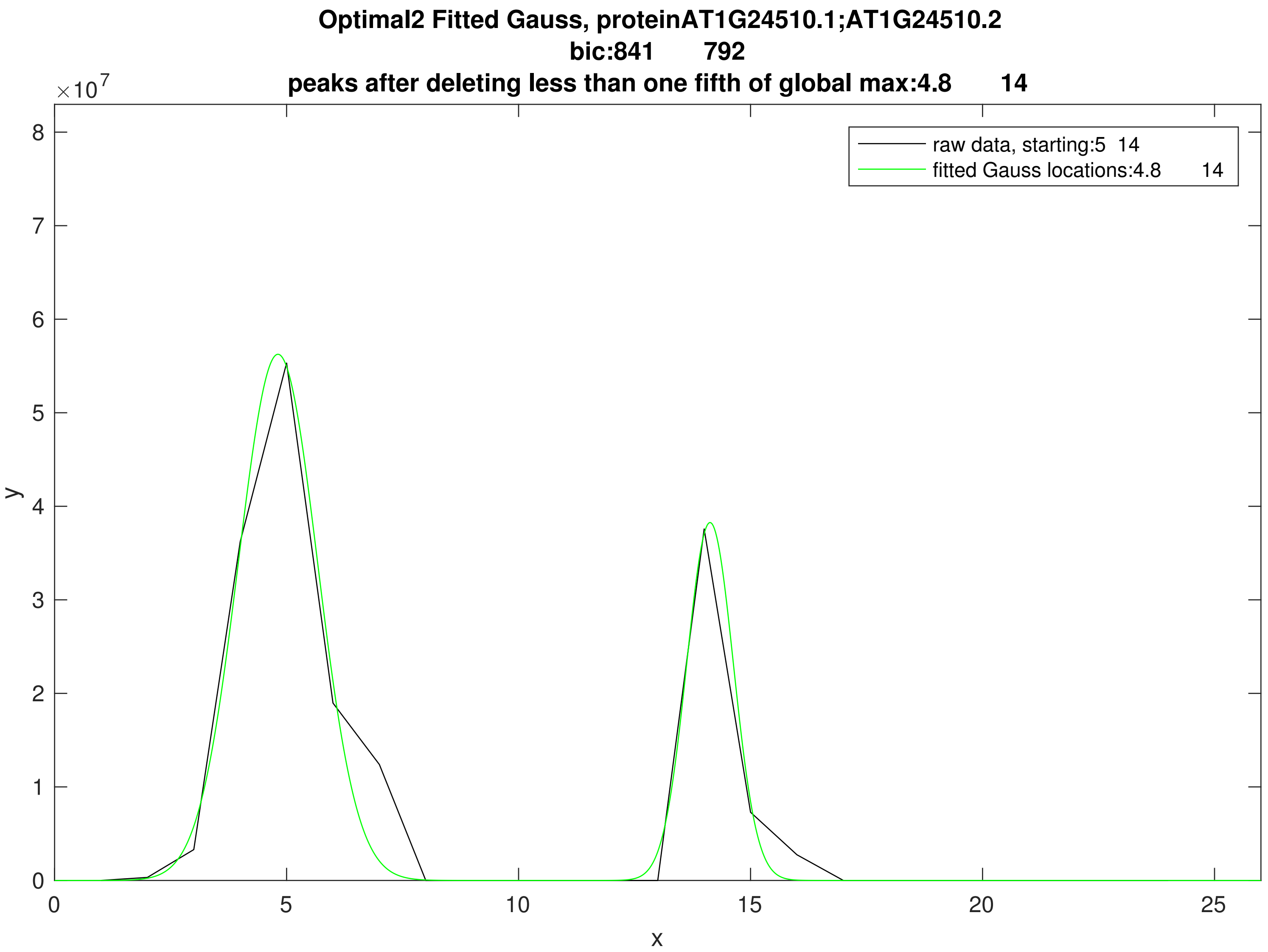
Optimal2 Fitted Gauss, proteinAT1G24510.1;AT1G24510.2
bic:841 792
peaks after deleting less than one fifth of global max:4.8 14
raw data, starting:5 14
fitted Gauss locations:4.8 14
×10^7
y
x

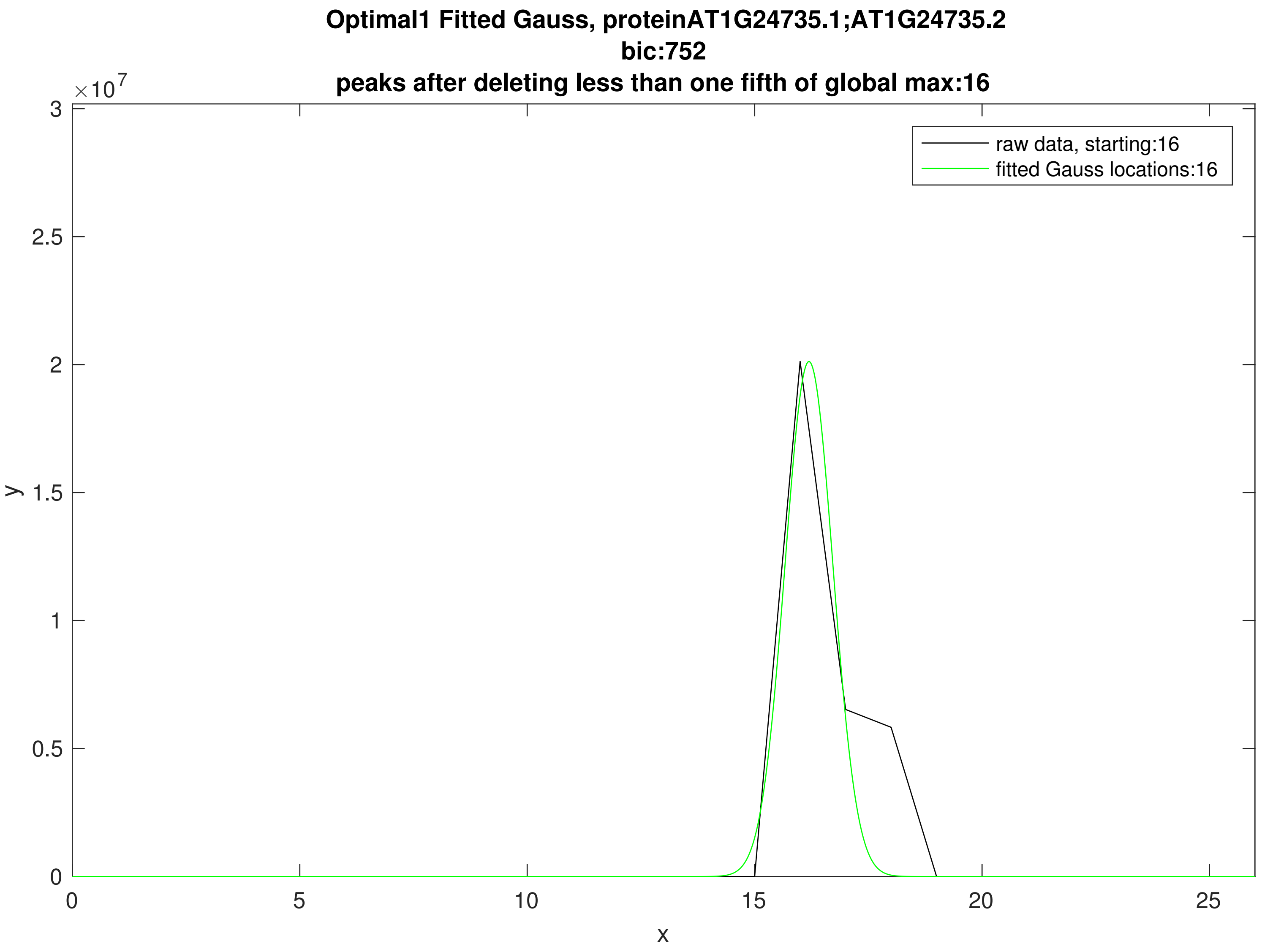
Optimal1 Fitted Gauss, proteinAT1G24735.1;AT1G24735.2
bic:752
peaks after deleting less than one fifth of global max:16
×10^7
3
2.5
2
1.5
1
0.5
0
y
0
5
10
15
20
25
x
raw data, starting:16
fitted Gauss locations:16

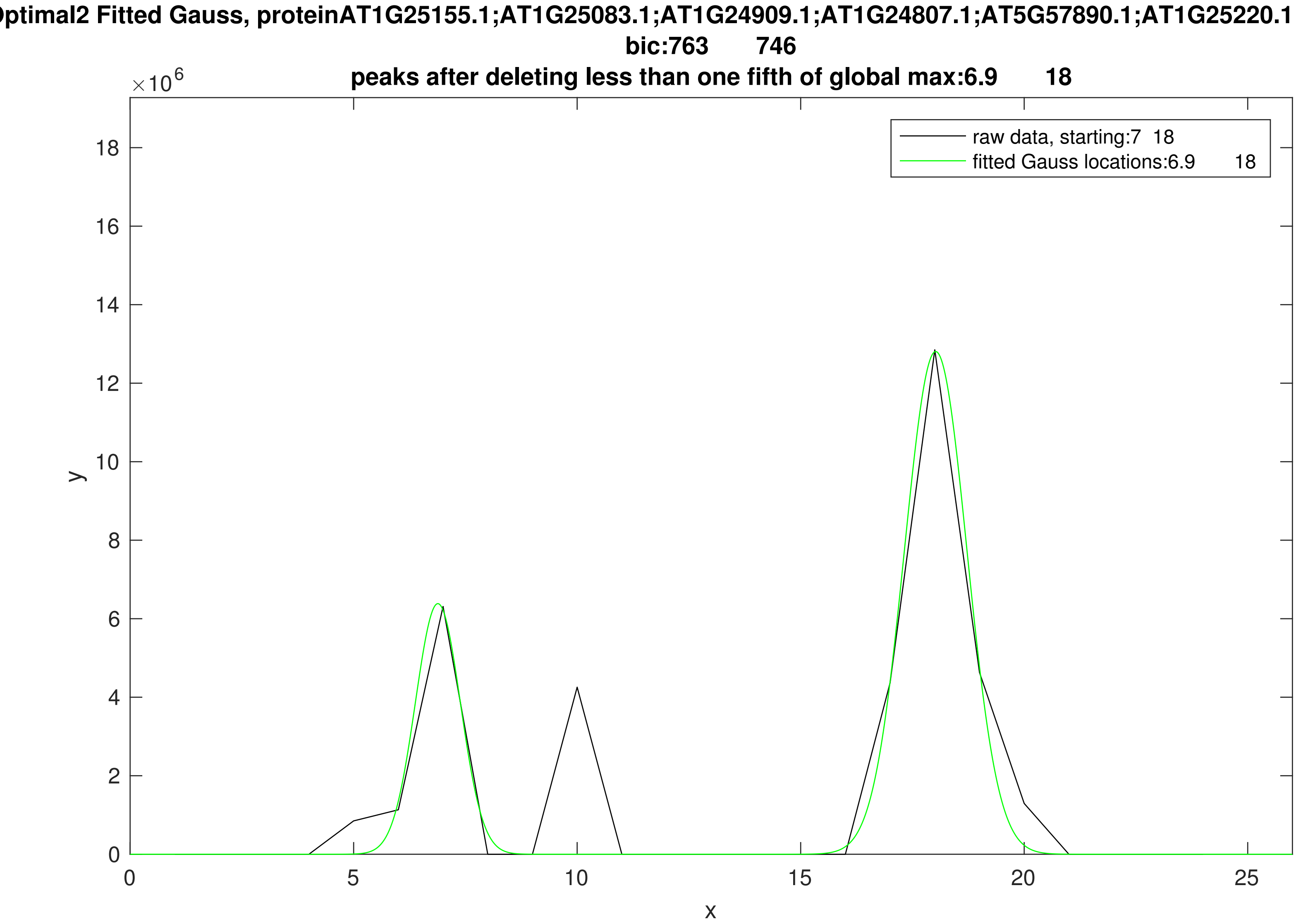
Optimal2 Fitted Gauss, proteinAT1G25155.1;AT1G25083.1;AT1G24909.1;AT1G24807.1;AT5G57890.1;AT1G25220.1;AT1
bic:763 746
peaks after deleting less than one fifth of global max:6.9 18
×10^6
raw data, starting:7 18
fitted Gauss locations:6.9 18
y
x
0
2
4
6
8
10
12
14
16
18
0
5
10
15
20
25

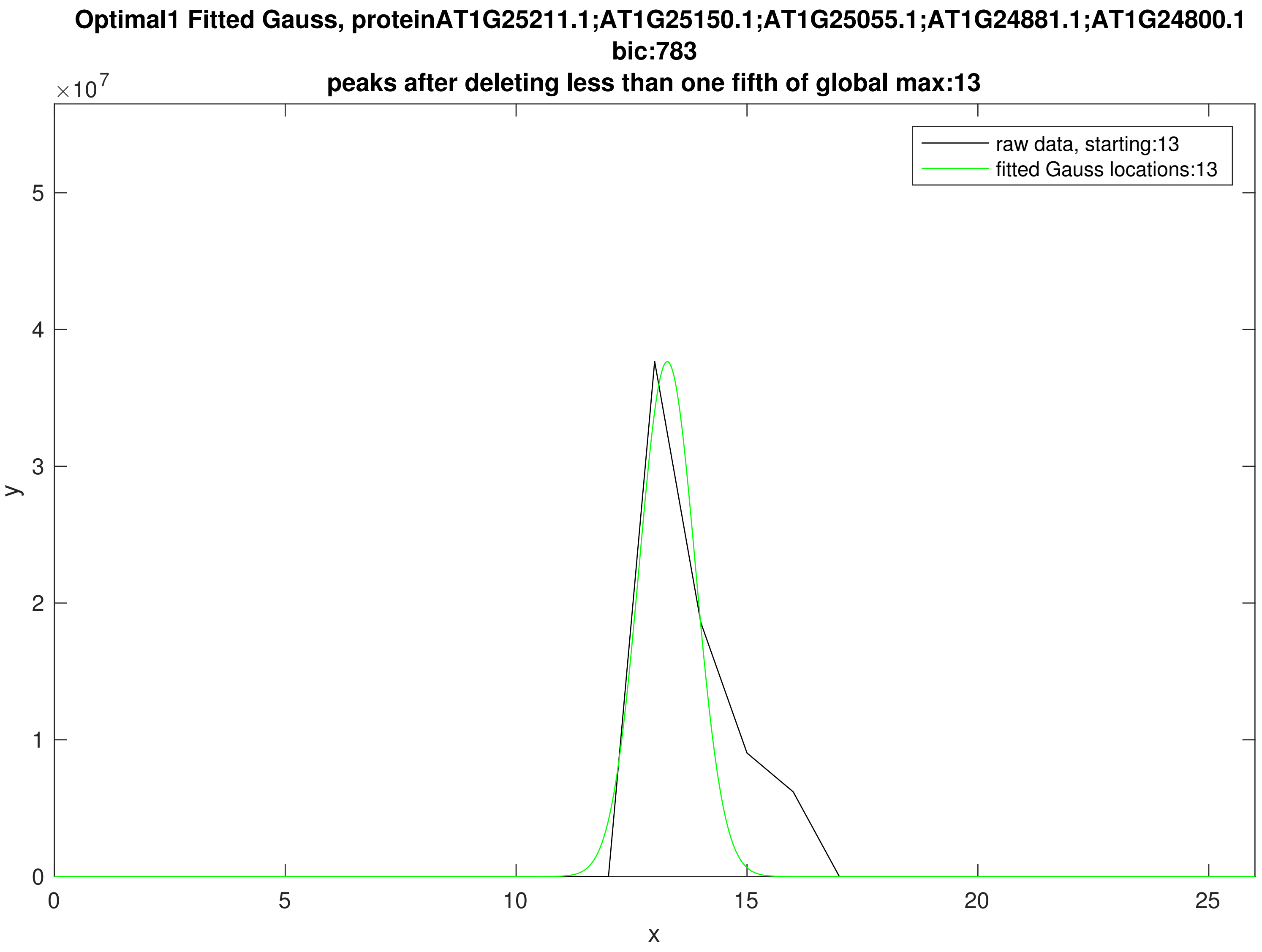
Optimal1 Fitted Gauss, proteinAT1G25211.1;AT1G25150.1;AT1G25055.1;AT1G24881.1;AT1G24800.1
bic:783
peaks after deleting less than one fifth of global max:13
×10^7
raw data, starting:13
fitted Gauss locations:13
y
x
0
1
2
3
4
5
0
5
10
15
20
25

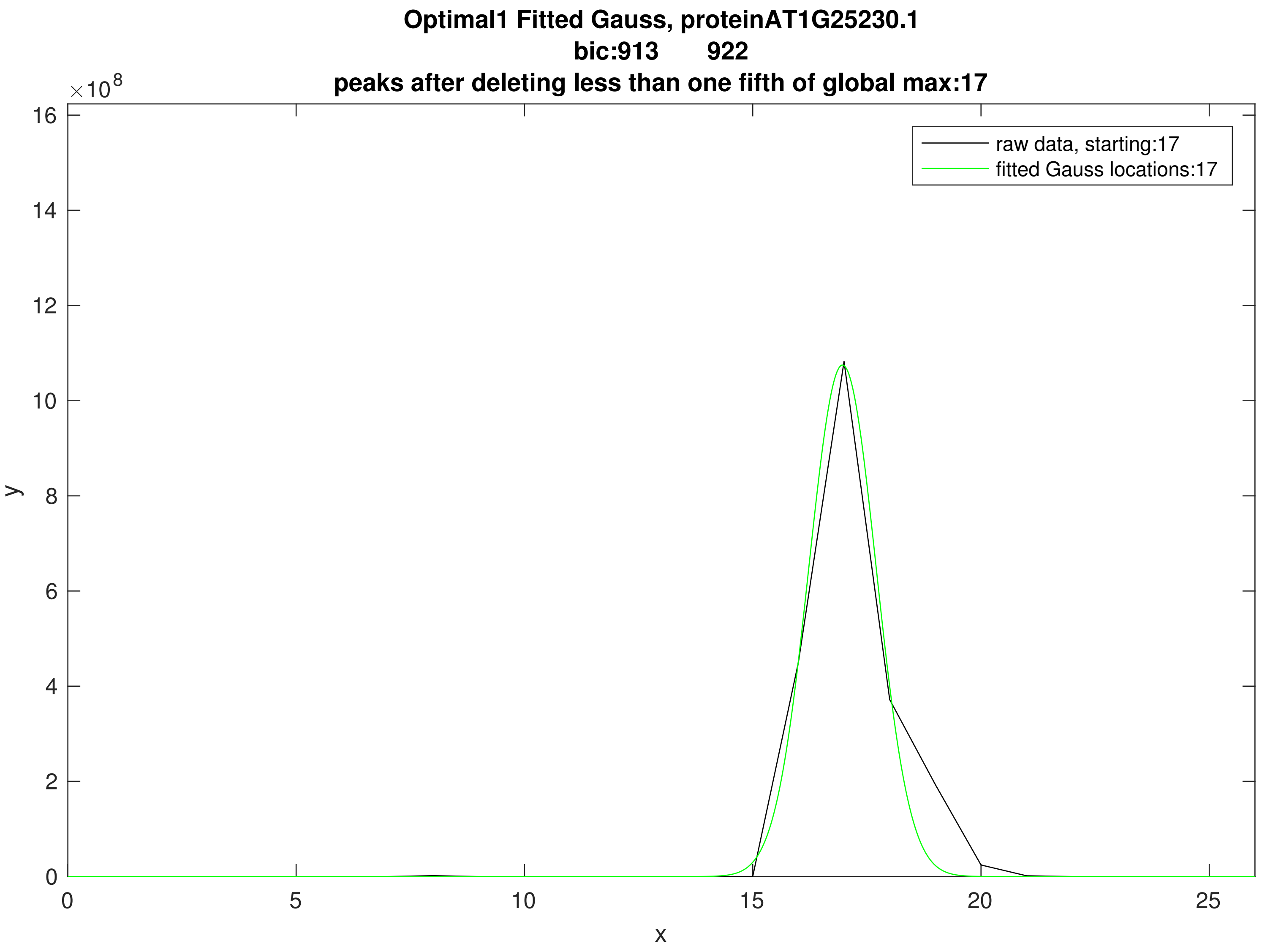
Optimal1 Fitted Gauss, proteinAT1G25230.1
bic:913 922
peaks after deleting less than one fifth of global max:17
×10^8
16
14
12
10
8
6
4
2
0
y
0
5
10
15
20
25
x
raw data, starting:17
fitted Gauss locations:17

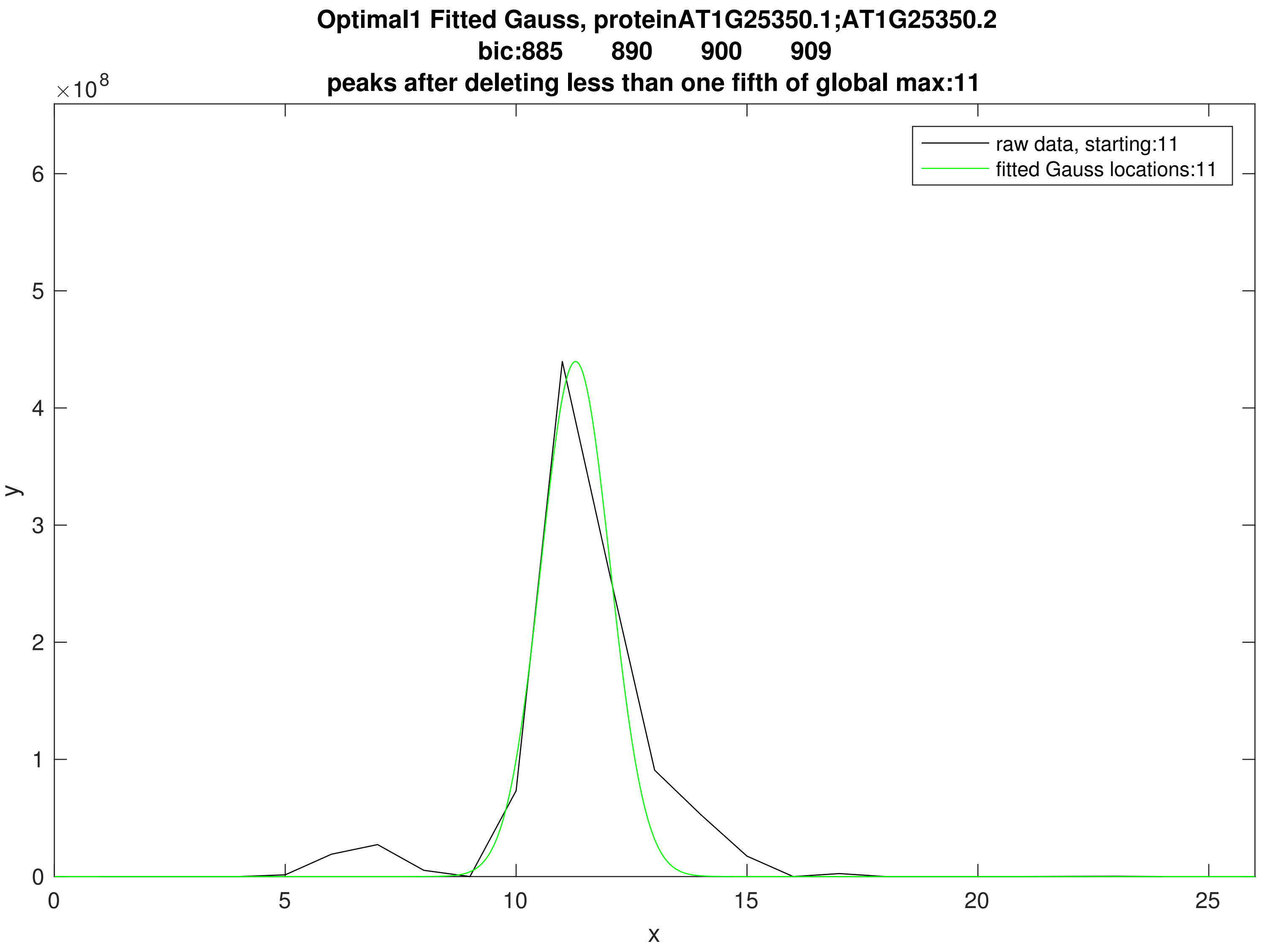
Optimal1 Fitted Gauss, proteinAT1G25350.1;AT1G25350.2
bic:885 890 900 909
peaks after deleting less than one fifth of global max:11
×10^8
raw data, starting:11
fitted Gauss locations:11
y
x
0
1
2
3
4
5
6
0
5
10
15
20
25

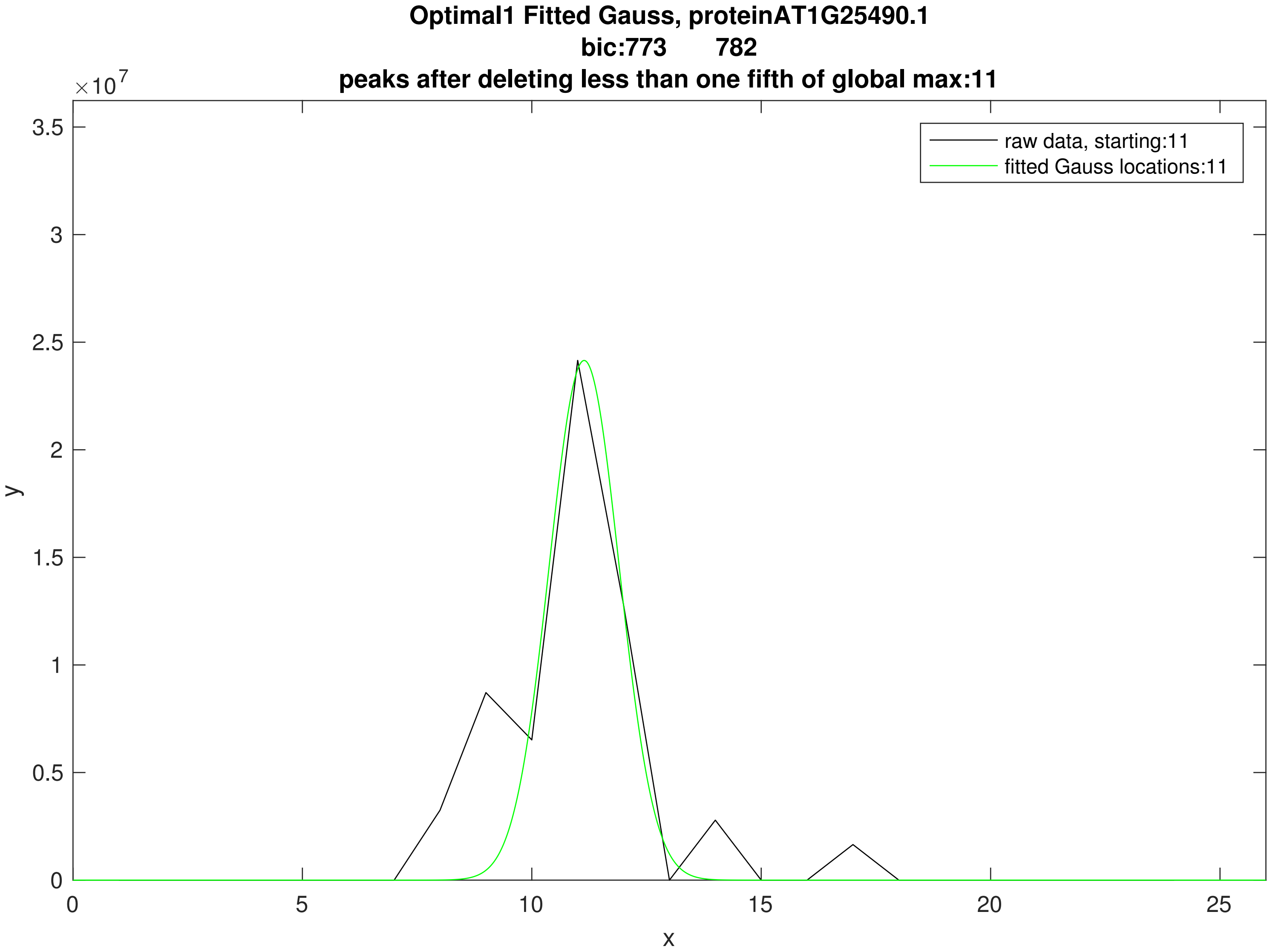

Optimal1 Fitted Gauss, proteinAT1G25490.1
bic:773 782
peaks after deleting less than one fifth of global max:11
×10^7
3.5
3
2.5
2
1.5
1
0.5
0
y
0
5
10
15
20
25
x
raw data, starting:11
fitted Gauss locations:11

**Optimal1 Fitted Gauss, proteinAT1G25570.1**
**bic:770**
**peaks after deleting less than one fifth of global max:15**

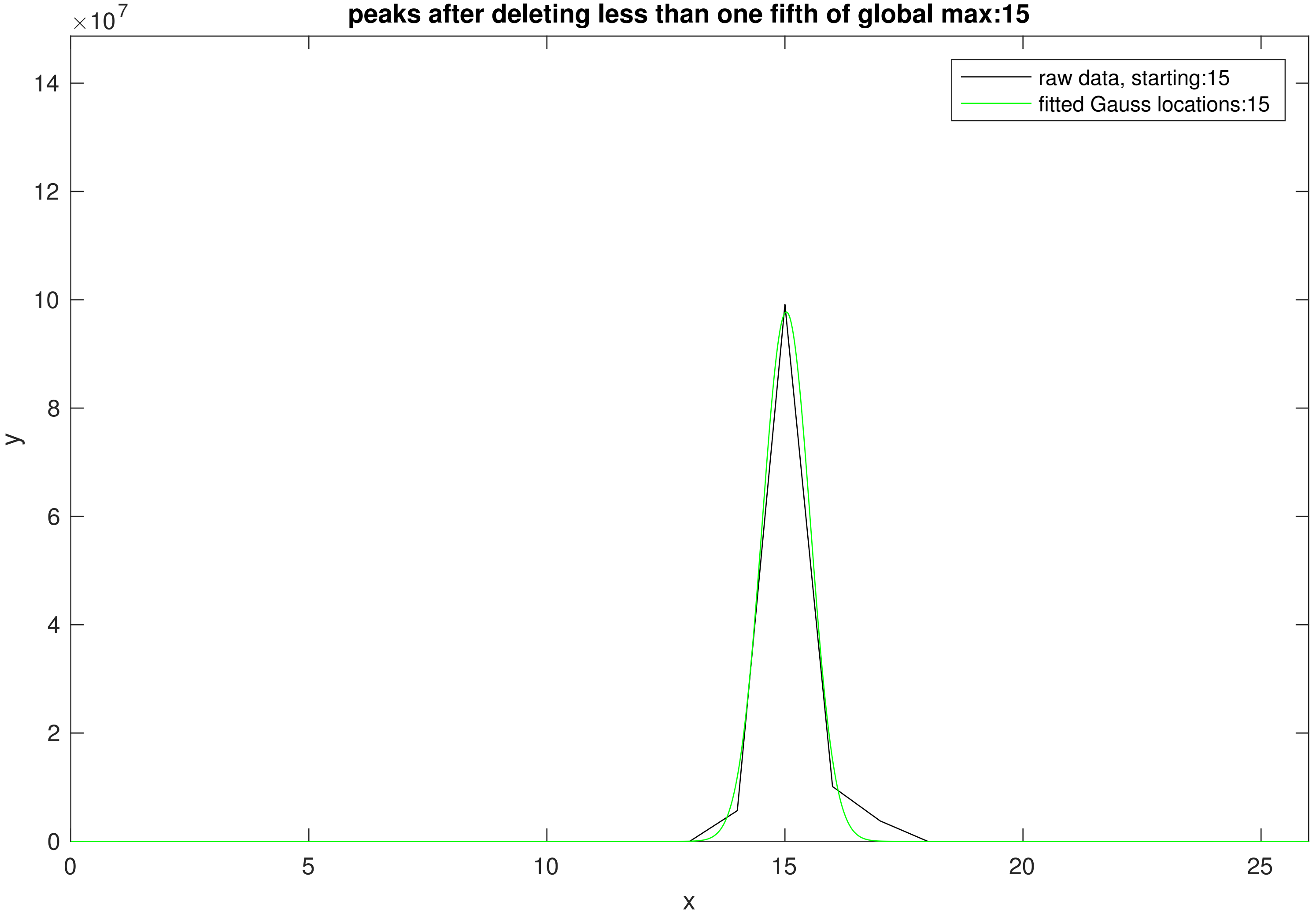

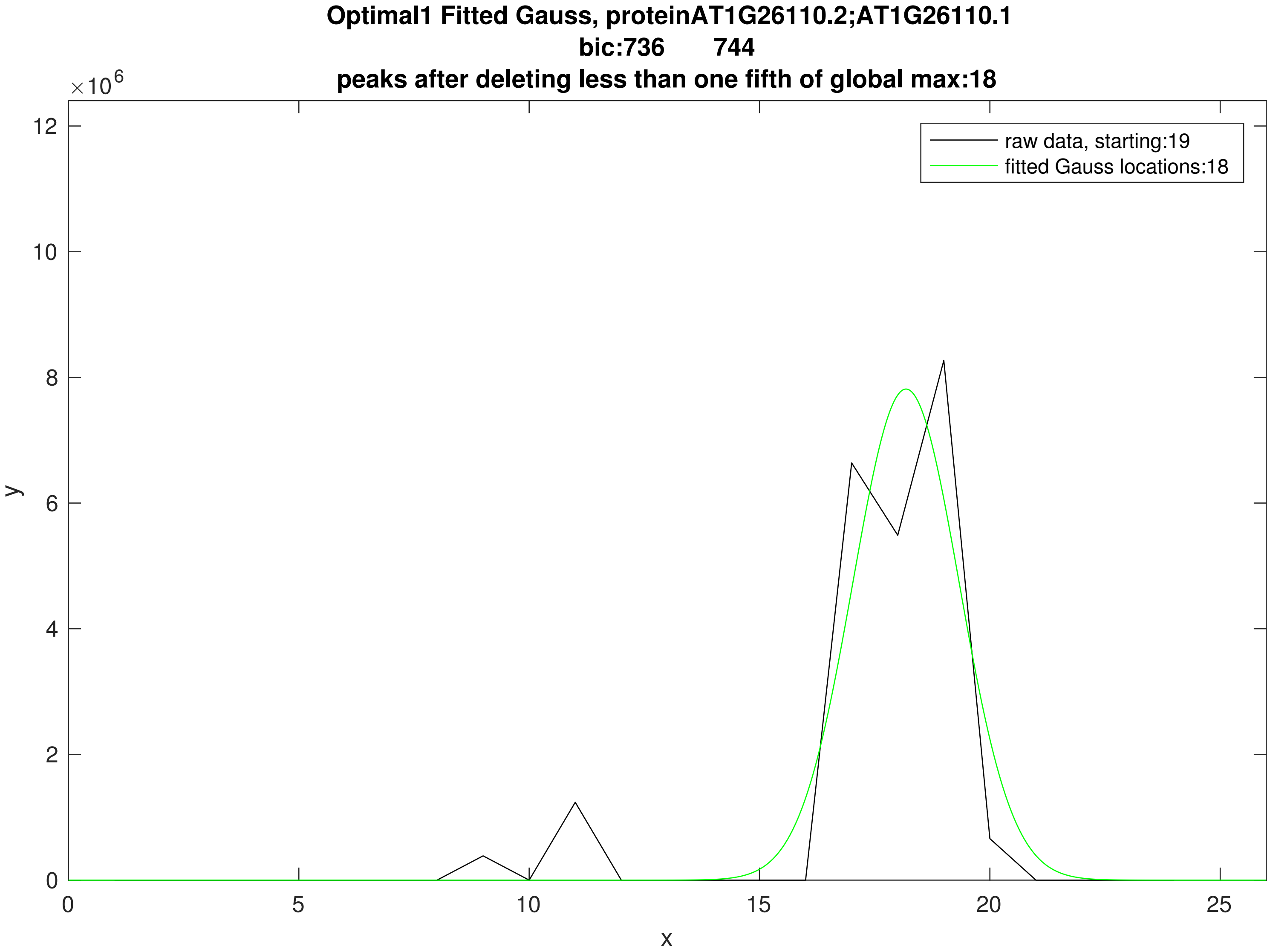

Optimal1 Fitted Gauss, proteinAT1G26110.2;AT1G26110.1
bic:736 744
peaks after deleting less than one fifth of global max:18
×10^6
raw data, starting:19
fitted Gauss locations:18
y
x
0
2
4
6
8
10
12
0
5
10
15
20
25

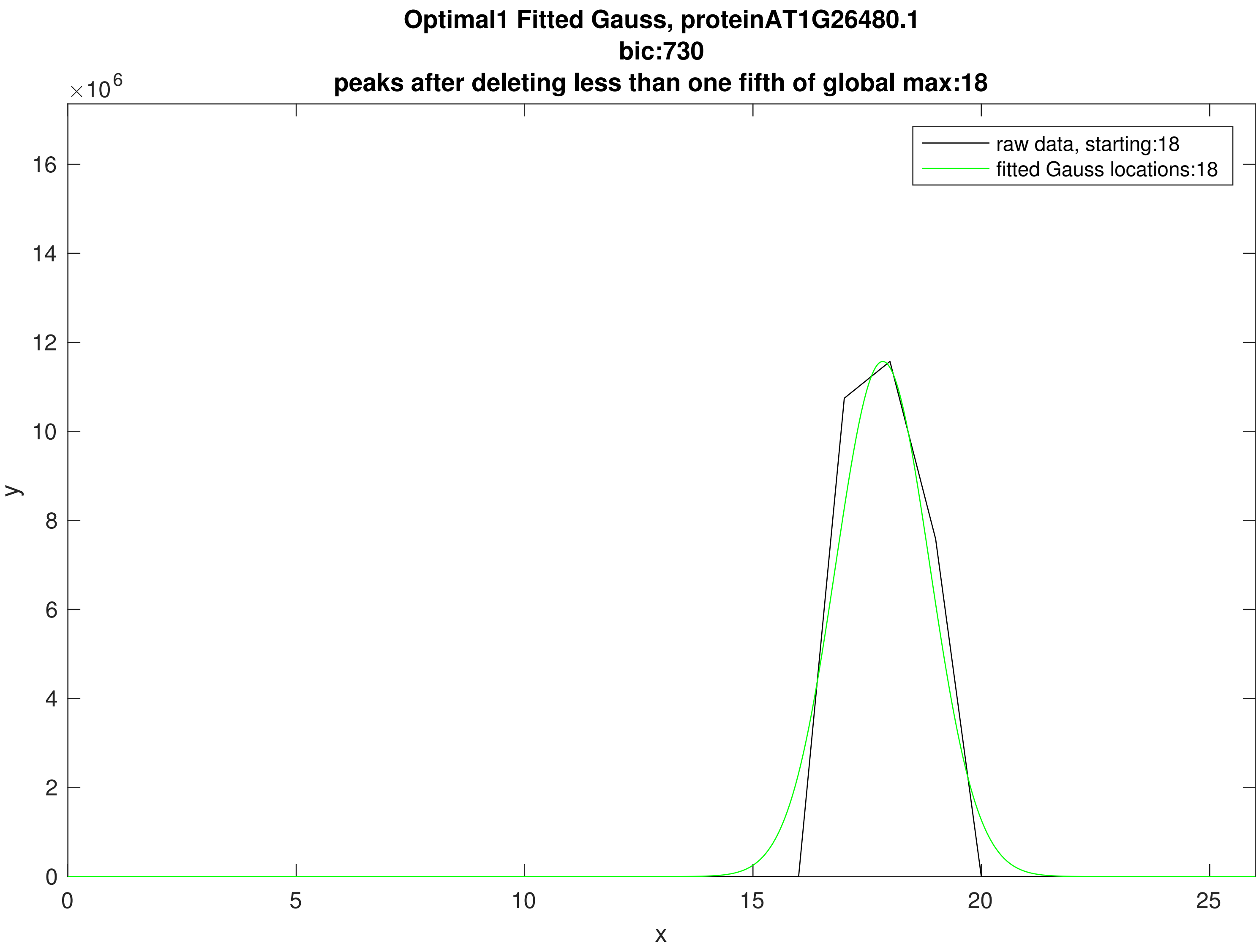
Optimal1 Fitted Gauss, proteinAT1G26480.1
bic:730
peaks after deleting less than one fifth of global max:18
×10^6
16
14
12
10
8
6
4
2
0
y
0
5
10
15
20
25
x
raw data, starting:18
fitted Gauss locations:18

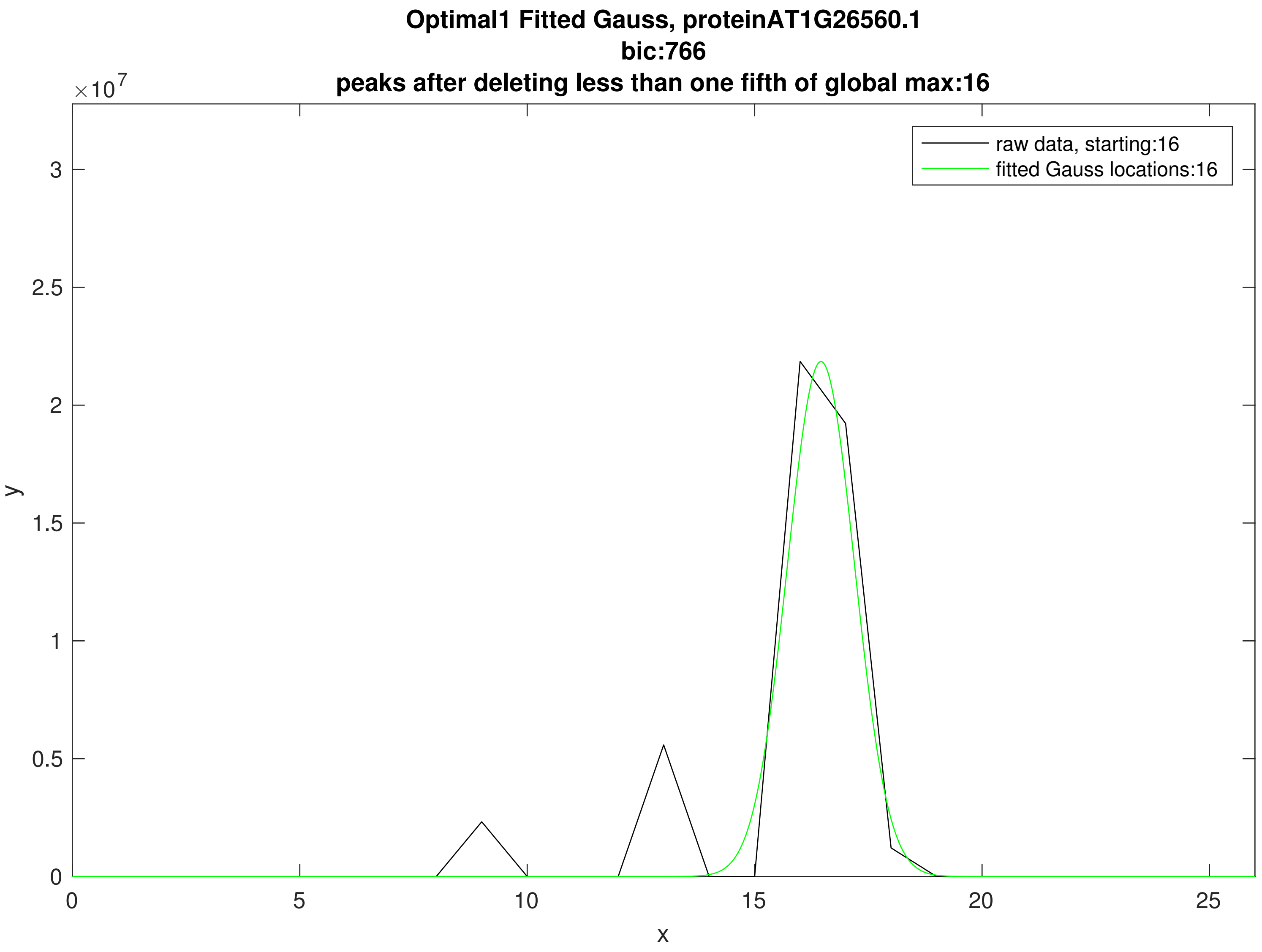

Optimal1 Fitted Gauss, proteinAT1G26560.1
bic:766
peaks after deleting less than one fifth of global max:16
×10^7
3
2.5
2
1.5
1
0.5
0
y
0
5
10
15
20
25
x
raw data, starting:16
fitted Gauss locations:16

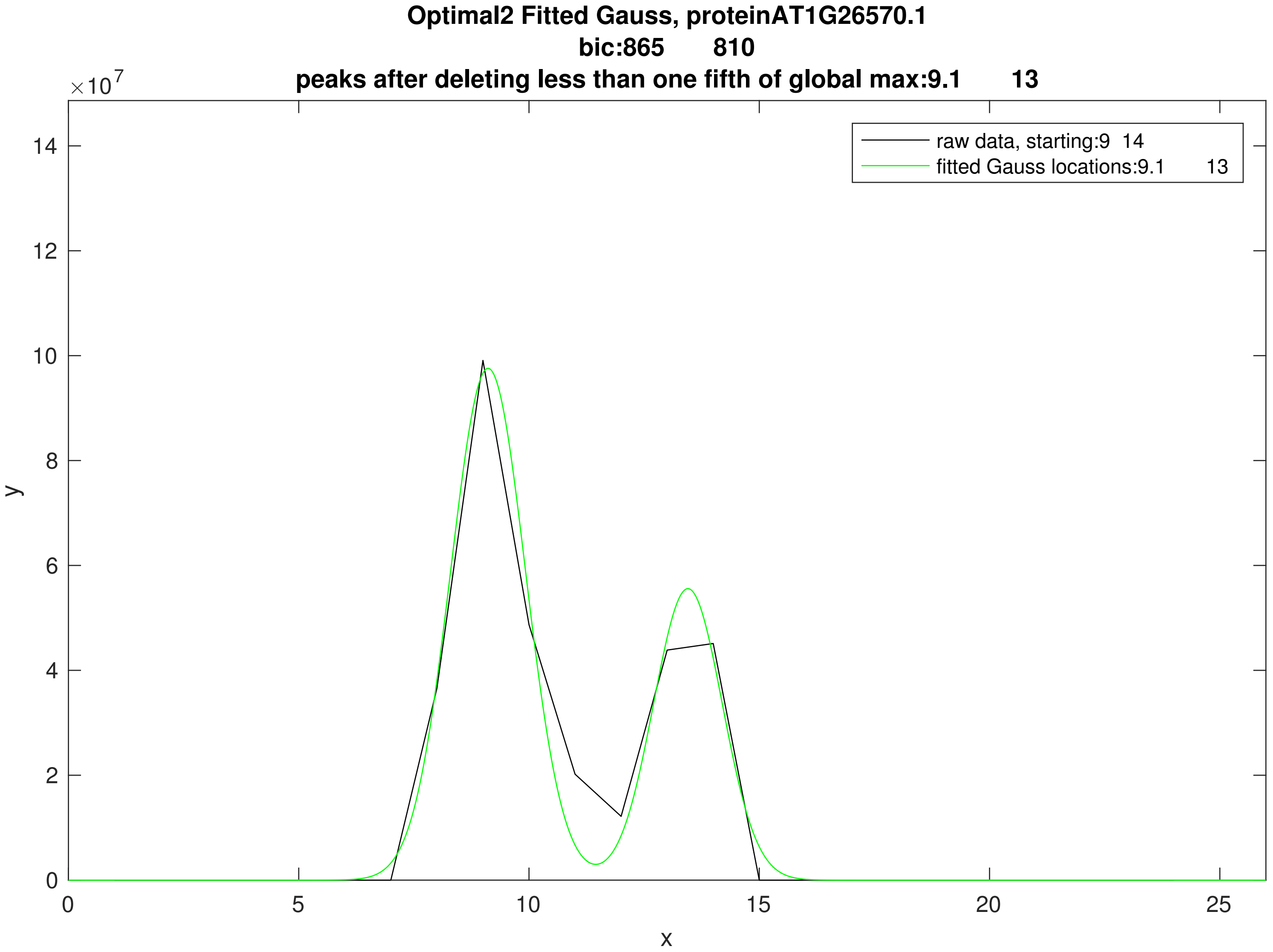
Optimal2 Fitted Gauss, proteinAT1G26570.1
bic:865 810
peaks after deleting less than one fifth of global max:9.1 13
×10^7
raw data, starting:9 14
fitted Gauss locations:9.1 13
y
x
0
2
4
6
8
10
12
14
0
5
10
15
20
25

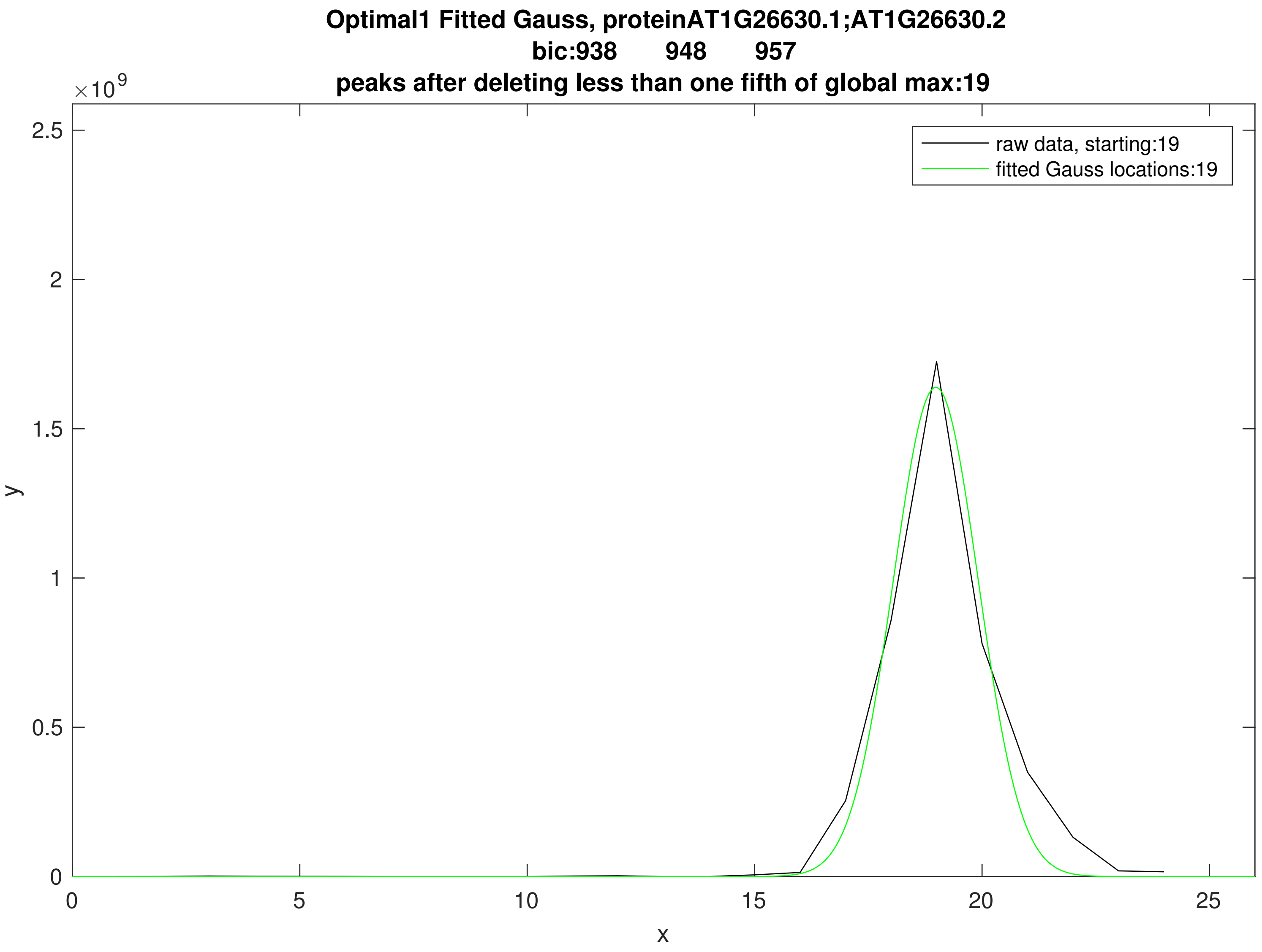

Optimal1 Fitted Gauss, proteinAT1G26630.1;AT1G26630.2
bic:938 948 957
peaks after deleting less than one fifth of global max:19
×10^9
raw data, starting:19
fitted Gauss locations:19
y
x
0
0.5
1
1.5
2
2.5
0
5
10
15
20
25

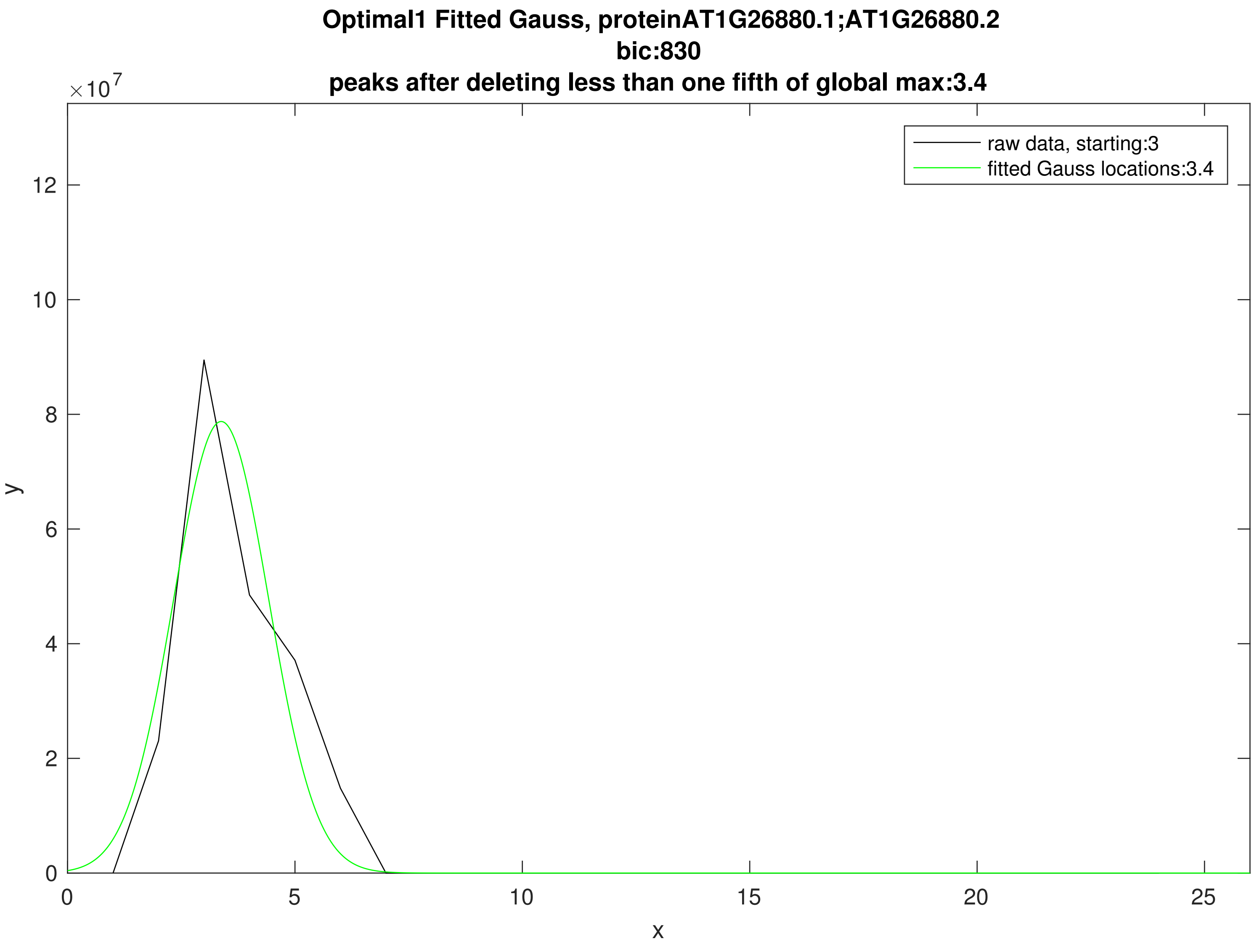

Optimal1 Fitted Gauss, proteinAT1G26880.1;AT1G26880.2
bic:830
peaks after deleting less than one fifth of global max:3.4
×10^7
raw data, starting:3
fitted Gauss locations:3.4
0
2
4
6
8
10
12
y
0
5
10
15
20
25
x

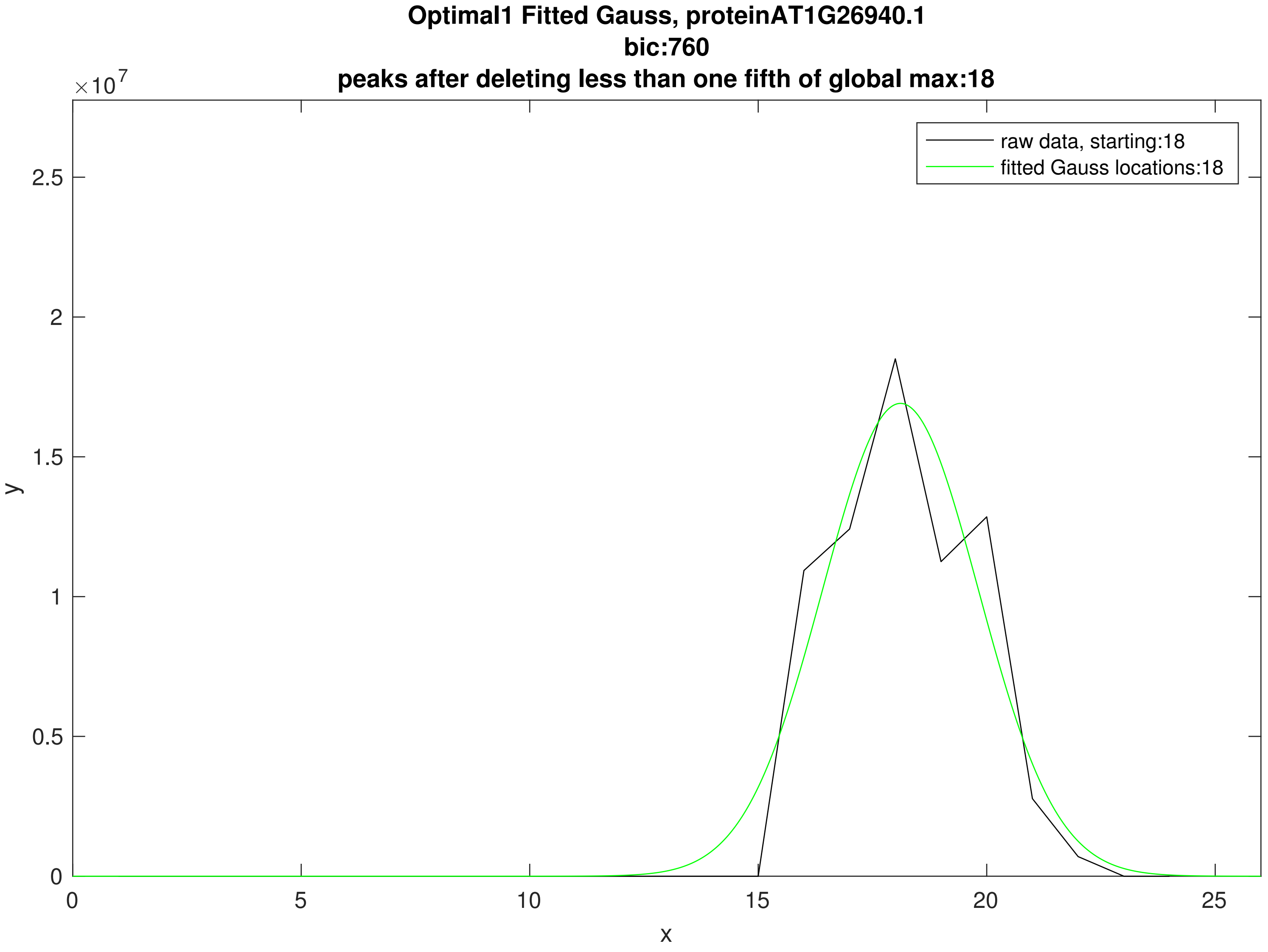
Optimal1 Fitted Gauss, proteinAT1G26940.1
bic:760
peaks after deleting less than one fifth of global max:18
×10^7
raw data, starting:18
fitted Gauss locations:18
2.5
2
1.5
1
0.5
0
y
0
5
10
15
20
25
x

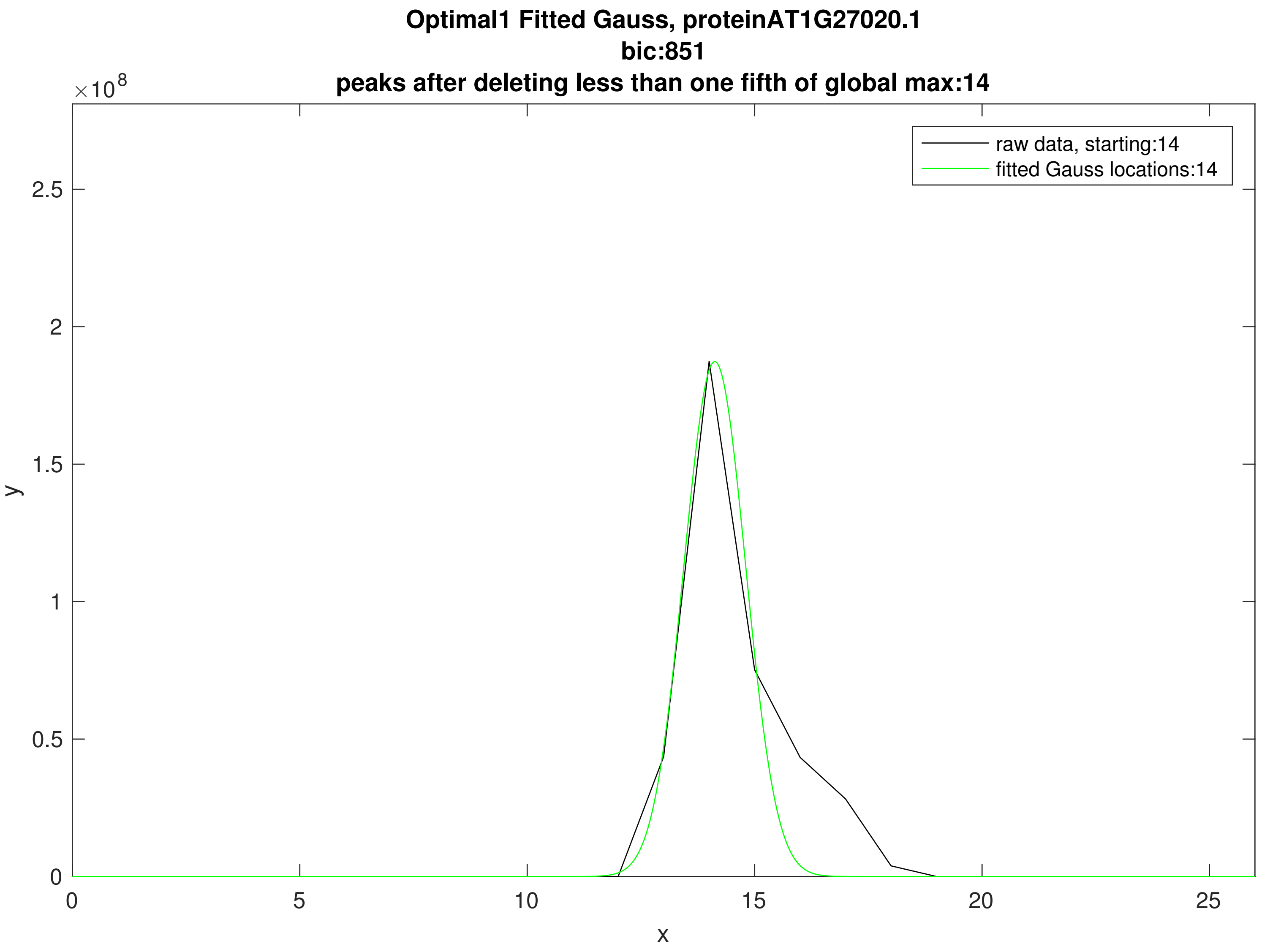
Optimal1 Fitted Gauss, proteinAT1G27020.1
bic:851
peaks after deleting less than one fifth of global max:14
×10^8
raw data, starting:14
fitted Gauss locations:14
0
0.5
1
1.5
2
2.5
y
0
5
10
15
20
25
x

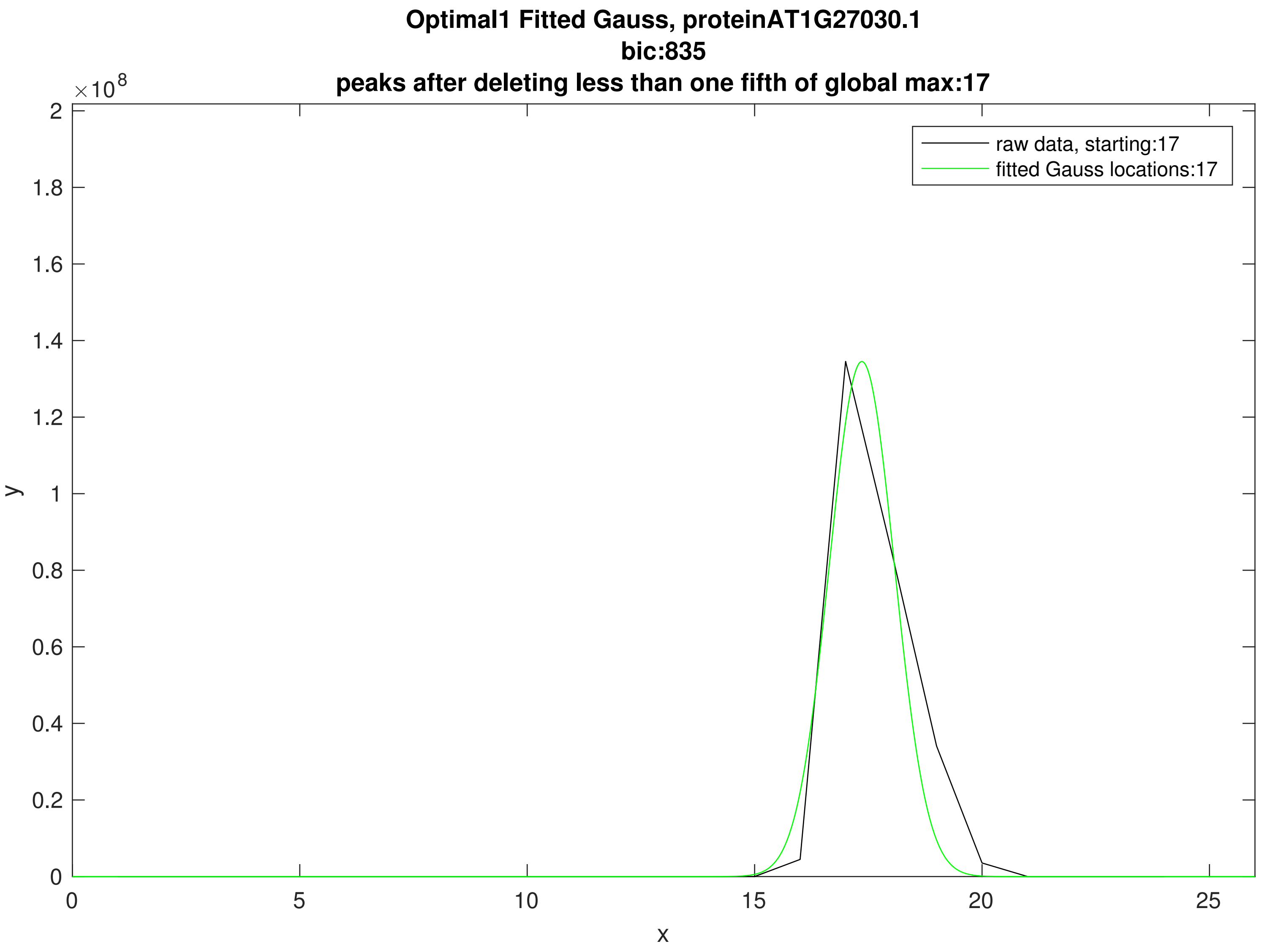
Optimal1 Fitted Gauss, proteinAT1G27030.1
bic:835
peaks after deleting less than one fifth of global max:17
×10^8
2
1.8
1.6
1.4
1.2
1
0.8
0.6
0.4
0.2
0
y
0
5
10
15
20
25
x
raw data, starting:17
fitted Gauss locations:17

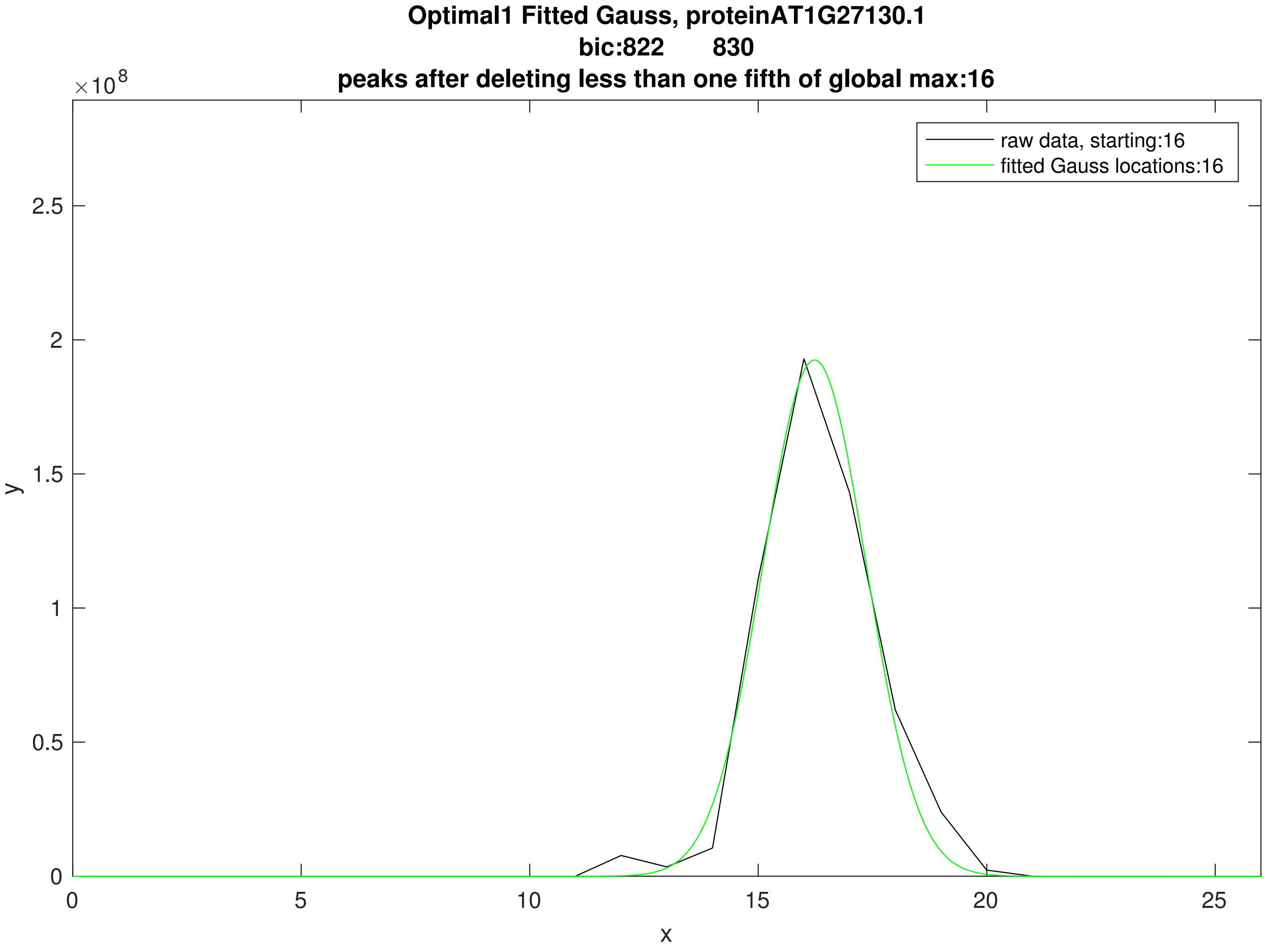
Optimal1 Fitted Gauss, proteinAT1G27130.1
bic:822 830
peaks after deleting less than one fifth of global max:16
×10^8
raw data, starting:16
fitted Gauss locations:16
2.5
2
1.5
1
0.5
0
y
0
5
10
15
20
25
x

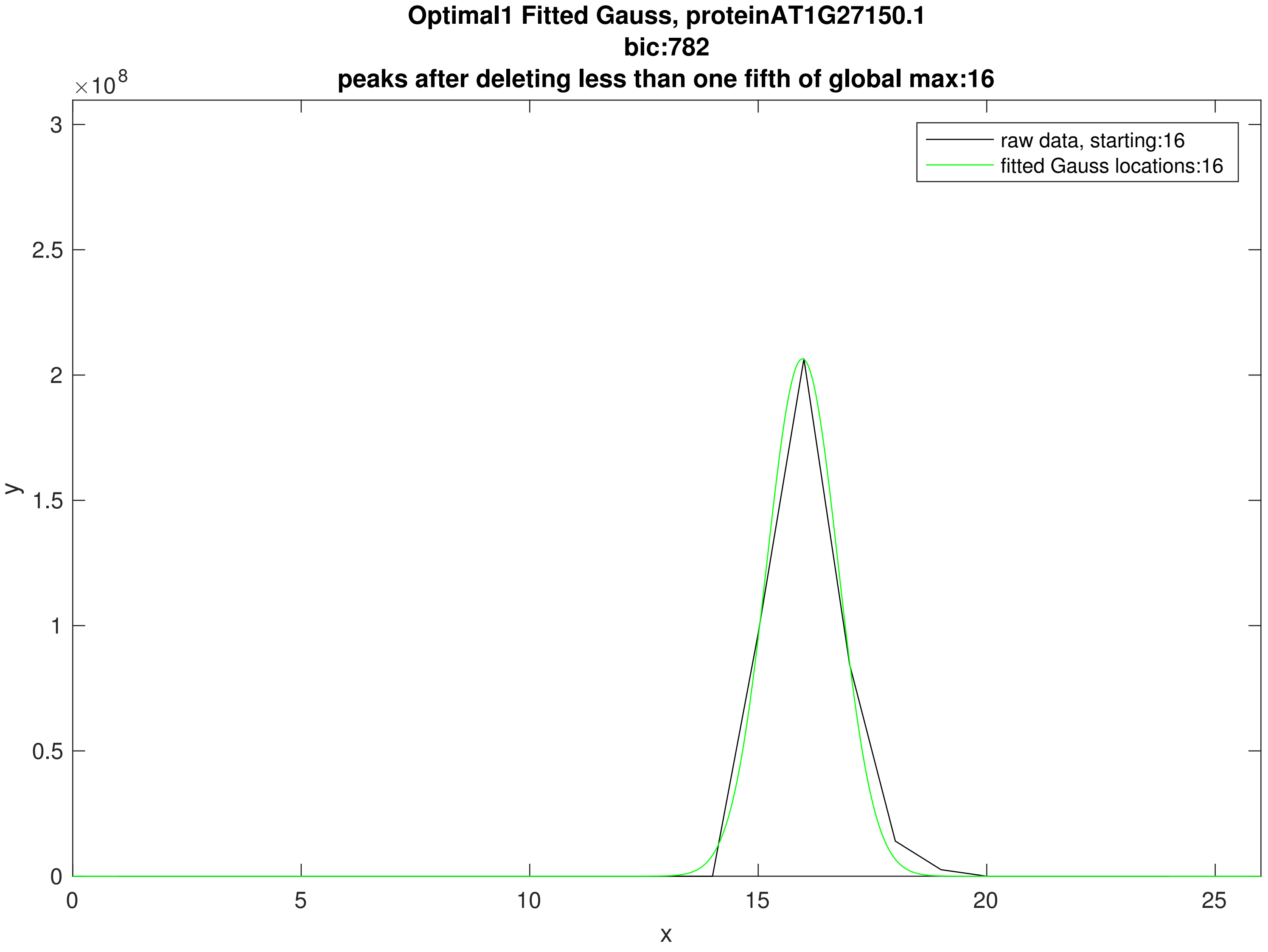
Optimal1 Fitted Gauss, proteinAT1G27150.1
bic:782
peaks after deleting less than one fifth of global max:16
×10^8
raw data, starting:16
fitted Gauss locations:16
y
x
0
0.5
1
1.5
2
2.5
3
0
5
10
15
20
25

**Optimal1 Fitted Gauss, proteinAT1G27310.1**
**bic:822**
**peaks after deleting less than one fifth of global max:19**

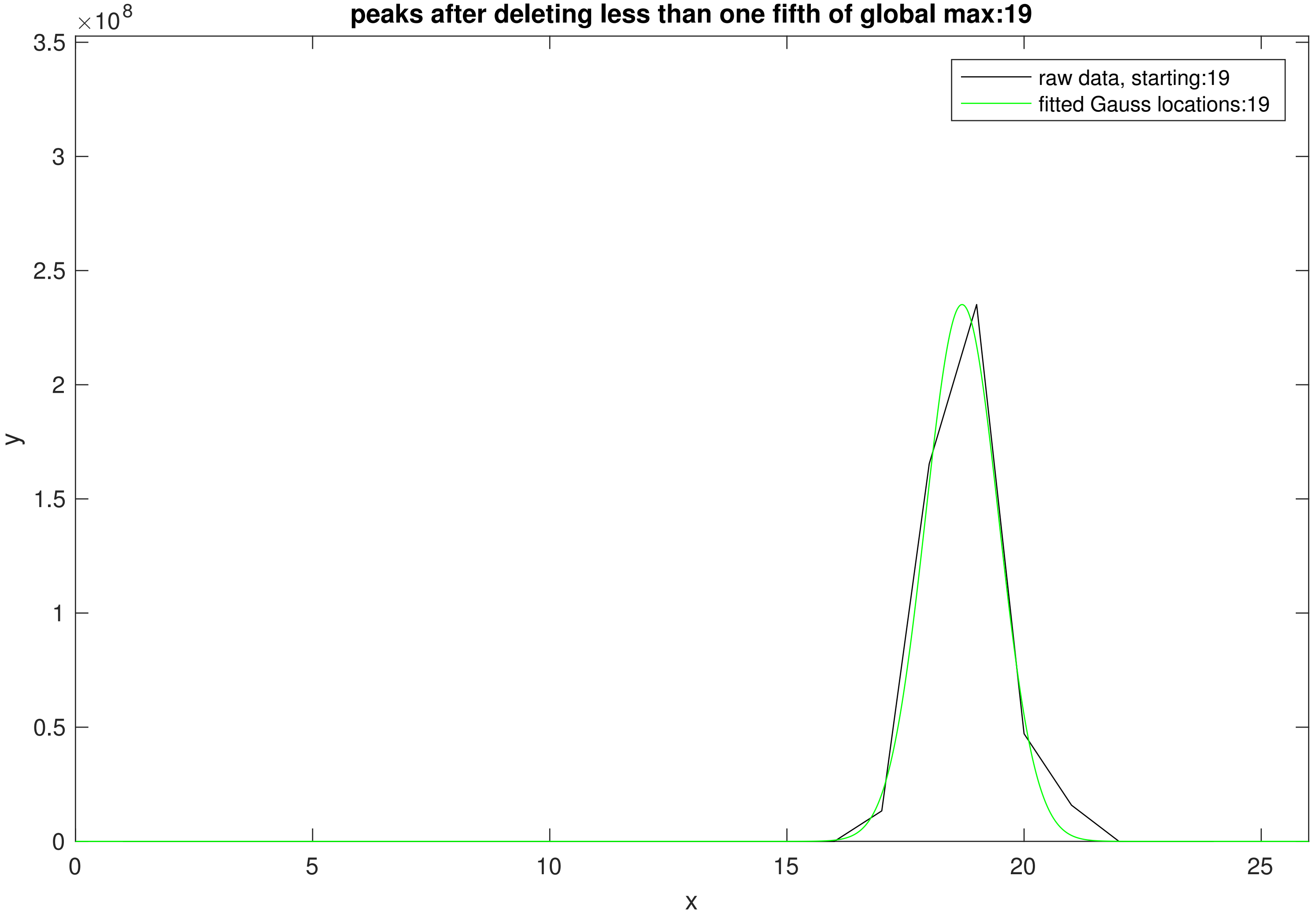

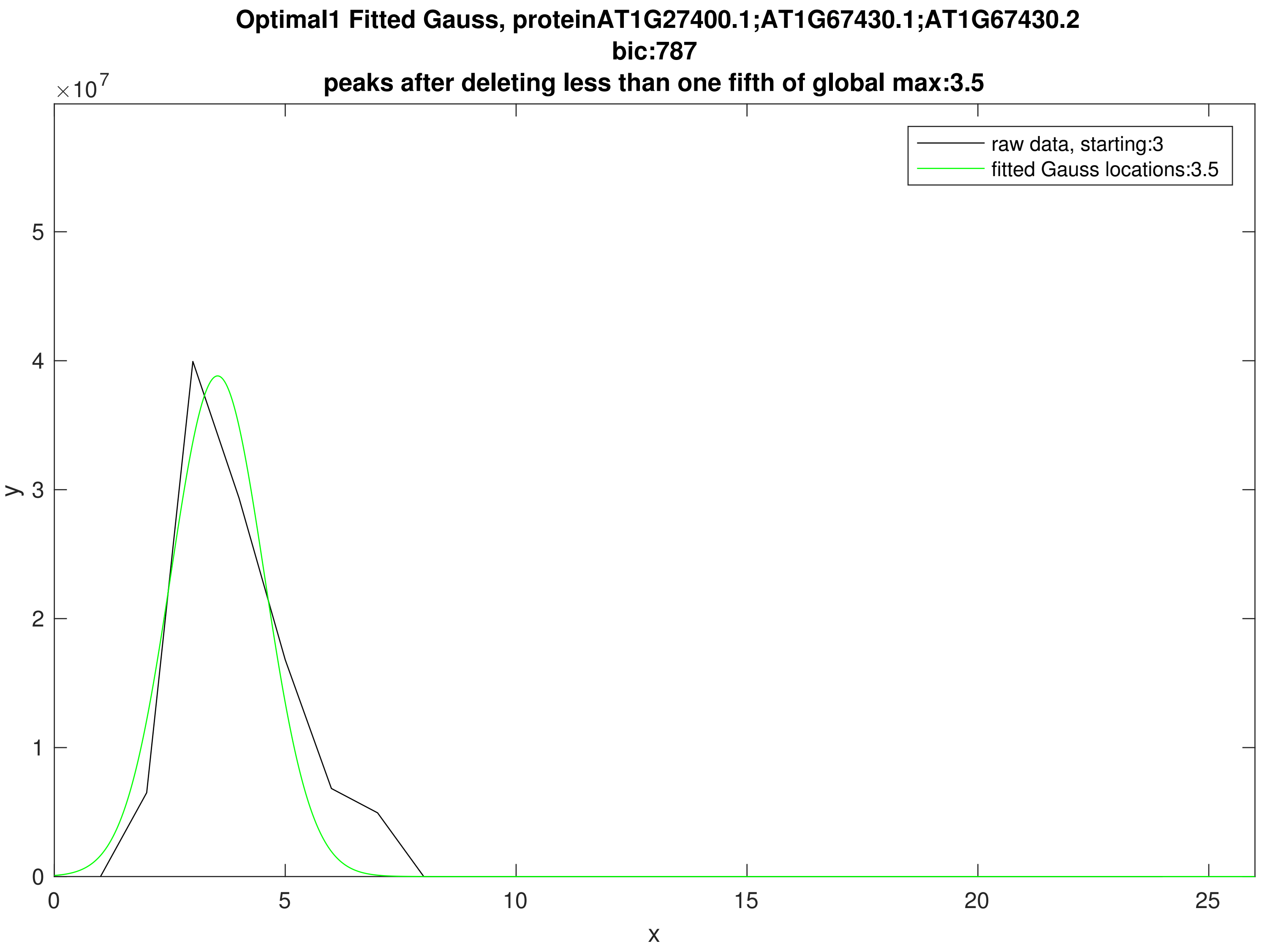
Optimal1 Fitted Gauss, proteinAT1G27400.1;AT1G67430.1;AT1G67430.2
bic:787
peaks after deleting less than one fifth of global max:3.5
×10^7
raw data, starting:3
fitted Gauss locations:3.5
y
x
0
1
2
3
4
5
0
5
10
15
20
25

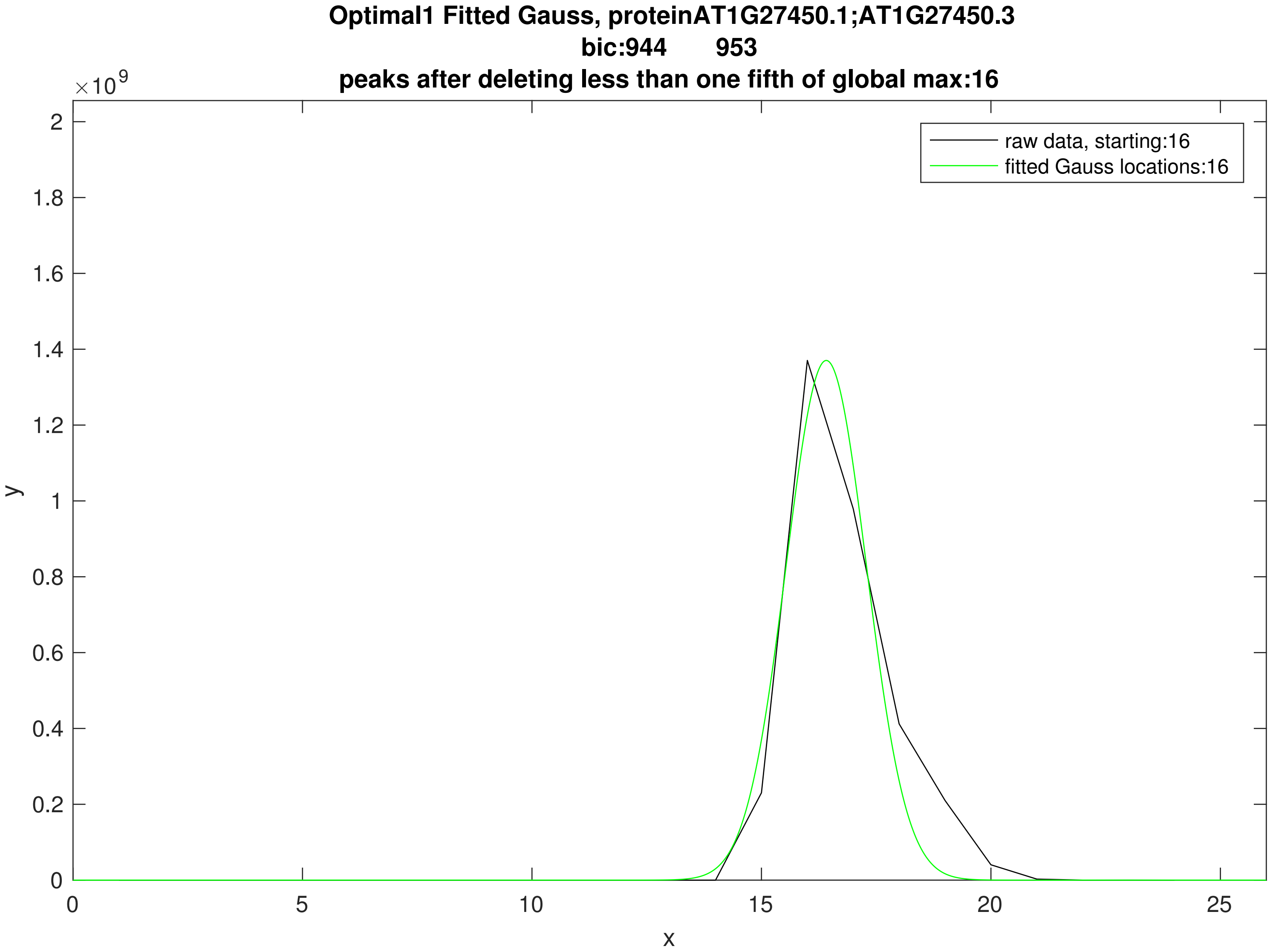
Optimal1 Fitted Gauss, proteinAT1G27450.1;AT1G27450.3
bic:944 953
peaks after deleting less than one fifth of global max:16
×10^9
raw data, starting:16
fitted Gauss locations:16
y
x
0
0.2
0.4
0.6
0.8
1
1.2
1.4
1.6
1.8
2
0
5
10
15
20
25

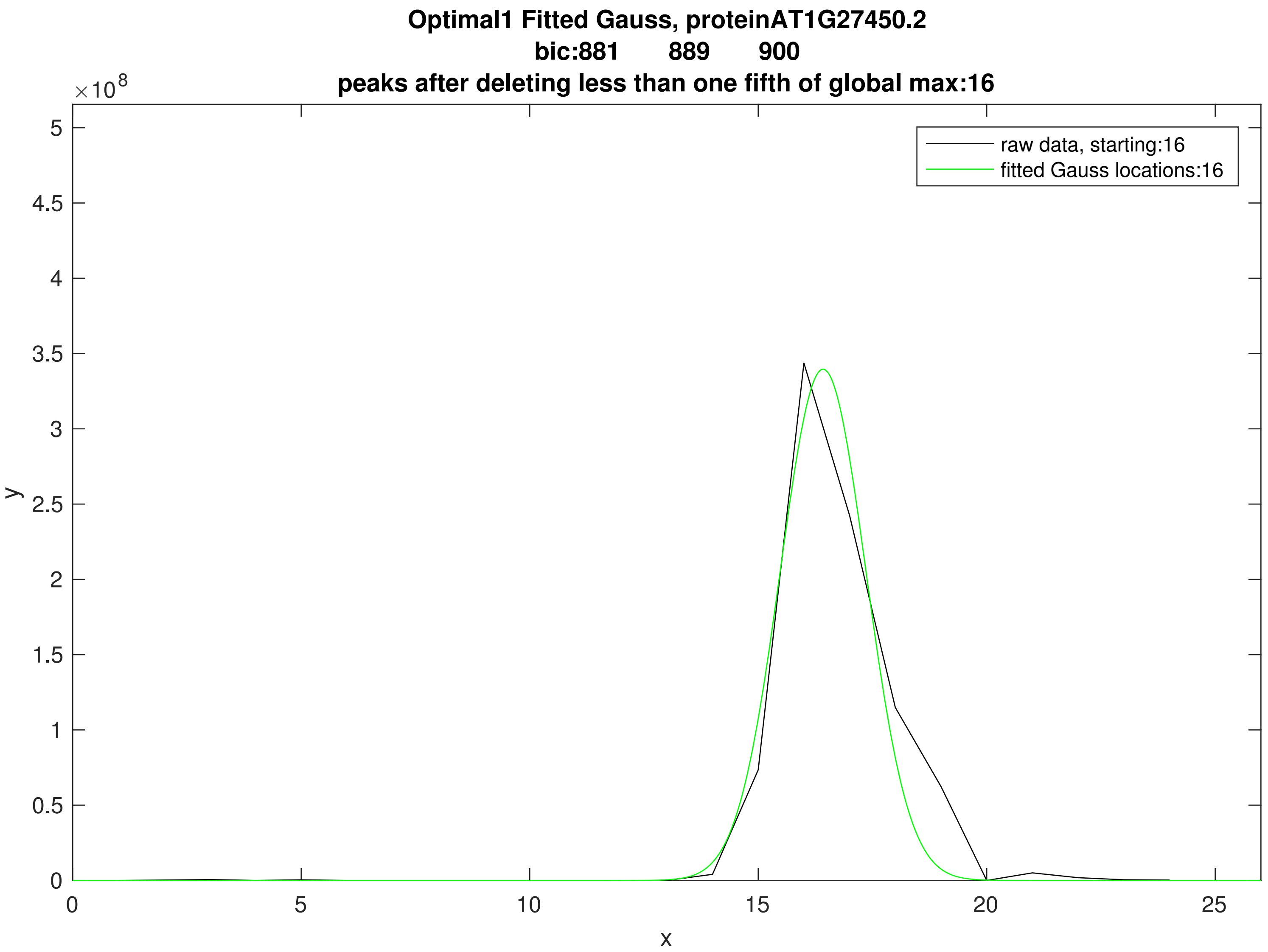
Optimal1 Fitted Gauss, proteinAT1G27450.2
bic:881 889 900
peaks after deleting less than one fifth of global max:16
×10^8
raw data, starting:16
fitted Gauss locations:16
y
x
0
0.5
1
1.5
2
2.5
3
3.5
4
4.5
5
0
5
10
15
20
25

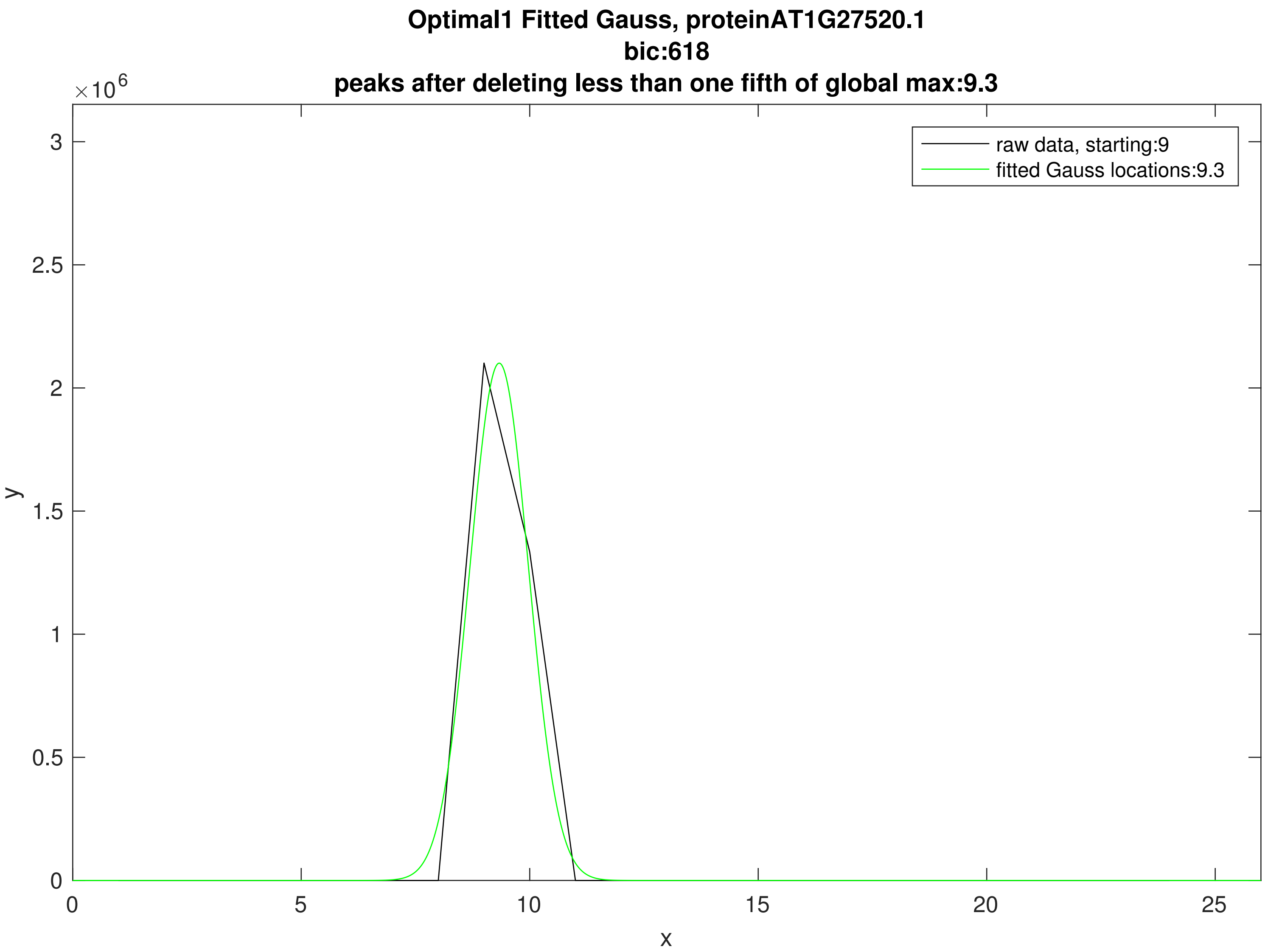
Optimal1 Fitted Gauss, proteinAT1G27520.1
bic:618
peaks after deleting less than one fifth of global max:9.3
×10^6
raw data, starting:9
fitted Gauss locations:9.3
y
x
0
0.5
1
1.5
2
2.5
3
0
5
10
15
20
25

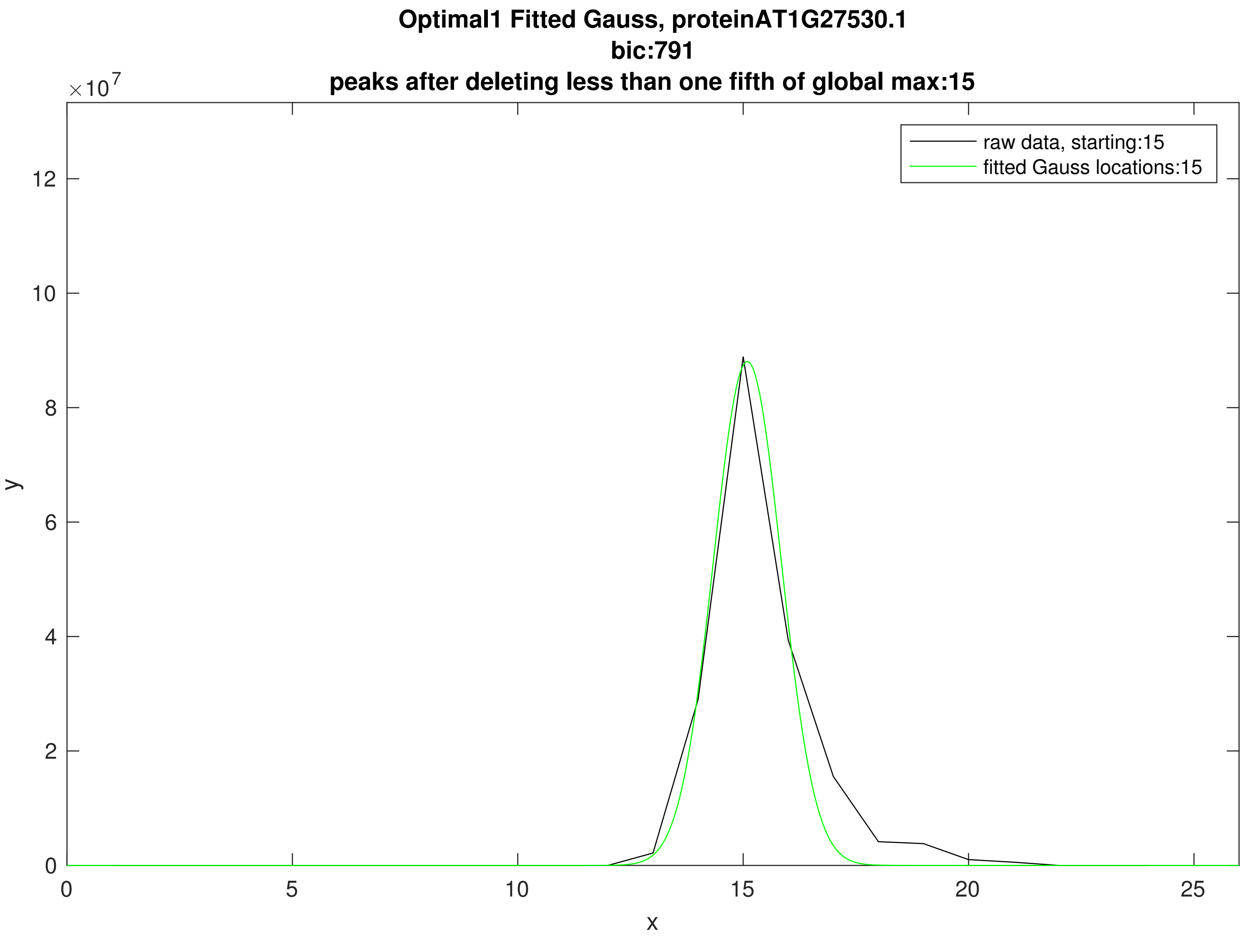
Optimal1 Fitted Gauss, proteinAT1G27530.1
bic:791
peaks after deleting less than one fifth of global max:15
×10^7
raw data, starting:15
fitted Gauss locations:15
y
x
0
2
4
6
8
10
12
0
5
10
15
20
25

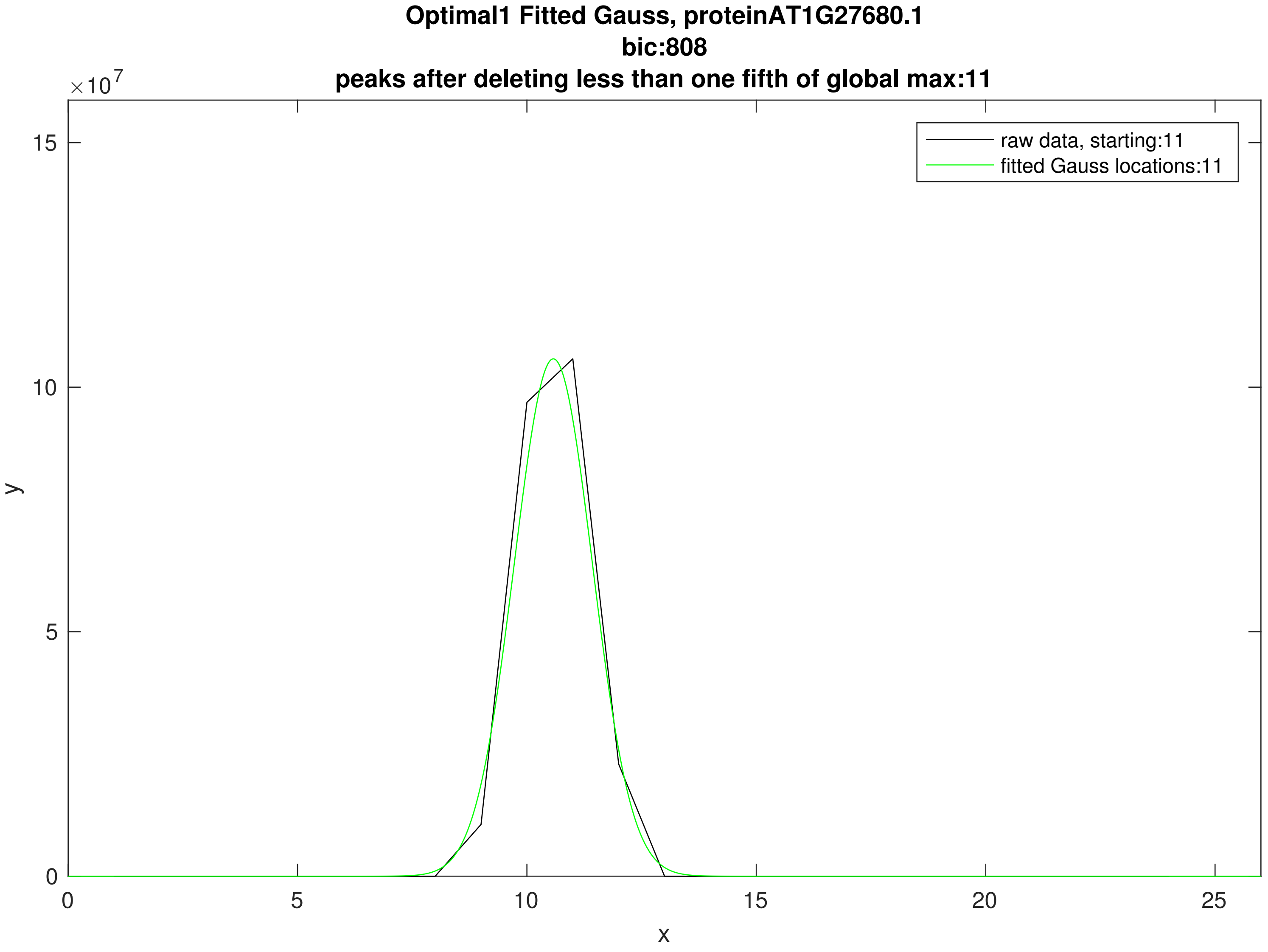

Optimal1 Fitted Gauss, proteinAT1G27680.1
bic:808
peaks after deleting less than one fifth of global max:11
×10^7
15
10
5
0
y
0
5
10
15
20
25
x
raw data, starting:11
fitted Gauss locations:11

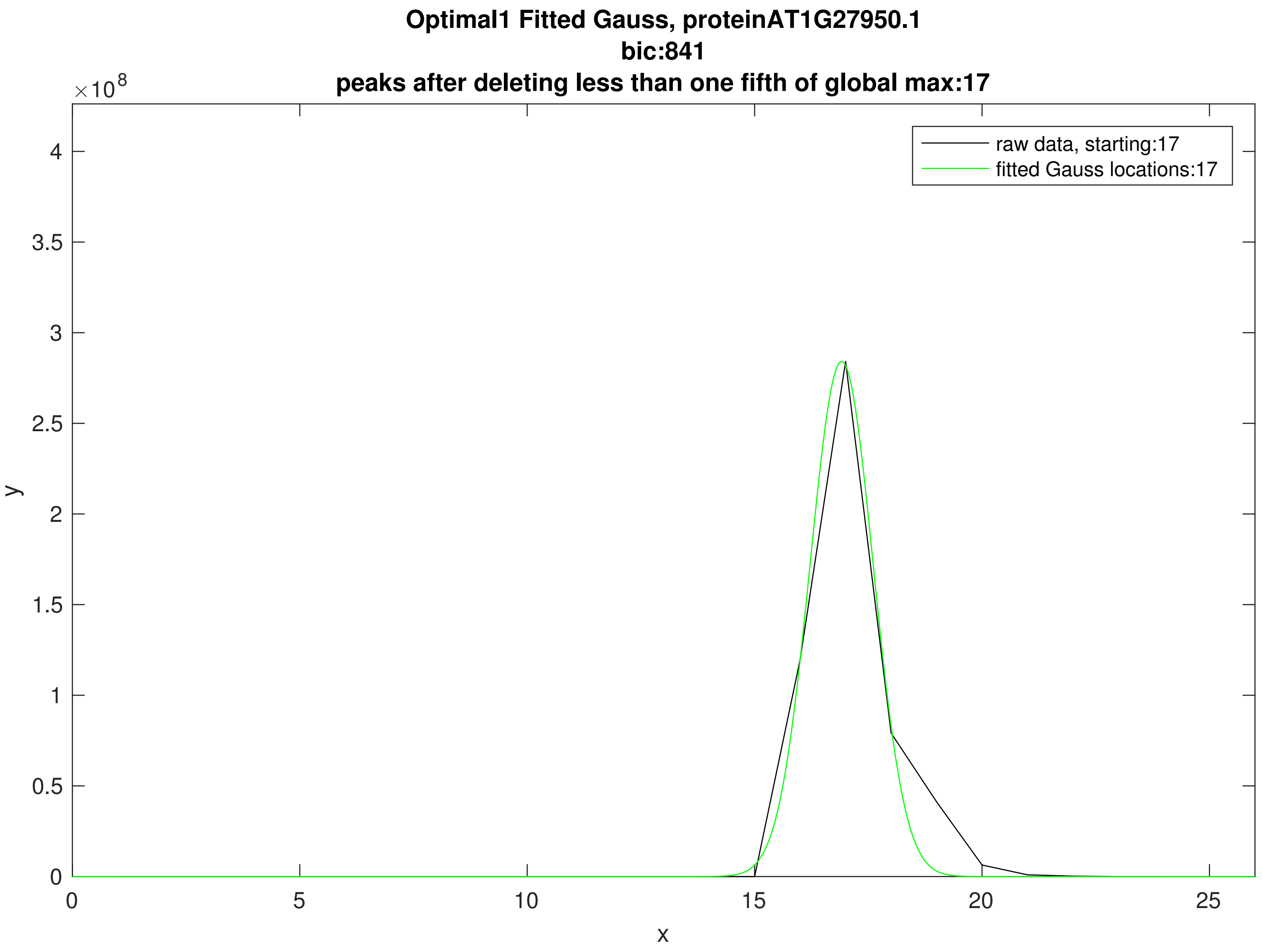
Optimal1 Fitted Gauss, proteinAT1G27950.1
bic:841
peaks after deleting less than one fifth of global max:17
×10^8
raw data, starting:17
fitted Gauss locations:17
y
x
0
0.5
1
1.5
2
2.5
3
3.5
4
0
5
10
15
20
25

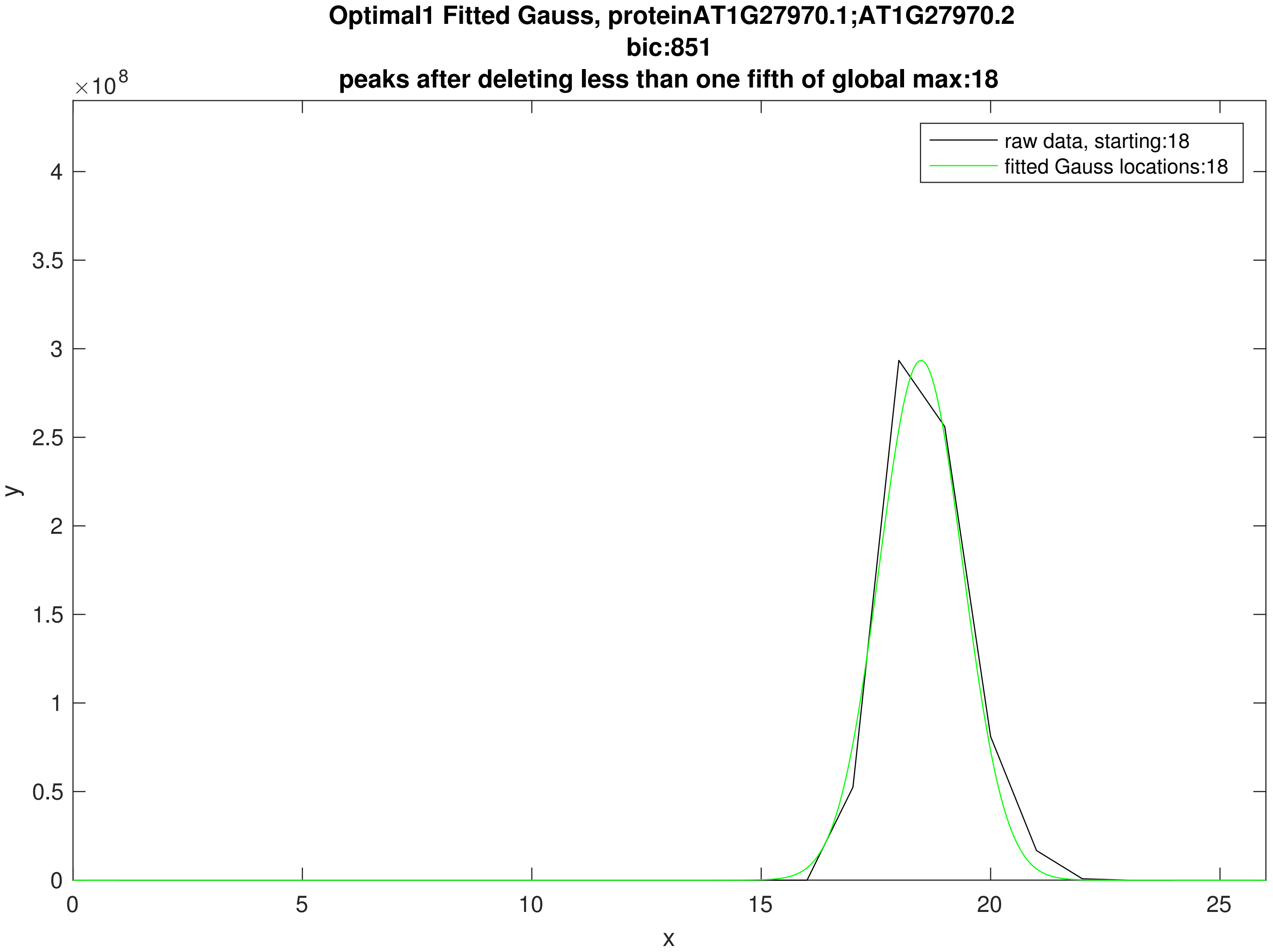
Optimal1 Fitted Gauss, proteinAT1G27970.1;AT1G27970.2
bic:851
peaks after deleting less than one fifth of global max:18
×10^8
raw data, starting:18
fitted Gauss locations:18
4
3.5
3
2.5
2
1.5
1
0.5
0
y
0
5
10
15
20
25
x

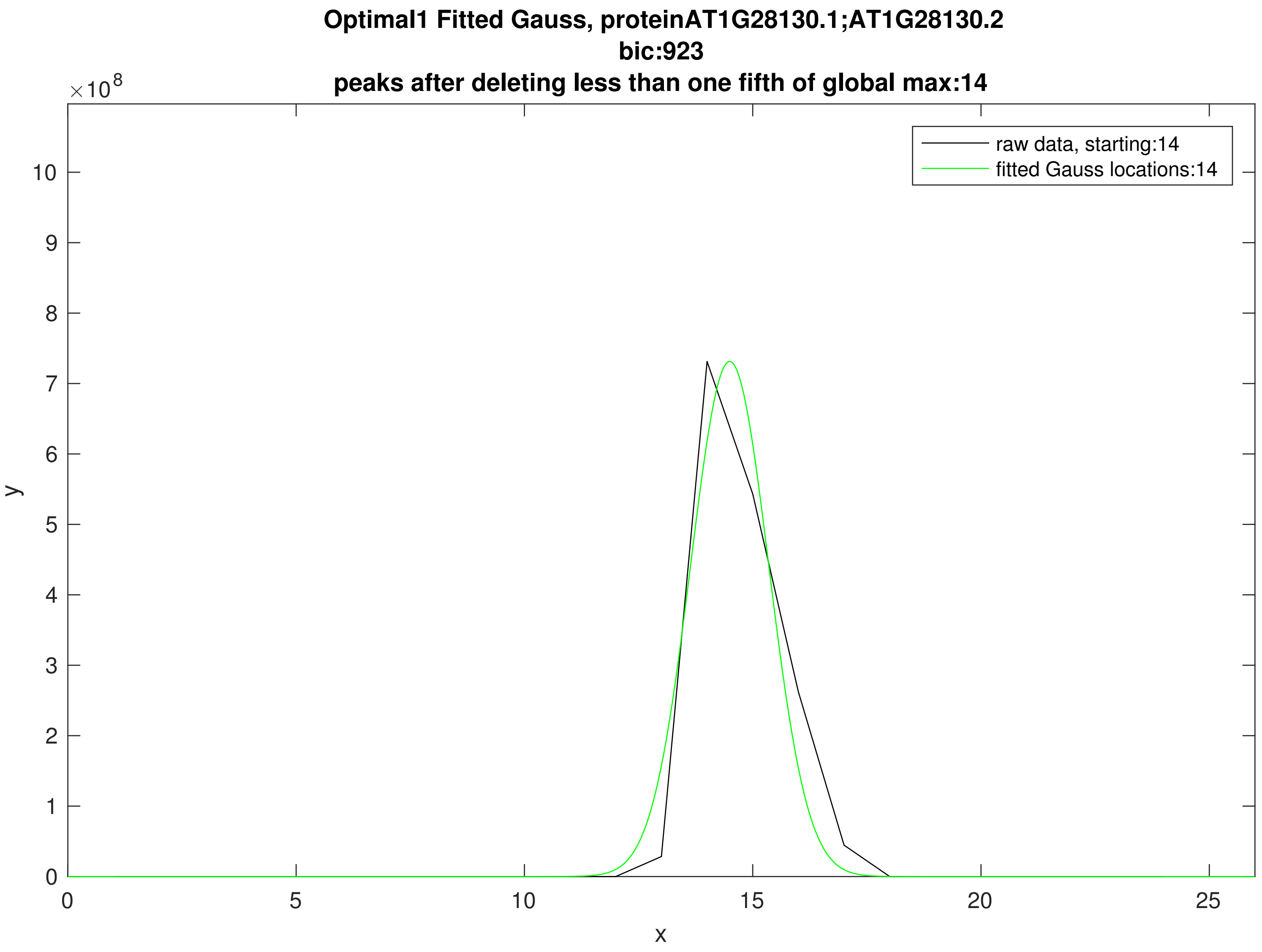
Optimal1 Fitted Gauss, proteinAT1G28130.1;AT1G28130.2
bic:923
peaks after deleting less than one fifth of global max:14
×10^8
raw data, starting:14
fitted Gauss locations:14
y
x
0
1
2
3
4
5
6
7
8
9
10
0
5
10
15
20
25

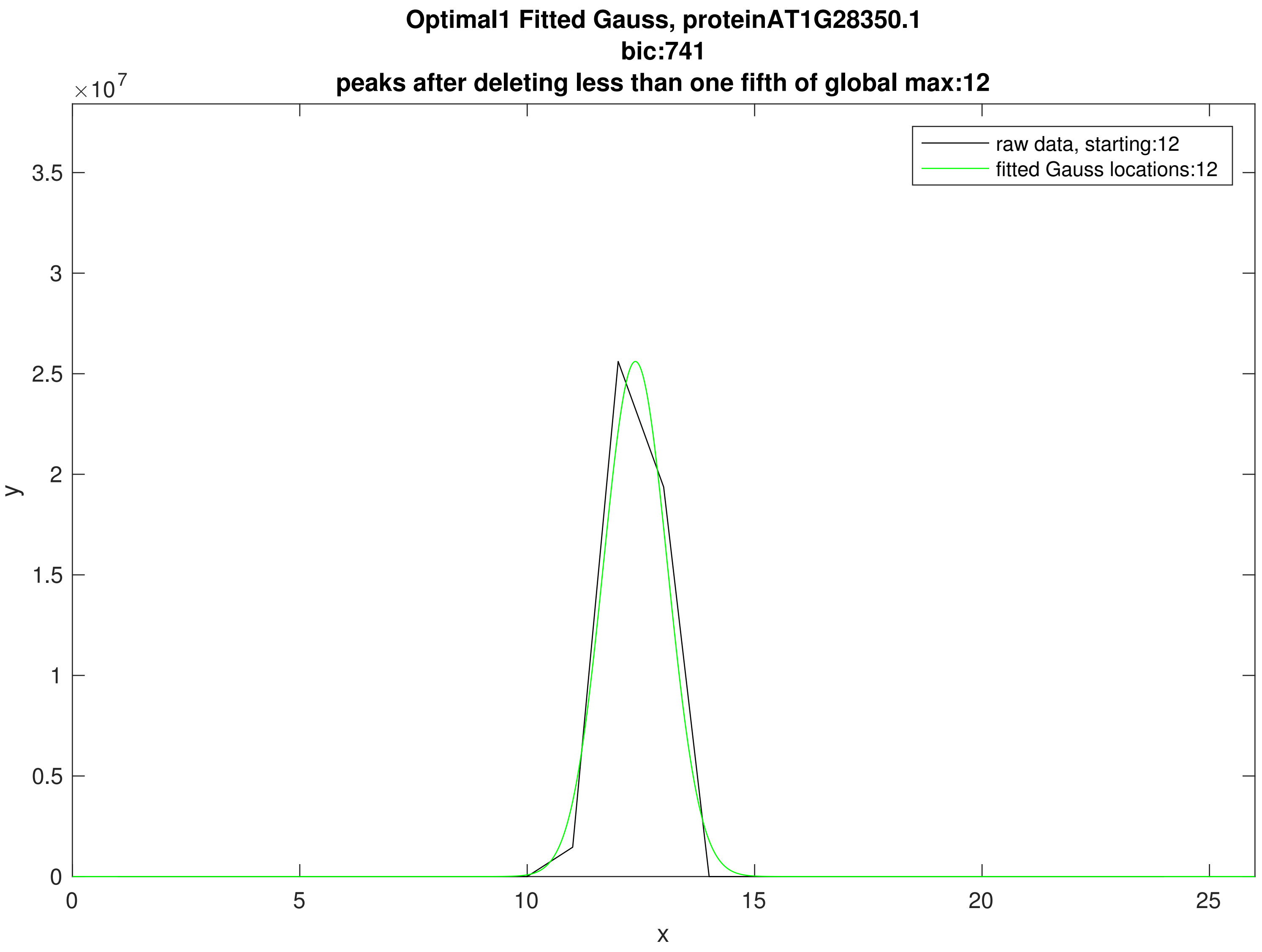
Optimal1 Fitted Gauss, proteinAT1G28350.1
bic:741
peaks after deleting less than one fifth of global max:12
×10^7
raw data, starting:12
fitted Gauss locations:12
3.5
3
2.5
2
1.5
1
0.5
0
y
0
5
10
15
20
25
x

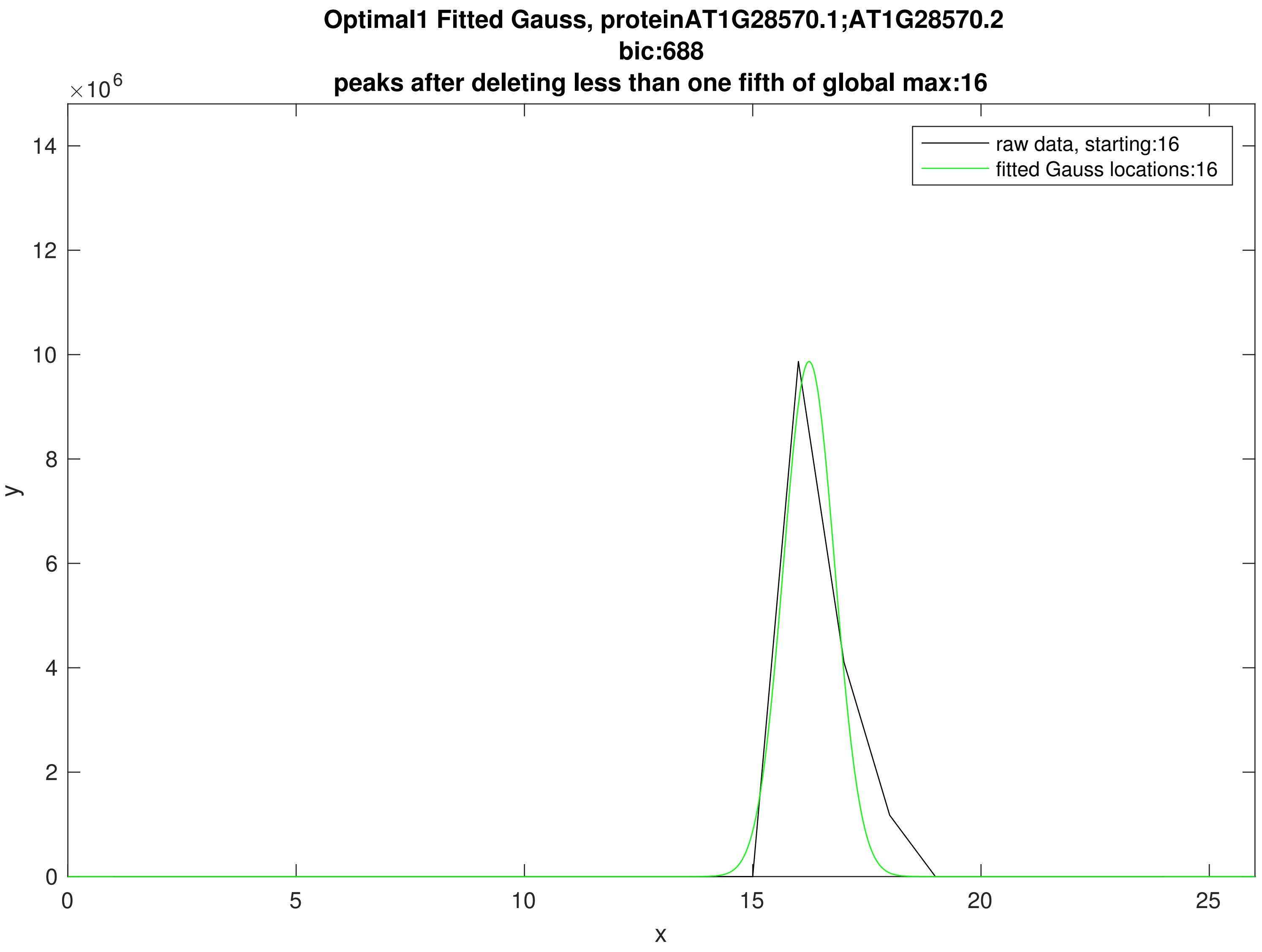
Optimal1 Fitted Gauss, proteinAT1G28570.1;AT1G28570.2
bic:688
peaks after deleting less than one fifth of global max:16
×10^6
raw data, starting:16
fitted Gauss locations:16
y
x
0
2
4
6
8
10
12
14
0
5
10
15
20
25

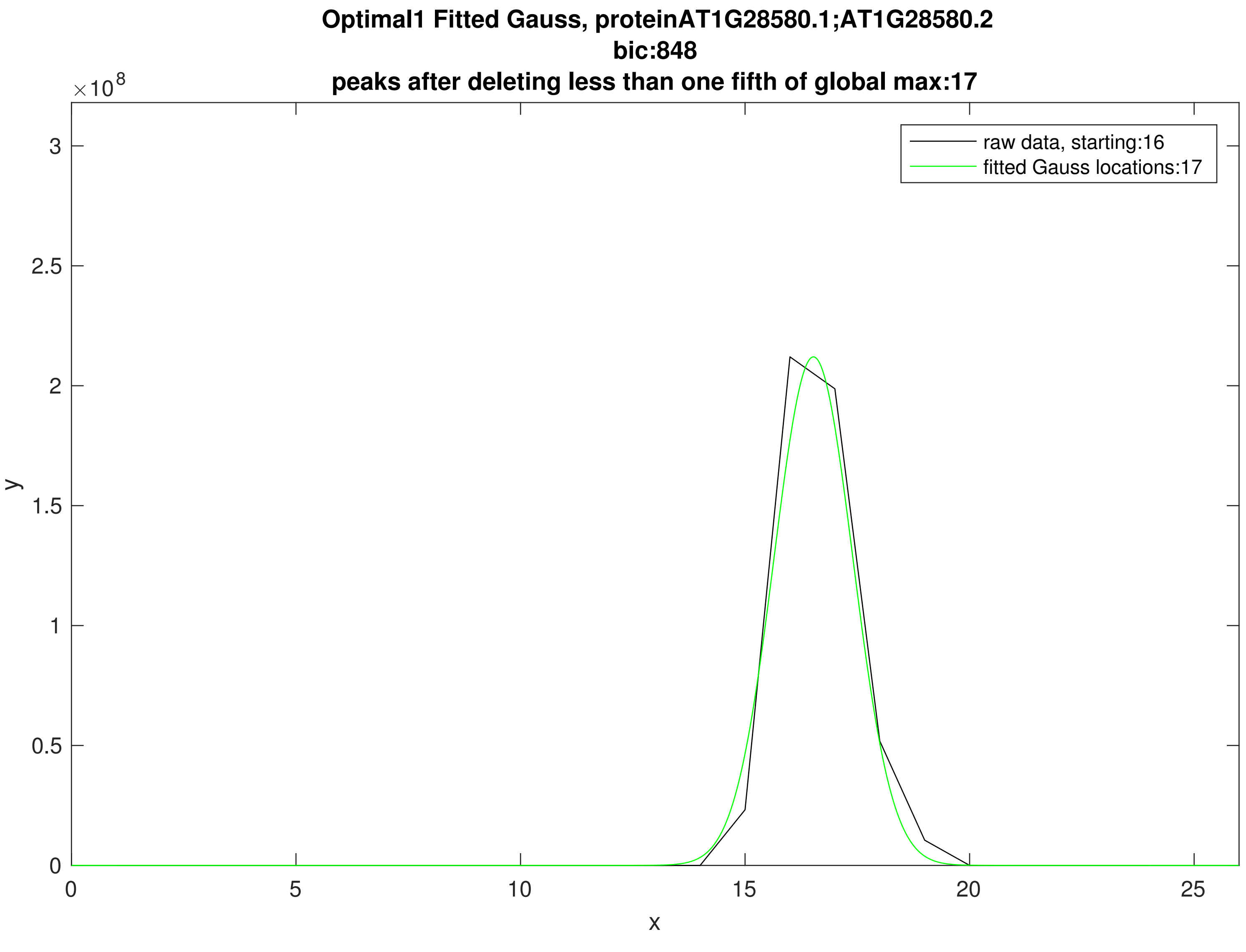

Optimal1 Fitted Gauss, proteinAT1G28580.1;AT1G28580.2
bic:848
peaks after deleting less than one fifth of global max:17
×10^8
raw data, starting:16
fitted Gauss locations:17
y
x
0
0.5
1
1.5
2
2.5
3
5
10
15
20
25

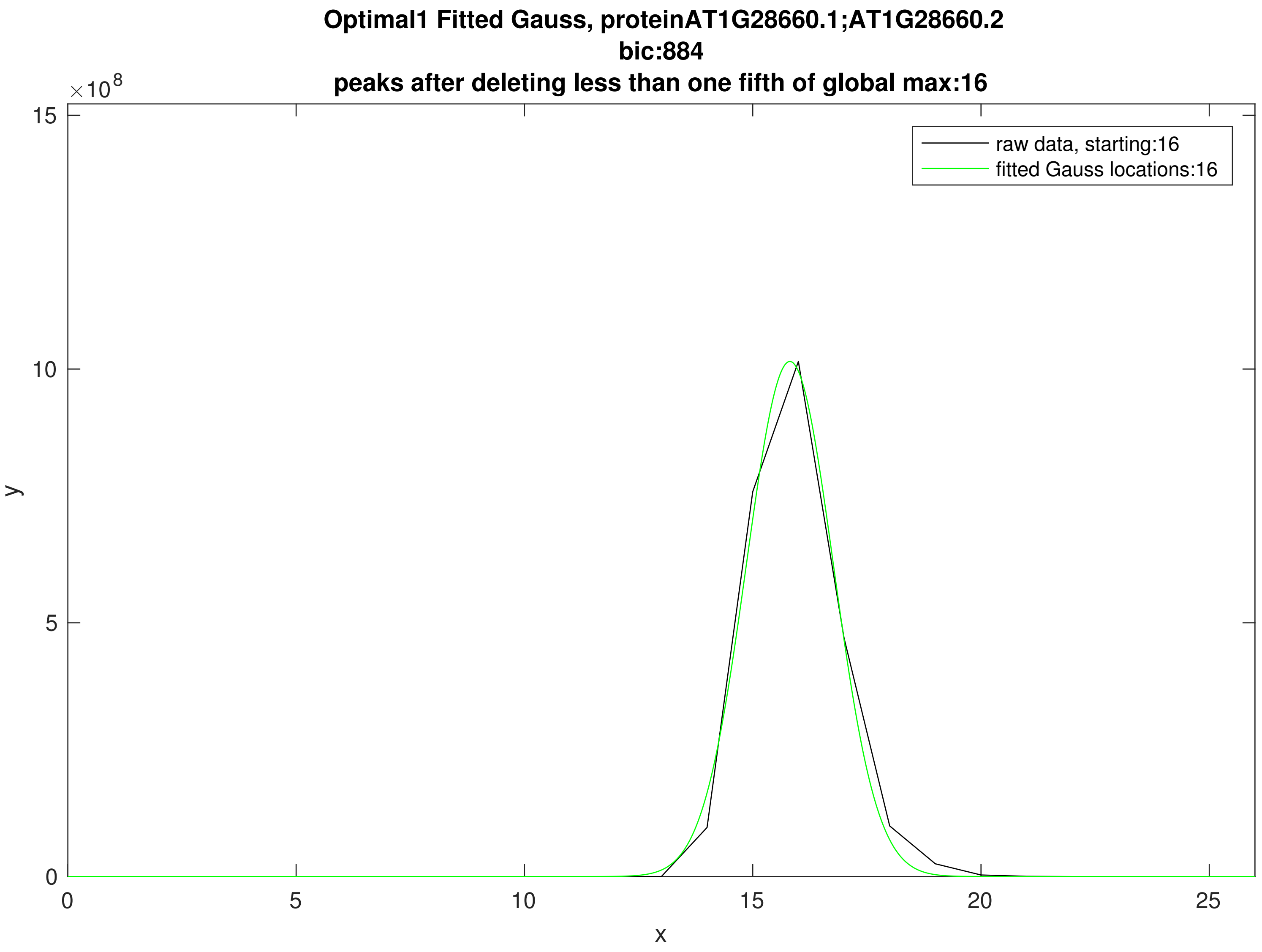

Optimal1 Fitted Gauss, proteinAT1G28660.1;AT1G28660.2
bic:884
peaks after deleting less than one fifth of global max:16
×10^8
15
10
5
0
y
0
5
10
15
20
25
x
raw data, starting:16
fitted Gauss locations:16

**Optimal1 Fitted Gauss, proteinAT1G28670.1**
**bic:874 880**
**peaks after deleting less than one fifth of global max:16**

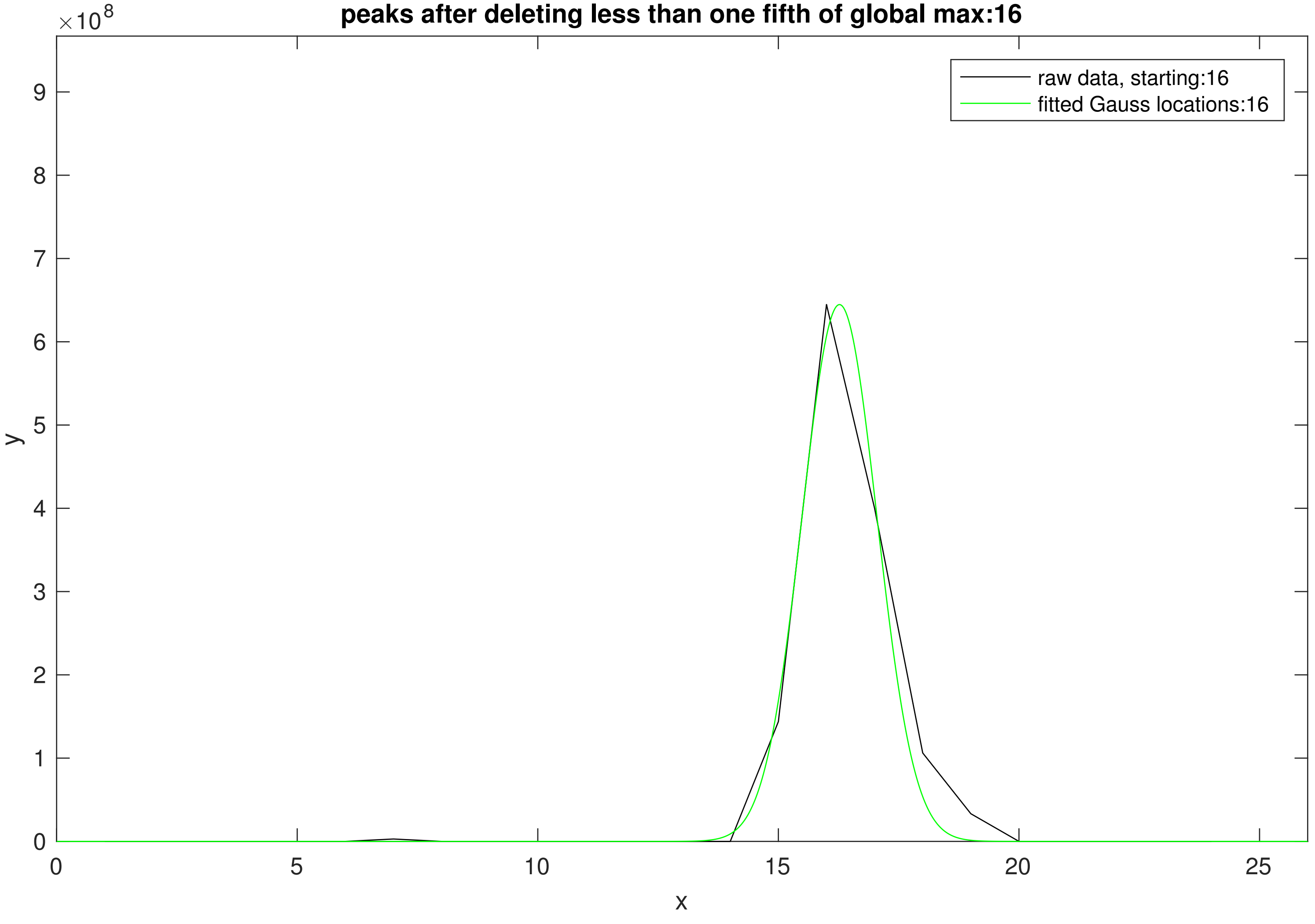

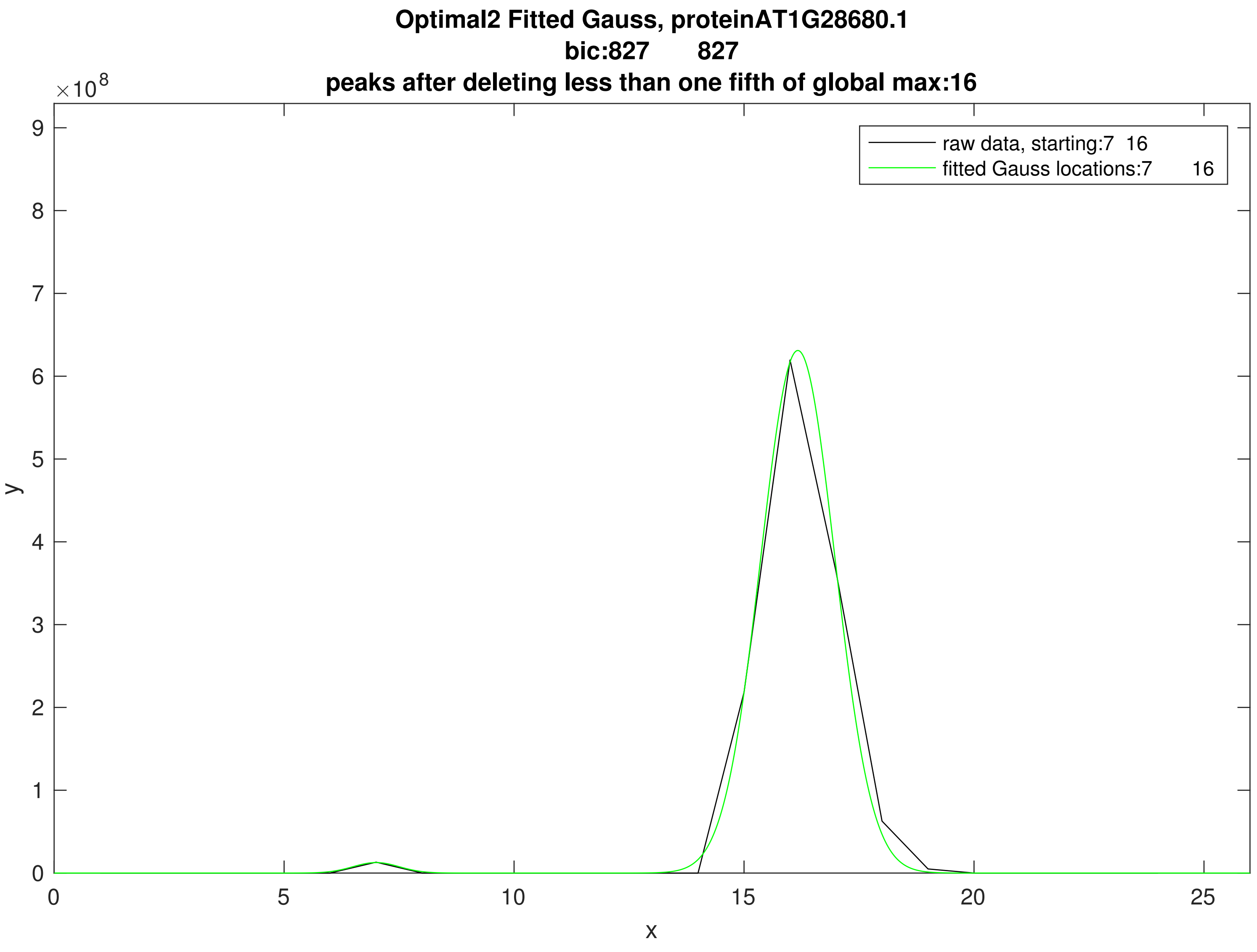

Optimal2 Fitted Gauss, proteinAT1G28680.1
bic:827 827
peaks after deleting less than one fifth of global max:16
×10^8
raw data, starting:7 16
fitted Gauss locations:7 16
y
x
0
1
2
3
4
5
6
7
8
9
0
5
10
15
20
25

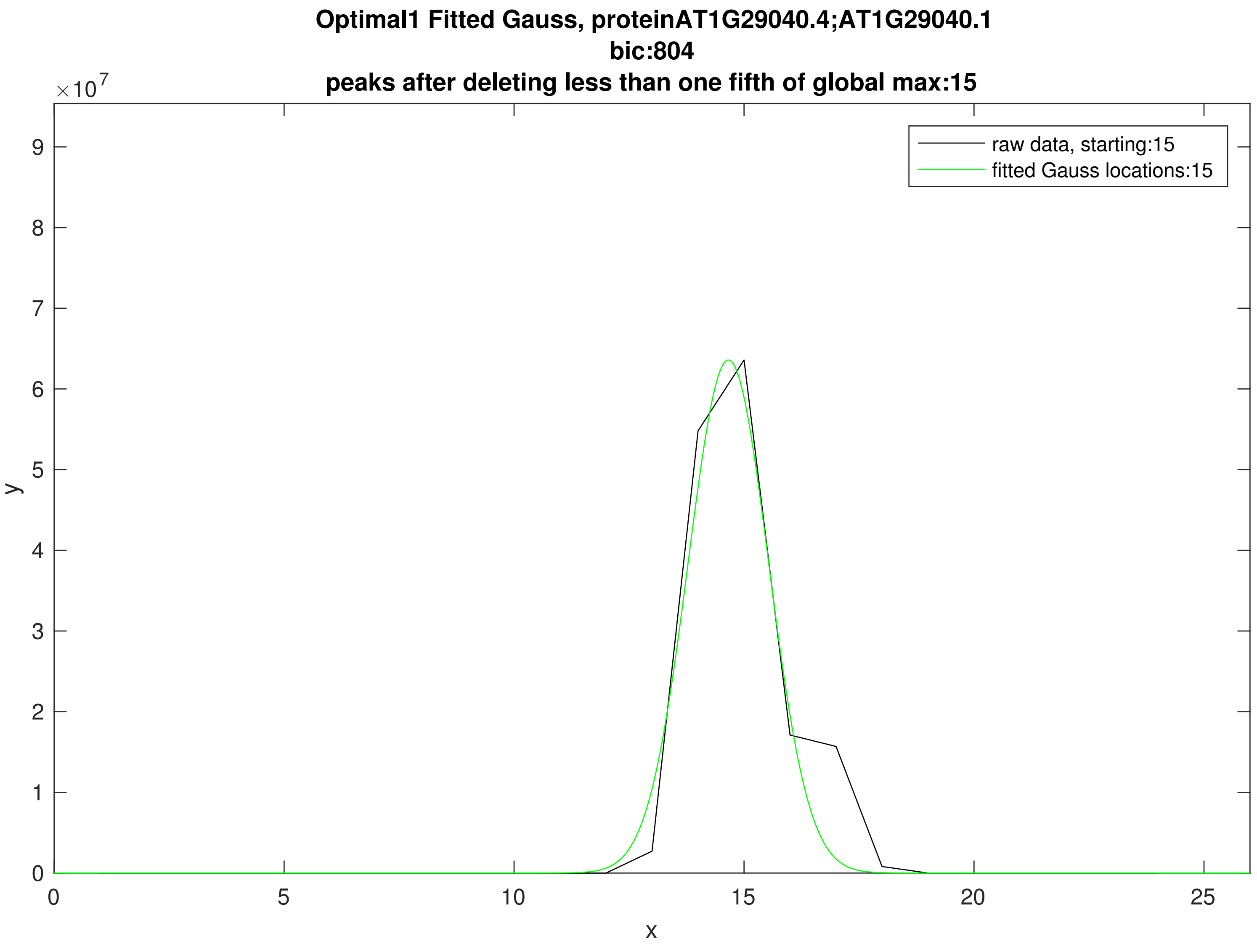

Optimal1 Fitted Gauss, proteinAT1G29040.4;AT1G29040.1
bic:804
peaks after deleting less than one fifth of global max:15
×10^7
raw data, starting:15
fitted Gauss locations:15
y
x
0
1
2
3
4
5
6
7
8
9
0
5
10
15
20
25

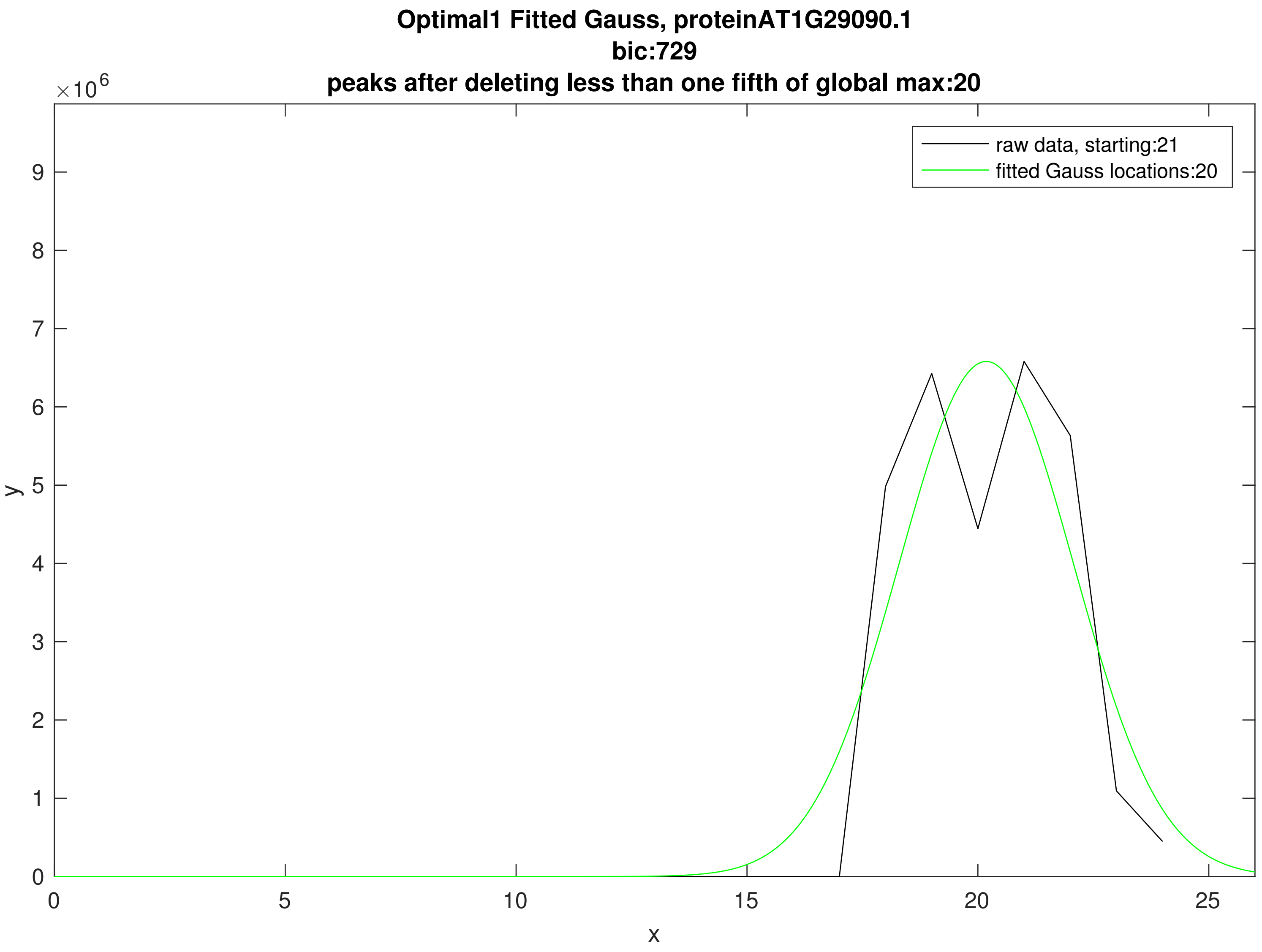

Optimal1 Fitted Gauss, proteinAT1G29090.1
bic:729
peaks after deleting less than one fifth of global max:20
×10^6
raw data, starting:21
fitted Gauss locations:20
y
0
1
2
3
4
5
6
7
8
9
0
5
10
15
20
25
x

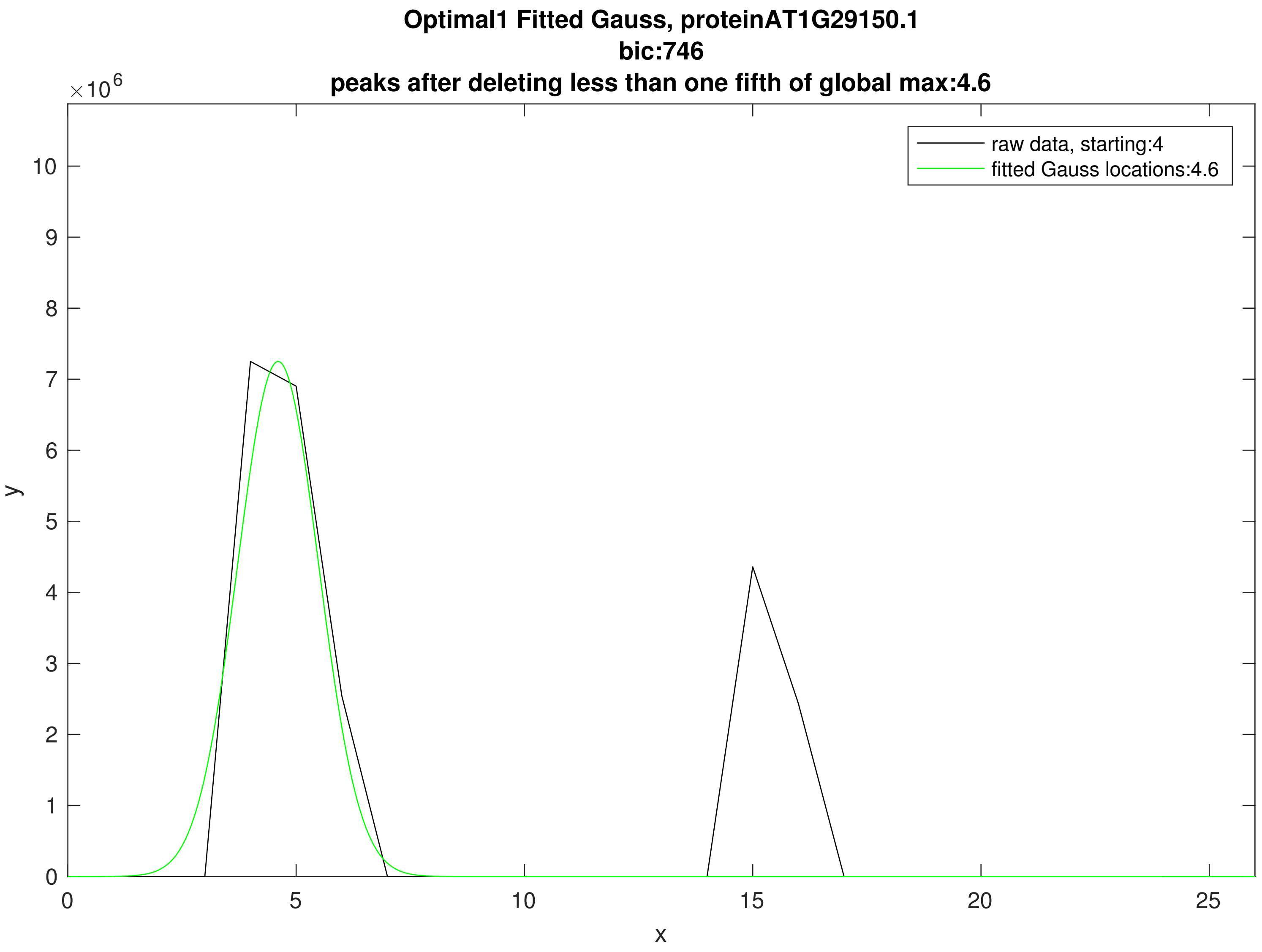
Optimal1 Fitted Gauss, proteinAT1G29150.1
bic:746
peaks after deleting less than one fifth of global max:4.6
×10^6
raw data, starting:4
fitted Gauss locations:4.6
y
x
0
1
2
3
4
5
6
7
8
9
10
0
5
10
15
20
25

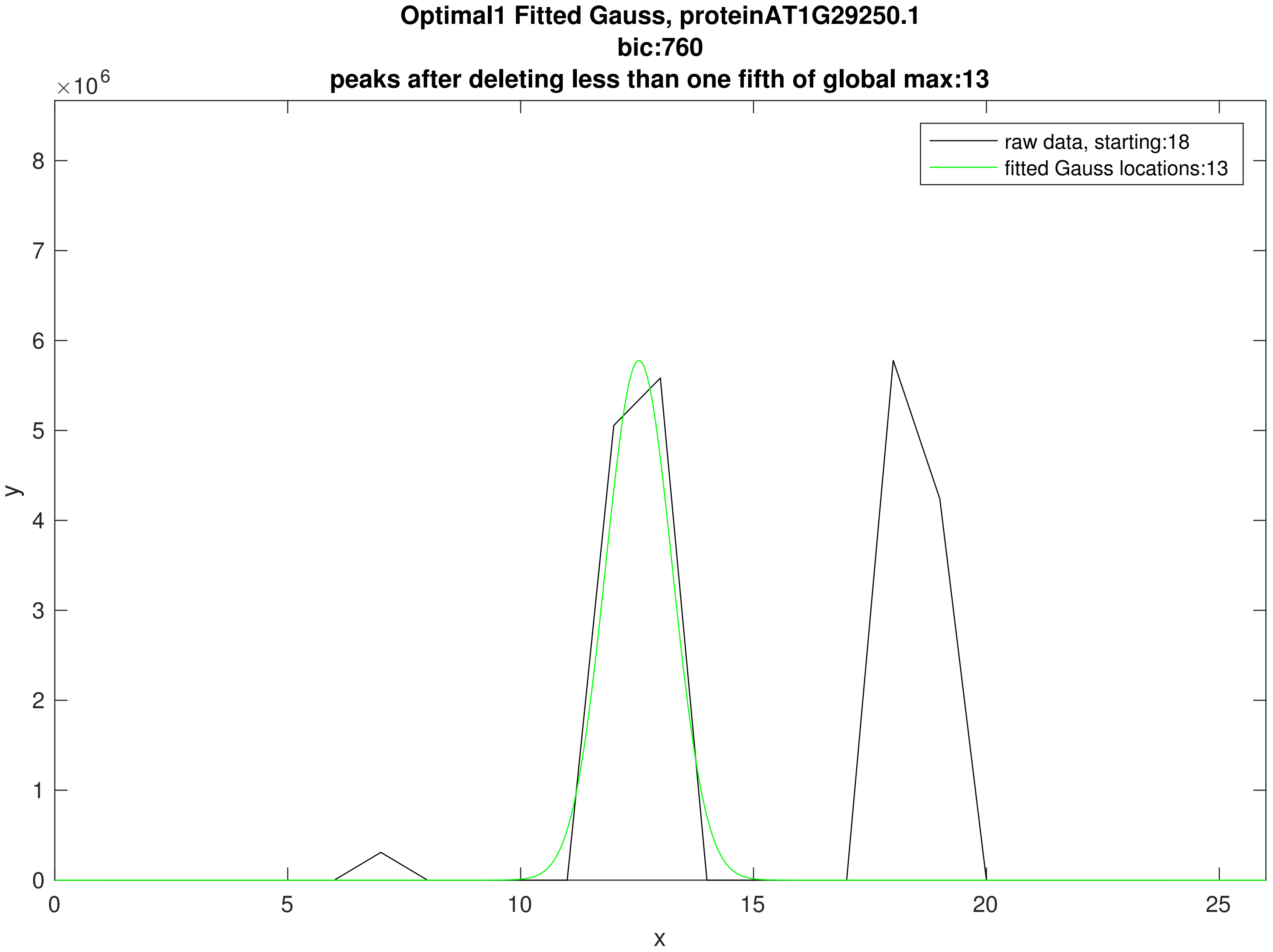
Optimal1 Fitted Gauss, proteinAT1G29250.1
bic:760
peaks after deleting less than one fifth of global max:13
×10^6
raw data, starting:18
fitted Gauss locations:13
y
x
0
1
2
3
4
5
6
7
8
0
5
10
15
20
25

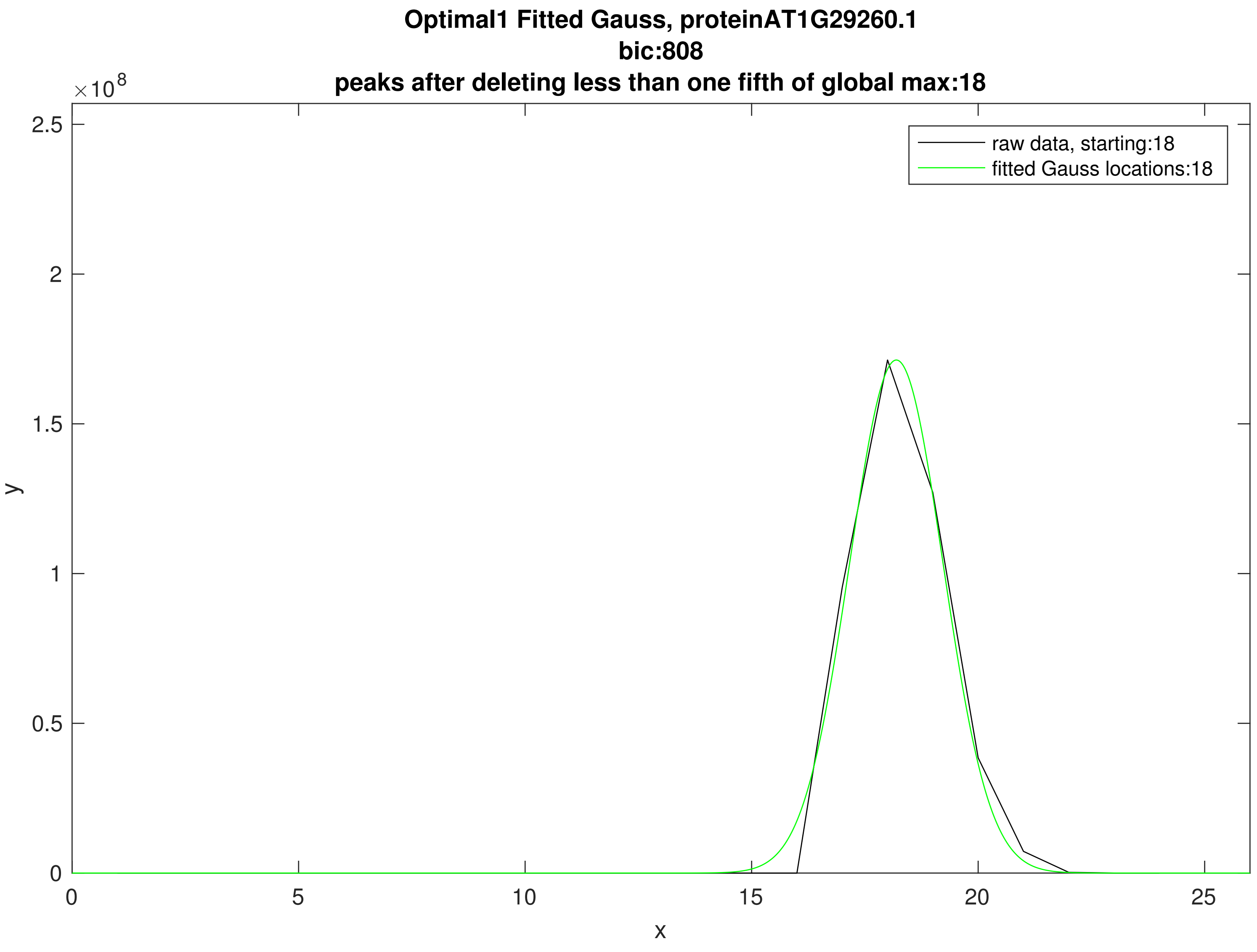
Optimal1 Fitted Gauss, proteinAT1G29260.1
bic:808
peaks after deleting less than one fifth of global max:18
×10^8
2.5
2
1.5
1
0.5
0
y
0
5
10
15
20
25
x
raw data, starting:18
fitted Gauss locations:18

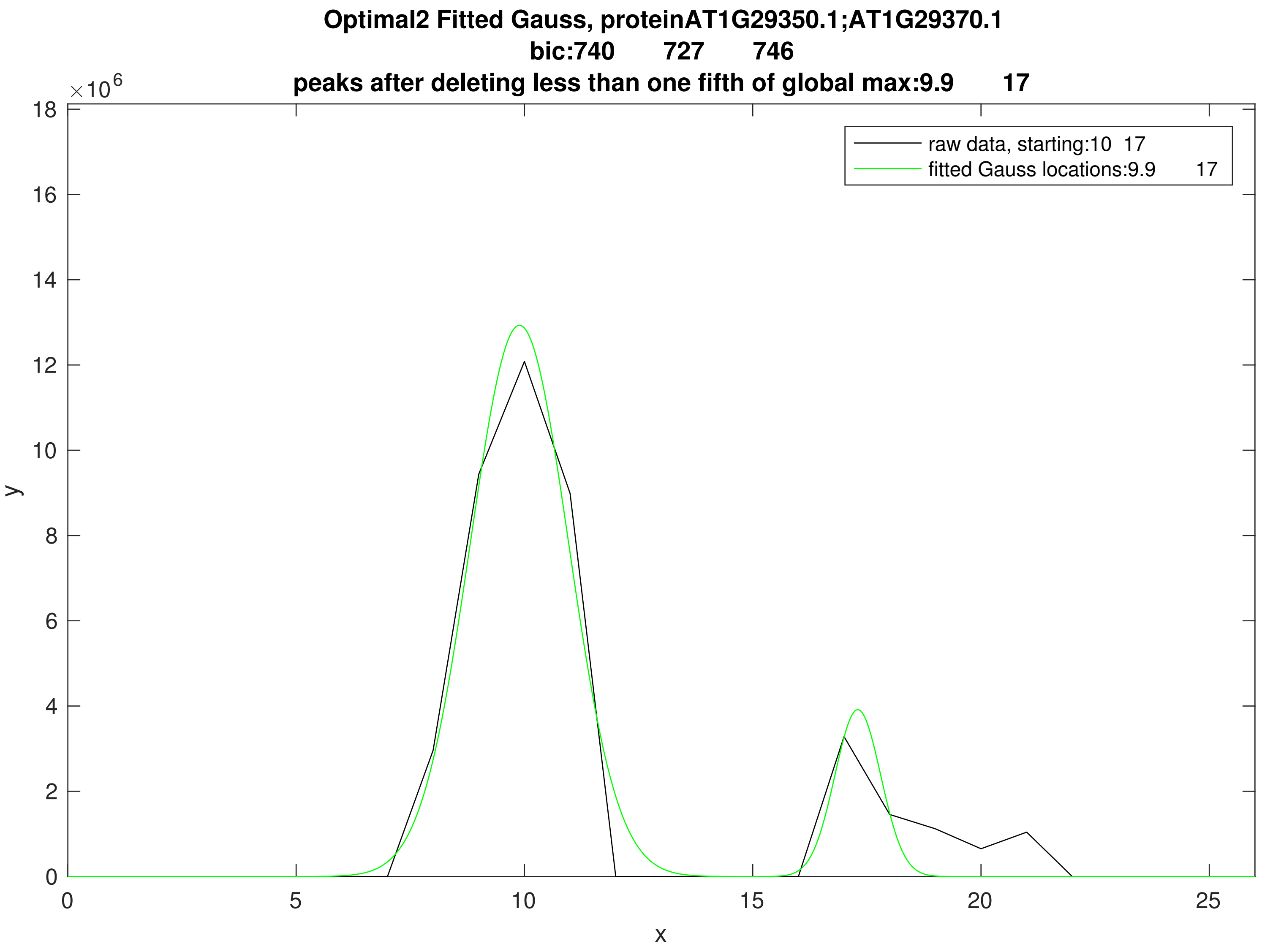
Optimal2 Fitted Gauss, proteinAT1G29350.1;AT1G29370.1
bic:740 727 746
peaks after deleting less than one fifth of global max:9.9 17
raw data, starting:10 17
fitted Gauss locations:9.9 17
×10^6
y
x

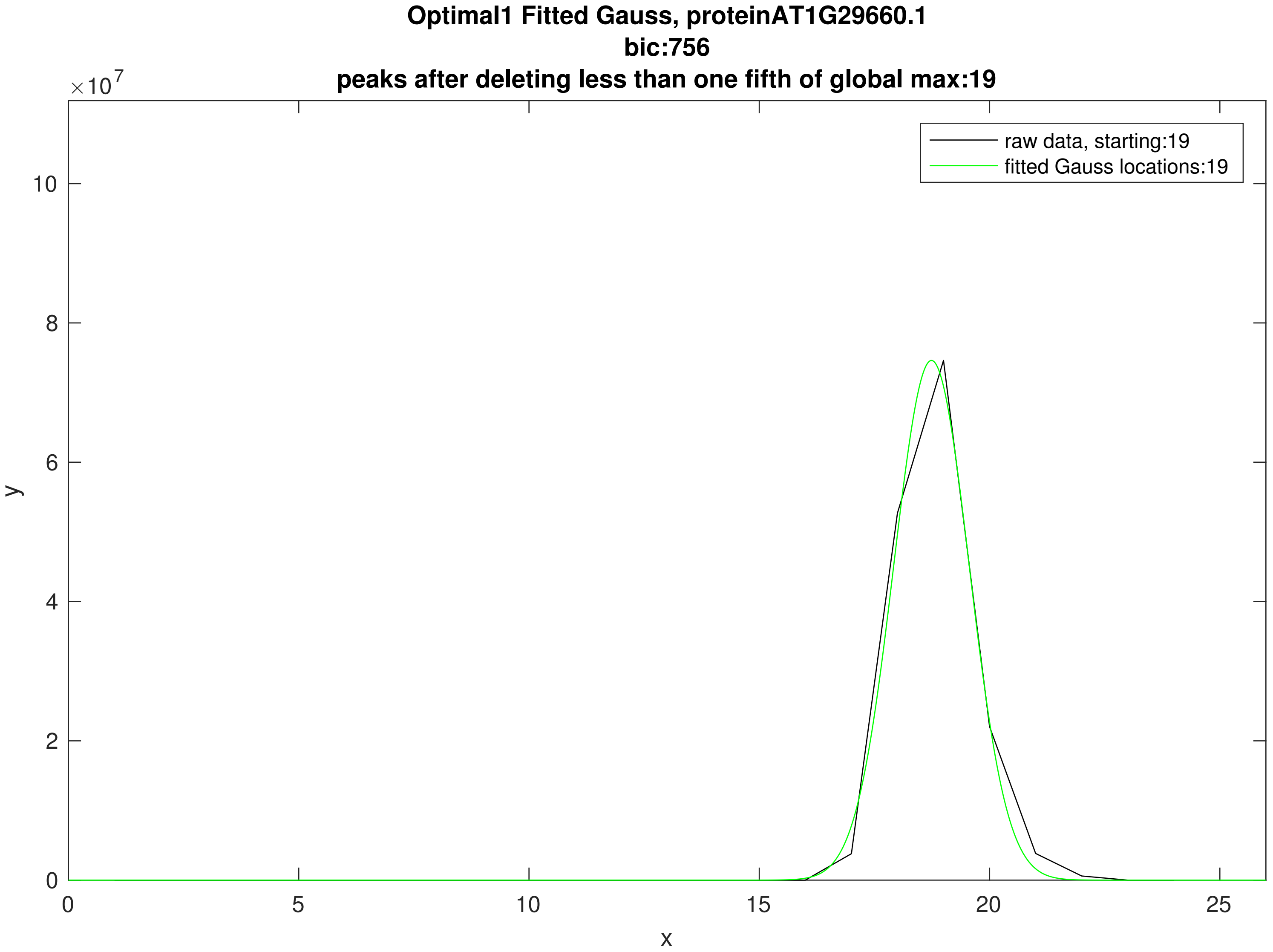
Optimal1 Fitted Gauss, proteinAT1G29660.1
bic:756
peaks after deleting less than one fifth of global max:19
×10^7
raw data, starting:19
fitted Gauss locations:19
10
8
6
4
2
0
y
0
5
10
15
20
25
x

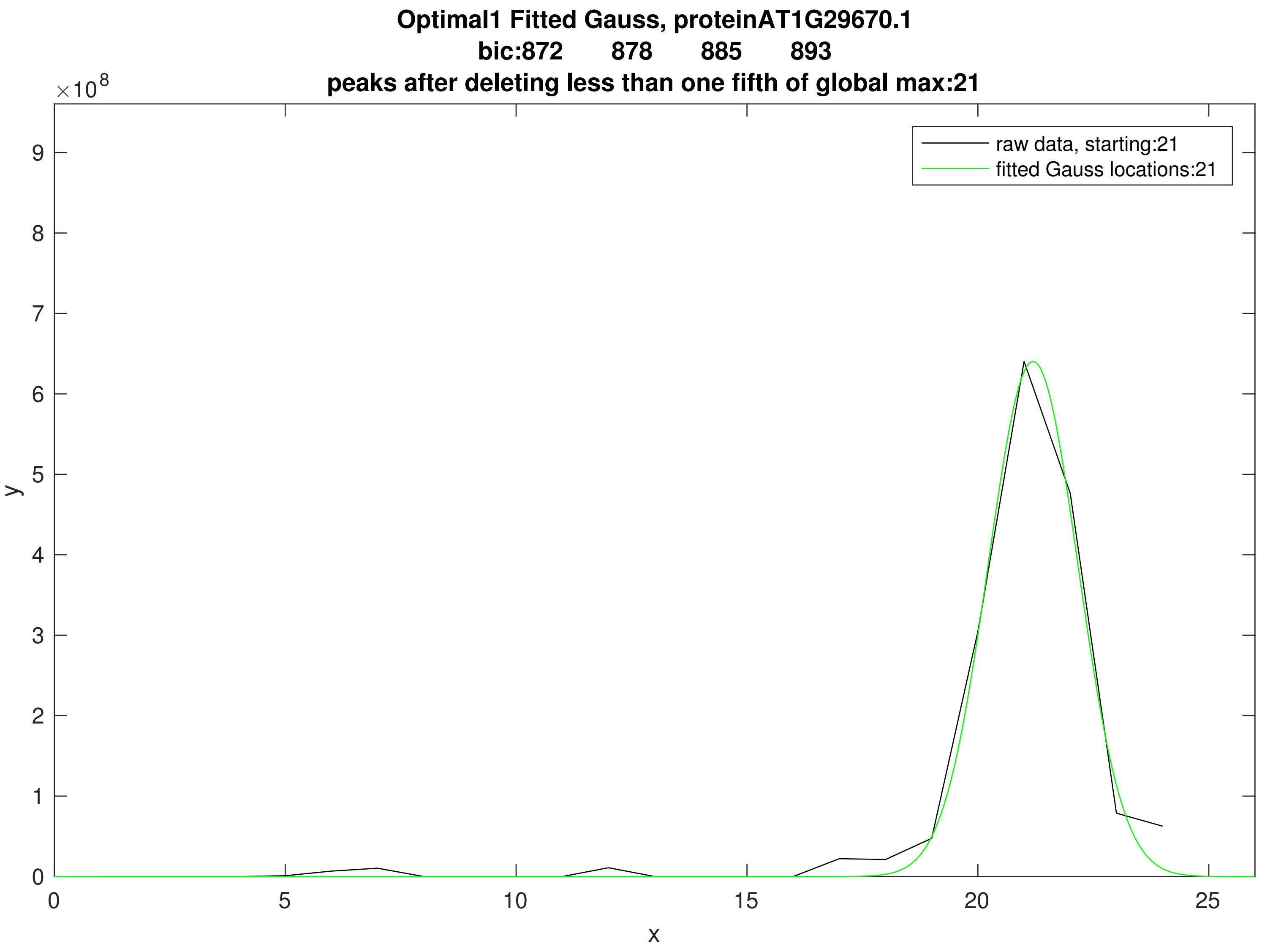

Optimal1 Fitted Gauss, proteinAT1G29670.1
bic:872 878 885 893
peaks after deleting less than one fifth of global max:21
×10^8
9
8
7
6
5
4
3
2
1
0
y
0
5
10
15
20
25
x
raw data, starting:21
fitted Gauss locations:21

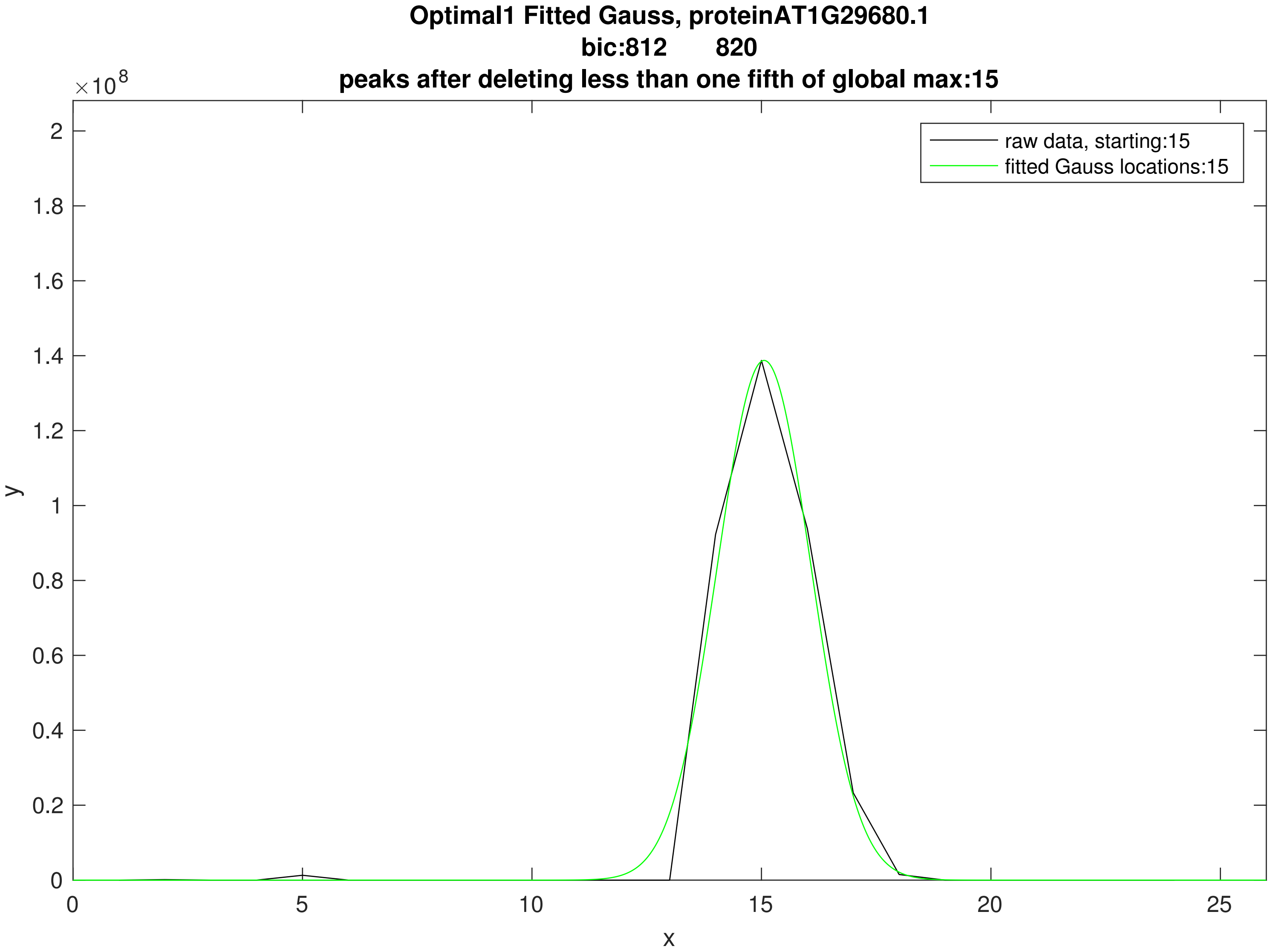

Optimal1 Fitted Gauss, proteinAT1G29680.1
bic:812 820
peaks after deleting less than one fifth of global max:15
×10^8
2
1.8
1.6
1.4
1.2
1
0.8
0.6
0.4
0.2
0
y
0
5
10
15
20
25
x
raw data, starting:15
fitted Gauss locations:15

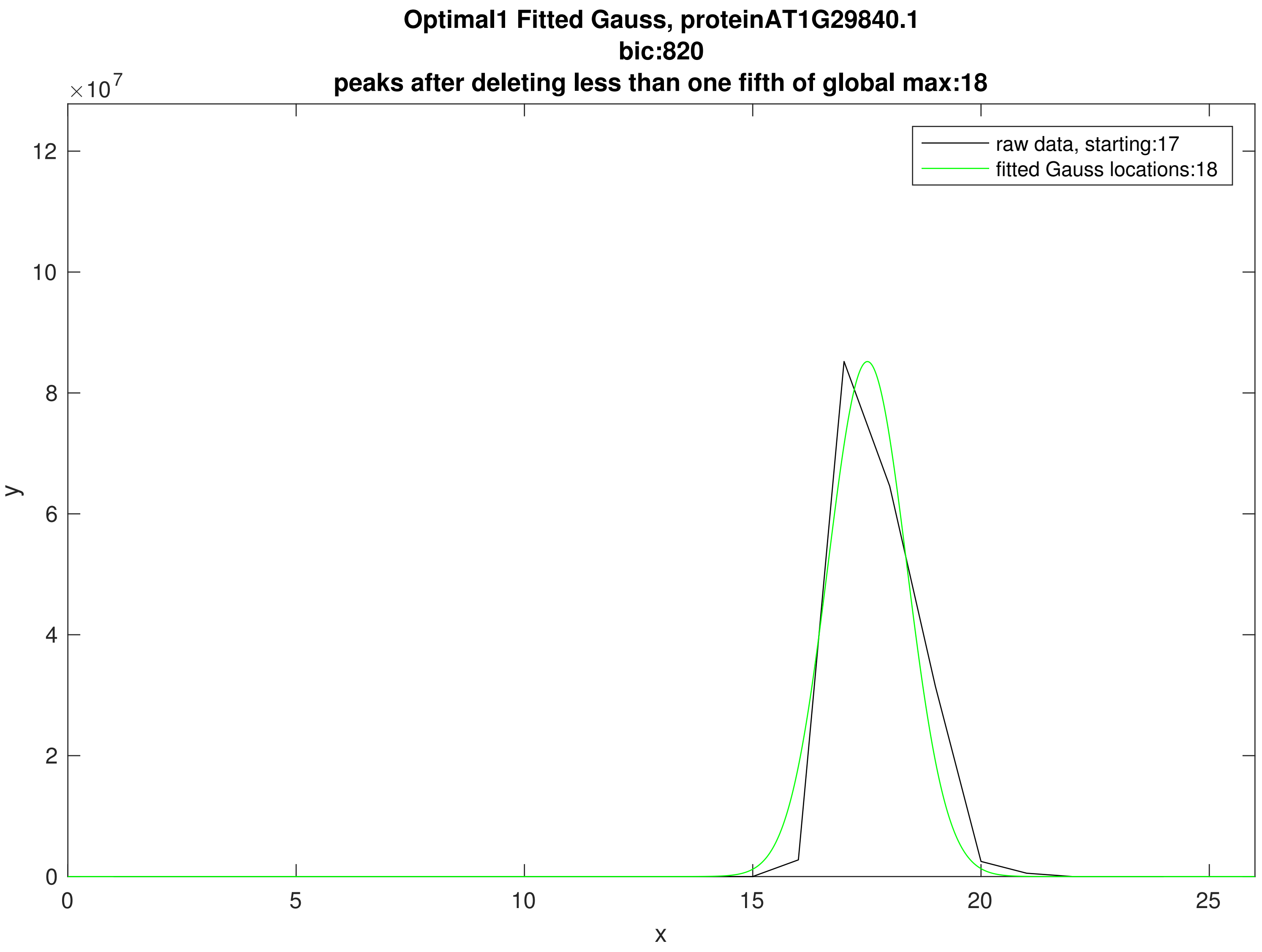

Optimal1 Fitted Gauss, proteinAT1G29840.1
bic:820
peaks after deleting less than one fifth of global max:18
×10^7
raw data, starting:17
fitted Gauss locations:18
y
x
0
2
4
6
8
10
12
0
5
10
15
20
25

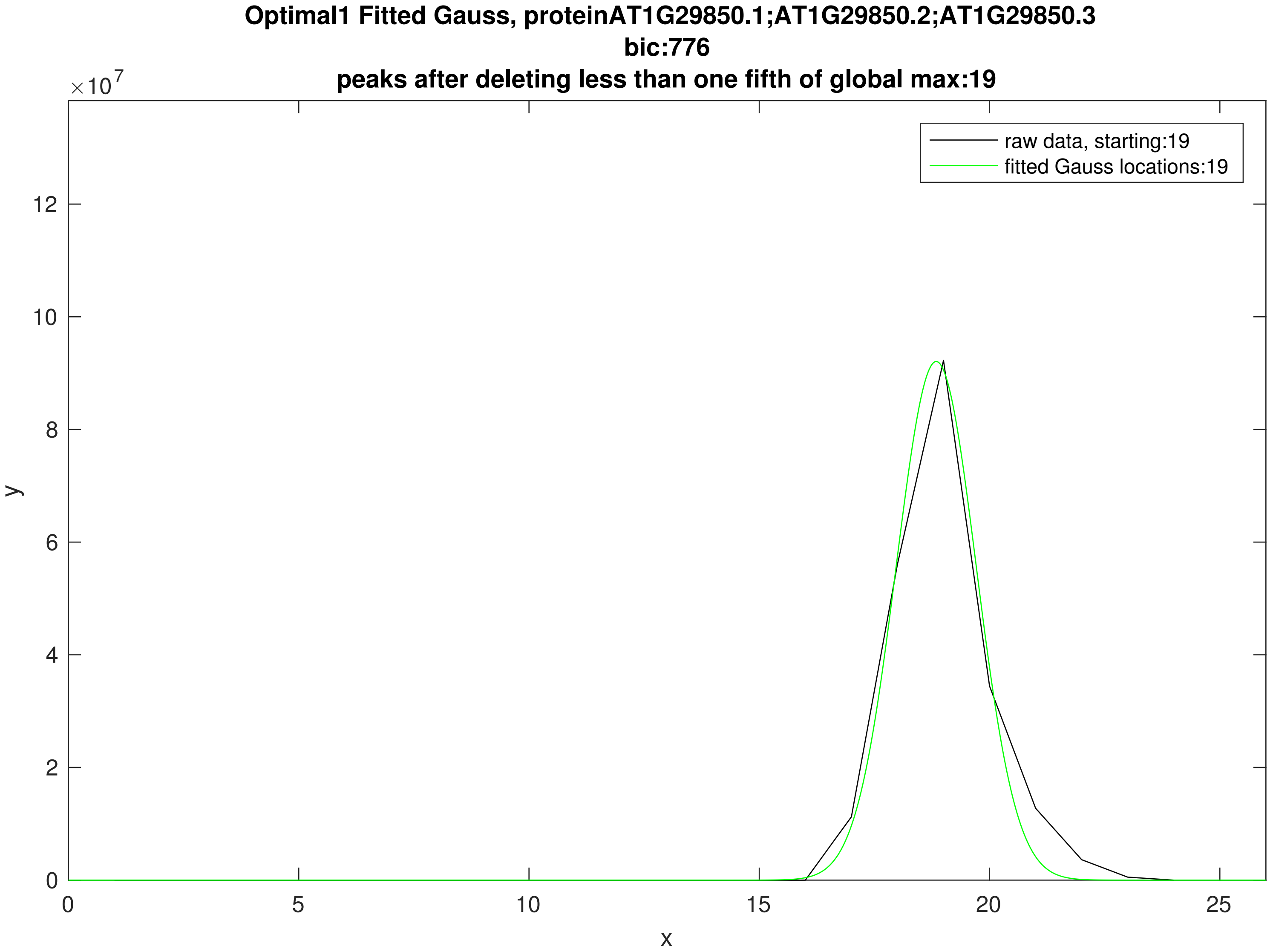

Optimal1 Fitted Gauss, proteinAT1G29850.1;AT1G29850.2;AT1G29850.3
bic:776
peaks after deleting less than one fifth of global max:19
×10^7
raw data, starting:19
fitted Gauss locations:19
y
x
0
2
4
6
8
10
12
0
5
10
15
20
25

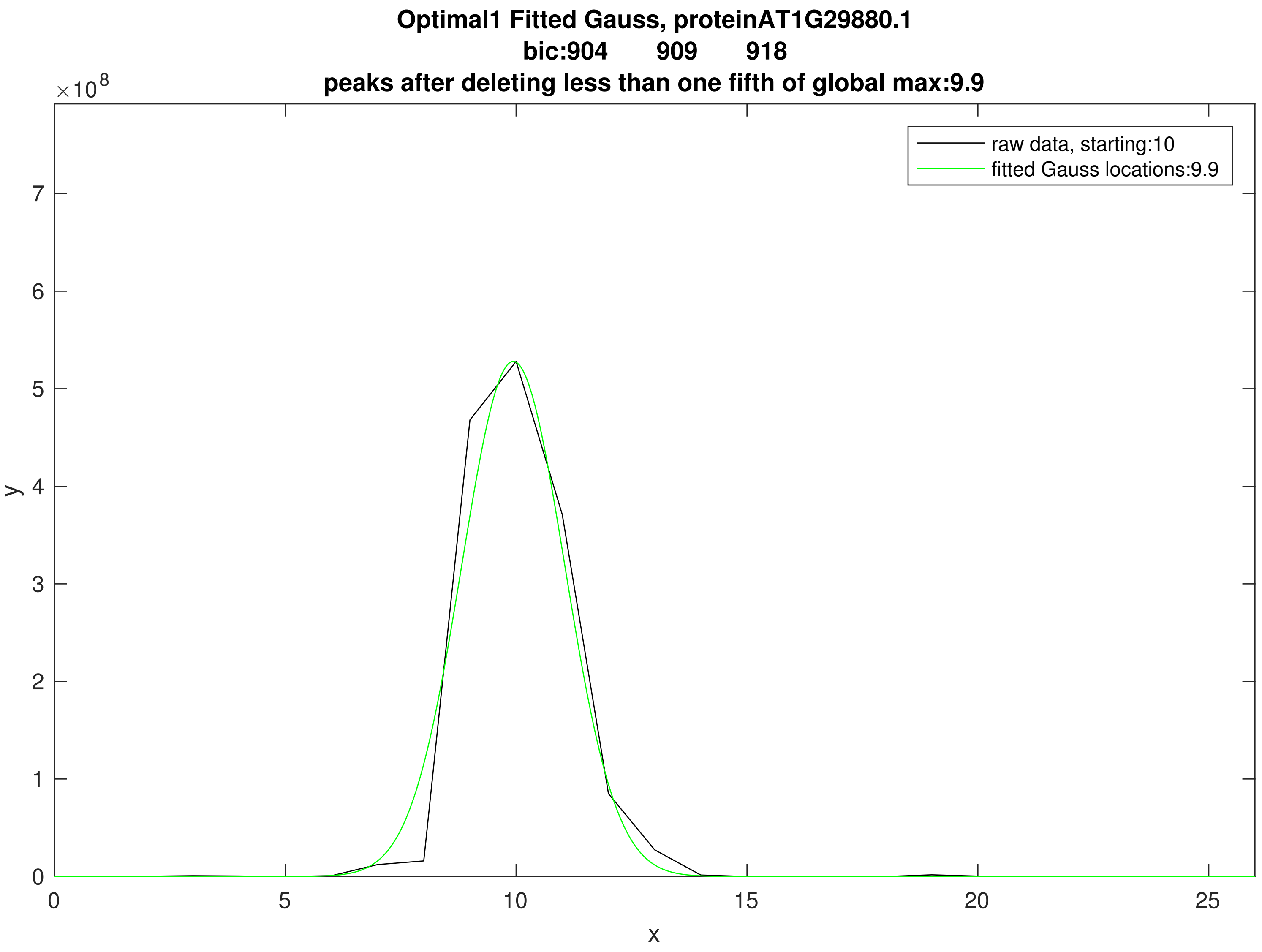
Optimal1 Fitted Gauss, proteinAT1G29880.1
bic:904 909 918
peaks after deleting less than one fifth of global max:9.9
×10^8
raw data, starting:10
fitted Gauss locations:9.9
y
x
0
1
2
3
4
5
6
7
0
5
10
15
20
25

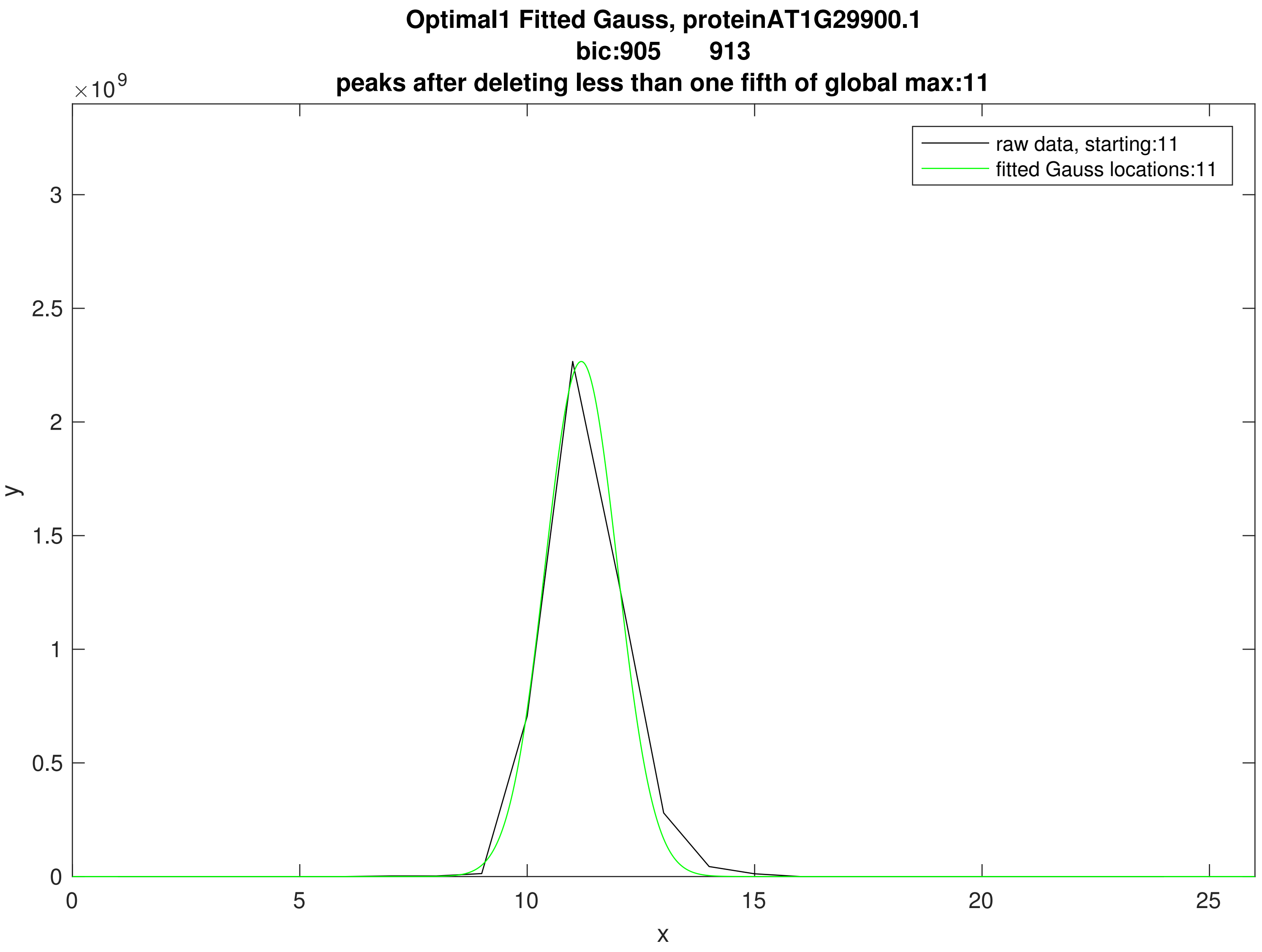
Optimal1 Fitted Gauss, proteinAT1G29900.1
bic:905 913
peaks after deleting less than one fifth of global max:11
×10^9
3
2.5
2
1.5
1
0.5
0
y
0
5
10
15
20
25
x
raw data, starting:11
fitted Gauss locations:11

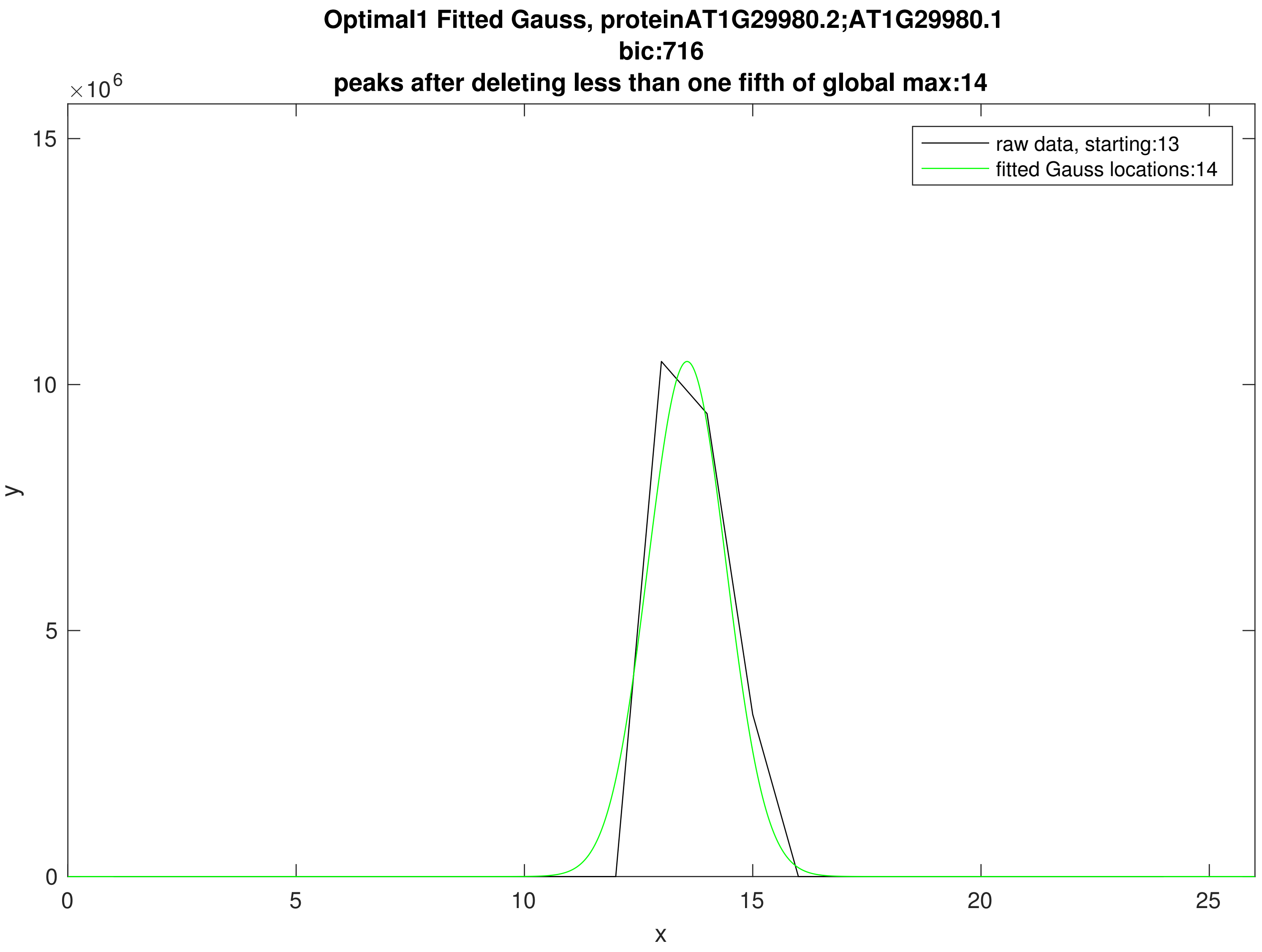

Optimal1 Fitted Gauss, proteinAT1G29980.2;AT1G29980.1
bic:716
peaks after deleting less than one fifth of global max:14
×10^6
15
10
5
0
y
0
5
10
15
20
25
x
raw data, starting:13
fitted Gauss locations:14

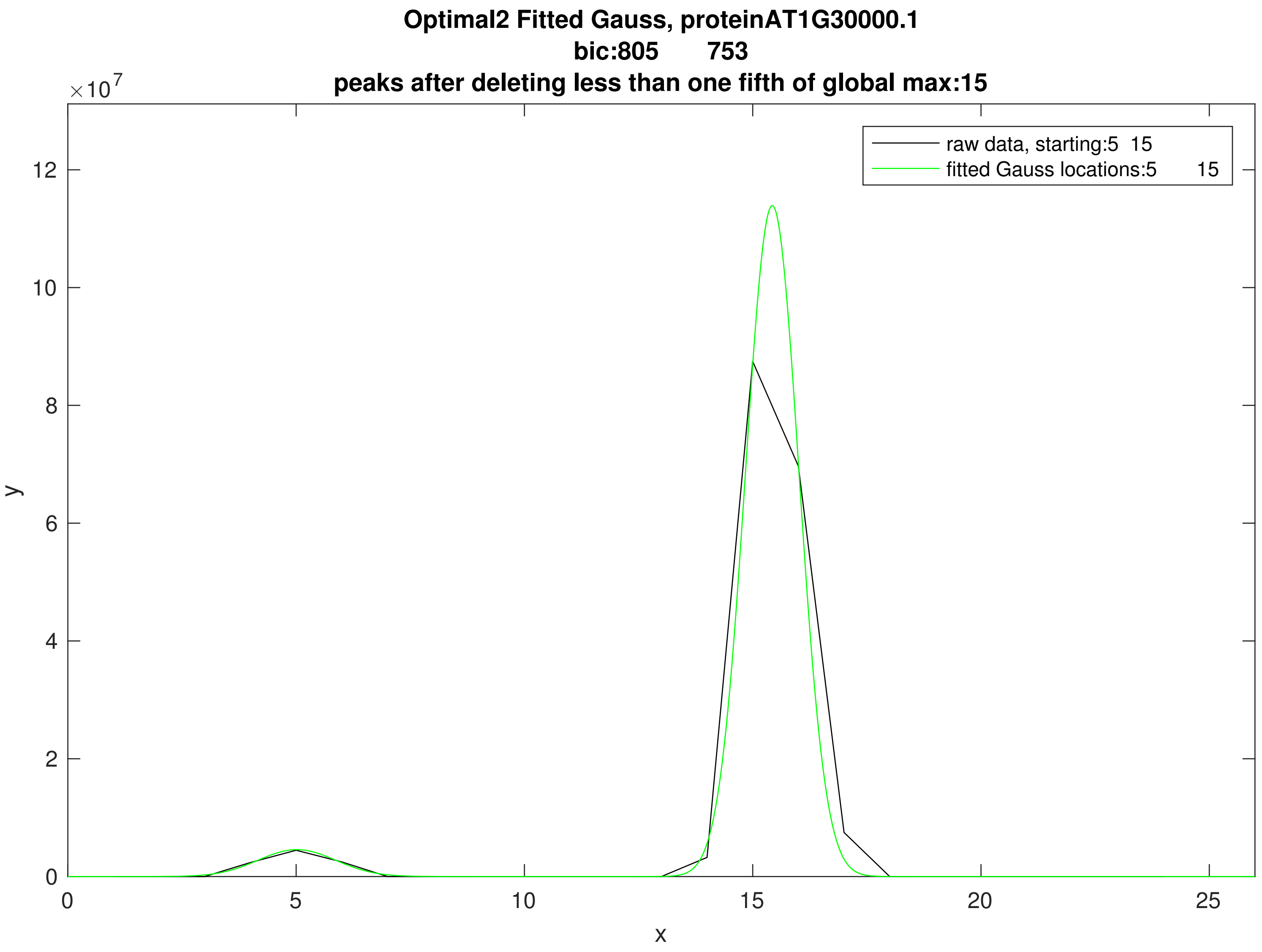
Optimal2 Fitted Gauss, proteinAT1G30000.1
bic:805 753
peaks after deleting less than one fifth of global max:15
×10^7
raw data, starting:5 15
fitted Gauss locations:5 15
y
x
0
2
4
6
8
10
12
0
5
10
15
20
25

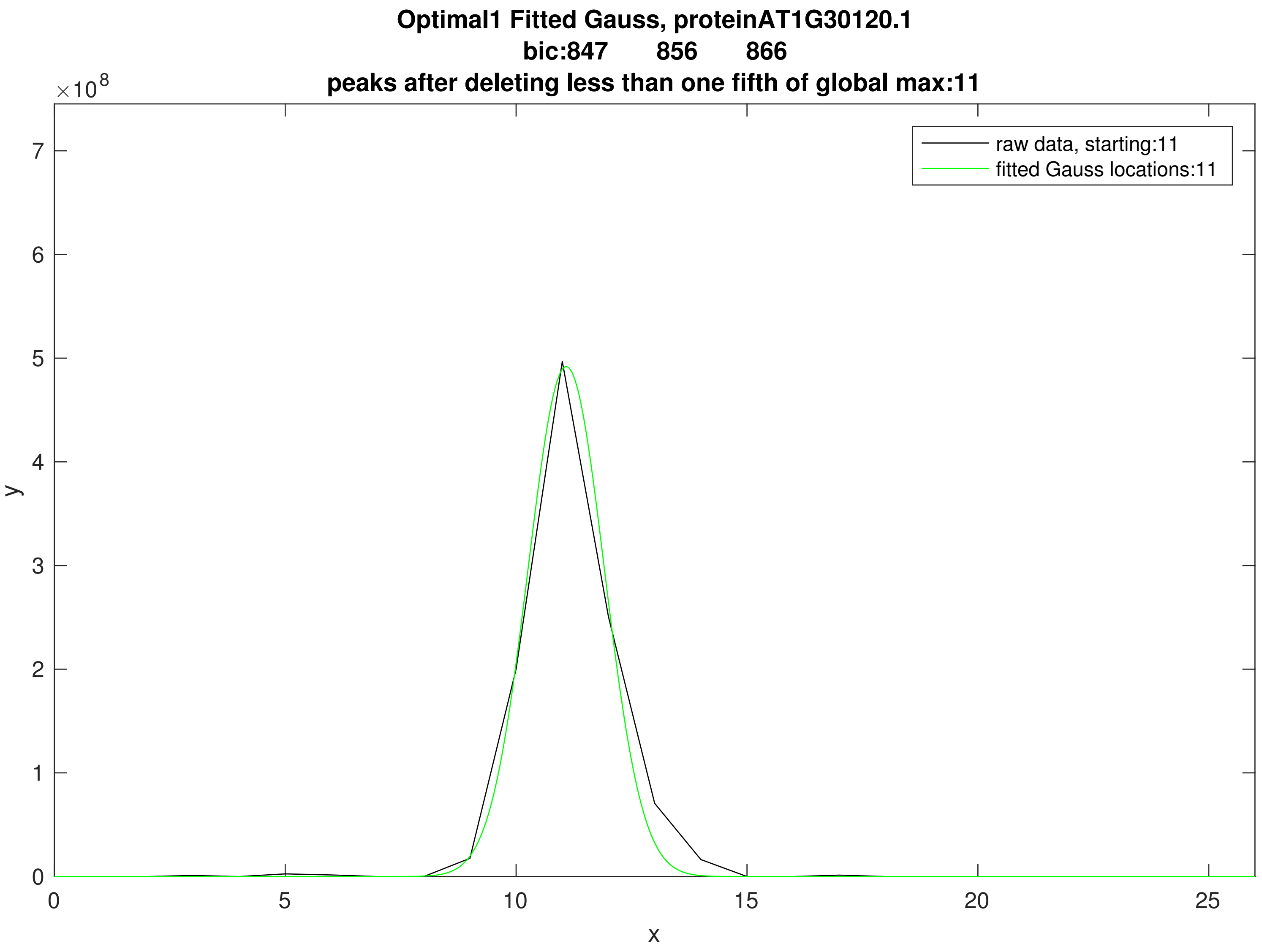
Optimal1 Fitted Gauss, proteinAT1G30120.1
bic:847 856 866
peaks after deleting less than one fifth of global max:11
×10^8
raw data, starting:11
fitted Gauss locations:11
y
x
0
1
2
3
4
5
6
7
0
5
10
15
20
25

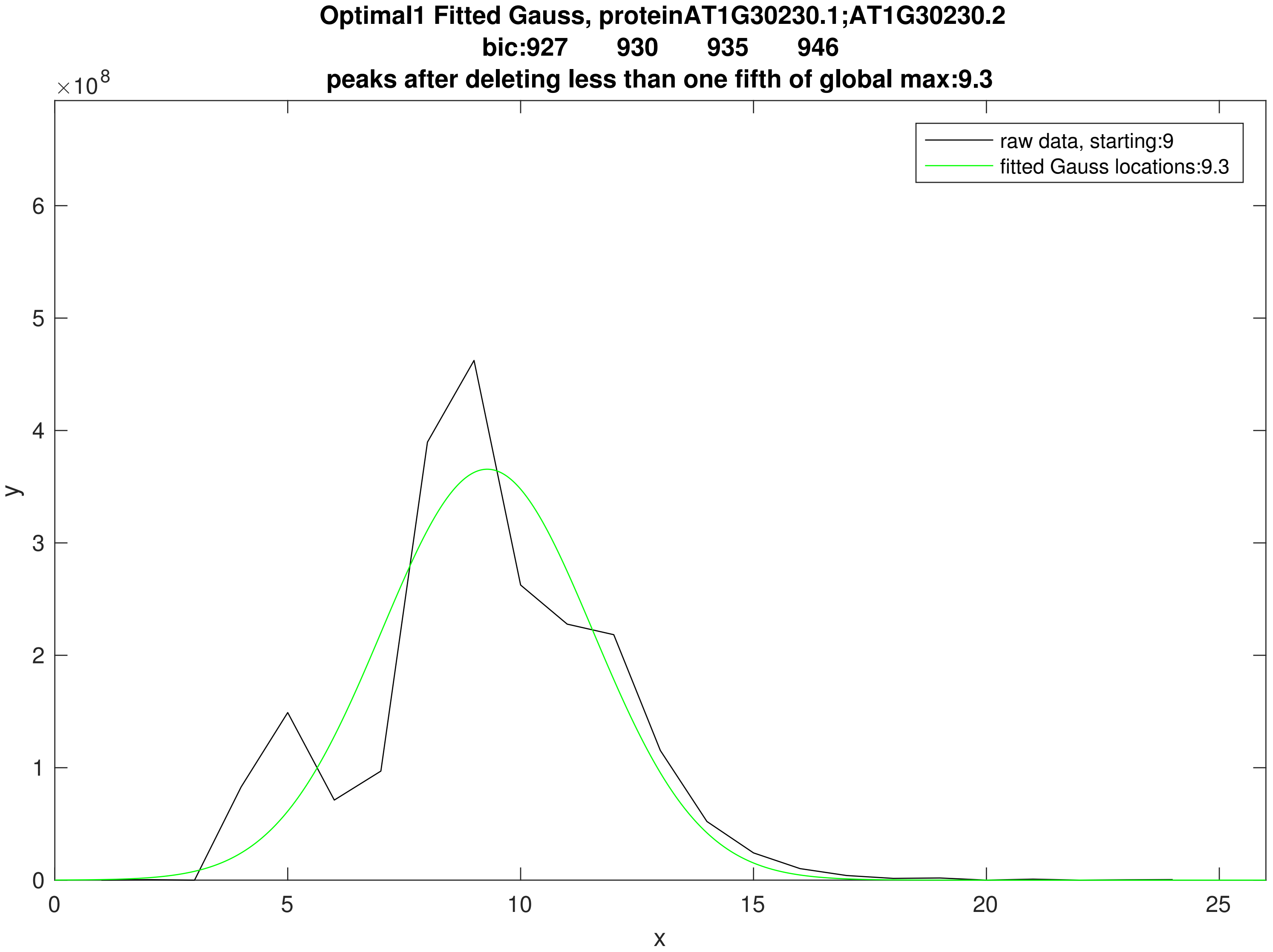
Optimal1 Fitted Gauss, proteinAT1G30230.1;AT1G30230.2
bic:927 930 935 946
peaks after deleting less than one fifth of global max:9.3
×10^8
raw data, starting:9
fitted Gauss locations:9.3
y
x
0
1
2
3
4
5
6
0
5
10
15
20
25

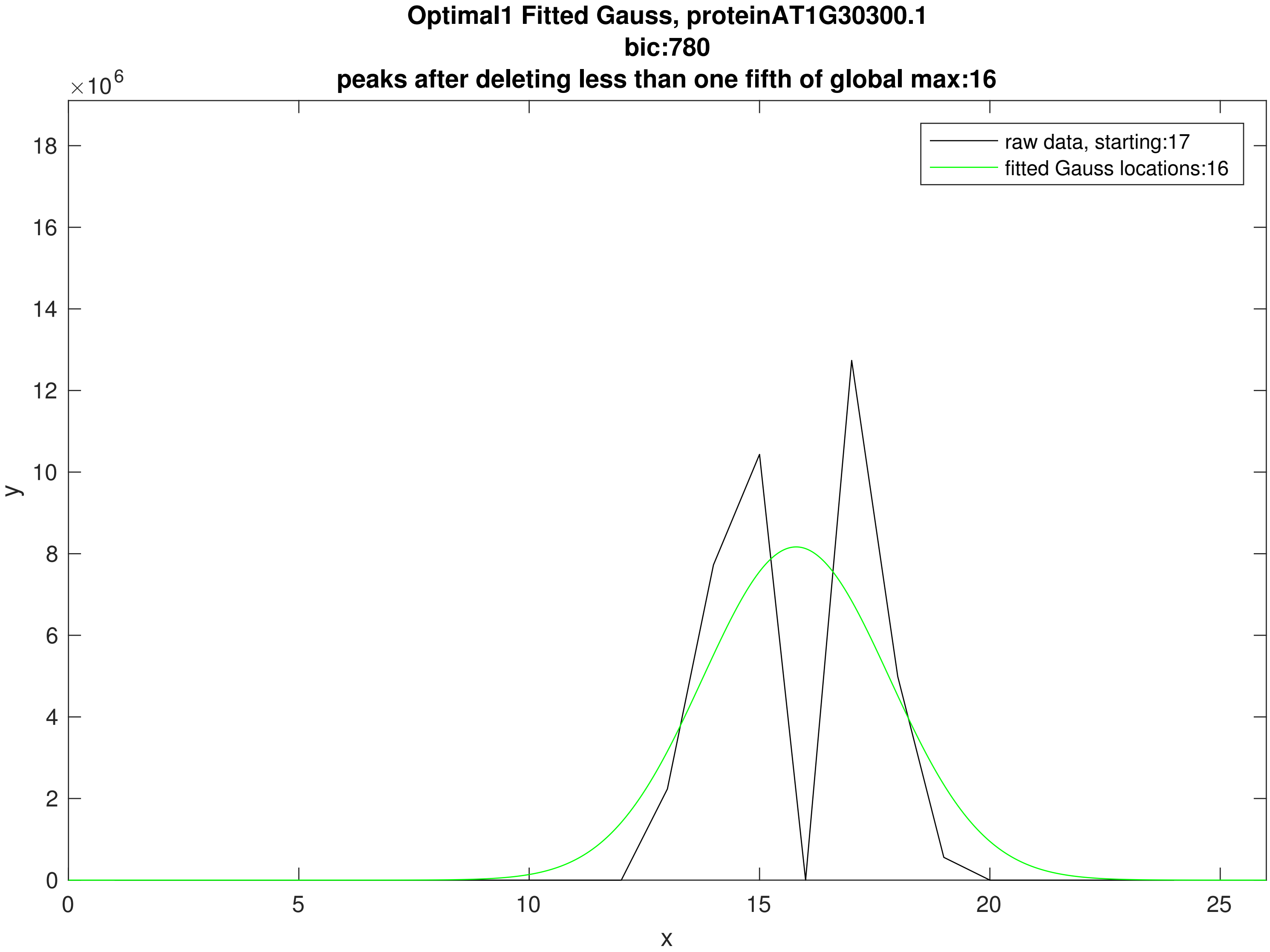

Optimal1 Fitted Gauss, proteinAT1G30300.1
bic:780
peaks after deleting less than one fifth of global max:16
×10^6
raw data, starting:17
fitted Gauss locations:16
y
x
0
2
4
6
8
10
12
14
16
18
0
5
10
15
20
25

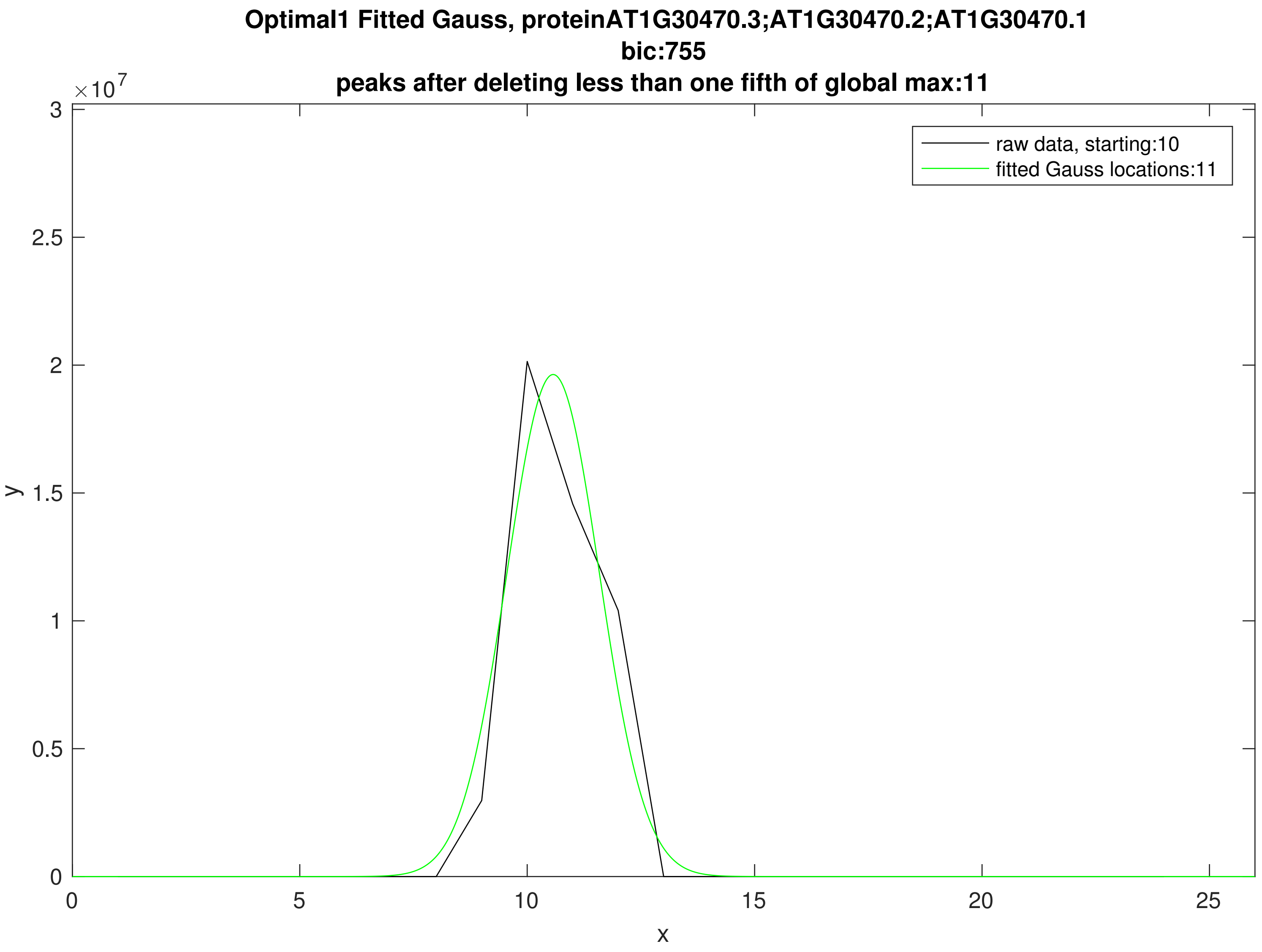
Optimal1 Fitted Gauss, proteinAT1G30470.3;AT1G30470.2;AT1G30470.1
bic:755
peaks after deleting less than one fifth of global max:11
×10^7
3
2.5
2
1.5
1
0.5
0
y
0
5
10
15
20
25
x
raw data, starting:10
fitted Gauss locations:11

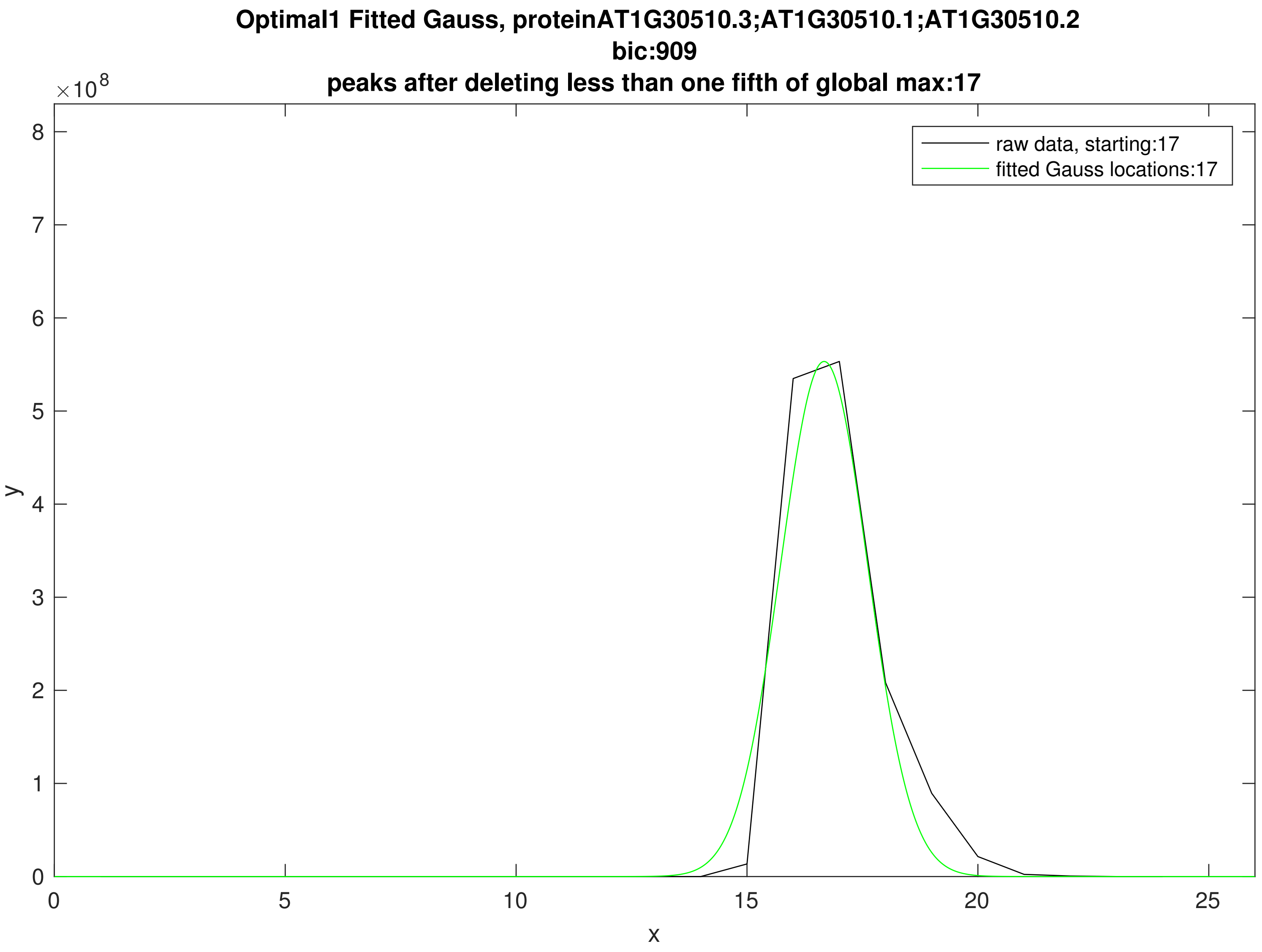
Optimal1 Fitted Gauss, proteinAT1G30510.3;AT1G30510.1;AT1G30510.2
bic:909
peaks after deleting less than one fifth of global max:17
×10^8
y
x
raw data, starting:17
fitted Gauss locations:17
0
1
2
3
4
5
6
7
8
0
5
10
15
20
25

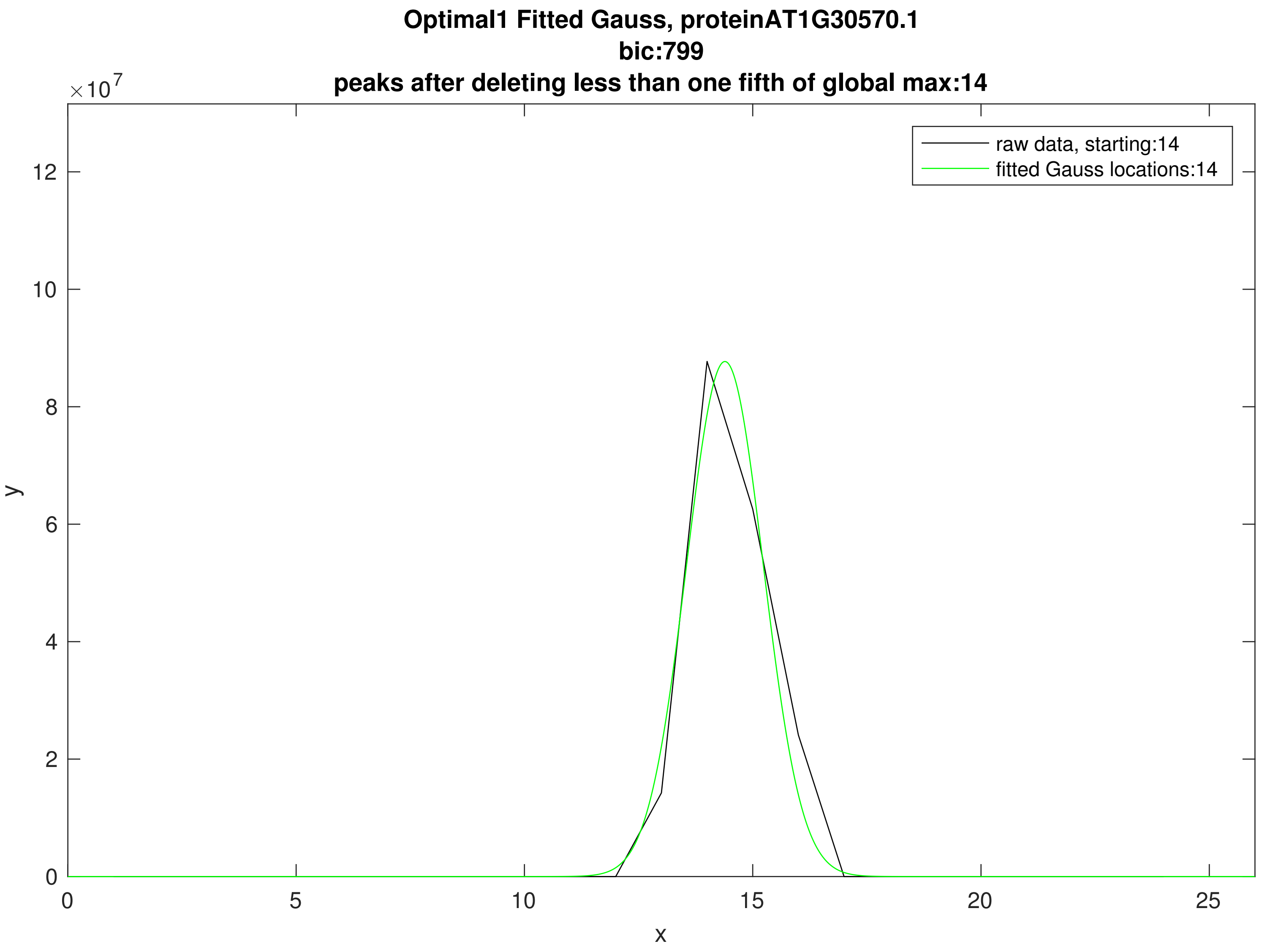
Optimal1 Fitted Gauss, proteinAT1G30570.1
bic:799
peaks after deleting less than one fifth of global max:14
×10^7
raw data, starting:14
fitted Gauss locations:14
y
x
0
2
4
6
8
10
12
5
15
20
25

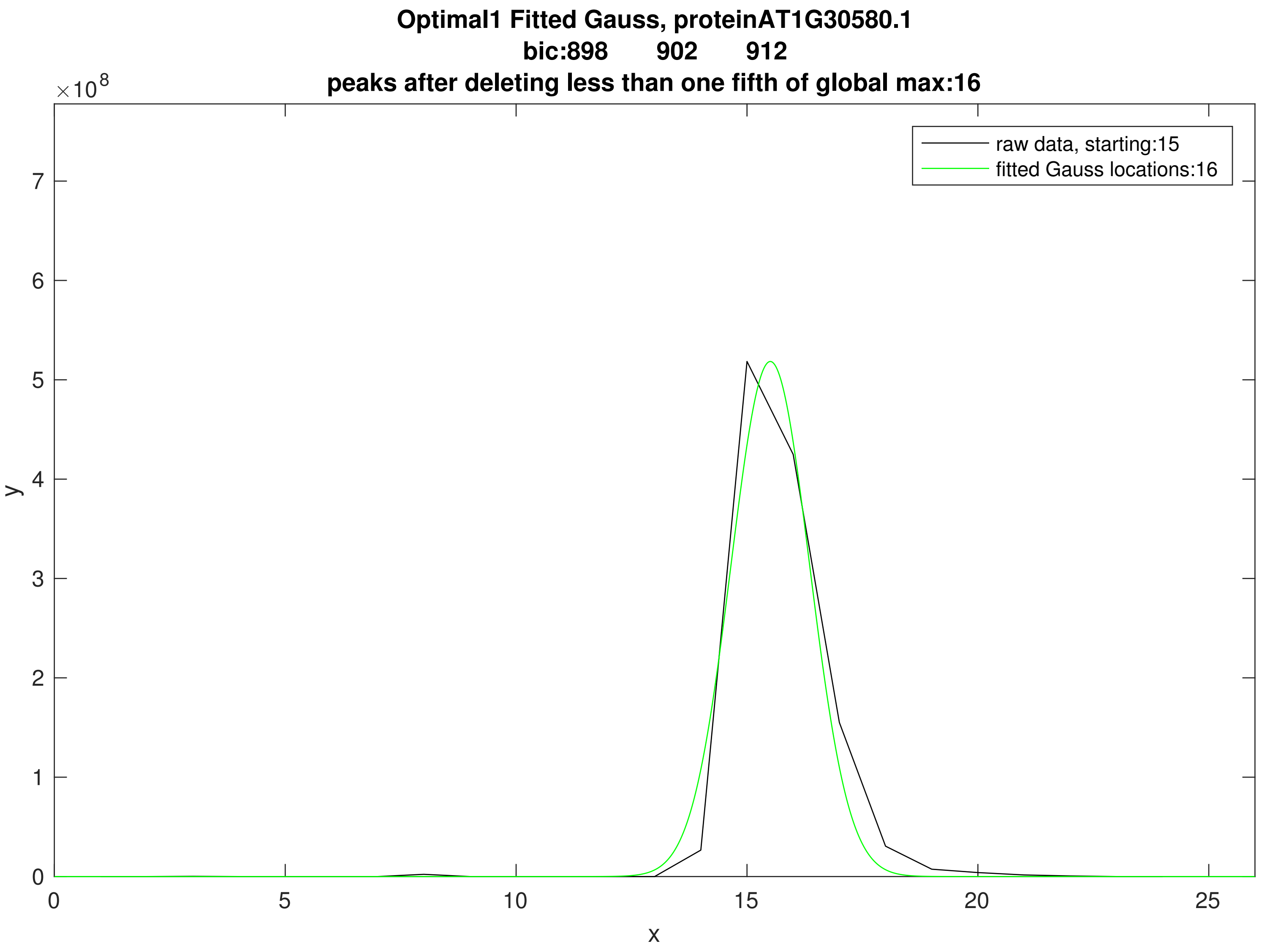

Optimal1 Fitted Gauss, proteinAT1G30580.1
bic:898 902 912
peaks after deleting less than one fifth of global max:16
×10^8
raw data, starting:15
fitted Gauss locations:16
y
x
0
1
2
3
4
5
6
7
0
5
10
15
20
25

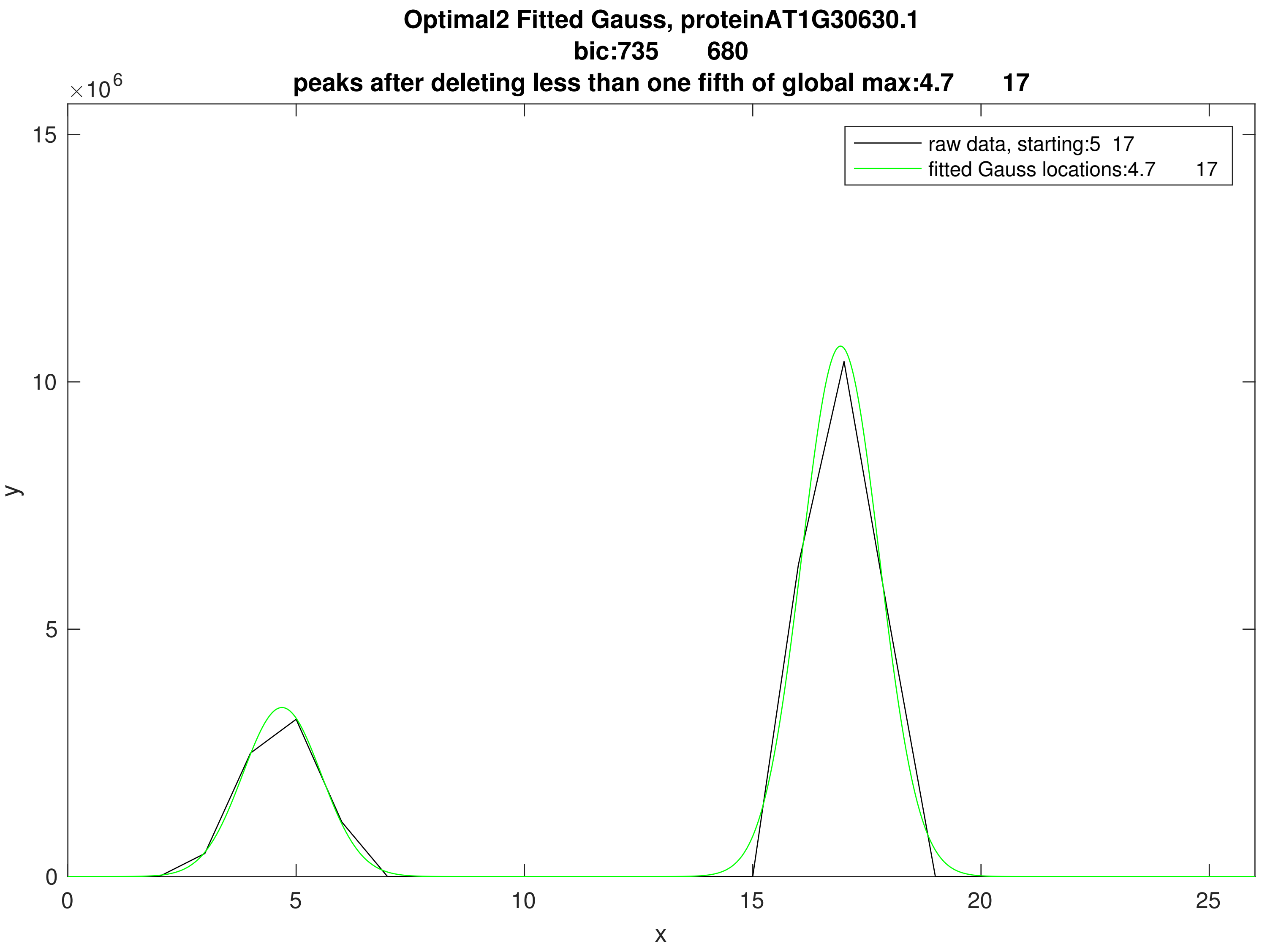
Optimal2 Fitted Gauss, proteinAT1G30630.1
bic:735 680
peaks after deleting less than one fifth of global max:4.7 17
×10^6
raw data, starting:5 17
fitted Gauss locations:4.7 17
y
x
0
5
10
15
20
25

Optimal1 Fitted Gauss, proteinAT1G30690.2;AT1G30690.1
bic:907 916 923
peaks after deleting less than one fifth of global max:11

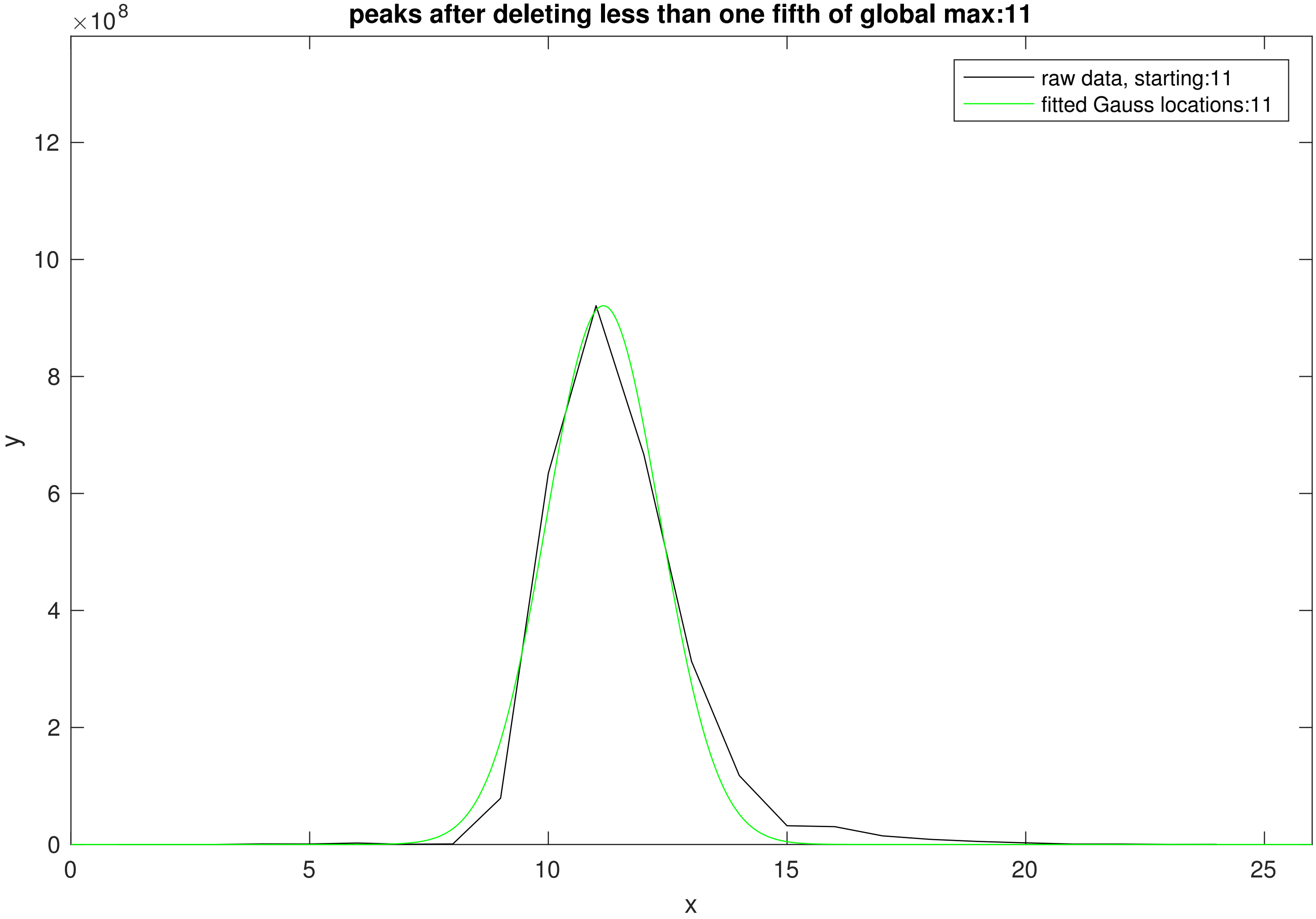

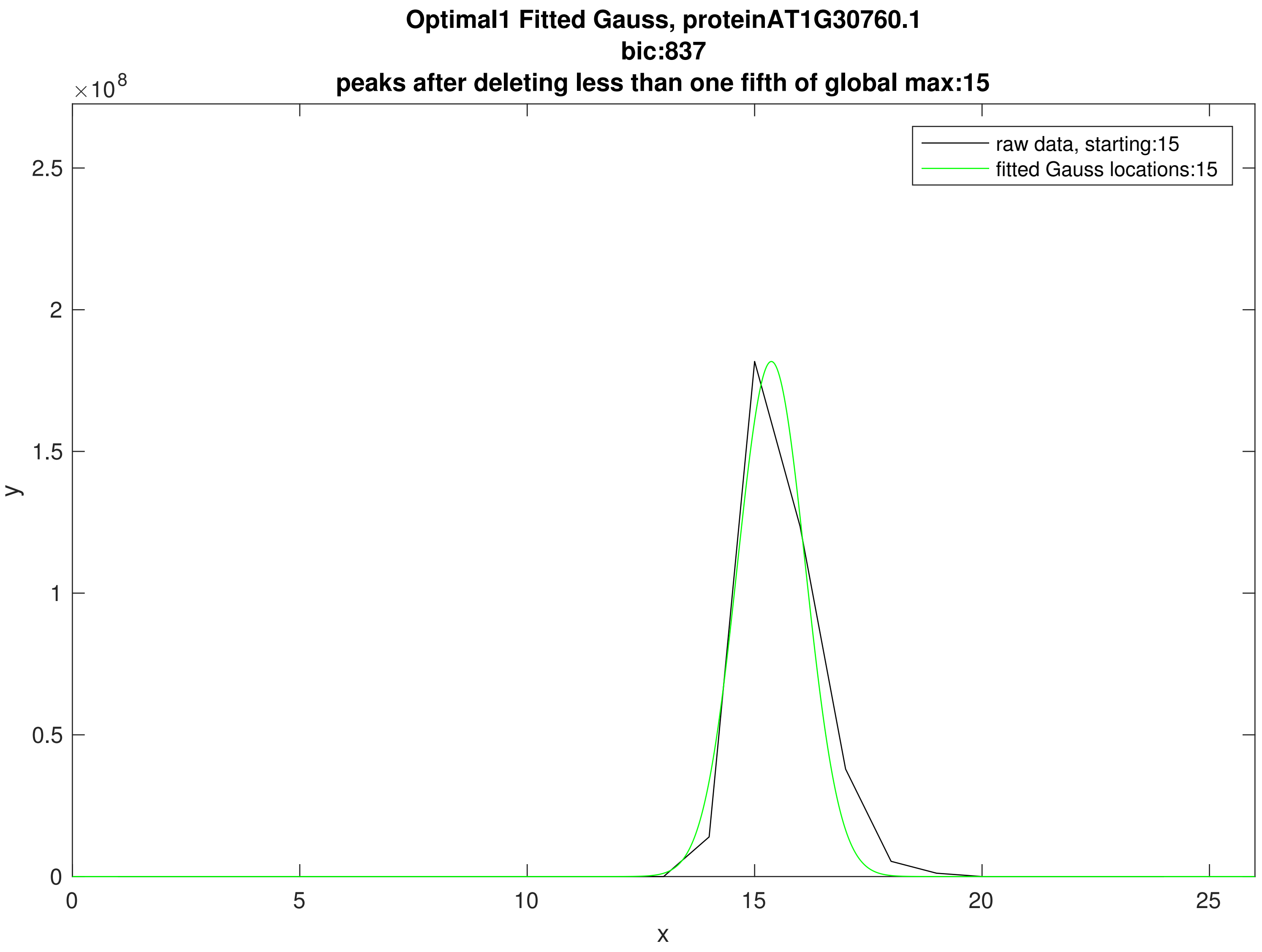

Optimal1 Fitted Gauss, proteinAT1G30760.1
bic:837
peaks after deleting less than one fifth of global max:15
×10^8
2.5
2
1.5
1
0.5
0
y
0
5
10
15
20
25
x
raw data, starting:15
fitted Gauss locations:15

**Optimal1 Fitted Gauss, proteinAT1G30910.1**
**bic:793**
**peaks after deleting less than one fifth of global max:17**

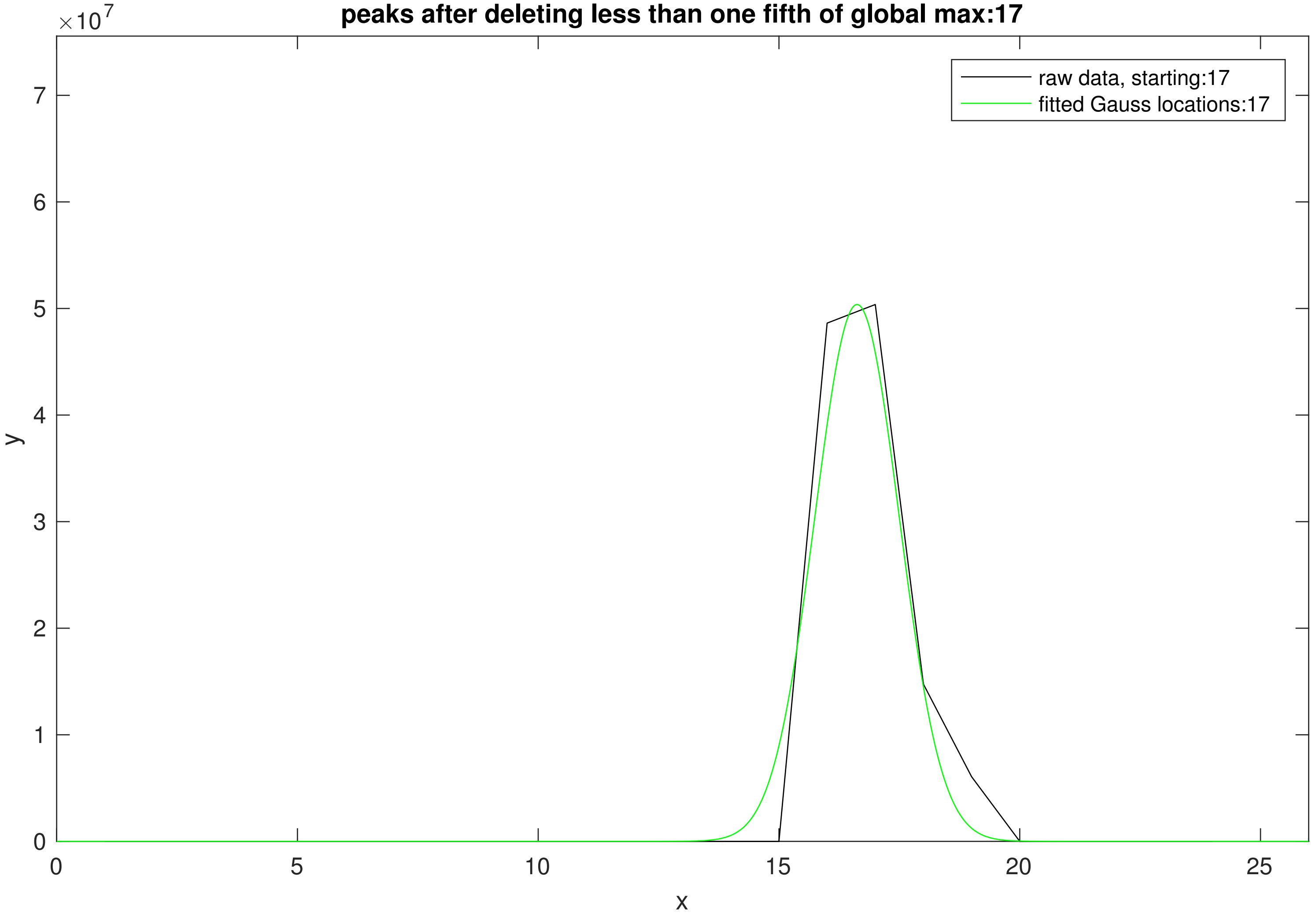

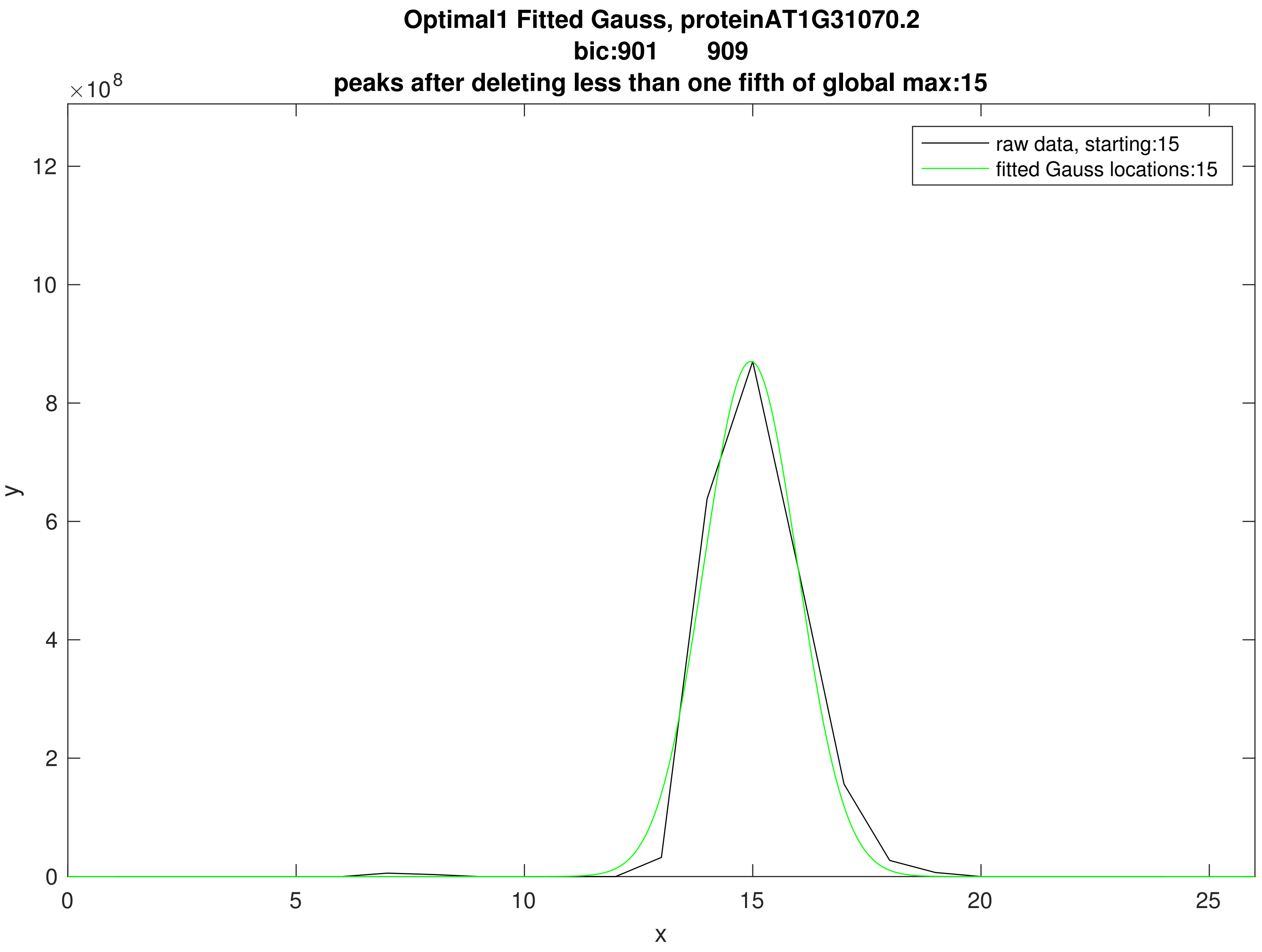

Optimal1 Fitted Gauss, proteinAT1G31070.2
bic:901 909
peaks after deleting less than one fifth of global max:15
×10^8
12
10
8
6
4
2
0
y
0
5
10
15
20
25
x
raw data, starting:15
fitted Gauss locations:15

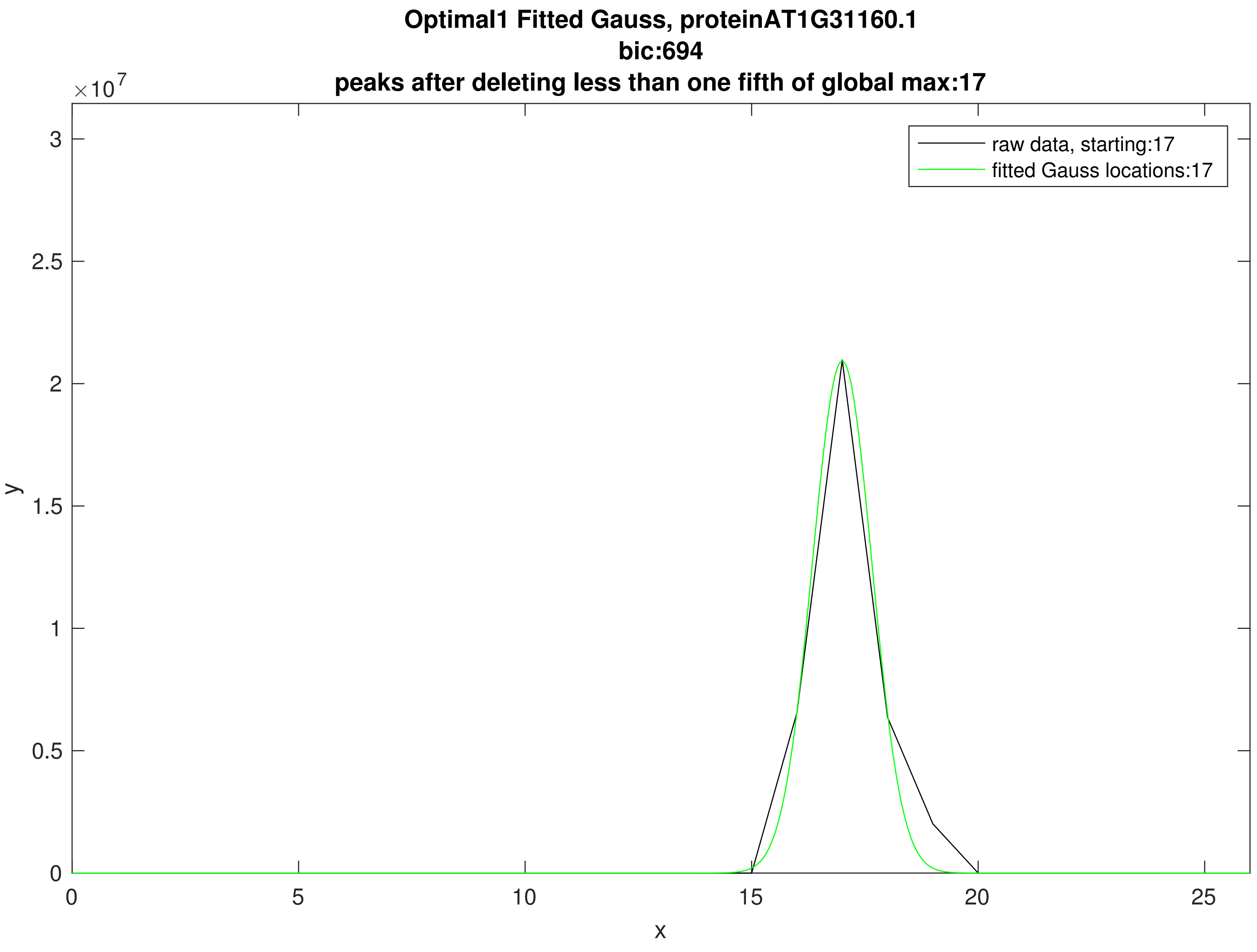
Optimal1 Fitted Gauss, proteinAT1G31160.1
bic:694
peaks after deleting less than one fifth of global max:17
×10^7
raw data, starting:17
fitted Gauss locations:17
y
x
0
0.5
1
1.5
2
2.5
3
0
5
10
15
20
25

Optimal1 Fitted Gauss, proteinAT1G31180.1
bic:847 857
peaks after deleting less than one fifth of global max:14

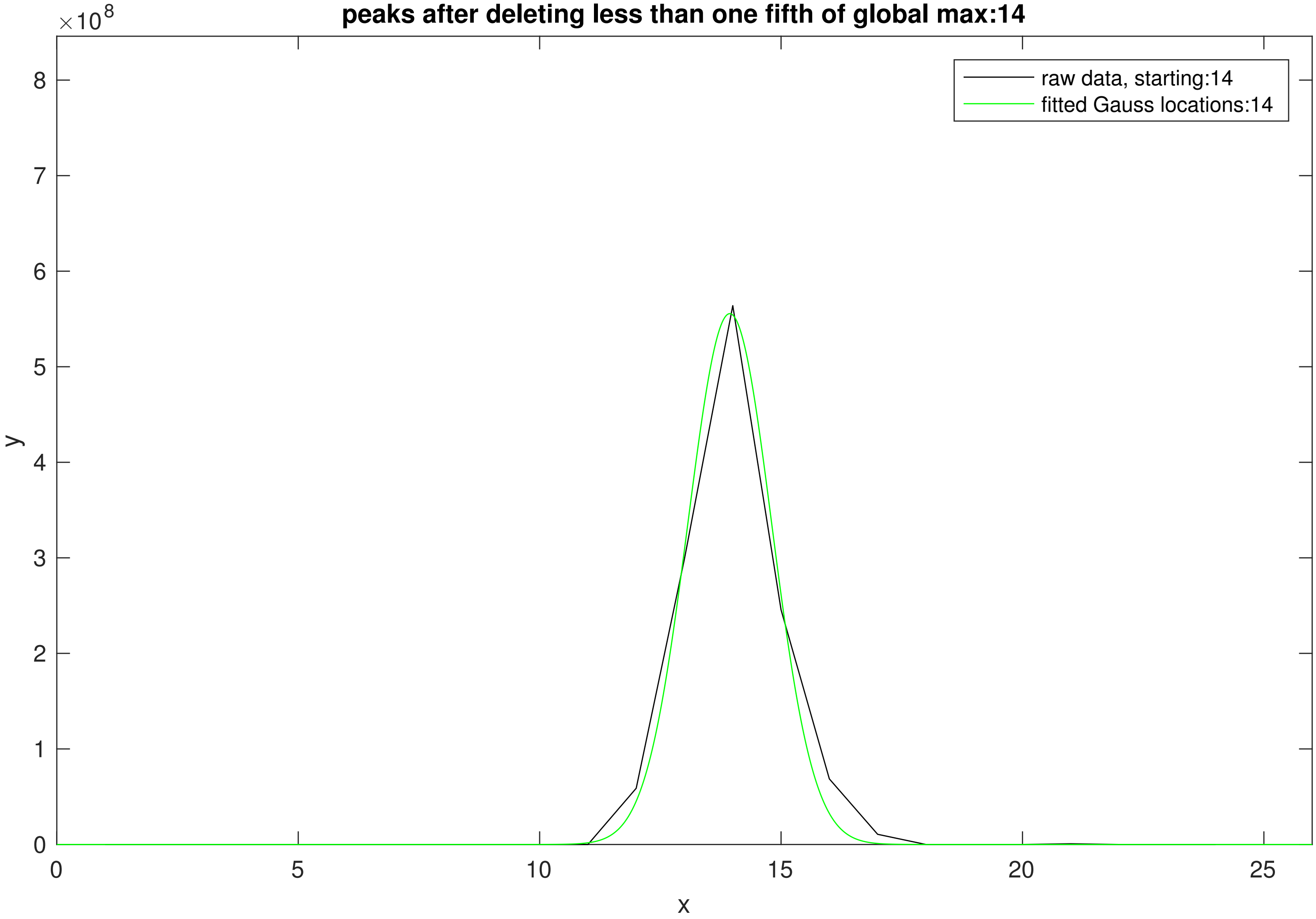

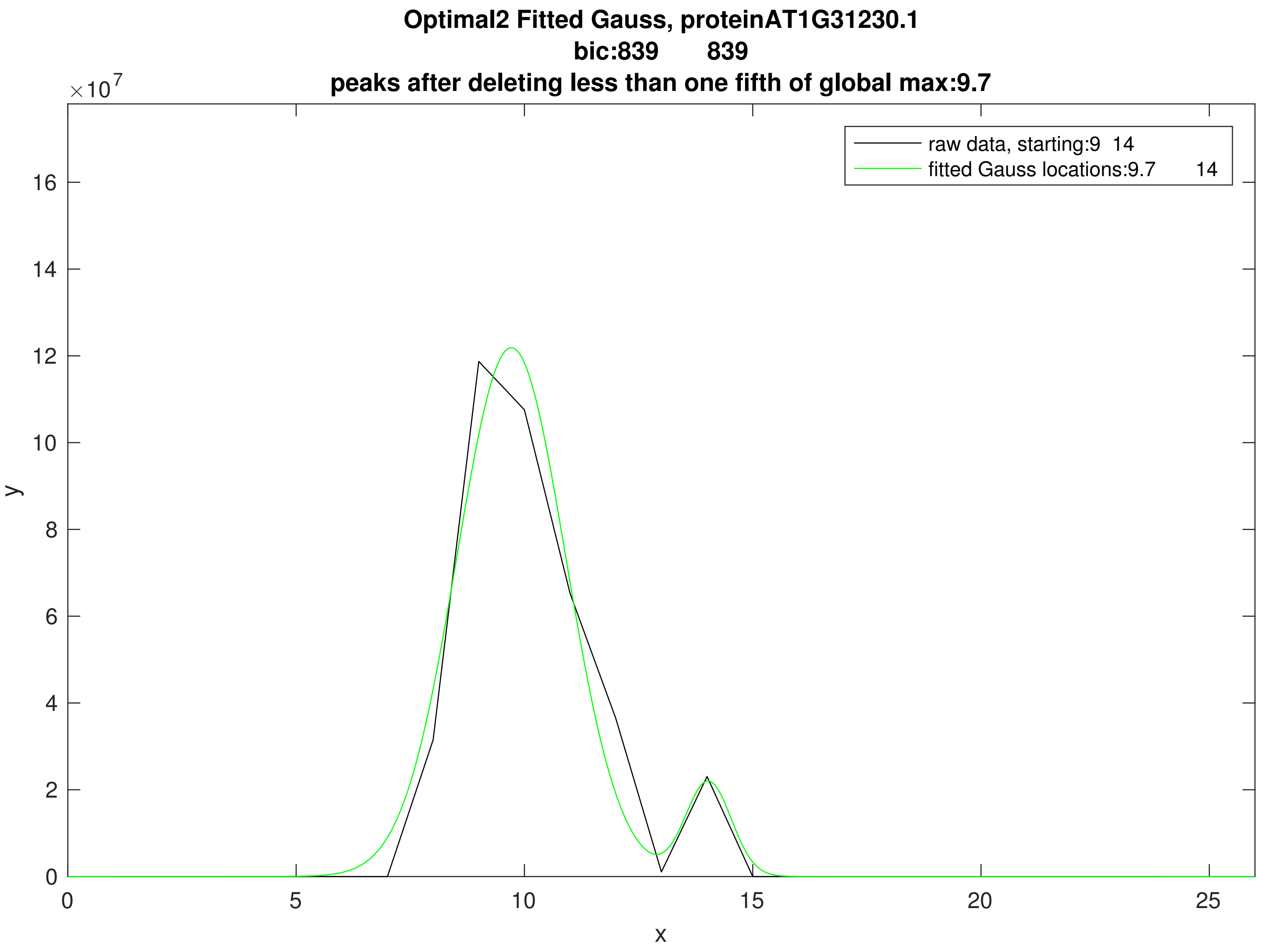

Optimal2 Fitted Gauss, proteinAT1G31230.1
bic:839 839
peaks after deleting less than one fifth of global max:9.7
×10^7
raw data, starting:9 14
fitted Gauss locations:9.7 14
16
14
12
10
8
6
4
2
0
y
0
5
10
15
20
25
x

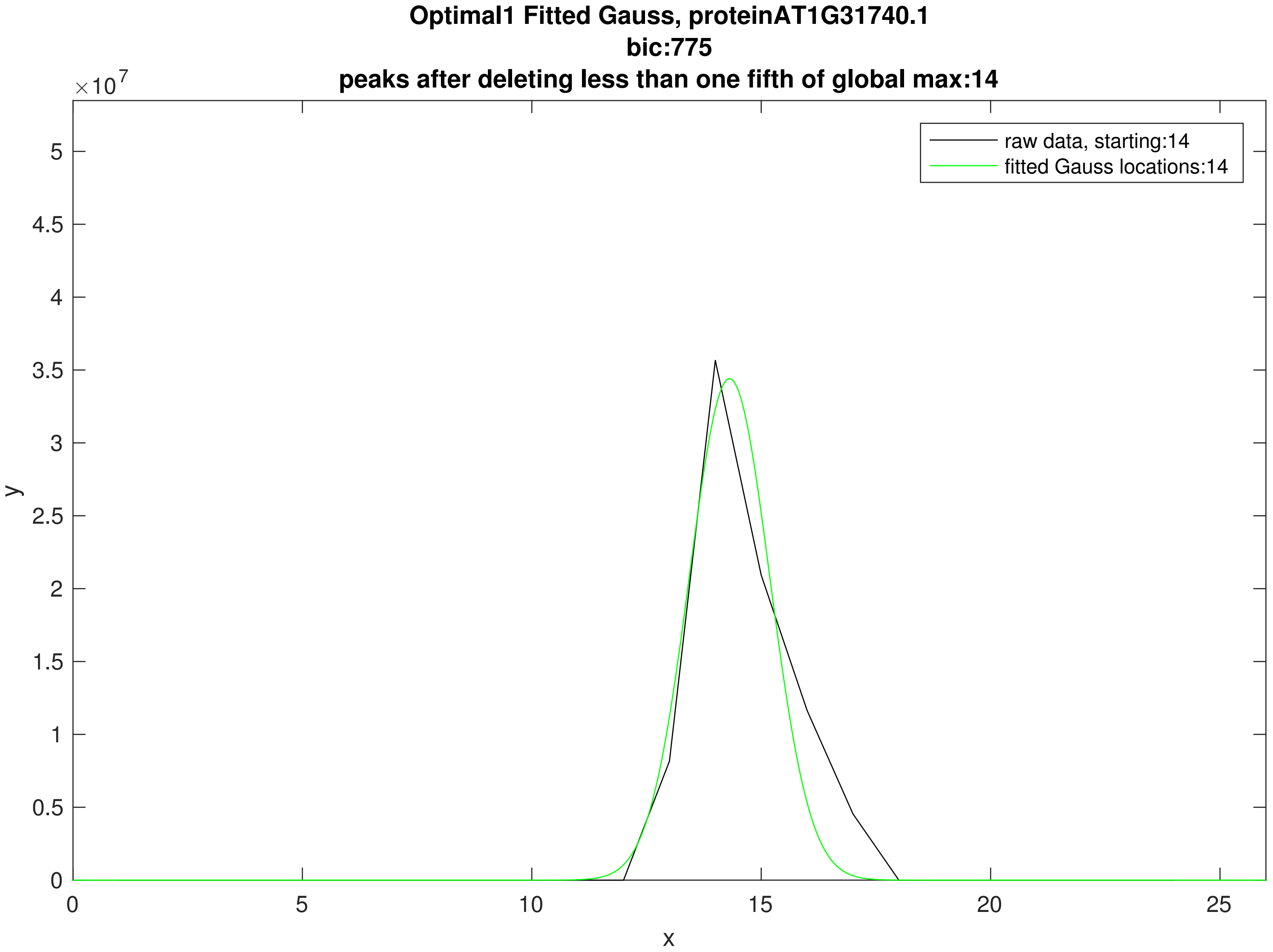
Optimal1 Fitted Gauss, proteinAT1G31740.1
bic:775
peaks after deleting less than one fifth of global max:14
×10^7
raw data, starting:14
fitted Gauss locations:14
y
x
0
0.5
1
1.5
2
2.5
3
3.5
4
4.5
5
0
5
10
15
20
25

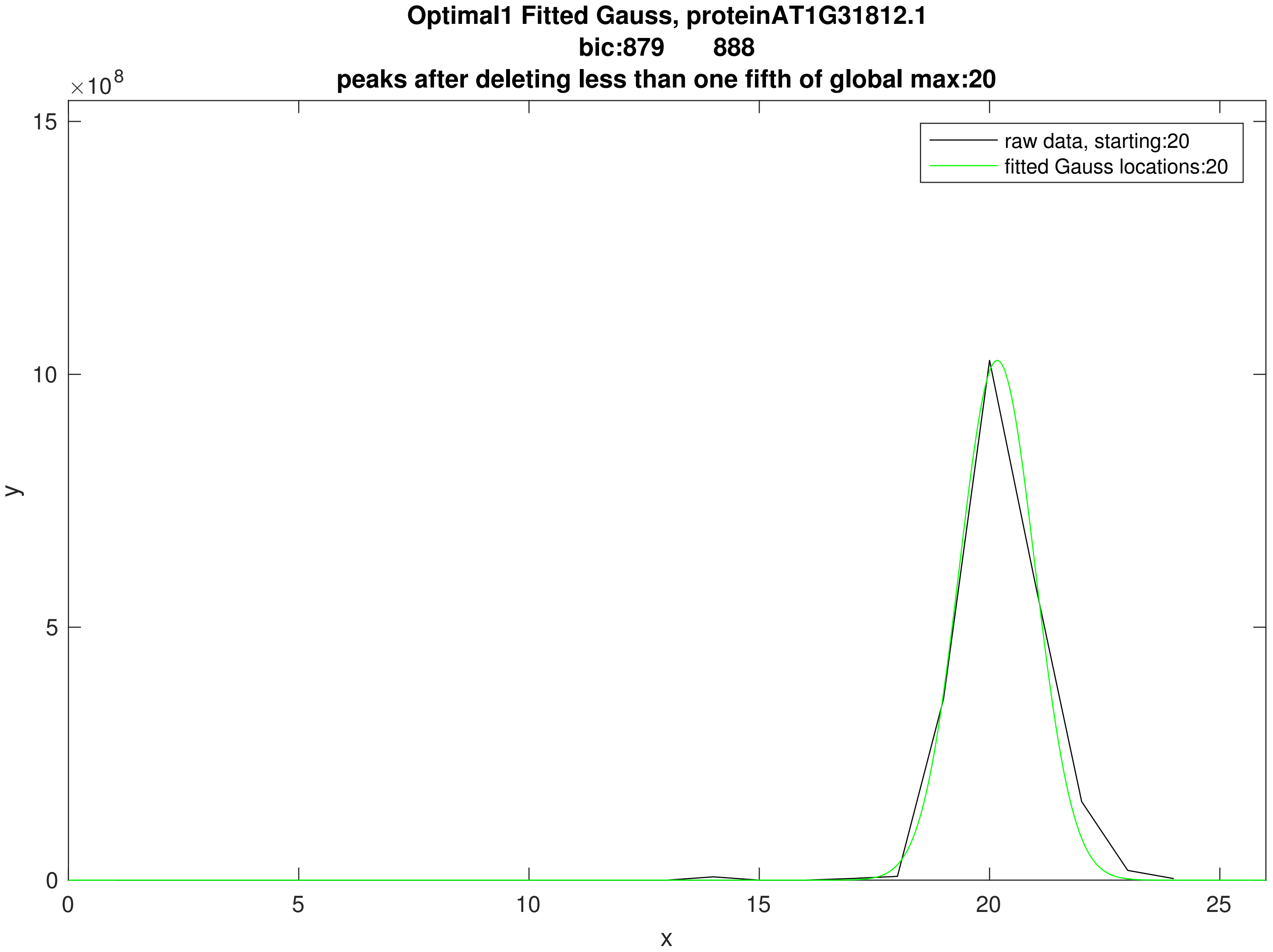
Optimal1 Fitted Gauss, proteinAT1G31812.1
bic:879 888
peaks after deleting less than one fifth of global max:20
×10^8
15
10
5
0
y
0
5
10
15
20
25
x
raw data, starting:20
fitted Gauss locations:20

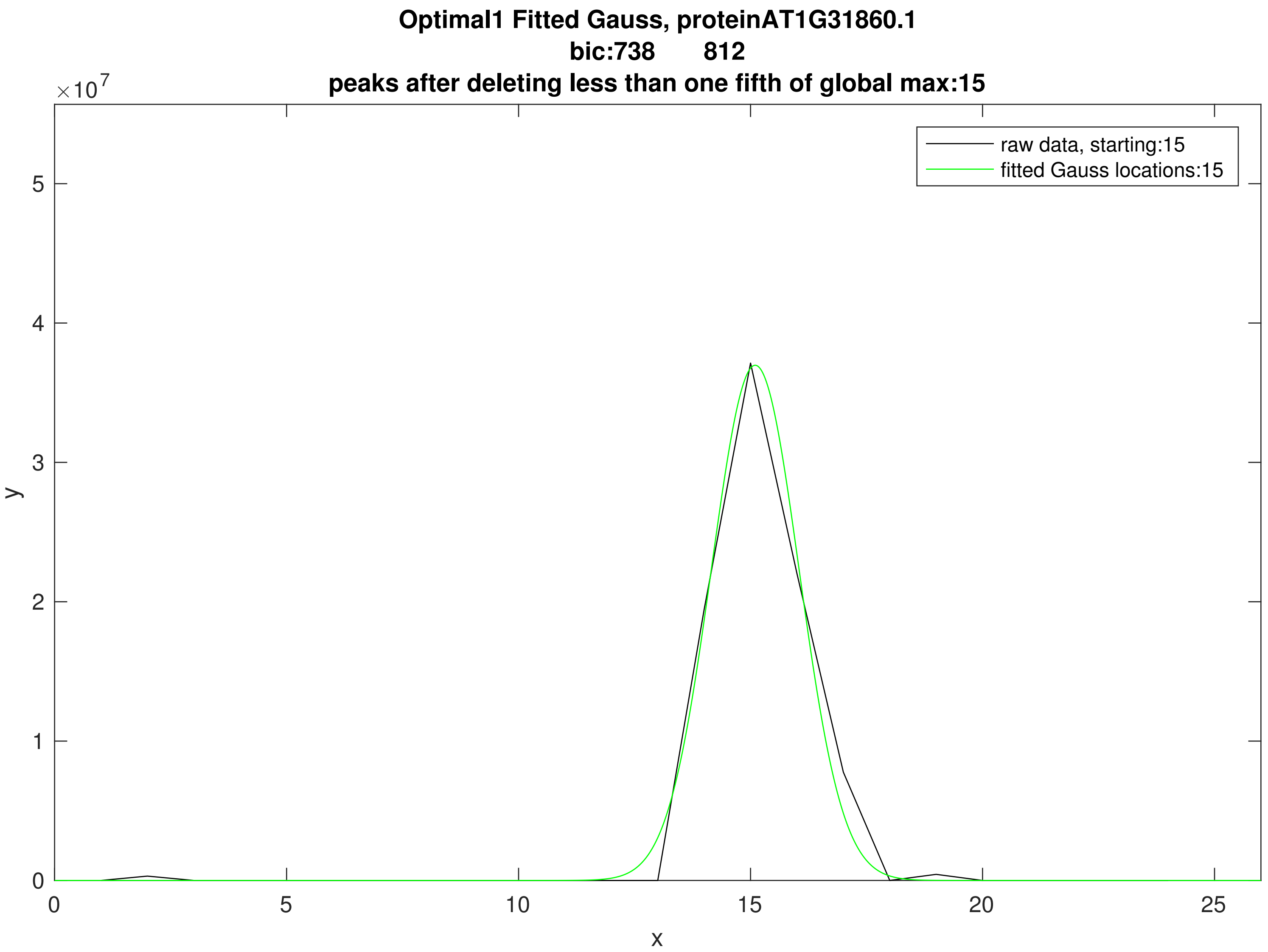
Optimal1 Fitted Gauss, proteinAT1G31860.1
bic:738 812
peaks after deleting less than one fifth of global max:15
×10^7
raw data, starting:15
fitted Gauss locations:15
y
x
0
1
2
3
4
5
0
5
10
15
20
25

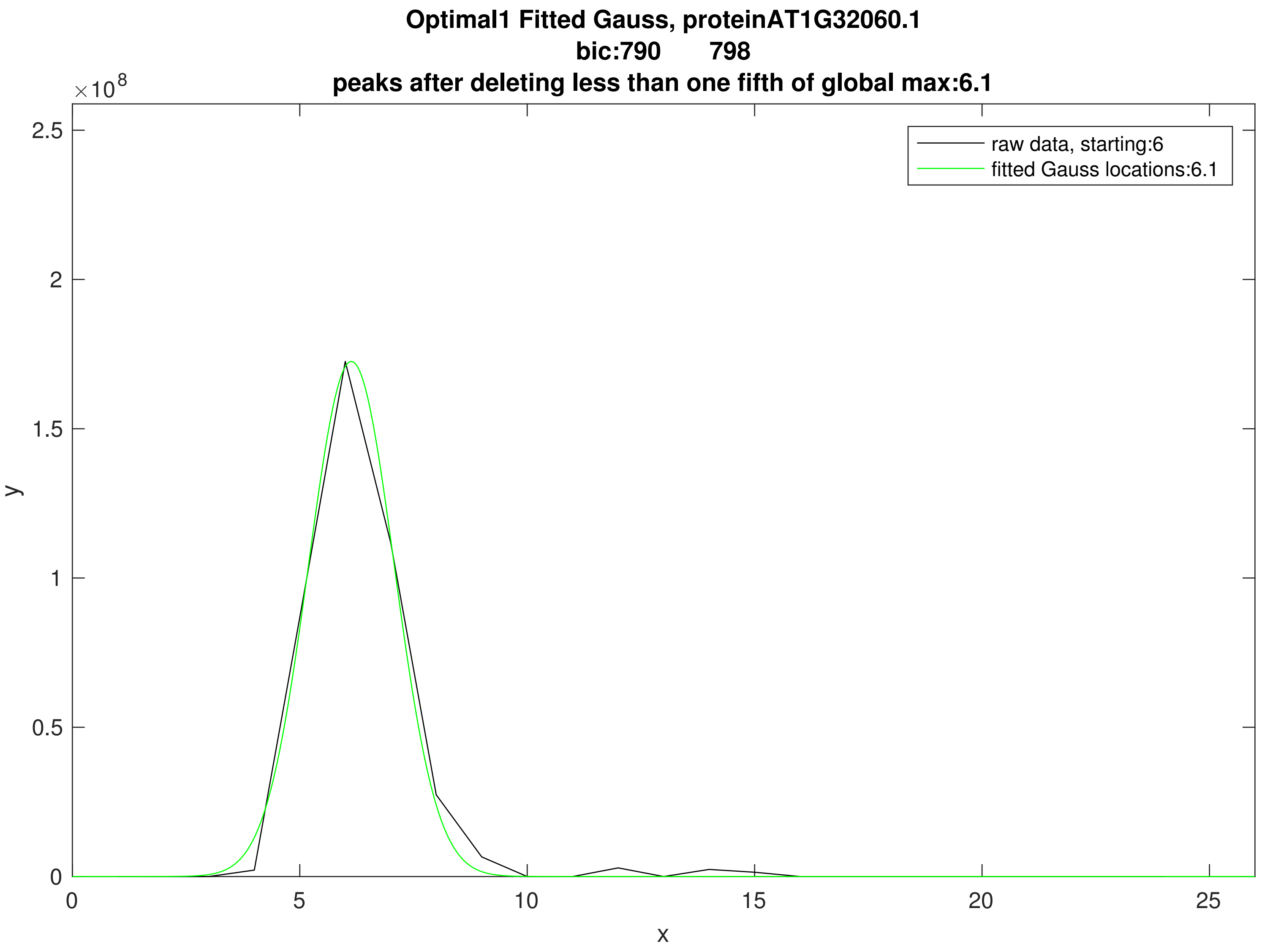

Optimal1 Fitted Gauss, proteinAT1G32060.1
bic:790 798
peaks after deleting less than one fifth of global max:6.1
×10^8
2.5
2
1.5
1
0.5
0
y
0
5
10
15
20
25
x
raw data, starting:6
fitted Gauss locations:6.1

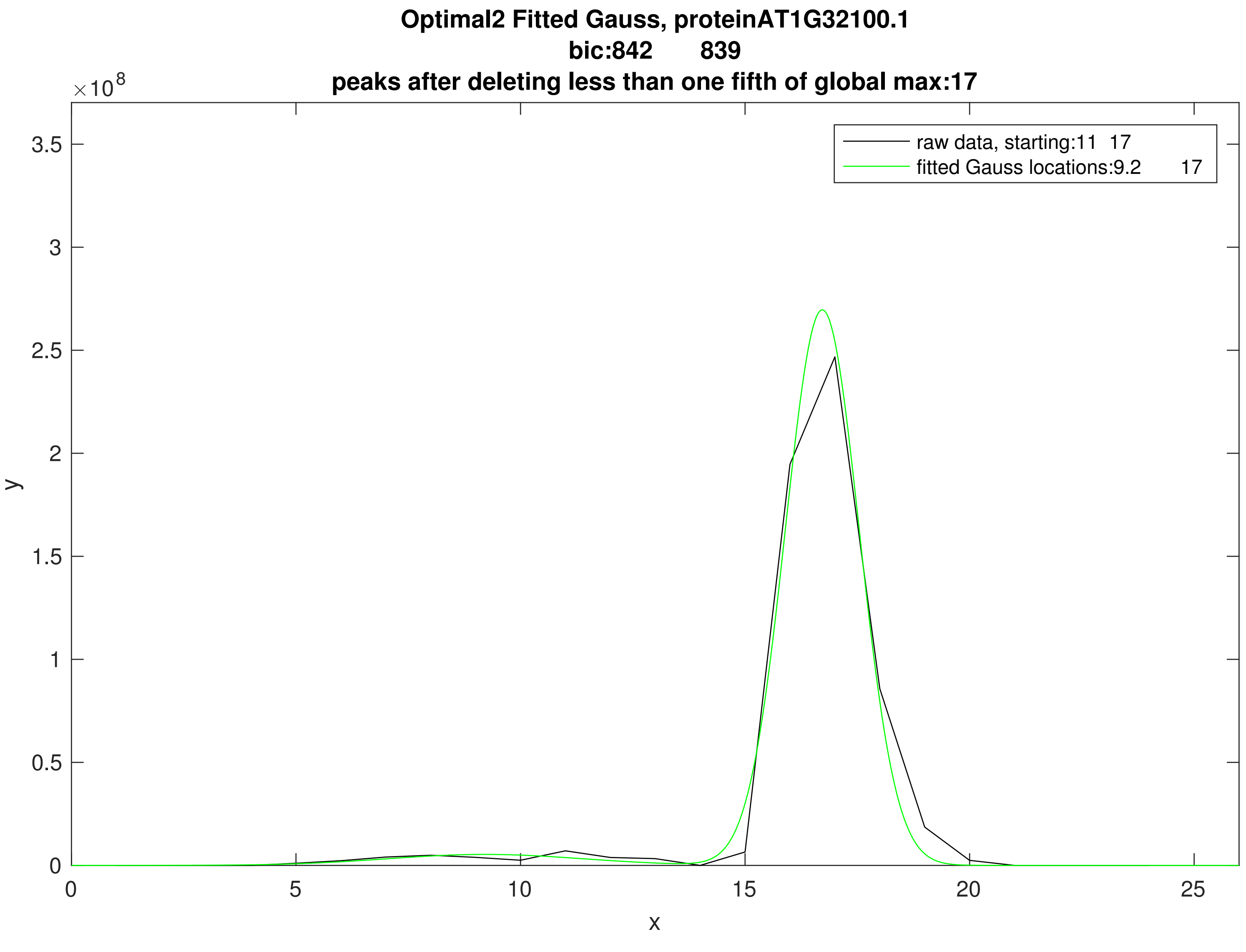

Optimal2 Fitted Gauss, proteinAT1G32100.1
bic:842 839
peaks after deleting less than one fifth of global max:17
×10^8
raw data, starting:11 17
fitted Gauss locations:9.2 17
y
x
0
0.5
1
1.5
2
2.5
3
3.5
0
5
10
15
20
25

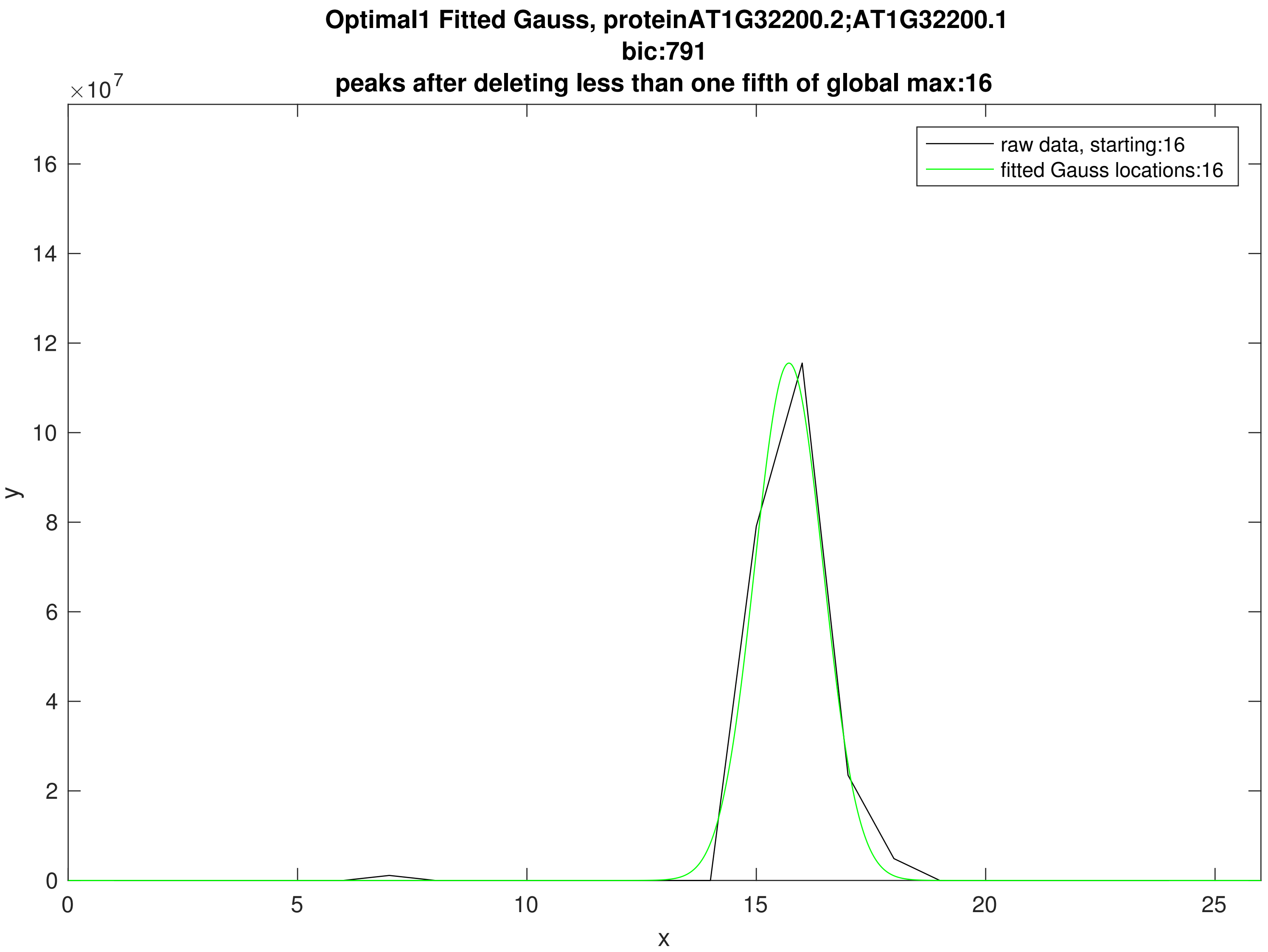
Optimal1 Fitted Gauss, proteinAT1G32200.2;AT1G32200.1
bic:791
peaks after deleting less than one fifth of global max:16
×10^7
raw data, starting:16
fitted Gauss locations:16
y
x
0
2
4
6
8
10
12
14
16
0
5
10
15
20
25

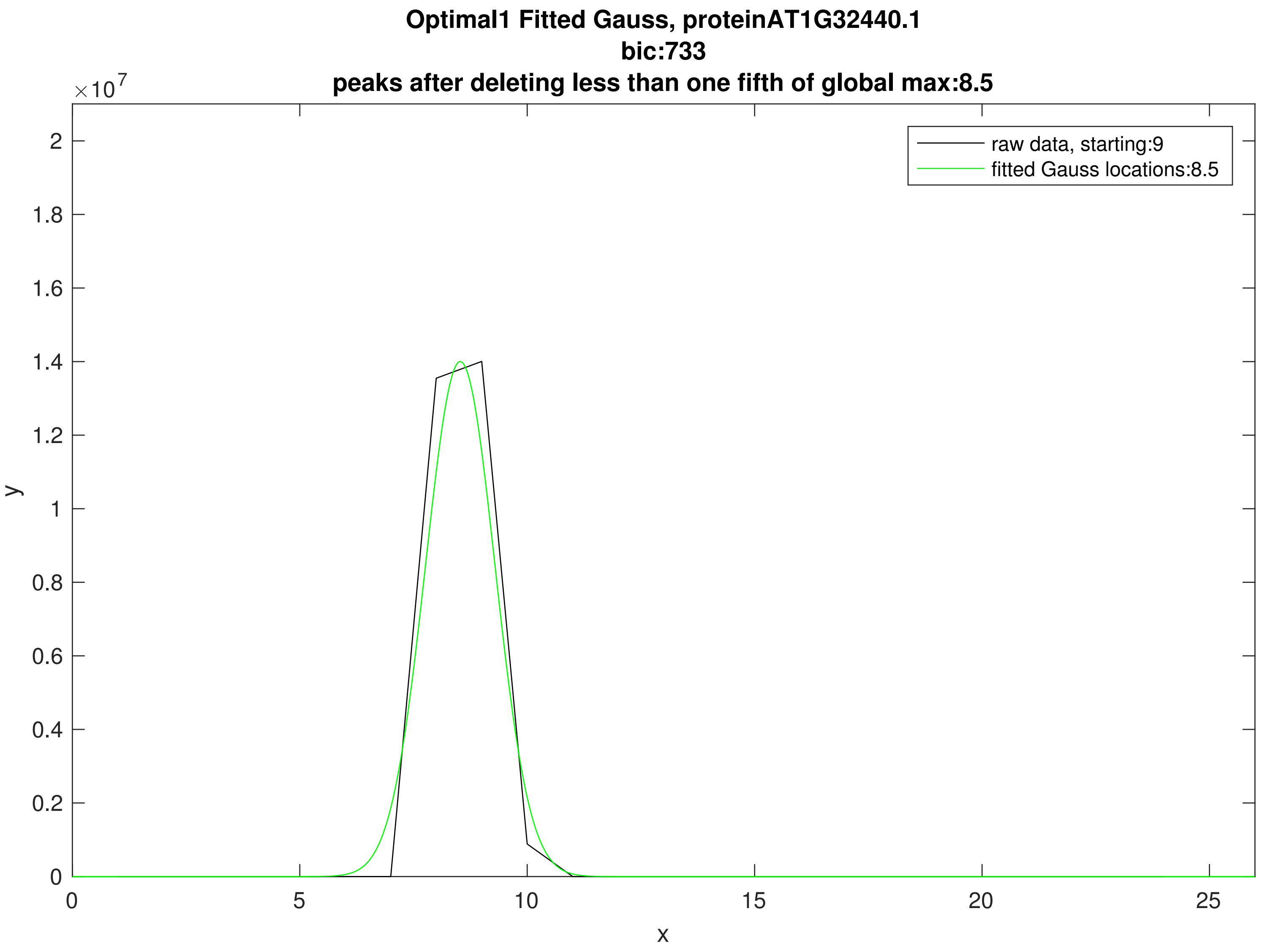
Optimal1 Fitted Gauss, proteinAT1G32440.1
bic:733
peaks after deleting less than one fifth of global max:8.5
×10^7
raw data, starting:9
fitted Gauss locations:8.5
y
x
0
0.2
0.4
0.6
0.8
1
1.2
1.4
1.6
1.8
2
0
5
10
15
20
25

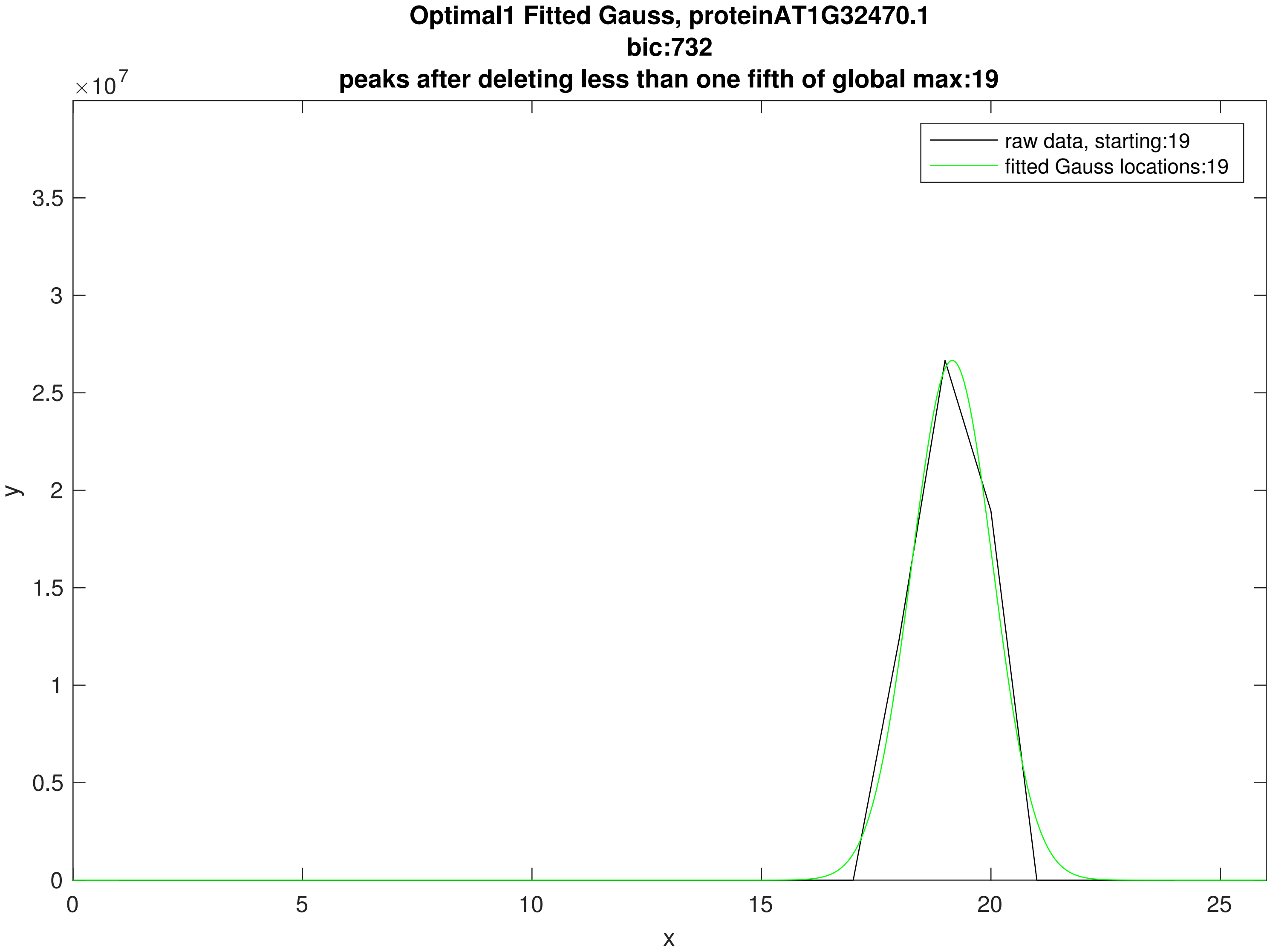
Optimal1 Fitted Gauss, proteinAT1G32470.1
bic:732
peaks after deleting less than one fifth of global max:19
×10^7
raw data, starting:19
fitted Gauss locations:19
y
x
0
0.5
1
1.5
2
2.5
3
3.5
0
5
10
15
20
25

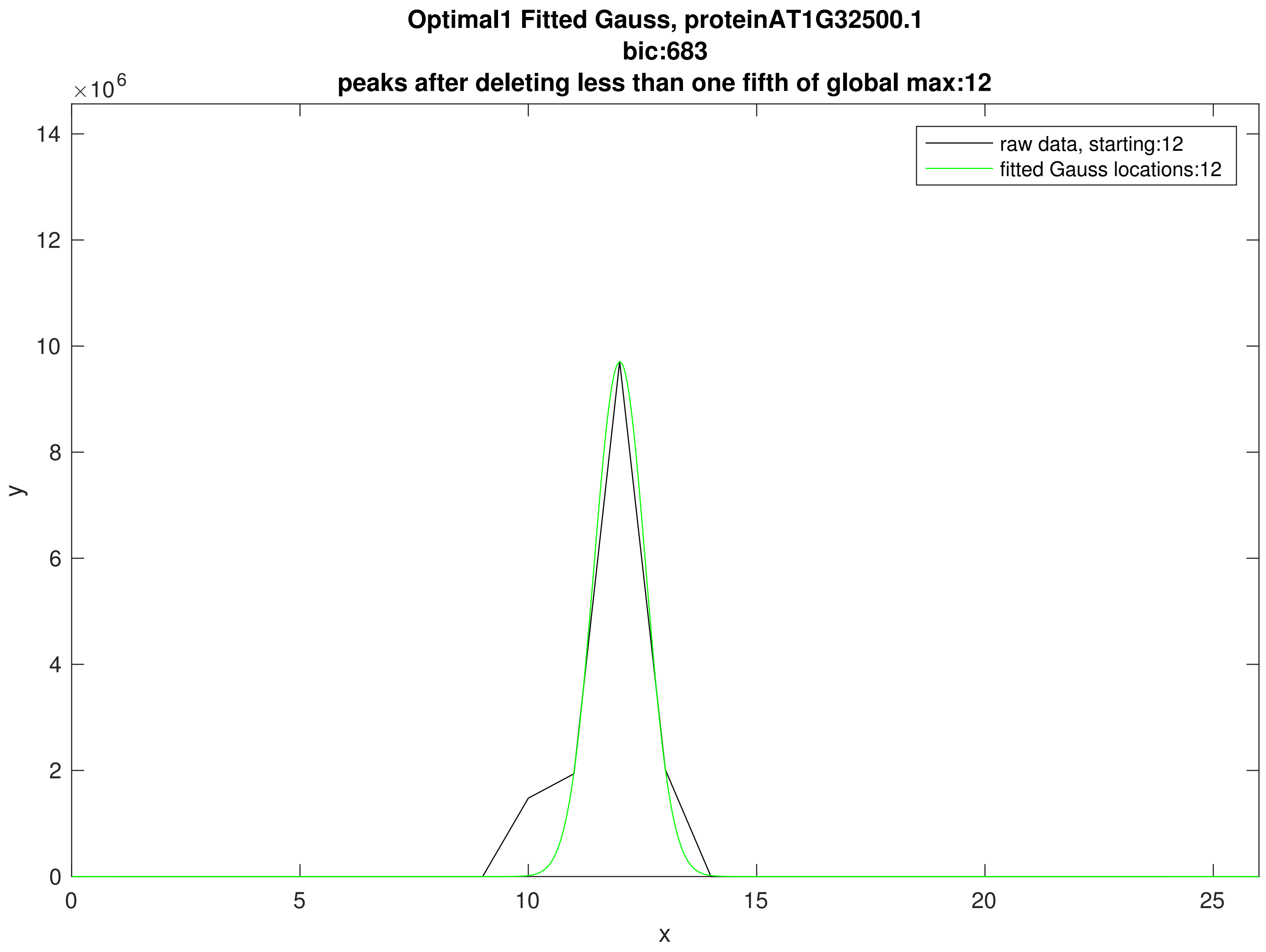
Optimal1 Fitted Gauss, proteinAT1G32500.1
bic:683
peaks after deleting less than one fifth of global max:12
×10^6
raw data, starting:12
fitted Gauss locations:12
y
x
0
2
4
6
8
10
12
14
0
5
10
15
20
25

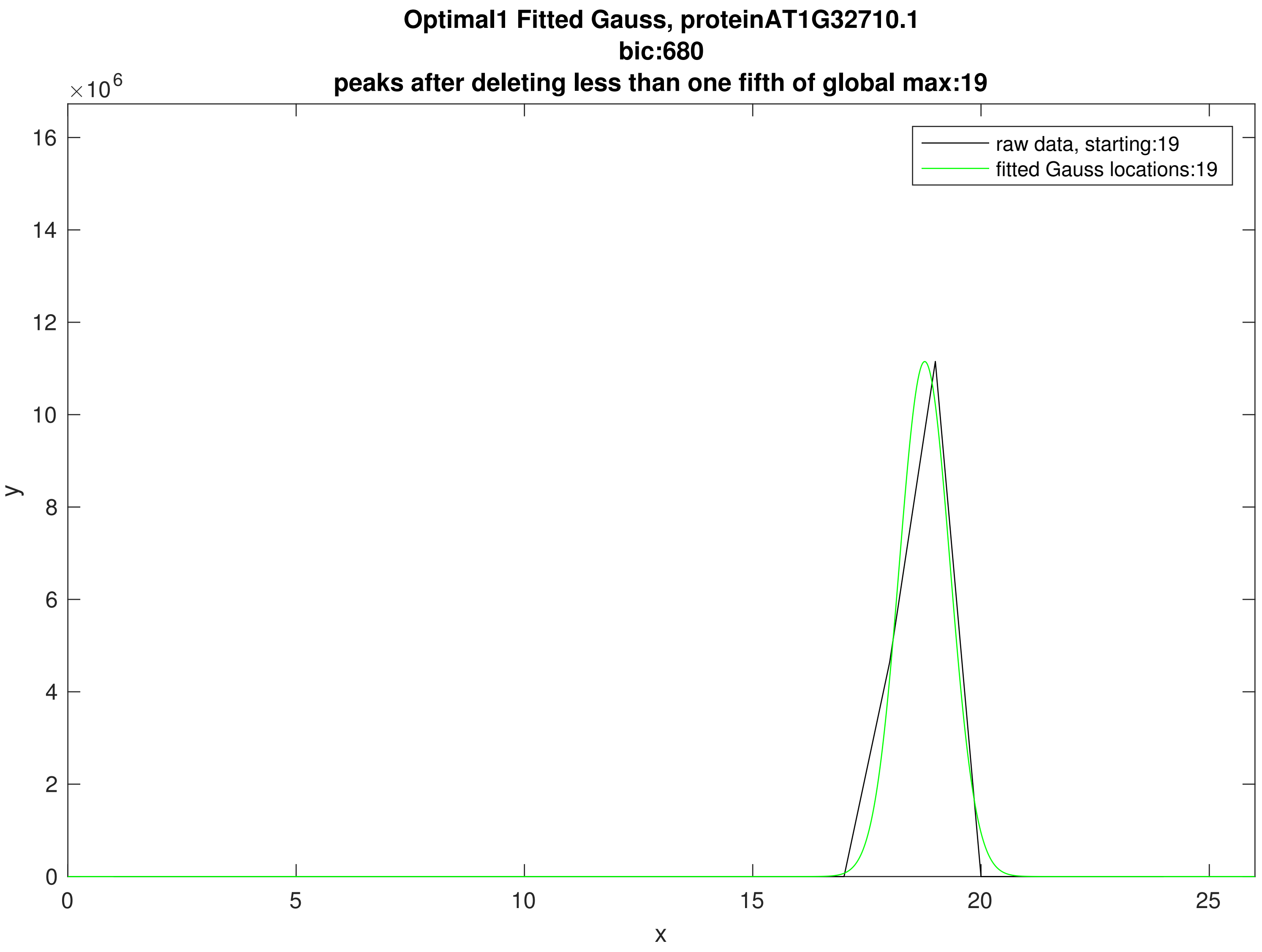

Optimal1 Fitted Gauss, proteinAT1G32710.1
bic:680
peaks after deleting less than one fifth of global max:19
×10^6
16
14
12
10
8
6
4
2
0
y
0
5
10
15
20
25
x
raw data, starting:19
fitted Gauss locations:19

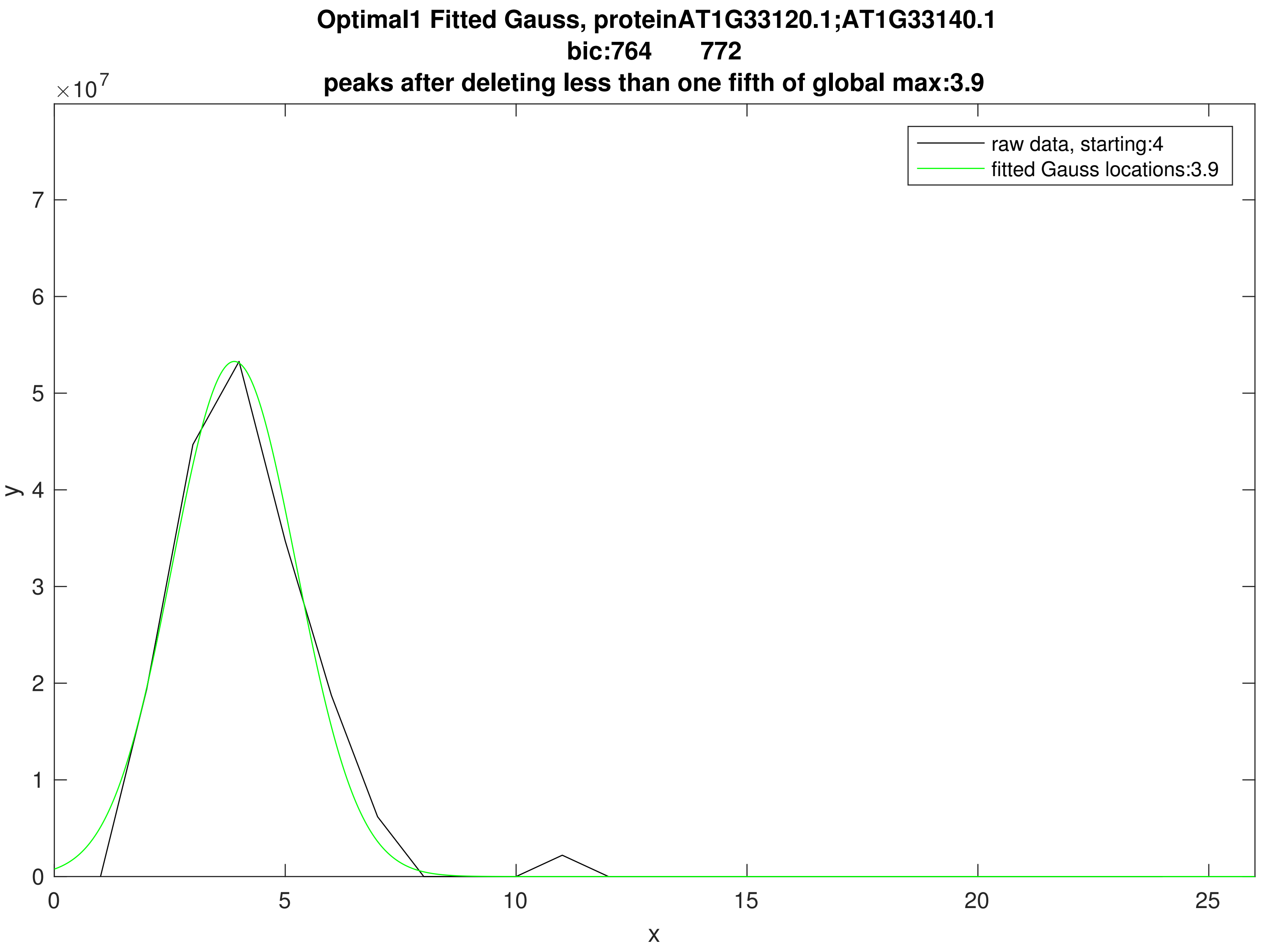
Optimal1 Fitted Gauss, proteinAT1G33120.1;AT1G33140.1
bic:764 772
peaks after deleting less than one fifth of global max:3.9
×10^7
raw data, starting:4
fitted Gauss locations:3.9
y
x
0
1
2
3
4
5
6
7
0
5
10
15
20
25

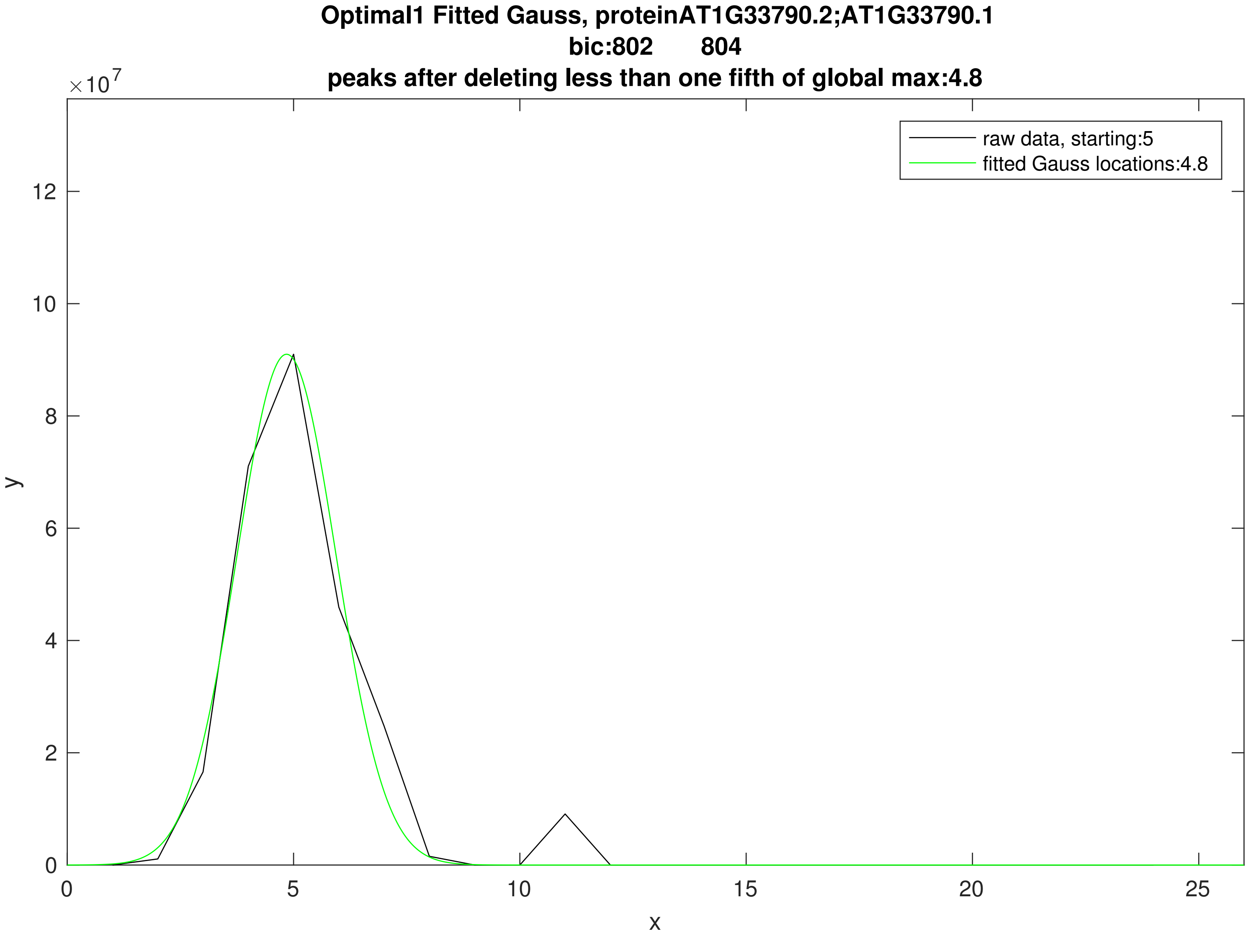
Optimal1 Fitted Gauss, proteinAT1G33790.2;AT1G33790.1
bic:802 804
peaks after deleting less than one fifth of global max:4.8
×10^7
raw data, starting:5
fitted Gauss locations:4.8
y
x
0
2
4
6
8
10
12
0
5
10
15
20
25

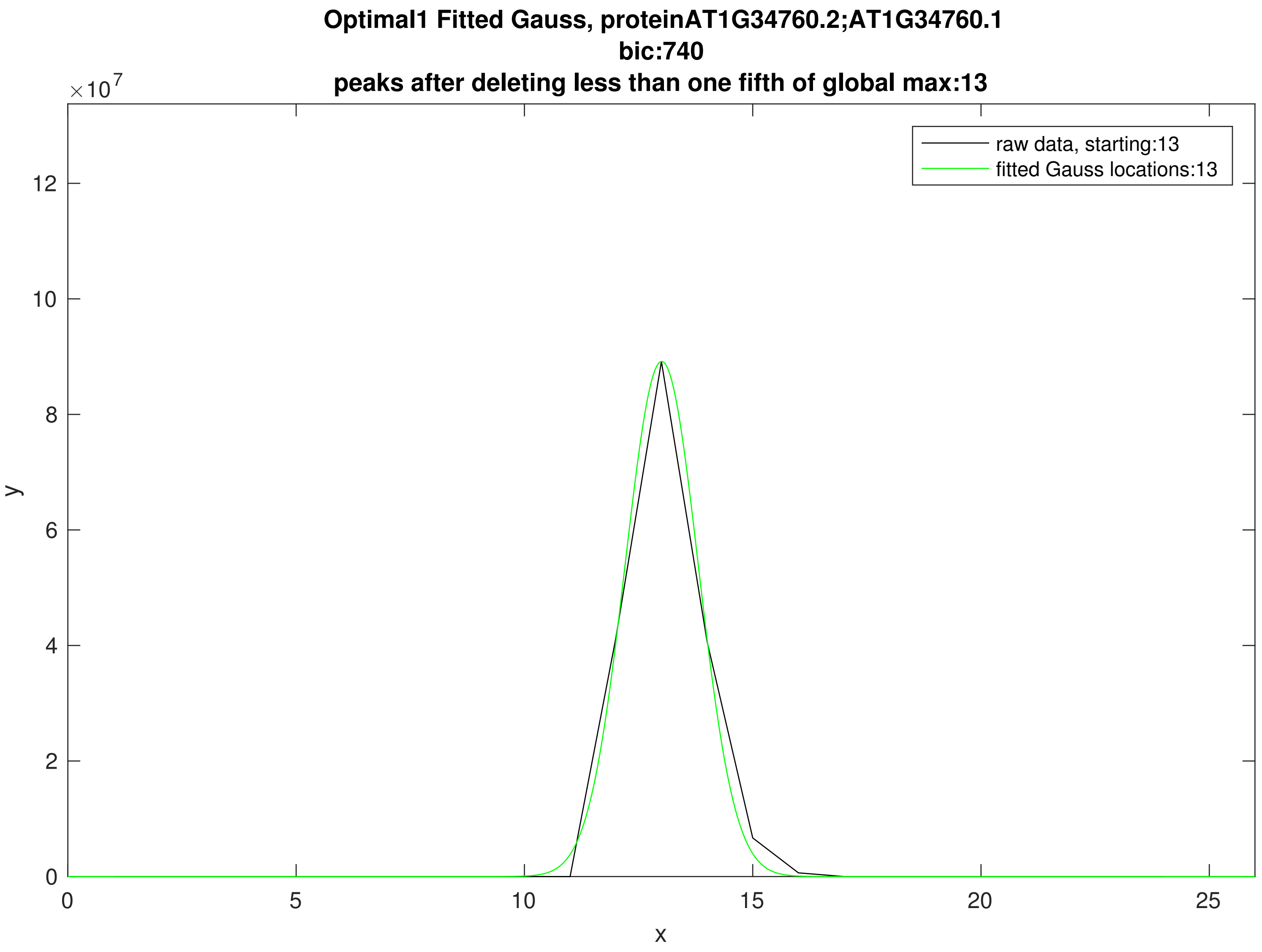

Optimal1 Fitted Gauss, proteinAT1G34760.2;AT1G34760.1
bic:740
peaks after deleting less than one fifth of global max:13
×10^7
raw data, starting:13
fitted Gauss locations:13
y
x
0
2
4
6
8
10
12
0
5
10
15
20
25

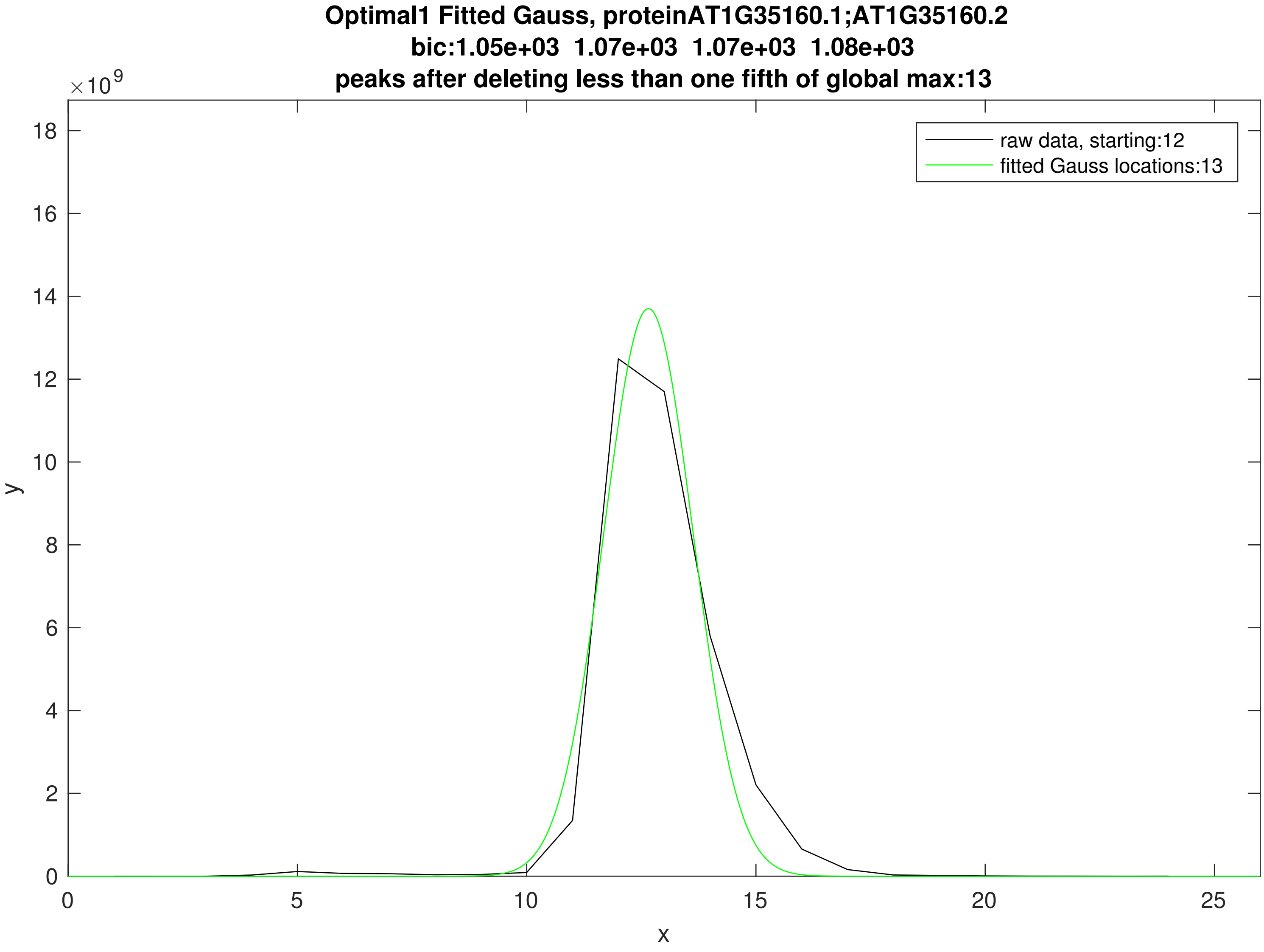
Optimal1 Fitted Gauss, proteinAT1G35160.1;AT1G35160.2
bic:1.05e+03  1.07e+03  1.07e+03  1.08e+03
peaks after deleting less than one fifth of global max:13
×10^9
raw data, starting:12
fitted Gauss locations:13
y
x
0
2
4
6
8
10
12
14
16
18
0
5
10
15
20
25

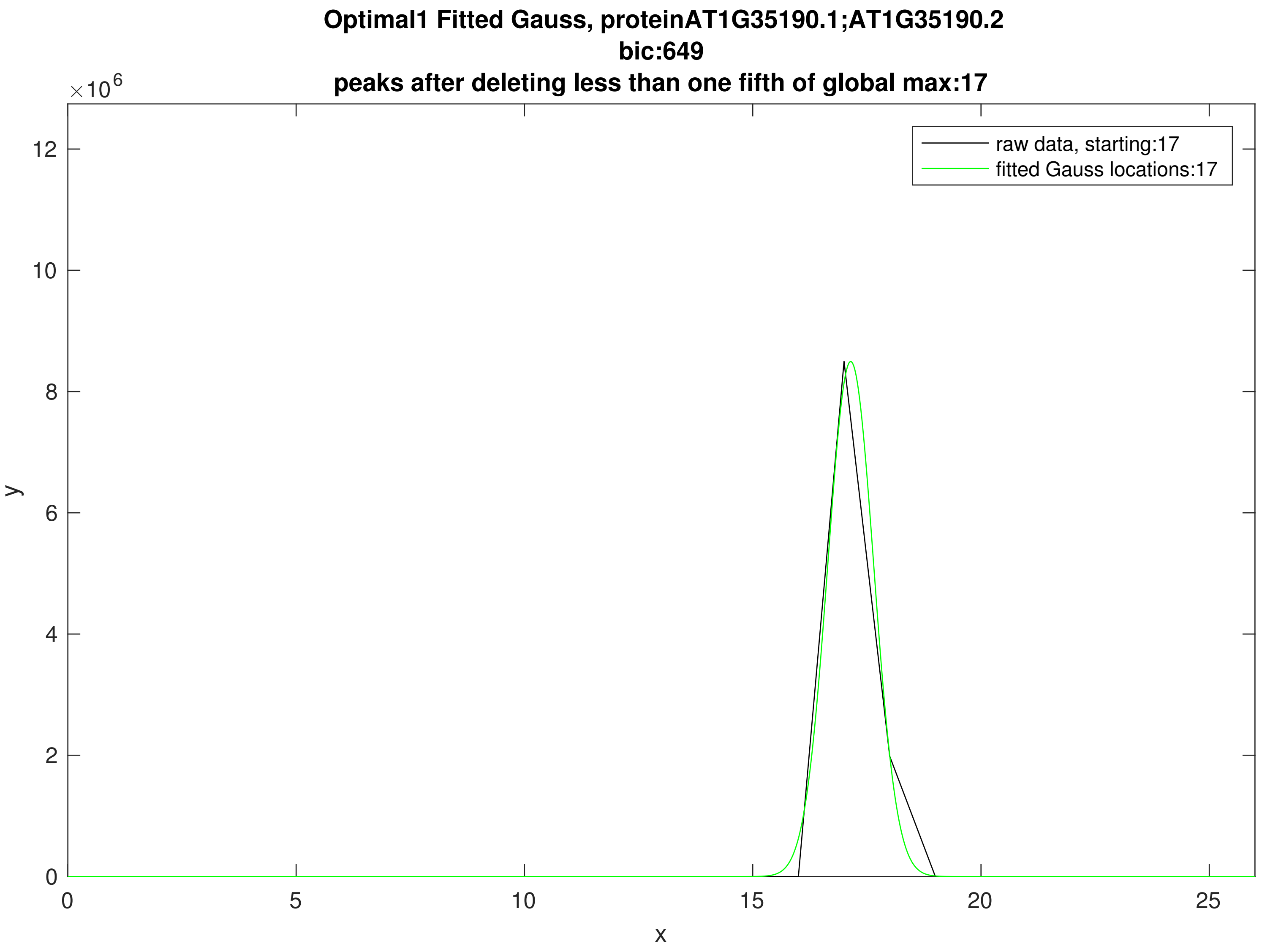
Optimal1 Fitted Gauss, proteinAT1G35190.1;AT1G35190.2
bic:649
peaks after deleting less than one fifth of global max:17
×10^6
raw data, starting:17
fitted Gauss locations:17
y
x
0
2
4
6
8
10
12
0
5
10
15
20
25

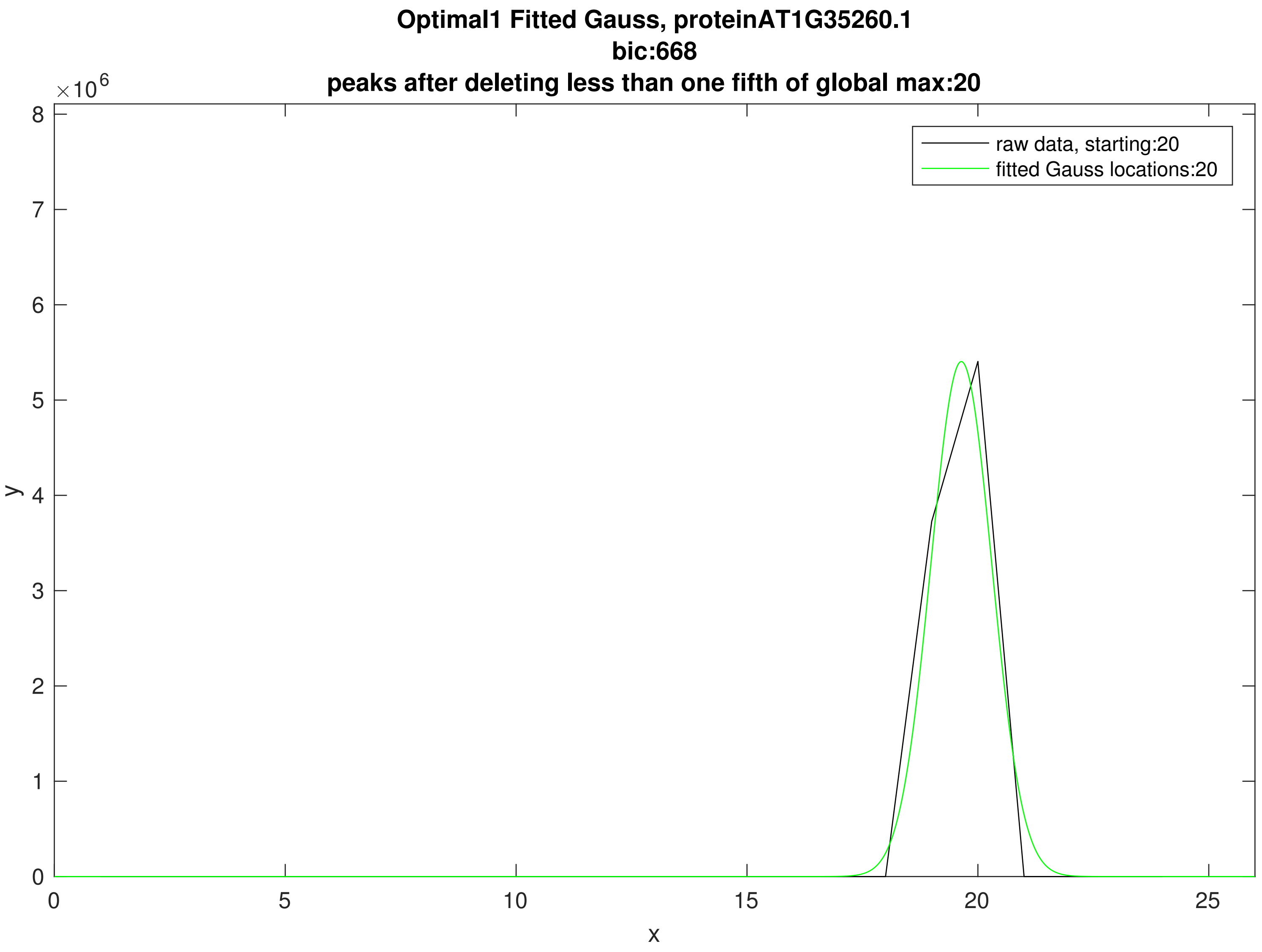

Optimal1 Fitted Gauss, proteinAT1G35260.1
bic:668
peaks after deleting less than one fifth of global max:20
×10^6
raw data, starting:20
fitted Gauss locations:20
y
x
0
1
2
3
4
5
6
7
8
0
5
10
15
20
25

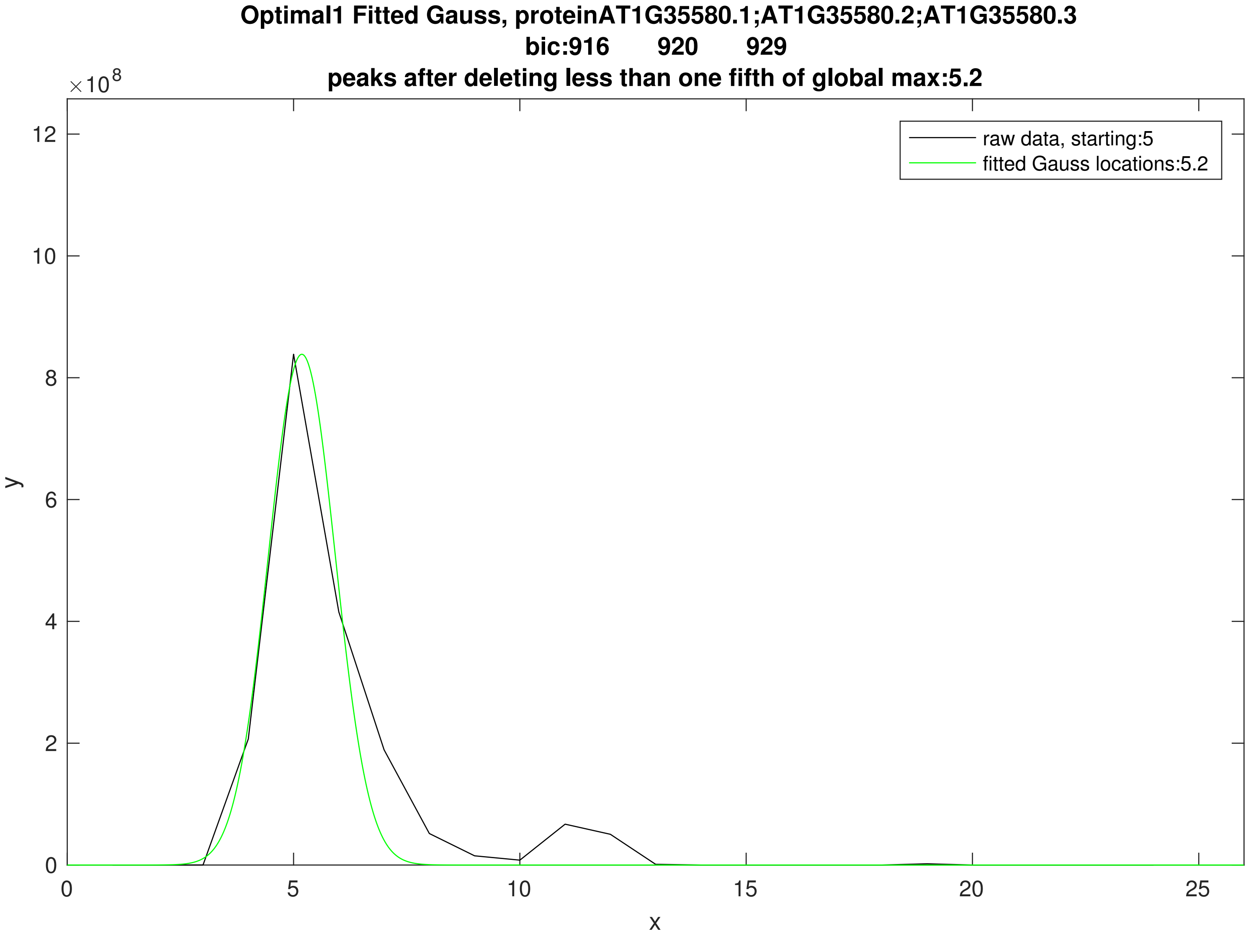
Optimal1 Fitted Gauss, proteinAT1G35580.1;AT1G35580.2;AT1G35580.3
bic:916 920 929
peaks after deleting less than one fifth of global max:5.2
×10^8
raw data, starting:5
fitted Gauss locations:5.2
y
x
0
2
4
6
8
10
12
0
5
10
15
20
25

**Optimal1 Fitted Gauss, proteinAT1G35720.1**
**bic:1.06e+03  1.07e+03  1.08e+03**
**peaks after deleting less than one fifth of global max:14**

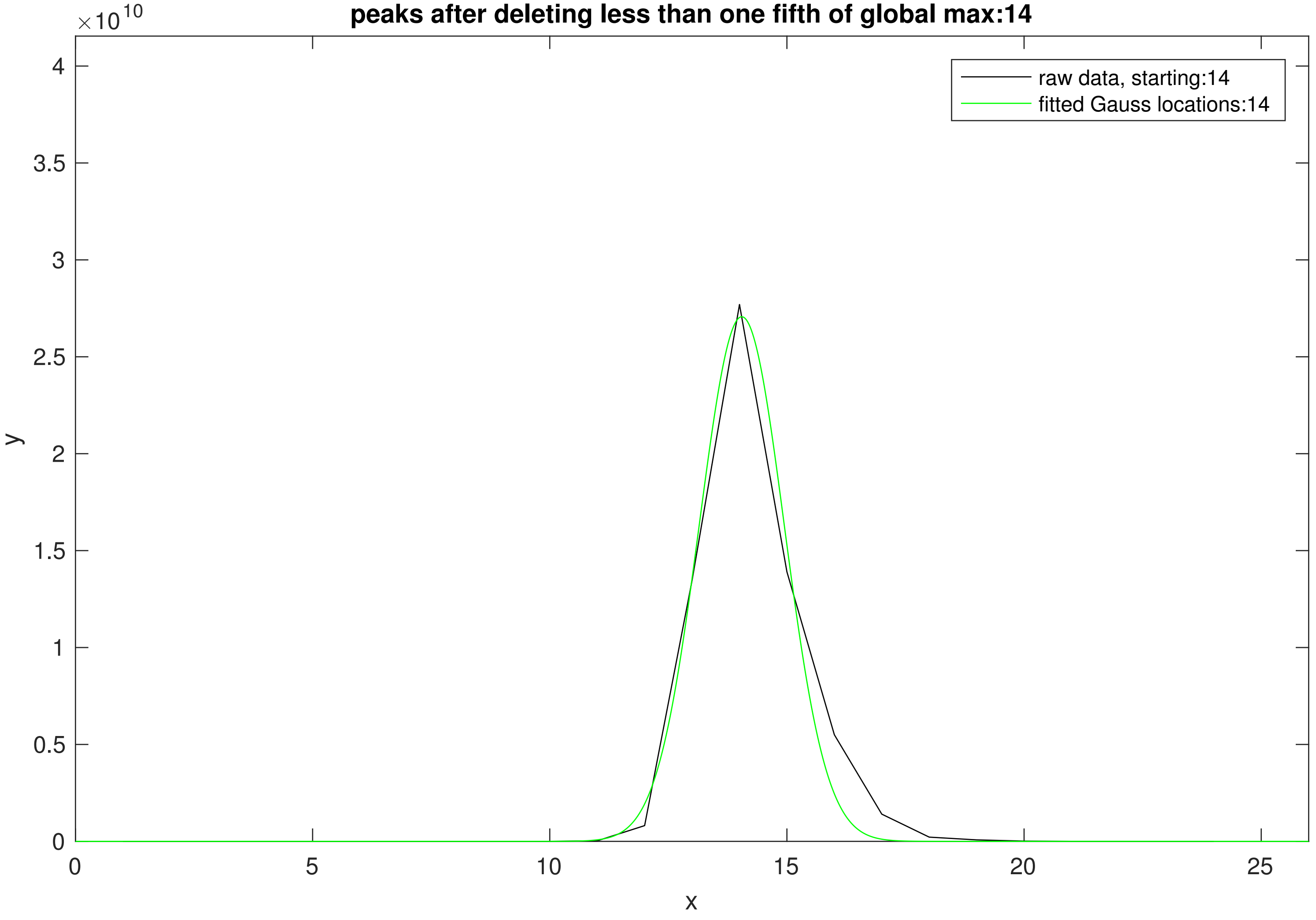

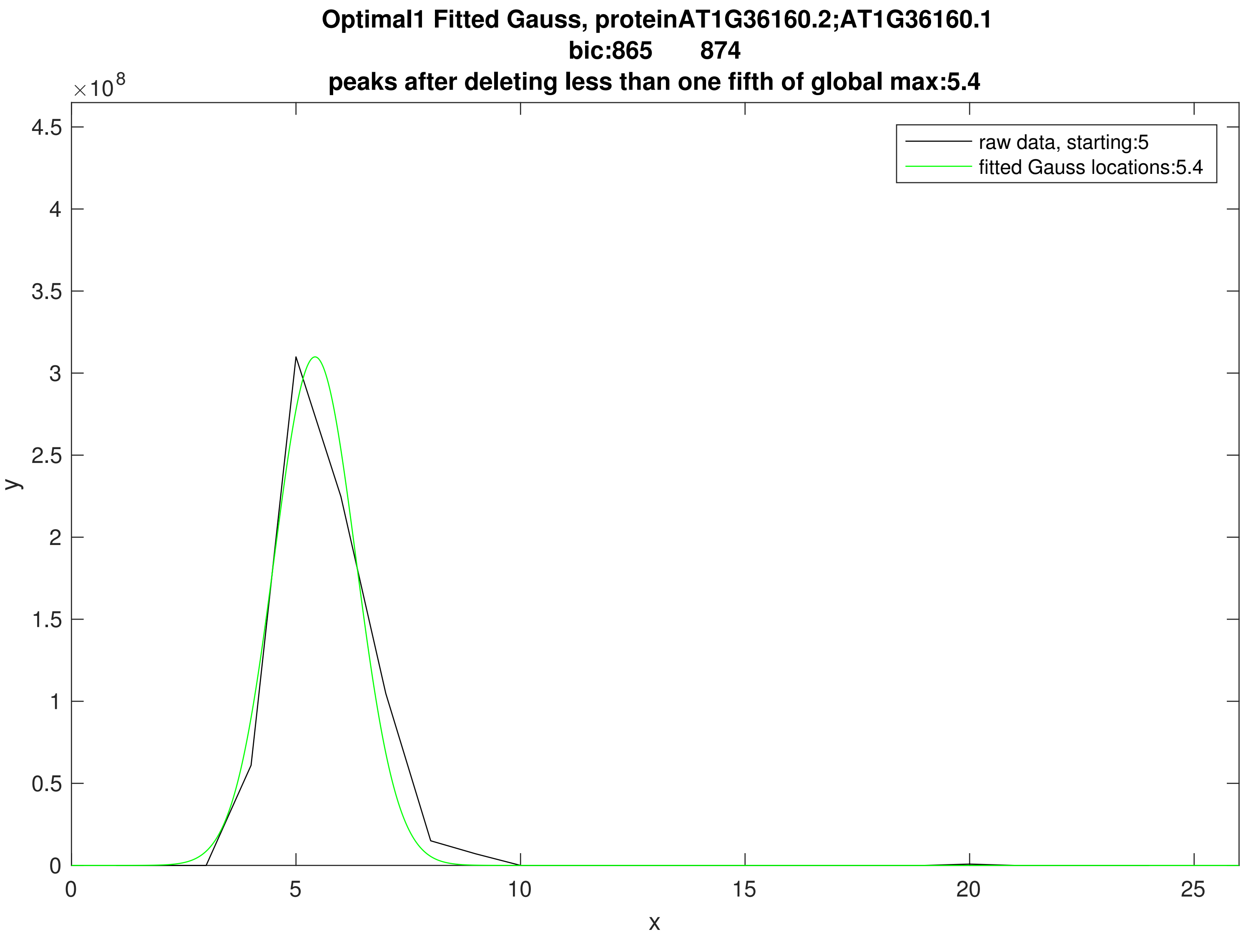
Optimal1 Fitted Gauss, proteinAT1G36160.2;AT1G36160.1
bic:865 874
peaks after deleting less than one fifth of global max:5.4
×10^8
4.5
4
3.5
3
2.5
2
1.5
1
0.5
0
y
0
5
10
15
20
25
x
raw data, starting:5
fitted Gauss locations:5.4

Optimal1 Fitted Gauss, proteinAT1G36390.2;AT1G36390.1
bic:680
peaks after deleting less than one fifth of global max:8.8

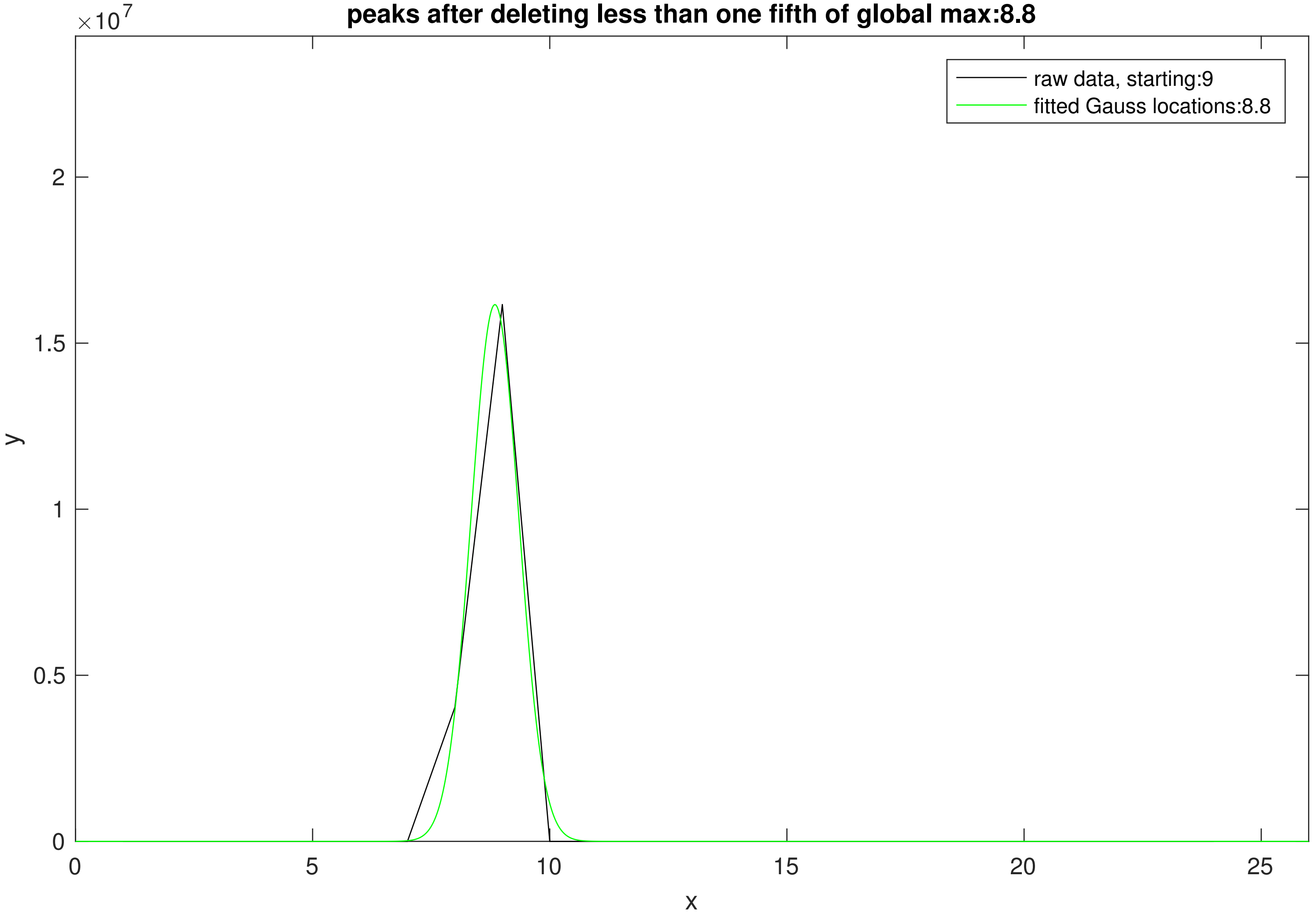

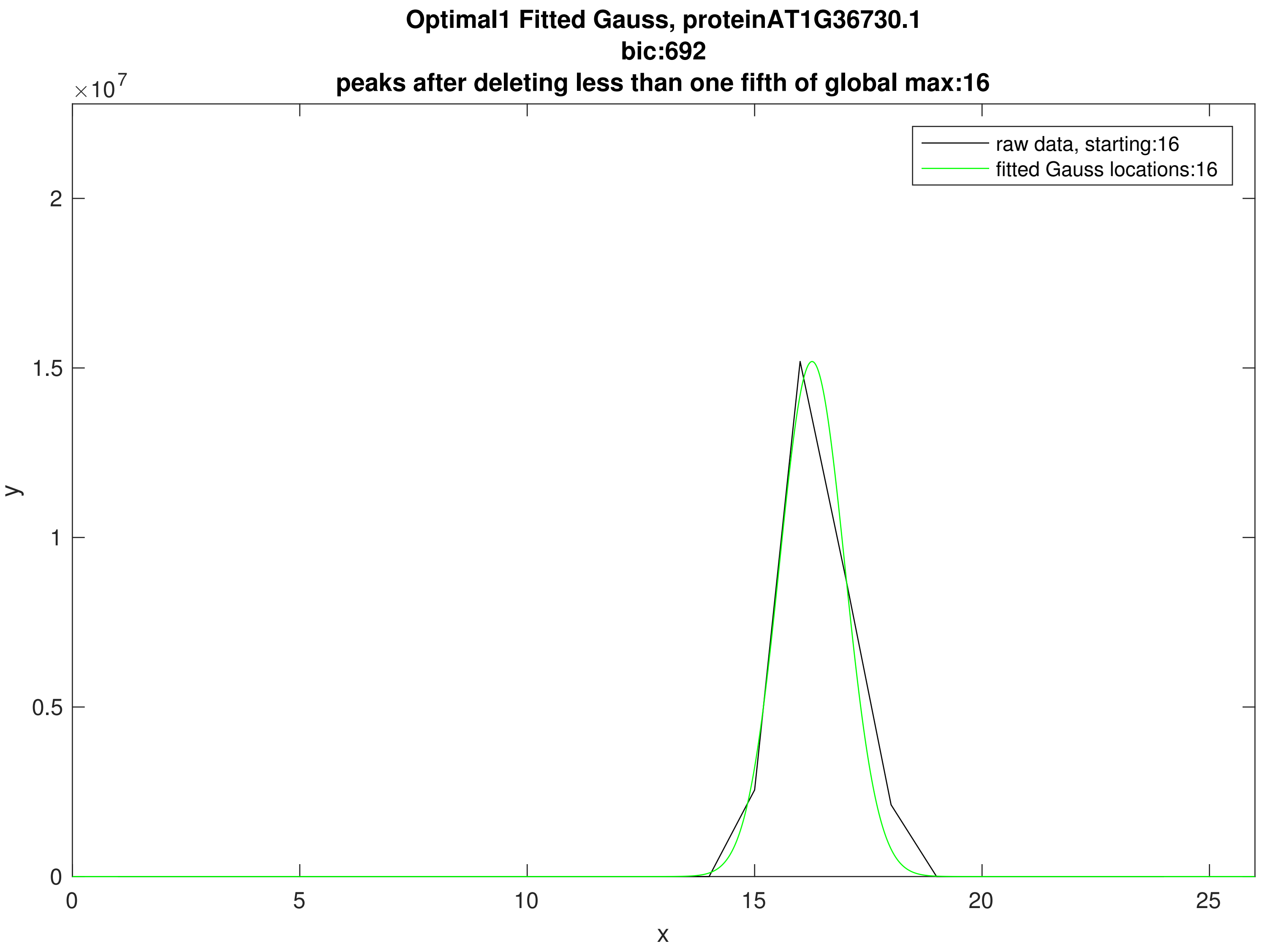
Optimal1 Fitted Gauss, proteinAT1G36730.1
bic:692
peaks after deleting less than one fifth of global max:16
×10^7
raw data, starting:16
fitted Gauss locations:16
y
x
0
0.5
1
1.5
2
0
5
10
15
20
25

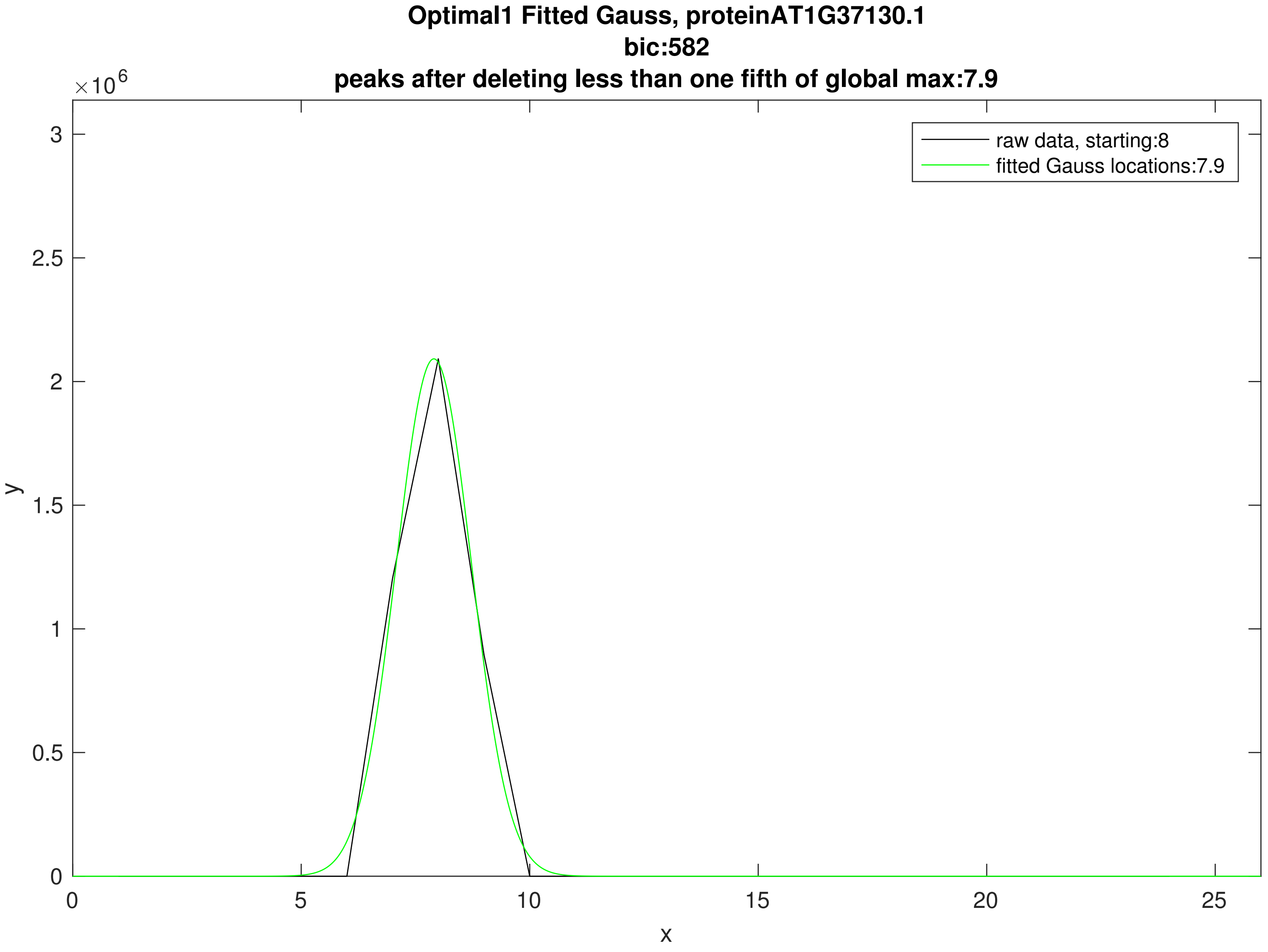
Optimal1 Fitted Gauss, proteinAT1G37130.1
bic:582
peaks after deleting less than one fifth of global max:7.9
×10^6
raw data, starting:8
fitted Gauss locations:7.9
y
0
0.5
1
1.5
2
2.5
3
x
0
5
10
15
20
25

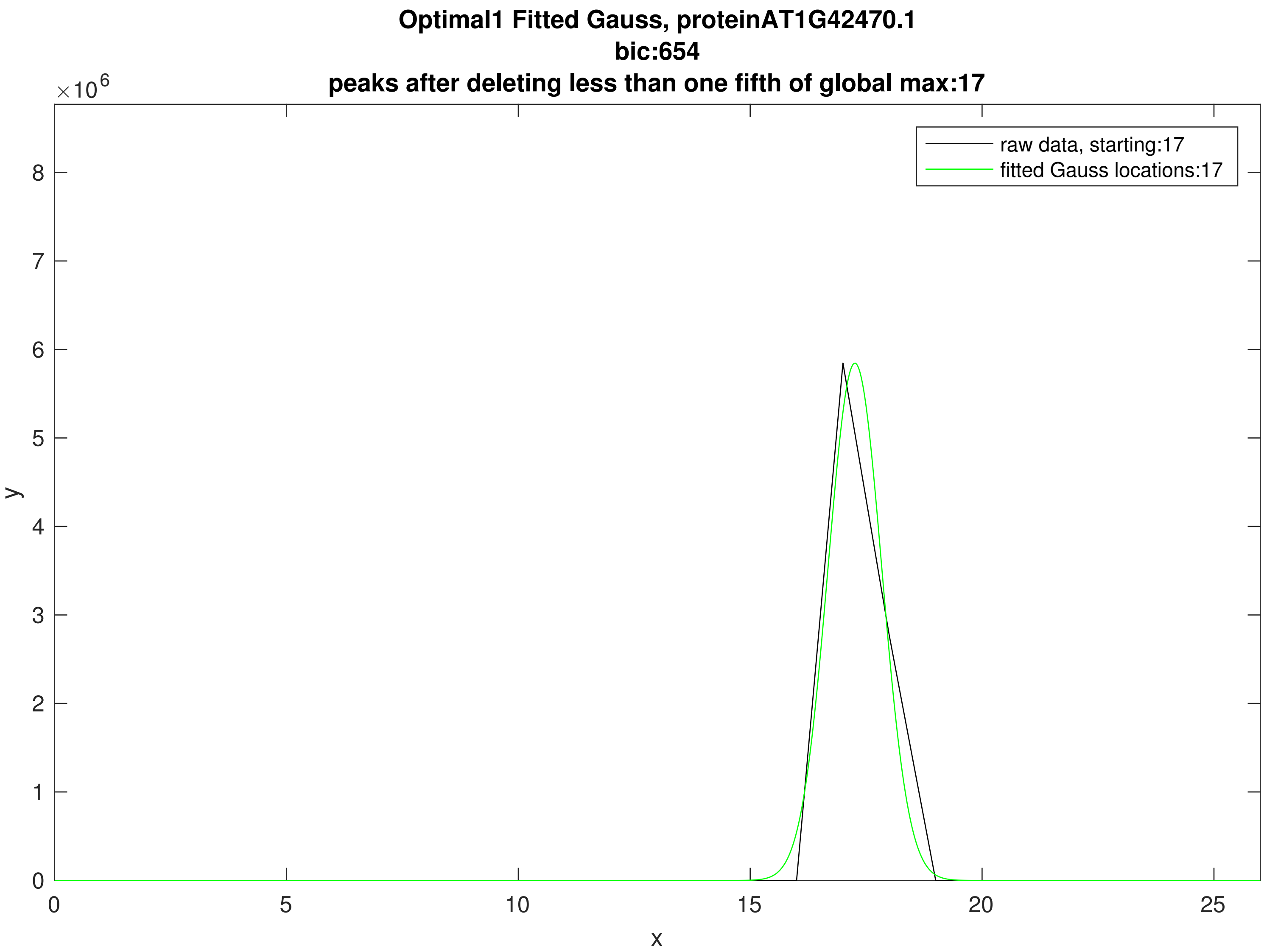

Optimal1 Fitted Gauss, proteinAT1G42470.1
bic:654
peaks after deleting less than one fifth of global max:17
×10^6
raw data, starting:17
fitted Gauss locations:17
y
x
0
1
2
3
4
5
6
7
8
0
5
10
15
20
25

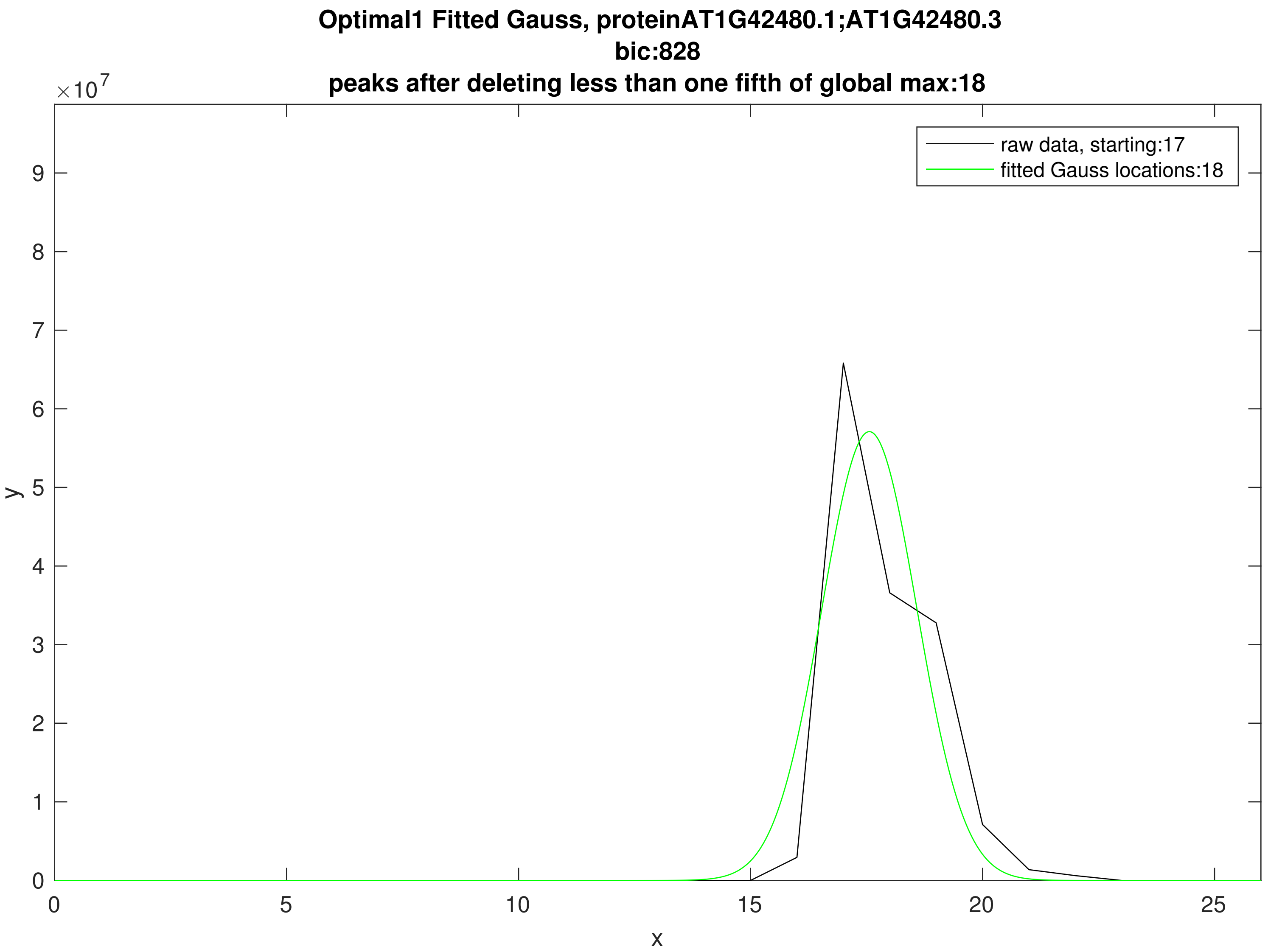

Optimal1 Fitted Gauss, proteinAT1G42480.1;AT1G42480.3
bic:828
peaks after deleting less than one fifth of global max:18
×10^7
raw data, starting:17
fitted Gauss locations:18
y
x
0
1
2
3
4
5
6
7
8
9
0
5
10
15
20
25

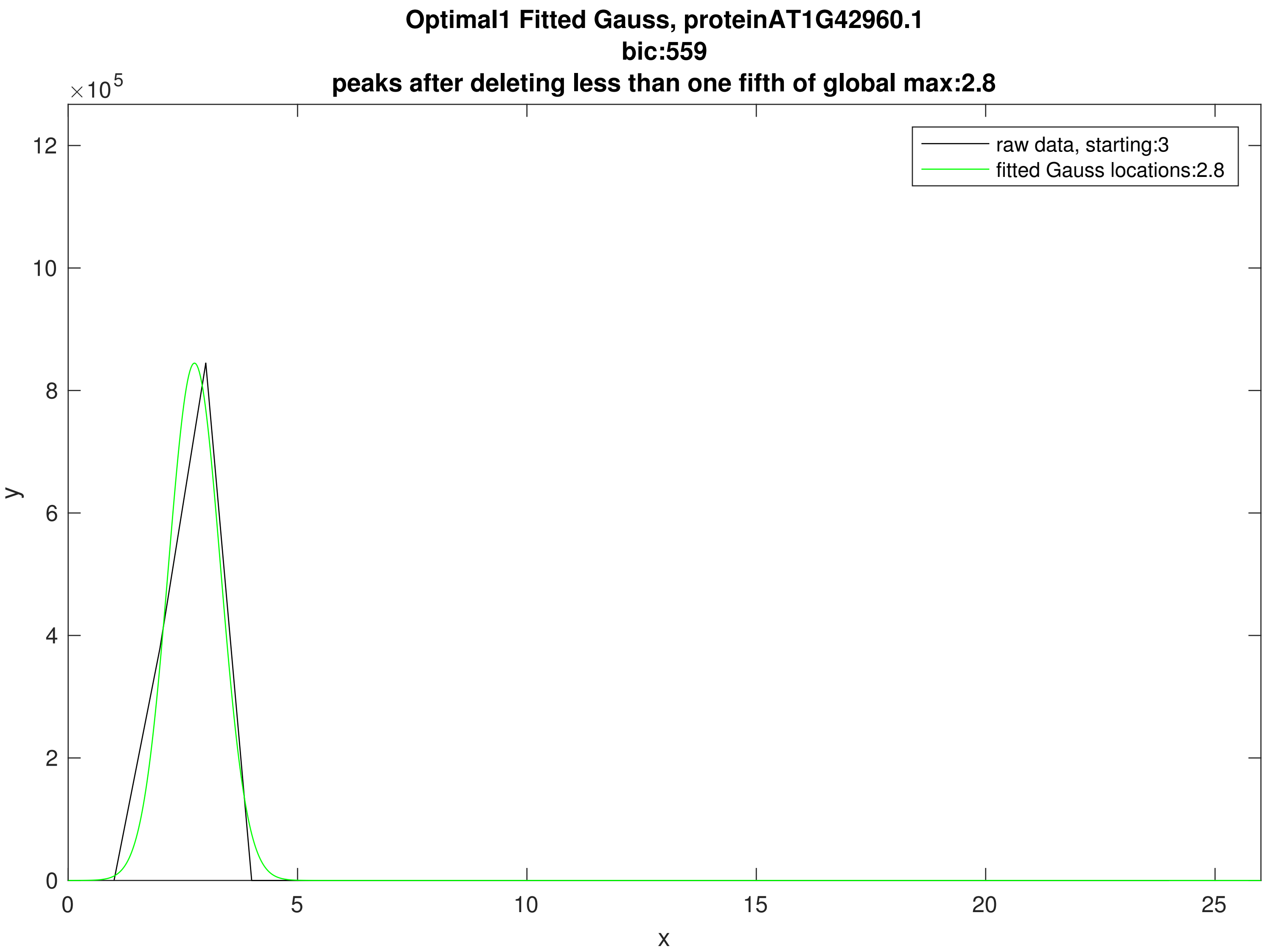

Optimal1 Fitted Gauss, proteinAT1G42960.1
bic:559
peaks after deleting less than one fifth of global max:2.8
×10^5
raw data, starting:3
fitted Gauss locations:2.8
y
x
0
2
4
6
8
10
12
0
5
10
15
20
25

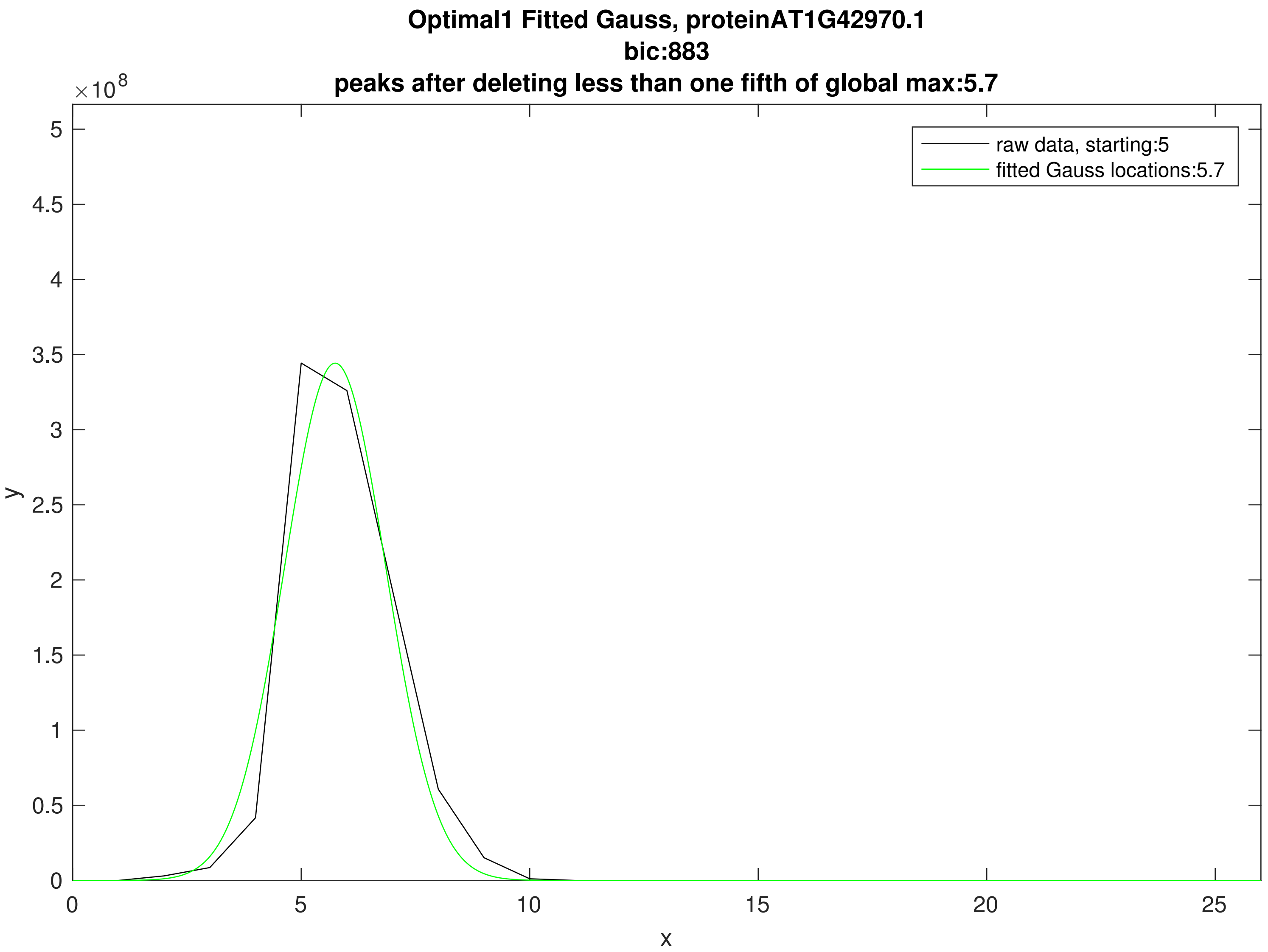
Optimal1 Fitted Gauss, proteinAT1G42970.1
bic:883
peaks after deleting less than one fifth of global max:5.7
×10^8
y
x
raw data, starting:5
fitted Gauss locations:5.7
0
0.5
1
1.5
2
2.5
3
3.5
4
4.5
5
0
5
10
15
20
25

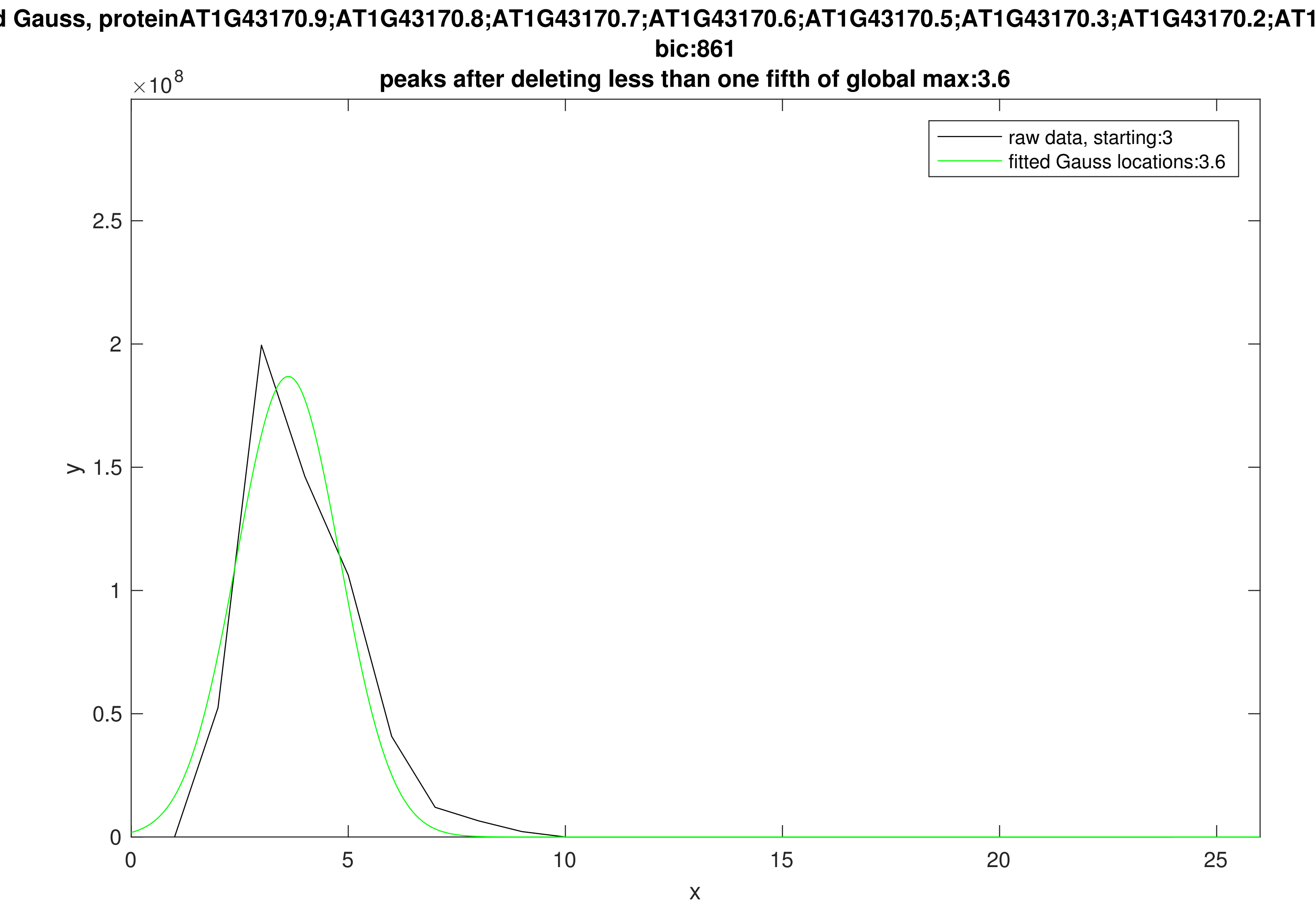

d Gauss, proteinAT1G43170.9;AT1G43170.8;AT1G43170.7;AT1G43170.6;AT1G43170.5;AT1G43170.3;AT1G43170.2;AT1
bic:861
peaks after deleting less than one fifth of global max:3.6
×10^8
raw data, starting:3
fitted Gauss locations:3.6
y
x

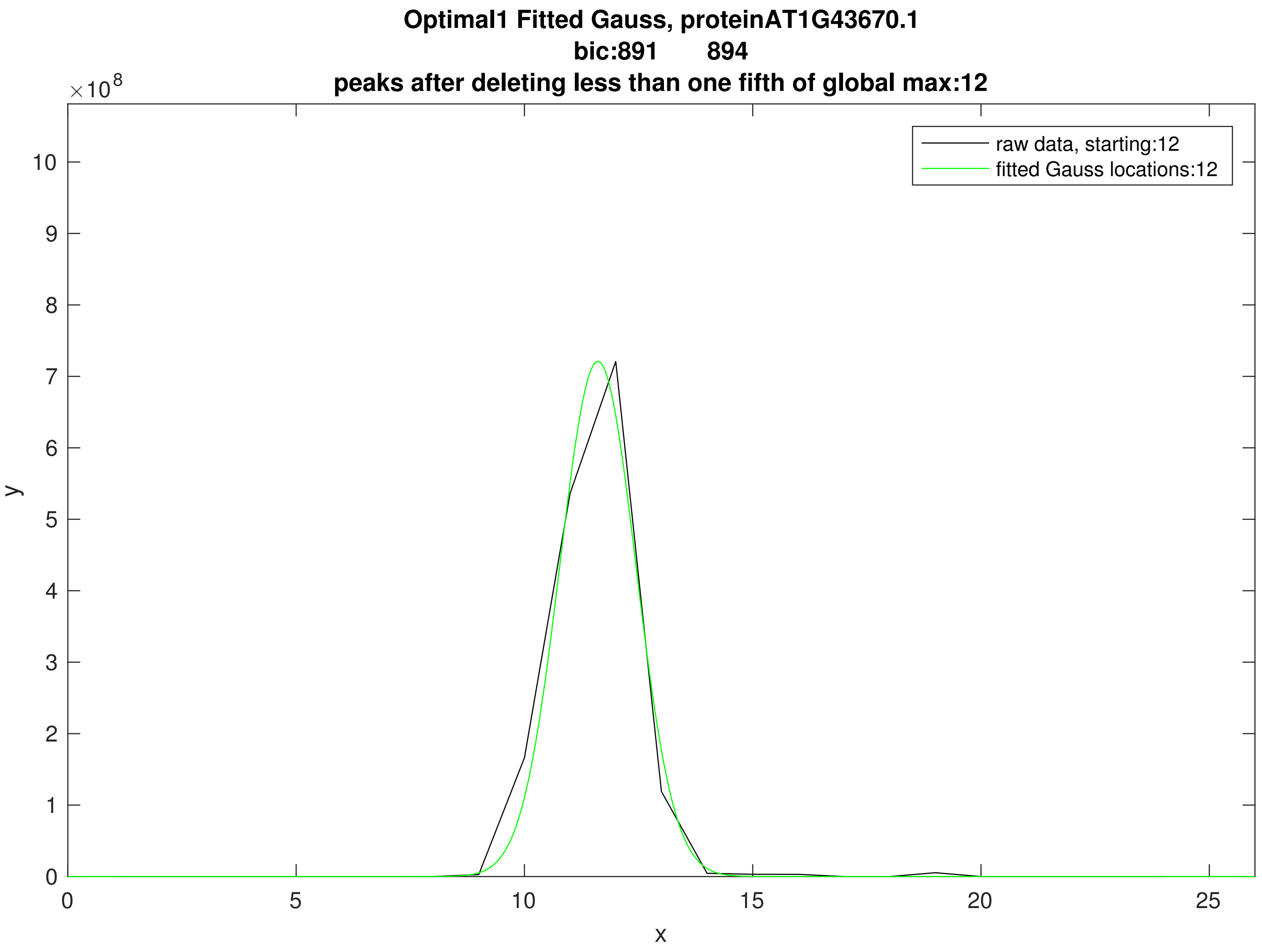

Optimal1 Fitted Gauss, proteinAT1G43670.1
bic:891 894
peaks after deleting less than one fifth of global max:12
×10^8
raw data, starting:12
fitted Gauss locations:12
y
x
0
1
2
3
4
5
6
7
8
9
10
0
5
10
15
20
25

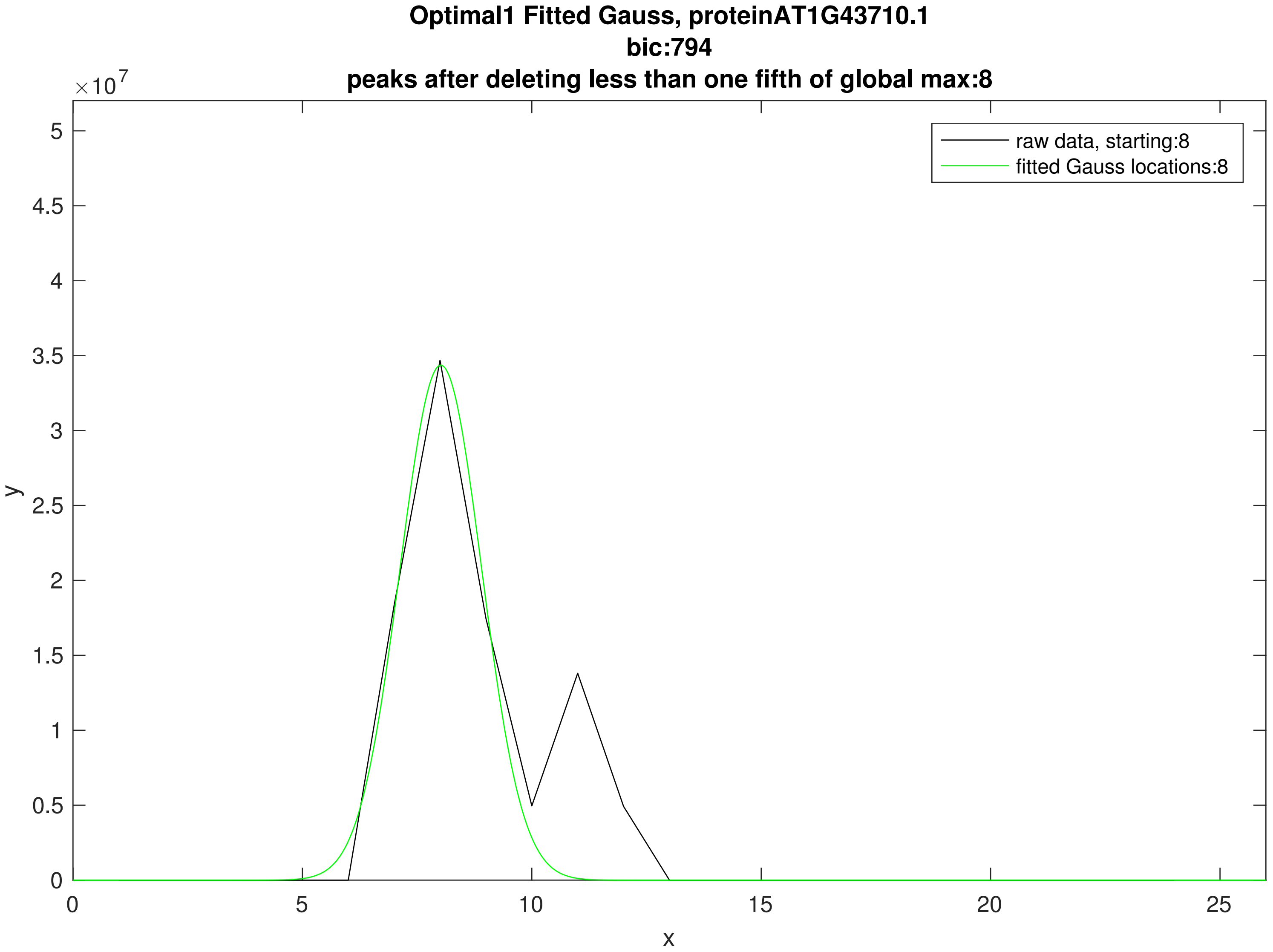
Optimal1 Fitted Gauss, proteinAT1G43710.1
bic:794
peaks after deleting less than one fifth of global max:8
×10^7
raw data, starting:8
fitted Gauss locations:8
y
x
0
0.5
1
1.5
2
2.5
3
3.5
4
4.5
5
0
5
10
15
20
25

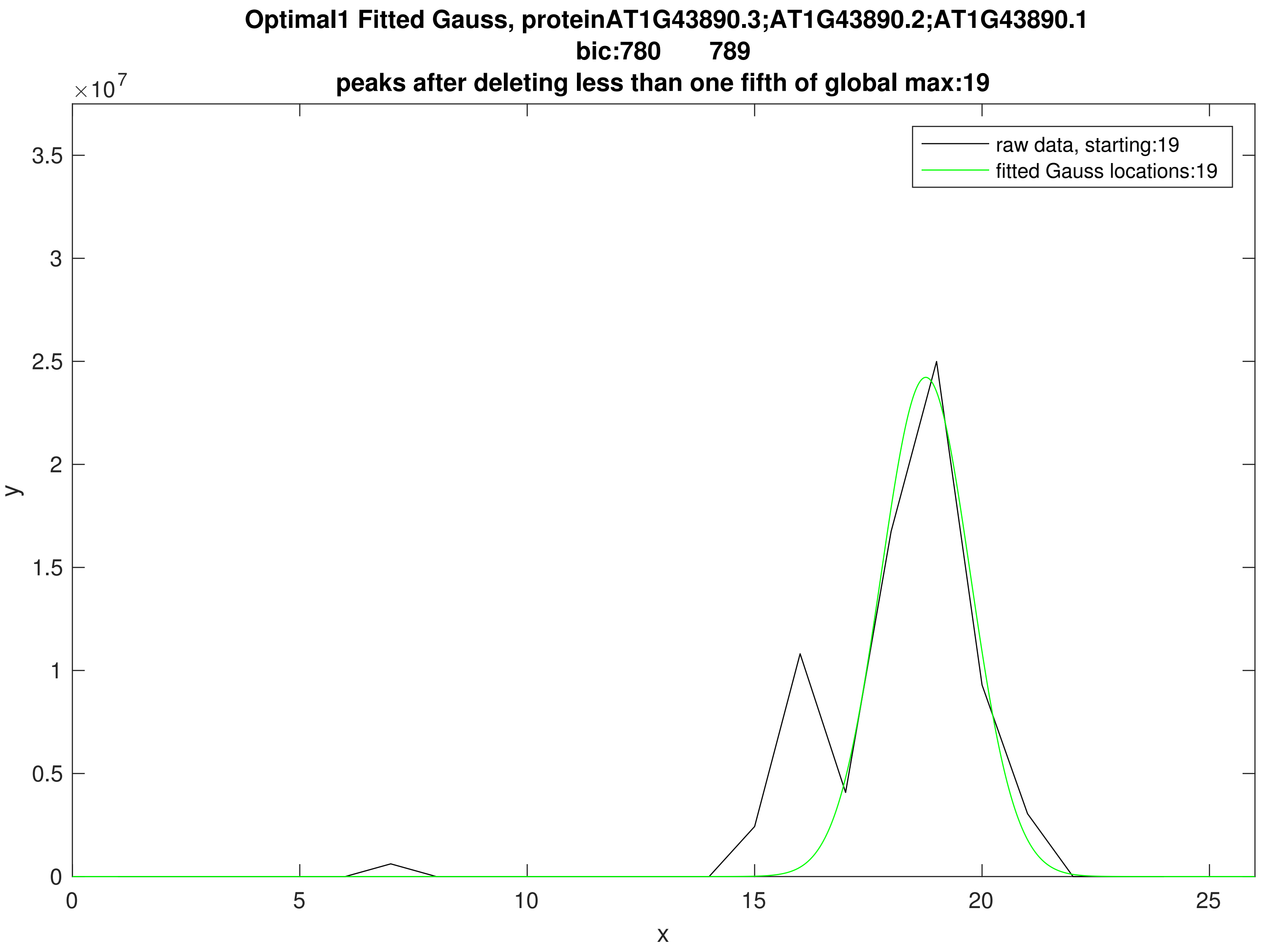
Optimal1 Fitted Gauss, proteinAT1G43890.3;AT1G43890.2;AT1G43890.1
bic:780 789
peaks after deleting less than one fifth of global max:19
×10^7
3.5
3
2.5
2
1.5
1
0.5
0
y
0
5
10
15
20
25
x
raw data, starting:19
fitted Gauss locations:19

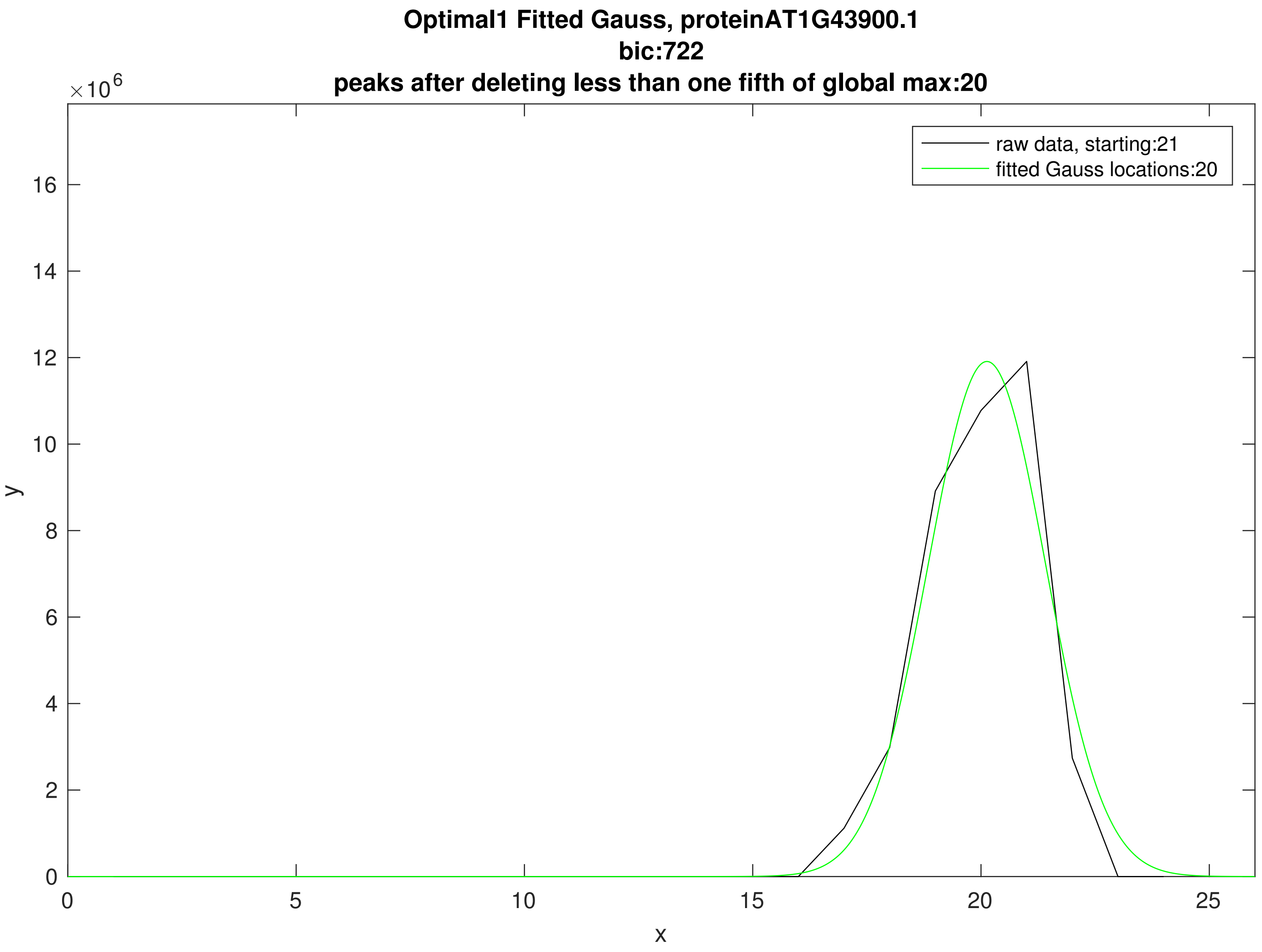

Optimal1 Fitted Gauss, proteinAT1G43900.1
bic:722
peaks after deleting less than one fifth of global max:20
×10^6
raw data, starting:21
fitted Gauss locations:20
16
14
12
10
8
6
4
2
0
y
0
5
10
15
20
25
x

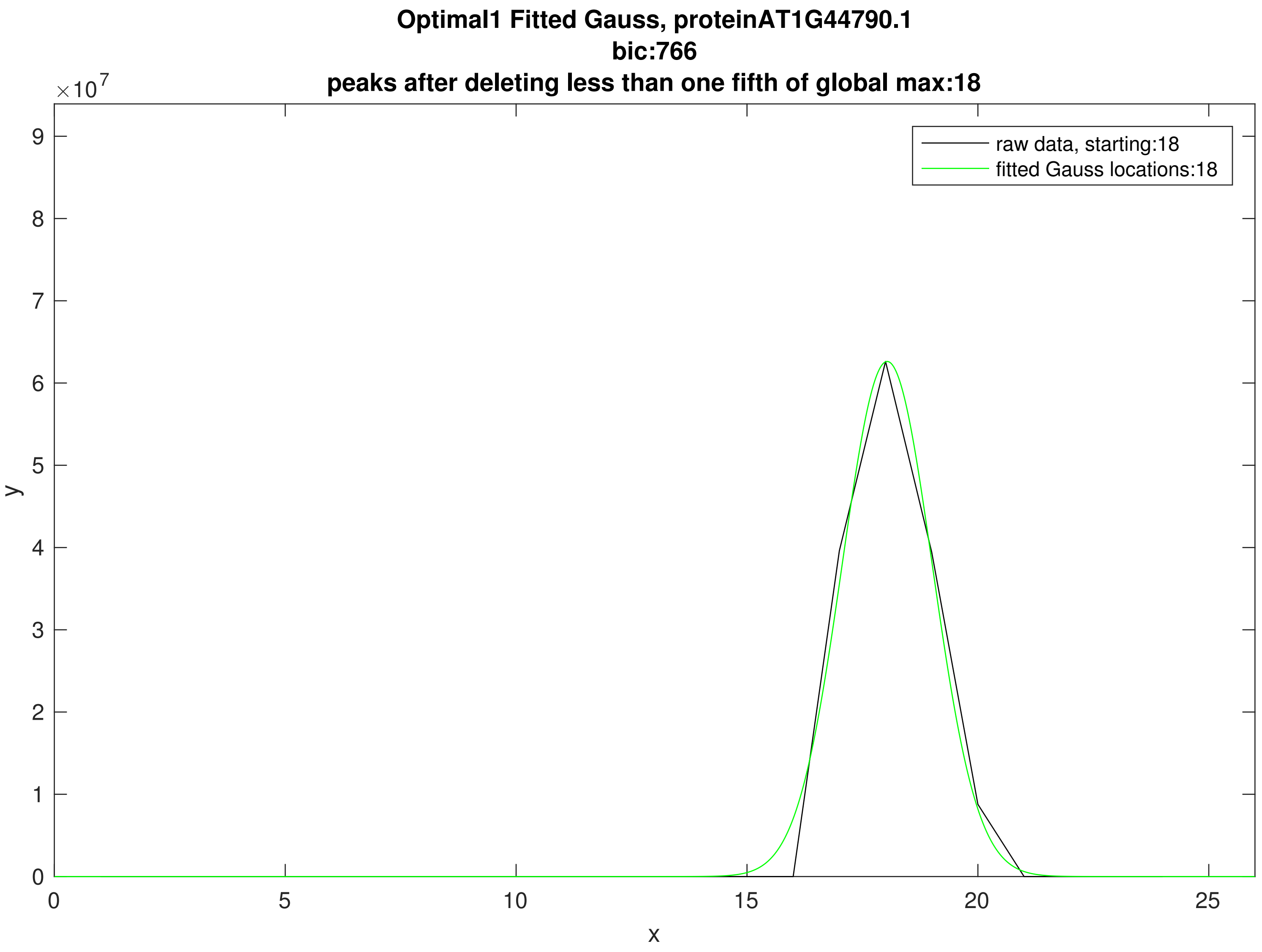

Optimal1 Fitted Gauss, proteinAT1G44790.1
bic:766
peaks after deleting less than one fifth of global max:18
×10^7
raw data, starting:18
fitted Gauss locations:18
y
x
0
1
2
3
4
5
6
7
8
9
0
5
10
15
20
25

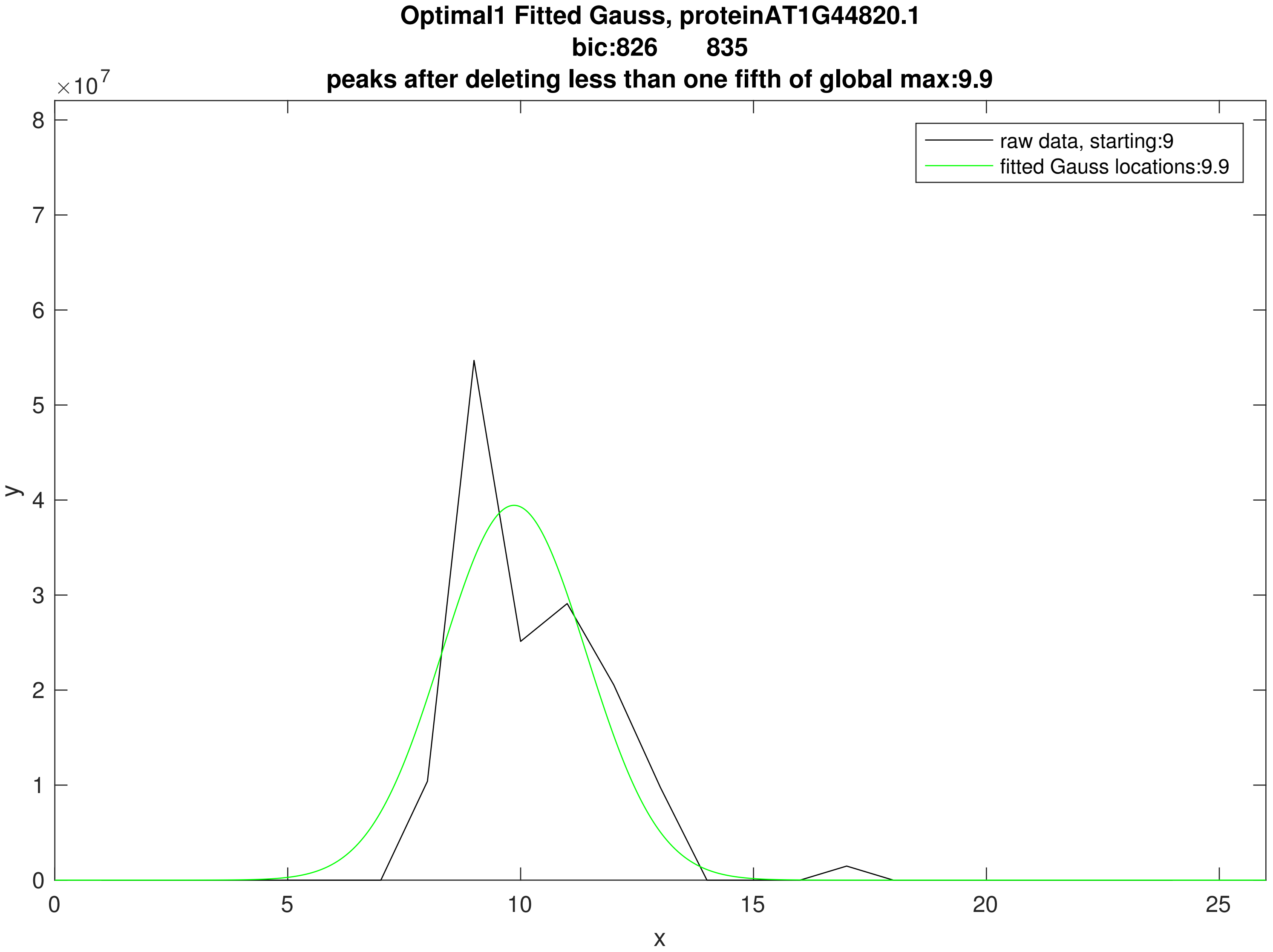
Optimal1 Fitted Gauss, proteinAT1G44820.1
bic:826 835
peaks after deleting less than one fifth of global max:9.9
×10^7
raw data, starting:9
fitted Gauss locations:9.9
y
x
0
1
2
3
4
5
6
7
8
0
5
10
15
20
25

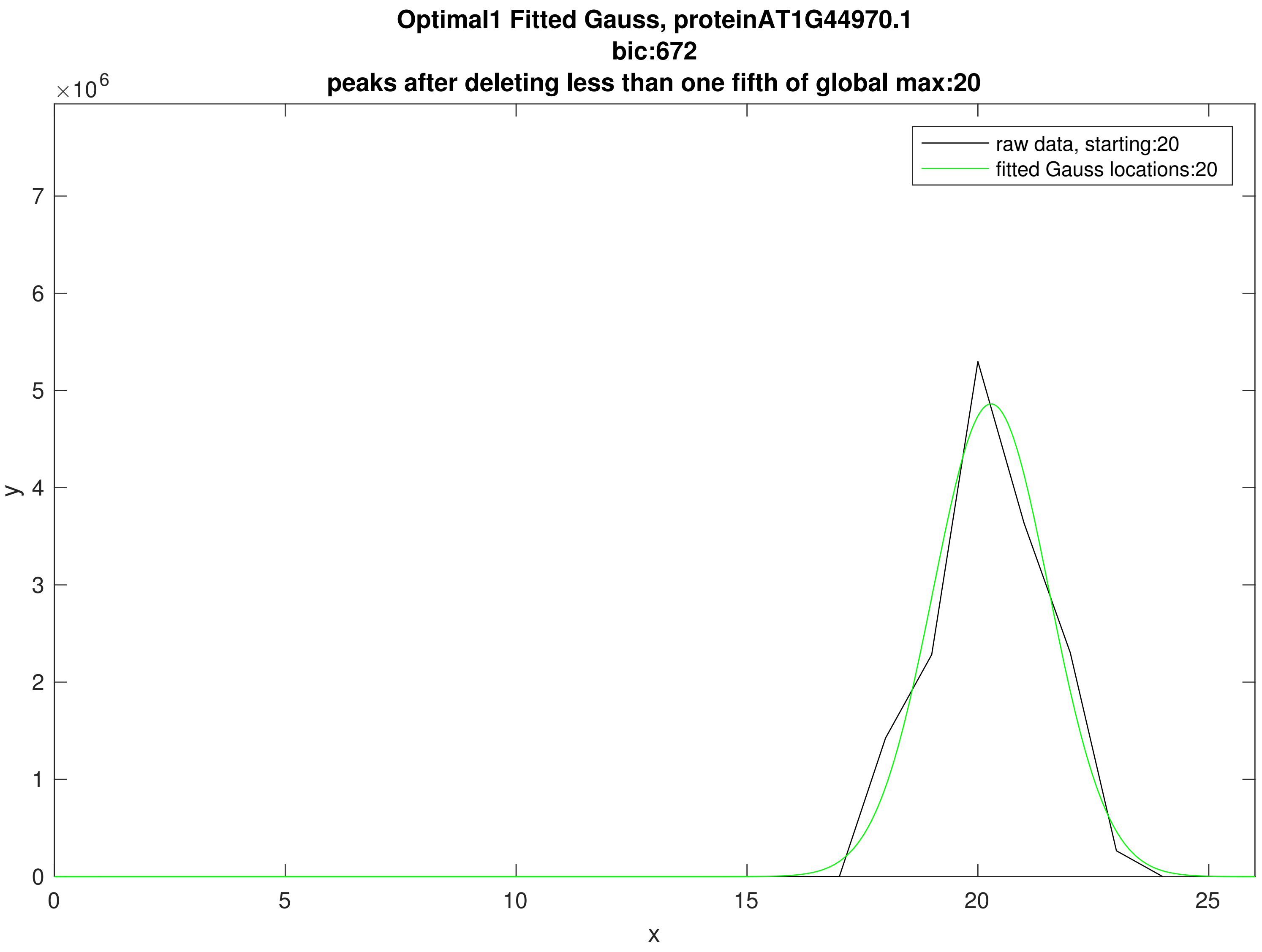
Optimal1 Fitted Gauss, proteinAT1G44970.1
bic:672
peaks after deleting less than one fifth of global max:20
×10^6
raw data, starting:20
fitted Gauss locations:20
y
x
0
1
2
3
4
5
6
7
0
5
10
15
20
25

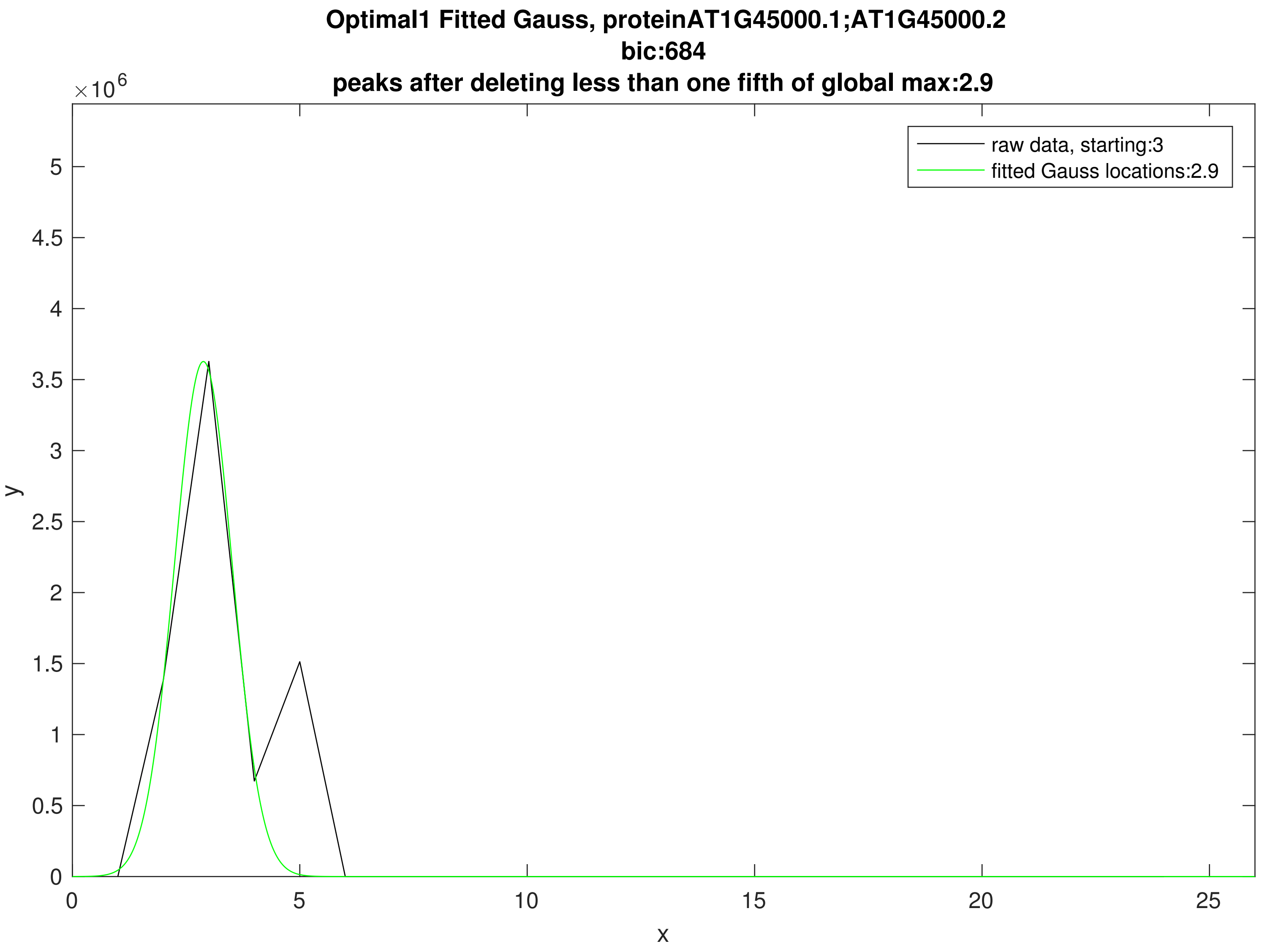
Optimal1 Fitted Gauss, proteinAT1G45000.1;AT1G45000.2
bic:684
peaks after deleting less than one fifth of global max:2.9
×10^6
raw data, starting:3
fitted Gauss locations:2.9
5
4.5
4
3.5
3
2.5
2
1.5
1
0.5
0
y
0
5
10
15
20
25
x

**Optimal1 Fitted Gauss, proteinAT1G45130.1**
**bic:869**
**peaks after deleting less than one fifth of global max:16**

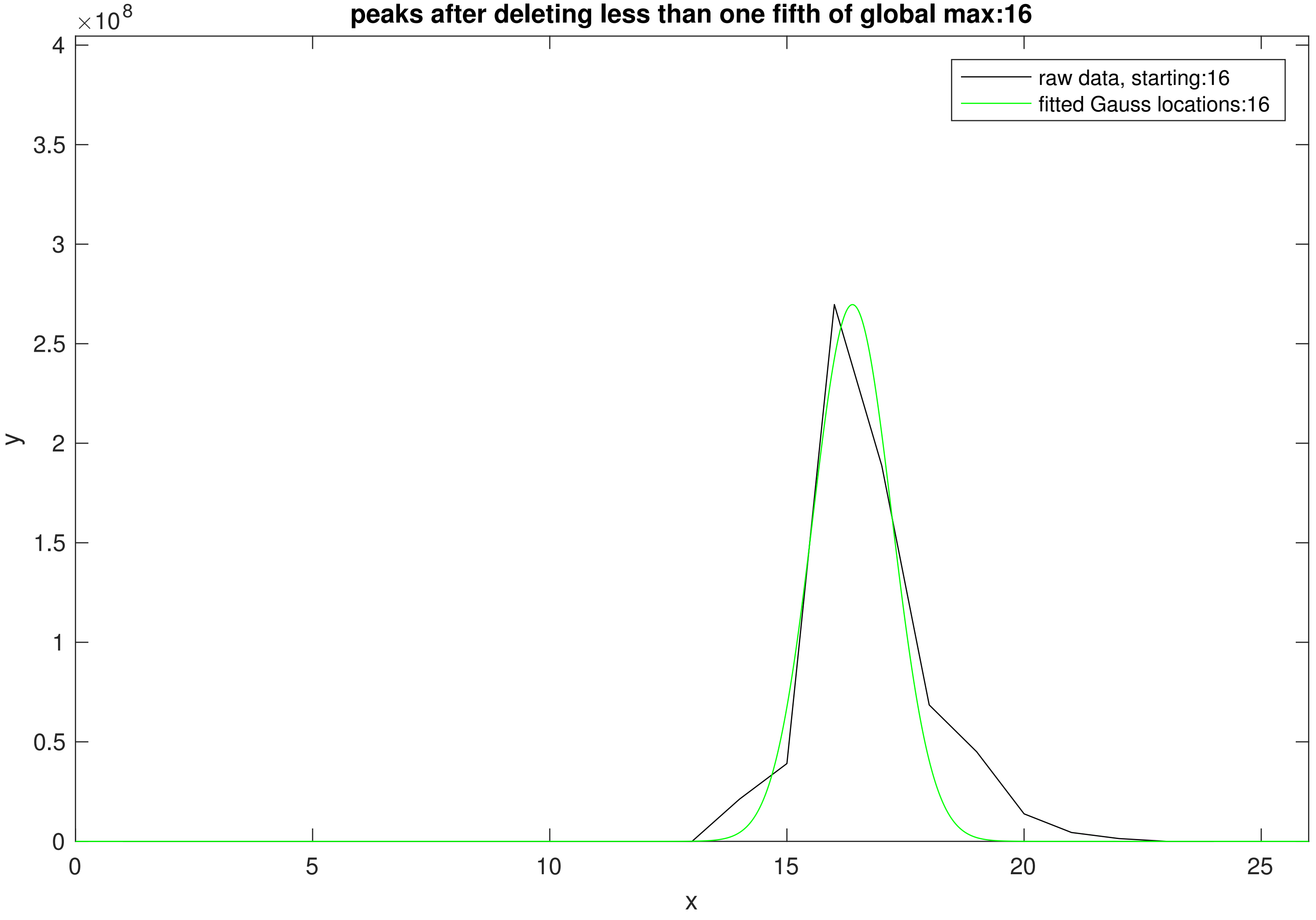

**Optimal1 Fitted Gauss, proteinAT1G45145.1**
**bic:681**
**peaks after deleting less than one fifth of global max:20**

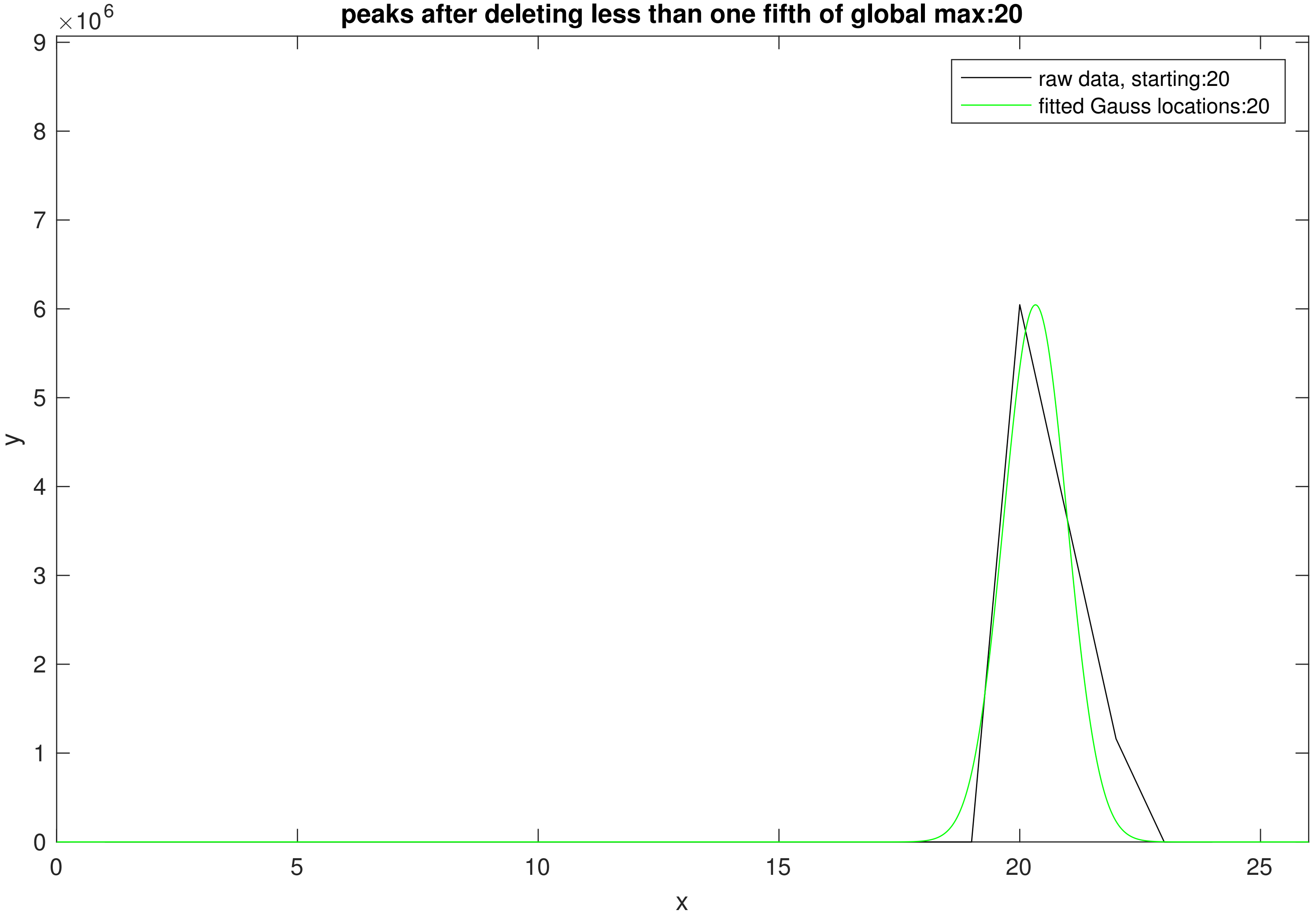

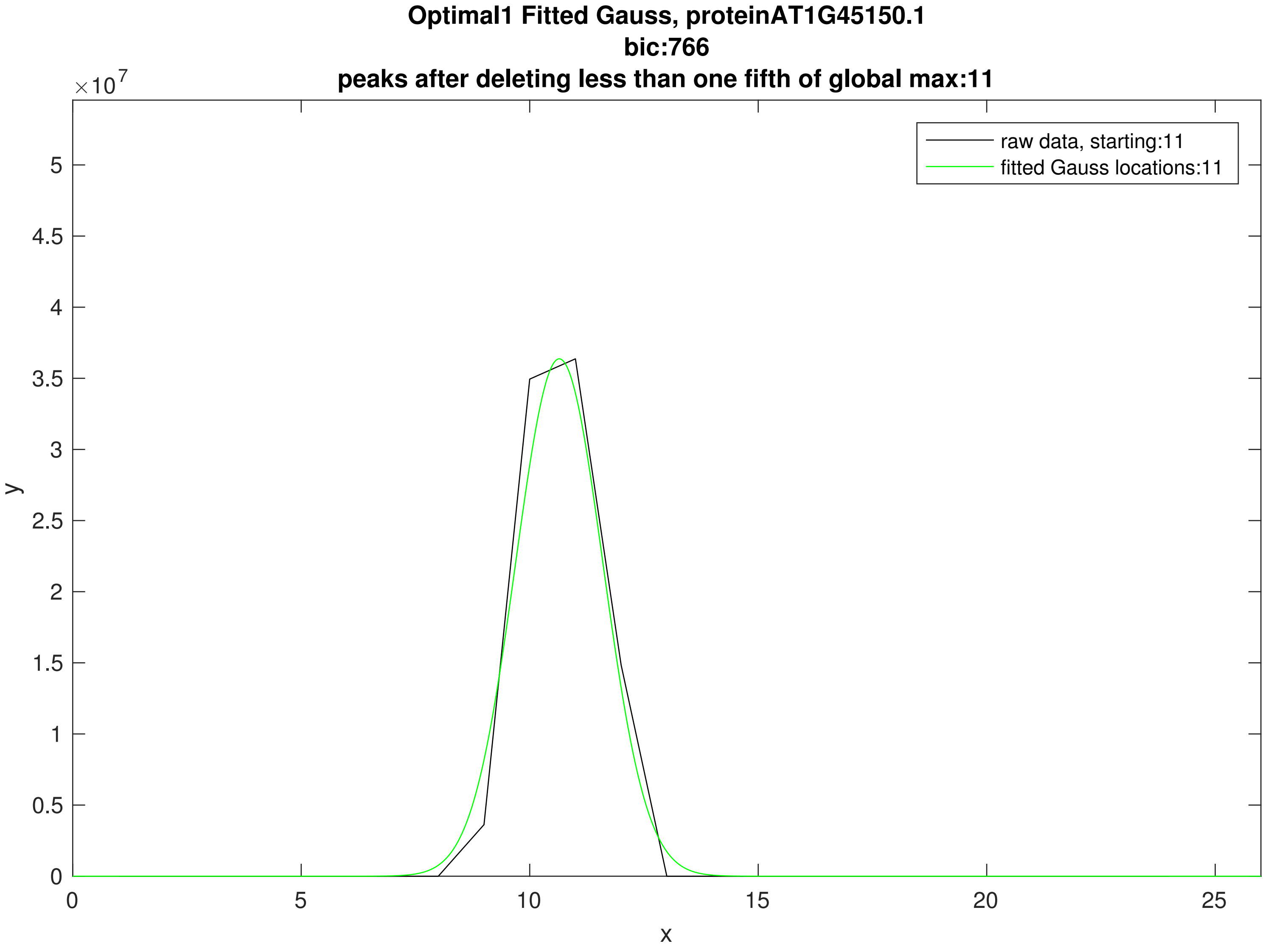
Optimal1 Fitted Gauss, proteinAT1G45150.1
bic:766
peaks after deleting less than one fifth of global max:11
×10^7
raw data, starting:11
fitted Gauss locations:11
y
x
0
0.5
1
1.5
2
2.5
3
3.5
4
4.5
5
0
5
10
15
20
25

**Optimal1 Fitted Gauss, proteinAT1G45201.2;AT1G45201.1**
**bic:702**
**peaks after deleting less than one fifth of global max:4.3**

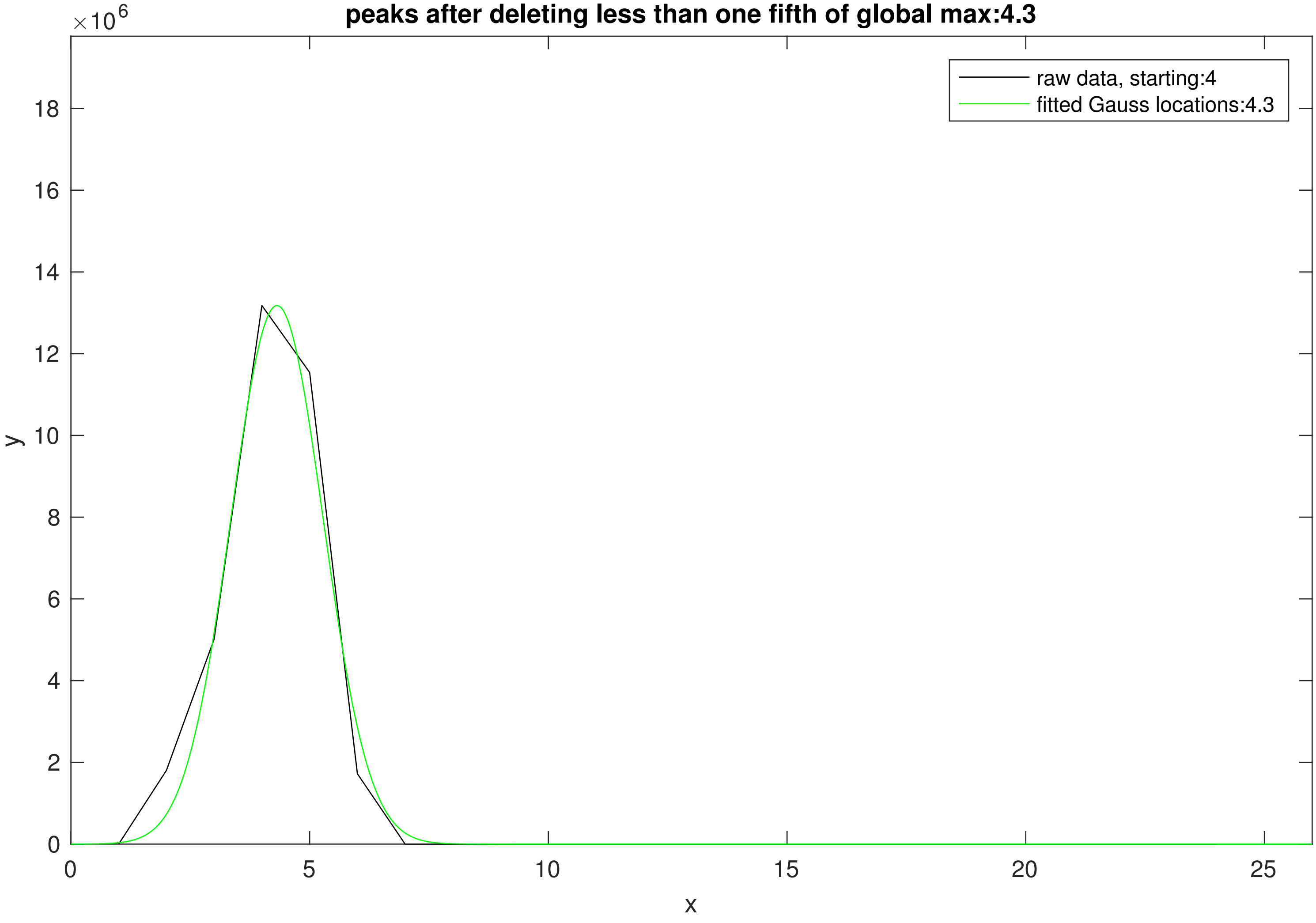

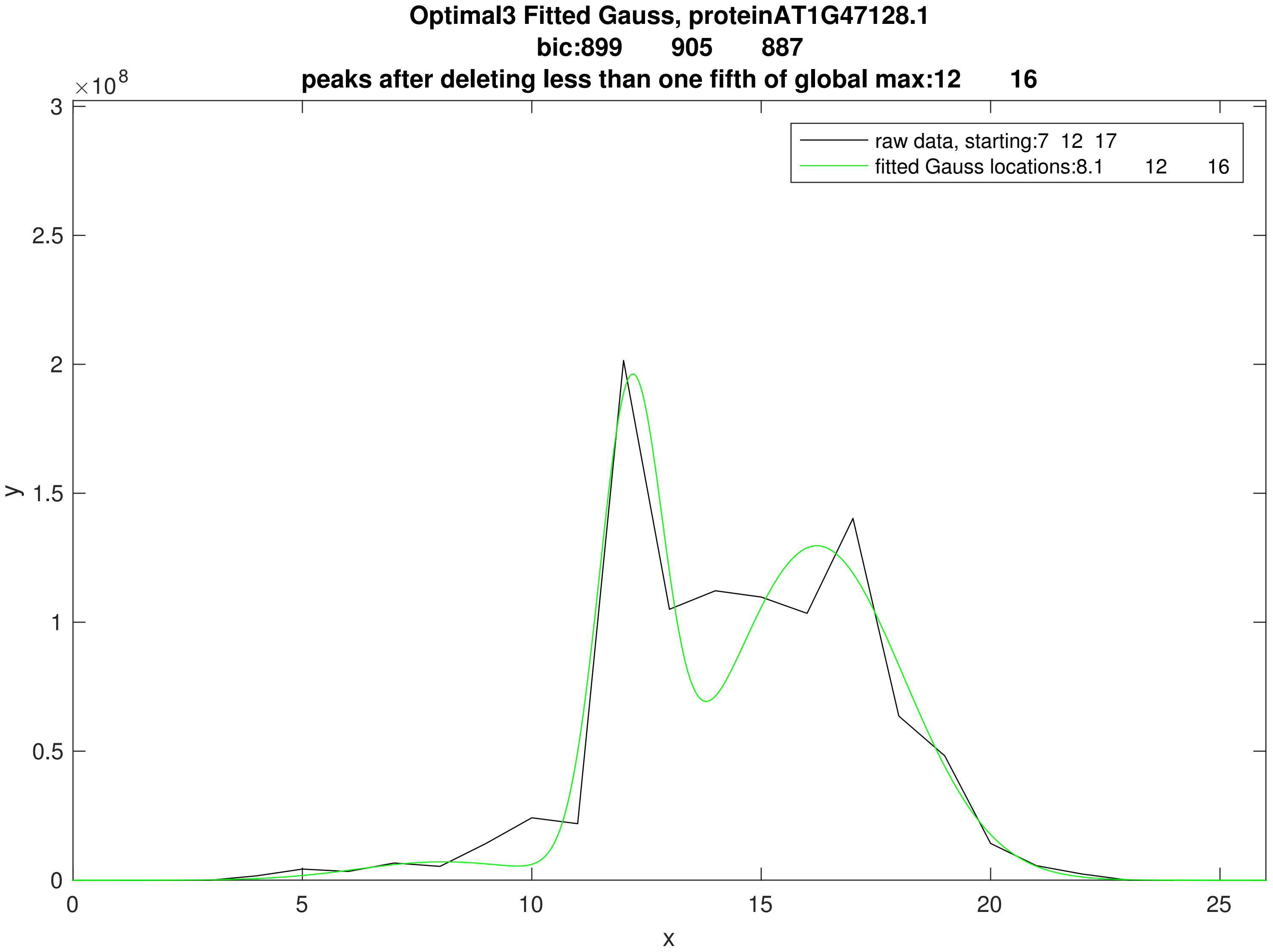
Optimal3 Fitted Gauss, proteinAT1G47128.1
bic:899 905 887
peaks after deleting less than one fifth of global max:12 16
×10^8
3
2.5
2
1.5
1
0.5
0
y
0
5
10
15
20
25
x
raw data, starting:7 12 17
fitted Gauss locations:8.1 12 16

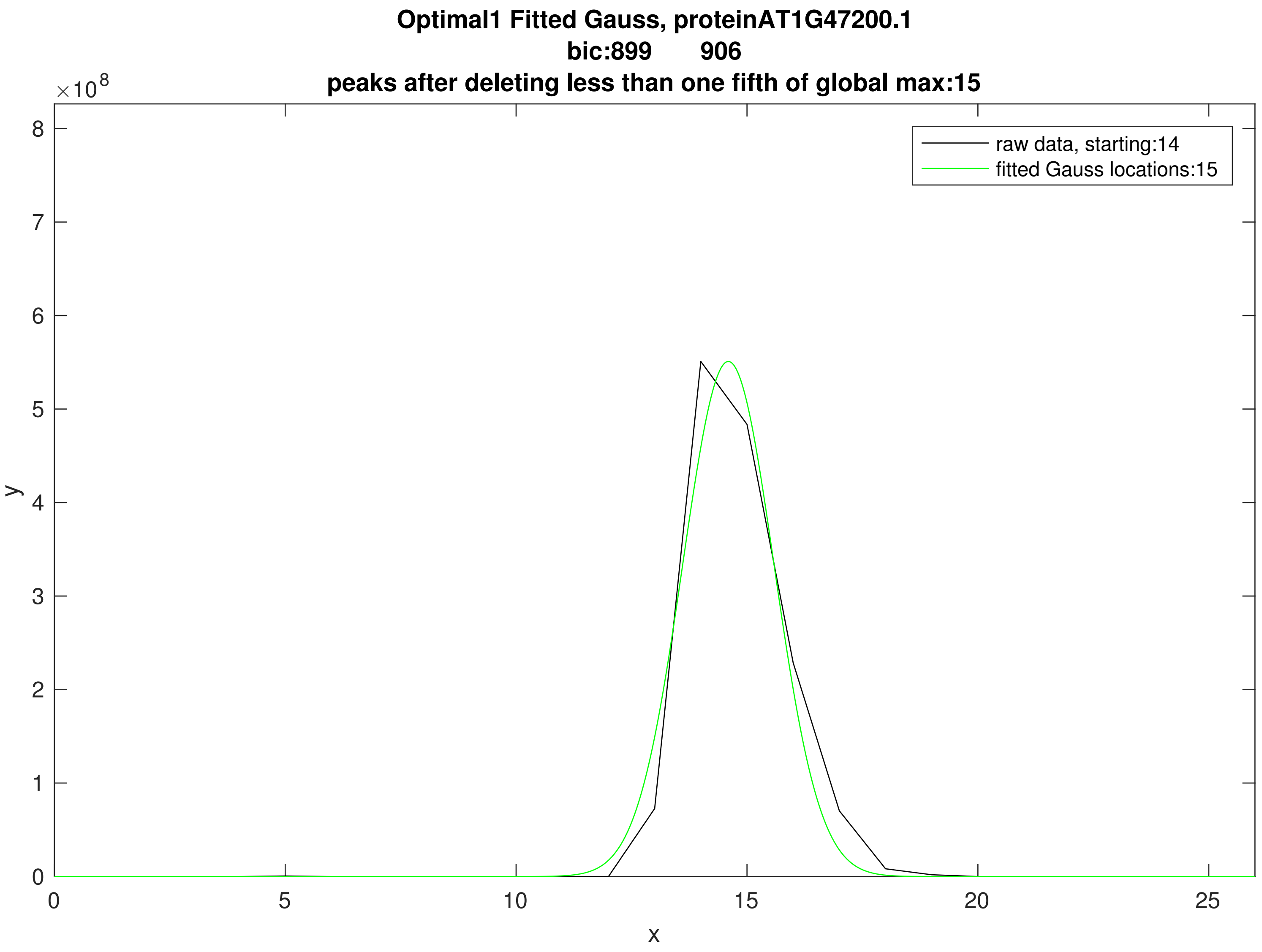

Optimal1 Fitted Gauss, proteinAT1G47200.1
bic:899 906
peaks after deleting less than one fifth of global max:15
×10^8
raw data, starting:14
fitted Gauss locations:15
y
x
0
1
2
3
4
5
6
7
8
0
5
10
15
20
25

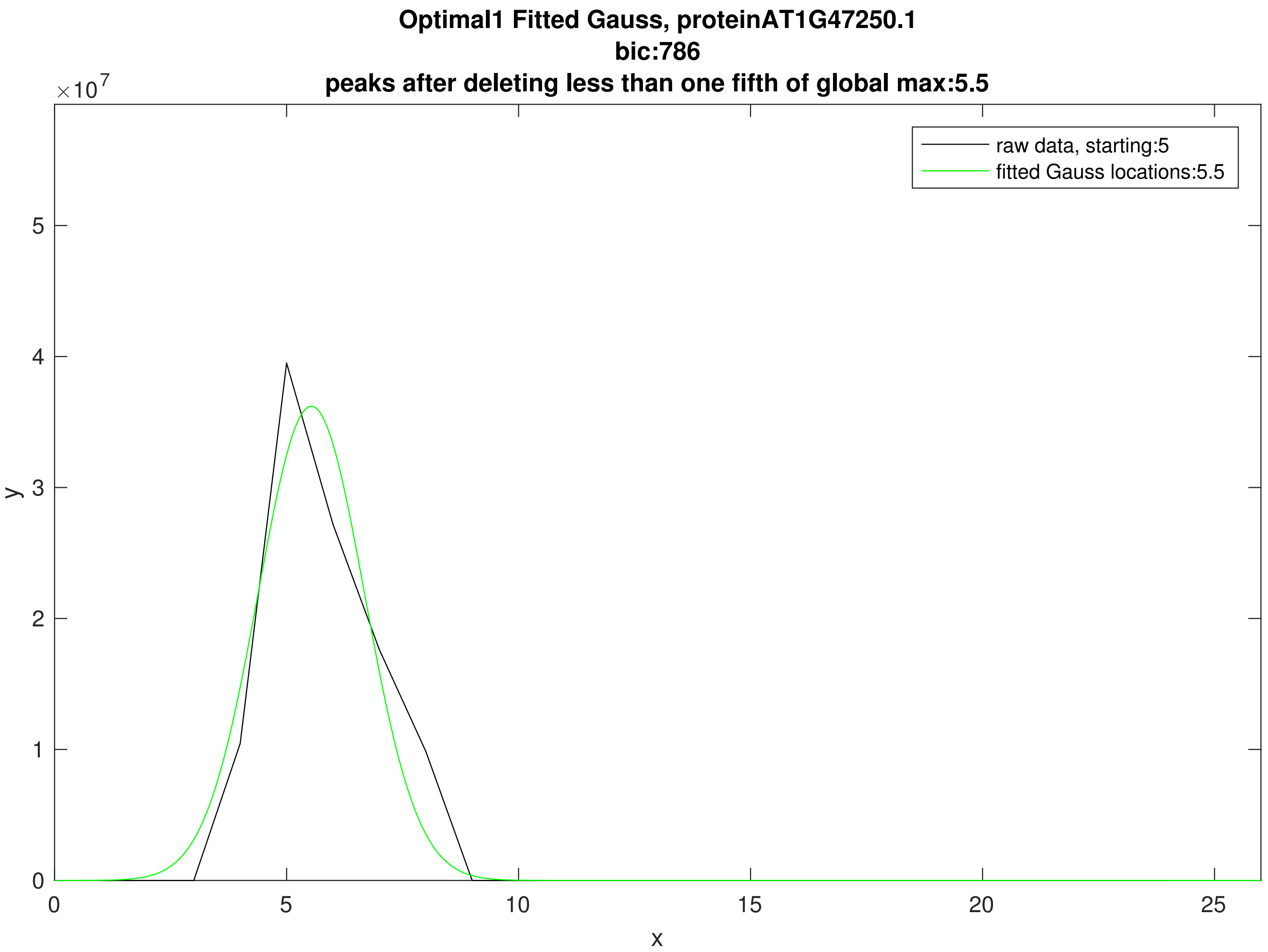
Optimal1 Fitted Gauss, proteinAT1G47250.1
bic:786
peaks after deleting less than one fifth of global max:5.5
×10^7
raw data, starting:5
fitted Gauss locations:5.5
y
x
0
1
2
3
4
5
0
5
10
15
20
25

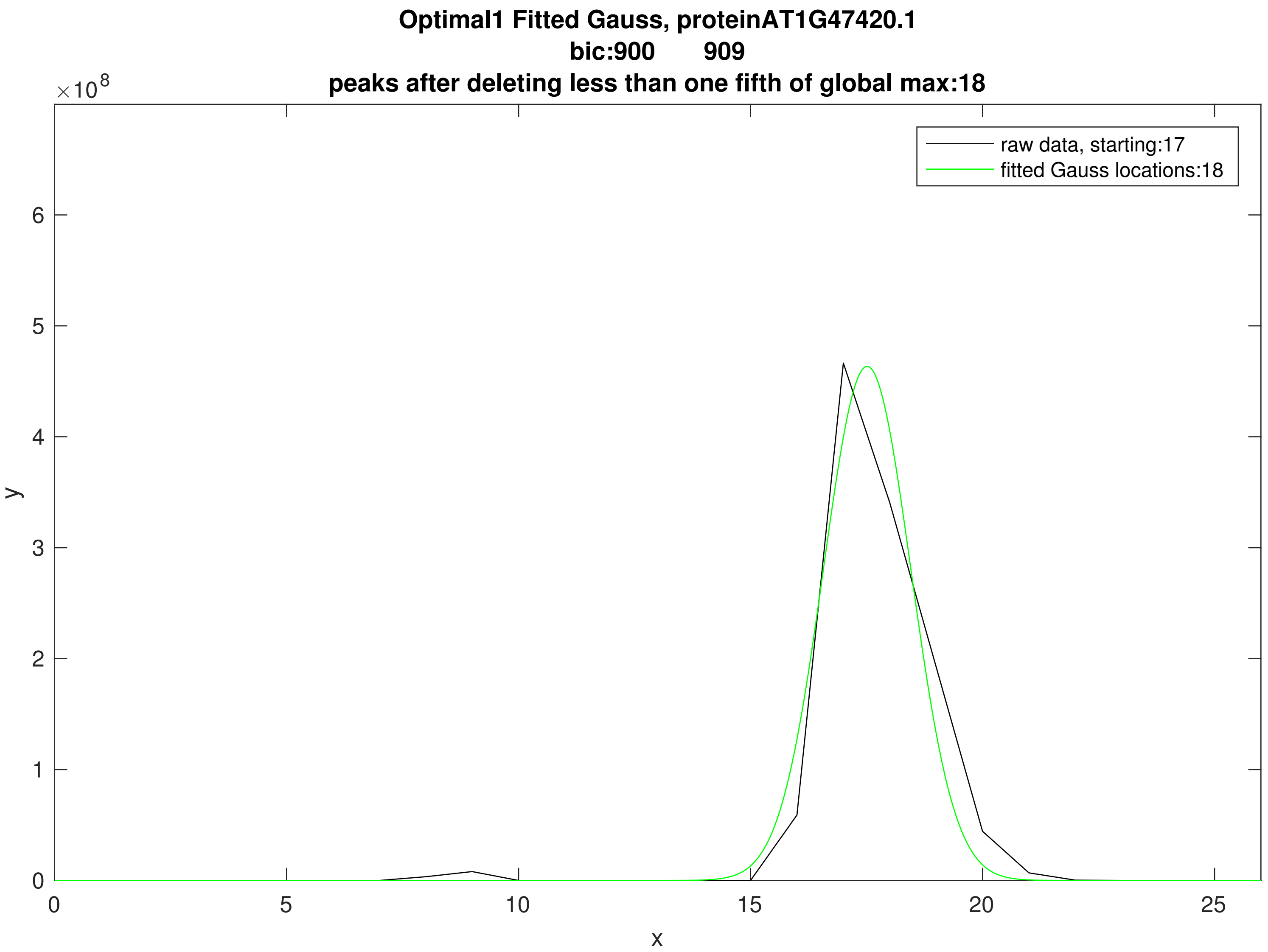
Optimal1 Fitted Gauss, proteinAT1G47420.1
bic:900 909
peaks after deleting less than one fifth of global max:18
×10^8
raw data, starting:17
fitted Gauss locations:18
y
x
0
1
2
3
4
5
6
0
5
10
15
20
25

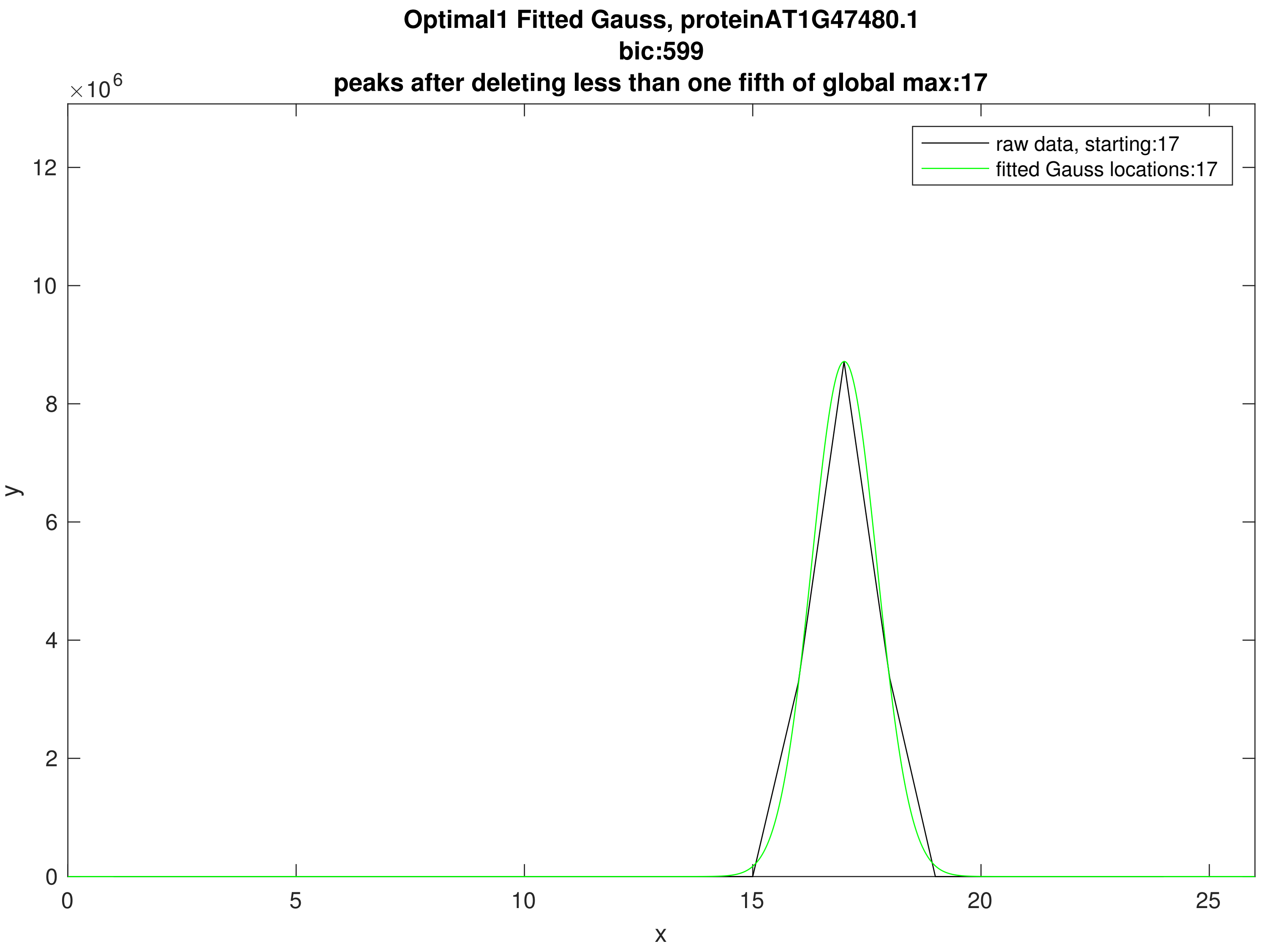
Optimal1 Fitted Gauss, proteinAT1G47480.1
bic:599
peaks after deleting less than one fifth of global max:17
×10^6
raw data, starting:17
fitted Gauss locations:17
12
10
8
6
4
2
0
y
0
5
10
15
20
25
x

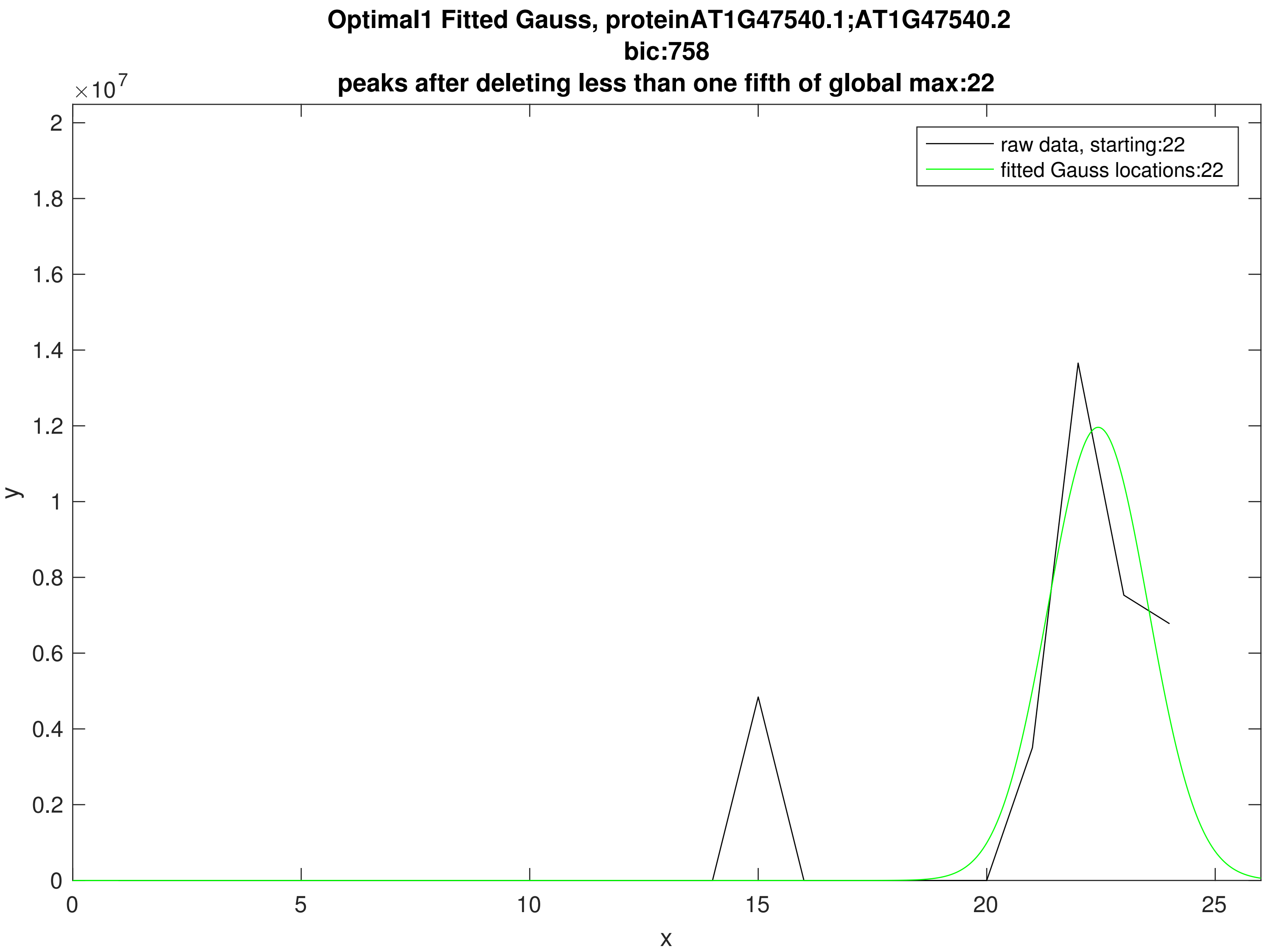
Optimal1 Fitted Gauss, proteinAT1G47540.1;AT1G47540.2
bic:758
peaks after deleting less than one fifth of global max:22
×10^7
raw data, starting:22
fitted Gauss locations:22
y
x
0
0.2
0.4
0.6
0.8
1
1.2
1.4
1.6
1.8
2
0
5
10
15
20
25

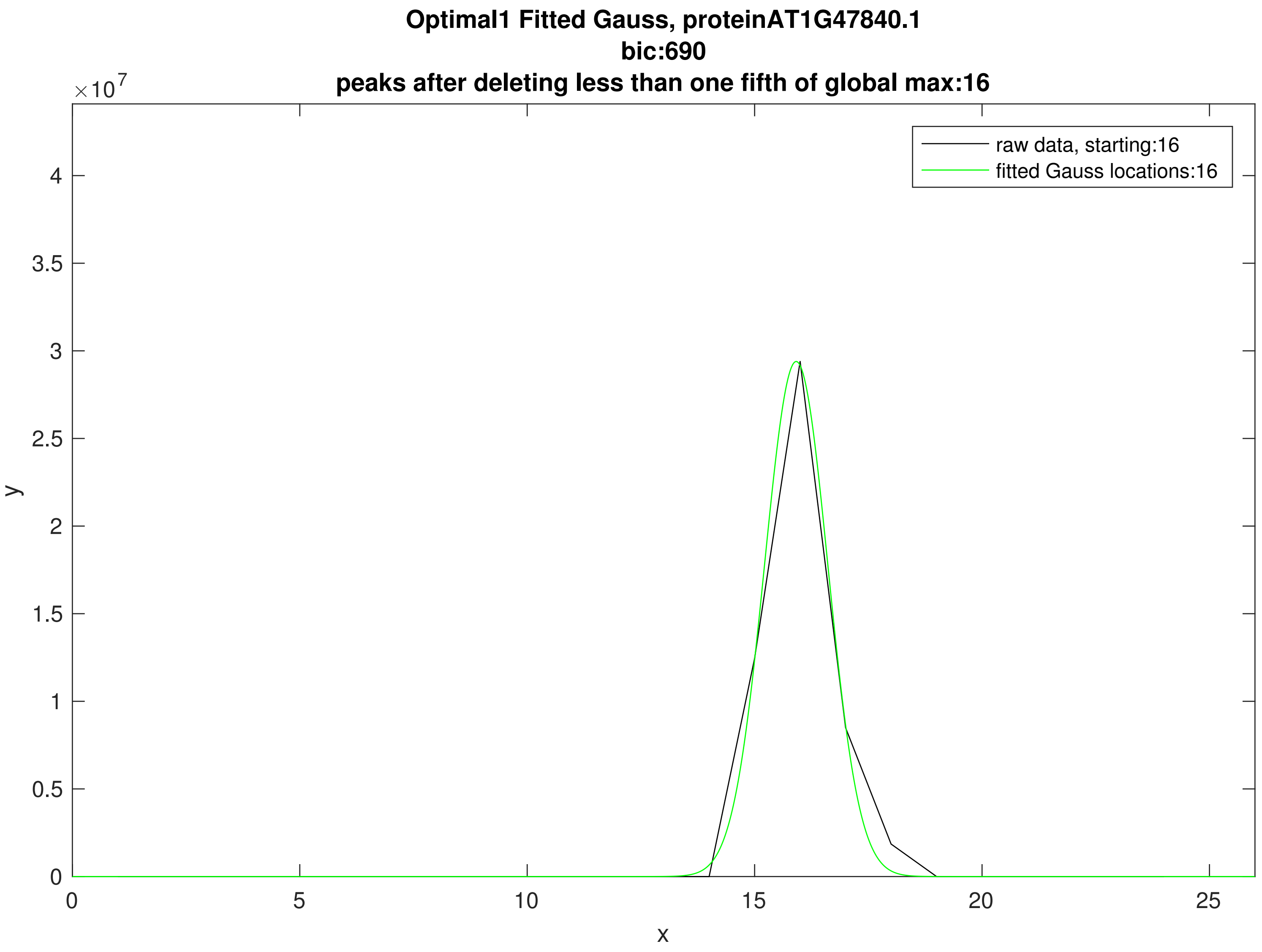
Optimal1 Fitted Gauss, proteinAT1G47840.1
bic:690
peaks after deleting less than one fifth of global max:16
×10^7
raw data, starting:16
fitted Gauss locations:16
y
x
0
0.5
1
1.5
2
2.5
3
3.5
4
0
5
10
15
20
25

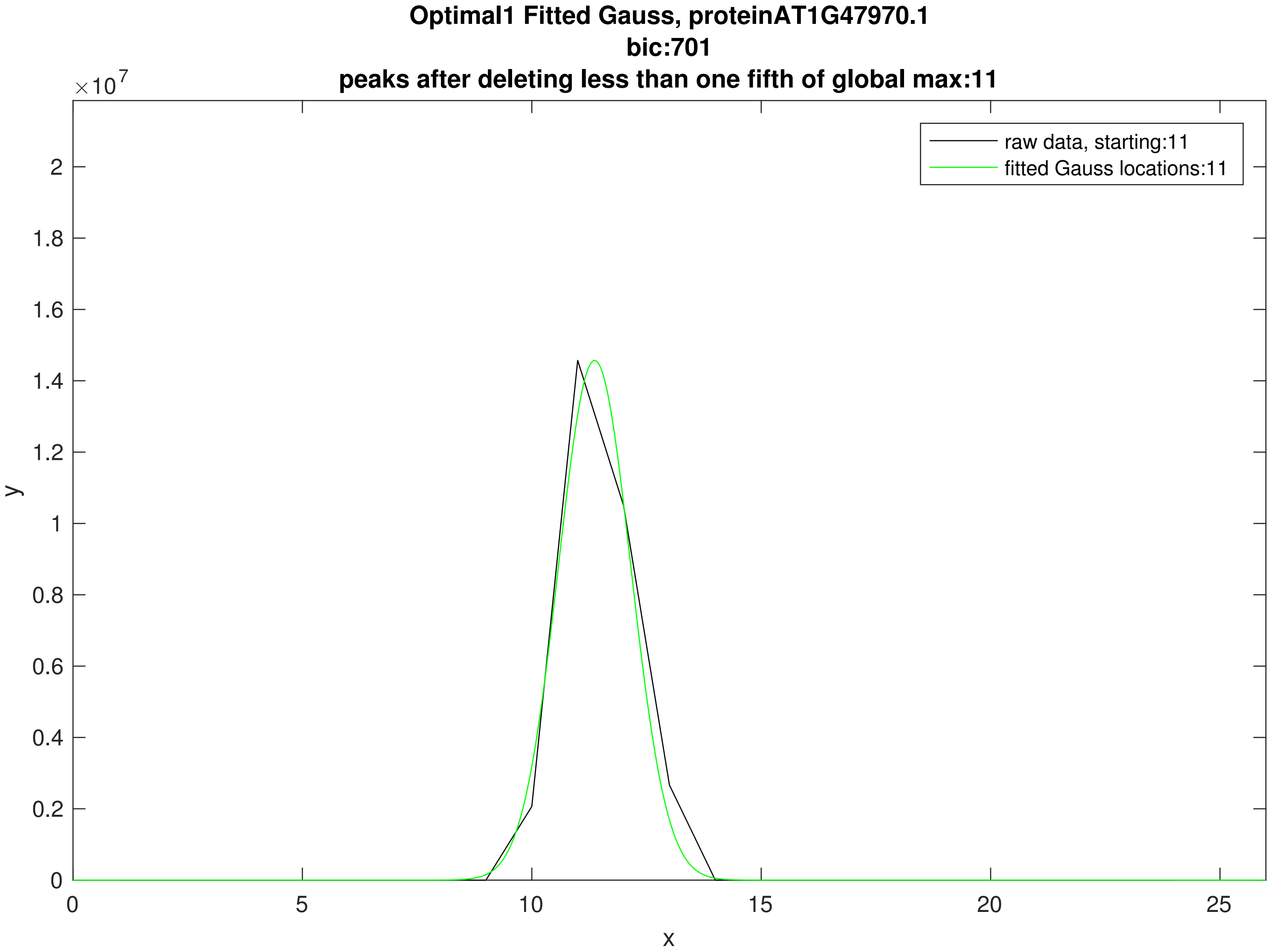
Optimal1 Fitted Gauss, proteinAT1G47970.1
bic:701
peaks after deleting less than one fifth of global max:11
×10^7
raw data, starting:11
fitted Gauss locations:11
2
1.8
1.6
1.4
1.2
1
0.8
0.6
0.4
0.2
0
y
0
5
10
15
20
25
x

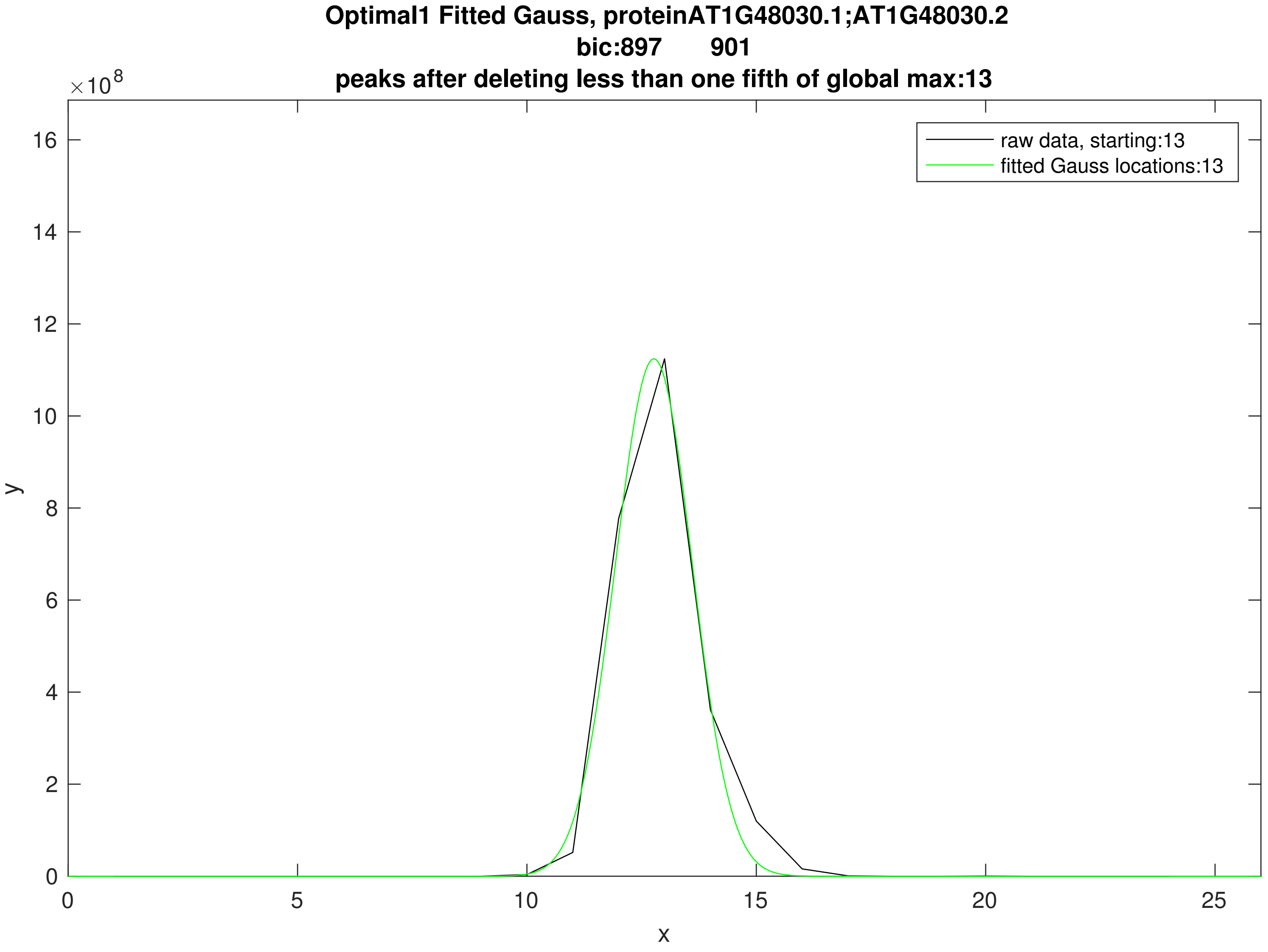

Optimal1 Fitted Gauss, proteinAT1G48030.1;AT1G48030.2
bic:897 901
peaks after deleting less than one fifth of global max:13
×10^8
16
14
12
10
8
6
4
2
0
y
0
5
10
15
20
25
x
raw data, starting:13
fitted Gauss locations:13

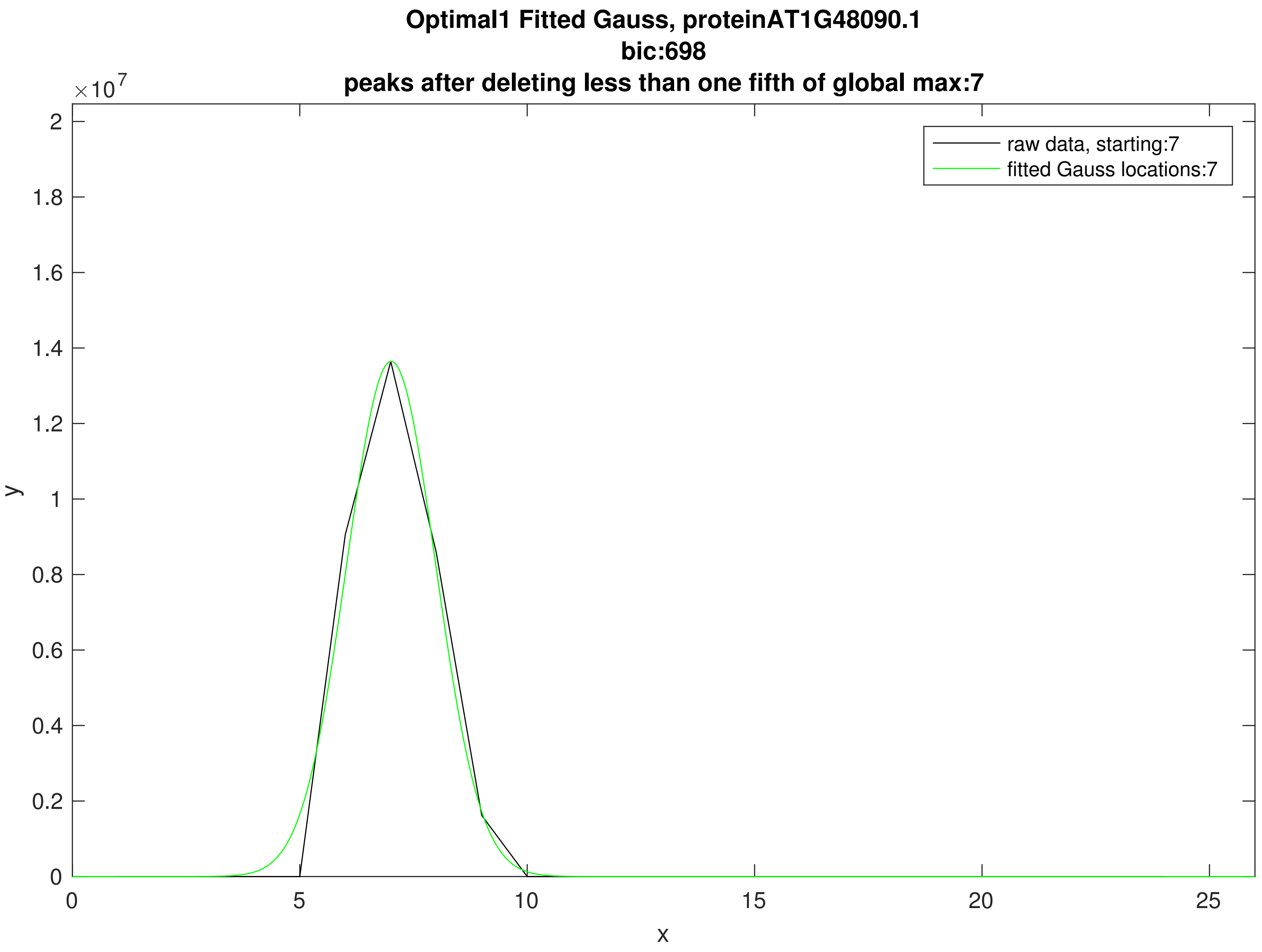
Optimal1 Fitted Gauss, proteinAT1G48090.1
bic:698
peaks after deleting less than one fifth of global max:7
×10^7
2
1.8
1.6
1.4
1.2
1
0.8
0.6
0.4
0.2
0
y
0
5
10
15
20
25
x
raw data, starting:7
fitted Gauss locations:7

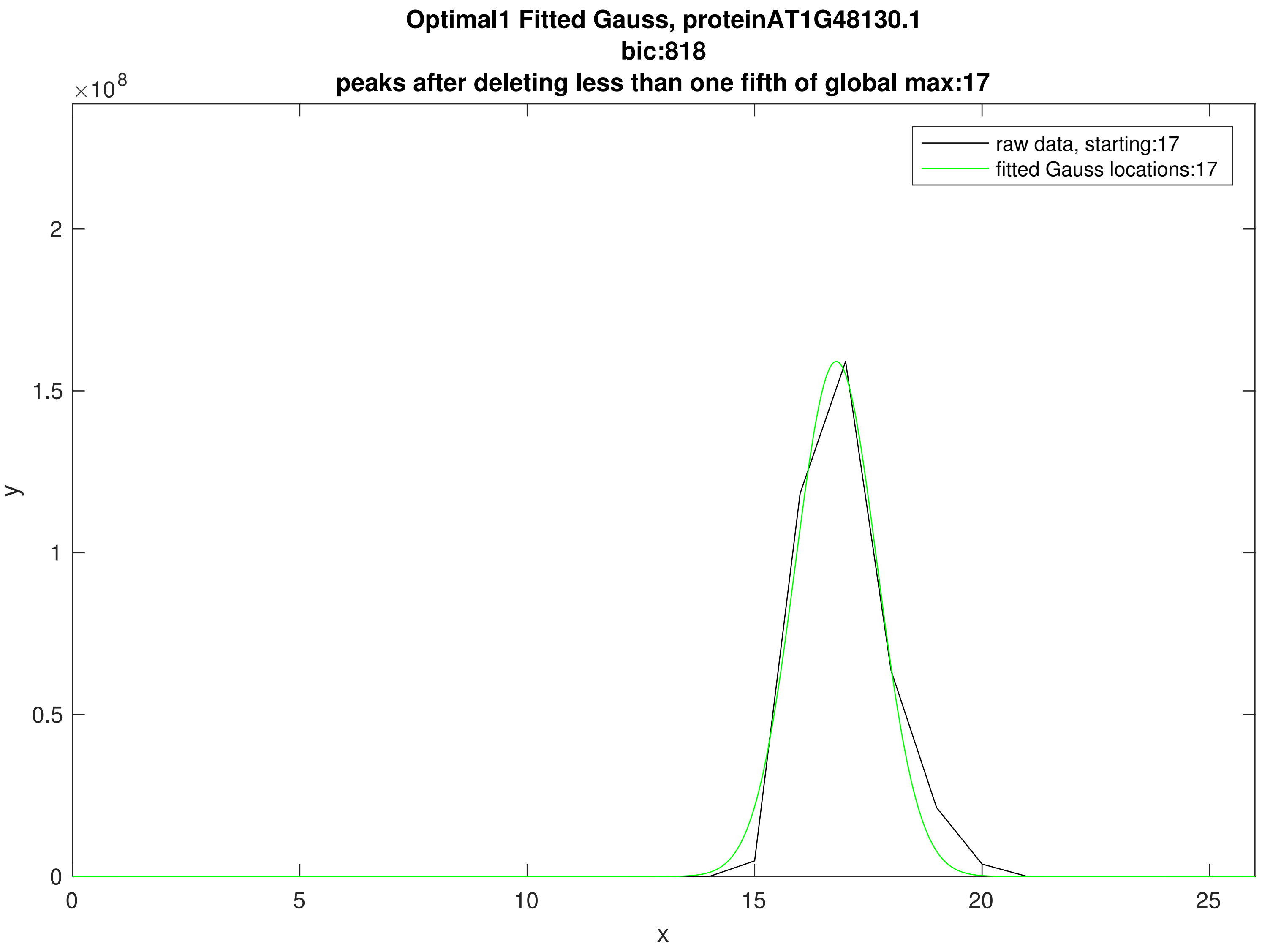
Optimal1 Fitted Gauss, proteinAT1G48130.1
bic:818
peaks after deleting less than one fifth of global max:17
×10^8
raw data, starting:17
fitted Gauss locations:17
y
x
0
0.5
1
1.5
2
0
5
10
15
20
25

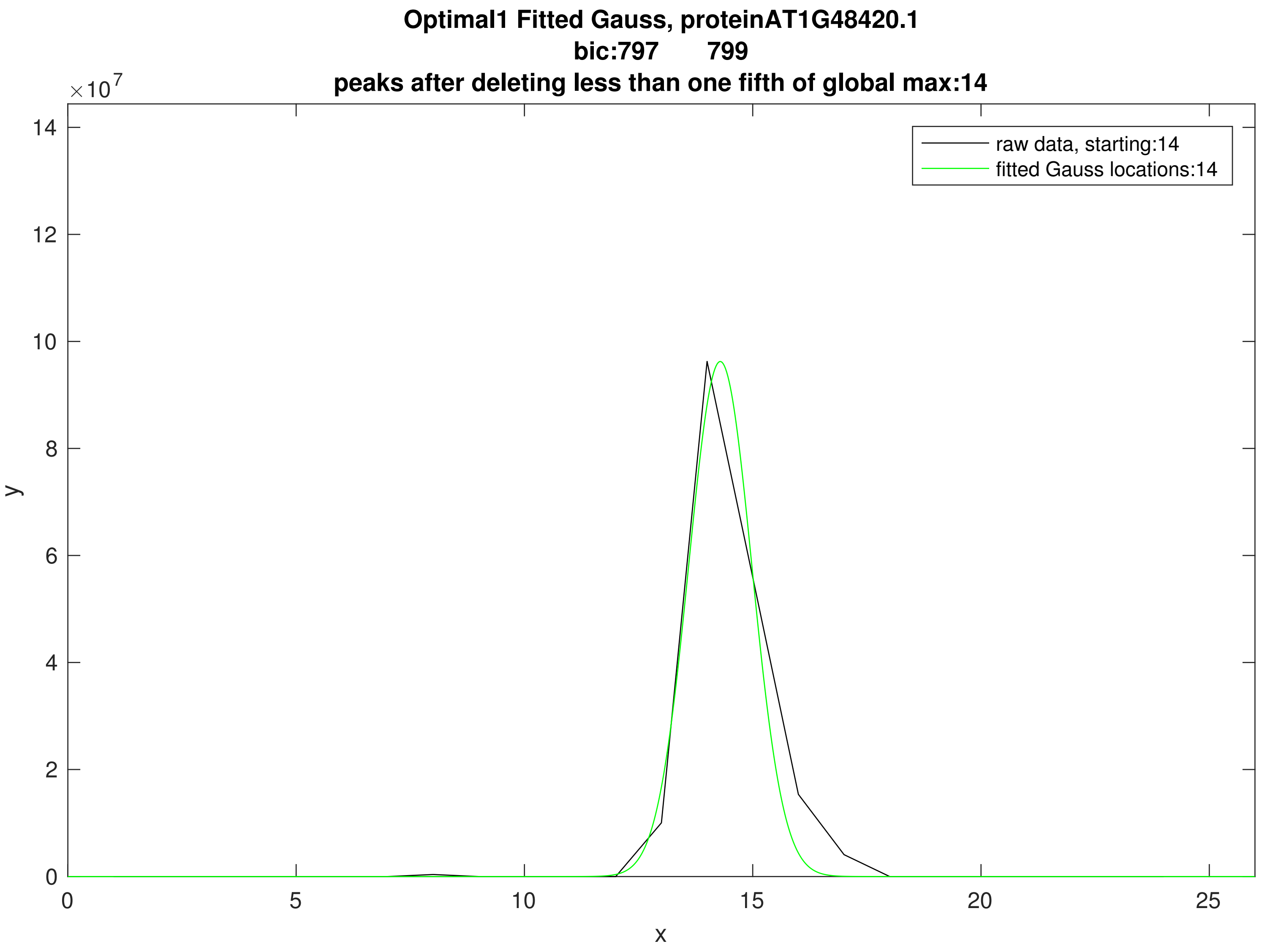
Optimal1 Fitted Gauss, proteinAT1G48420.1
bic:797 799
peaks after deleting less than one fifth of global max:14
×10^7
raw data, starting:14
fitted Gauss locations:14
y
x
0
2
4
6
8
10
12
14
0
5
10
15
20
25

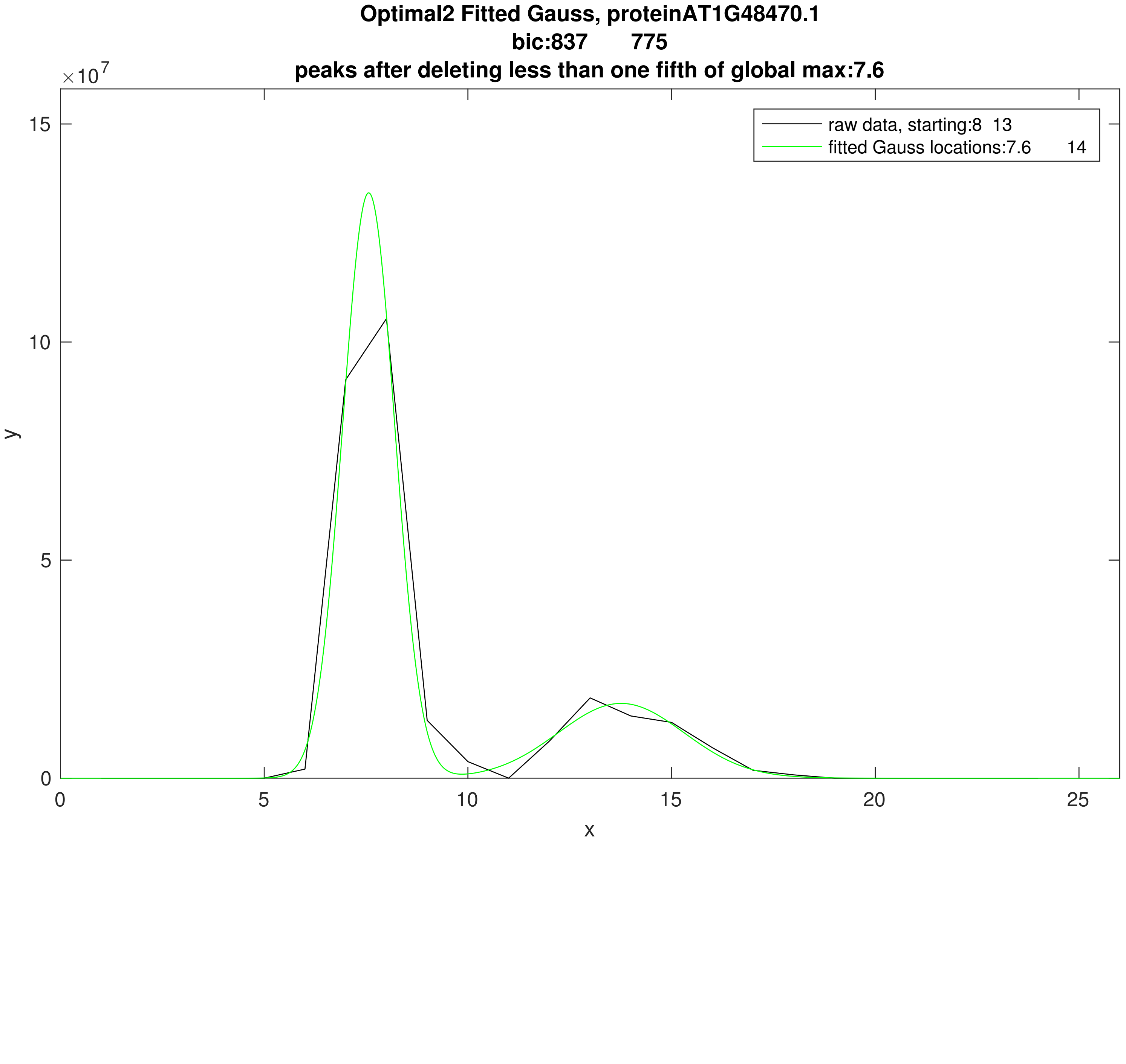

Optimal2 Fitted Gauss, proteinAT1G48470.1
bic:837 775
peaks after deleting less than one fifth of global max:7.6
×10^7
15
10
5
0
y
0
5
10
15
20
25
x
raw data, starting:8 13
fitted Gauss locations:7.6 14

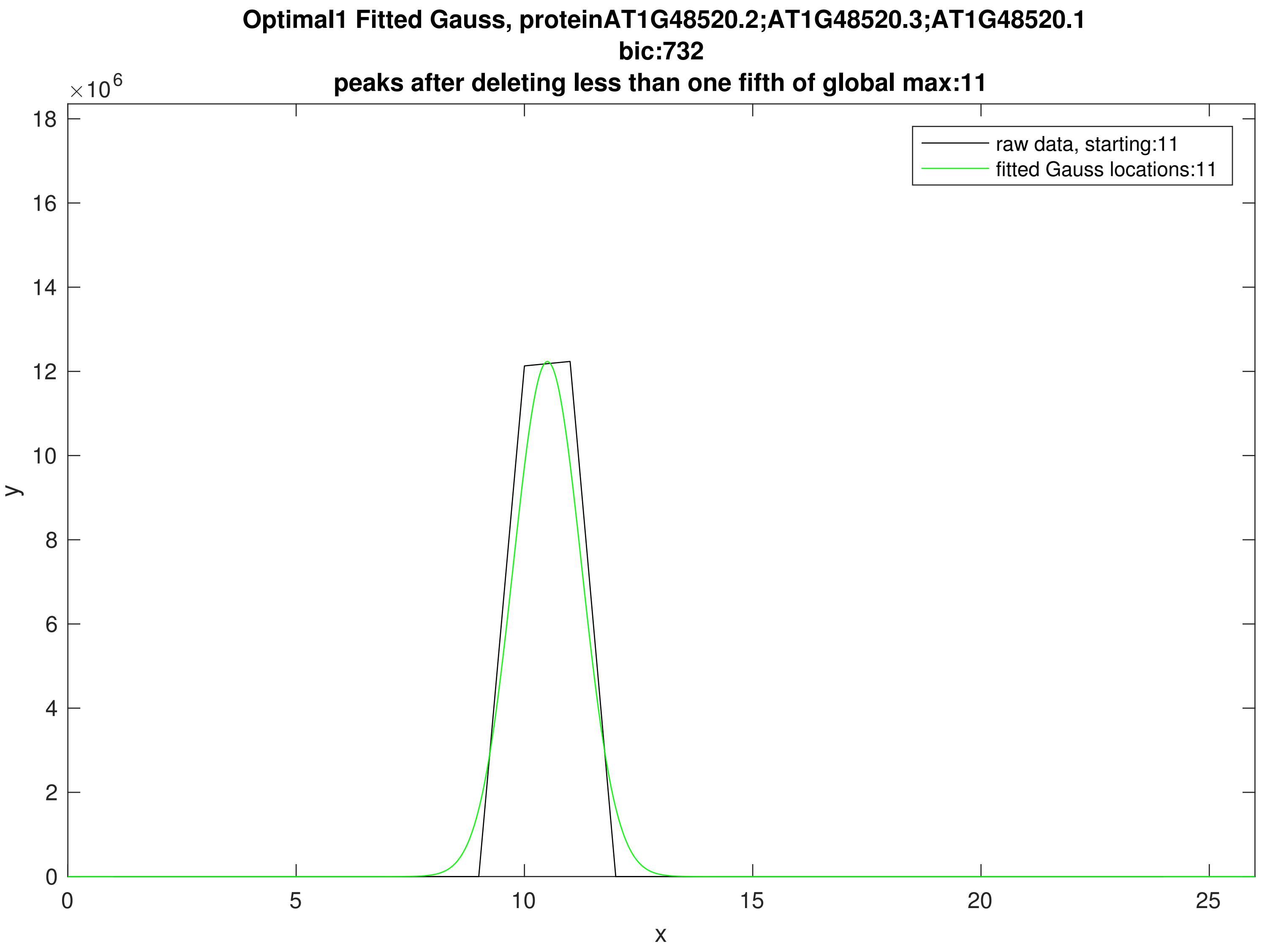
Optimal1 Fitted Gauss, proteinAT1G48520.2;AT1G48520.3;AT1G48520.1
bic:732
peaks after deleting less than one fifth of global max:11
×10^6
raw data, starting:11
fitted Gauss locations:11
y
x
0
2
4
6
8
10
12
14
16
18
0
5
10
15
20
25

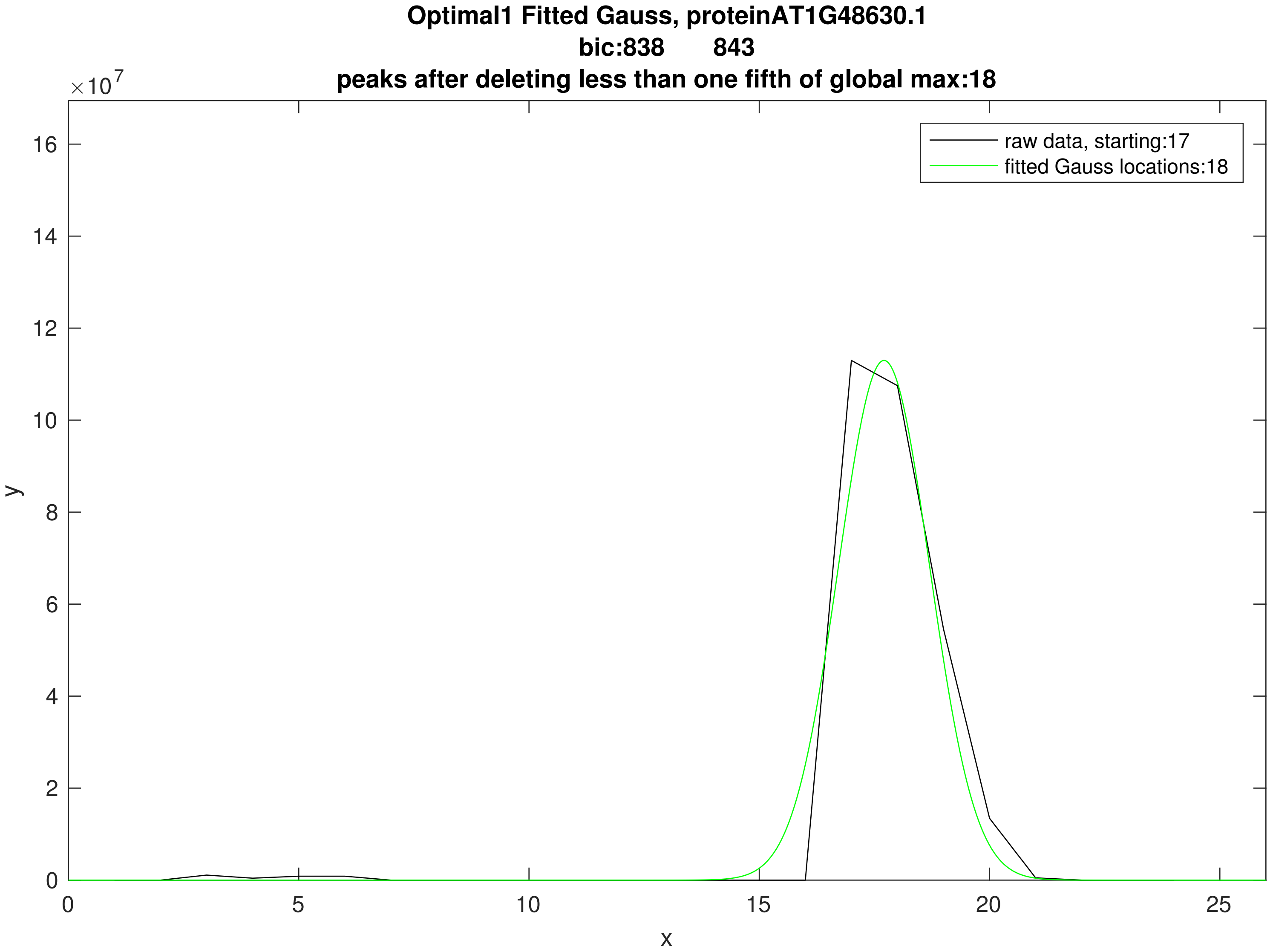
Optimal1 Fitted Gauss, proteinAT1G48630.1
bic:838 843
peaks after deleting less than one fifth of global max:18
×10^7
raw data, starting:17
fitted Gauss locations:18
y
x
0
2
4
6
8
10
12
14
16
0
5
10
15
20
25

Optimal1 Fitted Gauss, proteinAT1G48660.1
bic:740 744
peaks after deleting less than one fifth of global max:15

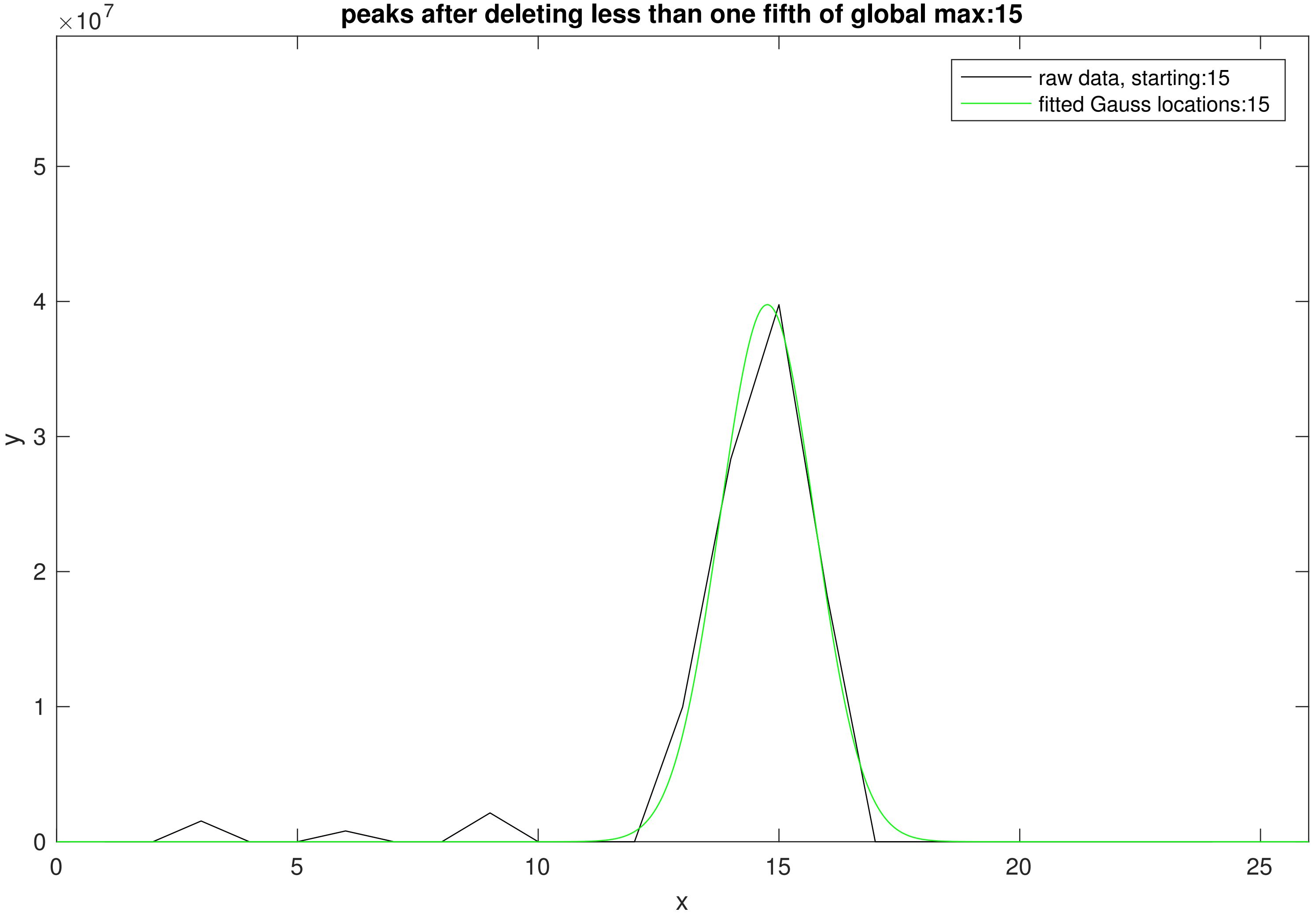

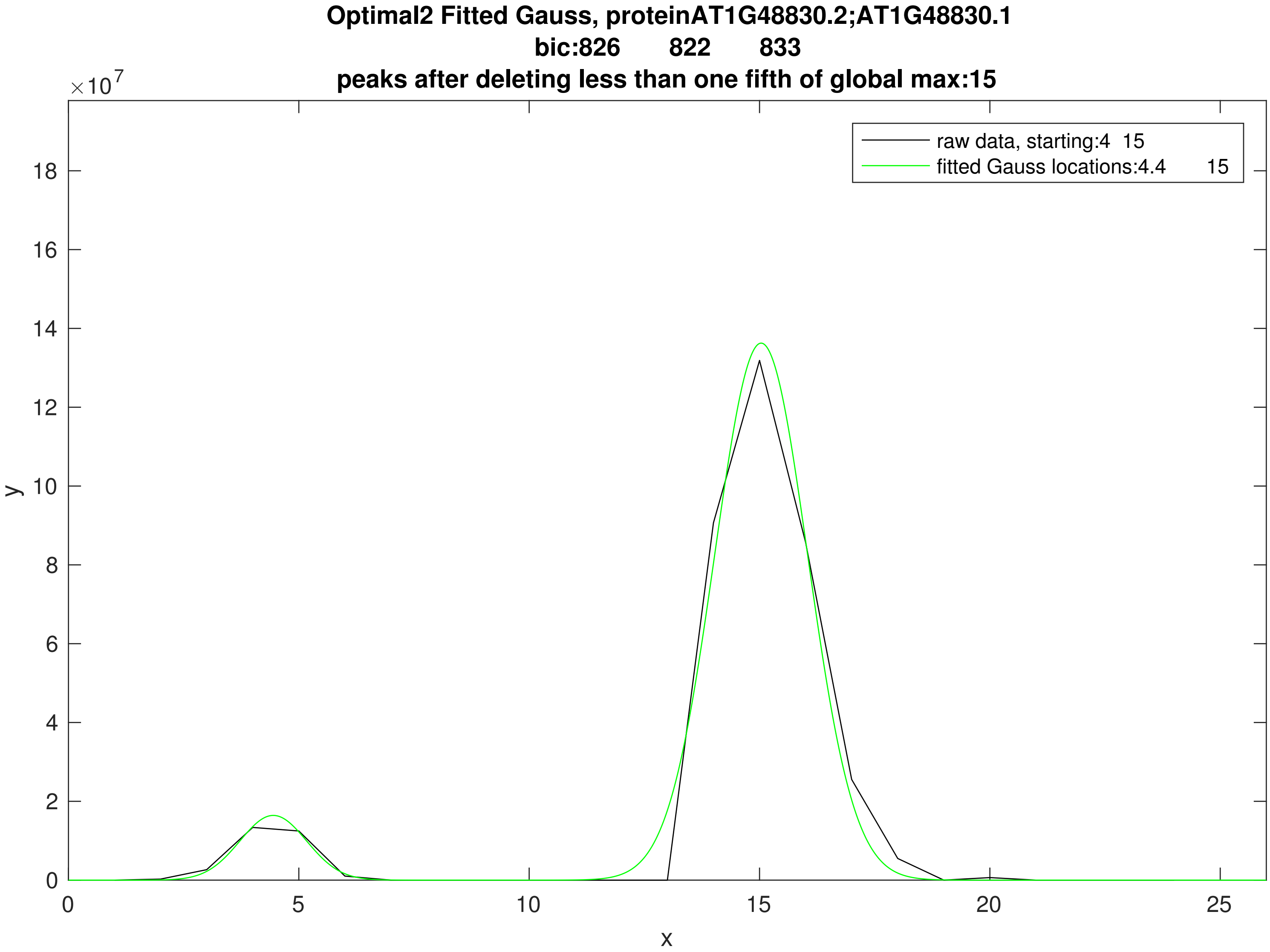
Optimal2 Fitted Gauss, proteinAT1G48830.2;AT1G48830.1
bic:826 822 833
peaks after deleting less than one fifth of global max:15
×10^7
raw data, starting:4 15
fitted Gauss locations:4.4 15
y
x
0
2
4
6
8
10
12
14
16
18
0
5
10
15
20
25

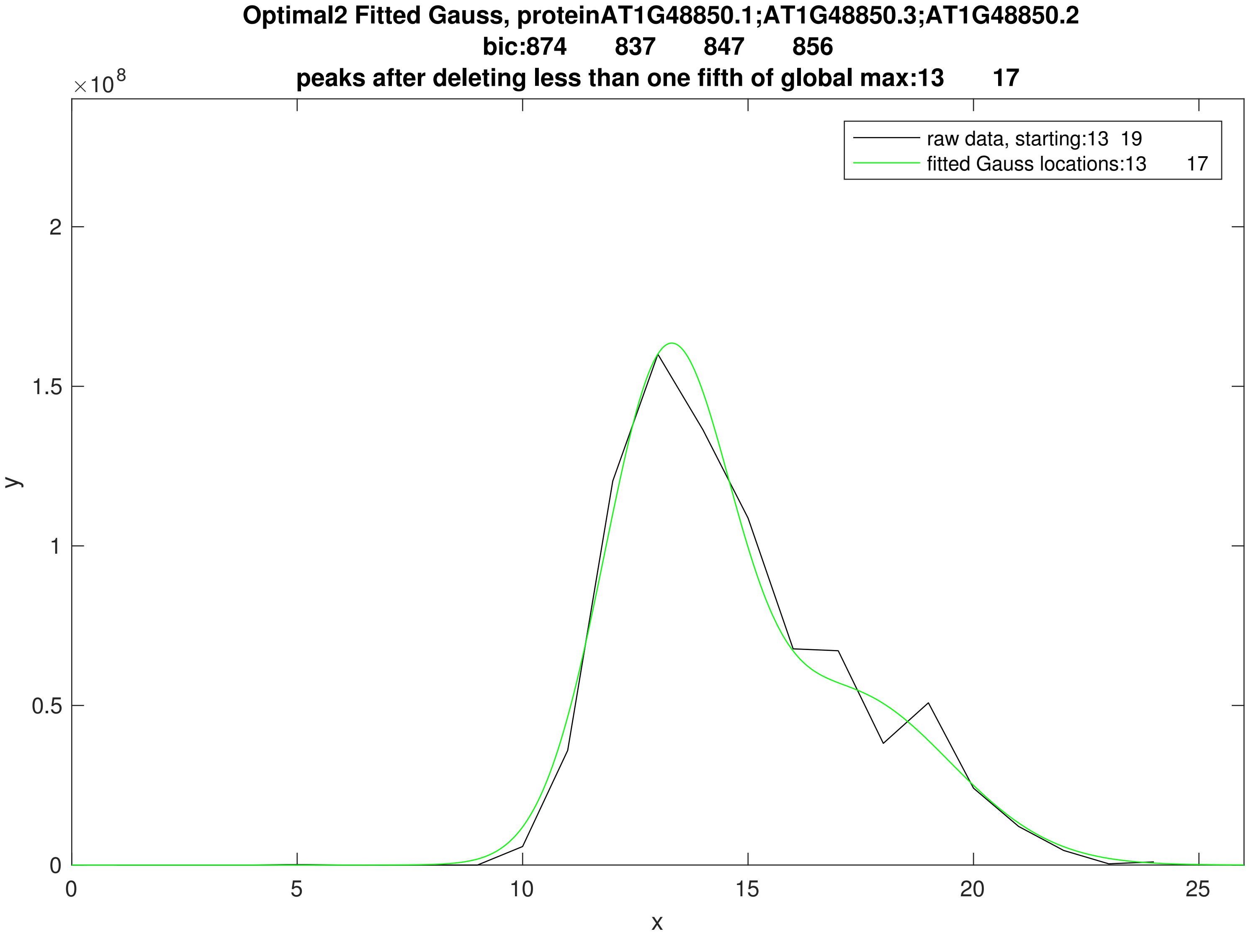

Optimal2 Fitted Gauss, proteinAT1G48850.1;AT1G48850.3;AT1G48850.2
bic:874 837 847 856
peaks after deleting less than one fifth of global max:13 17
raw data, starting:13 19
fitted Gauss locations:13 17
×10^8
y
x
0
0.5
1
1.5
2
5
10
15
20
25

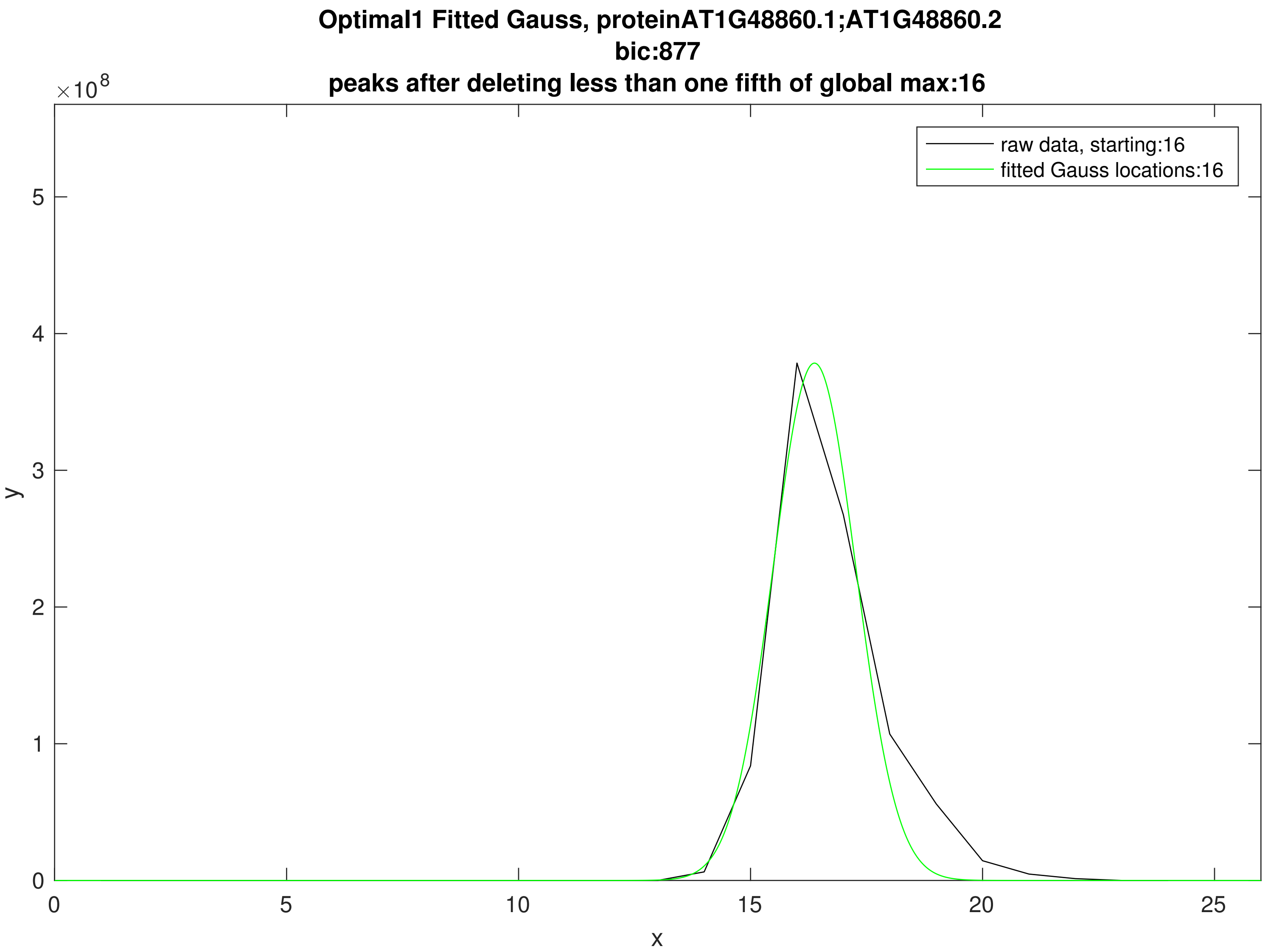

Optimal1 Fitted Gauss, proteinAT1G48860.1;AT1G48860.2
bic:877
peaks after deleting less than one fifth of global max:16
×10^8
raw data, starting:16
fitted Gauss locations:16
y
x
0
1
2
3
4
5
0
5
10
15
20
25

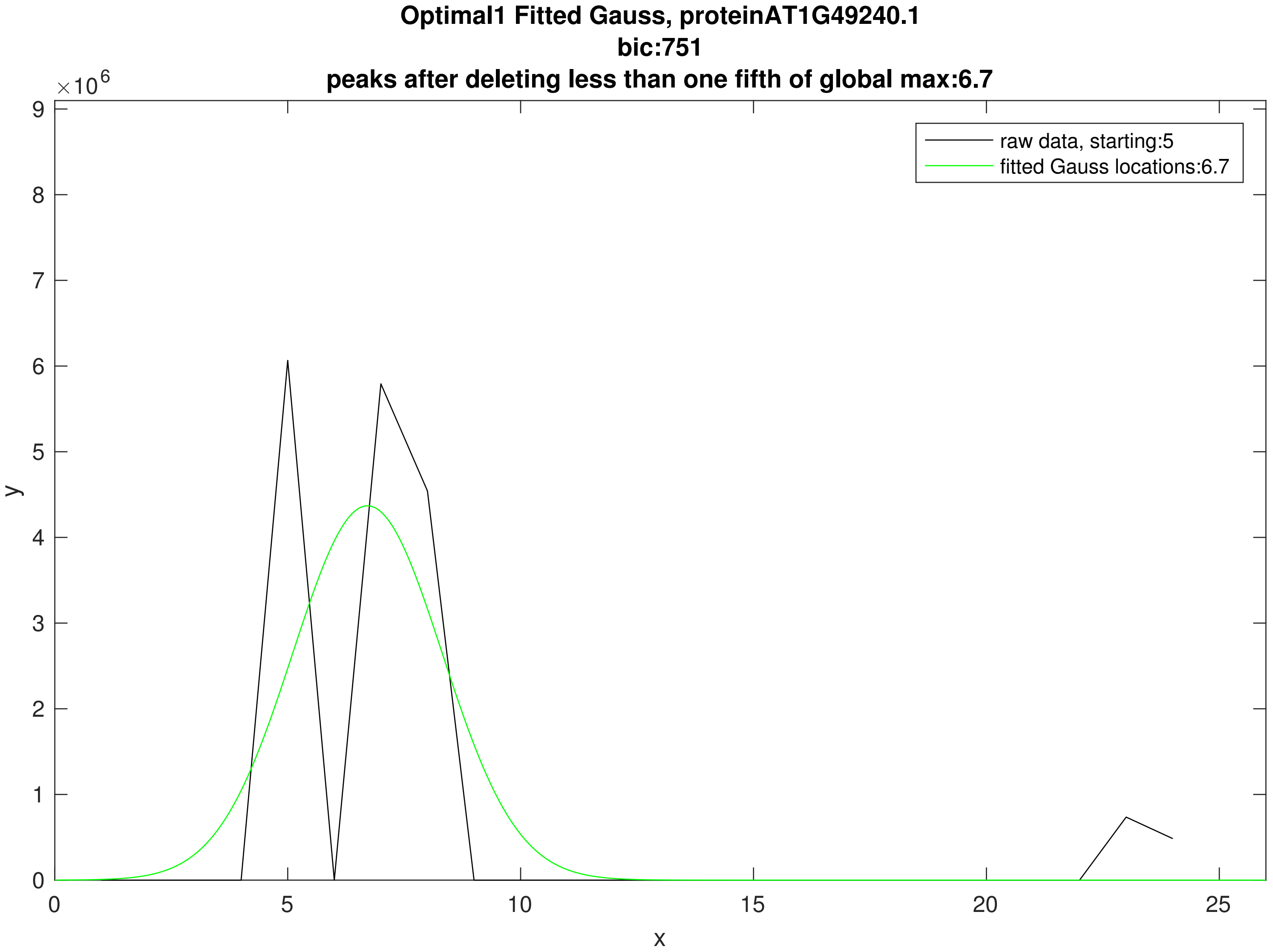
Optimal1 Fitted Gauss, proteinAT1G49240.1
bic:751
peaks after deleting less than one fifth of global max:6.7
×10^6
raw data, starting:5
fitted Gauss locations:6.7
y
x
0
1
2
3
4
5
6
7
8
9
0
5
10
15
20
25

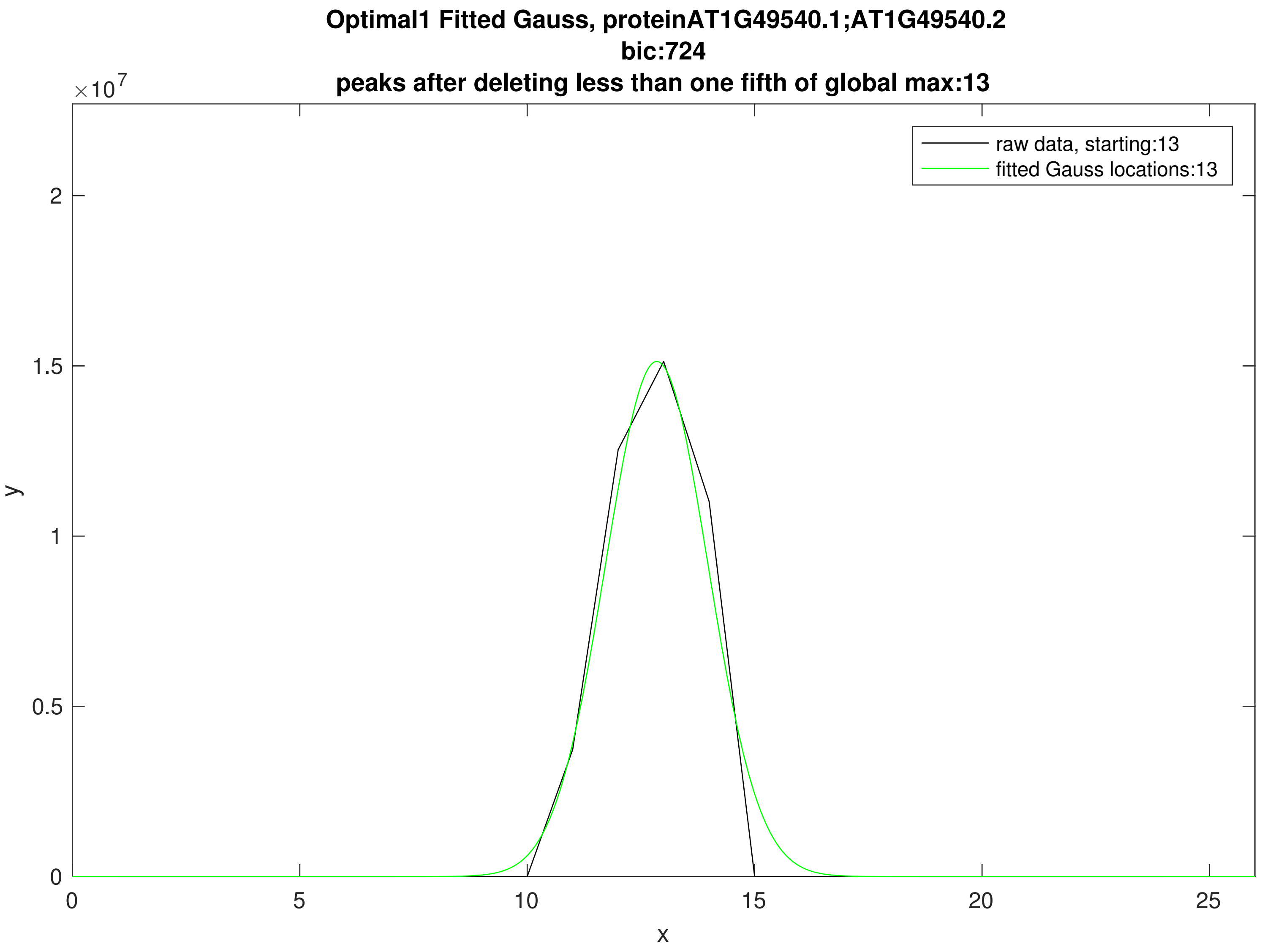
Optimal1 Fitted Gauss, proteinAT1G49540.1;AT1G49540.2
bic:724
peaks after deleting less than one fifth of global max:13
×10^7
raw data, starting:13
fitted Gauss locations:13
y
x
0
0.5
1
1.5
2
0
5
10
15
20
25

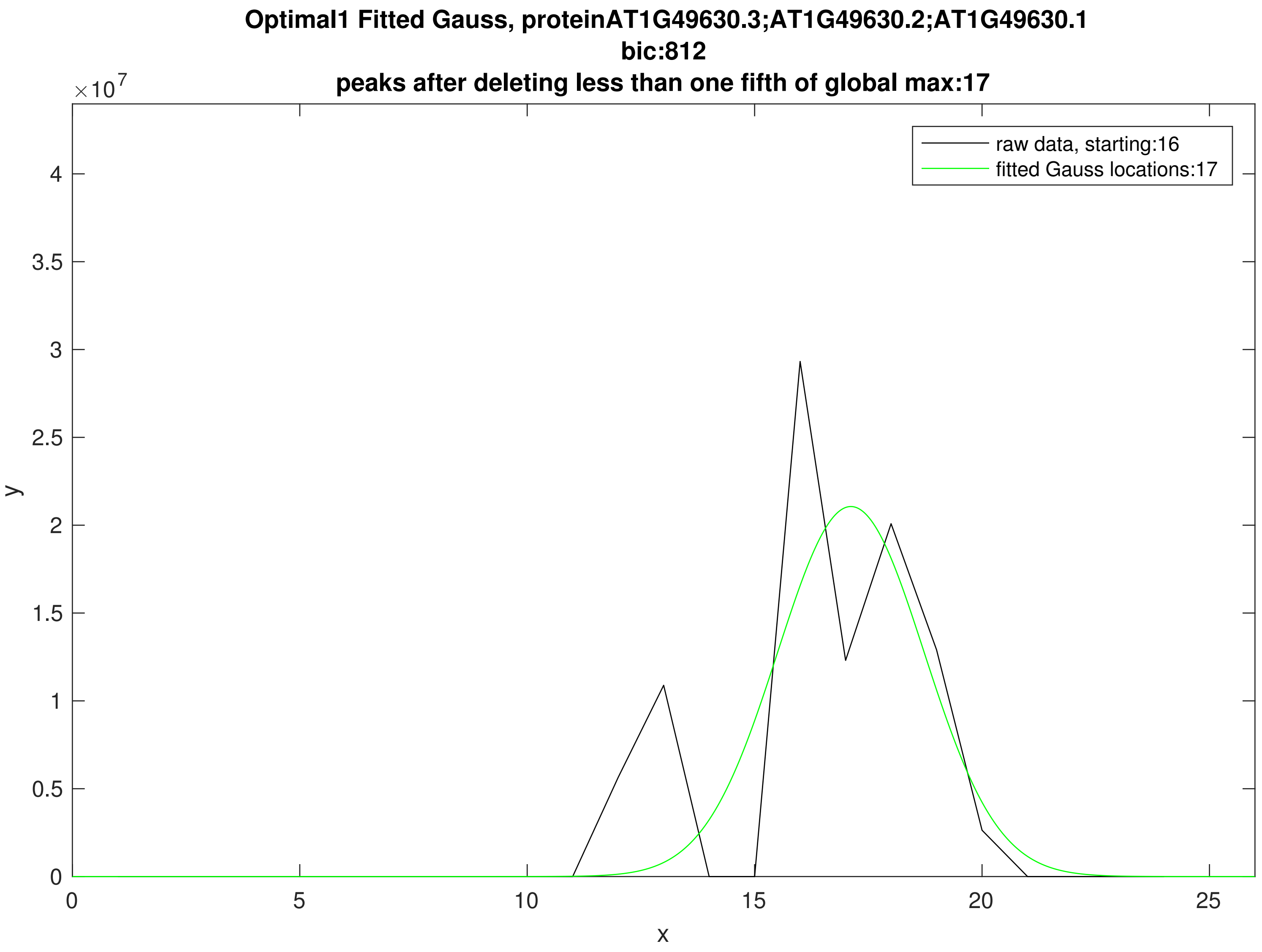
Optimal1 Fitted Gauss, proteinAT1G49630.3;AT1G49630.2;AT1G49630.1
bic:812
peaks after deleting less than one fifth of global max:17
×10^7
raw data, starting:16
fitted Gauss locations:17
y
x
0
0.5
1
1.5
2
2.5
3
3.5
4
0
5
10
15
20
25

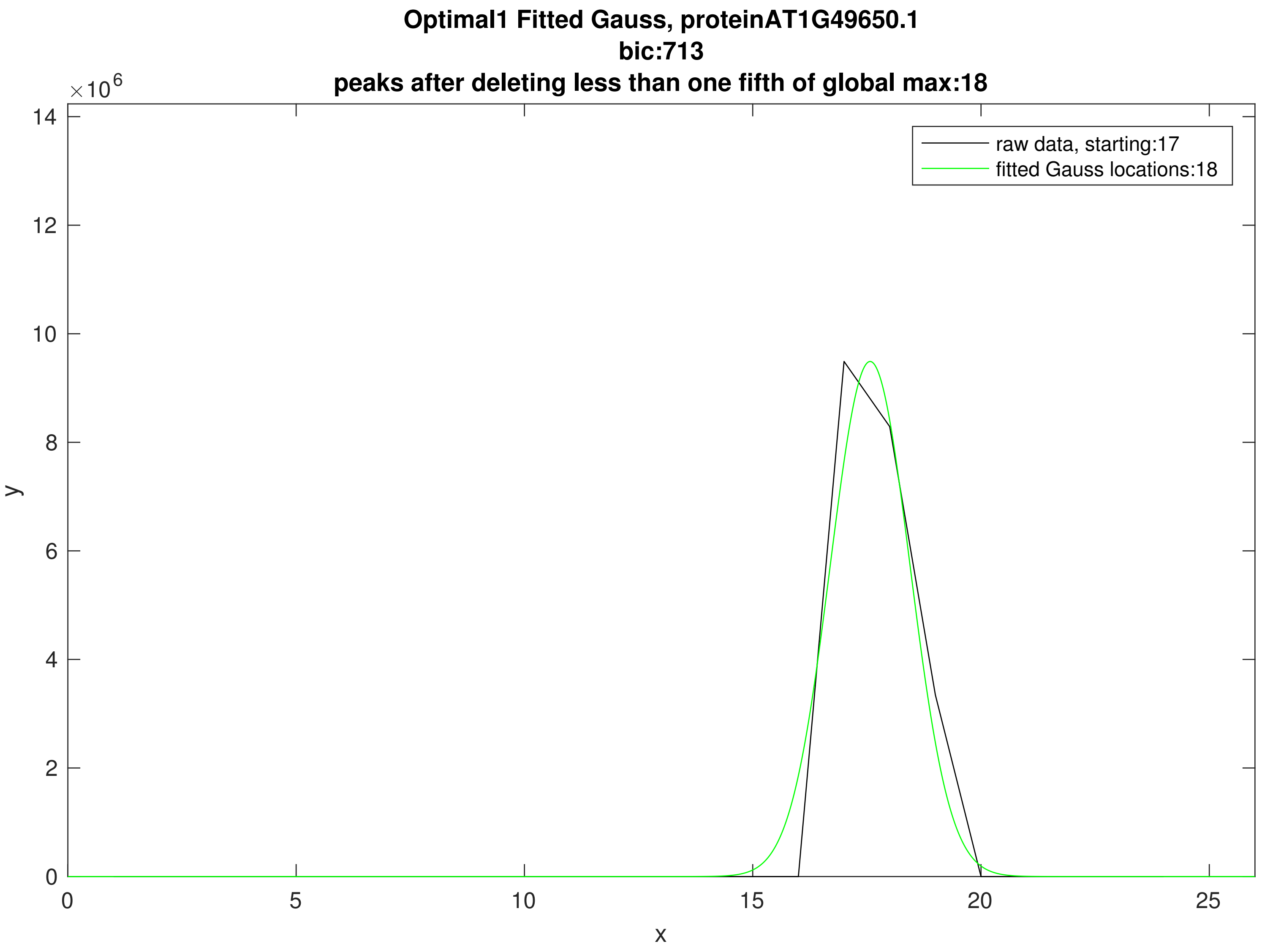
Optimal1 Fitted Gauss, proteinAT1G49650.1
bic:713
peaks after deleting less than one fifth of global max:18
×10^6
raw data, starting:17
fitted Gauss locations:18
y
x
0
2
4
6
8
10
12
14
0
5
10
15
20
25

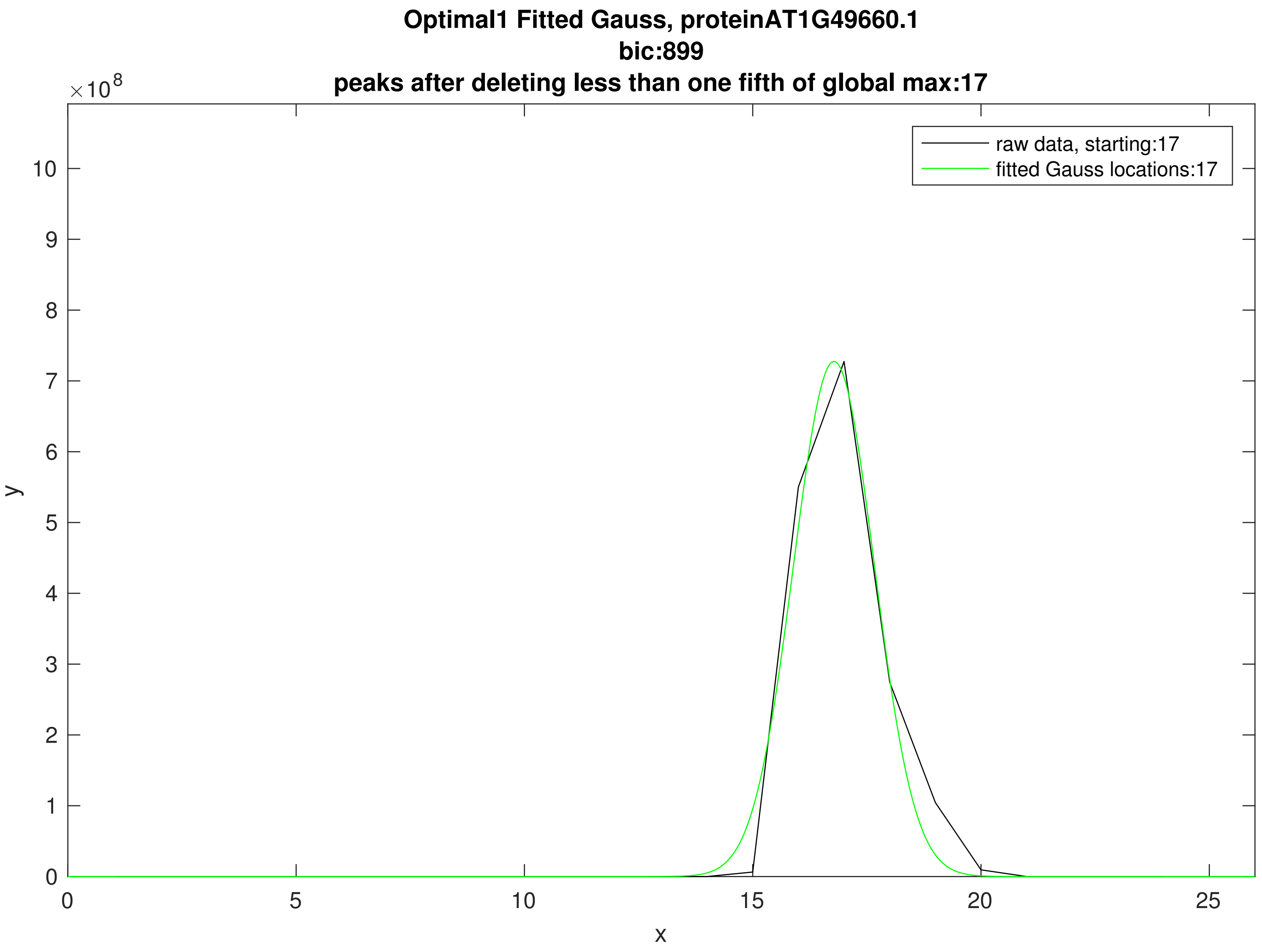

Optimal1 Fitted Gauss, proteinAT1G49660.1
bic:899
peaks after deleting less than one fifth of global max:17
×10^8
raw data, starting:17
fitted Gauss locations:17
y
x

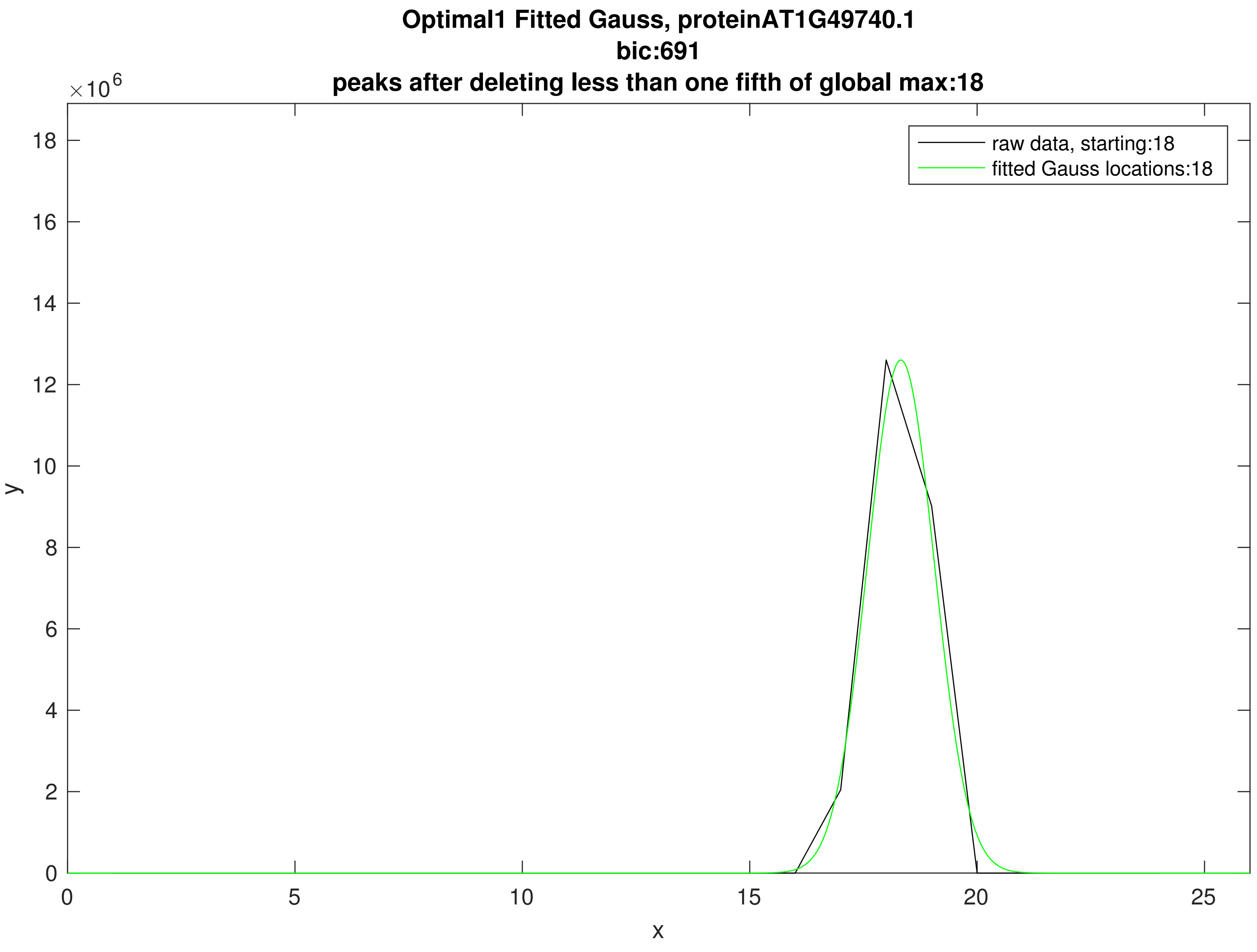

Optimal1 Fitted Gauss, proteinAT1G49740.1
bic:691
peaks after deleting less than one fifth of global max:18
×10^6
raw data, starting:18
fitted Gauss locations:18
18
16
14
12
10
8
6
4
2
0
y
0
5
10
15
20
25
x

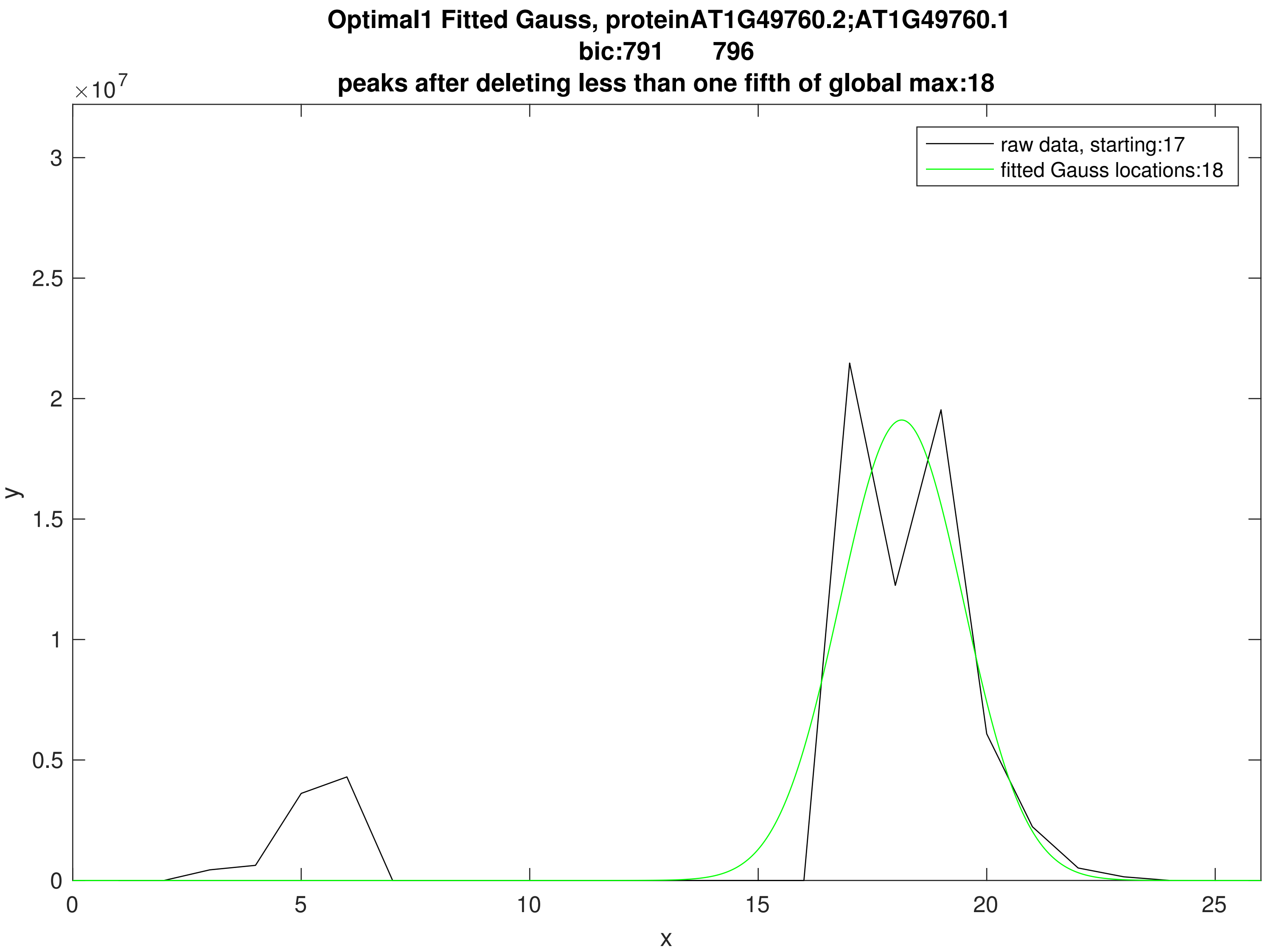

Optimal1 Fitted Gauss, proteinAT1G49760.2;AT1G49760.1
bic:791 796
peaks after deleting less than one fifth of global max:18
×10^7
raw data, starting:17
fitted Gauss locations:18
y
x
0
0.5
1
1.5
2
2.5
3
0
5
10
15
20
25

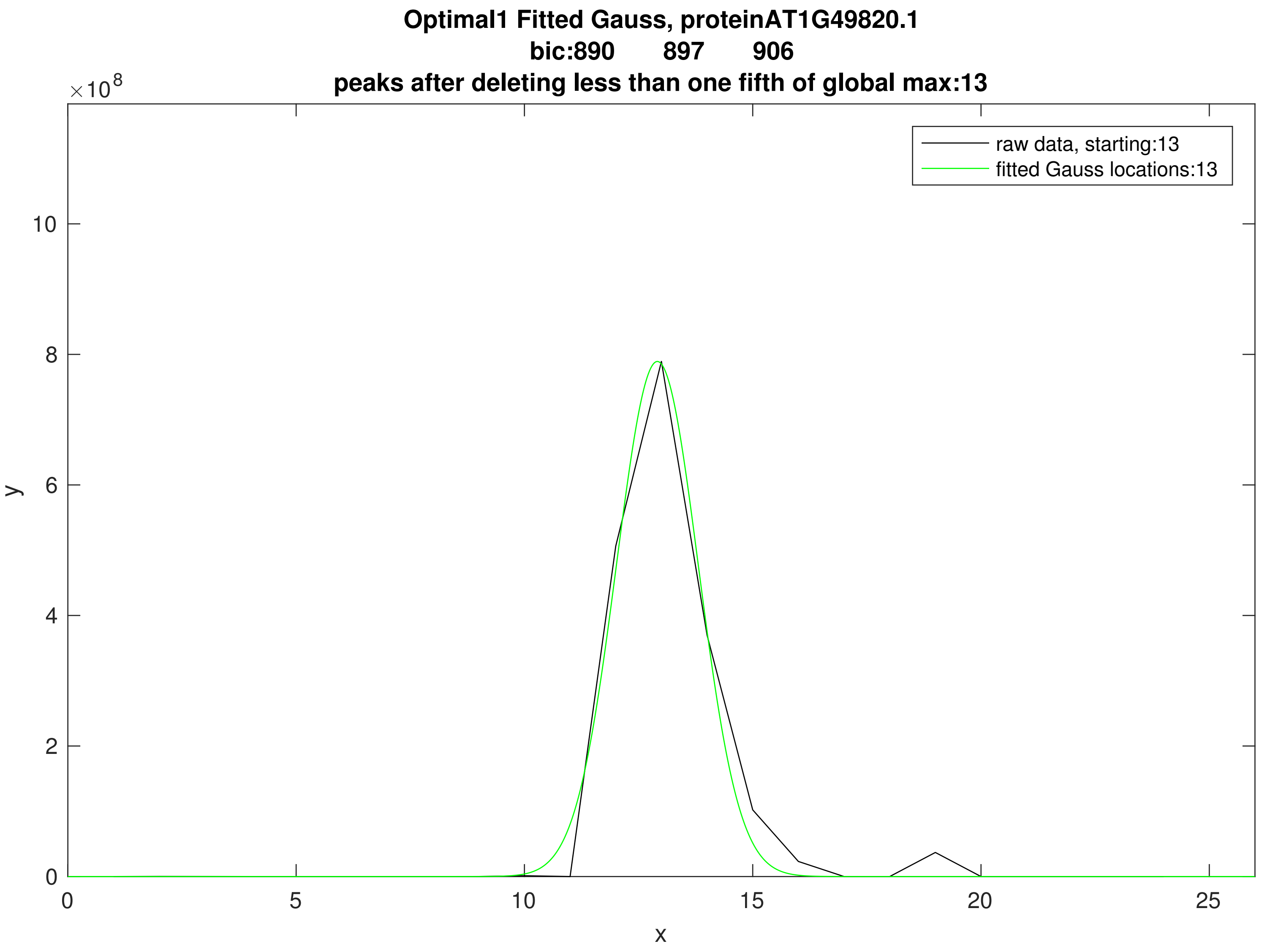
Optimal1 Fitted Gauss, proteinAT1G49820.1
bic:890 897 906
peaks after deleting less than one fifth of global max:13
×10^8
raw data, starting:13
fitted Gauss locations:13
y
x
0
2
4
6
8
10
0
5
10
15
20
25

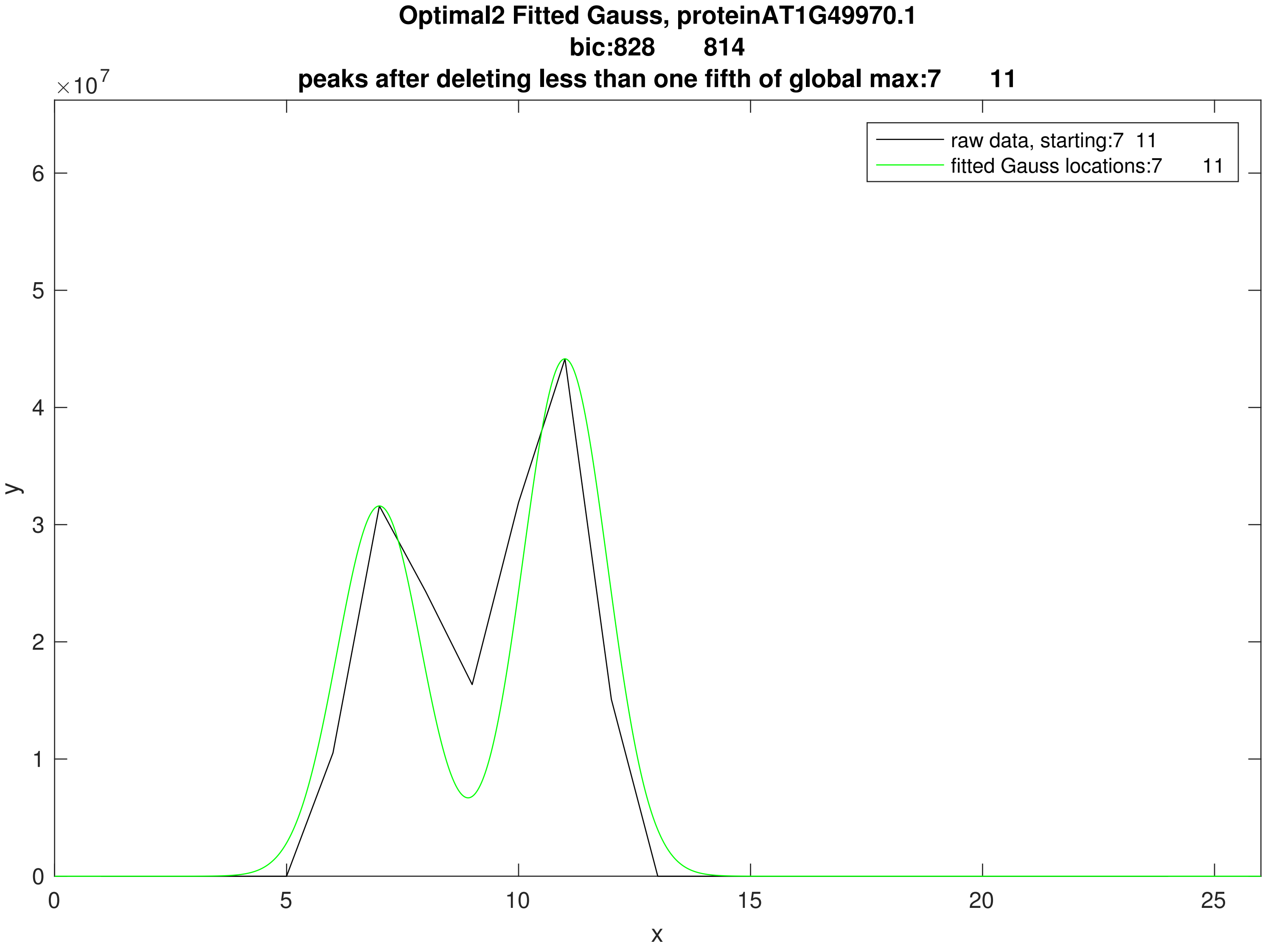
Optimal2 Fitted Gauss, proteinAT1G49970.1
bic:828 814
peaks after deleting less than one fifth of global max:7 11
×10^7
raw data, starting:7 11
fitted Gauss locations:7 11
y
x
0
1
2
3
4
5
6
0
5
10
15
20
25

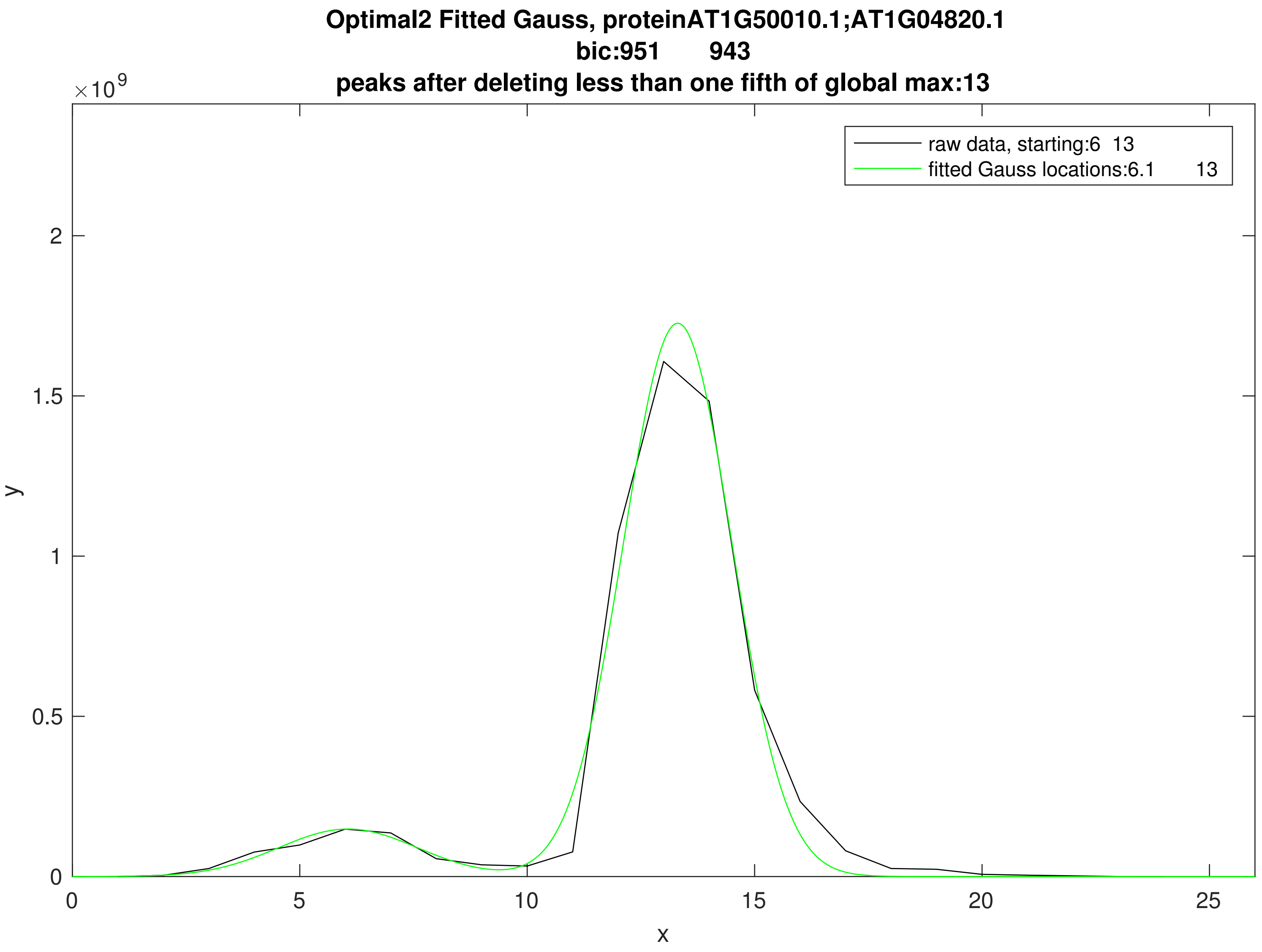
Optimal2 Fitted Gauss, proteinAT1G50010.1;AT1G04820.1
bic:951 943
peaks after deleting less than one fifth of global max:13
×10^9
raw data, starting:6 13
fitted Gauss locations:6.1 13
y
x
0
0.5
1
1.5
2
5
10
15
20
25

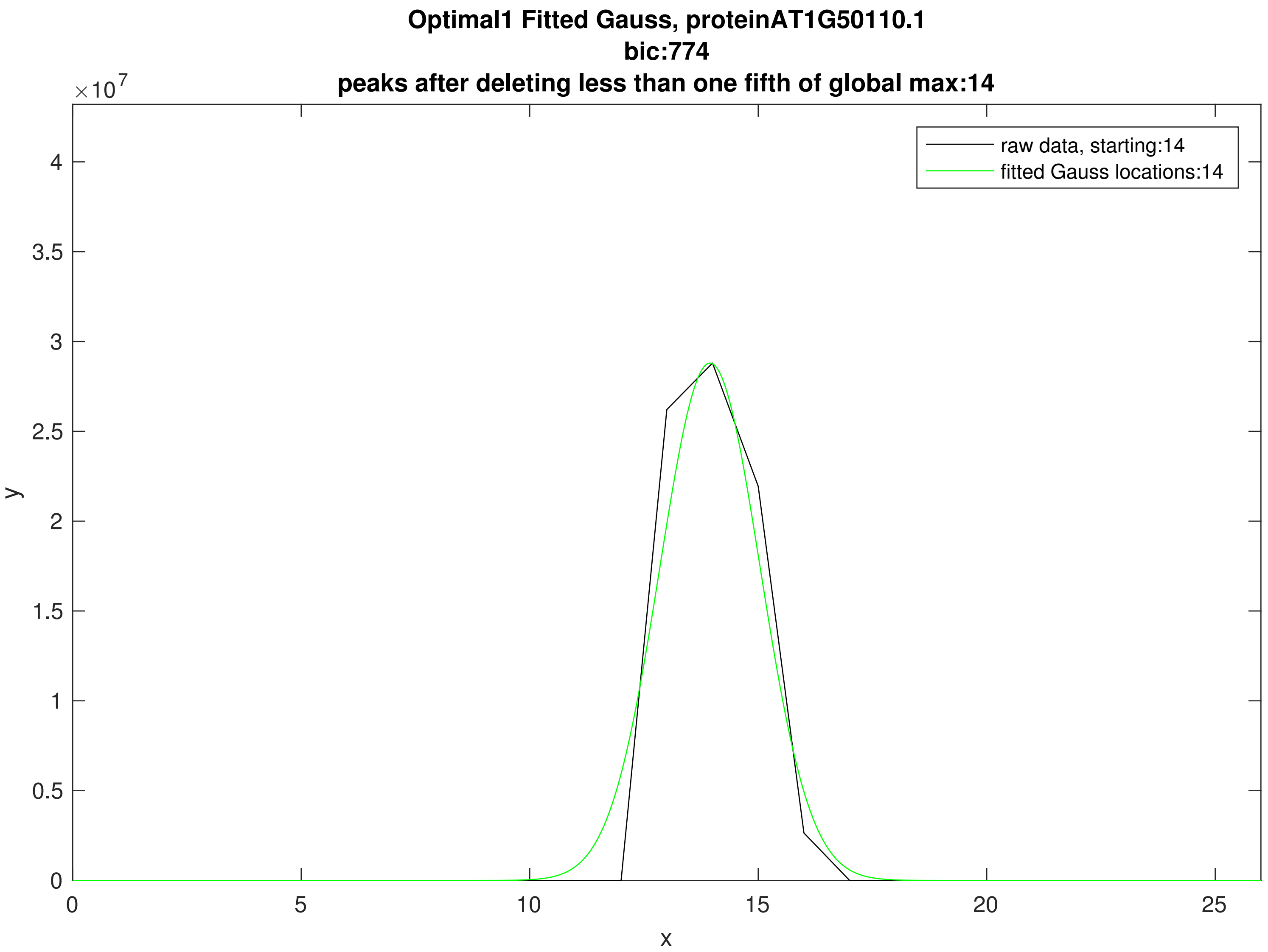

Optimal1 Fitted Gauss, proteinAT1G50110.1
bic:774
peaks after deleting less than one fifth of global max:14
×10^7
4
3.5
3
2.5
2
1.5
1
0.5
0
y
0
5
10
15
20
25
x
raw data, starting:14
fitted Gauss locations:14

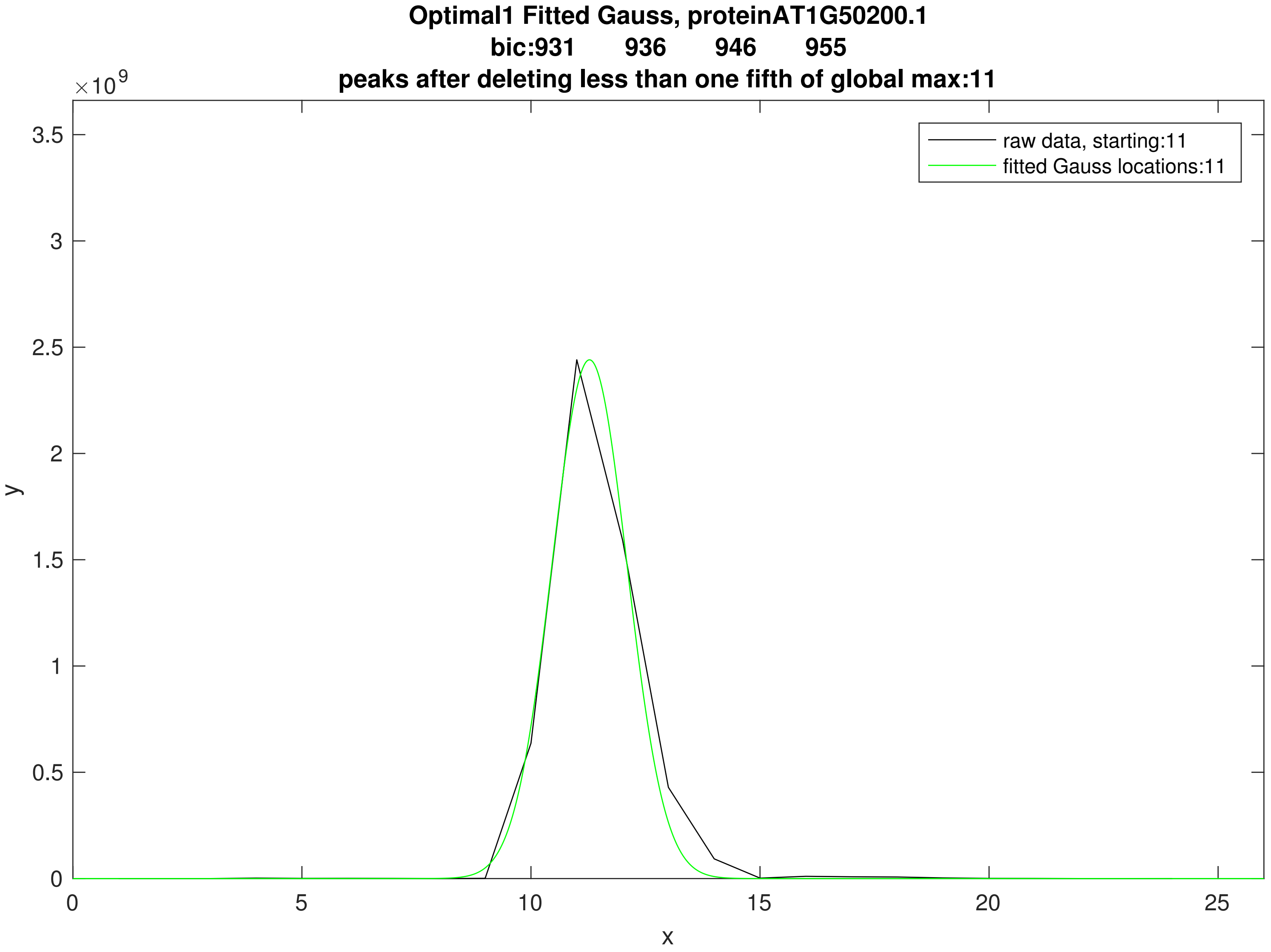
Optimal1 Fitted Gauss, proteinAT1G50200.1
bic:931 936 946 955
peaks after deleting less than one fifth of global max:11
×10^9
raw data, starting:11
fitted Gauss locations:11
y
x
0
0.5
1
1.5
2
2.5
3
3.5
0
5
10
15
20
25

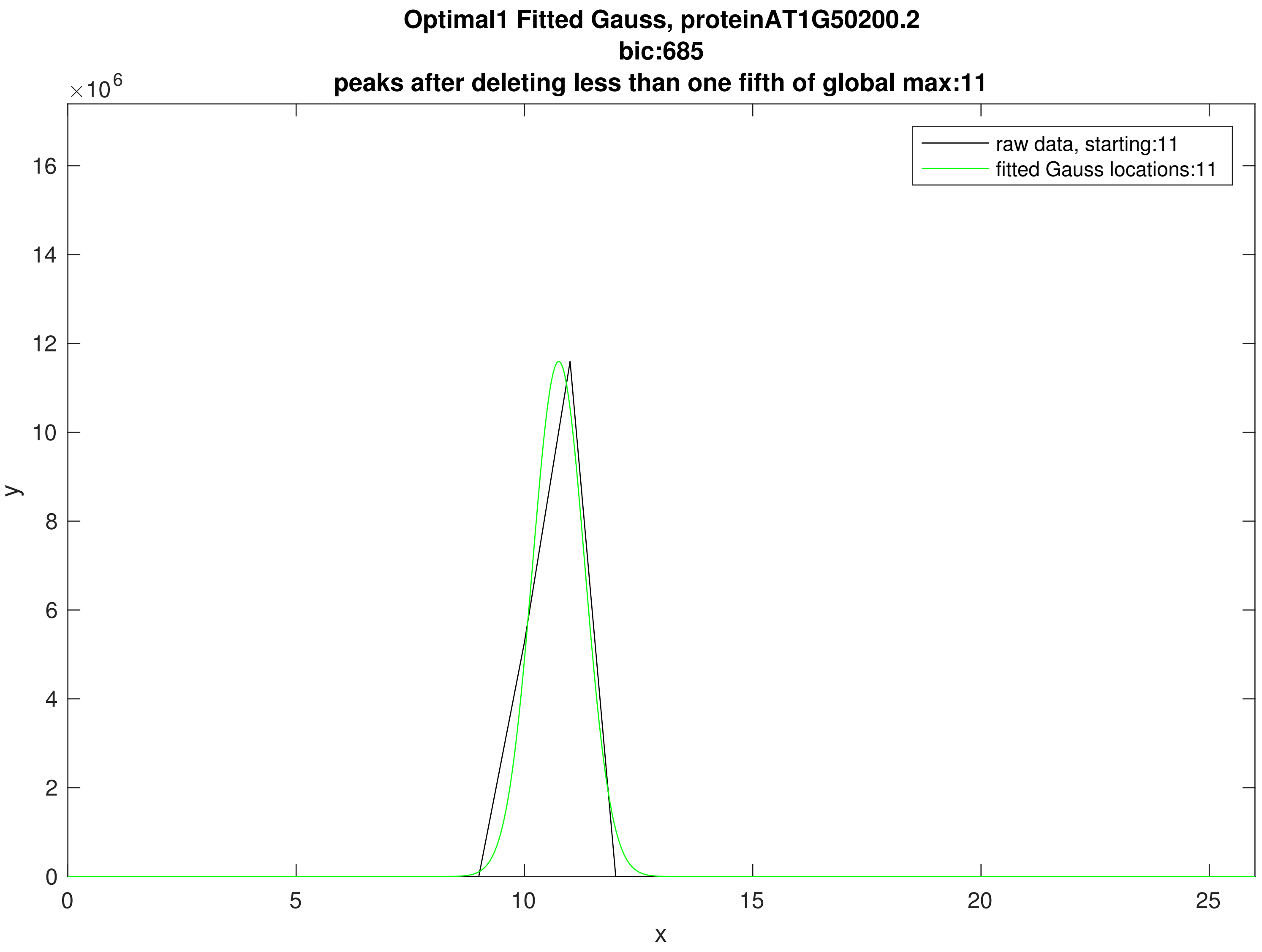
Optimal1 Fitted Gauss, proteinAT1G50200.2
bic:685
peaks after deleting less than one fifth of global max:11
×10^6
raw data, starting:11
fitted Gauss locations:11
y
x
0
2
4
6
8
10
12
14
16
5
15
20
25

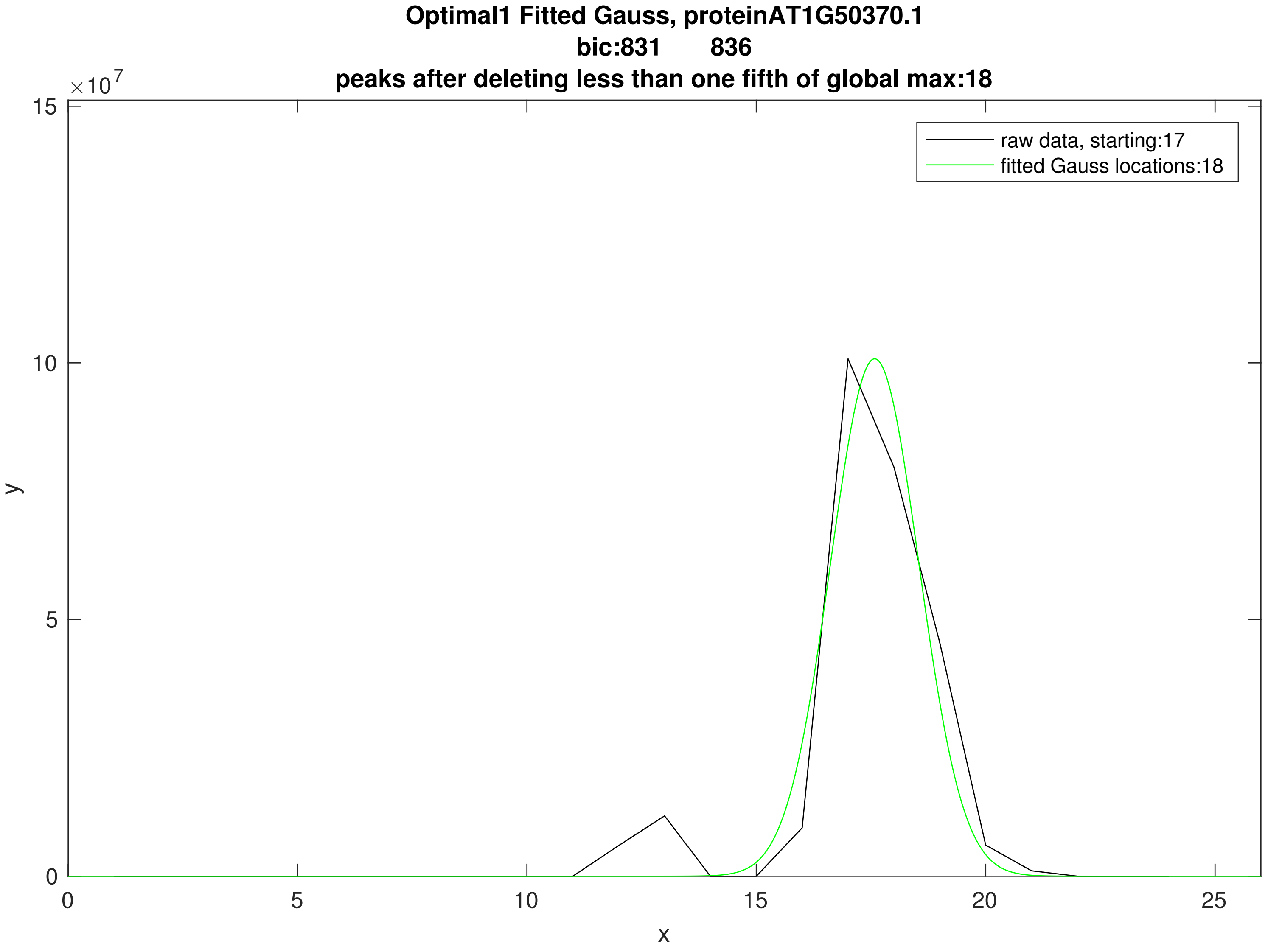
Optimal1 Fitted Gauss, proteinAT1G50370.1
bic:831 836
peaks after deleting less than one fifth of global max:18
×10^7
15
10
5
0
y
0
5
10
15
20
25
x
raw data, starting:17
fitted Gauss locations:18

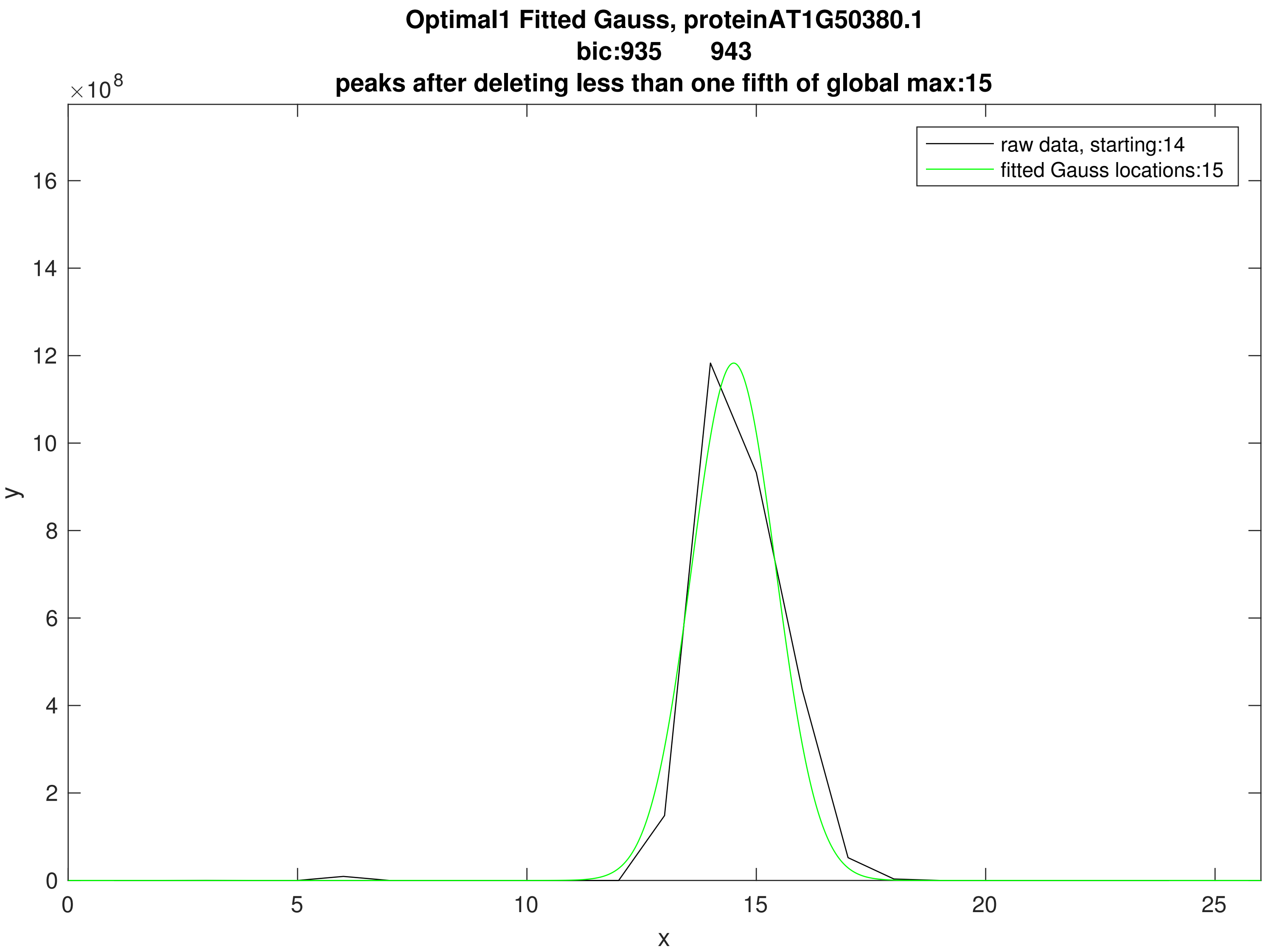
Optimal1 Fitted Gauss, proteinAT1G50380.1
bic:935 943
peaks after deleting less than one fifth of global max:15
×10^8
raw data, starting:14
fitted Gauss locations:15
y
x
0
2
4
6
8
10
12
14
16
0
5
10
15
20
25

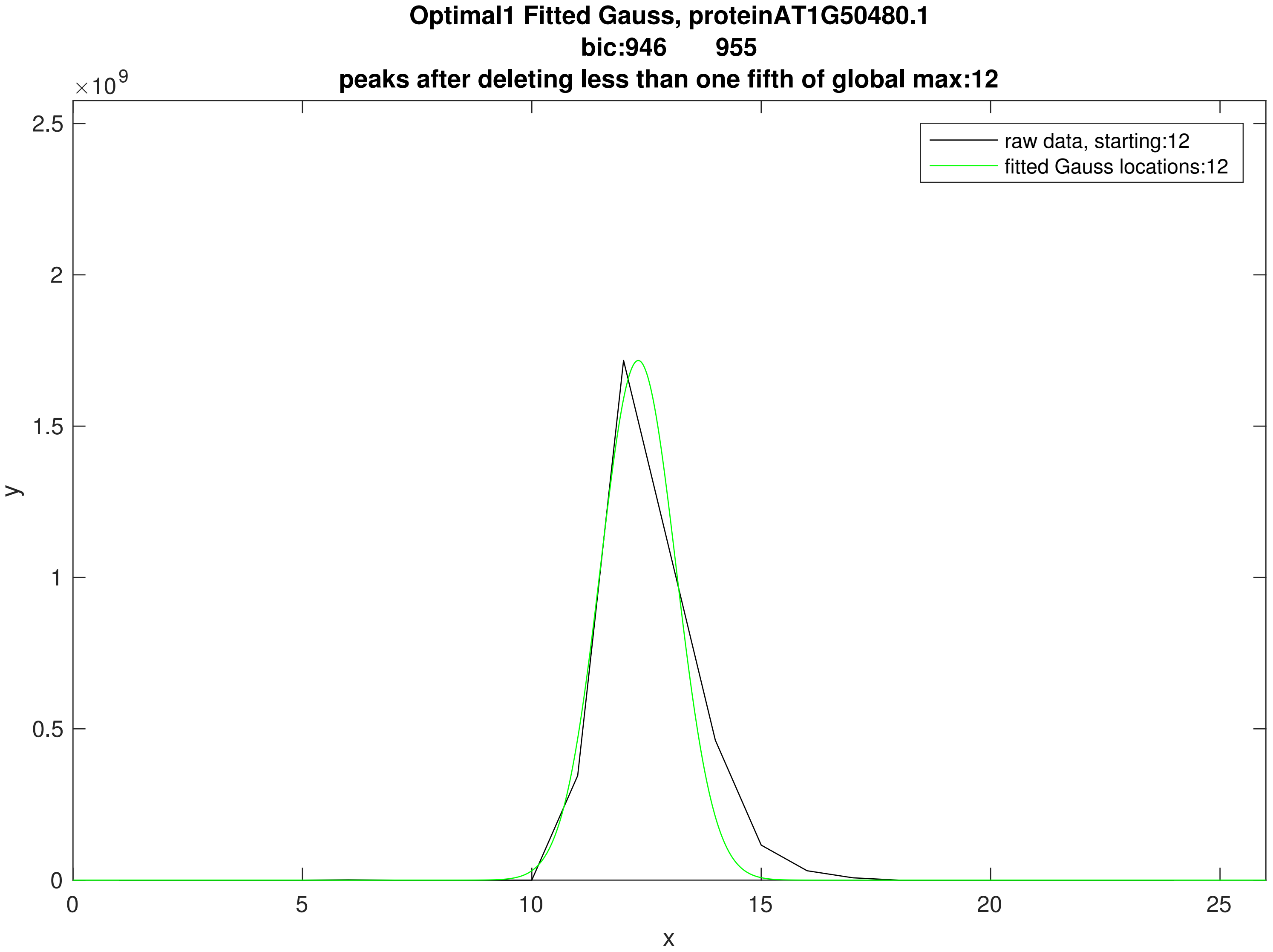
Optimal1 Fitted Gauss, proteinAT1G50480.1
bic:946 955
peaks after deleting less than one fifth of global max:12
×10^9
2.5
2
1.5
1
0.5
0
y
0
5
10
15
20
25
x
raw data, starting:12
fitted Gauss locations:12

Optimal1 Fitted Gauss, proteinAT1G50510.1
bic:783
peaks after deleting less than one fifth of global max:14

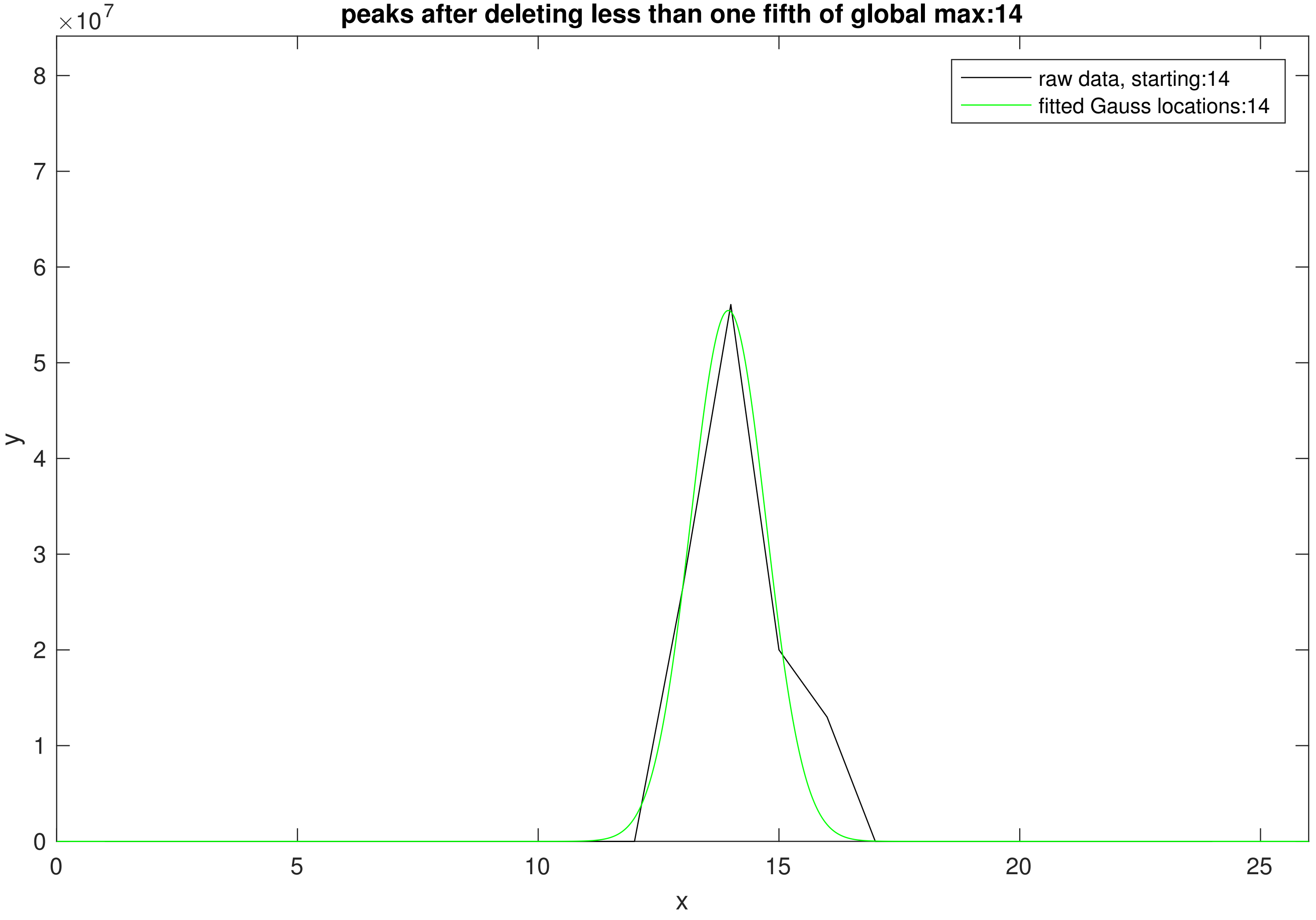

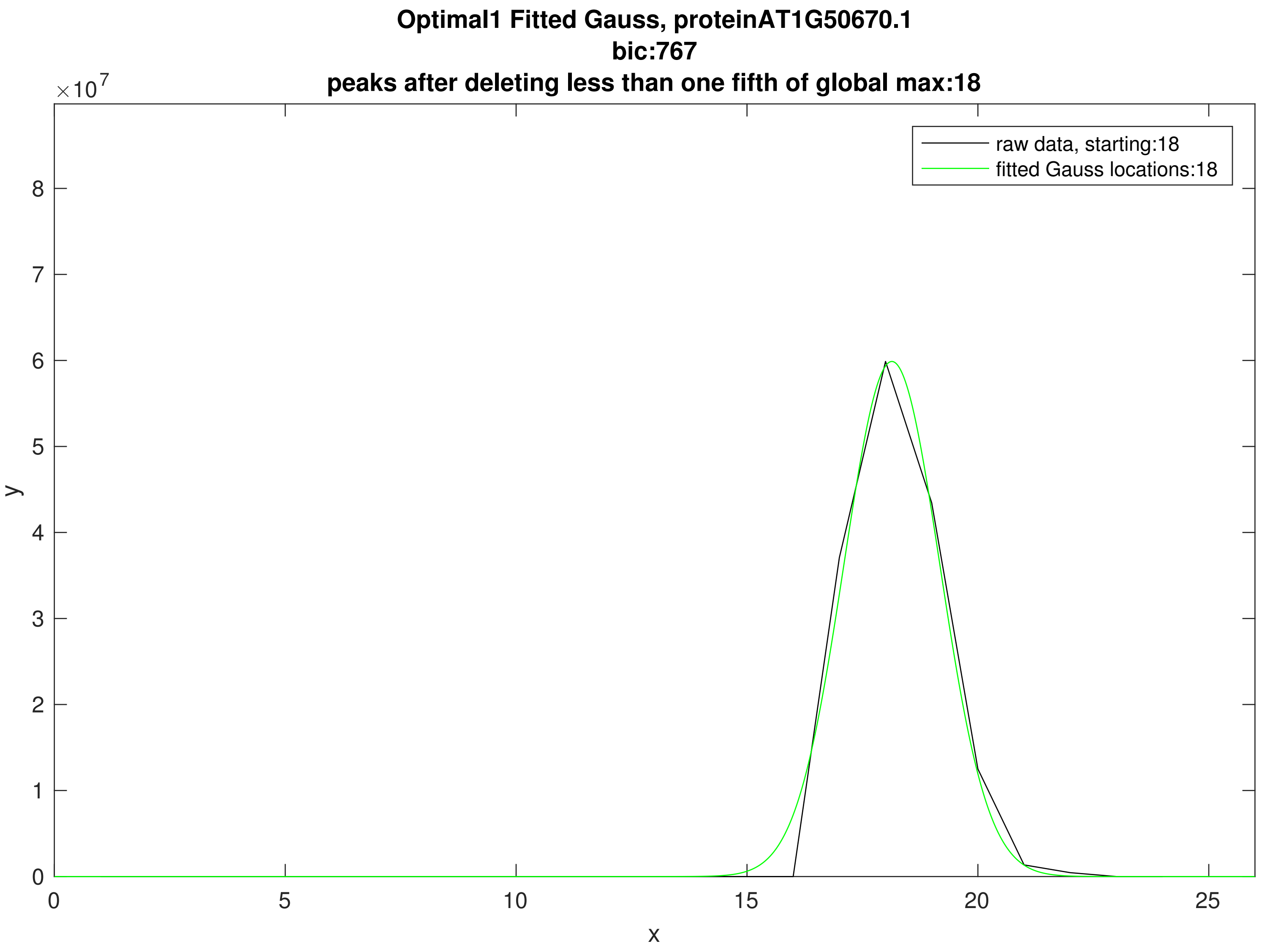
Optimal1 Fitted Gauss, proteinAT1G50670.1
bic:767
peaks after deleting less than one fifth of global max:18
×10^7
raw data, starting:18
fitted Gauss locations:18
y
x
0
1
2
3
4
5
6
7
8
0
5
10
15
20
25

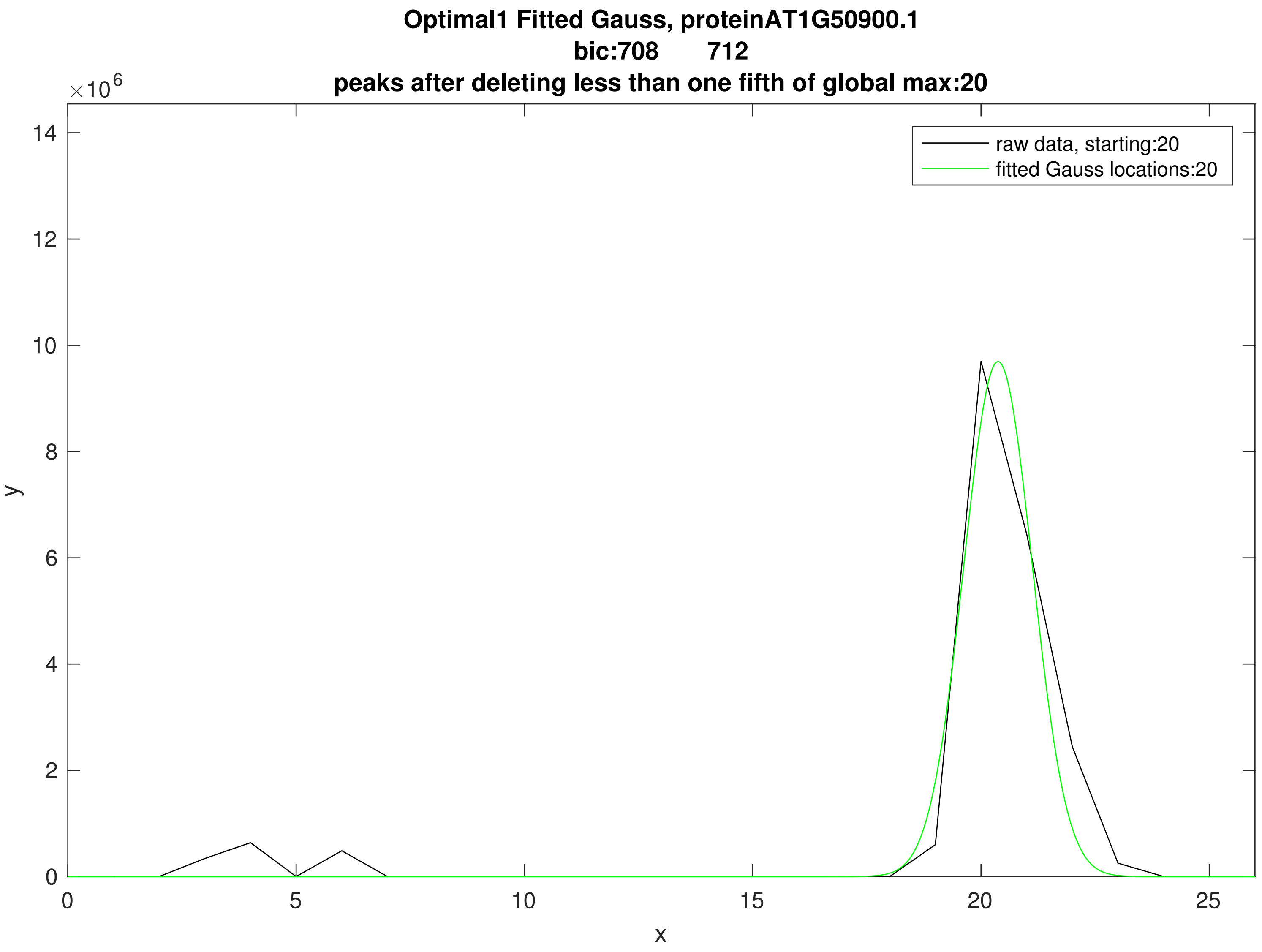

Optimal1 Fitted Gauss, proteinAT1G50900.1
bic:708 712
peaks after deleting less than one fifth of global max:20
×10^6
14
12
10
8
6
4
2
0
y
0
5
10
15
20
25
x
raw data, starting:20
fitted Gauss locations:20

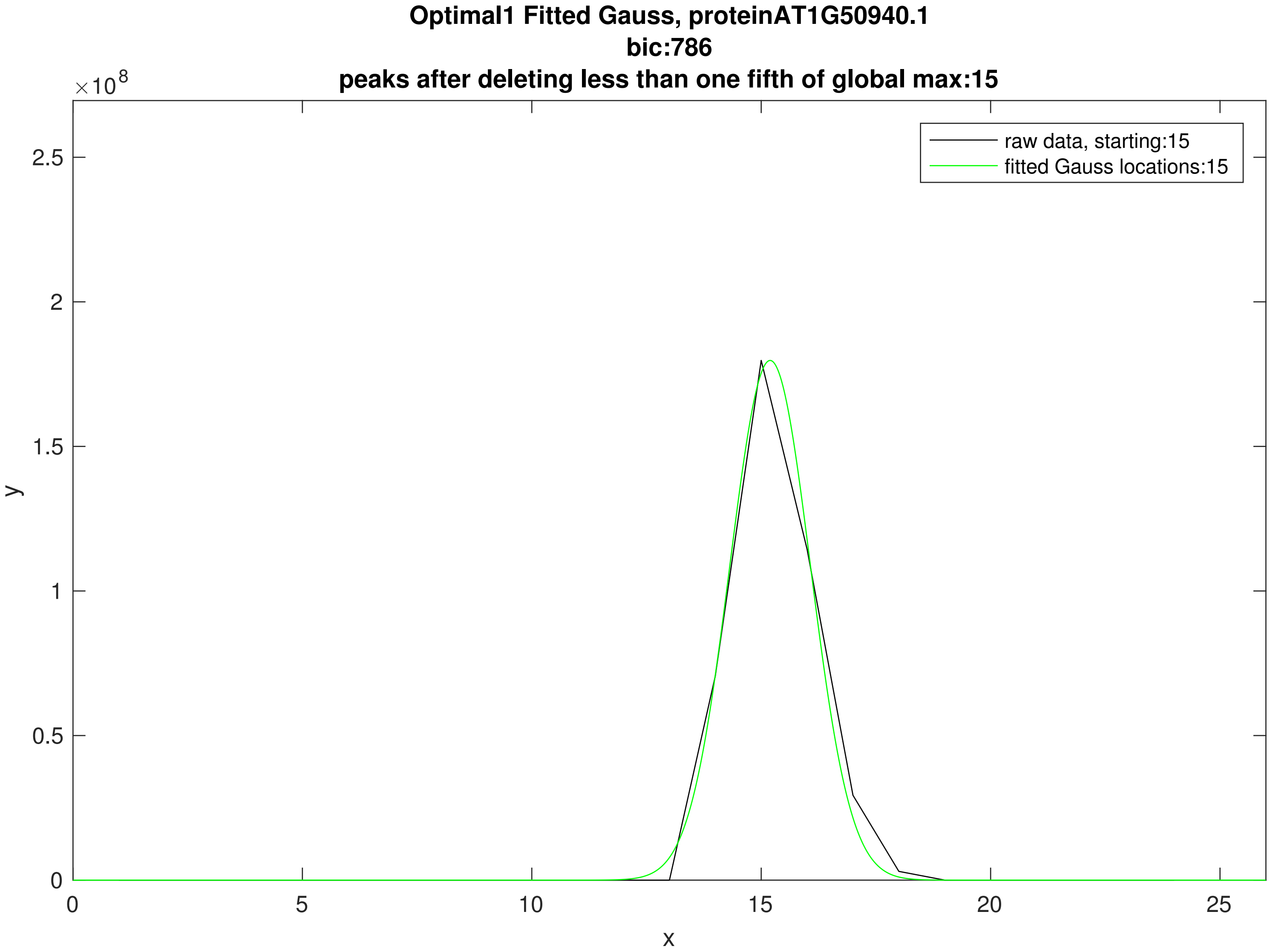
Optimal1 Fitted Gauss, proteinAT1G50940.1
bic:786
peaks after deleting less than one fifth of global max:15
×10^8
raw data, starting:15
fitted Gauss locations:15
y
x
0
0.5
1
1.5
2
2.5
0
5
10
15
20
25

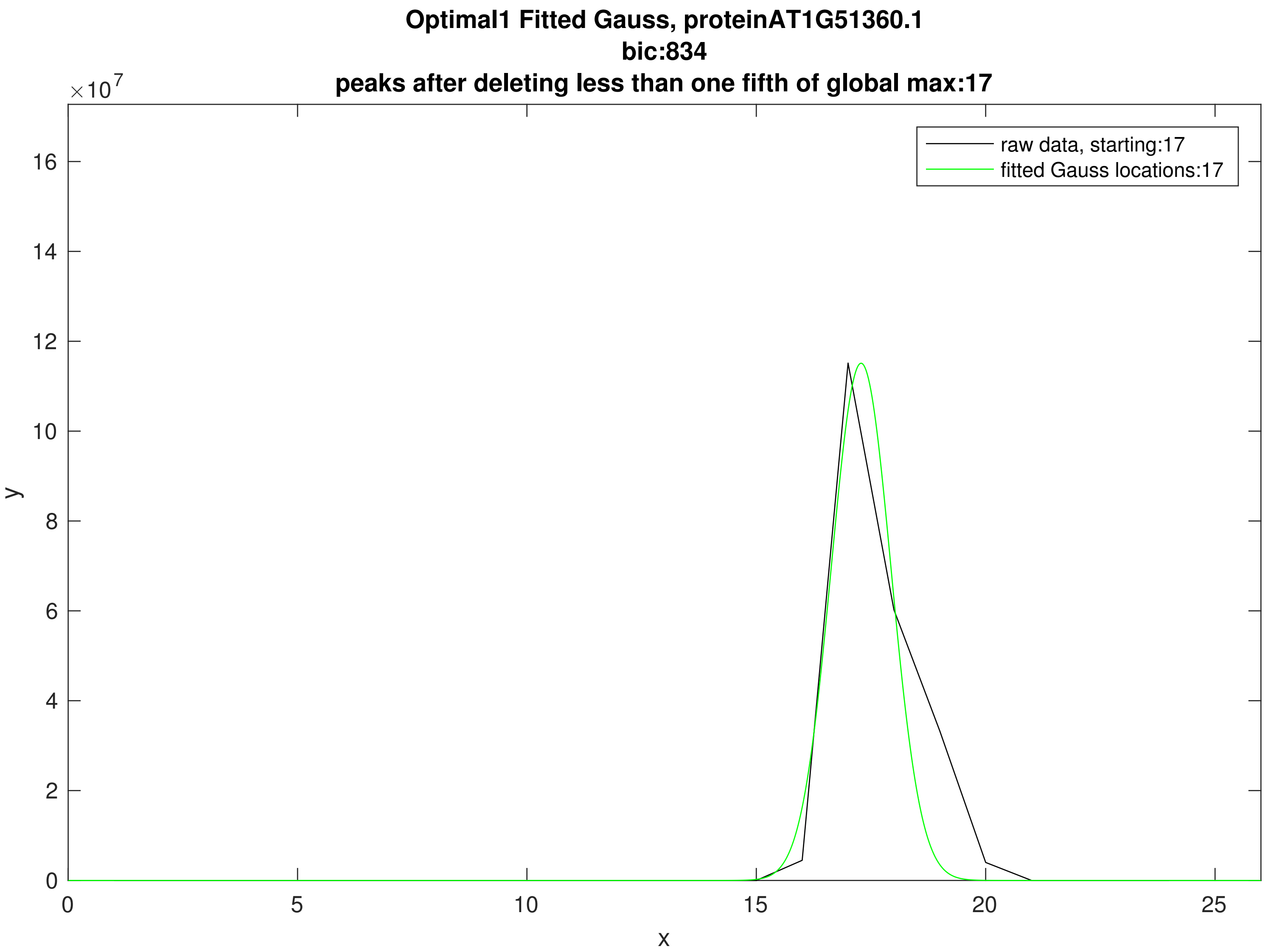
Optimal1 Fitted Gauss, proteinAT1G51360.1
bic:834
peaks after deleting less than one fifth of global max:17
×10^7
raw data, starting:17
fitted Gauss locations:17
y
x
0
2
4
6
8
10
12
14
16
0
5
10
15
20
25

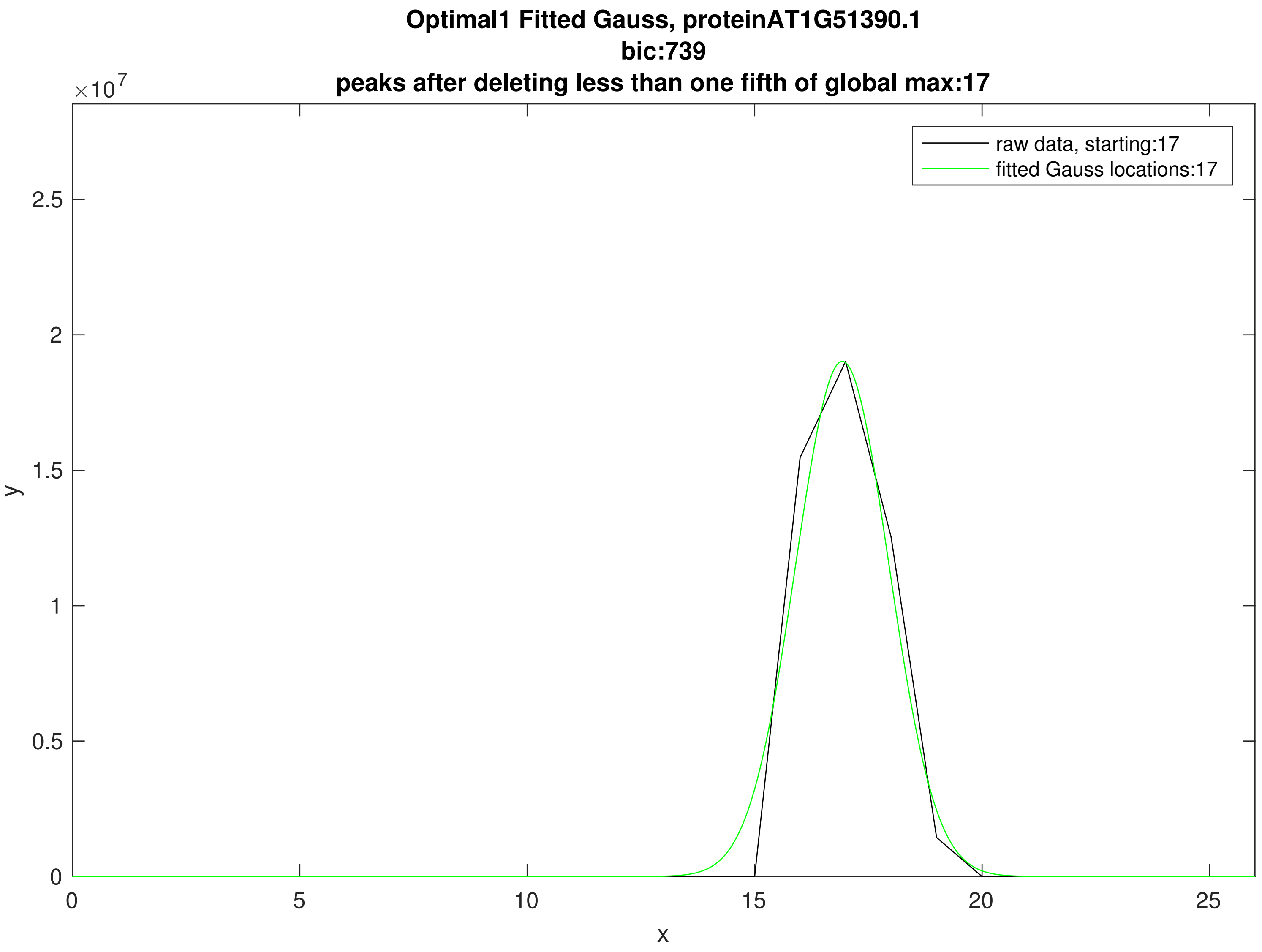
Optimal1 Fitted Gauss, proteinAT1G51390.1
bic:739
peaks after deleting less than one fifth of global max:17
×10^7
raw data, starting:17
fitted Gauss locations:17
y
2.5
2
1.5
1
0.5
0
0
5
10
15
20
25
x

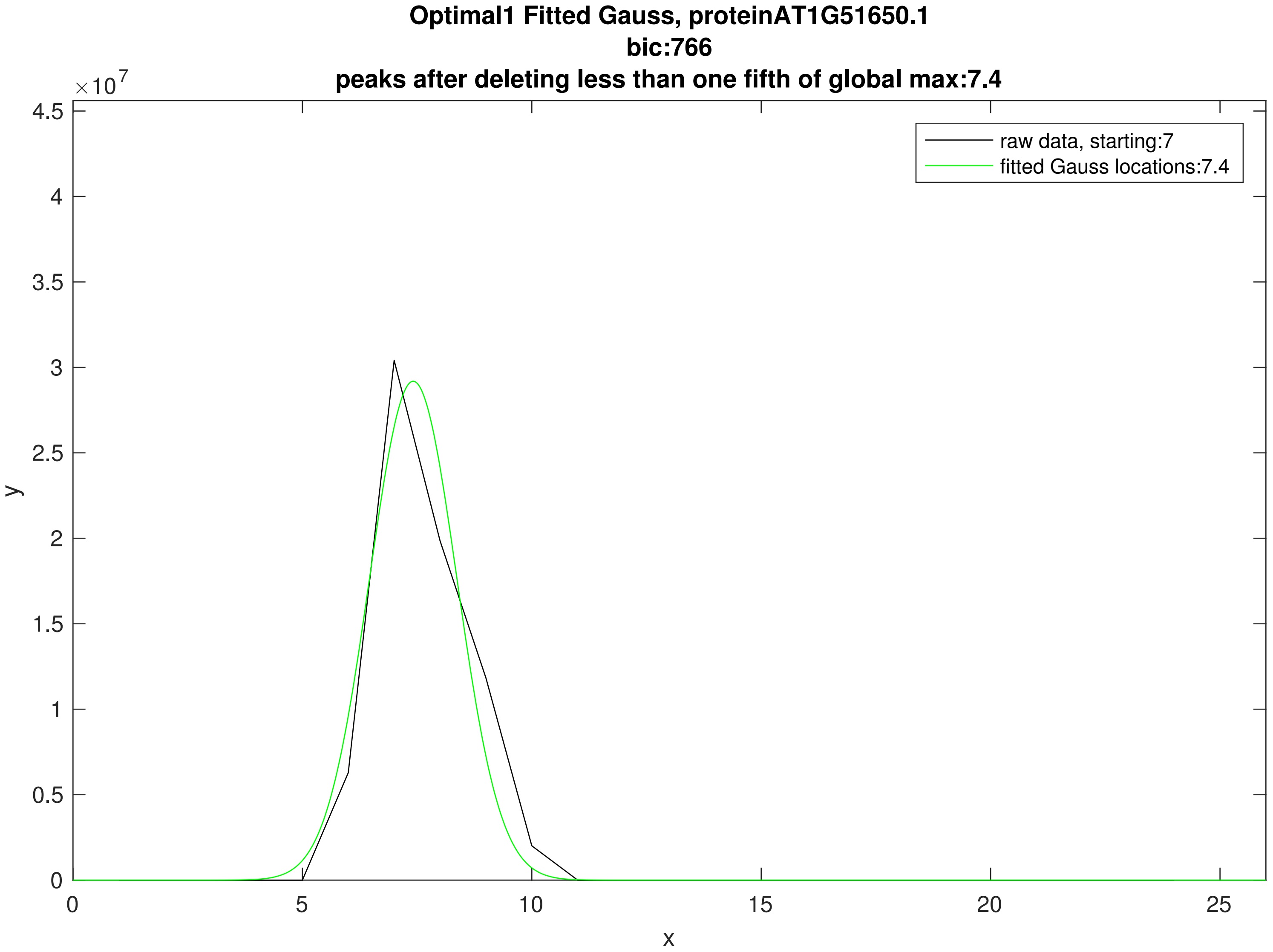

Optimal1 Fitted Gauss, proteinAT1G51650.1
bic:766
peaks after deleting less than one fifth of global max:7.4
×10^7
4.5
4
3.5
3
2.5
2
1.5
1
0.5
0
y
0
5
10
15
20
25
x
raw data, starting:7
fitted Gauss locations:7.4

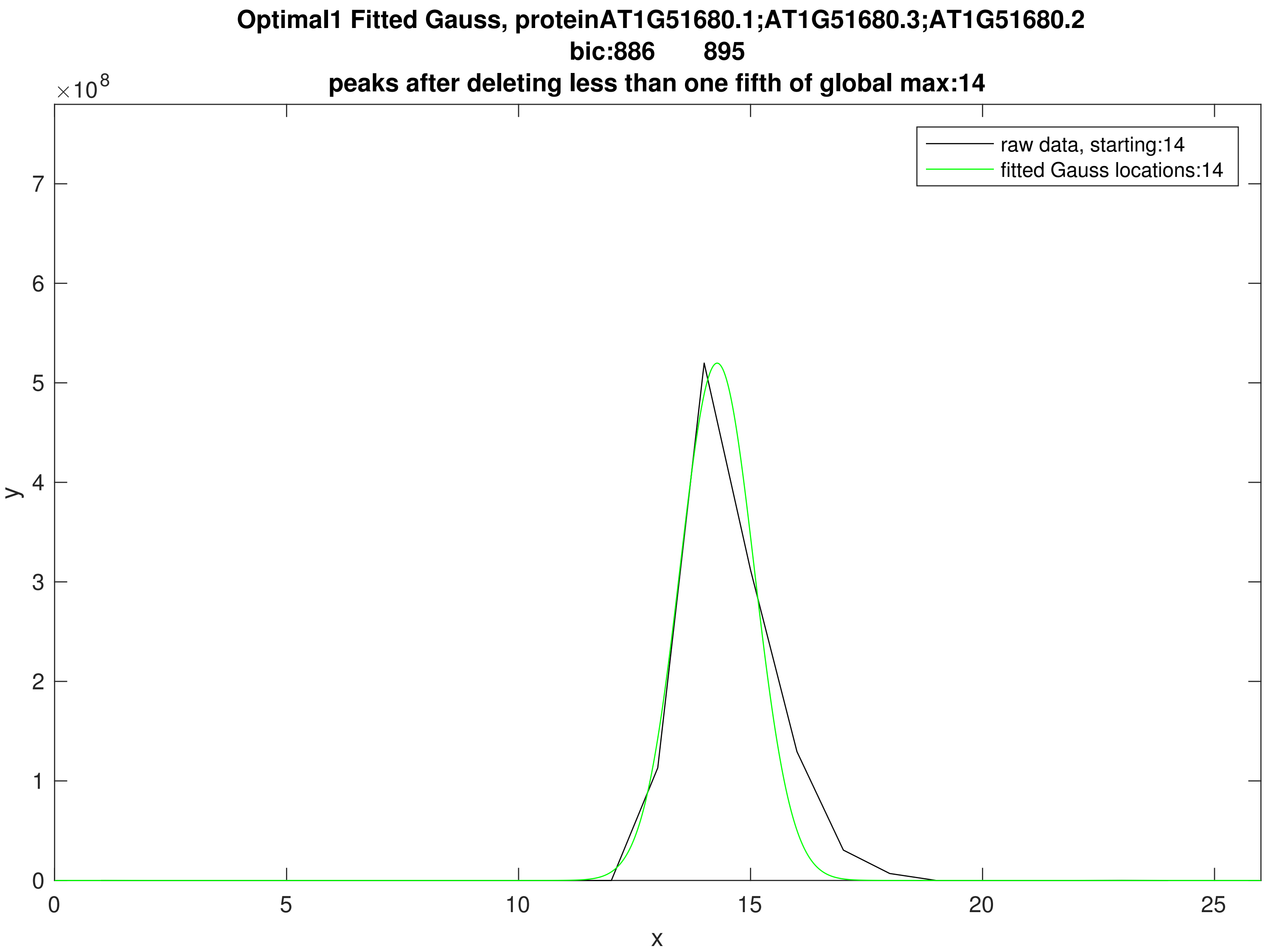

Optimal1 Fitted Gauss, proteinAT1G51680.1;AT1G51680.3;AT1G51680.2
bic:886 895
peaks after deleting less than one fifth of global max:14
×10^8
raw data, starting:14
fitted Gauss locations:14
y
x
0
1
2
3
4
5
6
7
0
5
10
15
20
25

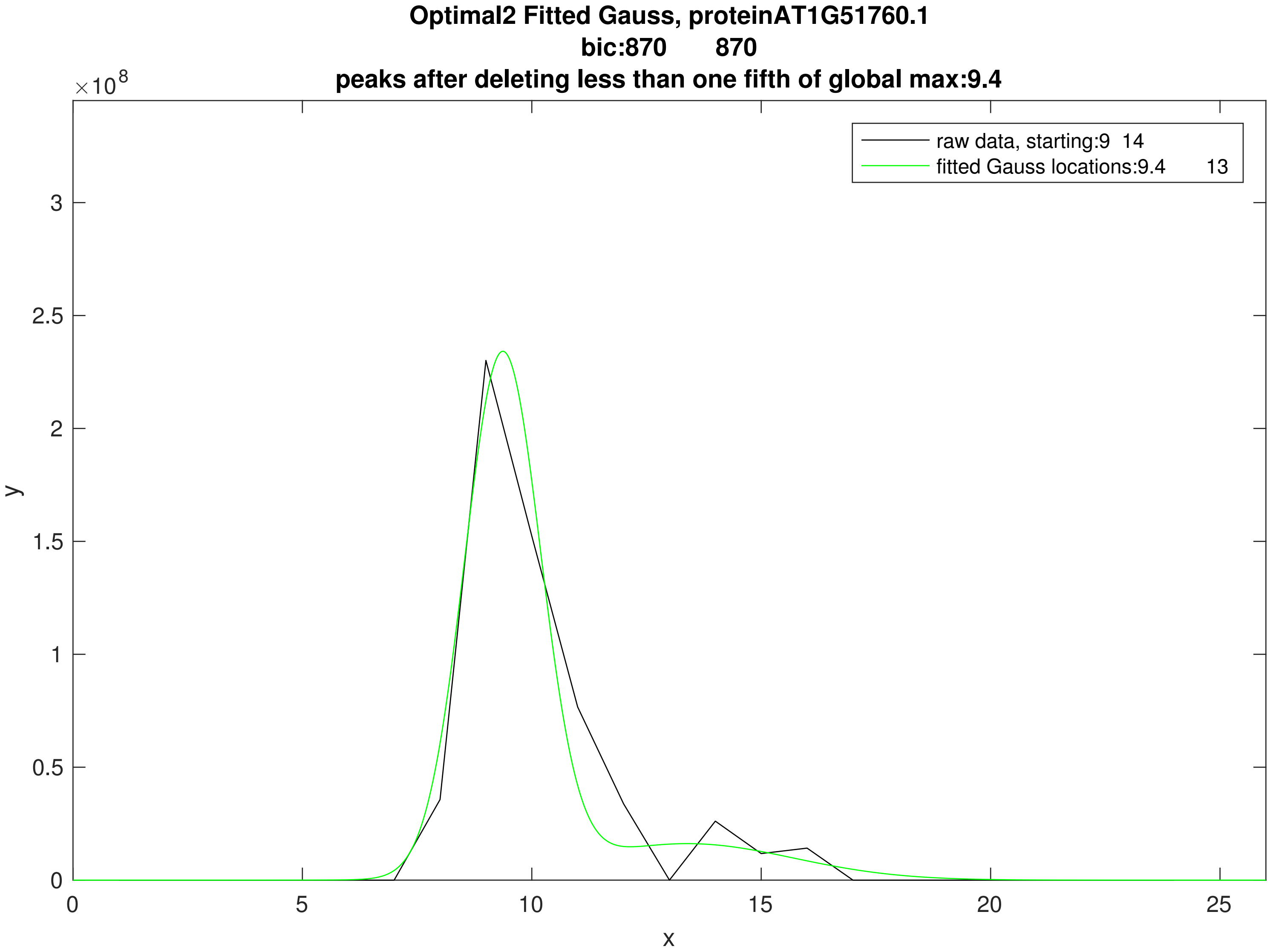

Optimal2 Fitted Gauss, proteinAT1G51760.1
bic:870 870
peaks after deleting less than one fifth of global max:9.4
×10^8
raw data, starting:9 14
fitted Gauss locations:9.4 13
y
x
0
0.5
1
1.5
2
2.5
3
0
5
10
15
20
25

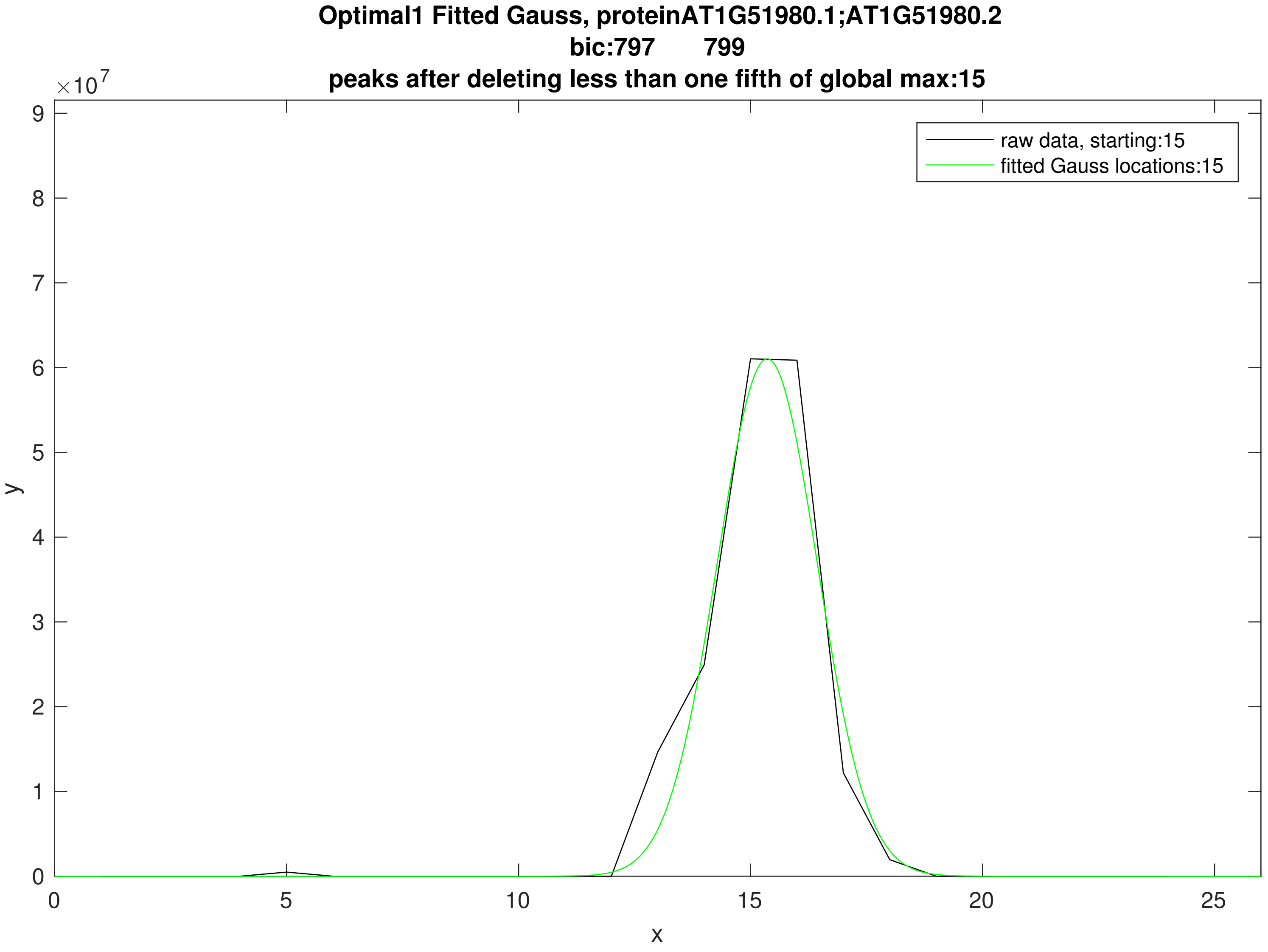
Optimal1 Fitted Gauss, proteinAT1G51980.1;AT1G51980.2
bic:797 799
peaks after deleting less than one fifth of global max:15
×10^7
raw data, starting:15
fitted Gauss locations:15
y
x
0
1
2
3
4
5
6
7
8
9
0
5
10
15
20
25

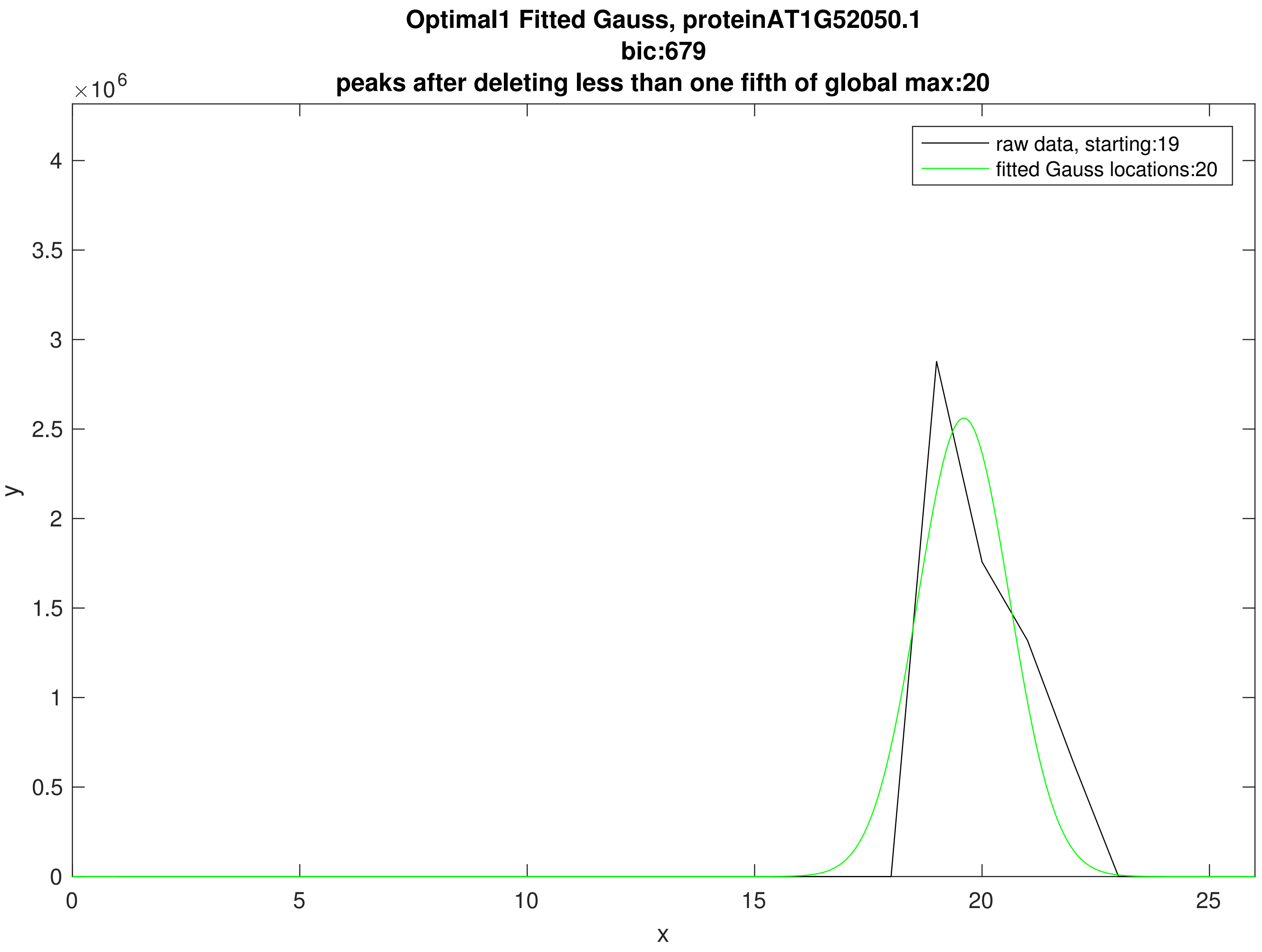

Optimal1 Fitted Gauss, proteinAT1G52050.1
bic:679
peaks after deleting less than one fifth of global max:20
×10^6
raw data, starting:19
fitted Gauss locations:20
4
3.5
3
2.5
2
1.5
1
0.5
0
y
0
5
10
15
20
25
x

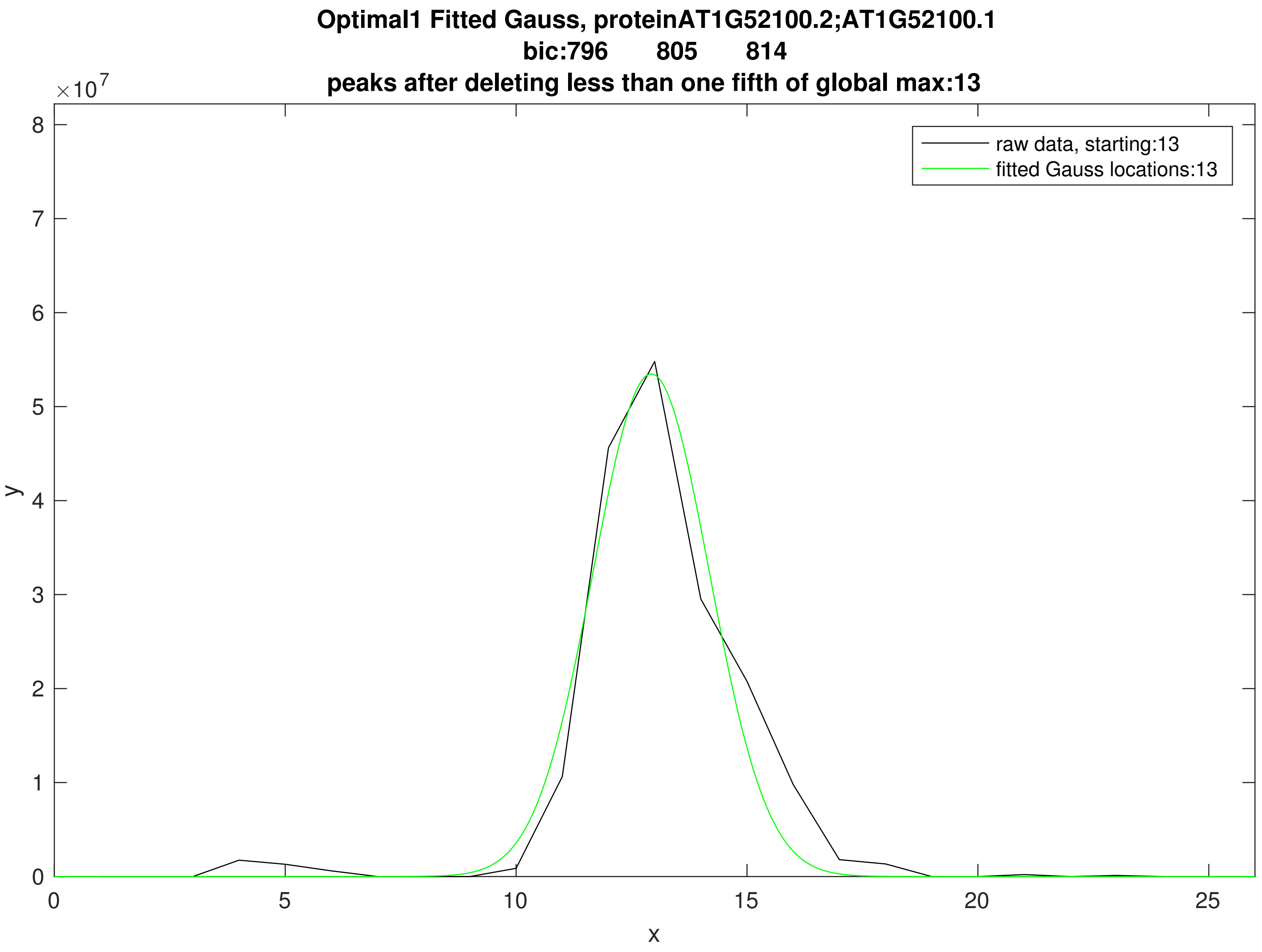
Optimal1 Fitted Gauss, proteinAT1G52100.2;AT1G52100.1
bic:796 805 814
peaks after deleting less than one fifth of global max:13
×10^7
raw data, starting:13
fitted Gauss locations:13
y
x
0
1
2
3
4
5
6
7
8
0
5
10
15
20
25

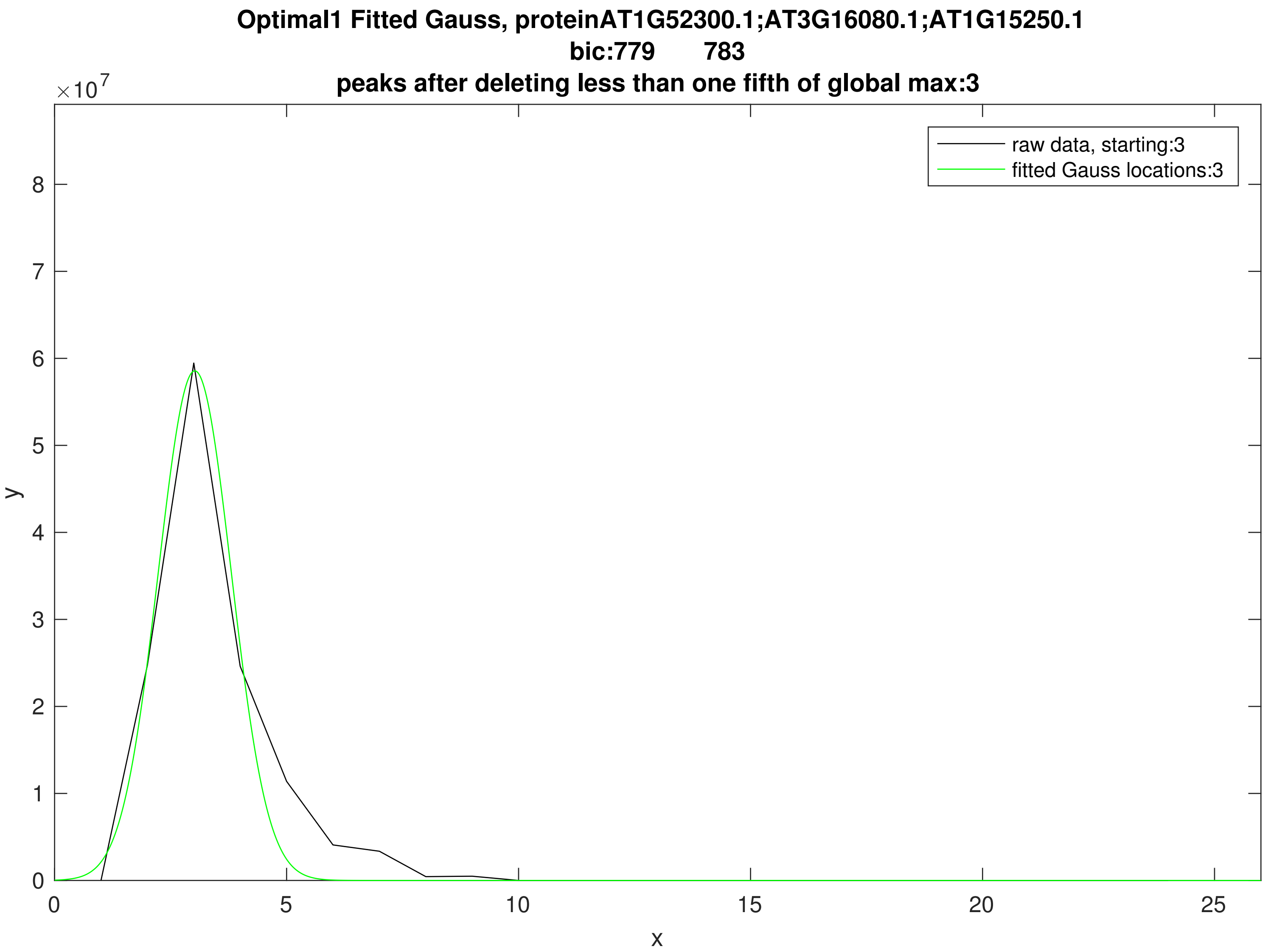

Optimal1 Fitted Gauss, proteinAT1G52300.1;AT3G16080.1;AT1G15250.1
bic:779 783
peaks after deleting less than one fifth of global max:3
×10^7
raw data, starting:3
fitted Gauss locations:3
y
x
0
1
2
3
4
5
6
7
8
0
5
10
15
20
25

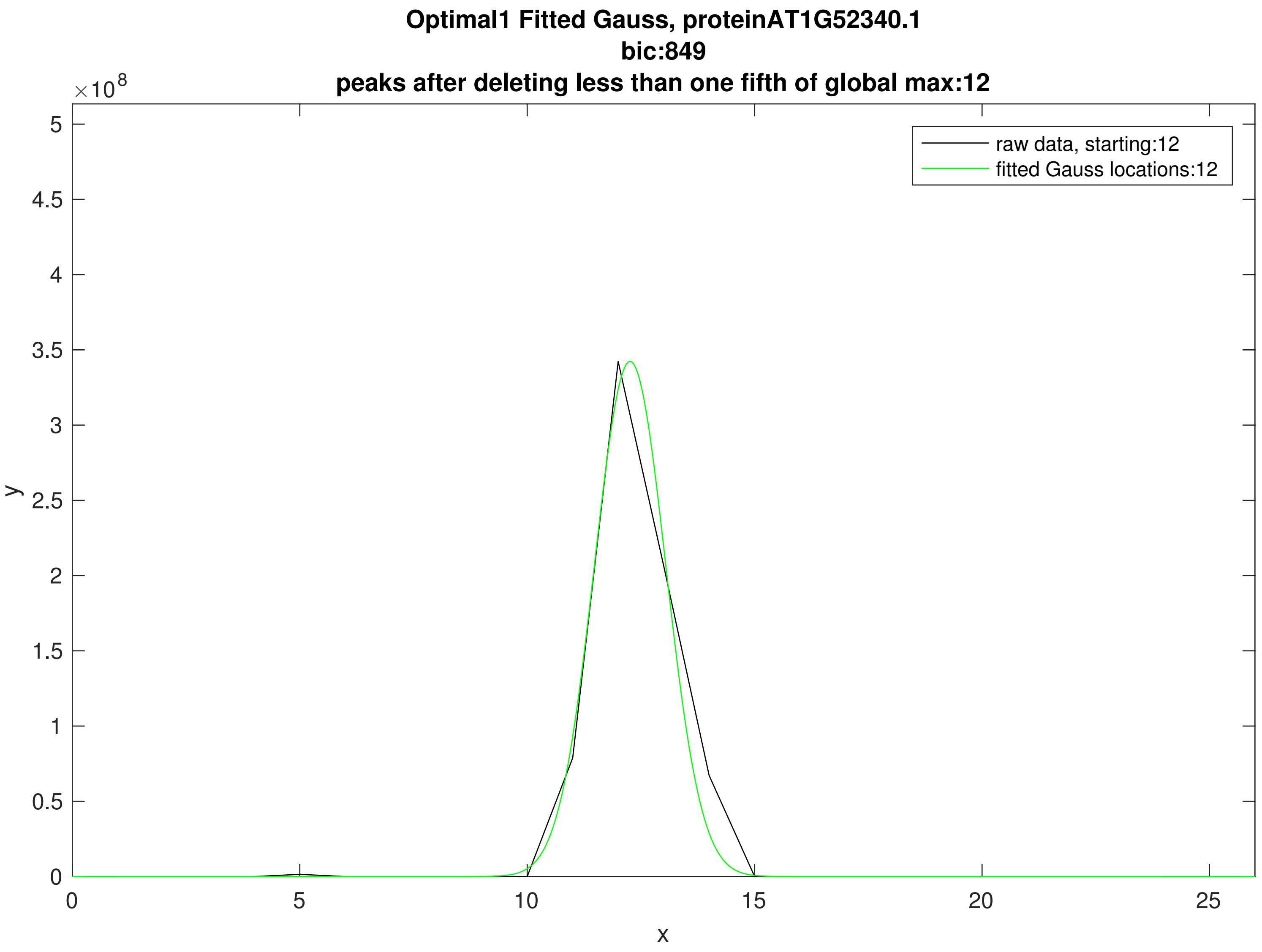
Optimal1 Fitted Gauss, proteinAT1G52340.1
bic:849
peaks after deleting less than one fifth of global max:12
×10^8
raw data, starting:12
fitted Gauss locations:12
y
x
0
0.5
1
1.5
2
2.5
3
3.5
4
4.5
5
0
5
10
15
20
25

#### Optimal1 Fitted Gauss, proteinAT1G52400.2;AT1G52400.3;AT1G52400.1
#### bic:773
#### peaks after deleting less than one fifth of global max:7.8

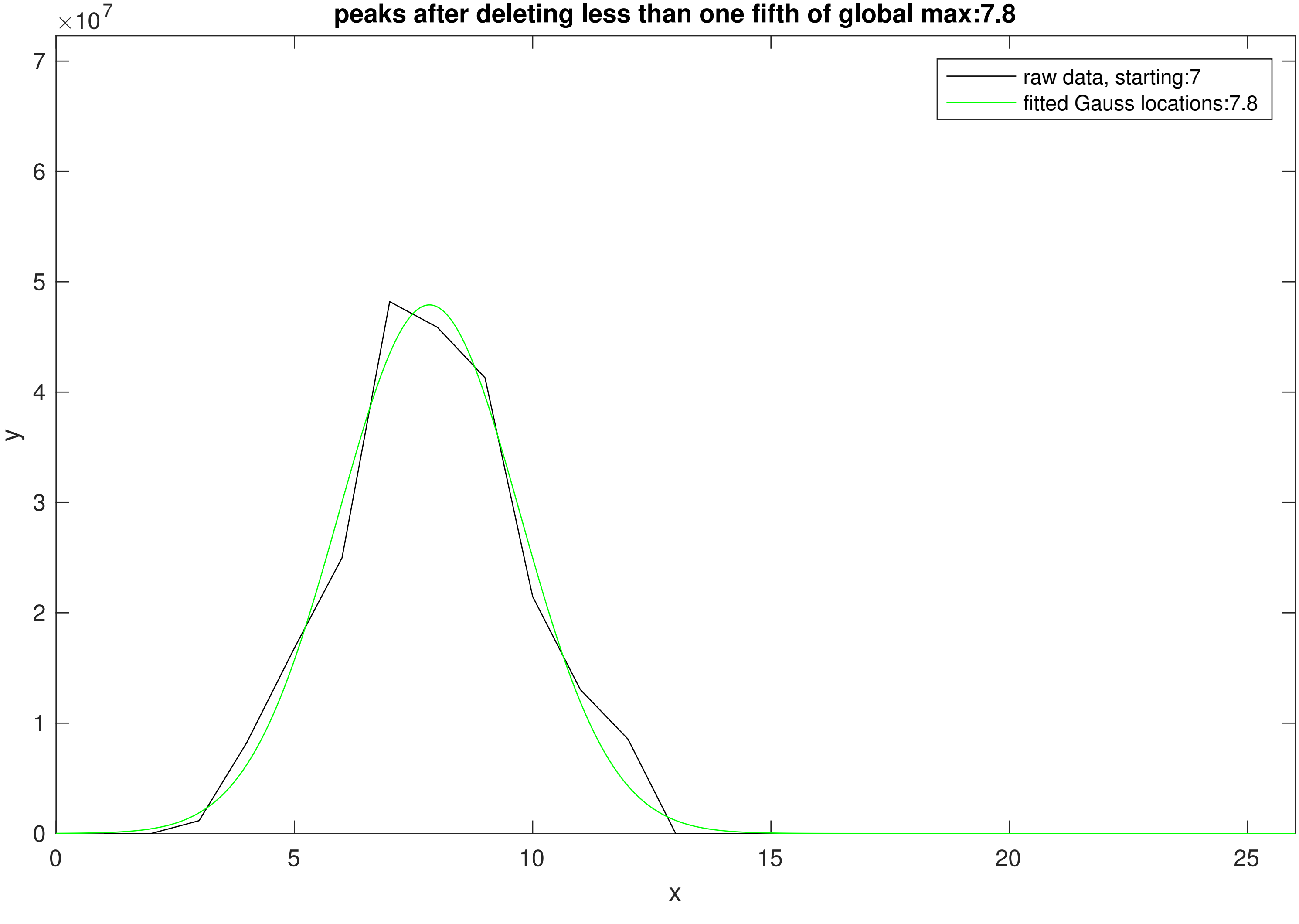

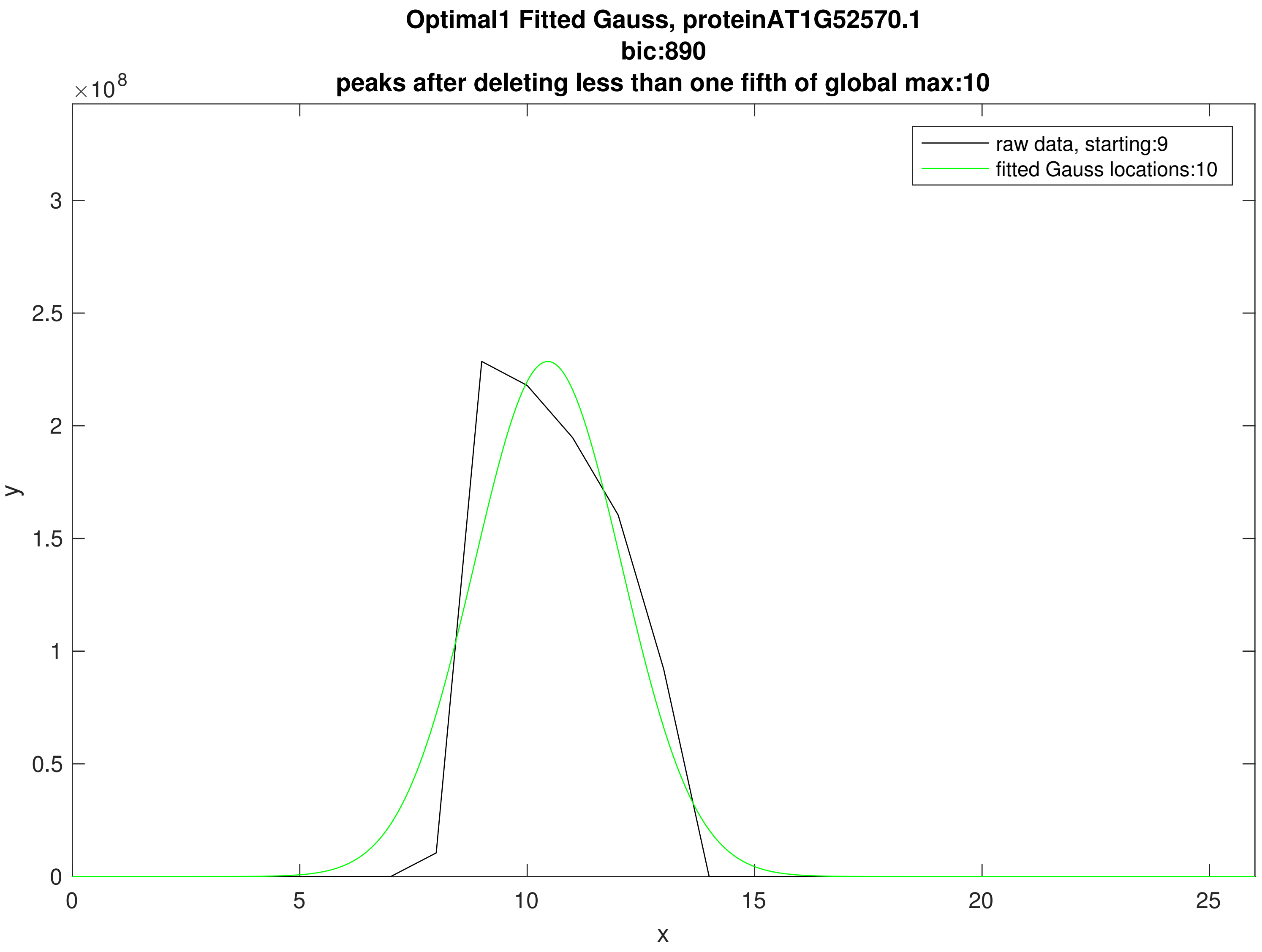
Optimal1 Fitted Gauss, proteinAT1G52570.1
bic:890
peaks after deleting less than one fifth of global max:10
×10^8
raw data, starting:9
fitted Gauss locations:10
y
x
0
0.5
1
1.5
2
2.5
3
0
5
10
15
20
25

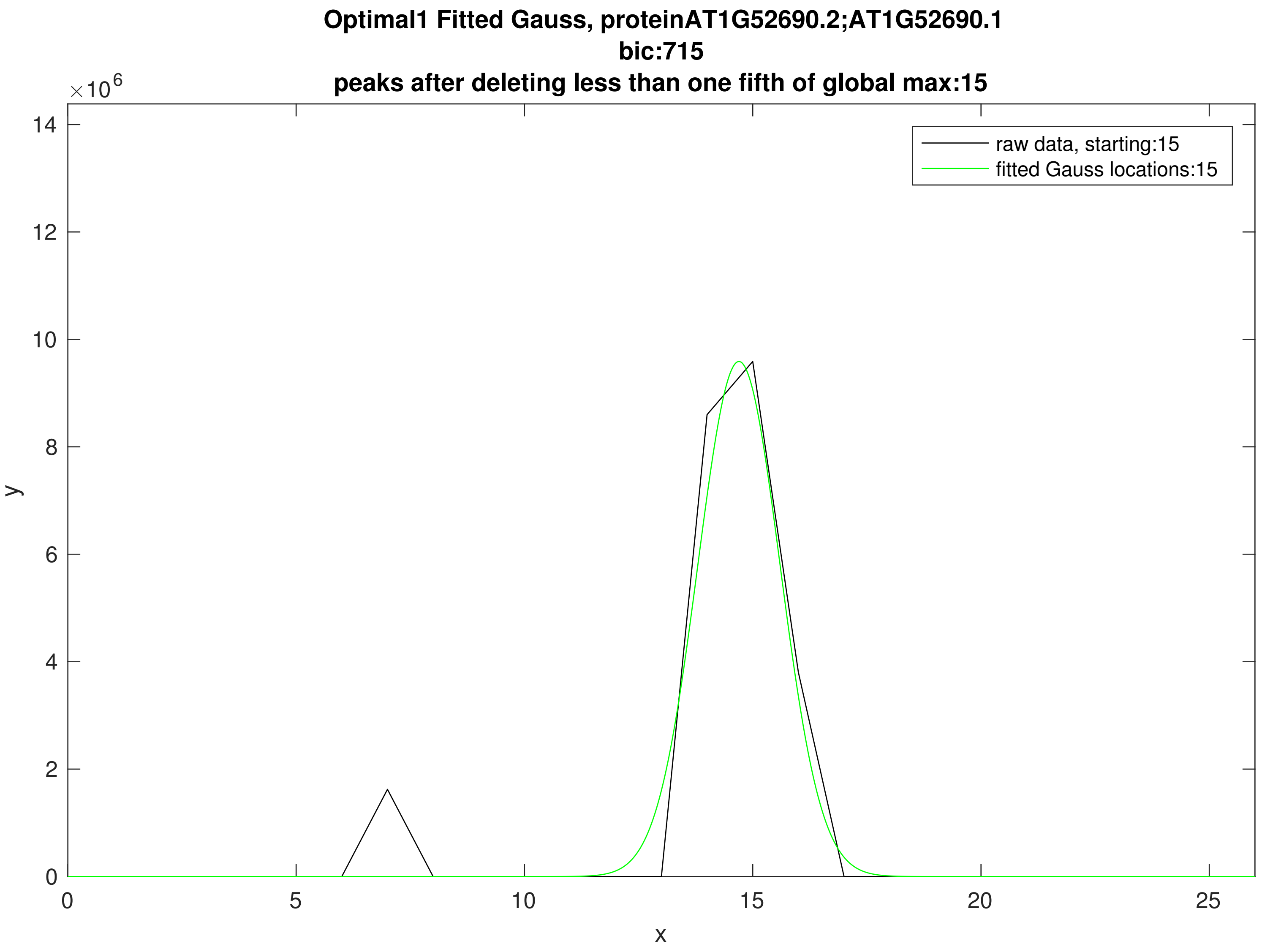
Optimal1 Fitted Gauss, proteinAT1G52690.2;AT1G52690.1
bic:715
peaks after deleting less than one fifth of global max:15
×10^6
raw data, starting:15
fitted Gauss locations:15
y
x
0
2
4
6
8
10
12
14
0
5
10
15
20
25

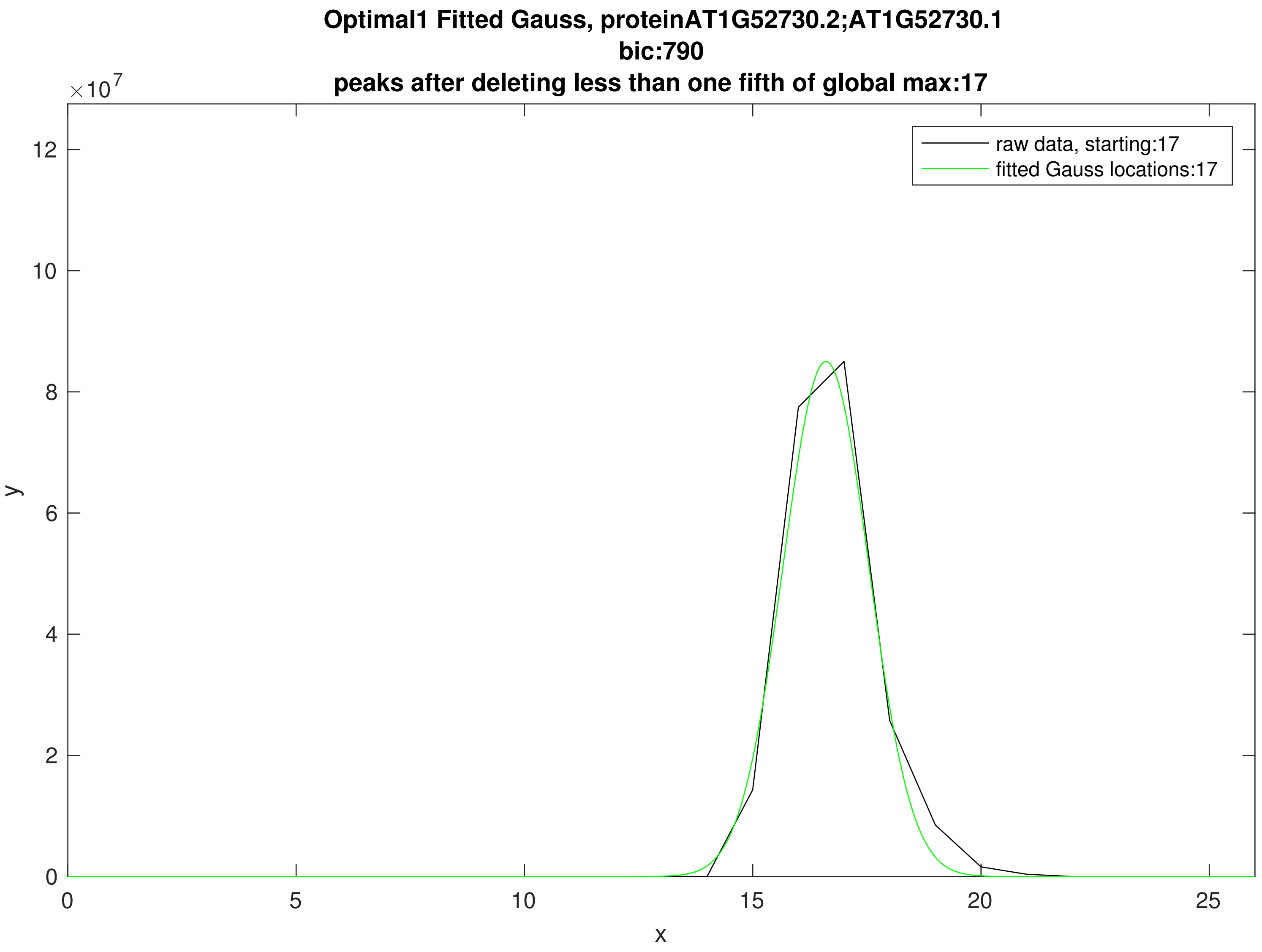
Optimal1 Fitted Gauss, proteinAT1G52730.2;AT1G52730.1
bic:790
peaks after deleting less than one fifth of global max:17
×10^7
12
10
8
6
4
2
0
y
0
5
10
15
20
25
x
raw data, starting:17
fitted Gauss locations:17

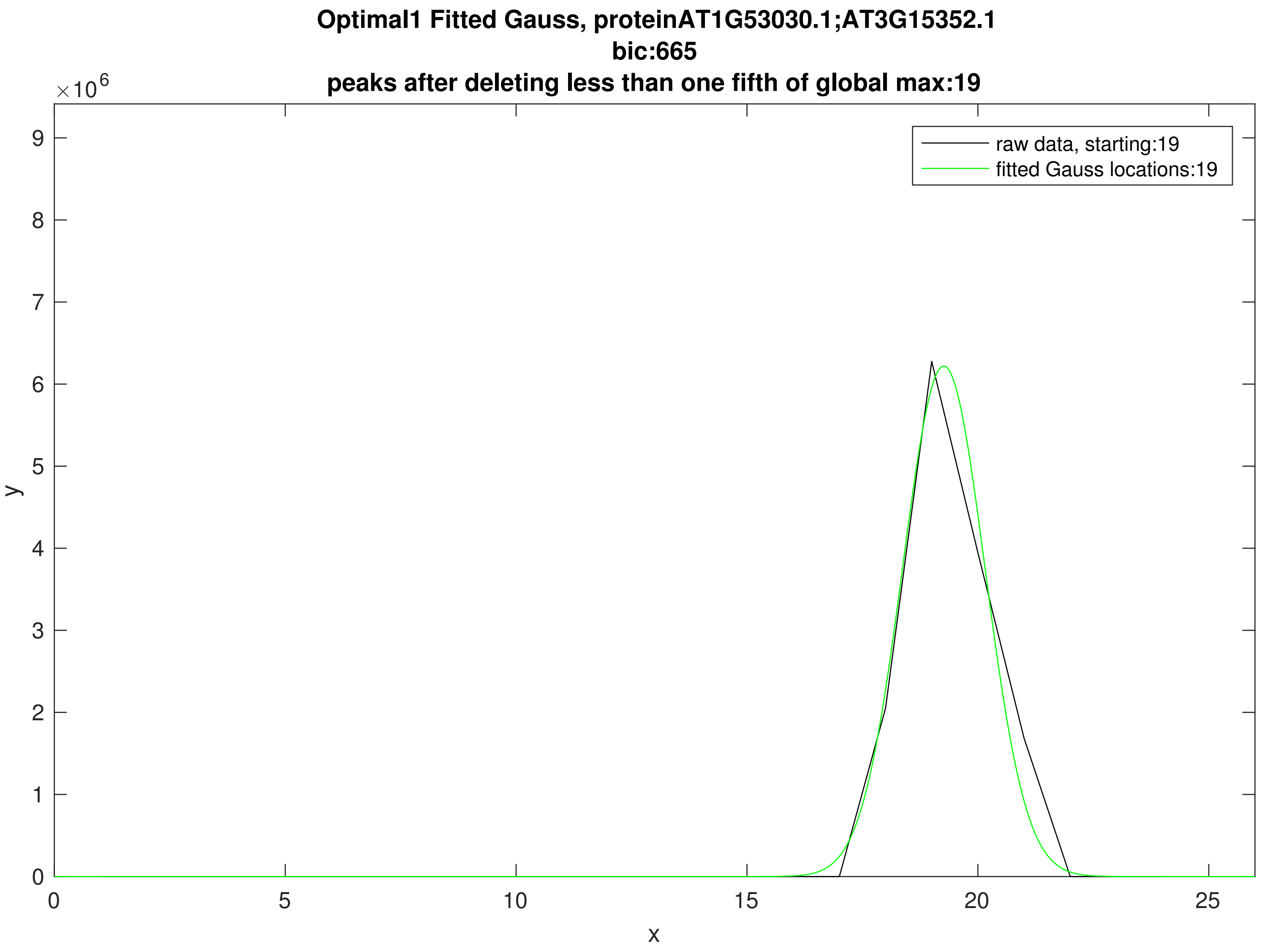
Optimal1 Fitted Gauss, proteinAT1G53030.1;AT3G15352.1
bic:665
peaks after deleting less than one fifth of global max:19
×10^6
raw data, starting:19
fitted Gauss locations:19
y
x
0
1
2
3
4
5
6
7
8
9
0
5
10
15
20
25

**Optimal1 Fitted Gauss, proteinAT1G53070.1**
**bic:688 780**
**peaks after deleting less than one fifth of global max:19**

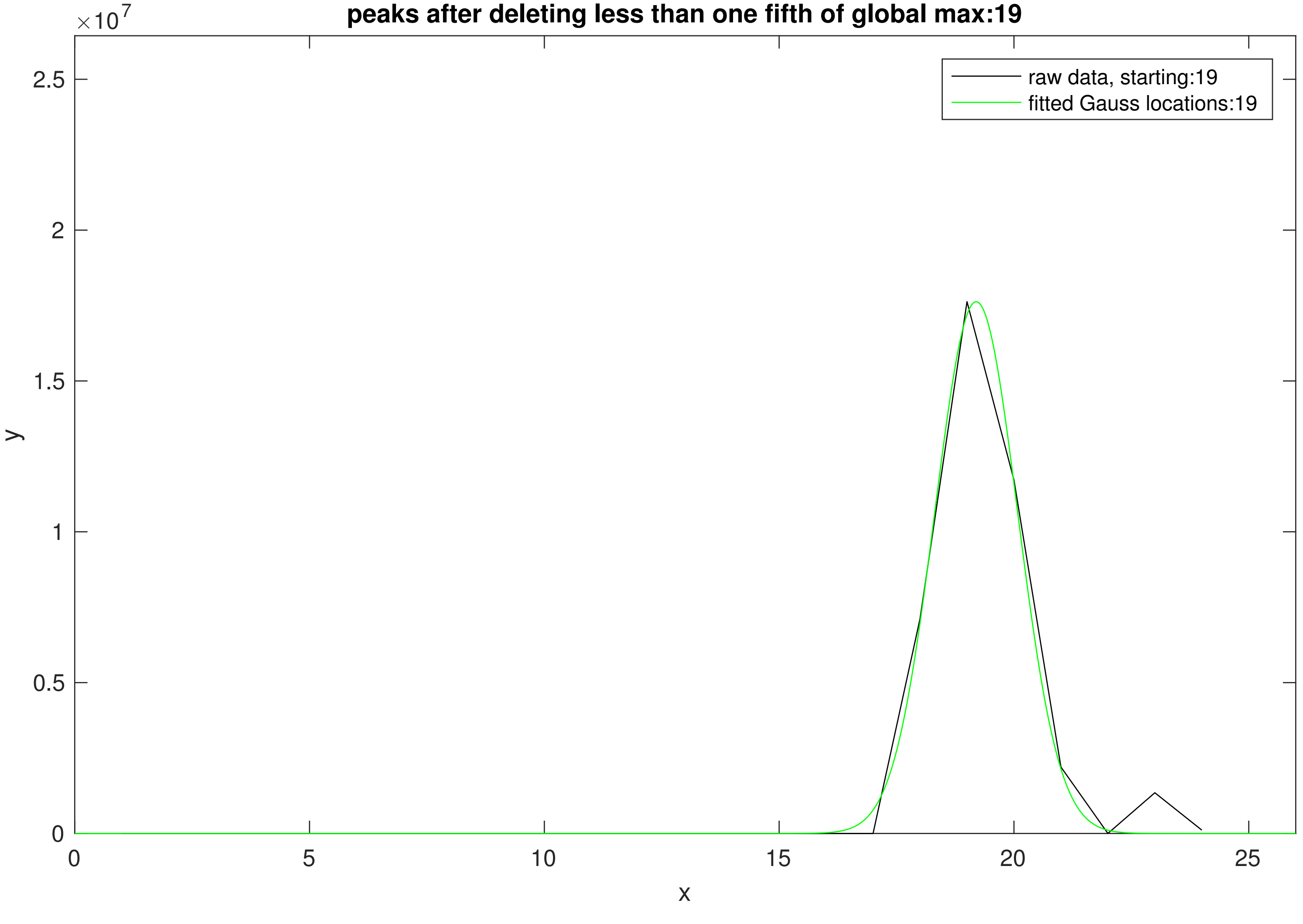

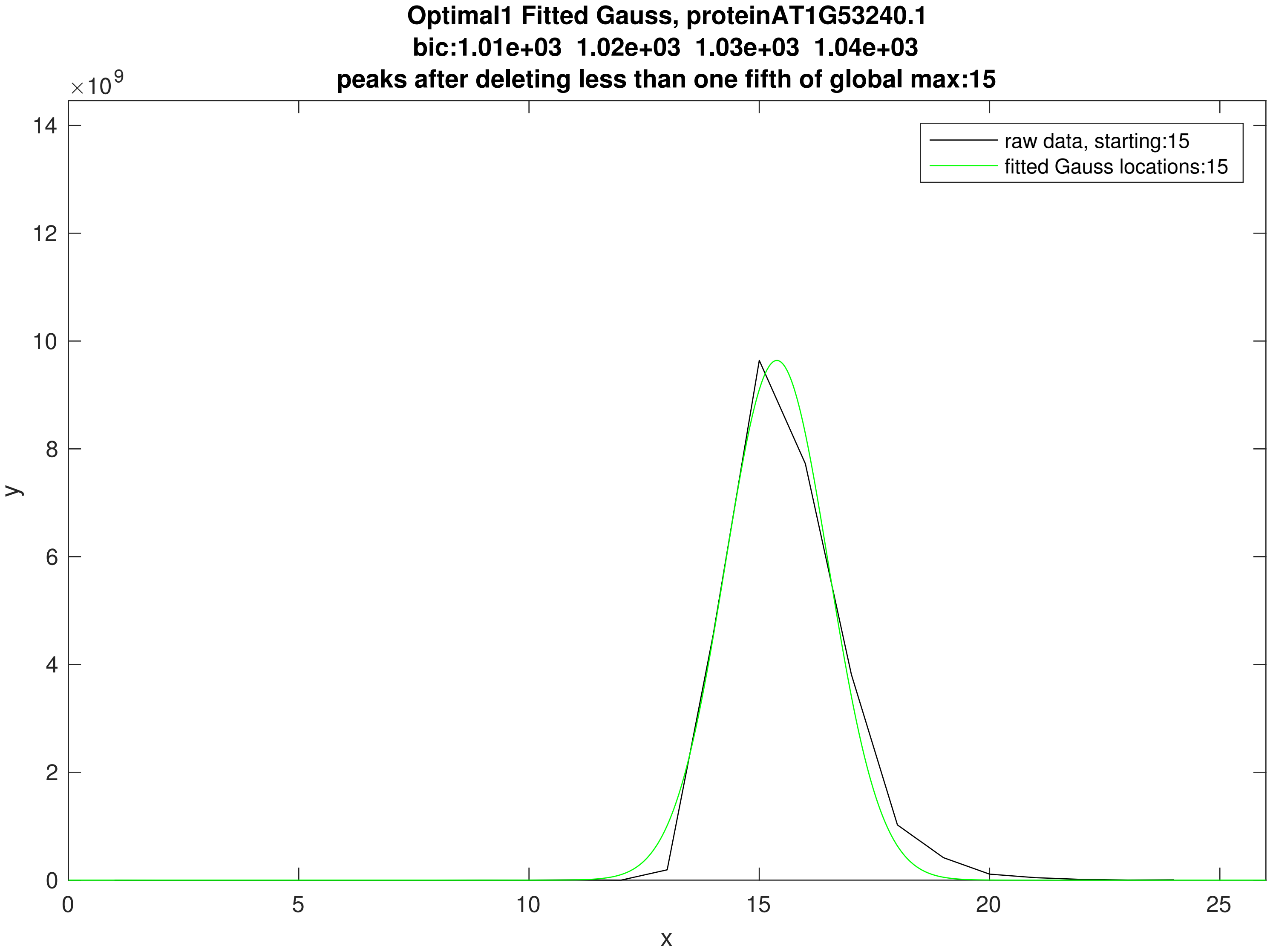

Optimal1 Fitted Gauss, proteinAT1G53240.1
bic:1.01e+03 1.02e+03 1.03e+03 1.04e+03
peaks after deleting less than one fifth of global max:15
×10^9
14
12
10
8
6
4
2
0
y
0
5
10
15
20
25
x
raw data, starting:15
fitted Gauss locations:15

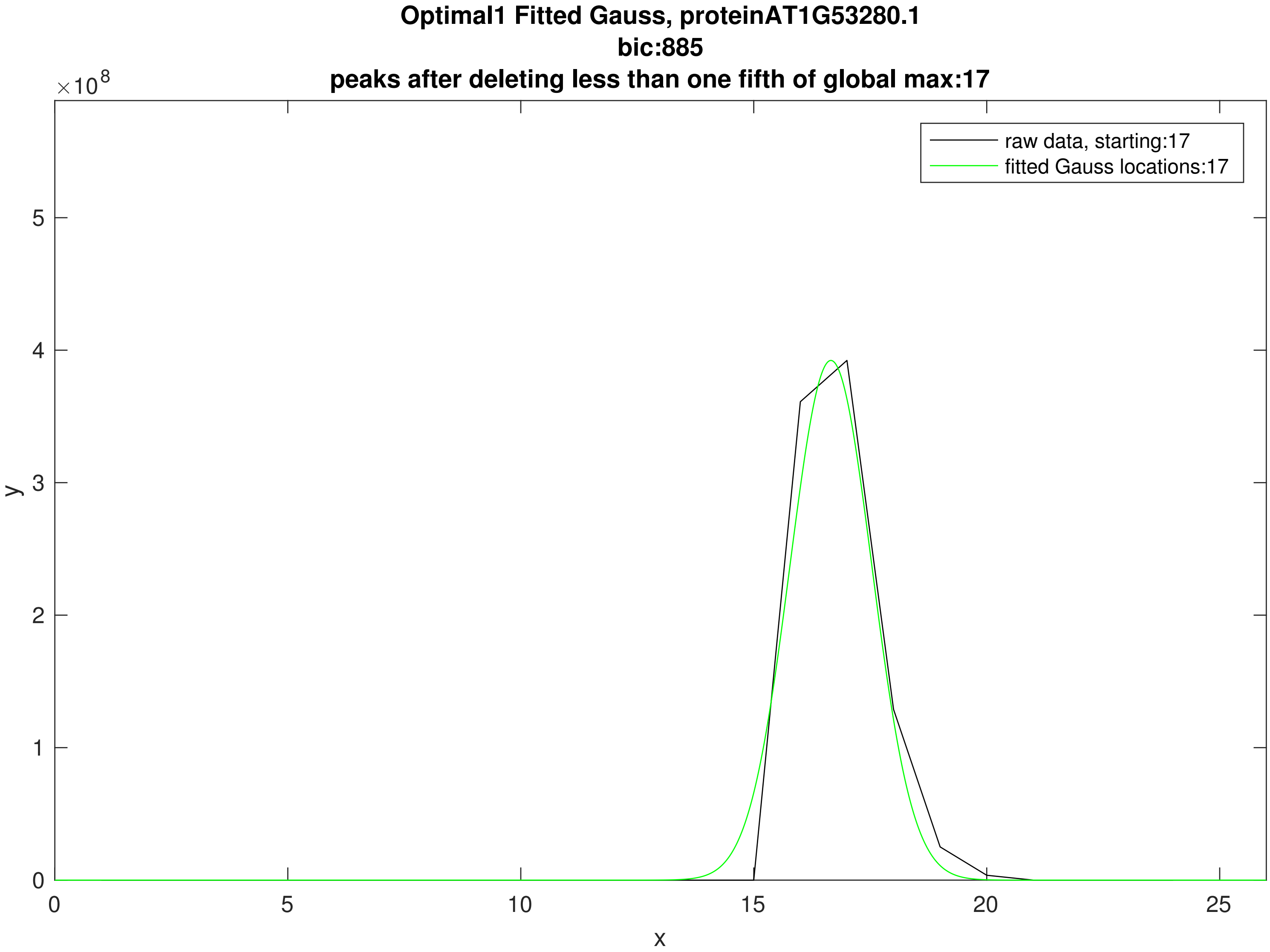

Optimal1 Fitted Gauss, proteinAT1G53280.1
bic:885
peaks after deleting less than one fifth of global max:17
×10^8
raw data, starting:17
fitted Gauss locations:17
y
x
0
1
2
3
4
5
0
5
10
15
20
25

#### Optimal2 Fitted Gauss, proteinAT1G53310.3;AT1G53310.2;AT1G53310.1
#### bic:1.07e+03 1.01e+03 1.02e+03 1.03e+03
#### peaks after deleting less than one fifth of global max:6.9 11

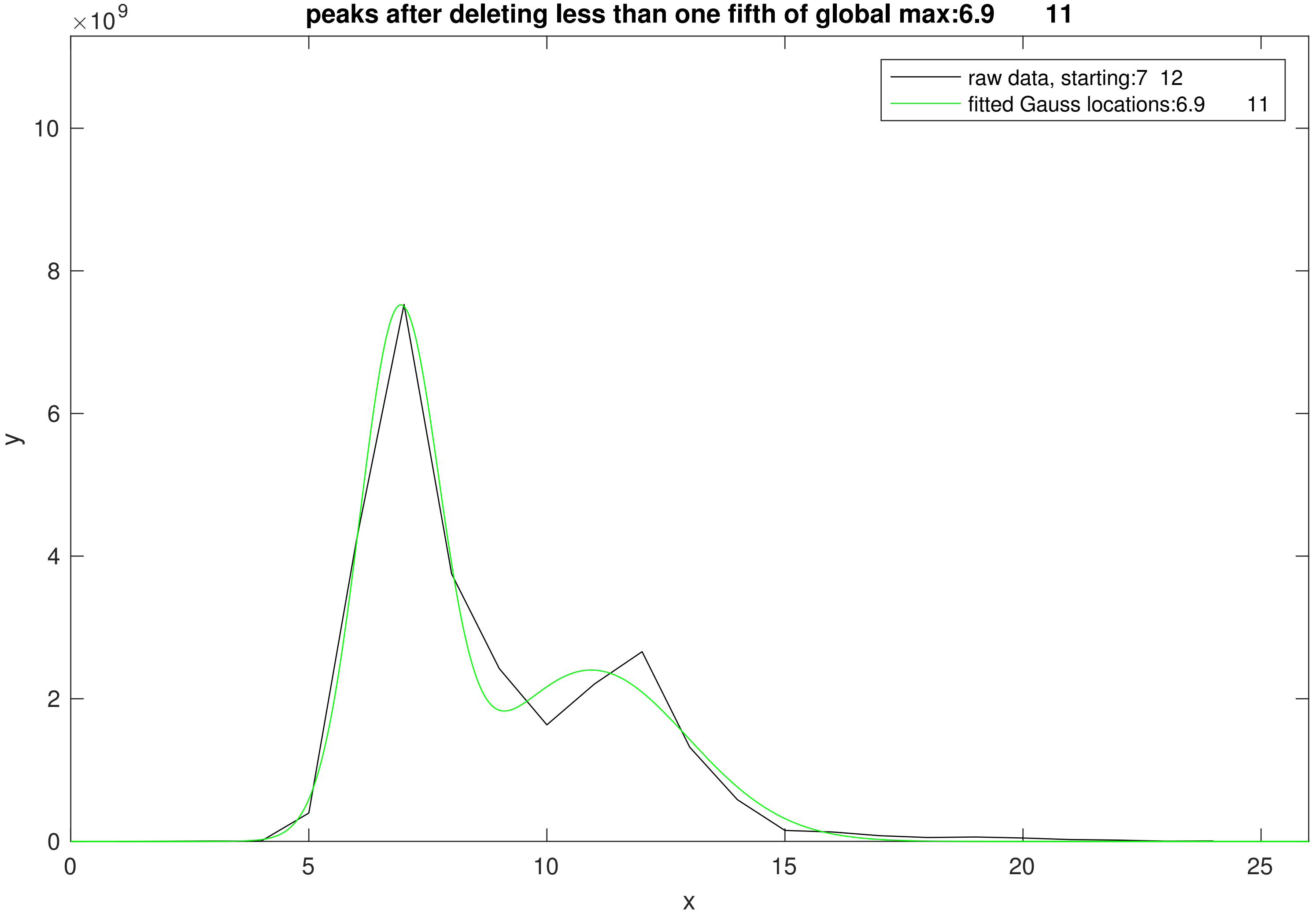

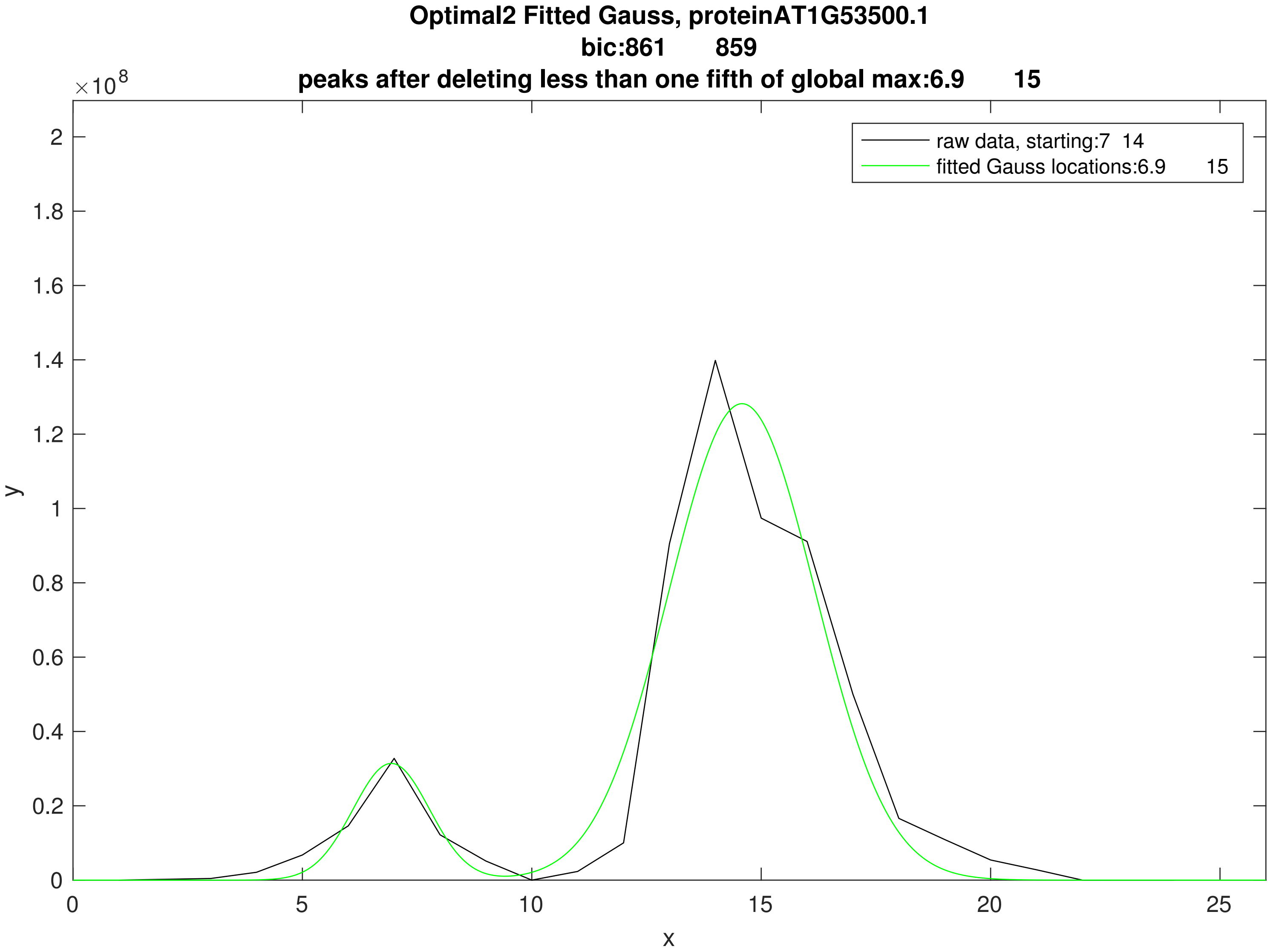

Optimal2 Fitted Gauss, proteinAT1G53500.1
bic:861 859
peaks after deleting less than one fifth of global max:6.9 15
×10^8
raw data, starting:7 14
fitted Gauss locations:6.9 15
y
x
0
0.2
0.4
0.6
0.8
1
1.2
1.4
1.6
1.8
2
0
5
10
15
20
25

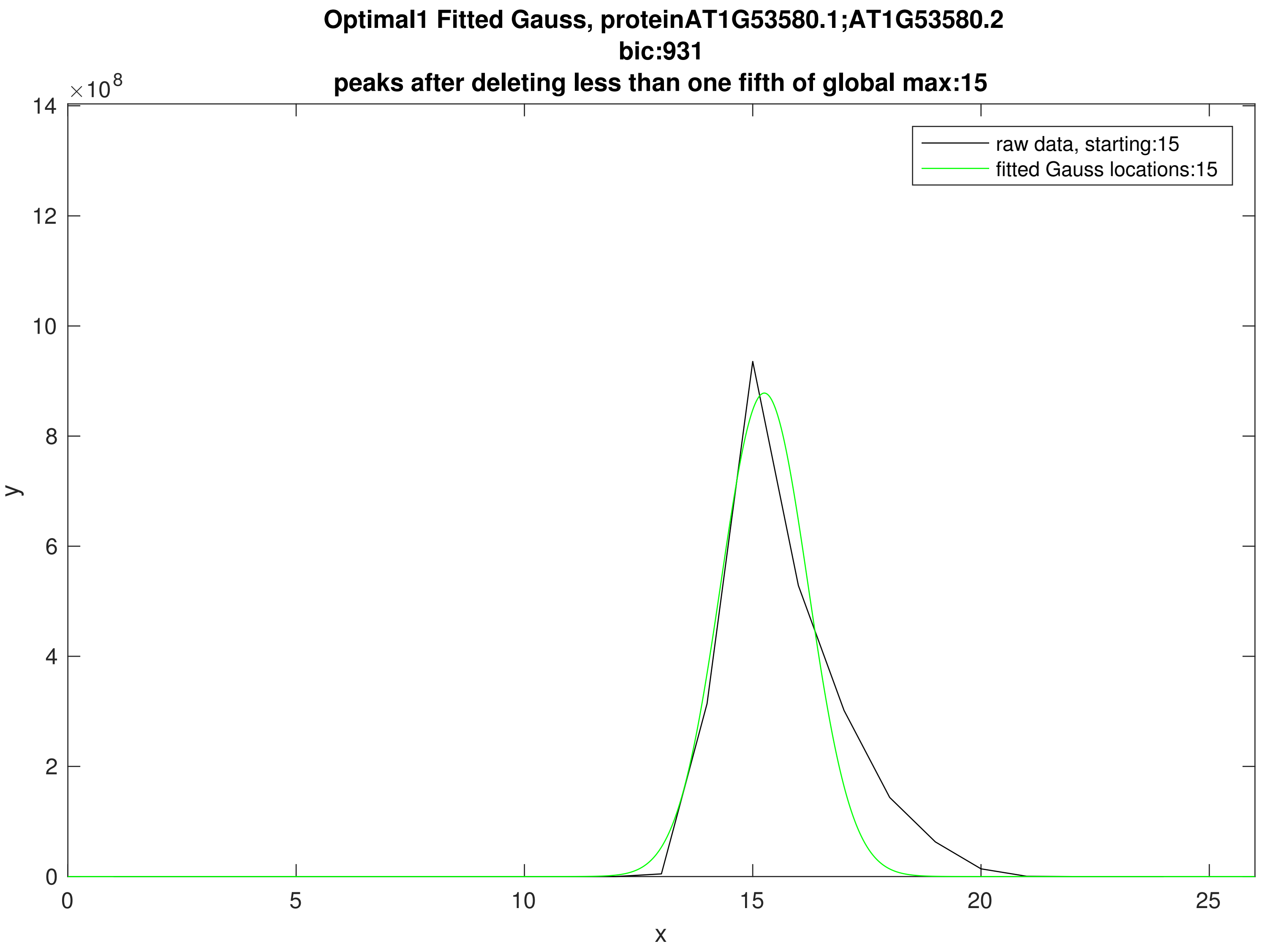
Optimal1 Fitted Gauss, proteinAT1G53580.1;AT1G53580.2
bic:931
peaks after deleting less than one fifth of global max:15
raw data, starting:15
fitted Gauss locations:15
×10^8
14
12
10
8
6
4
2
0
y
0
5
10
15
20
25
x

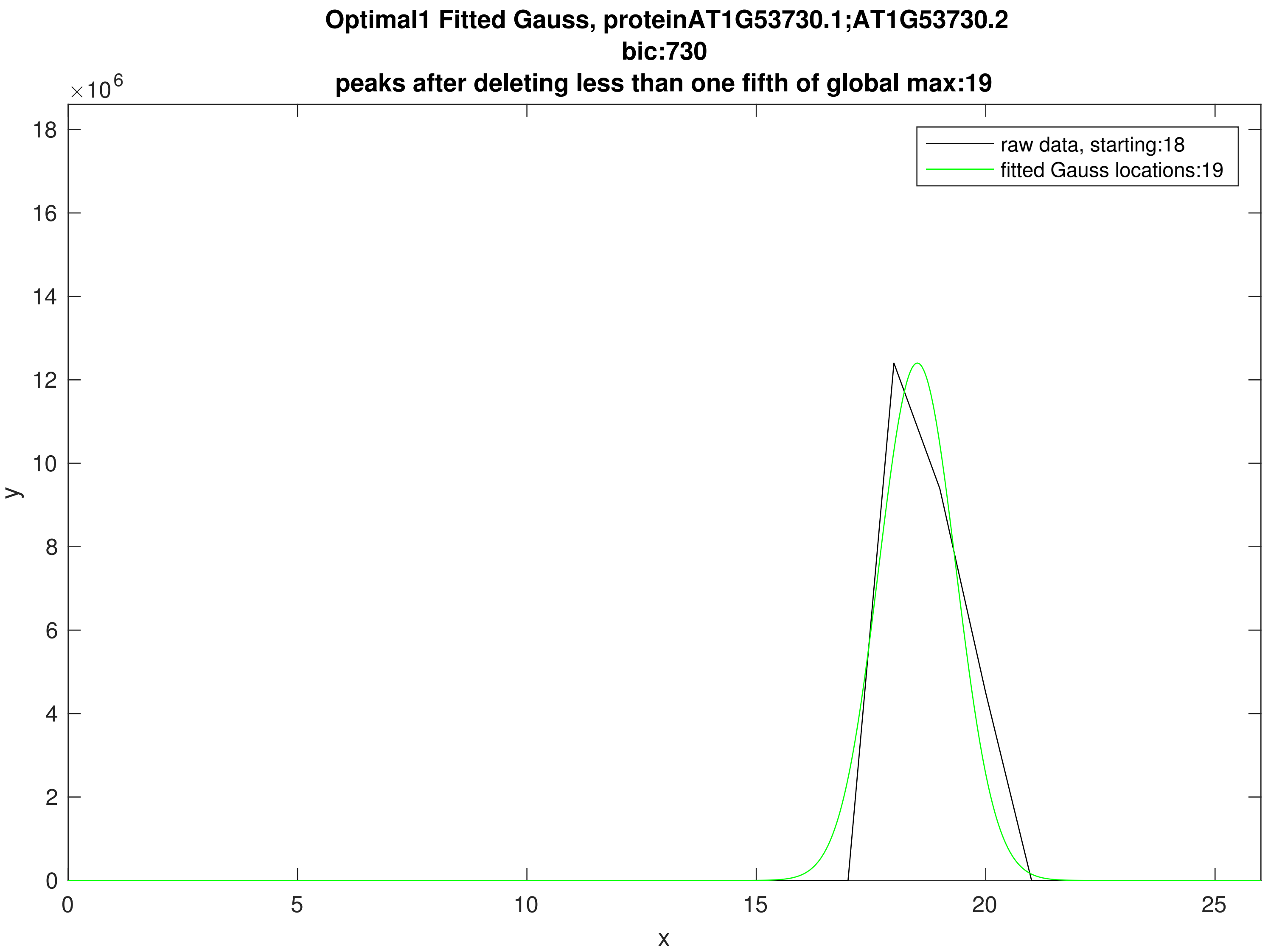
Optimal1 Fitted Gauss, proteinAT1G53730.1;AT1G53730.2
bic:730
peaks after deleting less than one fifth of global max:19
×10^6
raw data, starting:18
fitted Gauss locations:19
y
x
0
2
4
6
8
10
12
14
16
18
0
5
10
15
20
25

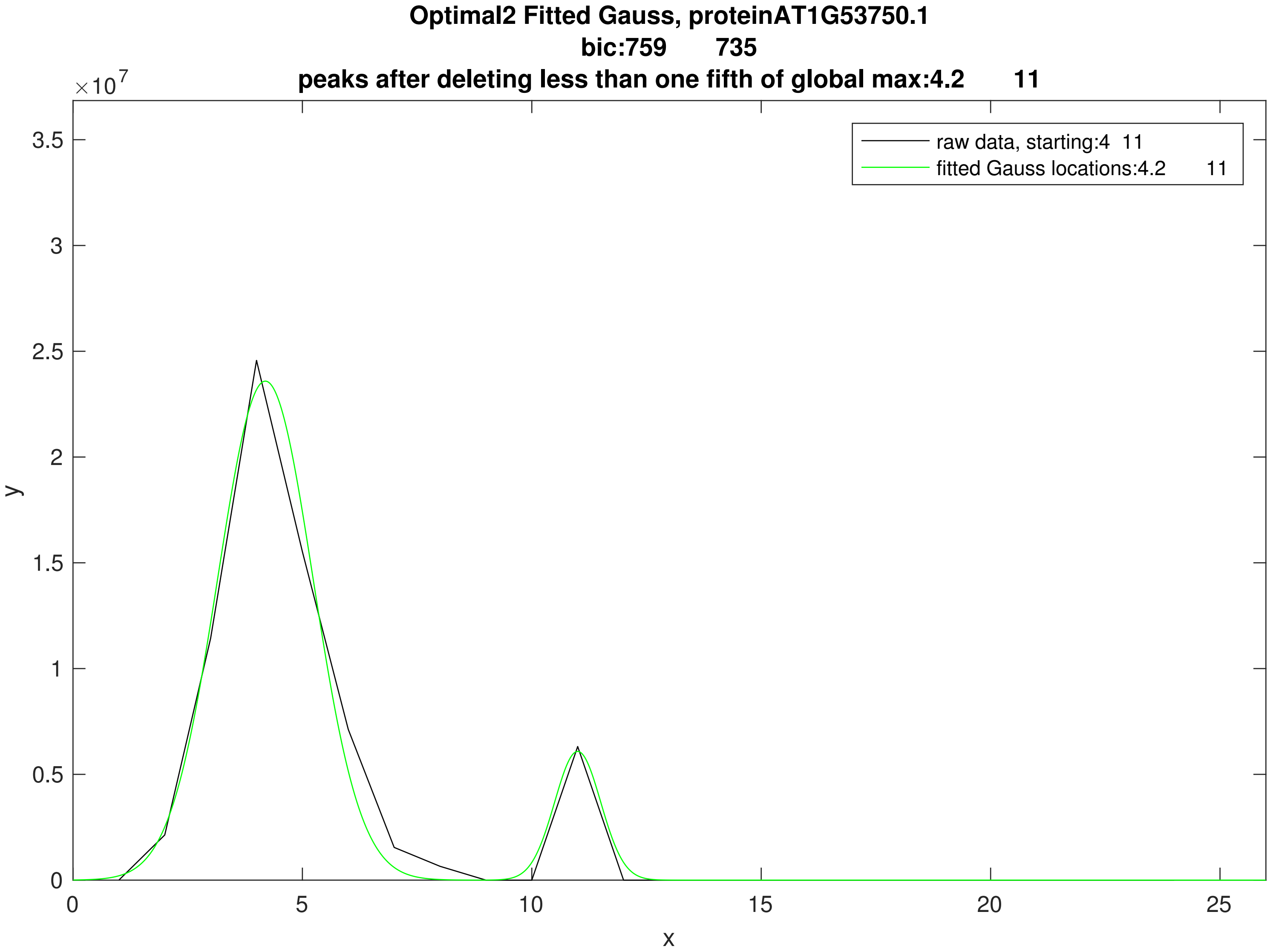
Optimal2 Fitted Gauss, proteinAT1G53750.1
bic:759 735
peaks after deleting less than one fifth of global max:4.2 11
×10^7
3.5
3
2.5
2
1.5
1
0.5
0
y
0
5
10
15
20
25
x
raw data, starting:4 11
fitted Gauss locations:4.2 11

Optimal1 Fitted Gauss, proteinAT1G53830.1
bic:820
peaks after deleting less than one fifth of global max:18

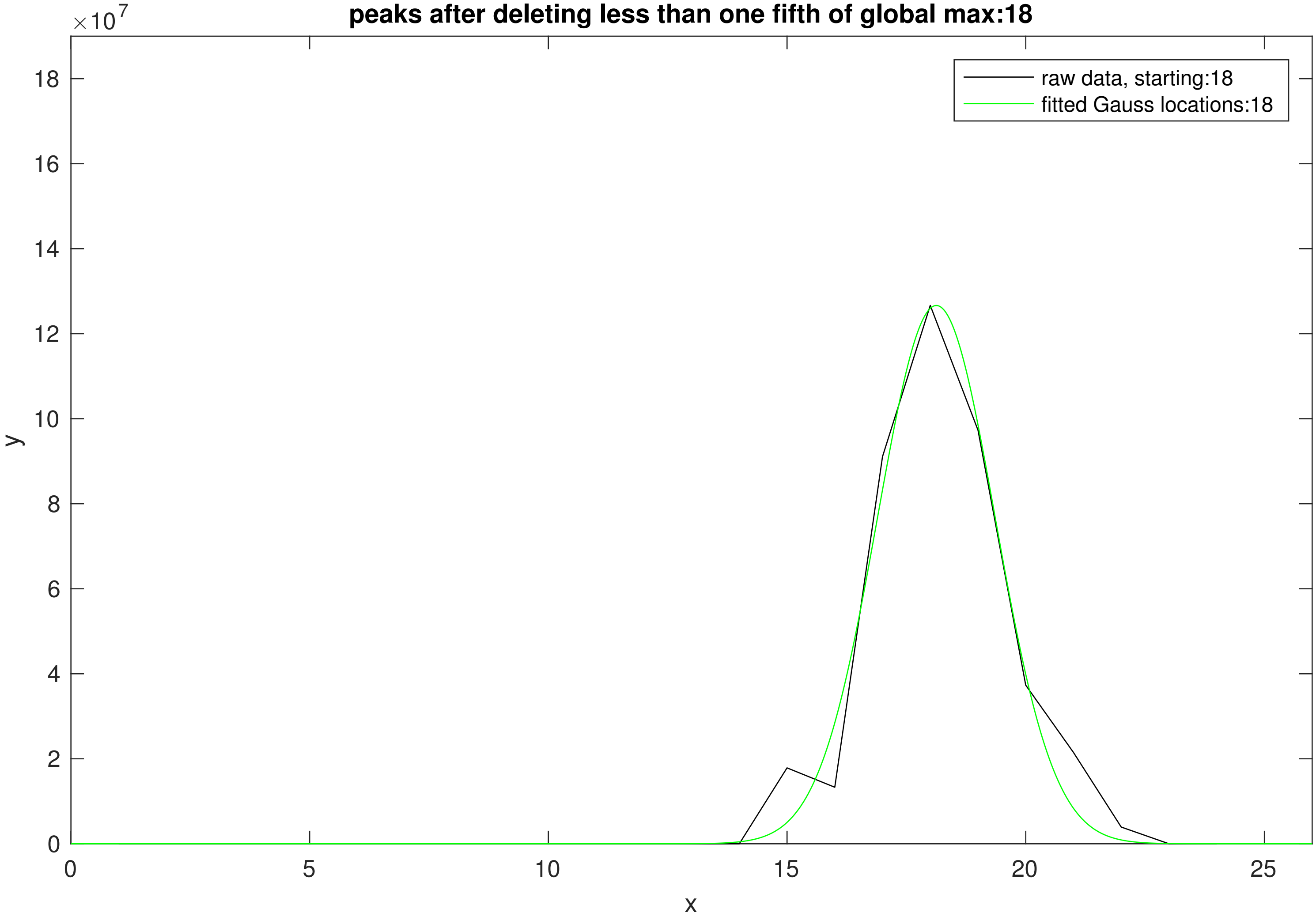

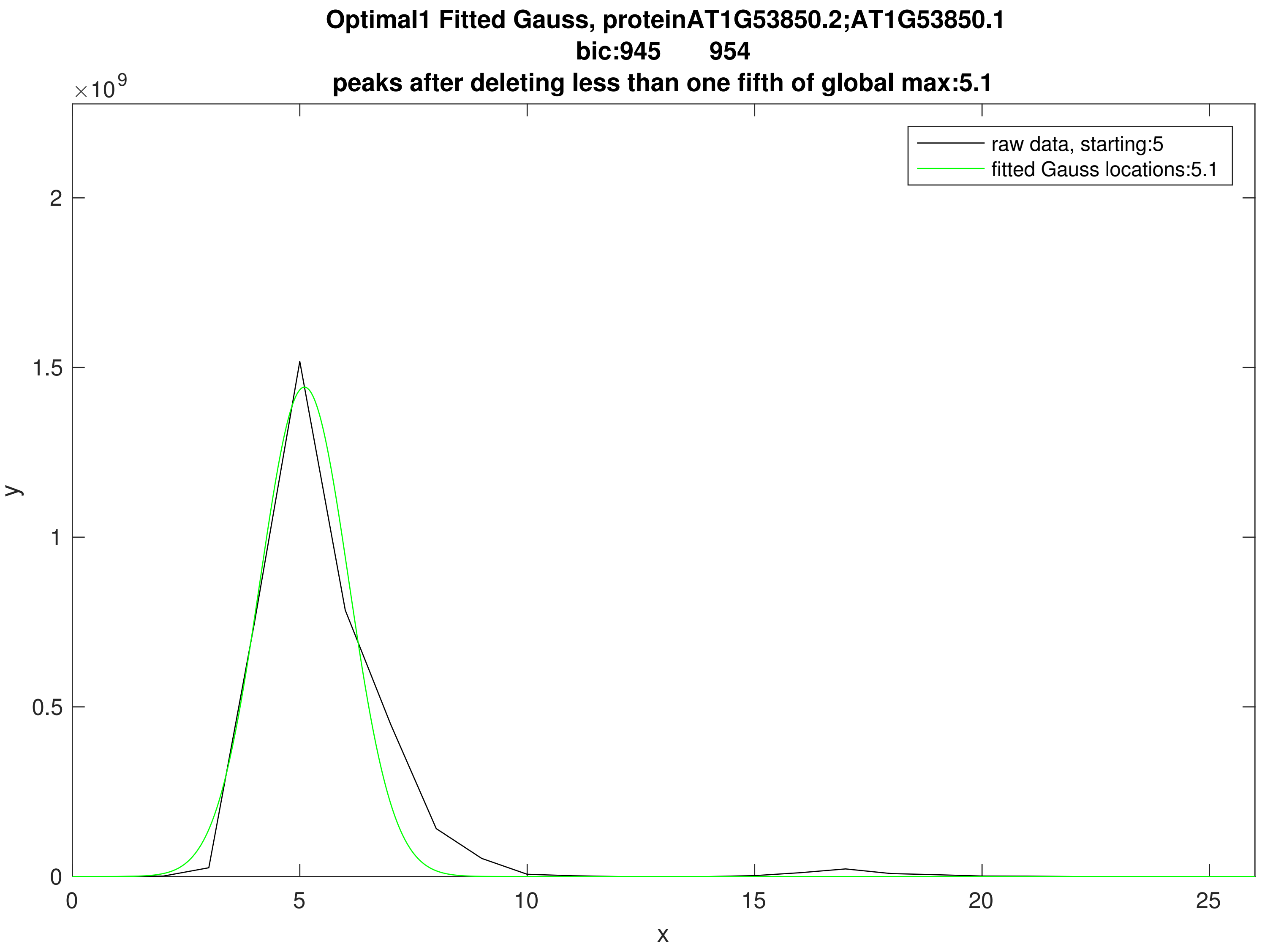
Optimal1 Fitted Gauss, proteinAT1G53850.2;AT1G53850.1
bic:945 954
peaks after deleting less than one fifth of global max:5.1
×10^9
2
1.5
1
0.5
0
y
0
5
10
15
20
25
x
raw data, starting:5
fitted Gauss locations:5.1

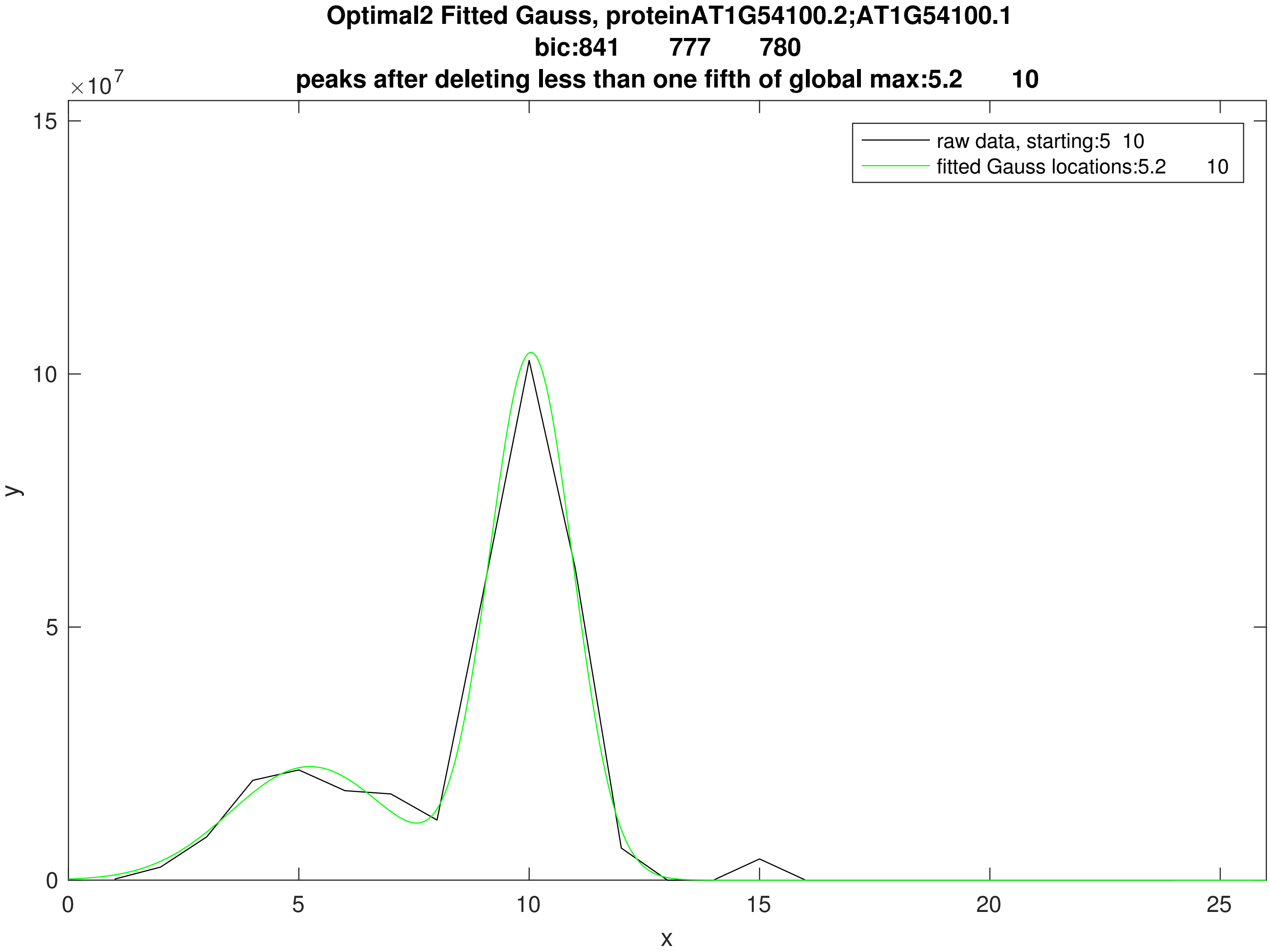

Optimal2 Fitted Gauss, proteinAT1G54100.2;AT1G54100.1
bic:841 777 780
peaks after deleting less than one fifth of global max:5.2 10
×10^7
raw data, starting:5 10
fitted Gauss locations:5.2 10
y
x
0
5
10
15
20
25

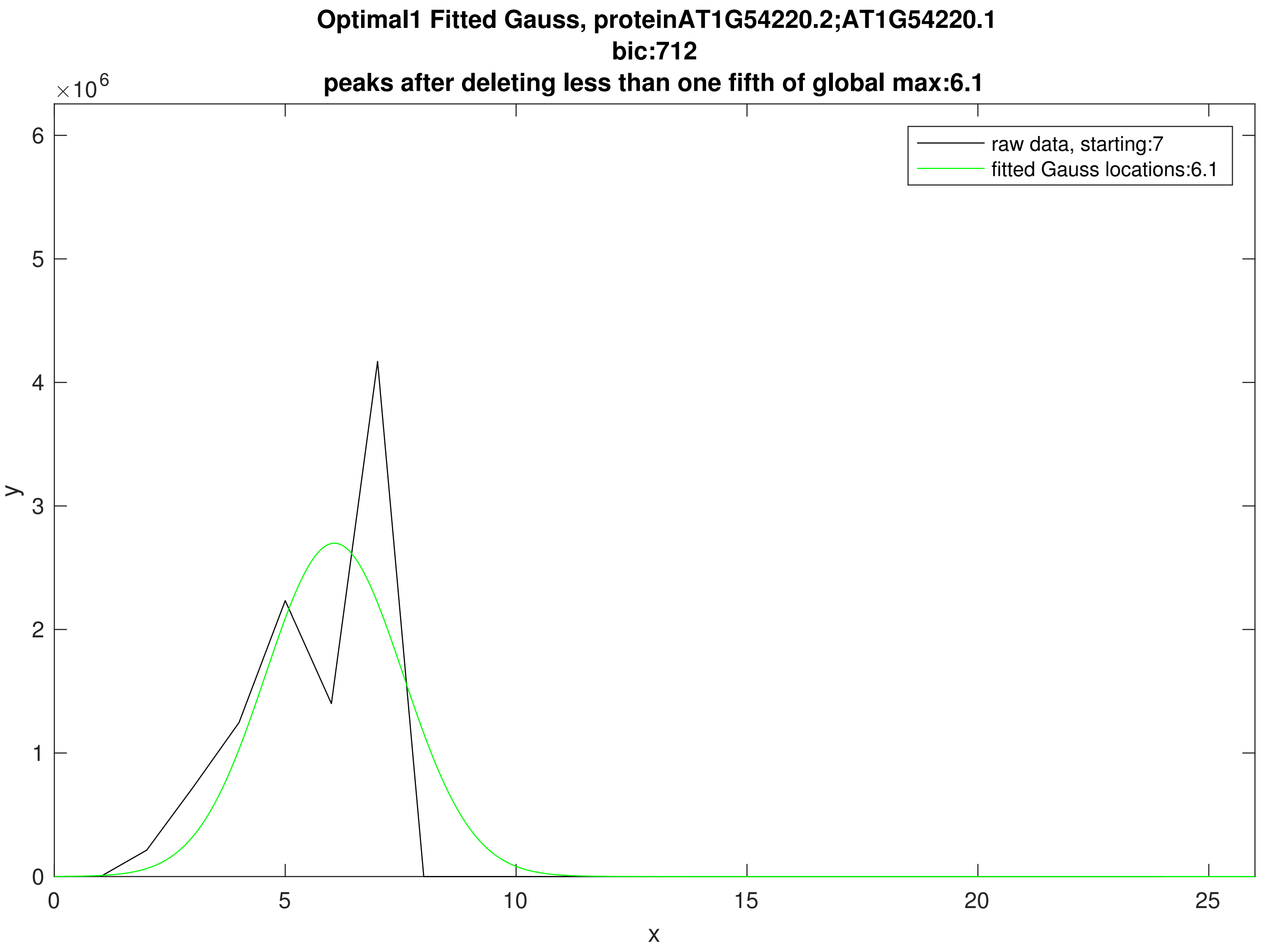
Optimal1 Fitted Gauss, proteinAT1G54220.2;AT1G54220.1
bic:712
peaks after deleting less than one fifth of global max:6.1
×10^6
raw data, starting:7
fitted Gauss locations:6.1
y
x
0
1
2
3
4
5
6
0
5
10
15
20
25

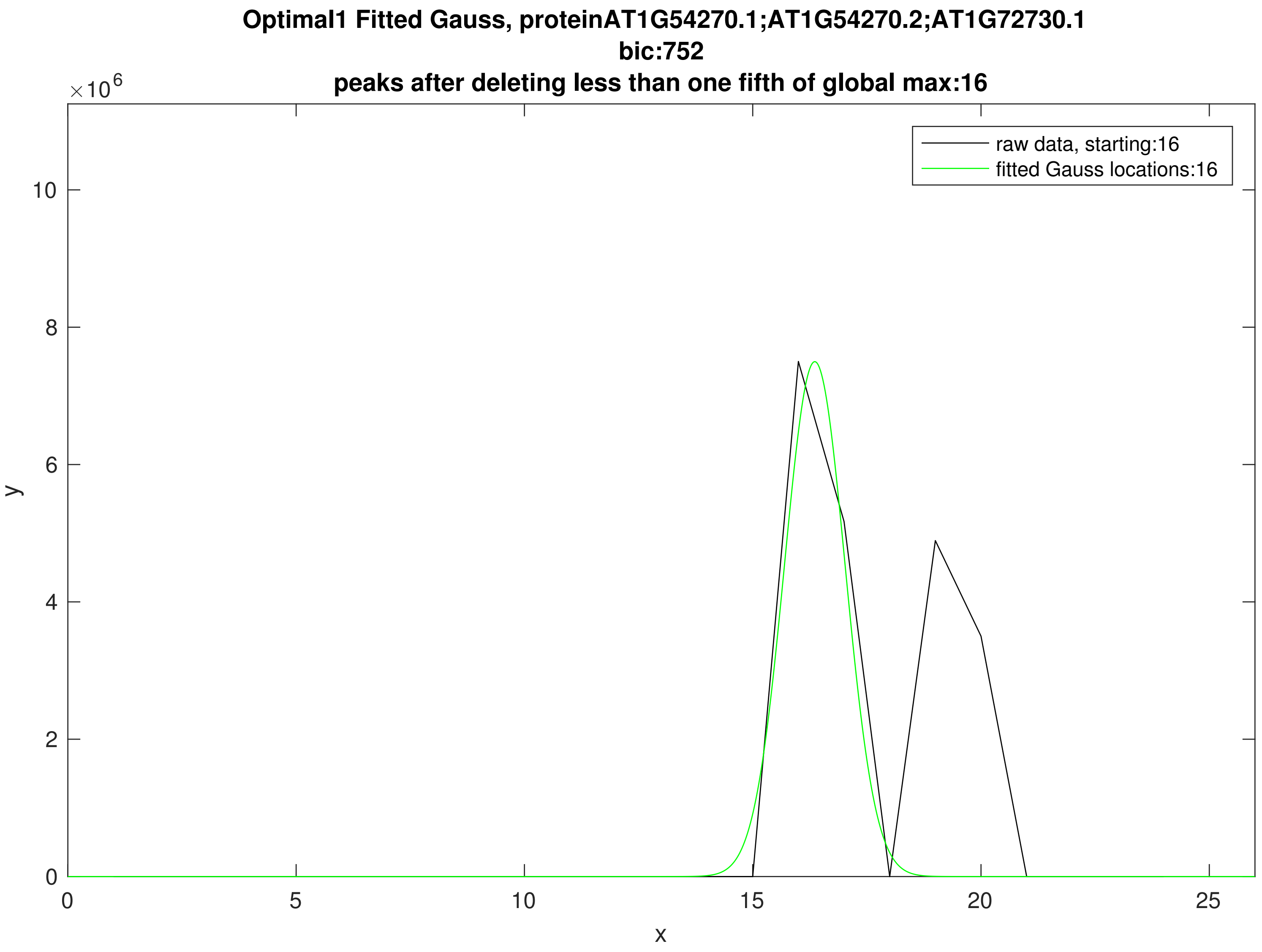
Optimal1 Fitted Gauss, proteinAT1G54270.1;AT1G54270.2;AT1G72730.1
bic:752
peaks after deleting less than one fifth of global max:16
×10^6
raw data, starting:16
fitted Gauss locations:16
10
8
6
4
2
0
y
0
5
10
15
20
25
x

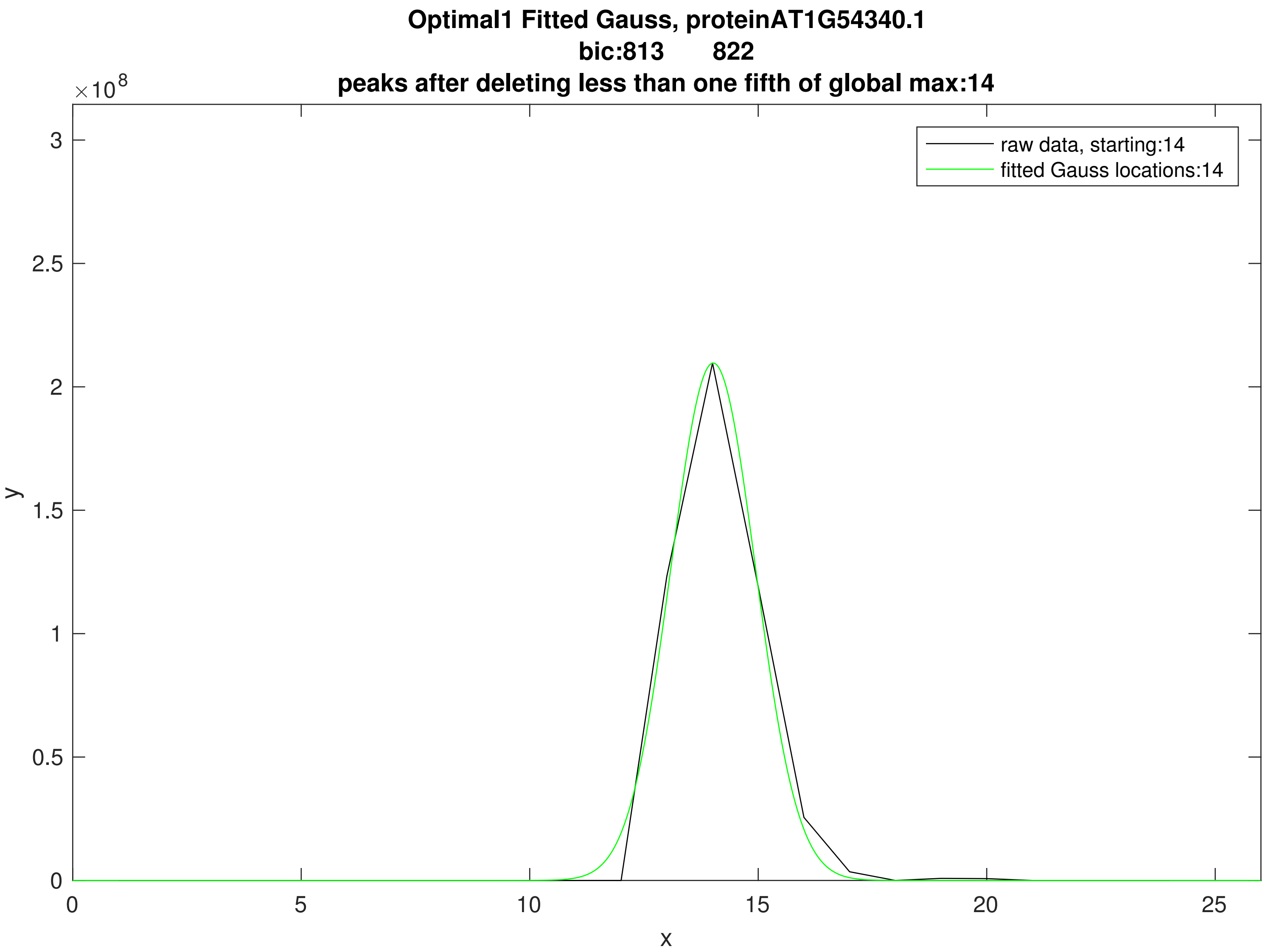
Optimal1 Fitted Gauss, proteinAT1G54340.1
bic:813 822
peaks after deleting less than one fifth of global max:14
×10^8
3
2.5
2
1.5
1
0.5
0
y
0
5
10
15
20
25
x
raw data, starting:14
fitted Gauss locations:14

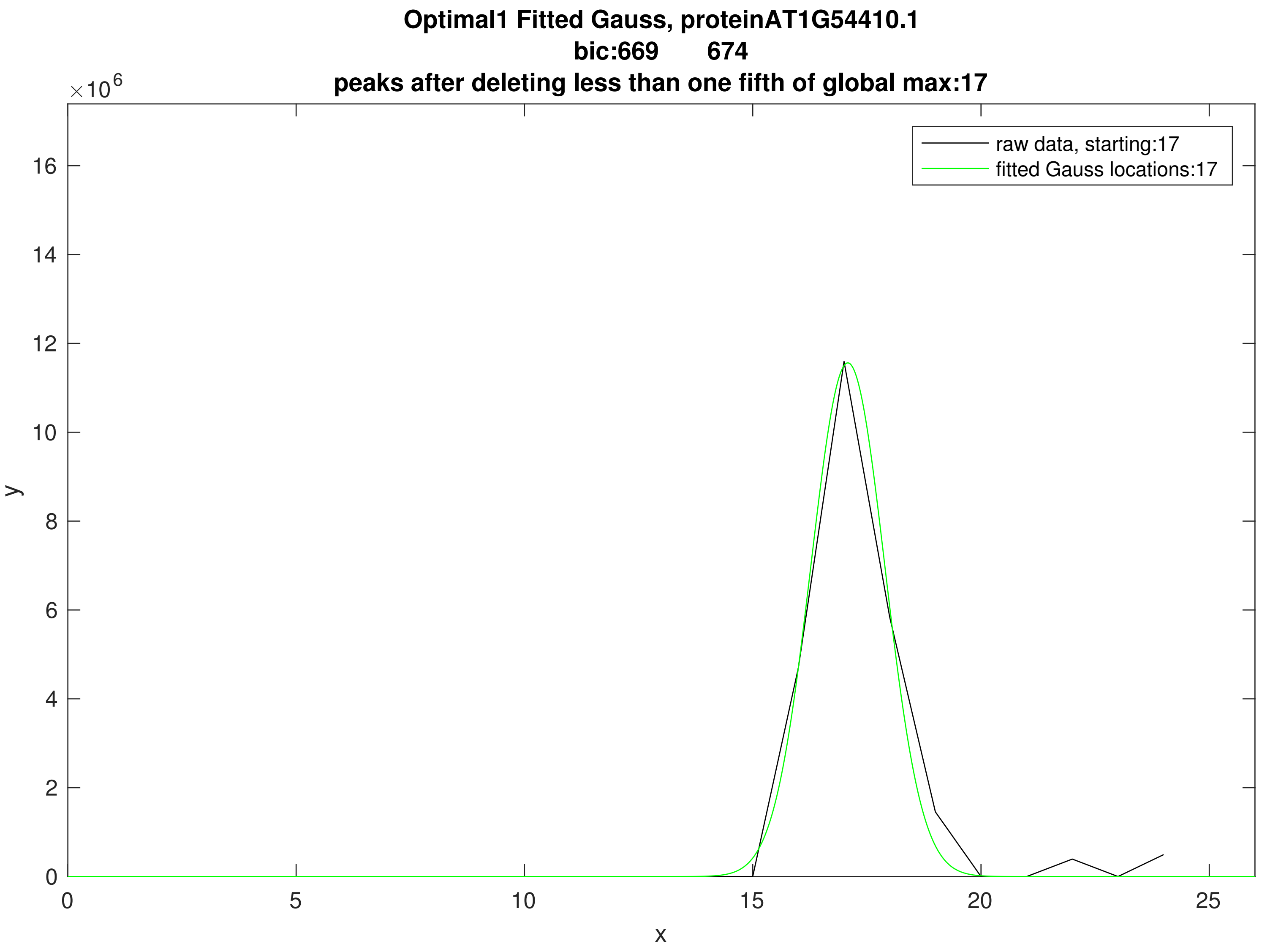
Optimal1 Fitted Gauss, proteinAT1G54410.1
bic:669 674
peaks after deleting less than one fifth of global max:17
×10^6
raw data, starting:17
fitted Gauss locations:17
y
x
0
2
4
6
8
10
12
14
16
0
5
10
15
20
25

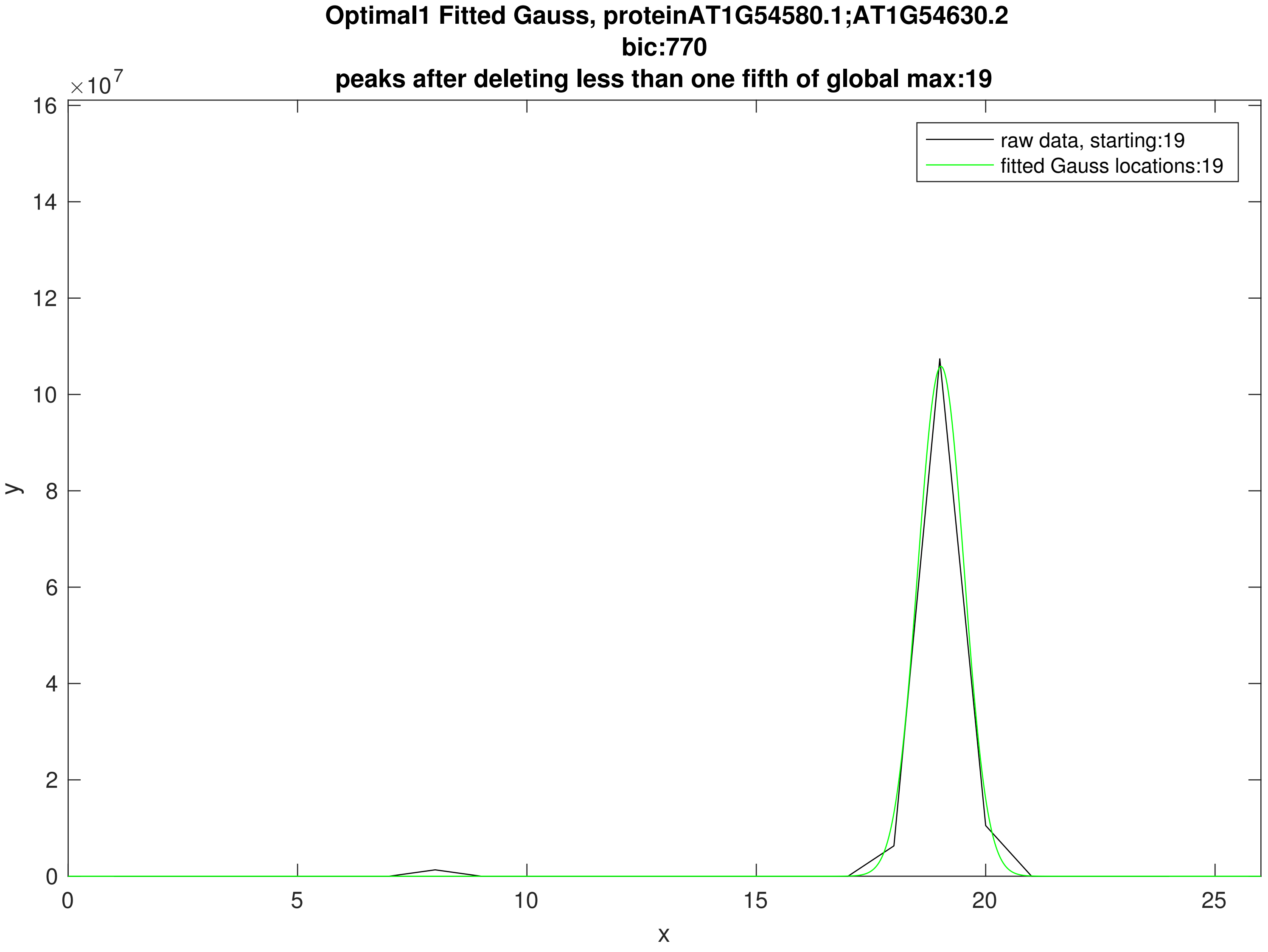

Optimal1 Fitted Gauss, proteinAT1G54580.1;AT1G54630.2
bic:770
peaks after deleting less than one fifth of global max:19
×10^7
16
14
12
10
8
6
4
2
0
y
0
5
10
15
20
25
x
raw data, starting:19
fitted Gauss locations:19

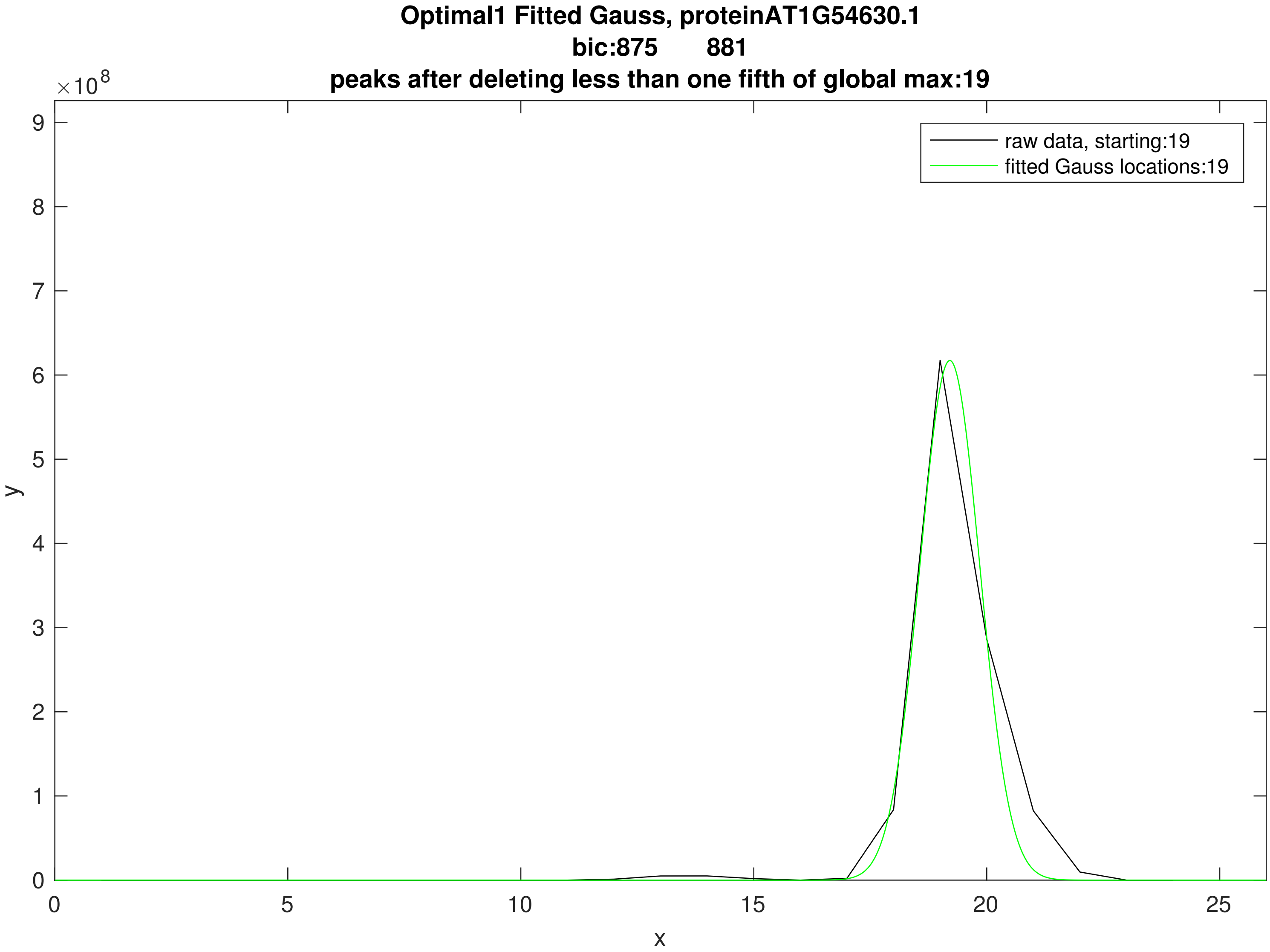

Optimal1 Fitted Gauss, proteinAT1G54630.1
bic:875 881
peaks after deleting less than one fifth of global max:19
×10^8
9
8
7
6
5
4
3
2
1
0
y
0
5
10
15
20
25
x
raw data, starting:19
fitted Gauss locations:19

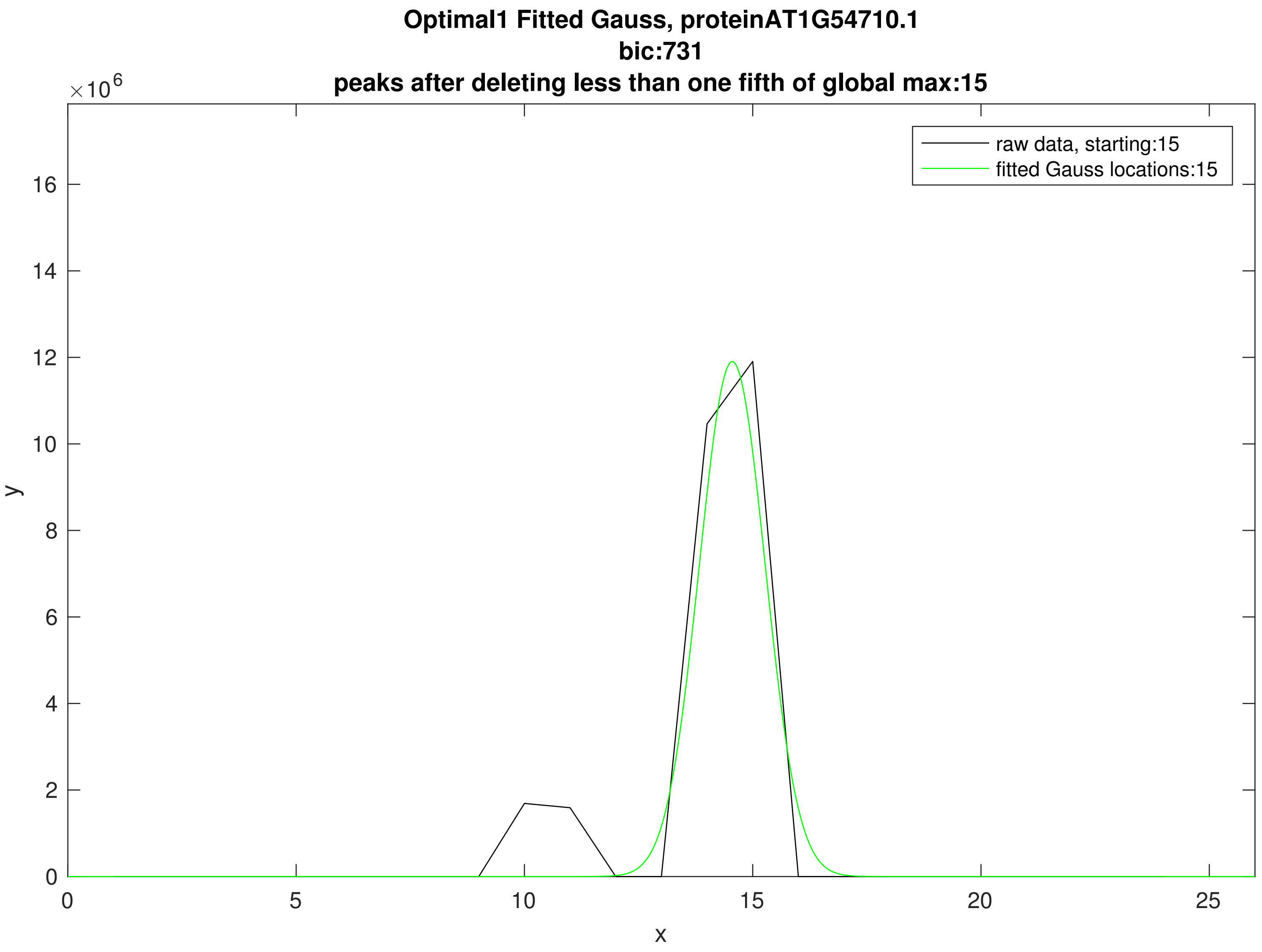
Optimal1 Fitted Gauss, proteinAT1G54710.1
bic:731
peaks after deleting less than one fifth of global max:15
×10^6
raw data, starting:15
fitted Gauss locations:15
16
14
12
10
8
6
4
2
0
y
0
5
10
15
20
25
x

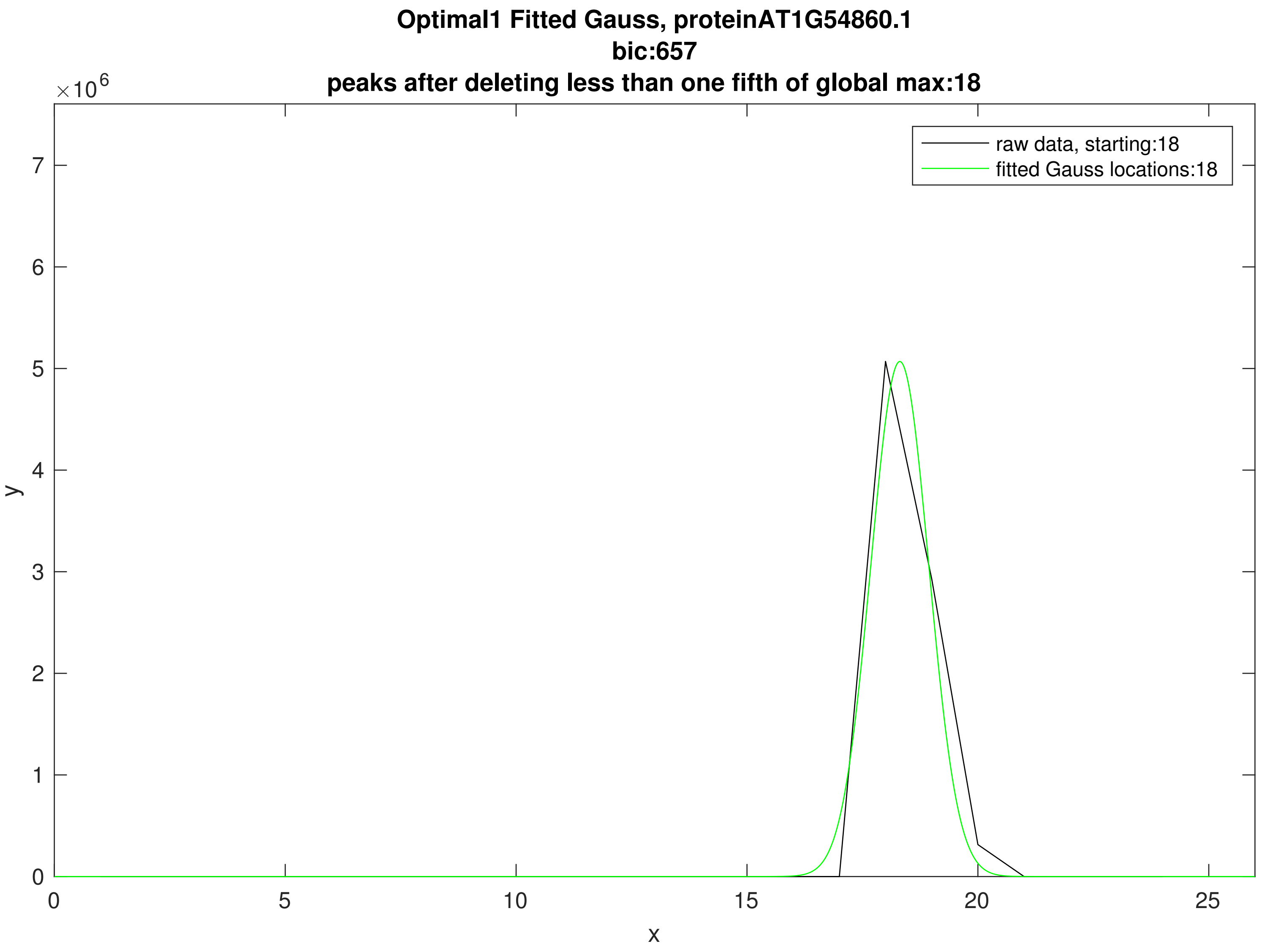
Optimal1 Fitted Gauss, proteinAT1G54860.1
bic:657
peaks after deleting less than one fifth of global max:18
×10^6
raw data, starting:18
fitted Gauss locations:18
y
x
0
1
2
3
4
5
6
7
0
5
10
15
20
25

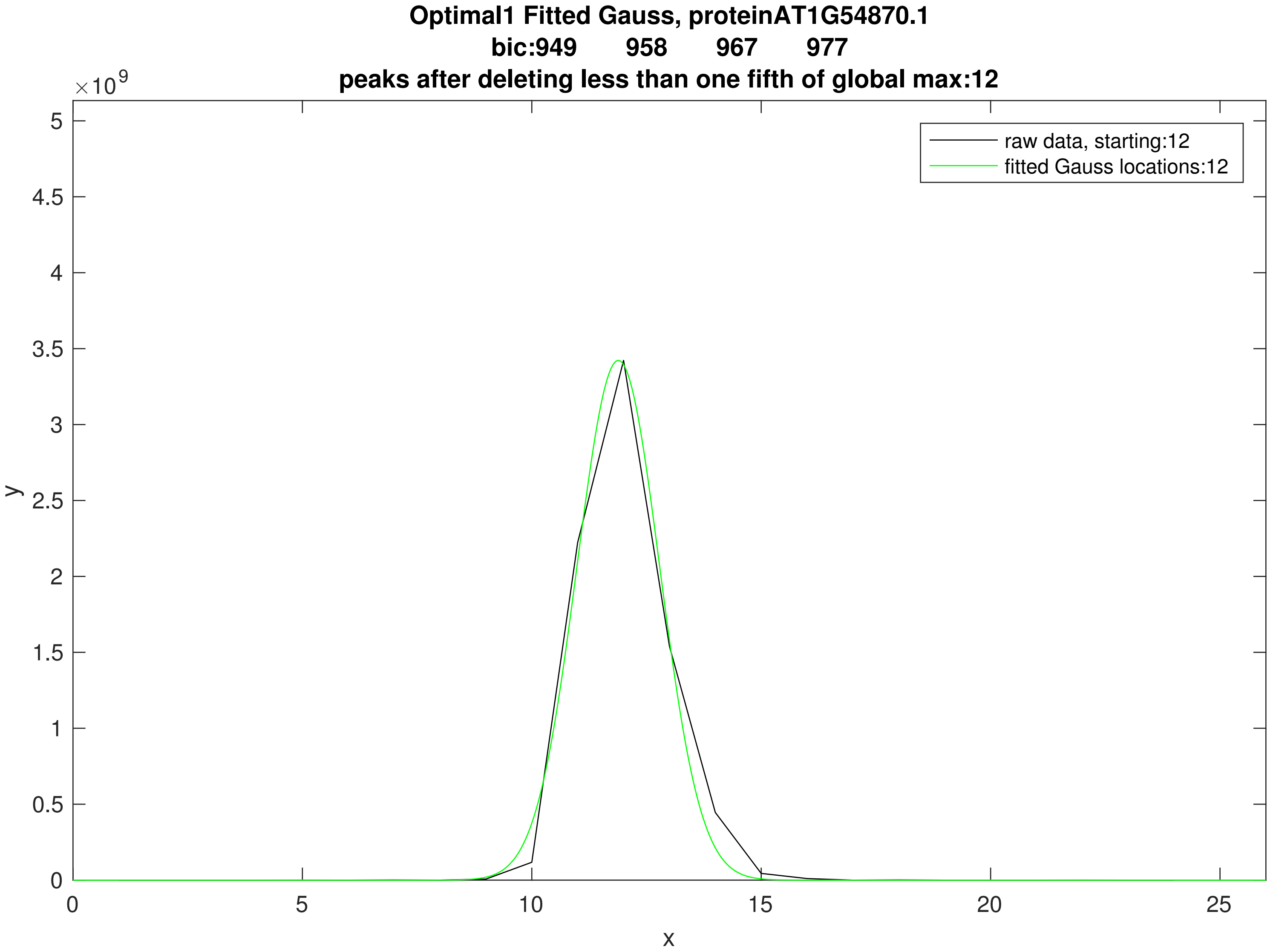
Optimal1 Fitted Gauss, proteinAT1G54870.1
bic:949 958 967 977
peaks after deleting less than one fifth of global max:12
×10^9
raw data, starting:12
fitted Gauss locations:12
y
x
0
0.5
1
1.5
2
2.5
3
3.5
4
4.5
5
0
5
10
15
20
25

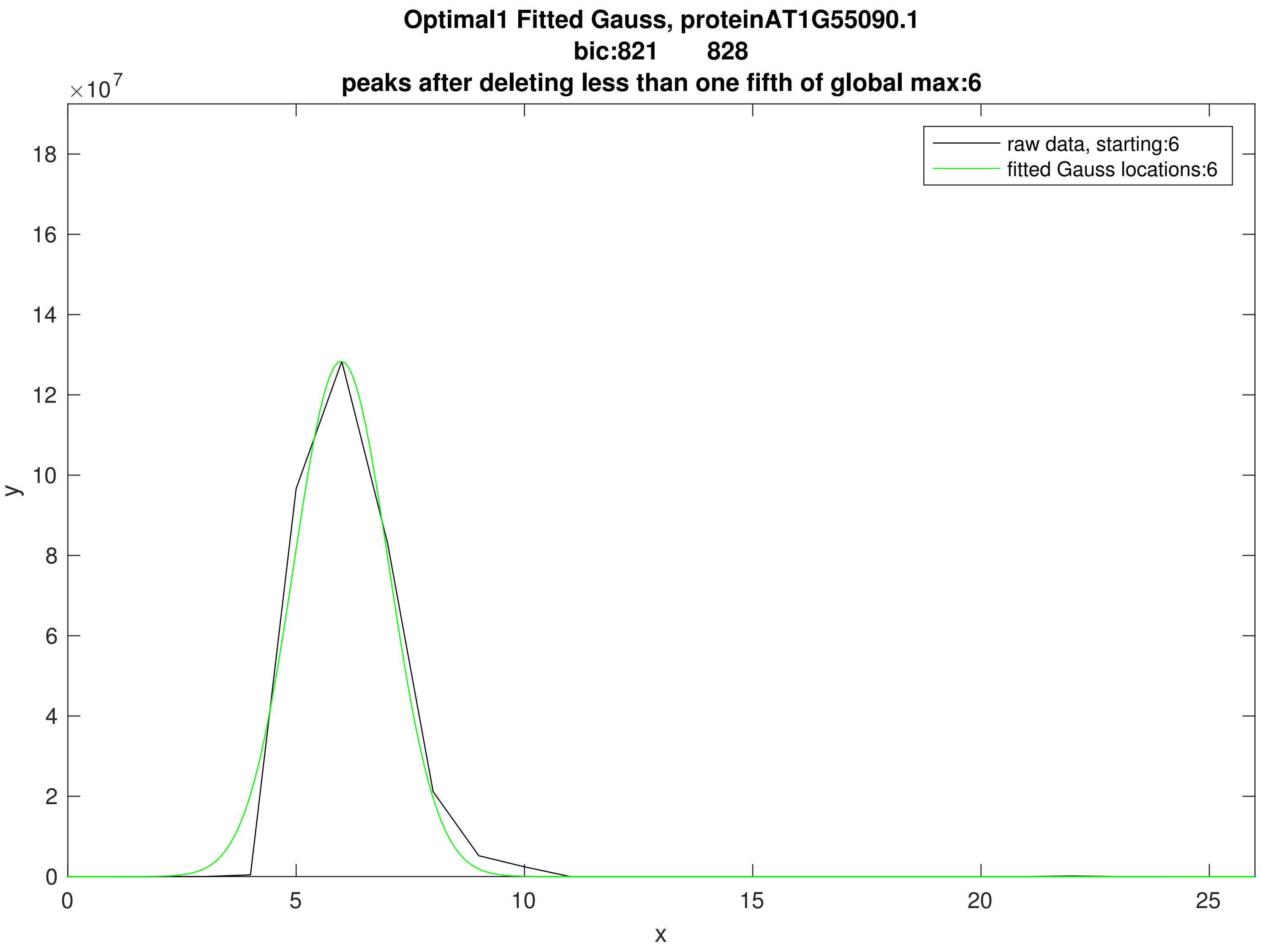
Optimal1 Fitted Gauss, proteinAT1G55090.1
bic:821 828
peaks after deleting less than one fifth of global max:6
×10^7
18
16
14
12
10
8
6
4
2
0
y
0
5
10
15
20
25
x
raw data, starting:6
fitted Gauss locations:6

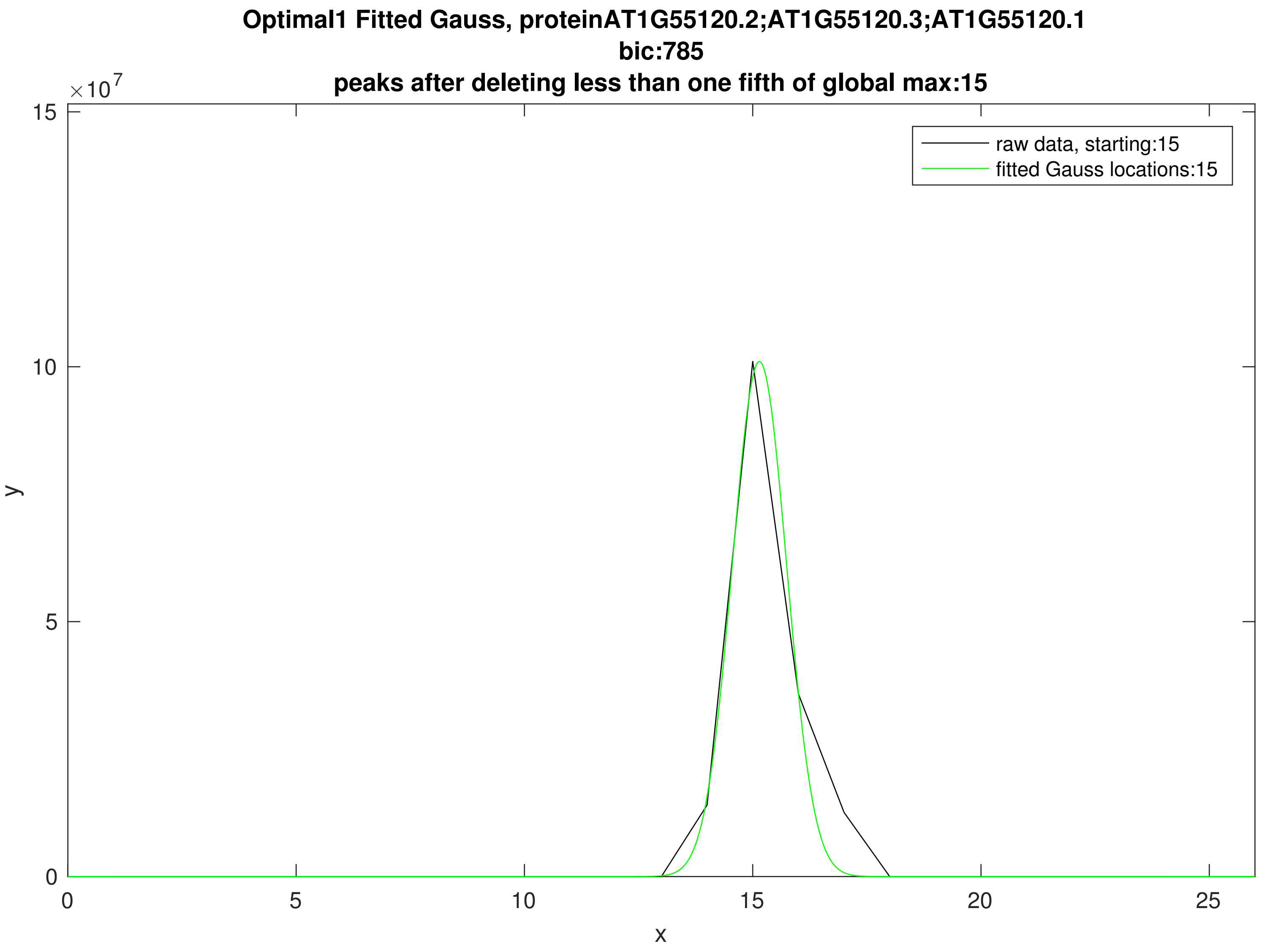
Optimal1 Fitted Gauss, proteinAT1G55120.2;AT1G55120.3;AT1G55120.1
bic:785
peaks after deleting less than one fifth of global max:15
×10^7
15
10
5
0
0
5
10
15
20
25
x
y
raw data, starting:15
fitted Gauss locations:15

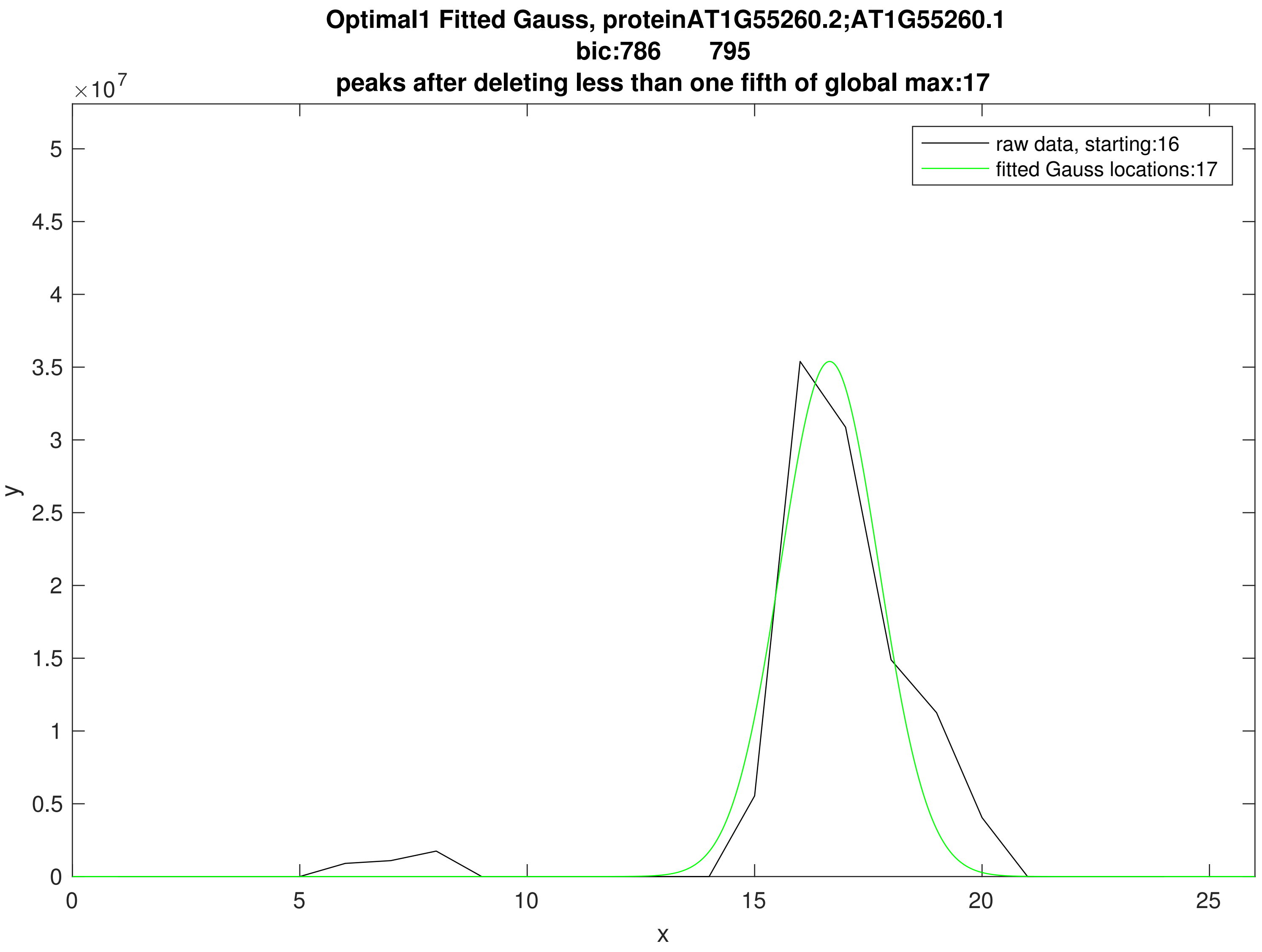

Optimal1 Fitted Gauss, proteinAT1G55260.2;AT1G55260.1
bic:786 795
peaks after deleting less than one fifth of global max:17
×10^7
raw data, starting:16
fitted Gauss locations:17
y
x
0
0.5
1
1.5
2
2.5
3
3.5
4
4.5
5
0
5
10
15
20
25

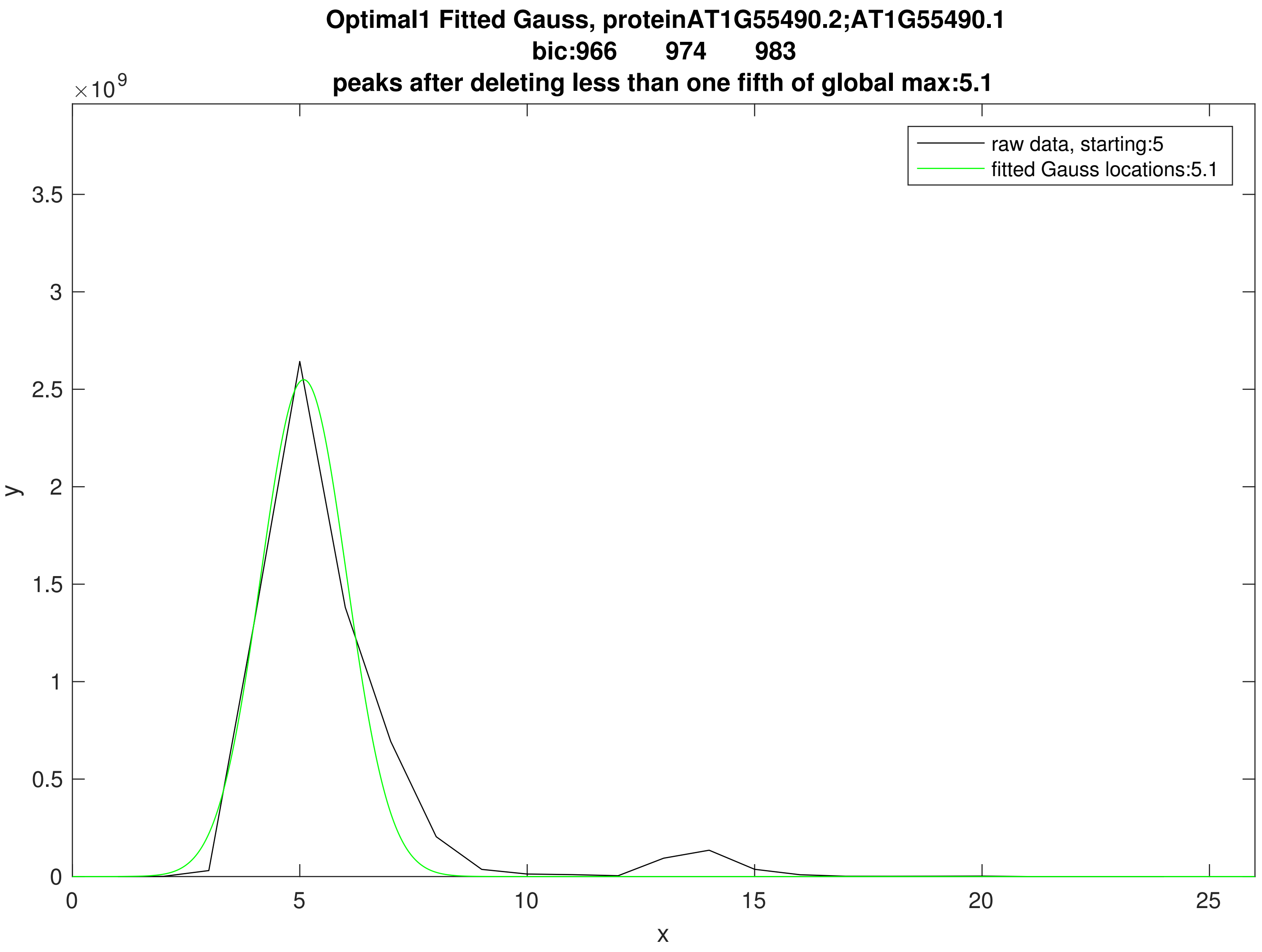

Optimal1 Fitted Gauss, proteinAT1G55490.2;AT1G55490.1
bic:966 974 983
peaks after deleting less than one fifth of global max:5.1
×10^9
3.5
3
2.5
2
1.5
1
0.5
0
y
0
5
10
15
20
25
x
raw data, starting:5
fitted Gauss locations:5.1

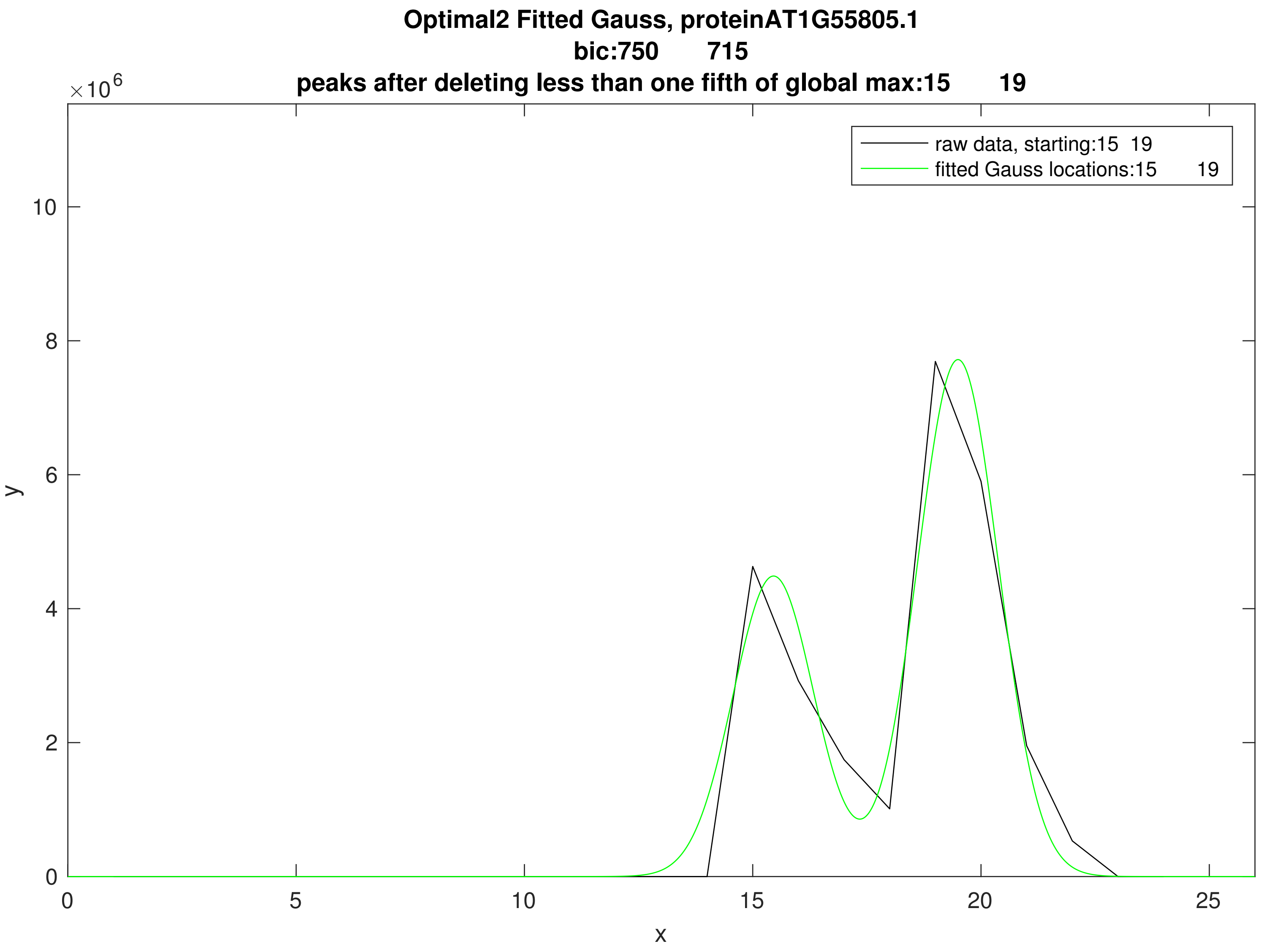
Optimal2 Fitted Gauss, proteinAT1G55805.1
bic:750 715
peaks after deleting less than one fifth of global max:15 19
×10^6
raw data, starting:15 19
fitted Gauss locations:15 19
y
x
0
2
4
6
8
10
0
5
10
15
20
25

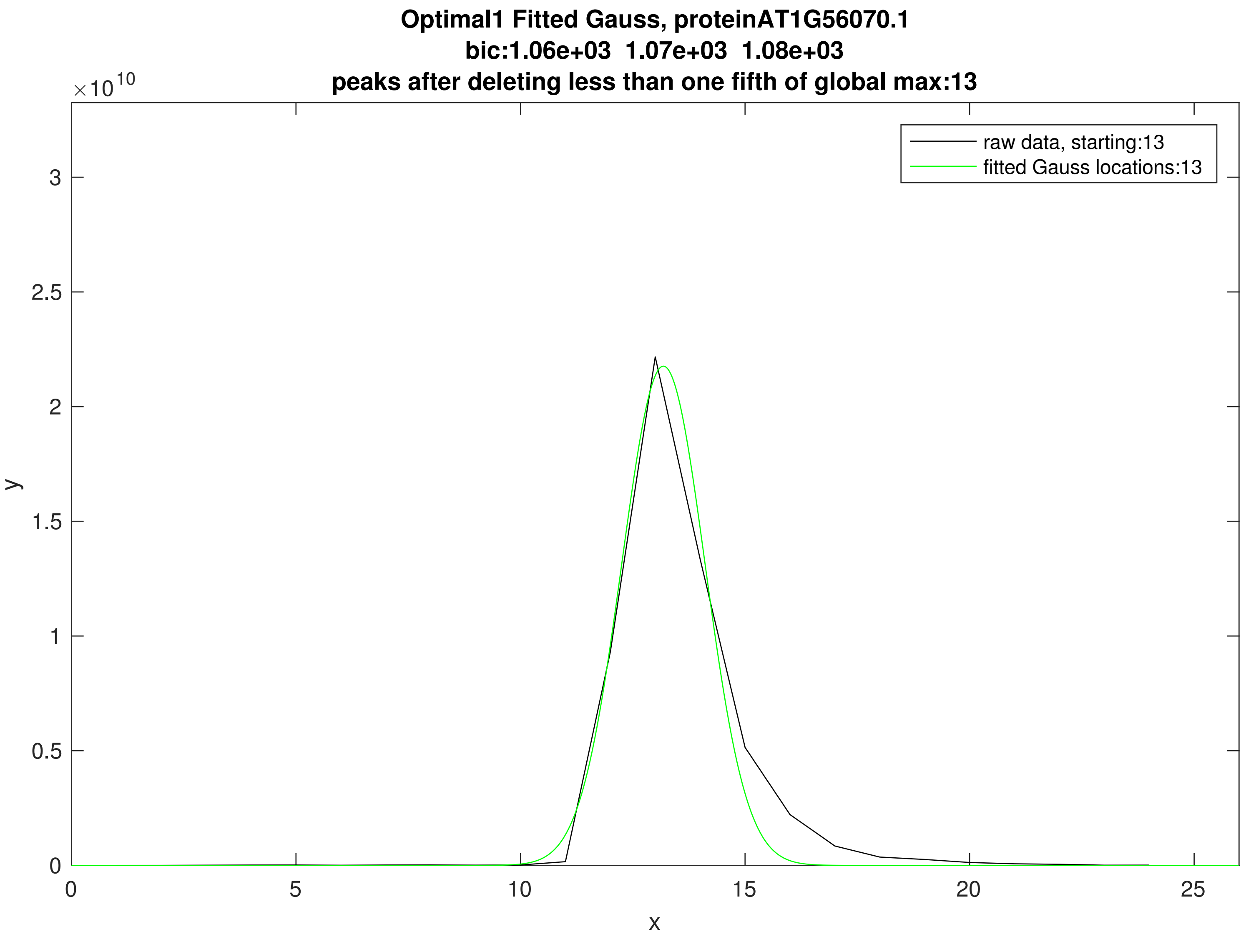

Optimal1 Fitted Gauss, proteinAT1G56070.1
bic:1.06e+03  1.07e+03  1.08e+03
peaks after deleting less than one fifth of global max:13
×10^10
raw data, starting:13
fitted Gauss locations:13
3
2.5
2
1.5
1
0.5
0
y
0
5
10
15
20
25
x

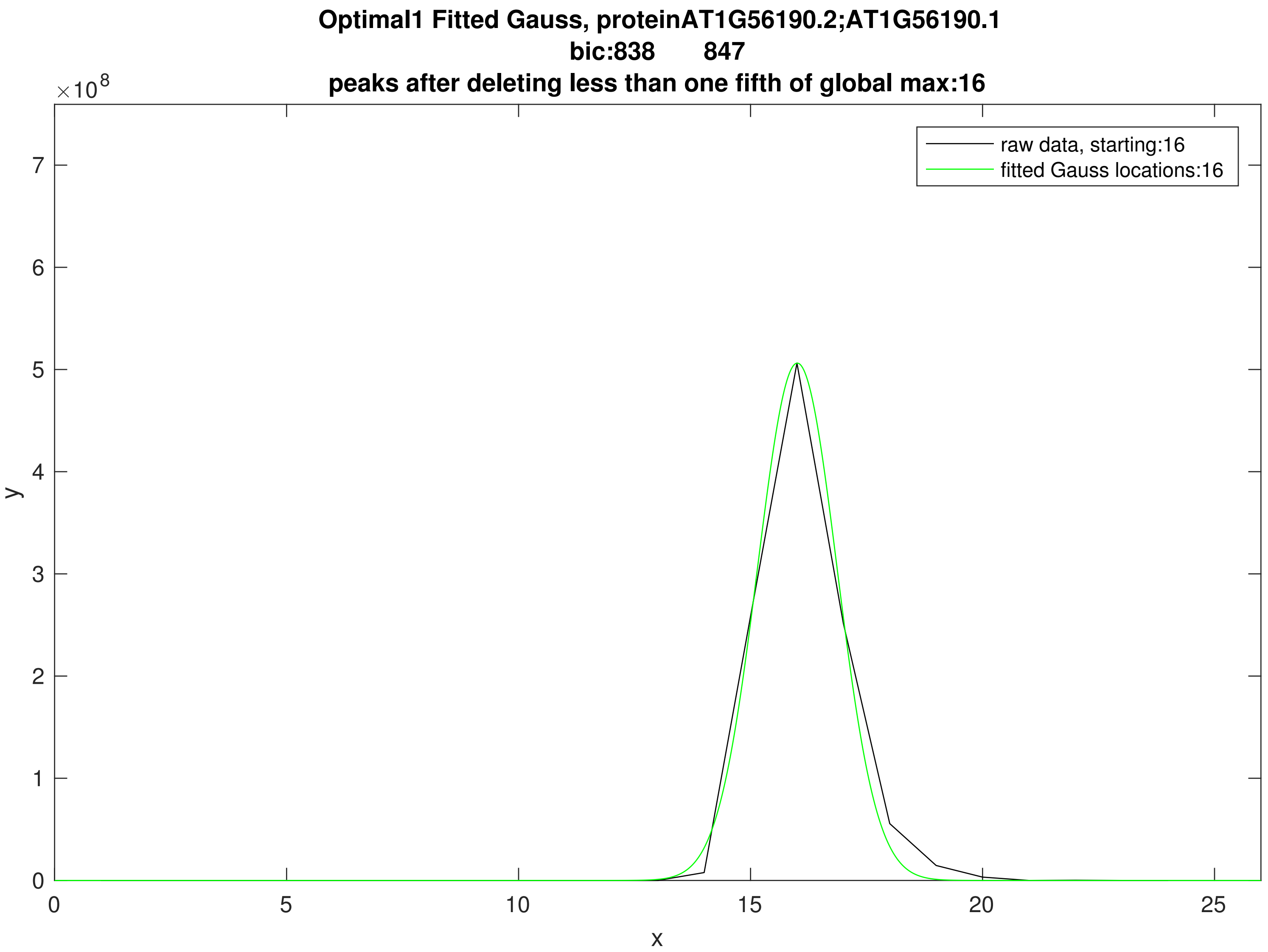

Optimal1 Fitted Gauss, proteinAT1G56190.2;AT1G56190.1
bic:838 847
peaks after deleting less than one fifth of global max:16
×10^8
raw data, starting:16
fitted Gauss locations:16
y
x
0
1
2
3
4
5
6
7
0
5
10
15
20
25

Optimal1 Fitted Gauss, proteinAT1G56220.4;AT1G56220.2;AT1G56220.1;AT1G56220.3
bic:683
peaks after deleting less than one fifth of global max:12

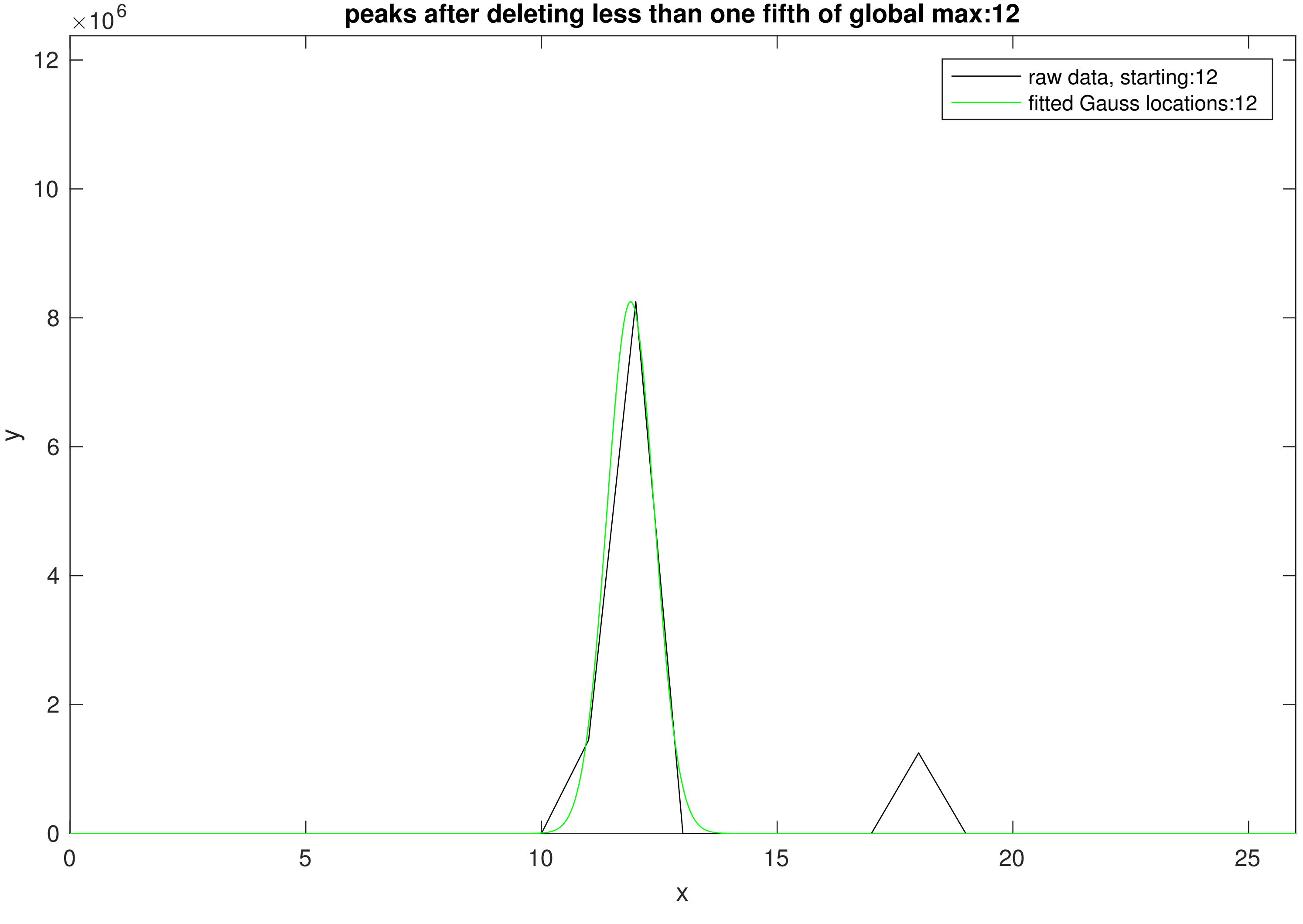

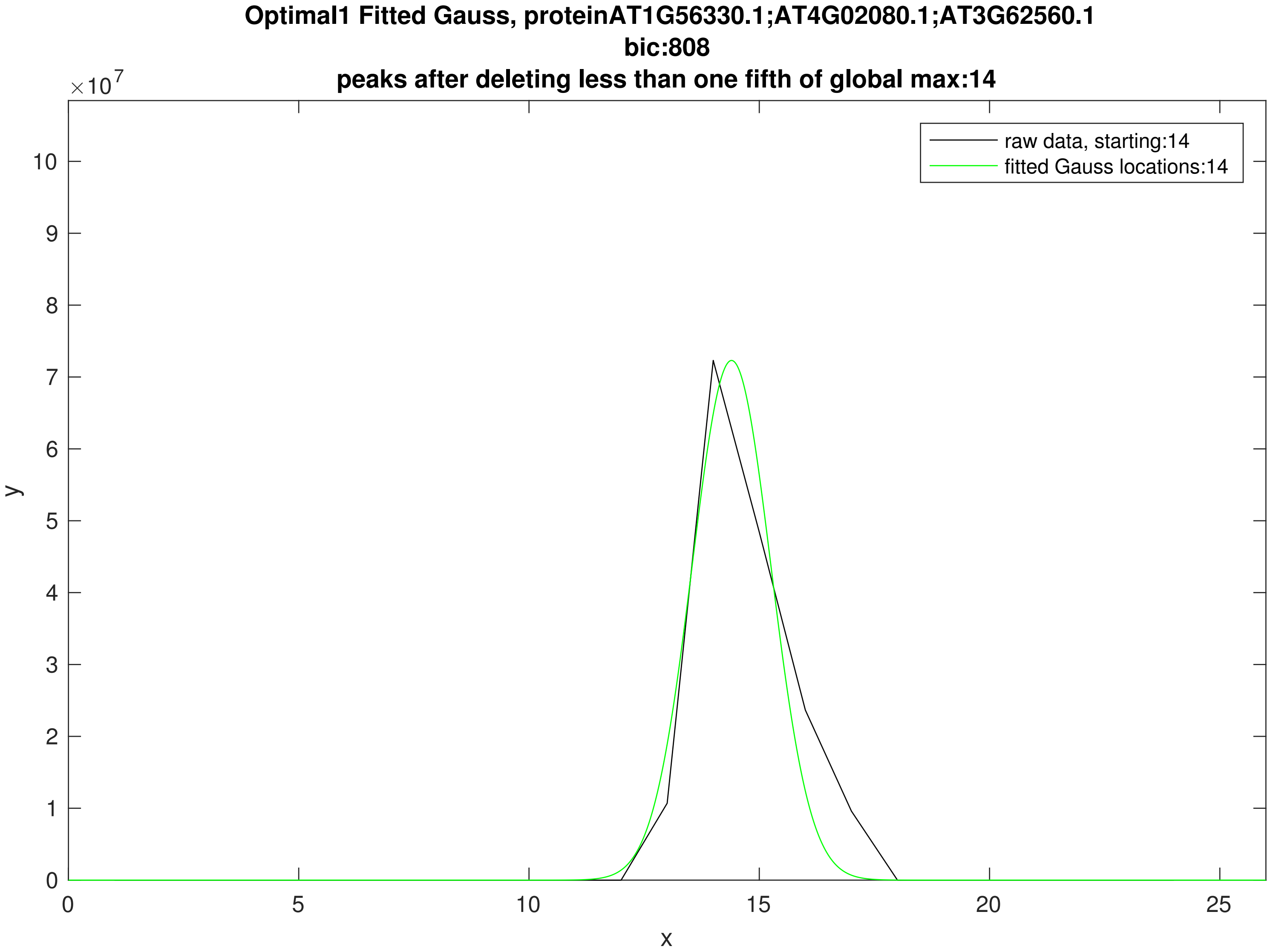

Optimal1 Fitted Gauss, proteinAT1G56330.1;AT4G02080.1;AT3G62560.1
bic:808
peaks after deleting less than one fifth of global max:14
×10^7
raw data, starting:14
fitted Gauss locations:14
y
x
0
1
2
3
4
5
6
7
8
9
10
0
5
10
15
20
25

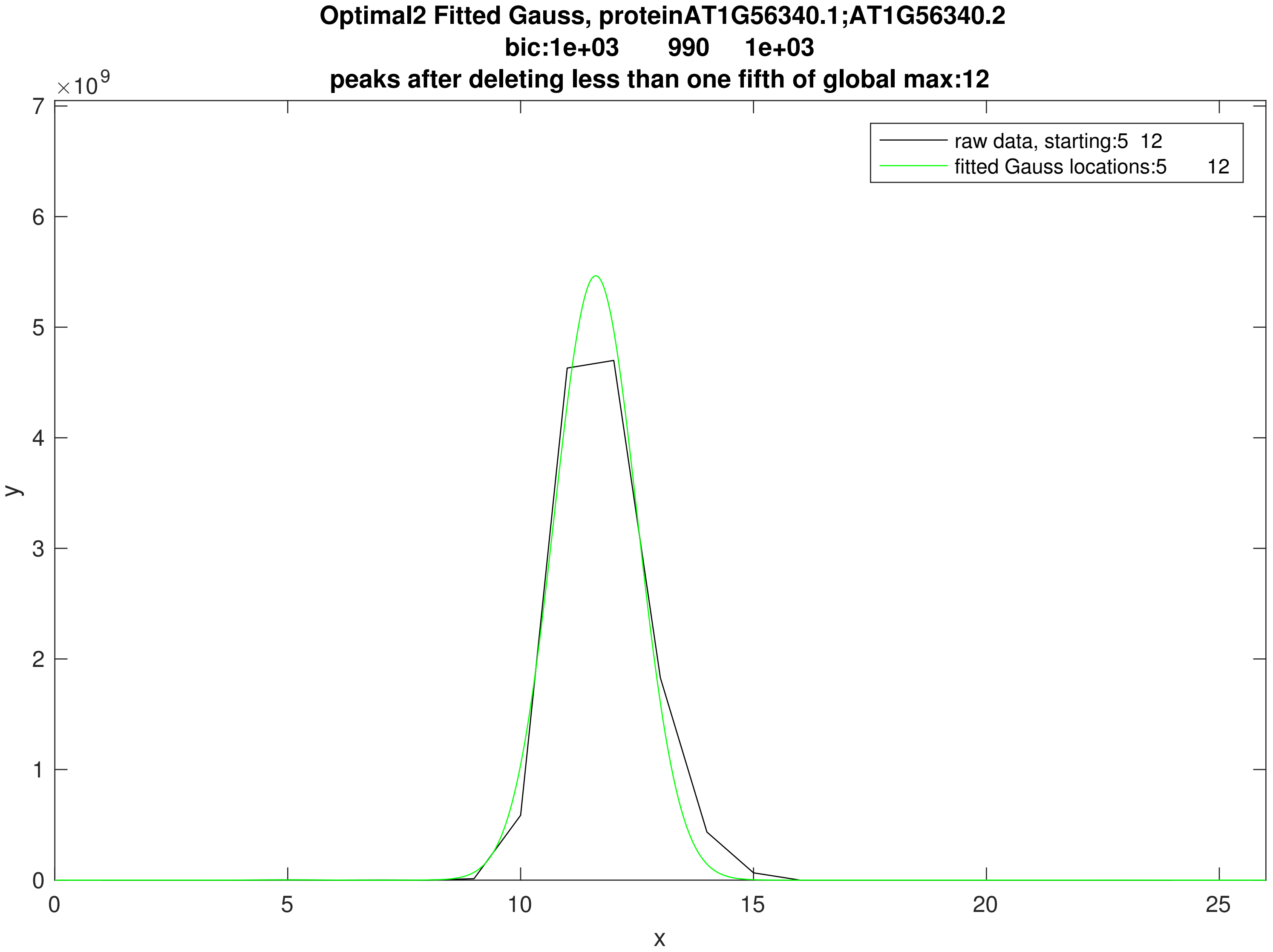

Optimal2 Fitted Gauss, proteinAT1G56340.1;AT1G56340.2
bic:1e+03 990 1e+03
peaks after deleting less than one fifth of global max:12
×10^9
raw data, starting:5 12
fitted Gauss locations:5 12
y
x
0
1
2
3
4
5
6
7
0
5
10
15
20
25

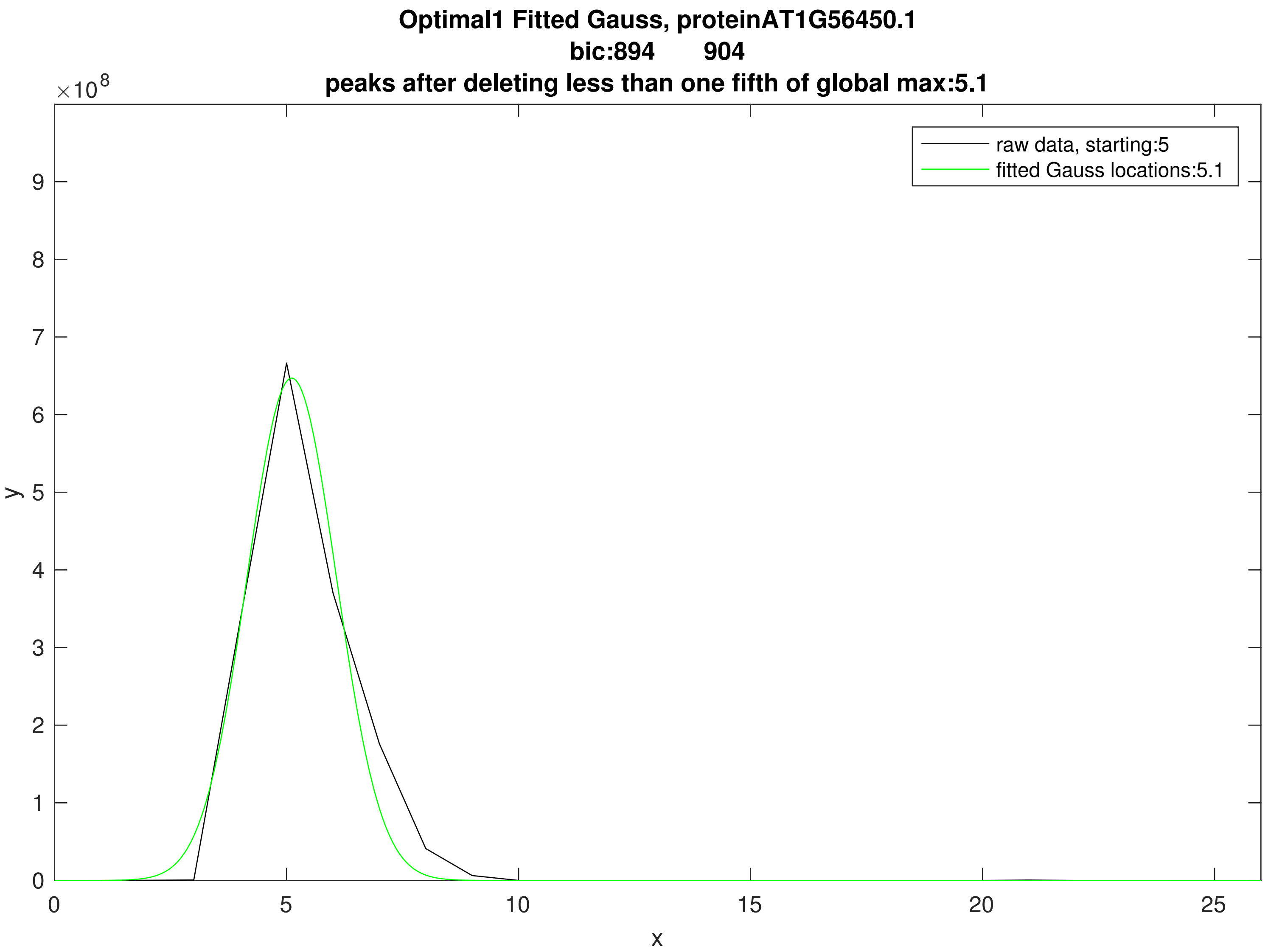

Optimal1 Fitted Gauss, proteinAT1G56450.1
bic:894 904
peaks after deleting less than one fifth of global max:5.1
×10^8
raw data, starting:5
fitted Gauss locations:5.1
y
x
0
1
2
3
4
5
6
7
8
9
10
15
20
25

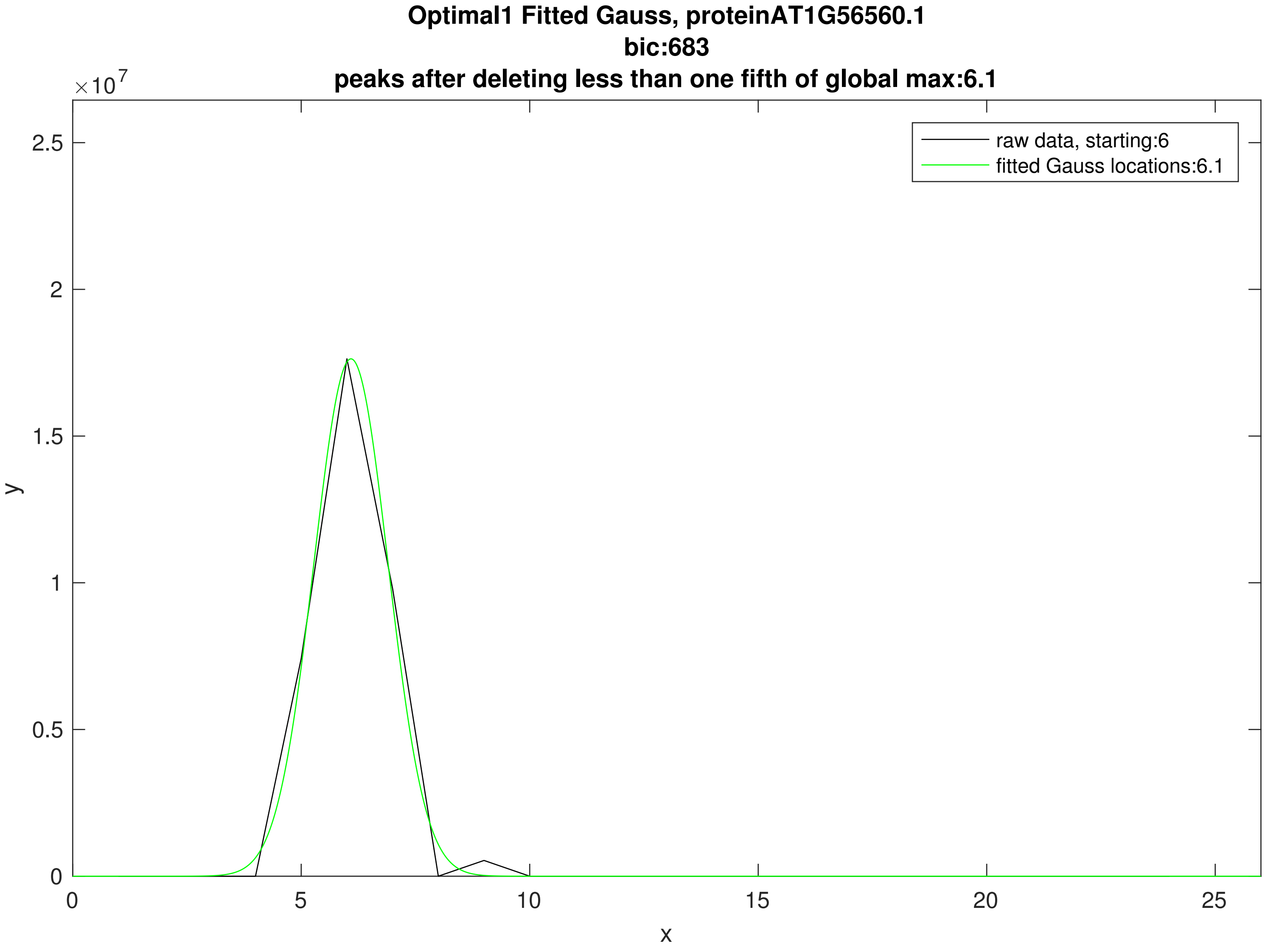
Optimal1 Fitted Gauss, proteinAT1G56560.1
bic:683
peaks after deleting less than one fifth of global max:6.1
×10^7
raw data, starting:6
fitted Gauss locations:6.1
y
x
0
0.5
1
1.5
2
2.5
5
10
15
20
25

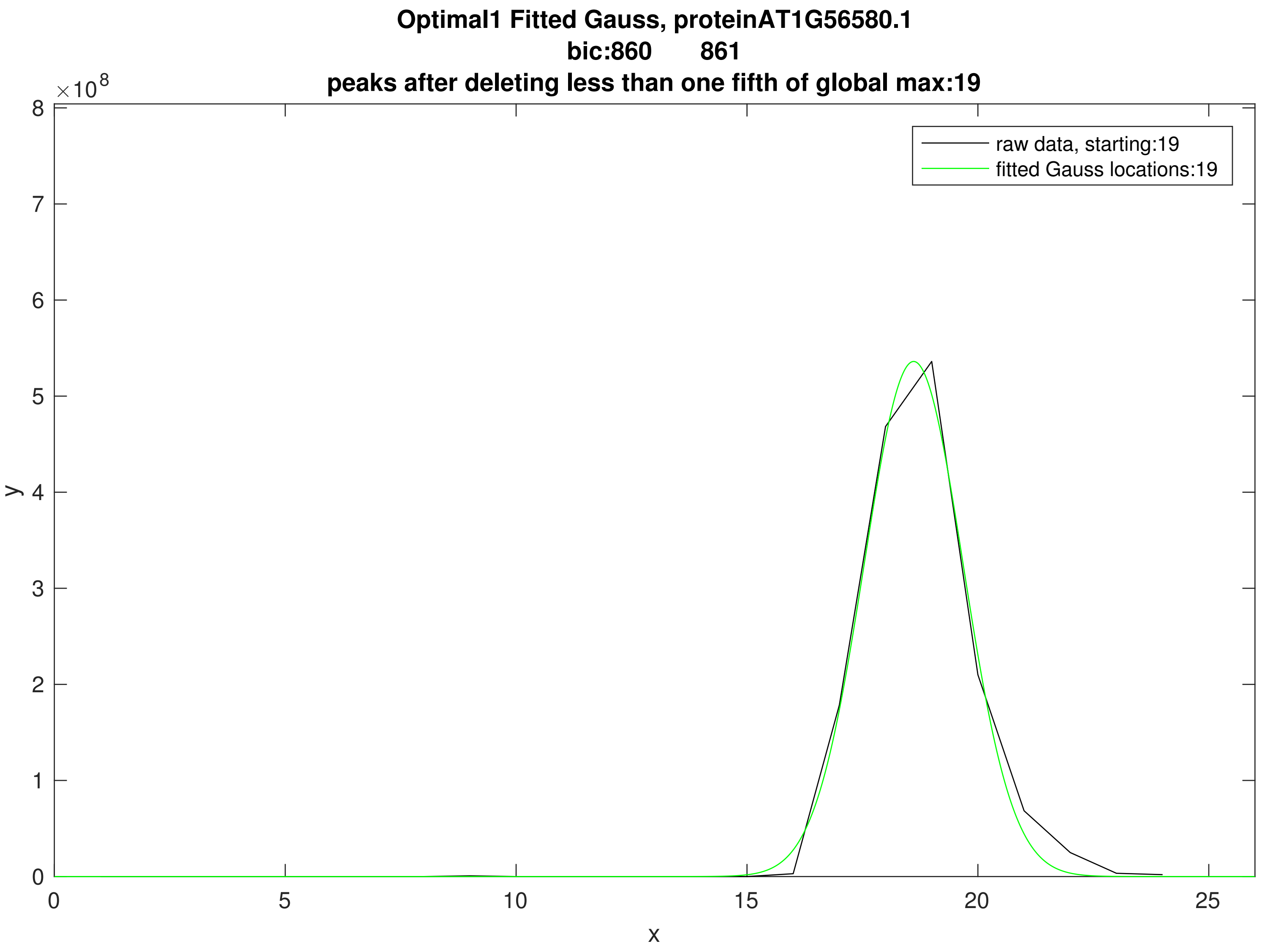
Optimal1 Fitted Gauss, proteinAT1G56580.1
bic:860 861
peaks after deleting less than one fifth of global max:19
×10^8
raw data, starting:19
fitted Gauss locations:19
y
x
0
1
2
3
4
5
6
7
8
0
5
10
15
20
25

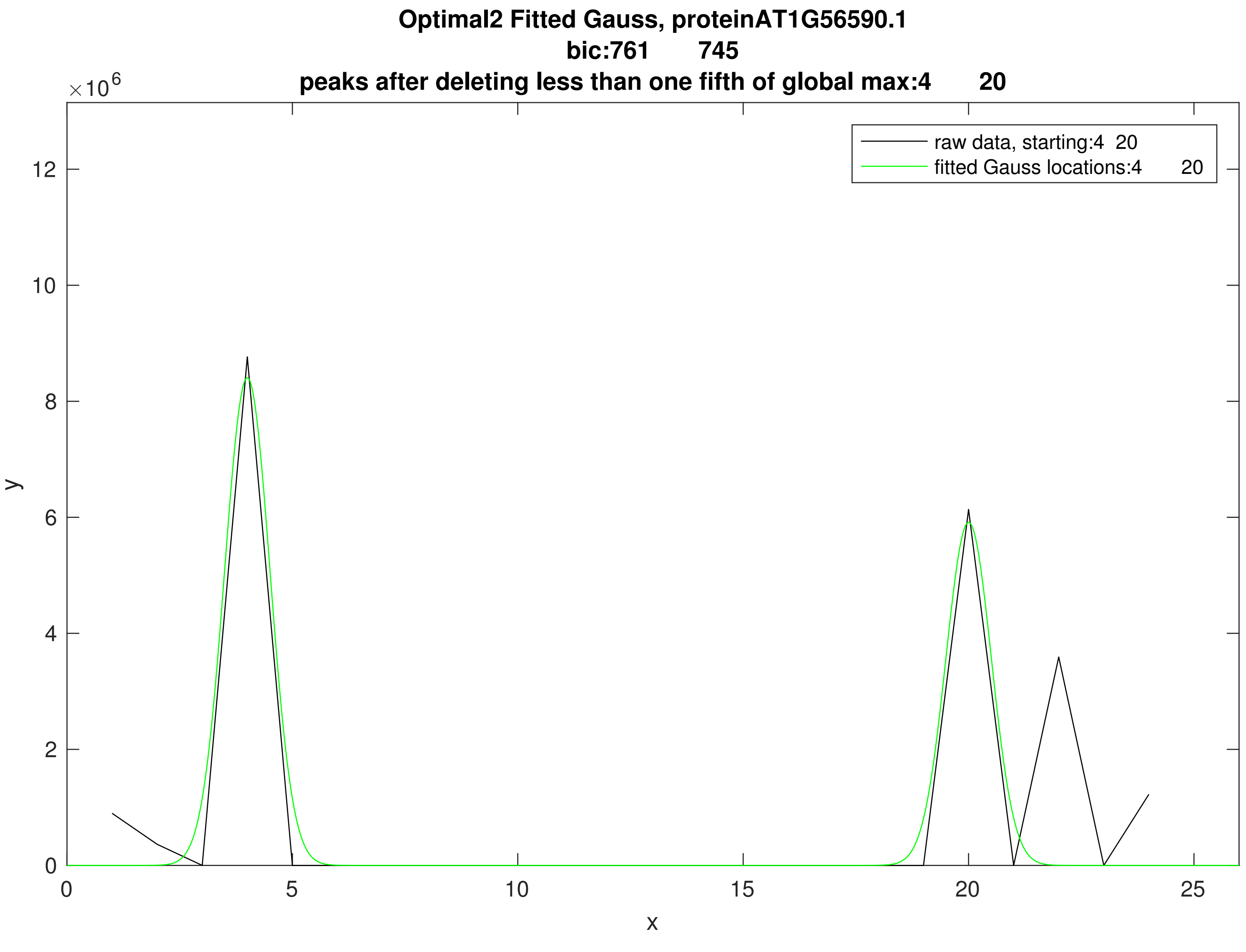
Optimal2 Fitted Gauss, proteinAT1G56590.1
bic:761 745
peaks after deleting less than one fifth of global max:4 20
×10^6
raw data, starting:4 20
fitted Gauss locations:4 20
12
10
8
6
4
2
0
y
0
5
10
15
20
25
x

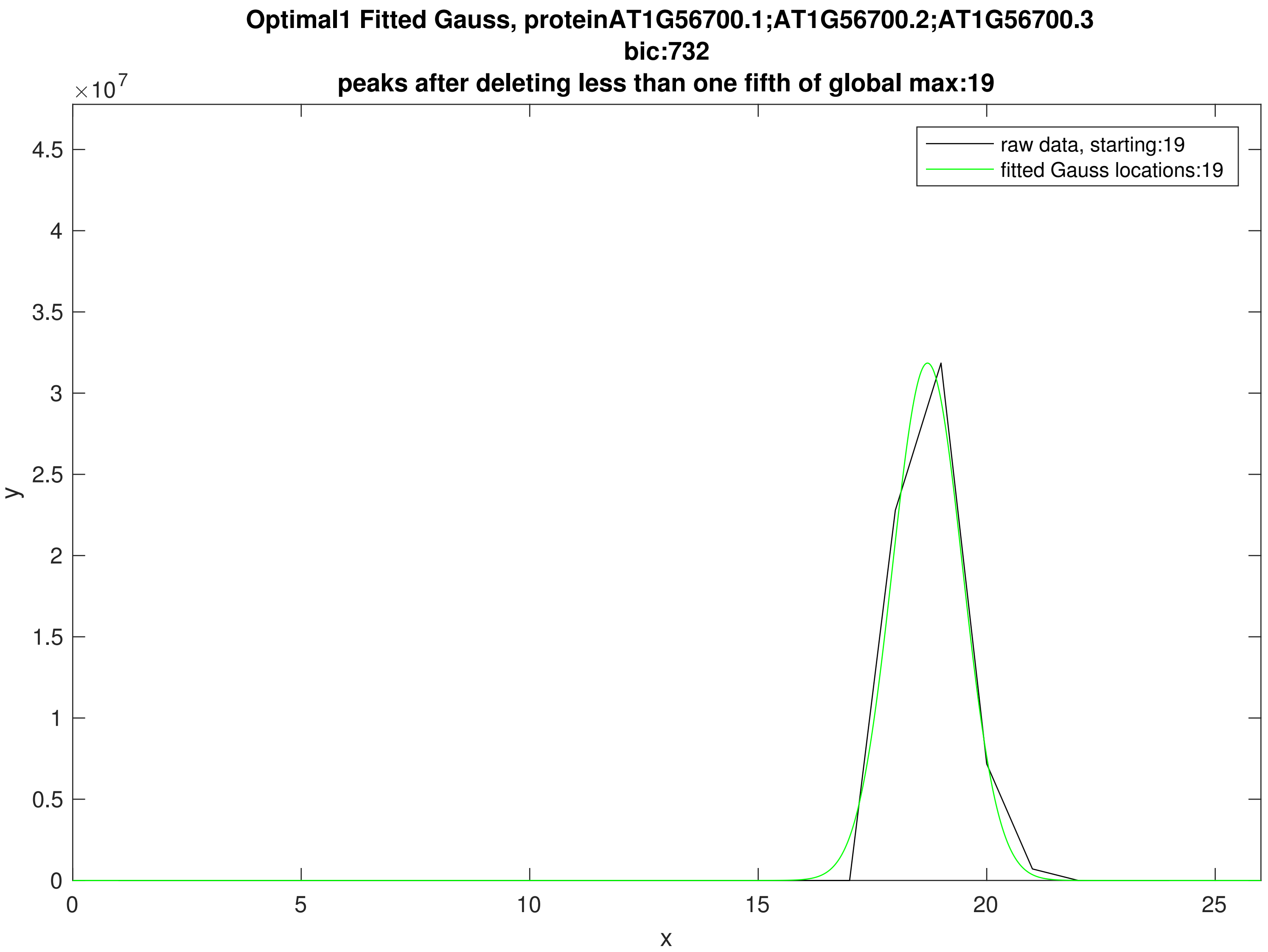
Optimal1 Fitted Gauss, proteinAT1G56700.1;AT1G56700.2;AT1G56700.3
bic:732
peaks after deleting less than one fifth of global max:19
×10^7
raw data, starting:19
fitted Gauss locations:19
y
x
0
0.5
1
1.5
2
2.5
3
3.5
4
4.5
5
10
15
20
25

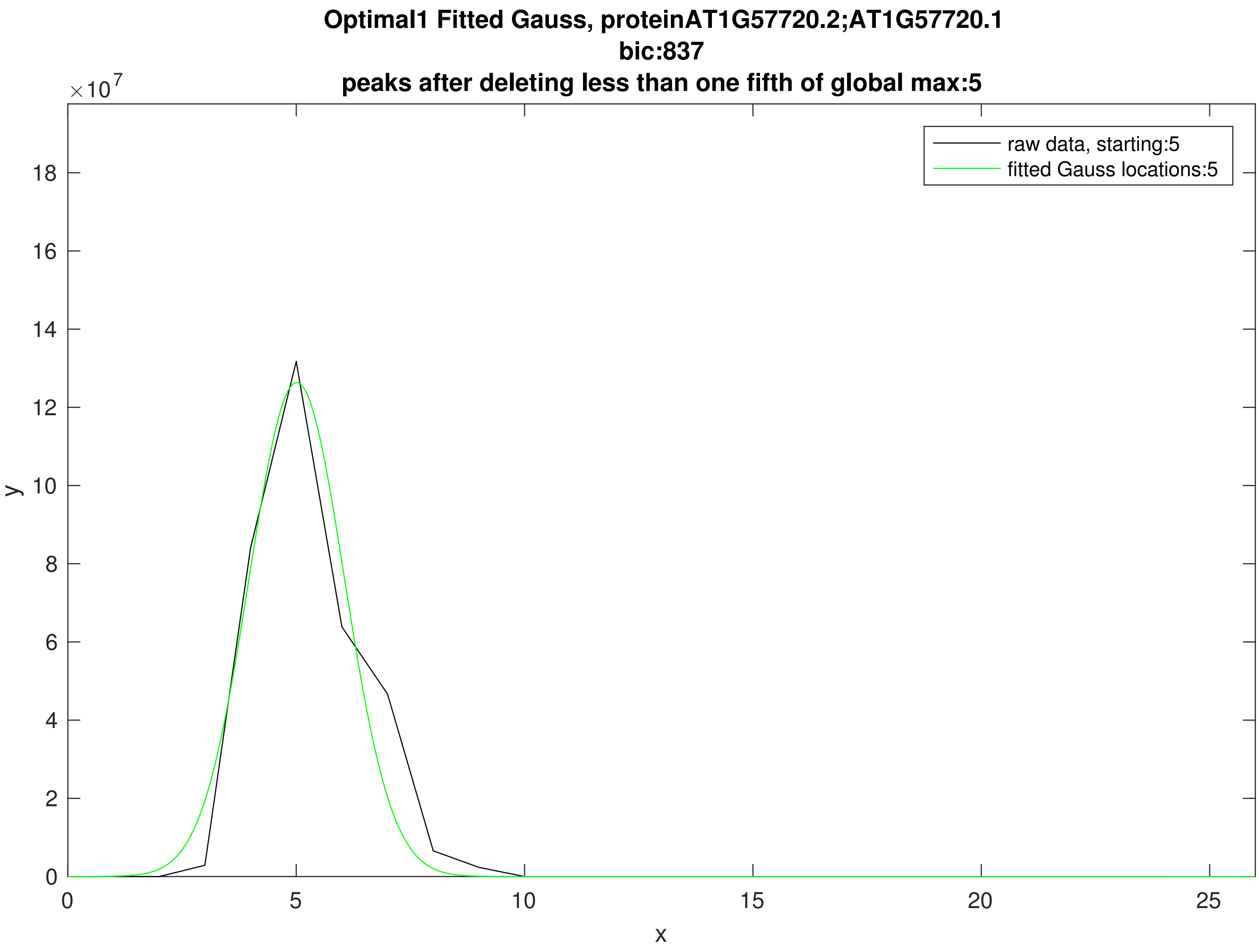
Optimal1 Fitted Gauss, proteinAT1G57720.2;AT1G57720.1
bic:837
peaks after deleting less than one fifth of global max:5
×10^7
raw data, starting:5
fitted Gauss locations:5
y
x
0
2
4
6
8
10
12
14
16
18
0
5
10
15
20
25

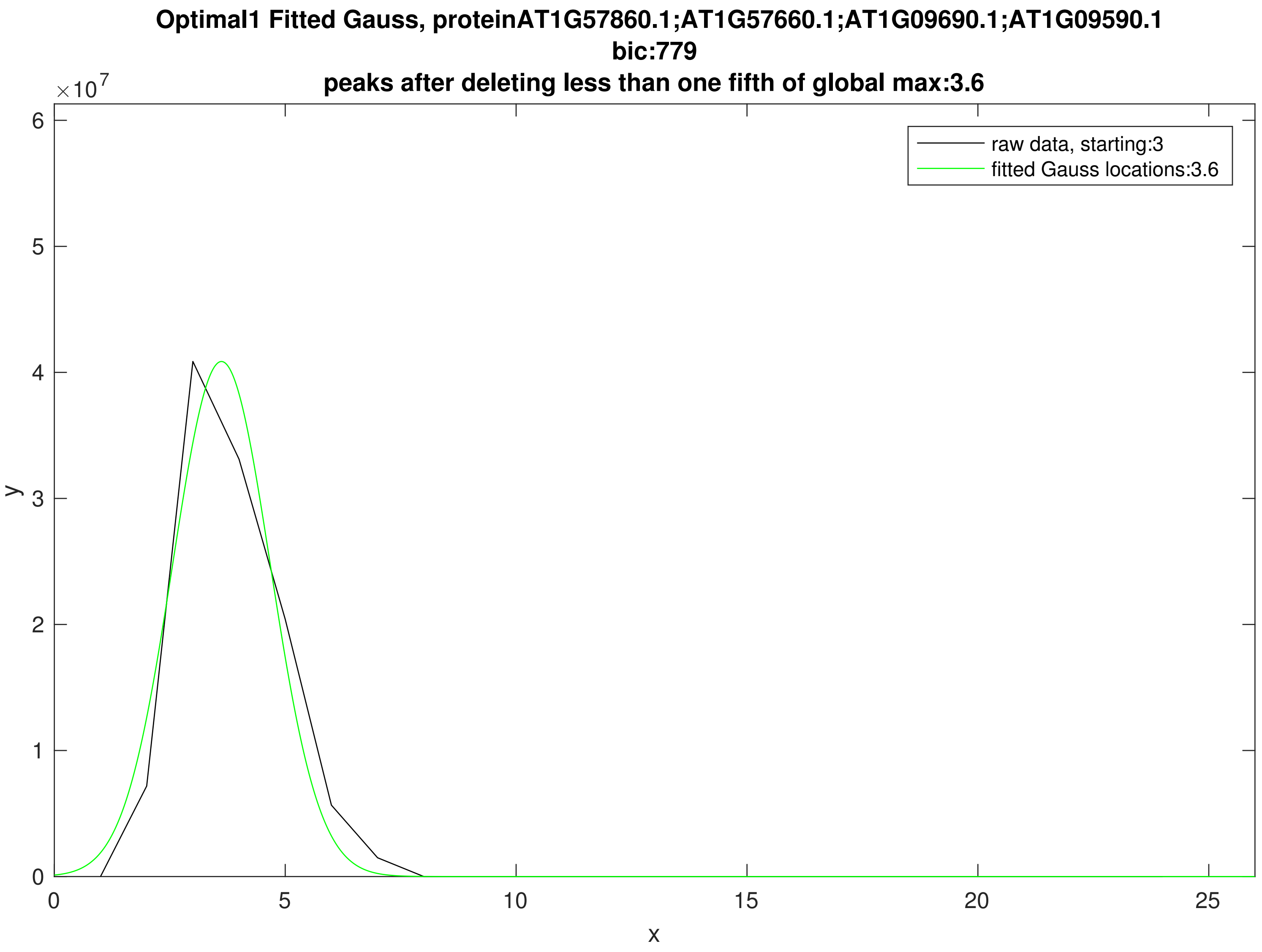

Optimal1 Fitted Gauss, proteinAT1G57860.1;AT1G57660.1;AT1G09690.1;AT1G09590.1
bic:779
peaks after deleting less than one fifth of global max:3.6
×10^7
raw data, starting:3
fitted Gauss locations:3.6
y
x
0
1
2
3
4
5
6
0
5
10
15
20
25

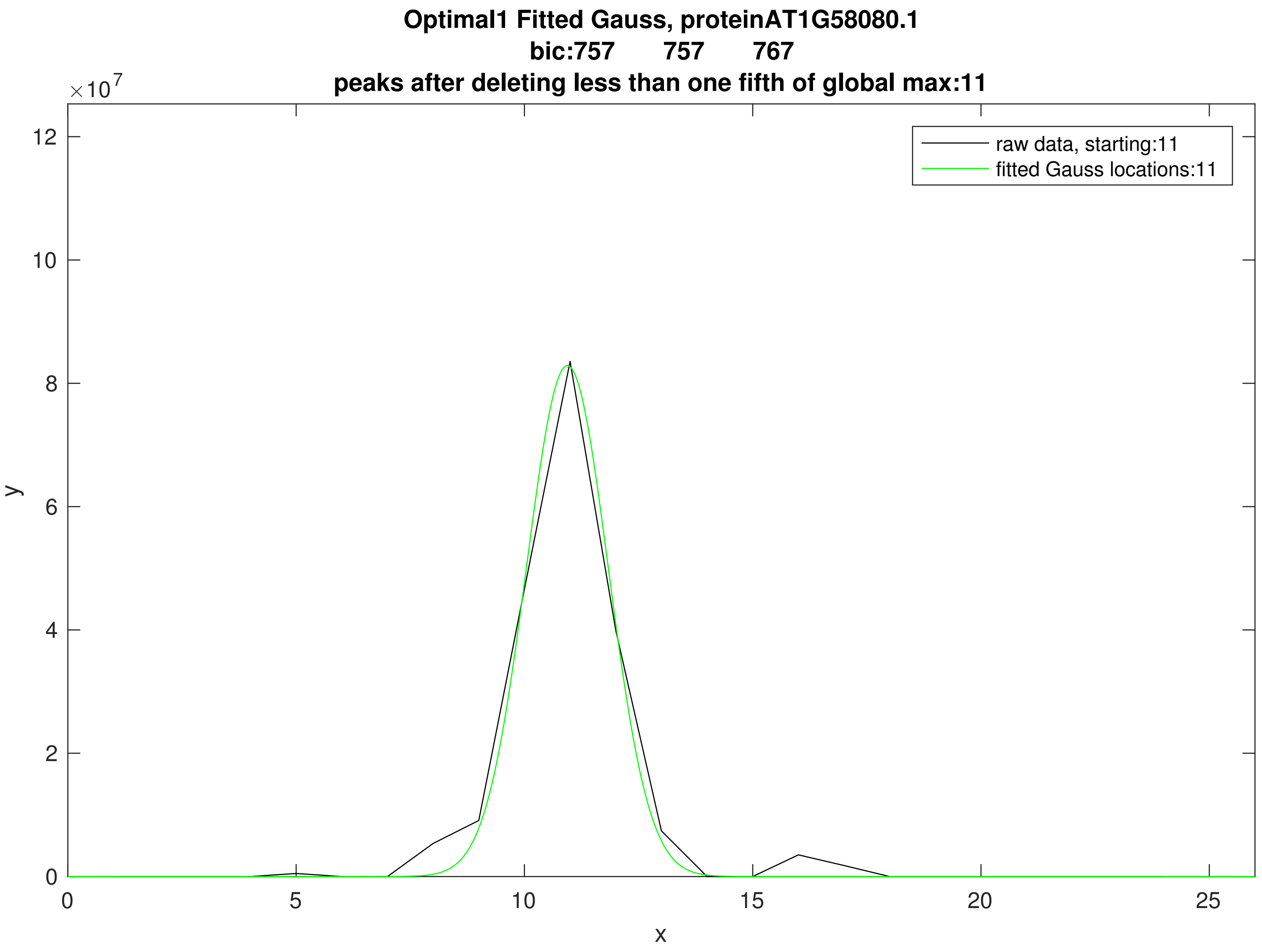

Optimal1 Fitted Gauss, proteinAT1G58080.1
bic:757 757 767
peaks after deleting less than one fifth of global max:11
×10^7
12
10
8
6
4
2
0
y
0
5
10
15
20
25
x
raw data, starting:11
fitted Gauss locations:11

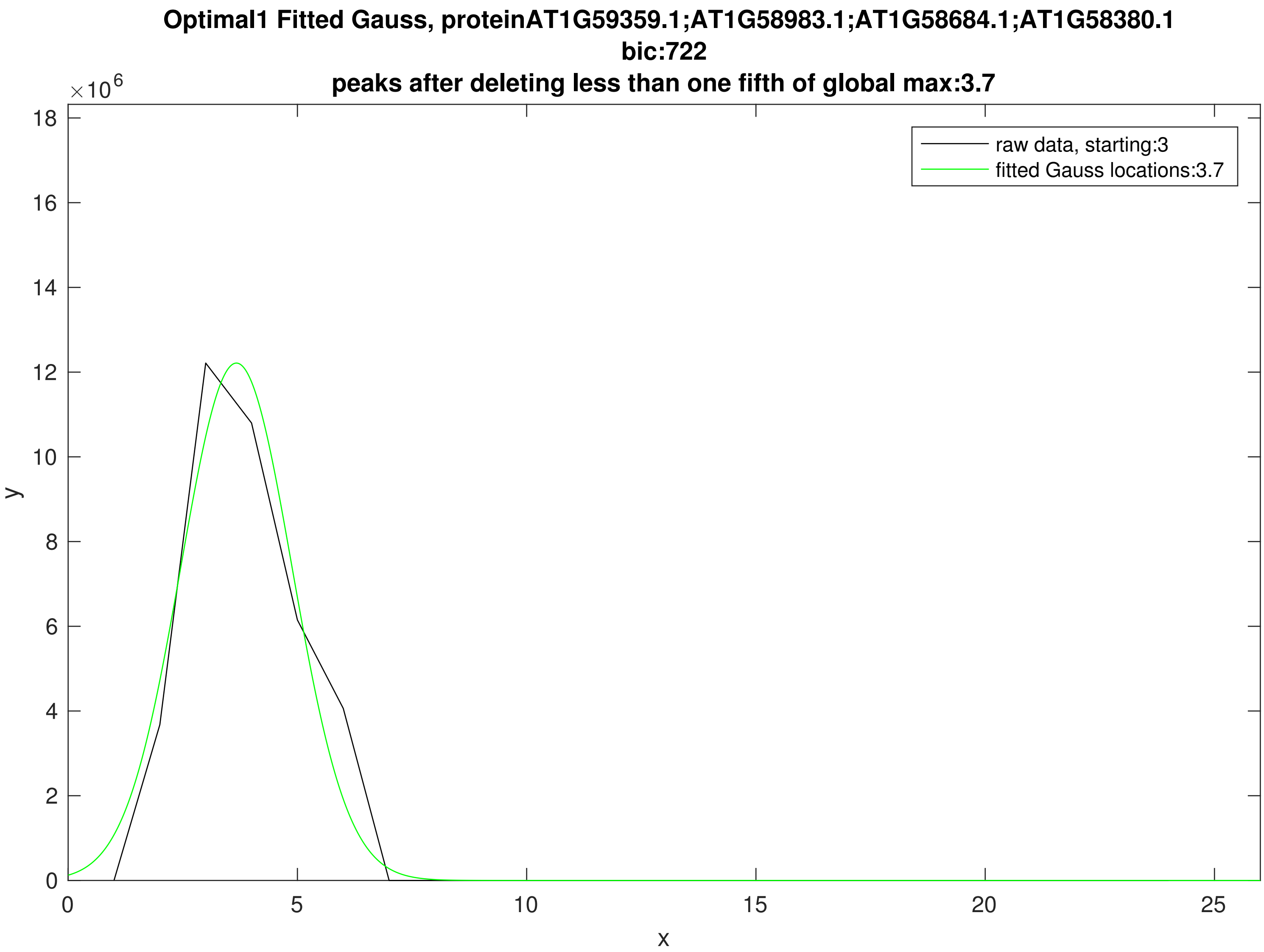

Optimal1 Fitted Gauss, proteinAT1G59359.1;AT1G58983.1;AT1G58684.1;AT1G58380.1
bic:722
peaks after deleting less than one fifth of global max:3.7
raw data, starting:3
fitted Gauss locations:3.7
y
x

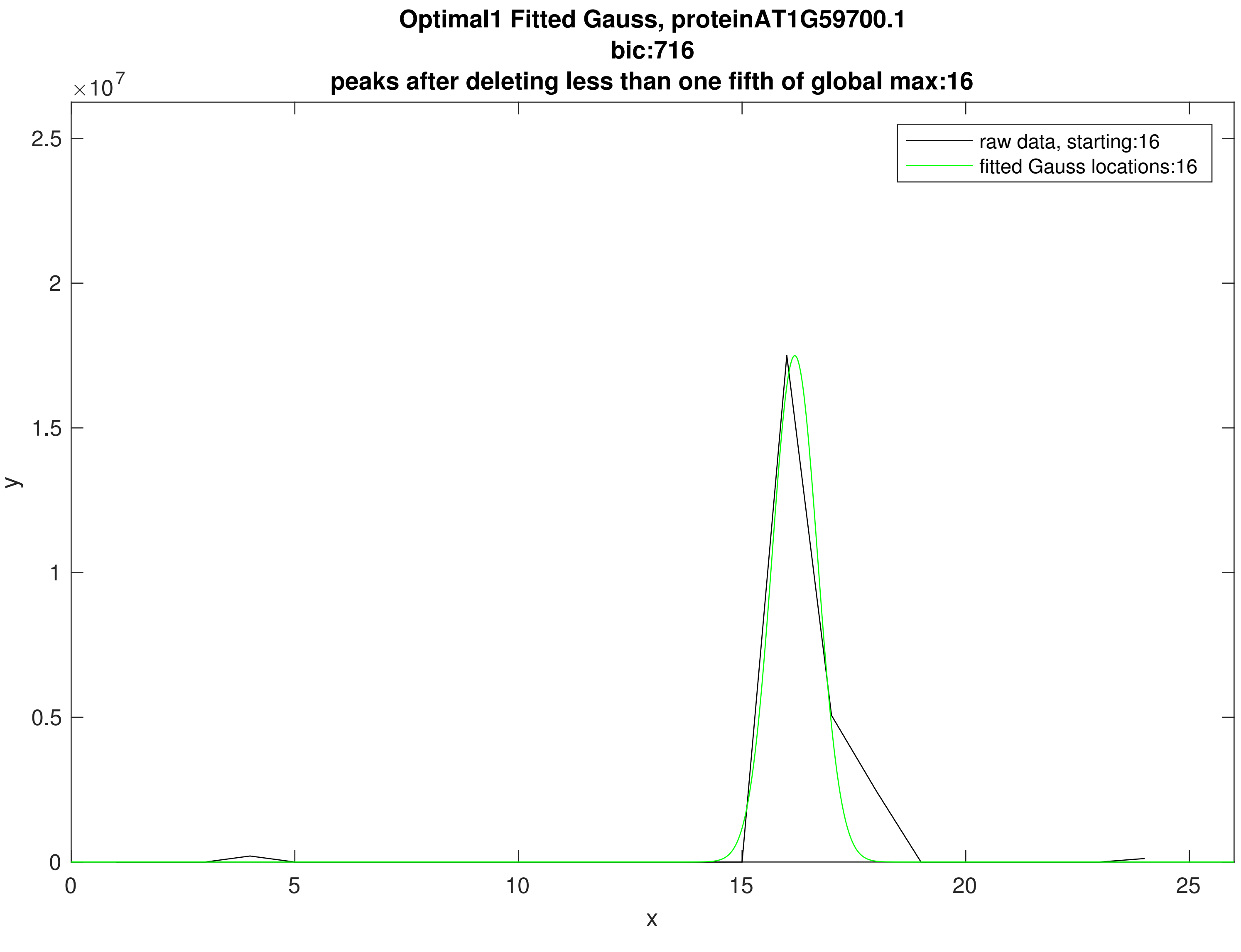
Optimal1 Fitted Gauss, proteinAT1G59700.1
bic:716
peaks after deleting less than one fifth of global max:16
×10^7
2.5
2
1.5
1
0.5
0
y
0
5
10
15
20
25
x
raw data, starting:16
fitted Gauss locations:16

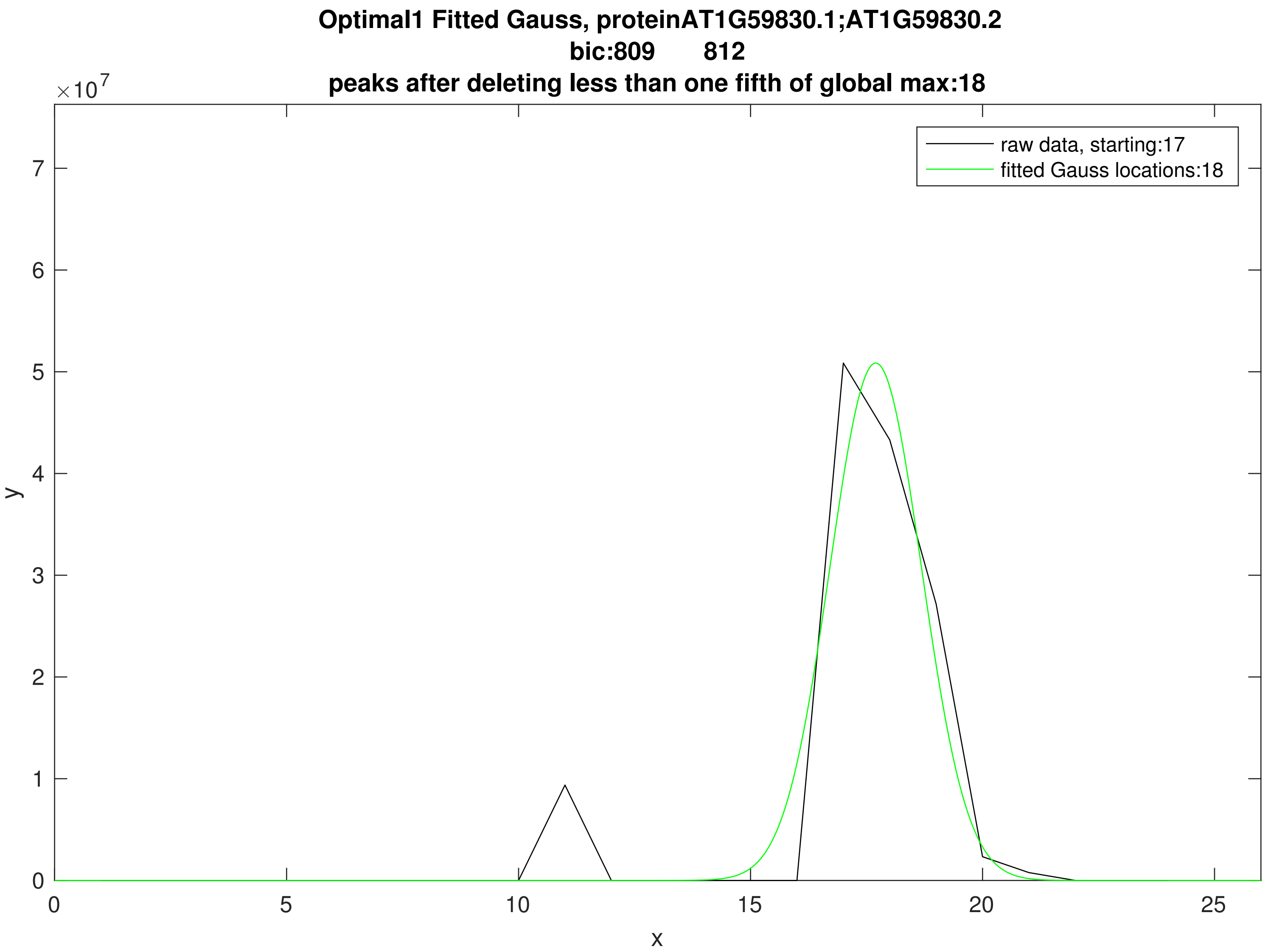
Optimal1 Fitted Gauss, proteinAT1G59830.1;AT1G59830.2
bic:809 812
peaks after deleting less than one fifth of global max:18
×10^7
raw data, starting:17
fitted Gauss locations:18
y
x
0
1
2
3
4
5
6
7
0
5
10
15
20
25

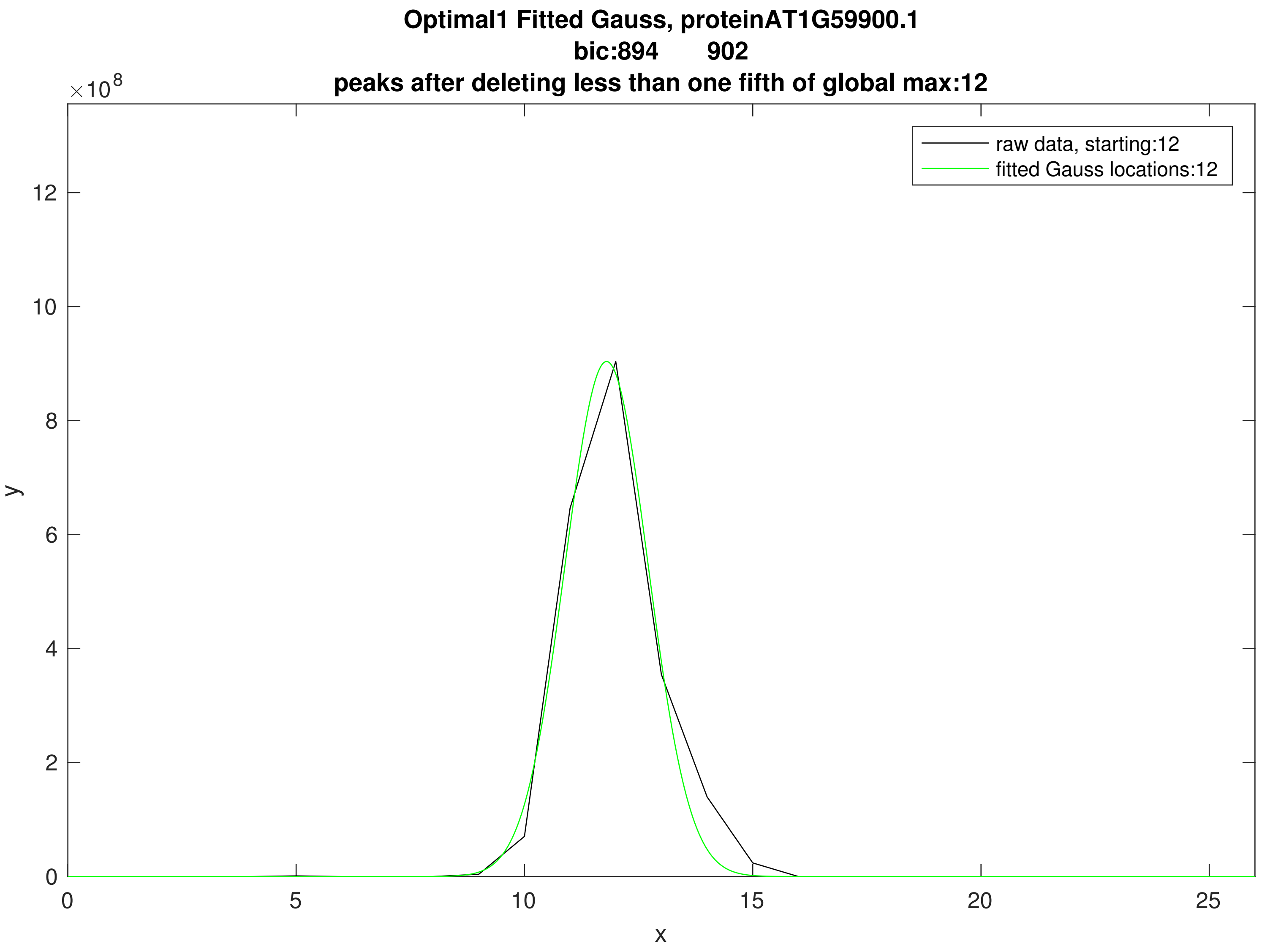

Optimal1 Fitted Gauss, proteinAT1G59900.1
bic:894 902
peaks after deleting less than one fifth of global max:12
×10^8
12
10
8
6
4
2
0
y
0
5
10
15
20
25
x
raw data, starting:12
fitted Gauss locations:12

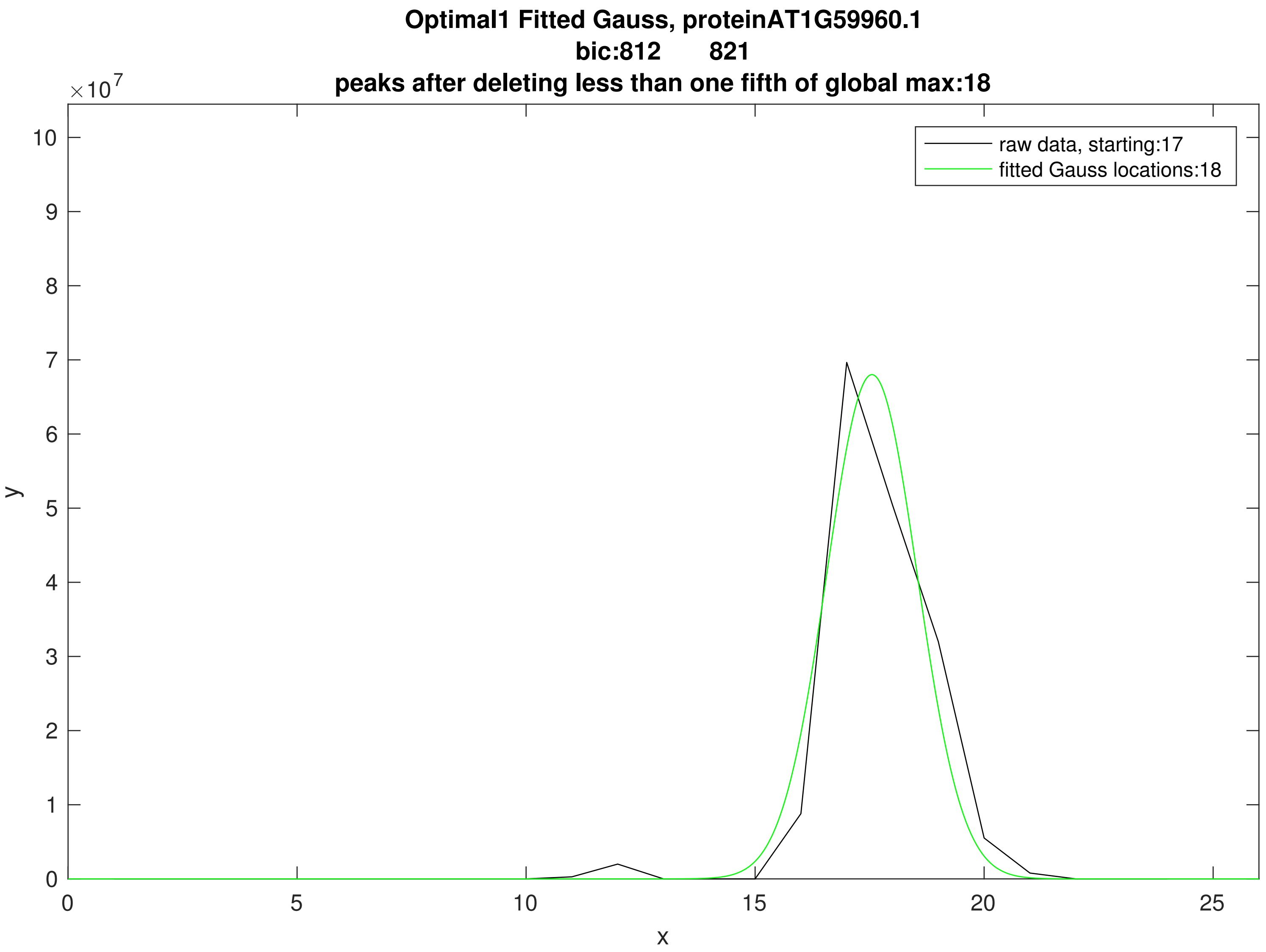

Optimal1 Fitted Gauss, proteinAT1G59960.1
bic:812 821
peaks after deleting less than one fifth of global max:18
×10^7
10
9
8
7
6
5
4
3
2
1
0
y
0
5
10
15
20
25
x
raw data, starting:17
fitted Gauss locations:18

#### Optimal1 Fitted Gauss, proteinAT1G60420.1
#### bic:877 950
#### peaks after deleting less than one fifth of global max:13

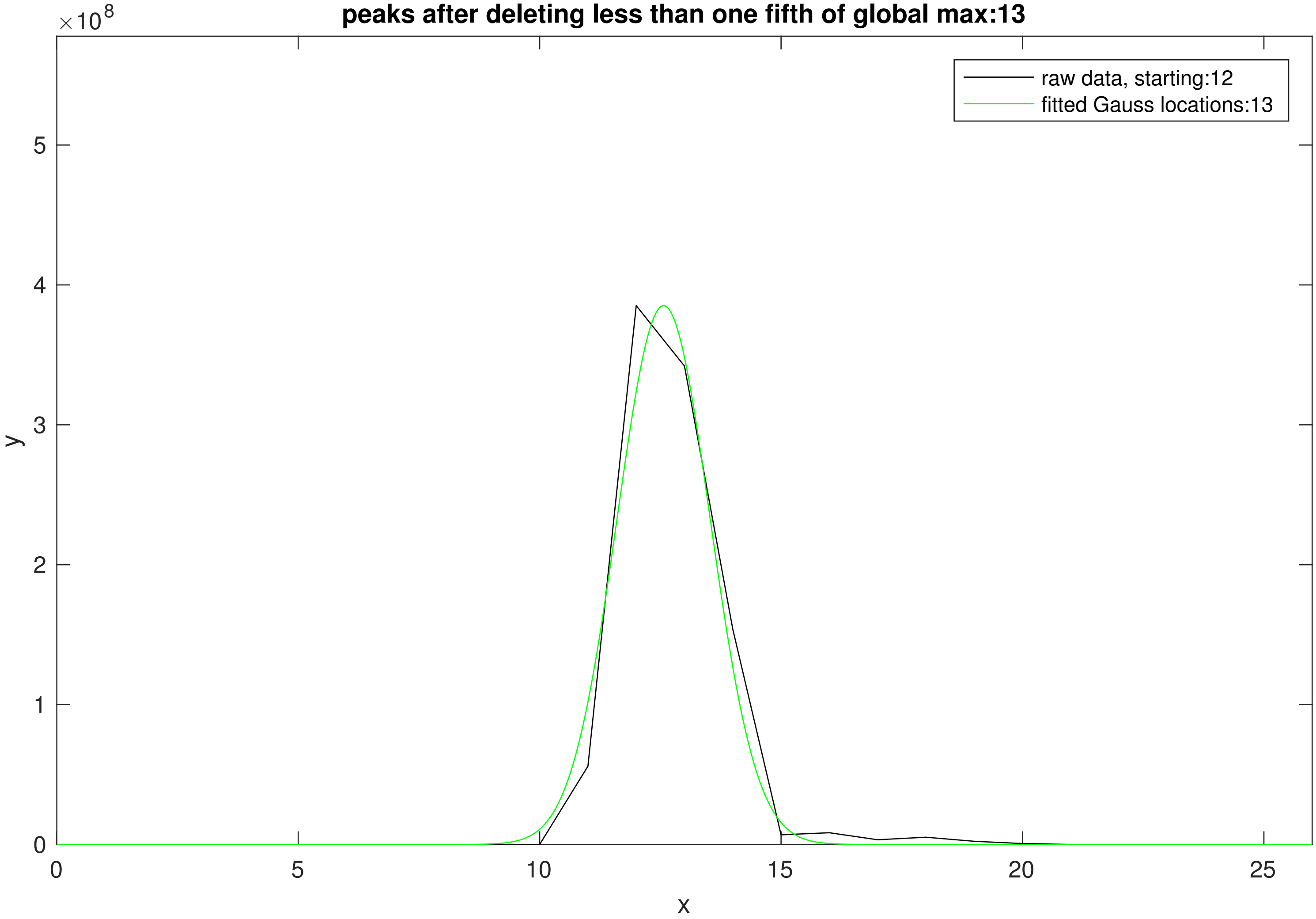

Optimal1 Fitted Gauss, proteinAT1G60680.1
bic:809
peaks after deleting less than one fifth of global max:16

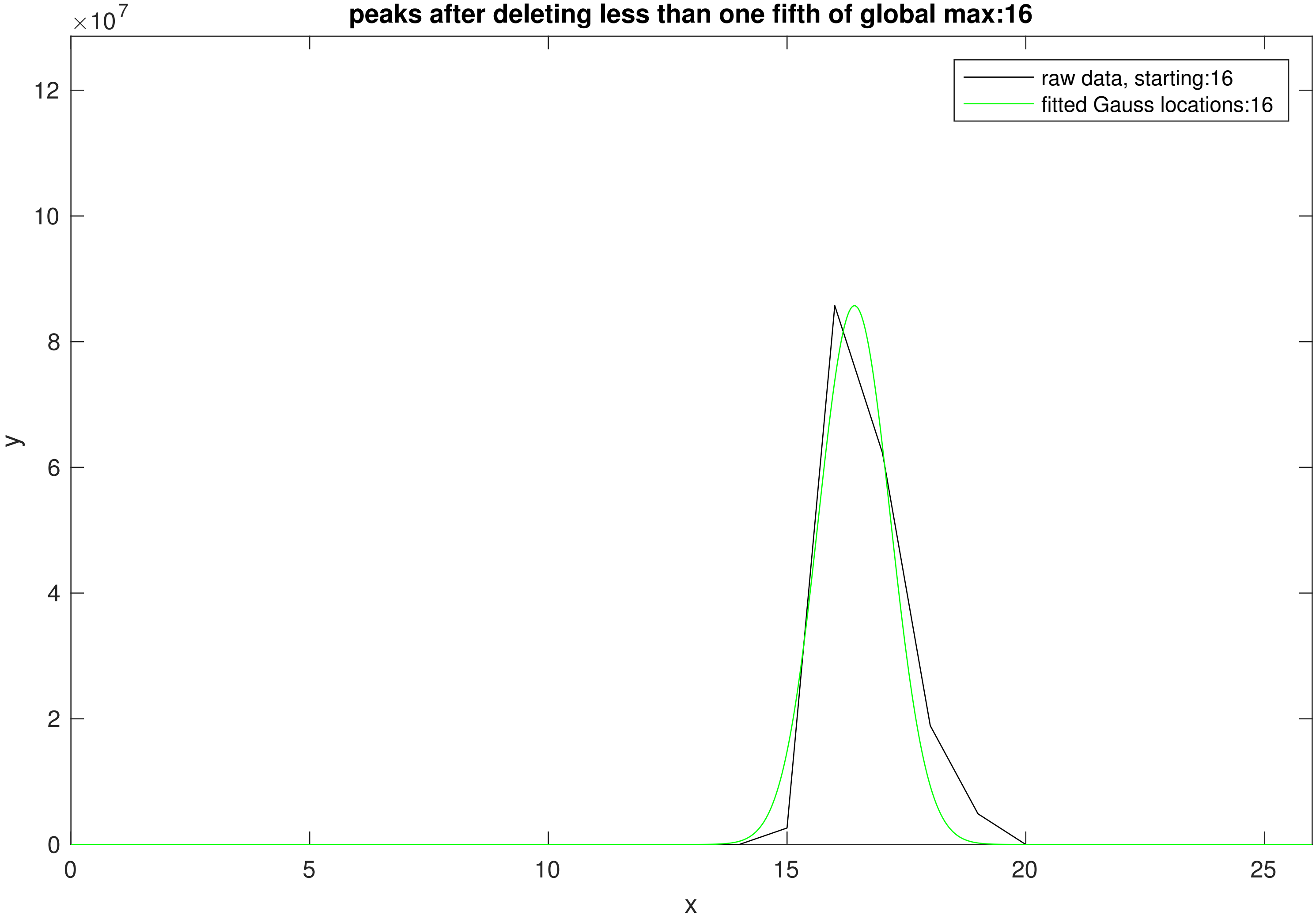

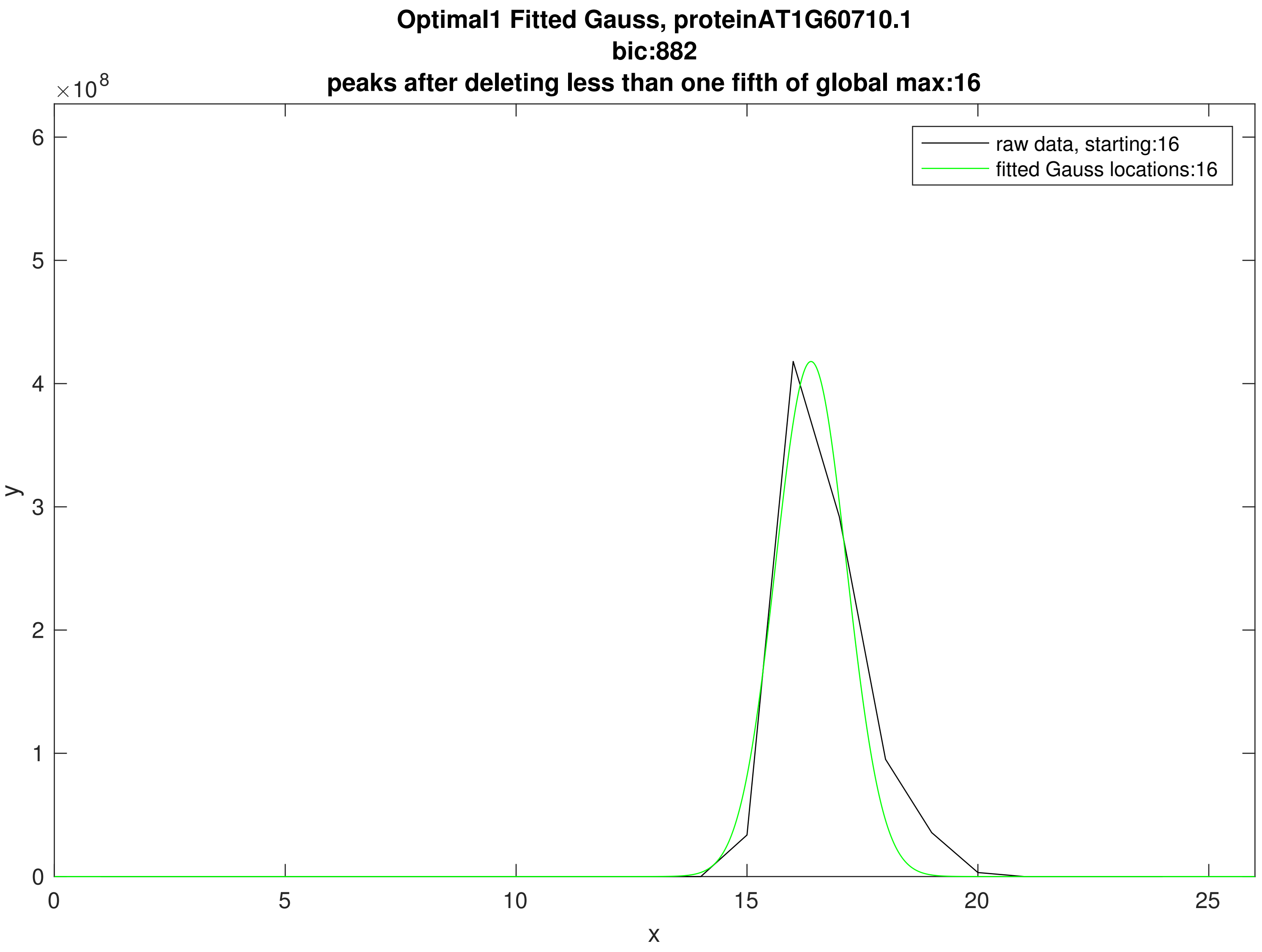
Optimal1 Fitted Gauss, proteinAT1G60710.1
bic:882
peaks after deleting less than one fifth of global max:16
×10^8
6
5
4
3
2
1
0
y
0
5
10
15
20
25
x
raw data, starting:16
fitted Gauss locations:16

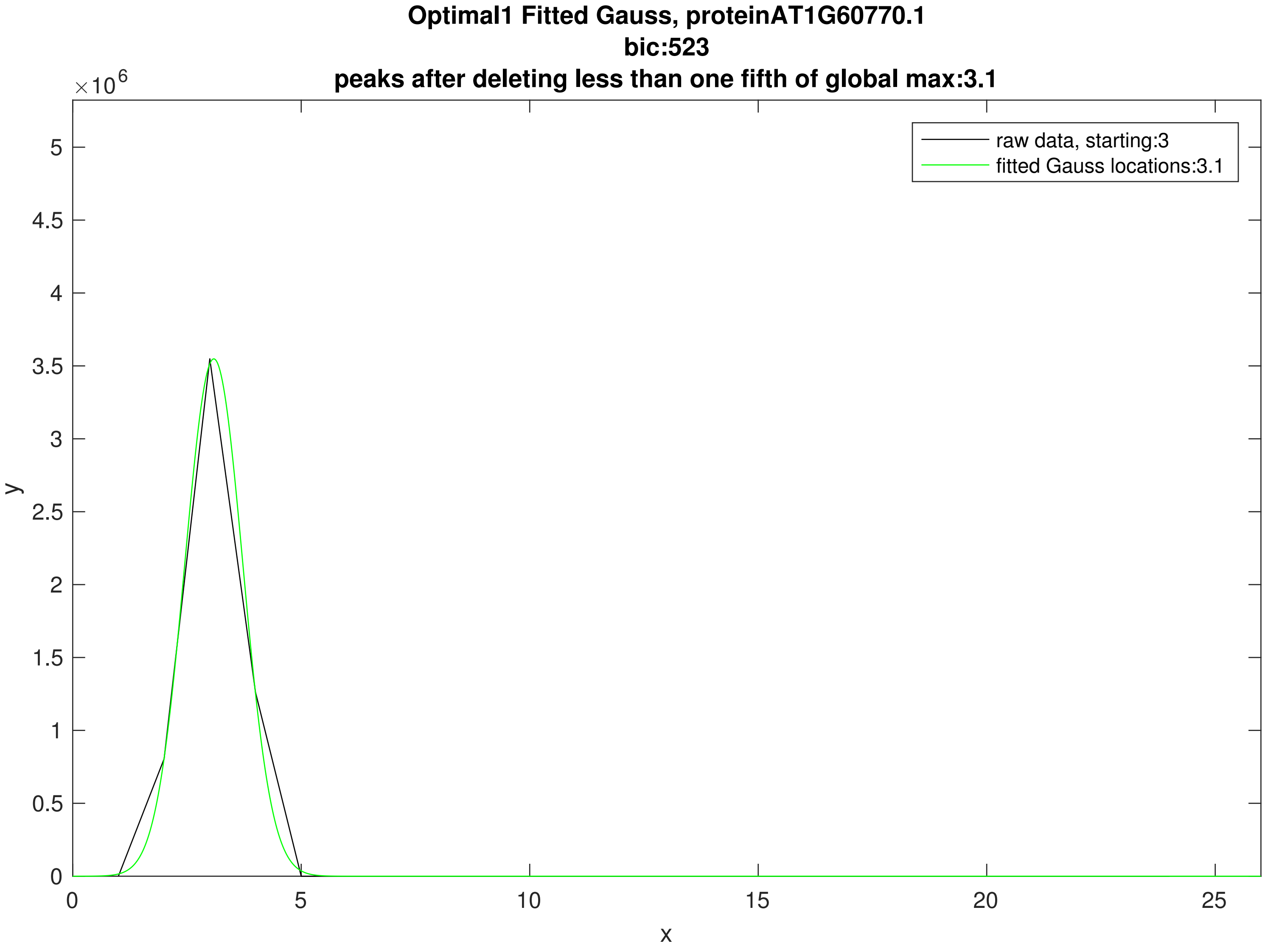

Optimal1 Fitted Gauss, proteinAT1G60770.1
bic:523
peaks after deleting less than one fifth of global max:3.1
×10^6
raw data, starting:3
fitted Gauss locations:3.1
y
x
0
0.5
1
1.5
2
2.5
3
3.5
4
4.5
5
0
5
10
15
20
25

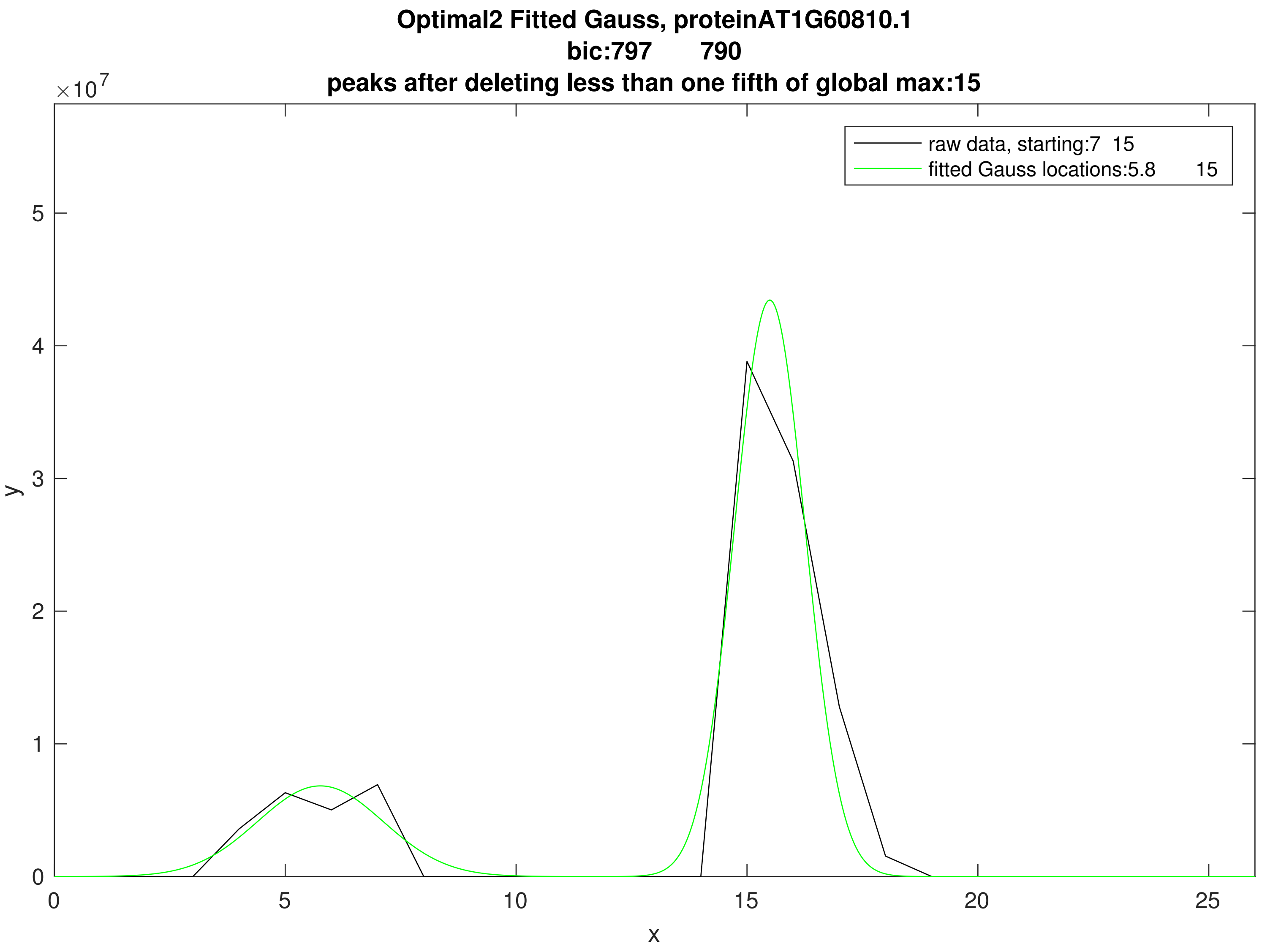

Optimal2 Fitted Gauss, proteinAT1G60810.1
bic:797 790
peaks after deleting less than one fifth of global max:15
×10^7
raw data, starting:7 15
fitted Gauss locations:5.8 15
y
x
0
1
2
3
4
5
0
5
10
15
20
25

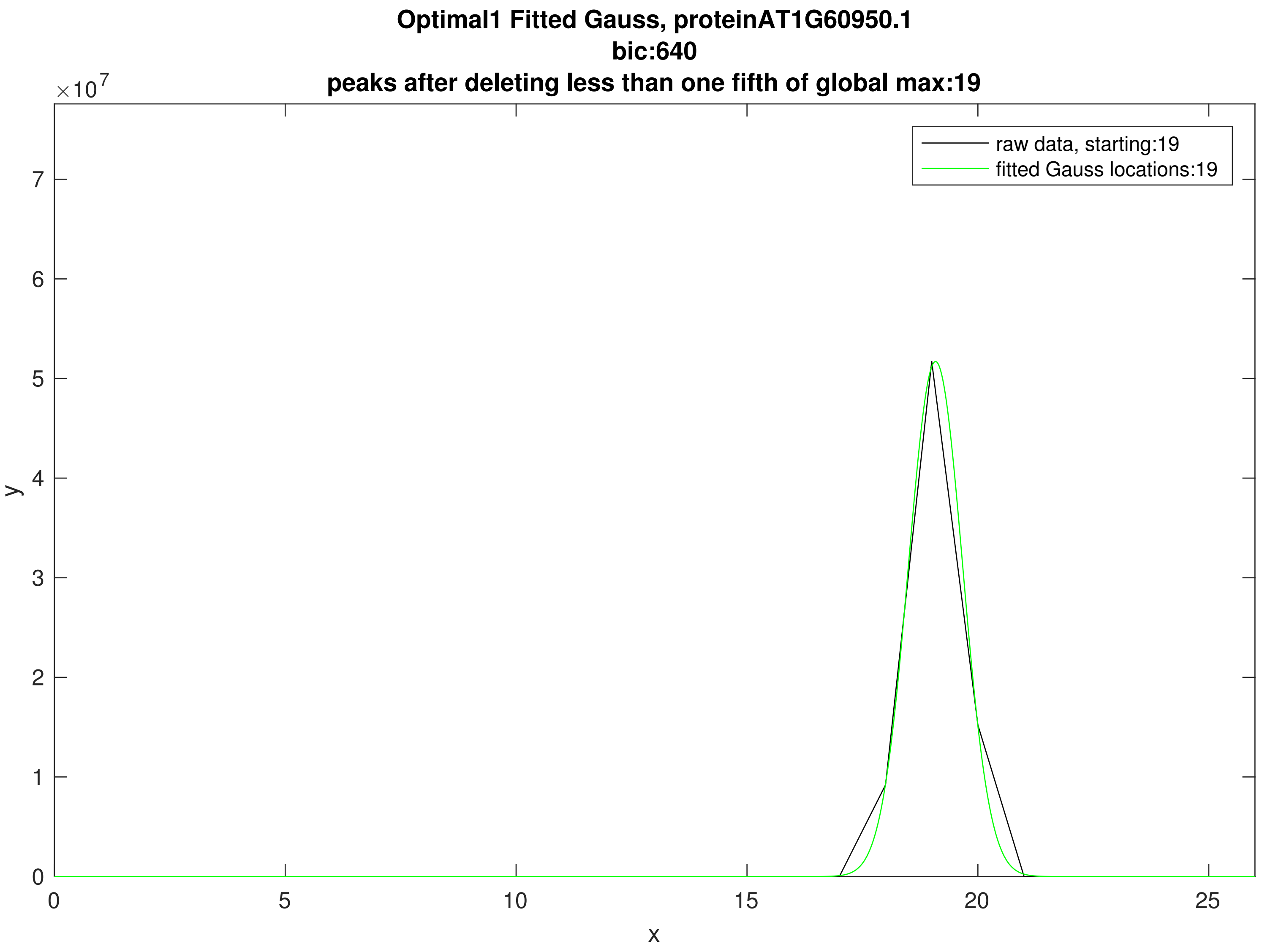

Optimal1 Fitted Gauss, proteinAT1G60950.1
bic:640
peaks after deleting less than one fifth of global max:19
×10^7
raw data, starting:19
fitted Gauss locations:19
y
x
0
1
2
3
4
5
6
7
0
5
10
15
20
25

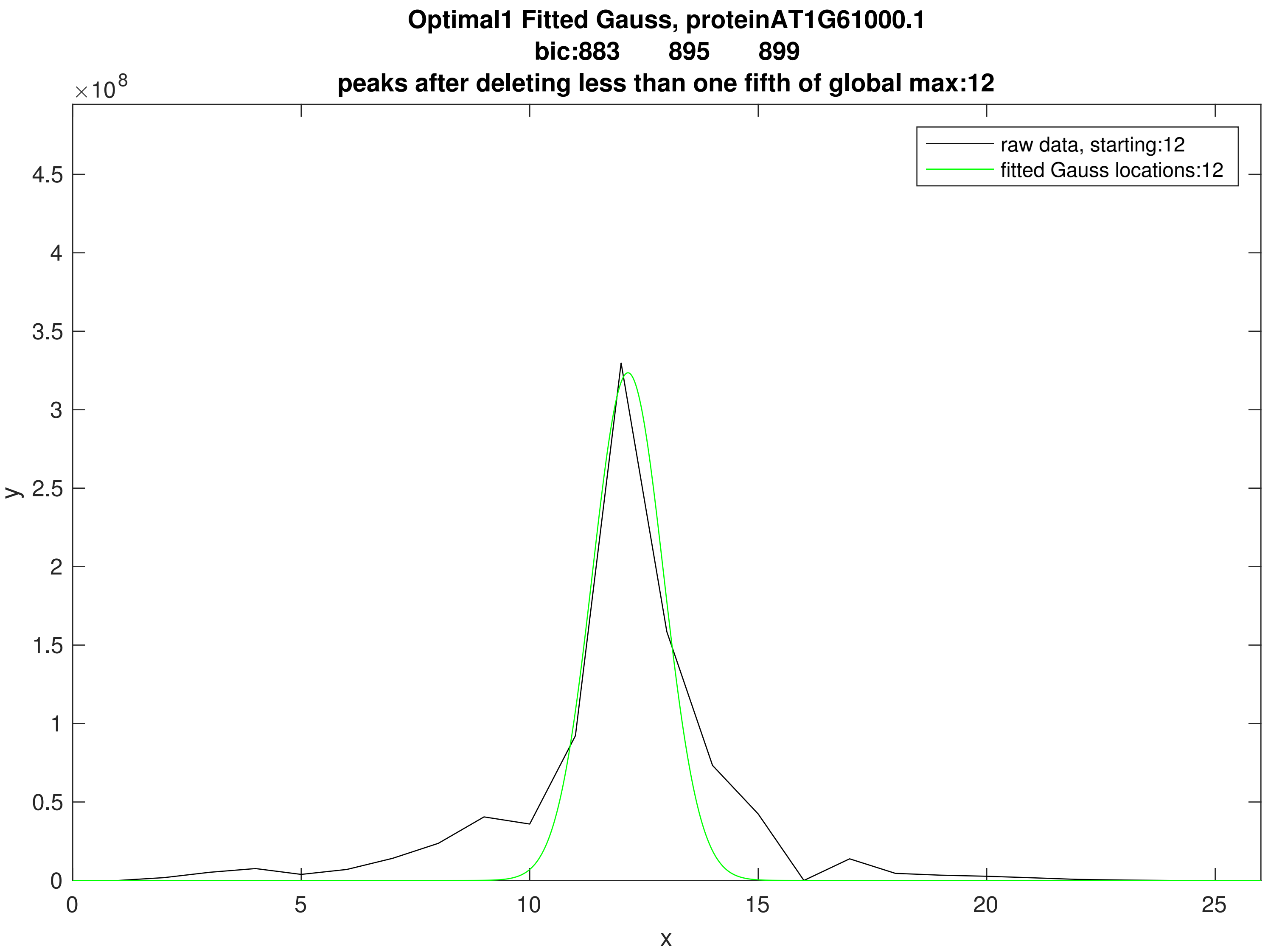

Optimal1 Fitted Gauss, proteinAT1G61000.1
bic:883 895 899
peaks after deleting less than one fifth of global max:12
×10^8
raw data, starting:12
fitted Gauss locations:12
y
x

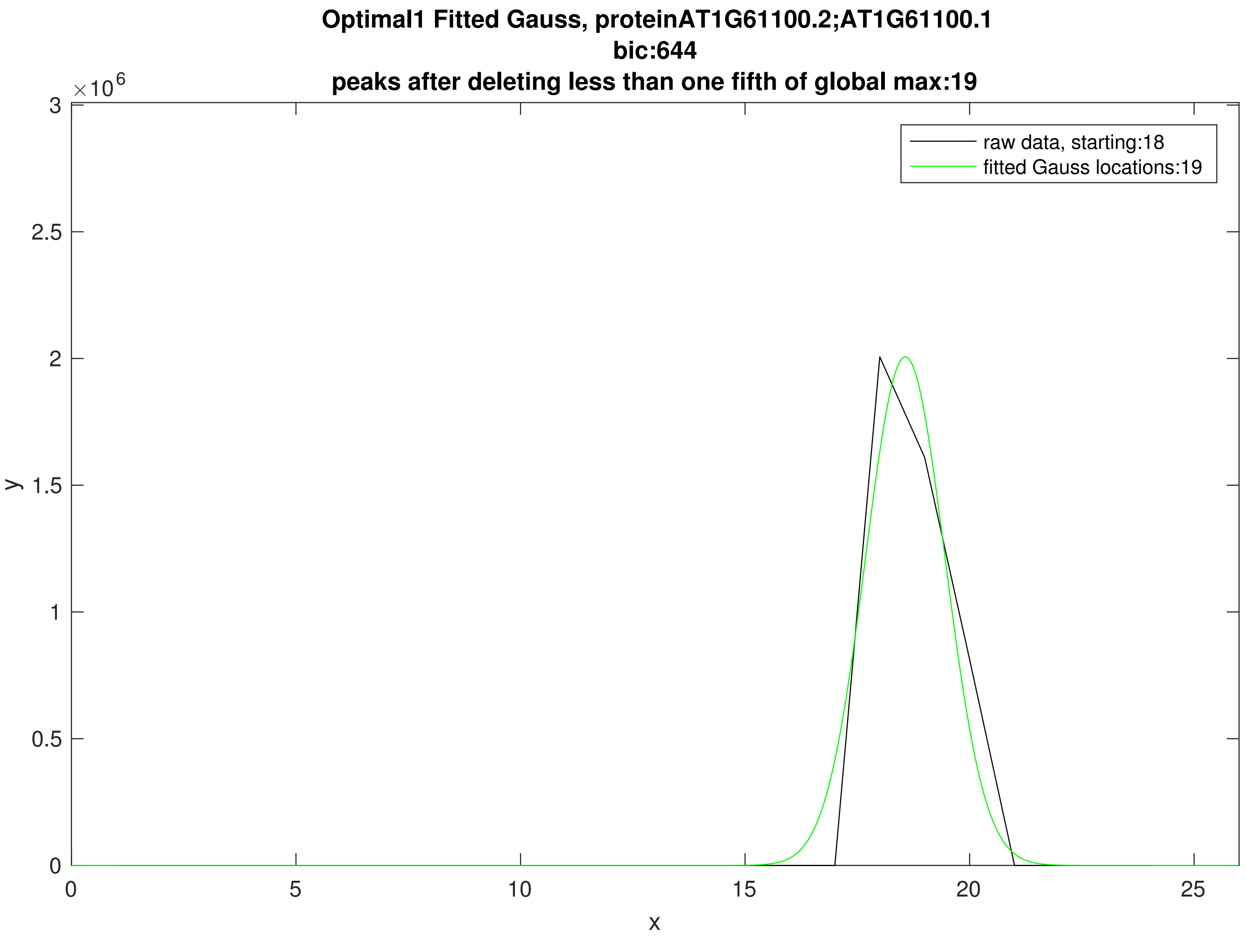

Optimal1 Fitted Gauss, proteinAT1G61100.2;AT1G61100.1
bic:644
peaks after deleting less than one fifth of global max:19
$\times 10^6$
3
2.5
2
1.5
1
0.5
0
y
0
5
10
15
20
25
x
raw data, starting:18
fitted Gauss locations:19

ptimal1 Fitted Gauss, proteinAT1G61150.5;AT1G61150.6;AT1G61150.4;AT1G61150.2;AT1G61150.1;AT1G61150.7;AT1G
bic:751 757 767
peaks after deleting less than one fifth of global max:16

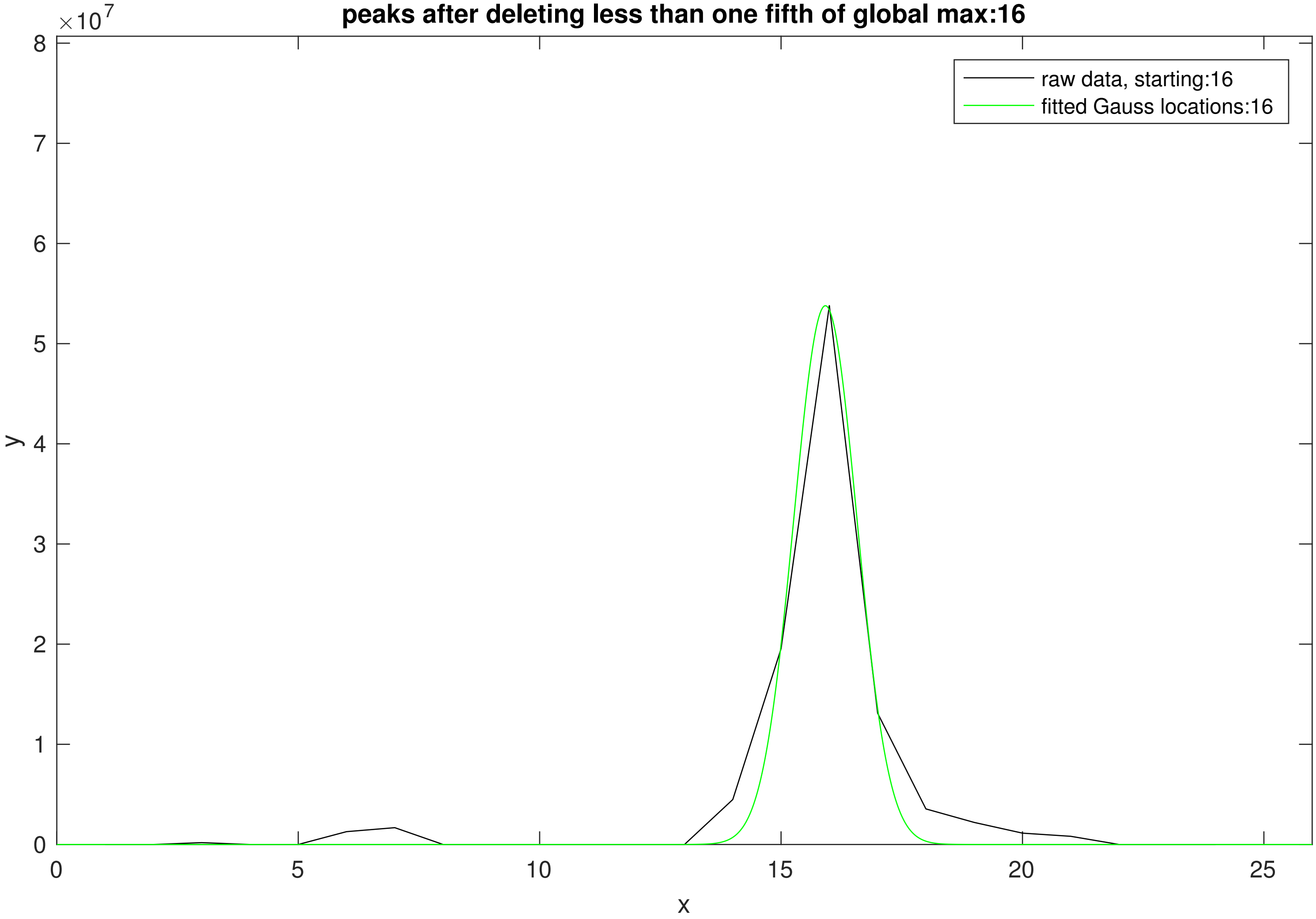

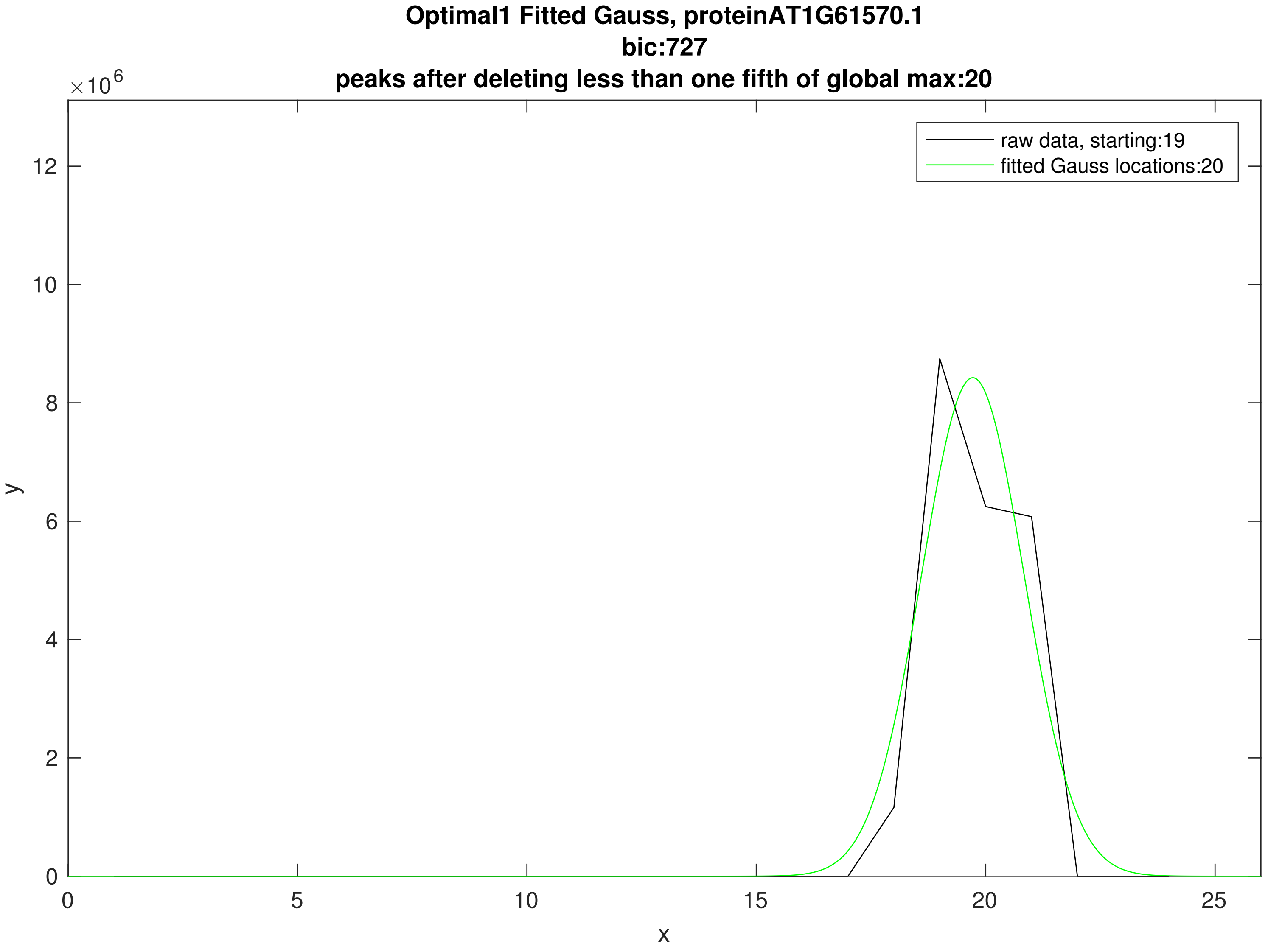

Optimal1 Fitted Gauss, proteinAT1G61570.1
bic:727
peaks after deleting less than one fifth of global max:20
×10^6
raw data, starting:19
fitted Gauss locations:20
12
10
8
6
4
2
0
y
0
5
10
15
20
25
x

#### Optimal1 Fitted Gauss, proteinAT1G61820.1
#### bic:787
#### peaks after deleting less than one fifth of global max:17

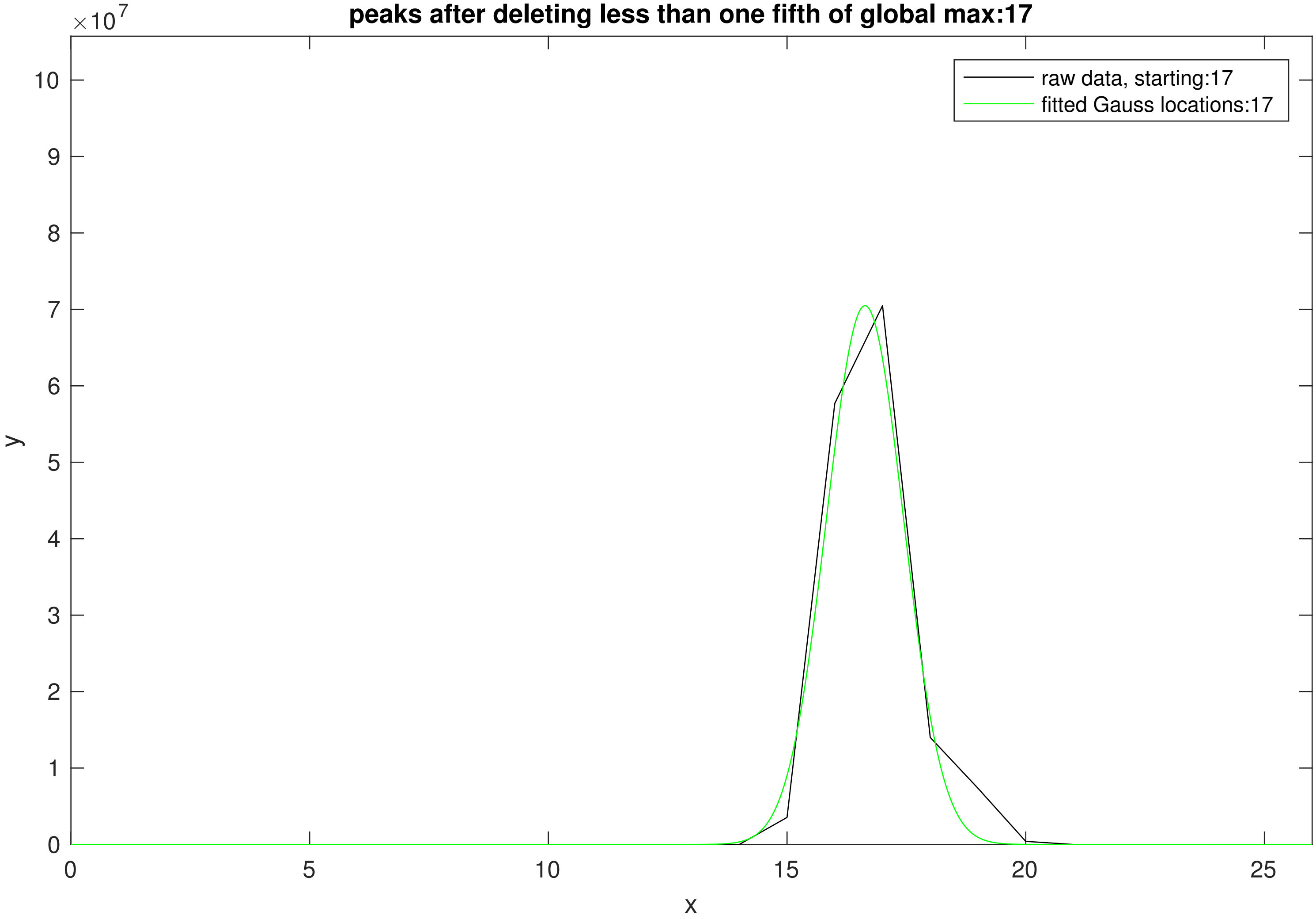

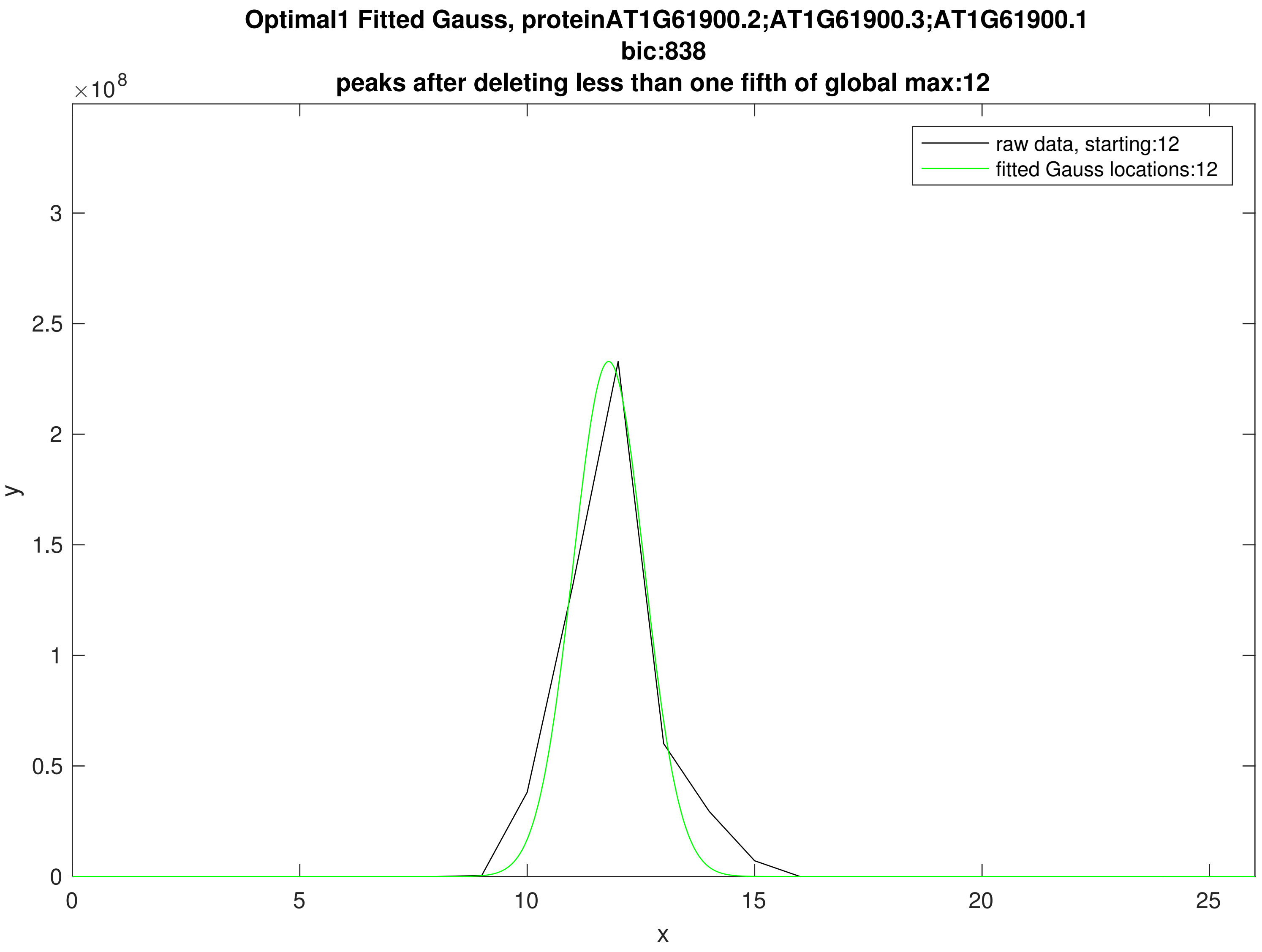
Optimal1 Fitted Gauss, proteinAT1G61900.2;AT1G61900.3;AT1G61900.1
bic:838
peaks after deleting less than one fifth of global max:12
×10^8
raw data, starting:12
fitted Gauss locations:12
y
x
0
0.5
1
1.5
2
2.5
3
5
10
15
20
25

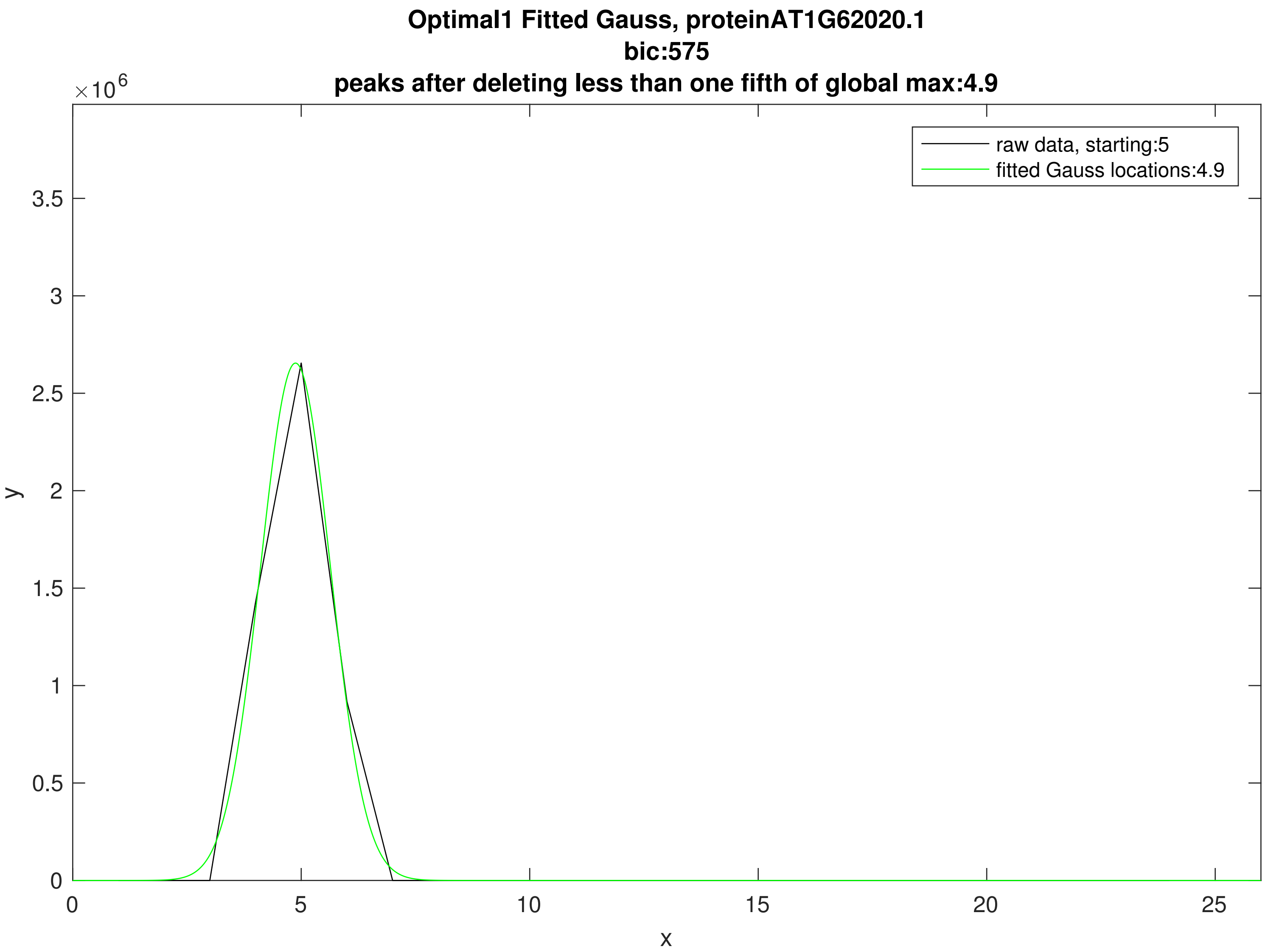
Optimal1 Fitted Gauss, proteinAT1G62020.1
bic:575
peaks after deleting less than one fifth of global max:4.9
×10^6
raw data, starting:5
fitted Gauss locations:4.9
3.5
3
2.5
2
1.5
1
0.5
0
y
0
5
10
15
20
25
x

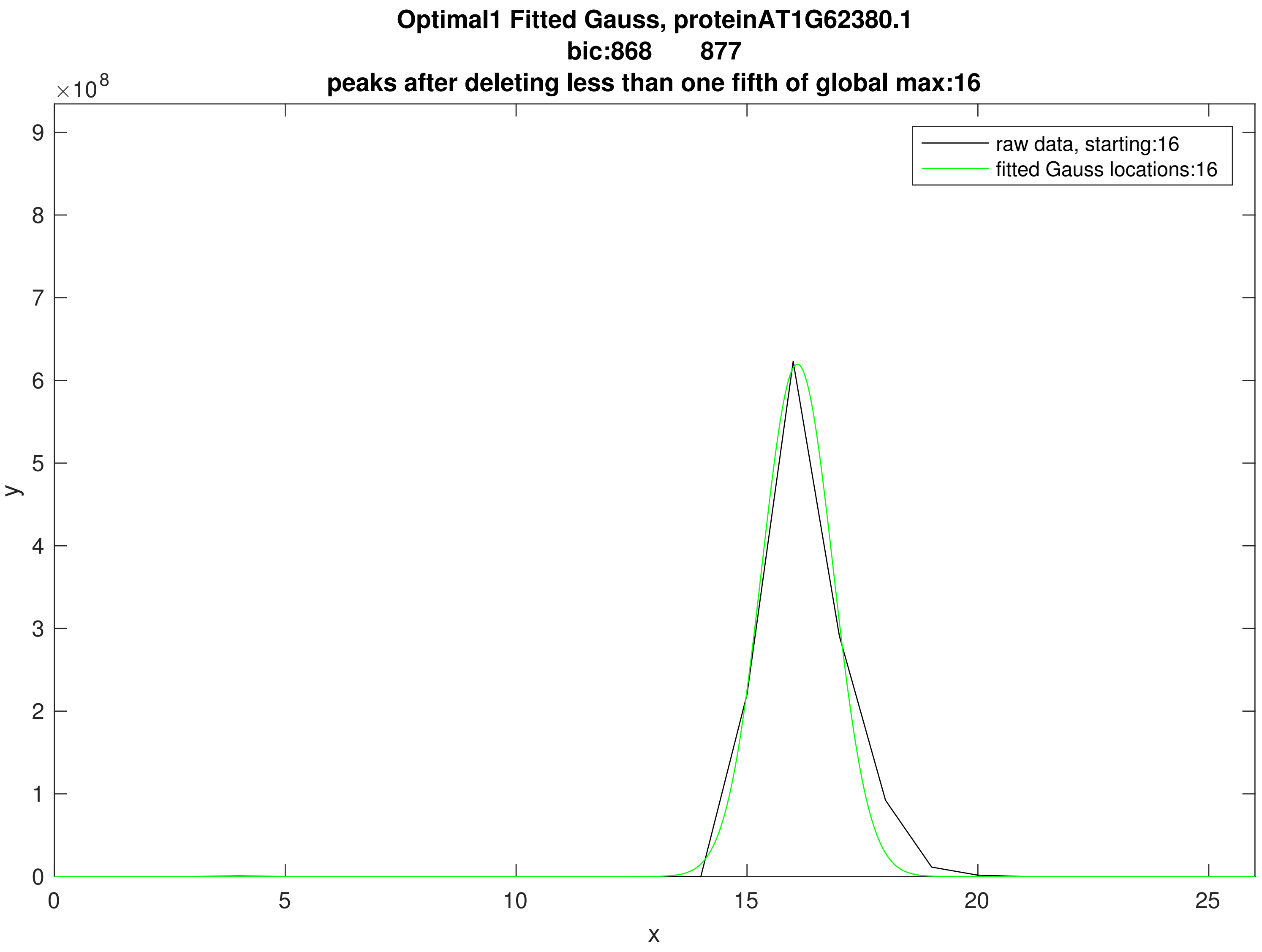

Optimal1 Fitted Gauss, proteinAT1G62380.1
bic:868 877
peaks after deleting less than one fifth of global max:16
×10^8
raw data, starting:16
fitted Gauss locations:16
y
x
0
1
2
3
4
5
6
7
8
9
0
5
10
15
20
25

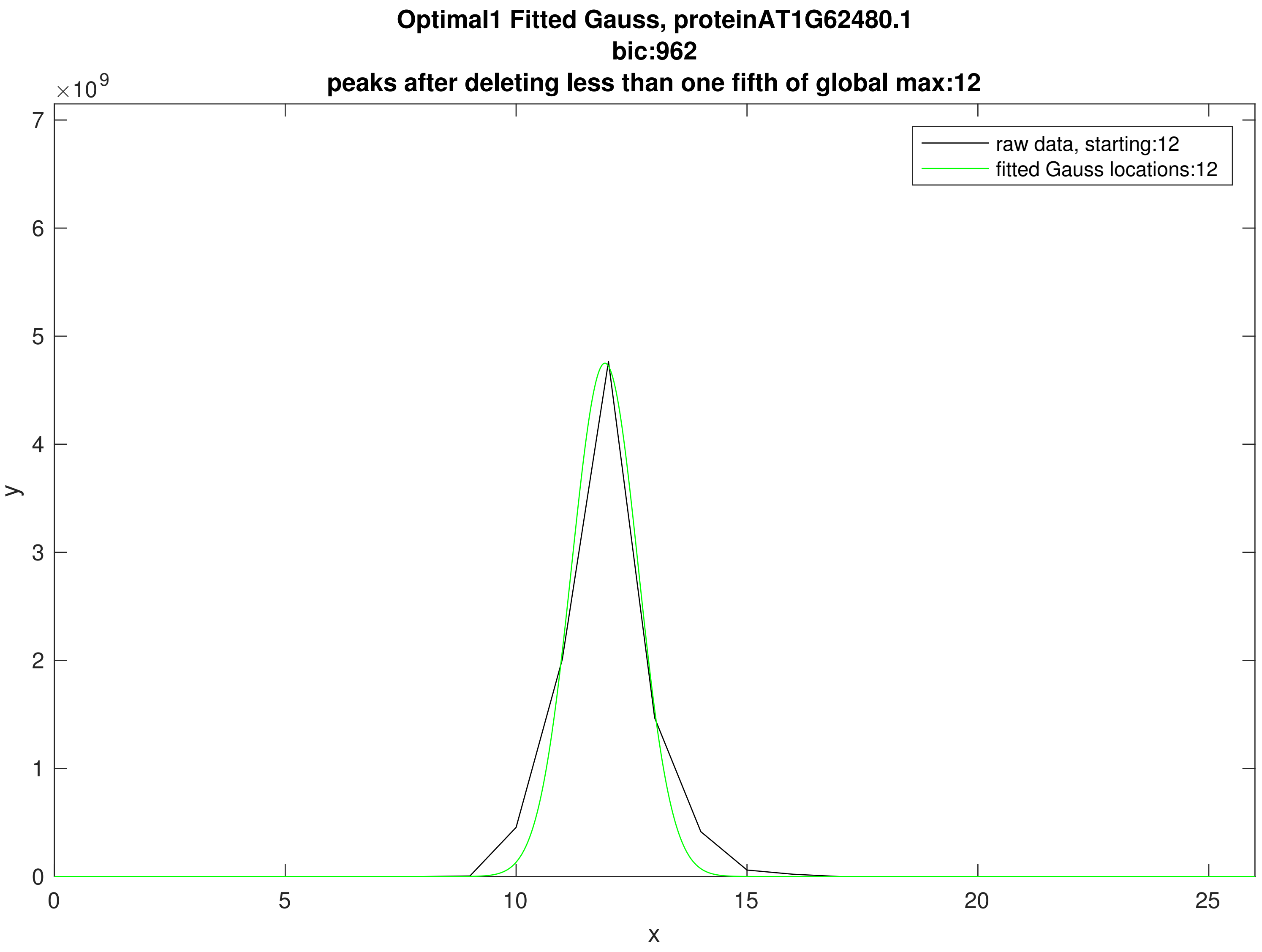

Optimal1 Fitted Gauss, proteinAT1G62480.1
bic:962
peaks after deleting less than one fifth of global max:12
×10^9
7
6
5
4
3
2
1
0
y
0
5
10
15
20
25
x
raw data, starting:12
fitted Gauss locations:12

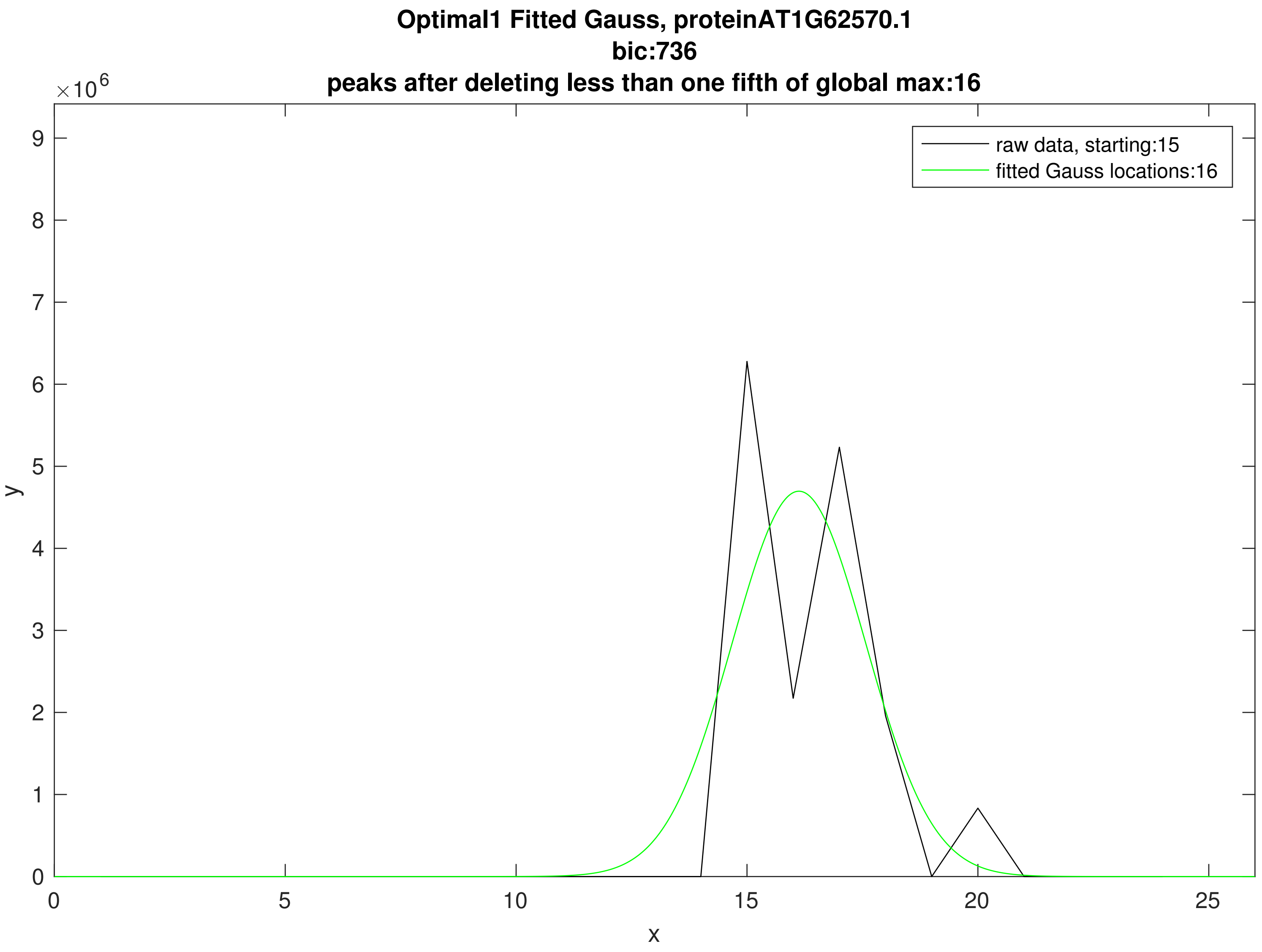

Optimal1 Fitted Gauss, proteinAT1G62570.1
bic:736
peaks after deleting less than one fifth of global max:16
×10^6
raw data, starting:15
fitted Gauss locations:16
y
x
0
1
2
3
4
5
6
7
8
9
0
5
10
15
20
25

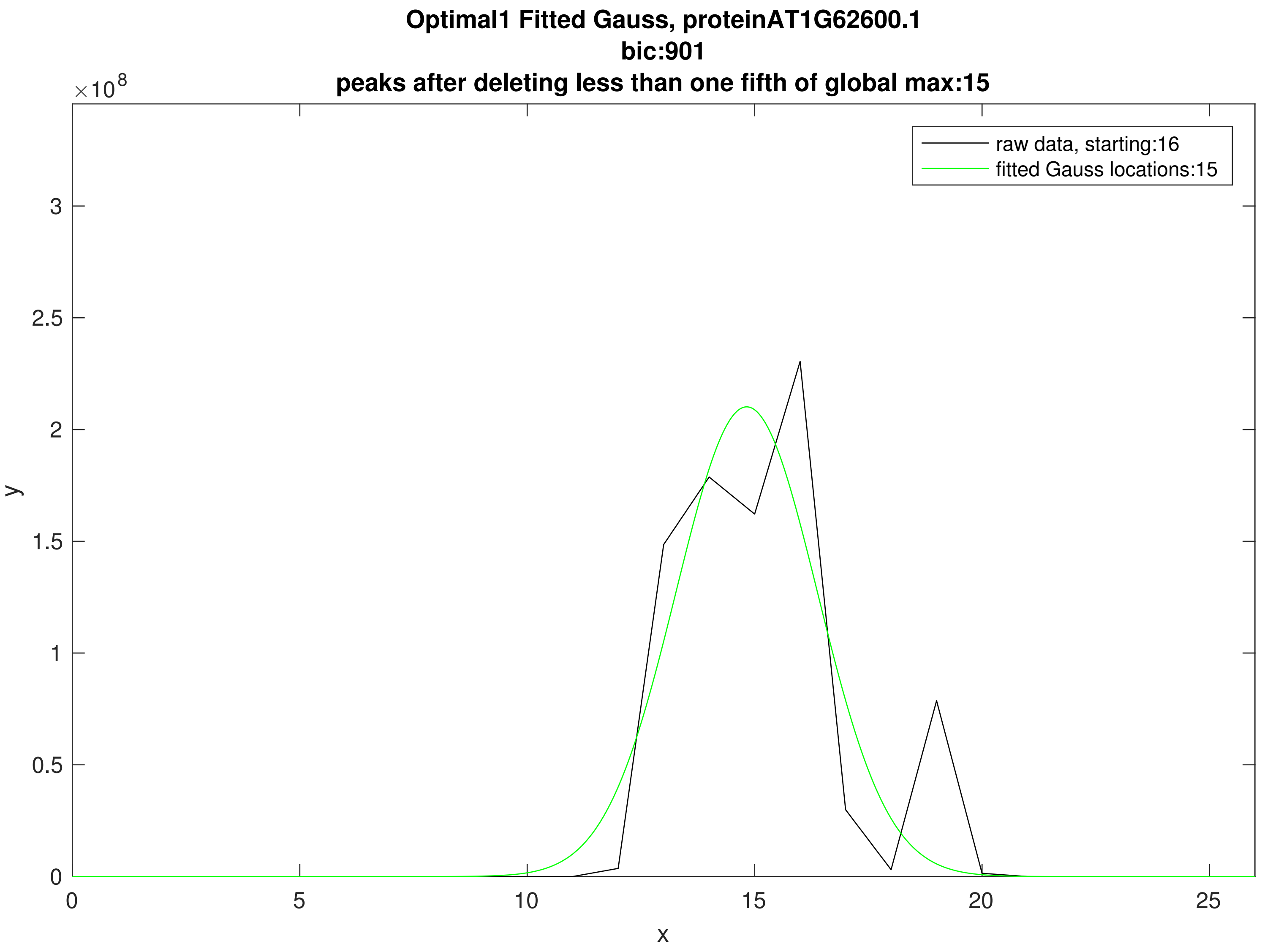
Optimal1 Fitted Gauss, proteinAT1G62600.1
bic:901
peaks after deleting less than one fifth of global max:15
×10^8
raw data, starting:16
fitted Gauss locations:15
y
x
0
0.5
1
1.5
2
2.5
3
0
5
10
15
20
25

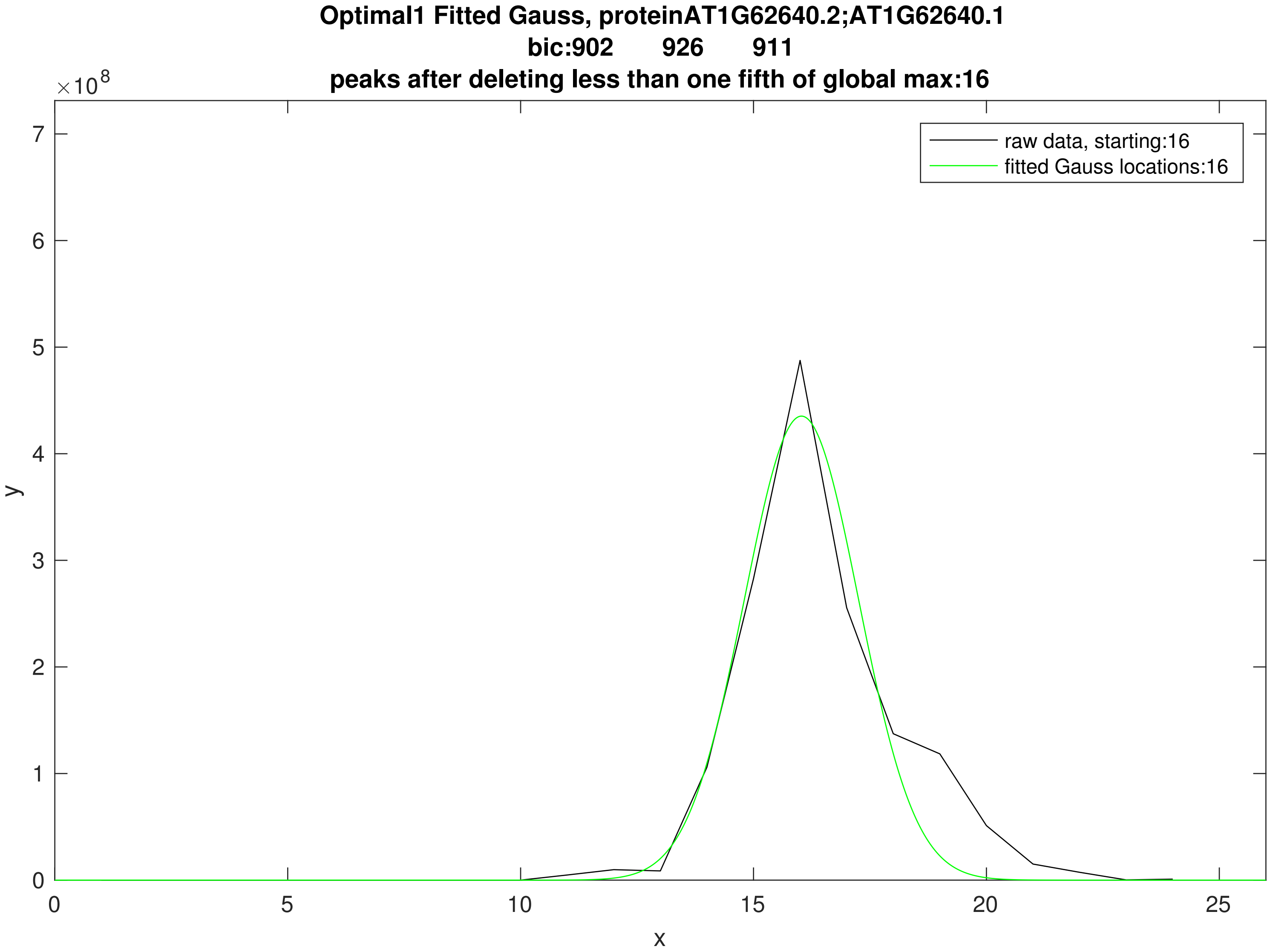
Optimal1 Fitted Gauss, proteinAT1G62640.2;AT1G62640.1
bic:902 926 911
peaks after deleting less than one fifth of global max:16
×10^8
raw data, starting:16
fitted Gauss locations:16
y
x
0
1
2
3
4
5
6
7
0
5
10
15
20
25

Optimal1 Fitted Gauss, proteinAT1G62660.1
bic:1.06e+03  1.07e+03
peaks after deleting less than one fifth of global max:14

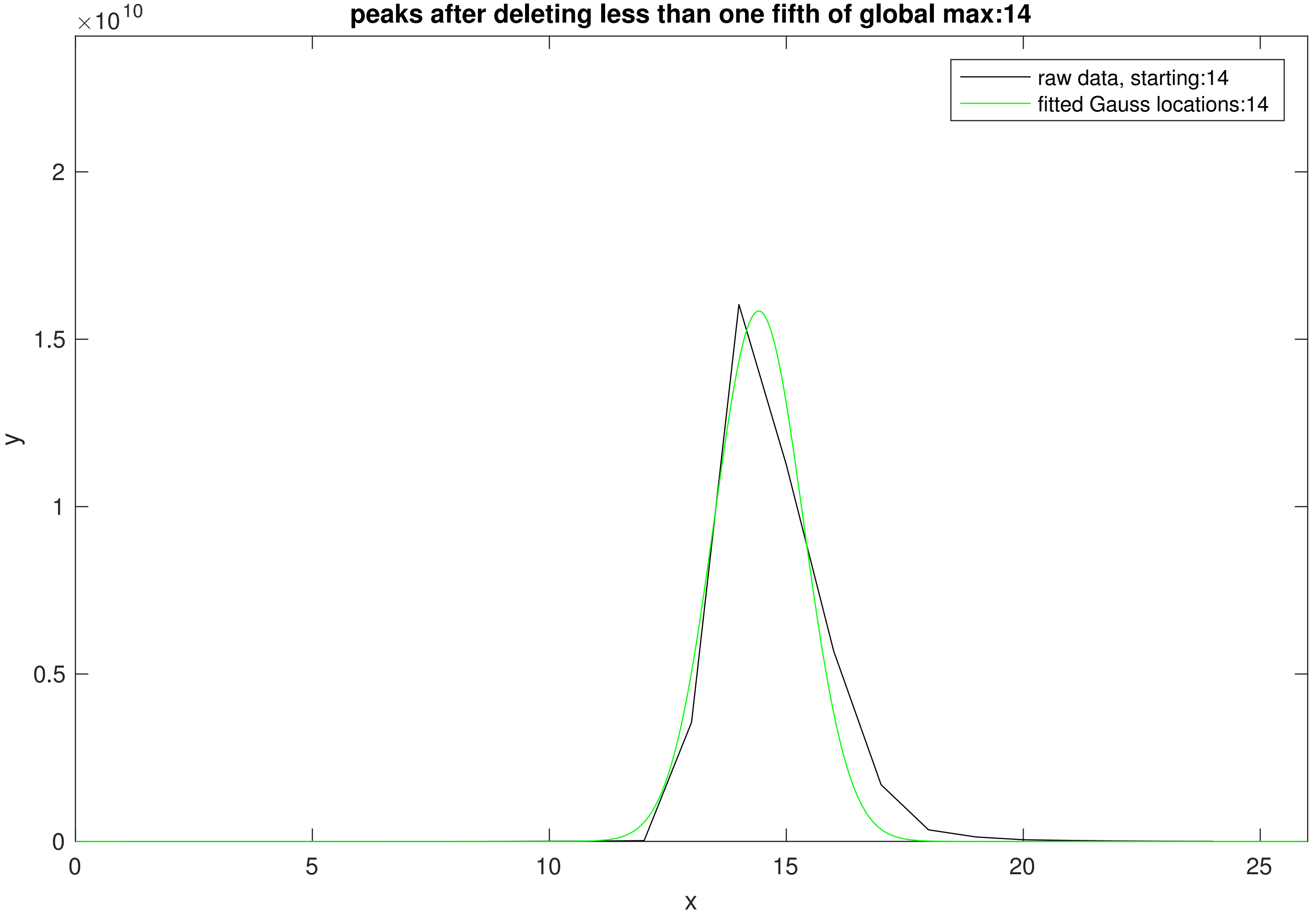

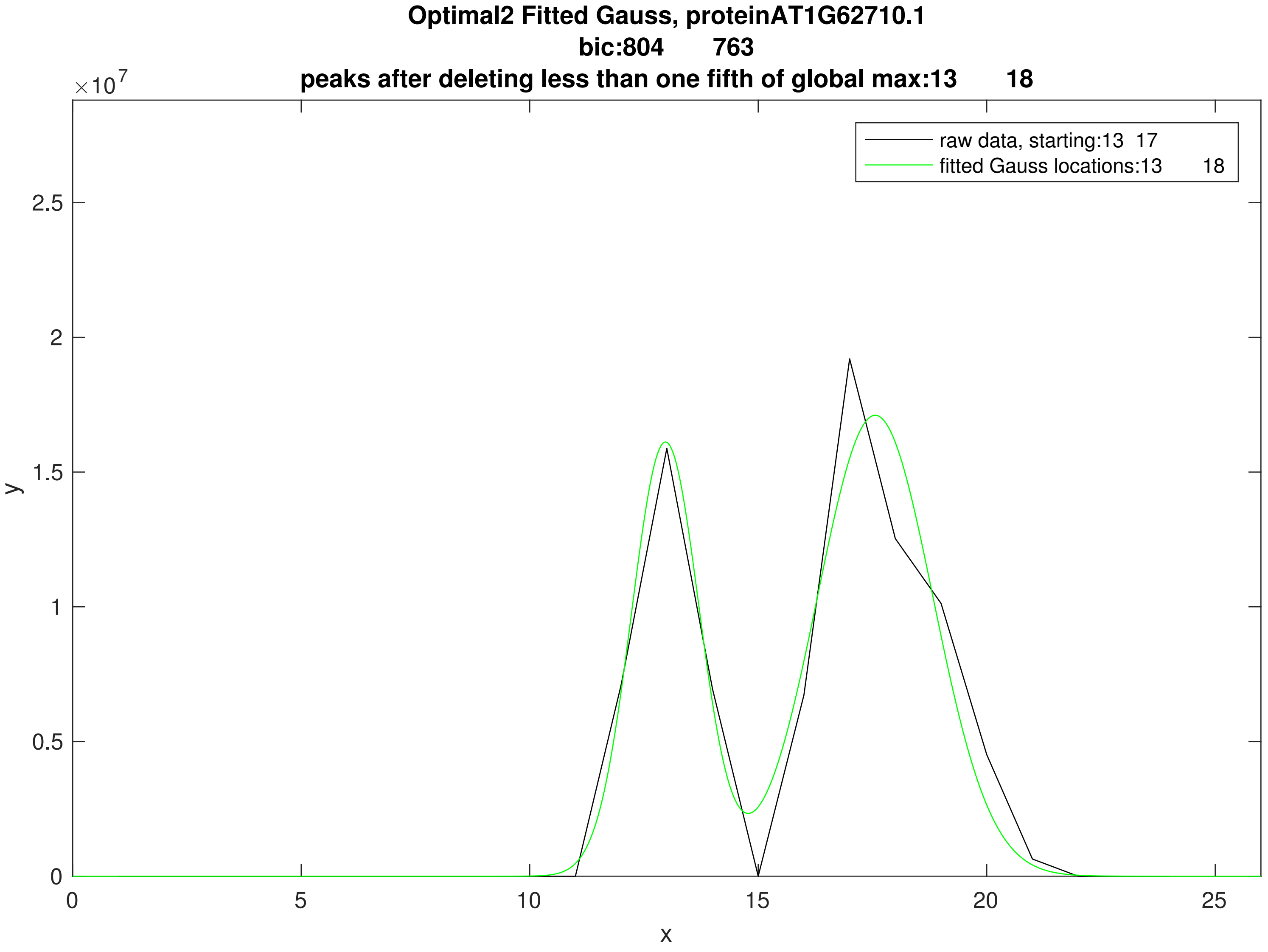
Optimal2 Fitted Gauss, proteinAT1G62710.1
bic:804 763
peaks after deleting less than one fifth of global max:13 18
×10^7
raw data, starting:13 17
fitted Gauss locations:13 18
y
x
0
0.5
1
1.5
2
2.5
5
10
15
20
25

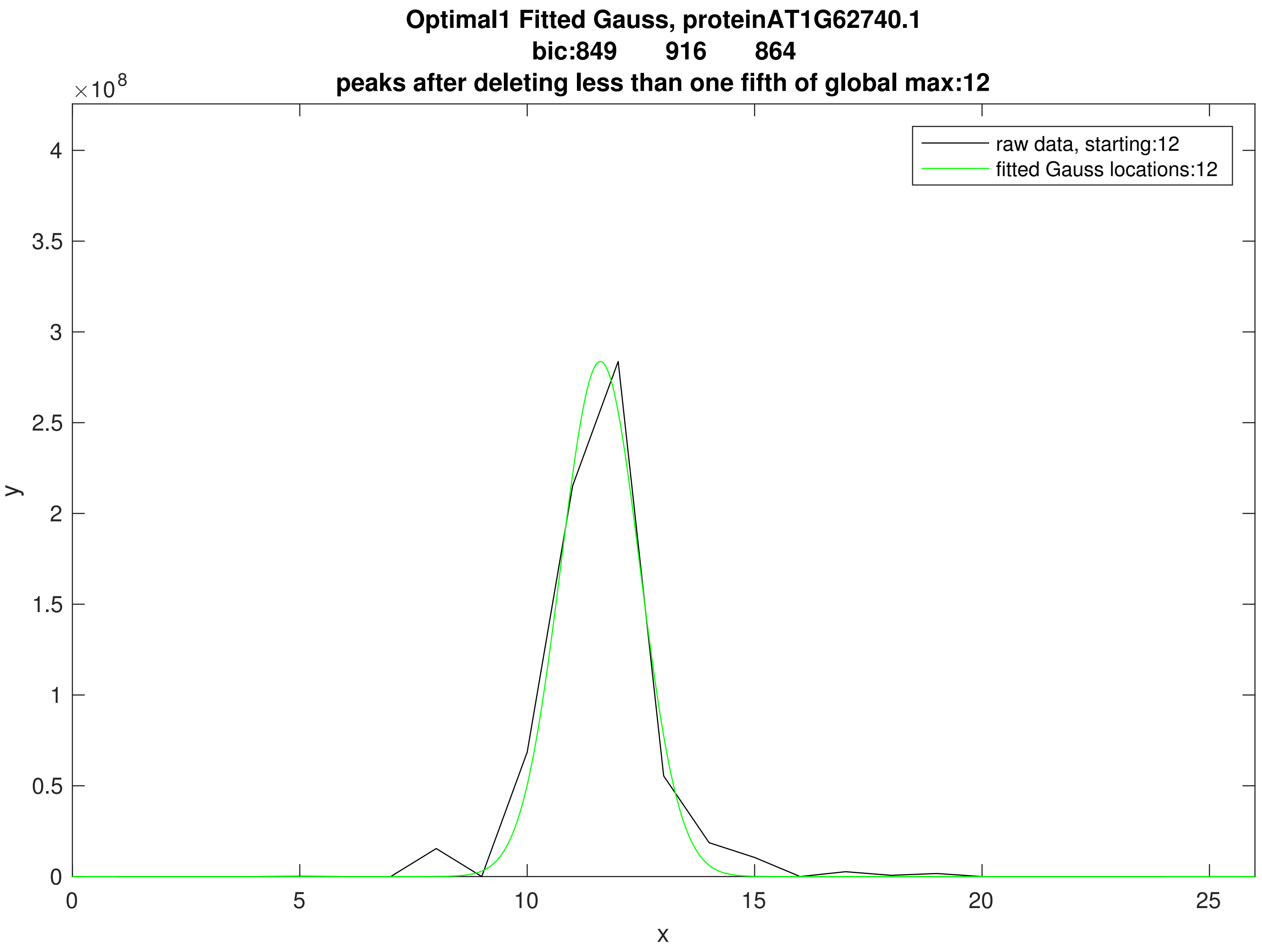
Optimal1 Fitted Gauss, proteinAT1G62740.1
bic:849 916 864
peaks after deleting less than one fifth of global max:12
×10^8
raw data, starting:12
fitted Gauss locations:12
y
x
0
0.5
1
1.5
2
2.5
3
3.5
4
0
5
10
15
20
25

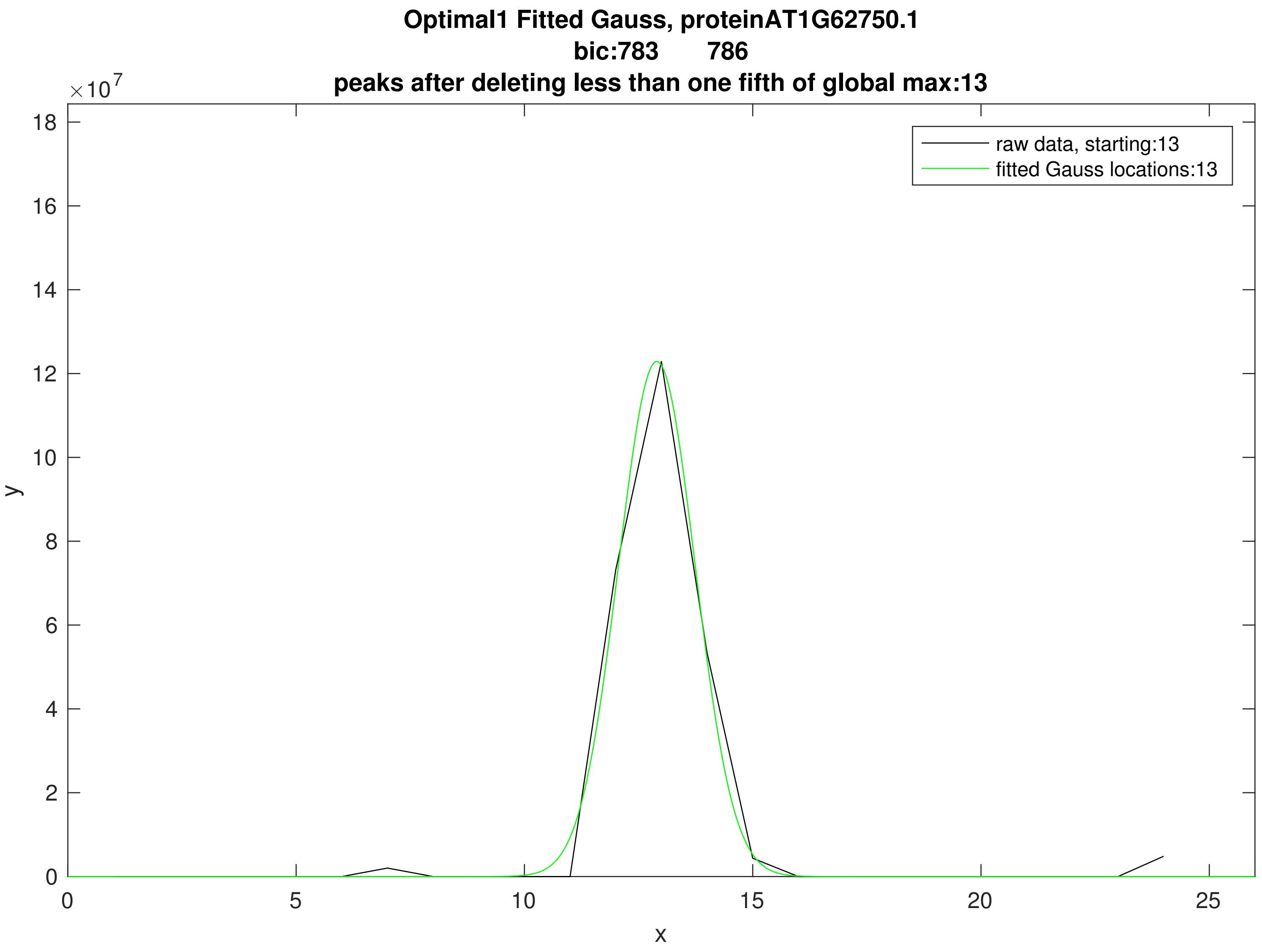
Optimal1 Fitted Gauss, proteinAT1G62750.1
bic:783 786
peaks after deleting less than one fifth of global max:13
×10^7
18
16
14
12
10
8
6
4
2
0
y
0
5
10
15
20
25
x
raw data, starting:13
fitted Gauss locations:13

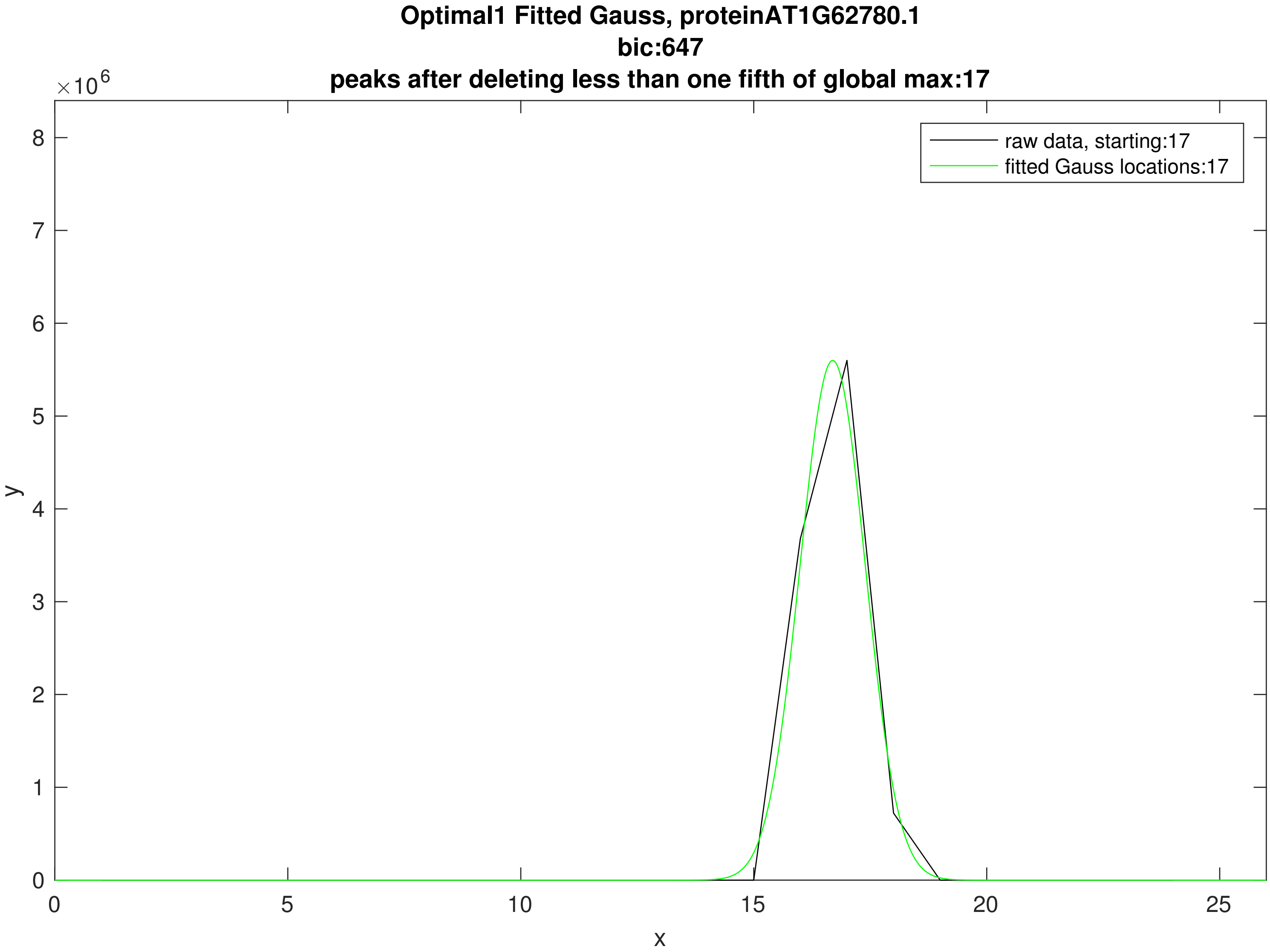

Optimal1 Fitted Gauss, proteinAT1G62780.1
bic:647
peaks after deleting less than one fifth of global max:17
×10^6
raw data, starting:17
fitted Gauss locations:17
y
x
0
1
2
3
4
5
6
7
8
0
5
10
15
20
25

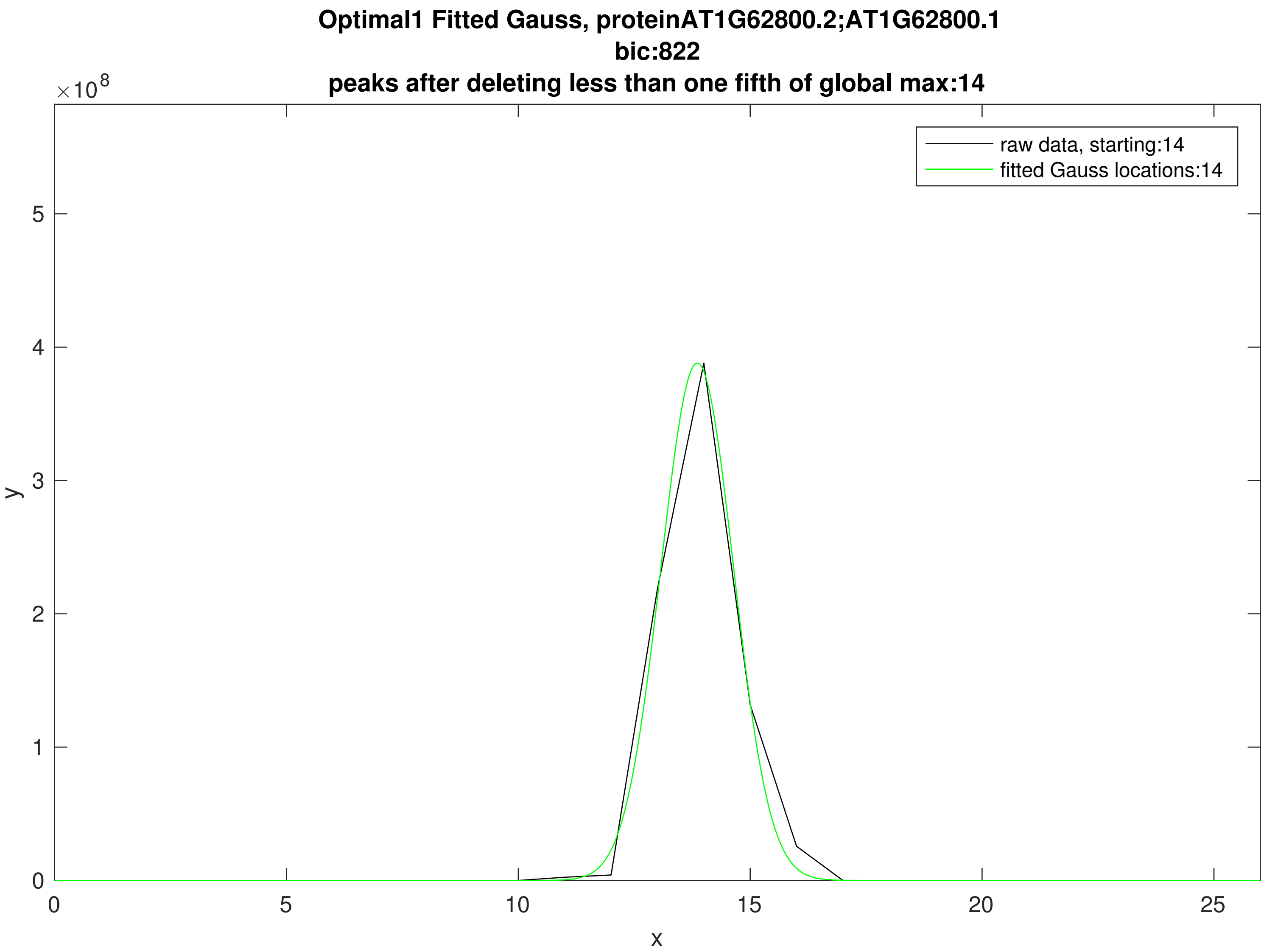
Optimal1 Fitted Gauss, proteinAT1G62800.2;AT1G62800.1
bic:822
peaks after deleting less than one fifth of global max:14
×10^8
raw data, starting:14
fitted Gauss locations:14
y
x
0
1
2
3
4
5
0
5
10
15
20
25

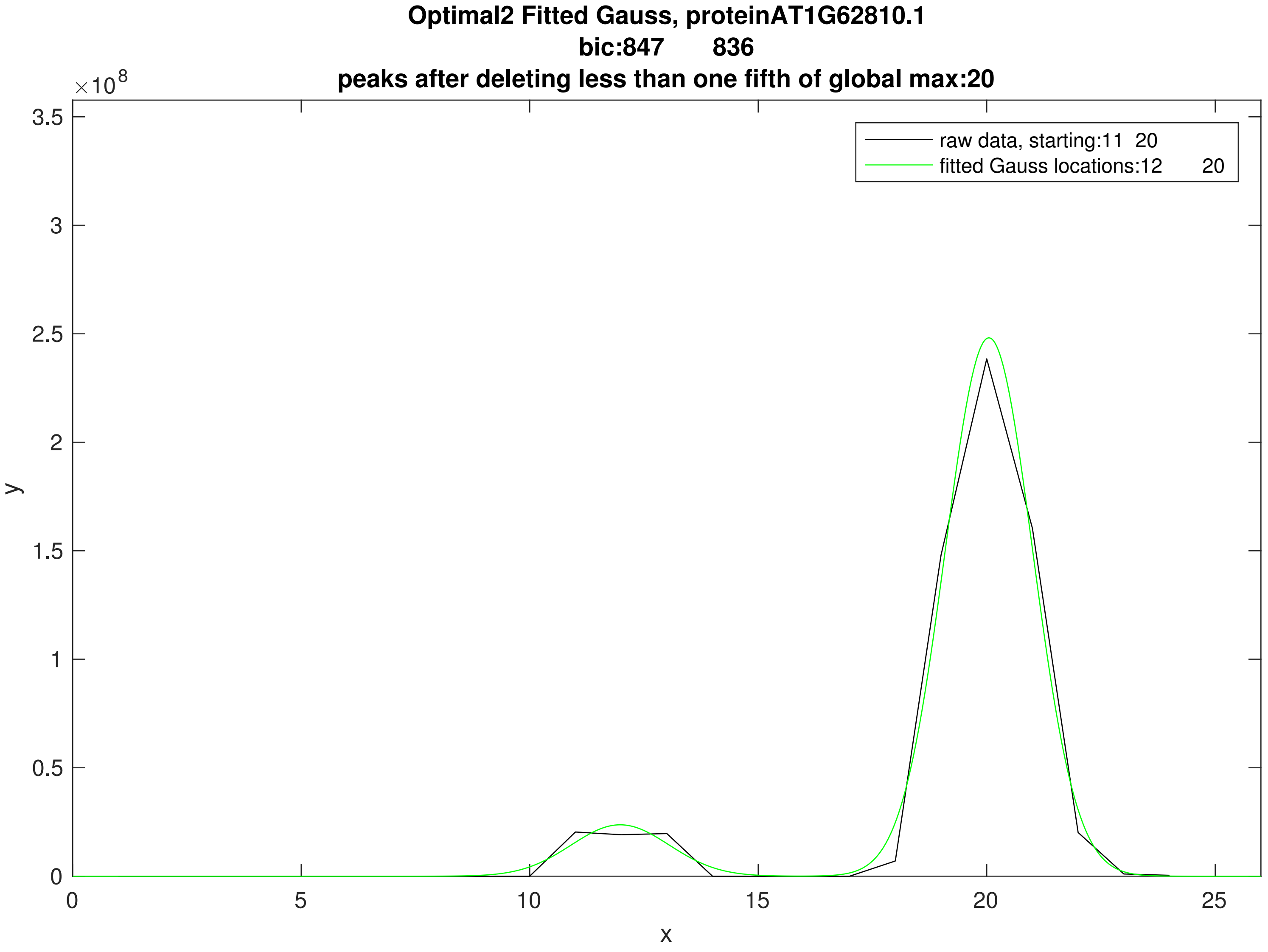
Optimal2 Fitted Gauss, proteinAT1G62810.1
bic:847 836
peaks after deleting less than one fifth of global max:20
×10^8
3.5
3
2.5
2
1.5
1
0.5
0
y
0
5
10
15
20
25
x
raw data, starting:11 20
fitted Gauss locations:12 20

**Optimal1 Fitted Gauss, proteinAT1G62820.1**
**bic:820**
**peaks after deleting less than one fifth of global max:19**

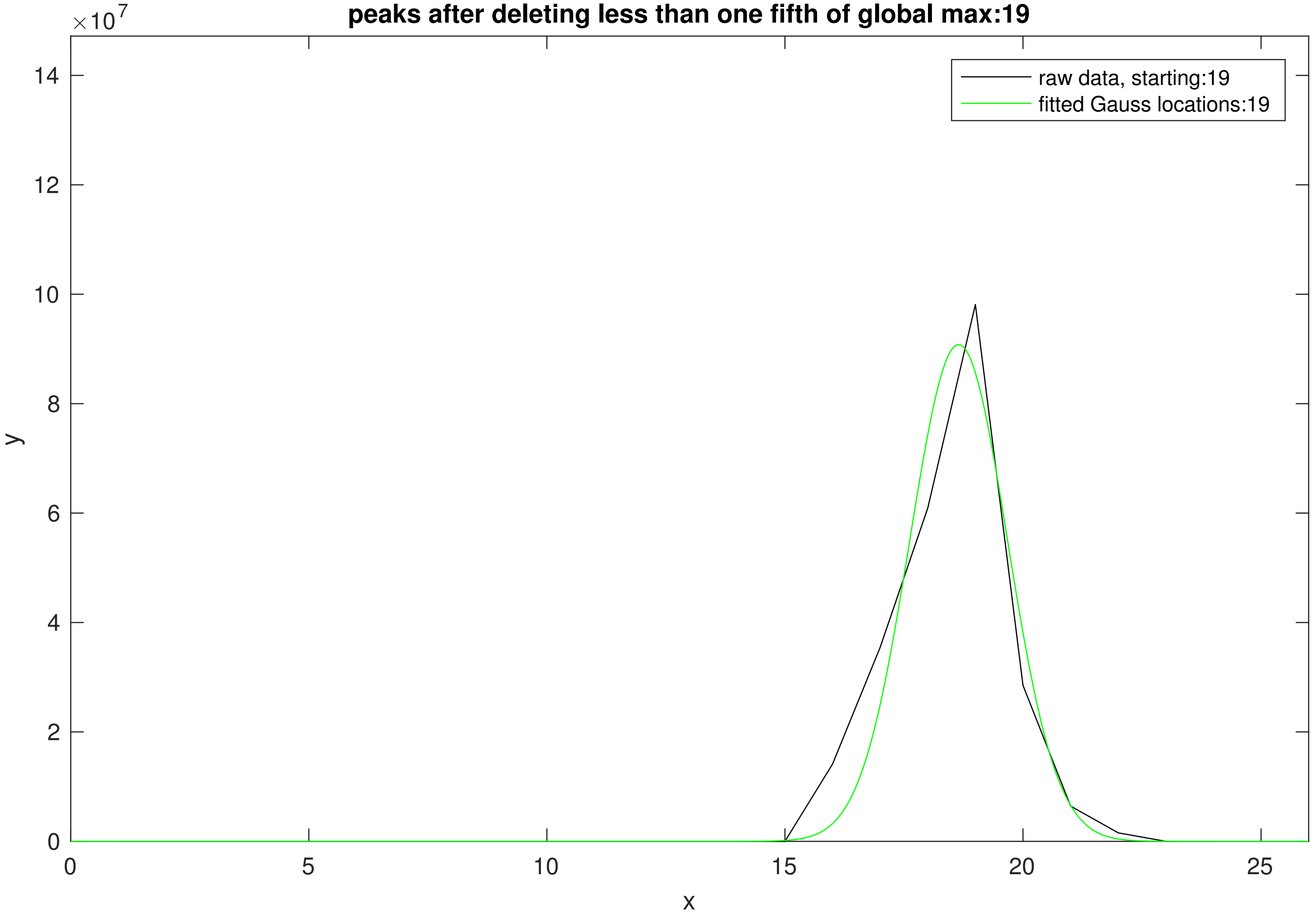

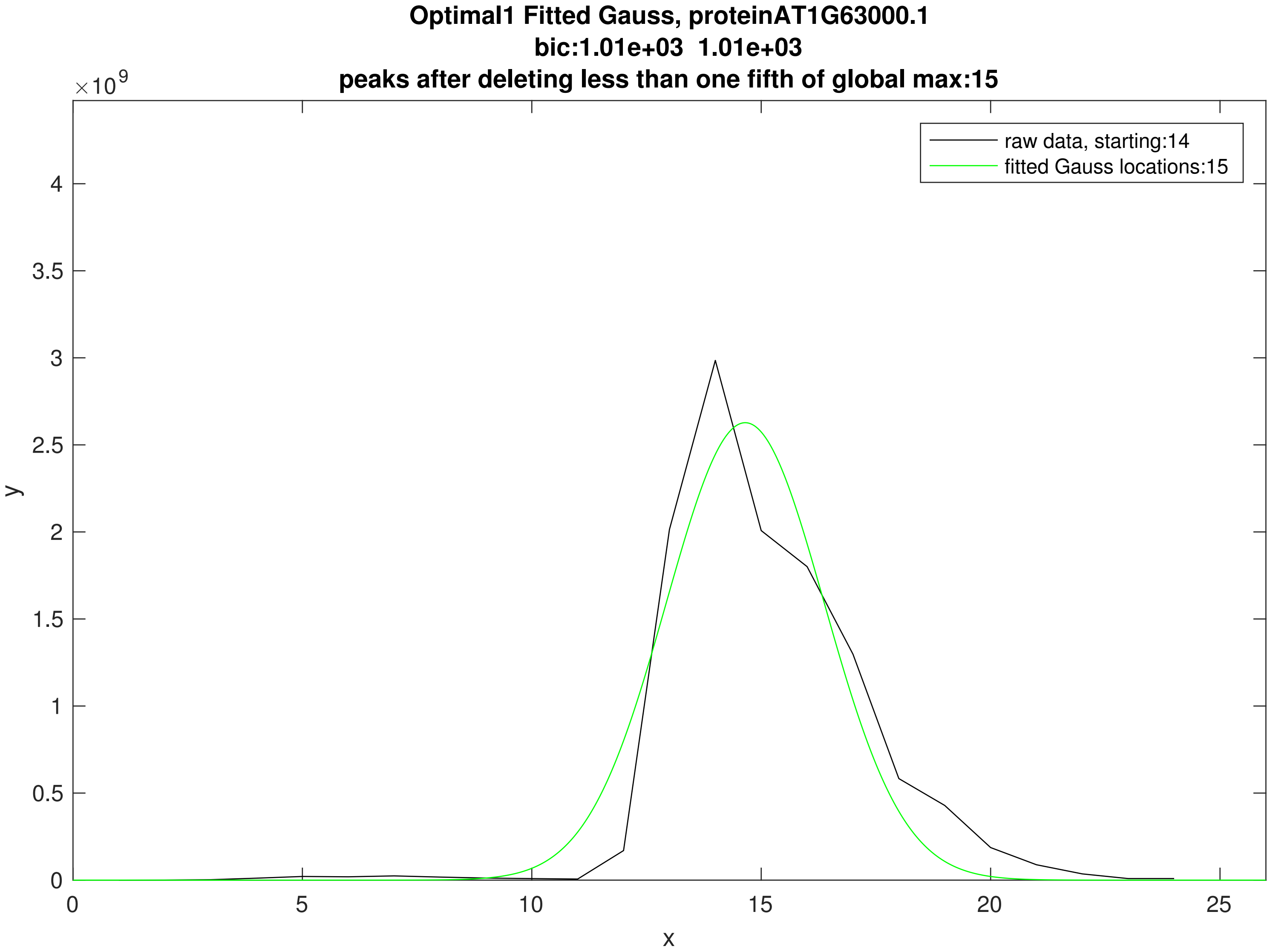

Optimal1 Fitted Gauss, proteinAT1G63000.1
bic:1.01e+03 1.01e+03
peaks after deleting less than one fifth of global max:15
×10^9
raw data, starting:14
fitted Gauss locations:15
y
x
0
0.5
1
1.5
2
2.5
3
3.5
4
5
10
15
20
25

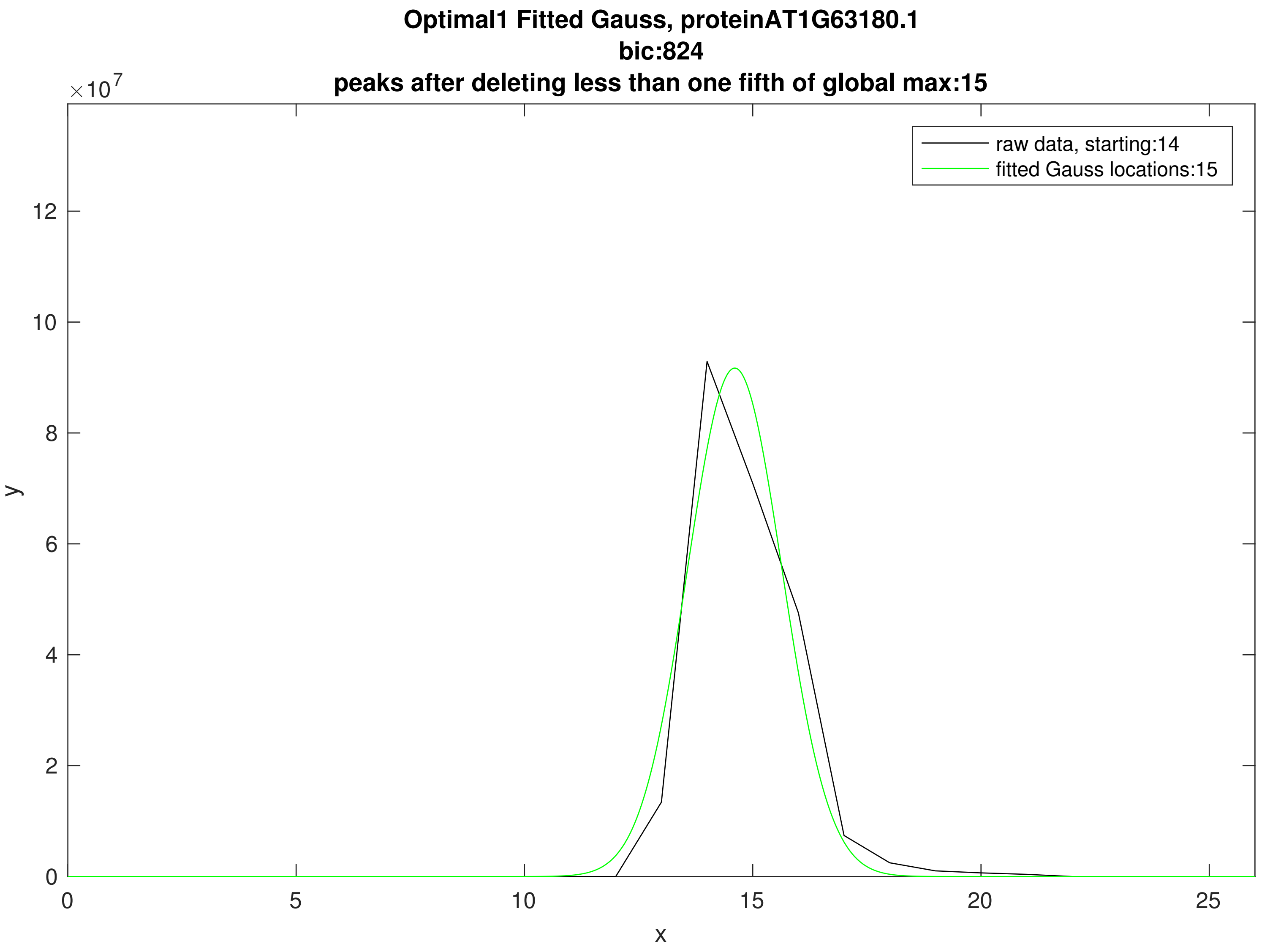

Optimal1 Fitted Gauss, proteinAT1G63180.1
bic:824
peaks after deleting less than one fifth of global max:15
×10^7
raw data, starting:14
fitted Gauss locations:15
y
x
0
2
4
6
8
10
12
0
5
10
15
20
25

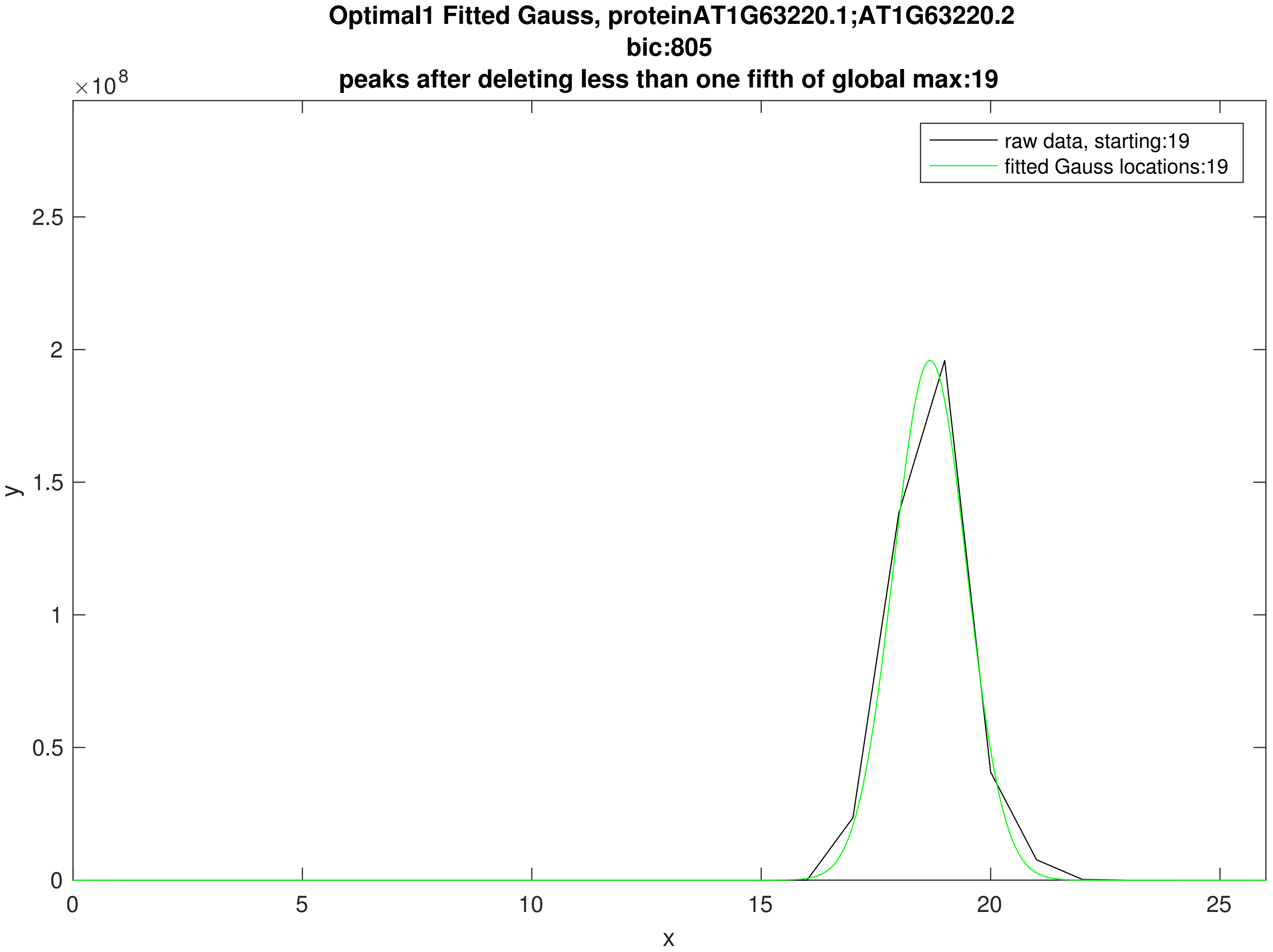
Optimal1 Fitted Gauss, proteinAT1G63220.1;AT1G63220.2
bic:805
peaks after deleting less than one fifth of global max:19
×10^8
raw data, starting:19
fitted Gauss locations:19
y
x
0
0.5
1
1.5
2
2.5
0
5
10
15
20
25

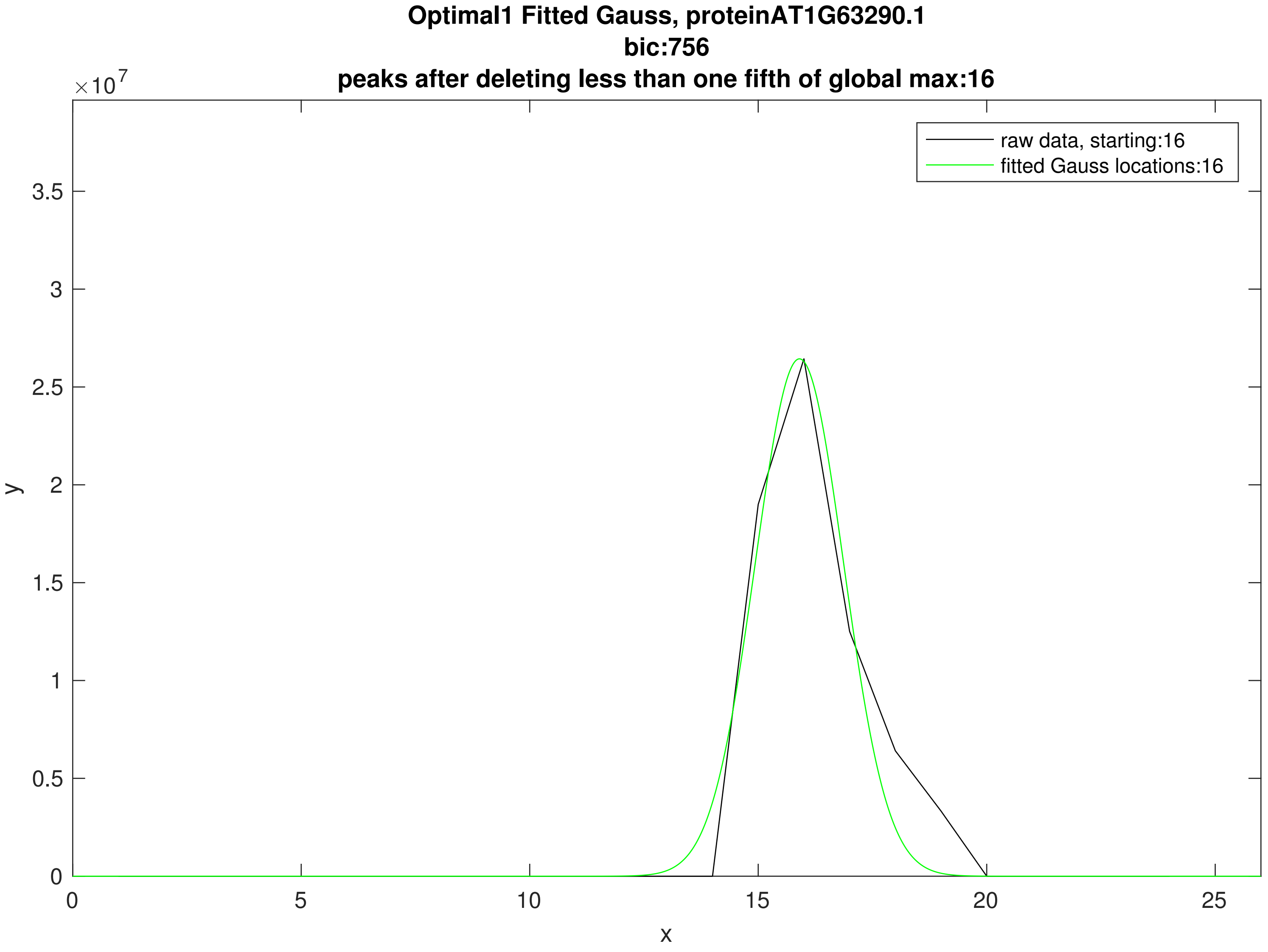

Optimal1 Fitted Gauss, proteinAT1G63290.1
bic:756
peaks after deleting less than one fifth of global max:16
×10^7
raw data, starting:16
fitted Gauss locations:16
y
x
0
0.5
1
1.5
2
2.5
3
3.5
0
5
10
15
20
25

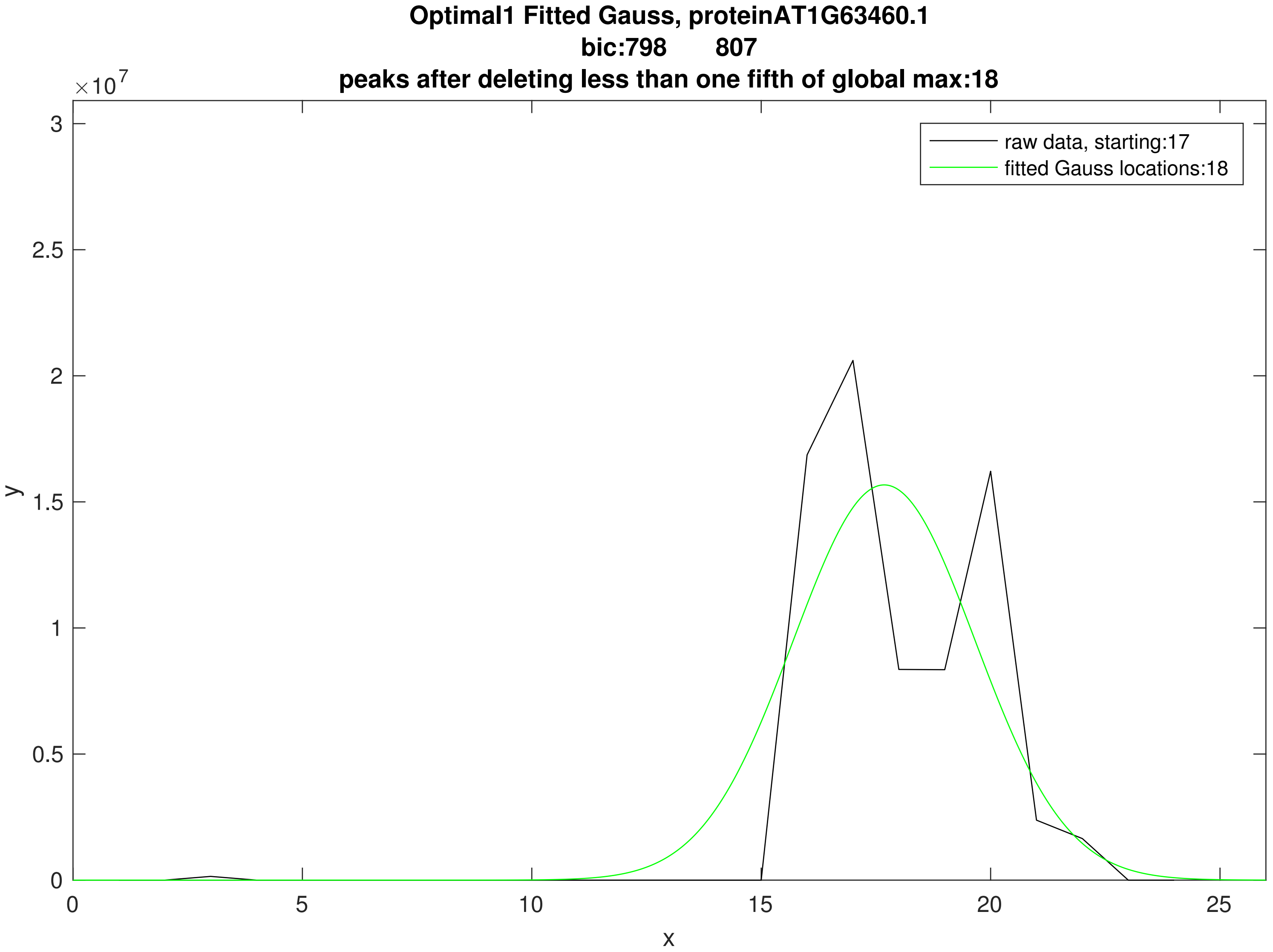

Optimal1 Fitted Gauss, proteinAT1G63460.1
bic:798 807
peaks after deleting less than one fifth of global max:18
×10^7
raw data, starting:17
fitted Gauss locations:18
y
x
0
0.5
1
1.5
2
2.5
3
0
5
10
15
20
25

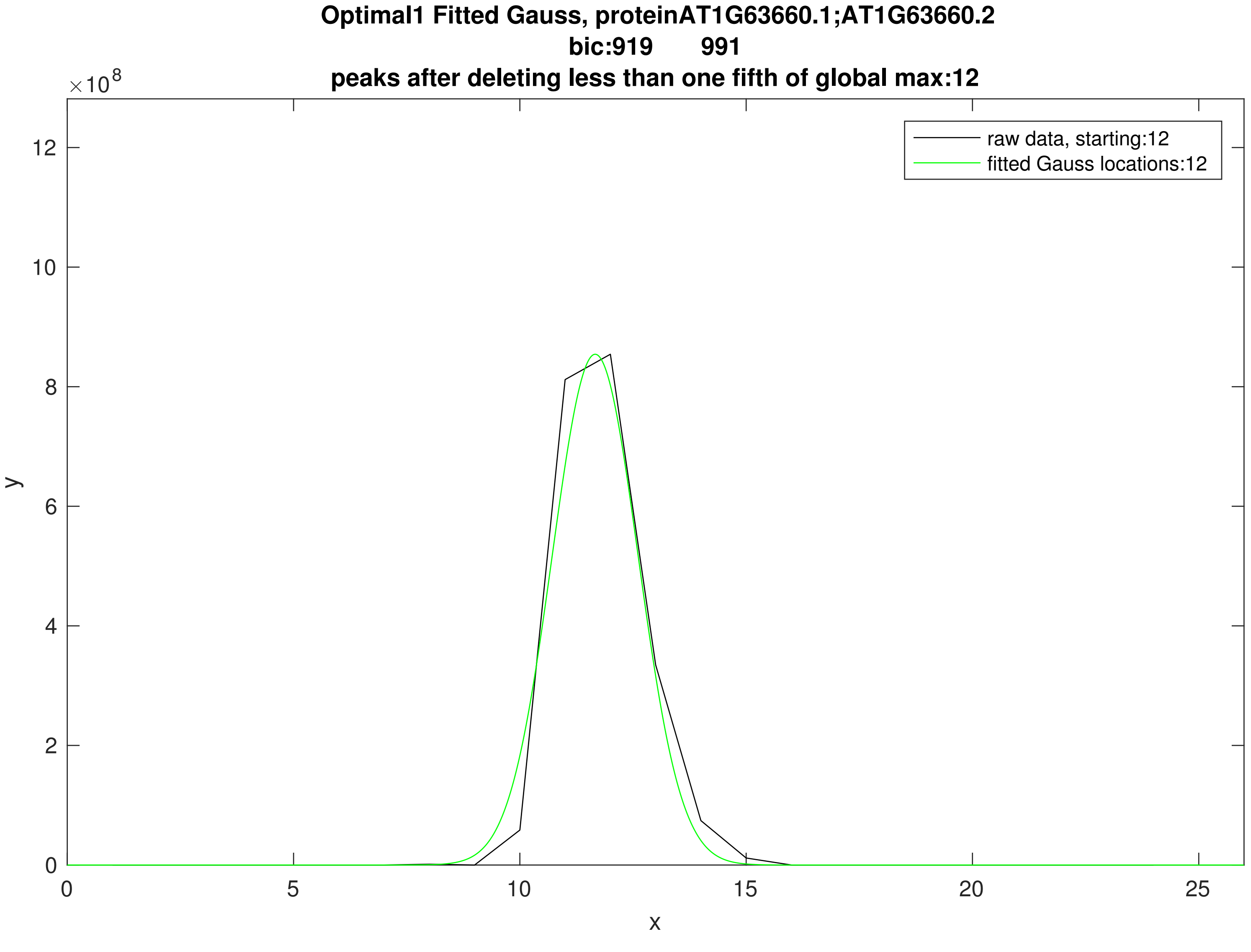
Optimal1 Fitted Gauss, proteinAT1G63660.1;AT1G63660.2
bic:919 991
peaks after deleting less than one fifth of global max:12
×10^8
raw data, starting:12
fitted Gauss locations:12
y
x
0
2
4
6
8
10
12
0
5
10
15
20
25

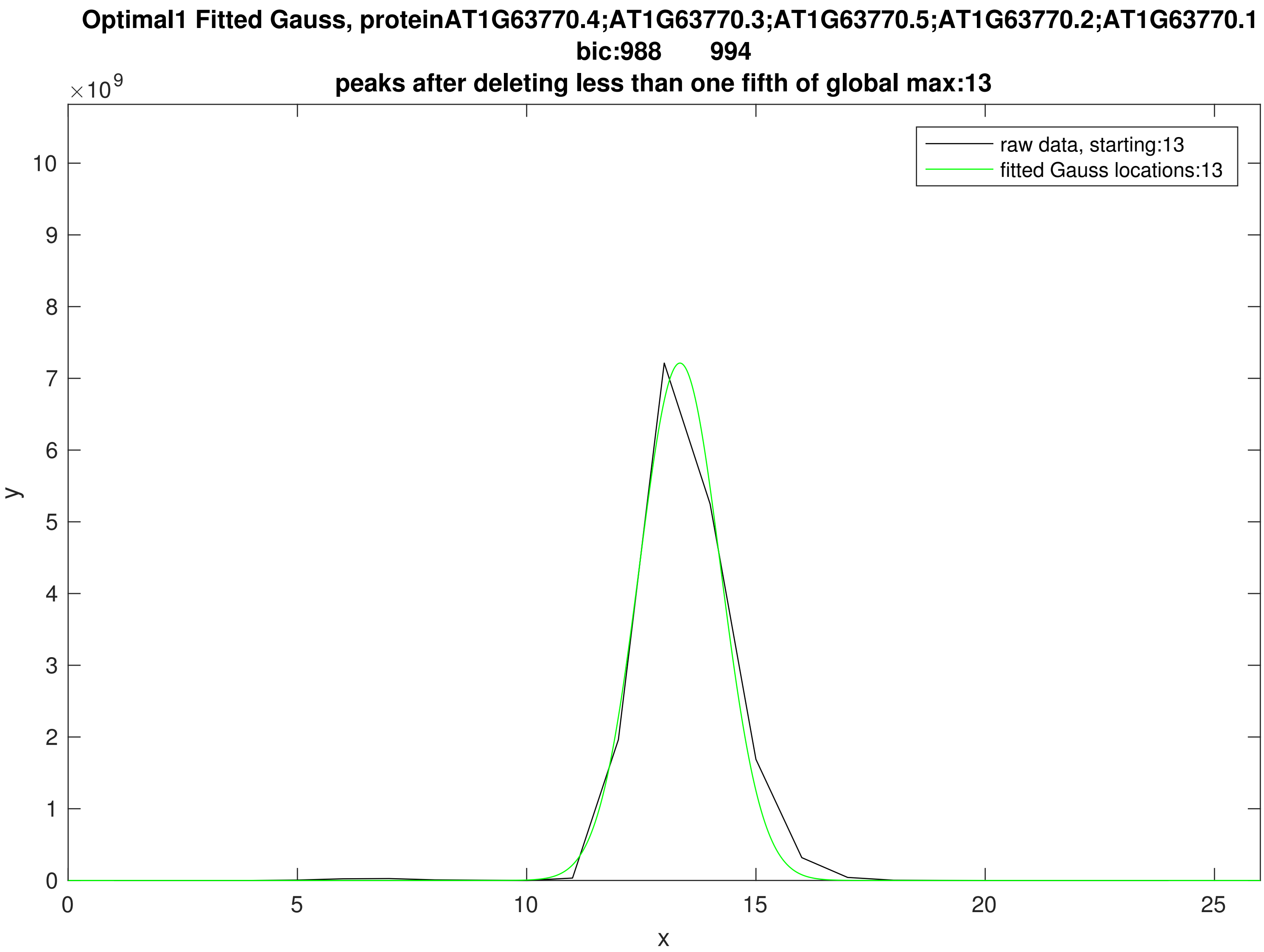

Optimal1 Fitted Gauss, proteinAT1G63770.4;AT1G63770.3;AT1G63770.5;AT1G63770.2;AT1G63770.1
bic:988 994
peaks after deleting less than one fifth of global max:13
×10^9
raw data, starting:13
fitted Gauss locations:13
y
x
0
5
10
15
20
25
1
2
3
4
5
6
7
8
9
10

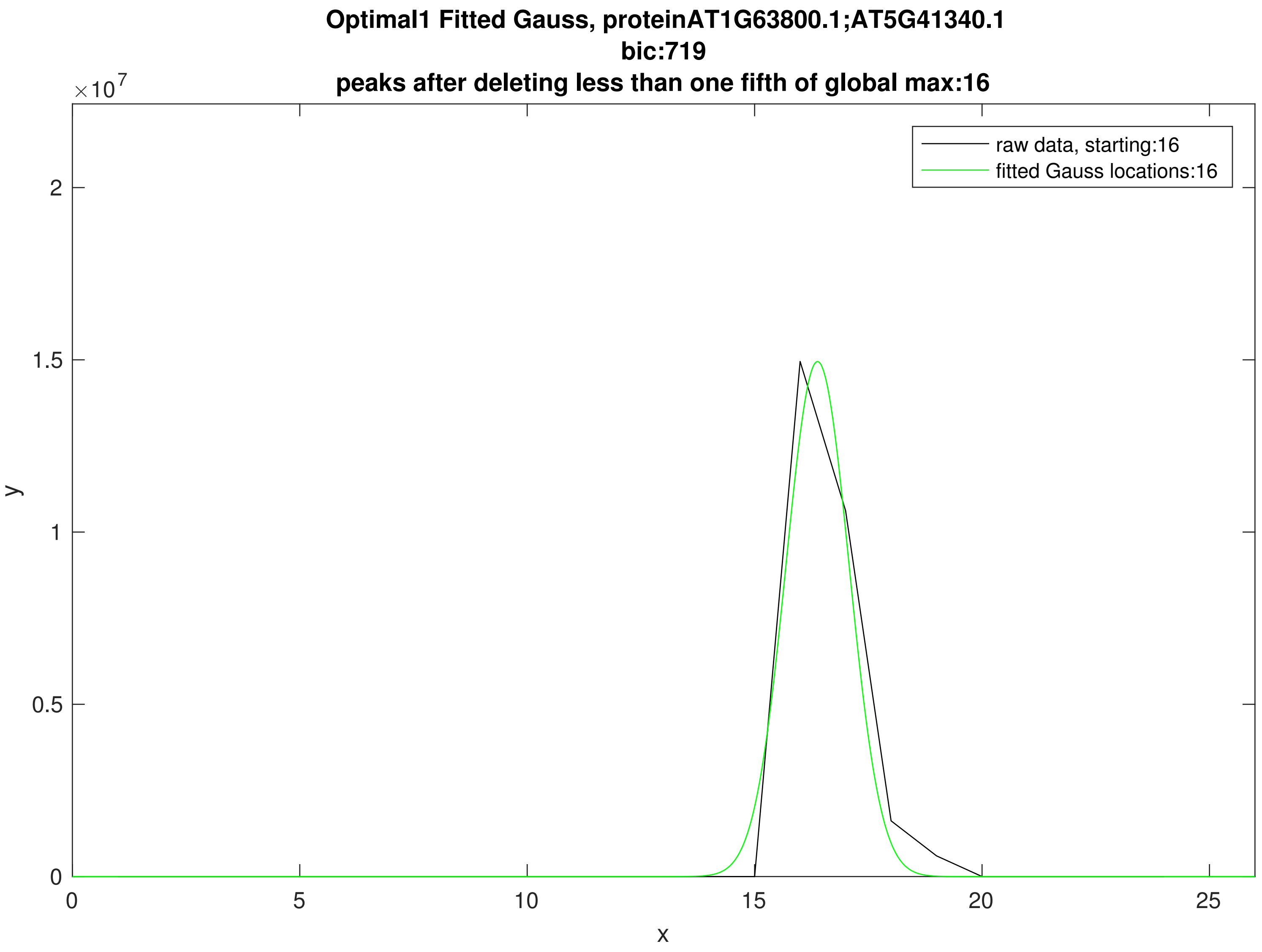
Optimal1 Fitted Gauss, proteinAT1G63800.1;AT5G41340.1
bic:719
peaks after deleting less than one fifth of global max:16
×10^7
raw data, starting:16
fitted Gauss locations:16
y
x
0
0.5
1
1.5
2
5
10
15
20
25

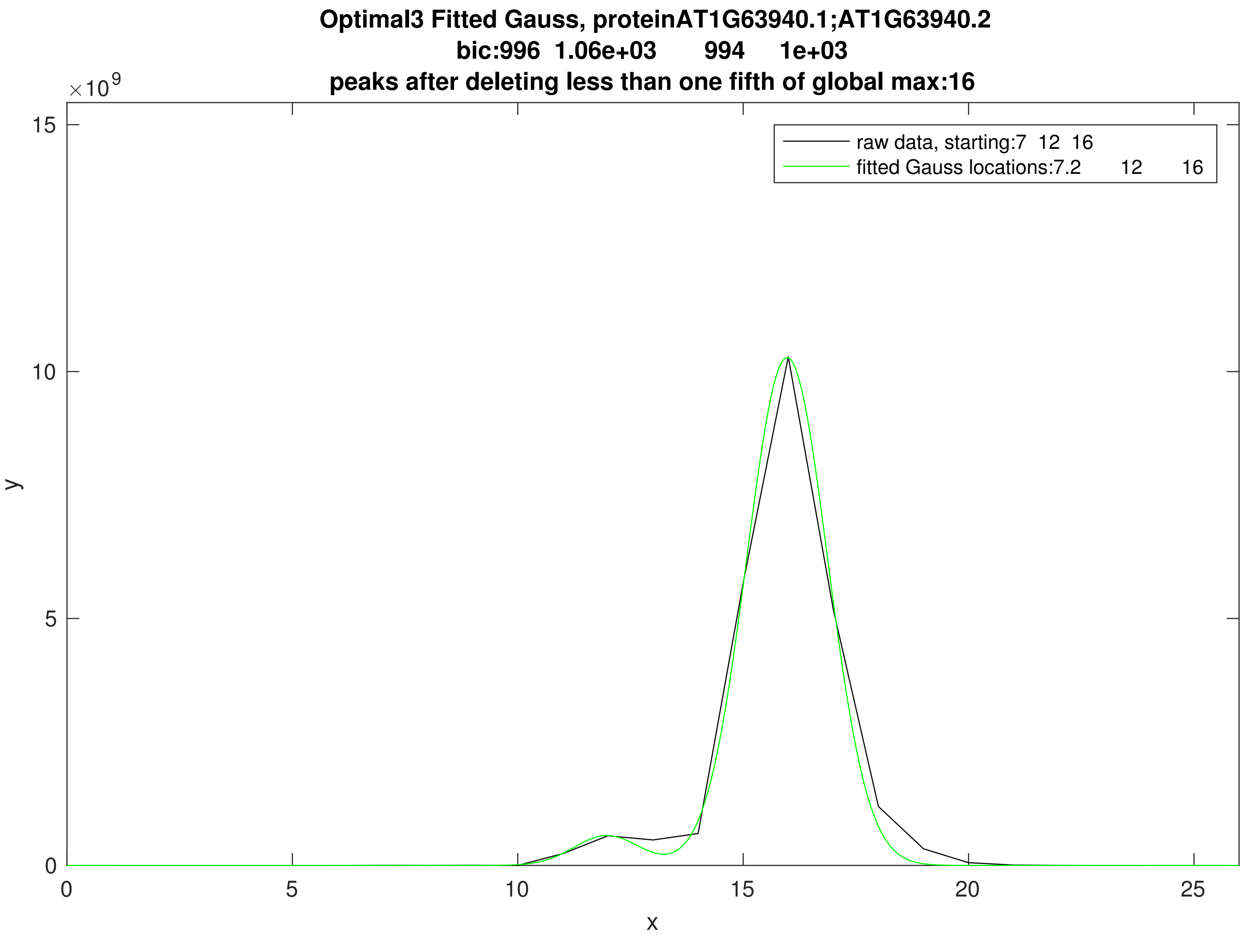

Optimal3 Fitted Gauss, proteinAT1G63940.1;AT1G63940.2
bic:996 1.06e+03 994 1e+03
peaks after deleting less than one fifth of global max:16
×10^9
15
10
5
0
y
0
5
10
15
20
25
x
raw data, starting:7 12 16
fitted Gauss locations:7.2 12 16

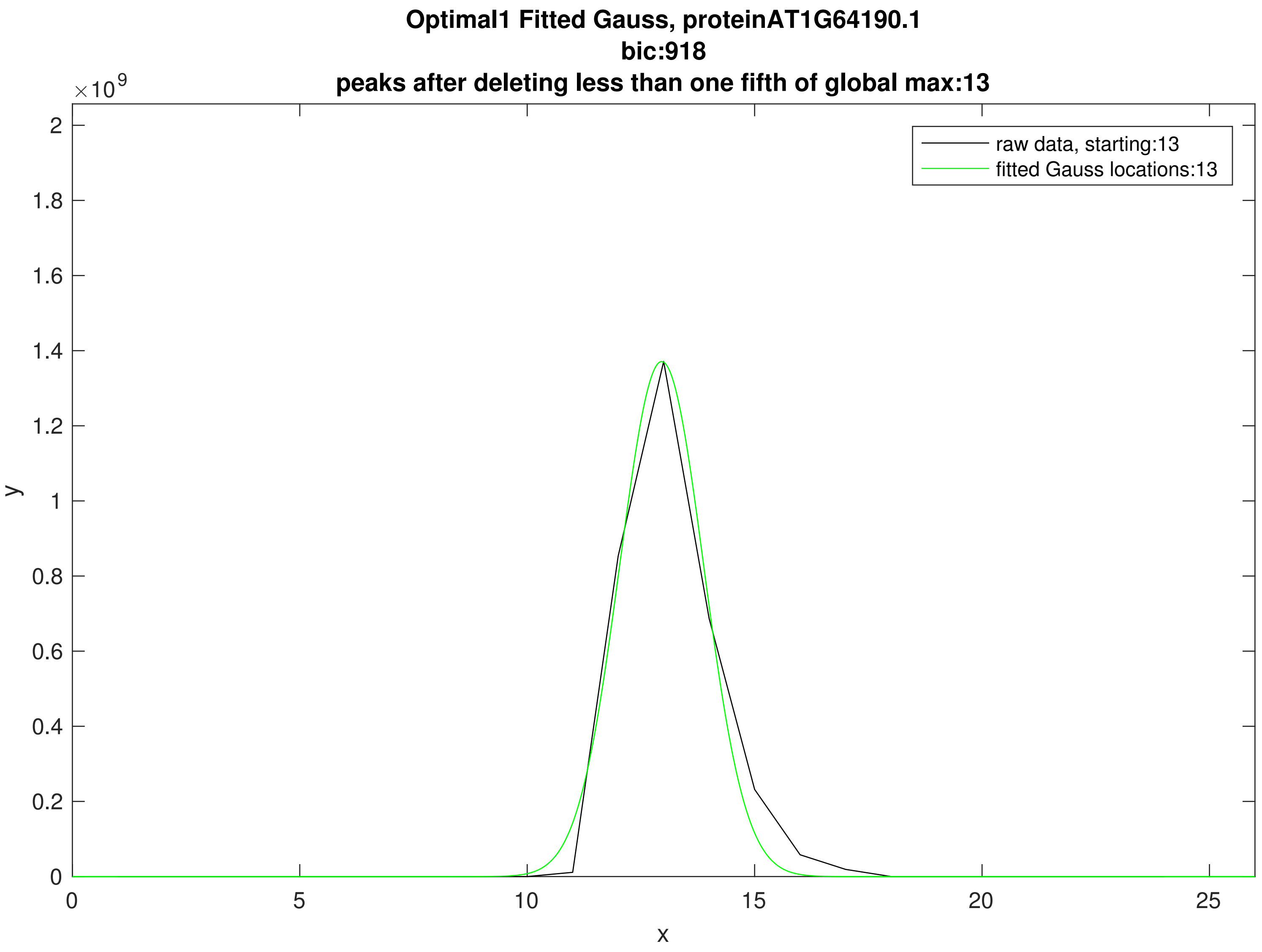
Optimal1 Fitted Gauss, proteinAT1G64190.1
bic:918
peaks after deleting less than one fifth of global max:13
×10^9
raw data, starting:13
fitted Gauss locations:13
y
x
0
0.2
0.4
0.6
0.8
1
1.2
1.4
1.6
1.8
2
0
5
10
15
20
25

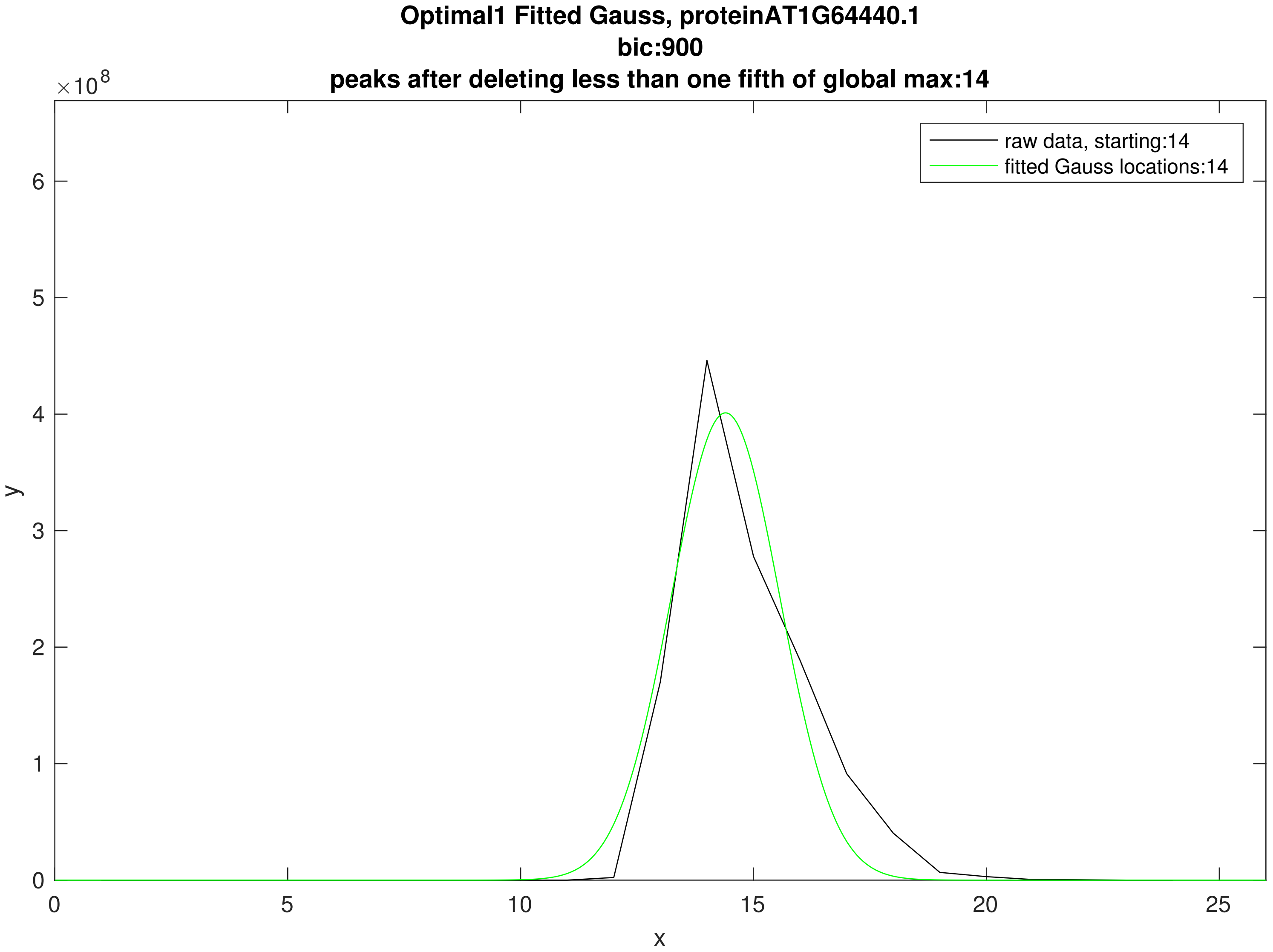
Optimal1 Fitted Gauss, proteinAT1G64440.1
bic:900
peaks after deleting less than one fifth of global max:14
×10^8
raw data, starting:14
fitted Gauss locations:14
y
x
0
1
2
3
4
5
6
0
5
10
15
20
25

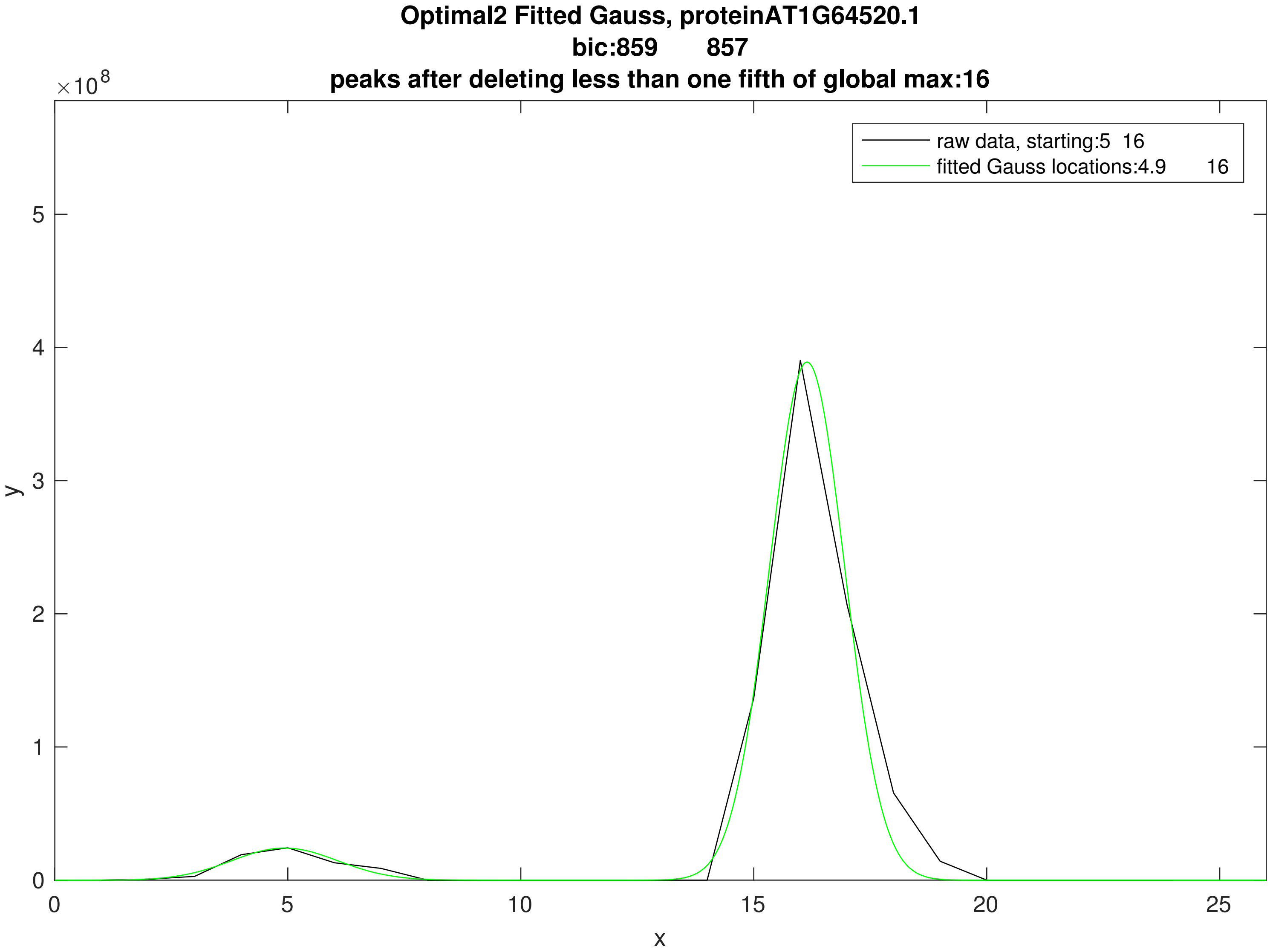
Optimal2 Fitted Gauss, proteinAT1G64520.1
bic:859 857
peaks after deleting less than one fifth of global max:16
×10^8
raw data, starting:5 16
fitted Gauss locations:4.9 16
y
x
0
1
2
3
4
5
0
5
10
15
20
25

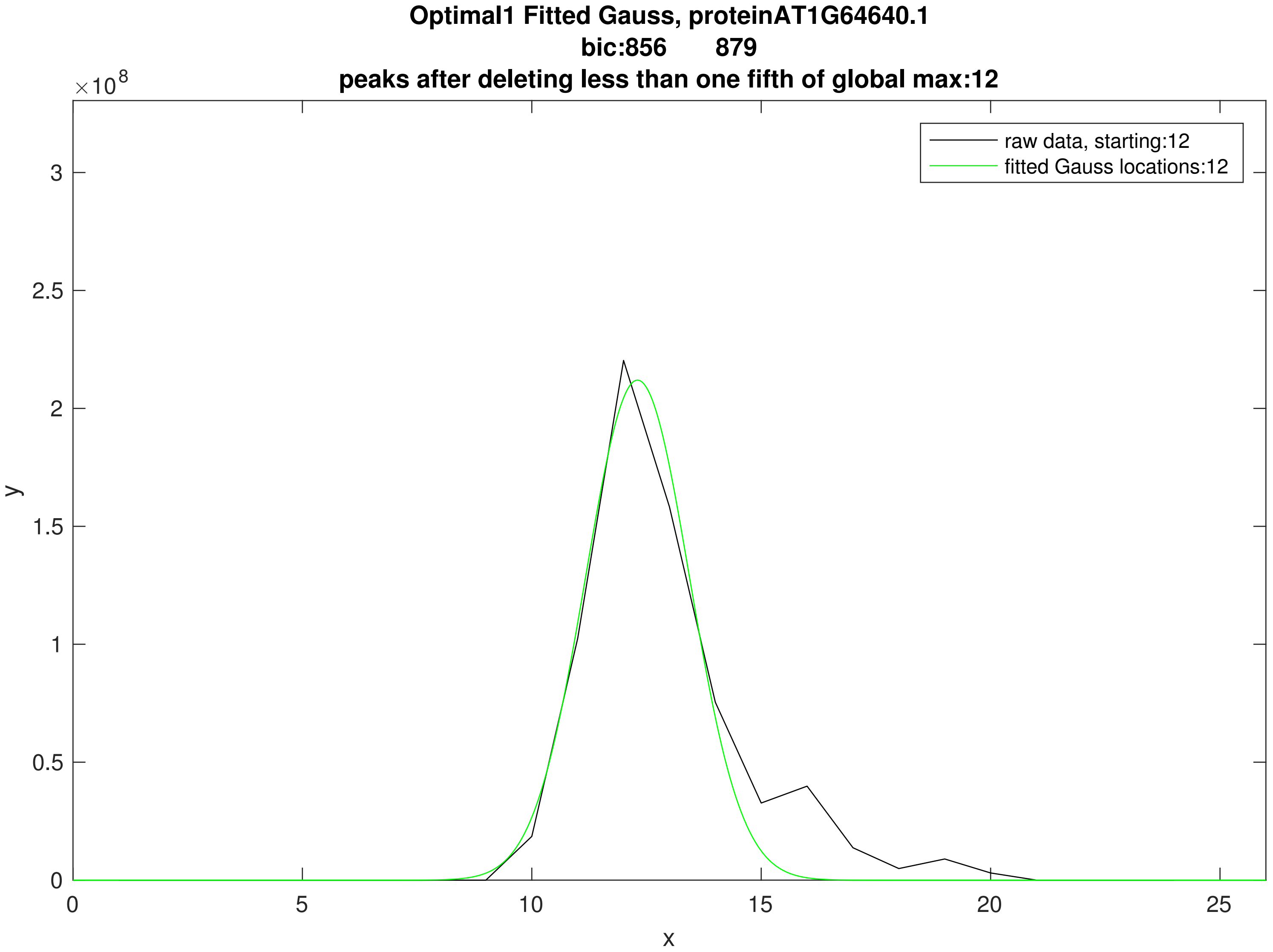

Optimal1 Fitted Gauss, proteinAT1G64640.1
bic:856 879
peaks after deleting less than one fifth of global max:12
×10^8
raw data, starting:12
fitted Gauss locations:12
y
x
0
0.5
1
1.5
2
2.5
3
0
5
10
15
20
25

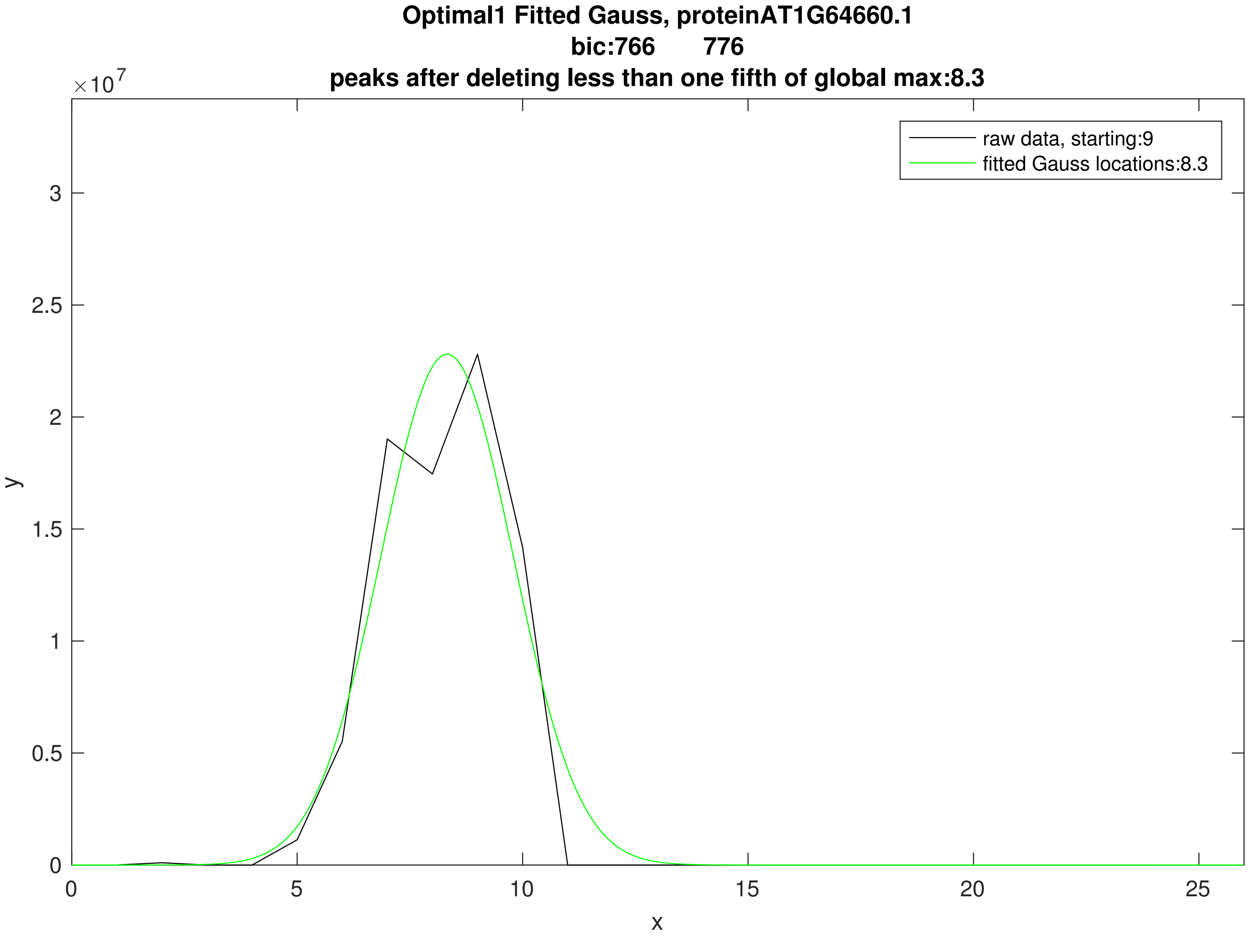

Optimal1 Fitted Gauss, proteinAT1G64660.1
bic:766 776
peaks after deleting less than one fifth of global max:8.3
×10^7
raw data, starting:9
fitted Gauss locations:8.3
y
x
0
0.5
1
1.5
2
2.5
3
0
5
10
15
20
25

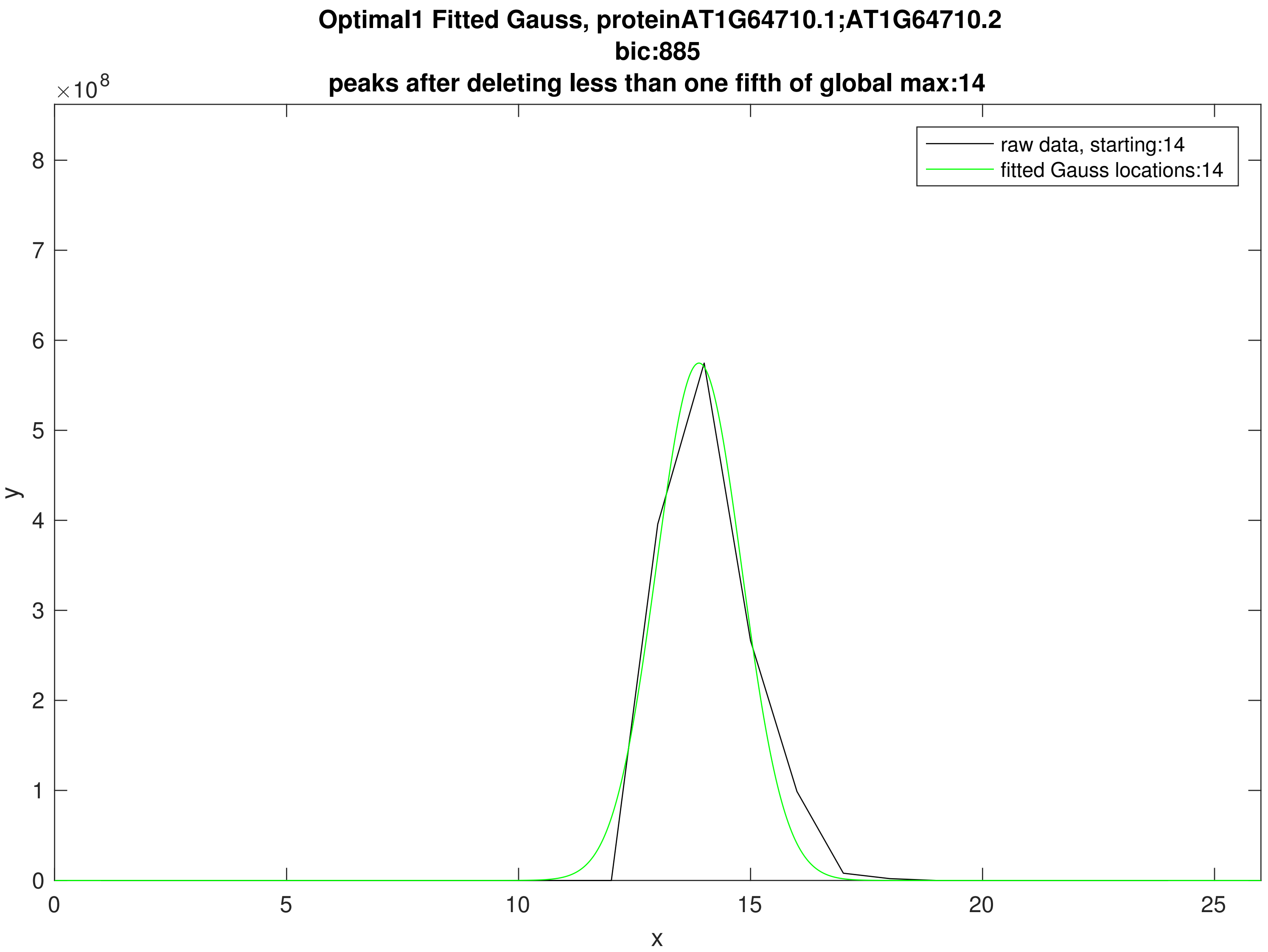
Optimal1 Fitted Gauss, proteinAT1G64710.1;AT1G64710.2
bic:885
peaks after deleting less than one fifth of global max:14
×10^8
raw data, starting:14
fitted Gauss locations:14
y
x
0
1
2
3
4
5
6
7
8
0
5
10
15
20
25

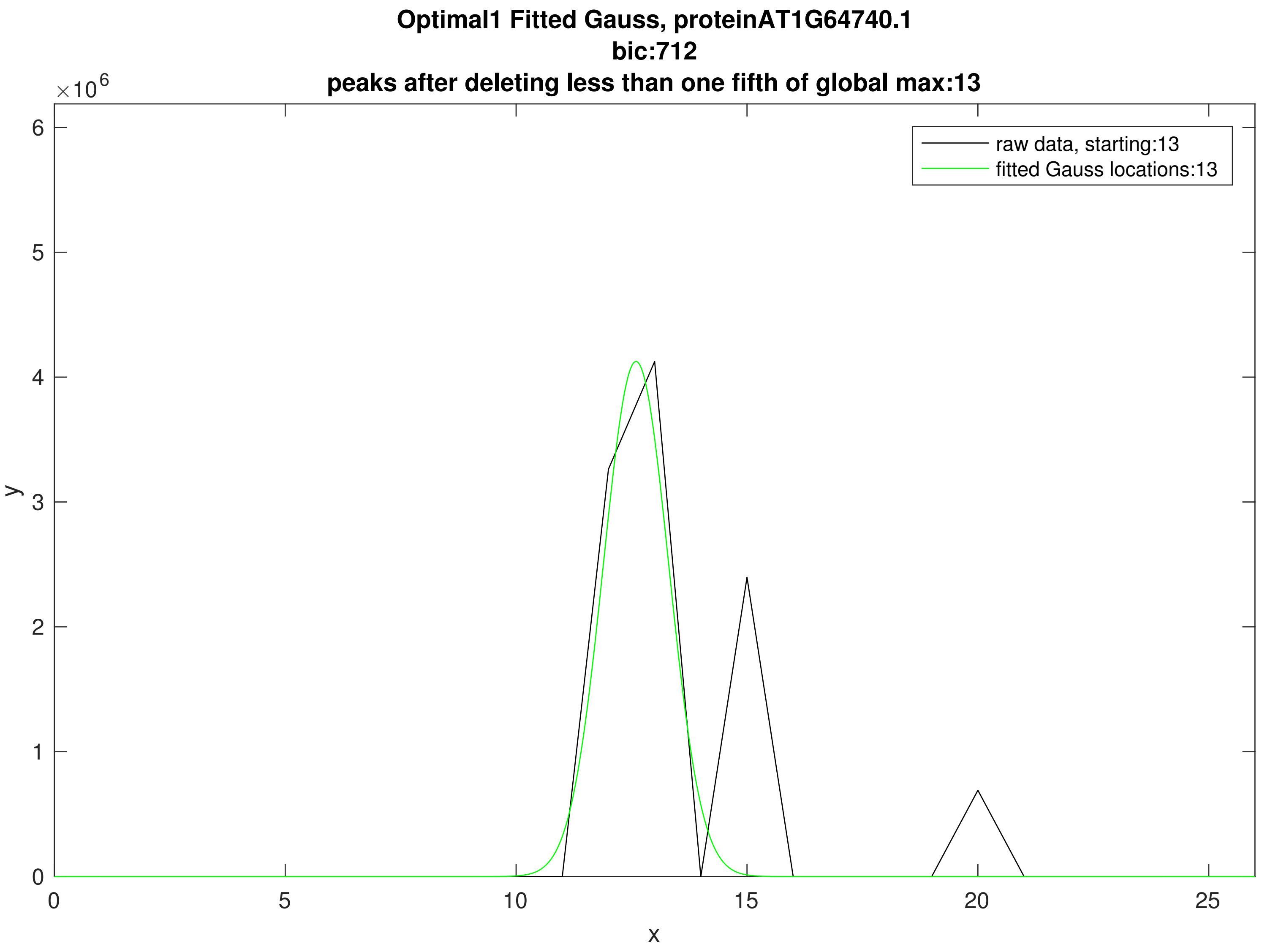

Optimal1 Fitted Gauss, proteinAT1G64740.1
bic:712
peaks after deleting less than one fifth of global max:13
×10^6
raw data, starting:13
fitted Gauss locations:13
y
x
0
1
2
3
4
5
6
0
5
10
15
20
25

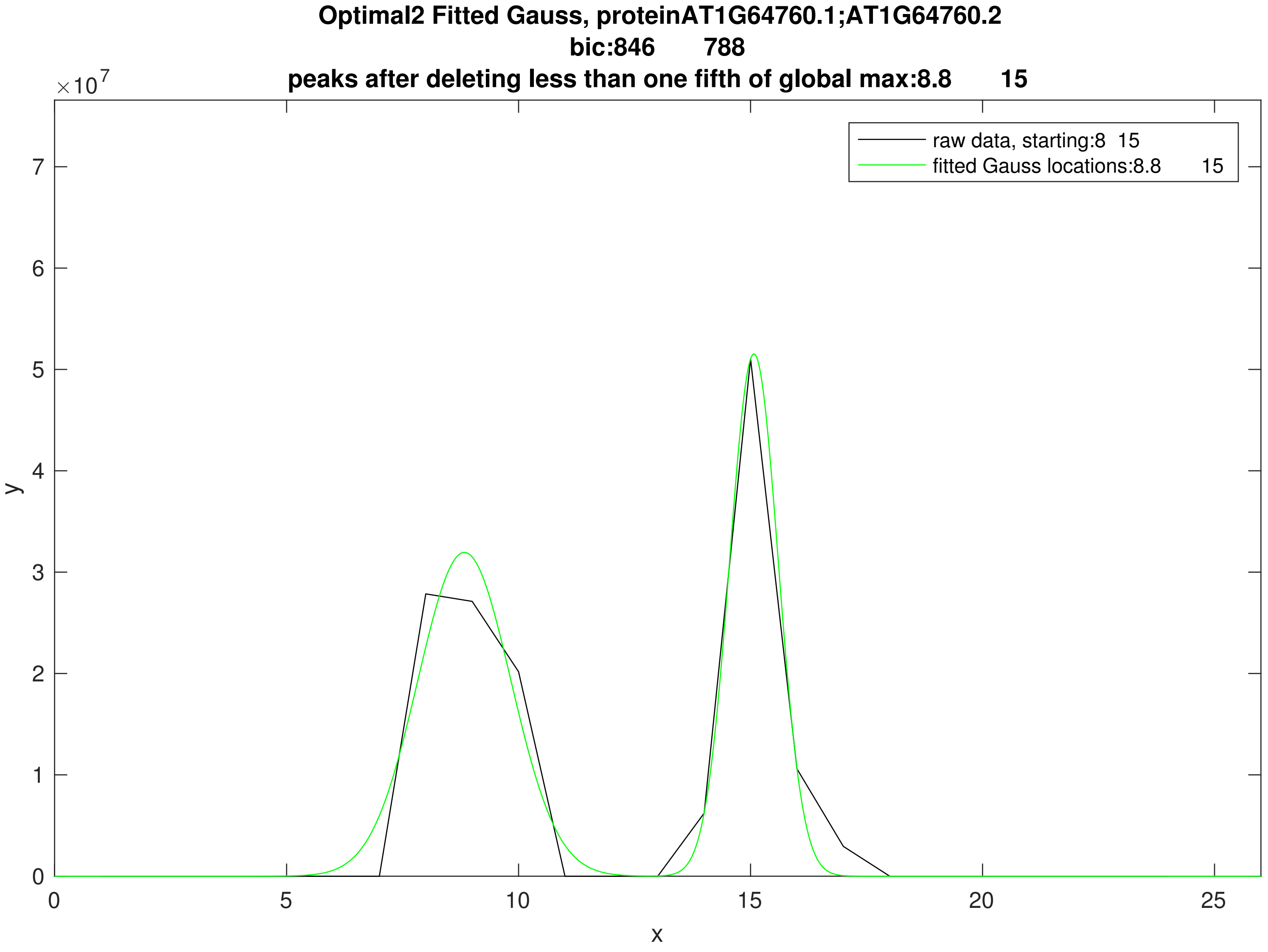
Optimal2 Fitted Gauss, proteinAT1G64760.1;AT1G64760.2
bic:846 788
peaks after deleting less than one fifth of global max:8.8 15
×10^7
raw data, starting:8 15
fitted Gauss locations:8.8 15
y
x
0
1
2
3
4
5
6
7
0
5
10
15
20
25

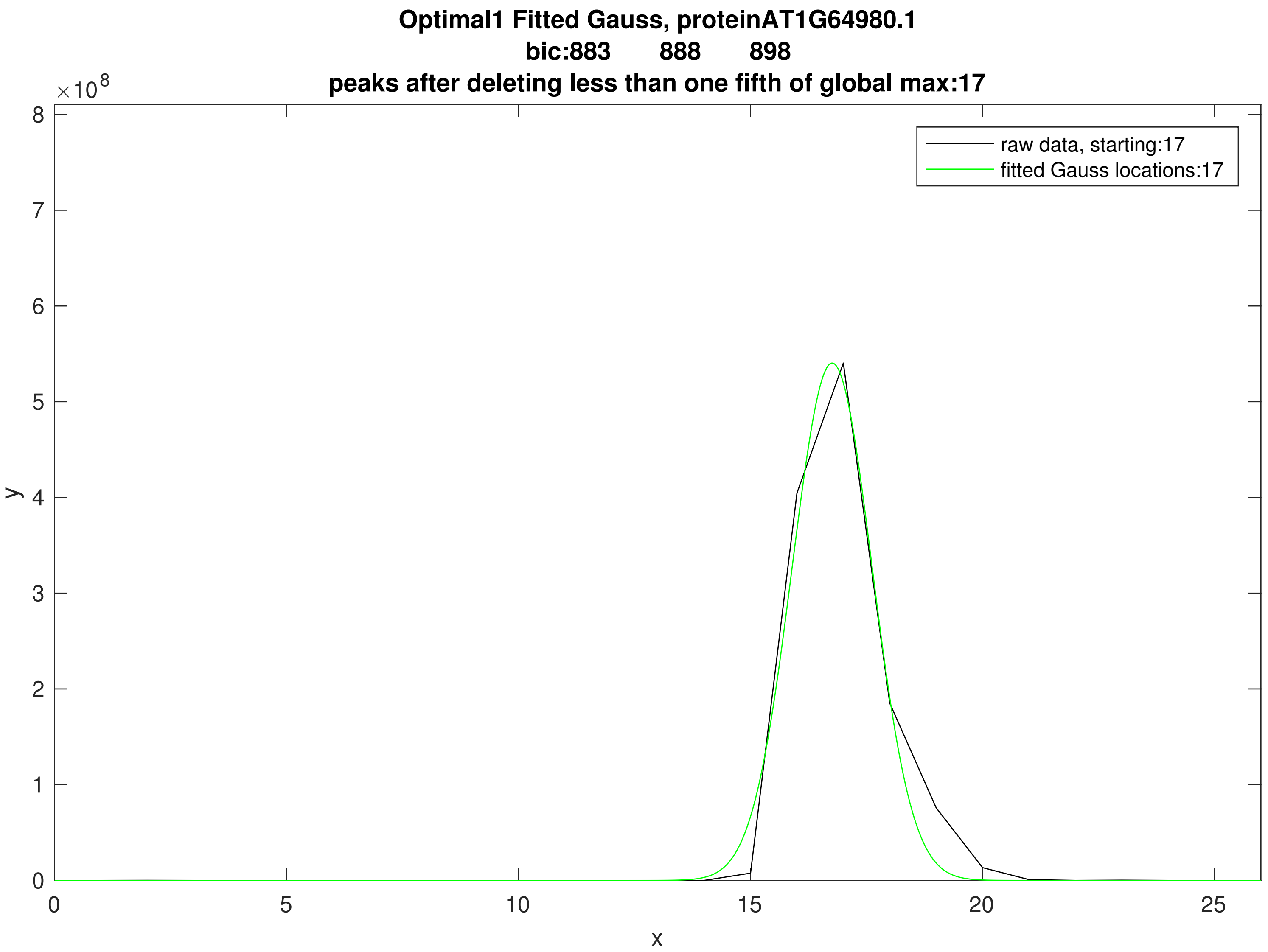
Optimal1 Fitted Gauss, proteinAT1G64980.1
bic:883 888 898
peaks after deleting less than one fifth of global max:17
×10^8
raw data, starting:17
fitted Gauss locations:17
y
x
0
1
2
3
4
5
6
7
8
0
5
10
15
20
25

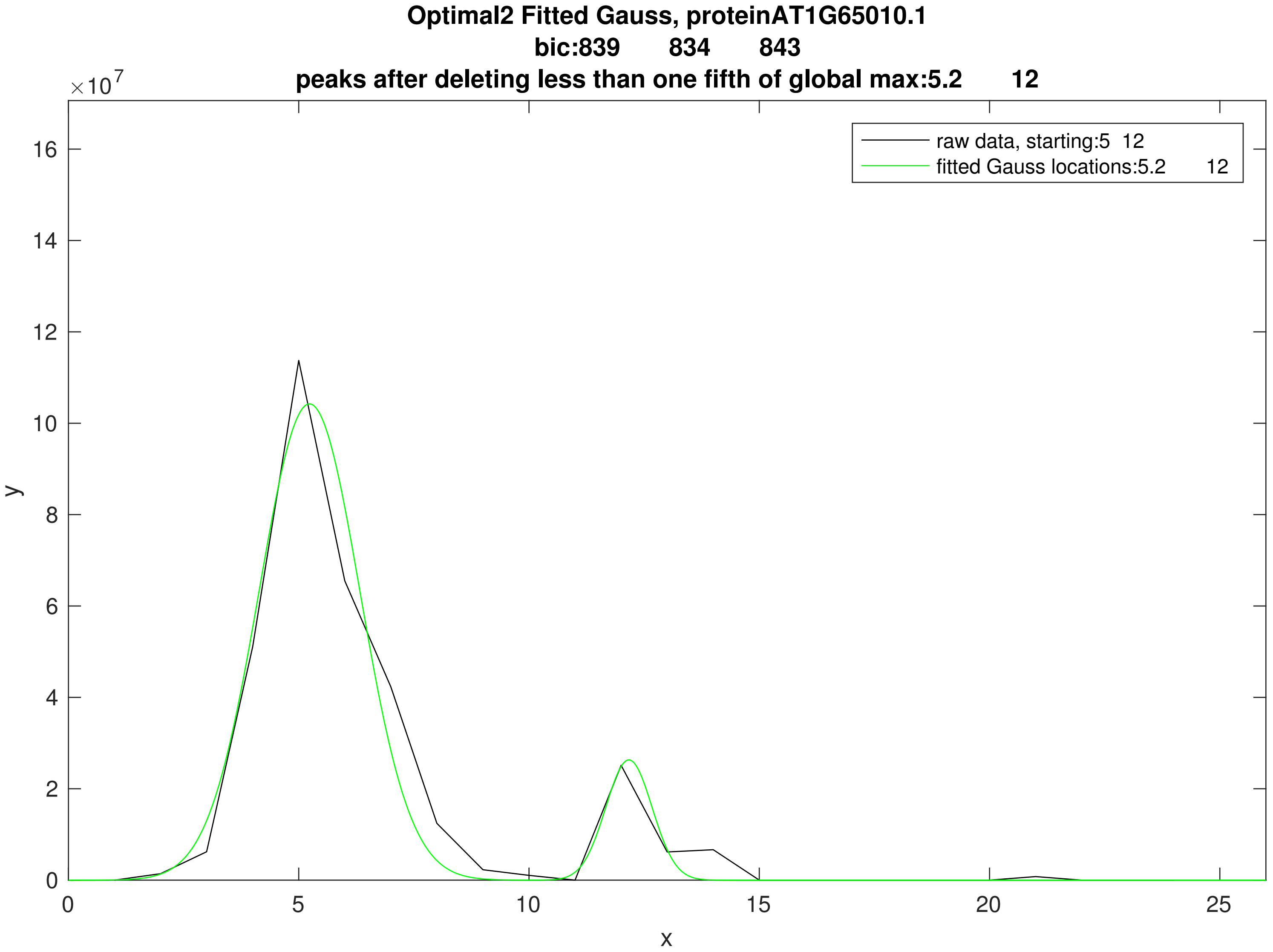
Optimal2 Fitted Gauss, proteinAT1G65010.1
bic:839 834 843
peaks after deleting less than one fifth of global max:5.2 12
×10^7
raw data, starting:5 12
fitted Gauss locations:5.2 12
y
x
0
2
4
6
8
10
12
14
16
0
5
10
15
20
25

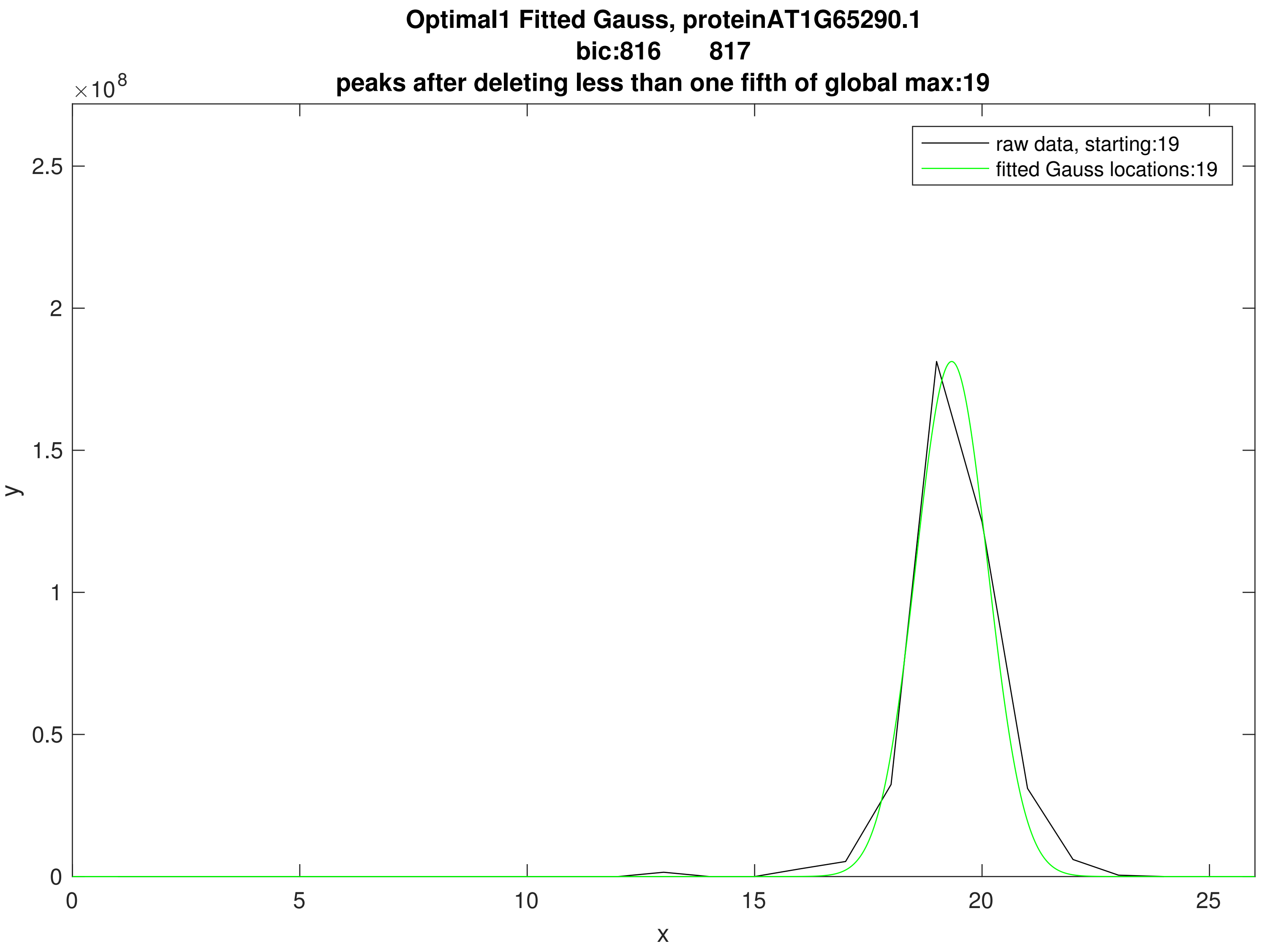

Optimal1 Fitted Gauss, proteinAT1G65290.1
bic:816 817
peaks after deleting less than one fifth of global max:19
×10^8
2.5
2
1.5
1
0.5
0
y
0
5
10
15
20
25
x
raw data, starting:19
fitted Gauss locations:19

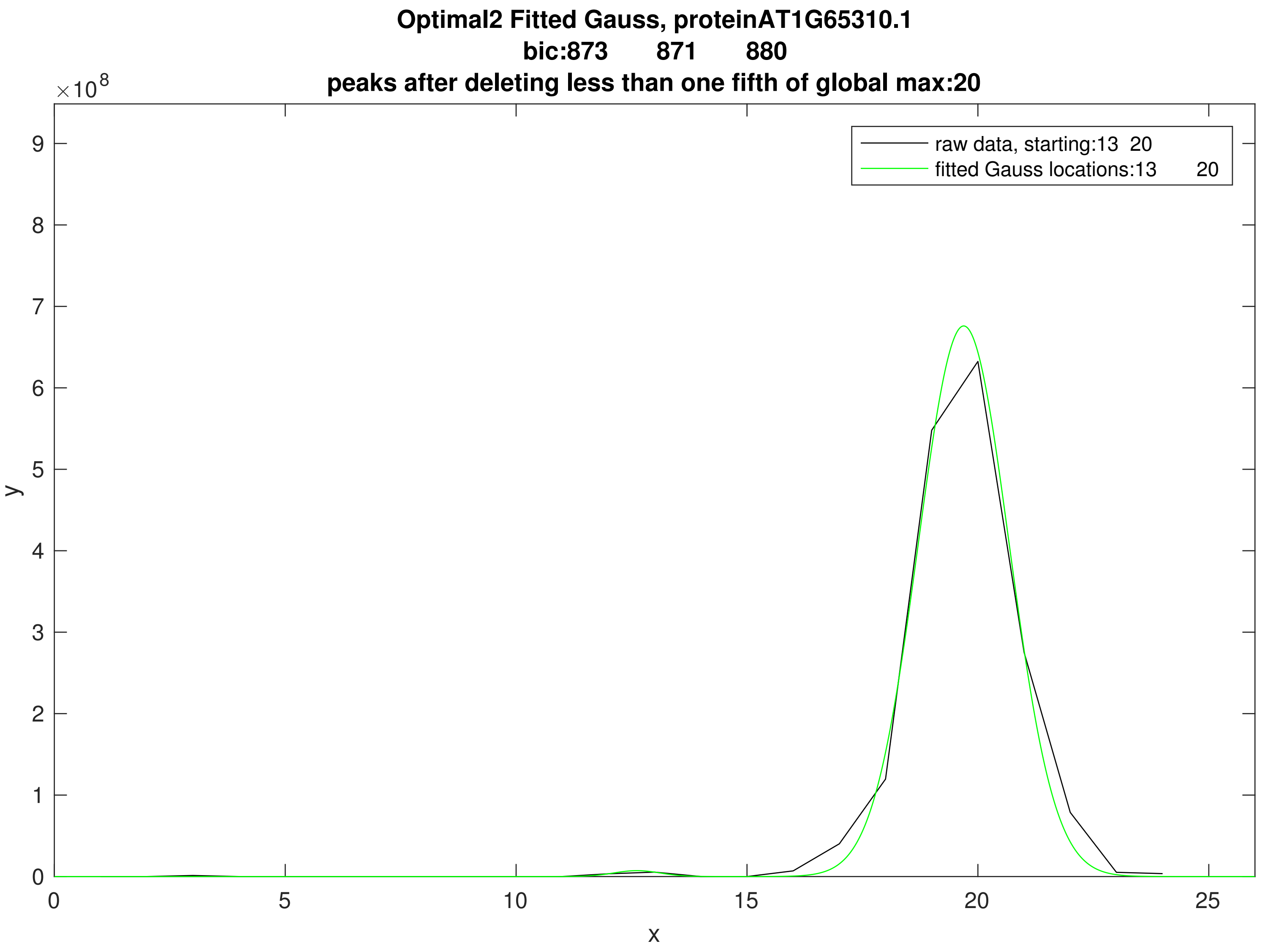
Optimal2 Fitted Gauss, proteinAT1G65310.1
bic:873 871 880
peaks after deleting less than one fifth of global max:20
×10^8
raw data, starting:13 20
fitted Gauss locations:13 20
y
x
0
1
2
3
4
5
6
7
8
9
0
5
10
15
20
25

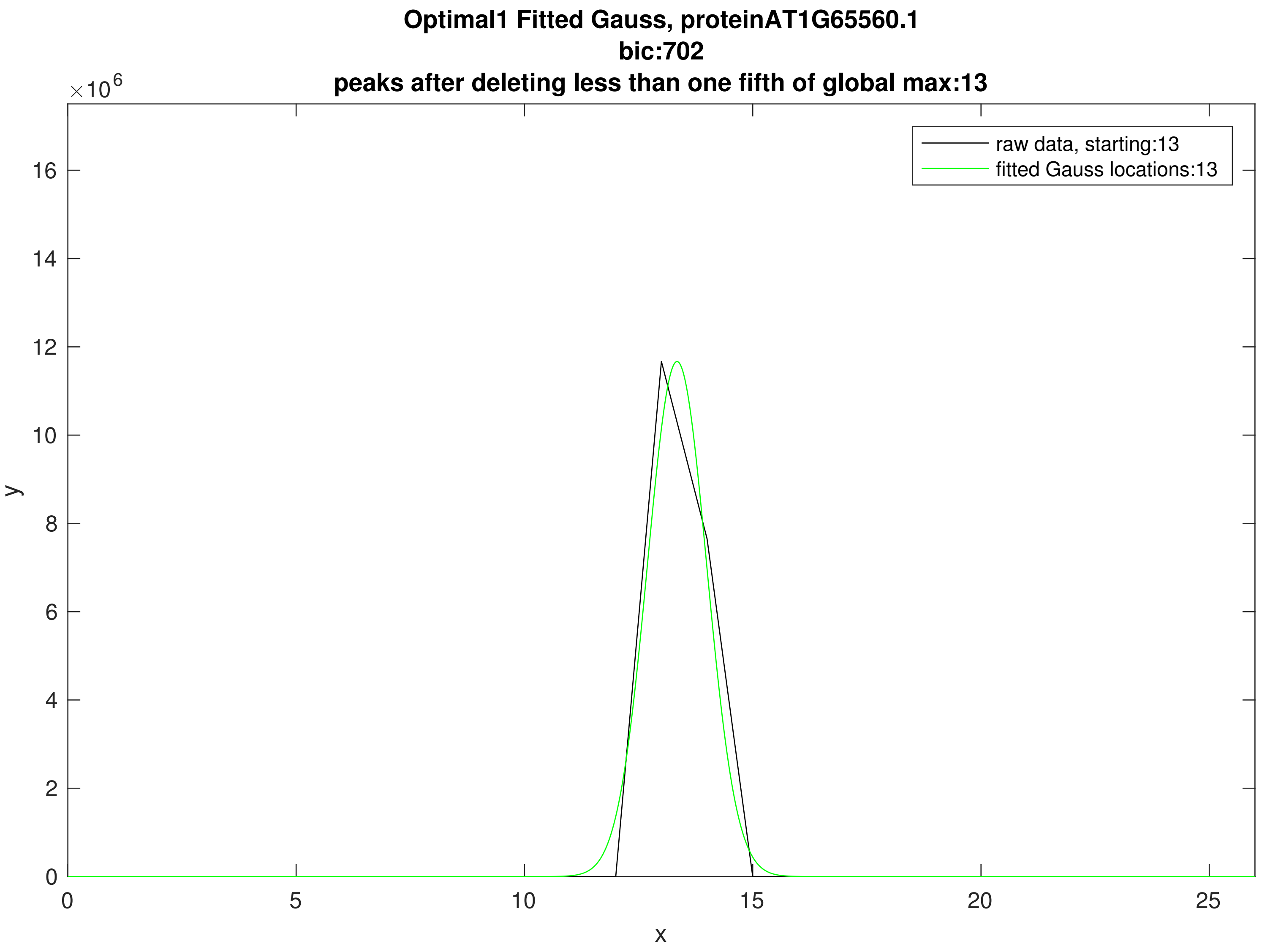
Optimal1 Fitted Gauss, proteinAT1G65560.1
bic:702
peaks after deleting less than one fifth of global max:13
×10^6
16
14
12
10
8
6
4
2
0
y
0
5
10
15
20
25
x
raw data, starting:13
fitted Gauss locations:13

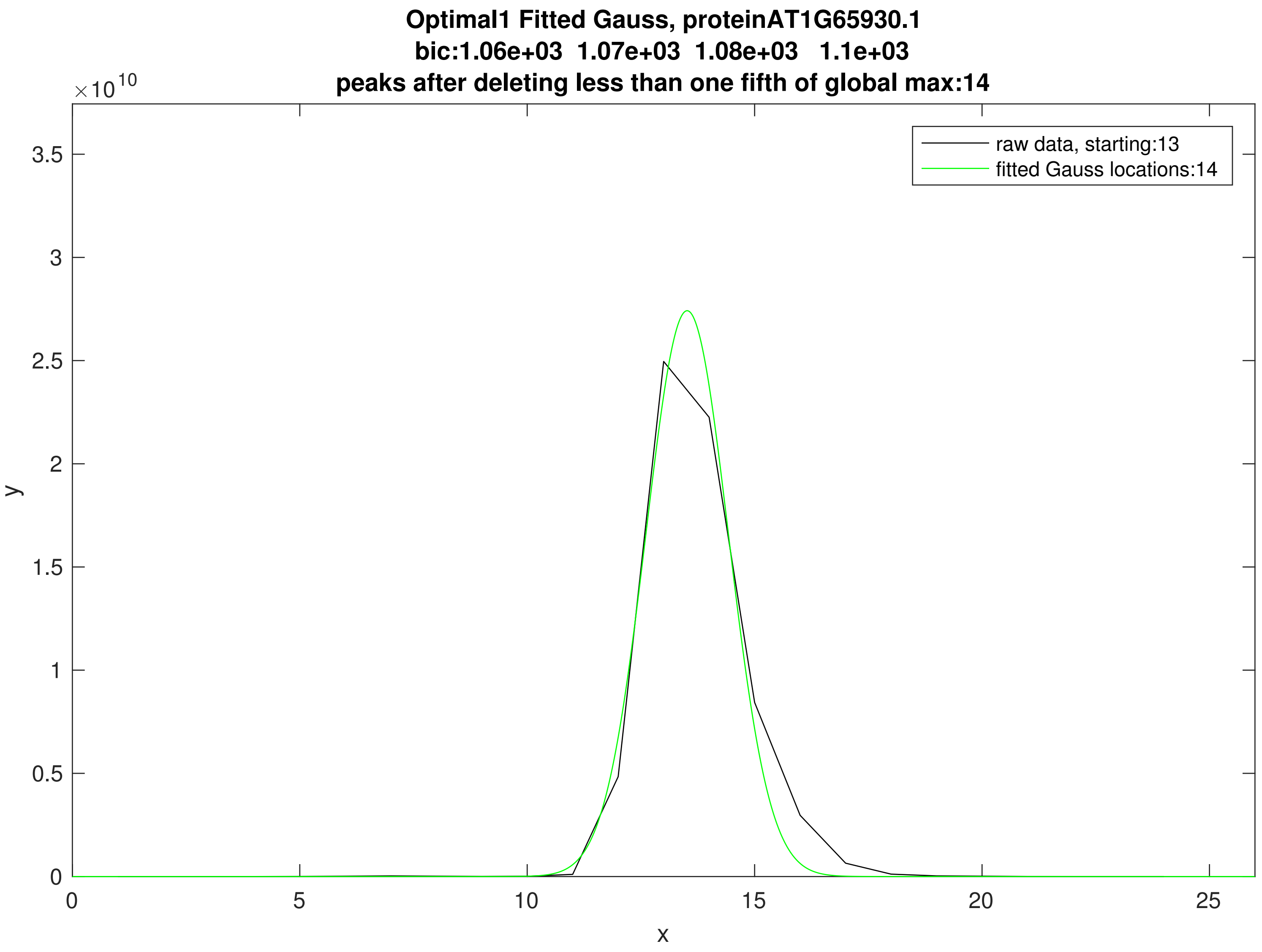
Optimal1 Fitted Gauss, proteinAT1G65930.1
bic:1.06e+03 1.07e+03 1.08e+03 1.1e+03
peaks after deleting less than one fifth of global max:14
×10^10
raw data, starting:13
fitted Gauss locations:14
3.5
3
2.5
2
1.5
1
0.5
0
y
0
5
10
15
20
25
x

Optimal2 Fitted Gauss, proteinAT1G65960.2;AT1G65960.1
bic:1.01e+03 957 967
peaks after deleting less than one fifth of global max:7.6 12

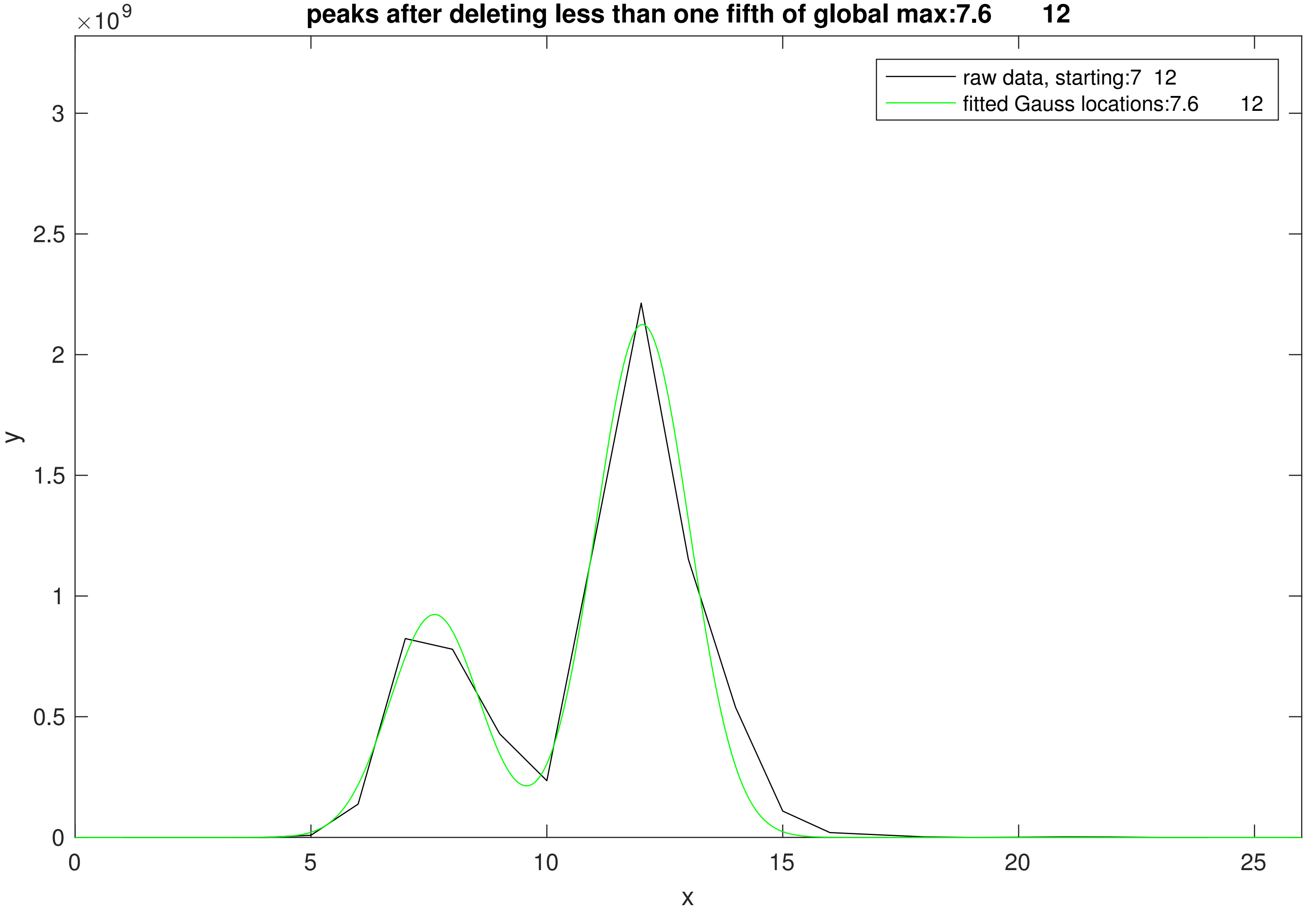

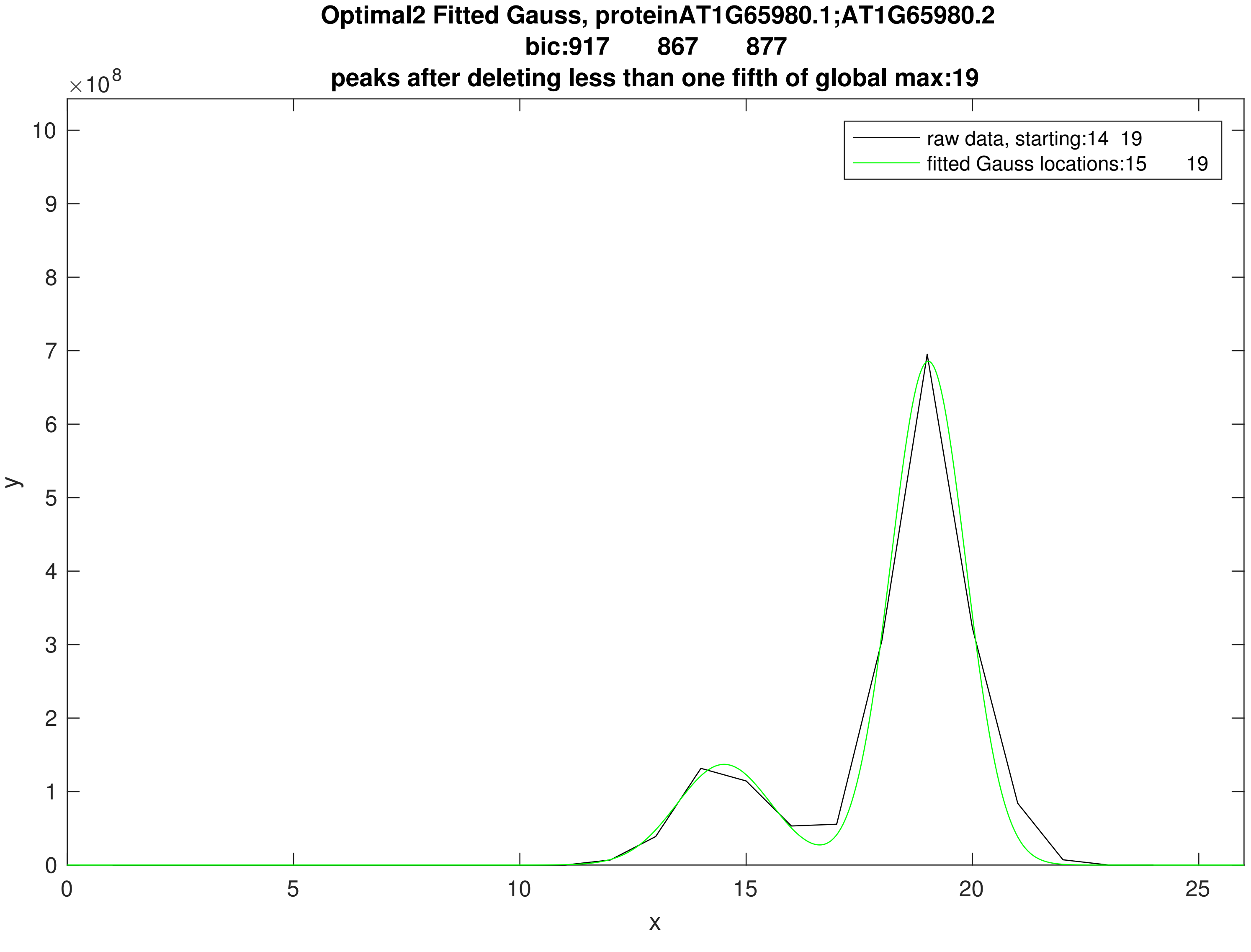
Optimal2 Fitted Gauss, proteinAT1G65980.1;AT1G65980.2
bic:917 867 877
peaks after deleting less than one fifth of global max:19
×10^8
raw data, starting:14 19
fitted Gauss locations:15 19
y
x
0
1
2
3
4
5
6
7
8
9
10
0
5
10
15
20
25

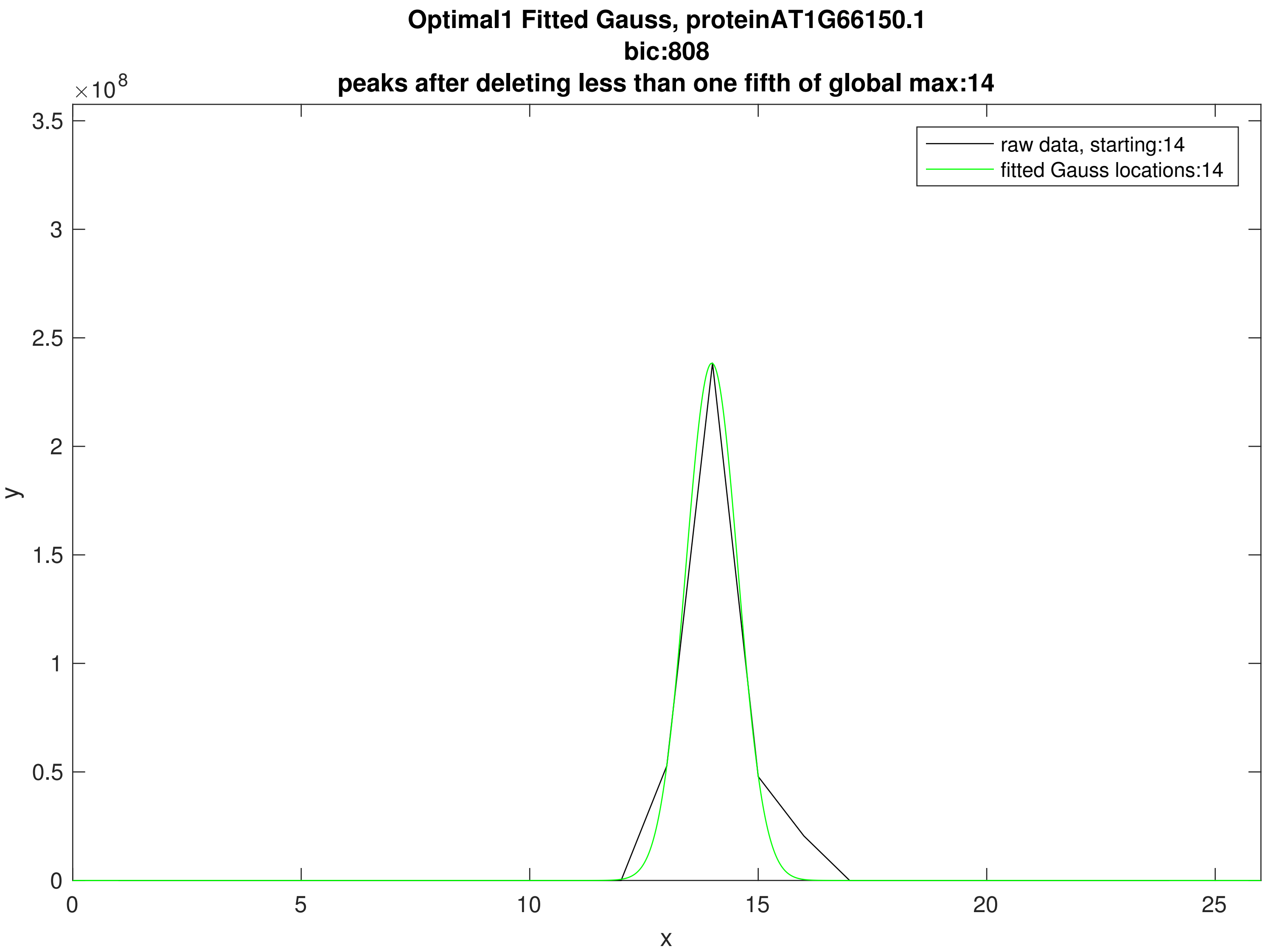
Optimal1 Fitted Gauss, proteinAT1G66150.1
bic:808
peaks after deleting less than one fifth of global max:14
×10^8
3.5
3
2.5
2
1.5
1
0.5
0
y
0
5
10
15
20
25
x
raw data, starting:14
fitted Gauss locations:14

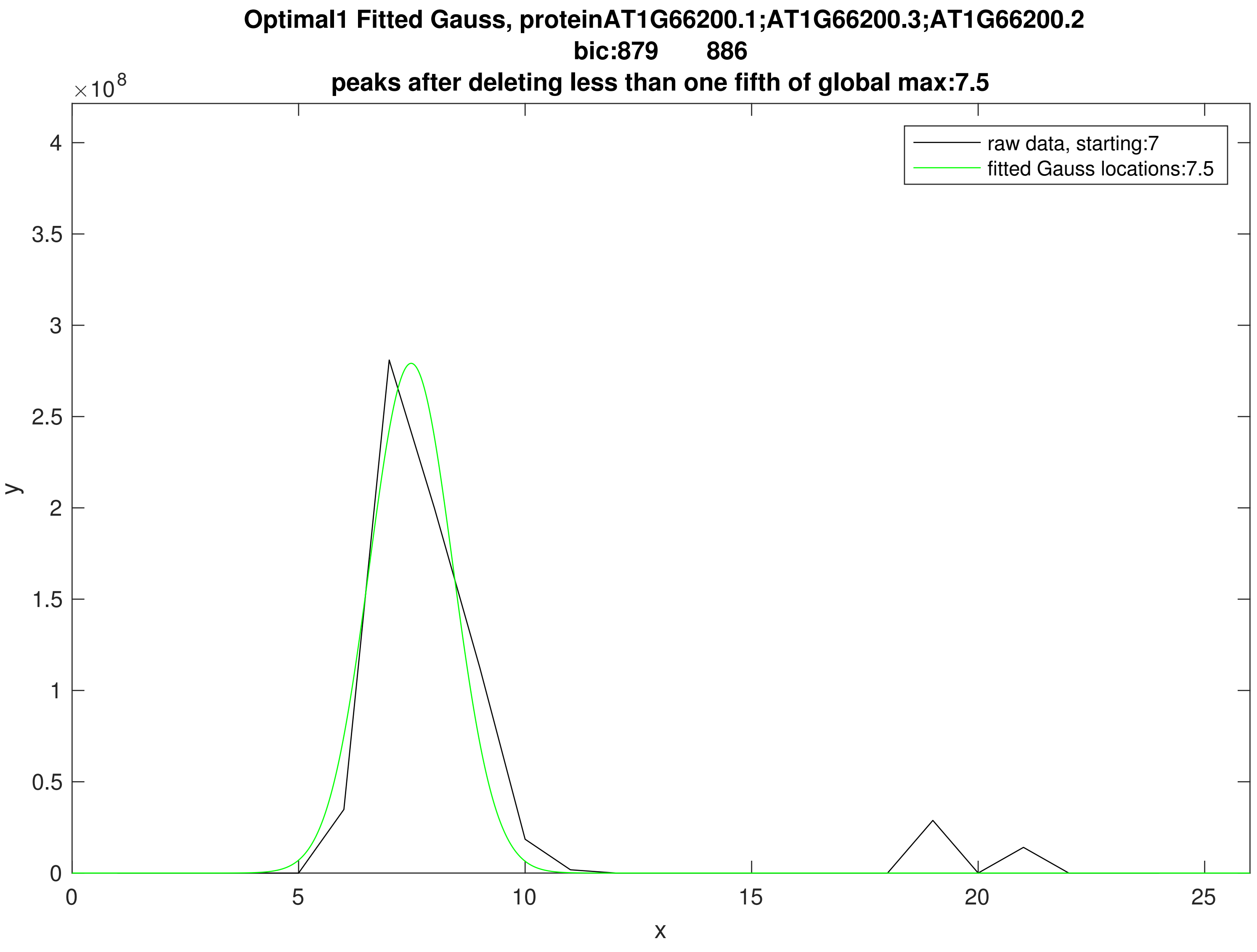
Optimal1 Fitted Gauss, proteinAT1G66200.1;AT1G66200.3;AT1G66200.2
bic:879 886
peaks after deleting less than one fifth of global max:7.5
×10^8
raw data, starting:7
fitted Gauss locations:7.5
y
x
0
0.5
1
1.5
2
2.5
3
3.5
4
0
5
10
15
20
25

Optimal2 Fitted Gauss, proteinAT1G66240.1;AT1G66240.2
bic:912 894
peaks after deleting less than one fifth of global max:21

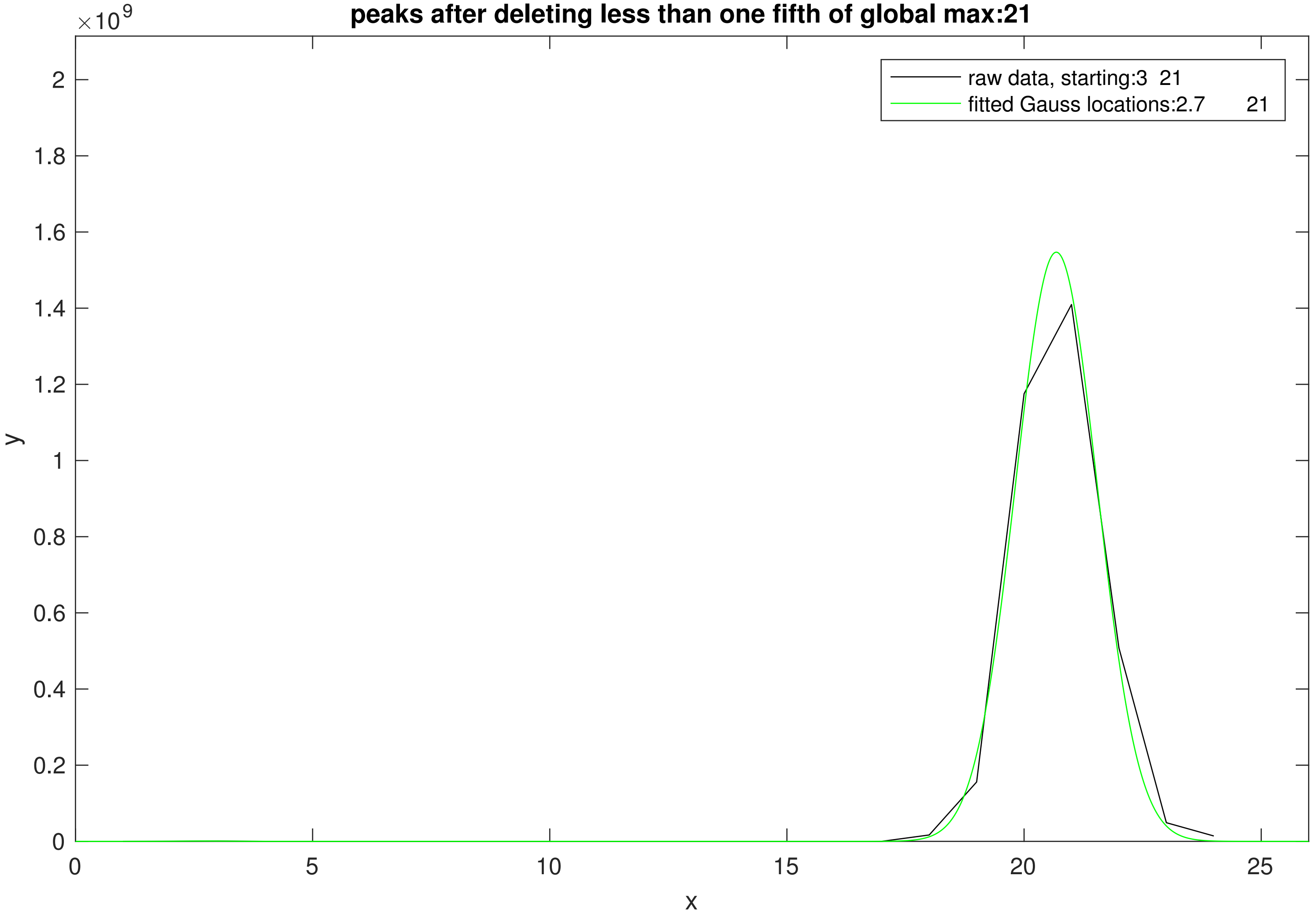

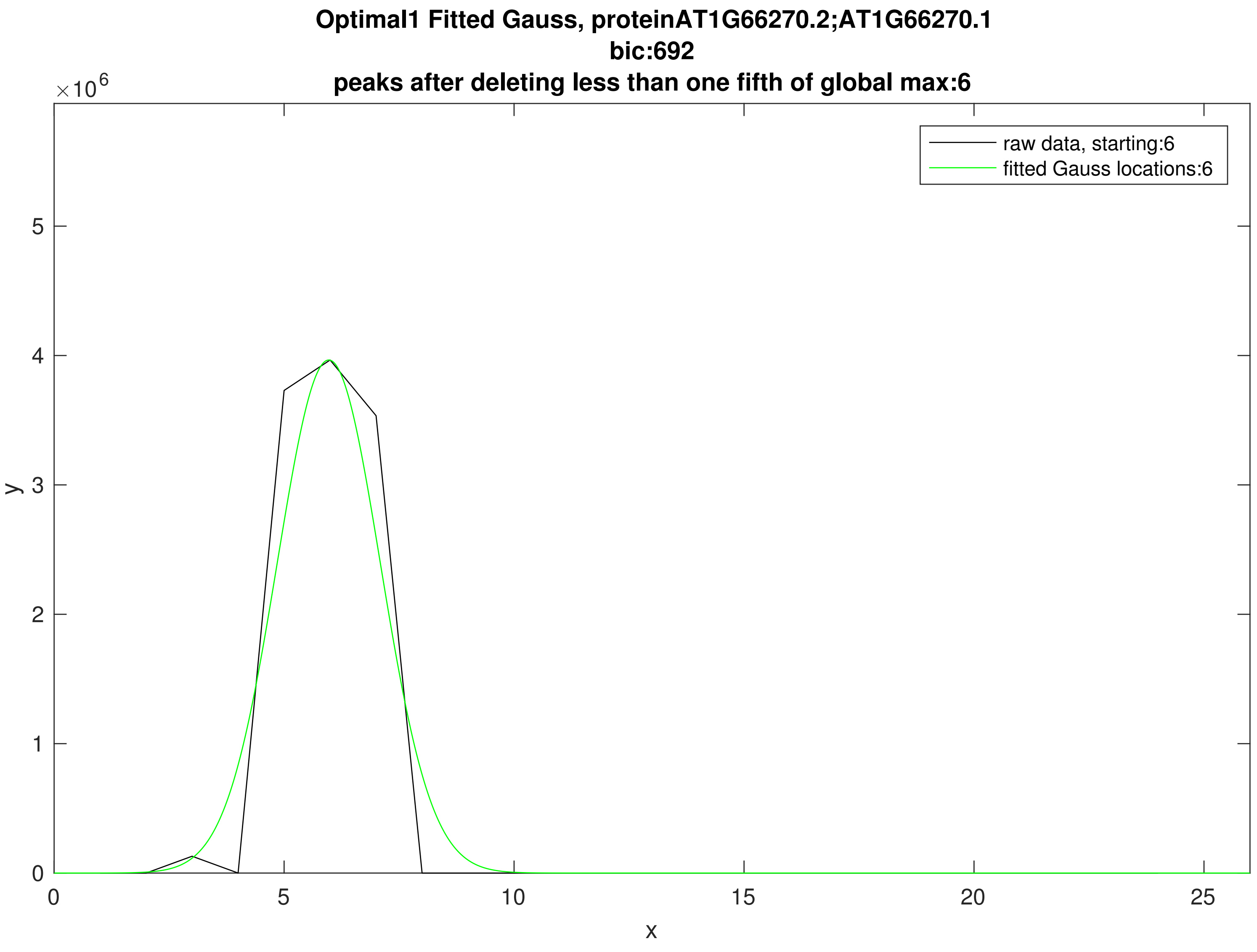

Optimal1 Fitted Gauss, proteinAT1G66270.2;AT1G66270.1
bic:692
peaks after deleting less than one fifth of global max:6
×10^6
raw data, starting:6
fitted Gauss locations:6
y
x
0
1
2
3
4
5
0
5
10
15
20
25

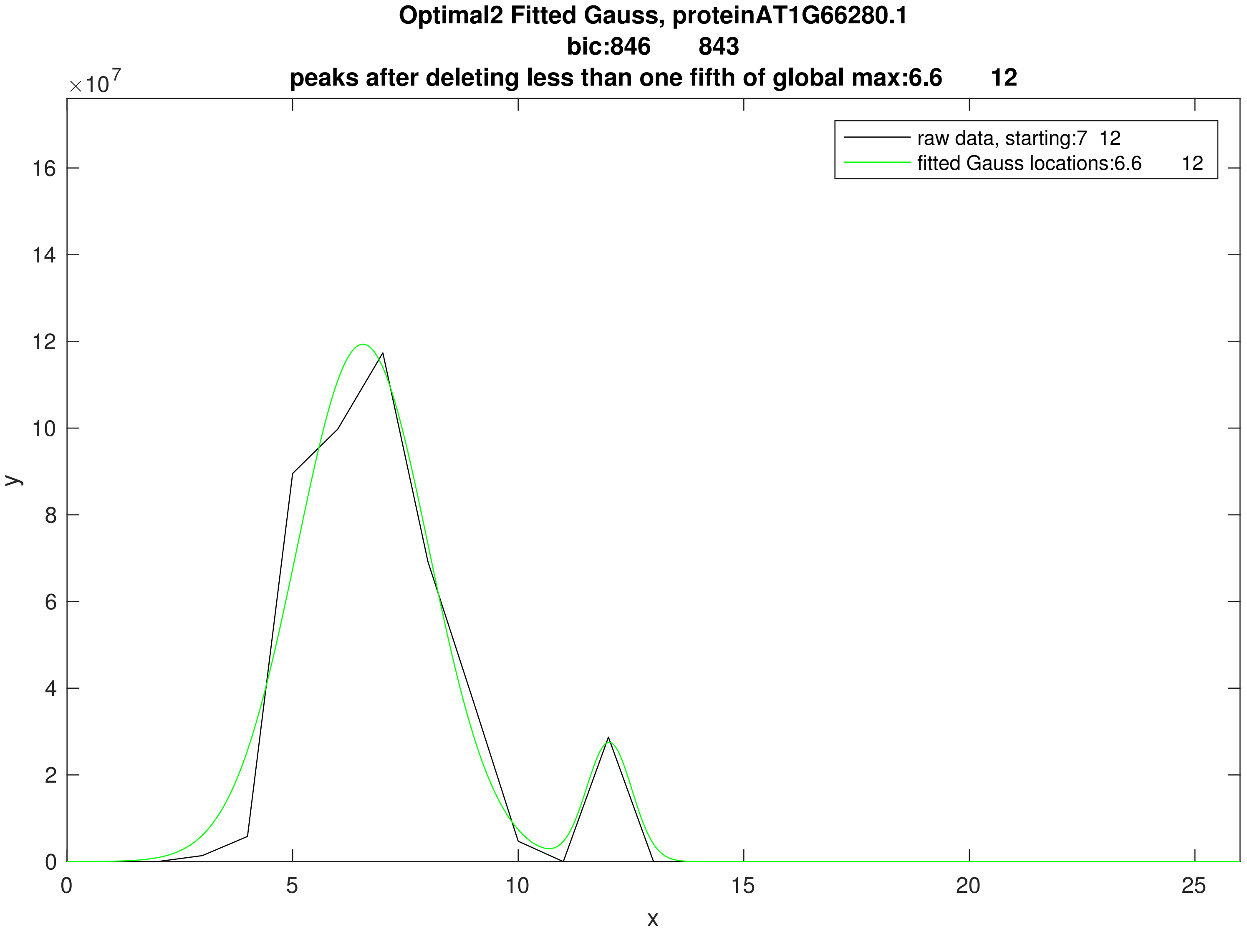
Optimal2 Fitted Gauss, proteinAT1G66280.1
bic:846 843
peaks after deleting less than one fifth of global max:6.6 12
×10^7
raw data, starting:7 12
fitted Gauss locations:6.6 12
y
x
0
2
4
6
8
10
12
14
16
0
5
10
15
20
25

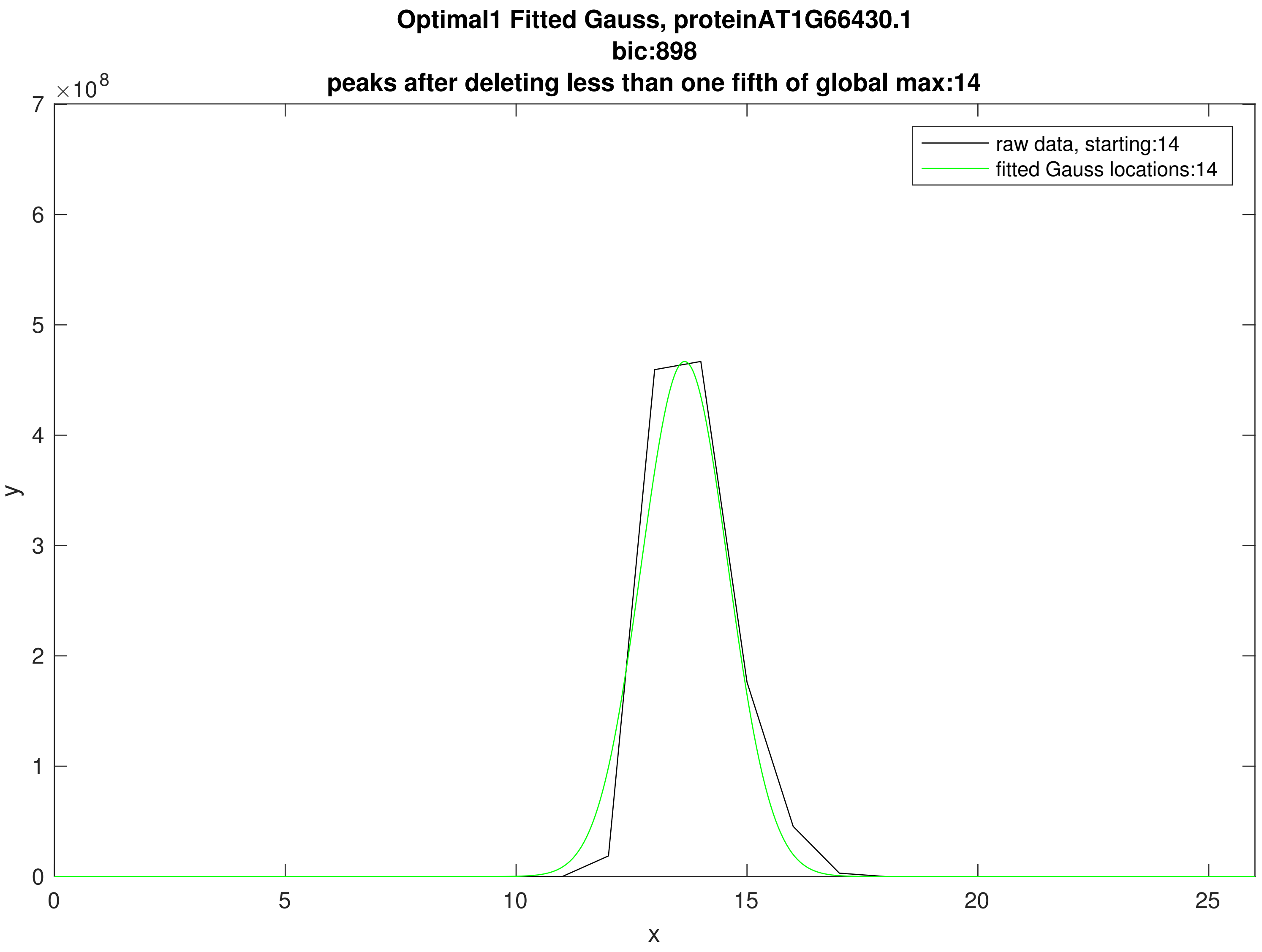
Optimal1 Fitted Gauss, proteinAT1G66430.1
bic:898
peaks after deleting less than one fifth of global max:14
$\times 10^8$
raw data, starting:14
fitted Gauss locations:14
y
x
0
1
2
3
4
5
6
7
0
5
10
15
20
25

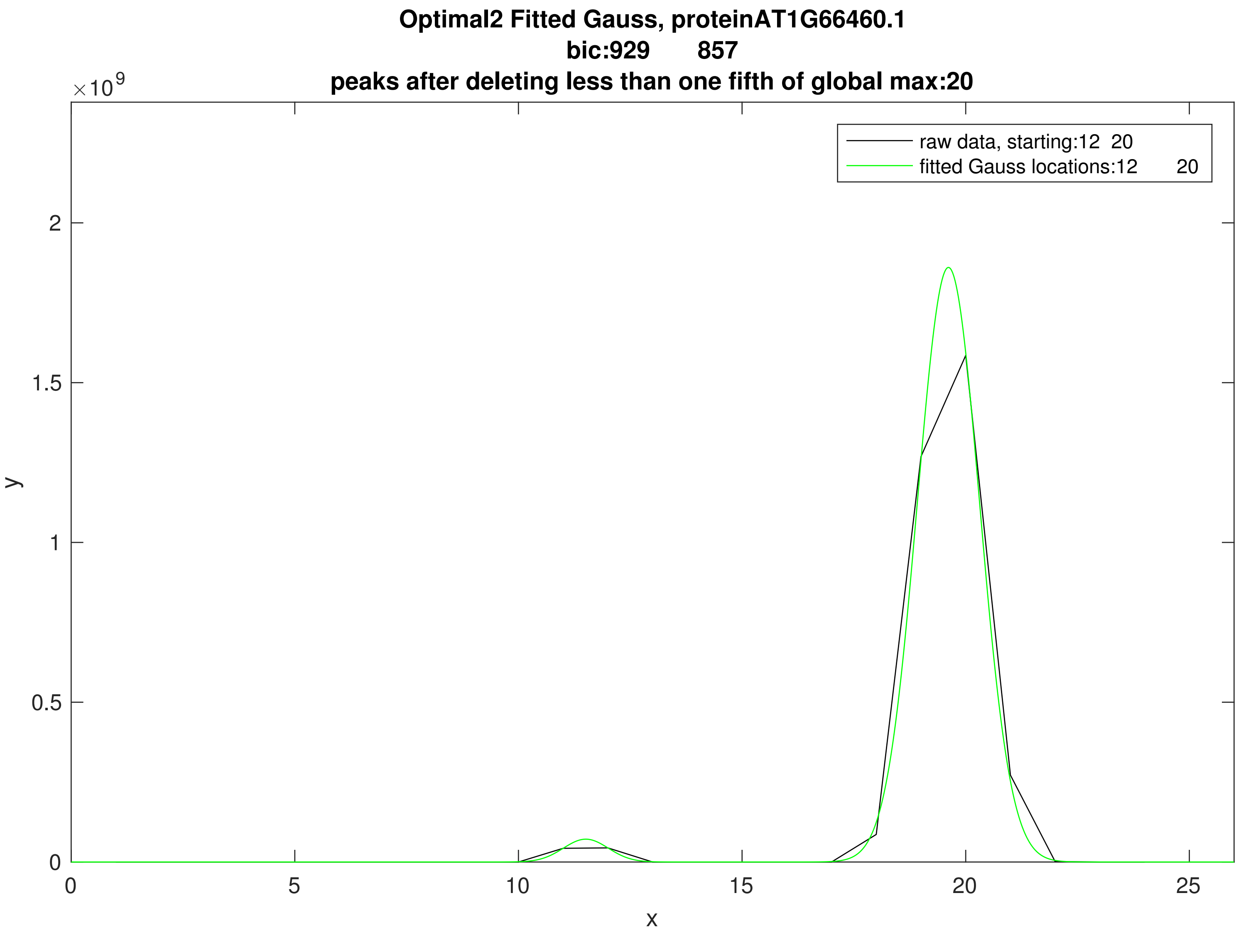
Optimal2 Fitted Gauss, proteinAT1G66460.1
bic:929 857
peaks after deleting less than one fifth of global max:20
×10^9
raw data, starting:12 20
fitted Gauss locations:12 20
2
1.5
1
0.5
0
y
0
5
10
15
20
25
x

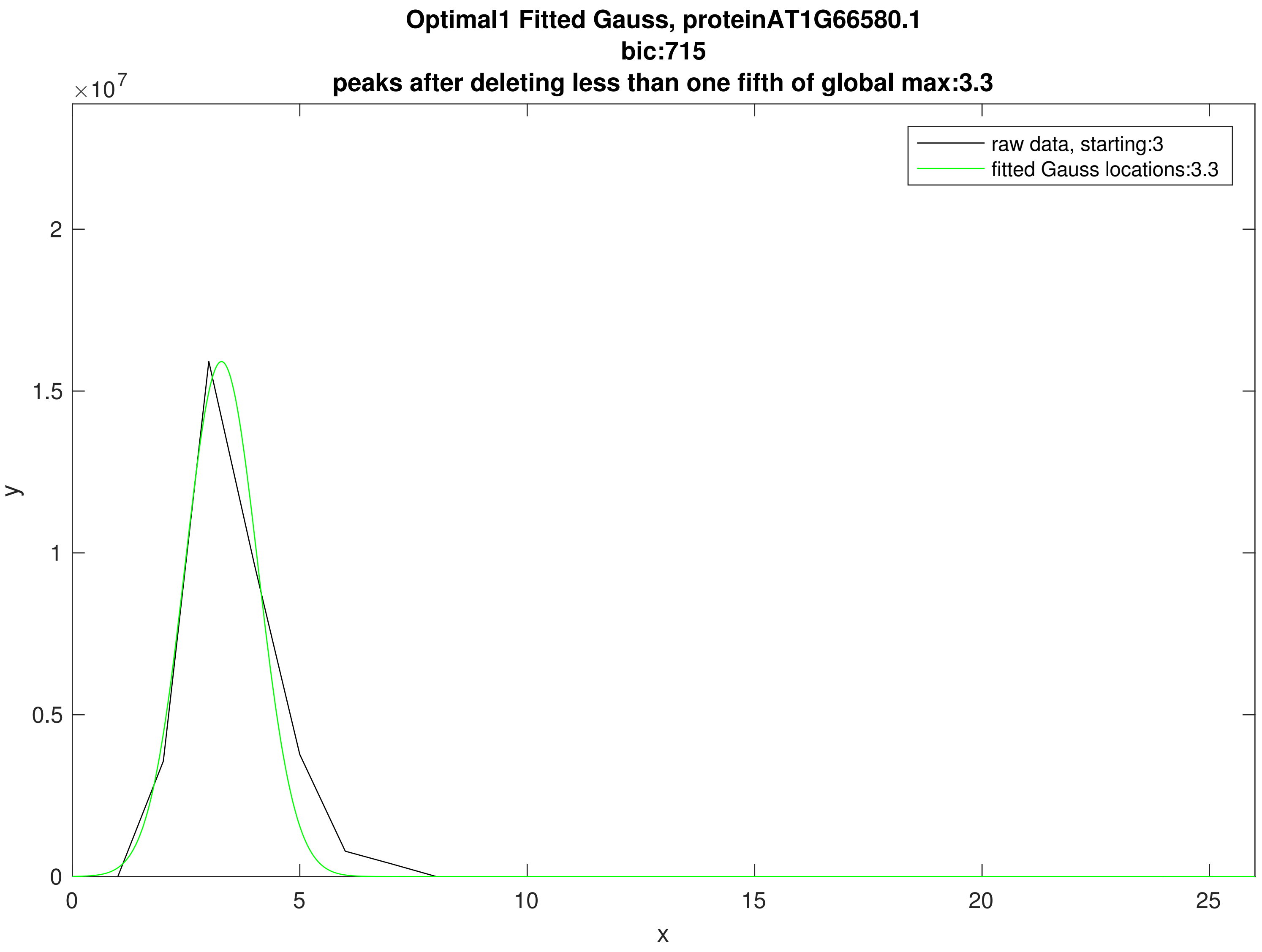
Optimal1 Fitted Gauss, proteinAT1G66580.1
bic:715
peaks after deleting less than one fifth of global max:3.3
×10^7
raw data, starting:3
fitted Gauss locations:3.3
y
x
0
0.5
1
1.5
2
5
10
15
20
25

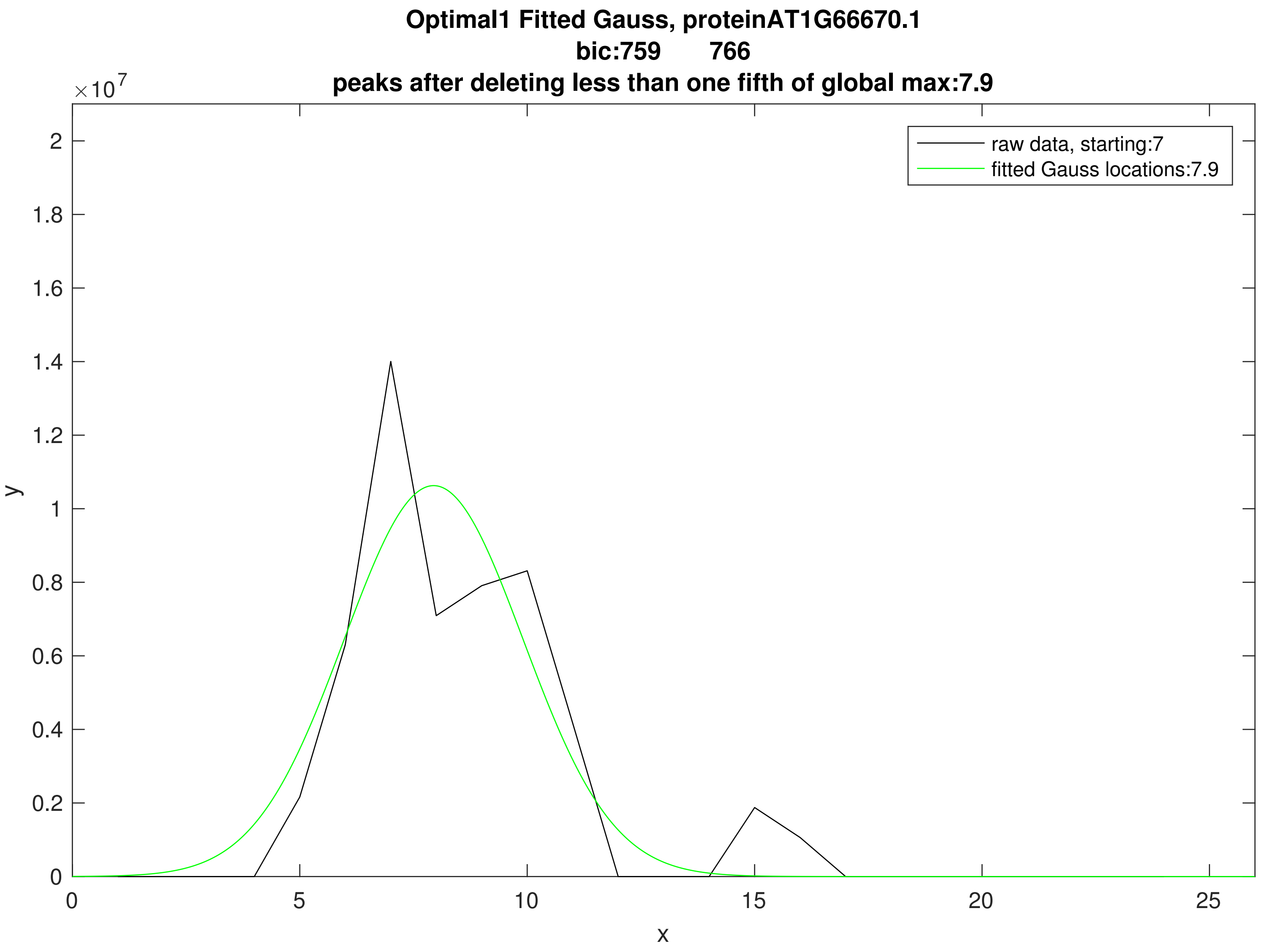

Optimal1 Fitted Gauss, proteinAT1G66670.1
bic:759 766
peaks after deleting less than one fifth of global max:7.9
×10^7
raw data, starting:7
fitted Gauss locations:7.9
y
x
0
0.2
0.4
0.6
0.8
1
1.2
1.4
1.6
1.8
2
0
5
10
15
20
25

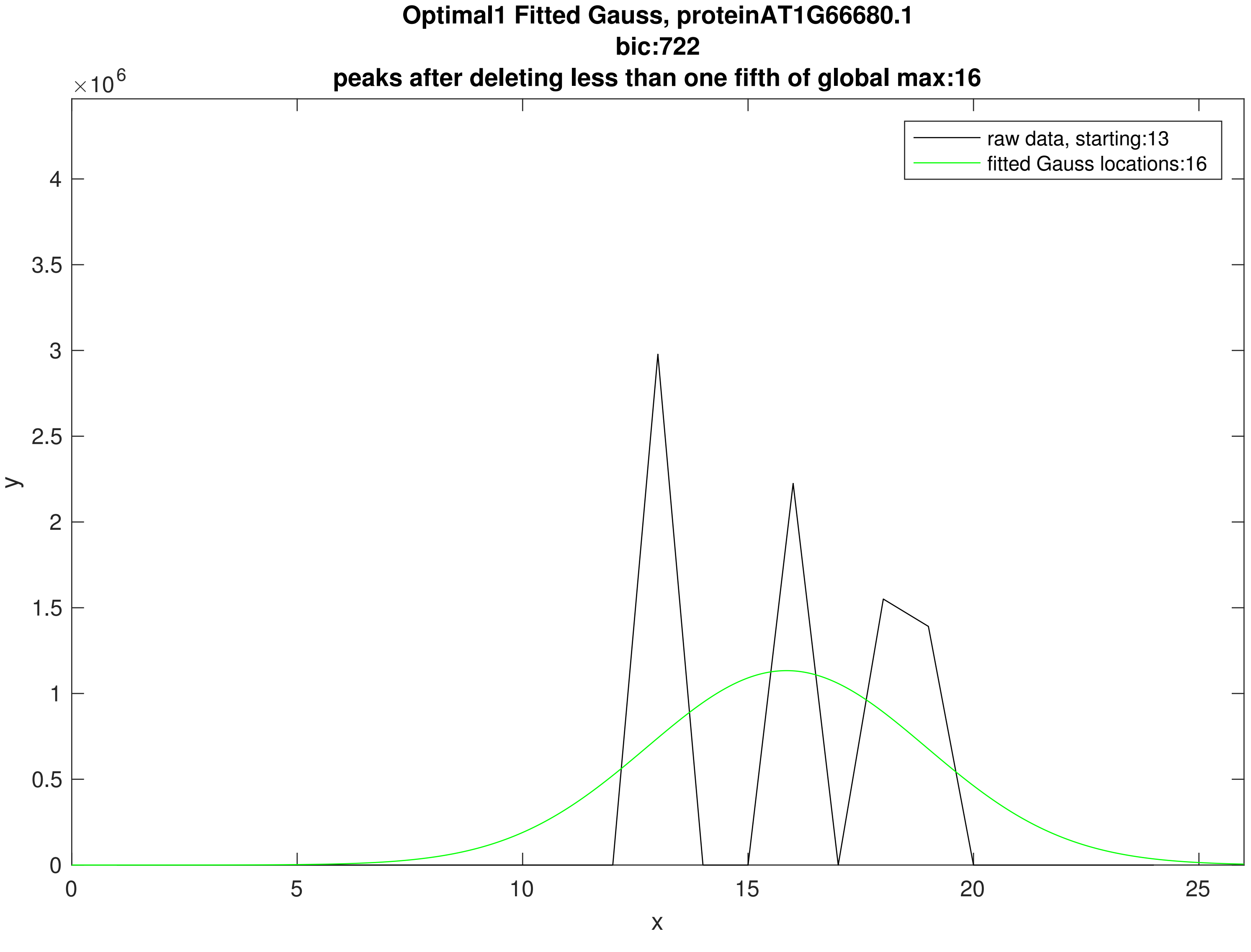
Optimal1 Fitted Gauss, proteinAT1G66680.1
bic:722
peaks after deleting less than one fifth of global max:16
×10^6
raw data, starting:13
fitted Gauss locations:16
y
x
0
0.5
1
1.5
2
2.5
3
3.5
4
0
5
10
15
20
25

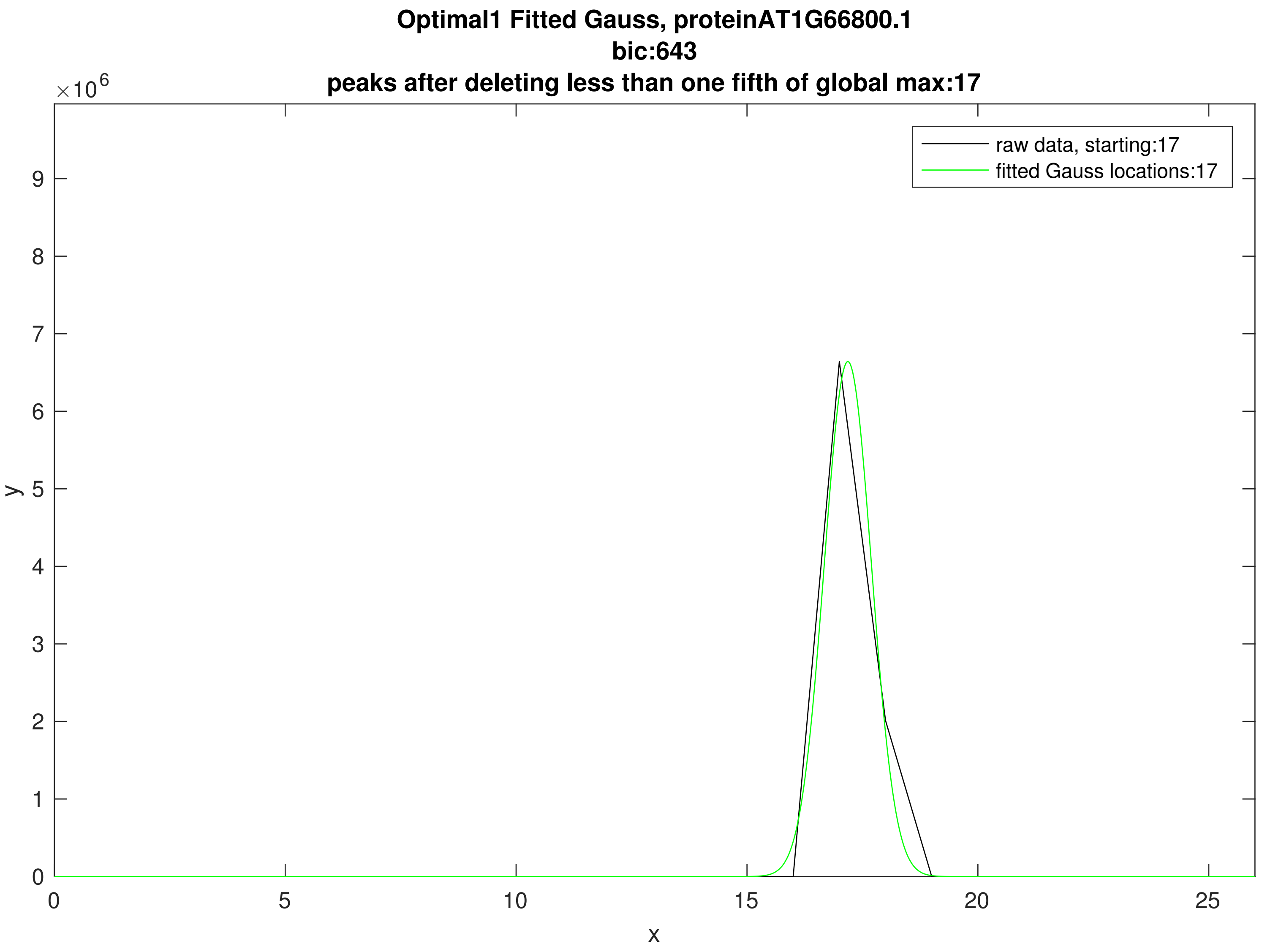

Optimal1 Fitted Gauss, proteinAT1G66800.1
bic:643
peaks after deleting less than one fifth of global max:17
×10^6
raw data, starting:17
fitted Gauss locations:17
y
x
0
1
2
3
4
5
6
7
8
9
0
5
10
15
20
25

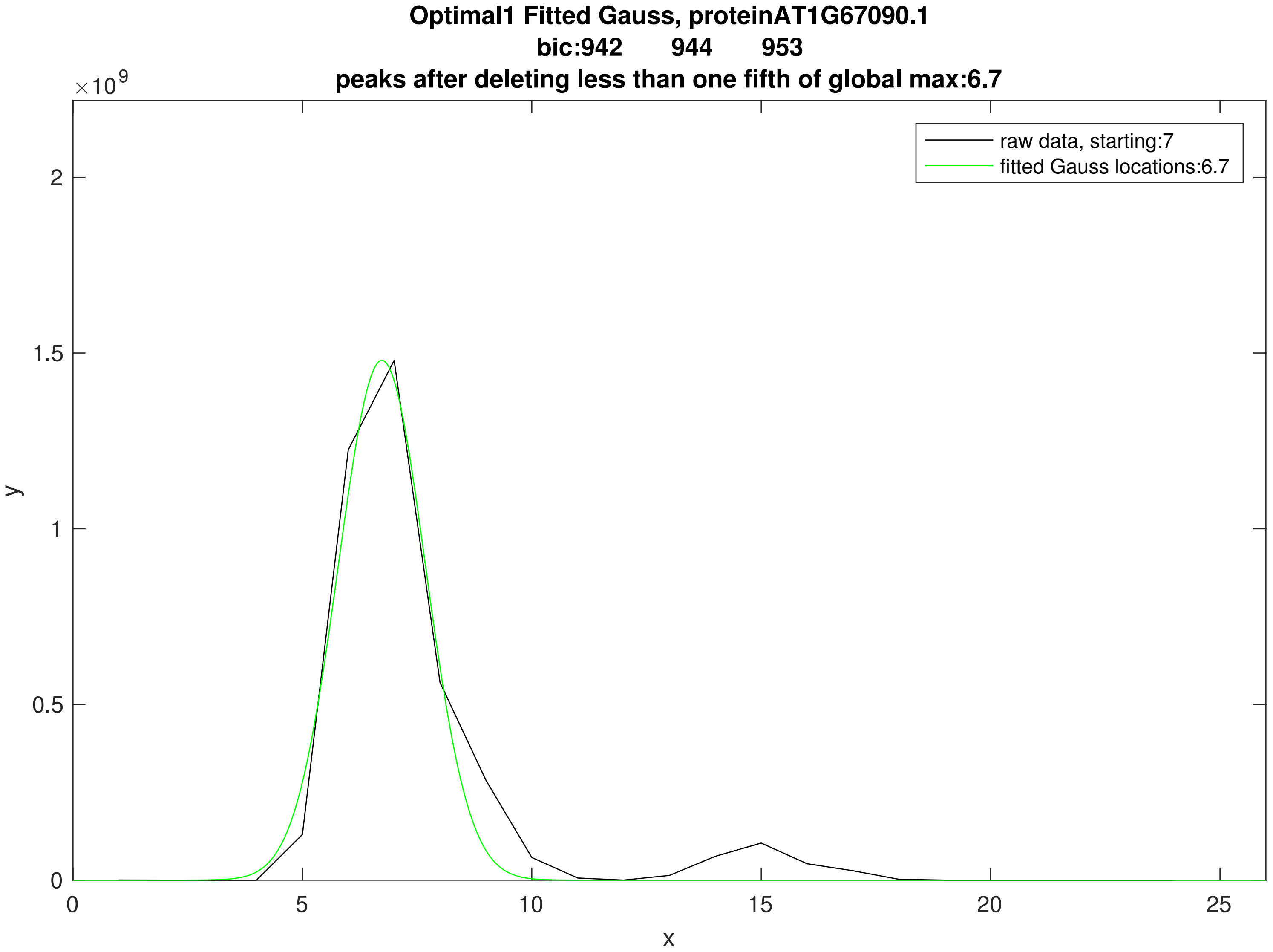

Optimal1 Fitted Gauss, proteinAT1G67090.1
bic:942 944 953
peaks after deleting less than one fifth of global max:6.7
×10^9
raw data, starting:7
fitted Gauss locations:6.7
y
x
0
0.5
1
1.5
2
0
5
10
15
20
25

**Optimal2 Fitted Gauss, proteinAT1G67280.1;AT1G67280.2**
**bic:824 821**
**peaks after deleting less than one fifth of global max:18**

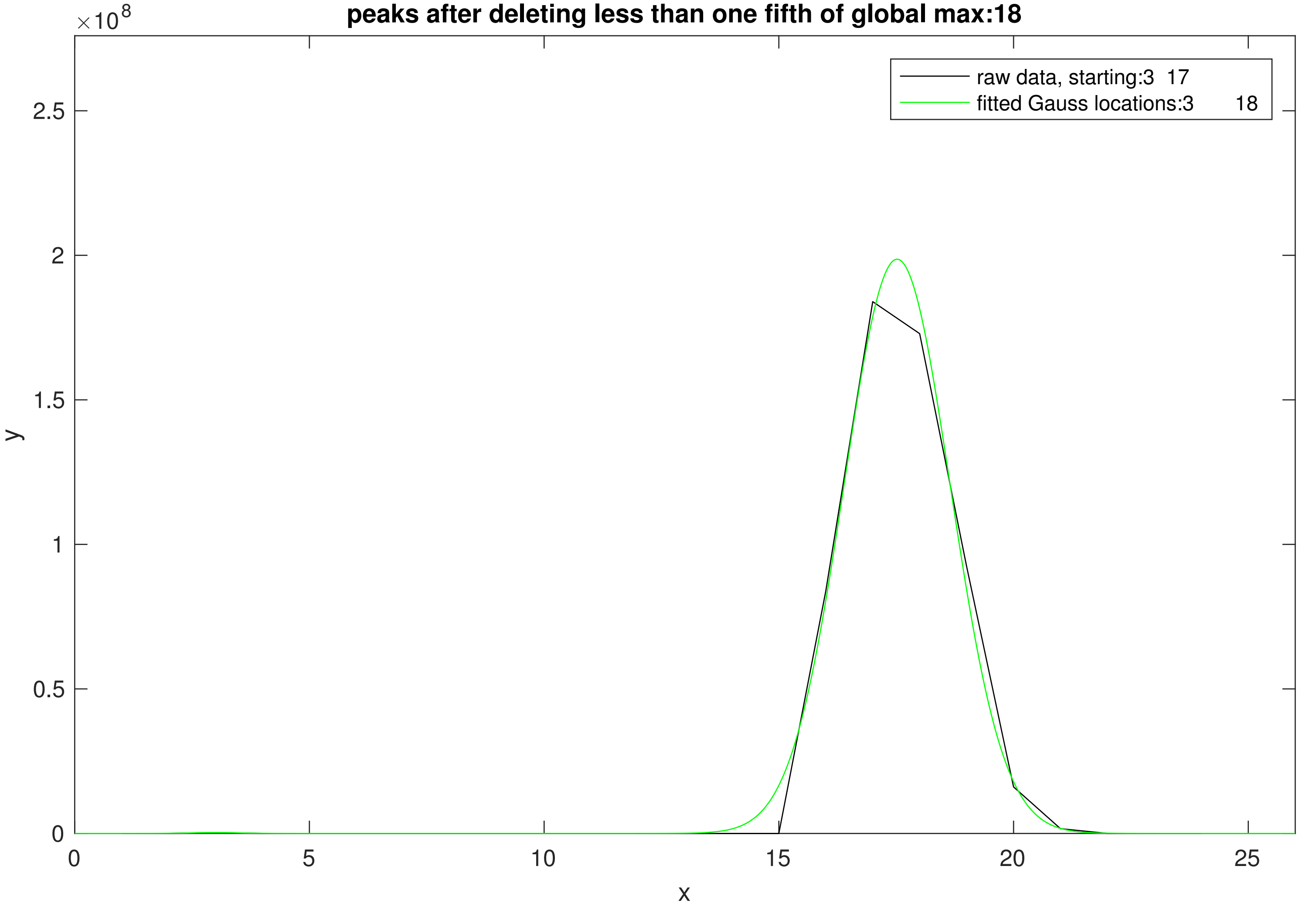

Optimal1 Fitted Gauss, proteinAT1G67550.1
bic:726
peaks after deleting less than one fifth of global max:9.6

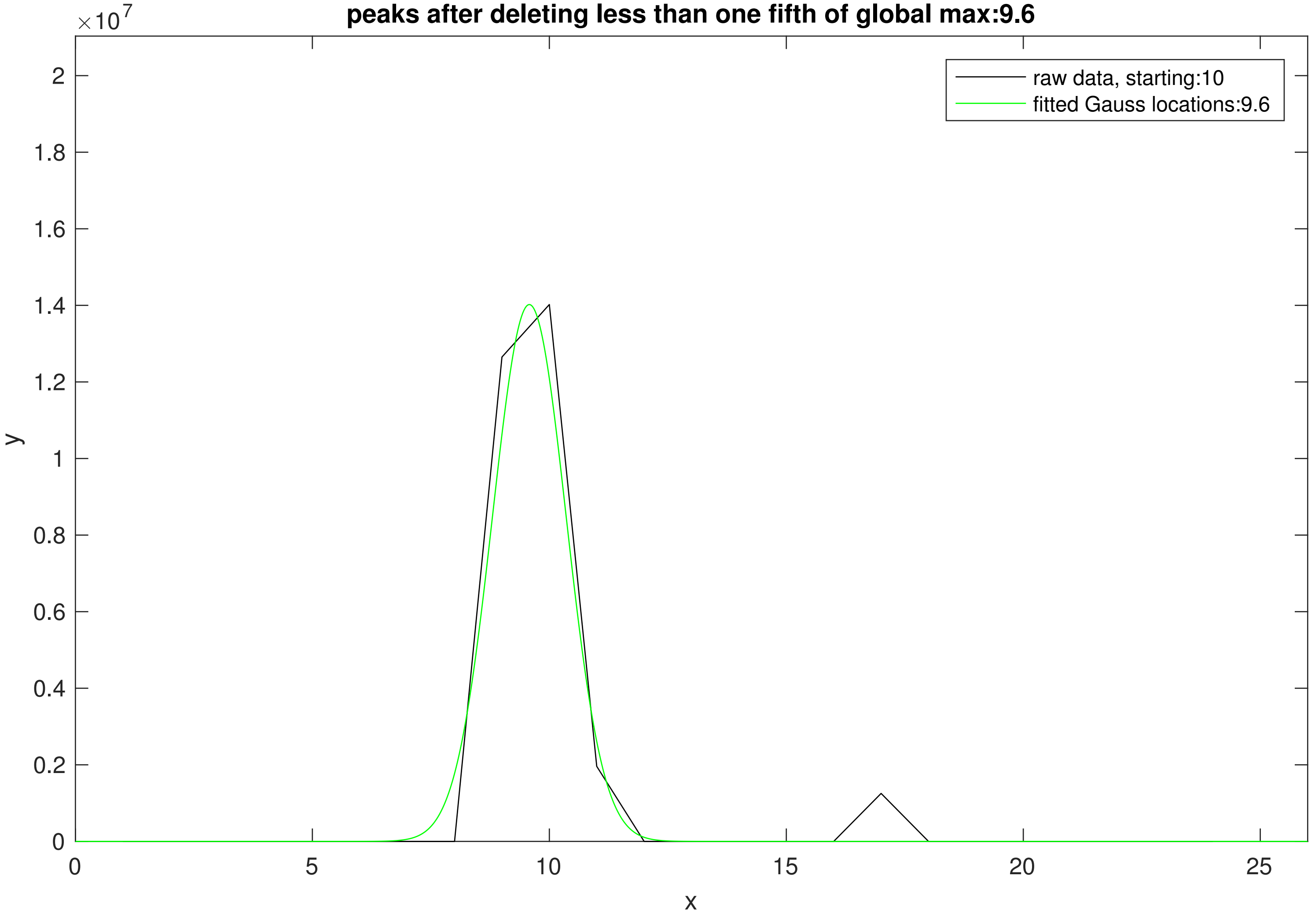

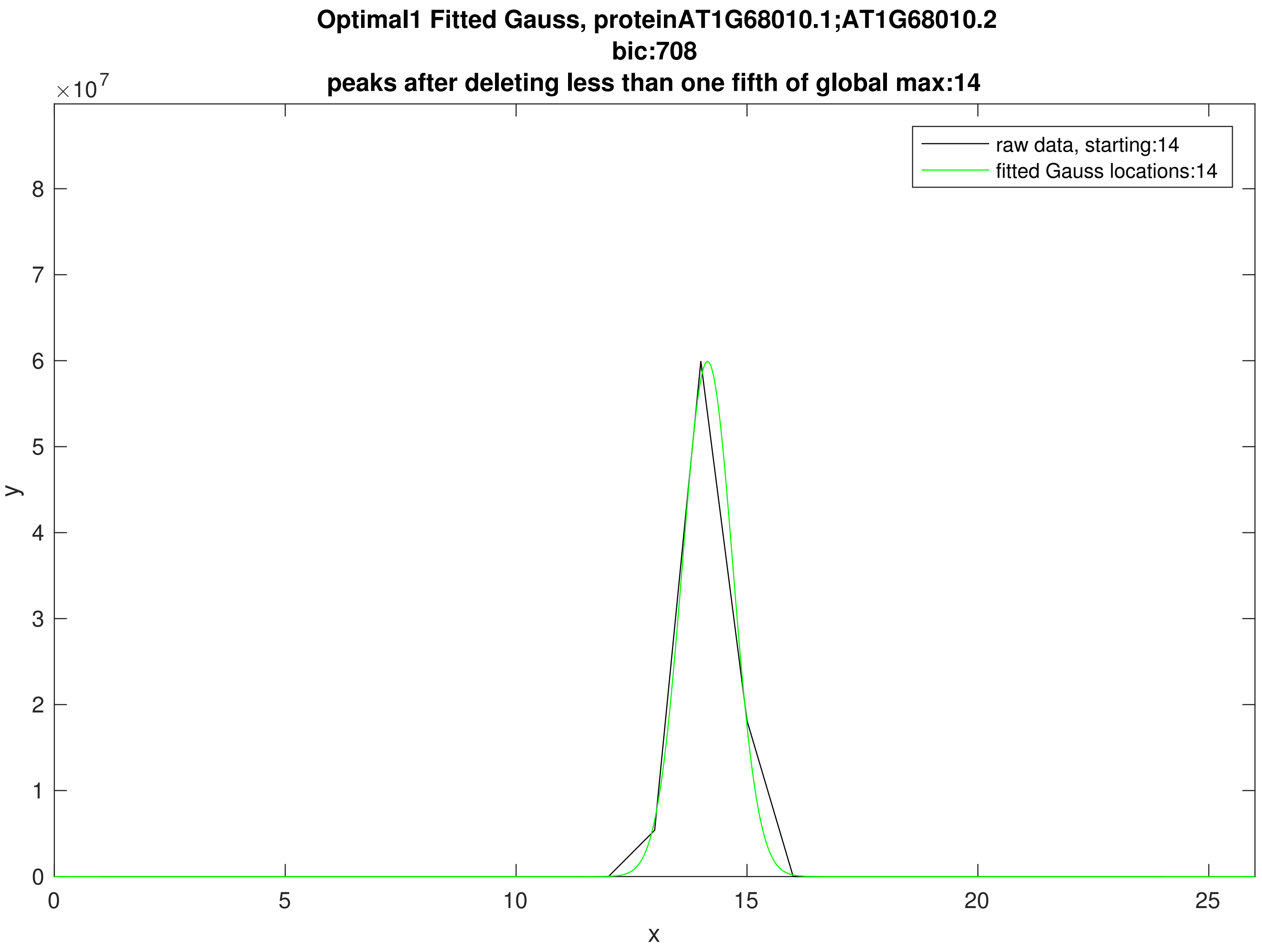
Optimal1 Fitted Gauss, proteinAT1G68010.1;AT1G68010.2
bic:708
peaks after deleting less than one fifth of global max:14
×10^7
raw data, starting:14
fitted Gauss locations:14
y
x
0
1
2
3
4
5
6
7
8
0
5
10
15
20
25

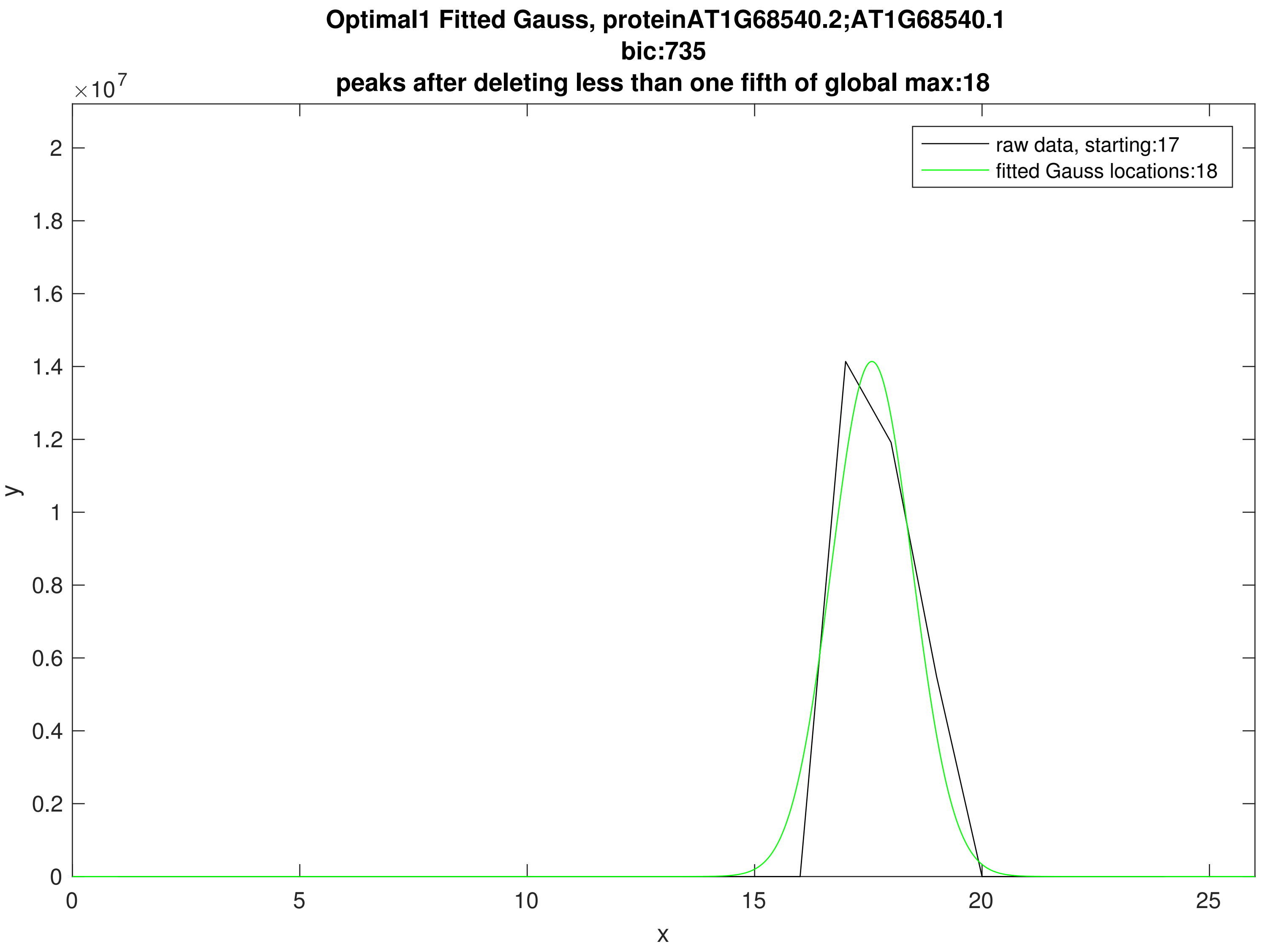

Optimal1 Fitted Gauss, proteinAT1G68540.2;AT1G68540.1
bic:735
peaks after deleting less than one fifth of global max:18
×10^7
raw data, starting:17
fitted Gauss locations:18
y
x
0
0.2
0.4
0.6
0.8
1
1.2
1.4
1.6
1.8
2
0
5
10
15
20
25

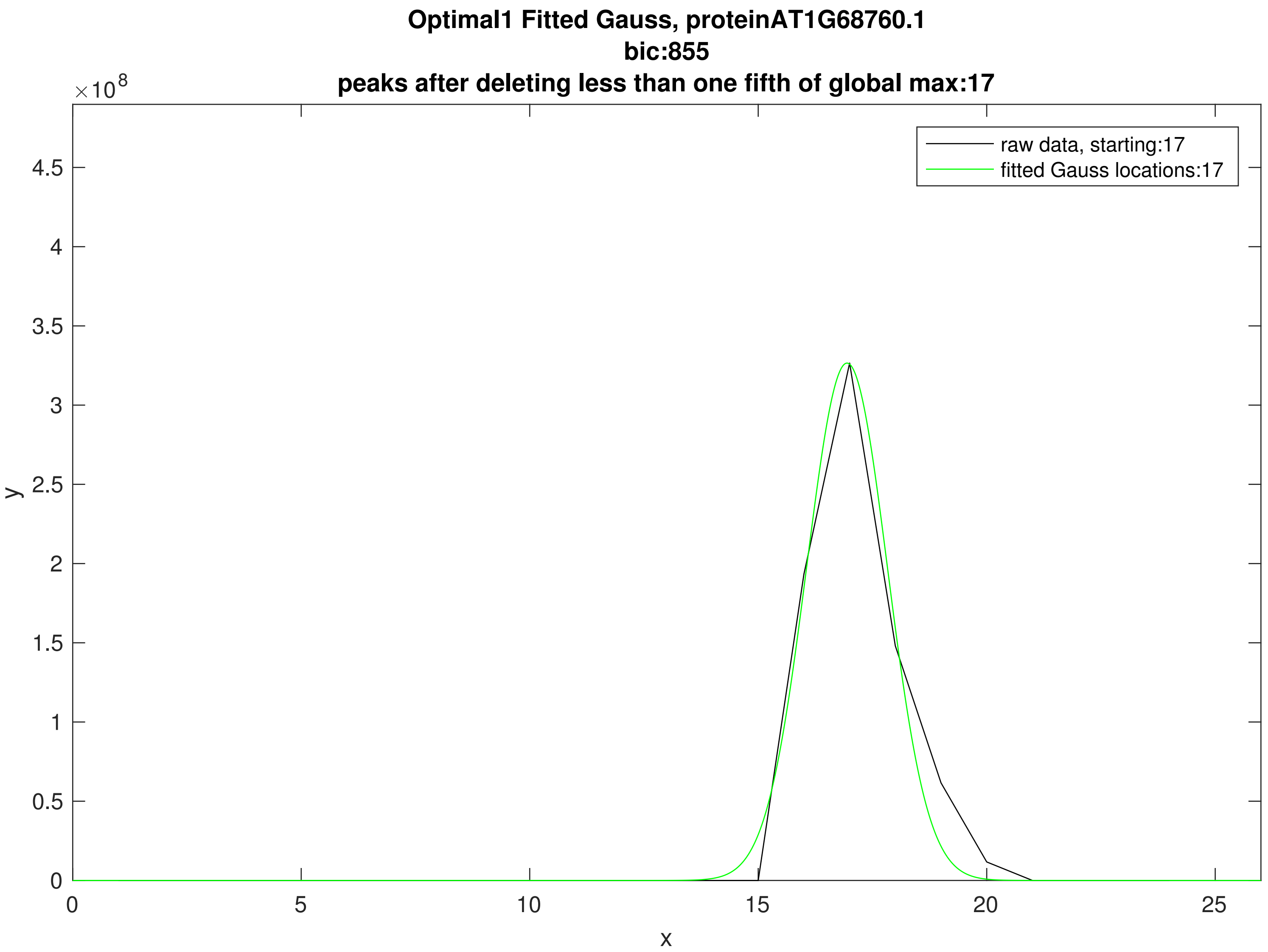
Optimal1 Fitted Gauss, proteinAT1G68760.1
bic:855
peaks after deleting less than one fifth of global max:17
×10^8
raw data, starting:17
fitted Gauss locations:17
y
x
0
0.5
1
1.5
2
2.5
3
3.5
4
4.5
5
10
15
20
25

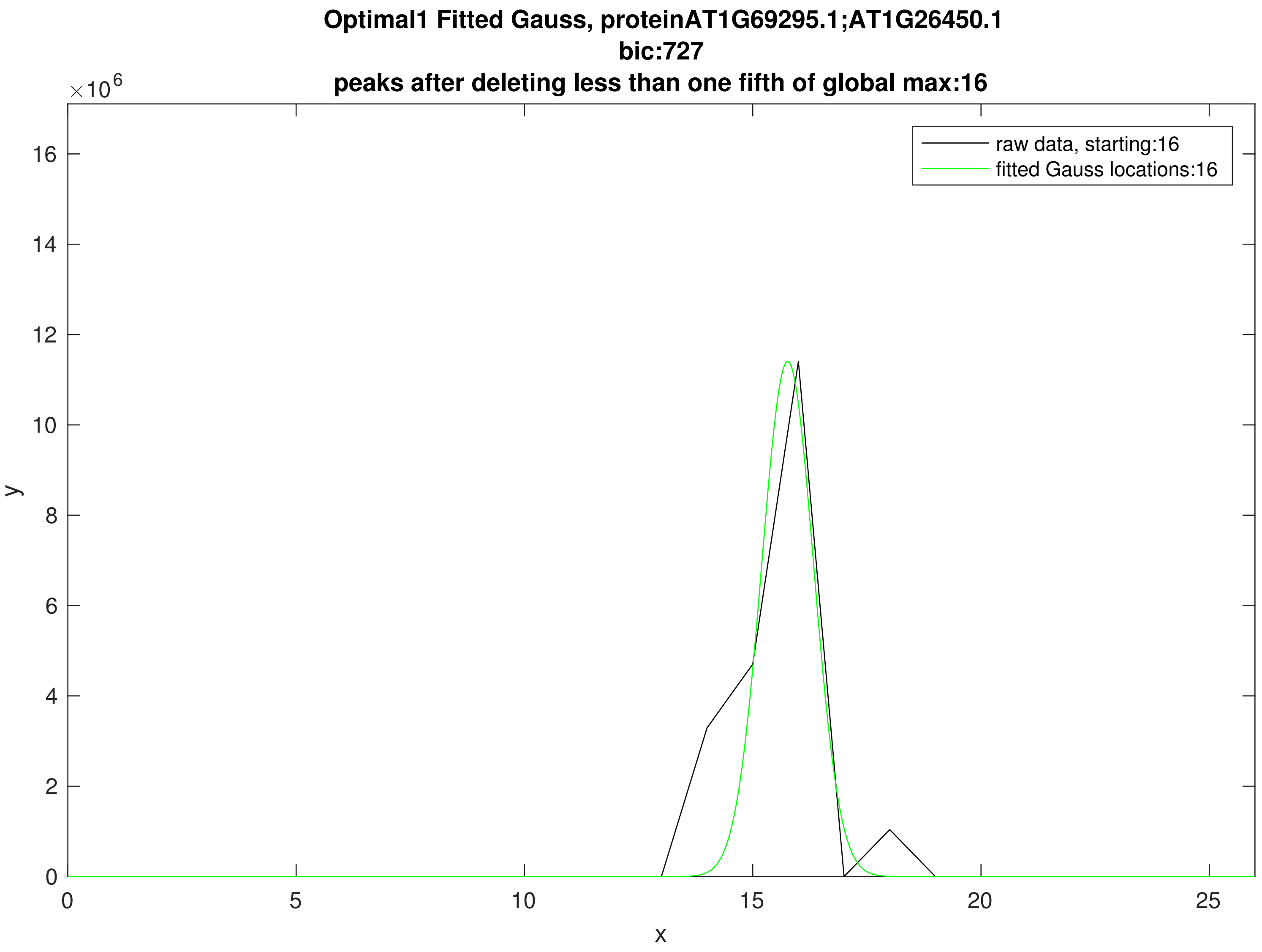
Optimal1 Fitted Gauss, proteinAT1G69295.1;AT1G26450.1
bic:727
peaks after deleting less than one fifth of global max:16
×10^6
raw data, starting:16
fitted Gauss locations:16
16
14
12
10
8
6
4
2
0
y
0
5
10
15
20
25
x

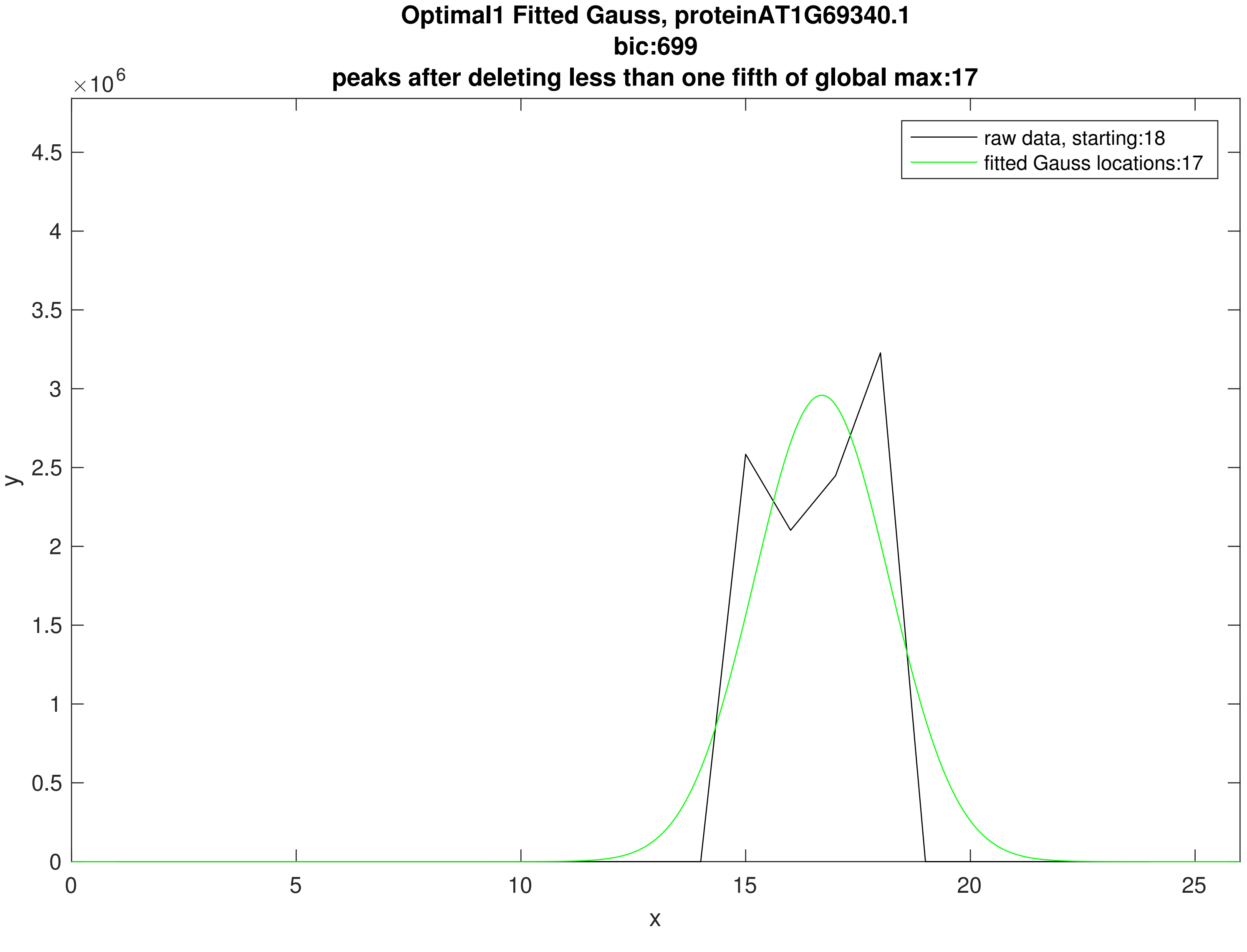
Optimal1 Fitted Gauss, proteinAT1G69340.1
bic:699
peaks after deleting less than one fifth of global max:17
×10^6
y
x
raw data, starting:18
fitted Gauss locations:17
0
0.5
1
1.5
2
2.5
3
3.5
4
4.5
0
5
10
15
20
25

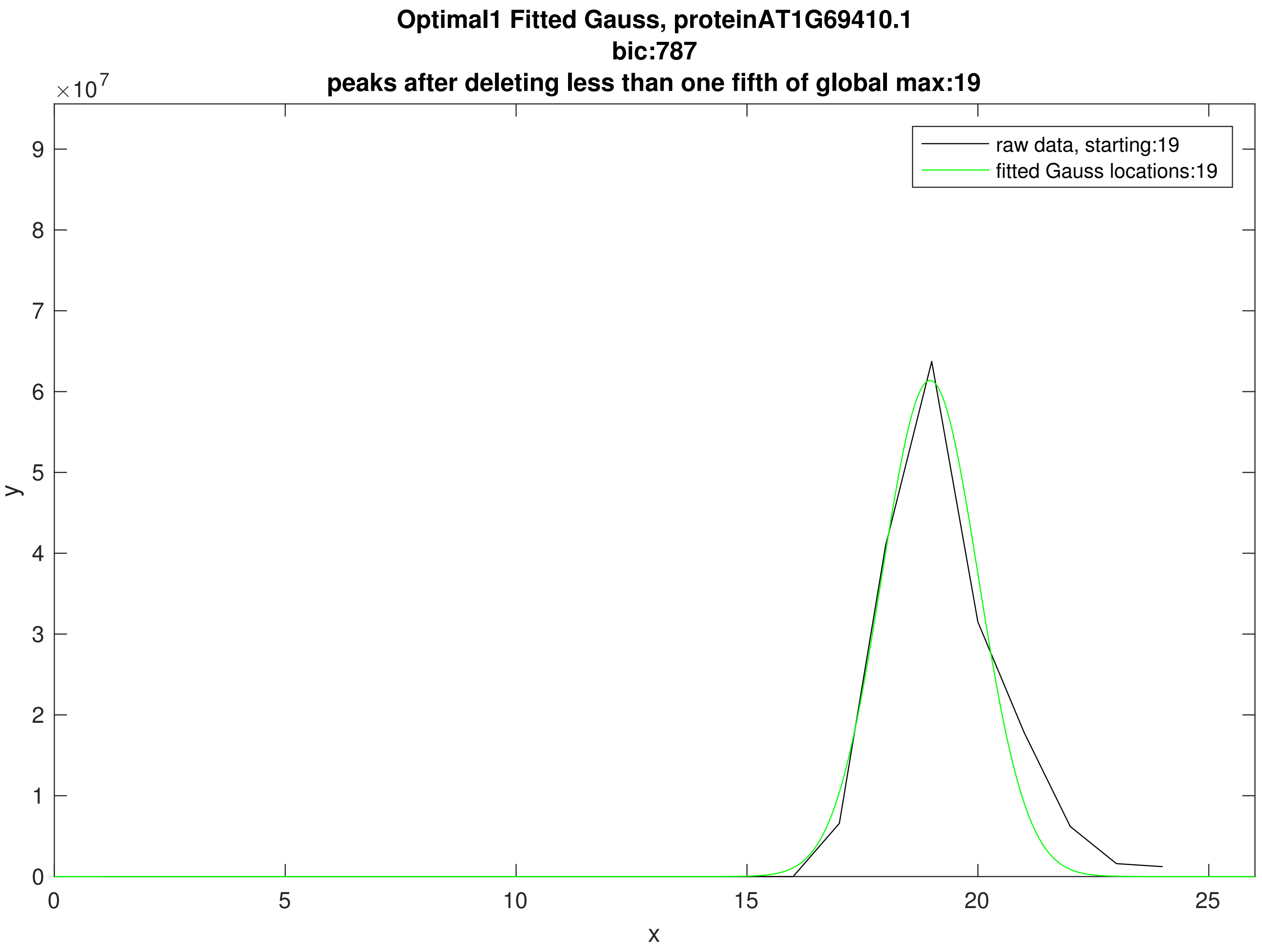

Optimal1 Fitted Gauss, proteinAT1G69410.1
bic:787
peaks after deleting less than one fifth of global max:19
×10^7
y
x
raw data, starting:19
fitted Gauss locations:19

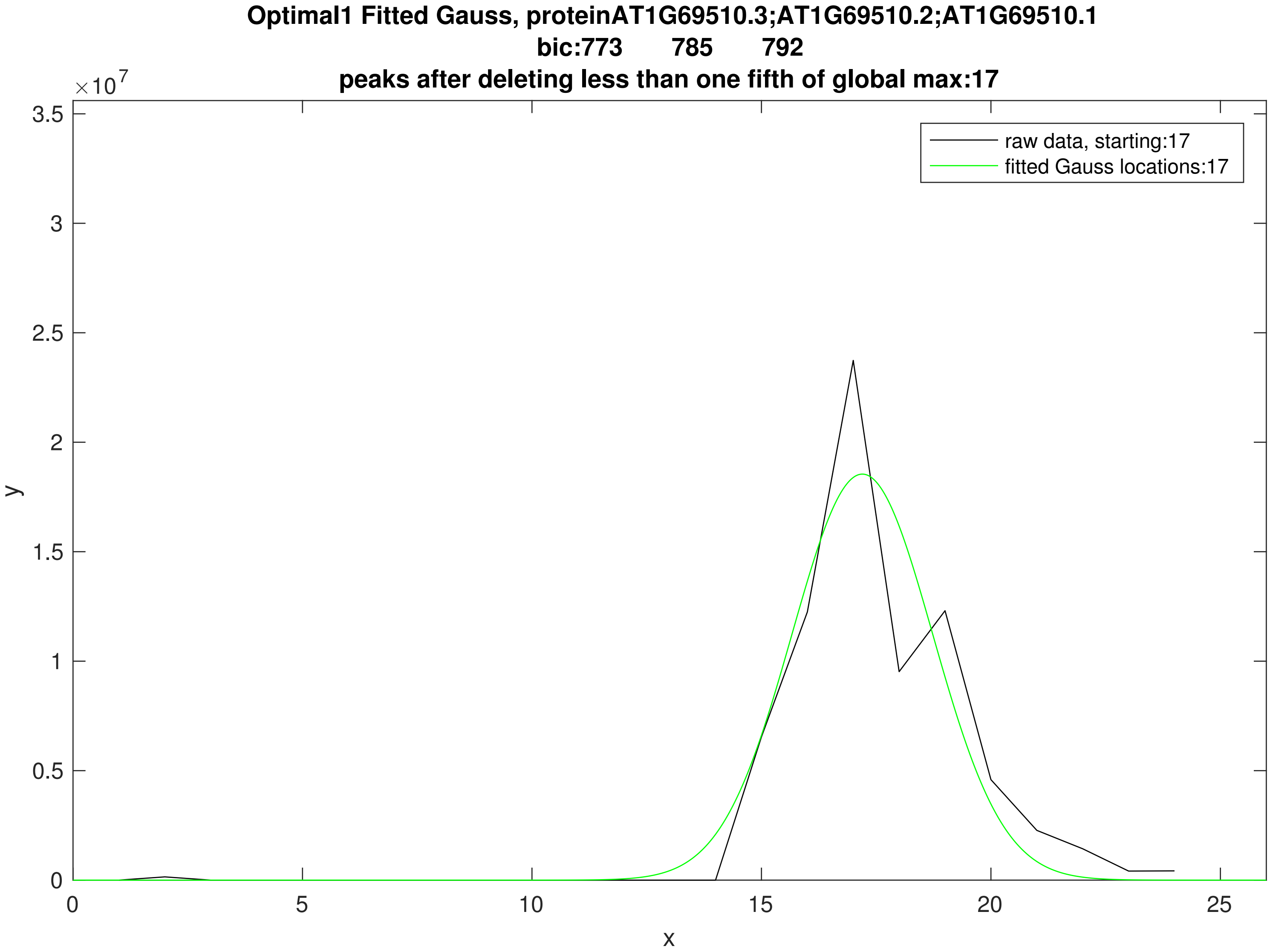
Optimal1 Fitted Gauss, proteinAT1G69510.3;AT1G69510.2;AT1G69510.1
bic:773 785 792
peaks after deleting less than one fifth of global max:17
×10^7
3.5
3
2.5
2
1.5
1
0.5
0
y
0
5
10
15
20
25
x
raw data, starting:17
fitted Gauss locations:17

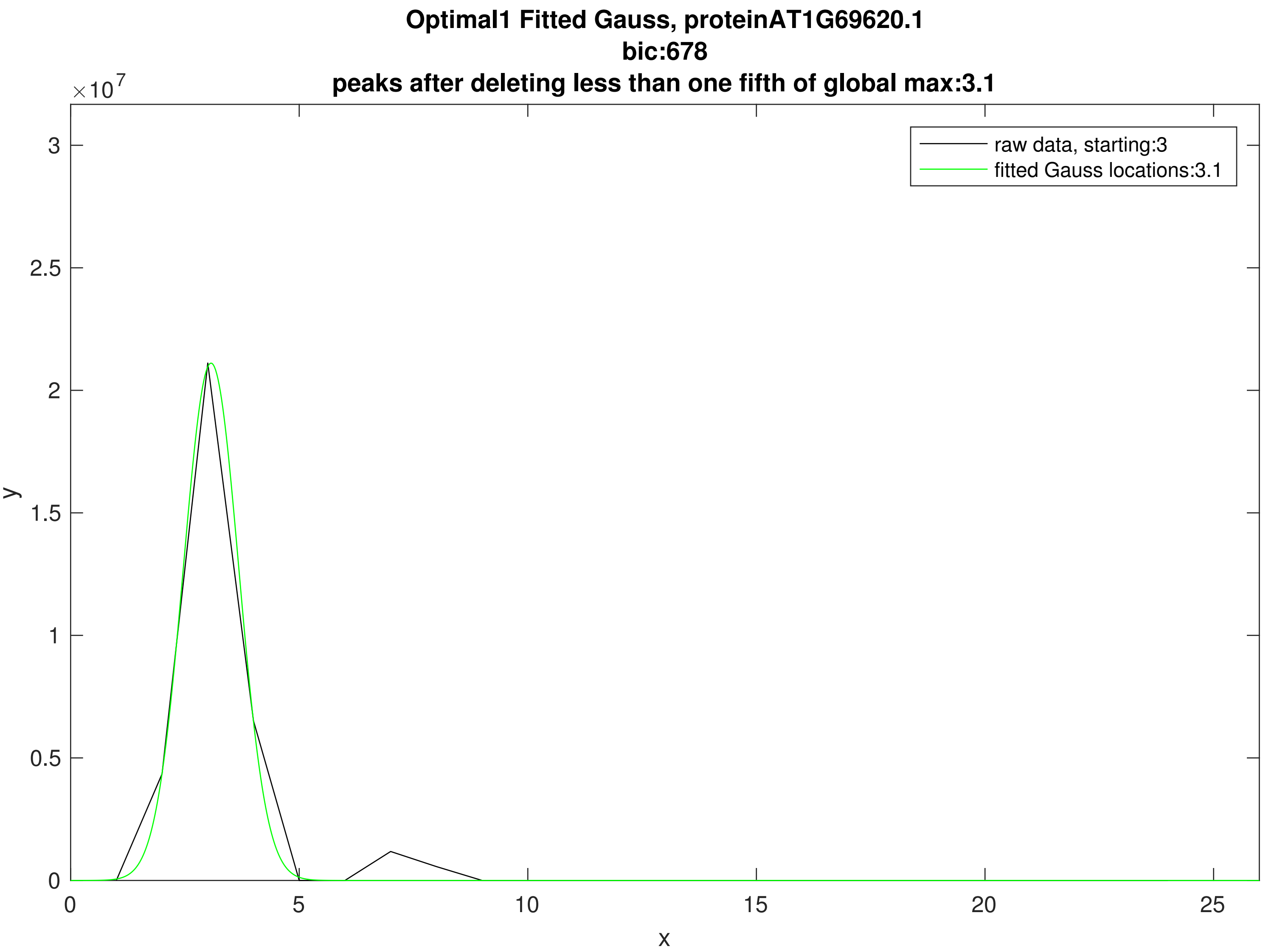

Optimal1 Fitted Gauss, proteinAT1G69620.1
bic:678
peaks after deleting less than one fifth of global max:3.1
×10^7
raw data, starting:3
fitted Gauss locations:3.1
y
x
0
0.5
1
1.5
2
2.5
3
0
5
10
15
20
25

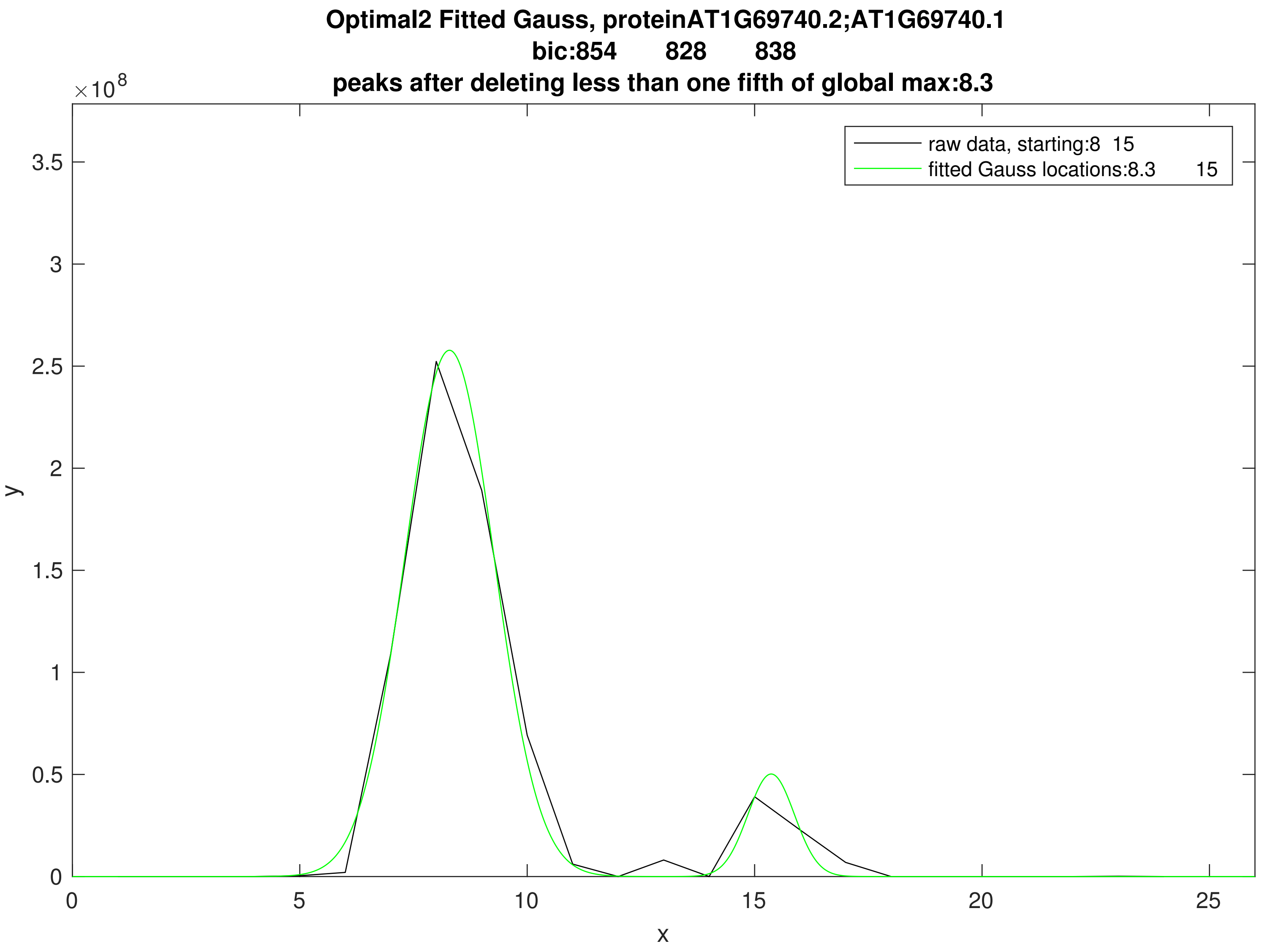

Optimal2 Fitted Gauss, proteinAT1G69740.2;AT1G69740.1
bic:854 828 838
peaks after deleting less than one fifth of global max:8.3
×10^8
raw data, starting:8 15
fitted Gauss locations:8.3 15
3.5
3
2.5
2
1.5
1
0.5
0
y
0
5
10
15
20
25
x

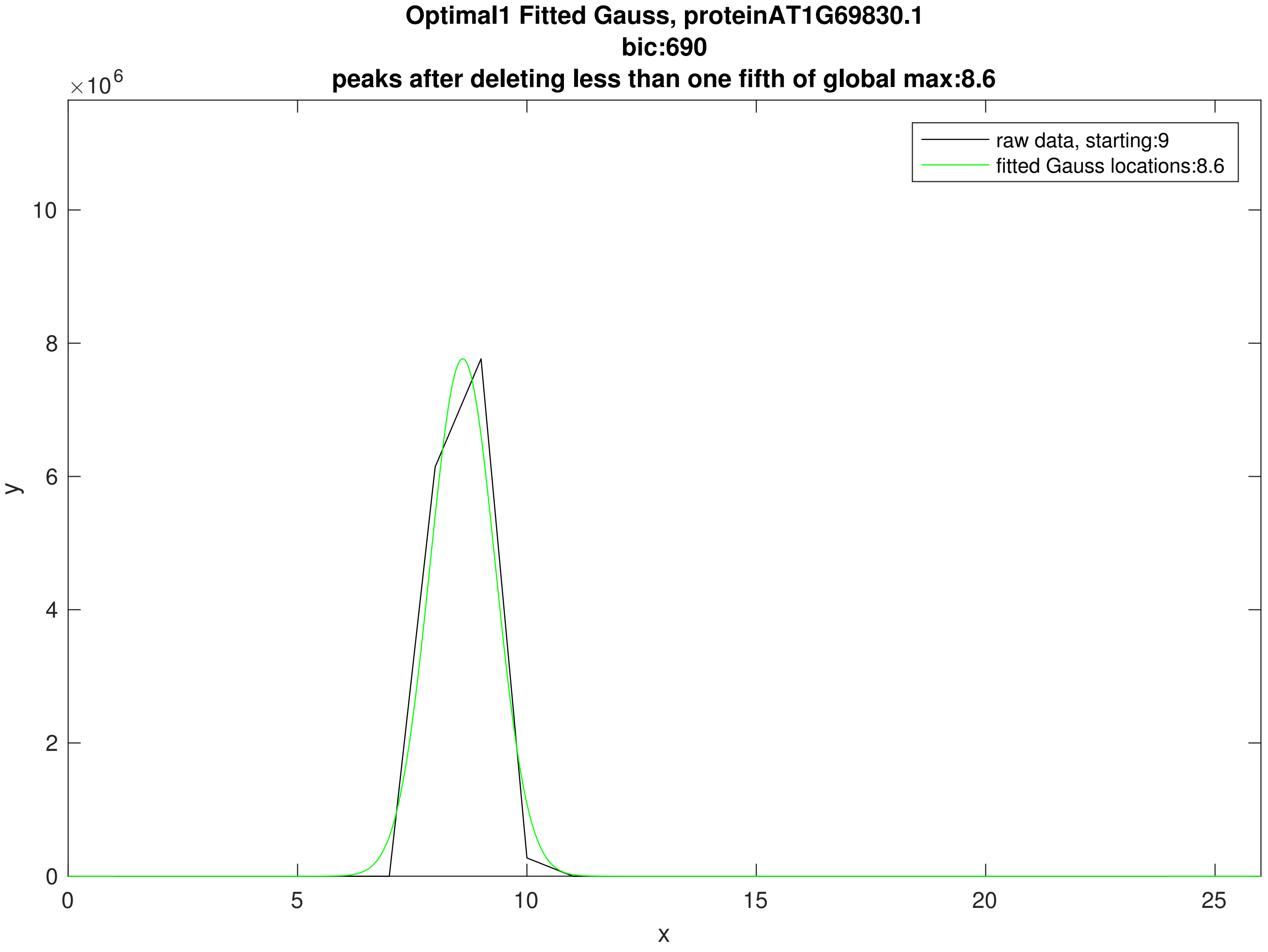
Optimal1 Fitted Gauss, proteinAT1G69830.1
bic:690
peaks after deleting less than one fifth of global max:8.6
×10^6
raw data, starting:9
fitted Gauss locations:8.6
10
8
6
4
2
0
y
0
5
10
15
20
25
x

#### Optimal1 Fitted Gauss, proteinAT1G69960.1
#### bic:698
#### peaks after deleting less than one fifth of global max:17

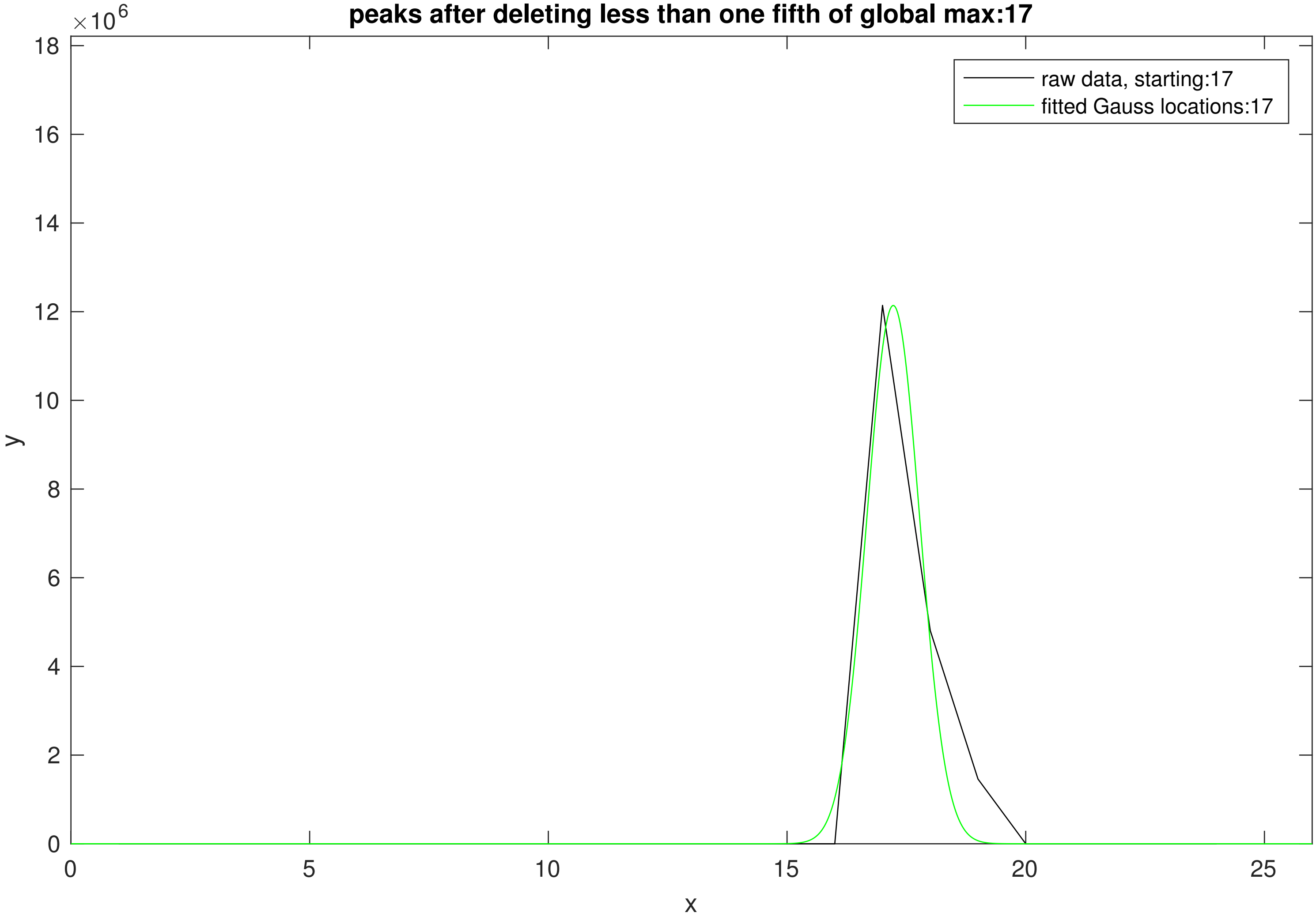

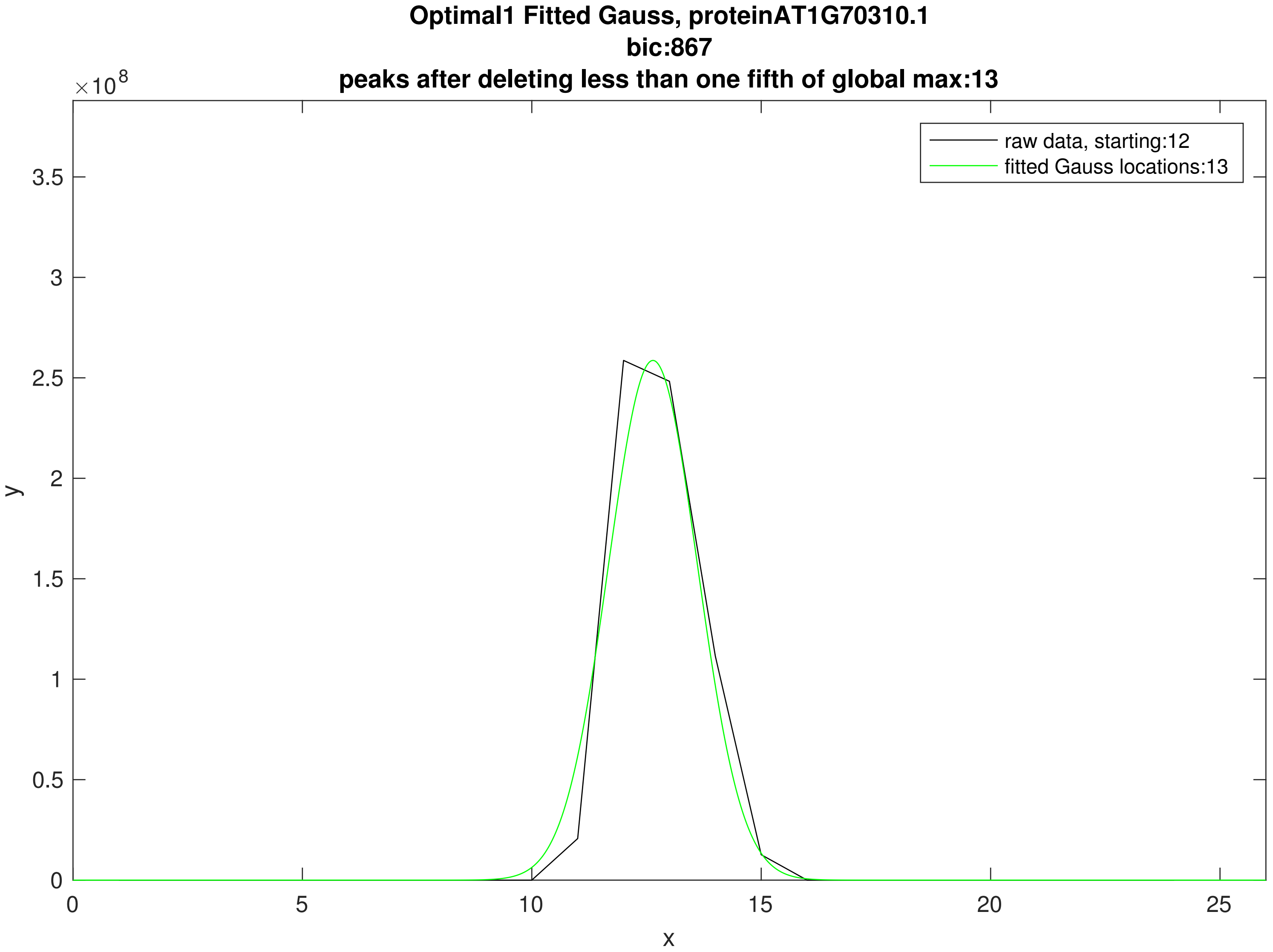

Optimal1 Fitted Gauss, proteinAT1G70310.1
bic:867
peaks after deleting less than one fifth of global max:13
×10^8
3.5
3
2.5
2
1.5
1
0.5
0
y
0
5
10
15
20
25
x
raw data, starting:12
fitted Gauss locations:13

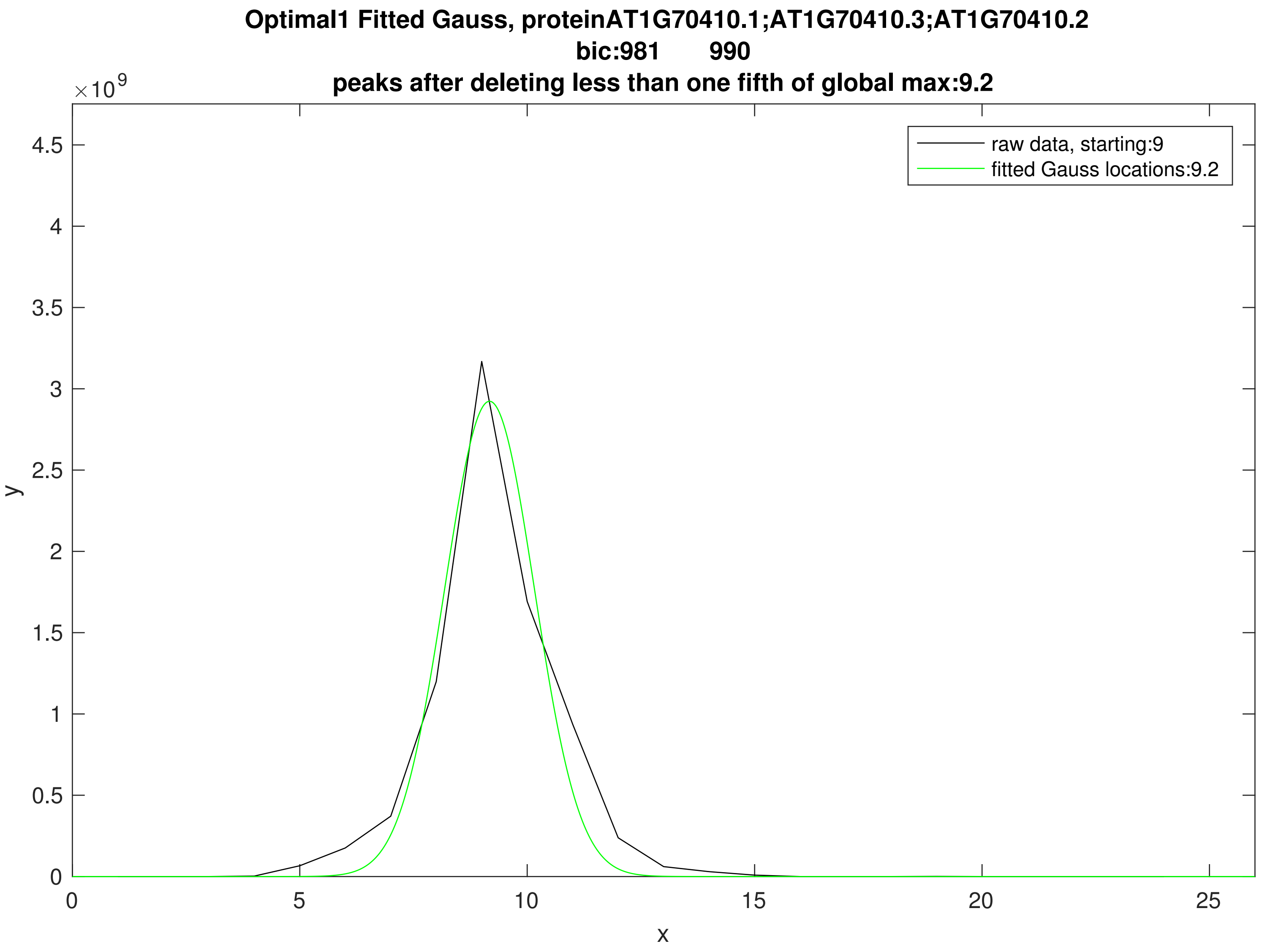
Optimal1 Fitted Gauss, proteinAT1G70410.1;AT1G70410.3;AT1G70410.2
bic:981 990
peaks after deleting less than one fifth of global max:9.2
×10^9
4.5
4
3.5
3
2.5
2
1.5
1
0.5
0
y
0
5
10
15
20
25
x
raw data, starting:9
fitted Gauss locations:9.2

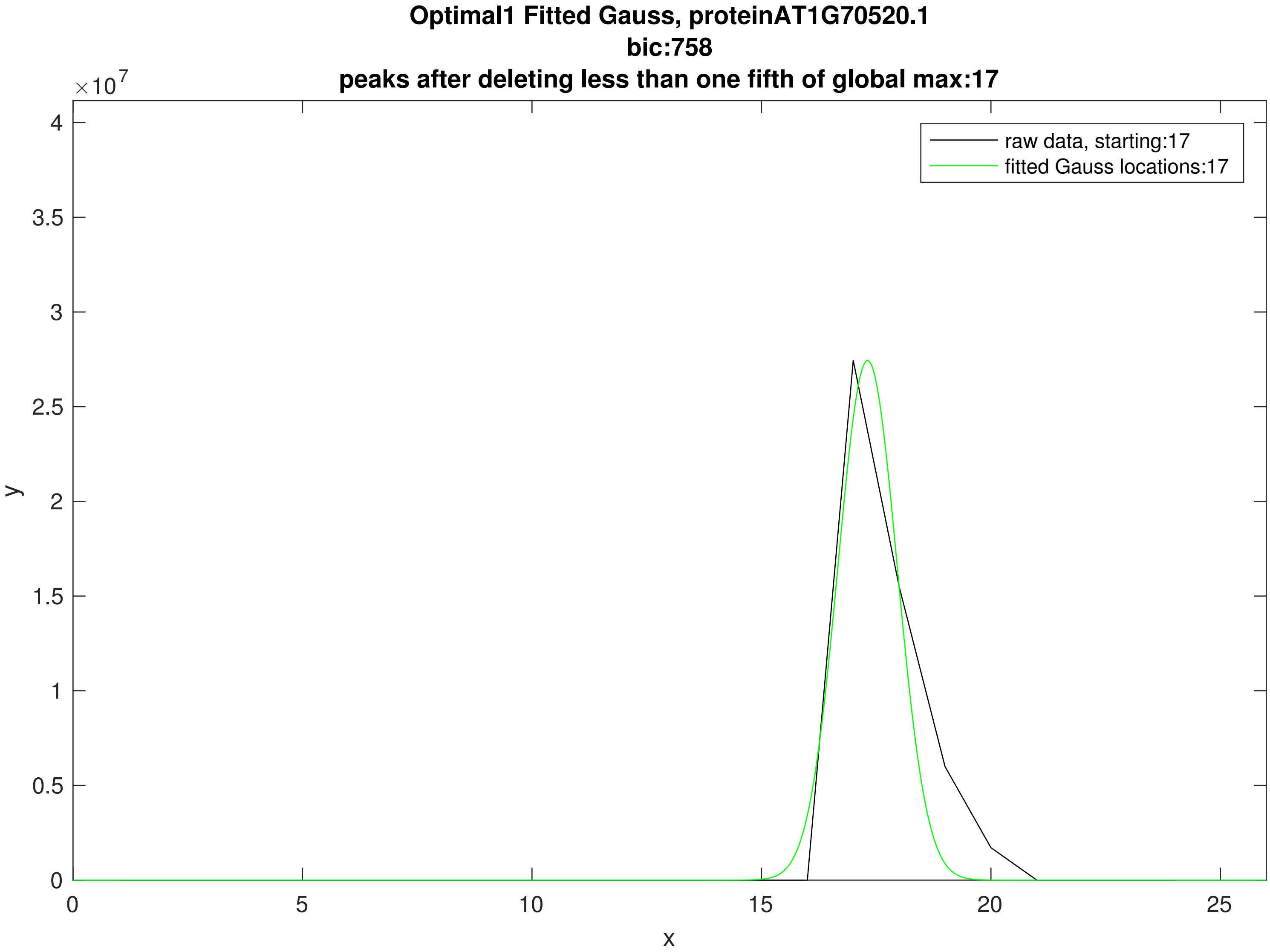
Optimal1 Fitted Gauss, proteinAT1G70520.1
bic:758
peaks after deleting less than one fifth of global max:17
×10^7
4
3.5
3
2.5
2
1.5
1
0.5
0
y
0
5
10
15
20
25
x
raw data, starting:17
fitted Gauss locations:17

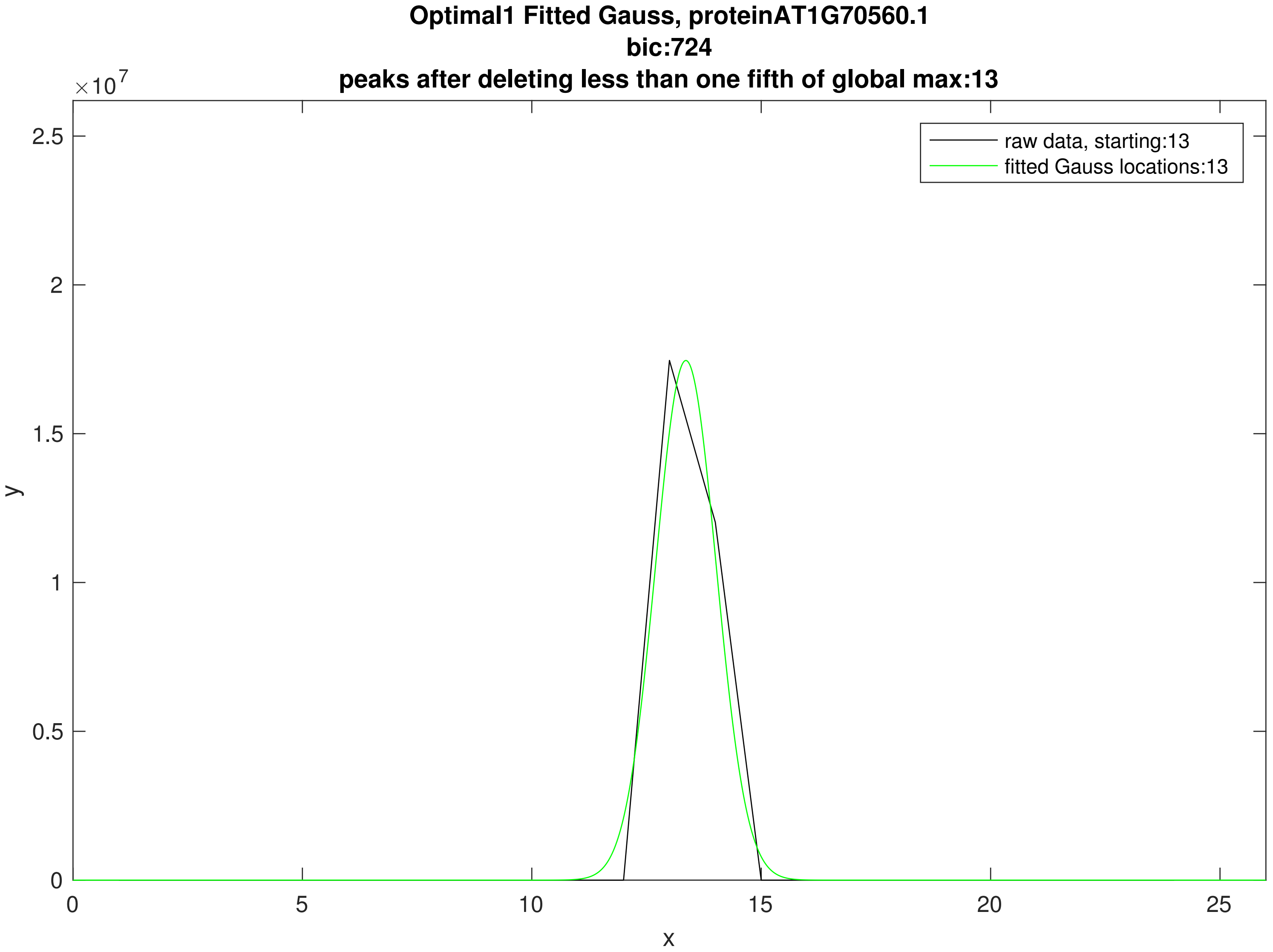
Optimal1 Fitted Gauss, proteinAT1G70560.1
bic:724
peaks after deleting less than one fifth of global max:13
×10^7
2.5
2
1.5
1
0.5
0
y
0
5
10
15
20
25
x
raw data, starting:13
fitted Gauss locations:13

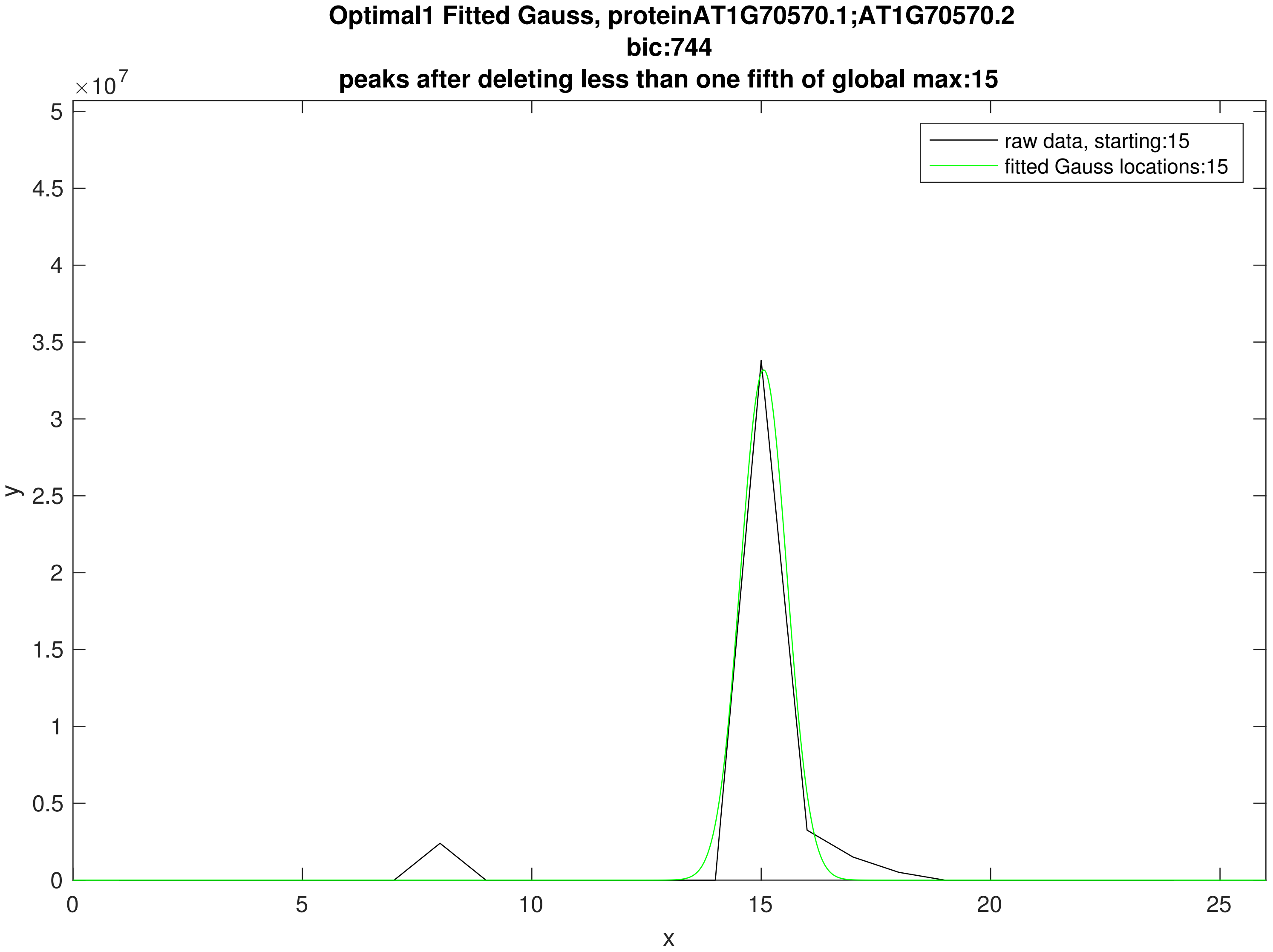

Optimal1 Fitted Gauss, proteinAT1G70570.1;AT1G70570.2
bic:744
peaks after deleting less than one fifth of global max:15
×10^7
raw data, starting:15
fitted Gauss locations:15
y
x
0
0.5
1
1.5
2
2.5
3
3.5
4
4.5
5
0
5
10
15
20
25

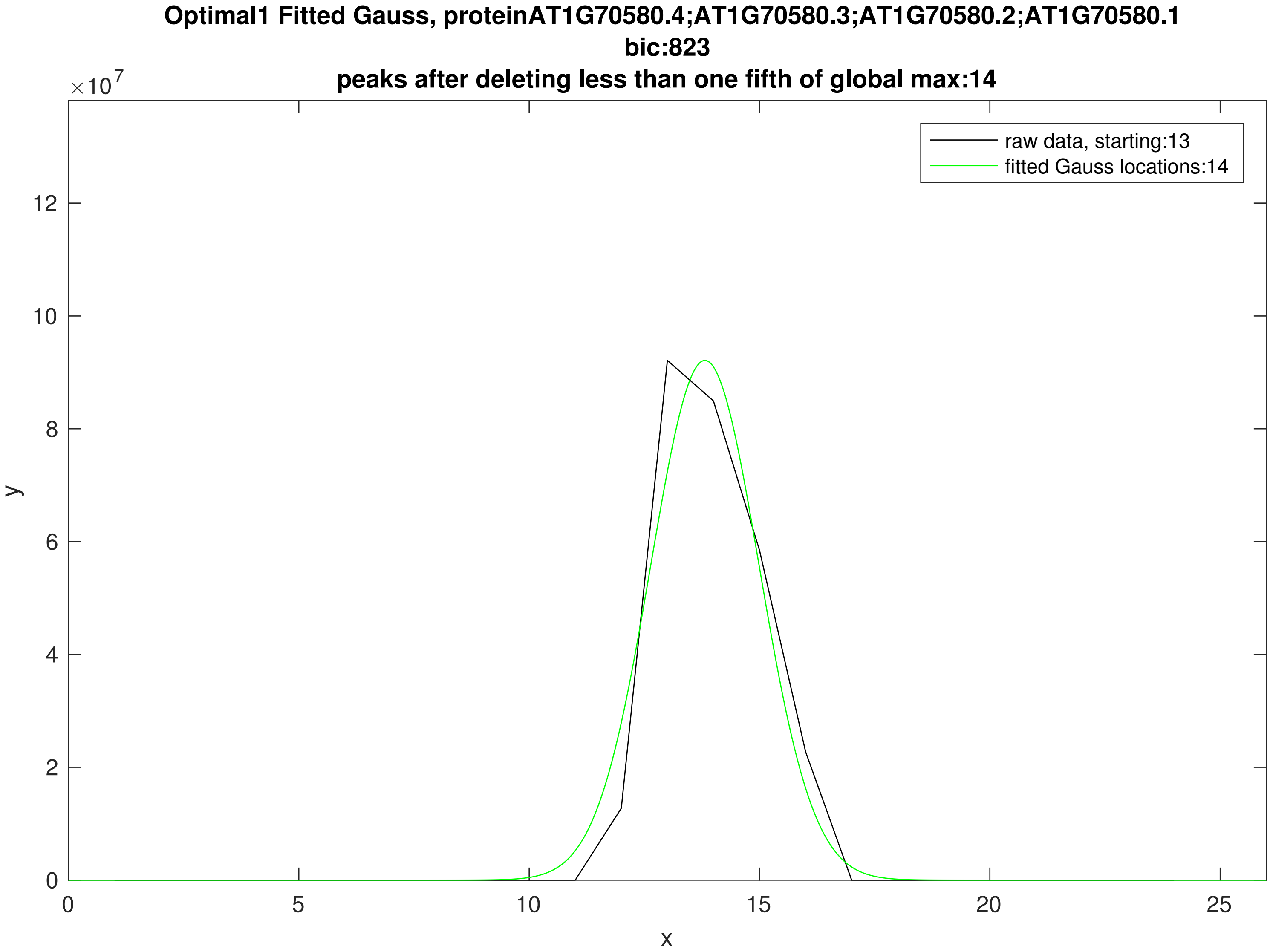
Optimal1 Fitted Gauss, proteinAT1G70580.4;AT1G70580.3;AT1G70580.2;AT1G70580.1
bic:823
peaks after deleting less than one fifth of global max:14
×10^7
raw data, starting:13
fitted Gauss locations:14
y
x
0
2
4
6
8
10
12
0
5
10
15
20
25

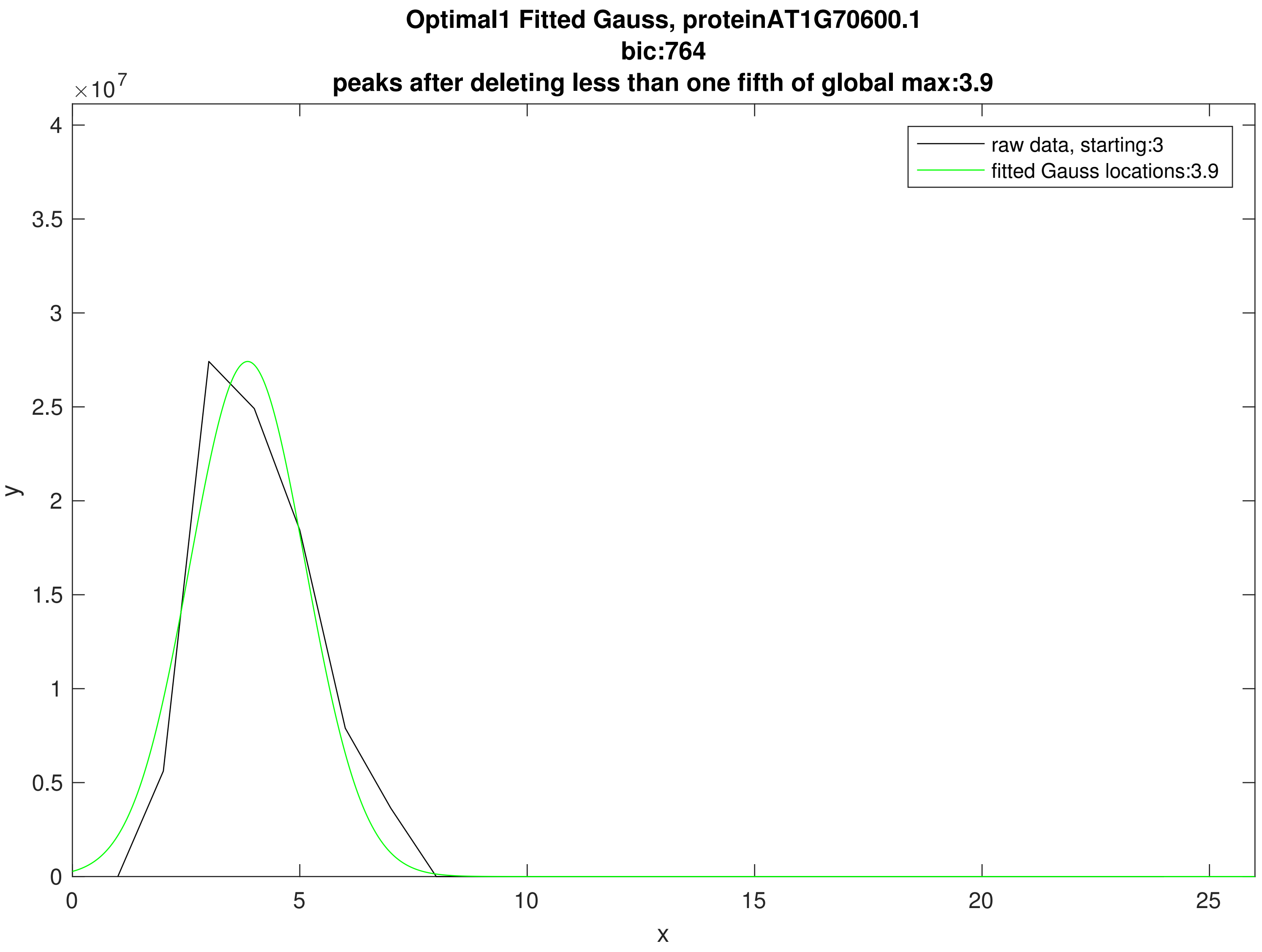
Optimal1 Fitted Gauss, proteinAT1G70600.1
bic:764
peaks after deleting less than one fifth of global max:3.9
×10^7
raw data, starting:3
fitted Gauss locations:3.9
y
x
0
0.5
1
1.5
2
2.5
3
3.5
4
0
5
10
15
20
25

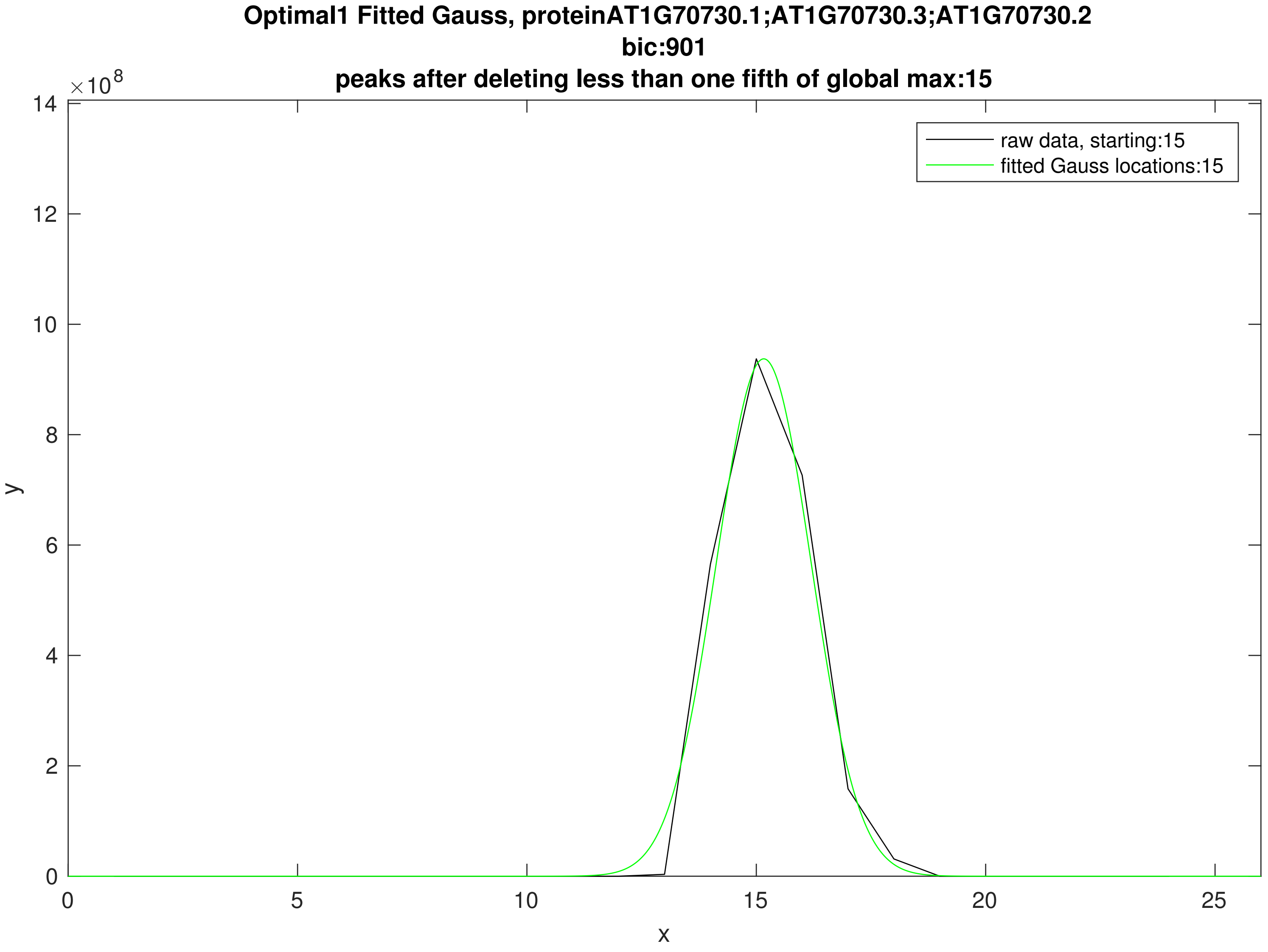

Optimal1 Fitted Gauss, proteinAT1G70730.1;AT1G70730.3;AT1G70730.2
bic:901
peaks after deleting less than one fifth of global max:15
×10^8
14
12
10
8
6
4
2
0
y
0
5
10
15
20
25
x
raw data, starting:15
fitted Gauss locations:15

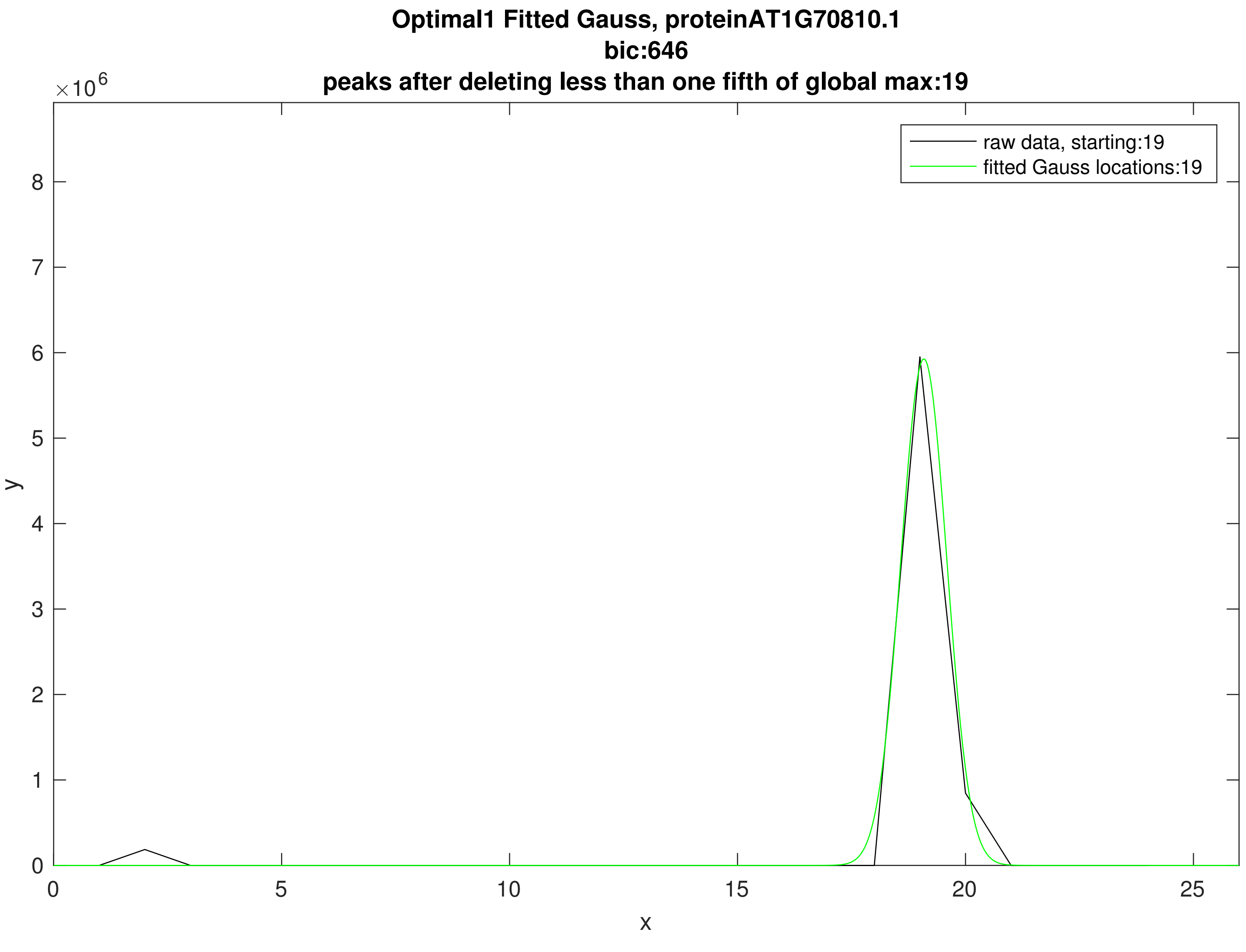
Optimal1 Fitted Gauss, proteinAT1G70810.1
bic:646
peaks after deleting less than one fifth of global max:19
×10^6
raw data, starting:19
fitted Gauss locations:19
y
x
0
1
2
3
4
5
6
7
8
0
5
10
15
20
25

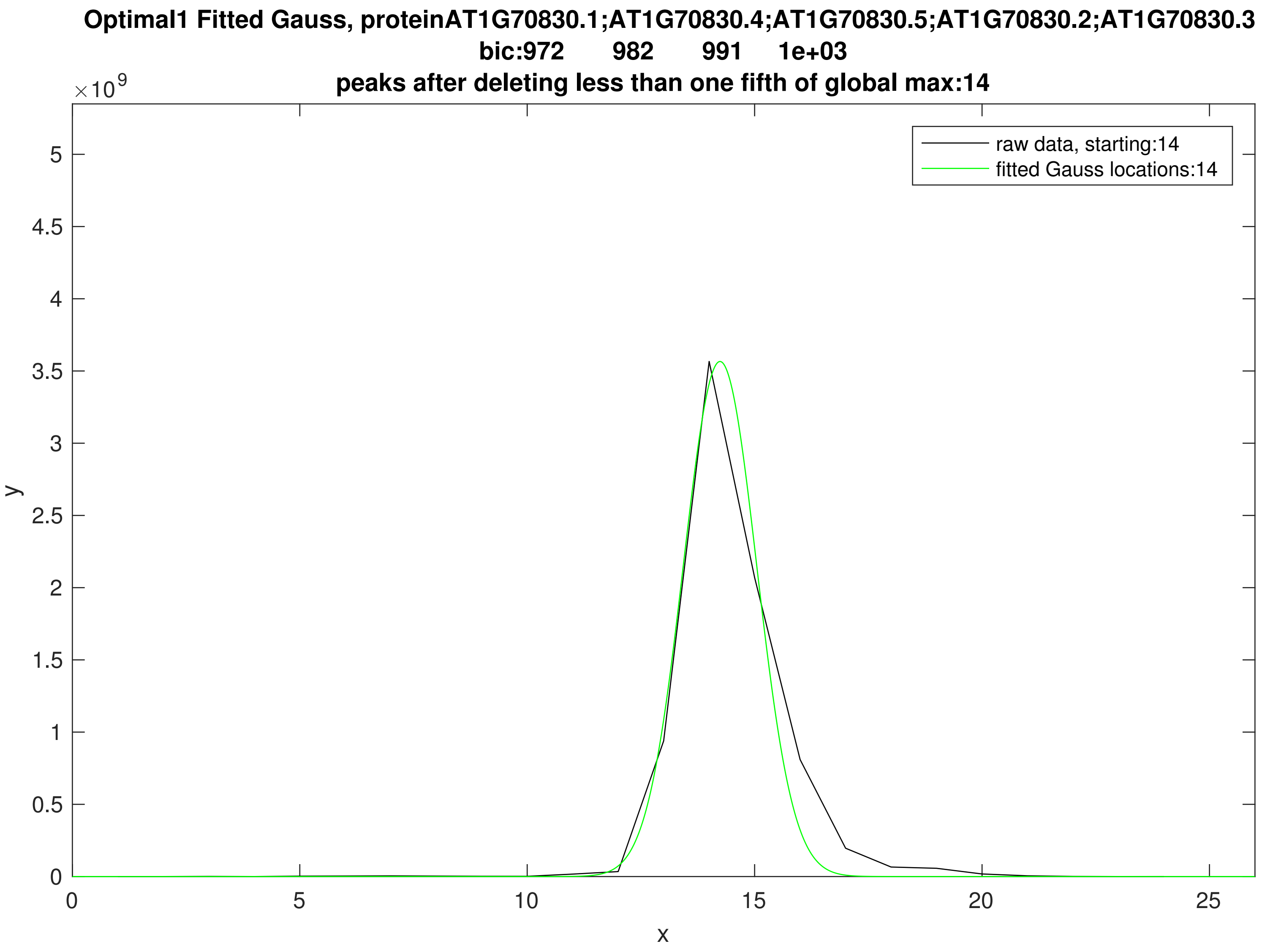

Optimal1 Fitted Gauss, proteinAT1G70830.1;AT1G70830.4;AT1G70830.5;AT1G70830.2;AT1G70830.3
bic:972 982 991 1e+03
peaks after deleting less than one fifth of global max:14
×10^9
raw data, starting:14
fitted Gauss locations:14
y
x
0
0.5
1
1.5
2
2.5
3
3.5
4
4.5
5
0
5
10
15
20
25

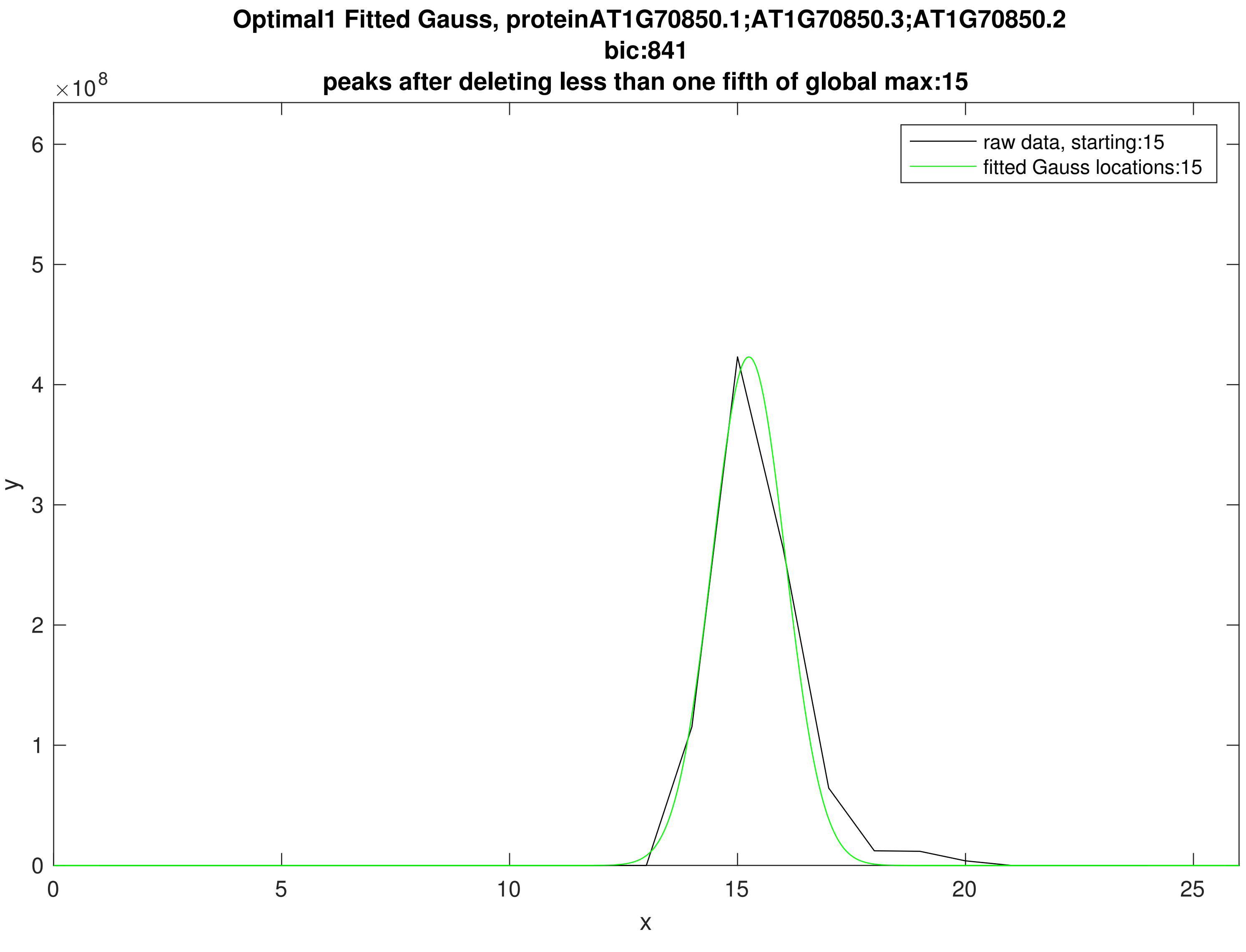
Optimal1 Fitted Gauss, proteinAT1G70850.1;AT1G70850.3;AT1G70850.2
bic:841
peaks after deleting less than one fifth of global max:15
×10^8
raw data, starting:15
fitted Gauss locations:15
y
x
0
1
2
3
4
5
6
0
5
10
15
20
25

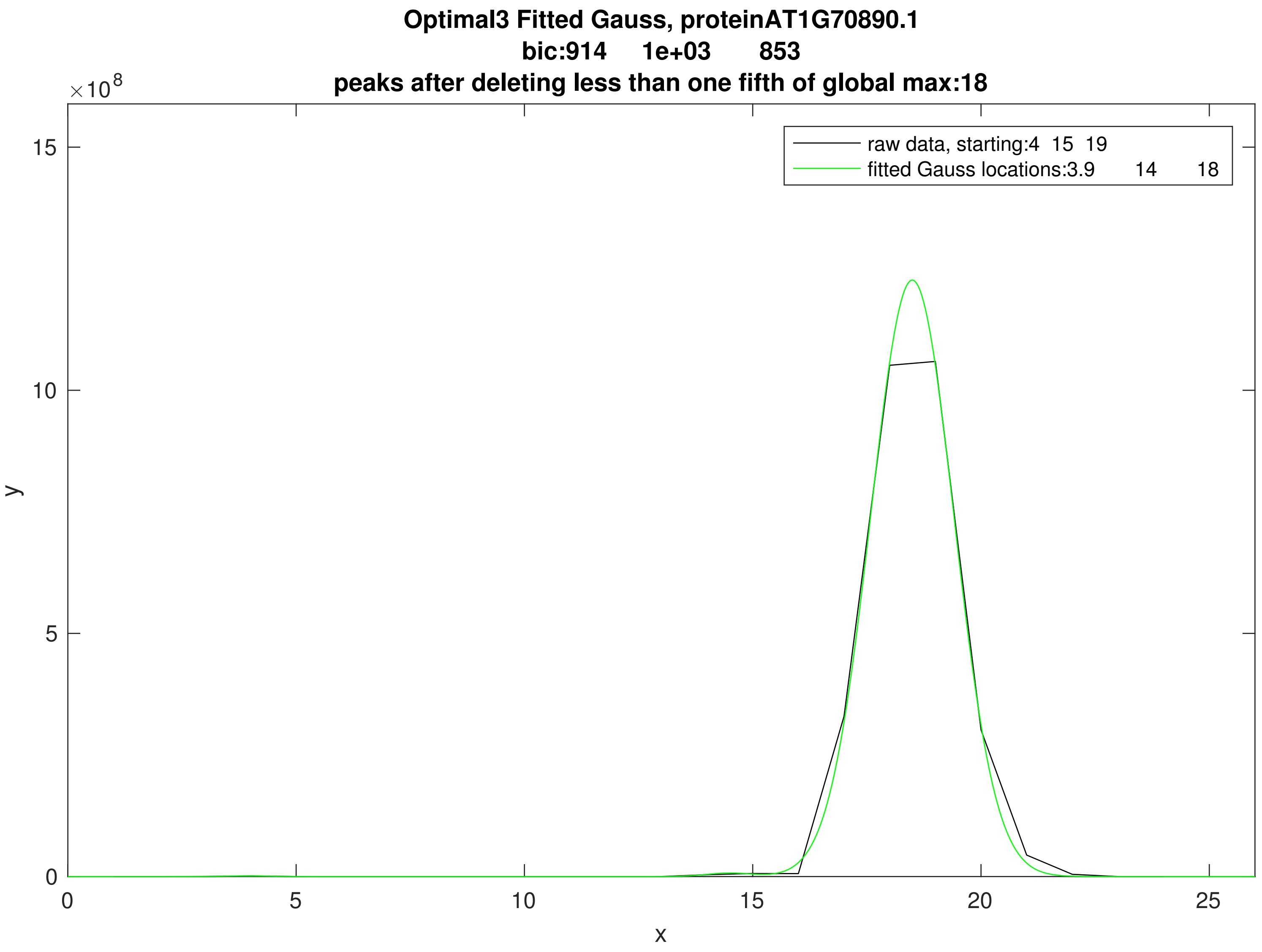

Optimal3 Fitted Gauss, proteinAT1G70890.1
bic:914 1e+03 853
peaks after deleting less than one fifth of global max:18
×10^8
15
10
5
0
0
5
10
15
20
25
y
x
raw data, starting:4 15 19
fitted Gauss locations:3.9 14 18

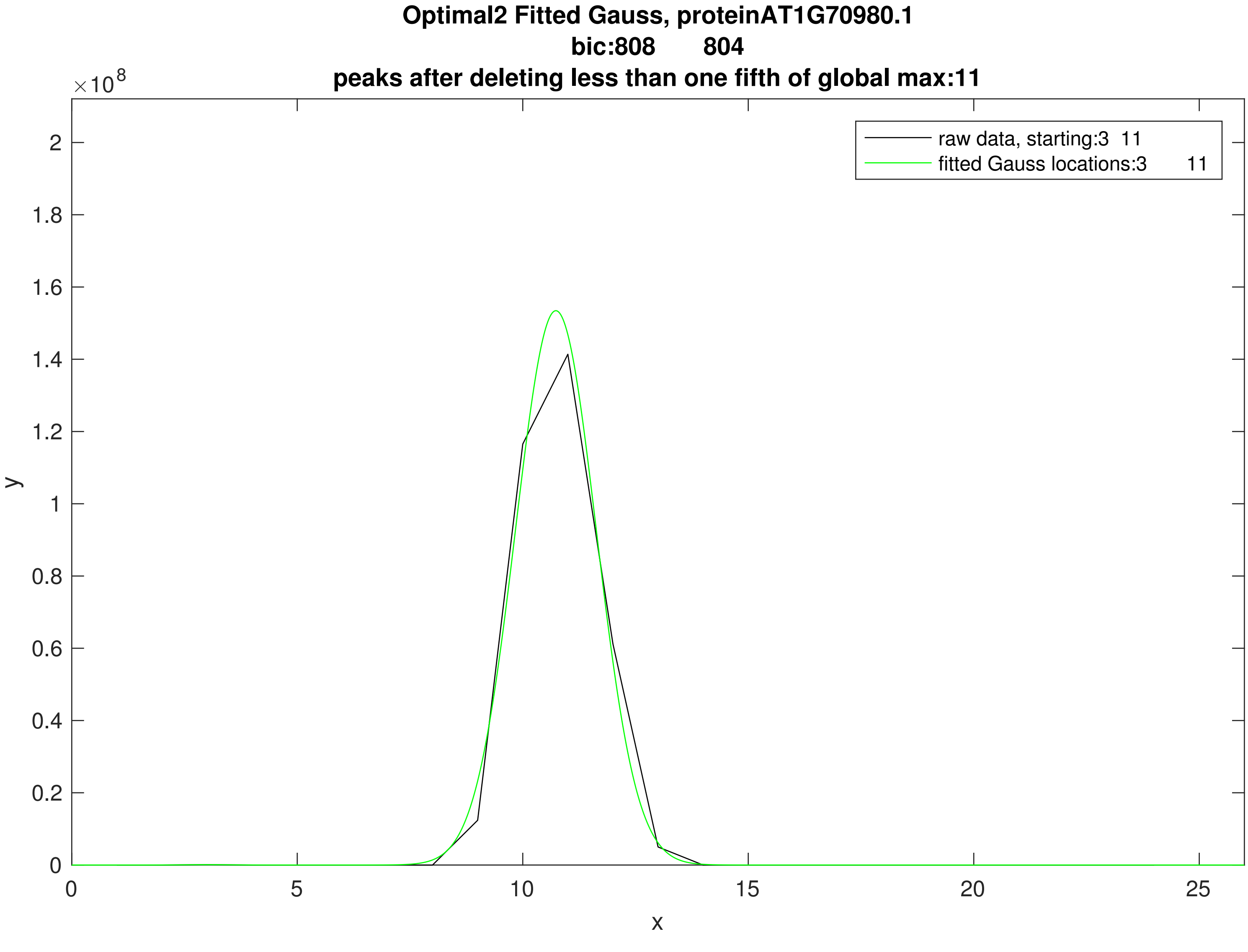
Optimal2 Fitted Gauss, proteinAT1G70980.1
bic:808 804
peaks after deleting less than one fifth of global max:11
×10^8
raw data, starting:3 11
fitted Gauss locations:3 11
y
x
0
0.2
0.4
0.6
0.8
1
1.2
1.4
1.6
1.8
2
0
5
10
15
20
25

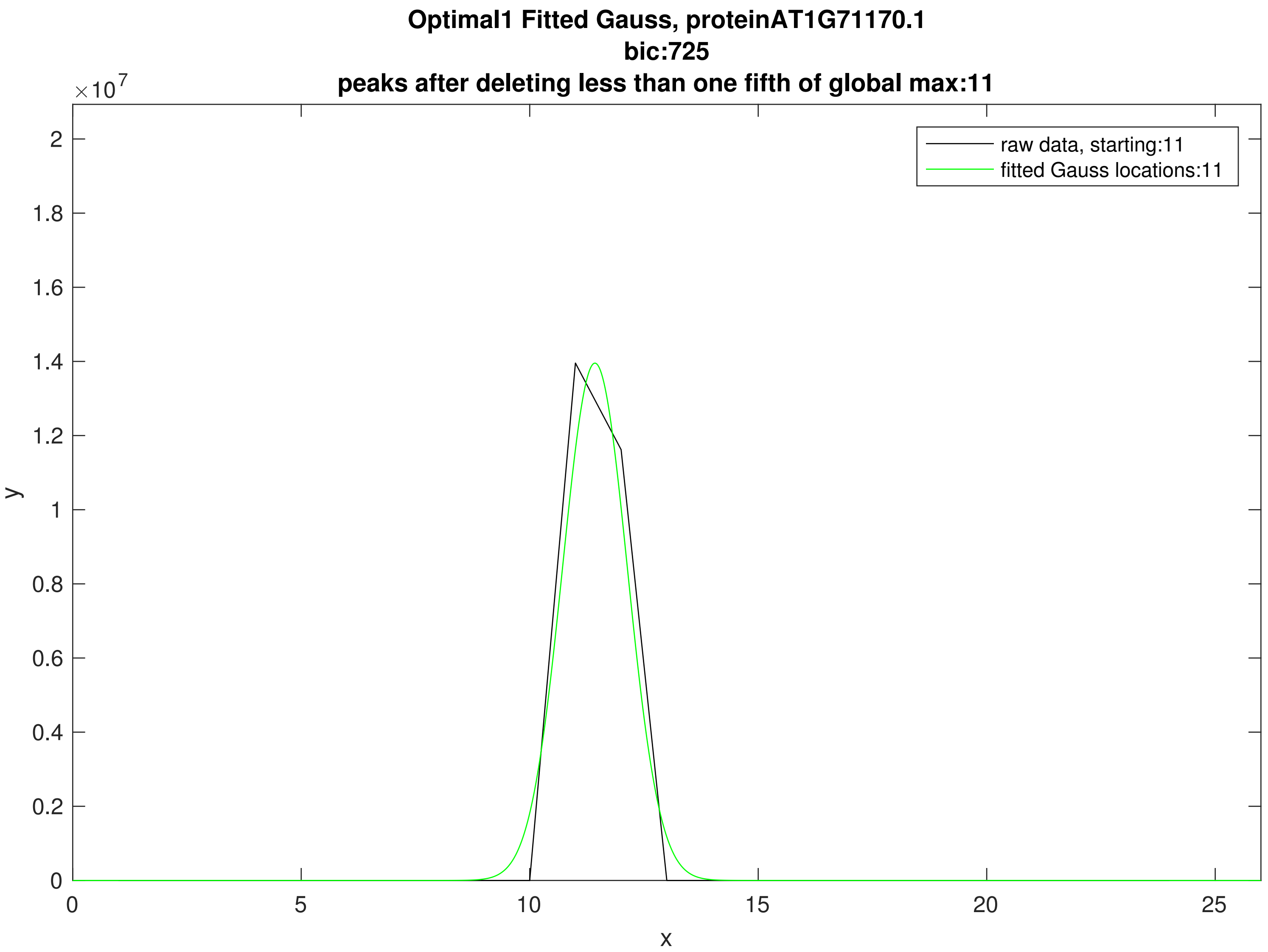
Optimal1 Fitted Gauss, proteinAT1G71170.1
bic:725
peaks after deleting less than one fifth of global max:11
×10^7
2
1.8
1.6
1.4
1.2
1
0.8
0.6
0.4
0.2
0
y
0
5
10
15
20
25
x
raw data, starting:11
fitted Gauss locations:11

Optimal1 Fitted Gauss, proteinAT1G71310.1;AT1G71310.2;AT1G71310.3
bic:698
peaks after deleting less than one fifth of global max:11

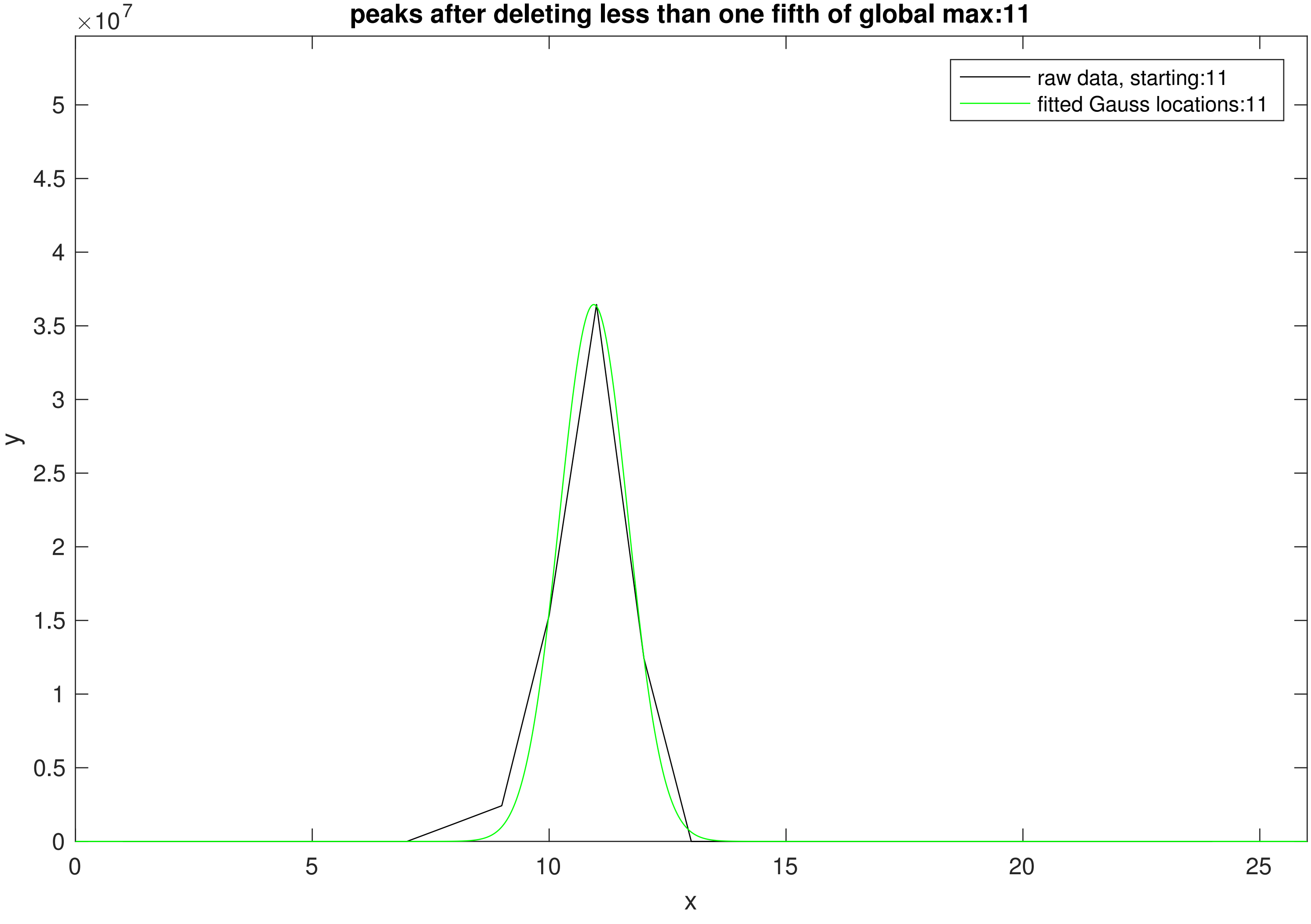

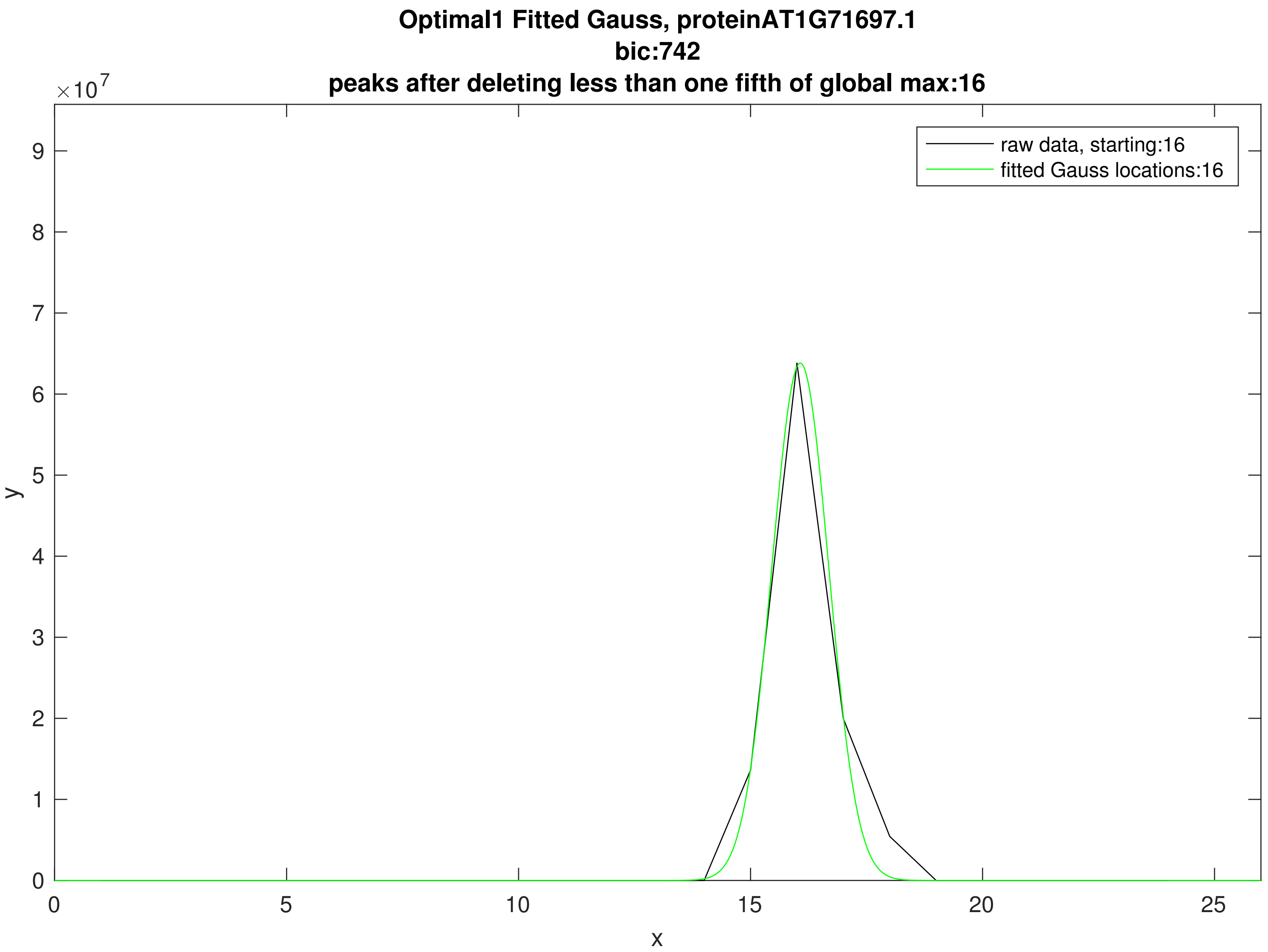
Optimal1 Fitted Gauss, proteinAT1G71697.1
bic:742
peaks after deleting less than one fifth of global max:16
×10^7
raw data, starting:16
fitted Gauss locations:16
y
x
0
1
2
3
4
5
6
7
8
9
0
5
10
15
20
25

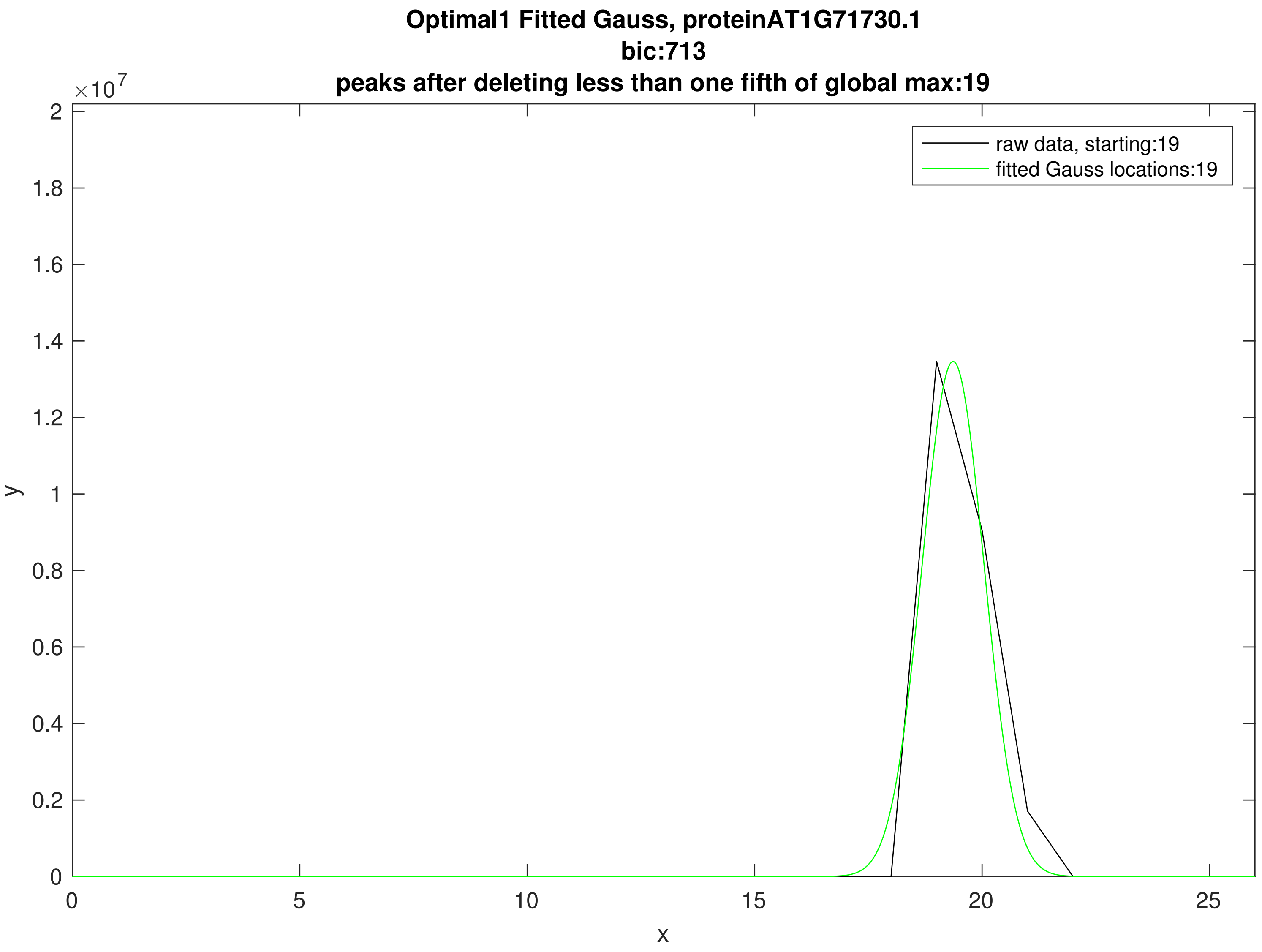

Optimal1 Fitted Gauss, proteinAT1G71730.1
bic:713
peaks after deleting less than one fifth of global max:19
×10^7
2
1.8
1.6
1.4
1.2
1
0.8
0.6
0.4
0.2
0
y
0
5
10
15
20
25
x
raw data, starting:19
fitted Gauss locations:19

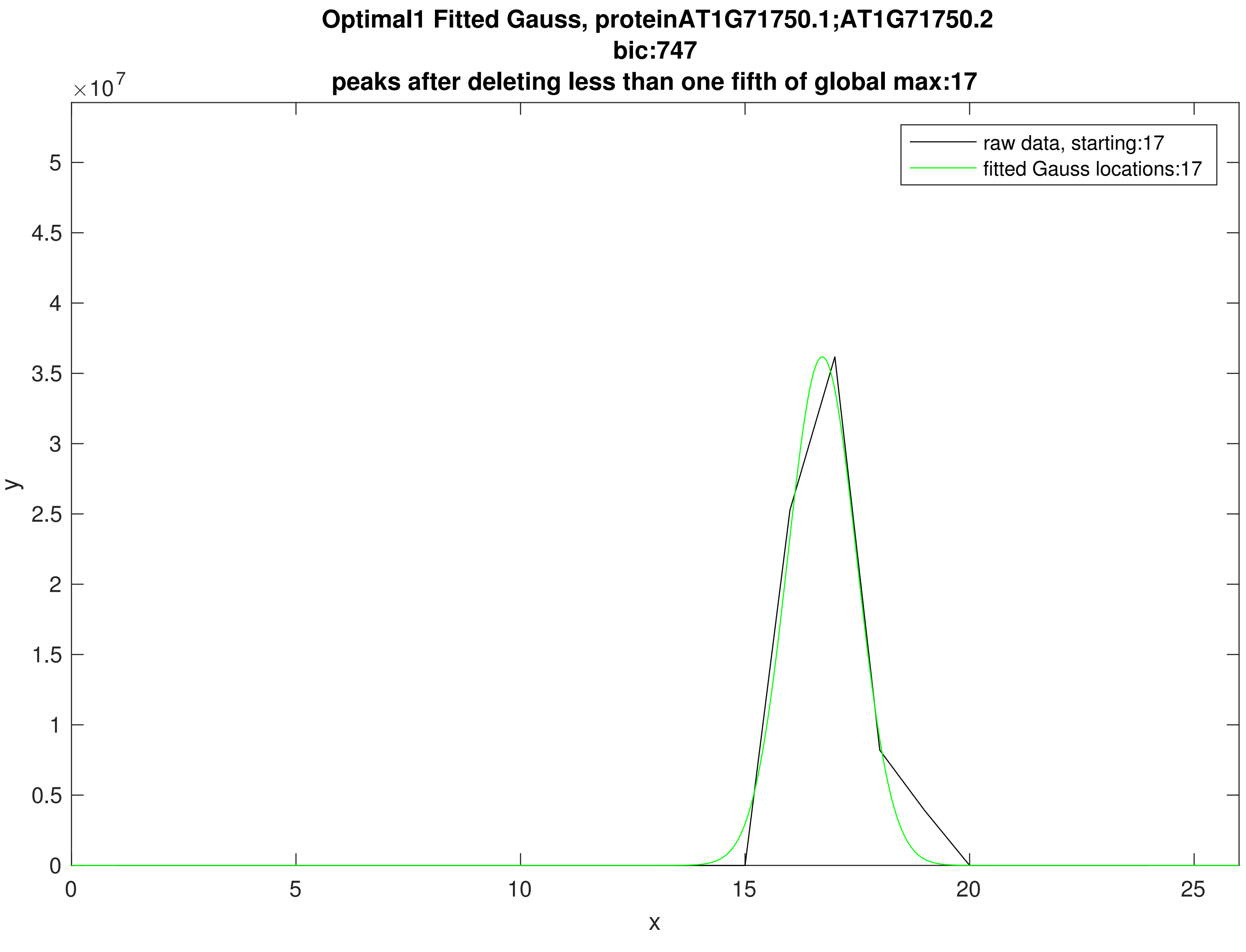
Optimal1 Fitted Gauss, proteinAT1G71750.1;AT1G71750.2
bic:747
peaks after deleting less than one fifth of global max:17
×10^7
5
4.5
4
3.5
3
2.5
2
1.5
1
0.5
0
y
0
5
10
15
20
25
x
raw data, starting:17
fitted Gauss locations:17

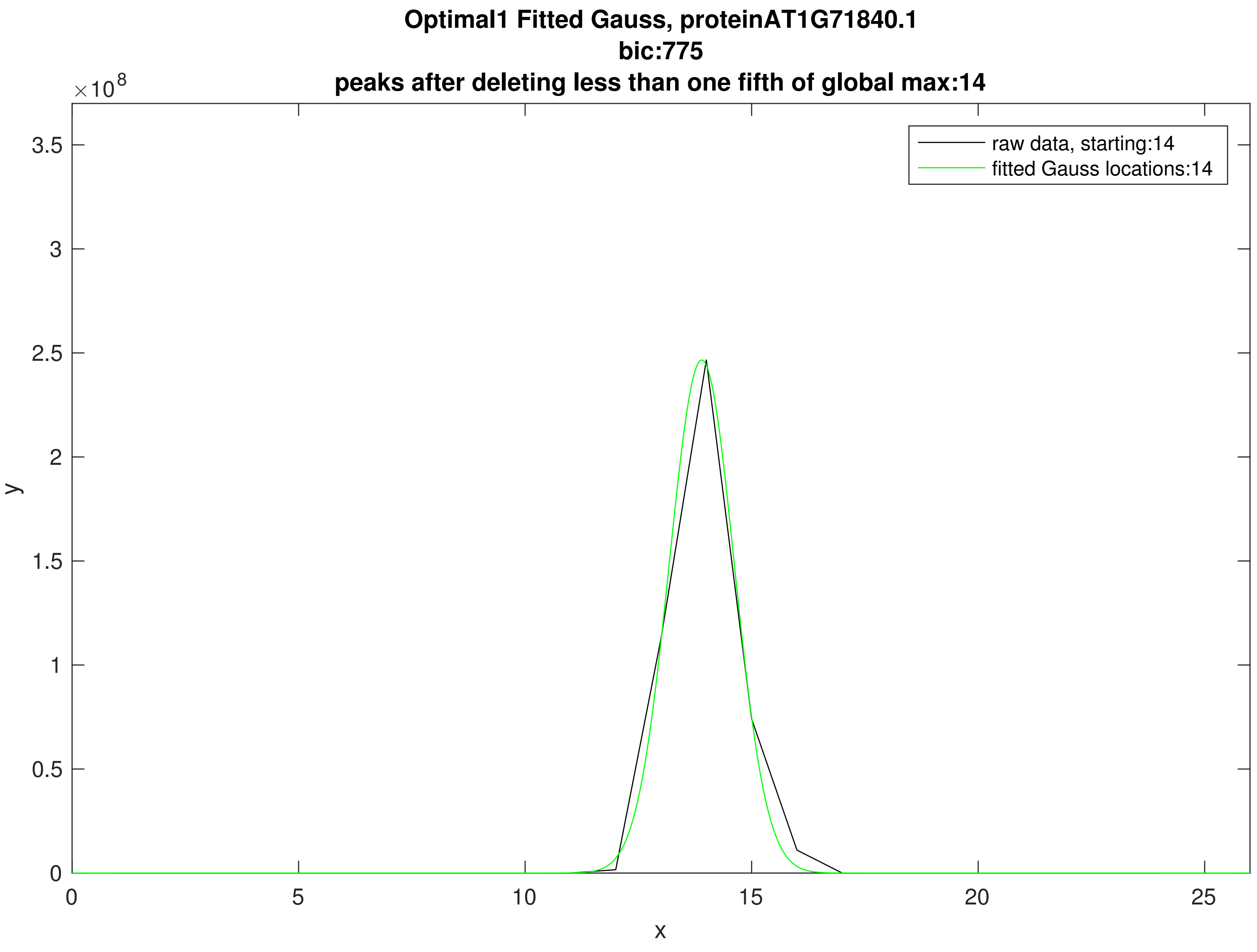
Optimal1 Fitted Gauss, proteinAT1G71840.1
bic:775
peaks after deleting less than one fifth of global max:14
×10^8
raw data, starting:14
fitted Gauss locations:14
y
x
0
0.5
1
1.5
2
2.5
3
3.5
0
5
10
15
20
25

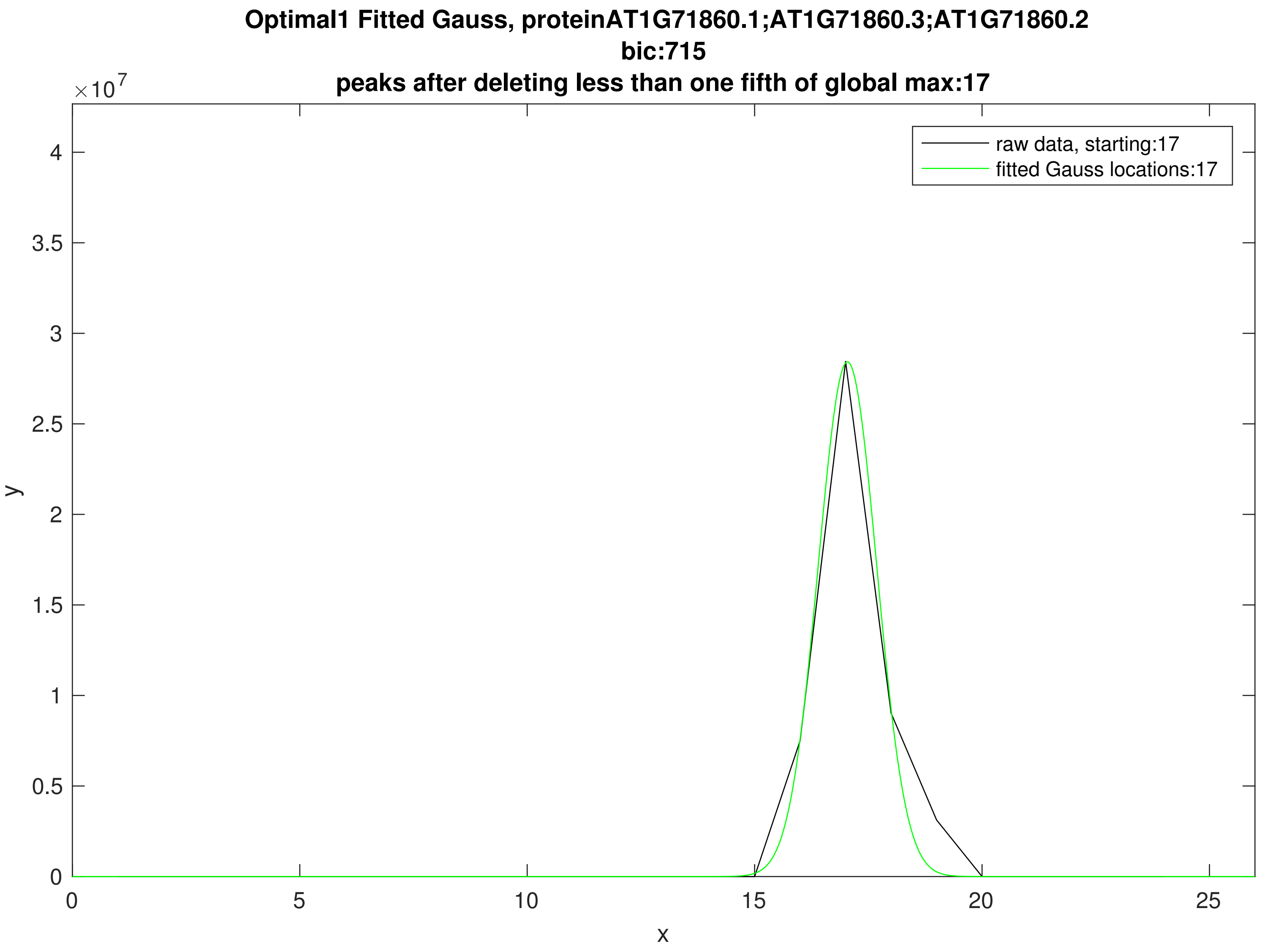

Optimal1 Fitted Gauss, proteinAT1G71860.1;AT1G71860.3;AT1G71860.2
bic:715
peaks after deleting less than one fifth of global max:17
×10^7
raw data, starting:17
fitted Gauss locations:17
y
x
0
0.5
1
1.5
2
2.5
3
3.5
4
5
10
15
20
25

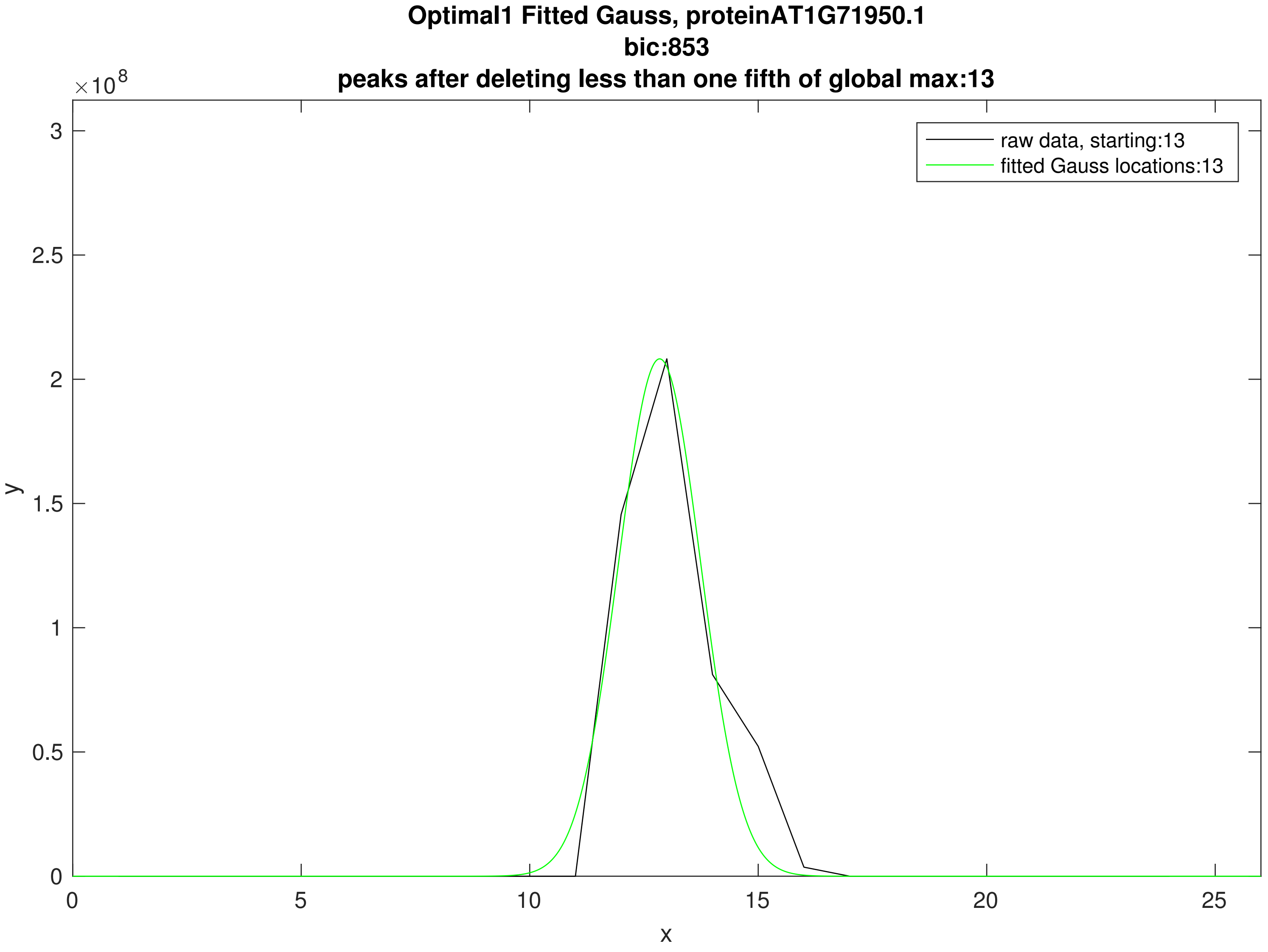

Optimal1 Fitted Gauss, proteinAT1G71950.1
bic:853
peaks after deleting less than one fifth of global max:13
×10^8
3
2.5
2
1.5
1
0.5
0
y
0
5
10
15
20
25
x
raw data, starting:13
fitted Gauss locations:13

**Optimal1 Fitted Gauss, proteinAT1G72150.1**
**bic:818 825 836 846**
**peaks after deleting less than one fifth of global max:9.8**

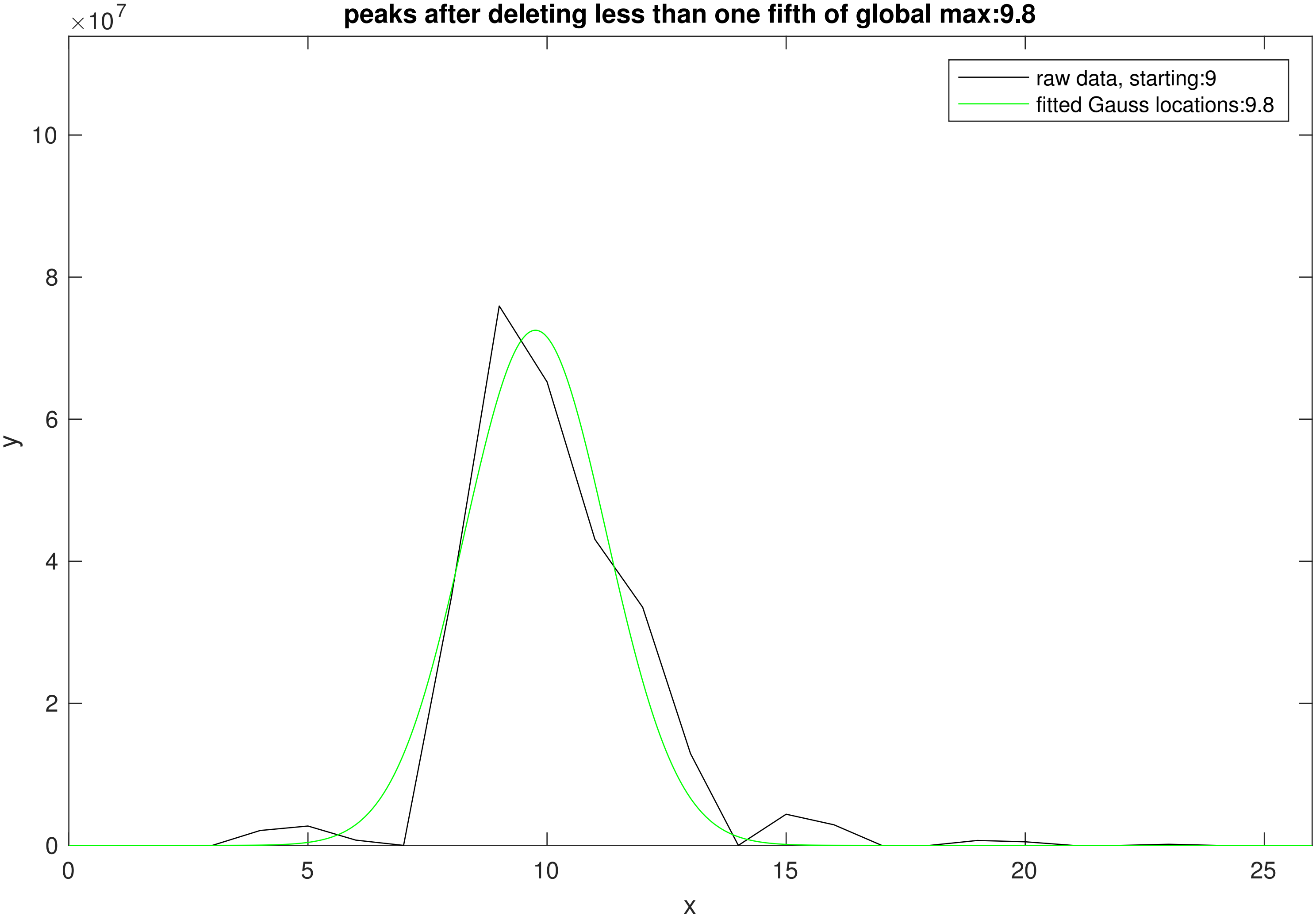

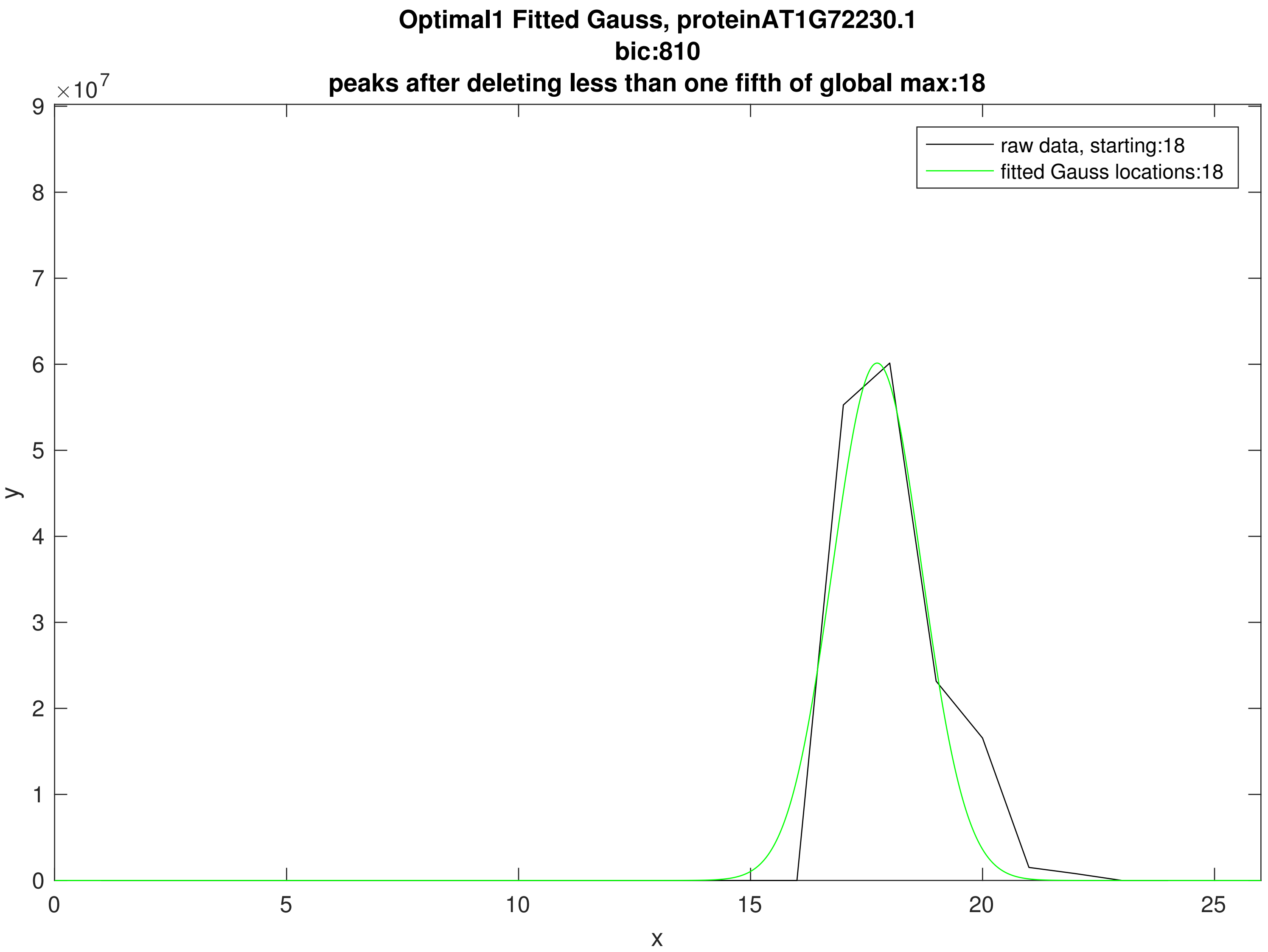
Optimal1 Fitted Gauss, proteinAT1G72230.1
bic:810
peaks after deleting less than one fifth of global max:18
×10^7
raw data, starting:18
fitted Gauss locations:18
y
x
0
5
10
15
20
25
0
1
2
3
4
5
6
7
8
9

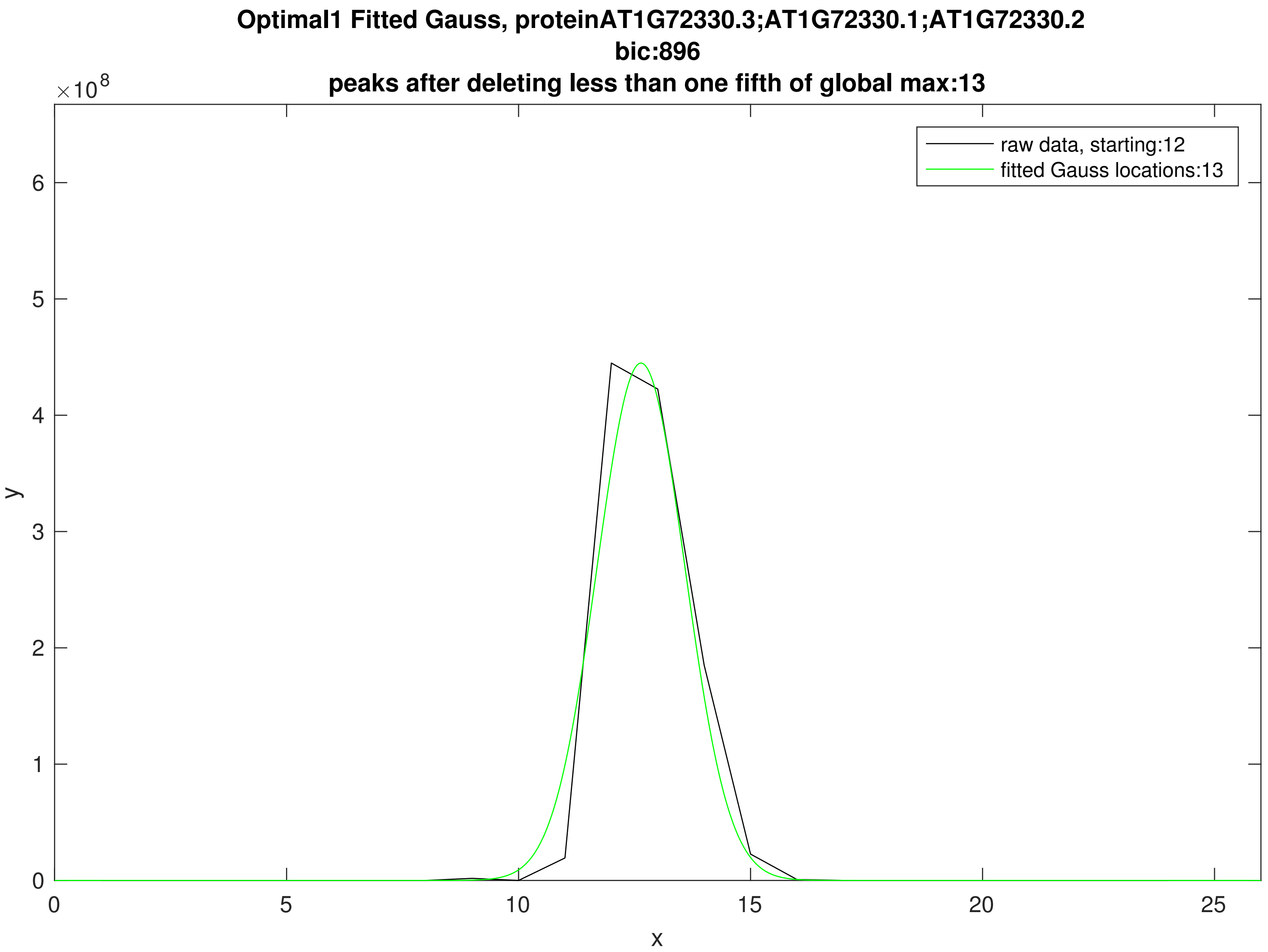

Optimal1 Fitted Gauss, proteinAT1G72330.3;AT1G72330.1;AT1G72330.2
bic:896
peaks after deleting less than one fifth of global max:13
×10^8
raw data, starting:12
fitted Gauss locations:13
y
x
0
1
2
3
4
5
6
0
5
10
15
20
25

Optimal2 Fitted Gauss, proteinAT1G72370.2;AT1G72370.1
bic:857 852 859 868
peaks after deleting less than one fifth of global max:4.2

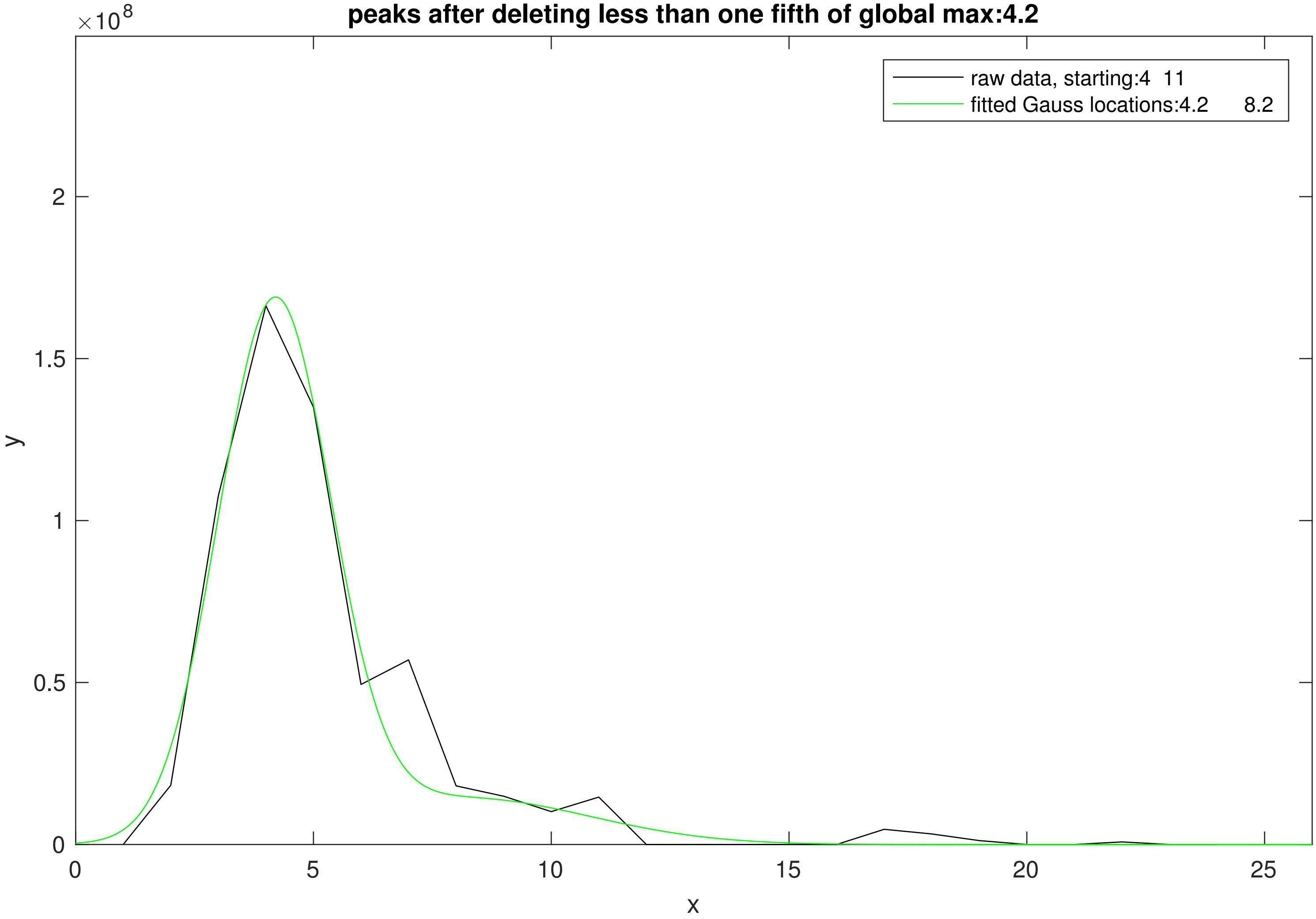

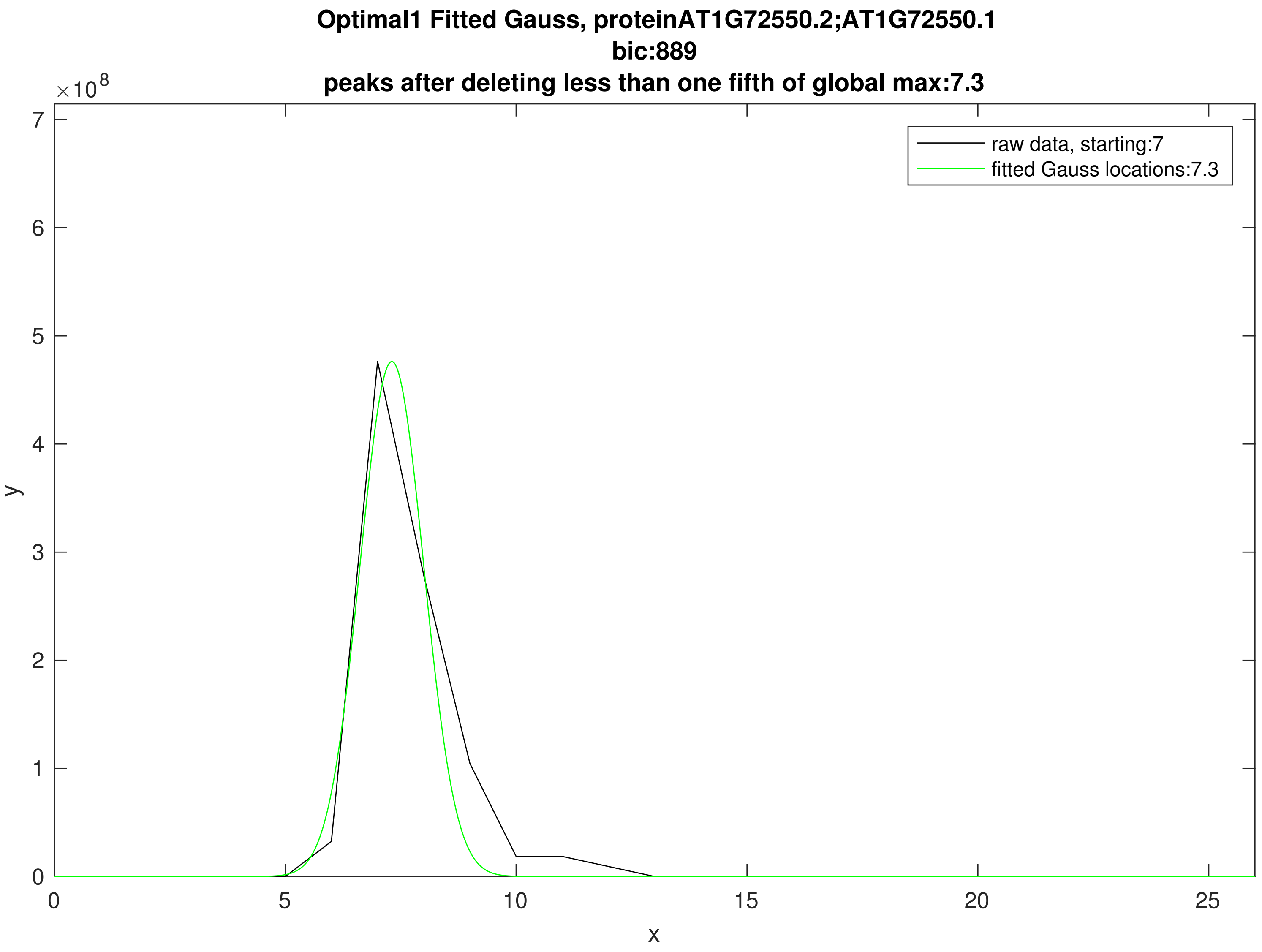
Optimal1 Fitted Gauss, proteinAT1G72550.2;AT1G72550.1
bic:889
peaks after deleting less than one fifth of global max:7.3
×10^8
7
6
5
4
3
2
1
0
y
0
5
10
15
20
25
x
raw data, starting:7
fitted Gauss locations:7.3

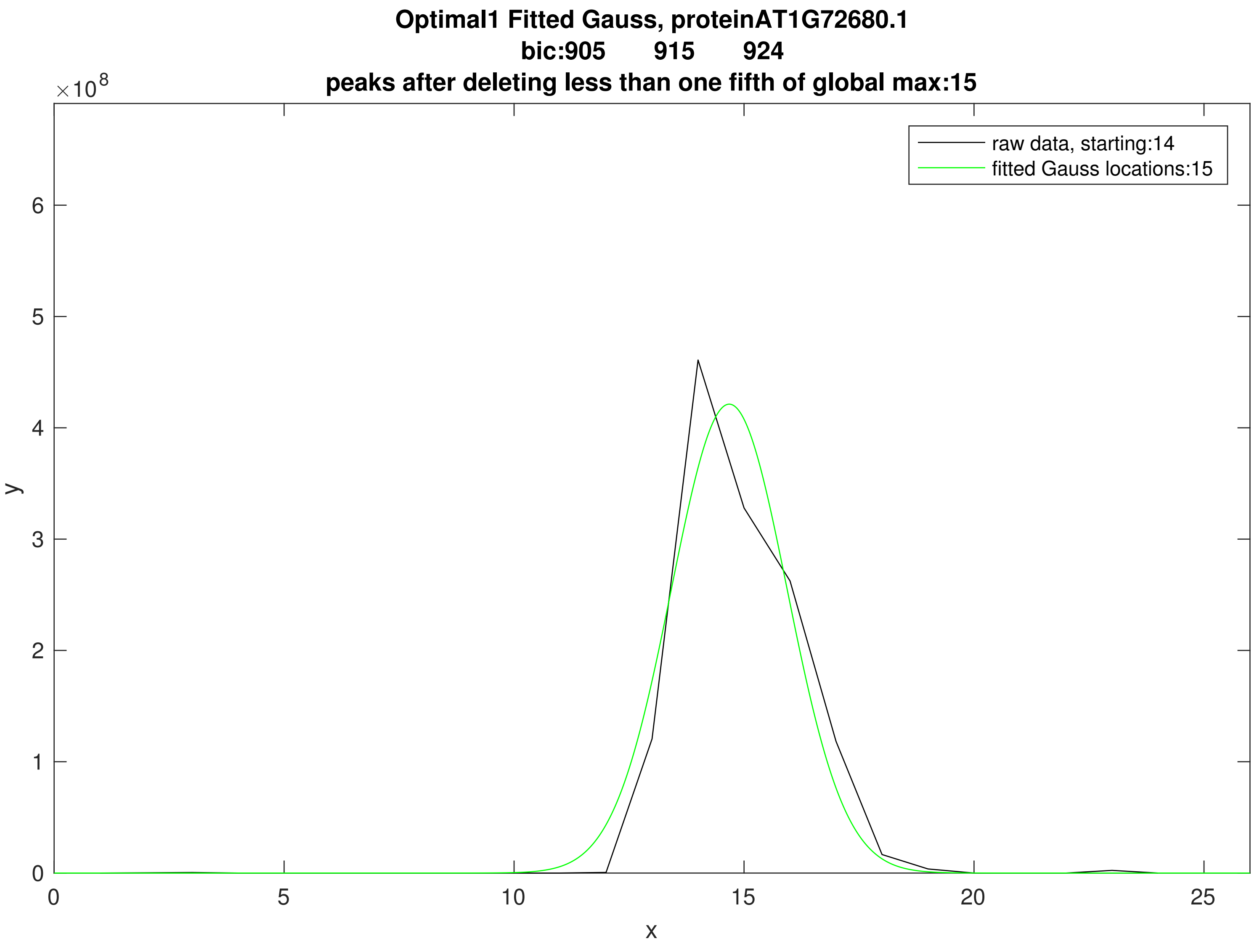
Optimal1 Fitted Gauss, proteinAT1G72680.1
bic:905 915 924
peaks after deleting less than one fifth of global max:15
×10^8
raw data, starting:14
fitted Gauss locations:15
y
x
0
1
2
3
4
5
6
0
5
10
15
20
25

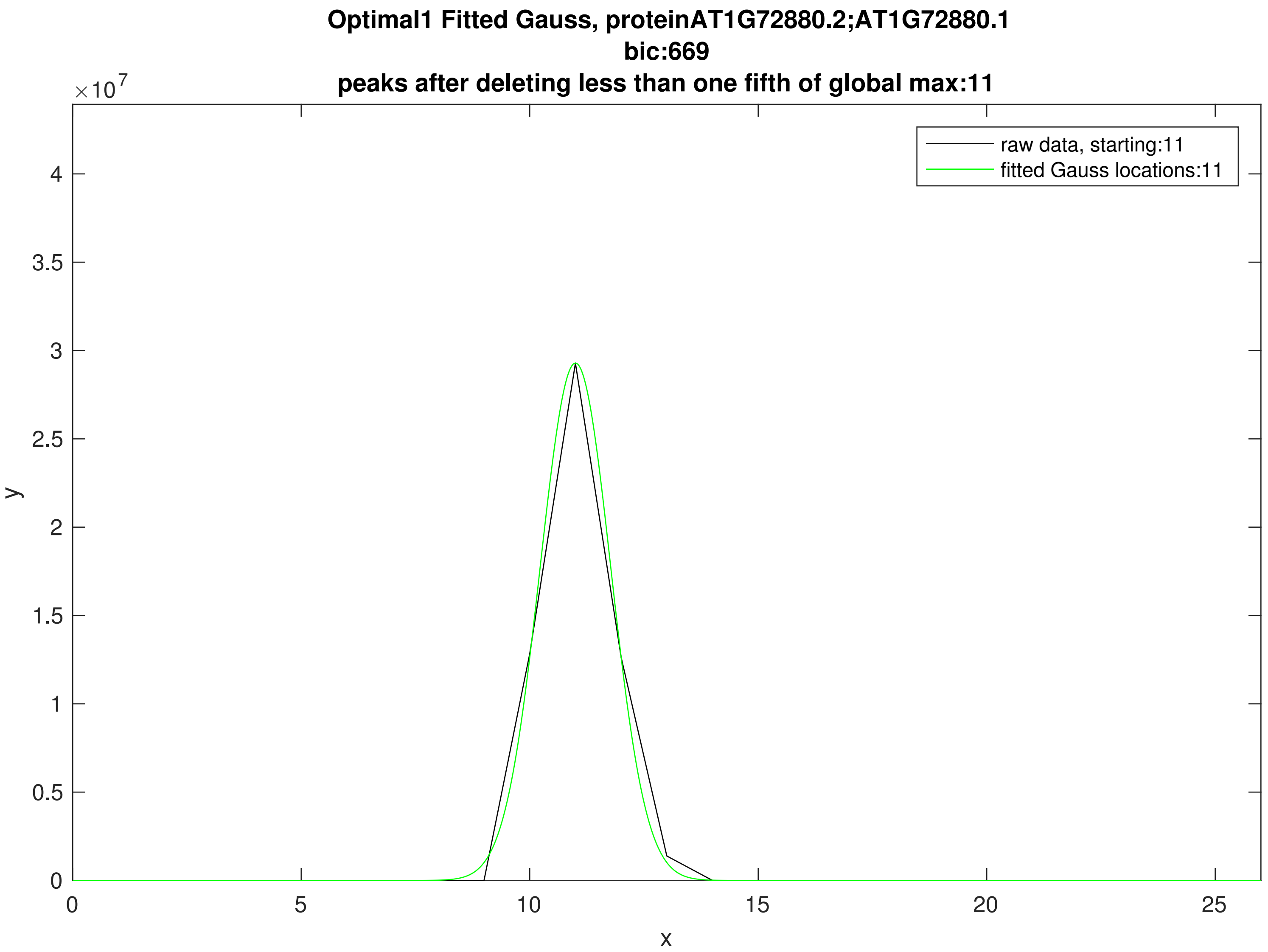
Optimal1 Fitted Gauss, proteinAT1G72880.2;AT1G72880.1
bic:669
peaks after deleting less than one fifth of global max:11
×10^7
raw data, starting:11
fitted Gauss locations:11
y
x
0
0.5
1
1.5
2
2.5
3
3.5
4
5
10
15
20
25

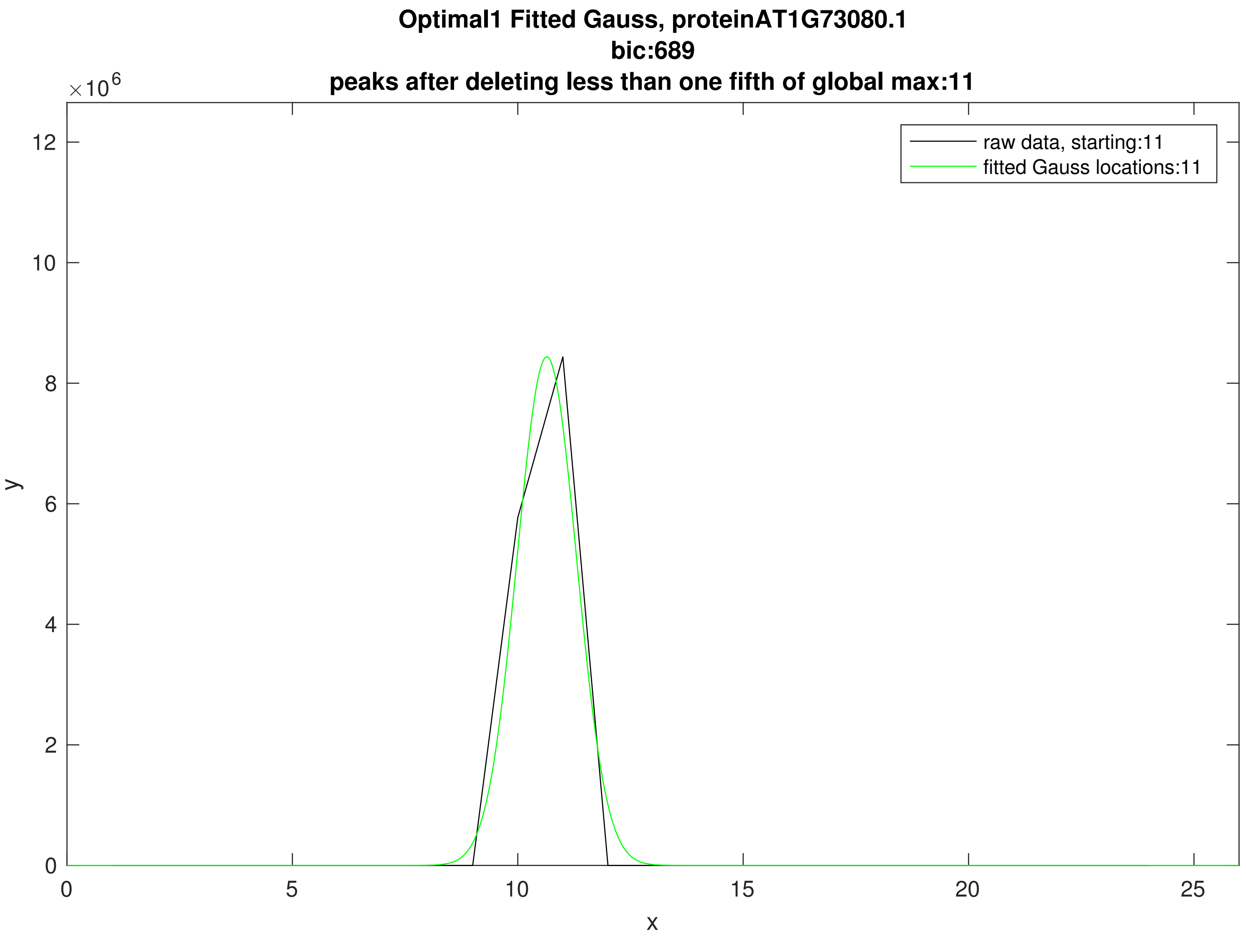
Optimal1 Fitted Gauss, proteinAT1G73080.1
bic:689
peaks after deleting less than one fifth of global max:11
×10^6
raw data, starting:11
fitted Gauss locations:11
y
x
0
2
4
6
8
10
12
0
5
10
15
20
25

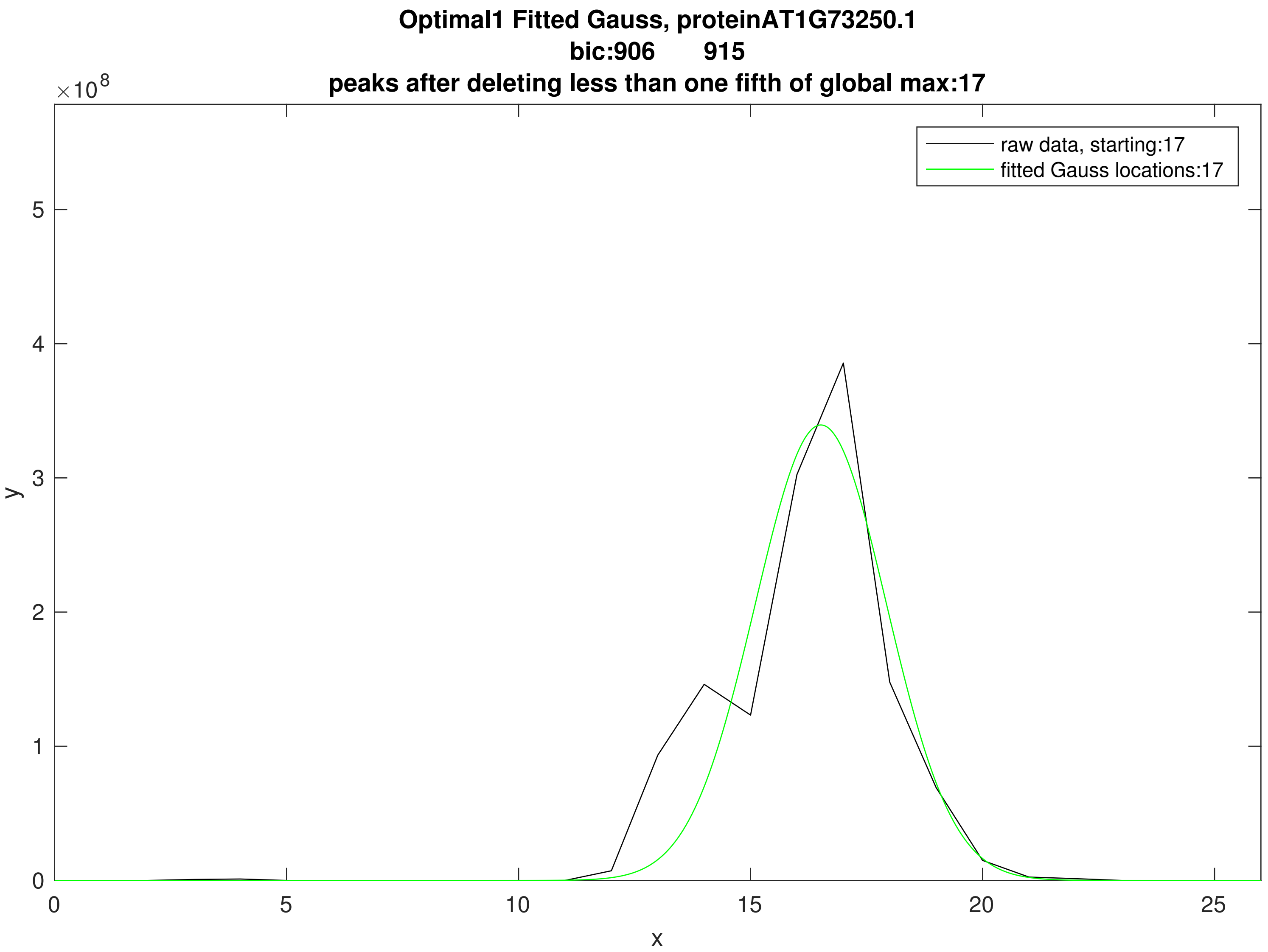
Optimal1 Fitted Gauss, proteinAT1G73250.1
bic:906 915
peaks after deleting less than one fifth of global max:17
raw data, starting:17
fitted Gauss locations:17
×10^8
5
4
3
2
1
0
y
0
5
10
15
20
25
x

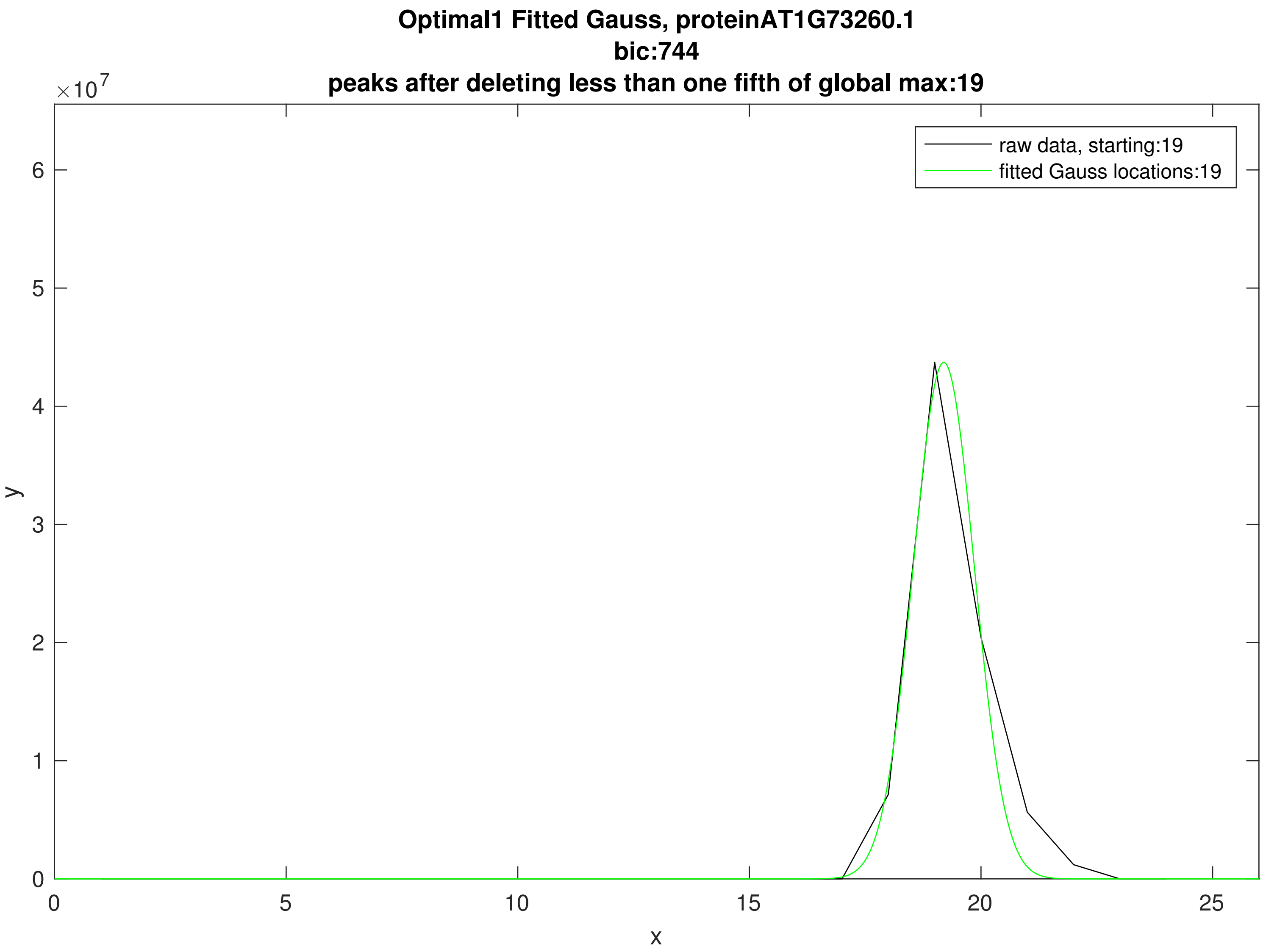
Optimal1 Fitted Gauss, proteinAT1G73260.1
bic:744
peaks after deleting less than one fifth of global max:19
×10^7
0
1
2
3
4
5
6
y
0
5
10
15
20
25
x
raw data, starting:19
fitted Gauss locations:19

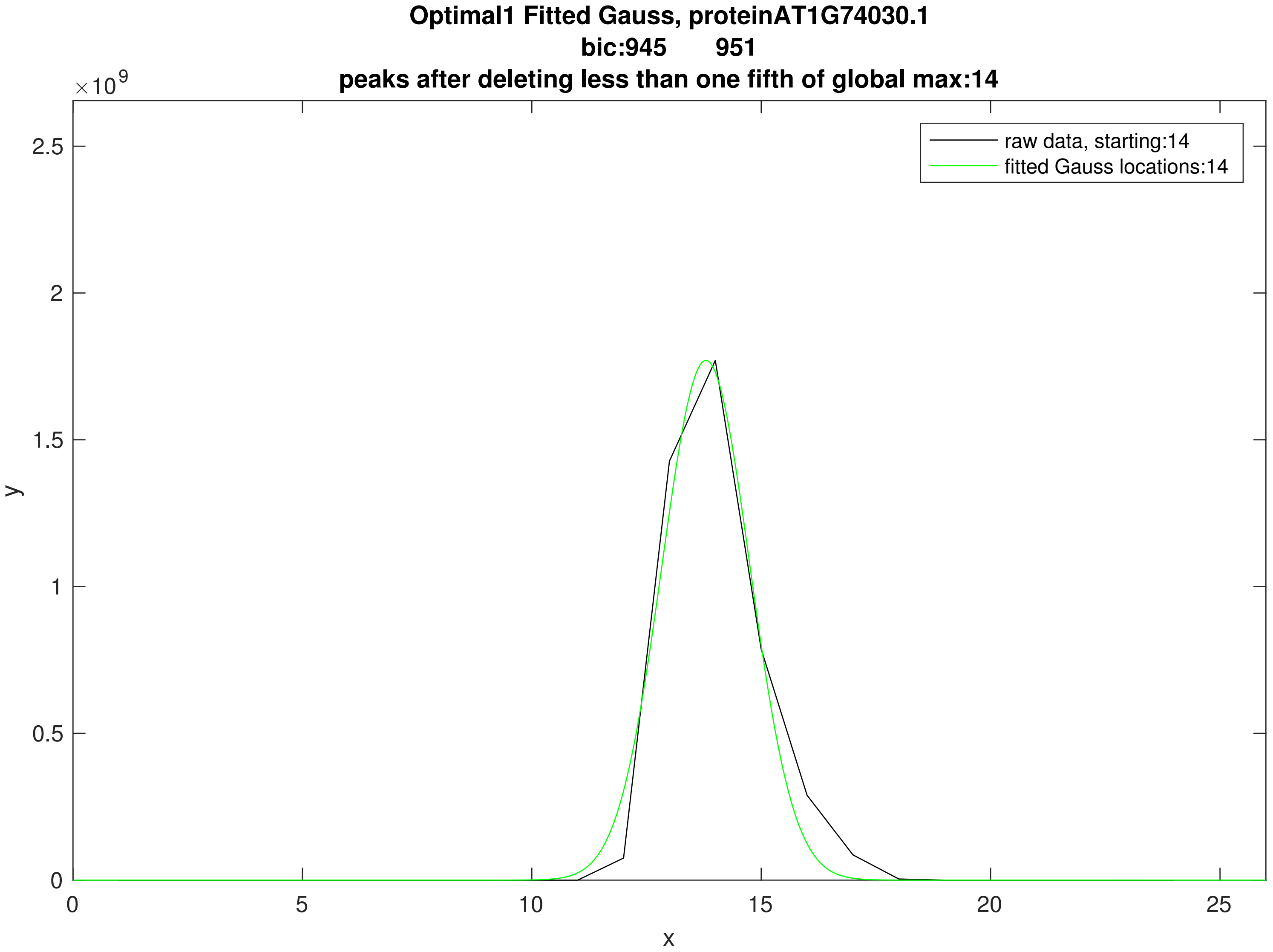

Optimal1 Fitted Gauss, proteinAT1G74030.1
bic:945 951
peaks after deleting less than one fifth of global max:14
×10^9
2.5
2
1.5
1
0.5
0
y
0
5
10
15
20
25
x
raw data, starting:14
fitted Gauss locations:14

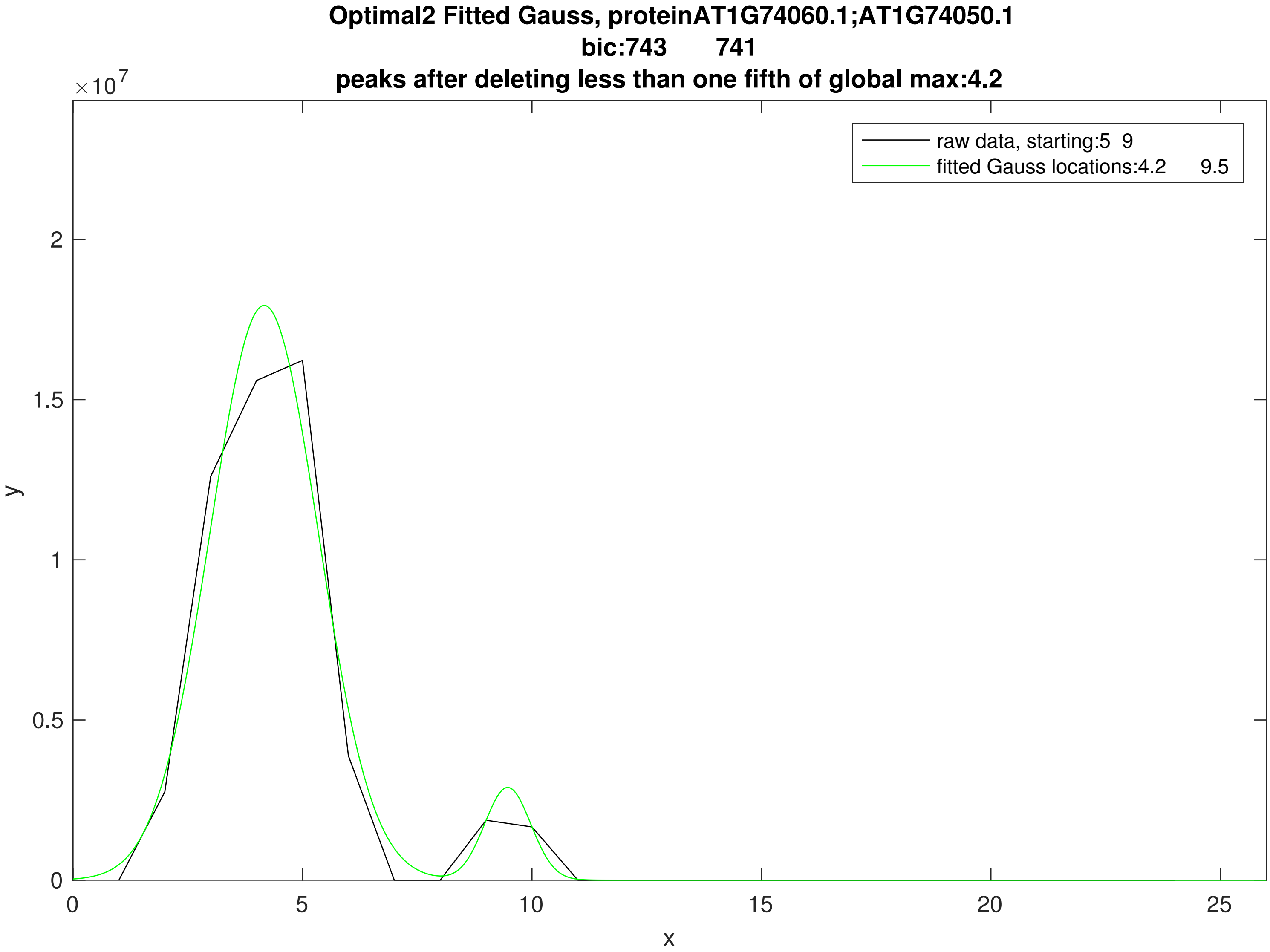
Optimal2 Fitted Gauss, proteinAT1G74060.1;AT1G74050.1
bic:743 741
peaks after deleting less than one fifth of global max:4.2
×10^7
raw data, starting:5 9
fitted Gauss locations:4.2 9.5
y
x
0
0.5
1
1.5
2
5
10
15
20
25

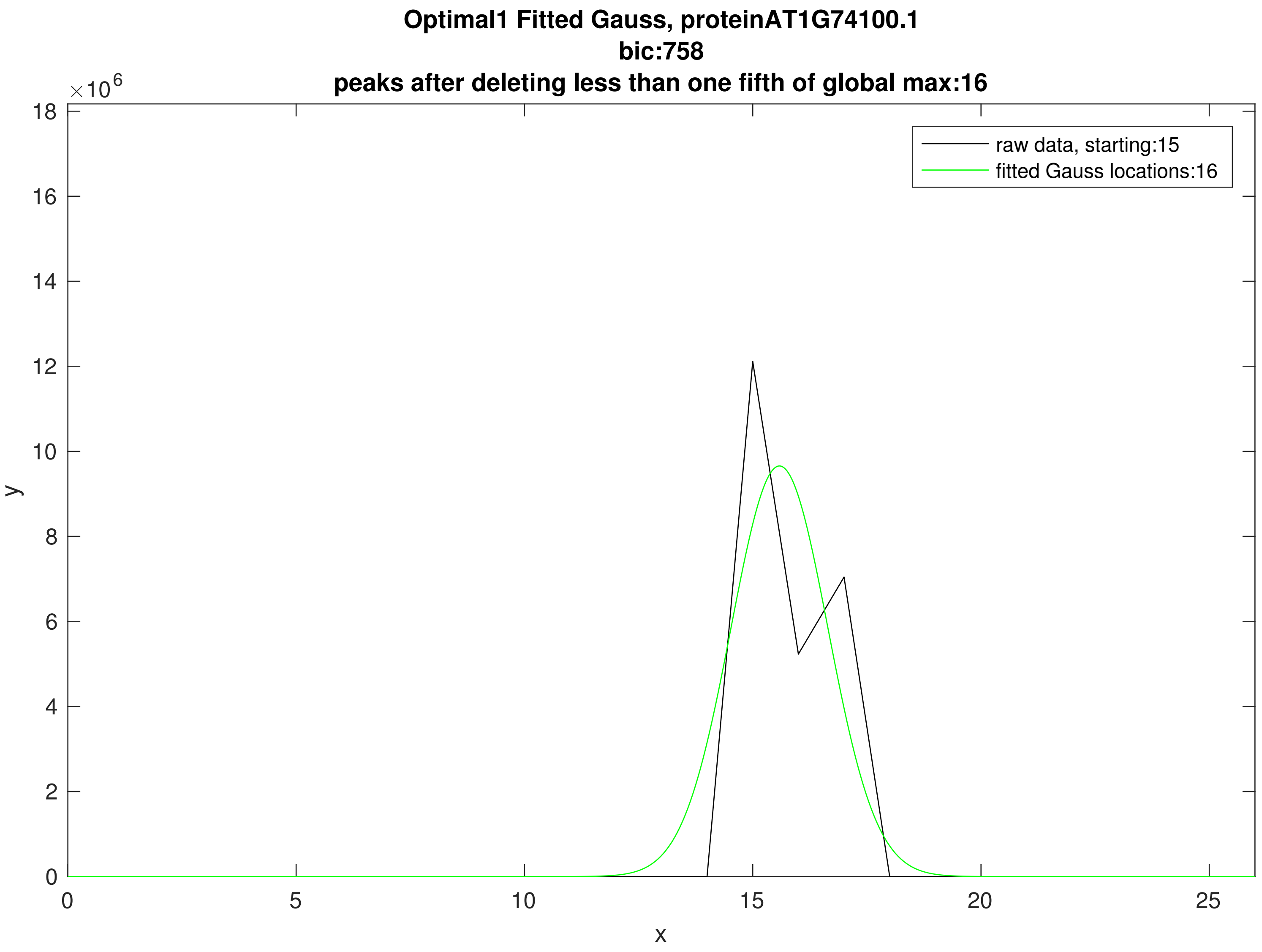
Optimal1 Fitted Gauss, proteinAT1G74100.1
bic:758
peaks after deleting less than one fifth of global max:16
×10^6
18
16
14
12
10
8
6
4
2
0
y
0
5
10
15
20
25
x
raw data, starting:15
fitted Gauss locations:16

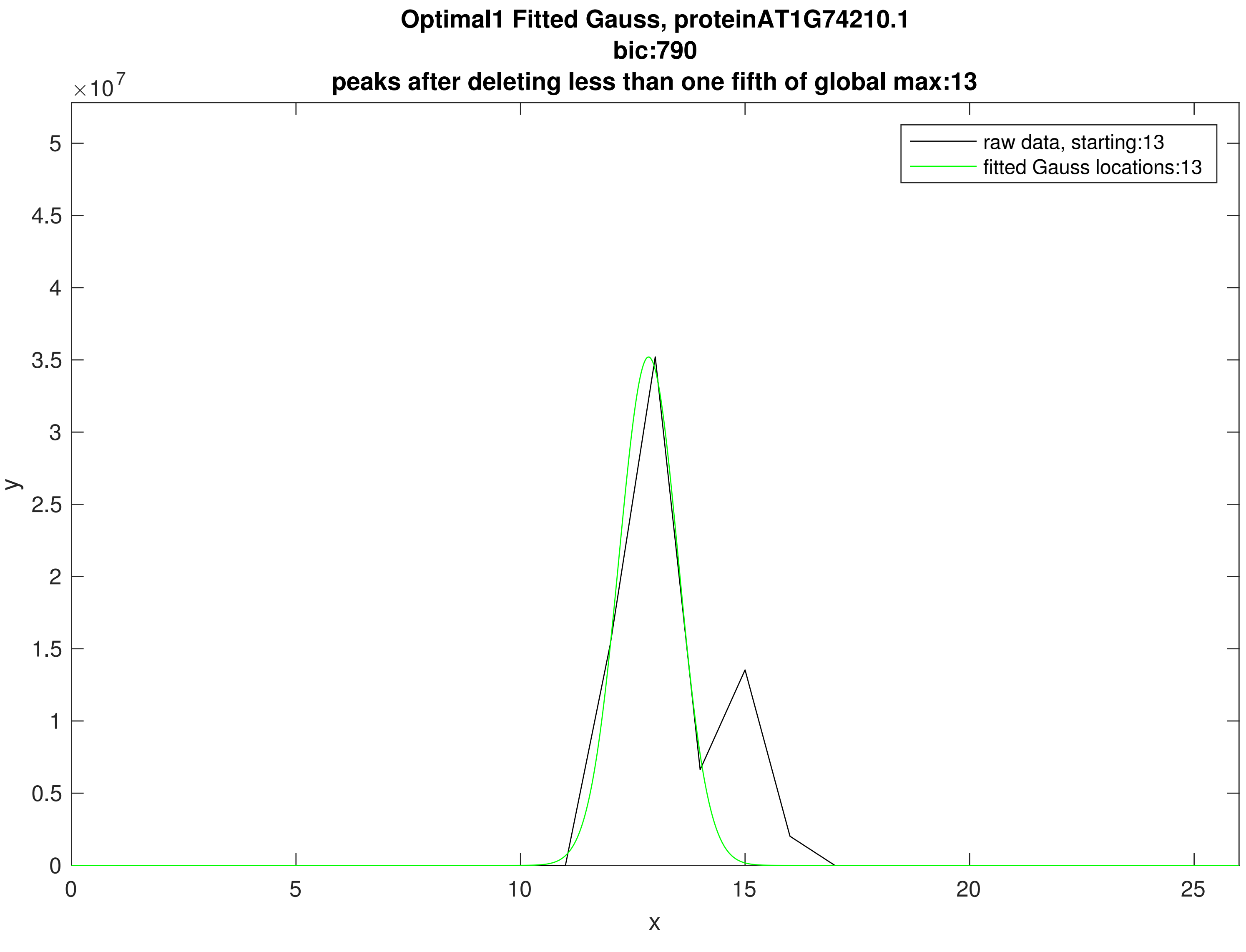
Optimal1 Fitted Gauss, proteinAT1G74210.1
bic:790
peaks after deleting less than one fifth of global max:13
×10^7
y
x
raw data, starting:13
fitted Gauss locations:13

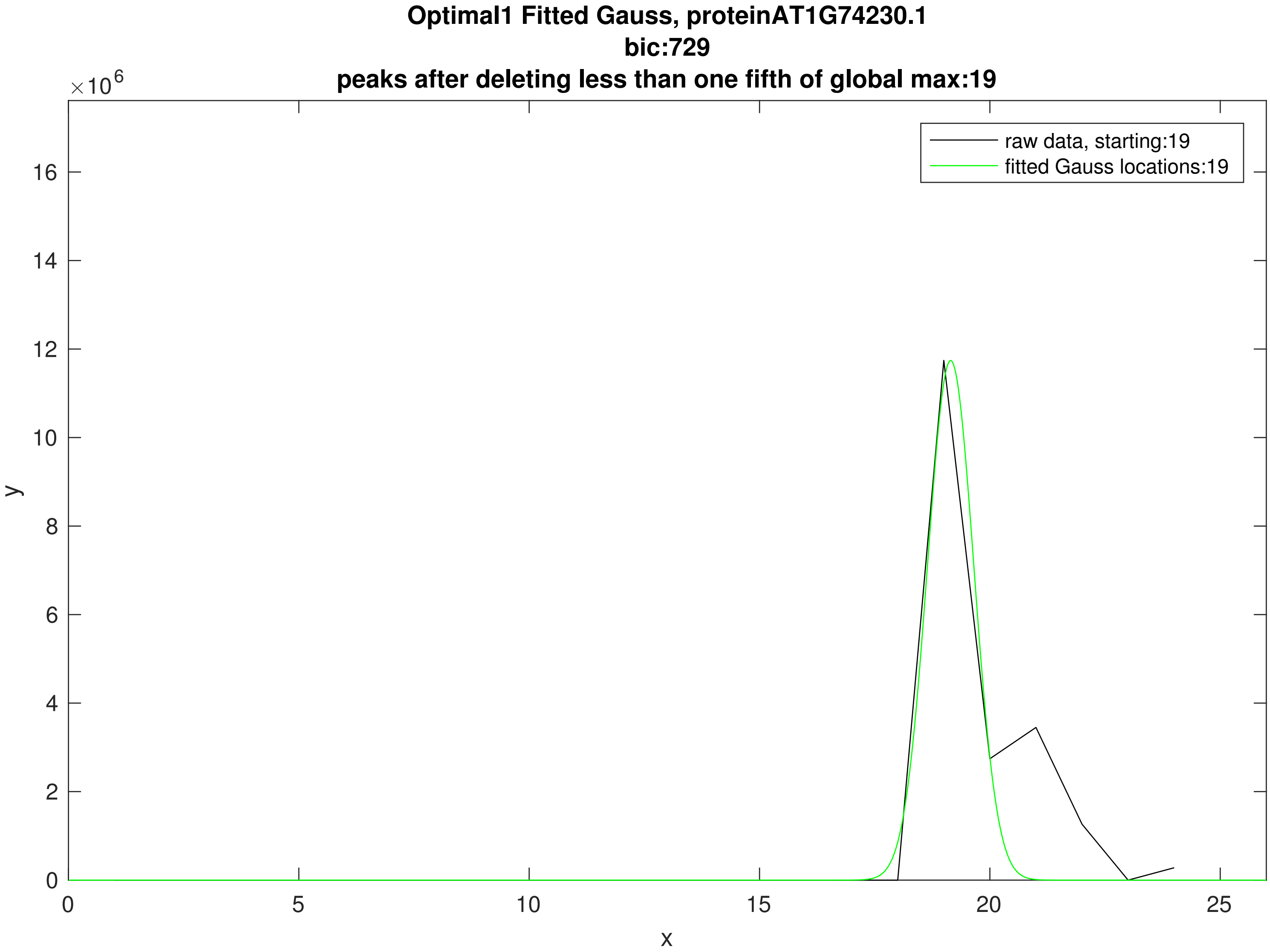
Optimal1 Fitted Gauss, proteinAT1G74230.1
bic:729
peaks after deleting less than one fifth of global max:19
×10^6
raw data, starting:19
fitted Gauss locations:19
y
x
0
2
4
6
8
10
12
14
16
0
5
10
15
20
25

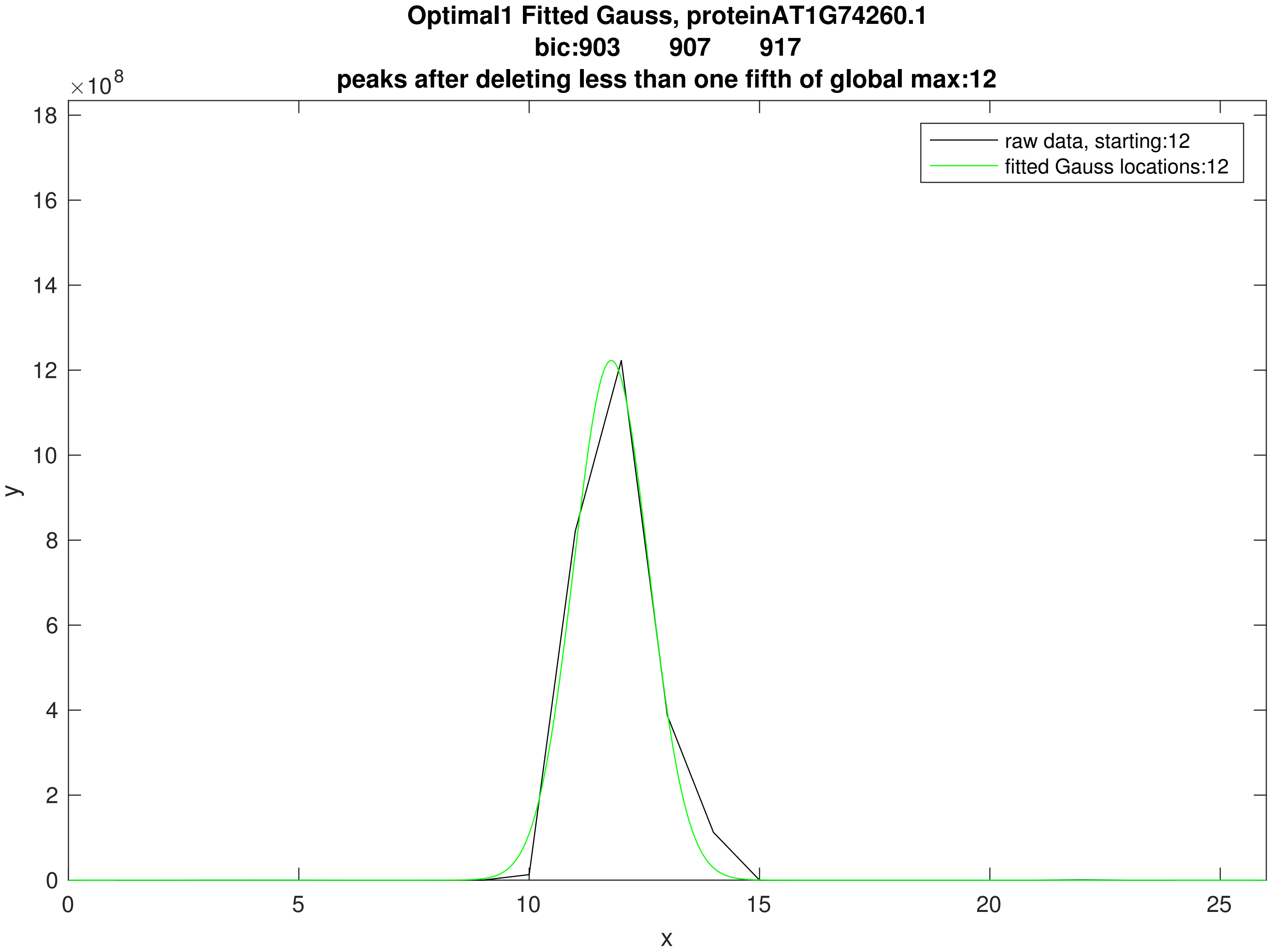

Optimal1 Fitted Gauss, proteinAT1G74260.1
bic:903 907 917
peaks after deleting less than one fifth of global max:12
×10^8
18
16
14
12
10
8
6
4
2
0
y
0
5
10
15
20
25
x
raw data, starting:12
fitted Gauss locations:12

**Optimal2 Fitted Gauss, proteinAT1G74320.1**
**bic:787 777**
**peaks after deleting less than one fifth of global max:16**

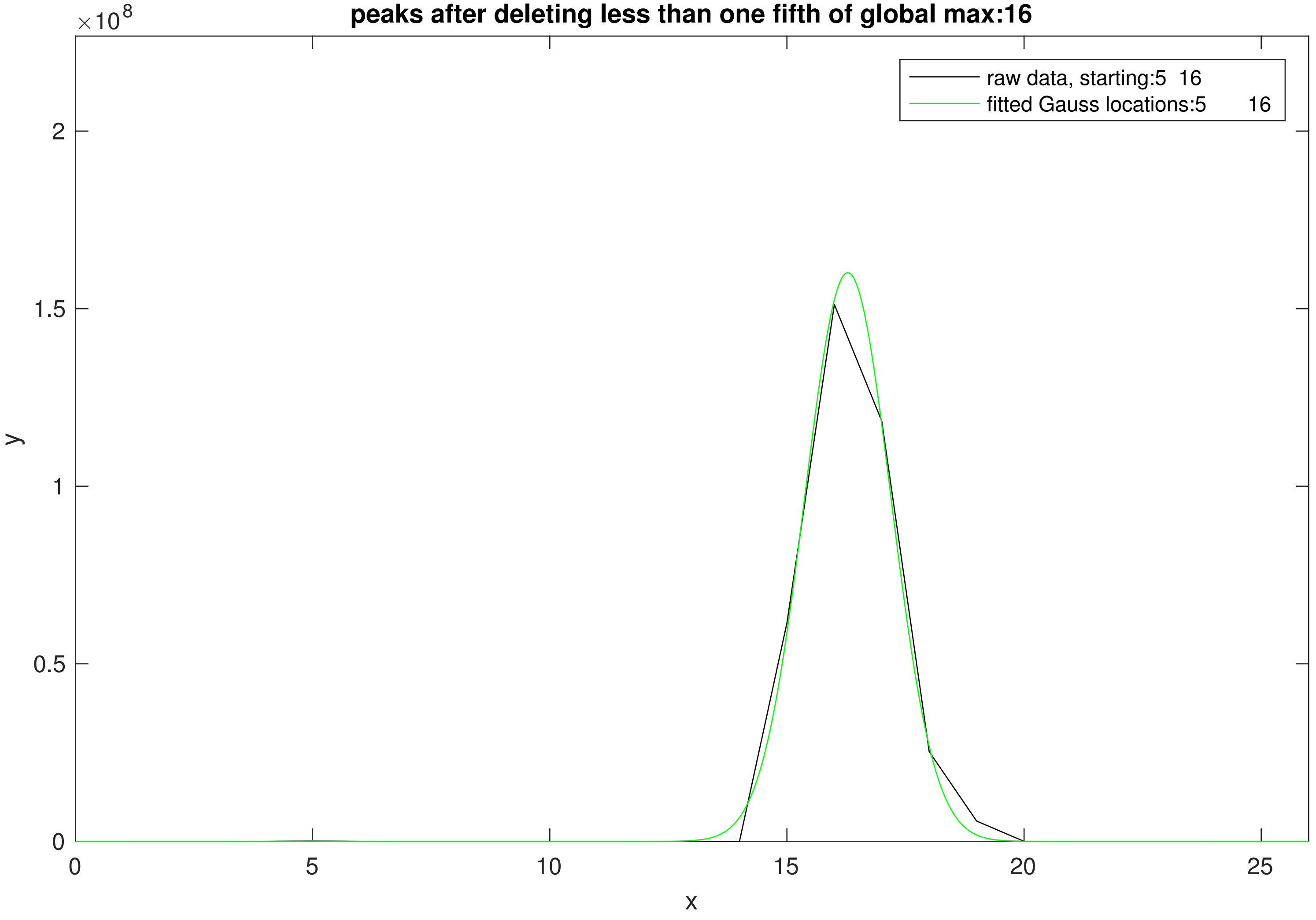

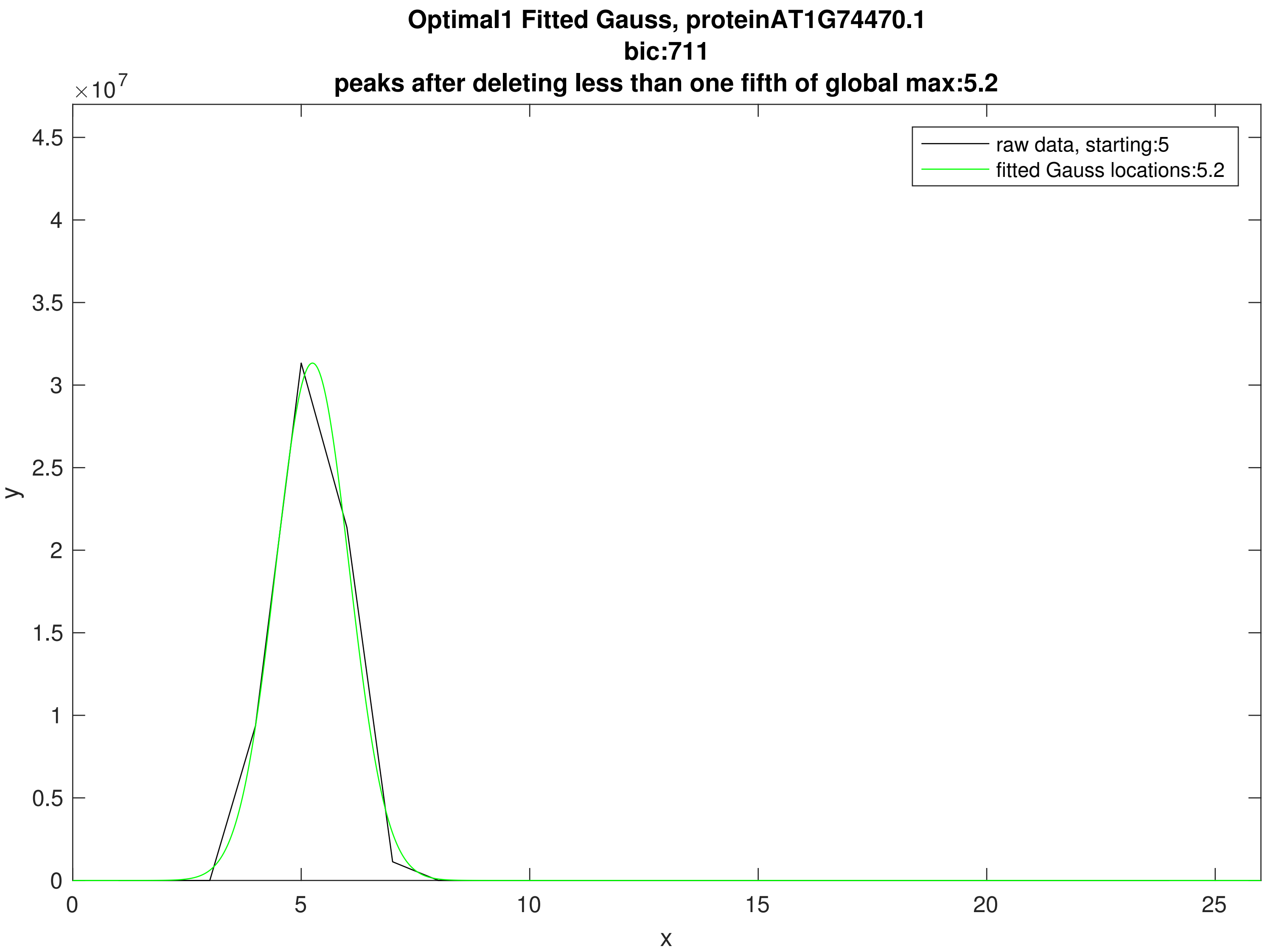
Optimal1 Fitted Gauss, proteinAT1G74470.1
bic:711
peaks after deleting less than one fifth of global max:5.2
×10^7
4.5
4
3.5
3
2.5
2
1.5
1
0.5
0
y
0
5
10
15
20
25
x
raw data, starting:5
fitted Gauss locations:5.2

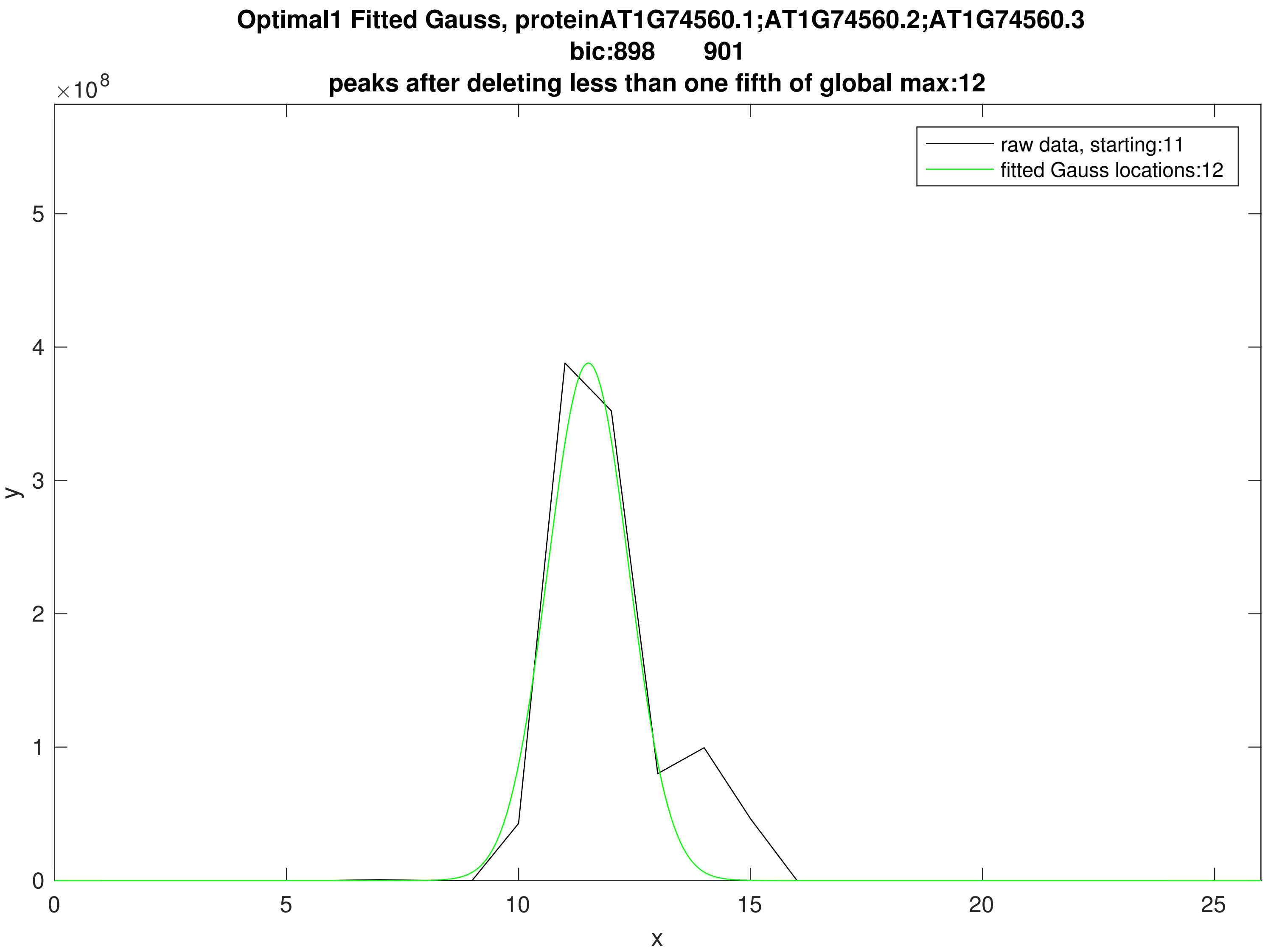
Optimal1 Fitted Gauss, proteinAT1G74560.1;AT1G74560.2;AT1G74560.3
bic:898 901
peaks after deleting less than one fifth of global max:12
×10^8
raw data, starting:11
fitted Gauss locations:12
y
x
0
1
2
3
4
5
0
5
10
15
20
25

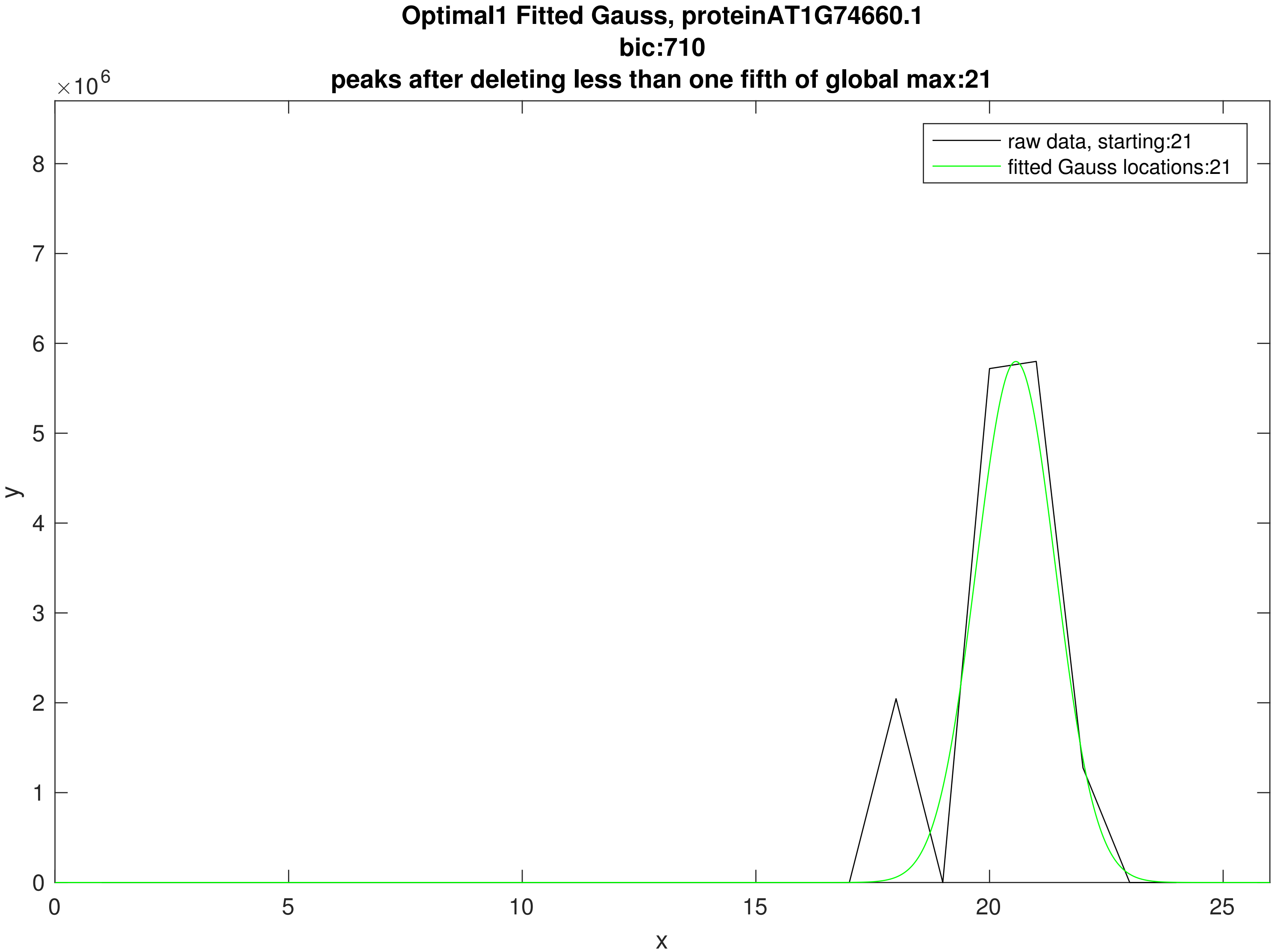
Optimal1 Fitted Gauss, proteinAT1G74660.1
bic:710
peaks after deleting less than one fifth of global max:21
×10^6
raw data, starting:21
fitted Gauss locations:21
y
x
0
1
2
3
4
5
6
7
8
0
5
10
15
20
25

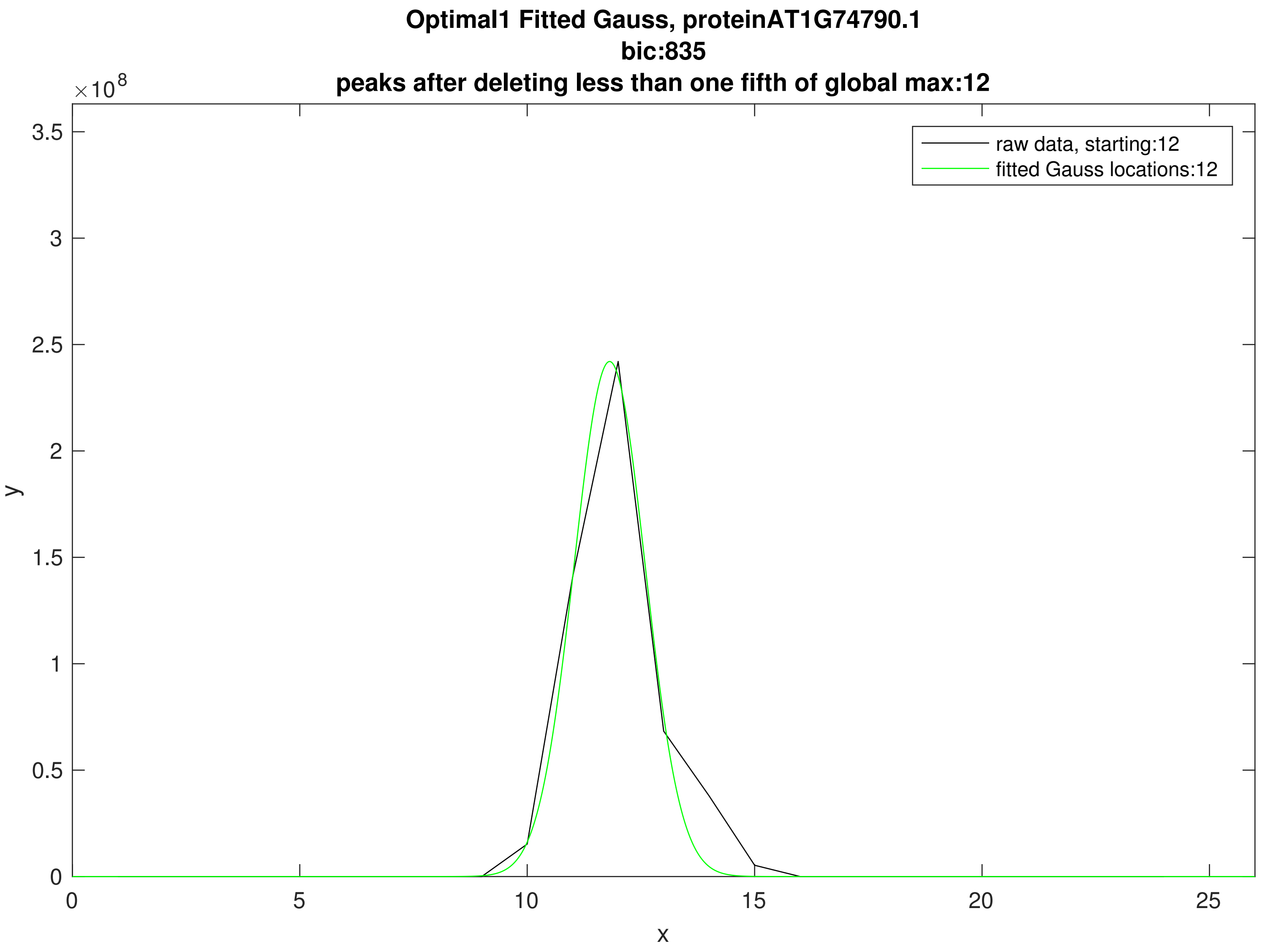
Optimal1 Fitted Gauss, proteinAT1G74790.1
bic:835
peaks after deleting less than one fifth of global max:12
×10^8
3.5
3
2.5
2
1.5
1
0.5
0
y
0
5
10
15
20
25
x
raw data, starting:12
fitted Gauss locations:12

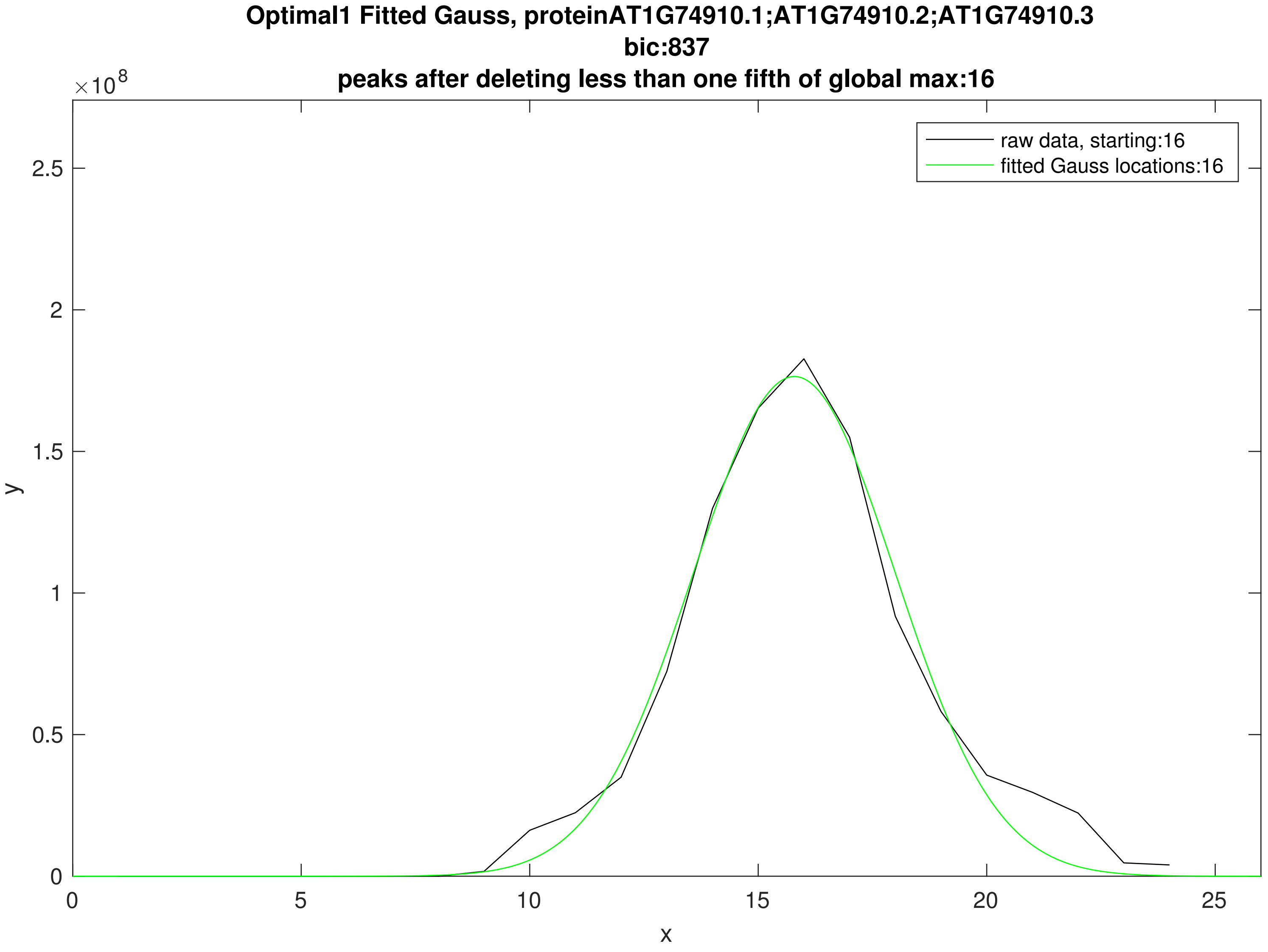

Optimal1 Fitted Gauss, proteinAT1G74910.1;AT1G74910.2;AT1G74910.3
bic:837
peaks after deleting less than one fifth of global max:16
×10^8
raw data, starting:16
fitted Gauss locations:16
y
x
0
0.5
1
1.5
2
2.5
0
5
10
15
20
25

**Optimal1 Fitted Gauss, proteinAT1G74920.1;AT1G74920.2**
**bic:961**
**peaks after deleting less than one fifth of global max:13**

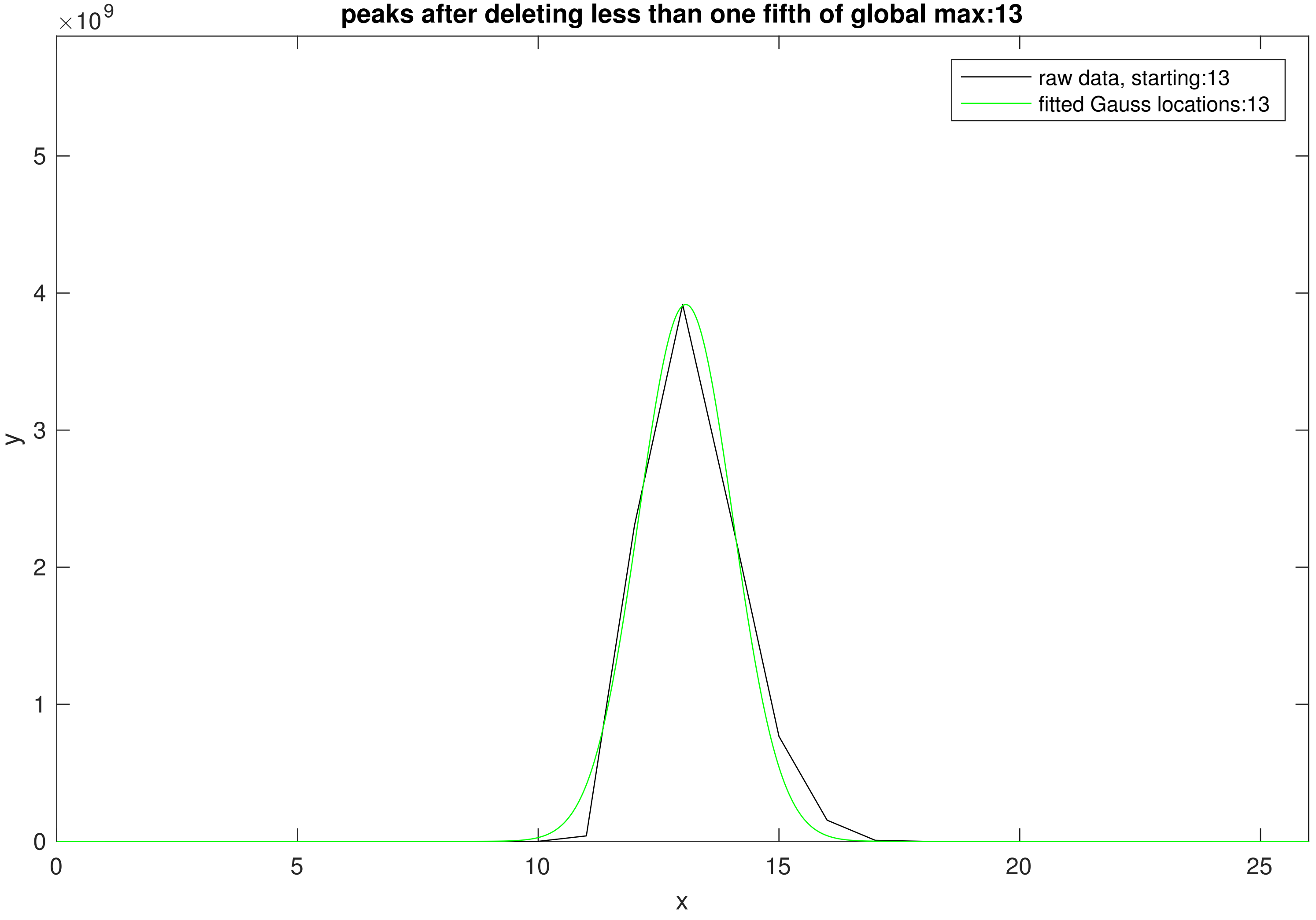

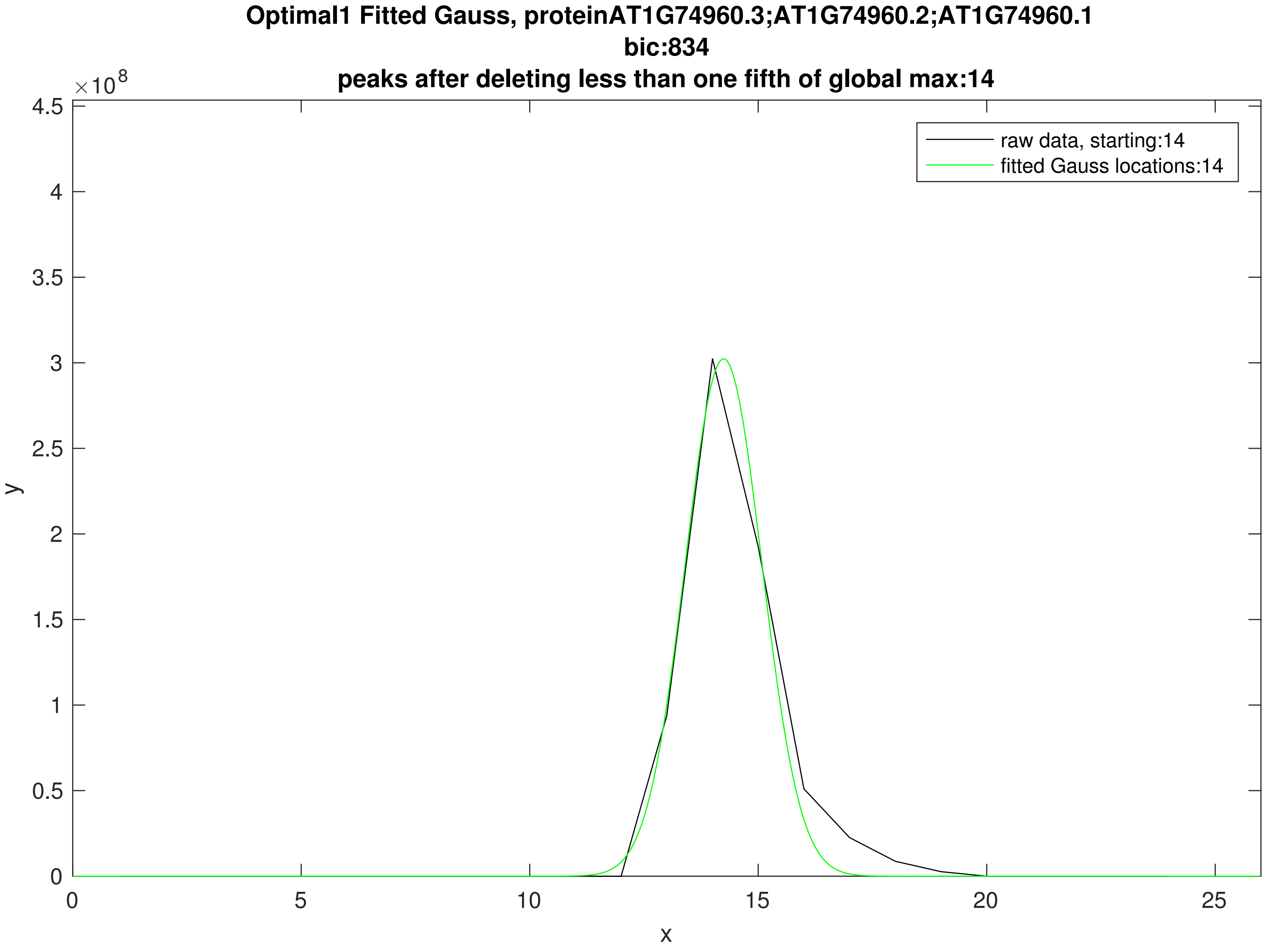

Optimal1 Fitted Gauss, proteinAT1G74960.3;AT1G74960.2;AT1G74960.1
bic:834
peaks after deleting less than one fifth of global max:14
×10^8
4.5
4
3.5
3
2.5
2
1.5
1
0.5
0
y
0
5
10
15
20
25
x
raw data, starting:14
fitted Gauss locations:14

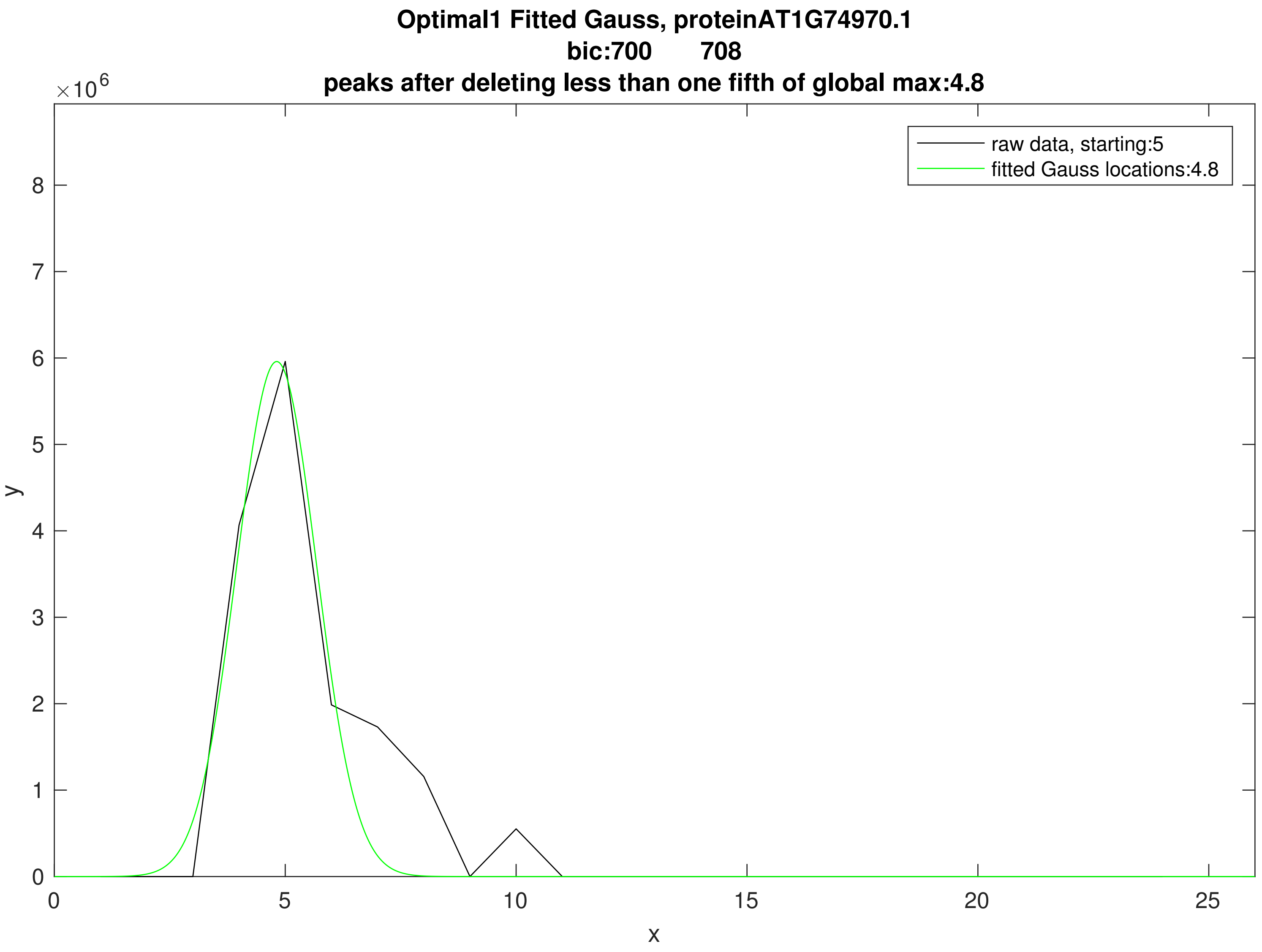

Optimal1 Fitted Gauss, proteinAT1G74970.1
bic:700 708
peaks after deleting less than one fifth of global max:4.8
×10^6
raw data, starting:5
fitted Gauss locations:4.8
y
x
0
1
2
3
4
5
6
7
8
0
5
10
15
20
25

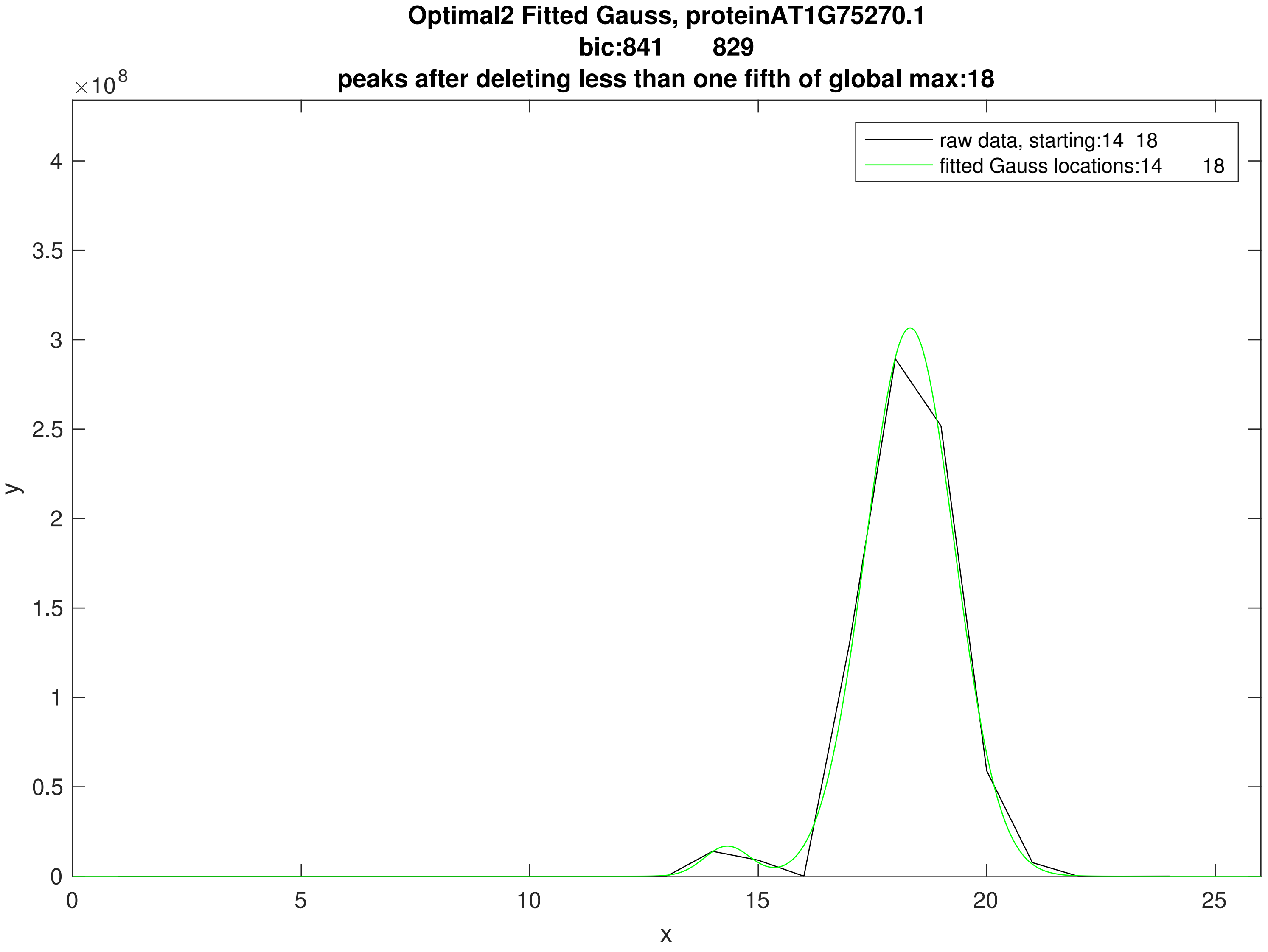

Optimal2 Fitted Gauss, proteinAT1G75270.1
bic:841 829
peaks after deleting less than one fifth of global max:18
×10^8
raw data, starting:14 18
fitted Gauss locations:14 18
y
x

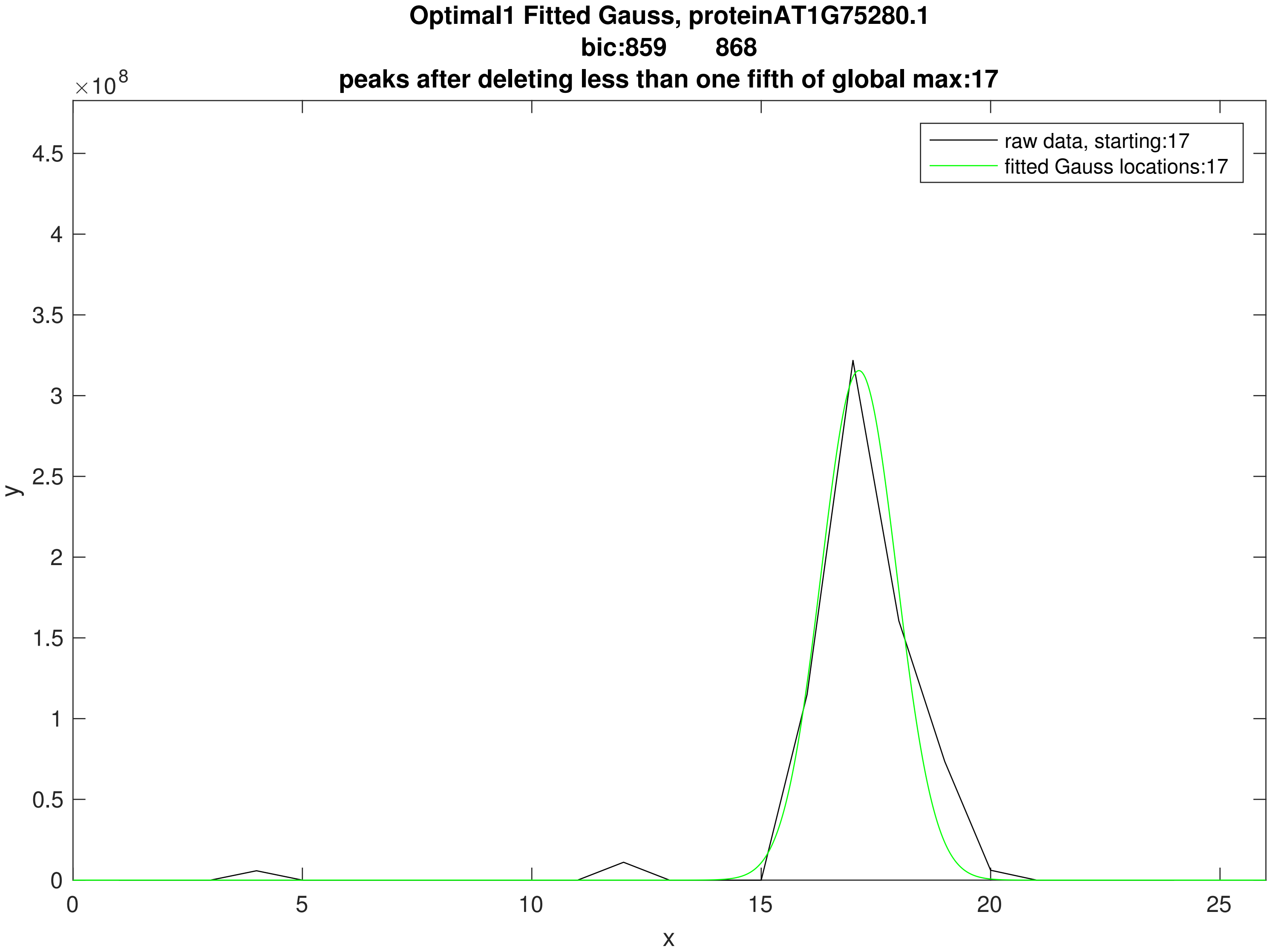
Optimal1 Fitted Gauss, proteinAT1G75280.1
bic:859 868
peaks after deleting less than one fifth of global max:17
×10^8
raw data, starting:17
fitted Gauss locations:17
y
x
0
0.5
1
1.5
2
2.5
3
3.5
4
4.5
0
5
10
15
20
25

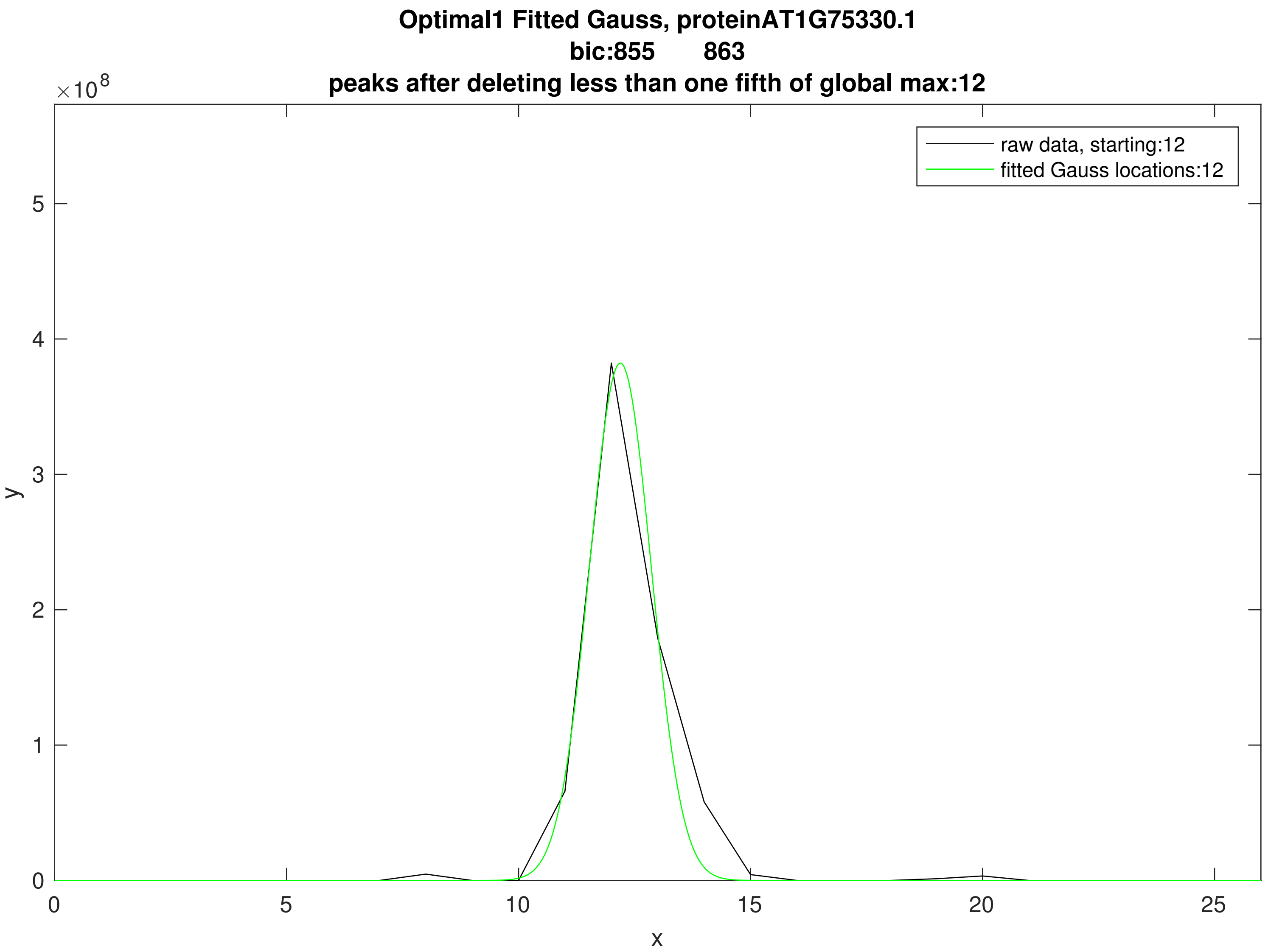

Optimal1 Fitted Gauss, proteinAT1G75330.1
bic:855 863
peaks after deleting less than one fifth of global max:12
×10^8
raw data, starting:12
fitted Gauss locations:12
y
x
0
1
2
3
4
5
0
5
10
15
20
25

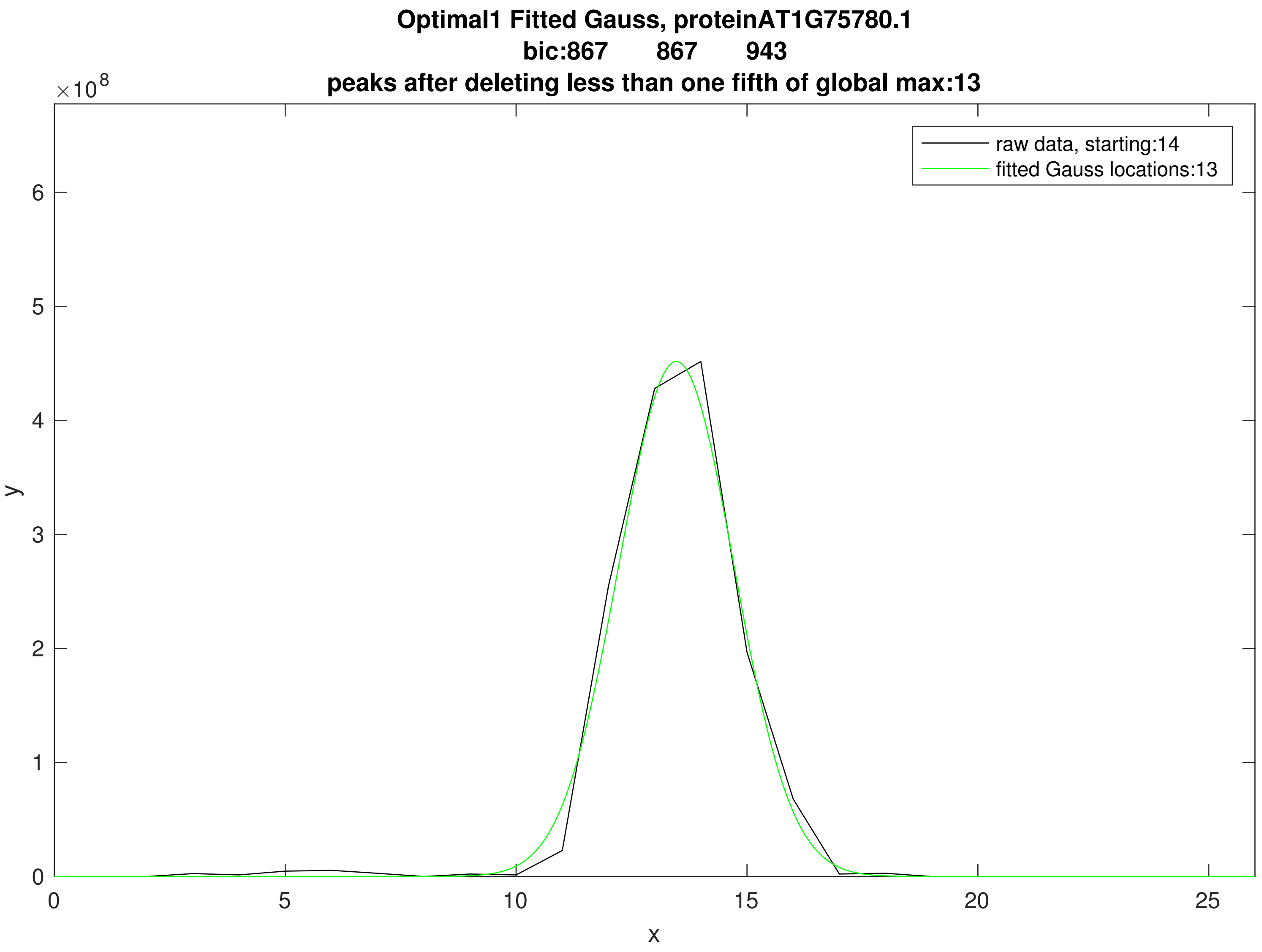
Optimal1 Fitted Gauss, proteinAT1G75780.1
bic:867 867 943
peaks after deleting less than one fifth of global max:13
×10^8
raw data, starting:14
fitted Gauss locations:13
y
x
0
1
2
3
4
5
6
0
5
10
15
20
25

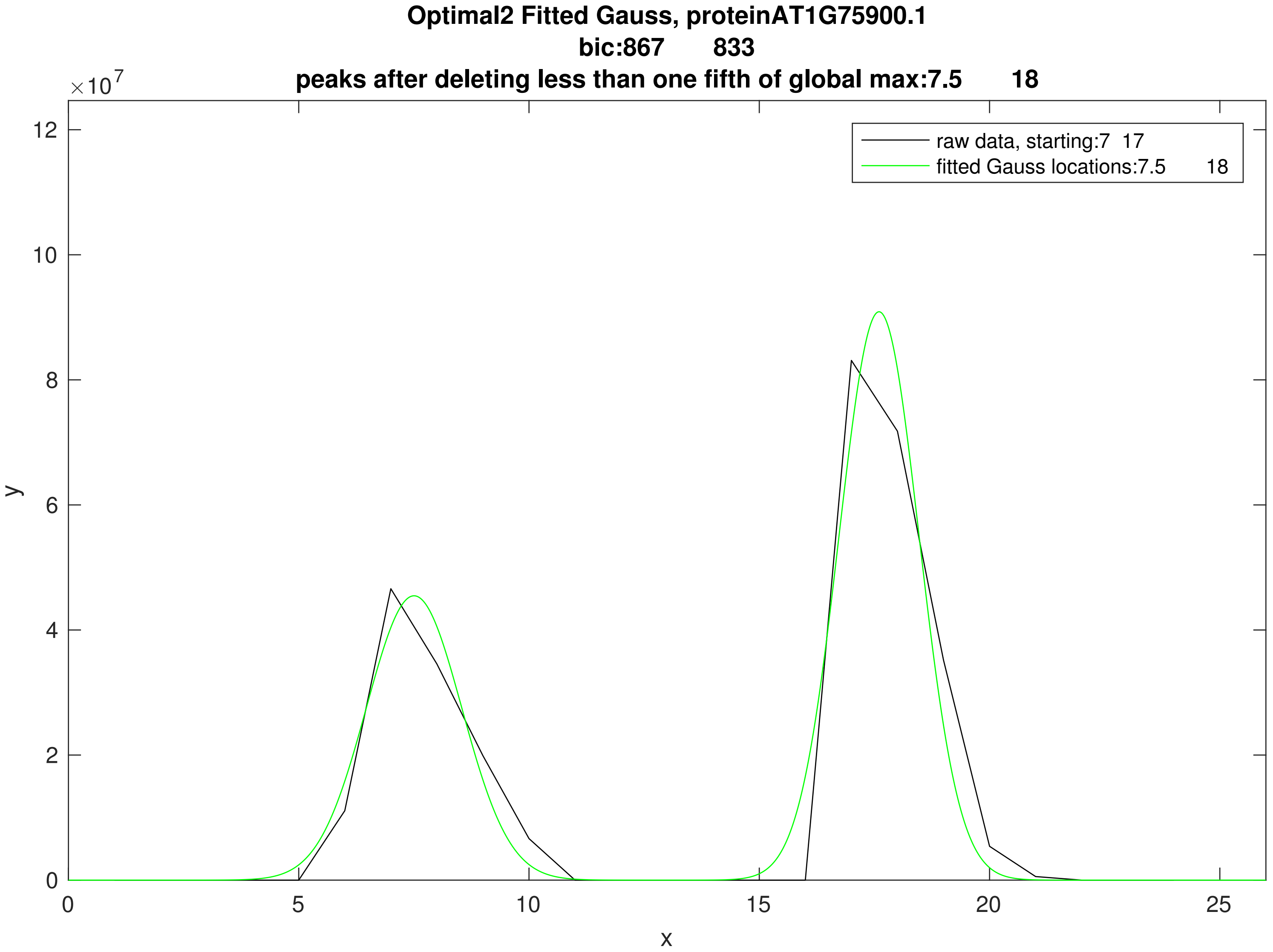
Optimal2 Fitted Gauss, proteinAT1G75900.1
bic:867 833
peaks after deleting less than one fifth of global max:7.5 18
×10^7
raw data, starting:7 17
fitted Gauss locations:7.5 18
y
x
0
2
4
6
8
10
12
5
15
20
25

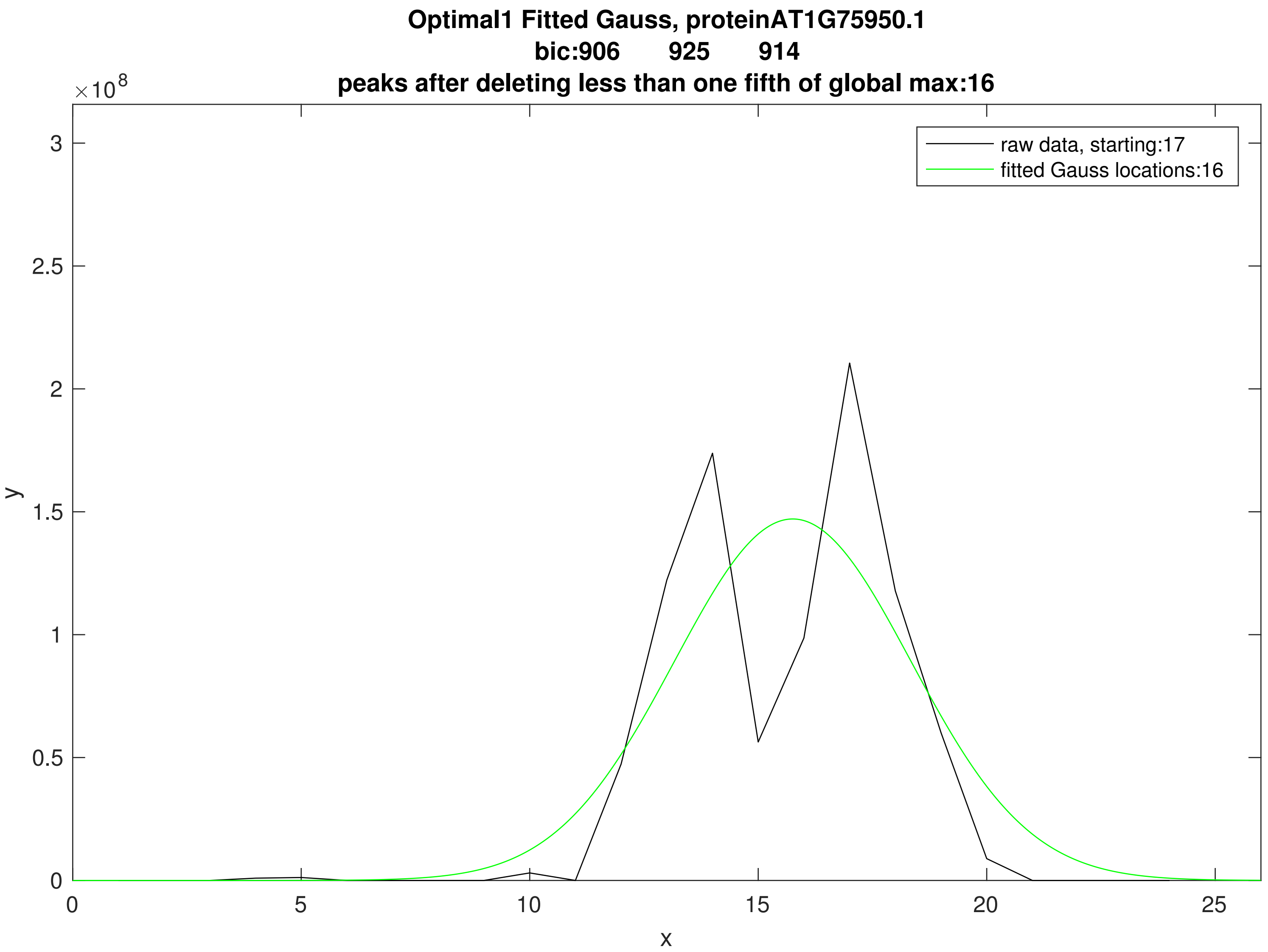
Optimal1 Fitted Gauss, proteinAT1G75950.1
bic:906 925 914
peaks after deleting less than one fifth of global max:16
×10^8
raw data, starting:17
fitted Gauss locations:16
y
x
0
0.5
1
1.5
2
2.5
3
0
5
10
15
20
25

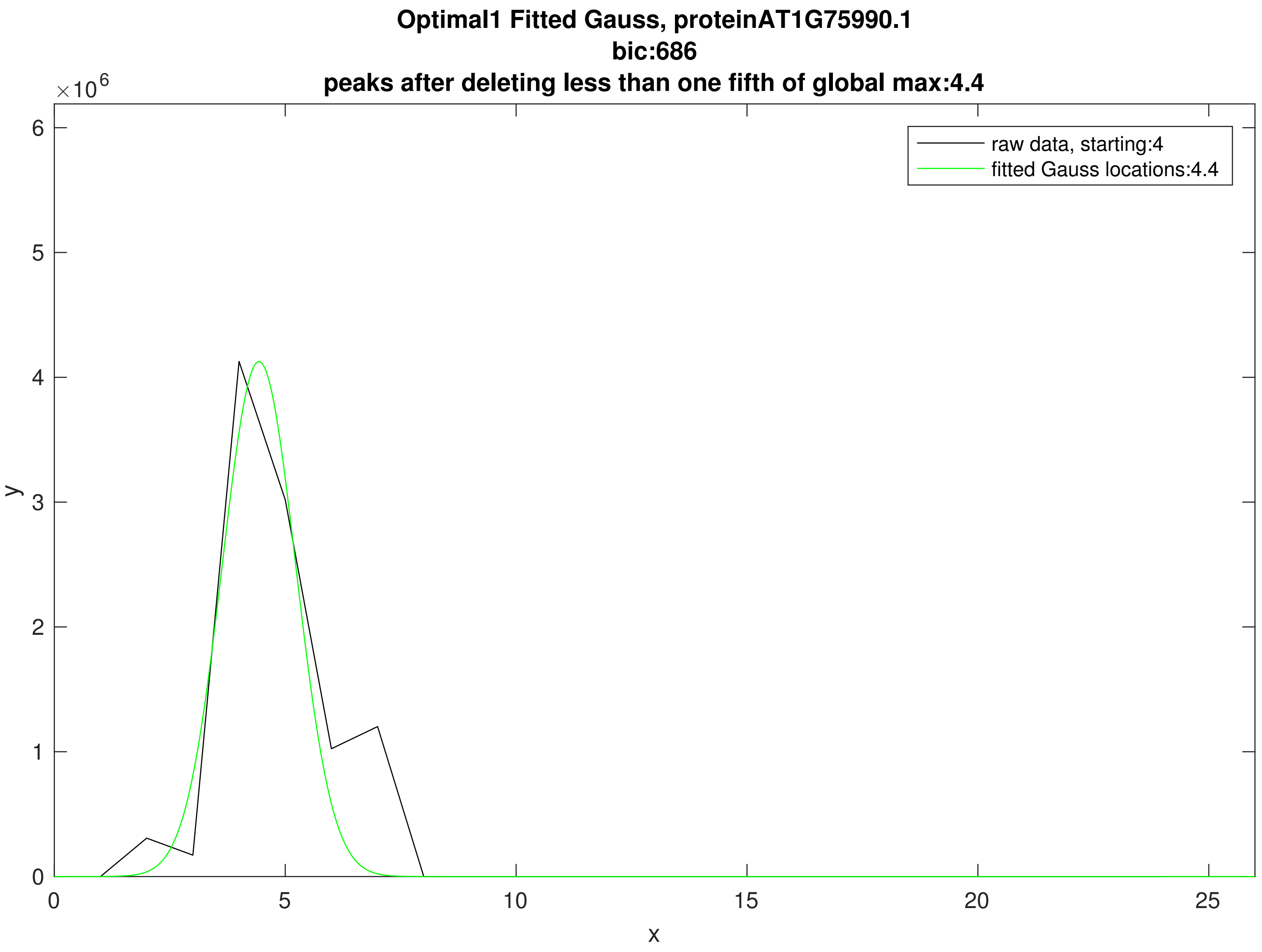
Optimal1 Fitted Gauss, proteinAT1G75990.1
bic:686
peaks after deleting less than one fifth of global max:4.4
×10^6
raw data, starting:4
fitted Gauss locations:4.4
y
x
0
1
2
3
4
5
6
0
5
10
15
20
25

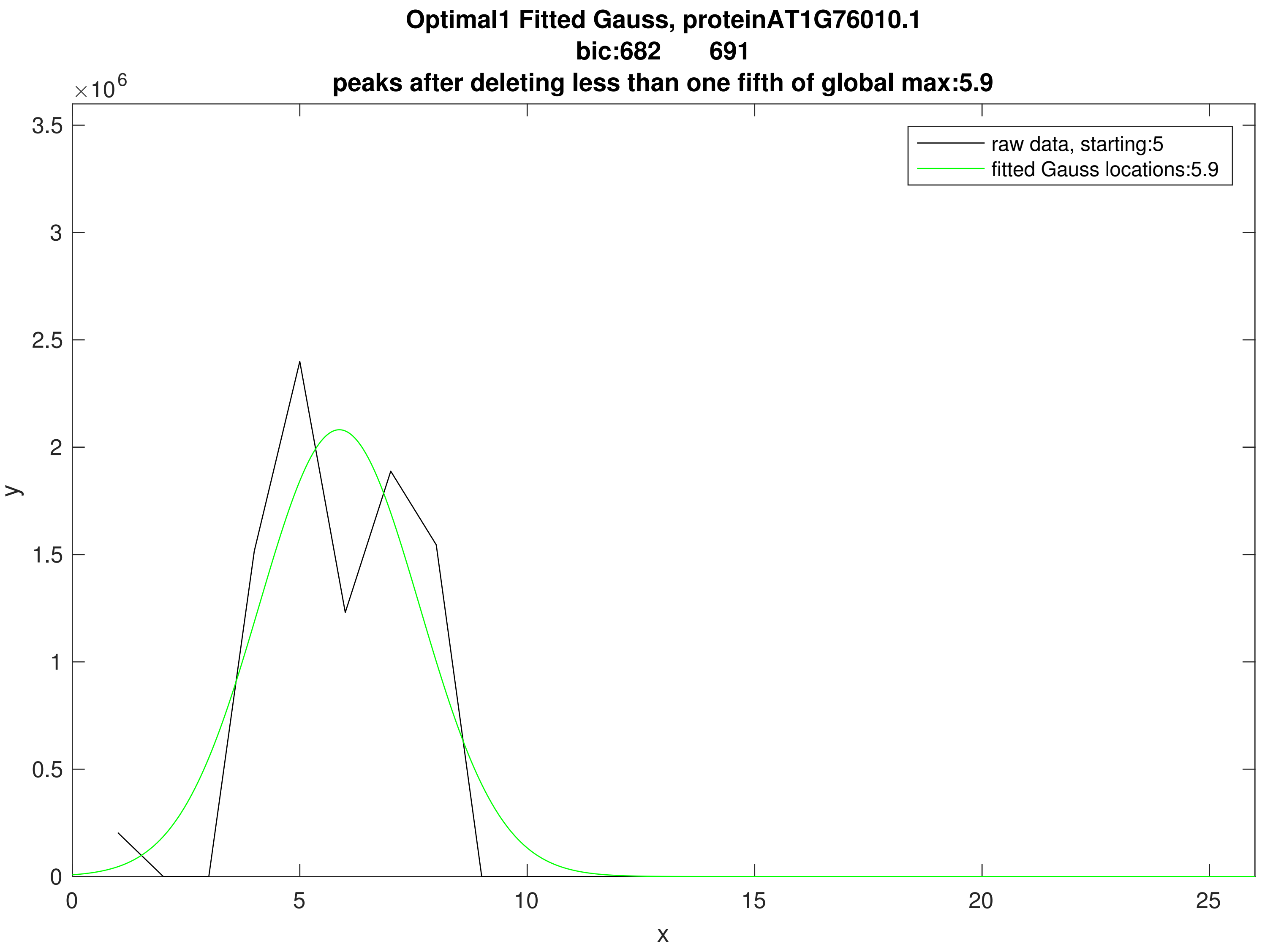
Optimal1 Fitted Gauss, proteinAT1G76010.1
bic:682 691
peaks after deleting less than one fifth of global max:5.9
×10^6
raw data, starting:5
fitted Gauss locations:5.9
3.5
3
2.5
2
1.5
1
0.5
0
y
0
5
10
15
20
25
x

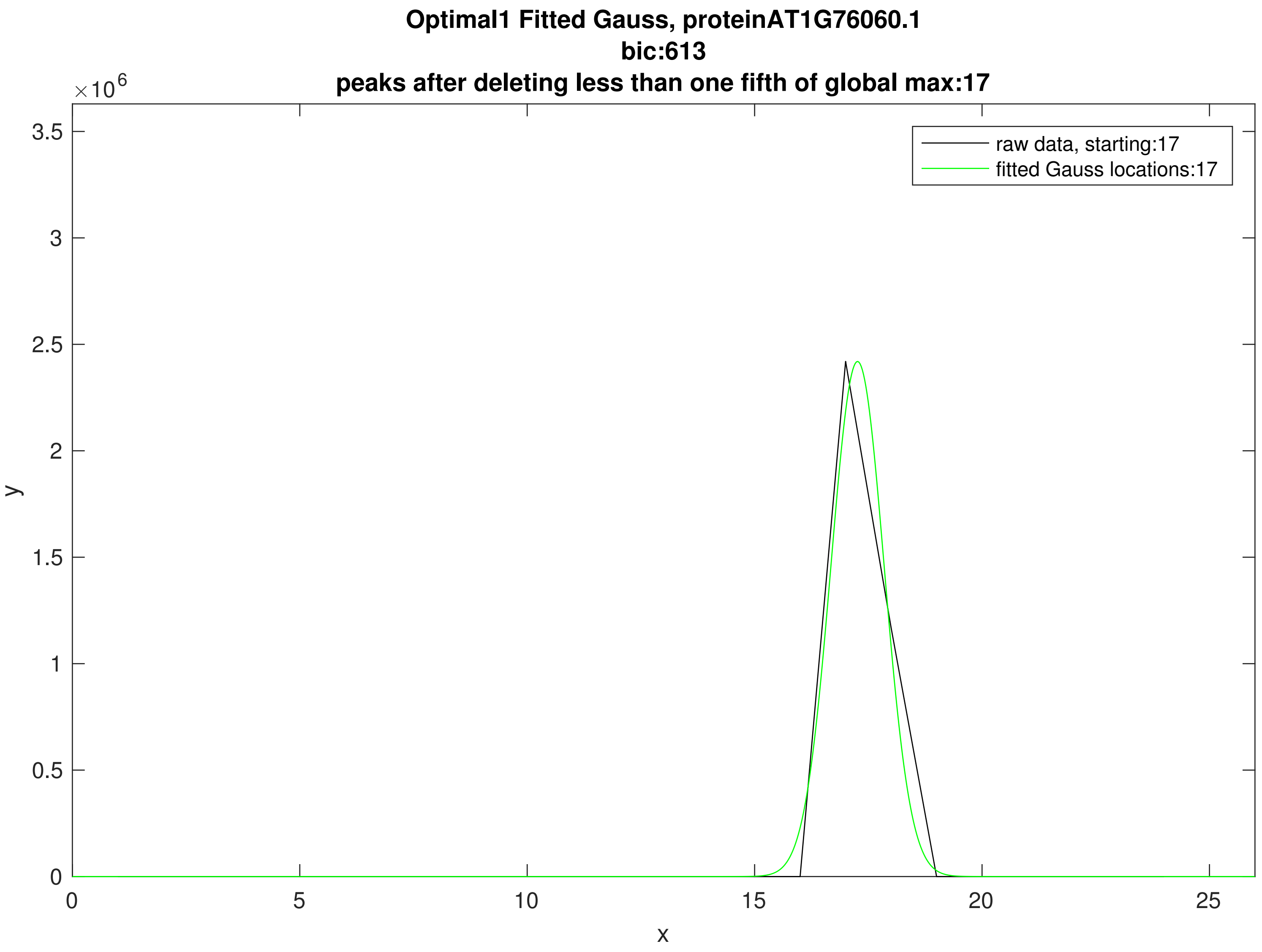
Optimal1 Fitted Gauss, proteinAT1G76060.1
bic:613
peaks after deleting less than one fifth of global max:17
×10^6
3.5
3
2.5
2
1.5
1
0.5
0
y
0
5
10
15
20
25
x
raw data, starting:17
fitted Gauss locations:17

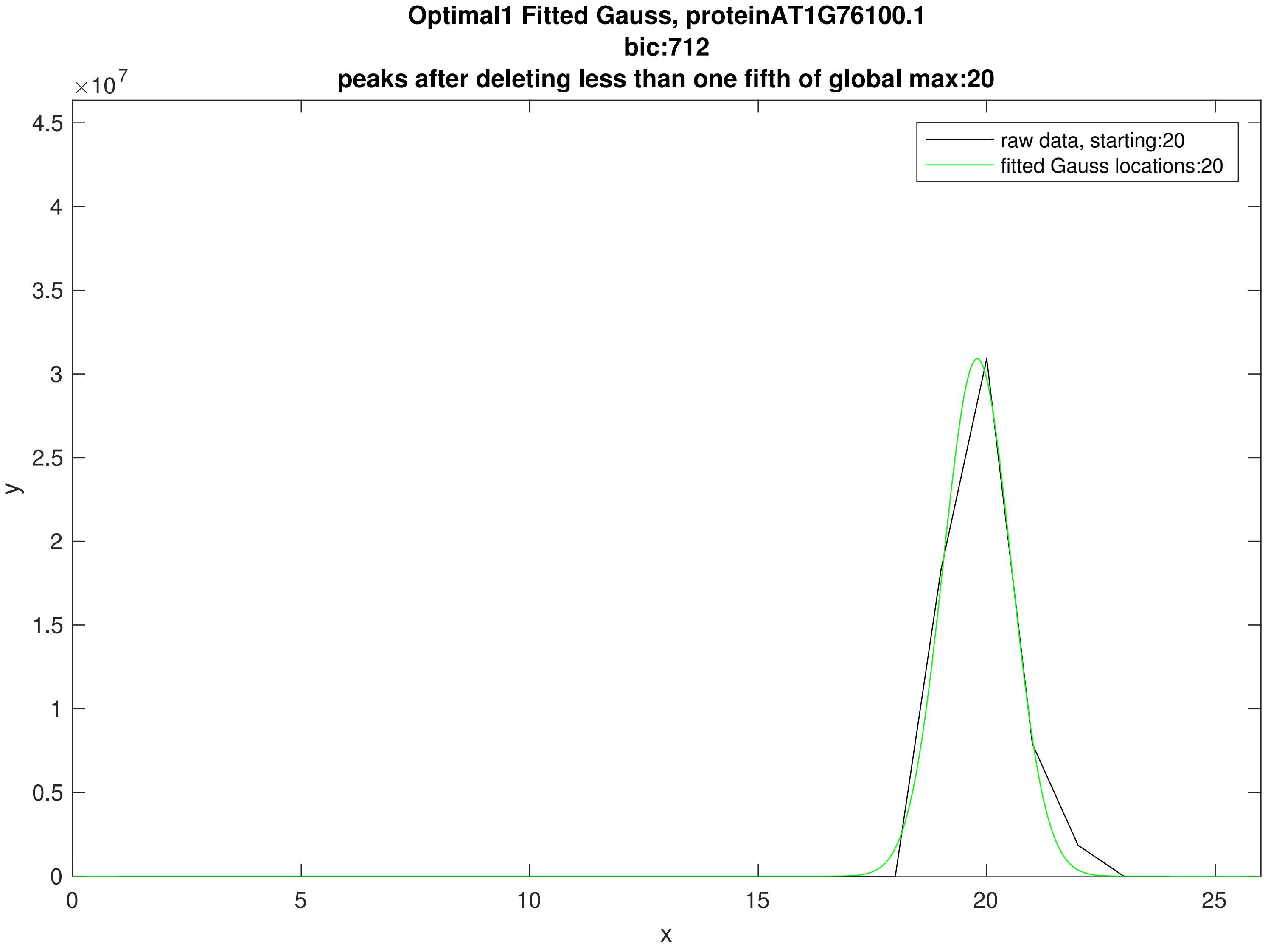
Optimal1 Fitted Gauss, proteinAT1G76100.1
bic:712
peaks after deleting less than one fifth of global max:20
×10^7
4.5
4
3.5
3
2.5
2
1.5
1
0.5
0
y
0
5
10
15
20
25
x
raw data, starting:20
fitted Gauss locations:20

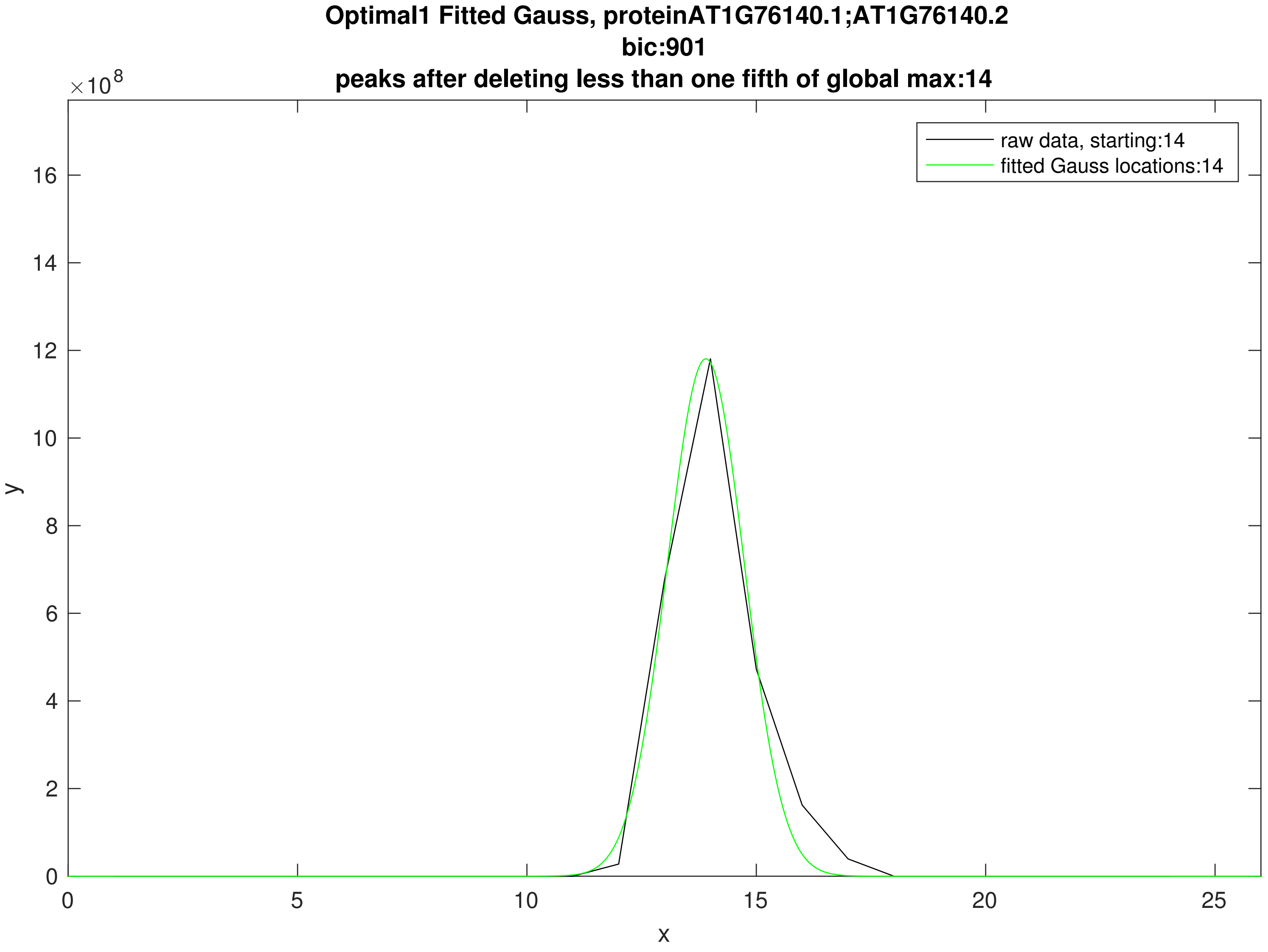
Optimal1 Fitted Gauss, proteinAT1G76140.1;AT1G76140.2
bic:901
peaks after deleting less than one fifth of global max:14
×10^8
raw data, starting:14
fitted Gauss locations:14
y
x
0
2
4
6
8
10
12
14
16
0
5
10
15
20
25

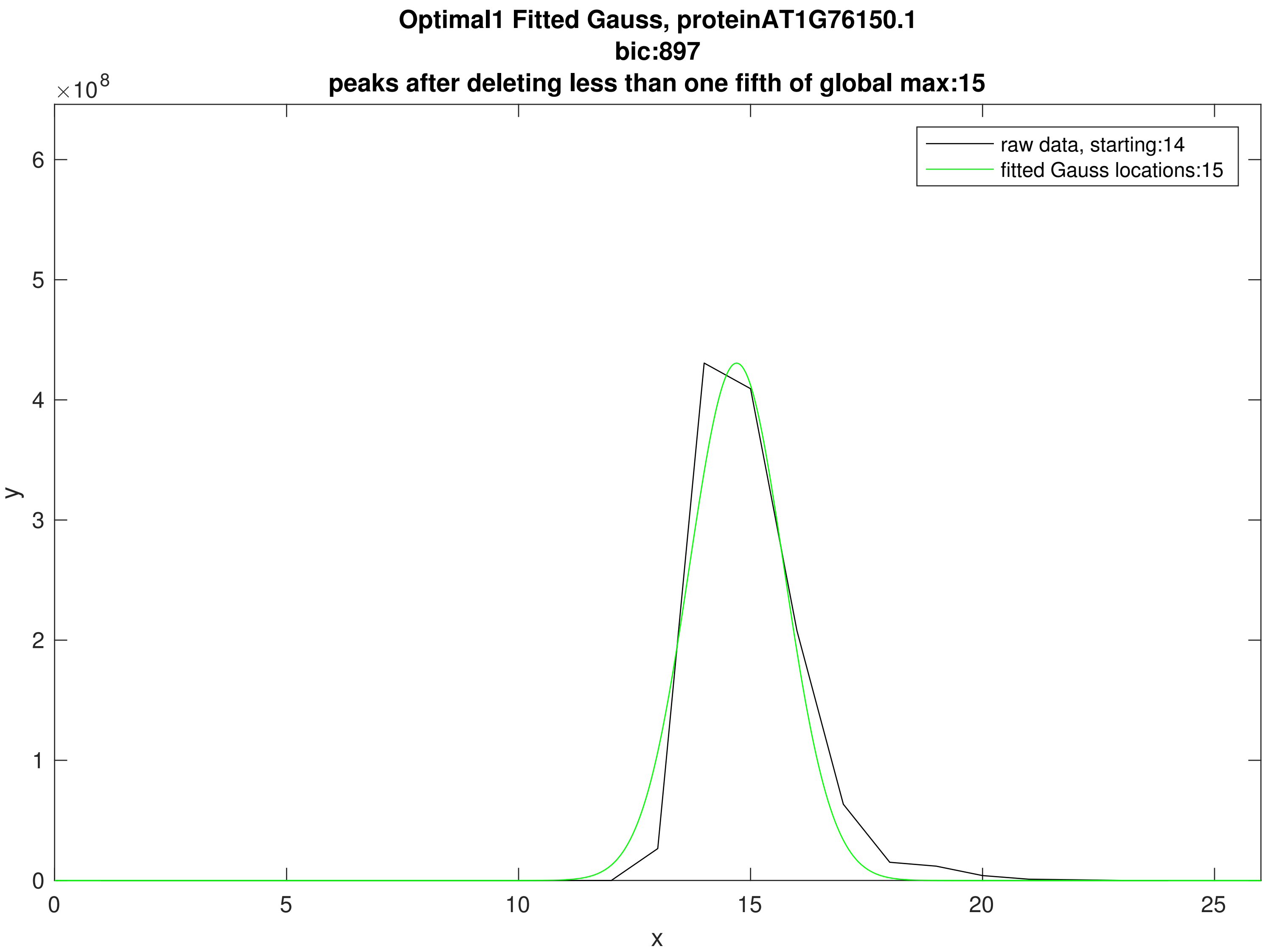
Optimal1 Fitted Gauss, proteinAT1G76150.1
bic:897
peaks after deleting less than one fifth of global max:15
×10^8
raw data, starting:14
fitted Gauss locations:15
y
x
0
1
2
3
4
5
6
0
5
10
15
20
25

#### Optimal1 Fitted Gauss, proteinAT1G76160.1
#### bic:995  1.01e+03  1.01e+03
#### peaks after deleting less than one fifth of global max:16

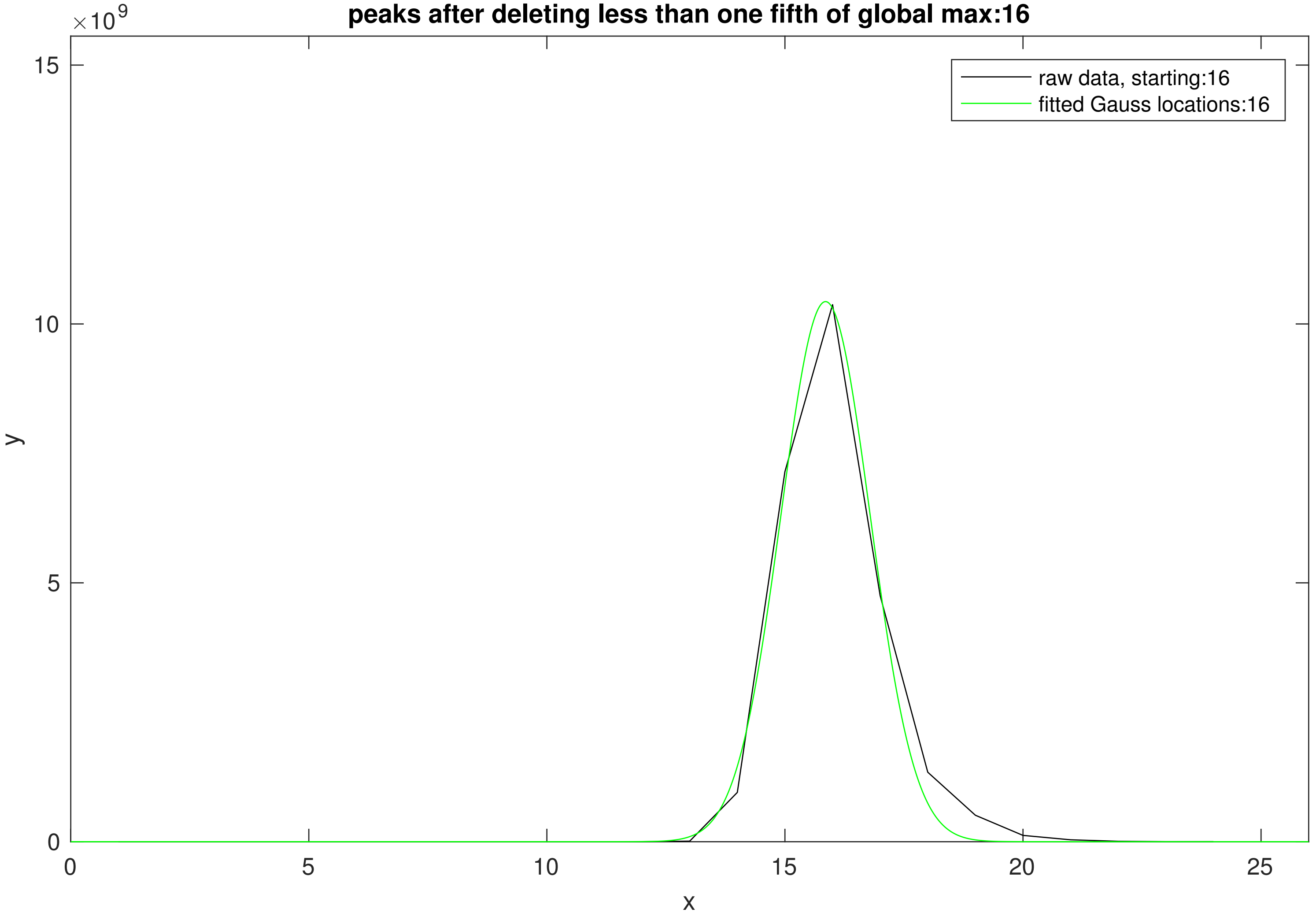

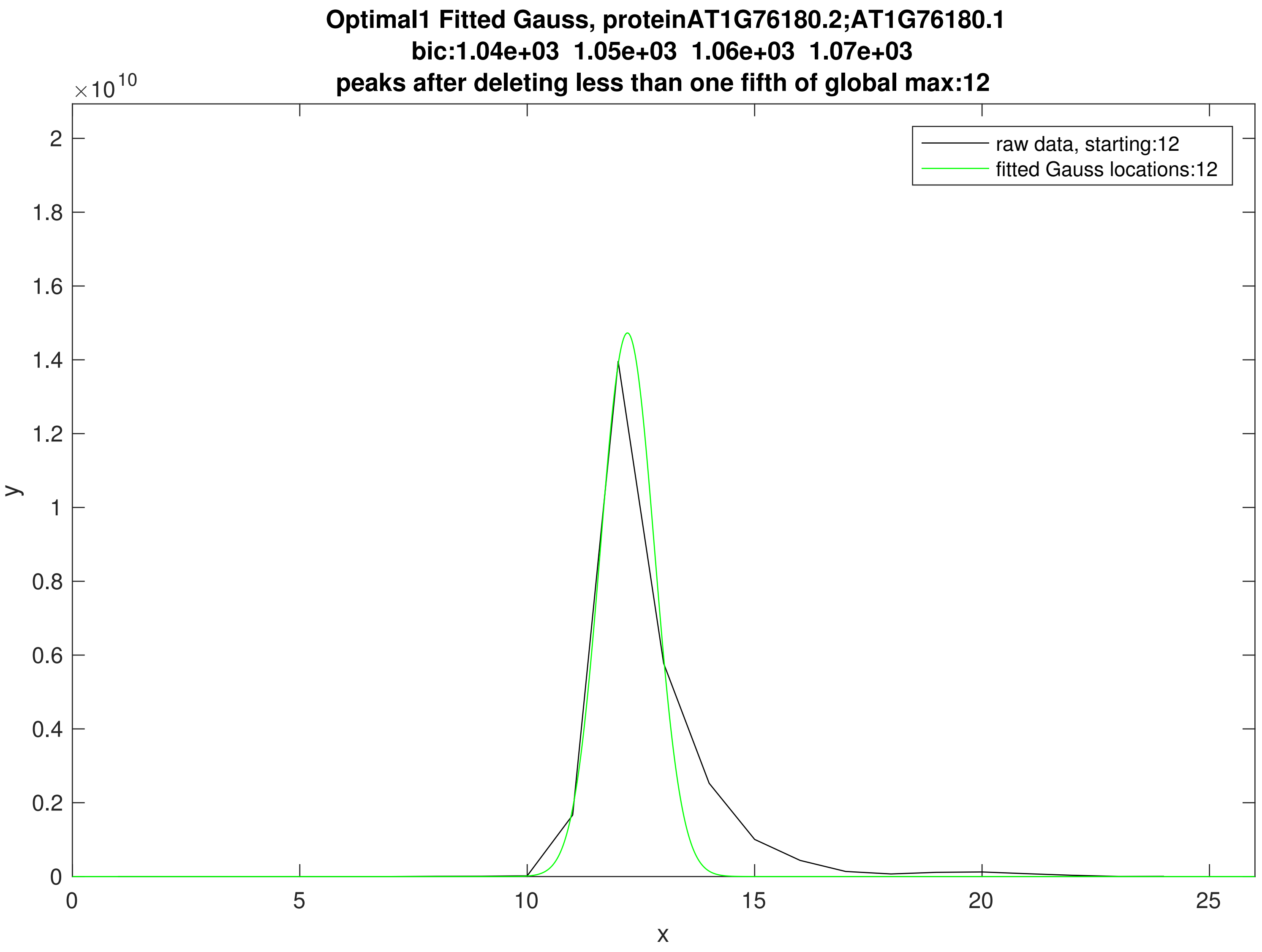

Optimal1 Fitted Gauss, proteinAT1G76180.2;AT1G76180.1
bic:1.04e+03  1.05e+03  1.06e+03  1.07e+03
peaks after deleting less than one fifth of global max:12
×10^10
raw data, starting:12
fitted Gauss locations:12
y
x
0
0.2
0.4
0.6
0.8
1
1.2
1.4
1.6
1.8
2
0
5
10
15
20
25

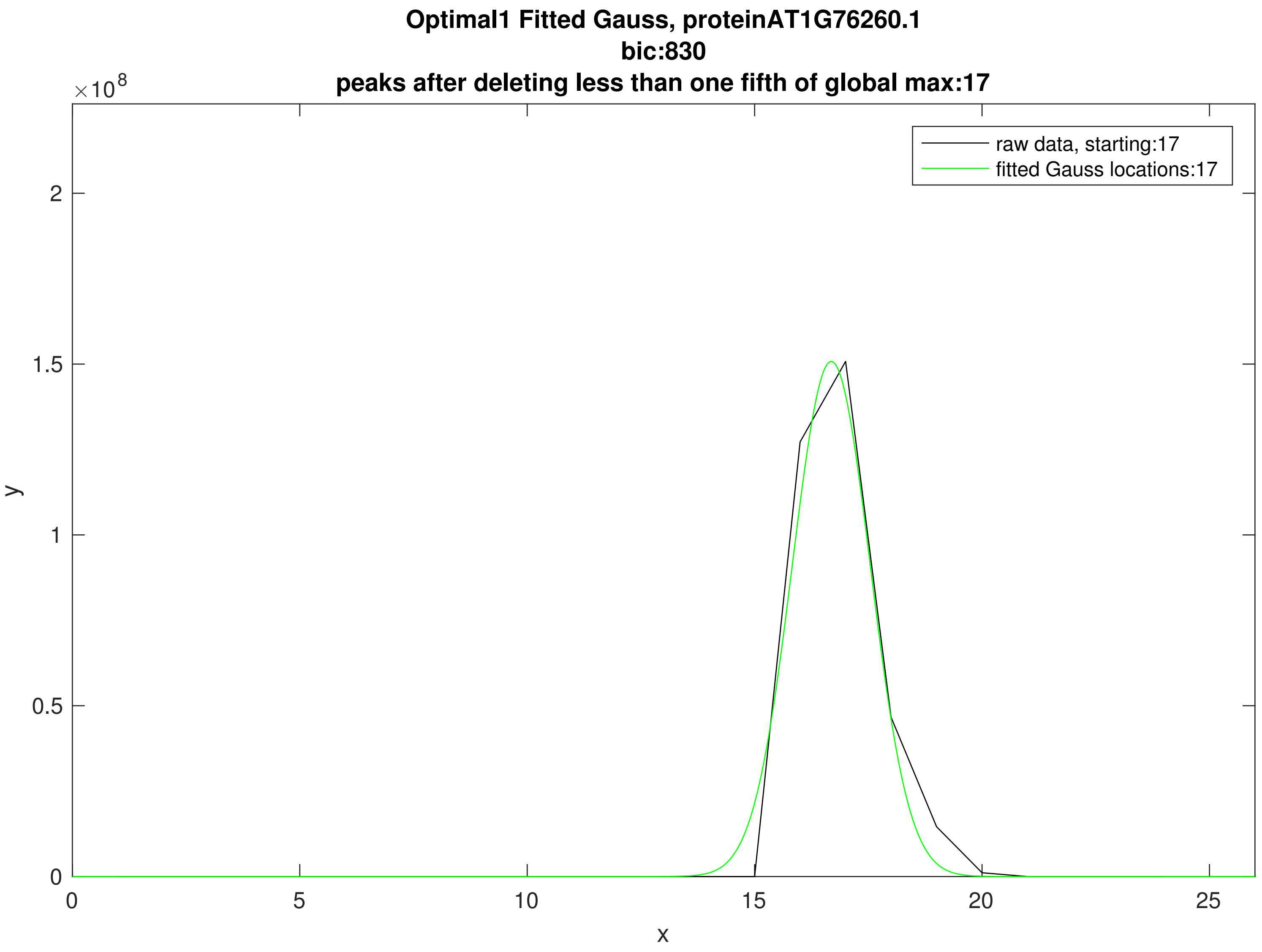
Optimal1 Fitted Gauss, proteinAT1G76260.1
bic:830
peaks after deleting less than one fifth of global max:17
×10^8
raw data, starting:17
fitted Gauss locations:17
y
x
0
0.5
1
1.5
2
0
5
10
15
20
25

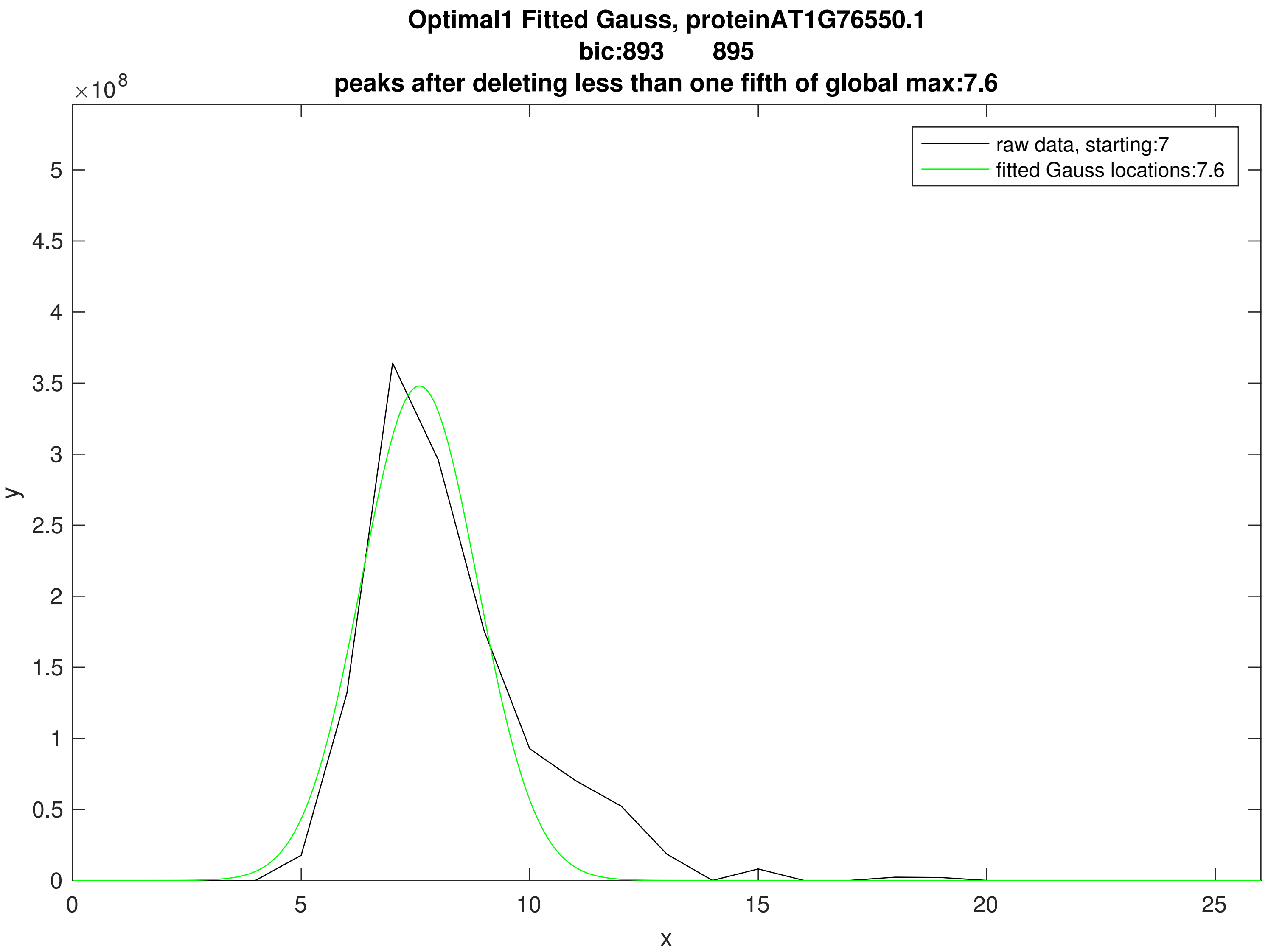
Optimal1 Fitted Gauss, proteinAT1G76550.1
bic:893 895
peaks after deleting less than one fifth of global max:7.6
×10^8
raw data, starting:7
fitted Gauss locations:7.6
y
x
0
0.5
1
1.5
2
2.5
3
3.5
4
4.5
5
0
5
10
15
20
25

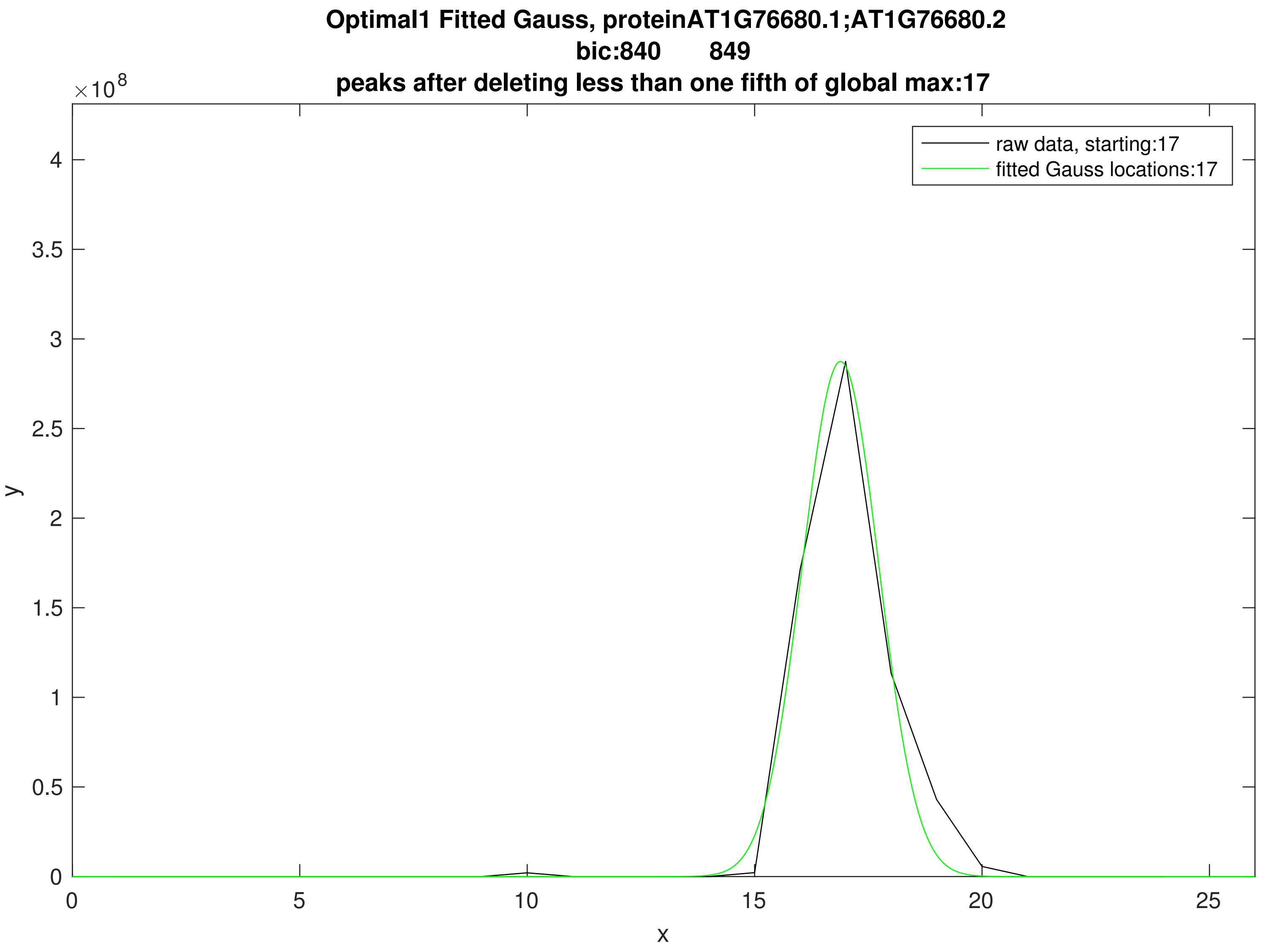
Optimal1 Fitted Gauss, proteinAT1G76680.1;AT1G76680.2
bic:840 849
peaks after deleting less than one fifth of global max:17
×10^8
raw data, starting:17
fitted Gauss locations:17
y
x
0
0.5
1
1.5
2
2.5
3
3.5
4
5
10
15
20
25

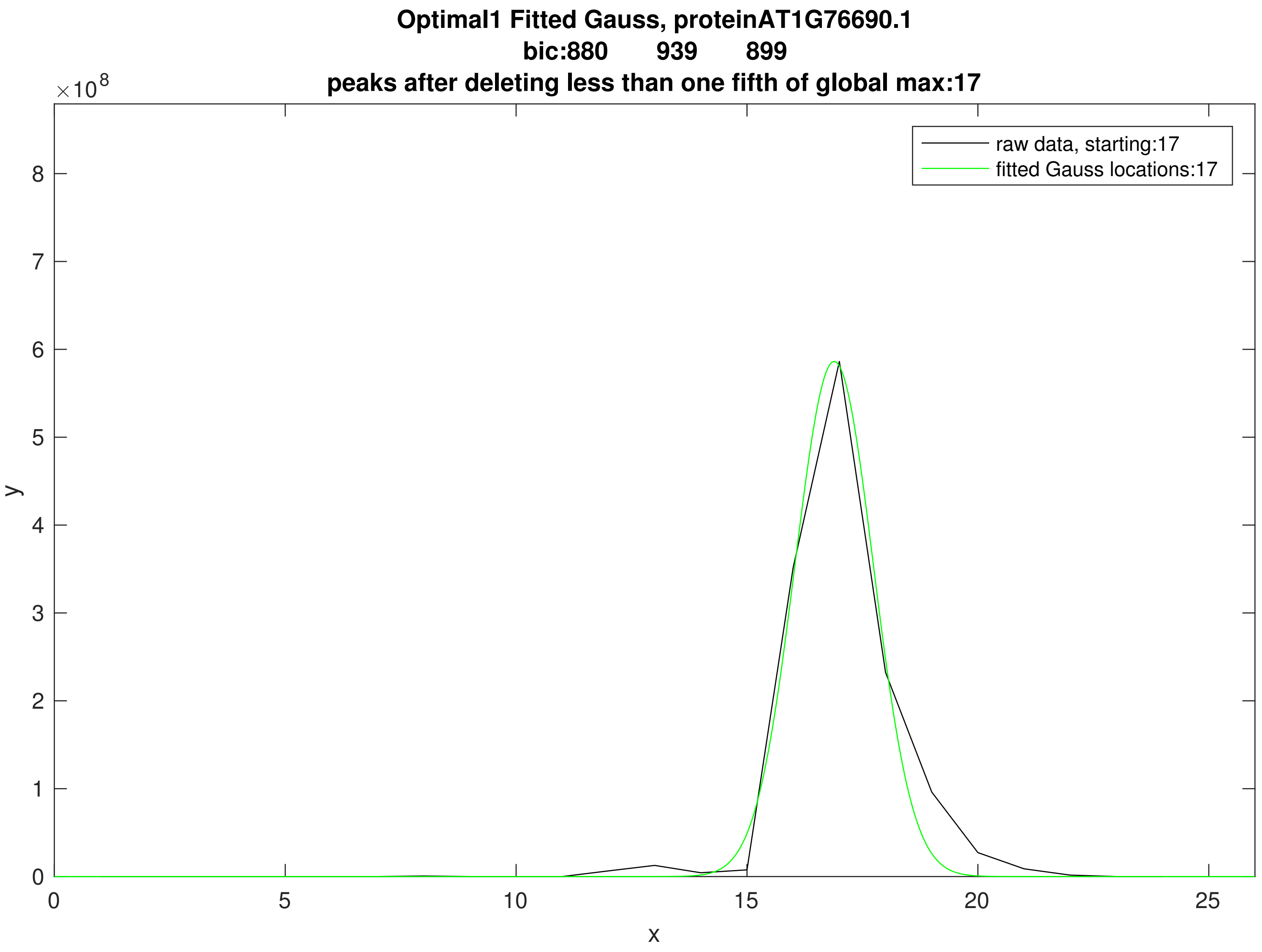

Optimal1 Fitted Gauss, proteinAT1G76690.1
bic:880 939 899
peaks after deleting less than one fifth of global max:17
×10^8
8
7
6
5
4
3
2
1
0
y
0
5
10
15
20
25
x
raw data, starting:17
fitted Gauss locations:17

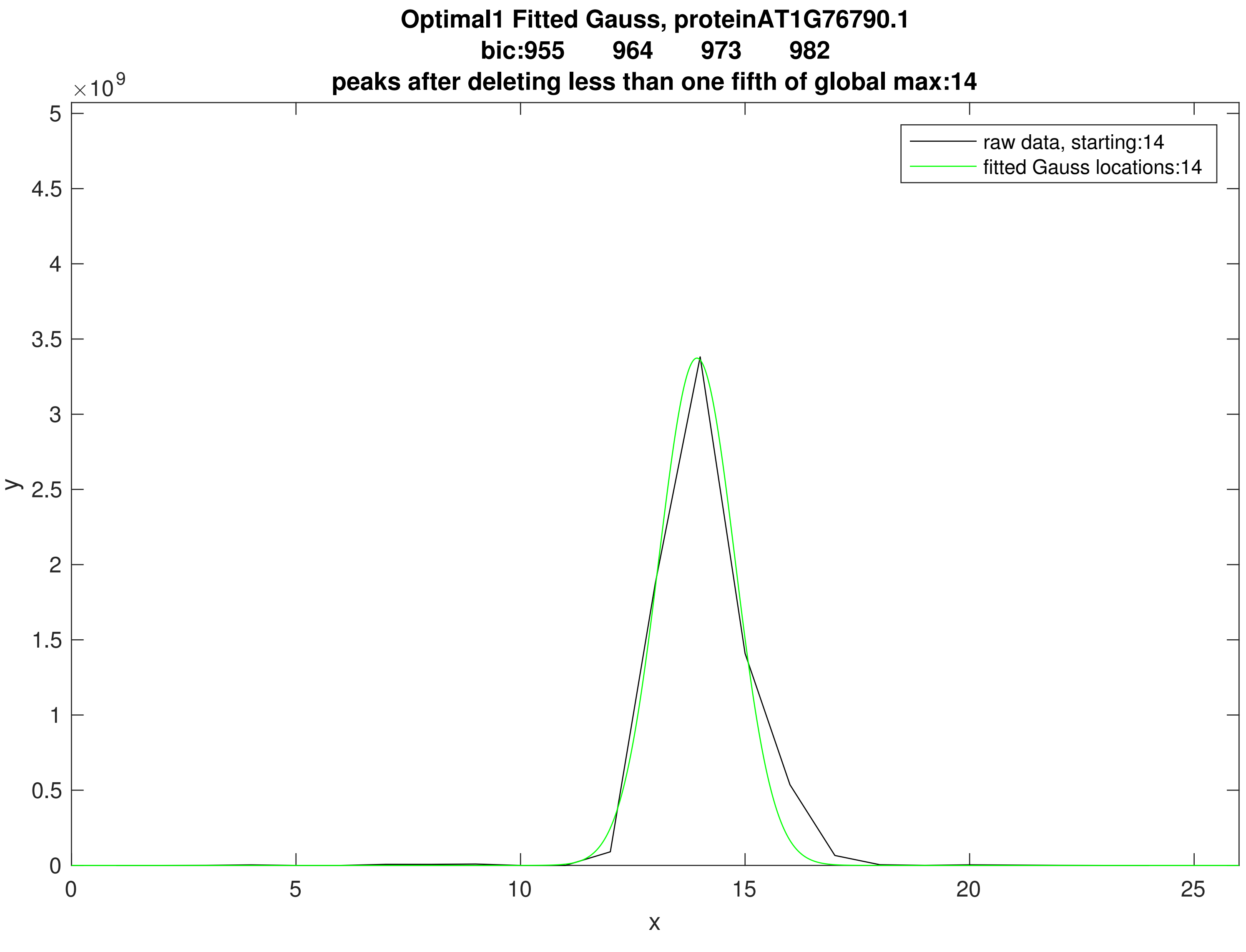
Optimal1 Fitted Gauss, proteinAT1G76790.1
bic:955 964 973 982
peaks after deleting less than one fifth of global max:14
×10^9
raw data, starting:14
fitted Gauss locations:14
y
x
0
0.5
1
1.5
2
2.5
3
3.5
4
4.5
5
0
5
10
15
20
25

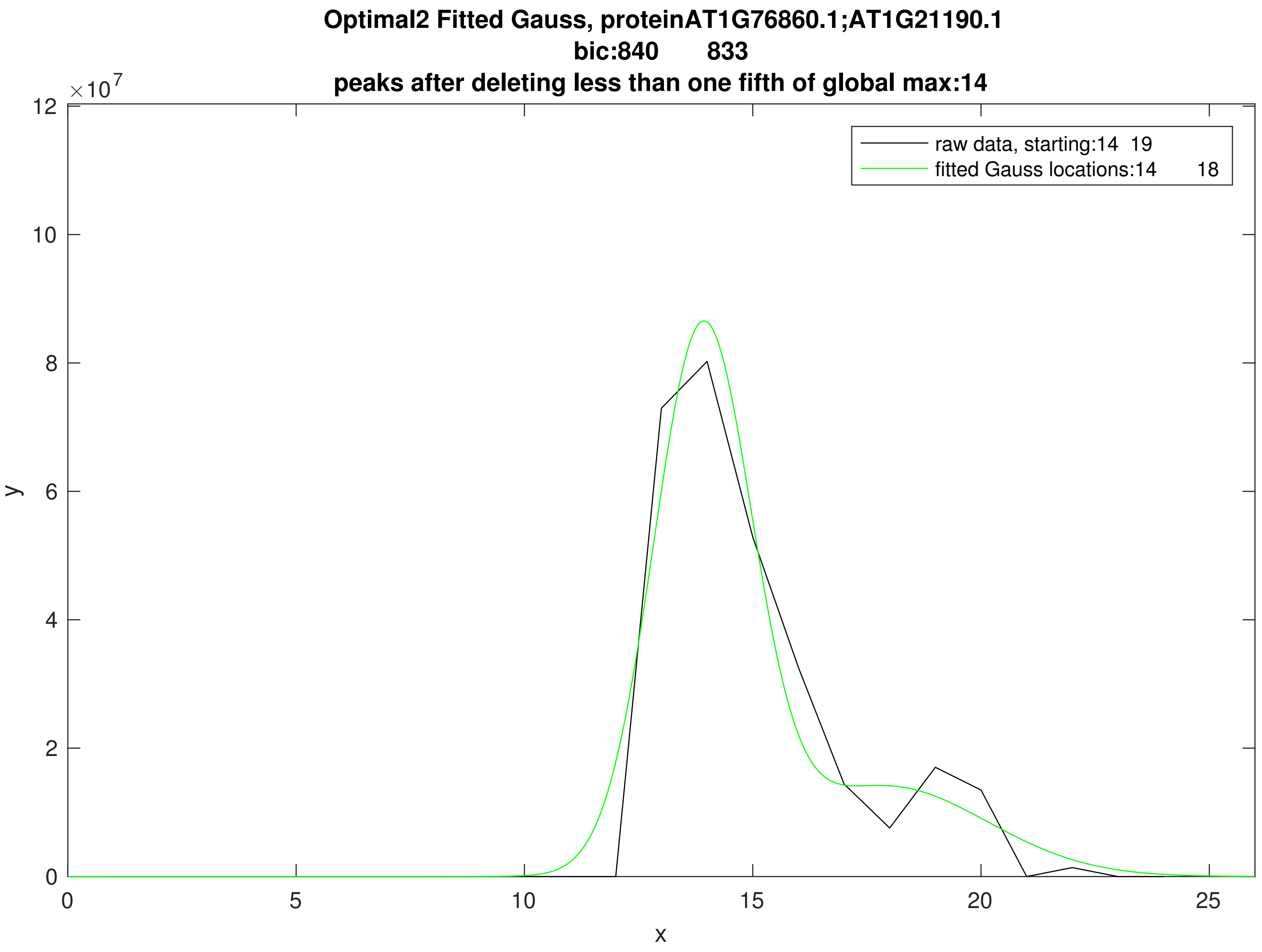

Optimal2 Fitted Gauss, proteinAT1G76860.1;AT1G21190.1
bic:840 833
peaks after deleting less than one fifth of global max:14
×10^7
12
10
8
6
4
2
0
y
0
5
10
15
20
25
x
raw data, starting:14 19
fitted Gauss locations:14 18

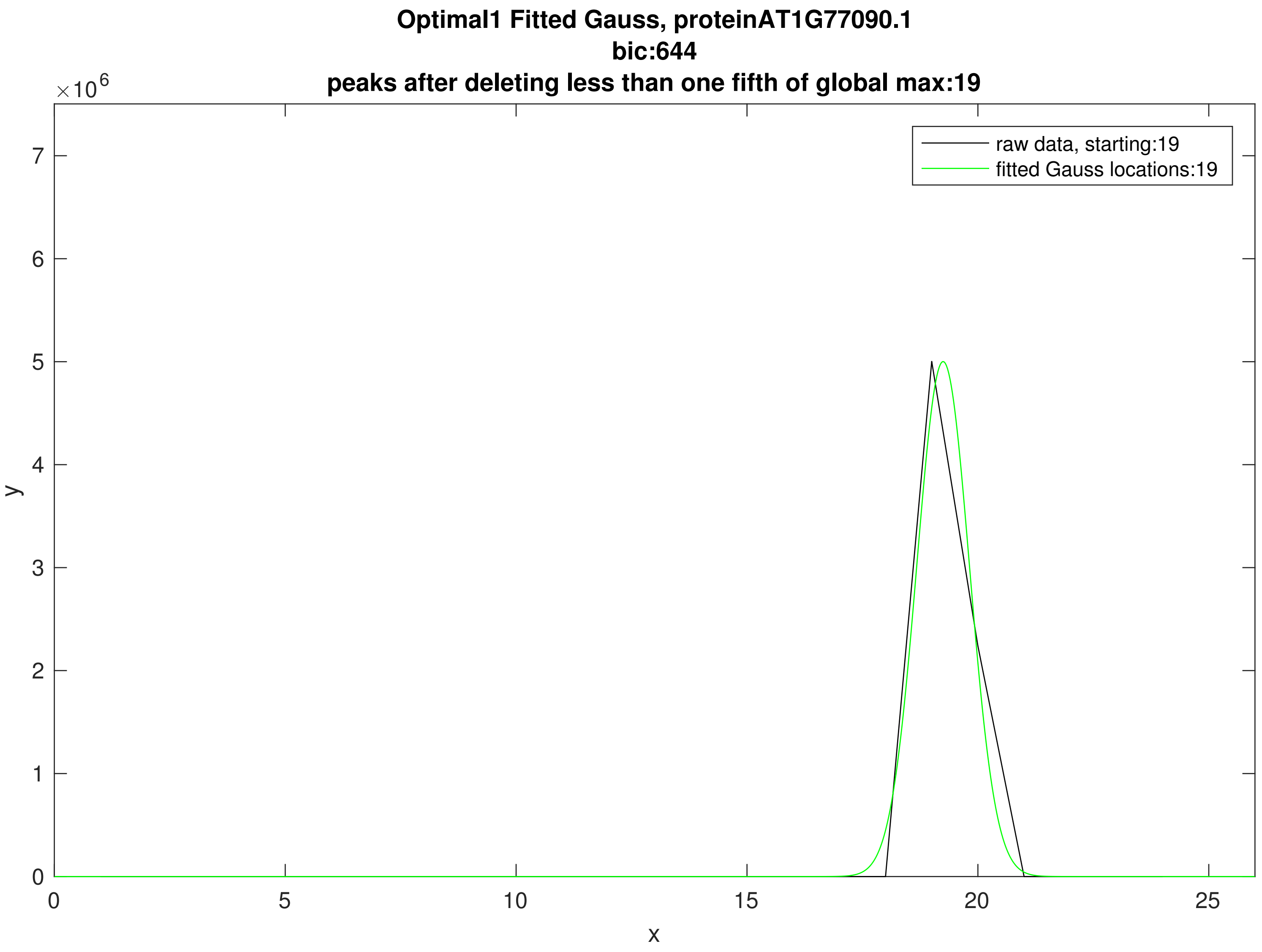
Optimal1 Fitted Gauss, proteinAT1G77090.1
bic:644
peaks after deleting less than one fifth of global max:19
×10^6
raw data, starting:19
fitted Gauss locations:19
y
x
0
1
2
3
4
5
6
7
0
5
10
15
20
25

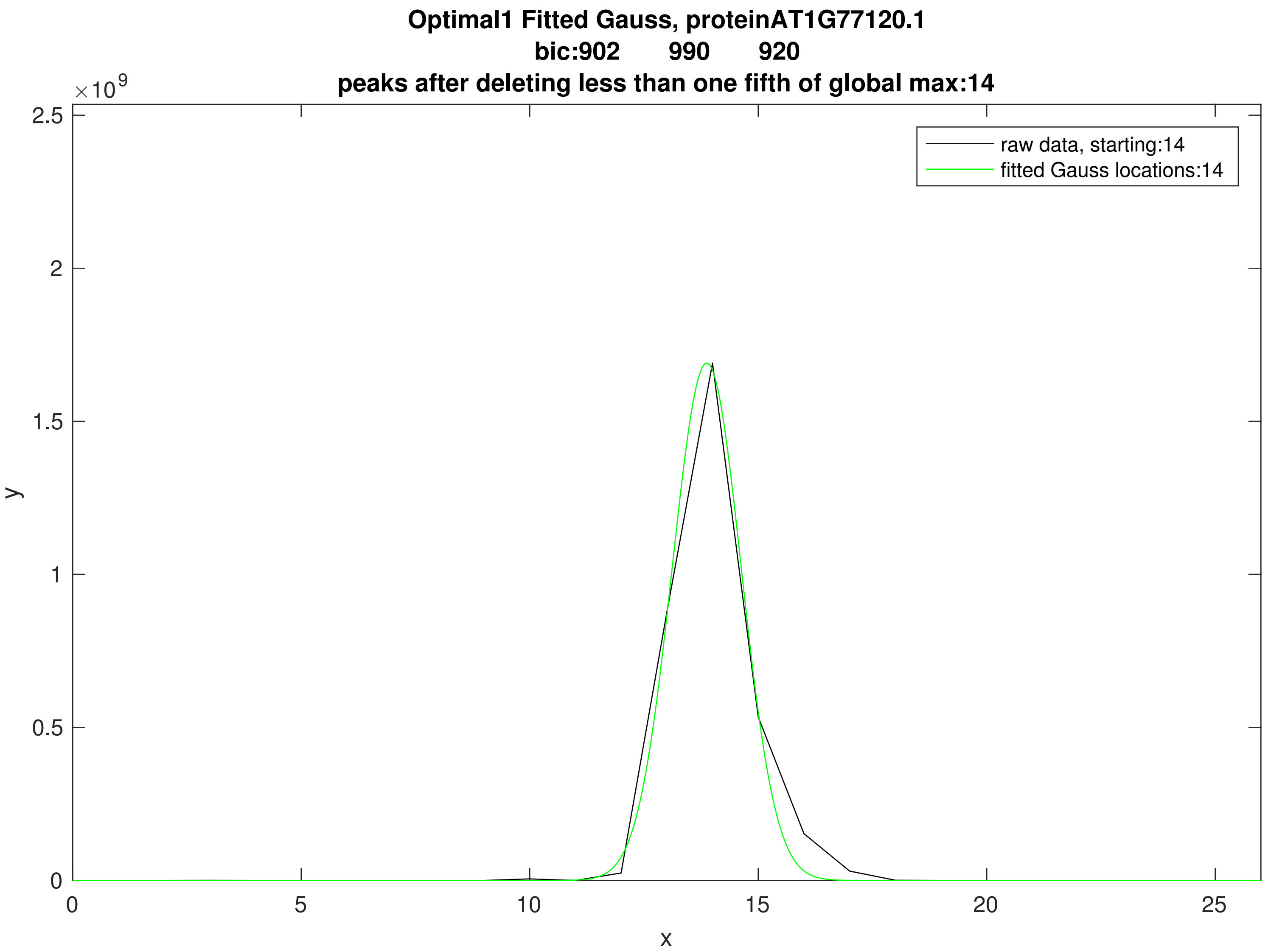
Optimal1 Fitted Gauss, proteinAT1G77120.1
bic:902 990 920
peaks after deleting less than one fifth of global max:14
×10^9
2.5
2
1.5
1
0.5
0
y
0
5
10
15
20
25
x
raw data, starting:14
fitted Gauss locations:14

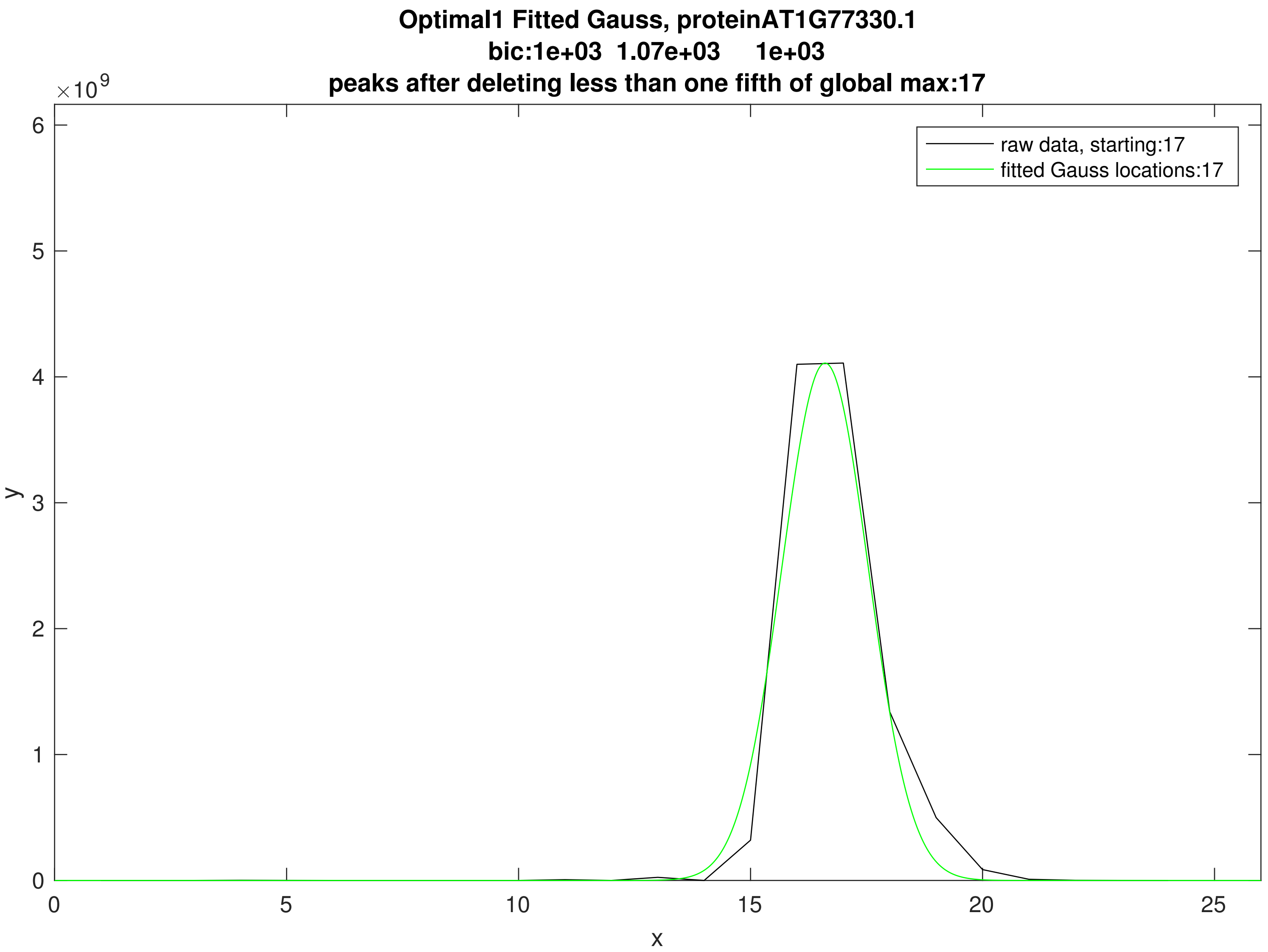
Optimal1 Fitted Gauss, proteinAT1G77330.1
bic:1e+03 1.07e+03 1e+03
peaks after deleting less than one fifth of global max:17
×10^9
raw data, starting:17
fitted Gauss locations:17
y
x
0
1
2
3
4
5
6
0
5
10
15
20
25

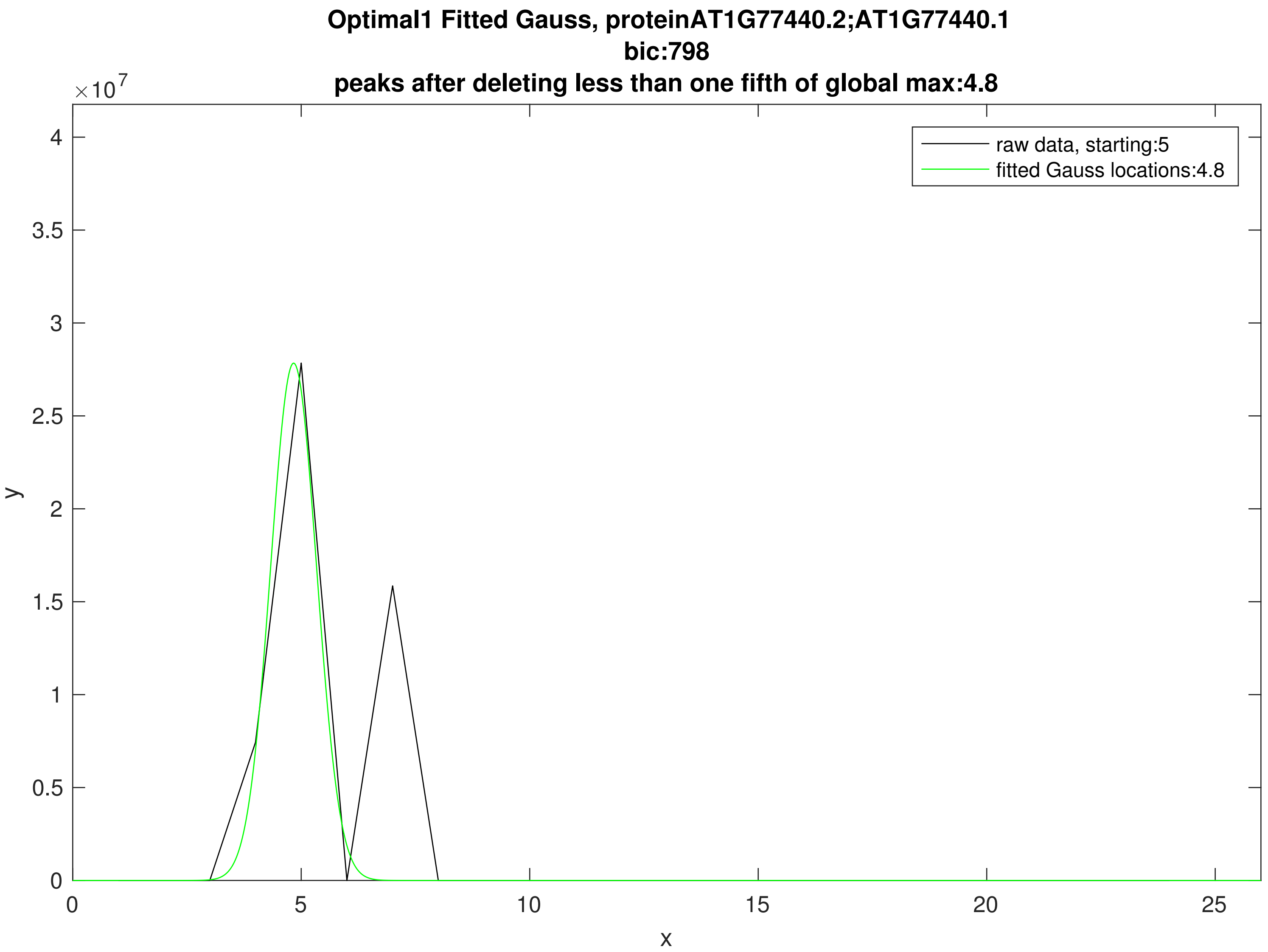
Optimal1 Fitted Gauss, proteinAT1G77440.2;AT1G77440.1
bic:798
peaks after deleting less than one fifth of global max:4.8
×10^7
raw data, starting:5
fitted Gauss locations:4.8
y
x
0
0.5
1
1.5
2
2.5
3
3.5
4
5
10
15
20
25

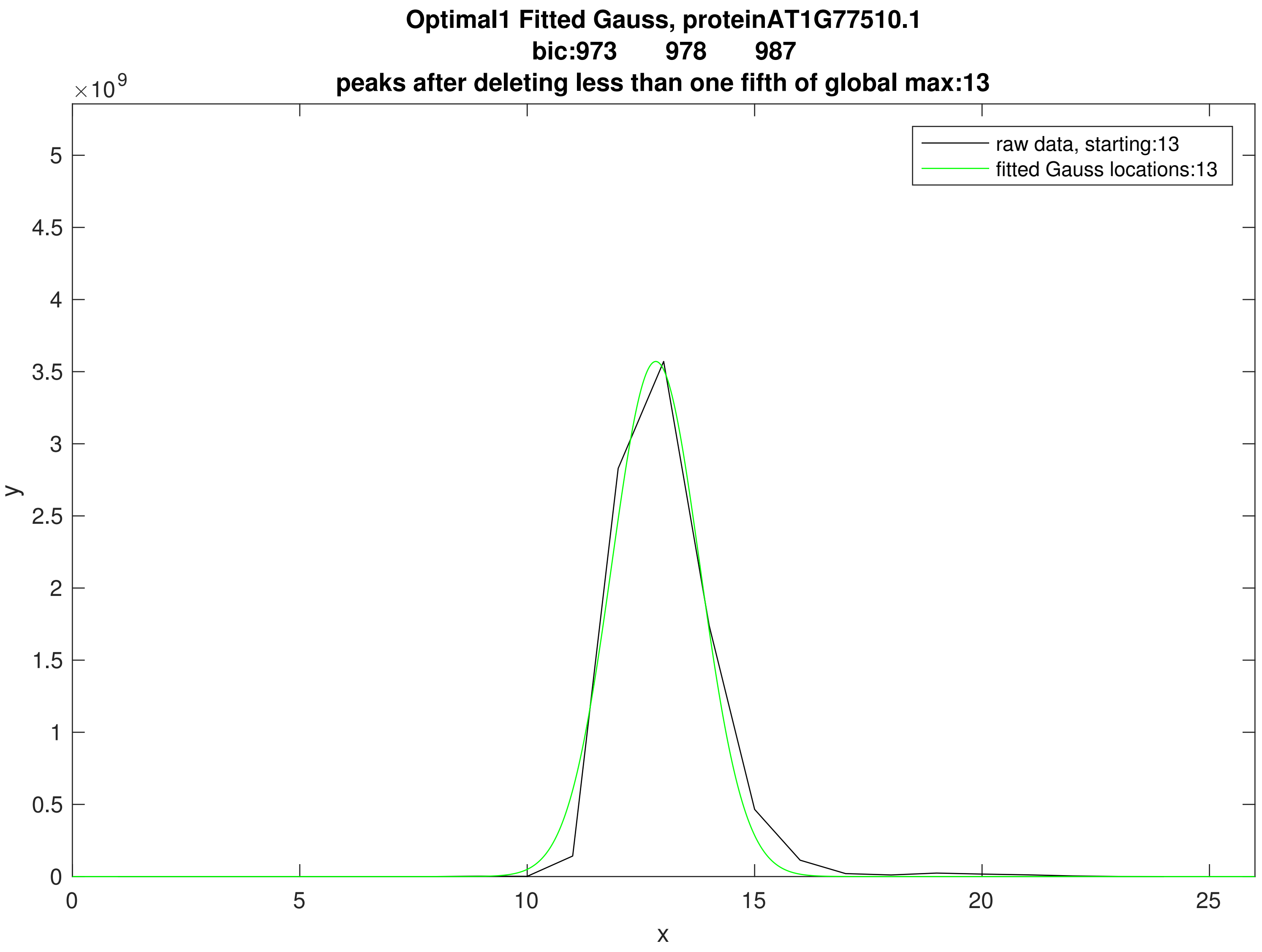
Optimal1 Fitted Gauss, proteinAT1G77510.1
bic:973 978 987
peaks after deleting less than one fifth of global max:13
×10^9
raw data, starting:13
fitted Gauss locations:13
y
x
0
0.5
1
1.5
2
2.5
3
3.5
4
4.5
5
0
5
10
15
20
25

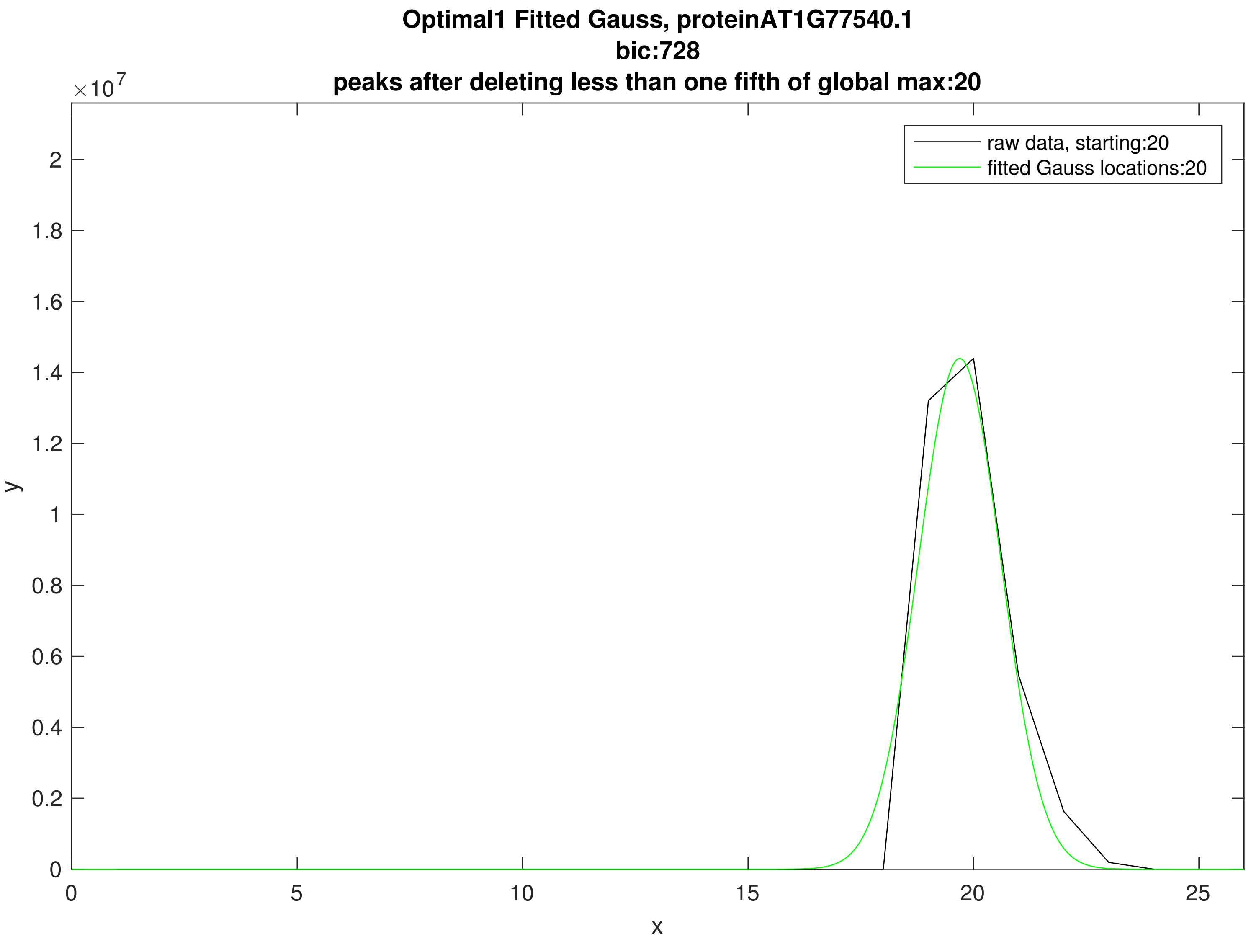

Optimal1 Fitted Gauss, proteinAT1G77540.1
bic:728
peaks after deleting less than one fifth of global max:20
×10^7
raw data, starting:20
fitted Gauss locations:20
y
x
0
0.2
0.4
0.6
0.8
1
1.2
1.4
1.6
1.8
2
5
10
15
20
25

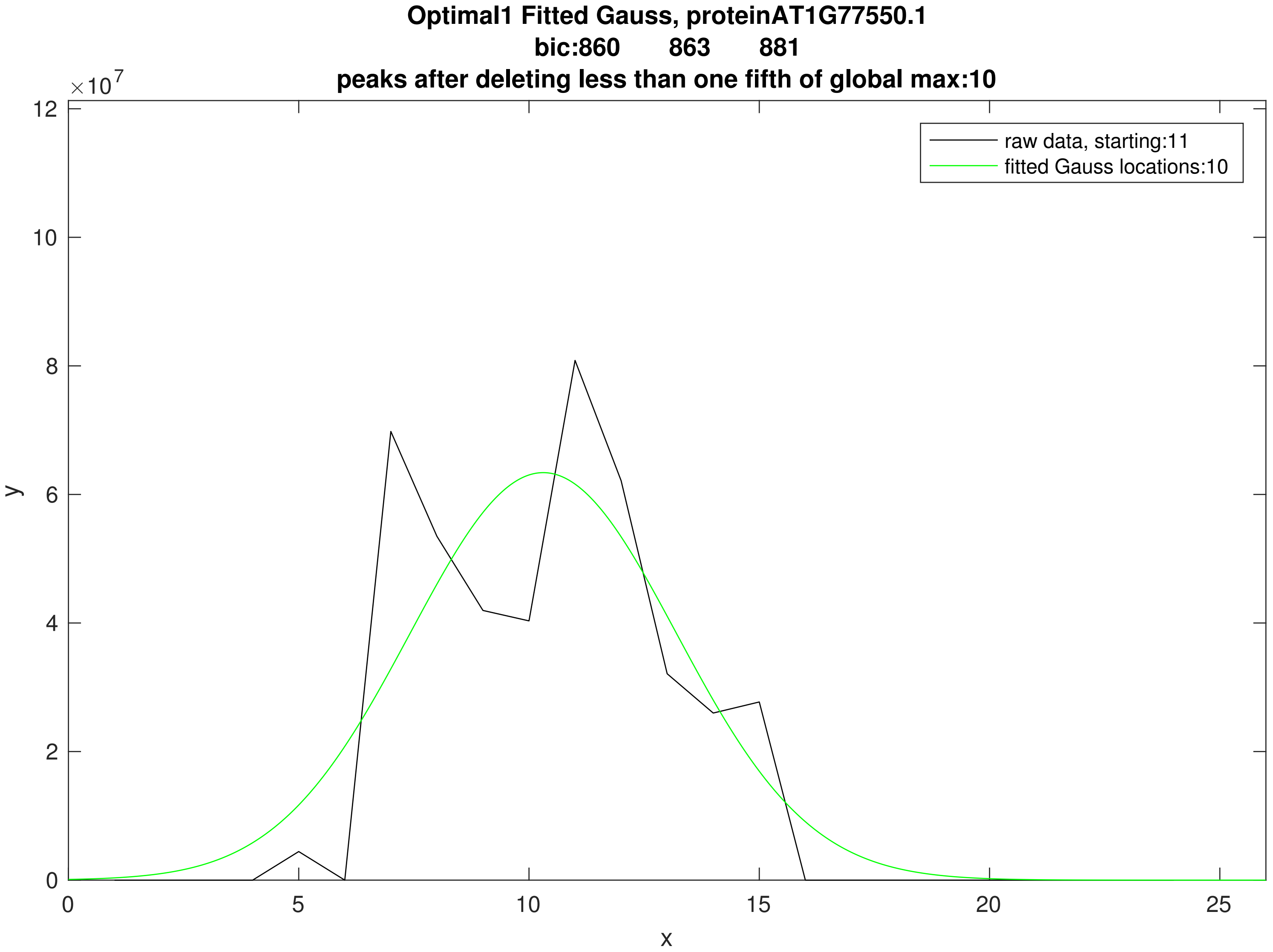
Optimal1 Fitted Gauss, proteinAT1G77550.1
bic:860 863 881
peaks after deleting less than one fifth of global max:10
×10^7
raw data, starting:11
fitted Gauss locations:10
y
x
0
2
4
6
8
10
12
0
5
10
15
20
25

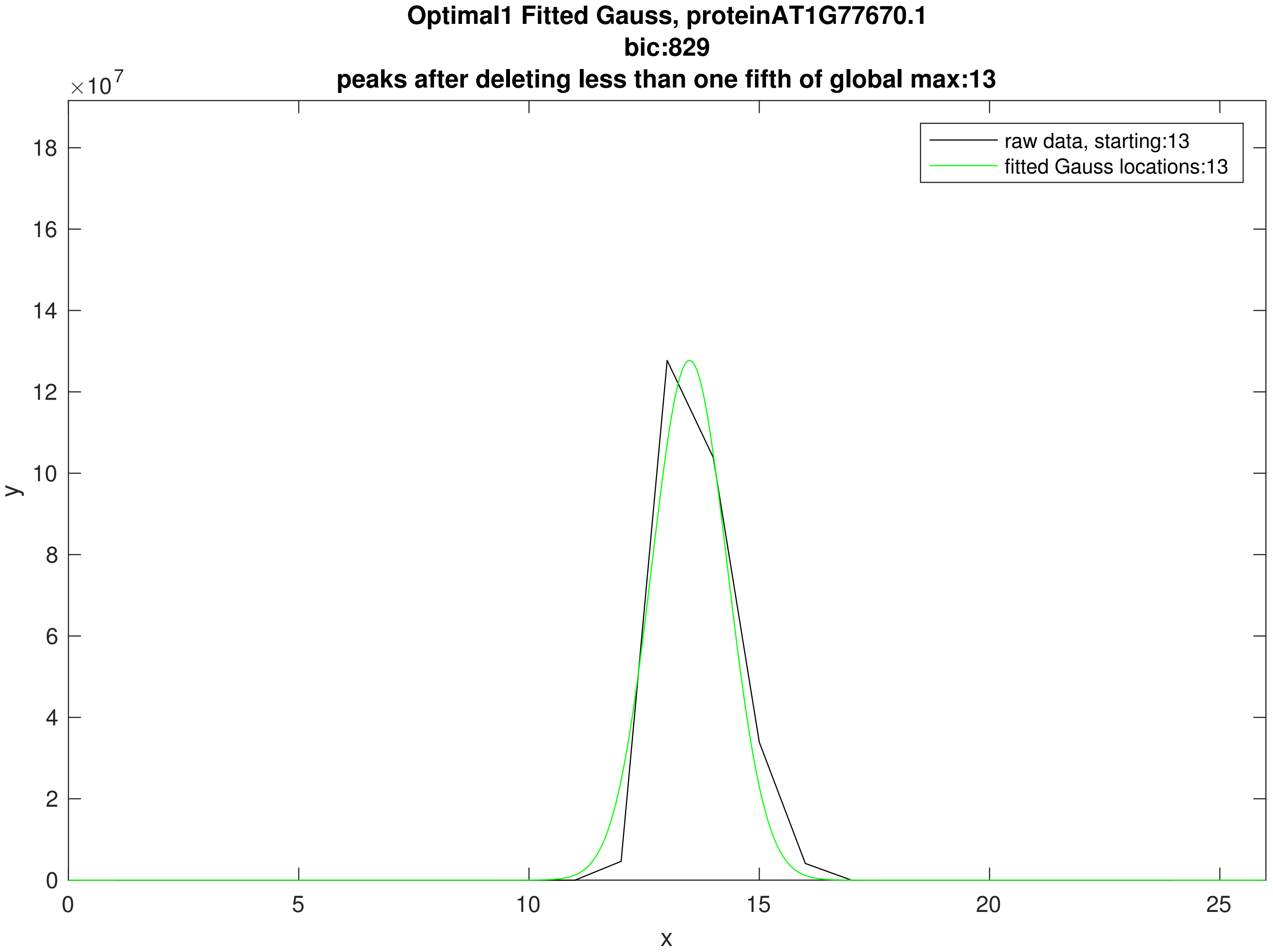

Optimal1 Fitted Gauss, proteinAT1G77670.1
bic:829
peaks after deleting less than one fifth of global max:13
×10^7
raw data, starting:13
fitted Gauss locations:13
y
x
0
2
4
6
8
10
12
14
16
18
0
5
10
15
20
25

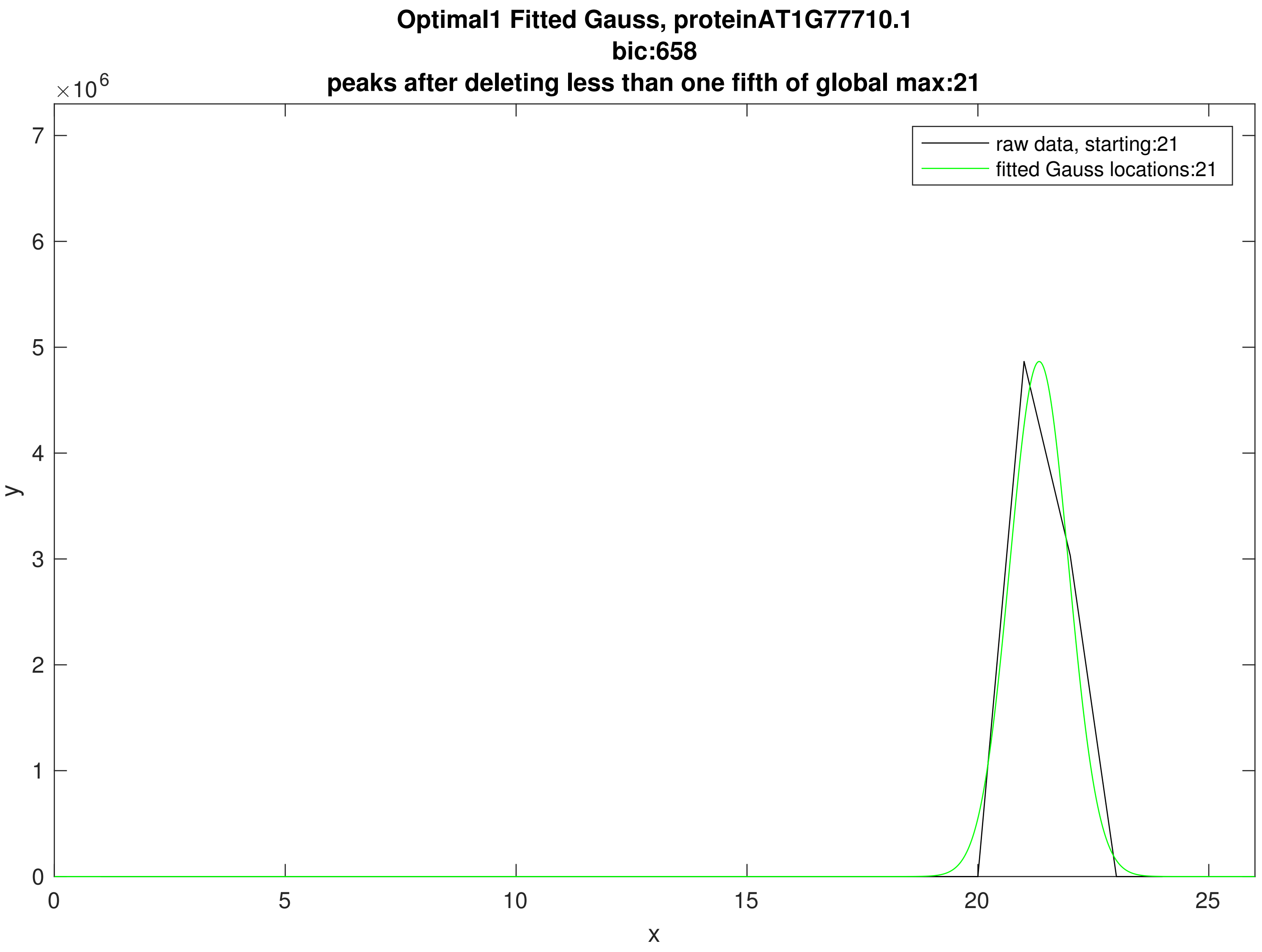

Optimal1 Fitted Gauss, proteinAT1G77710.1
bic:658
peaks after deleting less than one fifth of global max:21
×10^6
raw data, starting:21
fitted Gauss locations:21
y
x
0
1
2
3
4
5
6
7
0
5
10
15
20
25

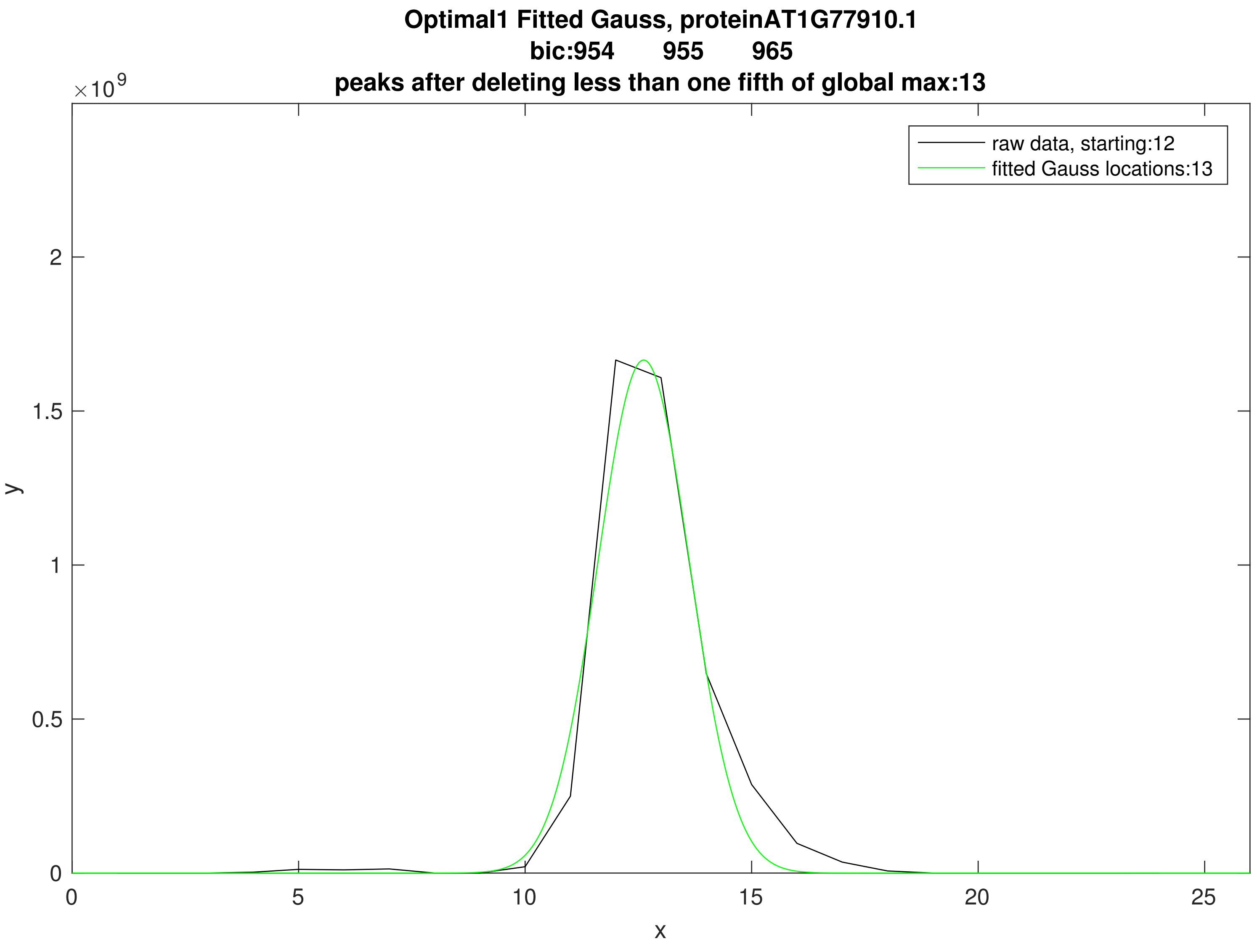
Optimal1 Fitted Gauss, proteinAT1G77910.1
bic:954 955 965
peaks after deleting less than one fifth of global max:13
×10^9
raw data, starting:12
fitted Gauss locations:13
y
x
0
0.5
1
1.5
2
0
5
10
15
20
25

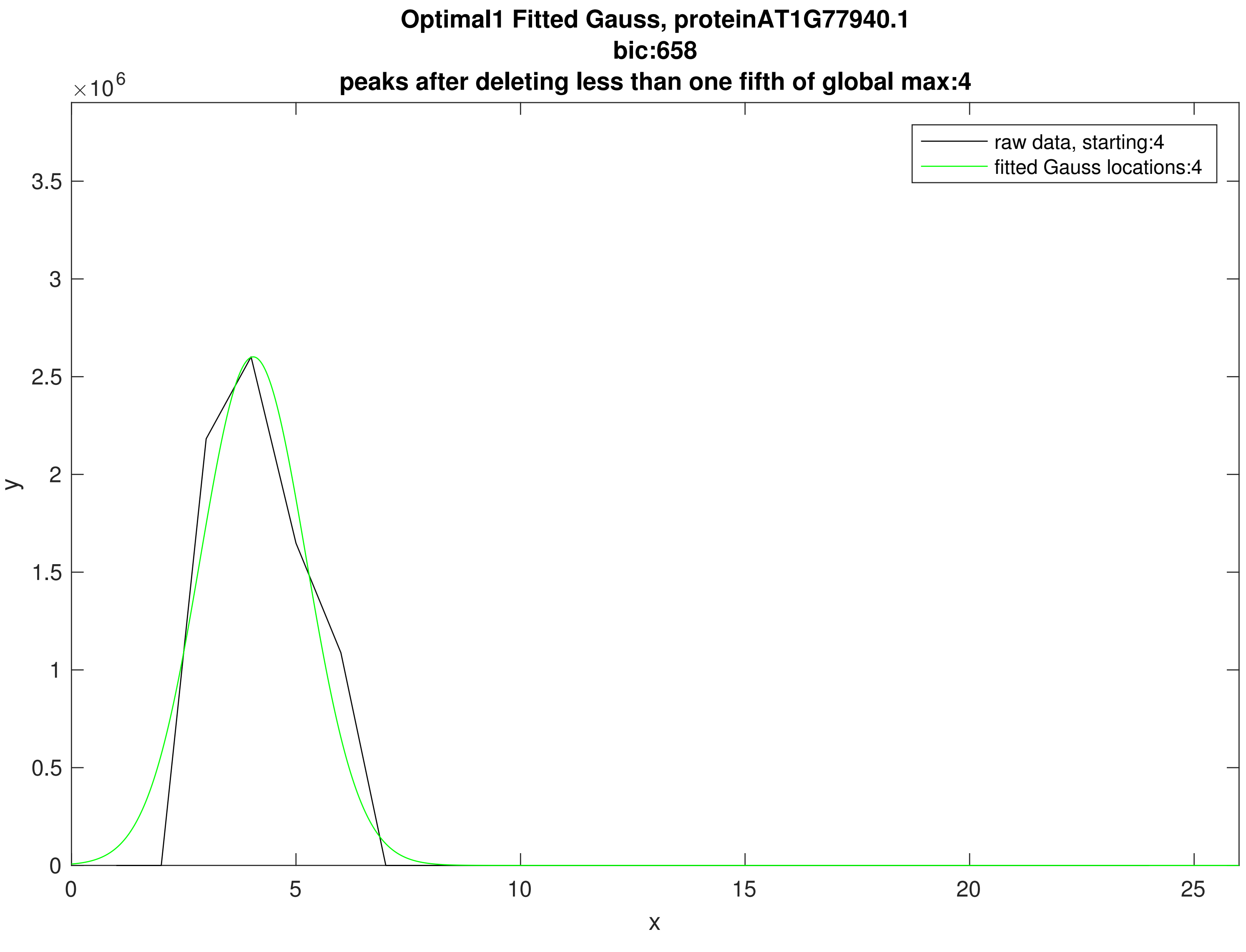
Optimal1 Fitted Gauss, proteinAT1G77940.1
bic:658
peaks after deleting less than one fifth of global max:4
×10^6
raw data, starting:4
fitted Gauss locations:4
y
x
0
0.5
1
1.5
2
2.5
3
3.5
0
5
10
15
20
25

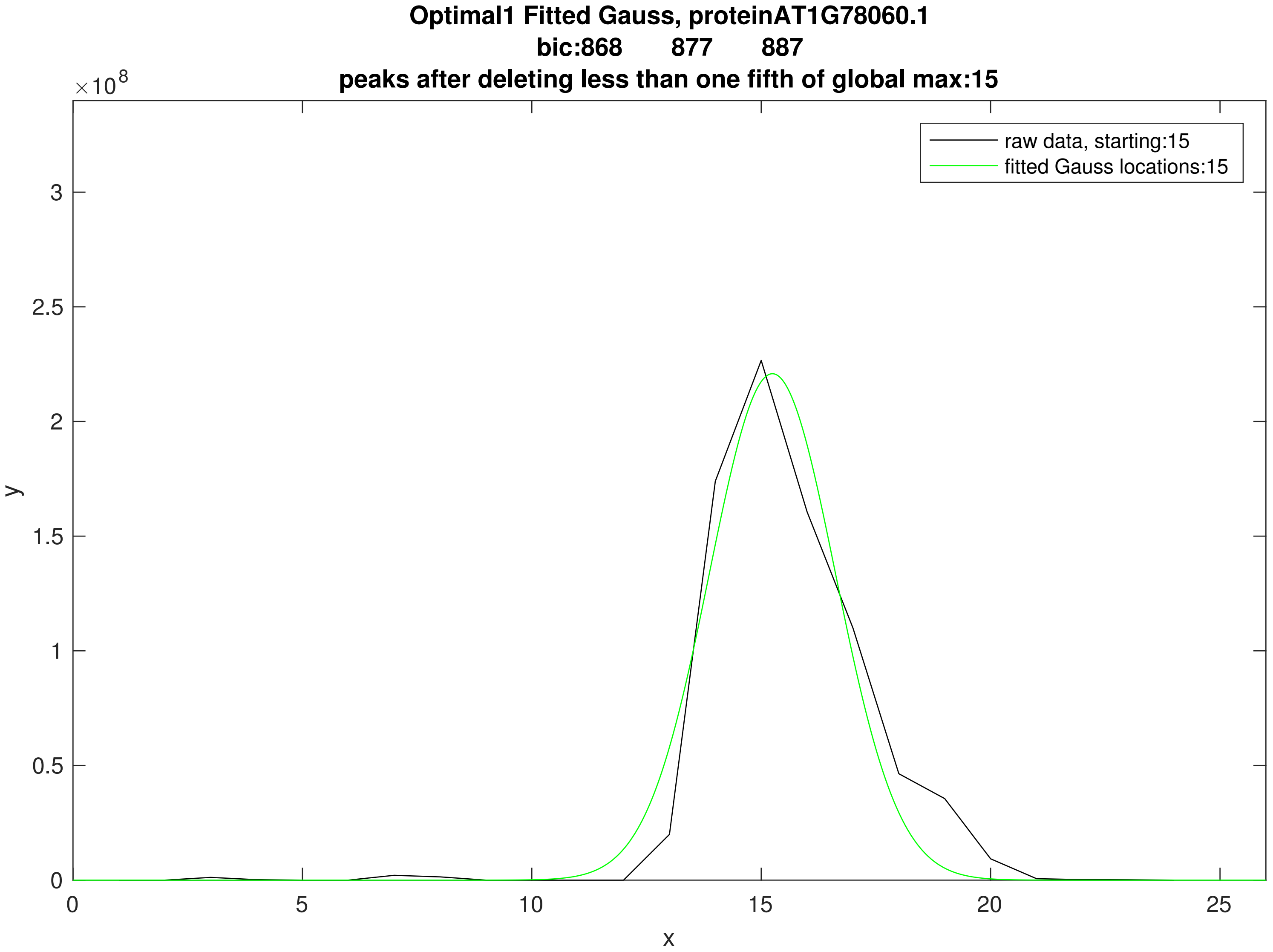
Optimal1 Fitted Gauss, proteinAT1G78060.1
bic:868 877 887
peaks after deleting less than one fifth of global max:15
×10^8
raw data, starting:15
fitted Gauss locations:15
y
x
0
0.5
1
1.5
2
2.5
3
0
5
10
15
20
25

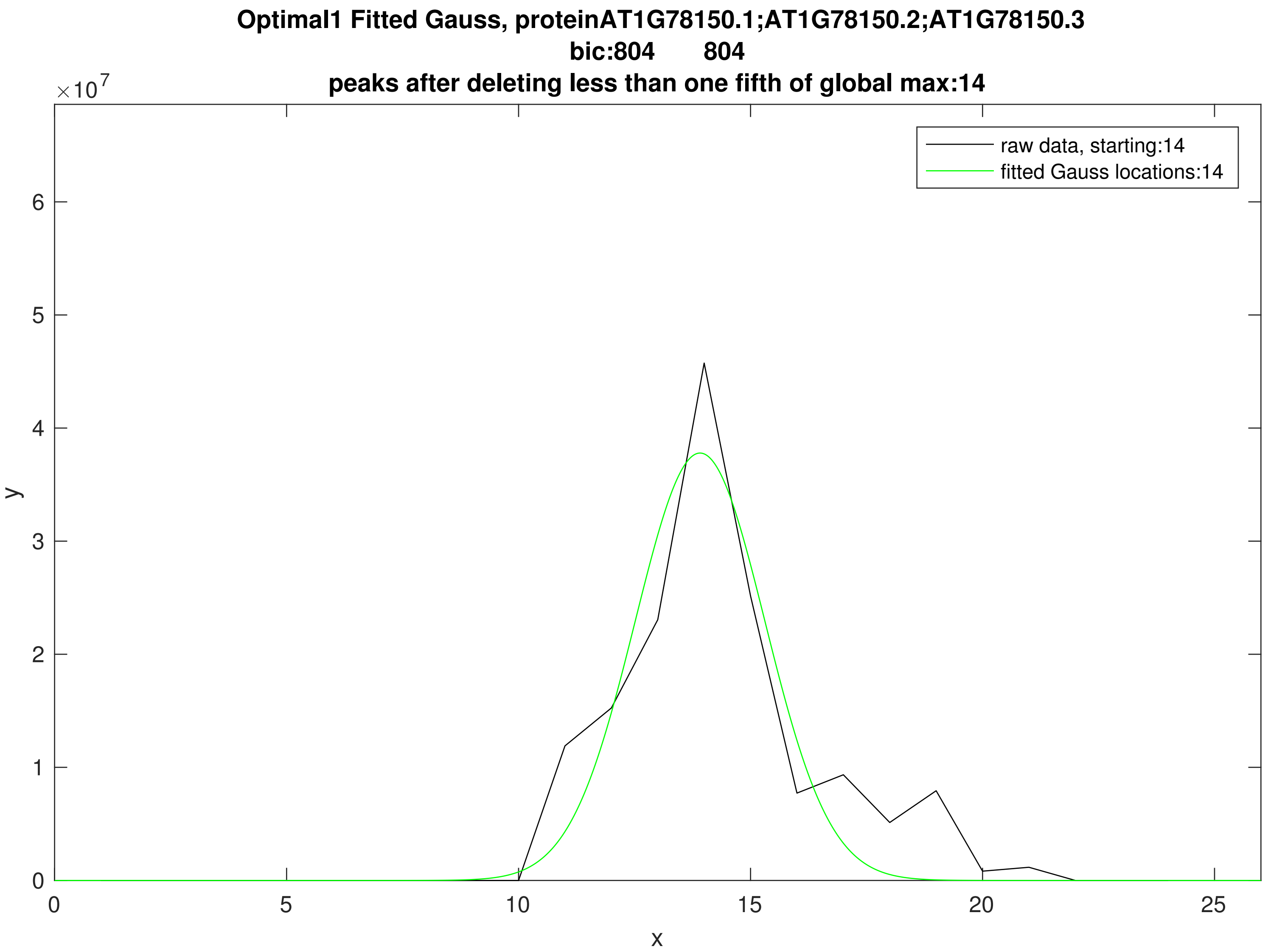

Optimal1 Fitted Gauss, proteinAT1G78150.1;AT1G78150.2;AT1G78150.3
bic:804 804
peaks after deleting less than one fifth of global max:14
×10^7
raw data, starting:14
fitted Gauss locations:14
y
x
0
1
2
3
4
5
6
0
5
10
15
20
25

**Optimal1 Fitted Gauss, proteinAT1G78260.2;AT1G78260.1;AT1G22330.1**
**bic:663**
**peaks after deleting less than one fifth of global max:21**

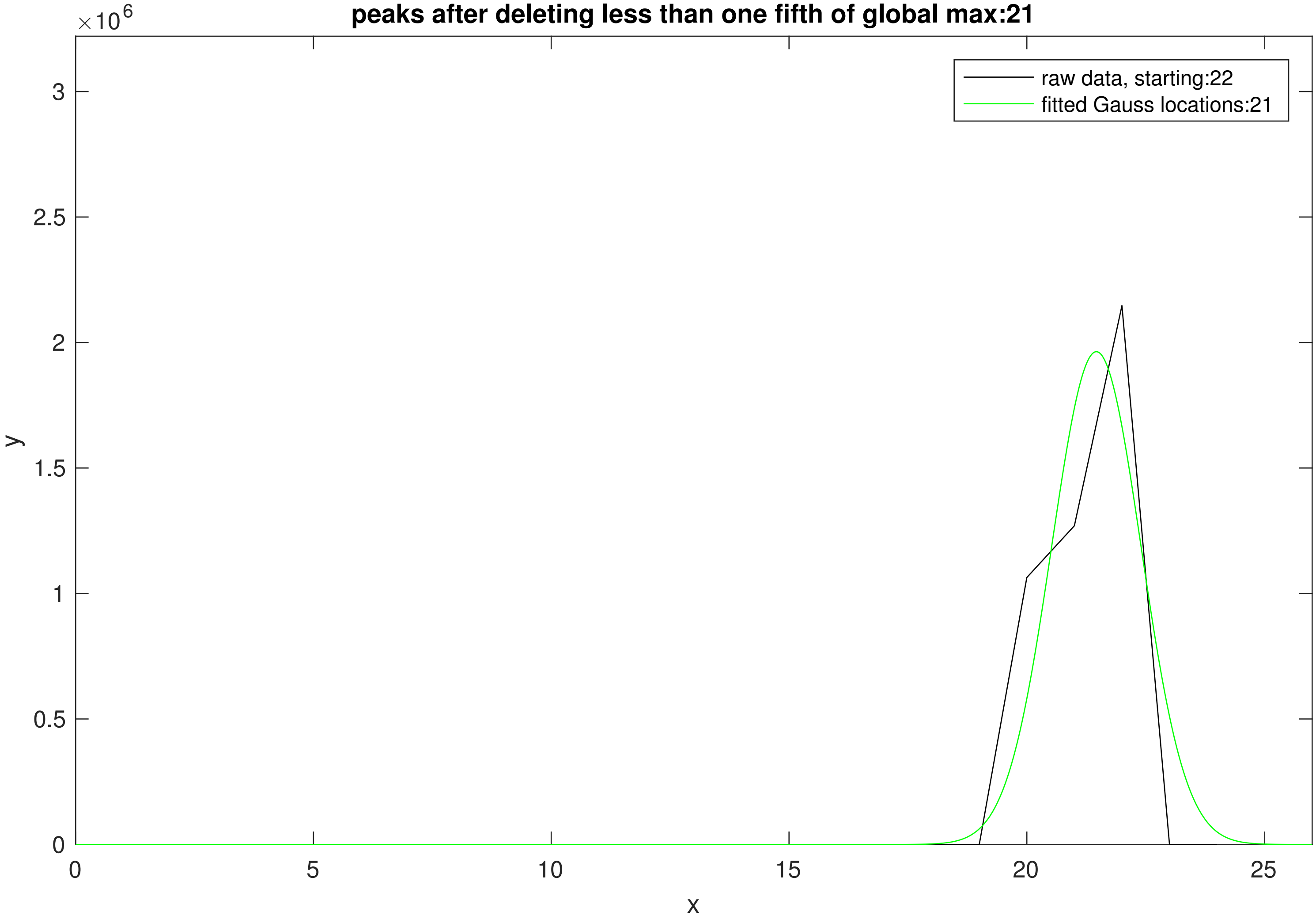

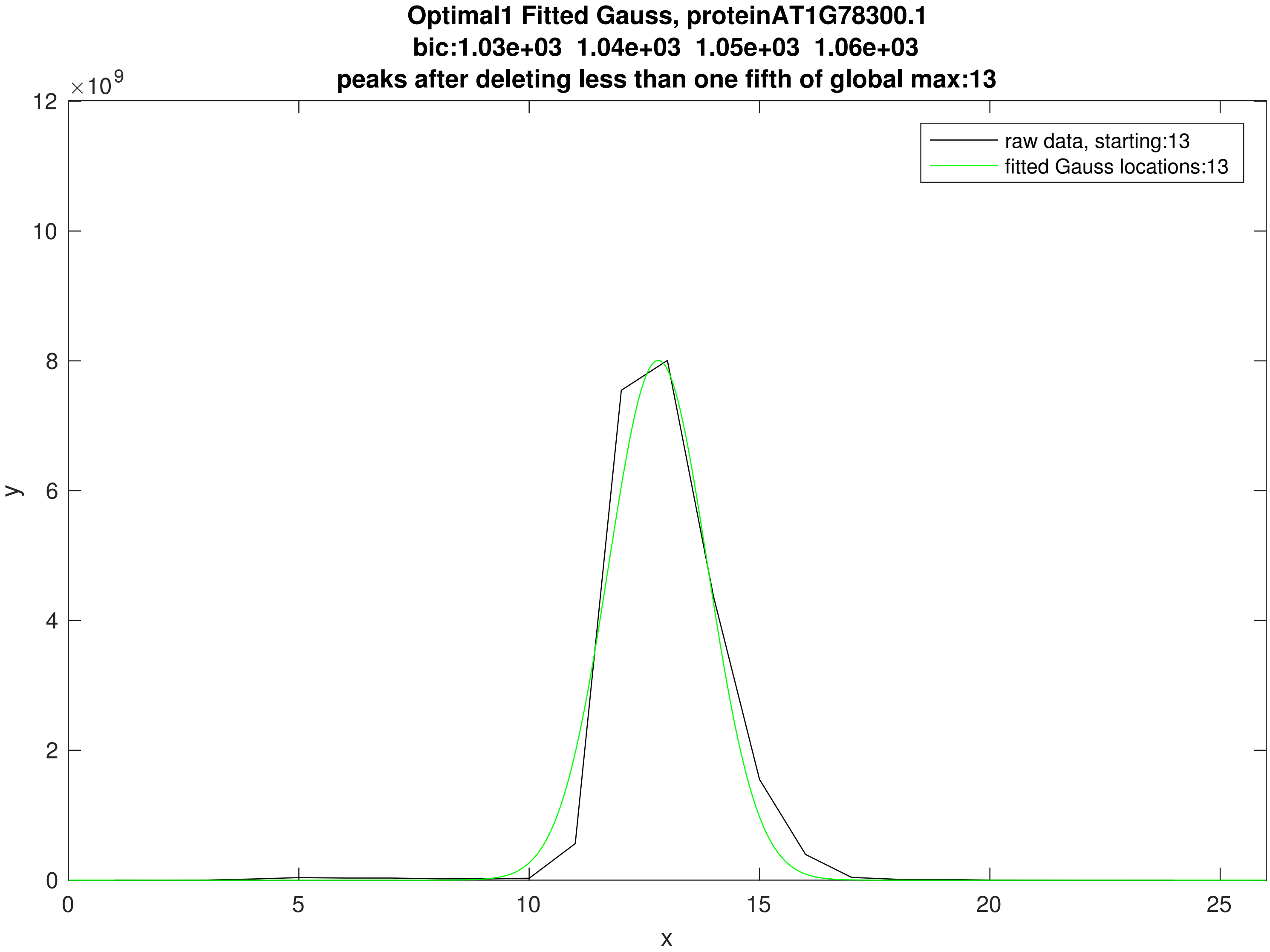
Optimal1 Fitted Gauss, proteinAT1G78300.1
bic:1.03e+03  1.04e+03  1.05e+03  1.06e+03
peaks after deleting less than one fifth of global max:13
×10^9
raw data, starting:13
fitted Gauss locations:13
y
x
0
2
4
6
8
10
12
0
5
10
15
20
25

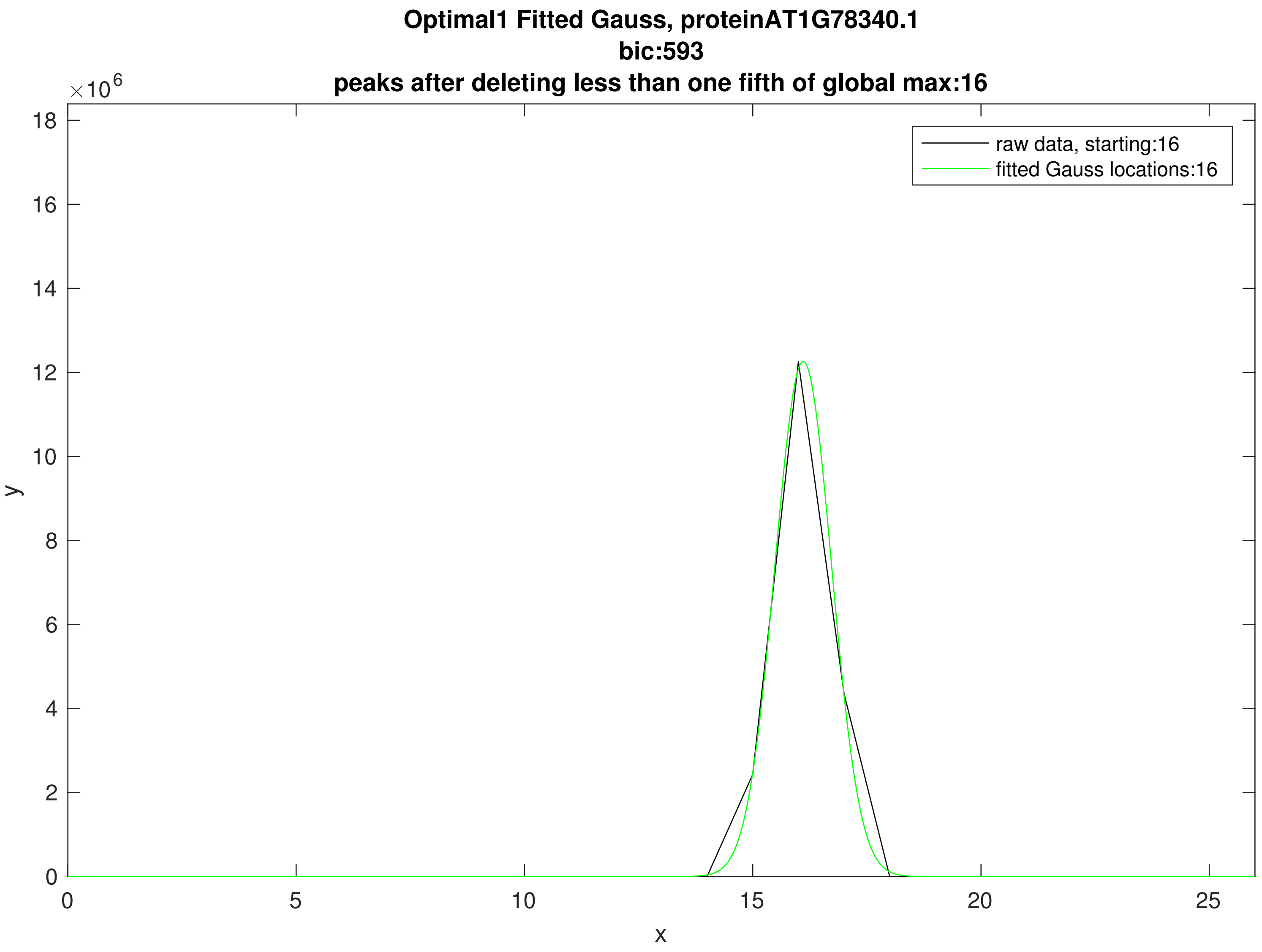
Optimal1 Fitted Gauss, proteinAT1G78340.1
bic:593
peaks after deleting less than one fifth of global max:16
×10^6
raw data, starting:16
fitted Gauss locations:16
18
16
14
12
10
8
6
4
2
0
y
0
5
10
15
20
25
x

**Optimal1 Fitted Gauss, proteinAT1G78370.1**
**bic:872 881**
**peaks after deleting less than one fifth of global max:16**

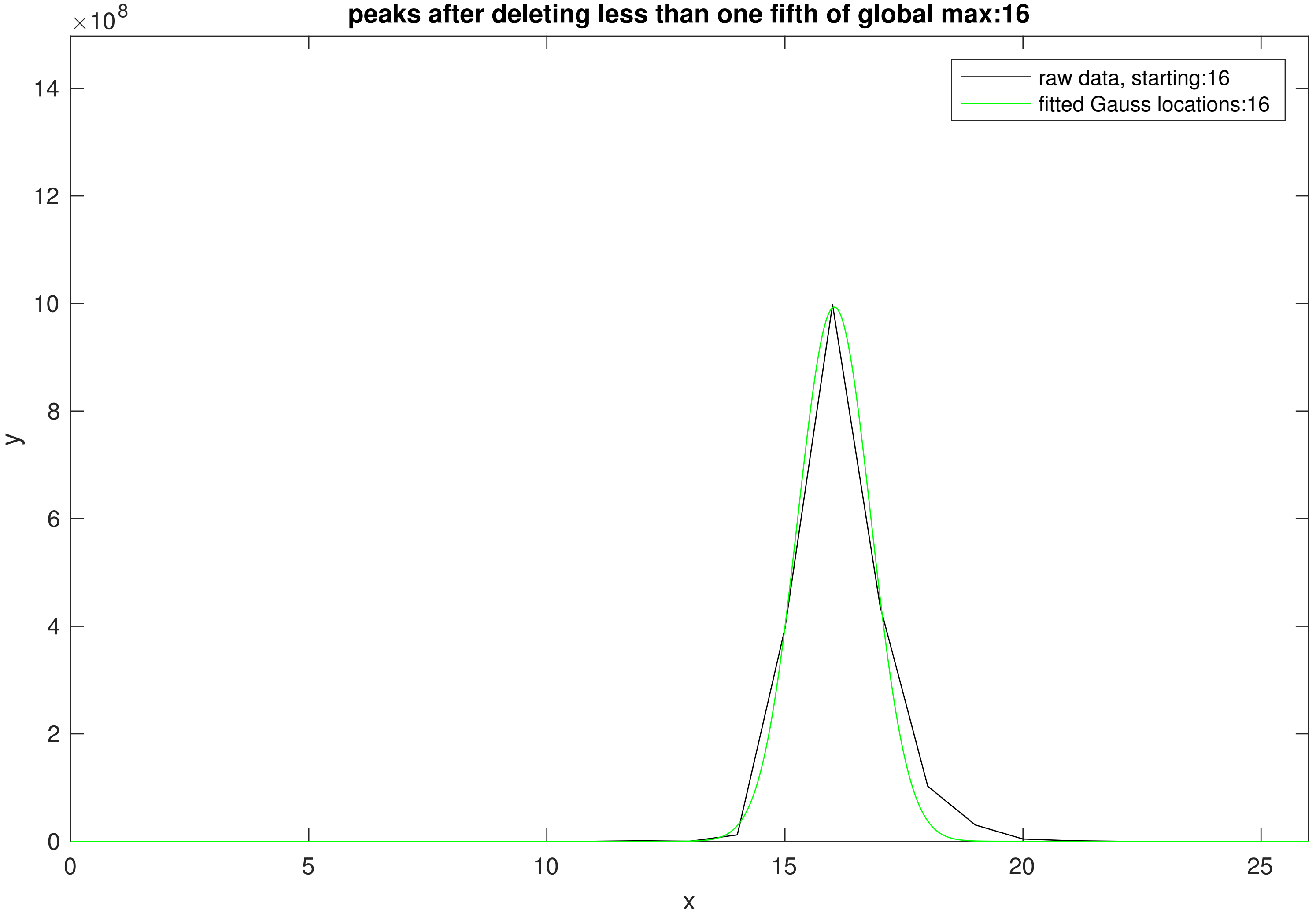

#### Optimal1 Fitted Gauss, proteinAT1G78380.1
#### bic:941 948 958
#### peaks after deleting less than one fifth of global max:16

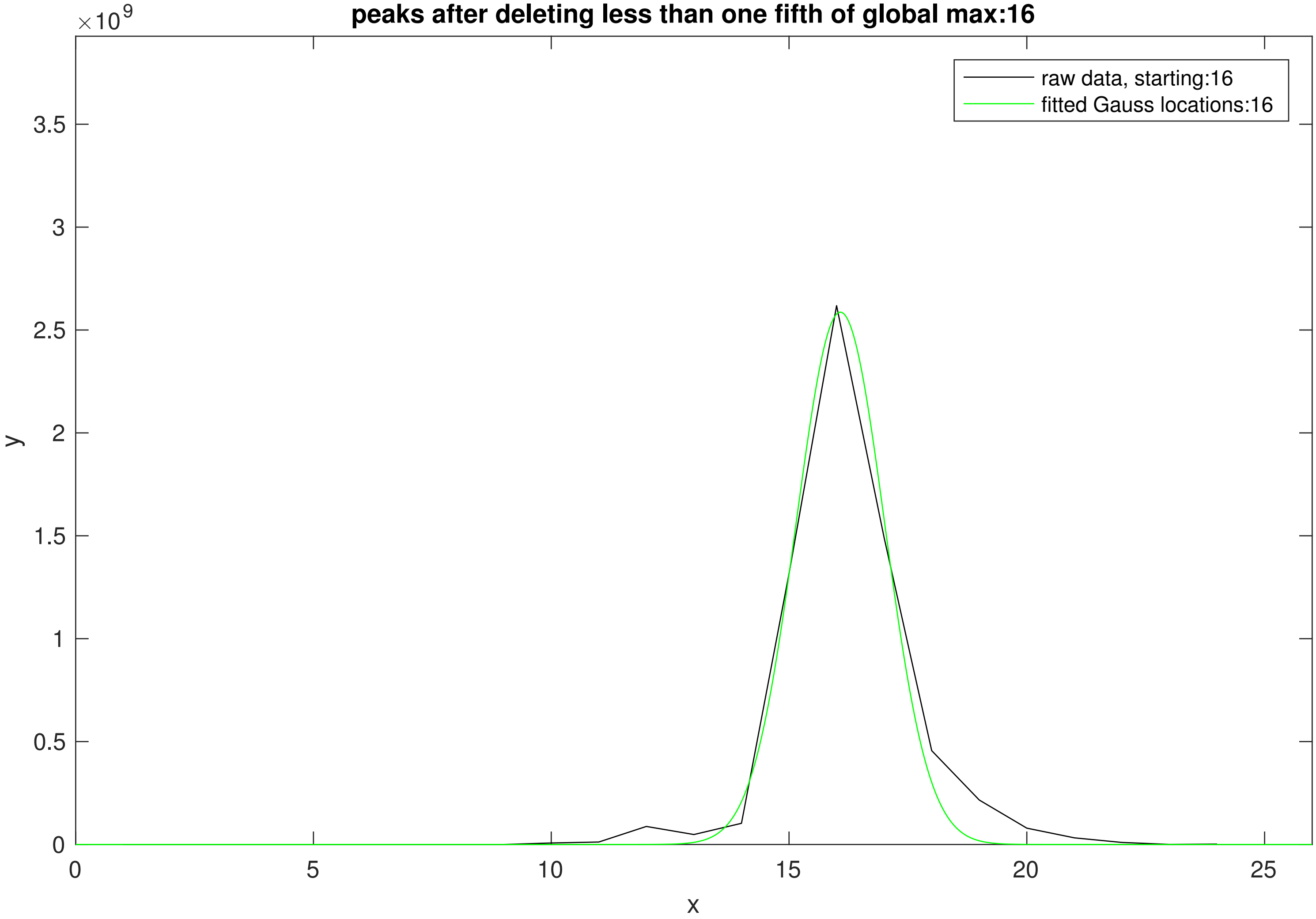

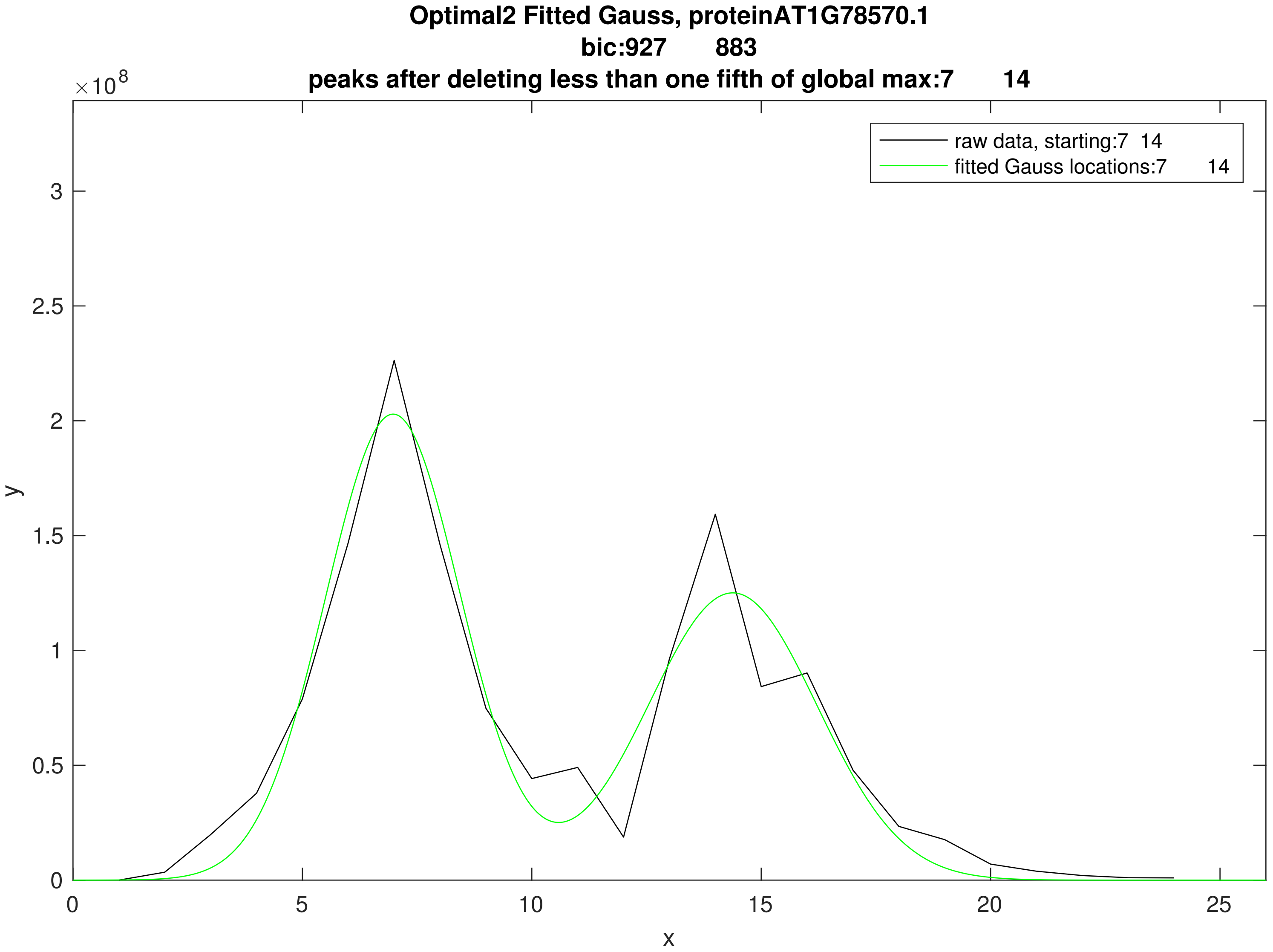

Optimal2 Fitted Gauss, proteinAT1G78570.1
bic:927 883
peaks after deleting less than one fifth of global max:7 14
×10^8
raw data, starting:7 14
fitted Gauss locations:7 14
y
x
0
0.5
1
1.5
2
2.5
3
5
10
15
20
25

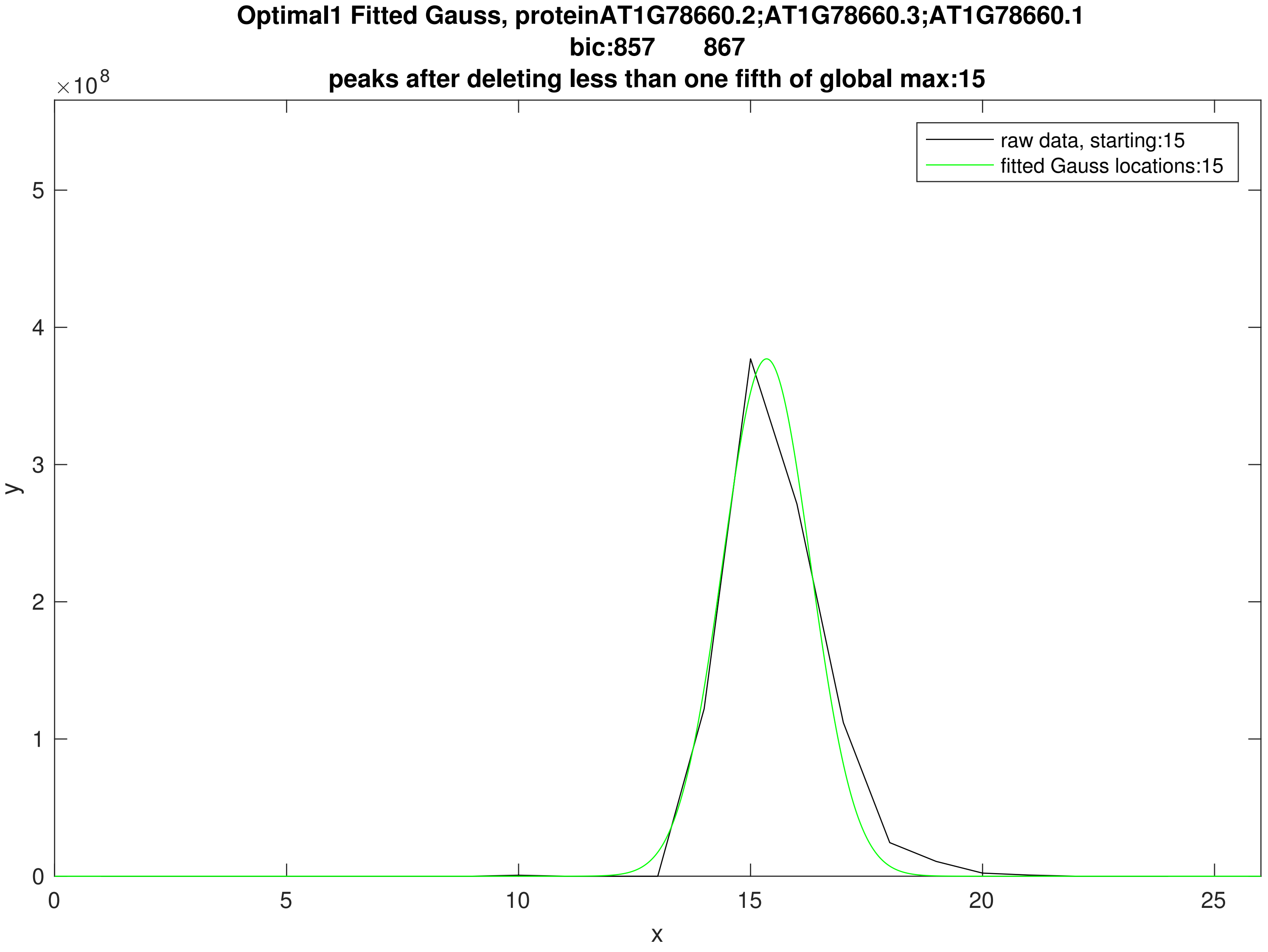
Optimal1 Fitted Gauss, proteinAT1G78660.2;AT1G78660.3;AT1G78660.1
bic:857 867
peaks after deleting less than one fifth of global max:15
×10^8
raw data, starting:15
fitted Gauss locations:15
y
x
0
1
2
3
4
5
0
5
10
15
20
25

#### Optimal1 Fitted Gauss, proteinAT1G78670.1
#### bic:804
#### peaks after deleting less than one fifth of global max:15

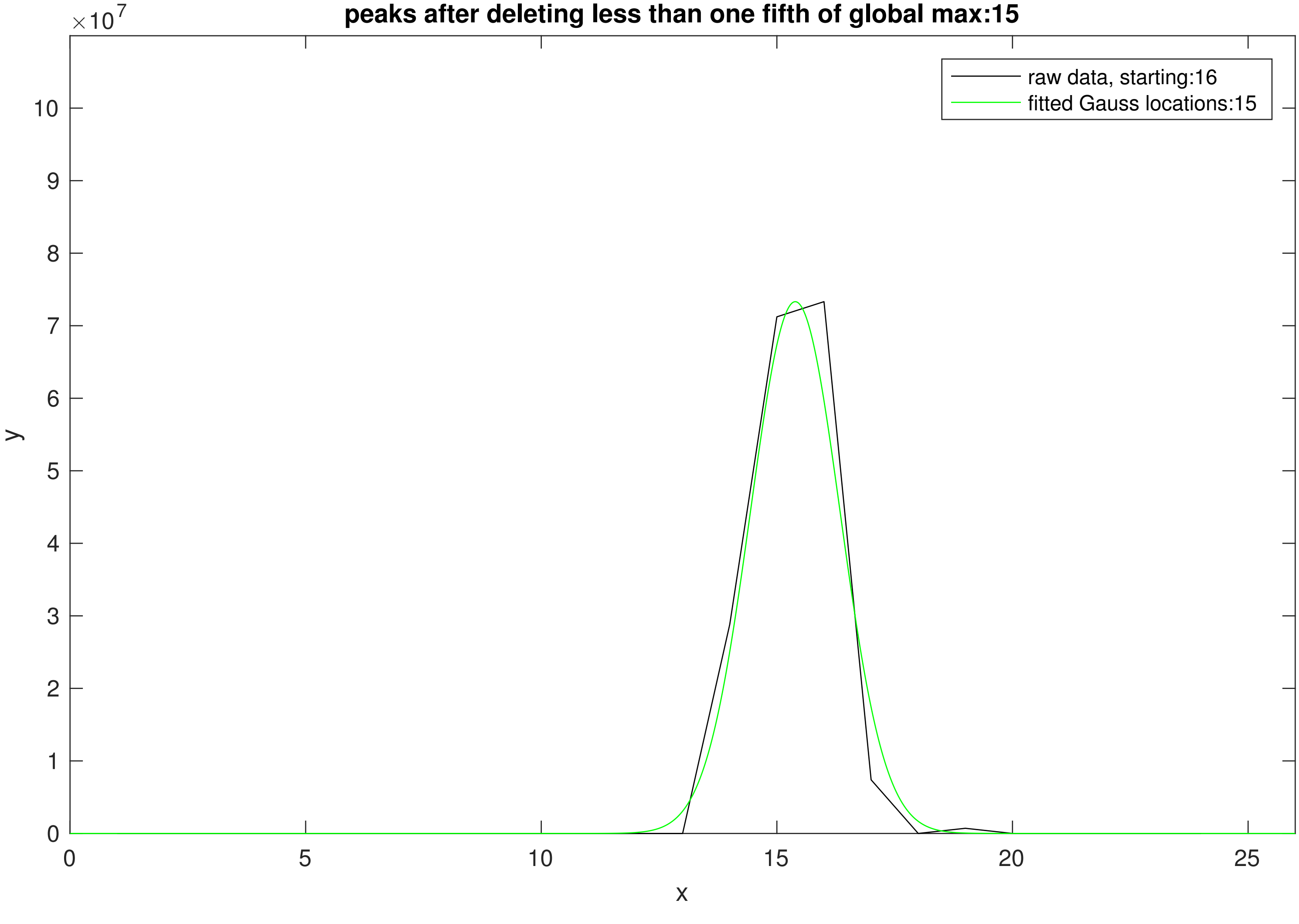

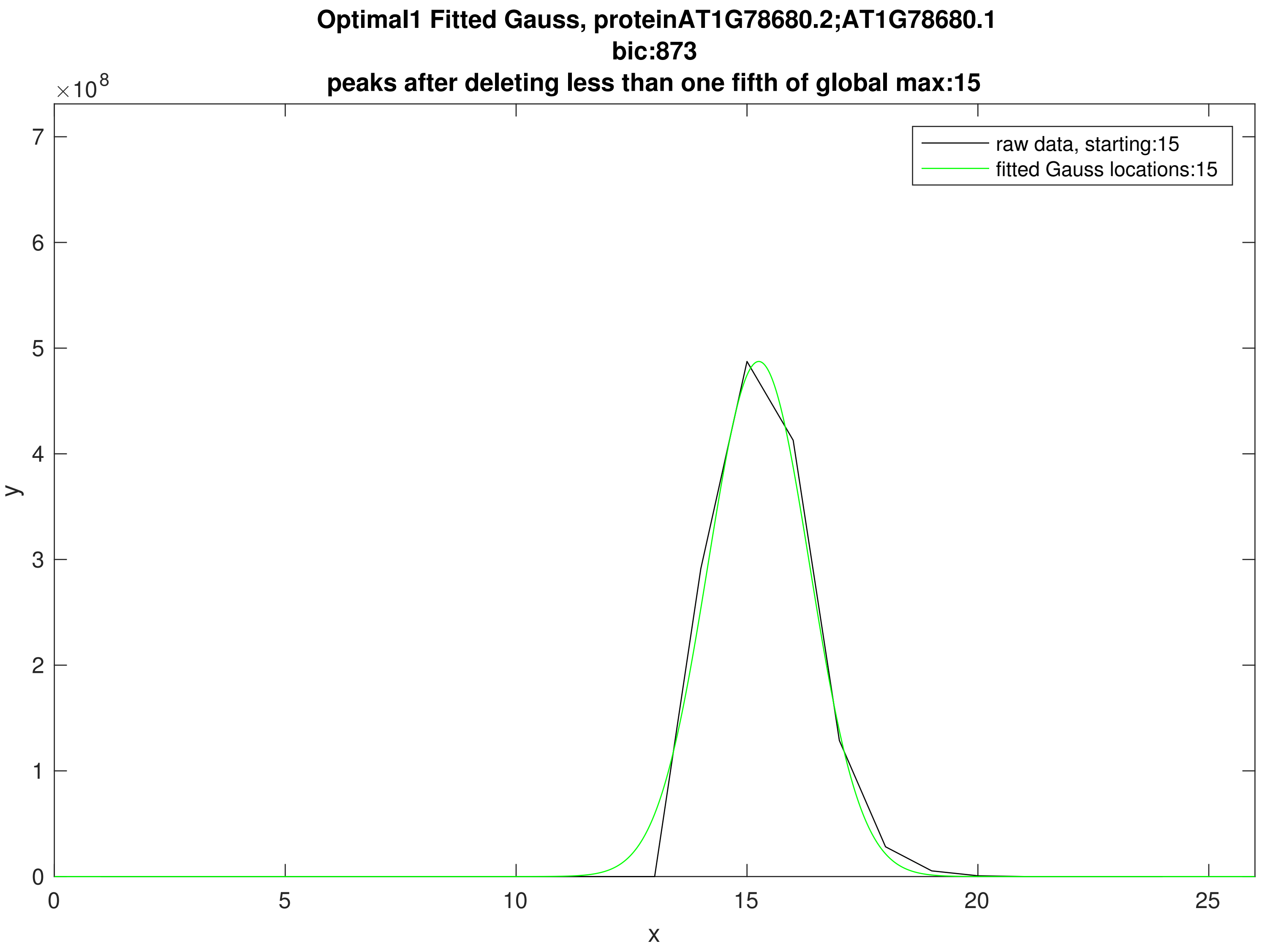

Optimal1 Fitted Gauss, proteinAT1G78680.2;AT1G78680.1
bic:873
peaks after deleting less than one fifth of global max:15
×10^8
raw data, starting:15
fitted Gauss locations:15
y
x
0
1
2
3
4
5
6
7
0
5
10
15
20
25

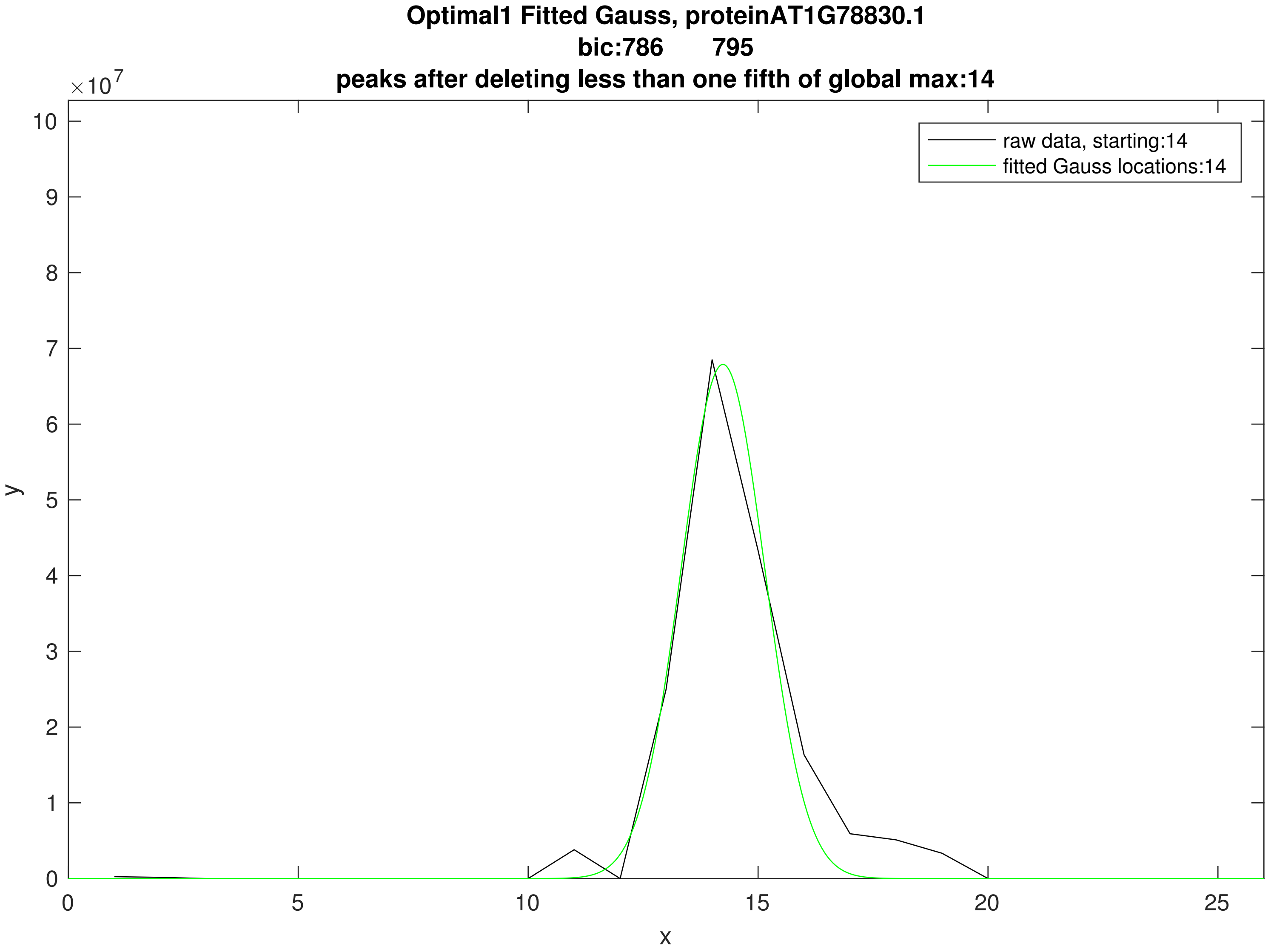

Optimal1 Fitted Gauss, proteinAT1G78830.1
bic:786 795
peaks after deleting less than one fifth of global max:14
×10^7
raw data, starting:14
fitted Gauss locations:14
y
x
0
1
2
3
4
5
6
7
8
9
10
0
5
10
15
20
25

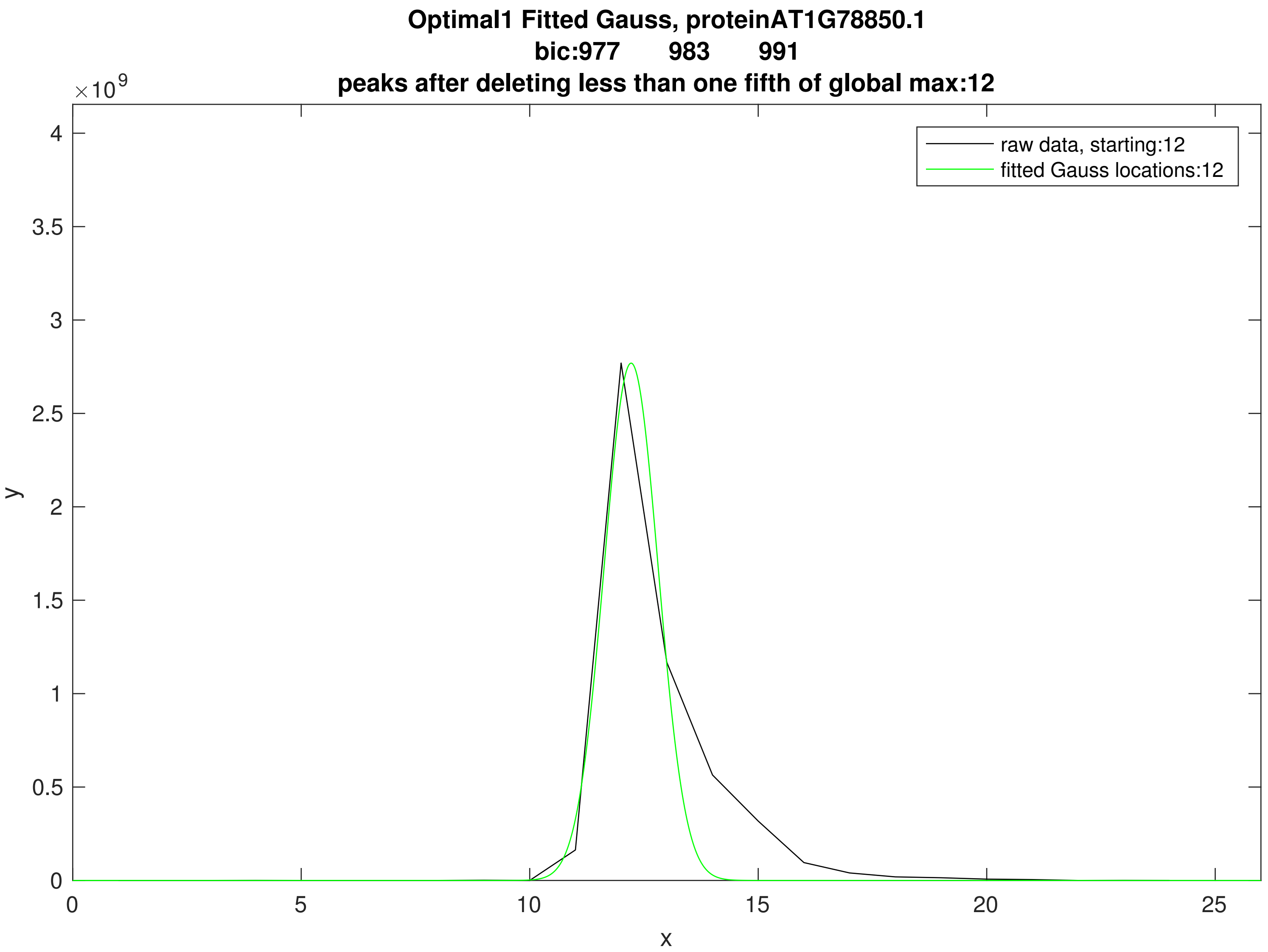
Optimal1 Fitted Gauss, proteinAT1G78850.1
bic:977 983 991
peaks after deleting less than one fifth of global max:12
×10^9
raw data, starting:12
fitted Gauss locations:12
y
x
0
0.5
1
1.5
2
2.5
3
3.5
4
5
10
15
20
25

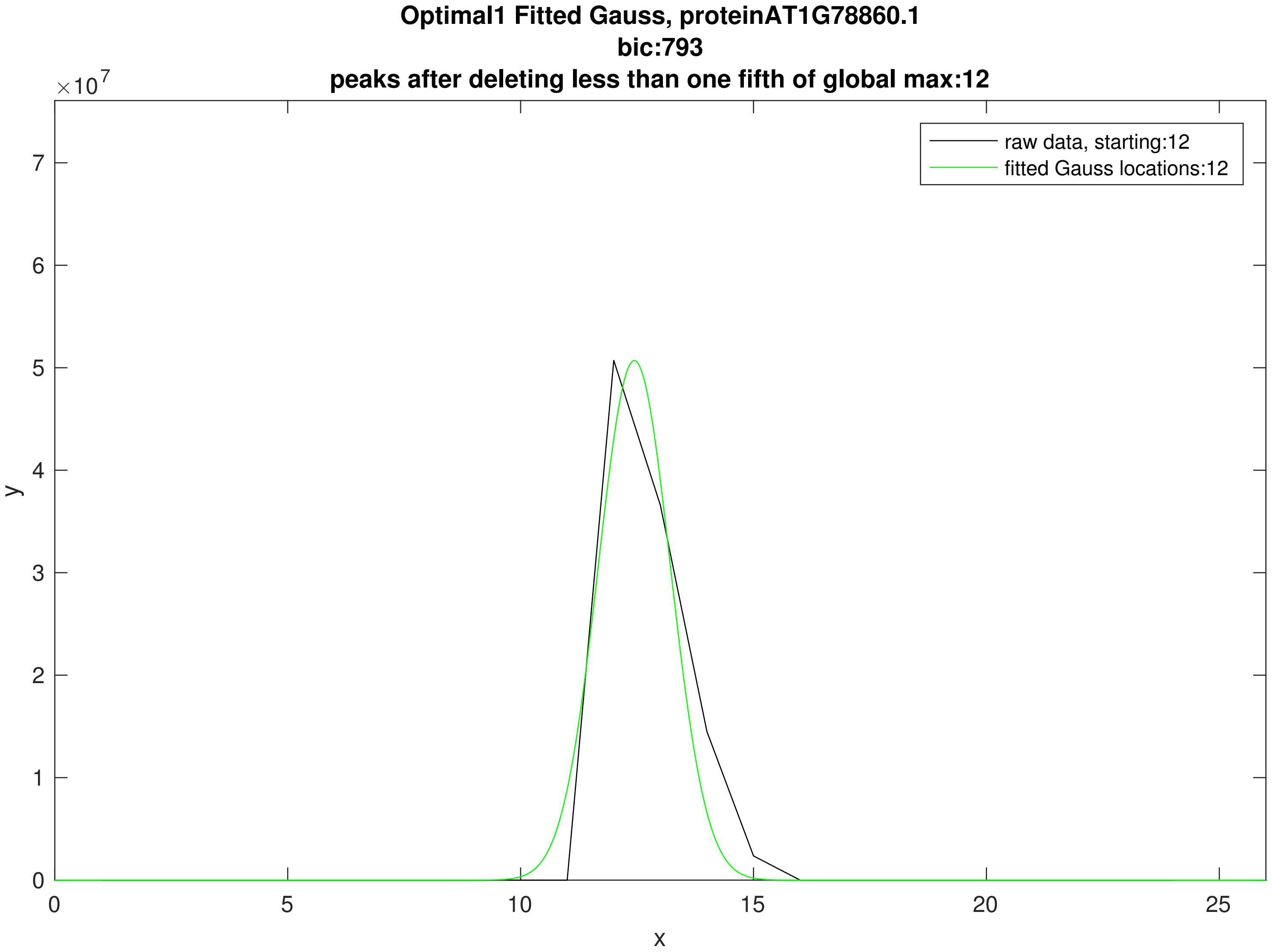
Optimal1 Fitted Gauss, proteinAT1G78860.1
bic:793
peaks after deleting less than one fifth of global max:12
×10^7
raw data, starting:12
fitted Gauss locations:12
y
x
0
1
2
3
4
5
6
7
0
5
10
15
20
25

**Optimal1 Fitted Gauss, proteinAT1G78870.1;AT1G78870.2;AT1G78870.3**
**bic:698**
**peaks after deleting less than one fifth of global max:19**

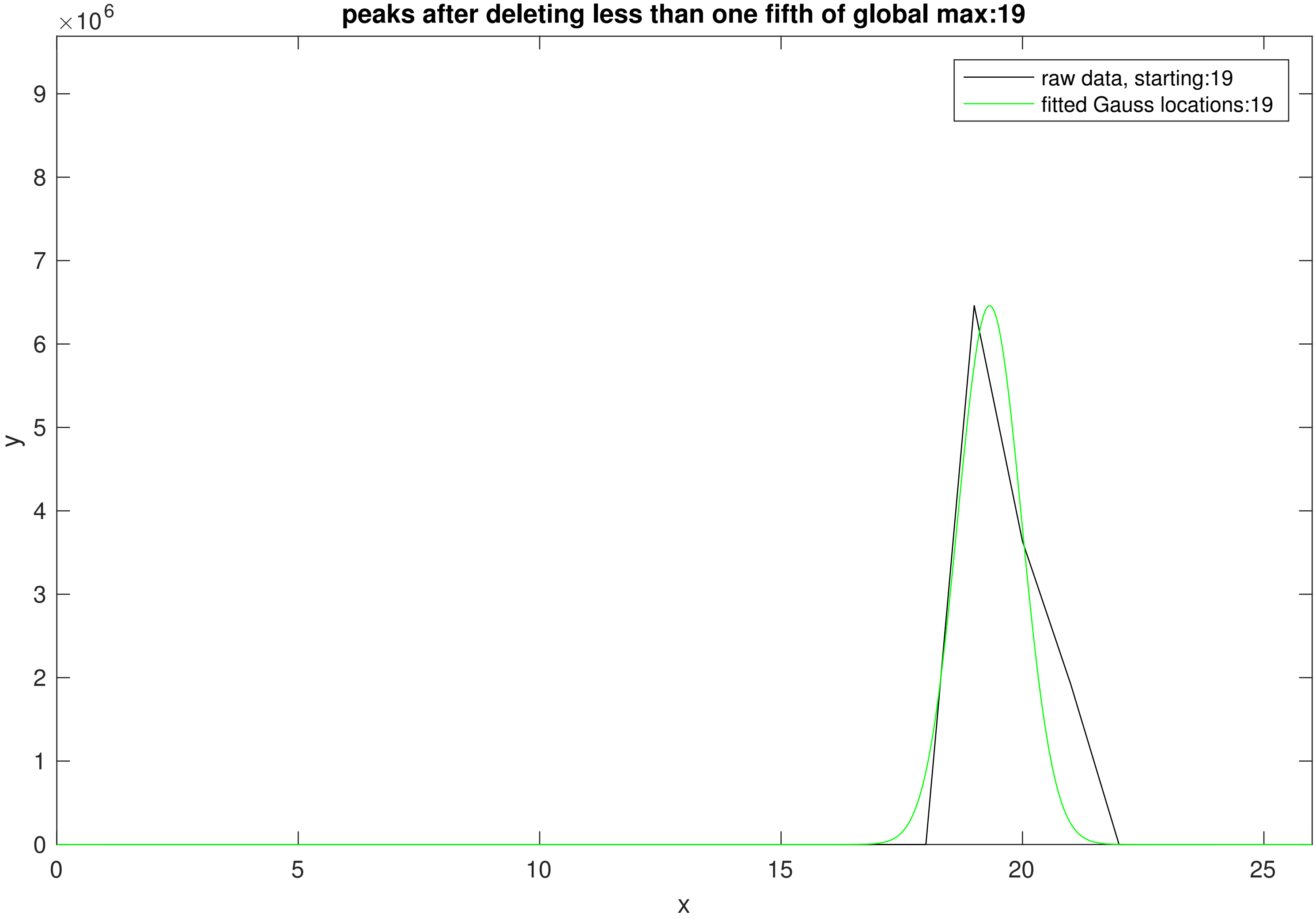

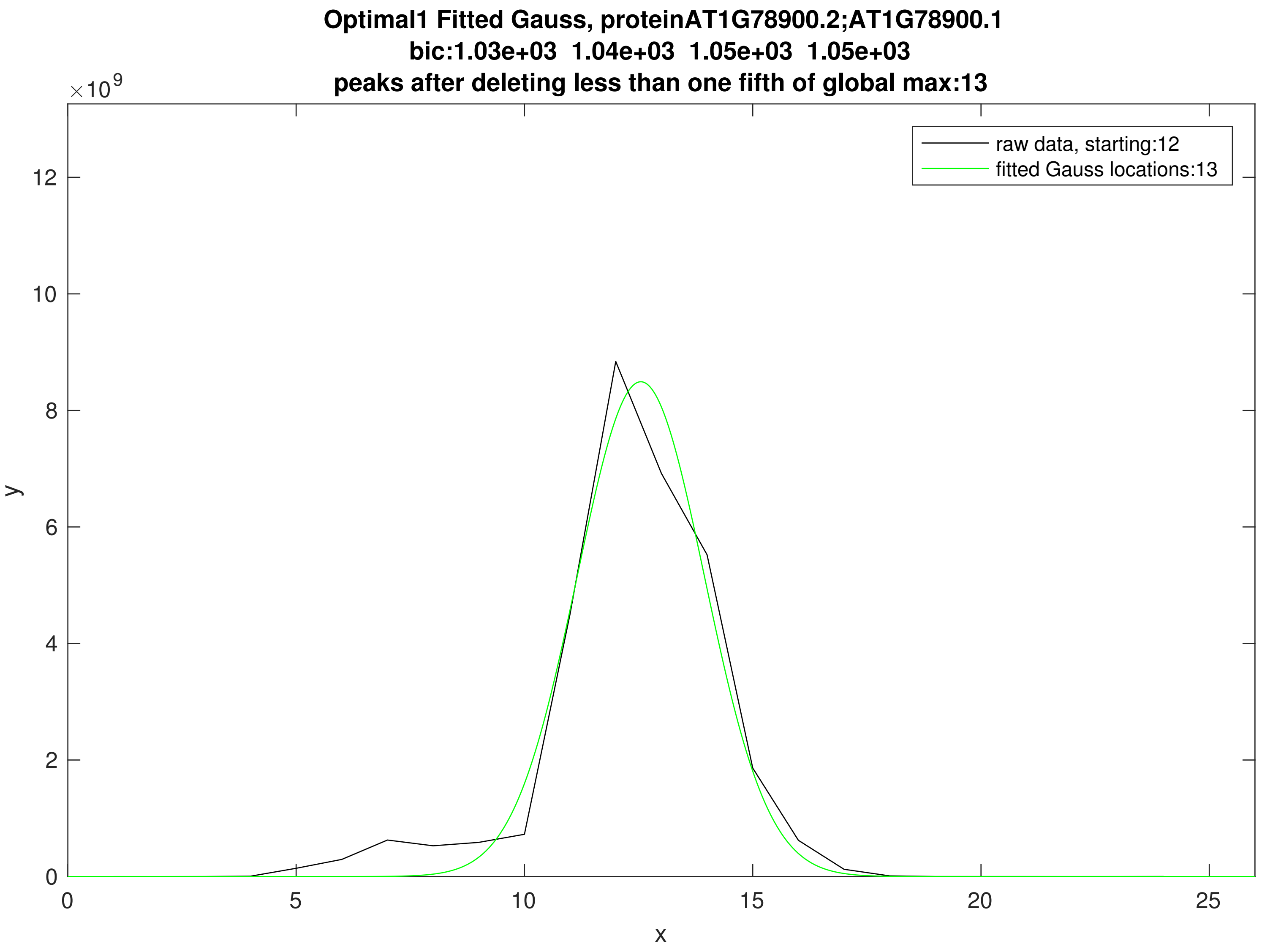
Optimal1 Fitted Gauss, proteinAT1G78900.2;AT1G78900.1
bic:1.03e+03  1.04e+03  1.05e+03  1.05e+03
peaks after deleting less than one fifth of global max:13
×10^9
raw data, starting:12
fitted Gauss locations:13
y
x
0
2
4
6
8
10
12
0
5
10
15
20
25

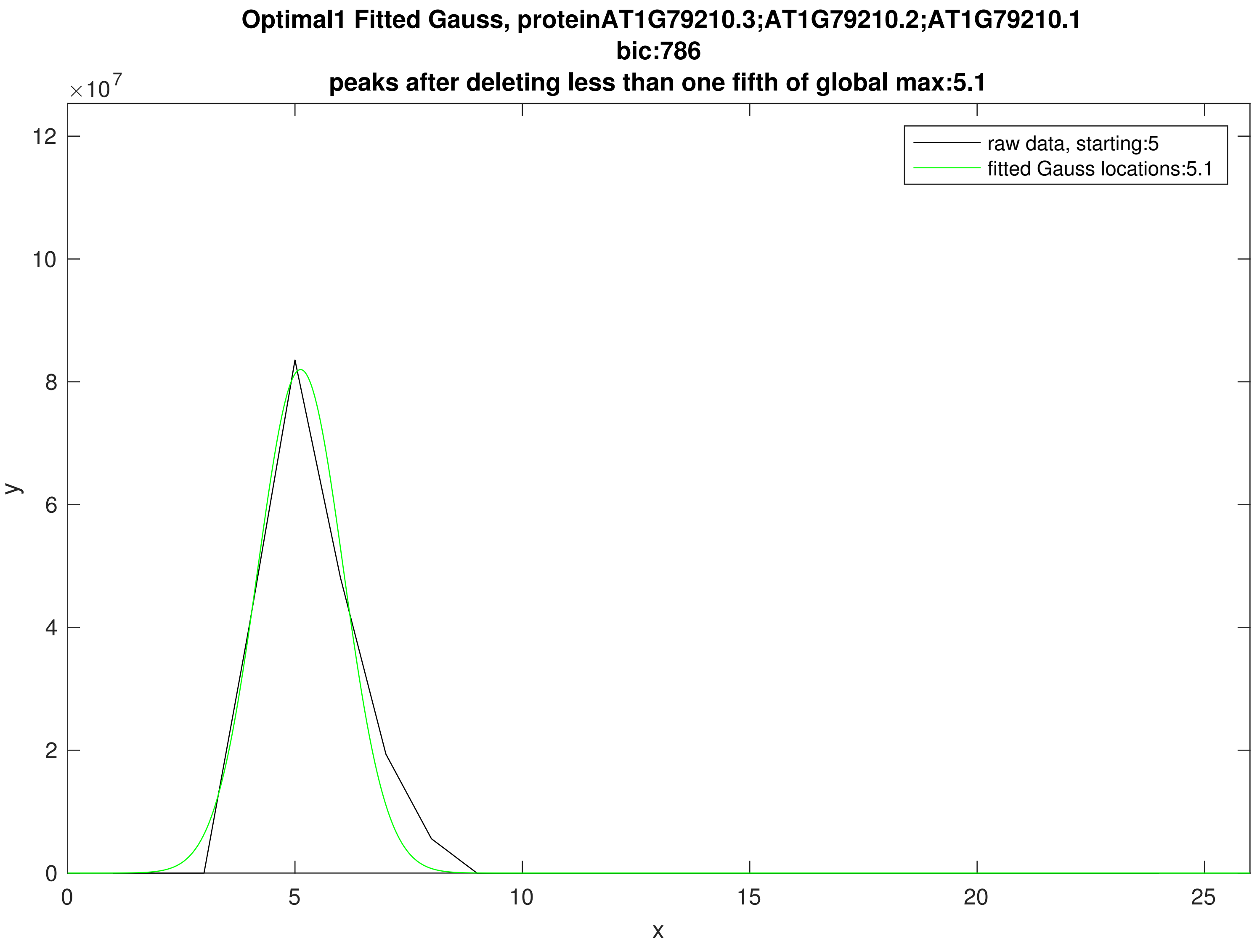

Optimal1 Fitted Gauss, proteinAT1G79210.3;AT1G79210.2;AT1G79210.1
bic:786
peaks after deleting less than one fifth of global max:5.1
×10^7
12
10
8
6
4
2
0
y
0
5
10
15
20
25
x
raw data, starting:5
fitted Gauss locations:5.1

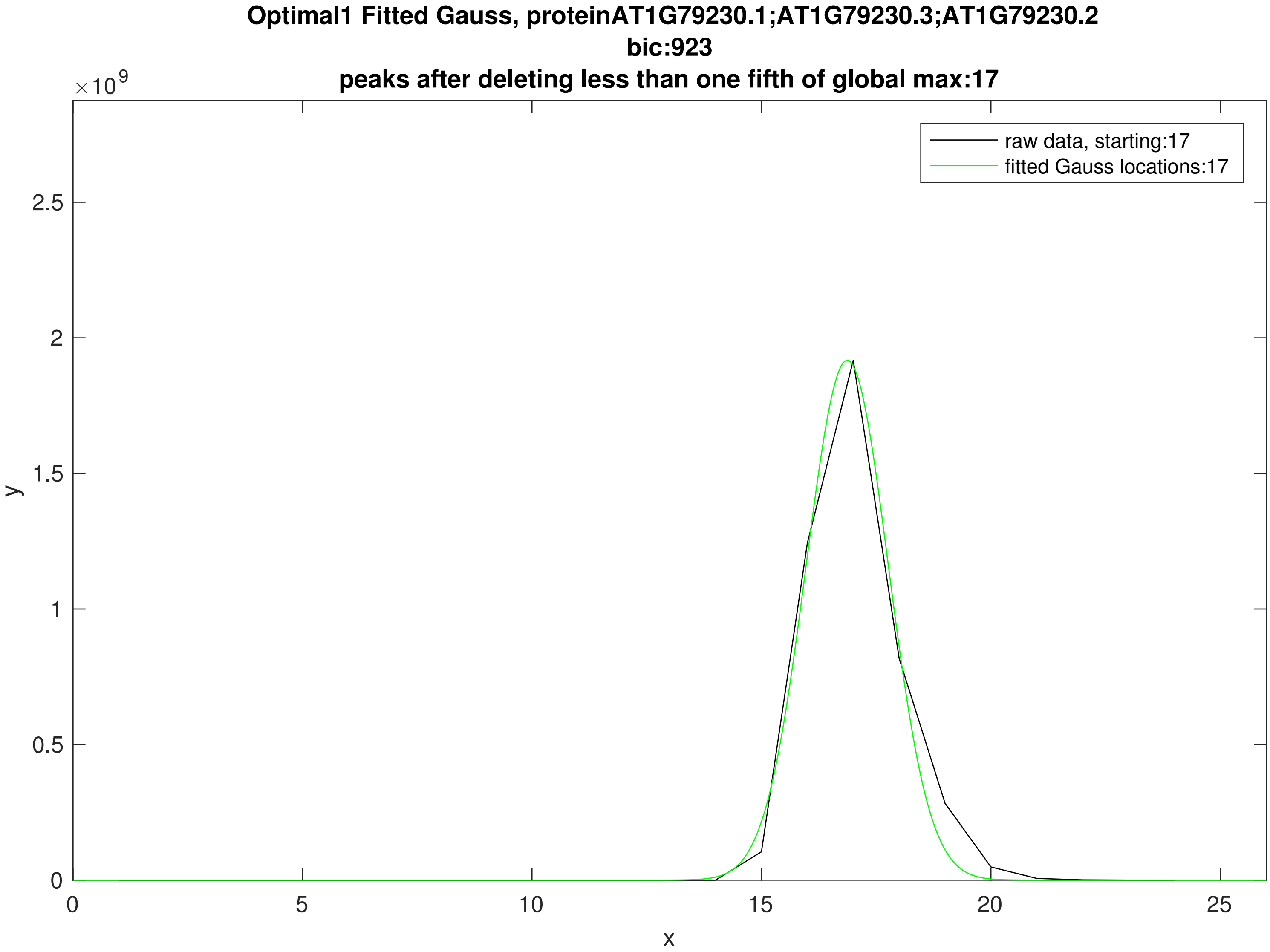

Optimal1 Fitted Gauss, proteinAT1G79230.1;AT1G79230.3;AT1G79230.2
bic:923
peaks after deleting less than one fifth of global max:17
×10^9
raw data, starting:17
fitted Gauss locations:17
2.5
2
1.5
1
0.5
0
y
0
5
10
15
20
25
x

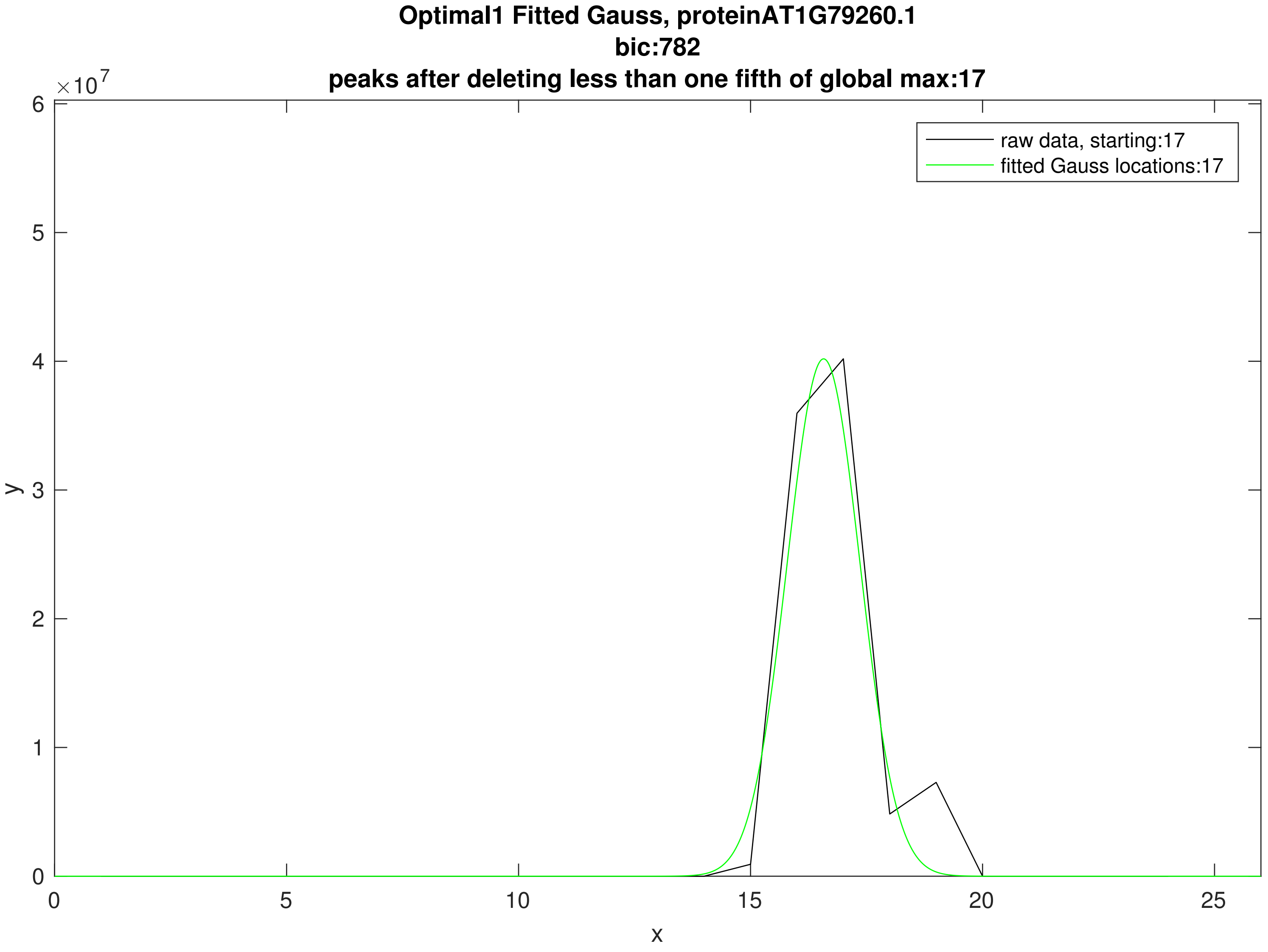

Optimal1 Fitted Gauss, proteinAT1G79260.1
bic:782
peaks after deleting less than one fifth of global max:17
×10^7
6
5
4
3
2
1
0
y
0
5
10
15
20
25
x
raw data, starting:17
fitted Gauss locations:17

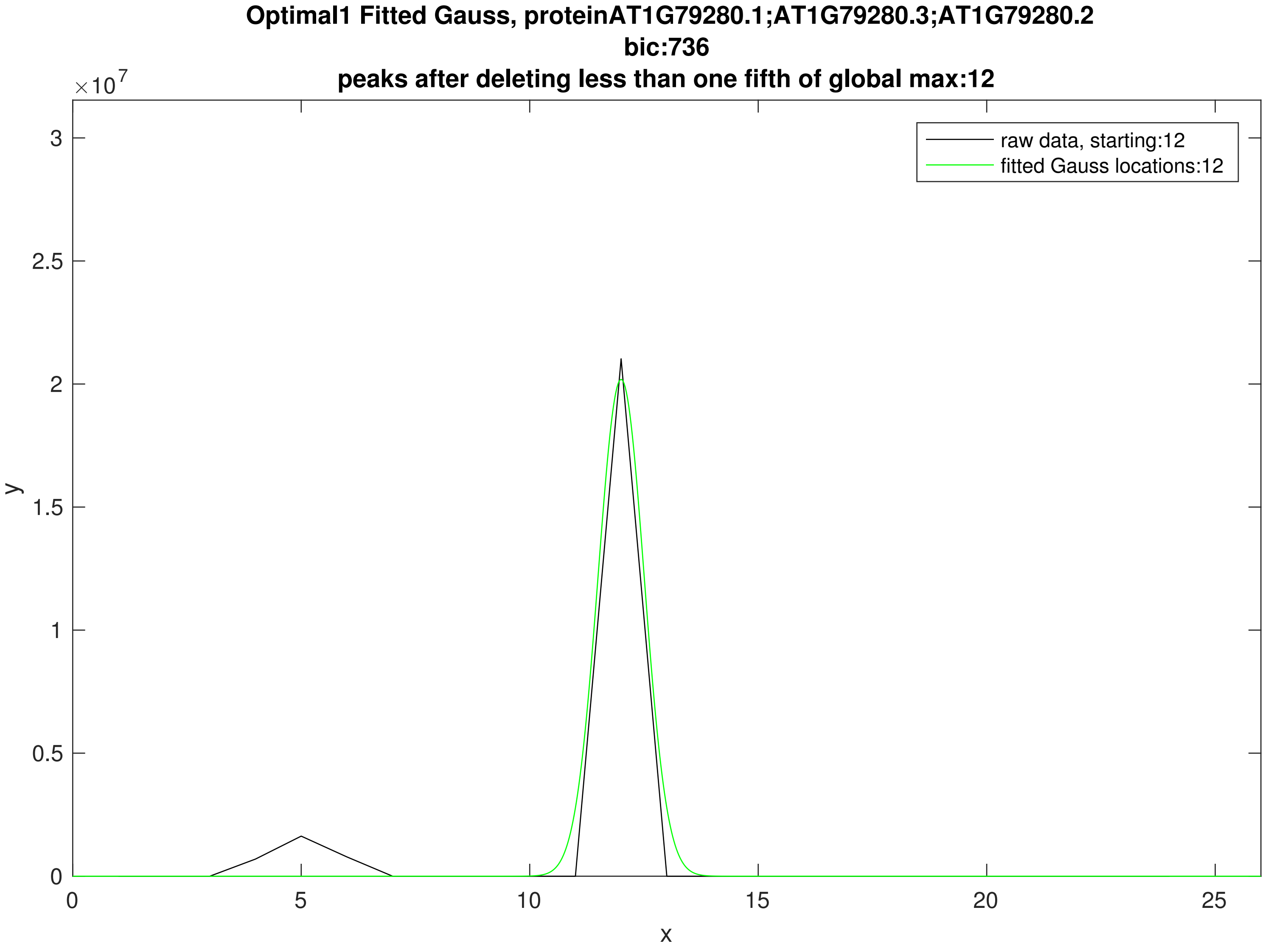
Optimal1 Fitted Gauss, proteinAT1G79280.1;AT1G79280.3;AT1G79280.2
bic:736
peaks after deleting less than one fifth of global max:12
×10^7
raw data, starting:12
fitted Gauss locations:12
y
x
0
0.5
1
1.5
2
2.5
3
0
5
10
15
20
25

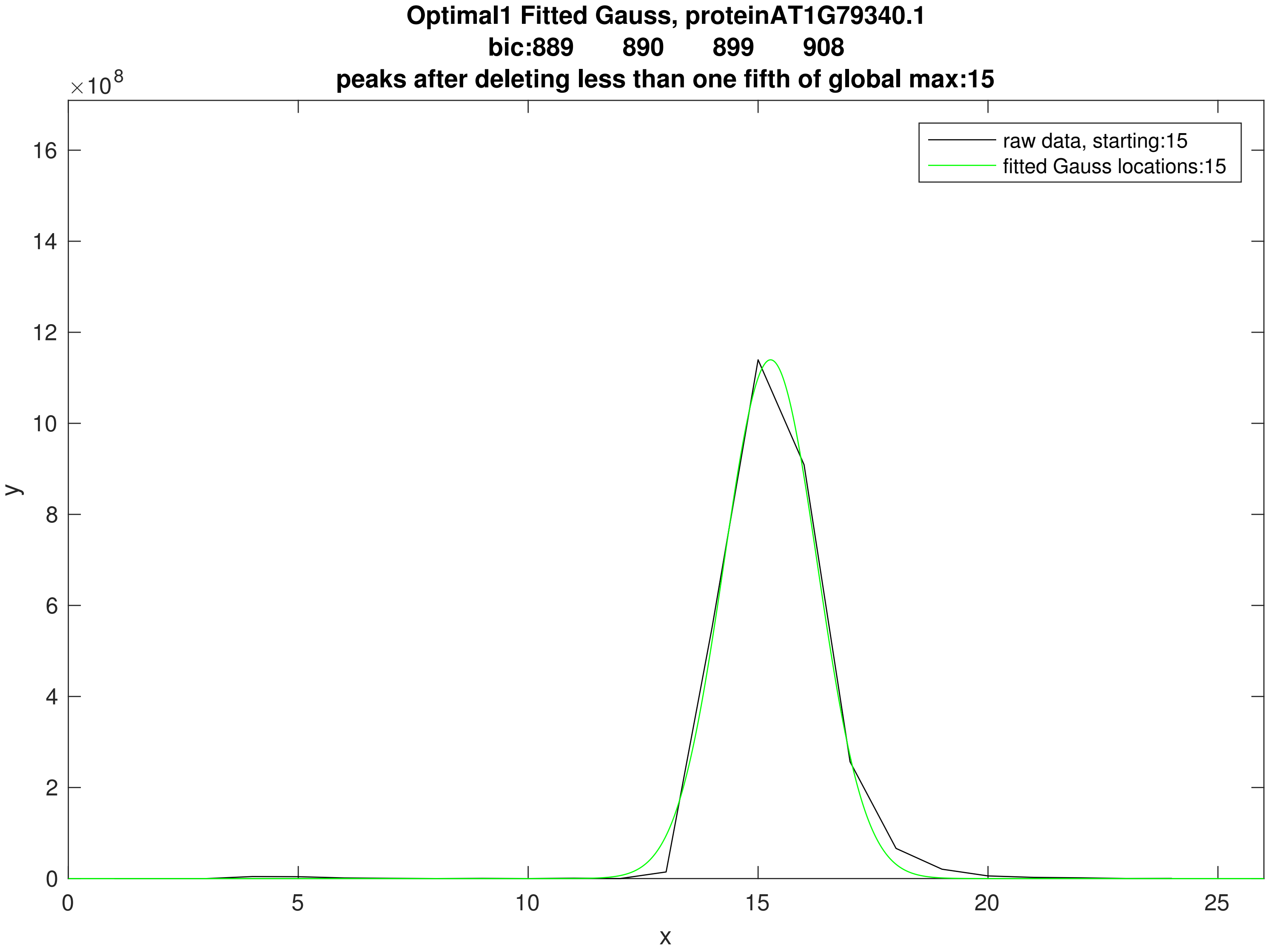

Optimal1 Fitted Gauss, proteinAT1G79340.1
bic:889 890 899 908
peaks after deleting less than one fifth of global max:15
×10^8
16
14
12
10
8
6
4
2
0
y
0
5
10
15
20
25
x
raw data, starting:15
fitted Gauss locations:15

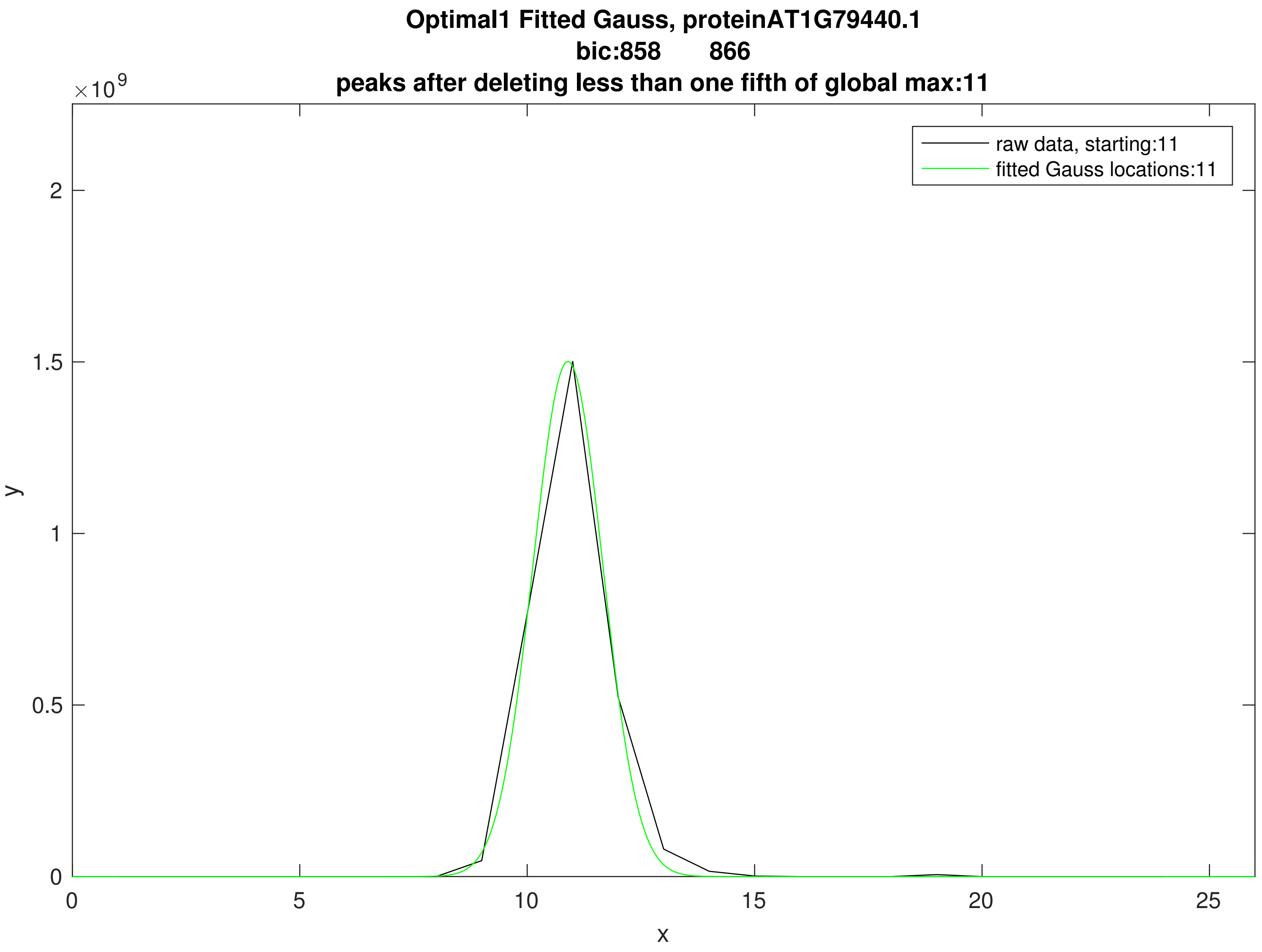
Optimal1 Fitted Gauss, proteinAT1G79440.1
bic:858 866
peaks after deleting less than one fifth of global max:11
×10^9
raw data, starting:11
fitted Gauss locations:11
y
x
0
0.5
1
1.5
2
0
5
10
15
20
25

**Optimal1 Fitted Gauss, proteinAT1G79470.1**
**bic:755**
**peaks after deleting less than one fifth of global max:10**

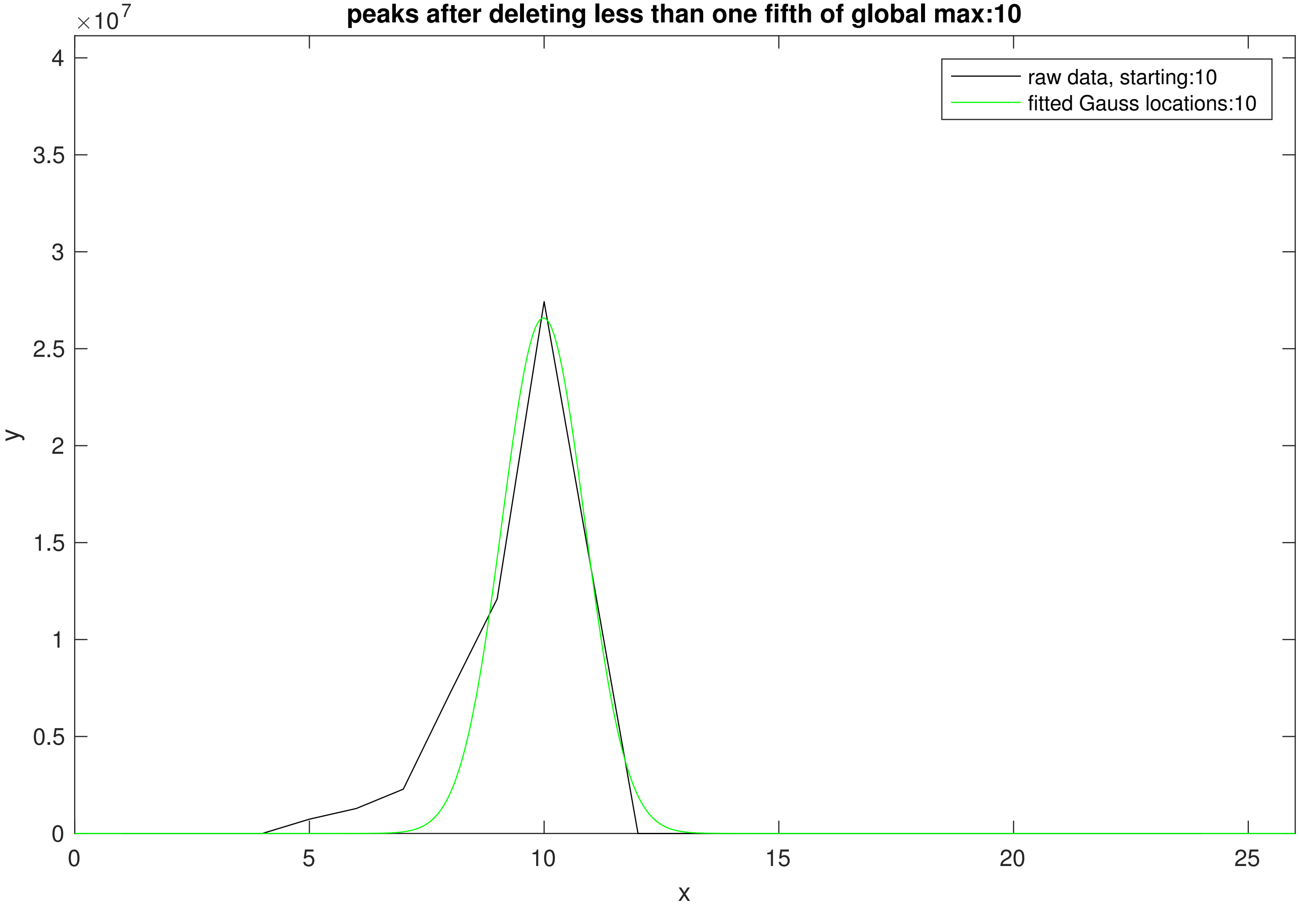

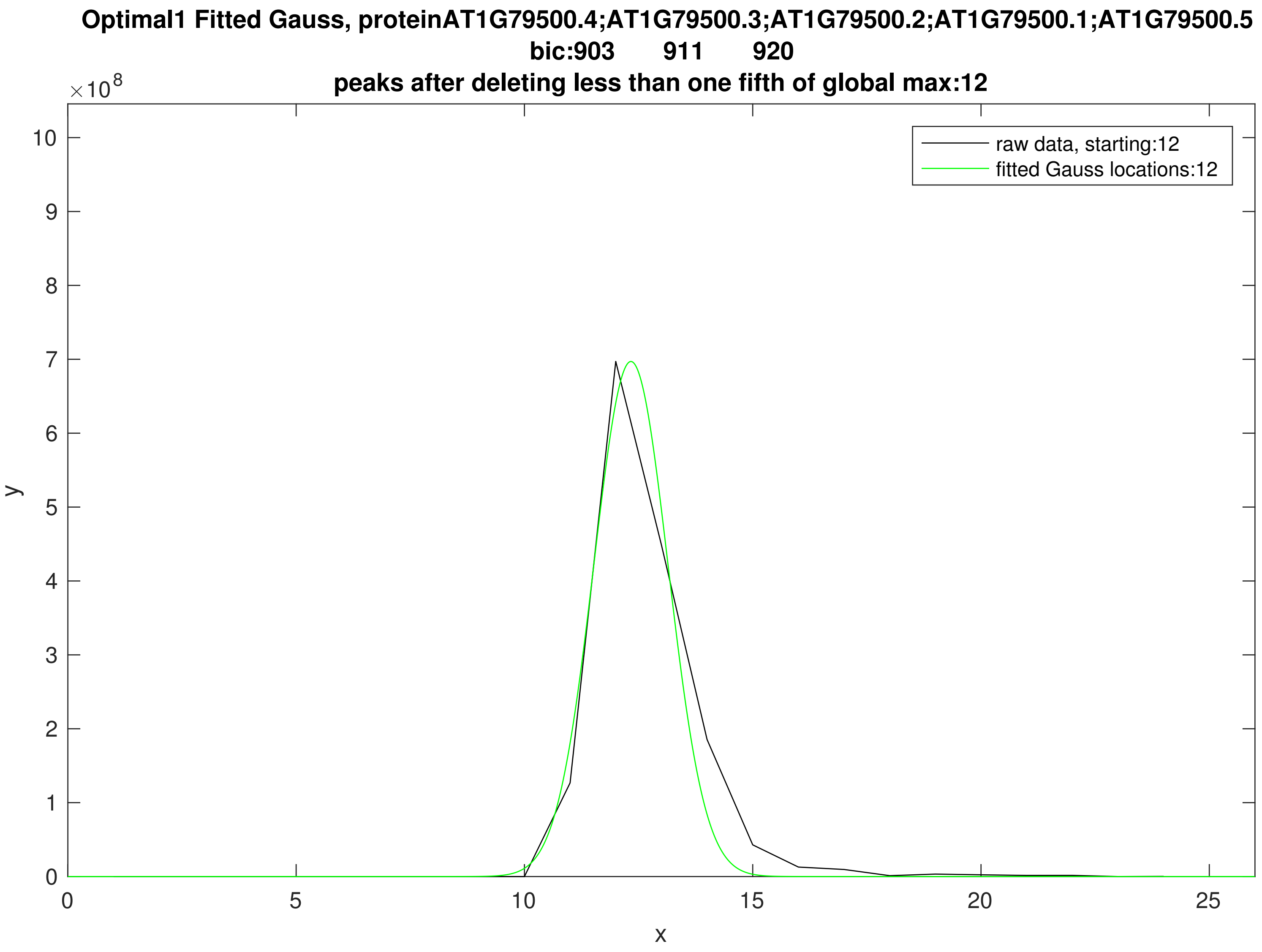
Optimal1 Fitted Gauss, proteinAT1G79500.4;AT1G79500.3;AT1G79500.2;AT1G79500.1;AT1G79500.5
bic:903 911 920
peaks after deleting less than one fifth of global max:12
×10^8
raw data, starting:12
fitted Gauss locations:12
y
x
0
1
2
3
4
5
6
7
8
9
10
0
5
10
15
20
25

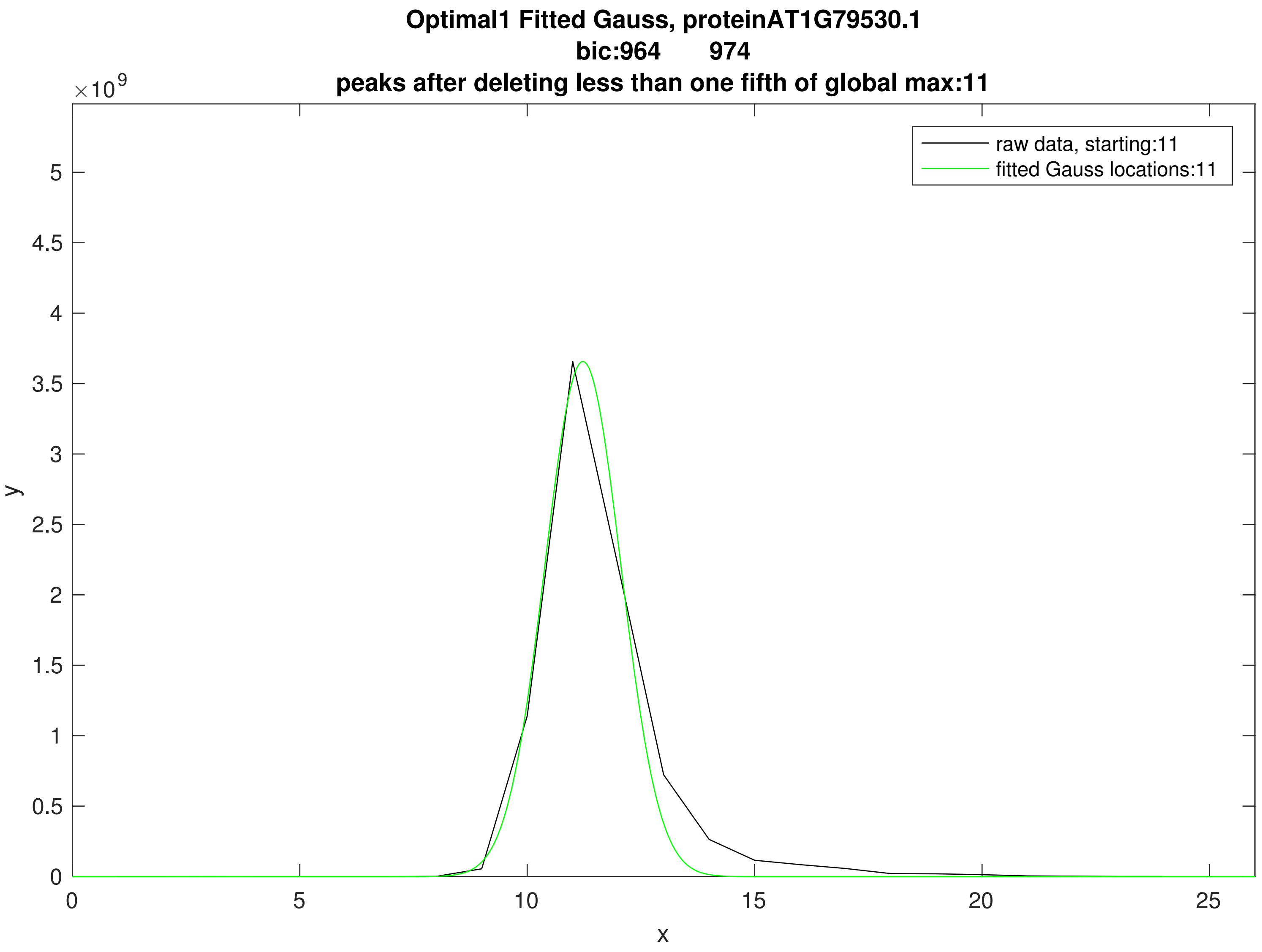

Optimal1 Fitted Gauss, proteinAT1G79530.1
bic:964 974
peaks after deleting less than one fifth of global max:11
×10^9
raw data, starting:11
fitted Gauss locations:11
y
x
0
0.5
1
1.5
2
2.5
3
3.5
4
4.5
5
0
5
10
15
20
25

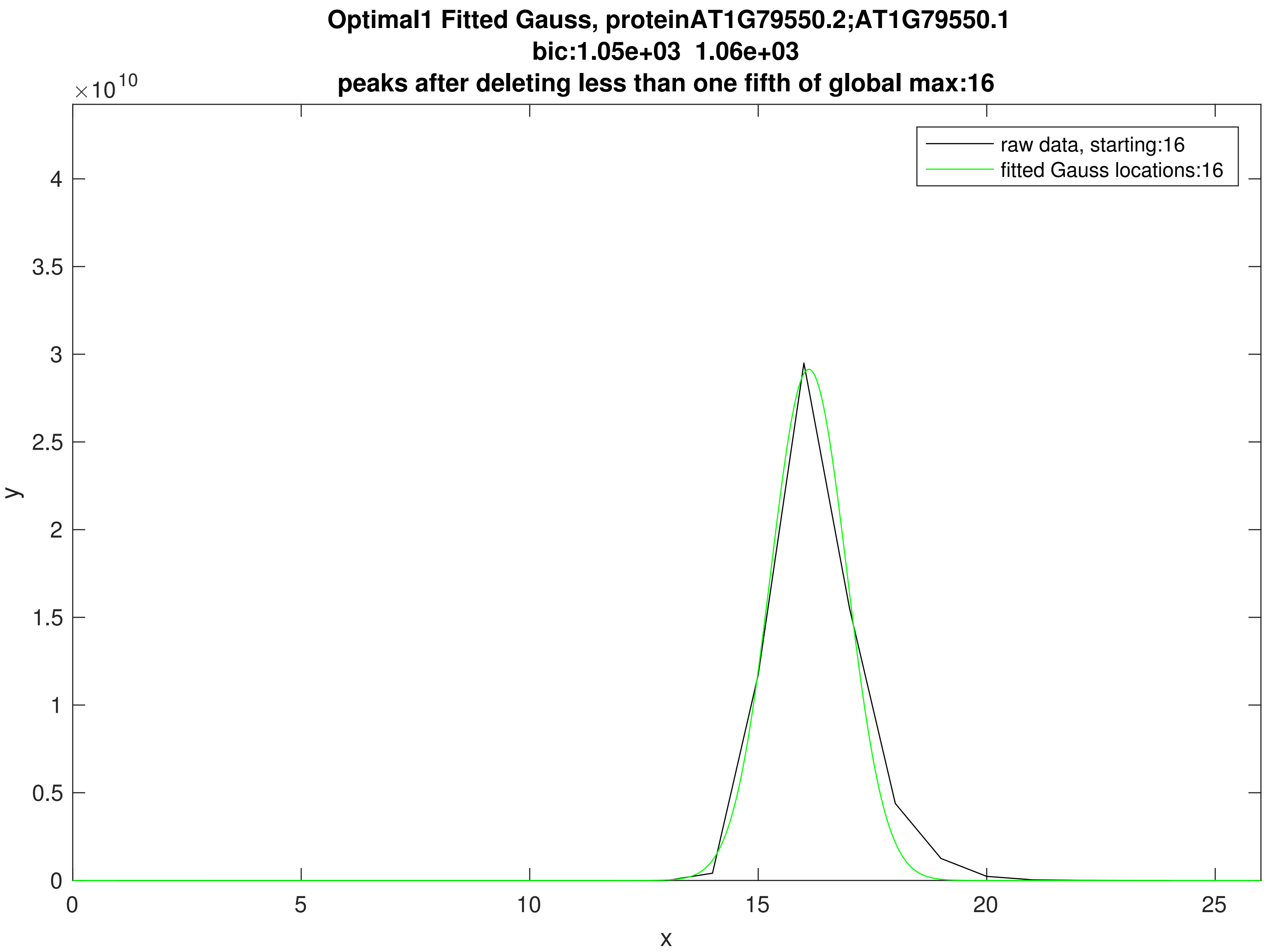
Optimal1 Fitted Gauss, proteinAT1G79550.2;AT1G79550.1
bic:1.05e+03  1.06e+03
peaks after deleting less than one fifth of global max:16
×10^10
raw data, starting:16
fitted Gauss locations:16
y
x
0
0.5
1
1.5
2
2.5
3
3.5
4
5
10
15
20
25

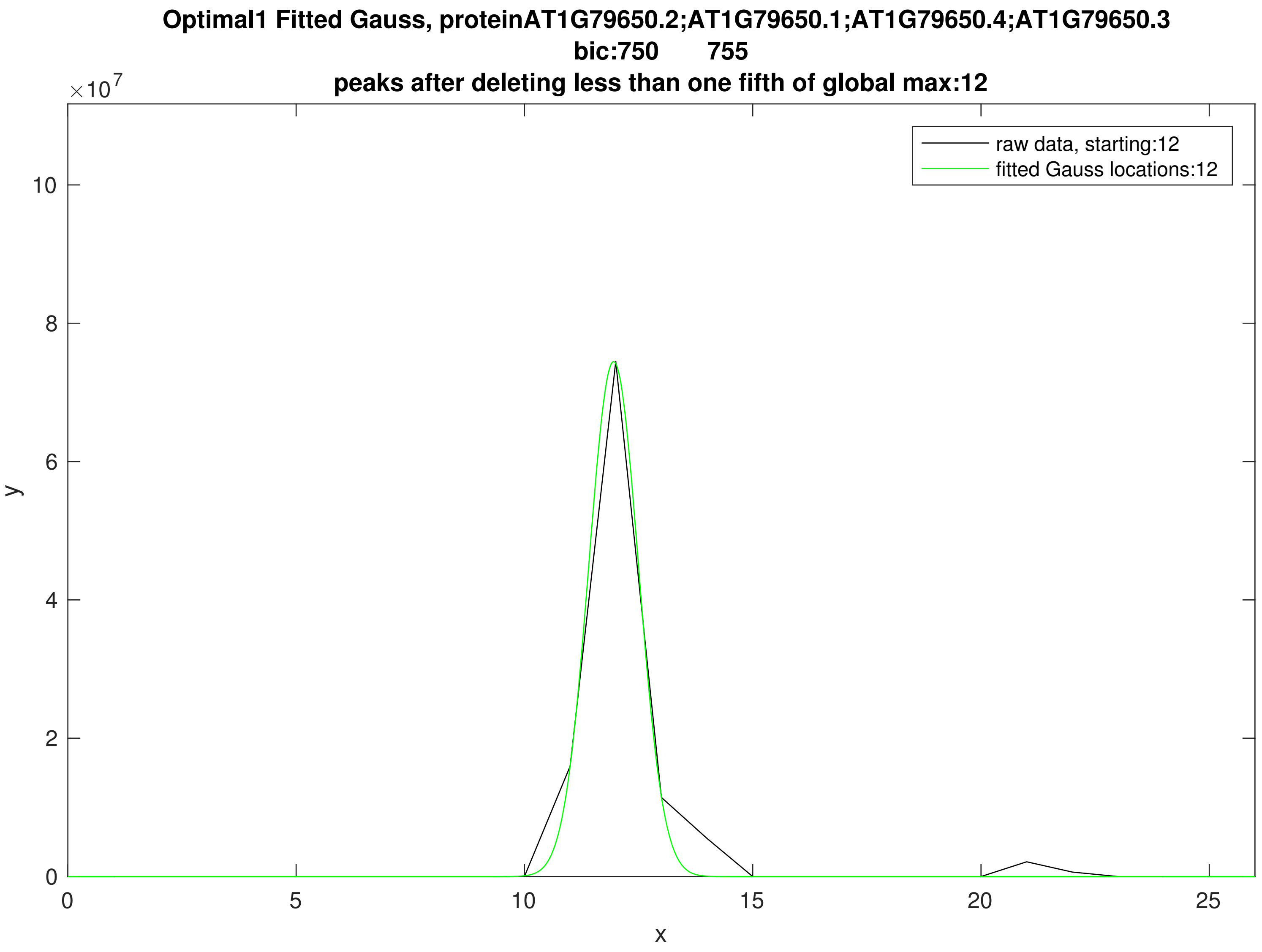
Optimal1 Fitted Gauss, proteinAT1G79650.2;AT1G79650.1;AT1G79650.4;AT1G79650.3
bic:750 755
peaks after deleting less than one fifth of global max:12
×10^7
raw data, starting:12
fitted Gauss locations:12
y
x
0
2
4
6
8
10
0
5
10
15
20
25

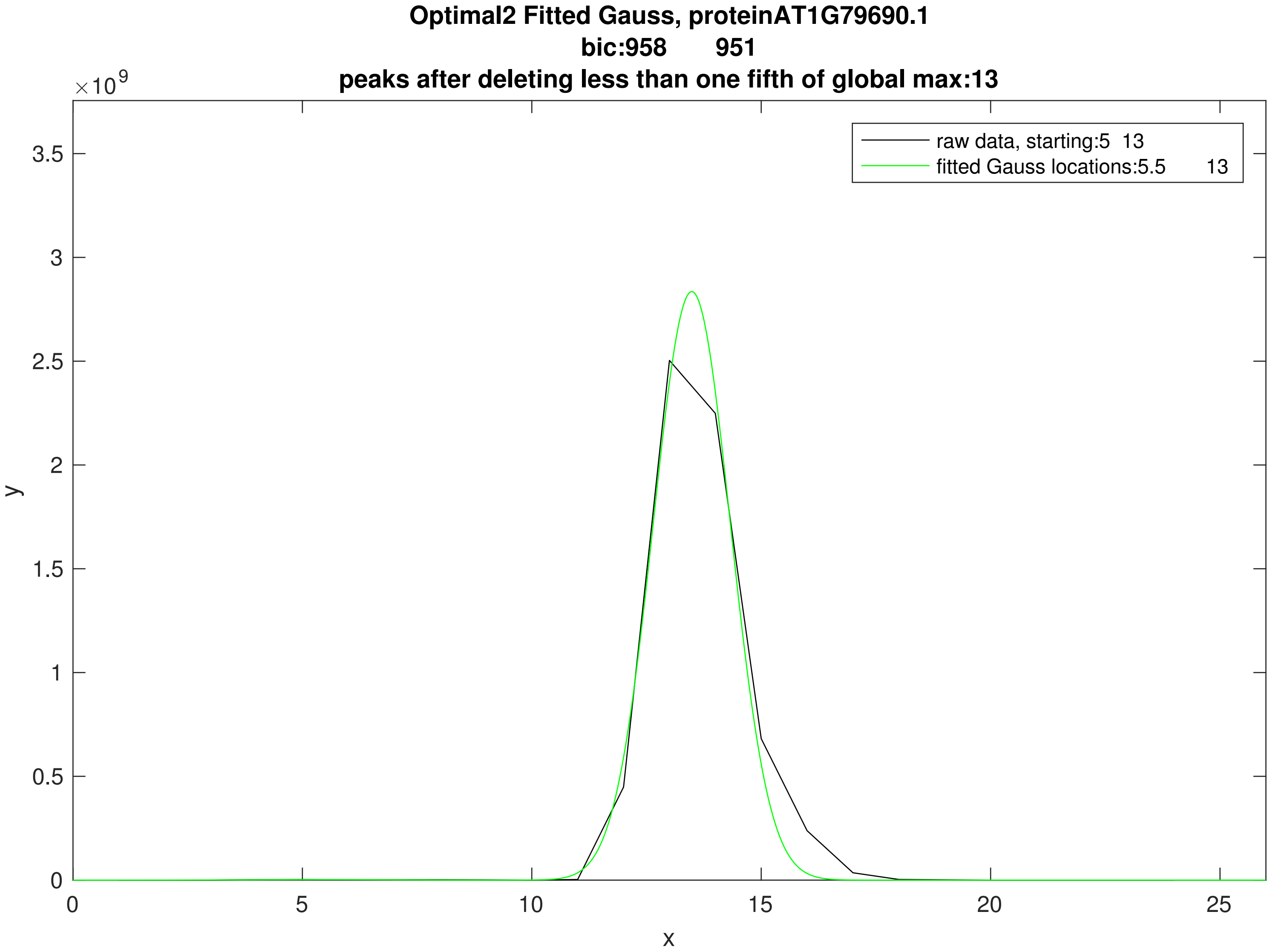
Optimal2 Fitted Gauss, proteinAT1G79690.1
bic:958 951
peaks after deleting less than one fifth of global max:13
×10^9
raw data, starting:5 13
fitted Gauss locations:5.5 13
3.5
3
2.5
2
1.5
1
0.5
0
y
0
5
10
15
20
25
x

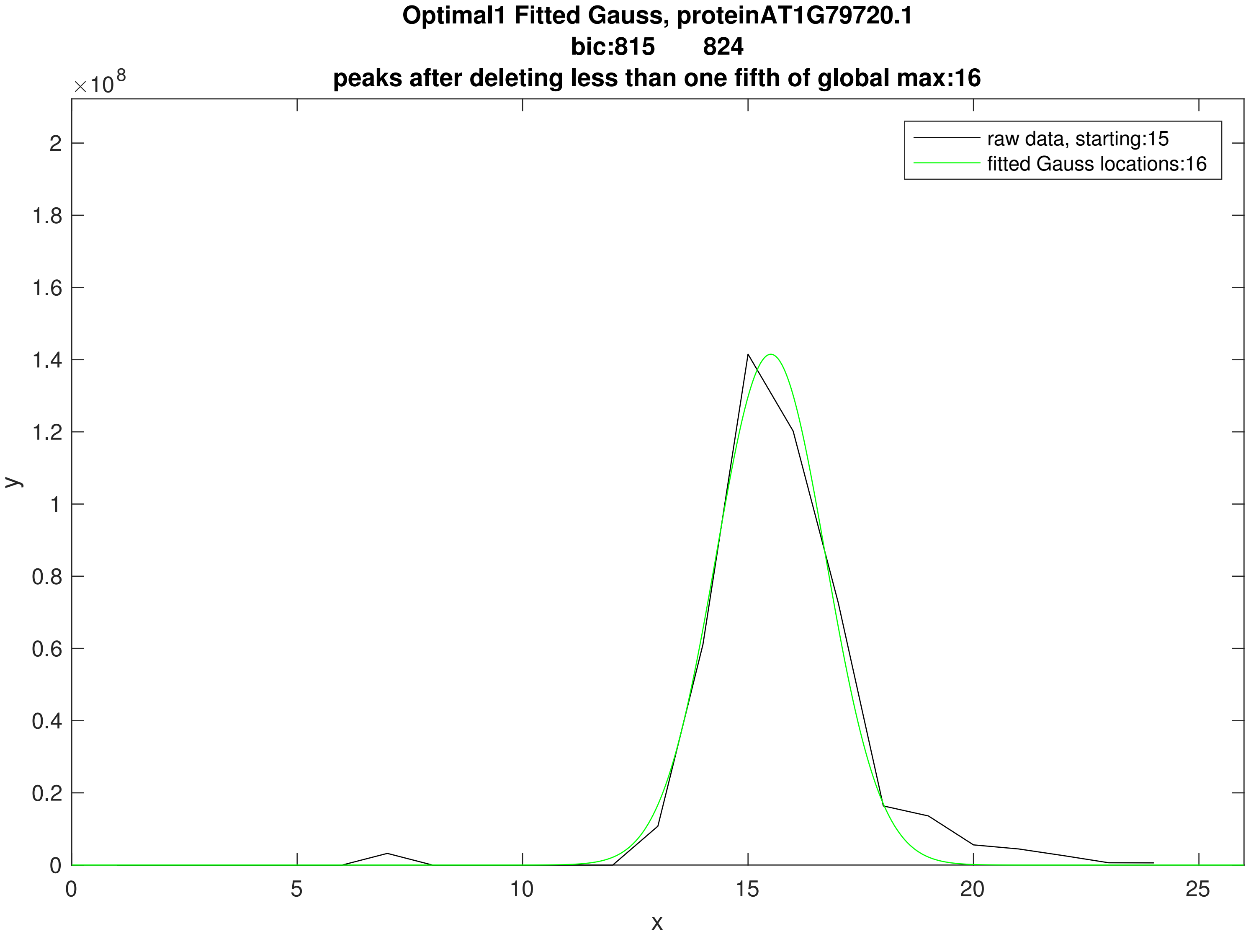

Optimal1 Fitted Gauss, proteinAT1G79720.1
bic:815 824
peaks after deleting less than one fifth of global max:16
×10^8
raw data, starting:15
fitted Gauss locations:16
y
x
0
0.2
0.4
0.6
0.8
1
1.2
1.4
1.6
1.8
2
0
5
10
15
20
25

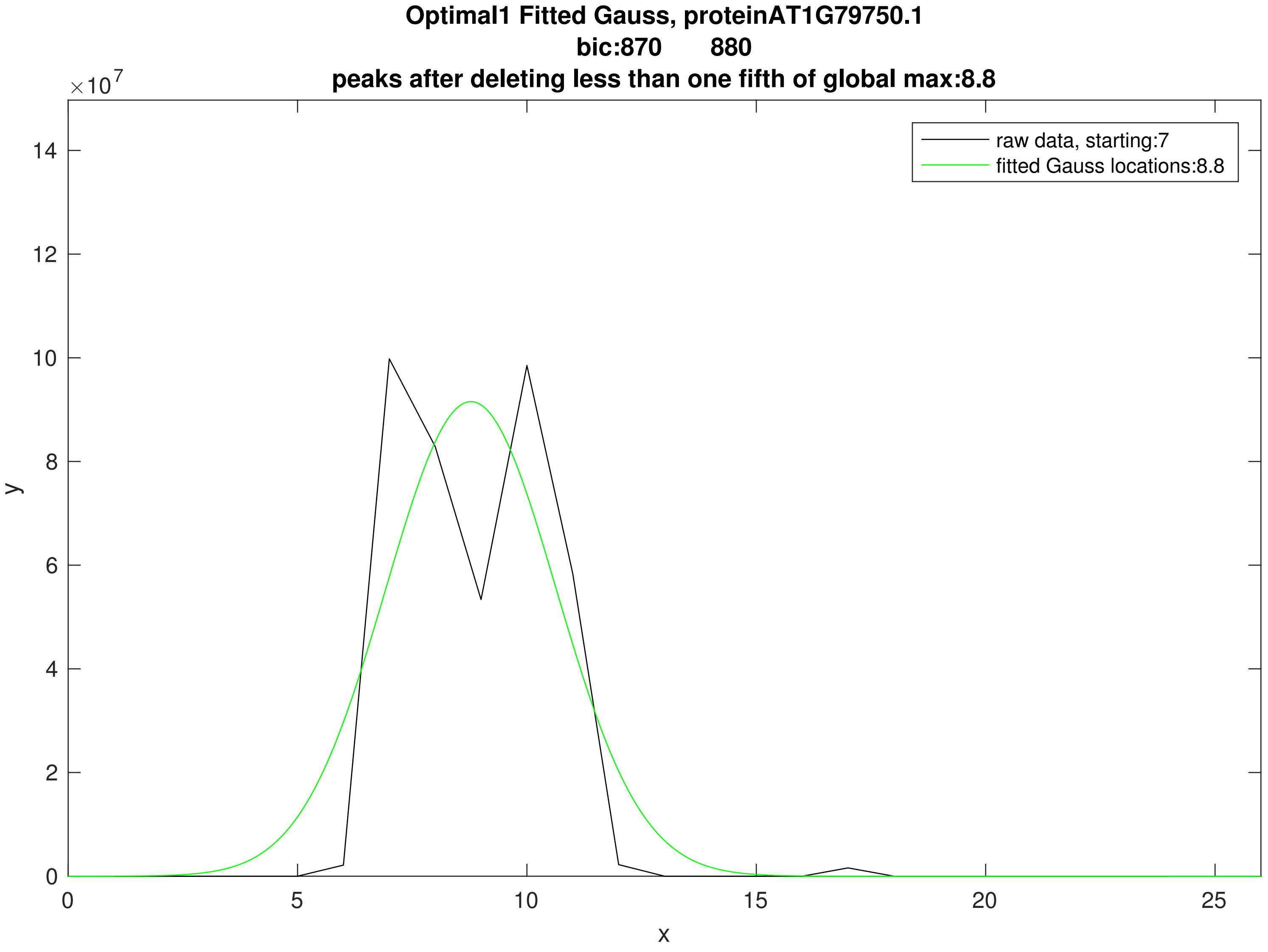

Optimal1 Fitted Gauss, proteinAT1G79750.1
bic:870 880
peaks after deleting less than one fifth of global max:8.8
×10^7
raw data, starting:7
fitted Gauss locations:8.8
14
12
10
8
6
4
2
0
y
0
5
10
15
20
25
x

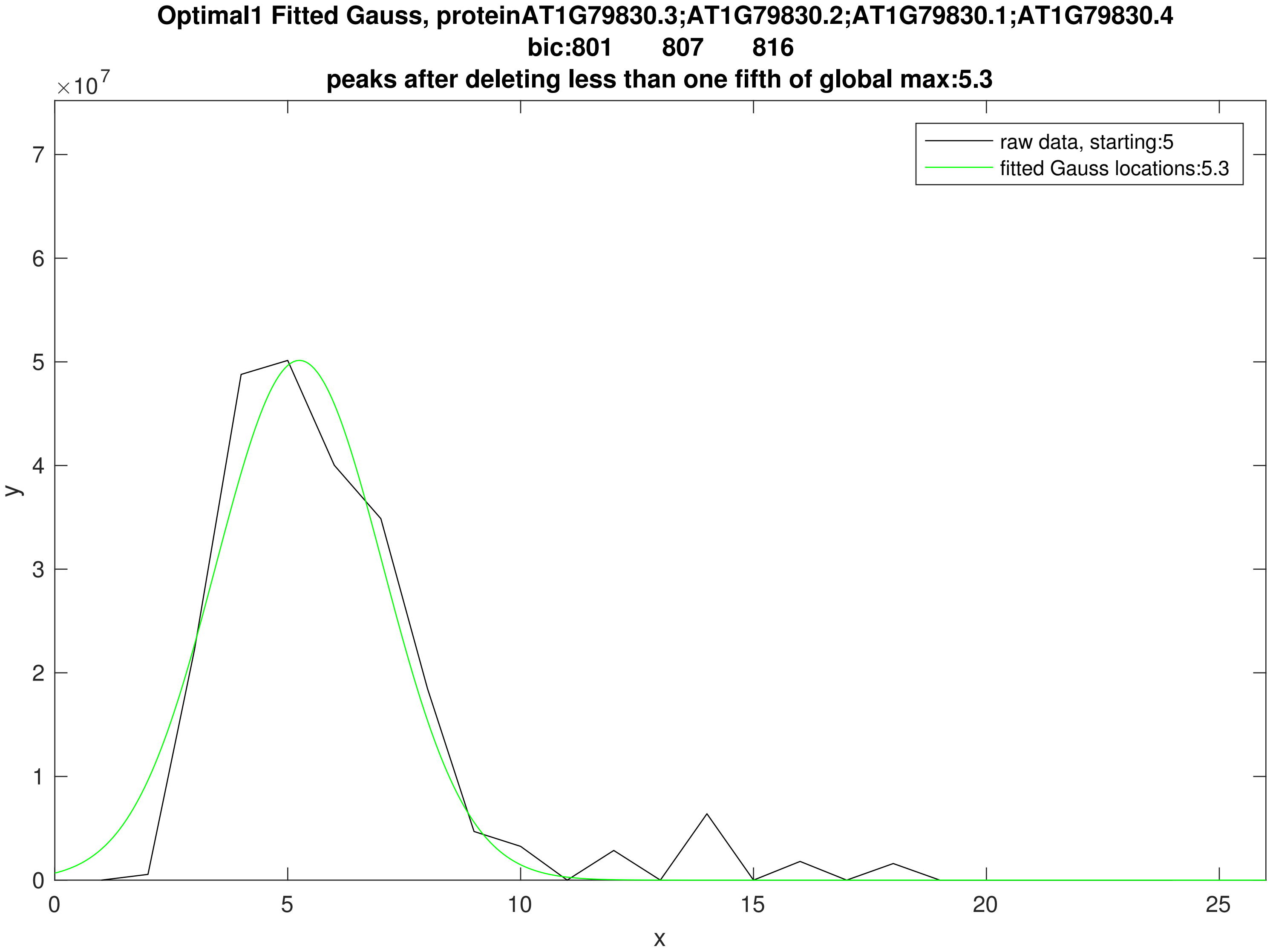

Optimal1 Fitted Gauss, proteinAT1G79830.3;AT1G79830.2;AT1G79830.1;AT1G79830.4
bic:801 807 816
peaks after deleting less than one fifth of global max:5.3
×10^7
raw data, starting:5
fitted Gauss locations:5.3
y
x

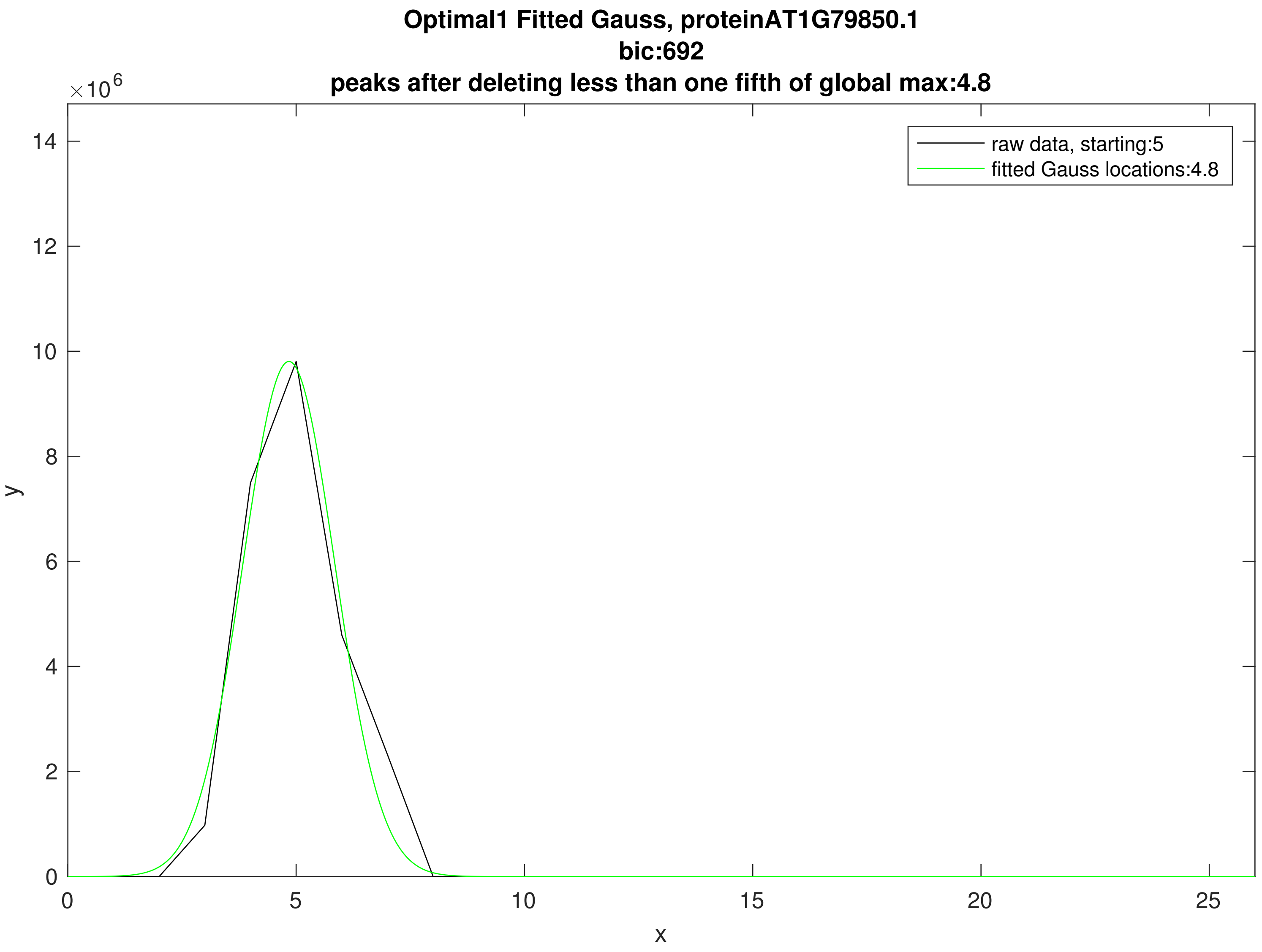

Optimal1 Fitted Gauss, proteinAT1G79850.1
bic:692
peaks after deleting less than one fifth of global max:4.8
×10^6
raw data, starting:5
fitted Gauss locations:4.8
y
x
0
2
4
6
8
10
12
14
0
5
10
15
20
25

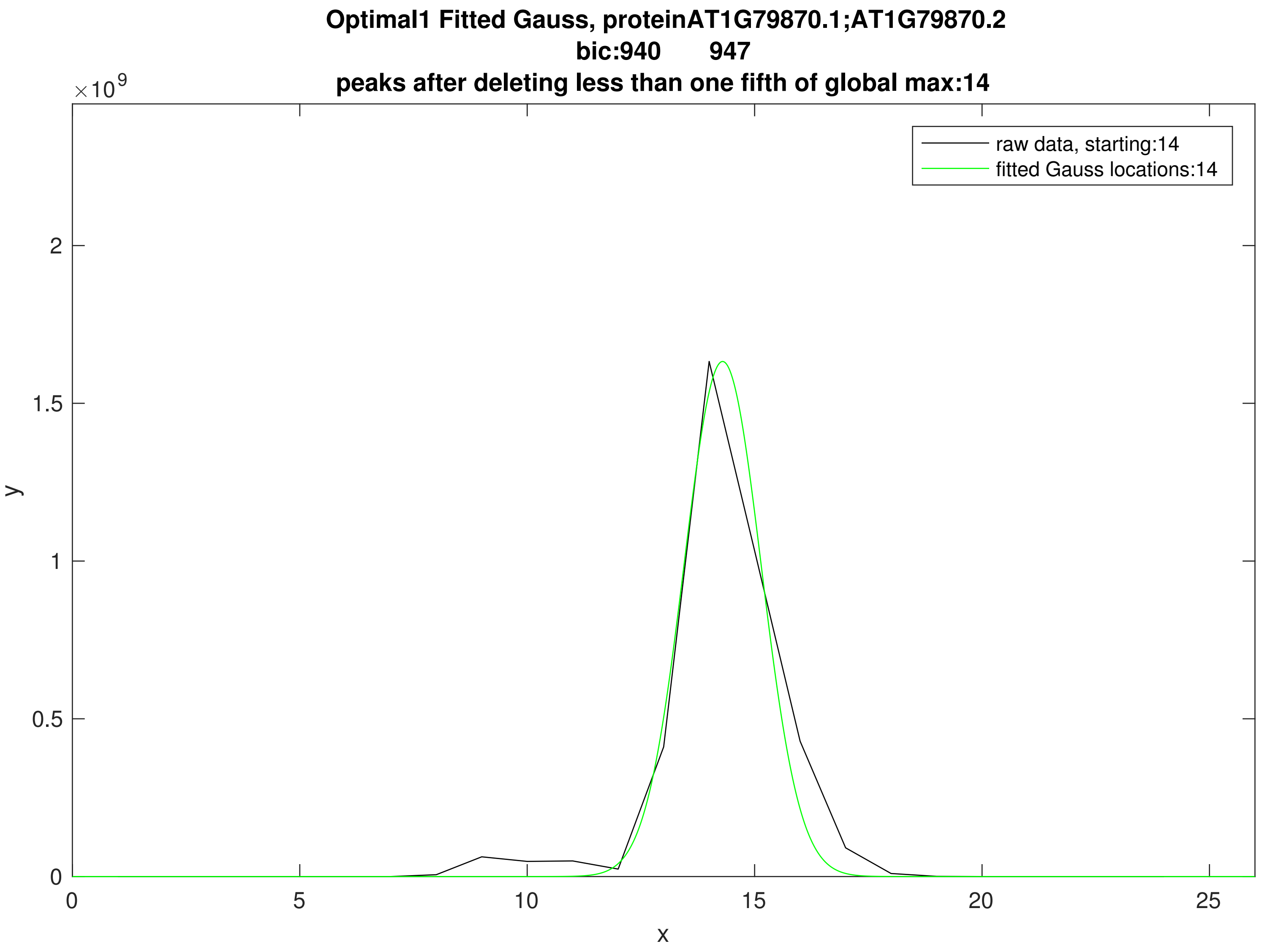

Optimal1 Fitted Gauss, proteinAT1G79870.1;AT1G79870.2
bic:940 947
peaks after deleting less than one fifth of global max:14
×10^9
raw data, starting:14
fitted Gauss locations:14
y
x
0
0.5
1
1.5
2
0
5
10
15
20
25

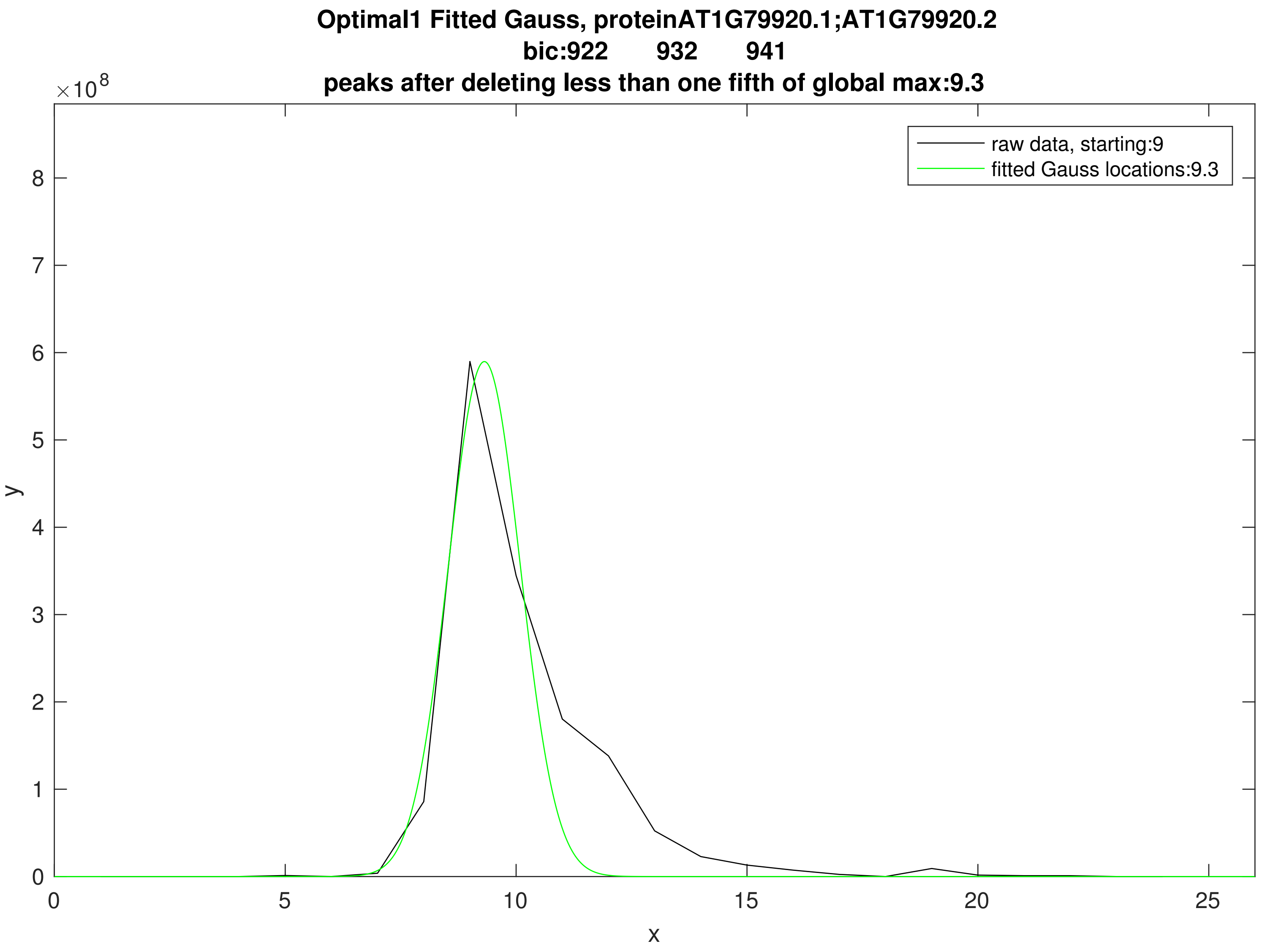

Optimal1 Fitted Gauss, proteinAT1G79920.1;AT1G79920.2
bic:922 932 941
peaks after deleting less than one fifth of global max:9.3
×10^8
raw data, starting:9
fitted Gauss locations:9.3
y
x
0
1
2
3
4
5
6
7
8
0
5
10
15
20
25

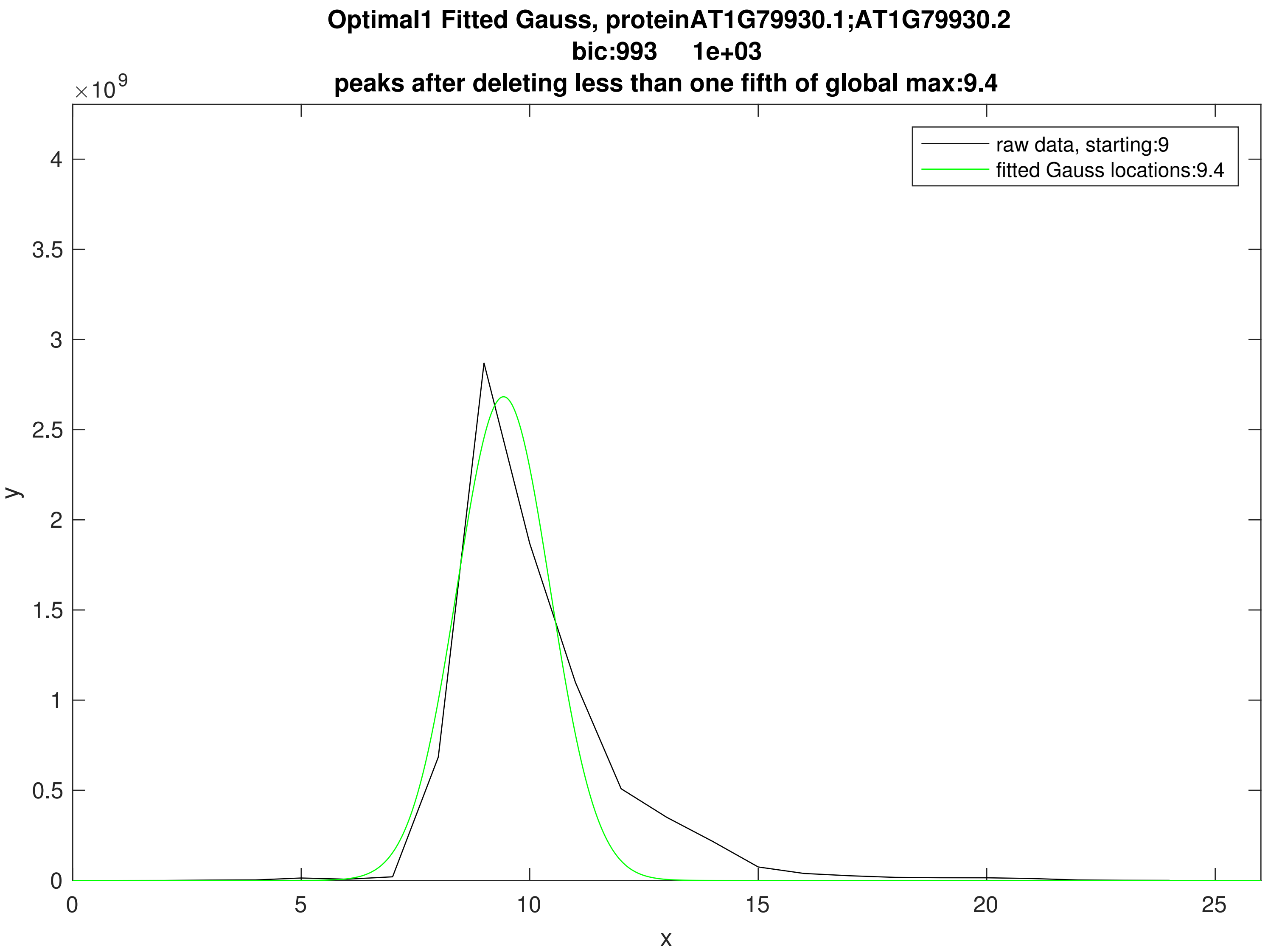
Optimal1 Fitted Gauss, proteinAT1G79930.1;AT1G79930.2
bic:993 1e+03
peaks after deleting less than one fifth of global max:9.4
×10^9
raw data, starting:9
fitted Gauss locations:9.4
y
x
0
0.5
1
1.5
2
2.5
3
3.5
4
5
10
15
20
25

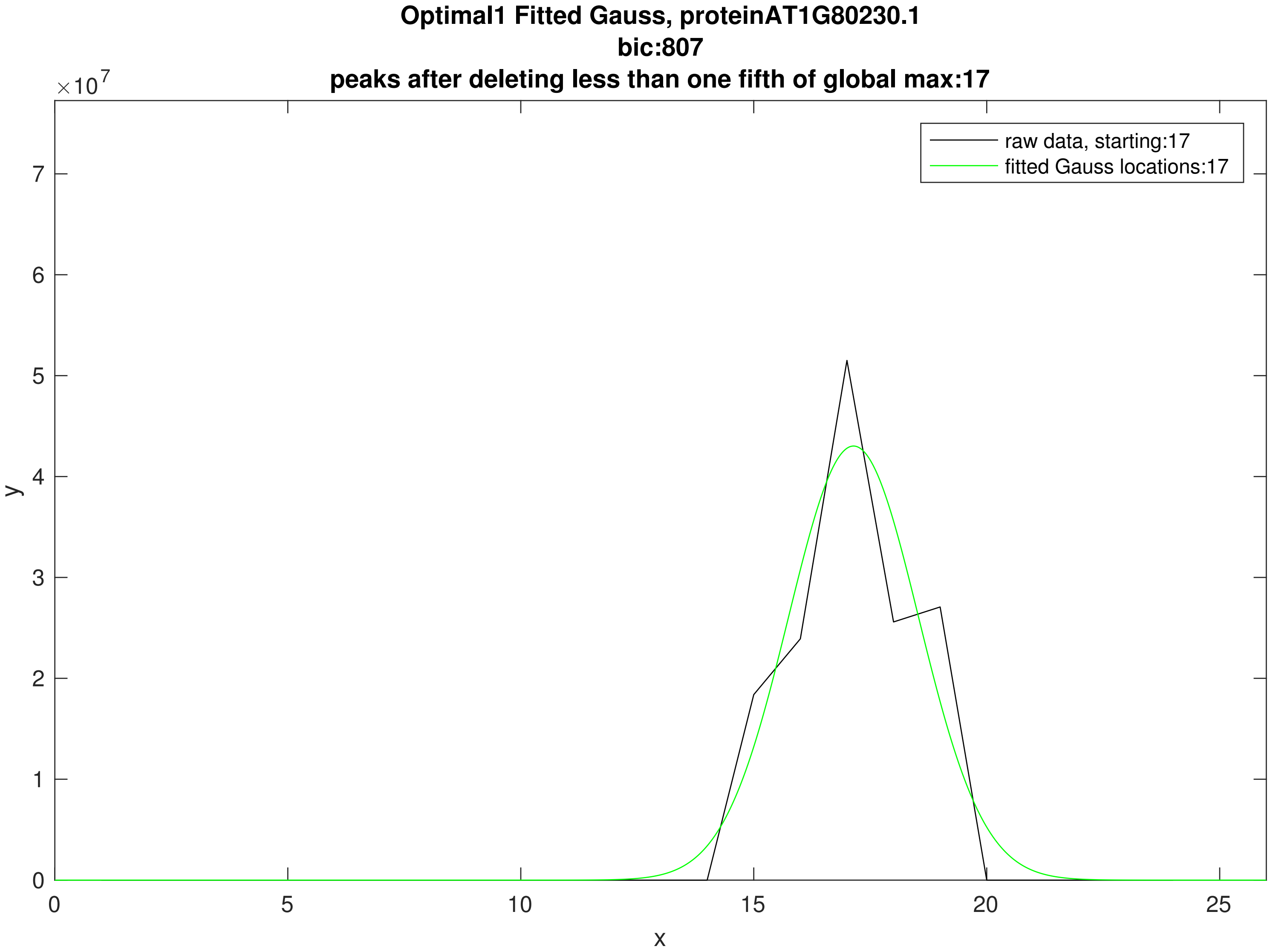

Optimal1 Fitted Gauss, proteinAT1G80230.1
bic:807
peaks after deleting less than one fifth of global max:17
×10^7
raw data, starting:17
fitted Gauss locations:17
y
x
0
1
2
3
4
5
6
7
0
5
10
15
20
25

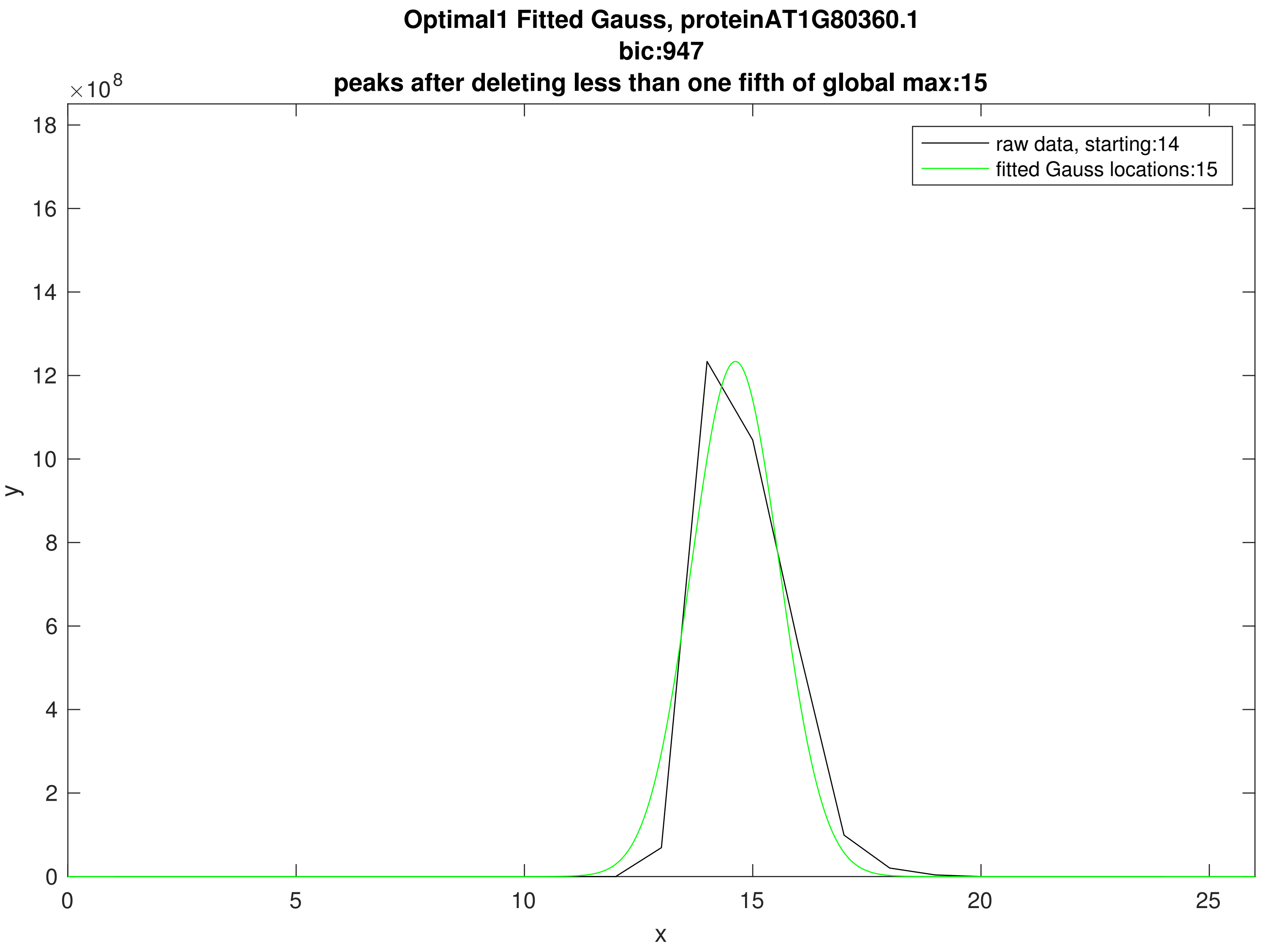
Optimal1 Fitted Gauss, proteinAT1G80360.1
bic:947
peaks after deleting less than one fifth of global max:15
×10^8
raw data, starting:14
fitted Gauss locations:15
y
x
0
2
4
6
8
10
12
14
16
18
0
5
10
15
20
25

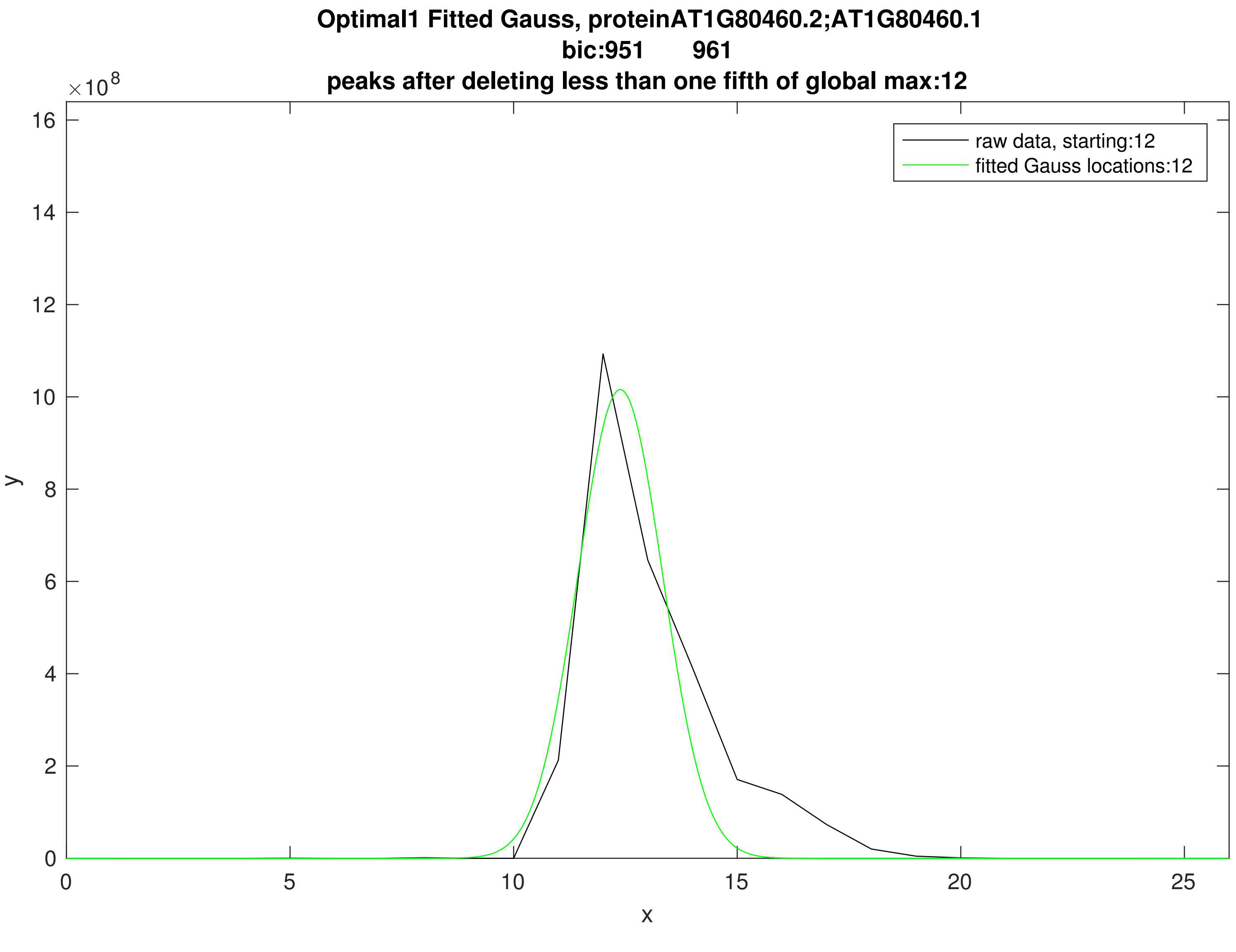

Optimal1 Fitted Gauss, proteinAT1G80460.2;AT1G80460.1
bic:951 961
peaks after deleting less than one fifth of global max:12
×10^8
raw data, starting:12
fitted Gauss locations:12
y
x
0
2
4
6
8
10
12
14
16
0
5
10
15
20
25

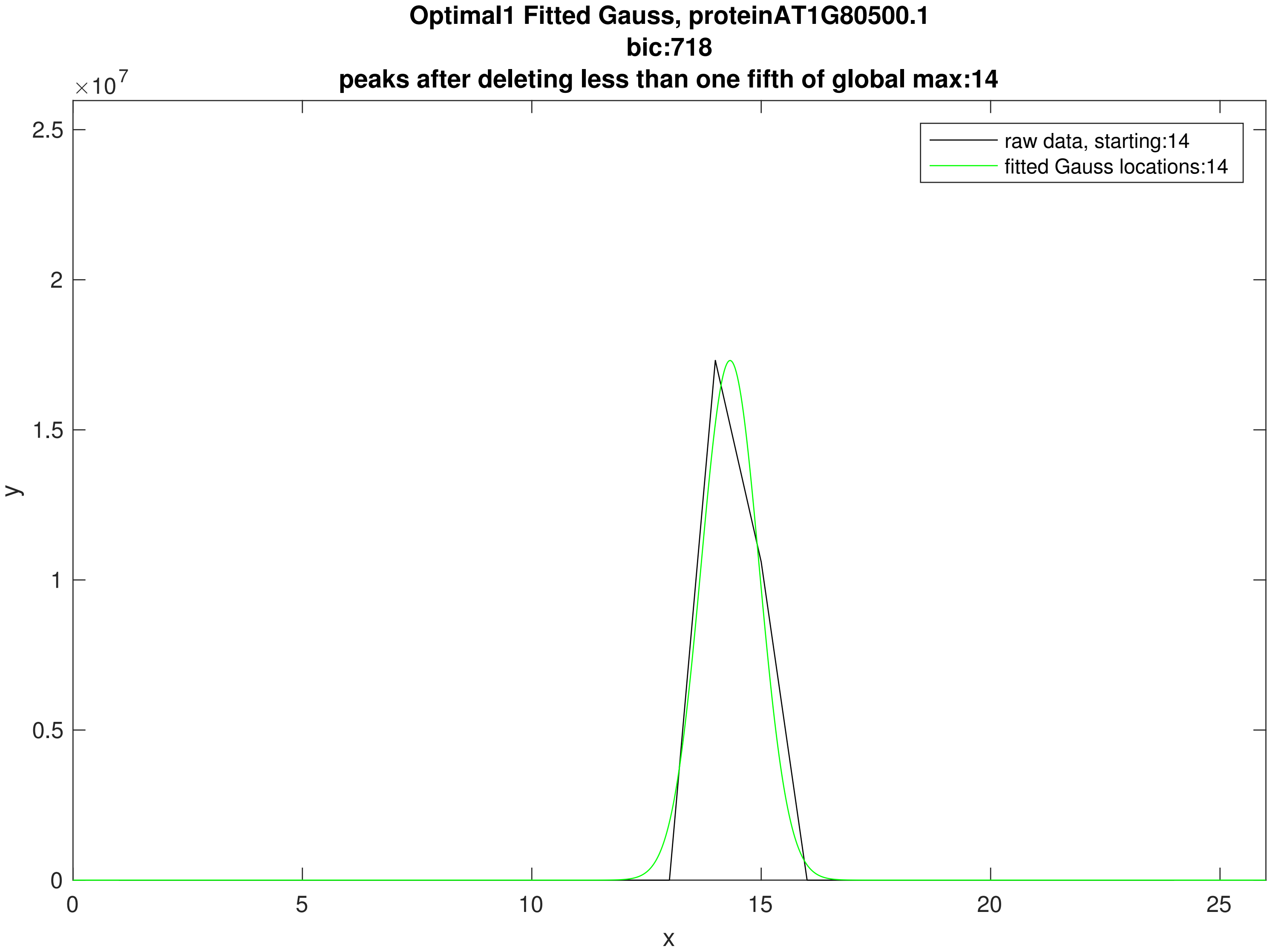
Optimal1 Fitted Gauss, proteinAT1G80500.1
bic:718
peaks after deleting less than one fifth of global max:14
×10^7
raw data, starting:14
fitted Gauss locations:14
y
x
0
0.5
1
1.5
2
2.5
0
5
10
15
20
25

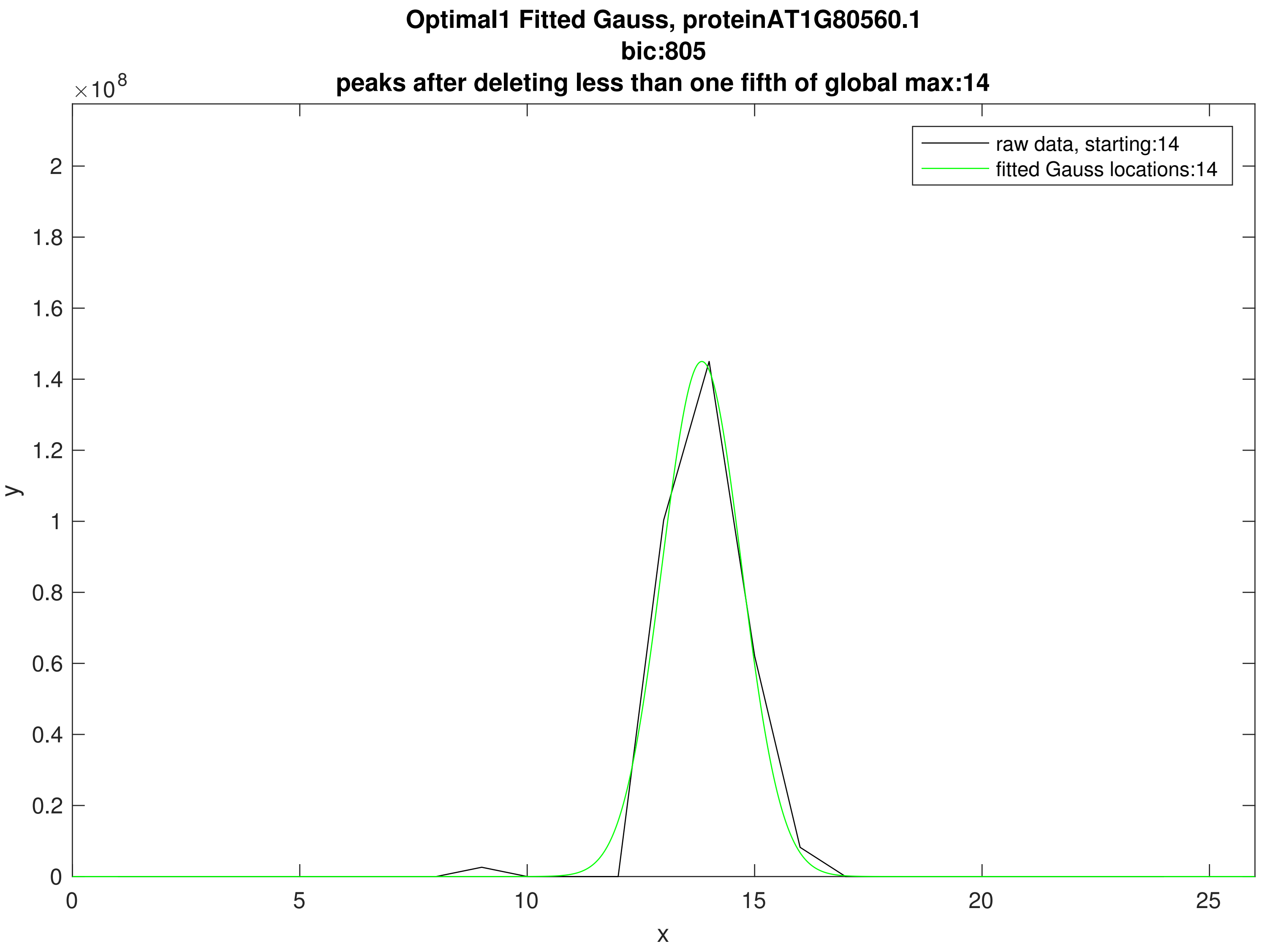
Optimal1 Fitted Gauss, proteinAT1G80560.1
bic:805
peaks after deleting less than one fifth of global max:14
×10^8
raw data, starting:14
fitted Gauss locations:14
2
1.8
1.6
1.4
1.2
1
0.8
0.6
0.4
0.2
0
y
0
5
10
15
20
25
x

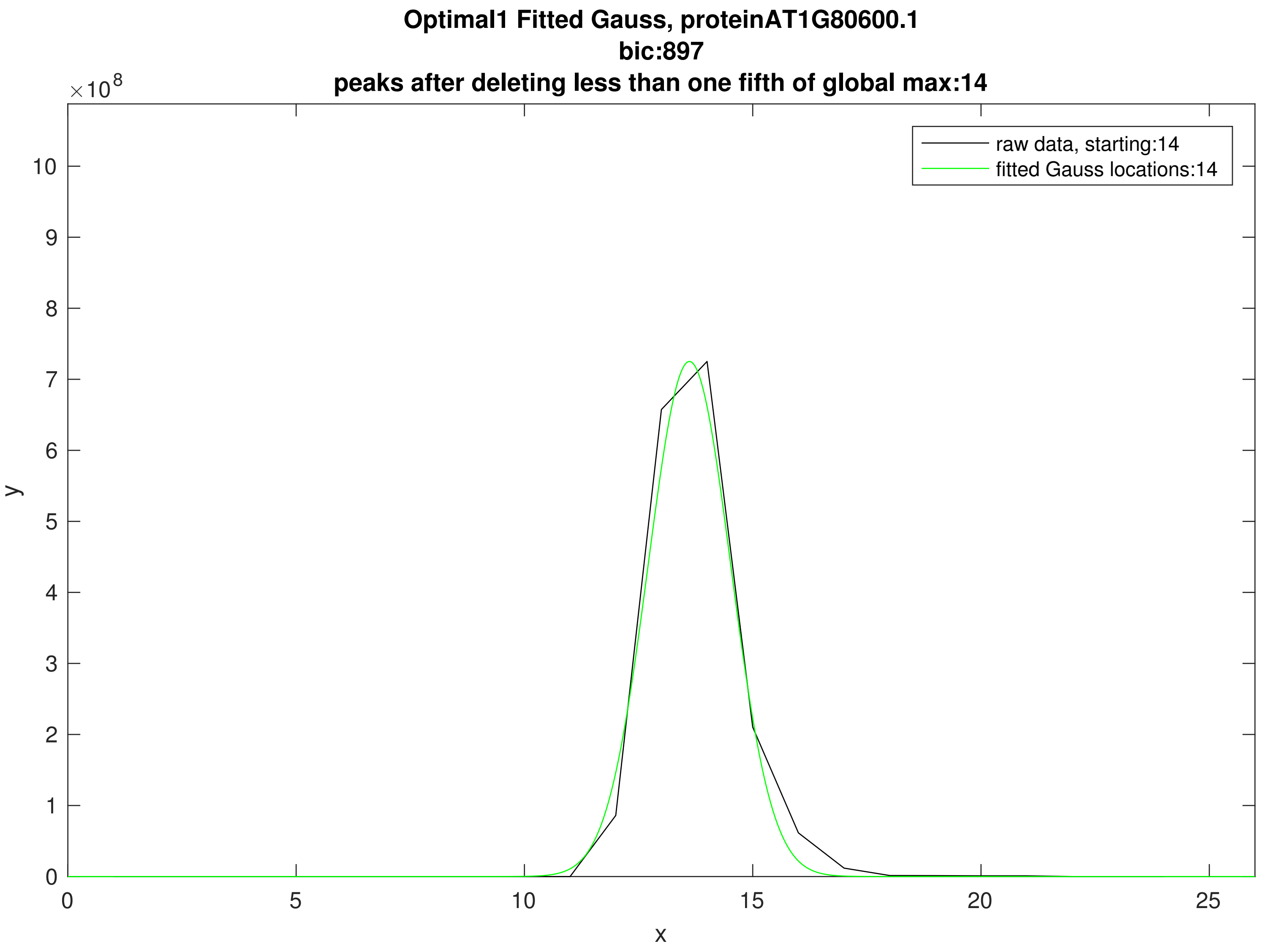
Optimal1 Fitted Gauss, proteinAT1G80600.1
bic:897
peaks after deleting less than one fifth of global max:14
×10^8
raw data, starting:14
fitted Gauss locations:14
y
x
0
1
2
3
4
5
6
7
8
9
10
0
5
10
15
20
25

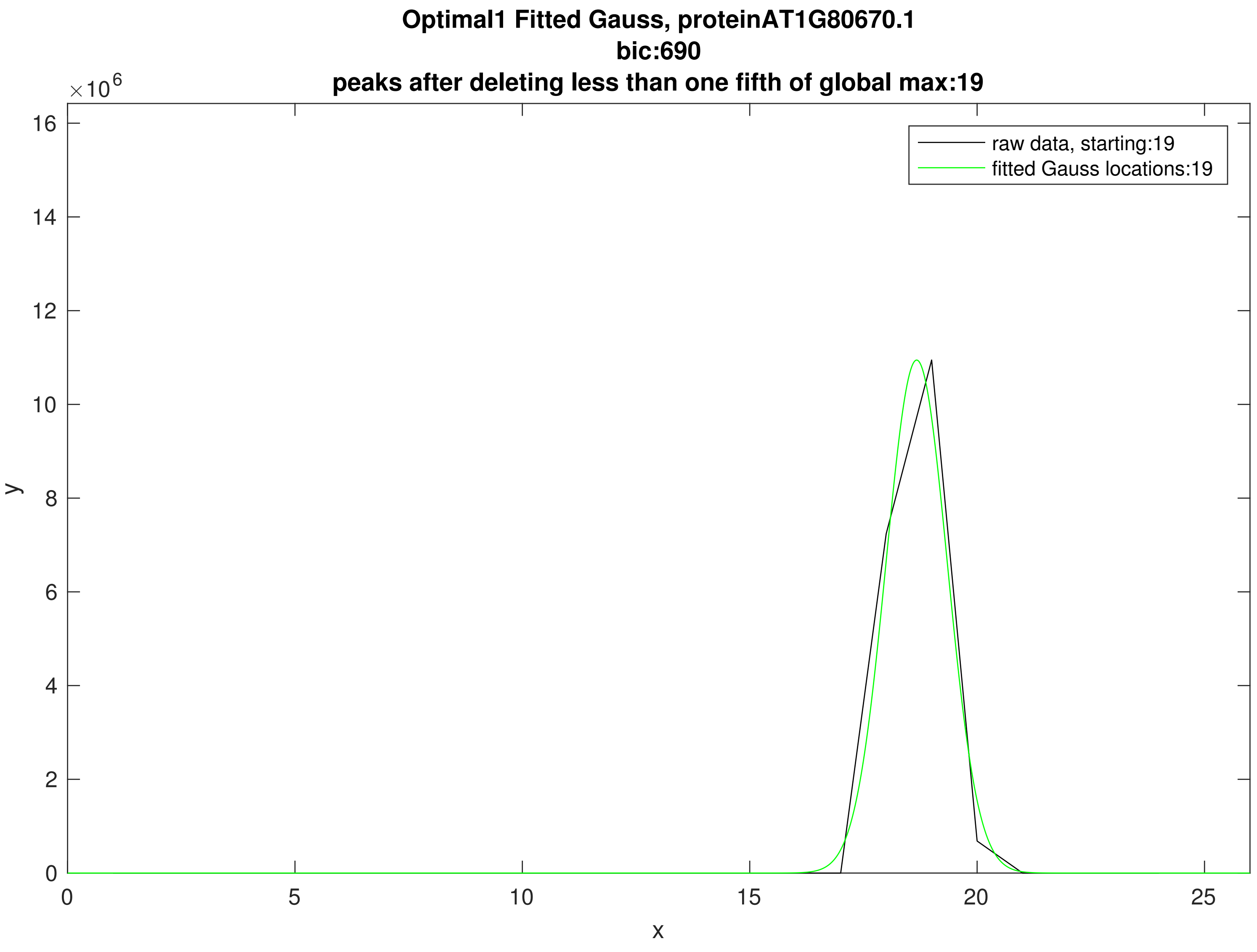
Optimal1 Fitted Gauss, proteinAT1G80670.1
bic:690
peaks after deleting less than one fifth of global max:19
×10^6
raw data, starting:19
fitted Gauss locations:19
y
x
0
2
4
6
8
10
12
14
16
0
5
10
15
20
25

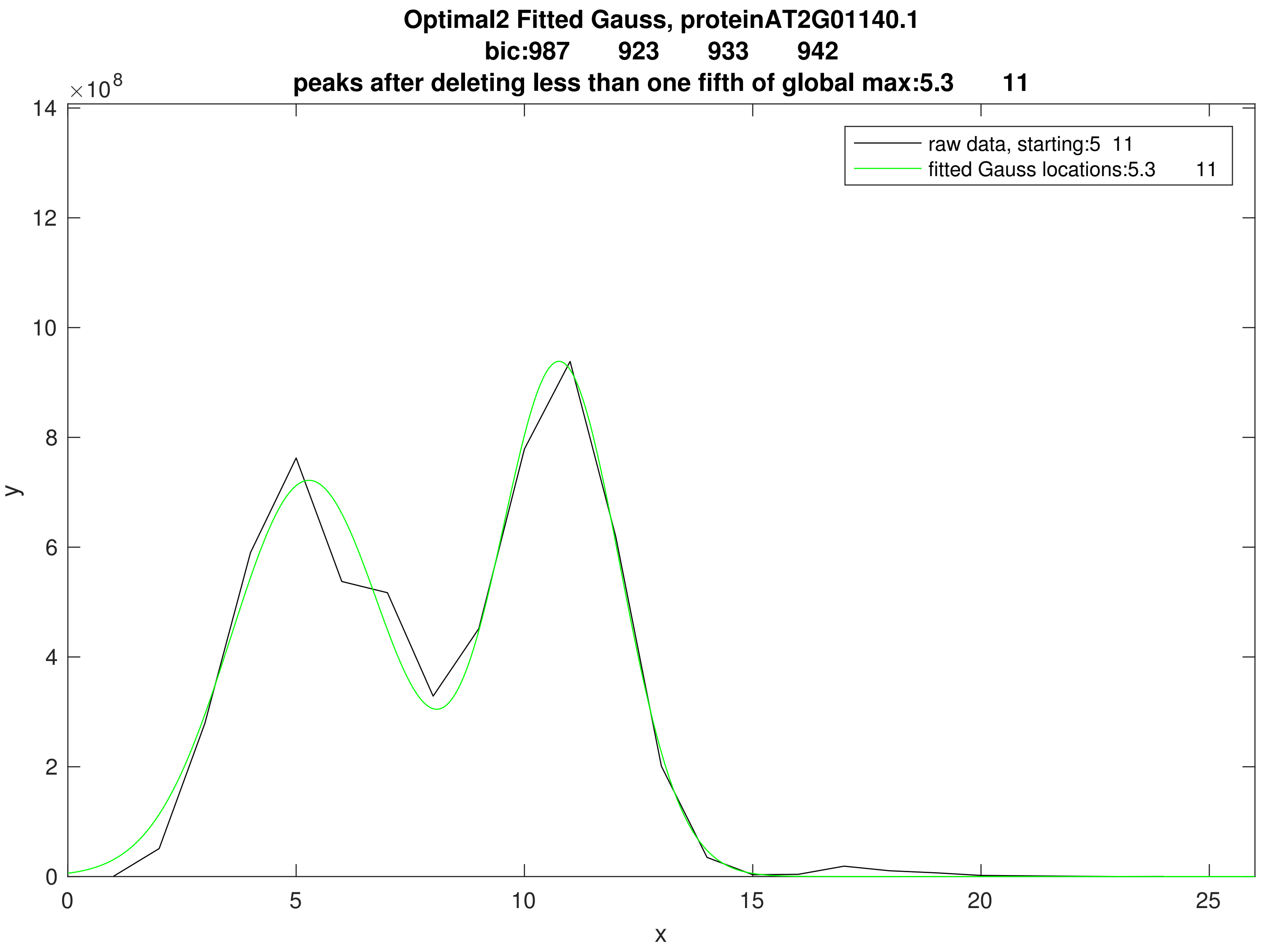
Optimal2 Fitted Gauss, proteinAT2G01140.1
bic:987 923 933 942
peaks after deleting less than one fifth of global max:5.3 11
×10^8
raw data, starting:5 11
fitted Gauss locations:5.3 11
y
x
0
2
4
6
8
10
12
14
0
5
10
15
20
25

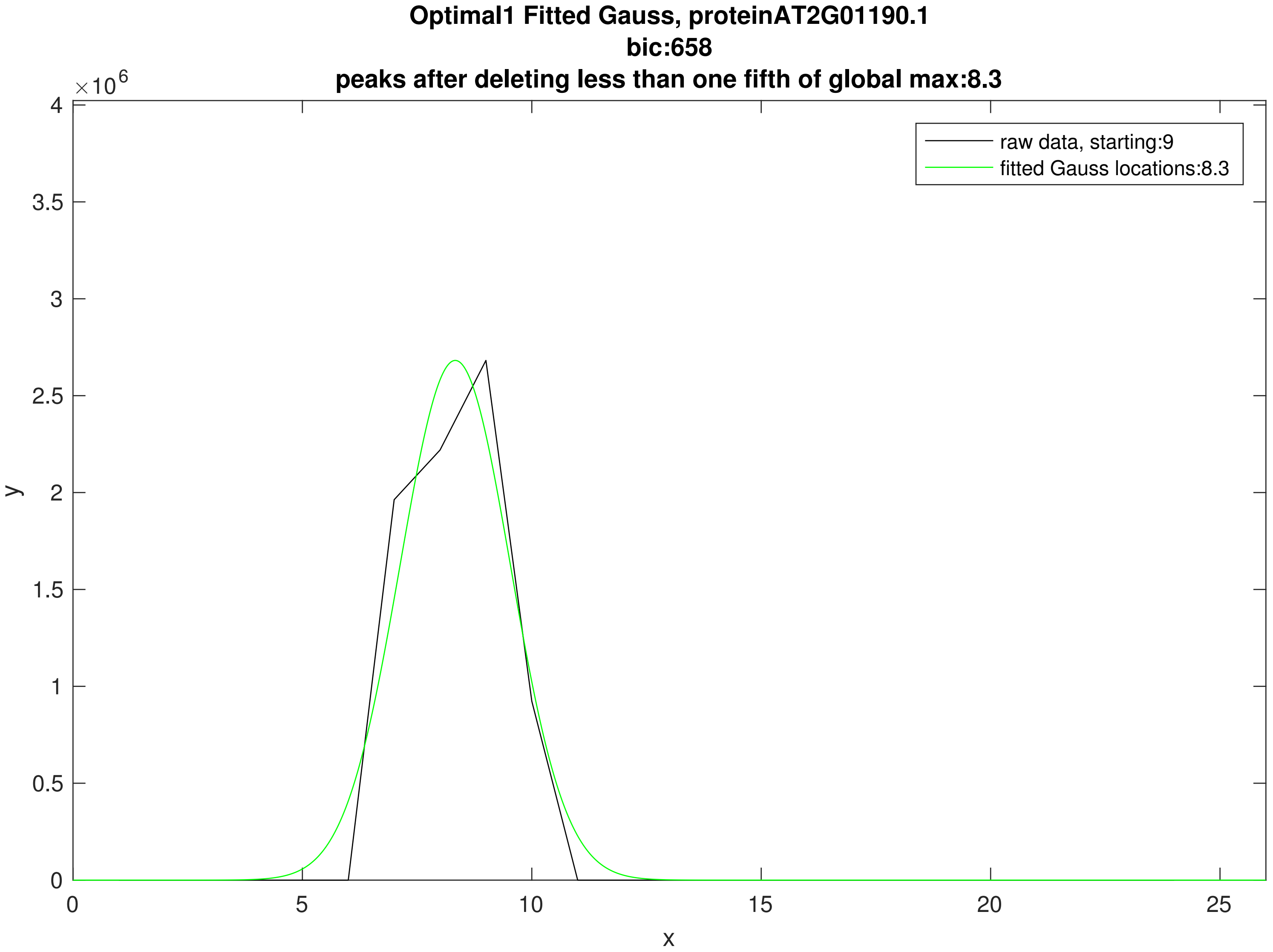
Optimal1 Fitted Gauss, proteinAT2G01190.1
bic:658
peaks after deleting less than one fifth of global max:8.3
raw data, starting:9
fitted Gauss locations:8.3
y
x
×10^6
0
0.5
1
1.5
2
2.5
3
3.5
4
5
10
15
20
25

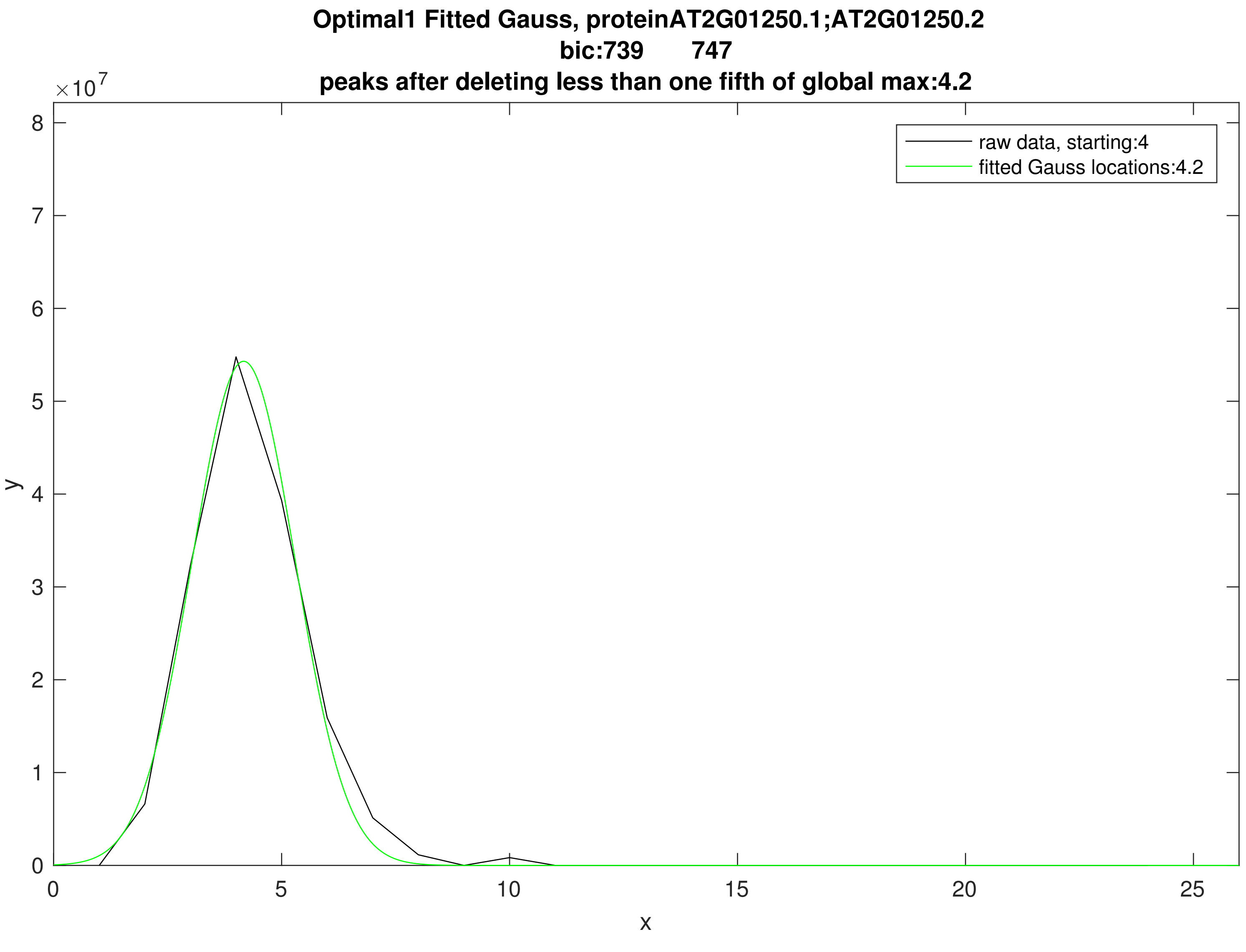

Optimal1 Fitted Gauss, proteinAT2G01250.1;AT2G01250.2
bic:739 747
peaks after deleting less than one fifth of global max:4.2
×10^7
raw data, starting:4
fitted Gauss locations:4.2
y
x
0
1
2
3
4
5
6
7
8
0
5
10
15
20
25

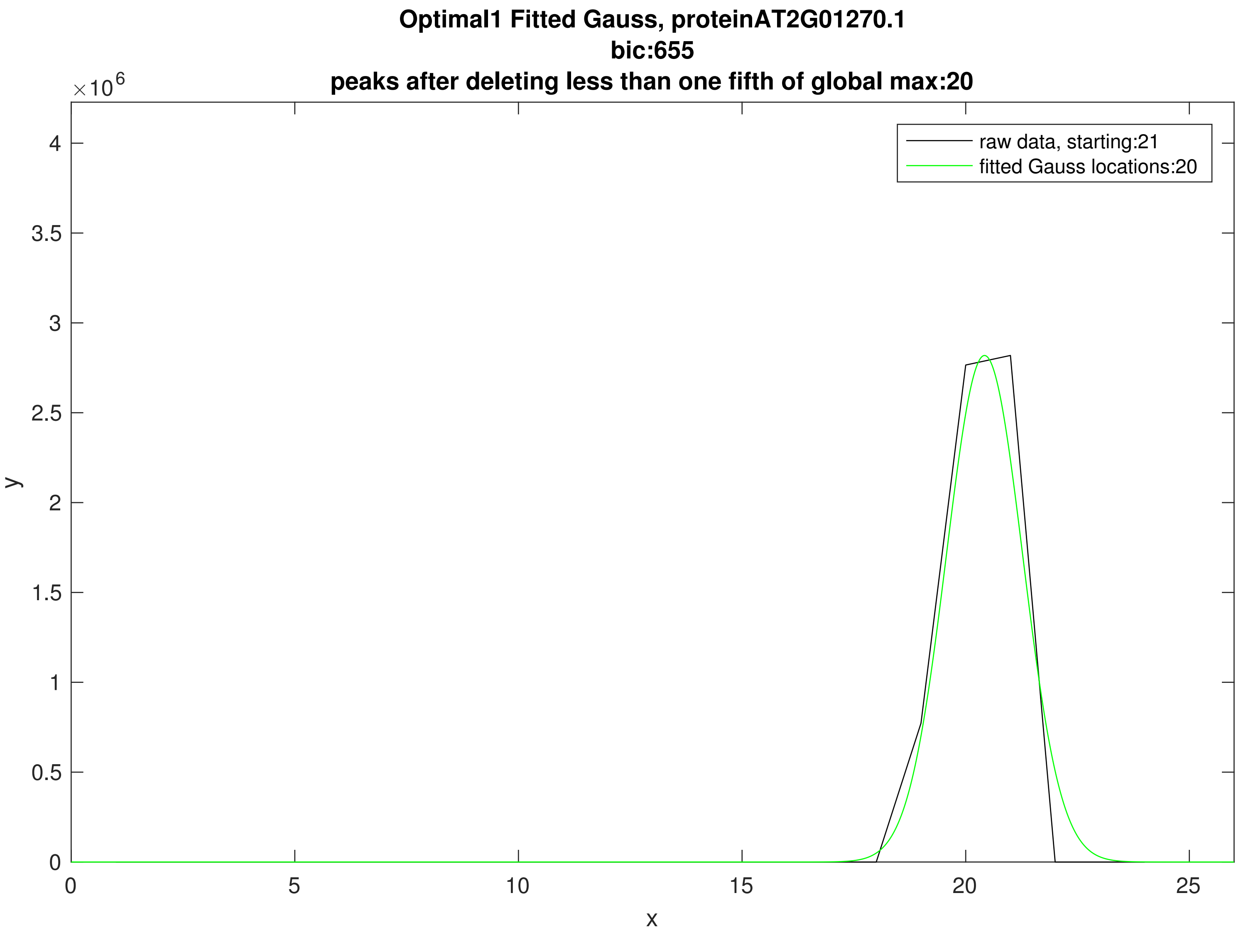

Optimal1 Fitted Gauss, proteinAT2G01270.1
bic:655
peaks after deleting less than one fifth of global max:20
×10^6
raw data, starting:21
fitted Gauss locations:20
y
x
0
0.5
1
1.5
2
2.5
3
3.5
4
0
5
10
15
20
25

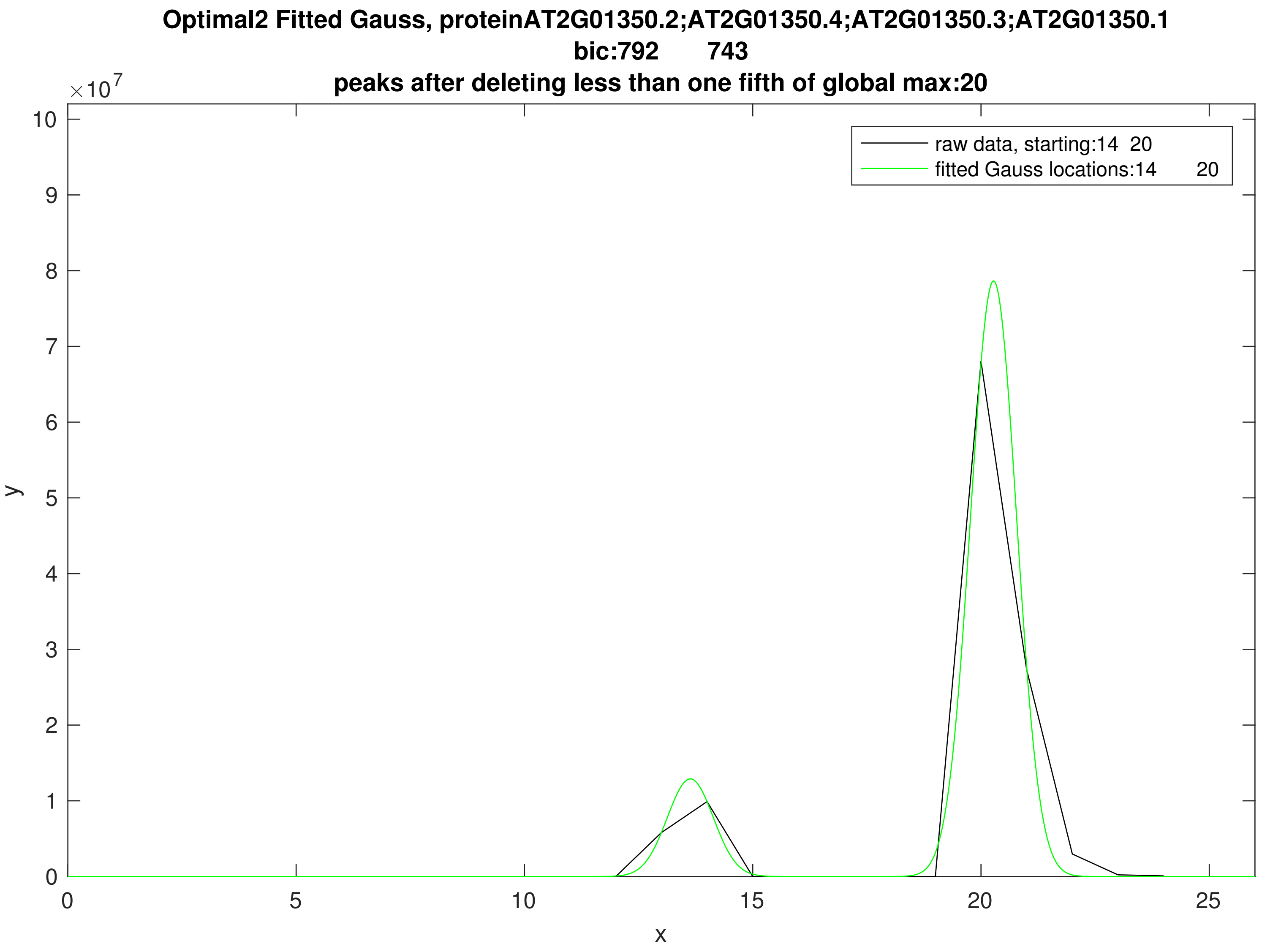
Optimal2 Fitted Gauss, proteinAT2G01350.2;AT2G01350.4;AT2G01350.3;AT2G01350.1
bic:792 743
peaks after deleting less than one fifth of global max:20
×10^7
raw data, starting:14 20
fitted Gauss locations:14 20
y
x
0
1
2
3
4
5
6
7
8
9
10
0
5
10
15
20
25

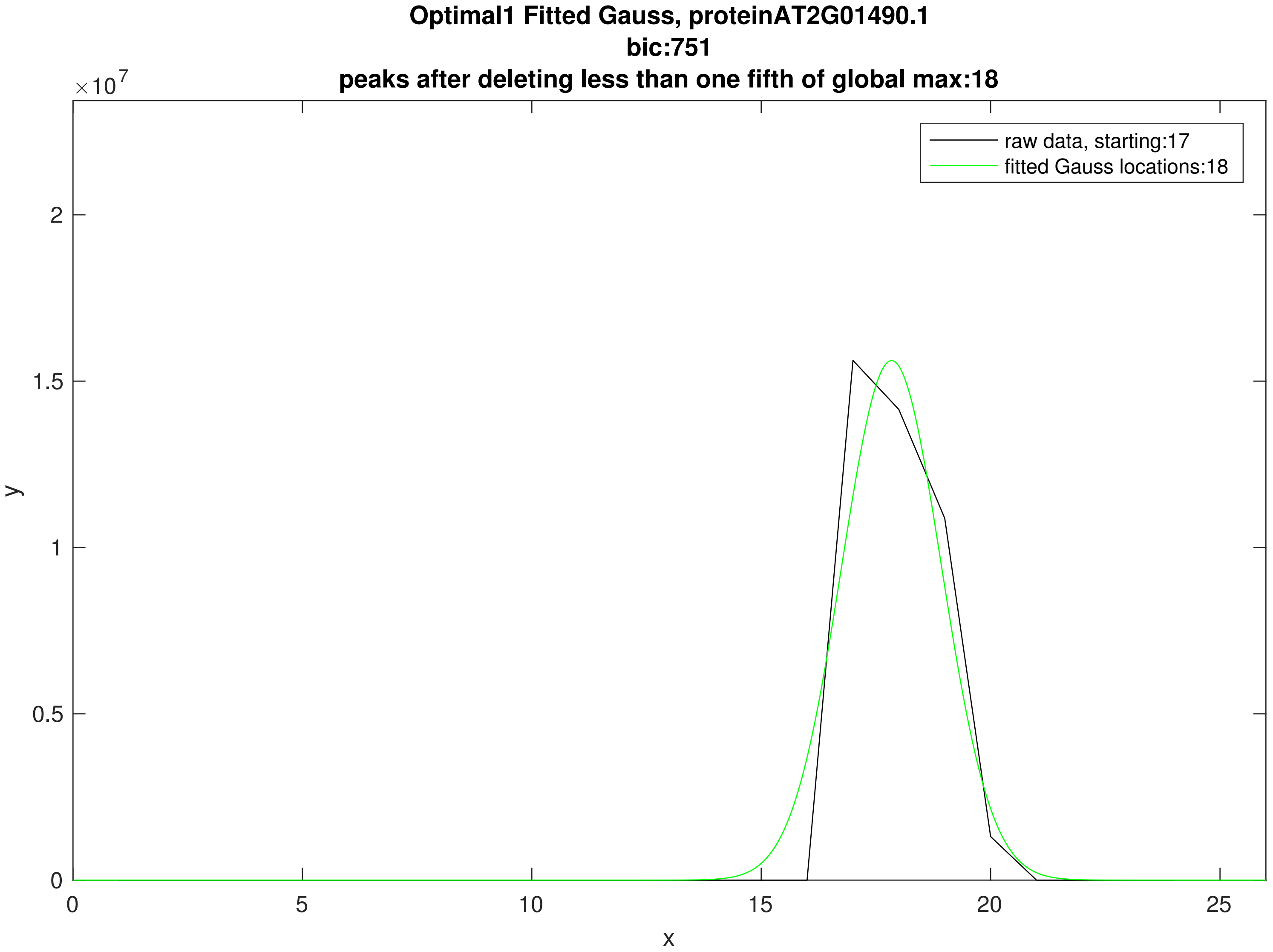
Optimal1 Fitted Gauss, proteinAT2G01490.1
bic:751
peaks after deleting less than one fifth of global max:18
×10^7
raw data, starting:17
fitted Gauss locations:18
y
x
0
0.5
1
1.5
2
5
10
15
20
25

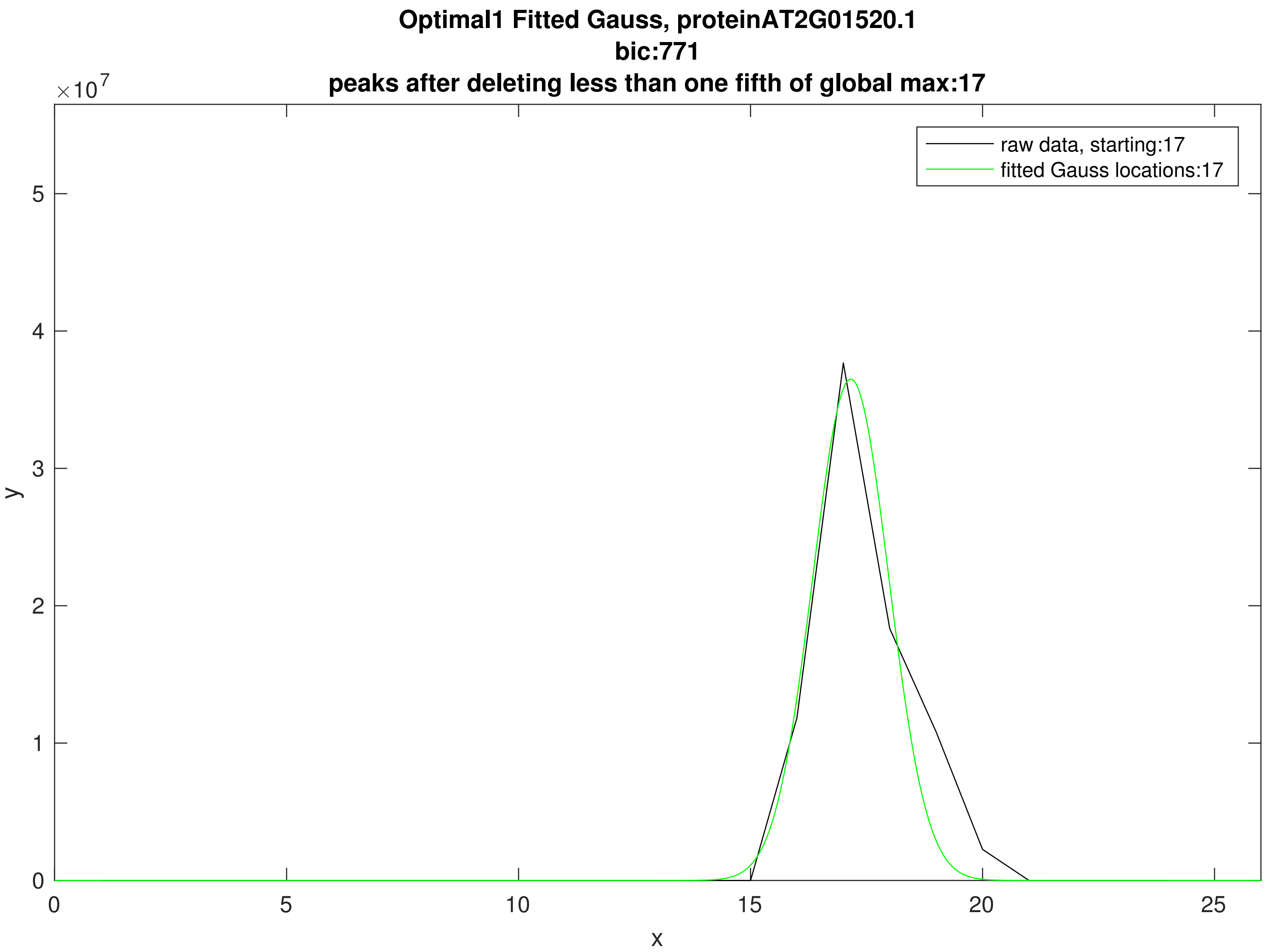

Optimal1 Fitted Gauss, proteinAT2G01520.1
bic:771
peaks after deleting less than one fifth of global max:17
×10^7
raw data, starting:17
fitted Gauss locations:17
y
x
0
1
2
3
4
5
0
5
10
15
20
25

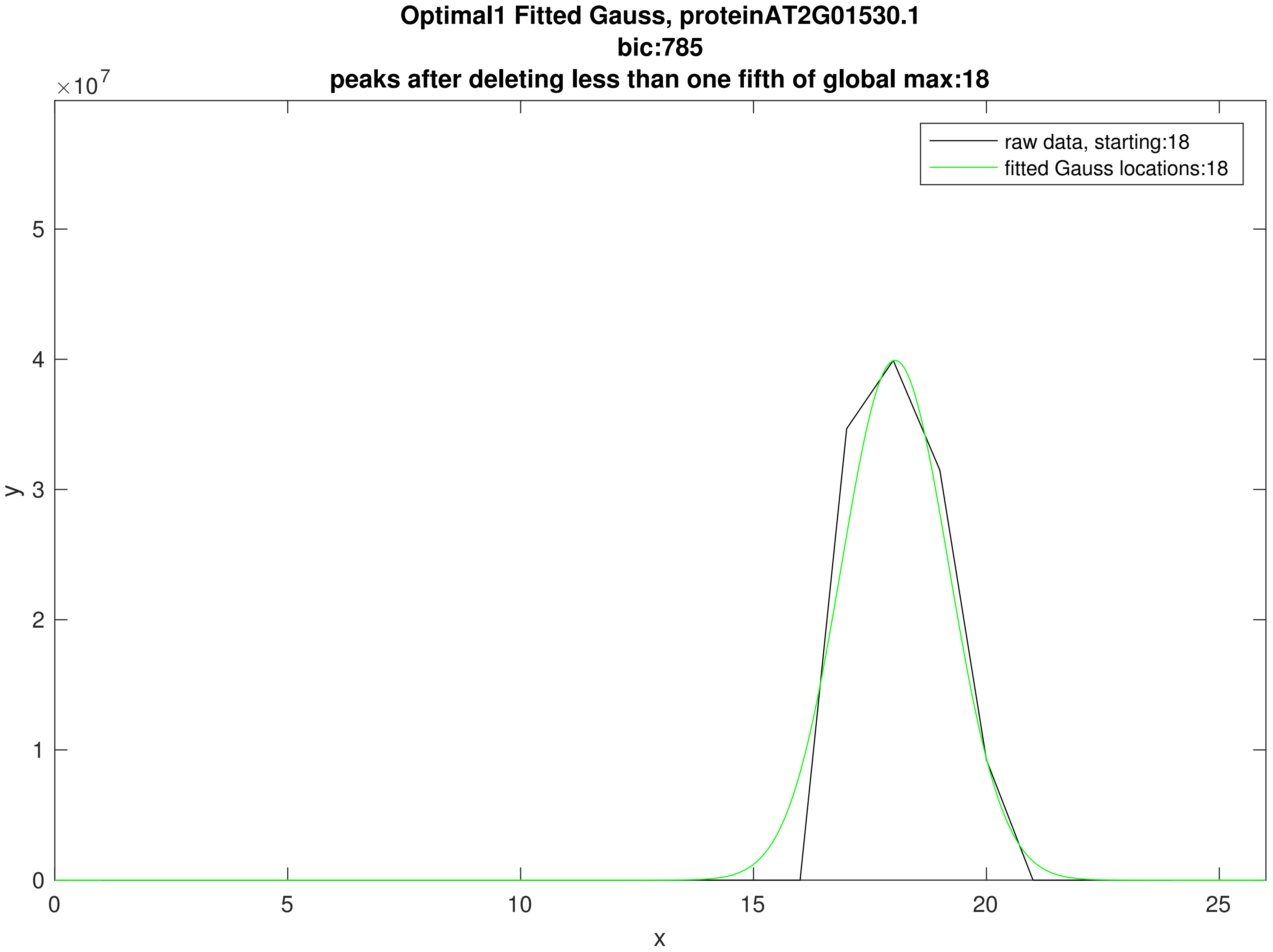

Optimal1 Fitted Gauss, proteinAT2G01530.1
bic:785
peaks after deleting less than one fifth of global max:18
×10^7
raw data, starting:18
fitted Gauss locations:18
y
x
0
1
2
3
4
5
0
5
10
15
20
25

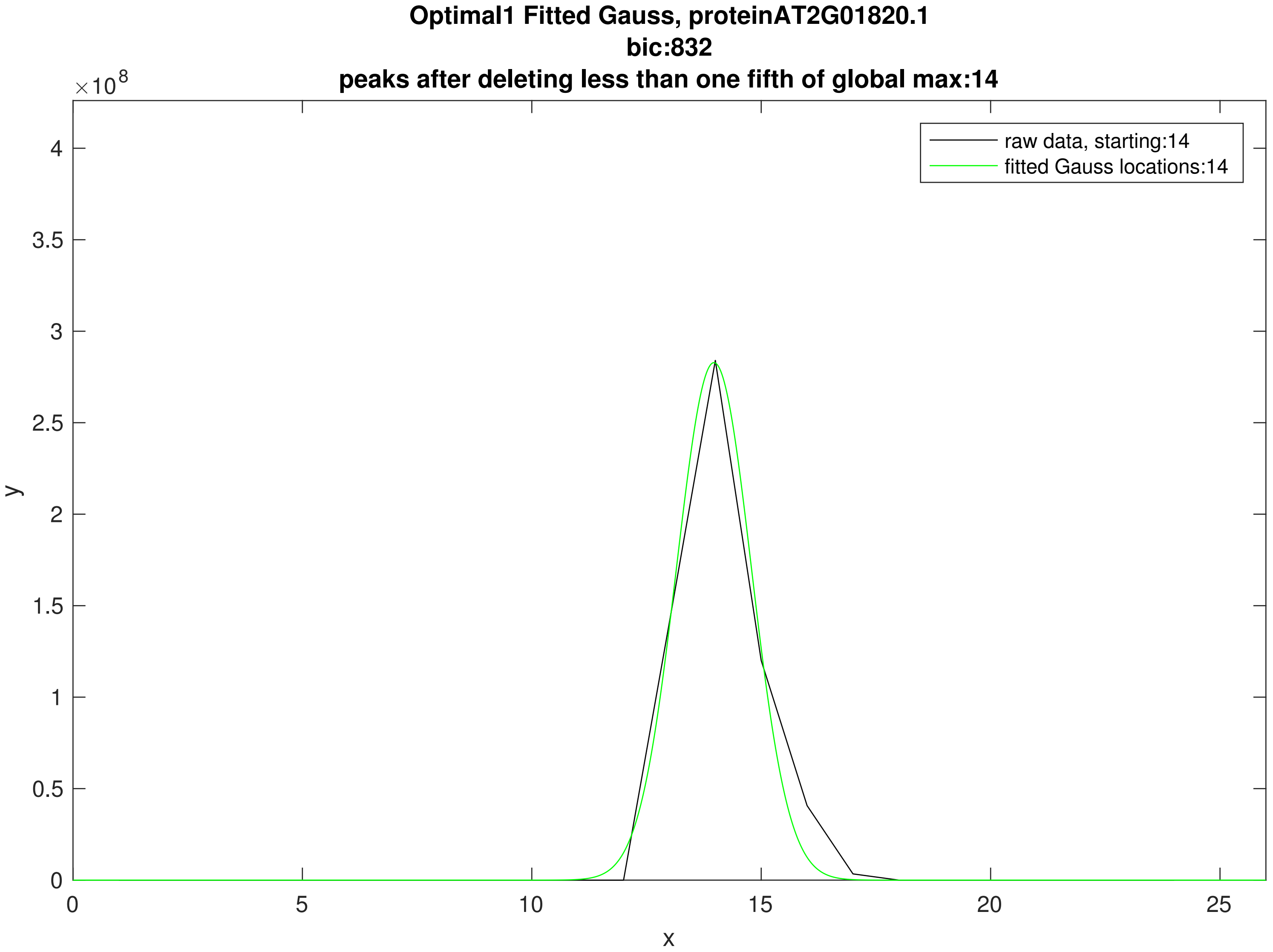
Optimal1 Fitted Gauss, proteinAT2G01820.1
bic:832
peaks after deleting less than one fifth of global max:14
×10^8
raw data, starting:14
fitted Gauss locations:14
y
x
0
0.5
1
1.5
2
2.5
3
3.5
4
0
5
10
15
20
25

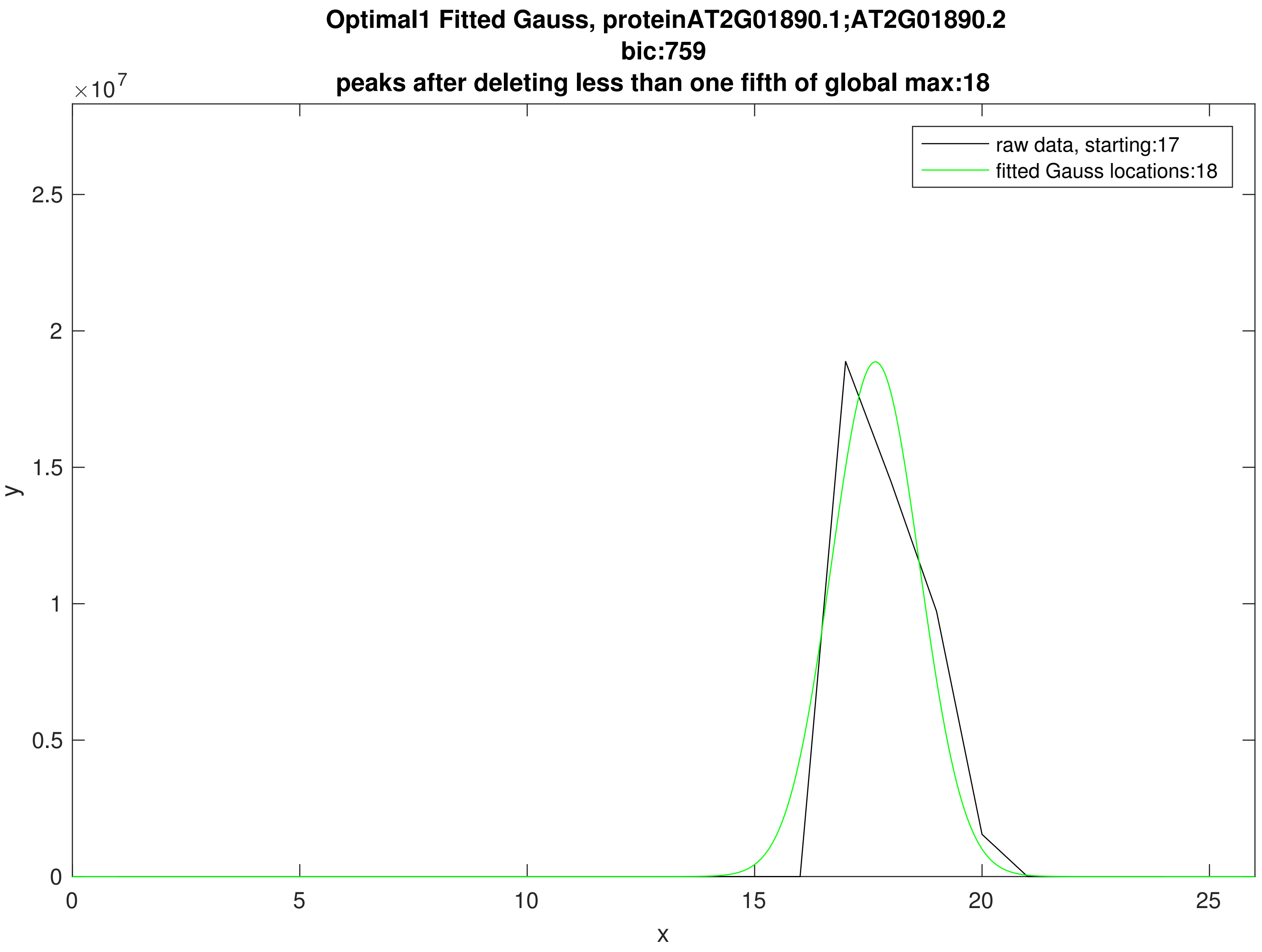
Optimal1 Fitted Gauss, proteinAT2G01890.1;AT2G01890.2
bic:759
peaks after deleting less than one fifth of global max:18
×10^7
raw data, starting:17
fitted Gauss locations:18
2.5
2
1.5
1
0.5
0
y
0
5
10
15
20
25
x

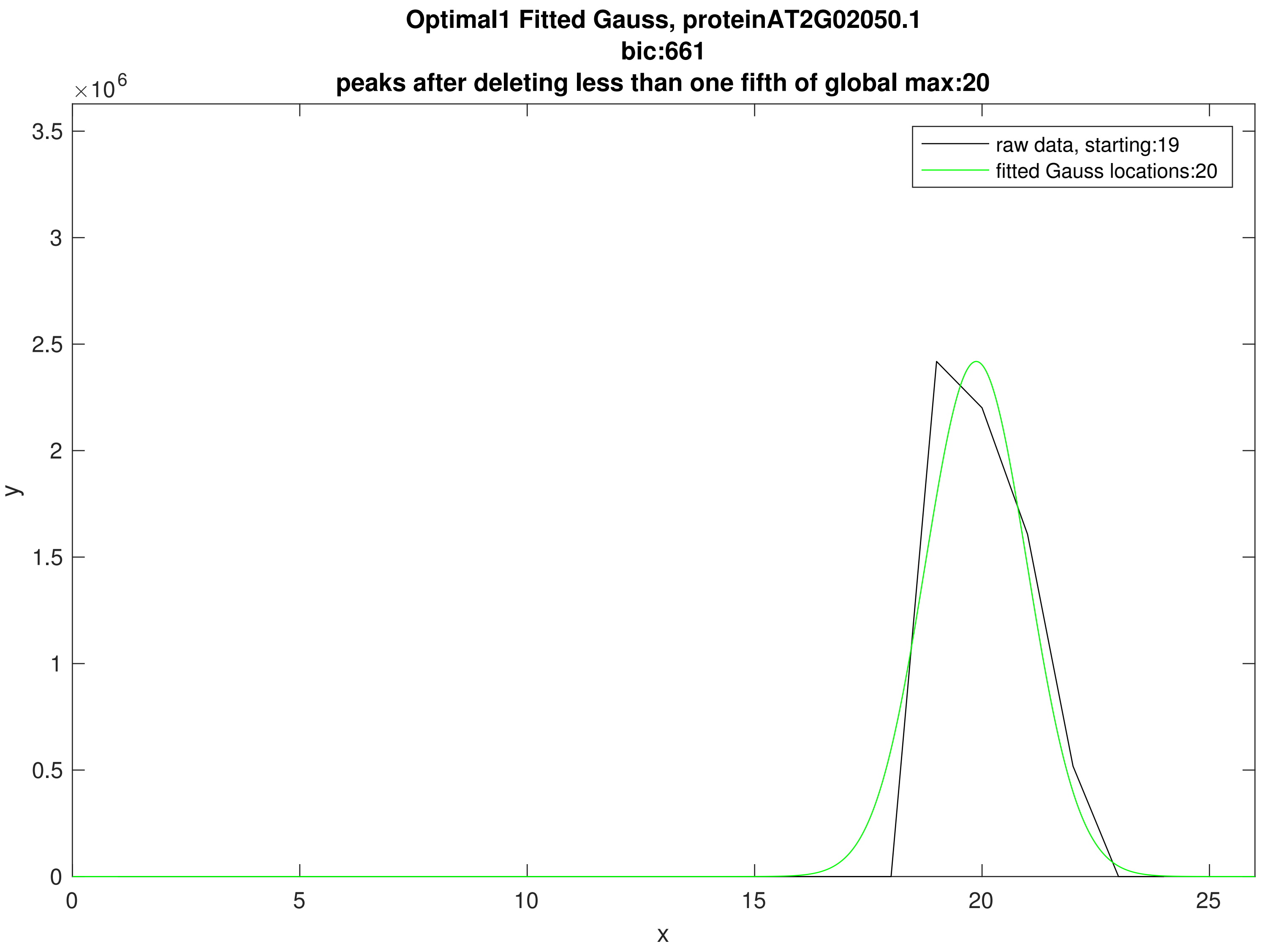
Optimal1 Fitted Gauss, proteinAT2G02050.1
bic:661
peaks after deleting less than one fifth of global max:20
×10^6
3.5
3
2.5
2
1.5
1
0.5
0
y
0
5
10
15
20
25
x
raw data, starting:19
fitted Gauss locations:20

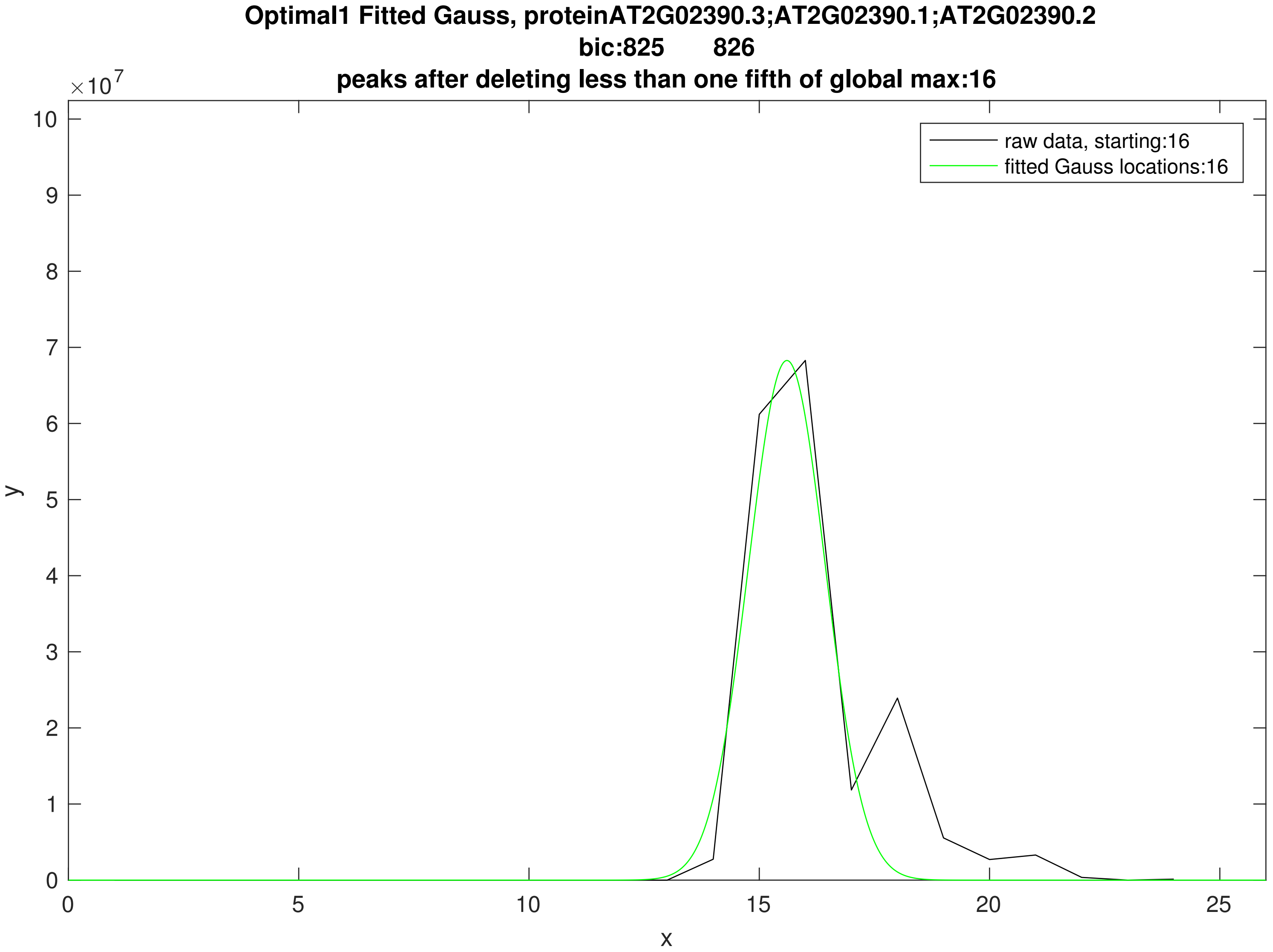

Optimal1 Fitted Gauss, proteinAT2G02390.3;AT2G02390.1;AT2G02390.2
bic:825 826
peaks after deleting less than one fifth of global max:16
×10^7
raw data, starting:16
fitted Gauss locations:16
y
x
0
1
2
3
4
5
6
7
8
9
10
0
5
10
15
20
25

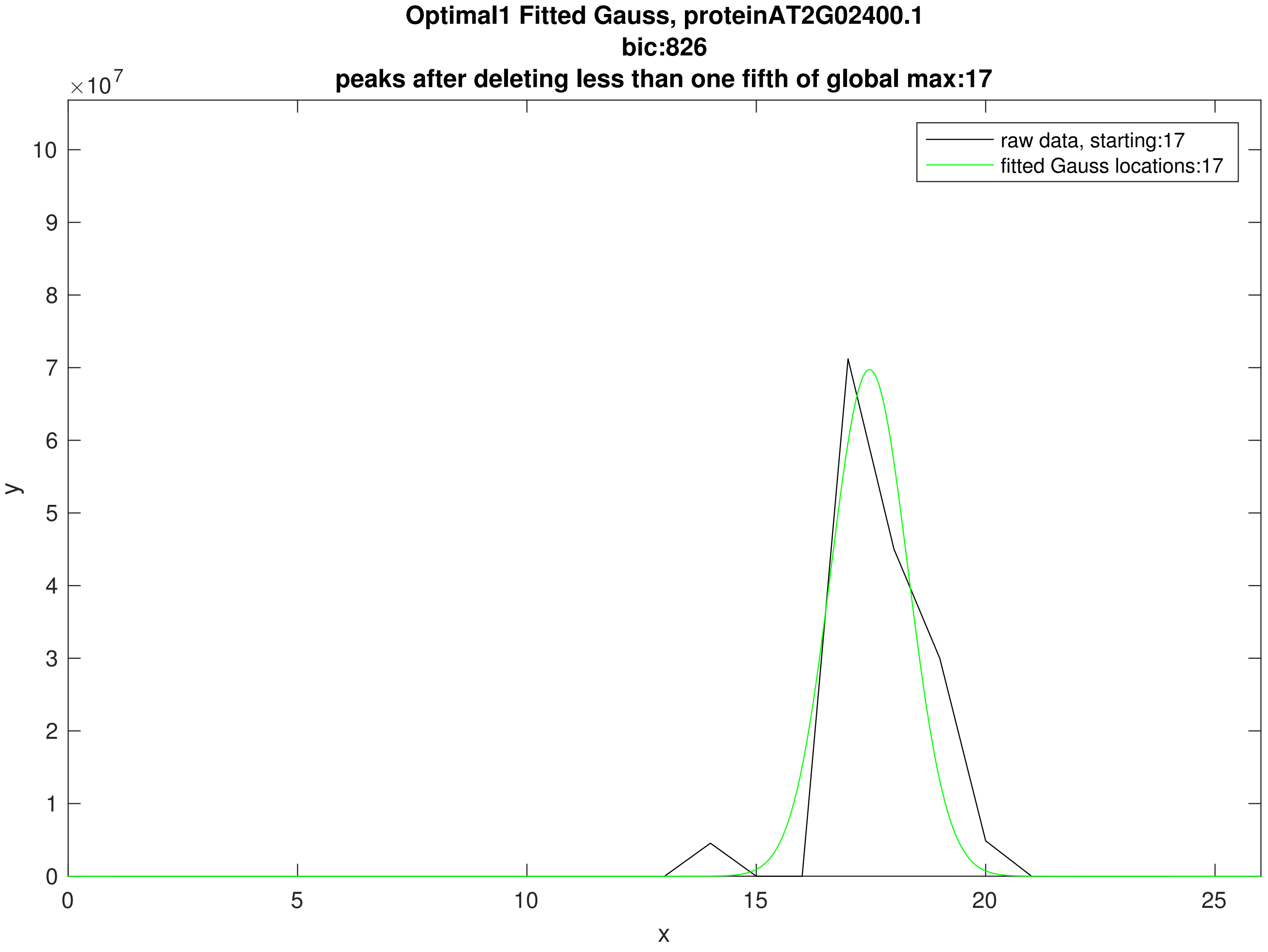

Optimal1 Fitted Gauss, proteinAT2G02400.1
bic:826
peaks after deleting less than one fifth of global max:17
×10^7
raw data, starting:17
fitted Gauss locations:17
y
x
0
1
2
3
4
5
6
7
8
9
10
0
5
10
15
20
25

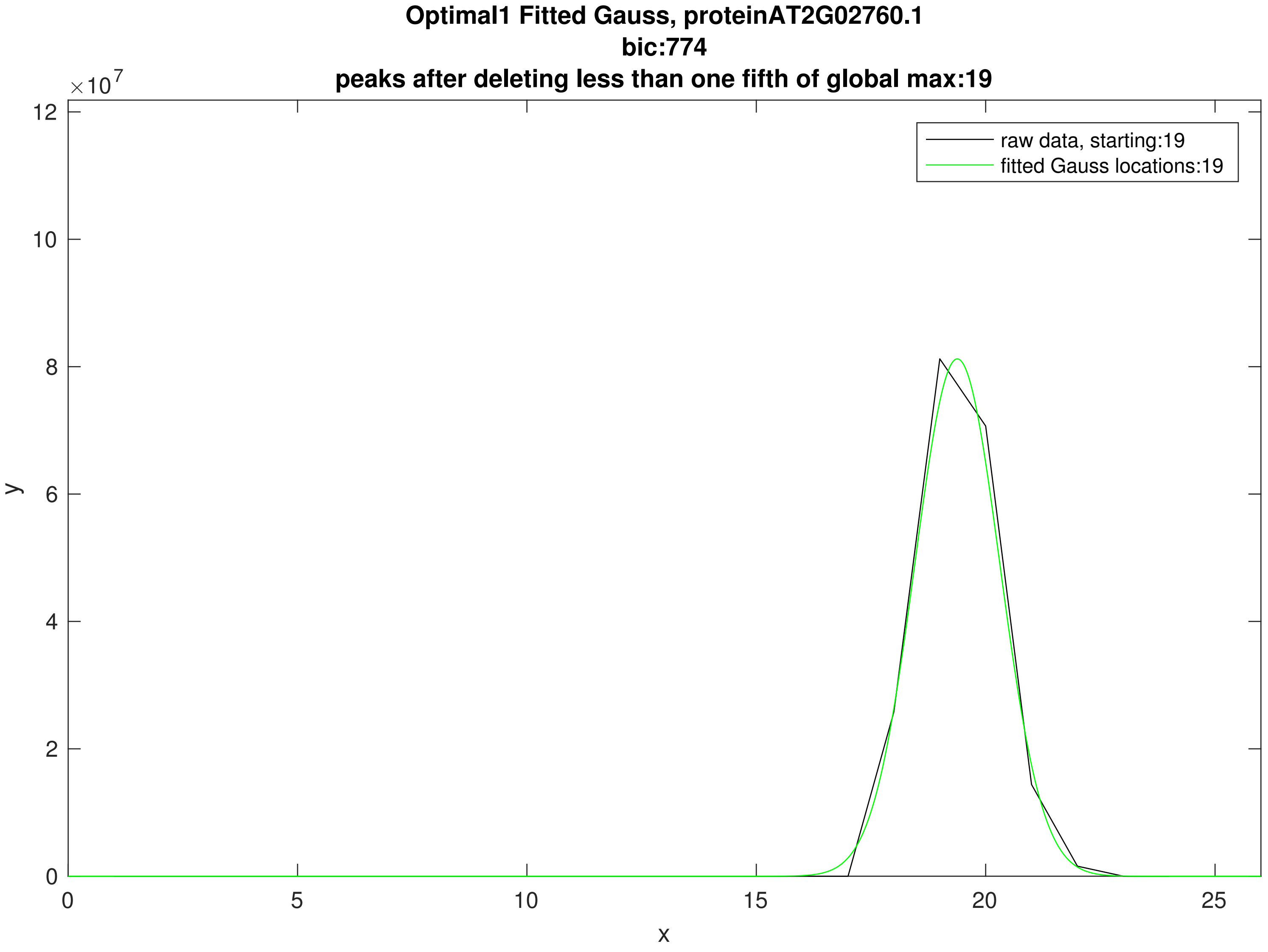
Optimal1 Fitted Gauss, proteinAT2G02760.1
bic:774
peaks after deleting less than one fifth of global max:19
×10^7
12
10
8
6
4
2
0
y
0
5
10
15
20
25
x
raw data, starting:19
fitted Gauss locations:19

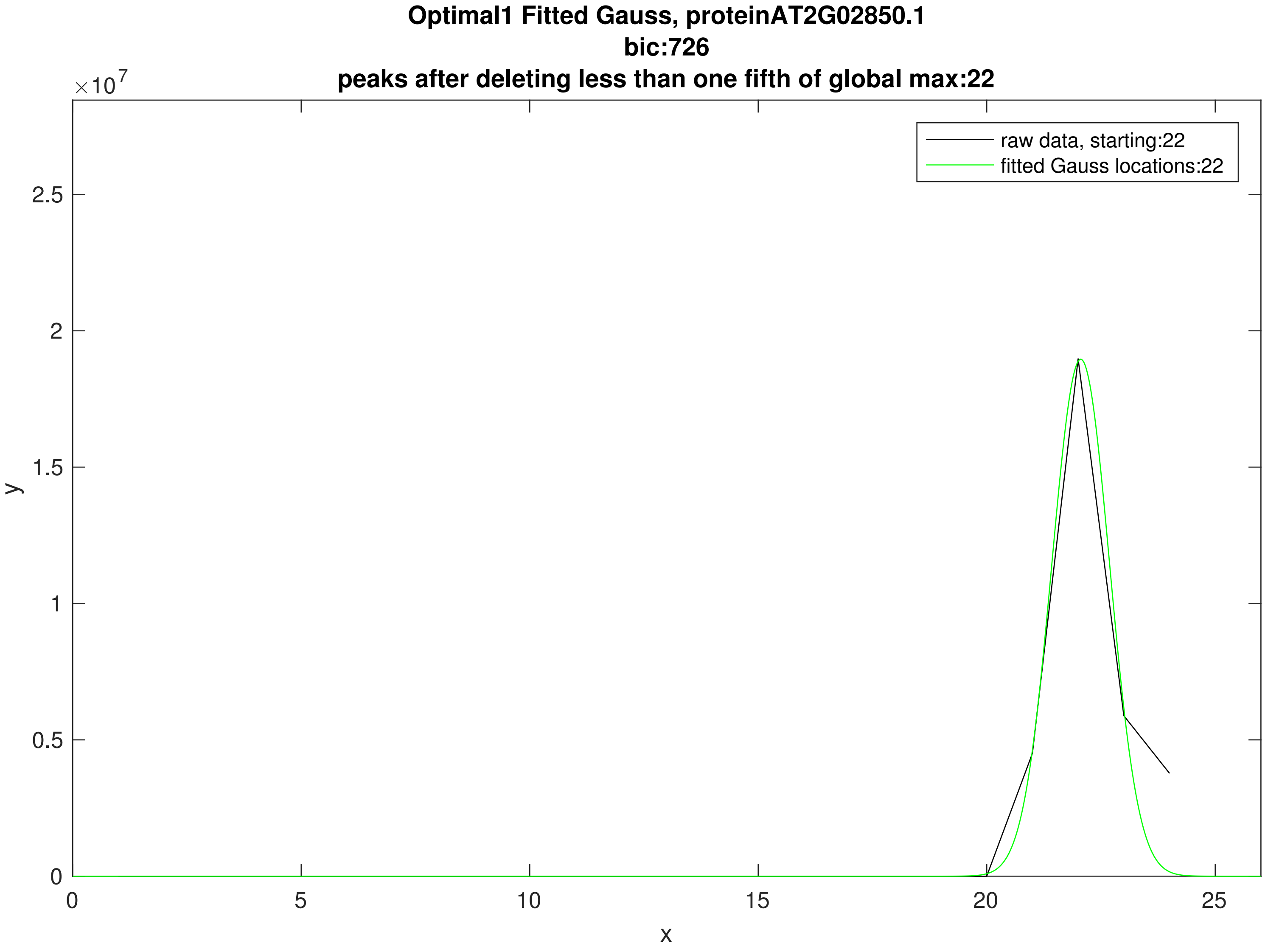

Optimal1 Fitted Gauss, proteinAT2G02850.1
bic:726
peaks after deleting less than one fifth of global max:22
×10^7
raw data, starting:22
fitted Gauss locations:22
2.5
2
1.5
1
0.5
0
y
0
5
10
15
20
25
x

**Optimal2 Fitted Gauss, proteinAT2G02990.1**
**bic:746 702**
**peaks after deleting less than one fifth of global max:18**

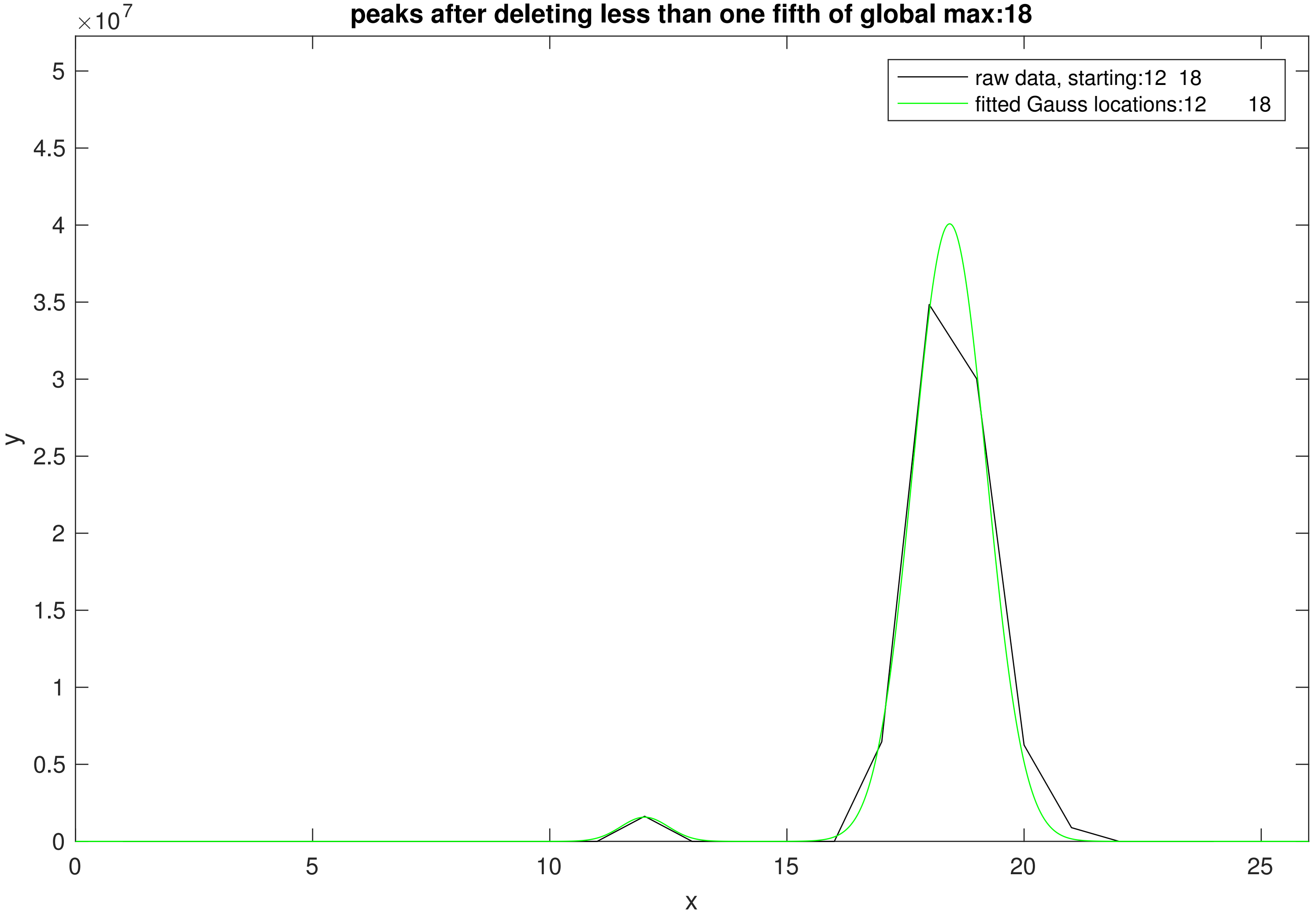

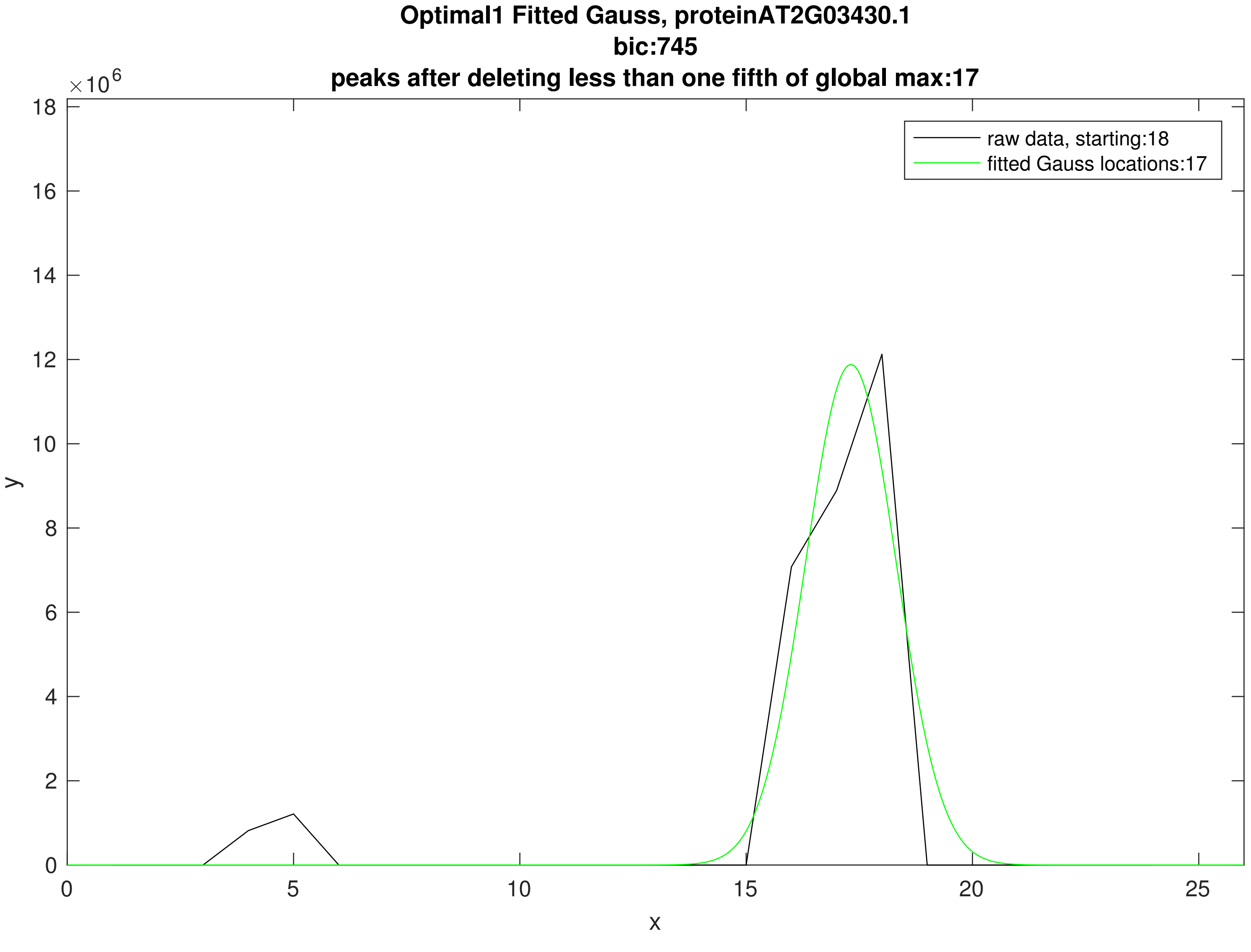
Optimal1 Fitted Gauss, proteinAT2G03430.1
bic:745
peaks after deleting less than one fifth of global max:17
×10^6
18
16
14
12
10
8
6
4
2
0
y
0
5
10
15
20
25
x
raw data, starting:18
fitted Gauss locations:17

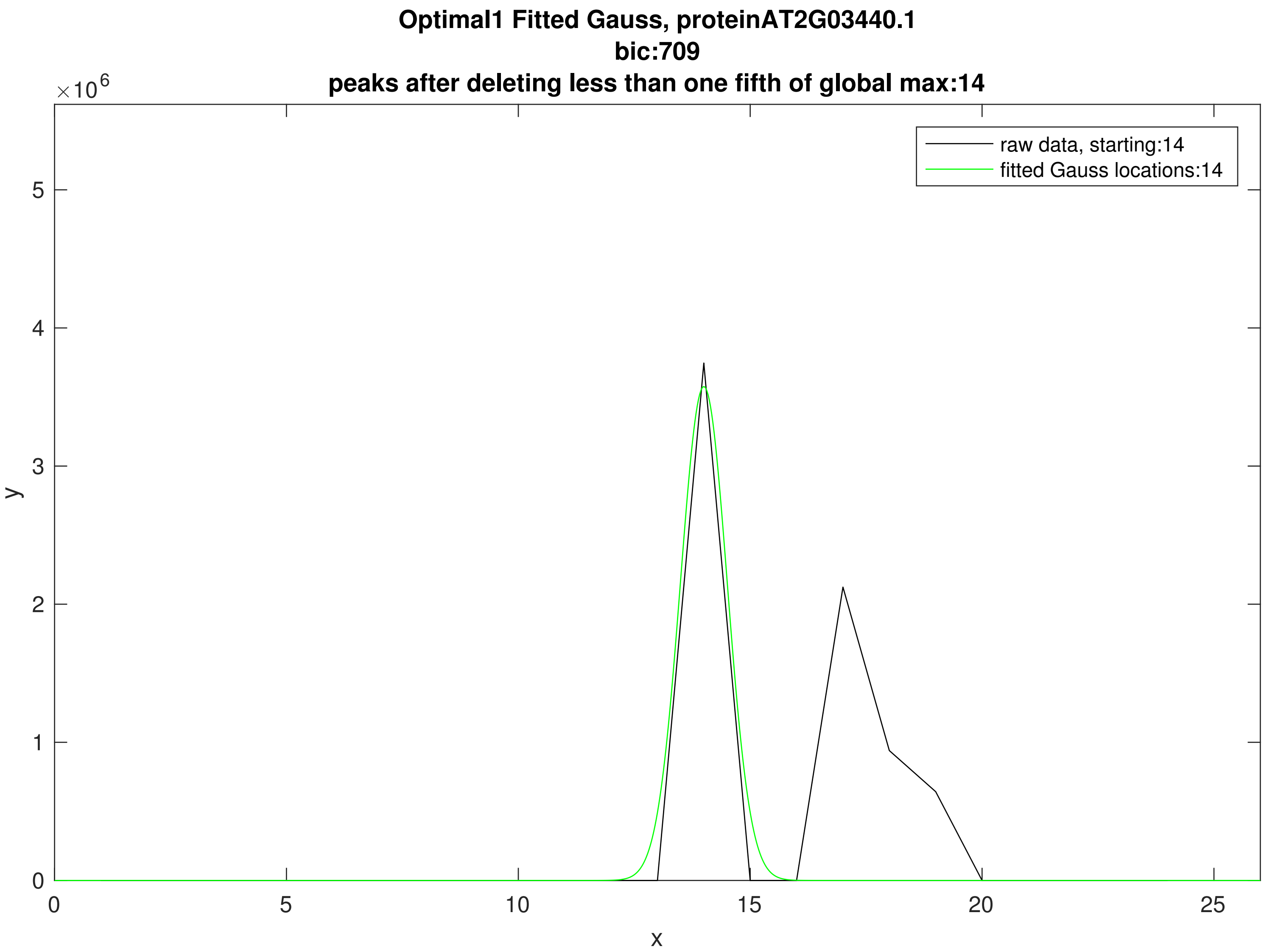

Optimal1 Fitted Gauss, proteinAT2G03440.1
bic:709
peaks after deleting less than one fifth of global max:14
×10^6
raw data, starting:14
fitted Gauss locations:14
y
x
0
1
2
3
4
5
0
5
10
15
20
25

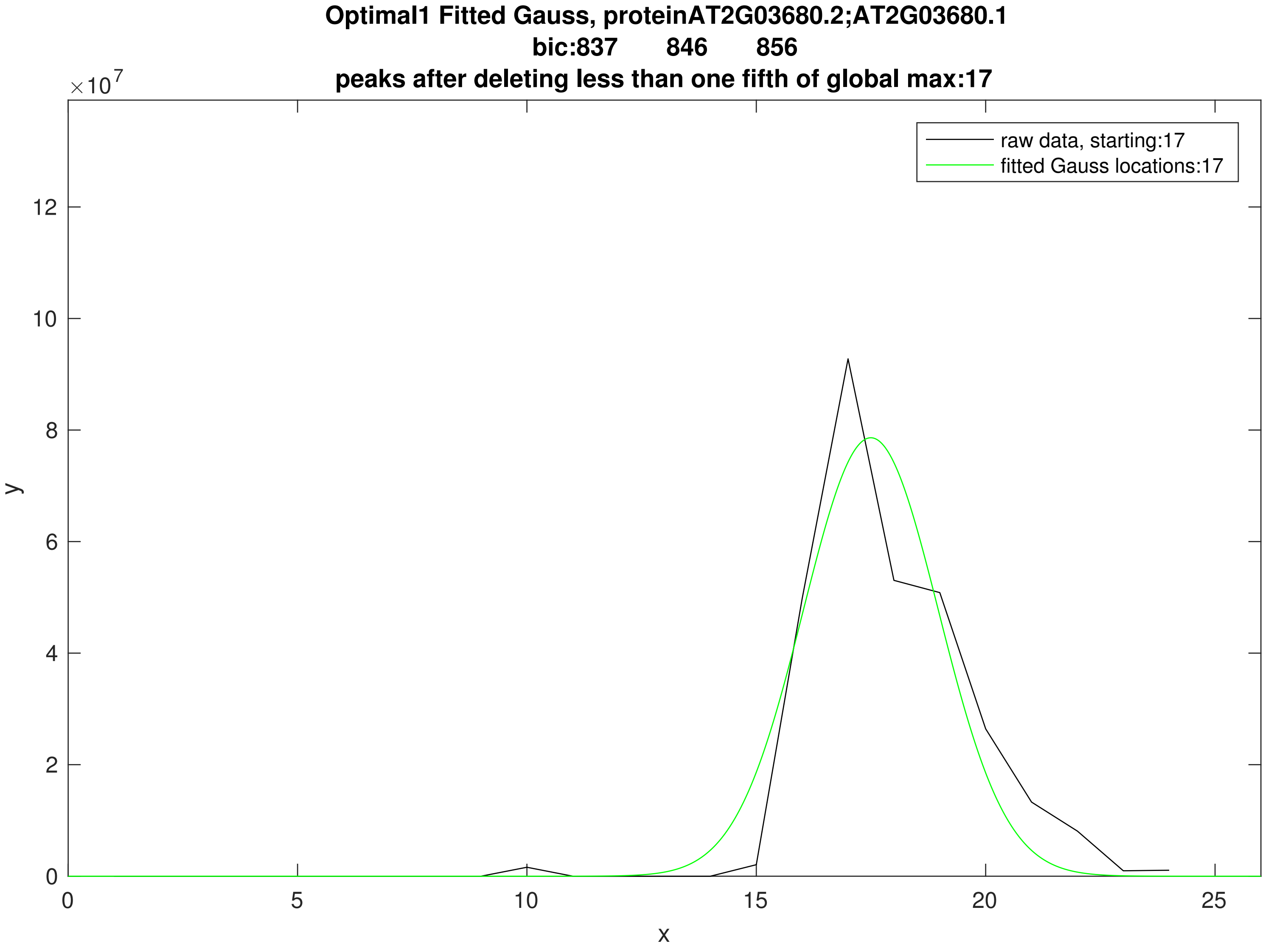
Optimal1 Fitted Gauss, proteinAT2G03680.2;AT2G03680.1
bic:837 846 856
peaks after deleting less than one fifth of global max:17
×10^7
raw data, starting:17
fitted Gauss locations:17
y
x
0
2
4
6
8
10
12
5
15
20
25

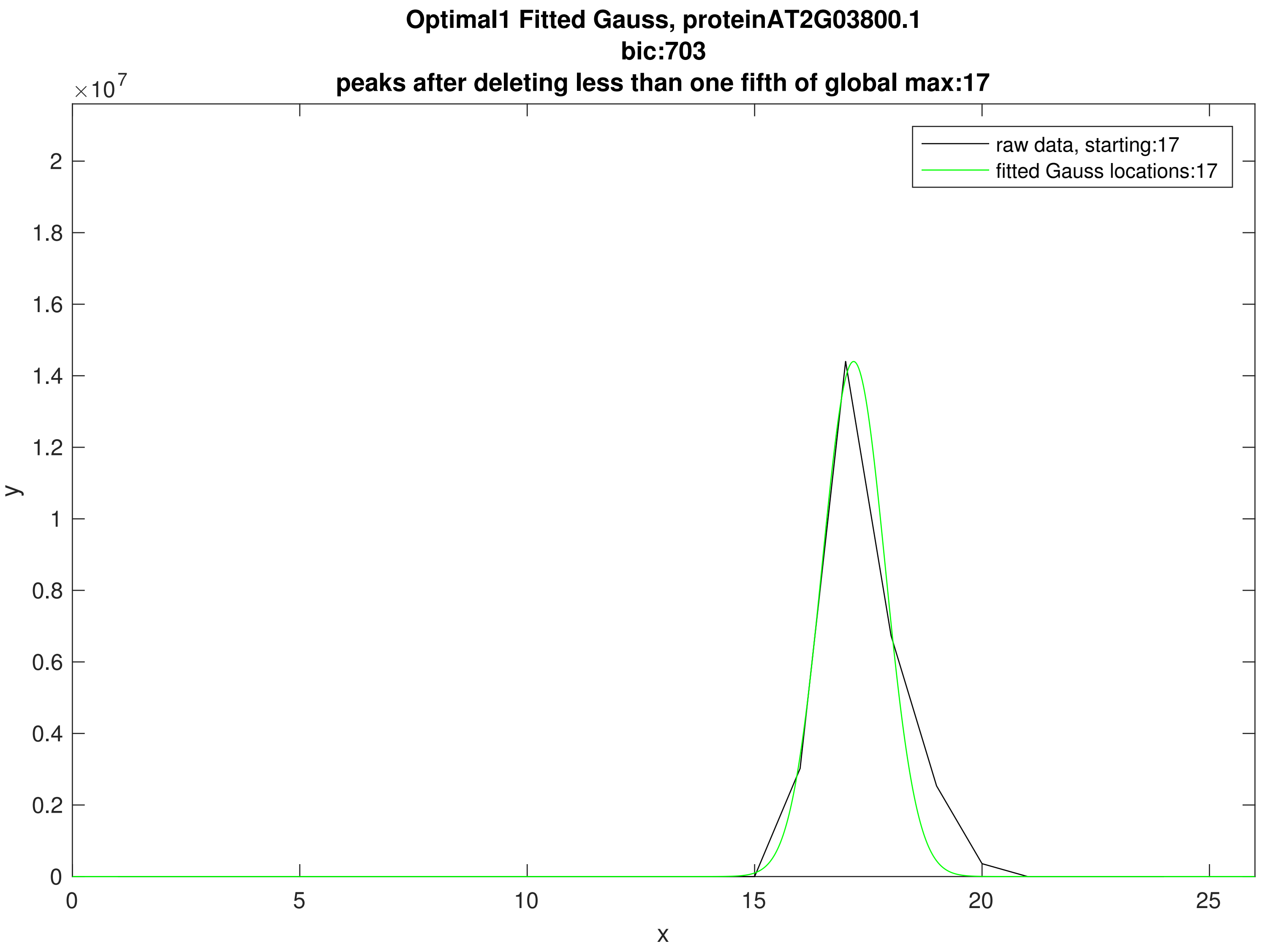

Optimal1 Fitted Gauss, proteinAT2G03800.1
bic:703
peaks after deleting less than one fifth of global max:17
×10^7
raw data, starting:17
fitted Gauss locations:17
y
x
0
0.2
0.4
0.6
0.8
1
1.2
1.4
1.6
1.8
2
0
5
10
15
20
25

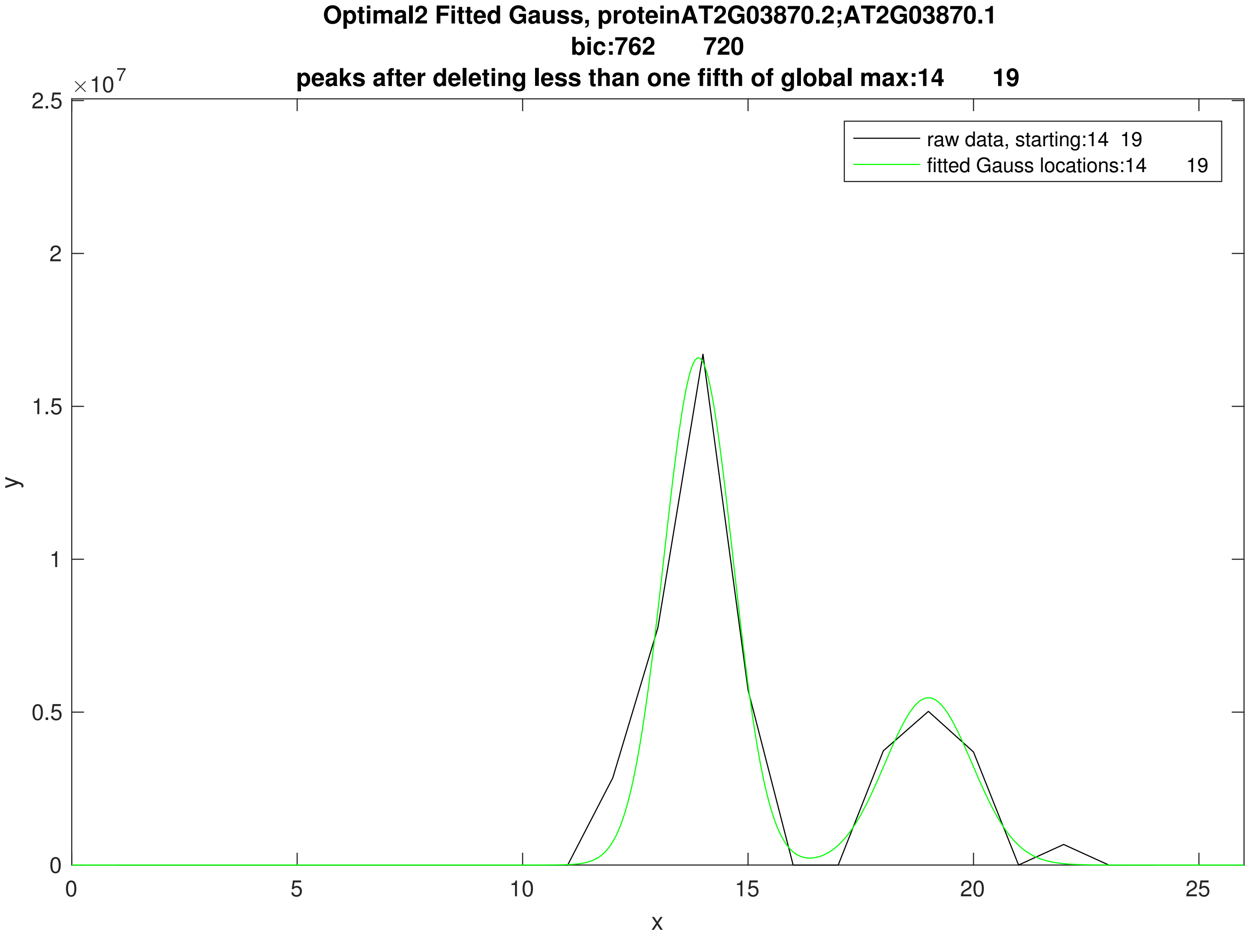

Optimal2 Fitted Gauss, proteinAT2G03870.2;AT2G03870.1
bic:762 720
peaks after deleting less than one fifth of global max:14 19
$\times 10^7$
raw data, starting:14 19
fitted Gauss locations:14 19
y
x
0
0.5
1
1.5
2
2.5
0
5
10
15
20
25

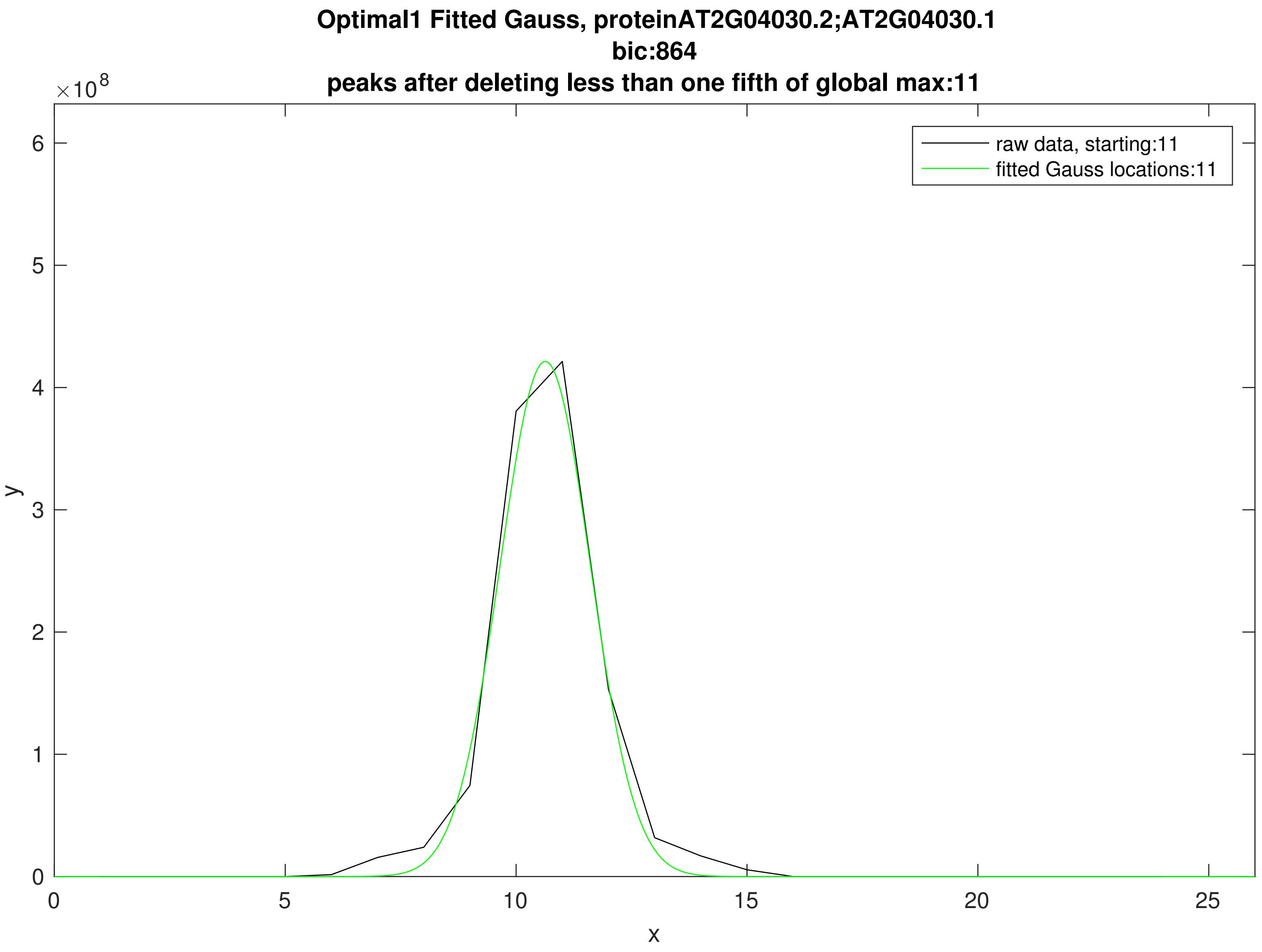
Optimal1 Fitted Gauss, proteinAT2G04030.2;AT2G04030.1
bic:864
peaks after deleting less than one fifth of global max:11
×10^8
raw data, starting:11
fitted Gauss locations:11
y
x
0
1
2
3
4
5
6
0
5
10
15
20
25

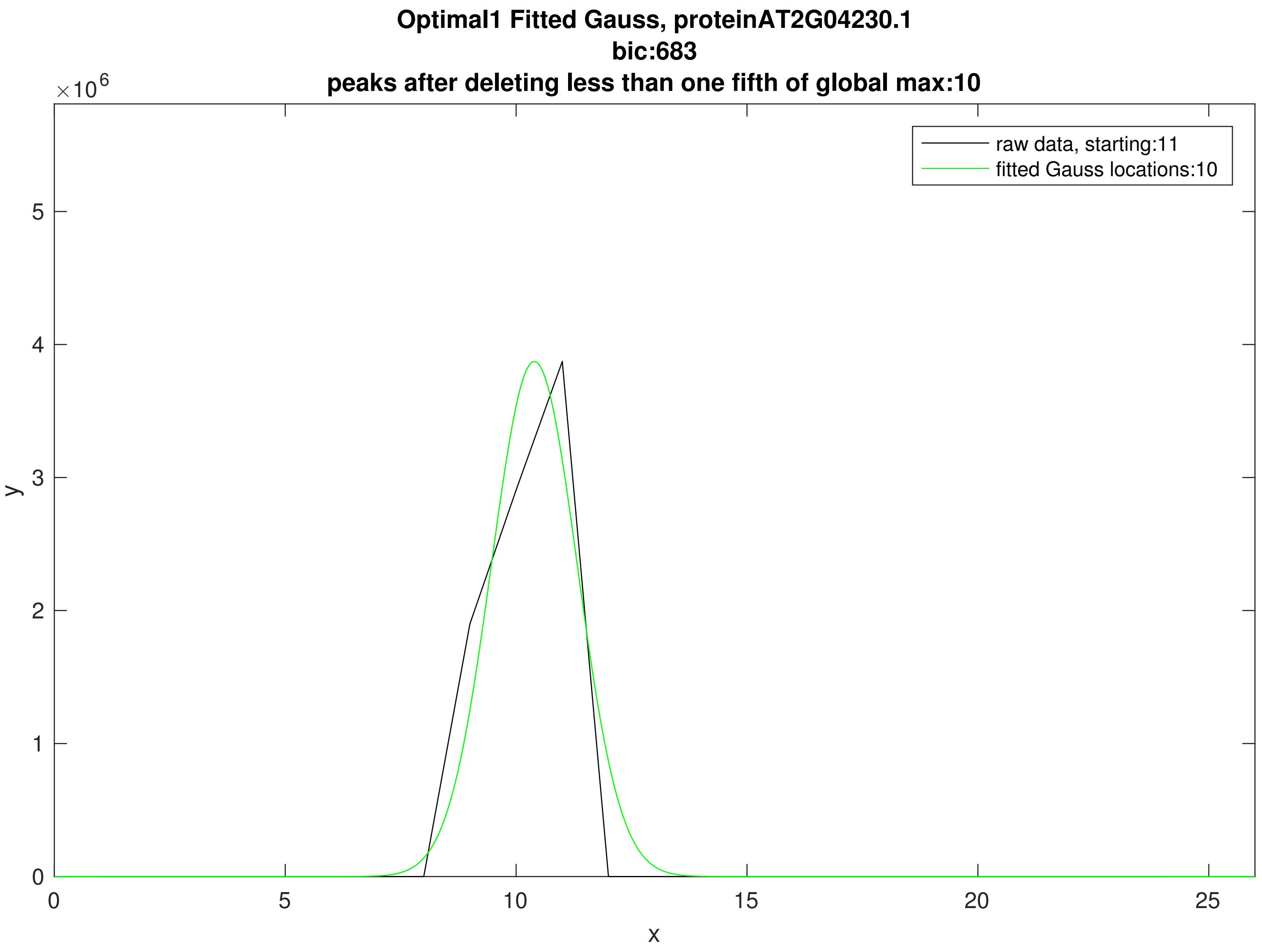

Optimal1 Fitted Gauss, proteinAT2G04230.1
bic:683
peaks after deleting less than one fifth of global max:10
×10^6
raw data, starting:11
fitted Gauss locations:10
y
x
0
1
2
3
4
5
0
5
10
15
20
25

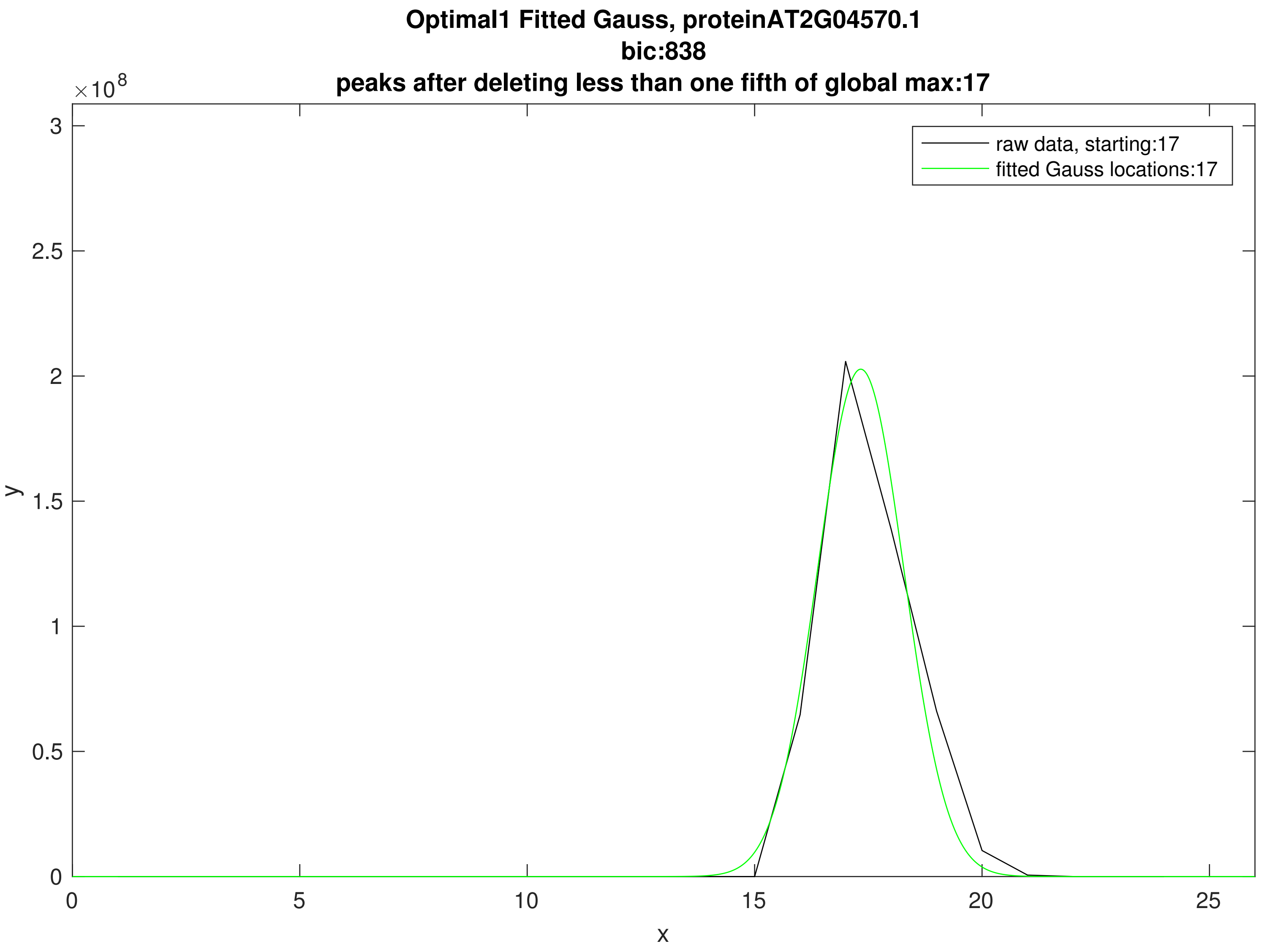

Optimal1 Fitted Gauss, proteinAT2G04570.1
bic:838
peaks after deleting less than one fifth of global max:17
×10^8
3
2.5
2
1.5
1
0.5
0
y
0
5
10
15
20
25
x
raw data, starting:17
fitted Gauss locations:17

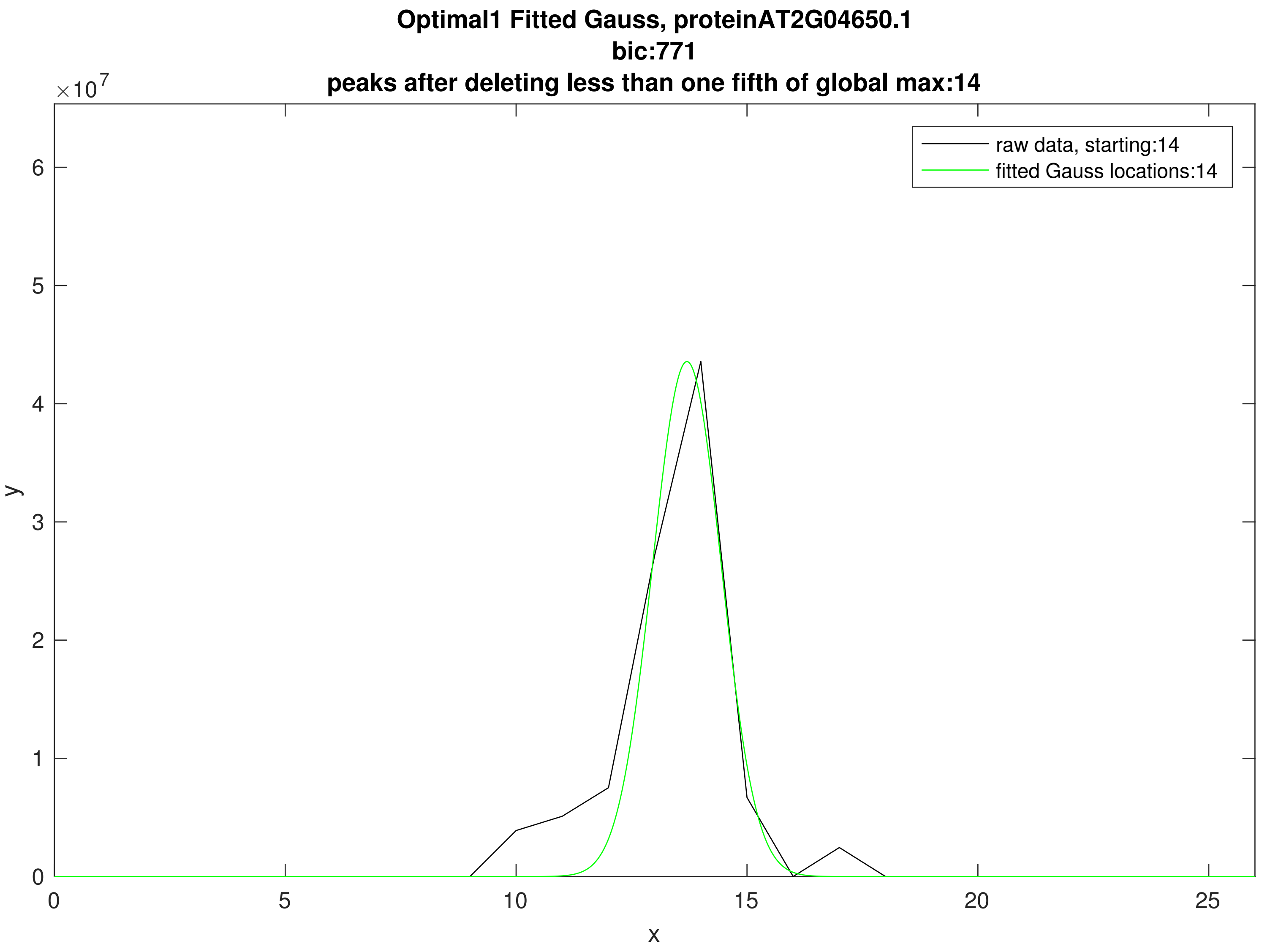
Optimal1 Fitted Gauss, proteinAT2G04650.1
bic:771
peaks after deleting less than one fifth of global max:14
×10^7
raw data, starting:14
fitted Gauss locations:14
y
x
0
1
2
3
4
5
6
0
5
10
15
20
25

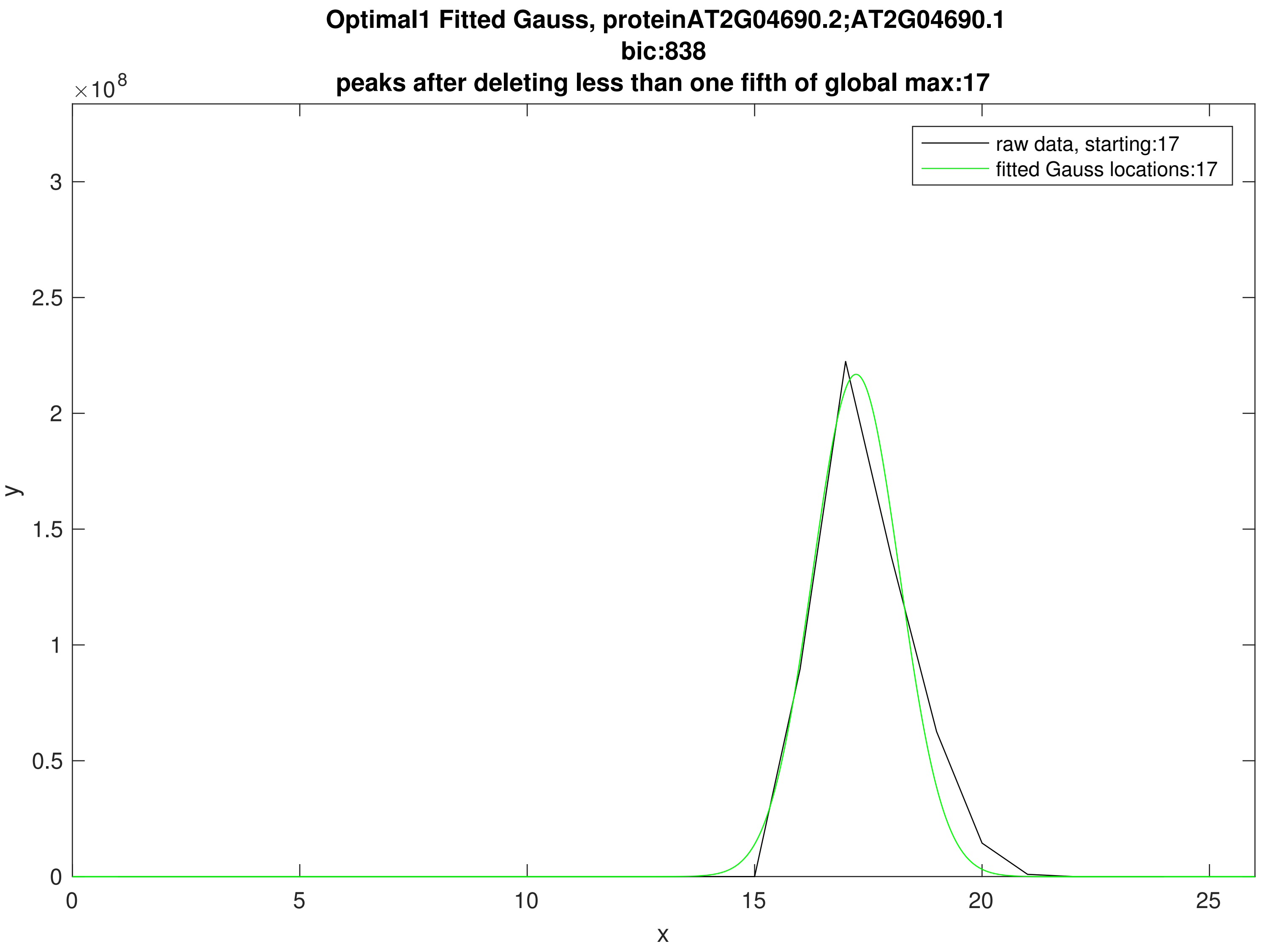

Optimal1 Fitted Gauss, proteinAT2G04690.2;AT2G04690.1
bic:838
peaks after deleting less than one fifth of global max:17
×10^8
raw data, starting:17
fitted Gauss locations:17
y
x
0
0.5
1
1.5
2
2.5
3
0
5
10
15
20
25

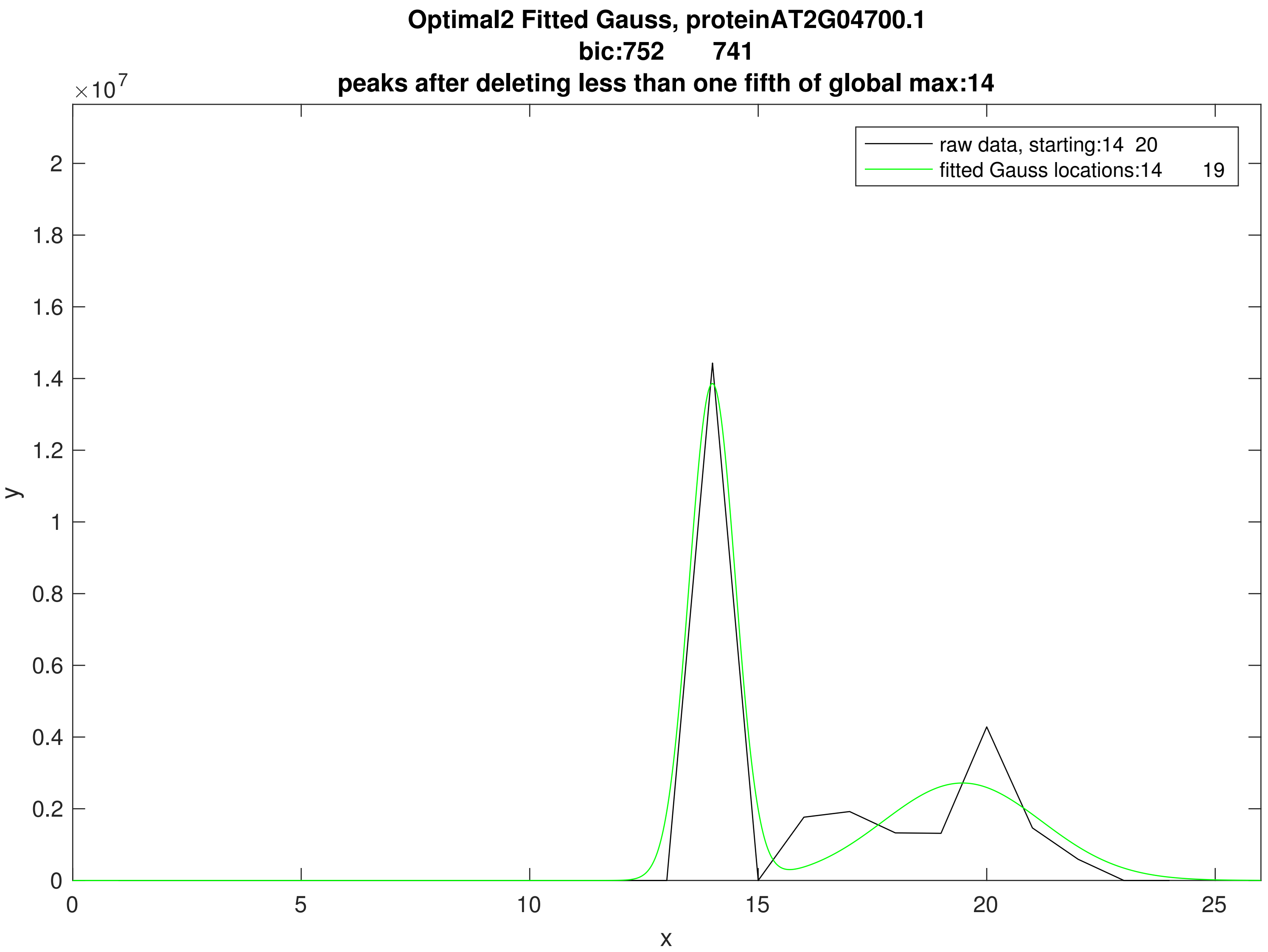

Optimal2 Fitted Gauss, proteinAT2G04700.1
bic:752 741
peaks after deleting less than one fifth of global max:14
×10^7
raw data, starting:14 20
fitted Gauss locations:14 19
y
x
0
0.2
0.4
0.6
0.8
1
1.2
1.4
1.6
1.8
2
0
5
10
15
20
25

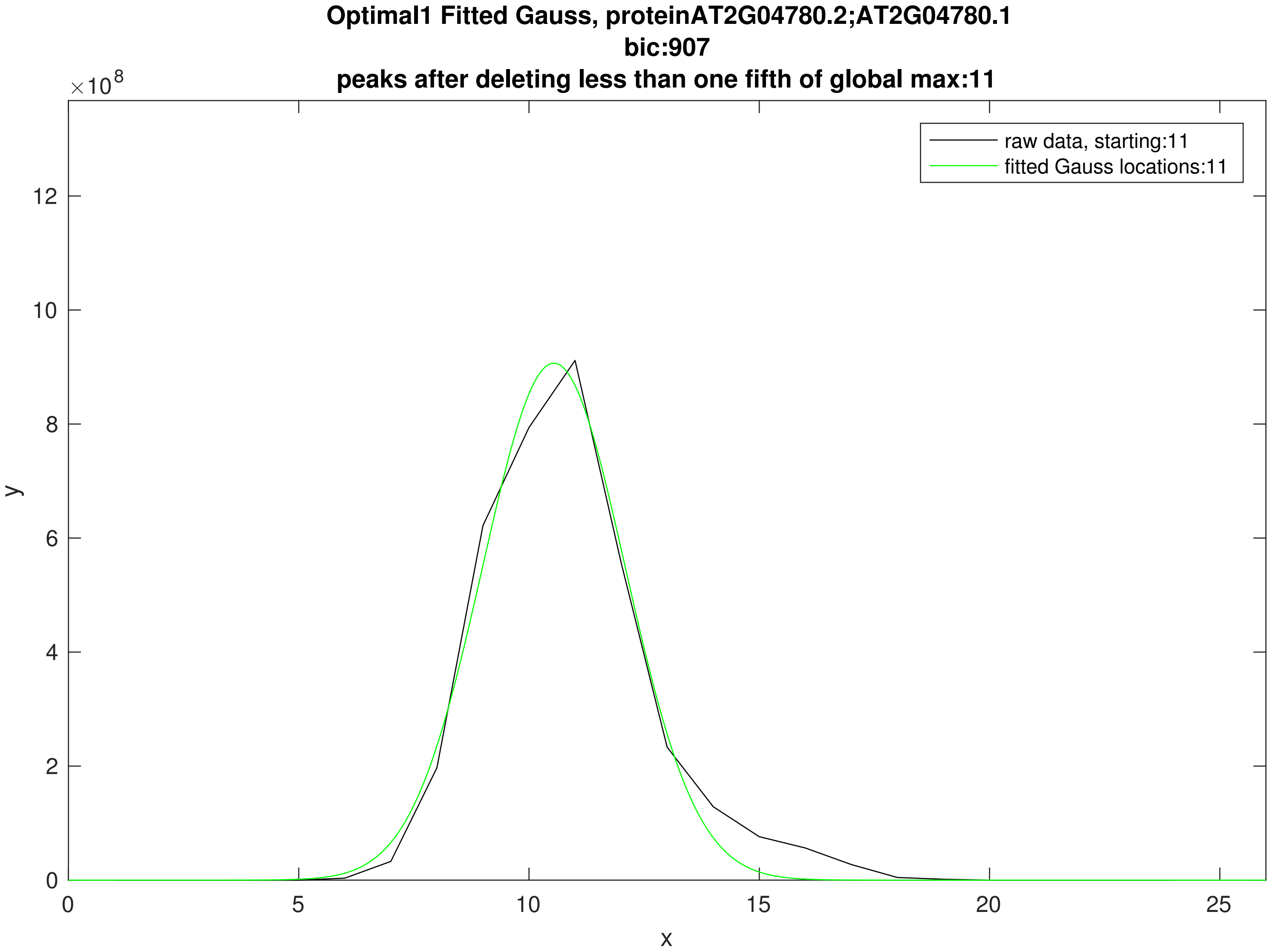

Optimal1 Fitted Gauss, proteinAT2G04780.2;AT2G04780.1
bic:907
peaks after deleting less than one fifth of global max:11
×10^8
raw data, starting:11
fitted Gauss locations:11
y
x
0
2
4
6
8
10
12
0
5
10
15
20
25

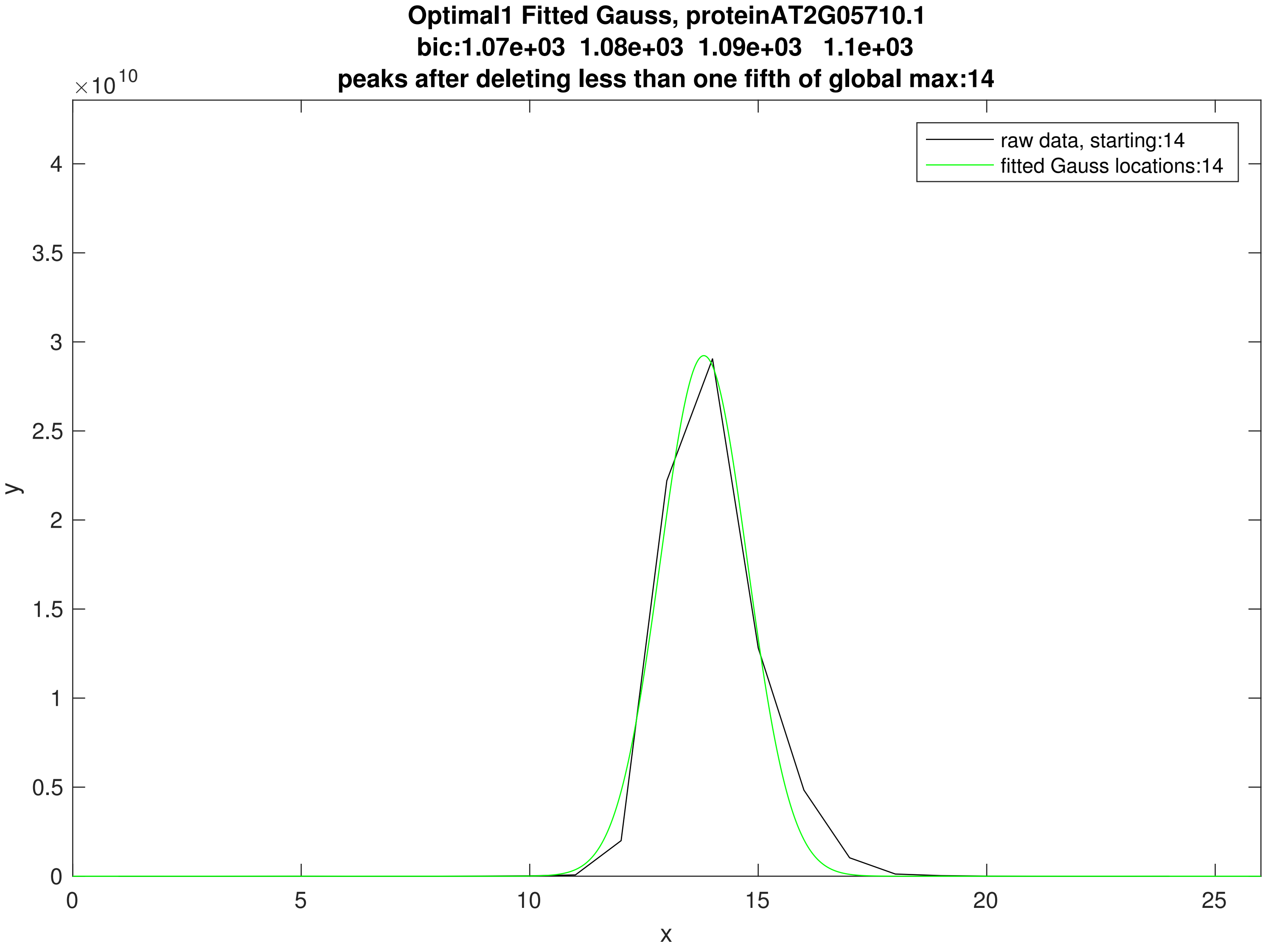
Optimal1 Fitted Gauss, proteinAT2G05710.1
bic:1.07e+03 1.08e+03 1.09e+03 1.1e+03
peaks after deleting less than one fifth of global max:14
×10^10
raw data, starting:14
fitted Gauss locations:14
y
x
0
0.5
1
1.5
2
2.5
3
3.5
4
0
5
10
15
20
25

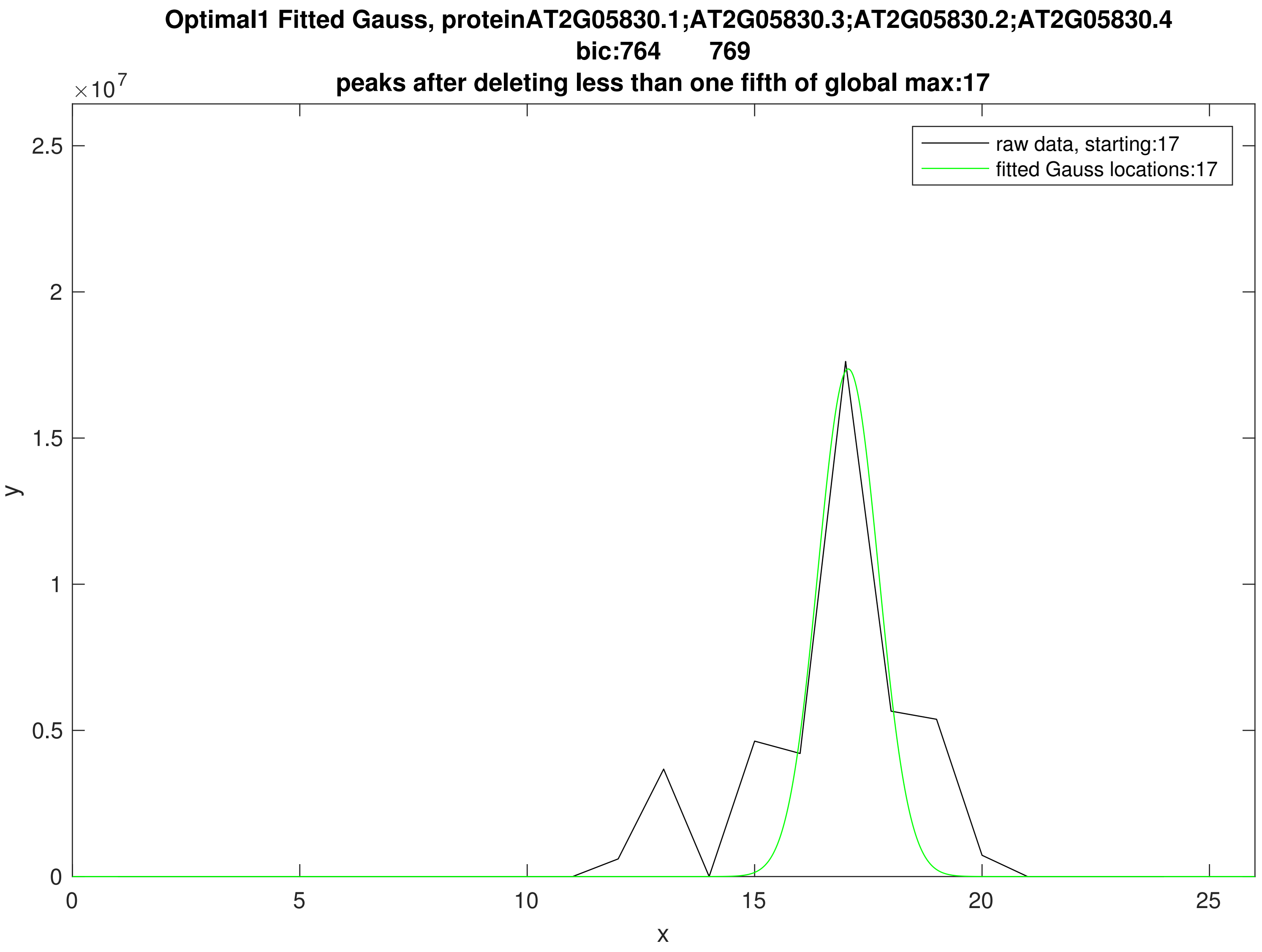

Optimal1 Fitted Gauss, proteinAT2G05830.1;AT2G05830.3;AT2G05830.2;AT2G05830.4
bic:764 769
peaks after deleting less than one fifth of global max:17
×10^7
2.5
2
1.5
1
0.5
0
y
0
5
10
15
20
25
x
raw data, starting:17
fitted Gauss locations:17

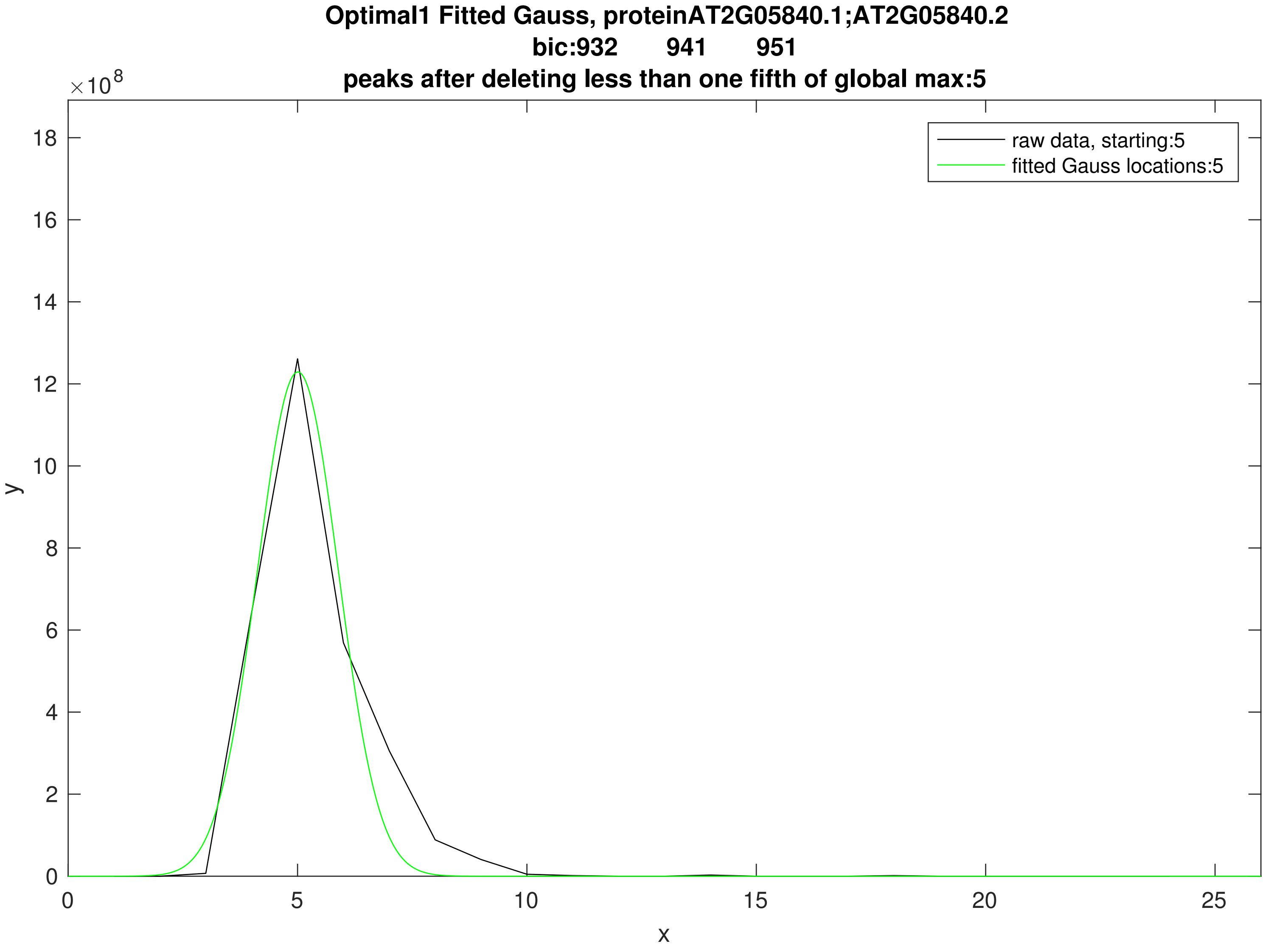

Optimal1 Fitted Gauss, proteinAT2G05840.1;AT2G05840.2
bic:932 941 951
peaks after deleting less than one fifth of global max:5
×10^8
18
16
14
12
10
8
6
4
2
0
y
0
5
10
15
20
25
x
raw data, starting:5
fitted Gauss locations:5

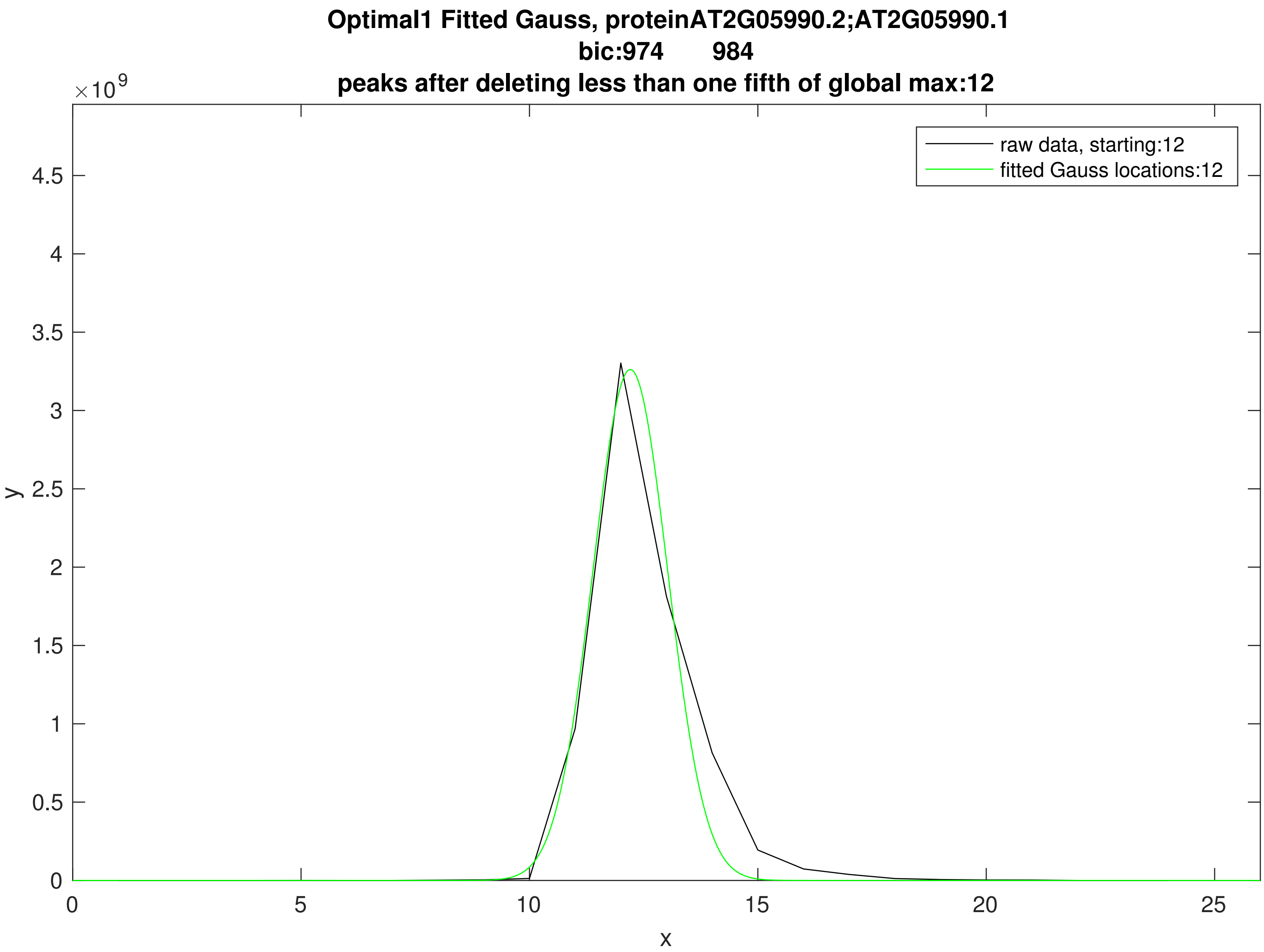
Optimal1 Fitted Gauss, proteinAT2G05990.2;AT2G05990.1
bic:974 984
peaks after deleting less than one fifth of global max:12
×10^9
4.5
4
3.5
3
2.5
2
1.5
1
0.5
0
y
0
5
10
15
20
25
x
raw data, starting:12
fitted Gauss locations:12

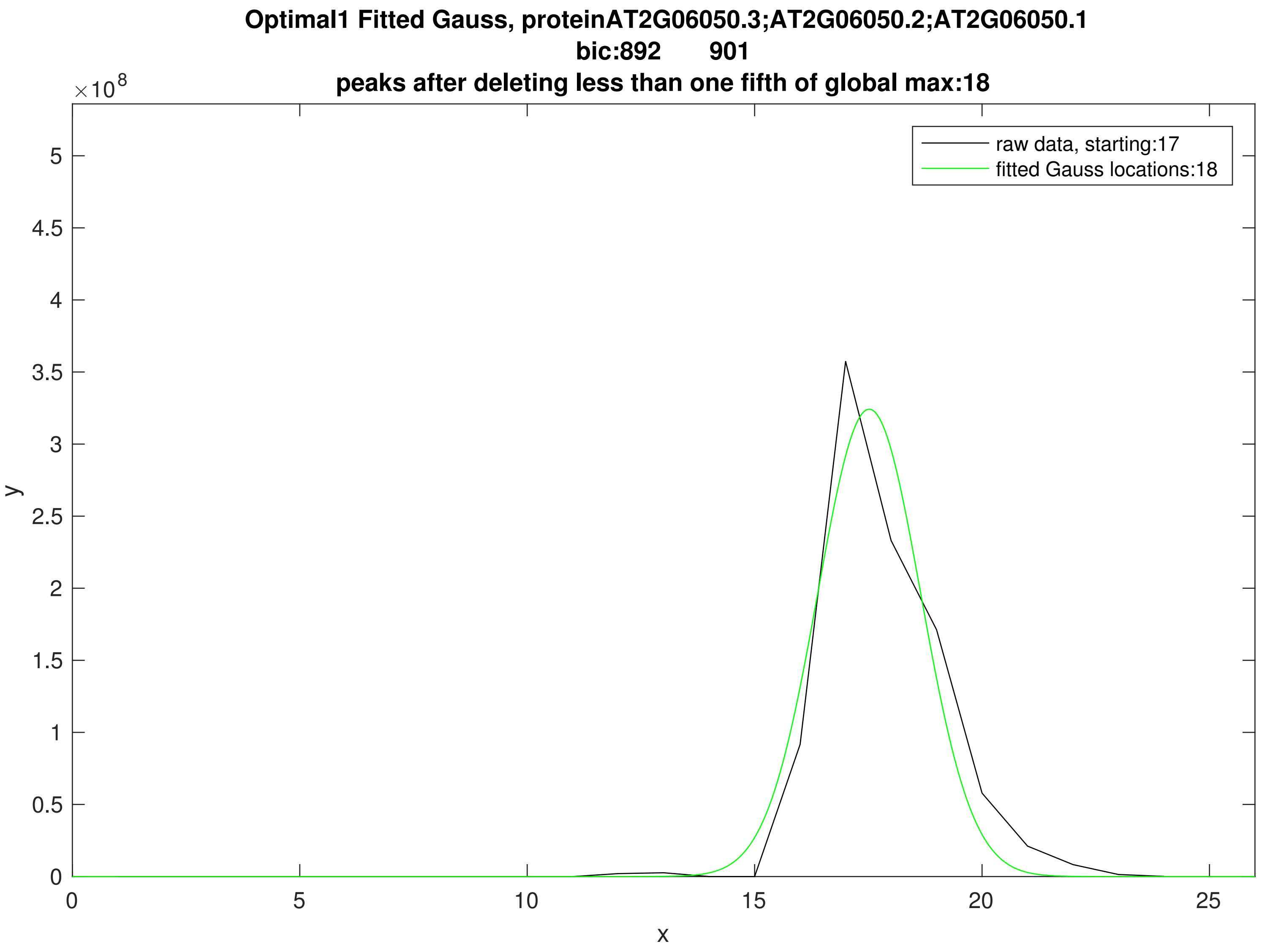

Optimal1 Fitted Gauss, proteinAT2G06050.3;AT2G06050.2;AT2G06050.1
bic:892 901
peaks after deleting less than one fifth of global max:18
×10^8
raw data, starting:17
fitted Gauss locations:18
y
x

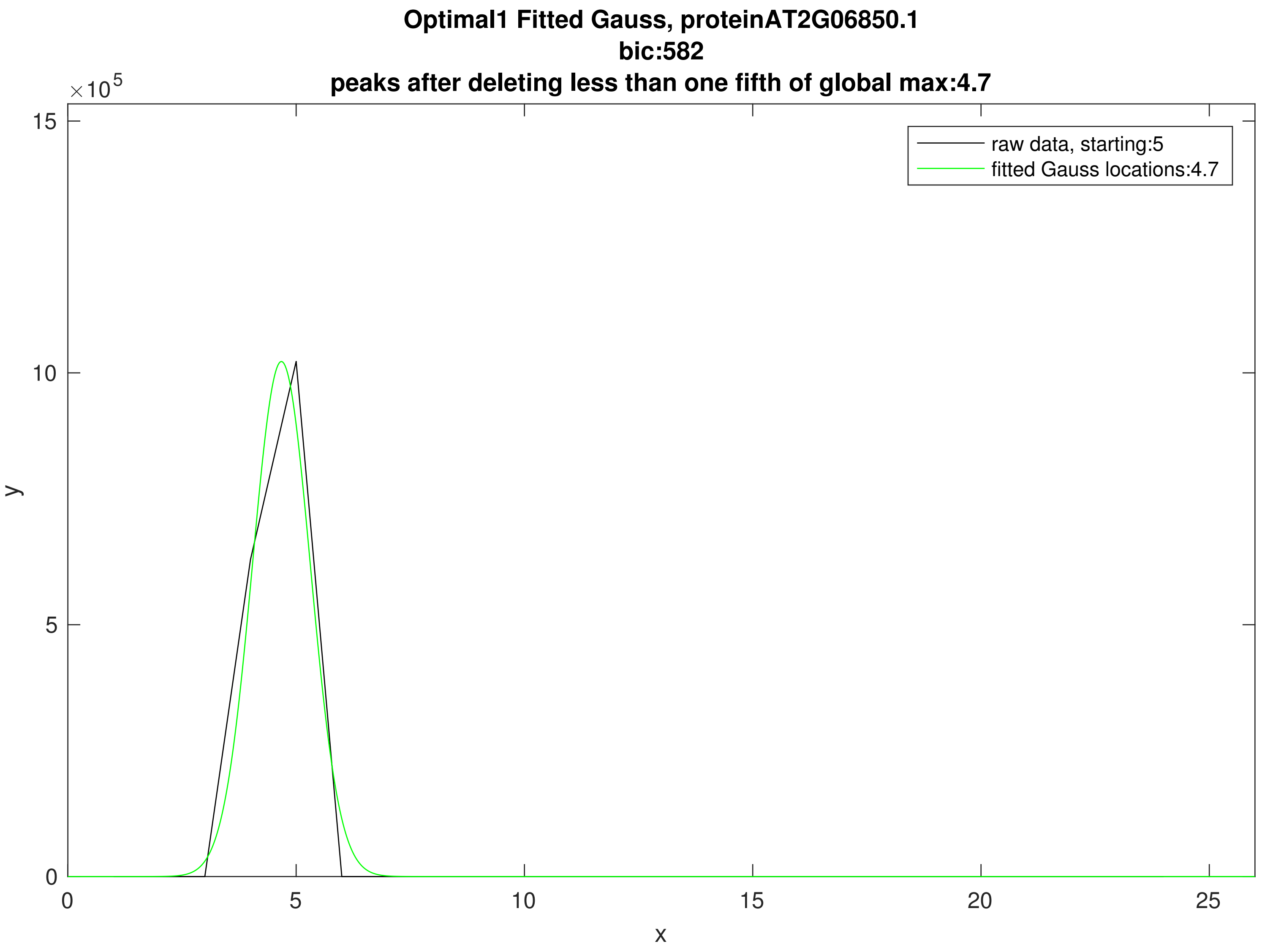

Optimal1 Fitted Gauss, proteinAT2G06850.1
bic:582
peaks after deleting less than one fifth of global max:4.7
×10^5
15
10
5
0
y
0
5
10
15
20
25
x
raw data, starting:5
fitted Gauss locations:4.7

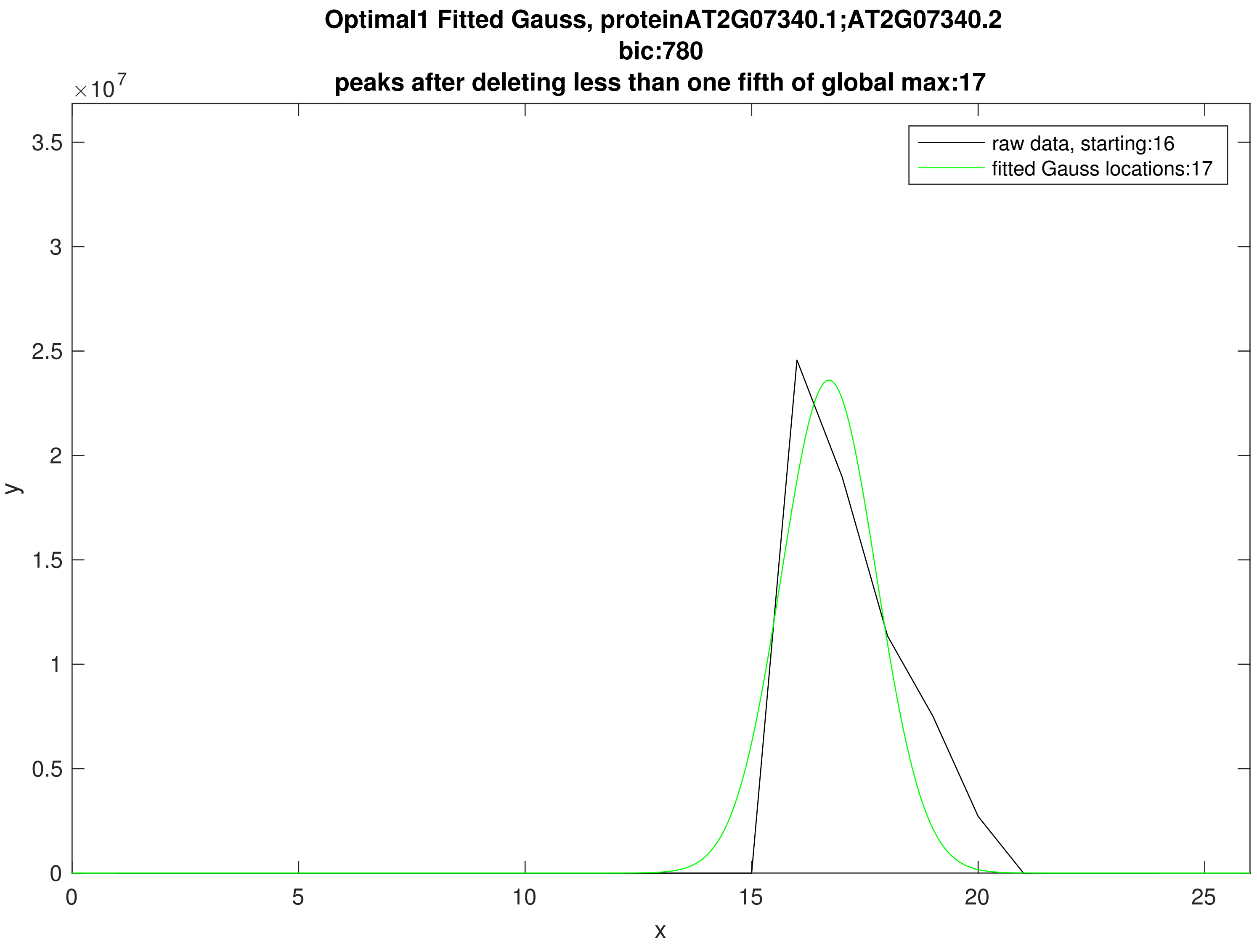

Optimal1 Fitted Gauss, proteinAT2G07340.1;AT2G07340.2
bic:780
peaks after deleting less than one fifth of global max:17
×10^7
3.5
3
2.5
2
1.5
1
0.5
0
y
0
5
10
15
20
25
x
raw data, starting:16
fitted Gauss locations:17

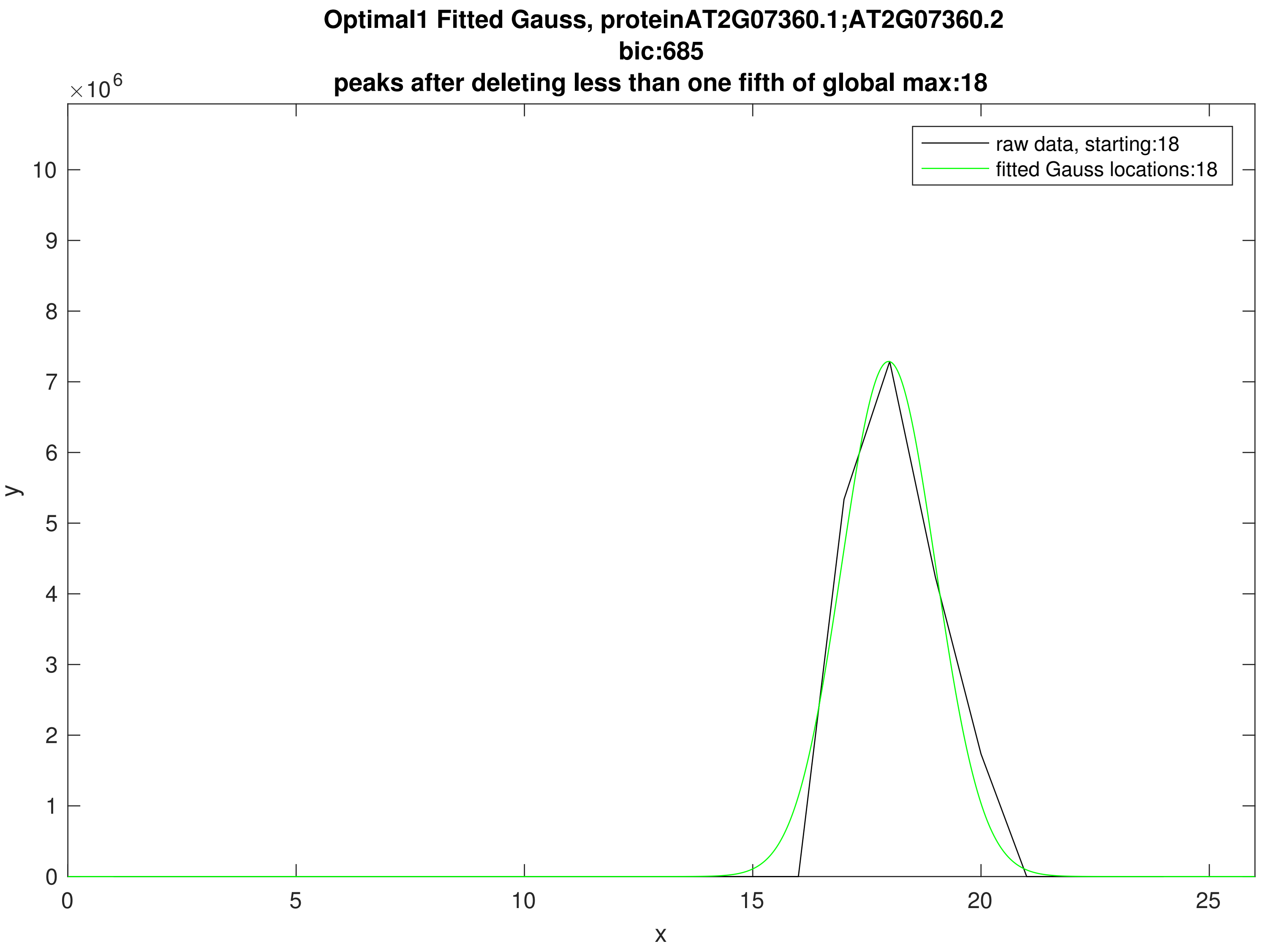
Optimal1 Fitted Gauss, proteinAT2G07360.1;AT2G07360.2
bic:685
peaks after deleting less than one fifth of global max:18
×10^6
raw data, starting:18
fitted Gauss locations:18
y
x
0
1
2
3
4
5
6
7
8
9
10
0
5
10
15
20
25

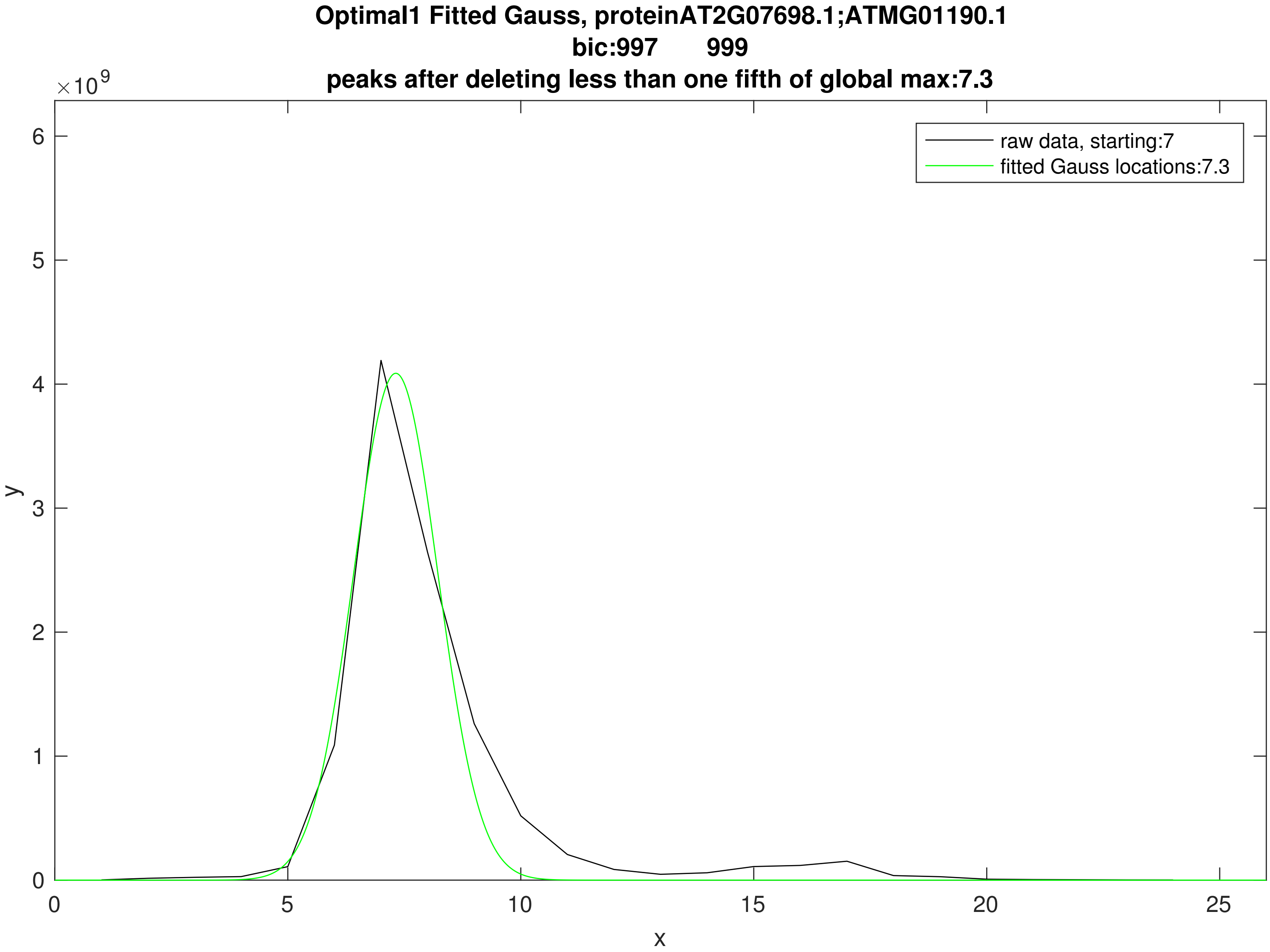

Optimal1 Fitted Gauss, proteinAT2G07698.1;ATMG01190.1
bic:997 999
peaks after deleting less than one fifth of global max:7.3
×10^9
raw data, starting:7
fitted Gauss locations:7.3
y
x
0
1
2
3
4
5
6
0
5
10
15
20
25

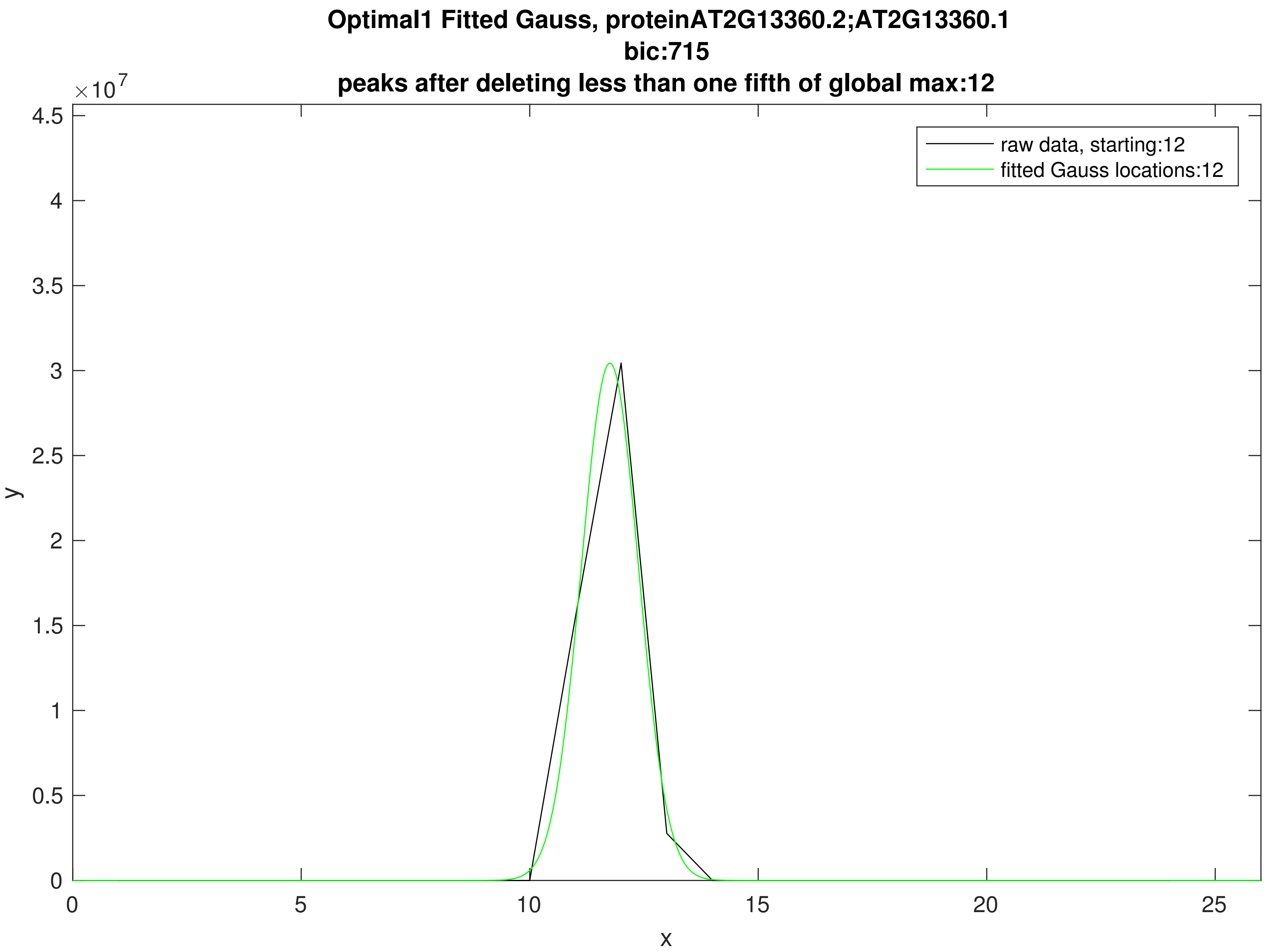

Optimal1 Fitted Gauss, proteinAT2G13360.2;AT2G13360.1
bic:715
peaks after deleting less than one fifth of global max:12
×10^7
4.5
4
3.5
3
2.5
2
1.5
1
0.5
0
y
0
5
10
15
20
25
x
raw data, starting:12
fitted Gauss locations:12

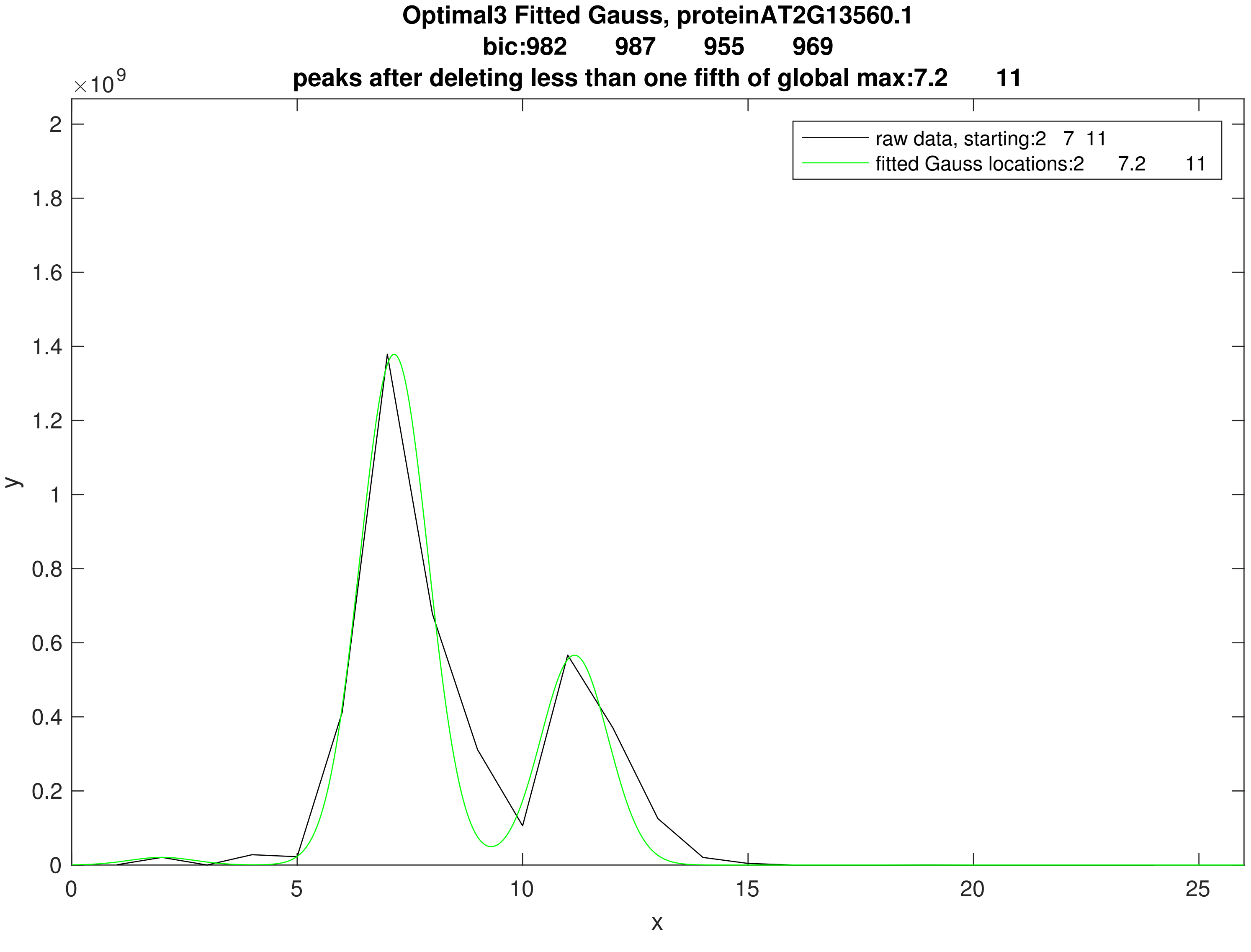

Optimal3 Fitted Gauss, proteinAT2G13560.1
bic:982 987 955 969
peaks after deleting less than one fifth of global max:7.2 11
×10^9
raw data, starting:2 7 11
fitted Gauss locations:2 7.2 11
y
x
0
0.2
0.4
0.6
0.8
1
1.2
1.4
1.6
1.8
2
0
5
10
15
20
25

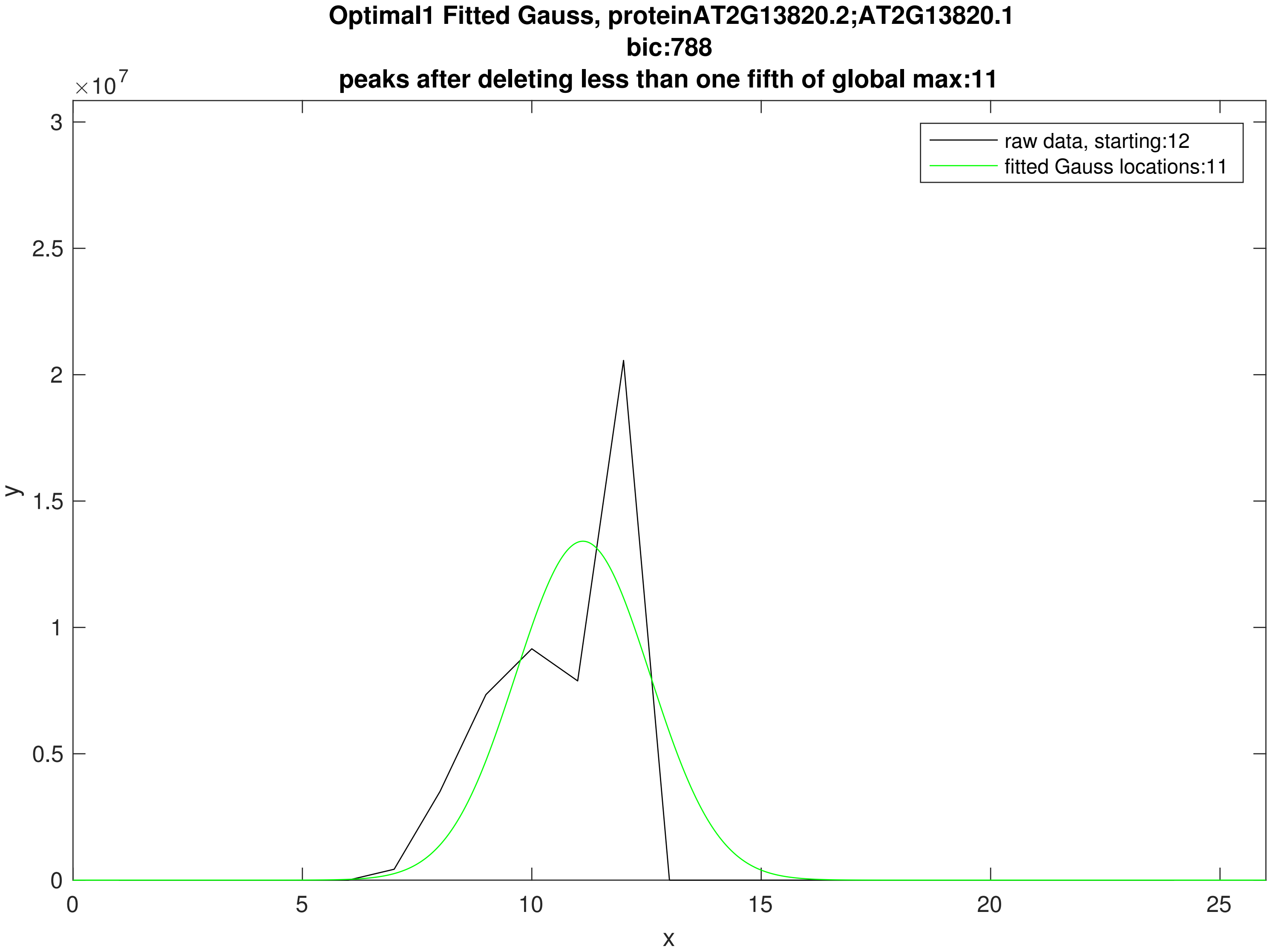
Optimal1 Fitted Gauss, proteinAT2G13820.2;AT2G13820.1
bic:788
peaks after deleting less than one fifth of global max:11
×10^7
3
2.5
2
1.5
1
0.5
0
y
0
5
10
15
20
25
x
raw data, starting:12
fitted Gauss locations:11

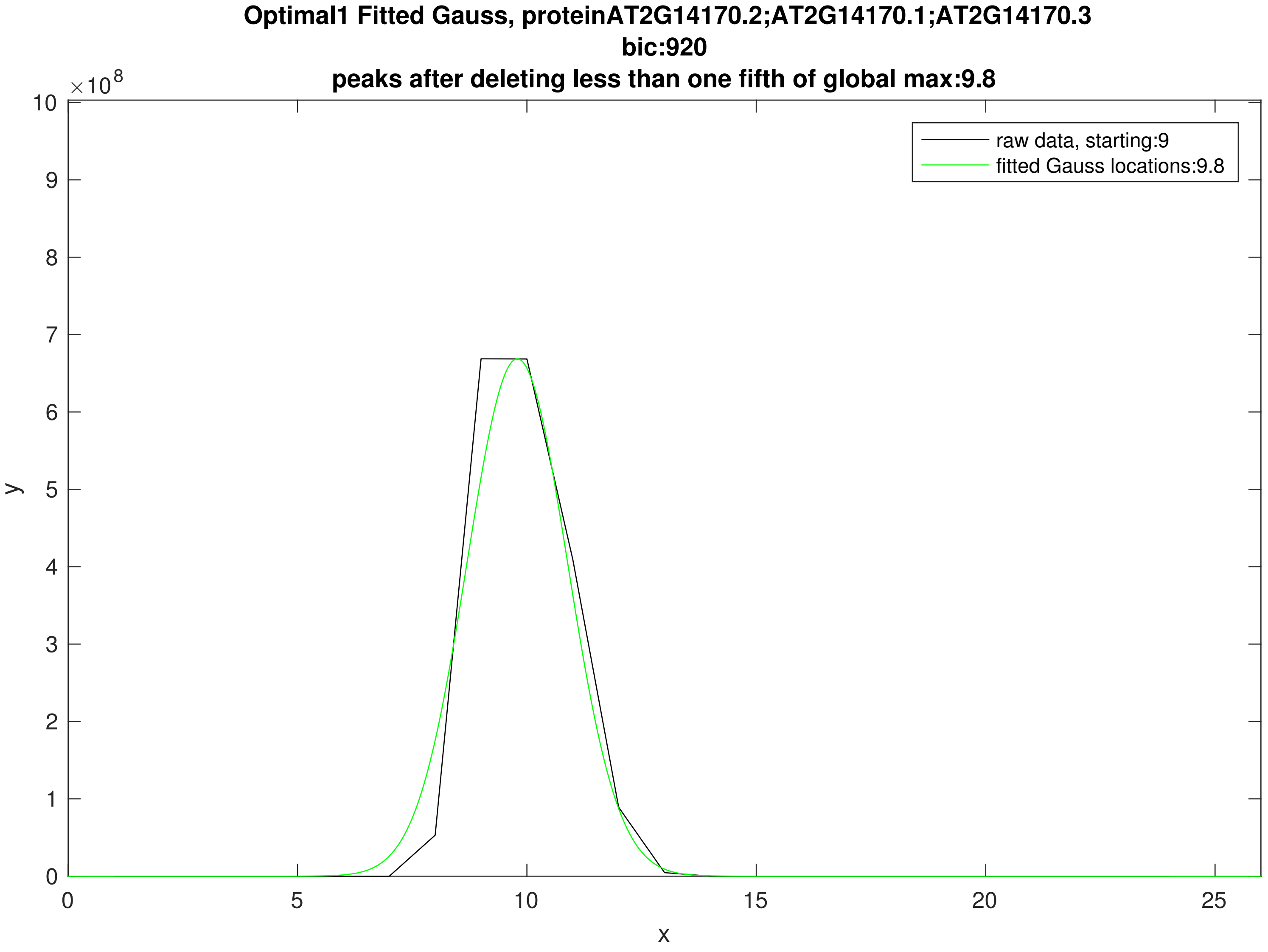

Optimal1 Fitted Gauss, proteinAT2G14170.2;AT2G14170.1;AT2G14170.3
bic:920
peaks after deleting less than one fifth of global max:9.8
raw data, starting:9
fitted Gauss locations:9.8
$\times 10^8$
10
9
8
7
6
5
4
3
2
1
0
y
0
5
10
15
20
25
x

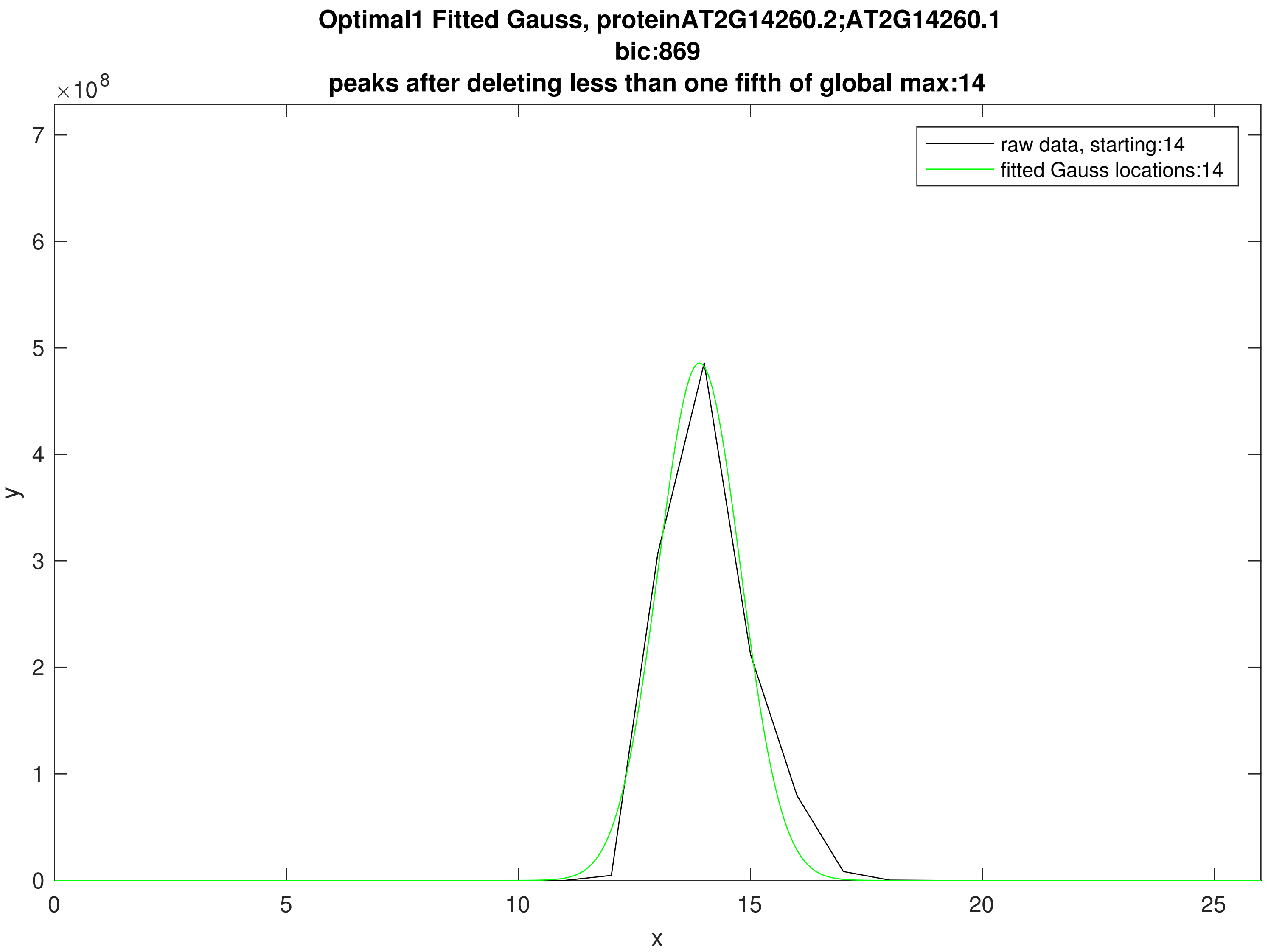
Optimal1 Fitted Gauss, proteinAT2G14260.2;AT2G14260.1
bic:869
peaks after deleting less than one fifth of global max:14
×10^8
raw data, starting:14
fitted Gauss locations:14
y
x
0
1
2
3
4
5
6
7
0
5
10
15
20
25

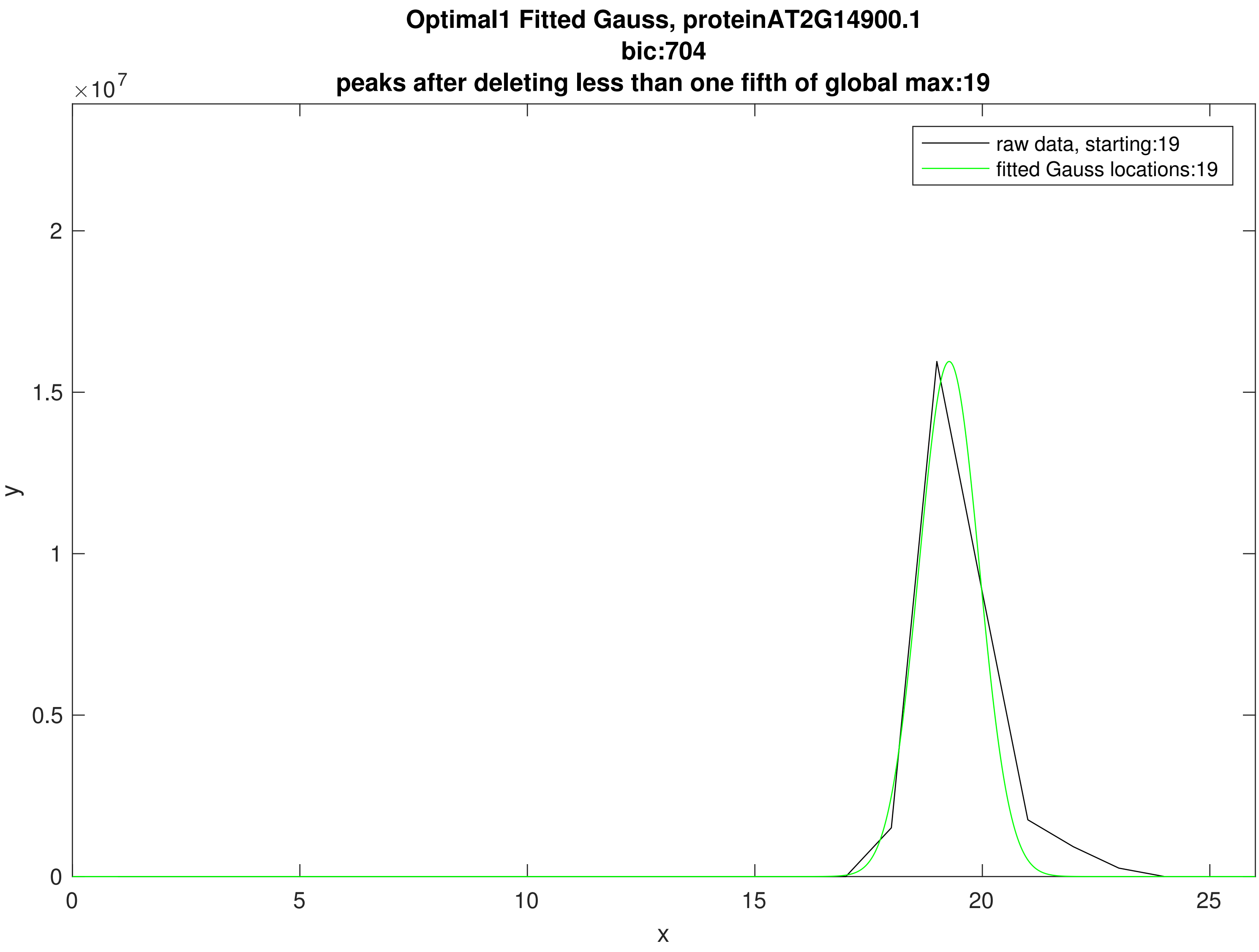
Optimal1 Fitted Gauss, proteinAT2G14900.1
bic:704
peaks after deleting less than one fifth of global max:19
×10^7
raw data, starting:19
fitted Gauss locations:19
y
x
0
0.5
1
1.5
2
0
5
10
15
20
25

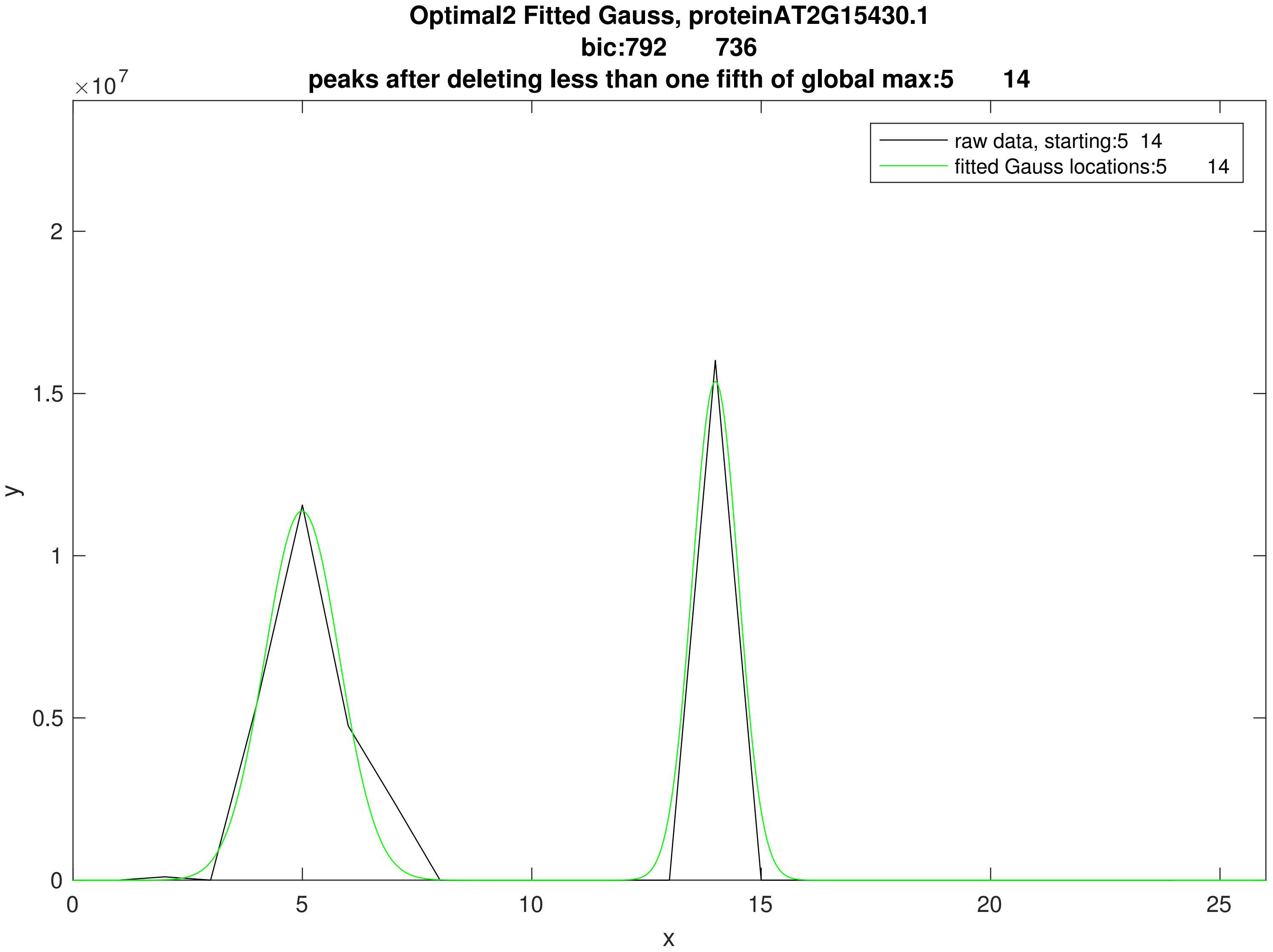
Optimal2 Fitted Gauss, proteinAT2G15430.1
bic:792 736
peaks after deleting less than one fifth of global max:5 14
×10^7
raw data, starting:5 14
fitted Gauss locations:5 14
2
1.5
1
0.5
0
y
0
5
10
15
20
25
x

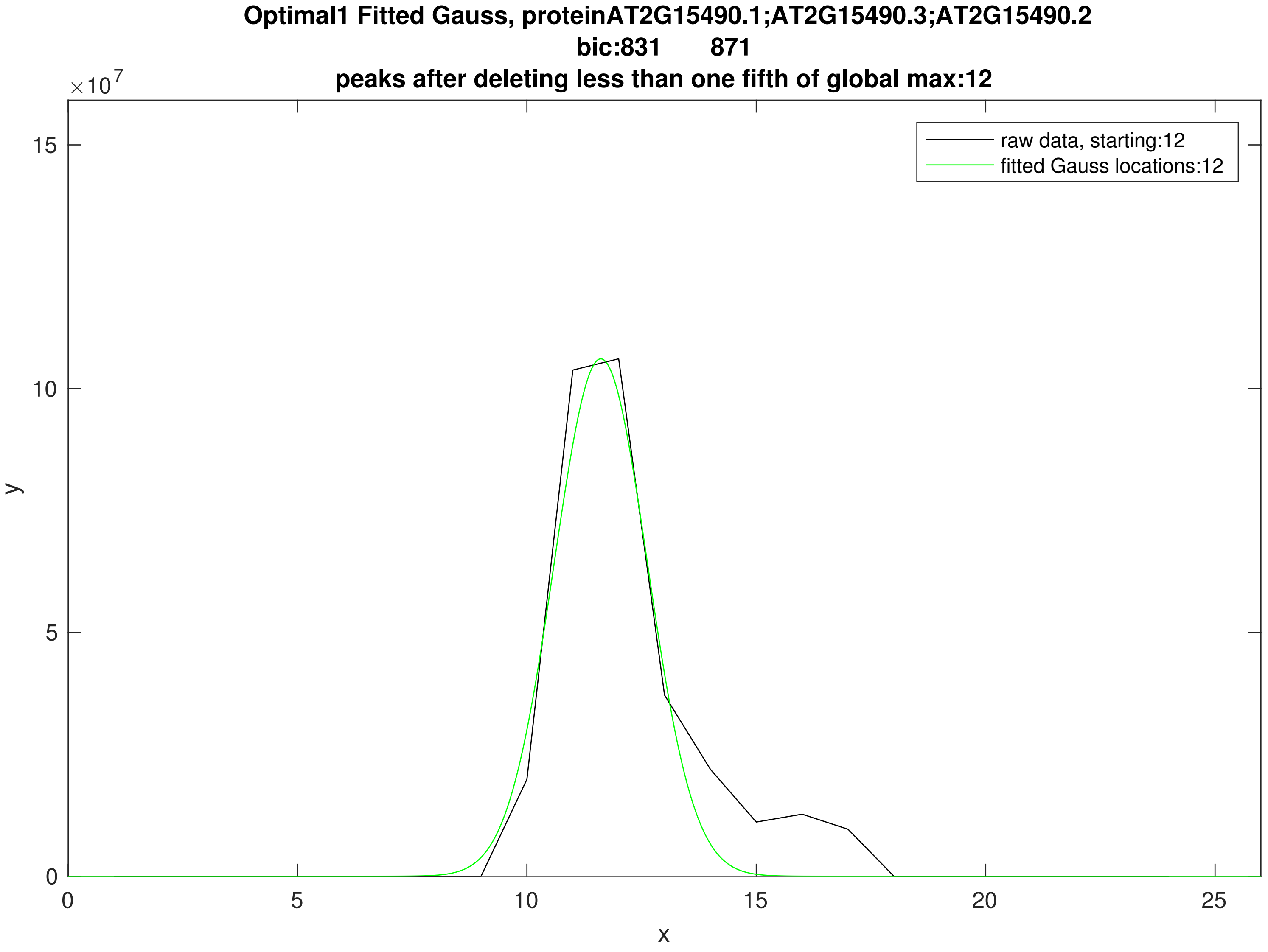
Optimal1 Fitted Gauss, proteinAT2G15490.1;AT2G15490.3;AT2G15490.2
bic:831 871
peaks after deleting less than one fifth of global max:12
×10^7
raw data, starting:12
fitted Gauss locations:12
15
10
5
0
y
0
5
10
15
20
25
x

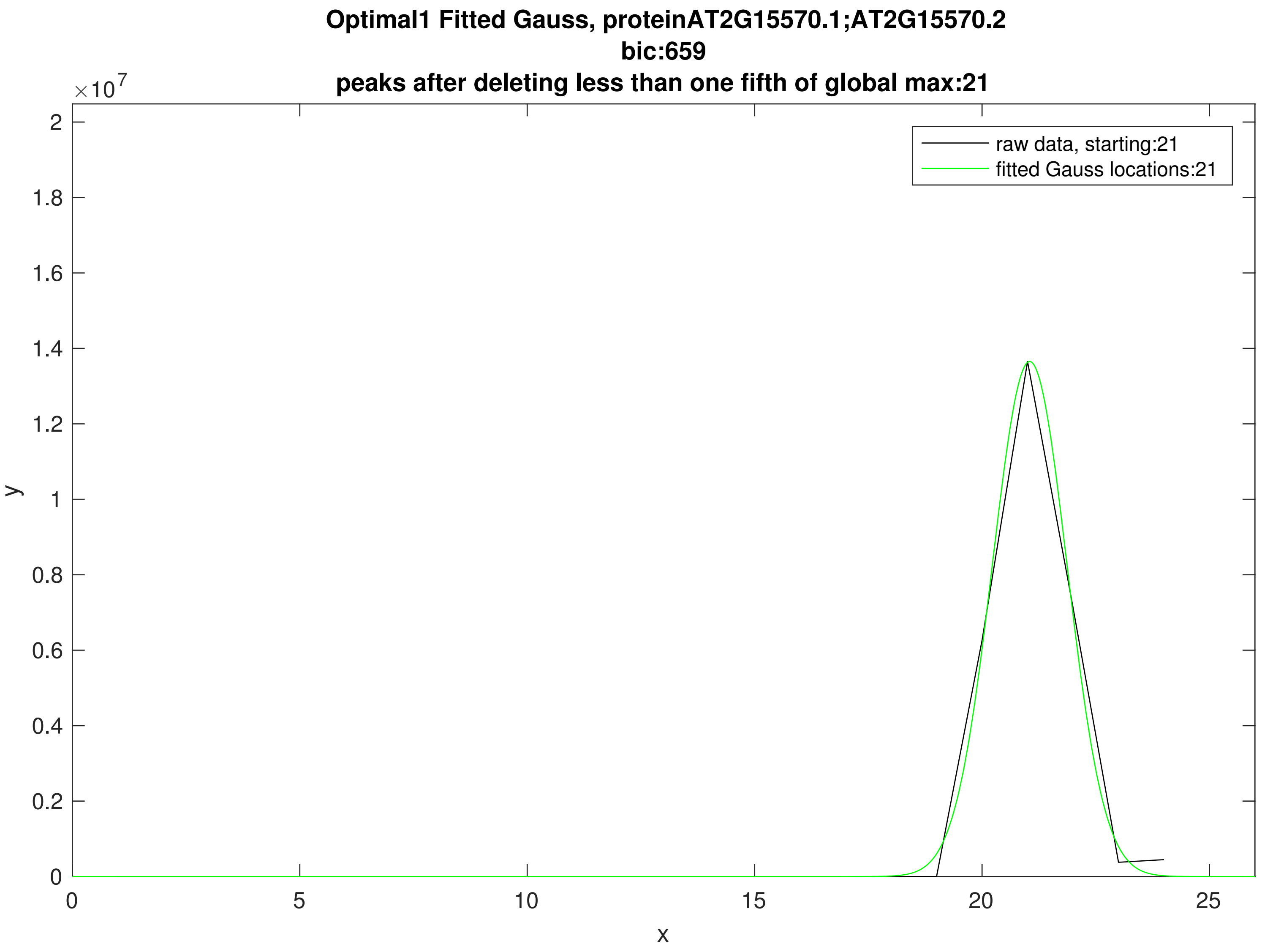
Optimal1 Fitted Gauss, proteinAT2G15570.1;AT2G15570.2
bic:659
peaks after deleting less than one fifth of global max:21
×10^7
2
1.8
1.6
1.4
1.2
1
0.8
0.6
0.4
0.2
0
y
0
5
10
15
20
25
x
raw data, starting:21
fitted Gauss locations:21

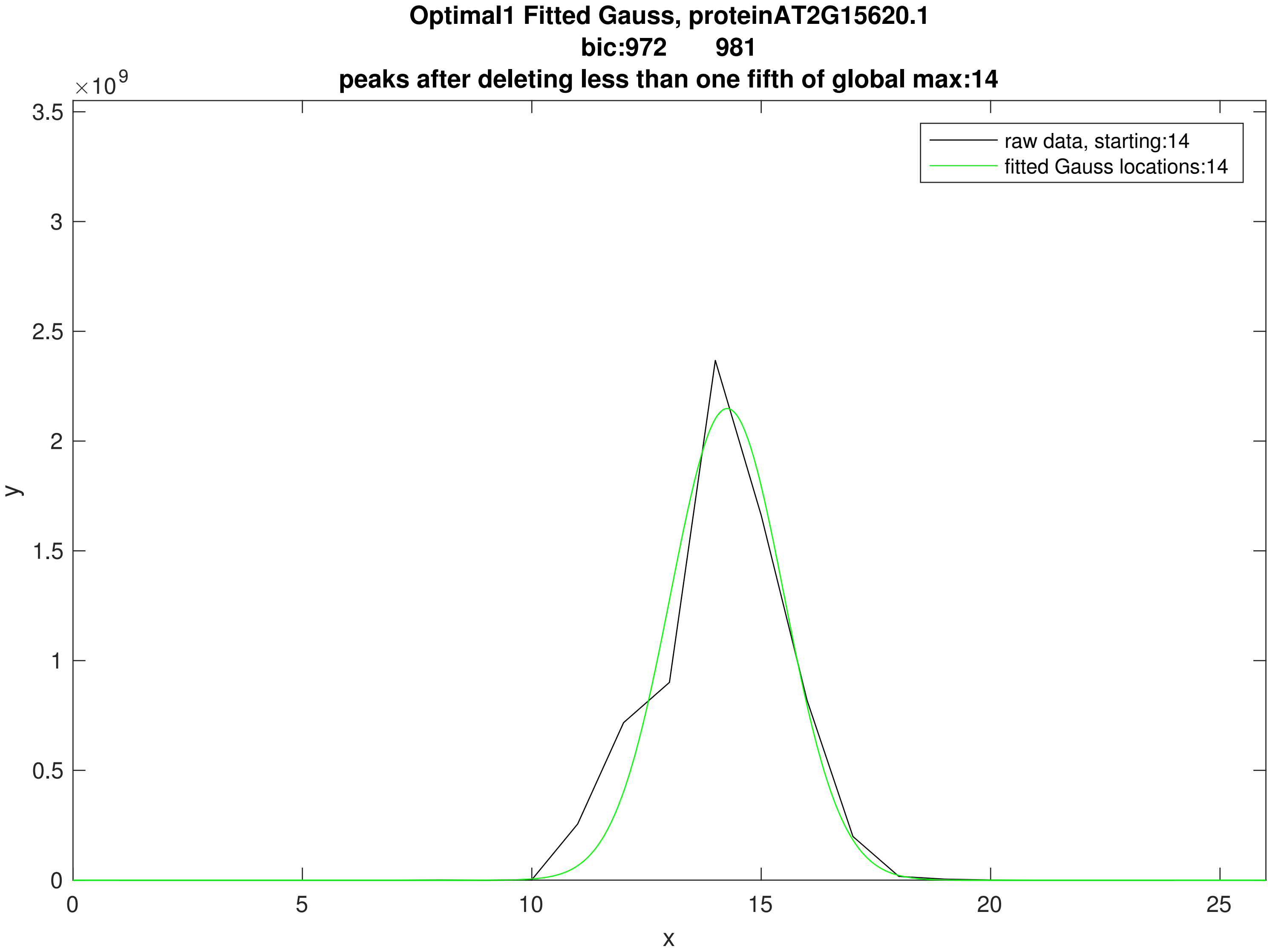

Optimal1 Fitted Gauss, proteinAT2G15620.1
bic:972 981
peaks after deleting less than one fifth of global max:14
×10^9
3.5
3
2.5
2
1.5
1
0.5
0
y
0
5
10
15
20
25
x
raw data, starting:14
fitted Gauss locations:14

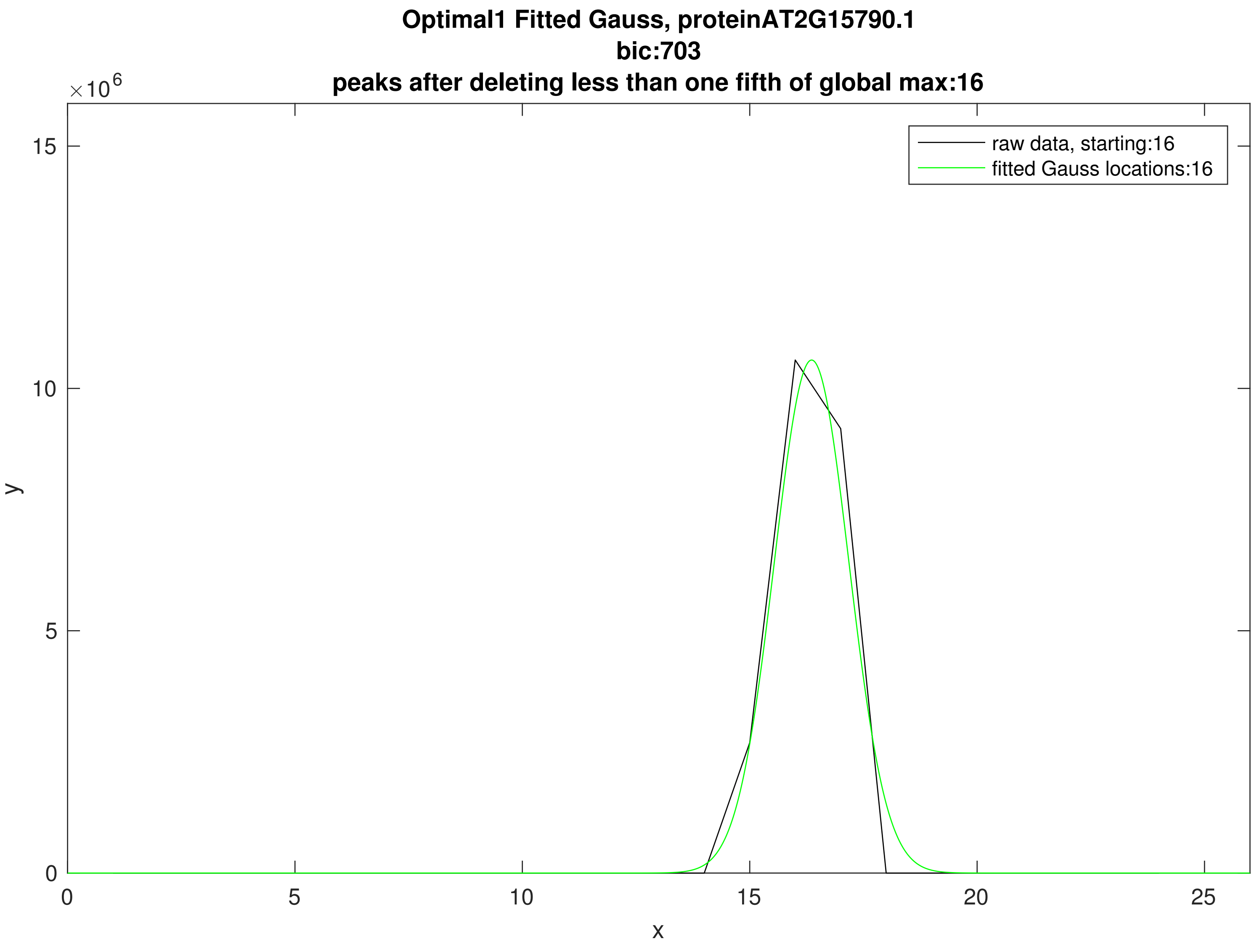

Optimal1 Fitted Gauss, proteinAT2G15790.1
bic:703
peaks after deleting less than one fifth of global max:16
×10^6
15
10
5
0
y
0
5
10
15
20
25
x
raw data, starting:16
fitted Gauss locations:16

**Optimal1 Fitted Gauss, proteinAT2G16060.1**
**bic:699**
**peaks after deleting less than one fifth of global max:19**

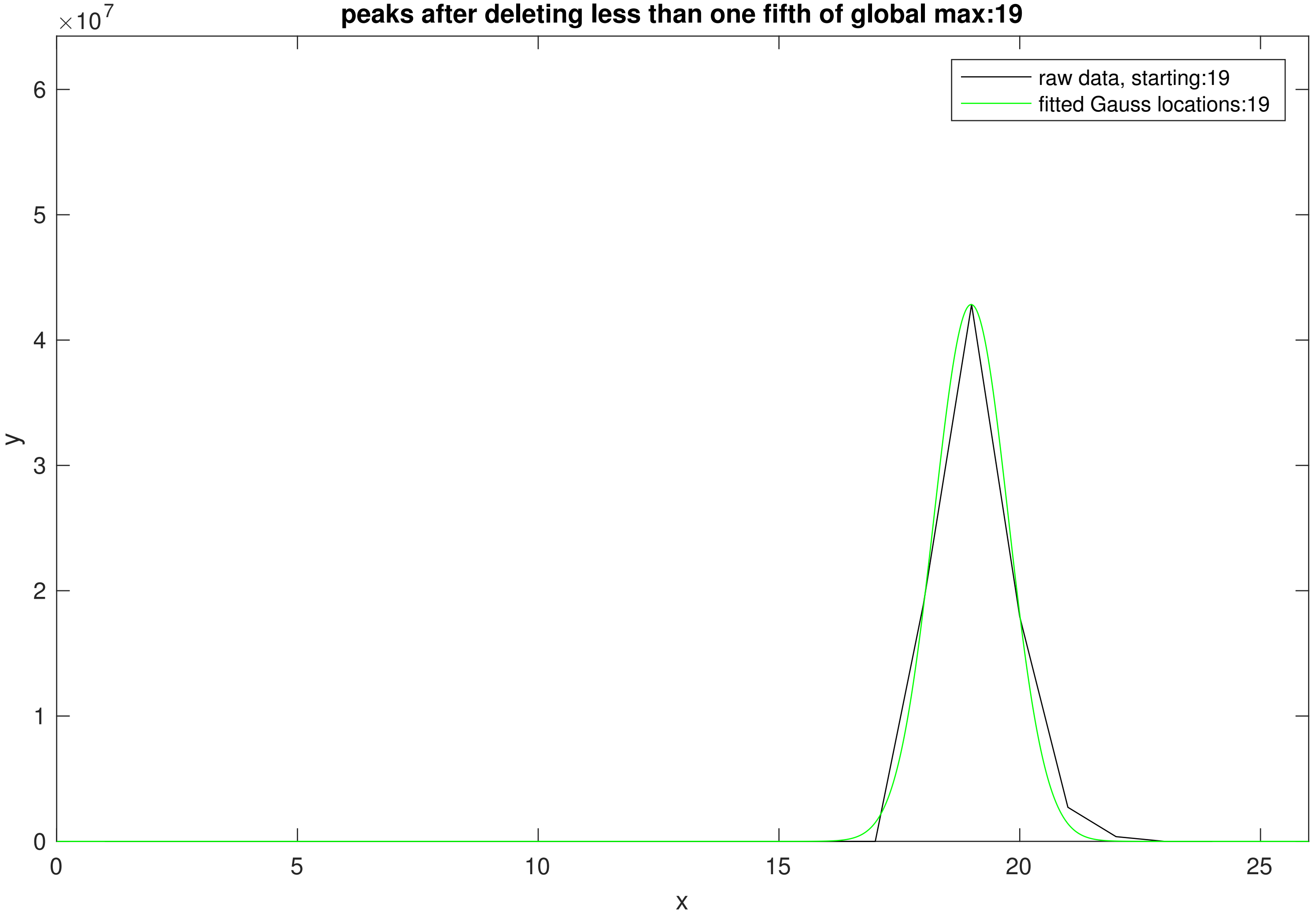

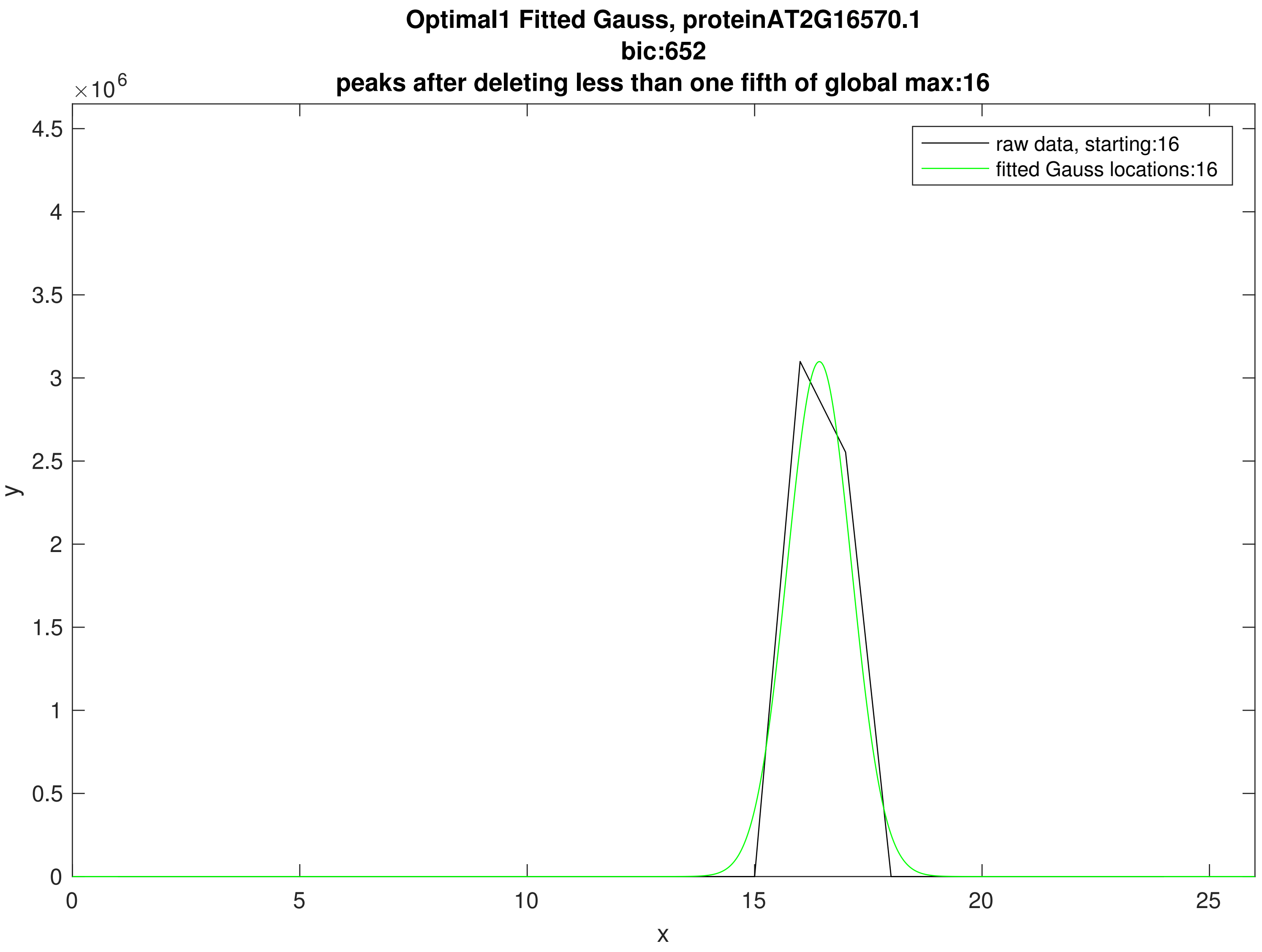

Optimal1 Fitted Gauss, proteinAT2G16570.1
bic:652
peaks after deleting less than one fifth of global max:16
×10^6
raw data, starting:16
fitted Gauss locations:16
y
x
0
0.5
1
1.5
2
2.5
3
3.5
4
4.5
5
10
15
20
25

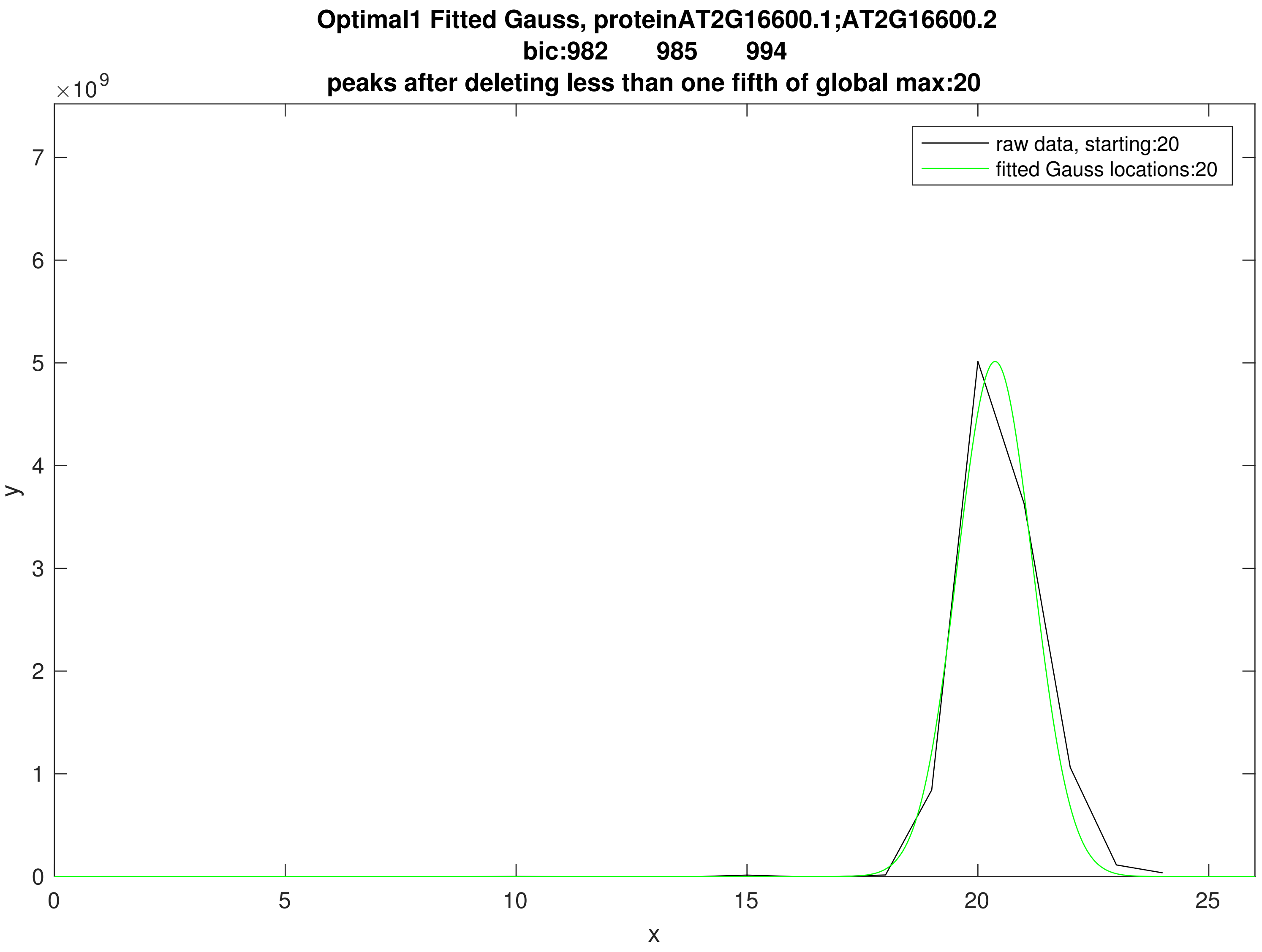
Optimal1 Fitted Gauss, proteinAT2G16600.1;AT2G16600.2
bic:982 985 994
peaks after deleting less than one fifth of global max:20
×10^9
raw data, starting:20
fitted Gauss locations:20
y
x
0
1
2
3
4
5
6
7
0
5
10
15
20
25

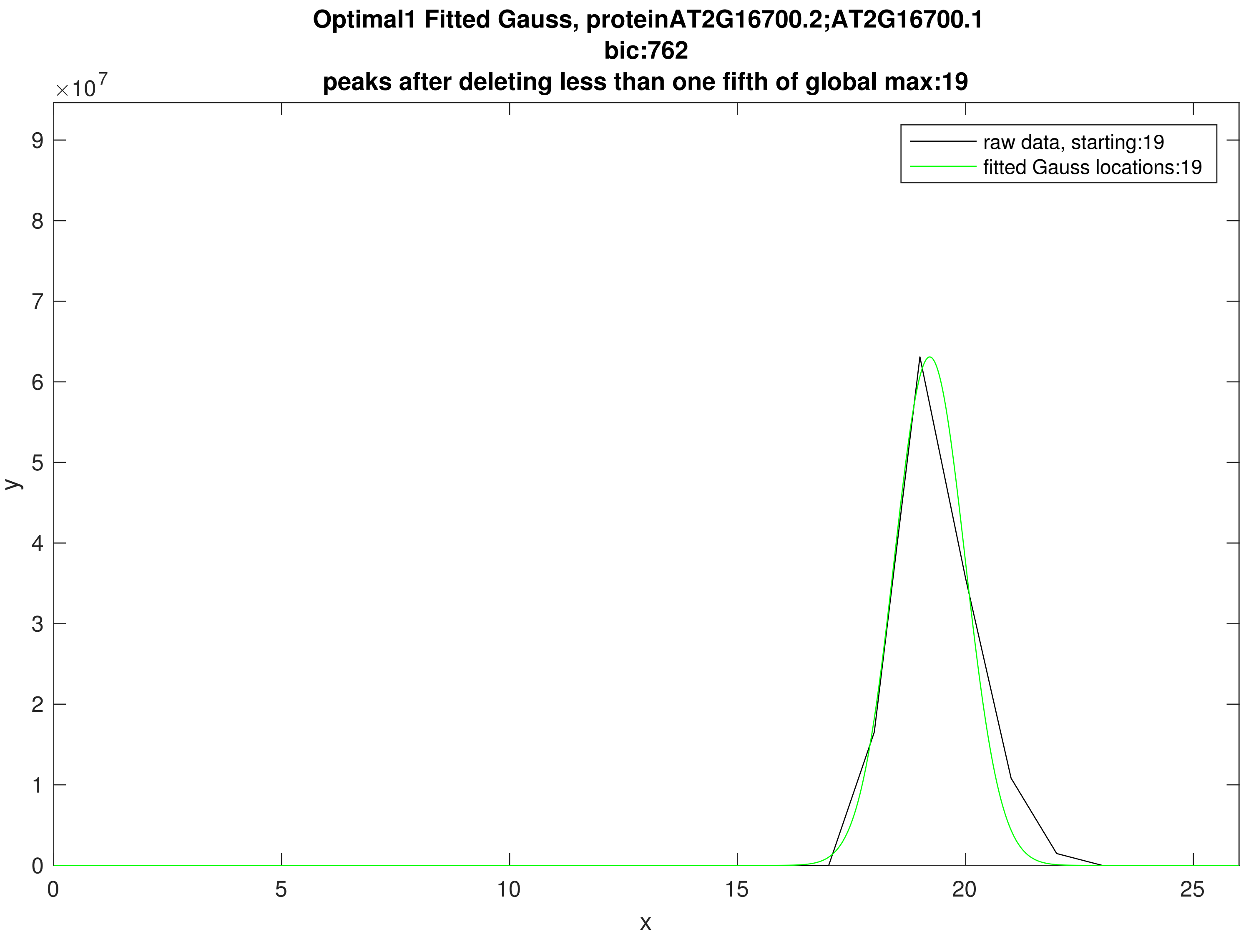
Optimal1 Fitted Gauss, proteinAT2G16700.2;AT2G16700.1
bic:762
peaks after deleting less than one fifth of global max:19
$\times 10^7$
9
8
7
6
5
4
3
2
1
0
y
0
5
10
15
20
25
x
raw data, starting:19
fitted Gauss locations:19

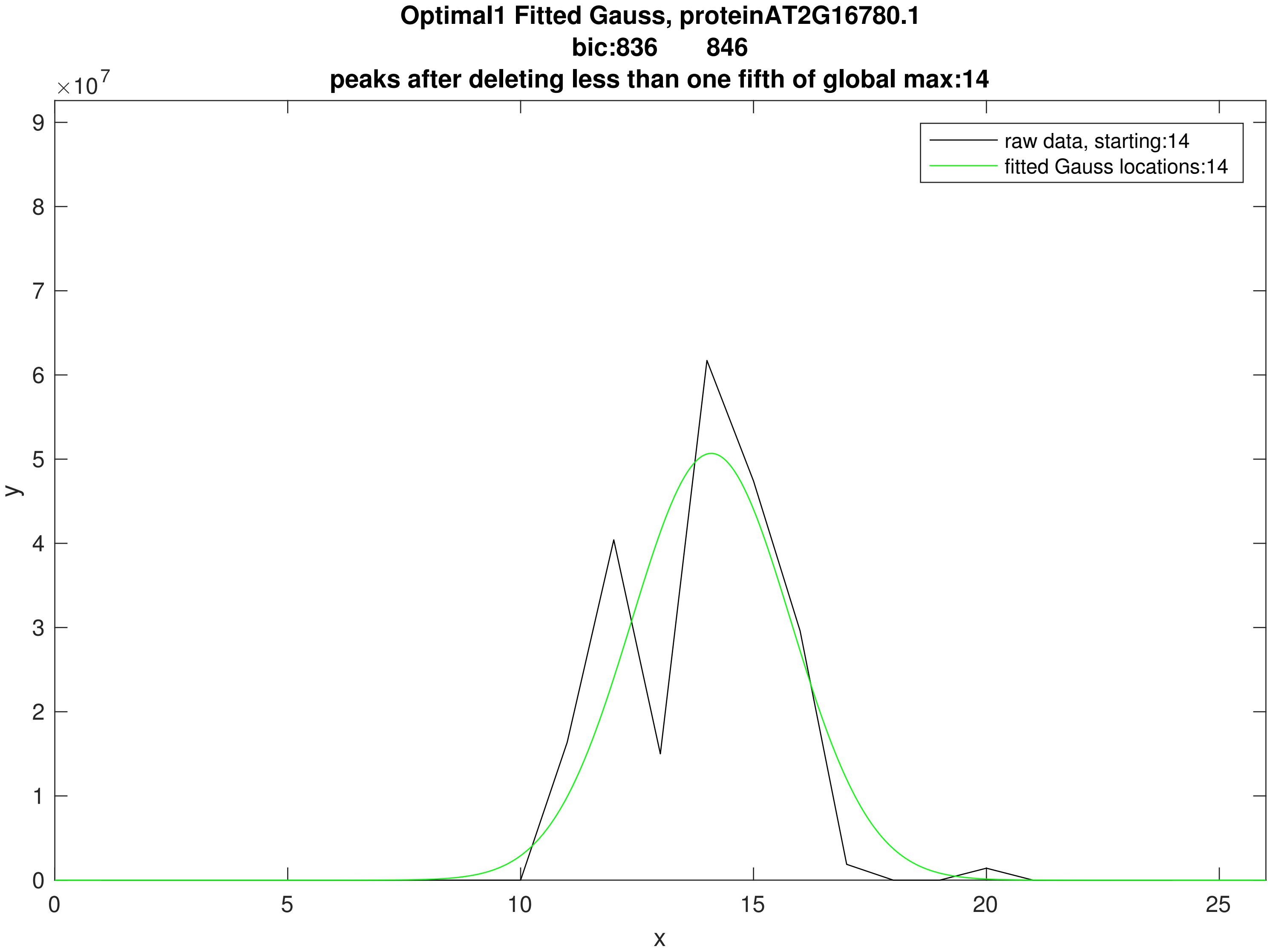

Optimal1 Fitted Gauss, proteinAT2G16780.1
bic:836 846
peaks after deleting less than one fifth of global max:14
×10^7
9
8
7
6
5
4
3
2
1
0
y
0
5
10
15
20
25
x
raw data, starting:14
fitted Gauss locations:14

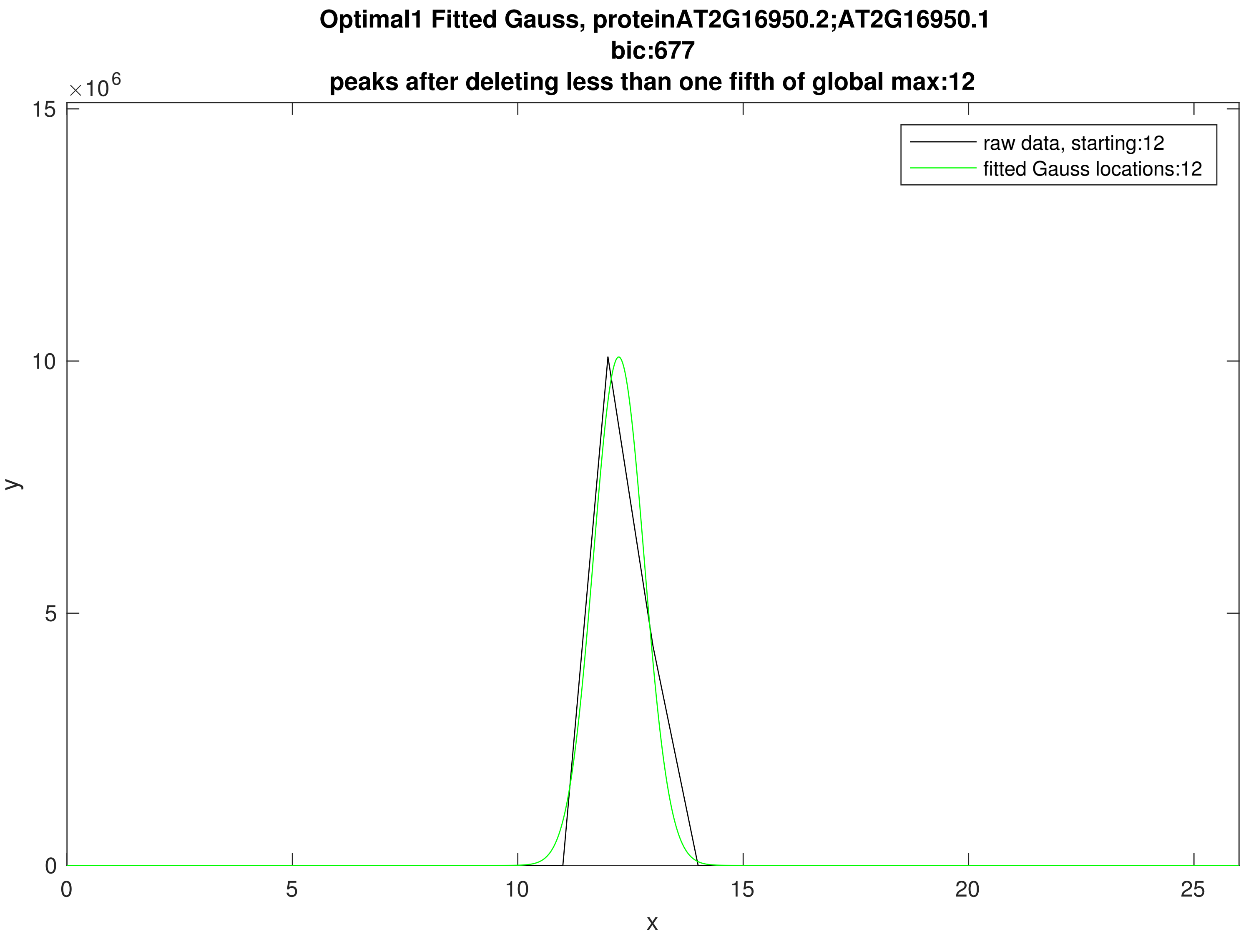
Optimal1 Fitted Gauss, proteinAT2G16950.2;AT2G16950.1
bic:677
peaks after deleting less than one fifth of global max:12
×10^6
15
10
5
0
y
0
5
10
15
20
25
x
raw data, starting:12
fitted Gauss locations:12

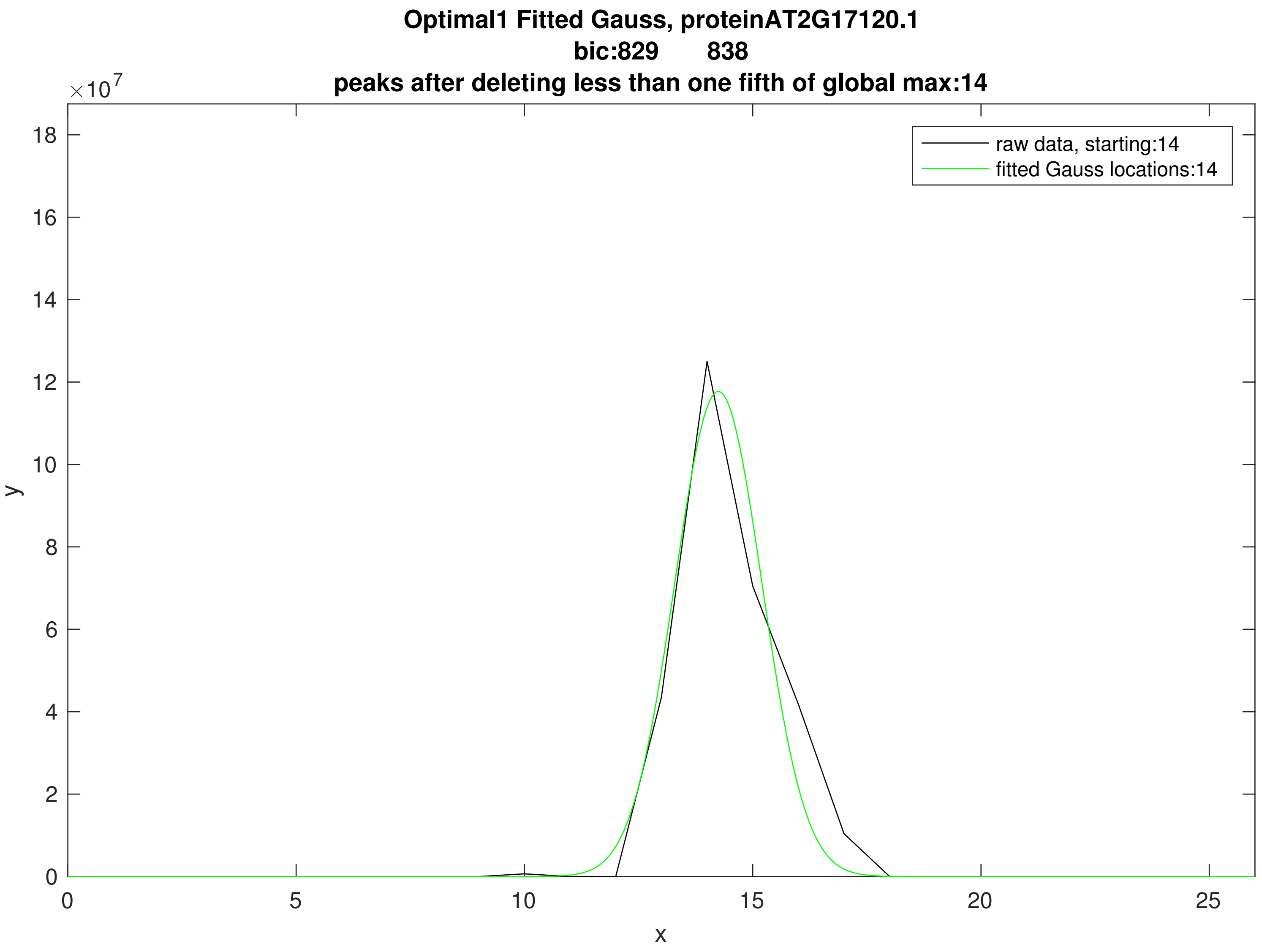

Optimal1 Fitted Gauss, proteinAT2G17120.1
bic:829 838
peaks after deleting less than one fifth of global max:14
×10^7
raw data, starting:14
fitted Gauss locations:14
y
x
0
2
4
6
8
10
12
14
16
18
0
5
10
15
20
25

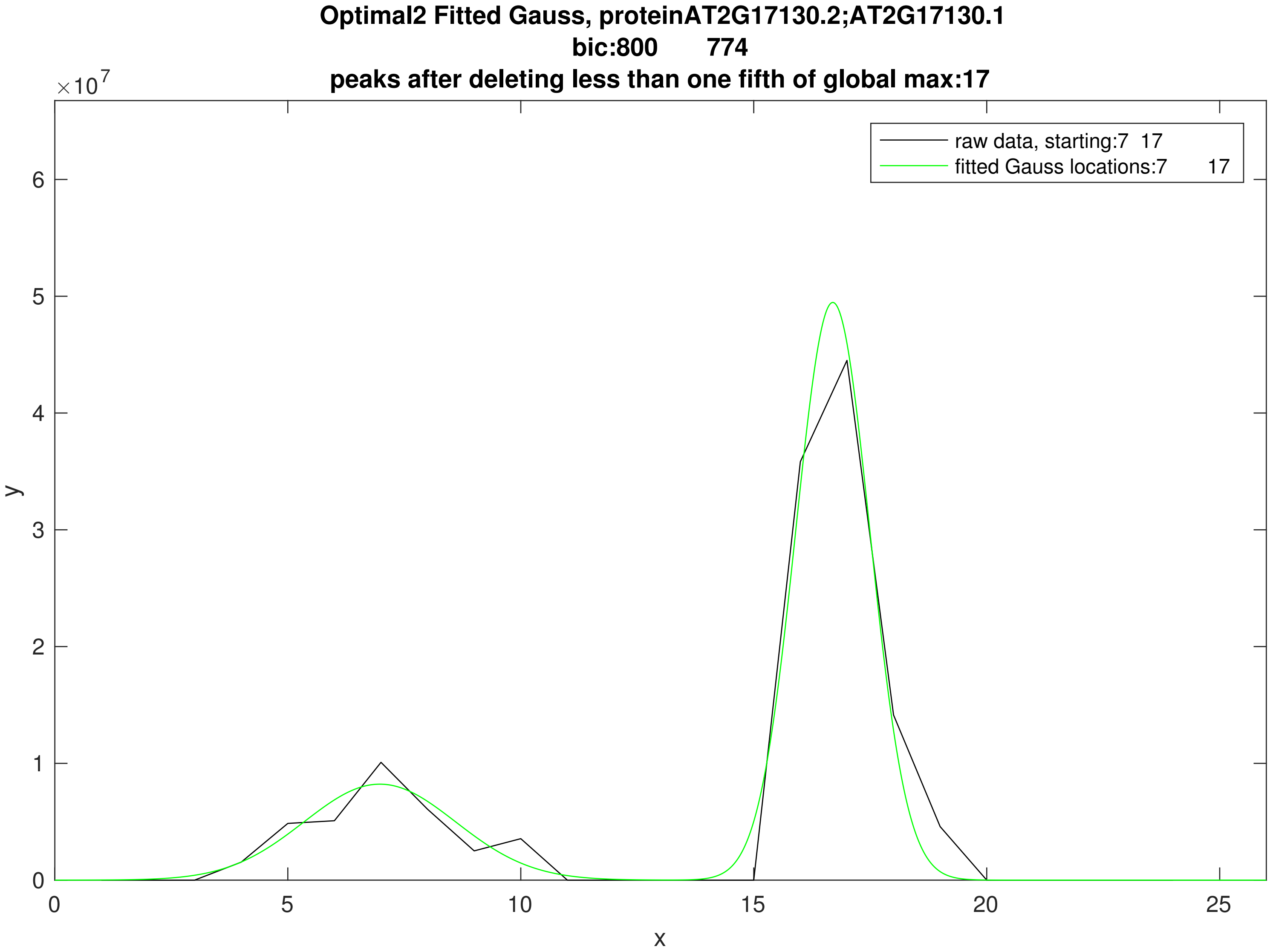
Optimal2 Fitted Gauss, proteinAT2G17130.2;AT2G17130.1
bic:800 774
peaks after deleting less than one fifth of global max:17
×10^7
raw data, starting:7 17
fitted Gauss locations:7 17
y
x
0
1
2
3
4
5
6
0
5
10
15
20
25

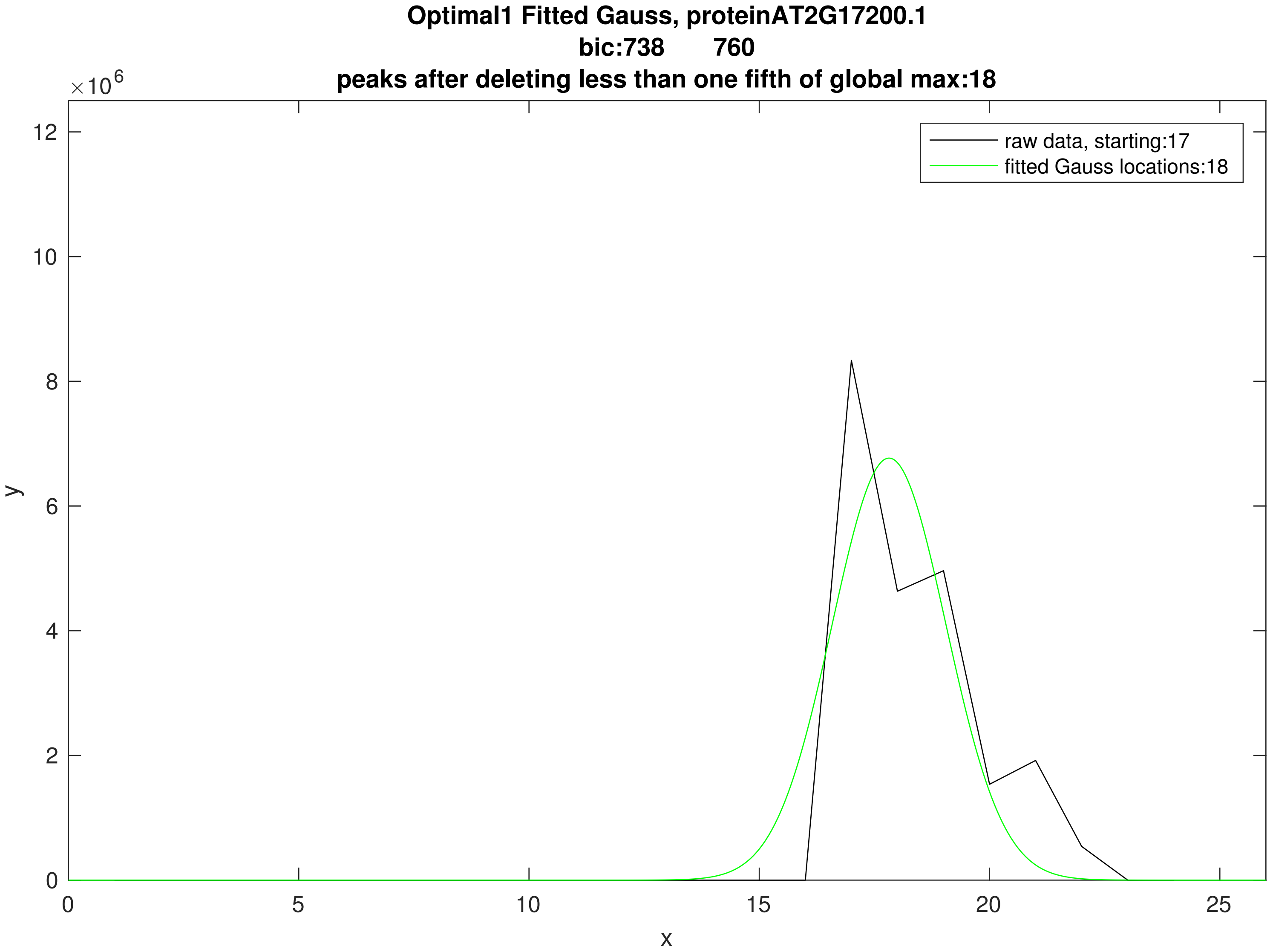

Optimal1 Fitted Gauss, proteinAT2G17200.1
bic:738 760
peaks after deleting less than one fifth of global max:18
×10^6
raw data, starting:17
fitted Gauss locations:18
y
x
0
2
4
6
8
10
12
0
5
10
15
20
25

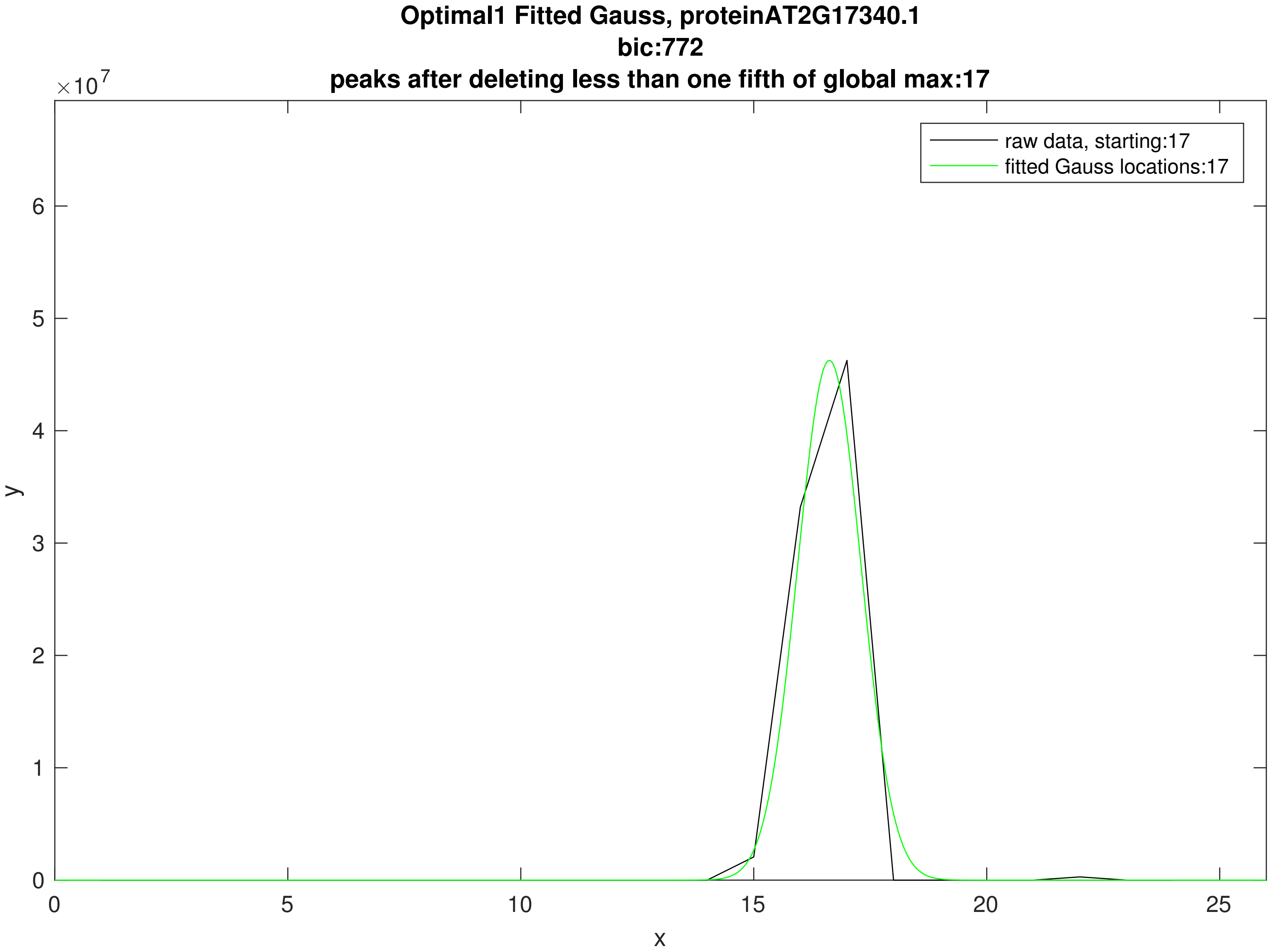

Optimal1 Fitted Gauss, proteinAT2G17340.1
bic:772
peaks after deleting less than one fifth of global max:17
×10^7
raw data, starting:17
fitted Gauss locations:17
y
x
0
1
2
3
4
5
6
0
5
10
15
20
25

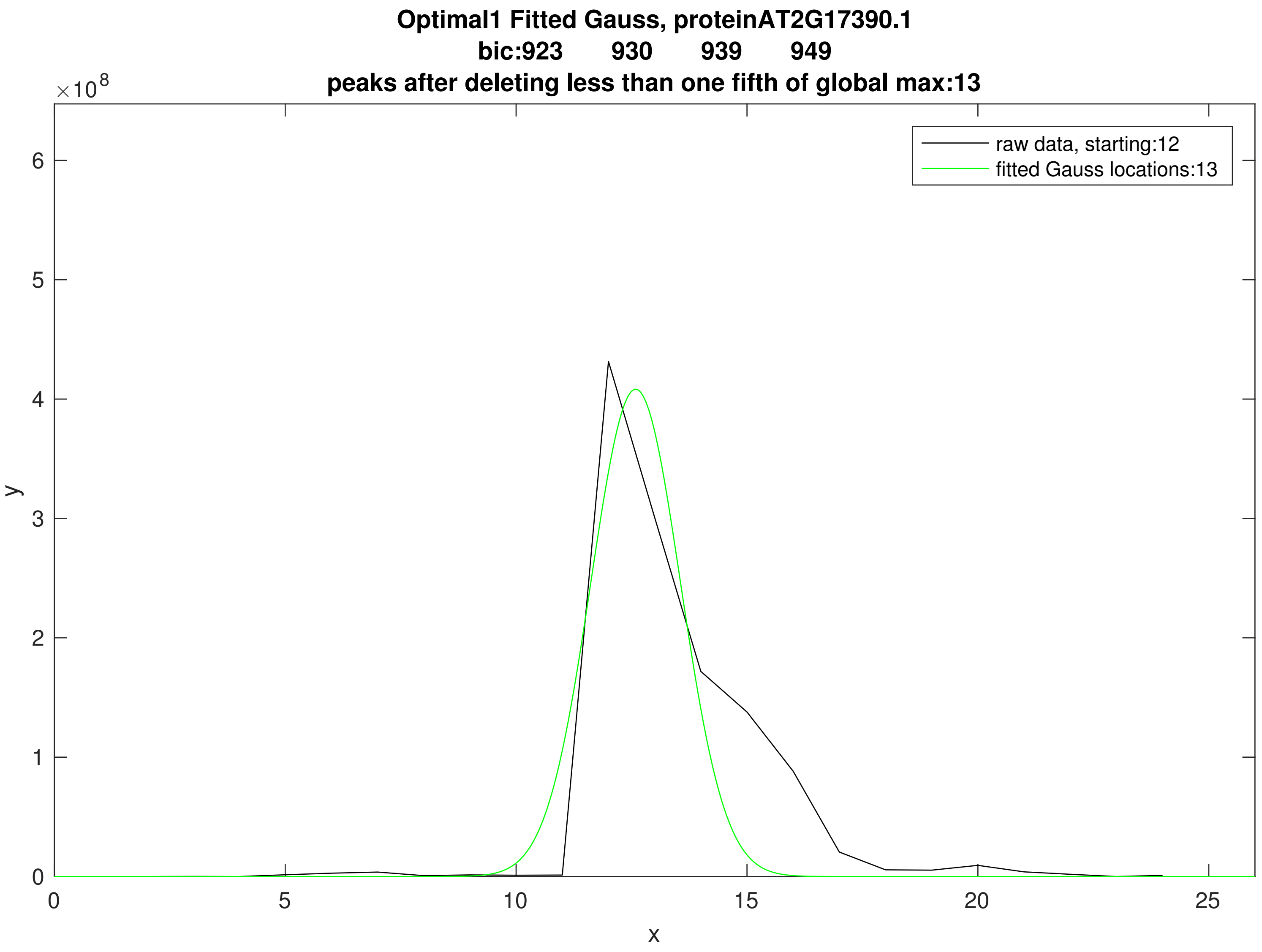
Optimal1 Fitted Gauss, proteinAT2G17390.1
bic:923 930 939 949
peaks after deleting less than one fifth of global max:13
raw data, starting:12
fitted Gauss locations:13
y
x

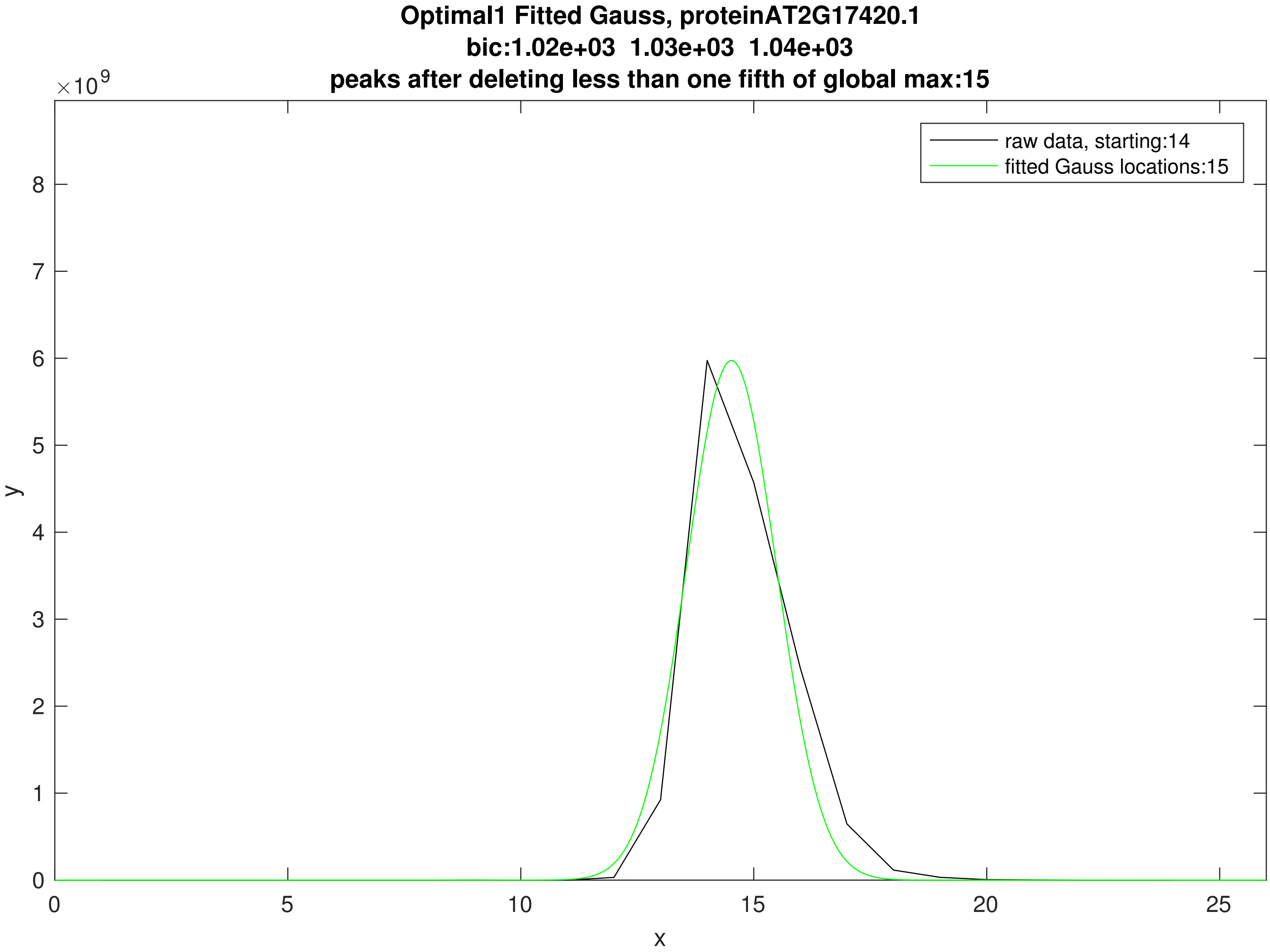
Optimal1 Fitted Gauss, proteinAT2G17420.1
bic:1.02e+03  1.03e+03  1.04e+03
peaks after deleting less than one fifth of global max:15
×10^9
raw data, starting:14
fitted Gauss locations:15
y
x
0
1
2
3
4
5
6
7
8
0
5
10
15
20
25

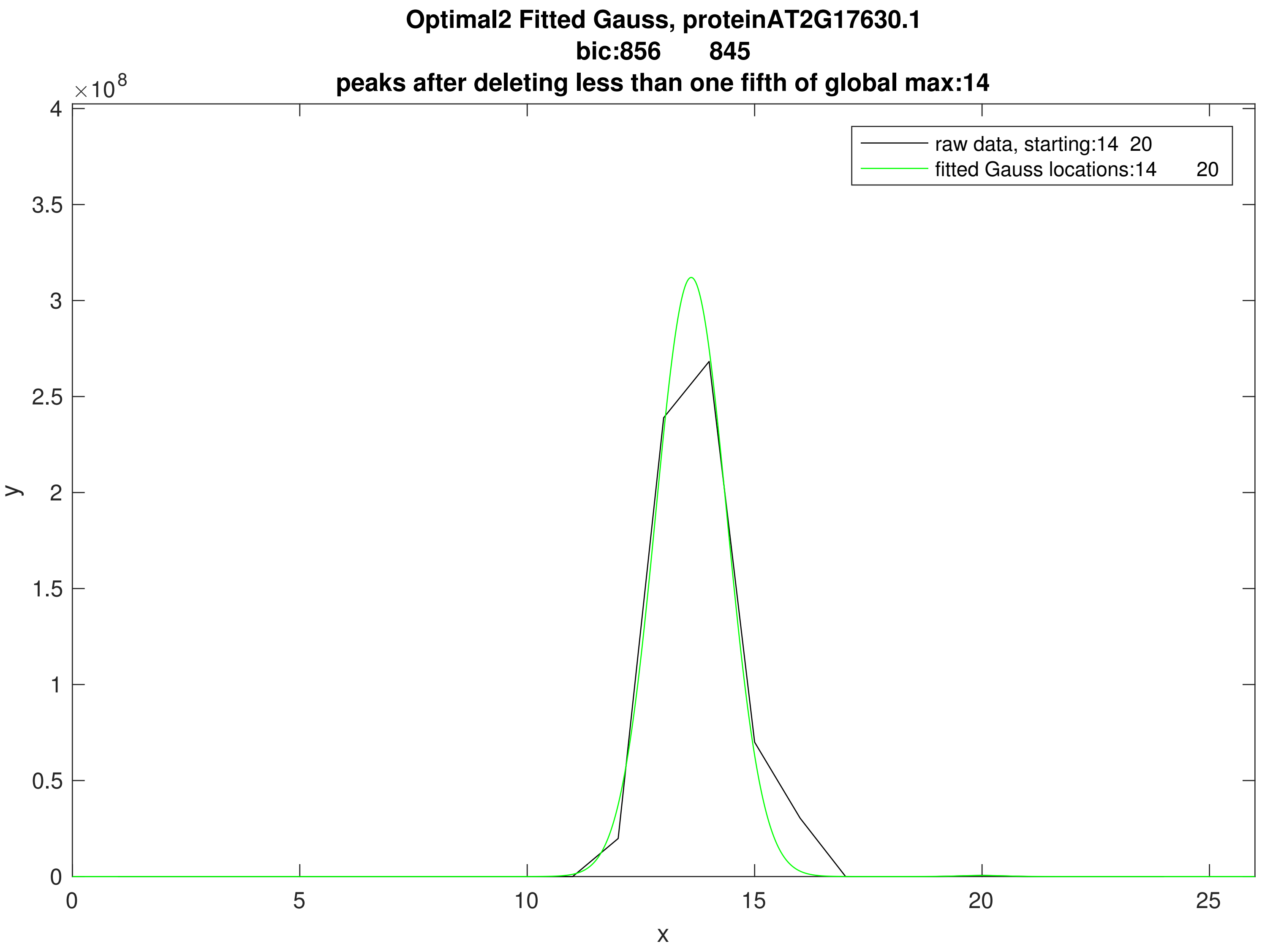

Optimal2 Fitted Gauss, proteinAT2G17630.1
bic:856 845
peaks after deleting less than one fifth of global max:14
×10^8
raw data, starting:14 20
fitted Gauss locations:14 20
y
x
0
0.5
1
1.5
2
2.5
3
3.5
4
5
10
15
20
25

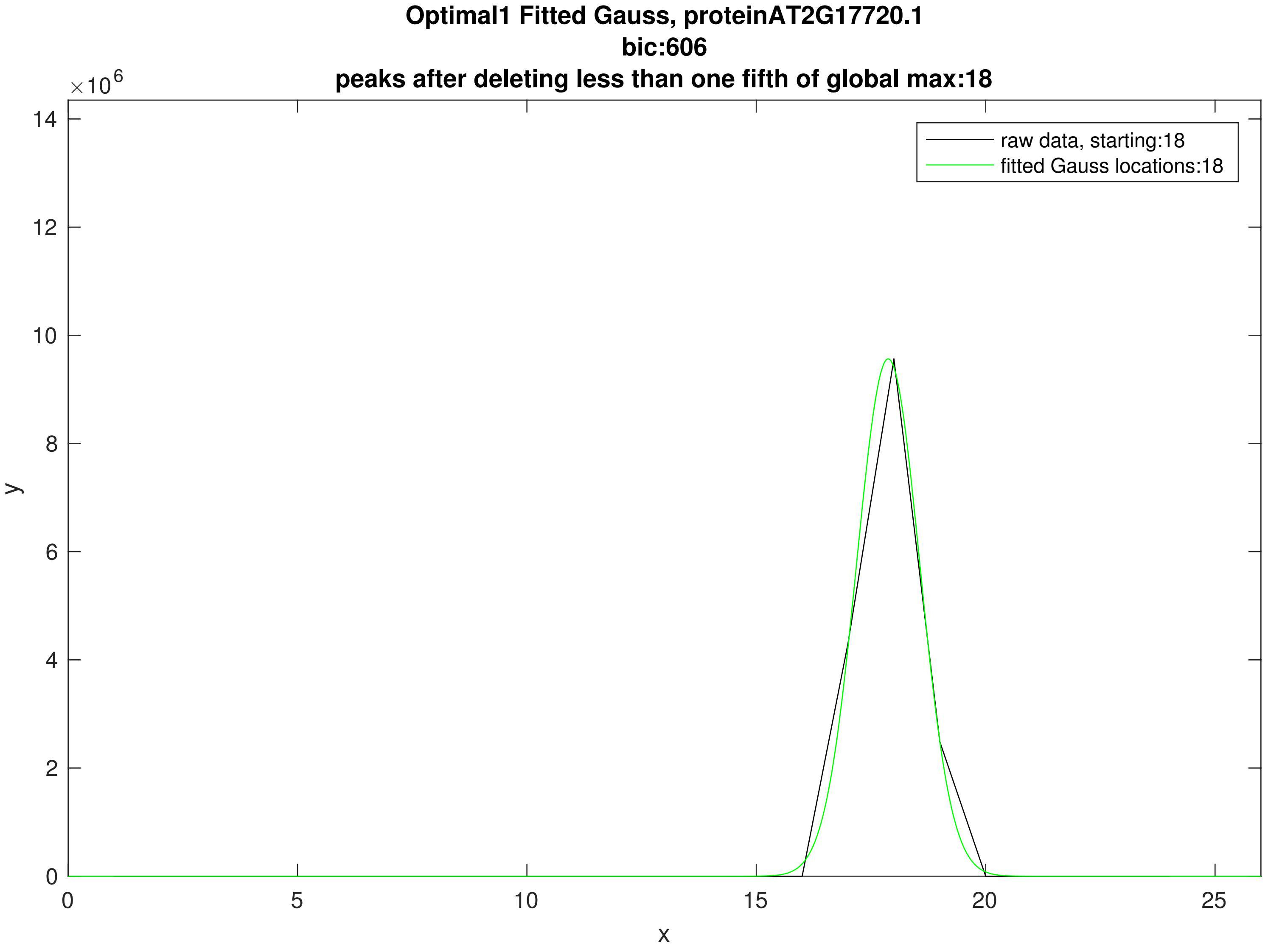
Optimal1 Fitted Gauss, proteinAT2G17720.1
bic:606
peaks after deleting less than one fifth of global max:18
×10^6
raw data, starting:18
fitted Gauss locations:18
y
x
0
2
4
6
8
10
12
14
0
5
10
15
20
25

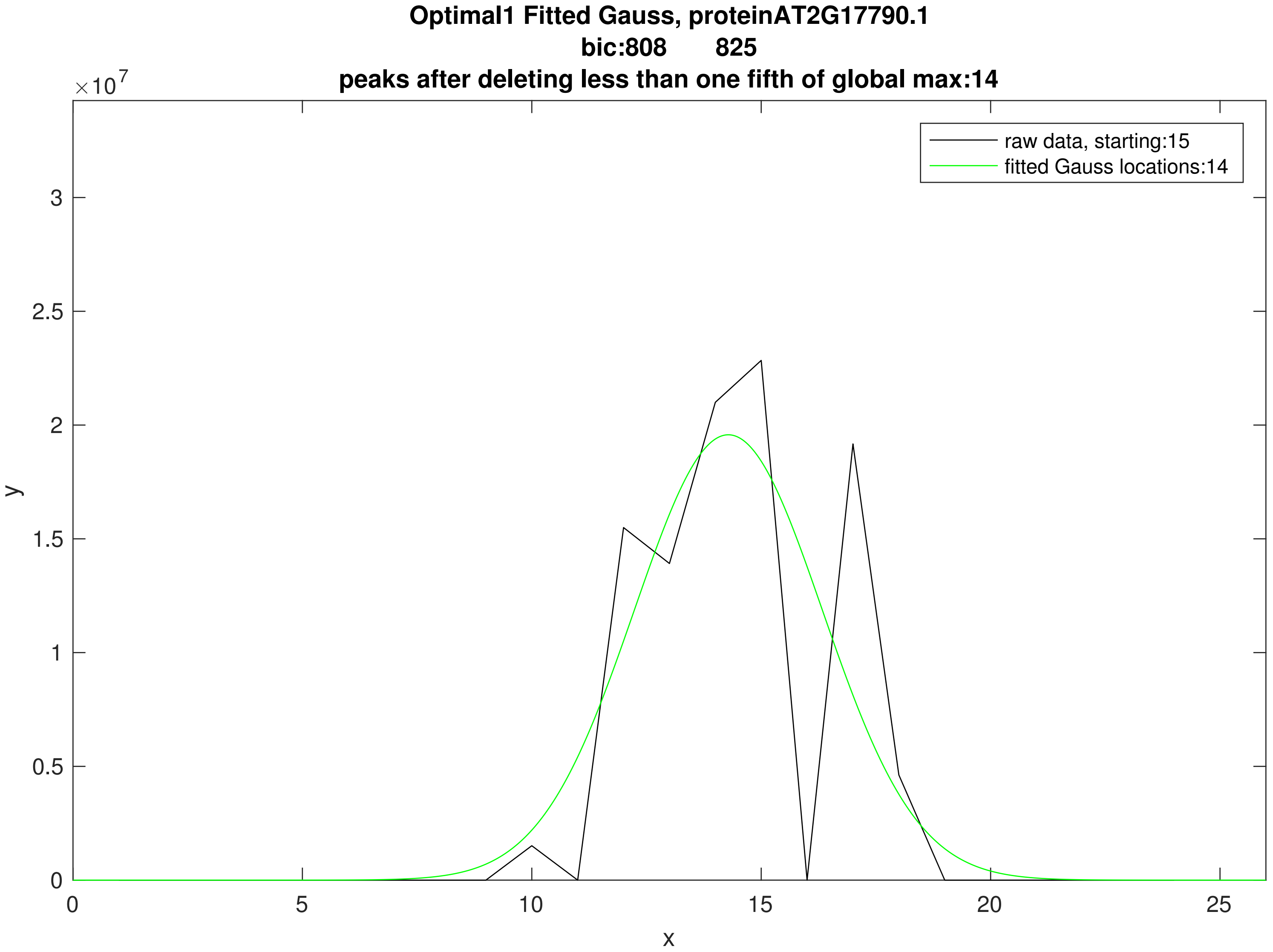

Optimal1 Fitted Gauss, proteinAT2G17790.1
bic:808 825
peaks after deleting less than one fifth of global max:14
×10^7
3
2.5
2
1.5
1
0.5
0
y
0
5
10
15
20
25
x
raw data, starting:15
fitted Gauss locations:14

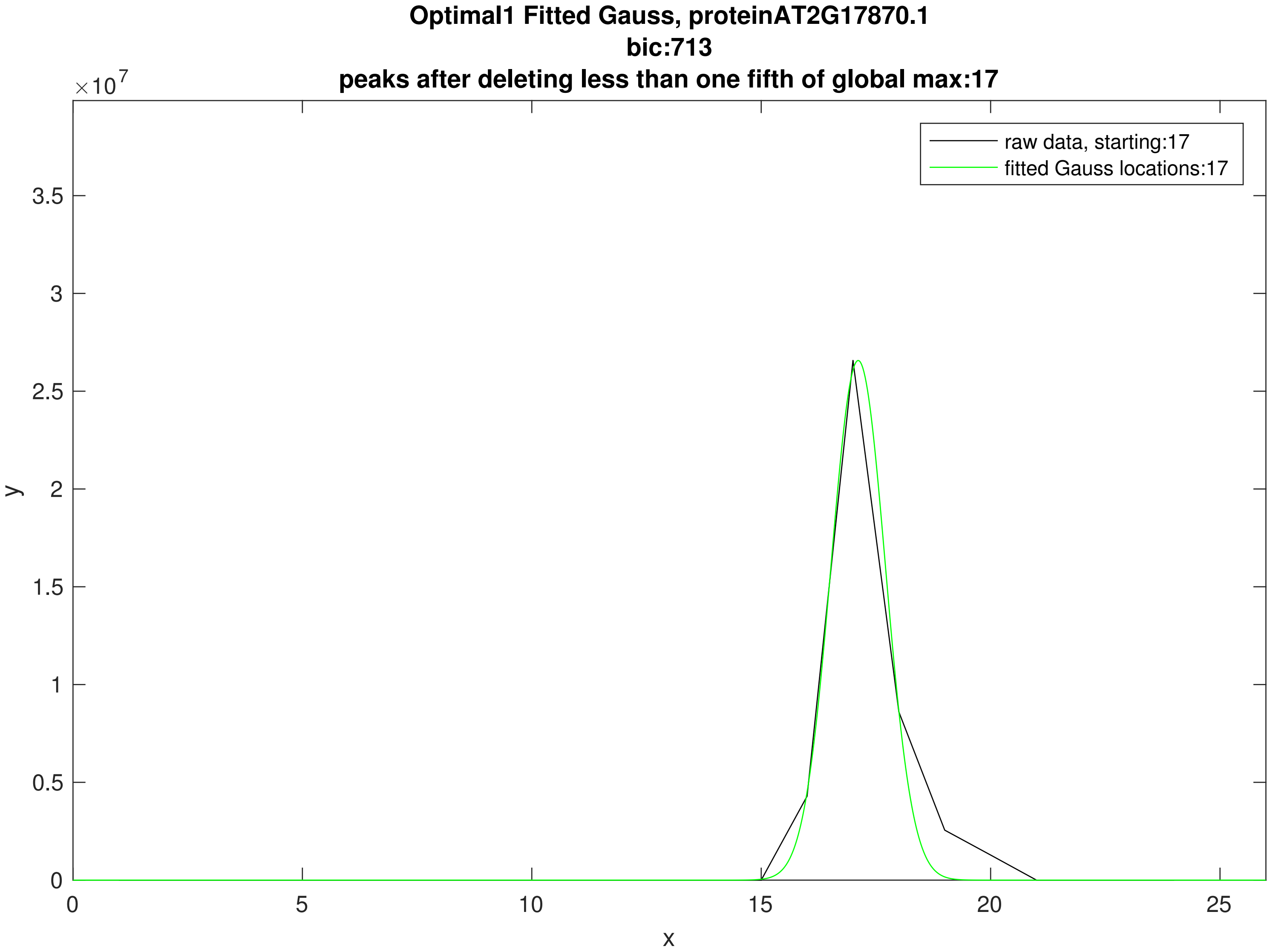
Optimal1 Fitted Gauss, proteinAT2G17870.1
bic:713
peaks after deleting less than one fifth of global max:17
×10^7
raw data, starting:17
fitted Gauss locations:17
y
x
0
0.5
1
1.5
2
2.5
3
3.5
0
5
10
15
20
25

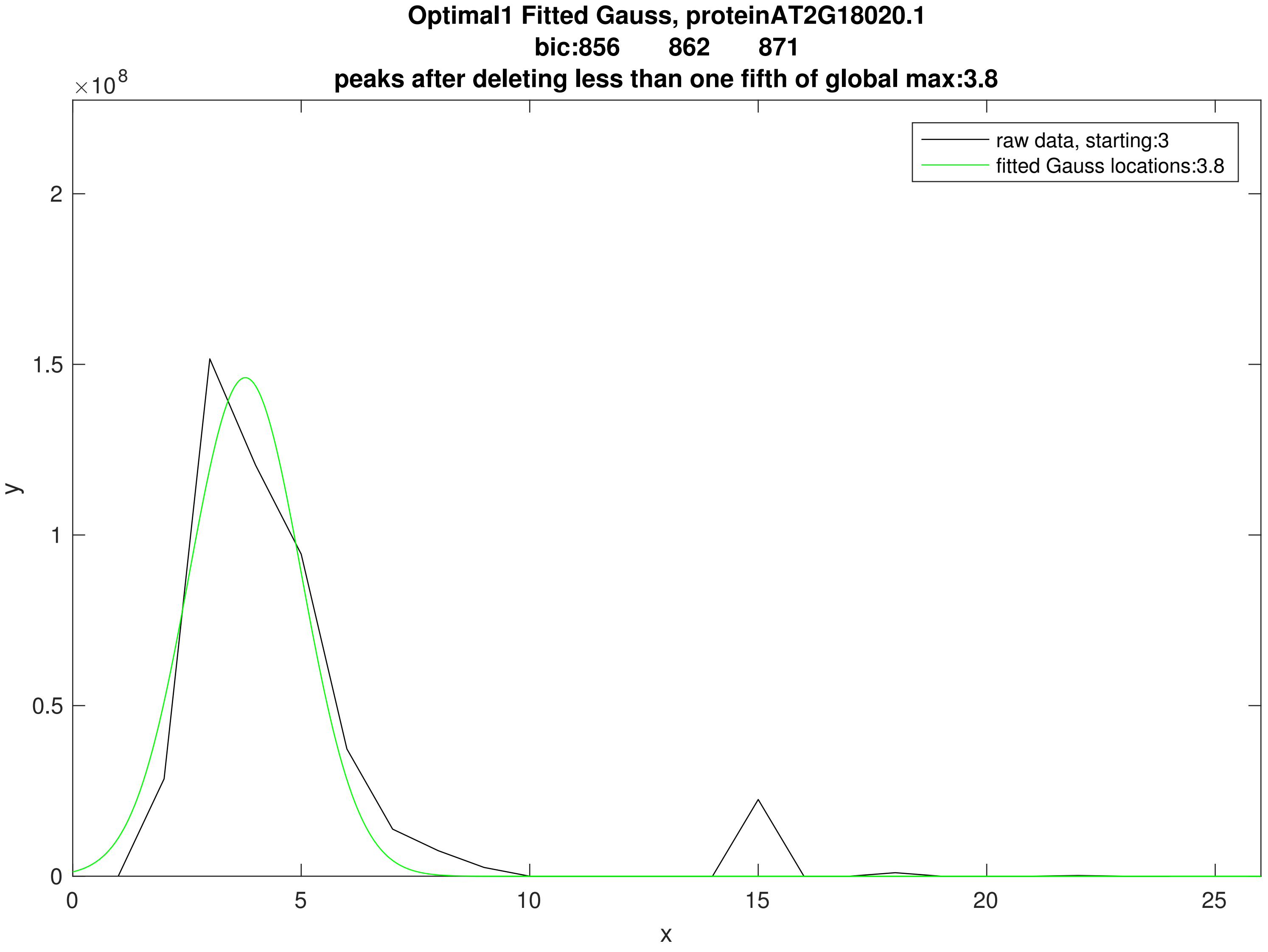

Optimal1 Fitted Gauss, proteinAT2G18020.1
bic:856 862 871
peaks after deleting less than one fifth of global max:3.8
×10^8
raw data, starting:3
fitted Gauss locations:3.8
y
x
0
0.5
1
1.5
2
0
5
10
15
20
25

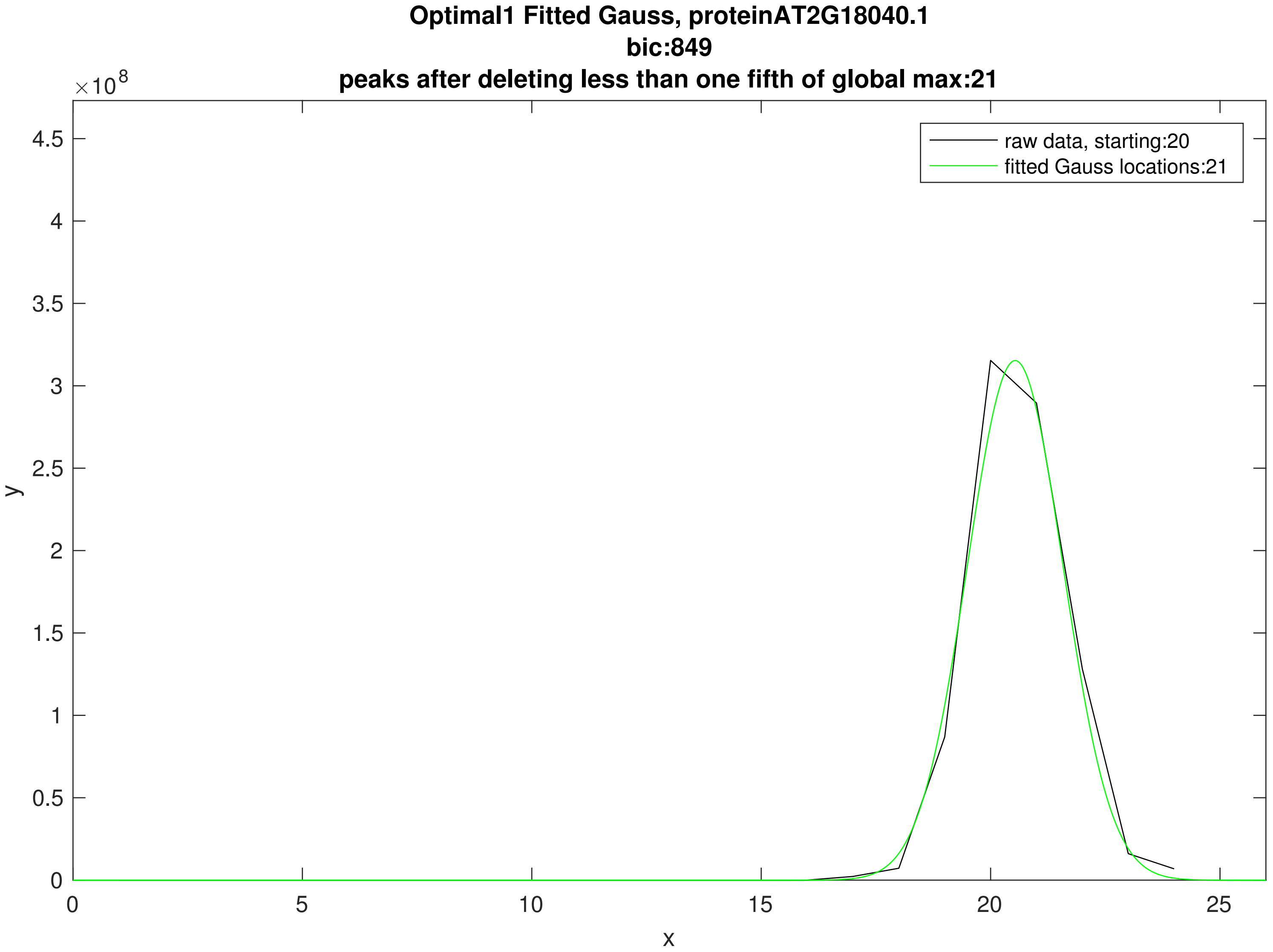
Optimal1 Fitted Gauss, proteinAT2G18040.1
bic:849
peaks after deleting less than one fifth of global max:21
×10^8
4.5
4
3.5
3
2.5
2
1.5
1
0.5
0
y
0
5
10
15
20
25
x
raw data, starting:20
fitted Gauss locations:21

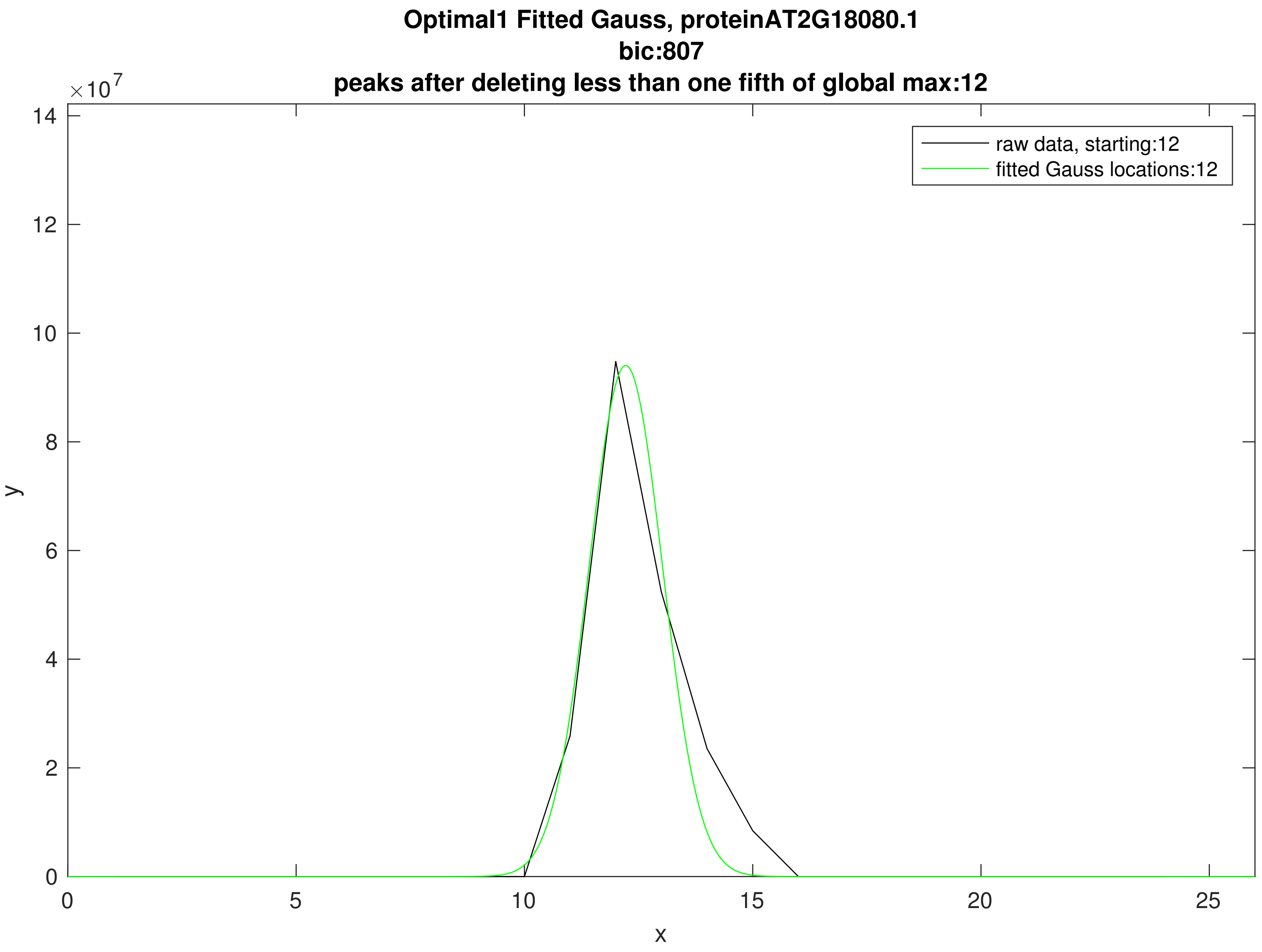

Optimal1 Fitted Gauss, proteinAT2G18080.1
bic:807
peaks after deleting less than one fifth of global max:12
×10^7
14
12
10
8
6
4
2
0
y
0
5
10
15
20
25
x
raw data, starting:12
fitted Gauss locations:12

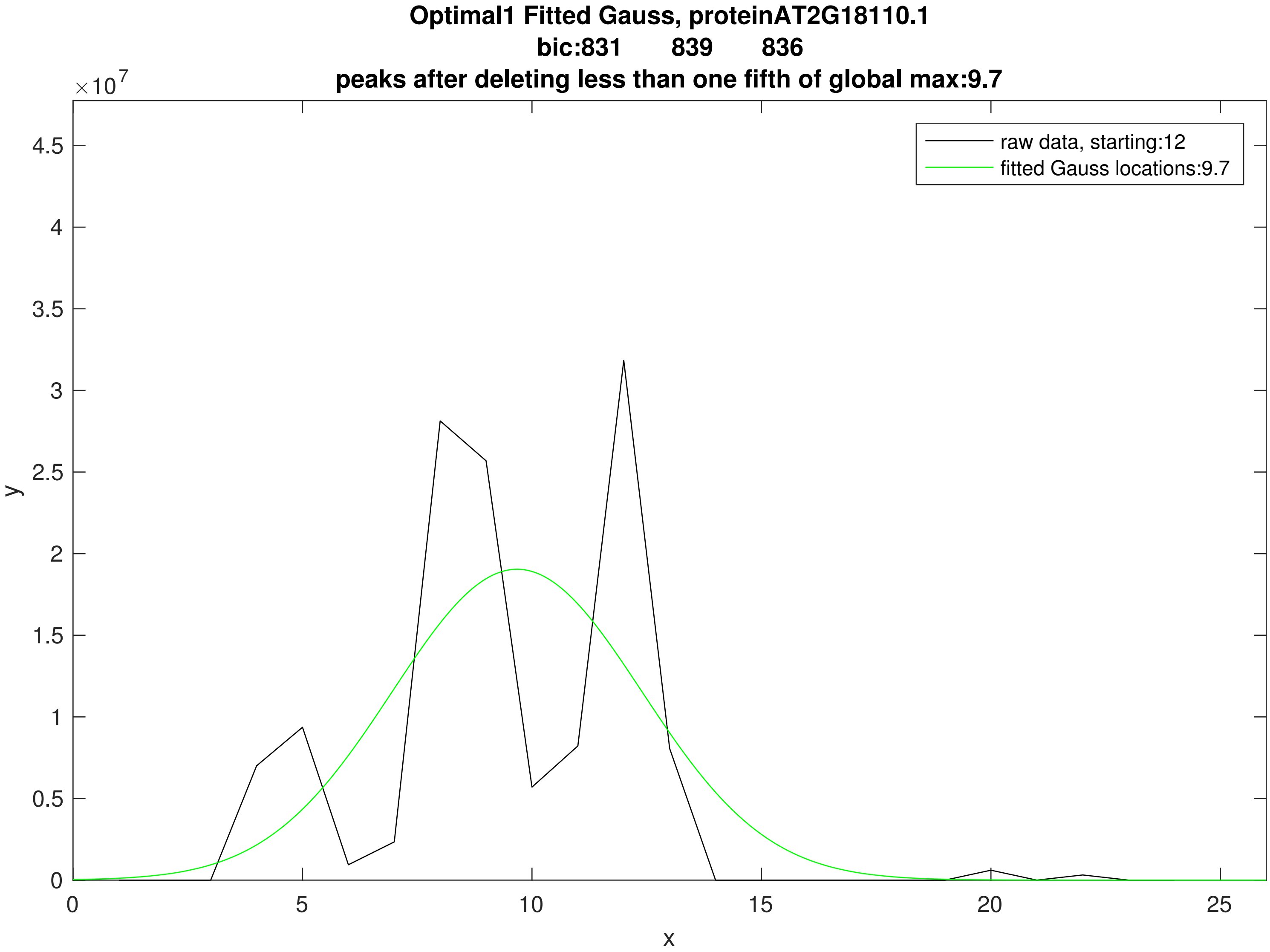
Optimal1 Fitted Gauss, proteinAT2G18110.1
bic:831 839 836
peaks after deleting less than one fifth of global max:9.7
×10^7
raw data, starting:12
fitted Gauss locations:9.7
y
x
0
0.5
1
1.5
2
2.5
3
3.5
4
4.5
5
10
15
20
25

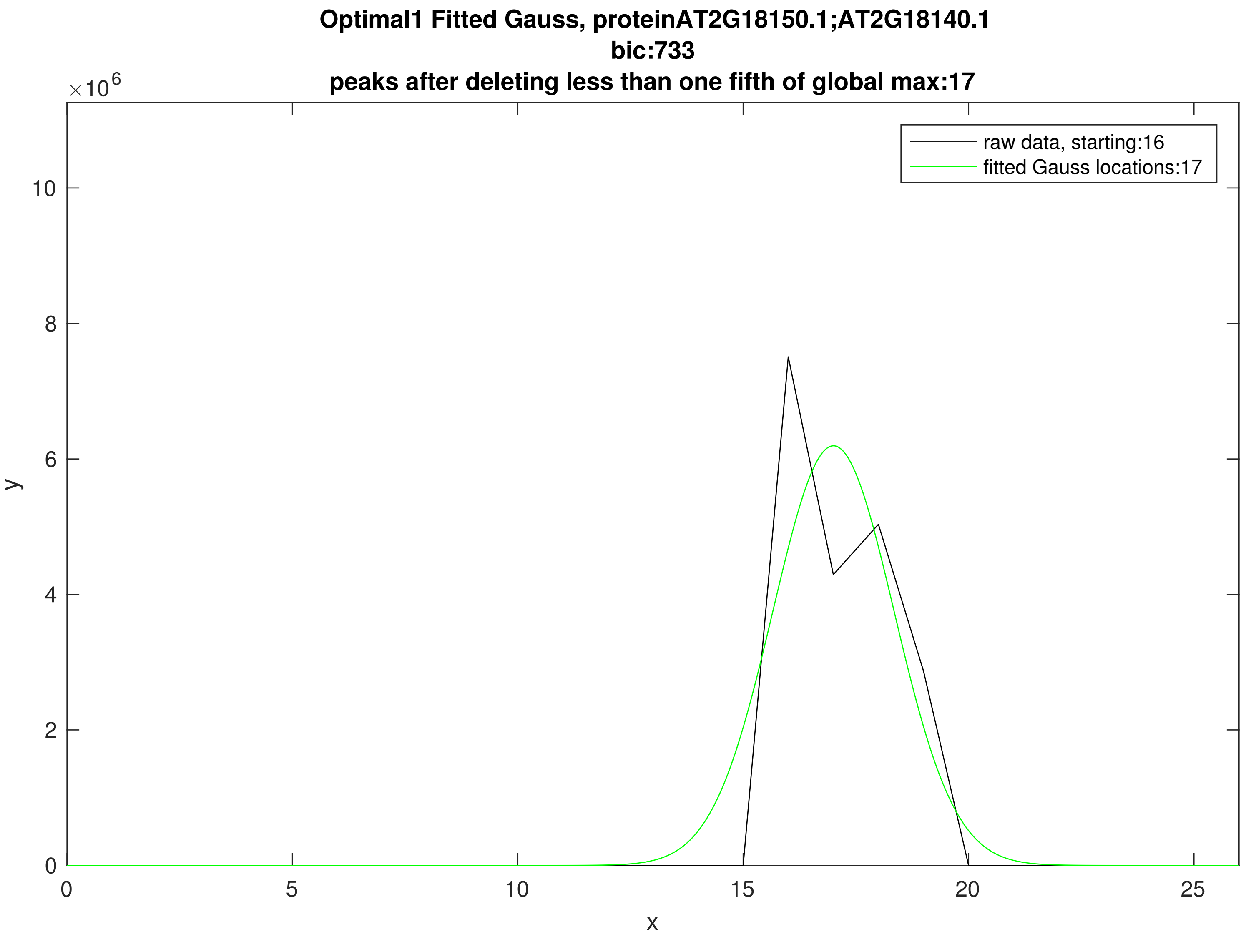
Optimal1 Fitted Gauss, proteinAT2G18150.1;AT2G18140.1
bic:733
peaks after deleting less than one fifth of global max:17
×10^6
10
8
6
4
2
0
y
0
5
10
15
20
25
x
raw data, starting:16
fitted Gauss locations:17

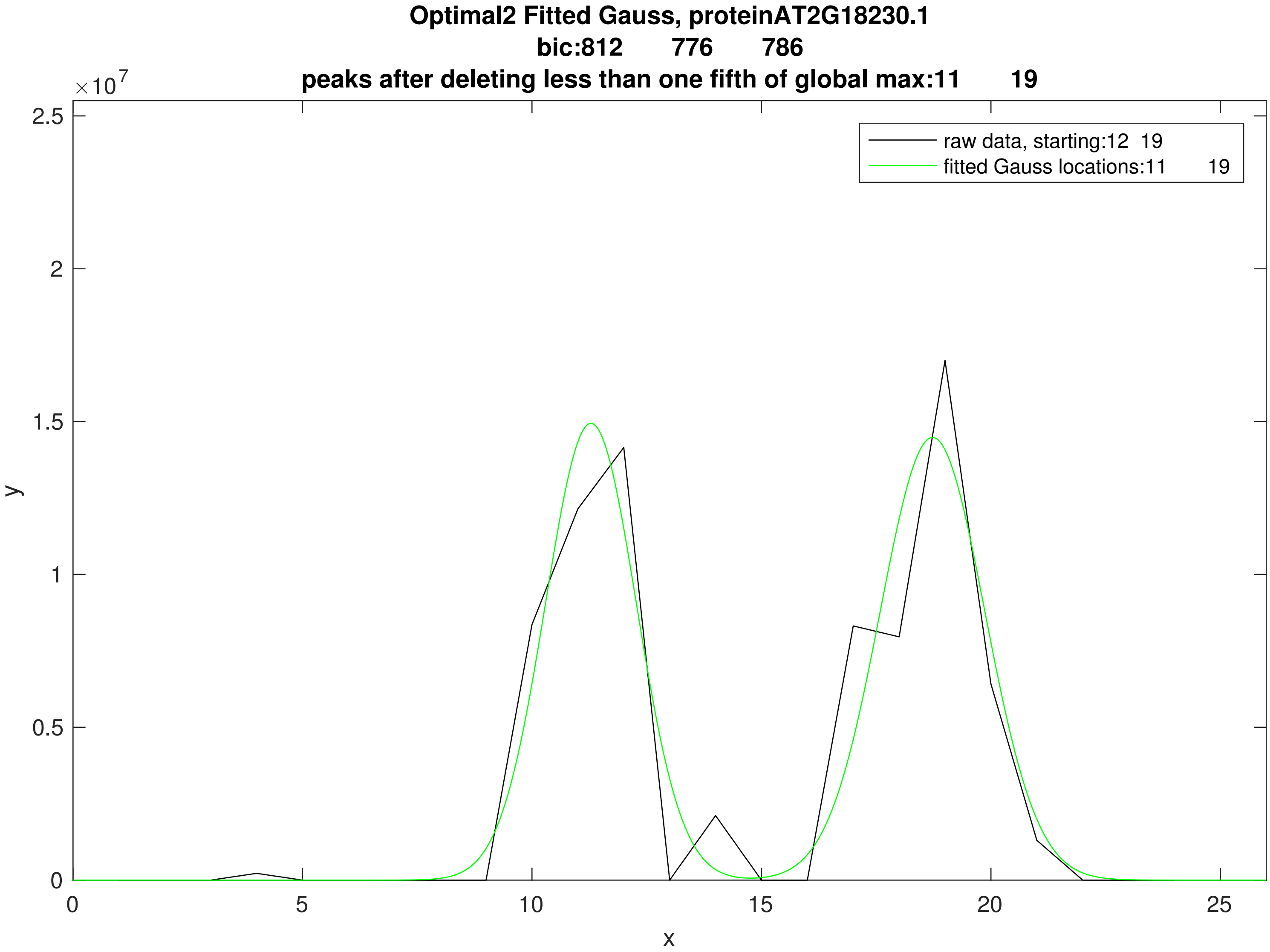

Optimal2 Fitted Gauss, proteinAT2G18230.1
bic:812 776 786
peaks after deleting less than one fifth of global max:11 19
×10^7
2.5
2
1.5
1
0.5
0
y
0
5
10
15
20
25
x
raw data, starting:12 19
fitted Gauss locations:11 19

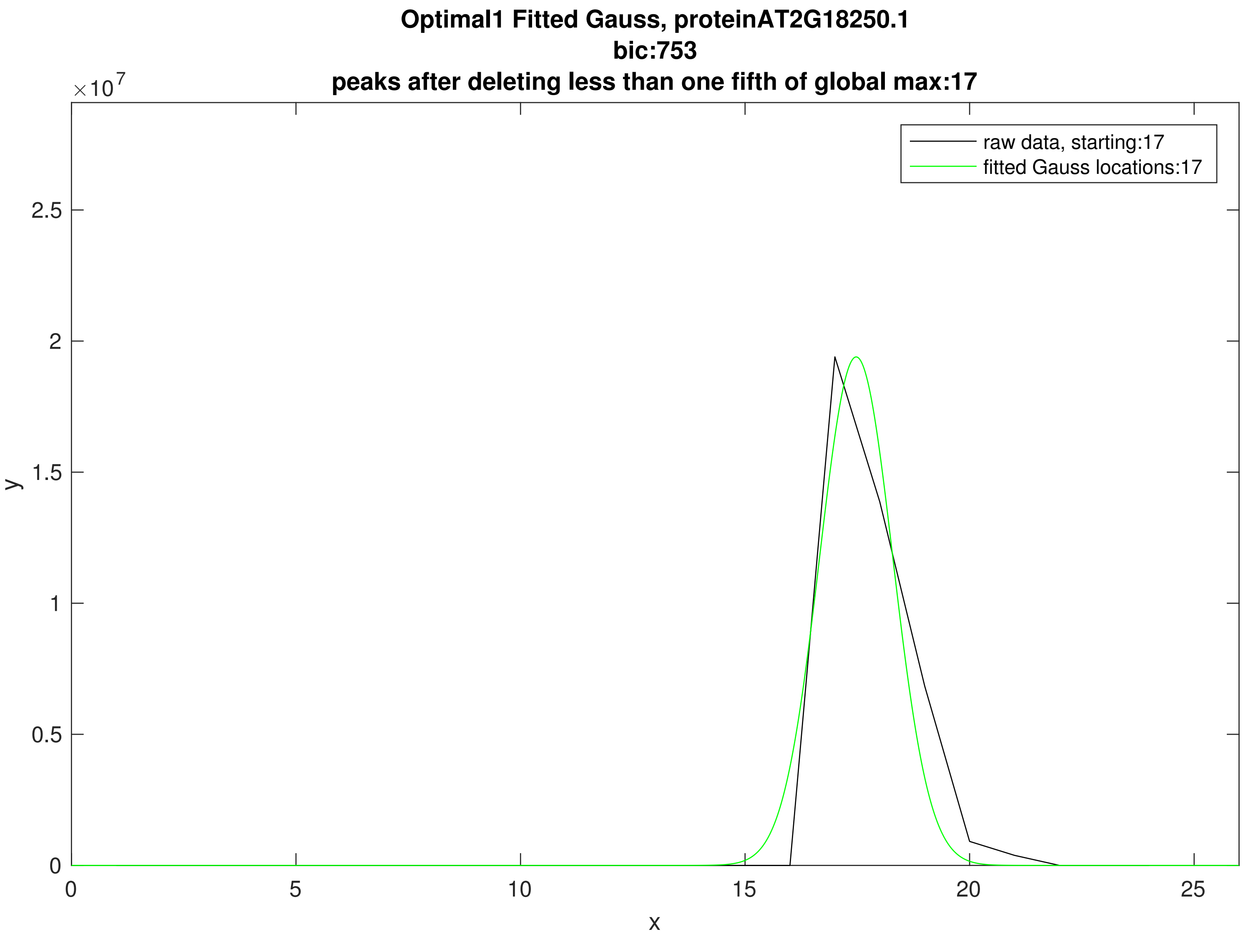
Optimal1 Fitted Gauss, proteinAT2G18250.1
bic:753
peaks after deleting less than one fifth of global max:17
×10^7
raw data, starting:17
fitted Gauss locations:17
y
x
0
0.5
1
1.5
2
2.5
5
10
15
20
25

**Optimal1 Fitted Gauss, proteinAT2G18370.1**
**bic:646**
**peaks after deleting less than one fifth of global max:21**

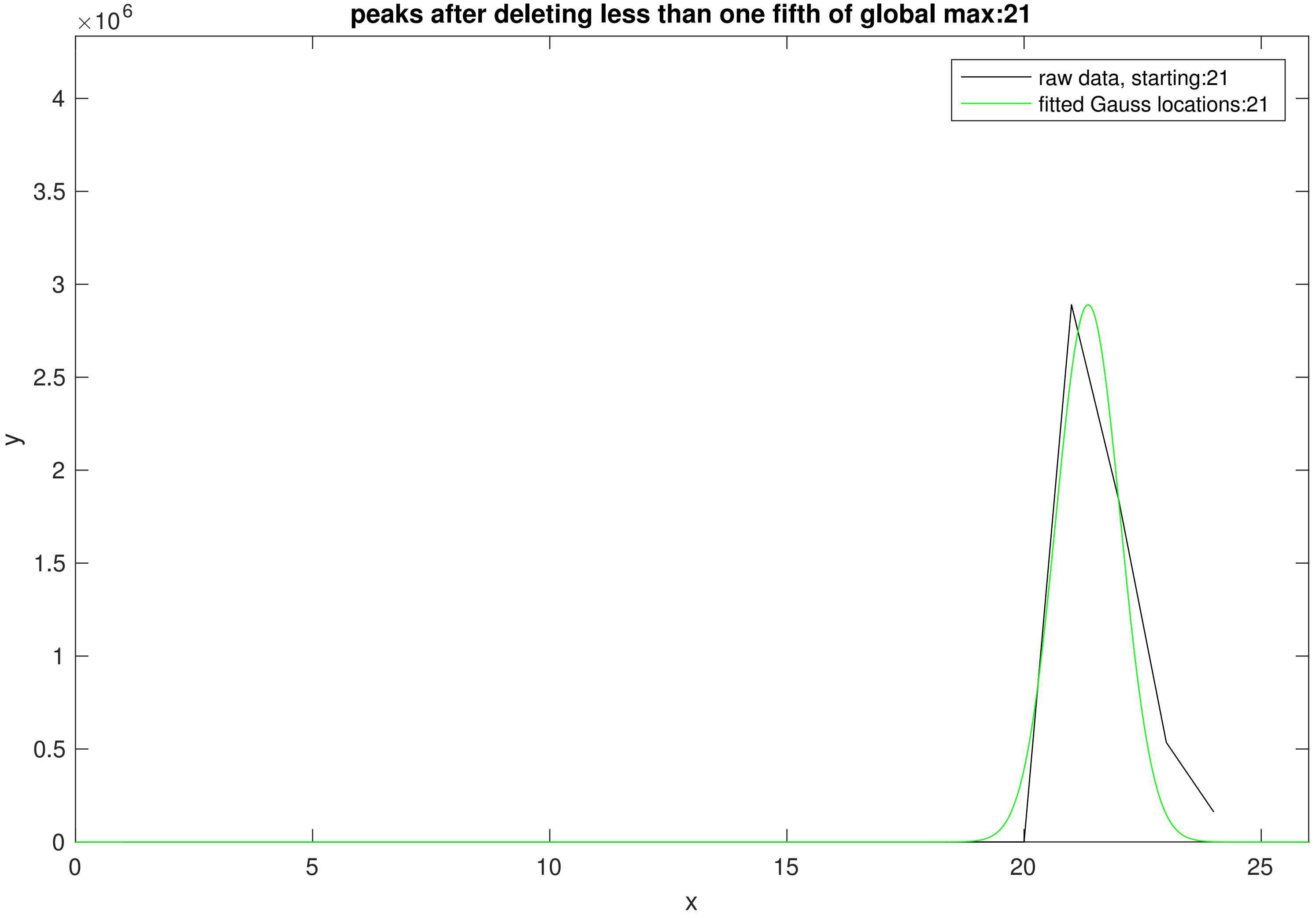

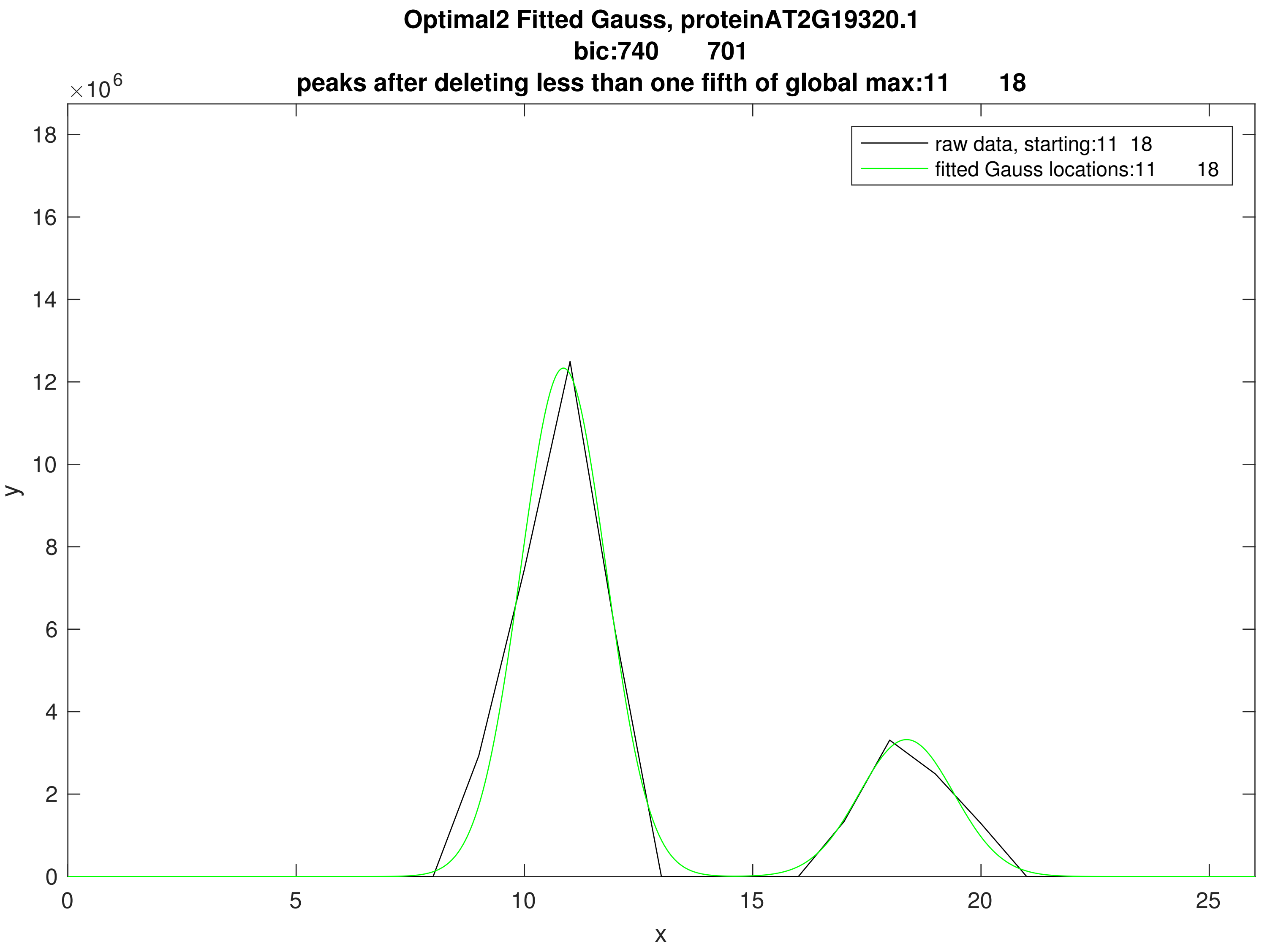

Optimal2 Fitted Gauss, proteinAT2G19320.1
bic:740 701
peaks after deleting less than one fifth of global max:11 18
×10^6
raw data, starting:11 18
fitted Gauss locations:11 18
y
x
0
2
4
6
8
10
12
14
16
18
0
5
10
15
20
25

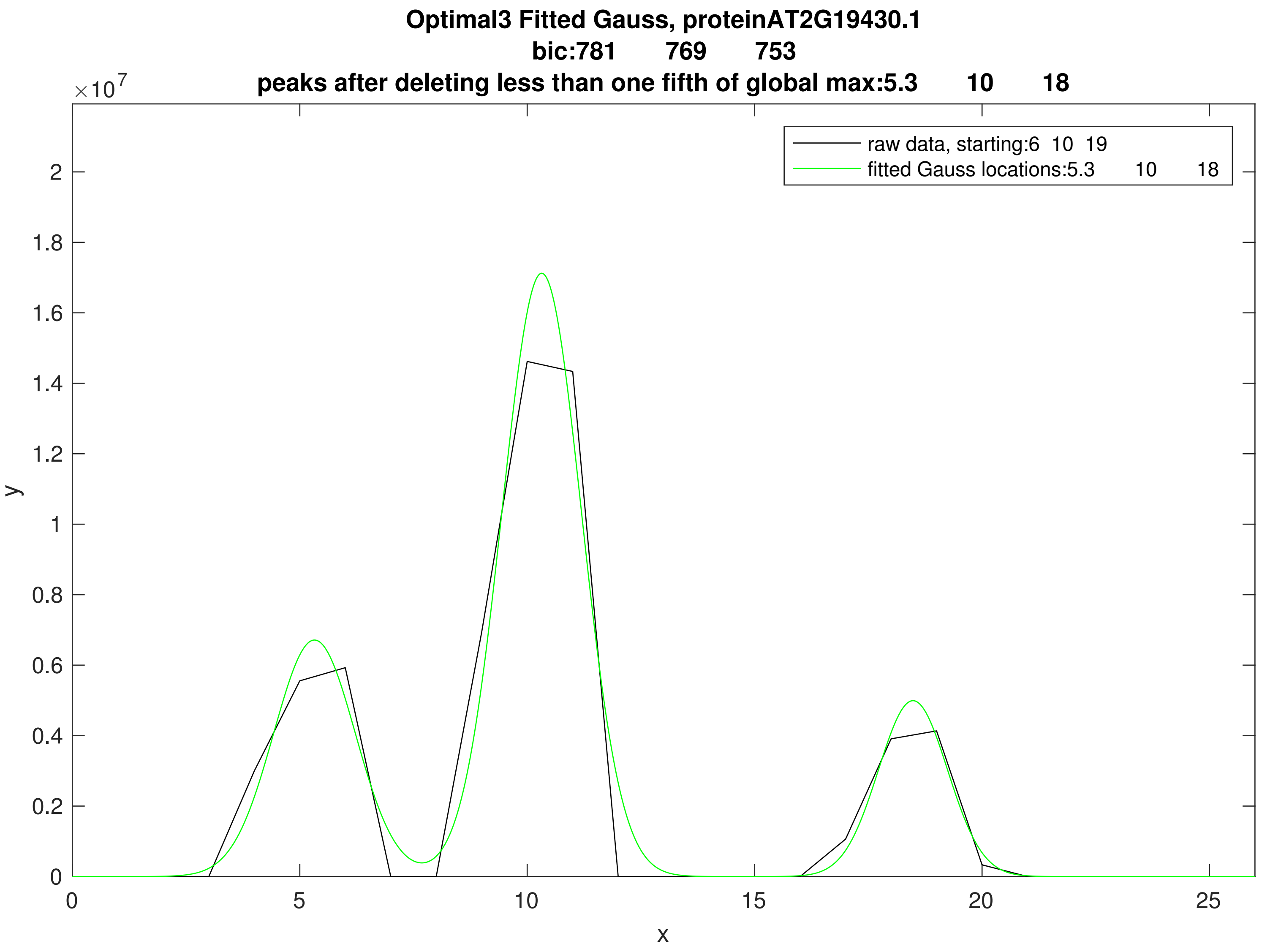

Optimal3 Fitted Gauss, proteinAT2G19430.1
bic:781 769 753
peaks after deleting less than one fifth of global max:5.3 10 18
×10^7
raw data, starting:6 10 19
fitted Gauss locations:5.3 10 18
y
x
0
0.2
0.4
0.6
0.8
1
1.2
1.4
1.6
1.8
2
0
5
10
15
20
25

**Optimal1 Fitted Gauss, proteinAT2G19480.3;AT2G19480.2;AT2G19480.1**
**bic:794 795 828**
**peaks after deleting less than one fifth of global max:13**

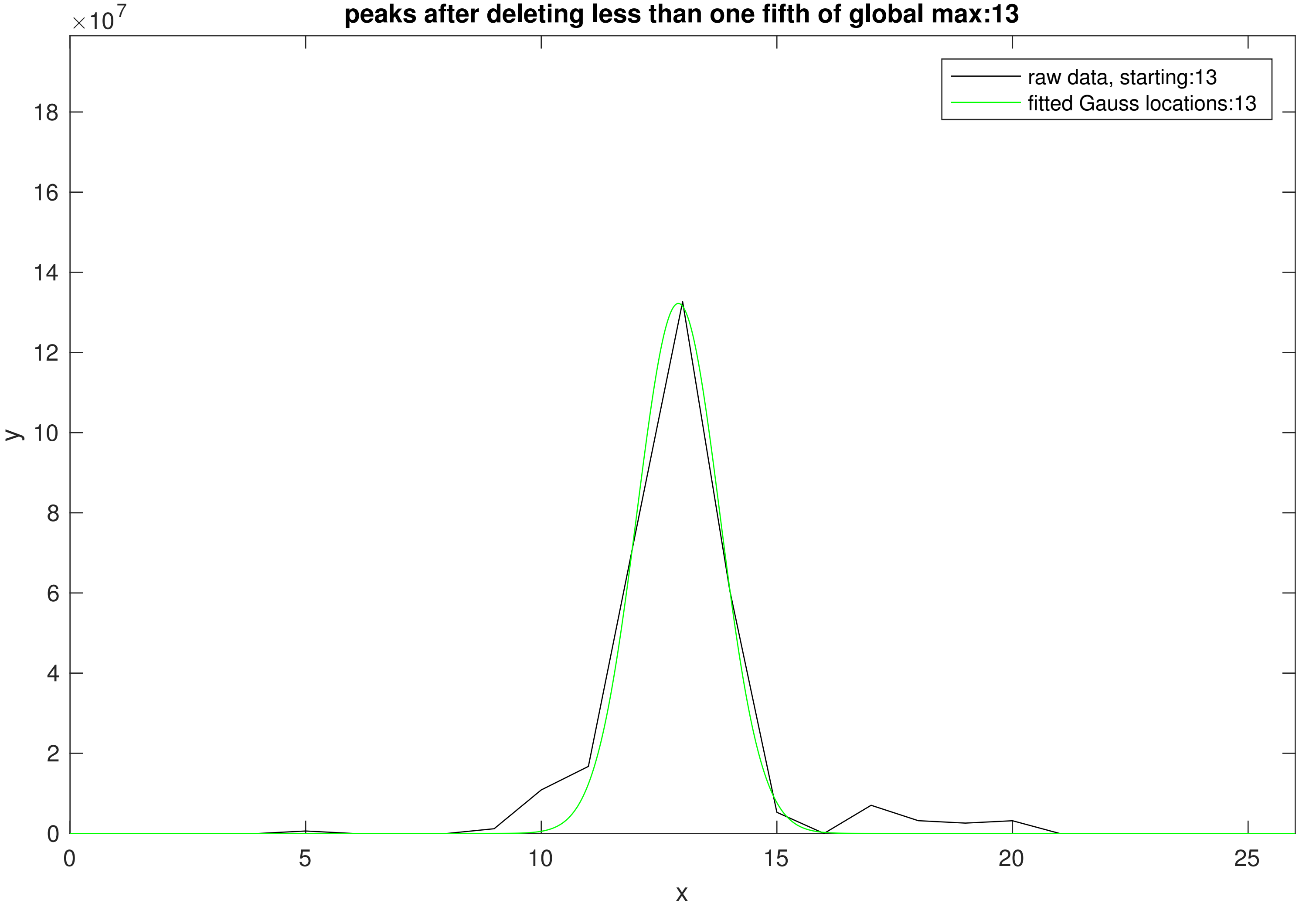

**Optimal1 Fitted Gauss, proteinAT2G19520.1**
**bic:870 880 889**
**peaks after deleting less than one fifth of global max:15**

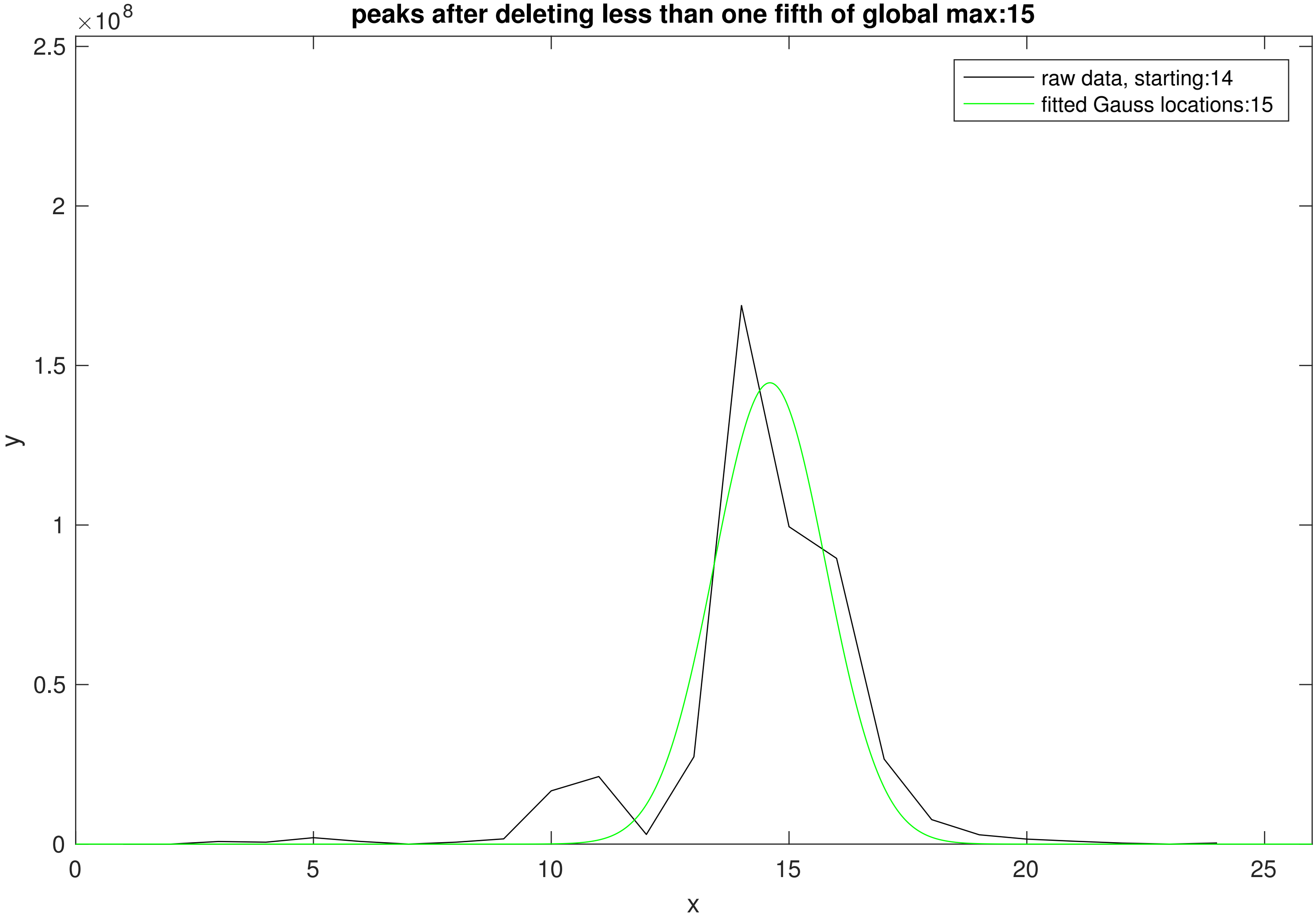

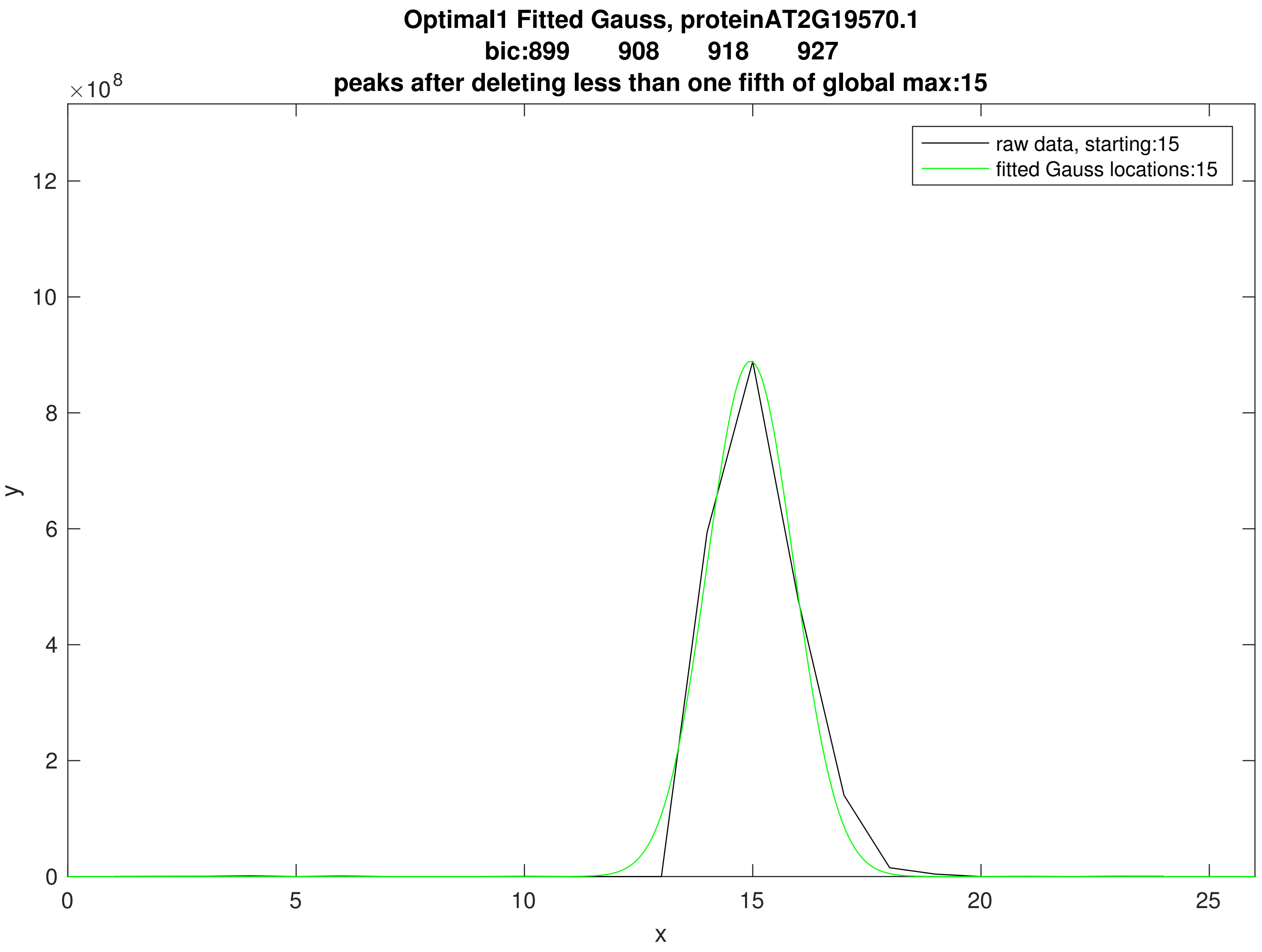

Optimal1 Fitted Gauss, proteinAT2G19570.1
bic:899 908 918 927
peaks after deleting less than one fifth of global max:15
×10^8
12
10
8
6
4
2
0
y
0
5
10
15
20
25
x
raw data, starting:15
fitted Gauss locations:15

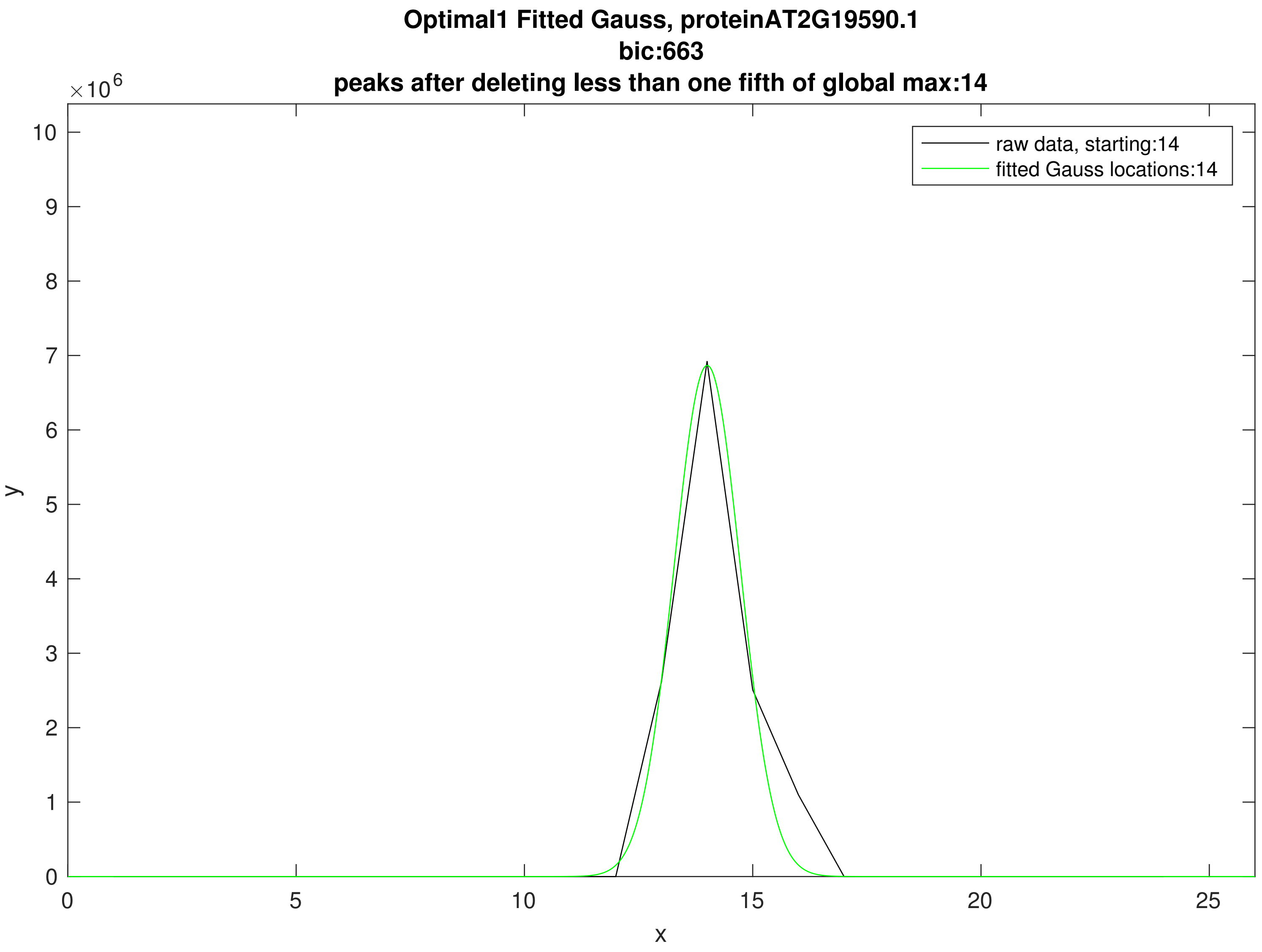
Optimal1 Fitted Gauss, proteinAT2G19590.1
bic:663
peaks after deleting less than one fifth of global max:14
×10^6
raw data, starting:14
fitted Gauss locations:14
y
x

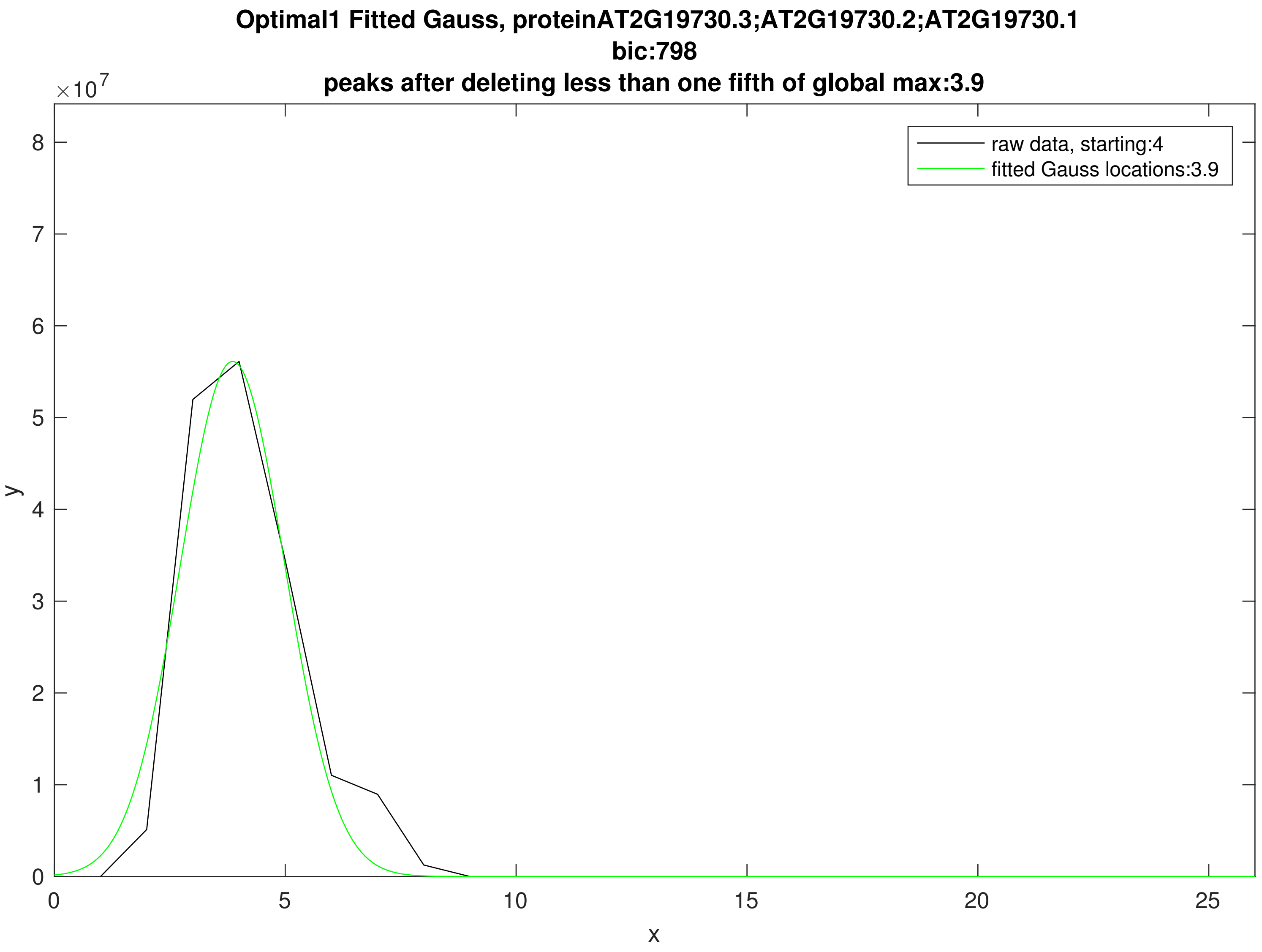

Optimal1 Fitted Gauss, proteinAT2G19730.3;AT2G19730.2;AT2G19730.1
bic:798
peaks after deleting less than one fifth of global max:3.9
×10^7
raw data, starting:4
fitted Gauss locations:3.9
y
x
0
1
2
3
4
5
6
7
8
0
5
10
15
20
25

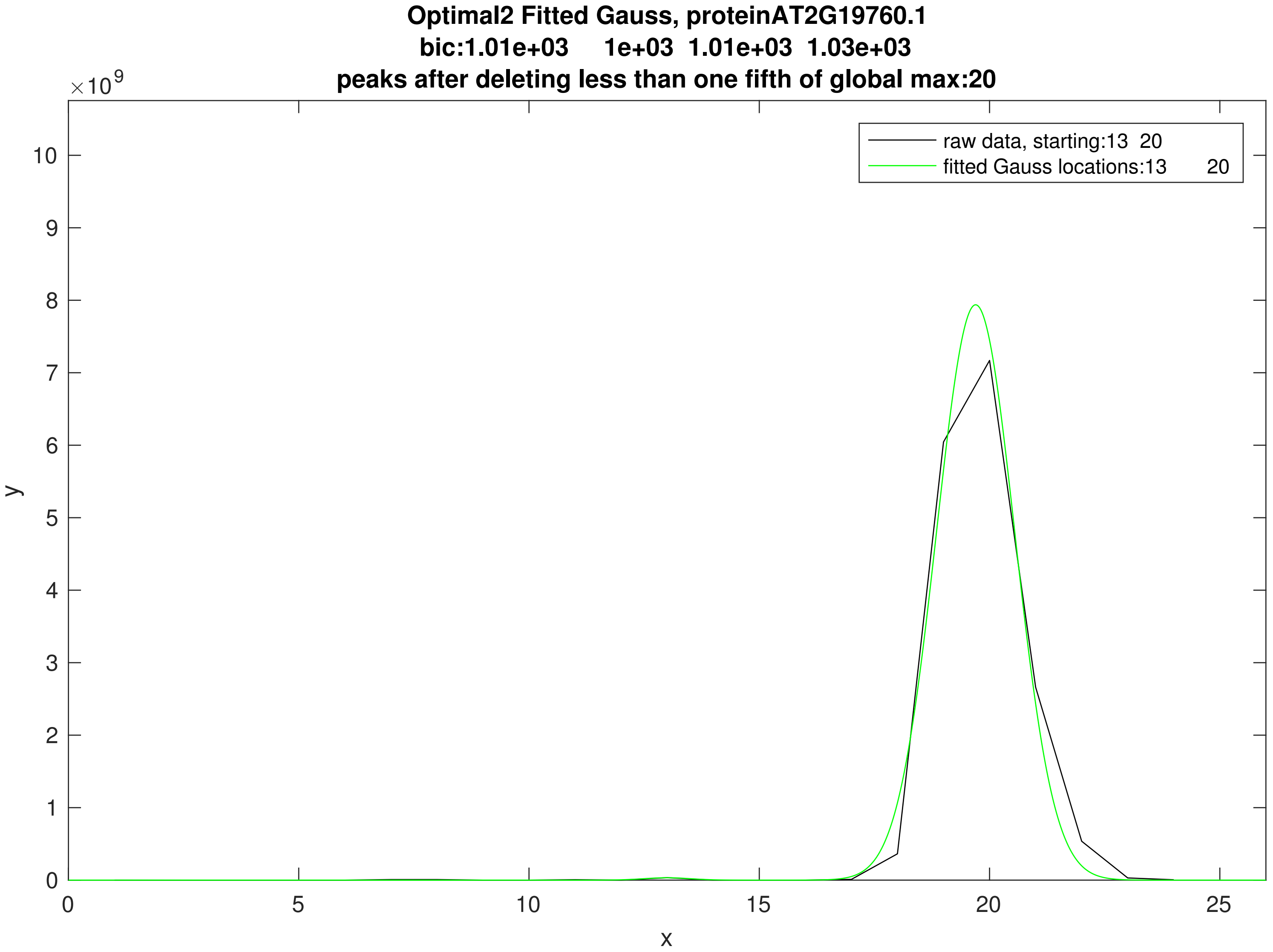
Optimal2 Fitted Gauss, proteinAT2G19760.1
bic:1.01e+03 1e+03 1.01e+03 1.03e+03
peaks after deleting less than one fifth of global max:20
×10^9
raw data, starting:13 20
fitted Gauss locations:13 20
y
x
0
1
2
3
4
5
6
7
8
9
10
0
5
10
15
20
25

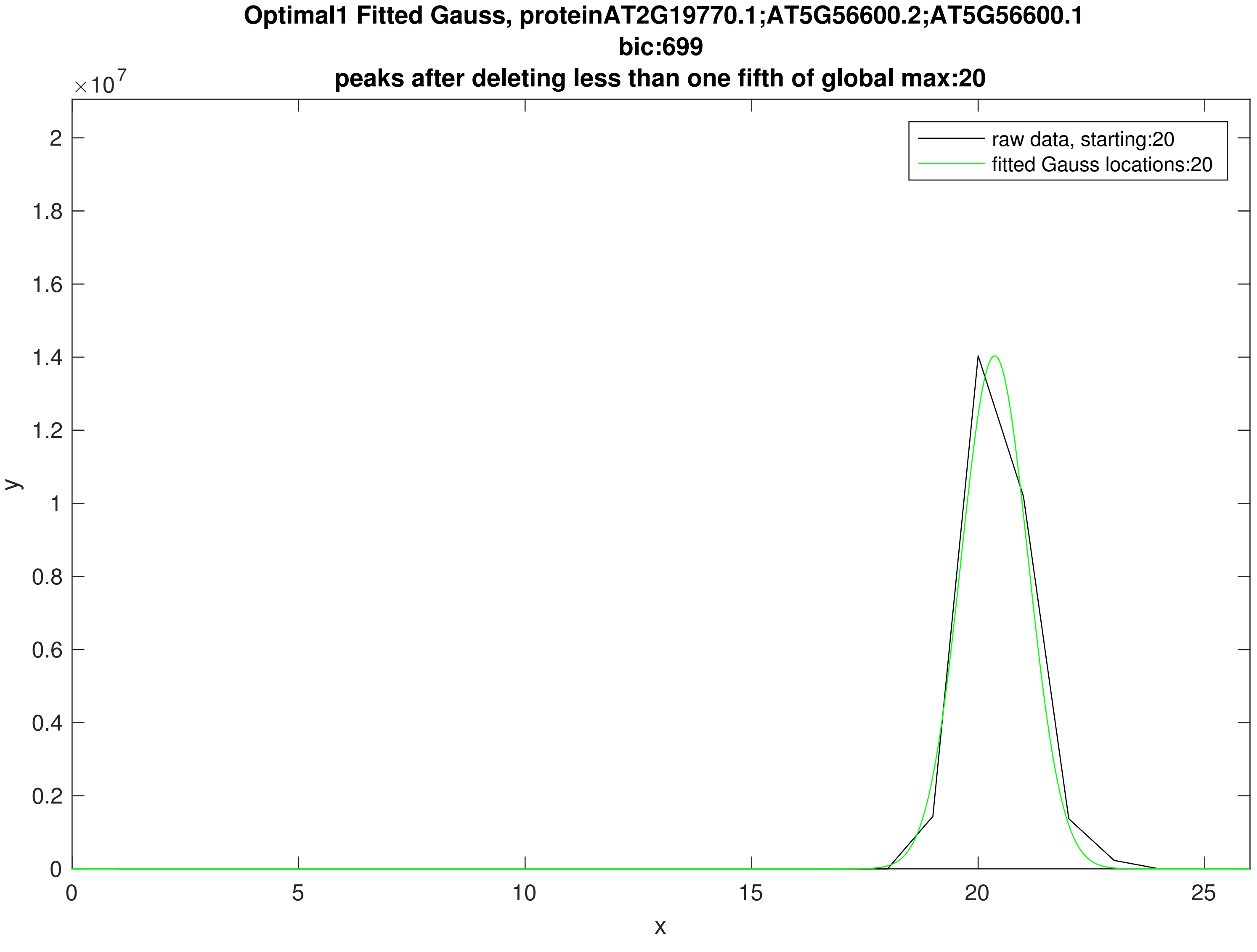

Optimal1 Fitted Gauss, proteinAT2G19770.1;AT5G56600.2;AT5G56600.1
bic:699
peaks after deleting less than one fifth of global max:20
×10^7
2
1.8
1.6
1.4
1.2
1
0.8
0.6
0.4
0.2
0
y
0
5
10
15
20
25
x
raw data, starting:20
fitted Gauss locations:20

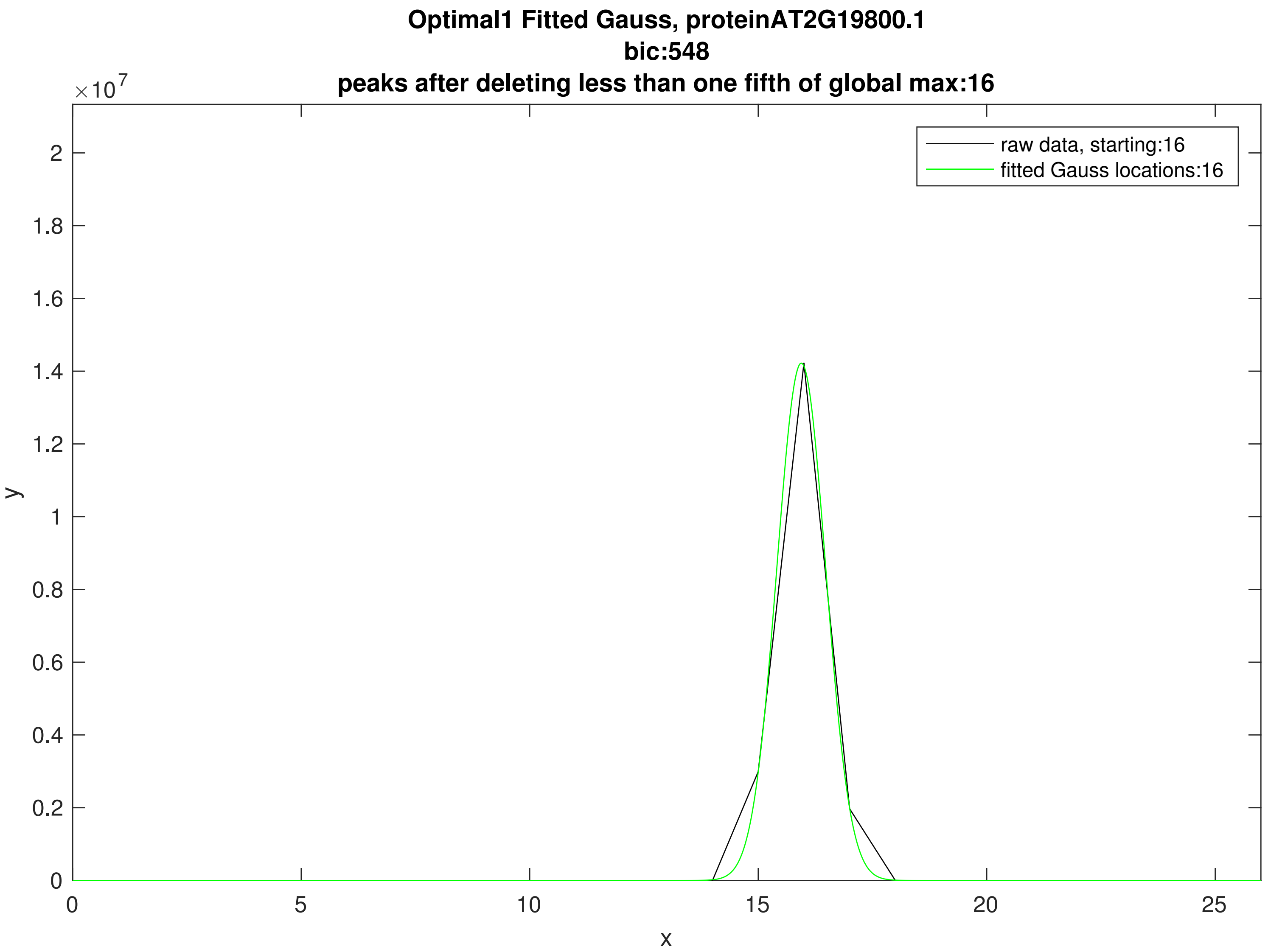
Optimal1 Fitted Gauss, proteinAT2G19800.1
bic:548
peaks after deleting less than one fifth of global max:16
×10^7
raw data, starting:16
fitted Gauss locations:16
y
x
0
0.2
0.4
0.6
0.8
1
1.2
1.4
1.6
1.8
2
0
5
10
15
20
25

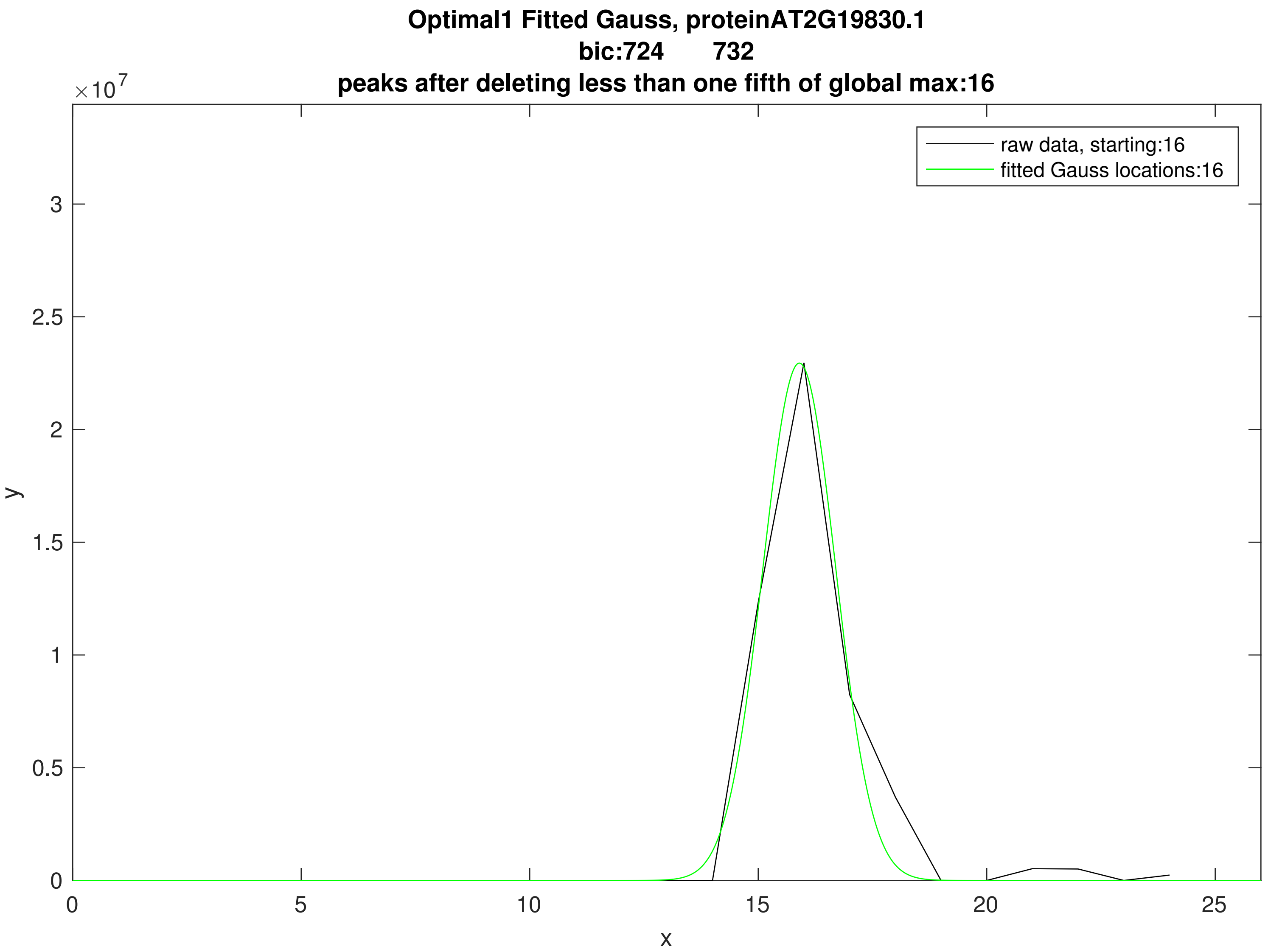

Optimal1 Fitted Gauss, proteinAT2G19830.1
bic:724 732
peaks after deleting less than one fifth of global max:16
×10^7
raw data, starting:16
fitted Gauss locations:16
0
0.5
1
1.5
2
2.5
3
y
0
5
10
15
20
25
x

**Optimal1 Fitted Gauss, proteinAT2G19900.1**
**bic:918 927 936 946**
**peaks after deleting less than one fifth of global max:9**

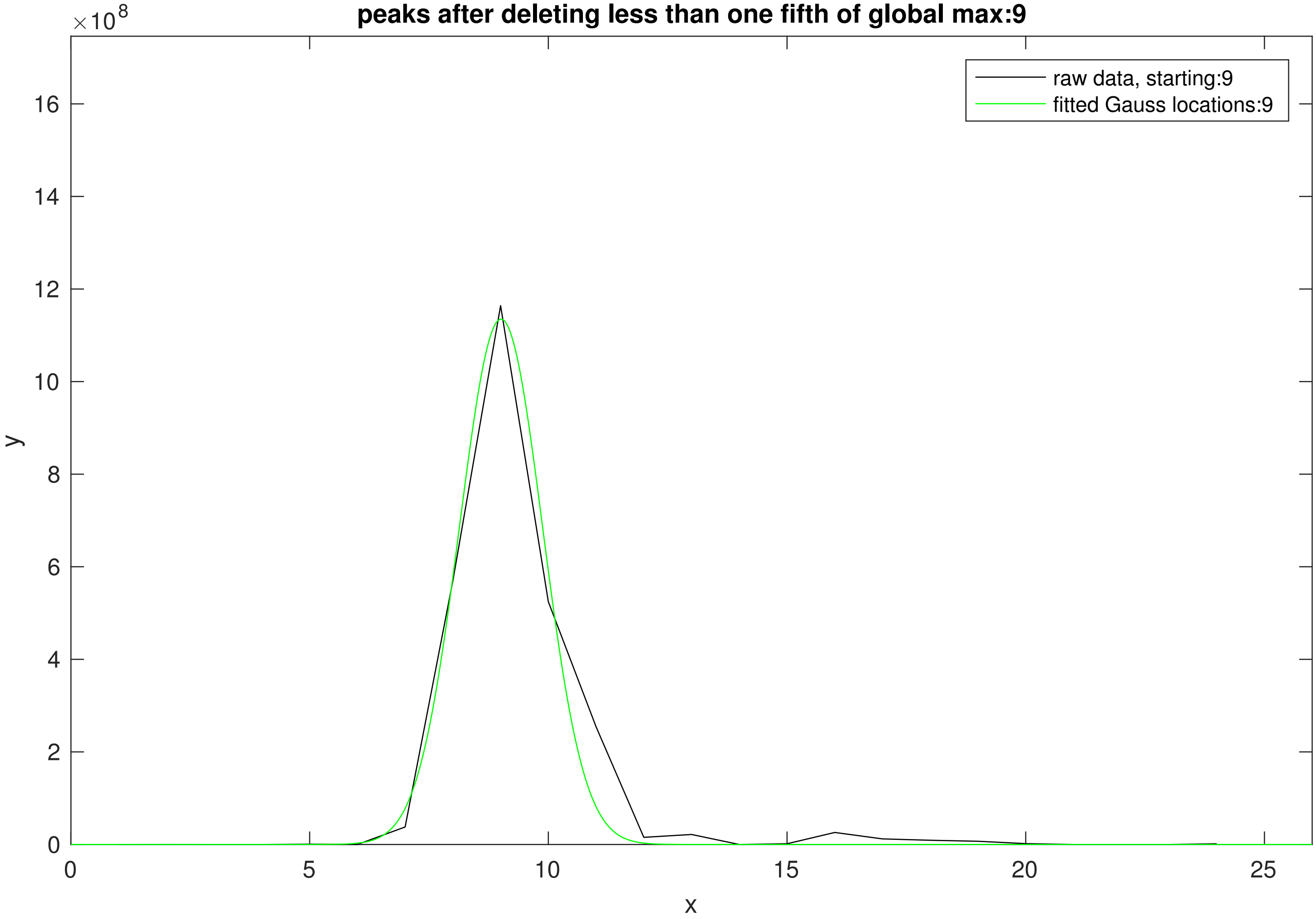

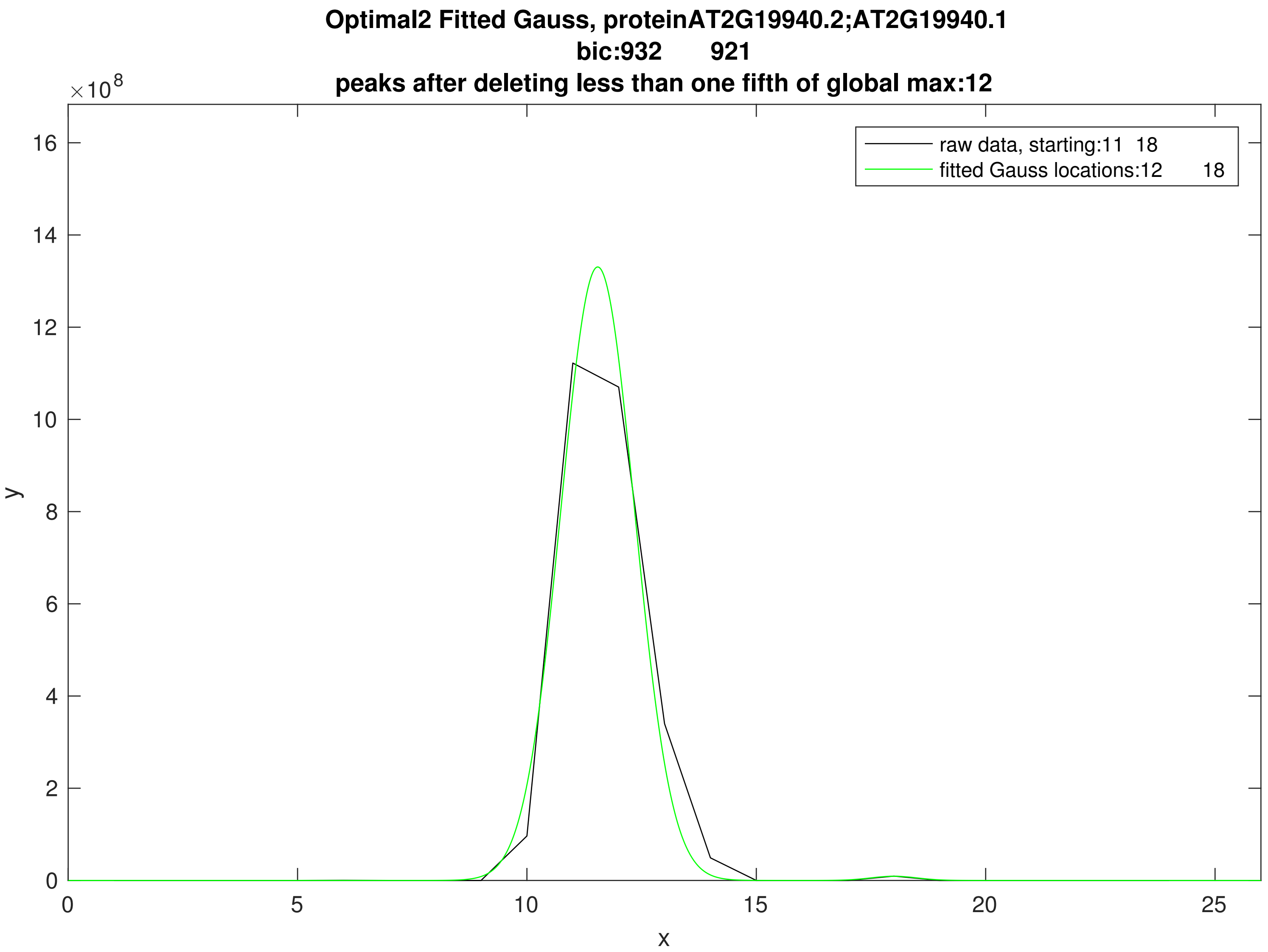
Optimal2 Fitted Gauss, proteinAT2G19940.2;AT2G19940.1
bic:932 921
peaks after deleting less than one fifth of global max:12
×10^8
raw data, starting:11 18
fitted Gauss locations:12 18
y
x
0
2
4
6
8
10
12
14
16
0
5
10
15
20
25

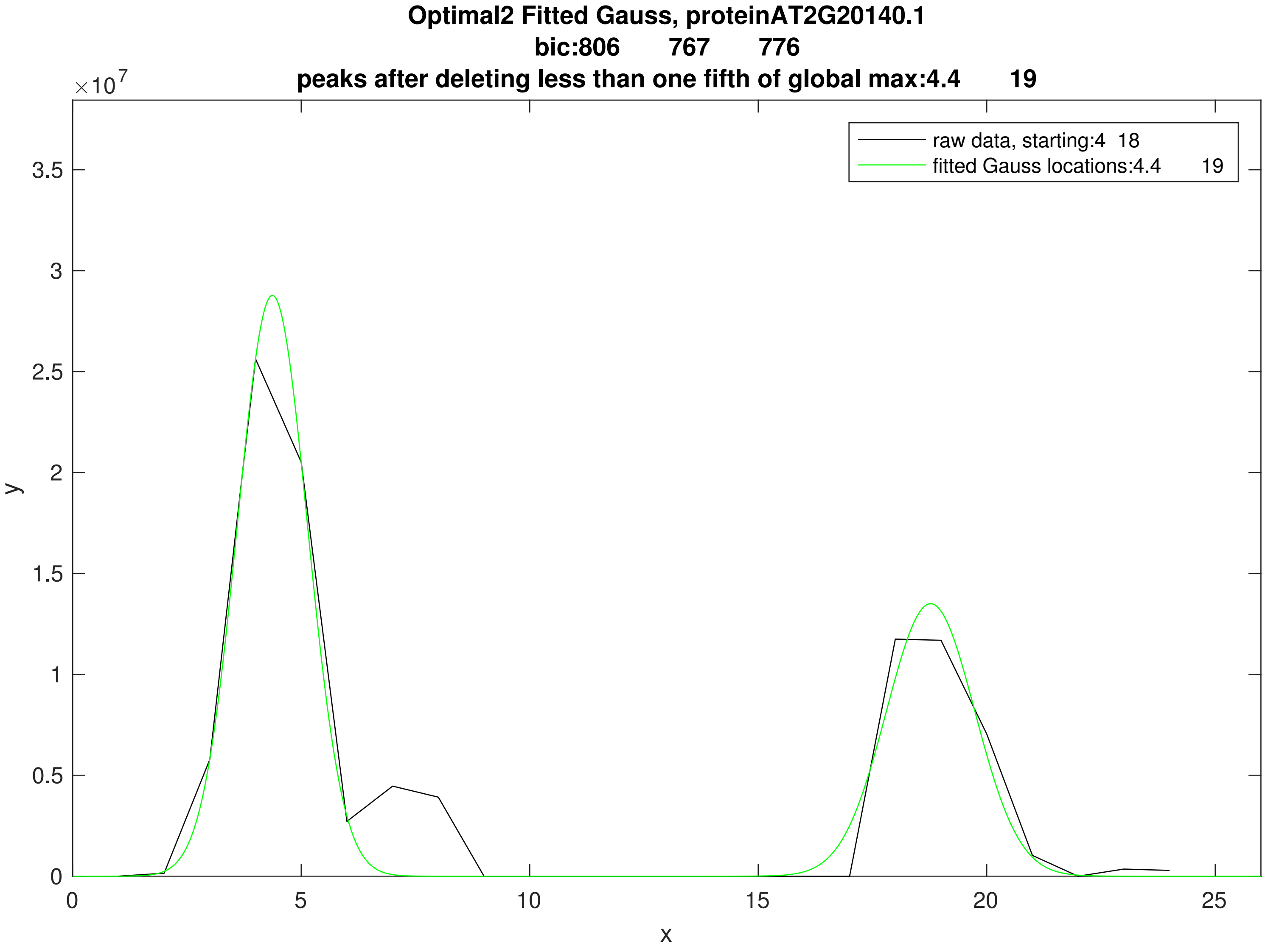
Optimal2 Fitted Gauss, proteinAT2G20140.1
bic:806 767 776
peaks after deleting less than one fifth of global max:4.4 19
×10^7
raw data, starting:4 18
fitted Gauss locations:4.4 19
y
x
0
0.5
1
1.5
2
2.5
3
3.5
5
10
15
20
25

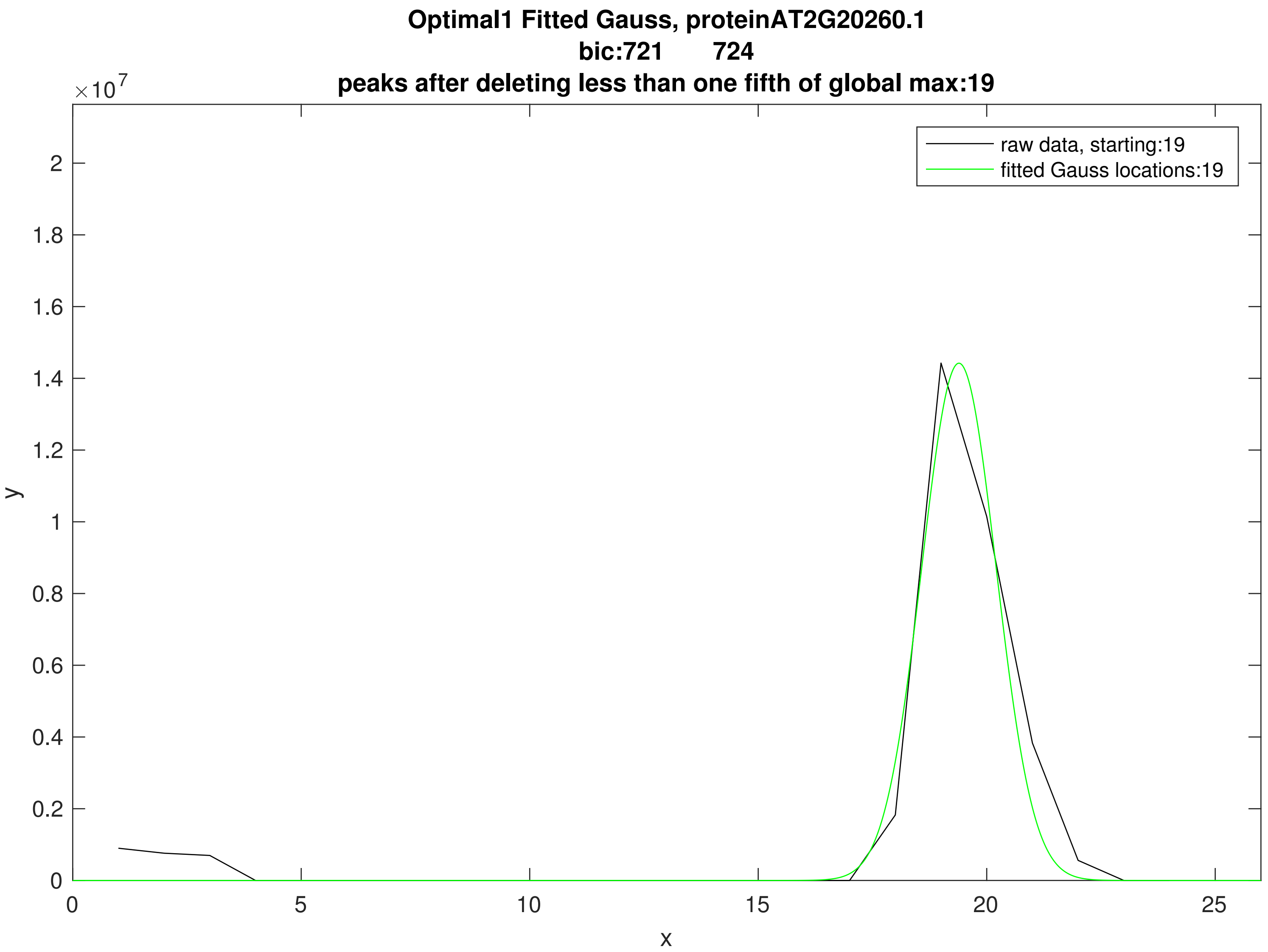

Optimal1 Fitted Gauss, proteinAT2G20260.1
bic:721 724
peaks after deleting less than one fifth of global max:19
×10^7
raw data, starting:19
fitted Gauss locations:19
y
x
0
0.2
0.4
0.6
0.8
1
1.2
1.4
1.6
1.8
2
0
5
10
15
20
25

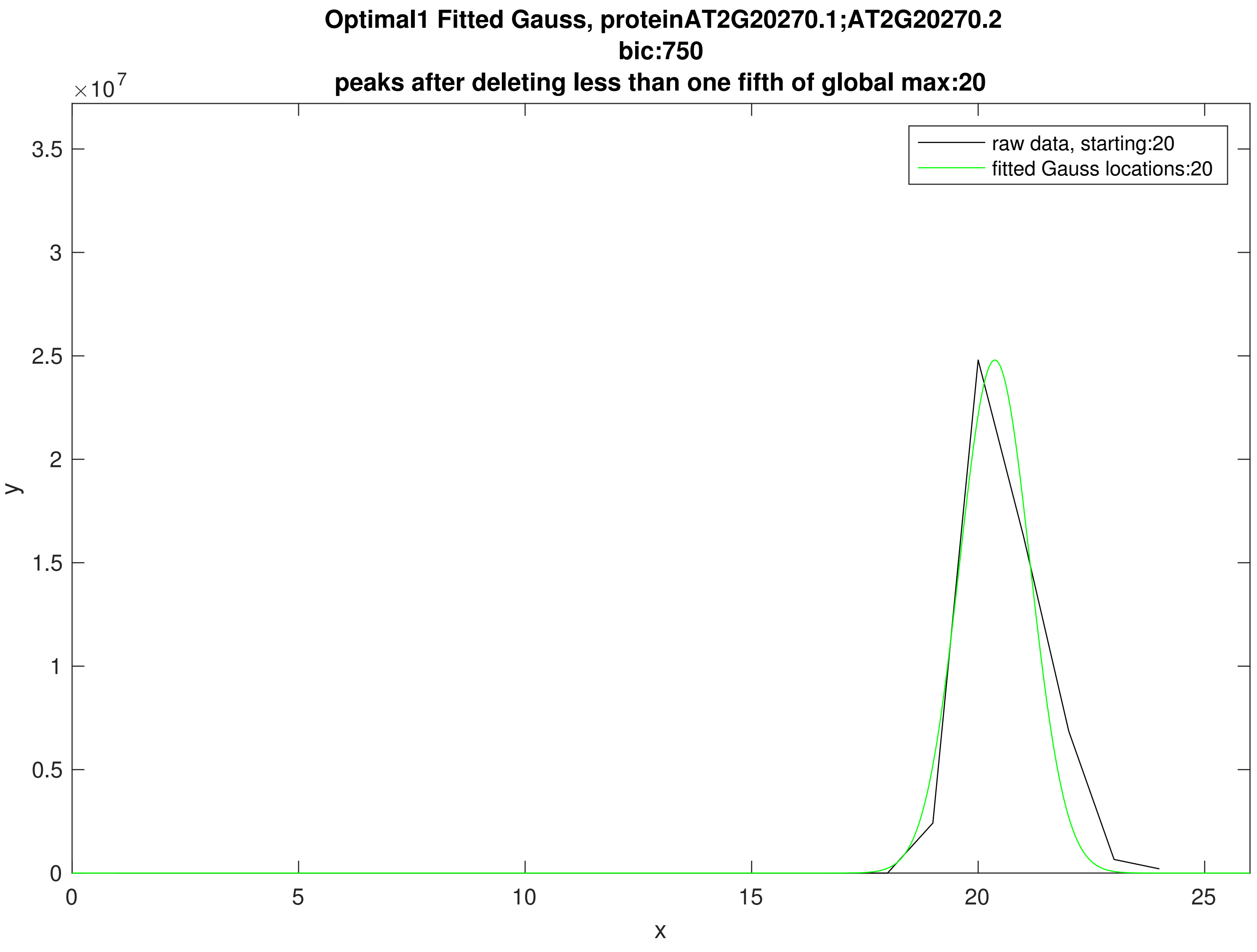

Optimal1 Fitted Gauss, proteinAT2G20270.1;AT2G20270.2
bic:750
peaks after deleting less than one fifth of global max:20
×10^7
3.5
3
2.5
2
1.5
1
0.5
0
y
0
5
10
15
20
25
x
raw data, starting:20
fitted Gauss locations:20

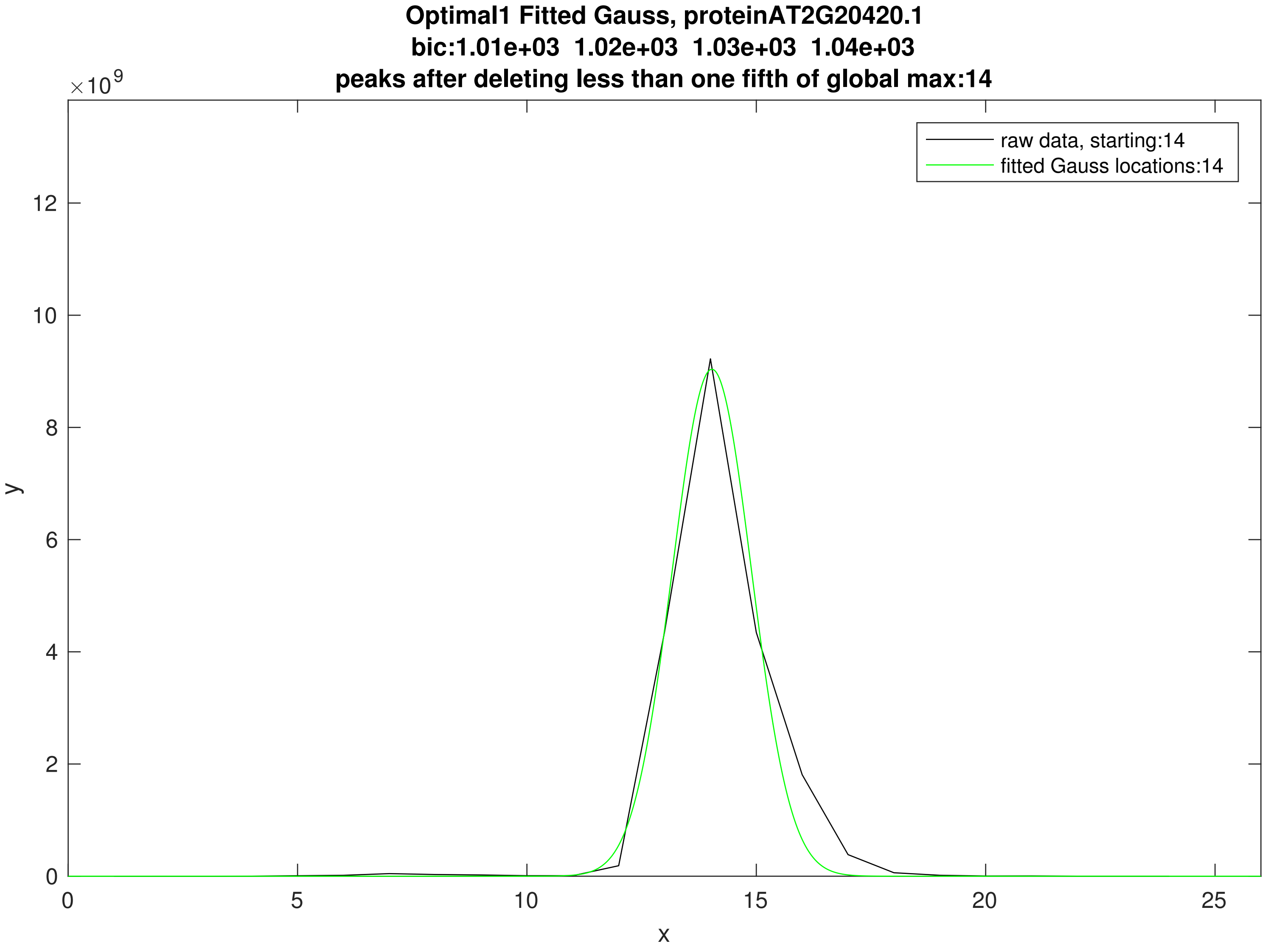

Optimal1 Fitted Gauss, proteinAT2G20420.1
bic:1.01e+03 1.02e+03 1.03e+03 1.04e+03
peaks after deleting less than one fifth of global max:14
×10^9
raw data, starting:14
fitted Gauss locations:14
y
x
0
2
4
6
8
10
12
0
5
10
15
20
25

**Optimal2 Fitted Gauss, proteinAT2G20580.1**
**bic:862 813 814**
**peaks after deleting less than one fifth of global max:4.2 12**

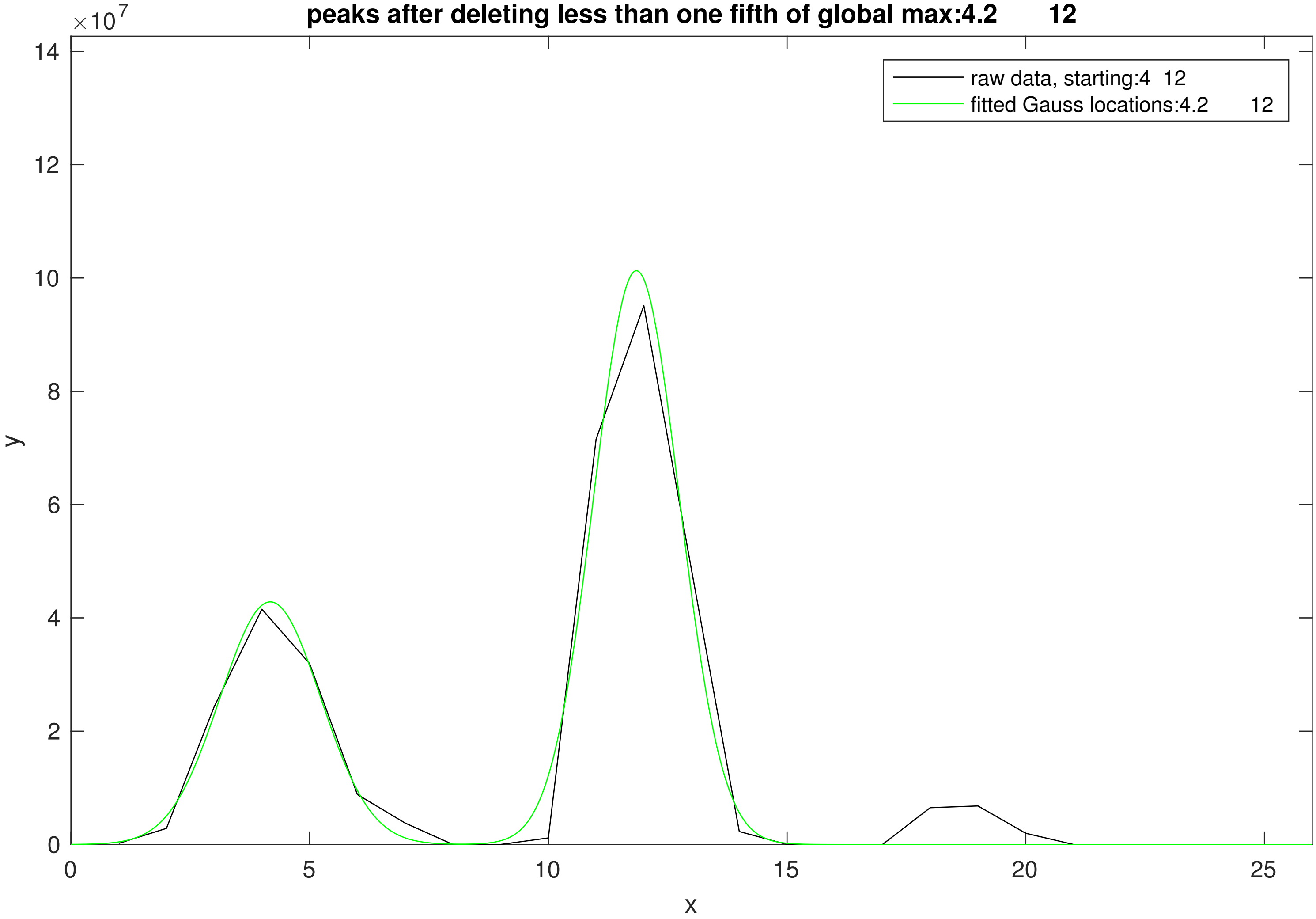

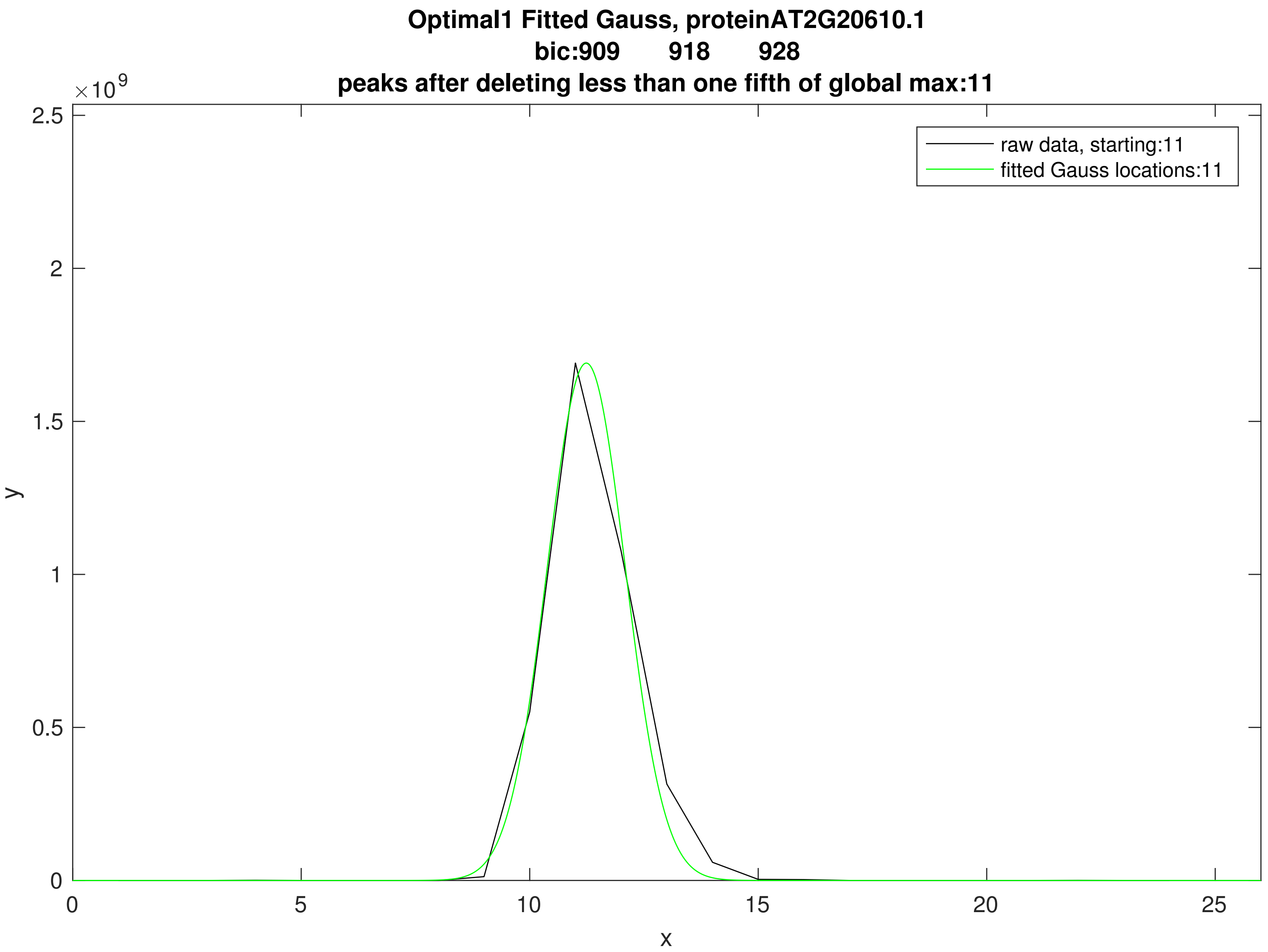
Optimal1 Fitted Gauss, proteinAT2G20610.1
bic:909 918 928
peaks after deleting less than one fifth of global max:11
×10^9
2.5
2
1.5
1
0.5
0
y
0
5
10
15
20
25
x
raw data, starting:11
fitted Gauss locations:11

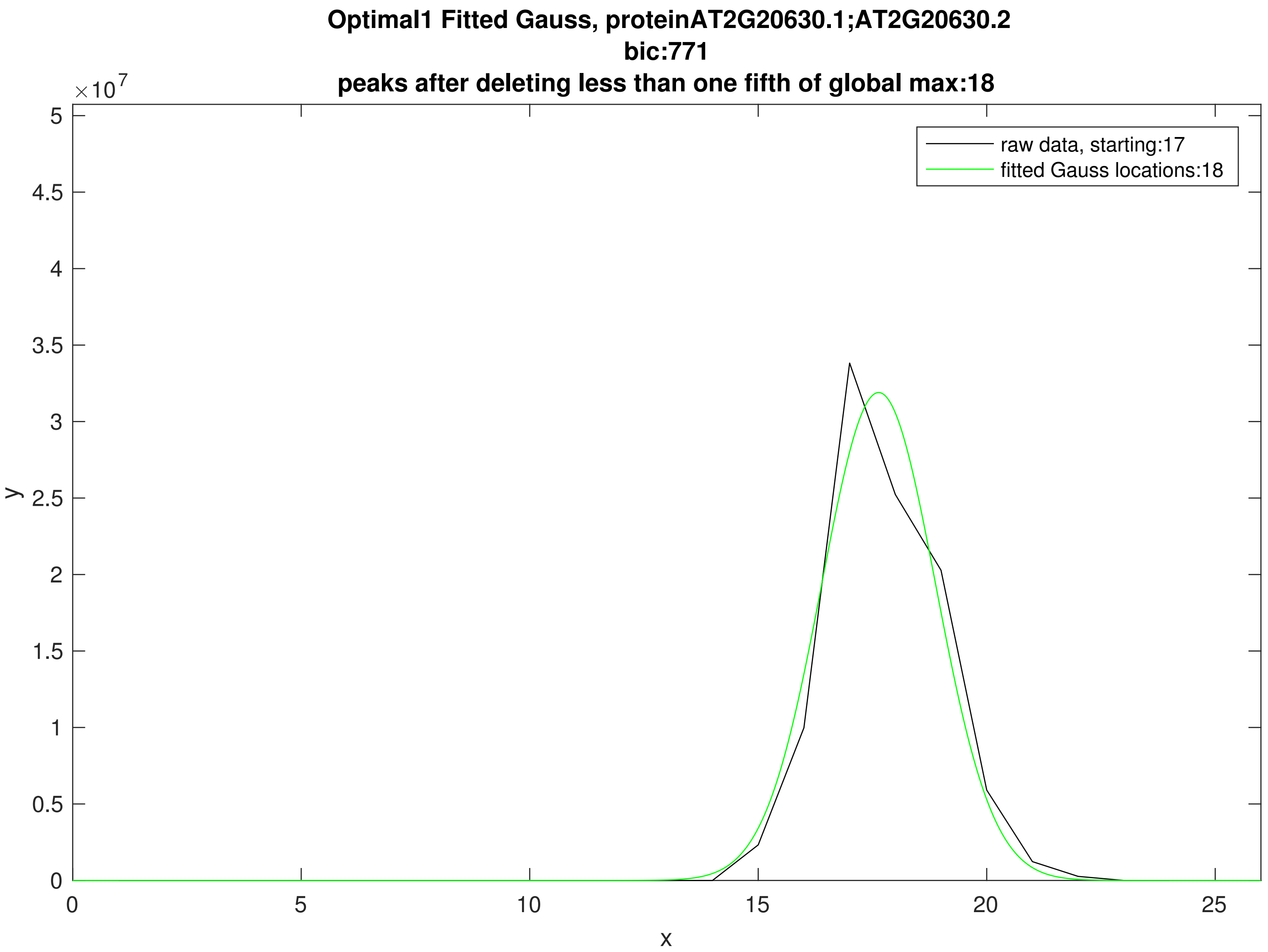

Optimal1 Fitted Gauss, proteinAT2G20630.1;AT2G20630.2
bic:771
peaks after deleting less than one fifth of global max:18
×10^7
raw data, starting:17
fitted Gauss locations:18
y
x
0
0.5
1
1.5
2
2.5
3
3.5
4
4.5
5
0
5
10
15
20
25

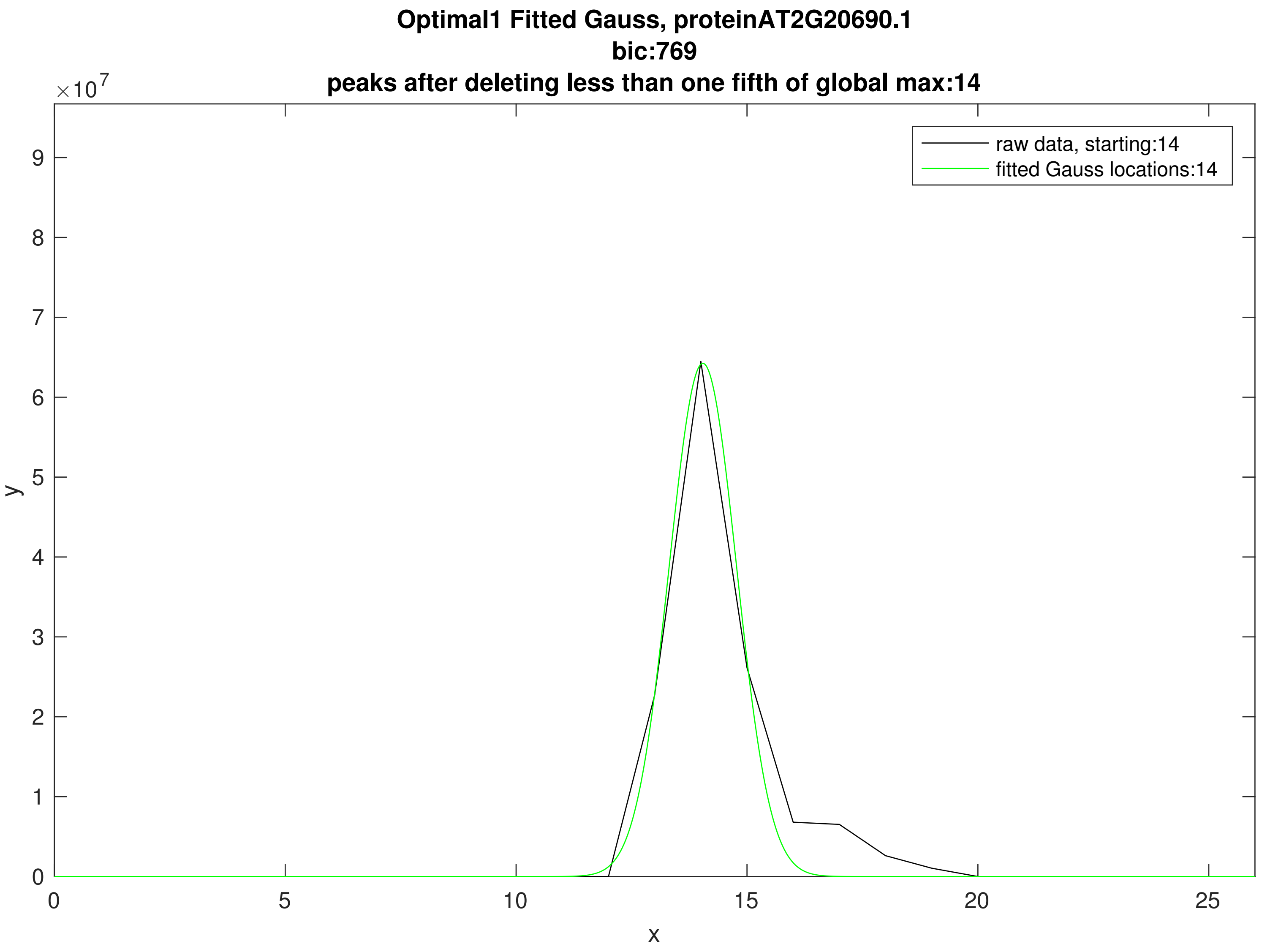

Optimal1 Fitted Gauss, proteinAT2G20690.1
bic:769
peaks after deleting less than one fifth of global max:14
×10^7
raw data, starting:14
fitted Gauss locations:14
y
x
0
1
2
3
4
5
6
7
8
9
0
5
10
15
20
25

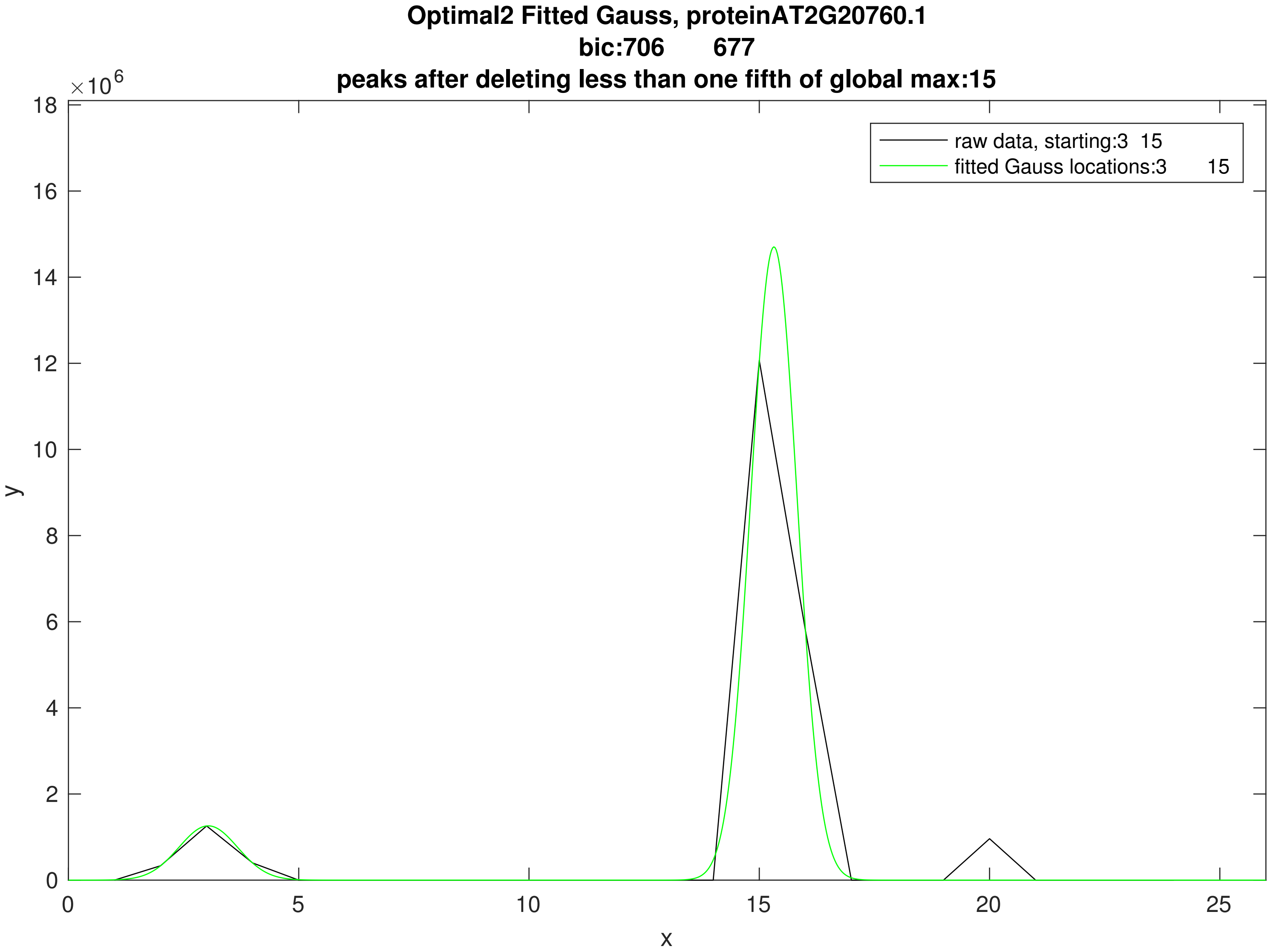
Optimal2 Fitted Gauss, proteinAT2G20760.1
bic:706 677
peaks after deleting less than one fifth of global max:15
×10^6
raw data, starting:3 15
fitted Gauss locations:3 15
y
x
0
2
4
6
8
10
12
14
16
18
0
5
10
15
20
25

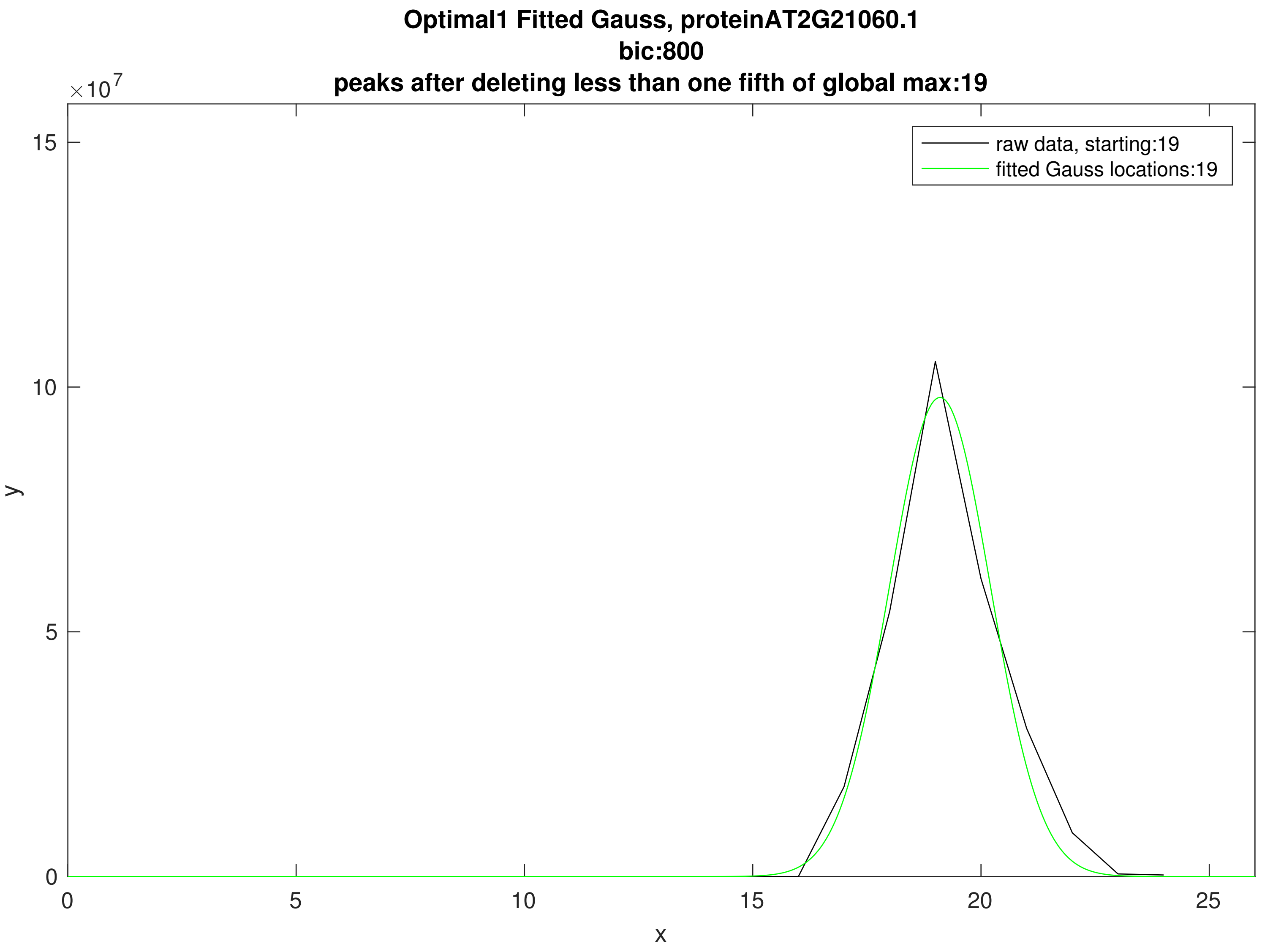
Optimal1 Fitted Gauss, proteinAT2G21060.1
bic:800
peaks after deleting less than one fifth of global max:19
×10^7
15
10
5
0
y
0
5
10
15
20
25
x
raw data, starting:19
fitted Gauss locations:19

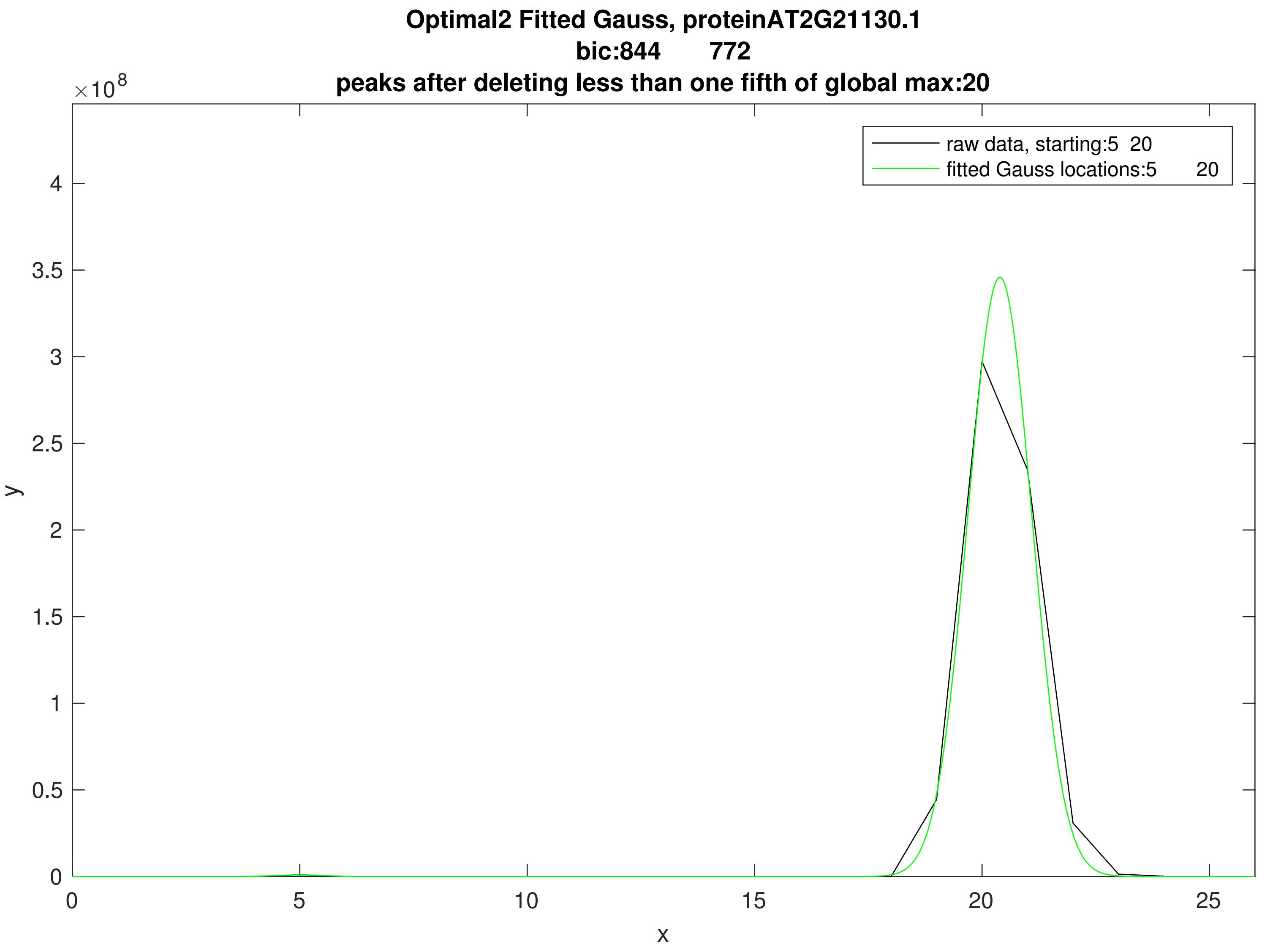
Optimal2 Fitted Gauss, proteinAT2G21130.1
bic:844 772
peaks after deleting less than one fifth of global max:20
×10^8
raw data, starting:5 20
fitted Gauss locations:5 20
y
x
0
0.5
1
1.5
2
2.5
3
3.5
4
0
5
10
15
20
25

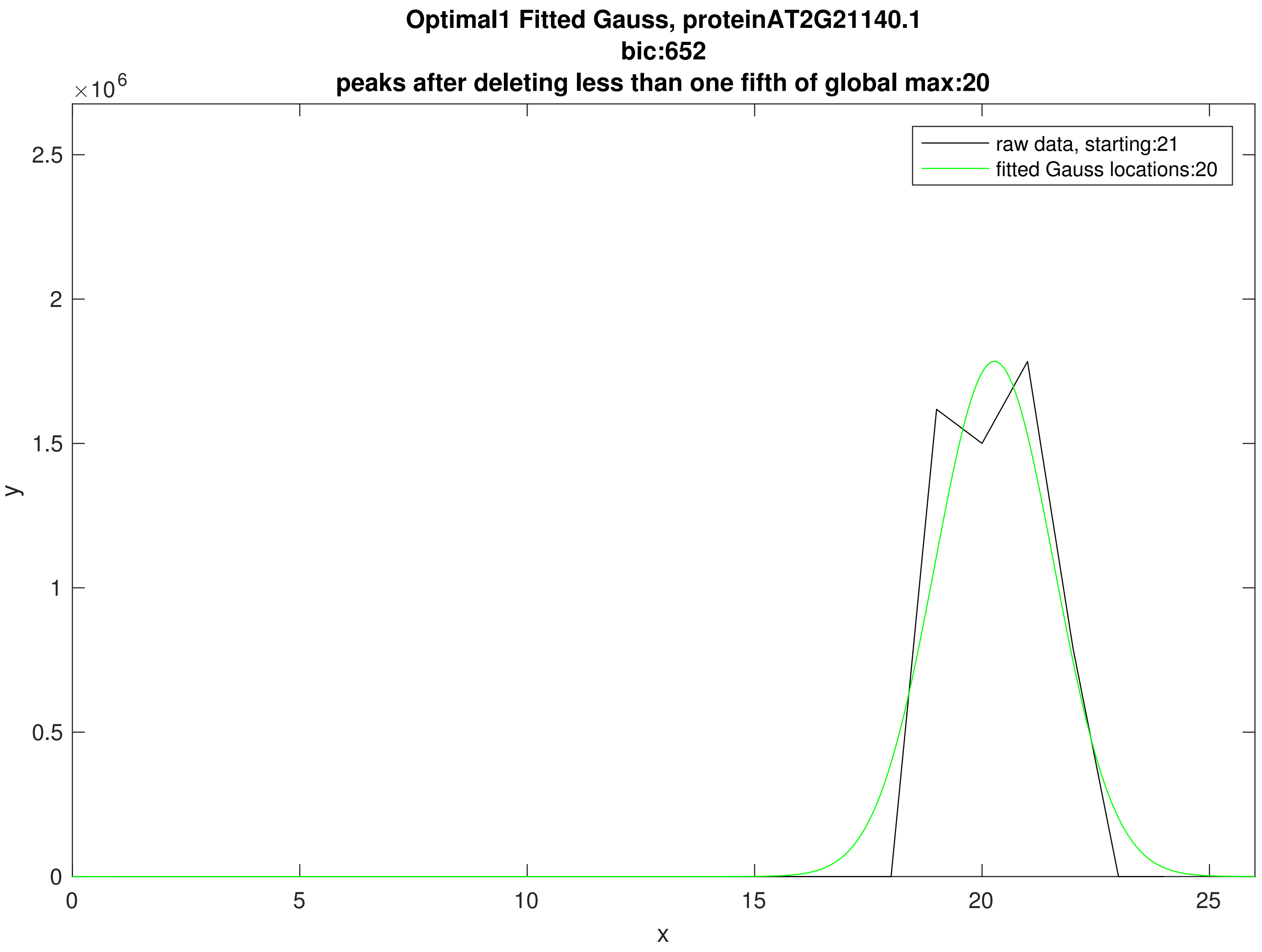

Optimal1 Fitted Gauss, proteinAT2G21140.1
bic:652
peaks after deleting less than one fifth of global max:20
×10^6
raw data, starting:21
fitted Gauss locations:20
2.5
2
1.5
1
0.5
0
y
0
5
10
15
20
25
x

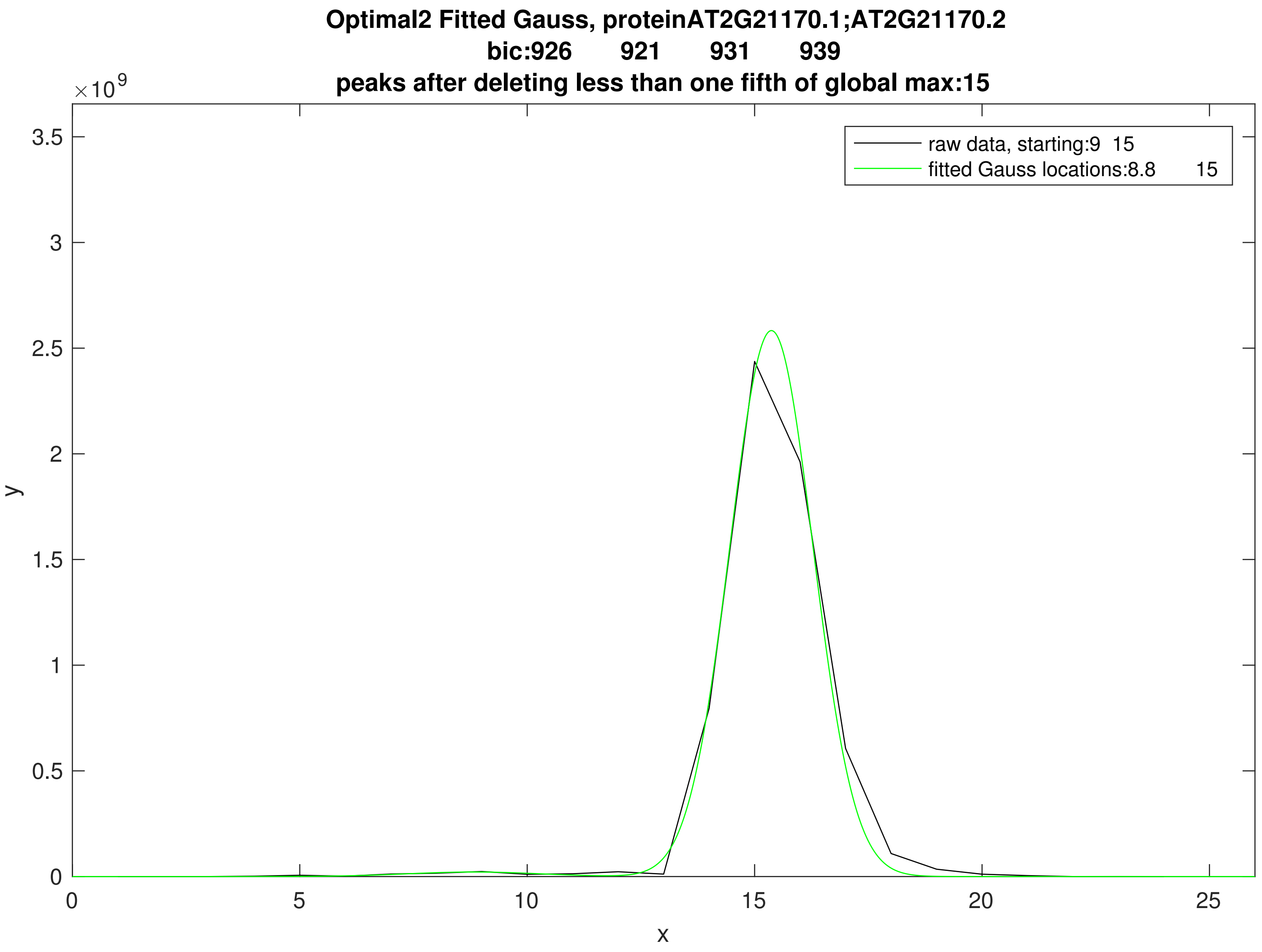

Optimal2 Fitted Gauss, proteinAT2G21170.1;AT2G21170.2
bic:926 921 931 939
peaks after deleting less than one fifth of global max:15
×10^9
raw data, starting:9 15
fitted Gauss locations:8.8 15
y
x
0
0.5
1
1.5
2
2.5
3
3.5
0
5
10
15
20
25

**Optimal1 Fitted Gauss, proteinAT2G21250.1;AT2G21250.2**
**bic:908**
**peaks after deleting less than one fifth of global max:15**

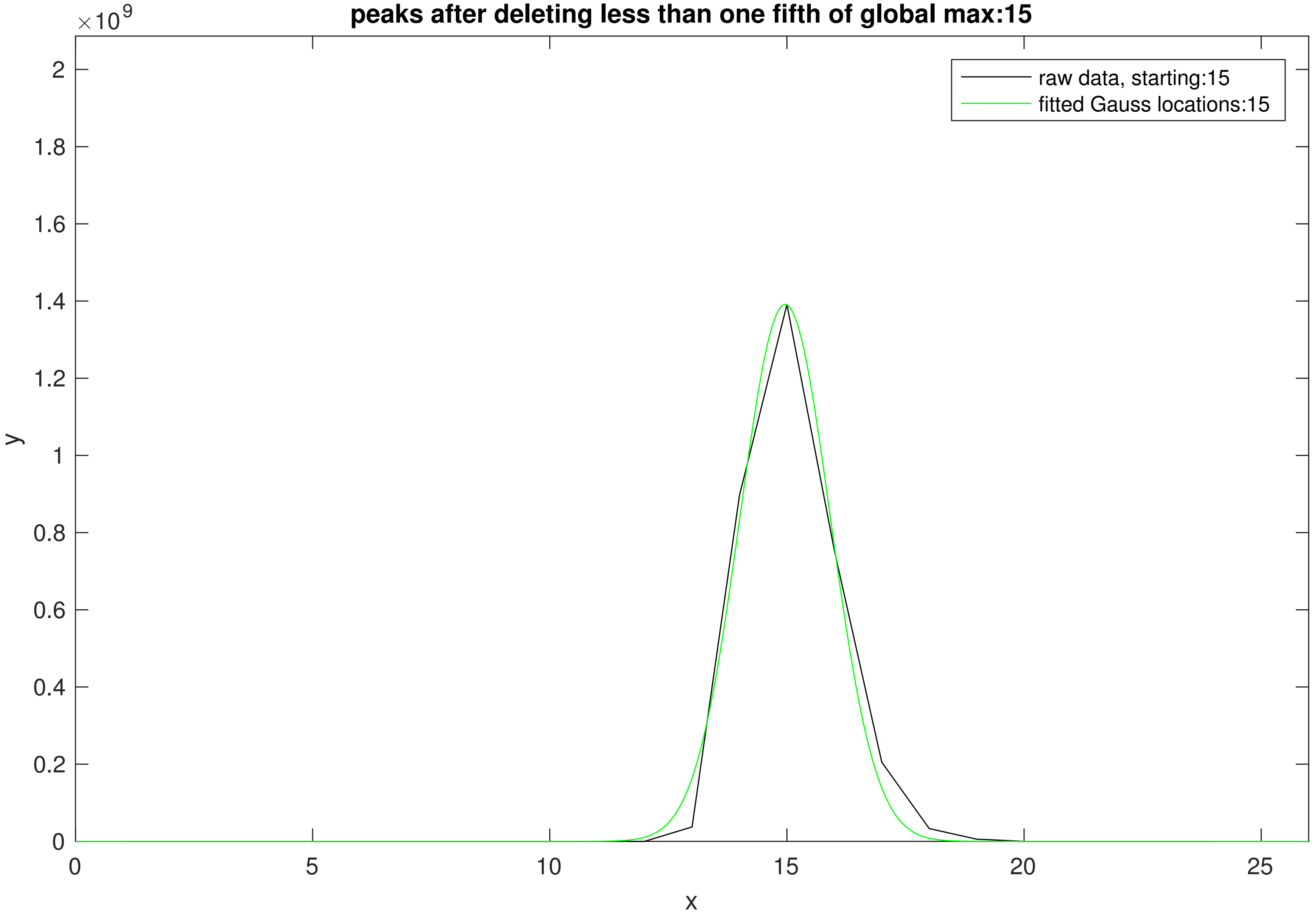

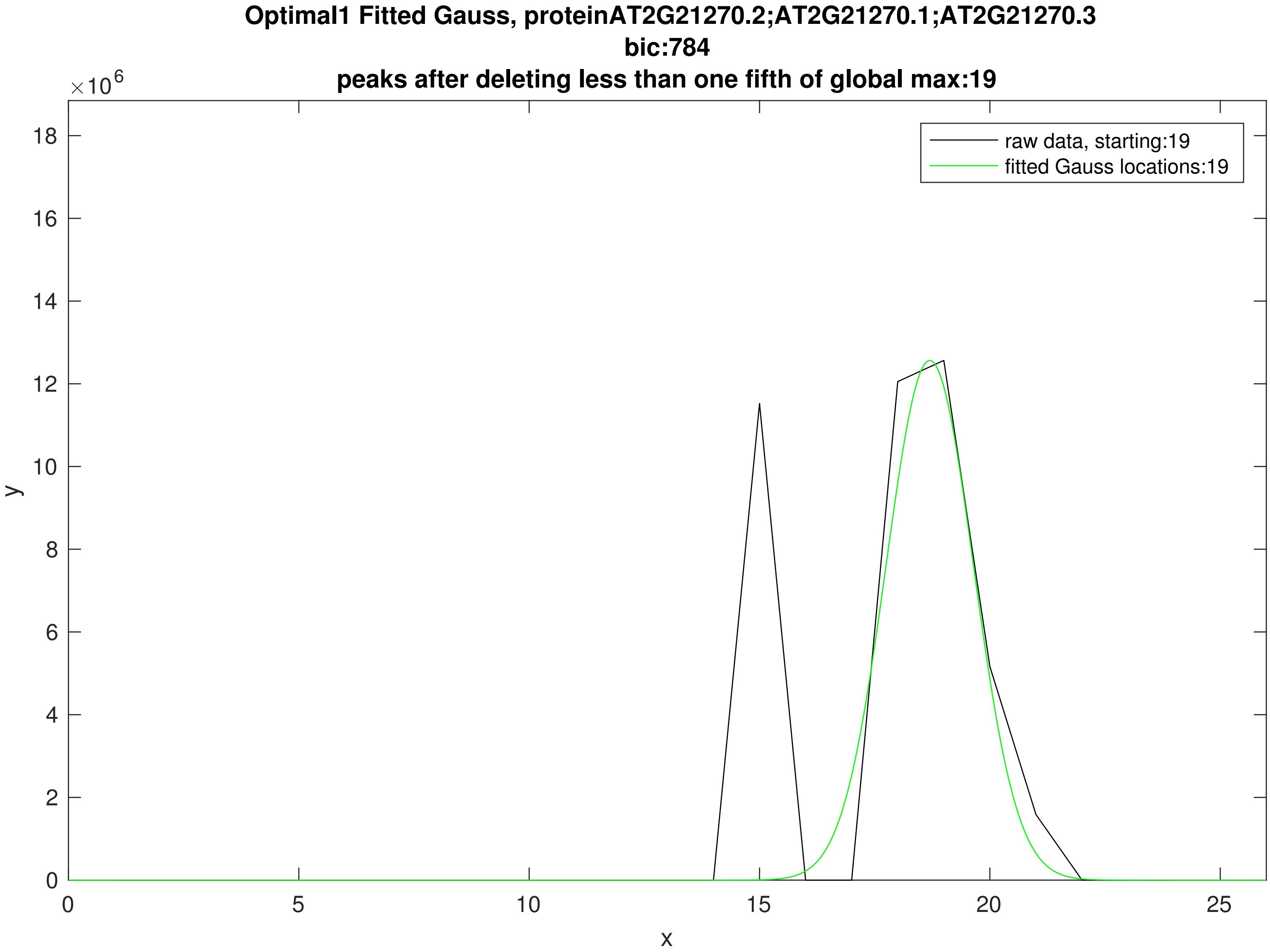
Optimal1 Fitted Gauss, proteinAT2G21270.2;AT2G21270.1;AT2G21270.3
bic:784
peaks after deleting less than one fifth of global max:19
×10^6
raw data, starting:19
fitted Gauss locations:19
y
x
0
2
4
6
8
10
12
14
16
18
0
5
10
15
20
25

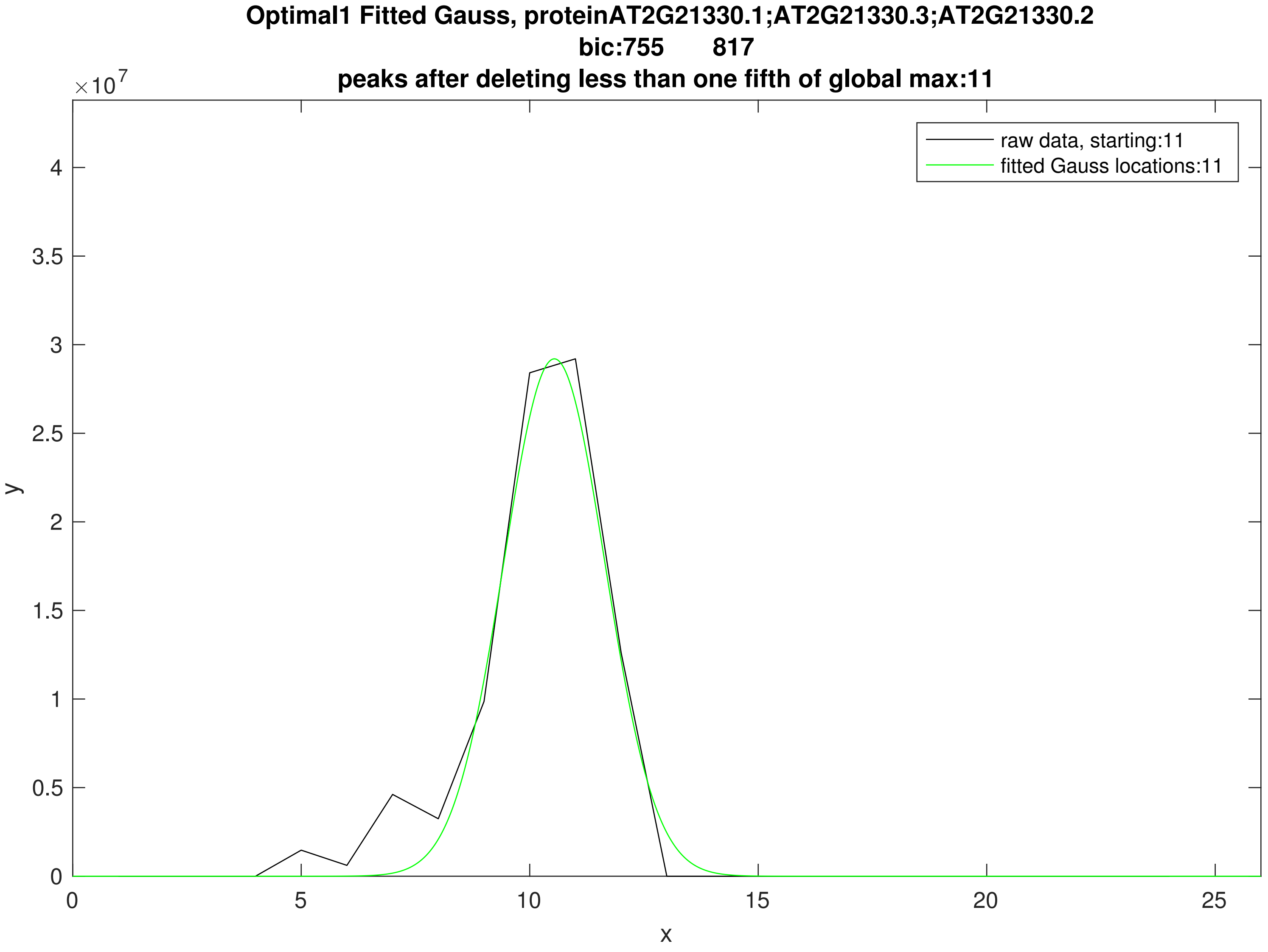

Optimal1 Fitted Gauss, proteinAT2G21330.1;AT2G21330.3;AT2G21330.2
bic:755 817
peaks after deleting less than one fifth of global max:11
×10^7
raw data, starting:11
fitted Gauss locations:11
y
x
0
0.5
1
1.5
2
2.5
3
3.5
4
5
10
15
20
25

**Optimal2 Fitted Gauss, proteinAT2G21390.1**
**bic:768 744**
**peaks after deleting less than one fifth of global max:4.7 14**

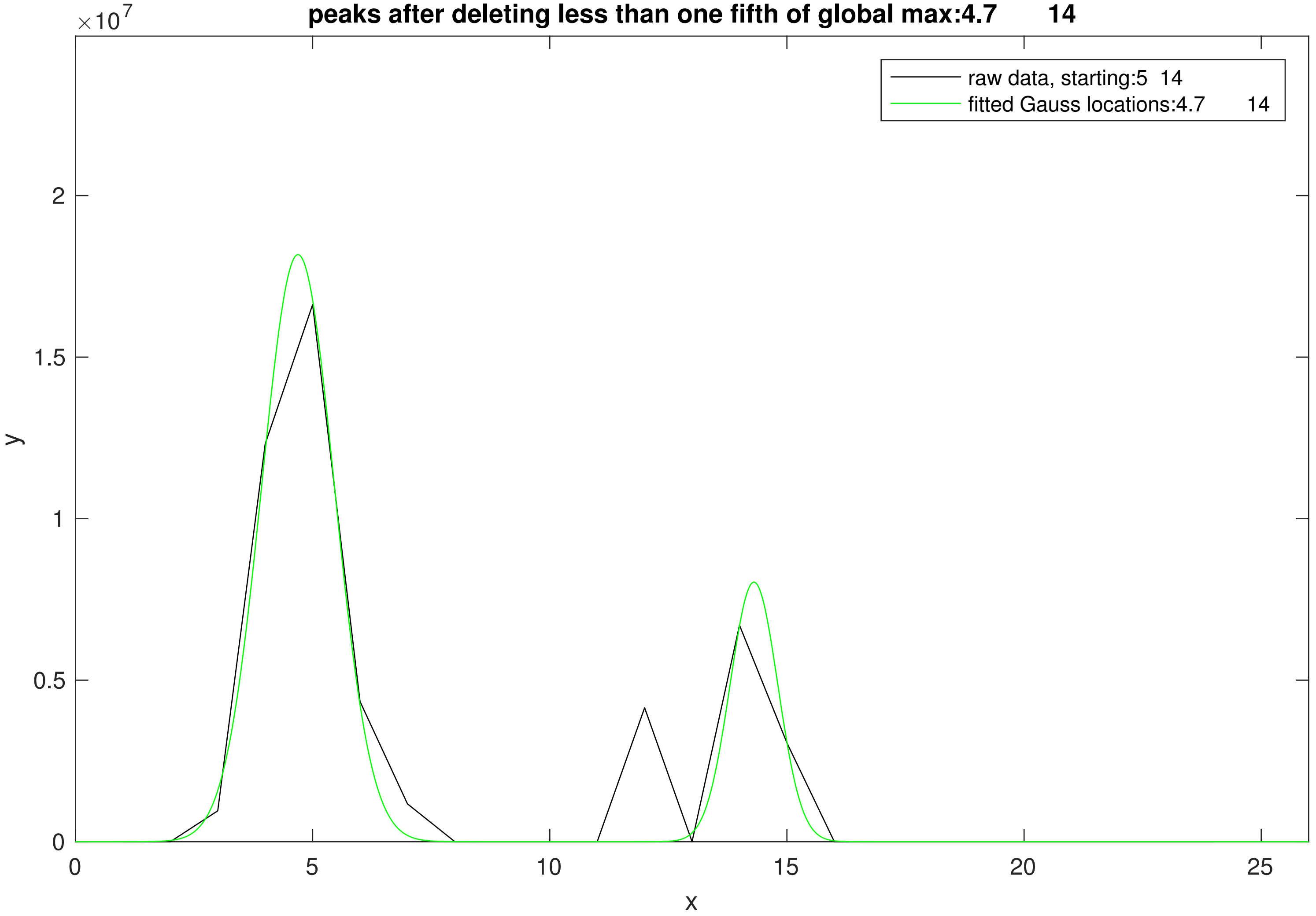

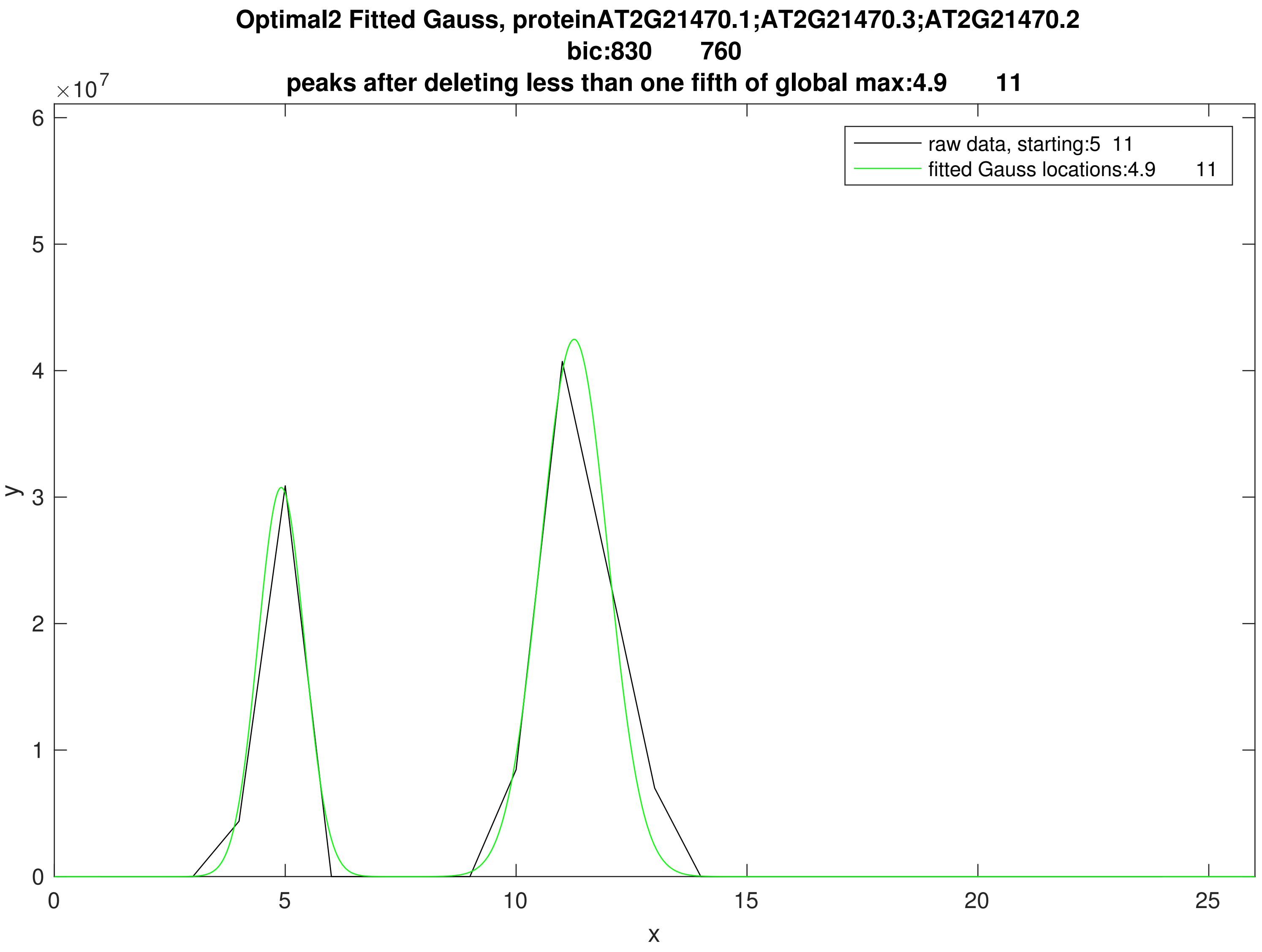

Optimal2 Fitted Gauss, proteinAT2G21470.1;AT2G21470.3;AT2G21470.2
bic:830 760
peaks after deleting less than one fifth of global max:4.9 11
×10^7
raw data, starting:5 11
fitted Gauss locations:4.9 11
y
x
0
1
2
3
4
5
6
0
5
10
15
20
25

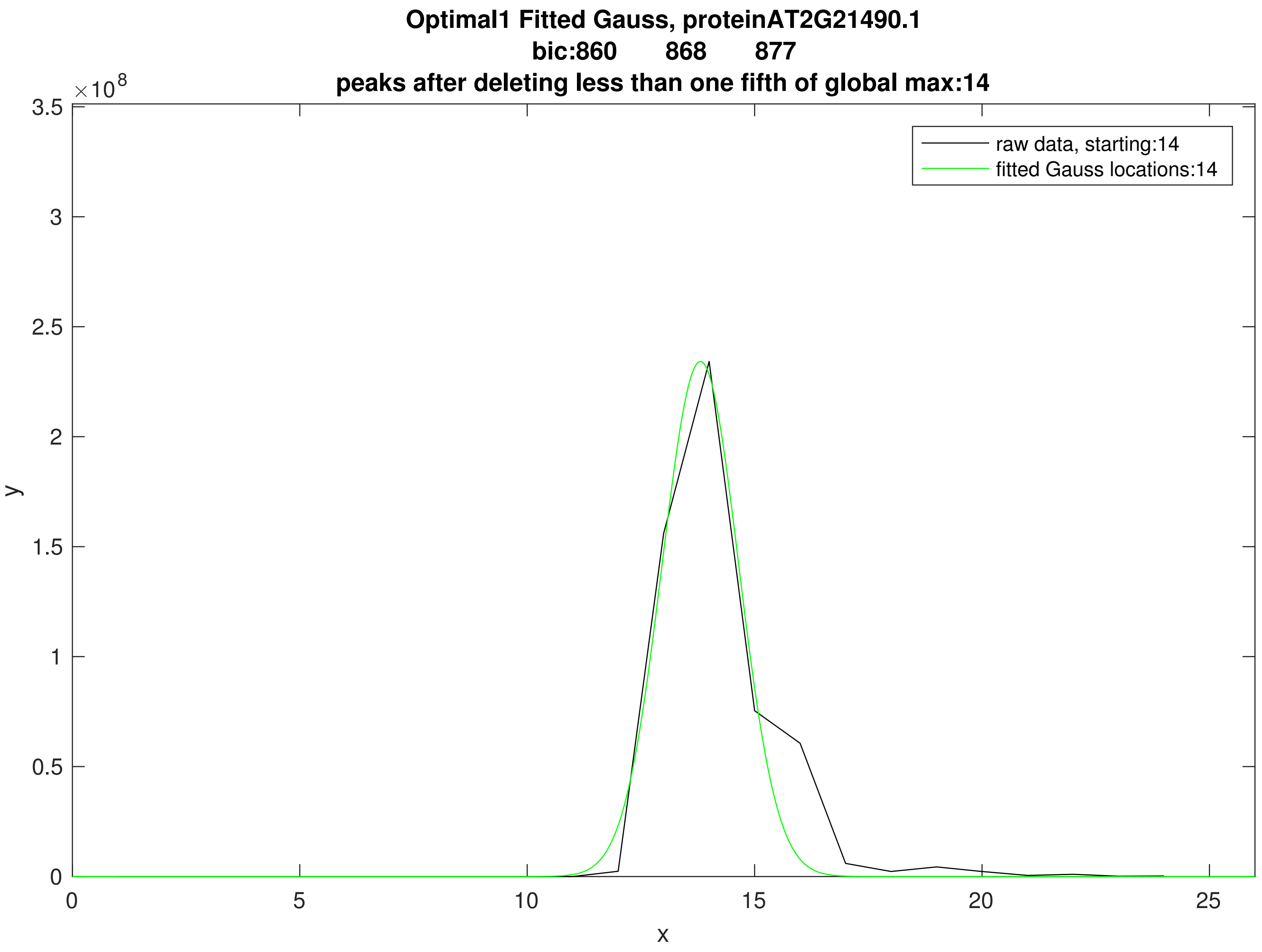

Optimal1 Fitted Gauss, proteinAT2G21490.1
bic:860 868 877
peaks after deleting less than one fifth of global max:14
×10^8
raw data, starting:14
fitted Gauss locations:14
y
x

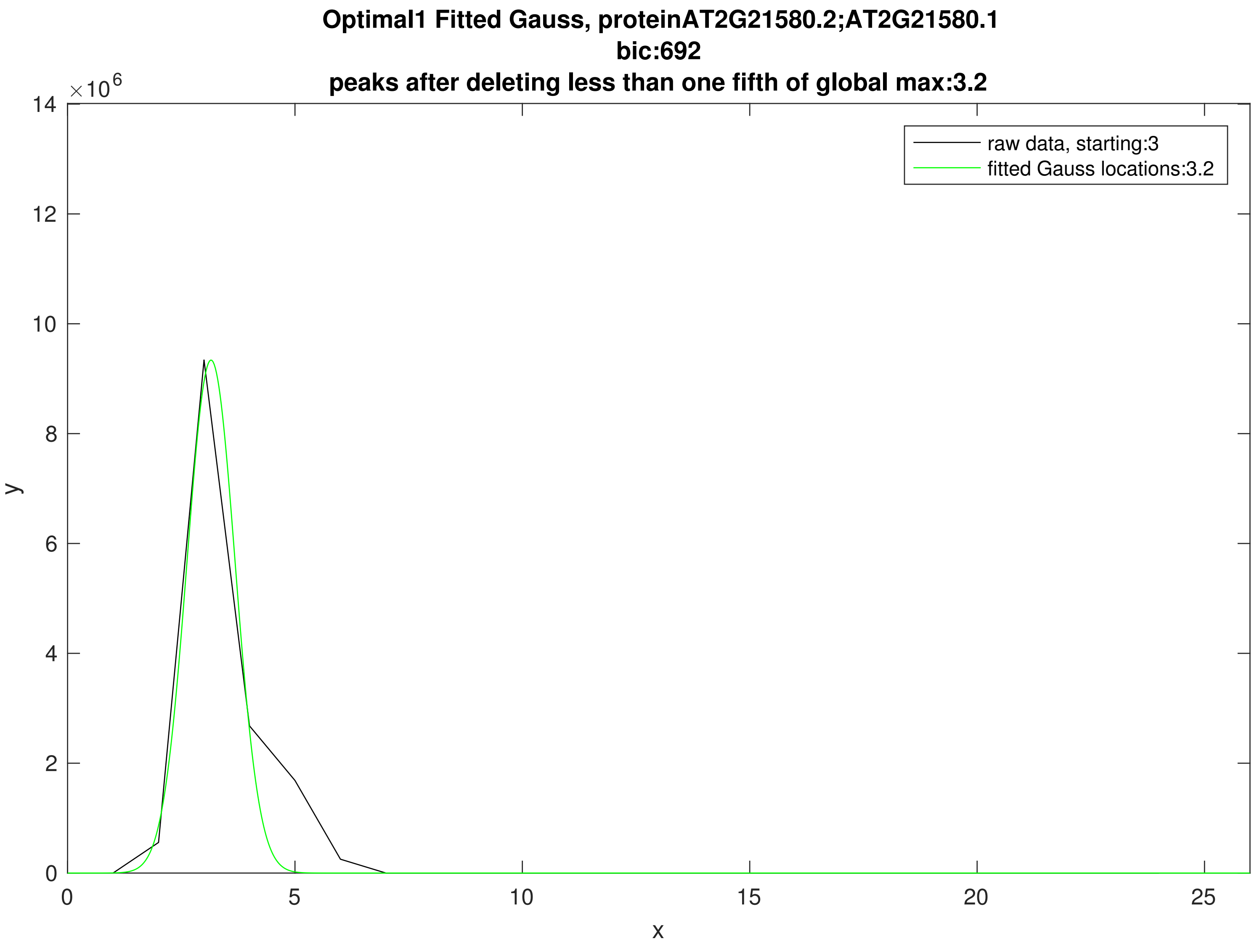
Optimal1 Fitted Gauss, proteinAT2G21580.2;AT2G21580.1
bic:692
peaks after deleting less than one fifth of global max:3.2
×10^6
14
12
10
8
6
4
2
0
y
0
5
10
15
20
25
x
raw data, starting:3
fitted Gauss locations:3.2

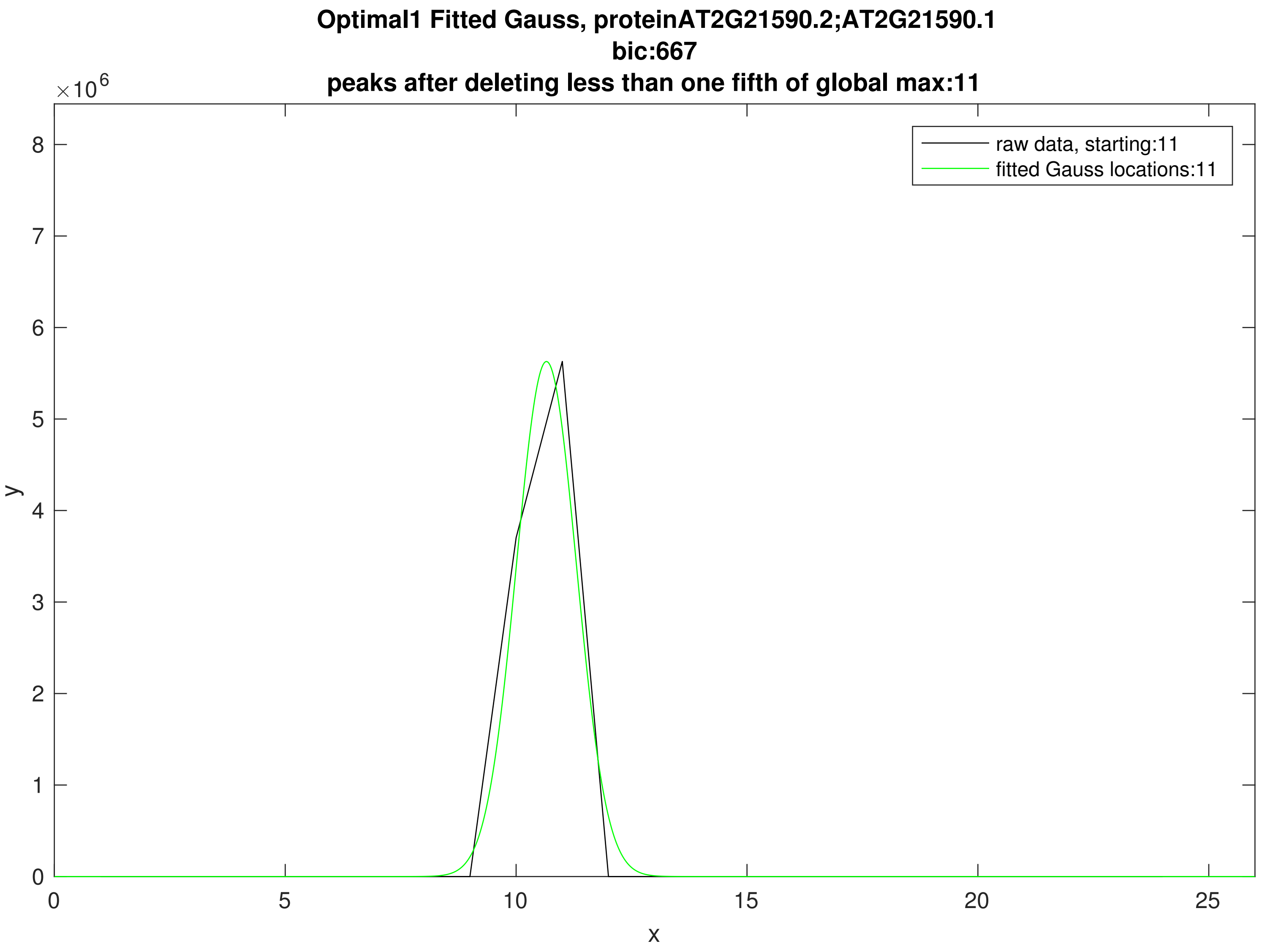
Optimal1 Fitted Gauss, proteinAT2G21590.2;AT2G21590.1
bic:667
peaks after deleting less than one fifth of global max:11
×10^6
raw data, starting:11
fitted Gauss locations:11
y
x
0
1
2
3
4
5
6
7
8
0
5
10
15
20
25

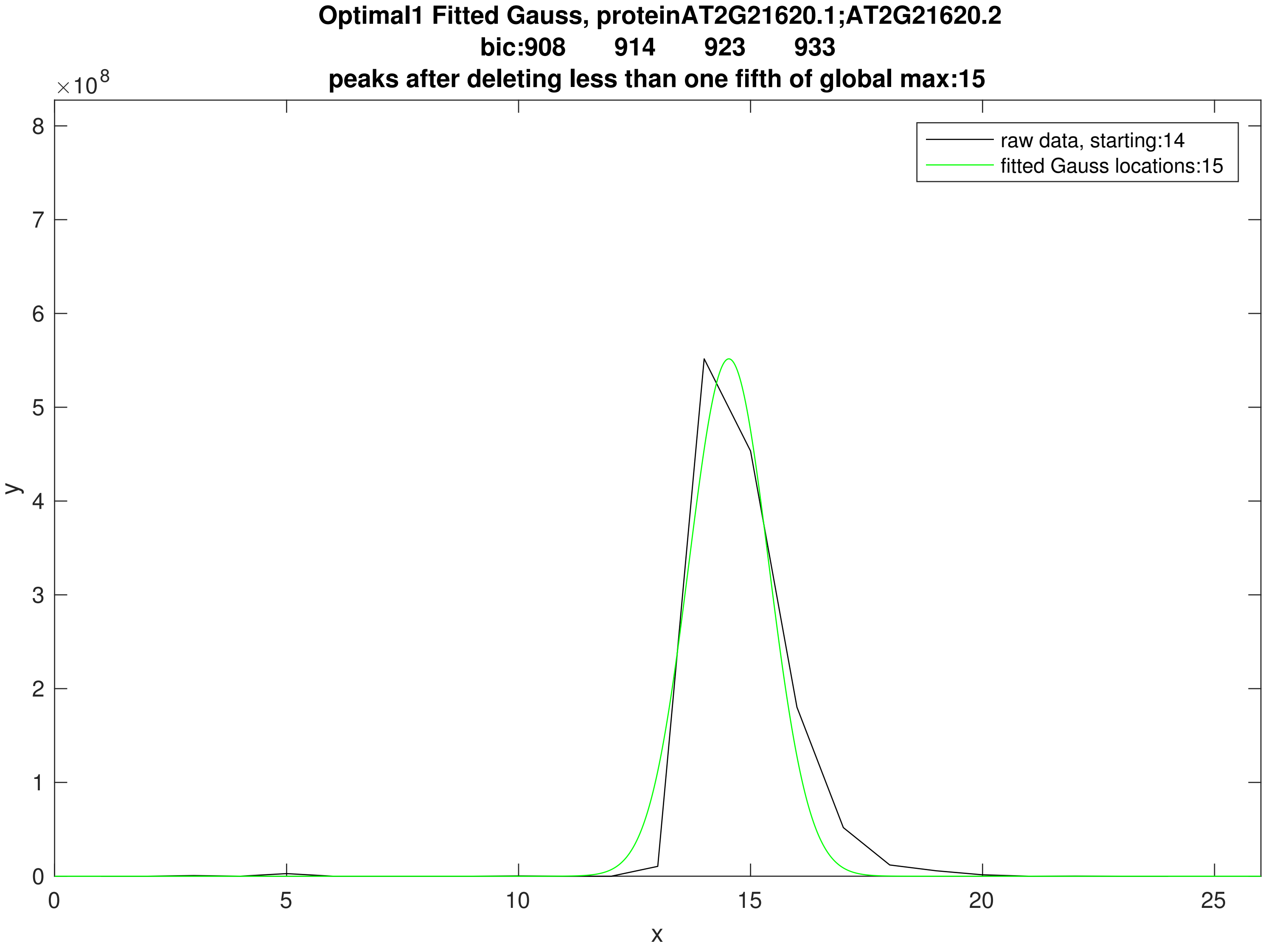
Optimal1 Fitted Gauss, proteinAT2G21620.1;AT2G21620.2
bic:908 914 923 933
peaks after deleting less than one fifth of global max:15
×10^8
raw data, starting:14
fitted Gauss locations:15
y
x
0
1
2
3
4
5
6
7
8
0
5
10
15
20
25

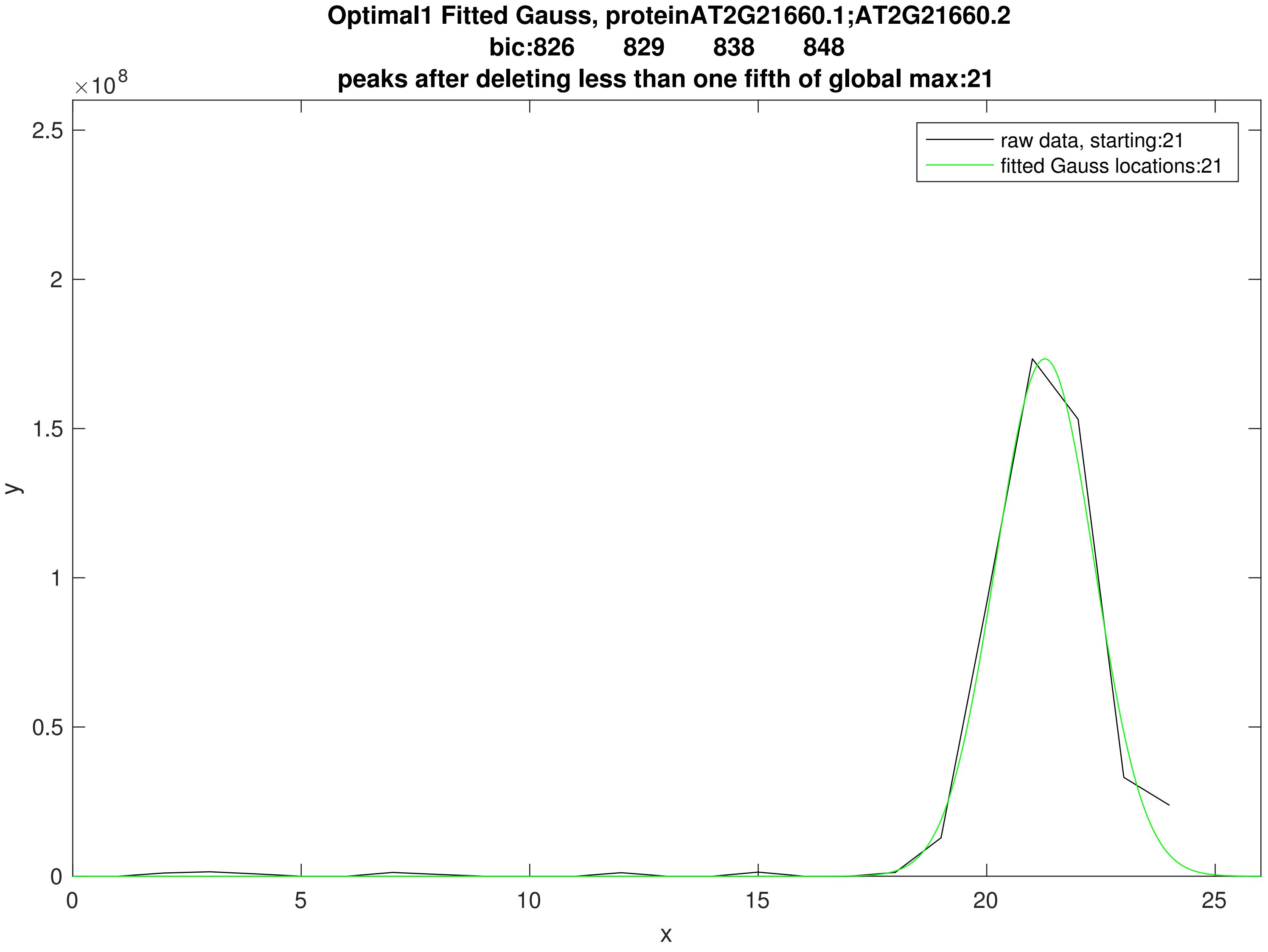
Optimal1 Fitted Gauss, proteinAT2G21660.1;AT2G21660.2
bic:826 829 838 848
peaks after deleting less than one fifth of global max:21
raw data, starting:21
fitted Gauss locations:21
$\times 10^8$
y
x

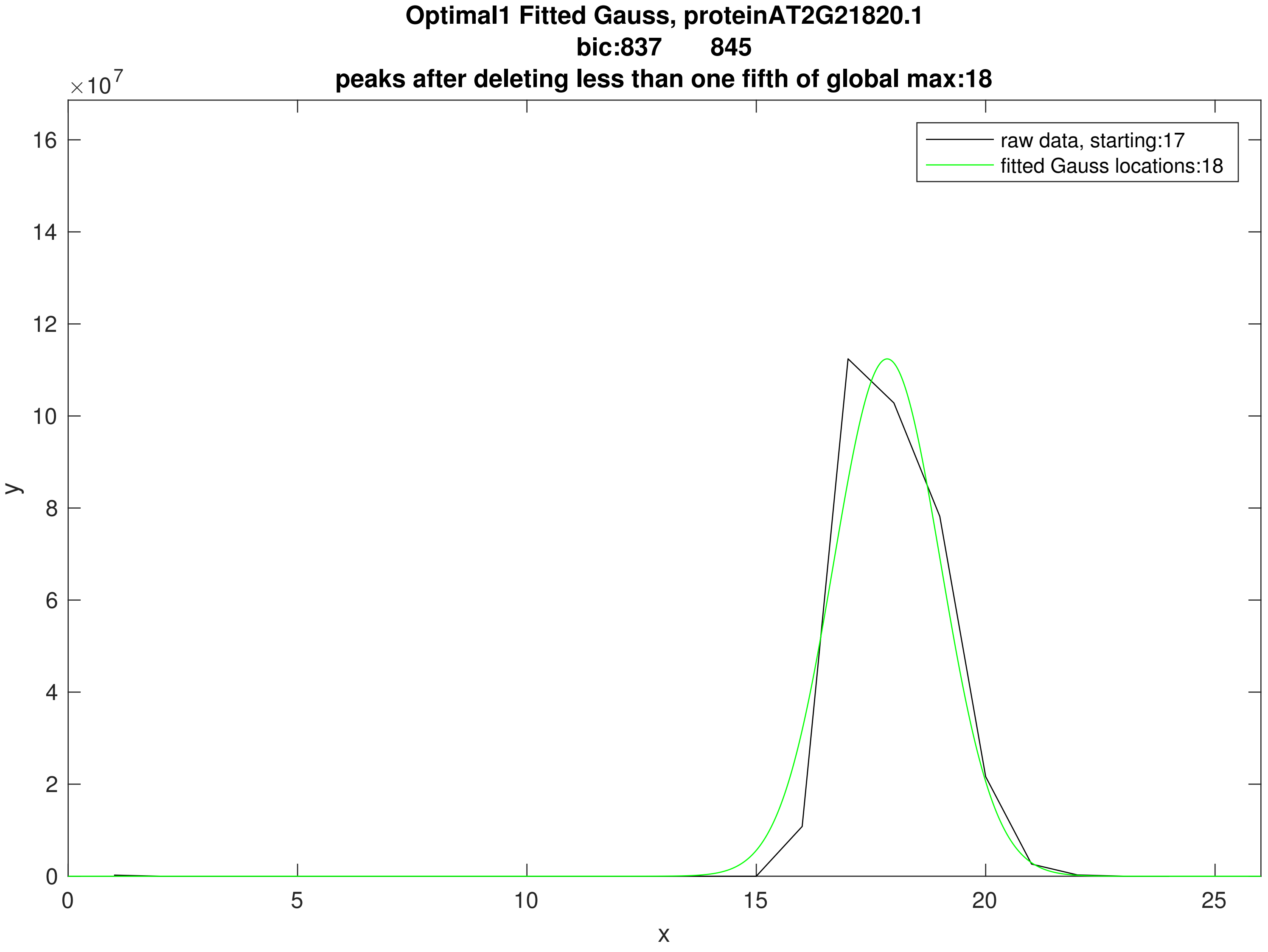
Optimal1 Fitted Gauss, proteinAT2G21820.1
bic:837 845
peaks after deleting less than one fifth of global max:18
×10^7
16
14
12
10
8
6
4
2
0
y
0
5
10
15
20
25
x
raw data, starting:17
fitted Gauss locations:18

#### Optimal1 Fitted Gauss, proteinAT2G21870.1;AT2G21870.2
#### bic:809 816 828
#### peaks after deleting less than one fifth of global max:16

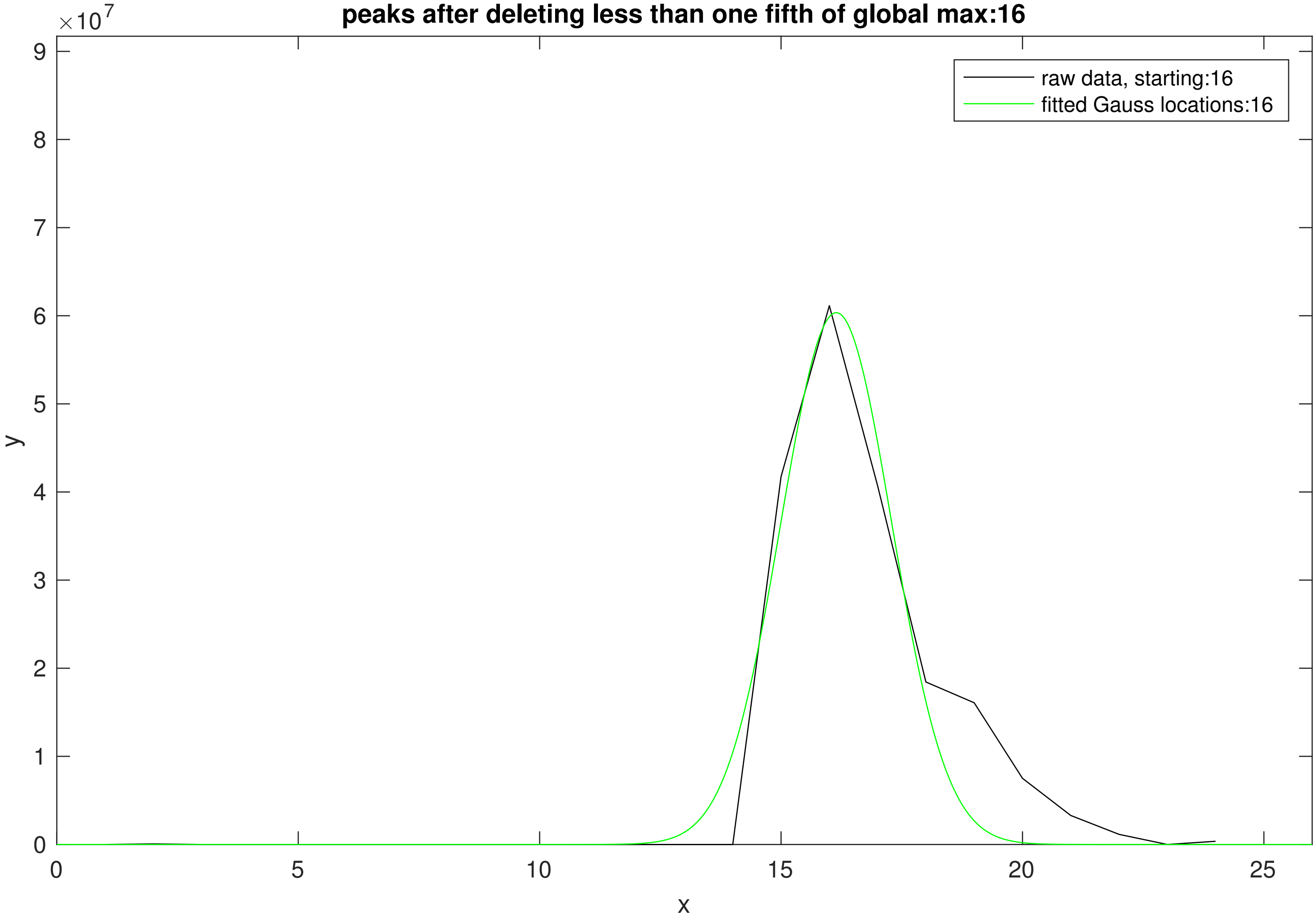

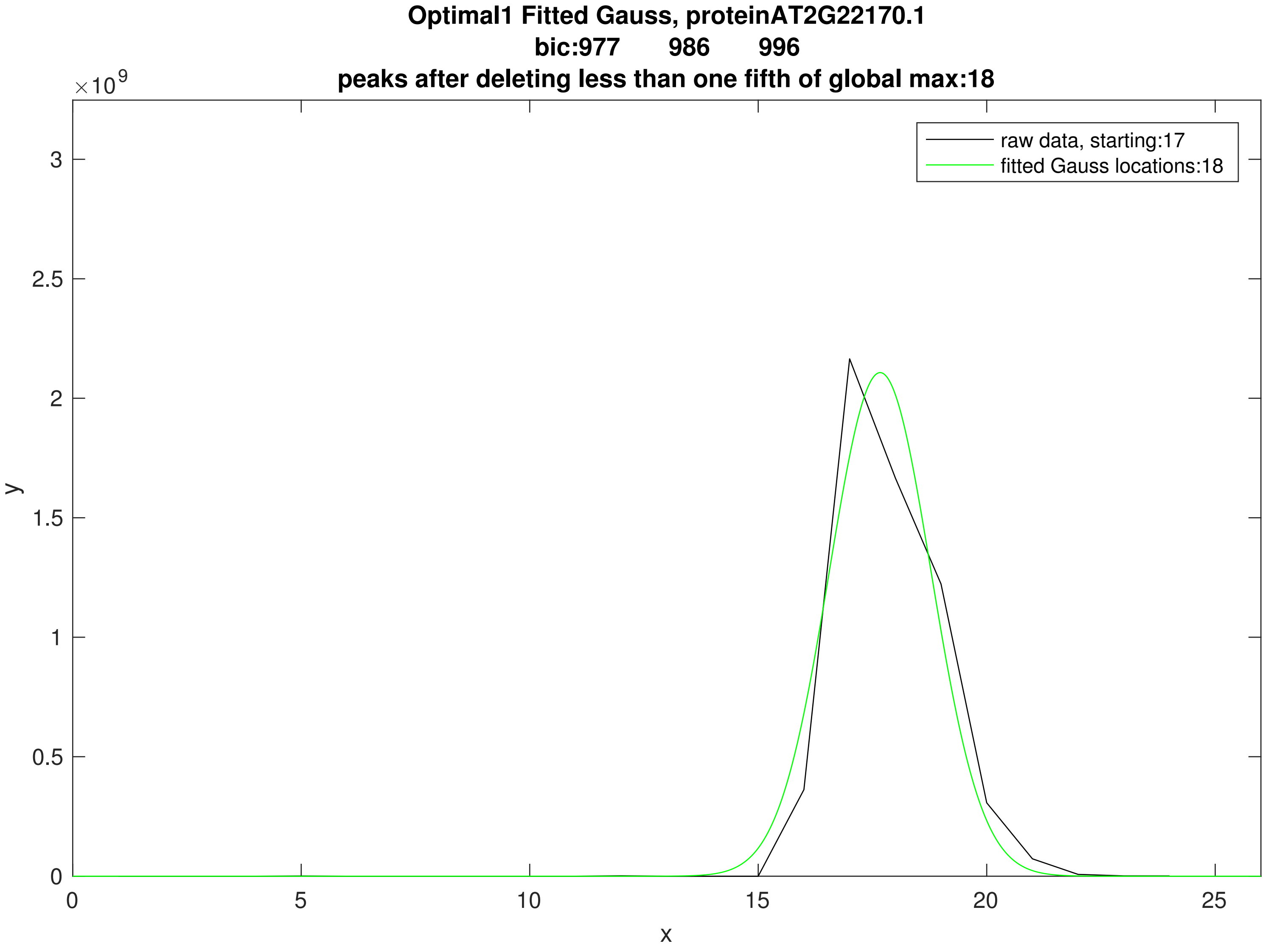
Optimal1 Fitted Gauss, proteinAT2G22170.1
bic:977 986 996
peaks after deleting less than one fifth of global max:18
×10^9
raw data, starting:17
fitted Gauss locations:18
y
x
0
0.5
1
1.5
2
2.5
3
5
10
15
20
25

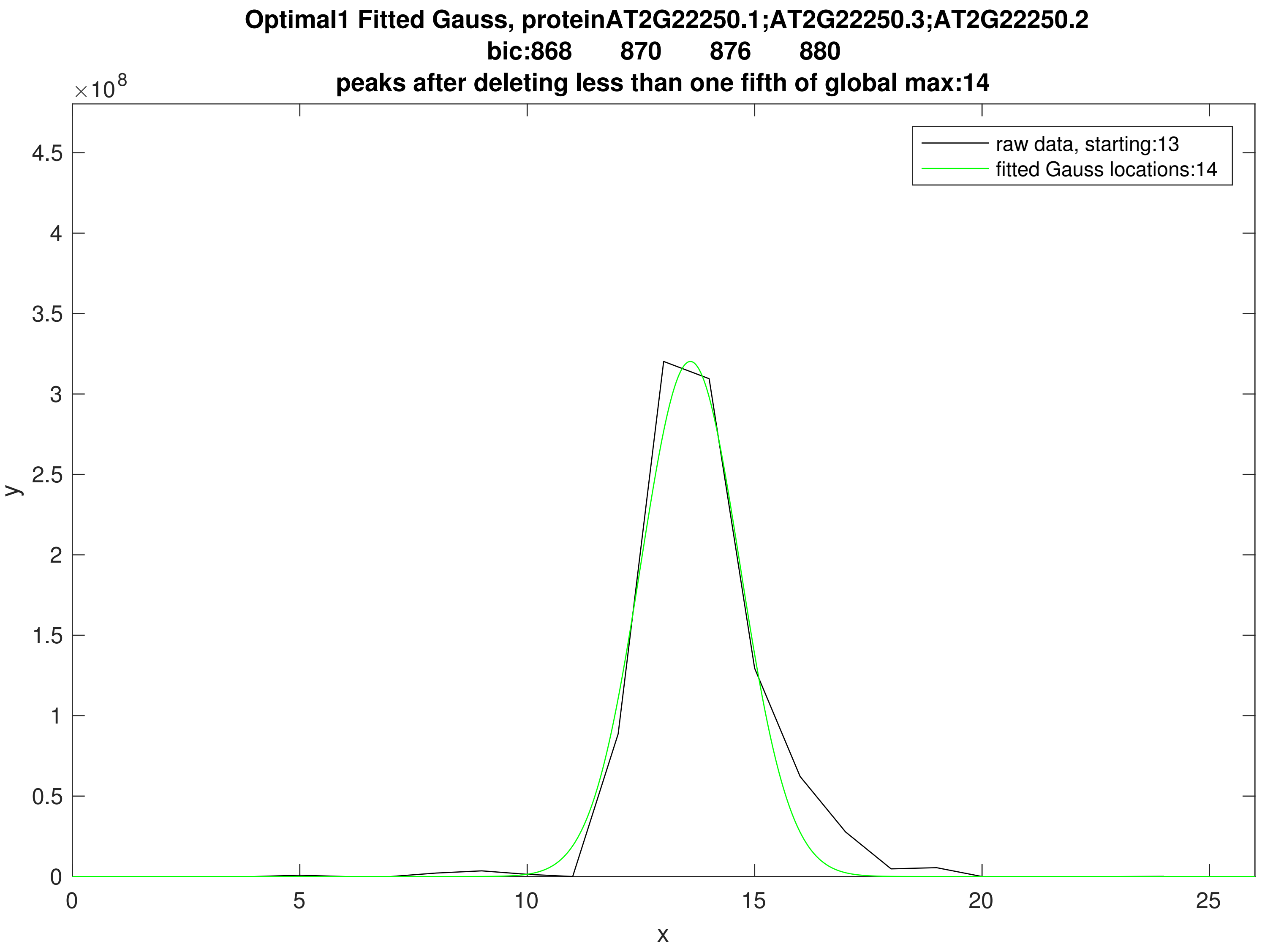
Optimal1 Fitted Gauss, proteinAT2G22250.1;AT2G22250.3;AT2G22250.2
bic:868 870 876 880
peaks after deleting less than one fifth of global max:14
×10^8
4.5
4
3.5
3
2.5
2
1.5
1
0.5
0
y
0
5
10
15
20
25
x
raw data, starting:13
fitted Gauss locations:14

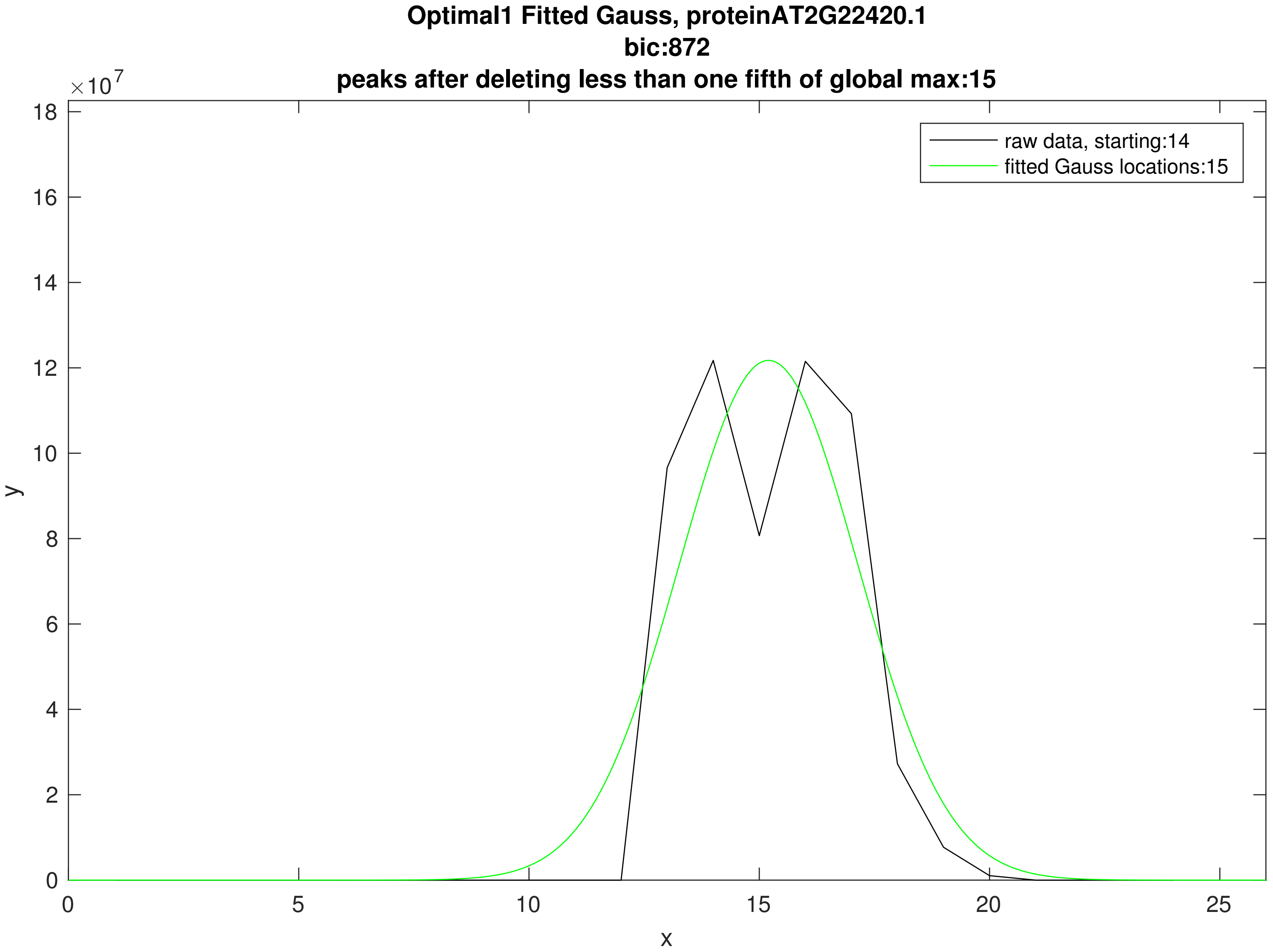
Optimal1 Fitted Gauss, proteinAT2G22420.1
bic:872
peaks after deleting less than one fifth of global max:15
×10^7
raw data, starting:14
fitted Gauss locations:15
y
x
0
2
4
6
8
10
12
14
16
18
0
5
10
15
20
25

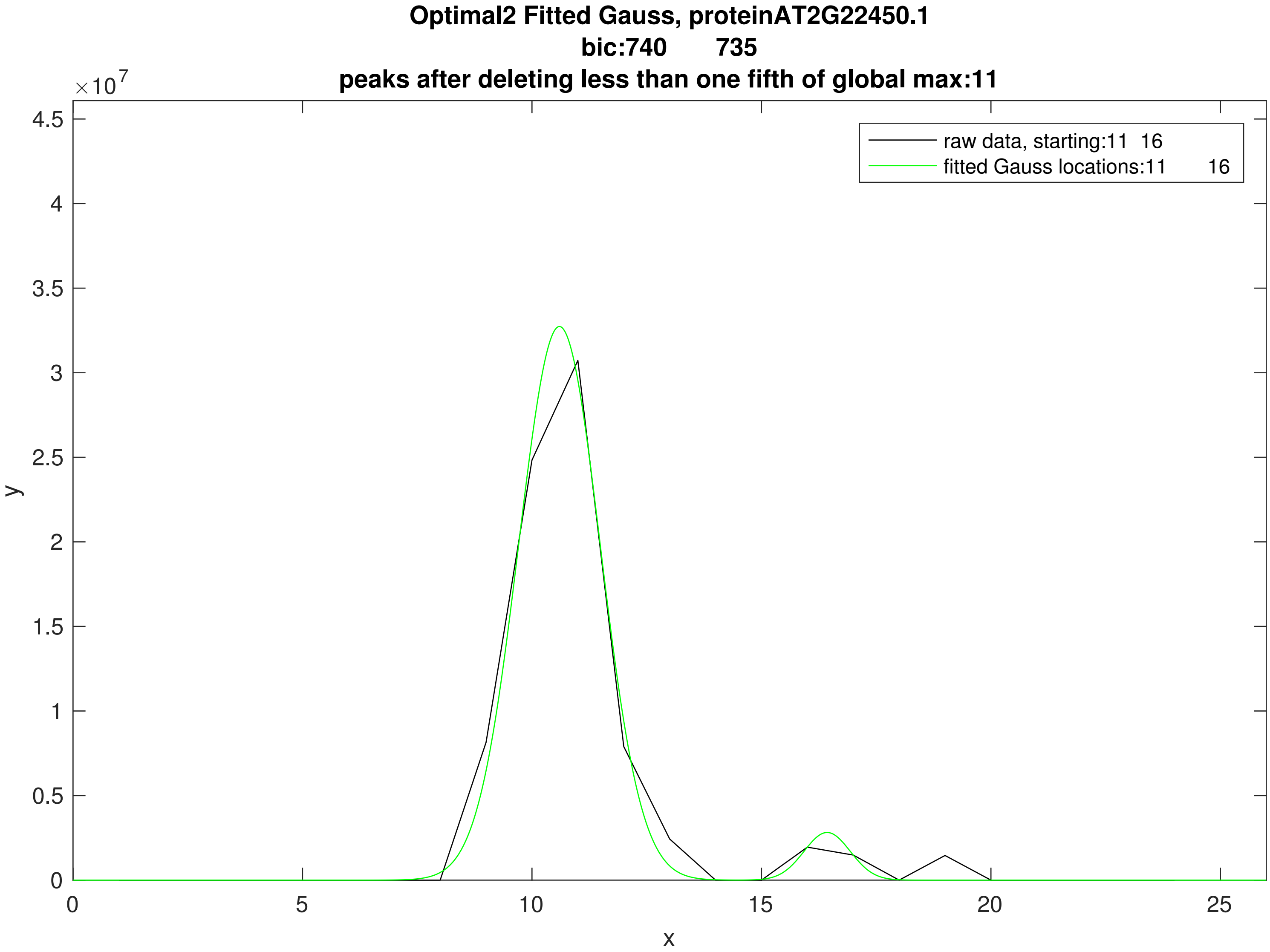
Optimal2 Fitted Gauss, proteinAT2G22450.1
bic:740 735
peaks after deleting less than one fifth of global max:11
×10^7
4.5
4
3.5
3
2.5
2
1.5
1
0.5
0
y
0
5
10
15
20
25
x
raw data, starting:11 16
fitted Gauss locations:11 16

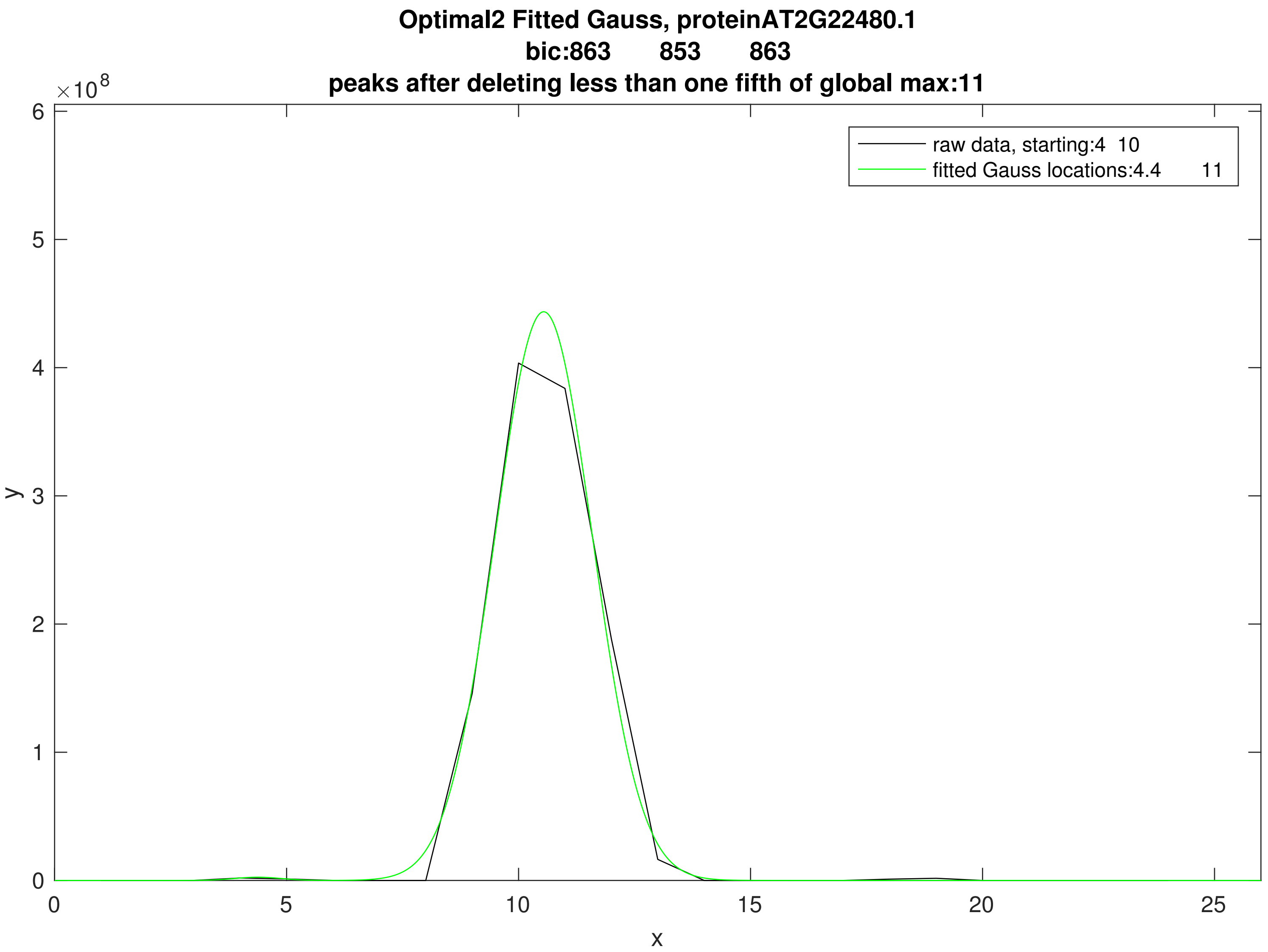
Optimal2 Fitted Gauss, proteinAT2G22480.1
bic:863 853 863
peaks after deleting less than one fifth of global max:11
×10^8
6
5
4
3
2
1
0
y
0
5
10
15
20
25
x
raw data, starting:4 10
fitted Gauss locations:4.4 11

Optimal1 Fitted Gauss, proteinAT2G22570.1;AT2G22570.2
bic:793
peaks after deleting less than one fifth of global max:15

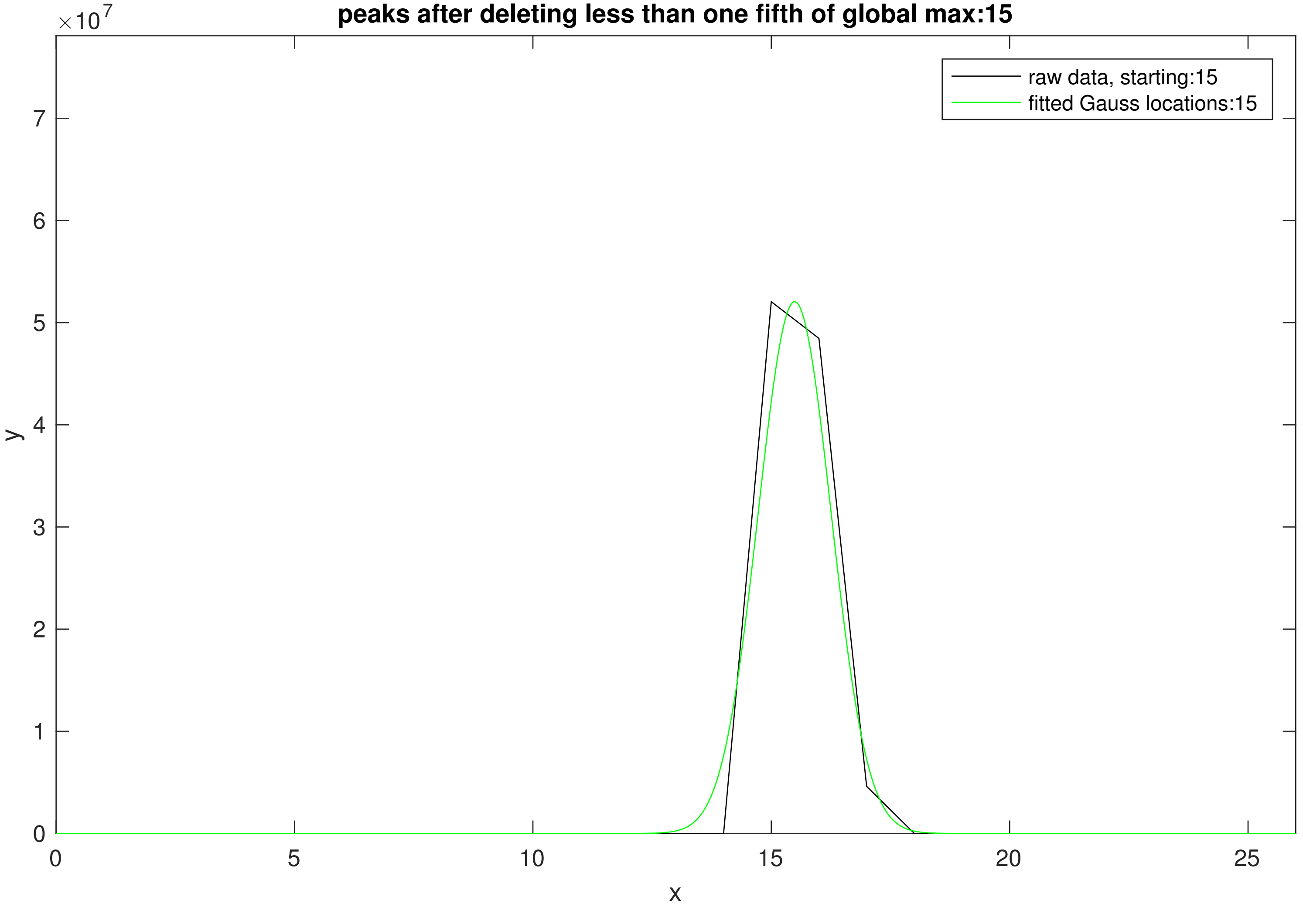

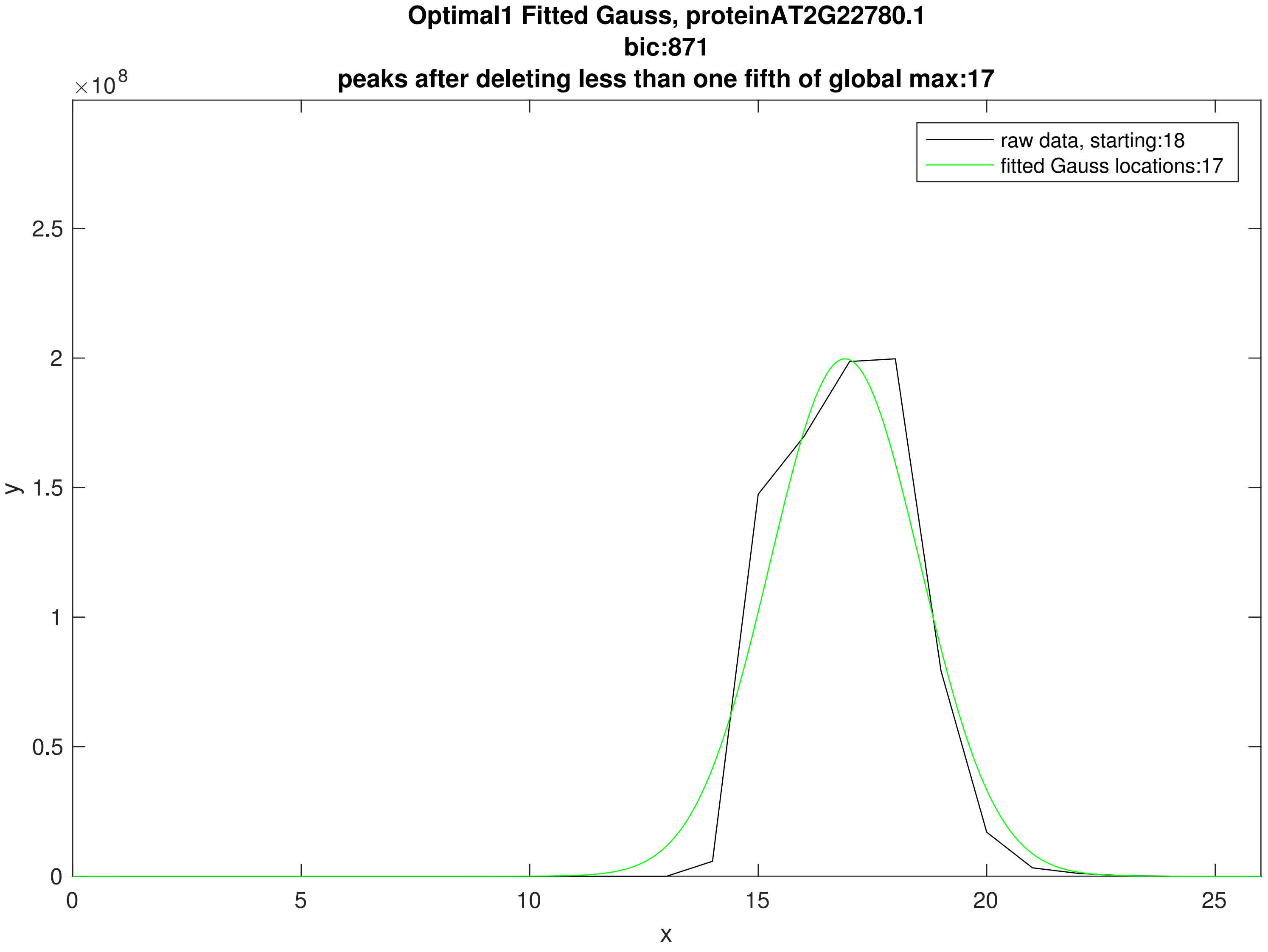

Optimal1 Fitted Gauss, proteinAT2G22780.1
bic:871
peaks after deleting less than one fifth of global max:17
×10^8
raw data, starting:18
fitted Gauss locations:17
y
x
2.5
2
1.5
1
0.5
0
5
10
15
20
25

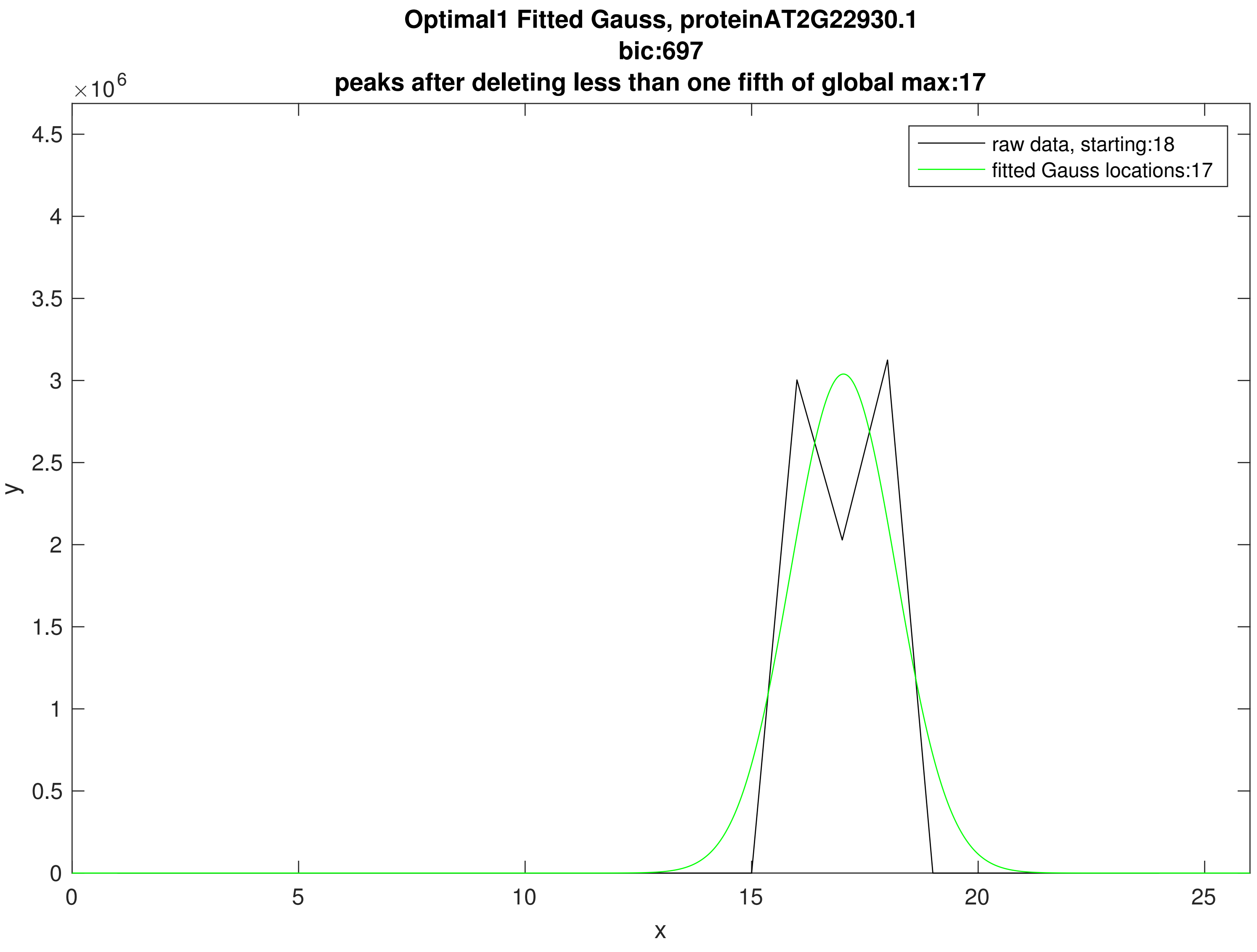

Optimal1 Fitted Gauss, proteinAT2G22930.1
bic:697
peaks after deleting less than one fifth of global max:17
×10^6
raw data, starting:18
fitted Gauss locations:17
y
x
0
0.5
1
1.5
2
2.5
3
3.5
4
4.5
0
5
10
15
20
25

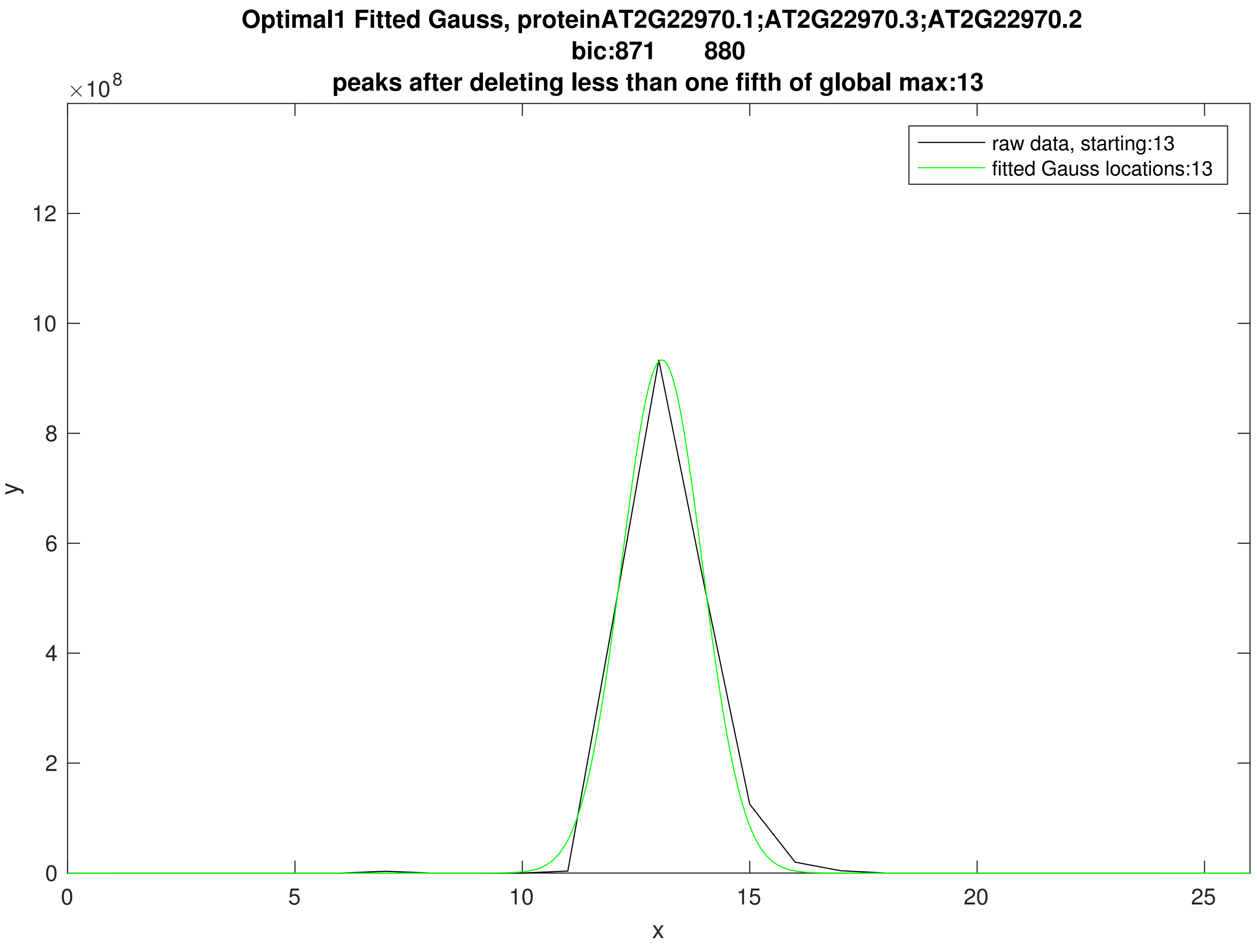

Optimal1 Fitted Gauss, proteinAT2G22970.1;AT2G22970.3;AT2G22970.2
bic:871 880
peaks after deleting less than one fifth of global max:13
×10^8
raw data, starting:13
fitted Gauss locations:13
y
x

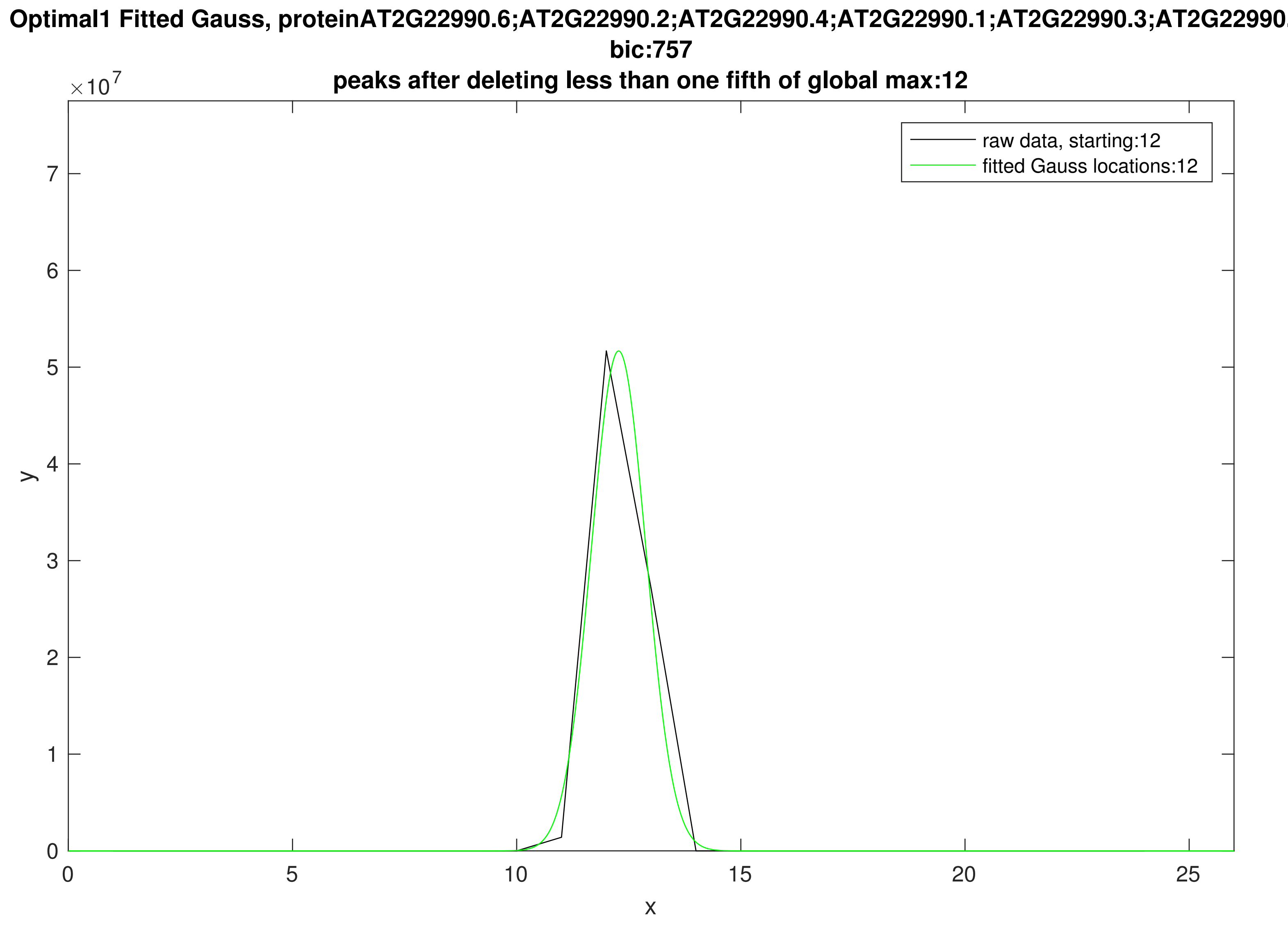
Optimal1 Fitted Gauss, proteinAT2G22990.6;AT2G22990.2;AT2G22990.4;AT2G22990.1;AT2G22990.3;AT2G22990.
bic:757
peaks after deleting less than one fifth of global max:12
×10^7
raw data, starting:12
fitted Gauss locations:12
y
x
0
1
2
3
4
5
6
7
0
5
10
15
20
25

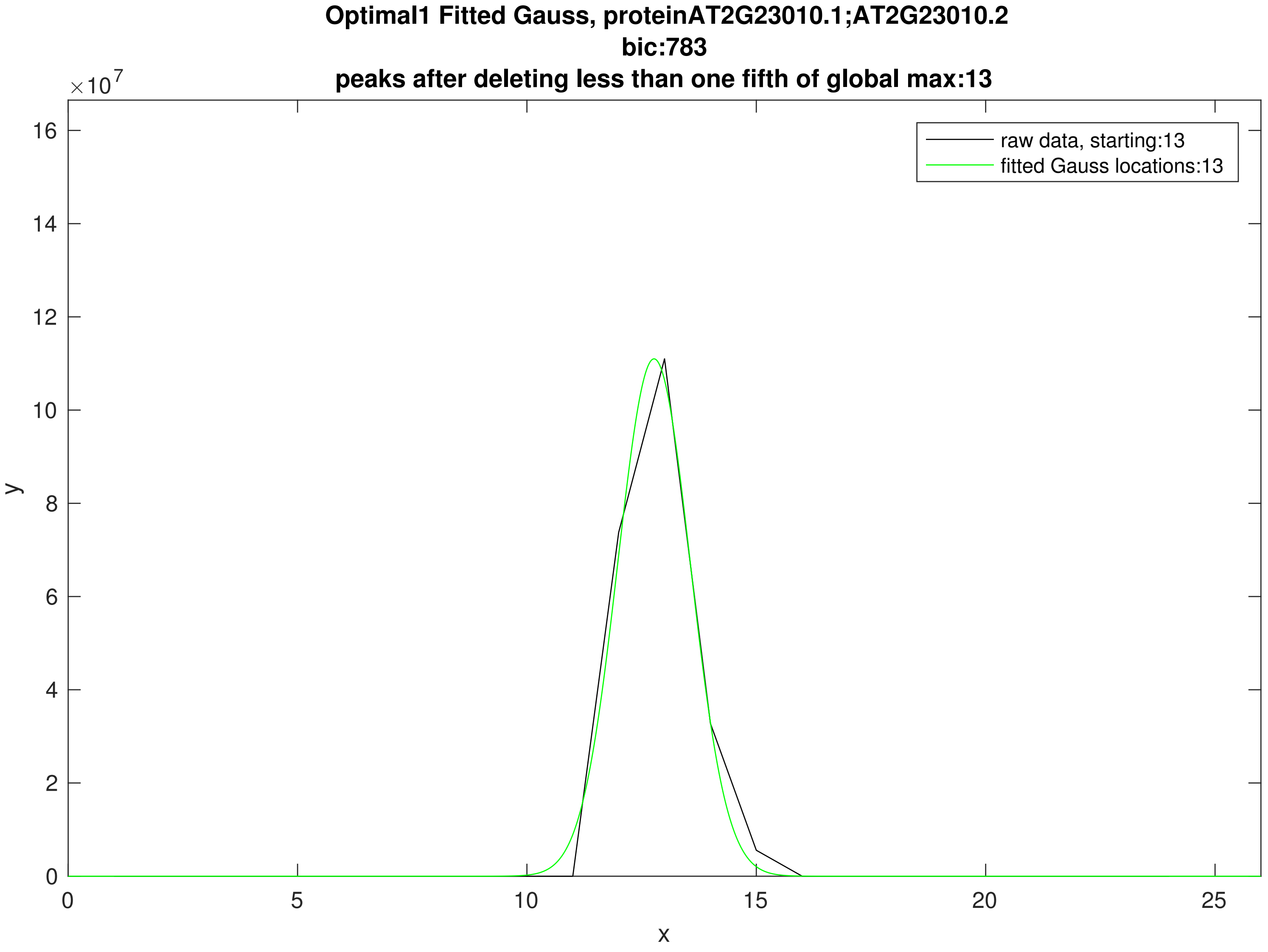
Optimal1 Fitted Gauss, proteinAT2G23010.1;AT2G23010.2
bic:783
peaks after deleting less than one fifth of global max:13
×10^7
raw data, starting:13
fitted Gauss locations:13
y
x
0
2
4
6
8
10
12
14
16
5
15
20
25

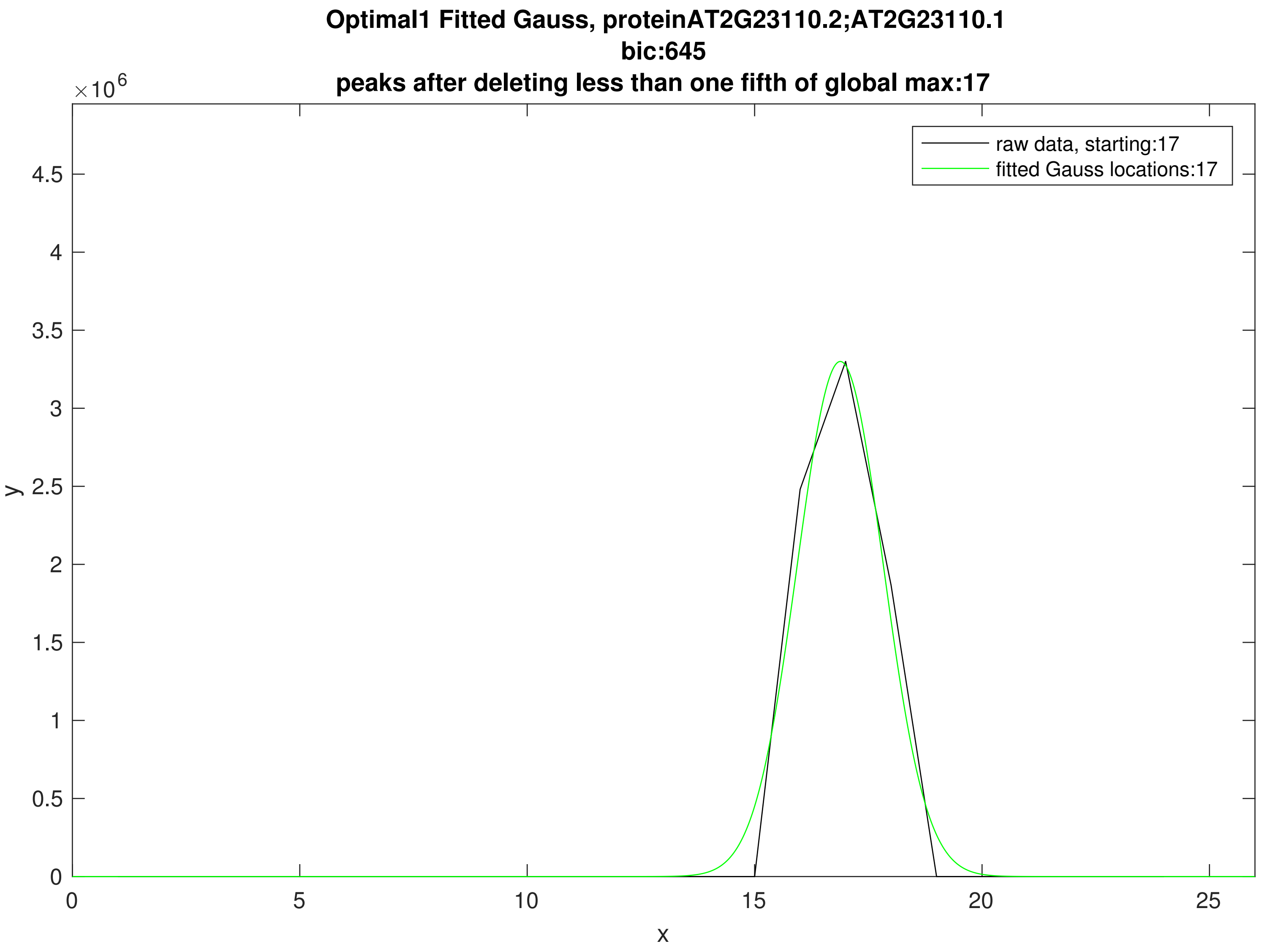

Optimal1 Fitted Gauss, proteinAT2G23110.2;AT2G23110.1
bic:645
peaks after deleting less than one fifth of global max:17
×10^6
raw data, starting:17
fitted Gauss locations:17
y
x
0
0.5
1
1.5
2
2.5
3
3.5
4
4.5
5
10
15
20
25

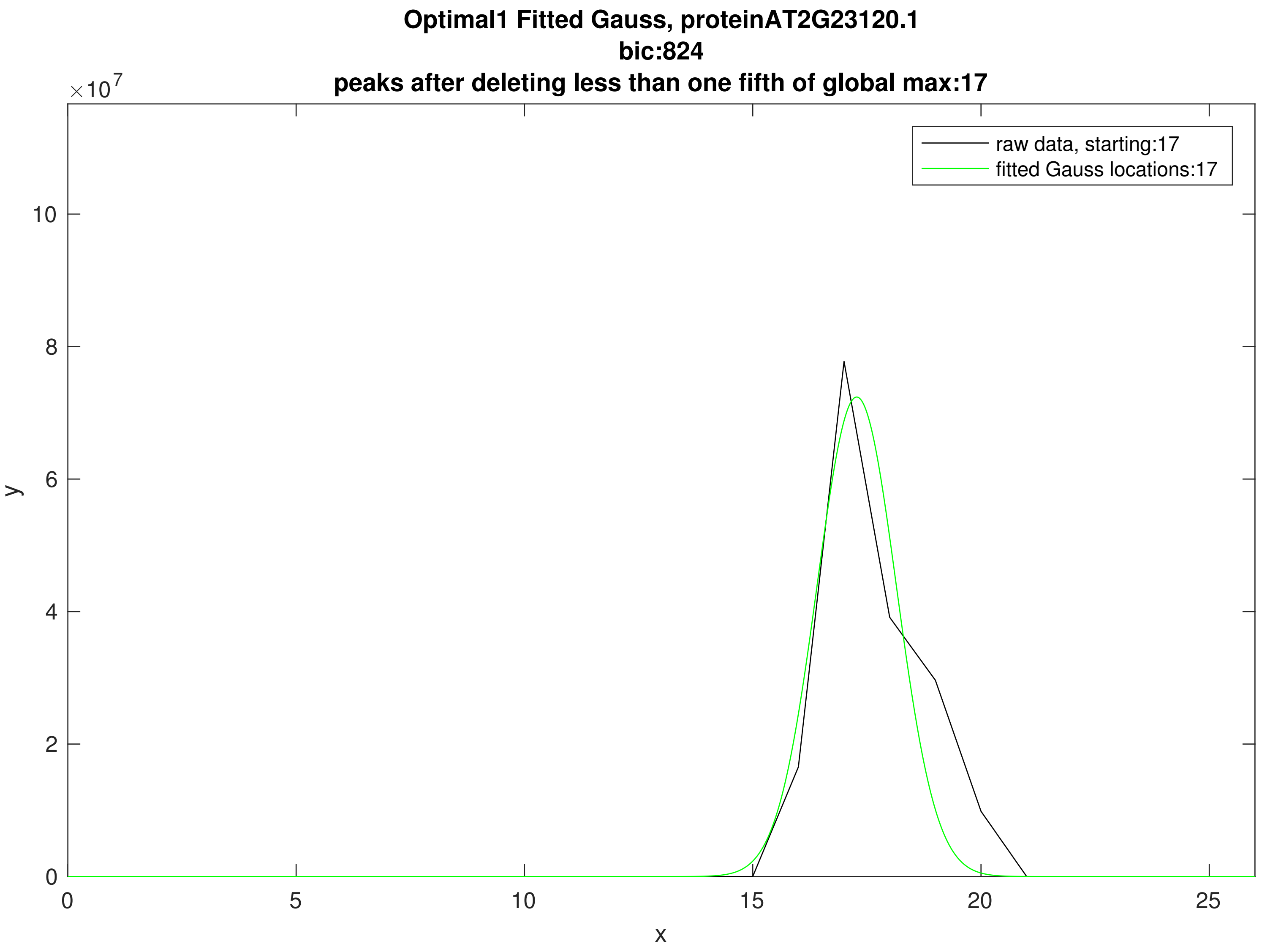

Optimal1 Fitted Gauss, proteinAT2G23120.1
bic:824
peaks after deleting less than one fifth of global max:17
×10^7
raw data, starting:17
fitted Gauss locations:17
10
8
6
4
2
0
y
0
5
10
15
20
25
x

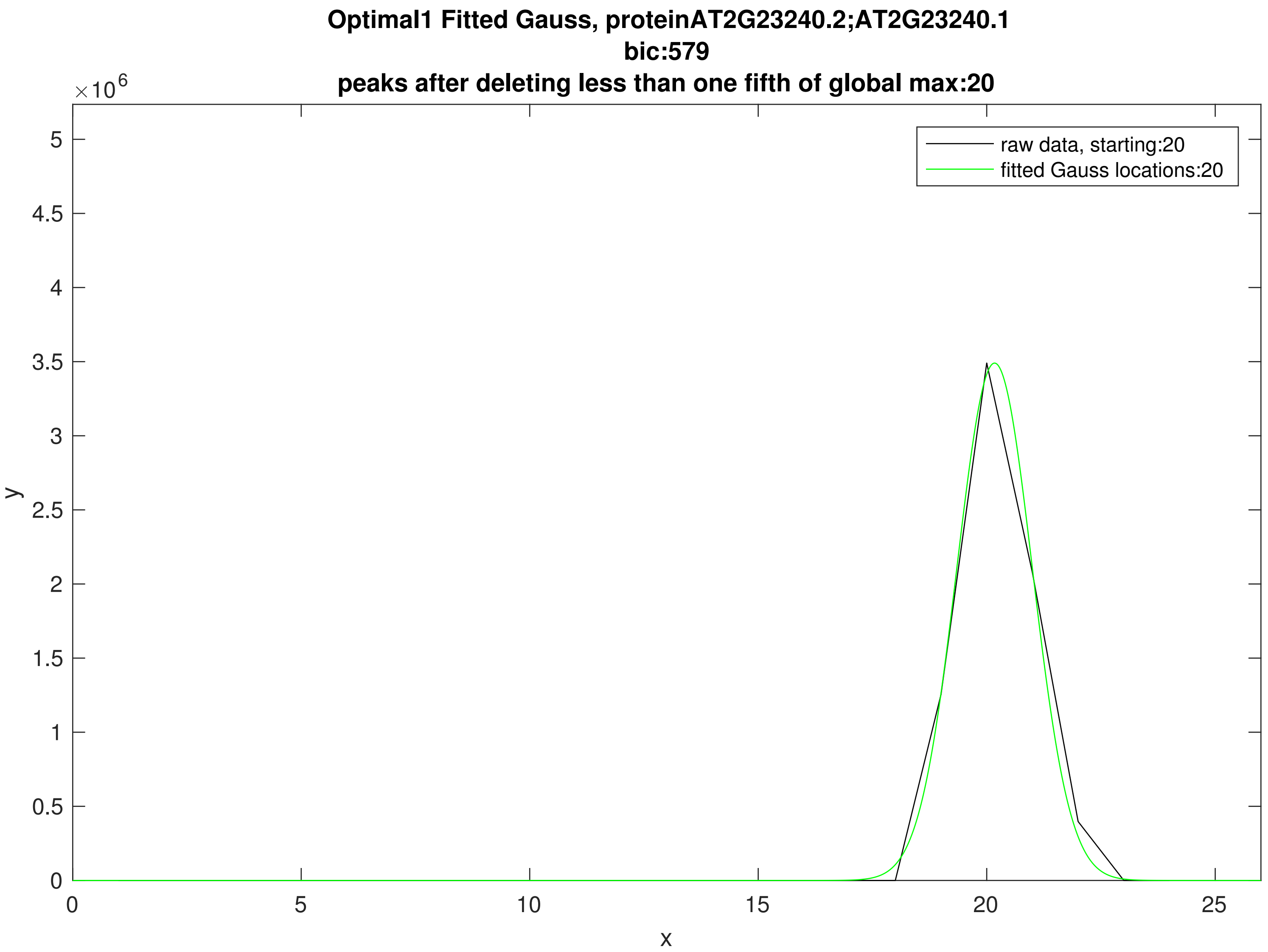
Optimal1 Fitted Gauss, proteinAT2G23240.2;AT2G23240.1
bic:579
peaks after deleting less than one fifth of global max:20
×10^6
raw data, starting:20
fitted Gauss locations:20
y
x
0
0.5
1
1.5
2
2.5
3
3.5
4
4.5
5
0
5
10
15
20
25

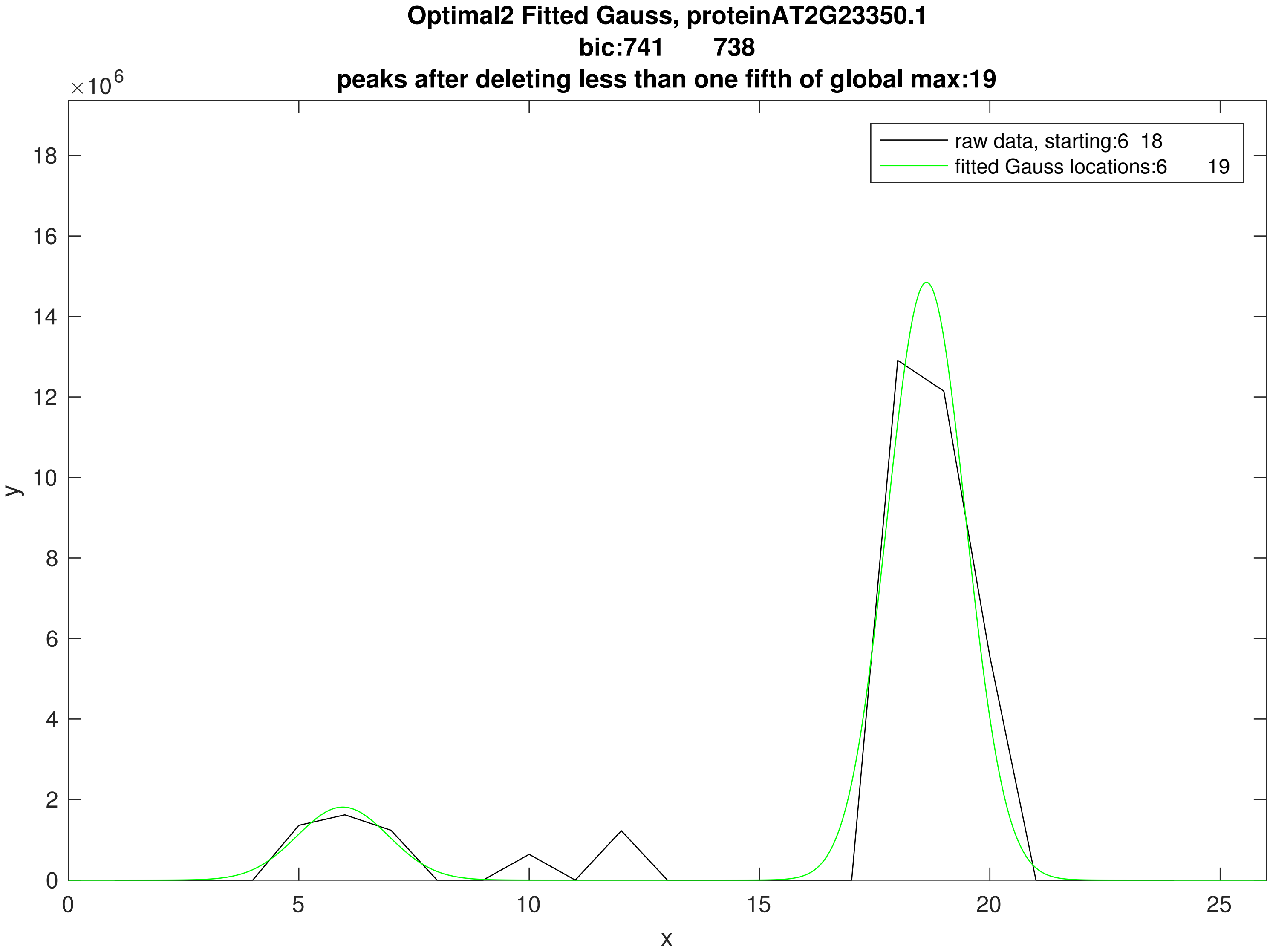
Optimal2 Fitted Gauss, proteinAT2G23350.1
bic:741 738
peaks after deleting less than one fifth of global max:19
×10^6
raw data, starting:6 18
fitted Gauss locations:6 19
y
x
0
2
4
6
8
10
12
14
16
18
0
5
10
15
20
25

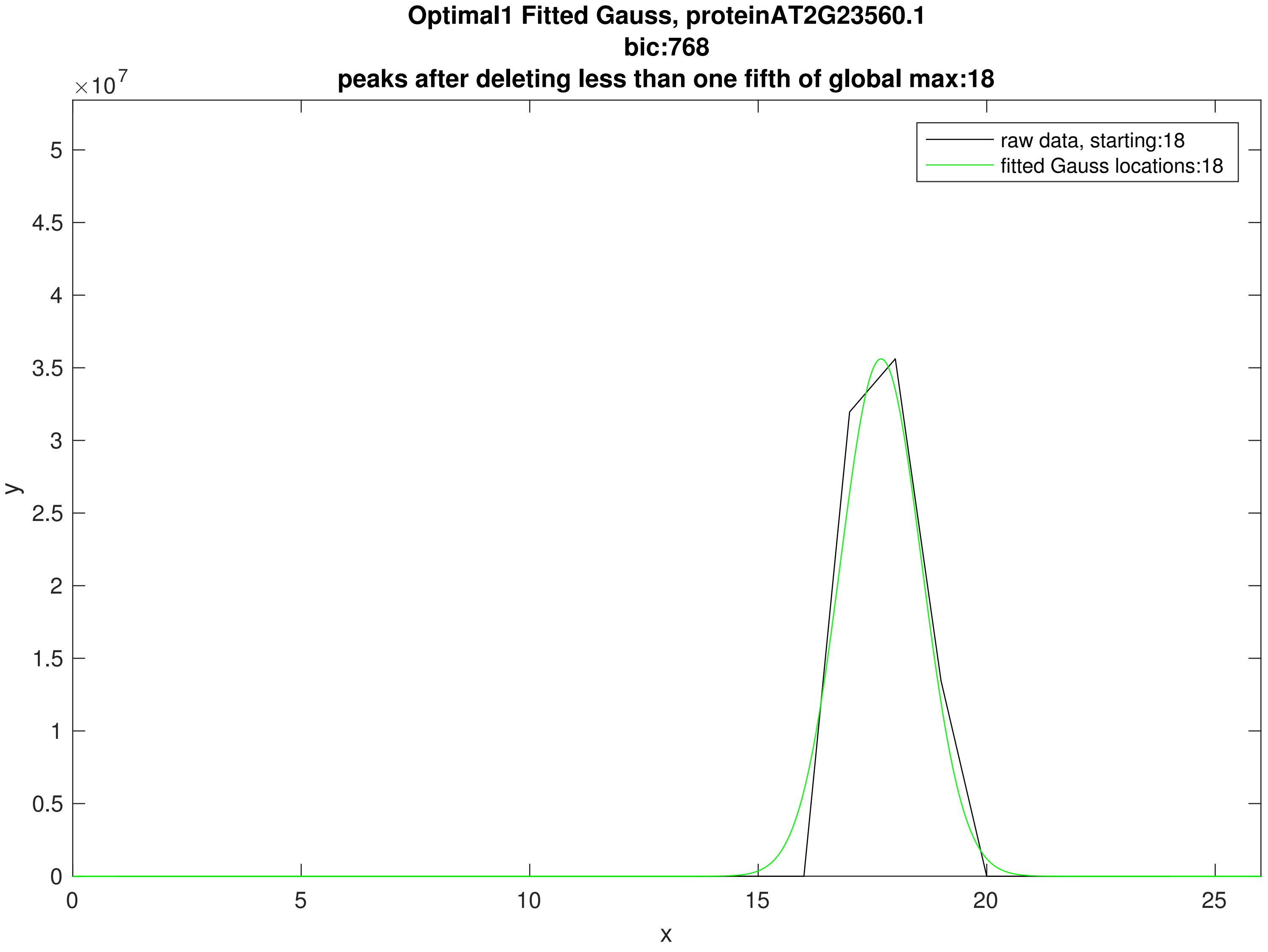

Optimal1 Fitted Gauss, proteinAT2G23560.1
bic:768
peaks after deleting less than one fifth of global max:18
×10^7
raw data, starting:18
fitted Gauss locations:18
y
x
0
0.5
1
1.5
2
2.5
3
3.5
4
4.5
5
0
5
10
15
20
25

**Optimal1 Fitted Gauss, proteinAT2G23590.1**
**bic:588**
**peaks after deleting less than one fifth of global max:16**

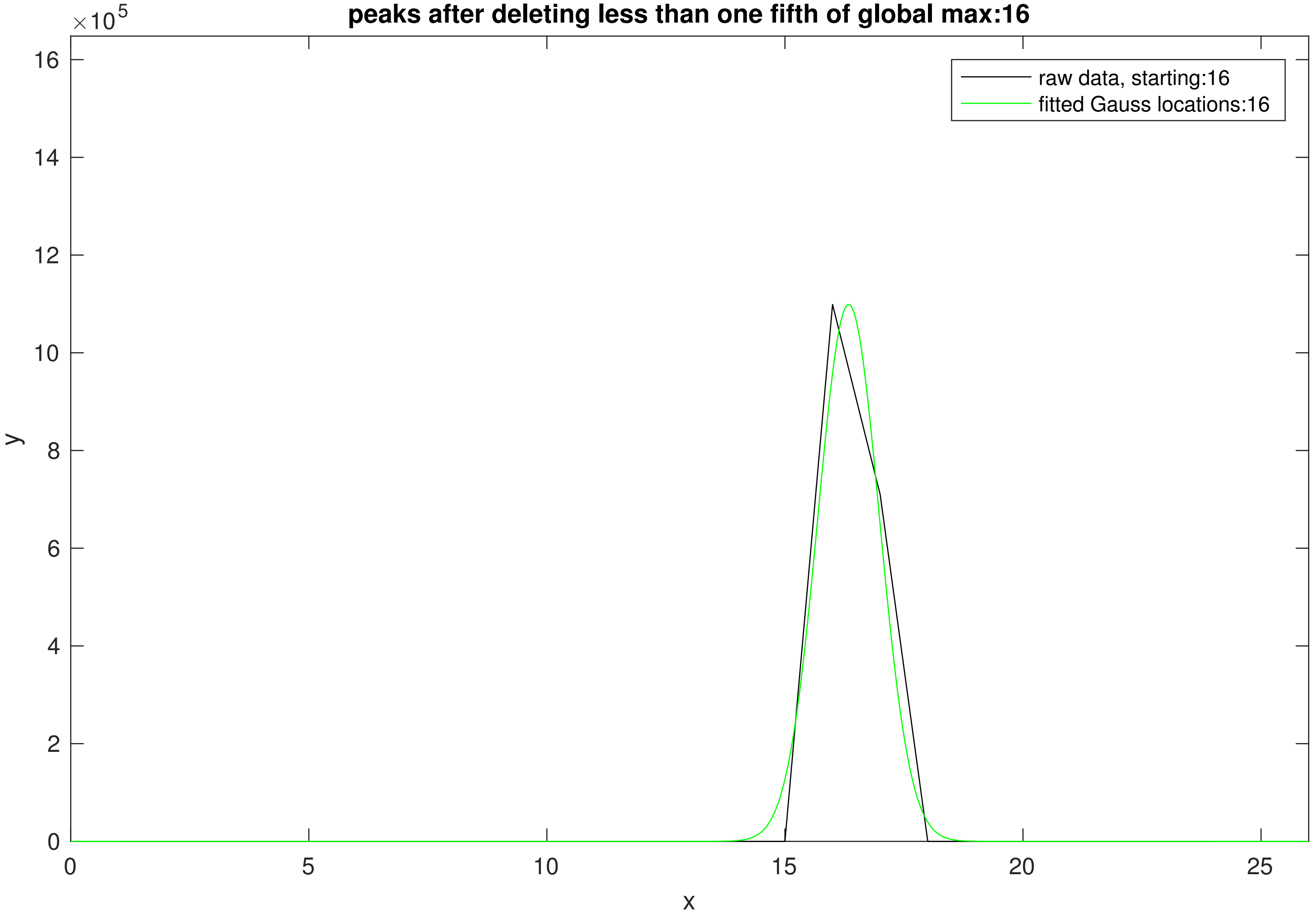

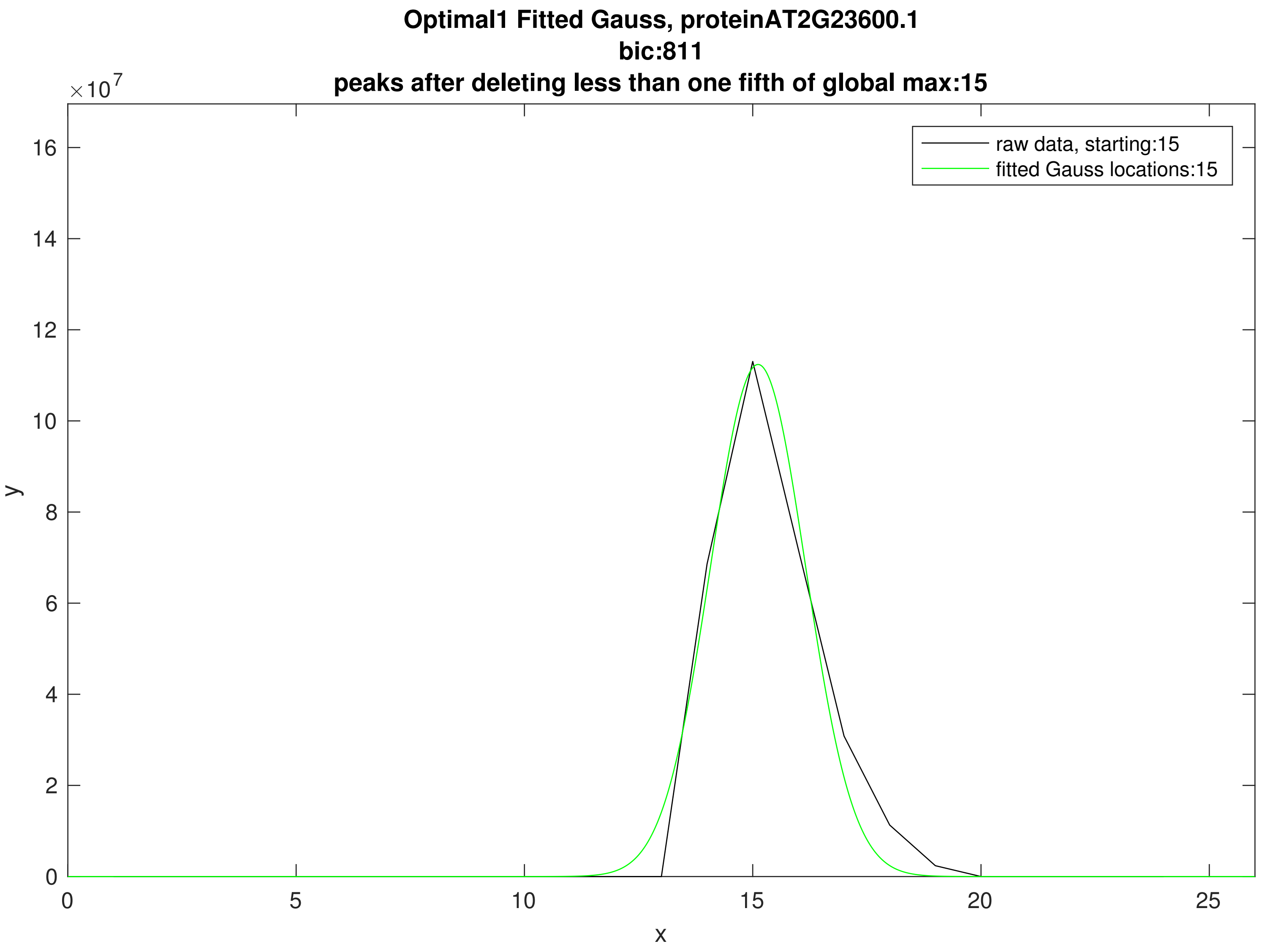

Optimal1 Fitted Gauss, proteinAT2G23600.1
bic:811
peaks after deleting less than one fifth of global max:15
×10^7
16
14
12
10
8
6
4
2
0
y
0
5
10
15
20
25
x
raw data, starting:15
fitted Gauss locations:15

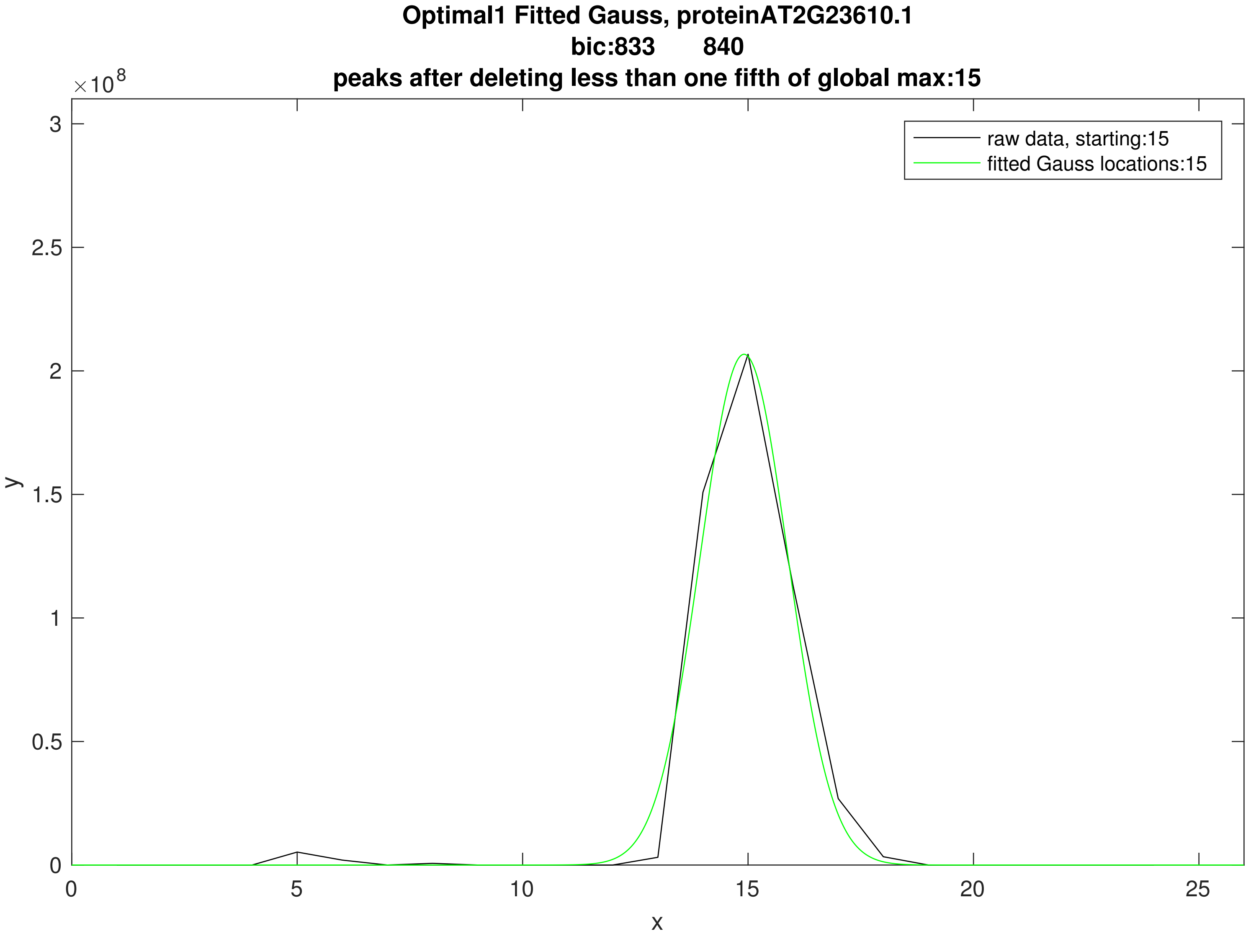

Optimal1 Fitted Gauss, proteinAT2G23610.1
bic:833 840
peaks after deleting less than one fifth of global max:15
×10^8
raw data, starting:15
fitted Gauss locations:15
y
3
2.5
2
1.5
1
0.5
0
0
5
10
15
20
25
x

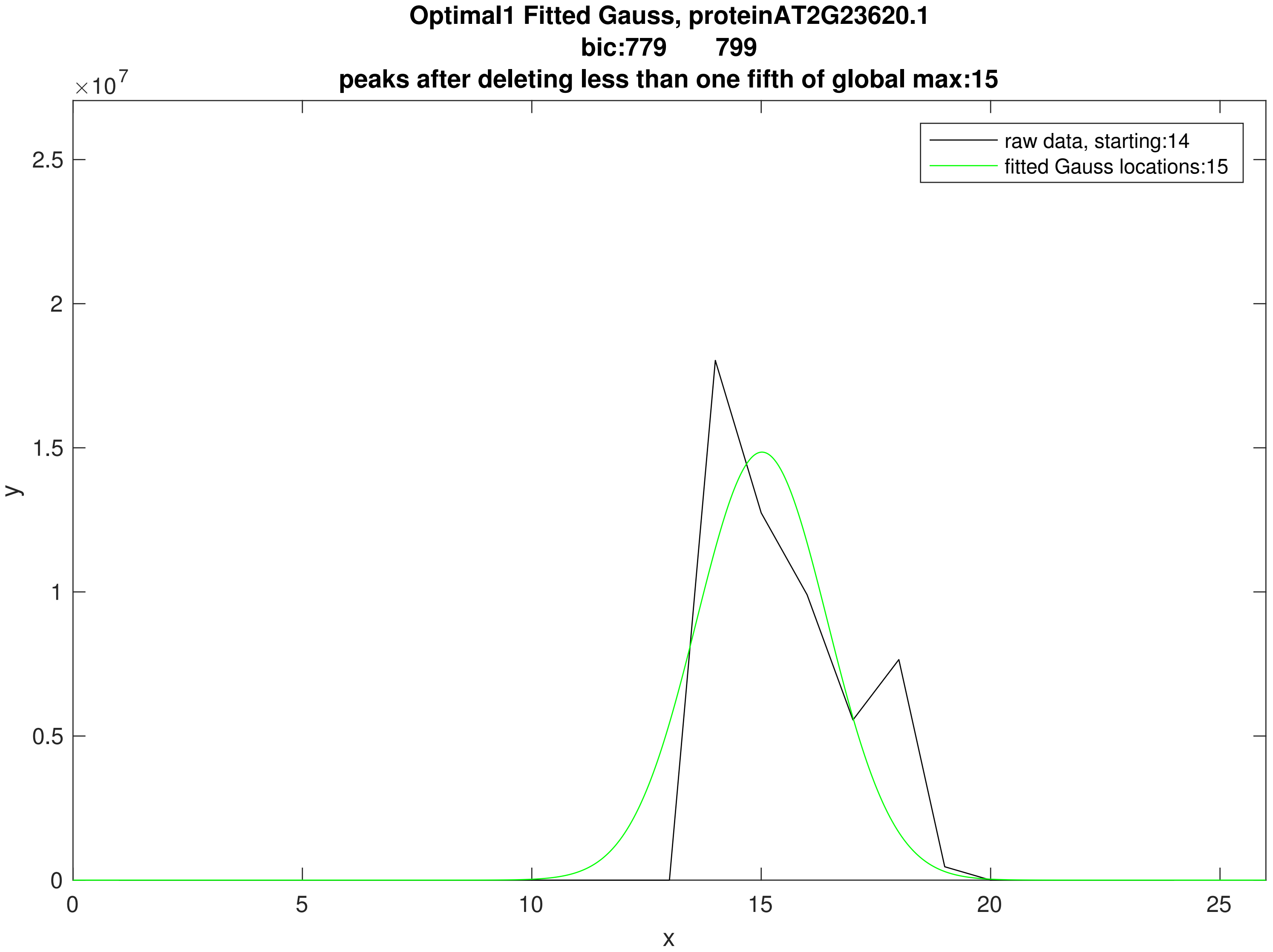

Optimal1 Fitted Gauss, proteinAT2G23620.1
bic:779 799
peaks after deleting less than one fifth of global max:15
×10^7
2.5
2
1.5
1
0.5
0
y
0
5
10
15
20
25
x
raw data, starting:14
fitted Gauss locations:15

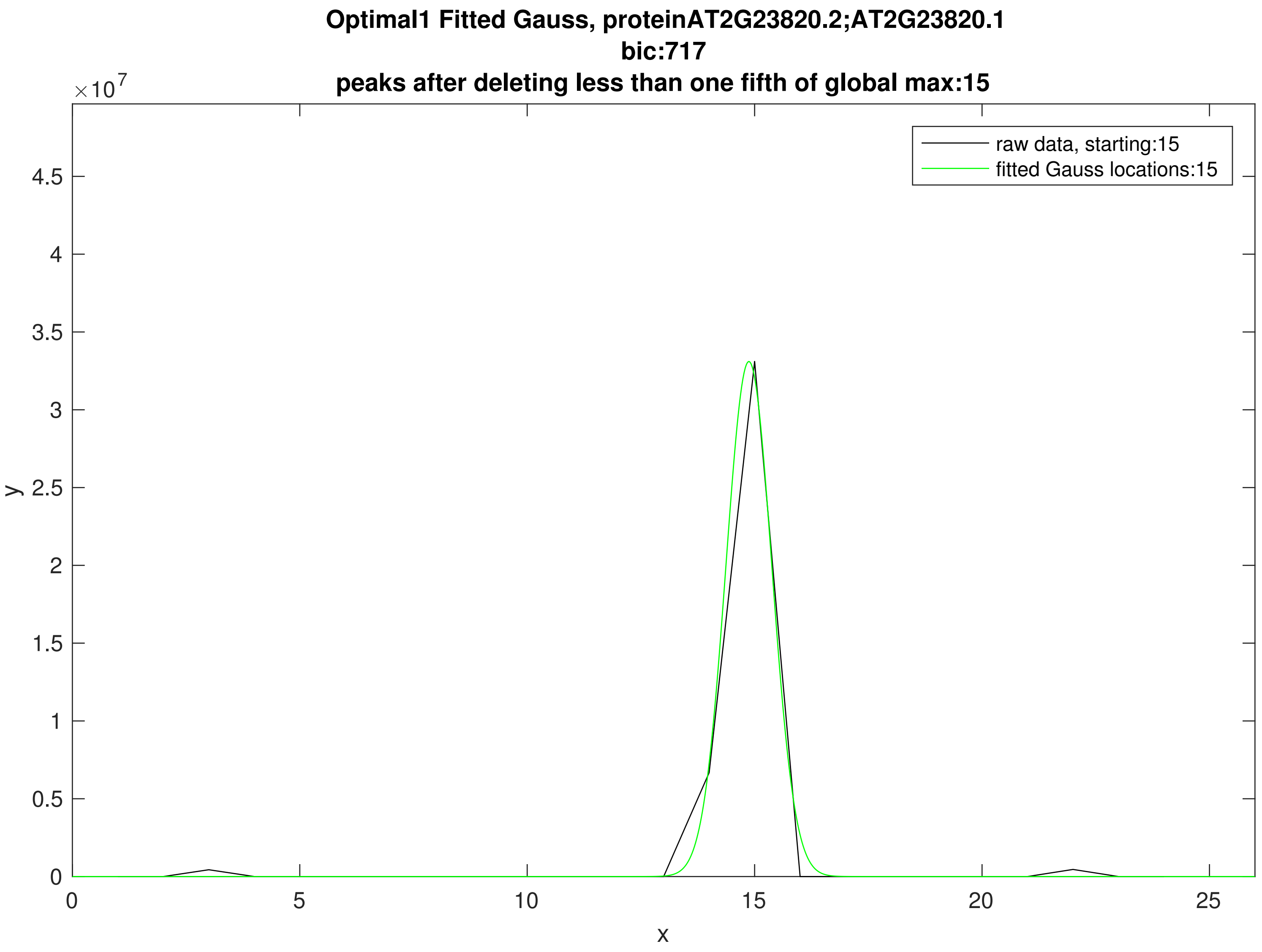
Optimal1 Fitted Gauss, proteinAT2G23820.2;AT2G23820.1
bic:717
peaks after deleting less than one fifth of global max:15
×10^7
raw data, starting:15
fitted Gauss locations:15
y
x
0
0.5
1
1.5
2
2.5
3
3.5
4
4.5
5
10
15
20
25

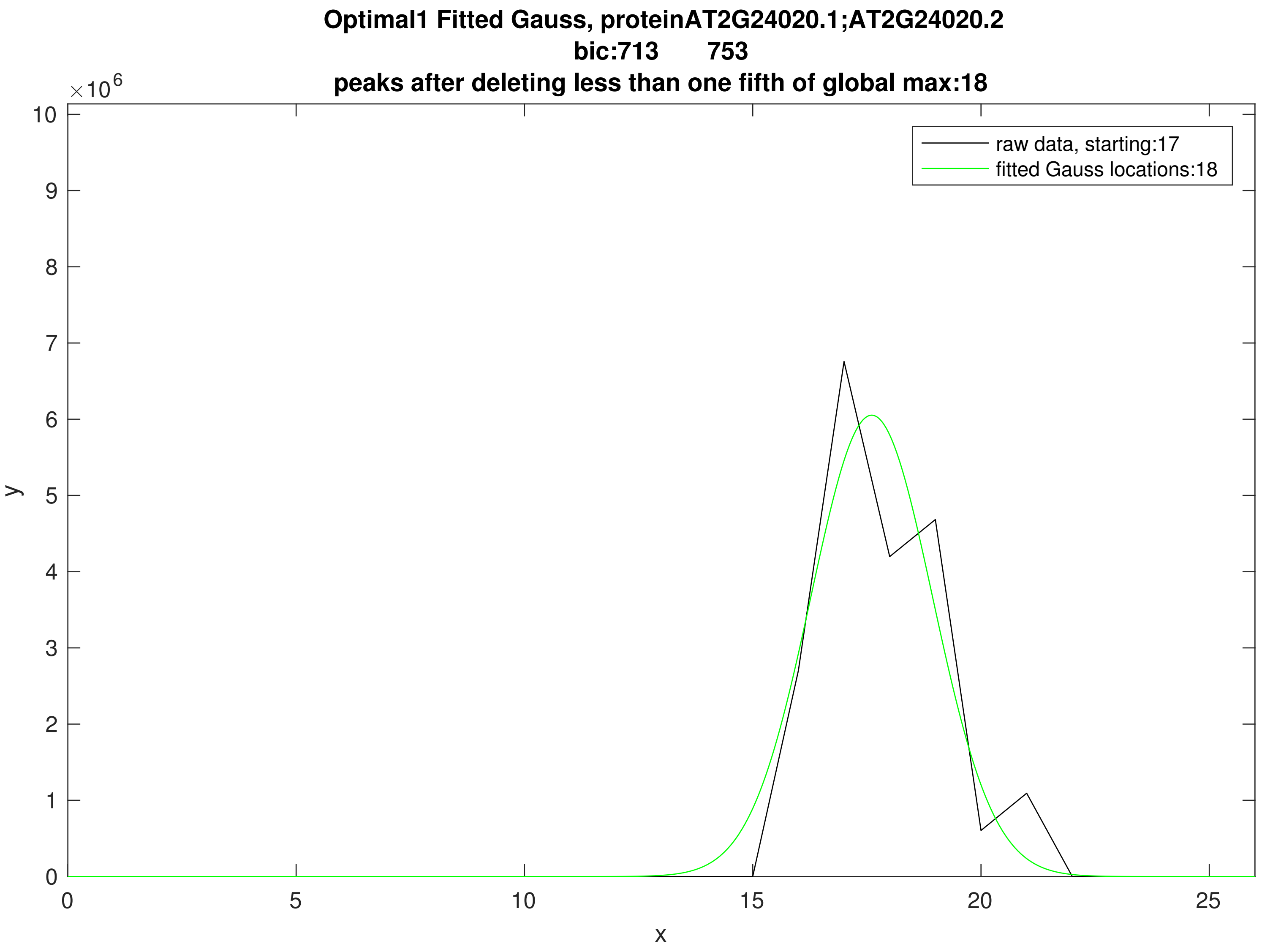
Optimal1 Fitted Gauss, proteinAT2G24020.1;AT2G24020.2
bic:713 753
peaks after deleting less than one fifth of global max:18
×10^6
raw data, starting:17
fitted Gauss locations:18
y
x
0
5
10
15
20
25
1
2
3
4
6
7
8
9

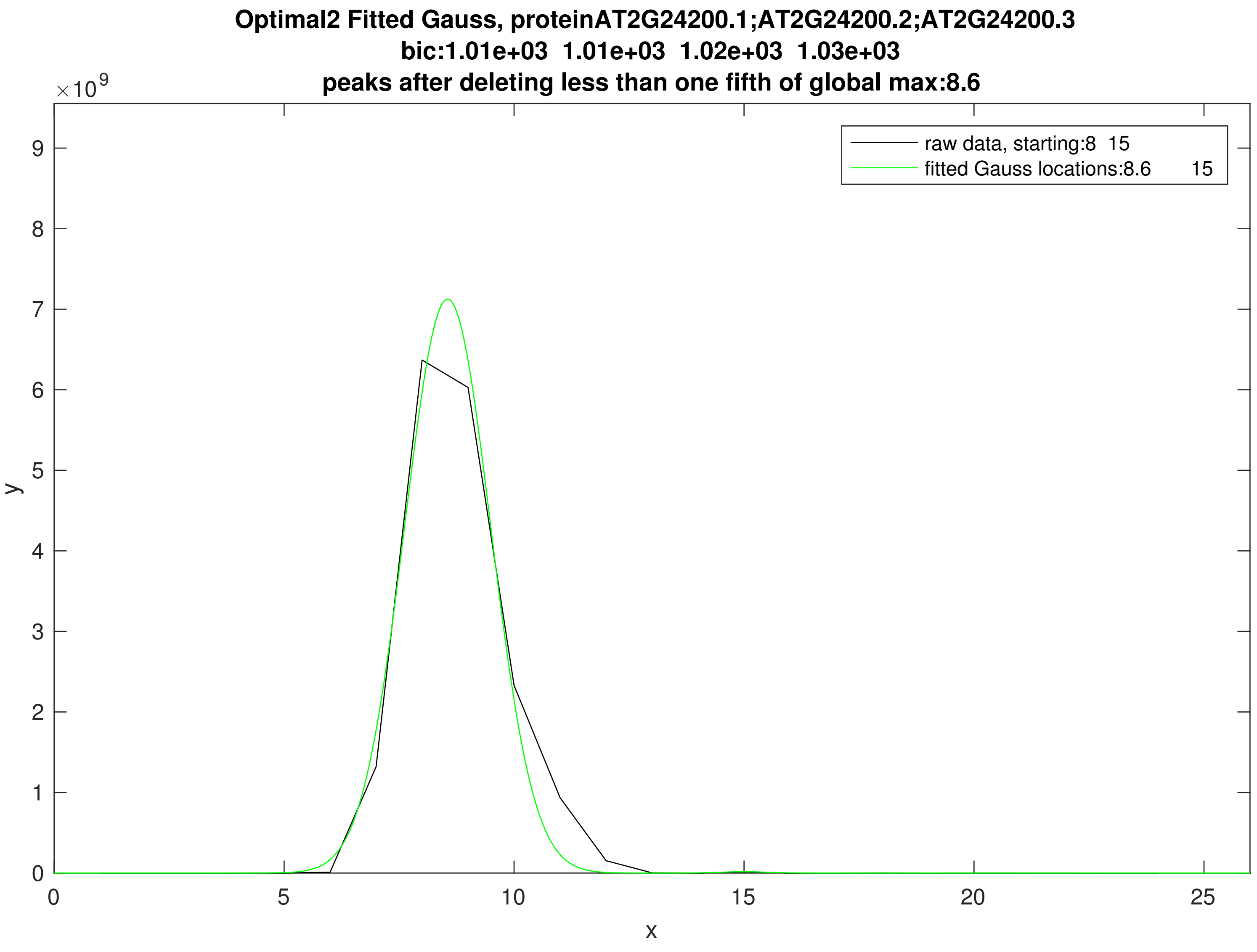
Optimal2 Fitted Gauss, proteinAT2G24200.1;AT2G24200.2;AT2G24200.3
bic:1.01e+03 1.01e+03 1.02e+03 1.03e+03
peaks after deleting less than one fifth of global max:8.6
×10^9
raw data, starting:8 15
fitted Gauss locations:8.6 15
y
x
0
1
2
3
4
5
6
7
8
9
0
5
10
15
20
25

**Optimal1 Fitted Gauss, proteinAT2G24270.3;AT2G24270.2;AT2G24270.1;AT2G24270.4**
**bic:835**
**peaks after deleting less than one fifth of global max:11**

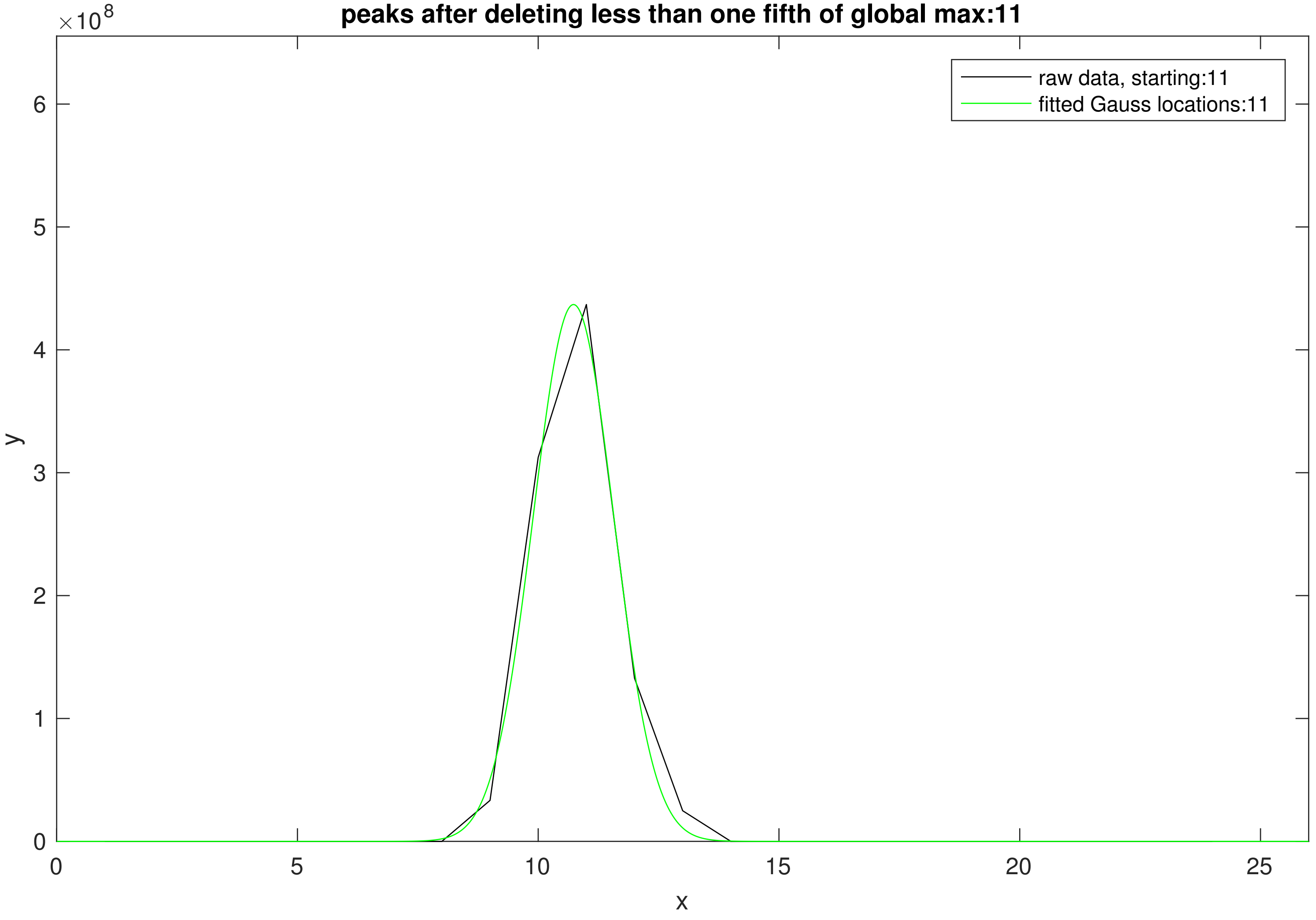

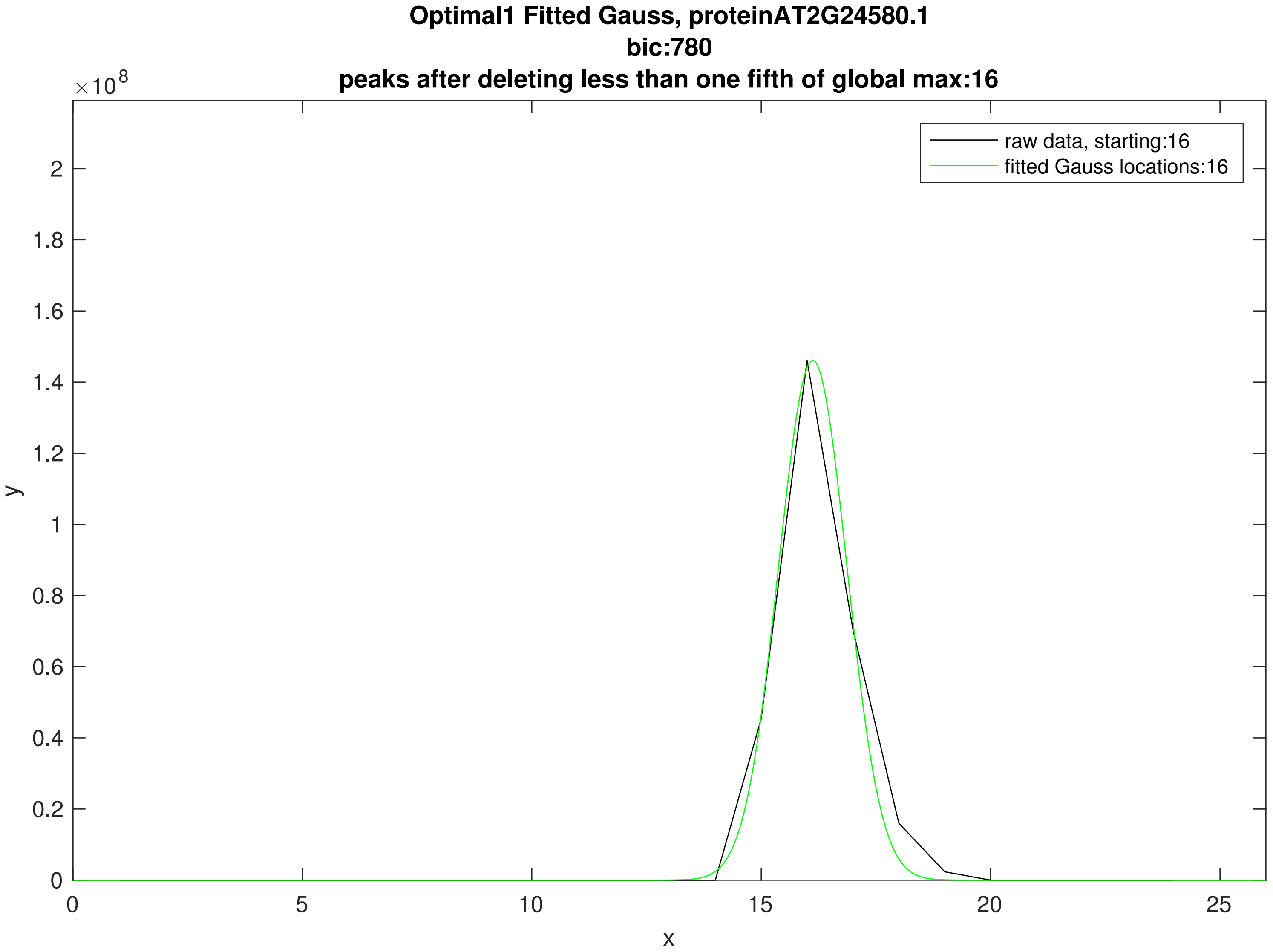
Optimal1 Fitted Gauss, proteinAT2G24580.1
bic:780
peaks after deleting less than one fifth of global max:16
×10^8
raw data, starting:16
fitted Gauss locations:16
y
x
0
0.2
0.4
0.6
0.8
1
1.2
1.4
1.6
1.8
2
0
5
10
15
20
25

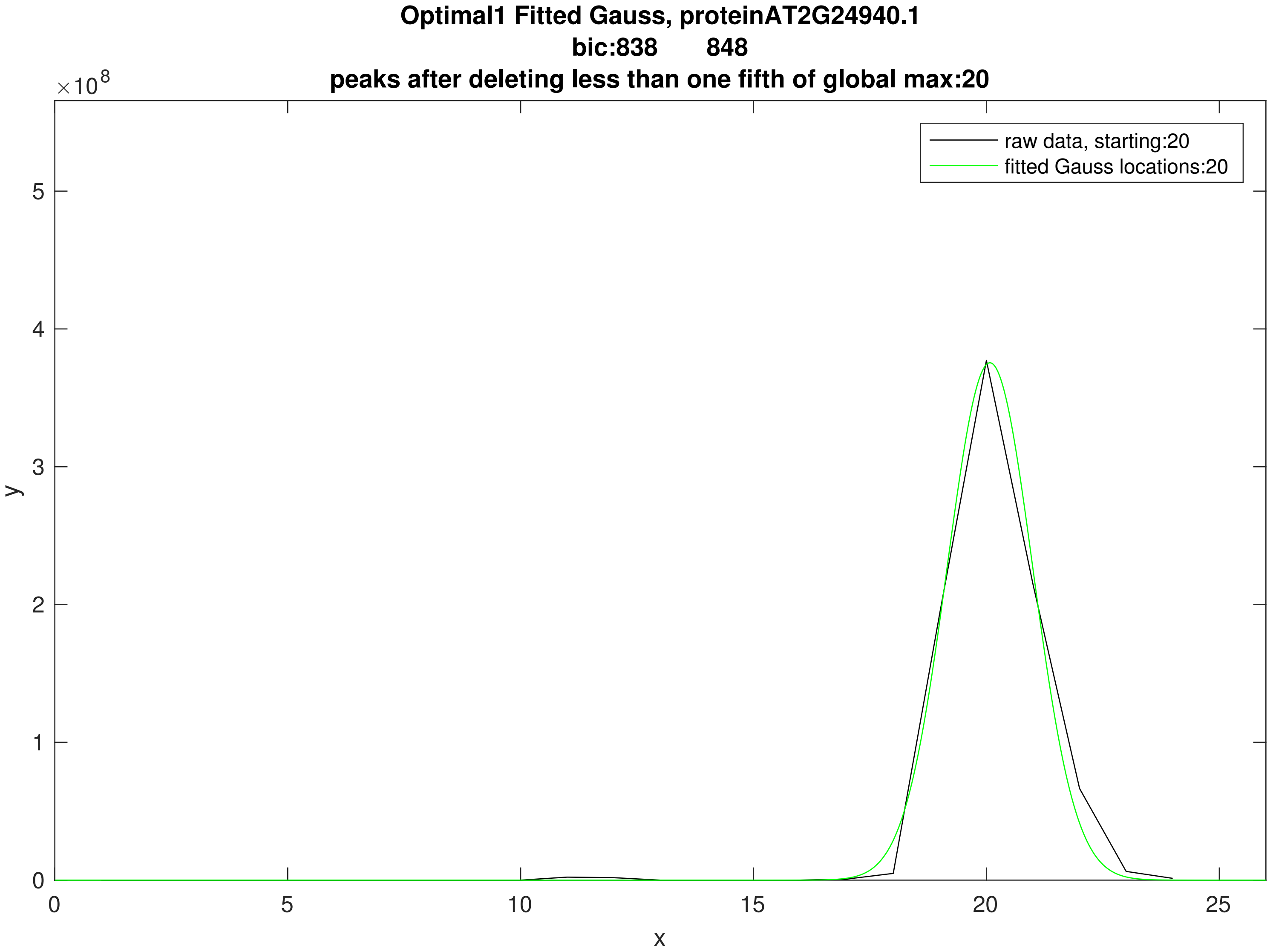

Optimal1 Fitted Gauss, proteinAT2G24940.1
bic:838 848
peaks after deleting less than one fifth of global max:20
×10^8
raw data, starting:20
fitted Gauss locations:20
5
4
3
2
1
0
y
0
5
10
15
20
25
x

**Optimal1 Fitted Gauss, proteinAT2G25060.1**
**bic:762**
**peaks after deleting less than one fifth of global max:8.8**

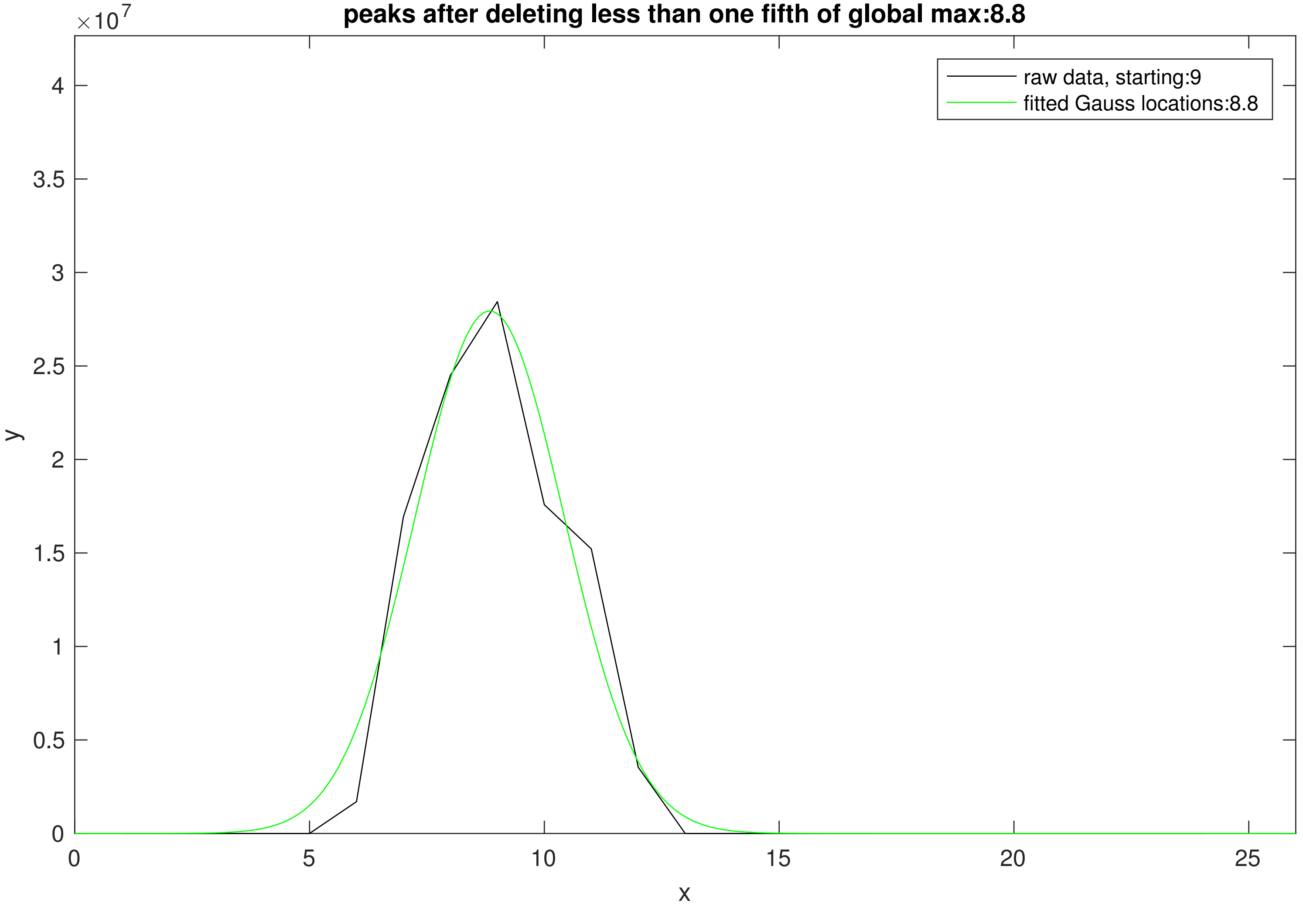

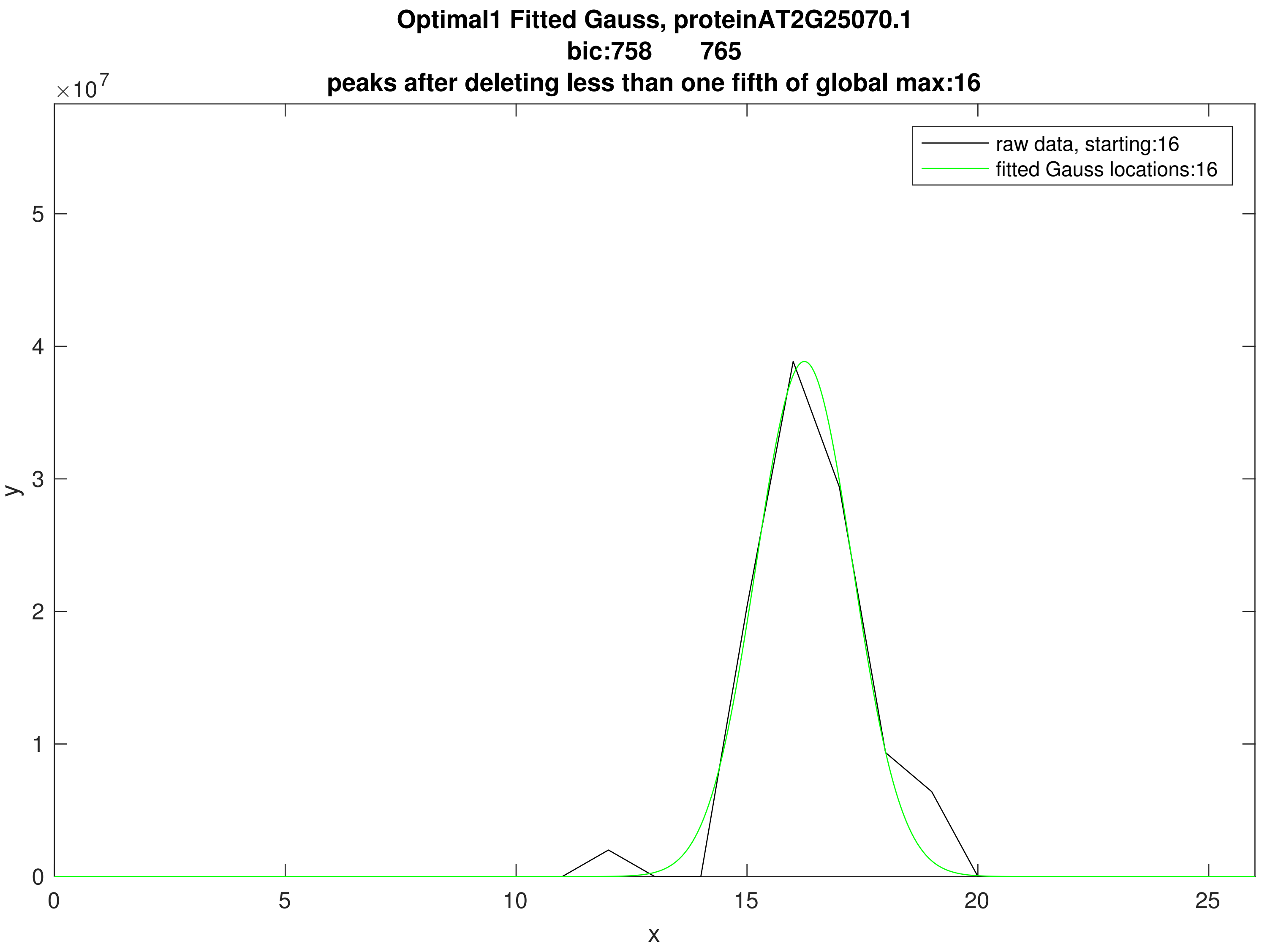

Optimal1 Fitted Gauss, proteinAT2G25070.1
bic:758 765
peaks after deleting less than one fifth of global max:16
×10^7
raw data, starting:16
fitted Gauss locations:16
y
x
0
1
2
3
4
5
0
5
10
15
20
25

**Optimal1 Fitted Gauss, proteinAT2G25280.1**
**bic:840 843 853**
**peaks after deleting less than one fifth of global max:19**

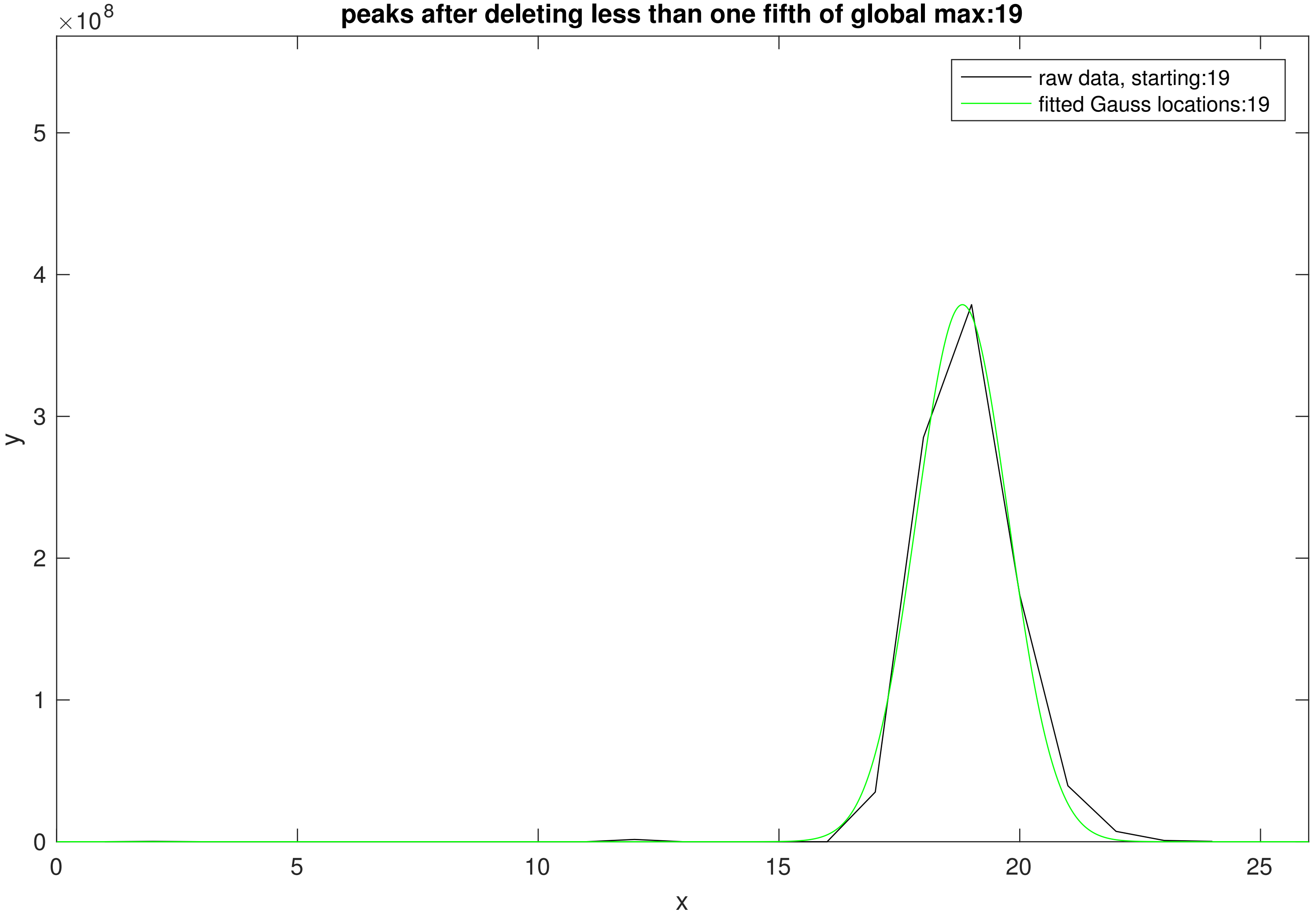

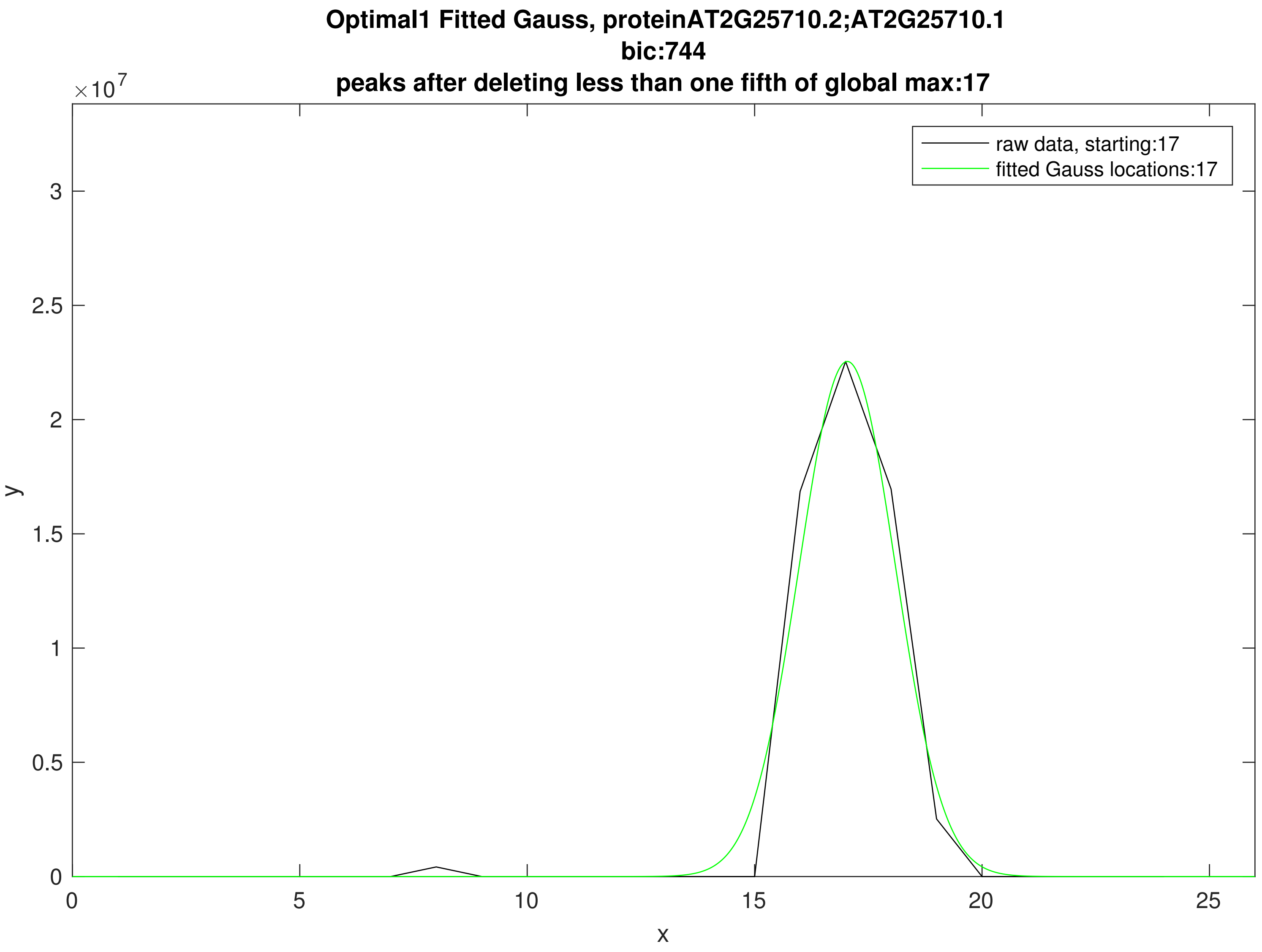
Optimal1 Fitted Gauss, proteinAT2G25710.2;AT2G25710.1
bic:744
peaks after deleting less than one fifth of global max:17
×10^7
raw data, starting:17
fitted Gauss locations:17
y
x
0
0.5
1
1.5
2
2.5
3
0
5
10
15
20
25

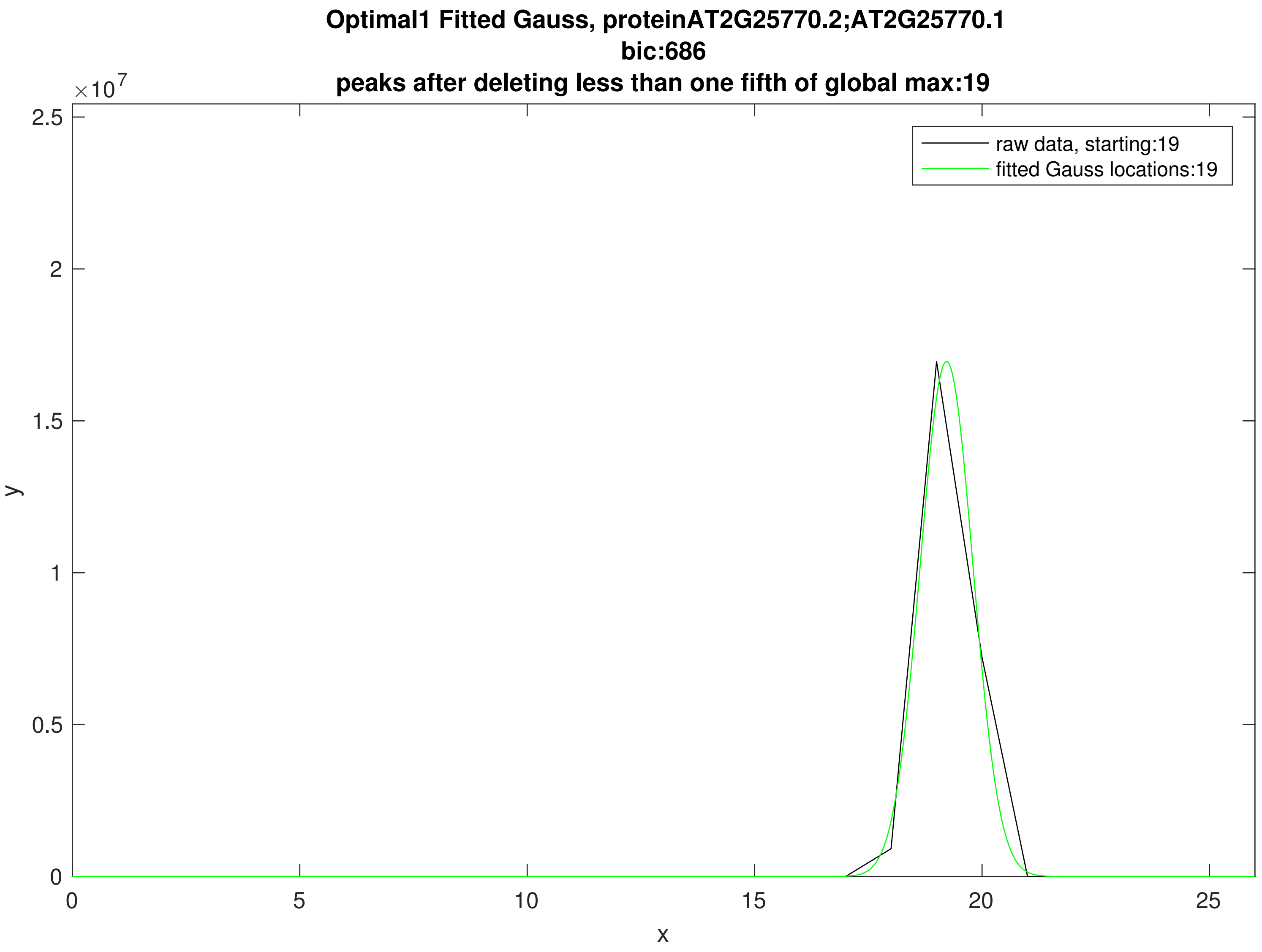

Optimal1 Fitted Gauss, proteinAT2G25770.2;AT2G25770.1
bic:686
peaks after deleting less than one fifth of global max:19
×10^7
2.5
2
1.5
1
0.5
0
y
0
5
10
15
20
25
x
raw data, starting:19
fitted Gauss locations:19

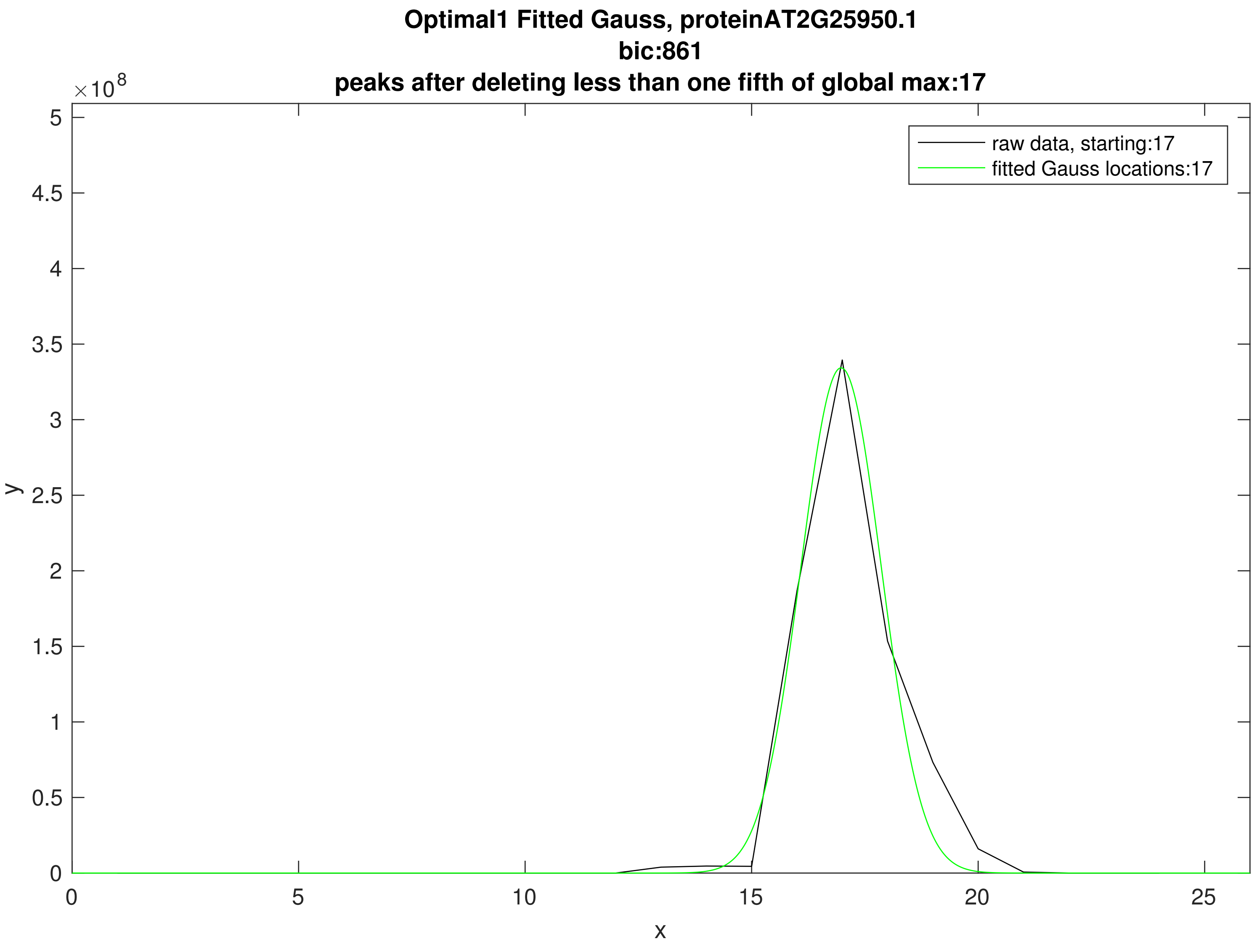

Optimal1 Fitted Gauss, proteinAT2G25950.1
bic:861
peaks after deleting less than one fifth of global max:17
×10^8
raw data, starting:17
fitted Gauss locations:17
y
x
0
0.5
1
1.5
2
2.5
3
3.5
4
4.5
5
0
5
10
15
20
25

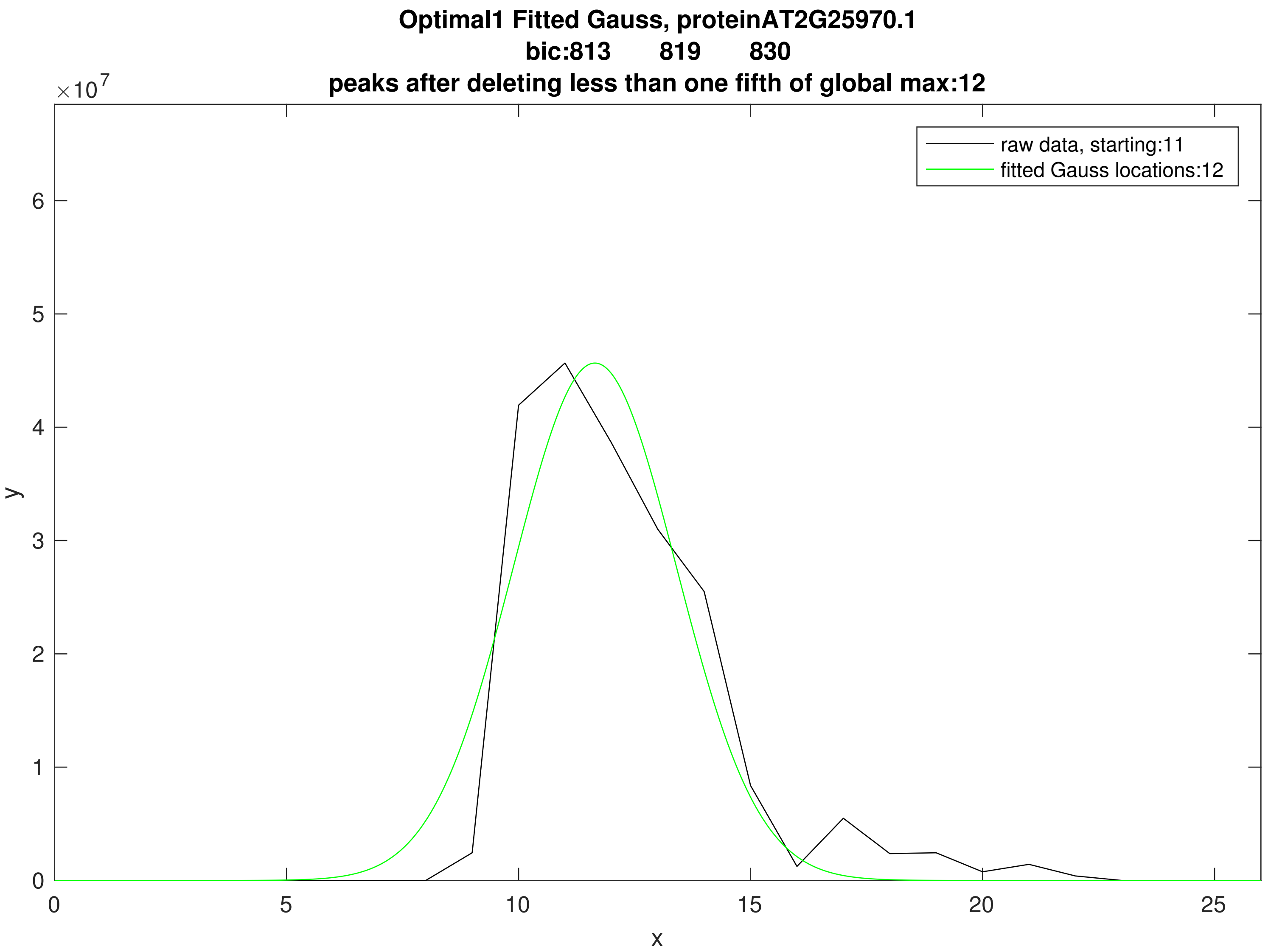

Optimal1 Fitted Gauss, proteinAT2G25970.1
bic:813 819 830
peaks after deleting less than one fifth of global max:12
×10^7
raw data, starting:11
fitted Gauss locations:12
y
x
0
1
2
3
4
5
6
0
5
10
15
20
25

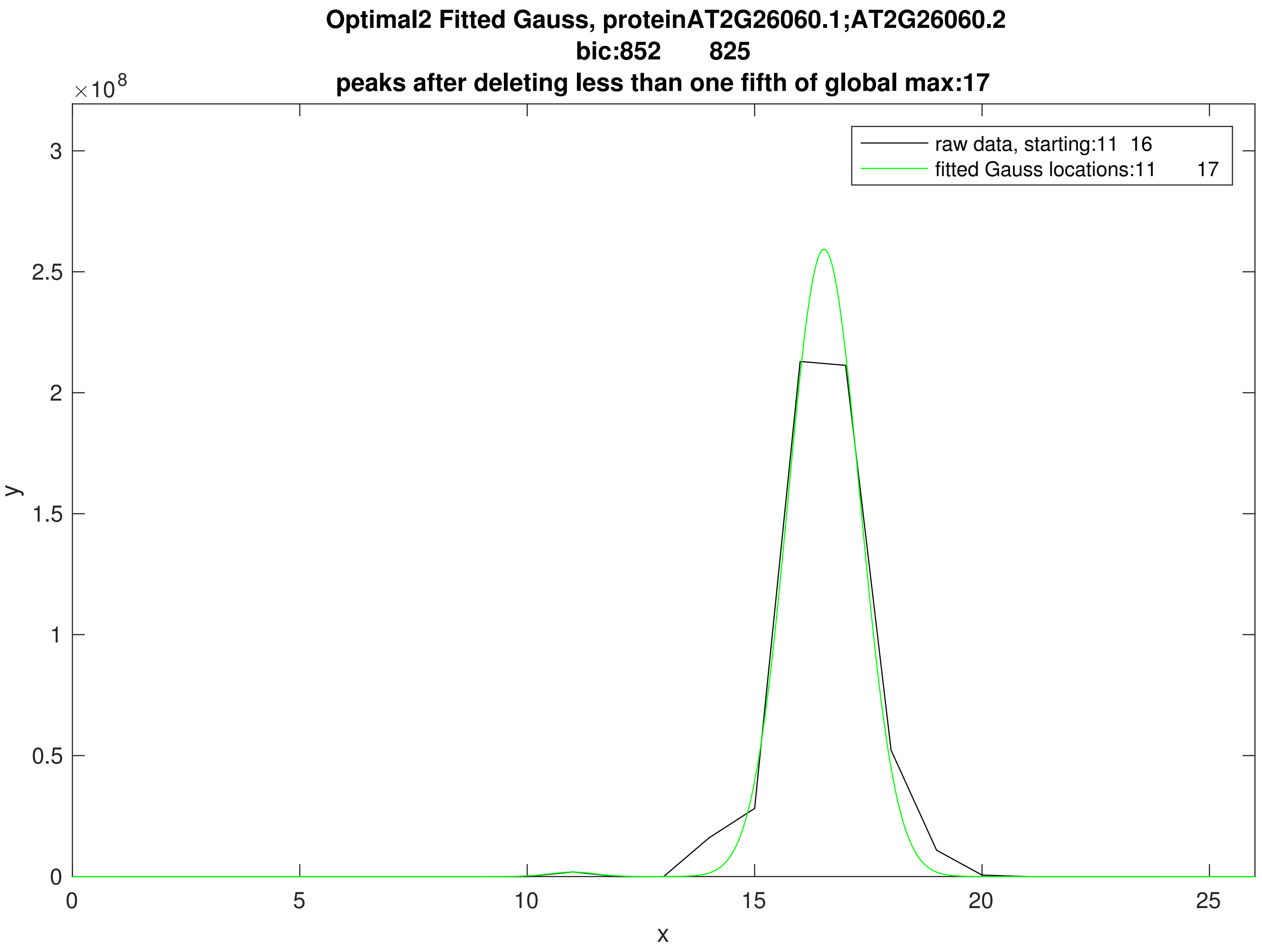
Optimal2 Fitted Gauss, proteinAT2G26060.1;AT2G26060.2
bic:852 825
peaks after deleting less than one fifth of global max:17
×10^8
raw data, starting:11 16
fitted Gauss locations:11 17
y
x
0
0.5
1
1.5
2
2.5
3
0
5
10
15
20
25

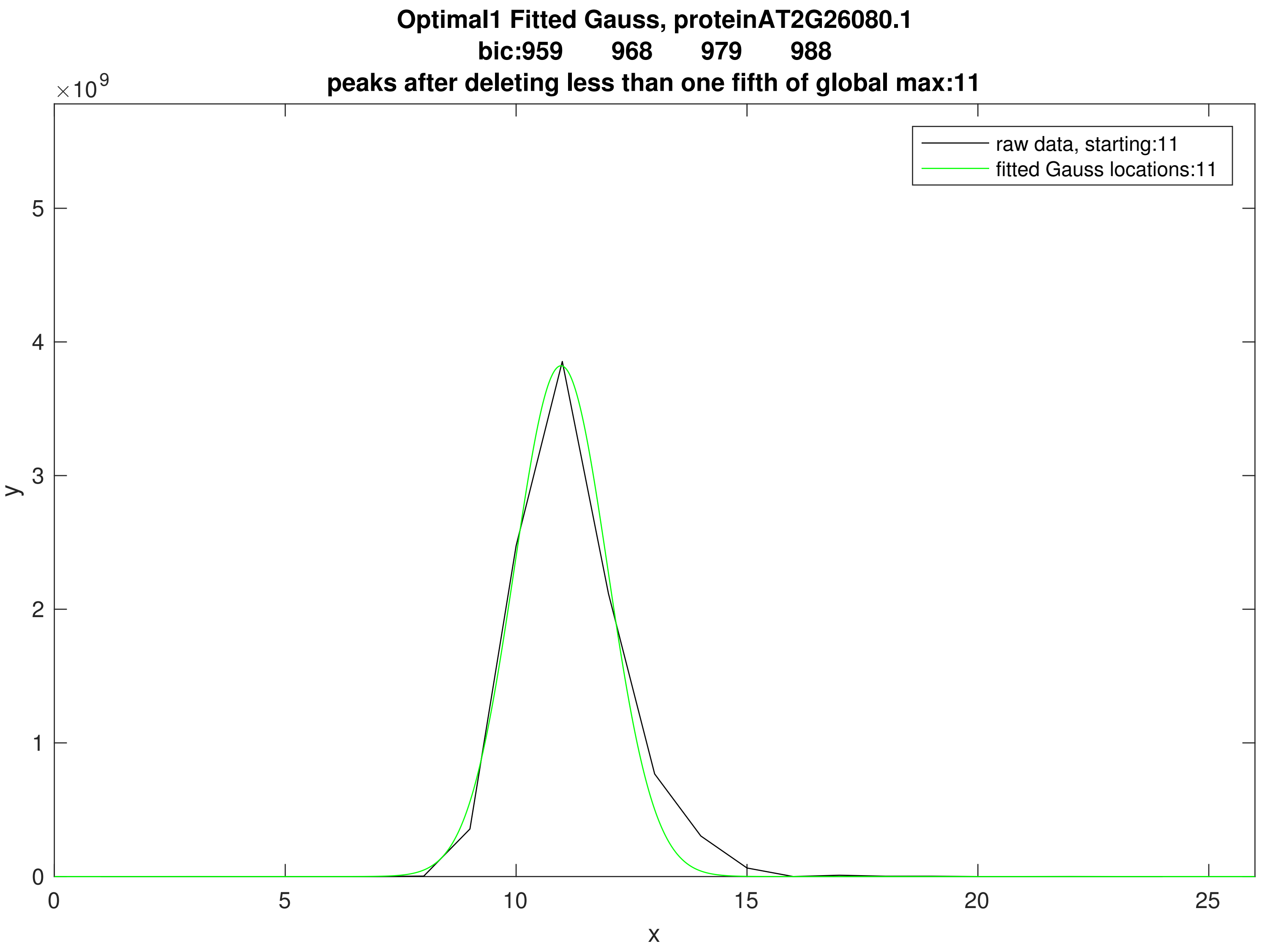
Optimal1 Fitted Gauss, proteinAT2G26080.1
bic:959 968 979 988
peaks after deleting less than one fifth of global max:11
×10^9
raw data, starting:11
fitted Gauss locations:11
y
x
0
1
2
3
4
5
0
5
10
15
20
25

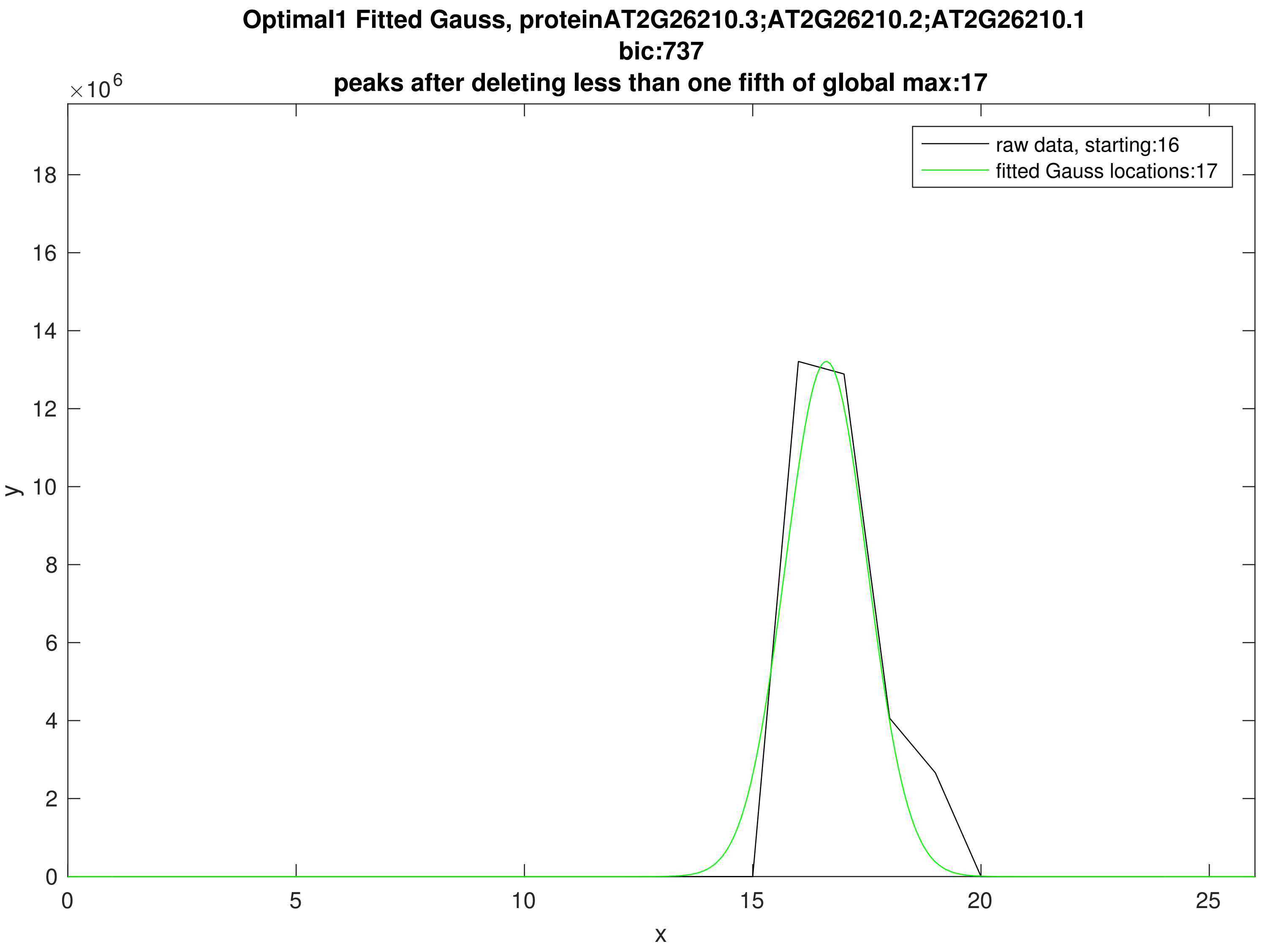
Optimal1 Fitted Gauss, proteinAT2G26210.3;AT2G26210.2;AT2G26210.1
bic:737
peaks after deleting less than one fifth of global max:17
×10^6
raw data, starting:16
fitted Gauss locations:17
y
x
0
2
4
6
8
10
12
14
16
18
0
5
10
15
20
25

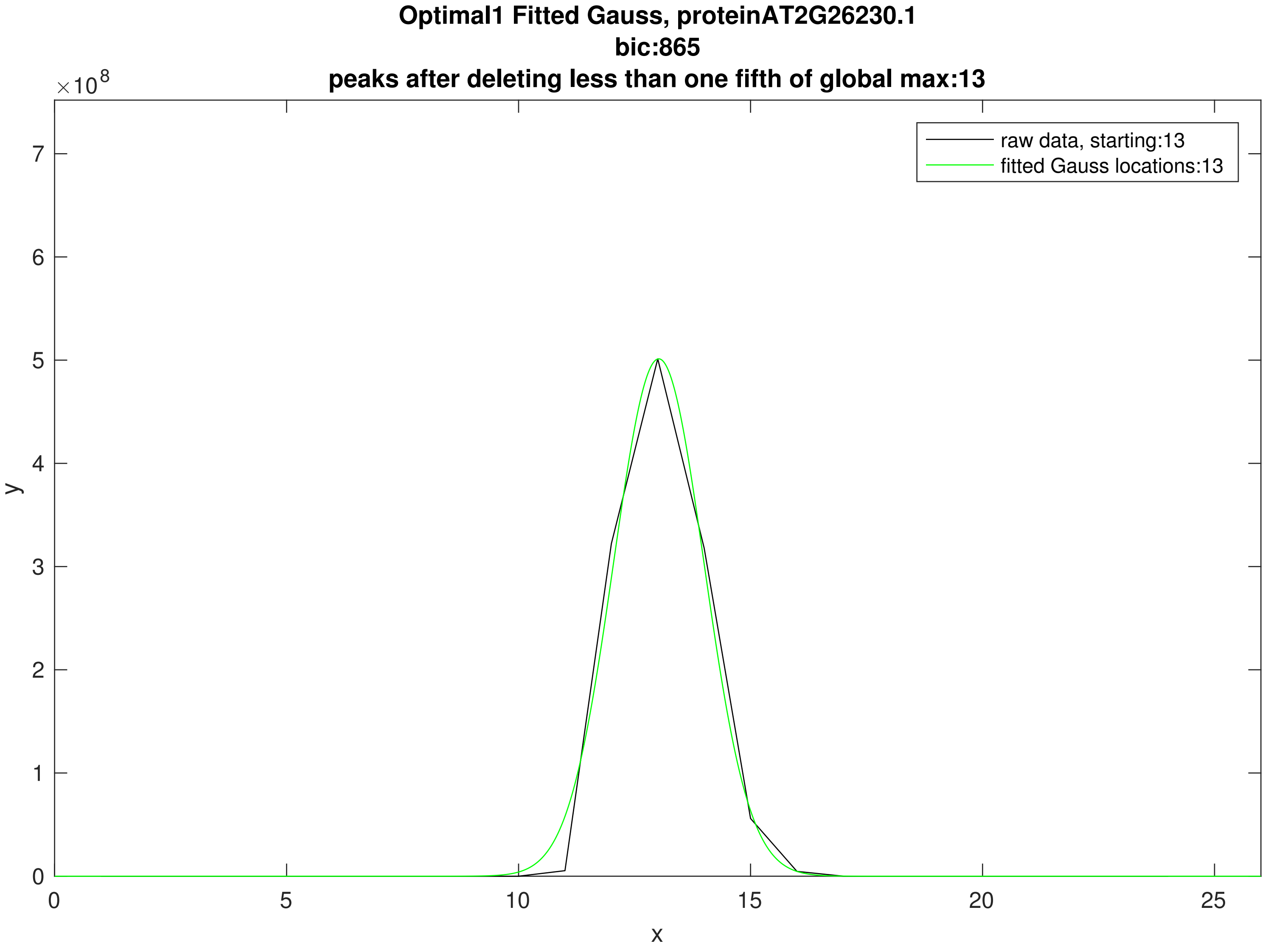
Optimal1 Fitted Gauss, proteinAT2G26230.1
bic:865
peaks after deleting less than one fifth of global max:13
×10^8
raw data, starting:13
fitted Gauss locations:13
y
x
0
1
2
3
4
5
6
7
0
5
10
15
20
25

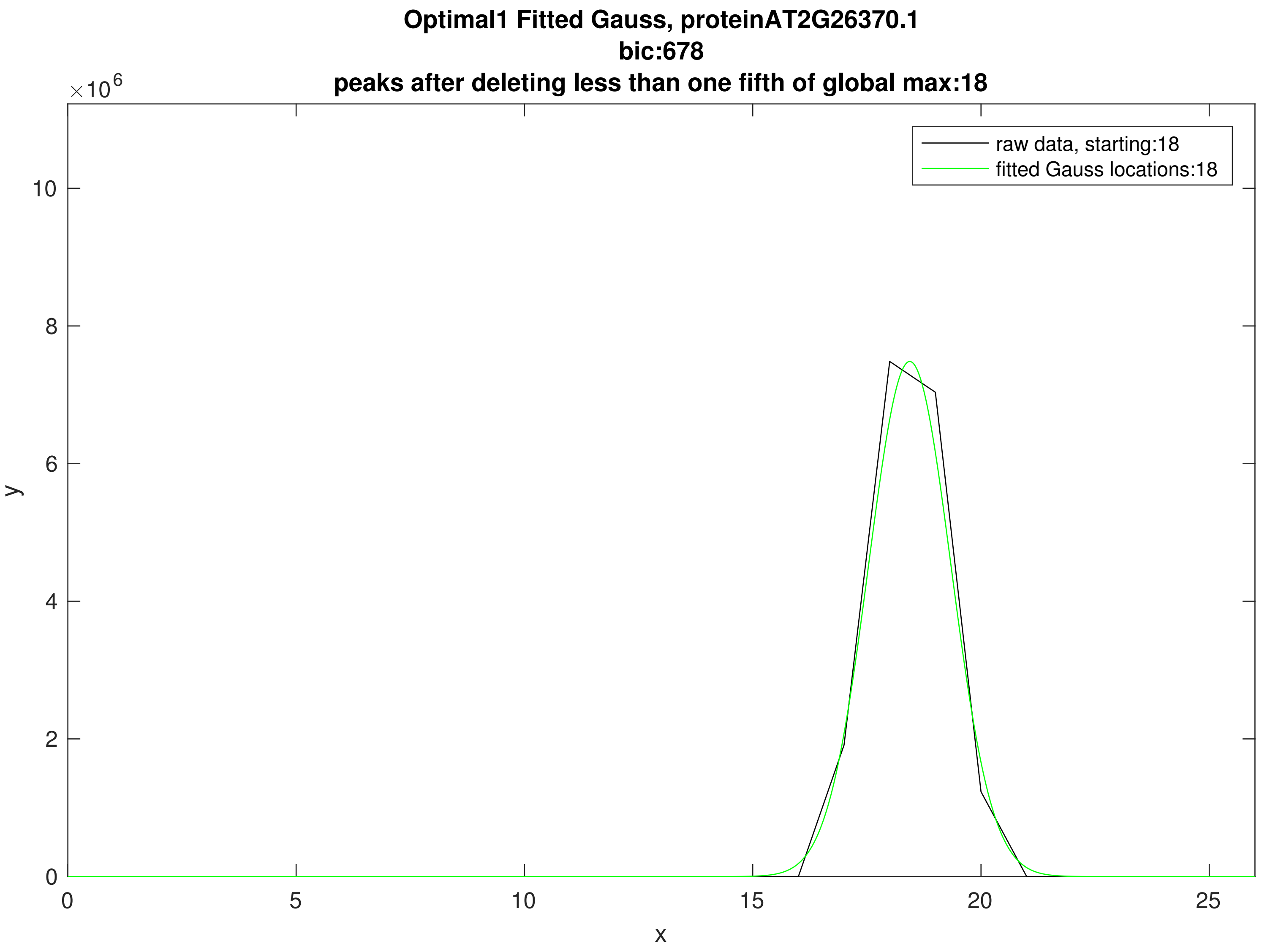
Optimal1 Fitted Gauss, proteinAT2G26370.1
bic:678
peaks after deleting less than one fifth of global max:18
×10^6
raw data, starting:18
fitted Gauss locations:18
y
x
0
2
4
6
8
10
0
5
10
15
20
25

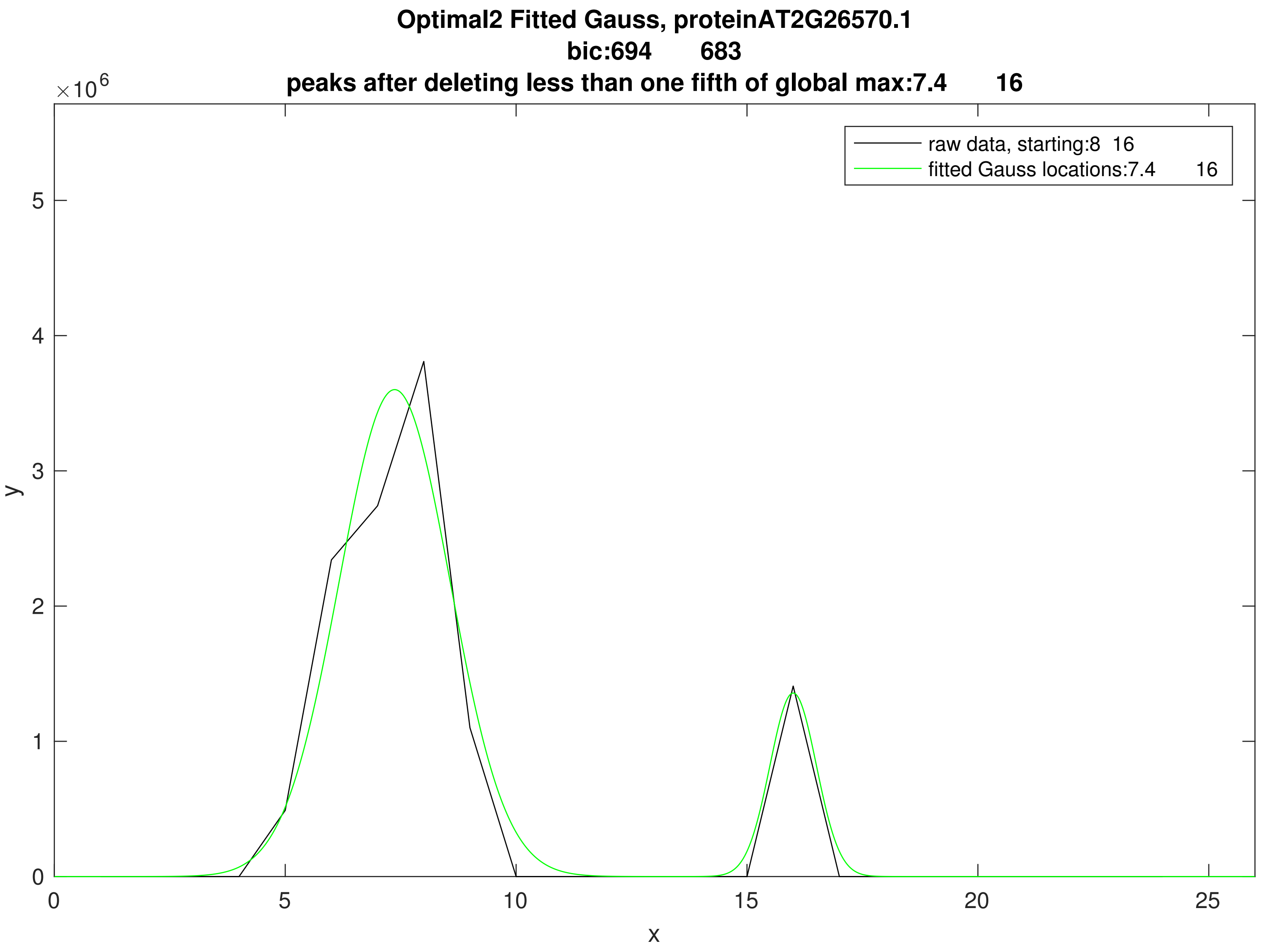
Optimal2 Fitted Gauss, proteinAT2G26570.1
bic:694 683
peaks after deleting less than one fifth of global max:7.4 16
×10^6
raw data, starting:8 16
fitted Gauss locations:7.4 16
y
x
0
1
2
3
4
5
0
5
10
15
20
25

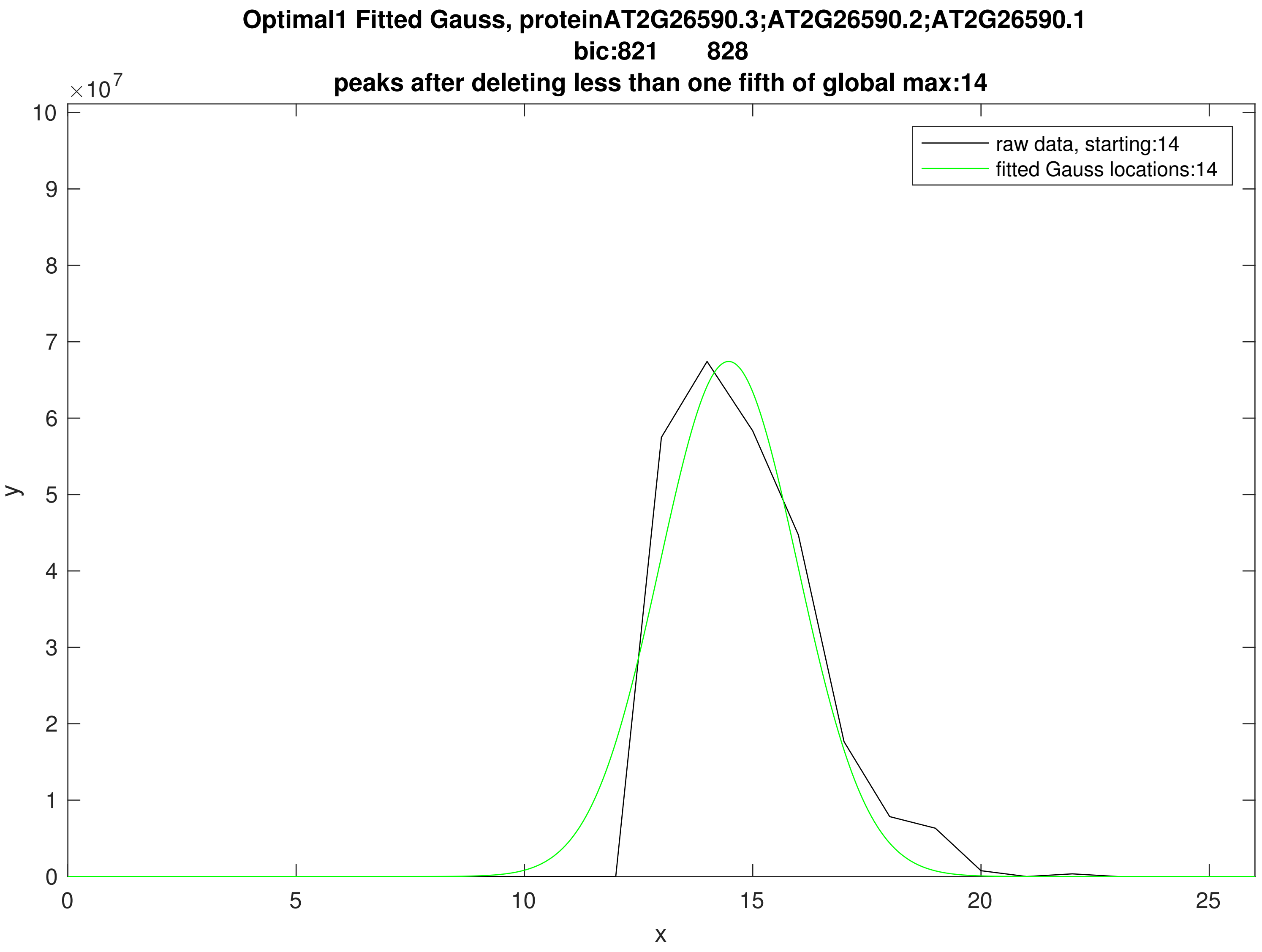

Optimal1 Fitted Gauss, proteinAT2G26590.3;AT2G26590.2;AT2G26590.1
bic:821 828
peaks after deleting less than one fifth of global max:14
×10^7
raw data, starting:14
fitted Gauss locations:14
y
x
0
1
2
3
4
5
6
7
8
9
10
0
5
10
15
20
25

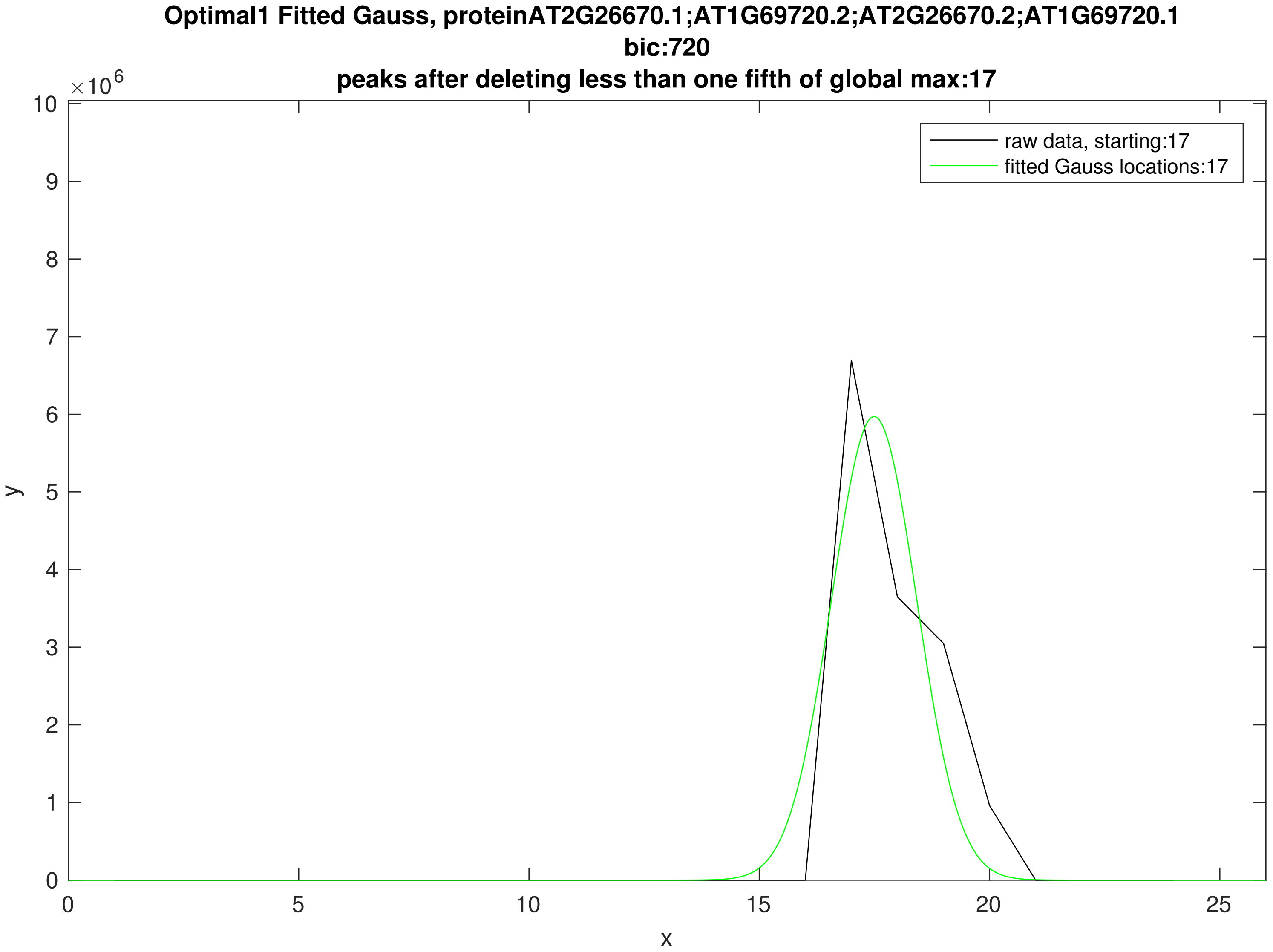

Optimal1 Fitted Gauss, proteinAT2G26670.1;AT1G69720.2;AT2G26670.2;AT1G69720.1
bic:720
peaks after deleting less than one fifth of global max:17
×10^6
raw data, starting:17
fitted Gauss locations:17
y
x

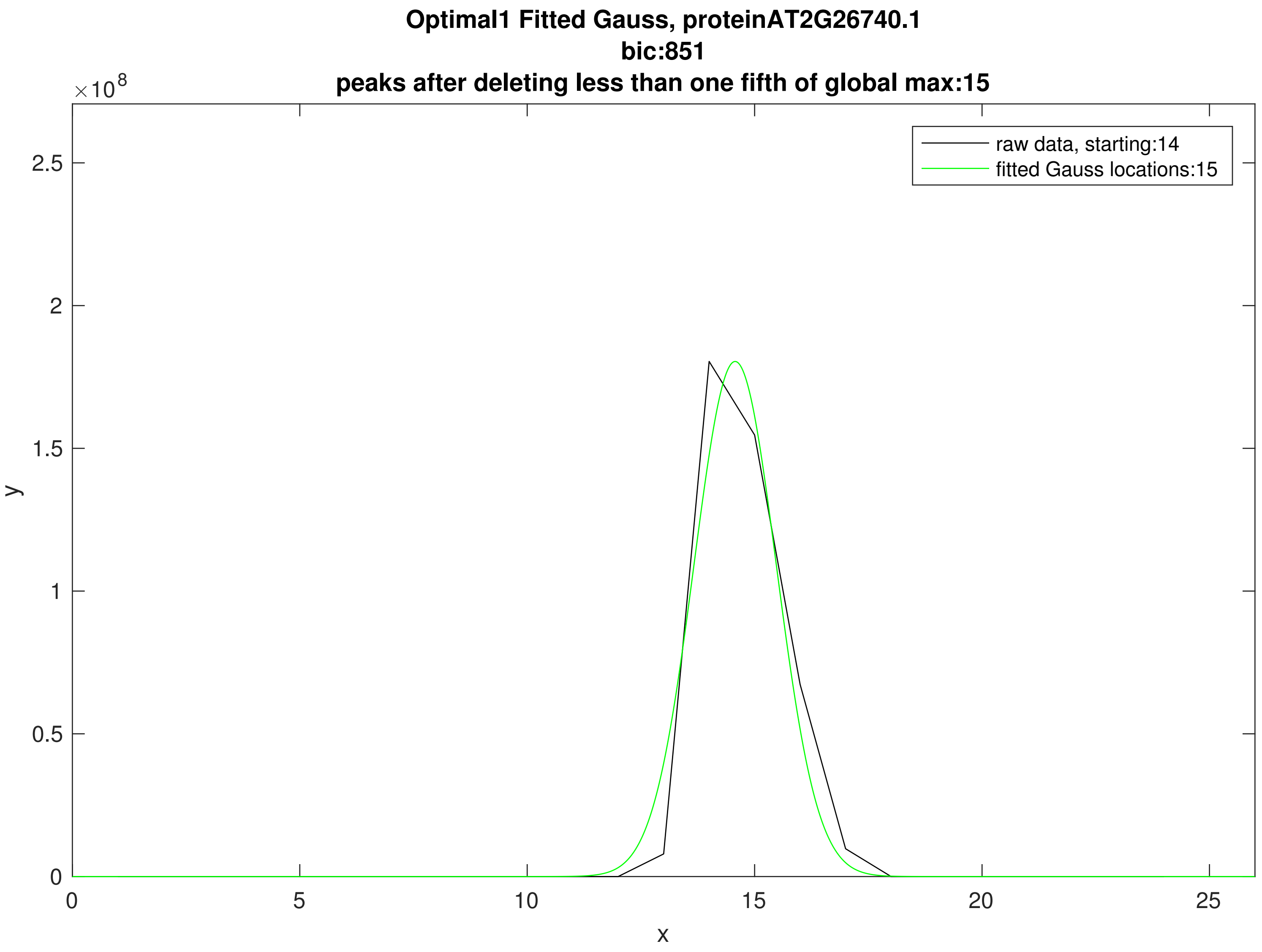
Optimal1 Fitted Gauss, proteinAT2G26740.1
bic:851
peaks after deleting less than one fifth of global max:15
×10^8
2.5
2
1.5
1
0.5
0
y
0
5
10
15
20
25
x
raw data, starting:14
fitted Gauss locations:15

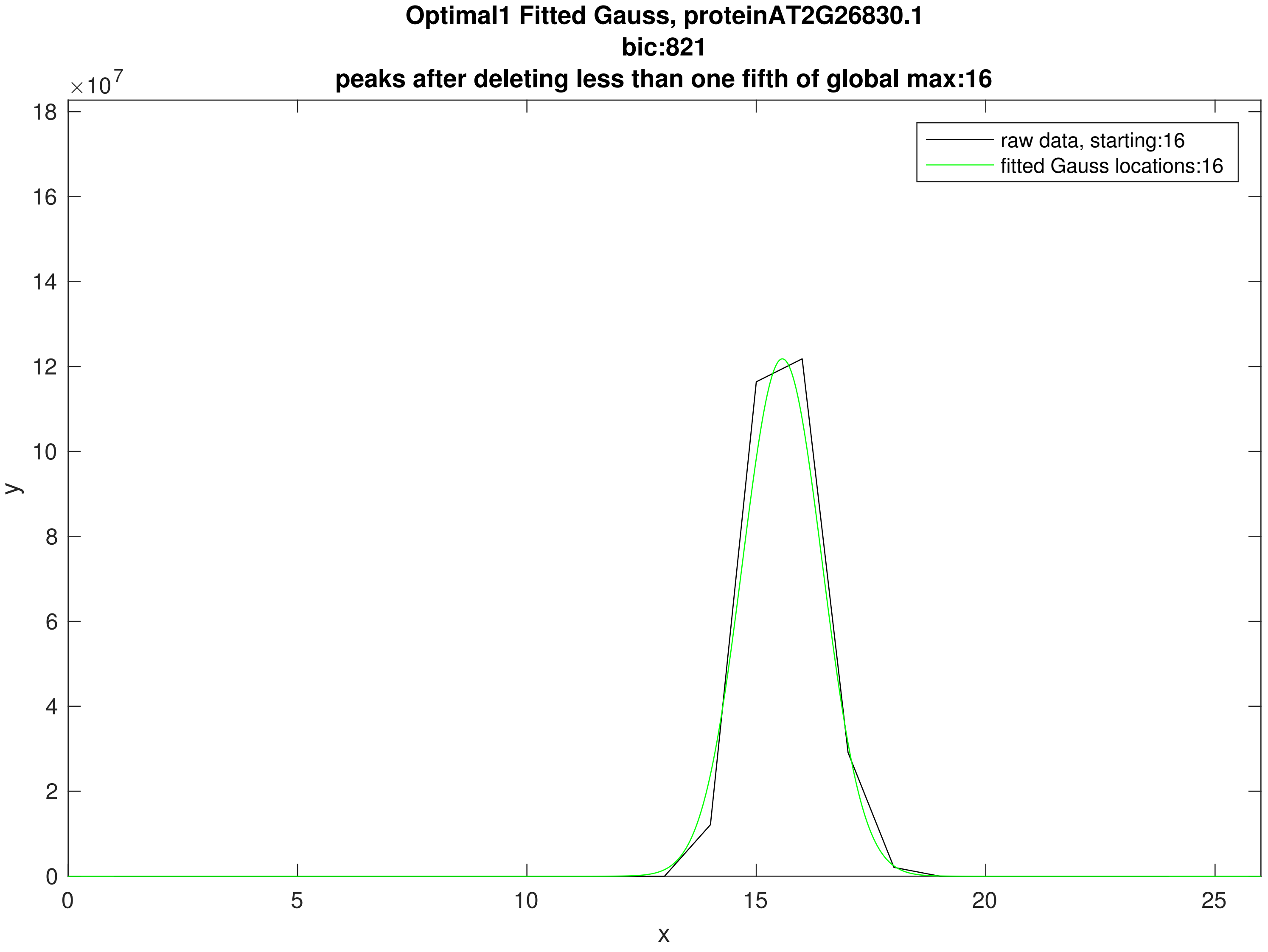
Optimal1 Fitted Gauss, proteinAT2G26830.1
bic:821
peaks after deleting less than one fifth of global max:16
×10^7
raw data, starting:16
fitted Gauss locations:16
y
x
0
2
4
6
8
10
12
14
16
18
0
5
10
15
20
25

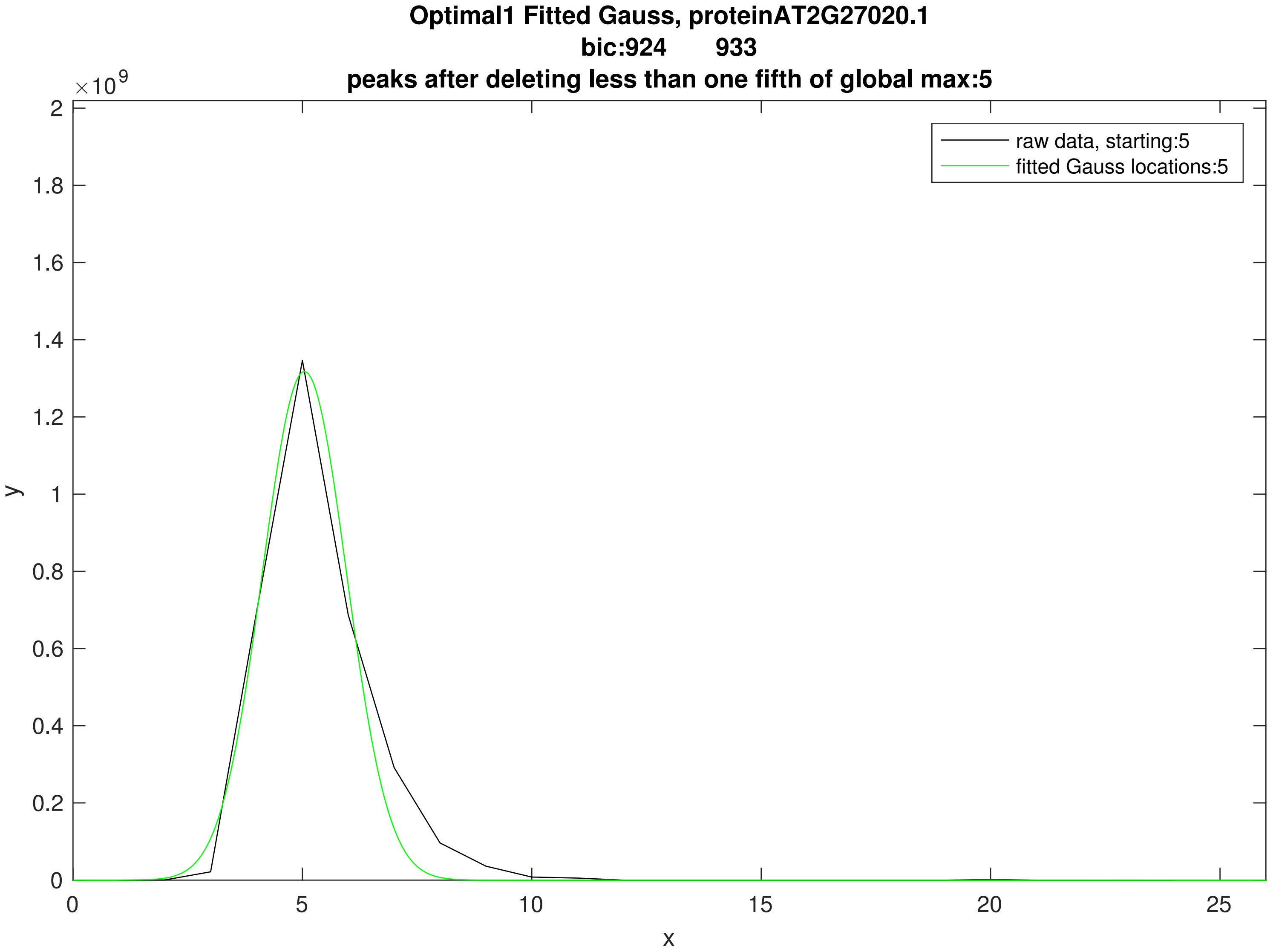

Optimal1 Fitted Gauss, proteinAT2G27020.1
bic:924 933
peaks after deleting less than one fifth of global max:5
×10^9
2
1.8
1.6
1.4
1.2
1
0.8
0.6
0.4
0.2
0
y
0
5
10
15
20
25
x
raw data, starting:5
fitted Gauss locations:5

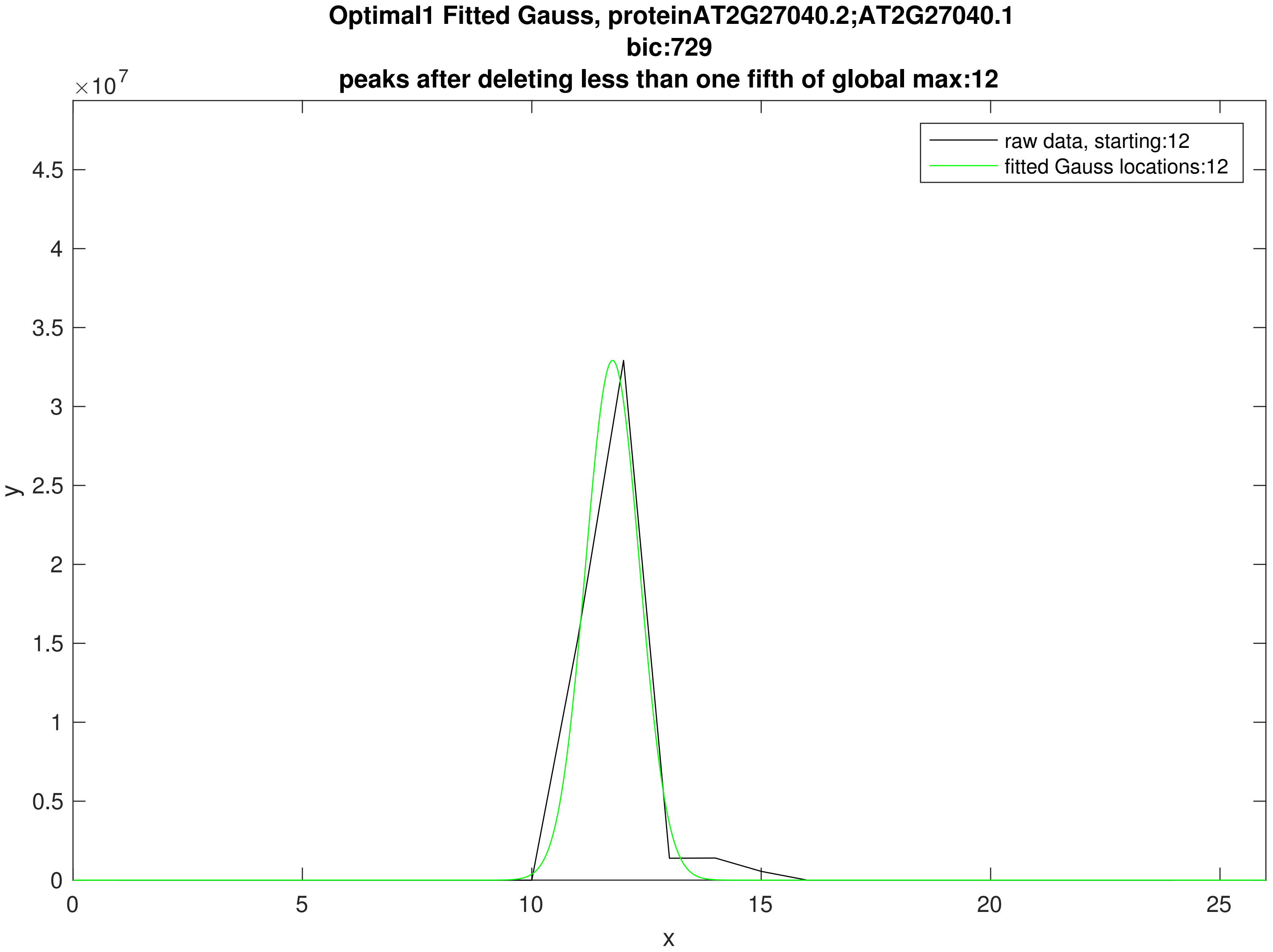
Optimal1 Fitted Gauss, proteinAT2G27040.2;AT2G27040.1
bic:729
peaks after deleting less than one fifth of global max:12
×10^7
4.5
4
3.5
3
2.5
2
1.5
1
0.5
0
y
0
5
10
15
20
25
x
raw data, starting:12
fitted Gauss locations:12

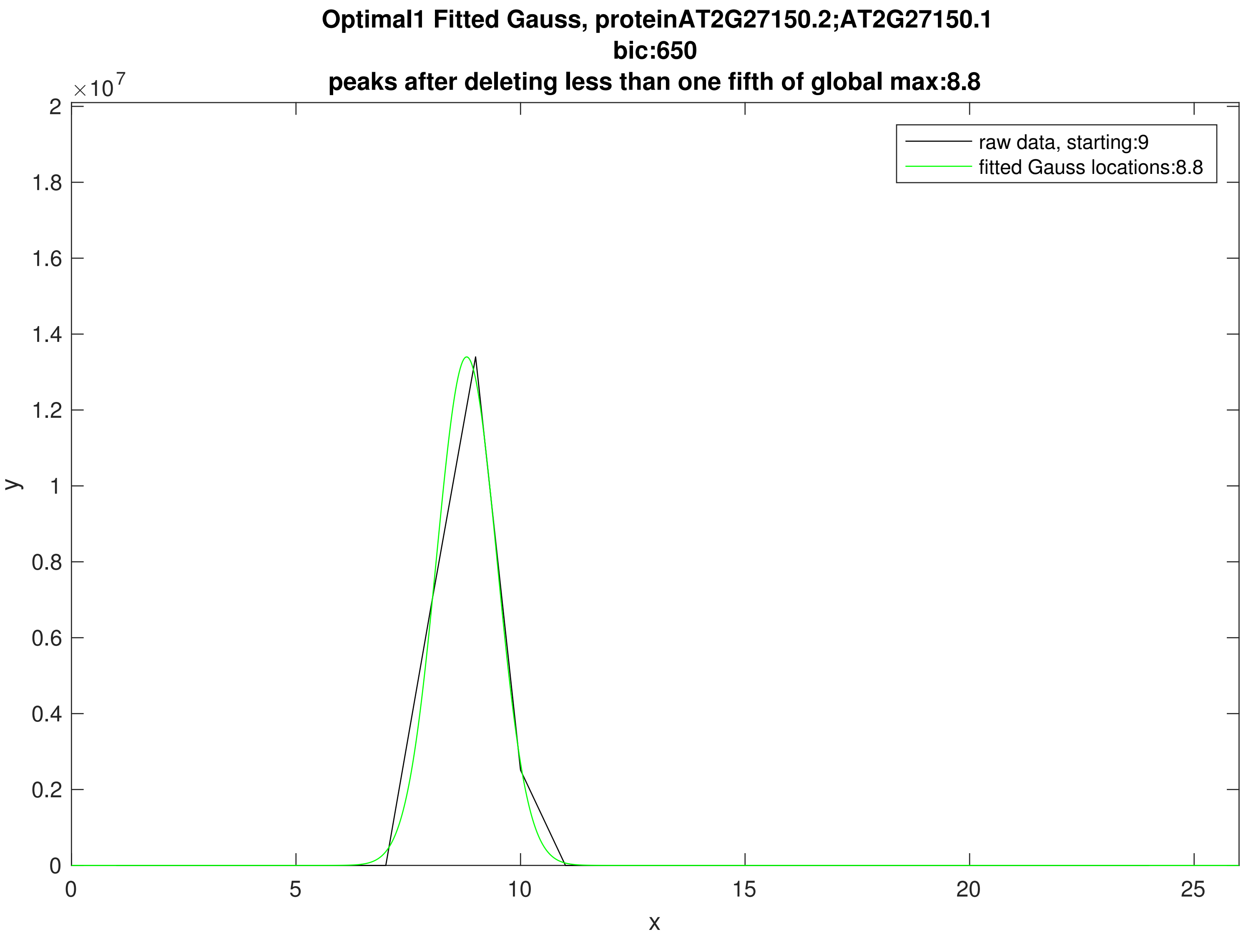
Optimal1 Fitted Gauss, proteinAT2G27150.2;AT2G27150.1
bic:650
peaks after deleting less than one fifth of global max:8.8
×10^7
2
1.8
1.6
1.4
1.2
1
0.8
0.6
0.4
0.2
0
y
0
5
10
15
20
25
x
raw data, starting:9
fitted Gauss locations:8.8

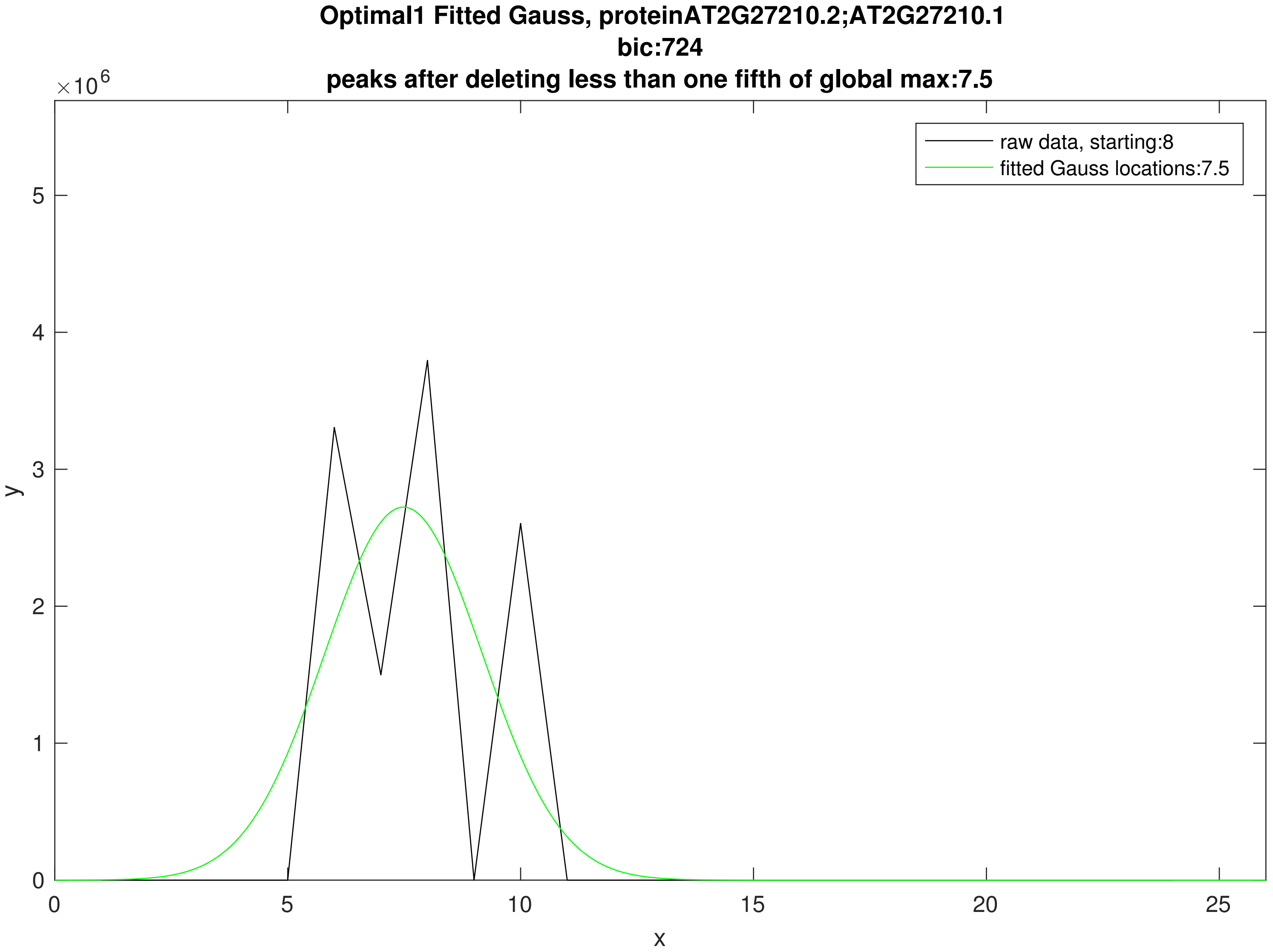
Optimal1 Fitted Gauss, proteinAT2G27210.2;AT2G27210.1
bic:724
peaks after deleting less than one fifth of global max:7.5
×10^6
raw data, starting:8
fitted Gauss locations:7.5
y
x
0
1
2
3
4
5
0
5
10
15
20
25

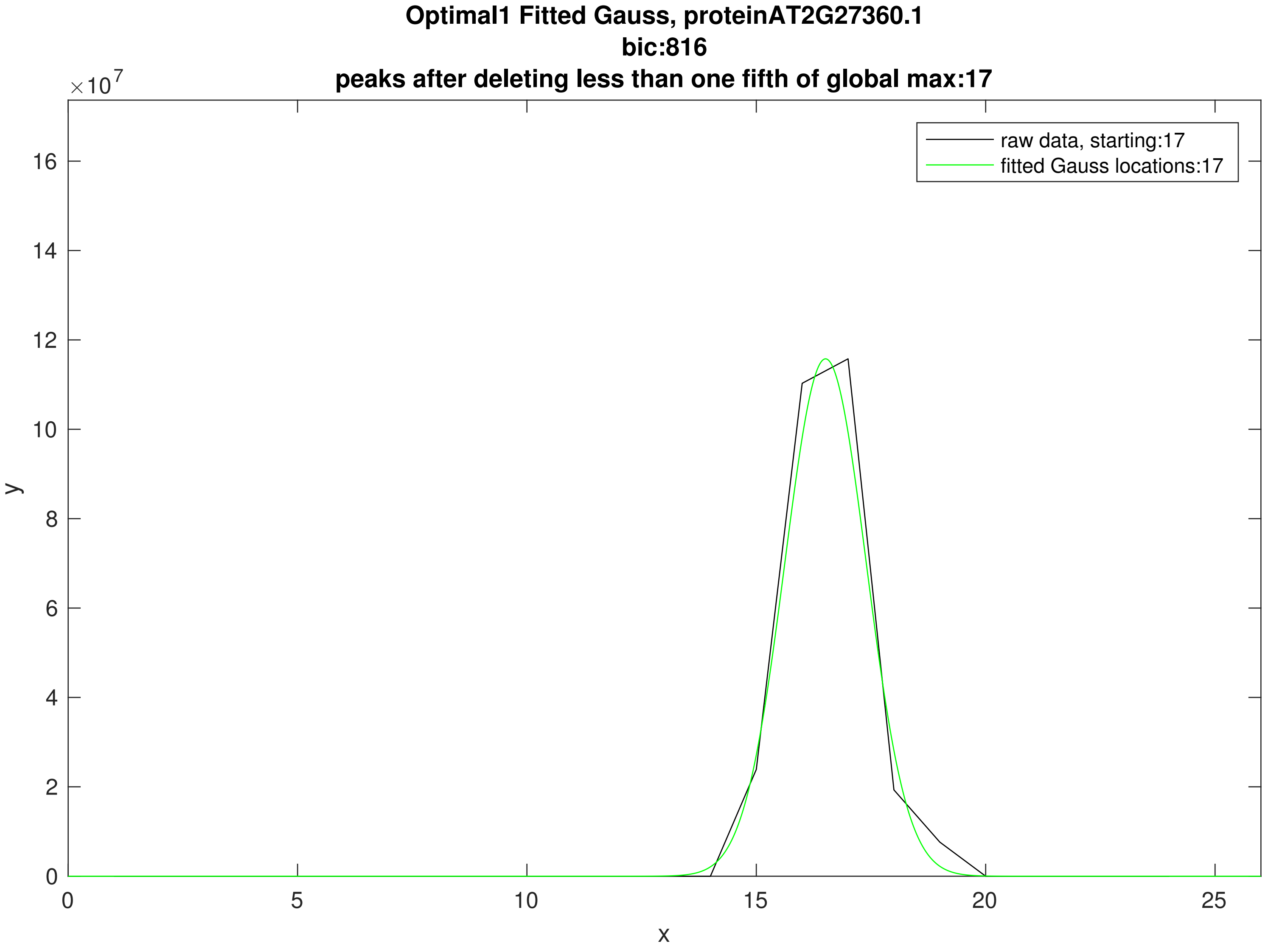
Optimal1 Fitted Gauss, proteinAT2G27360.1
bic:816
peaks after deleting less than one fifth of global max:17
×10^7
raw data, starting:17
fitted Gauss locations:17
y
x
0
2
4
6
8
10
12
14
16
0
5
10
15
20
25

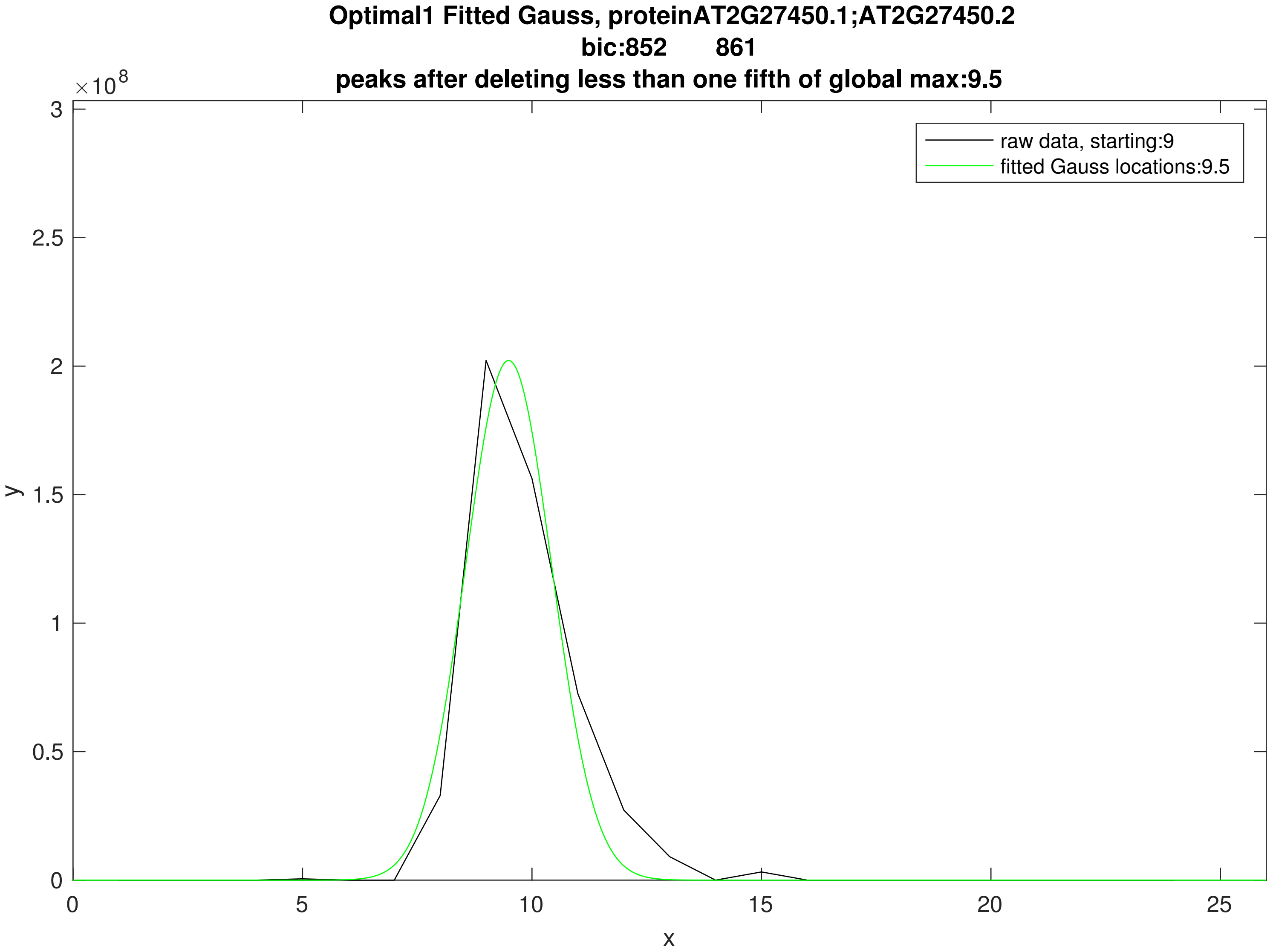
Optimal1 Fitted Gauss, proteinAT2G27450.1;AT2G27450.2
bic:852 861
peaks after deleting less than one fifth of global max:9.5
×10^8
raw data, starting:9
fitted Gauss locations:9.5
y
x
0
0.5
1
1.5
2
2.5
3
5
10
15
20
25

**Optimal1 Fitted Gauss, proteinAT2G27510.1**
**bic:876**
**peaks after deleting less than one fifth of global max:19**

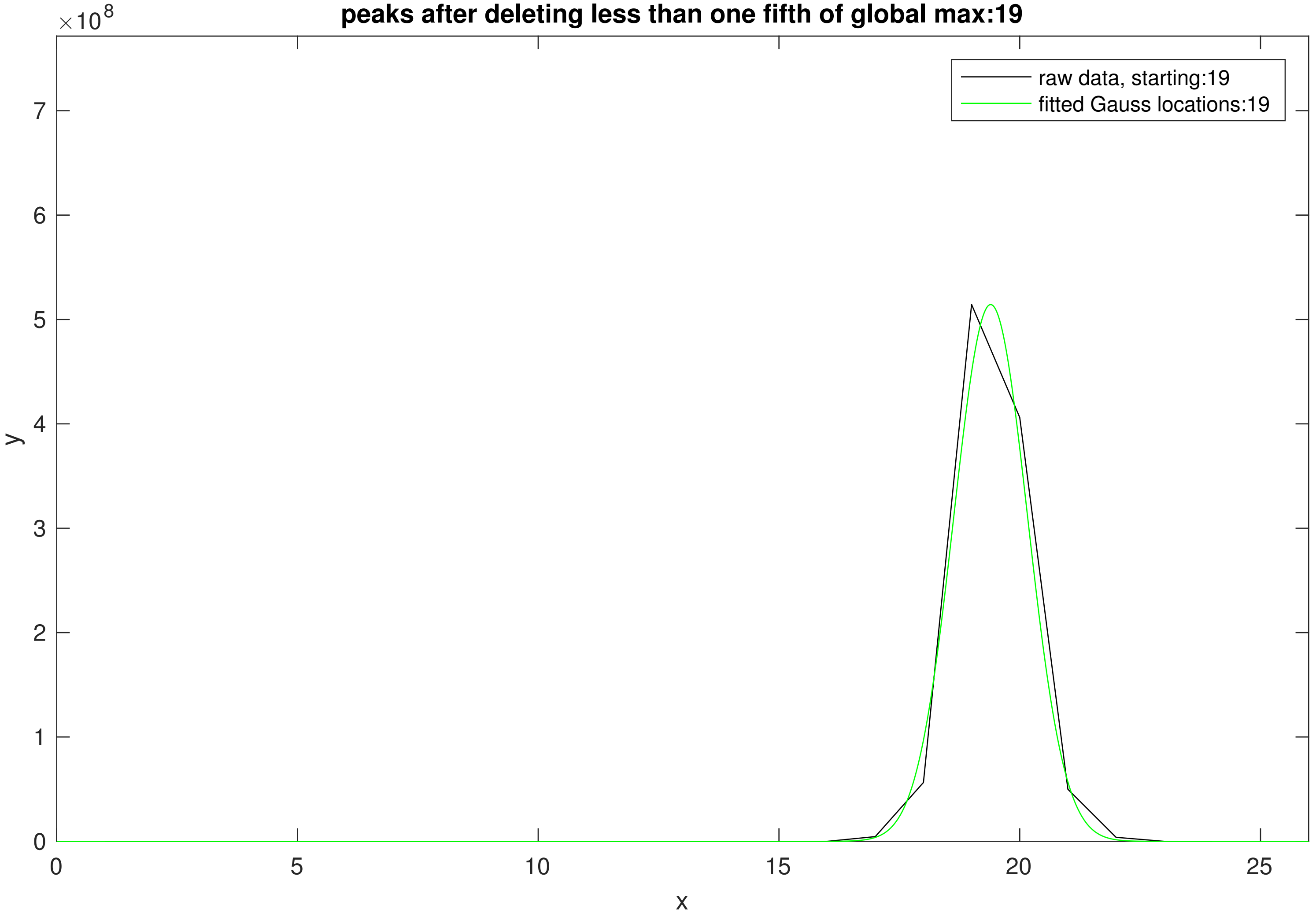

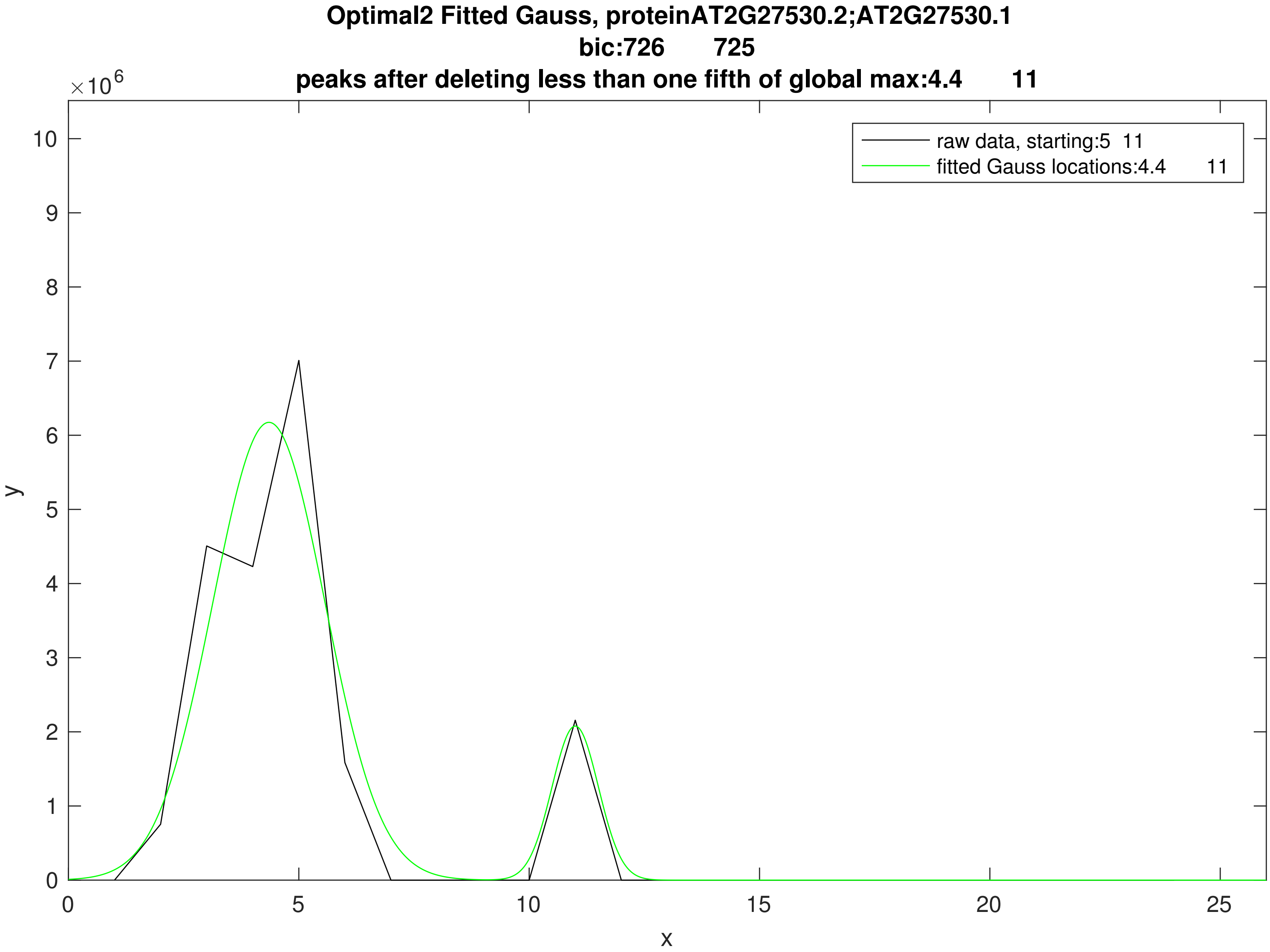

Optimal2 Fitted Gauss, proteinAT2G27530.2;AT2G27530.1
bic:726 725
peaks after deleting less than one fifth of global max:4.4 11
×10^6
raw data, starting:5 11
fitted Gauss locations:4.4 11
y
x
0
1
2
3
4
5
6
7
8
9
10
0
5
10
15
20
25

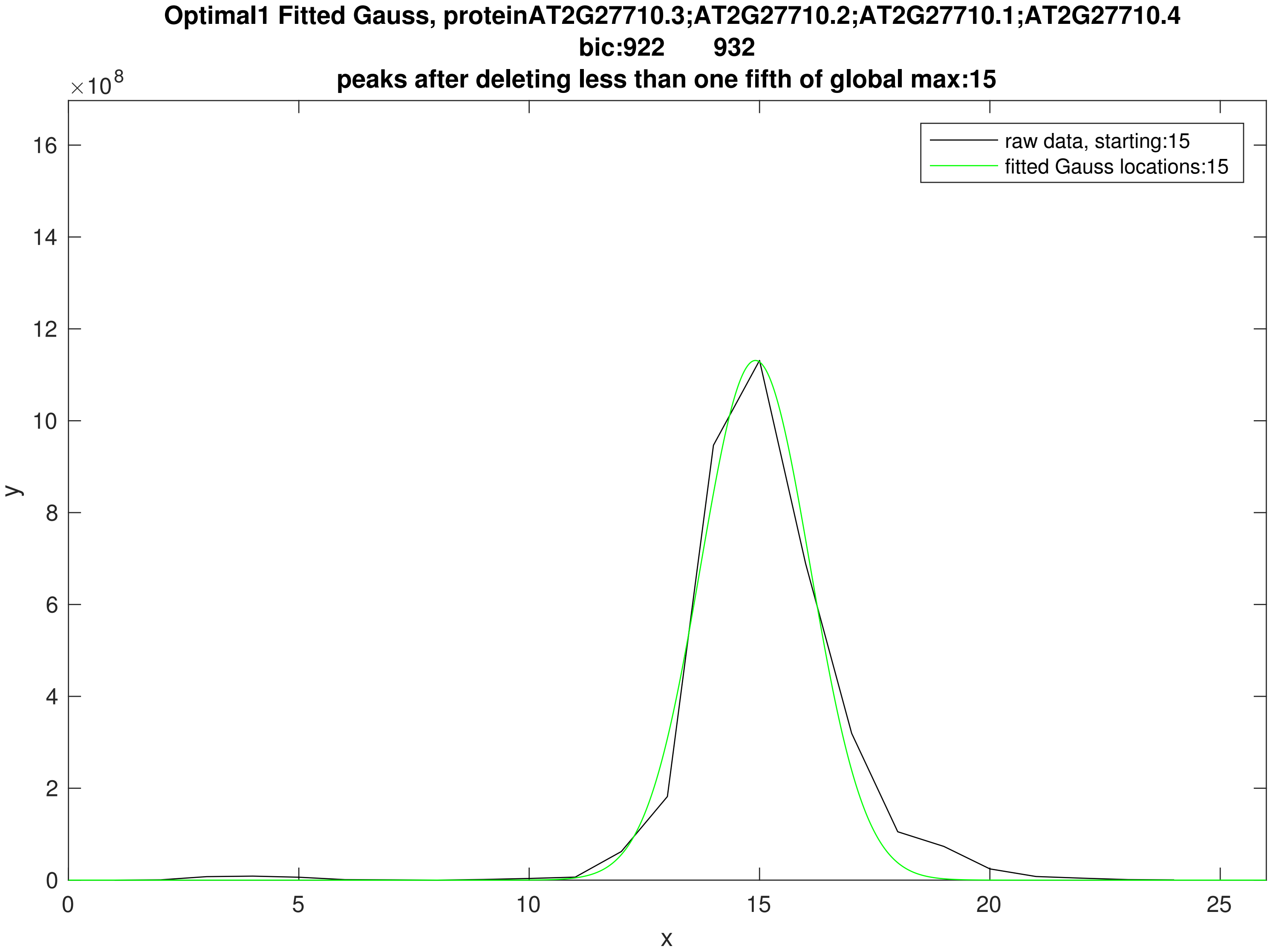

Optimal1 Fitted Gauss, proteinAT2G27710.3;AT2G27710.2;AT2G27710.1;AT2G27710.4
bic:922 932
peaks after deleting less than one fifth of global max:15
×10^8
16
14
12
10
8
6
4
2
0
y
0
5
10
15
20
25
x
raw data, starting:15
fitted Gauss locations:15

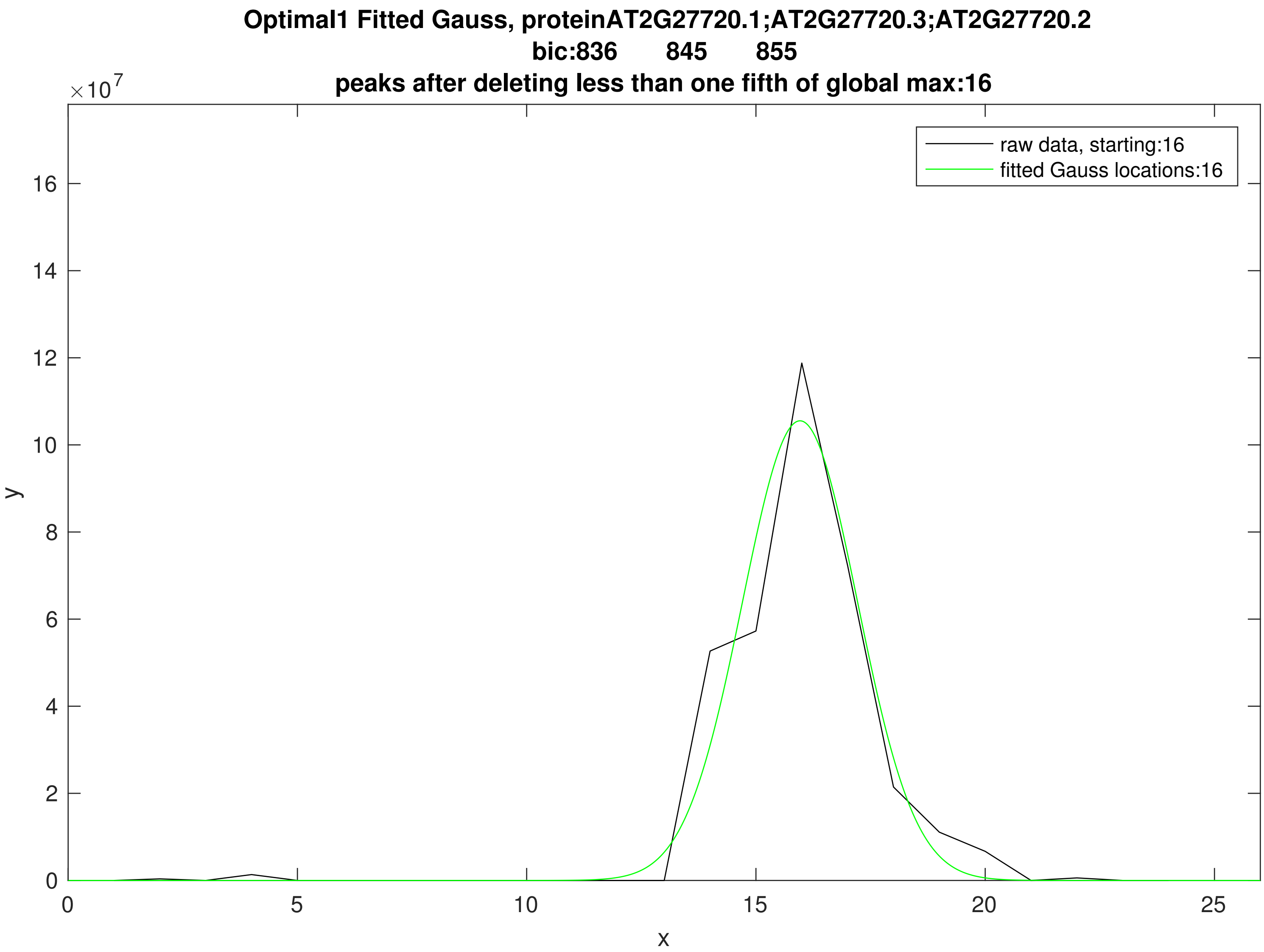

Optimal1 Fitted Gauss, proteinAT2G27720.1;AT2G27720.3;AT2G27720.2
bic:836 845 855
peaks after deleting less than one fifth of global max:16
×10^7
raw data, starting:16
fitted Gauss locations:16
16
14
12
10
8
6
4
2
0
y
0
5
10
15
20
25
x

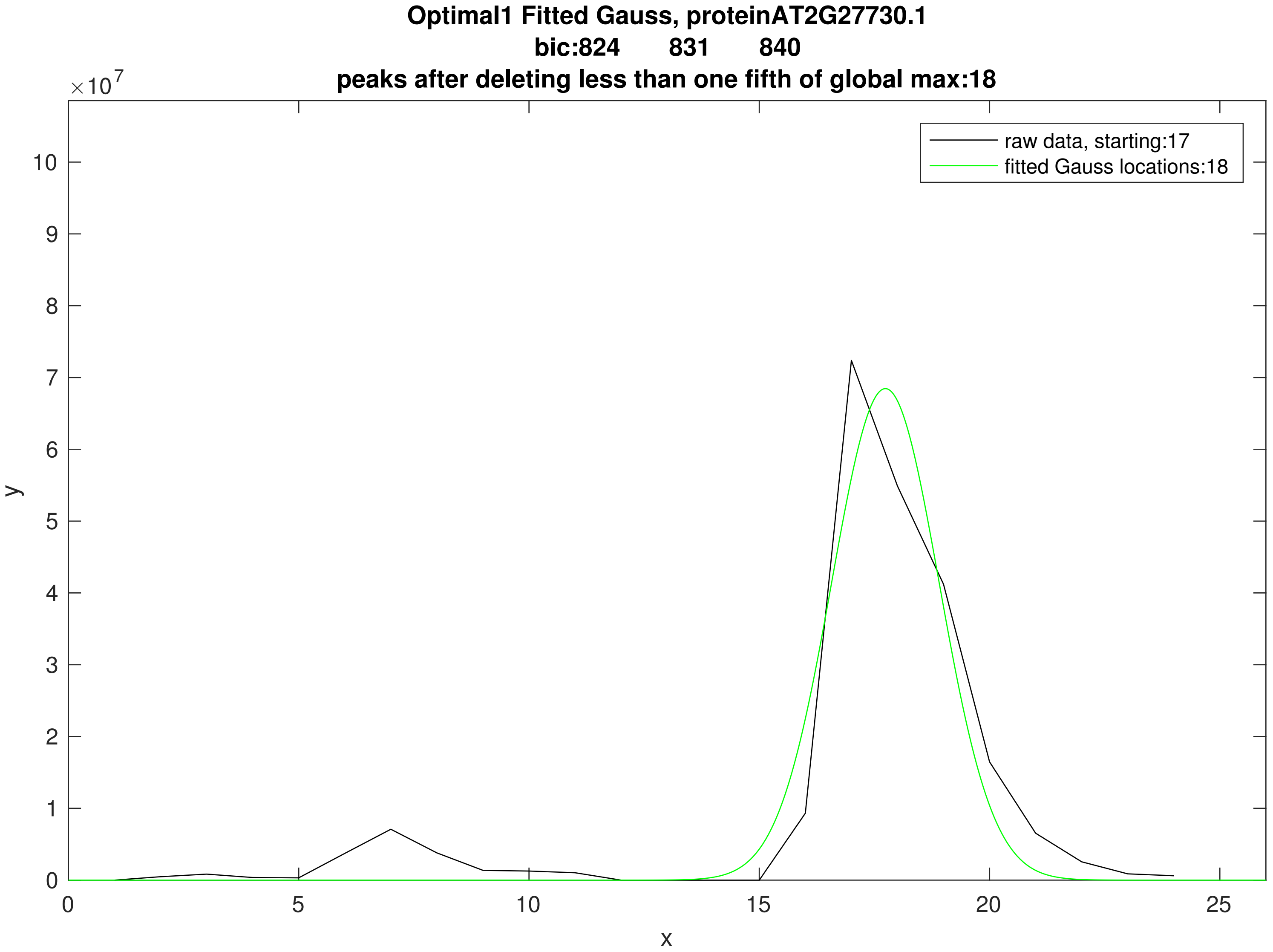
Optimal1 Fitted Gauss, proteinAT2G27730.1
bic:824 831 840
peaks after deleting less than one fifth of global max:18
×10^7
raw data, starting:17
fitted Gauss locations:18
y
x
0
5
10
15
20
25
1
2
3
4
5
6
7
8
9
10

**Optimal1 Fitted Gauss, proteinAT2G27860.1**
**bic:930**
**peaks after deleting less than one fifth of global max:13**

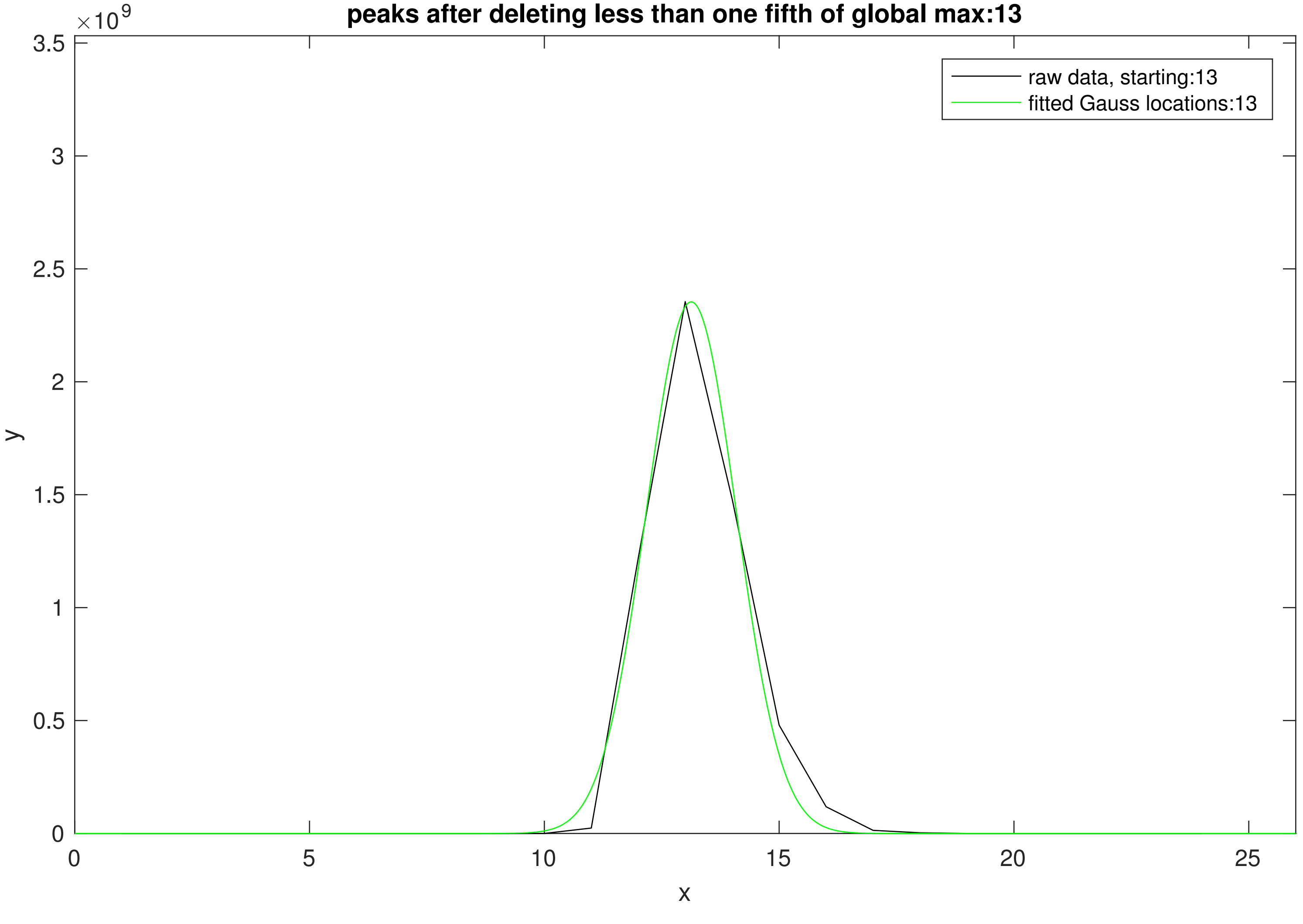

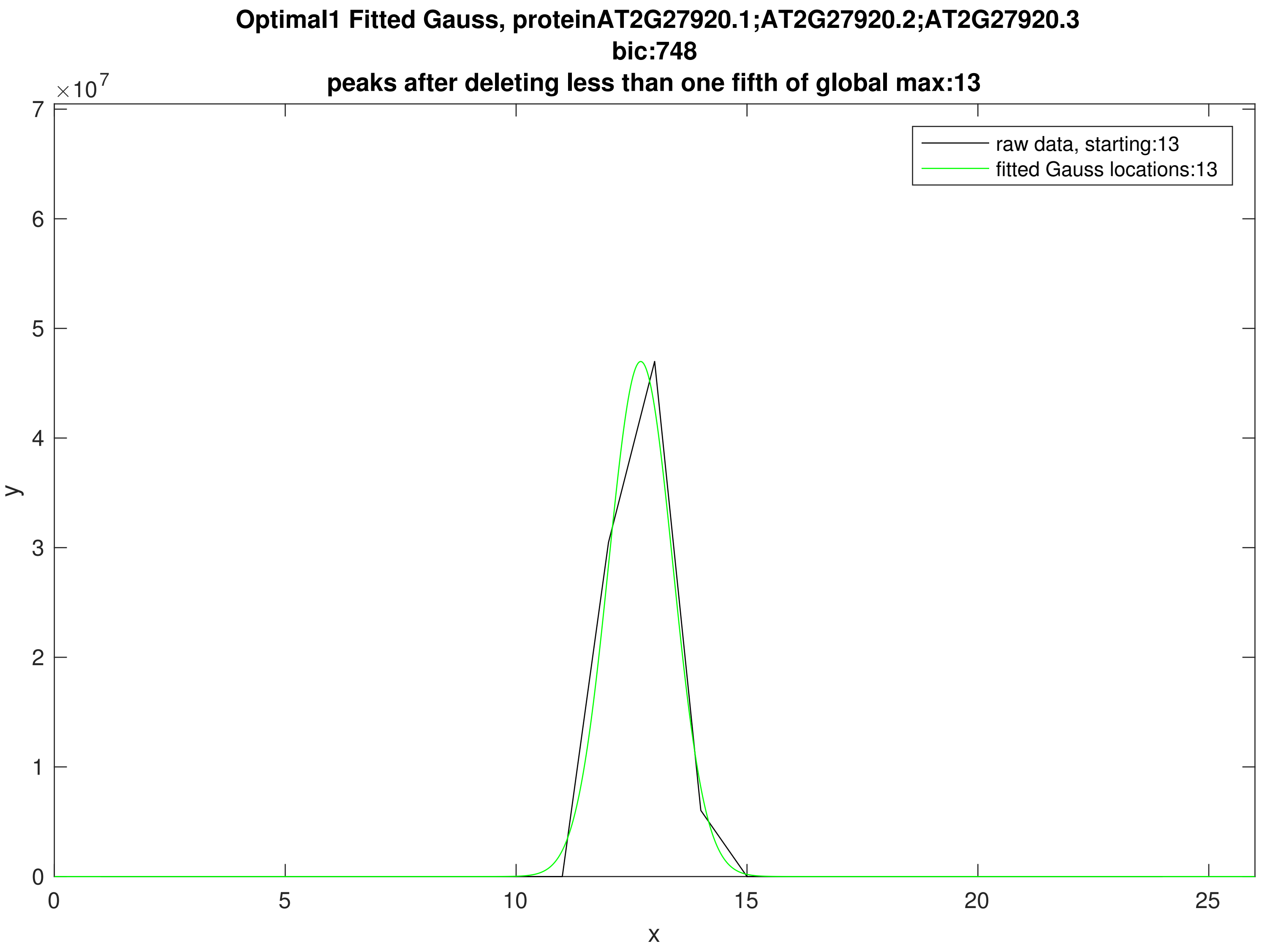

Optimal1 Fitted Gauss, proteinAT2G27920.1;AT2G27920.2;AT2G27920.3
bic:748
peaks after deleting less than one fifth of global max:13
×10^7
raw data, starting:13
fitted Gauss locations:13
y
x
0
1
2
3
4
5
6
7
0
5
10
15
20
25

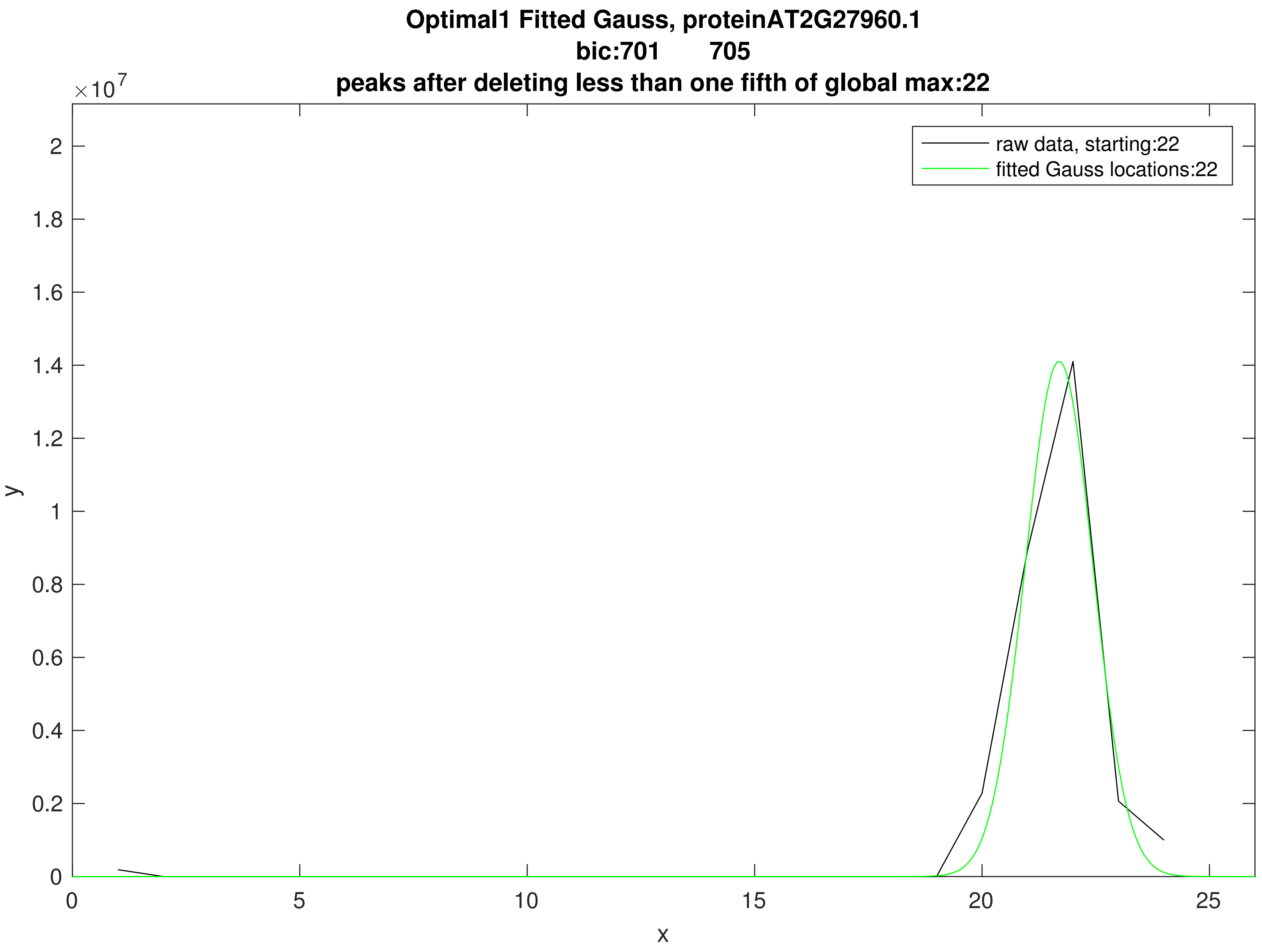
Optimal1 Fitted Gauss, proteinAT2G27960.1
bic:701 705
peaks after deleting less than one fifth of global max:22
×10^7
raw data, starting:22
fitted Gauss locations:22
y
x
0
0.2
0.4
0.6
0.8
1
1.2
1.4
1.6
1.8
2
0
5
10
15
20
25

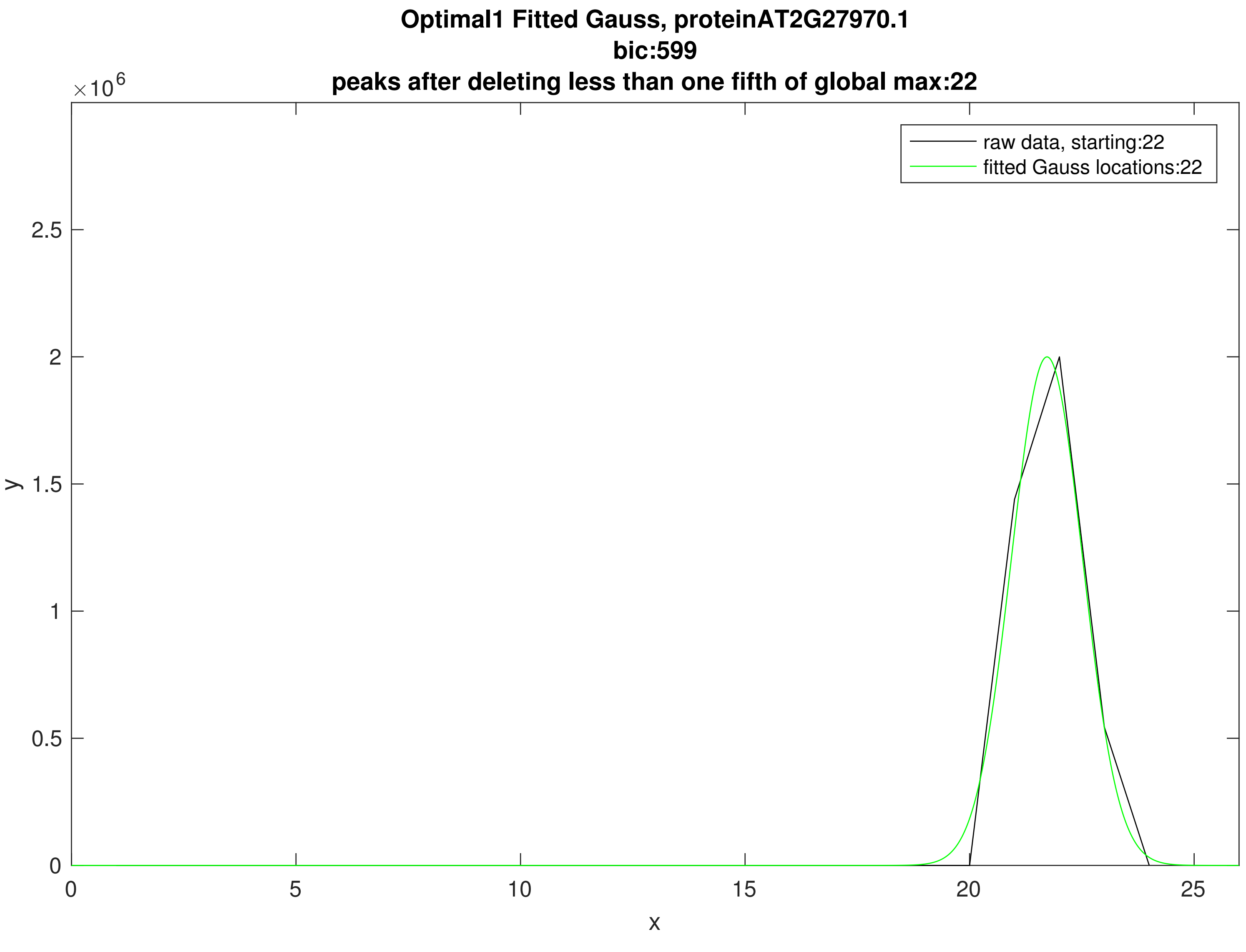
Optimal1 Fitted Gauss, proteinAT2G27970.1
bic:599
peaks after deleting less than one fifth of global max:22
×10^6
raw data, starting:22
fitted Gauss locations:22
2.5
2
1.5
1
0.5
0
y
0
5
10
15
20
25
x

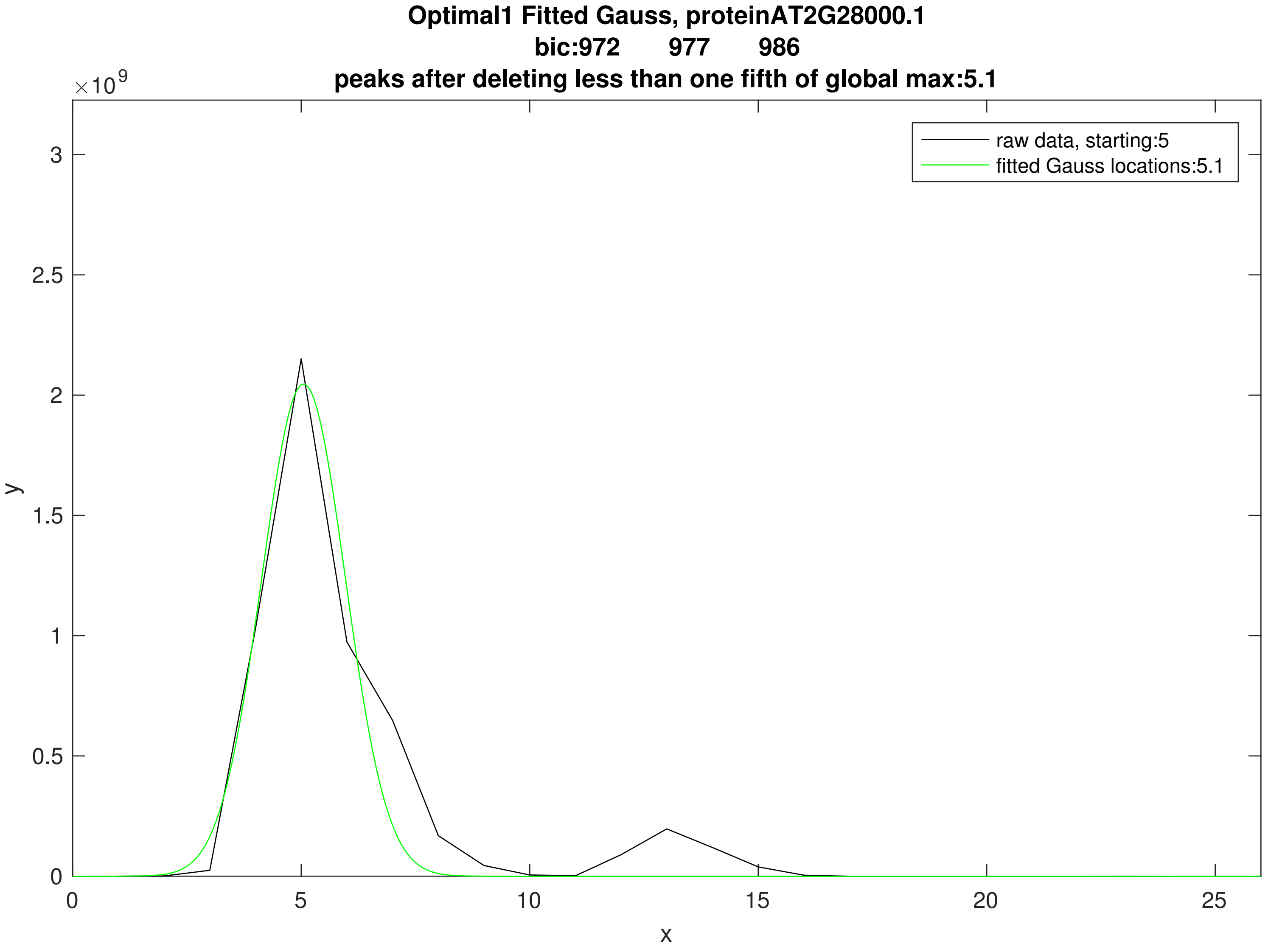
Optimal1 Fitted Gauss, proteinAT2G28000.1
bic:972 977 986
peaks after deleting less than one fifth of global max:5.1
×10^9
raw data, starting:5
fitted Gauss locations:5.1
y
x
0
0.5
1
1.5
2
2.5
3
0
5
10
15
20
25

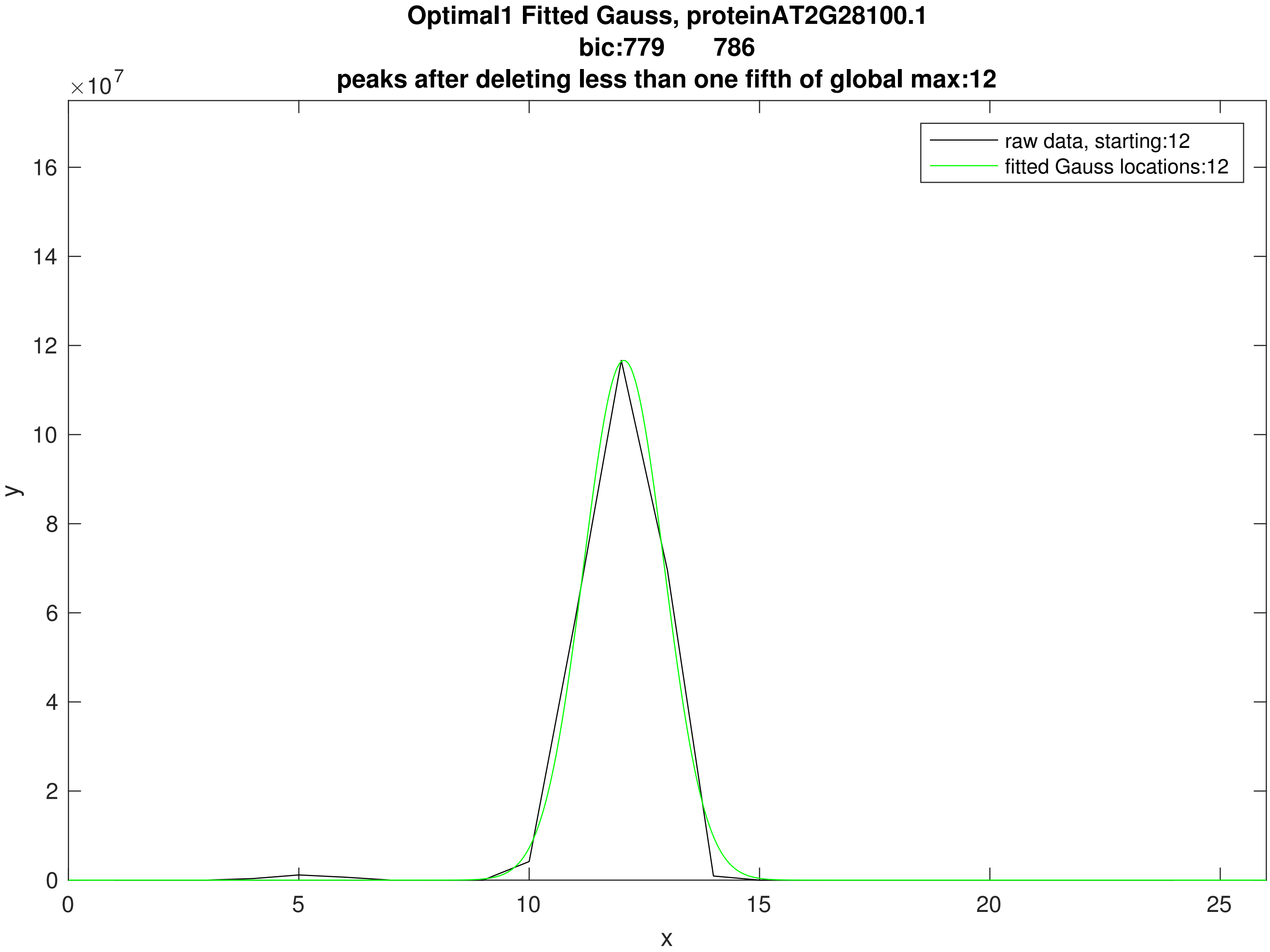

Optimal1 Fitted Gauss, proteinAT2G28100.1
bic:779 786
peaks after deleting less than one fifth of global max:12
×10^7
raw data, starting:12
fitted Gauss locations:12
16
14
12
10
8
6
4
2
0
y
0
5
10
15
20
25
x

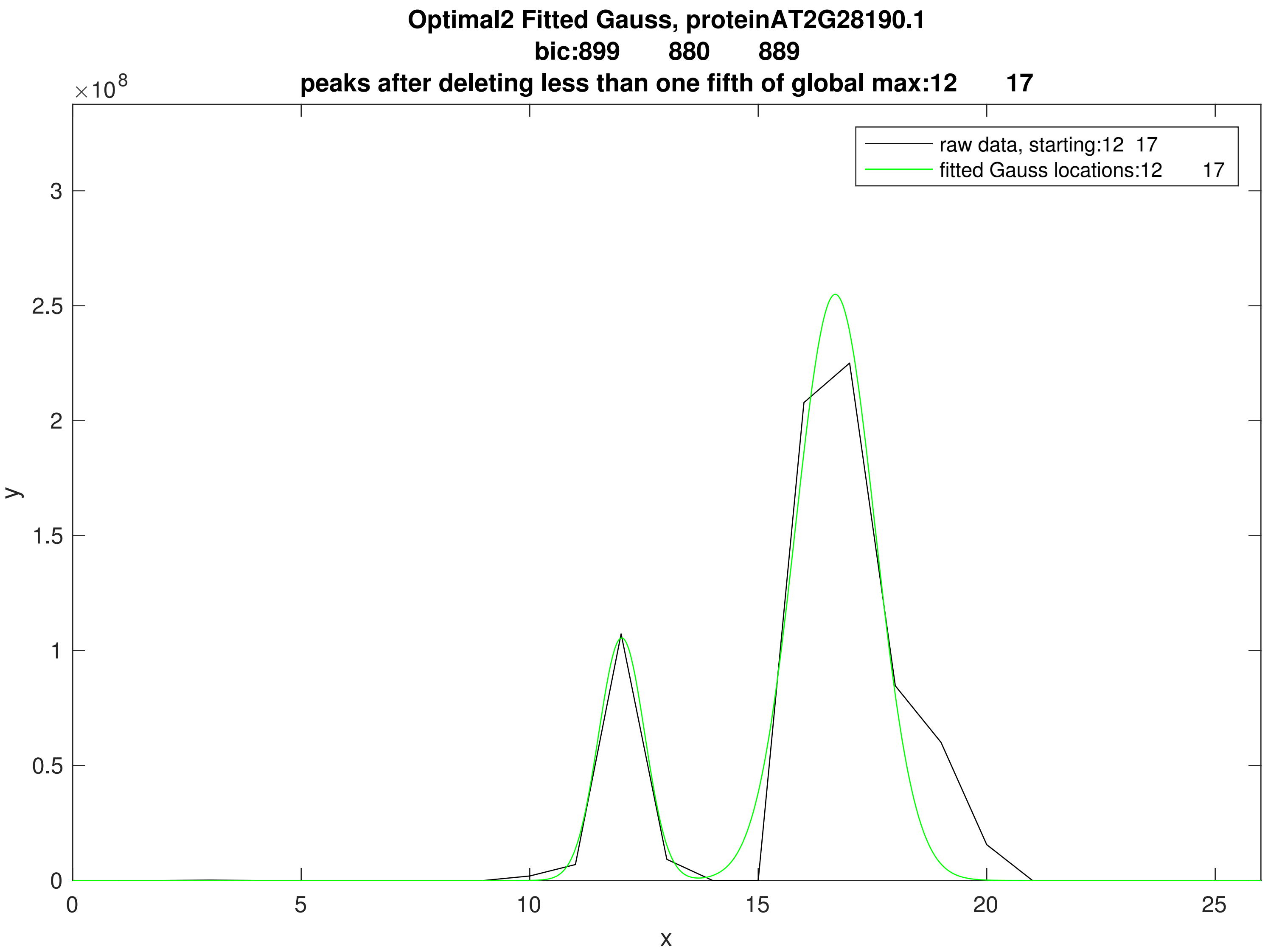
Optimal2 Fitted Gauss, proteinAT2G28190.1
bic:899 880 889
peaks after deleting less than one fifth of global max:12 17
×10^8
raw data, starting:12 17
fitted Gauss locations:12 17
y
x
0
0.5
1
1.5
2
2.5
3
0
5
10
15
20
25

**Optimal1 Fitted Gauss, proteinAT2G28290.2;AT2G28290.3;AT2G28290.1**
**bic:818**
**peaks after deleting less than one fifth of global max:10**

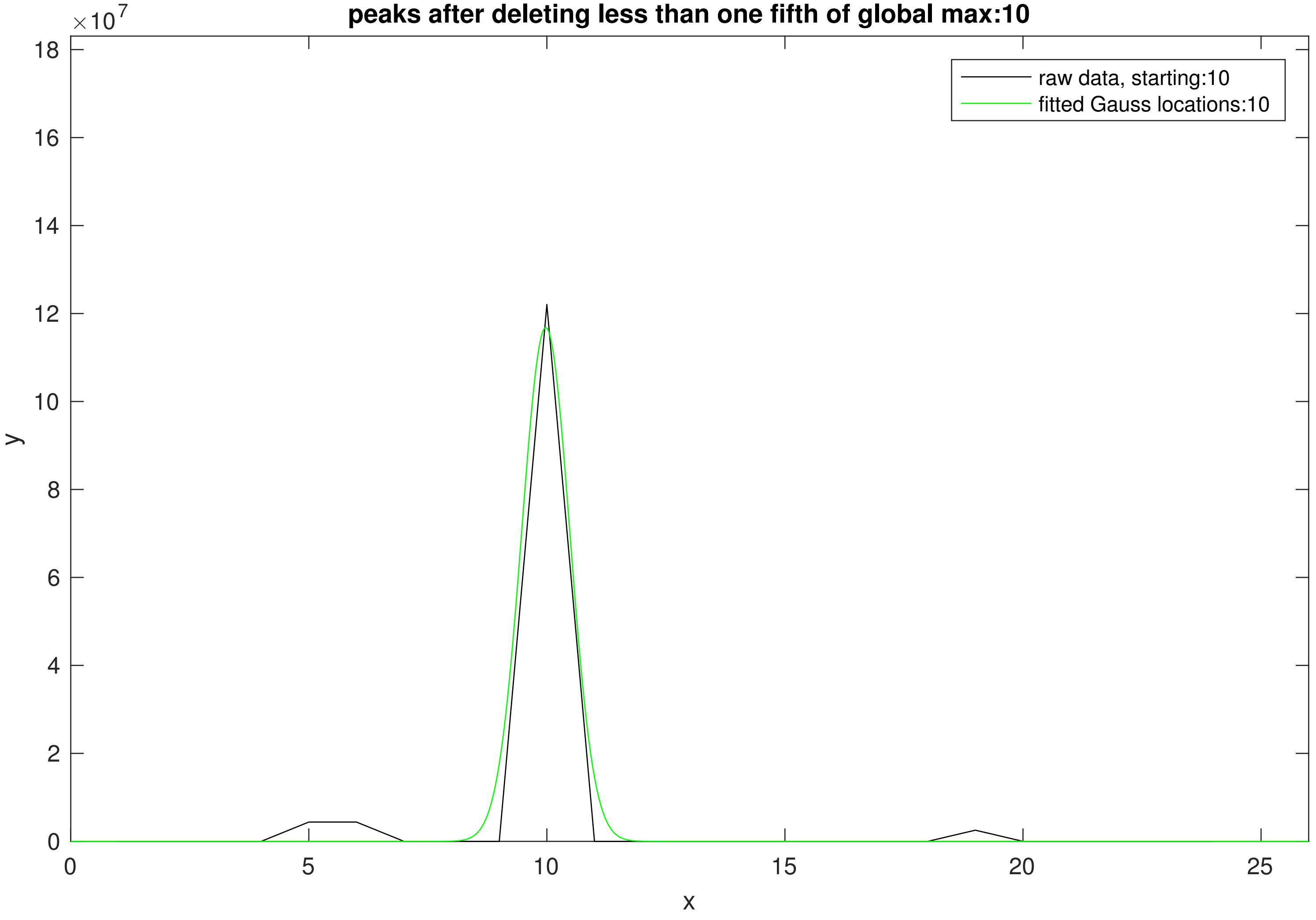

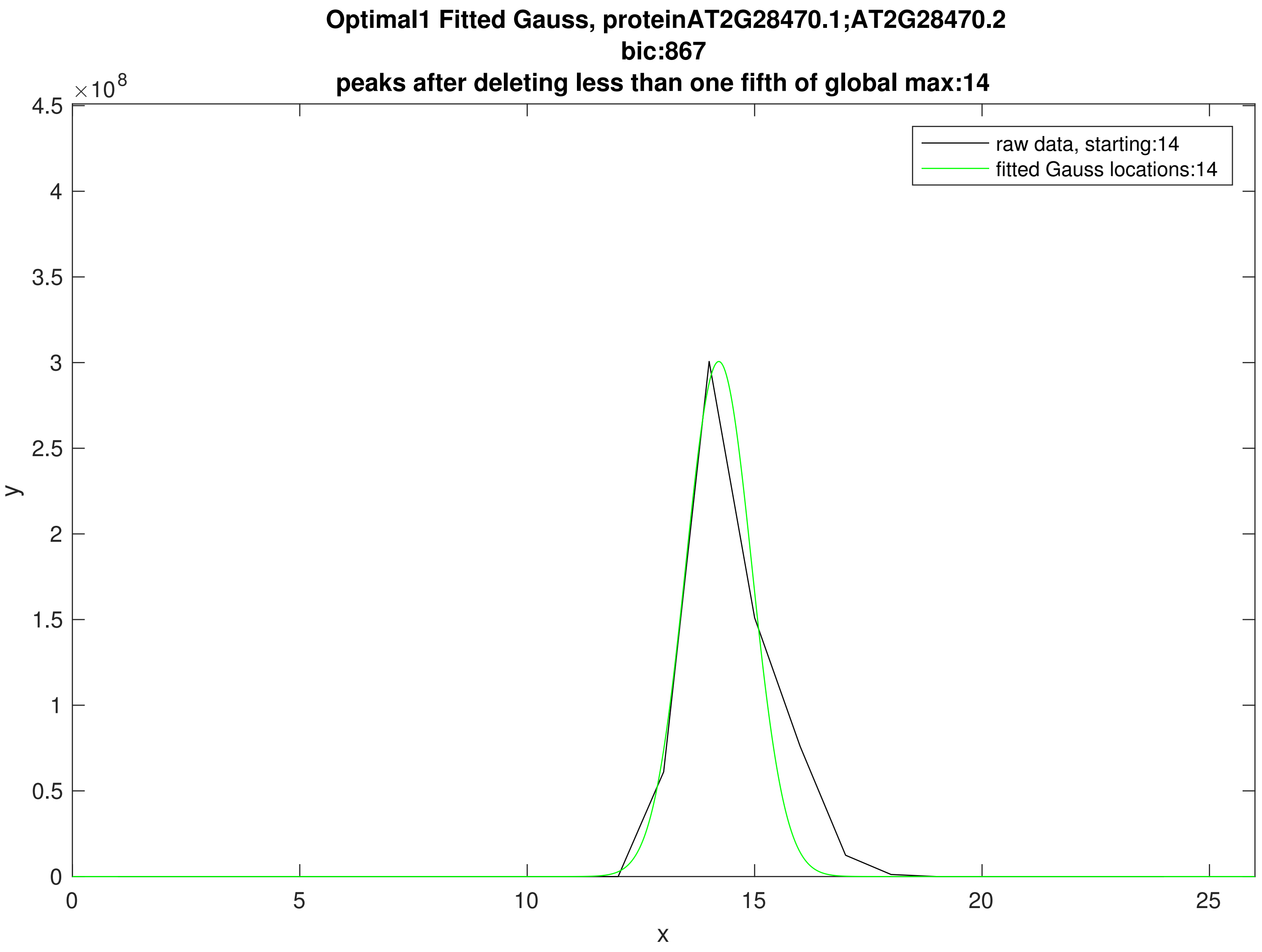

Optimal1 Fitted Gauss, proteinAT2G28470.1;AT2G28470.2
bic:867
peaks after deleting less than one fifth of global max:14
×10^8
4.5
4
3.5
3
2.5
2
1.5
1
0.5
0
y
0
5
10
15
20
25
x
raw data, starting:14
fitted Gauss locations:14

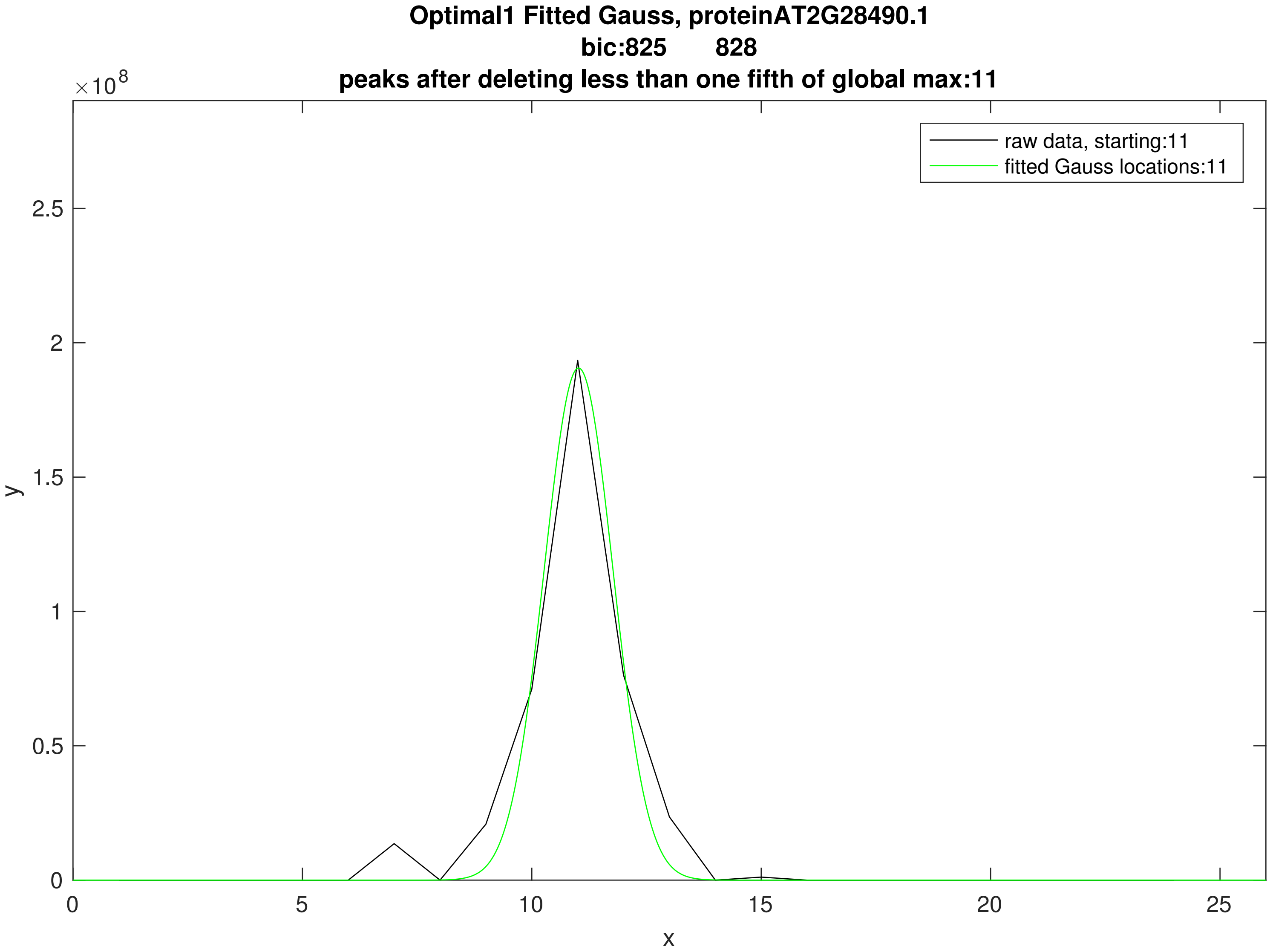

Optimal1 Fitted Gauss, proteinAT2G28490.1
bic:825 828
peaks after deleting less than one fifth of global max:11
×10^8
raw data, starting:11
fitted Gauss locations:11
2.5
2
1.5
1
0.5
0
y
0
5
10
15
20
25
x

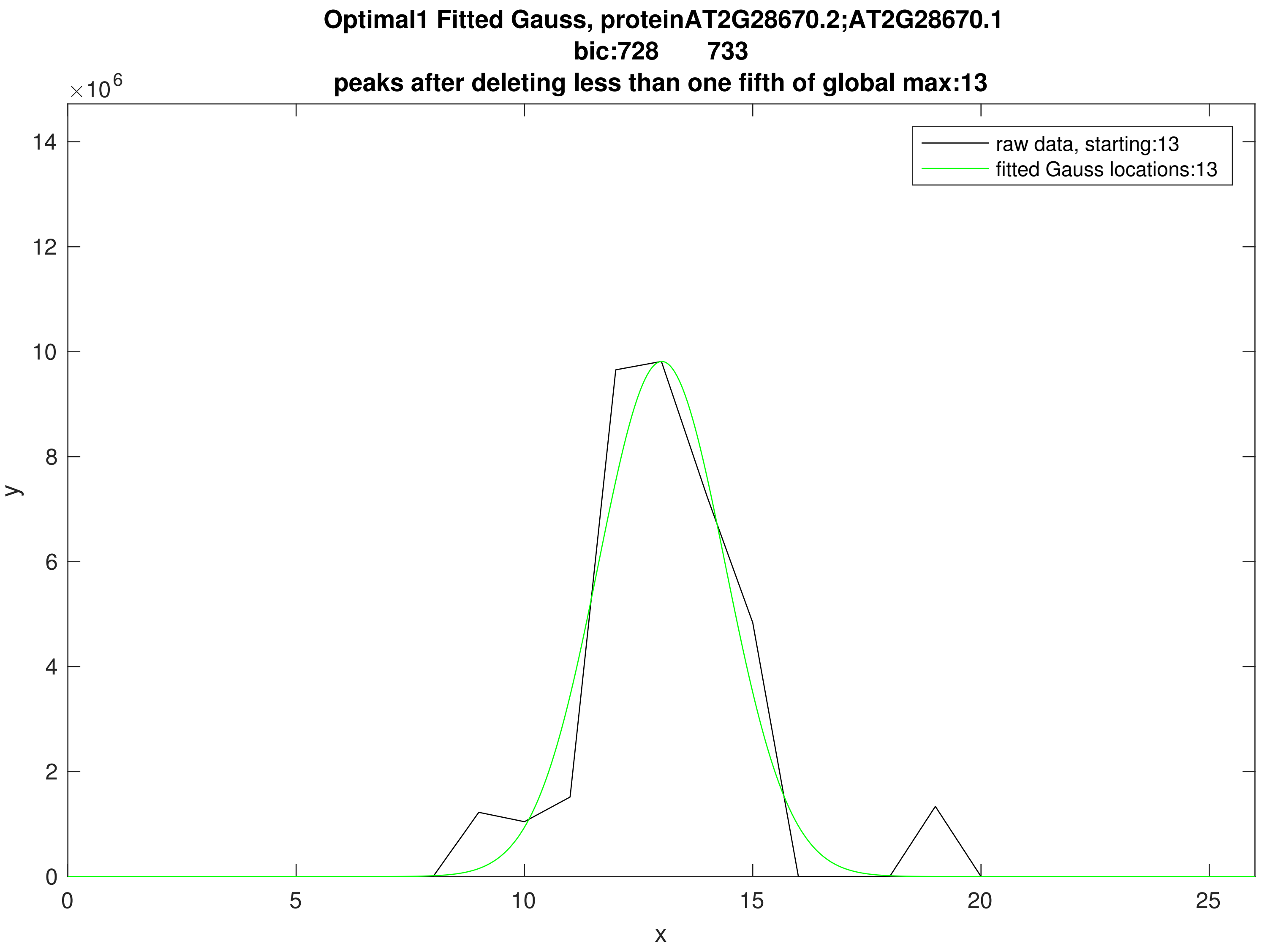
Optimal1 Fitted Gauss, proteinAT2G28670.2;AT2G28670.1
bic:728 733
peaks after deleting less than one fifth of global max:13
×10^6
raw data, starting:13
fitted Gauss locations:13
y
x
0
2
4
6
8
10
12
14
0
5
10
15
20
25

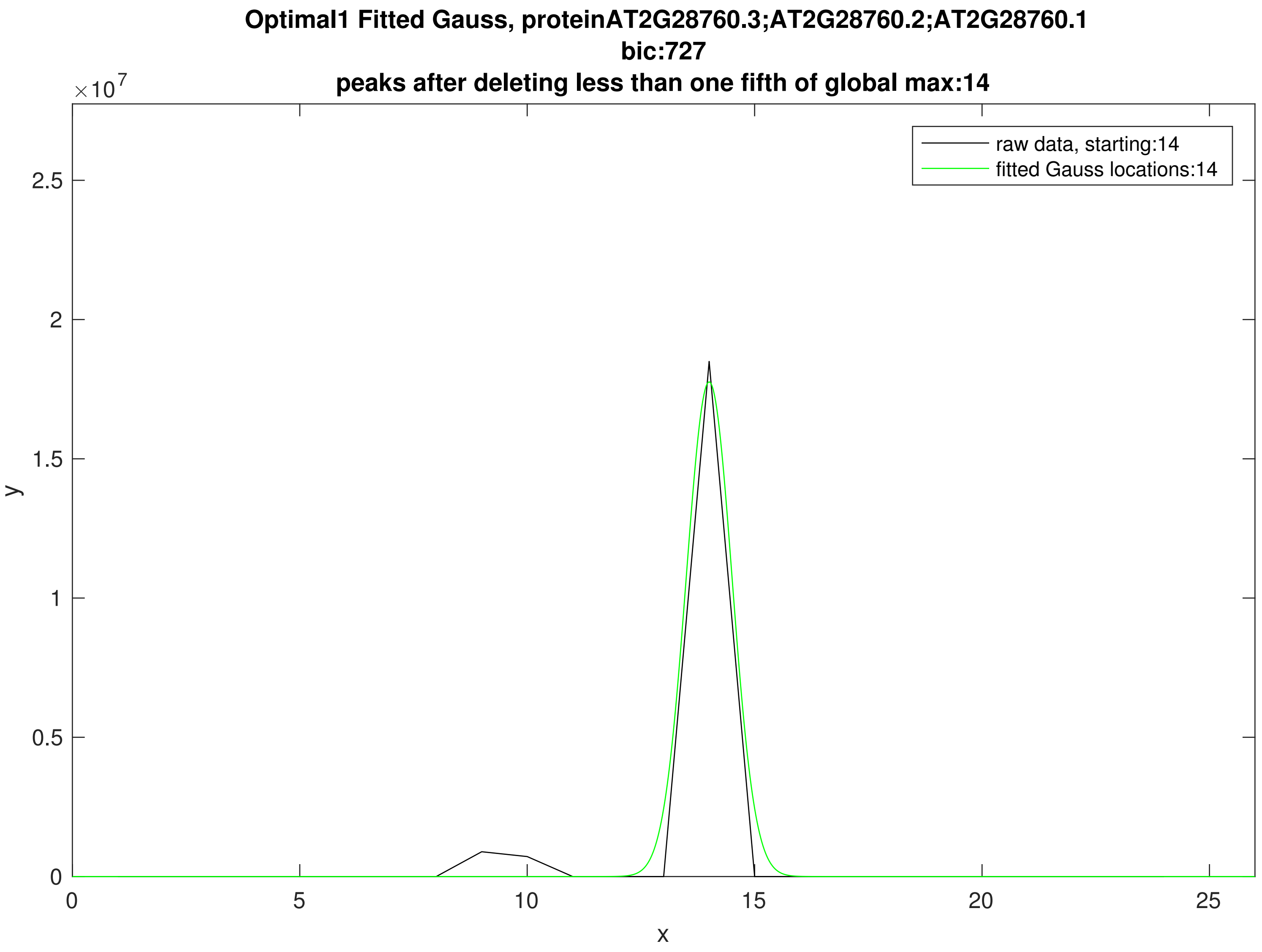
Optimal1 Fitted Gauss, proteinAT2G28760.3;AT2G28760.2;AT2G28760.1
bic:727
peaks after deleting less than one fifth of global max:14
×10^7
2.5
2
1.5
1
0.5
0
y
0
5
10
15
20
25
x
raw data, starting:14
fitted Gauss locations:14

**Optimal1 Fitted Gauss, proteinAT2G29300.1;AT2G29300.2;AT2G29320.1**
**bic:758**
**peaks after deleting less than one fifth of global max:13**

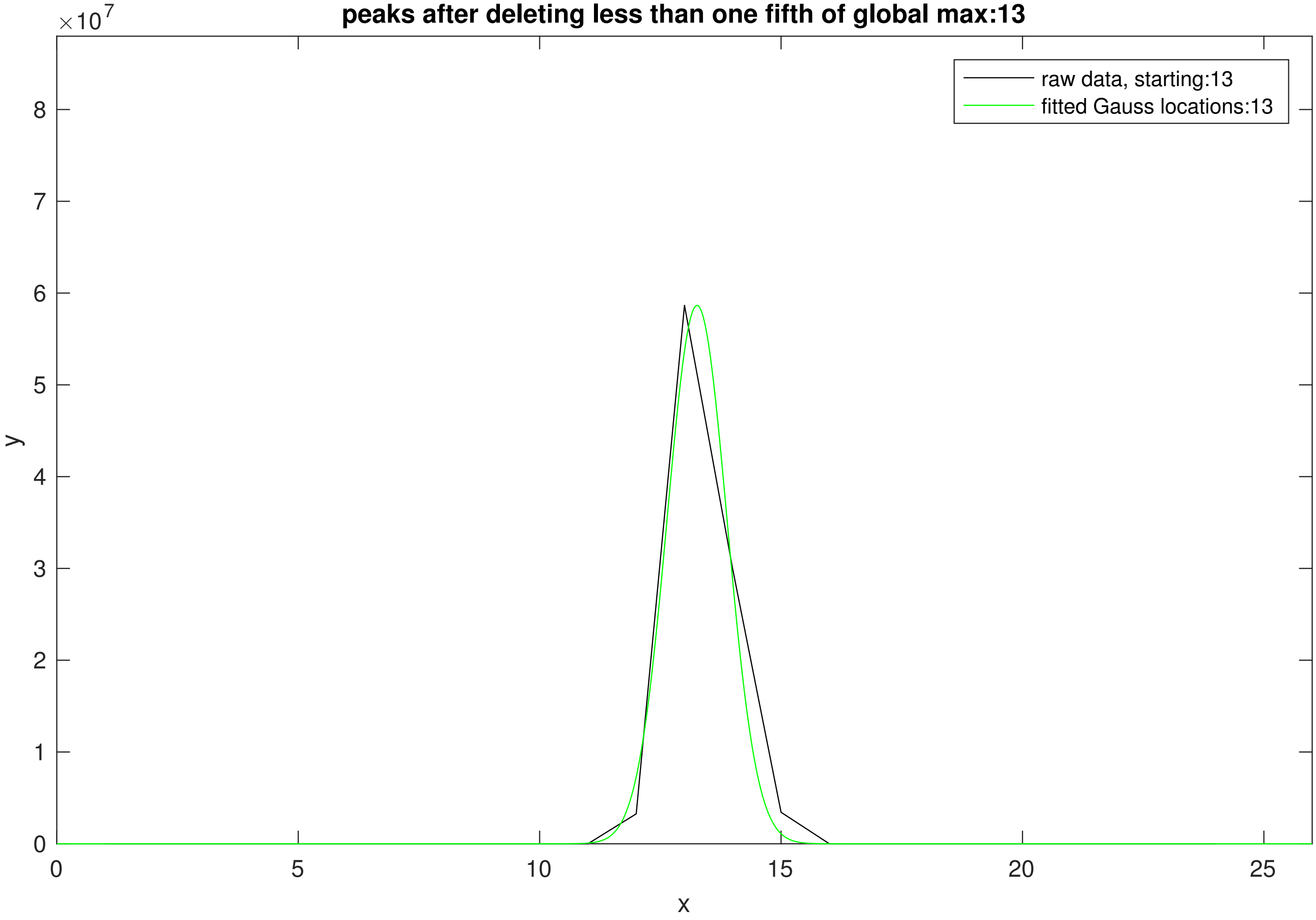

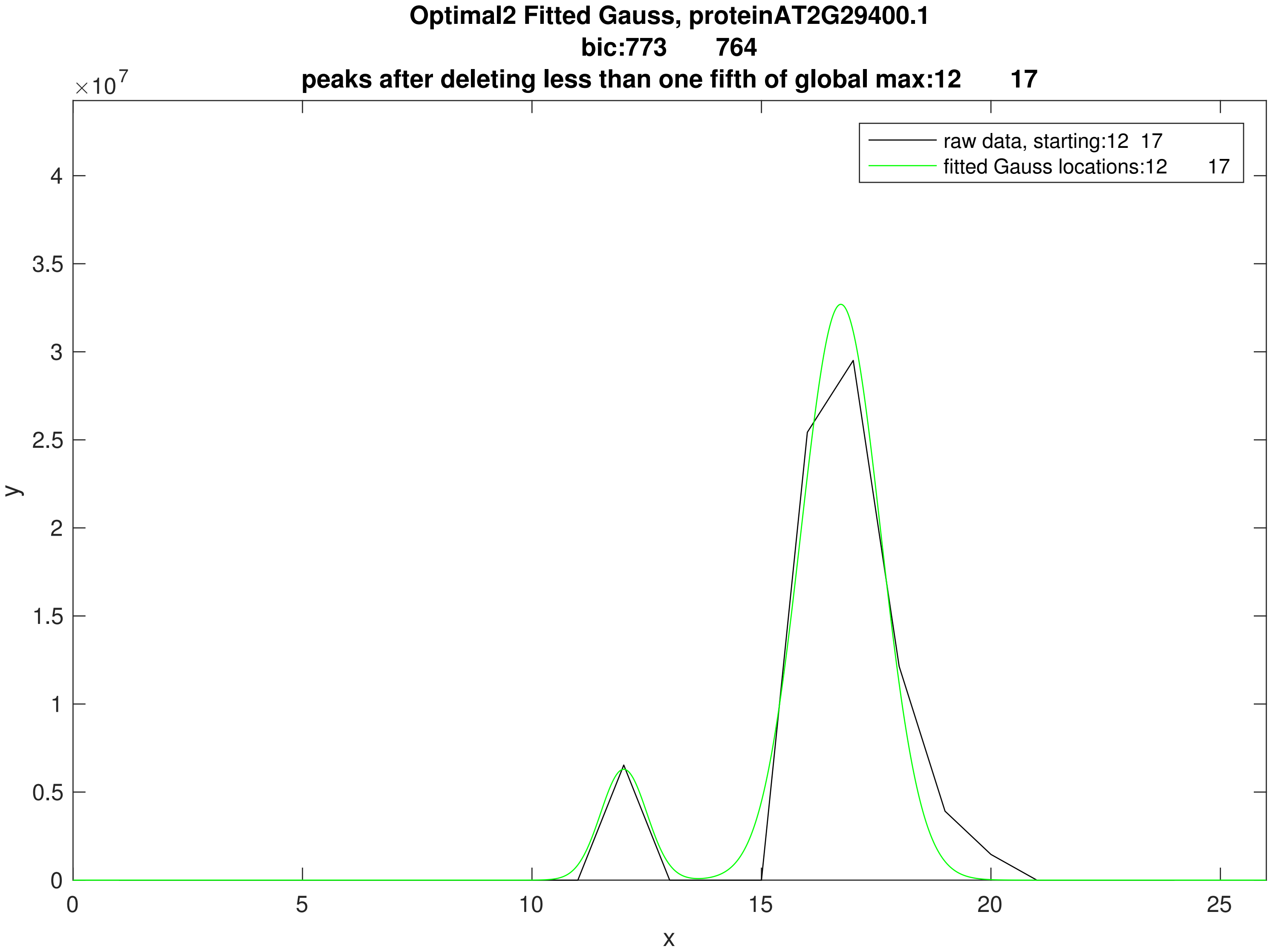

Optimal2 Fitted Gauss, proteinAT2G29400.1
bic:773 764
peaks after deleting less than one fifth of global max:12 17
×10^7
raw data, starting:12 17
fitted Gauss locations:12 17
y
x
0
0.5
1
1.5
2
2.5
3
3.5
4
5
10
15
20
25

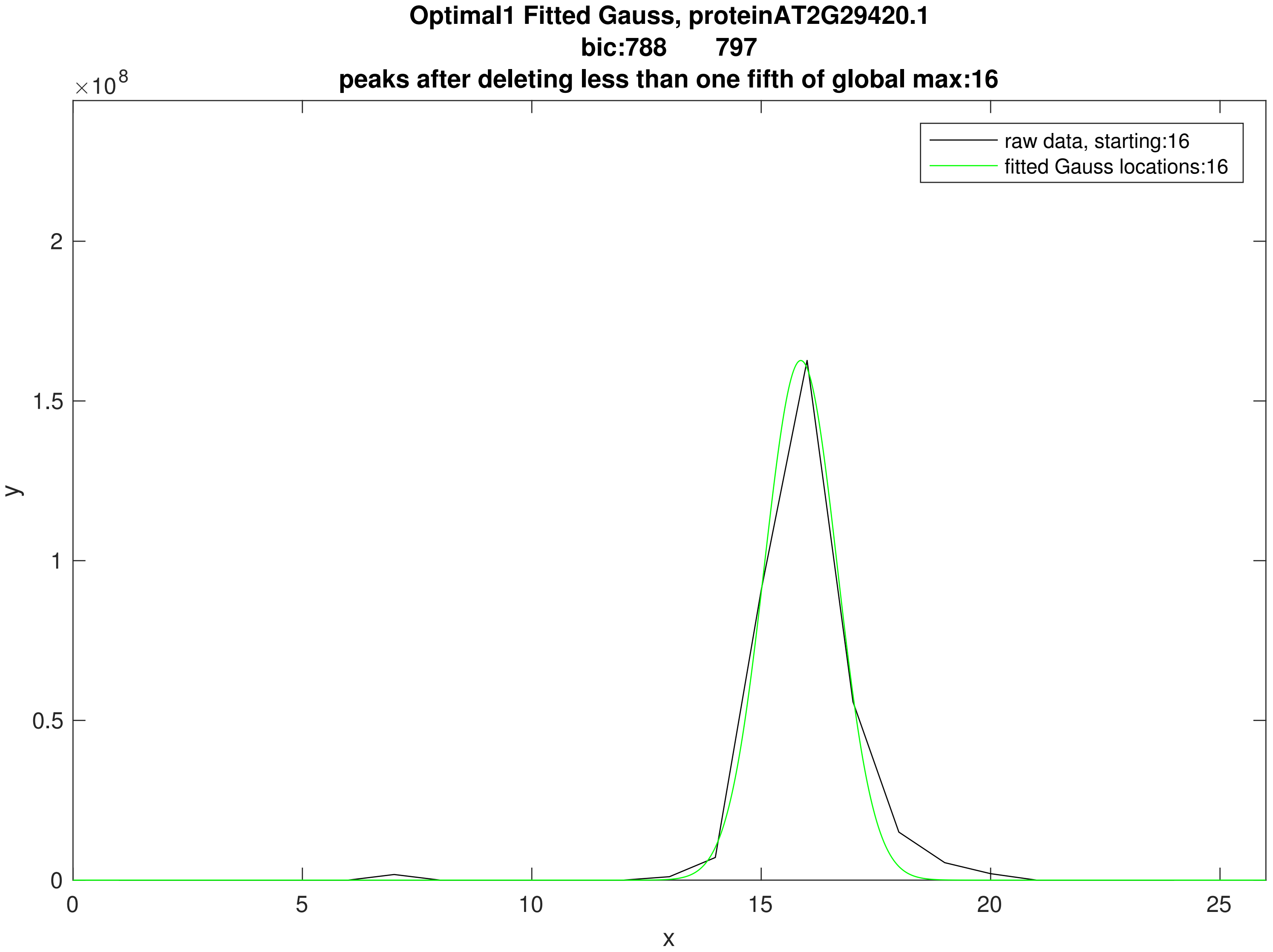

Optimal1 Fitted Gauss, proteinAT2G29420.1
bic:788 797
peaks after deleting less than one fifth of global max:16
×10^8
raw data, starting:16
fitted Gauss locations:16
y
x
0
0.5
1
1.5
2
5
10
15
20
25

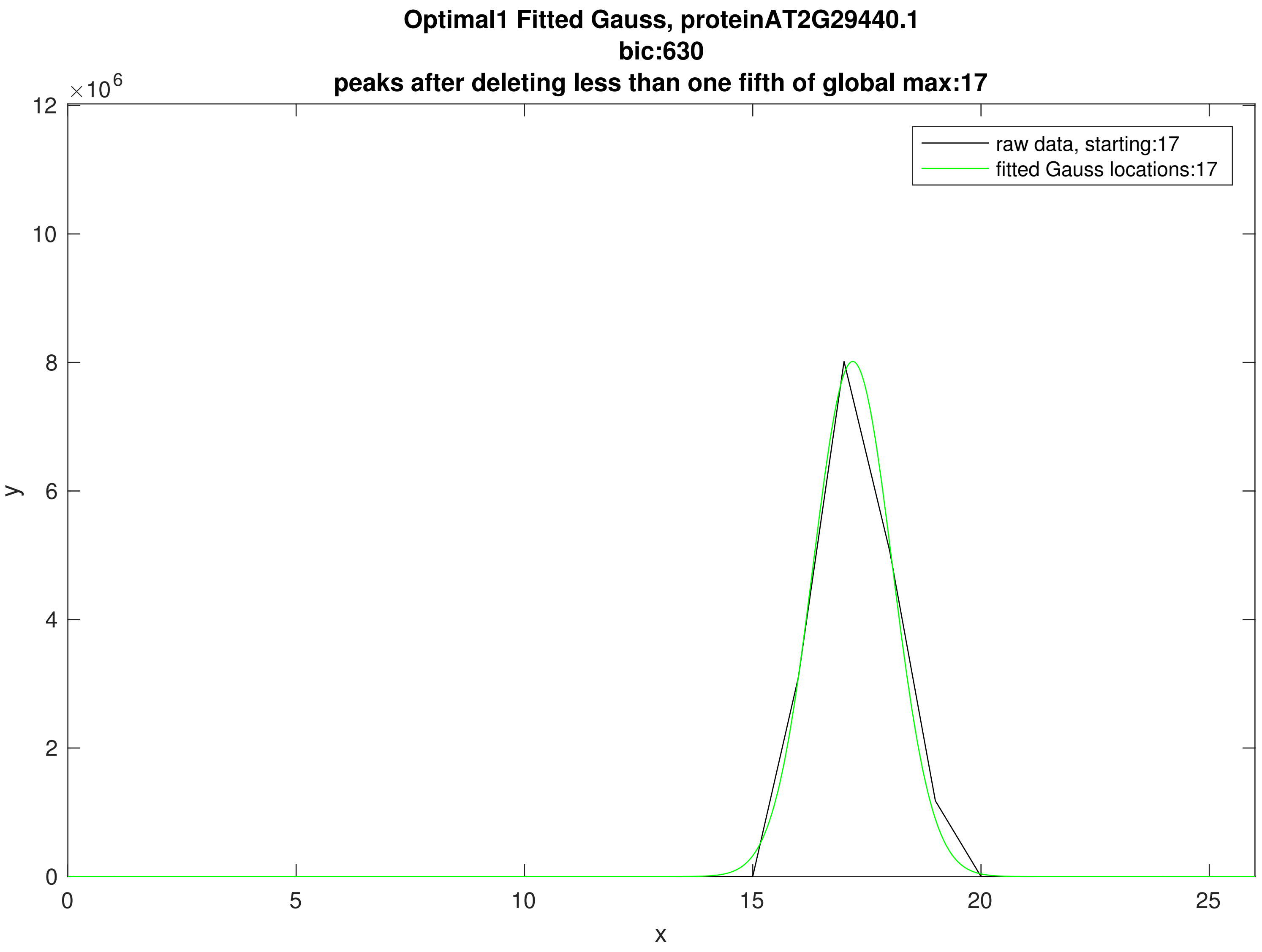

Optimal1 Fitted Gauss, proteinAT2G29440.1
bic:630
peaks after deleting less than one fifth of global max:17
×10^6
raw data, starting:17
fitted Gauss locations:17
y
x
0
2
4
6
8
10
12
0
5
10
15
20
25

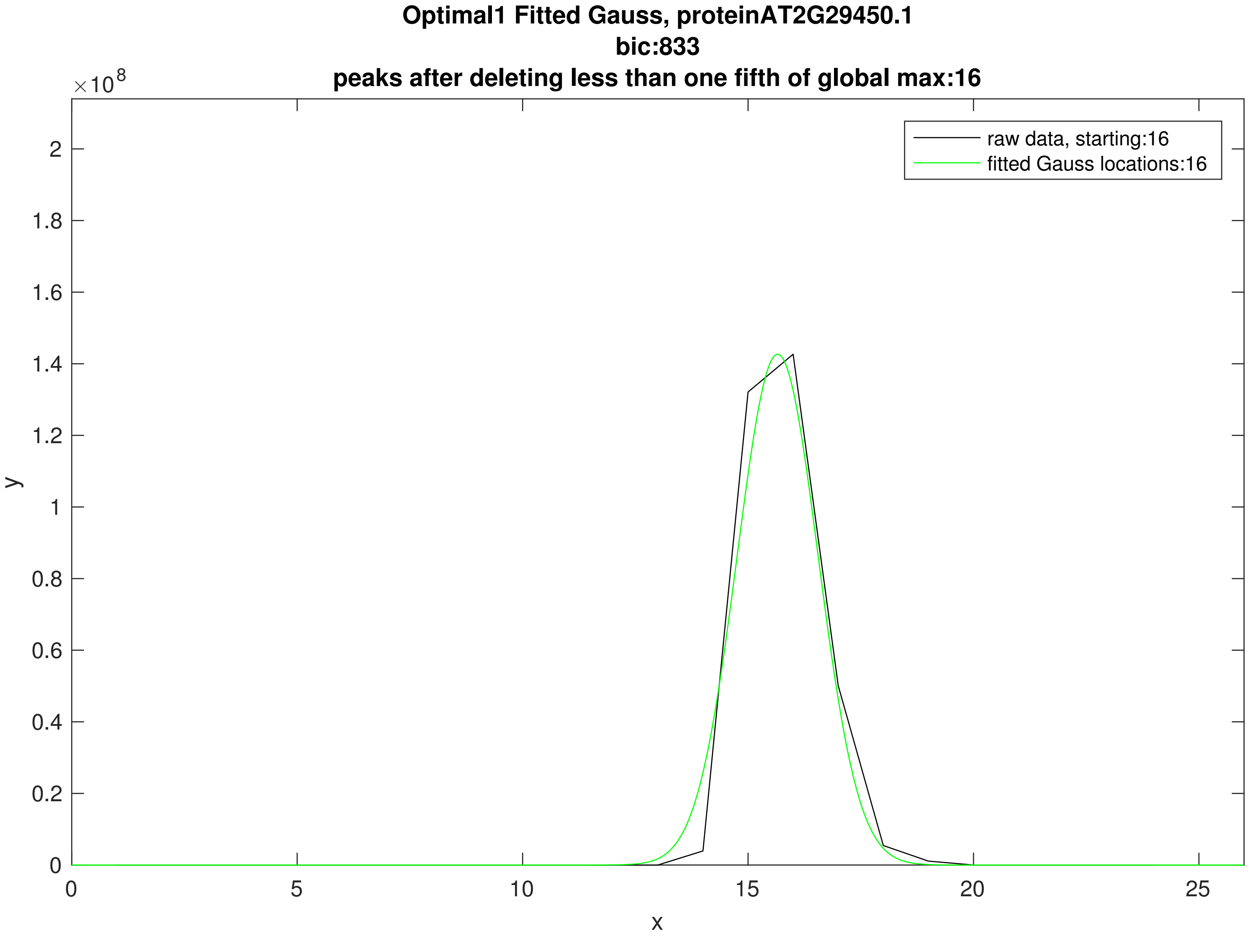
Optimal1 Fitted Gauss, proteinAT2G29450.1
bic:833
peaks after deleting less than one fifth of global max:16
×10^8
raw data, starting:16
fitted Gauss locations:16
y
x
0
0.2
0.4
0.6
0.8
1
1.2
1.4
1.6
1.8
2
5
10
15
20
25

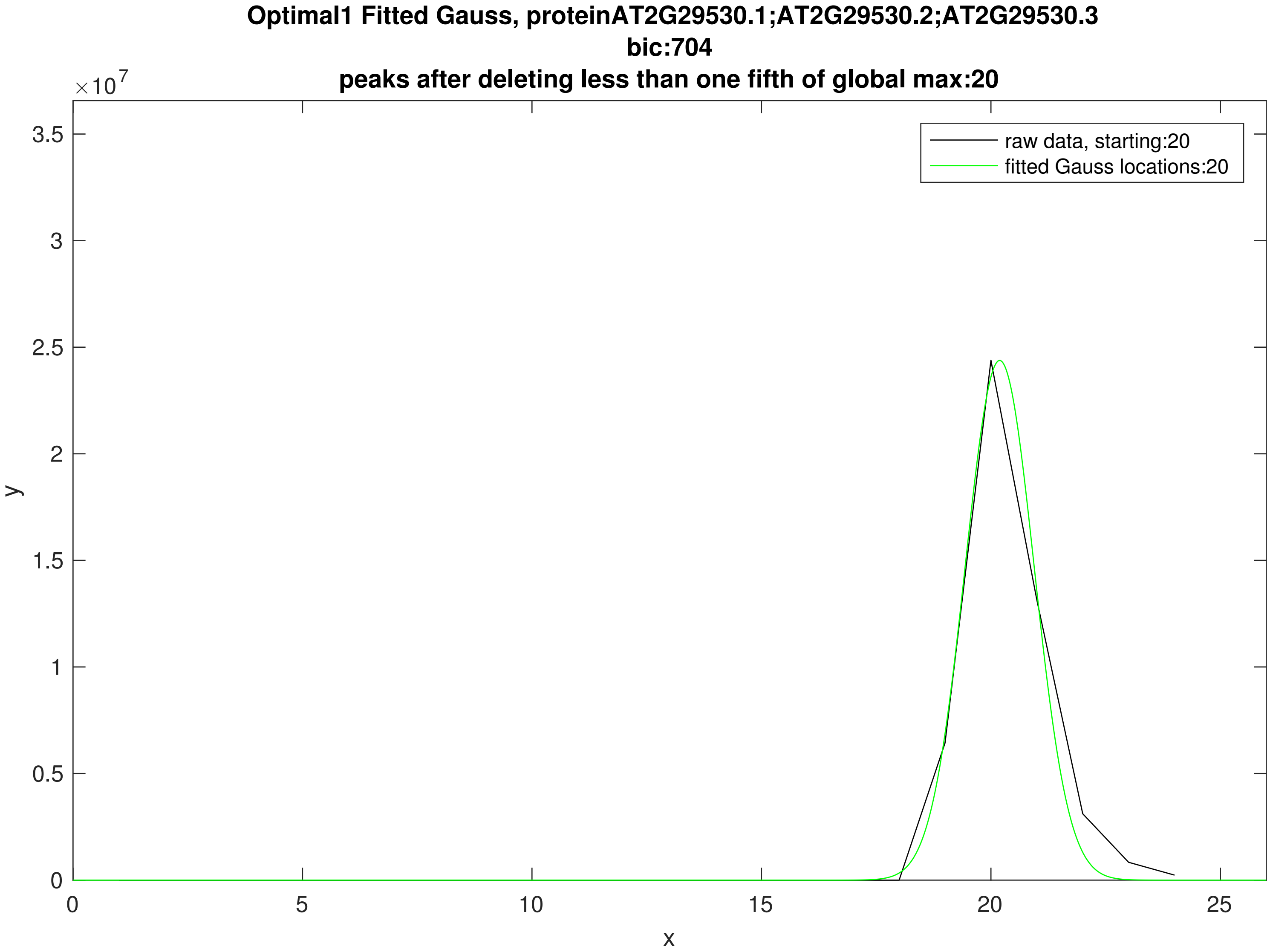
Optimal1 Fitted Gauss, proteinAT2G29530.1;AT2G29530.2;AT2G29530.3
bic:704
peaks after deleting less than one fifth of global max:20
×10^7
raw data, starting:20
fitted Gauss locations:20
3.5
3
2.5
2
1.5
1
0.5
0
y
0
5
10
15
20
25
x

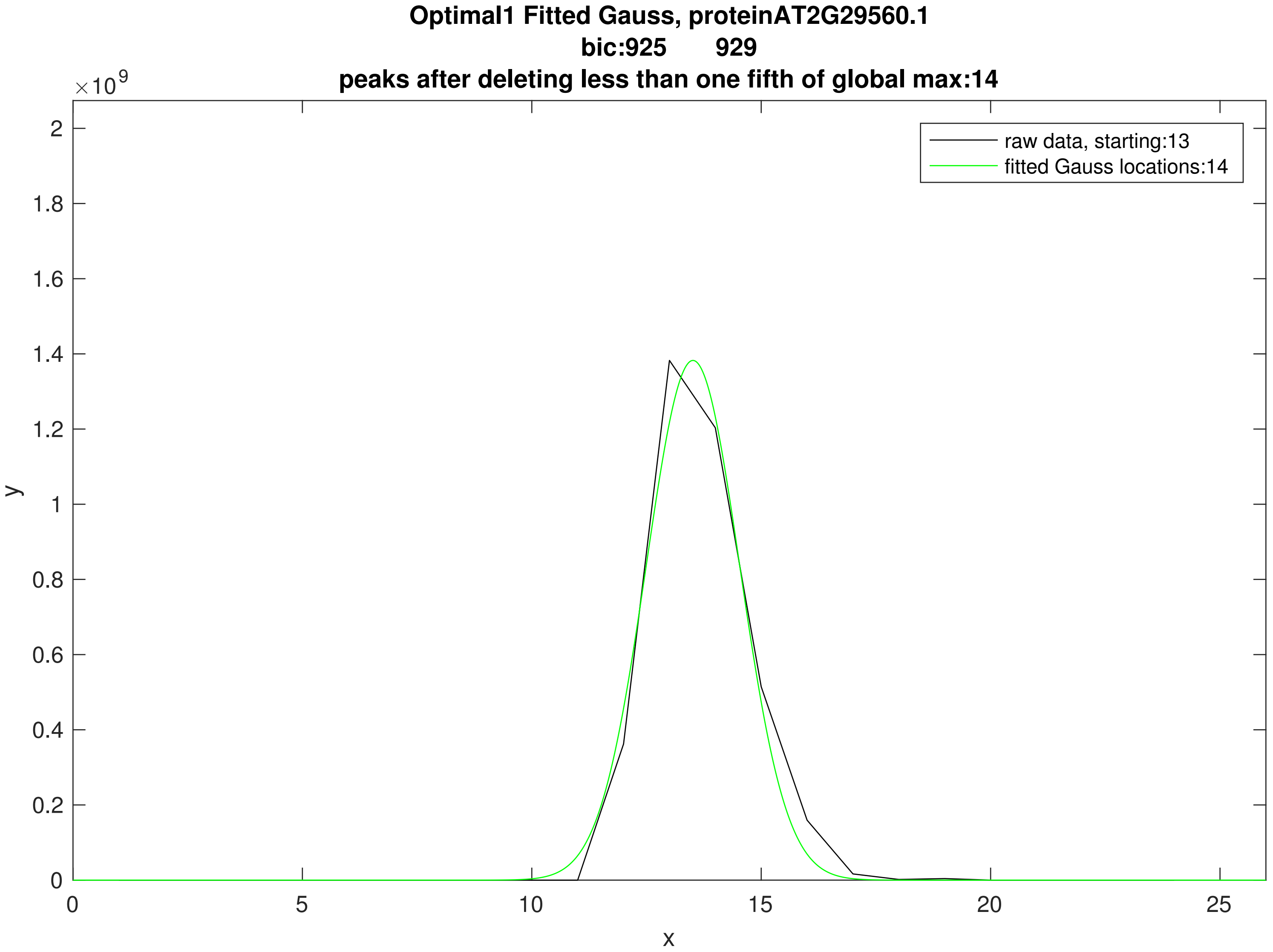
Optimal1 Fitted Gauss, proteinAT2G29560.1
bic:925 929
peaks after deleting less than one fifth of global max:14
×10^9
raw data, starting:13
fitted Gauss locations:14
y
x
0
0.2
0.4
0.6
0.8
1
1.2
1.4
1.6
1.8
2
5
10
15
20
25

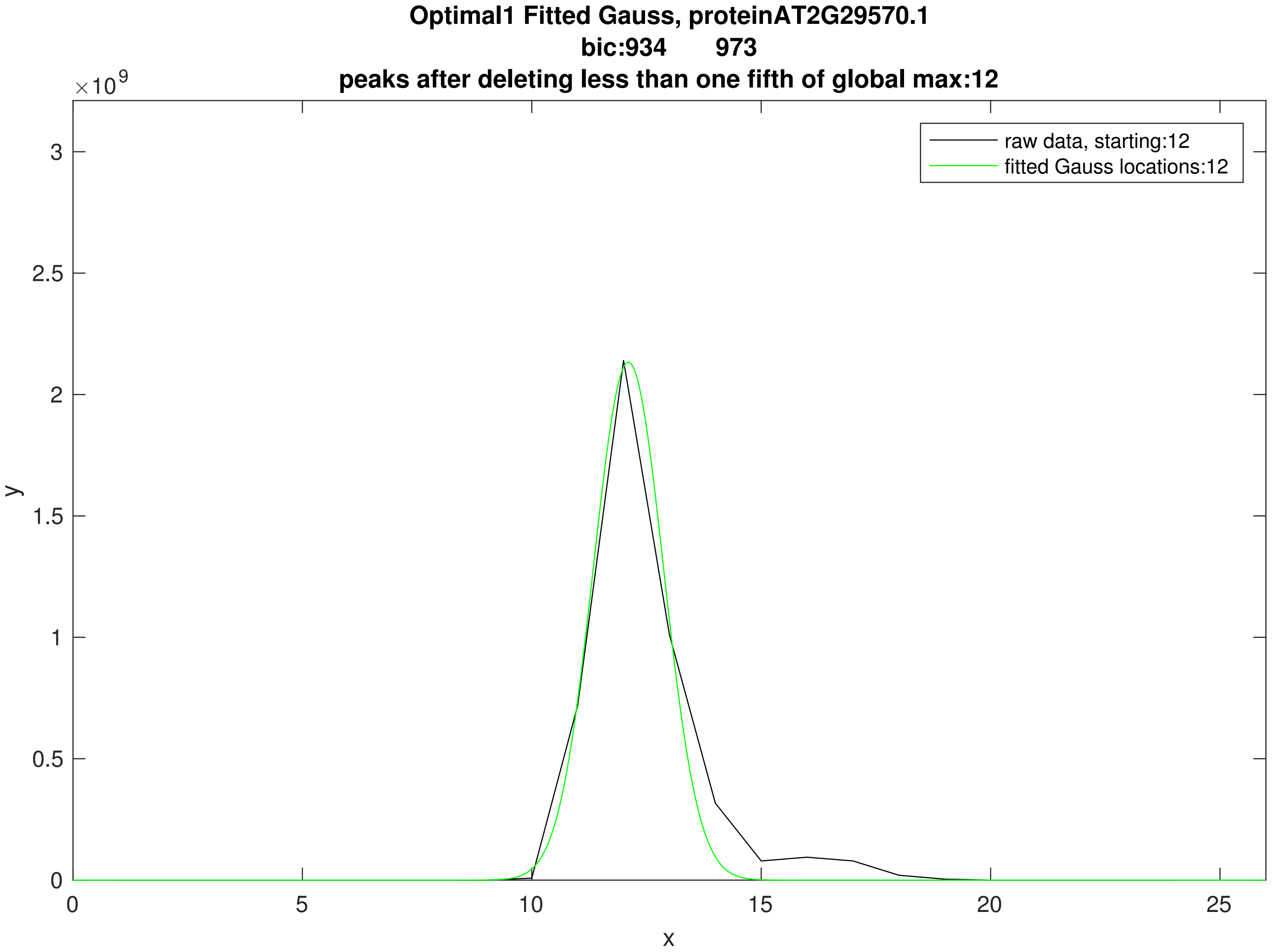

Optimal1 Fitted Gauss, proteinAT2G29570.1
bic:934 973
peaks after deleting less than one fifth of global max:12
×10^9
3
2.5
2
1.5
1
0.5
0
y
0
5
10
15
20
25
x
raw data, starting:12
fitted Gauss locations:12

**Optimal3 Fitted Gauss, proteinAT2G29690.1**
**bic:831 806 770**
**peaks after deleting less than one fifth of global max:6.4 11 16**

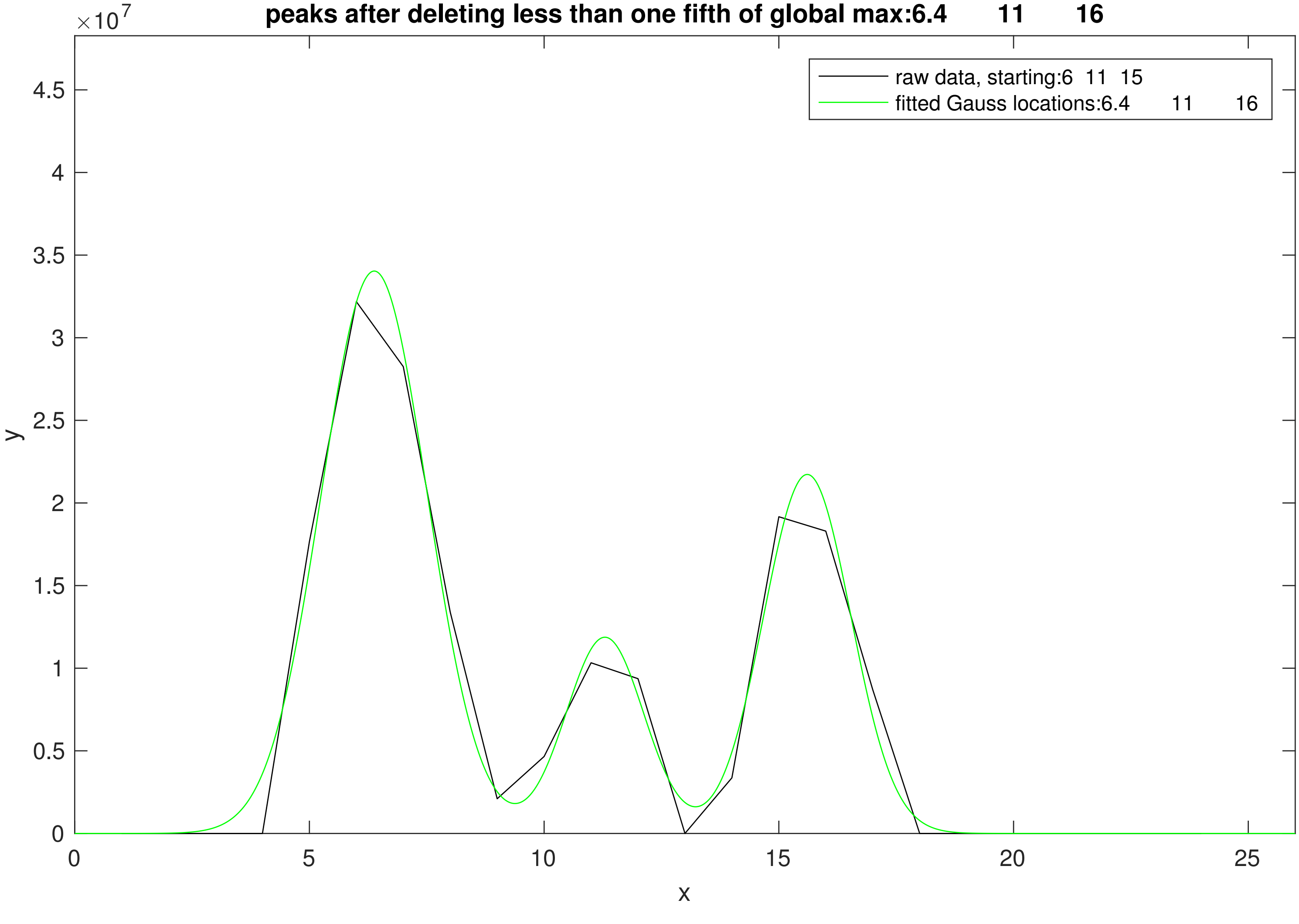

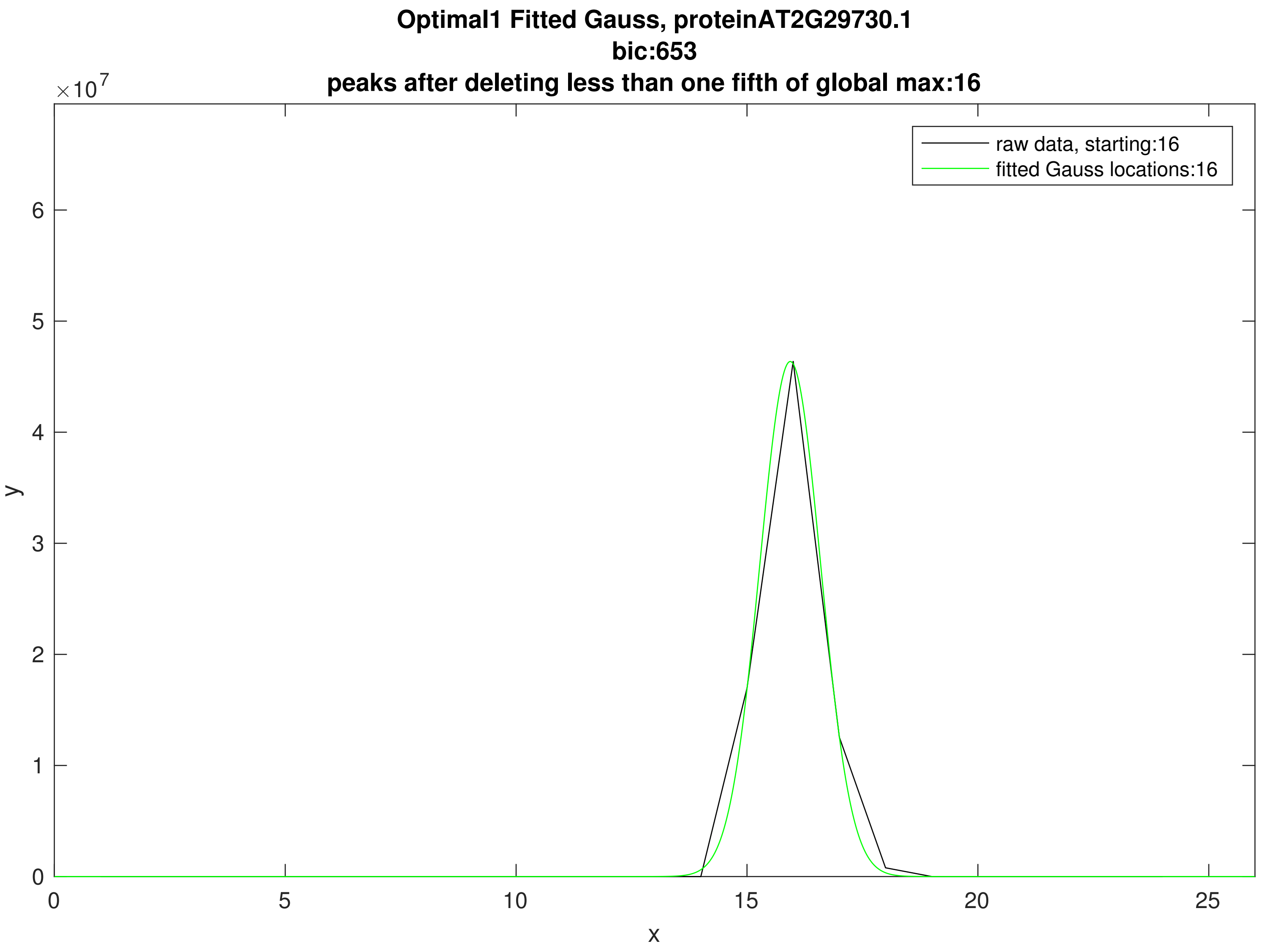
Optimal1 Fitted Gauss, proteinAT2G29730.1
bic:653
peaks after deleting less than one fifth of global max:16
×10^7
raw data, starting:16
fitted Gauss locations:16
y
x
0
1
2
3
4
5
6
0
5
10
15
20
25

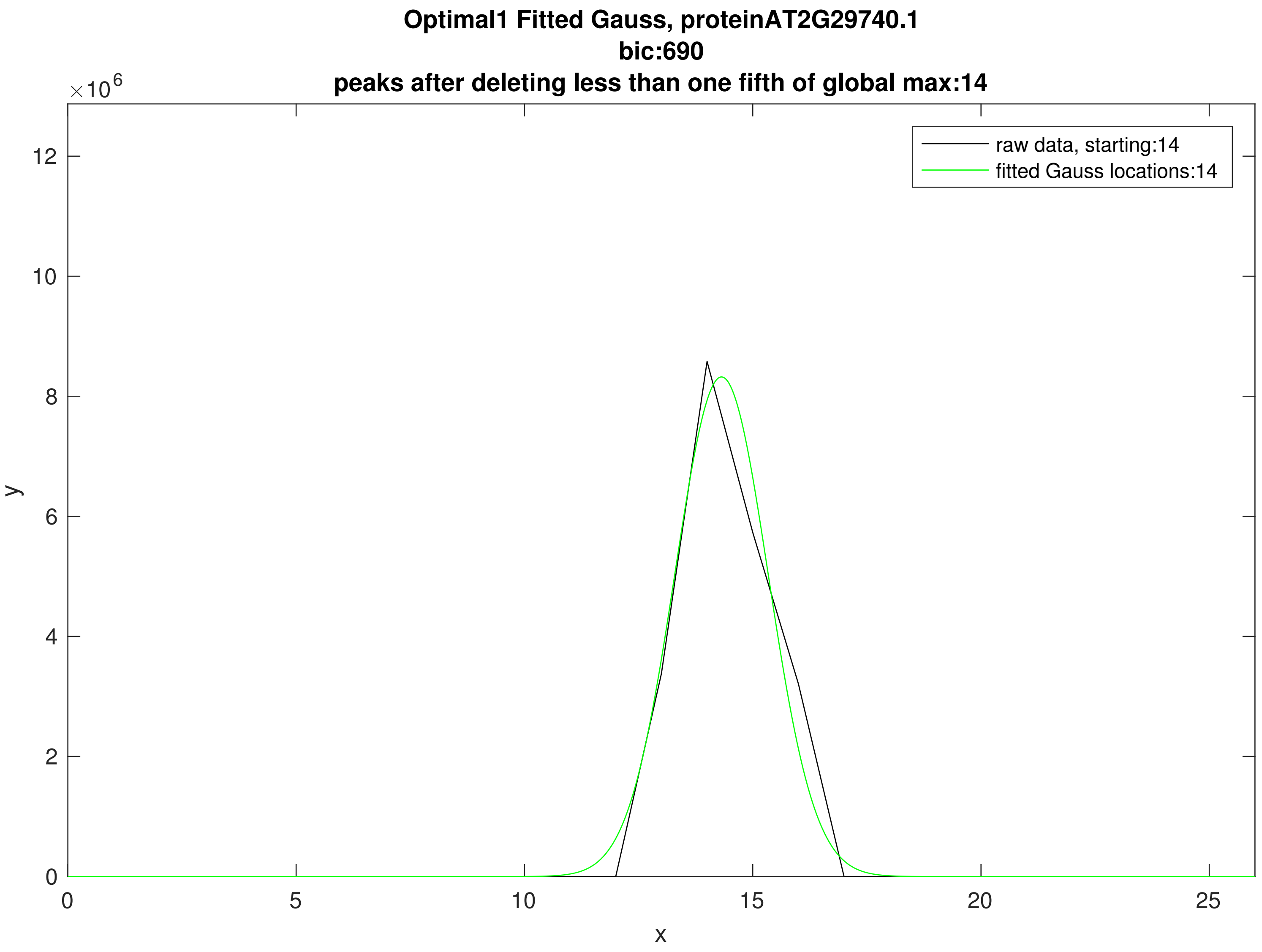
Optimal1 Fitted Gauss, proteinAT2G29740.1
bic:690
peaks after deleting less than one fifth of global max:14
×10^6
12
10
8
6
4
2
0
y
0
5
10
15
20
25
x
raw data, starting:14
fitted Gauss locations:14

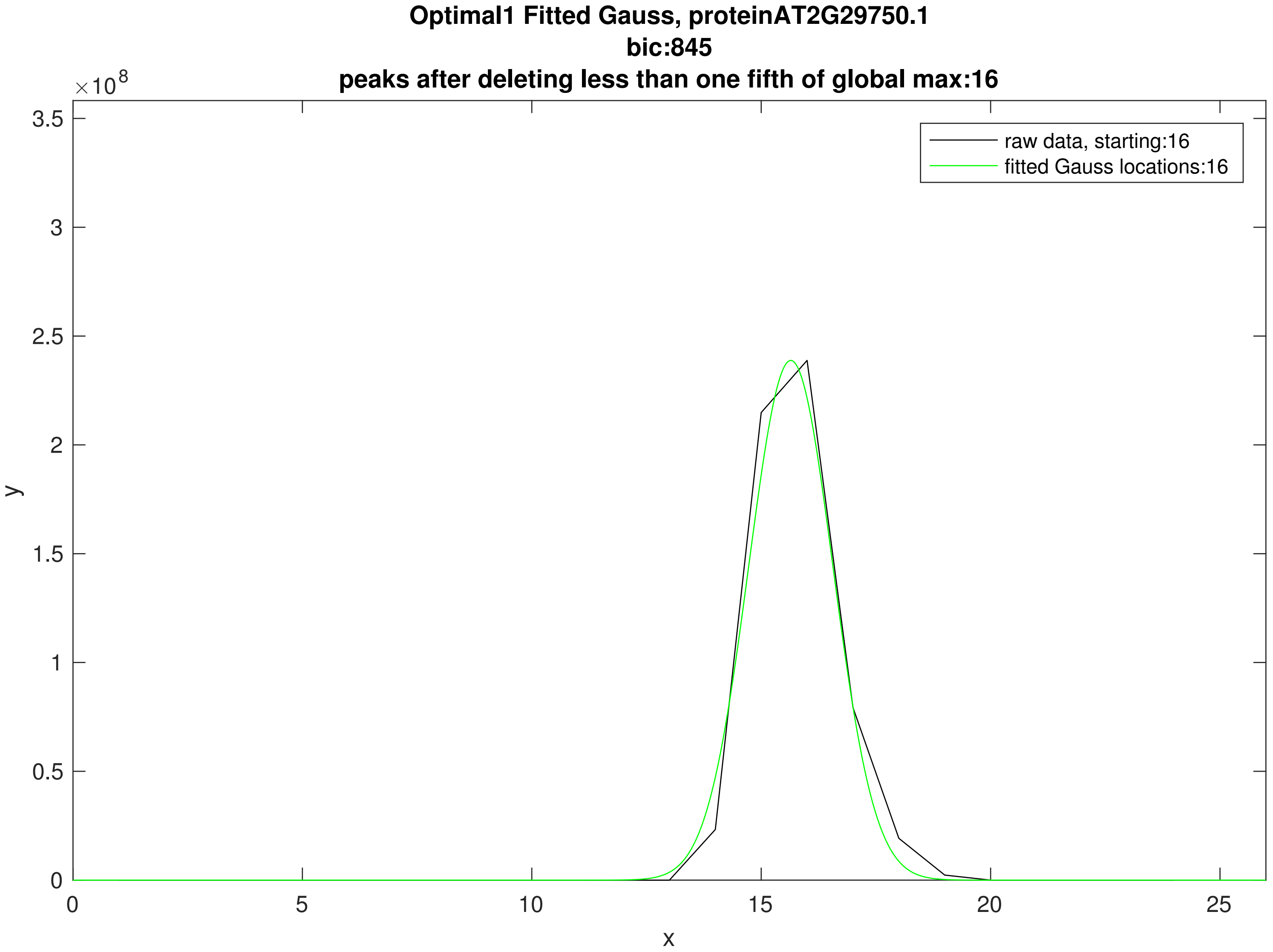

Optimal1 Fitted Gauss, proteinAT2G29750.1
bic:845
peaks after deleting less than one fifth of global max:16
×10^8
3.5
3
2.5
2
1.5
1
0.5
0
y
0
5
10
15
20
25
x
raw data, starting:16
fitted Gauss locations:16

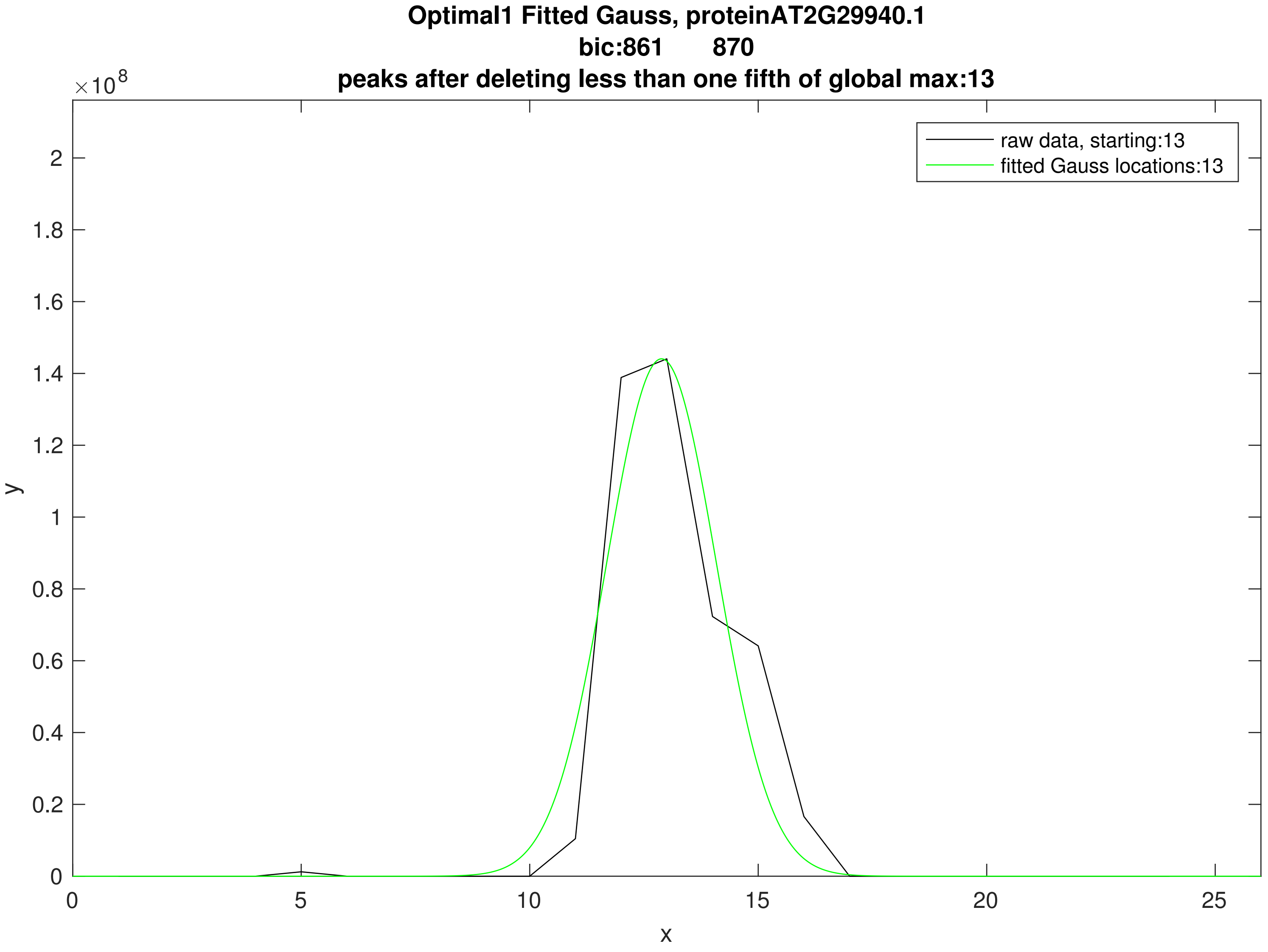
Optimal1 Fitted Gauss, proteinAT2G29940.1
bic:861 870
peaks after deleting less than one fifth of global max:13
×10^8
raw data, starting:13
fitted Gauss locations:13
y
x
0
0.2
0.4
0.6
0.8
1
1.2
1.4
1.6
1.8
2
5
10
15
20
25

#### Optimal1 Fitted Gauss, proteinAT2G29960.1;AT2G29960.2
#### bic:854 859
#### peaks after deleting less than one fifth of global max:20

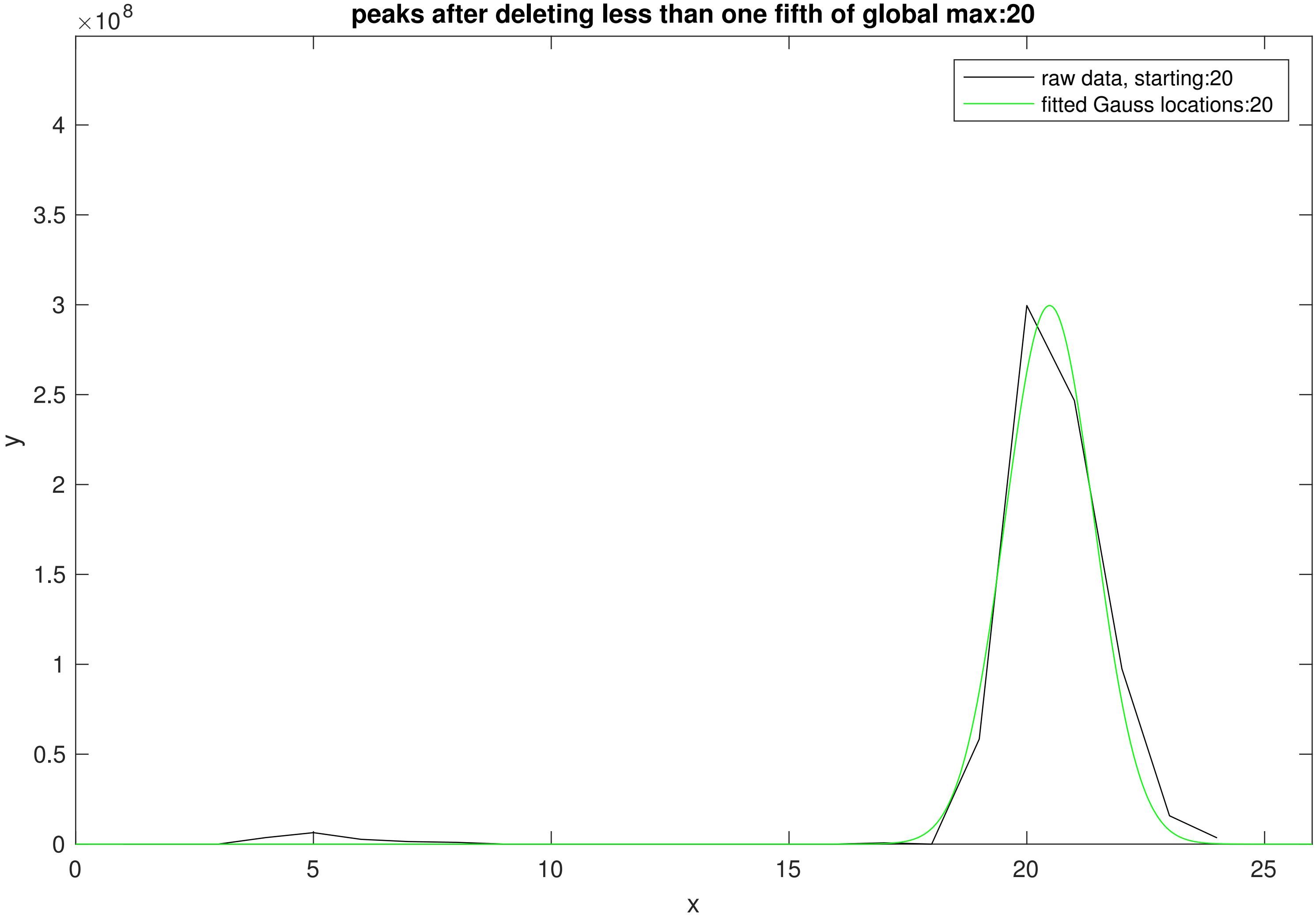

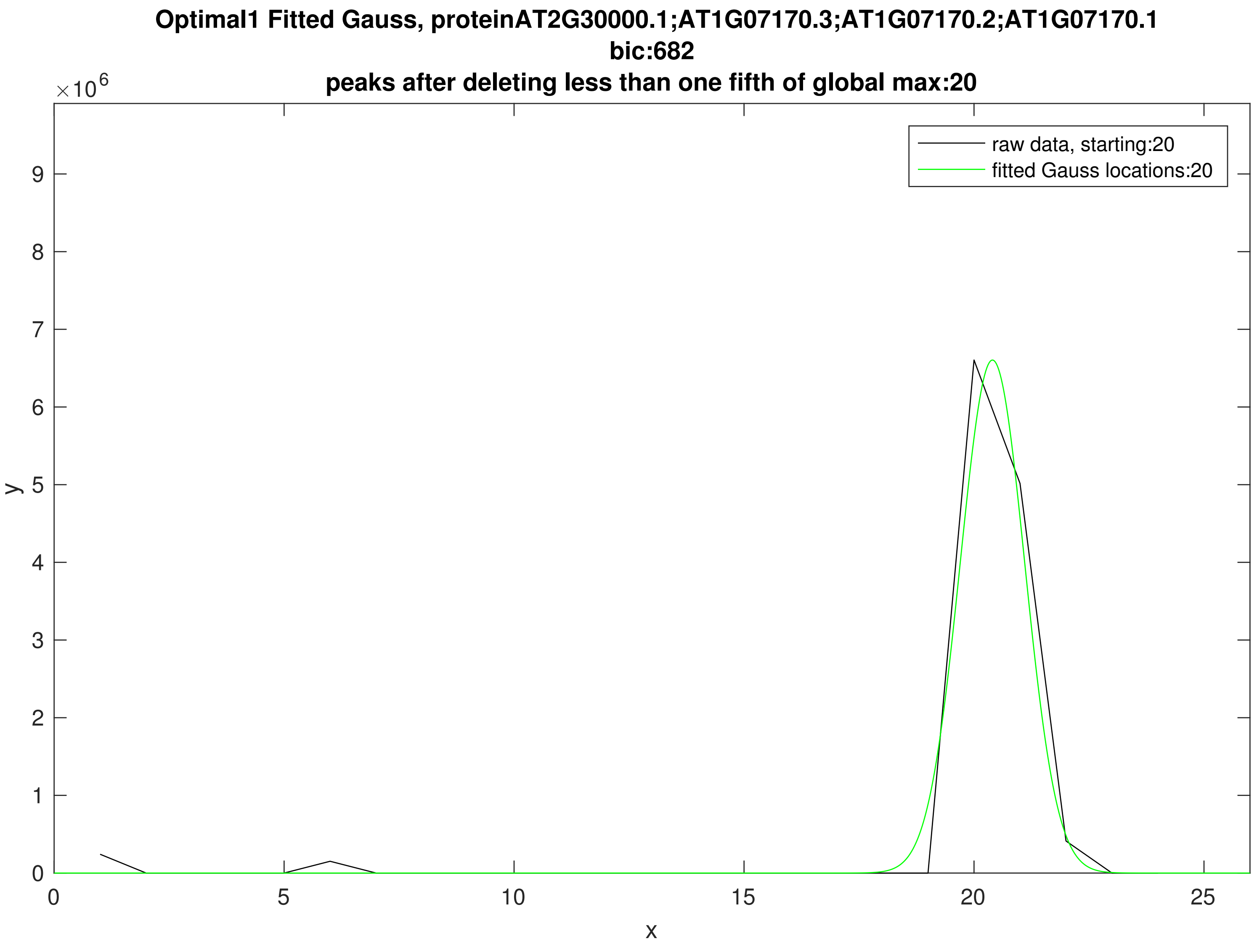
Optimal1 Fitted Gauss, proteinAT2G30000.1;AT1G07170.3;AT1G07170.2;AT1G07170.1
bic:682
peaks after deleting less than one fifth of global max:20
×10^6
raw data, starting:20
fitted Gauss locations:20
y
x
0
1
2
3
4
5
6
7
8
9
0
5
10
15
20
25

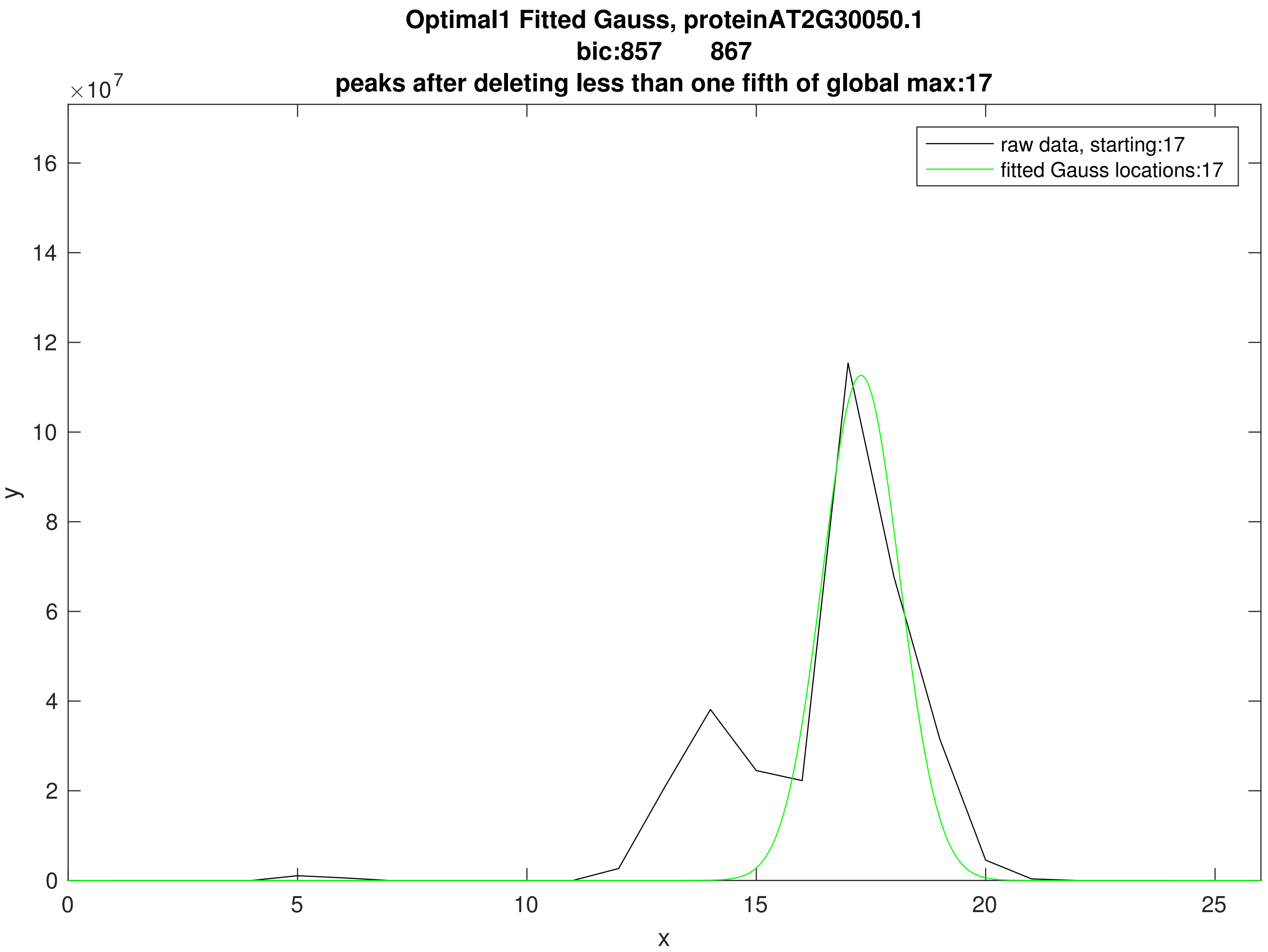
Optimal1 Fitted Gauss, proteinAT2G30050.1
bic:857 867
peaks after deleting less than one fifth of global max:17
×10^7
raw data, starting:17
fitted Gauss locations:17
y
x
0
2
4
6
8
10
12
14
16
0
5
10
15
20
25

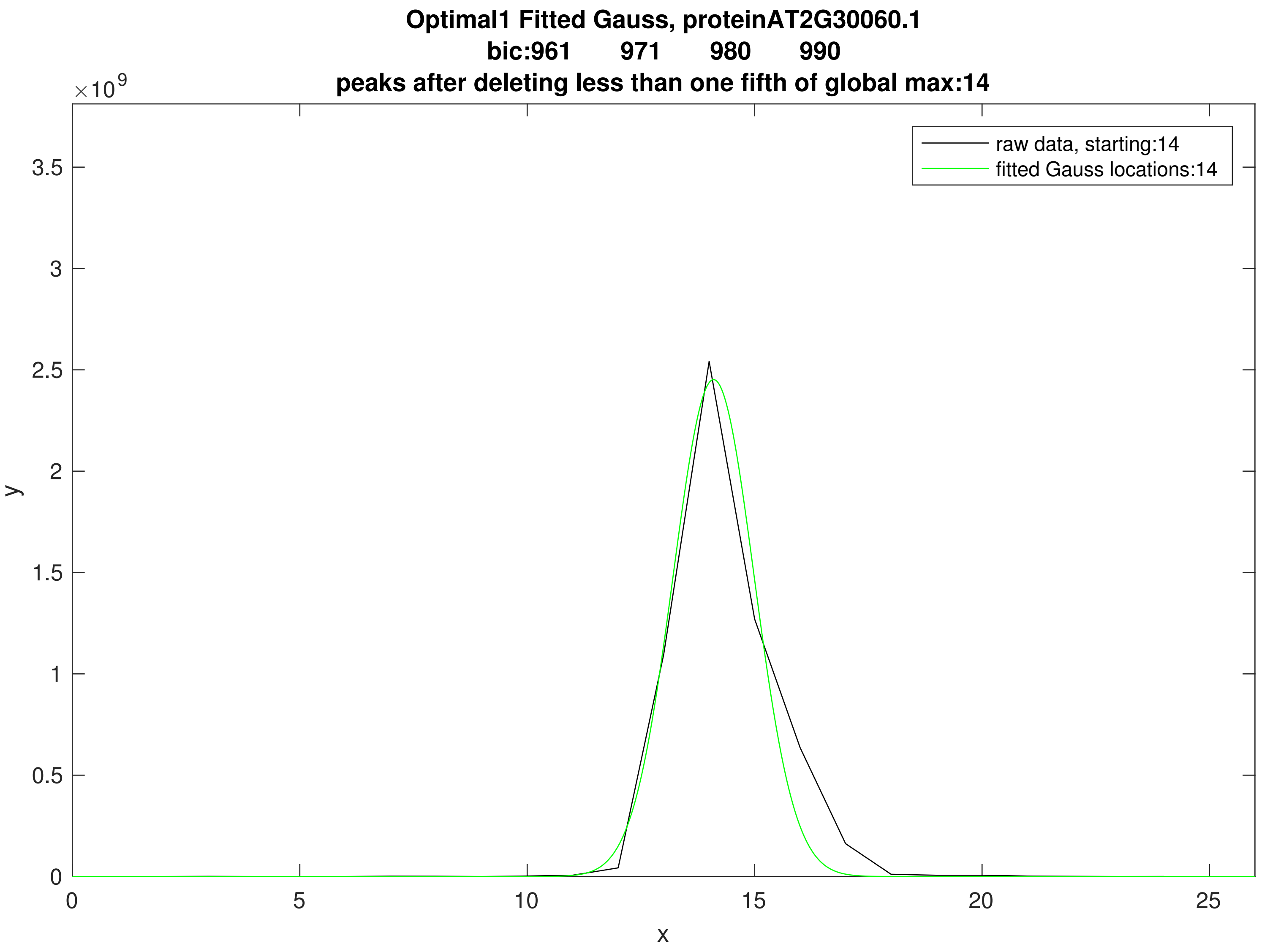
Optimal1 Fitted Gauss, proteinAT2G30060.1
bic:961 971 980 990
peaks after deleting less than one fifth of global max:14
×10^9
3.5
3
2.5
2
1.5
1
0.5
0
y
0
5
10
15
20
25
x
raw data, starting:14
fitted Gauss locations:14

**Optimal1 Fitted Gauss, proteinAT2G30110.1**
**bic:956 966 975**
**peaks after deleting less than one fifth of global max:12**

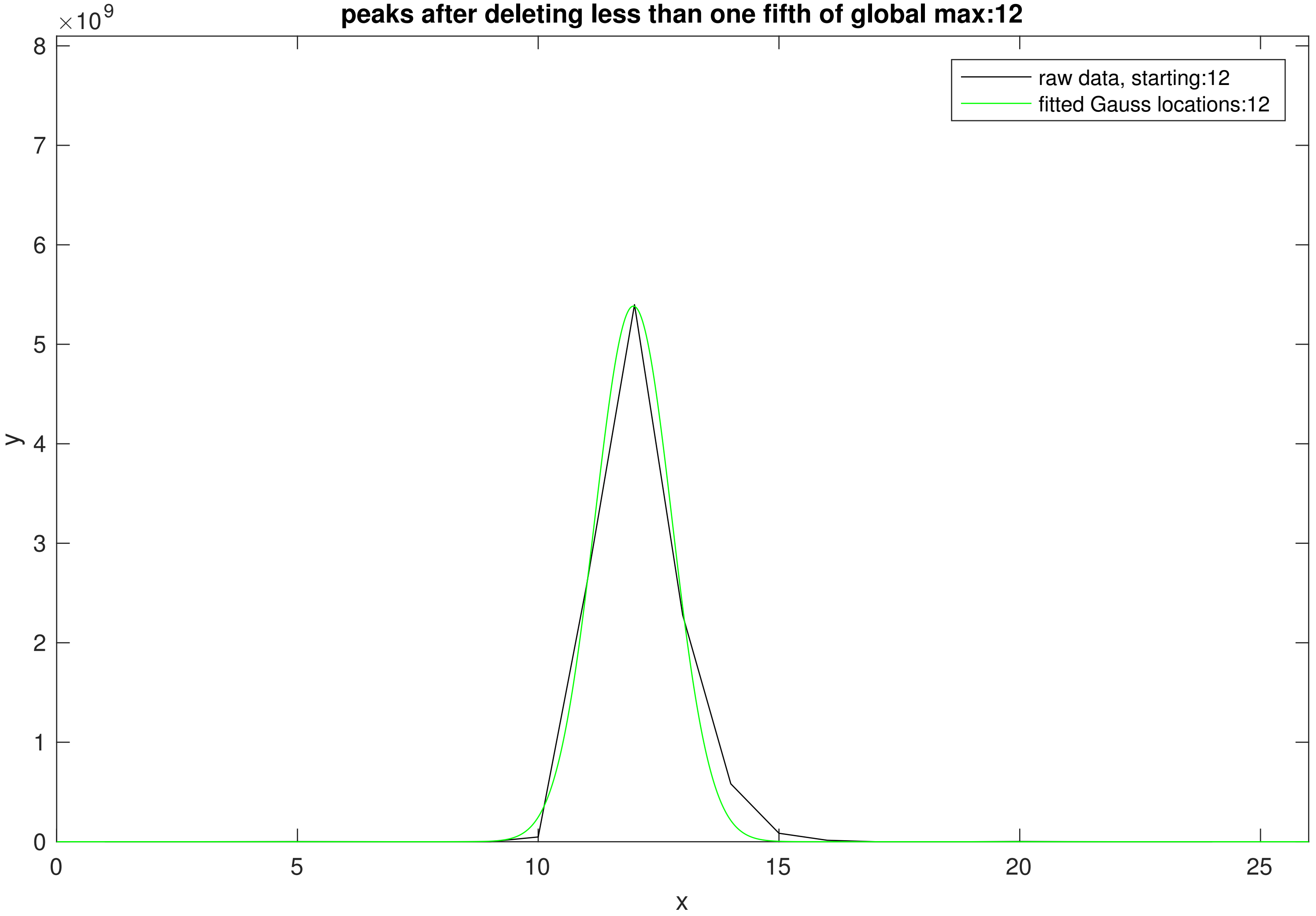

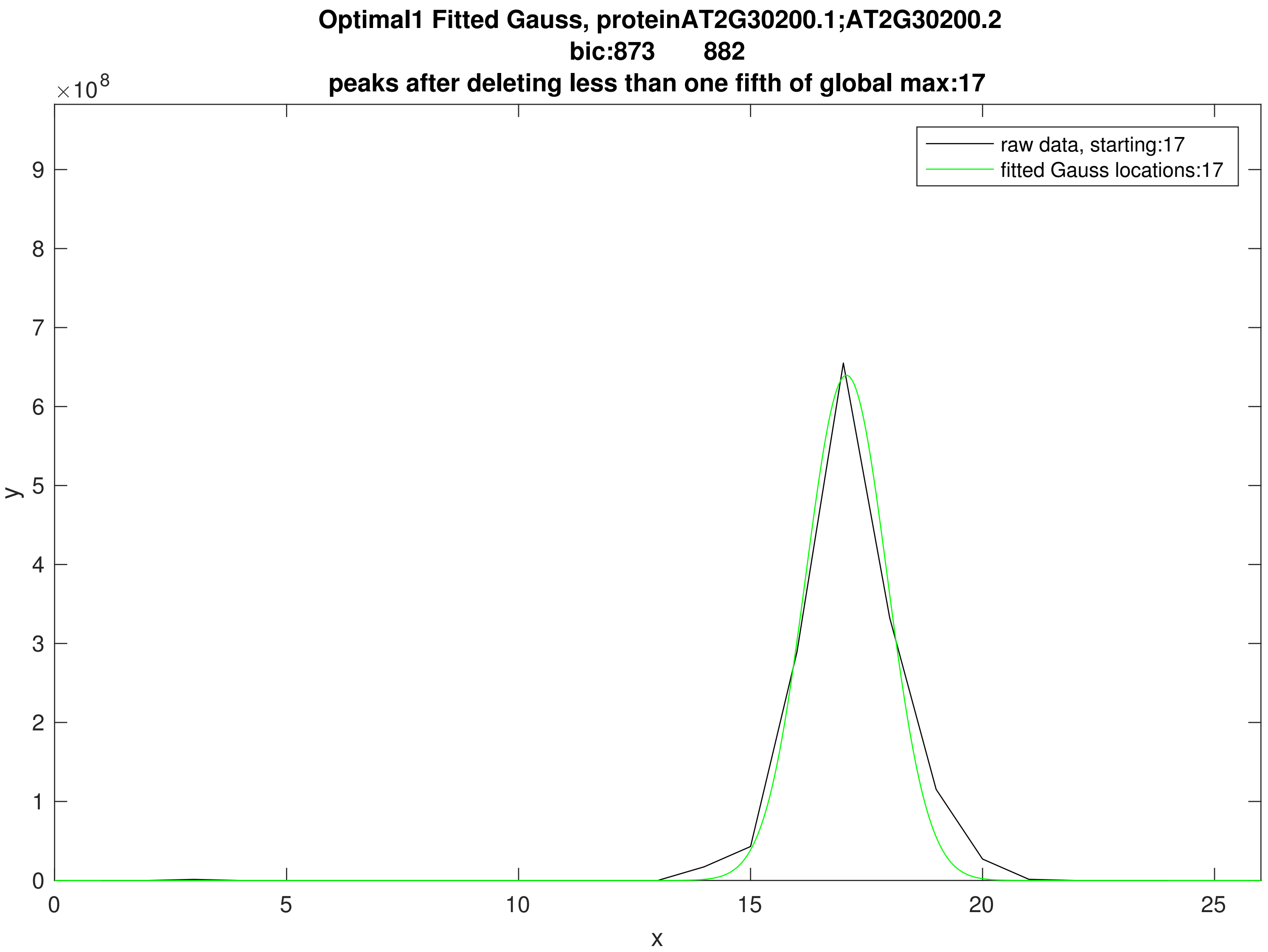
Optimal1 Fitted Gauss, proteinAT2G30200.1;AT2G30200.2
bic:873 882
peaks after deleting less than one fifth of global max:17
×10^8
raw data, starting:17
fitted Gauss locations:17
y
x
0
1
2
3
4
5
6
7
8
9
0
5
10
15
20
25

**Optimal2 Fitted Gauss, proteinAT2G30410.2;AT2G30410.1**
**bic:917 844**
**peaks after deleting less than one fifth of global max:14 19**

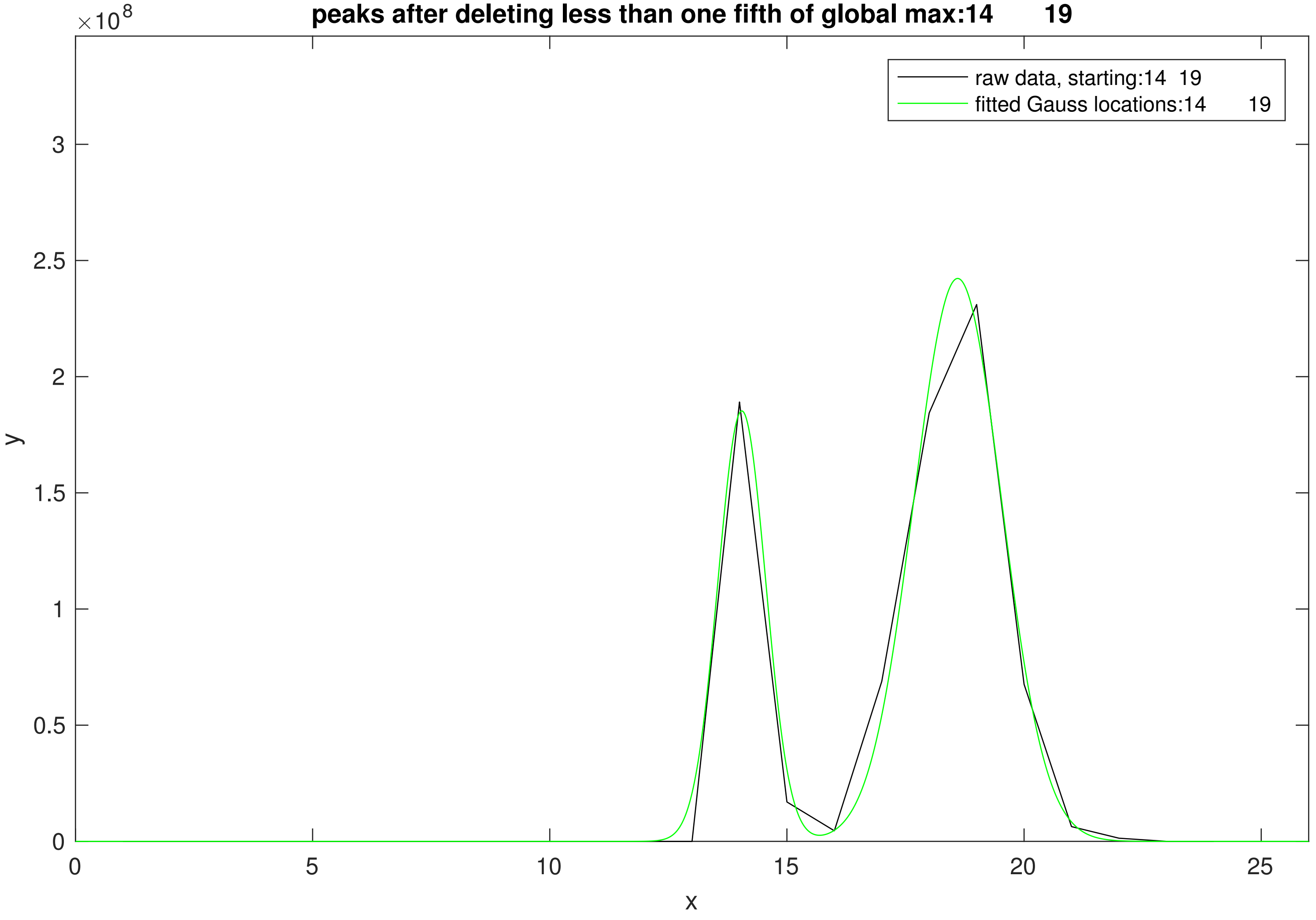

**Optimal1 Fitted Gauss, proteinAT2G30860.1;AT2G30860.2**
**bic:995 999 1.01e+03**
**peaks after deleting less than one fifth of global max:17**

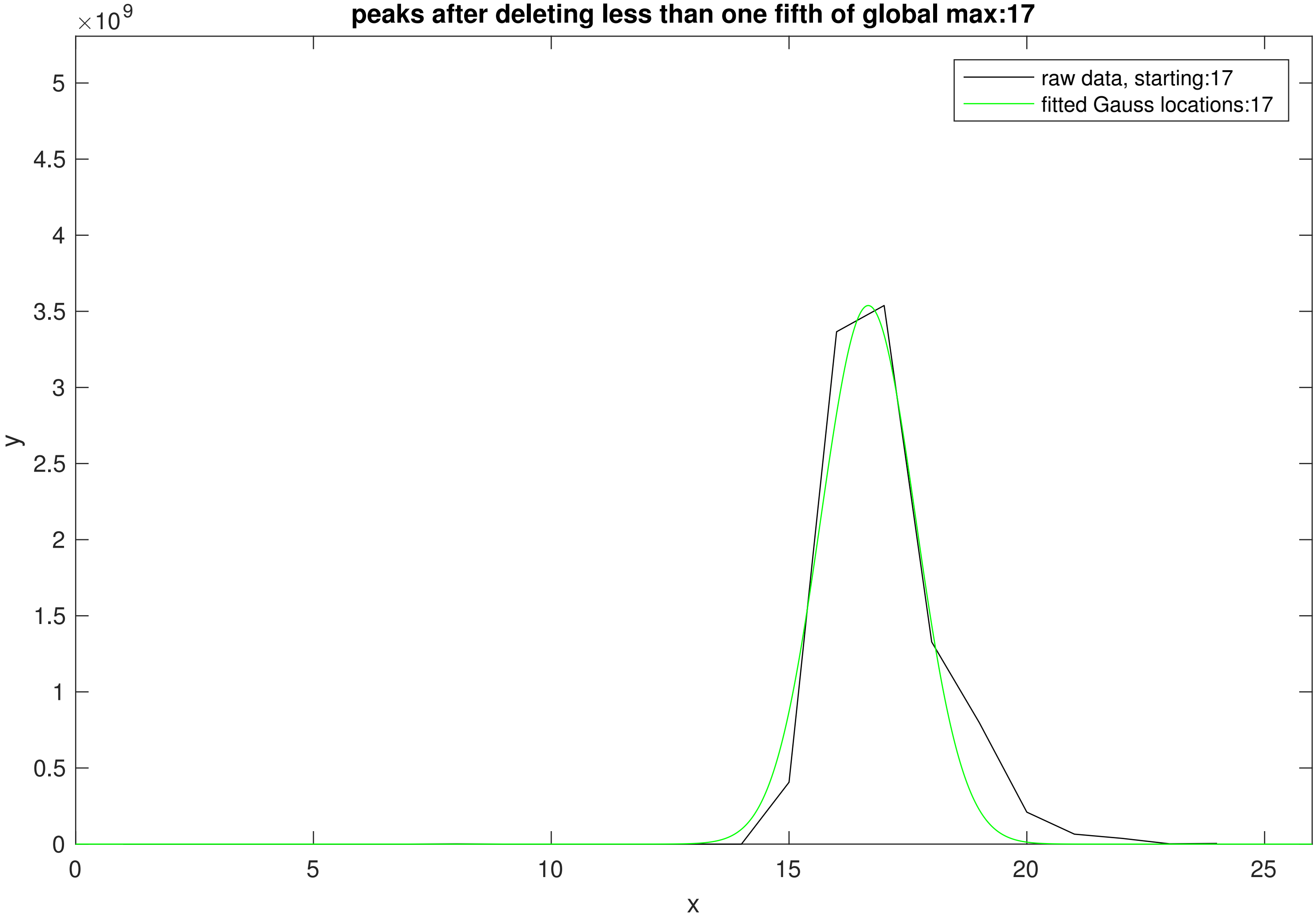

**Optimal1 Fitted Gauss, proteinAT2G30870.1**
**bic:1.01e+03  1.02e+03  1.03e+03  1.04e+03**
**peaks after deleting less than one fifth of global max:16**

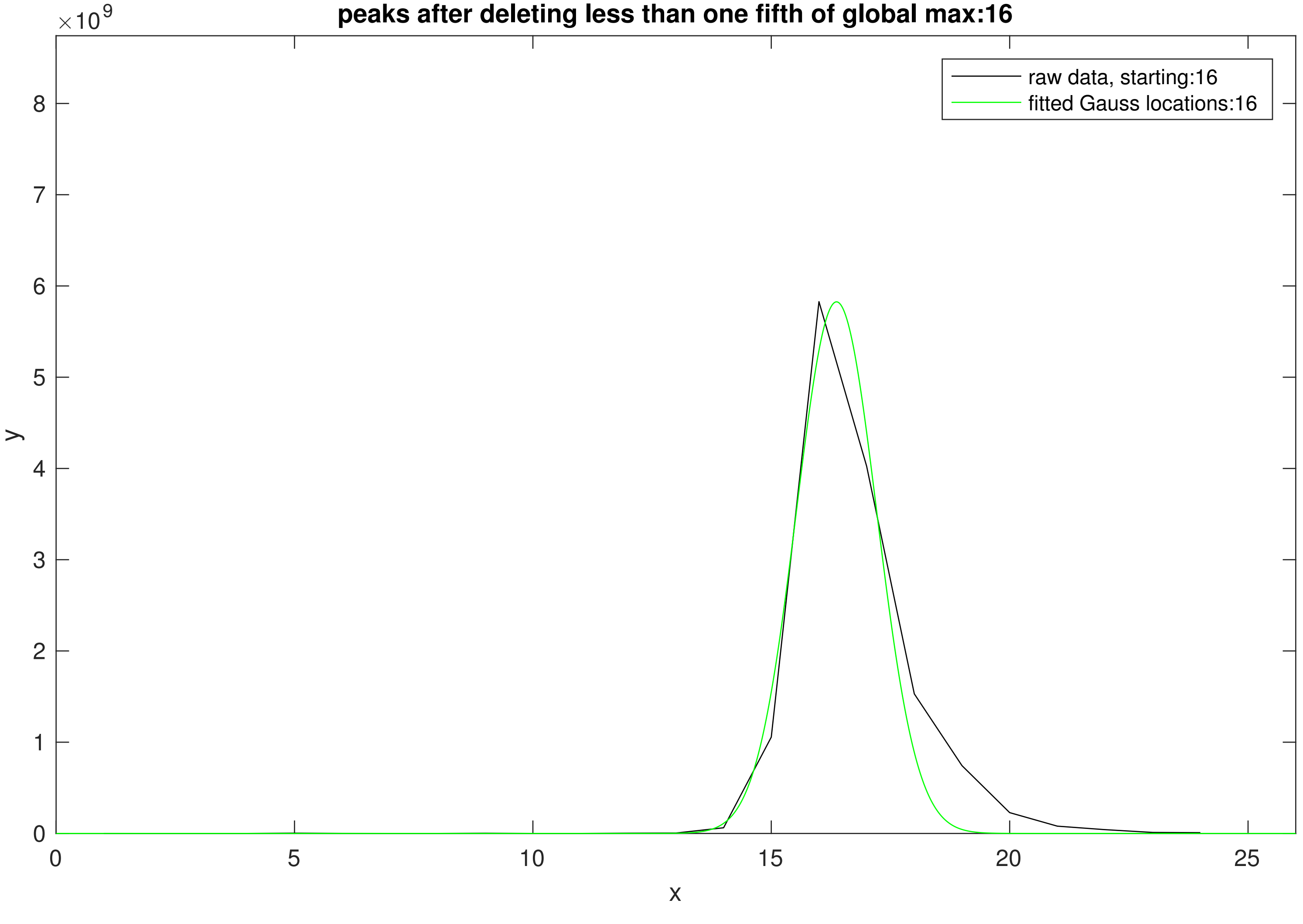

**Optimal1 Fitted Gauss, proteinAT2G30930.1**
**bic:841 850**
**peaks after deleting less than one fifth of global max:16**

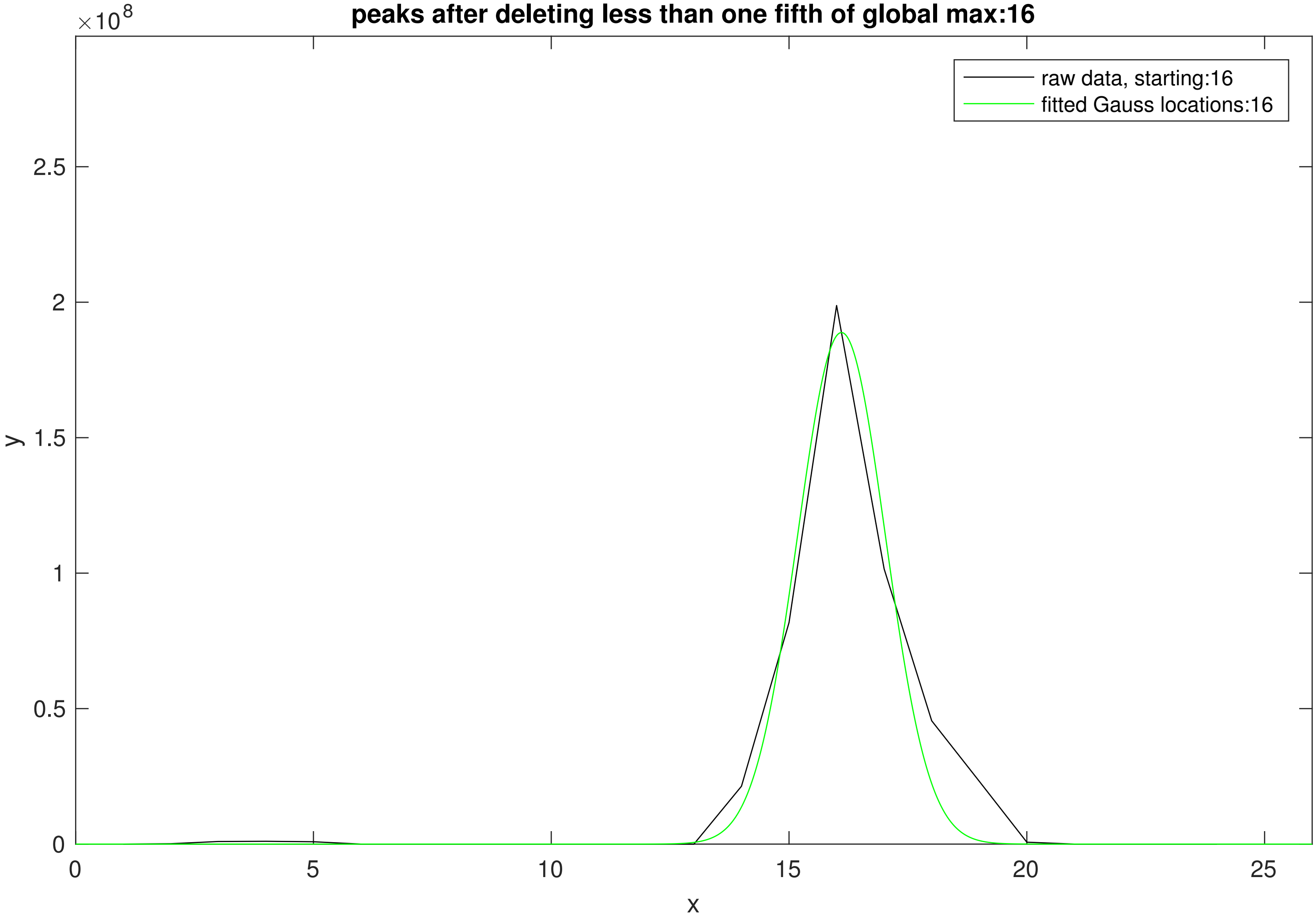

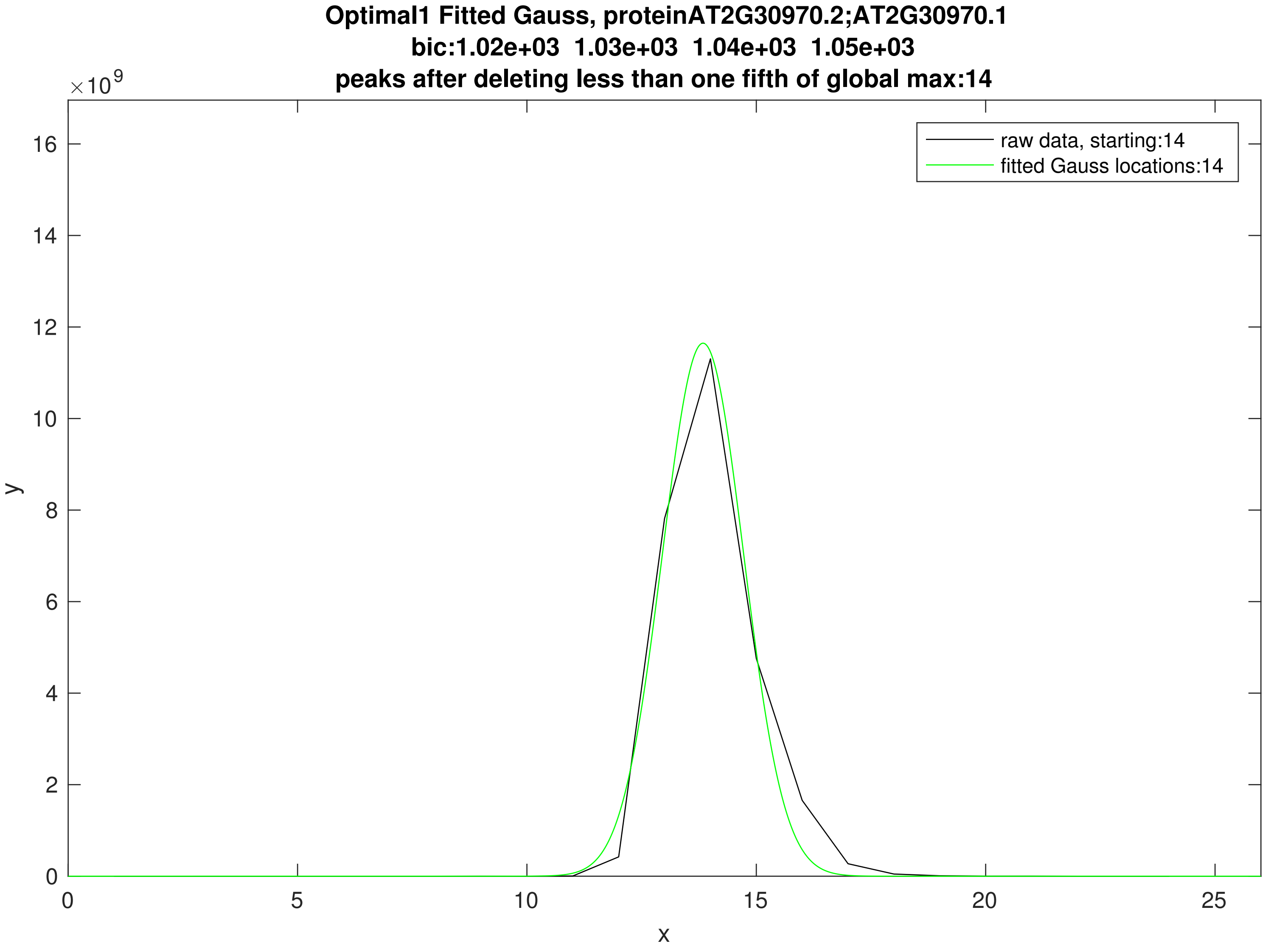
Optimal1 Fitted Gauss, proteinAT2G30970.2;AT2G30970.1
bic:1.02e+03  1.03e+03  1.04e+03  1.05e+03
peaks after deleting less than one fifth of global max:14
×10^9
16
14
12
10
8
6
4
2
0
y
0
5
10
15
20
25
x
raw data, starting:14
fitted Gauss locations:14

#### Optimal1 Fitted Gauss, proteinAT2G31390.1
#### bic:1e+03  1.01e+03  1.02e+03
#### peaks after deleting less than one fifth of global max:14

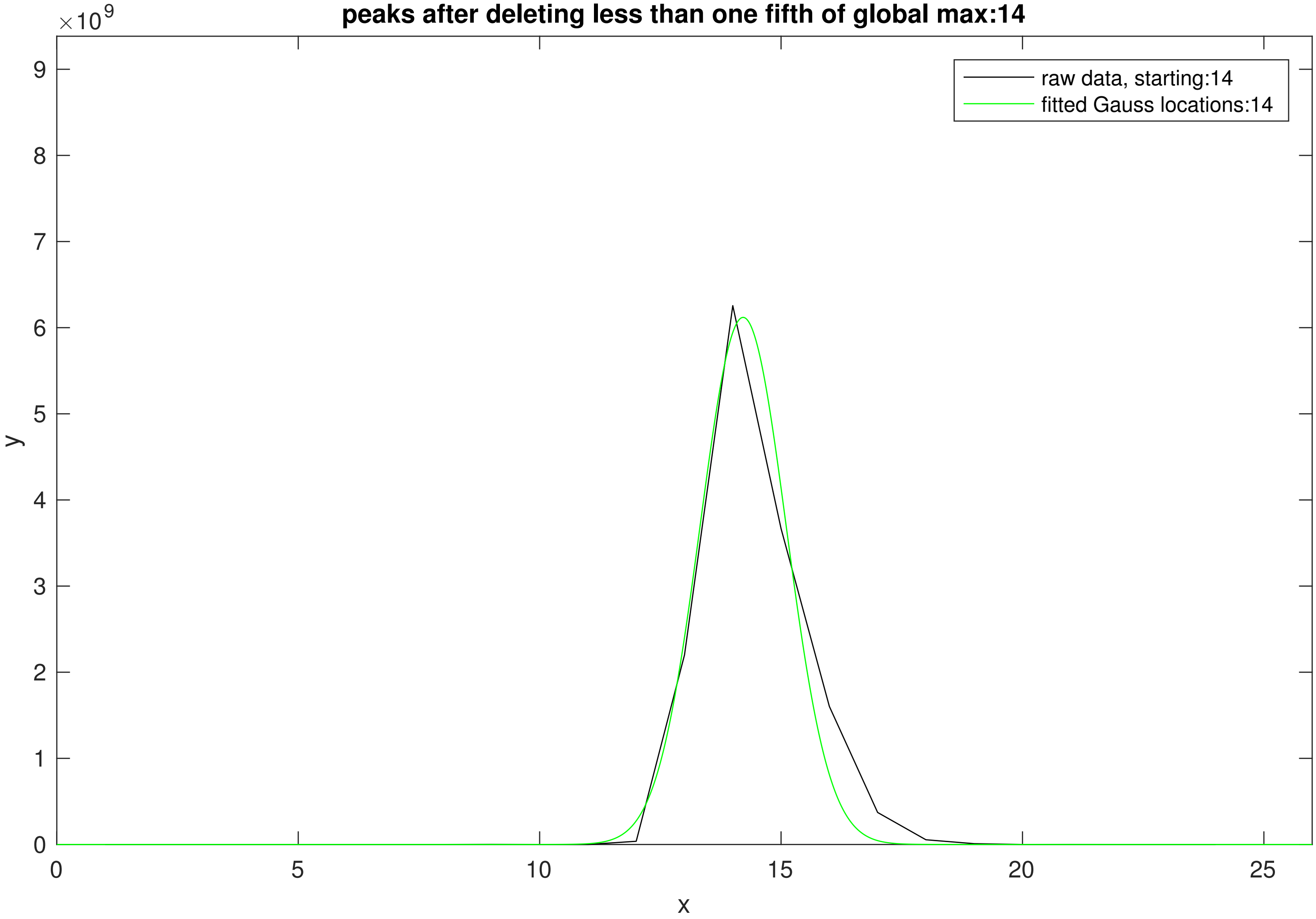

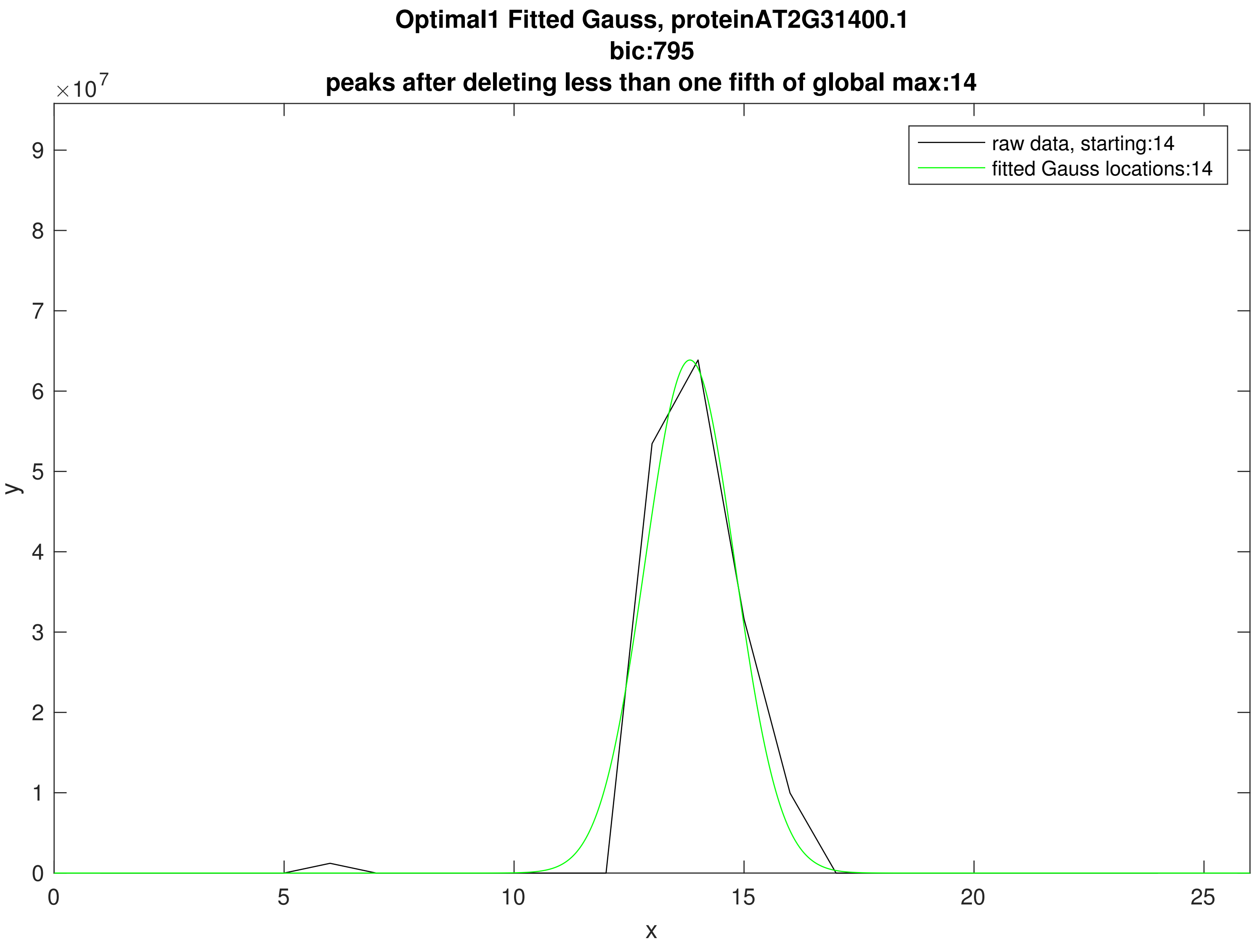
Optimal1 Fitted Gauss, proteinAT2G31400.1
bic:795
peaks after deleting less than one fifth of global max:14
×10^7
raw data, starting:14
fitted Gauss locations:14
y
x
0
1
2
3
4
5
6
7
8
9
0
5
10
15
20
25

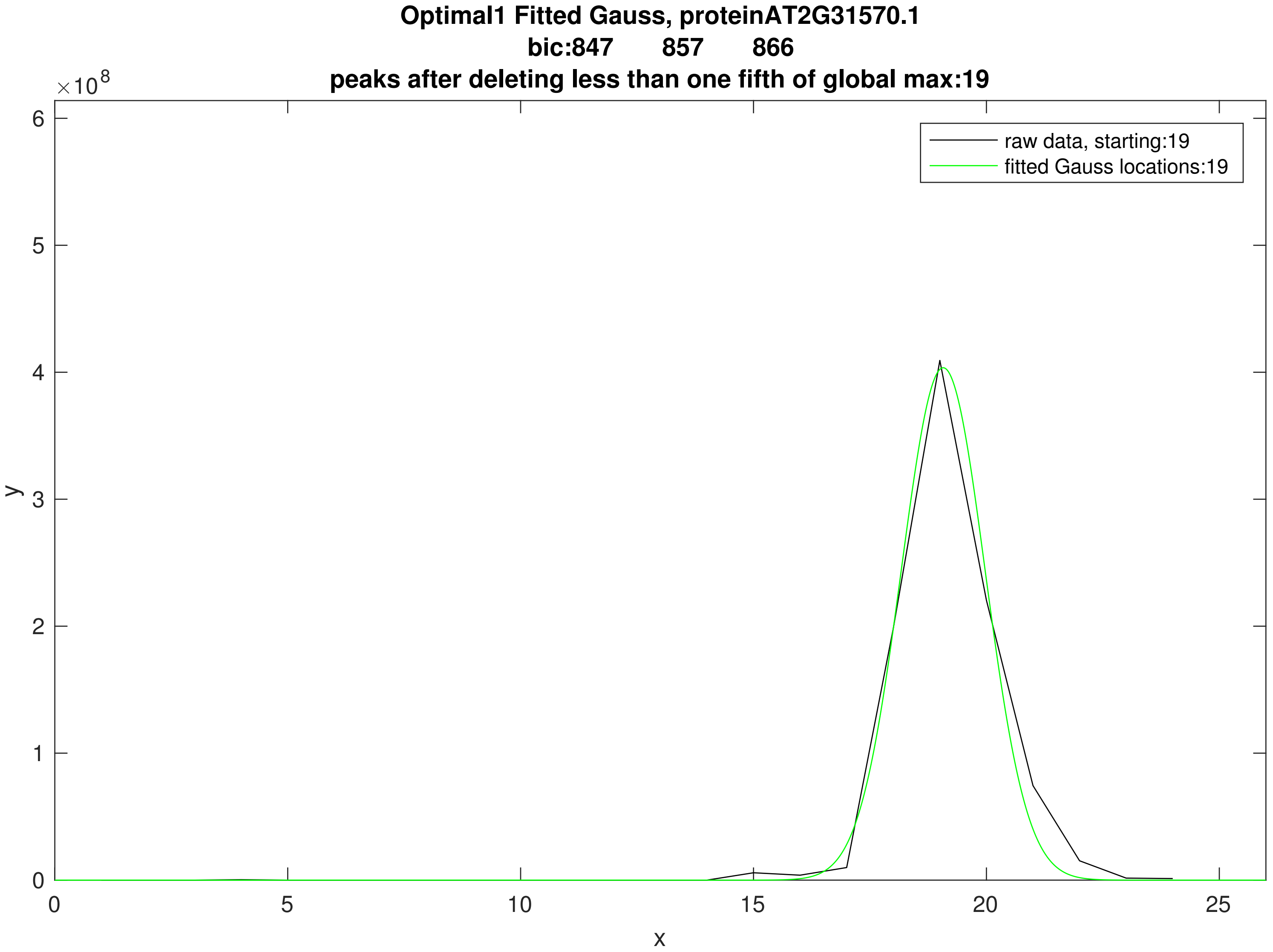
Optimal1 Fitted Gauss, proteinAT2G31570.1
bic:847 857 866
peaks after deleting less than one fifth of global max:19
×10^8
raw data, starting:19
fitted Gauss locations:19
y
x
0
1
2
3
4
5
6
0
5
10
15
20
25

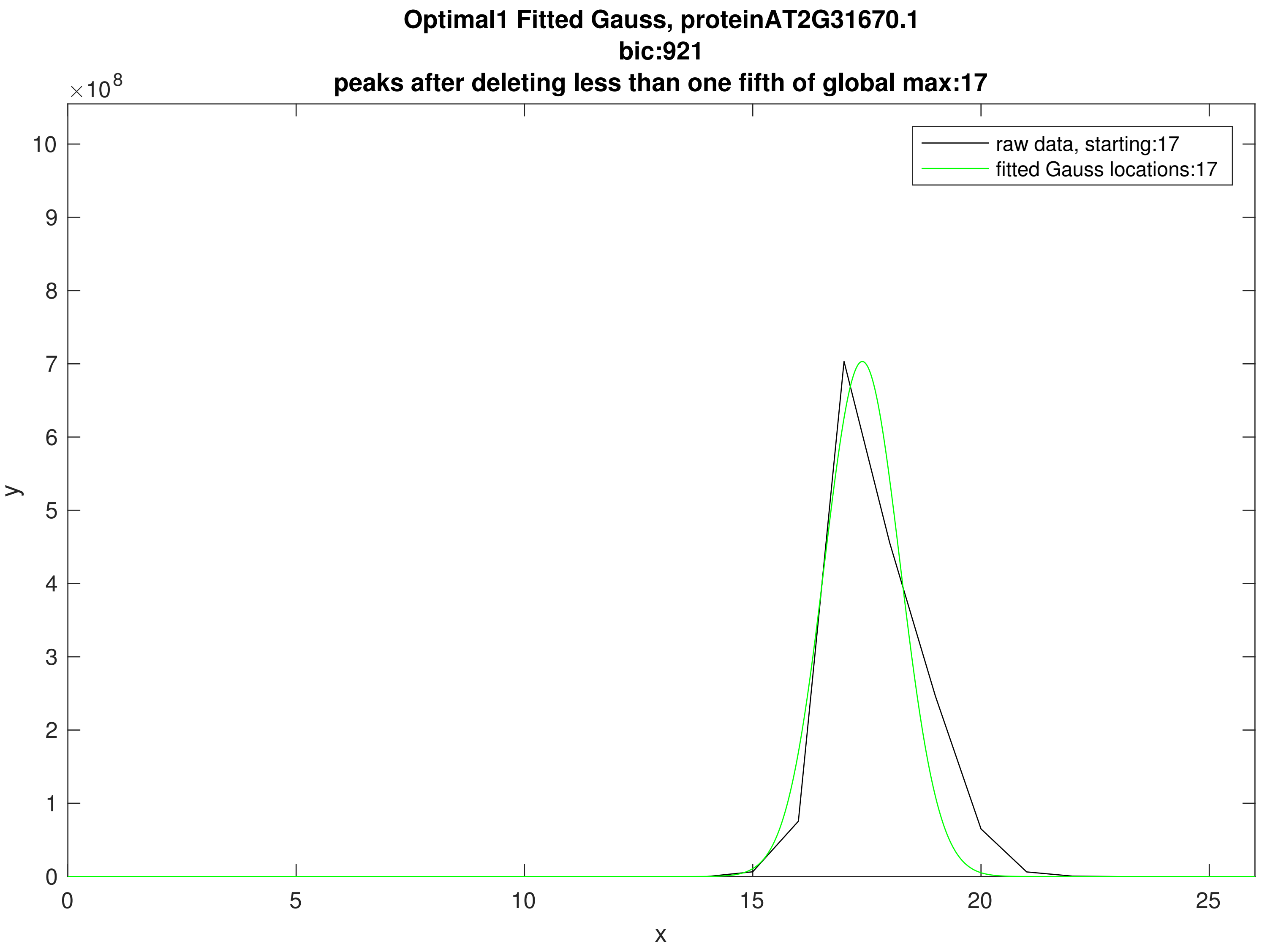

Optimal1 Fitted Gauss, proteinAT2G31670.1
bic:921
peaks after deleting less than one fifth of global max:17
×10^8
raw data, starting:17
fitted Gauss locations:17
y
x
0
1
2
3
4
5
6
7
8
9
10
0
5
10
15
20
25

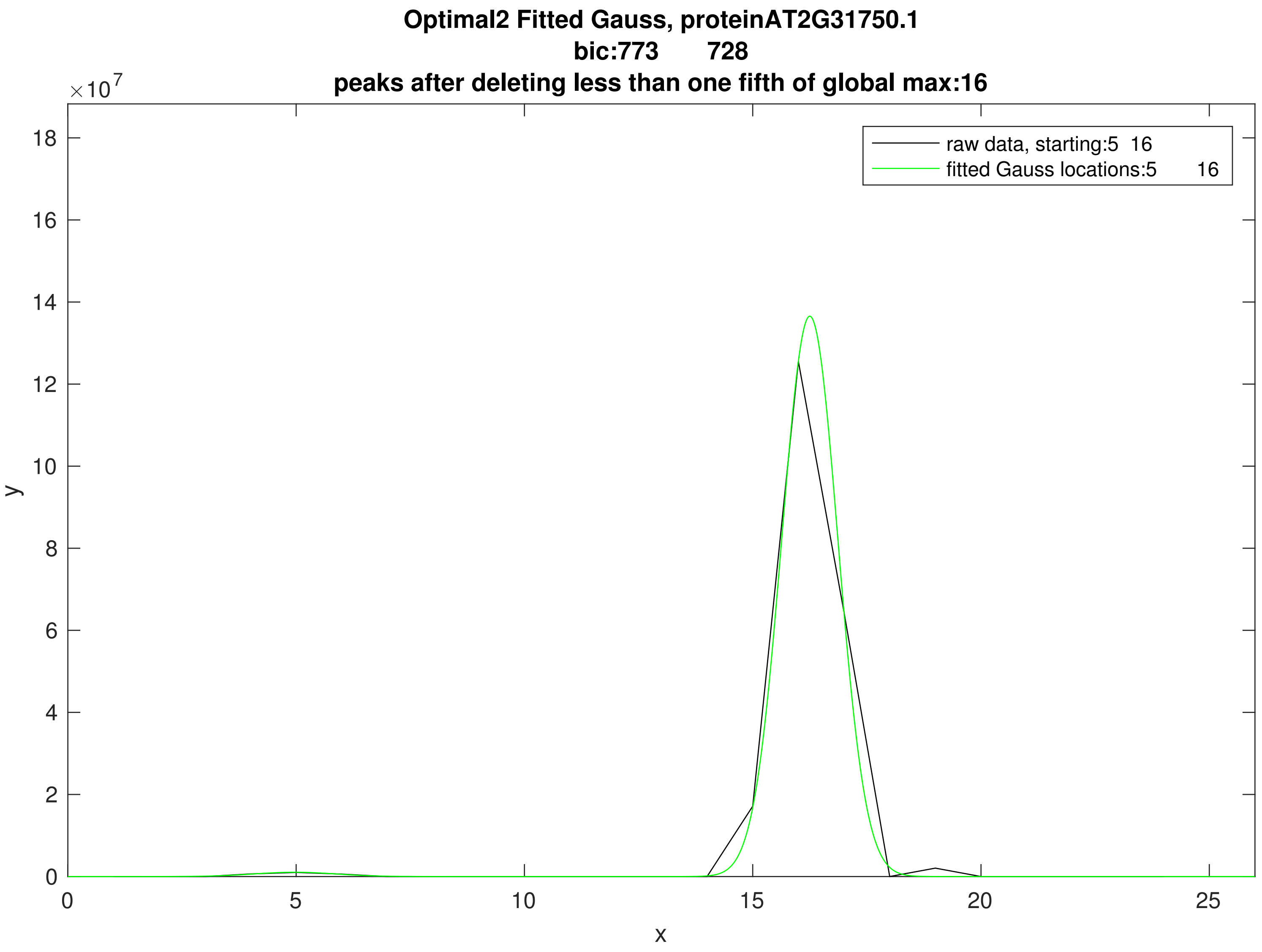

Optimal2 Fitted Gauss, proteinAT2G31750.1
bic:773 728
peaks after deleting less than one fifth of global max:16
×10^7
18
16
14
12
10
8
6
4
2
0
y
0
5
10
15
20
25
x
raw data, starting:5 16
fitted Gauss locations:5 16

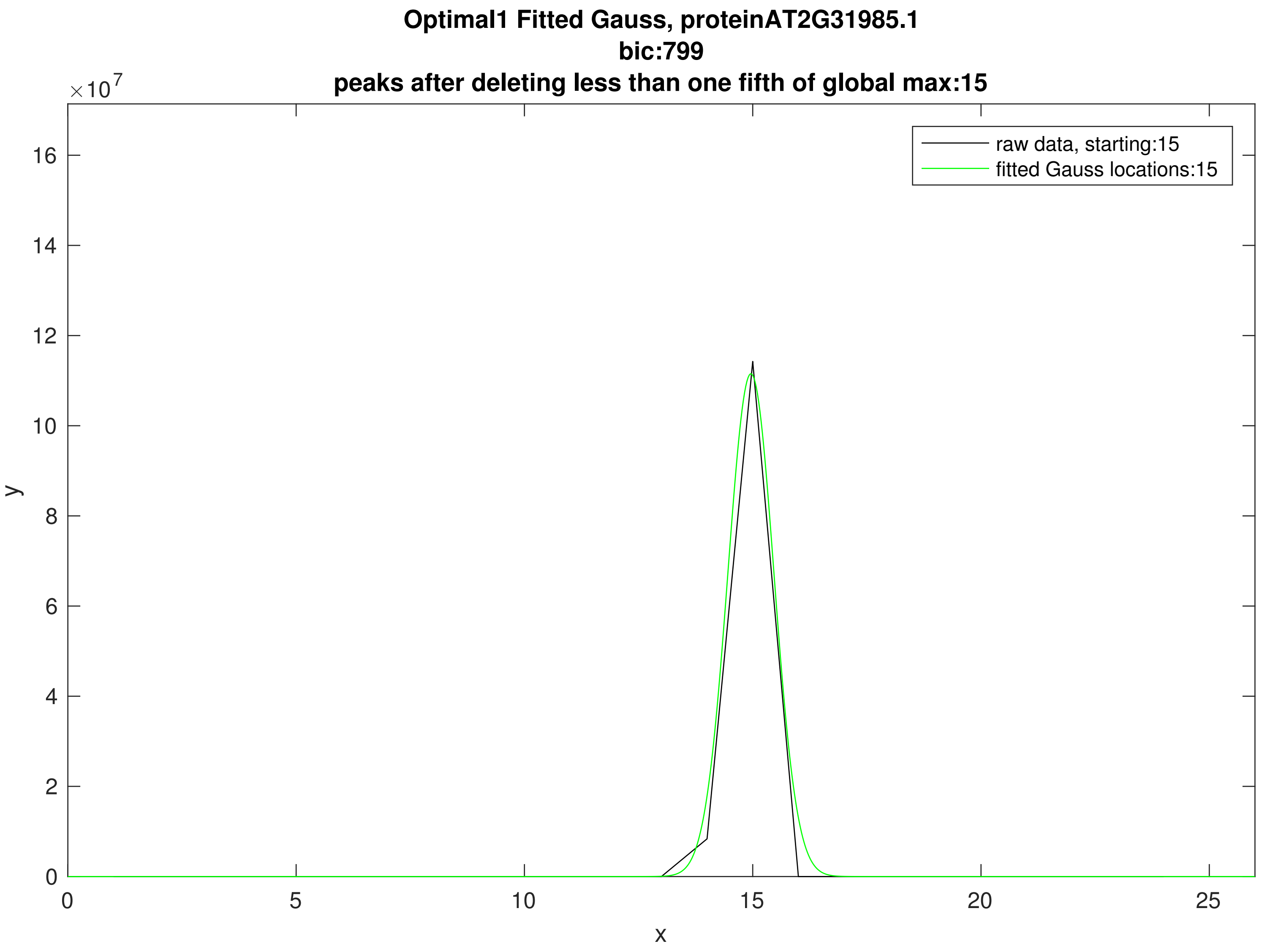
Optimal1 Fitted Gauss, proteinAT2G31985.1
bic:799
peaks after deleting less than one fifth of global max:15
×10^7
16
14
12
10
8
6
4
2
0
y
0
5
10
15
20
25
x
raw data, starting:15
fitted Gauss locations:15

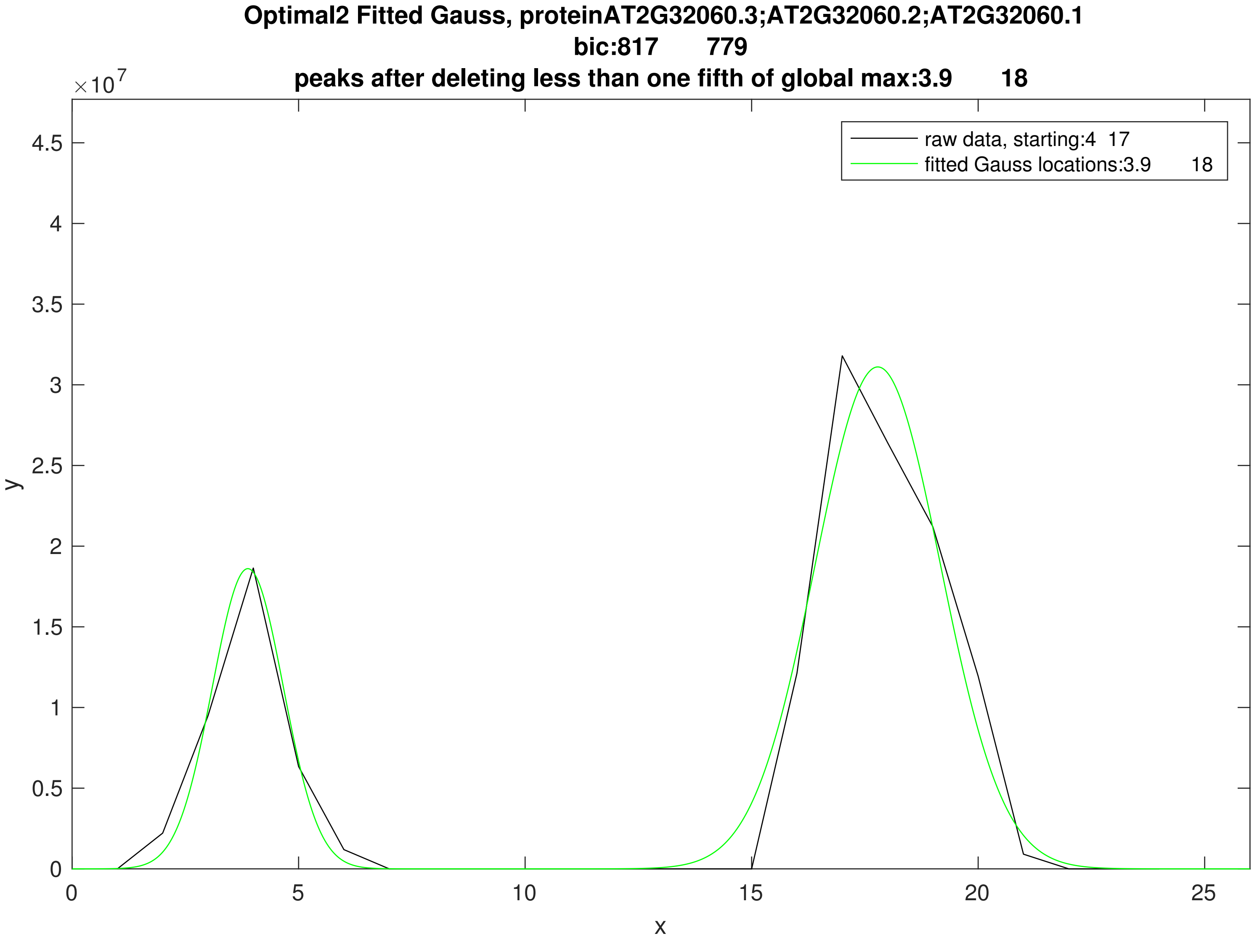

Optimal2 Fitted Gauss, proteinAT2G32060.3;AT2G32060.2;AT2G32060.1
bic:817 779
peaks after deleting less than one fifth of global max:3.9 18
×10^7
raw data, starting:4 17
fitted Gauss locations:3.9 18
y
x
0
0.5
1
1.5
2
2.5
3
3.5
4
4.5
5
10
15
20
25

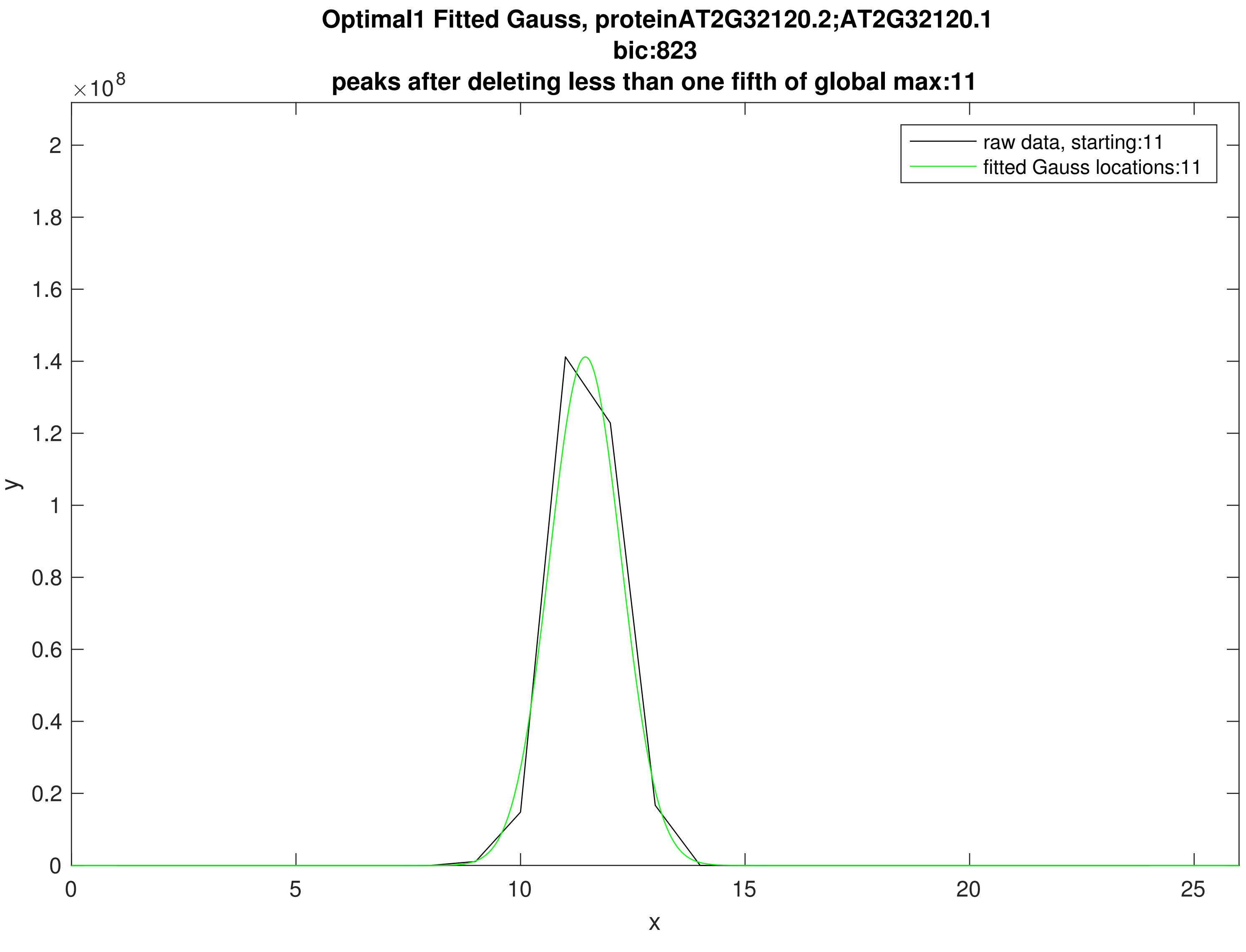
Optimal1 Fitted Gauss, proteinAT2G32120.2;AT2G32120.1
bic:823
peaks after deleting less than one fifth of global max:11
×10^8
raw data, starting:11
fitted Gauss locations:11
y
x
0
0.2
0.4
0.6
0.8
1
1.2
1.4
1.6
1.8
2
5
10
15
20
25

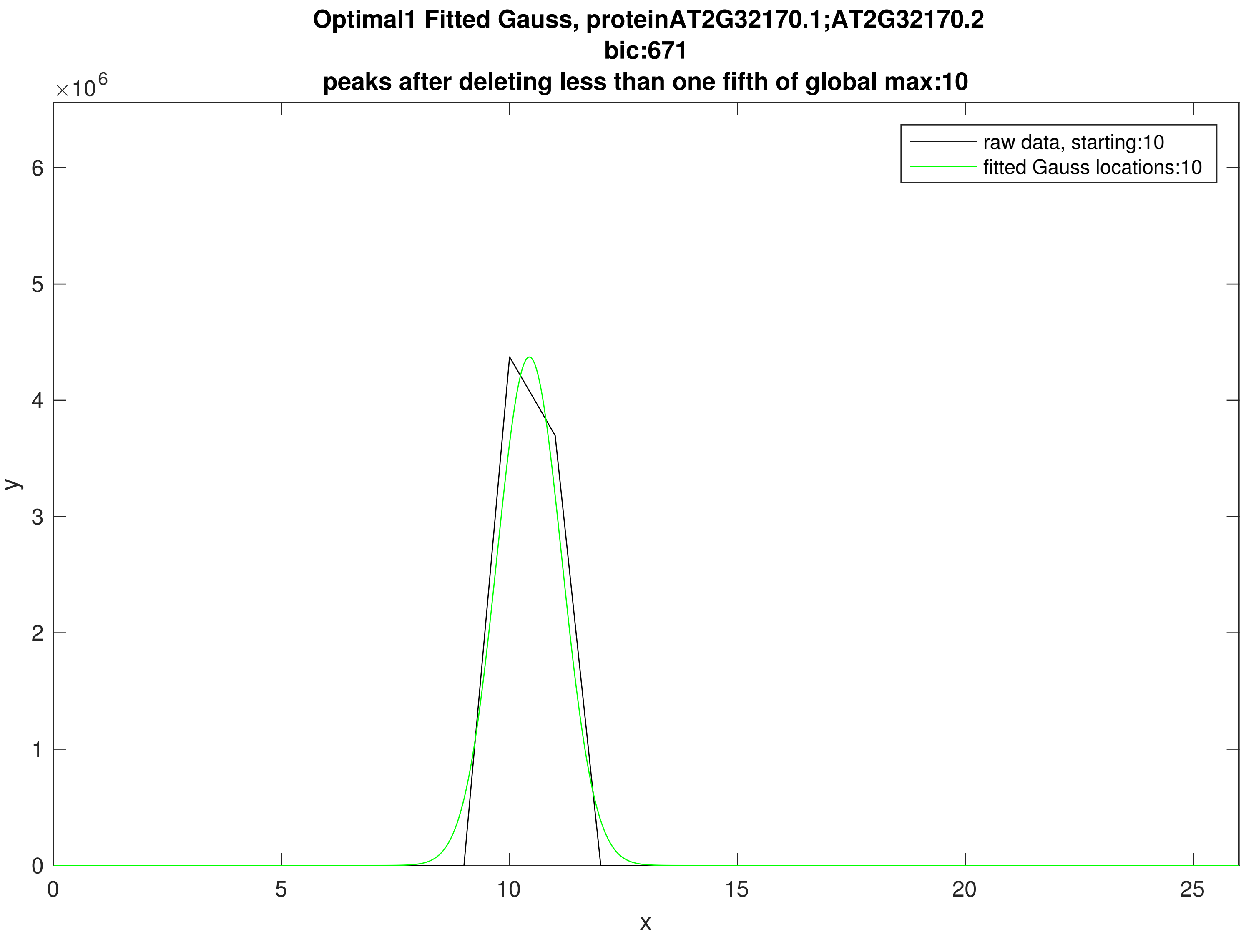
Optimal1 Fitted Gauss, proteinAT2G32170.1;AT2G32170.2
bic:671
peaks after deleting less than one fifth of global max:10
×10^6
raw data, starting:10
fitted Gauss locations:10
y
x
0
1
2
3
4
5
6
0
5
10
15
20
25

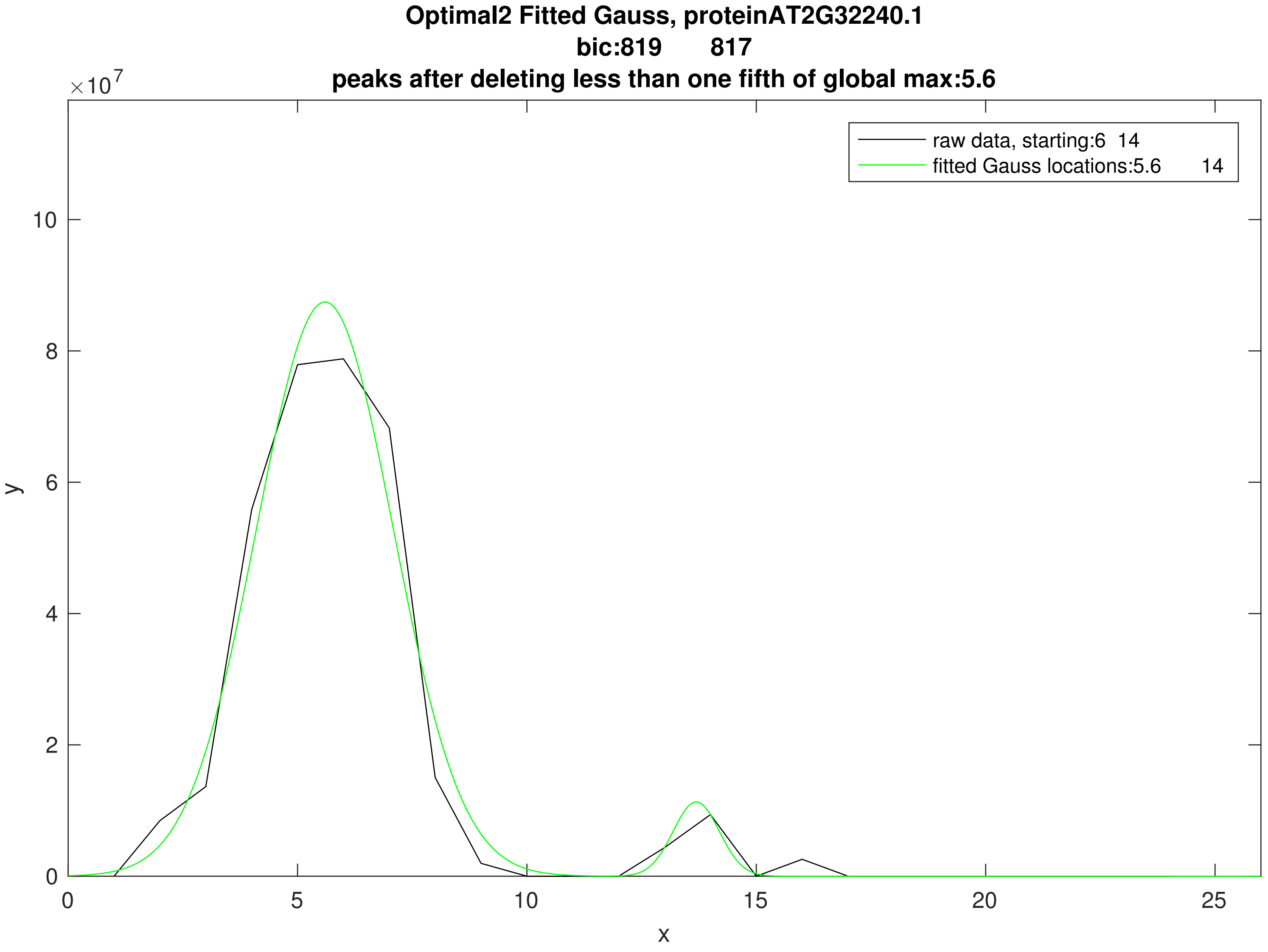
Optimal2 Fitted Gauss, proteinAT2G32240.1
bic:819 817
peaks after deleting less than one fifth of global max:5.6
×10^7
raw data, starting:6 14
fitted Gauss locations:5.6 14
y
x
0
2
4
6
8
10
5
15
20
25

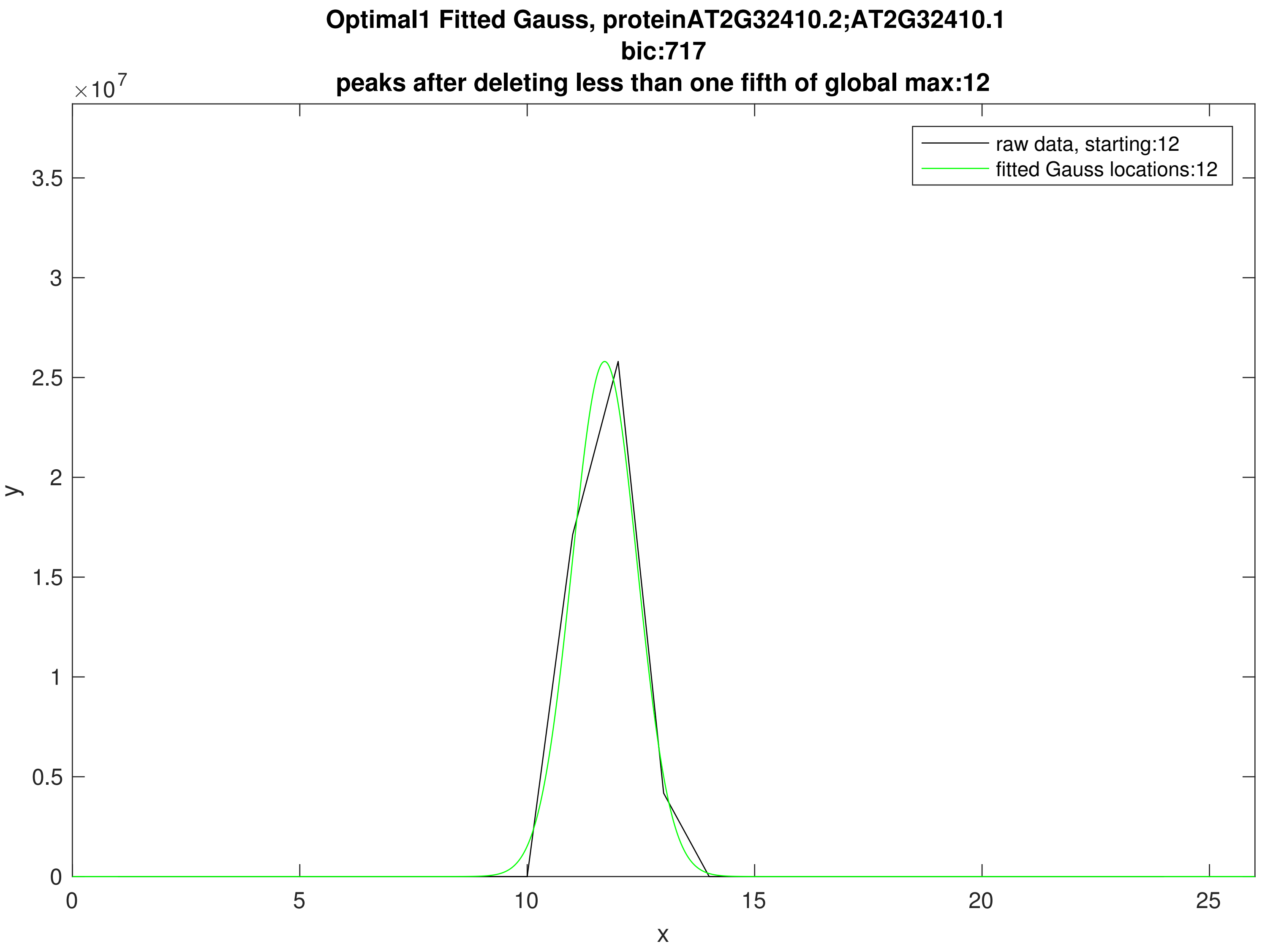
Optimal1 Fitted Gauss, proteinAT2G32410.2;AT2G32410.1
bic:717
peaks after deleting less than one fifth of global max:12
×10^7
3.5
3
2.5
2
1.5
1
0.5
0
y
0
5
10
15
20
25
x
raw data, starting:12
fitted Gauss locations:12

#### Optimal1 Fitted Gauss, proteinAT2G32520.1
#### bic:831
#### peaks after deleting less than one fifth of global max:18

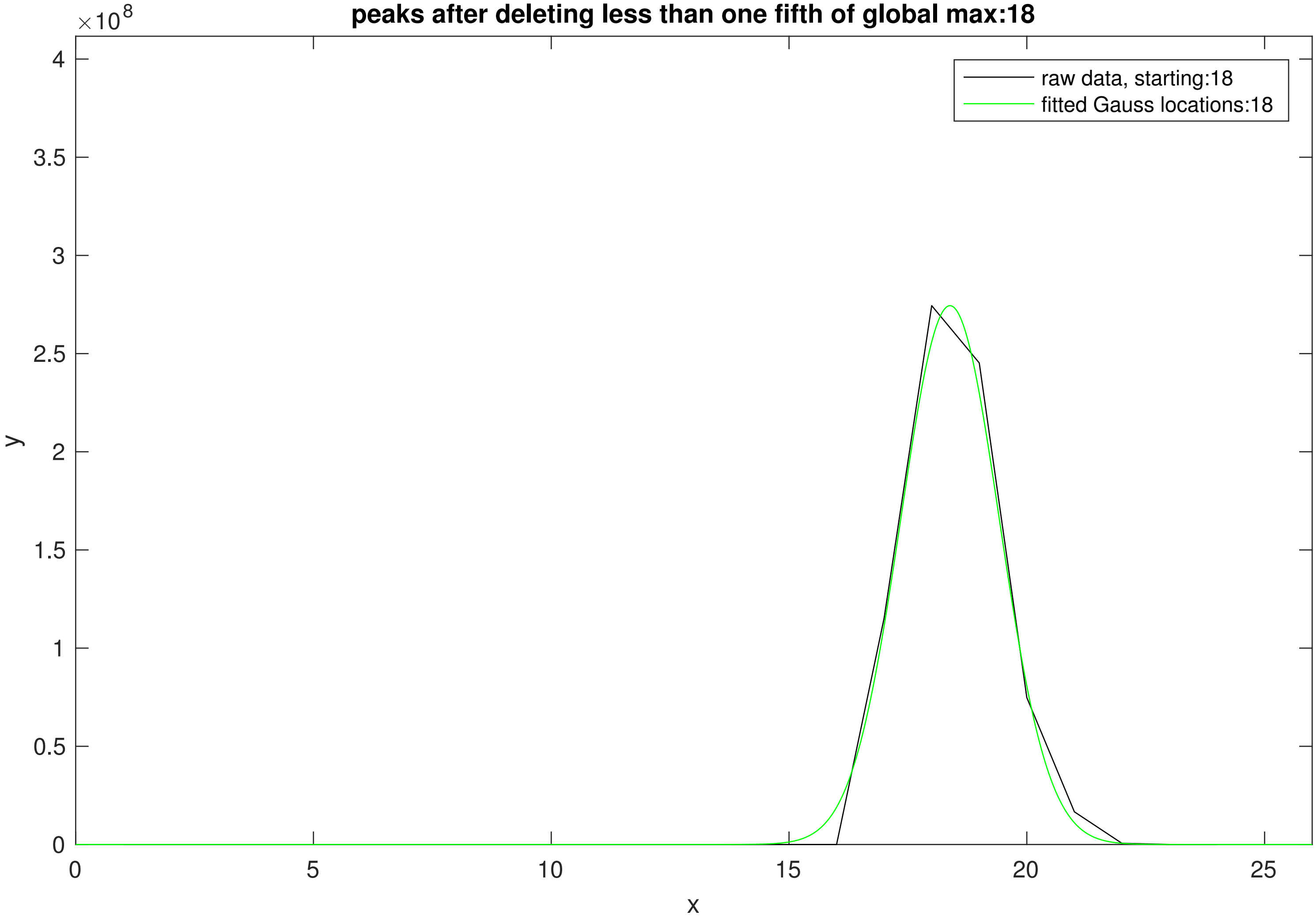

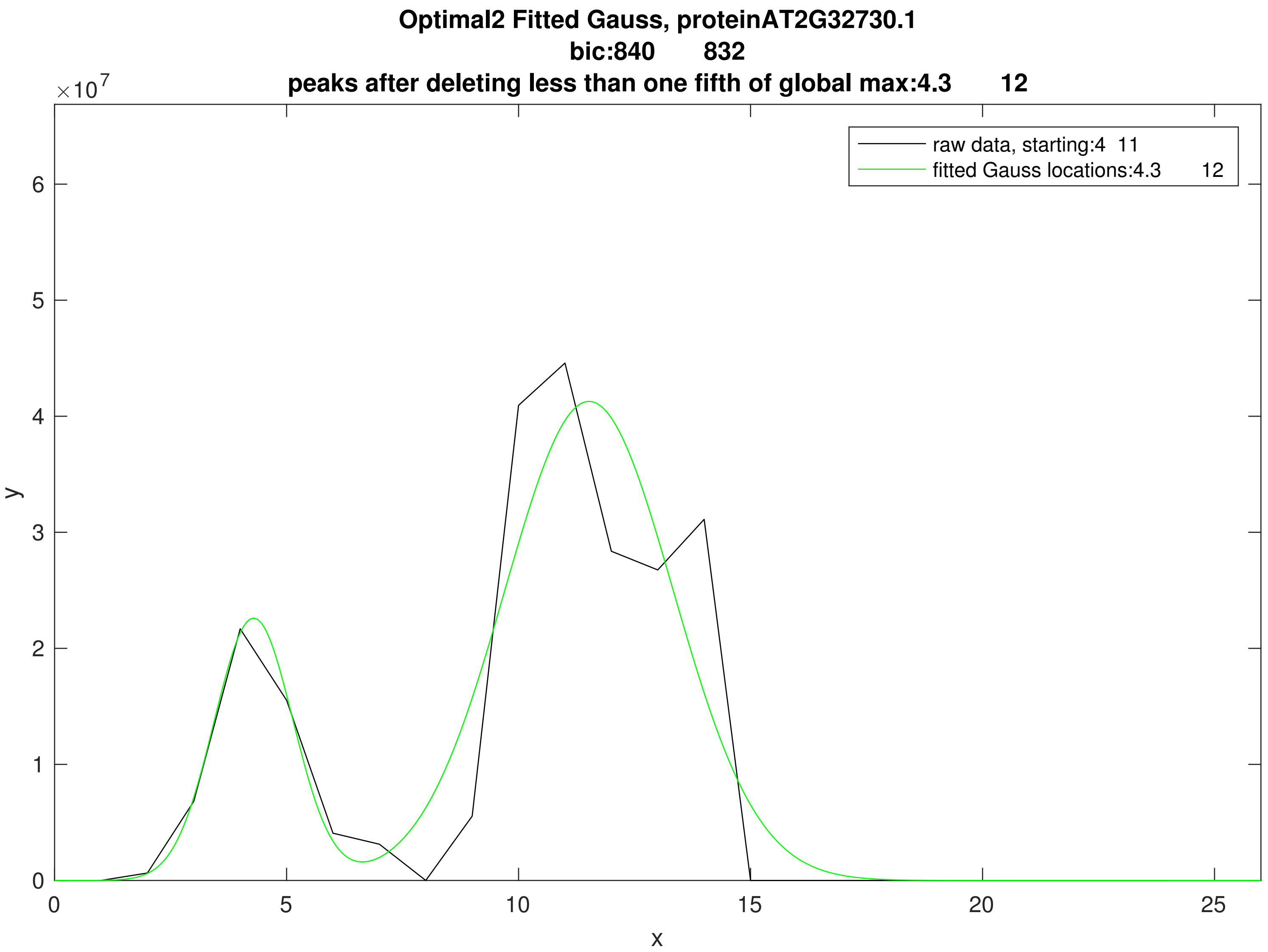
Optimal2 Fitted Gauss, proteinAT2G32730.1
bic:840 832
peaks after deleting less than one fifth of global max:4.3 12
×10^7
raw data, starting:4 11
fitted Gauss locations:4.3 12
y
x
0
1
2
3
4
5
6
0
5
10
15
20
25

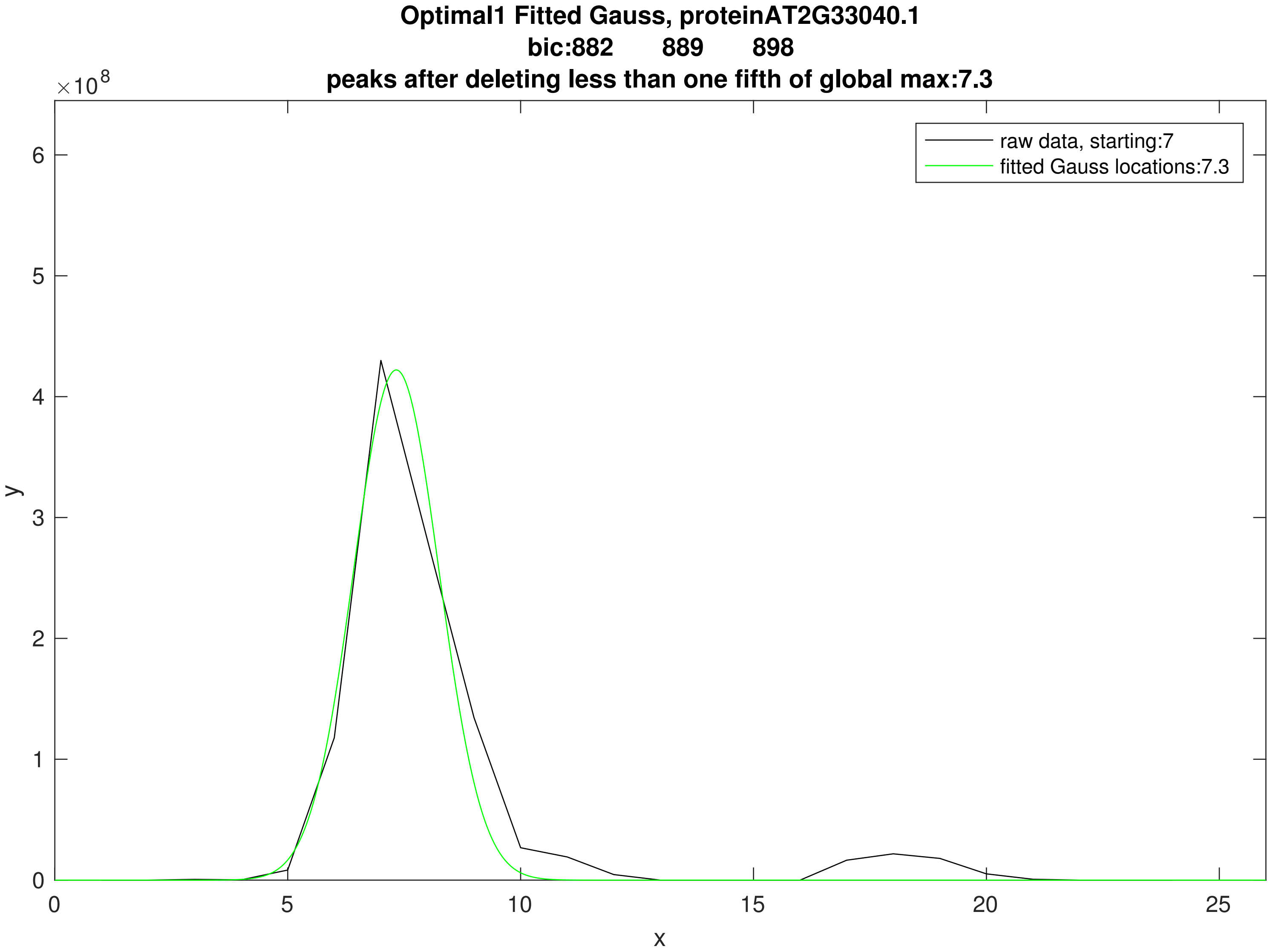

Optimal1 Fitted Gauss, proteinAT2G33040.1
bic:882 889 898
peaks after deleting less than one fifth of global max:7.3
×10^8
raw data, starting:7
fitted Gauss locations:7.3
y
x
0
1
2
3
4
5
6
0
5
10
15
20
25

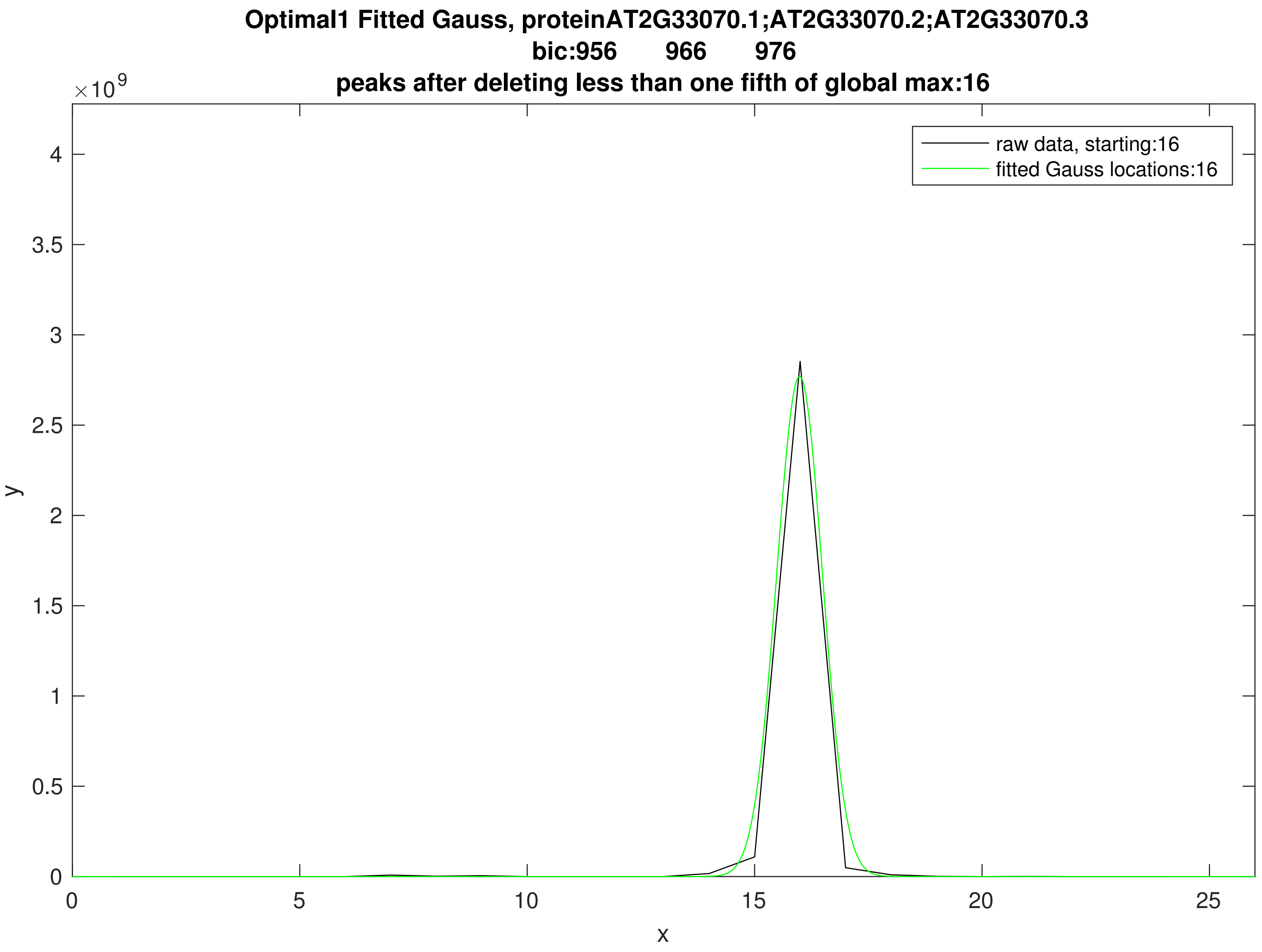
Optimal1 Fitted Gauss, proteinAT2G33070.1;AT2G33070.2;AT2G33070.3
bic:956 966 976
peaks after deleting less than one fifth of global max:16
×10^9
raw data, starting:16
fitted Gauss locations:16
y
x
0
0.5
1
1.5
2
2.5
3
3.5
4
0
5
10
15
20
25

**Optimal1 Fitted Gauss, proteinAT2G33150.1**
**bic:966 976 985**
**peaks after deleting less than one fifth of global max:14**

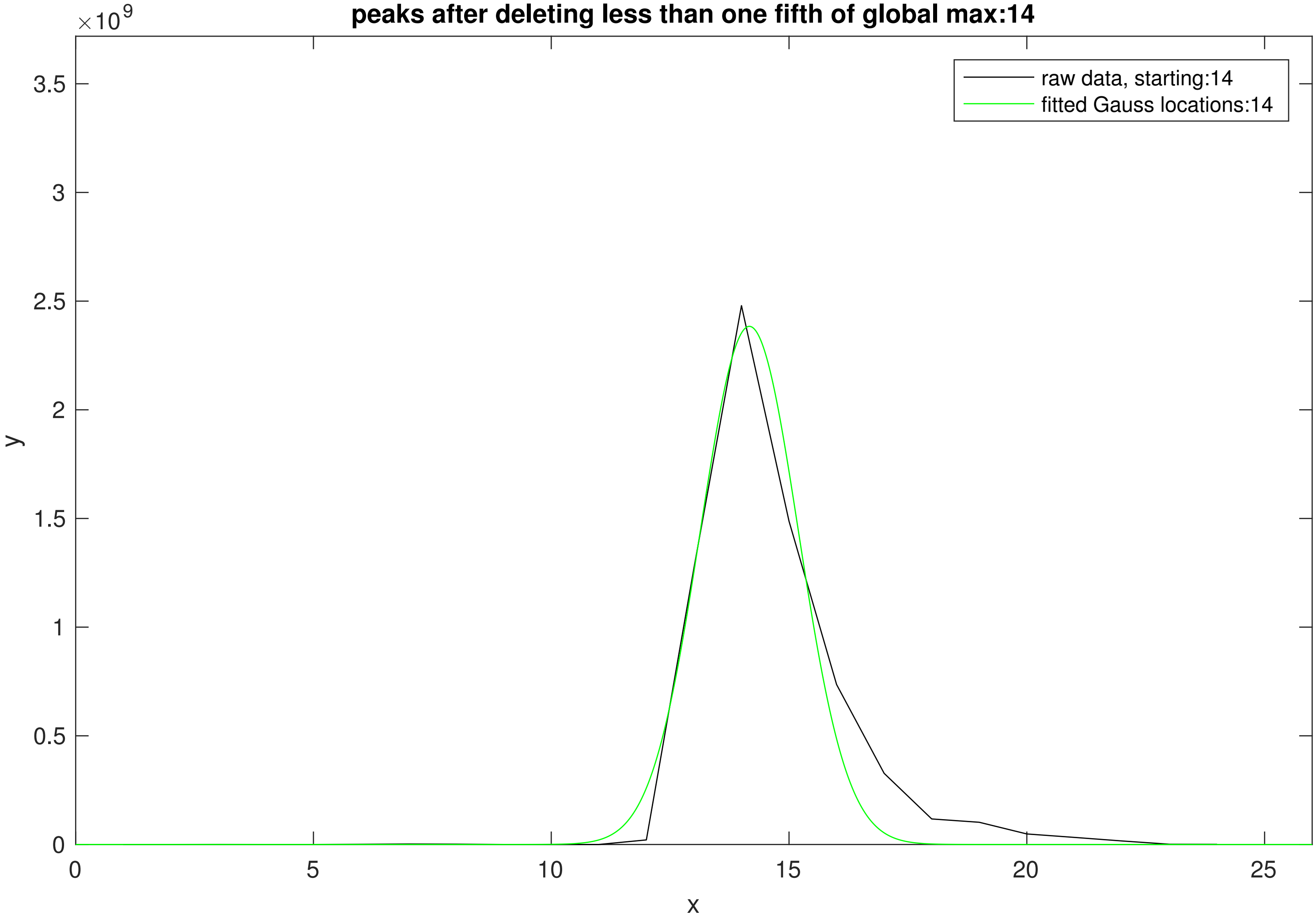

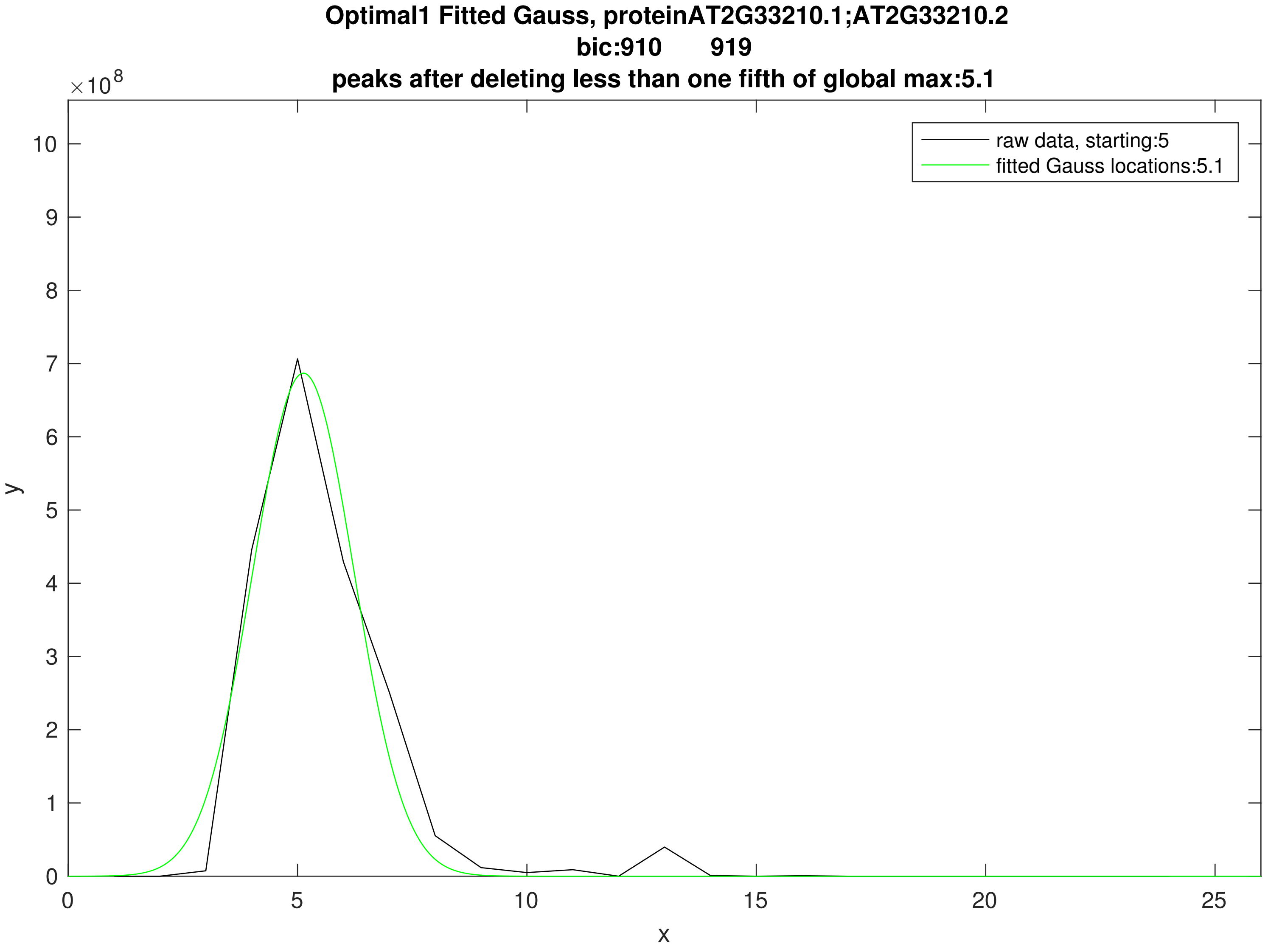

Optimal1 Fitted Gauss, proteinAT2G33210.1;AT2G33210.2
bic:910 919
peaks after deleting less than one fifth of global max:5.1
×10^8
raw data, starting:5
fitted Gauss locations:5.1
y
x
0
1
2
3
4
5
6
7
8
9
10
0
5
10
15
20
25

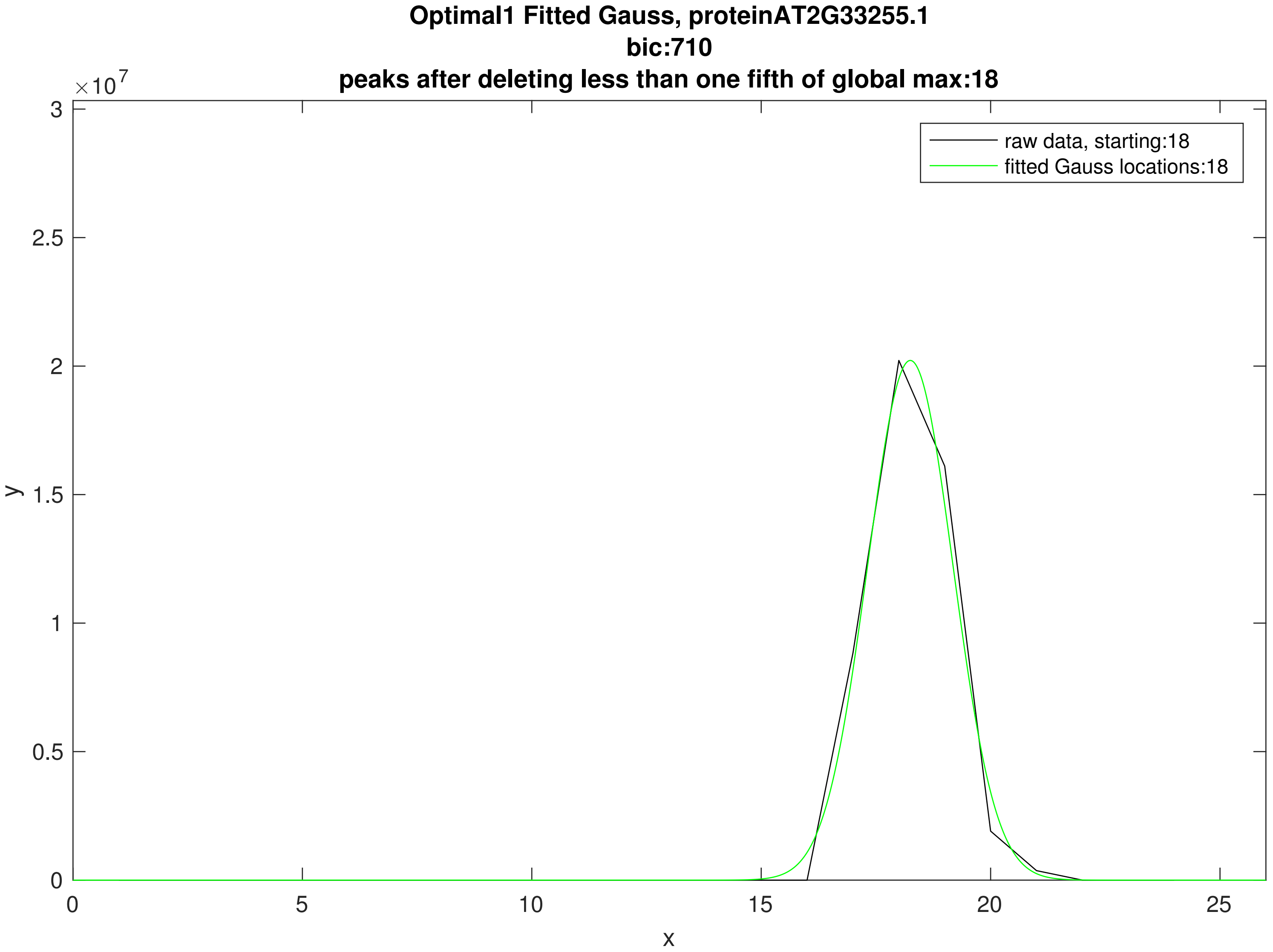

Optimal1 Fitted Gauss, proteinAT2G33255.1
bic:710
peaks after deleting less than one fifth of global max:18
×10^7
3
2.5
2
1.5
1
0.5
0
y
0
5
10
15
20
25
x
raw data, starting:18
fitted Gauss locations:18

#### Optimal1 Fitted Gauss, proteinAT2G33410.1
#### bic:756
#### peaks after deleting less than one fifth of global max:19

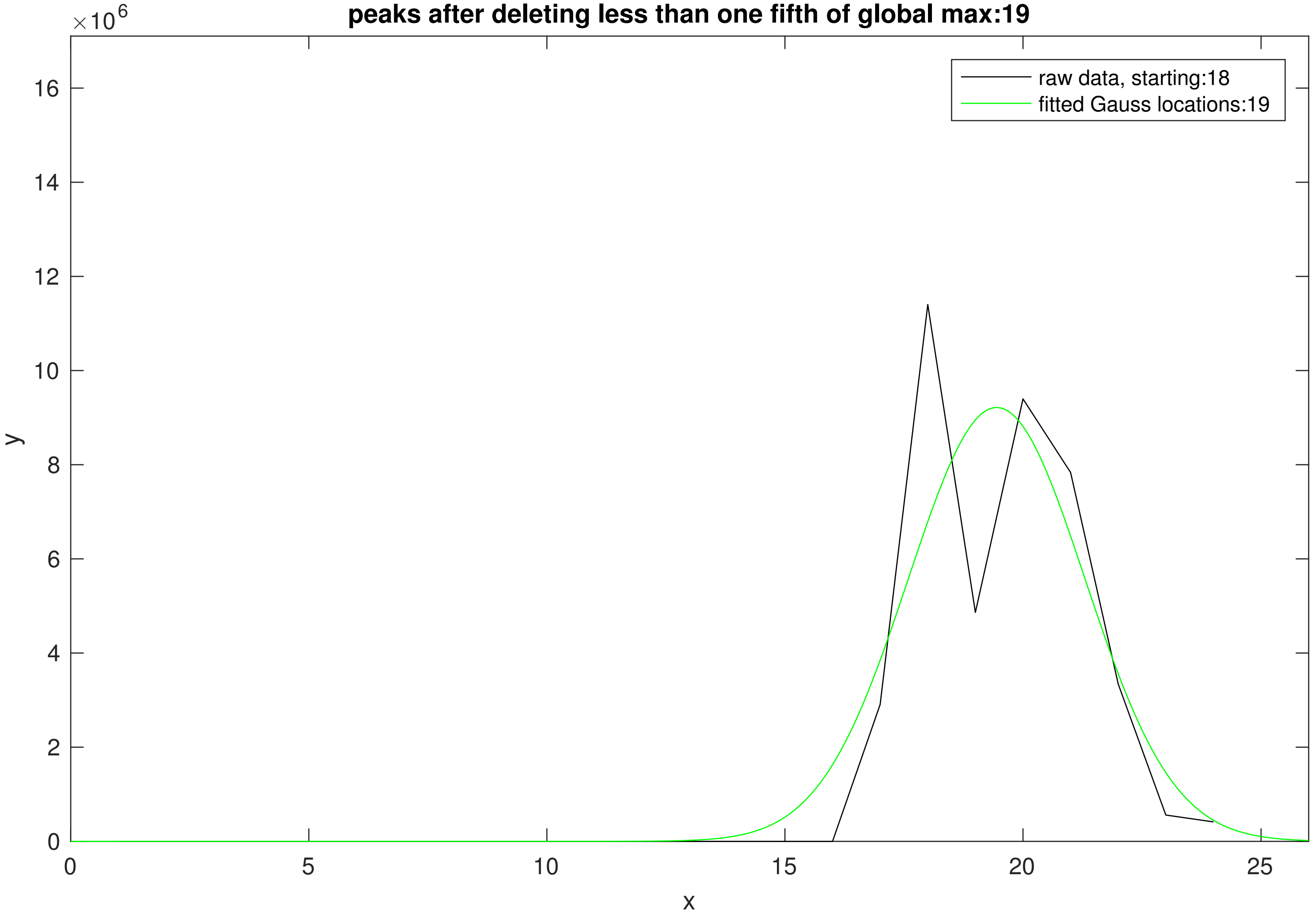

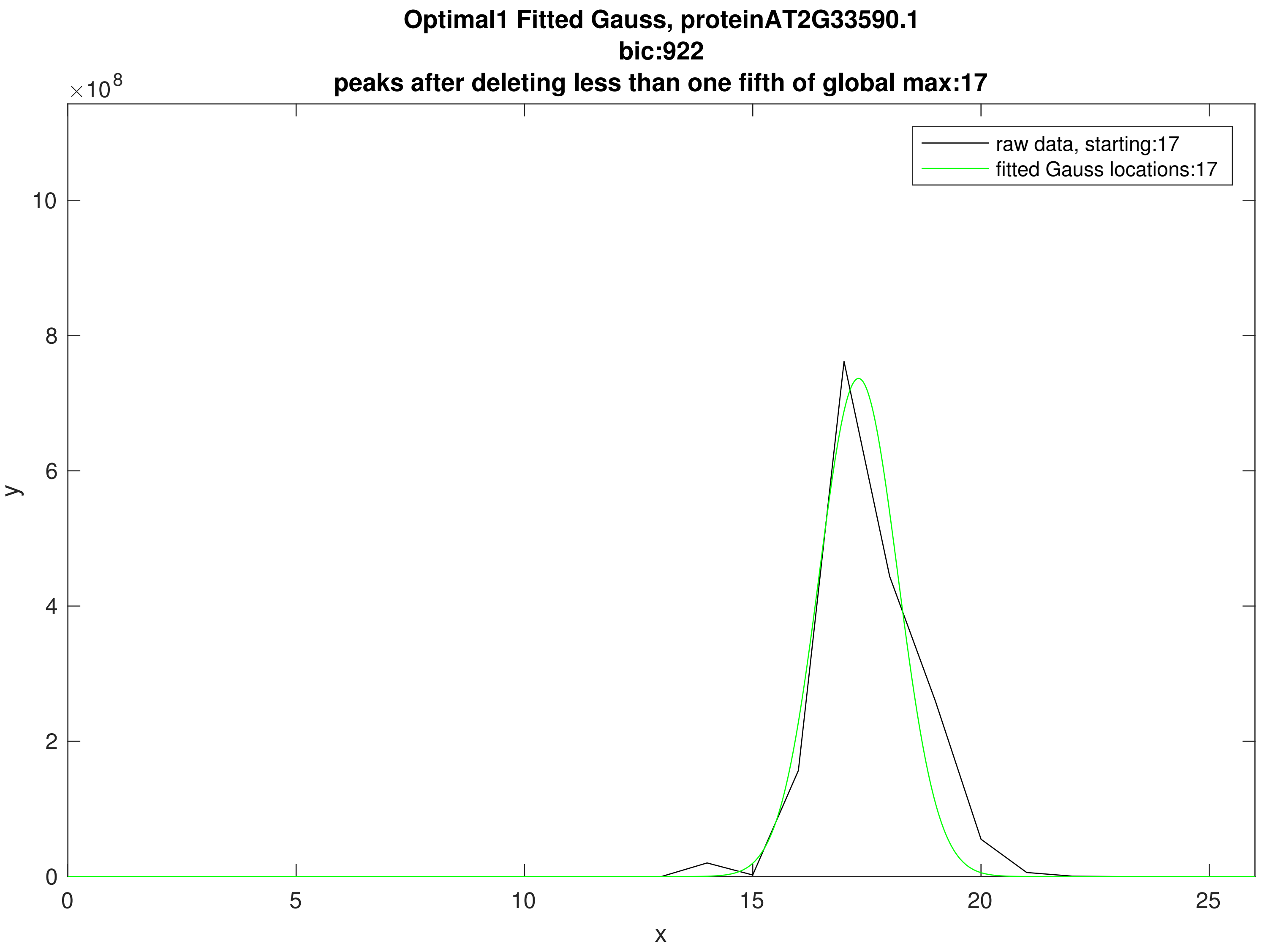
Optimal1 Fitted Gauss, proteinAT2G33590.1
bic:922
peaks after deleting less than one fifth of global max:17
$\times 10^8$
raw data, starting:17
fitted Gauss locations:17
10
8
6
4
2
0
y
0
5
10
15
20
25
x

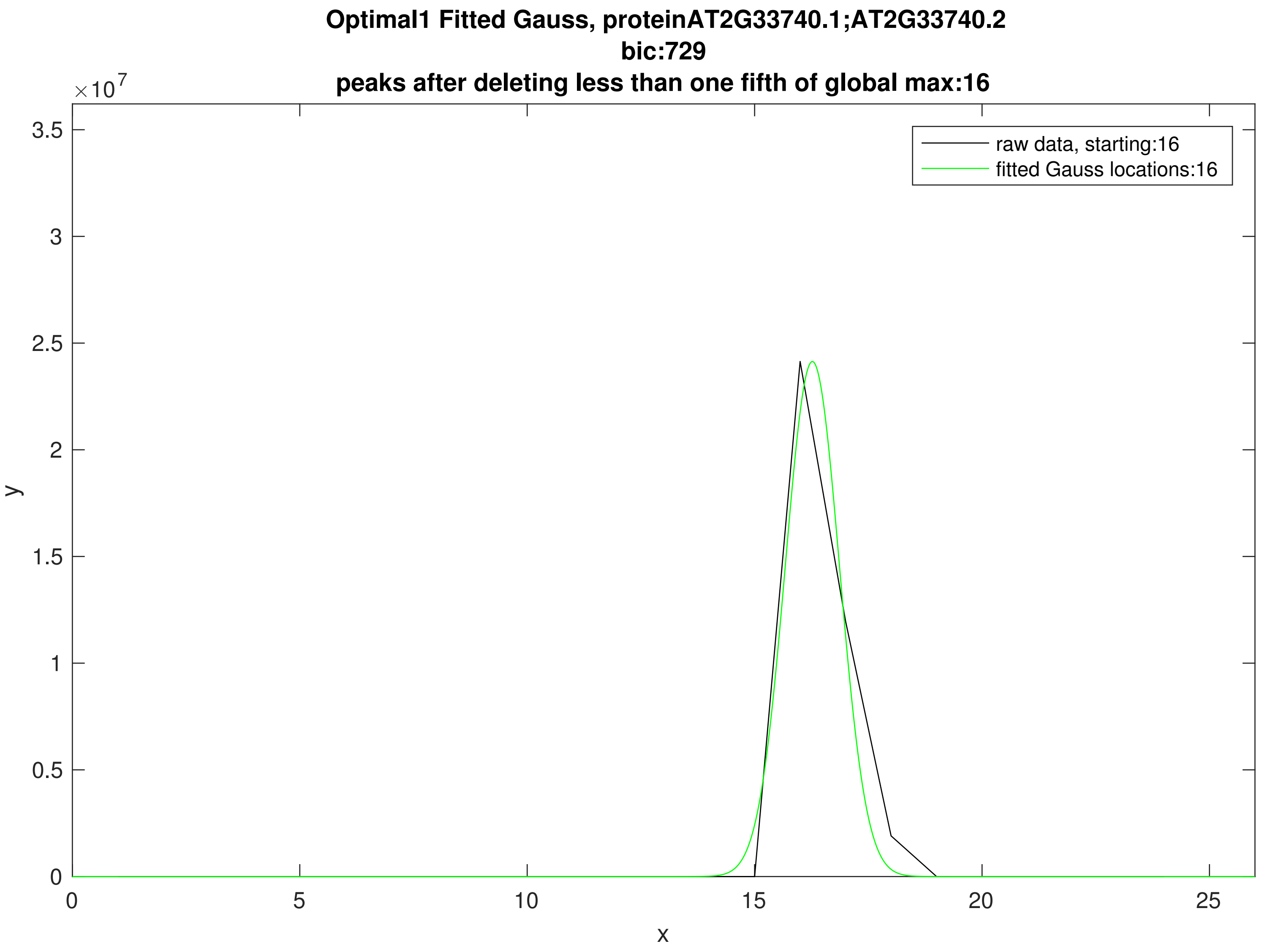
Optimal1 Fitted Gauss, proteinAT2G33740.1;AT2G33740.2
bic:729
peaks after deleting less than one fifth of global max:16
×10^7
3.5
3
2.5
2
1.5
1
0.5
0
y
0
5
10
15
20
25
x
raw data, starting:16
fitted Gauss locations:16

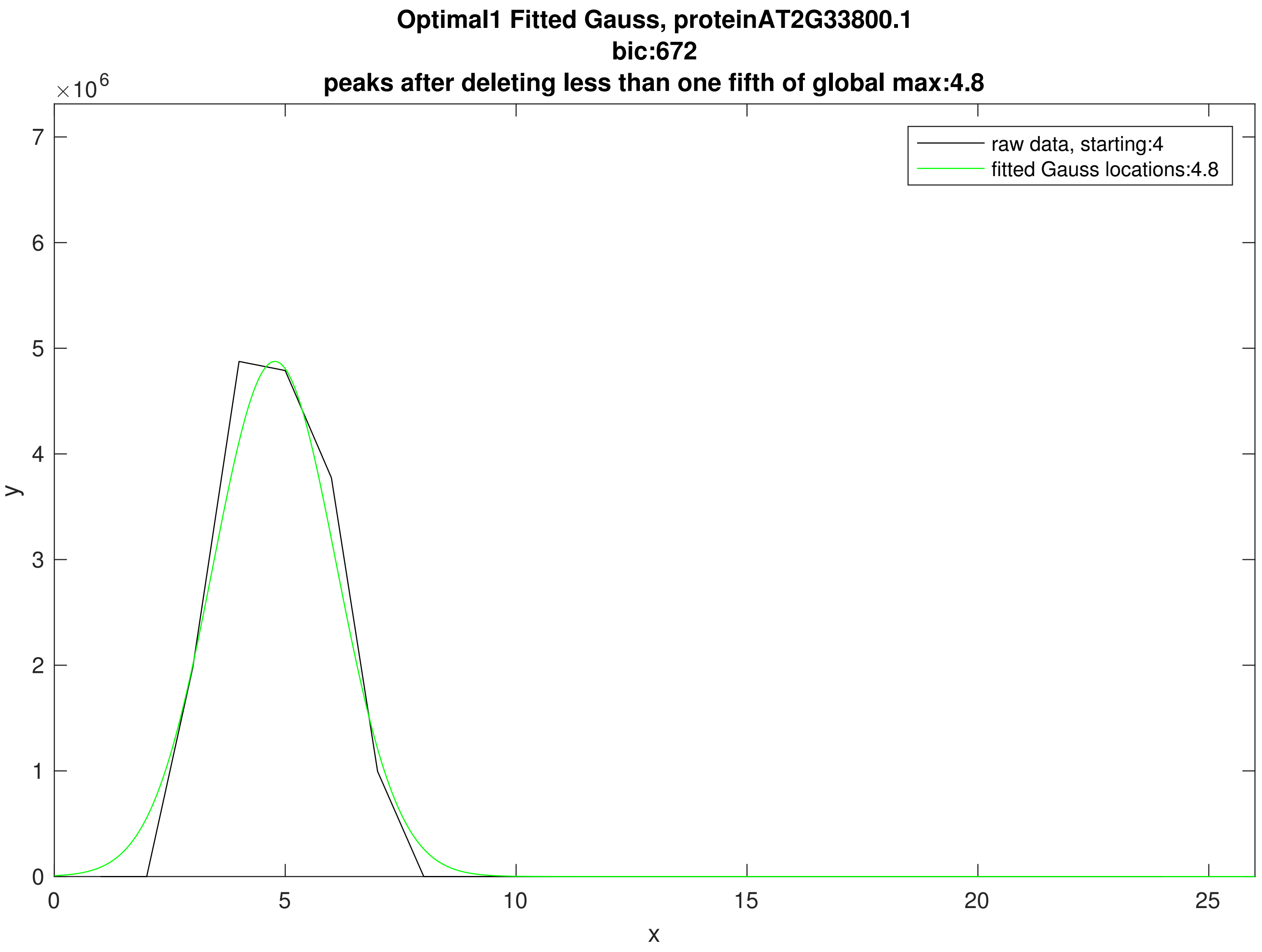
Optimal1 Fitted Gauss, proteinAT2G33800.1
bic:672
peaks after deleting less than one fifth of global max:4.8
raw data, starting:4
fitted Gauss locations:4.8
$\times 10^6$
y
x
0
1
2
3
4
5
6
7
0
5
10
15
20
25

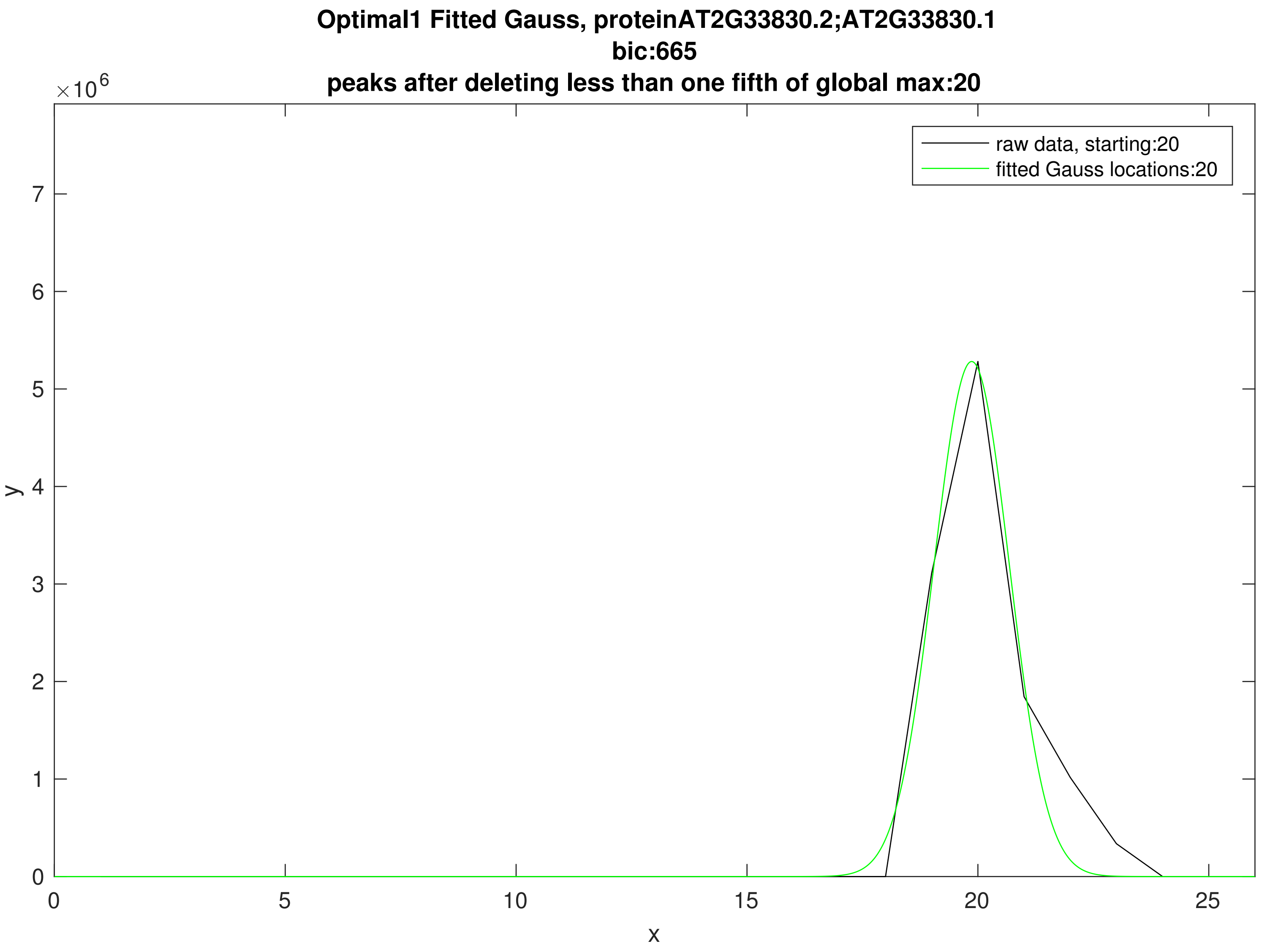

Optimal1 Fitted Gauss, proteinAT2G33830.2;AT2G33830.1
bic:665
peaks after deleting less than one fifth of global max:20
×10^6
raw data, starting:20
fitted Gauss locations:20
y
x
0
1
2
3
4
5
6
7
0
5
10
15
20
25

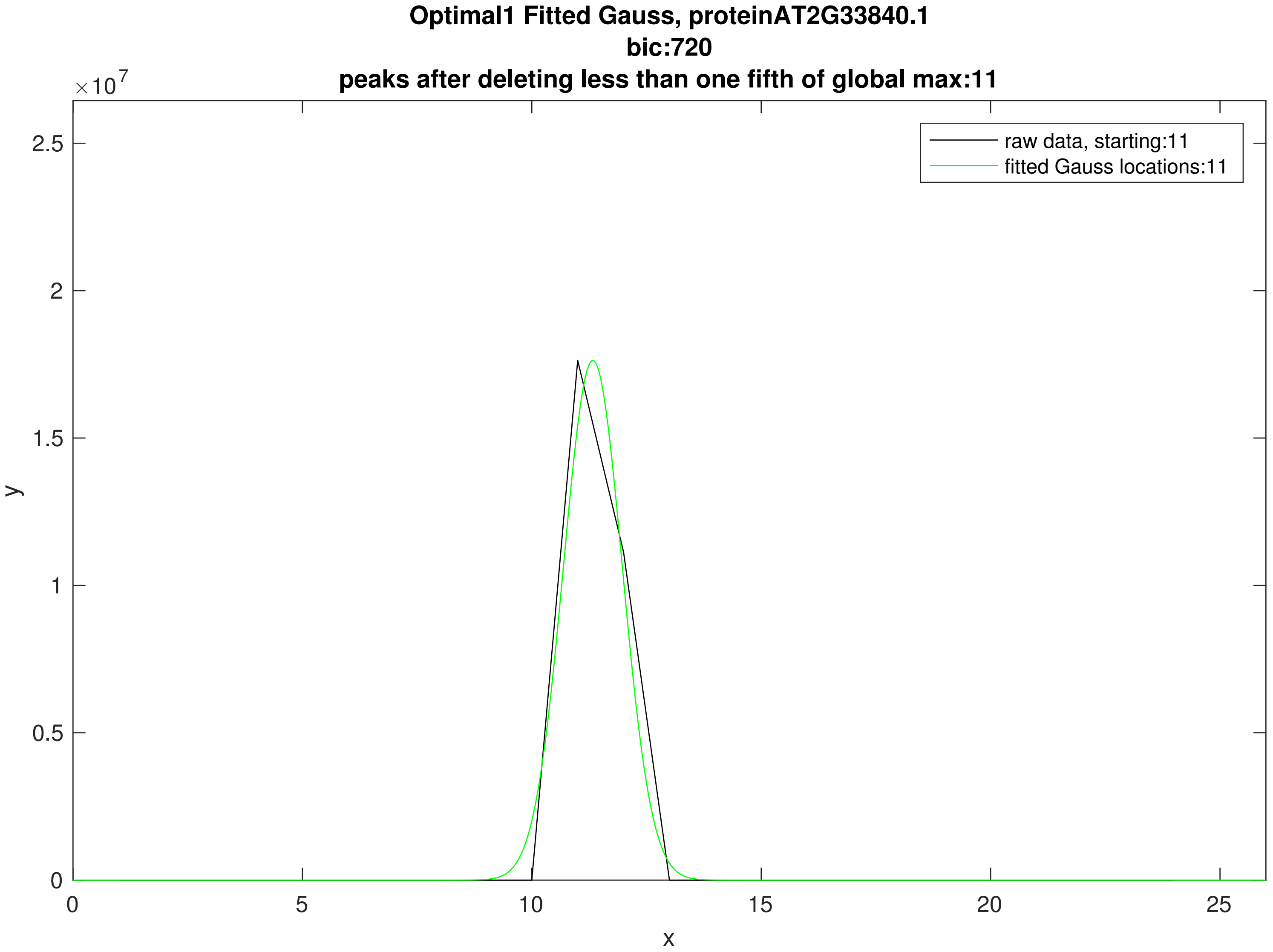

Optimal1 Fitted Gauss, proteinAT2G33840.1
bic:720
peaks after deleting less than one fifth of global max:11
×10^7
2.5
2
1.5
1
0.5
0
y
0
5
10
15
20
25
x
raw data, starting:11
fitted Gauss locations:11

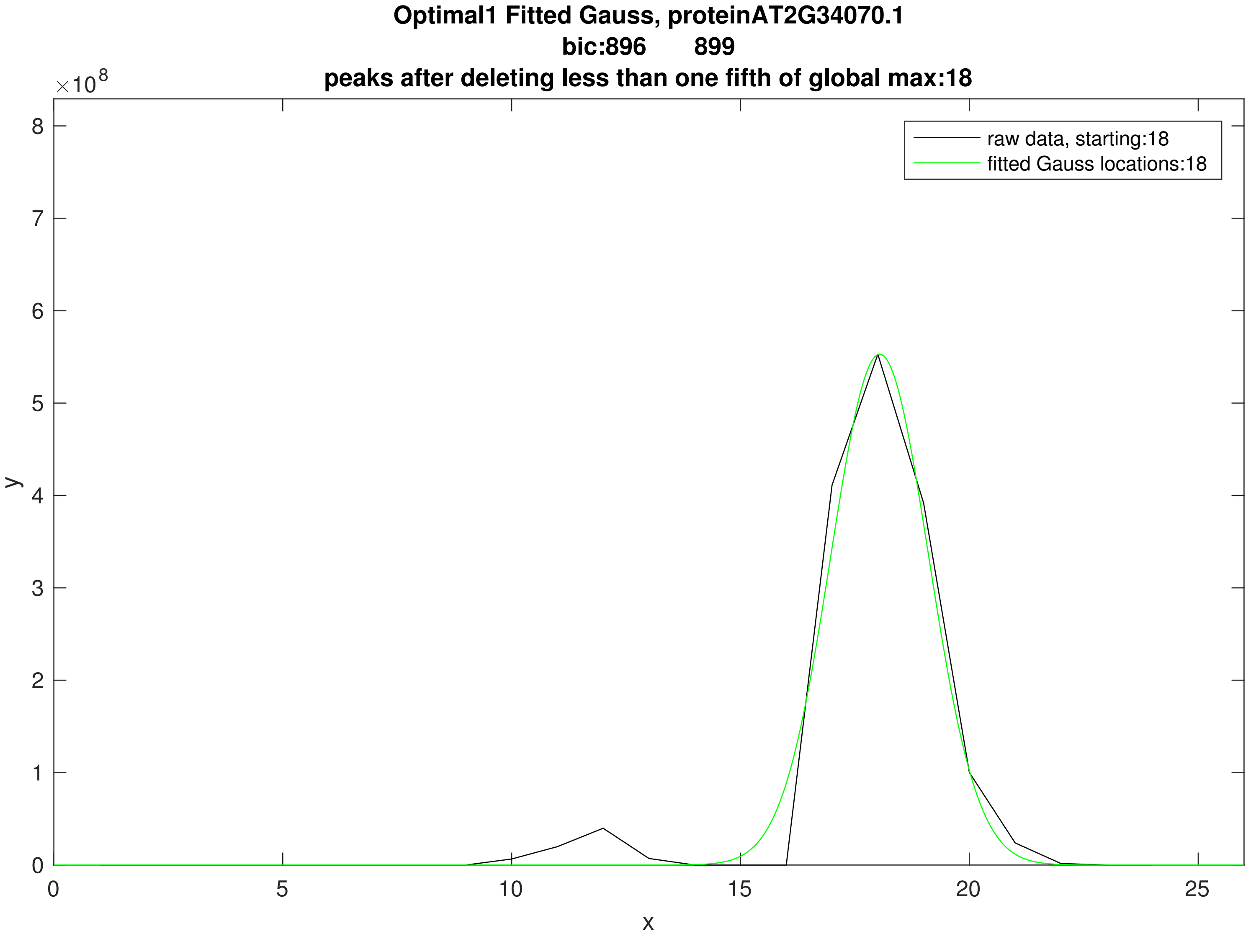

Optimal1 Fitted Gauss, proteinAT2G34070.1
bic:896 899
peaks after deleting less than one fifth of global max:18
×10^8
raw data, starting:18
fitted Gauss locations:18
y
x
0
1
2
3
4
5
6
7
8
0
5
10
15
20
25

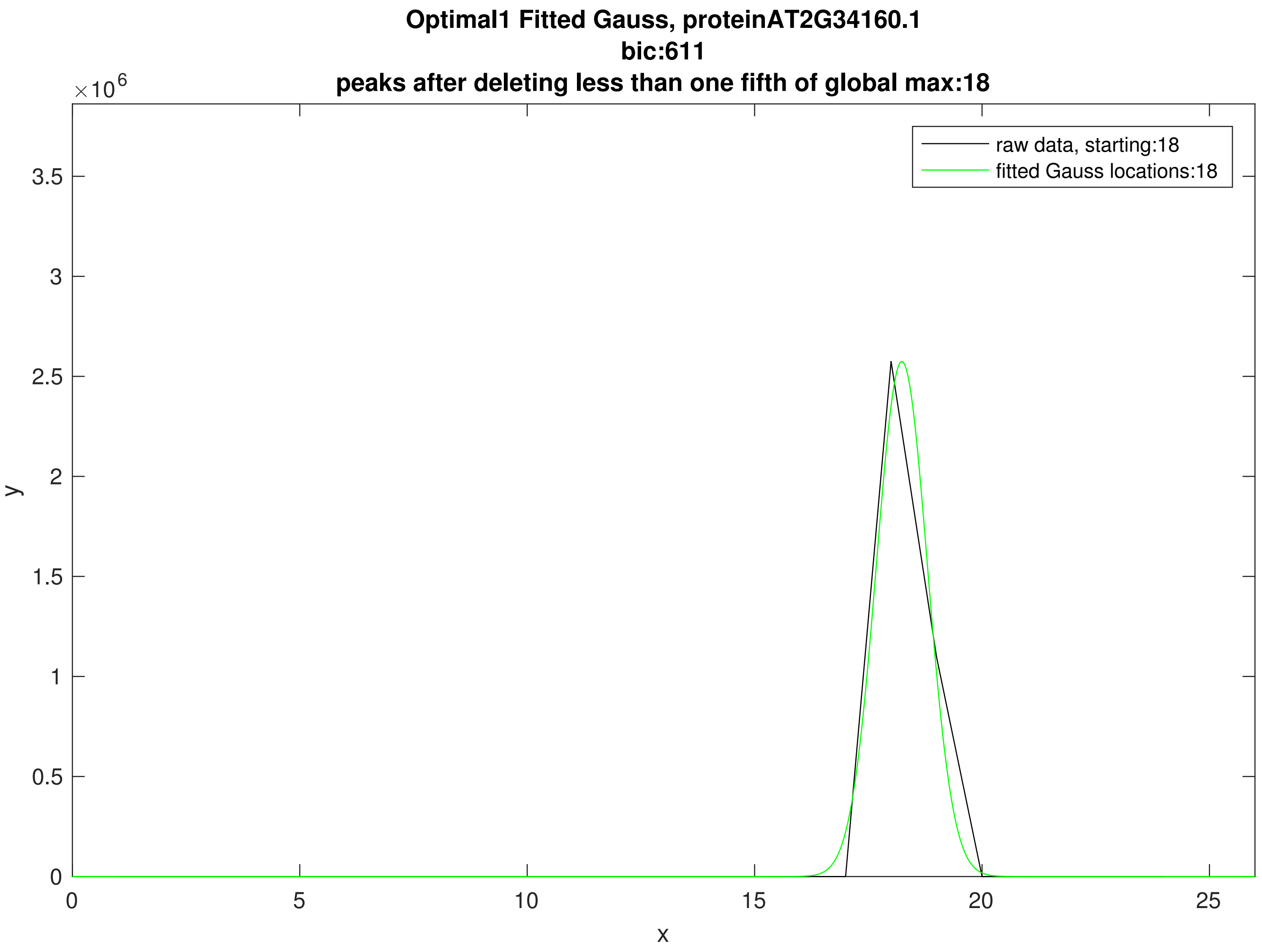

Optimal1 Fitted Gauss, proteinAT2G34160.1
bic:611
peaks after deleting less than one fifth of global max:18
×10^6
3.5
3
2.5
2
1.5
1
0.5
0
y
0
5
10
15
20
25
x
raw data, starting:18
fitted Gauss locations:18

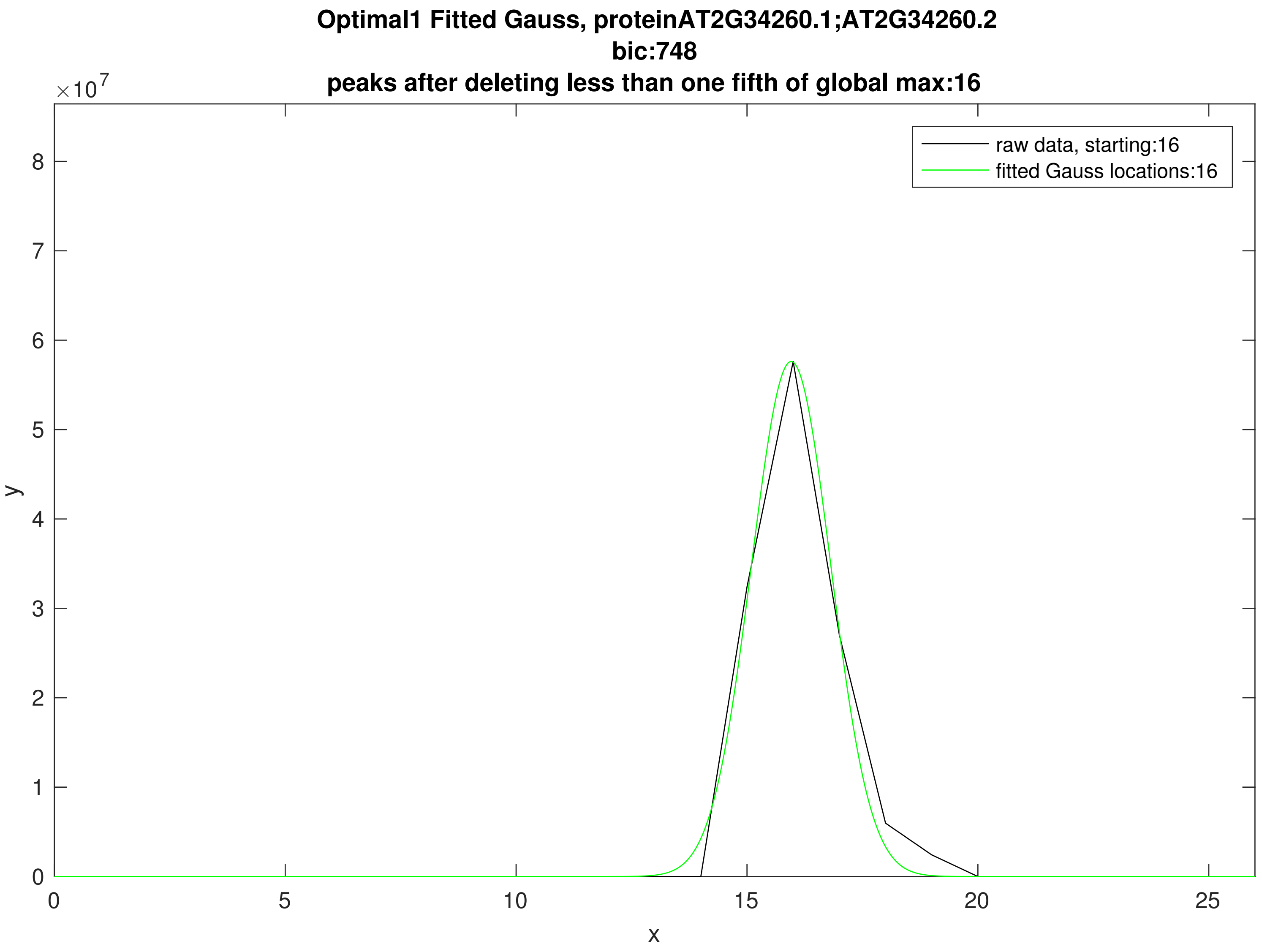

Optimal1 Fitted Gauss, proteinAT2G34260.1;AT2G34260.2
bic:748
peaks after deleting less than one fifth of global max:16
×10^7
raw data, starting:16
fitted Gauss locations:16
y
x
0
1
2
3
4
5
6
7
8
0
5
10
15
20
25

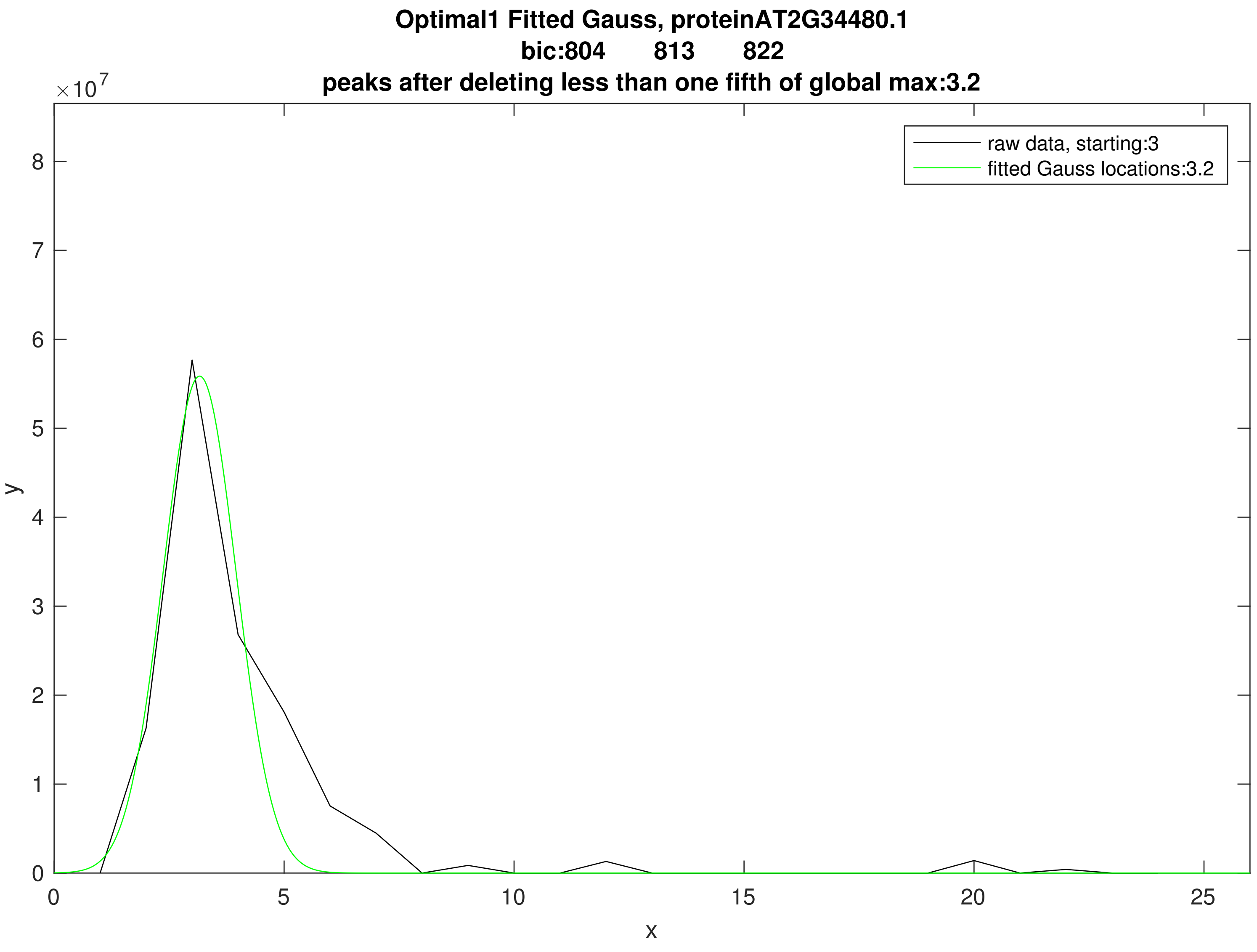

Optimal1 Fitted Gauss, proteinAT2G34480.1
bic:804 813 822
peaks after deleting less than one fifth of global max:3.2
×10^7
raw data, starting:3
fitted Gauss locations:3.2
y
x
0
1
2
3
4
5
6
7
8
0
5
10
15
20
25

**Optimal1 Fitted Gauss, proteinAT2G34590.1**
**bic:798 799**
**peaks after deleting less than one fifth of global max:11**

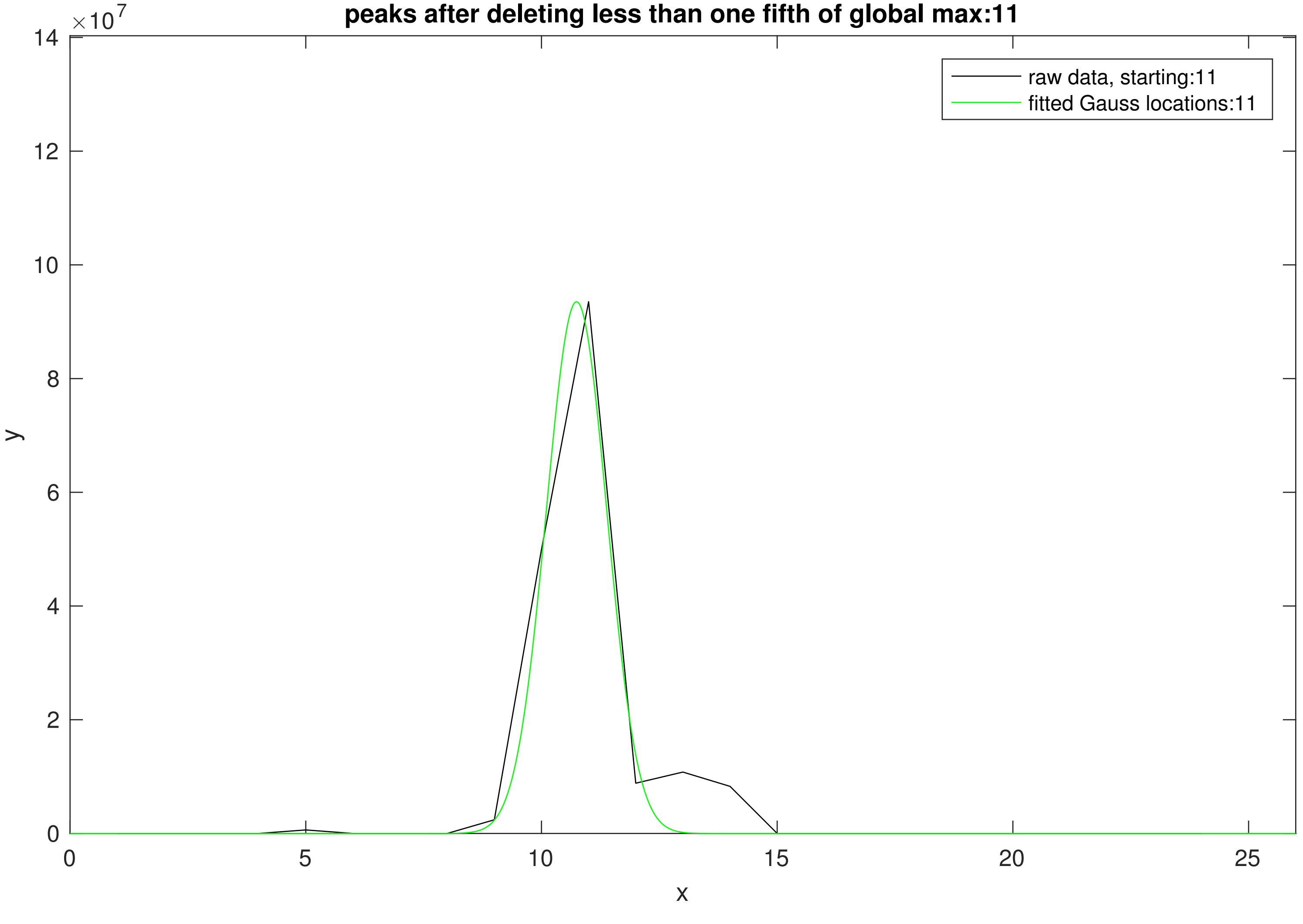

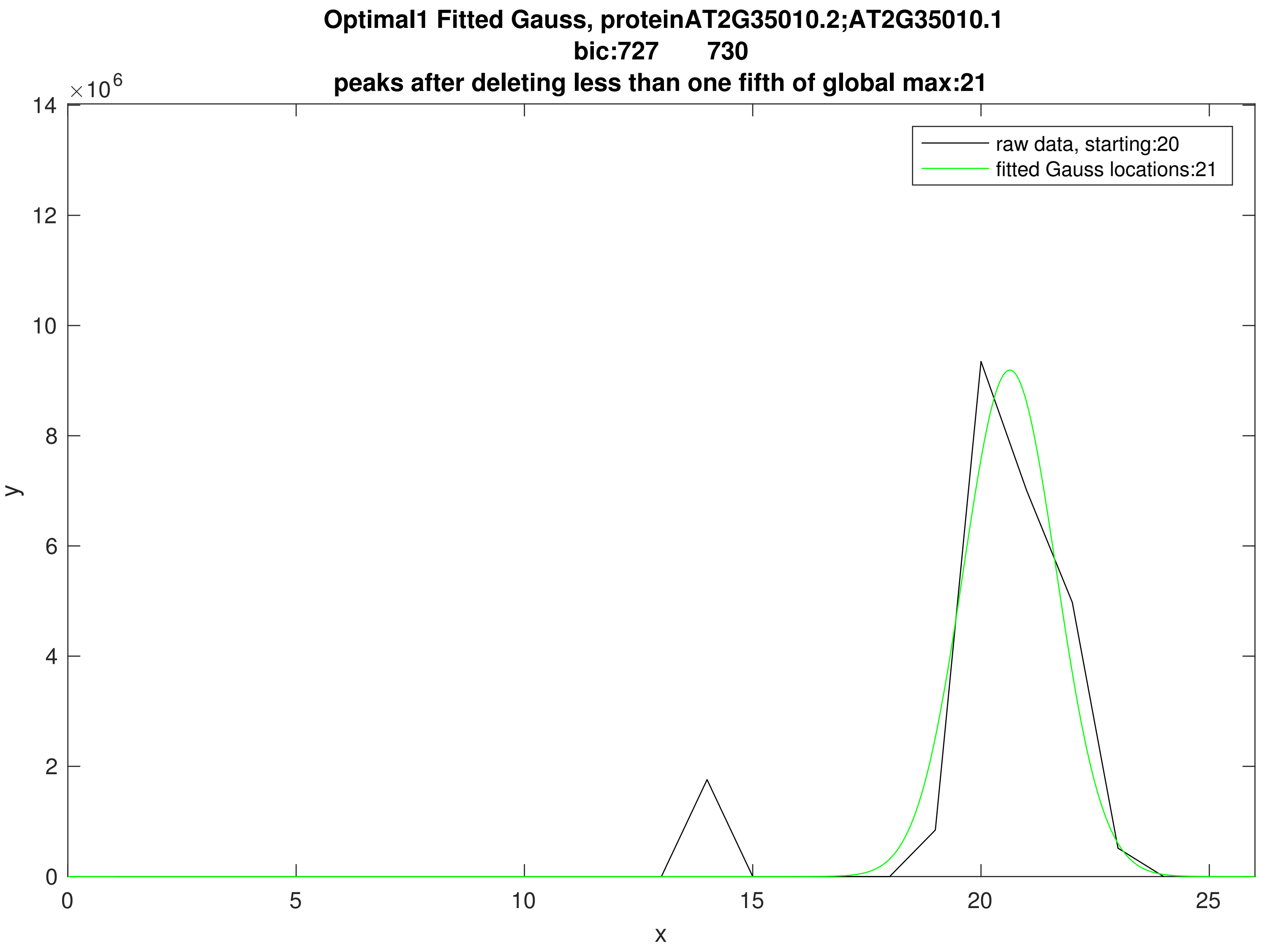
Optimal1 Fitted Gauss, proteinAT2G35010.2;AT2G35010.1
bic:727 730
peaks after deleting less than one fifth of global max:21
×10^6
raw data, starting:20
fitted Gauss locations:21
y
x
0
2
4
6
8
10
12
14
0
5
10
15
20
25

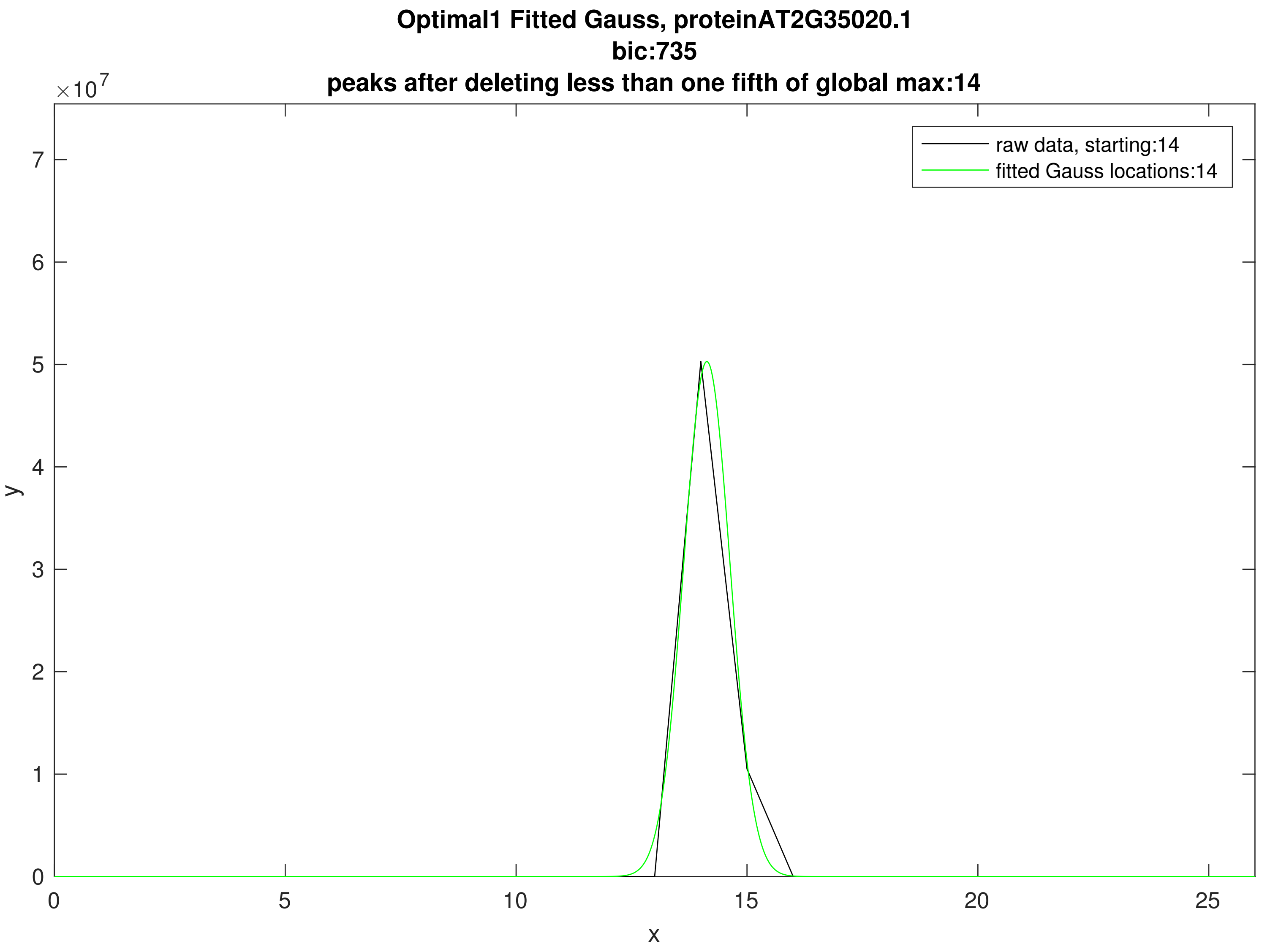

Optimal1 Fitted Gauss, proteinAT2G35020.1
bic:735
peaks after deleting less than one fifth of global max:14
×10^7
7
6
5
4
3
2
1
0
y
0
5
10
15
20
25
x
raw data, starting:14
fitted Gauss locations:14

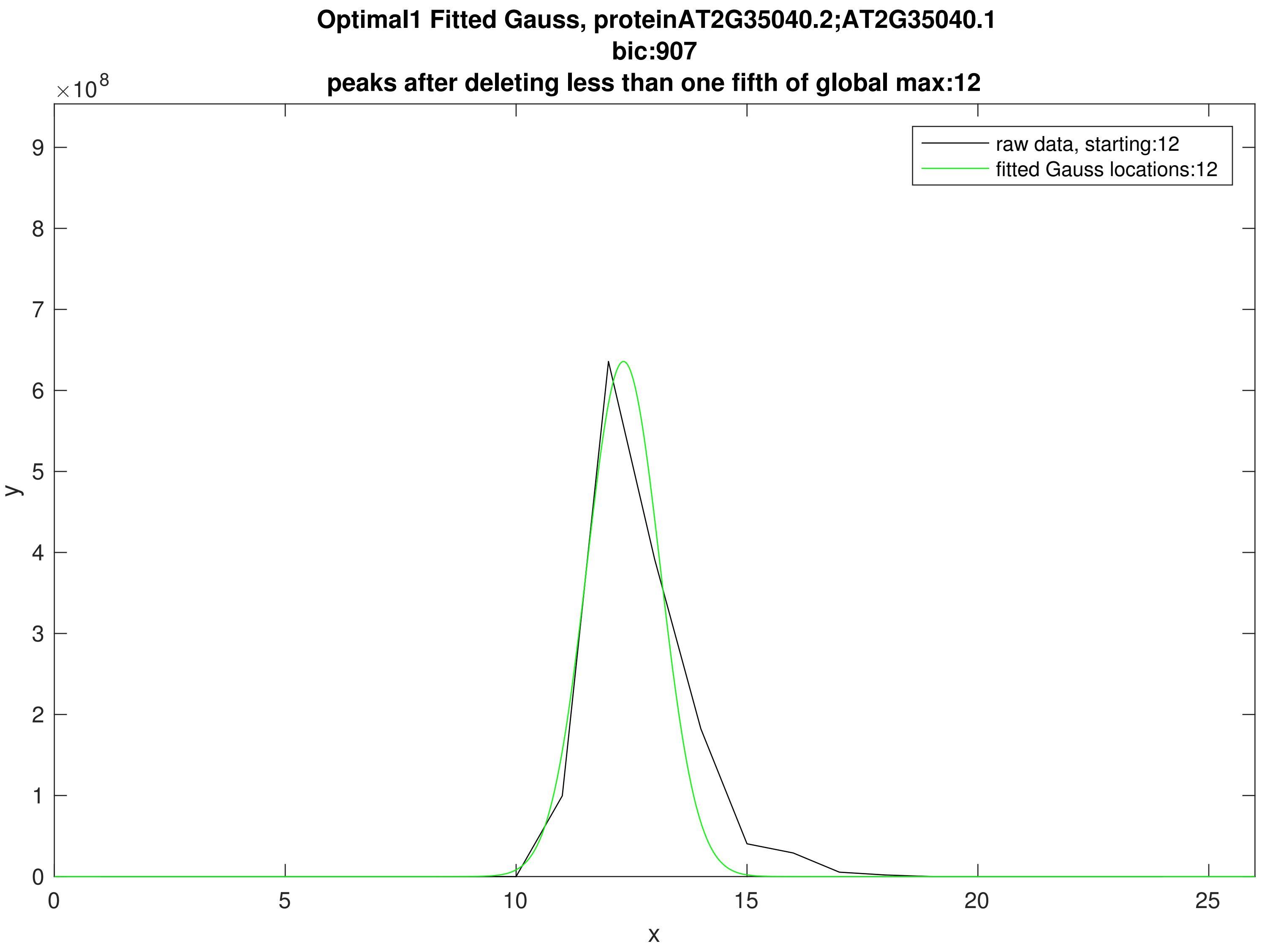

Optimal1 Fitted Gauss, proteinAT2G35040.2;AT2G35040.1
bic:907
peaks after deleting less than one fifth of global max:12
×10^8
raw data, starting:12
fitted Gauss locations:12
y
x
0
1
2
3
4
5
6
7
8
9
0
5
10
15
20
25

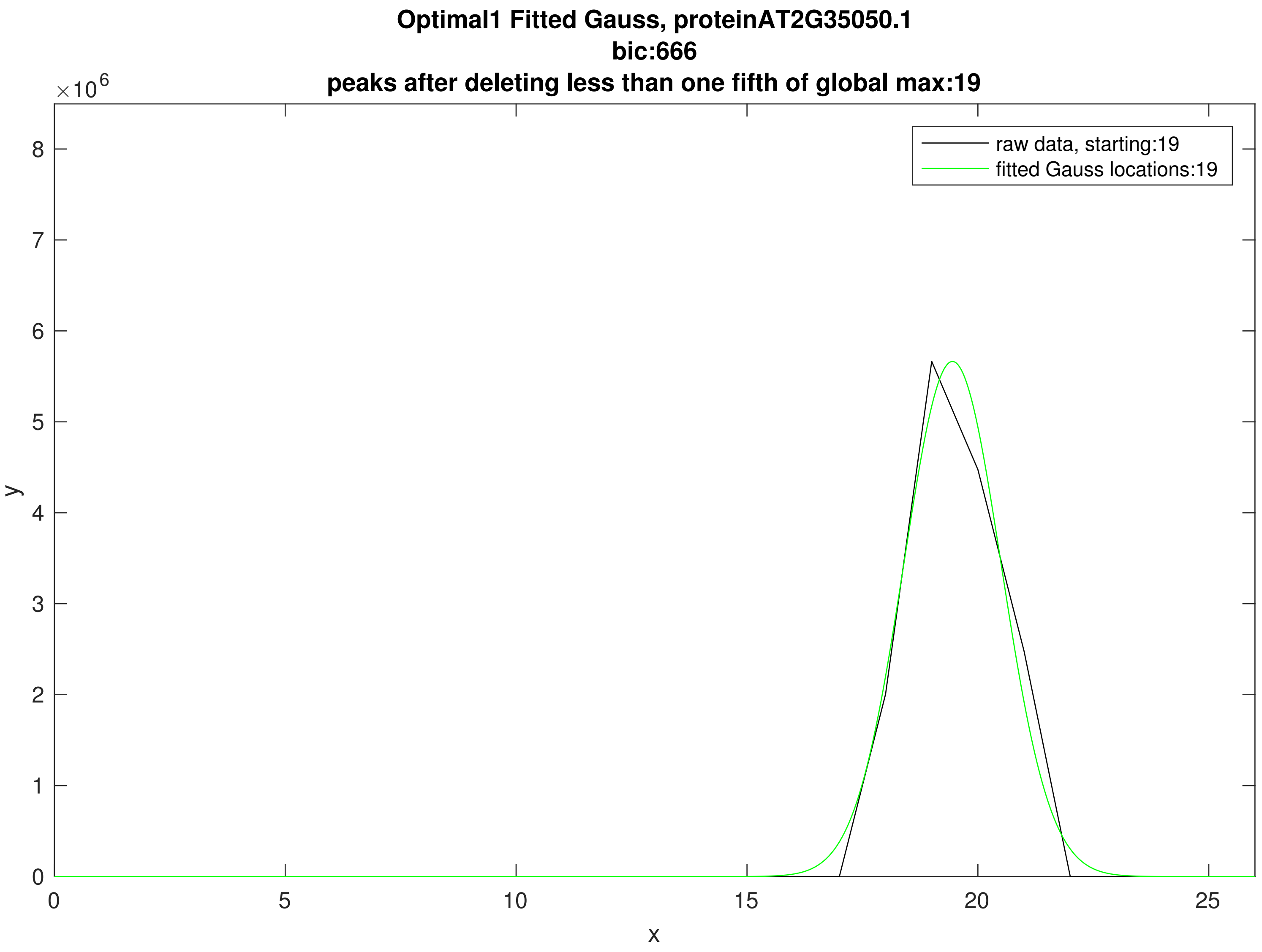
Optimal1 Fitted Gauss, proteinAT2G35050.1
bic:666
peaks after deleting less than one fifth of global max:19
×10^6
raw data, starting:19
fitted Gauss locations:19
y
x
0
1
2
3
4
5
6
7
8
0
5
10
15
20
25

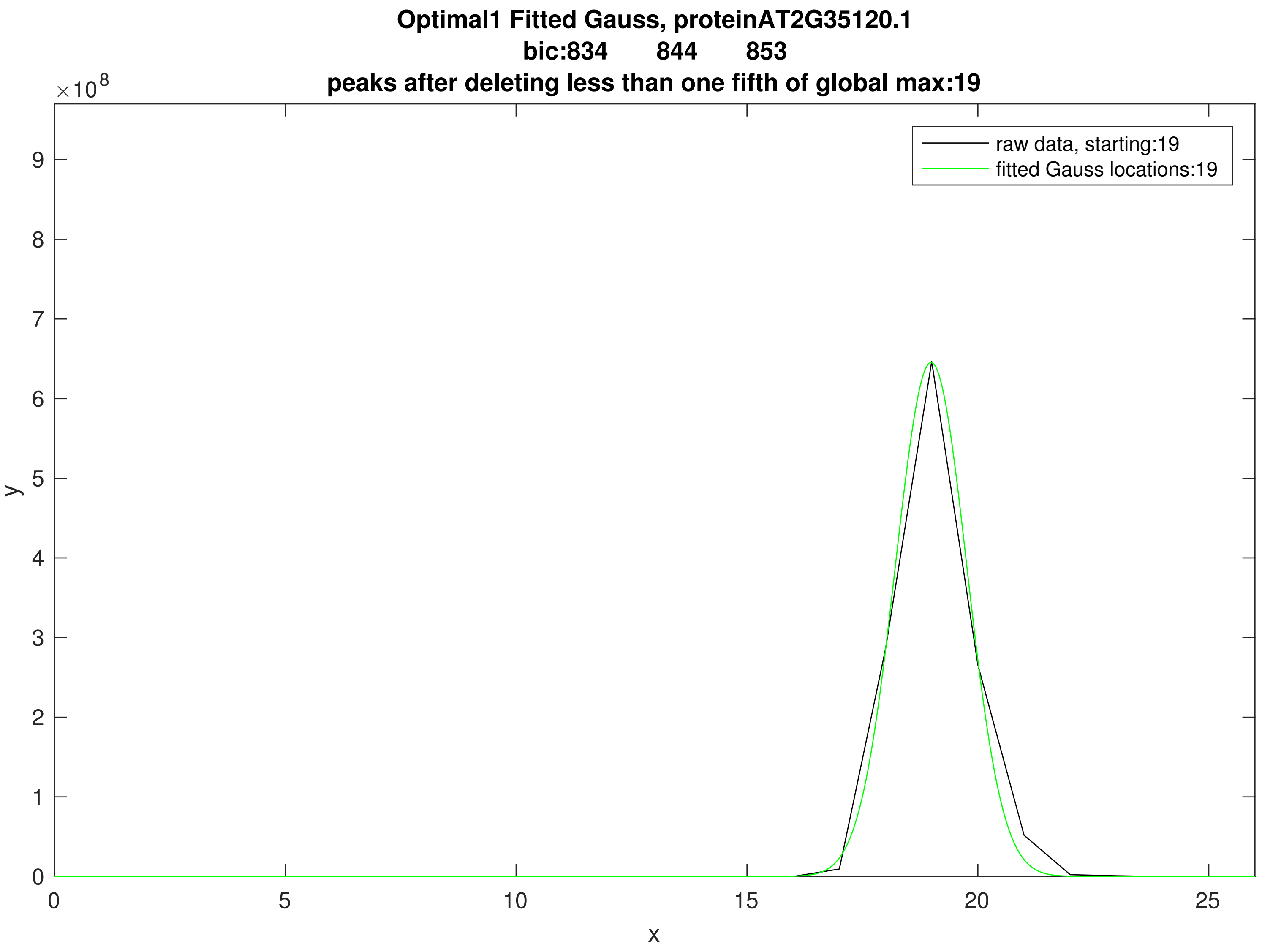
Optimal1 Fitted Gauss, proteinAT2G35120.1
bic:834 844 853
peaks after deleting less than one fifth of global max:19
×10^8
raw data, starting:19
fitted Gauss locations:19
y
x
0
1
2
3
4
5
6
7
8
9
0
5
10
15
20
25

#### Optimal1 Fitted Gauss, proteinAT2G35200.1
#### bic:748
#### peaks after deleting less than one fifth of global max:8.3

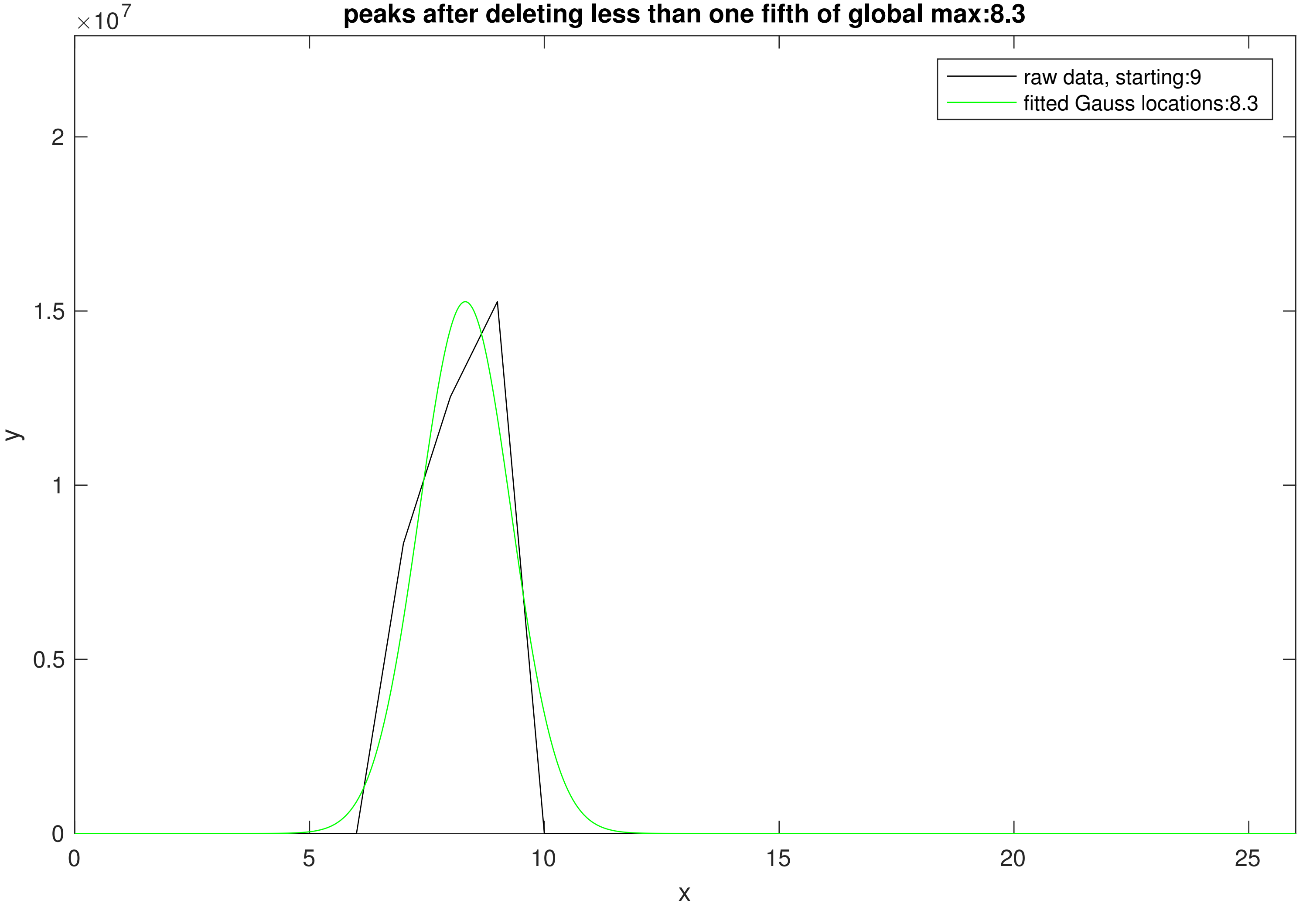

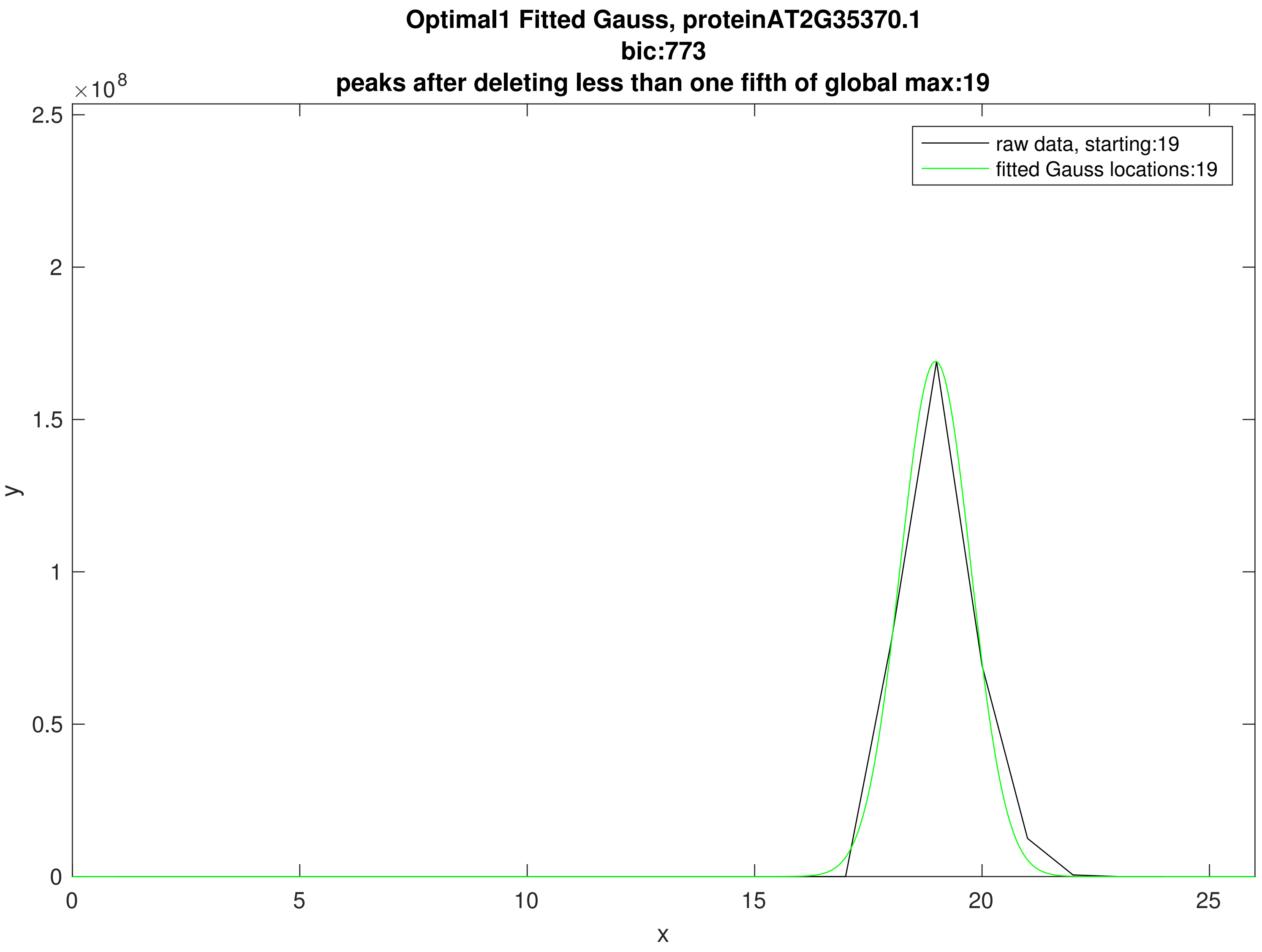

Optimal1 Fitted Gauss, proteinAT2G35370.1
bic:773
peaks after deleting less than one fifth of global max:19
×10^8
2.5
2
1.5
1
0.5
0
y
0
5
10
15
20
25
x
raw data, starting:19
fitted Gauss locations:19

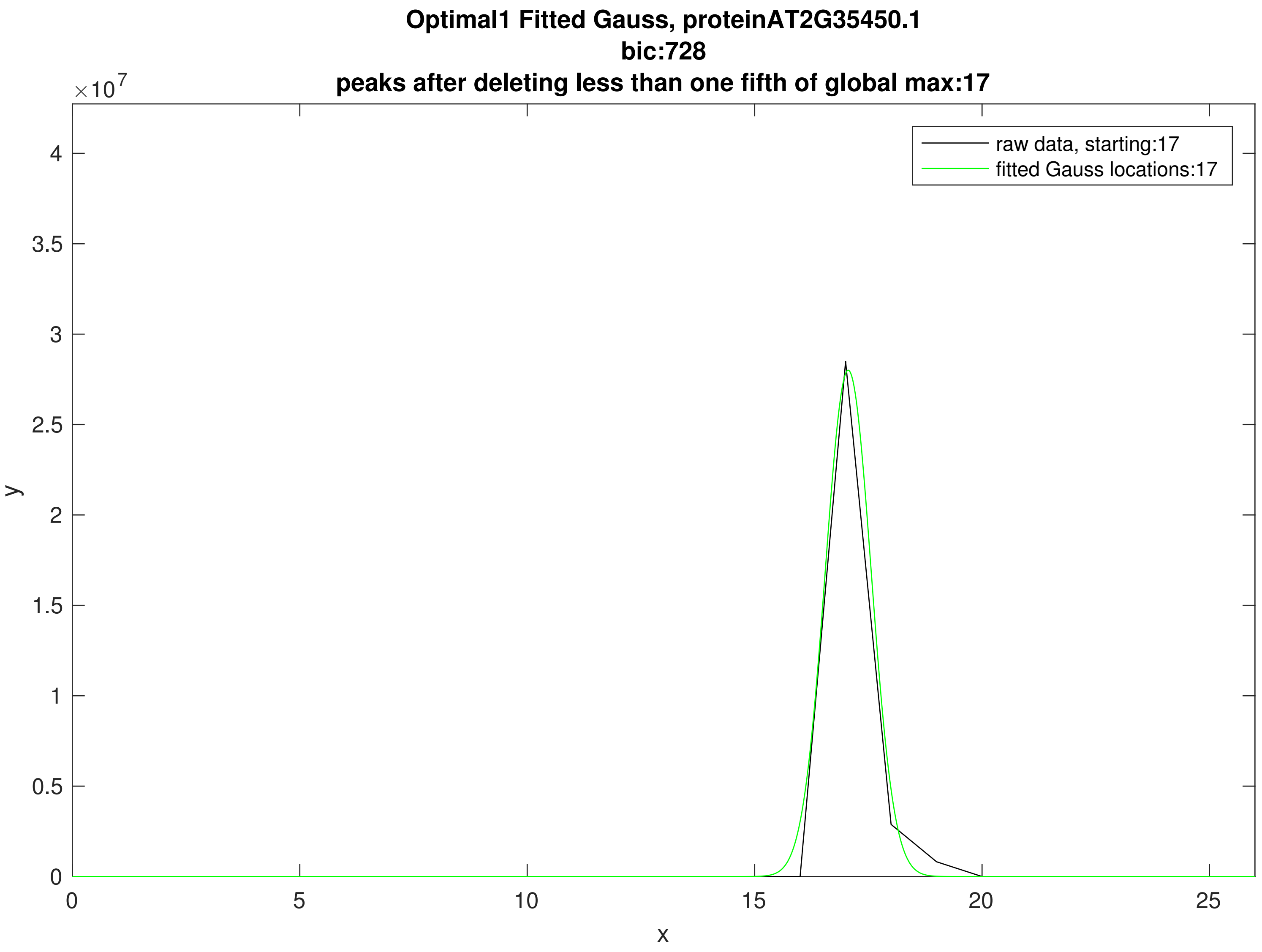
Optimal1 Fitted Gauss, proteinAT2G35450.1
bic:728
peaks after deleting less than one fifth of global max:17
×10^7
raw data, starting:17
fitted Gauss locations:17
4
3.5
3
2.5
2
1.5
1
0.5
0
y
0
5
10
15
20
25
x

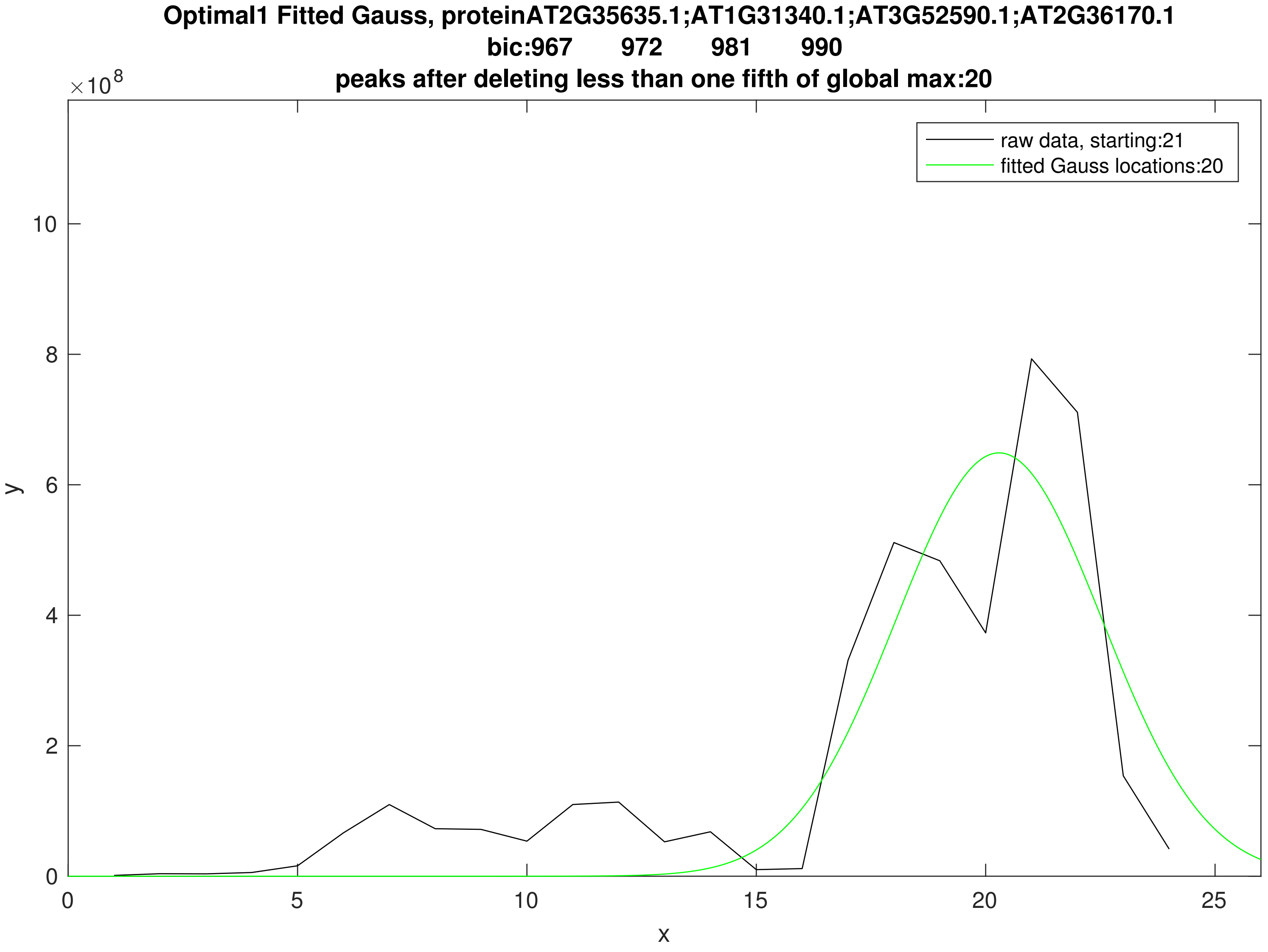
Optimal1 Fitted Gauss, proteinAT2G35635.1;AT1G31340.1;AT3G52590.1;AT2G36170.1
bic:967 972 981 990
peaks after deleting less than one fifth of global max:20
×10^8
raw data, starting:21
fitted Gauss locations:20
y
x
0
2
4
6
8
10
0
5
10
15
20
25

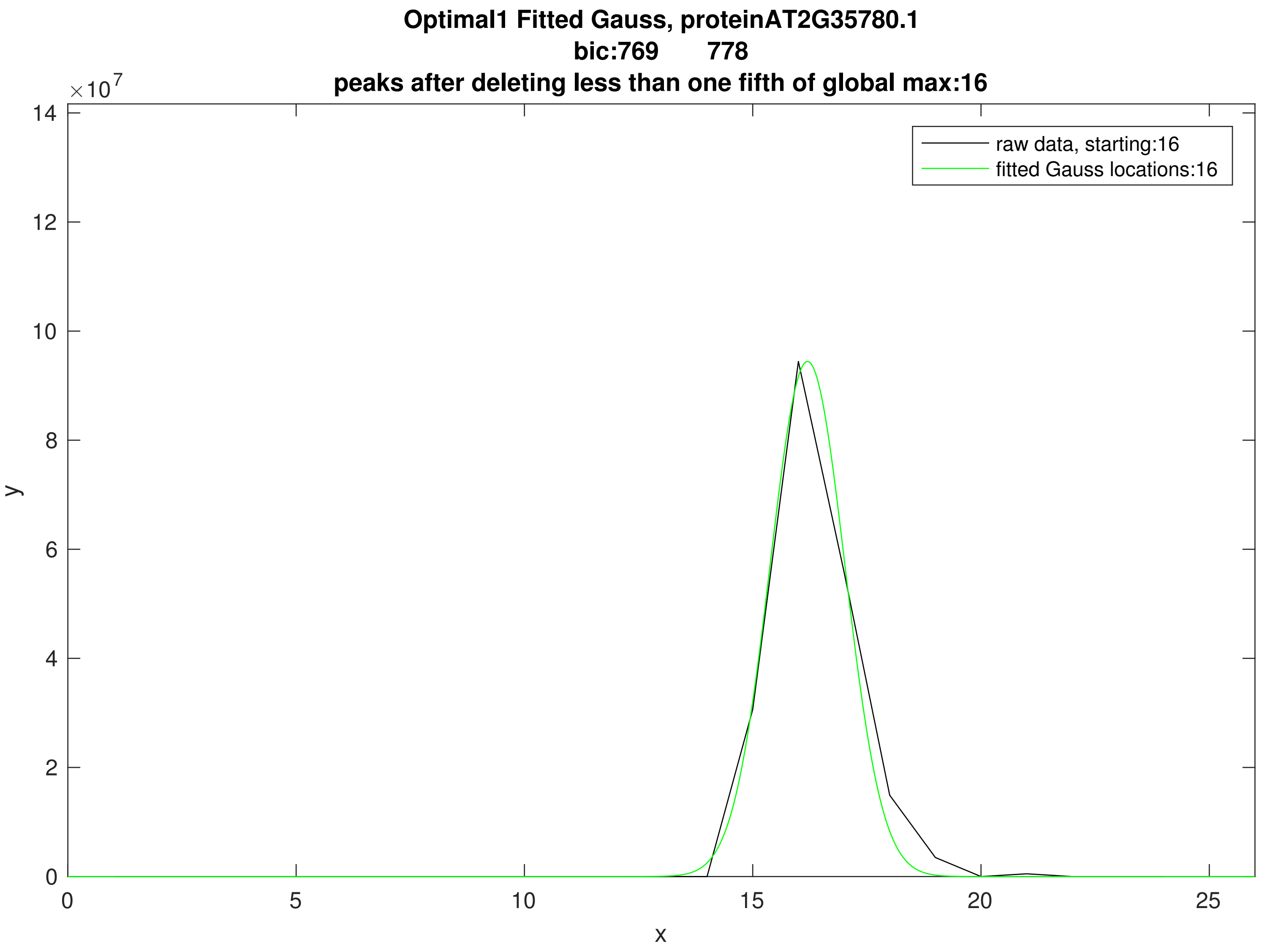
Optimal1 Fitted Gauss, proteinAT2G35780.1
bic:769 778
peaks after deleting less than one fifth of global max:16
×10^7
14
12
10
8
6
4
2
0
y
0
5
10
15
20
25
x
raw data, starting:16
fitted Gauss locations:16

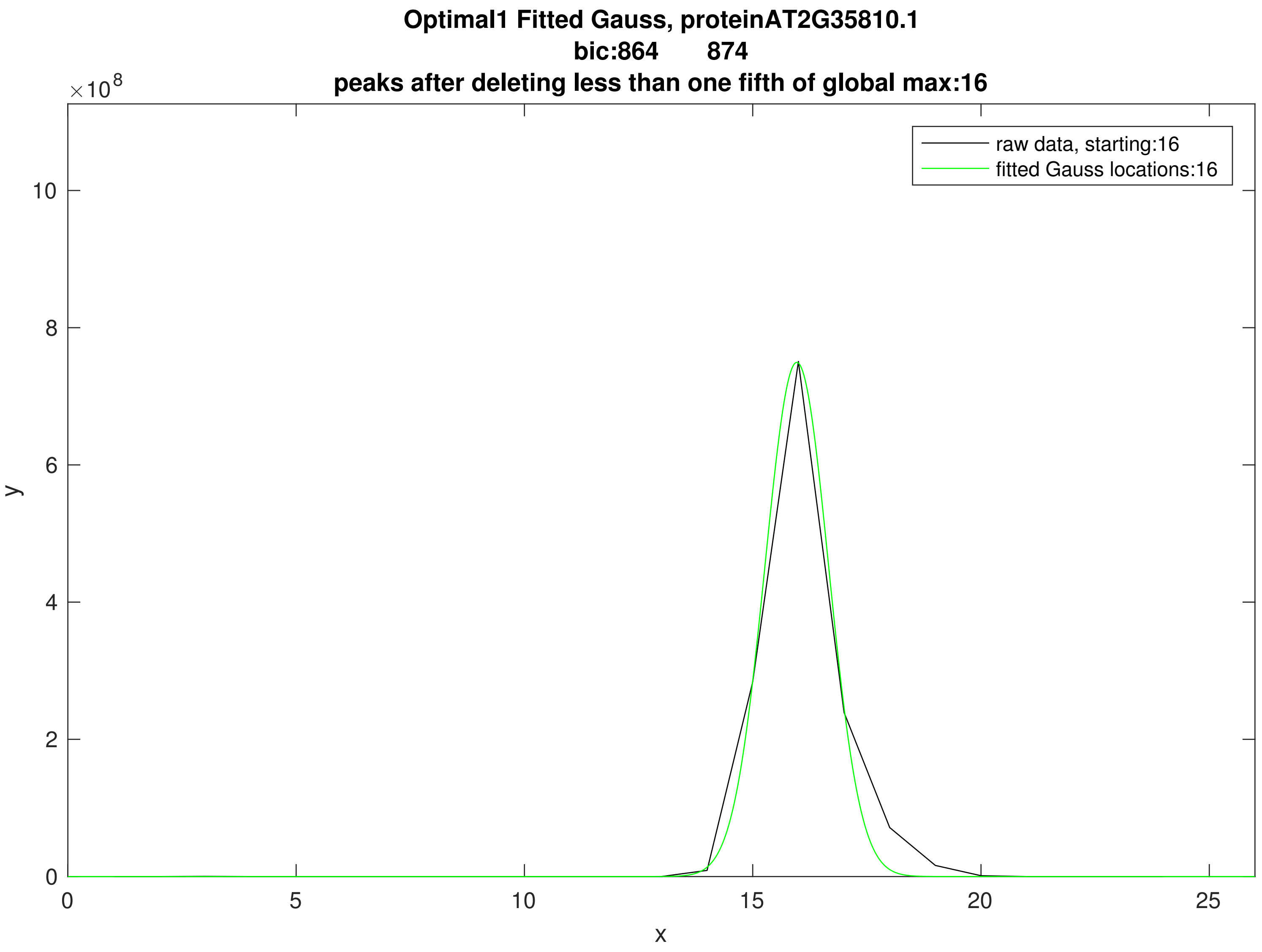
Optimal1 Fitted Gauss, proteinAT2G35810.1
bic:864 874
peaks after deleting less than one fifth of global max:16
×10^8
10
8
6
4
2
0
y
0
5
10
15
20
25
x
raw data, starting:16
fitted Gauss locations:16

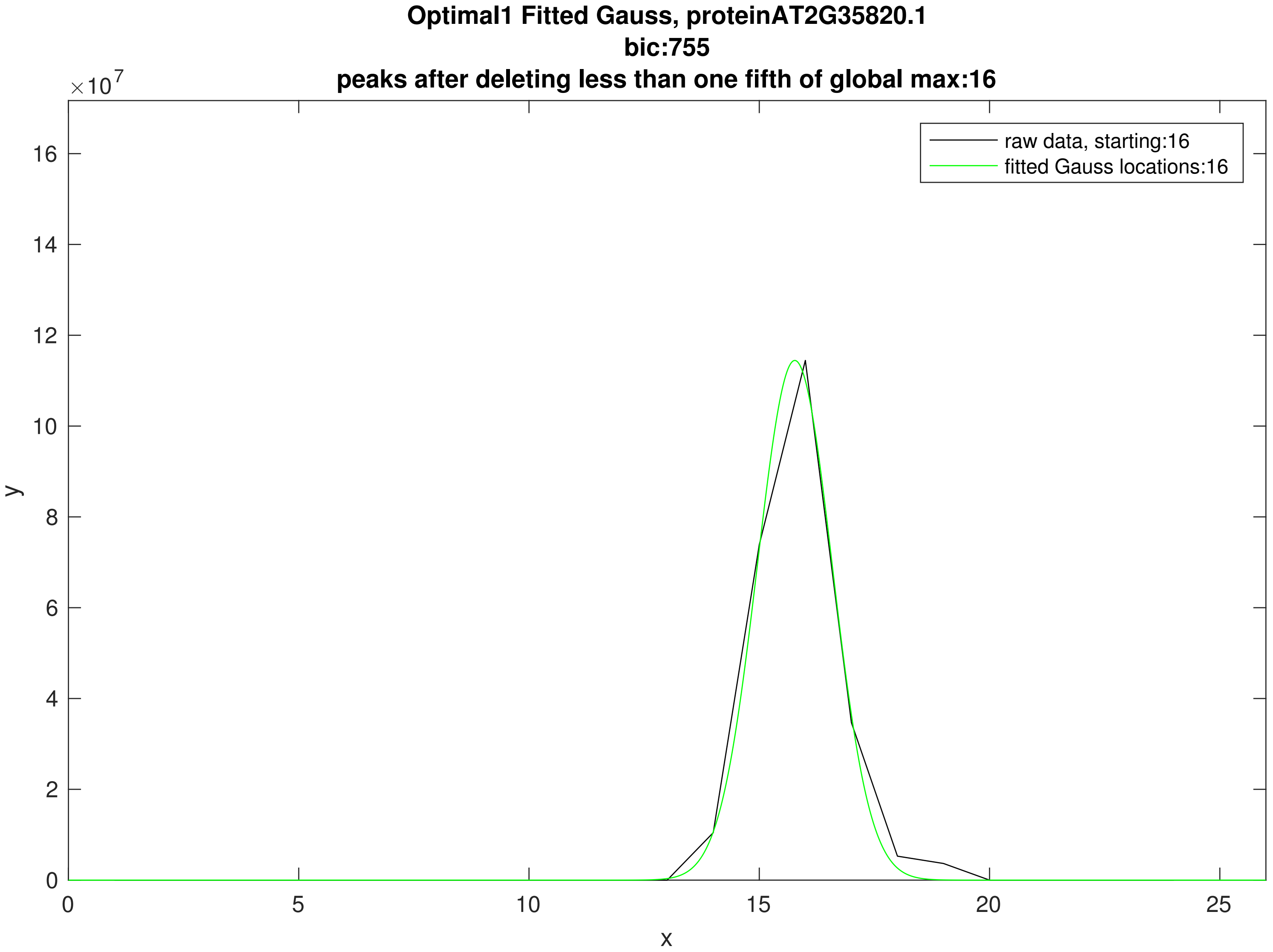
Optimal1 Fitted Gauss, proteinAT2G35820.1
bic:755
peaks after deleting less than one fifth of global max:16
×10^7
raw data, starting:16
fitted Gauss locations:16
y
x
0
2
4
6
8
10
12
14
16
0
5
10
15
20
25

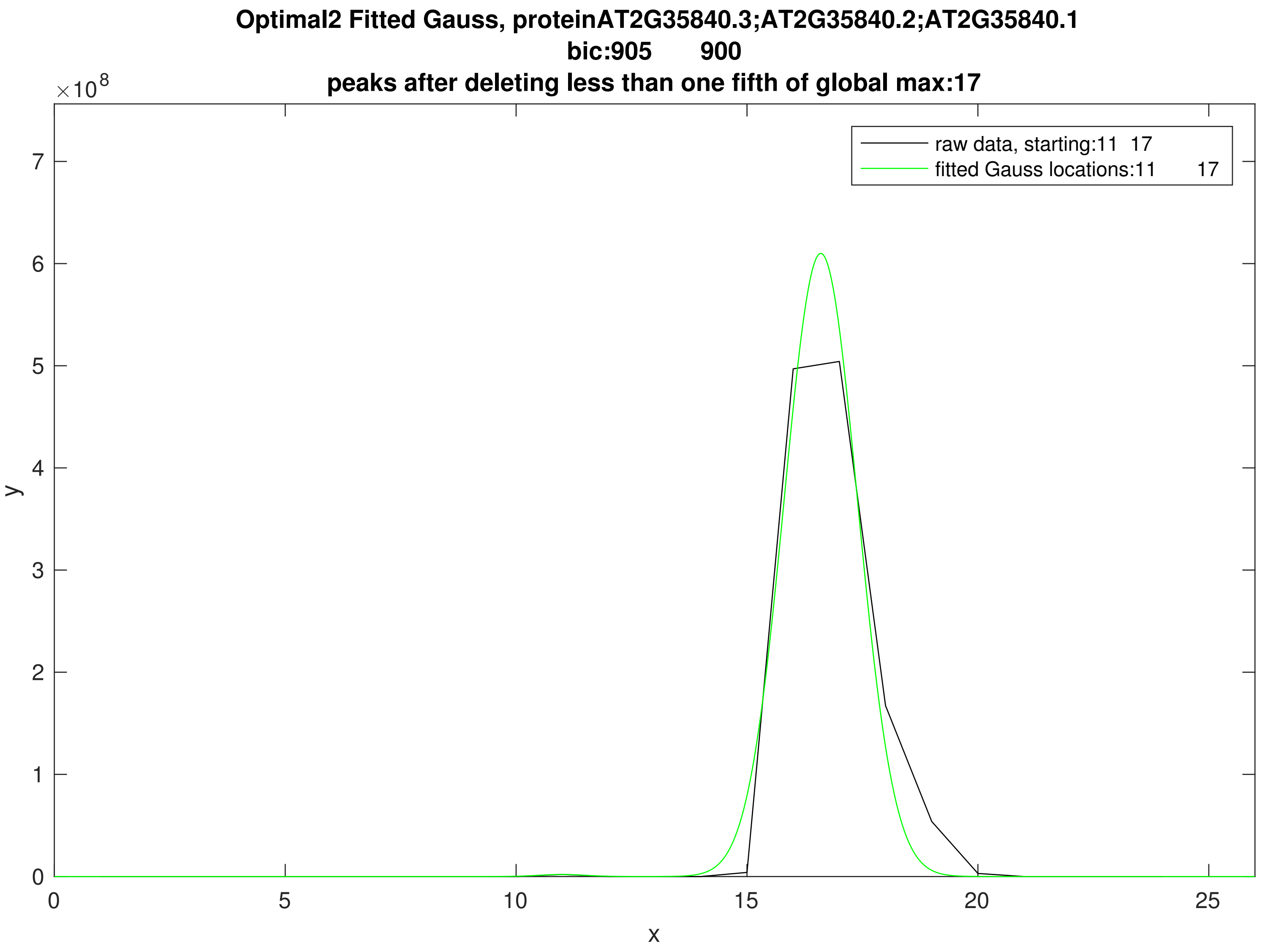
Optimal2 Fitted Gauss, proteinAT2G35840.3;AT2G35840.2;AT2G35840.1
bic:905 900
peaks after deleting less than one fifth of global max:17
×10^8
raw data, starting:11 17
fitted Gauss locations:11 17
y
x
0
1
2
3
4
5
6
7
0
5
10
15
20
25

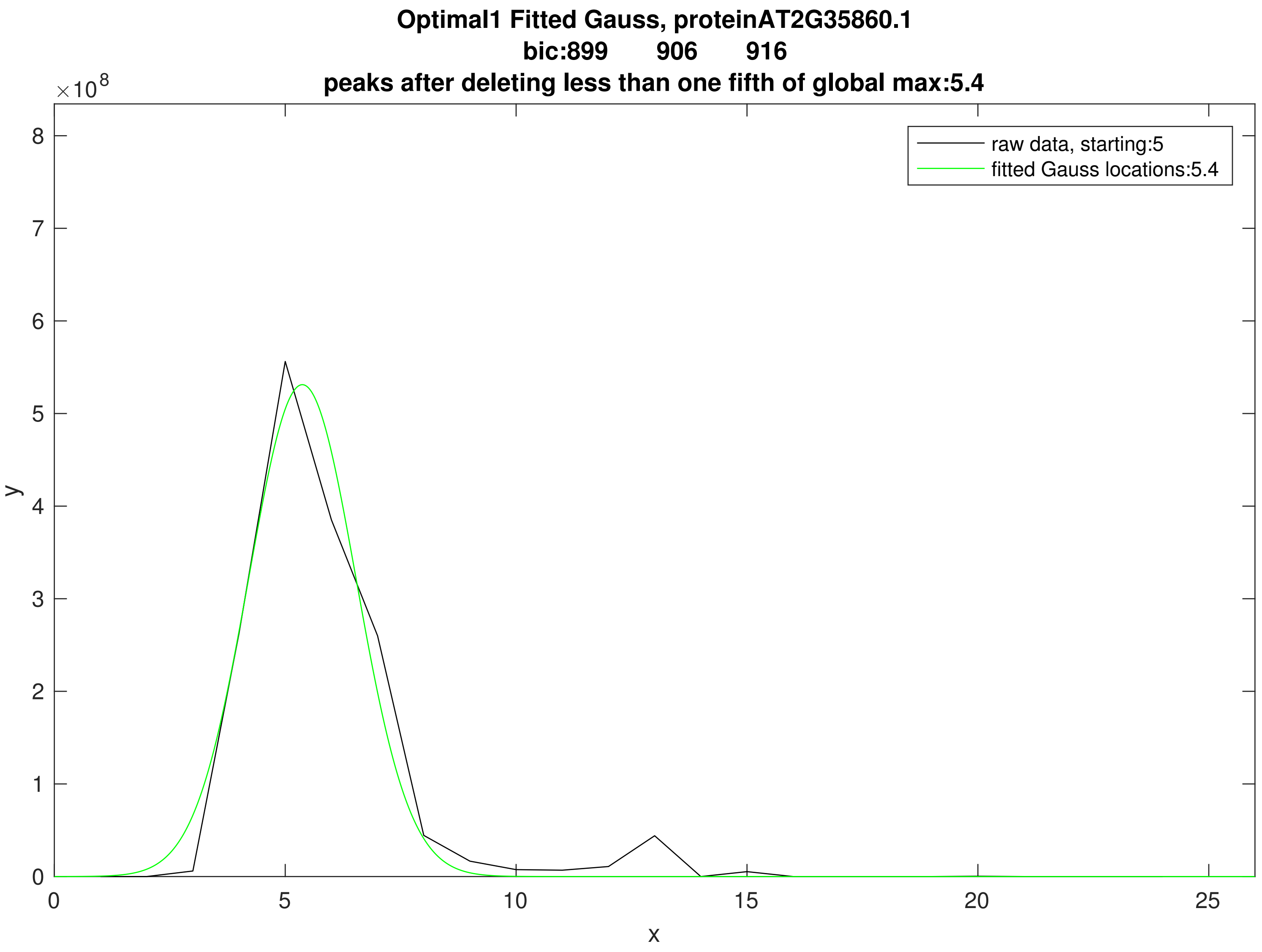

Optimal1 Fitted Gauss, proteinAT2G35860.1
bic:899 906 916
peaks after deleting less than one fifth of global max:5.4
×10^8
raw data, starting:5
fitted Gauss locations:5.4
y
x
0
1
2
3
4
5
6
7
8
10
15
20
25

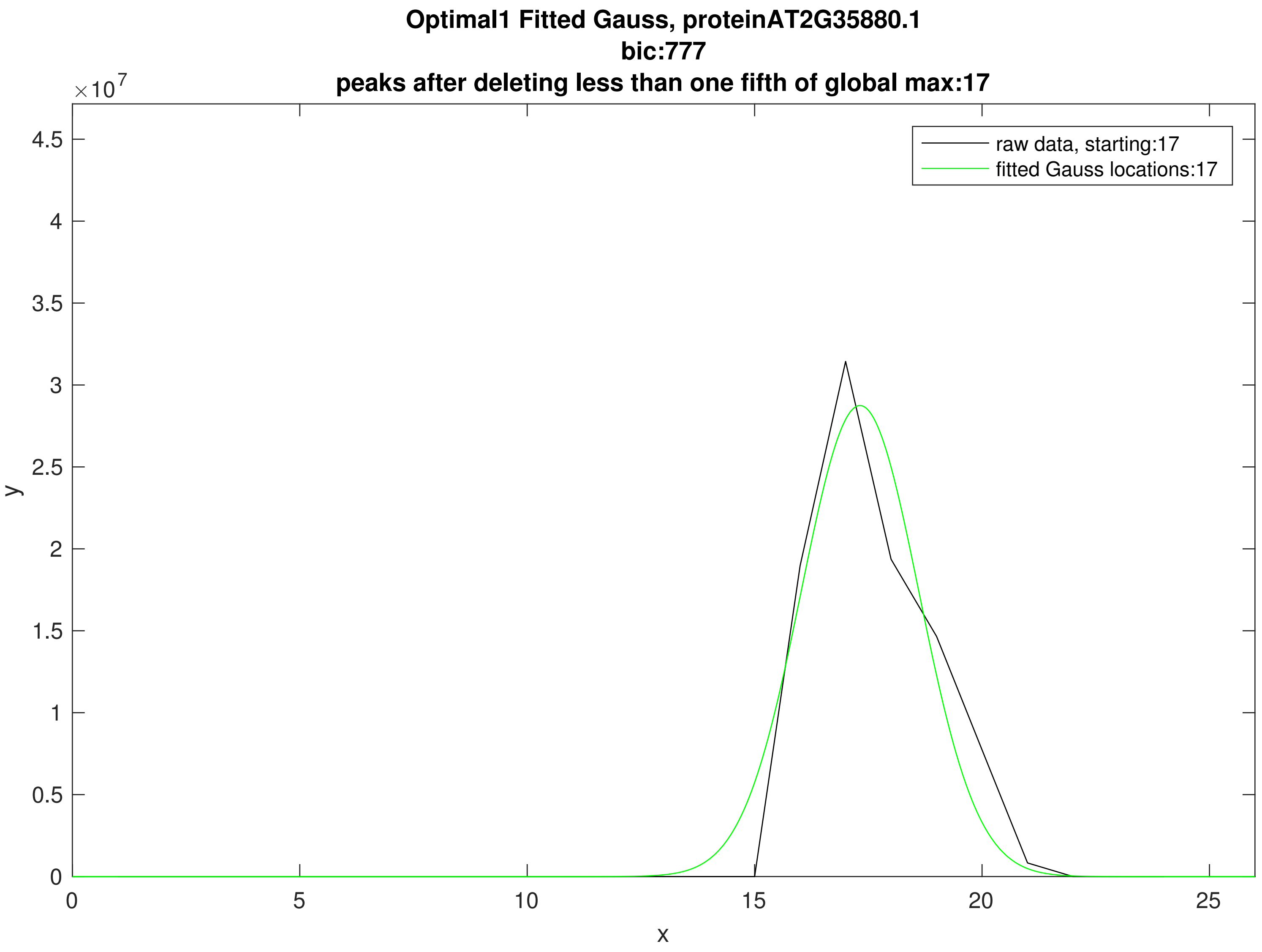
Optimal1 Fitted Gauss, proteinAT2G35880.1
bic:777
peaks after deleting less than one fifth of global max:17
$\times 10^7$
4.5
4
3.5
3
2.5
2
1.5
1
0.5
0
y
0
5
10
15
20
25
x
raw data, starting:17
fitted Gauss locations:17

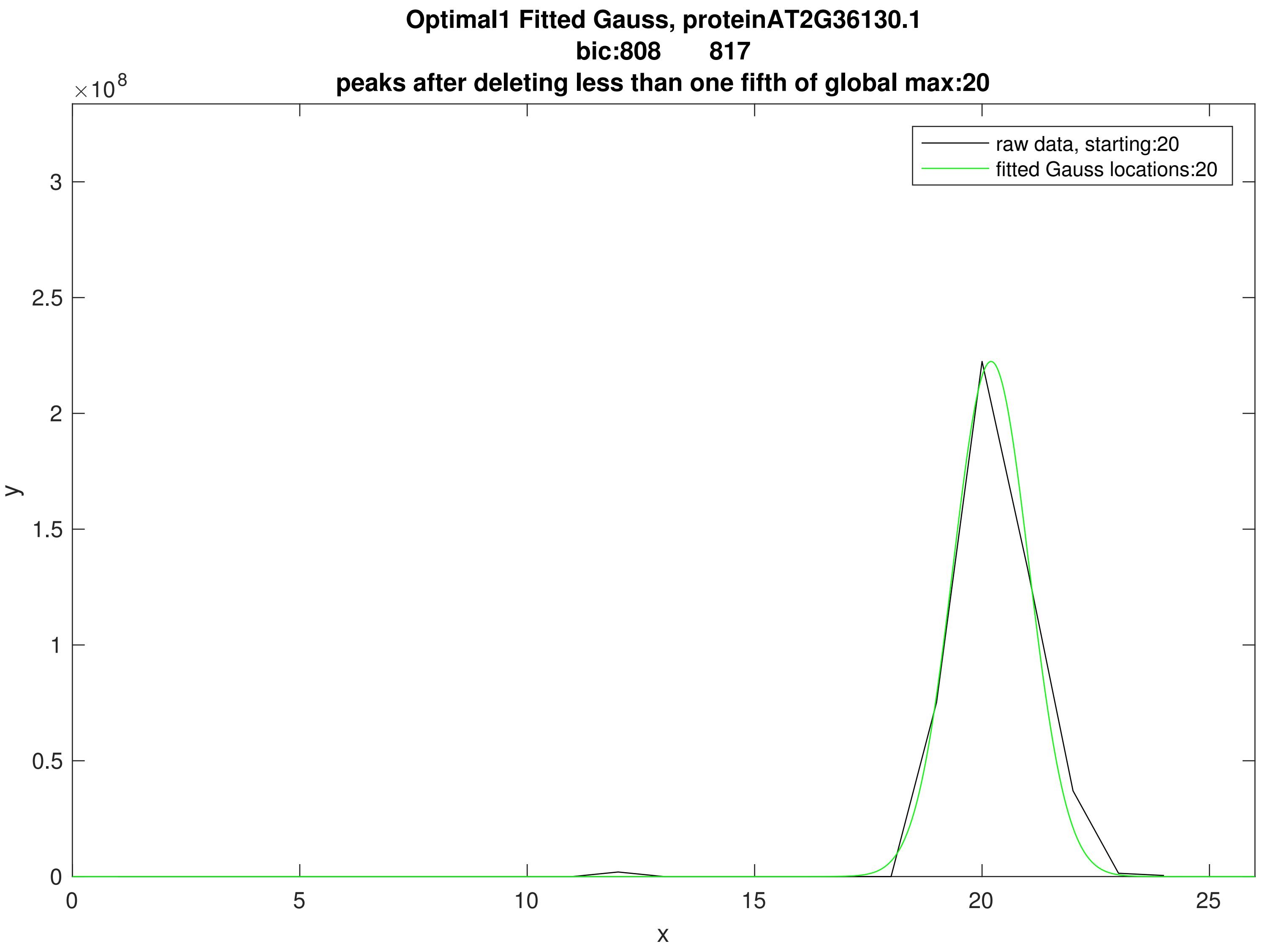

Optimal1 Fitted Gauss, proteinAT2G36130.1
bic:808 817
peaks after deleting less than one fifth of global max:20
×10^8
raw data, starting:20
fitted Gauss locations:20
y
x
0
0.5
1
1.5
2
2.5
3
0
5
10
15
20
25

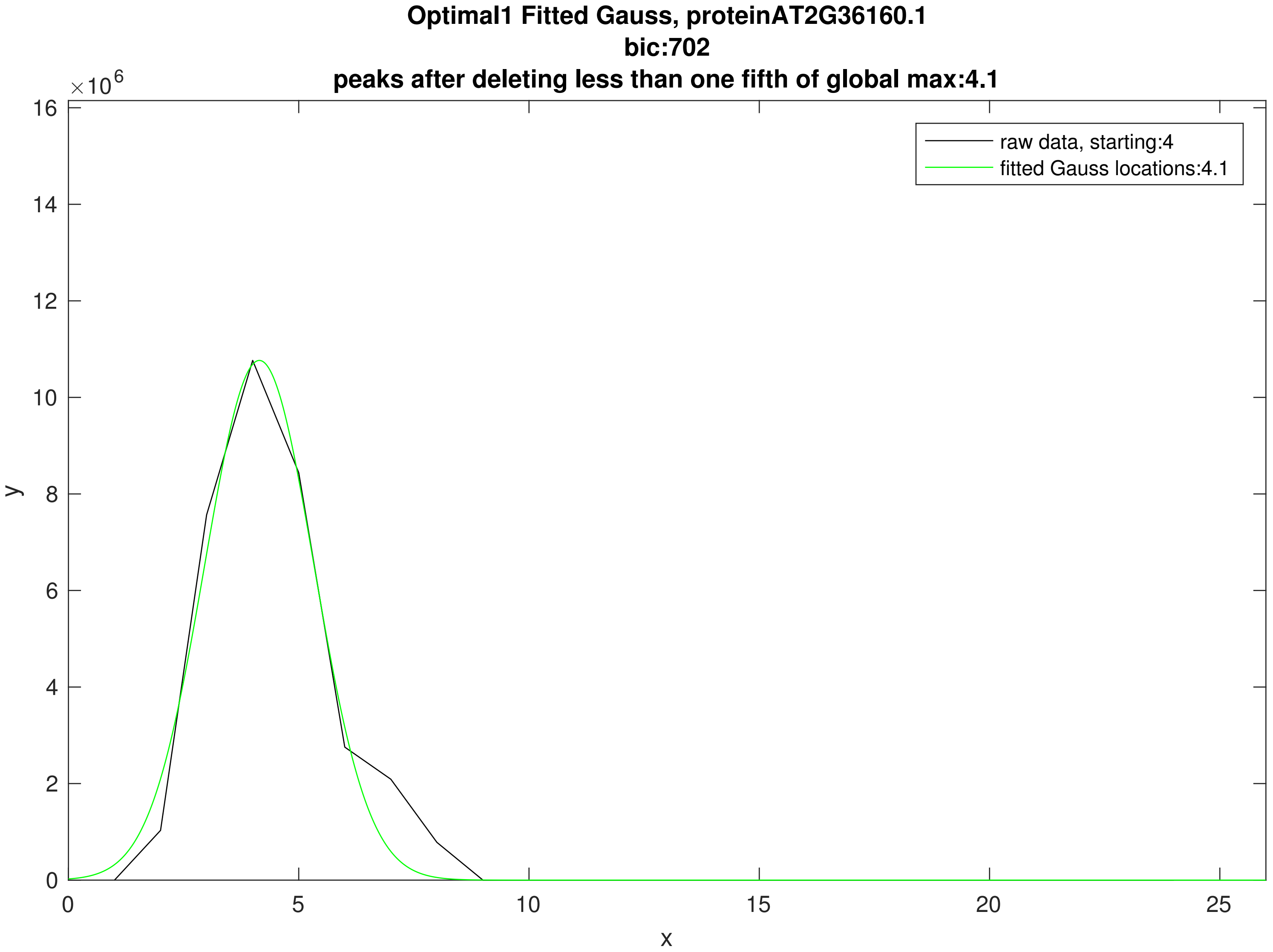
Optimal1 Fitted Gauss, proteinAT2G36160.1
bic:702
peaks after deleting less than one fifth of global max:4.1
×10^6
16
14
12
10
8
6
4
2
0
y
0
5
10
15
20
25
x
raw data, starting:4
fitted Gauss locations:4.1

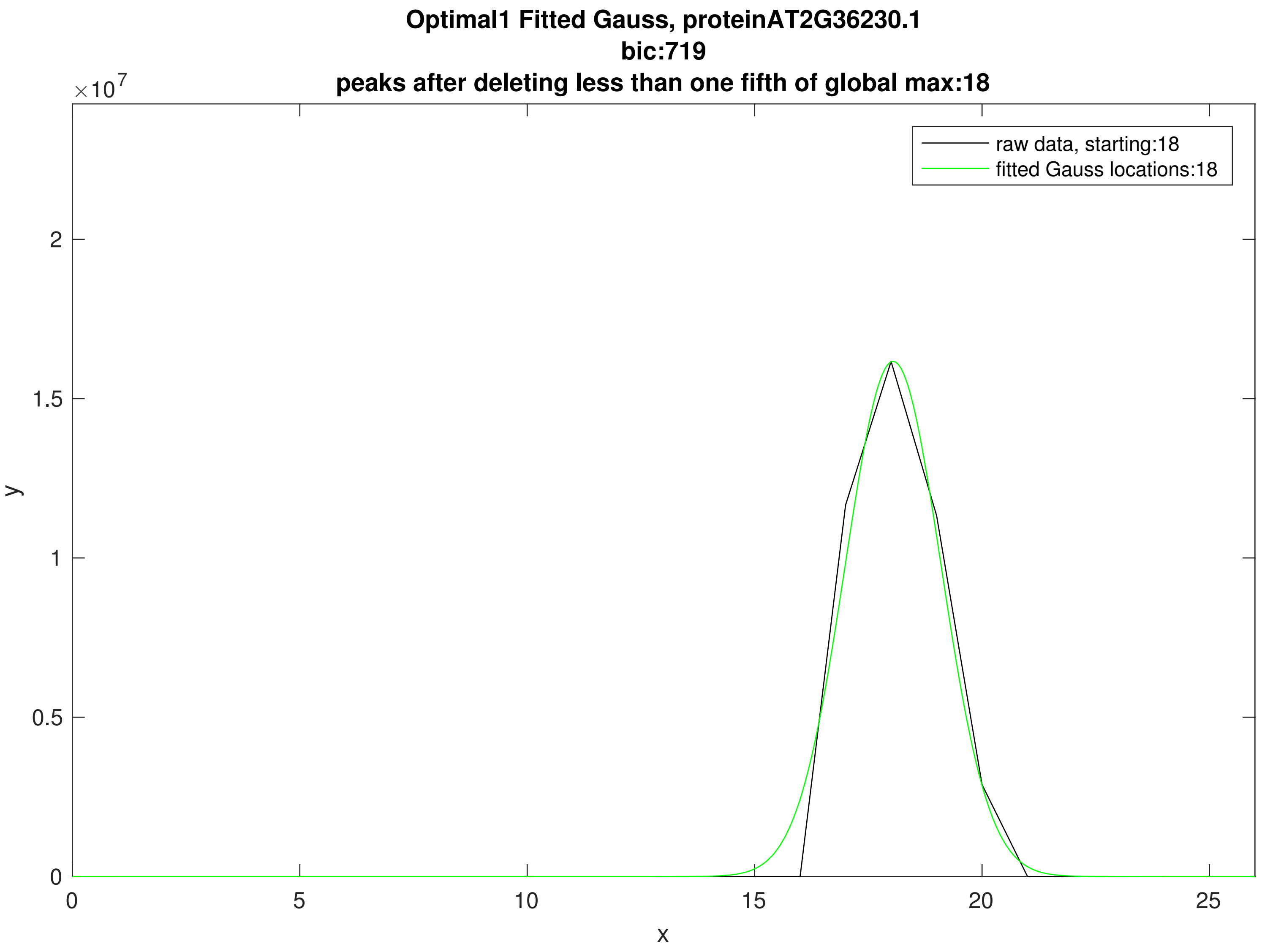
Optimal1 Fitted Gauss, proteinAT2G36230.1
bic:719
peaks after deleting less than one fifth of global max:18
×10^7
raw data, starting:18
fitted Gauss locations:18
2
1.5
1
0.5
0
y
0
5
10
15
20
25
x

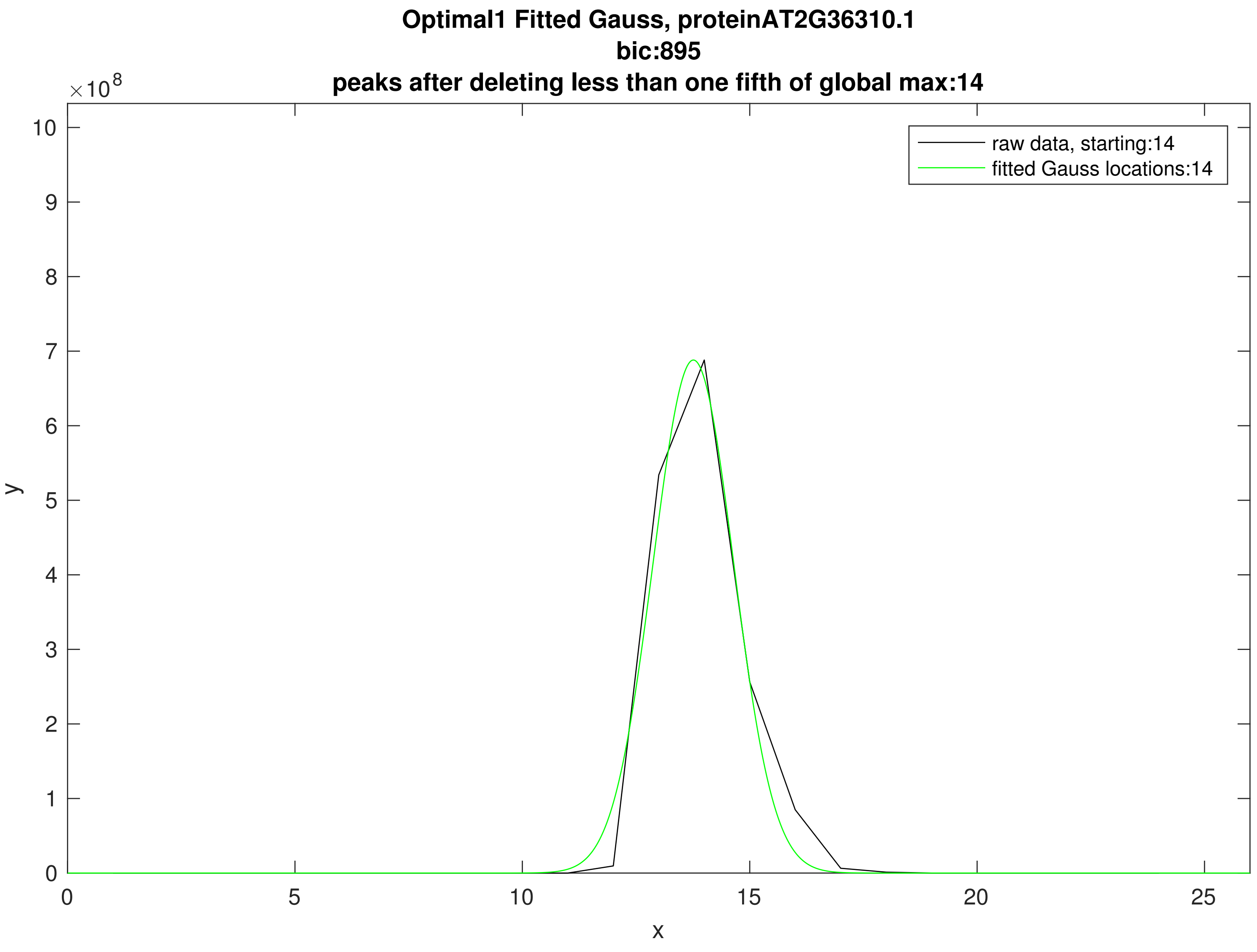

Optimal1 Fitted Gauss, proteinAT2G36310.1
bic:895
peaks after deleting less than one fifth of global max:14
×10^8
raw data, starting:14
fitted Gauss locations:14
y
x
0
1
2
3
4
5
6
7
8
9
10
0
5
10
15
20
25

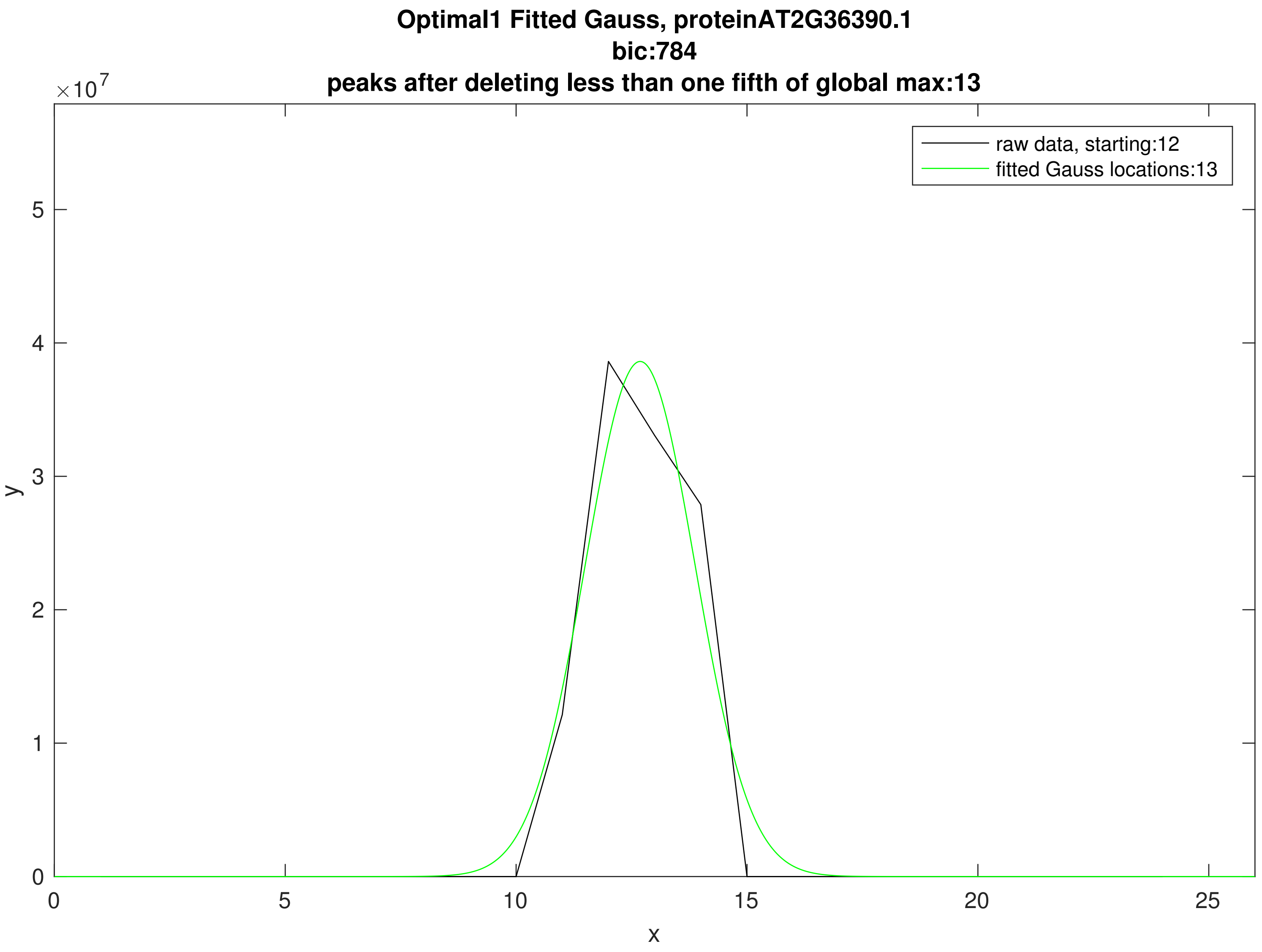

Optimal1 Fitted Gauss, proteinAT2G36390.1
bic:784
peaks after deleting less than one fifth of global max:13
×10^7
raw data, starting:12
fitted Gauss locations:13
y
x
0
1
2
3
4
5
0
5
10
15
20
25

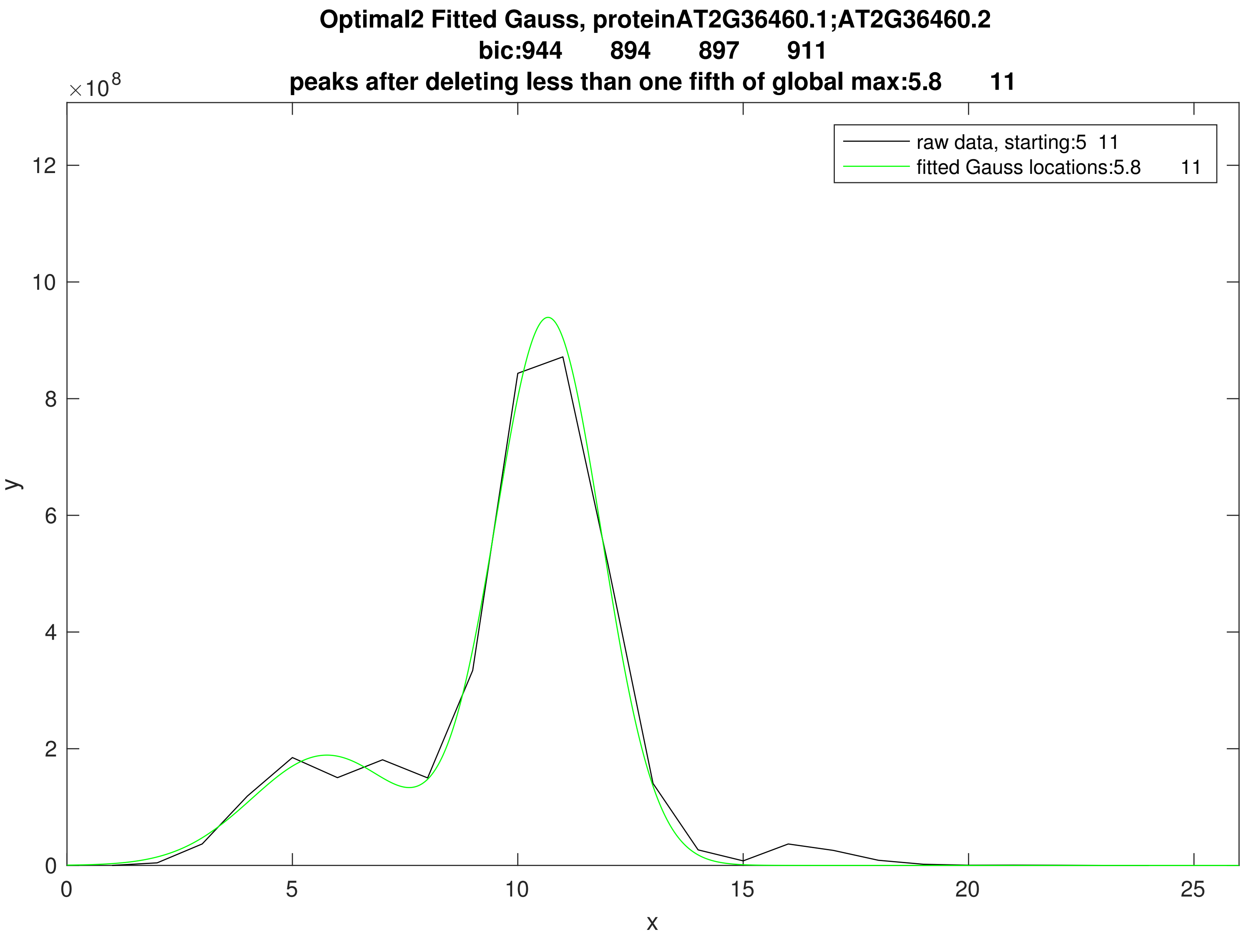
Optimal2 Fitted Gauss, proteinAT2G36460.1;AT2G36460.2
bic:944 894 897 911
peaks after deleting less than one fifth of global max:5.8 11
×10^8
raw data, starting:5 11
fitted Gauss locations:5.8 11
y
x
0
2
4
6
8
10
12
0
5
10
15
20
25

**Optimal1 Fitted Gauss, proteinAT2G36530.1**
**bic:1.08e+03  1.09e+03   1.1e+03  1.11e+03**
**peaks after deleting less than one fifth of global max:14**

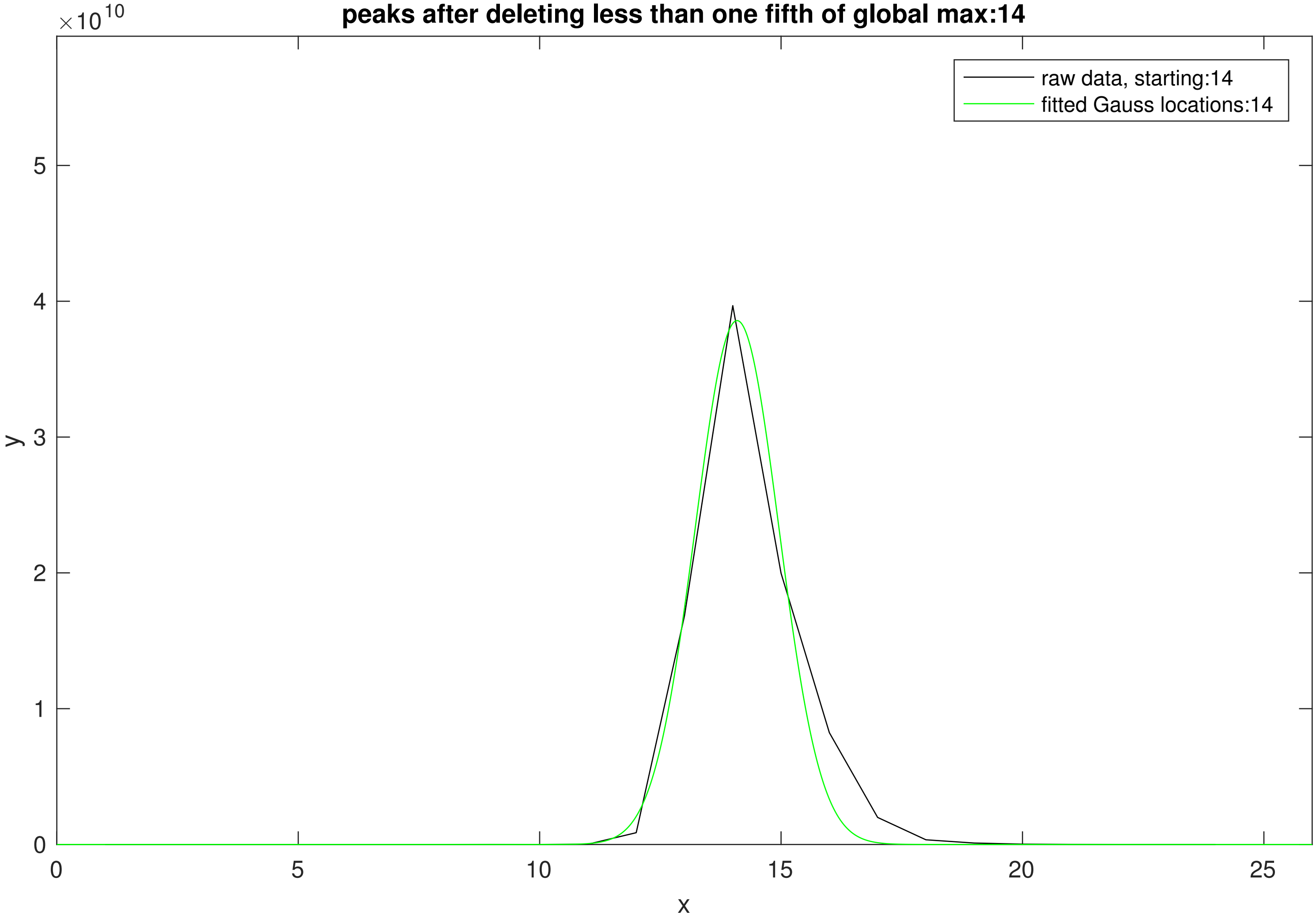

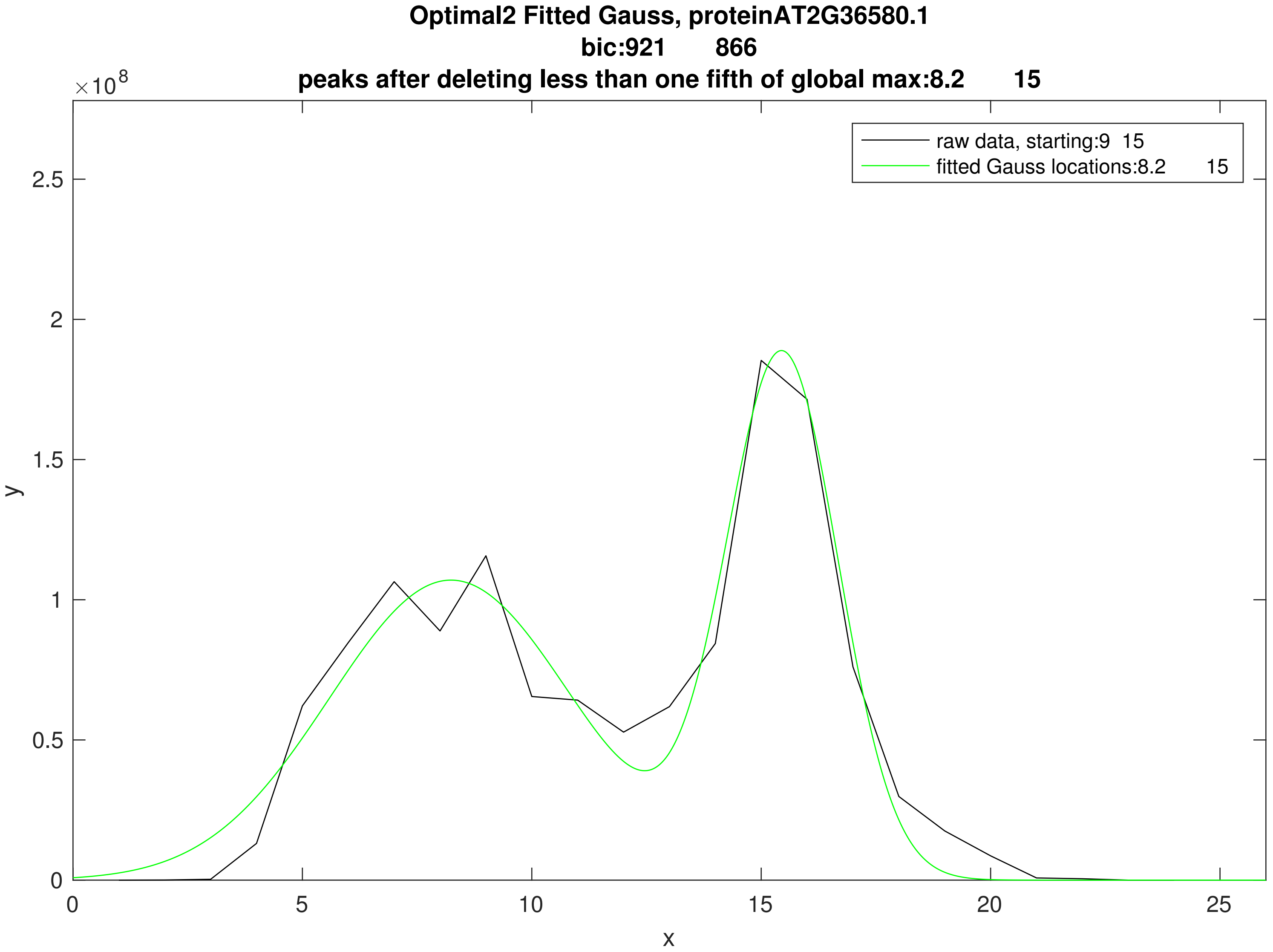
Optimal2 Fitted Gauss, proteinAT2G36580.1
bic:921 866
peaks after deleting less than one fifth of global max:8.2 15
×10^8
raw data, starting:9 15
fitted Gauss locations:8.2 15
y
x
0
0.5
1
1.5
2
2.5
5
10
15
20
25

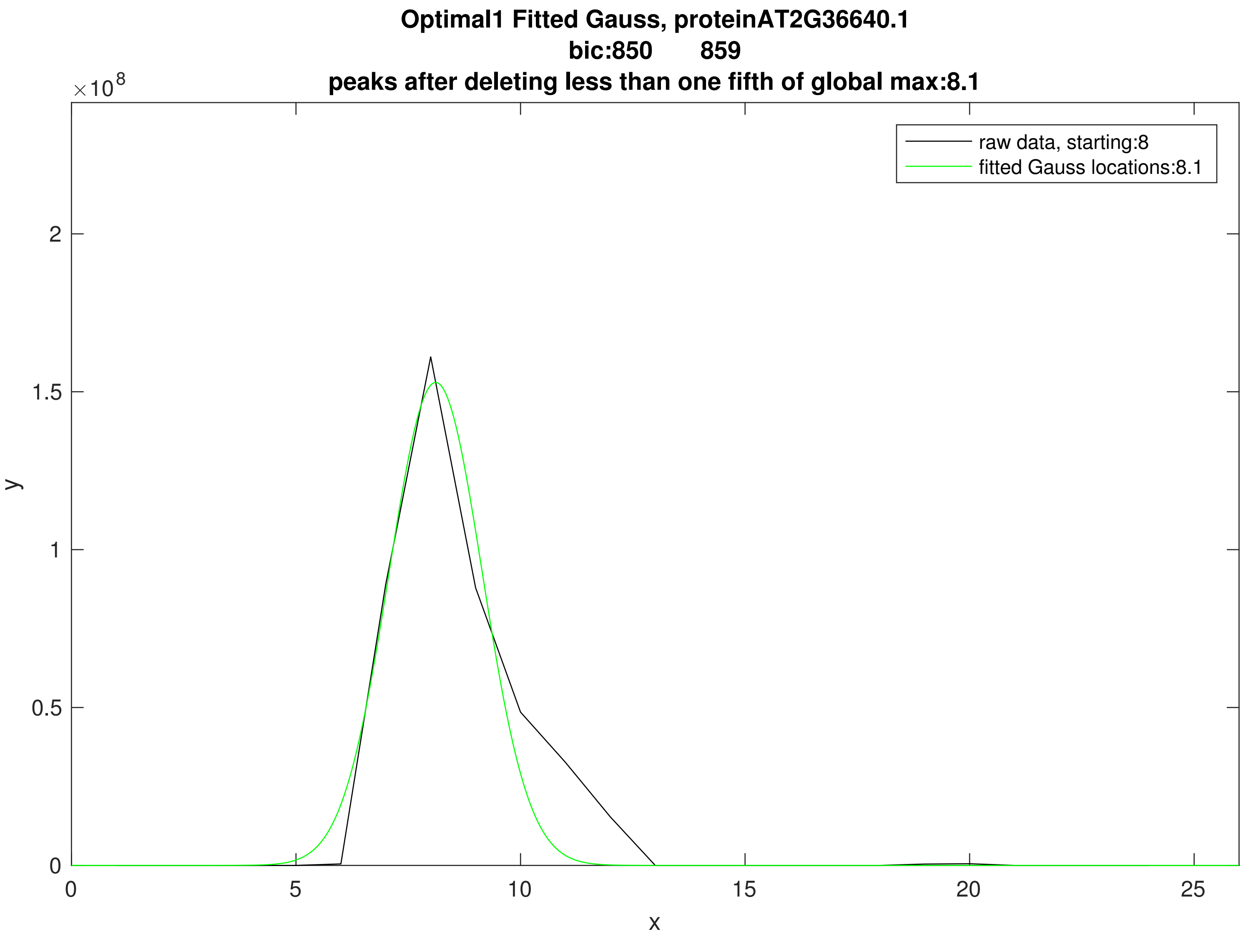

Optimal1 Fitted Gauss, proteinAT2G36640.1
bic:850 859
peaks after deleting less than one fifth of global max:8.1
×10^8
raw data, starting:8
fitted Gauss locations:8.1
y
x
0
0.5
1
1.5
2
0
5
10
15
20
25

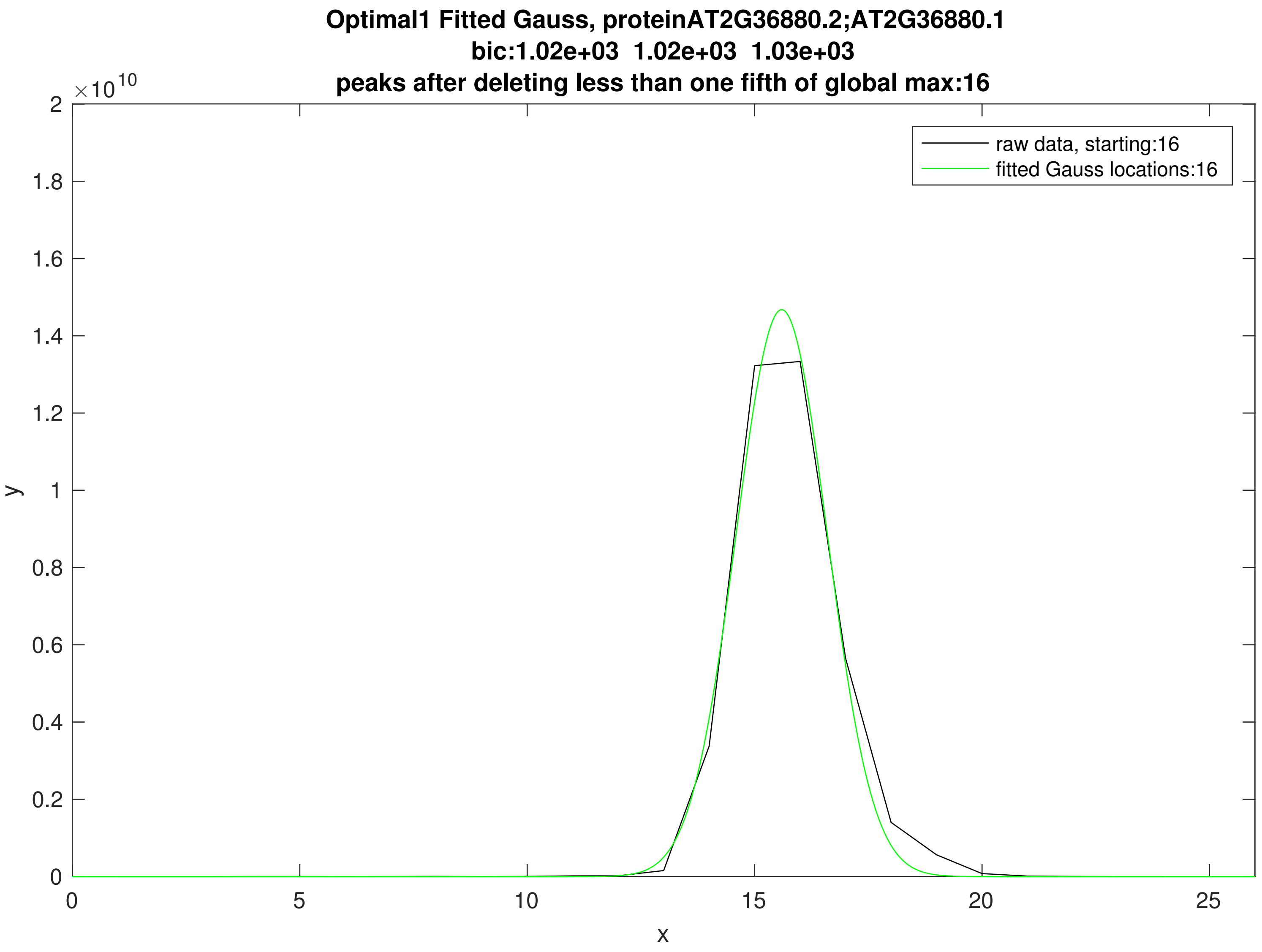
Optimal1 Fitted Gauss, proteinAT2G36880.2;AT2G36880.1
bic:1.02e+03 1.02e+03 1.03e+03
peaks after deleting less than one fifth of global max:16
×10^10
raw data, starting:16
fitted Gauss locations:16
y
x
0
0.2
0.4
0.6
0.8
1
1.2
1.4
1.6
1.8
2
0
5
10
15
20
25

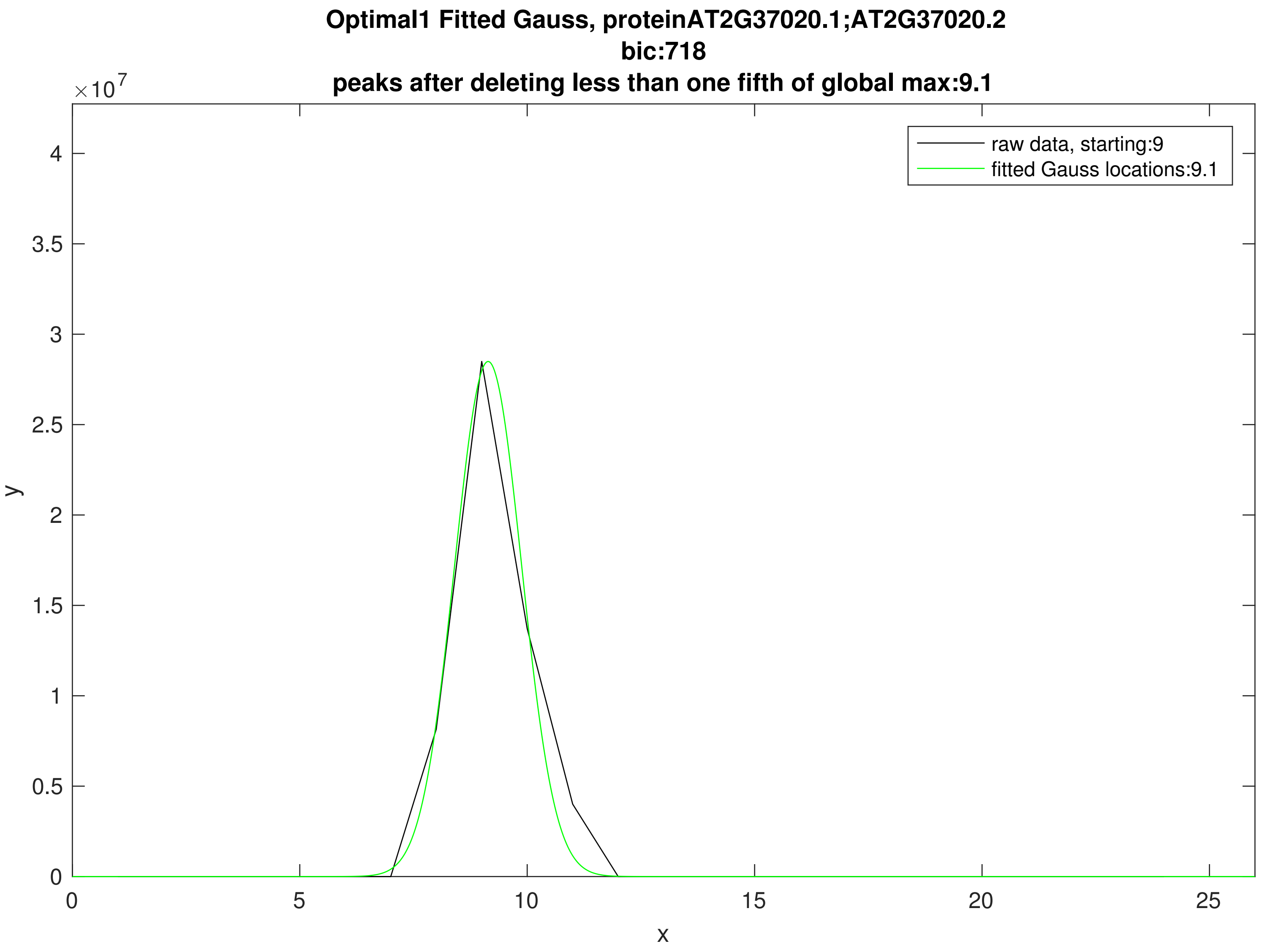

Optimal1 Fitted Gauss, proteinAT2G37020.1;AT2G37020.2
bic:718
peaks after deleting less than one fifth of global max:9.1
×10^7
raw data, starting:9
fitted Gauss locations:9.1
y
x
0
0.5
1
1.5
2
2.5
3
3.5
4
0
5
10
15
20
25

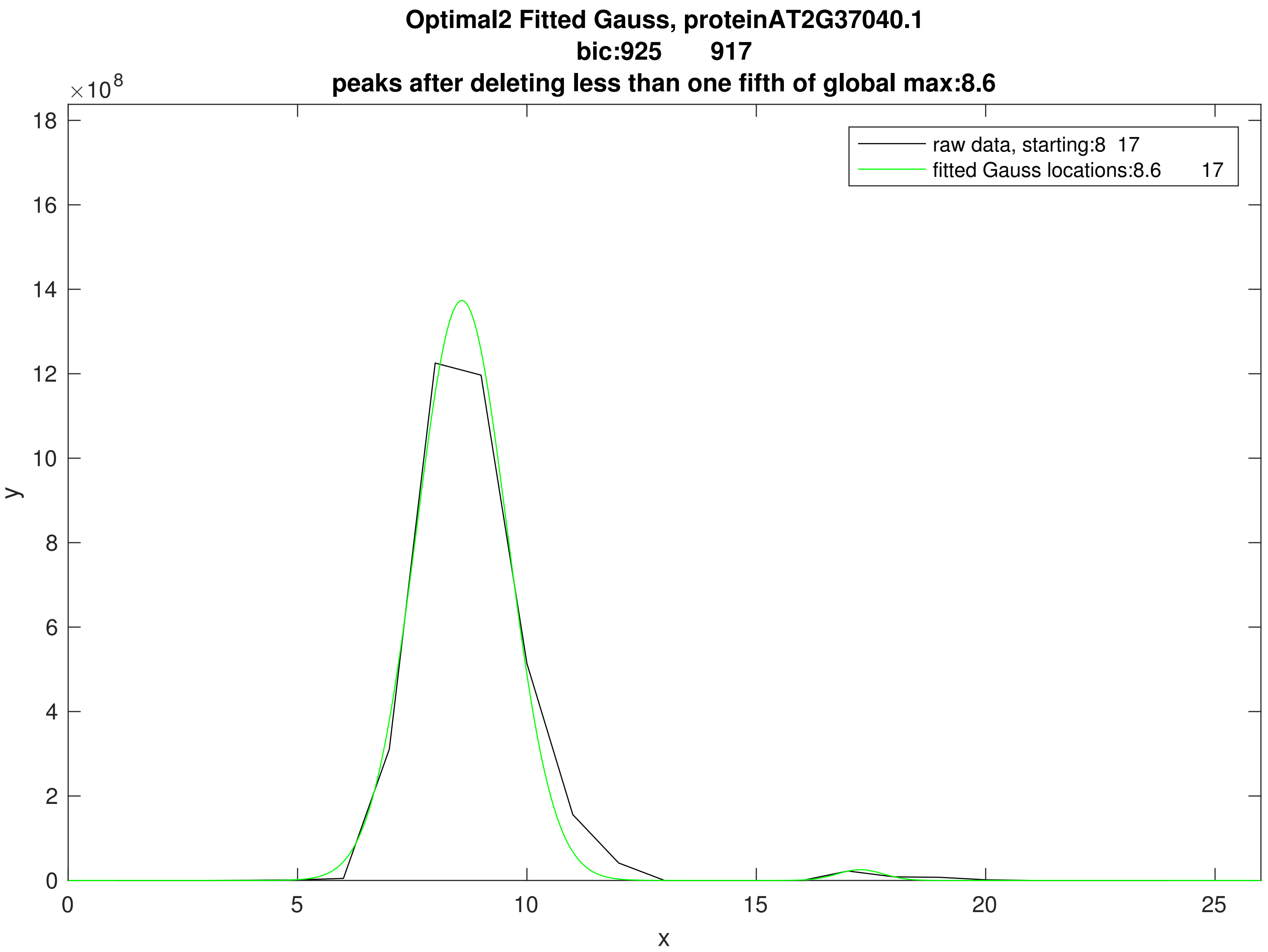
Optimal2 Fitted Gauss, proteinAT2G37040.1
bic:925 917
peaks after deleting less than one fifth of global max:8.6
×10^8
raw data, starting:8 17
fitted Gauss locations:8.6 17
y
x
0
2
4
6
8
10
12
14
16
18
0
5
10
15
20
25

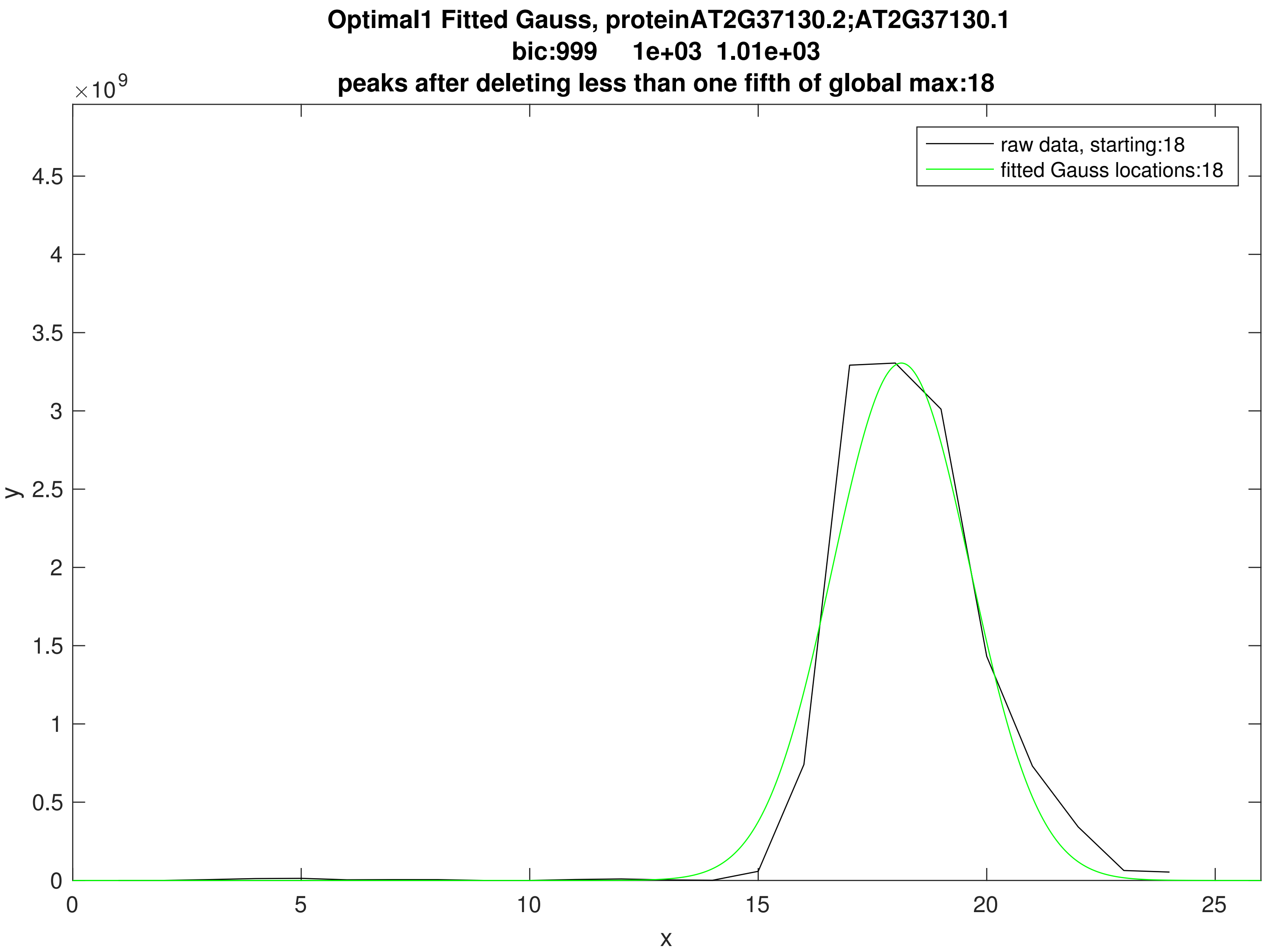
Optimal1 Fitted Gauss, proteinAT2G37130.2;AT2G37130.1
bic:999 1e+03 1.01e+03
peaks after deleting less than one fifth of global max:18
×10^9
raw data, starting:18
fitted Gauss locations:18
y
x
0
0.5
1
1.5
2
2.5
3
3.5
4
4.5
5
10
15
20
25

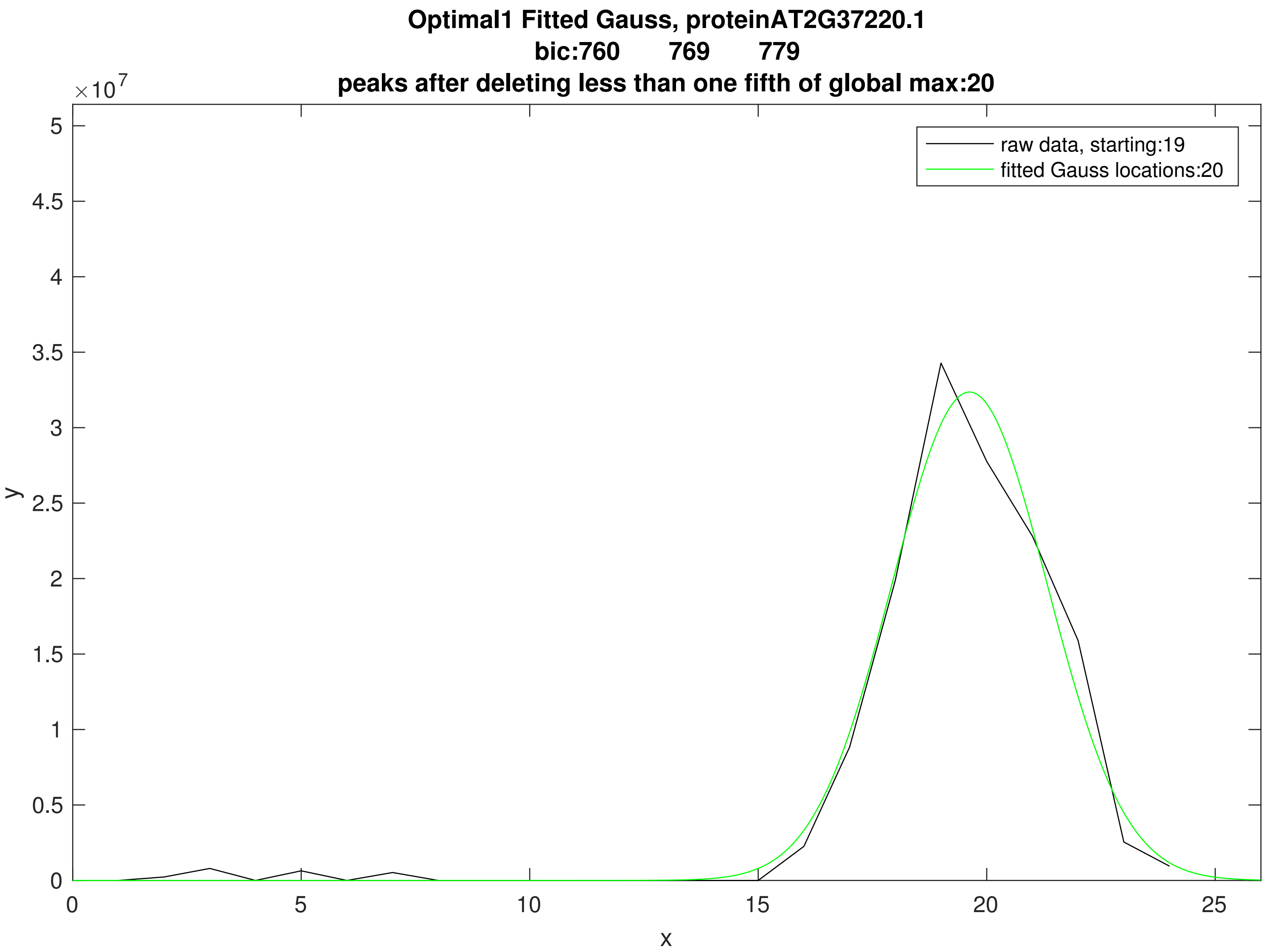

Optimal1 Fitted Gauss, proteinAT2G37220.1
bic:760 769 779
peaks after deleting less than one fifth of global max:20
×10^7
raw data, starting:19
fitted Gauss locations:20
y
x
0
0.5
1
1.5
2
2.5
3
3.5
4
4.5
5
0
5
10
15
20
25

**Optimal1 Fitted Gauss, proteinAT2G37230.1**
**bic:643**
**peaks after deleting less than one fifth of global max:3.6**

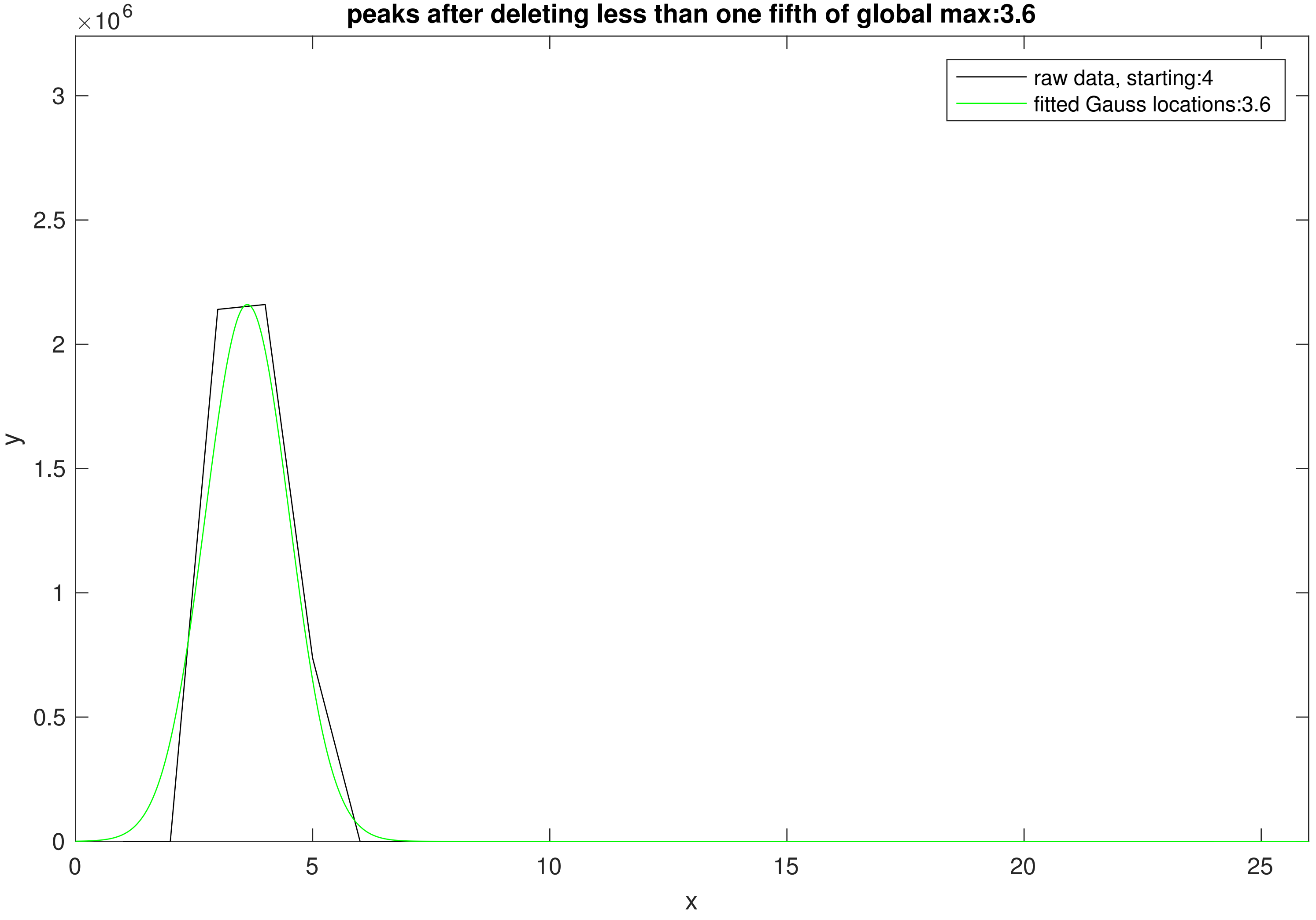

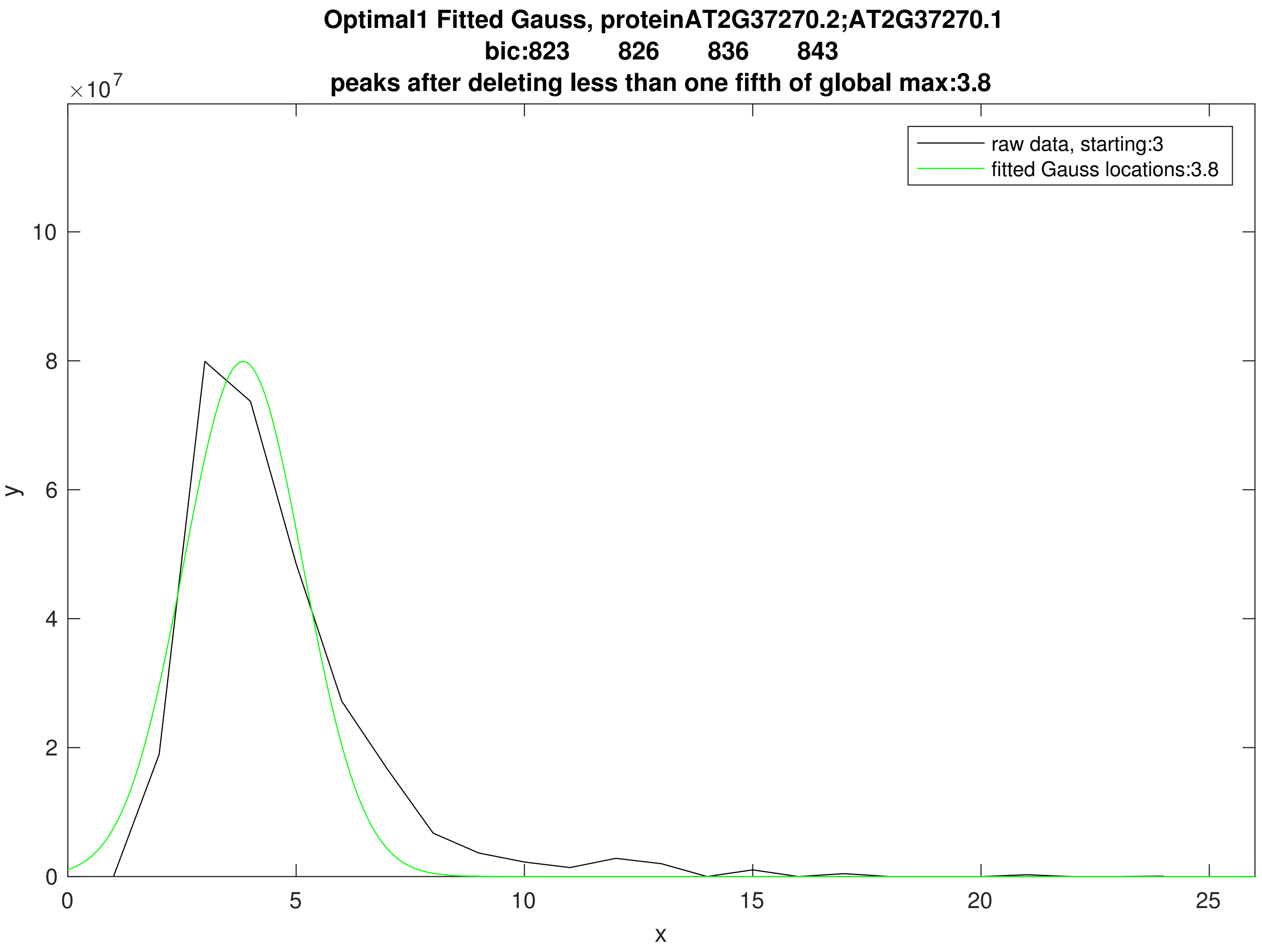

Optimal1 Fitted Gauss, proteinAT2G37270.2;AT2G37270.1
bic:823 826 836 843
peaks after deleting less than one fifth of global max:3.8
×10^7
raw data, starting:3
fitted Gauss locations:3.8
y
x
0
2
4
6
8
10
5
15
20
25

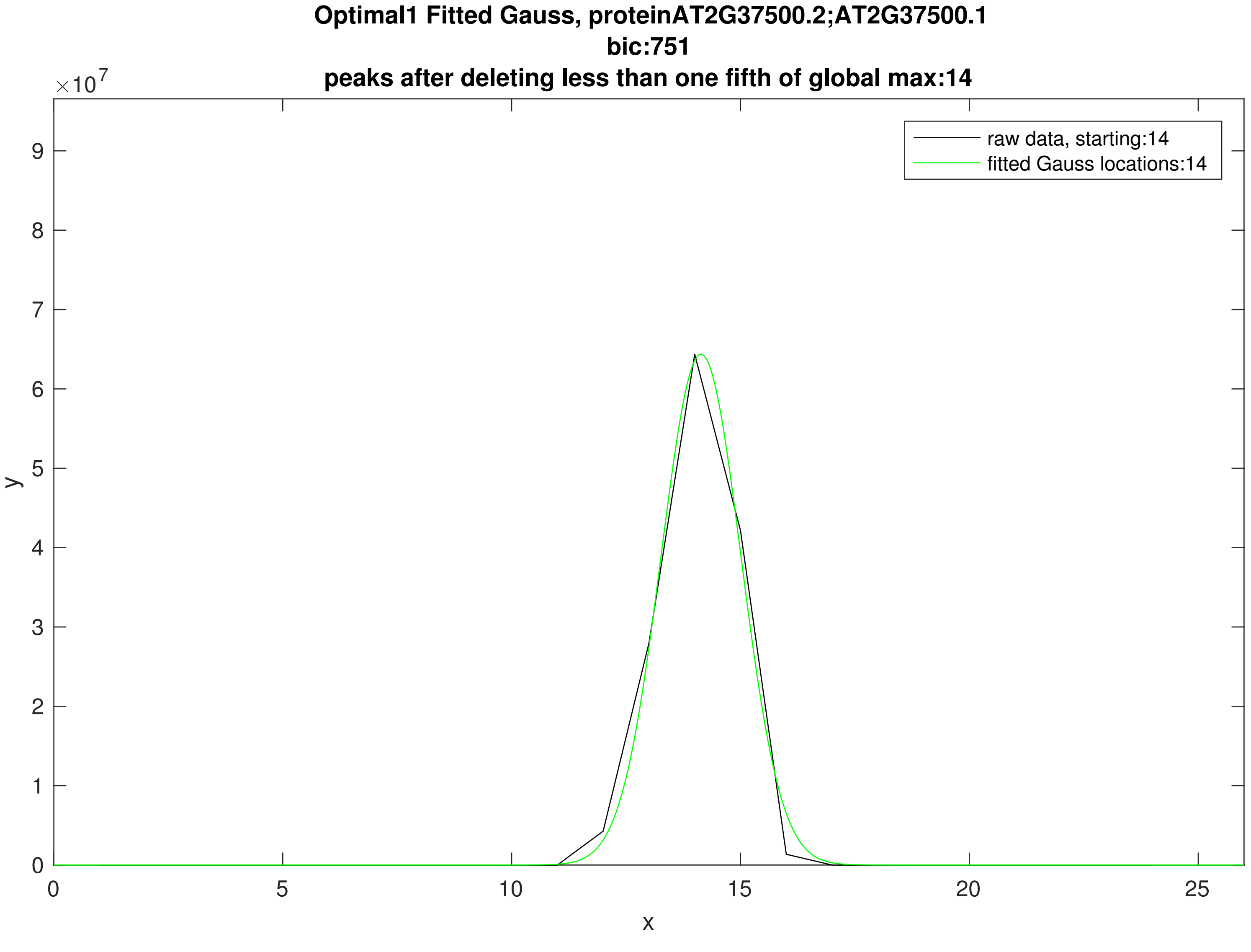
Optimal1 Fitted Gauss, proteinAT2G37500.2;AT2G37500.1
bic:751
peaks after deleting less than one fifth of global max:14
×10^7
raw data, starting:14
fitted Gauss locations:14
y
x
0
1
2
3
4
5
6
7
8
9
0
5
10
15
20
25

Optimal1 Fitted Gauss, proteinAT2G37550.2;AT2G37550.1
bic:674
peaks after deleting less than one fifth of global max:20

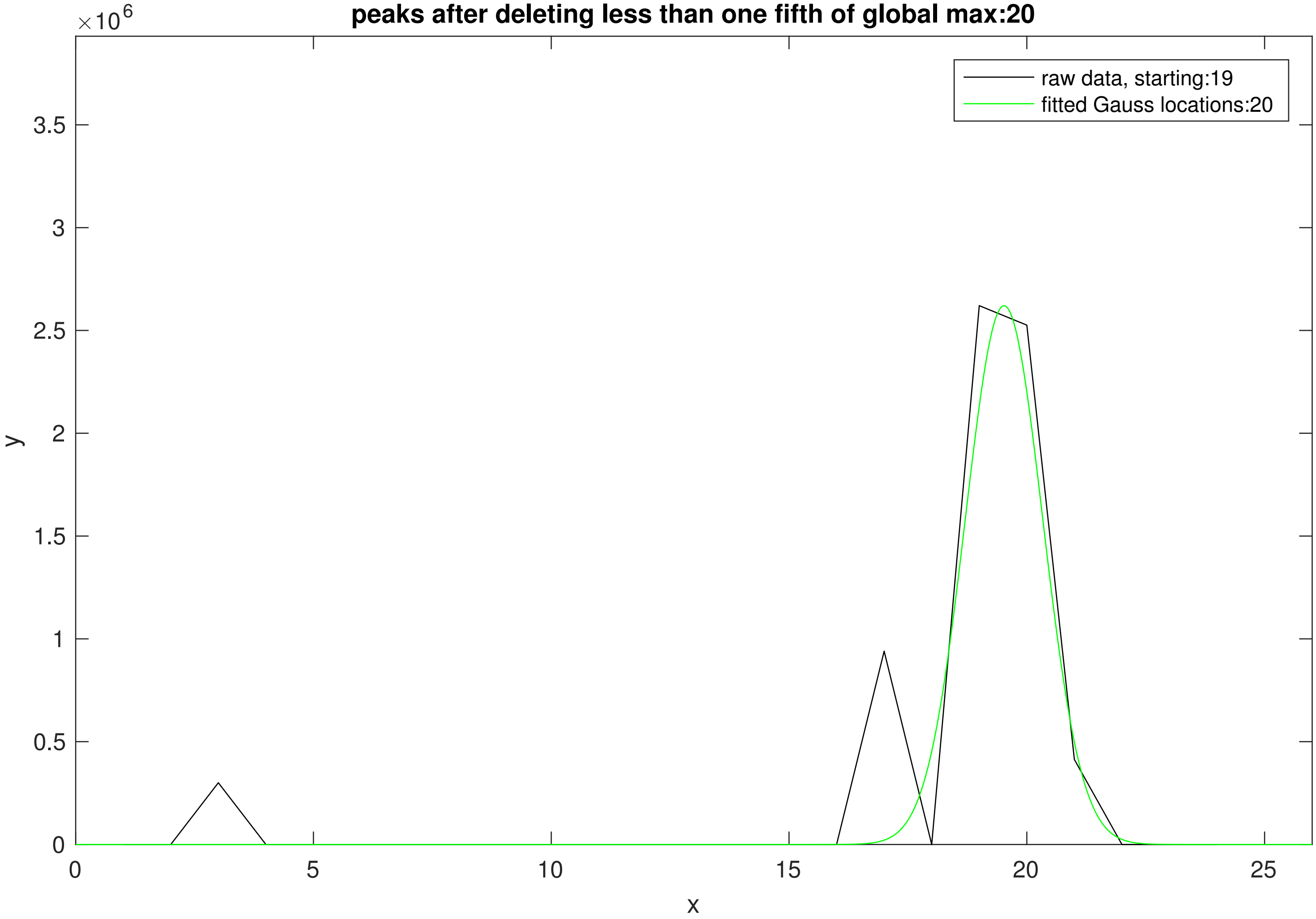

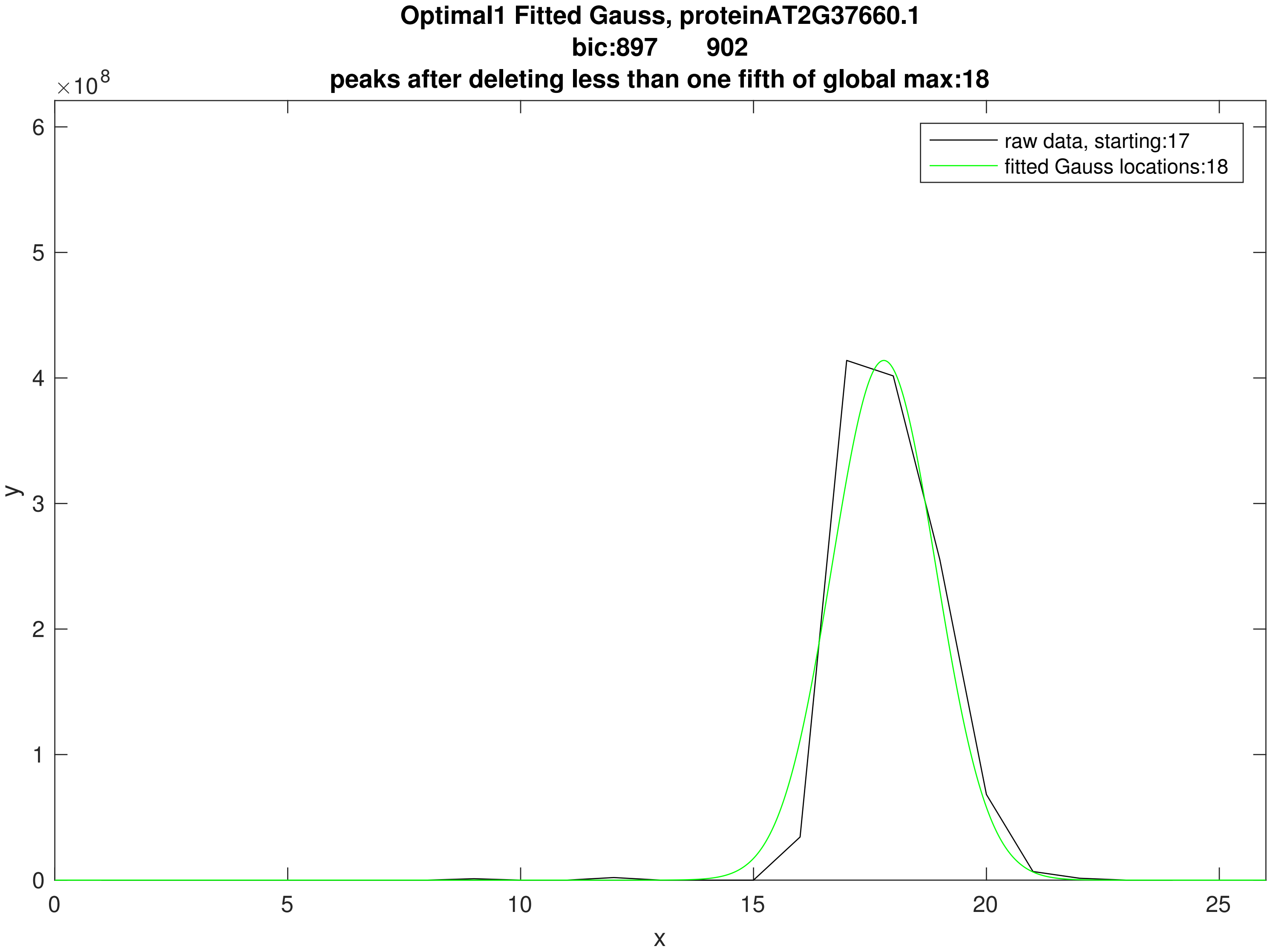
Optimal1 Fitted Gauss, proteinAT2G37660.1
bic:897 902
peaks after deleting less than one fifth of global max:18
×10^8
raw data, starting:17
fitted Gauss locations:18
6
5
4
3
2
1
0
y
0
5
10
15
20
25
x

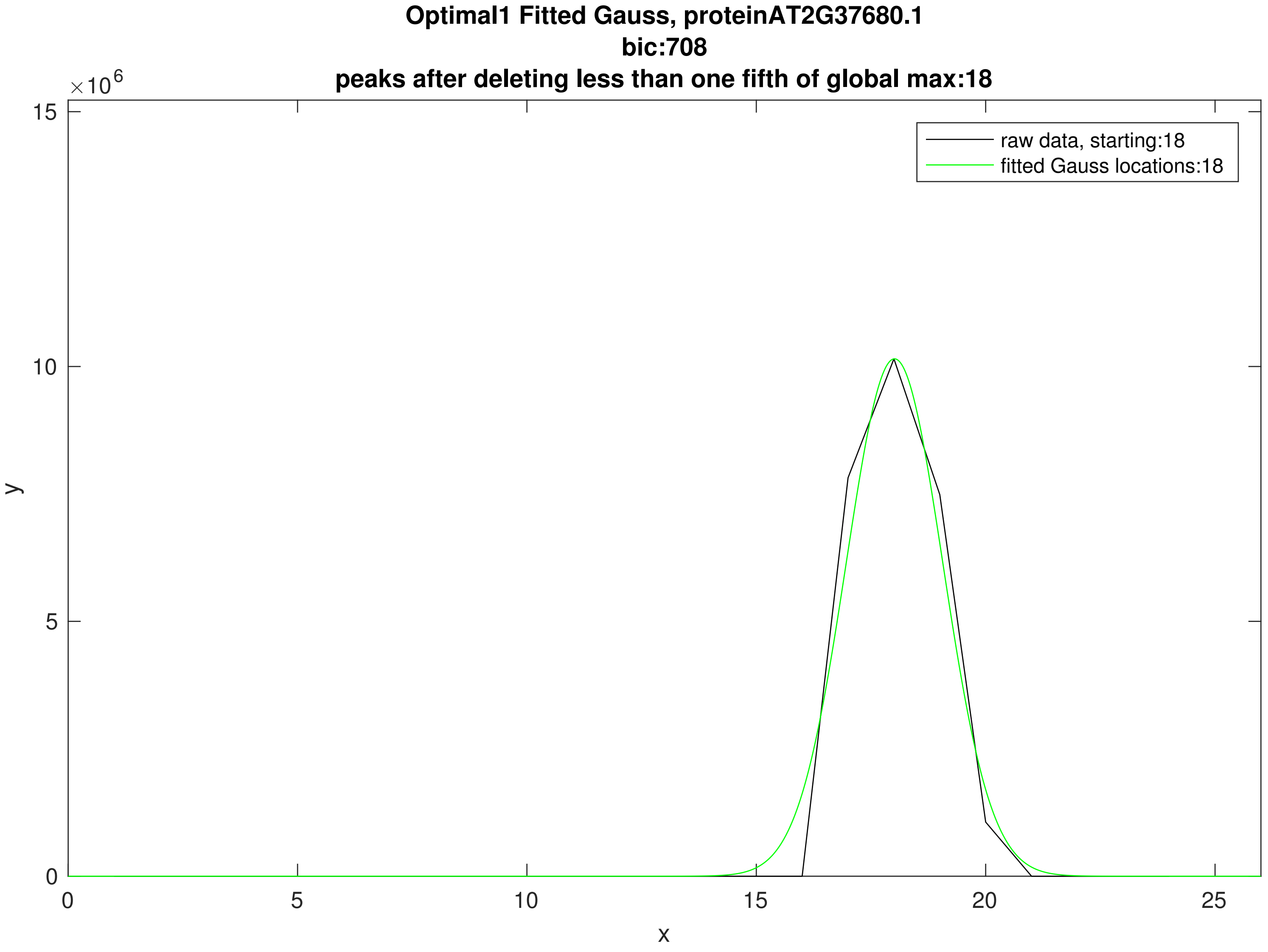
Optimal1 Fitted Gauss, proteinAT2G37680.1
bic:708
peaks after deleting less than one fifth of global max:18
×10^6
15
10
5
0
y
0
5
10
15
20
25
x
raw data, starting:18
fitted Gauss locations:18

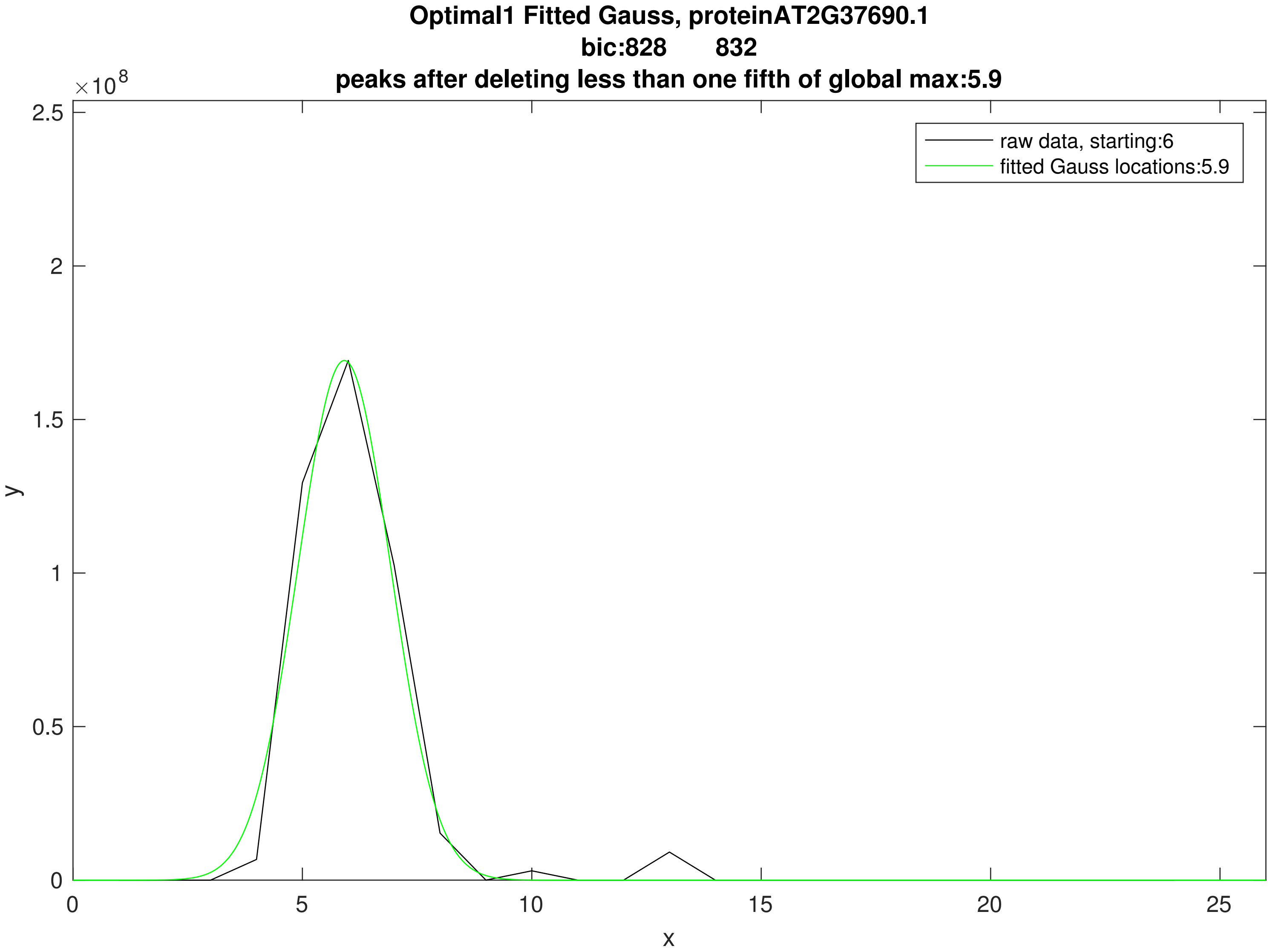
Optimal1 Fitted Gauss, proteinAT2G37690.1
bic:828 832
peaks after deleting less than one fifth of global max:5.9
×10^8
2.5
2
1.5
1
0.5
0
y
0
5
10
15
20
25
x
raw data, starting:6
fitted Gauss locations:5.9

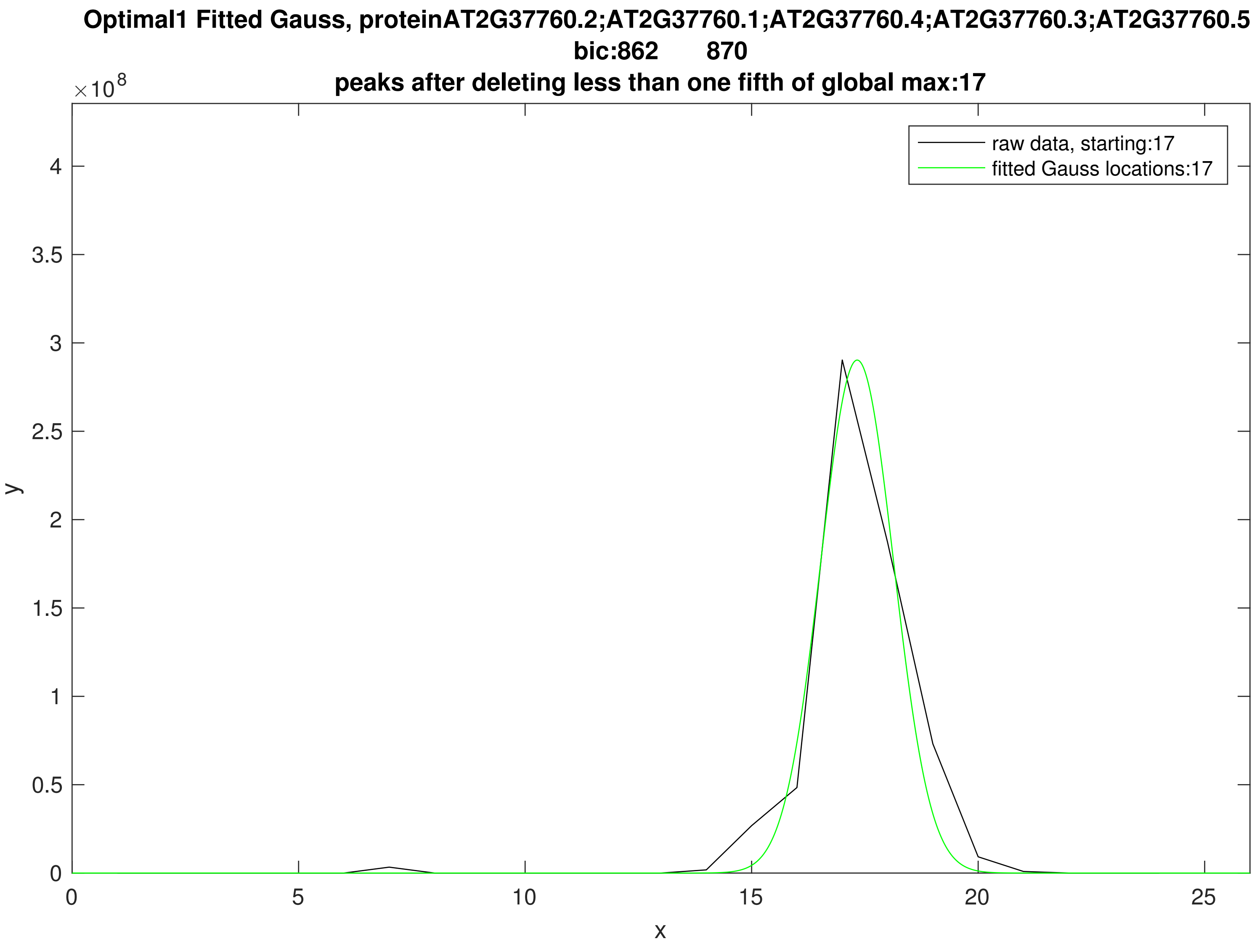

Optimal1 Fitted Gauss, proteinAT2G37760.2;AT2G37760.1;AT2G37760.4;AT2G37760.3;AT2G37760.5
bic:862 870
peaks after deleting less than one fifth of global max:17
×10^8
4
3.5
3
2.5
2
1.5
1
0.5
0
y
0
5
10
15
20
25
x
raw data, starting:17
fitted Gauss locations:17

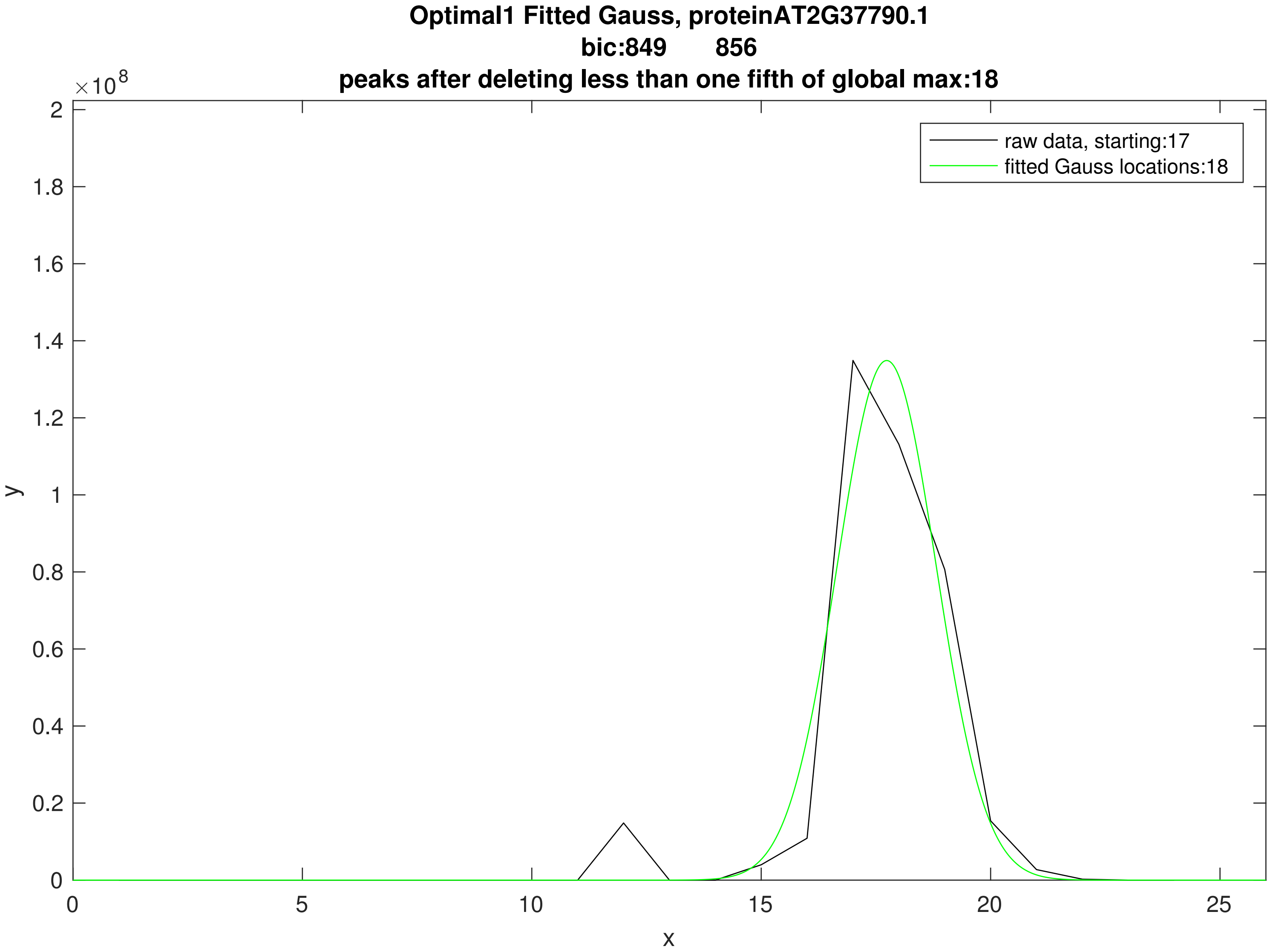

Optimal1 Fitted Gauss, proteinAT2G37790.1
bic:849 856
peaks after deleting less than one fifth of global max:18
×10^8
raw data, starting:17
fitted Gauss locations:18
y
x
0
0.2
0.4
0.6
0.8
1
1.2
1.4
1.6
1.8
2
5
10
15
20
25

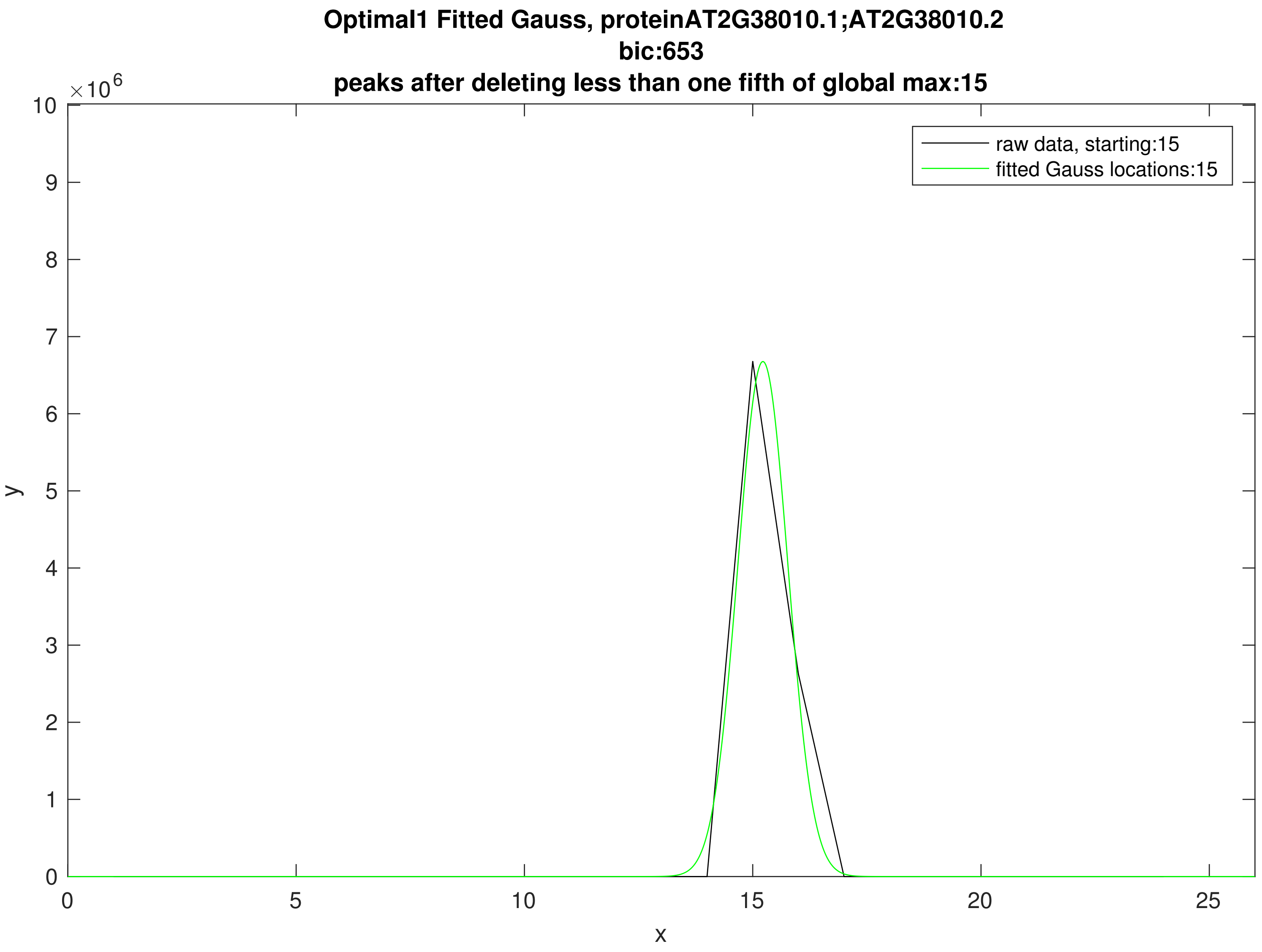
Optimal1 Fitted Gauss, proteinAT2G38010.1;AT2G38010.2
bic:653
peaks after deleting less than one fifth of global max:15
×10^6
raw data, starting:15
fitted Gauss locations:15
y
x
0
1
2
3
4
5
6
7
8
9
10
0
5
10
15
20
25

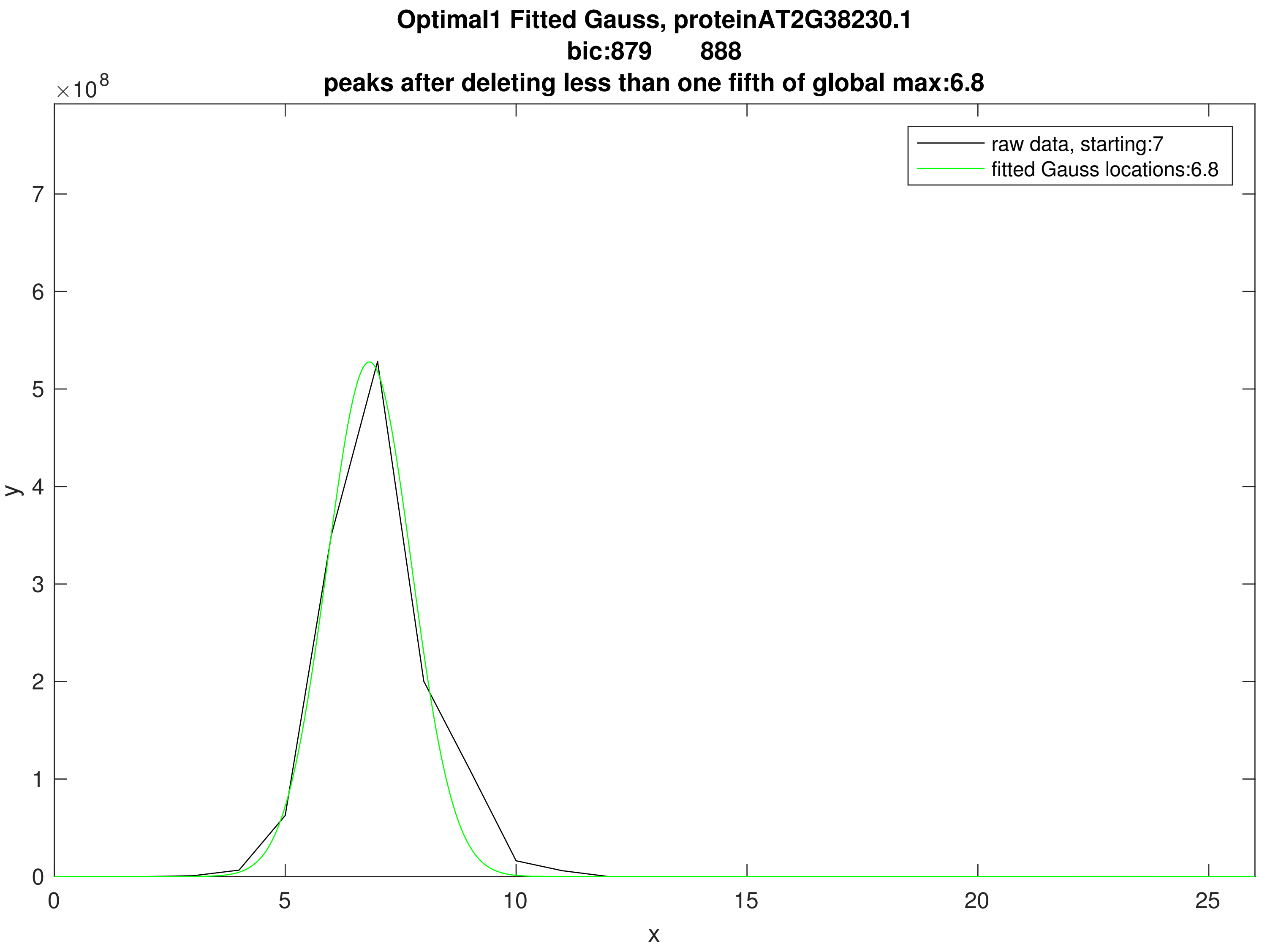

Optimal1 Fitted Gauss, proteinAT2G38230.1
bic:879 888
peaks after deleting less than one fifth of global max:6.8
×10^8
raw data, starting:7
fitted Gauss locations:6.8
y
x
0
1
2
3
4
5
6
7
0
5
10
15
20
25

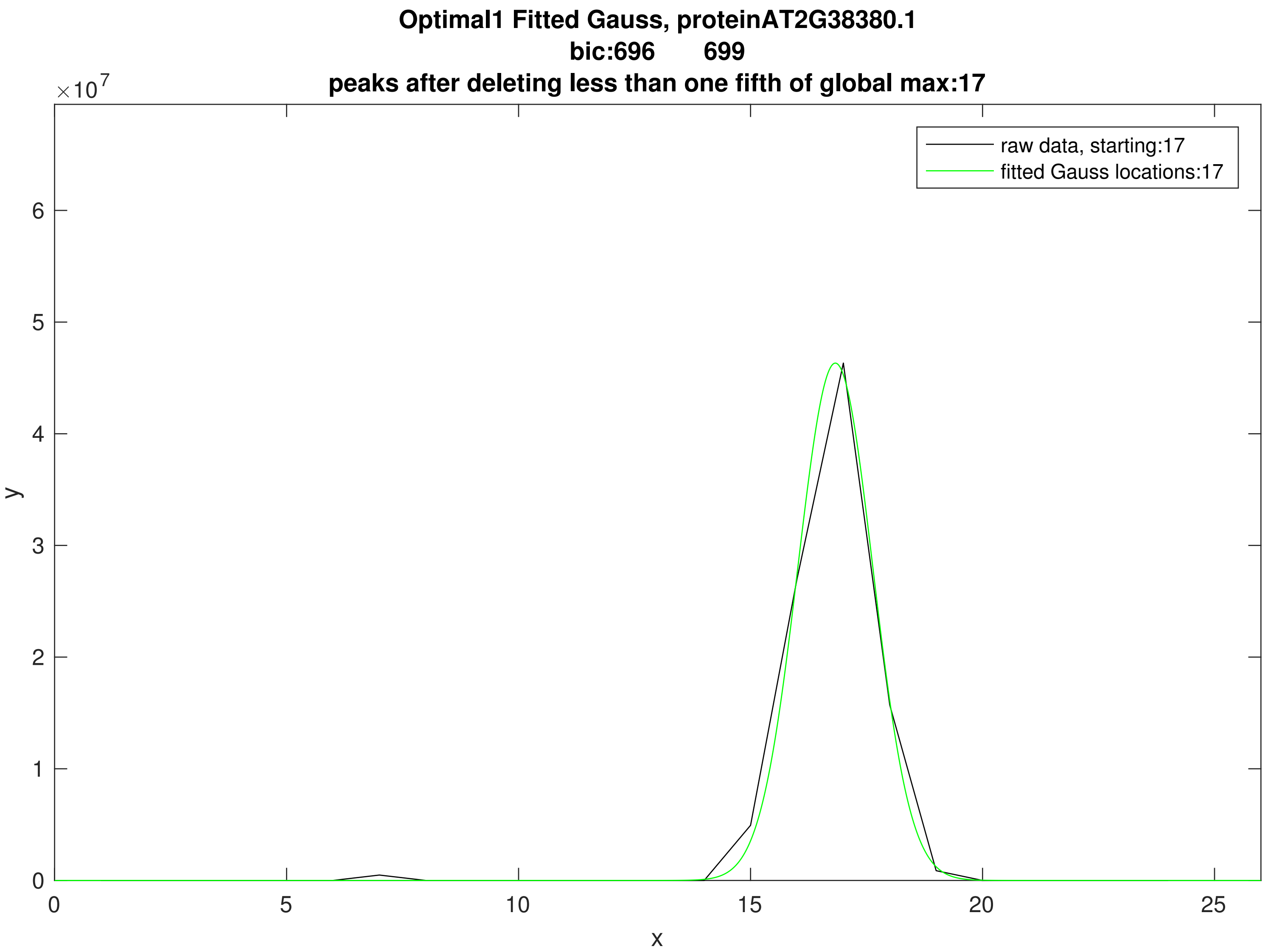

Optimal1 Fitted Gauss, proteinAT2G38380.1
bic:696 699
peaks after deleting less than one fifth of global max:17
×10^7
raw data, starting:17
fitted Gauss locations:17
y
x
0
1
2
3
4
5
6
0
5
10
15
20
25

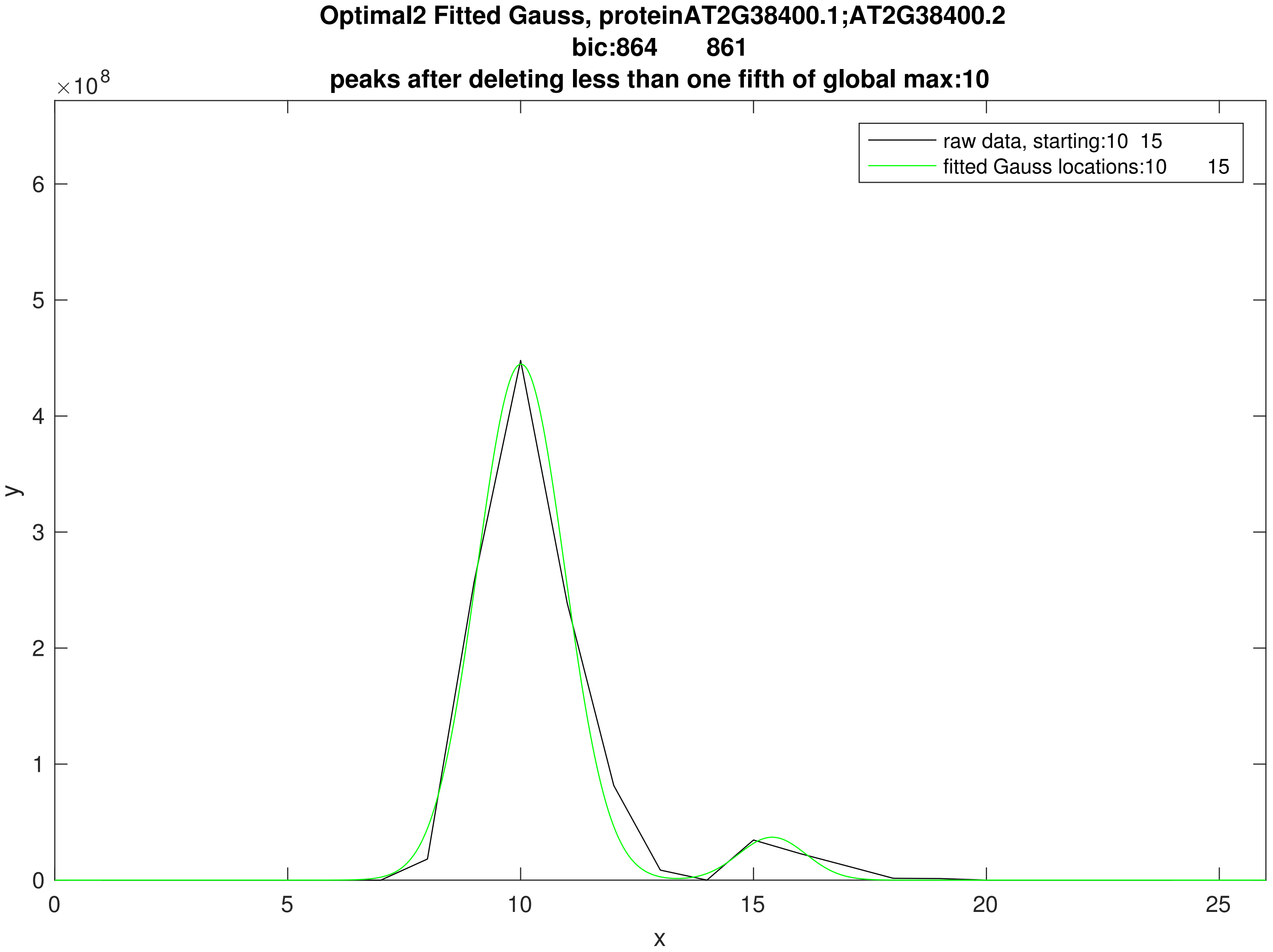
Optimal2 Fitted Gauss, proteinAT2G38400.1;AT2G38400.2
bic:864 861
peaks after deleting less than one fifth of global max:10
×10^8
raw data, starting:10 15
fitted Gauss locations:10 15
y
x
0
1
2
3
4
5
6
0
5
10
15
20
25

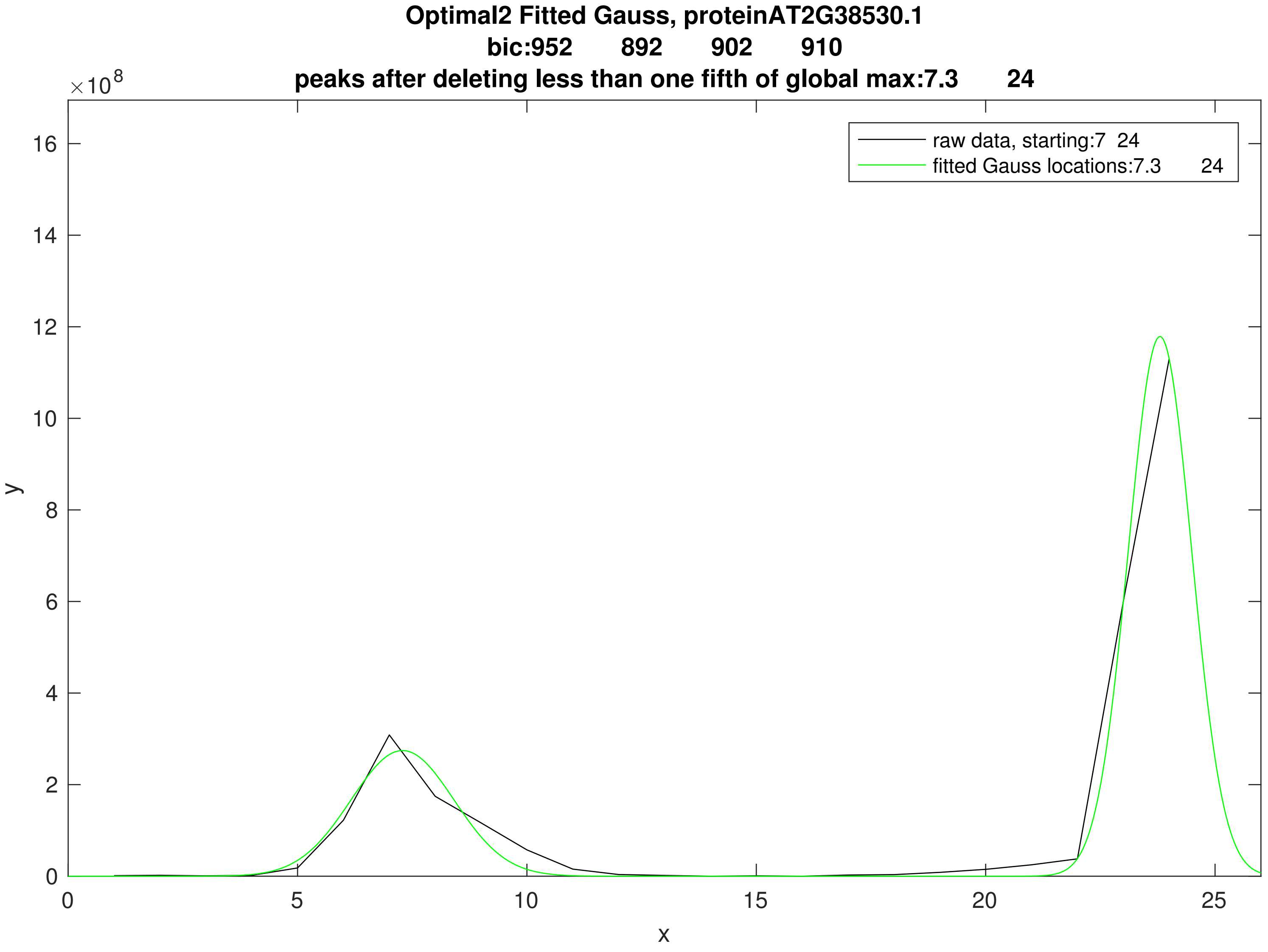
Optimal2 Fitted Gauss, proteinAT2G38530.1
bic:952 892 902 910
peaks after deleting less than one fifth of global max:7.3 24
×10^8
raw data, starting:7 24
fitted Gauss locations:7.3 24
y
x
0
2
4
6
8
10
12
14
16
0
5
10
15
20
25

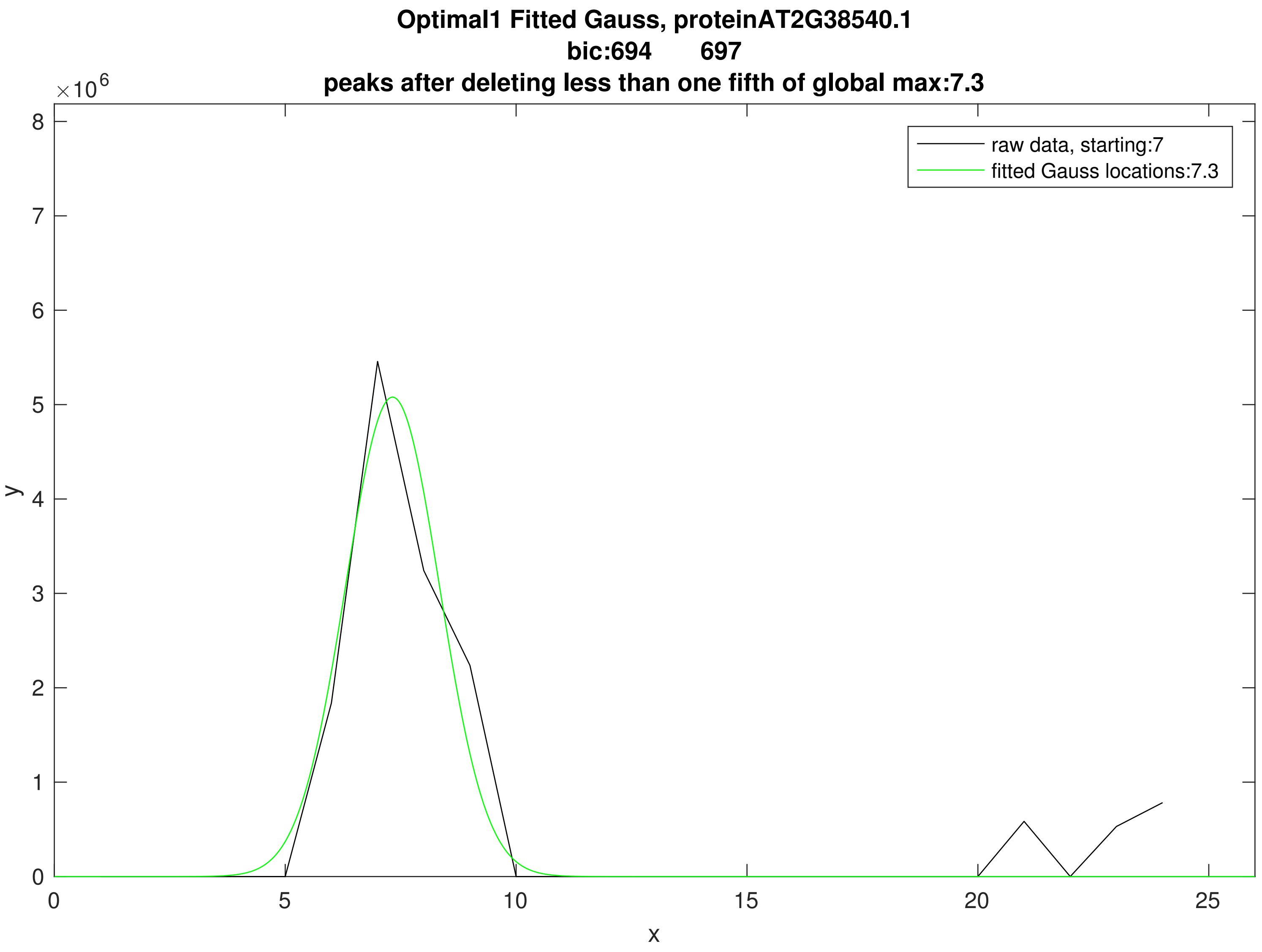

Optimal1 Fitted Gauss, proteinAT2G38540.1
bic:694 697
peaks after deleting less than one fifth of global max:7.3
×10^6
raw data, starting:7
fitted Gauss locations:7.3
y
x
0
1
2
3
4
5
6
7
8
0
5
10
15
20
25

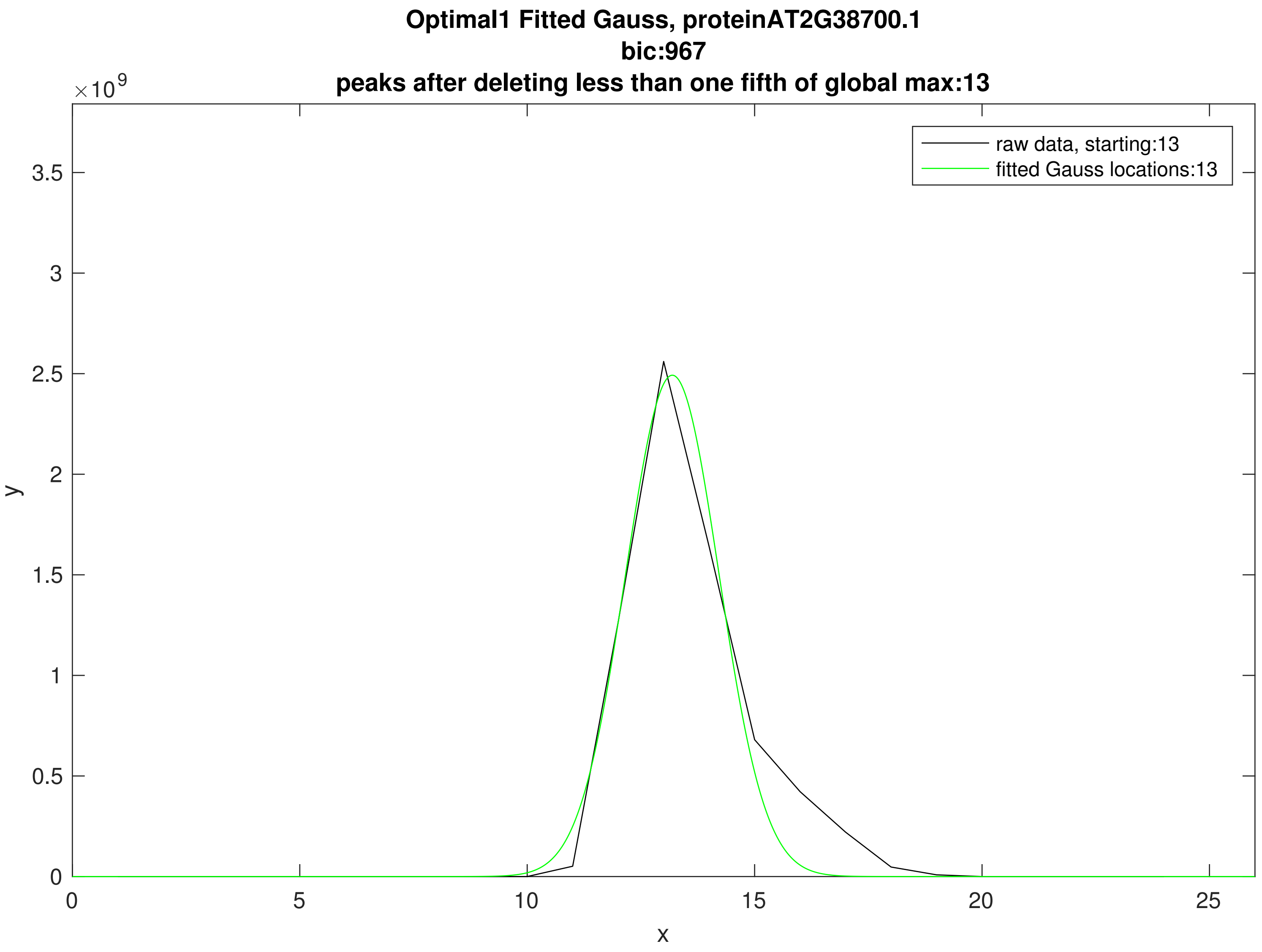

Optimal1 Fitted Gauss, proteinAT2G38700.1
bic:967
peaks after deleting less than one fifth of global max:13
×10^9
raw data, starting:13
fitted Gauss locations:13
y
x
0
0.5
1
1.5
2
2.5
3
3.5
0
5
10
15
20
25

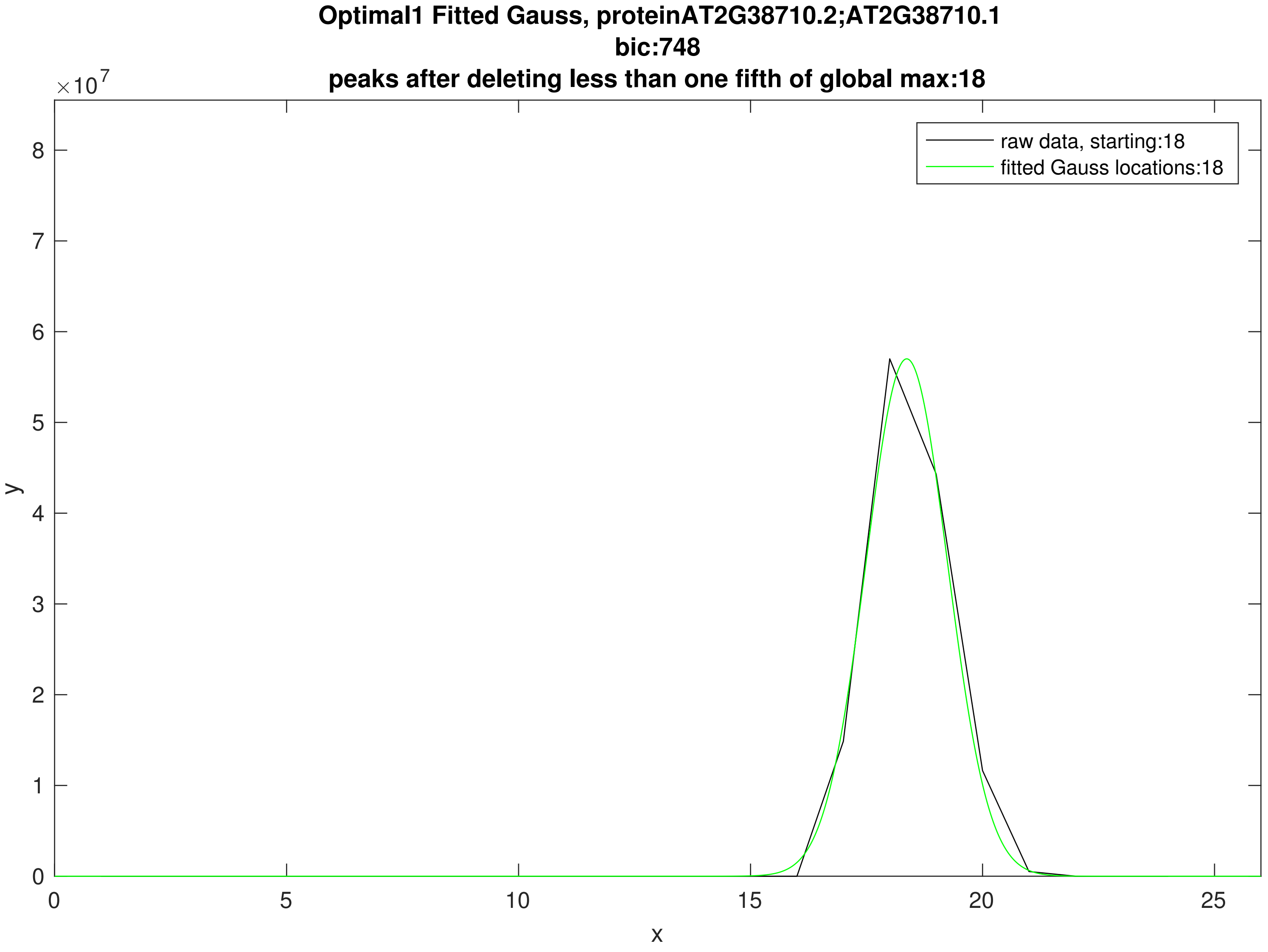
Optimal1 Fitted Gauss, proteinAT2G38710.2;AT2G38710.1
bic:748
peaks after deleting less than one fifth of global max:18
×10^7
raw data, starting:18
fitted Gauss locations:18
y
x
0
1
2
3
4
5
6
7
8
0
5
10
15
20
25

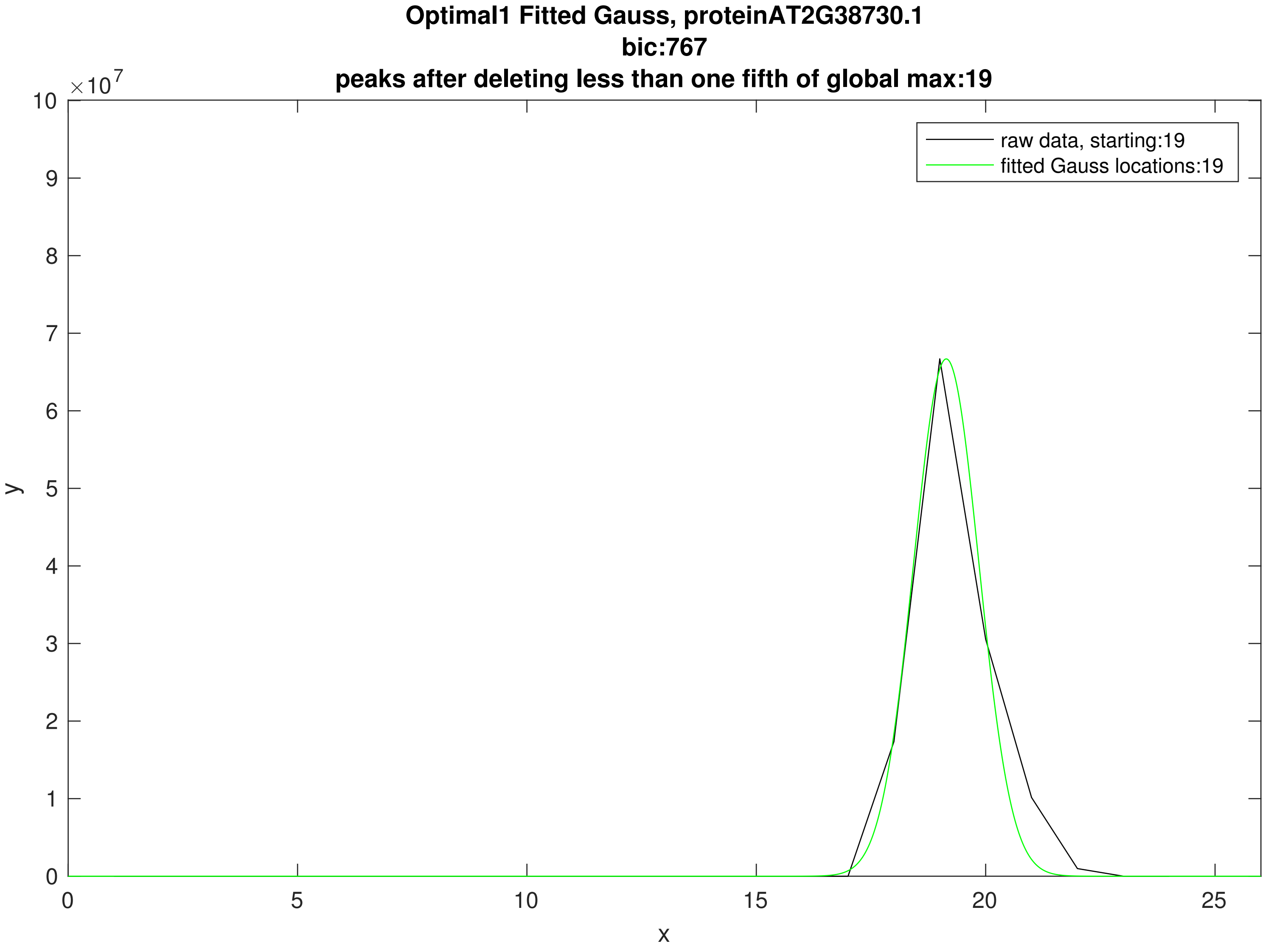
Optimal1 Fitted Gauss, proteinAT2G38730.1
bic:767
peaks after deleting less than one fifth of global max:19
×10^7
raw data, starting:19
fitted Gauss locations:19
y
x
0
1
2
3
4
5
6
7
8
9
10
0
5
10
15
20
25

**Optimal2 Fitted Gauss, proteinAT2G38750.1**
**bic:777 771**
**peaks after deleting less than one fifth of global max:5.7 11**

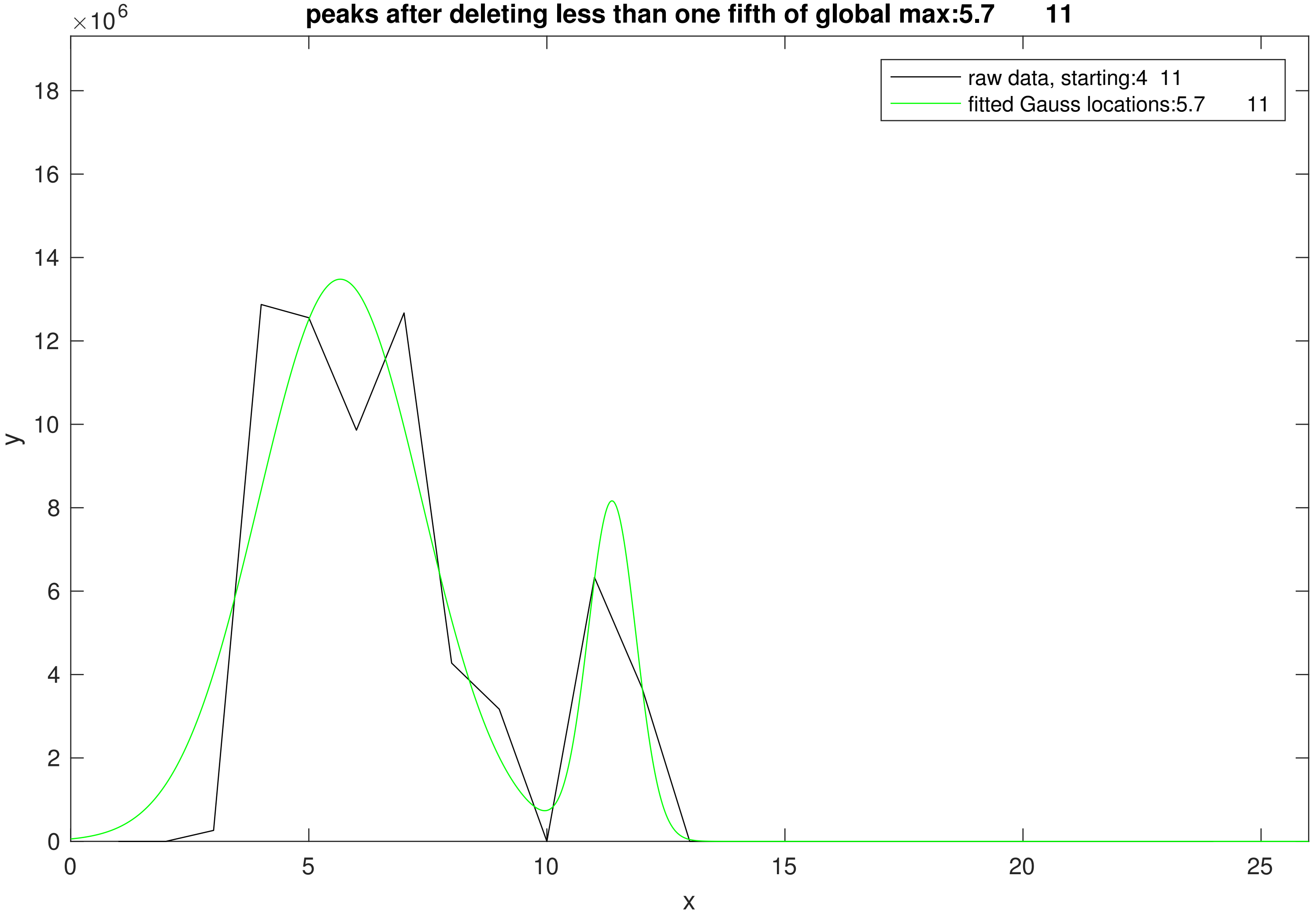

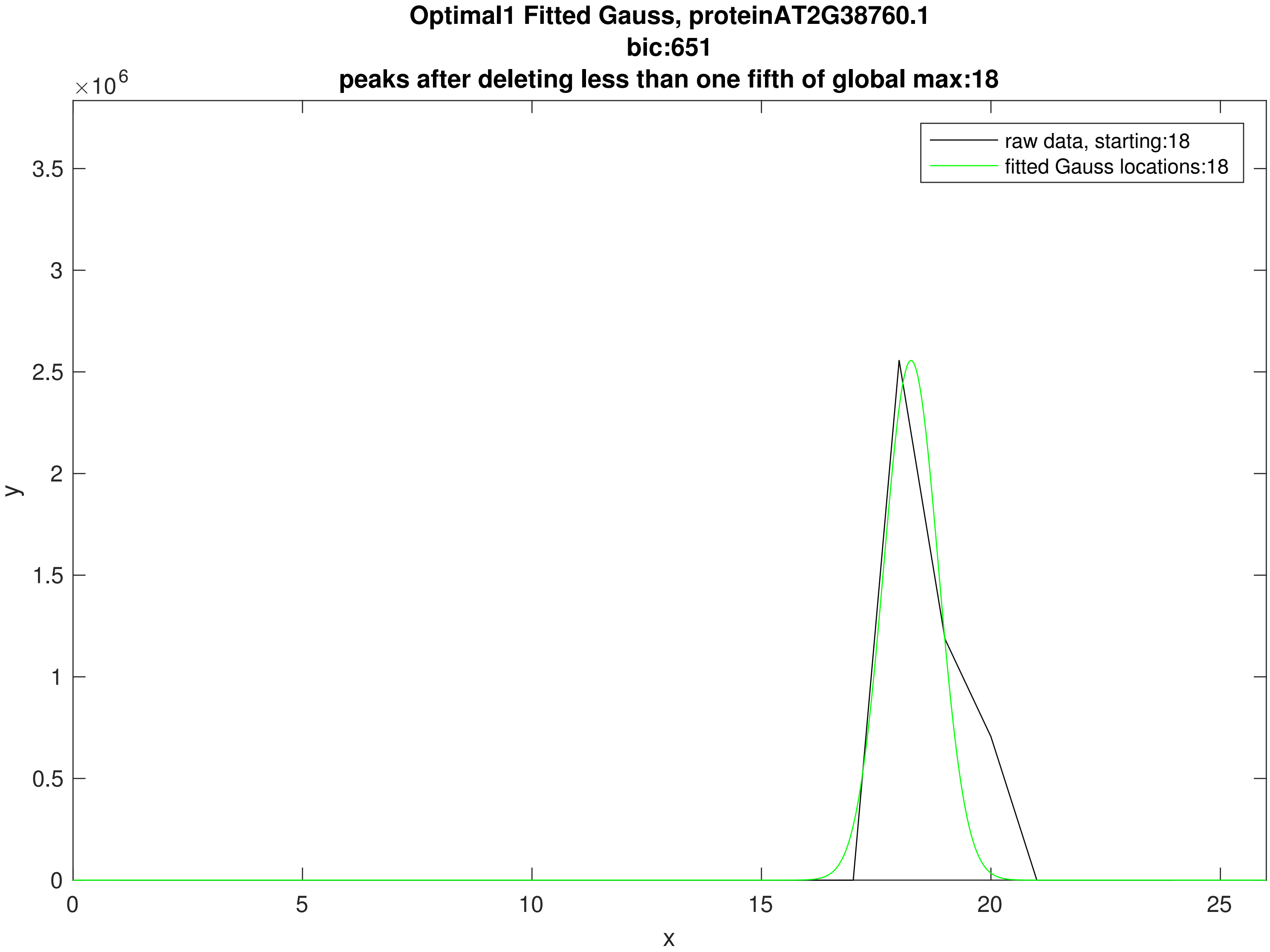
Optimal1 Fitted Gauss, proteinAT2G38760.1
bic:651
peaks after deleting less than one fifth of global max:18
×10^6
raw data, starting:18
fitted Gauss locations:18
y
x
0
0.5
1
1.5
2
2.5
3
3.5
0
5
10
15
20
25

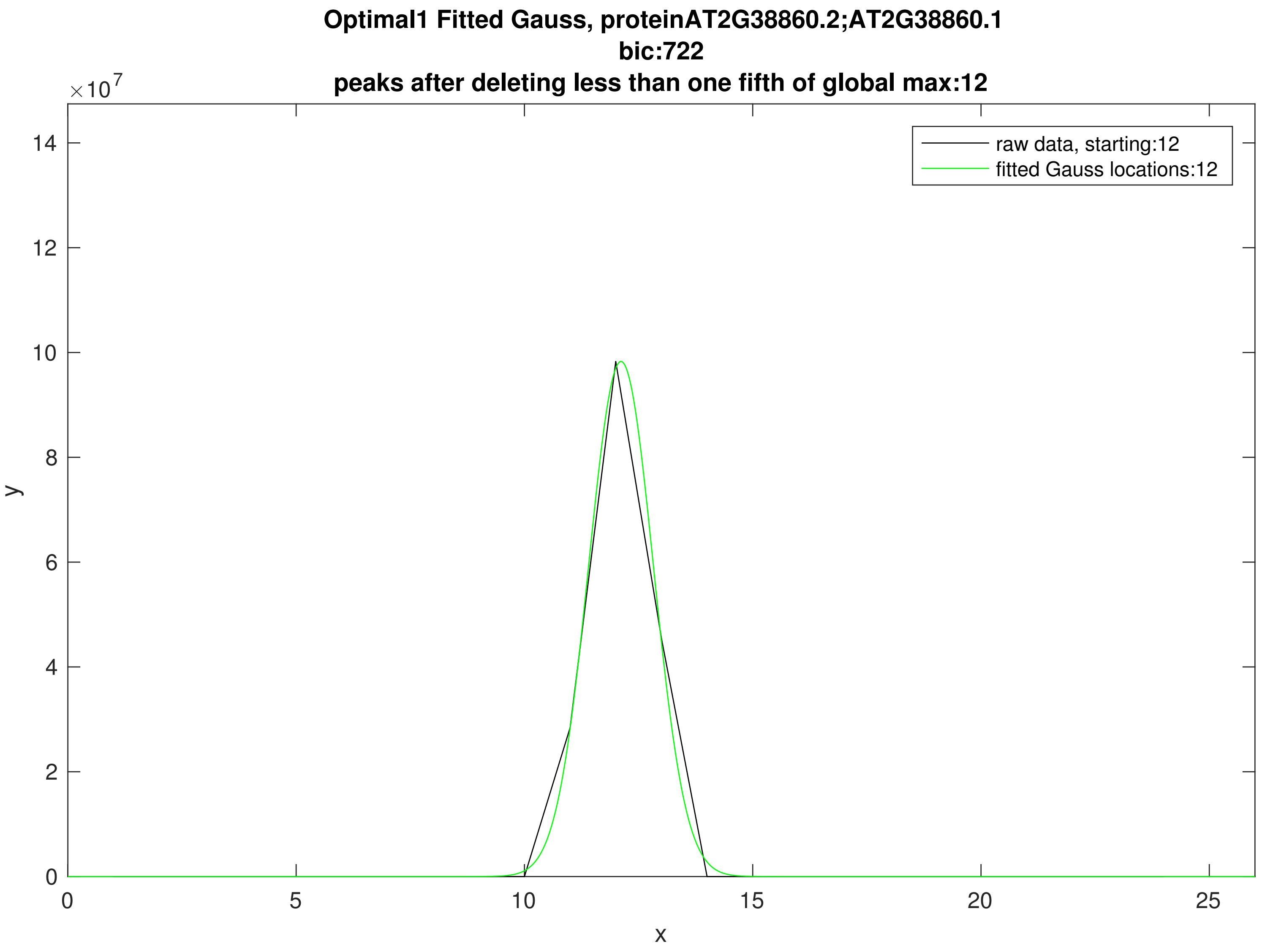

Optimal1 Fitted Gauss, proteinAT2G38860.2;AT2G38860.1
bic:722
peaks after deleting less than one fifth of global max:12
×10^7
14
12
10
8
6
4
2
0
y
0
5
10
15
20
25
x
raw data, starting:12
fitted Gauss locations:12

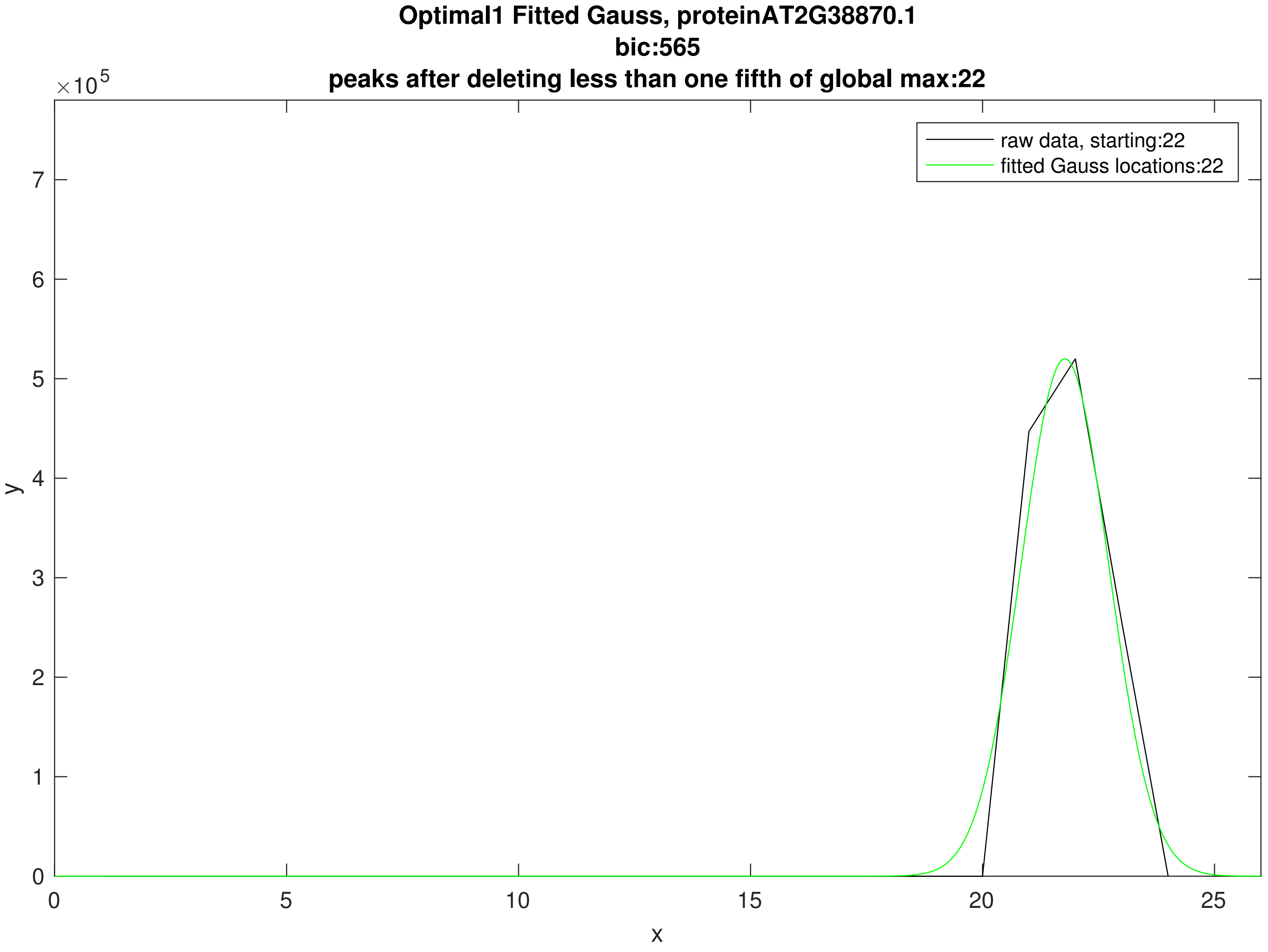

Optimal1 Fitted Gauss, proteinAT2G38870.1
bic:565
peaks after deleting less than one fifth of global max:22
×10^5
raw data, starting:22
fitted Gauss locations:22
y
x
0
1
2
3
4
5
6
7
0
5
10
15
20
25

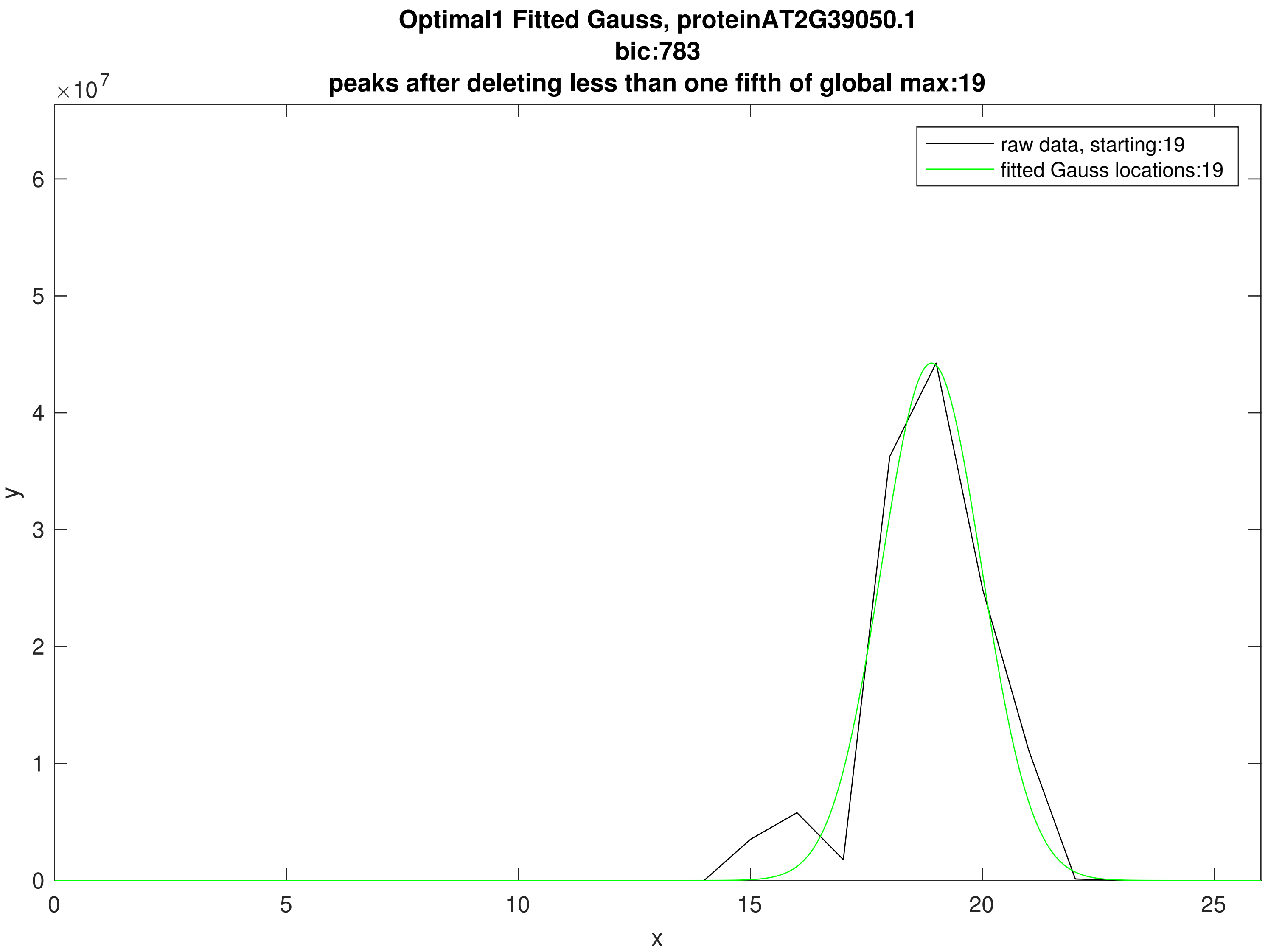
Optimal1 Fitted Gauss, proteinAT2G39050.1
bic:783
peaks after deleting less than one fifth of global max:19
×10^7
raw data, starting:19
fitted Gauss locations:19
y
x
0
1
2
3
4
5
6
0
5
10
15
20
25

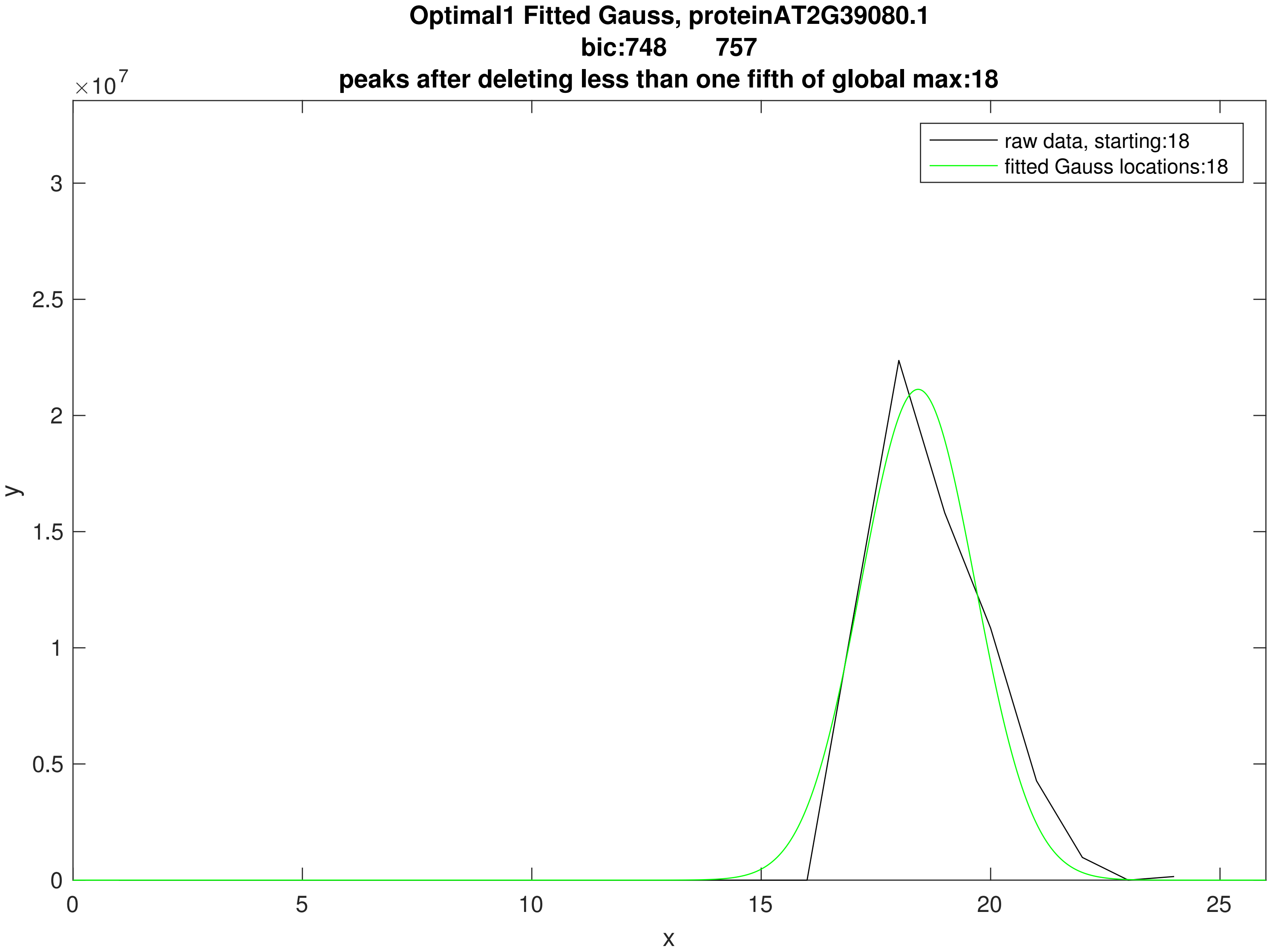

Optimal1 Fitted Gauss, proteinAT2G39080.1
bic:748 757
peaks after deleting less than one fifth of global max:18
×10^7
raw data, starting:18
fitted Gauss locations:18
y
x
0
0.5
1
1.5
2
2.5
3
0
5
10
15
20
25

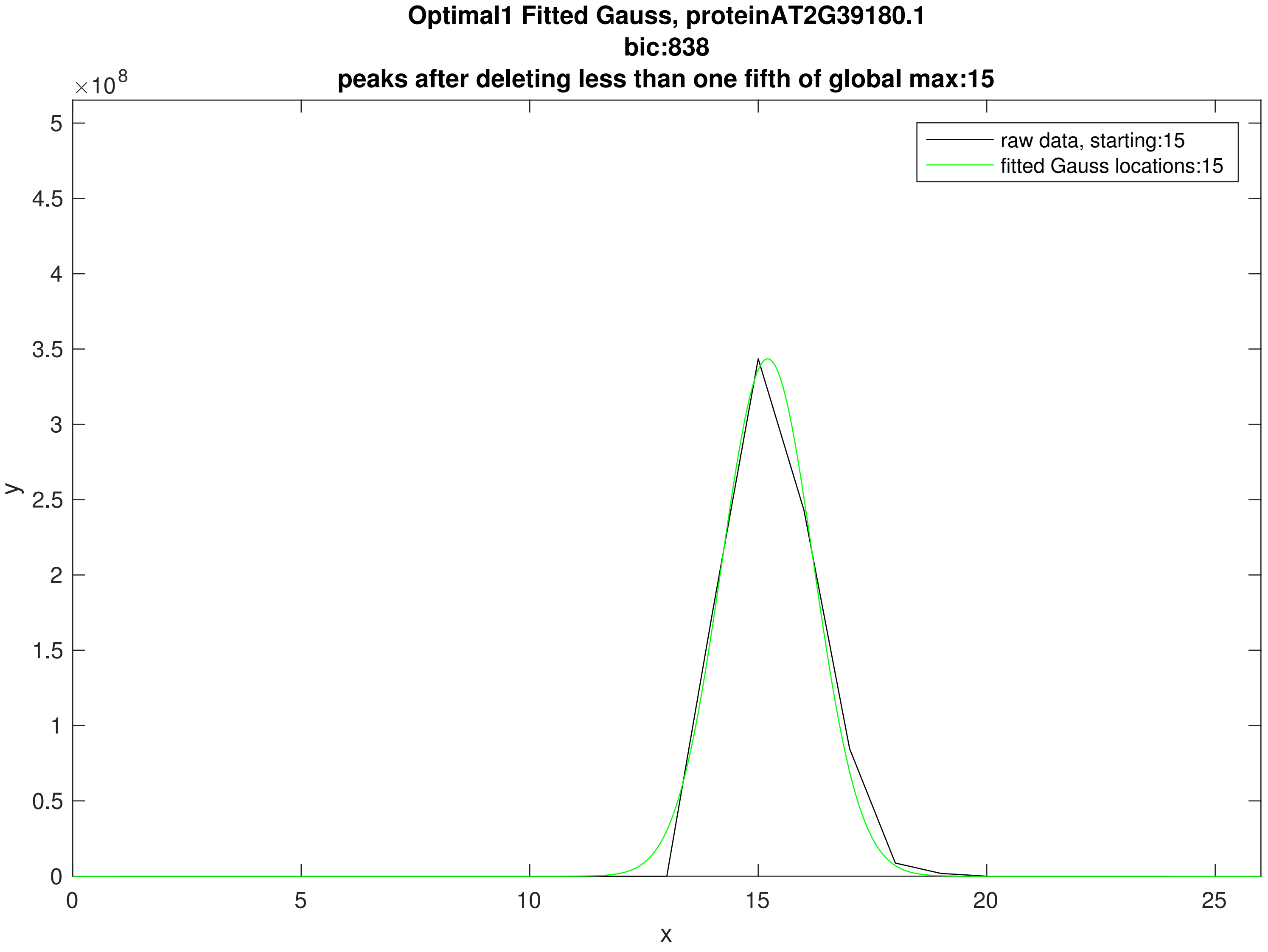
Optimal1 Fitted Gauss, proteinAT2G39180.1
bic:838
peaks after deleting less than one fifth of global max:15
×10^8
raw data, starting:15
fitted Gauss locations:15
y
x
0
0.5
1
1.5
2
2.5
3
3.5
4
4.5
5
0
5
10
15
20
25

Optimal2 Fitted Gauss, proteinAT2G39310.3;AT2G39310.1;AT2G39310.2
bic:1e+03 984 1e+03 1.01e+03
peaks after deleting less than one fifth of global max:15

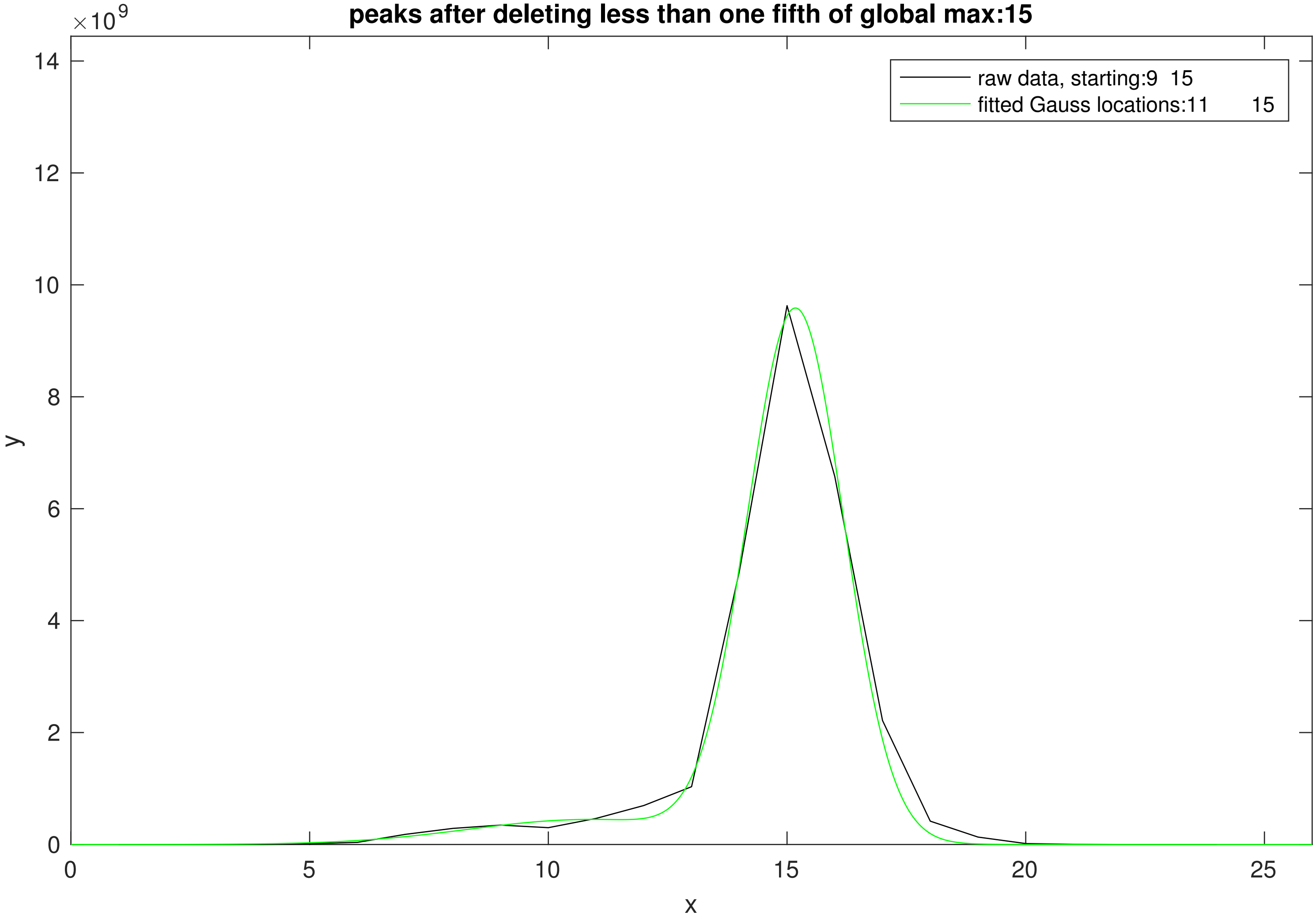

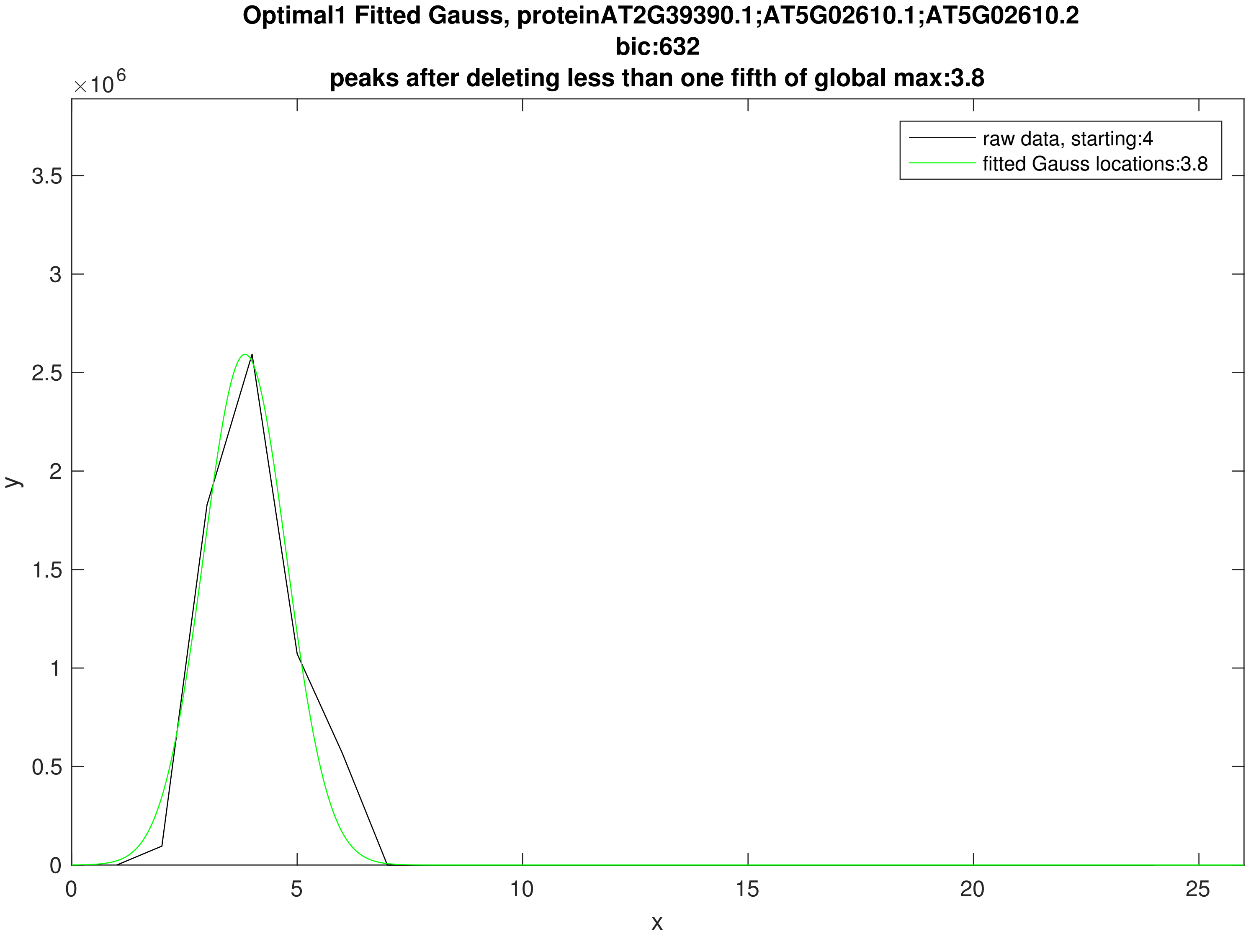

Optimal1 Fitted Gauss, proteinAT2G39390.1;AT5G02610.1;AT5G02610.2
bic:632
peaks after deleting less than one fifth of global max:3.8
×10^6
3.5
3
2.5
2
1.5
1
0.5
0
y
0
5
10
15
20
25
x
raw data, starting:4
fitted Gauss locations:3.8

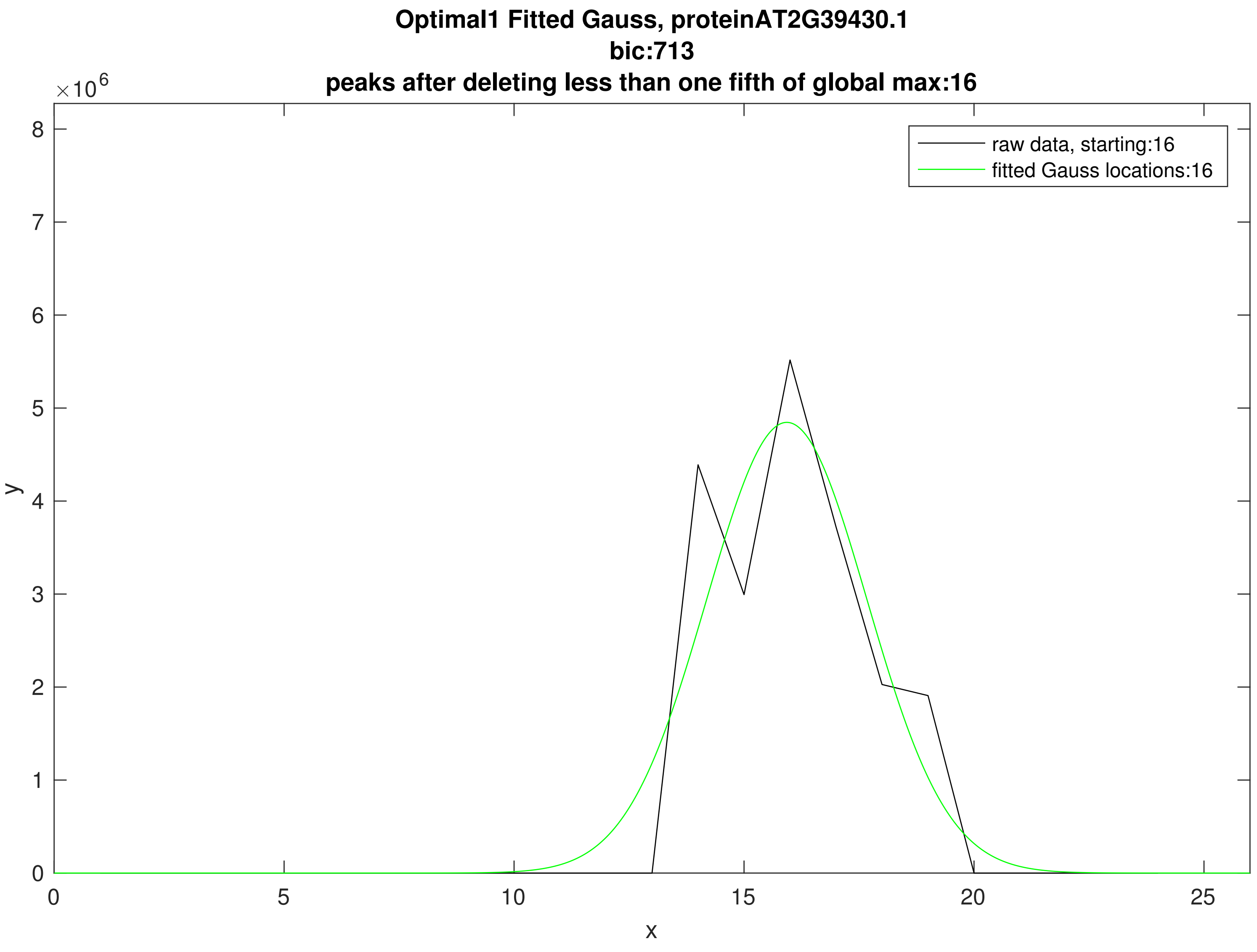

Optimal1 Fitted Gauss, proteinAT2G39430.1
bic:713
peaks after deleting less than one fifth of global max:16
×10^6
8
7
6
5
4
3
2
1
0
y
0
5
10
15
20
25
x
raw data, starting:16
fitted Gauss locations:16

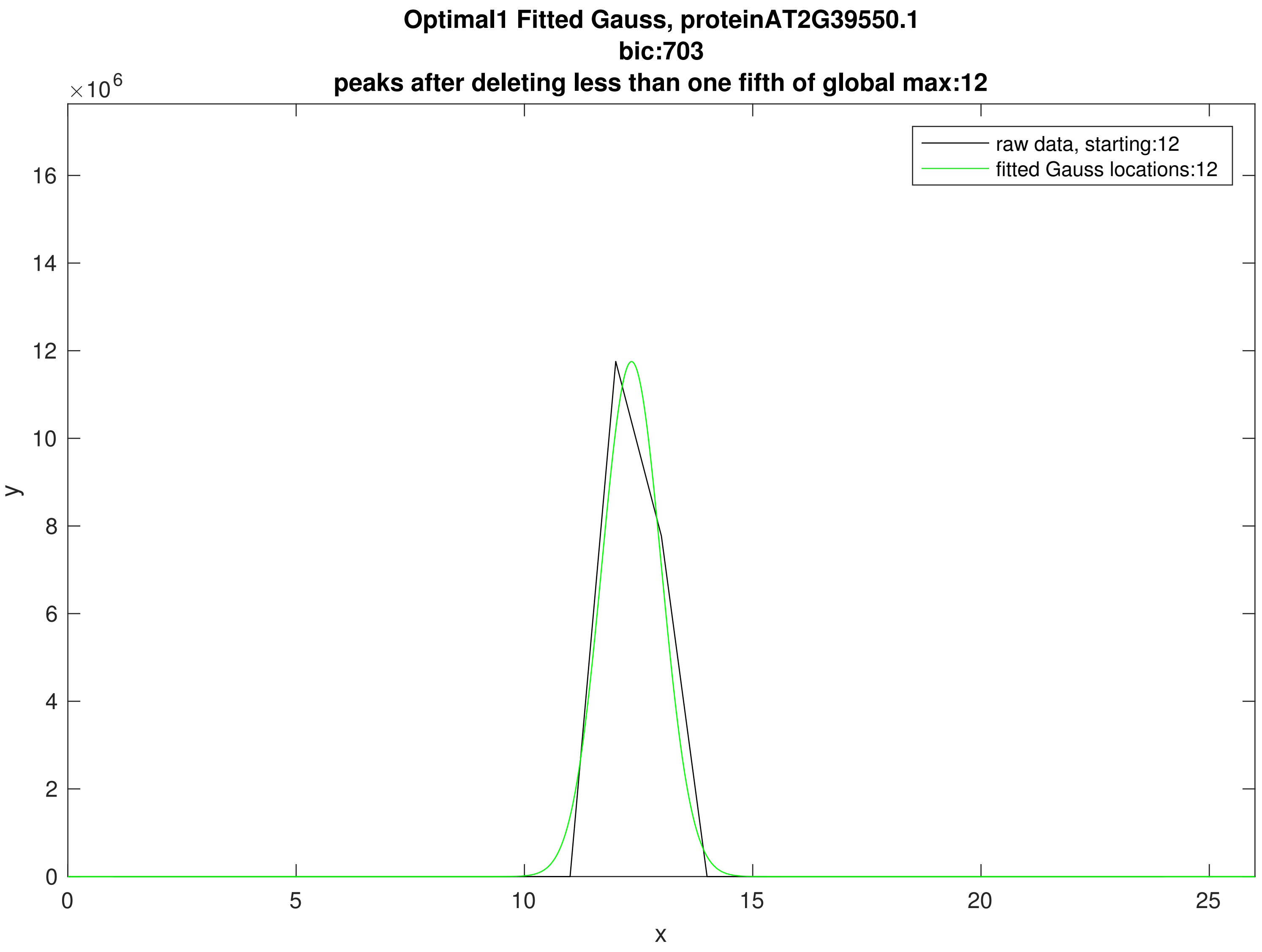

Optimal1 Fitted Gauss, proteinAT2G39550.1
bic:703
peaks after deleting less than one fifth of global max:12
×10^6
raw data, starting:12
fitted Gauss locations:12
0
2
4
6
8
10
12
14
16
y
0
5
10
15
20
25
x

**Optimal1 Fitted Gauss, proteinAT2G39730.3;AT2G39730.2;AT2G39730.1**
**bic:787 796**
**peaks after deleting less than one fifth of global max:14**

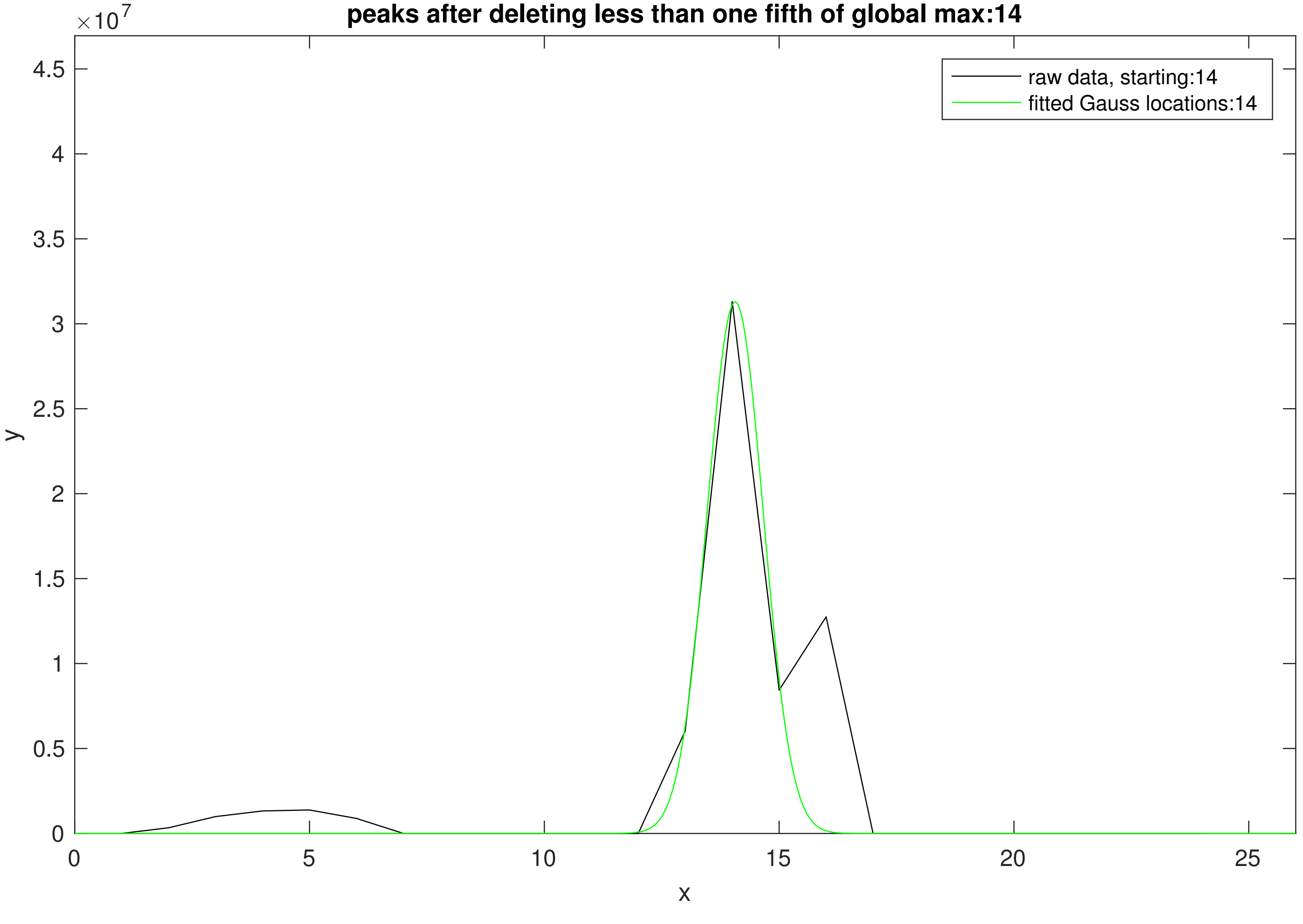

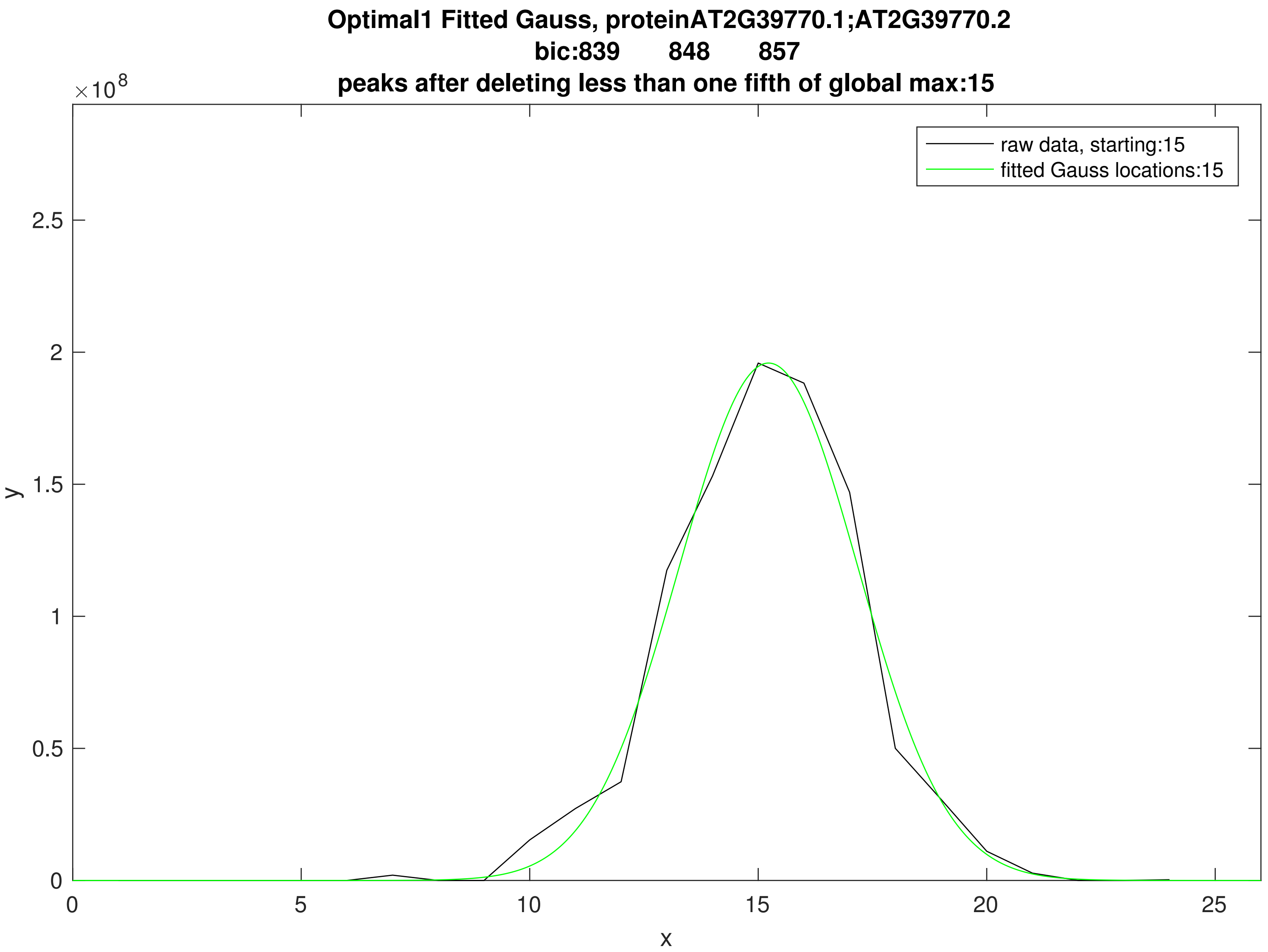

Optimal1 Fitted Gauss, proteinAT2G39770.1;AT2G39770.2
bic:839 848 857
peaks after deleting less than one fifth of global max:15
×10^8
raw data, starting:15
fitted Gauss locations:15
y
x
0
0.5
1
1.5
2
2.5
0
5
10
15
20
25

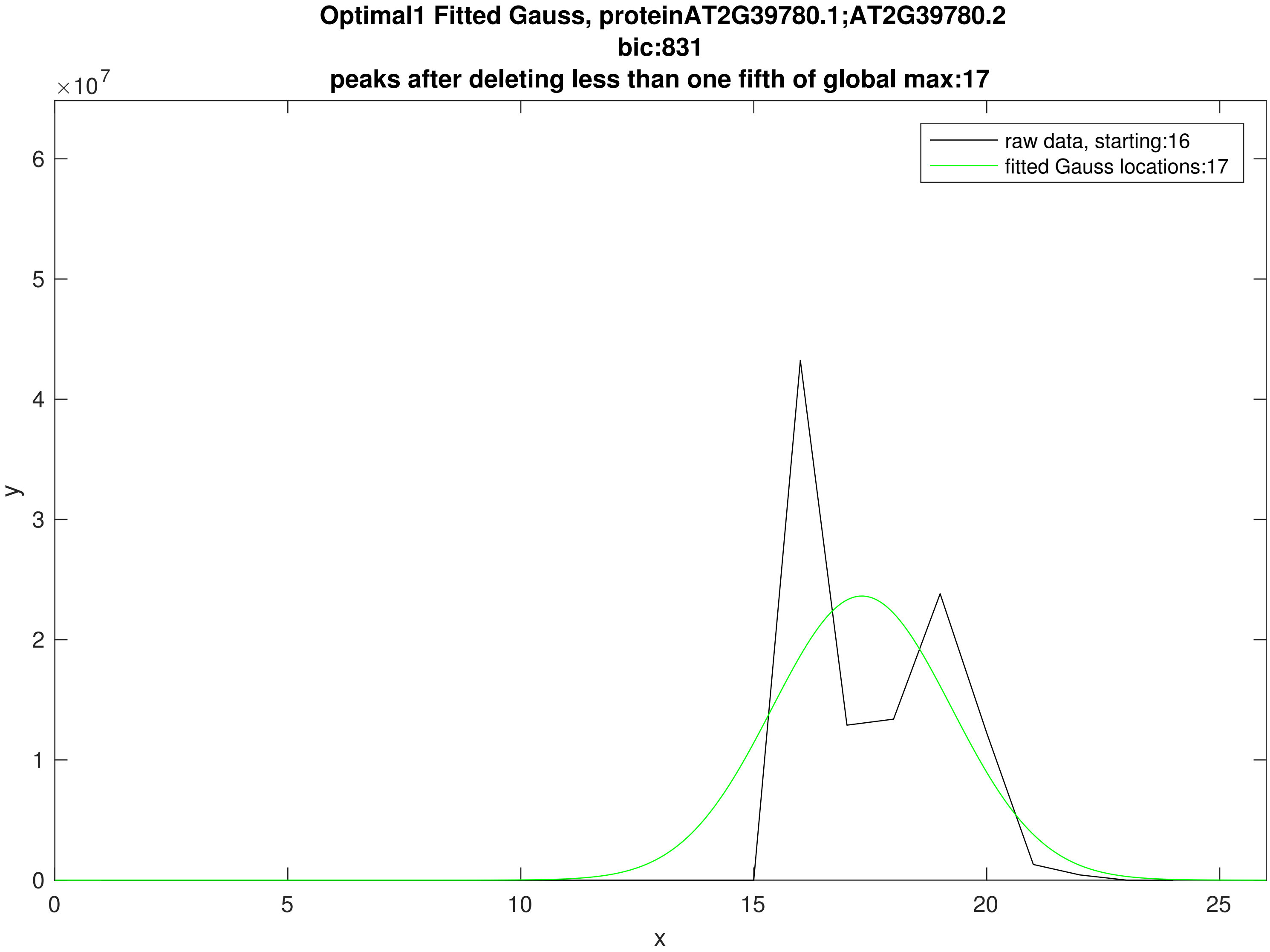
Optimal1 Fitted Gauss, proteinAT2G39780.1;AT2G39780.2
bic:831
peaks after deleting less than one fifth of global max:17
×10^7
raw data, starting:16
fitted Gauss locations:17
y
x
0
1
2
3
4
5
6
0
5
10
15
20
25

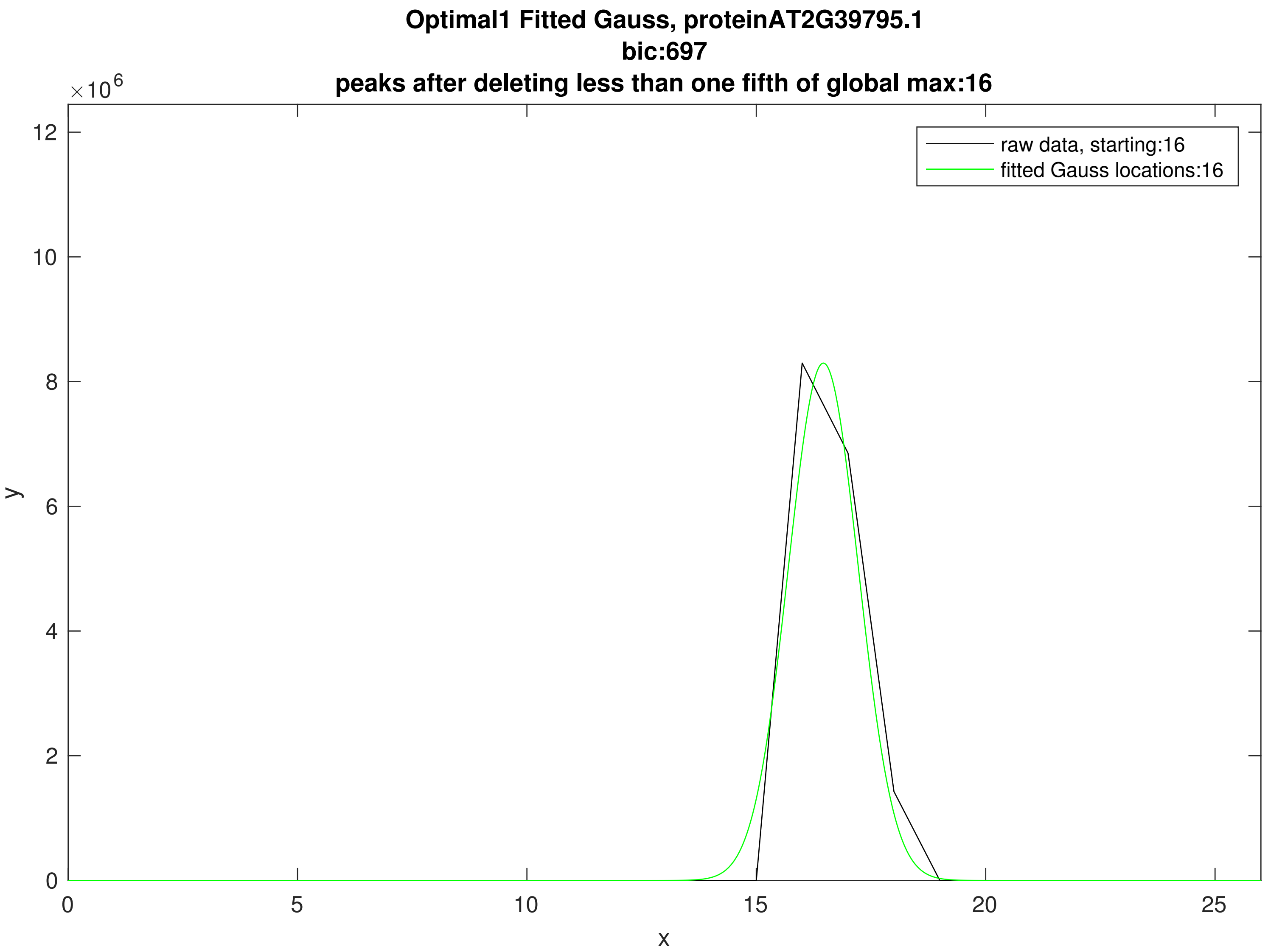
Optimal1 Fitted Gauss, proteinAT2G39795.1
bic:697
peaks after deleting less than one fifth of global max:16
×10^6
raw data, starting:16
fitted Gauss locations:16
y
x
0
2
4
6
8
10
12
0
5
10
15
20
25

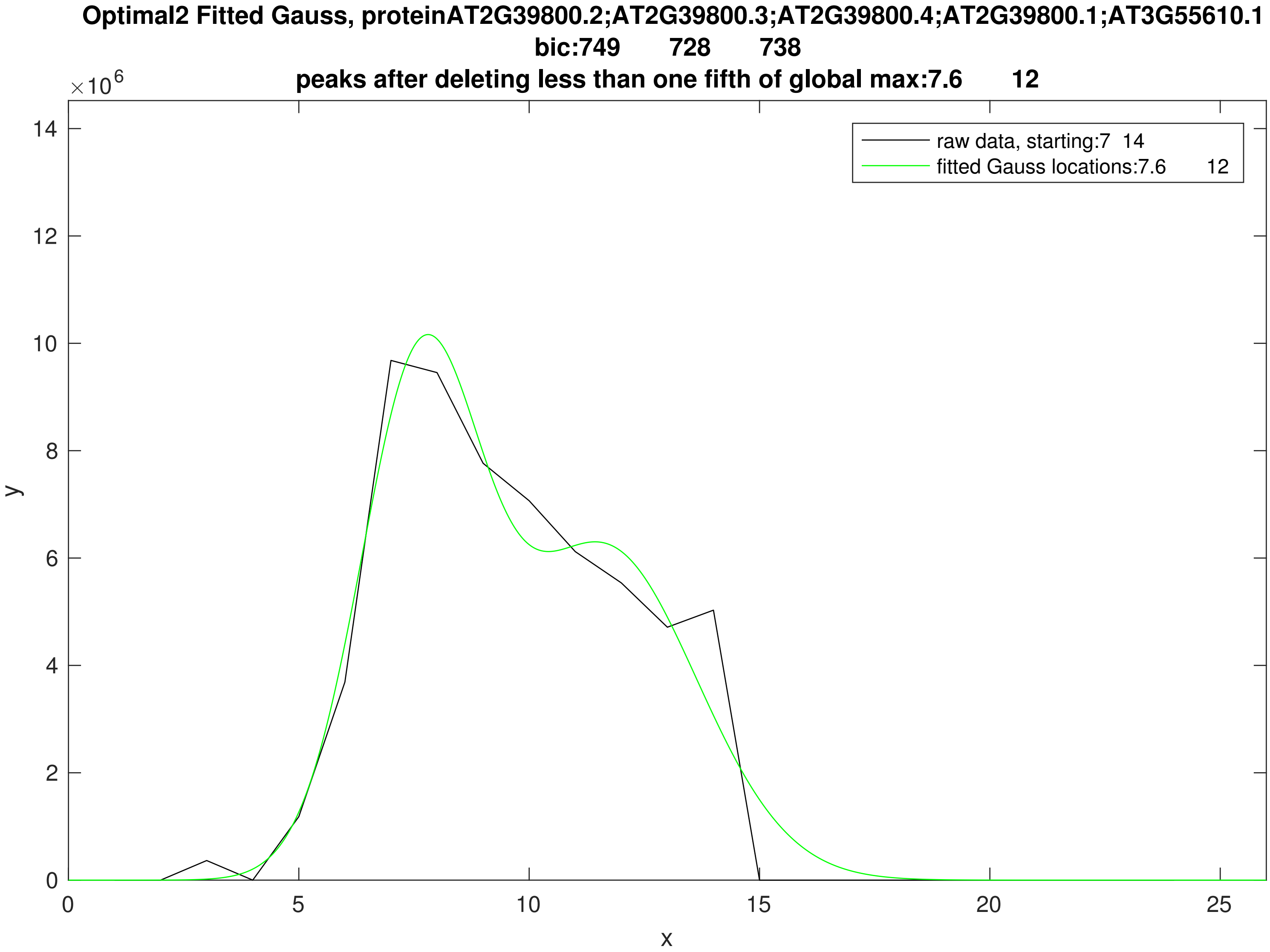
Optimal2 Fitted Gauss, proteinAT2G39800.2;AT2G39800.3;AT2G39800.4;AT2G39800.1;AT3G55610.1
bic:749 728 738
peaks after deleting less than one fifth of global max:7.6 12
raw data, starting:7 14
fitted Gauss locations:7.6 12
×10^6
y
x
0
2
4
6
8
10
12
14
0
5
10
15
20
25

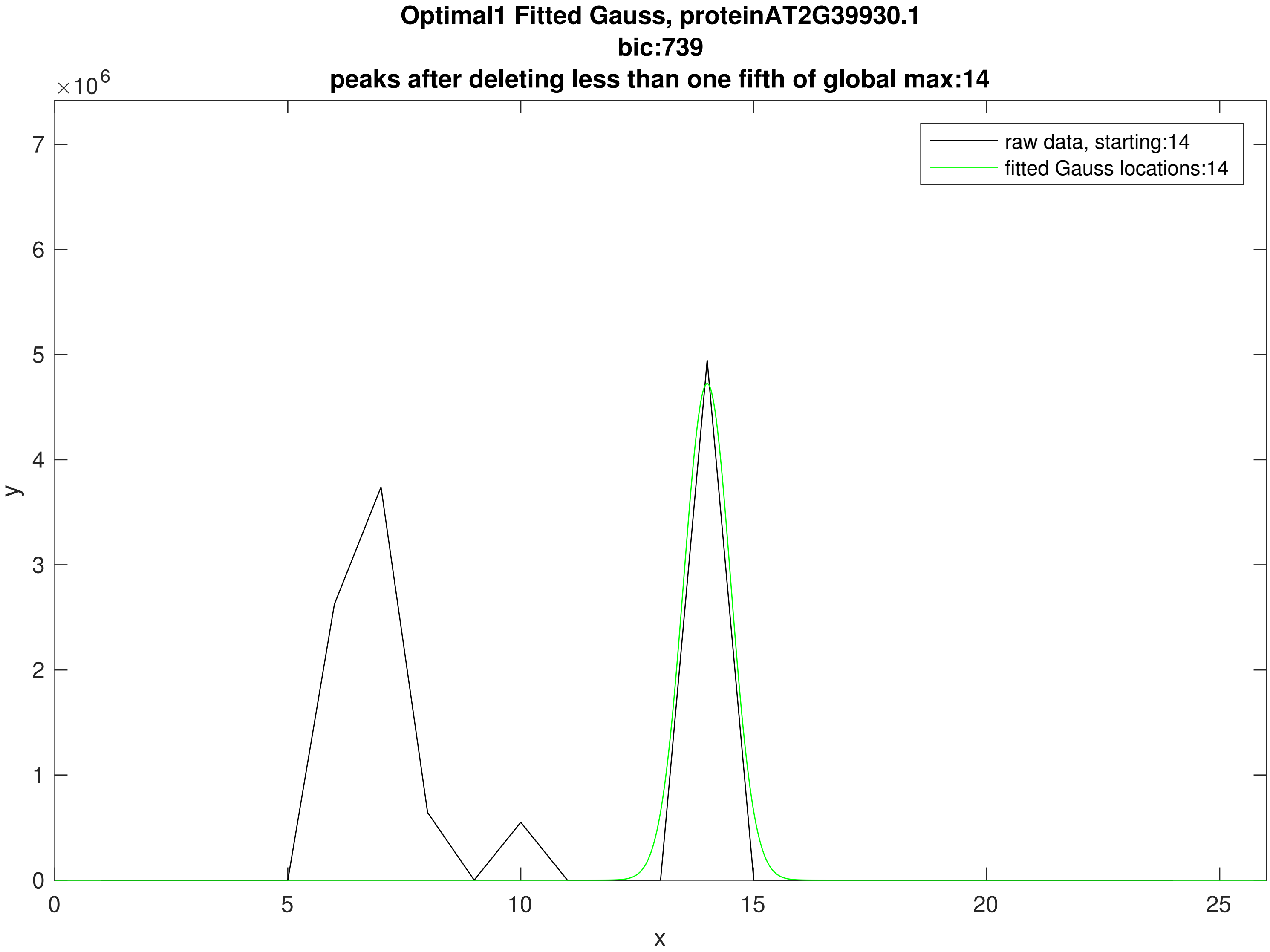

Optimal1 Fitted Gauss, proteinAT2G39930.1
bic:739
peaks after deleting less than one fifth of global max:14
×10^6
raw data, starting:14
fitted Gauss locations:14
y
x
0
1
2
3
4
5
6
7
0
5
10
15
20
25

**Optimal1 Fitted Gauss, proteinAT2G39940.1**
**bic:685**
**peaks after deleting less than one fifth of global max:13**

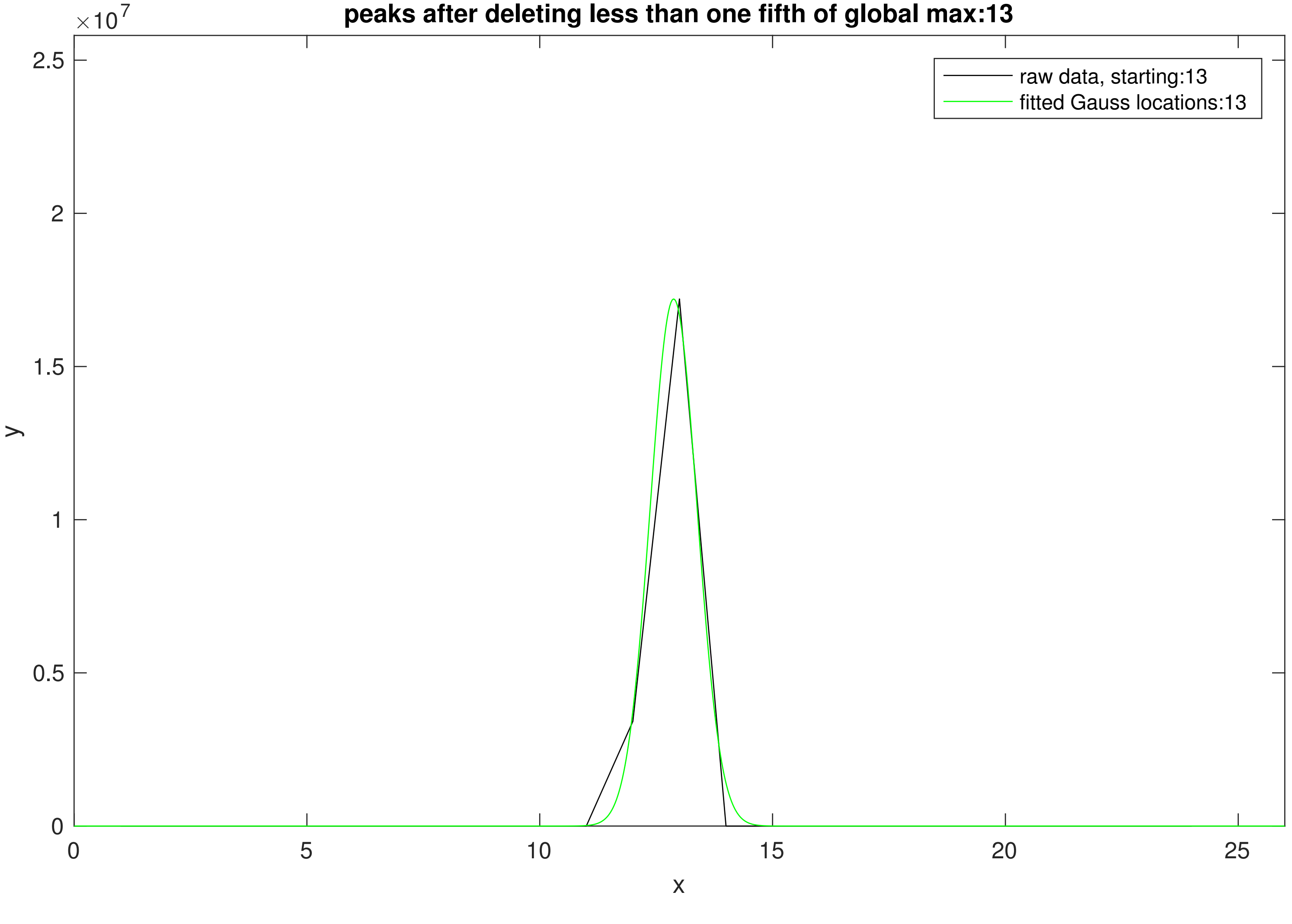

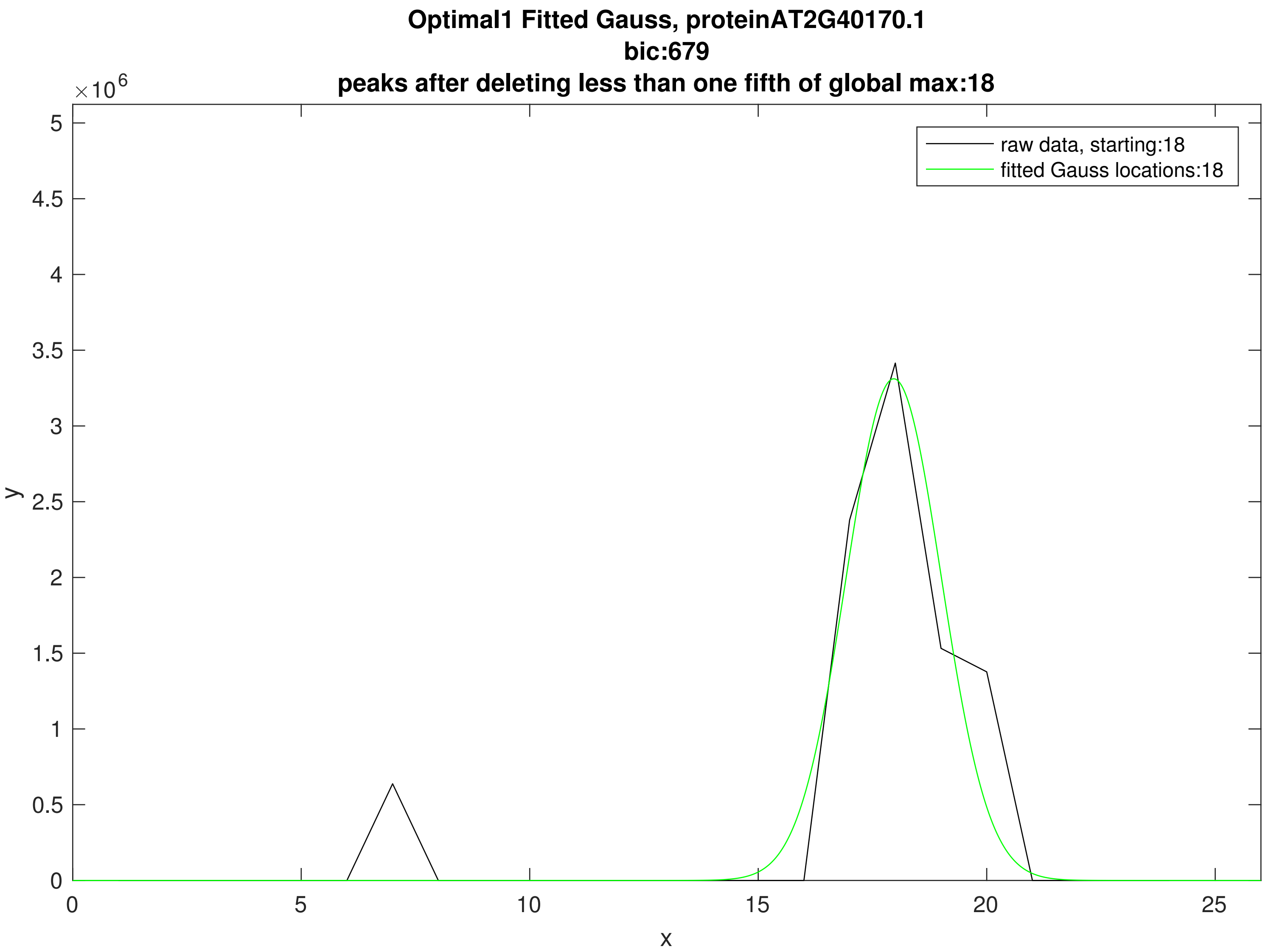

Optimal1 Fitted Gauss, proteinAT2G40170.1
bic:679
peaks after deleting less than one fifth of global max:18
×10^6
raw data, starting:18
fitted Gauss locations:18
y
x
0
0.5
1
1.5
2
2.5
3
3.5
4
4.5
5
0
5
10
15
20
25

#### Optimal1 Fitted Gauss, proteinAT2G40290.1
#### bic:730
#### peaks after deleting less than one fifth of global max:14

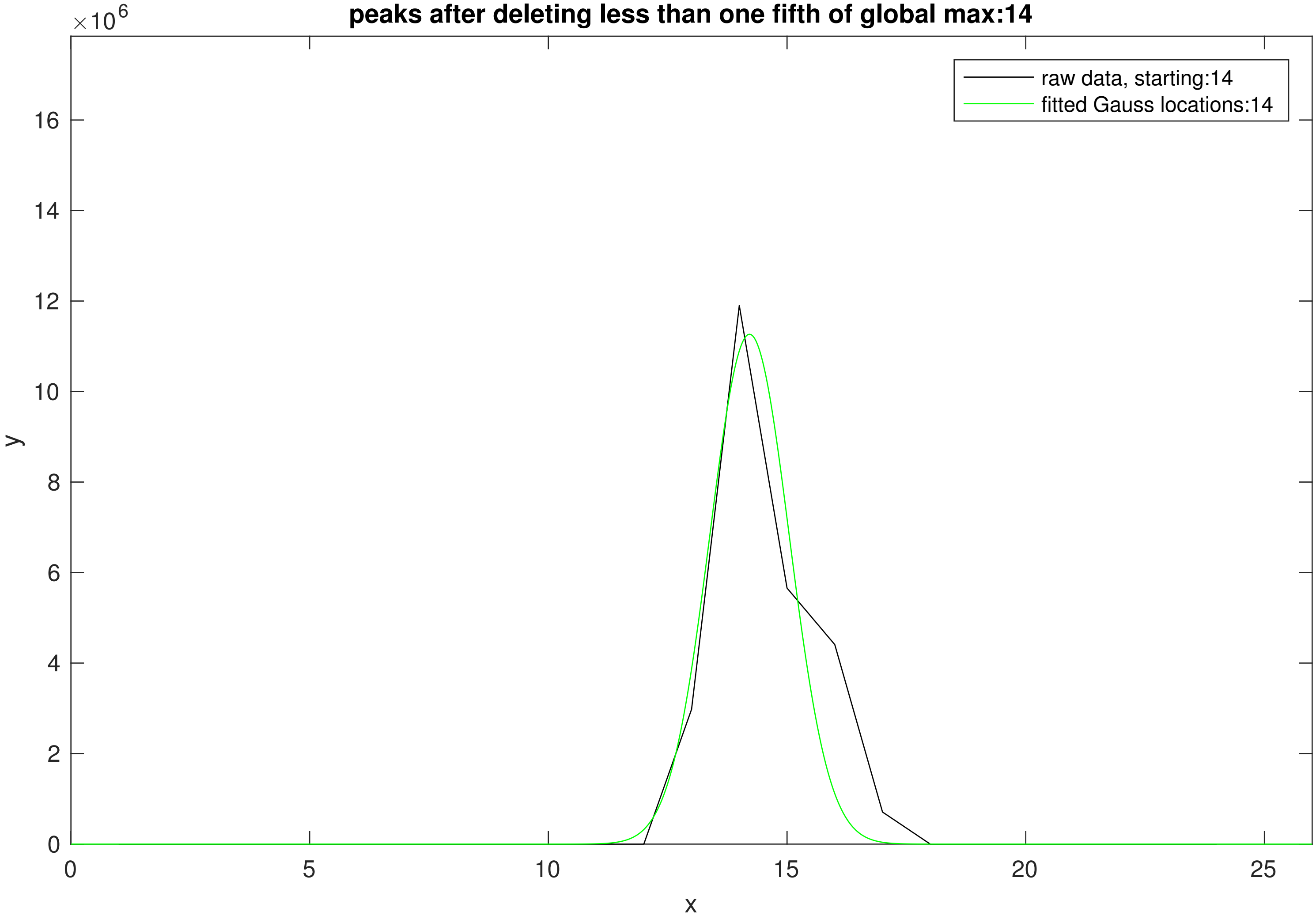

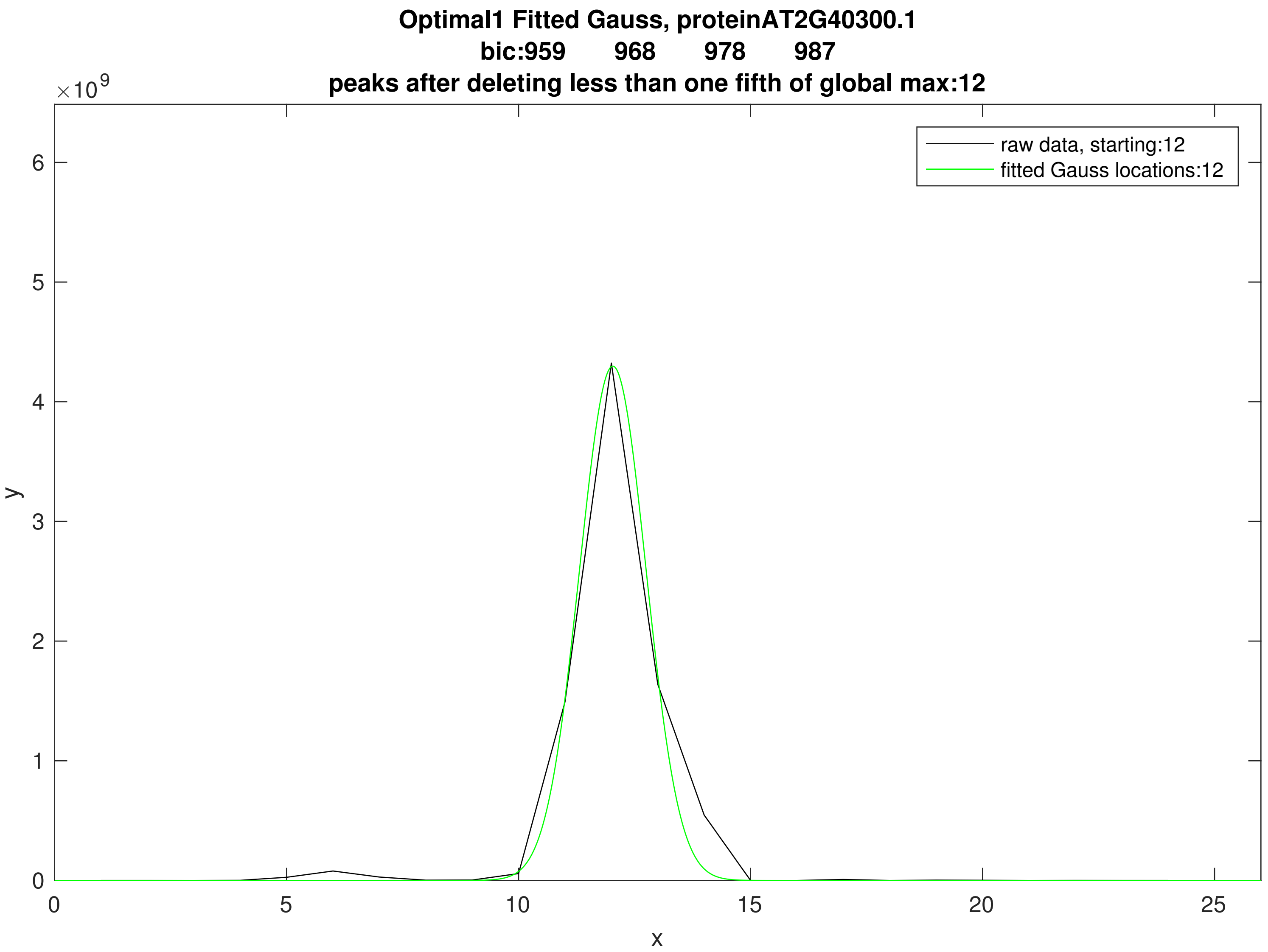

Optimal1 Fitted Gauss, proteinAT2G40300.1
bic:959 968 978 987
peaks after deleting less than one fifth of global max:12
×10^9
raw data, starting:12
fitted Gauss locations:12
y
x
0
1
2
3
4
5
6
0
5
10
15
20
25

**Optimal1 Fitted Gauss, proteinAT2G40490.1**
**bic:778 782**
**peaks after deleting less than one fifth of global max:16**

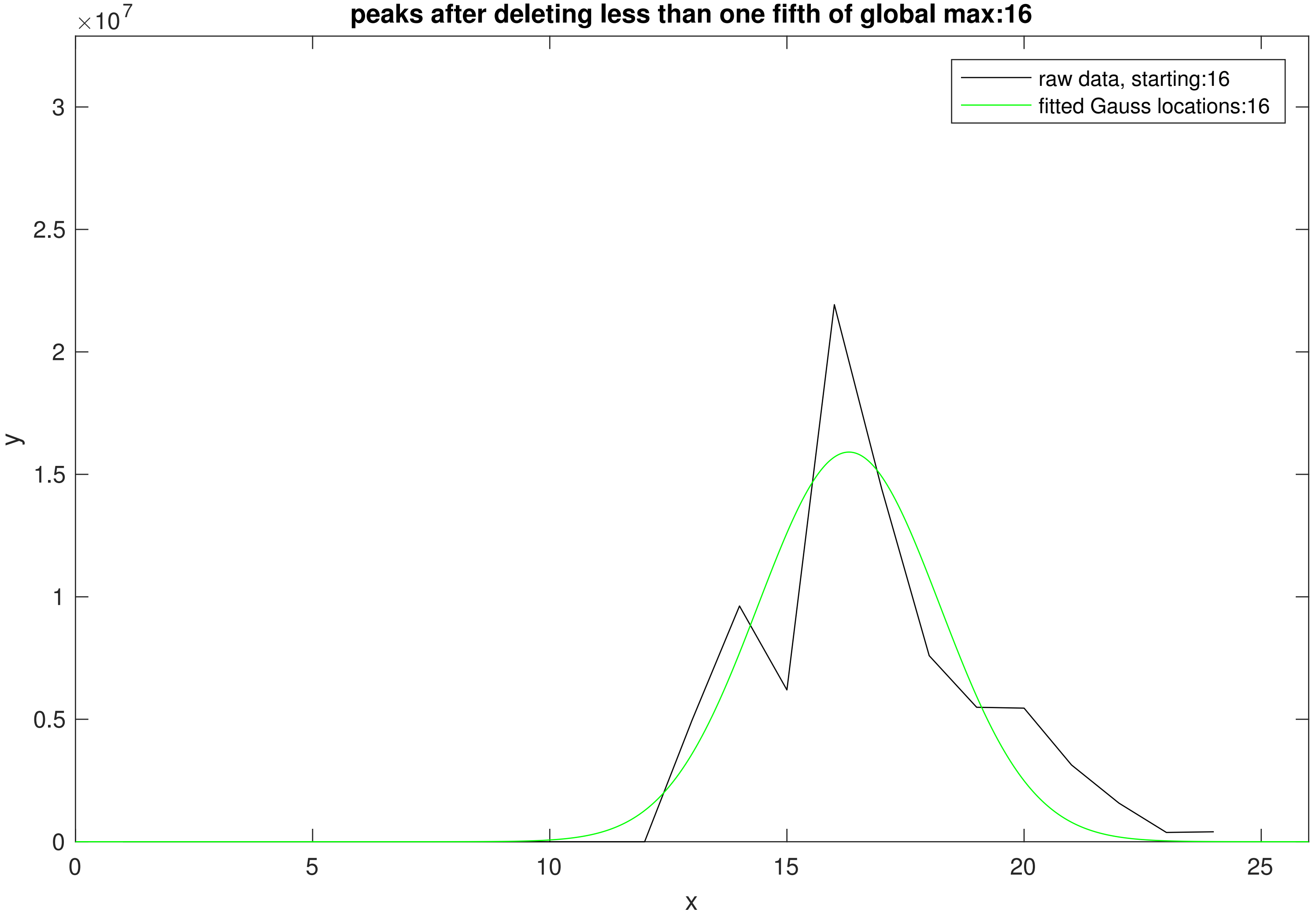

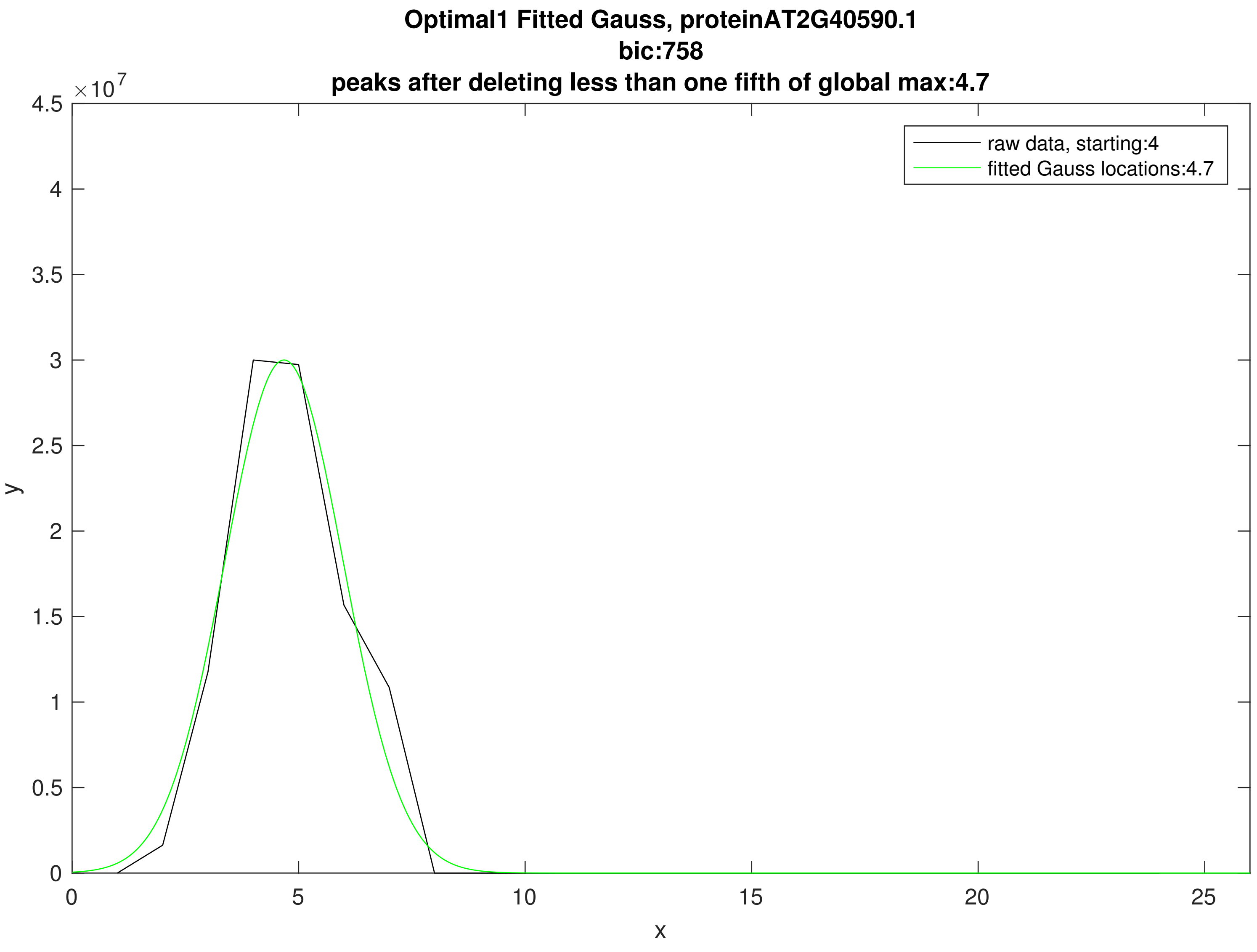
Optimal1 Fitted Gauss, proteinAT2G40590.1
bic:758
peaks after deleting less than one fifth of global max:4.7
×10^7
4.5
4
3.5
3
2.5
2
1.5
1
0.5
0
y
0
5
10
15
20
25
x
raw data, starting:4
fitted Gauss locations:4.7

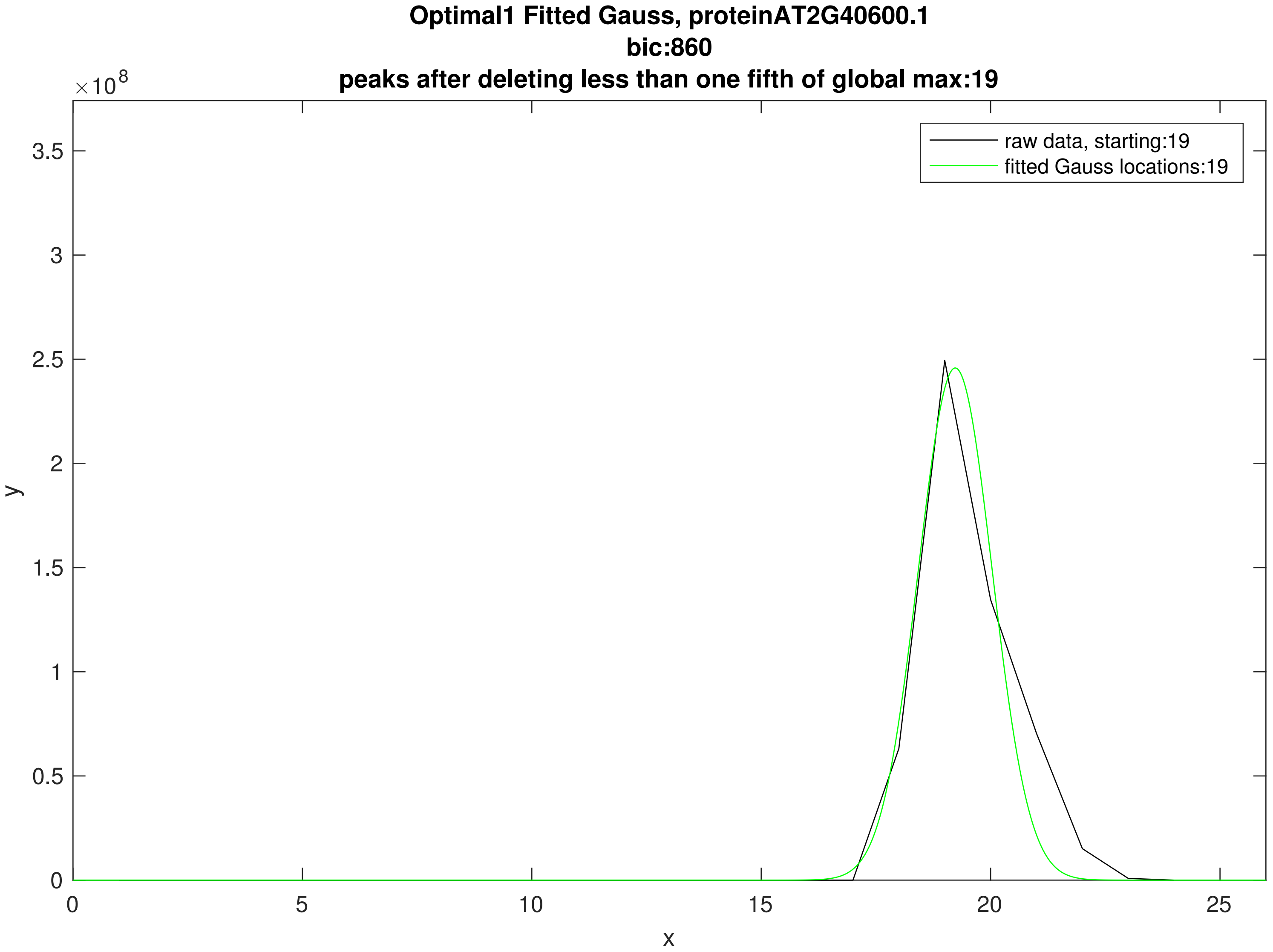
Optimal1 Fitted Gauss, proteinAT2G40600.1
bic:860
peaks after deleting less than one fifth of global max:19
×10^8
3.5
3
2.5
2
1.5
1
0.5
0
y
0
5
10
15
20
25
x
raw data, starting:19
fitted Gauss locations:19

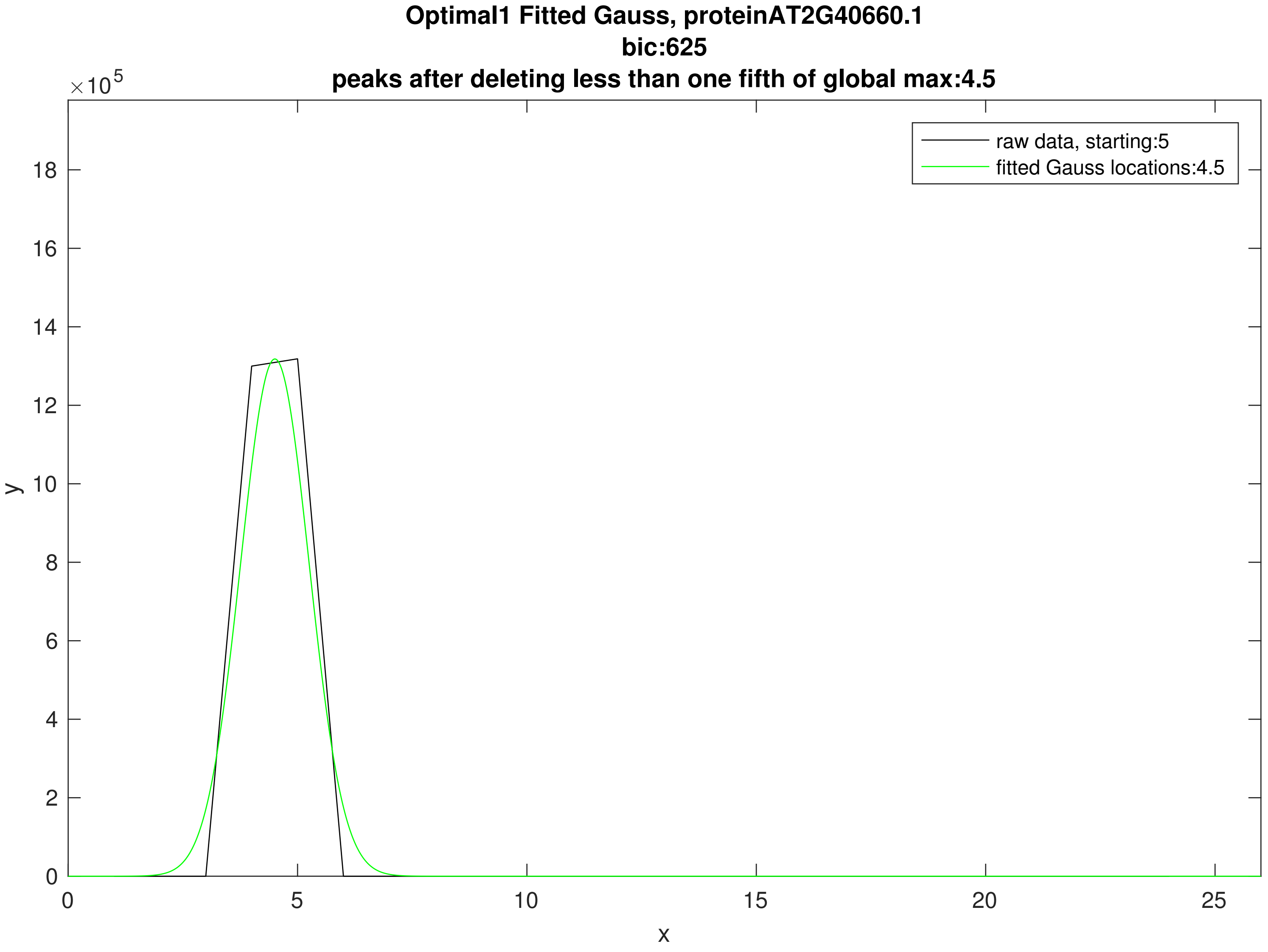
Optimal1 Fitted Gauss, proteinAT2G40660.1
bic:625
peaks after deleting less than one fifth of global max:4.5
×10^5
raw data, starting:5
fitted Gauss locations:4.5
y
x
0
2
4
6
8
10
12
14
16
18
0
5
10
15
20
25

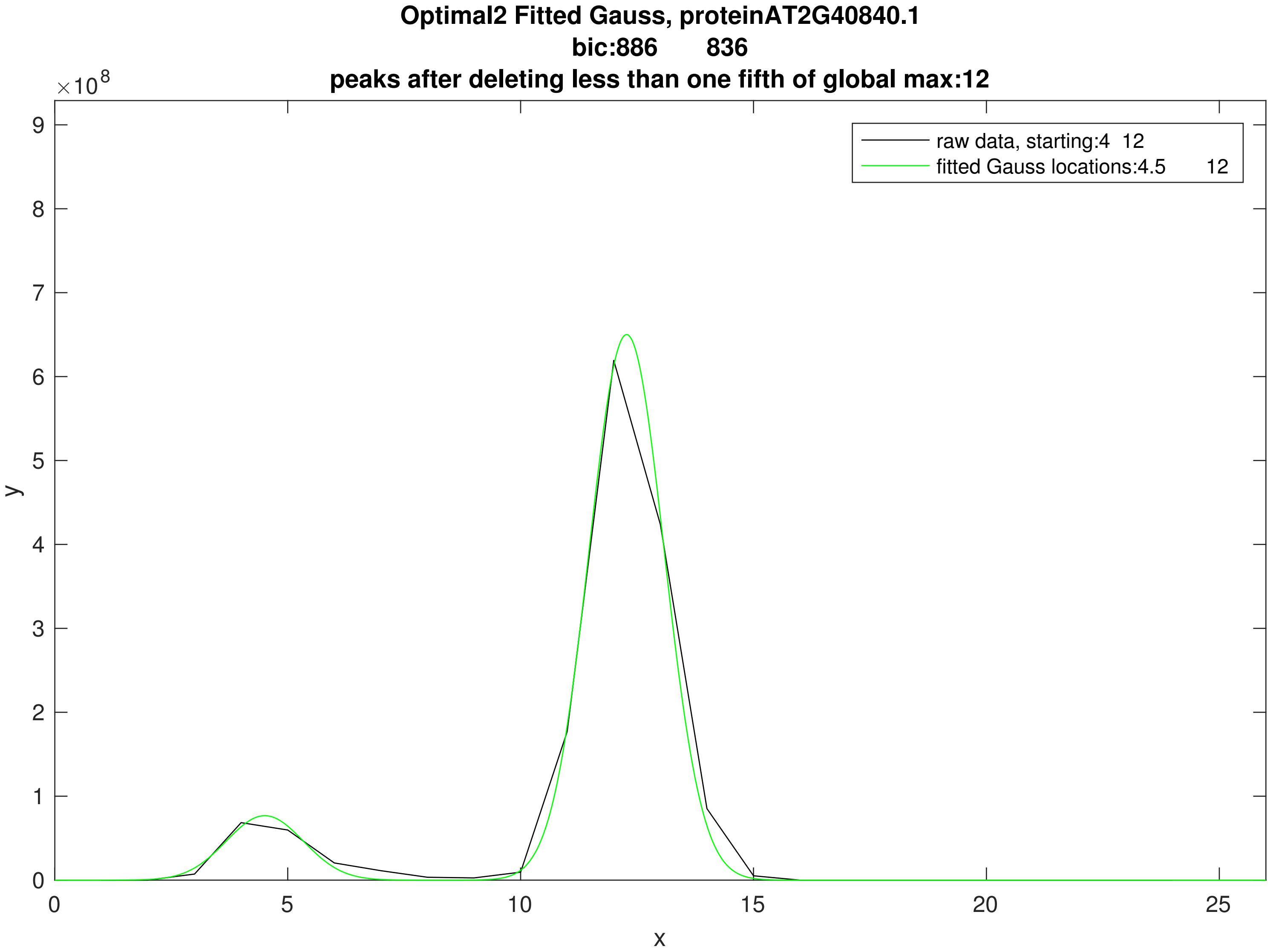

Optimal2 Fitted Gauss, proteinAT2G40840.1
bic:886 836
peaks after deleting less than one fifth of global max:12
×10^8
raw data, starting:4 12
fitted Gauss locations:4.5 12
y
x
0
1
2
3
4
5
6
7
8
9
0
5
10
15
20
25

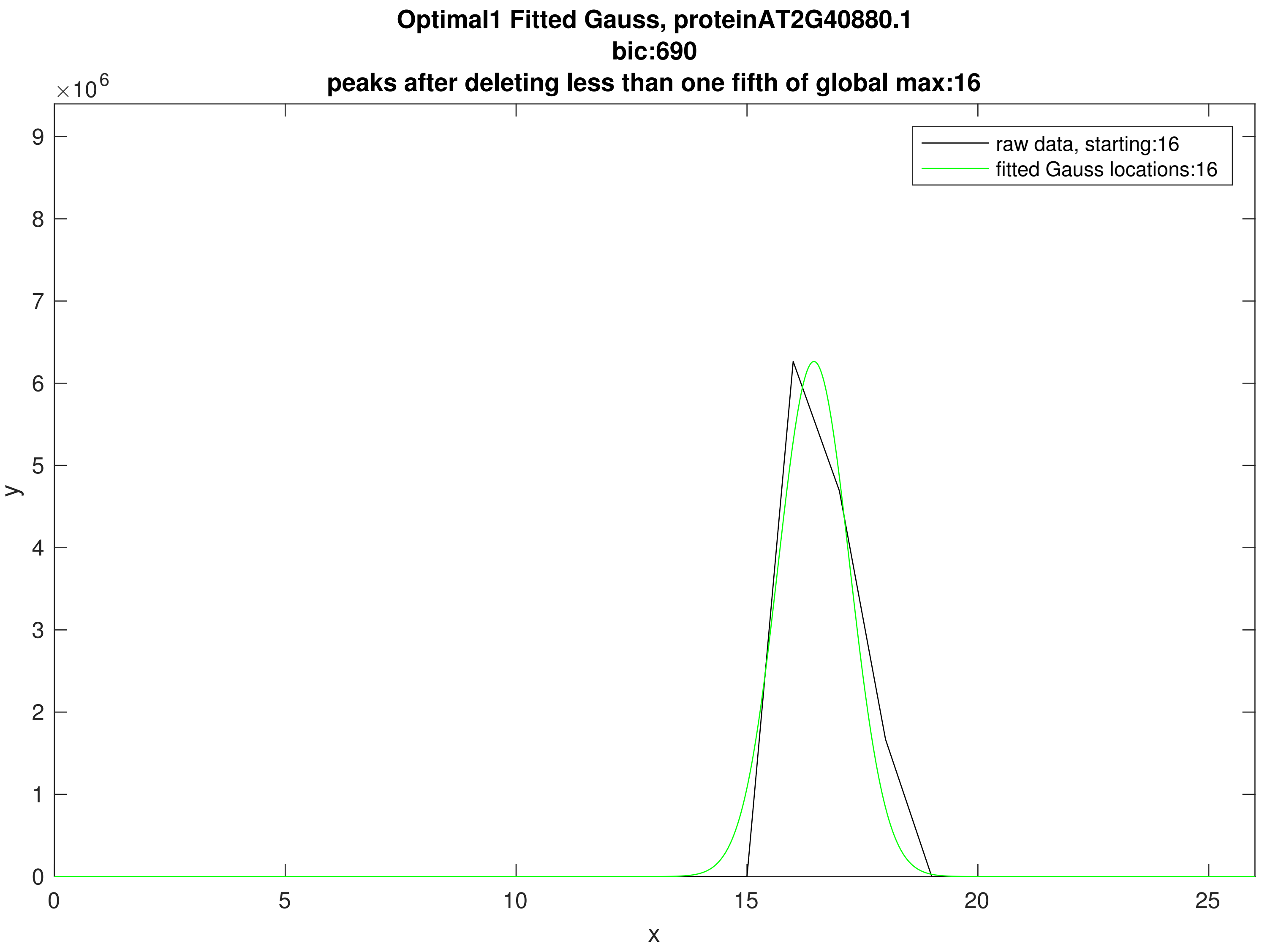
Optimal1 Fitted Gauss, proteinAT2G40880.1
bic:690
peaks after deleting less than one fifth of global max:16
×10^6
raw data, starting:16
fitted Gauss locations:16
y
x
0
1
2
3
4
5
6
7
8
9
0
5
10
15
20
25

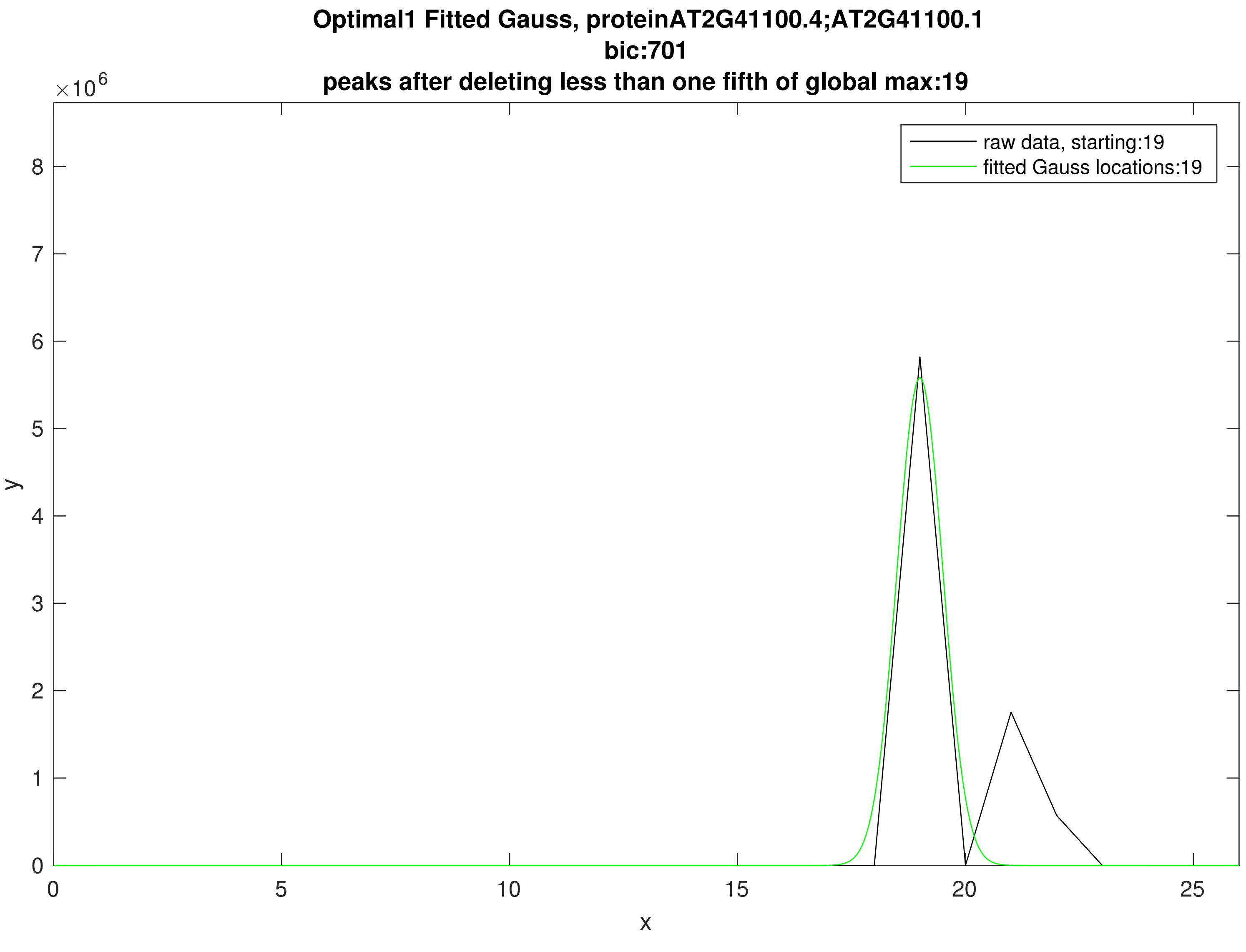

Optimal1 Fitted Gauss, proteinAT2G41100.4;AT2G41100.1
bic:701
peaks after deleting less than one fifth of global max:19
×10^6
raw data, starting:19
fitted Gauss locations:19
y
x
0
1
2
3
4
5
6
7
8
0
5
10
15
20
25

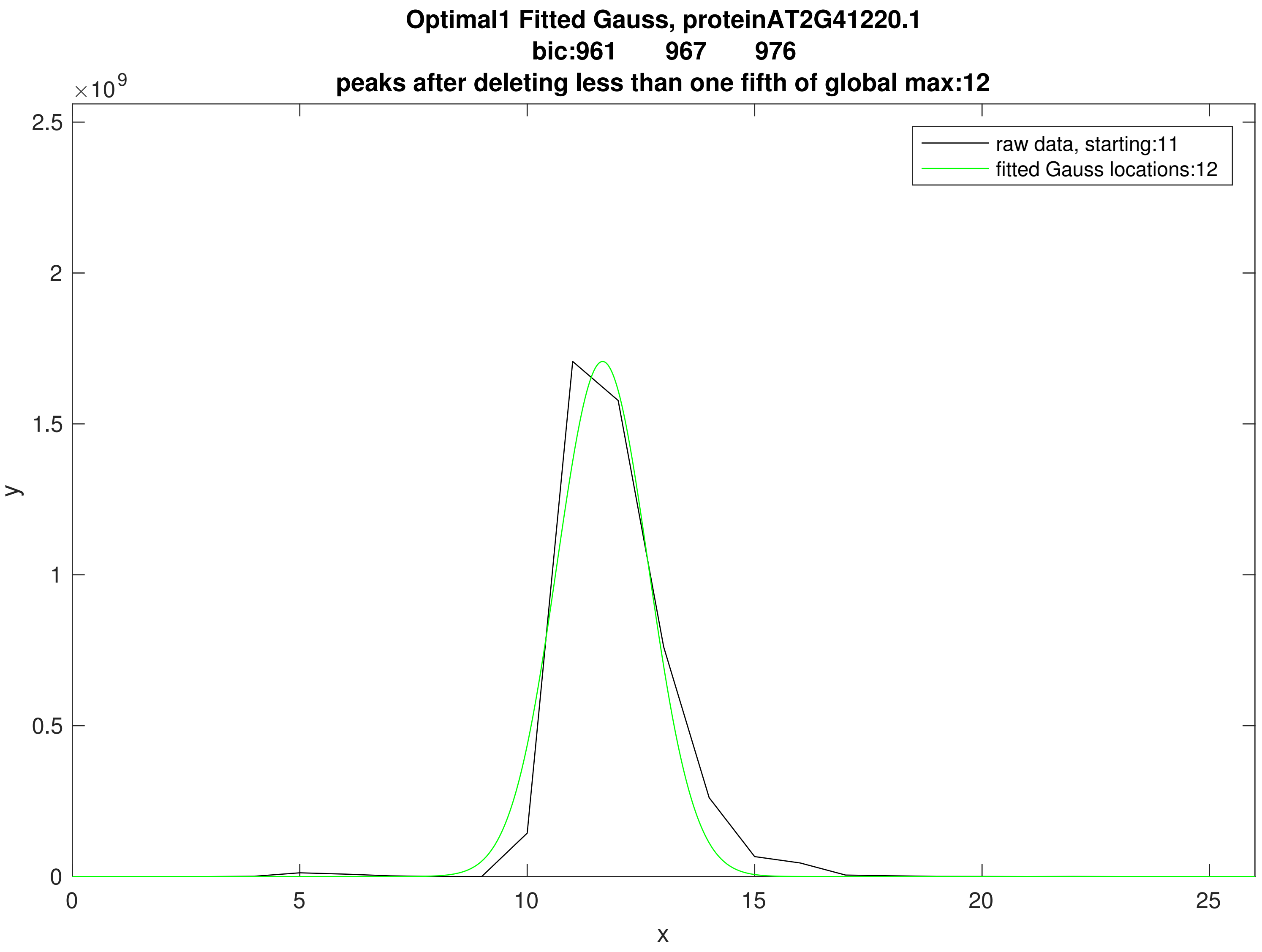

Optimal1 Fitted Gauss, proteinAT2G41220.1
bic:961 967 976
peaks after deleting less than one fifth of global max:12
×10^9
raw data, starting:11
fitted Gauss locations:12
y
x
0
0.5
1
1.5
2
2.5
0
5
10
15
20
25

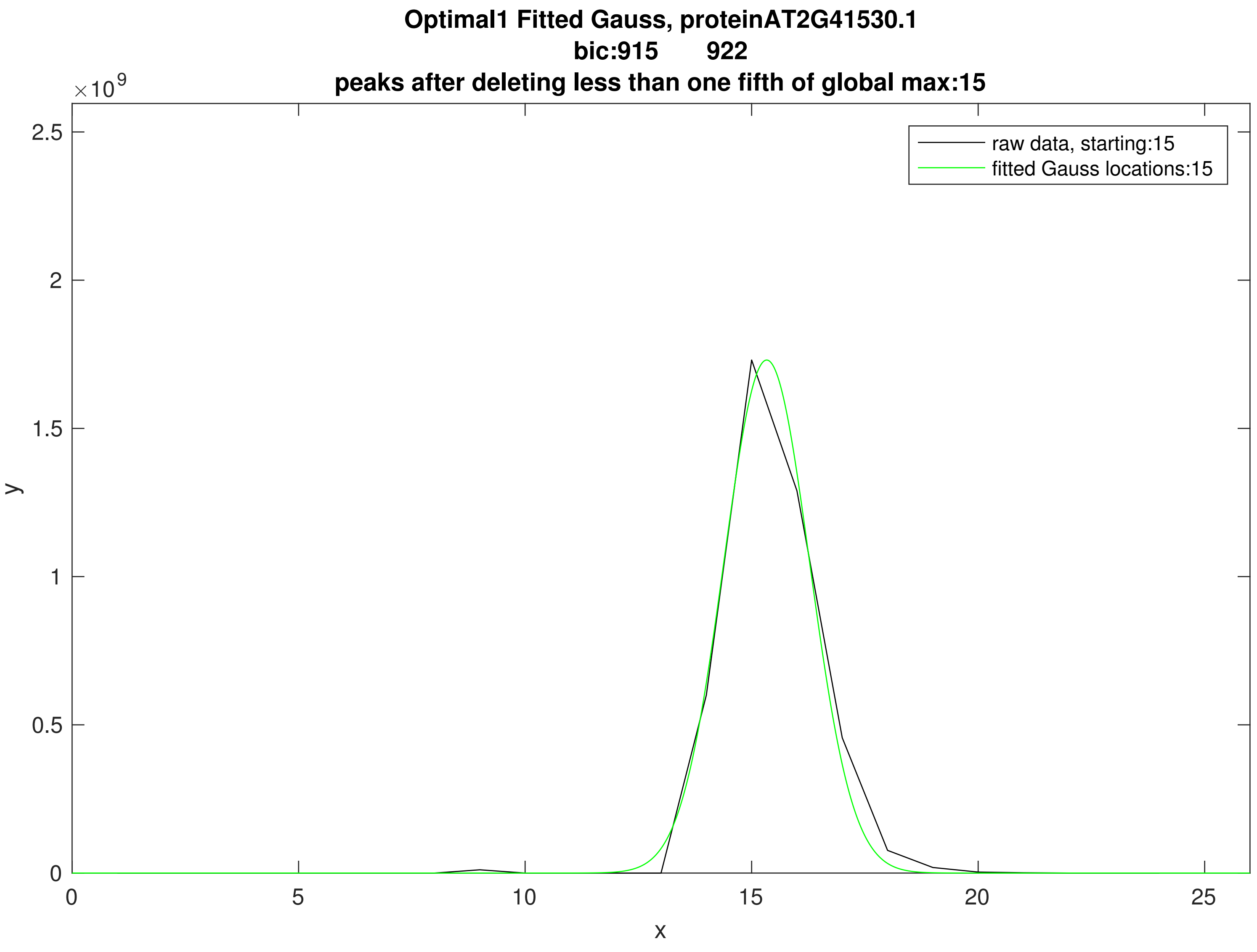
Optimal1 Fitted Gauss, proteinAT2G41530.1
bic:915 922
peaks after deleting less than one fifth of global max:15
×10^9
2.5
2
1.5
1
0.5
0
y
0
5
10
15
20
25
x
raw data, starting:15
fitted Gauss locations:15

Optimal1 Fitted Gauss, proteinAT2G41680.1
bic:805 824
peaks after deleting less than one fifth of global max:12

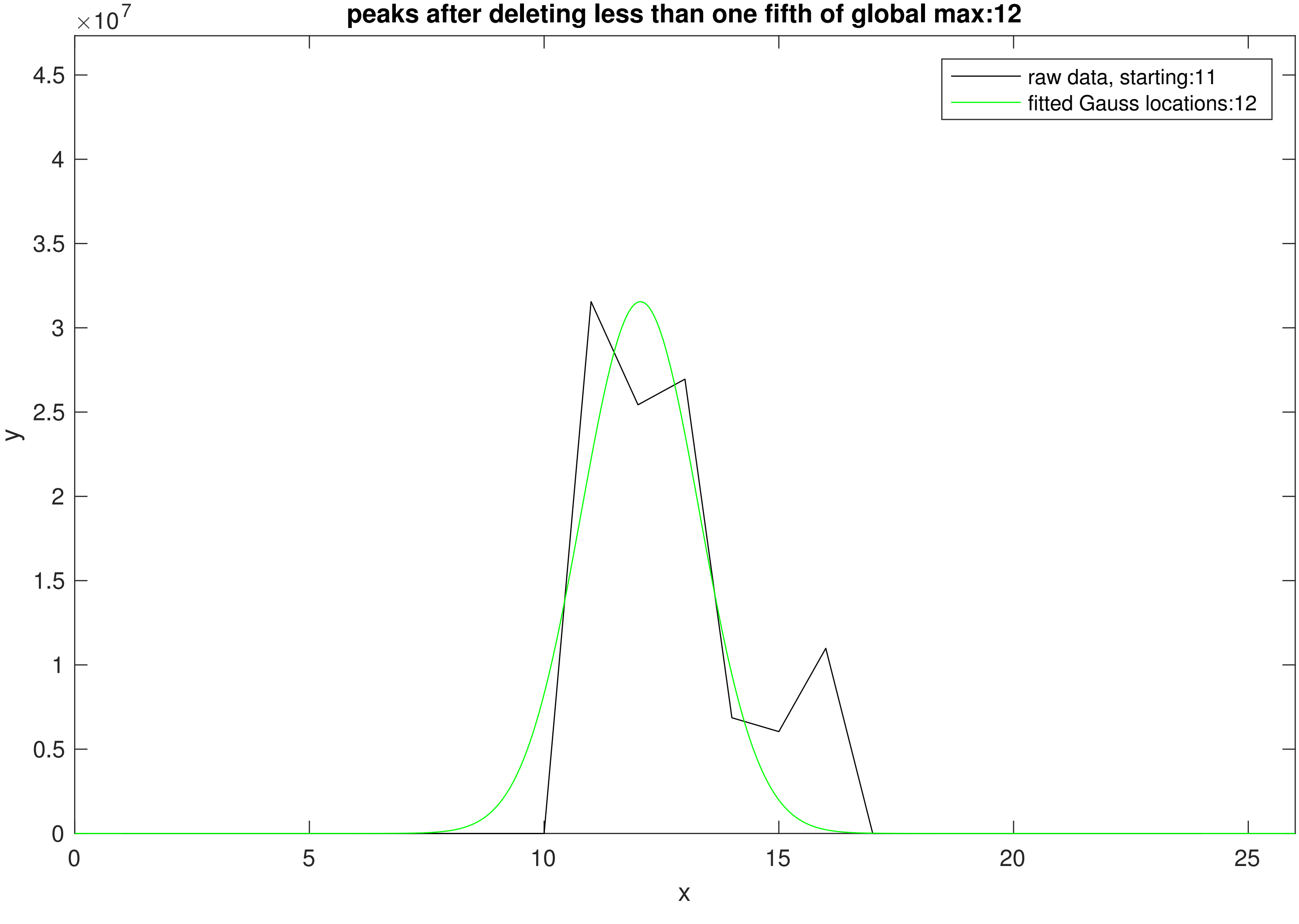

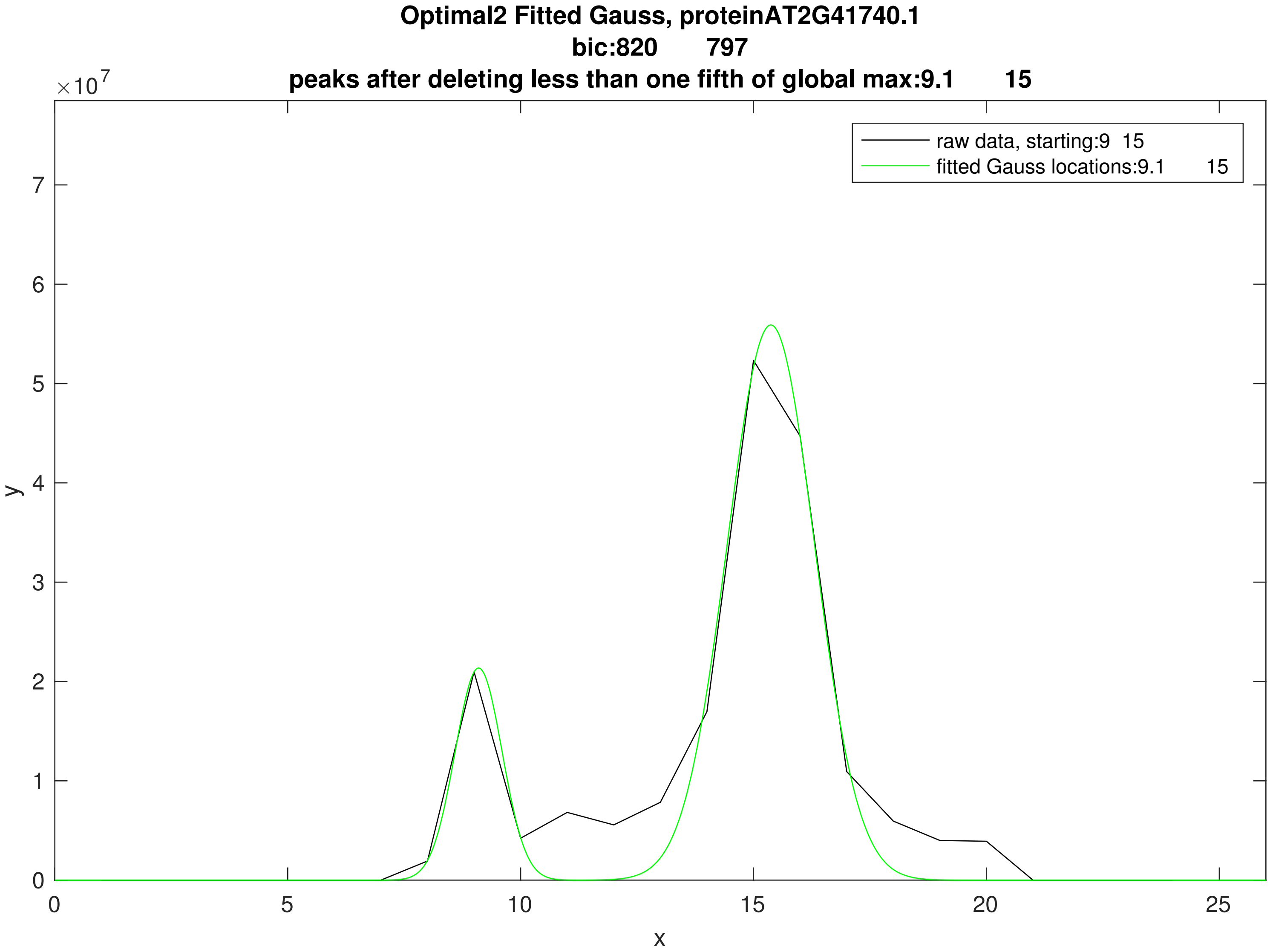
Optimal2 Fitted Gauss, proteinAT2G41740.1
bic:820 797
peaks after deleting less than one fifth of global max:9.1 15
×10^7
raw data, starting:9 15
fitted Gauss locations:9.1 15
y
x
0
1
2
3
4
5
6
7
0
5
10
15
20
25

**Optimal1 Fitted Gauss, proteinAT2G41790.1**
**bic:908 917**
**peaks after deleting less than one fifth of global max:9.3**

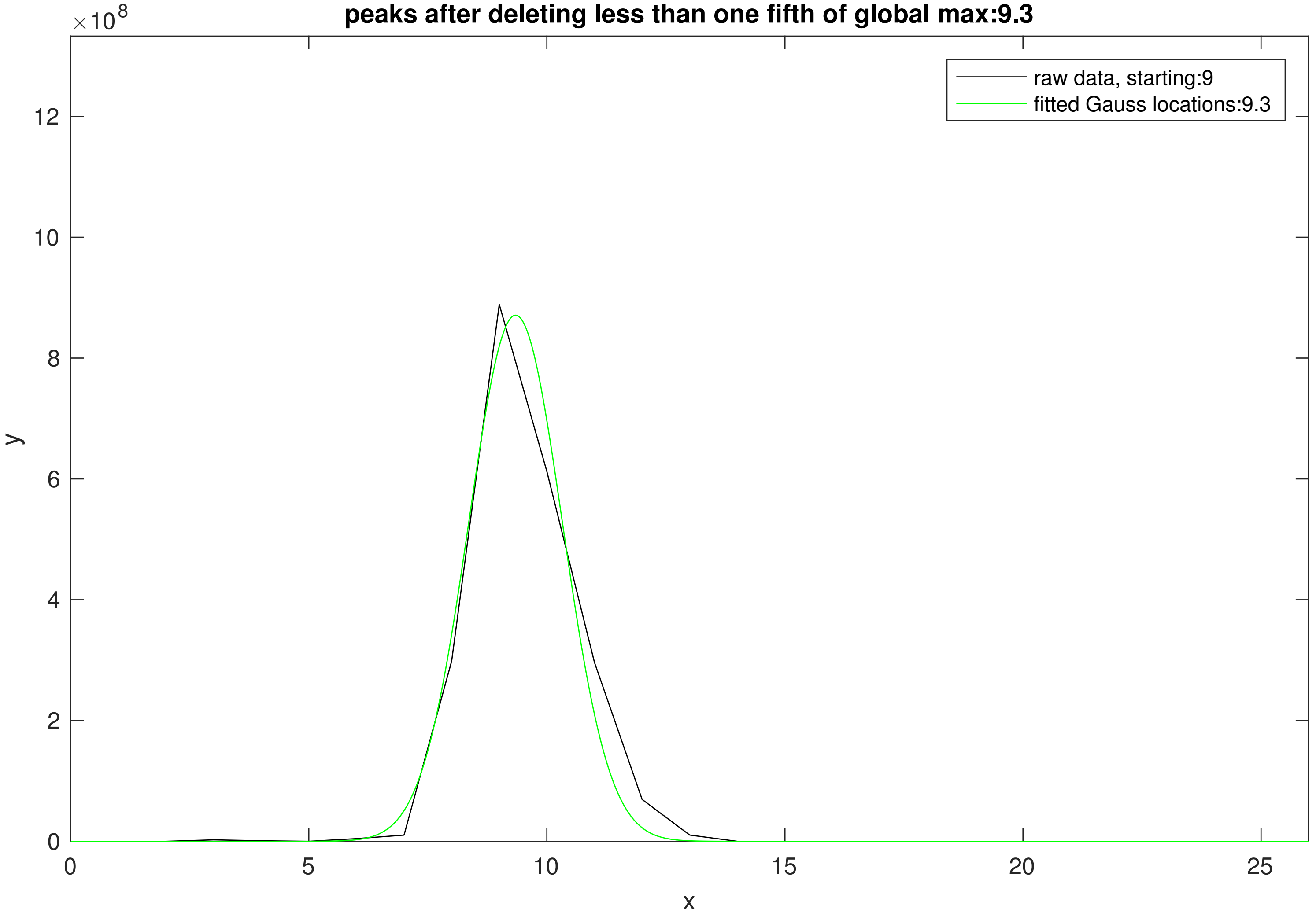

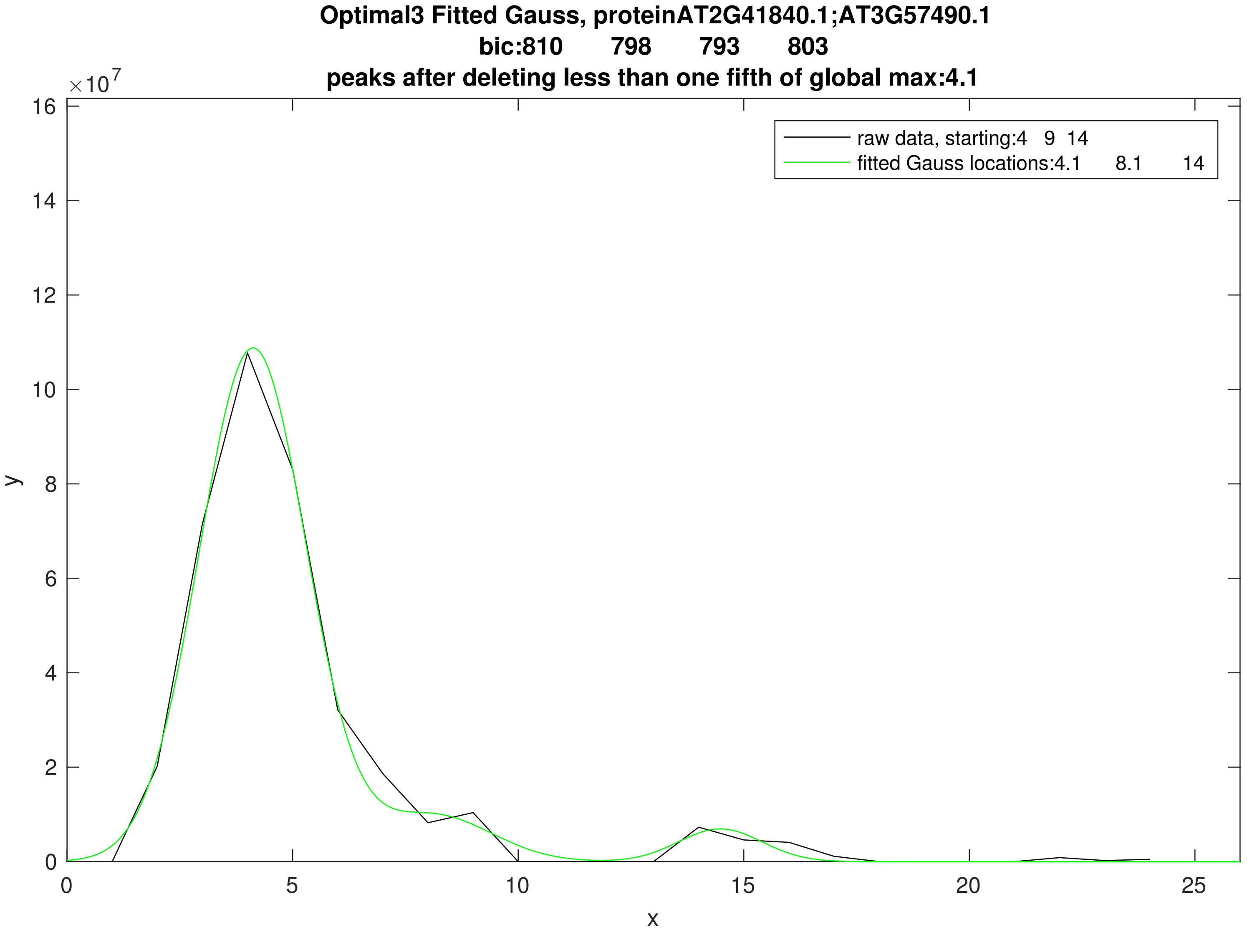

Optimal3 Fitted Gauss, proteinAT2G41840.1;AT3G57490.1
bic:810 798 793 803
peaks after deleting less than one fifth of global max:4.1
×10^7
raw data, starting:4 9 14
fitted Gauss locations:4.1 8.1 14
y
x

#### Optimal1 Fitted Gauss, proteinAT2G42560.1
#### bic:747 751
#### peaks after deleting less than one fifth of global max:8.2

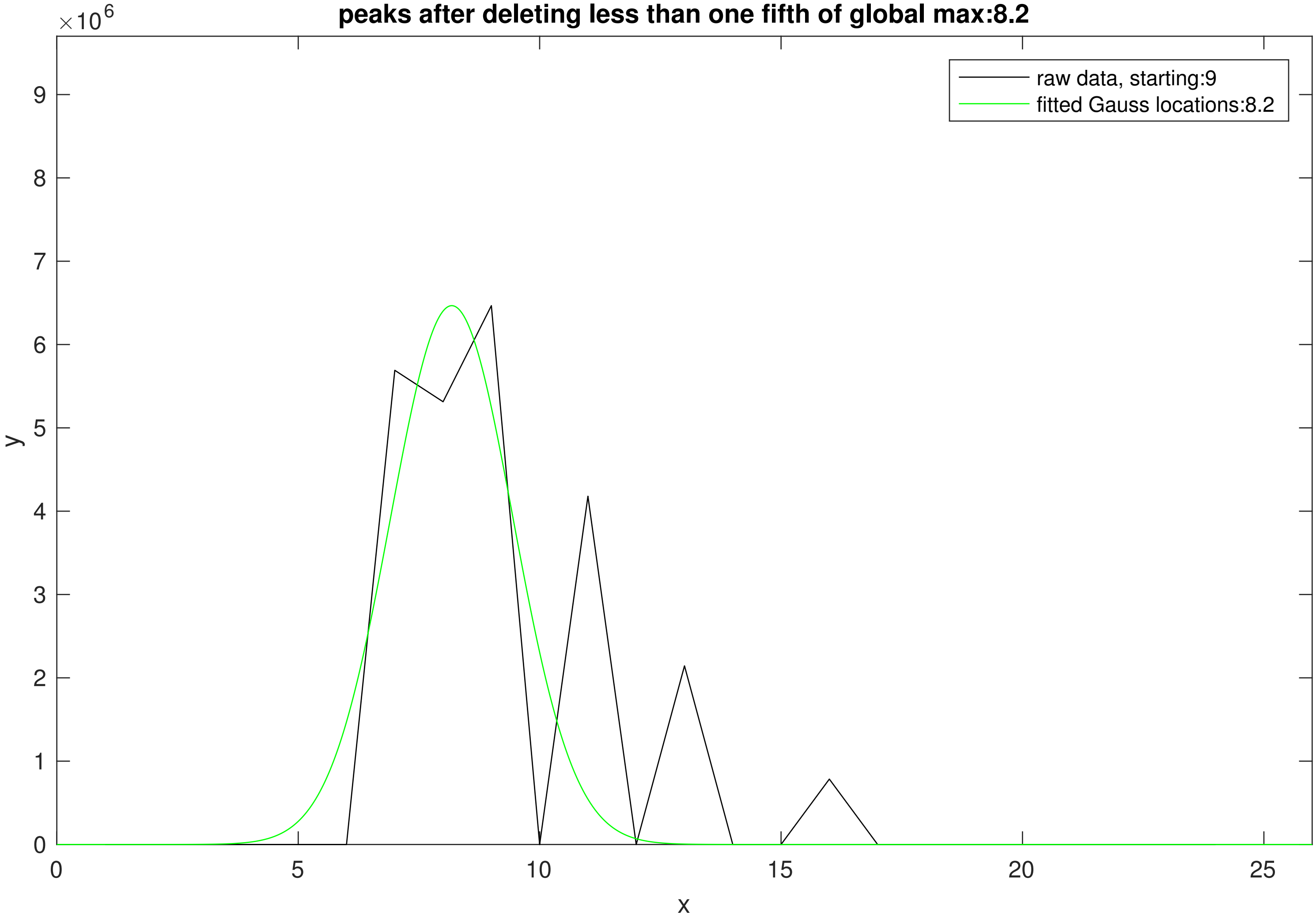

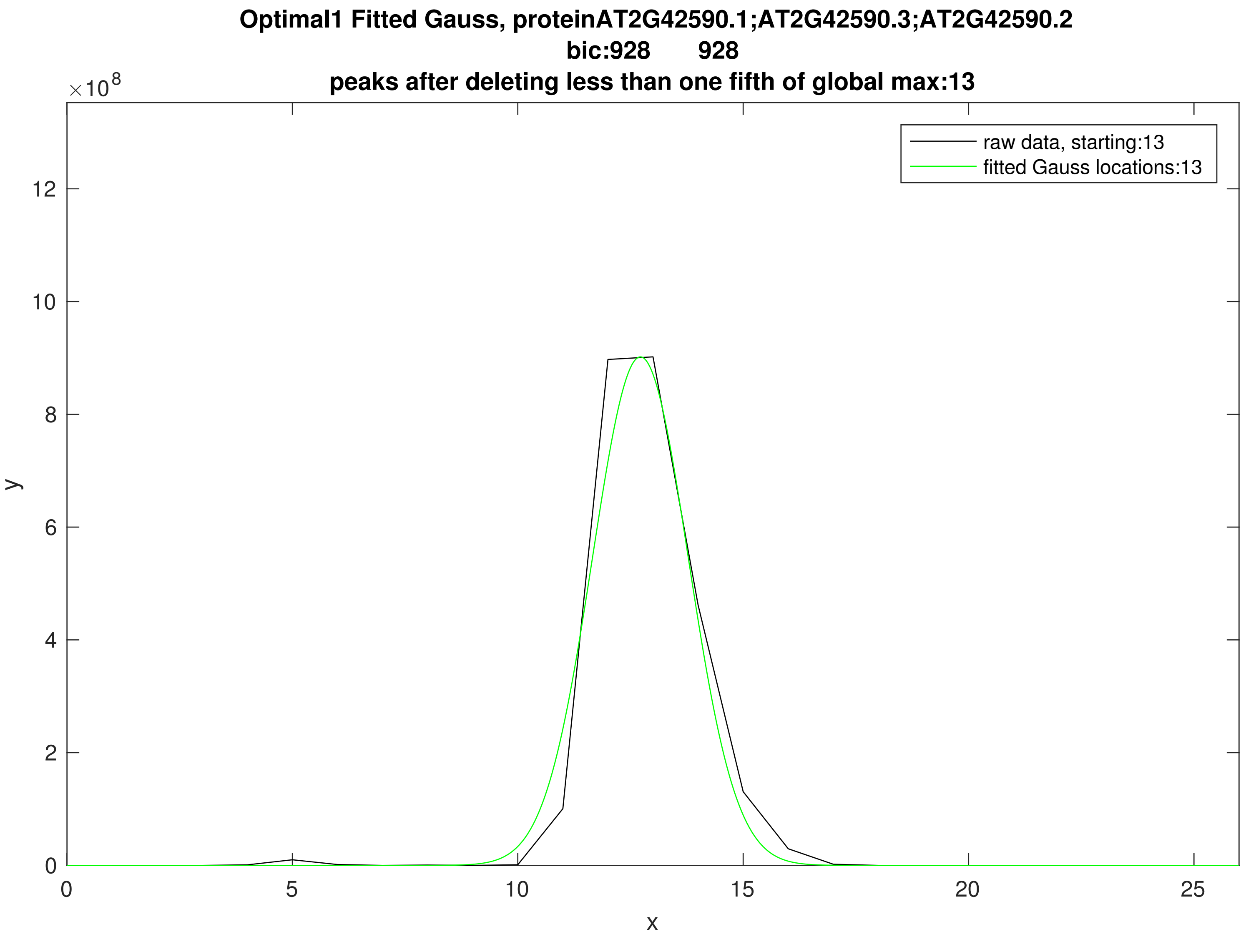
Optimal1 Fitted Gauss, proteinAT2G42590.1;AT2G42590.3;AT2G42590.2
bic:928 928
peaks after deleting less than one fifth of global max:13
$\times 10^8$
raw data, starting:13
fitted Gauss locations:13
y
x
0
2
4
6
8
10
12
5
15
20
25

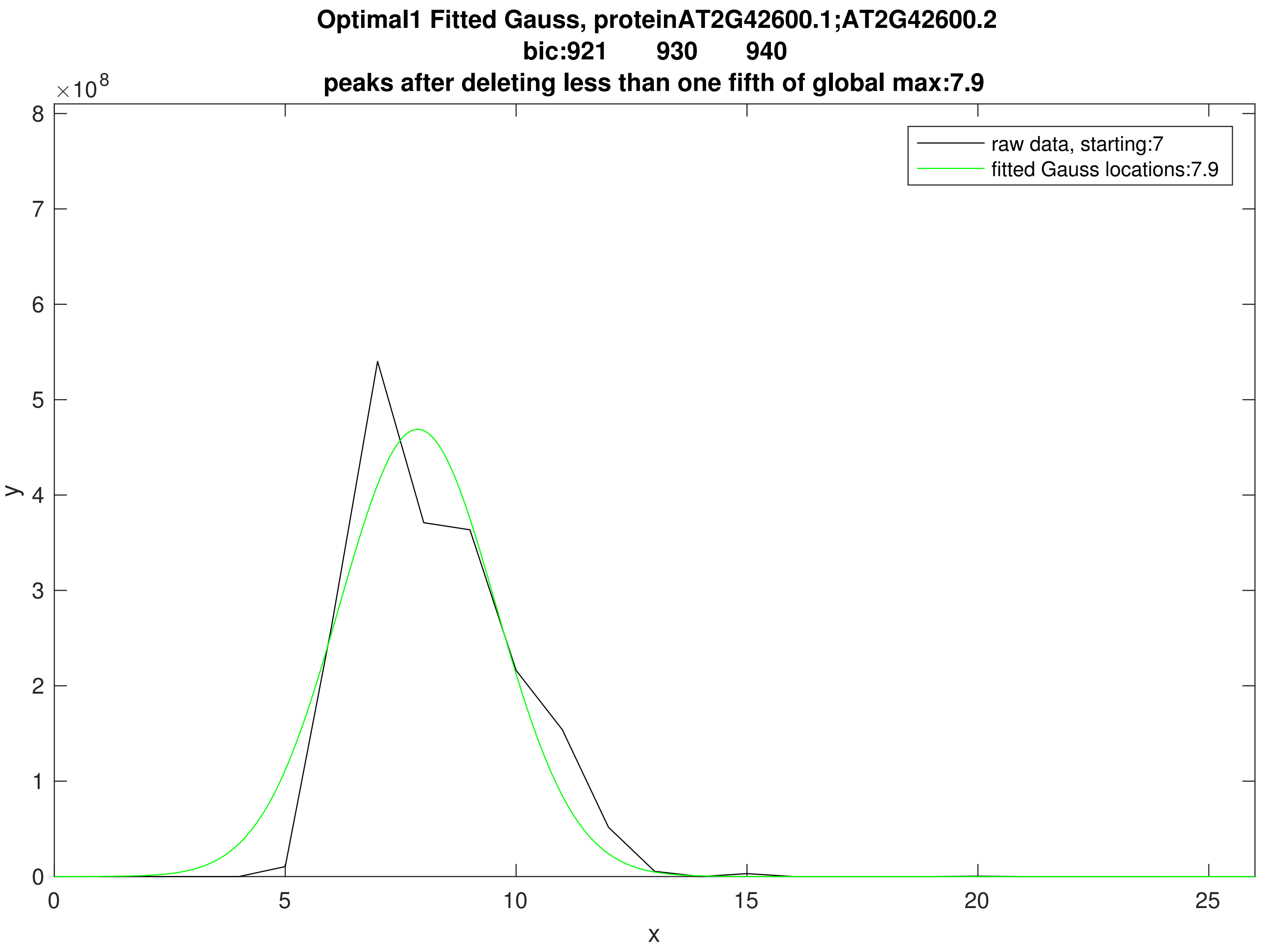
Optimal1 Fitted Gauss, proteinAT2G42600.1;AT2G42600.2
bic:921 930 940
peaks after deleting less than one fifth of global max:7.9
×10^8
raw data, starting:7
fitted Gauss locations:7.9
y
x
0
1
2
3
4
5
6
7
8
0
5
10
15
20
25

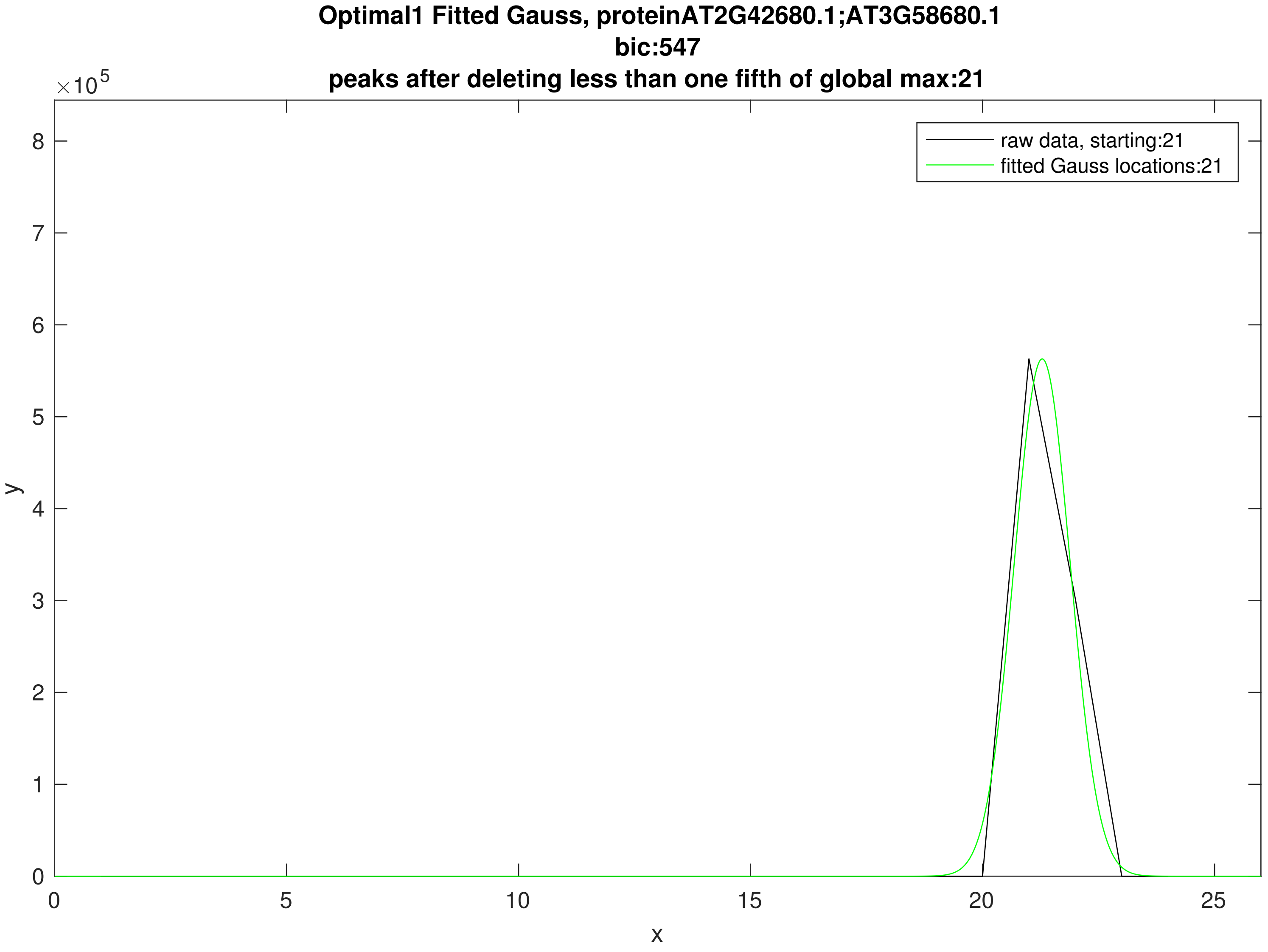
Optimal1 Fitted Gauss, proteinAT2G42680.1;AT3G58680.1
bic:547
peaks after deleting less than one fifth of global max:21
×10^5
8
7
6
5
4
3
2
1
0
y
0
5
10
15
20
25
x
raw data, starting:21
fitted Gauss locations:21

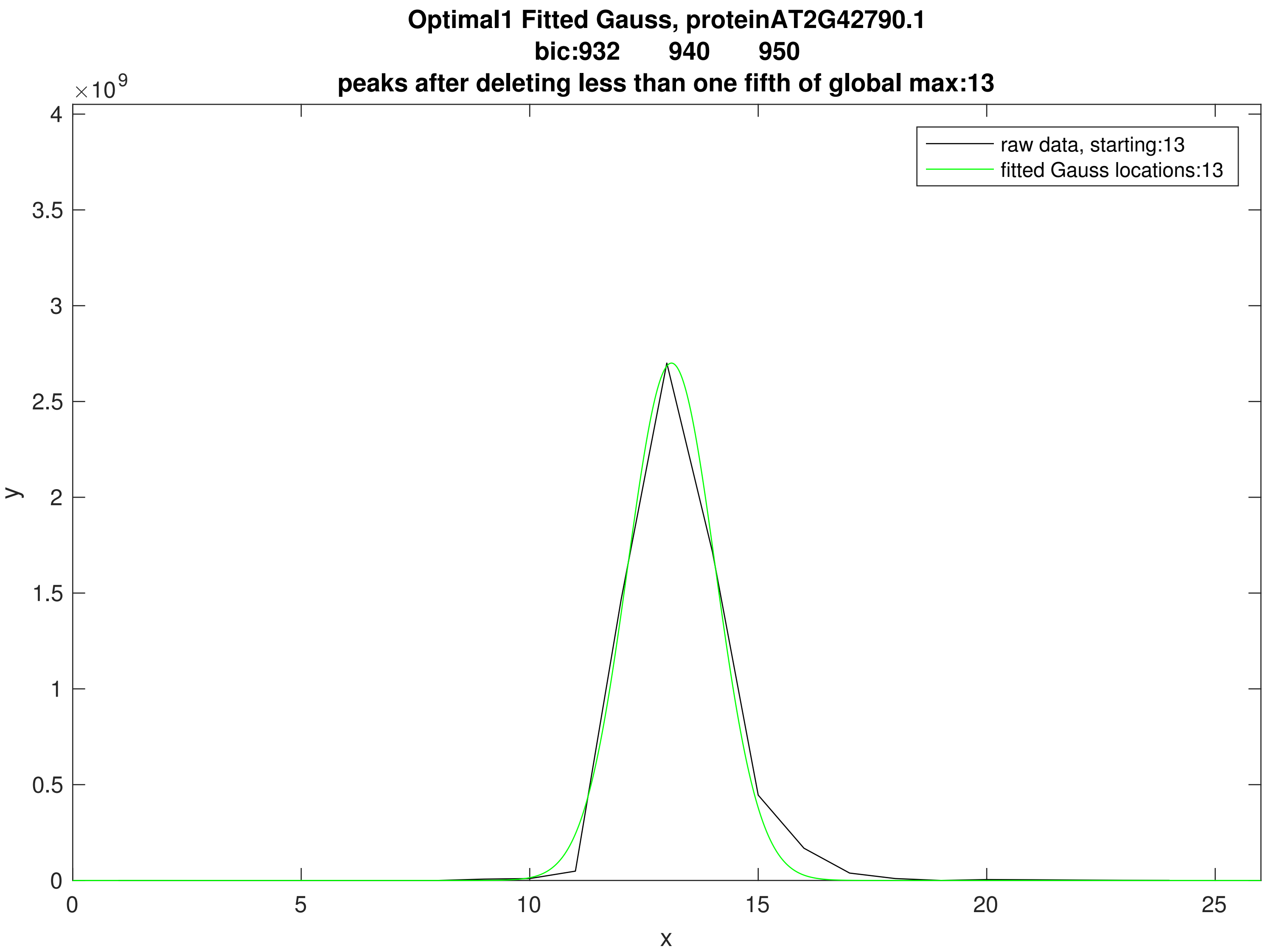
Optimal1 Fitted Gauss, proteinAT2G42790.1
bic:932 940 950
peaks after deleting less than one fifth of global max:13
×10^9
4
3.5
3
2.5
2
1.5
1
0.5
0
y
0
5
10
15
20
25
x
raw data, starting:13
fitted Gauss locations:13

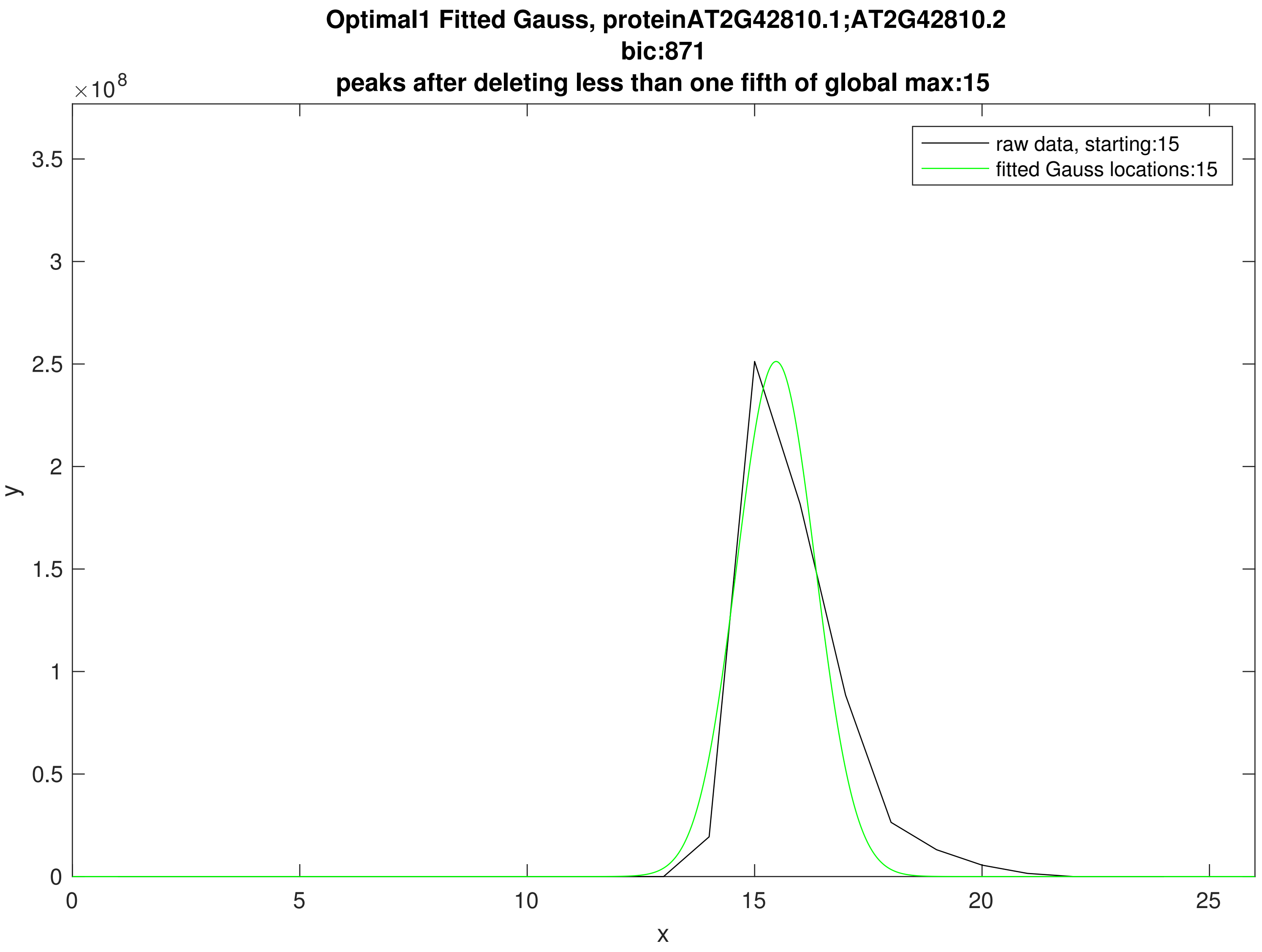
Optimal1 Fitted Gauss, proteinAT2G42810.1;AT2G42810.2
bic:871
peaks after deleting less than one fifth of global max:15
×10^8
raw data, starting:15
fitted Gauss locations:15
3.5
3
2.5
2
1.5
1
0.5
0
y
0
5
10
15
20
25
x

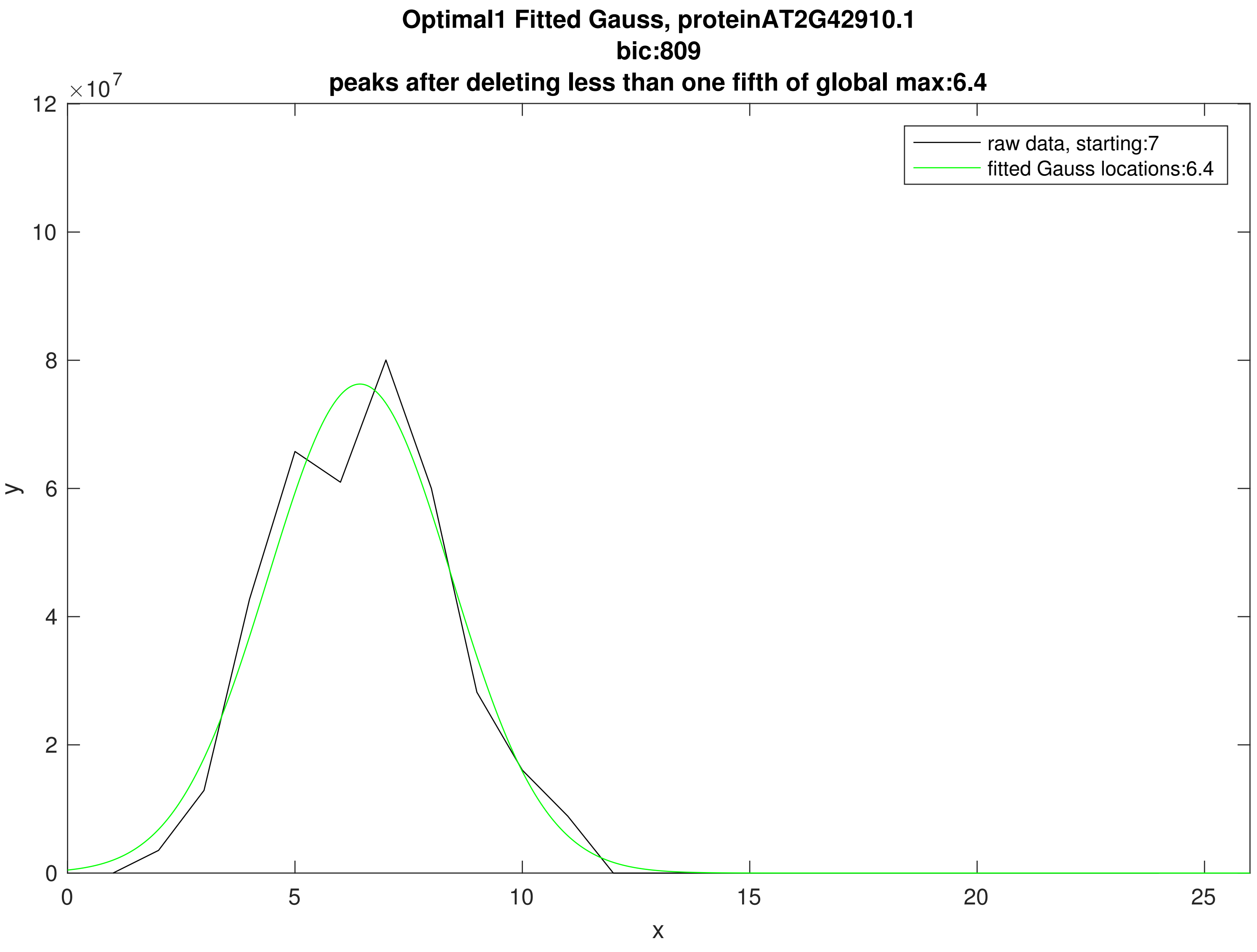
Optimal1 Fitted Gauss, proteinAT2G42910.1
bic:809
peaks after deleting less than one fifth of global max:6.4
×10^7
raw data, starting:7
fitted Gauss locations:6.4
y
x
0
2
4
6
8
10
12
0
5
10
15
20
25

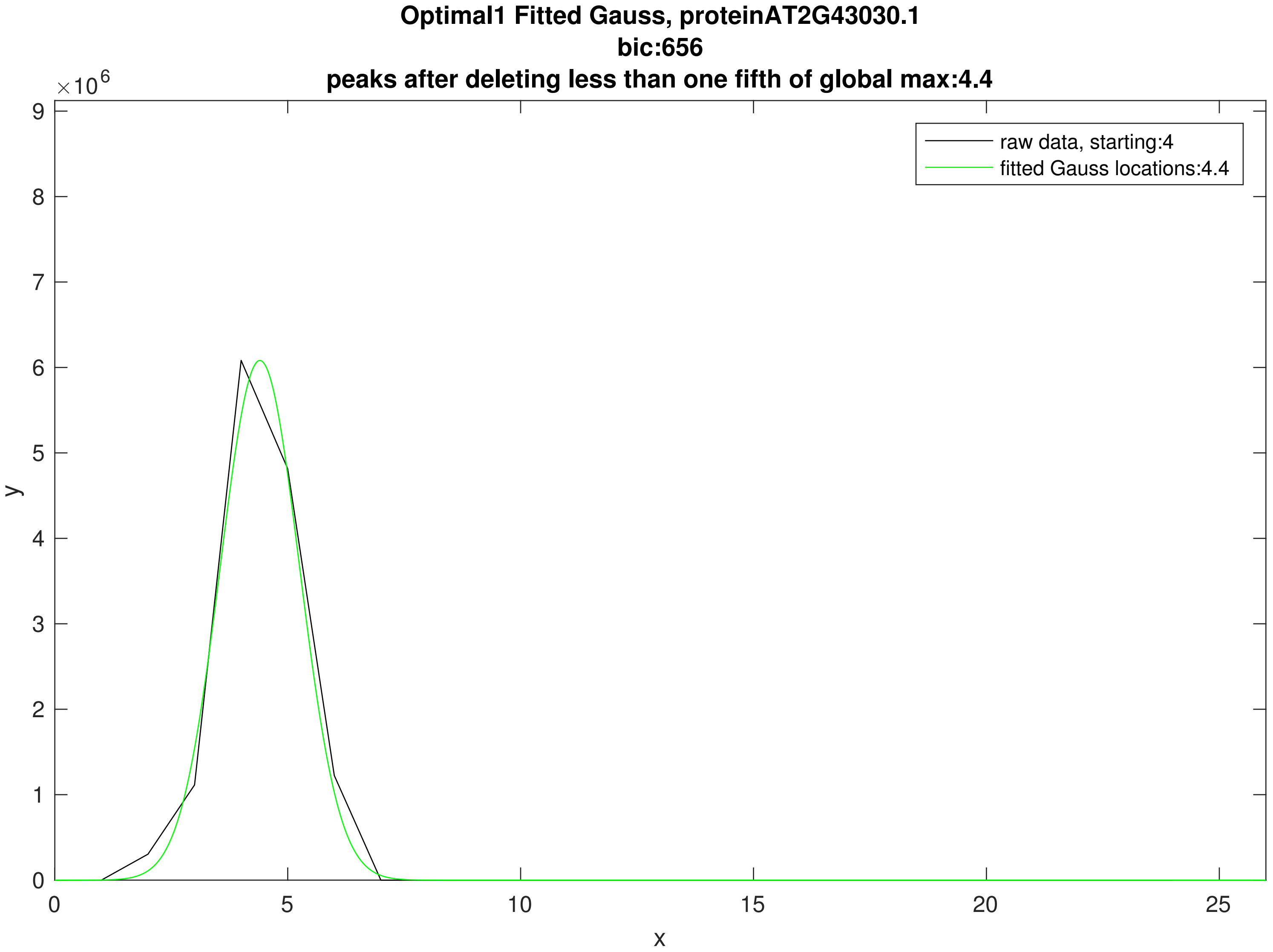
Optimal1 Fitted Gauss, proteinAT2G43030.1
bic:656
peaks after deleting less than one fifth of global max:4.4
×10^6
raw data, starting:4
fitted Gauss locations:4.4
y
x
0
1
2
3
4
5
6
7
8
9
0
5
10
15
20
25

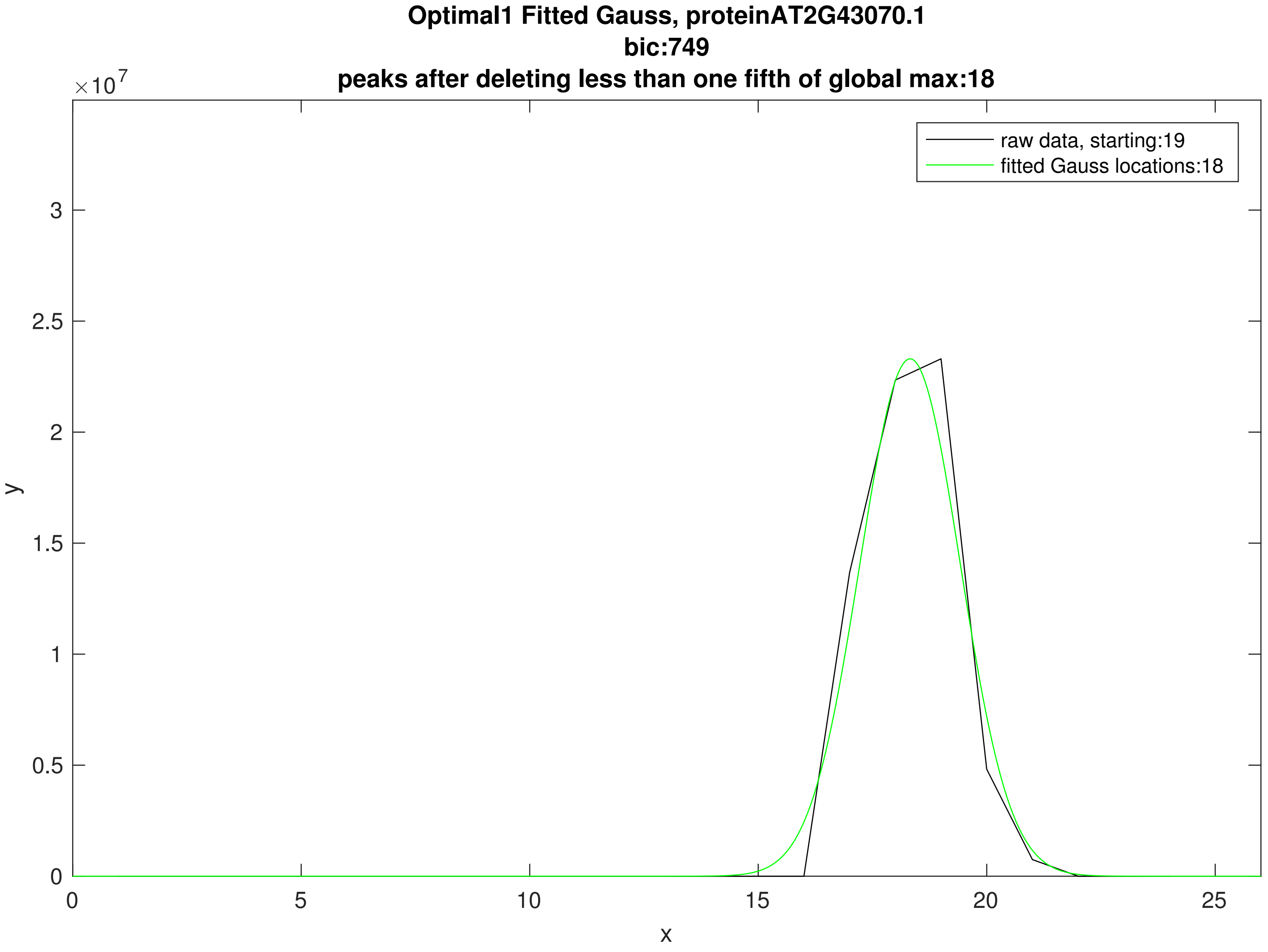
Optimal1 Fitted Gauss, proteinAT2G43070.1
bic:749
peaks after deleting less than one fifth of global max:18
×10^7
raw data, starting:19
fitted Gauss locations:18
y
x
0
0.5
1
1.5
2
2.5
3
5
10
15
20
25

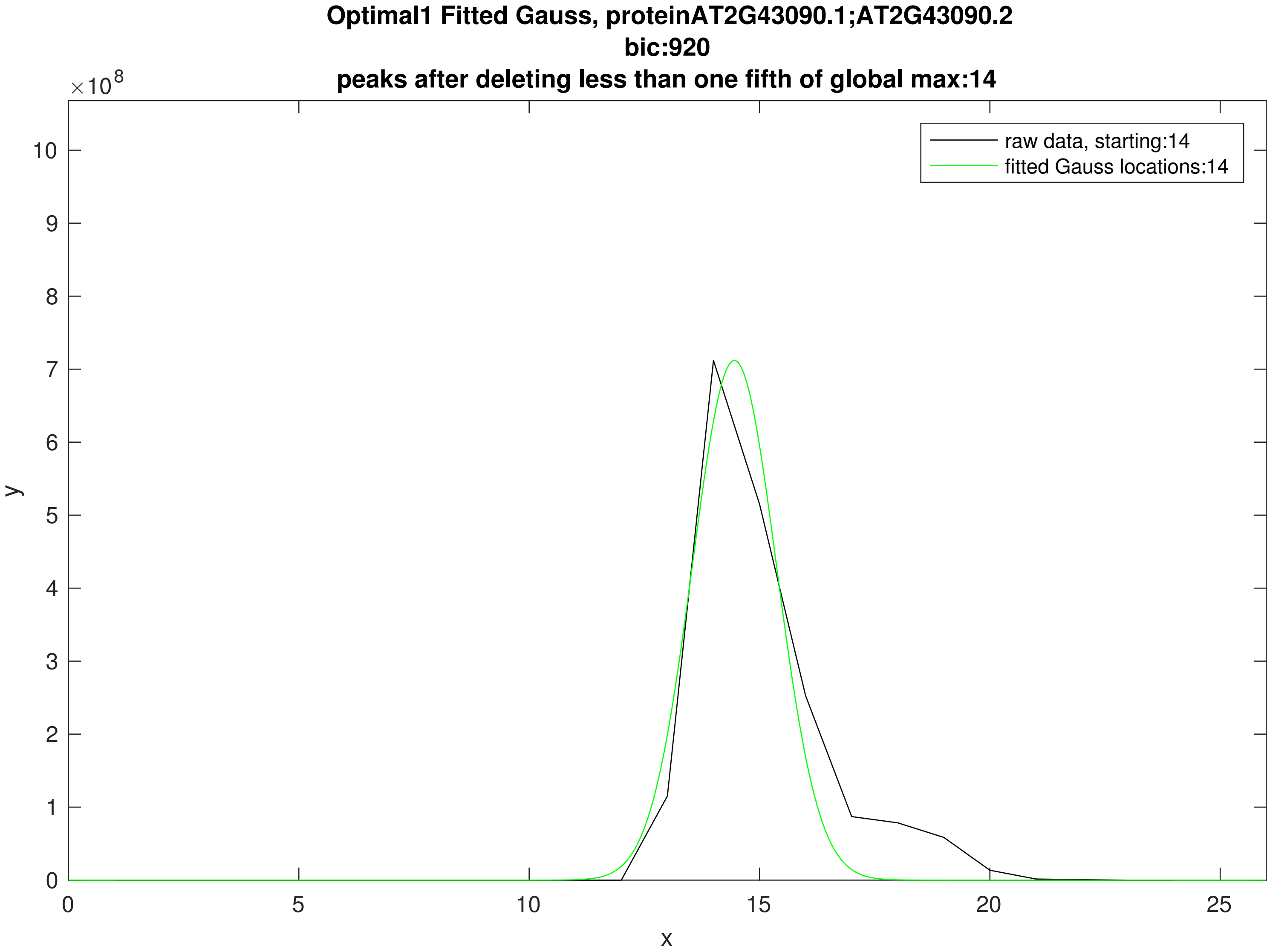
Optimal1 Fitted Gauss, proteinAT2G43090.1;AT2G43090.2
bic:920
peaks after deleting less than one fifth of global max:14
×10^8
raw data, starting:14
fitted Gauss locations:14
y
x
0
1
2
3
4
5
6
7
8
9
10
0
5
10
15
20
25

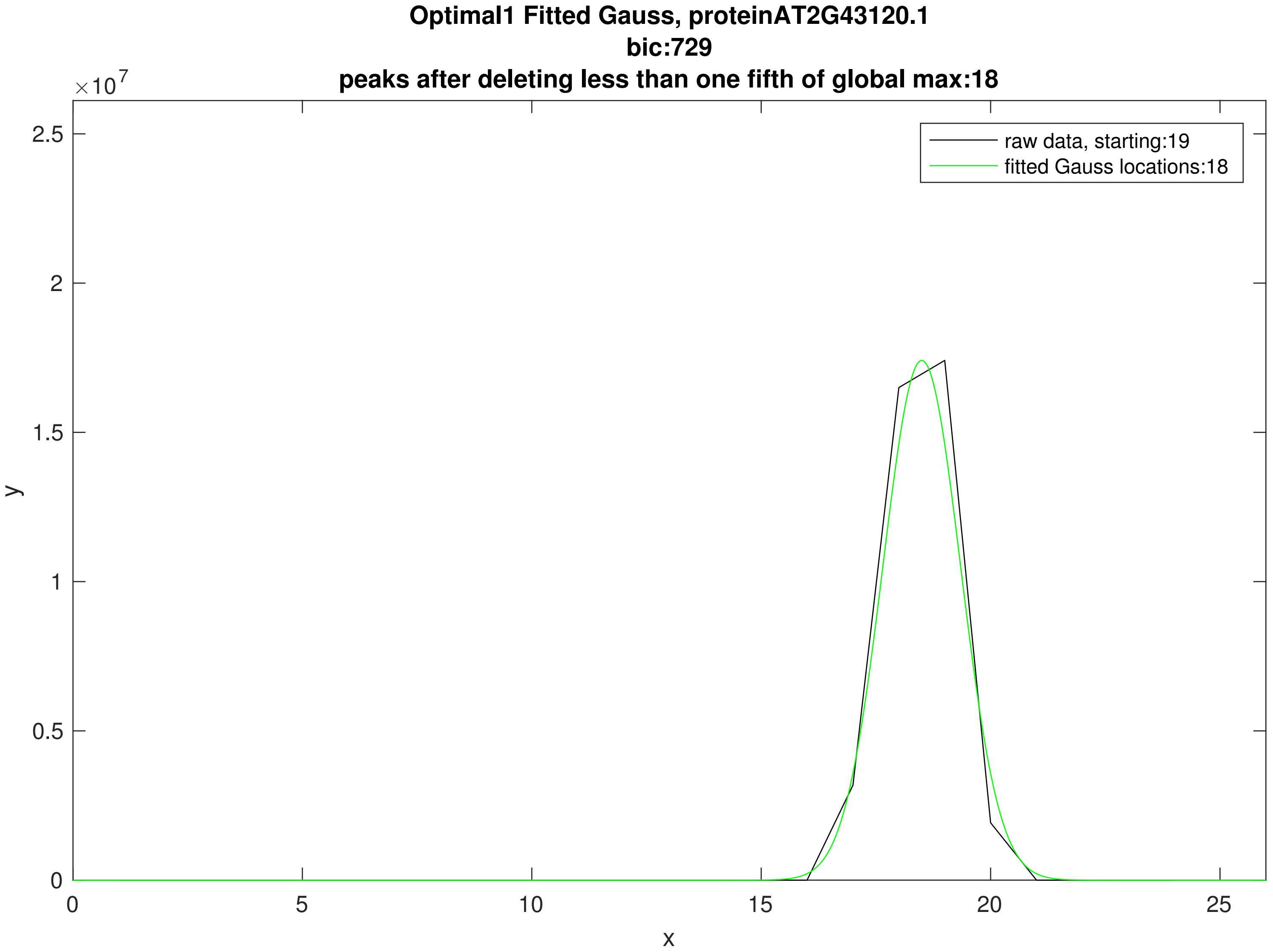
Optimal1 Fitted Gauss, proteinAT2G43120.1
bic:729
peaks after deleting less than one fifth of global max:18
×10^7
raw data, starting:19
fitted Gauss locations:18
y
x
0
0.5
1
1.5
2
2.5
0
5
10
15
20
25

**Optimal1 Fitted Gauss, proteinAT2G43410.1;AT2G43410.4;AT2G43410.3;AT2G43410.2**
**bic:669**
**peaks after deleting less than one fifth of global max:11**

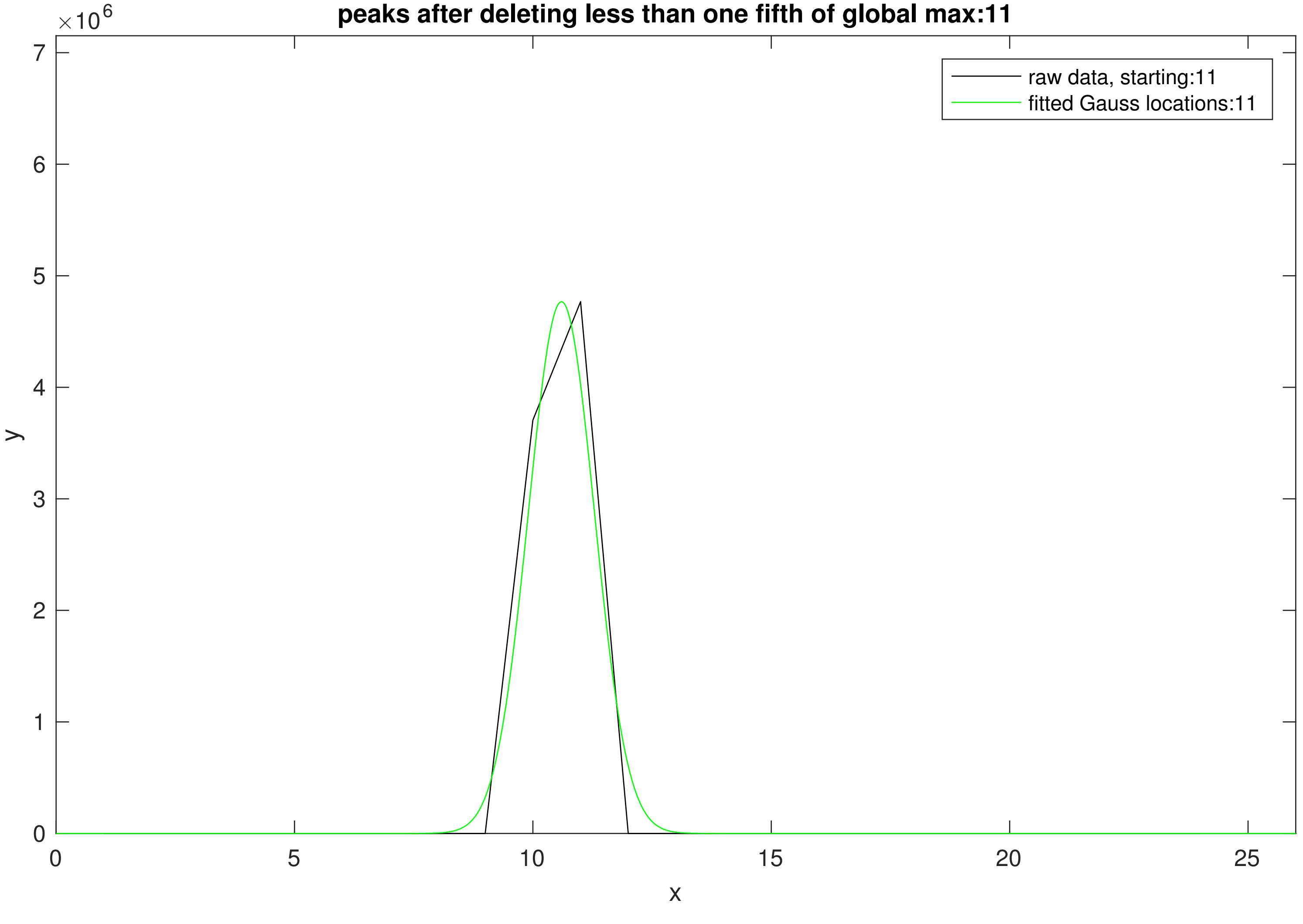

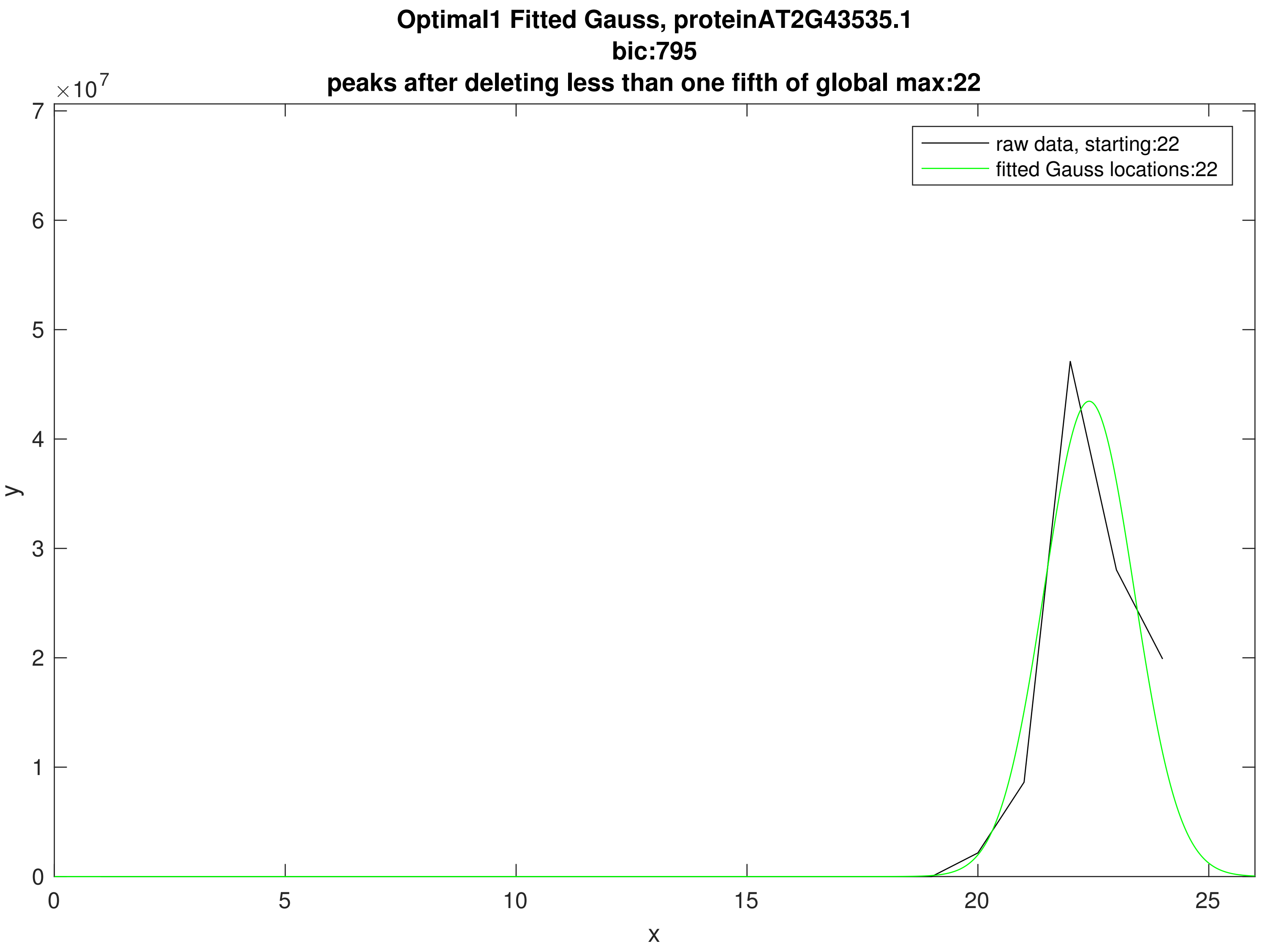

Optimal1 Fitted Gauss, proteinAT2G43535.1
bic:795
peaks after deleting less than one fifth of global max:22
×10^7
raw data, starting:22
fitted Gauss locations:22
y
x
0
1
2
3
4
5
6
7
0
5
10
15
20
25

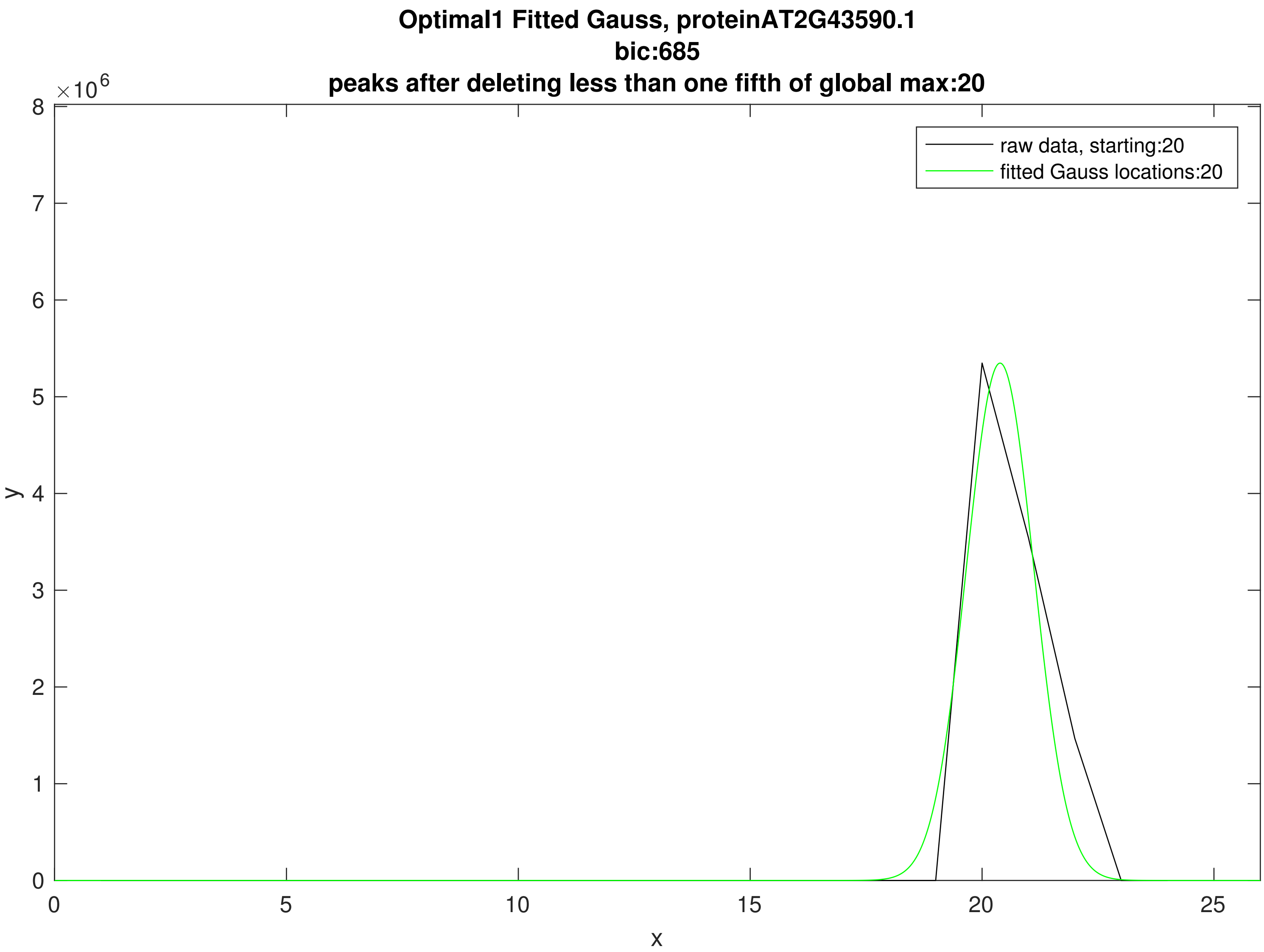

Optimal1 Fitted Gauss, proteinAT2G43590.1
bic:685
peaks after deleting less than one fifth of global max:20
×10^6
8
7
6
5
4
3
2
1
0
y
0
5
10
15
20
25
x
raw data, starting:20
fitted Gauss locations:20

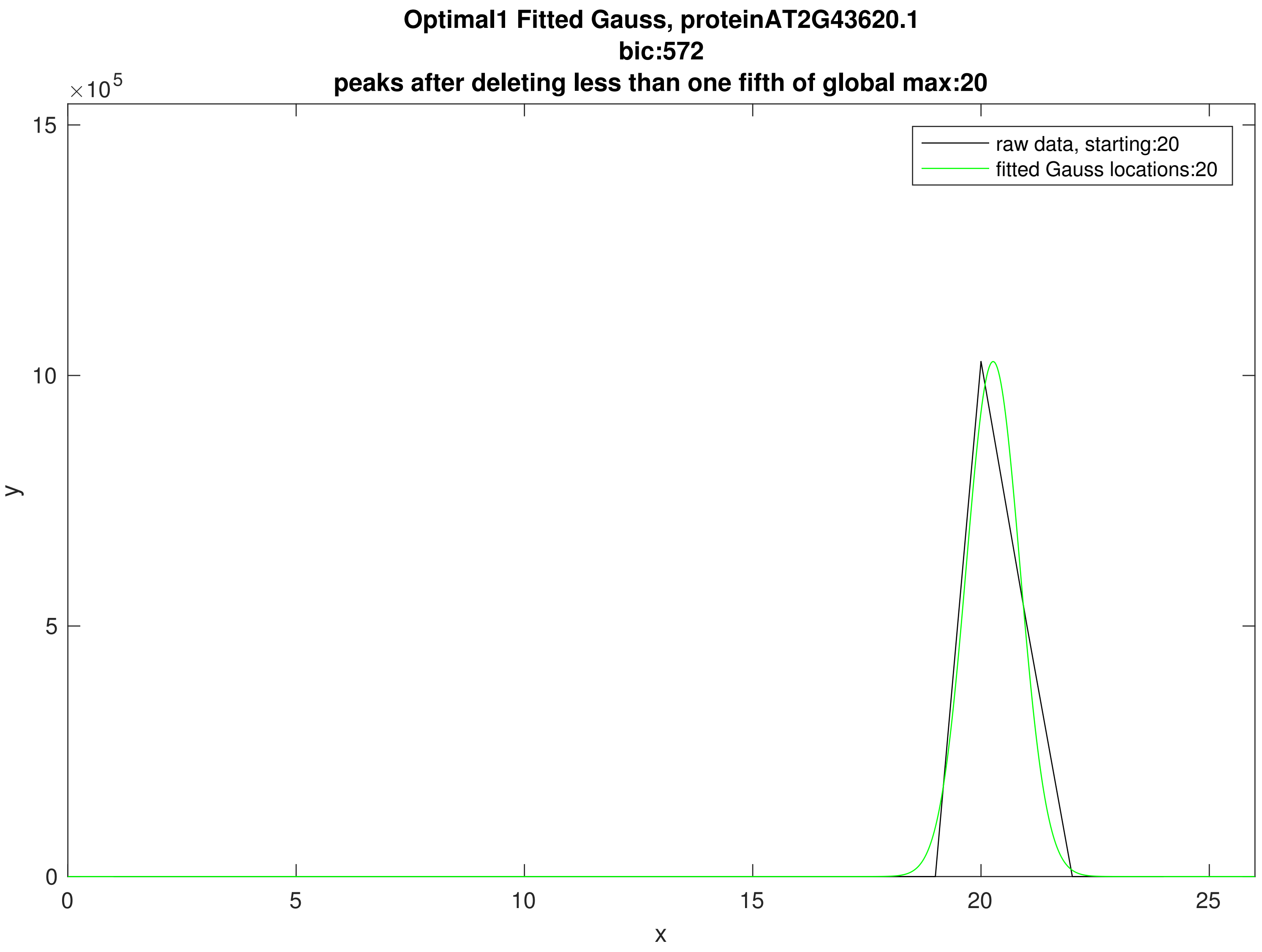
Optimal1 Fitted Gauss, proteinAT2G43620.1
bic:572
peaks after deleting less than one fifth of global max:20
×10^5
15
10
5
0
0
5
10
15
20
25
x
y
raw data, starting:20
fitted Gauss locations:20

#### Optimal1 Fitted Gauss, proteinAT2G43640.3;AT2G43640.2;AT2G43640.1
#### bic:677
#### peaks after deleting less than one fifth of global max:12

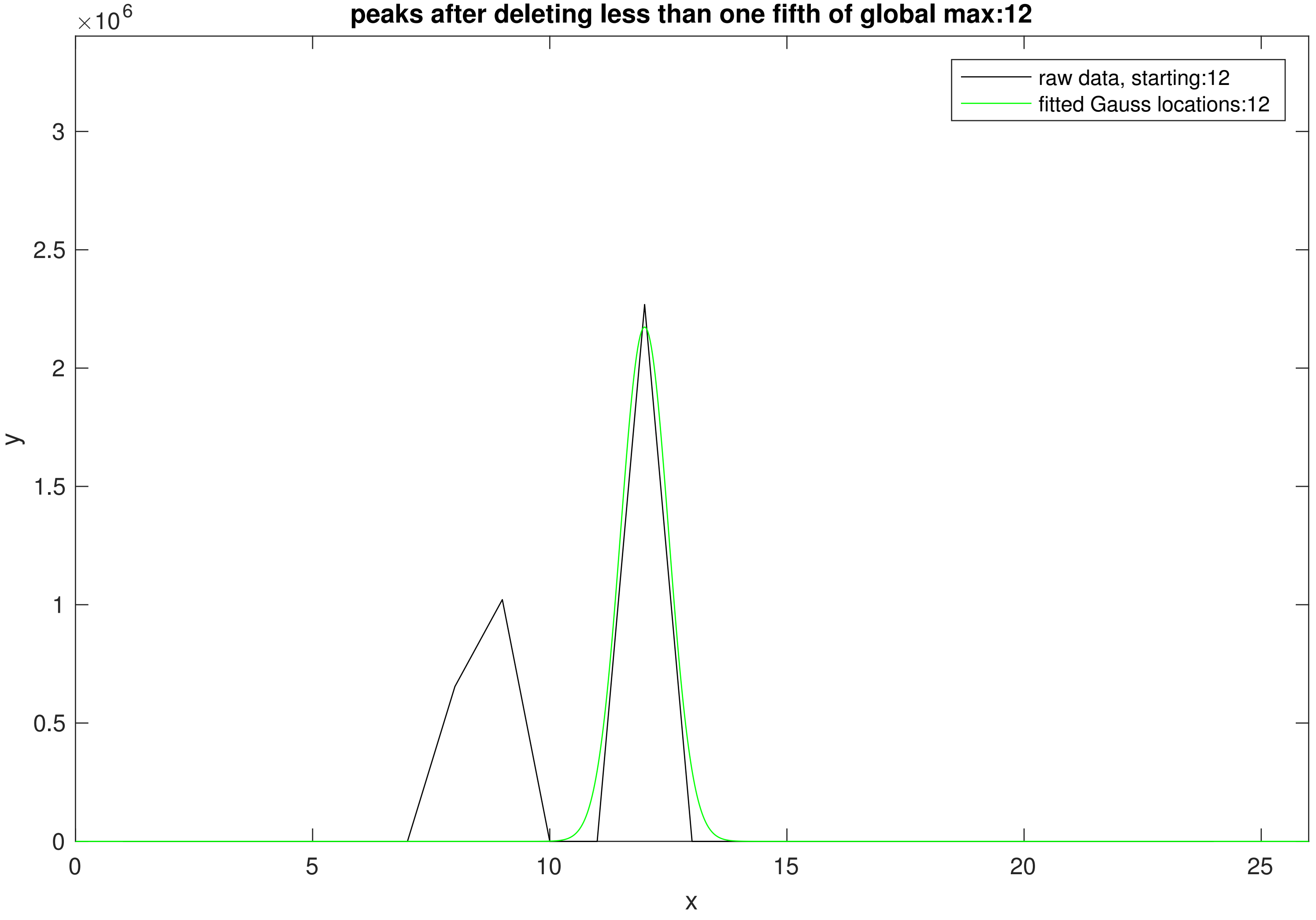

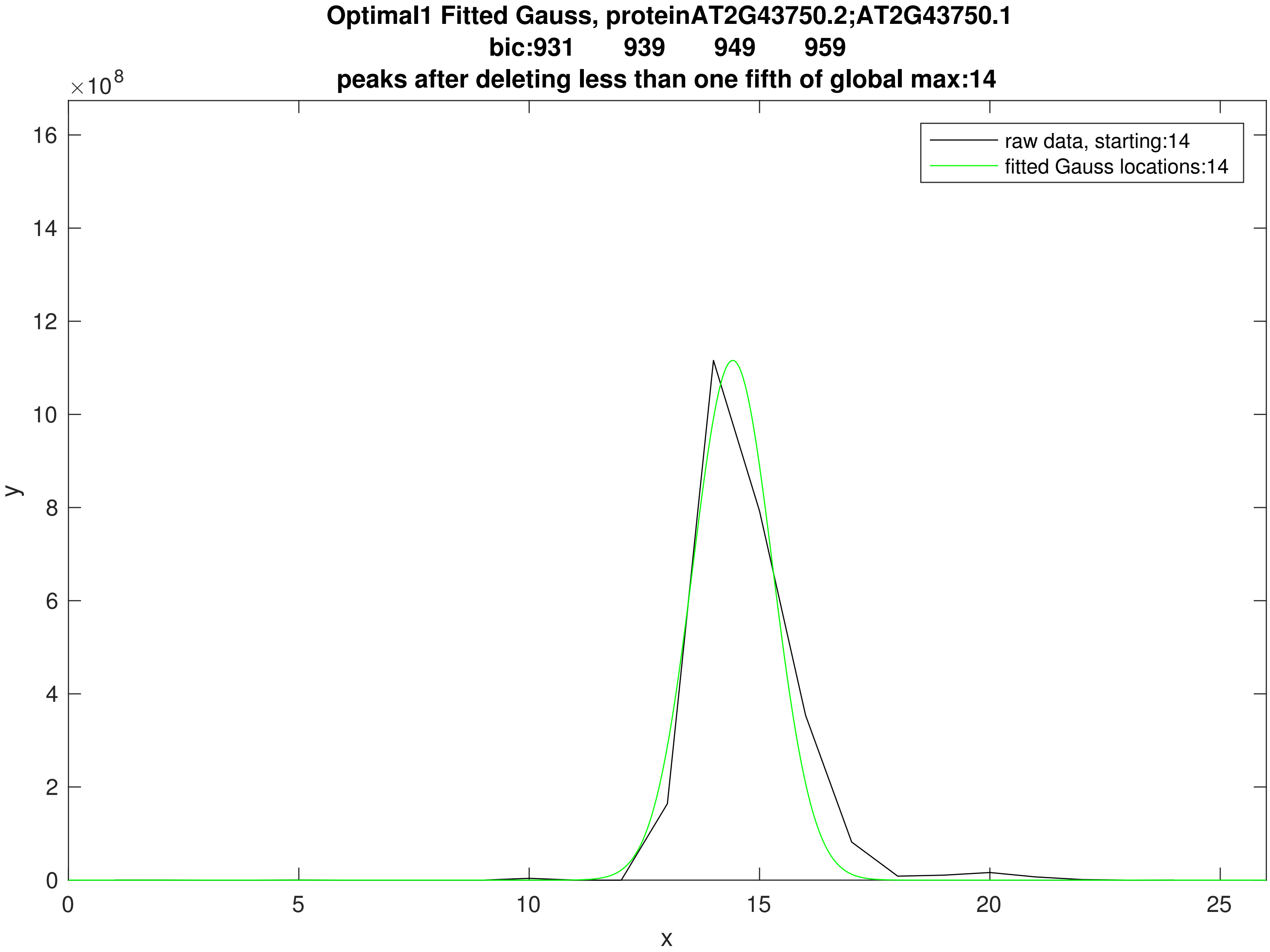
Optimal1 Fitted Gauss, proteinAT2G43750.2;AT2G43750.1
bic:931 939 949 959
peaks after deleting less than one fifth of global max:14
×10^8
raw data, starting:14
fitted Gauss locations:14
y
x
0
2
4
6
8
10
12
14
16
0
5
10
15
20
25

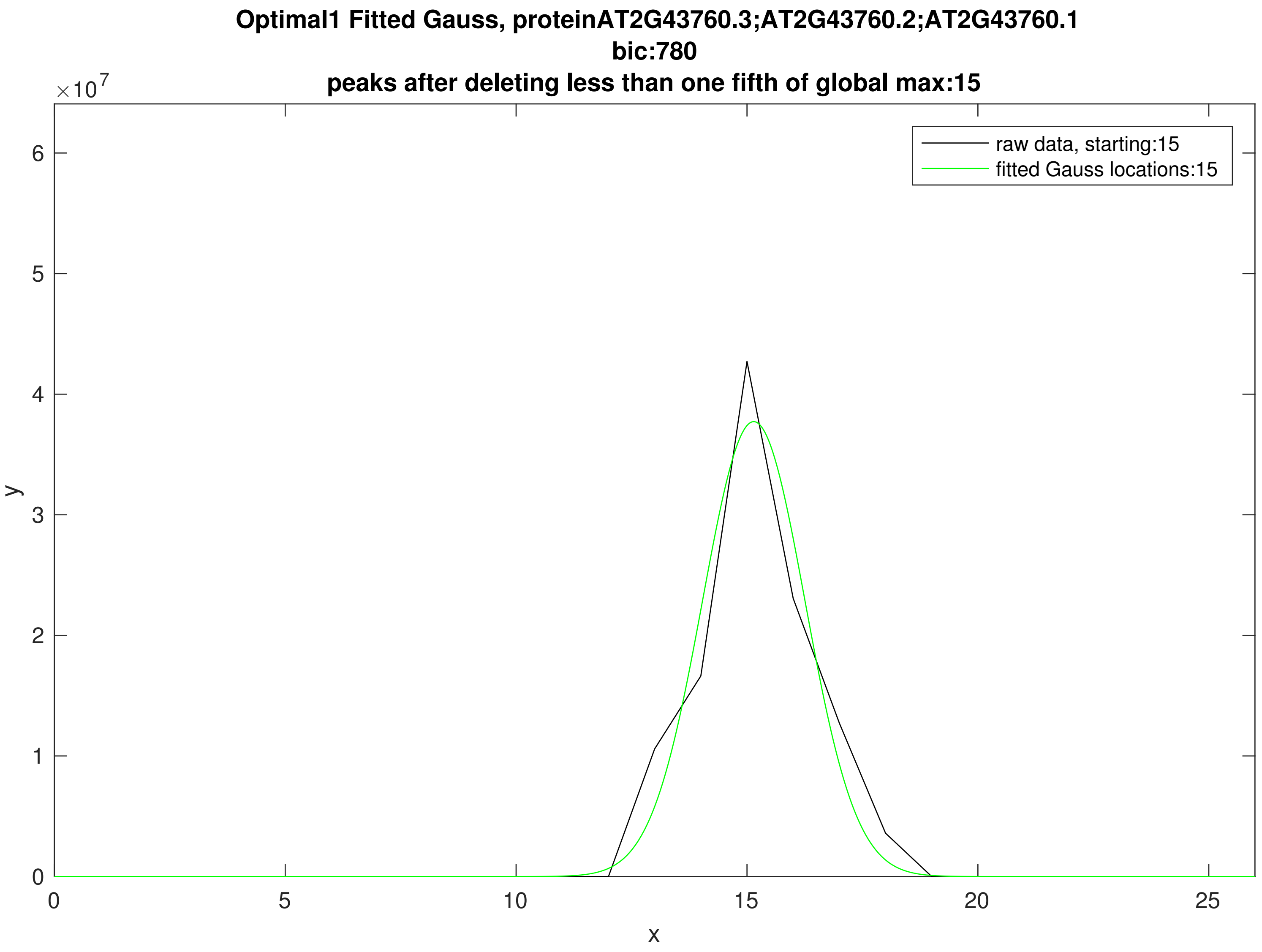
Optimal1 Fitted Gauss, proteinAT2G43760.3;AT2G43760.2;AT2G43760.1
bic:780
peaks after deleting less than one fifth of global max:15
×10^7
raw data, starting:15
fitted Gauss locations:15
y
x
0
1
2
3
4
5
6
0
5
10
15
20
25

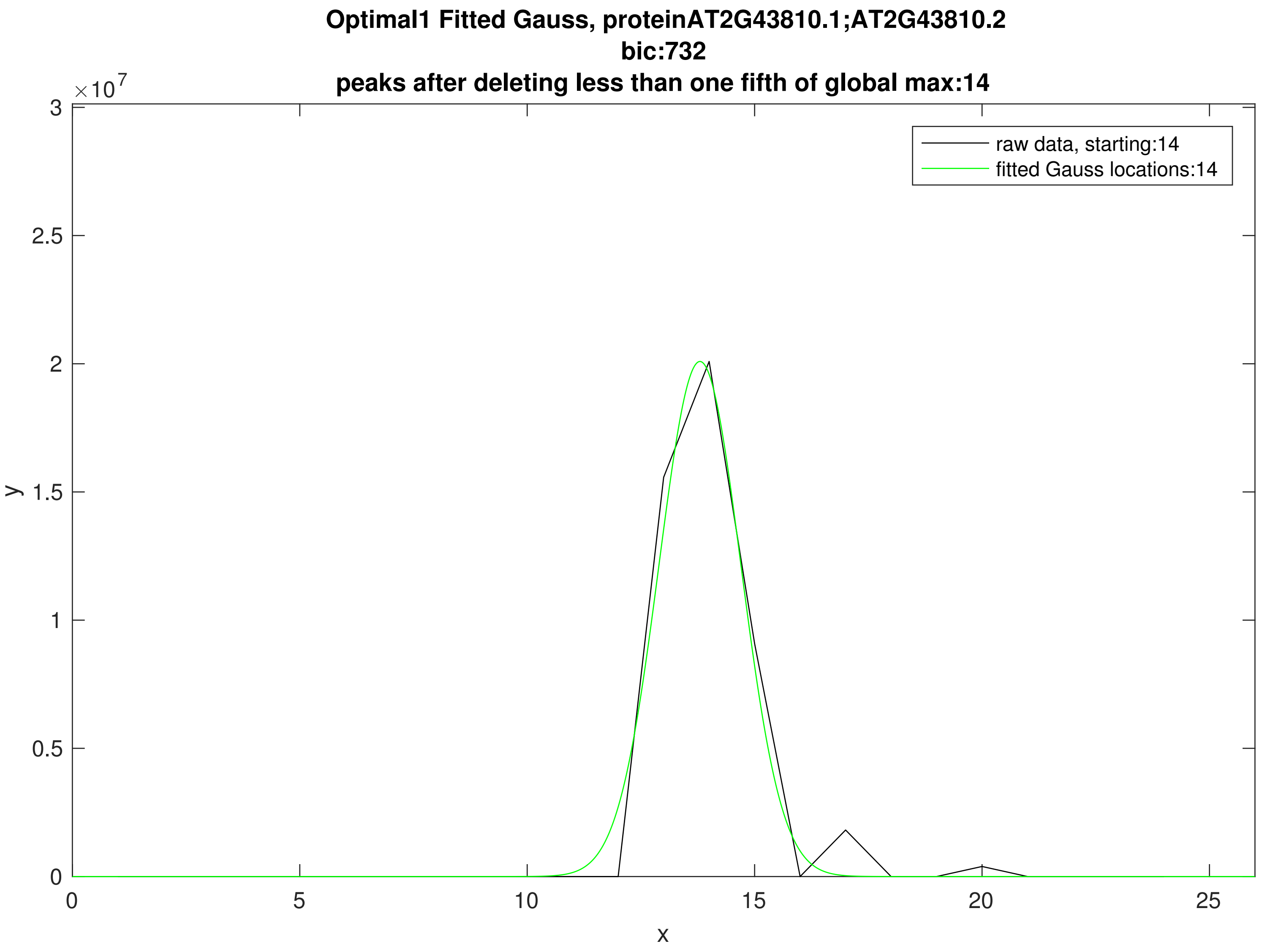

Optimal1 Fitted Gauss, proteinAT2G43810.1;AT2G43810.2
bic:732
peaks after deleting less than one fifth of global max:14
×10^7
3
2.5
2
1.5
1
0.5
0
y
0
5
10
15
20
25
x
raw data, starting:14
fitted Gauss locations:14

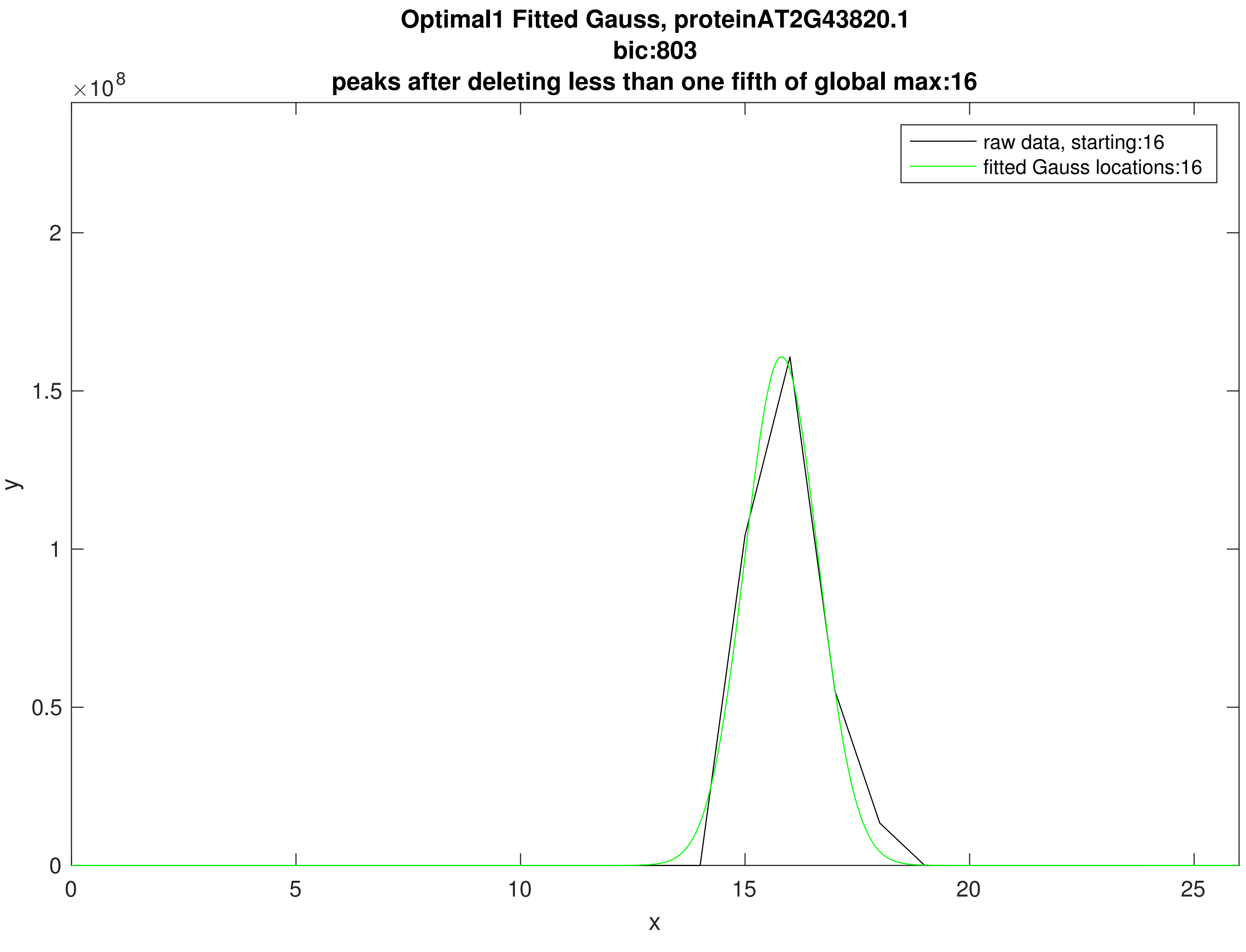
Optimal1 Fitted Gauss, proteinAT2G43820.1
bic:803
peaks after deleting less than one fifth of global max:16
×10^8
raw data, starting:16
fitted Gauss locations:16
y
x
0
0.5
1
1.5
2
0
5
10
15
20
25

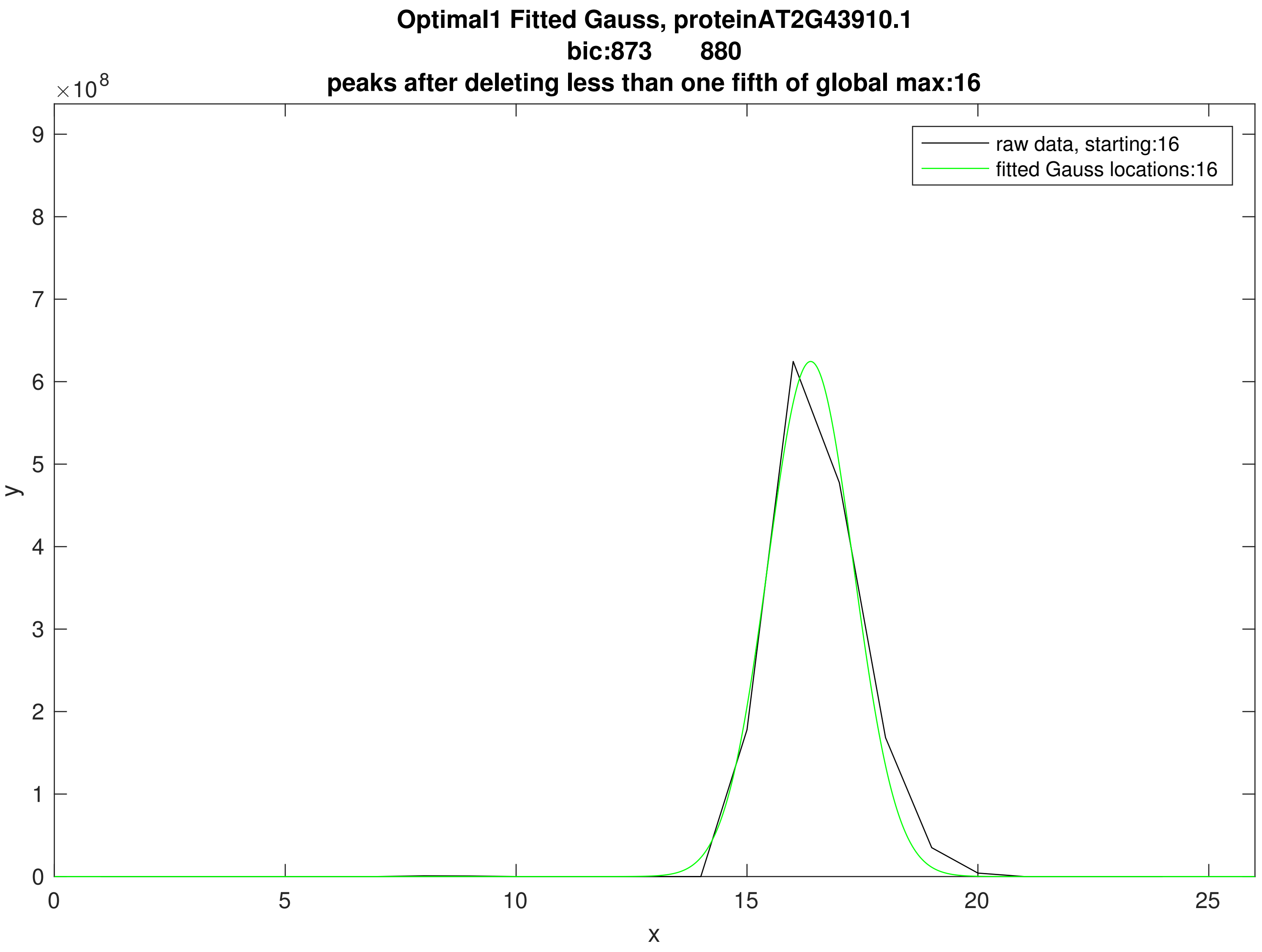
Optimal1 Fitted Gauss, proteinAT2G43910.1
bic:873 880
peaks after deleting less than one fifth of global max:16
×10^8
9
8
7
6
5
4
3
2
1
0
y
raw data, starting:16
fitted Gauss locations:16
0
5
10
15
20
25
x

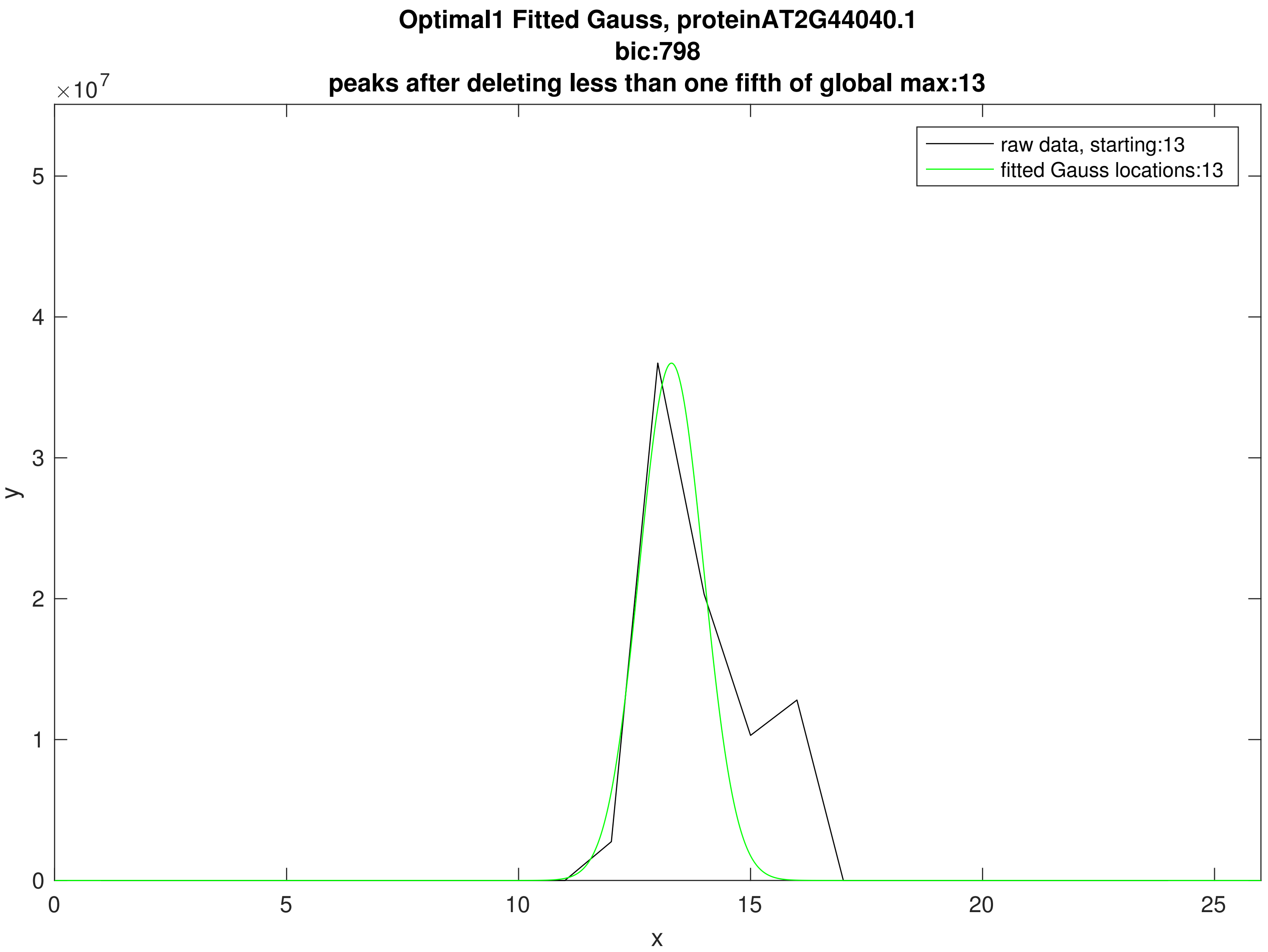

Optimal1 Fitted Gauss, proteinAT2G44040.1
bic:798
peaks after deleting less than one fifth of global max:13
×10^7
raw data, starting:13
fitted Gauss locations:13
y
x
0
1
2
3
4
5
0
5
10
15
20
25

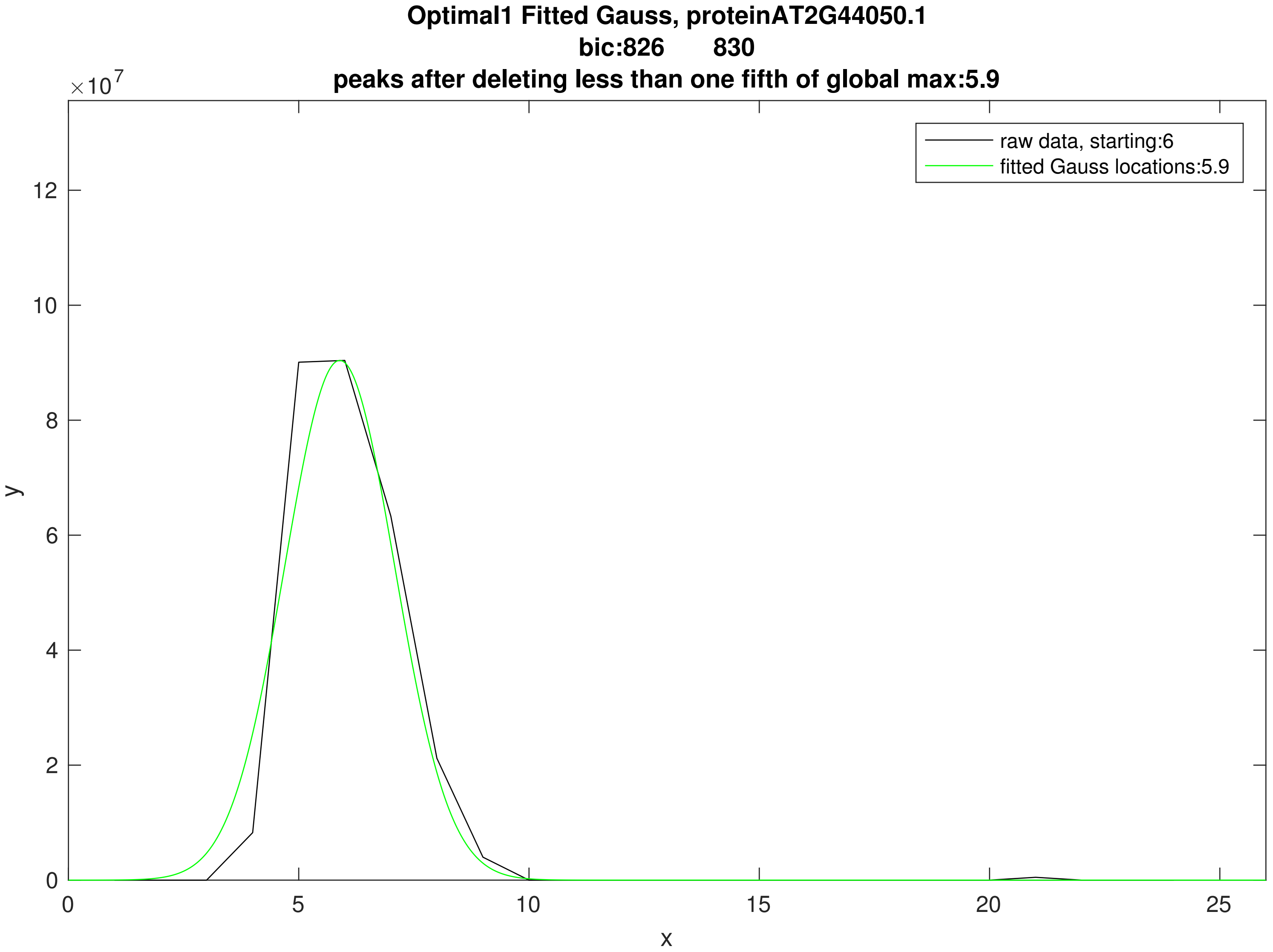
Optimal1 Fitted Gauss, proteinAT2G44050.1
bic:826 830
peaks after deleting less than one fifth of global max:5.9
×10^7
raw data, starting:6
fitted Gauss locations:5.9
y
x
0
2
4
6
8
10
12
0
5
10
15
20
25

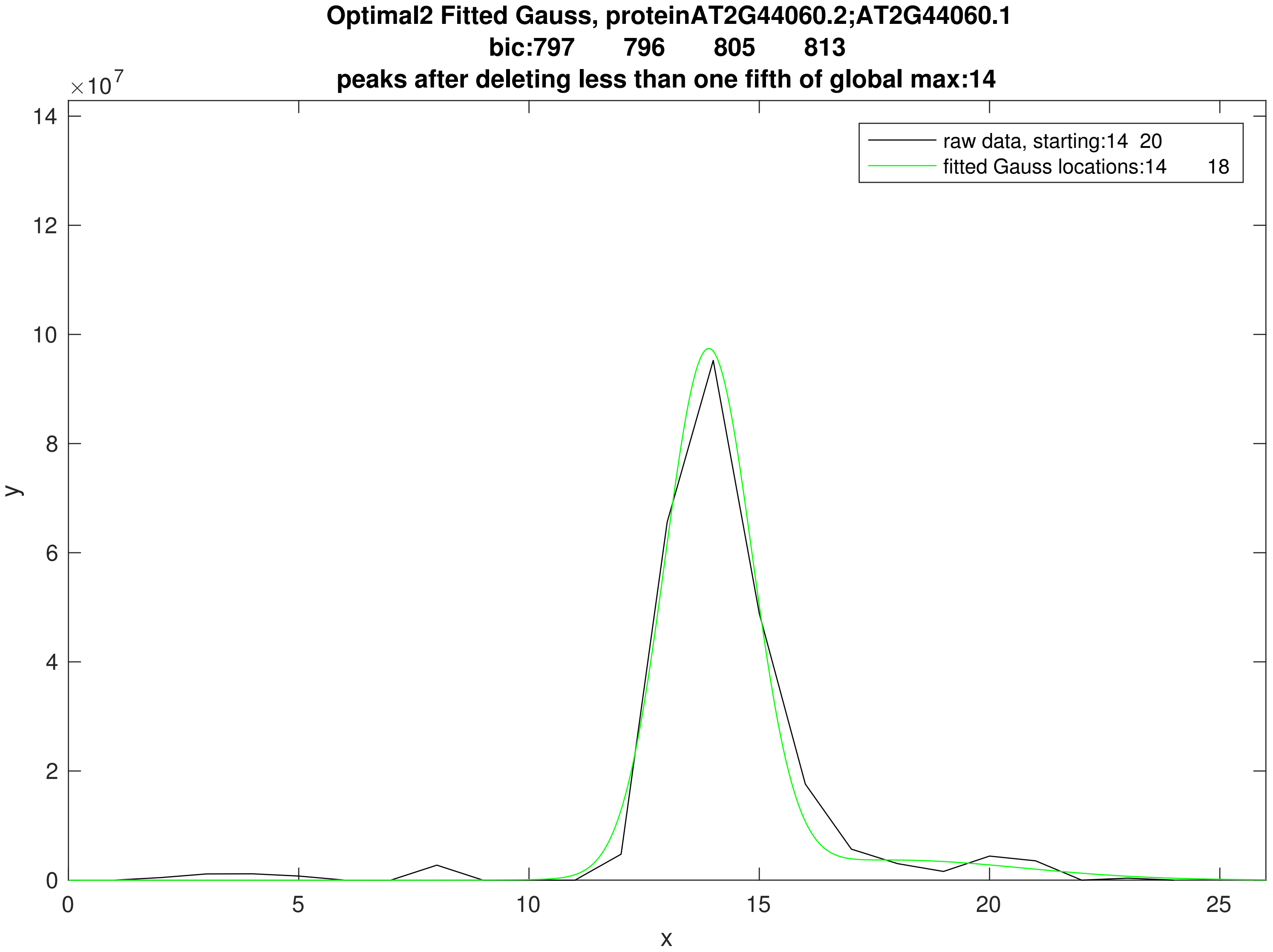

Optimal2 Fitted Gauss, proteinAT2G44060.2;AT2G44060.1
bic:797 796 805 813
peaks after deleting less than one fifth of global max:14
×10^7
raw data, starting:14 20
fitted Gauss locations:14 18
y
x
0
2
4
6
8
10
12
14
0
5
10
15
20
25

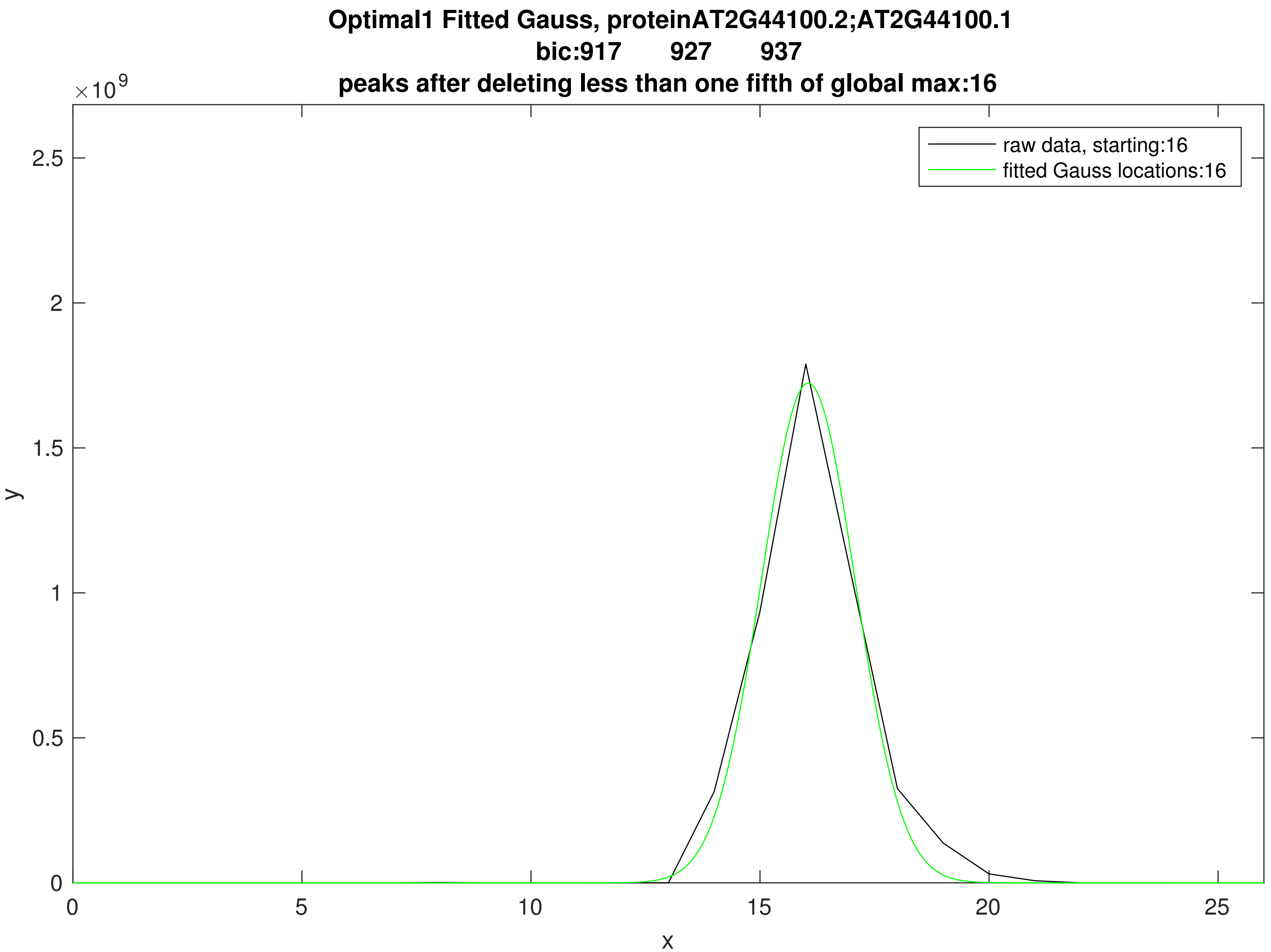

Optimal1 Fitted Gauss, proteinAT2G44100.2;AT2G44100.1
bic:917 927 937
peaks after deleting less than one fifth of global max:16
×10^9
2.5
2
1.5
1
0.5
0
y
0
5
10
15
20
25
x
raw data, starting:16
fitted Gauss locations:16

**Optimal1 Fitted Gauss, proteinAT2G44120.1;AT2G44120.2**
**bic:741**
**peaks after deleting less than one fifth of global max:4.6**

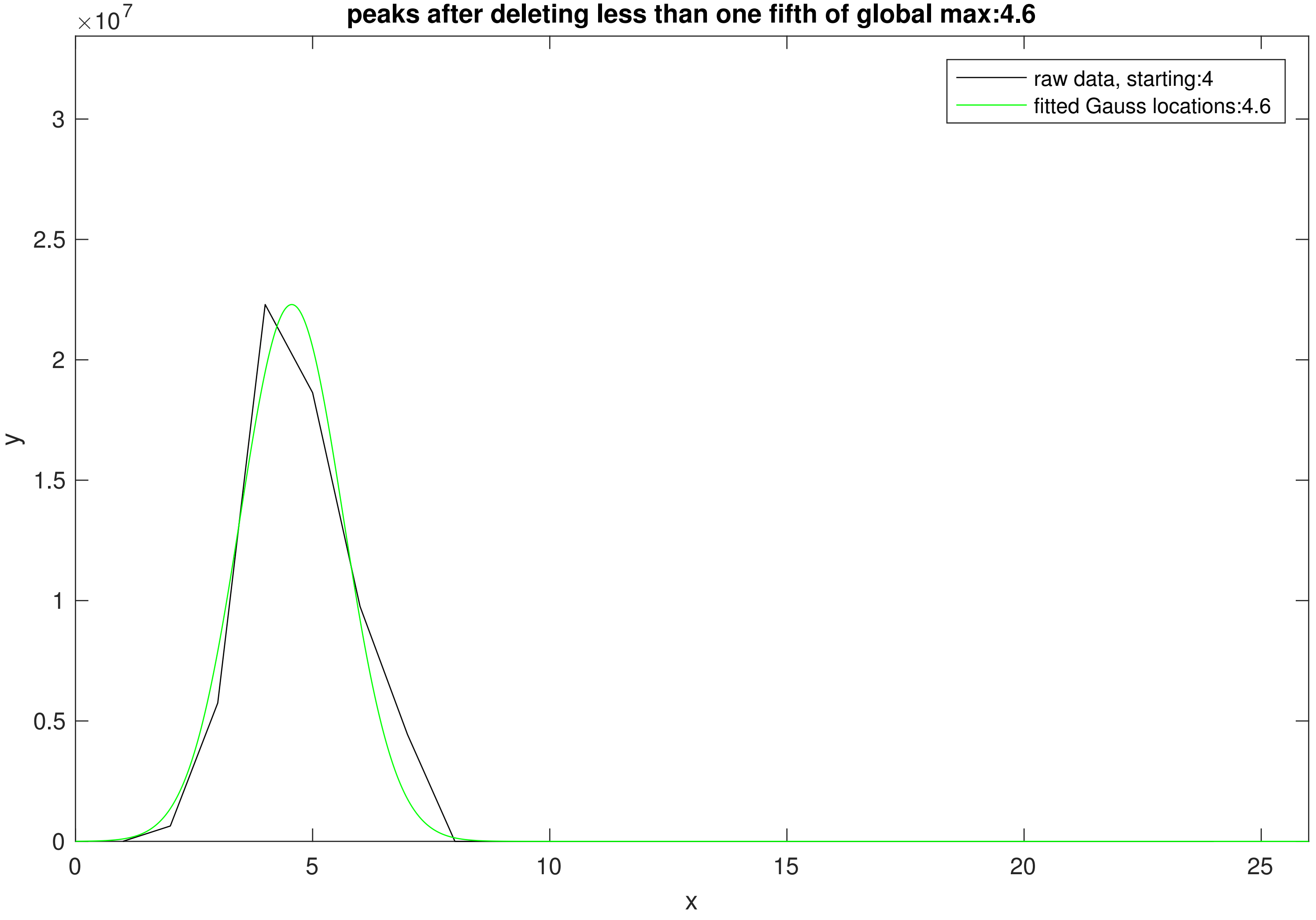

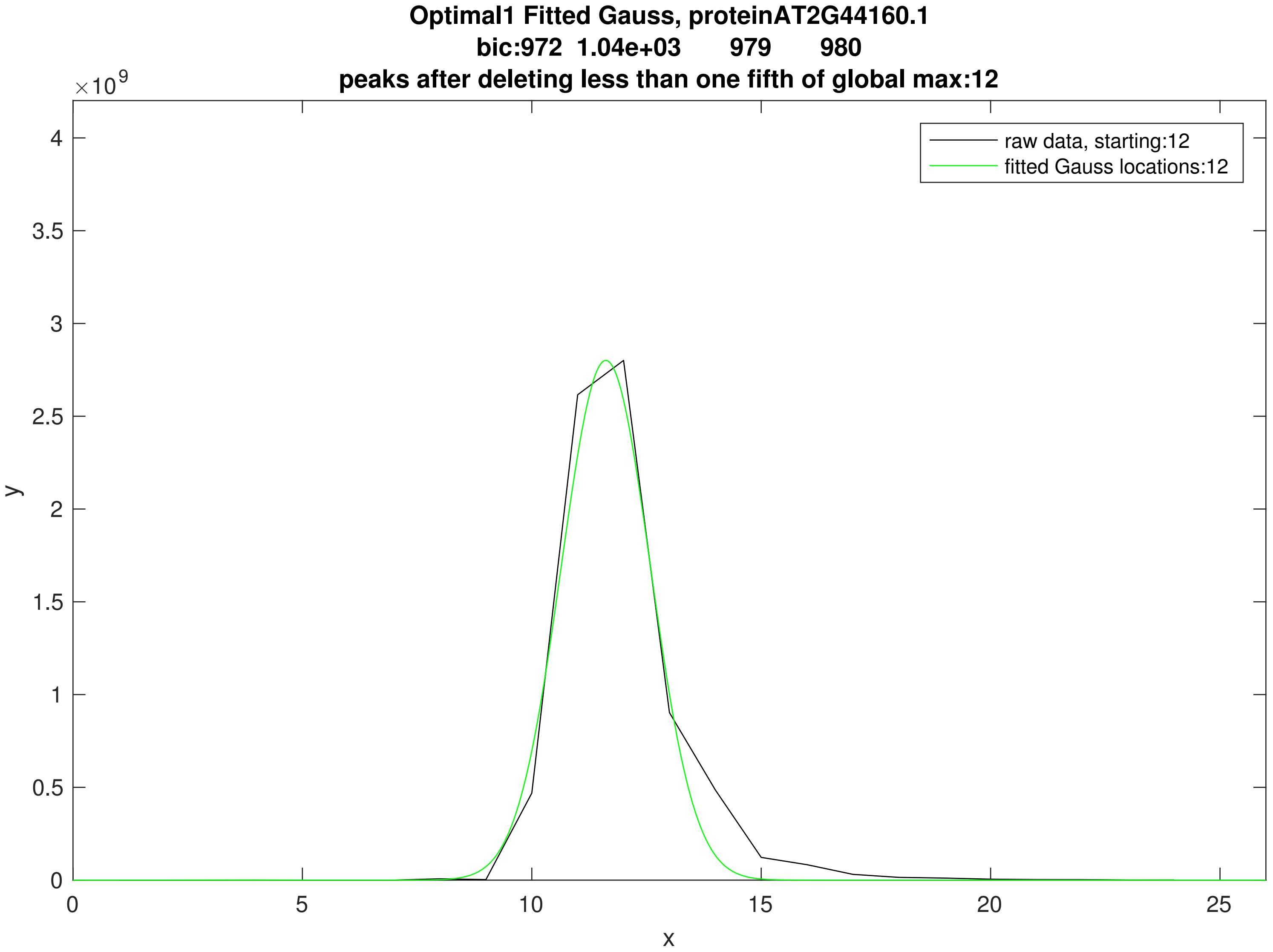

Optimal1 Fitted Gauss, proteinAT2G44160.1
bic:972 1.04e+03 979 980
peaks after deleting less than one fifth of global max:12
×10^9
raw data, starting:12
fitted Gauss locations:12
y
x
0
0.5
1
1.5
2
2.5
3
3.5
4
0
5
10
15
20
25

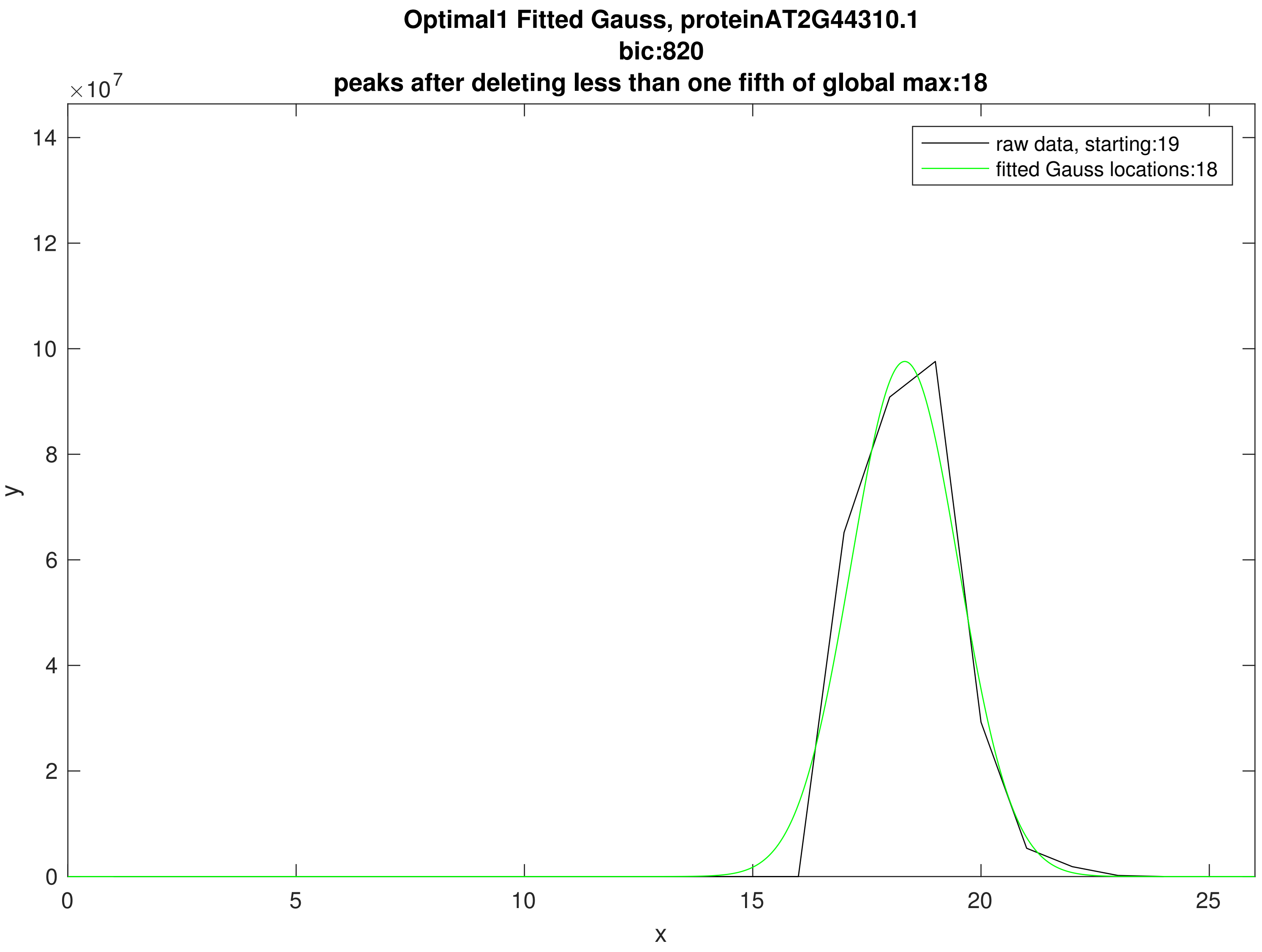

Optimal1 Fitted Gauss, proteinAT2G44310.1
bic:820
peaks after deleting less than one fifth of global max:18
×10^7
14
12
10
8
6
4
2
0
y
0
5
10
15
20
25
x
raw data, starting:19
fitted Gauss locations:18

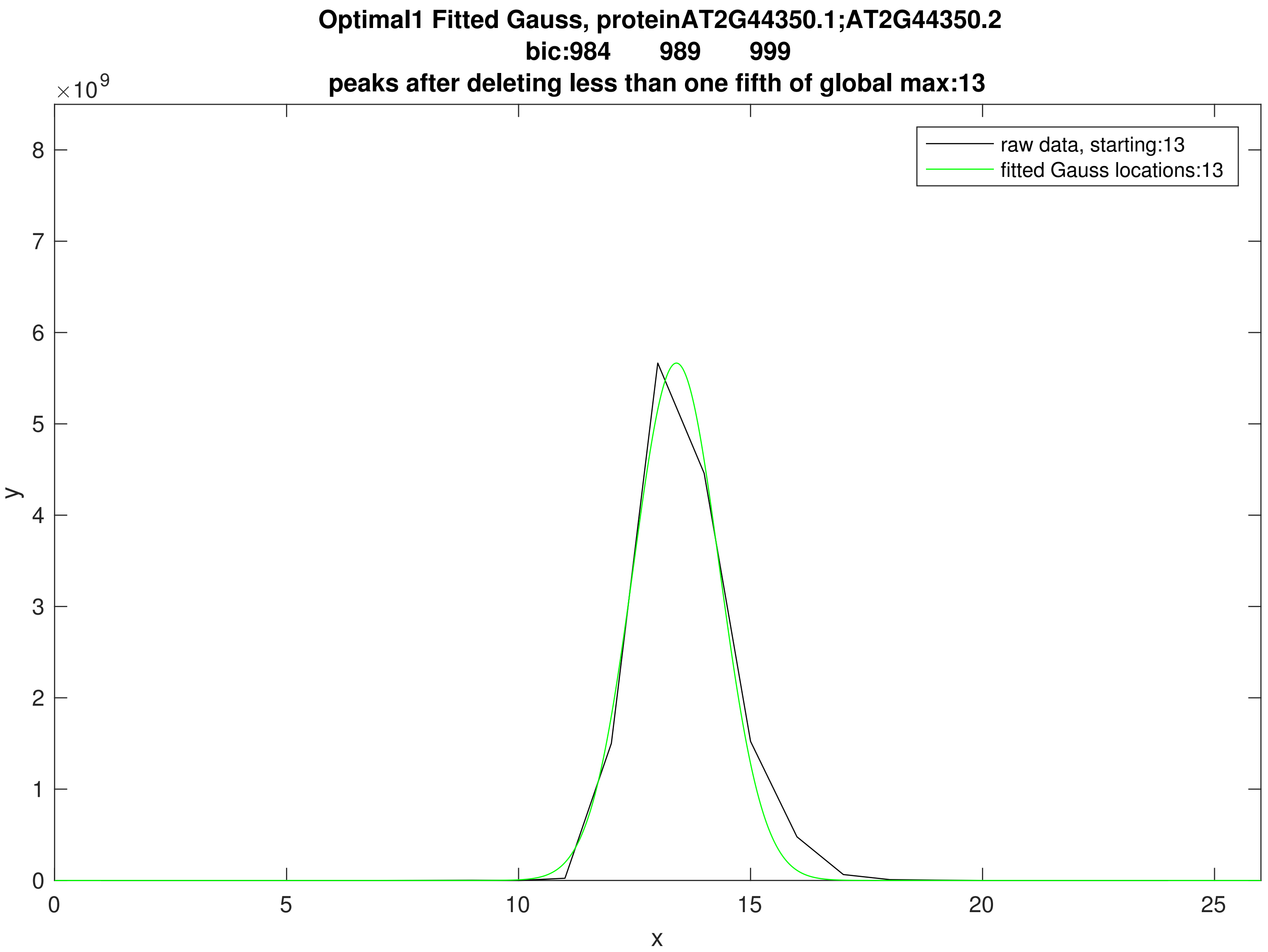
Optimal1 Fitted Gauss, proteinAT2G44350.1;AT2G44350.2
bic:984 989 999
peaks after deleting less than one fifth of global max:13
×10^9
raw data, starting:13
fitted Gauss locations:13
y
x
0
1
2
3
4
5
6
7
8
0
5
10
15
20
25

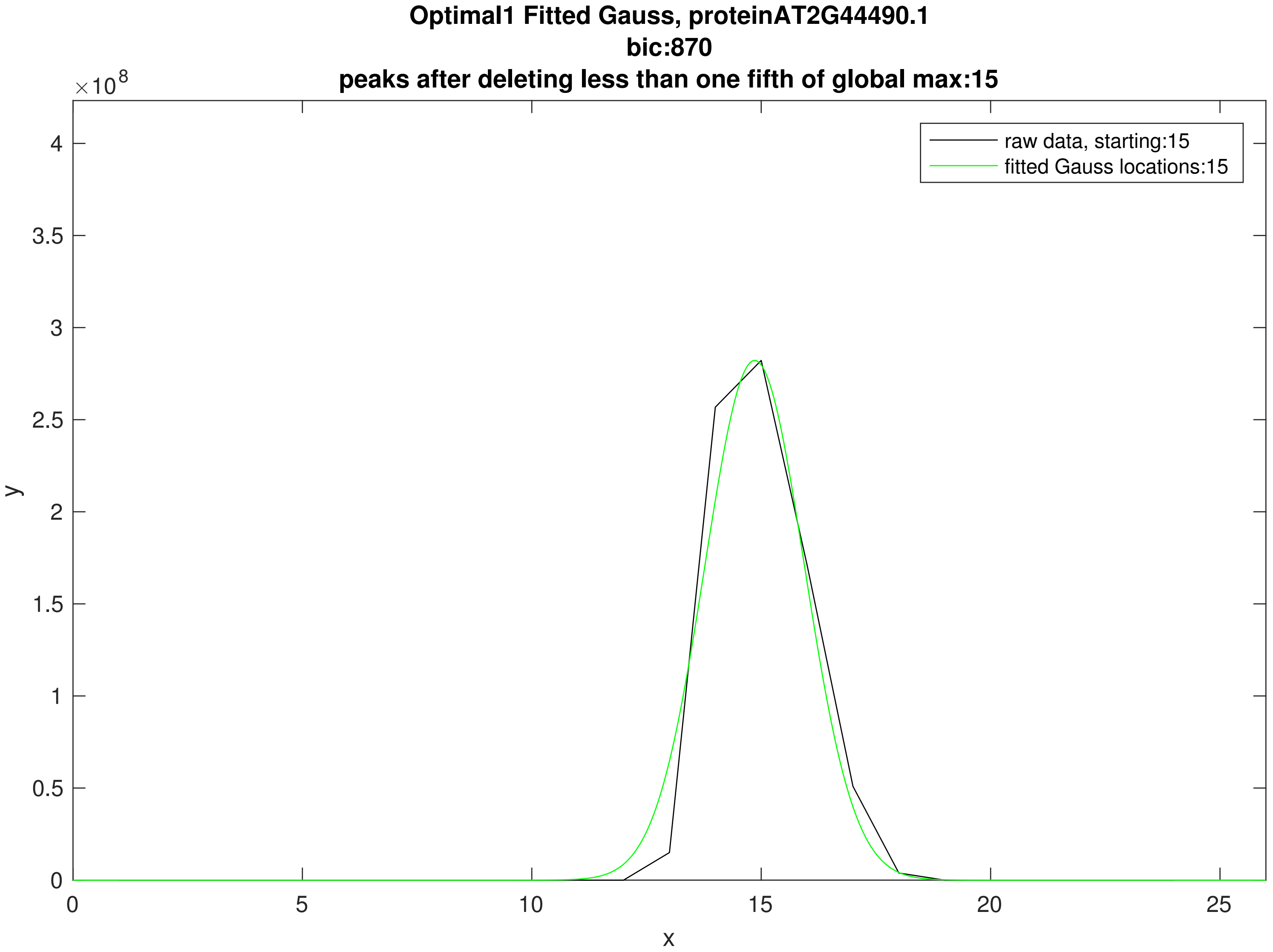
Optimal1 Fitted Gauss, proteinAT2G44490.1
bic:870
peaks after deleting less than one fifth of global max:15
×10^8
raw data, starting:15
fitted Gauss locations:15
y
x
0
0.5
1
1.5
2
2.5
3
3.5
4
0
5
10
15
20
25

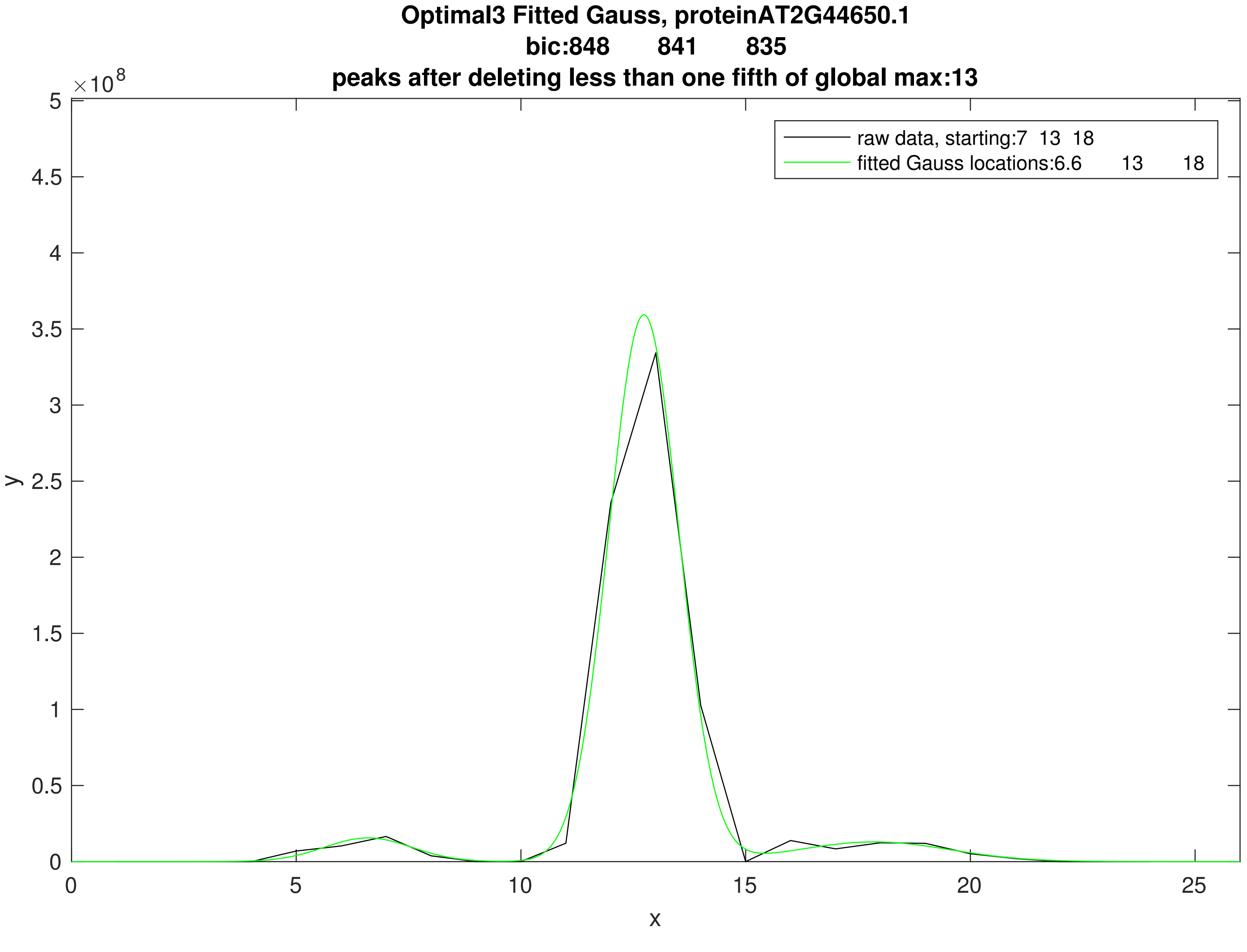
Optimal3 Fitted Gauss, proteinAT2G44650.1
bic:848 841 835
peaks after deleting less than one fifth of global max:13
×10^8
raw data, starting:7 13 18
fitted Gauss locations:6.6 13 18
y
x
0
0.5
1
1.5
2
2.5
3
3.5
4
4.5
5
0
5
10
15
20
25

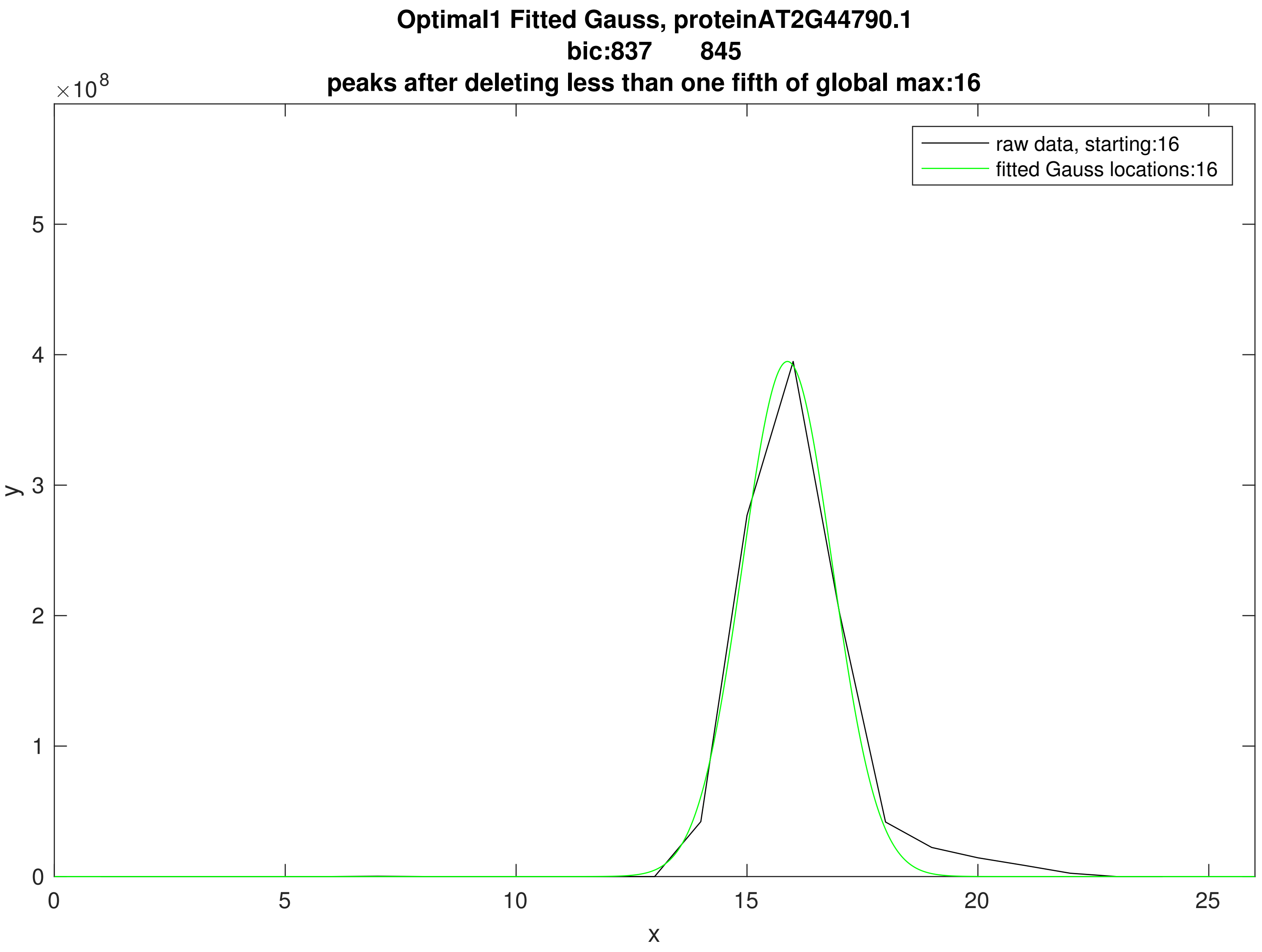
Optimal1 Fitted Gauss, proteinAT2G44790.1
bic:837 845
peaks after deleting less than one fifth of global max:16
×10^8
raw data, starting:16
fitted Gauss locations:16
y
x
0
1
2
3
4
5
0
5
10
15
20
25

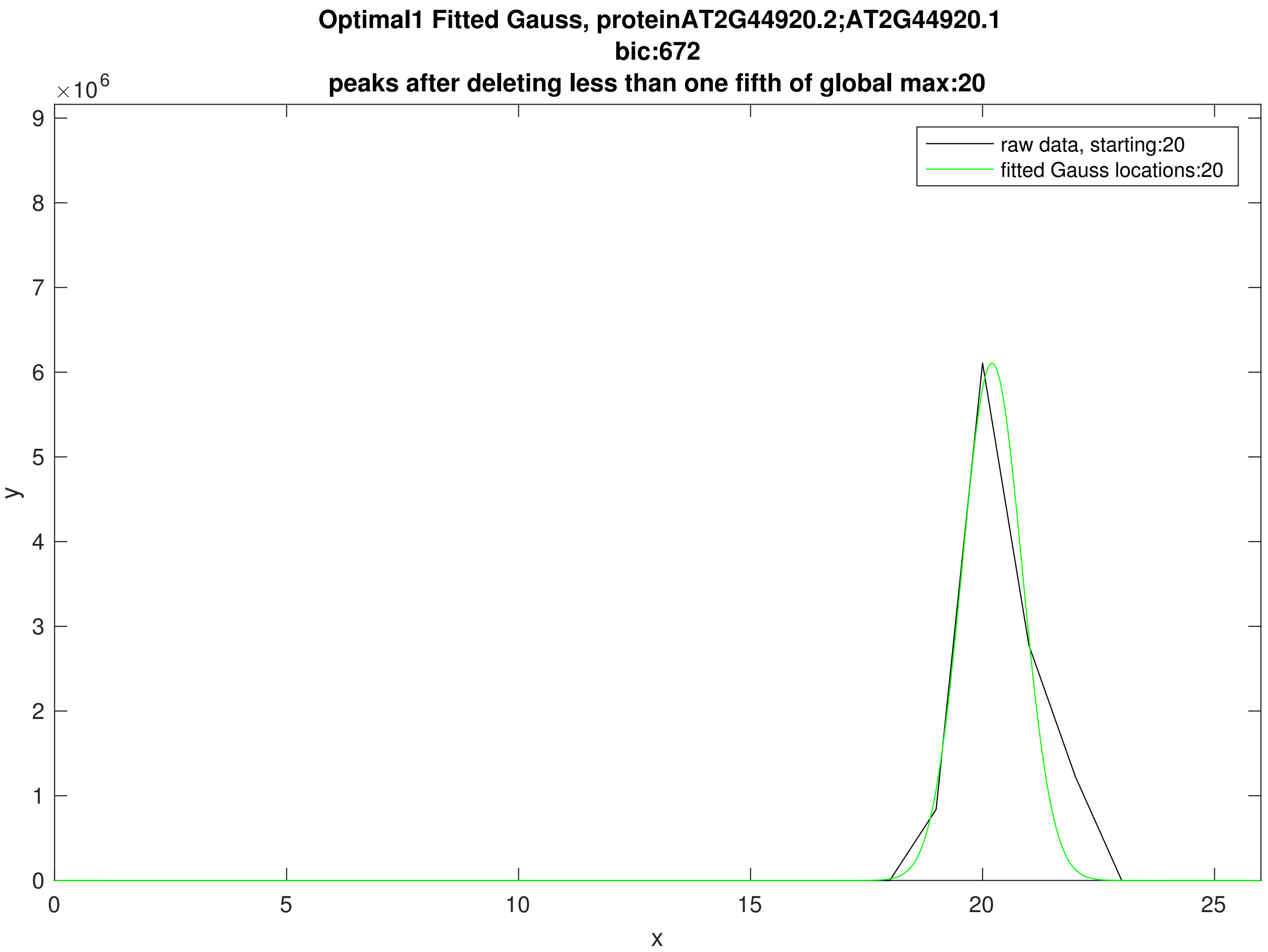

Optimal1 Fitted Gauss, proteinAT2G44920.2;AT2G44920.1
bic:672
peaks after deleting less than one fifth of global max:20
×10^6
9
8
7
6
5
4
3
2
1
0
y
0
5
10
15
20
25
x
raw data, starting:20
fitted Gauss locations:20

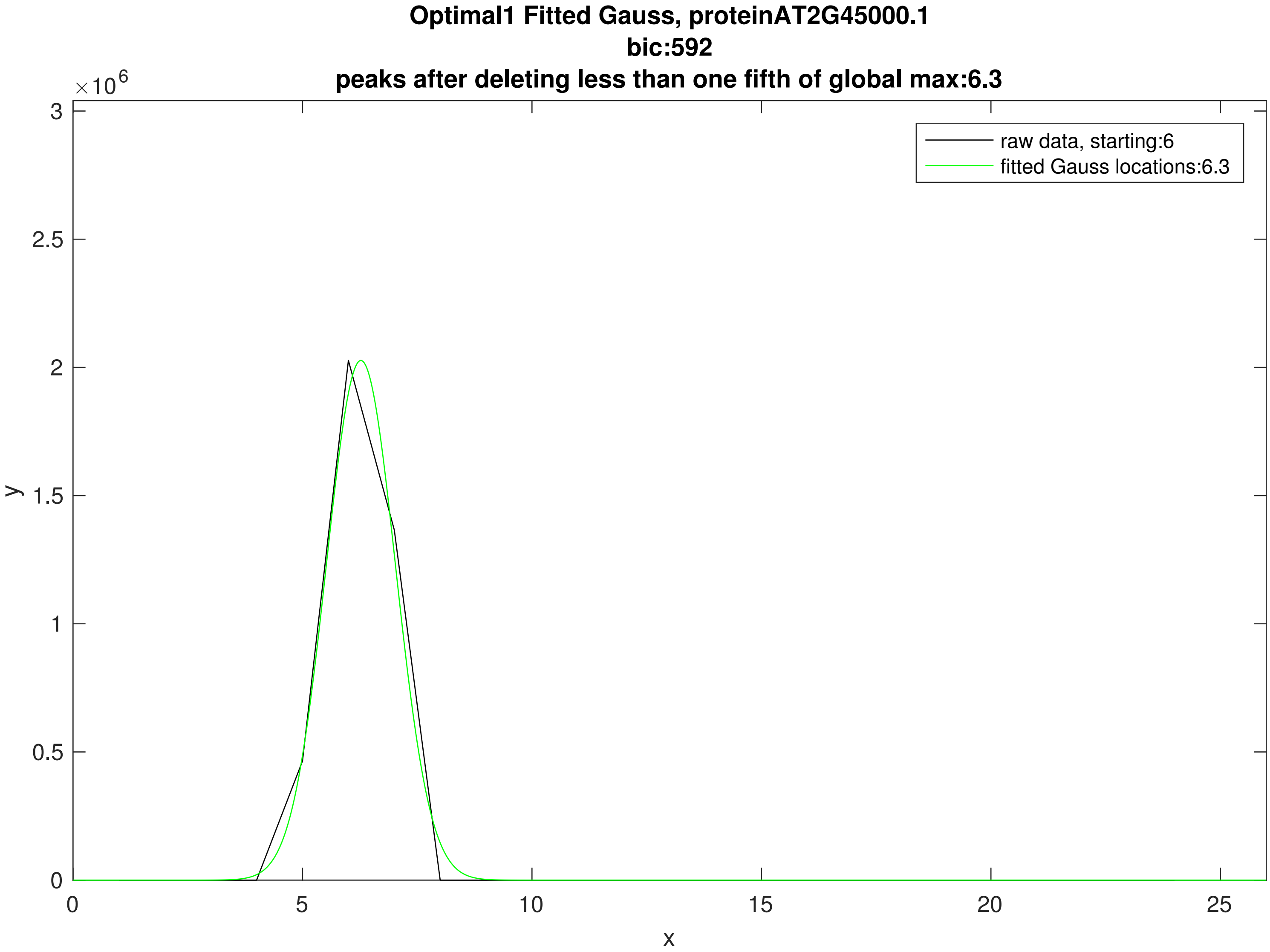
Optimal1 Fitted Gauss, proteinAT2G45000.1
bic:592
peaks after deleting less than one fifth of global max:6.3
×10^6
3
2.5
2
1.5
1
0.5
0
y
0
5
10
15
20
25
x
raw data, starting:6
fitted Gauss locations:6.3

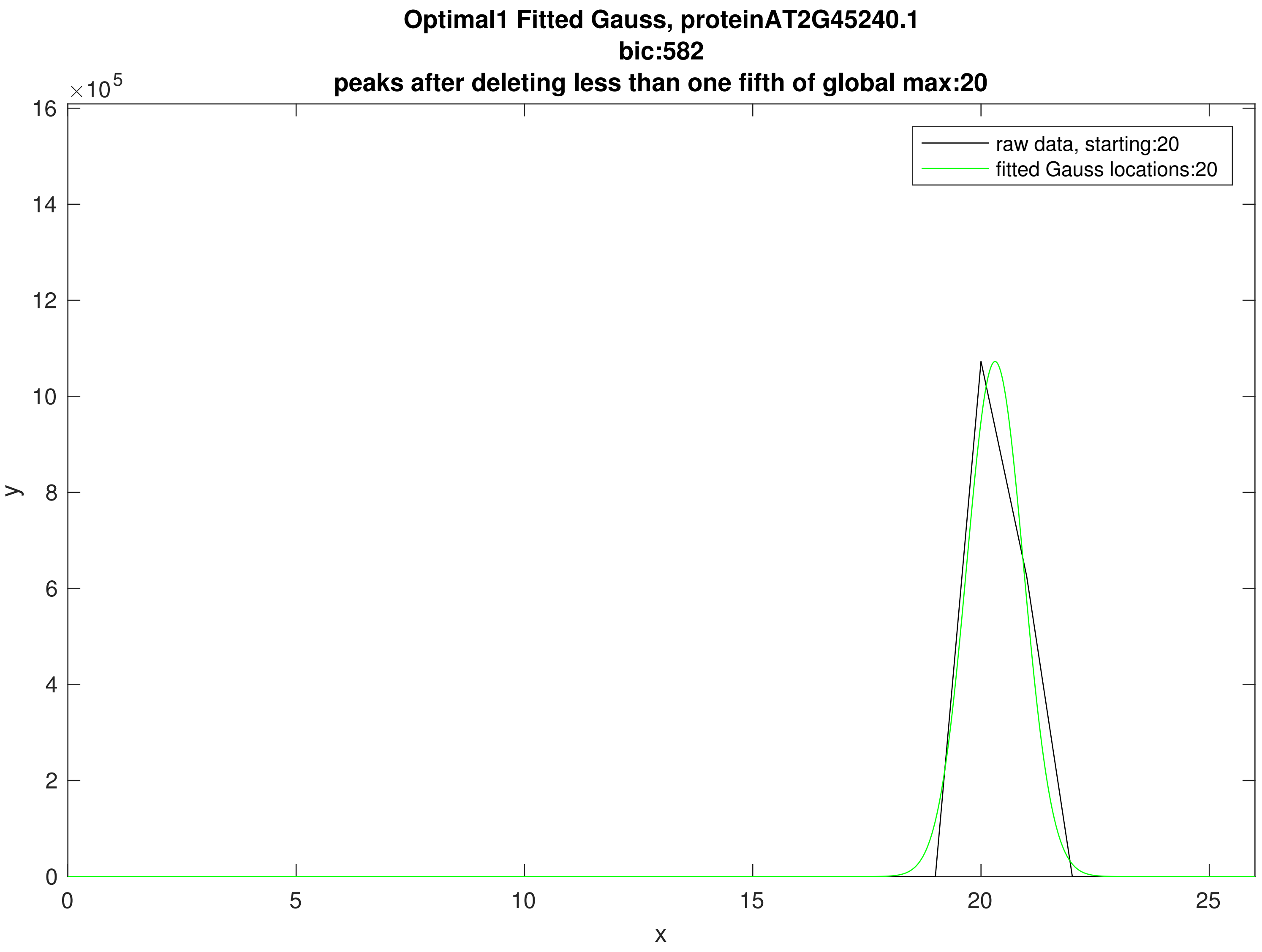
Optimal1 Fitted Gauss, proteinAT2G45240.1
bic:582
peaks after deleting less than one fifth of global max:20
×10^5
y
x
raw data, starting:20
fitted Gauss locations:20
0
2
4
6
8
10
12
14
16
0
5
10
15
20
25

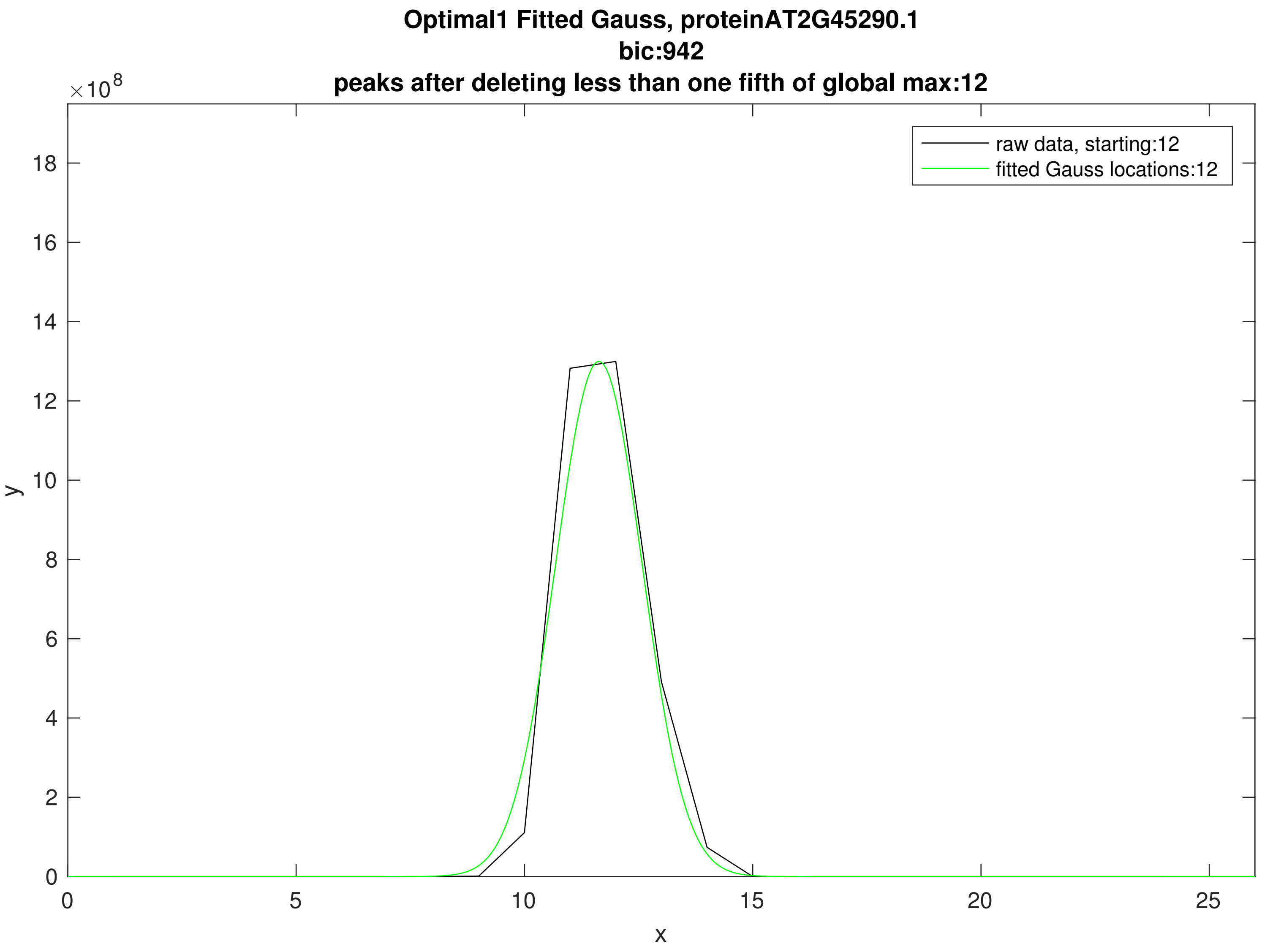

Optimal1 Fitted Gauss, proteinAT2G45290.1
bic:942
peaks after deleting less than one fifth of global max:12
×10^8
raw data, starting:12
fitted Gauss locations:12
y
x
0
2
4
6
8
10
12
14
16
18
0
5
10
15
20
25

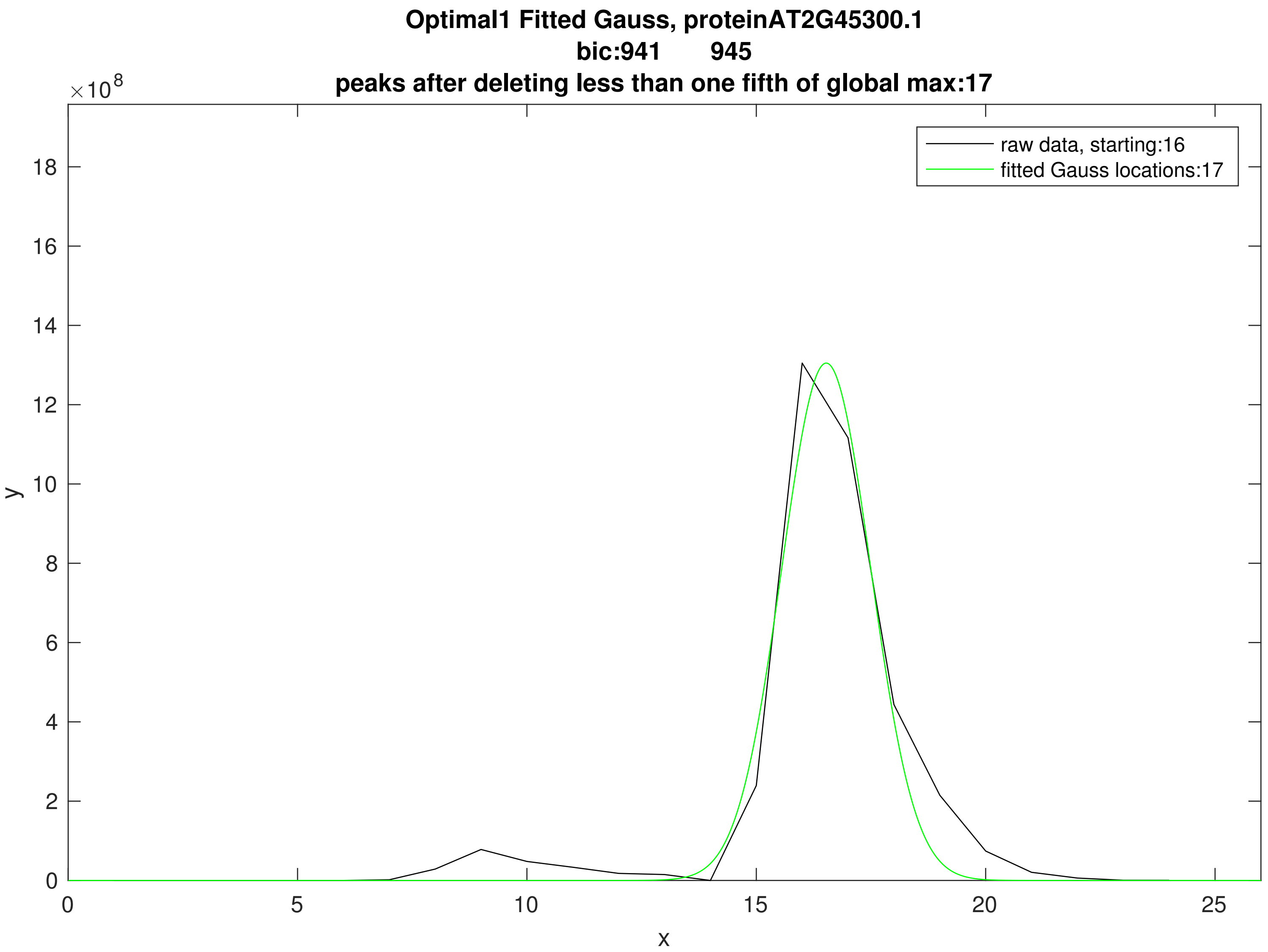
Optimal1 Fitted Gauss, proteinAT2G45300.1
bic:941 945
peaks after deleting less than one fifth of global max:17
×10^8
raw data, starting:16
fitted Gauss locations:17
y
x
0
2
4
6
8
10
12
14
16
18
0
5
10
15
20
25

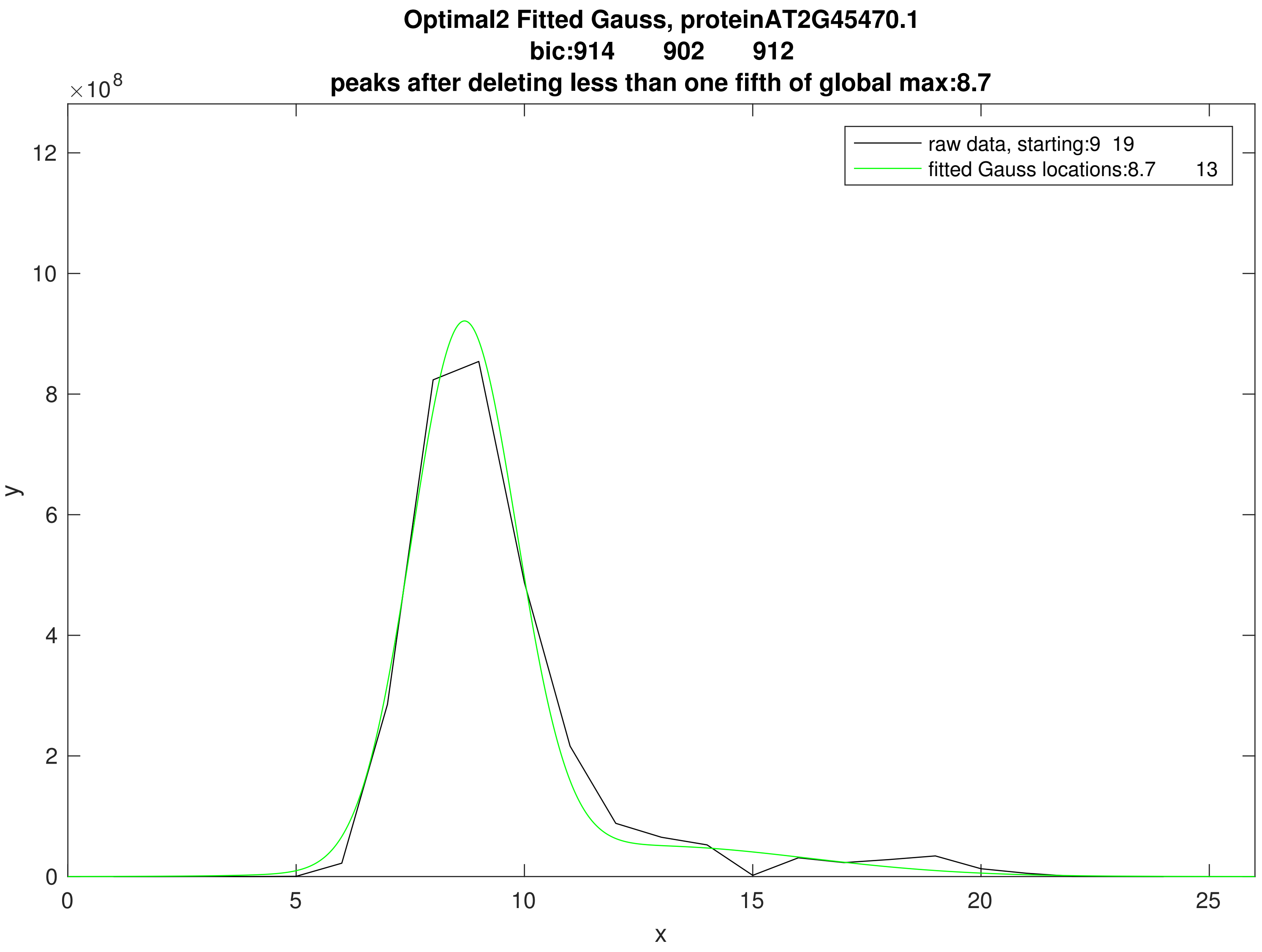

Optimal2 Fitted Gauss, proteinAT2G45470.1
bic:914 902 912
peaks after deleting less than one fifth of global max:8.7
×10^8
raw data, starting:9 19
fitted Gauss locations:8.7 13
y
x
0
2
4
6
8
10
12
0
5
10
15
20
25

**Optimal1 Fitted Gauss, proteinAT2G45710.1;AT3G61110.1;AT3G61111.1**
**bic:733 733**
**peaks after deleting less than one fifth of global max:4**

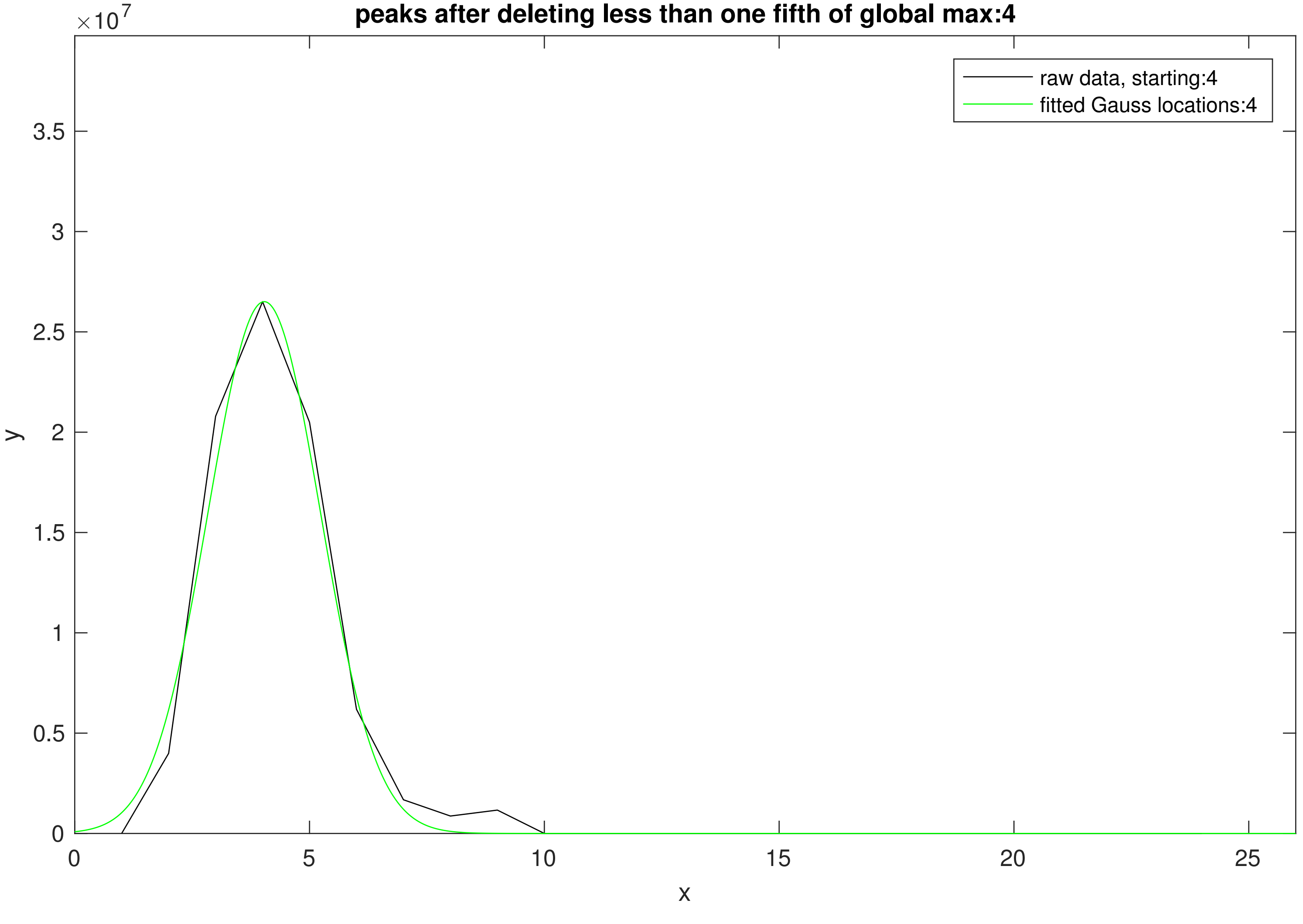

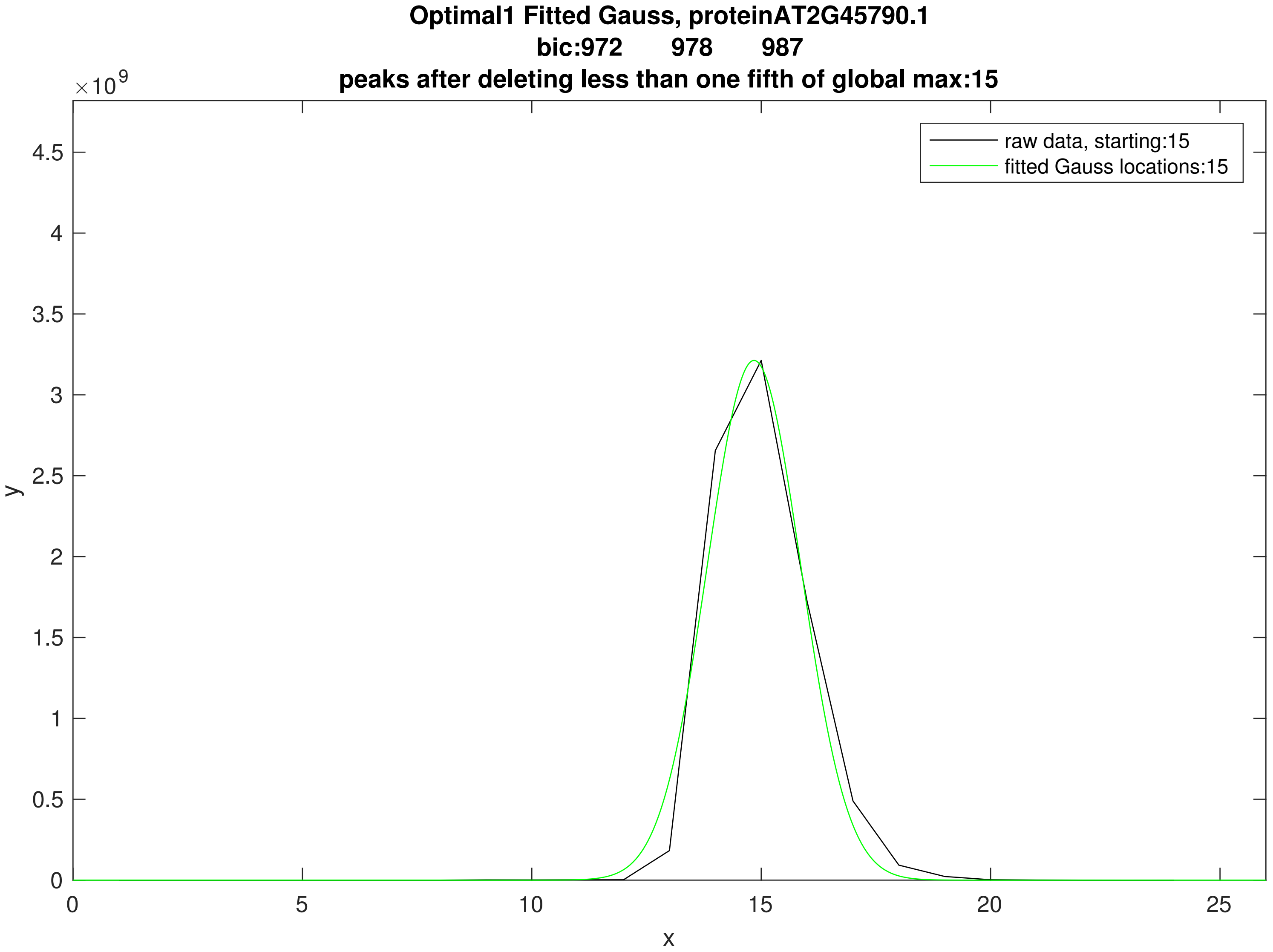

Optimal1 Fitted Gauss, proteinAT2G45790.1
bic:972 978 987
peaks after deleting less than one fifth of global max:15
×10^9
4.5
4
3.5
3
2.5
2
1.5
1
0.5
0
y
0
5
10
15
20
25
x
raw data, starting:15
fitted Gauss locations:15

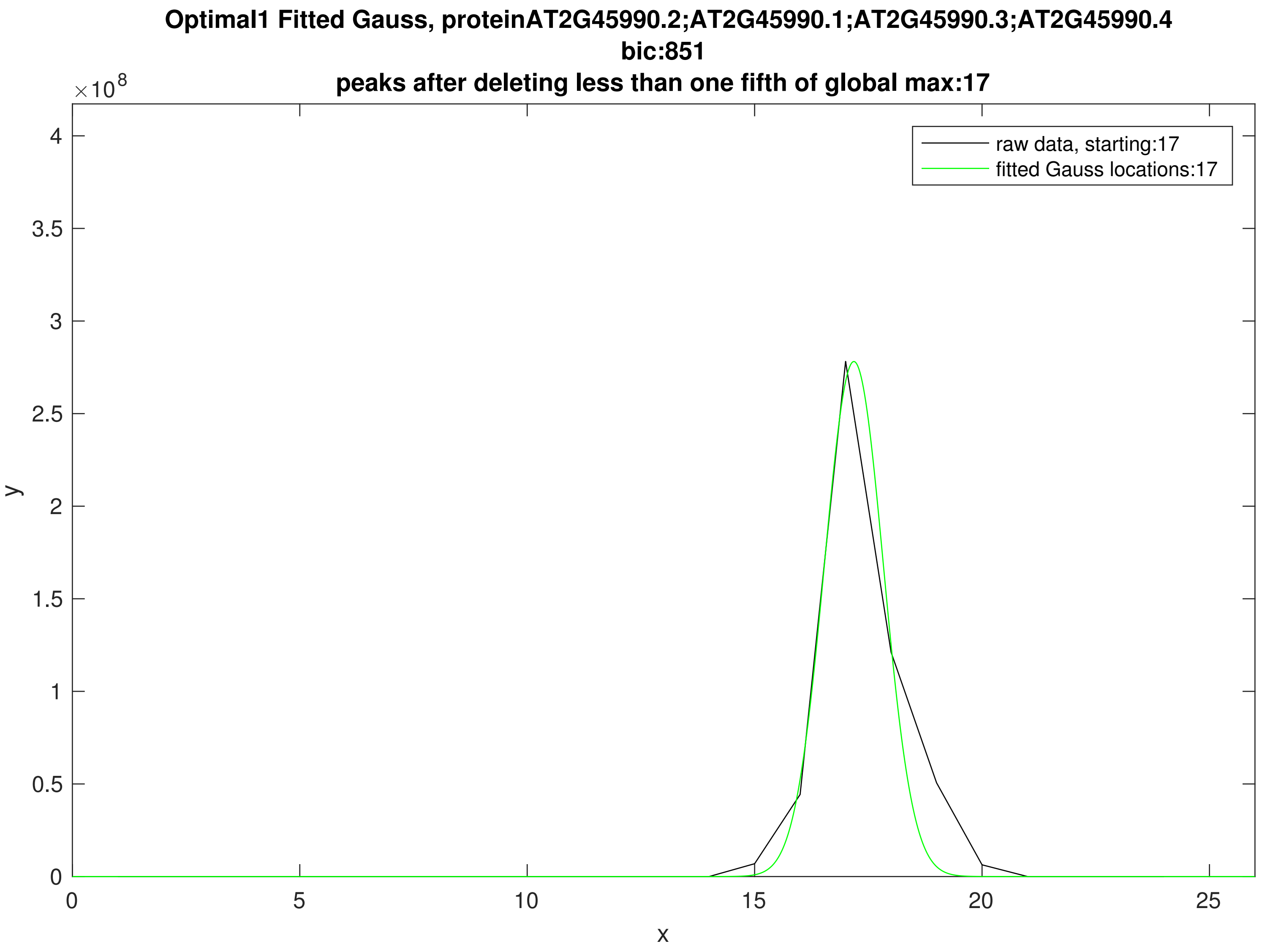
Optimal1 Fitted Gauss, proteinAT2G45990.2;AT2G45990.1;AT2G45990.3;AT2G45990.4
bic:851
peaks after deleting less than one fifth of global max:17
×10^8
raw data, starting:17
fitted Gauss locations:17
y
x
0
0.5
1
1.5
2
2.5
3
3.5
4
5
10
15
20
25

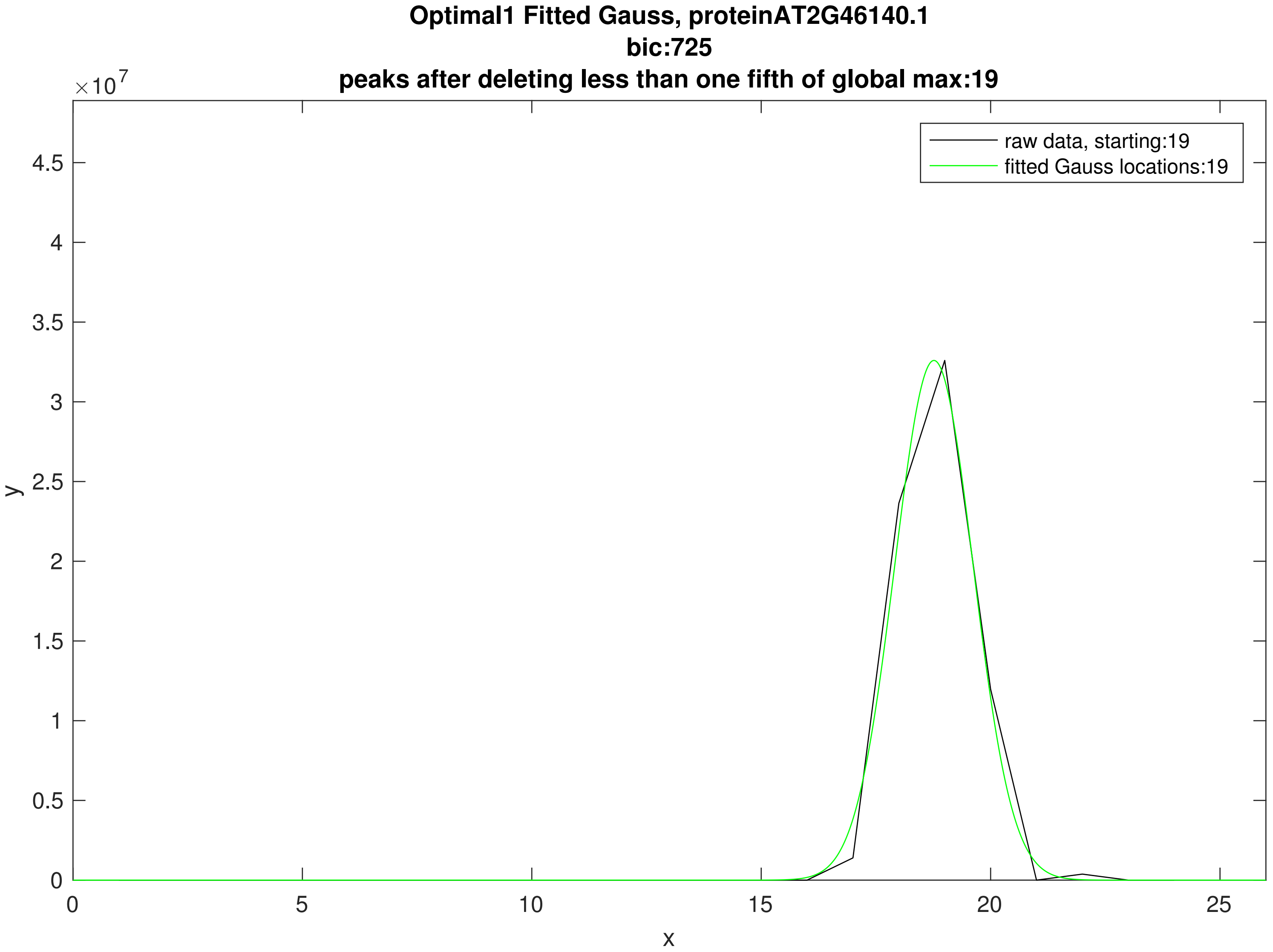
Optimal1 Fitted Gauss, proteinAT2G46140.1
bic:725
peaks after deleting less than one fifth of global max:19
×10^7
raw data, starting:19
fitted Gauss locations:19
y
x
0
0.5
1
1.5
2
2.5
3
3.5
4
4.5
0
5
10
15
20
25

**Optimal1 Fitted Gauss, proteinAT2G46280.2;AT2G46280.1;AT2G46280.3**
**bic:925 934 944**
**peaks after deleting less than one fifth of global max:18**

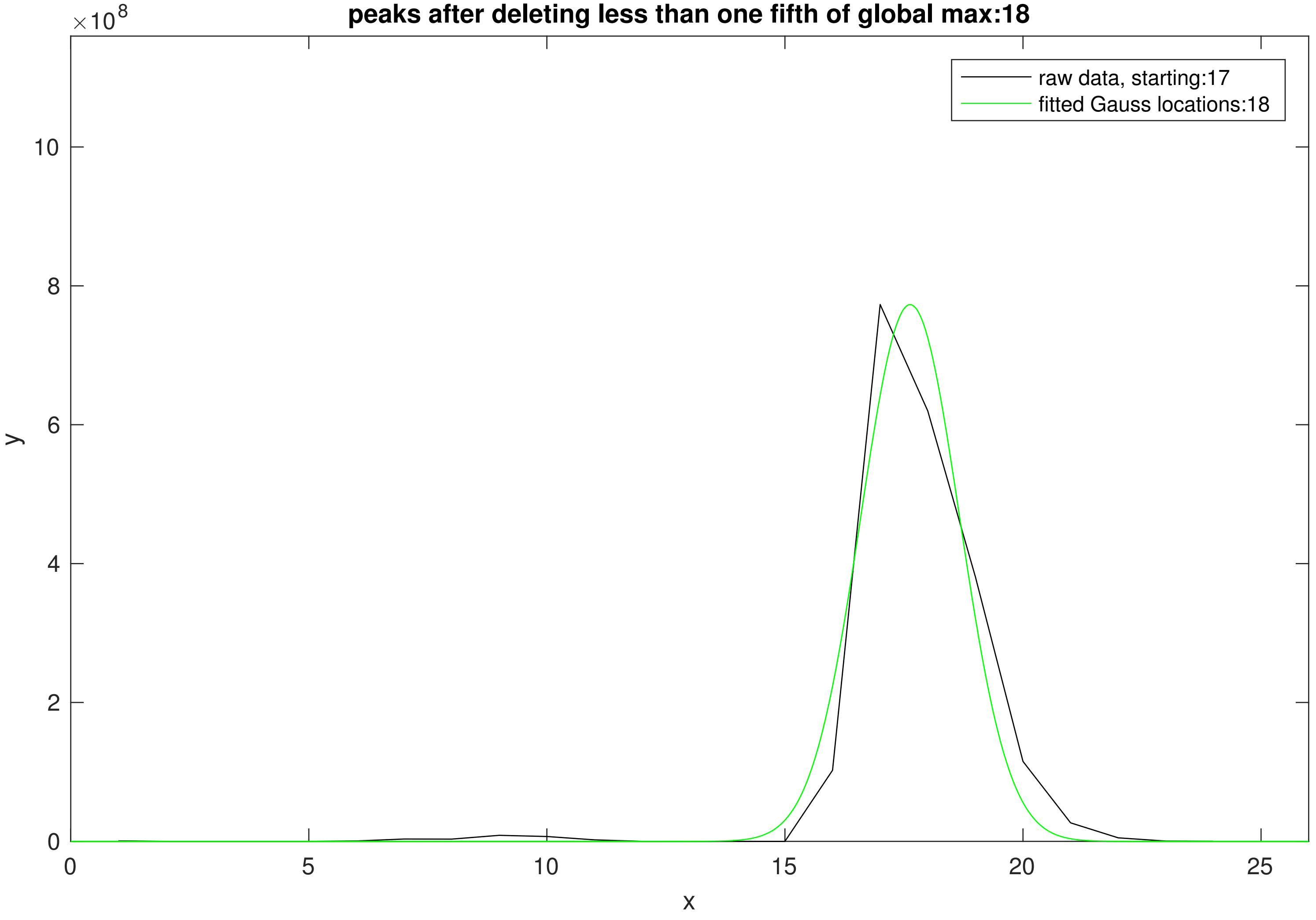

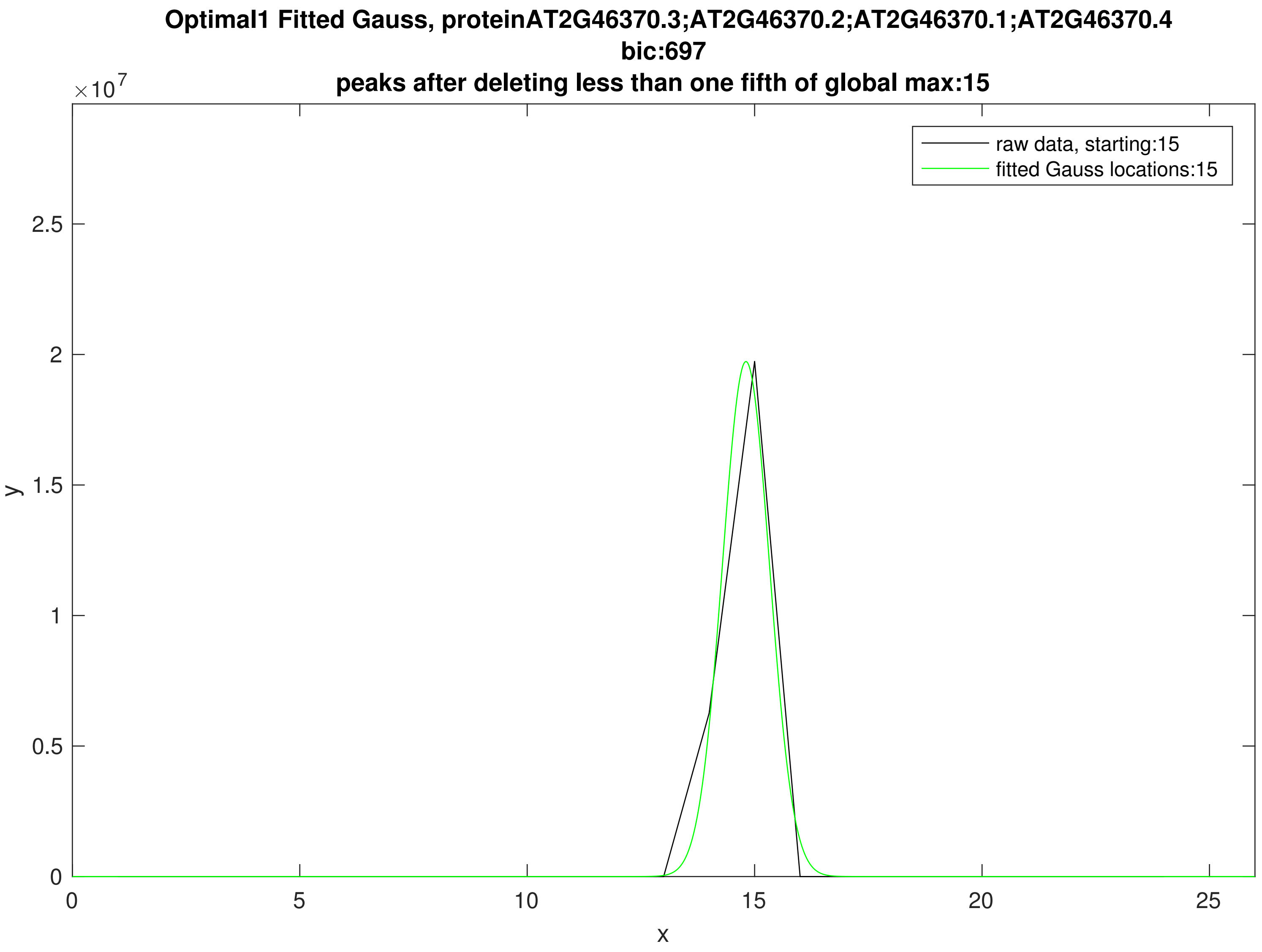
Optimal1 Fitted Gauss, proteinAT2G46370.3;AT2G46370.2;AT2G46370.1;AT2G46370.4
bic:697
peaks after deleting less than one fifth of global max:15
×10^7
raw data, starting:15
fitted Gauss locations:15
2.5
2
1.5
1
0.5
0
y
0
5
10
15
20
25
x

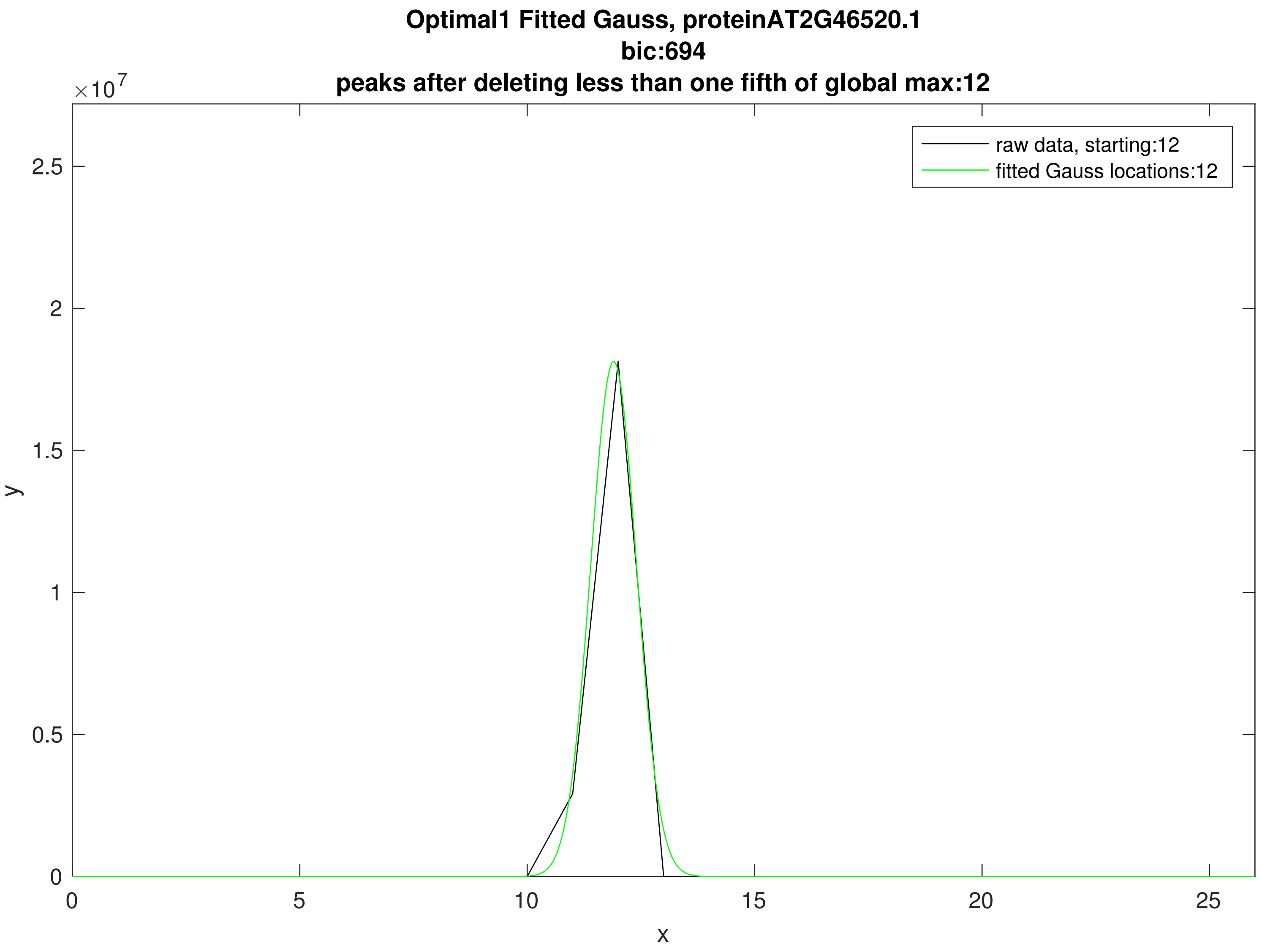

Optimal1 Fitted Gauss, proteinAT2G46520.1
bic:694
peaks after deleting less than one fifth of global max:12
×10^7
2.5
2
1.5
1
0.5
0
y
0
5
10
15
20
25
x
raw data, starting:12
fitted Gauss locations:12

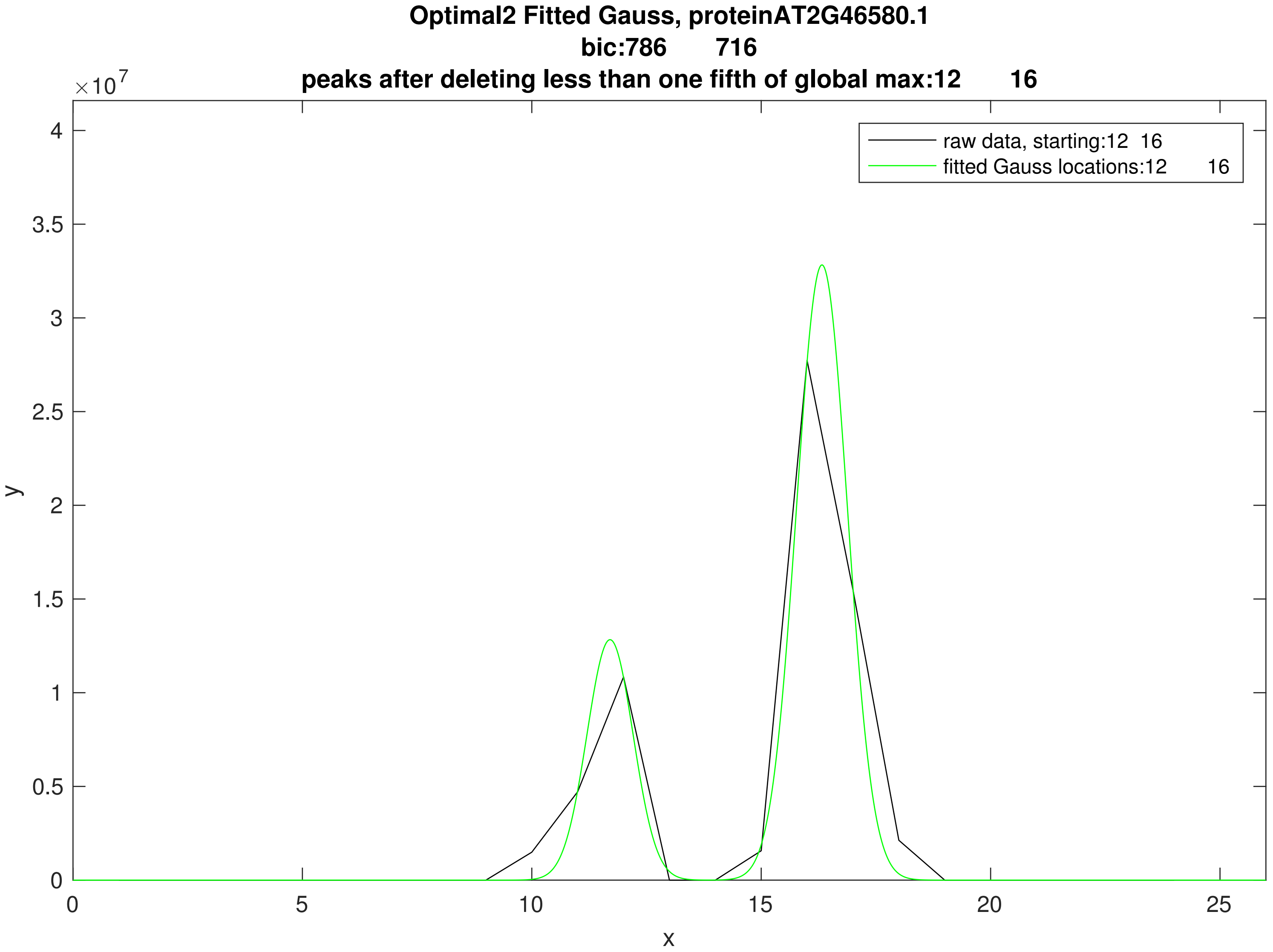

Optimal2 Fitted Gauss, proteinAT2G46580.1
bic:786 716
peaks after deleting less than one fifth of global max:12 16
×10^7
raw data, starting:12 16
fitted Gauss locations:12 16
y
x
0
0.5
1
1.5
2
2.5
3
3.5
4
5
10
15
20
25

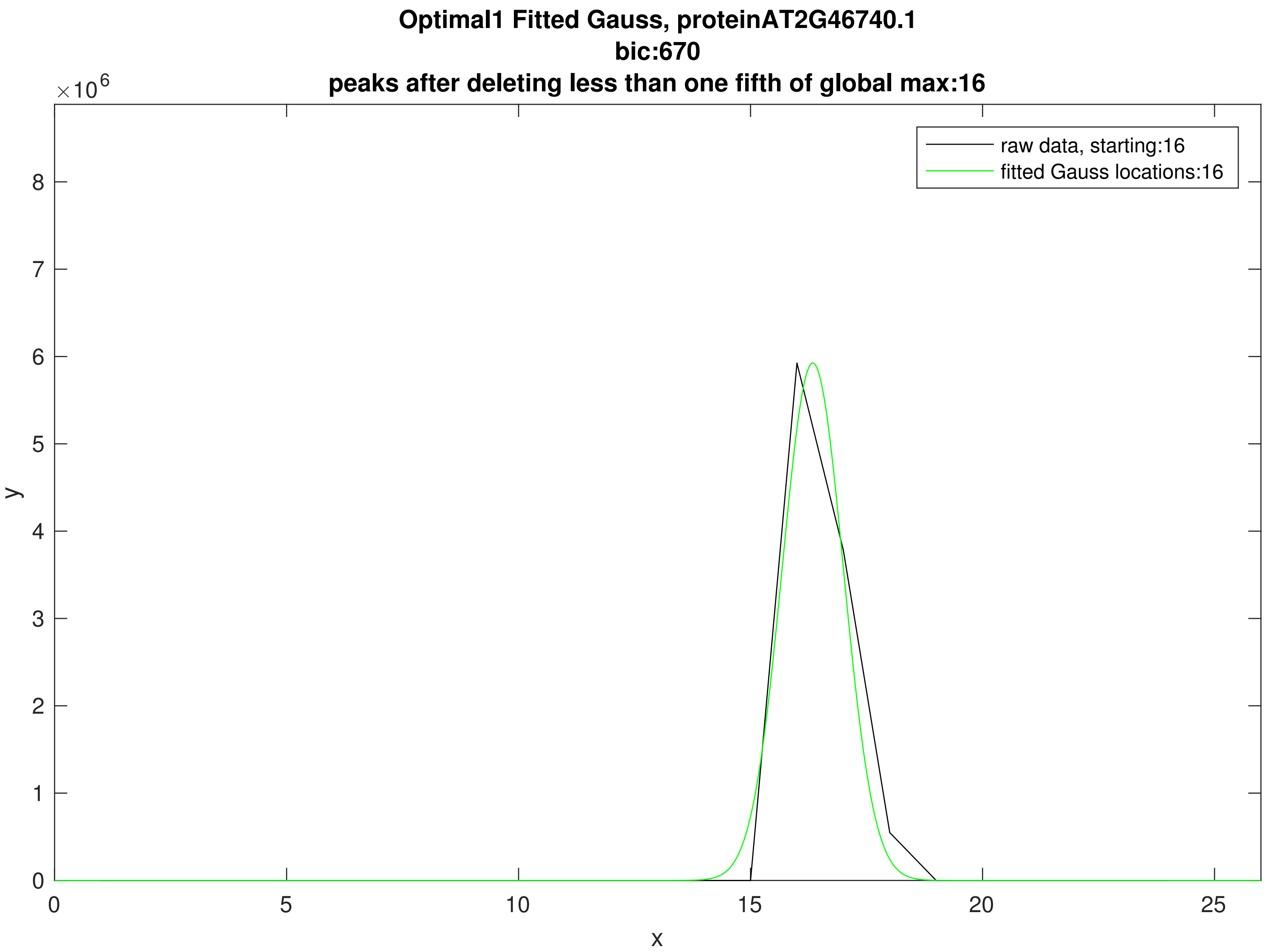
Optimal1 Fitted Gauss, proteinAT2G46740.1
bic:670
peaks after deleting less than one fifth of global max:16
×10^6
raw data, starting:16
fitted Gauss locations:16
y
x
0
1
2
3
4
5
6
7
8
0
5
10
15
20
25

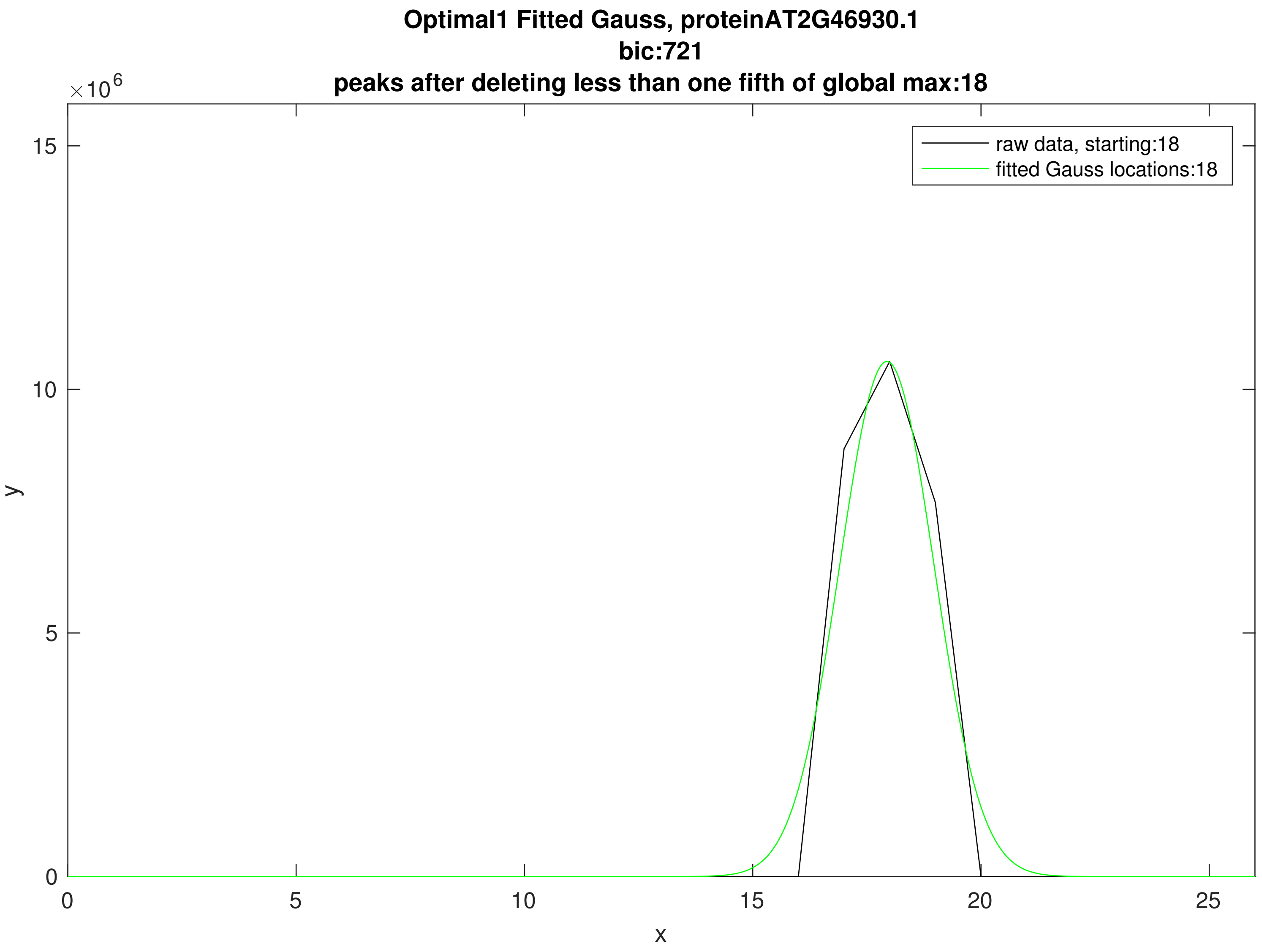

Optimal1 Fitted Gauss, proteinAT2G46930.1
bic:721
peaks after deleting less than one fifth of global max:18
×10^6
15
10
5
0
y
0
5
10
15
20
25
x
raw data, starting:18
fitted Gauss locations:18

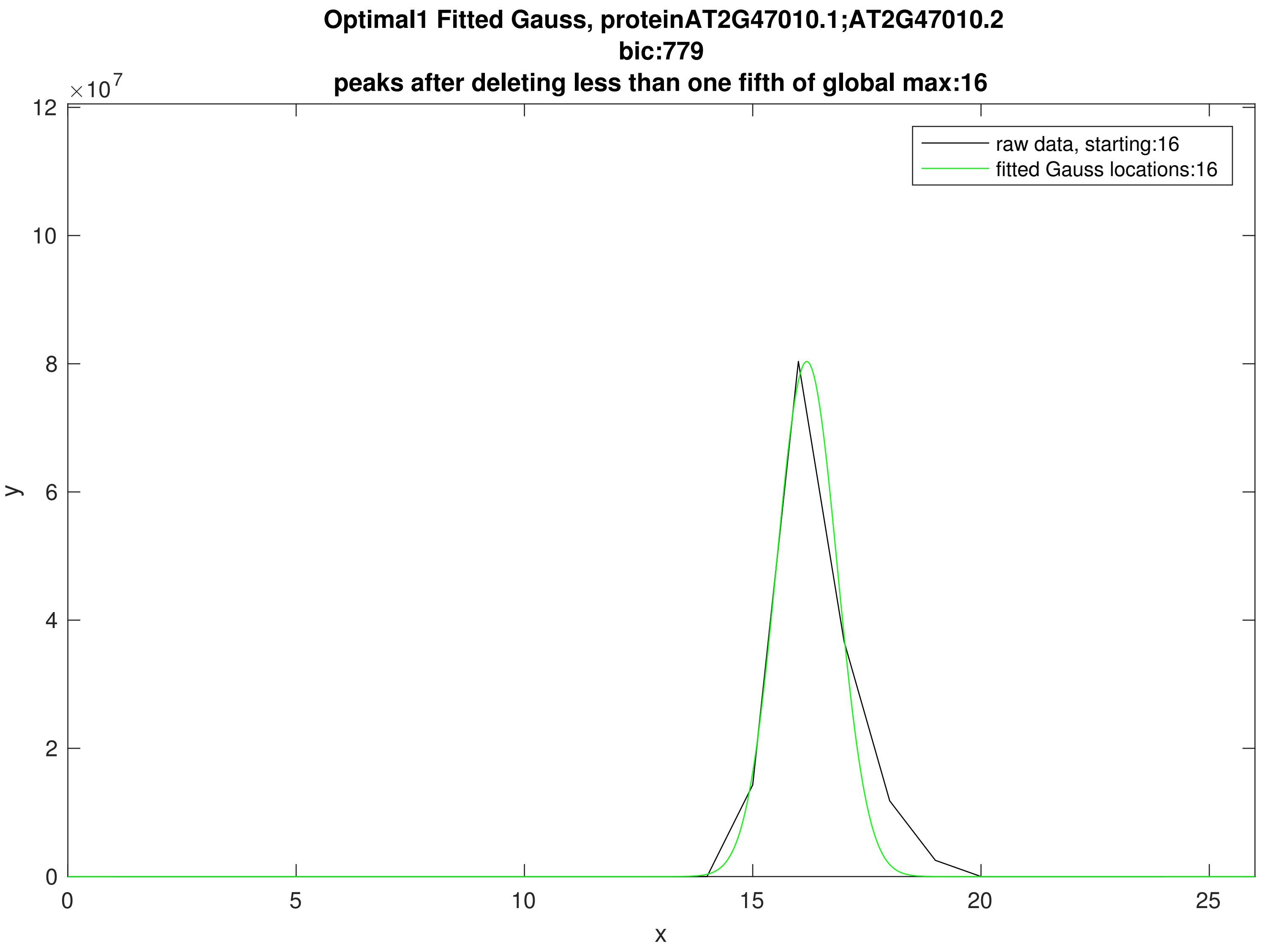
Optimal1 Fitted Gauss, proteinAT2G47010.1;AT2G47010.2
bic:779
peaks after deleting less than one fifth of global max:16
×10^7
12
10
8
6
4
2
0
y
0
5
10
15
20
25
x
raw data, starting:16
fitted Gauss locations:16

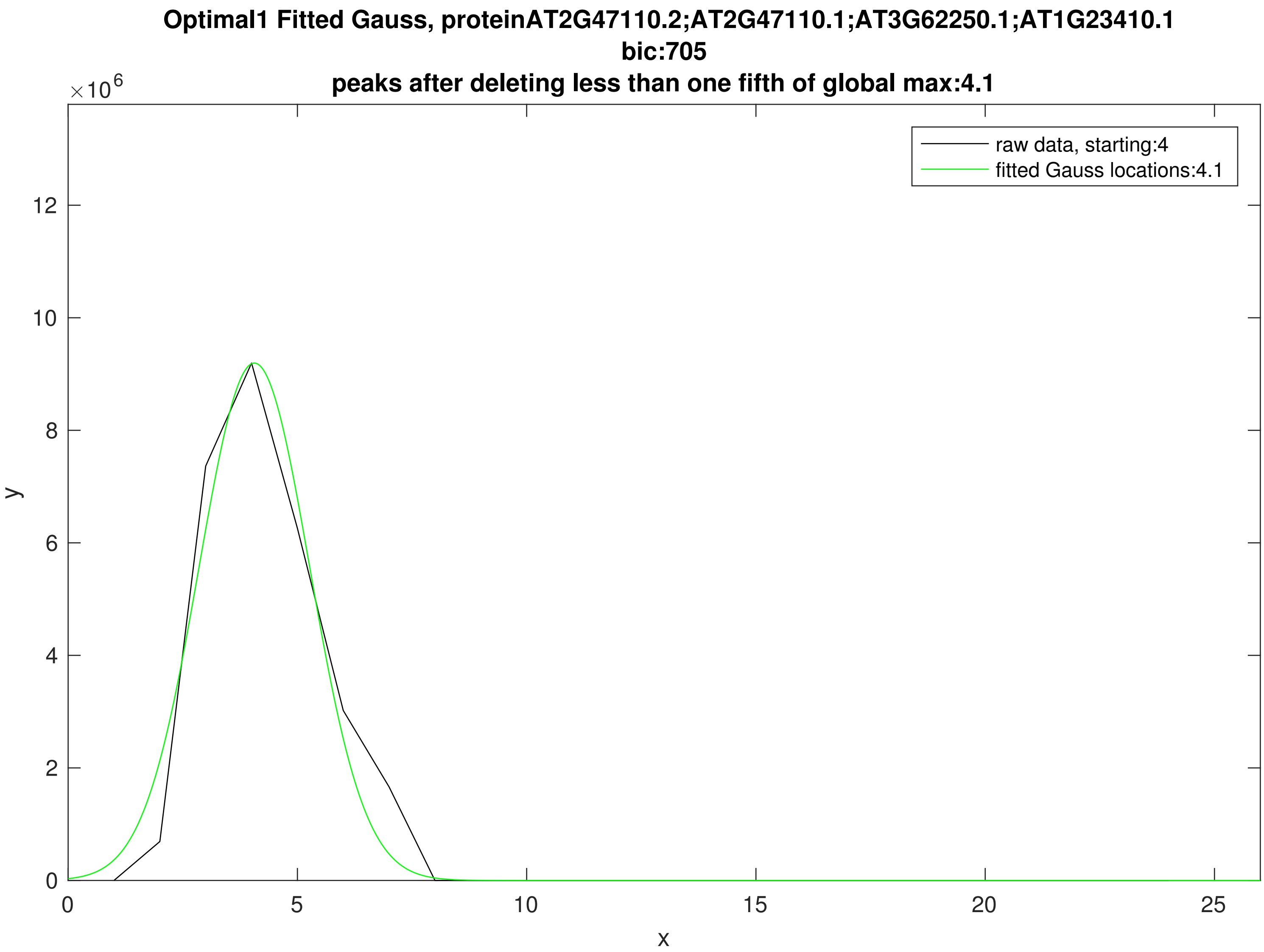

Optimal1 Fitted Gauss, proteinAT2G47110.2;AT2G47110.1;AT3G62250.1;AT1G23410.1
bic:705
peaks after deleting less than one fifth of global max:4.1
×10^6
raw data, starting:4
fitted Gauss locations:4.1
y
x
0
2
4
6
8
10
12
0
5
10
15
20
25

**Optimal1 Fitted Gauss, proteinAT2G47120.1**
**bic:701**
**peaks after deleting less than one fifth of global max:13**

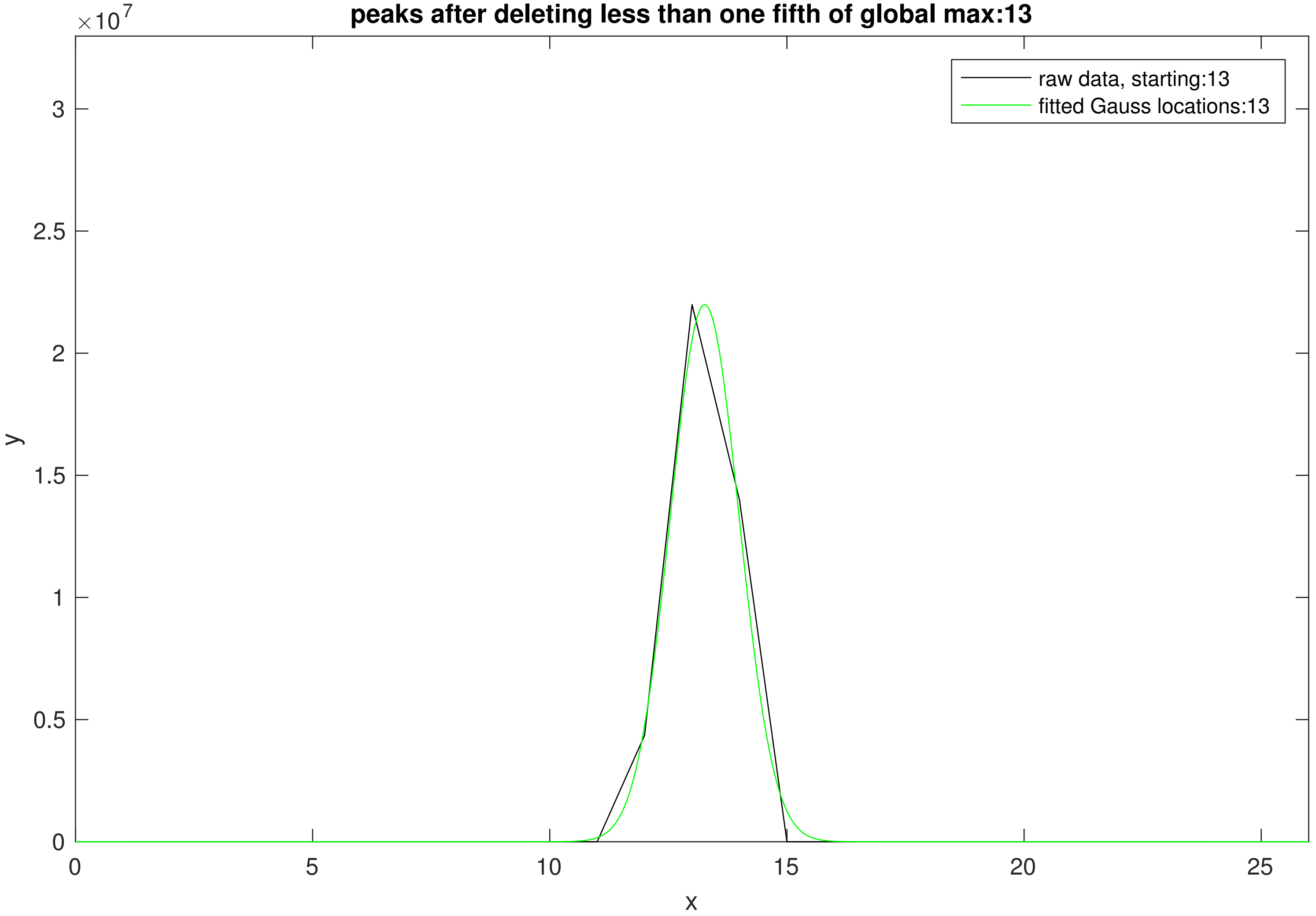

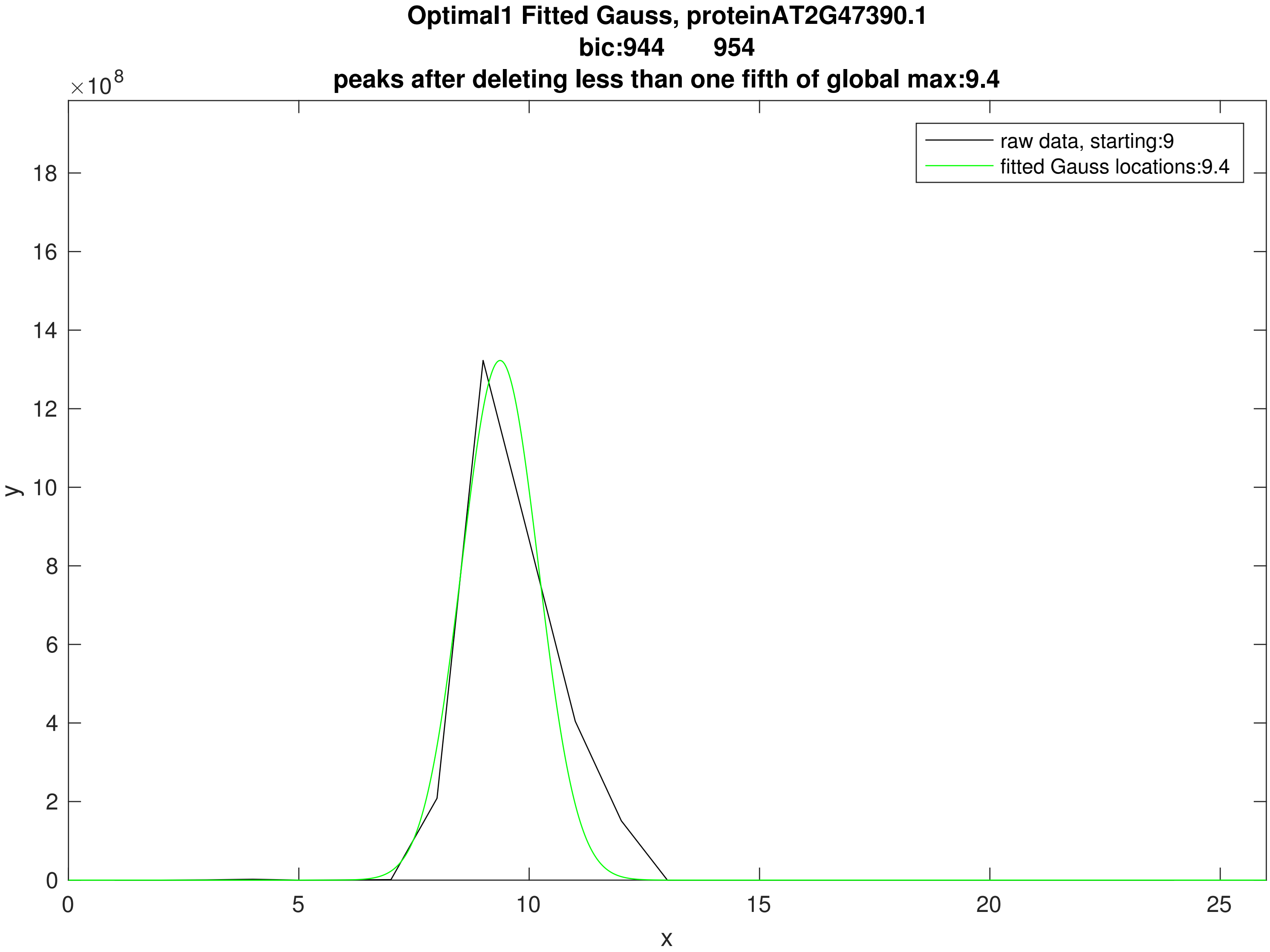
Optimal1 Fitted Gauss, proteinAT2G47390.1
bic:944 954
peaks after deleting less than one fifth of global max:9.4
×10^8
raw data, starting:9
fitted Gauss locations:9.4
y
x
0
2
4
6
8
10
12
14
16
18
0
5
10
15
20
25

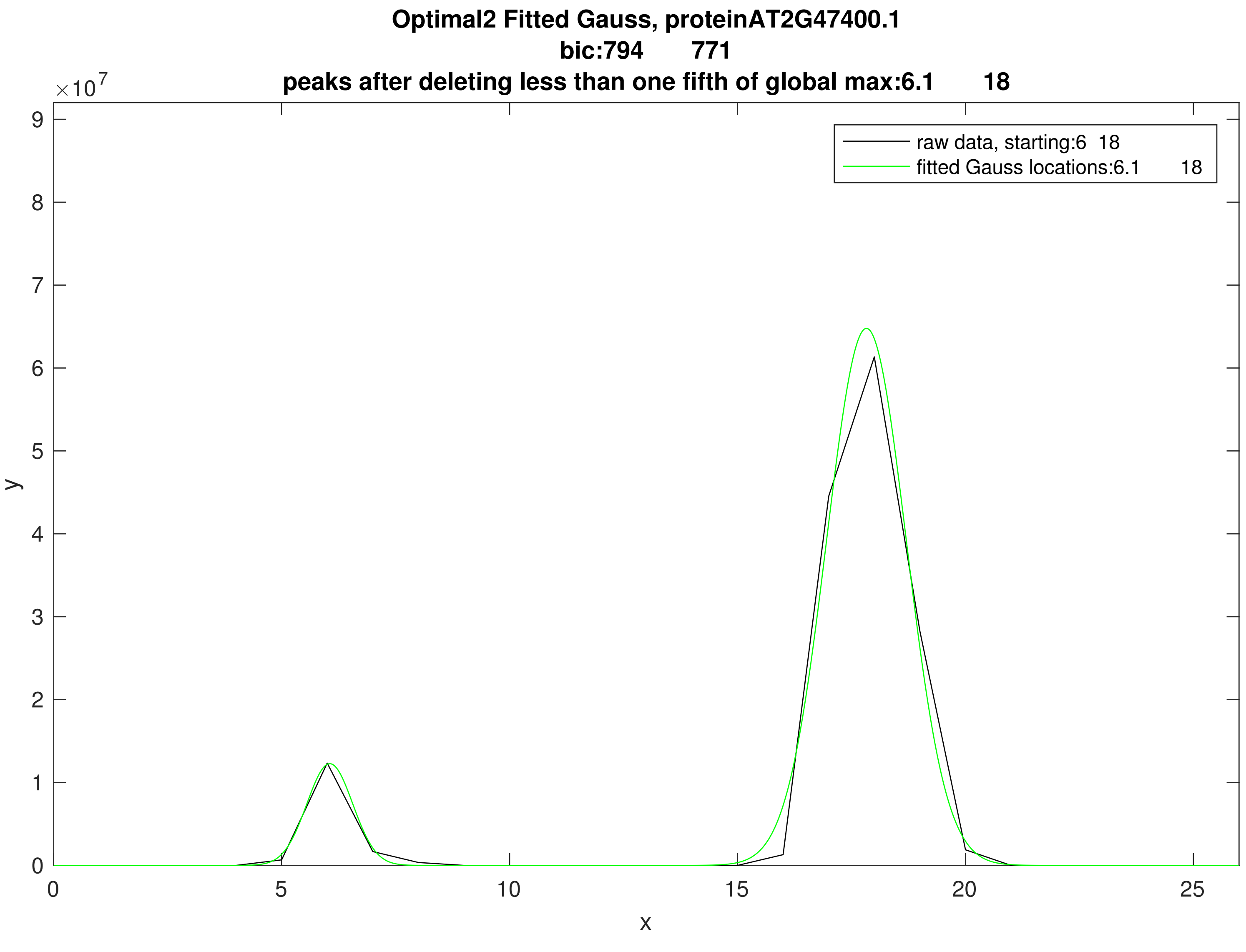

Optimal2 Fitted Gauss, proteinAT2G47400.1
bic:794 771
peaks after deleting less than one fifth of global max:6.1 18
×10^7
raw data, starting:6 18
fitted Gauss locations:6.1 18
y
x
0
1
2
3
4
5
6
7
8
9
0
5
10
15
20
25

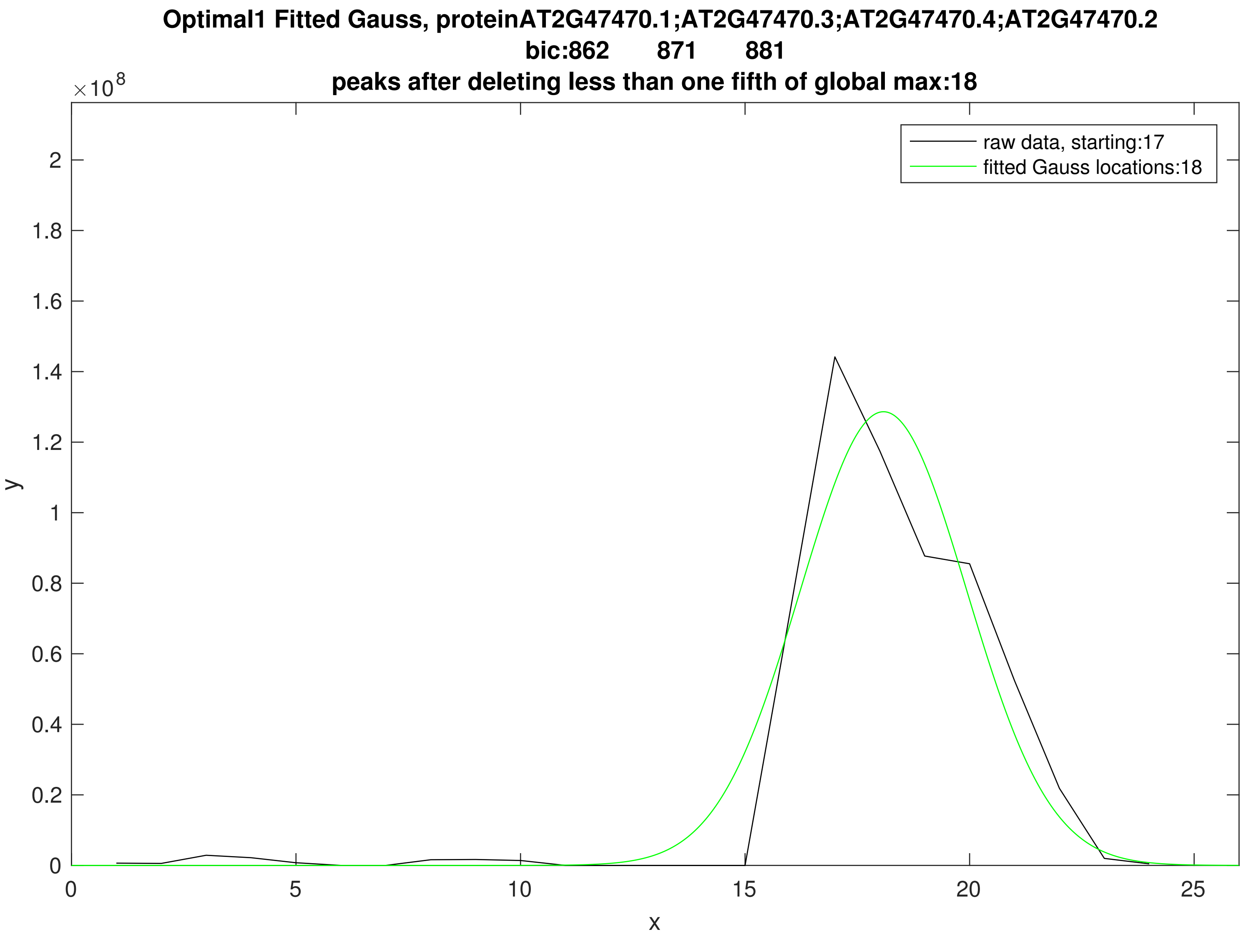

Optimal1 Fitted Gauss, proteinAT2G47470.1;AT2G47470.3;AT2G47470.4;AT2G47470.2
bic:862 871 881
peaks after deleting less than one fifth of global max:18
×10^8
2
1.8
1.6
1.4
1.2
1
0.8
0.6
0.4
0.2
0
y
0
5
10
15
20
25
x
raw data, starting:17
fitted Gauss locations:18

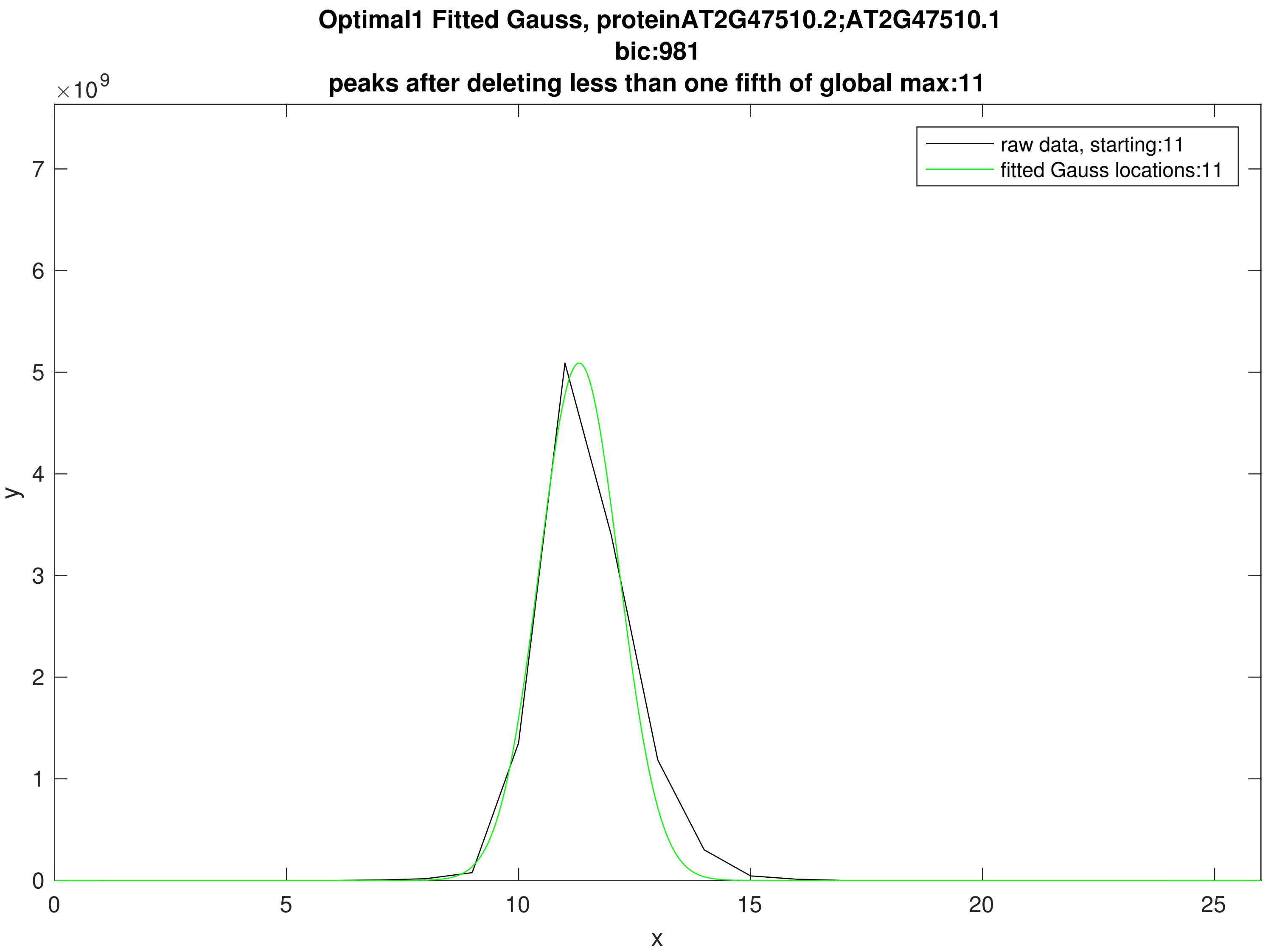

Optimal1 Fitted Gauss, proteinAT2G47510.2;AT2G47510.1
bic:981
peaks after deleting less than one fifth of global max:11
×10^9
raw data, starting:11
fitted Gauss locations:11
y
x
0
1
2
3
4
5
6
7
0
5
10
15
20
25

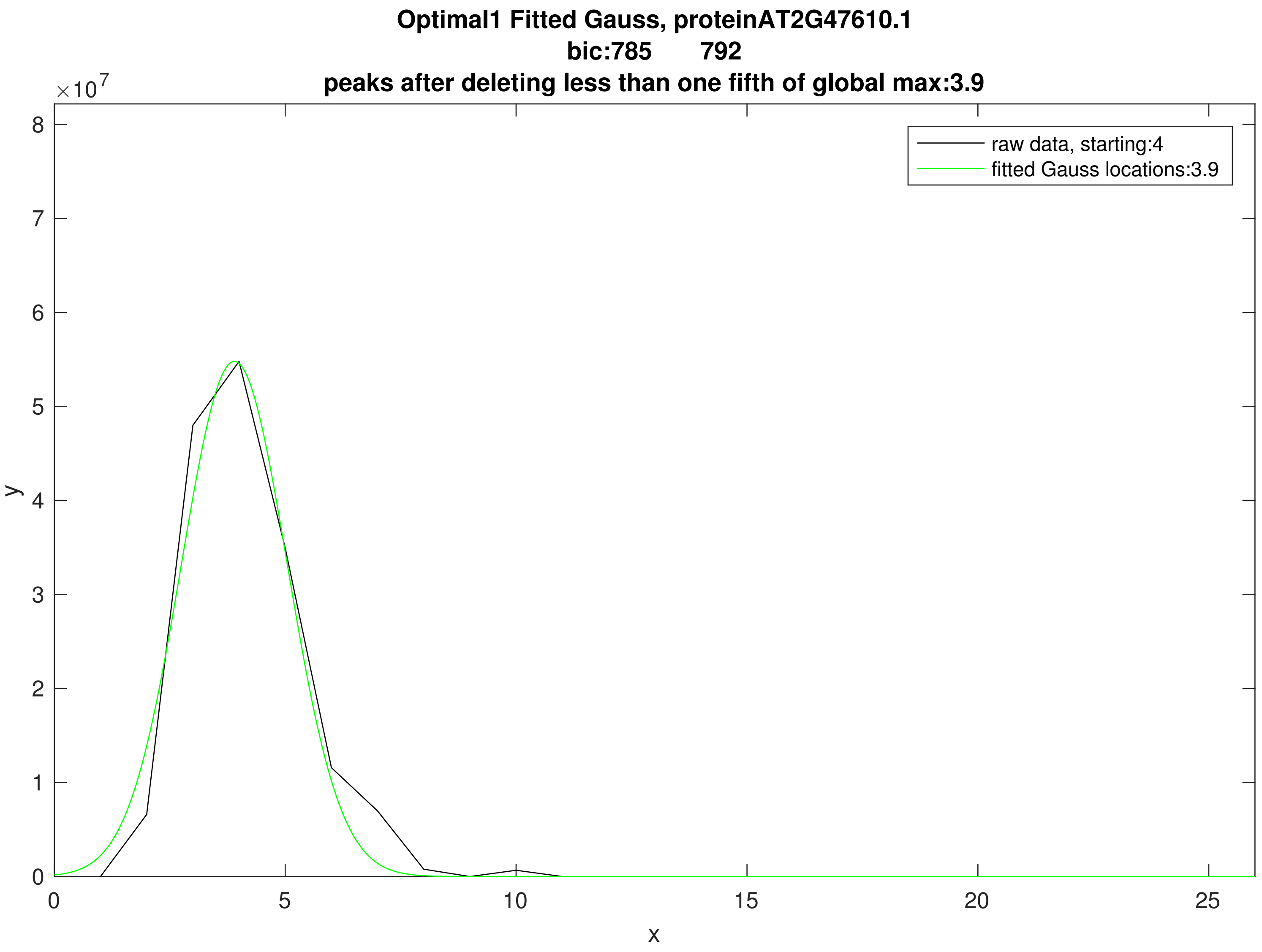

Optimal1 Fitted Gauss, proteinAT2G47610.1
bic:785 792
peaks after deleting less than one fifth of global max:3.9
×10^7
raw data, starting:4
fitted Gauss locations:3.9
y
x
0
1
2
3
4
5
6
7
8
0
5
10
15
20
25

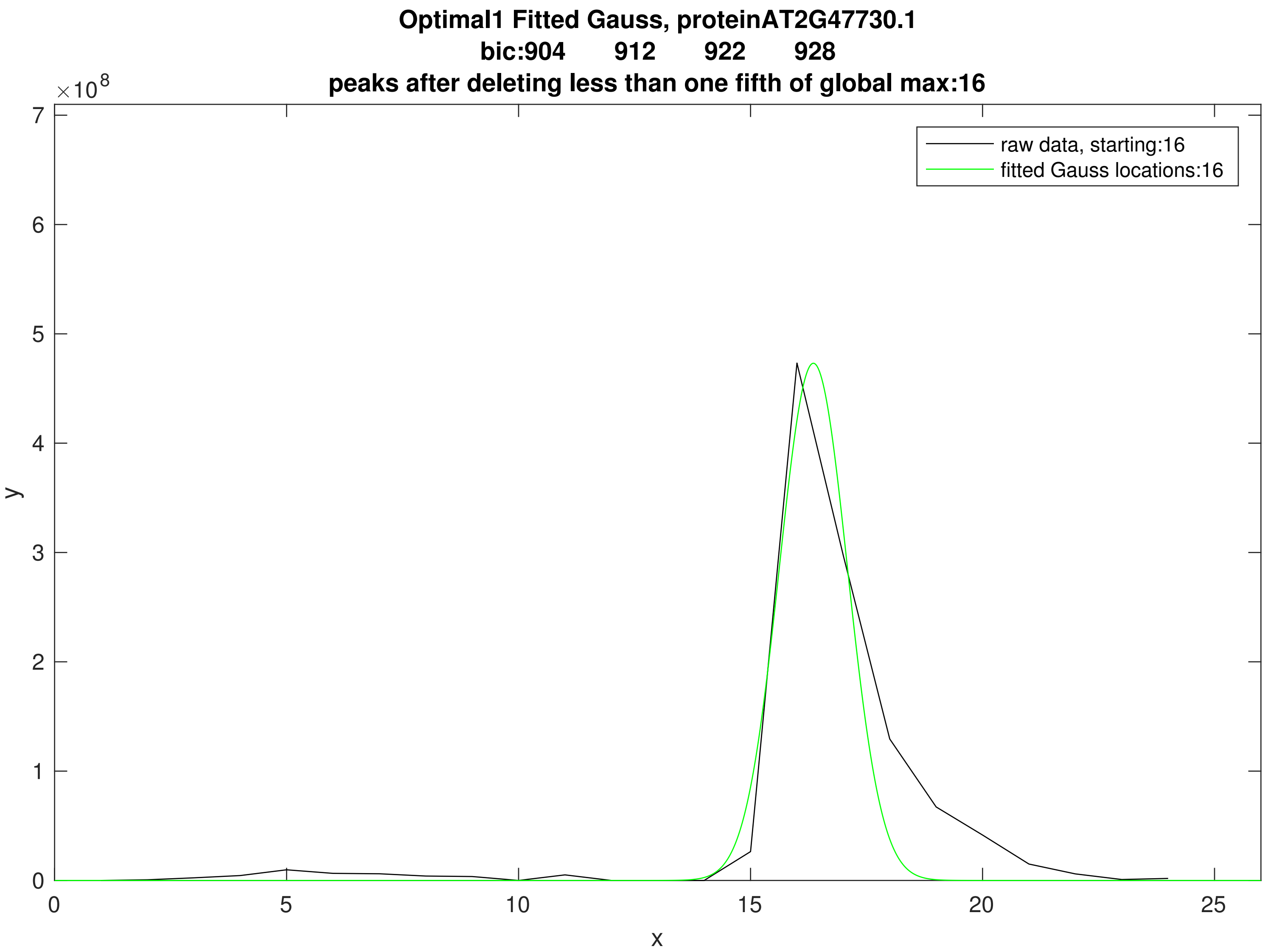
Optimal1 Fitted Gauss, proteinAT2G47730.1
bic:904 912 922 928
peaks after deleting less than one fifth of global max:16
raw data, starting:16
fitted Gauss locations:16
$\times 10^8$
y
x
0
1
2
3
4
5
6
7
10
15
20
25

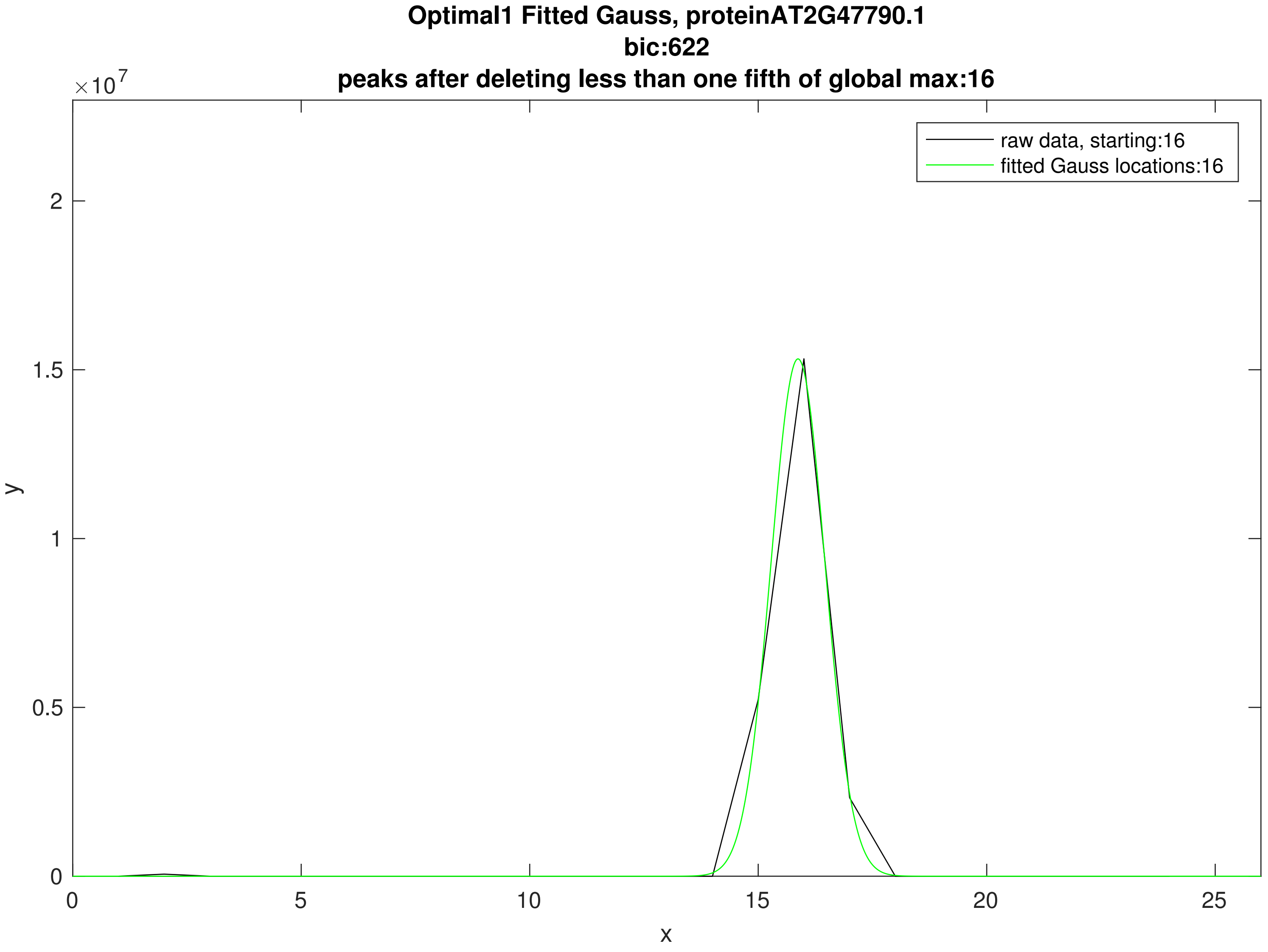

Optimal1 Fitted Gauss, proteinAT2G47790.1
bic:622
peaks after deleting less than one fifth of global max:16
×10^7
raw data, starting:16
fitted Gauss locations:16
2
1.5
1
0.5
0
y
0
5
10
15
20
25
x

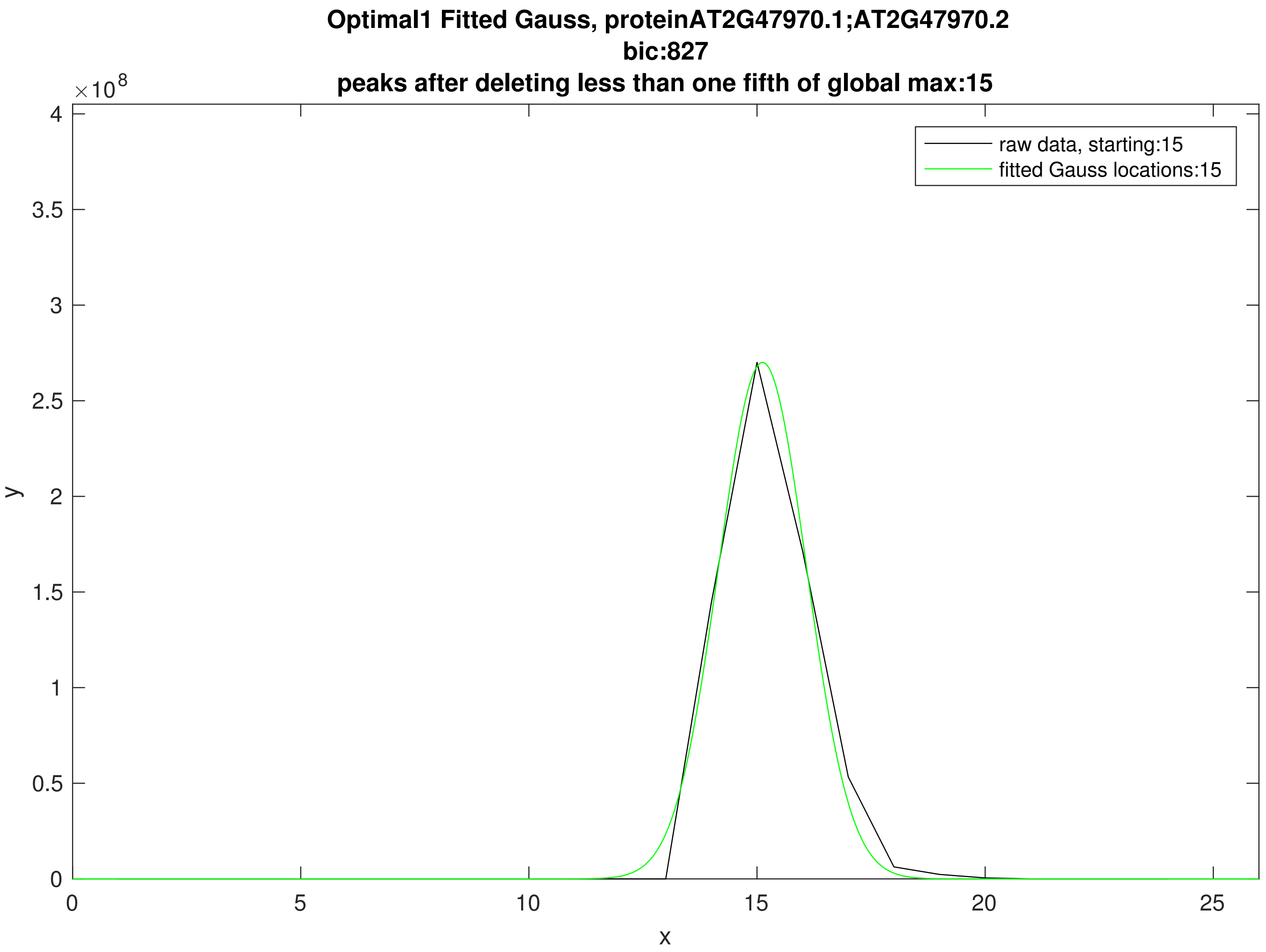
Optimal1 Fitted Gauss, proteinAT2G47970.1;AT2G47970.2
bic:827
peaks after deleting less than one fifth of global max:15
×10^8
4
3.5
3
2.5
2
1.5
1
0.5
0
y
0
5
10
15
20
25
x
raw data, starting:15
fitted Gauss locations:15

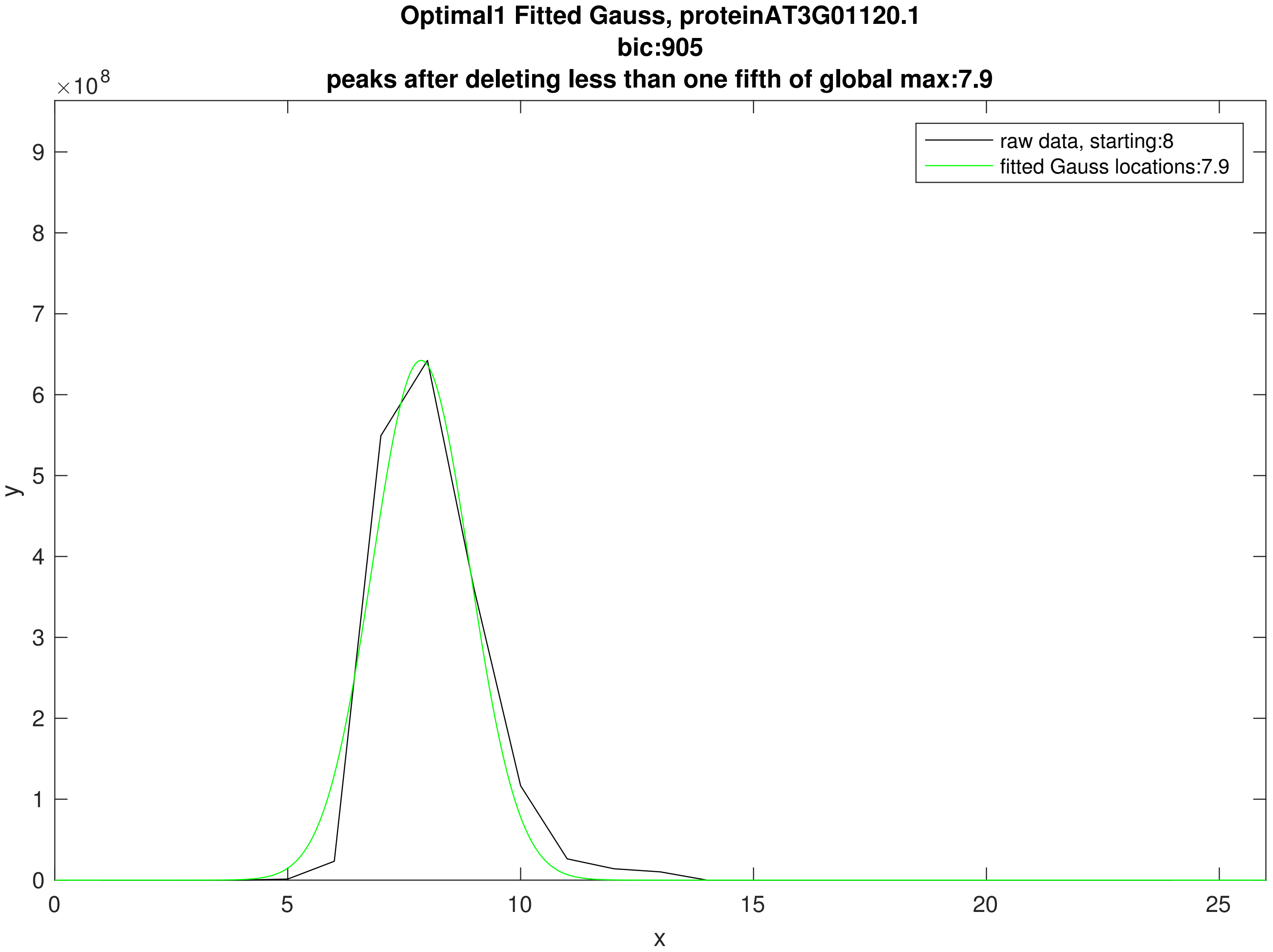
Optimal1 Fitted Gauss, proteinAT3G01120.1
bic:905
peaks after deleting less than one fifth of global max:7.9
×10^8
raw data, starting:8
fitted Gauss locations:7.9
y
x
0
1
2
3
4
5
6
7
8
9
0
5
10
15
20
25

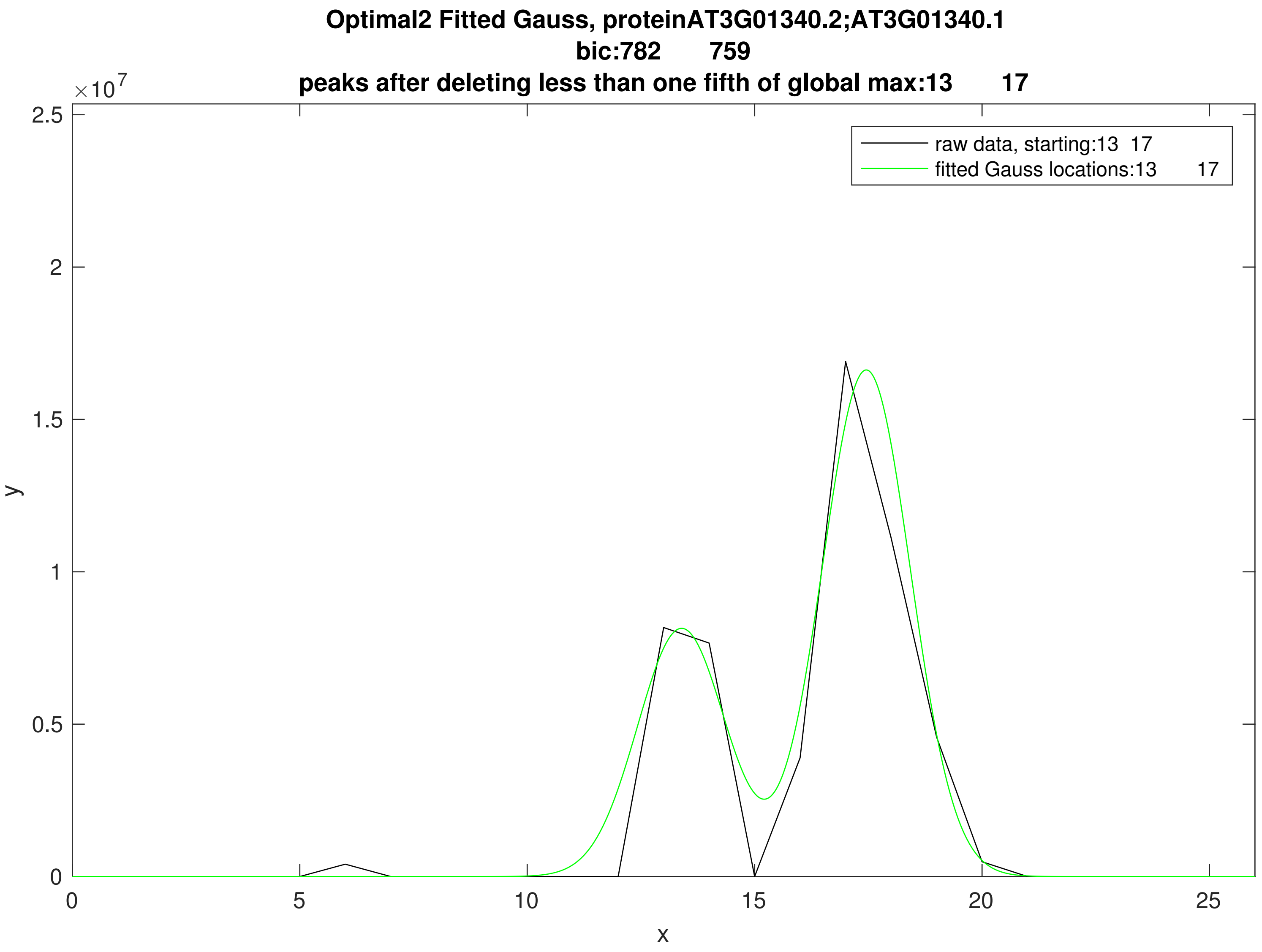
Optimal2 Fitted Gauss, proteinAT3G01340.2;AT3G01340.1
bic:782 759
peaks after deleting less than one fifth of global max:13 17
×10^7
2.5
2
1.5
1
0.5
0
y
0
5
10
15
20
25
x
raw data, starting:13 17
fitted Gauss locations:13 17

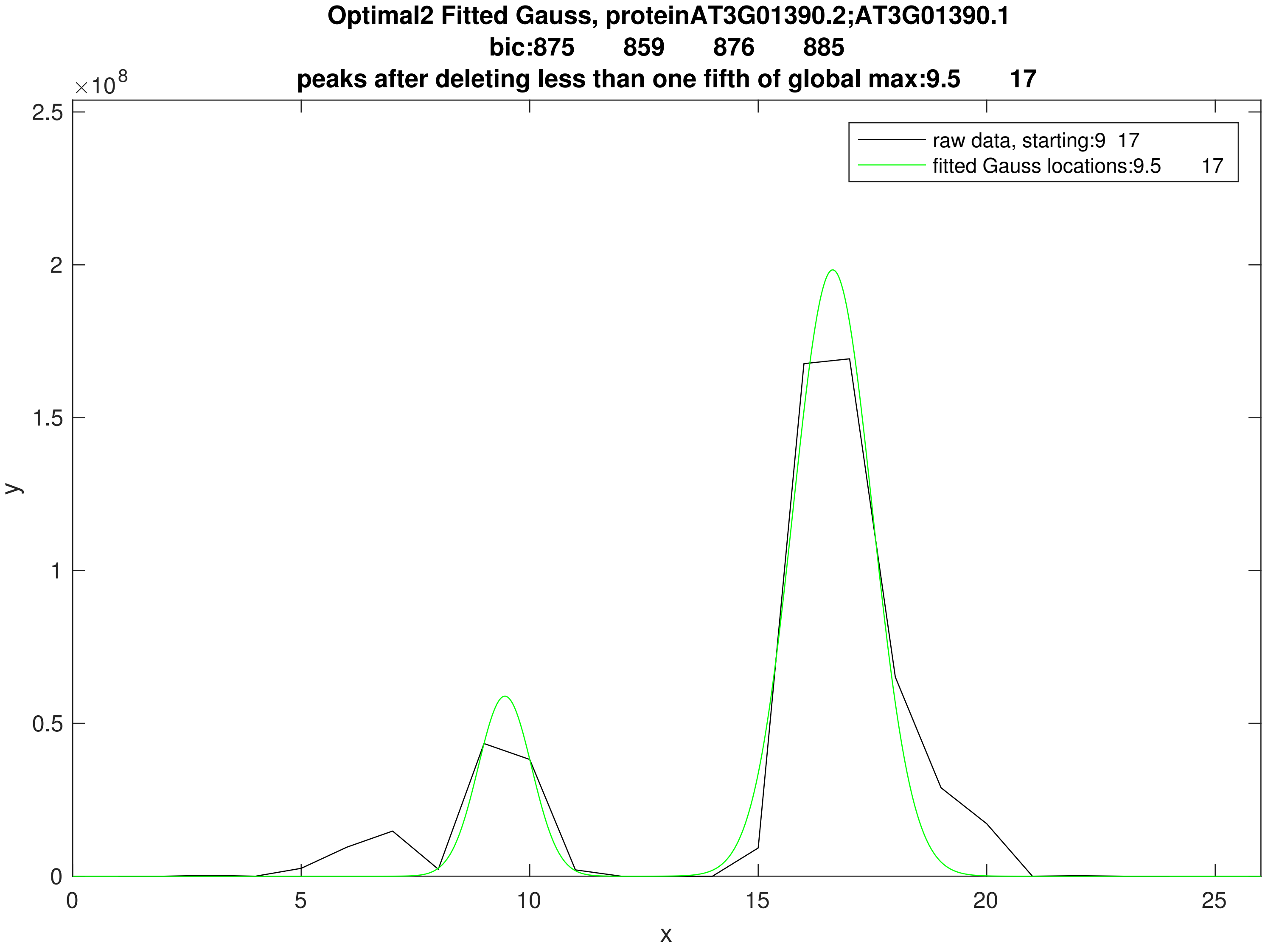
Optimal2 Fitted Gauss, proteinAT3G01390.2;AT3G01390.1
bic:875 859 876 885
peaks after deleting less than one fifth of global max:9.5 17
×10^8
2.5
2
1.5
1
0.5
0
y
0
5
10
15
20
25
x
raw data, starting:9 17
fitted Gauss locations:9.5 17

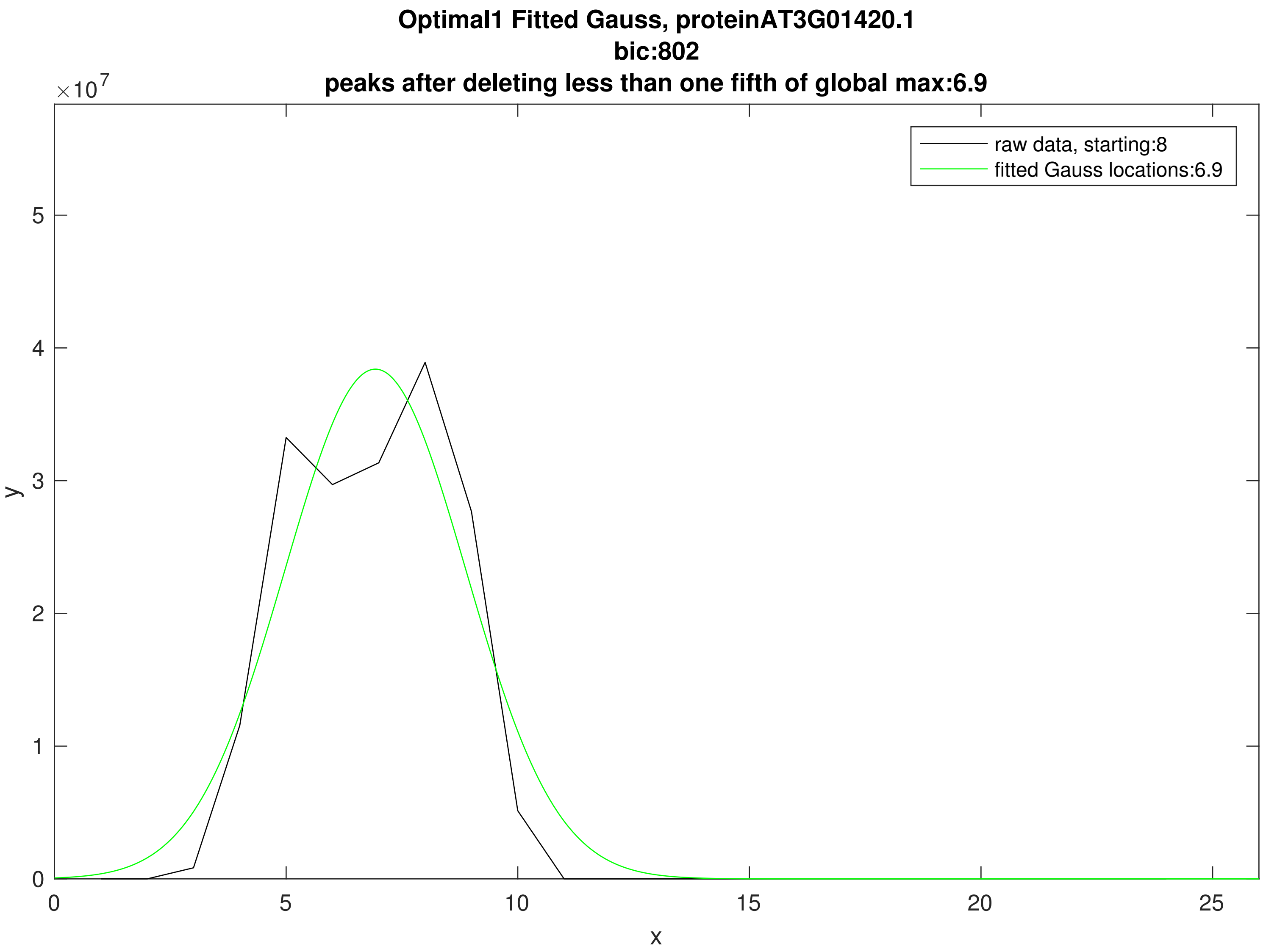
Optimal1 Fitted Gauss, proteinAT3G01420.1
bic:802
peaks after deleting less than one fifth of global max:6.9
×10^7
raw data, starting:8
fitted Gauss locations:6.9
y
x
0
1
2
3
4
5
10
15
20
25

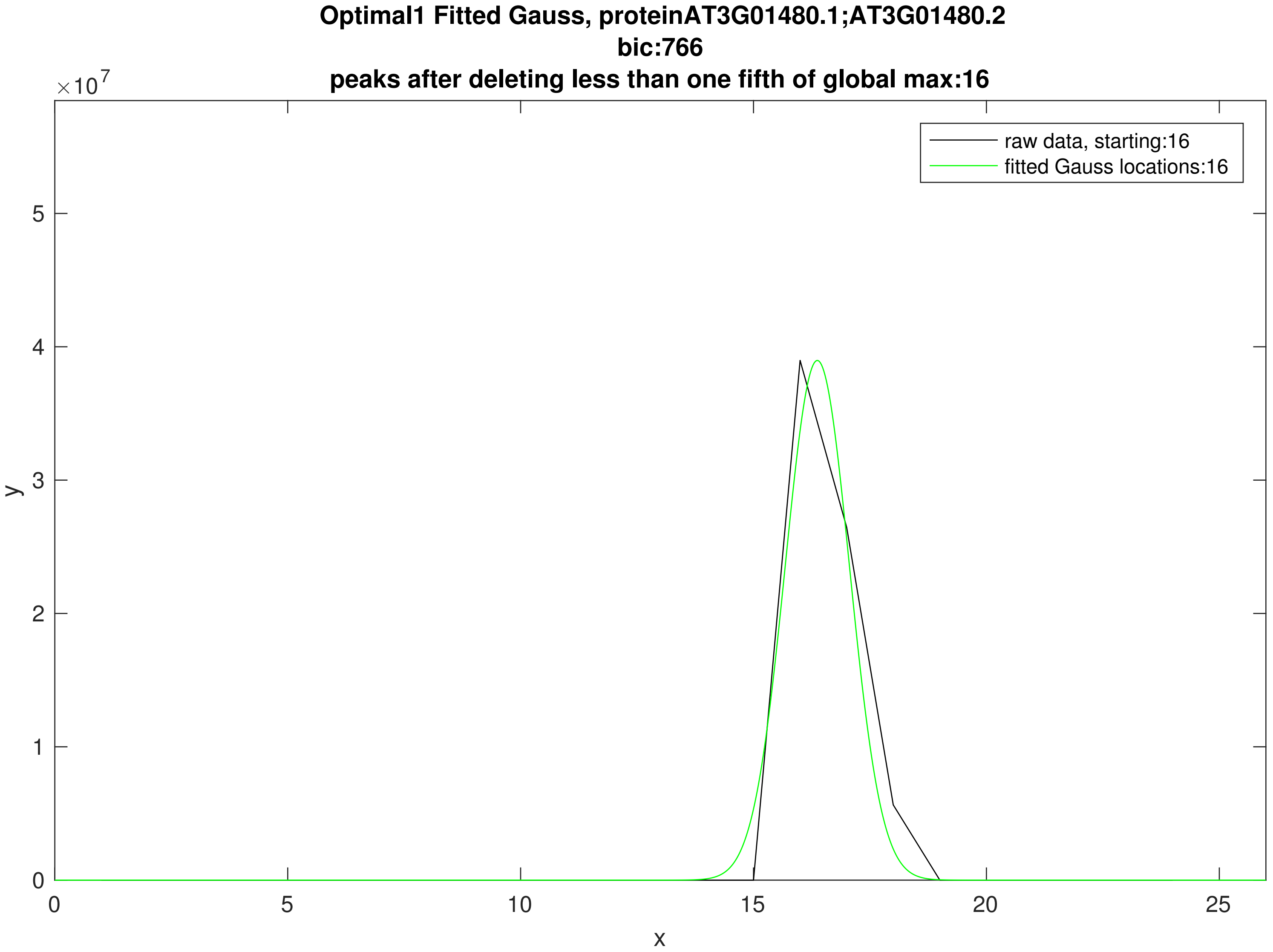
Optimal1 Fitted Gauss, proteinAT3G01480.1;AT3G01480.2
bic:766
peaks after deleting less than one fifth of global max:16
×10^7
raw data, starting:16
fitted Gauss locations:16
y
x
0
1
2
3
4
5
0
5
10
15
20
25

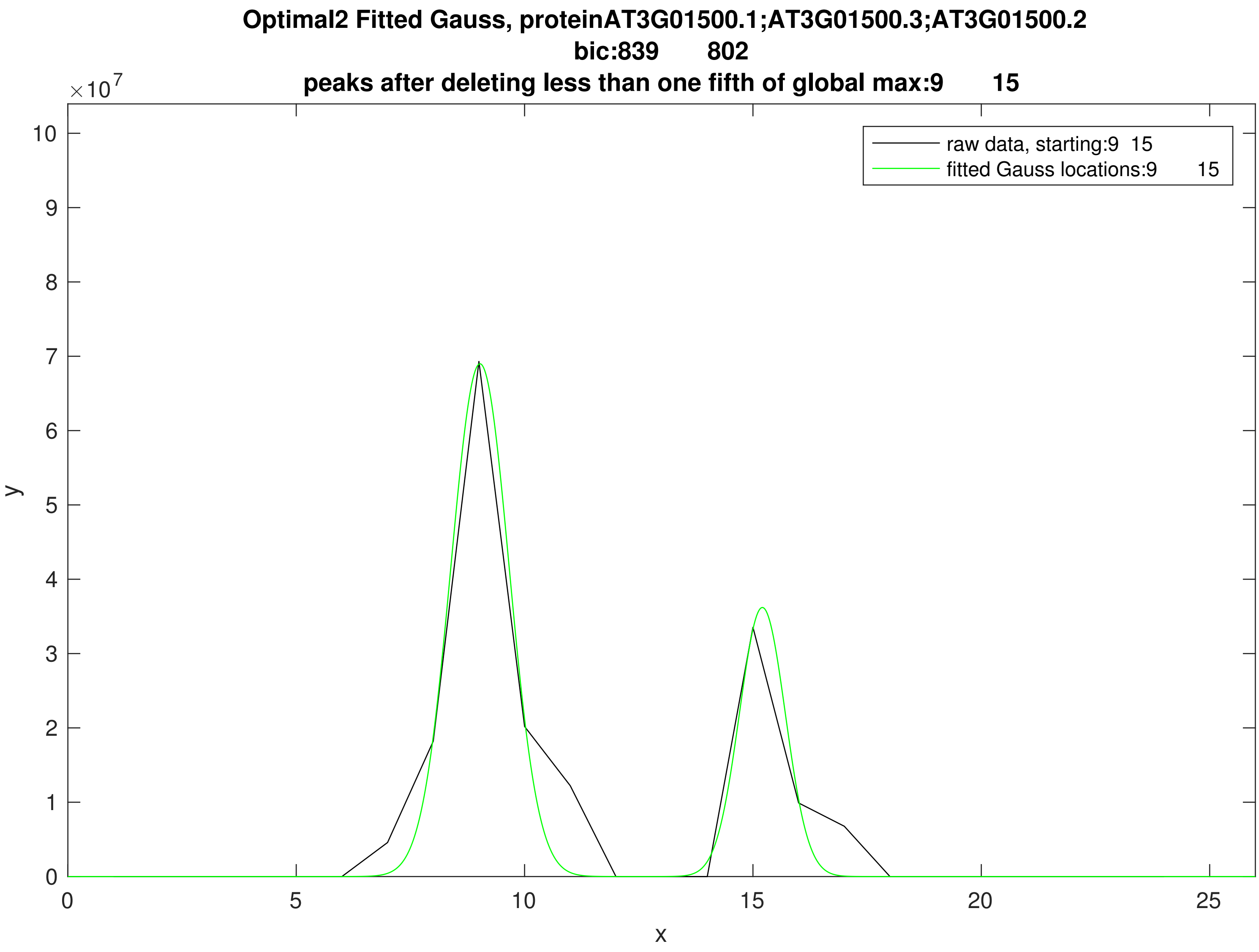
Optimal2 Fitted Gauss, proteinAT3G01500.1;AT3G01500.3;AT3G01500.2
bic:839 802
peaks after deleting less than one fifth of global max:9 15
×10^7
raw data, starting:9 15
fitted Gauss locations:9 15
y
x
0
1
2
3
4
5
6
7
8
9
10
0
5
10
15
20
25

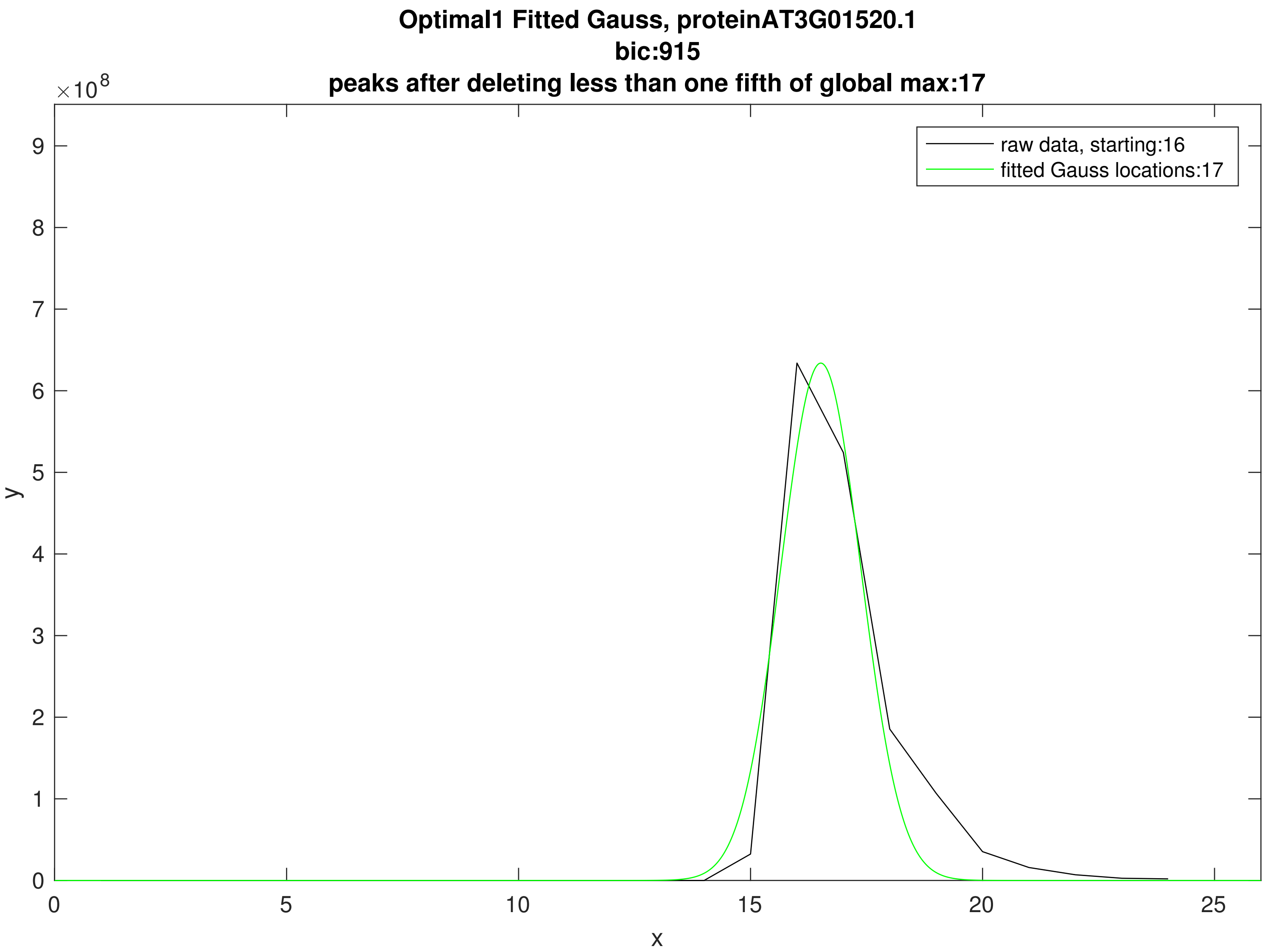

Optimal1 Fitted Gauss, proteinAT3G01520.1
bic:915
peaks after deleting less than one fifth of global max:17
×10^8
raw data, starting:16
fitted Gauss locations:17
y
x
0
1
2
3
4
5
6
7
8
9
0
5
10
15
20
25

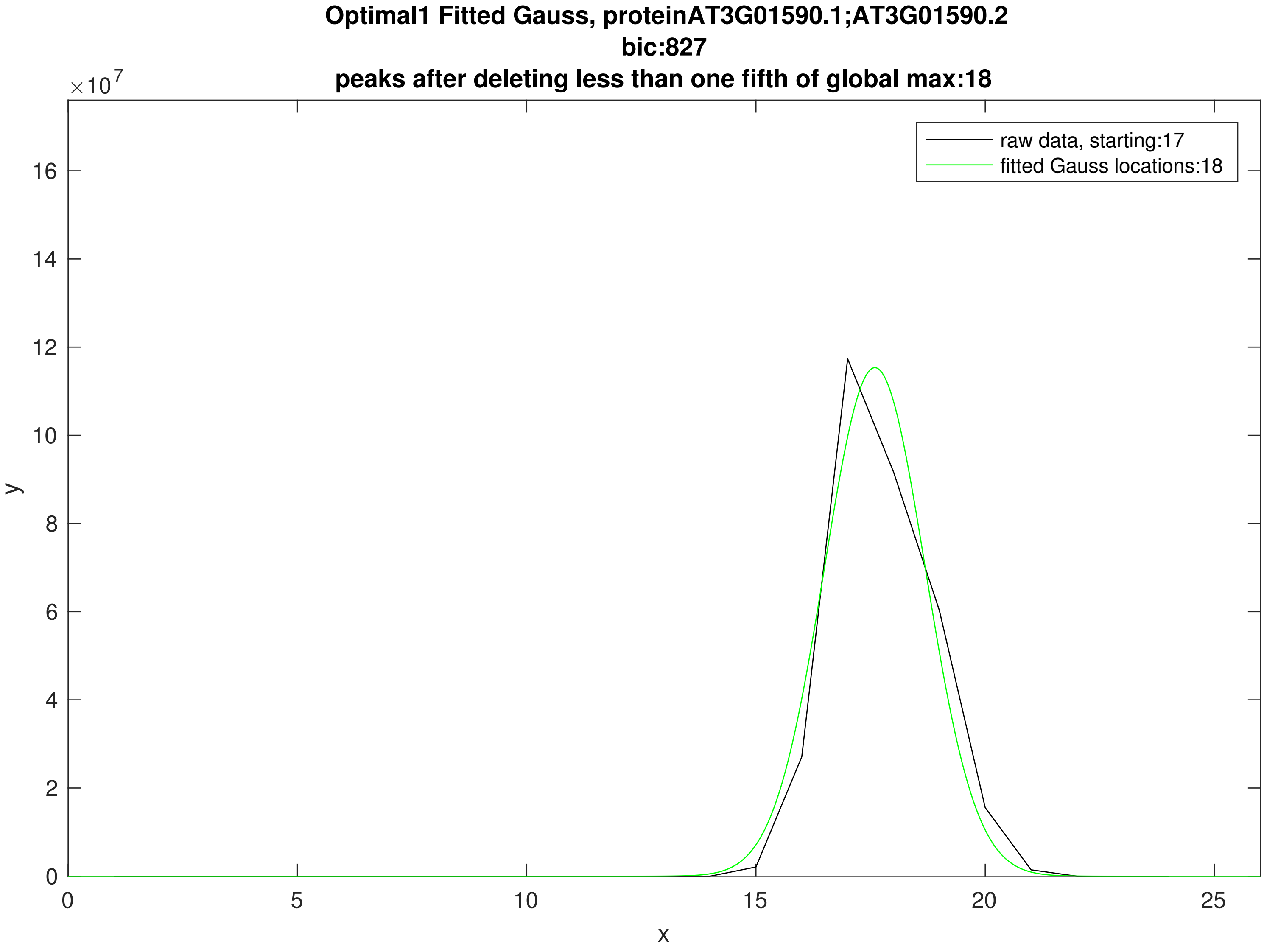
Optimal1 Fitted Gauss, proteinAT3G01590.1;AT3G01590.2
bic:827
peaks after deleting less than one fifth of global max:18
×10^7
raw data, starting:17
fitted Gauss locations:18
y
x
0
2
4
6
8
10
12
14
16
5
15
20
25

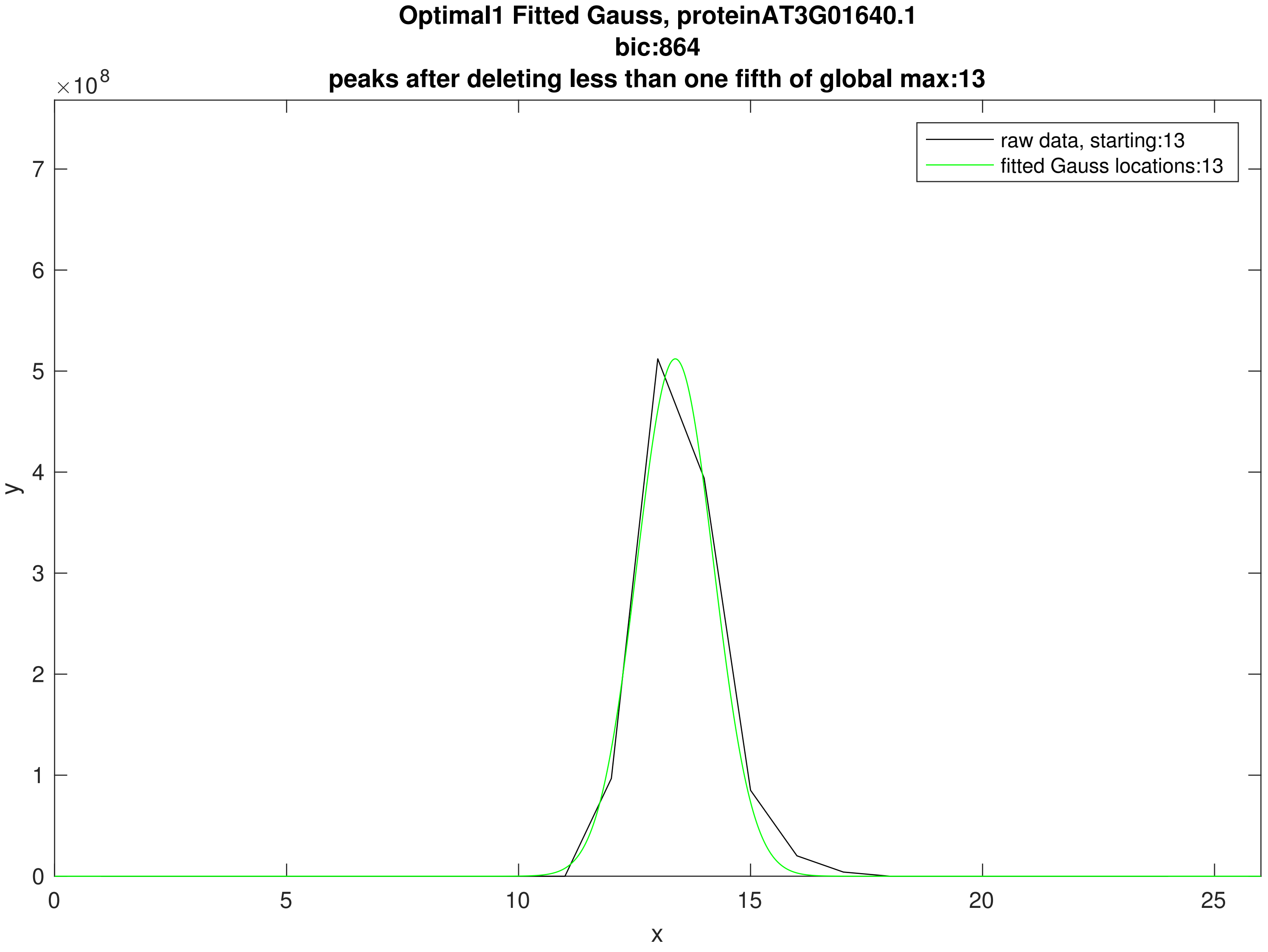
Optimal1 Fitted Gauss, proteinAT3G01640.1
bic:864
peaks after deleting less than one fifth of global max:13
×10^8
raw data, starting:13
fitted Gauss locations:13
y
x
0
1
2
3
4
5
6
7
0
5
10
15
20
25

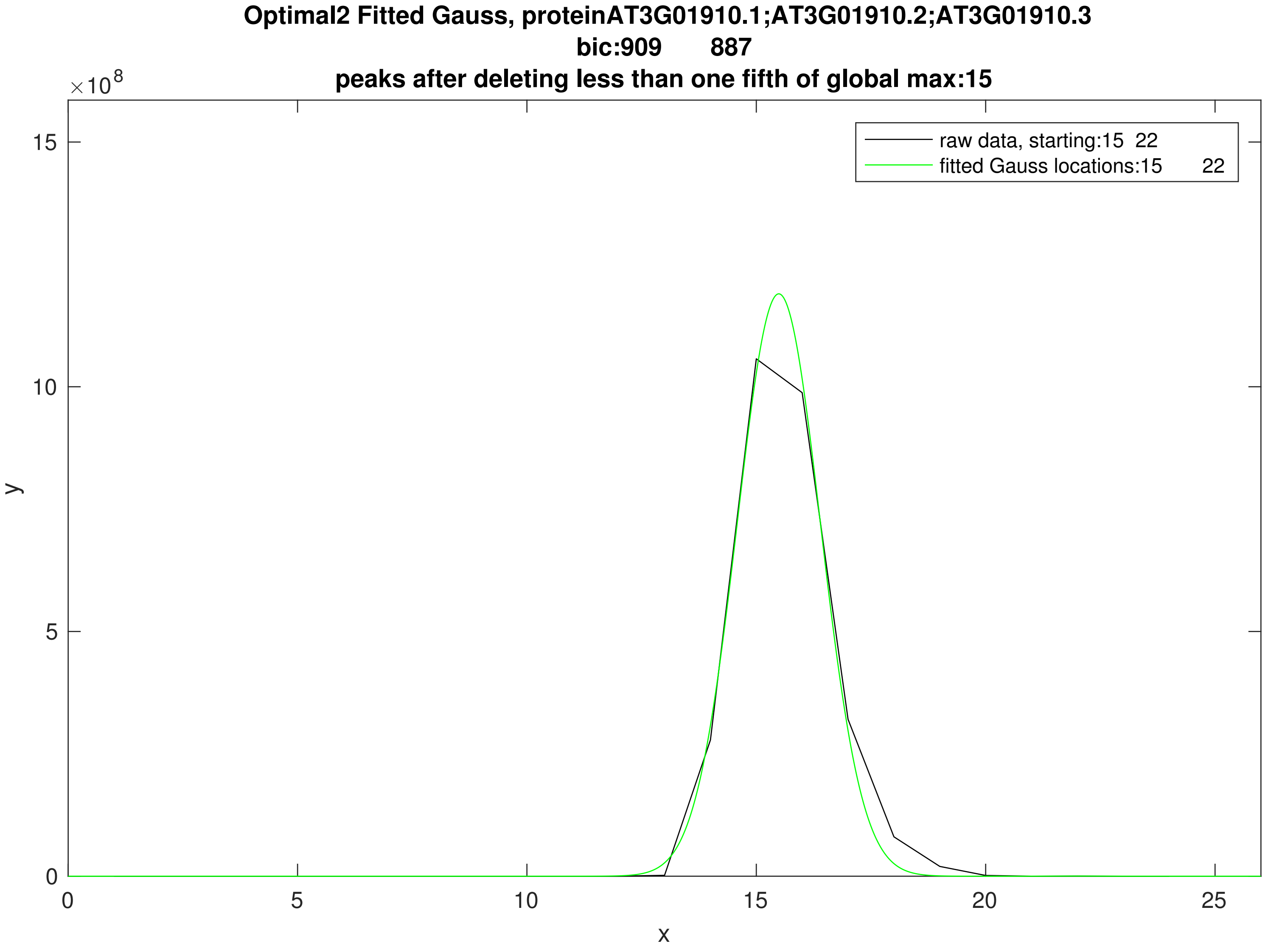

Optimal2 Fitted Gauss, proteinAT3G01910.1;AT3G01910.2;AT3G01910.3
bic:909 887
peaks after deleting less than one fifth of global max:15
×10^8
raw data, starting:15 22
fitted Gauss locations:15 22
y
x
0
5
10
15
20
25

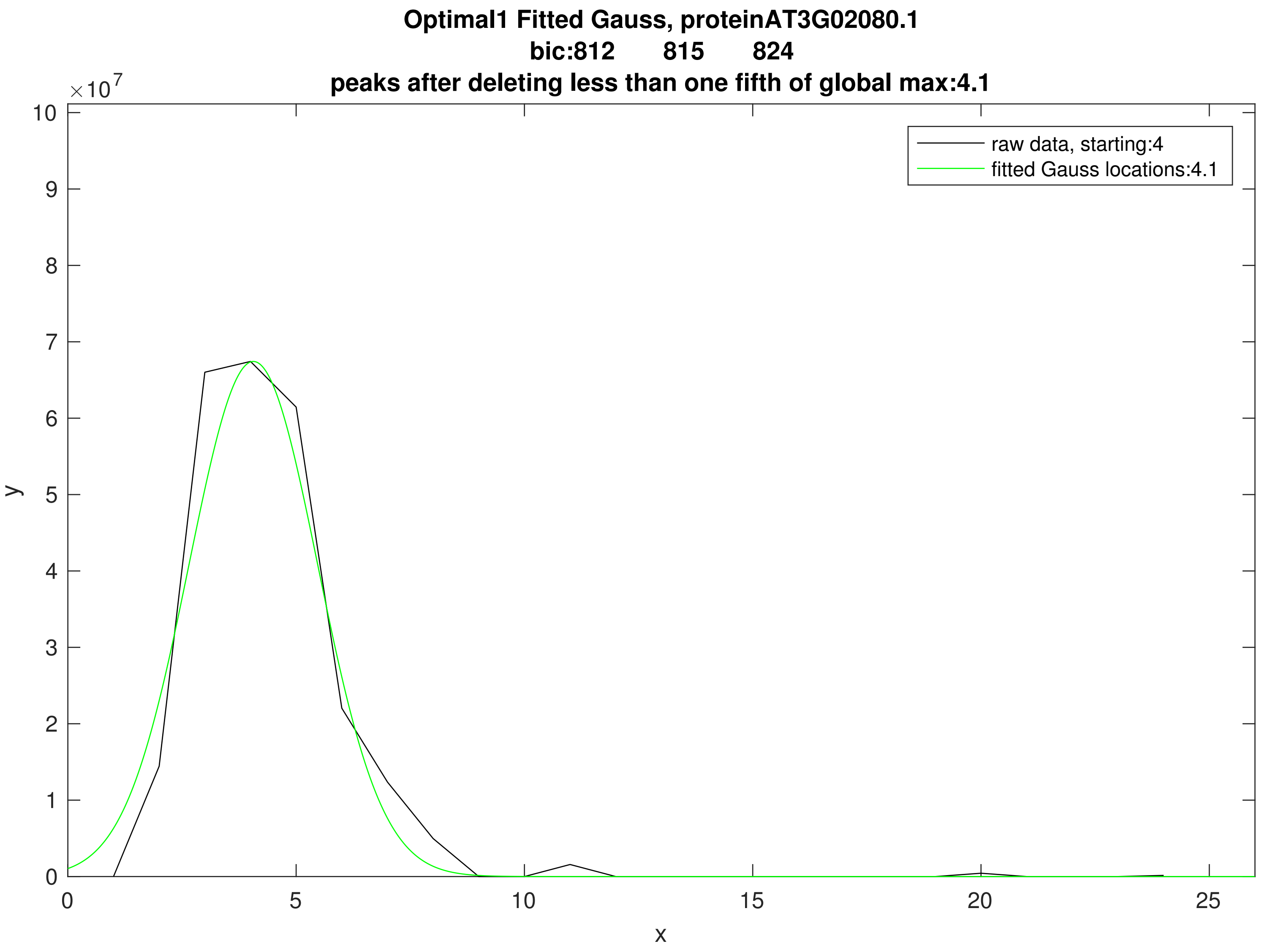
Optimal1 Fitted Gauss, proteinAT3G02080.1
bic:812 815 824
peaks after deleting less than one fifth of global max:4.1
×10^7
raw data, starting:4
fitted Gauss locations:4.1
y
x
0
1
2
3
4
5
6
7
8
9
10
0
5
10
15
20
25

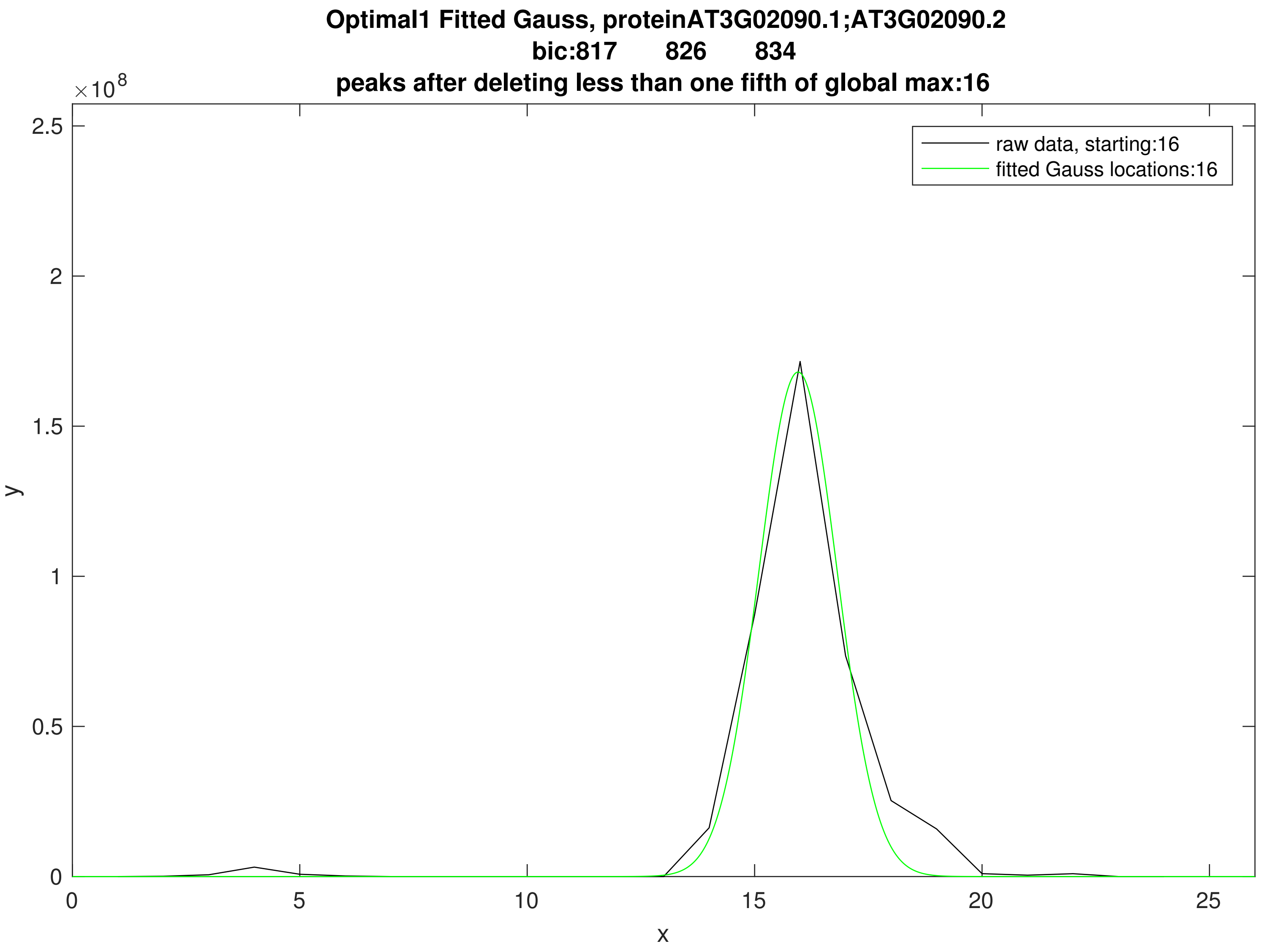
Optimal1 Fitted Gauss, proteinAT3G02090.1;AT3G02090.2
bic:817 826 834
peaks after deleting less than one fifth of global max:16
×10^8
2.5
2
1.5
1
0.5
0
y
0
5
10
15
20
25
x
raw data, starting:16
fitted Gauss locations:16

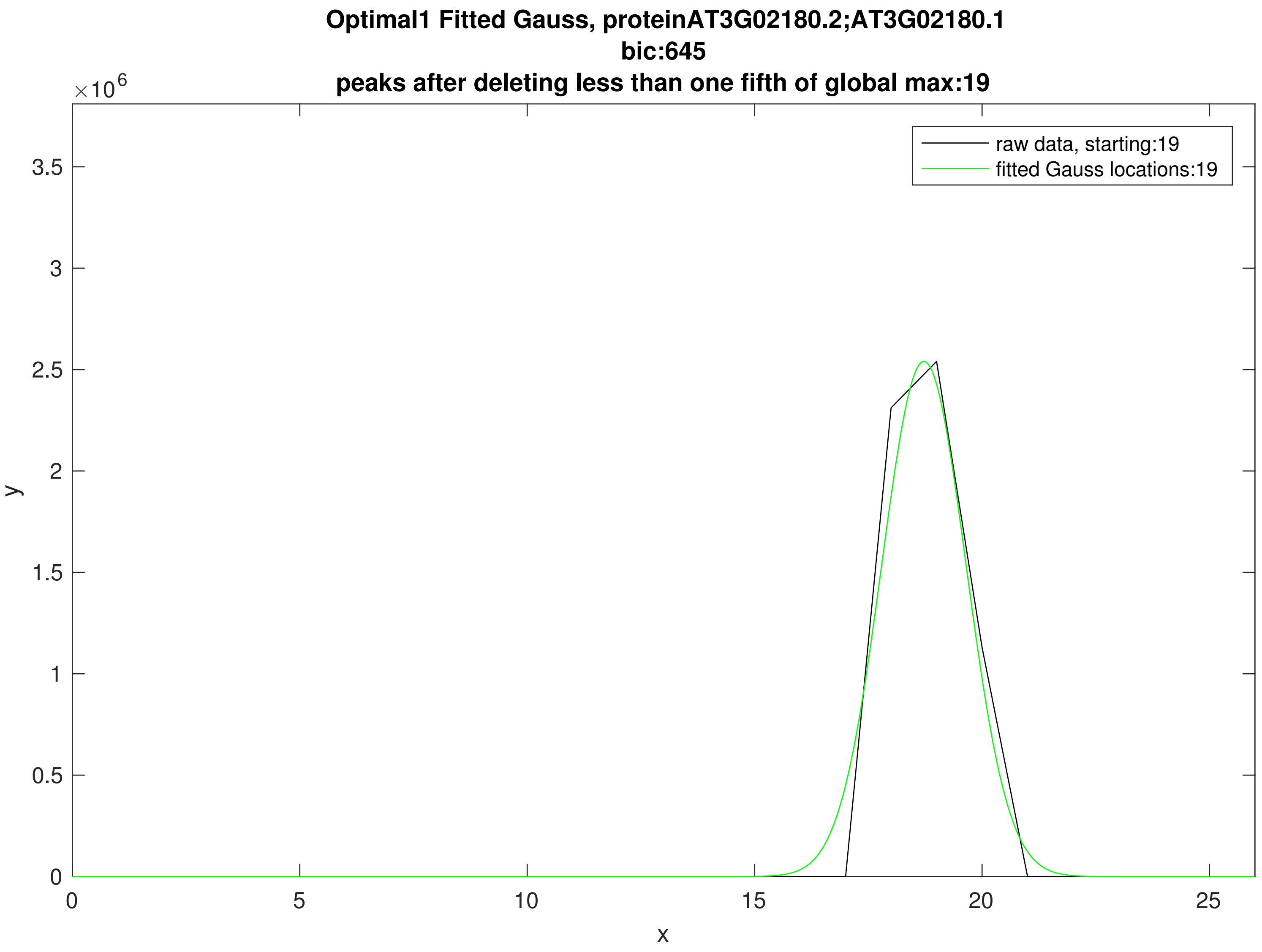

Optimal1 Fitted Gauss, proteinAT3G02180.2;AT3G02180.1
bic:645
peaks after deleting less than one fifth of global max:19
×10^6
raw data, starting:19
fitted Gauss locations:19
3.5
3
2.5
2
1.5
1
0.5
0
y
0
5
10
15
20
25
x

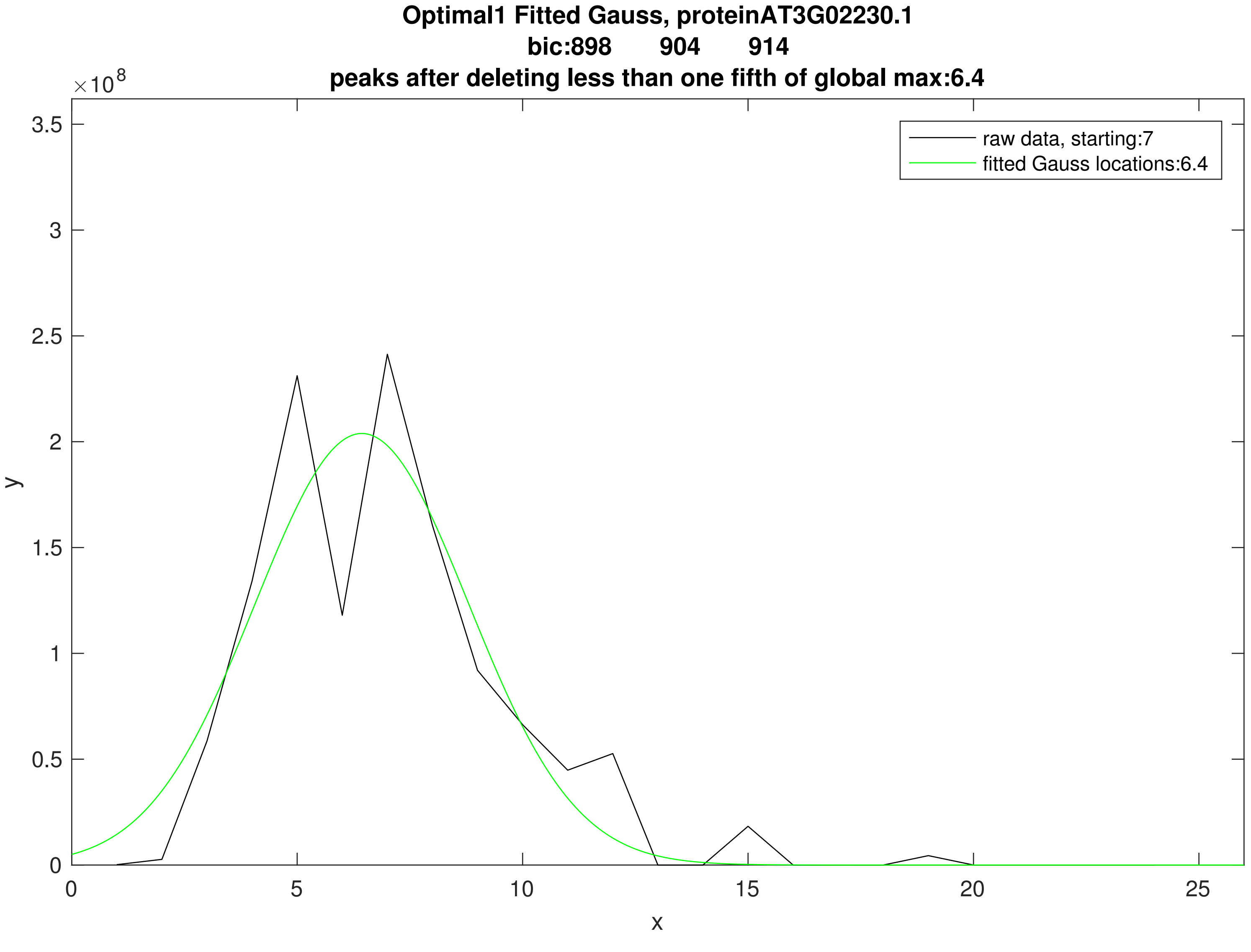

Optimal1 Fitted Gauss, proteinAT3G02230.1
bic:898 904 914
peaks after deleting less than one fifth of global max:6.4
×10^8
raw data, starting:7
fitted Gauss locations:6.4
y
x
0
0.5
1
1.5
2
2.5
3
3.5
5
10
15
20
25

**Optimal1 Fitted Gauss, proteinAT3G02360.2;AT3G02360.1**
**bic:983 990 1e+03**
**peaks after deleting less than one fifth of global max:13**

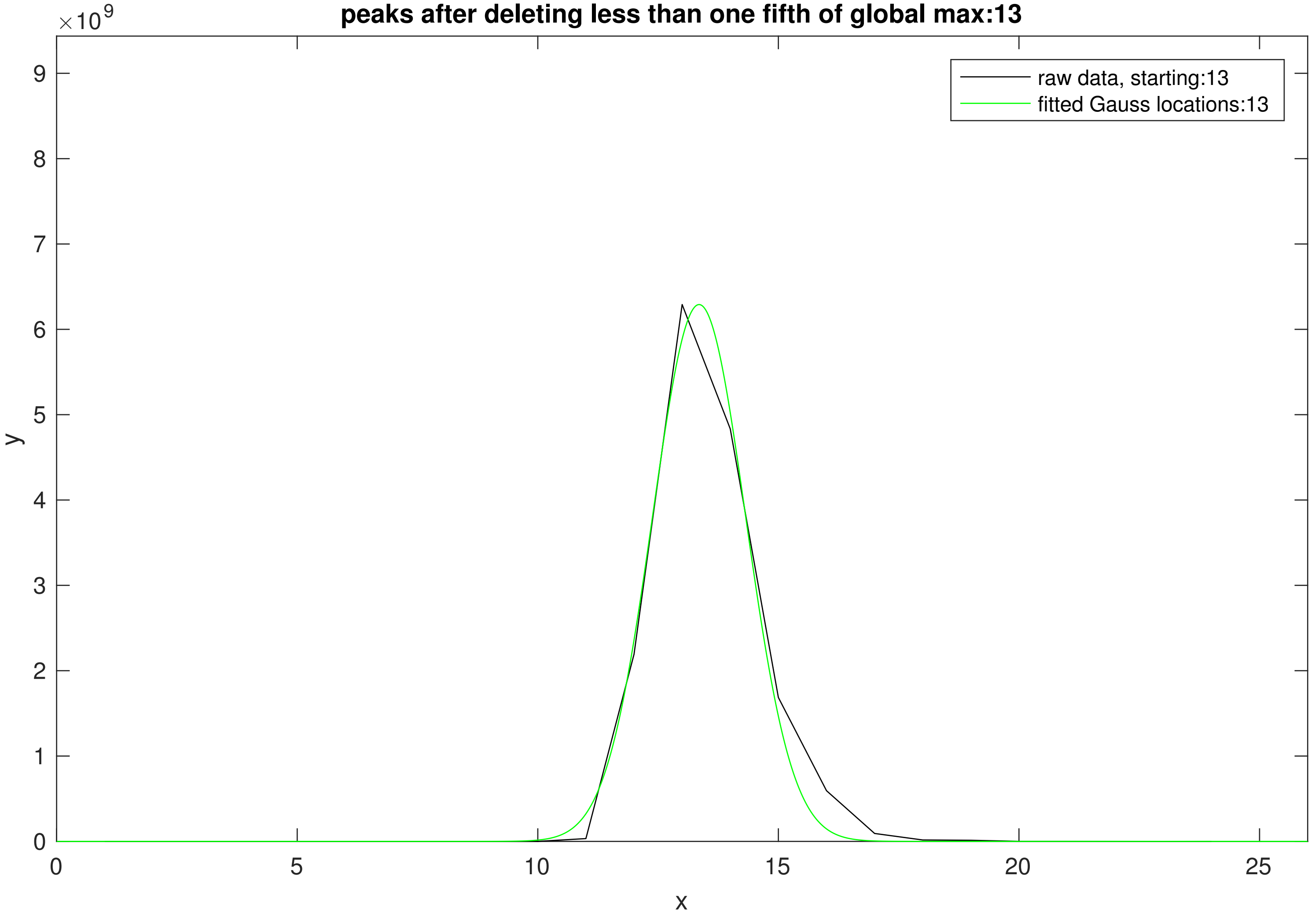

**Optimal1 Fitted Gauss, proteinAT3G02480.1**
**bic:791 799**
**peaks after deleting less than one fifth of global max:19**

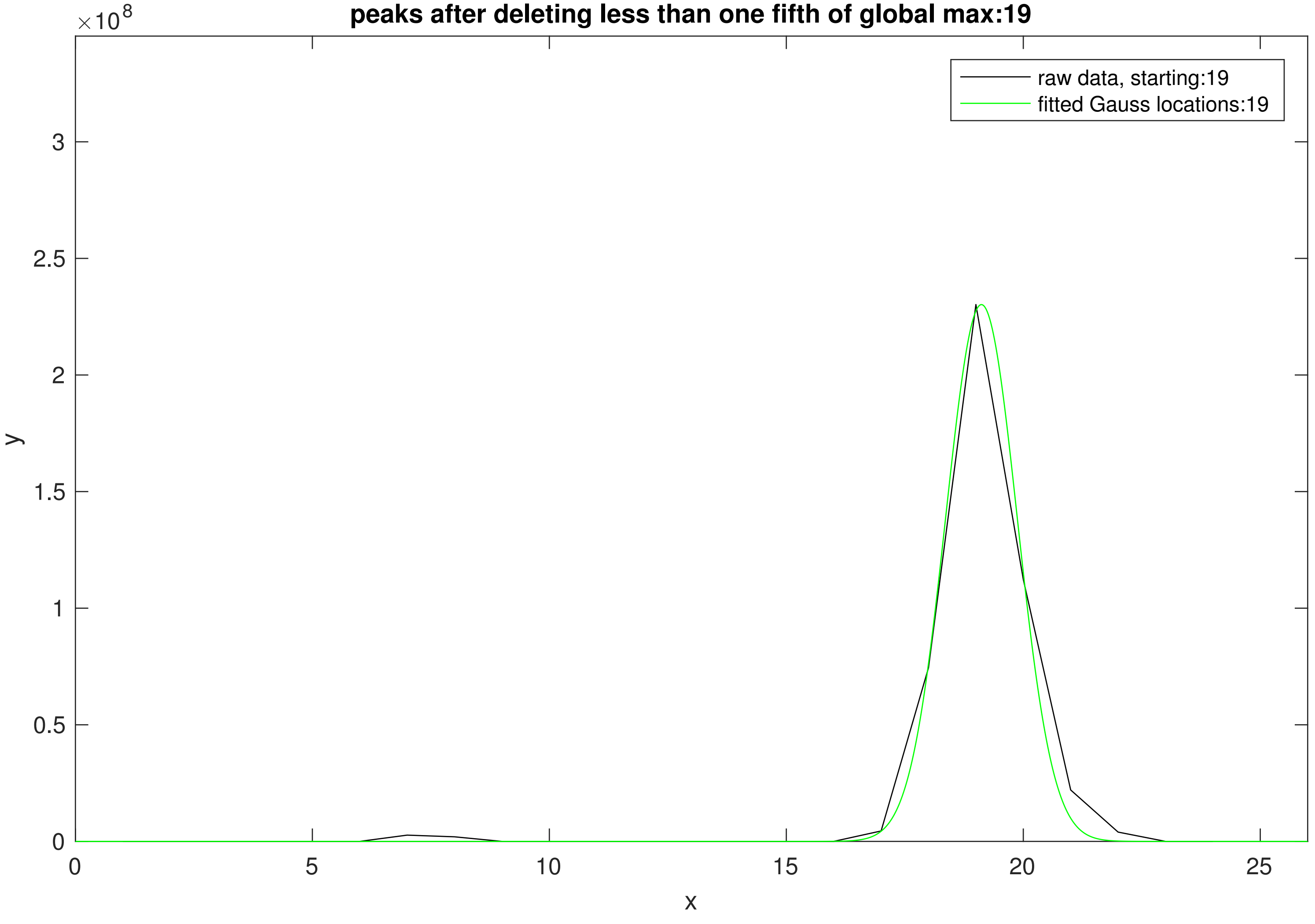

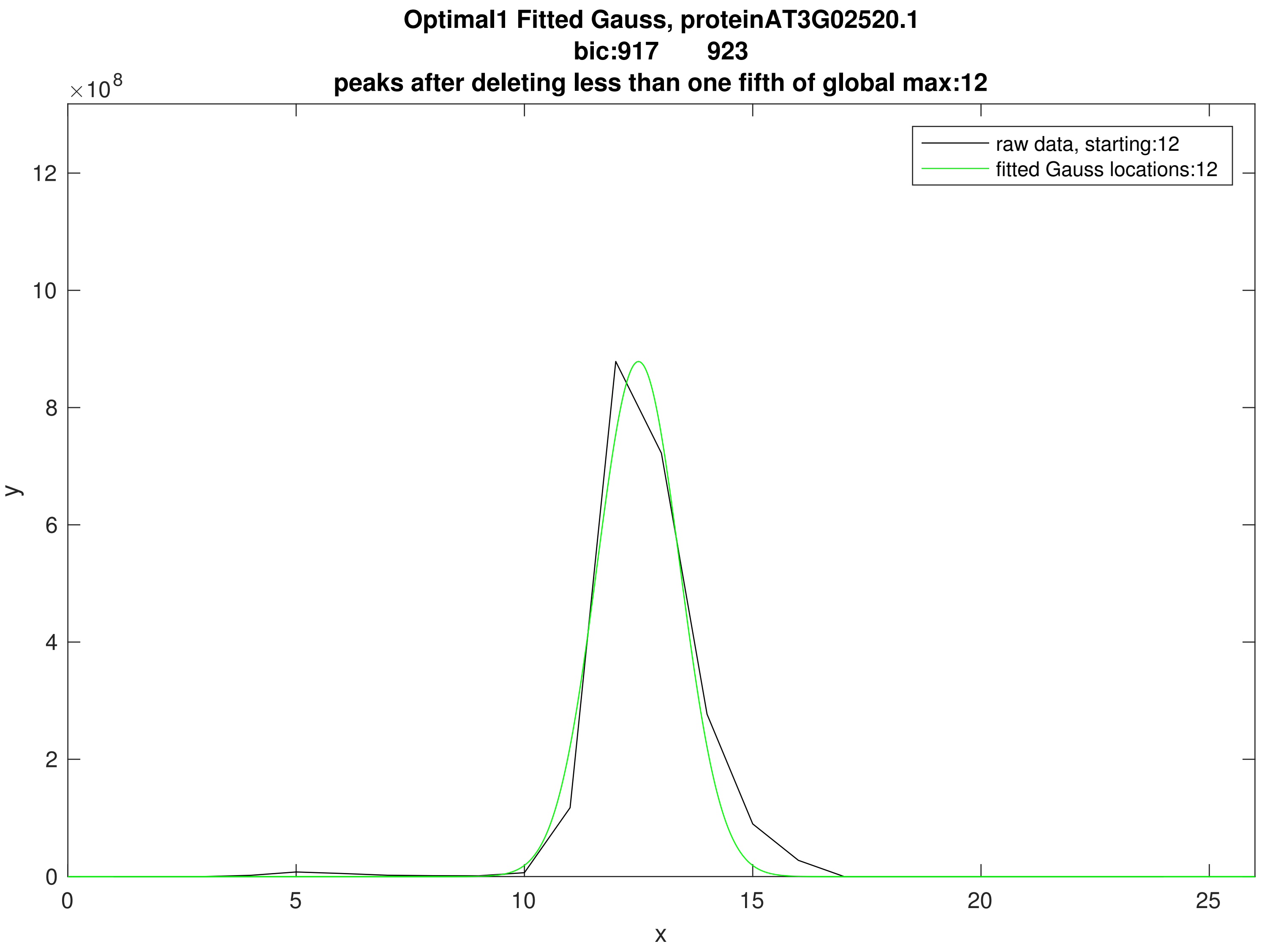

Optimal1 Fitted Gauss, proteinAT3G02520.1
bic:917 923
peaks after deleting less than one fifth of global max:12
×10^8
raw data, starting:12
fitted Gauss locations:12
y
x
0
2
4
6
8
10
12
0
5
10
15
20
25

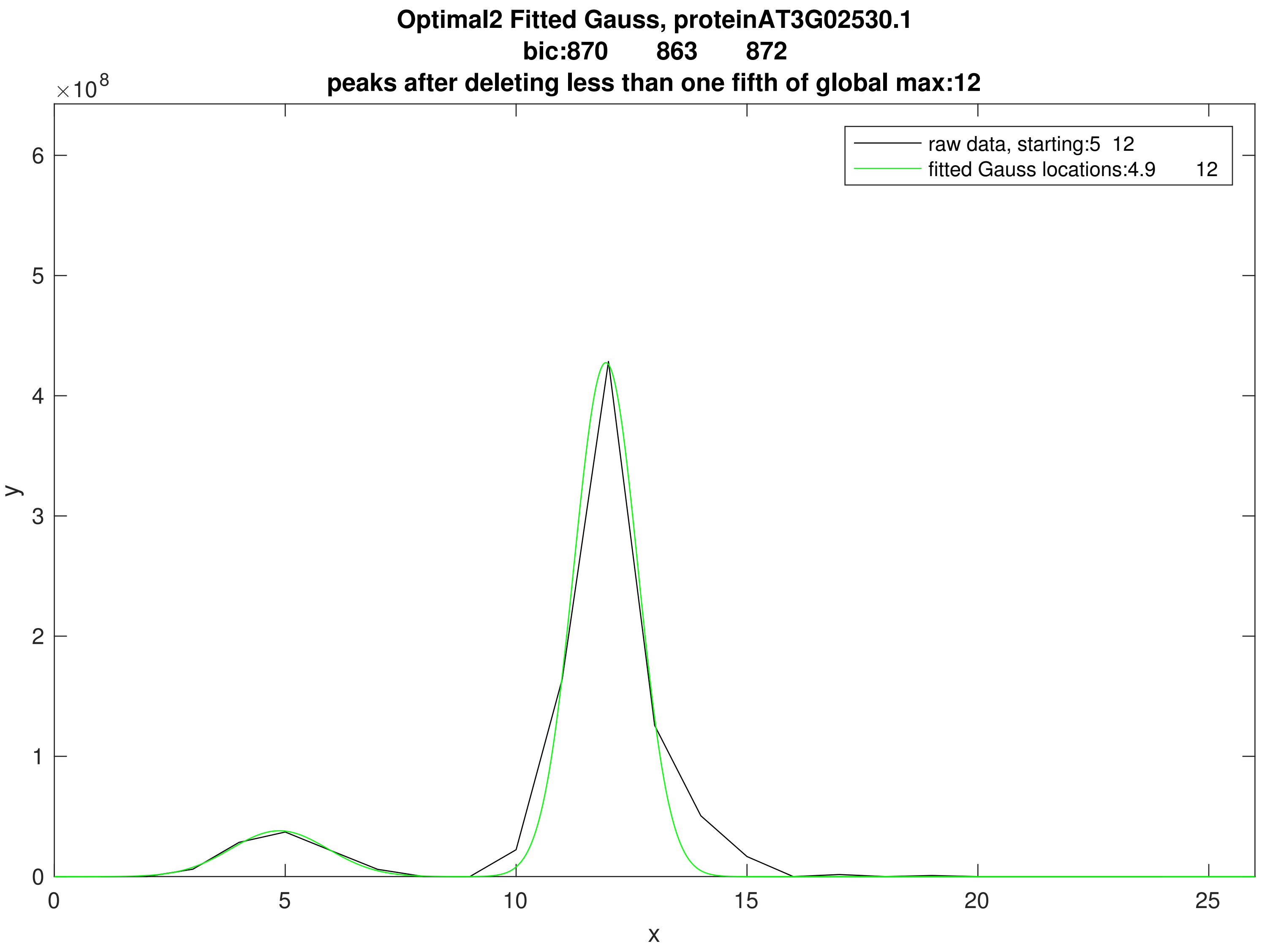

Optimal2 Fitted Gauss, proteinAT3G02530.1
bic:870 863 872
peaks after deleting less than one fifth of global max:12
×10^8
raw data, starting:5 12
fitted Gauss locations:4.9 12
y
x
0
1
2
3
4
5
6
0
5
10
15
20
25

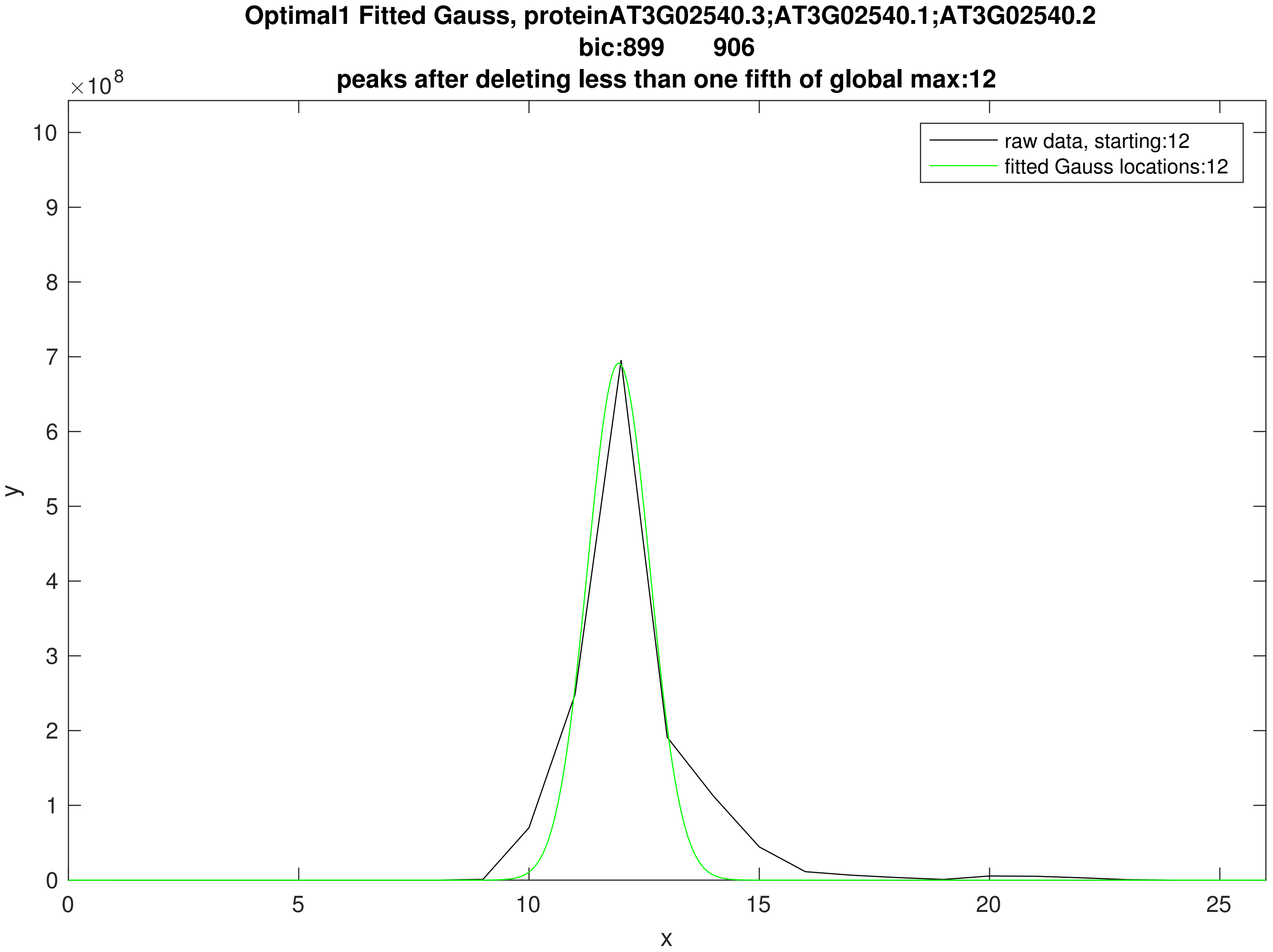
Optimal1 Fitted Gauss, proteinAT3G02540.3;AT3G02540.1;AT3G02540.2
bic:899 906
peaks after deleting less than one fifth of global max:12
×10^8
raw data, starting:12
fitted Gauss locations:12
y
x

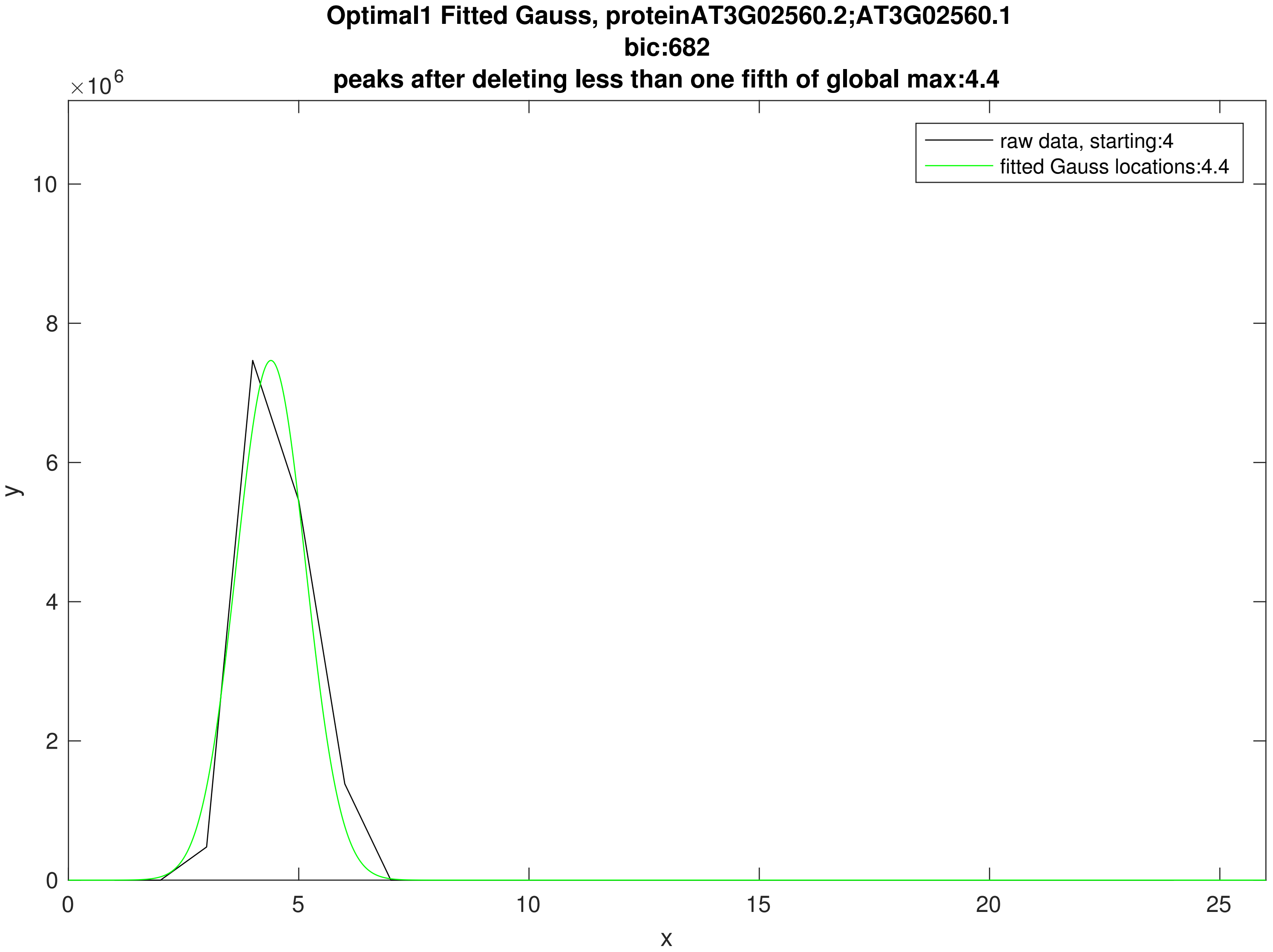

Optimal1 Fitted Gauss, proteinAT3G02560.2;AT3G02560.1
bic:682
peaks after deleting less than one fifth of global max:4.4
×10^6
raw data, starting:4
fitted Gauss locations:4.4
y
x
0
2
4
6
8
10
0
5
10
15
20
25

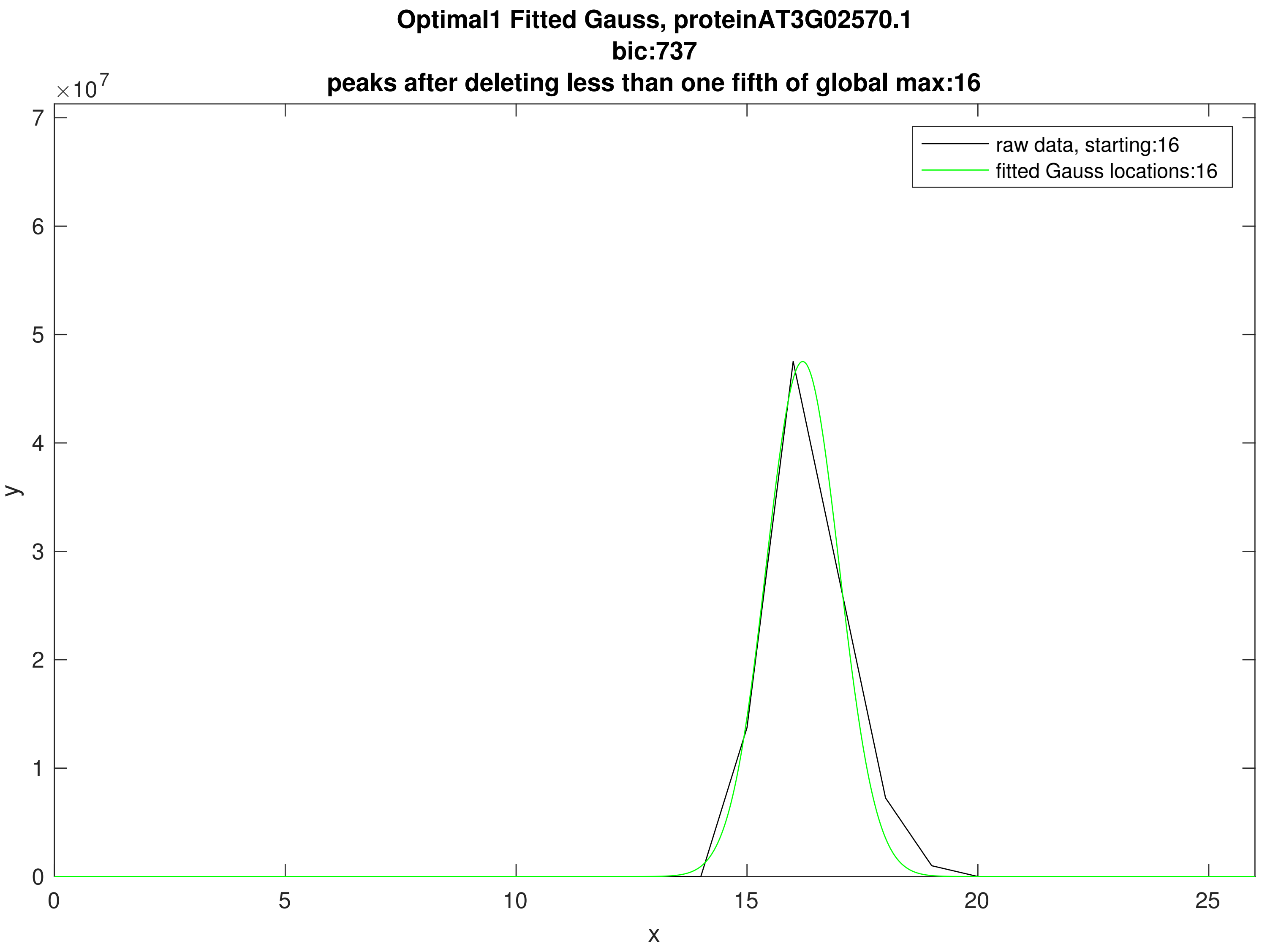

Optimal1 Fitted Gauss, proteinAT3G02570.1
bic:737
peaks after deleting less than one fifth of global max:16
×10^7
raw data, starting:16
fitted Gauss locations:16
y
x
0
1
2
3
4
5
6
7
0
5
10
15
20
25

Optimal1 Fitted Gauss, proteinAT3G02660.1
bic:809 826
peaks after deleting less than one fifth of global max:15

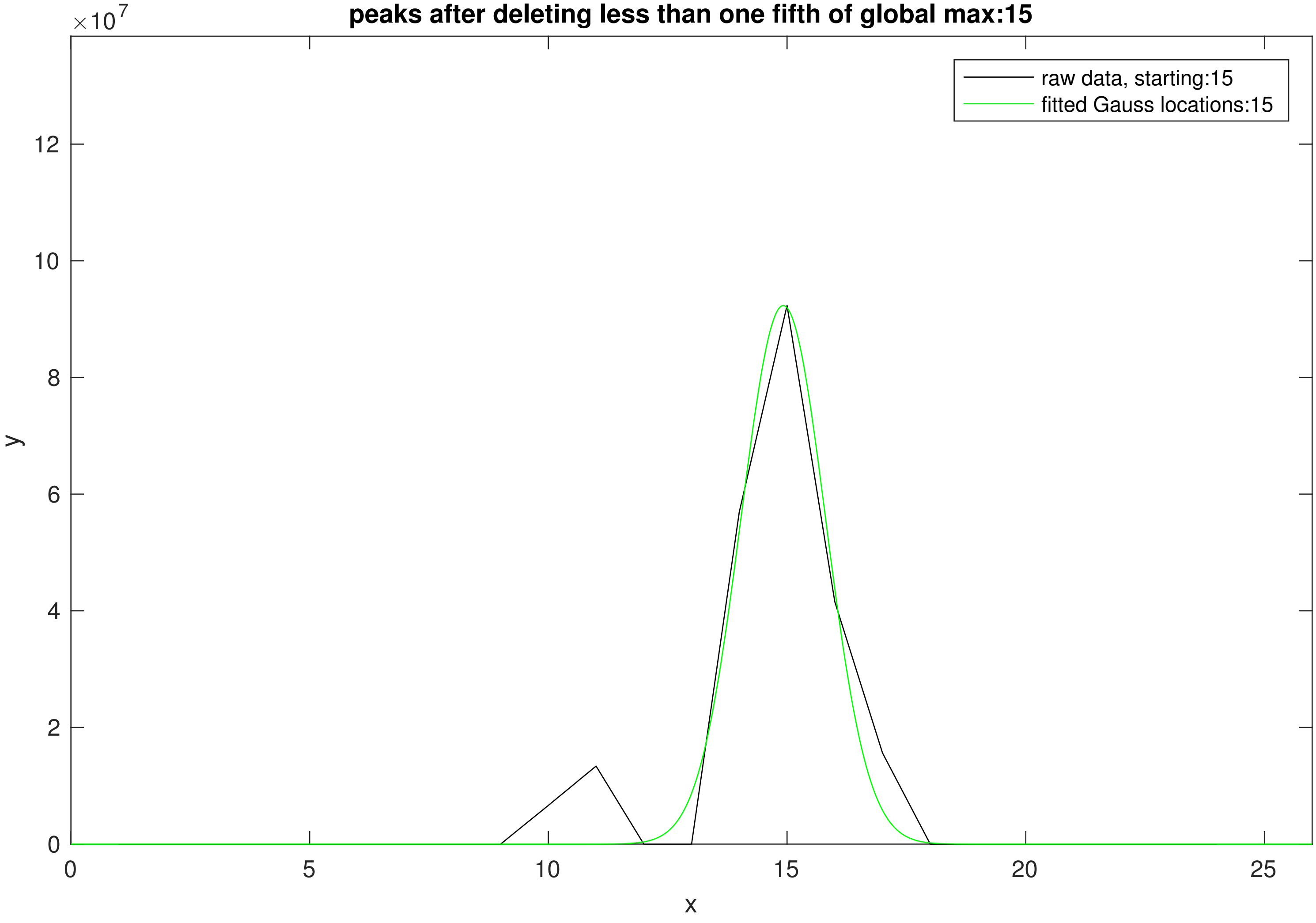

Optimal1 Fitted Gauss, proteinAT3G02720.1
bic:864 868
peaks after deleting less than one fifth of global max:12

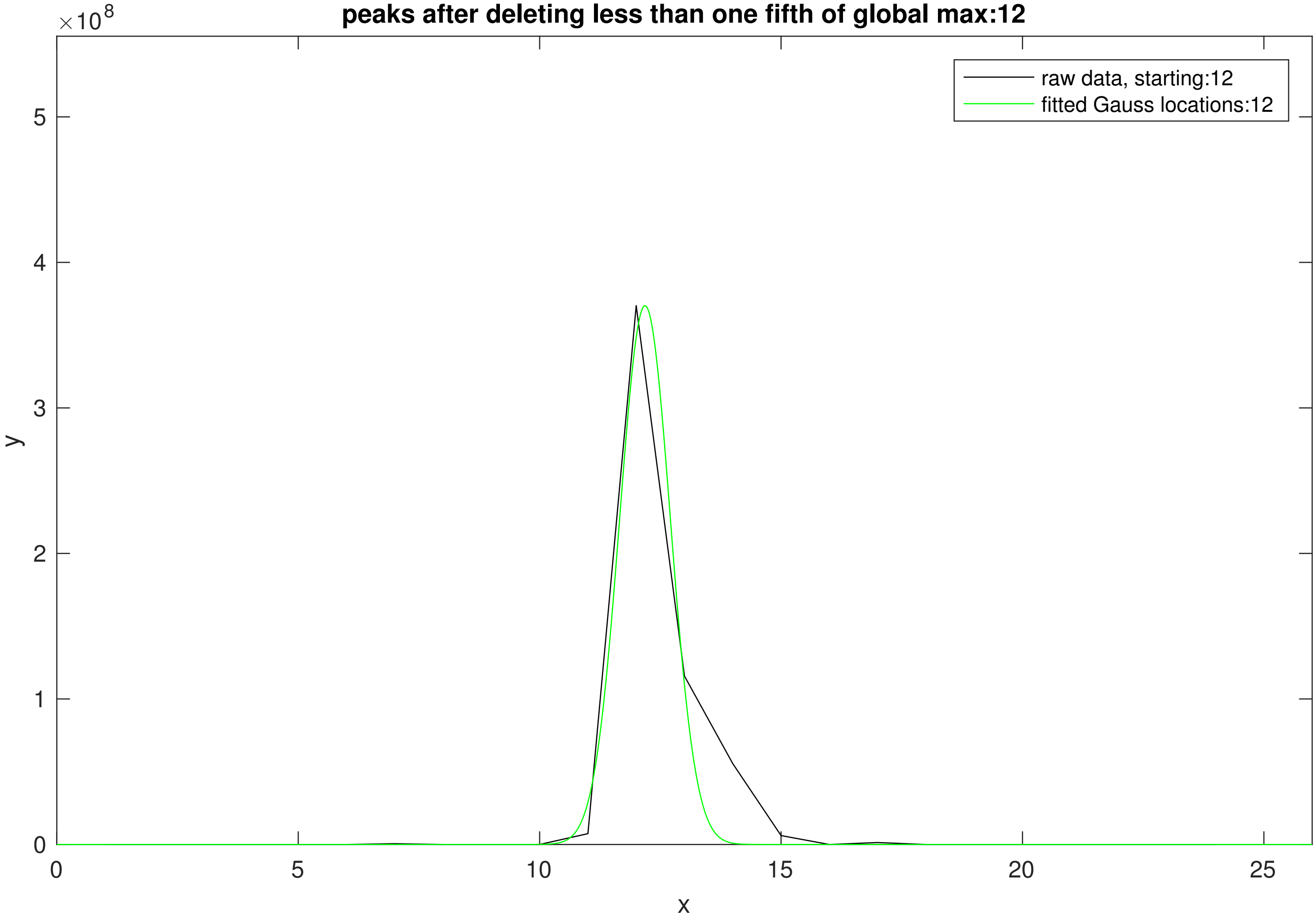

**Optimal1 Fitted Gauss, proteinAT3G02740.1**
**bic:821 831**
**peaks after deleting less than one fifth of global max:12**

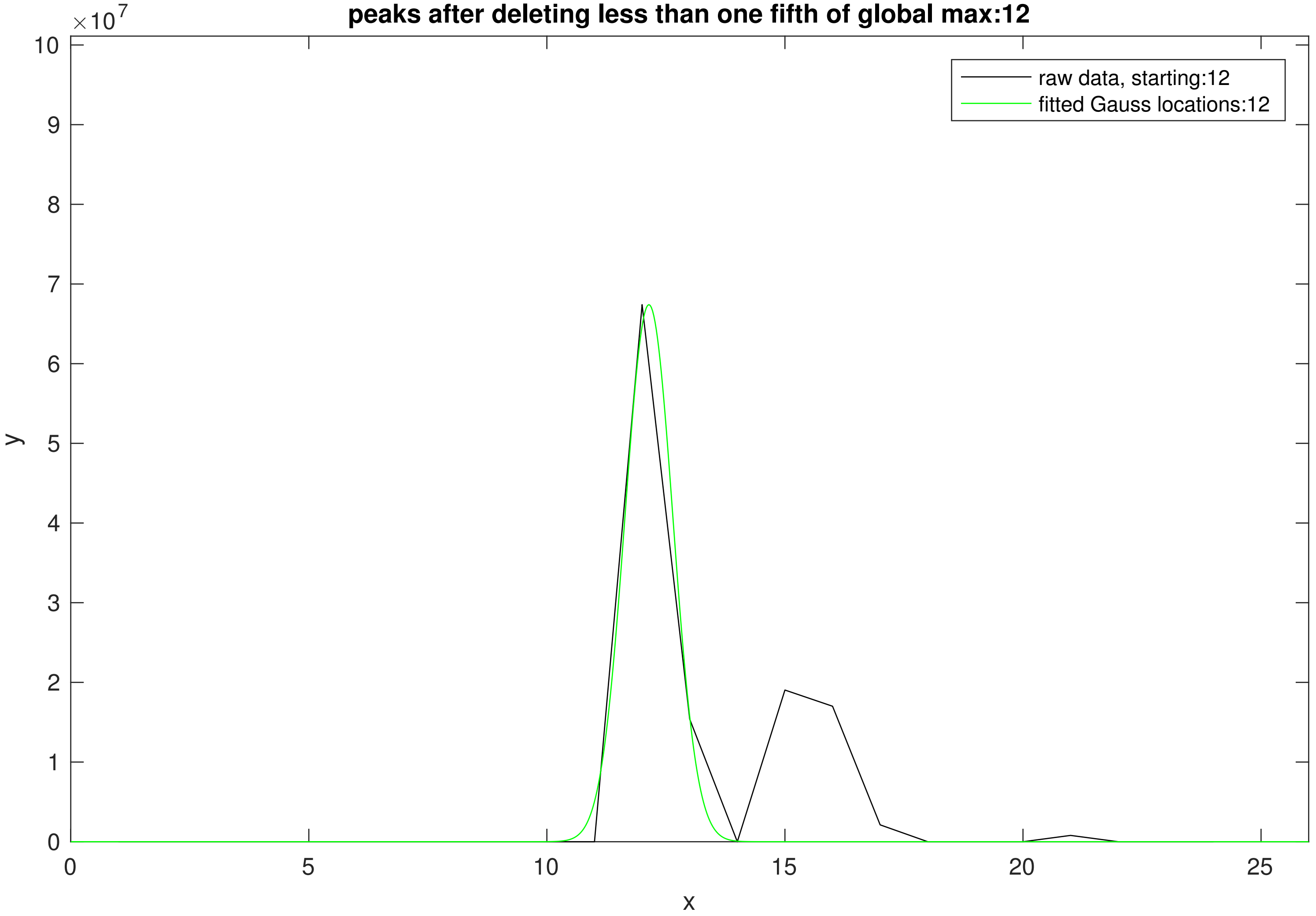

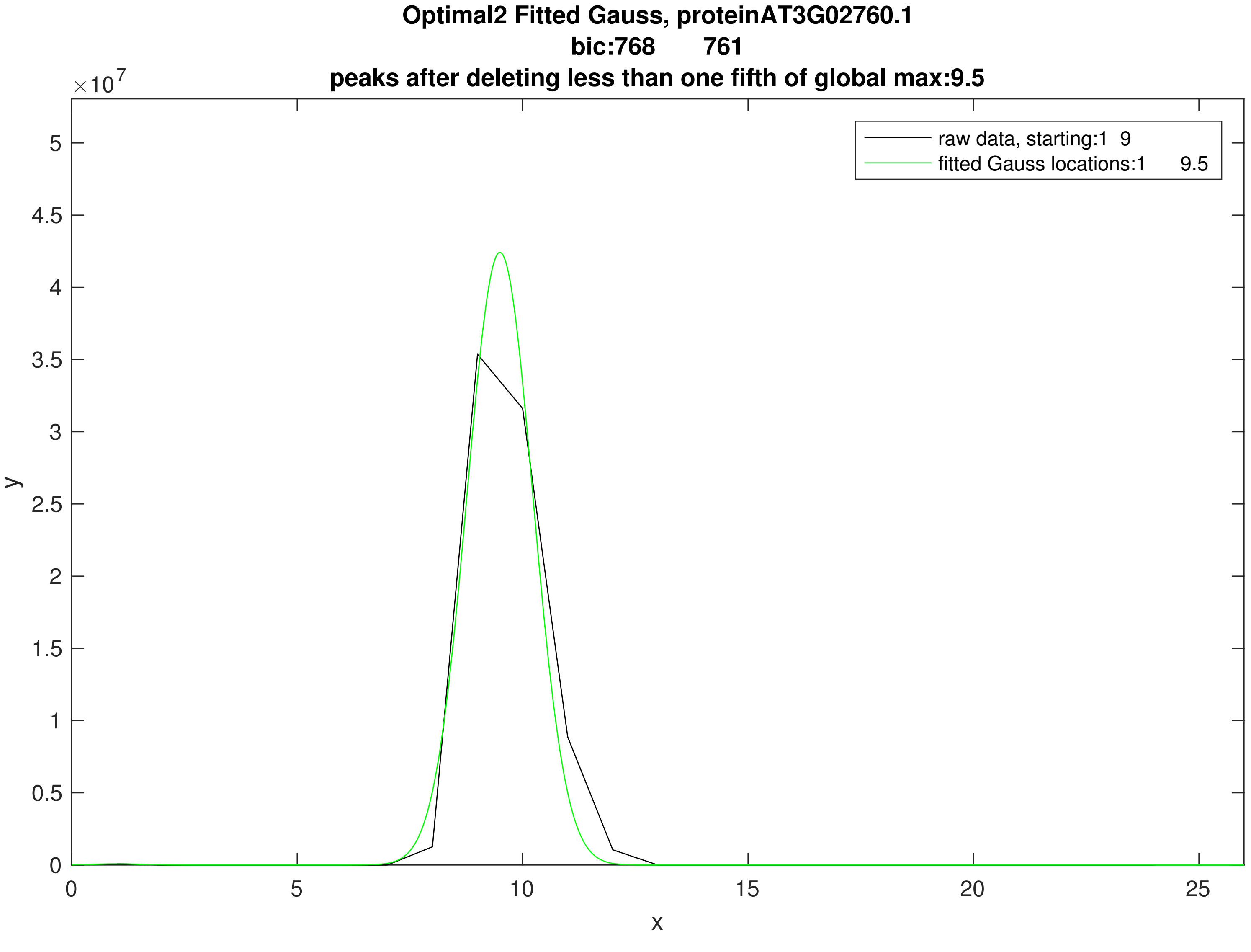

Optimal2 Fitted Gauss, proteinAT3G02760.1
bic:768 761
peaks after deleting less than one fifth of global max:9.5
×10^7
raw data, starting:1 9
fitted Gauss locations:1 9.5
y
x
0
0.5
1
1.5
2
2.5
3
3.5
4
4.5
5
0
5
10
15
20
25

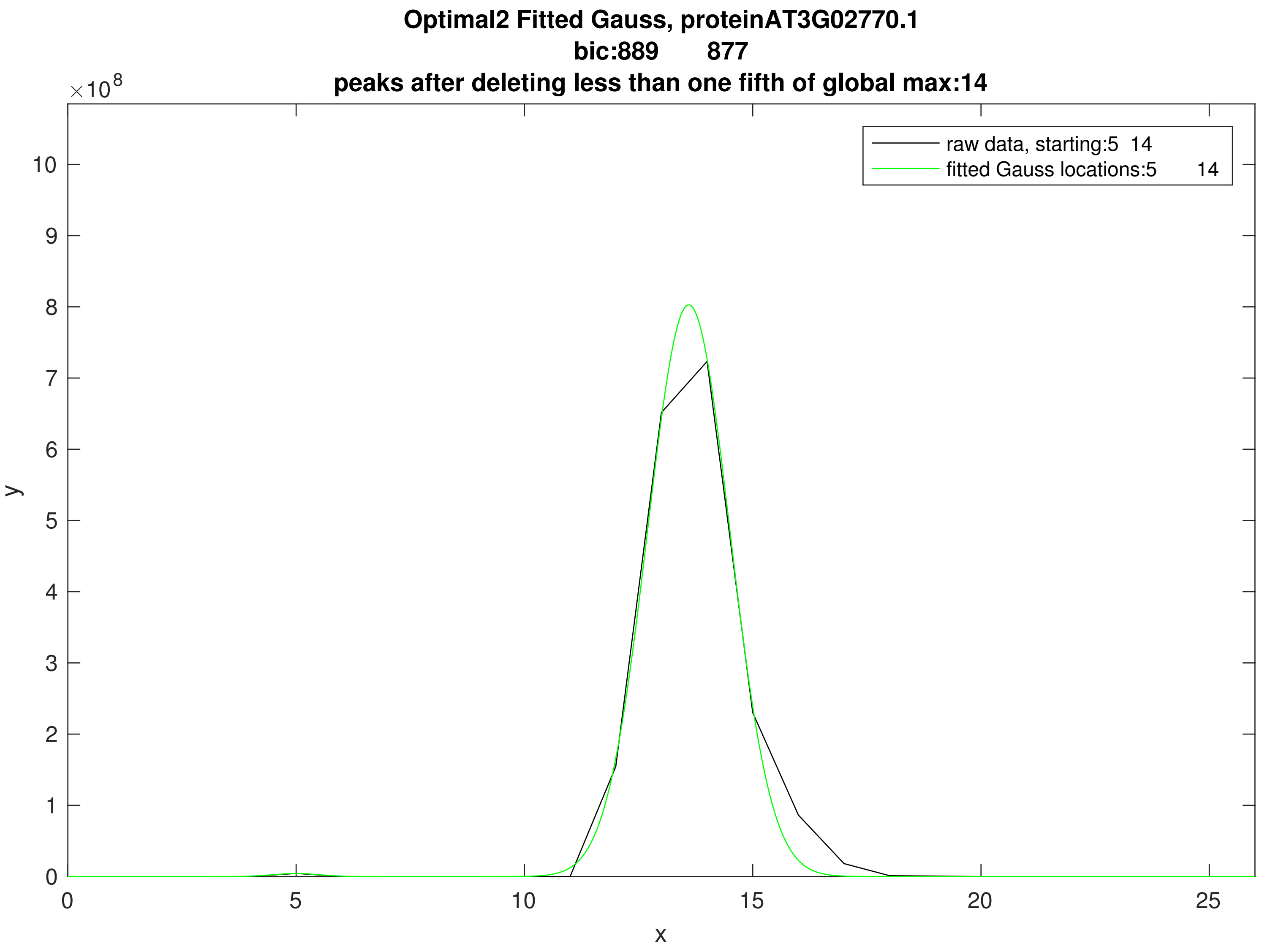

Optimal2 Fitted Gauss, proteinAT3G02770.1
bic:889 877
peaks after deleting less than one fifth of global max:14
×10^8
raw data, starting:5 14
fitted Gauss locations:5 14
y
x
0
1
2
3
4
5
6
7
8
9
10
0
5
10
15
20
25

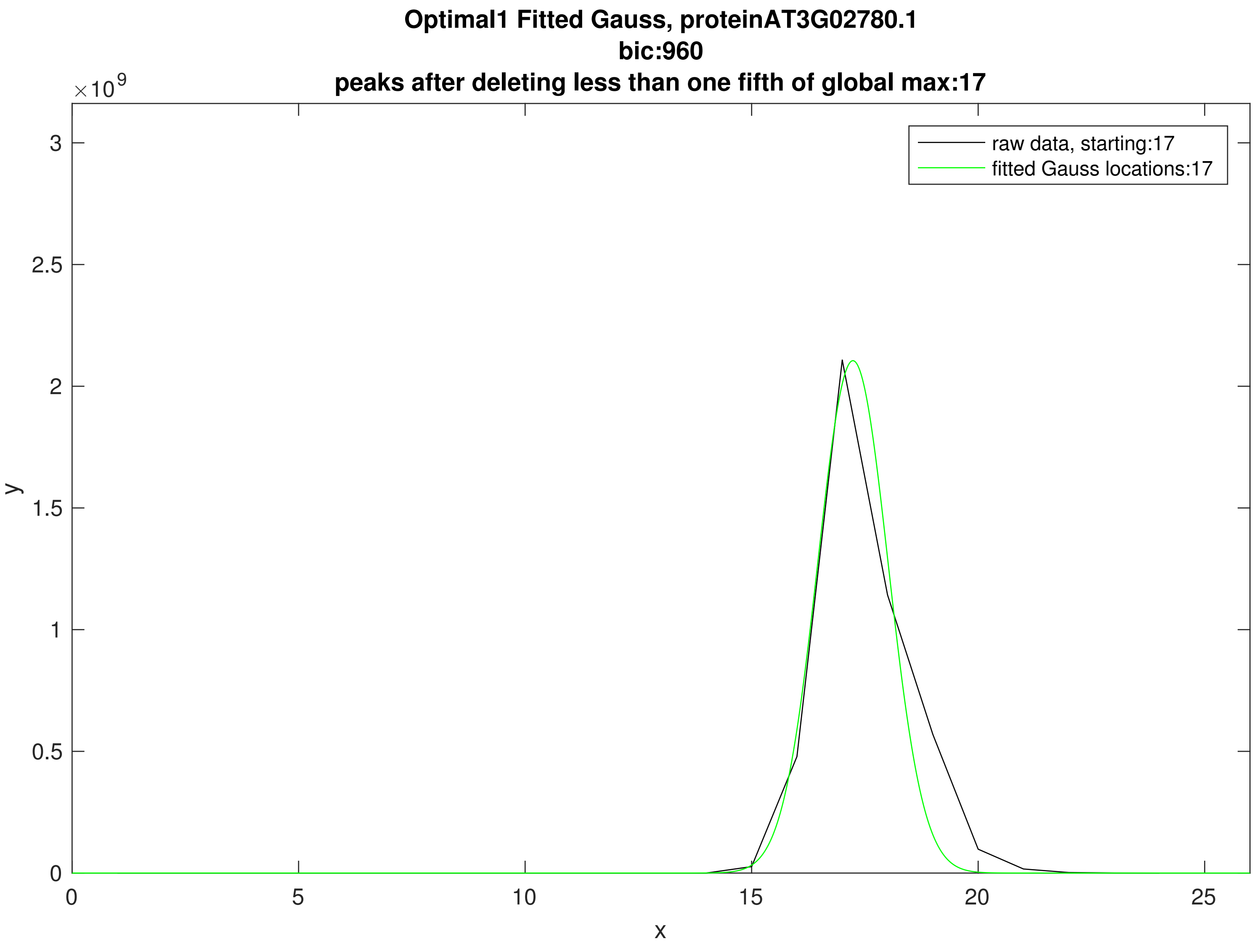

Optimal1 Fitted Gauss, proteinAT3G02780.1
bic:960
peaks after deleting less than one fifth of global max:17
×10^9
3
2.5
2
1.5
1
0.5
0
y
0
5
10
15
20
25
x
raw data, starting:17
fitted Gauss locations:17

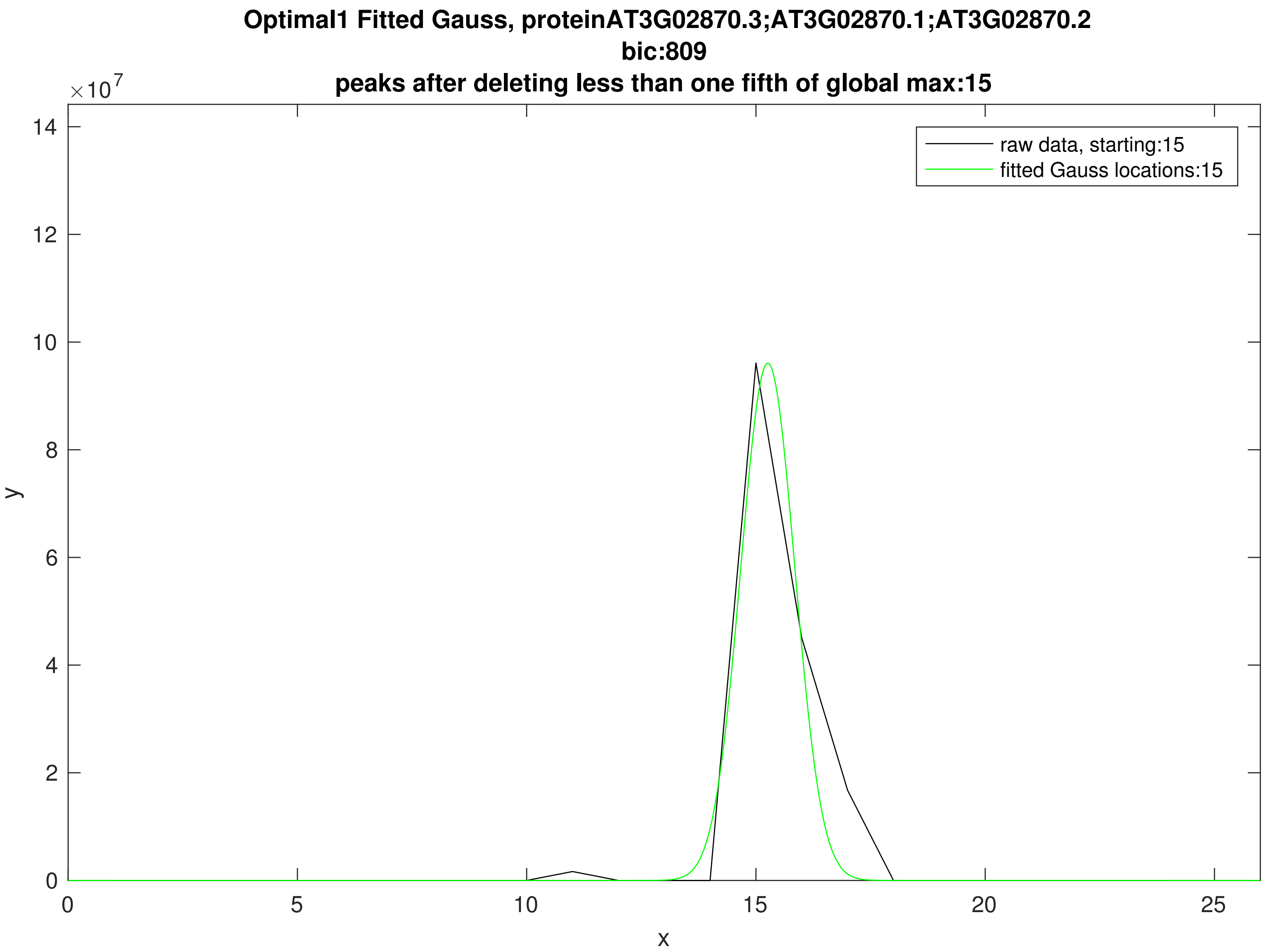

Optimal1 Fitted Gauss, proteinAT3G02870.3;AT3G02870.1;AT3G02870.2
bic:809
peaks after deleting less than one fifth of global max:15
×10^7
raw data, starting:15
fitted Gauss locations:15
y
x
0
2
4
6
8
10
12
14
0
5
10
15
20
25

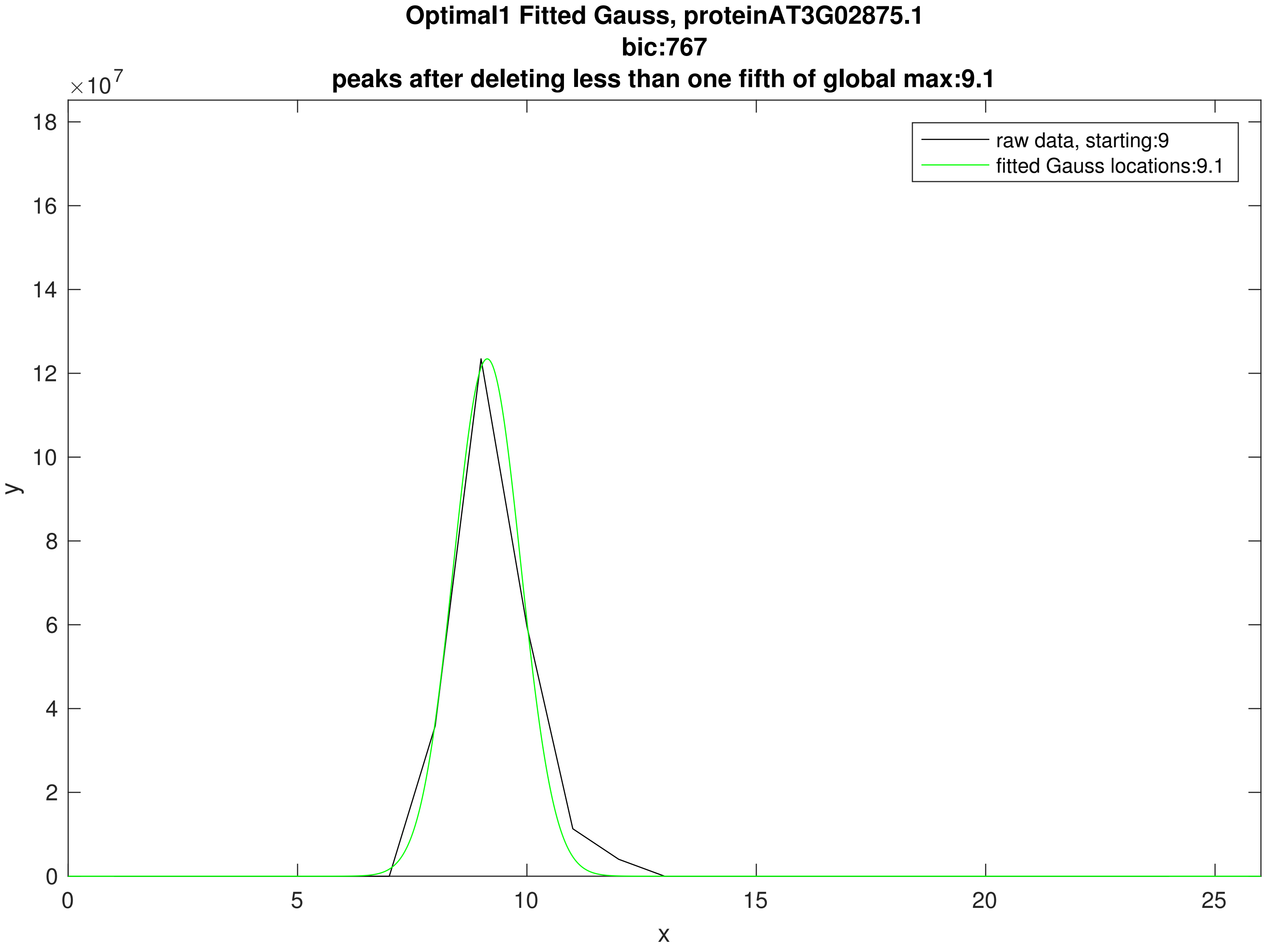

Optimal1 Fitted Gauss, proteinAT3G02875.1
bic:767
peaks after deleting less than one fifth of global max:9.1
×10^7
raw data, starting:9
fitted Gauss locations:9.1
y
x
0
2
4
6
8
10
12
14
16
18
0
5
10
15
20
25

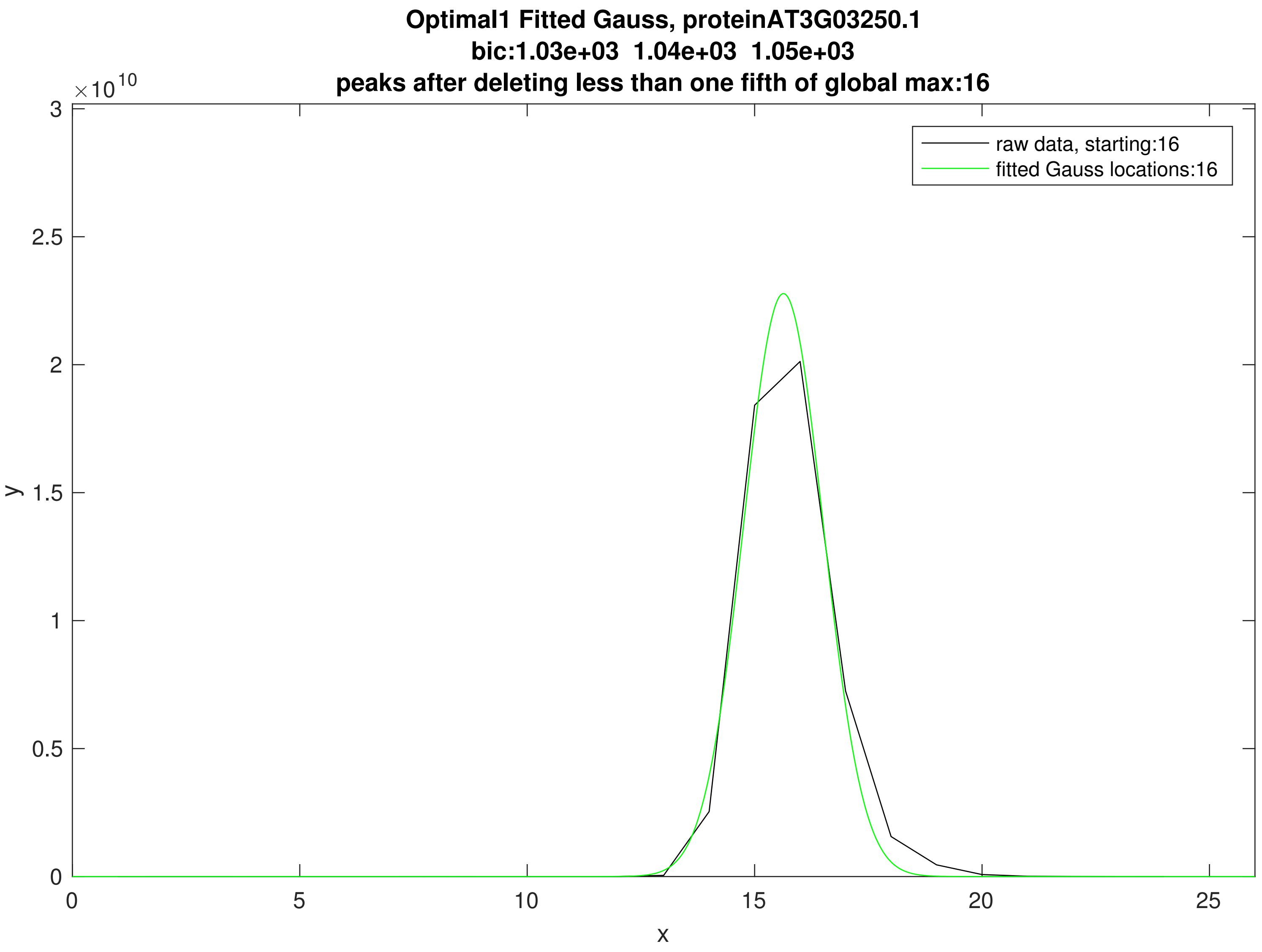
Optimal1 Fitted Gauss, proteinAT3G03250.1
bic:1.03e+03 1.04e+03 1.05e+03
peaks after deleting less than one fifth of global max:16
×10^10
raw data, starting:16
fitted Gauss locations:16
3
2.5
2
1.5
1
0.5
0
y
0
5
10
15
20
25
x

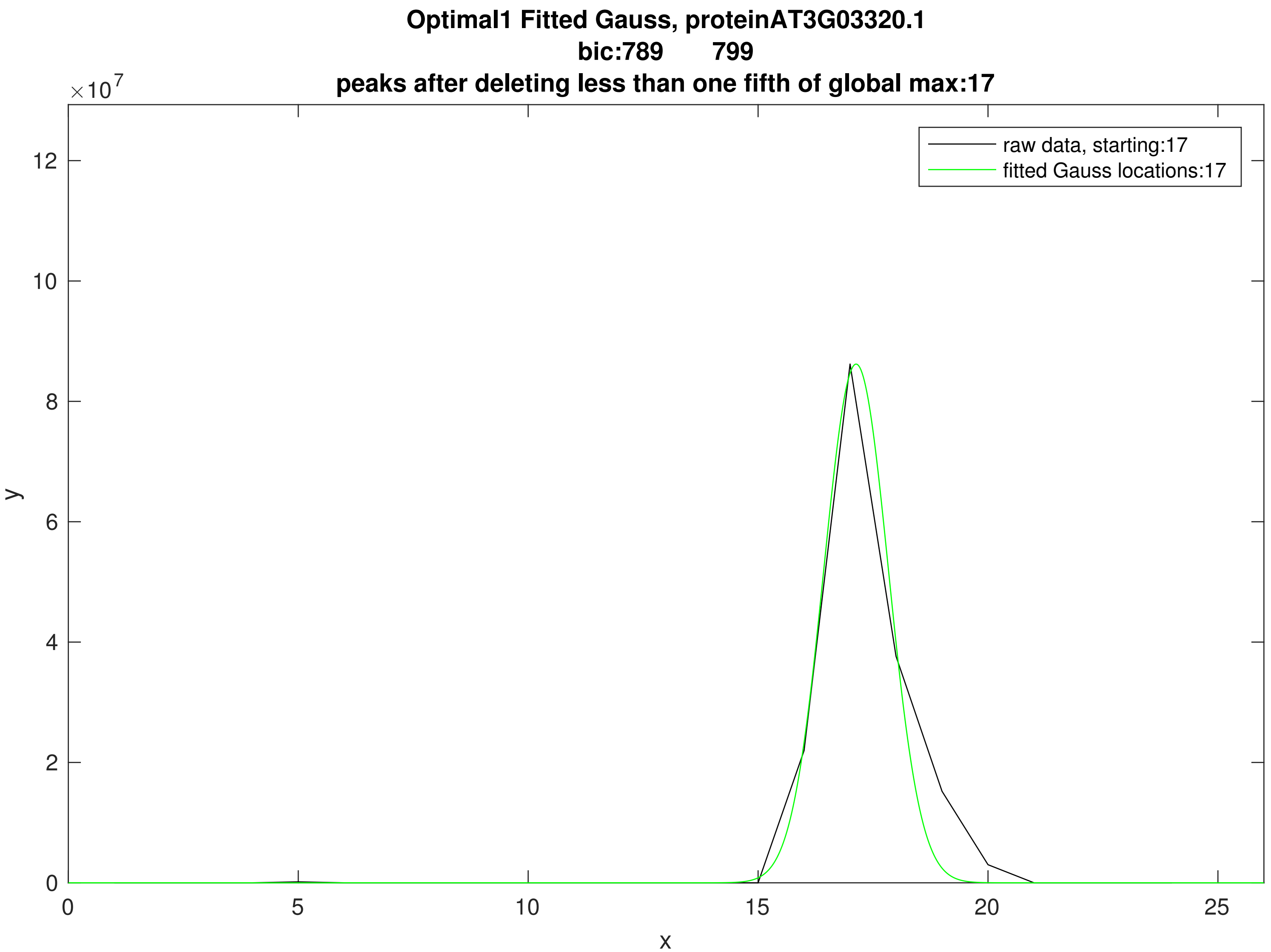

Optimal1 Fitted Gauss, proteinAT3G03320.1
bic:789 799
peaks after deleting less than one fifth of global max:17
×10^7
raw data, starting:17
fitted Gauss locations:17
y
x
0
2
4
6
8
10
12
0
5
10
15
20
25

Optimal1 Fitted Gauss, proteinAT3G03380.1
bic:865 873 883
peaks after deleting less than one fifth of global max:6.8

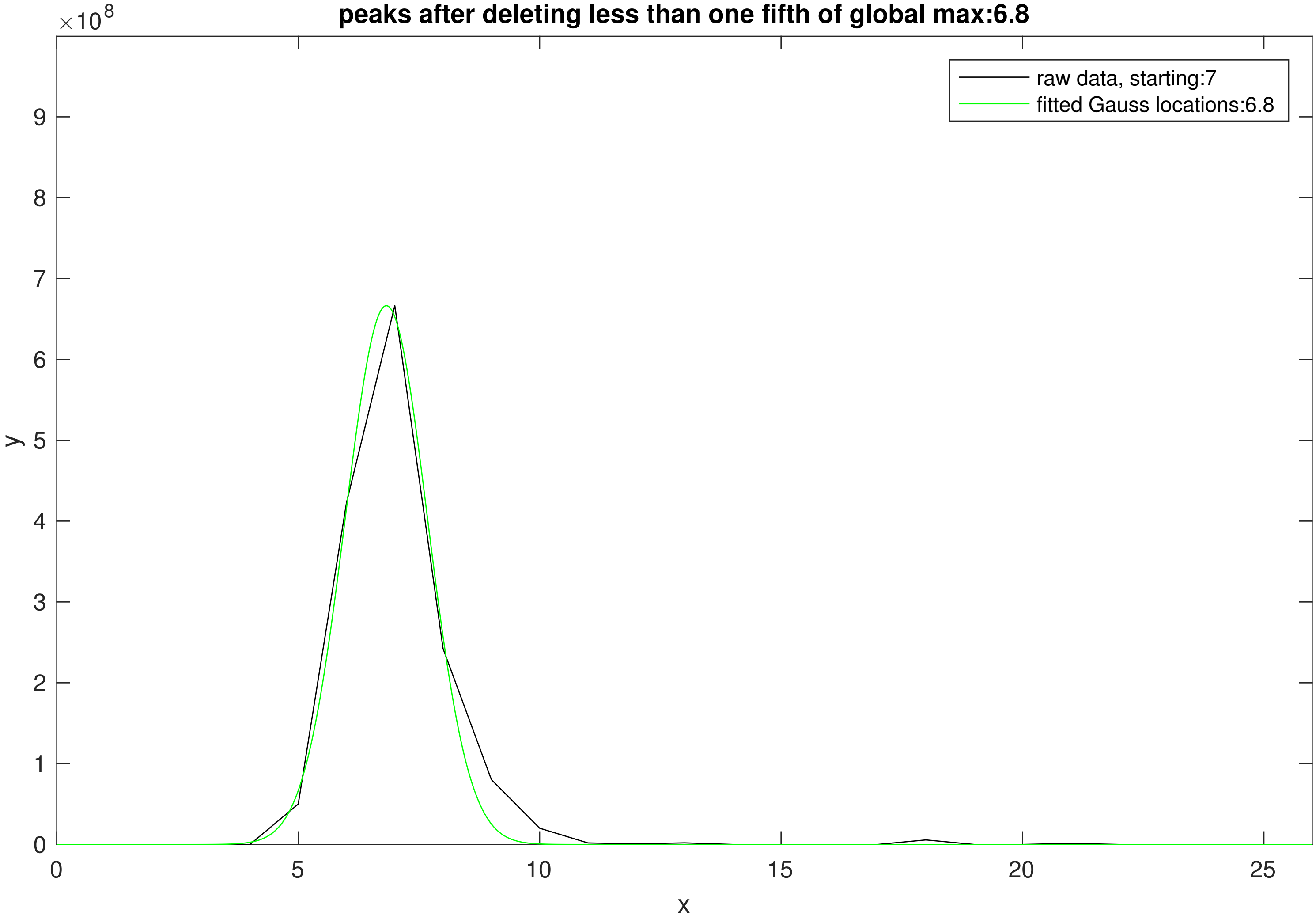

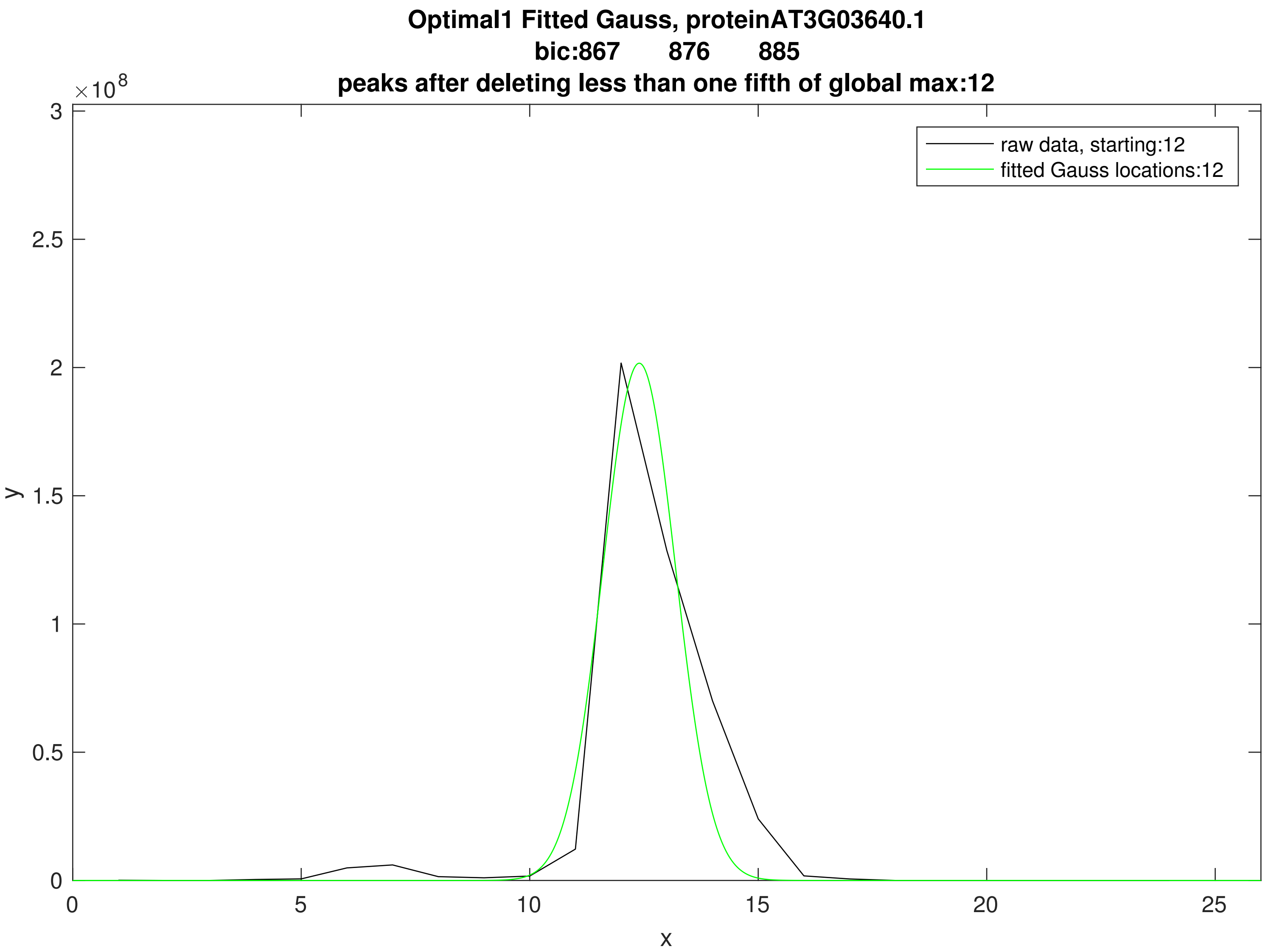
Optimal1 Fitted Gauss, proteinAT3G03640.1
bic:867 876 885
peaks after deleting less than one fifth of global max:12
raw data, starting:12
fitted Gauss locations:12
×10^8
3
2.5
2
1.5
1
0.5
0
y
0
5
10
15
20
25
x

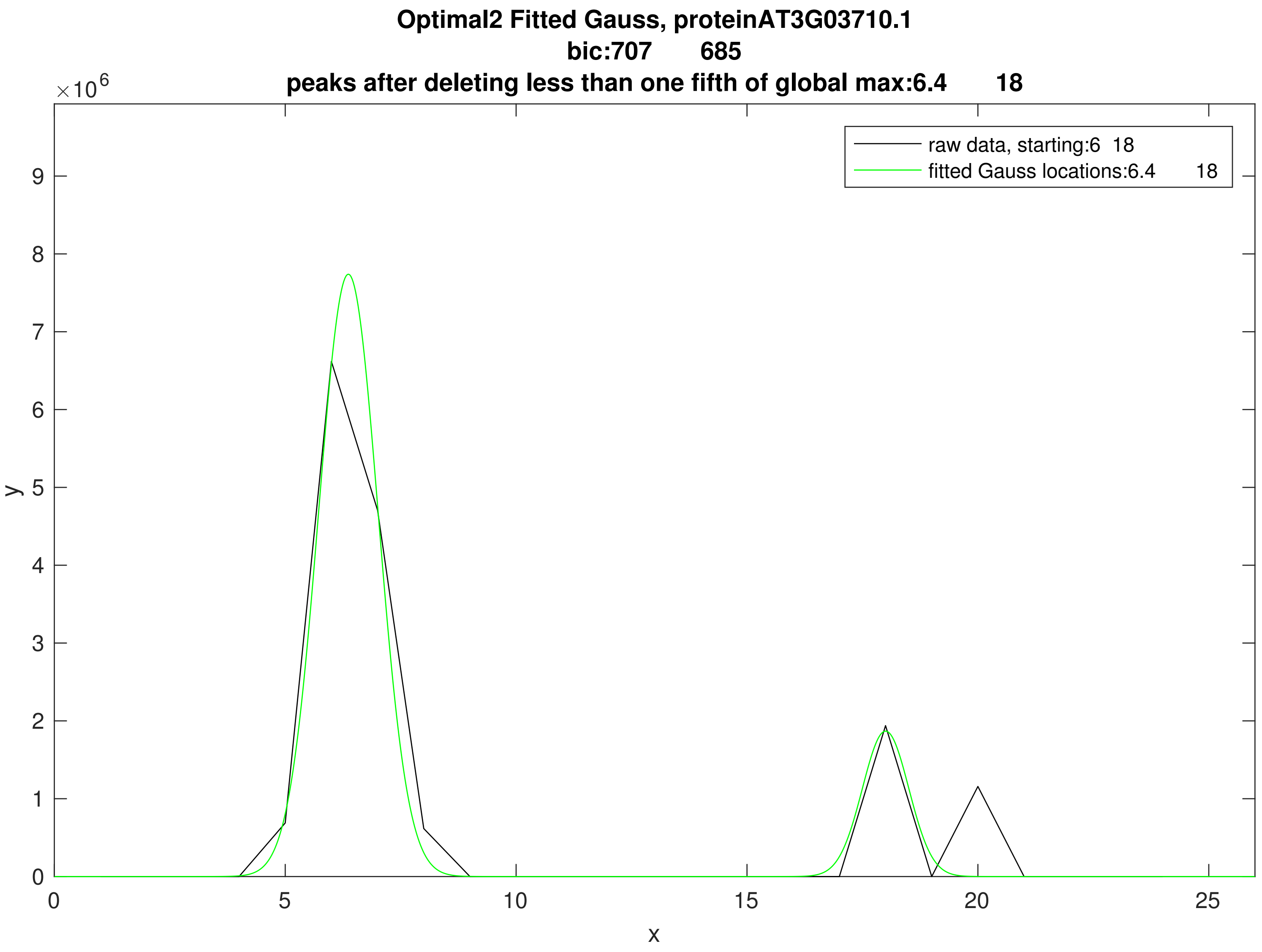
Optimal2 Fitted Gauss, proteinAT3G03710.1
bic:707 685
peaks after deleting less than one fifth of global max:6.4 18
×10^6
raw data, starting:6 18
fitted Gauss locations:6.4 18
y
x
0
1
2
3
4
5
6
7
8
9
0
5
10
15
20
25

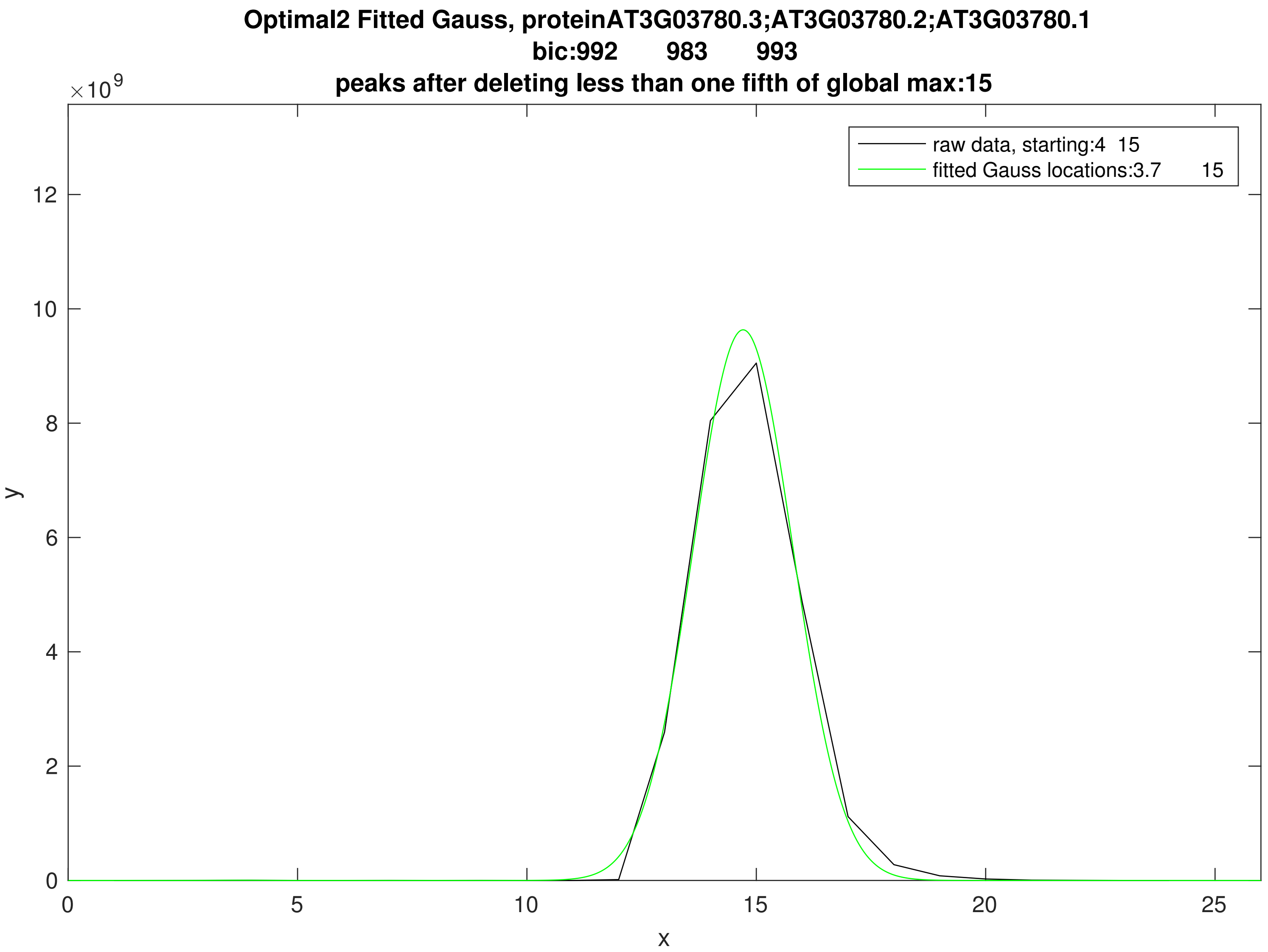
Optimal2 Fitted Gauss, proteinAT3G03780.3;AT3G03780.2;AT3G03780.1
bic:992 983 993
peaks after deleting less than one fifth of global max:15
×10^9
raw data, starting:4 15
fitted Gauss locations:3.7 15
12
10
8
6
4
2
0
y
0
5
10
15
20
25
x

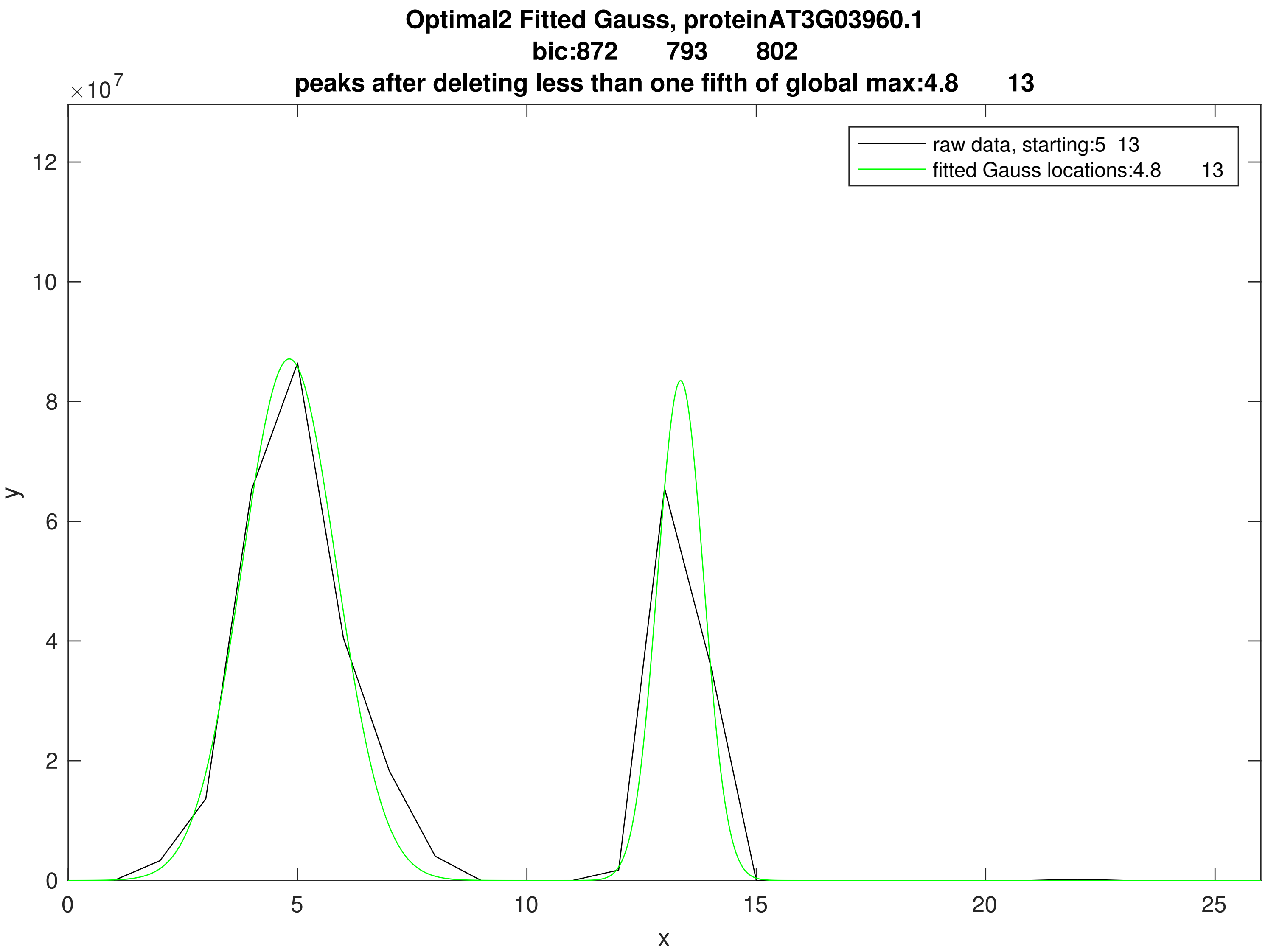

Optimal2 Fitted Gauss, proteinAT3G03960.1
bic:872 793 802
peaks after deleting less than one fifth of global max:4.8 13
×10^7
raw data, starting:5 13
fitted Gauss locations:4.8 13
y
x
0
2
4
6
8
10
12
0
5
10
15
20
25

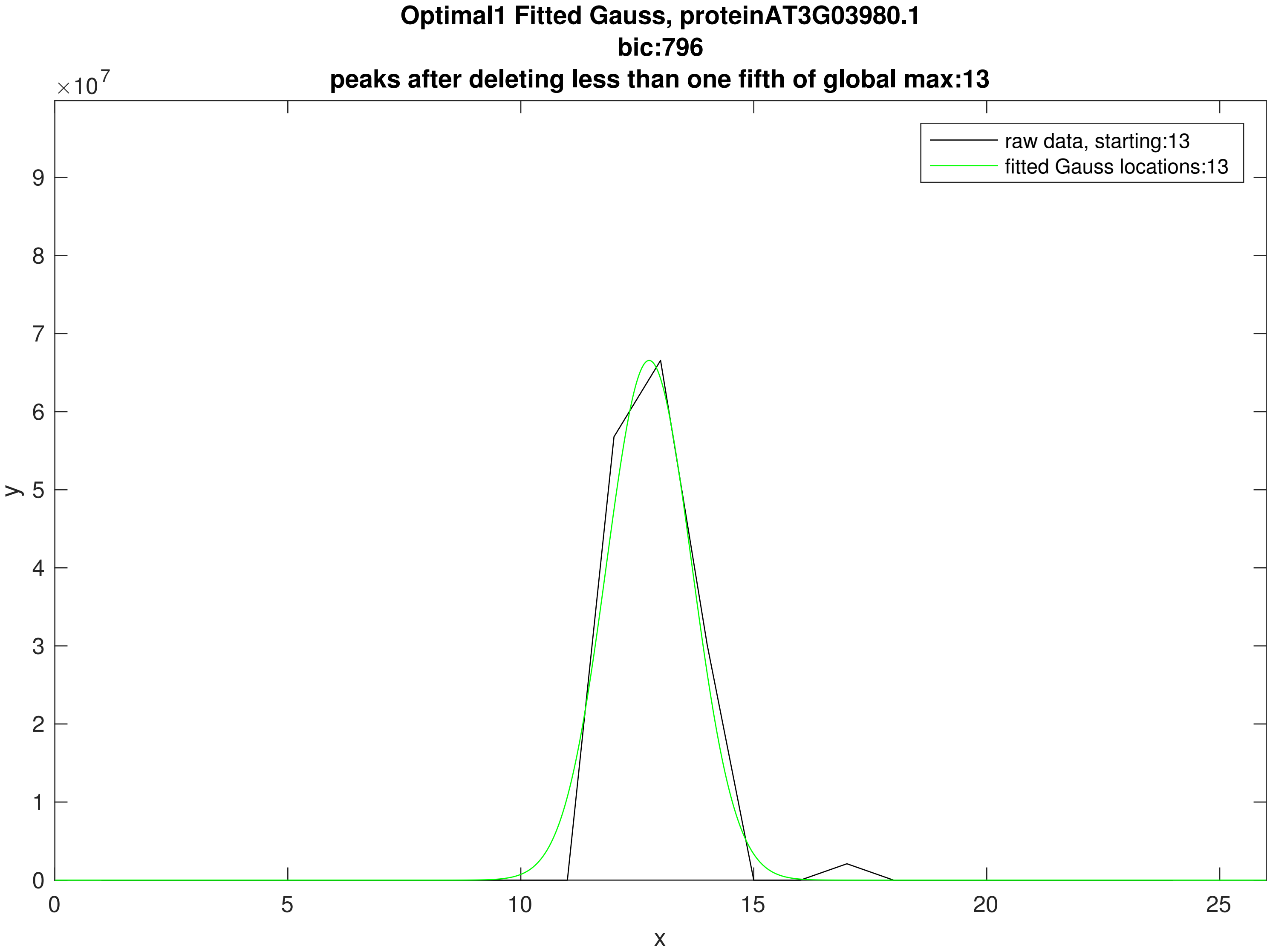

Optimal1 Fitted Gauss, proteinAT3G03980.1
bic:796
peaks after deleting less than one fifth of global max:13
×10^7
raw data, starting:13
fitted Gauss locations:13
y
x
0
1
2
3
4
5
6
7
8
9
0
5
10
15
20
25

**Optimal1 Fitted Gauss, proteinAT3G03990.1**
**bic:814**
**peaks after deleting less than one fifth of global max:18**

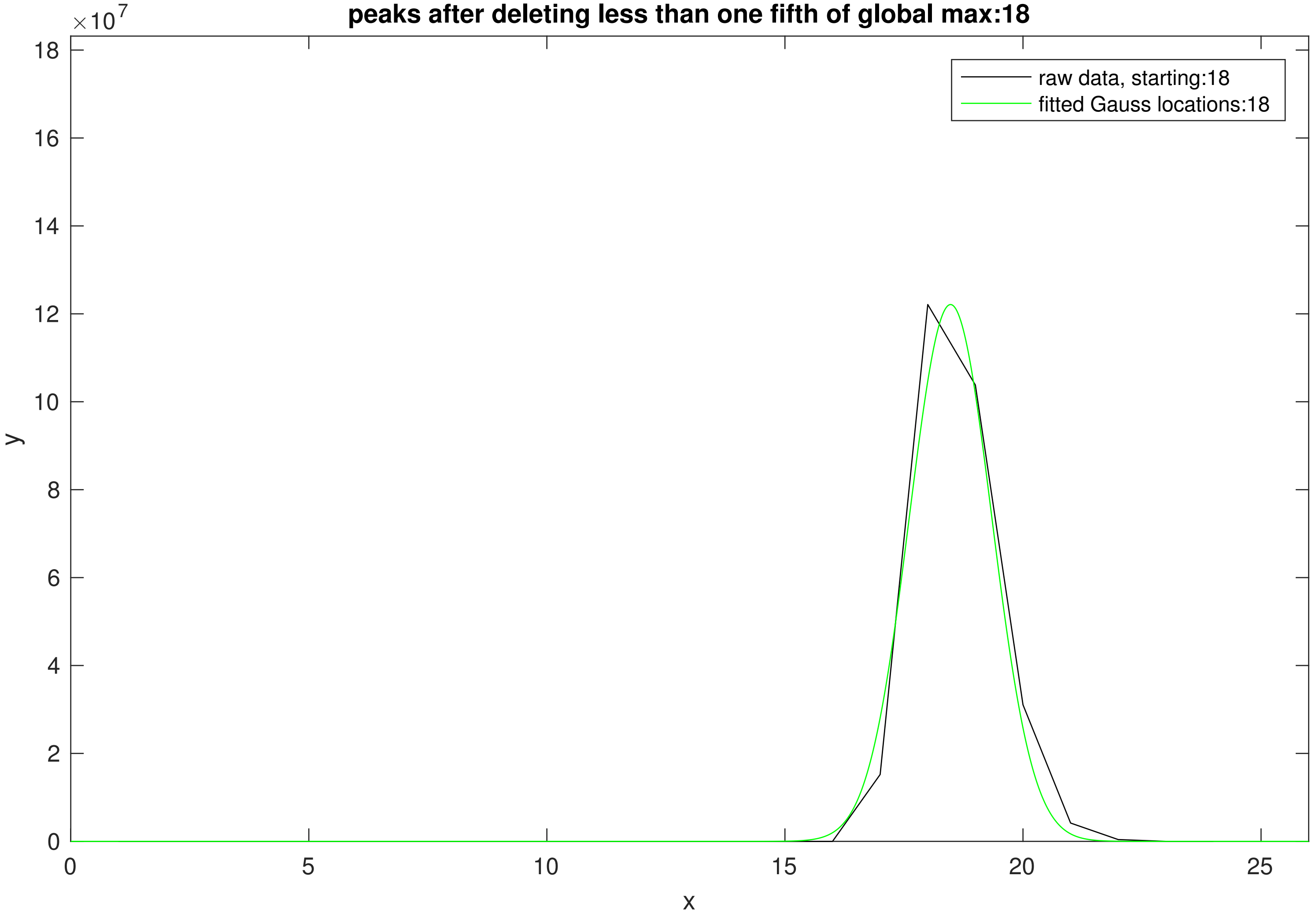

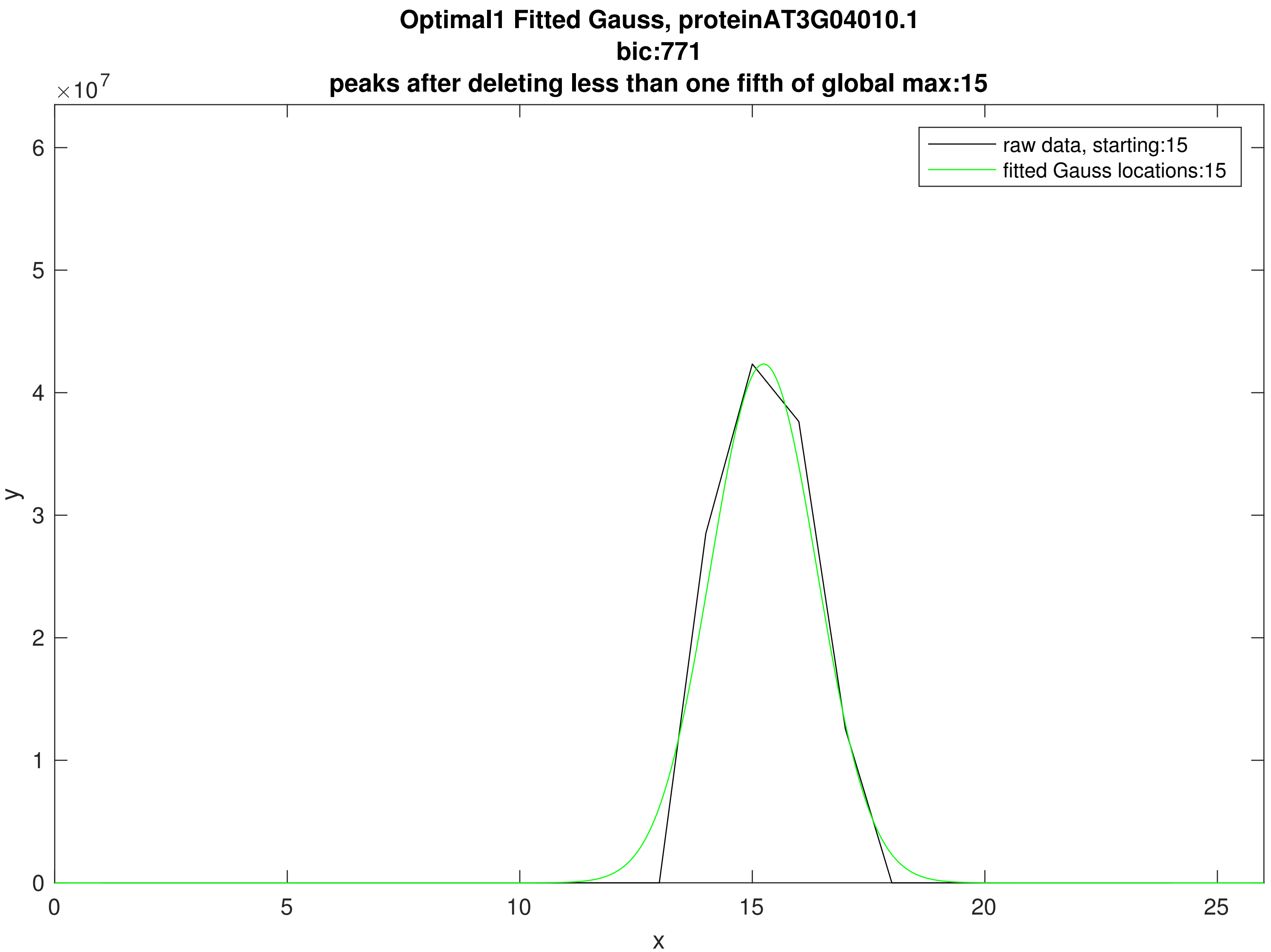

Optimal1 Fitted Gauss, proteinAT3G04010.1
bic:771
peaks after deleting less than one fifth of global max:15
×10^7
raw data, starting:15
fitted Gauss locations:15
y
x
0
1
2
3
4
5
6
0
5
10
15
20
25

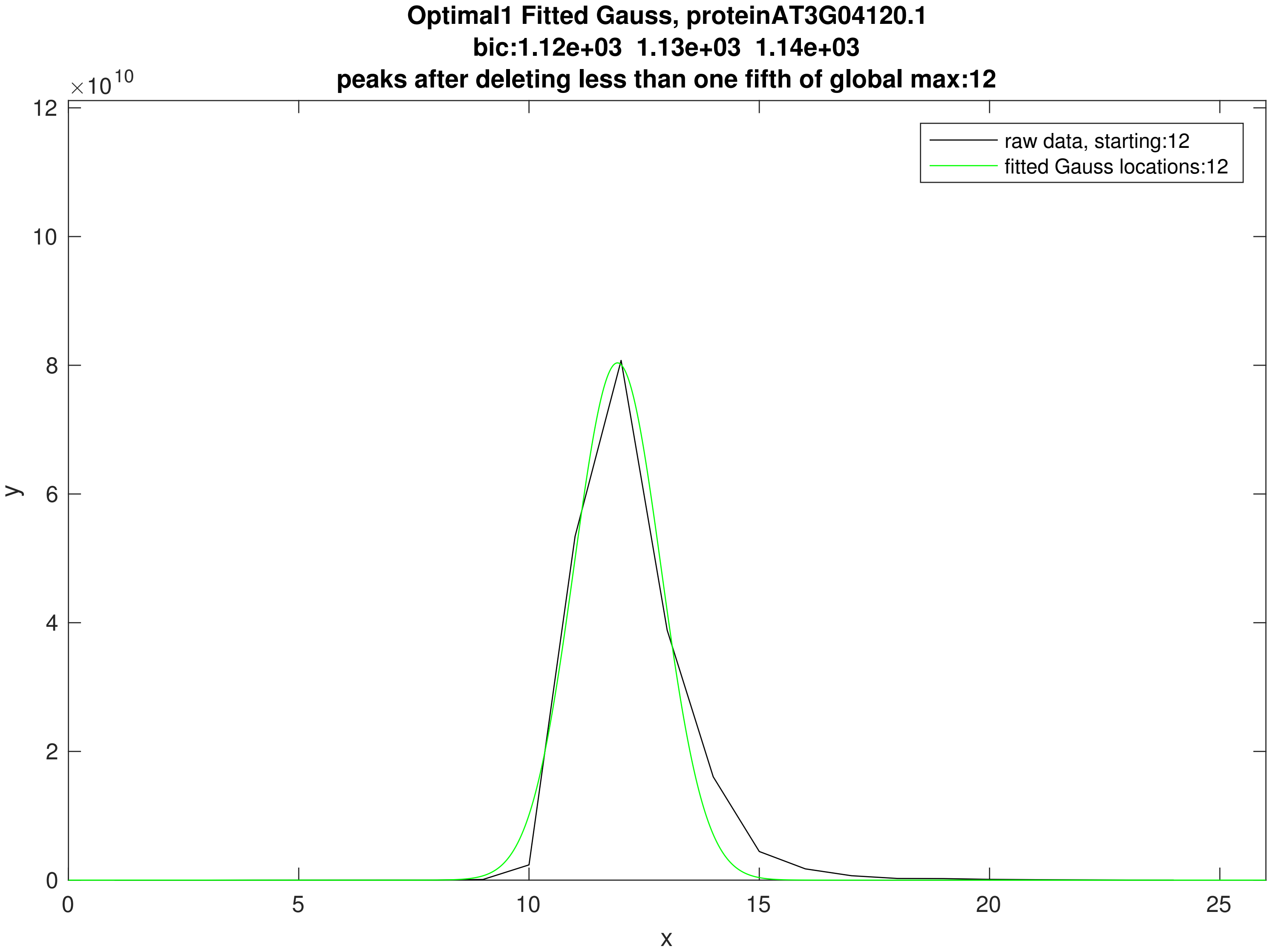

Optimal1 Fitted Gauss, proteinAT3G04120.1
bic:1.12e+03 1.13e+03 1.14e+03
peaks after deleting less than one fifth of global max:12
×10^10
raw data, starting:12
fitted Gauss locations:12
y
x
0
2
4
6
8
10
12
0
5
10
15
20
25

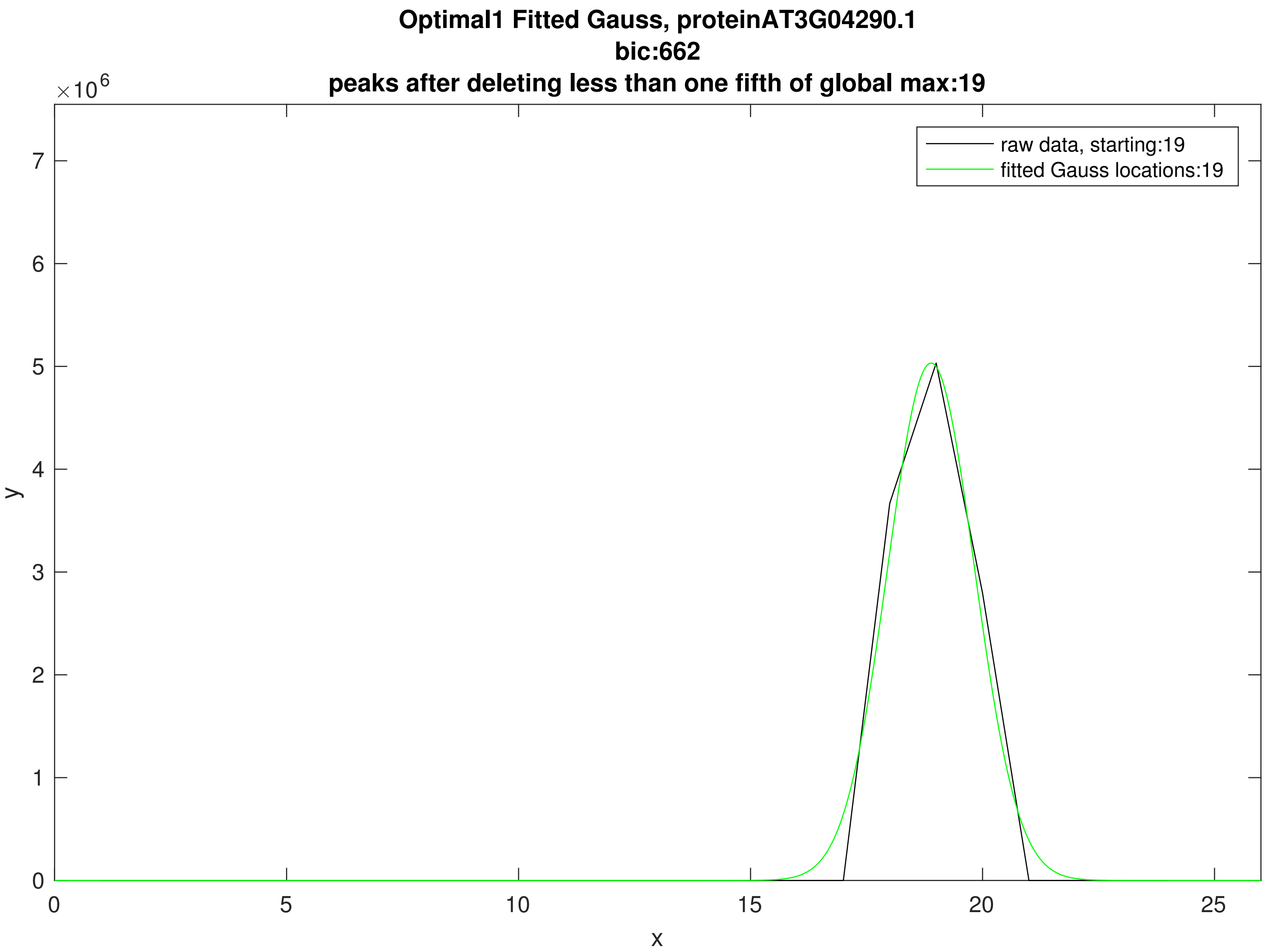

Optimal1 Fitted Gauss, proteinAT3G04290.1
bic:662
peaks after deleting less than one fifth of global max:19
×10^6
raw data, starting:19
fitted Gauss locations:19
y
x
0
1
2
3
4
5
6
7
0
5
10
15
20
25

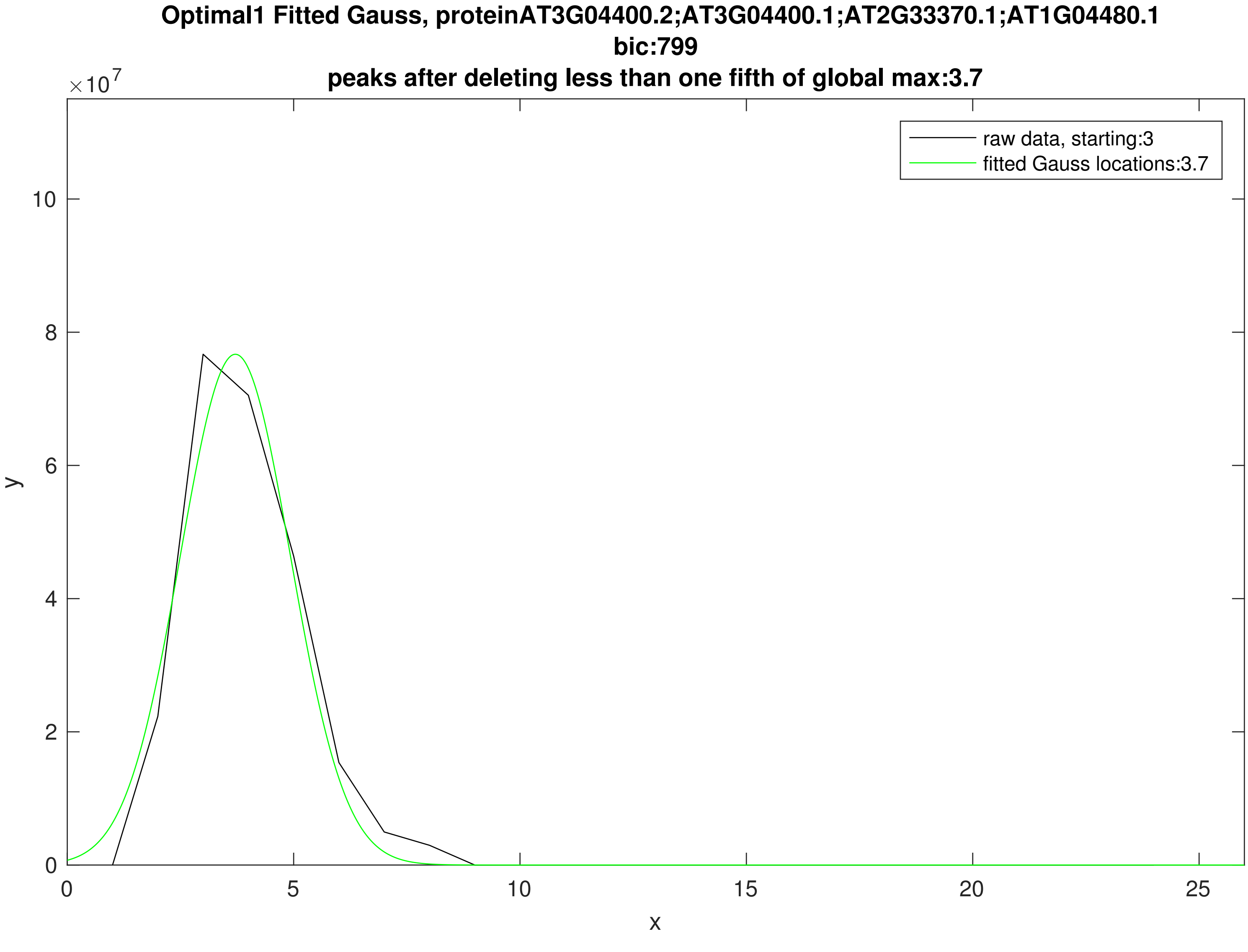
Optimal1 Fitted Gauss, proteinAT3G04400.2;AT3G04400.1;AT2G33370.1;AT1G04480.1
bic:799
peaks after deleting less than one fifth of global max:3.7
×10^7
raw data, starting:3
fitted Gauss locations:3.7
y
x
0
2
4
6
8
10
0
5
10
15
20
25

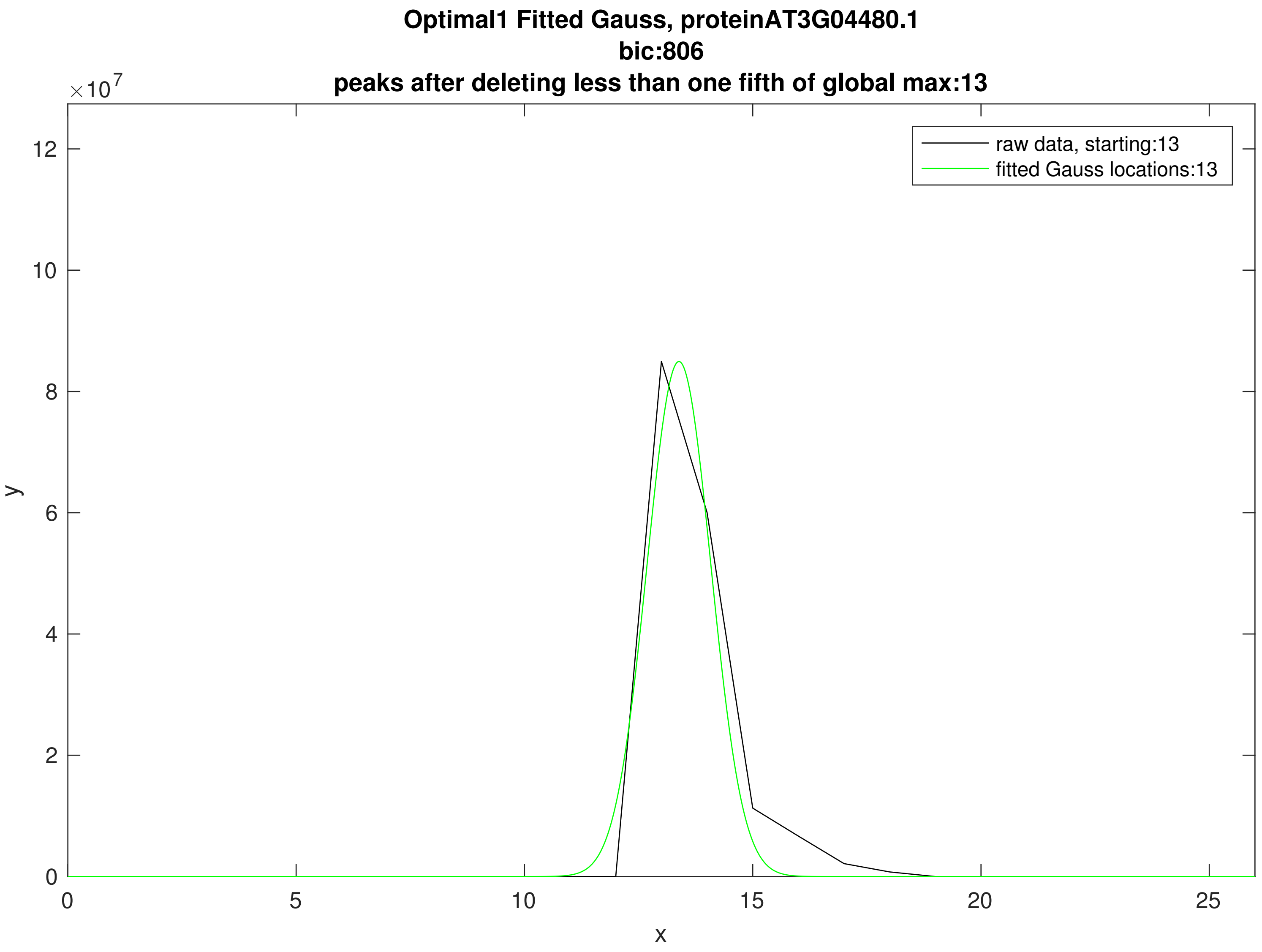
Optimal1 Fitted Gauss, proteinAT3G04480.1
bic:806
peaks after deleting less than one fifth of global max:13
×10^7
raw data, starting:13
fitted Gauss locations:13
y
x
0
2
4
6
8
10
12
0
5
10
15
20
25

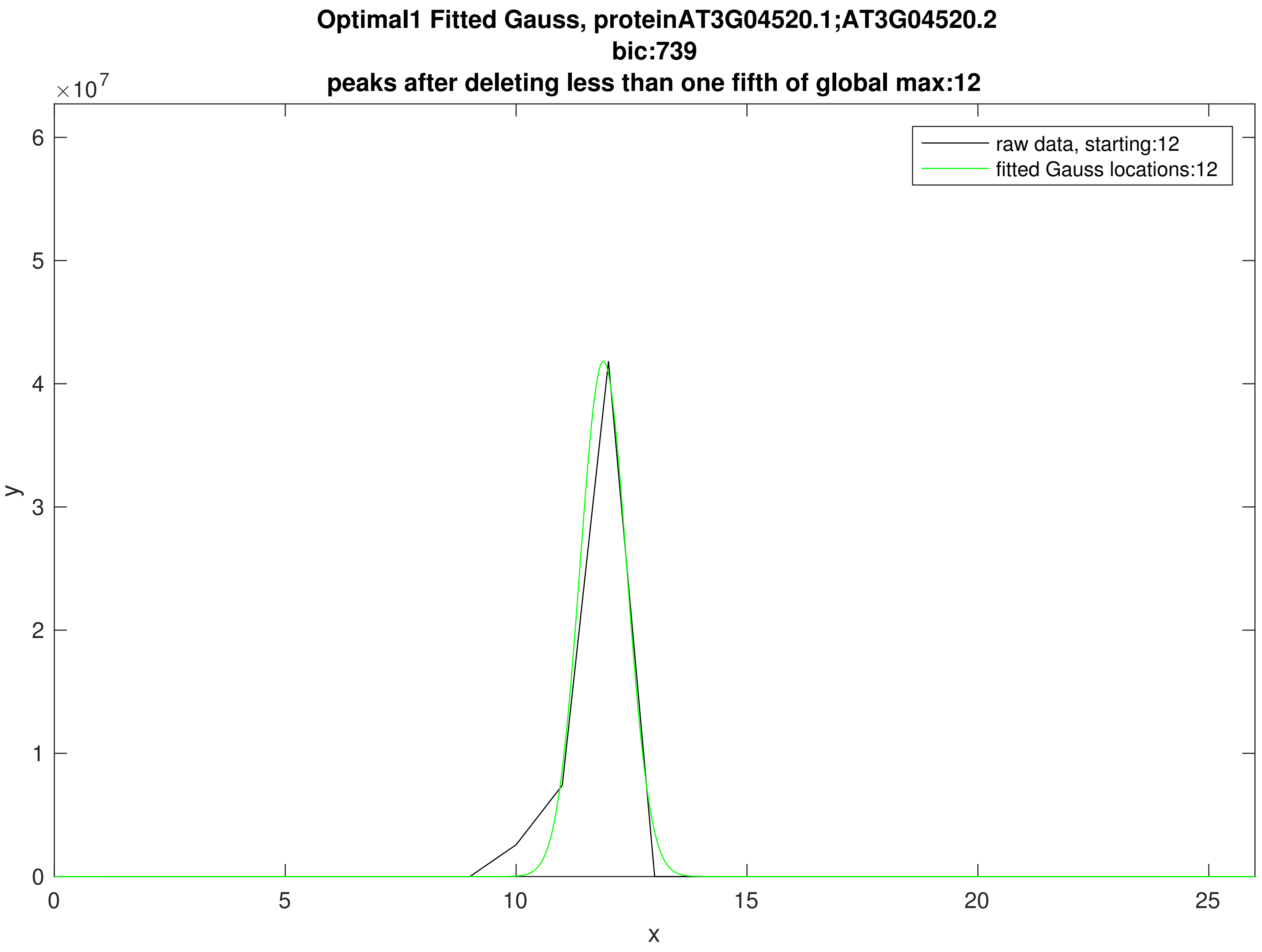
Optimal1 Fitted Gauss, proteinAT3G04520.1;AT3G04520.2
bic:739
peaks after deleting less than one fifth of global max:12
×10^7
6
5
4
3
2
1
0
y
0
5
10
15
20
25
x
raw data, starting:12
fitted Gauss locations:12

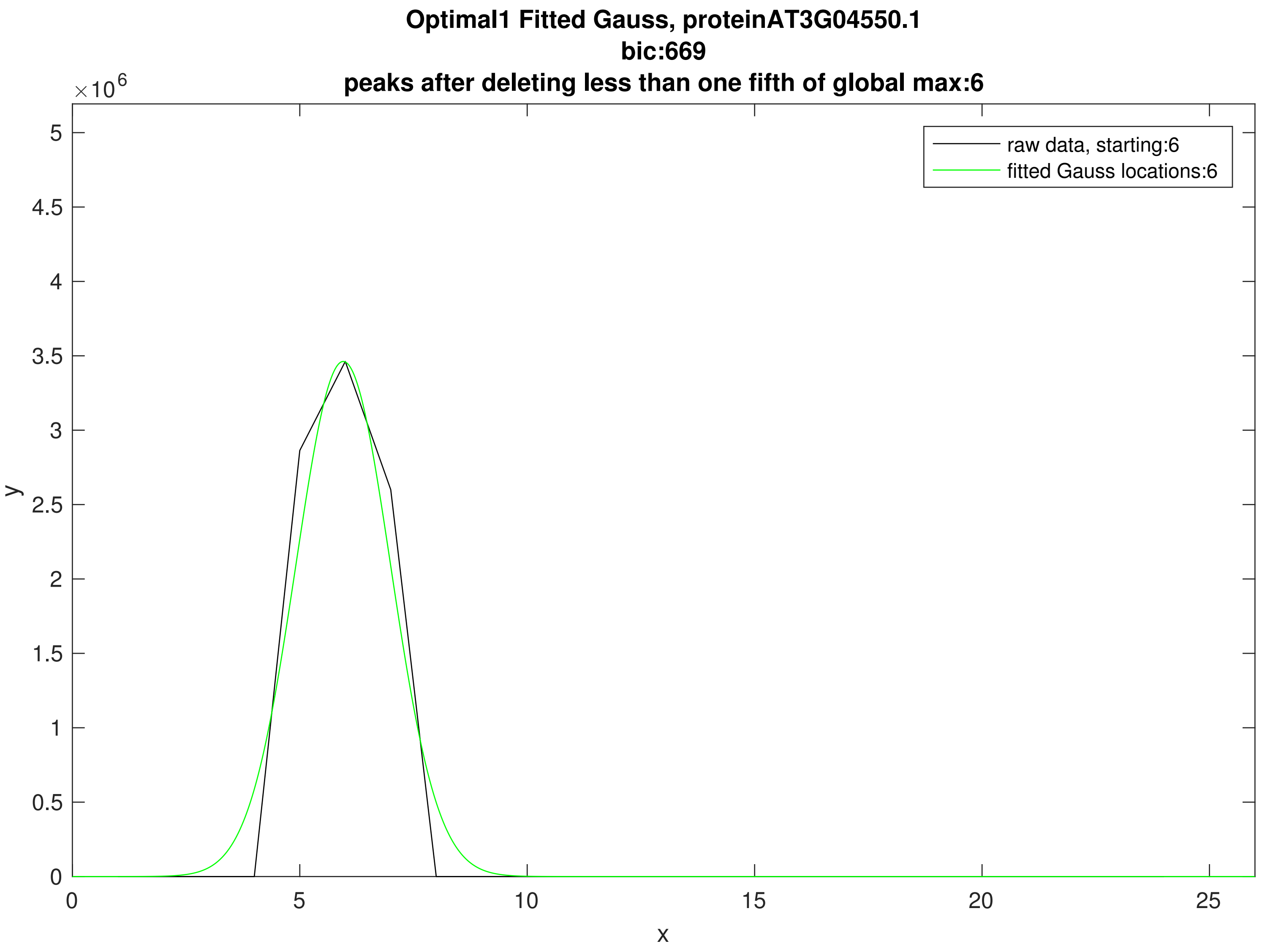

Optimal1 Fitted Gauss, proteinAT3G04550.1
bic:669
peaks after deleting less than one fifth of global max:6
raw data, starting:6
fitted Gauss locations:6
y
x

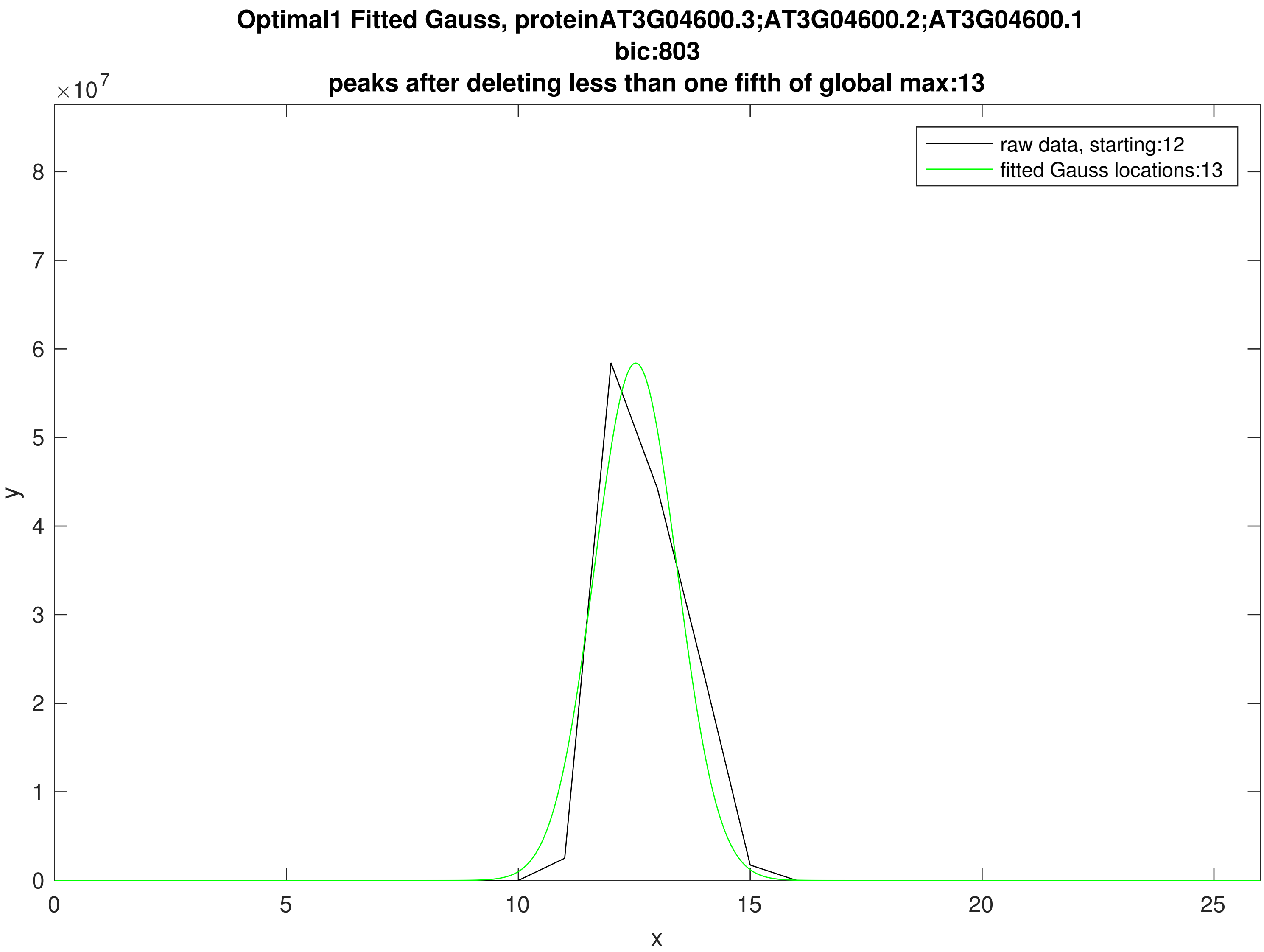
Optimal1 Fitted Gauss, proteinAT3G04600.3;AT3G04600.2;AT3G04600.1
bic:803
peaks after deleting less than one fifth of global max:13
×10^7
8
7
6
5
4
3
2
1
0
y
0
5
10
15
20
25
x
raw data, starting:12
fitted Gauss locations:13

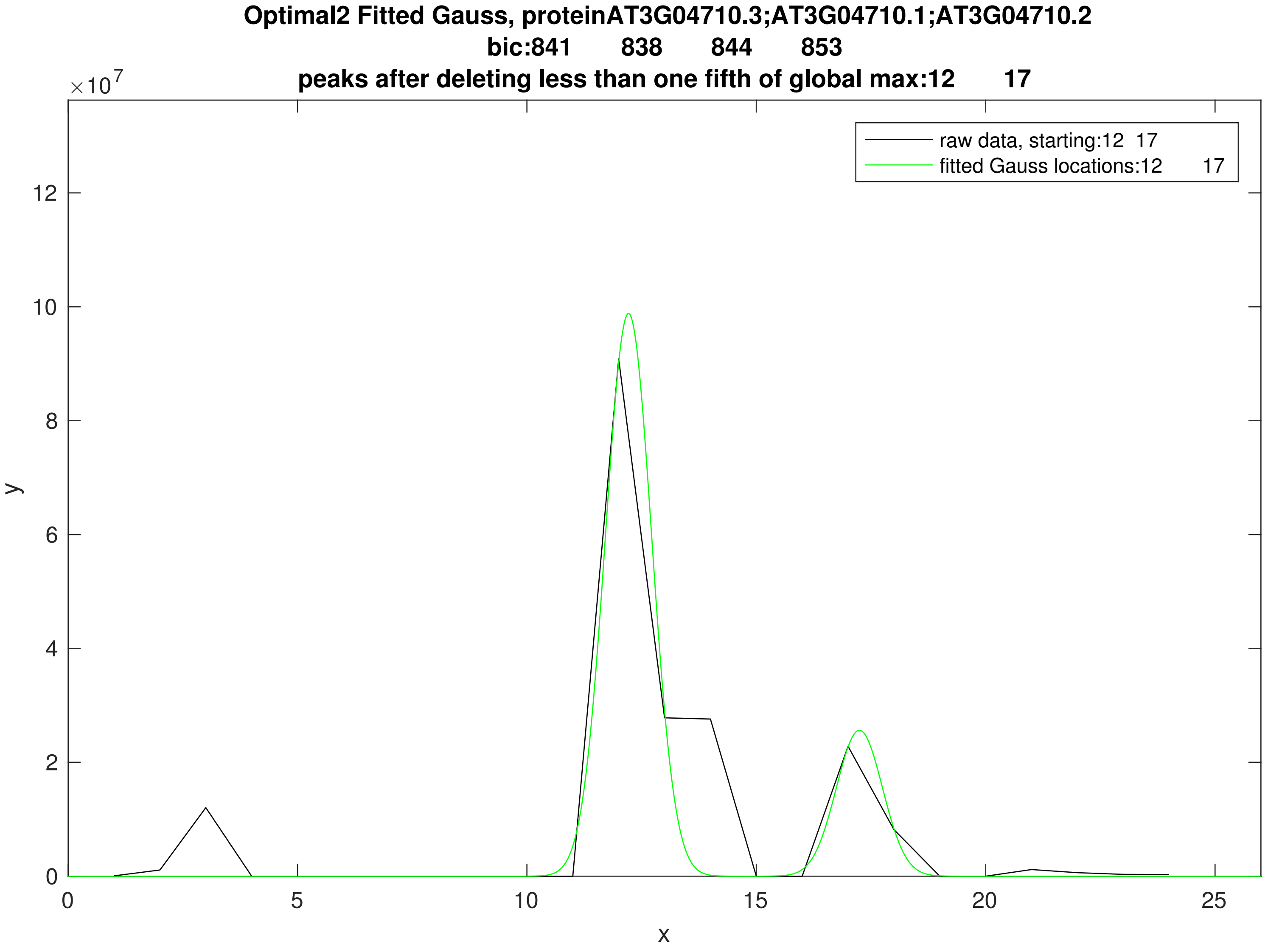

Optimal2 Fitted Gauss, proteinAT3G04710.3;AT3G04710.1;AT3G04710.2
bic:841 838 844 853
peaks after deleting less than one fifth of global max:12 17
×10^7
raw data, starting:12 17
fitted Gauss locations:12 17
y
x
0
2
4
6
8
10
12
0
5
10
15
20
25

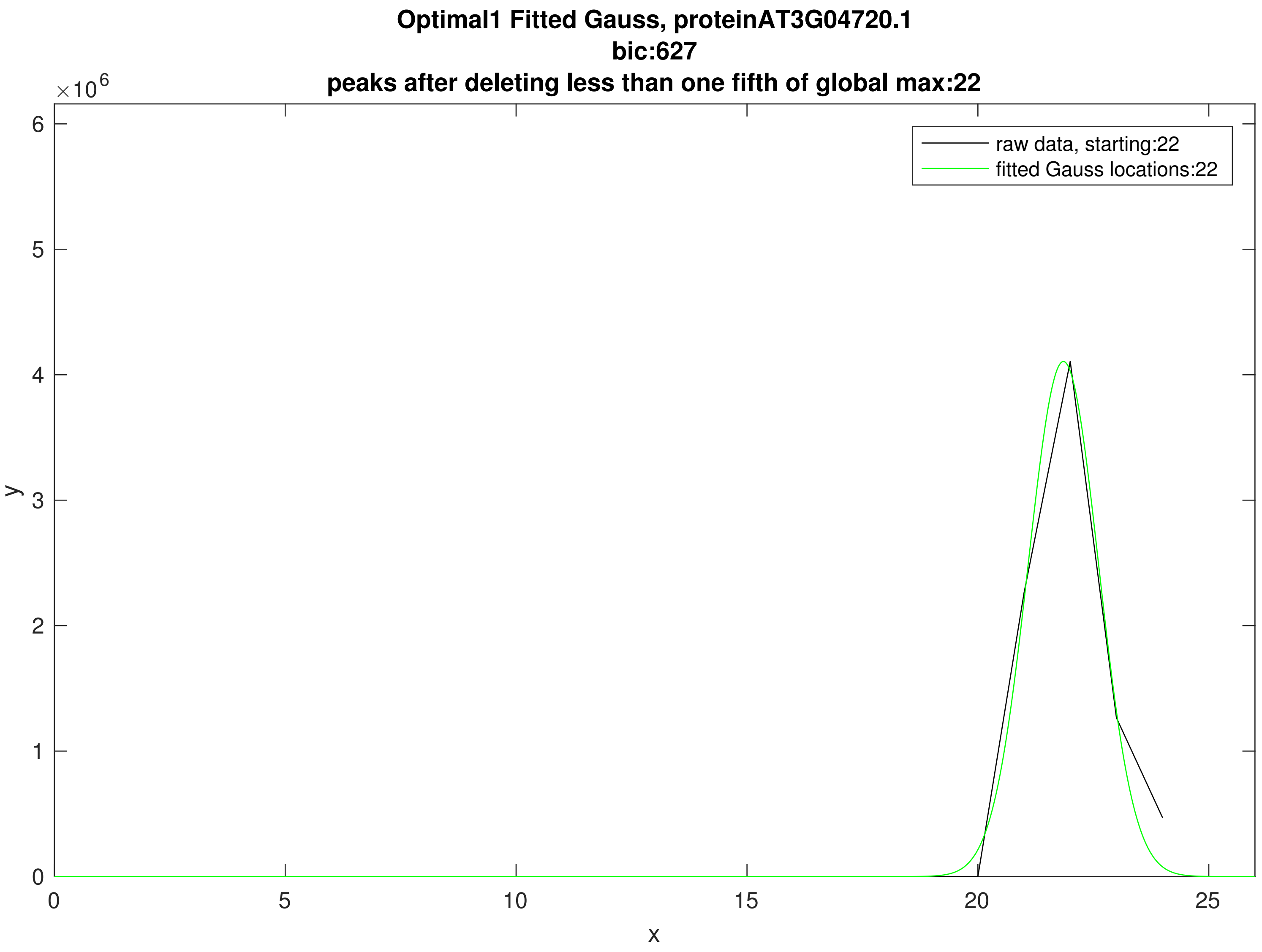

Optimal1 Fitted Gauss, proteinAT3G04720.1
bic:627
peaks after deleting less than one fifth of global max:22
×10^6
raw data, starting:22
fitted Gauss locations:22
y
x
0
1
2
3
4
5
6
0
5
10
15
20
25

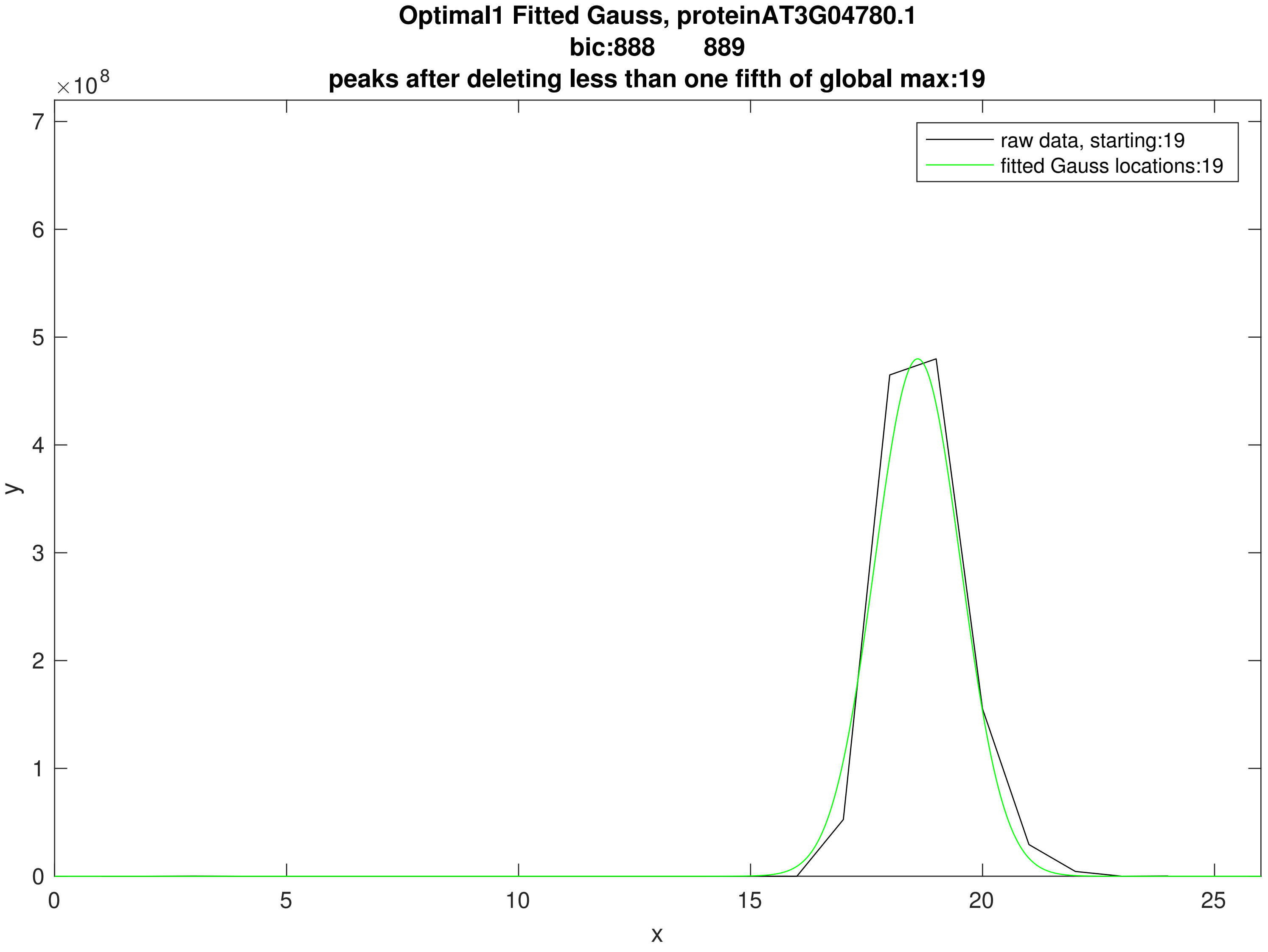

Optimal1 Fitted Gauss, proteinAT3G04780.1
bic:888 889
peaks after deleting less than one fifth of global max:19
×10^8
raw data, starting:19
fitted Gauss locations:19
y
x
0
1
2
3
4
5
6
7
0
5
10
15
20
25

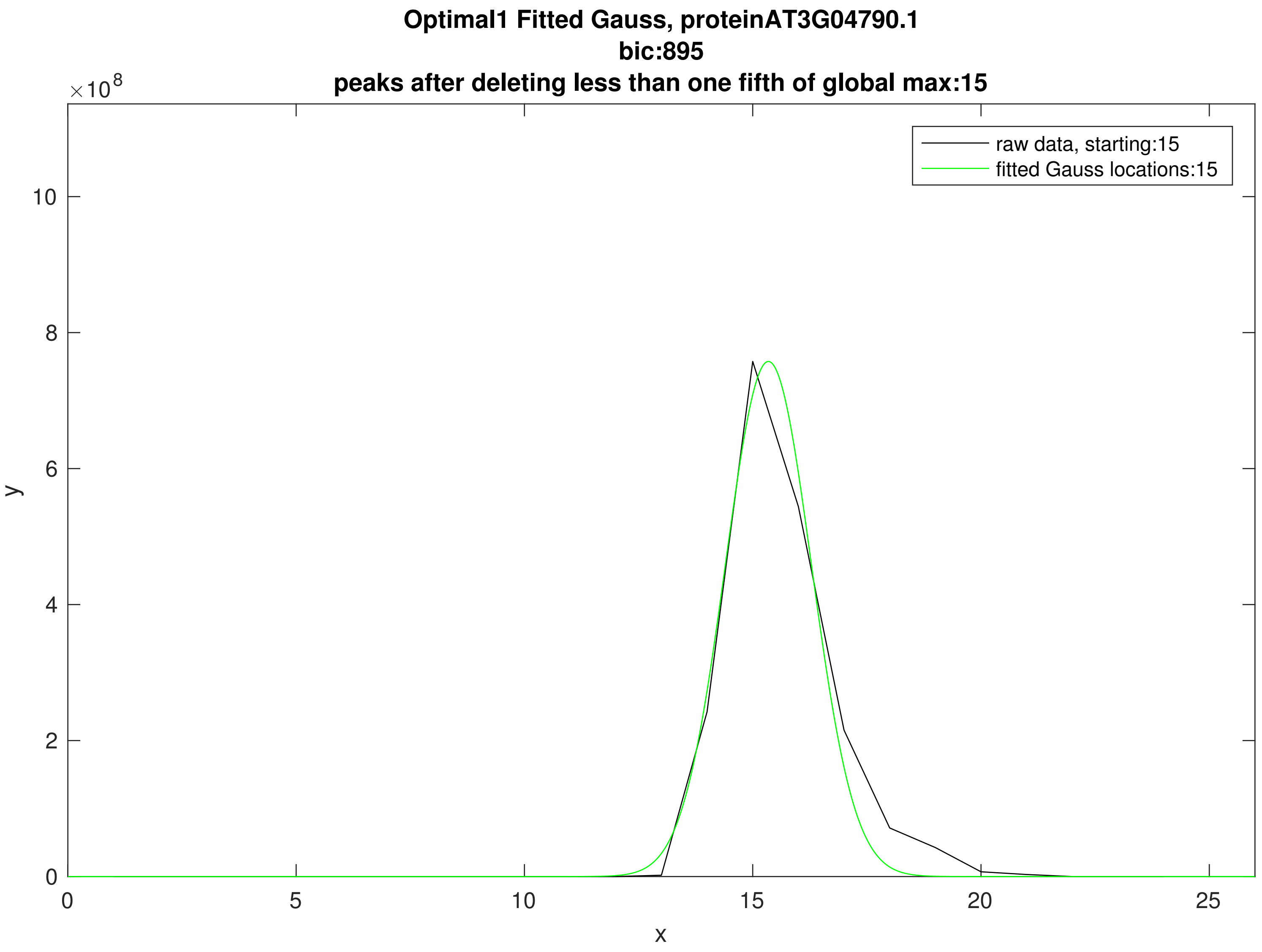

Optimal1 Fitted Gauss, proteinAT3G04790.1
bic:895
peaks after deleting less than one fifth of global max:15
×10^8
raw data, starting:15
fitted Gauss locations:15
10
8
6
4
2
0
y
0
5
10
15
20
25
x

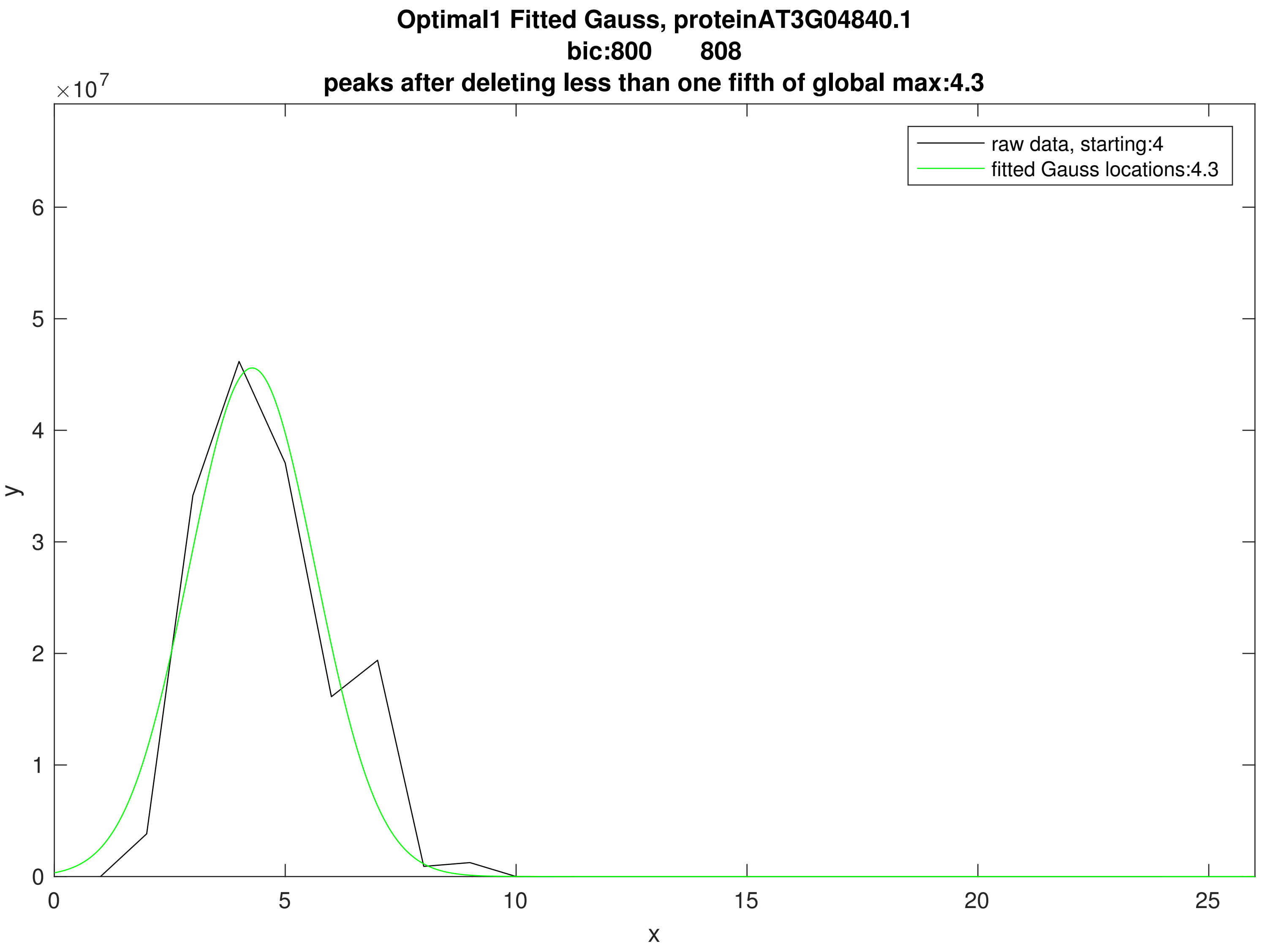

Optimal1 Fitted Gauss, proteinAT3G04840.1
bic:800 808
peaks after deleting less than one fifth of global max:4.3
×10^7
raw data, starting:4
fitted Gauss locations:4.3
y
x
0
1
2
3
4
5
6
0
5
10
15
20
25

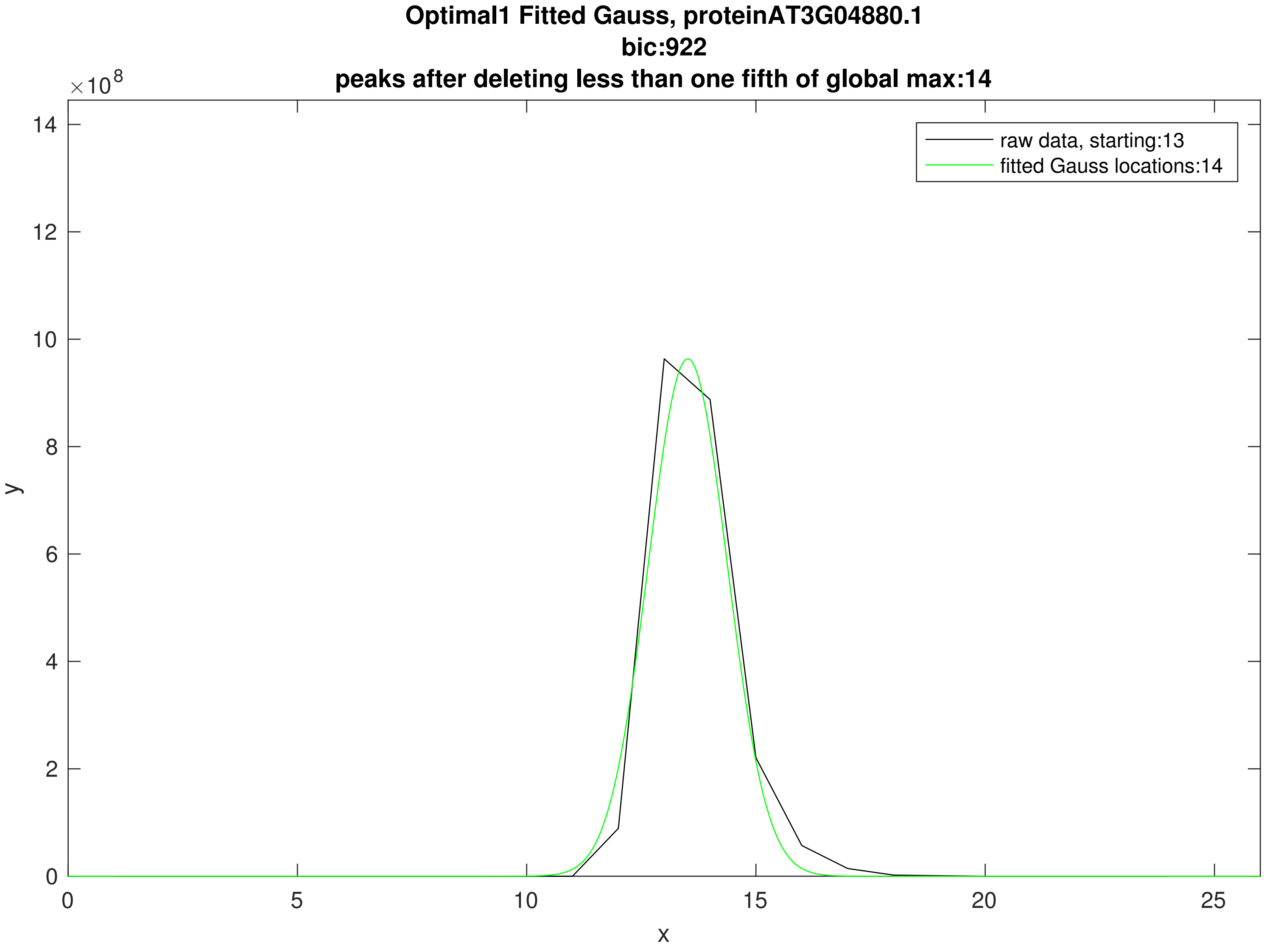

Optimal1 Fitted Gauss, proteinAT3G04880.1
bic:922
peaks after deleting less than one fifth of global max:14
×10^8
raw data, starting:13
fitted Gauss locations:14
y
x
0
2
4
6
8
10
12
14
0
5
10
15
20
25

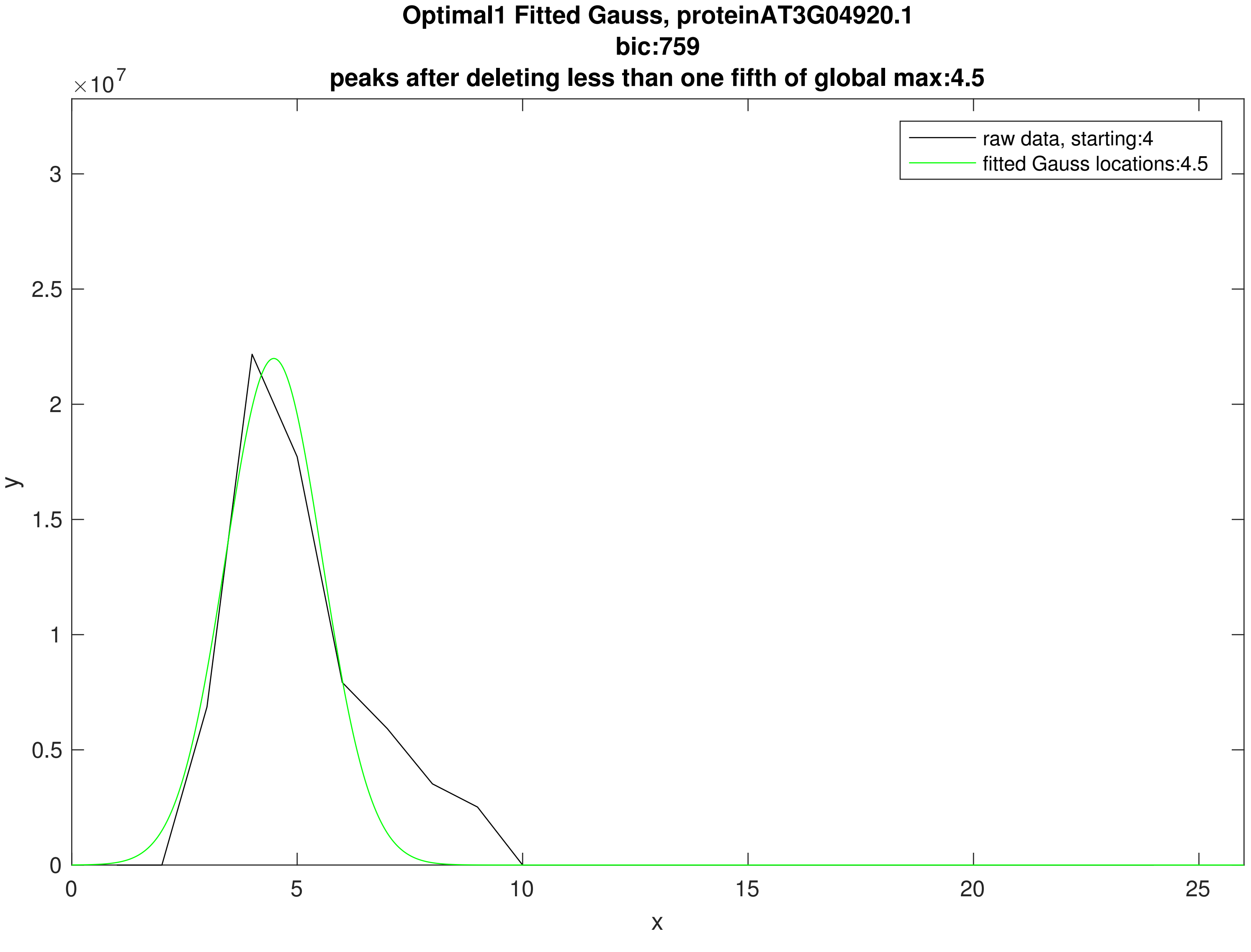
Optimal1 Fitted Gauss, proteinAT3G04920.1
bic:759
peaks after deleting less than one fifth of global max:4.5
×10^7
raw data, starting:4
fitted Gauss locations:4.5
y
x
0
0.5
1
1.5
2
2.5
3
0
5
10
15
20
25

**Optimal1 Fitted Gauss, proteinAT3G04940.1**
**bic:740**
**peaks after deleting less than one fifth of global max:15**

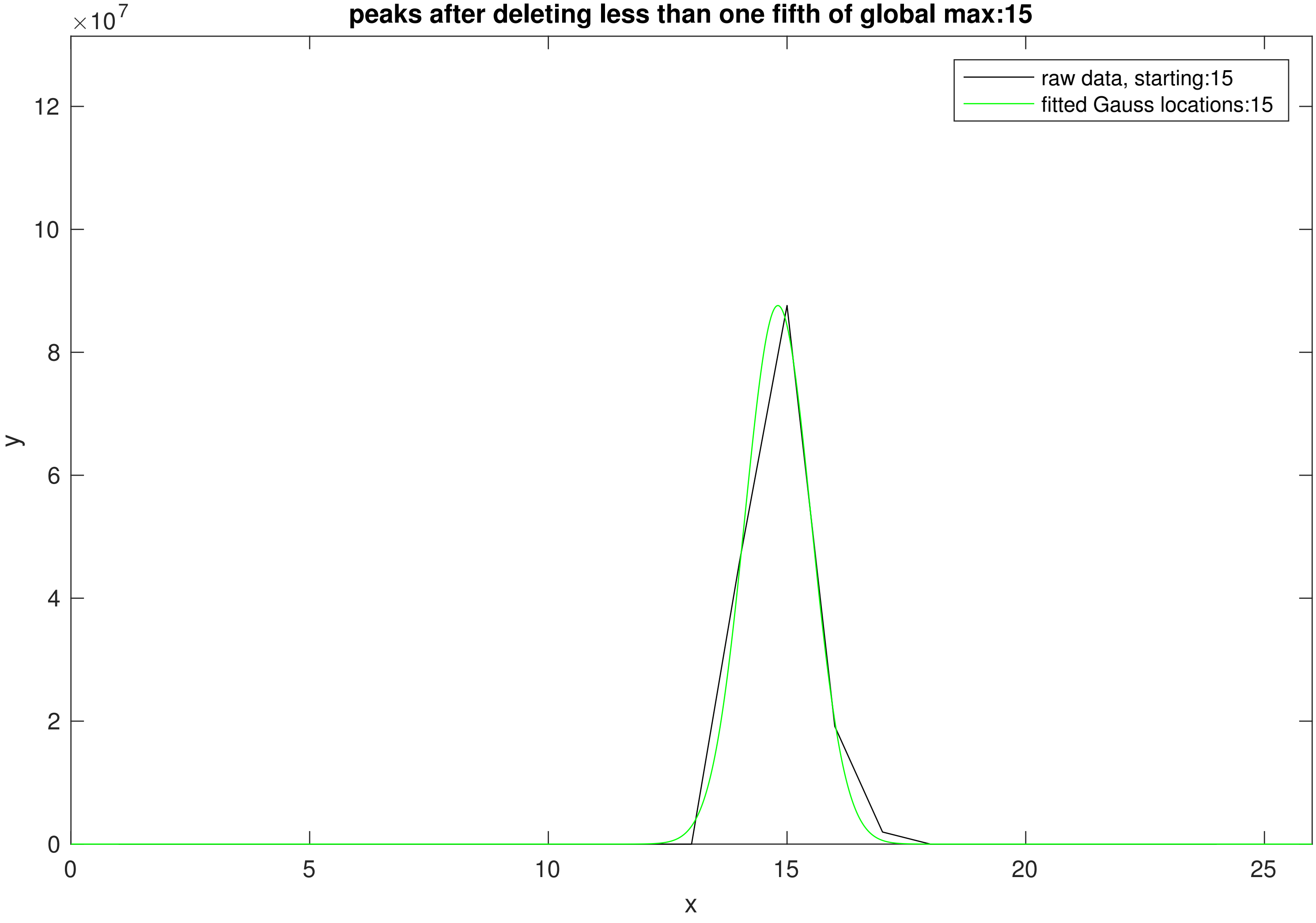

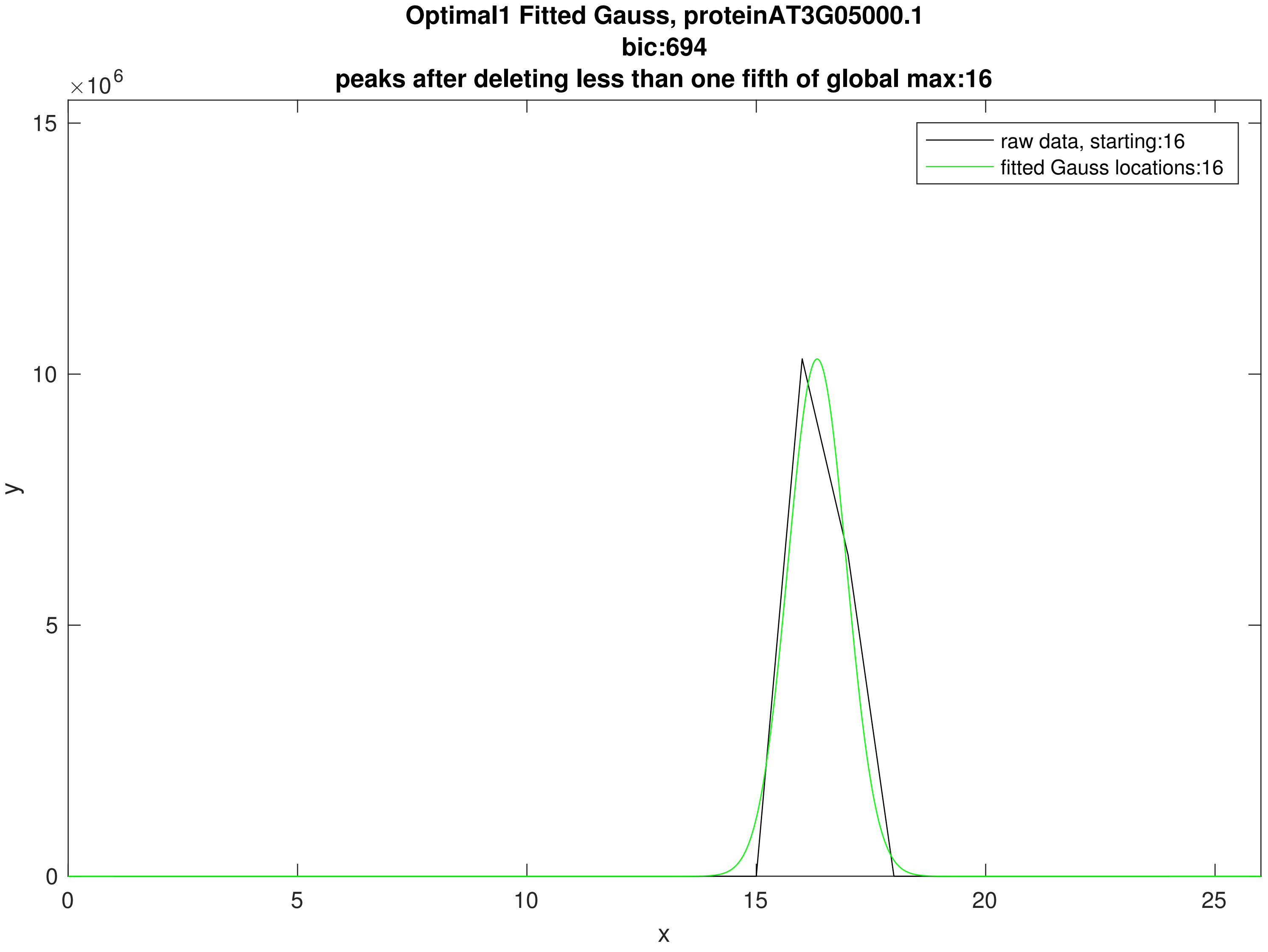

Optimal1 Fitted Gauss, proteinAT3G05000.1
bic:694
peaks after deleting less than one fifth of global max:16
×10^6
15
10
5
0
y
0
5
10
15
20
25
x
raw data, starting:16
fitted Gauss locations:16

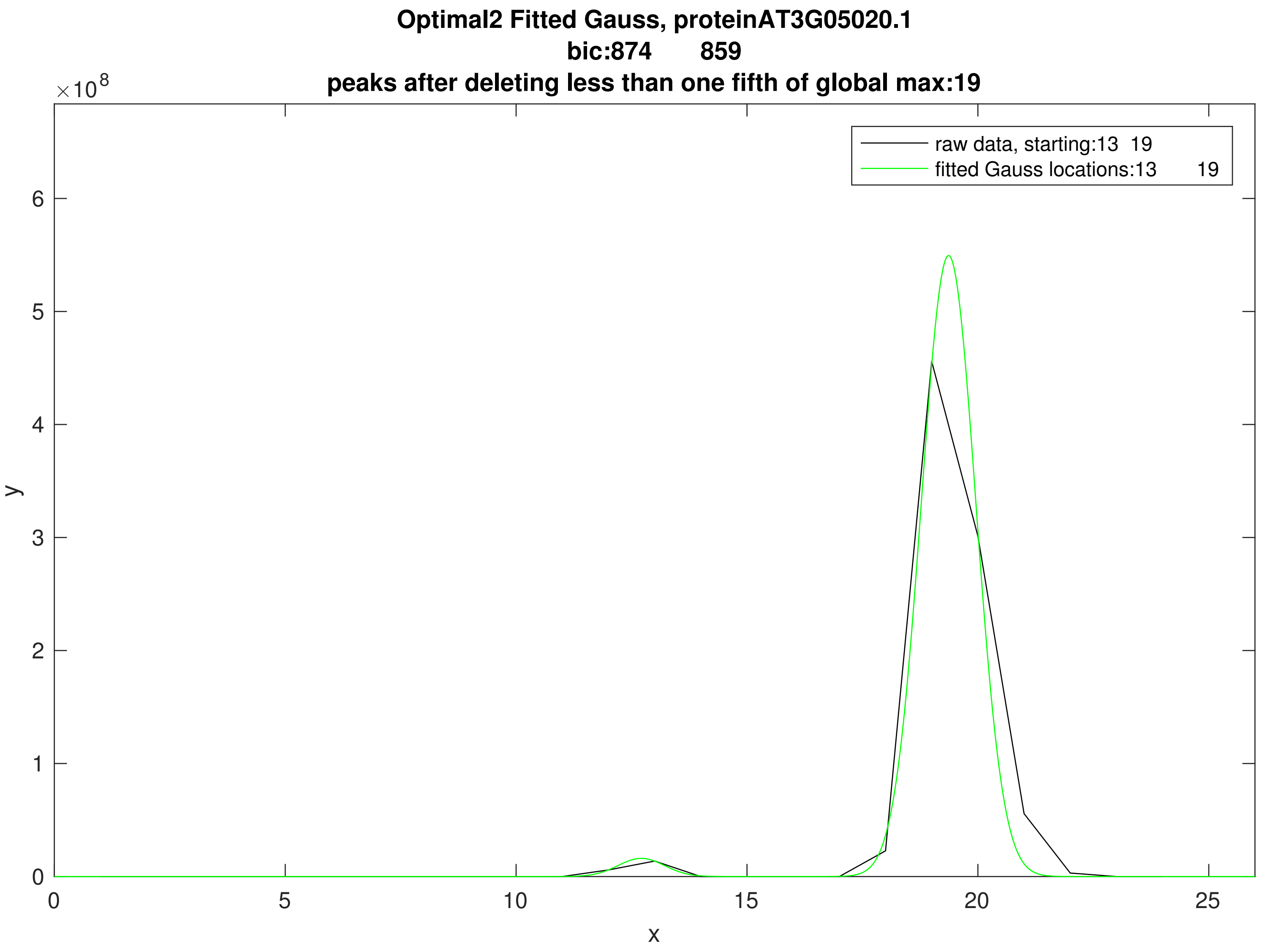
Optimal2 Fitted Gauss, proteinAT3G05020.1
bic:874 859
peaks after deleting less than one fifth of global max:19
×10^8
y
x
raw data, starting:13 19
fitted Gauss locations:13 19
0
1
2
3
4
5
6
0
5
10
15
20
25

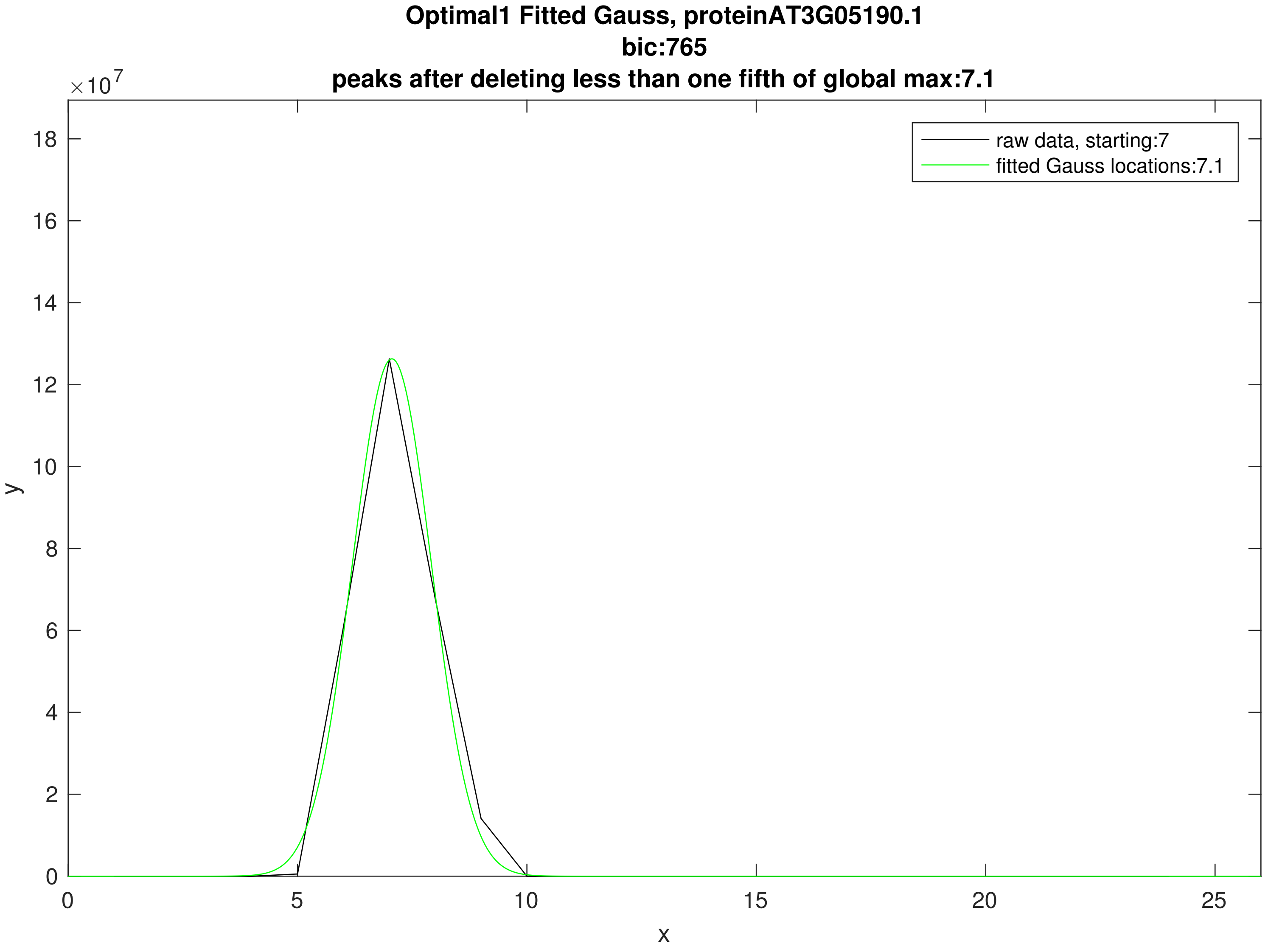

Optimal1 Fitted Gauss, proteinAT3G05190.1
bic:765
peaks after deleting less than one fifth of global max:7.1
×10^7
18
16
14
12
10
8
6
4
2
0
y
0
5
10
15
20
25
x
raw data, starting:7
fitted Gauss locations:7.1

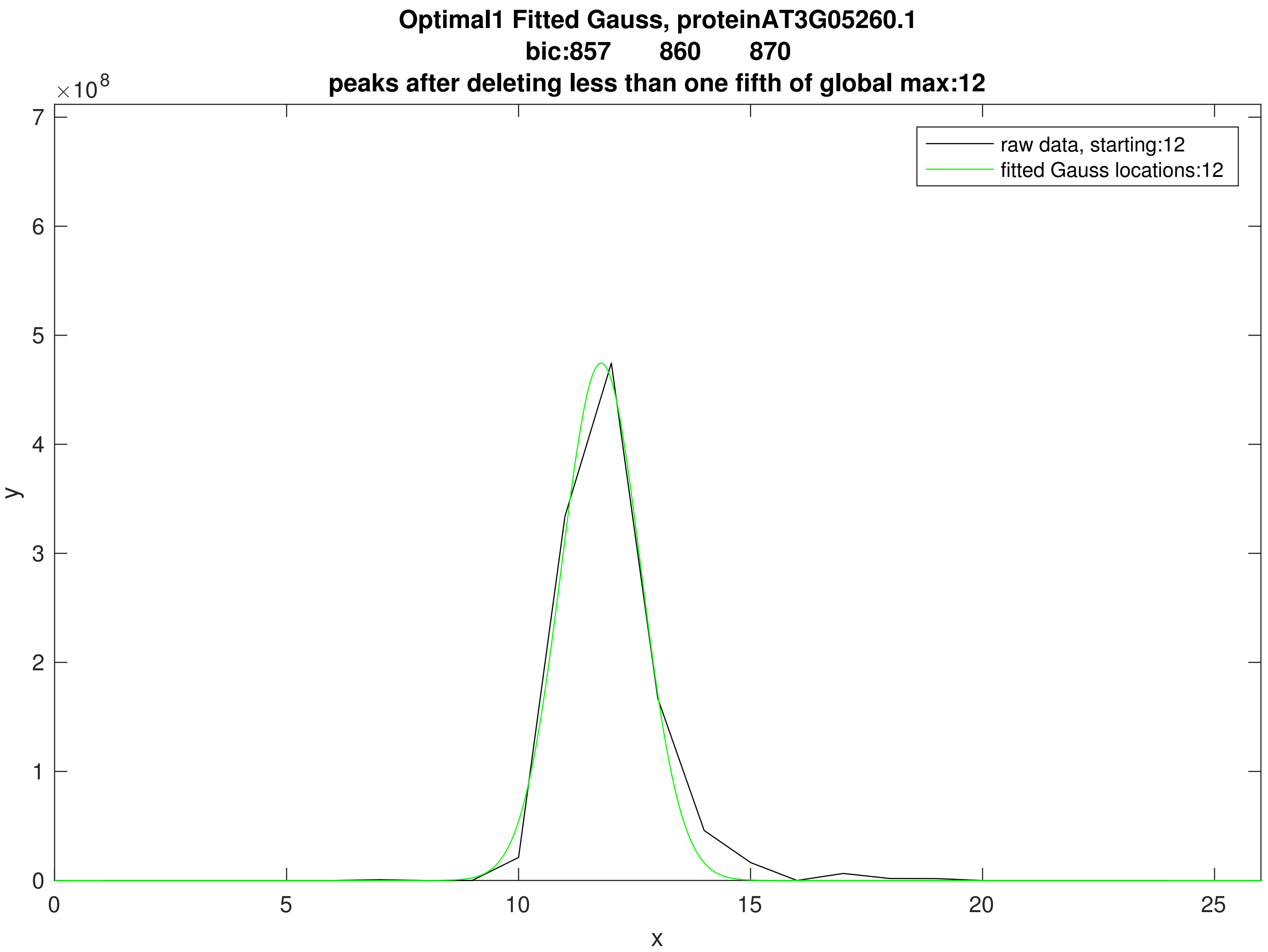
Optimal1 Fitted Gauss, proteinAT3G05260.1
bic:857 860 870
peaks after deleting less than one fifth of global max:12
×10^8
raw data, starting:12
fitted Gauss locations:12
y
x
0
1
2
3
4
5
6
7
0
5
10
15
20
25

#### Optimal1 Fitted Gauss, proteinAT3G05350.1
#### bic:735
#### peaks after deleting less than one fifth of global max:11

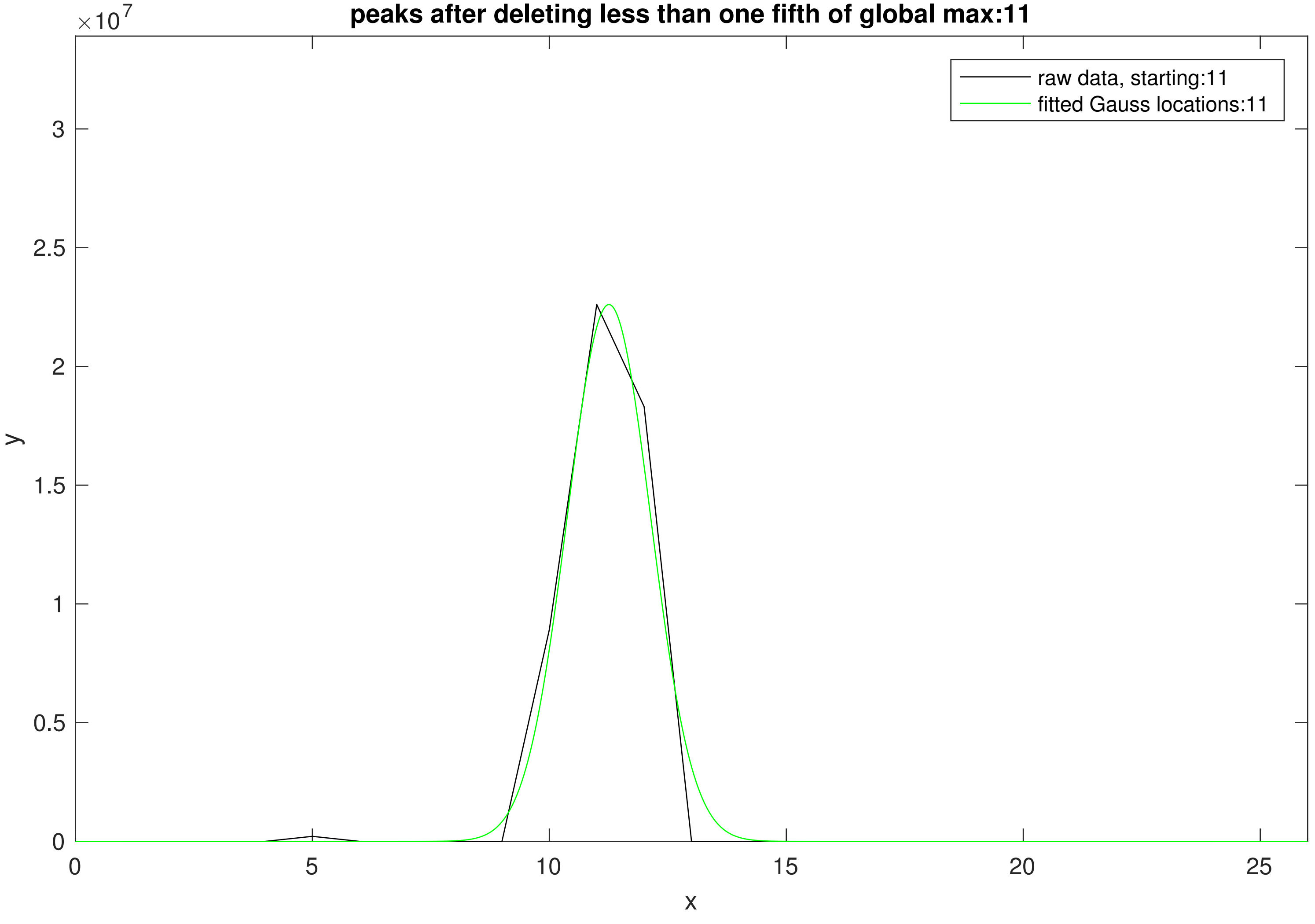

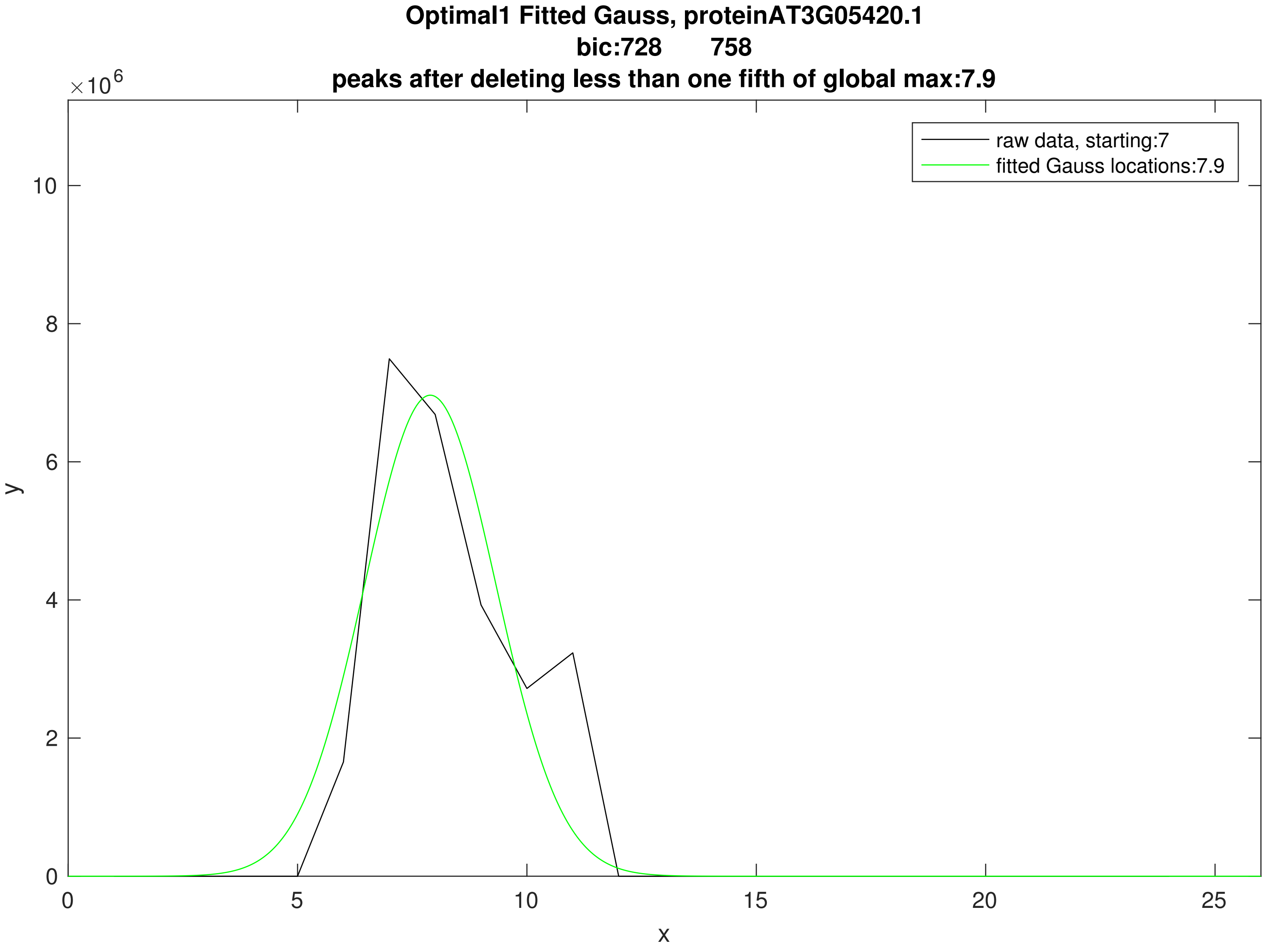
Optimal1 Fitted Gauss, proteinAT3G05420.1
bic:728 758
peaks after deleting less than one fifth of global max:7.9
×10^6
raw data, starting:7
fitted Gauss locations:7.9
y
x
0
2
4
6
8
10
0
5
10
15
20
25

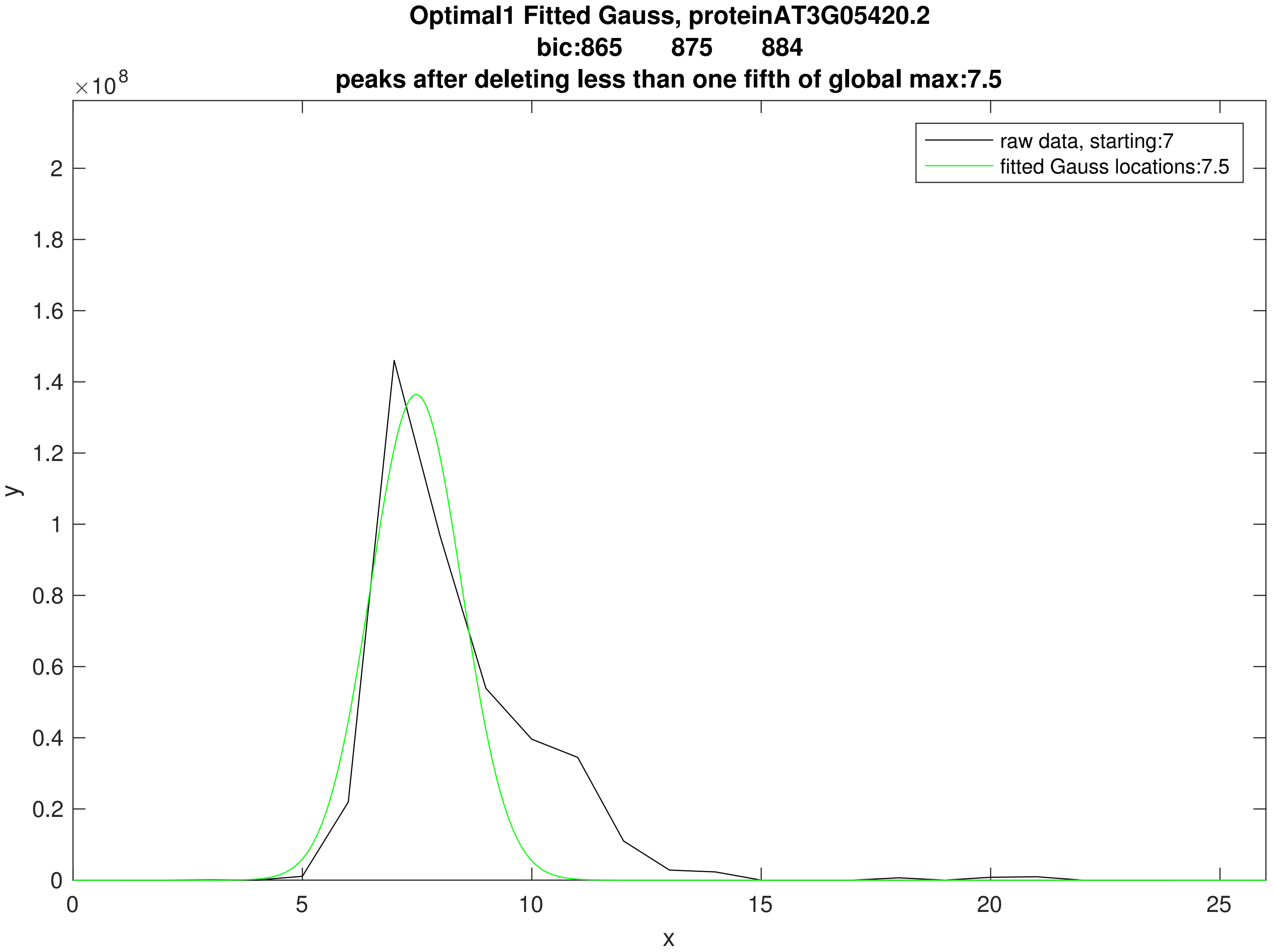
Optimal1 Fitted Gauss, proteinAT3G05420.2
bic:865 875 884
peaks after deleting less than one fifth of global max:7.5
×10^8
raw data, starting:7
fitted Gauss locations:7.5
y
x
0
0.2
0.4
0.6
0.8
1
1.2
1.4
1.6
1.8
2
0
5
10
15
20
25

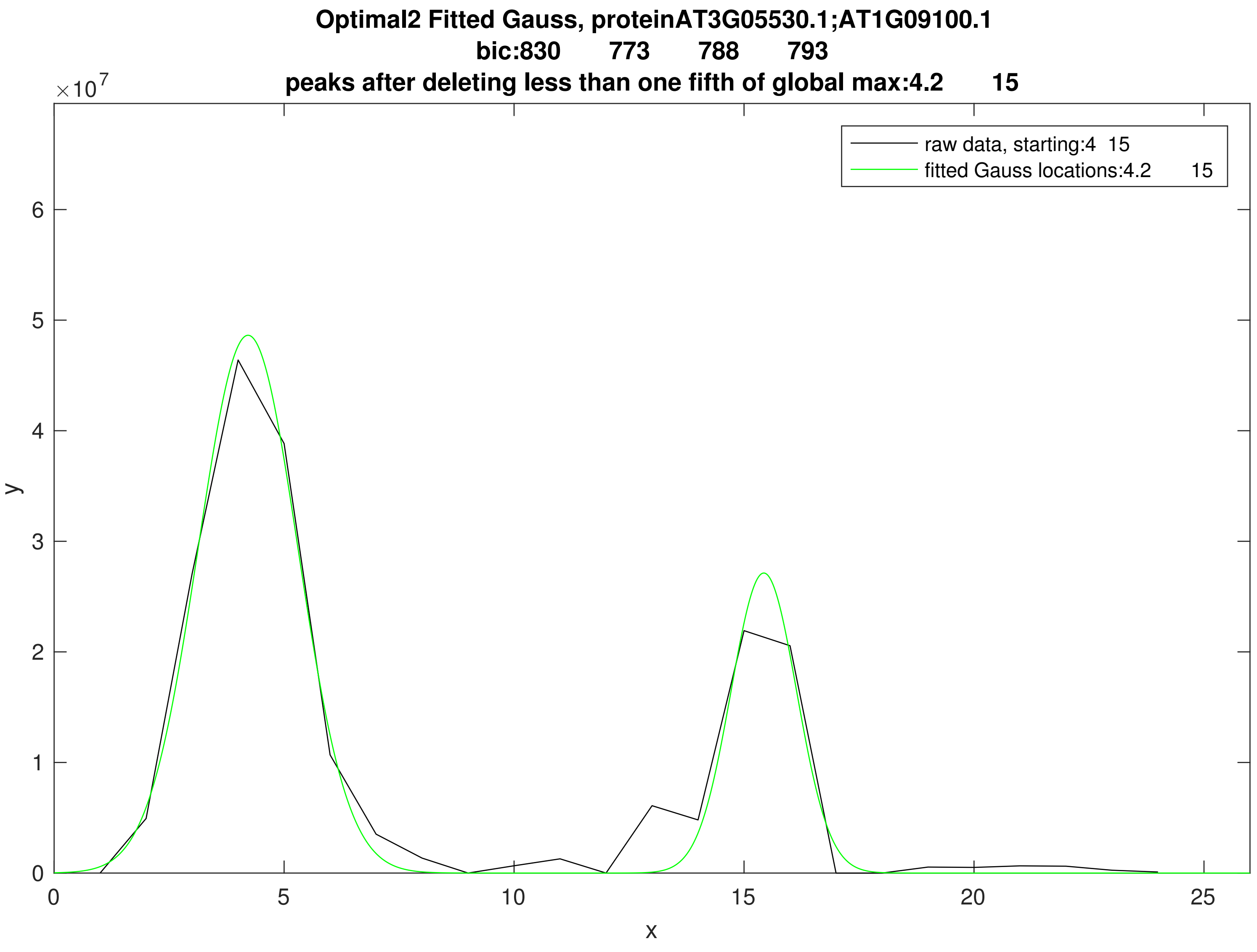

Optimal2 Fitted Gauss, proteinAT3G05530.1;AT1G09100.1
bic:830 773 788 793
peaks after deleting less than one fifth of global max:4.2 15
×10^7
raw data, starting:4 15
fitted Gauss locations:4.2 15
y
x
0
1
2
3
4
5
6
0
5
10
15
20
25

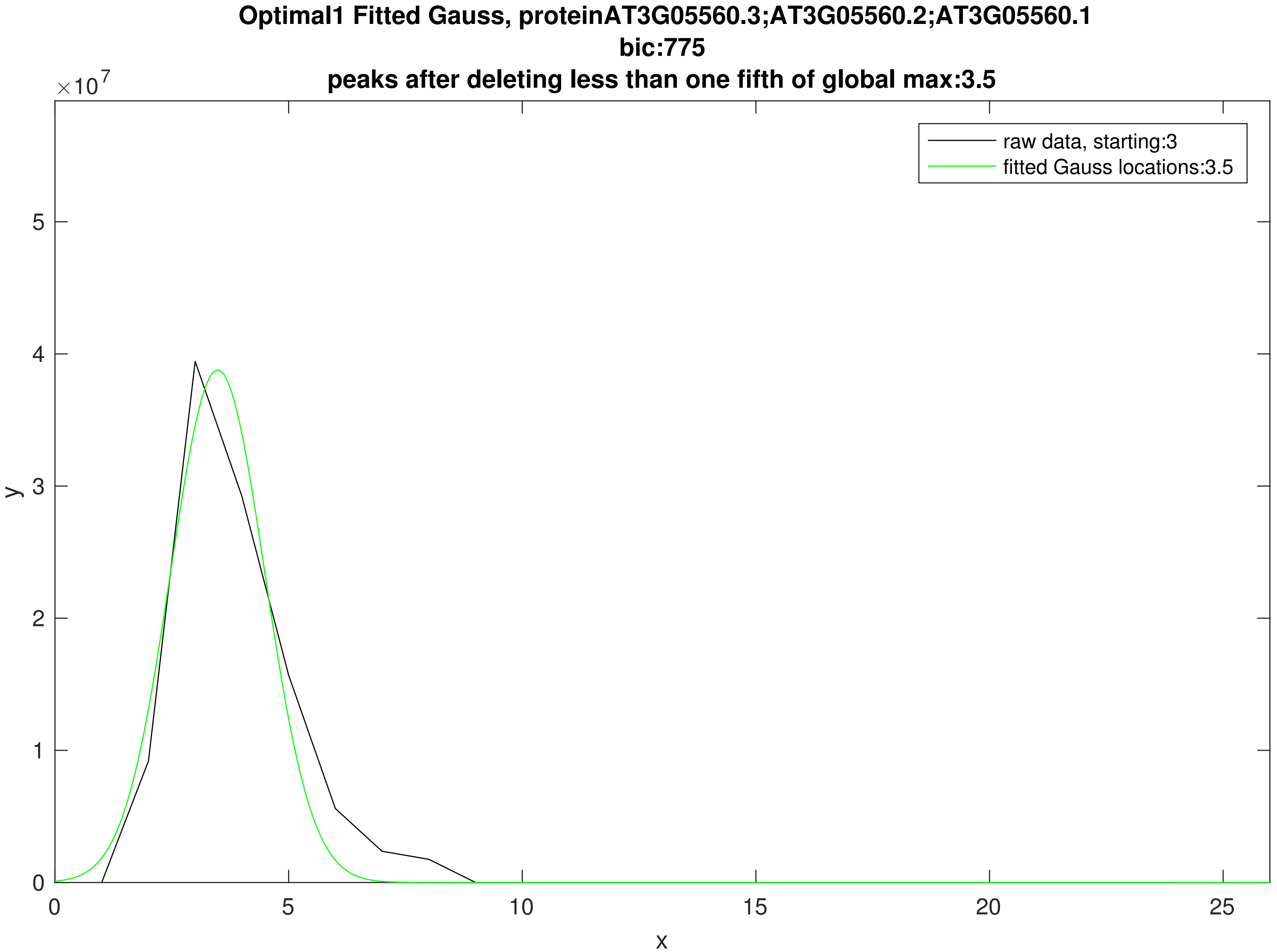
Optimal1 Fitted Gauss, proteinAT3G05560.3;AT3G05560.2;AT3G05560.1
bic:775
peaks after deleting less than one fifth of global max:3.5
×10^7
raw data, starting:3
fitted Gauss locations:3.5
y
x
0
1
2
3
4
5
0
5
10
15
20
25

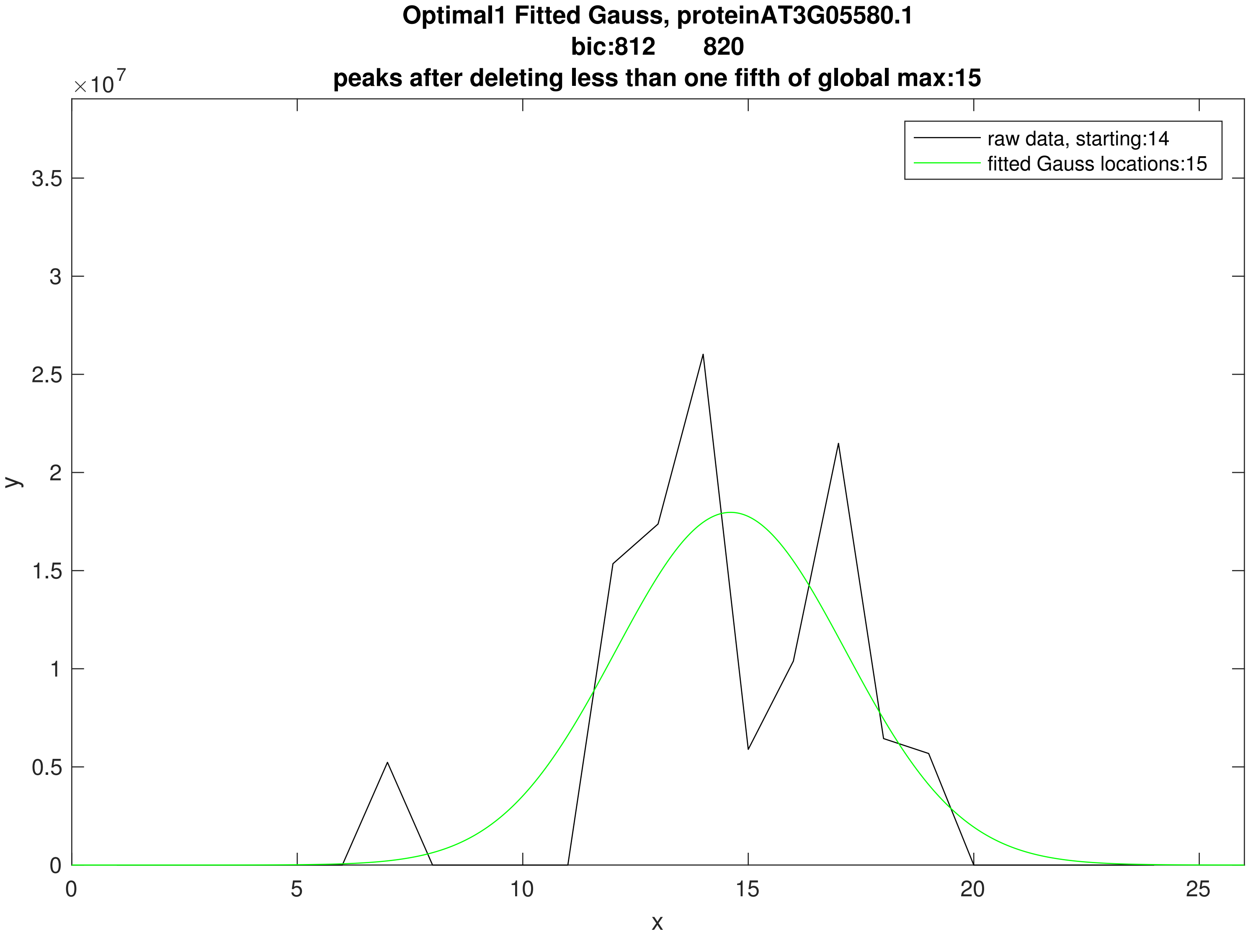
Optimal1 Fitted Gauss, proteinAT3G05580.1
bic:812 820
peaks after deleting less than one fifth of global max:15
×10^7
raw data, starting:14
fitted Gauss locations:15
y
x
0
0.5
1
1.5
2
2.5
3
3.5
5
10
15
20
25

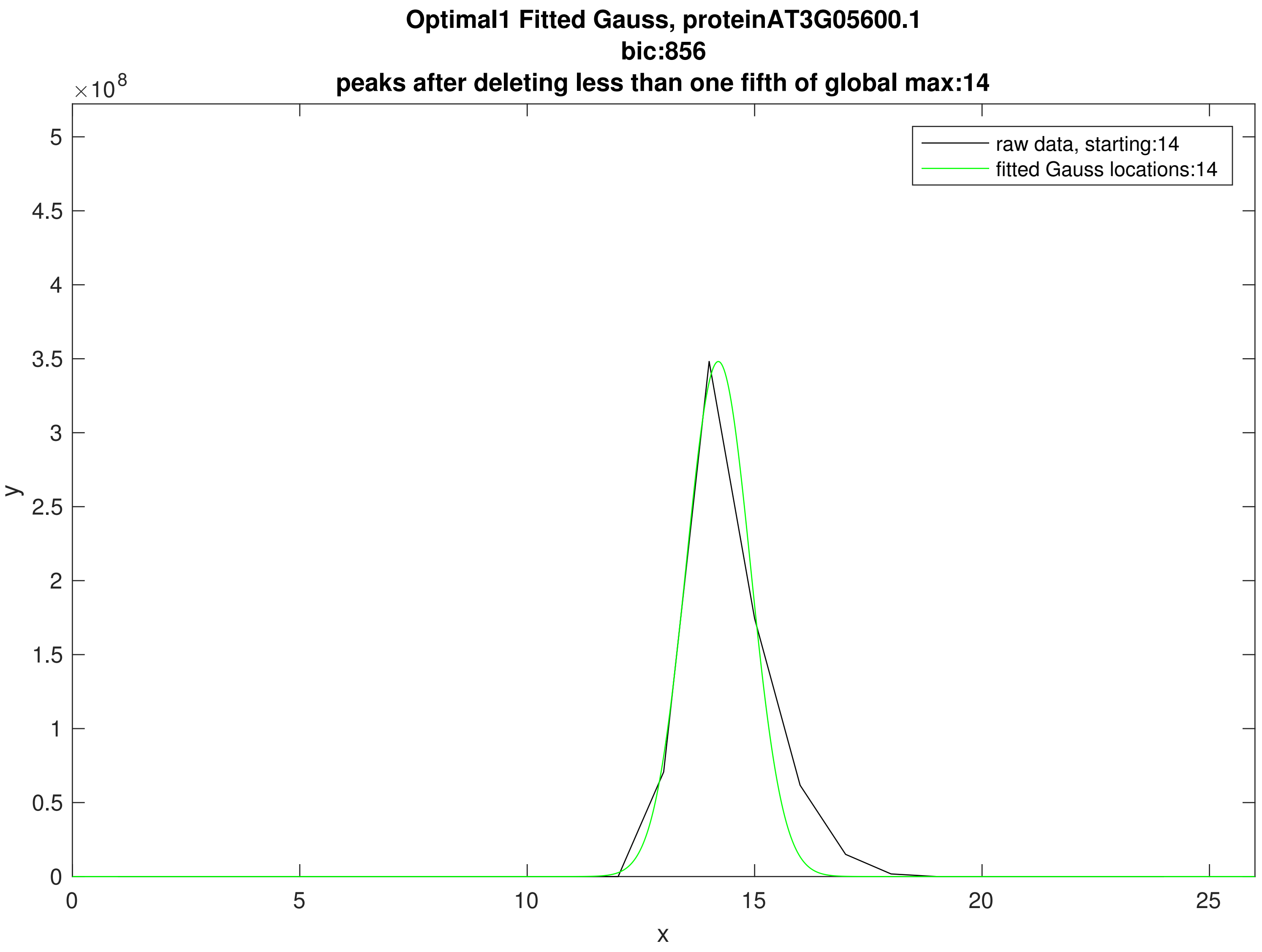

Optimal1 Fitted Gauss, proteinAT3G05600.1
bic:856
peaks after deleting less than one fifth of global max:14
×10^8
raw data, starting:14
fitted Gauss locations:14
y
x
0
0.5
1
1.5
2
2.5
3
3.5
4
4.5
5
0
5
10
15
20
25

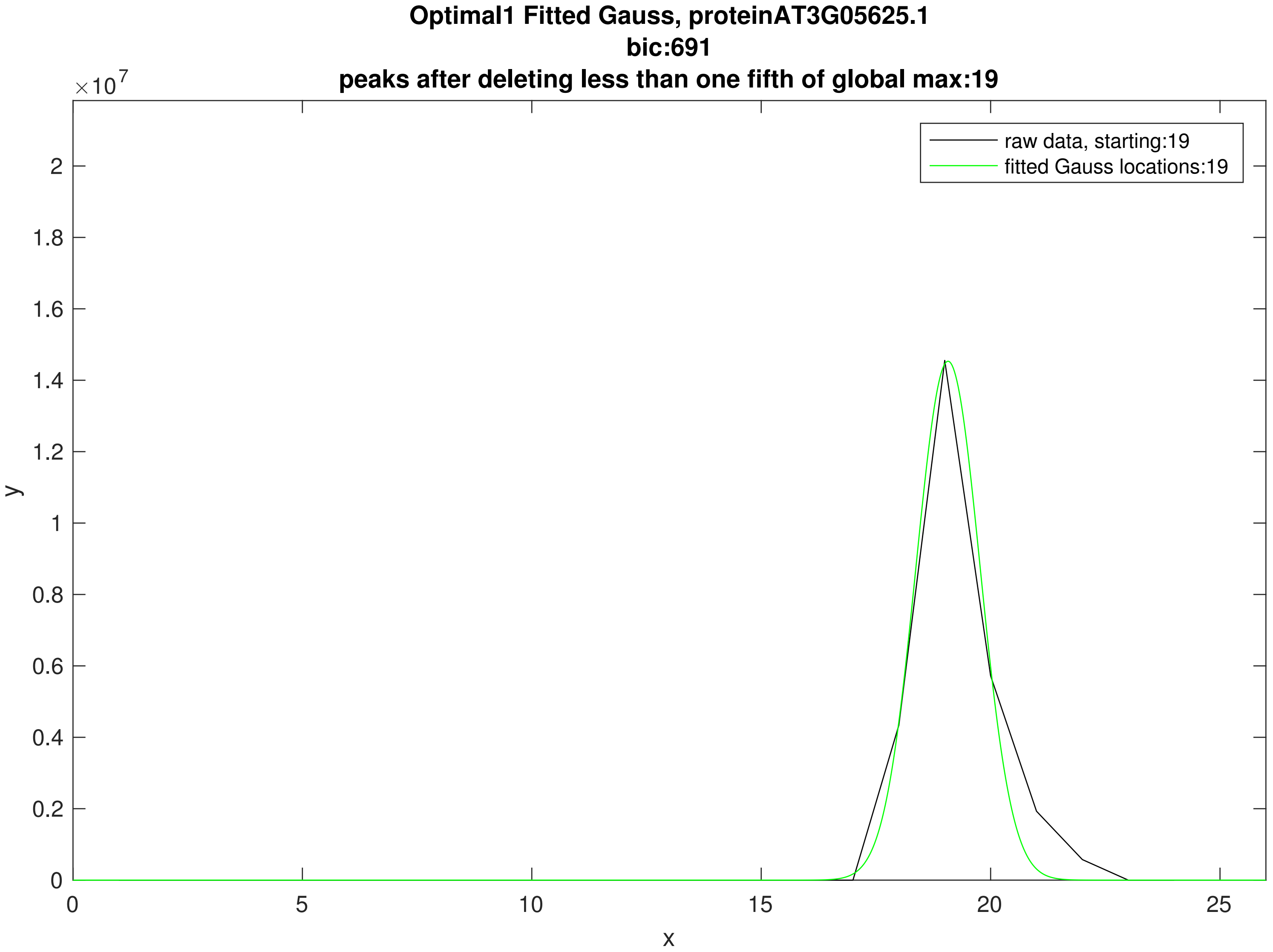

Optimal1 Fitted Gauss, proteinAT3G05625.1
bic:691
peaks after deleting less than one fifth of global max:19
×10^7
2
1.8
1.6
1.4
1.2
1
0.8
0.6
0.4
0.2
0
y
0
5
10
15
20
25
x
raw data, starting:19
fitted Gauss locations:19

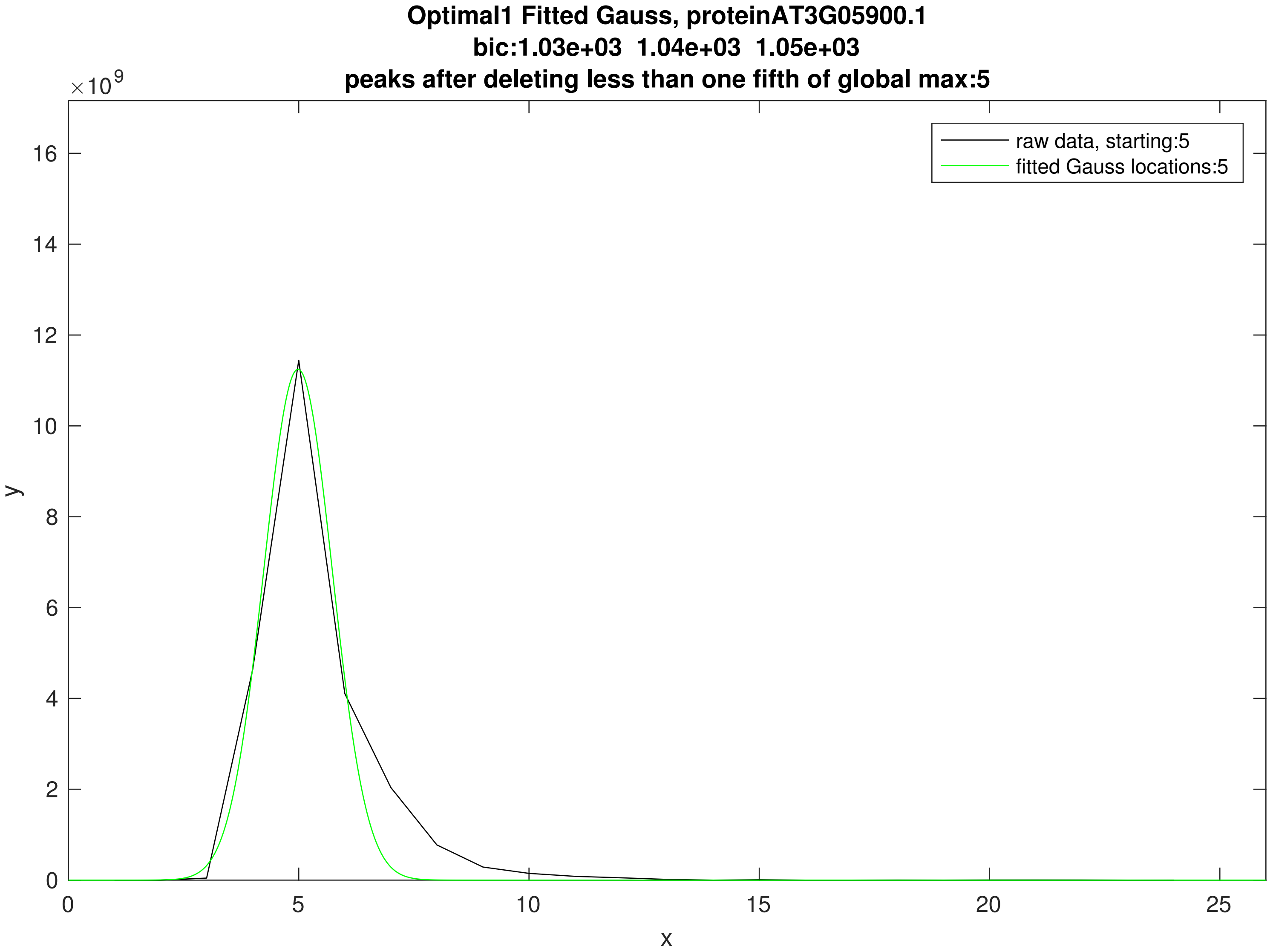
Optimal1 Fitted Gauss, proteinAT3G05900.1
bic:1.03e+03 1.04e+03 1.05e+03
peaks after deleting less than one fifth of global max:5
×10^9
raw data, starting:5
fitted Gauss locations:5
16
14
12
10
8
6
4
2
0
y
0
5
10
15
20
25
x

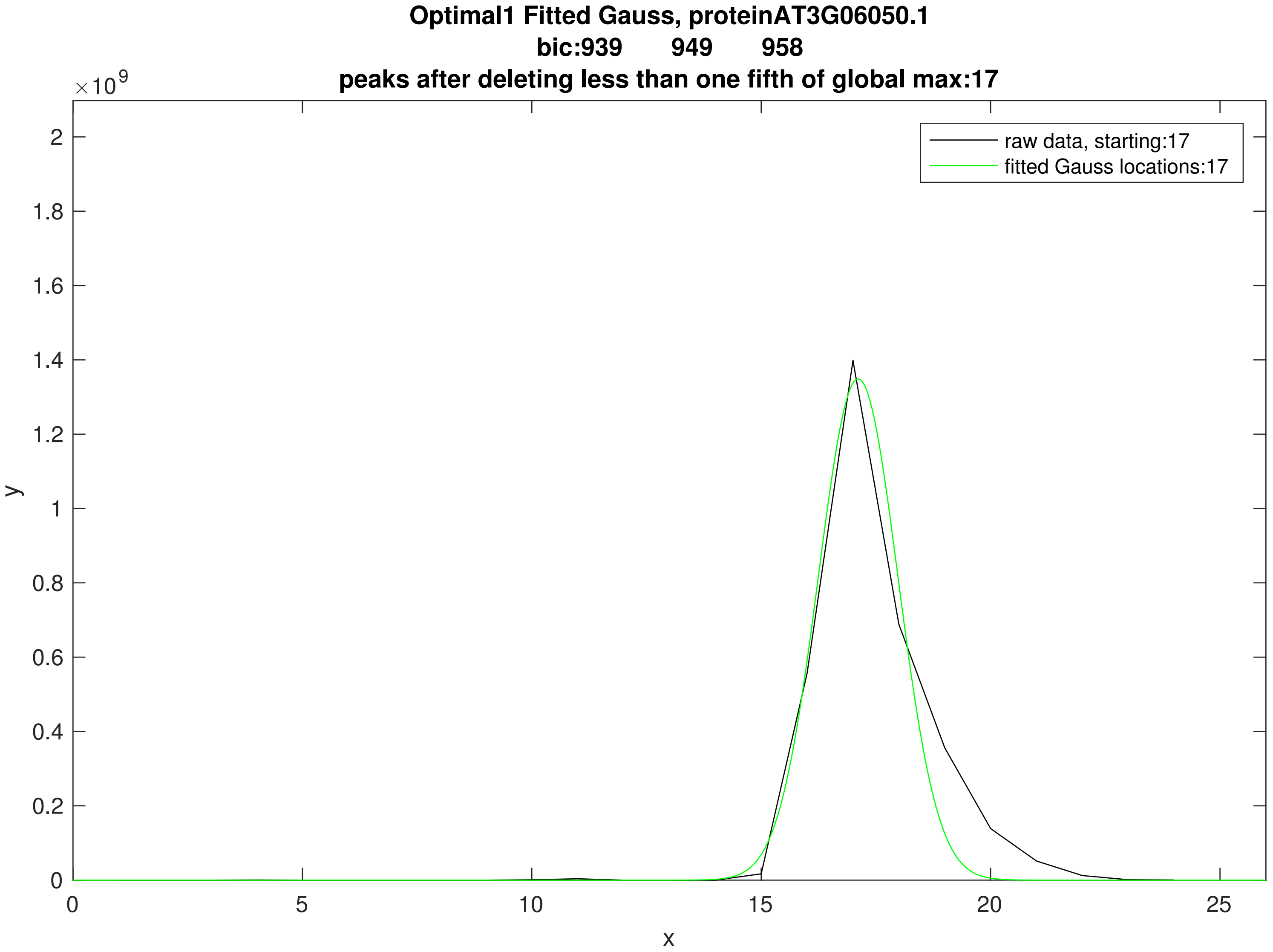
Optimal1 Fitted Gauss, proteinAT3G06050.1
bic:939 949 958
peaks after deleting less than one fifth of global max:17
×10^9
raw data, starting:17
fitted Gauss locations:17
y
x
0
0.2
0.4
0.6
0.8
1
1.2
1.4
1.6
1.8
2
0
5
10
15
20
25

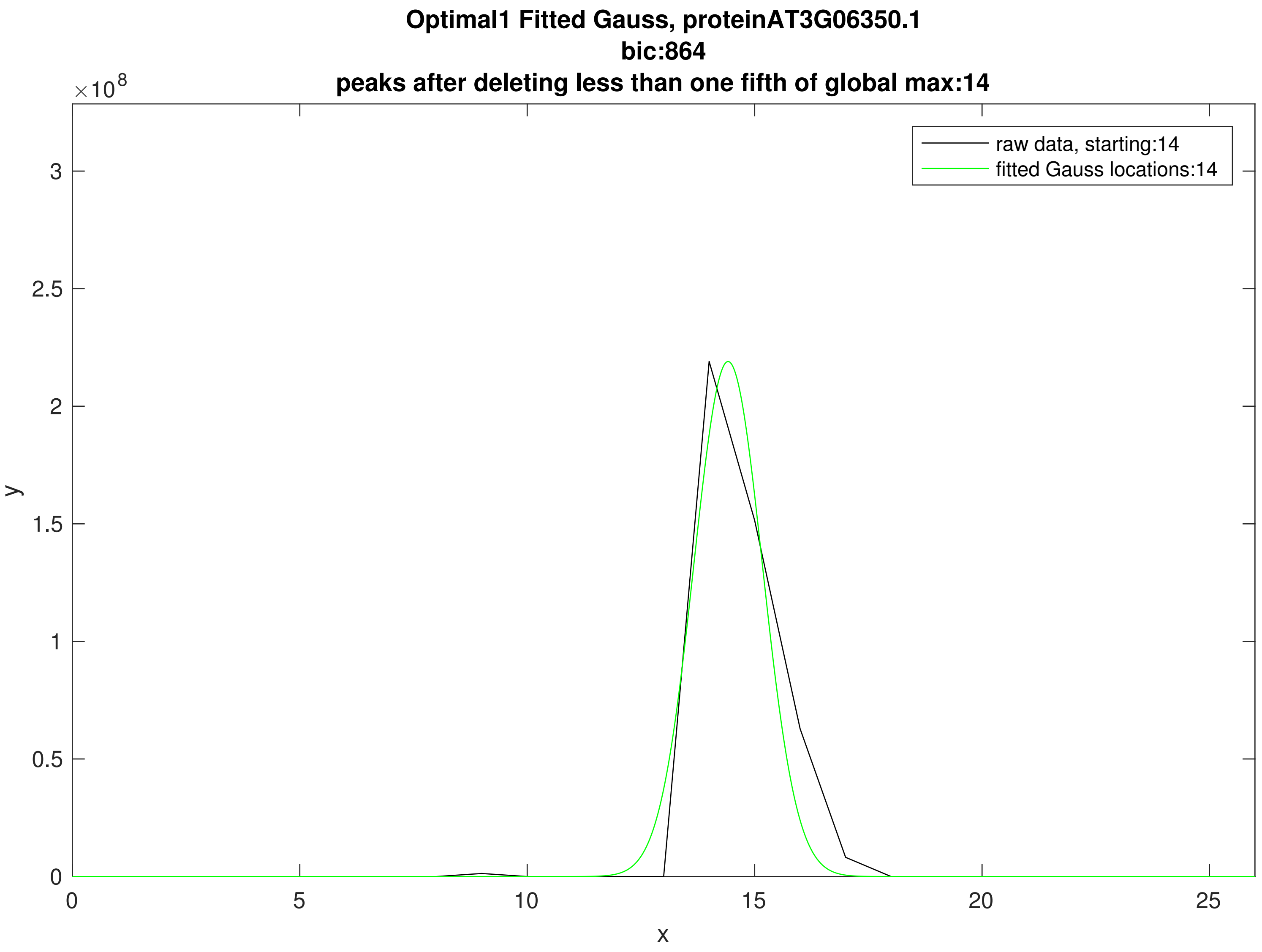
Optimal1 Fitted Gauss, proteinAT3G06350.1
bic:864
peaks after deleting less than one fifth of global max:14
×10^8
raw data, starting:14
fitted Gauss locations:14
y
x
0
0.5
1
1.5
2
2.5
3
0
5
10
15
20
25

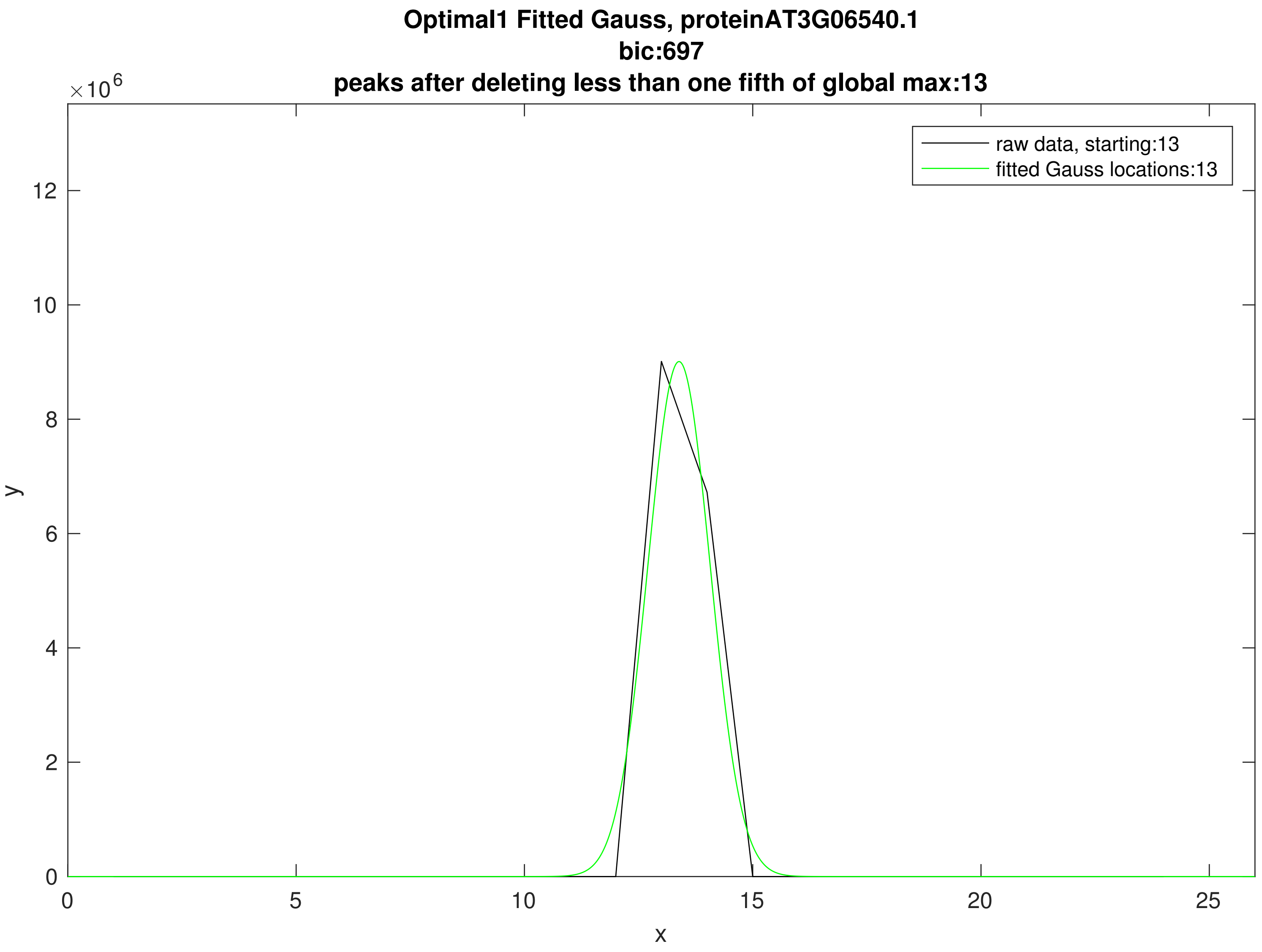

Optimal1 Fitted Gauss, proteinAT3G06540.1
bic:697
peaks after deleting less than one fifth of global max:13
×10^6
raw data, starting:13
fitted Gauss locations:13
y
x
0
2
4
6
8
10
12
0
5
10
15
20
25

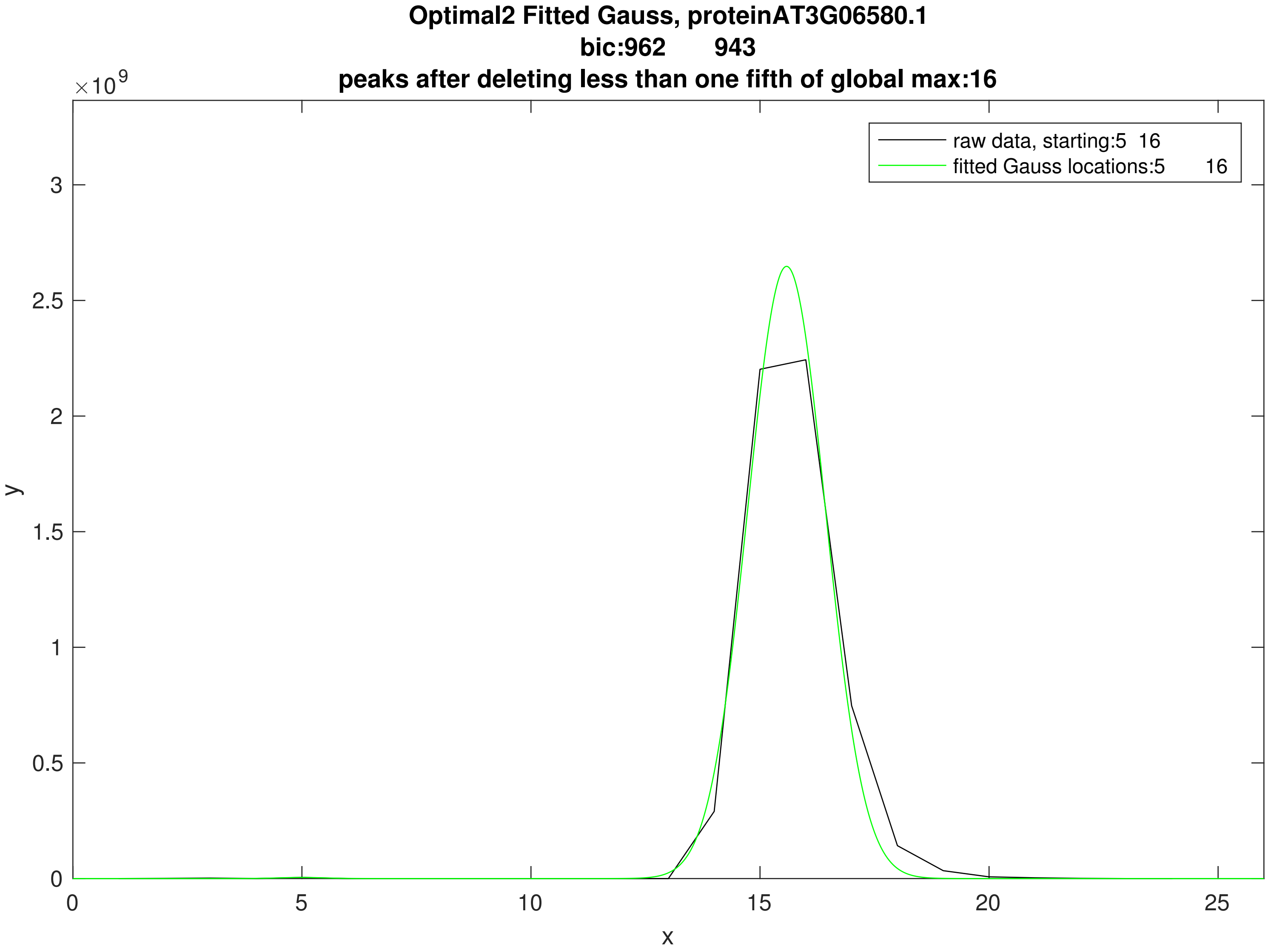

Optimal2 Fitted Gauss, proteinAT3G06580.1
bic:962 943
peaks after deleting less than one fifth of global max:16
×10^9
raw data, starting:5 16
fitted Gauss locations:5 16
3
2.5
2
1.5
1
0.5
0
y
0
5
10
15
20
25
x

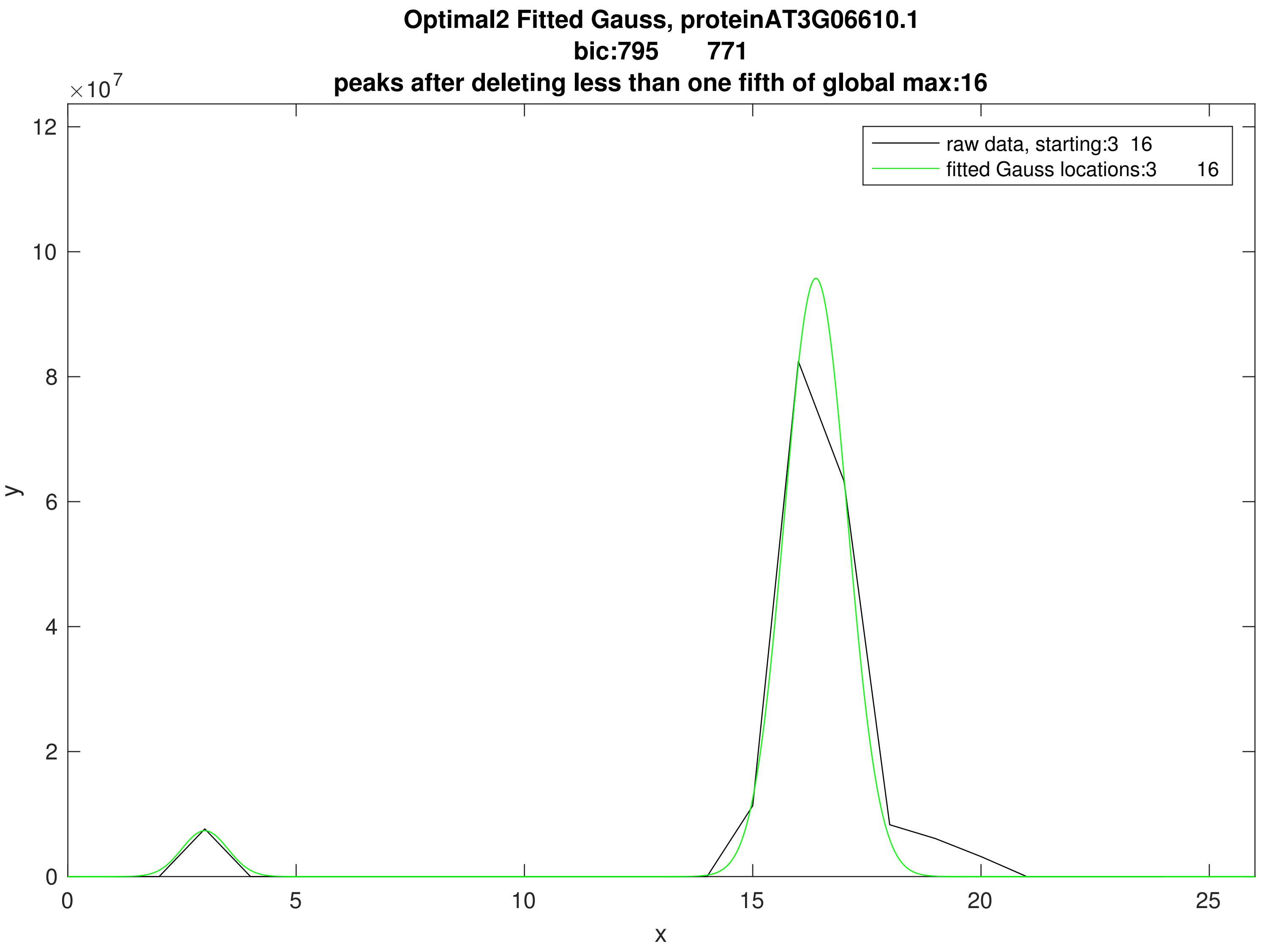
Optimal2 Fitted Gauss, proteinAT3G06610.1
bic:795 771
peaks after deleting less than one fifth of global max:16
×10^7
12
10
8
6
4
2
0
y
0
5
10
15
20
25
x
raw data, starting:3 16
fitted Gauss locations:3 16

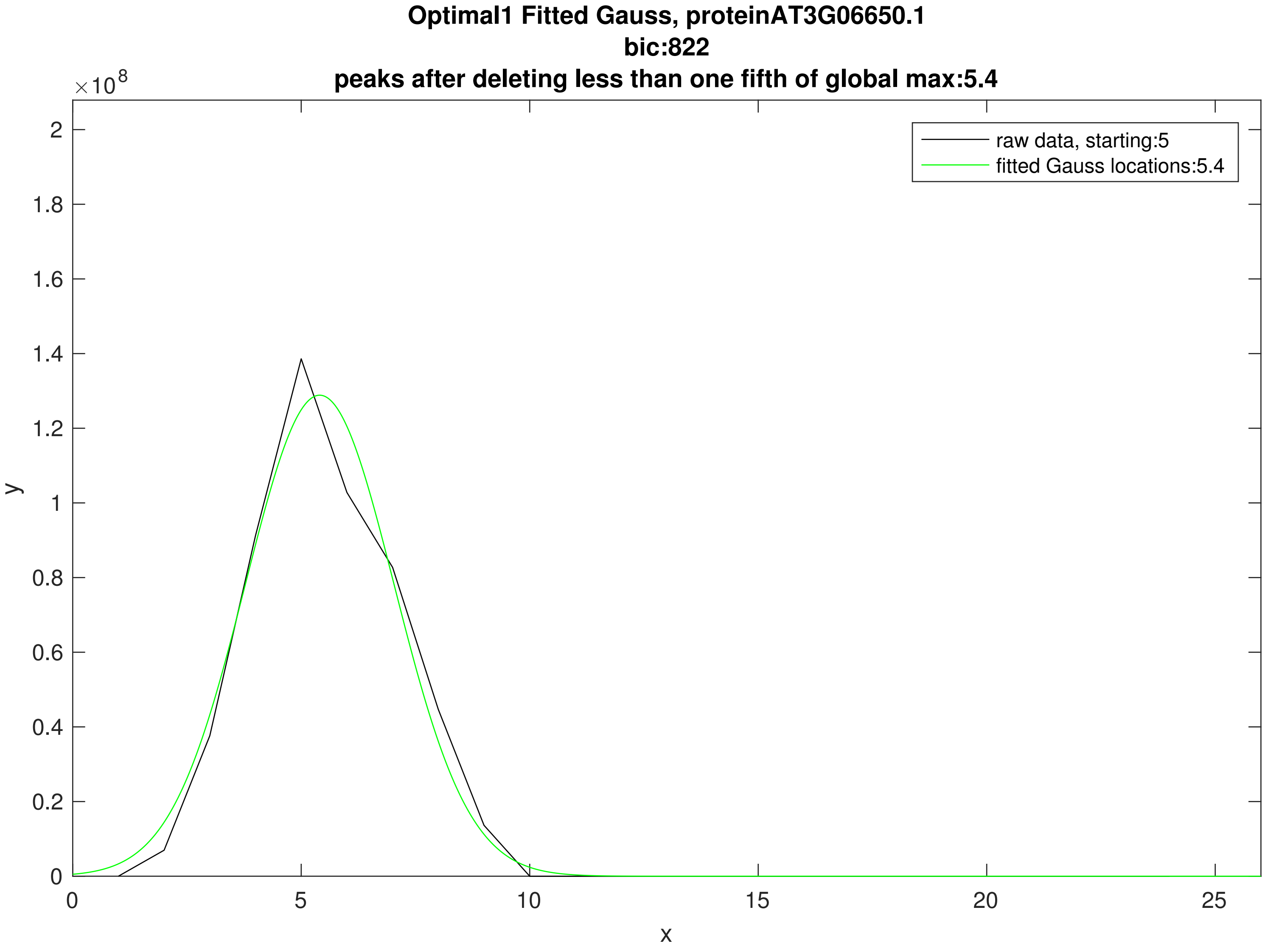

Optimal1 Fitted Gauss, proteinAT3G06650.1
bic:822
peaks after deleting less than one fifth of global max:5.4
×10^8
raw data, starting:5
fitted Gauss locations:5.4
y
x
0
0.2
0.4
0.6
0.8
1
1.2
1.4
1.6
1.8
2
0
5
10
15
20
25

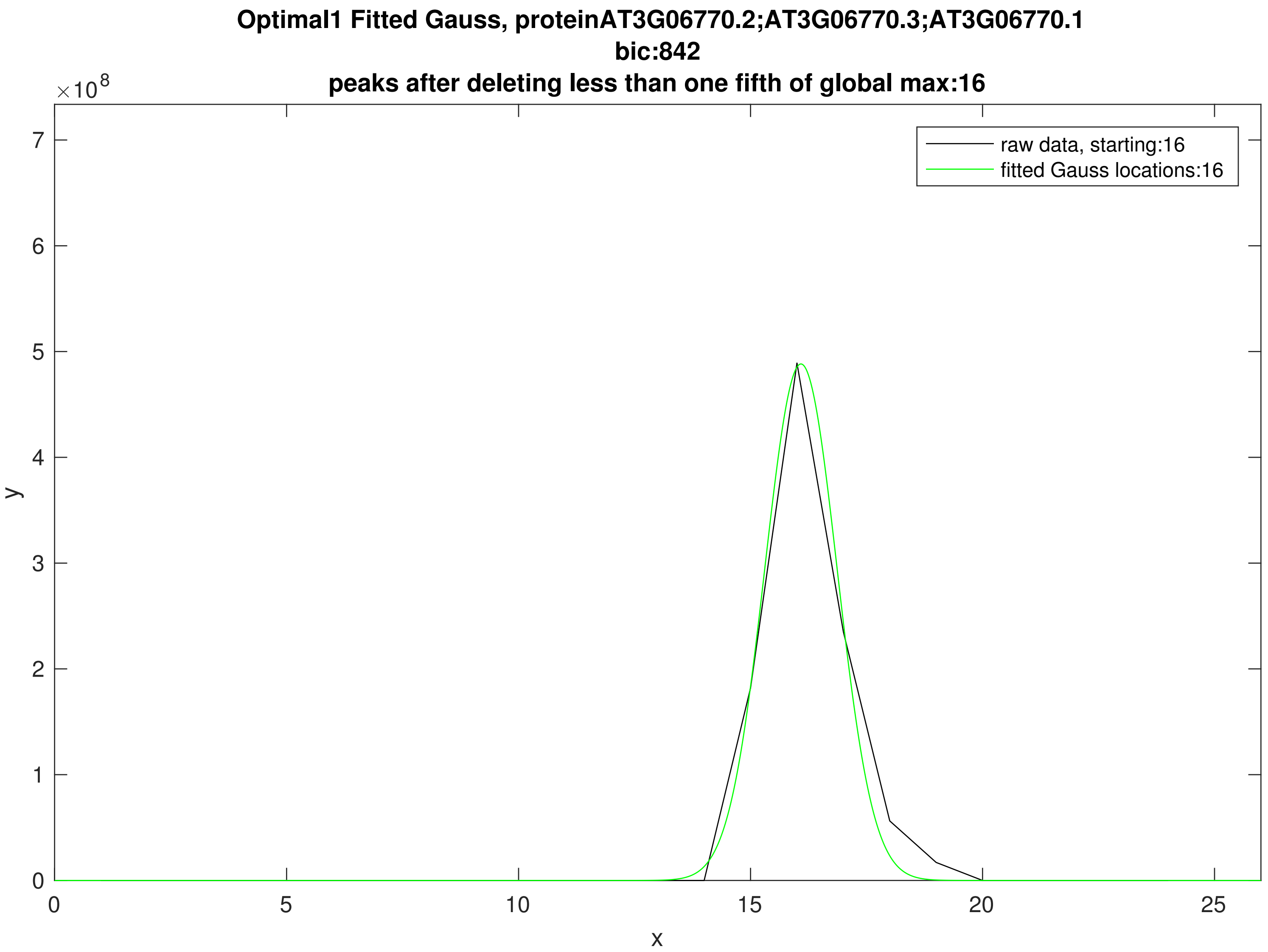

Optimal1 Fitted Gauss, proteinAT3G06770.2;AT3G06770.3;AT3G06770.1
bic:842
peaks after deleting less than one fifth of global max:16
×10^8
raw data, starting:16
fitted Gauss locations:16
0
1
2
3
4
5
6
7
y
0
5
10
15
20
25
x

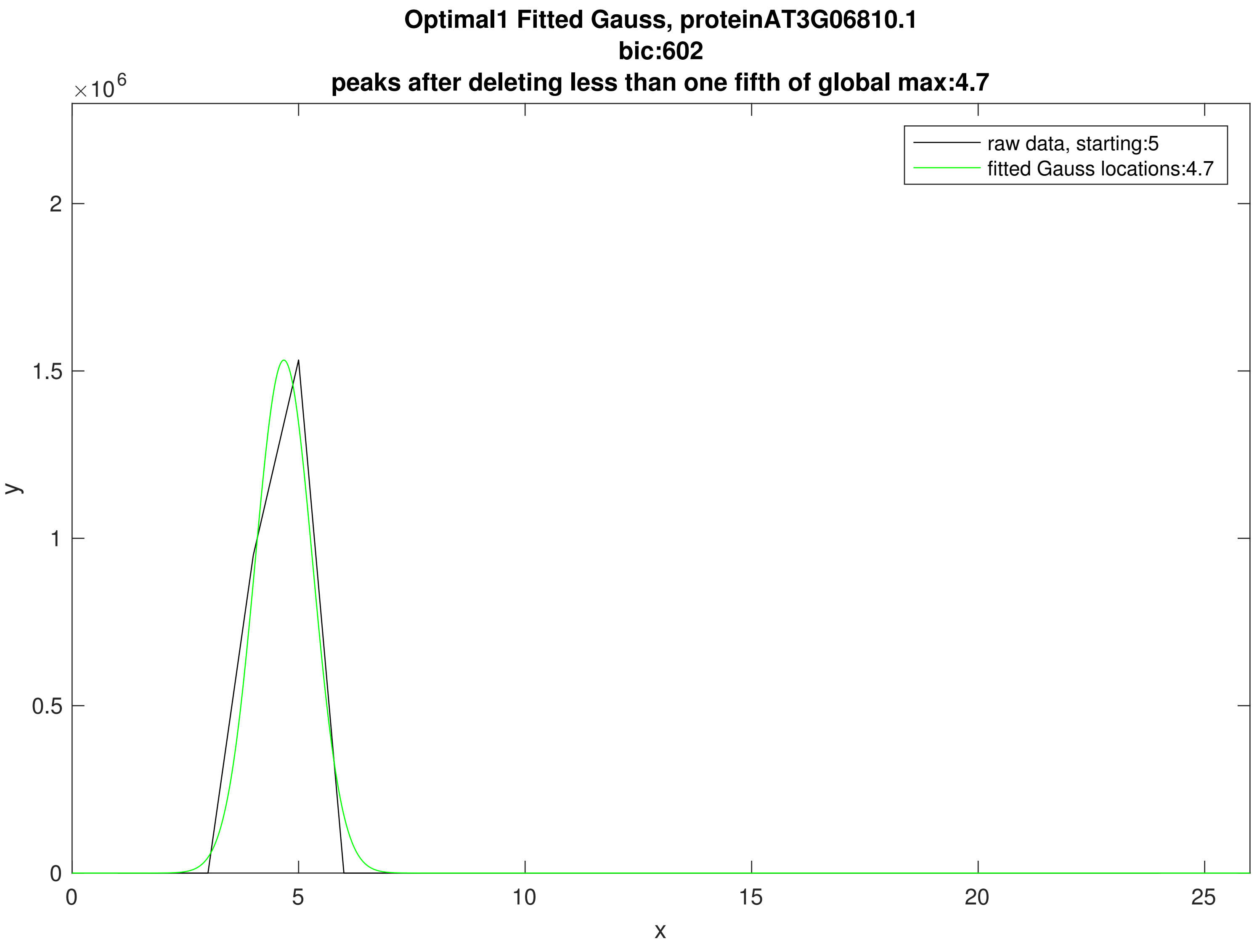
Optimal1 Fitted Gauss, proteinAT3G06810.1
bic:602
peaks after deleting less than one fifth of global max:4.7
×10^6
raw data, starting:5
fitted Gauss locations:4.7
2
1.5
1
0.5
0
y
0
5
10
15
20
25
x

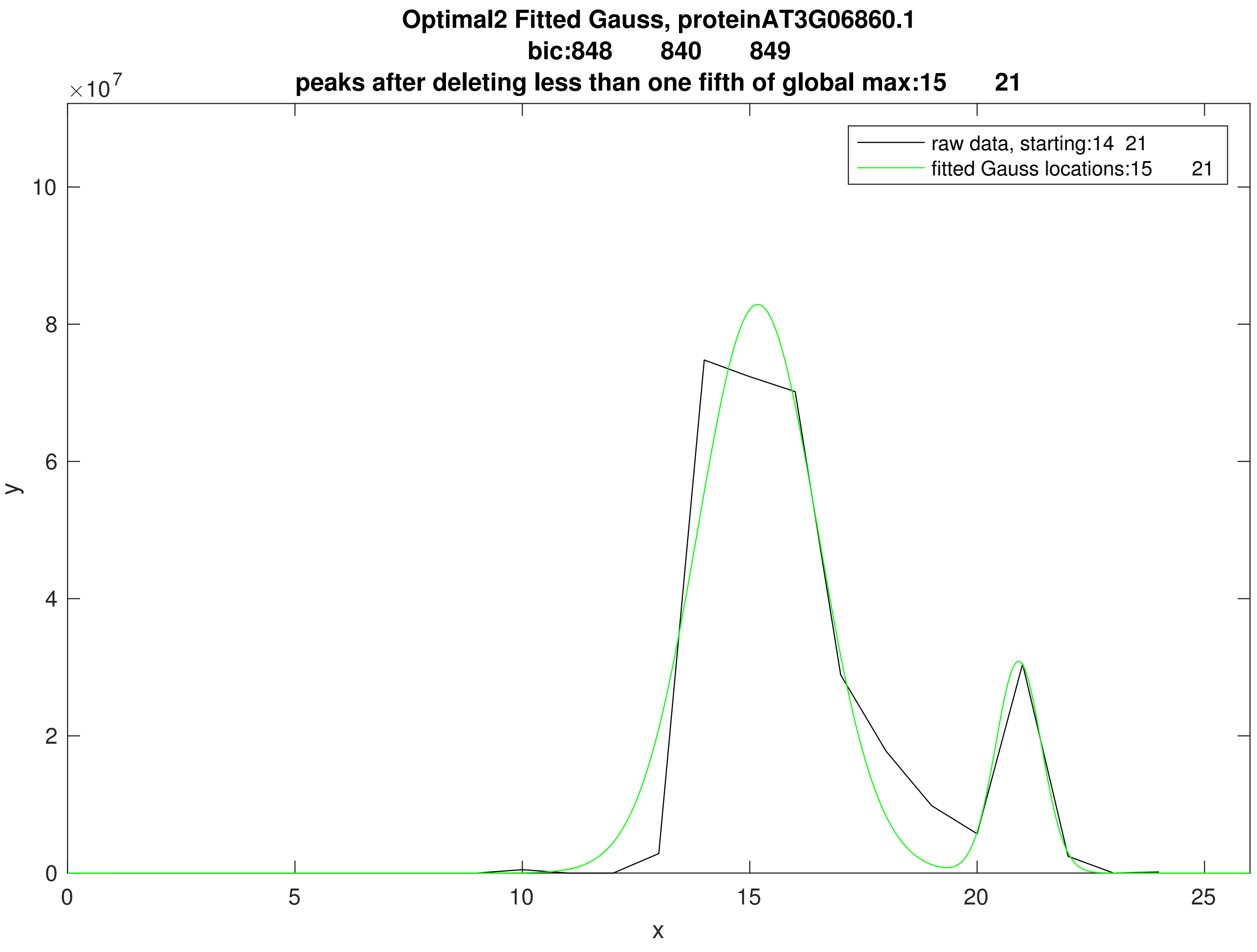
Optimal2 Fitted Gauss, proteinAT3G06860.1
bic:848 840 849
peaks after deleting less than one fifth of global max:15 21
×10^7
raw data, starting:14 21
fitted Gauss locations:15 21
10
8
6
4
2
0
y
0
5
10
15
20
25
x

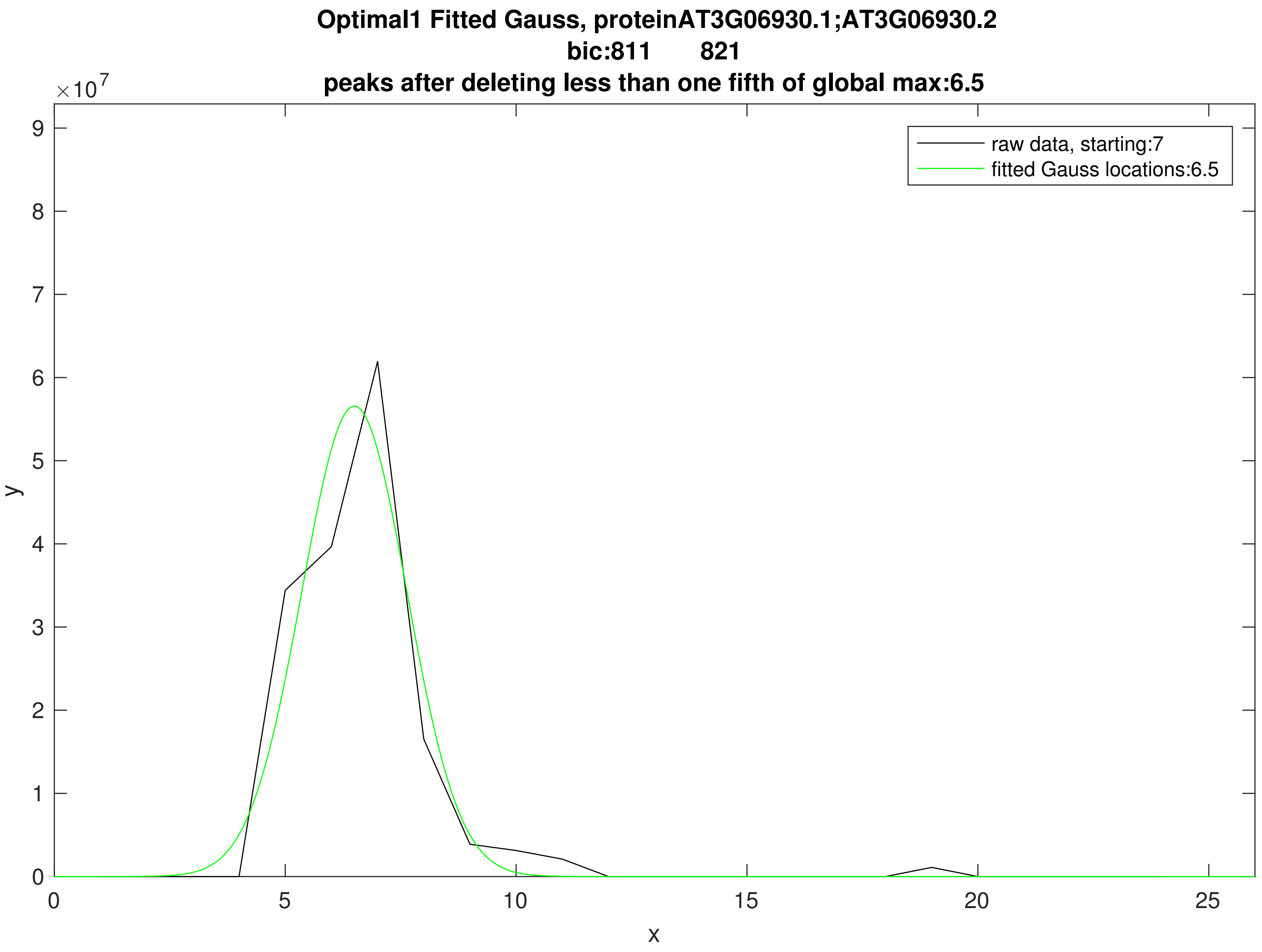
Optimal1 Fitted Gauss, proteinAT3G06930.1;AT3G06930.2
bic:811 821
peaks after deleting less than one fifth of global max:6.5
×10^7
raw data, starting:7
fitted Gauss locations:6.5
y
x
0
1
2
3
4
5
6
7
8
9
0
5
10
15
20
25

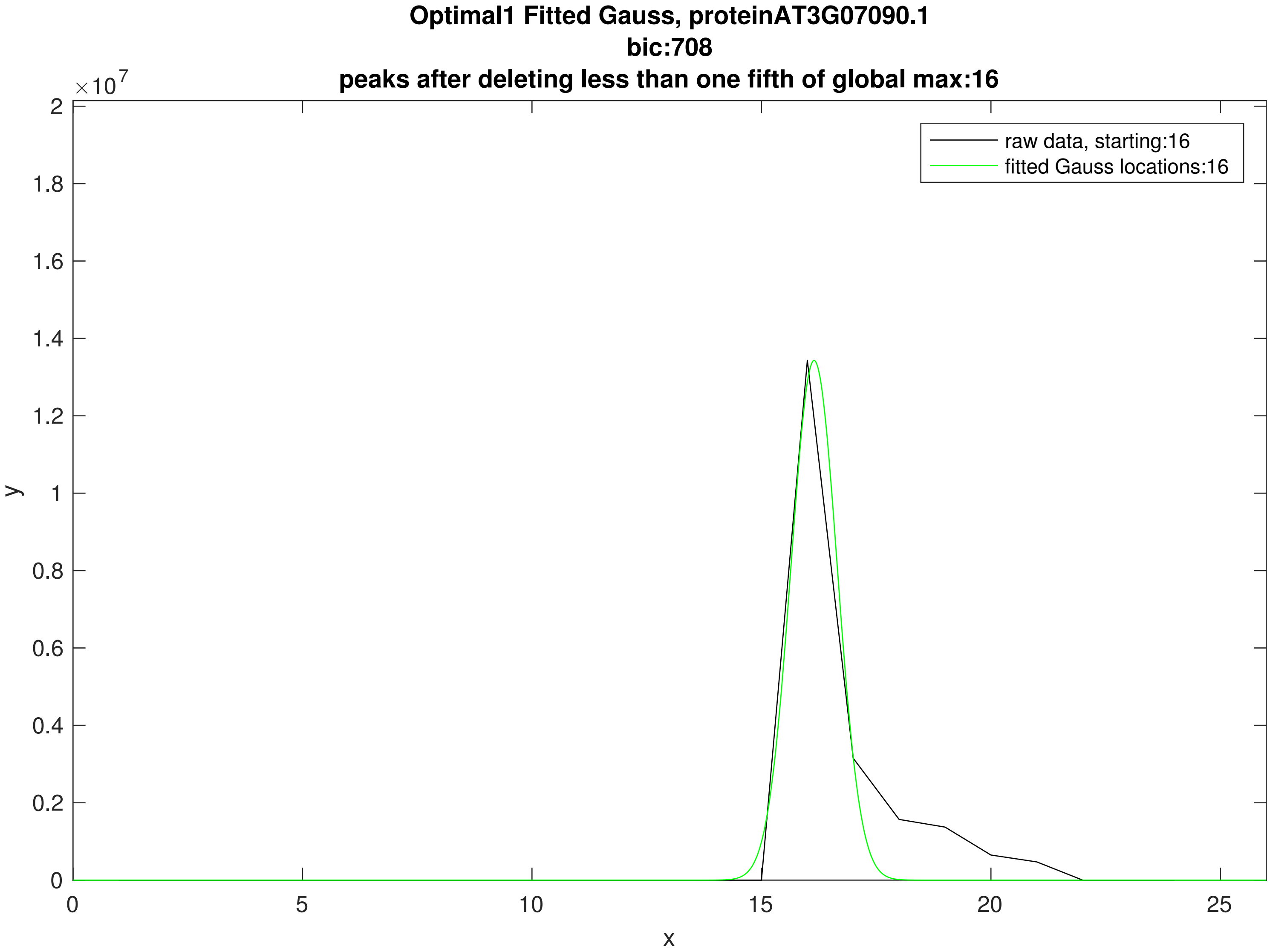
Optimal1 Fitted Gauss, proteinAT3G07090.1
bic:708
peaks after deleting less than one fifth of global max:16
×10^7
raw data, starting:16
fitted Gauss locations:16
y
x
0
0.2
0.4
0.6
0.8
1
1.2
1.4
1.6
1.8
2
0
5
10
15
20
25

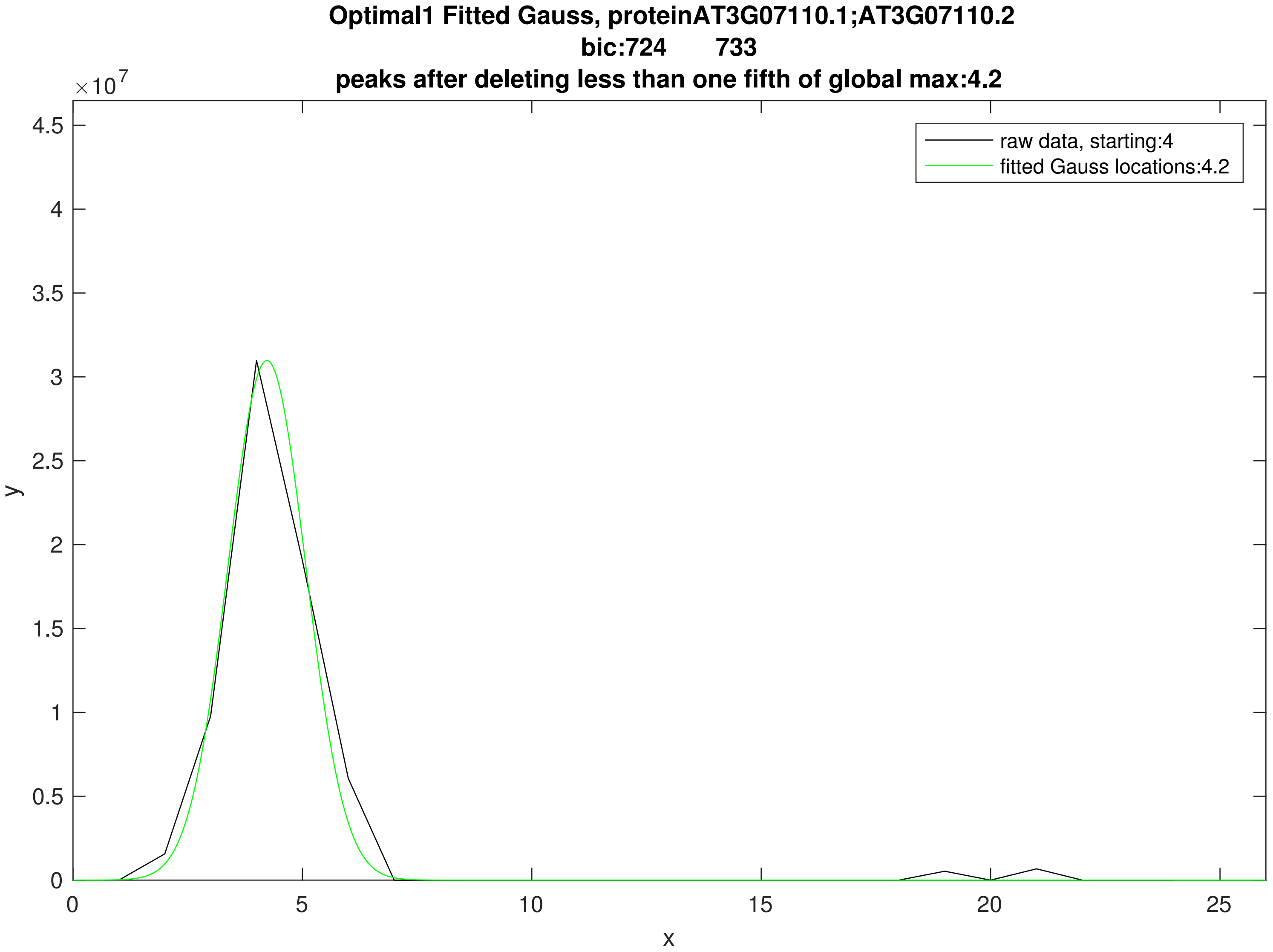

Optimal1 Fitted Gauss, proteinAT3G07110.1;AT3G07110.2
bic:724 733
peaks after deleting less than one fifth of global max:4.2
×10^7
4.5
4
3.5
3
2.5
2
1.5
1
0.5
0
y
0
5
10
15
20
25
x
raw data, starting:4
fitted Gauss locations:4.2

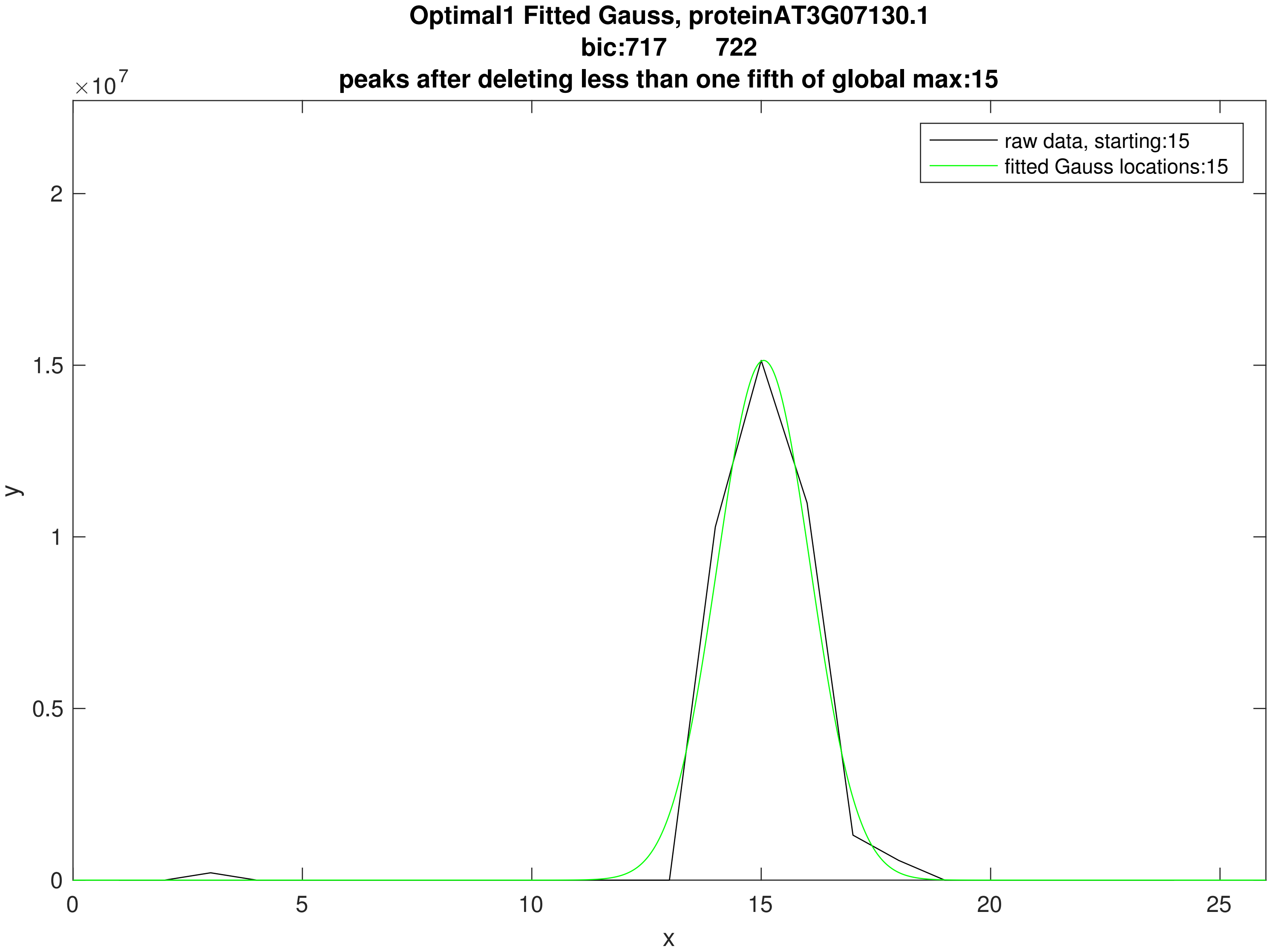

Optimal1 Fitted Gauss, proteinAT3G07130.1
bic:717 722
peaks after deleting less than one fifth of global max:15
×10^7
2
1.5
1
0.5
0
y
0
5
10
15
20
25
x
raw data, starting:15
fitted Gauss locations:15

**Optimal1 Fitted Gauss, proteinAT3G07390.1**
**bic:787 791**
**peaks after deleting less than one fifth of global max:12**

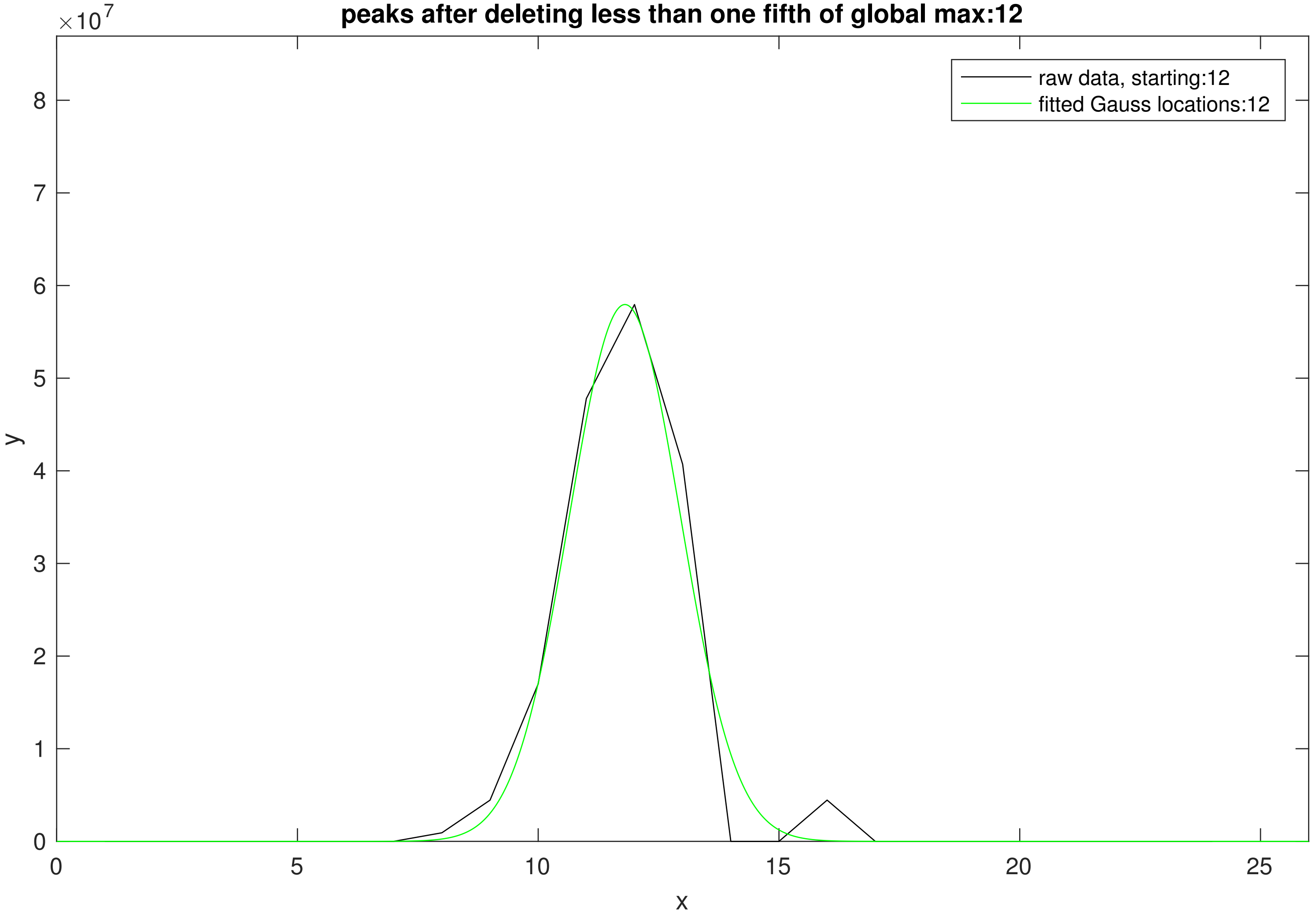

**Optimal2 Fitted Gauss, proteinAT3G07460.1;AT3G07460.2**
**bic:845 824**
**peaks after deleting less than one fifth of global max:20**

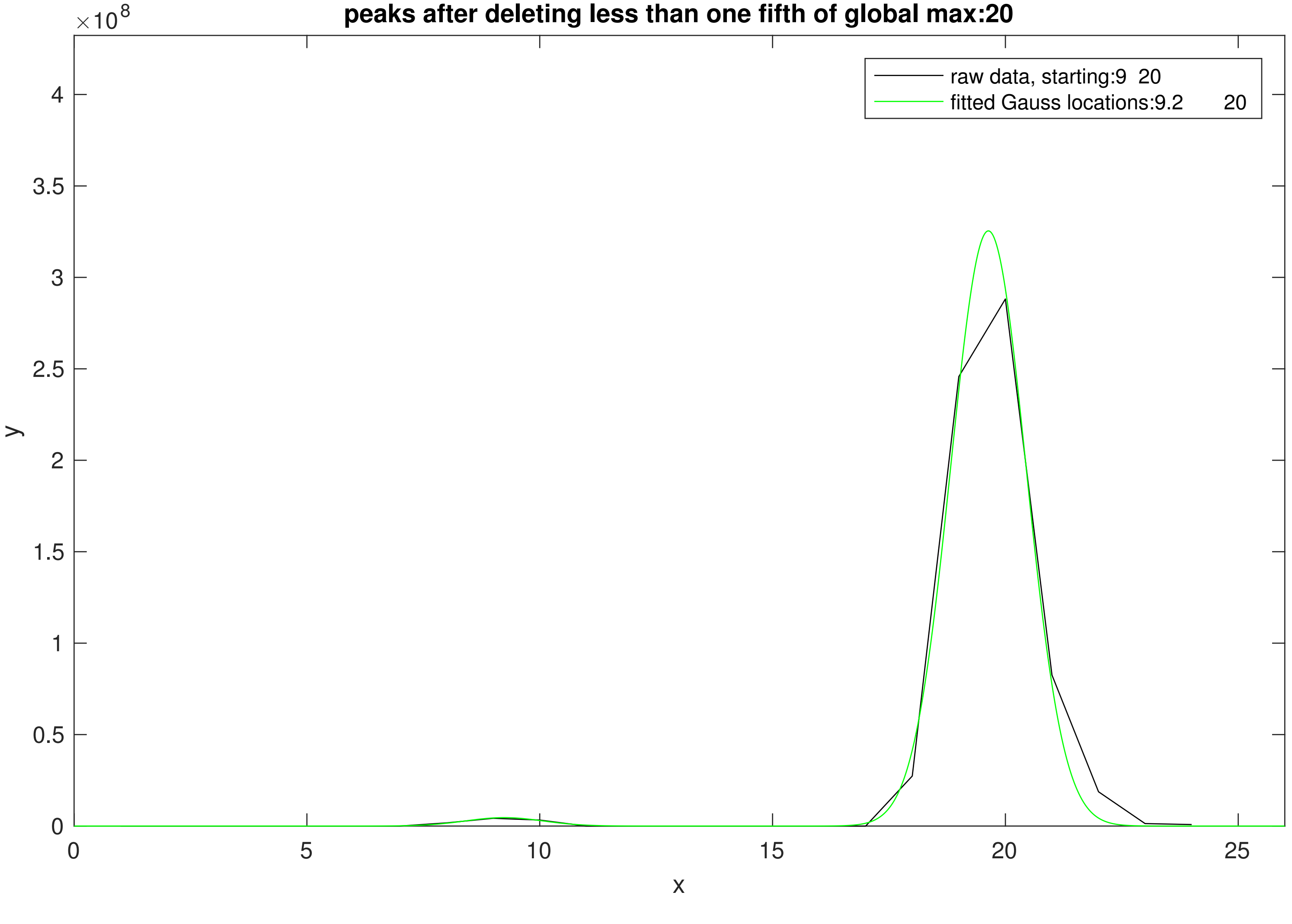

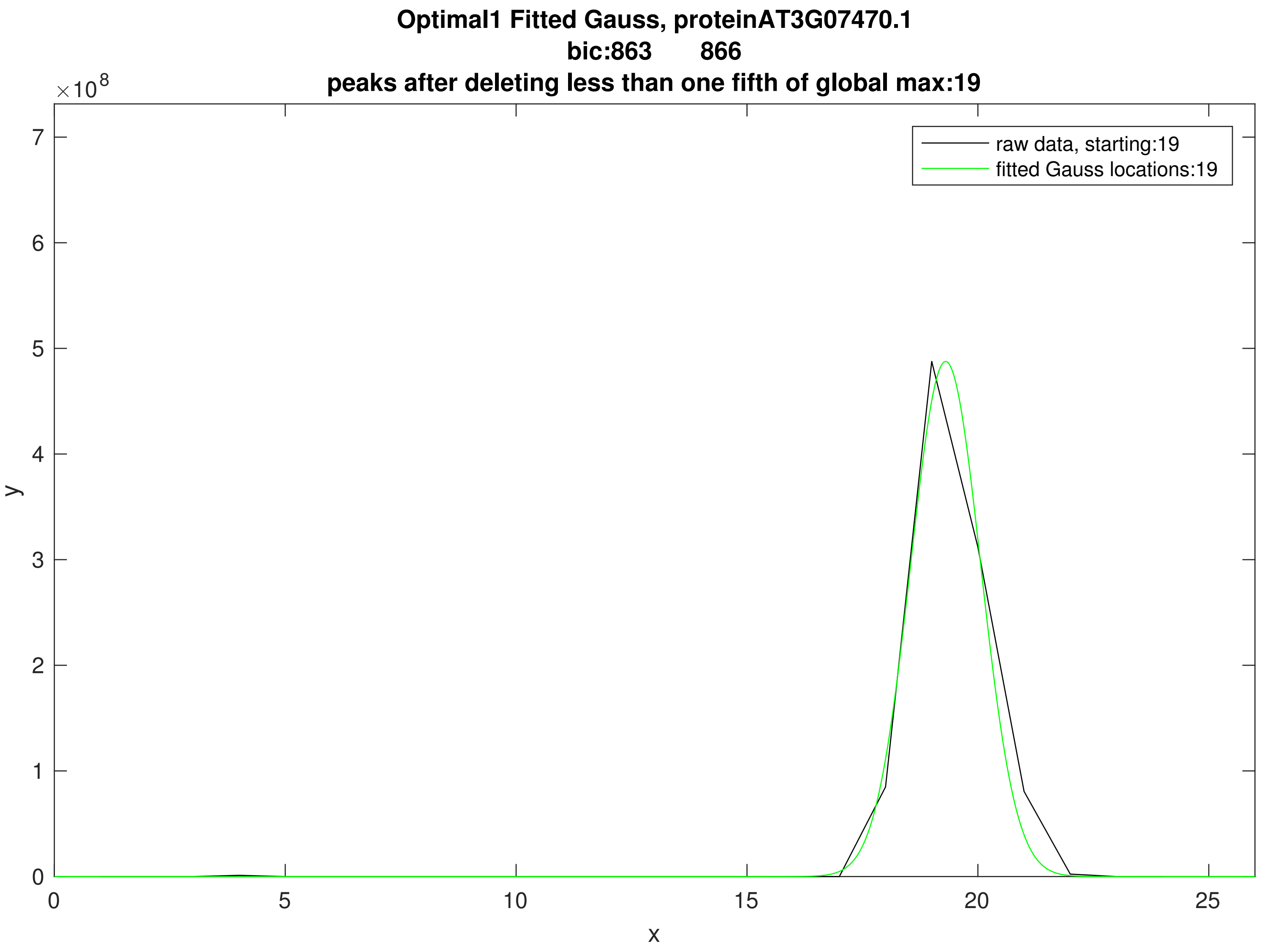
Optimal1 Fitted Gauss, proteinAT3G07470.1
bic:863 866
peaks after deleting less than one fifth of global max:19
×10^8
raw data, starting:19
fitted Gauss locations:19
y
x
0
1
2
3
4
5
6
7
0
5
10
15
20
25

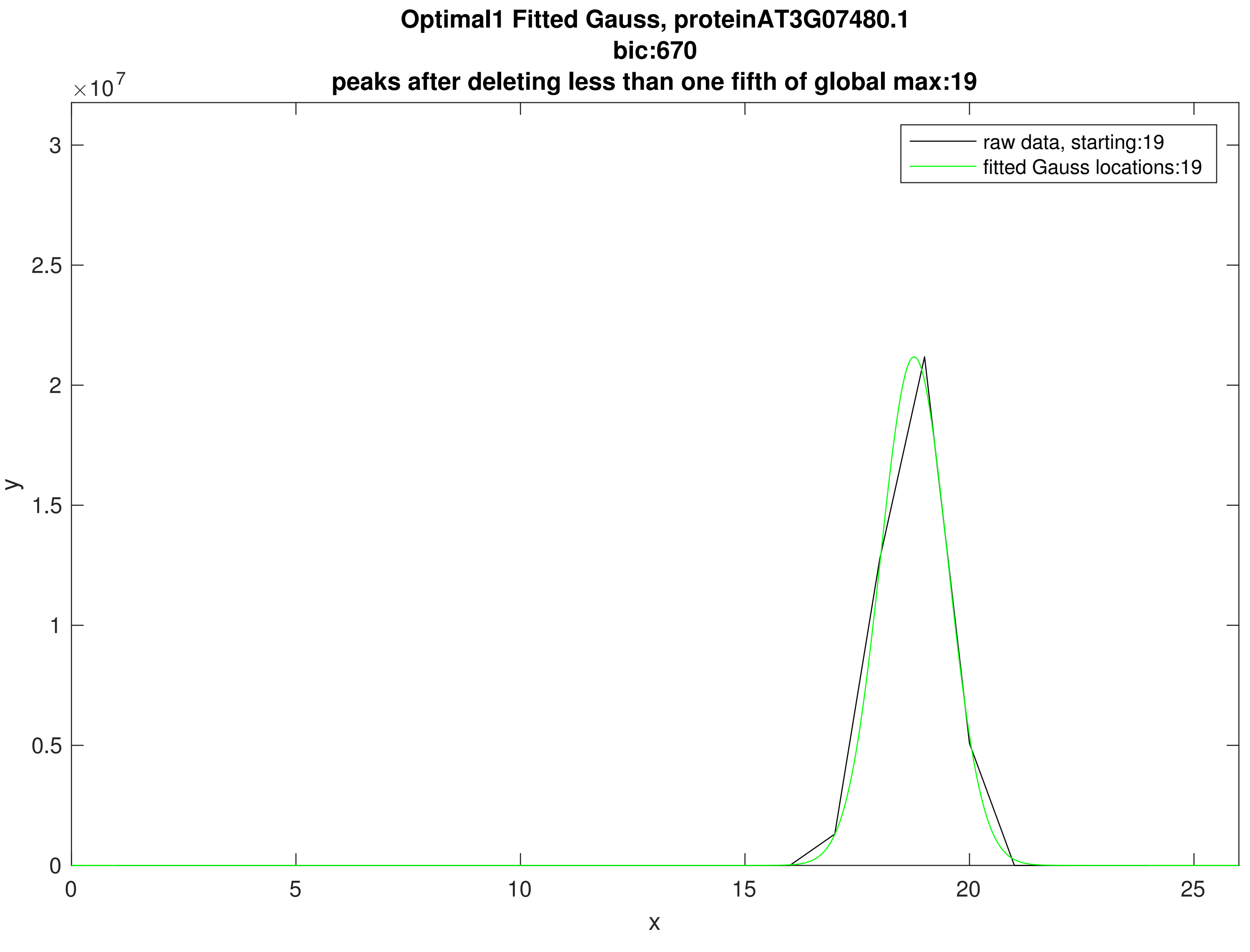
Optimal1 Fitted Gauss, proteinAT3G07480.1
bic:670
peaks after deleting less than one fifth of global max:19
×10^7
raw data, starting:19
fitted Gauss locations:19
y
x
0
0.5
1
1.5
2
2.5
3
0
5
10
15
20
25

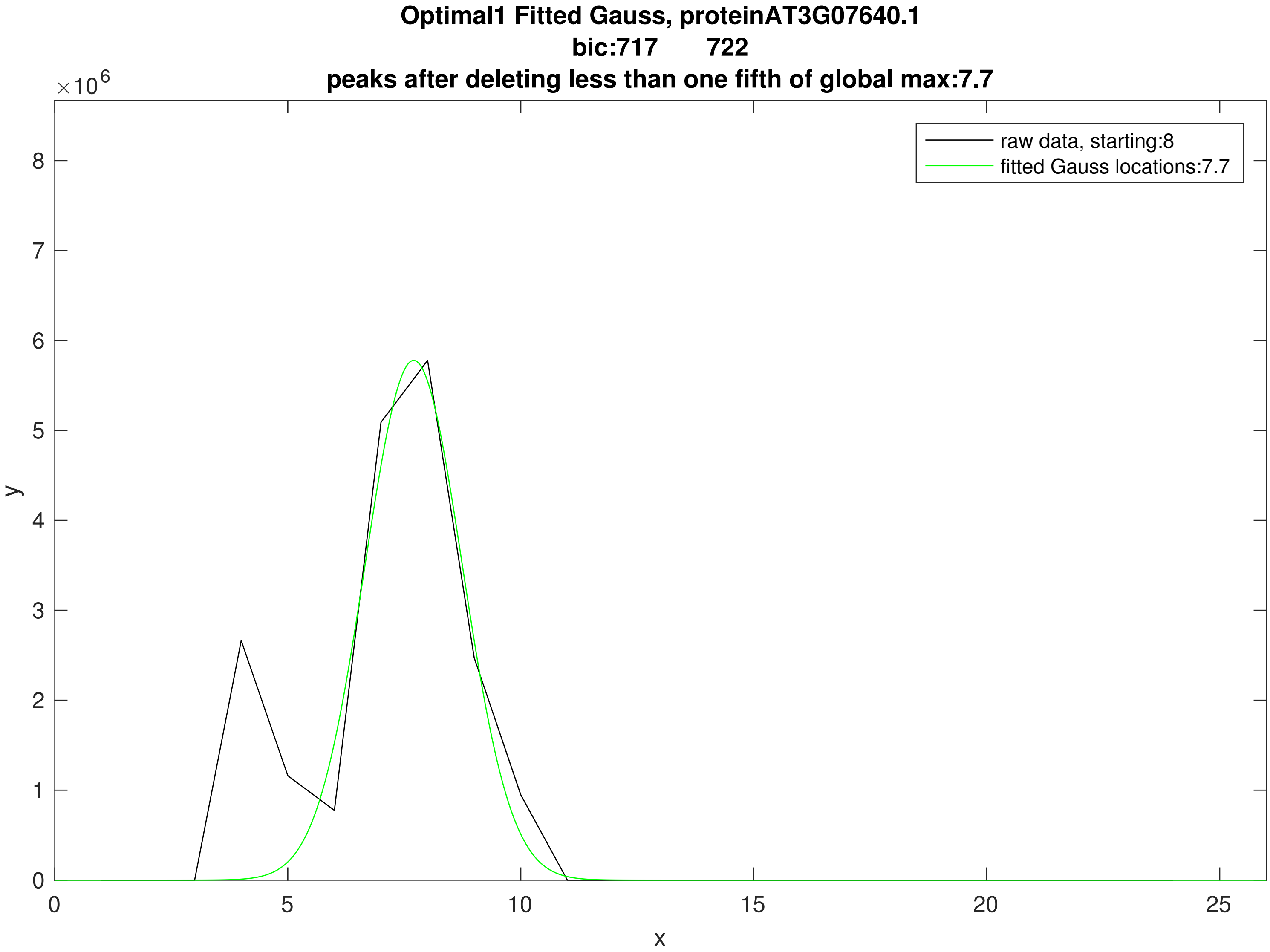
Optimal1 Fitted Gauss, proteinAT3G07640.1
bic:717 722
peaks after deleting less than one fifth of global max:7.7
raw data, starting:8
fitted Gauss locations:7.7
x
y

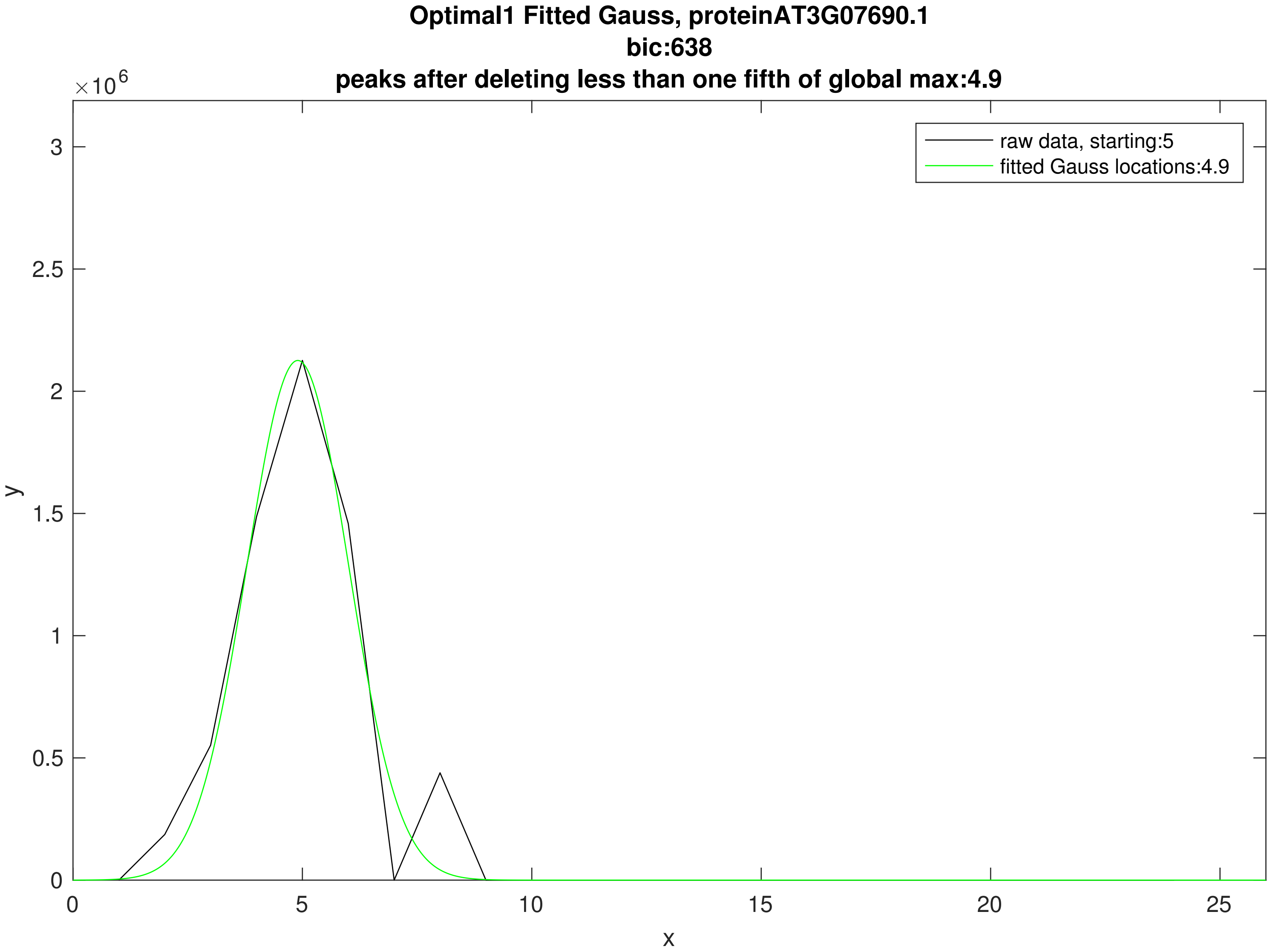
Optimal1 Fitted Gauss, proteinAT3G07690.1
bic:638
peaks after deleting less than one fifth of global max:4.9
×10^6
raw data, starting:5
fitted Gauss locations:4.9
y
x
0
0.5
1
1.5
2
2.5
3
0
5
10
15
20
25

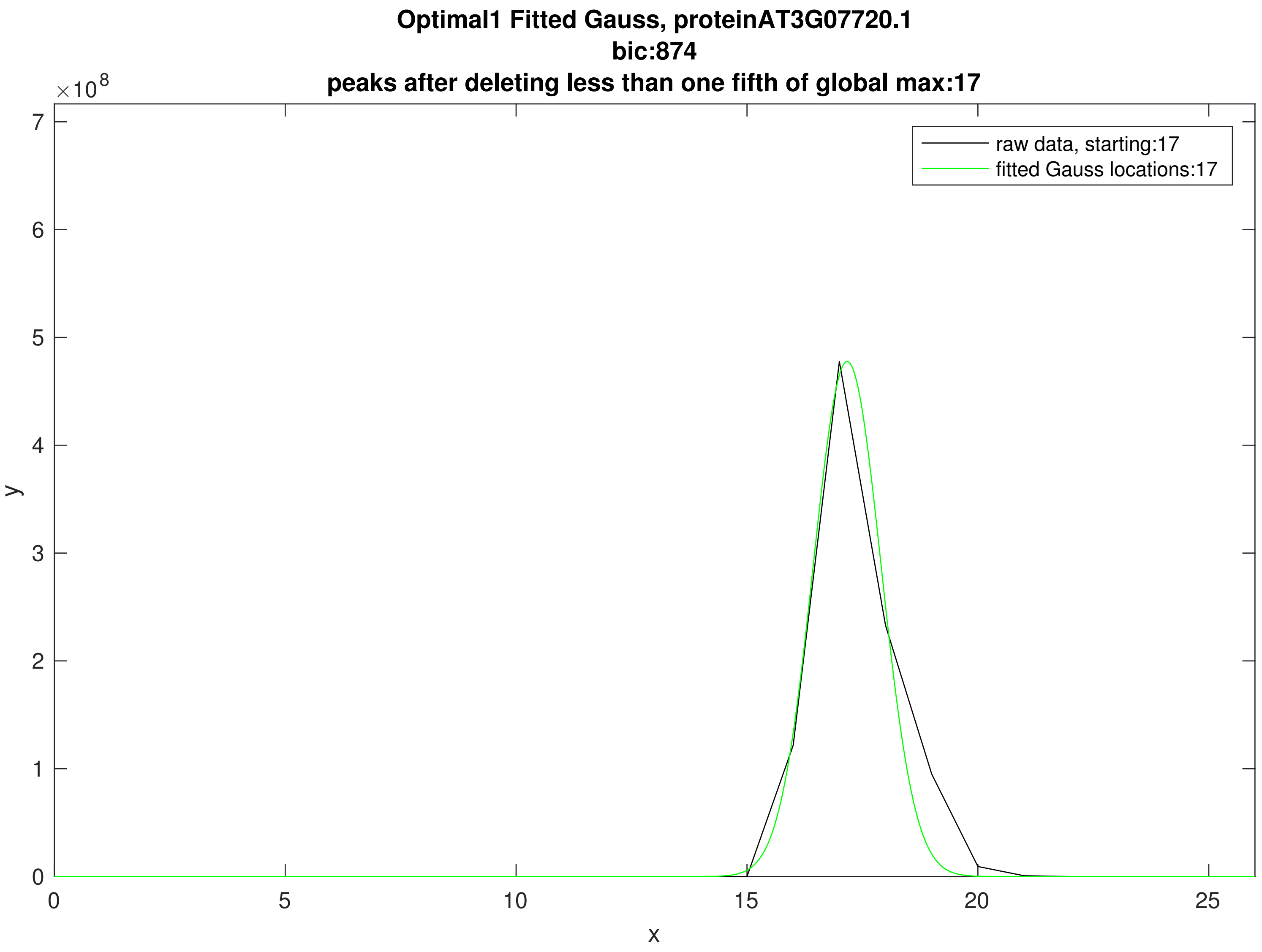
Optimal1 Fitted Gauss, proteinAT3G07720.1
bic:874
peaks after deleting less than one fifth of global max:17
×10^8
raw data, starting:17
fitted Gauss locations:17
y
x
0
1
2
3
4
5
6
7
0
5
10
15
20
25

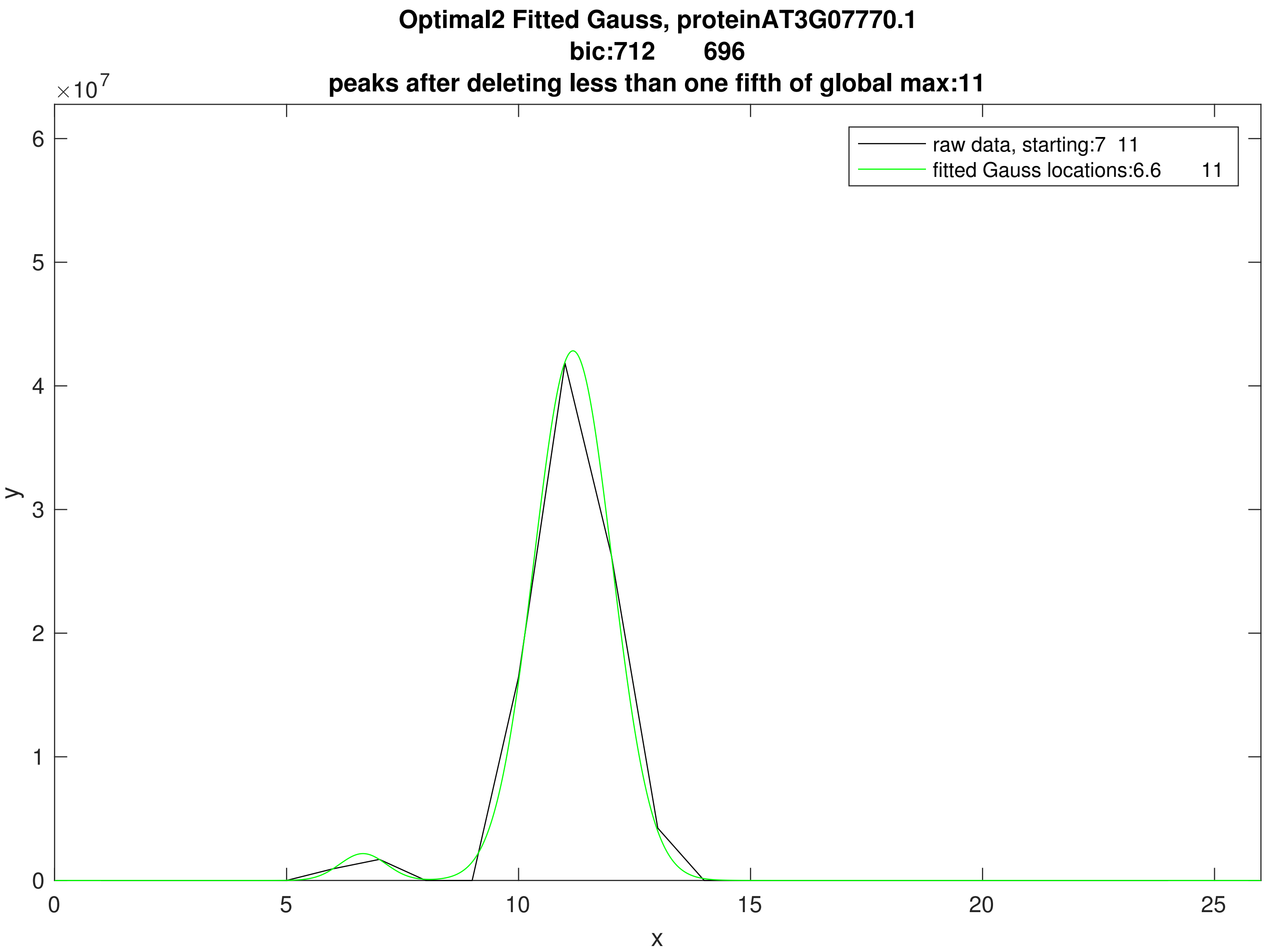
Optimal2 Fitted Gauss, proteinAT3G07770.1
bic:712 696
peaks after deleting less than one fifth of global max:11
×10^7
raw data, starting:7 11
fitted Gauss locations:6.6 11
y
x
0
1
2
3
4
5
6
0
5
10
15
20
25

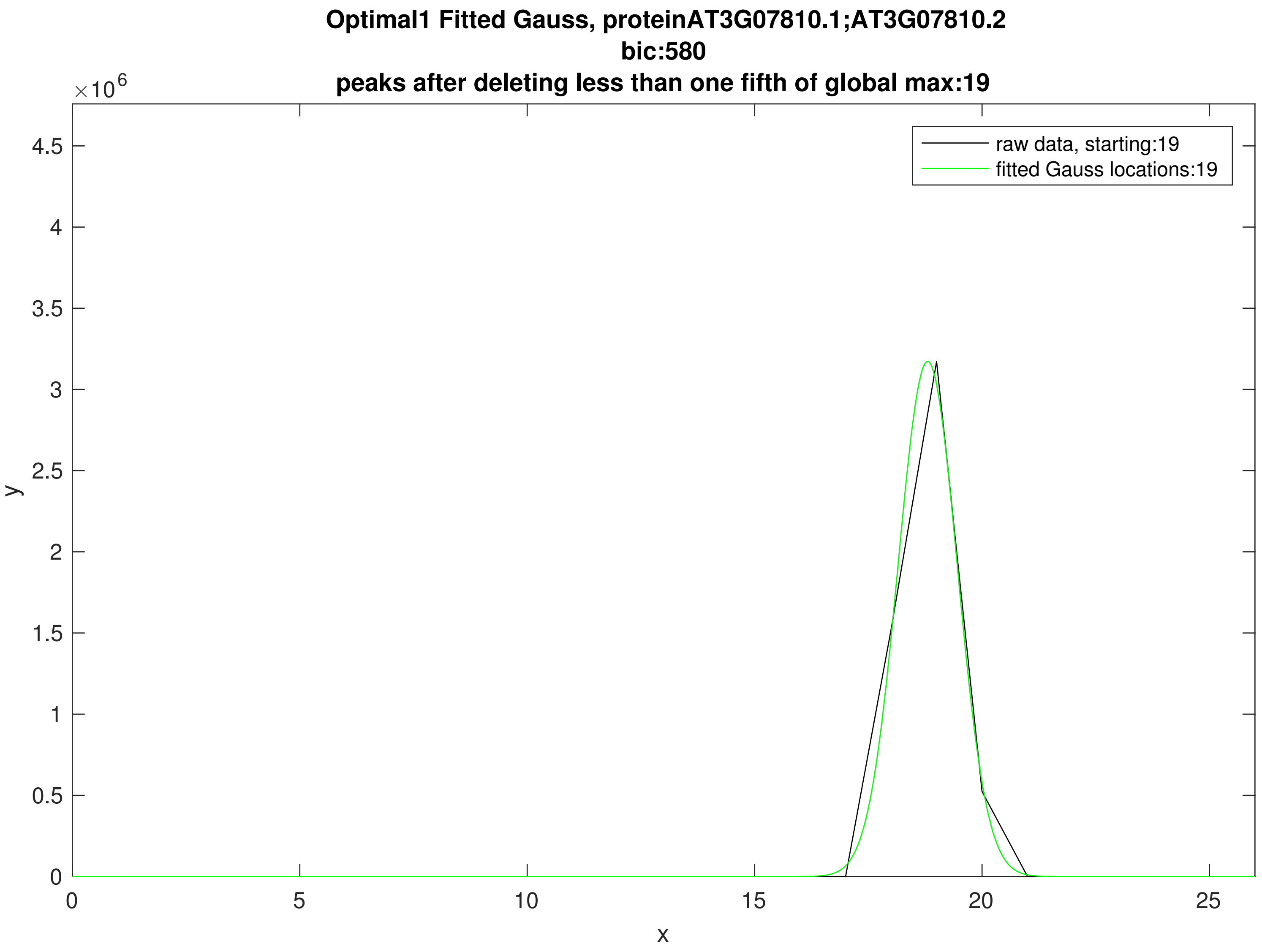
Optimal1 Fitted Gauss, proteinAT3G07810.1;AT3G07810.2
bic:580
peaks after deleting less than one fifth of global max:19
×10^6
raw data, starting:19
fitted Gauss locations:19
4.5
4
3.5
3
2.5
2
1.5
1
0.5
0
y
0
5
10
15
20
25
x

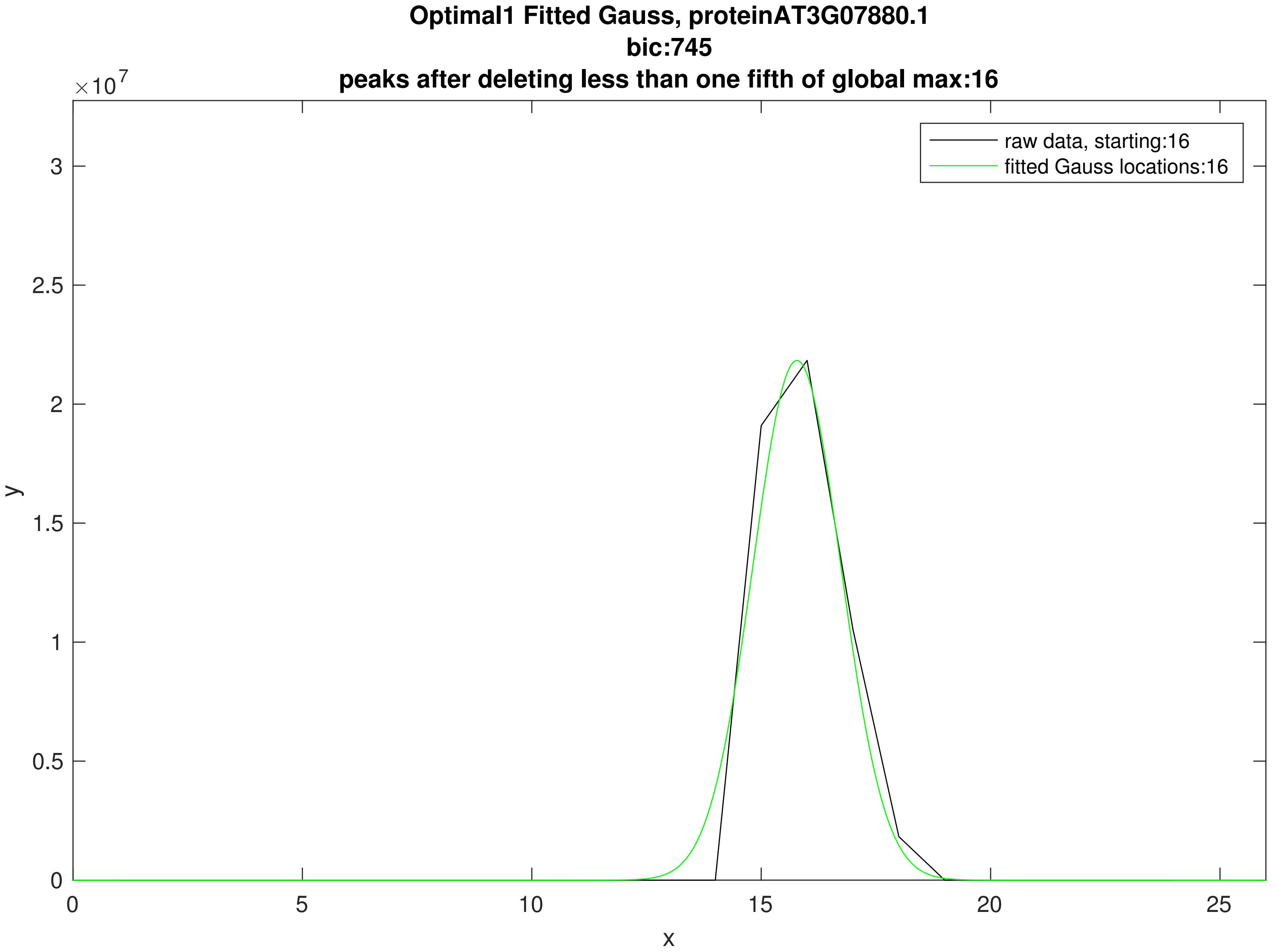
Optimal1 Fitted Gauss, proteinAT3G07880.1
bic:745
peaks after deleting less than one fifth of global max:16
×10^7
3
2.5
2
1.5
1
0.5
0
y
0
5
10
15
20
25
x
raw data, starting:16
fitted Gauss locations:16

**Optimal1 Fitted Gauss, proteinAT3G08030.1;AT3G08030.2**
**bic:1.03e+03  1.04e+03**
**peaks after deleting less than one fifth of global max:17**

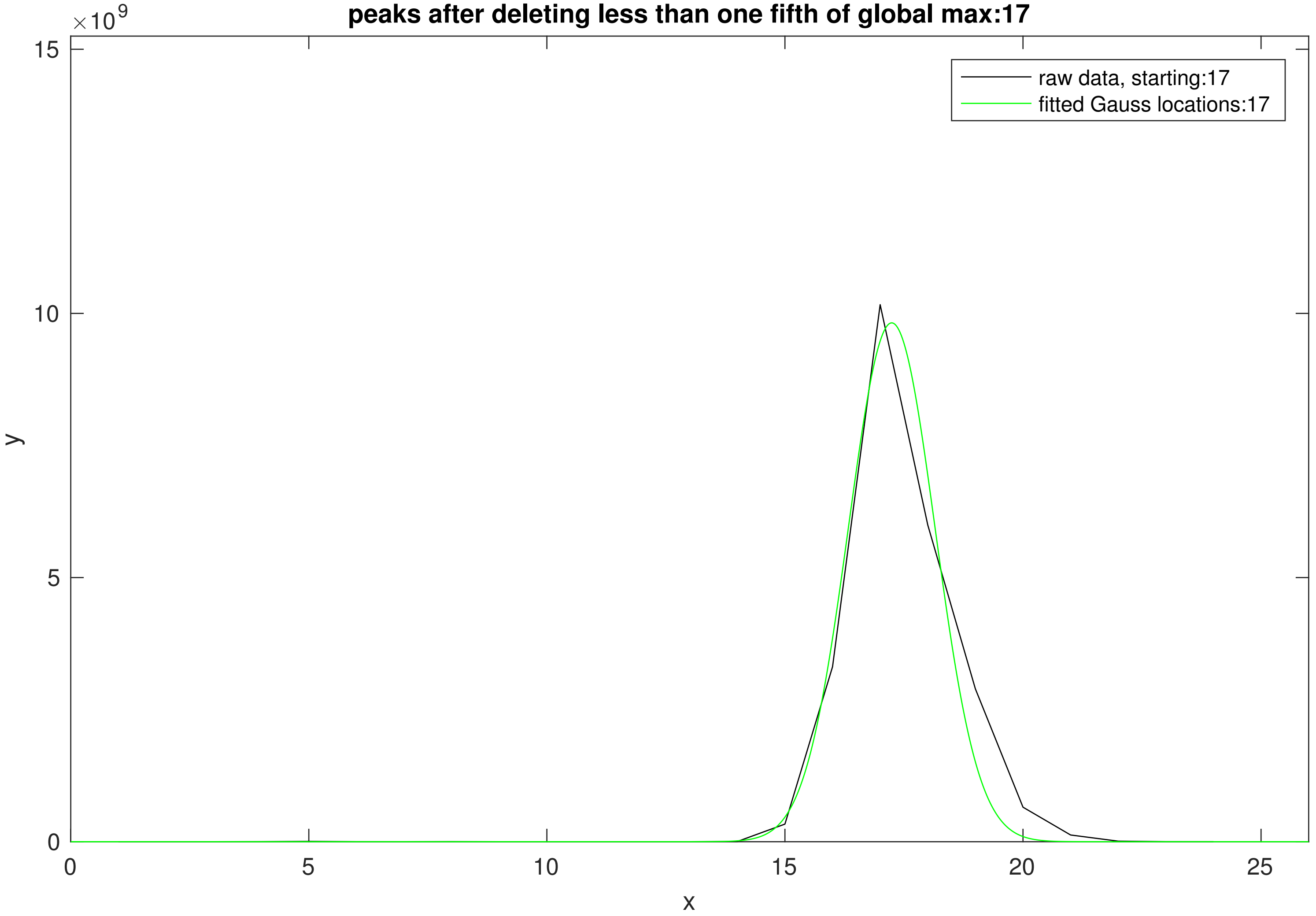

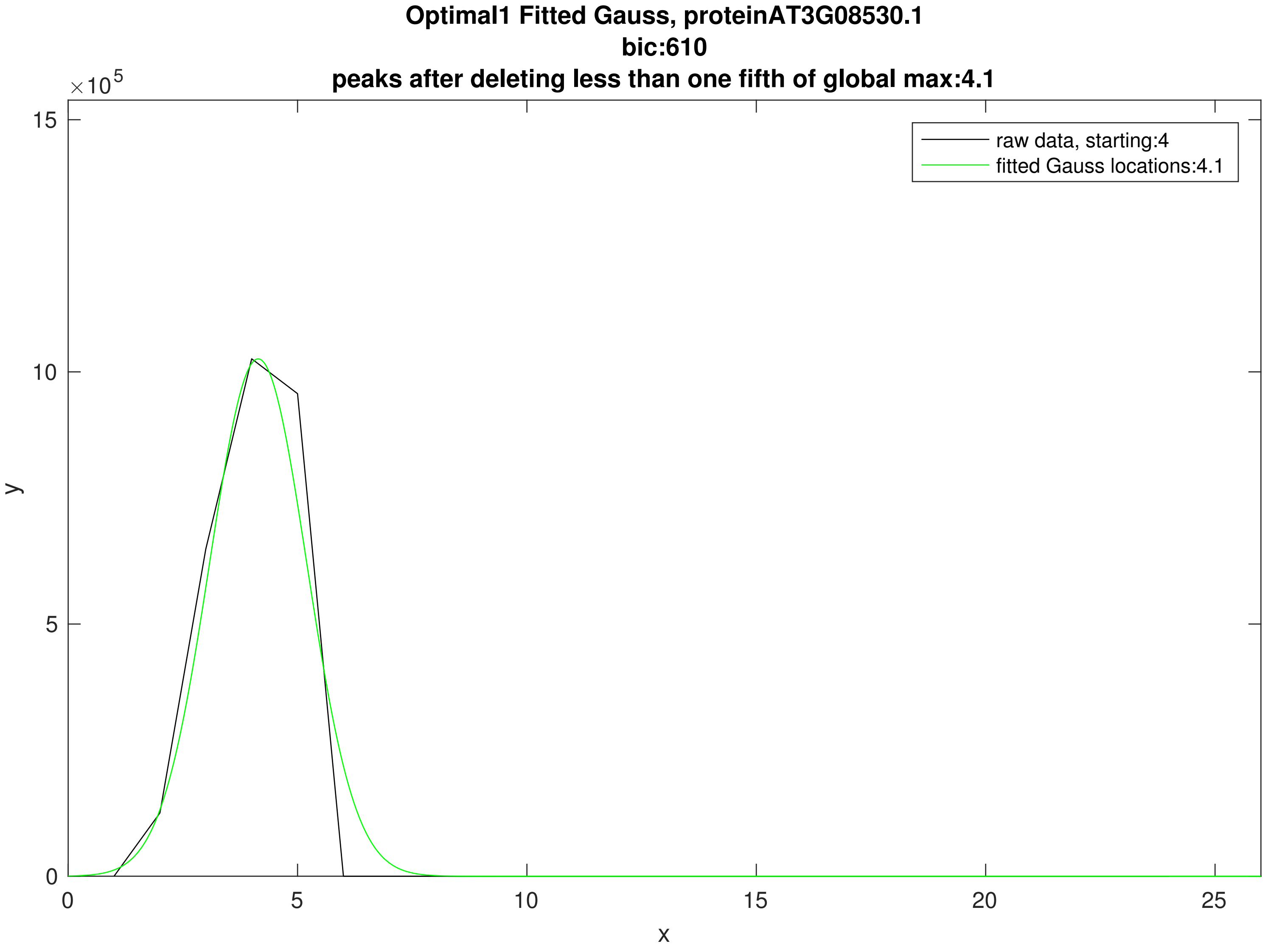
Optimal1 Fitted Gauss, proteinAT3G08530.1
bic:610
peaks after deleting less than one fifth of global max:4.1
×10^5
raw data, starting:4
fitted Gauss locations:4.1
y
x
0
5
10
15
20
25

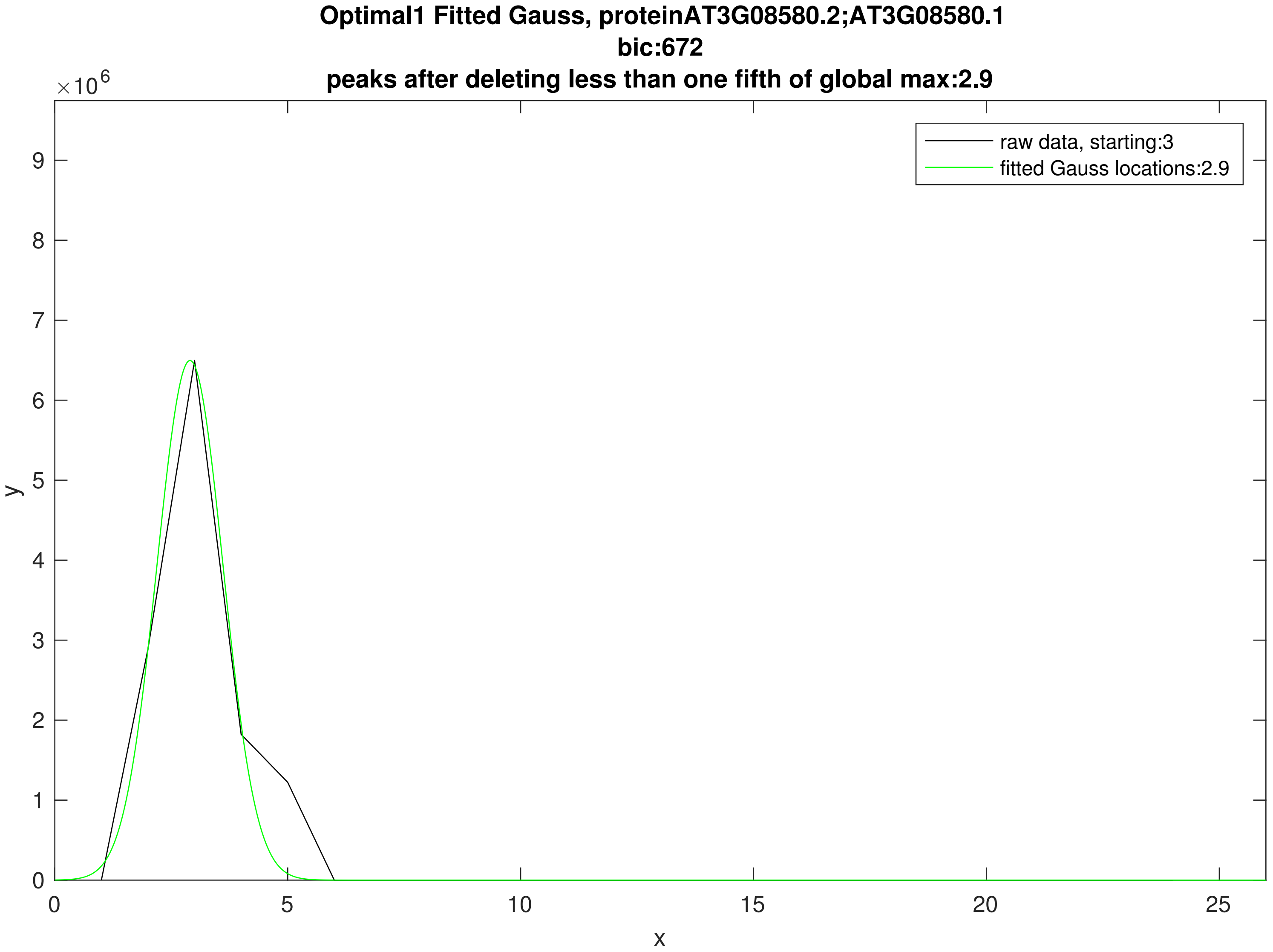

Optimal1 Fitted Gauss, proteinAT3G08580.2;AT3G08580.1
bic:672
peaks after deleting less than one fifth of global max:2.9
×10^6
raw data, starting:3
fitted Gauss locations:2.9
y
x
0
1
2
3
4
5
6
7
8
9
0
5
10
15
20
25

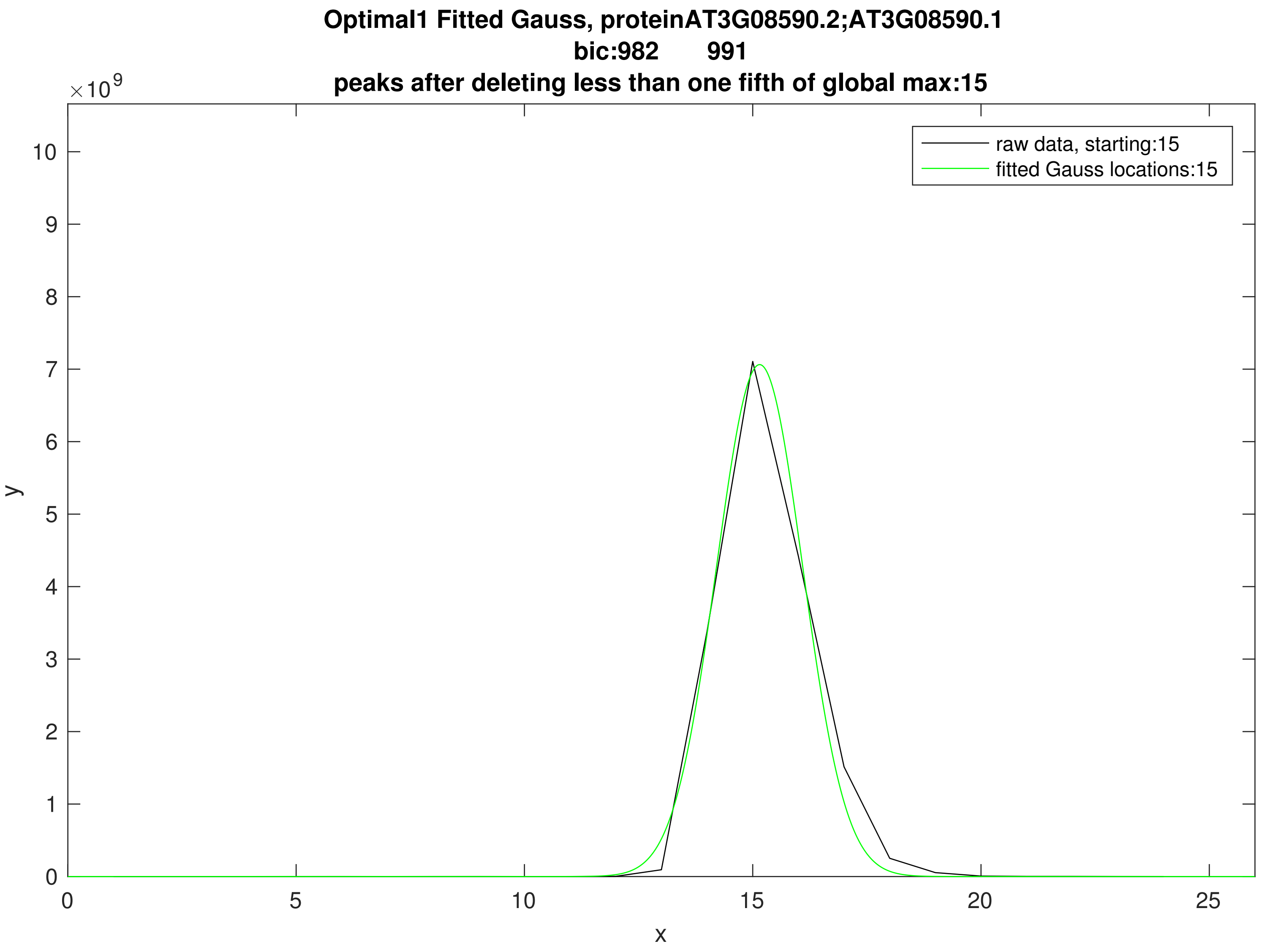

Optimal1 Fitted Gauss, proteinAT3G08590.2;AT3G08590.1
bic:982 991
peaks after deleting less than one fifth of global max:15
×10^9
raw data, starting:15
fitted Gauss locations:15
y
x
0
1
2
3
4
5
6
7
8
9
10
0
5
10
15
20
25

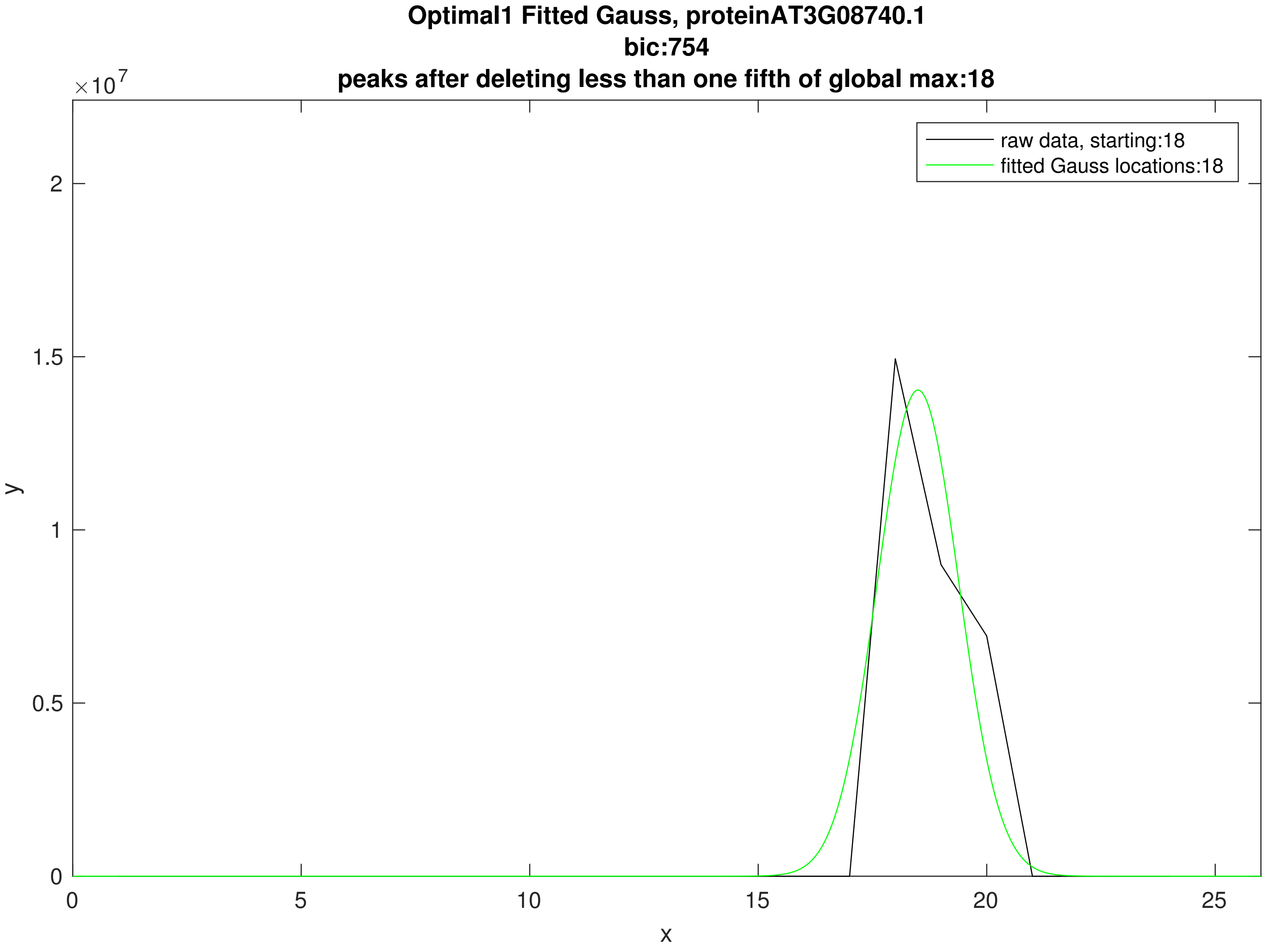
Optimal1 Fitted Gauss, proteinAT3G08740.1
bic:754
peaks after deleting less than one fifth of global max:18
×10^7
raw data, starting:18
fitted Gauss locations:18
y
x
0
0.5
1
1.5
2
0
5
10
15
20
25

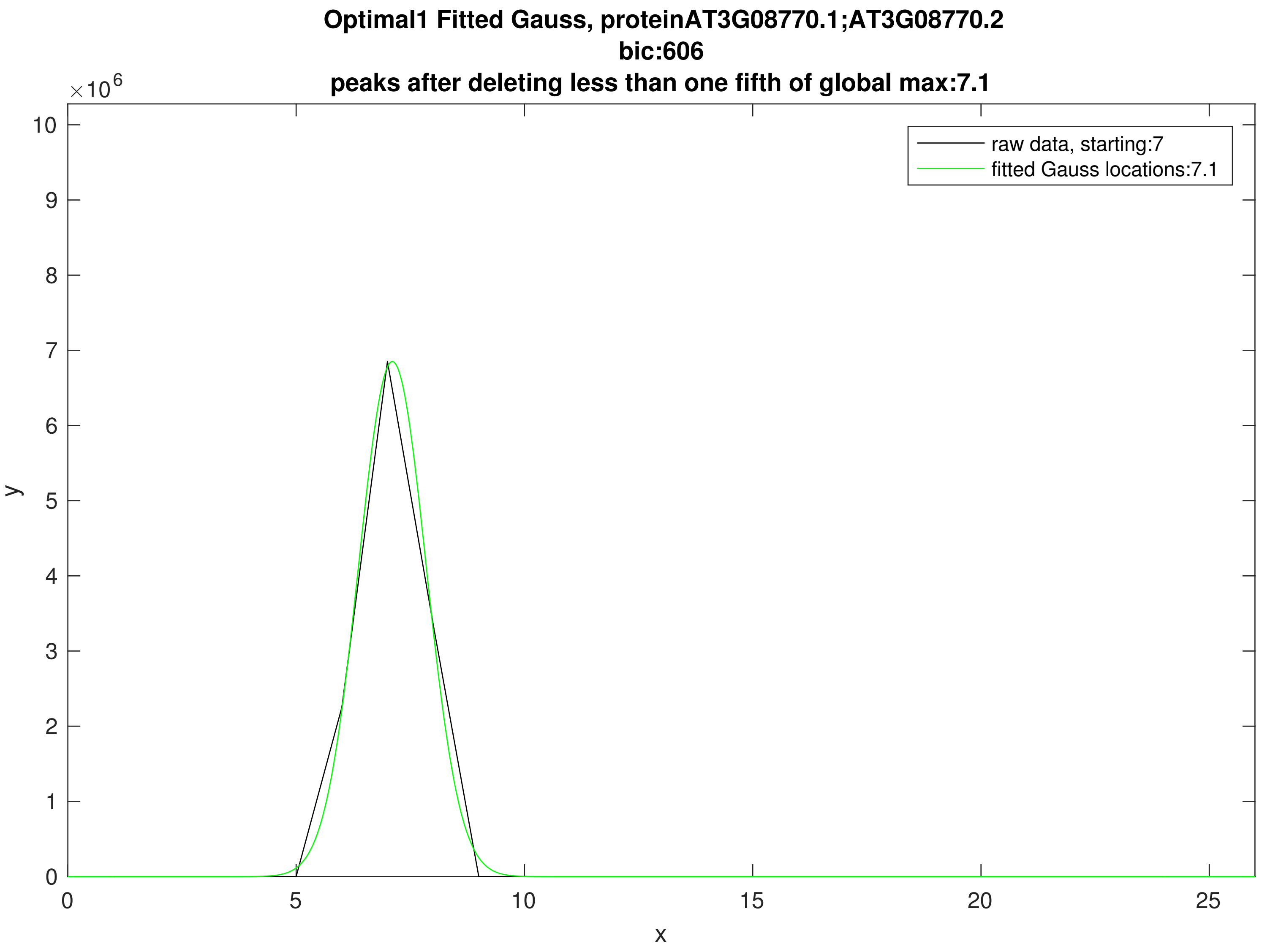
Optimal1 Fitted Gauss, proteinAT3G08770.1;AT3G08770.2
bic:606
peaks after deleting less than one fifth of global max:7.1
×10^6
raw data, starting:7
fitted Gauss locations:7.1
y
x
0
1
2
3
4
5
6
7
8
9
10
0
5
10
15
20
25

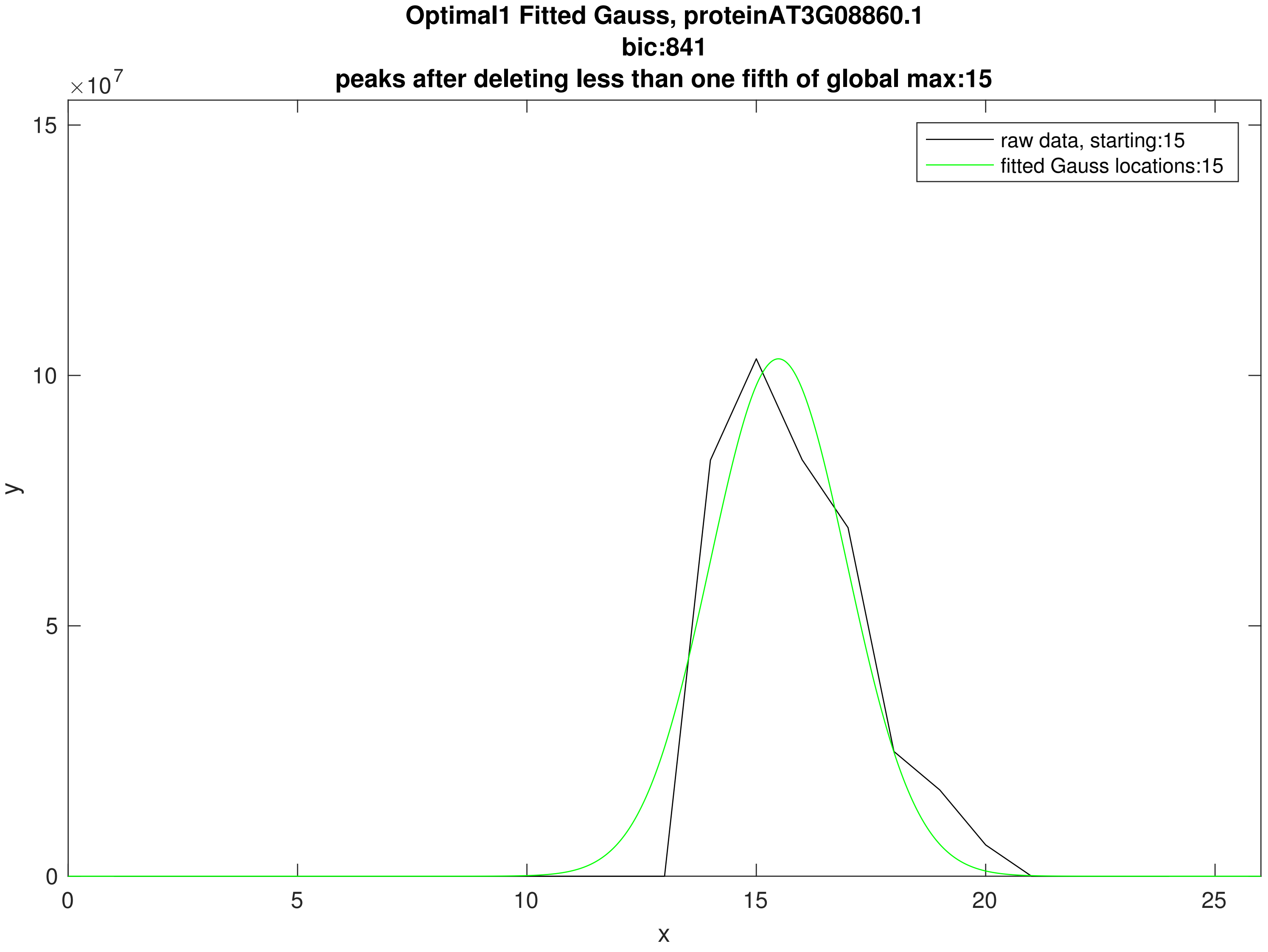

Optimal1 Fitted Gauss, proteinAT3G08860.1
bic:841
peaks after deleting less than one fifth of global max:15
×10^7
15
10
5
0
y
0
5
10
15
20
25
x
raw data, starting:15
fitted Gauss locations:15

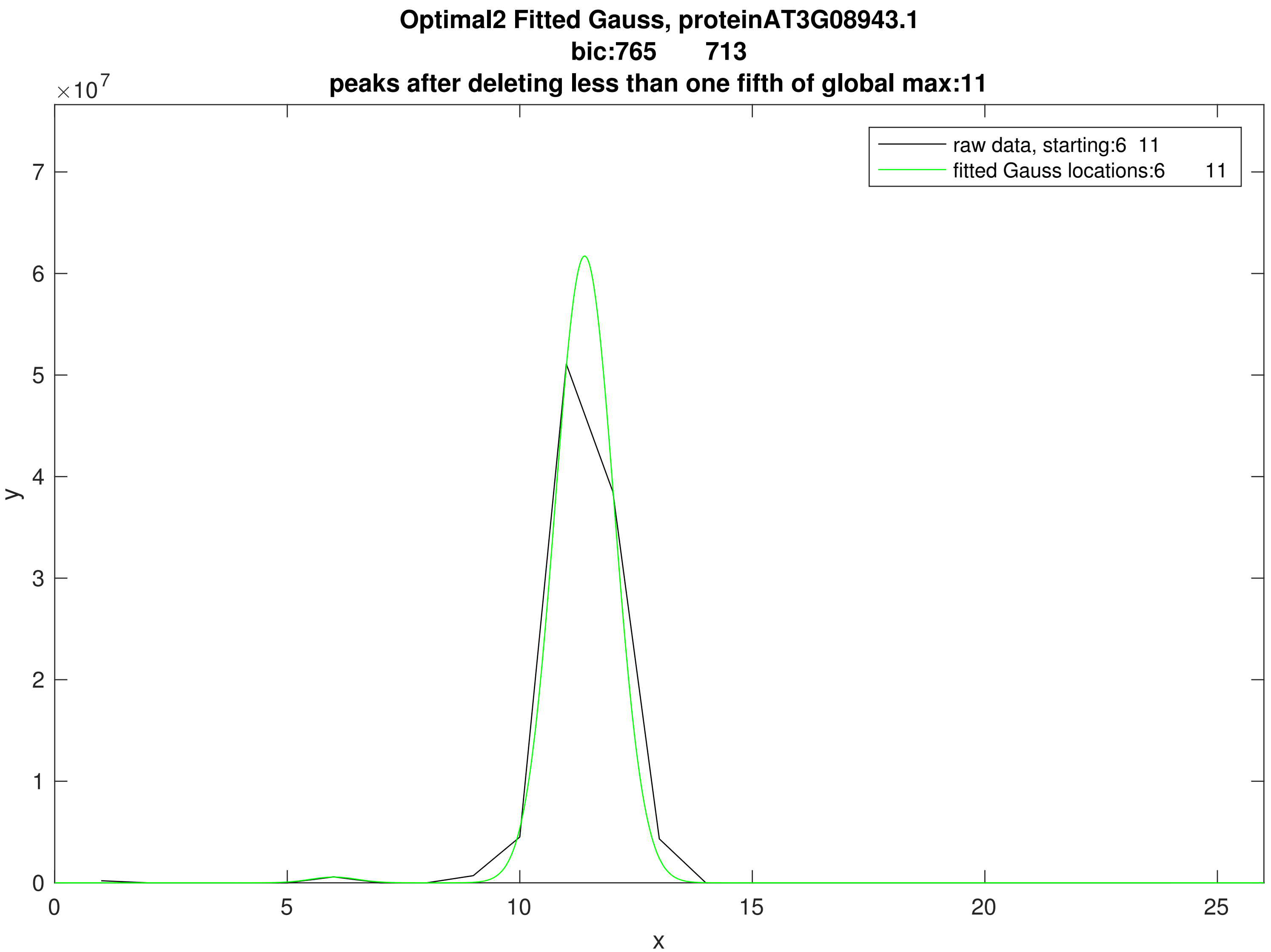

Optimal2 Fitted Gauss, proteinAT3G08943.1
bic:765 713
peaks after deleting less than one fifth of global max:11
×10^7
raw data, starting:6 11
fitted Gauss locations:6 11
y
x
0
1
2
3
4
5
6
7
0
5
10
15
20
25

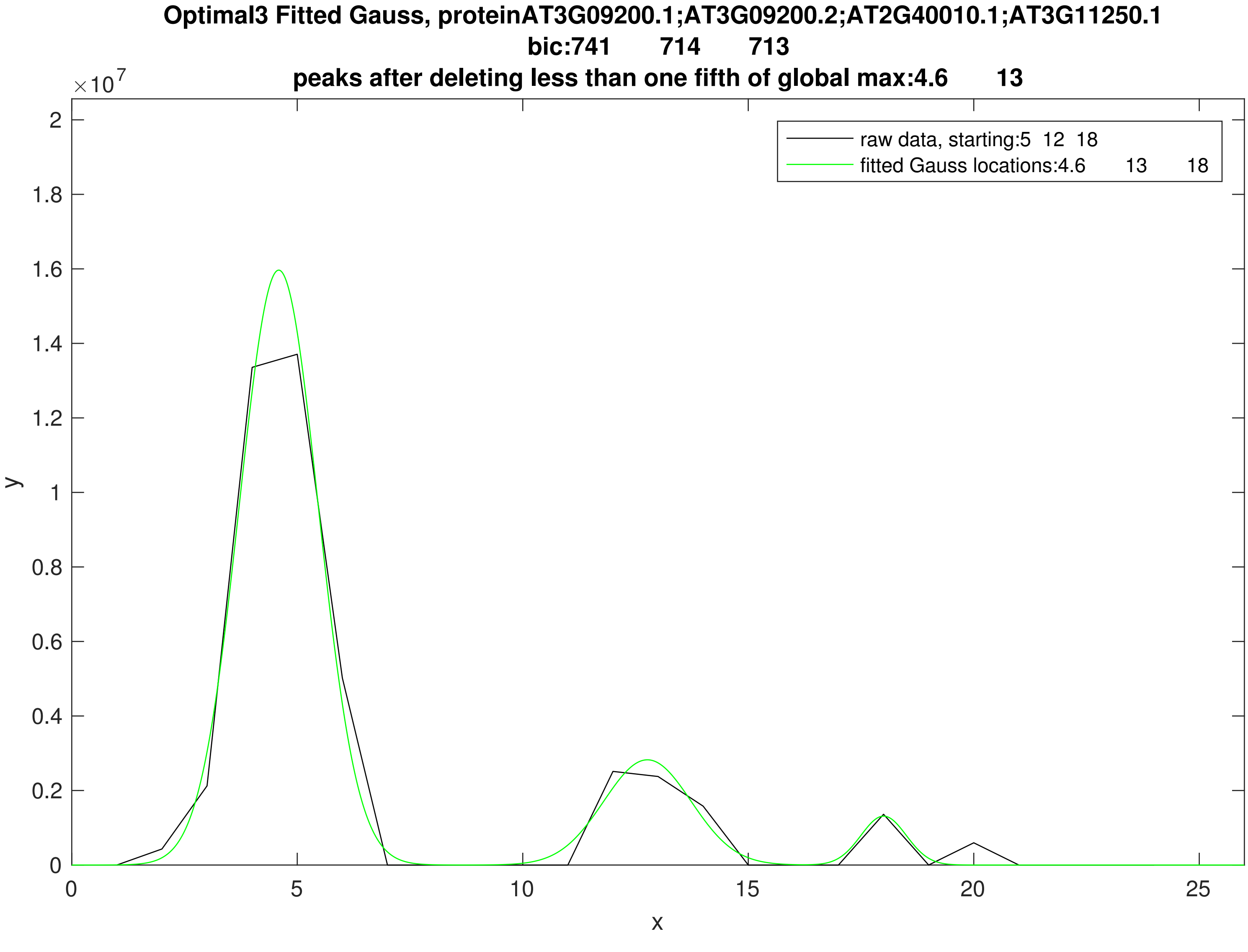
Optimal3 Fitted Gauss, proteinAT3G09200.1;AT3G09200.2;AT2G40010.1;AT3G11250.1
bic:741 714 713
peaks after deleting less than one fifth of global max:4.6 13
×10^7
raw data, starting:5 12 18
fitted Gauss locations:4.6 13 18
y
x
0
0.2
0.4
0.6
0.8
1
1.2
1.4
1.6
1.8
2
5
10
15
20
25

#### Optimal1 Fitted Gauss, proteinAT3G09260.1
#### bic:1.1e+03
#### peaks after deleting less than one fifth of global max:7.8

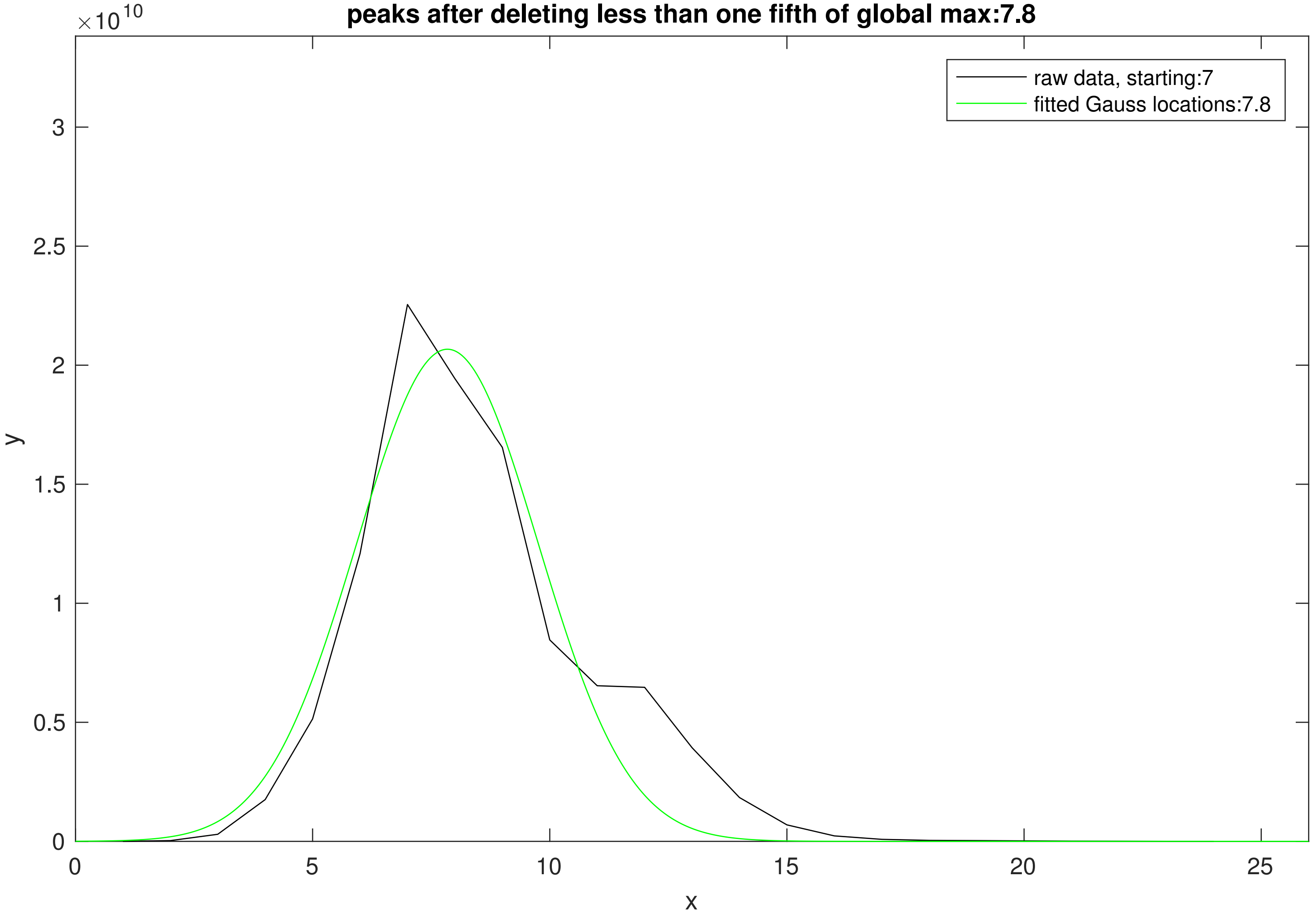

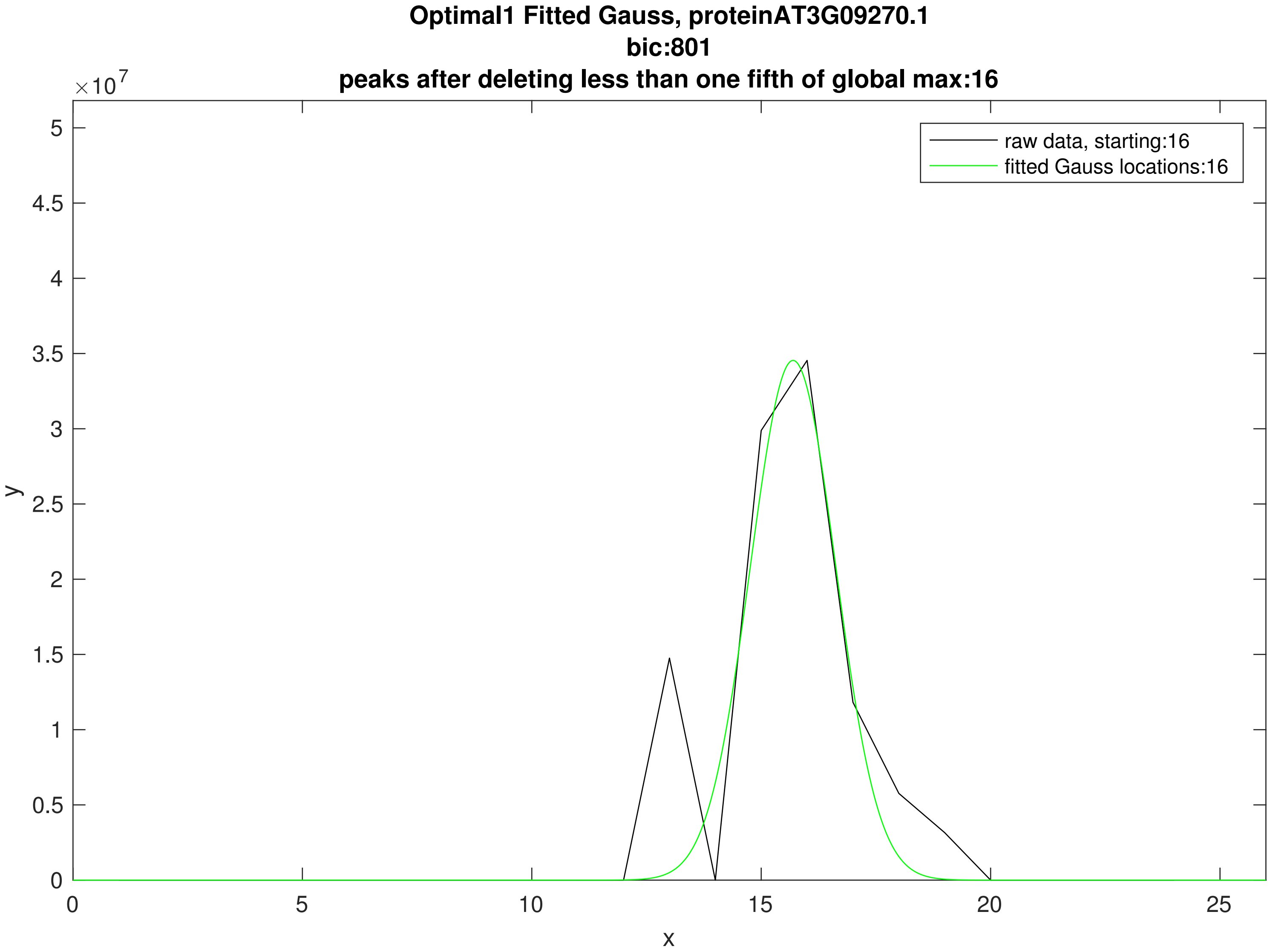

Optimal1 Fitted Gauss, proteinAT3G09270.1
bic:801
peaks after deleting less than one fifth of global max:16
×10^7
raw data, starting:16
fitted Gauss locations:16
y
x
0
0.5
1
1.5
2
2.5
3
3.5
4
4.5
5
0
5
10
15
20
25

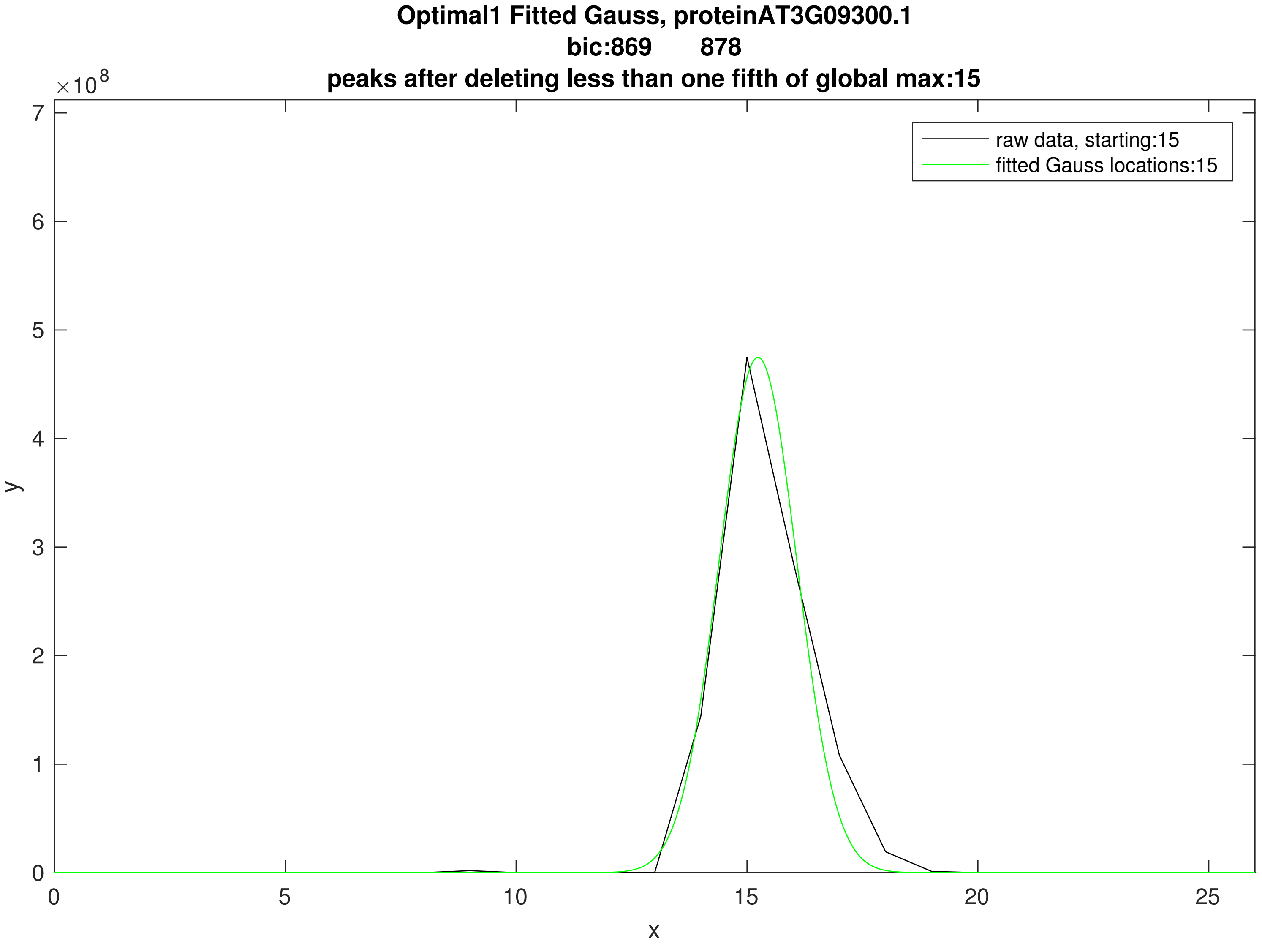

Optimal1 Fitted Gauss, proteinAT3G09300.1
bic:869 878
peaks after deleting less than one fifth of global max:15
×10^8
7
6
5
4
3
2
1
0
y
0
5
10
15
20
25
x
raw data, starting:15
fitted Gauss locations:15

**Optimal1 Fitted Gauss, proteinAT3G09350.1;AT3G09350.3;AT3G09350.2**
**bic:720**
**peaks after deleting less than one fifth of global max:11**

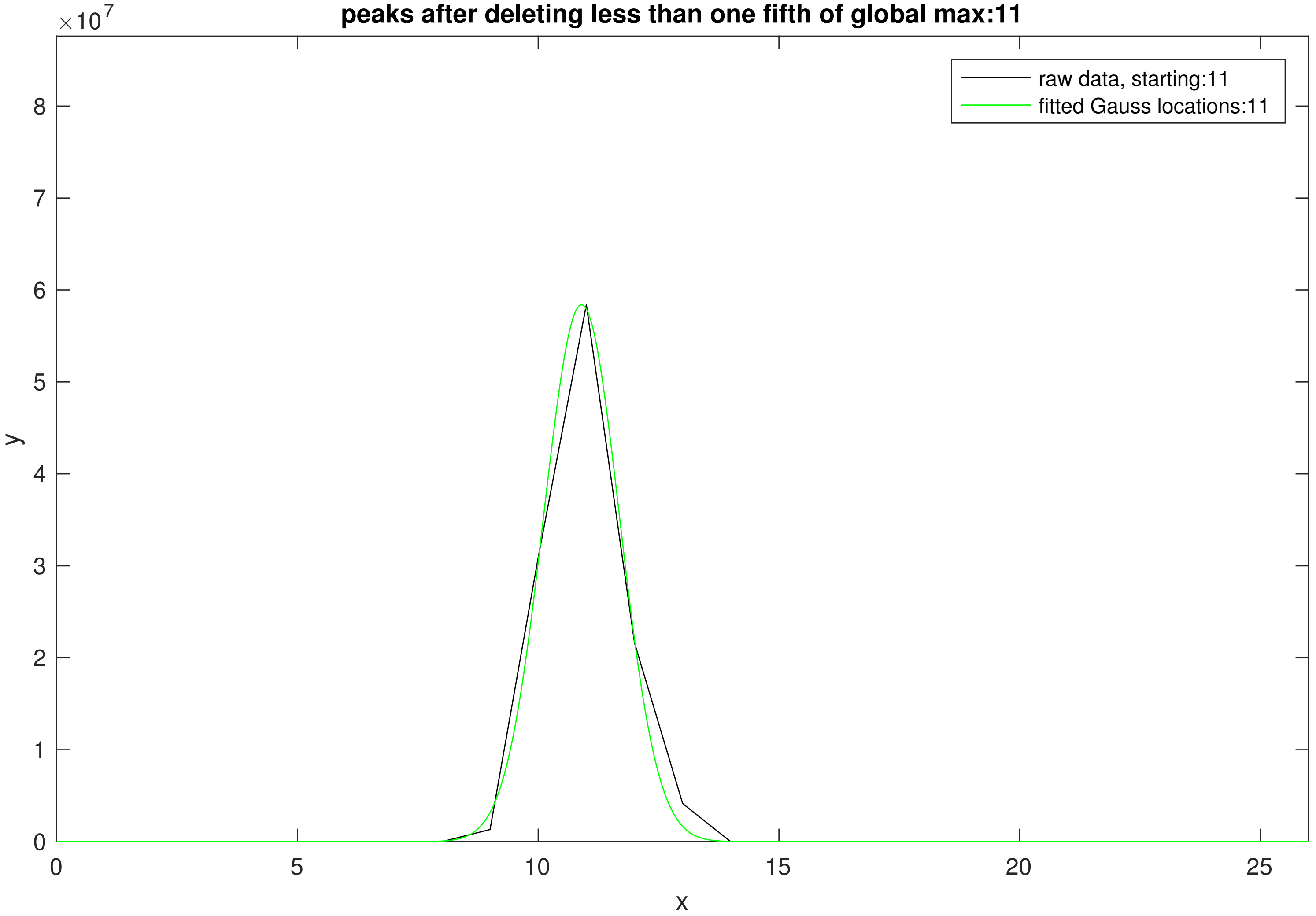

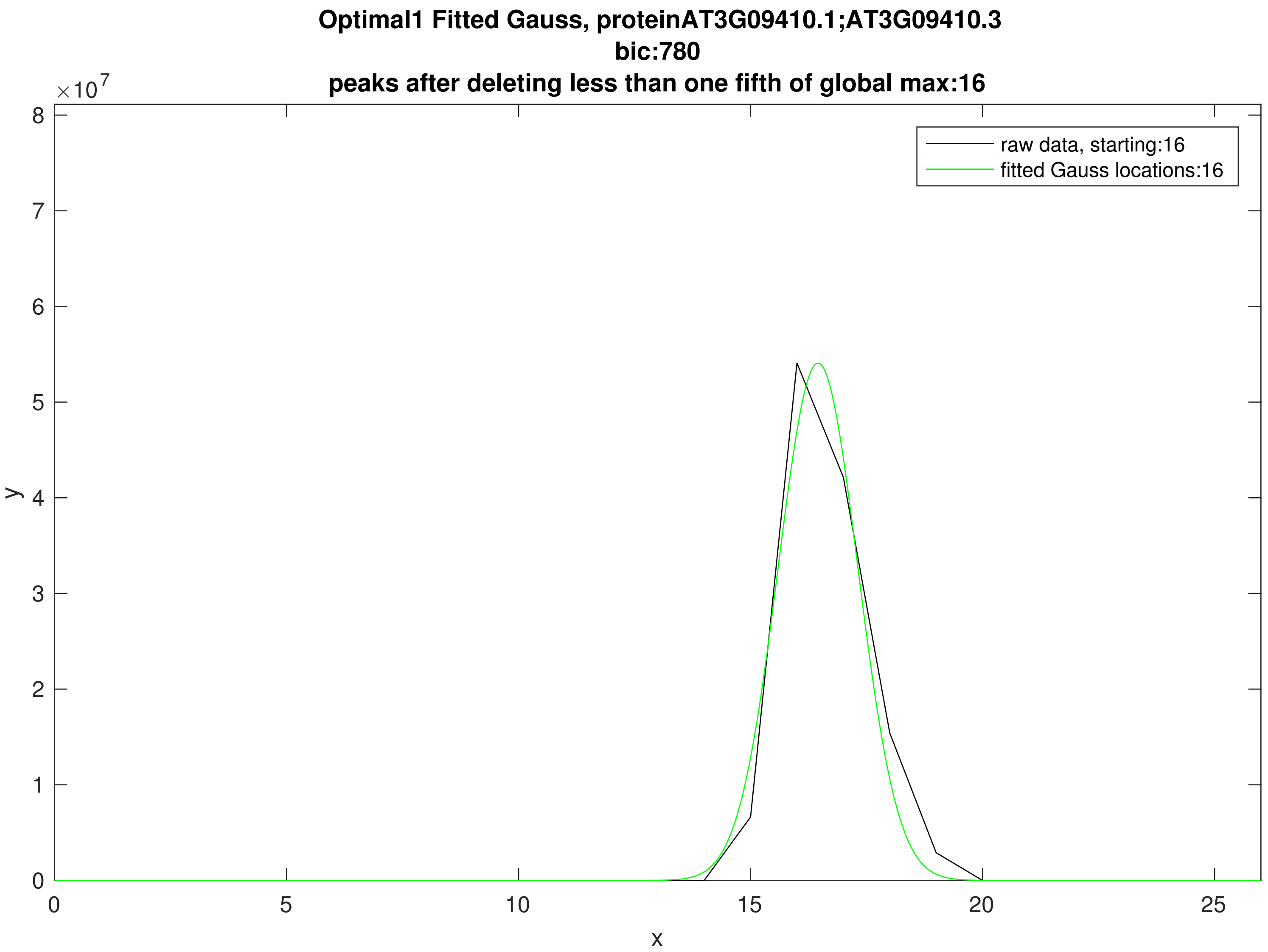
Optimal1 Fitted Gauss, proteinAT3G09410.1;AT3G09410.3
bic:780
peaks after deleting less than one fifth of global max:16
×10^7
raw data, starting:16
fitted Gauss locations:16
y
x
0
1
2
3
4
5
6
7
8
0
5
10
15
20
25

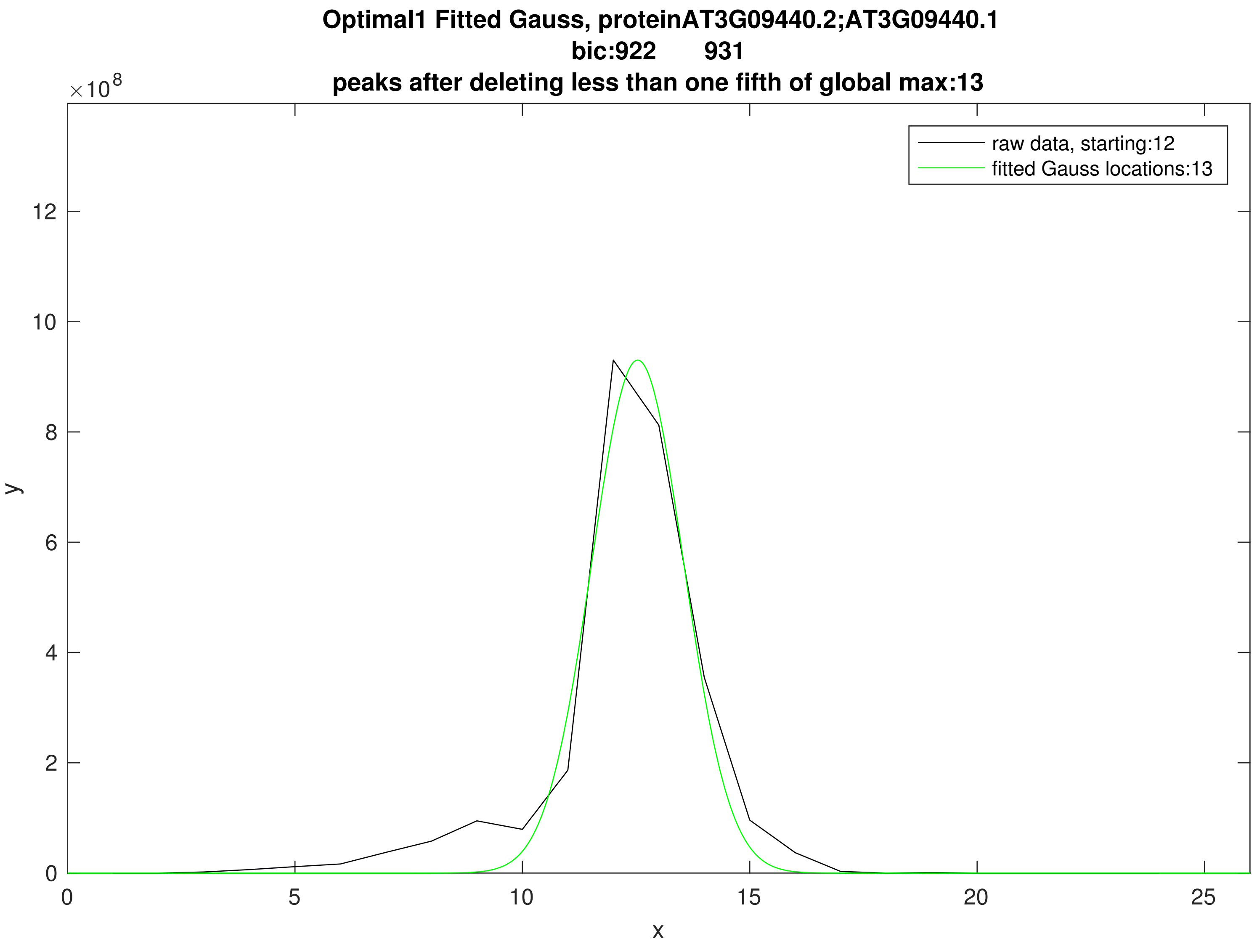

Optimal1 Fitted Gauss, proteinAT3G09440.2;AT3G09440.1
bic:922 931
peaks after deleting less than one fifth of global max:13
×10^8
raw data, starting:12
fitted Gauss locations:13
y
x
0
2
4
6
8
10
12
0
5
10
15
20
25

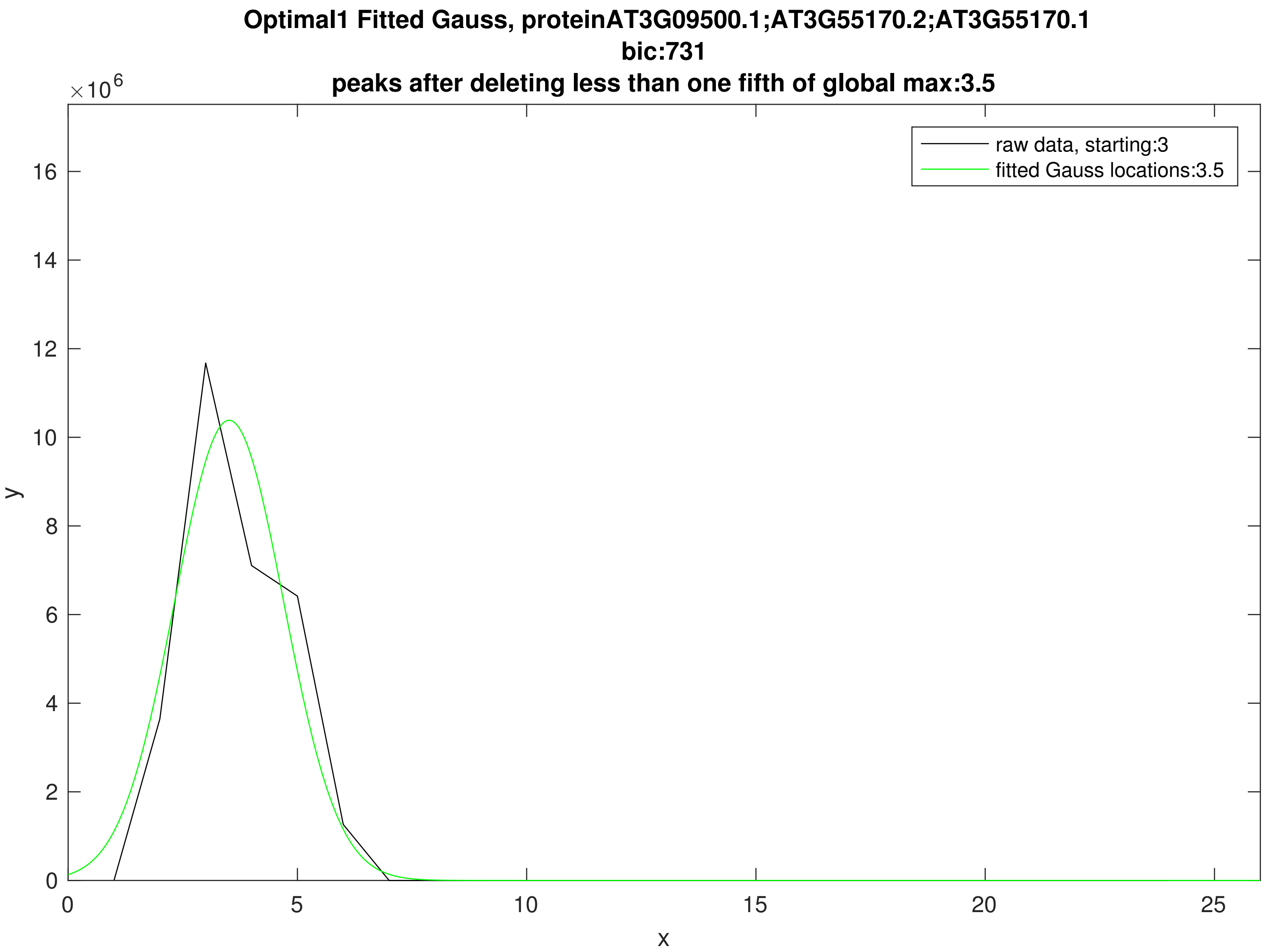
Optimal1 Fitted Gauss, proteinAT3G09500.1;AT3G55170.2;AT3G55170.1
bic:731
peaks after deleting less than one fifth of global max:3.5
×10^6
raw data, starting:3
fitted Gauss locations:3.5
y
x
0
2
4
6
8
10
12
14
16
0
5
10
15
20
25

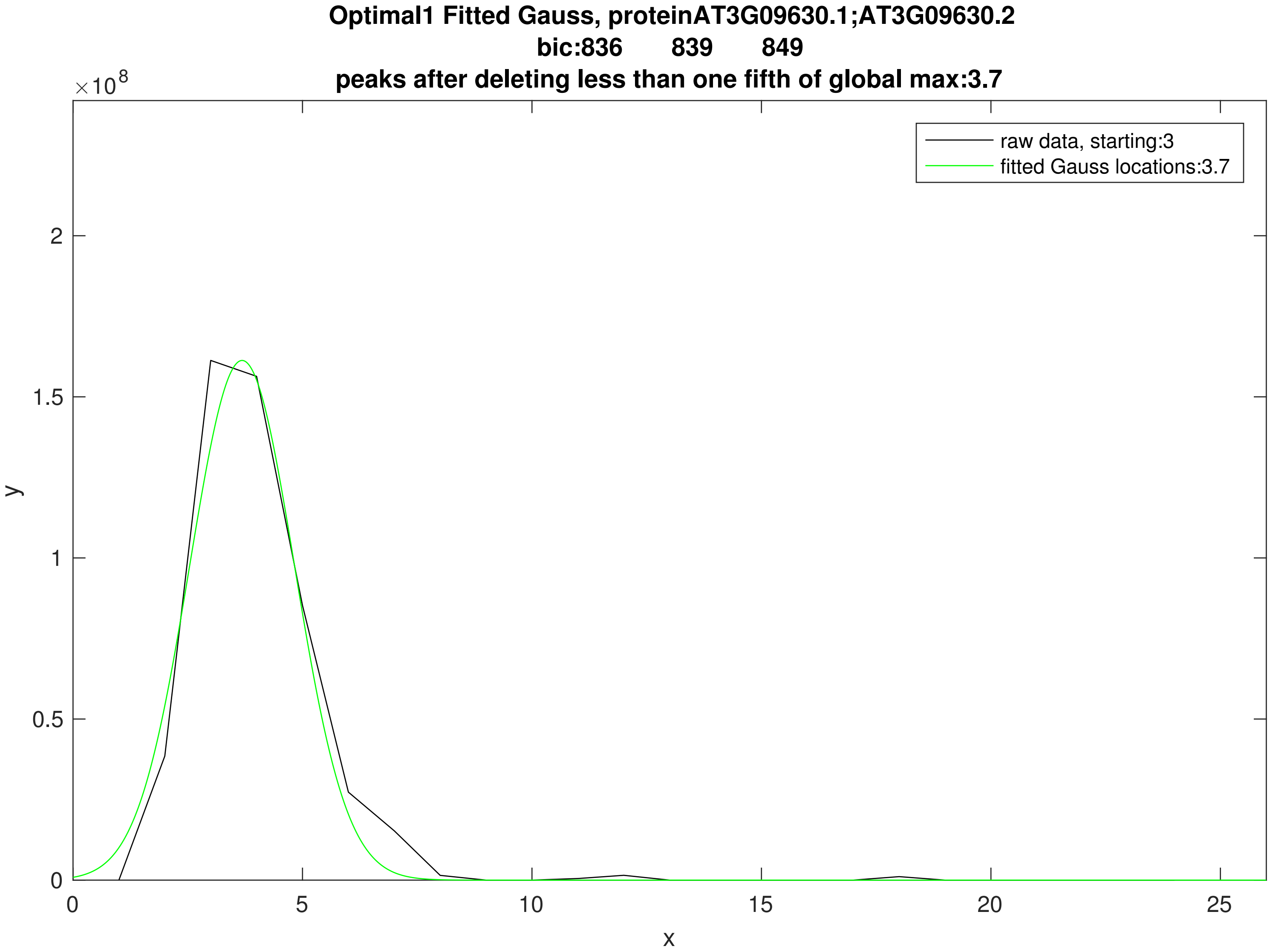

Optimal1 Fitted Gauss, proteinAT3G09630.1;AT3G09630.2
bic:836 839 849
peaks after deleting less than one fifth of global max:3.7
×10^8
raw data, starting:3
fitted Gauss locations:3.7
y
x
0
0.5
1
1.5
2
5
10
15
20
25

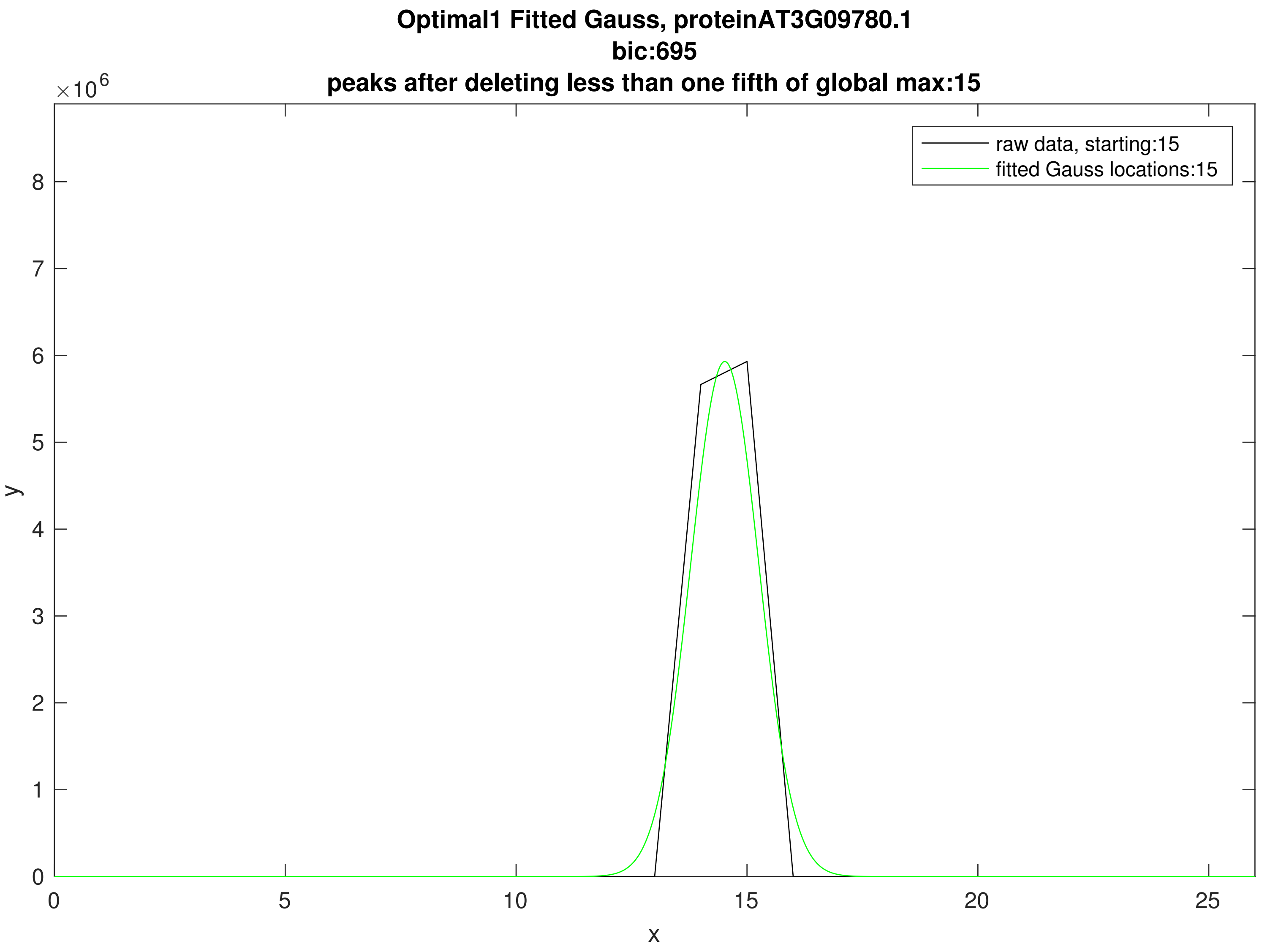
Optimal1 Fitted Gauss, proteinAT3G09780.1
bic:695
peaks after deleting less than one fifth of global max:15
×10^6
raw data, starting:15
fitted Gauss locations:15
y
x
0
1
2
3
4
5
6
7
8
0
5
10
15
20
25

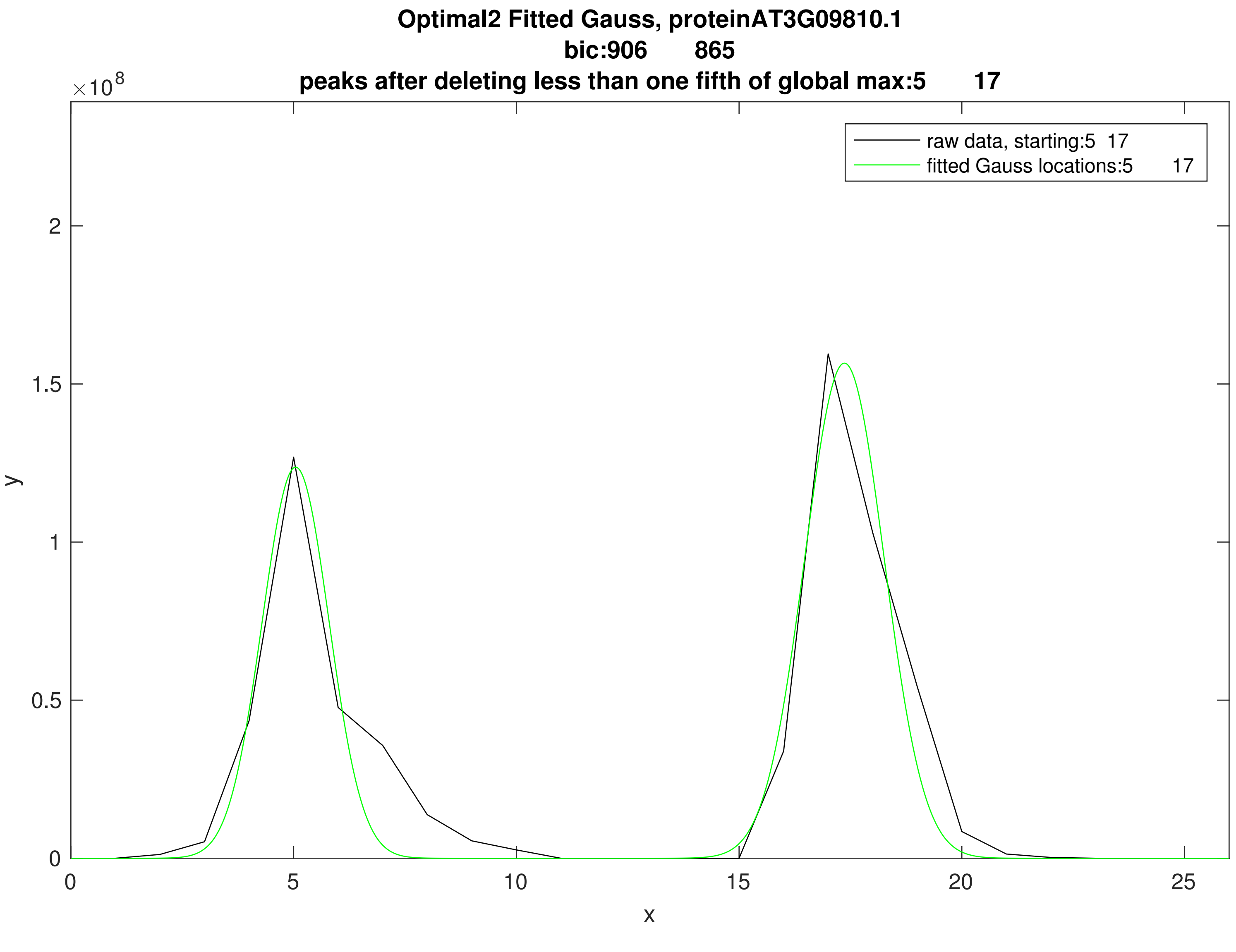
Optimal2 Fitted Gauss, proteinAT3G09810.1
bic:906 865
peaks after deleting less than one fifth of global max:5 17
×10^8
raw data, starting:5 17
fitted Gauss locations:5 17
y
x
0
0.5
1
1.5
2
0
5
10
15
20
25

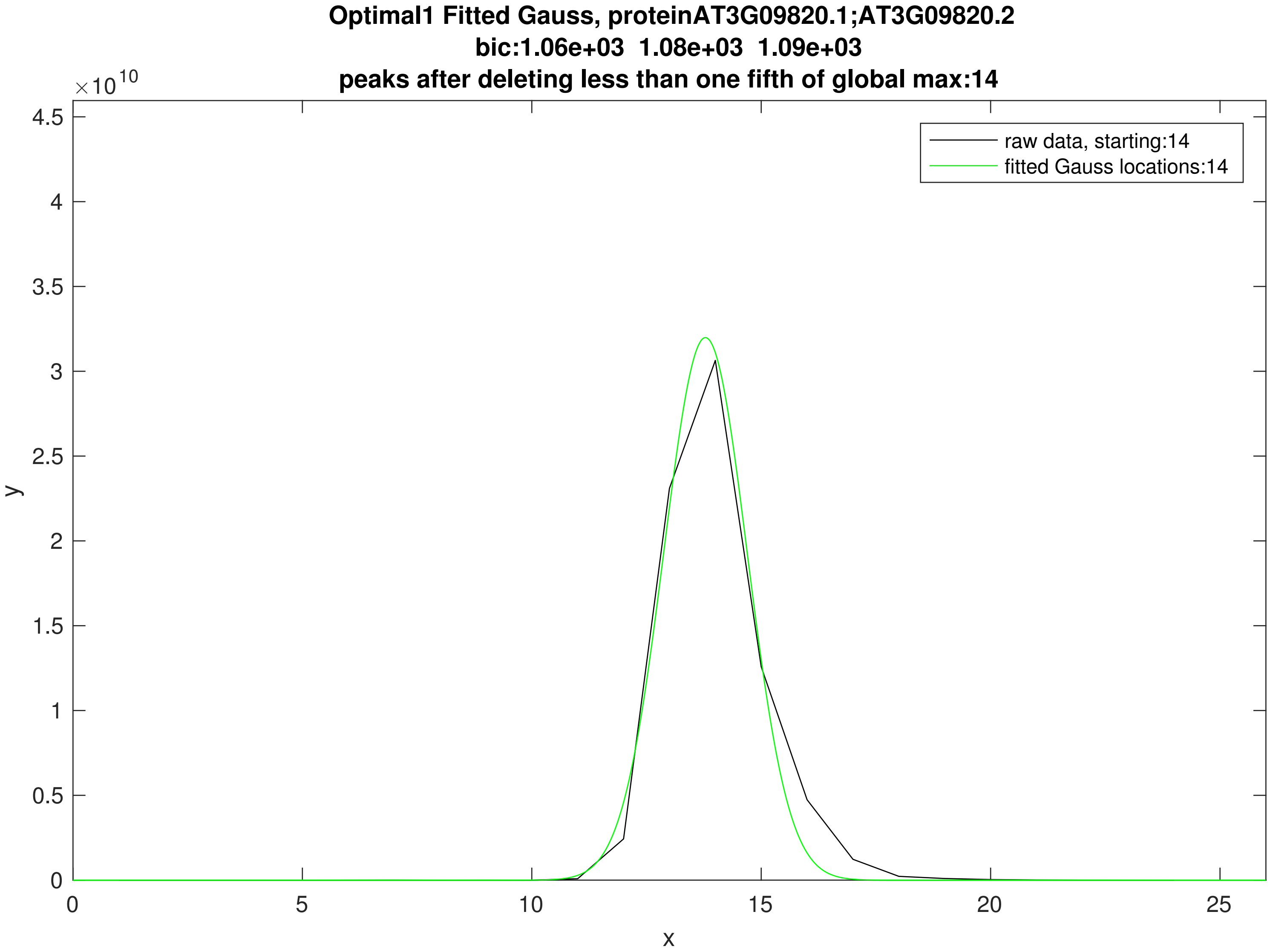
Optimal1 Fitted Gauss, proteinAT3G09820.1;AT3G09820.2
bic:1.06e+03  1.08e+03  1.09e+03
peaks after deleting less than one fifth of global max:14
×10^10
4.5
4
3.5
3
2.5
2
1.5
1
0.5
0
y
0
5
10
15
20
25
x
raw data, starting:14
fitted Gauss locations:14

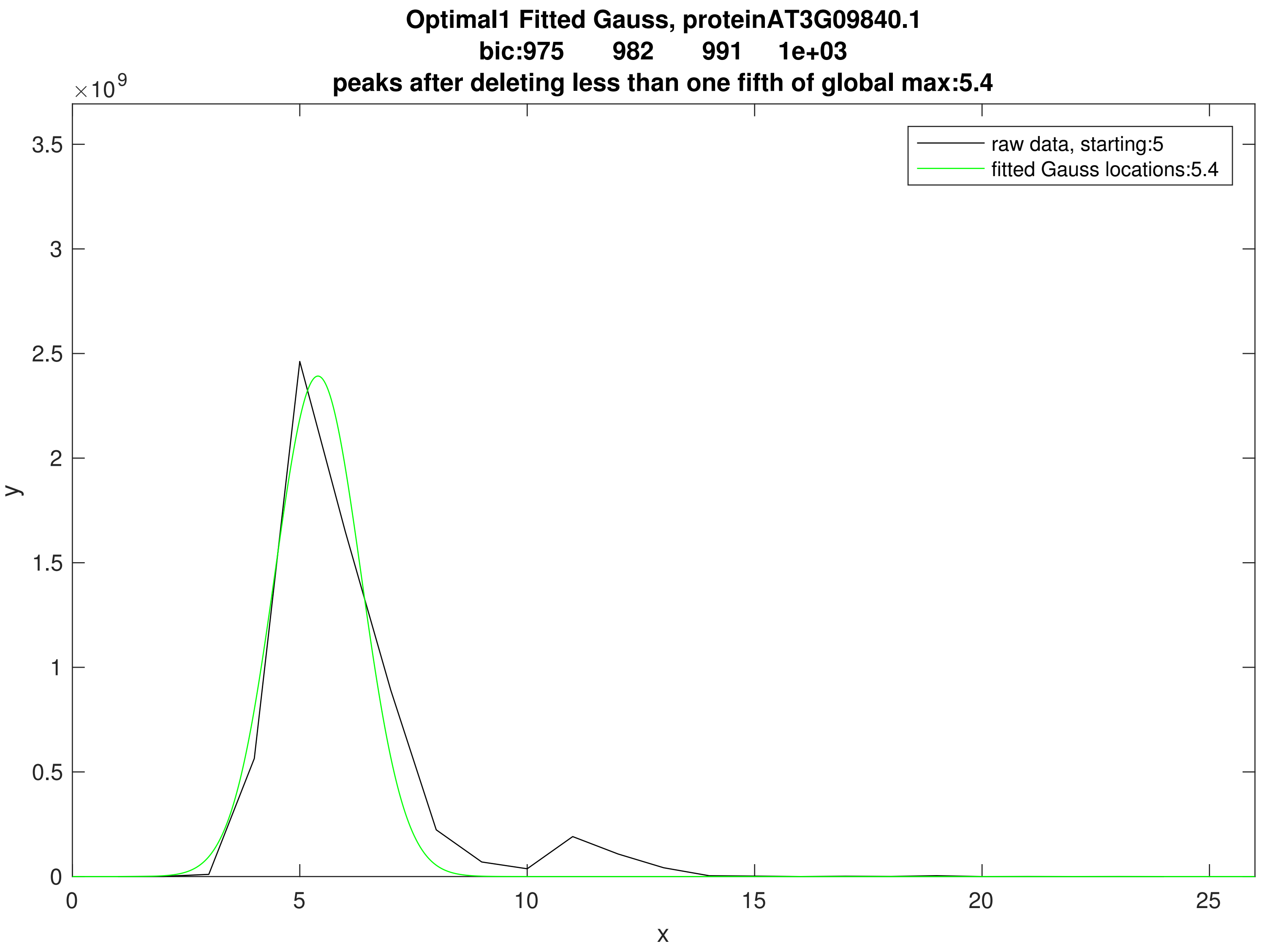

Optimal1 Fitted Gauss, proteinAT3G09840.1
bic:975 982 991 1e+03
peaks after deleting less than one fifth of global max:5.4
×10^9
3.5
3
2.5
2
1.5
1
0.5
0
y
0
5
10
15
20
25
x
raw data, starting:5
fitted Gauss locations:5.4

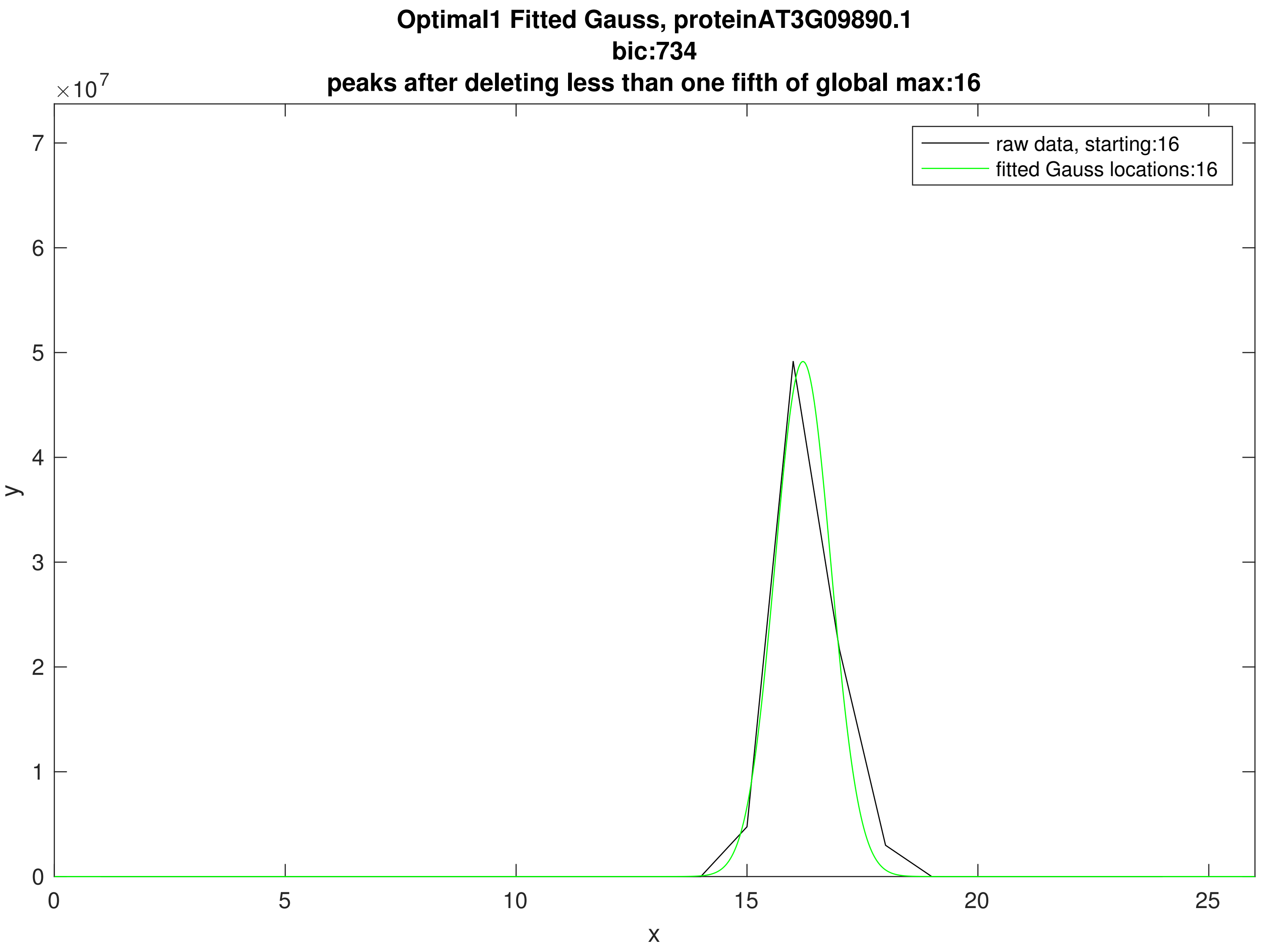
Optimal1 Fitted Gauss, proteinAT3G09890.1
bic:734
peaks after deleting less than one fifth of global max:16
×10^7
raw data, starting:16
fitted Gauss locations:16
y
x
0
1
2
3
4
5
6
7
0
5
10
15
20
25

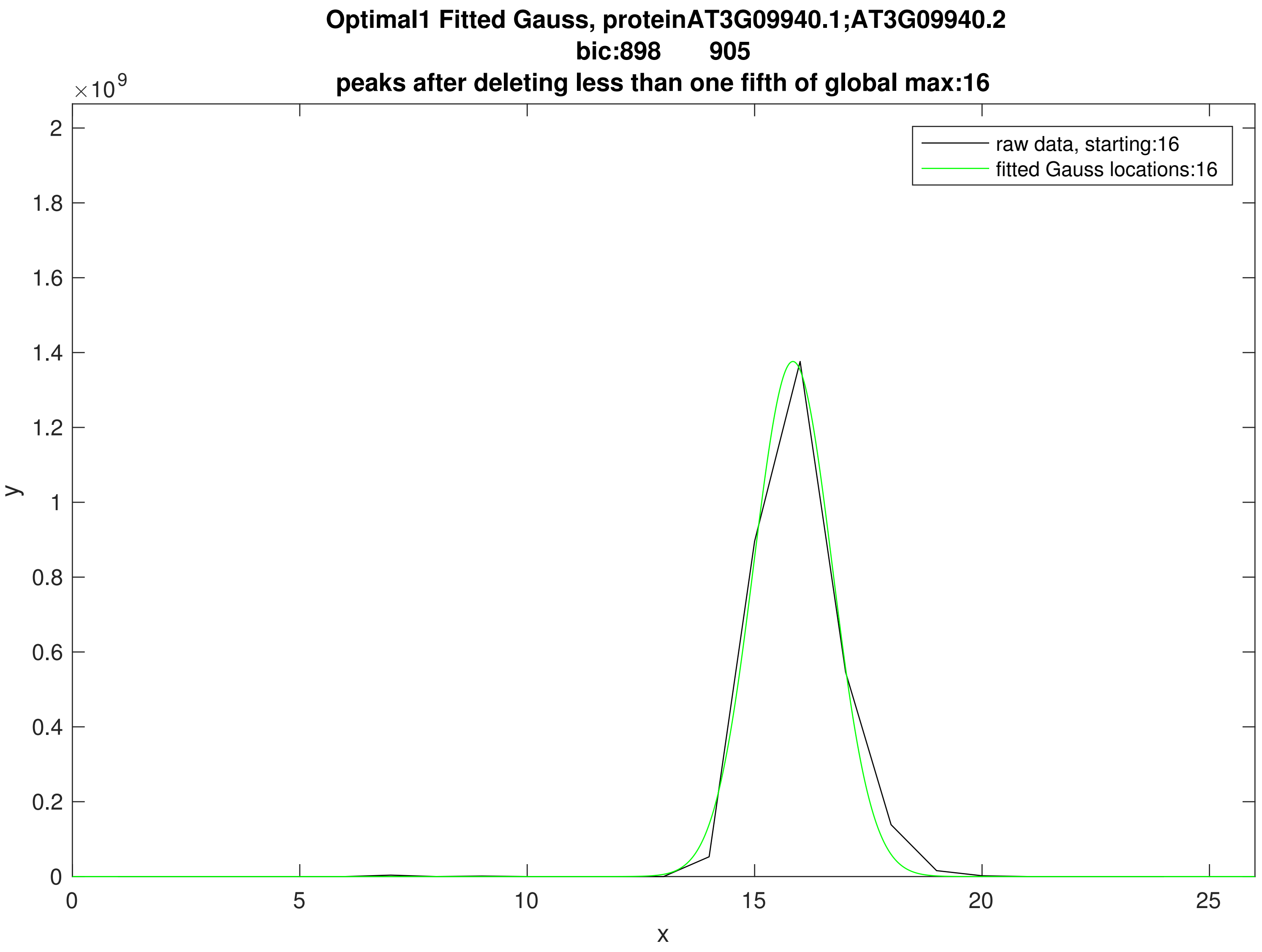

Optimal1 Fitted Gauss, proteinAT3G09940.1;AT3G09940.2
bic:898 905
peaks after deleting less than one fifth of global max:16
×10^9
raw data, starting:16
fitted Gauss locations:16
y
x
0
0.2
0.4
0.6
0.8
1
1.2
1.4
1.6
1.8
2
5
10
15
20
25

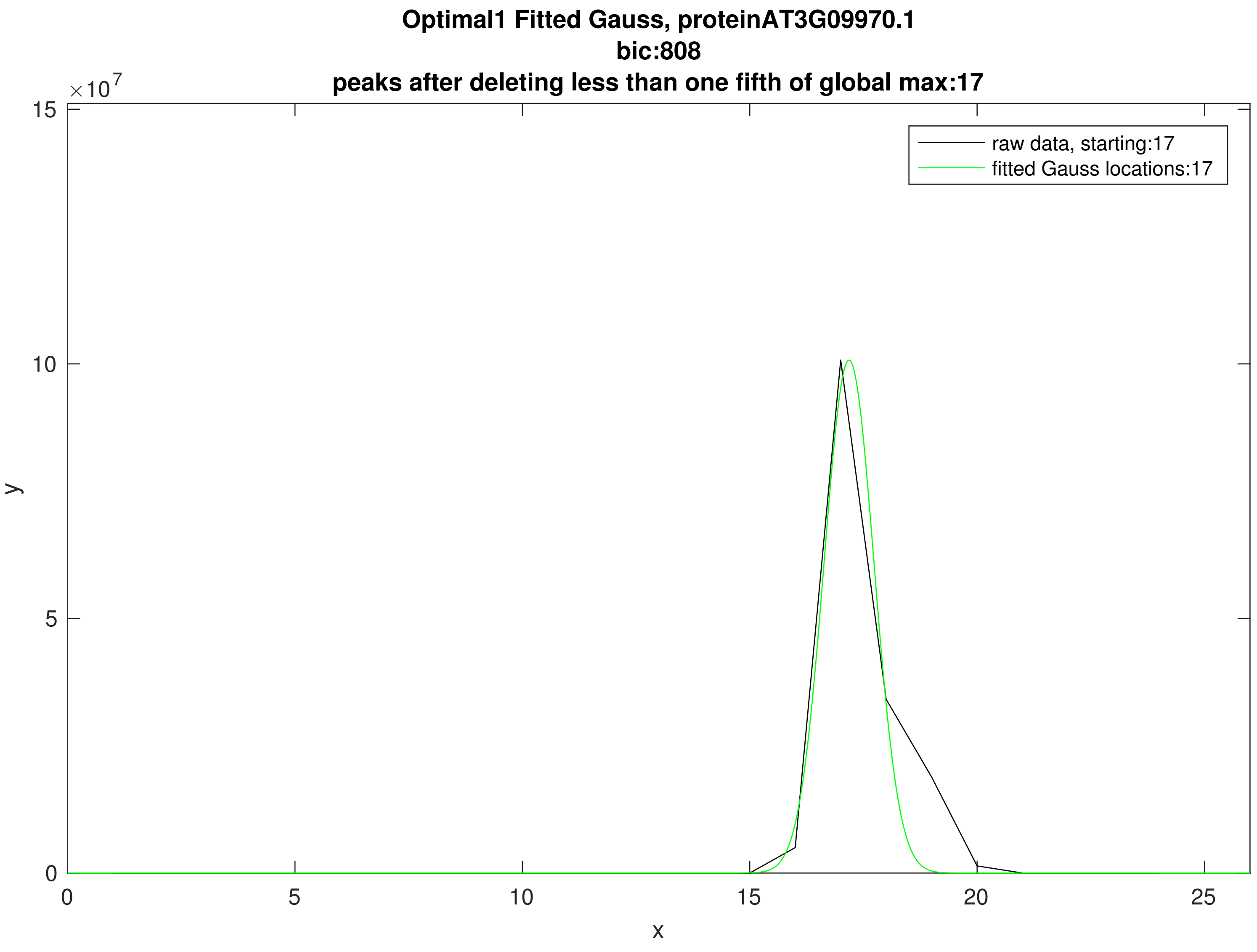
Optimal1 Fitted Gauss, proteinAT3G09970.1
bic:808
peaks after deleting less than one fifth of global max:17
×10^7
15
10
5
0
y
0
5
10
15
20
25
x
raw data, starting:17
fitted Gauss locations:17

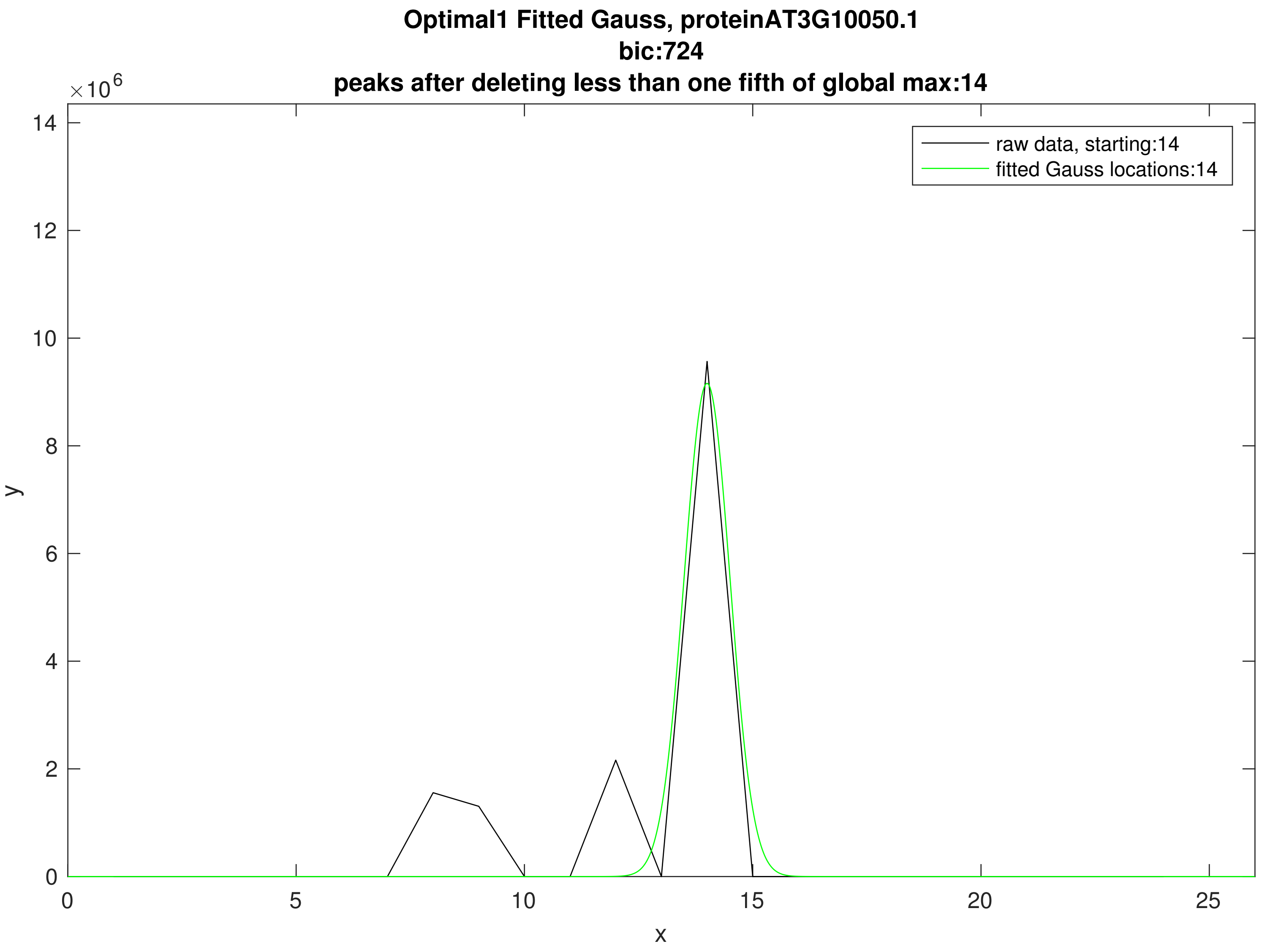
Optimal1 Fitted Gauss, proteinAT3G10050.1
bic:724
peaks after deleting less than one fifth of global max:14
×10^6
raw data, starting:14
fitted Gauss locations:14
y
x
0
2
4
6
8
10
12
14
0
5
10
15
20
25

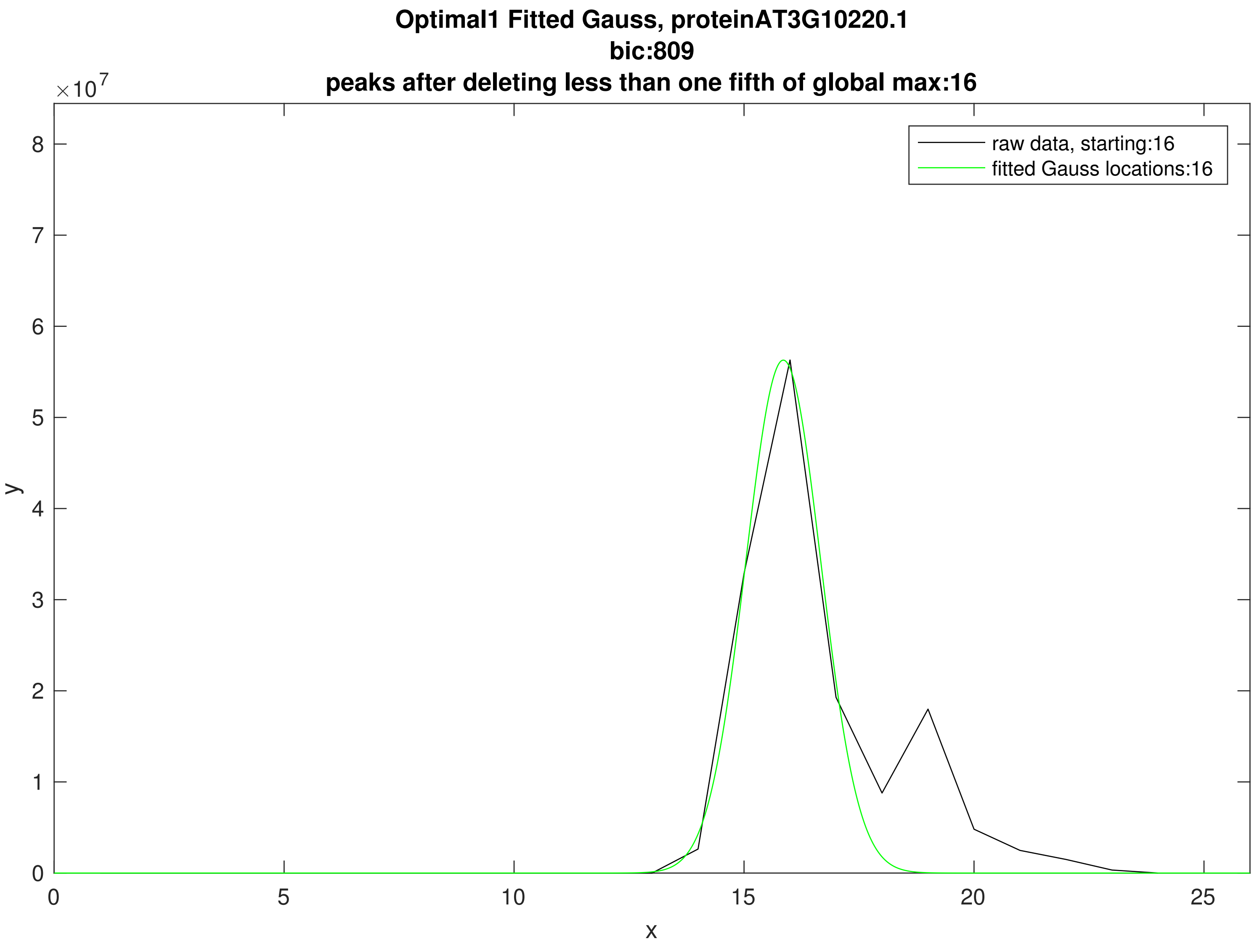

Optimal1 Fitted Gauss, proteinAT3G10220.1
bic:809
peaks after deleting less than one fifth of global max:16
×10^7
raw data, starting:16
fitted Gauss locations:16
y
x
0
1
2
3
4
5
6
7
8
0
5
10
15
20
25

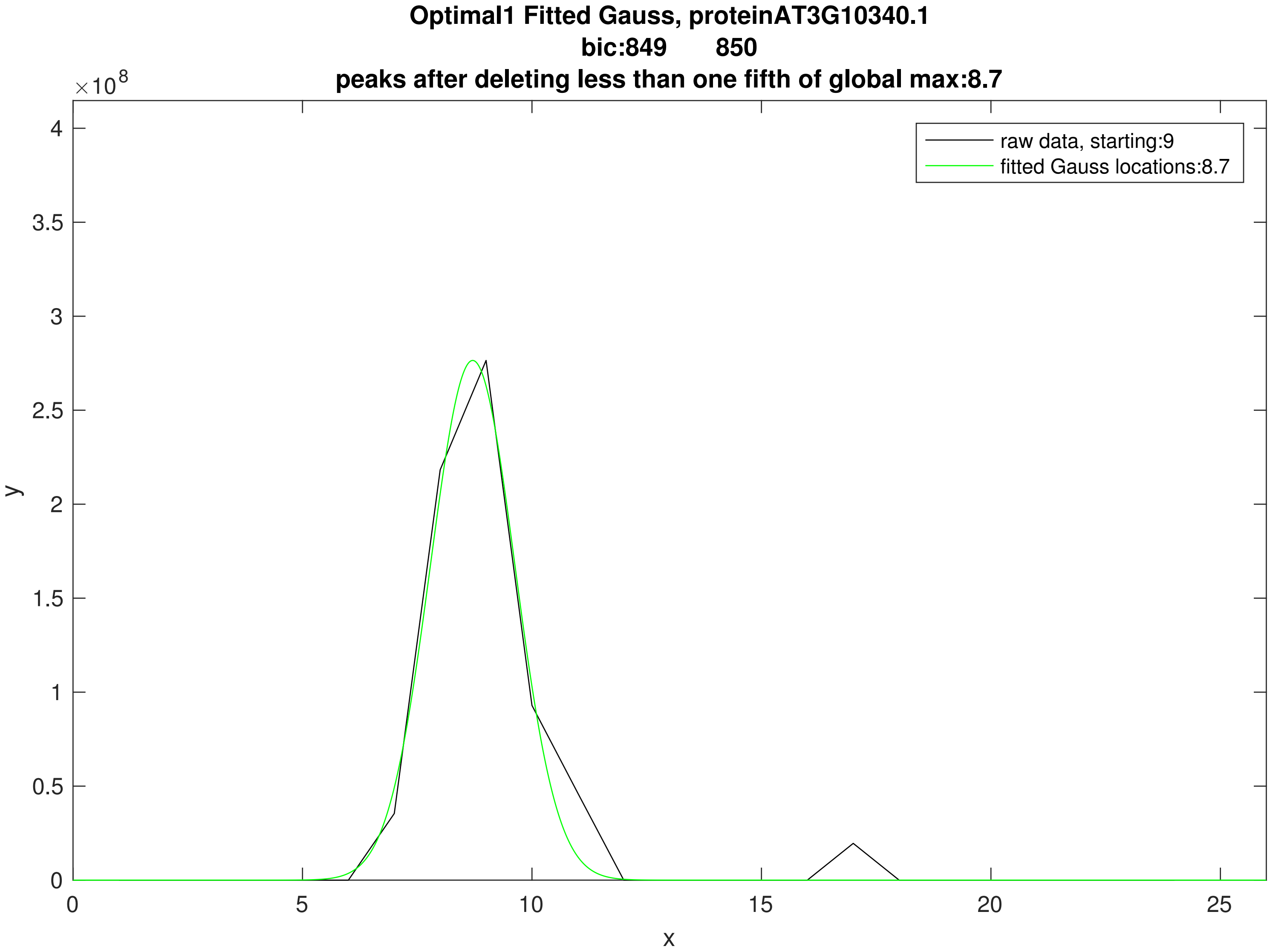
Optimal1 Fitted Gauss, proteinAT3G10340.1
bic:849 850
peaks after deleting less than one fifth of global max:8.7
×10^8
raw data, starting:9
fitted Gauss locations:8.7
y
x
0
0.5
1
1.5
2
2.5
3
3.5
4
0
5
10
15
20
25

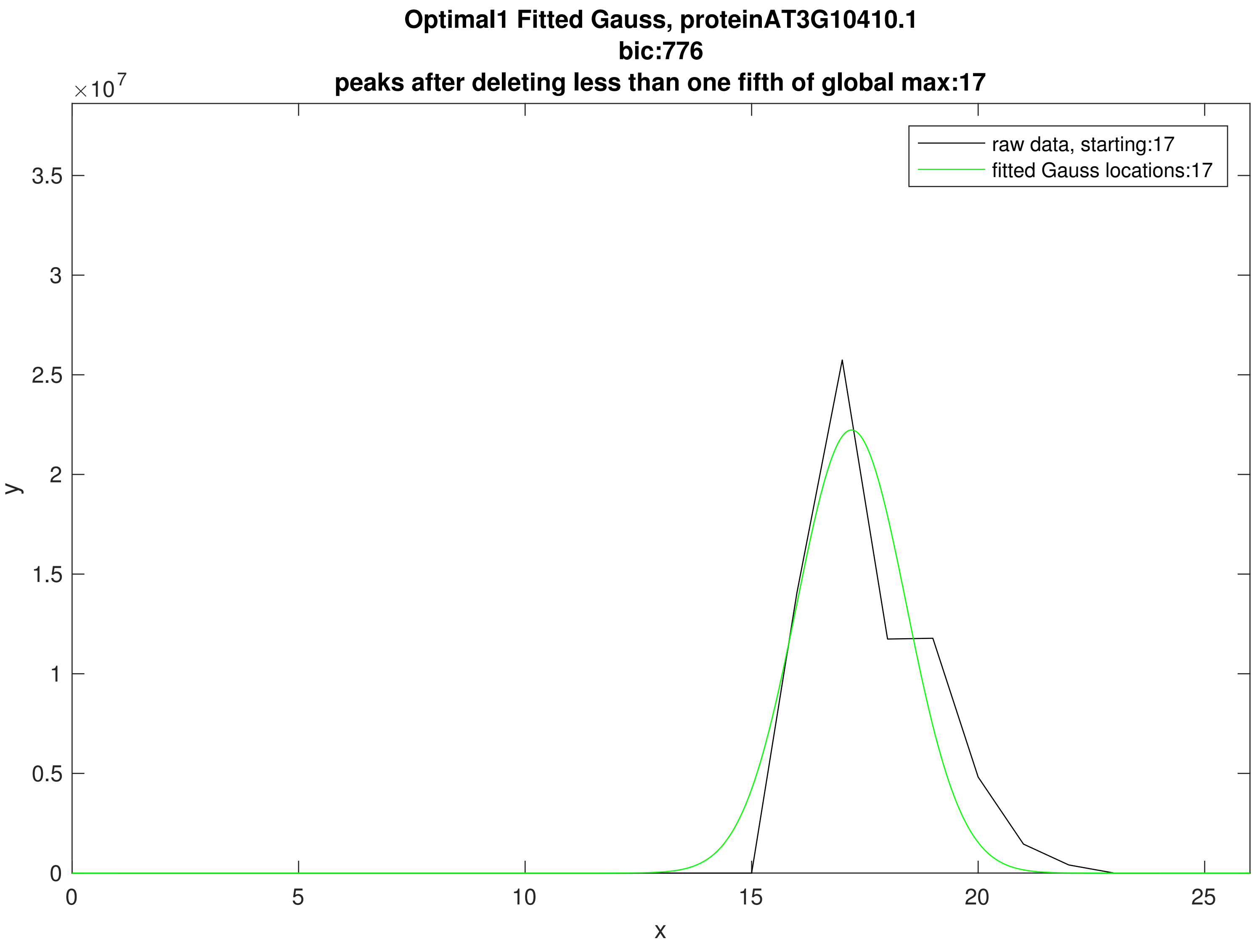

Optimal1 Fitted Gauss, proteinAT3G10410.1
bic:776
peaks after deleting less than one fifth of global max:17
×10^7
3.5
3
2.5
2
1.5
1
0.5
0
y
0
5
10
15
20
25
x
raw data, starting:17
fitted Gauss locations:17

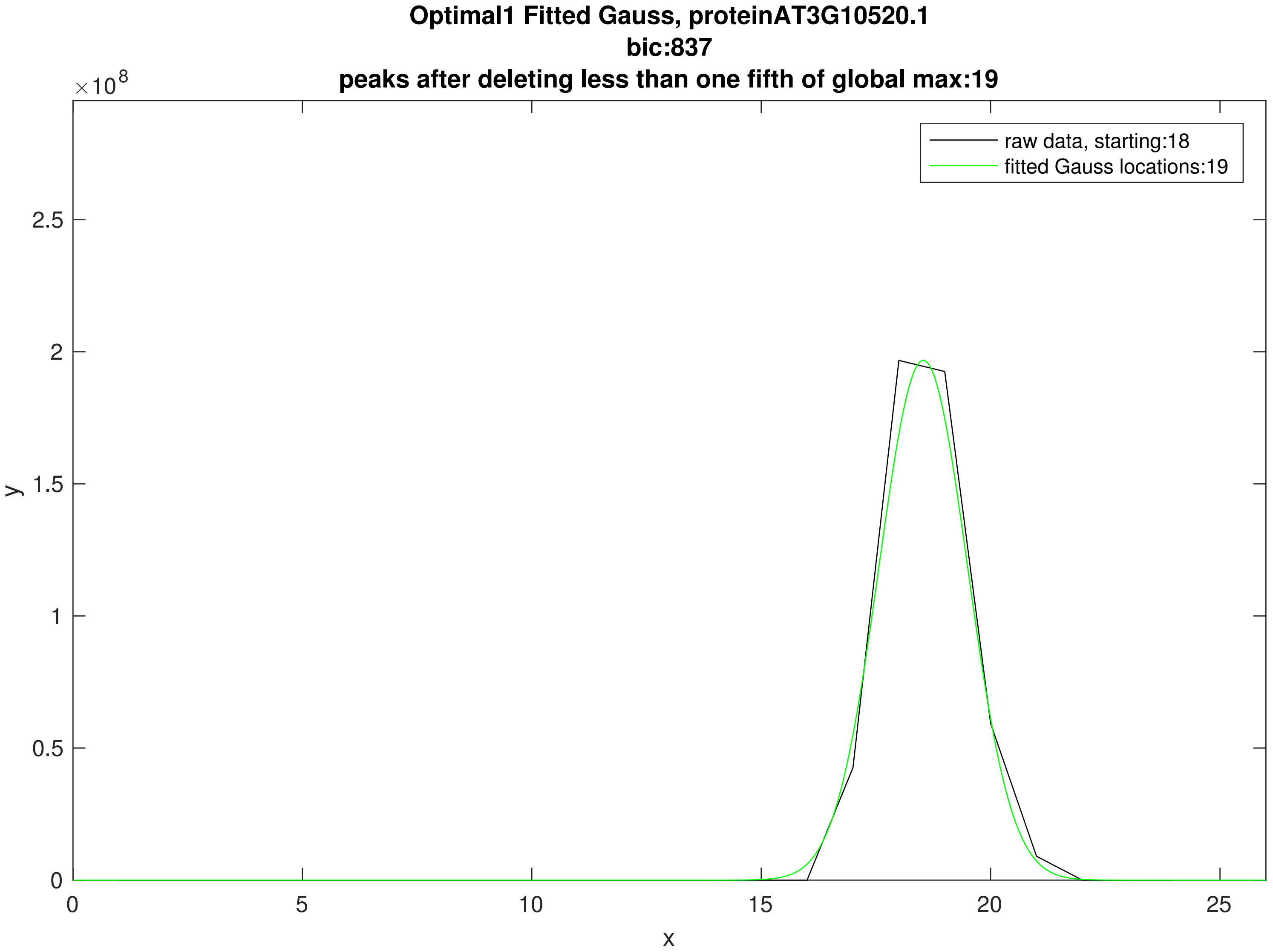
Optimal1 Fitted Gauss, proteinAT3G10520.1
bic:837
peaks after deleting less than one fifth of global max:19
×10^8
raw data, starting:18
fitted Gauss locations:19
2.5
2
1.5
1
0.5
0
y
0
5
10
15
20
25
x

1 Fitted Gauss, proteinAT3G10610.1;AT2G05220.2;AT2G05220.1;AT5G04800.4;AT5G04800.3;AT5G04800.2;AT5G0480
bic:782 786
peaks after deleting less than one fifth of global max:4

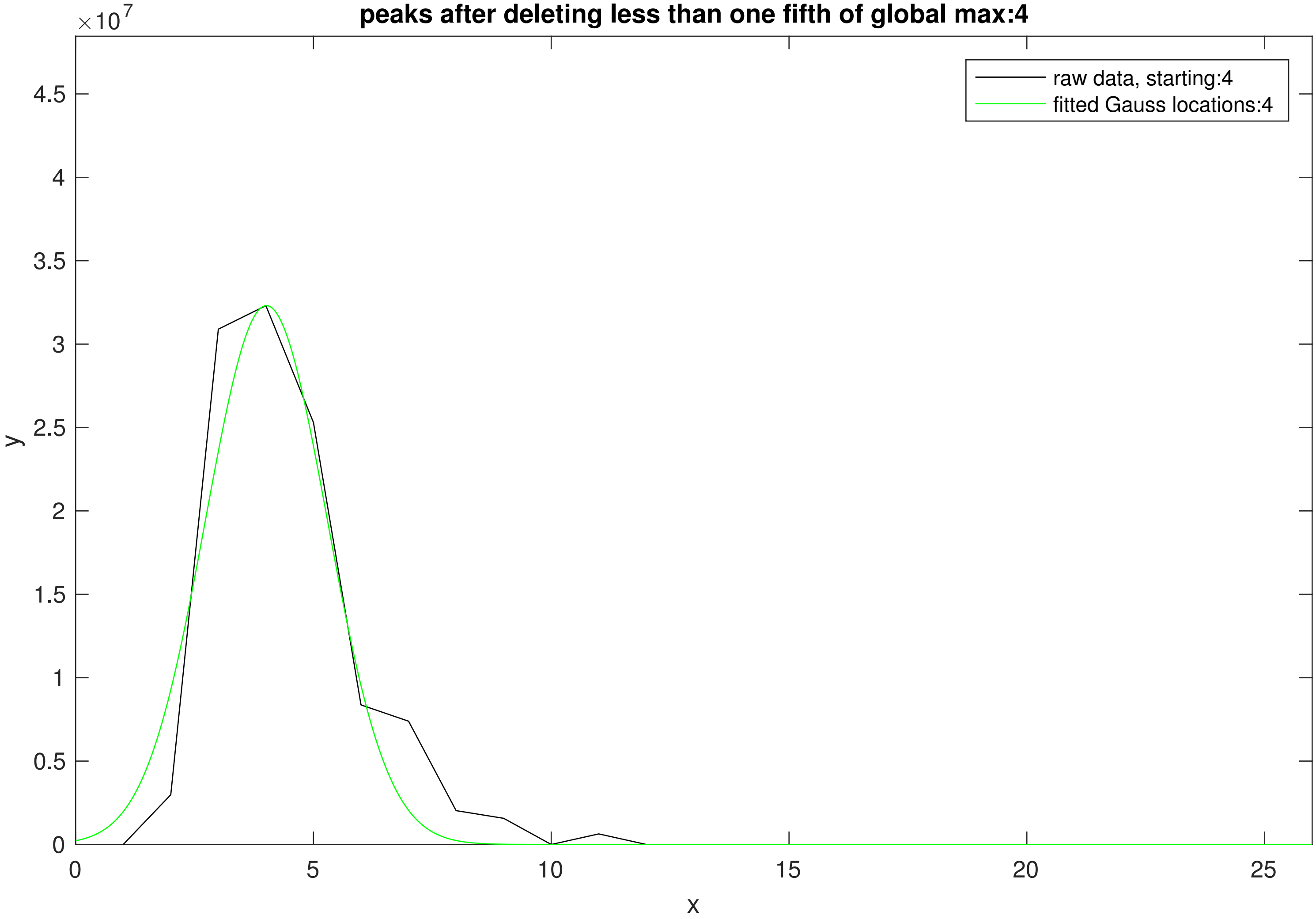

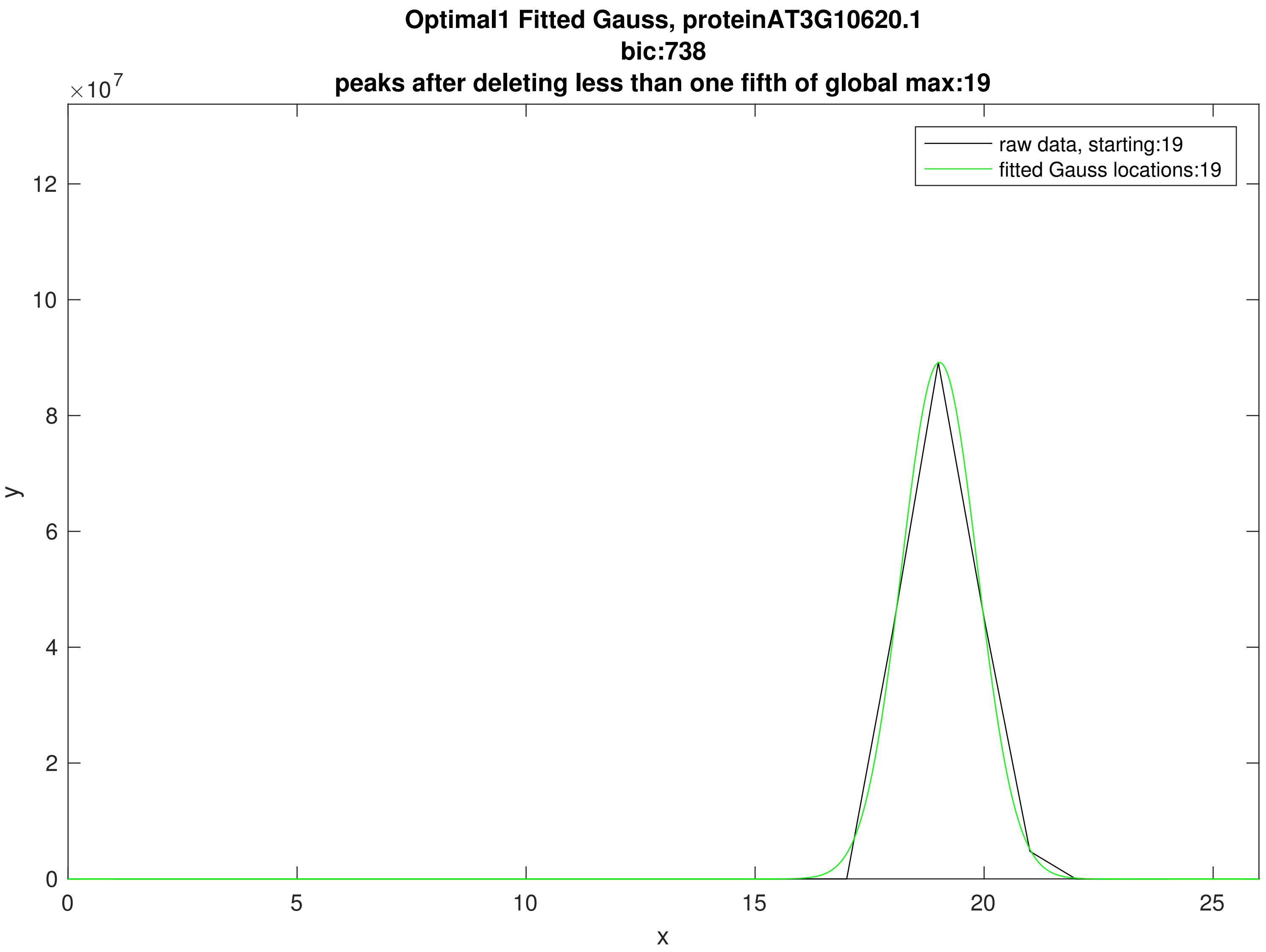

Optimal1 Fitted Gauss, proteinAT3G10620.1
bic:738
peaks after deleting less than one fifth of global max:19
×10^7
raw data, starting:19
fitted Gauss locations:19
0
2
4
6
8
10
12
y
0
5
10
15
20
25
x

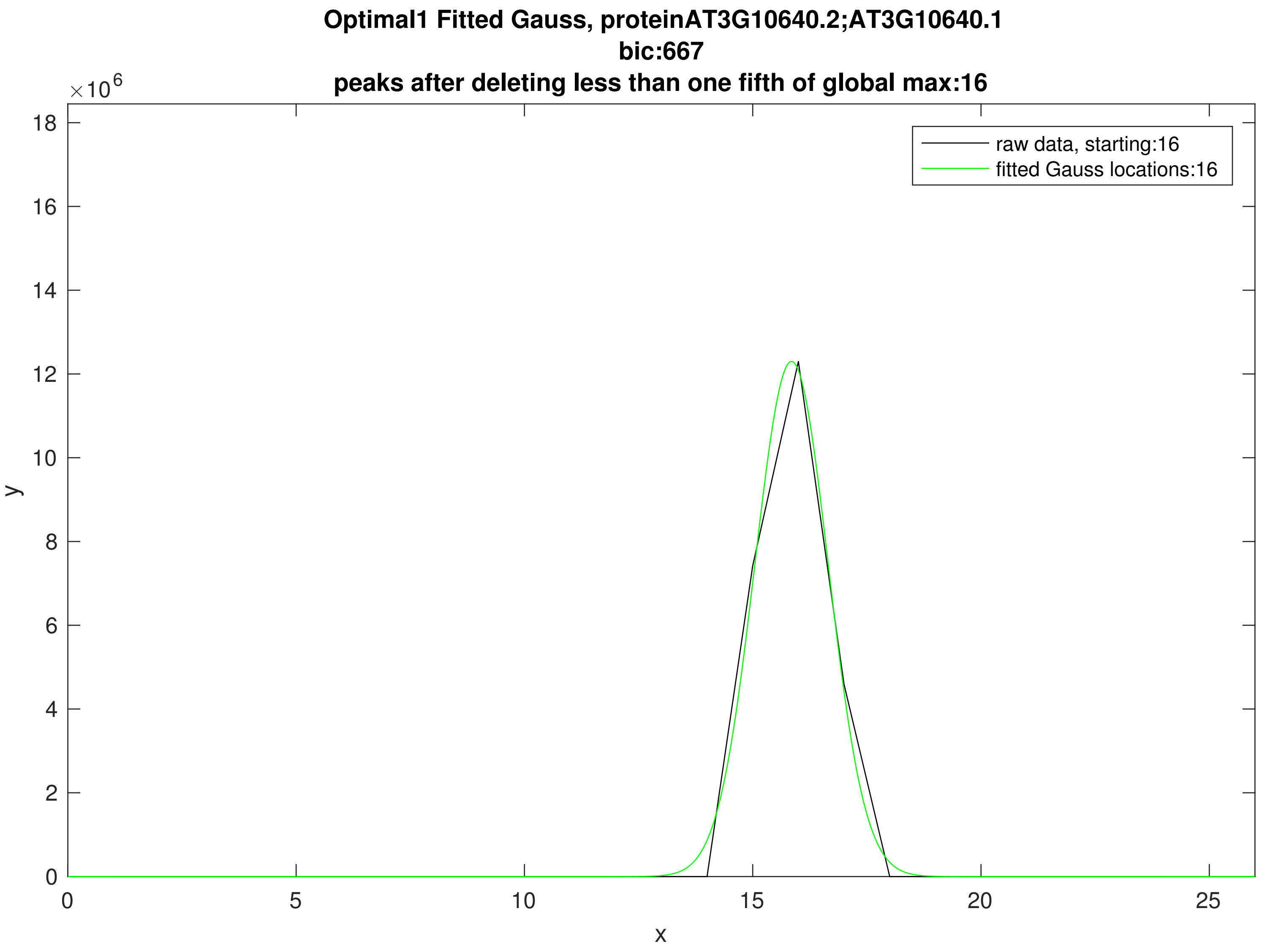
Optimal1 Fitted Gauss, proteinAT3G10640.2;AT3G10640.1
bic:667
peaks after deleting less than one fifth of global max:16
×10^6
raw data, starting:16
fitted Gauss locations:16
y
x
0
2
4
6
8
10
12
14
16
18
0
5
10
15
20
25

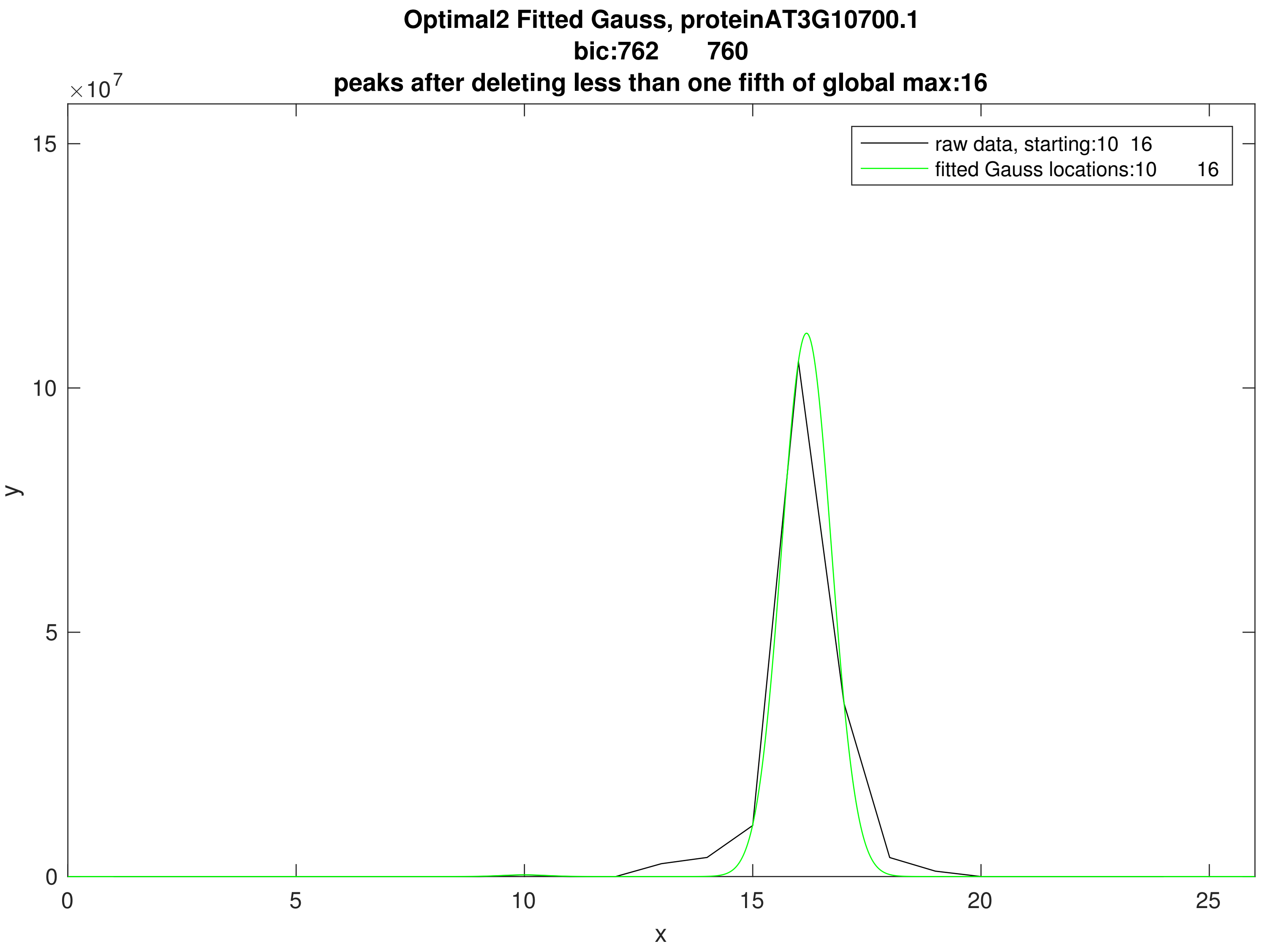
Optimal2 Fitted Gauss, proteinAT3G10700.1
bic:762 760
peaks after deleting less than one fifth of global max:16
×10^7
raw data, starting:10 16
fitted Gauss locations:10 16
y
x
0
5
10
15
20
25

**Optimal1 Fitted Gauss, proteinAT3G10720.1;AT1G02810.1;AT3G10720.2**
**bic:683**
**peaks after deleting less than one fifth of global max:14**

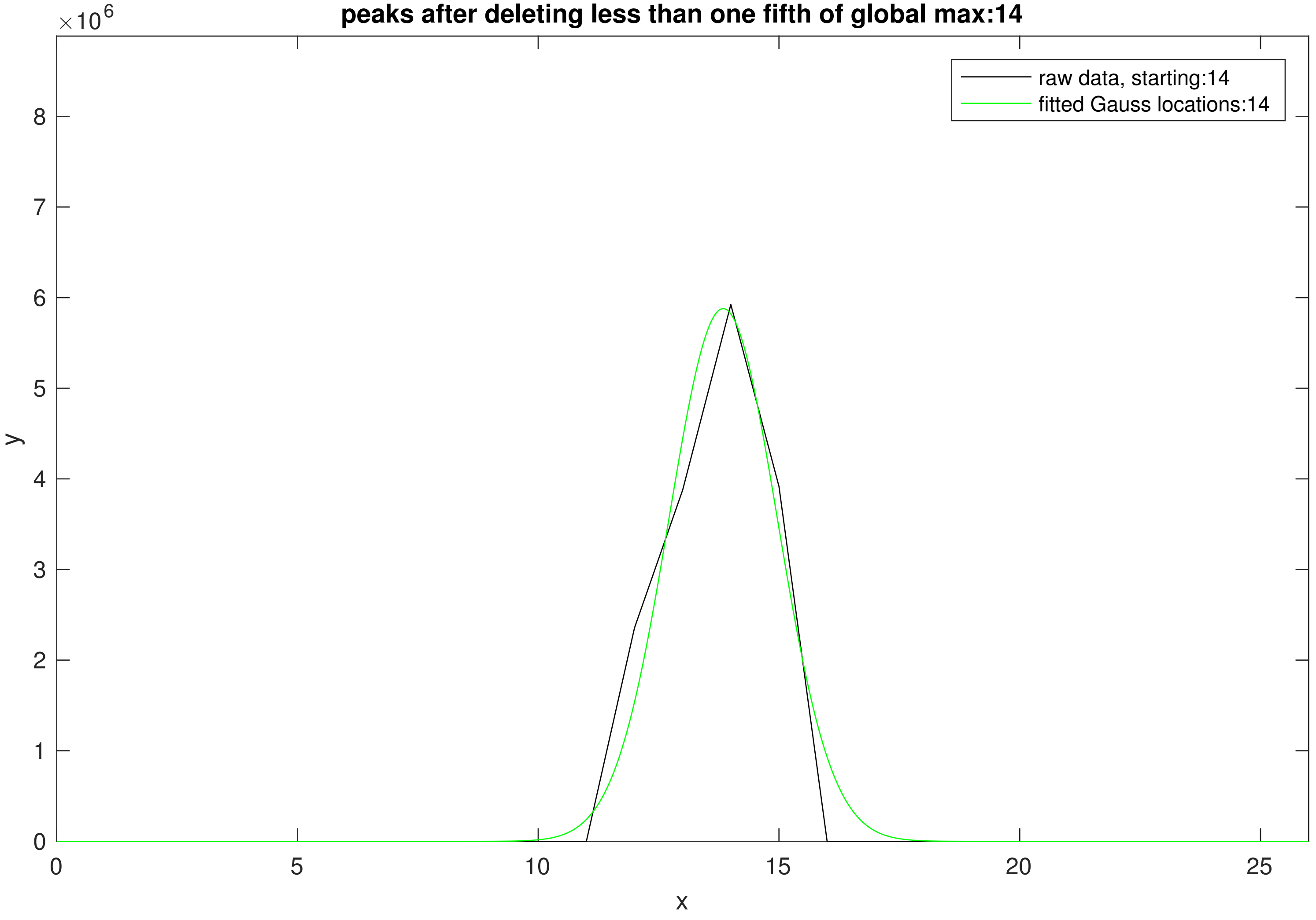

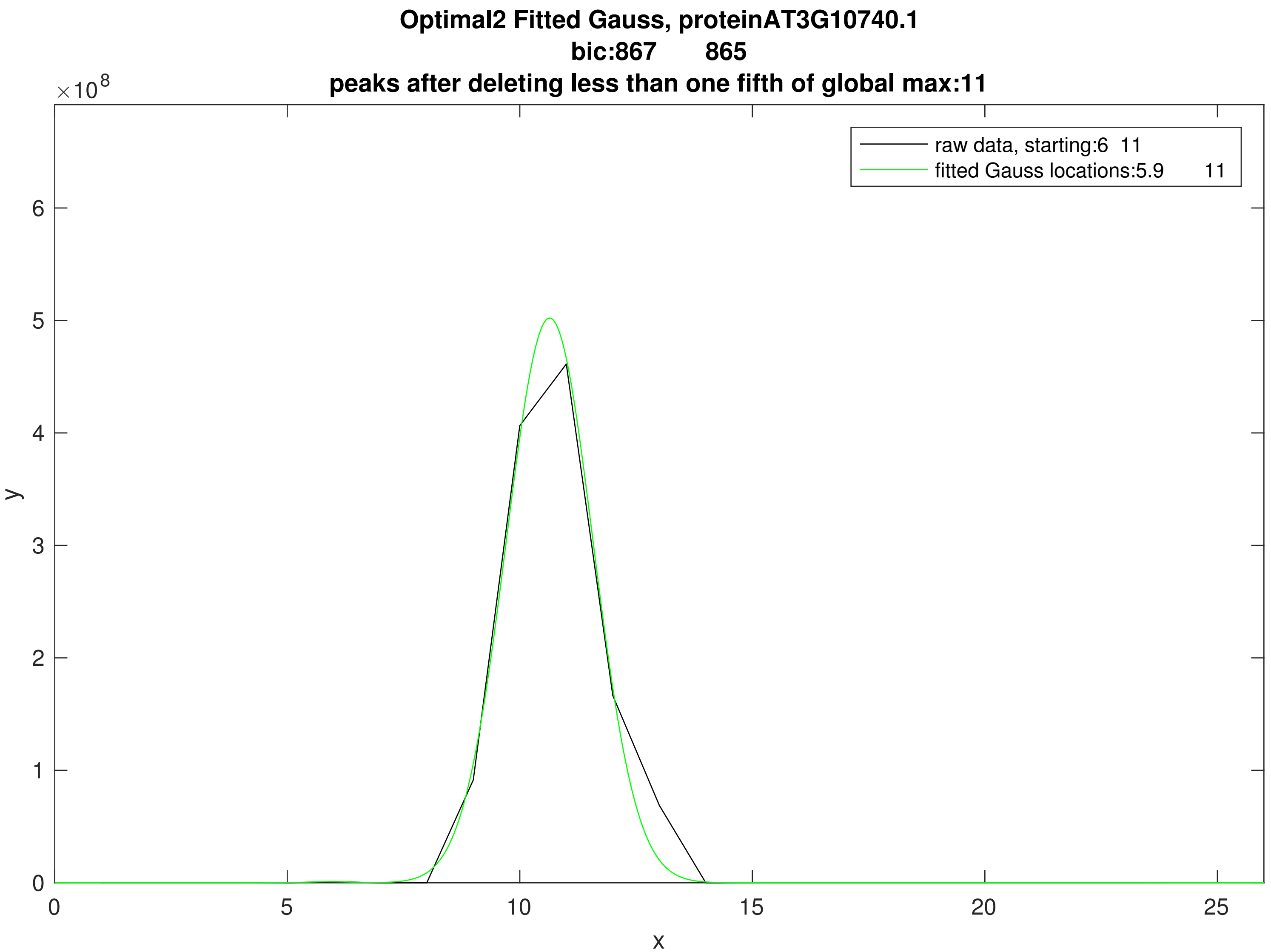

Optimal2 Fitted Gauss, proteinAT3G10740.1
bic:867 865
peaks after deleting less than one fifth of global max:11
×10^8
raw data, starting:6 11
fitted Gauss locations:5.9 11
y
x
0
1
2
3
4
5
6
0
5
10
15
20
25

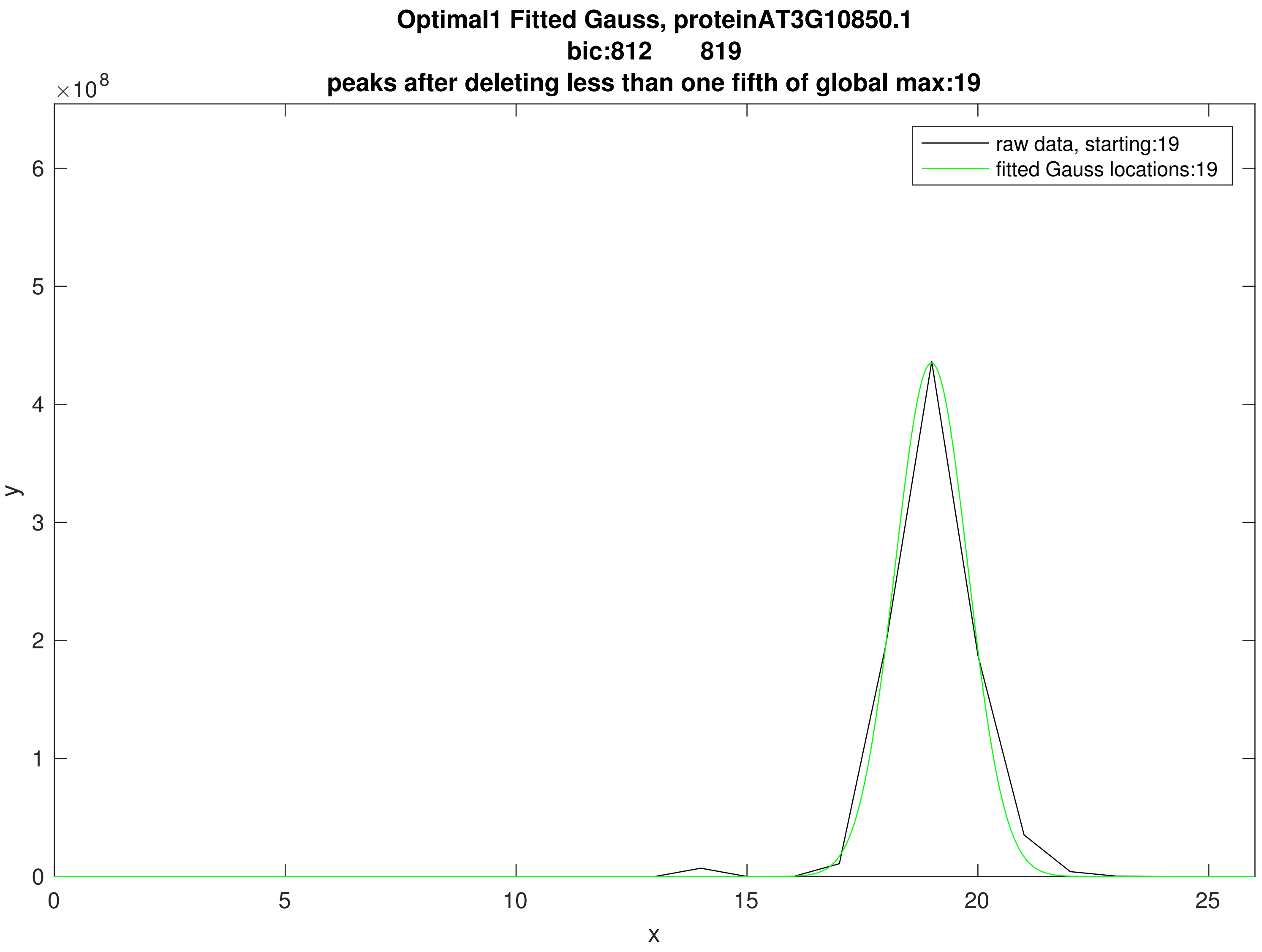

Optimal1 Fitted Gauss, proteinAT3G10850.1
bic:812 819
peaks after deleting less than one fifth of global max:19
×10^8
raw data, starting:19
fitted Gauss locations:19
y
x
0
1
2
3
4
5
6
0
5
10
15
20
25

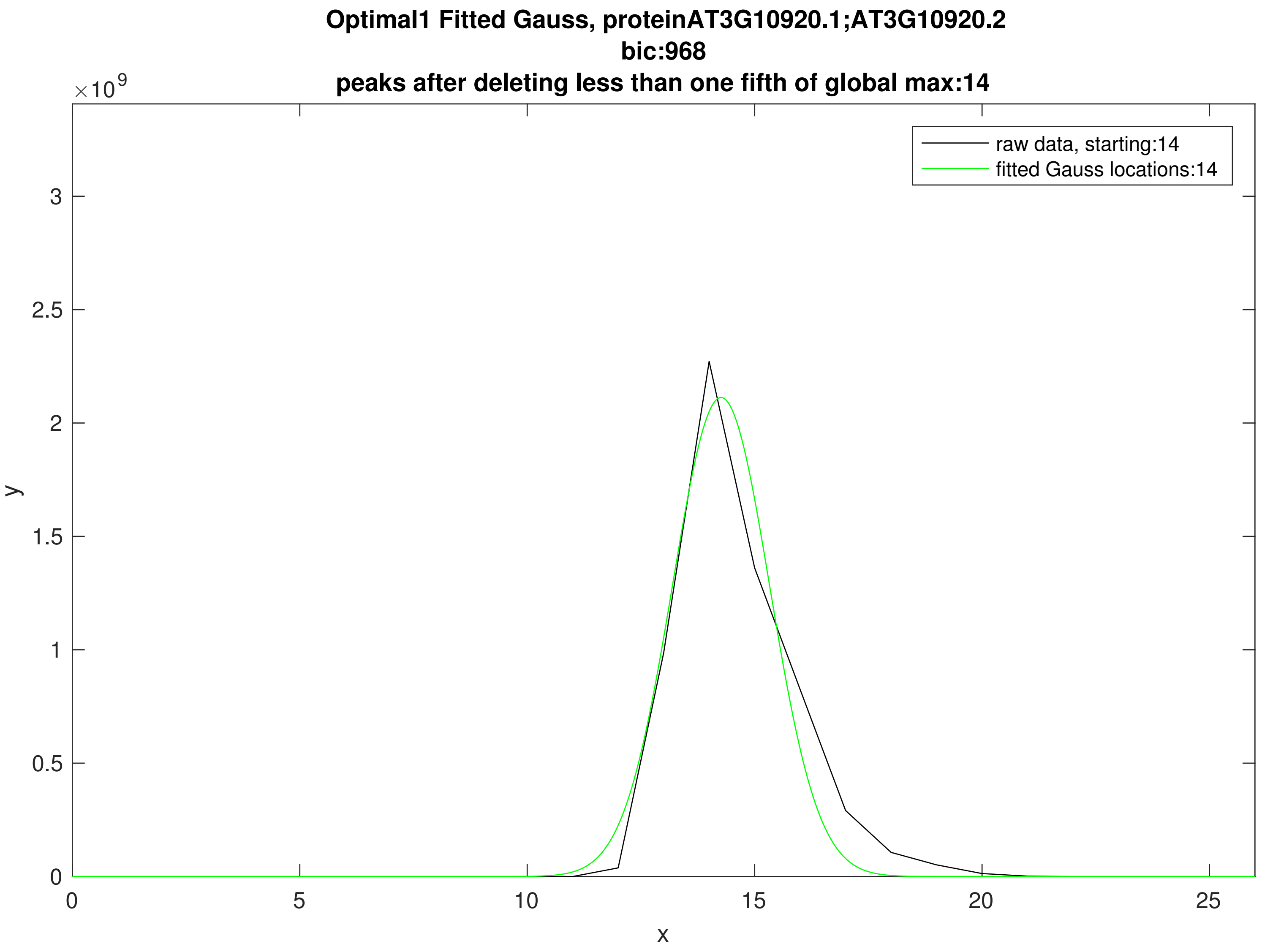
Optimal1 Fitted Gauss, proteinAT3G10920.1;AT3G10920.2
bic:968
peaks after deleting less than one fifth of global max:14
×10^9
raw data, starting:14
fitted Gauss locations:14
y
x
0
0.5
1
1.5
2
2.5
3
0
5
10
15
20
25

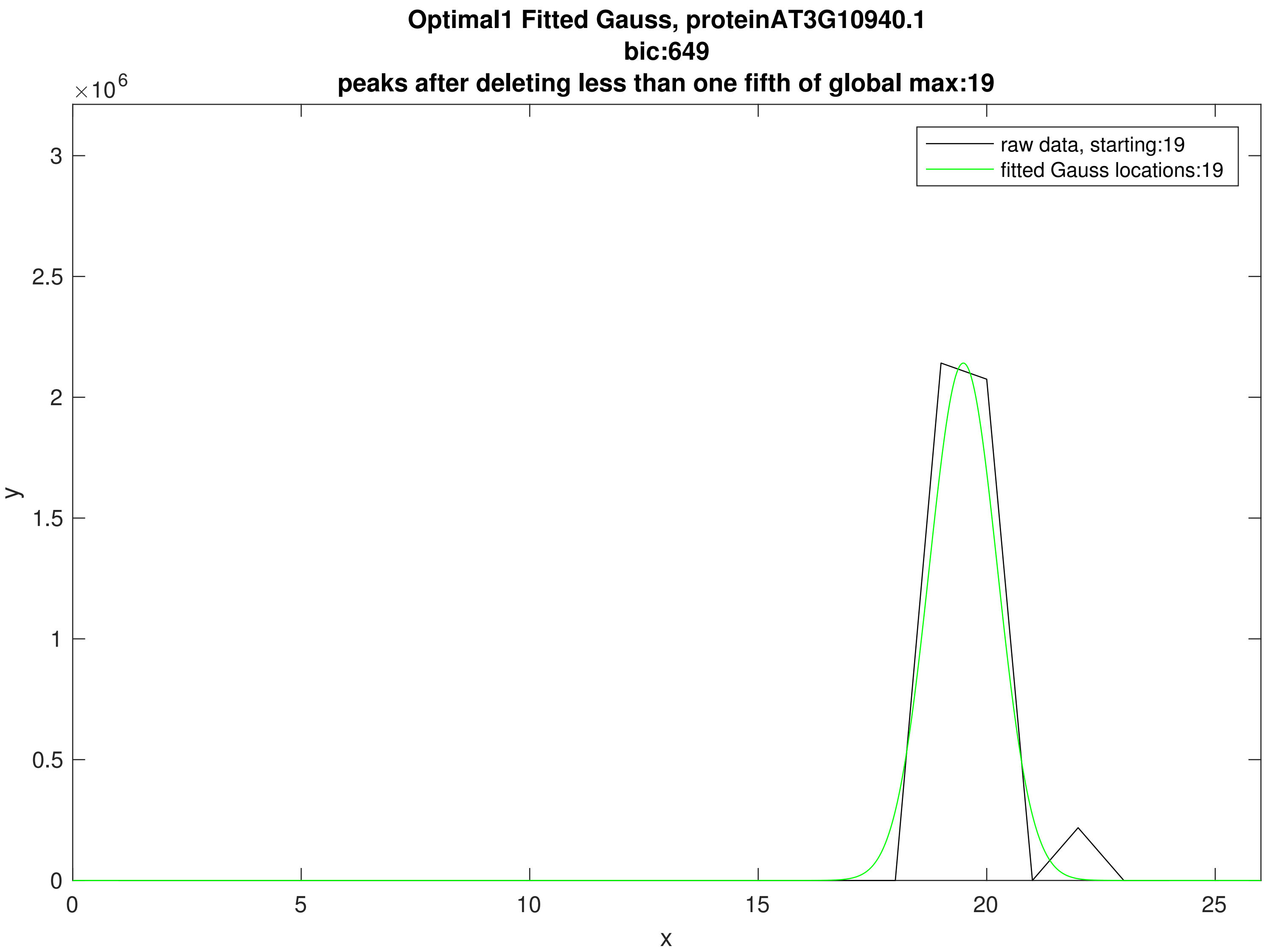

Optimal1 Fitted Gauss, proteinAT3G10940.1
bic:649
peaks after deleting less than one fifth of global max:19
×10^6
3
2.5
2
1.5
1
0.5
0
y
0
5
10
15
20
25
x
raw data, starting:19
fitted Gauss locations:19

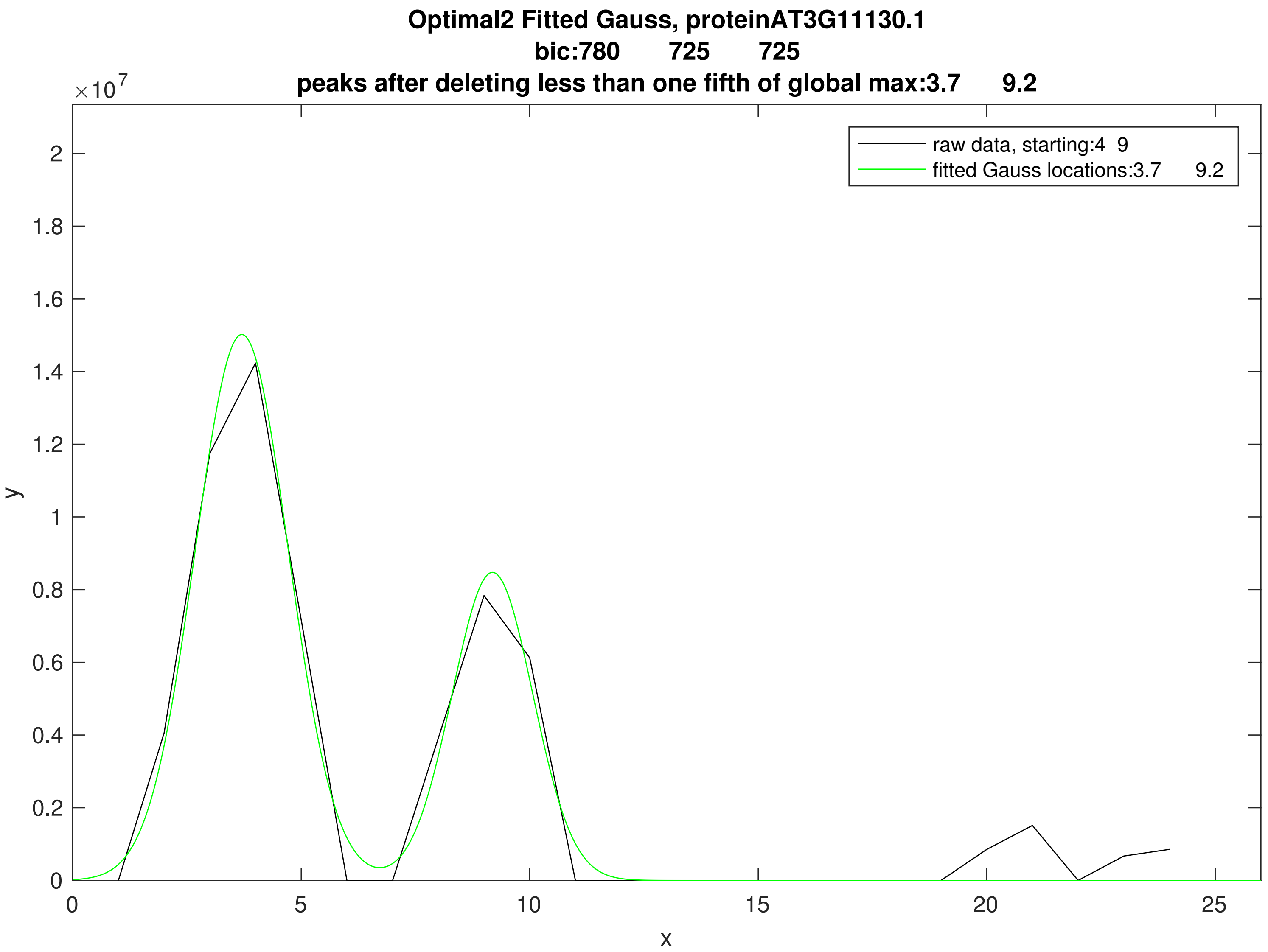

Optimal2 Fitted Gauss, proteinAT3G11130.1
bic:780 725 725
peaks after deleting less than one fifth of global max:3.7 9.2
×10^7
raw data, starting:4 9
fitted Gauss locations:3.7 9.2
y
x
0
0.2
0.4
0.6
0.8
1
1.2
1.4
1.6
1.8
2
0
5
10
15
20
25

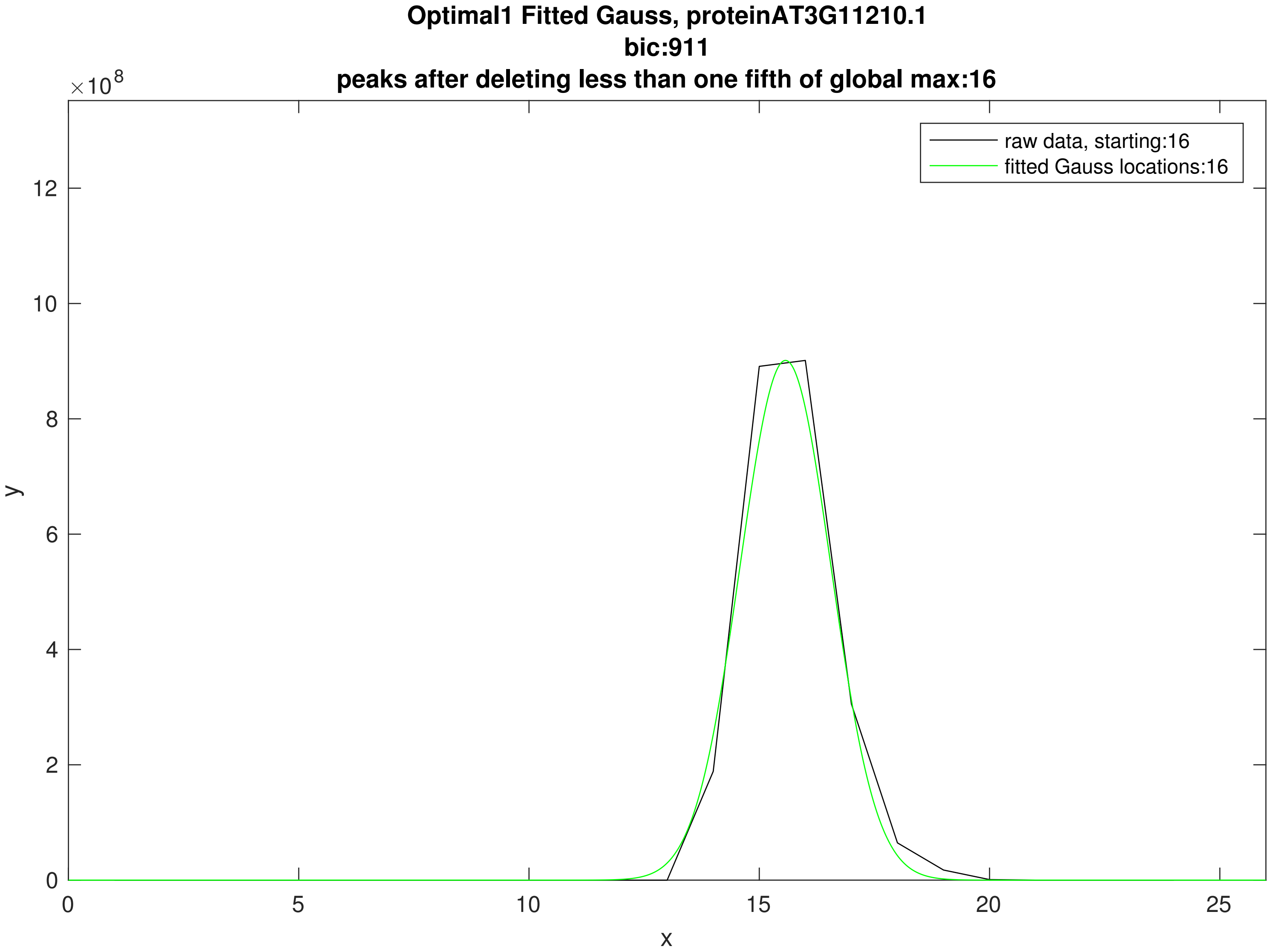
Optimal1 Fitted Gauss, proteinAT3G11210.1
bic:911
peaks after deleting less than one fifth of global max:16
×10^8
raw data, starting:16
fitted Gauss locations:16
y
x
0
2
4
6
8
10
12
0
5
10
15
20
25

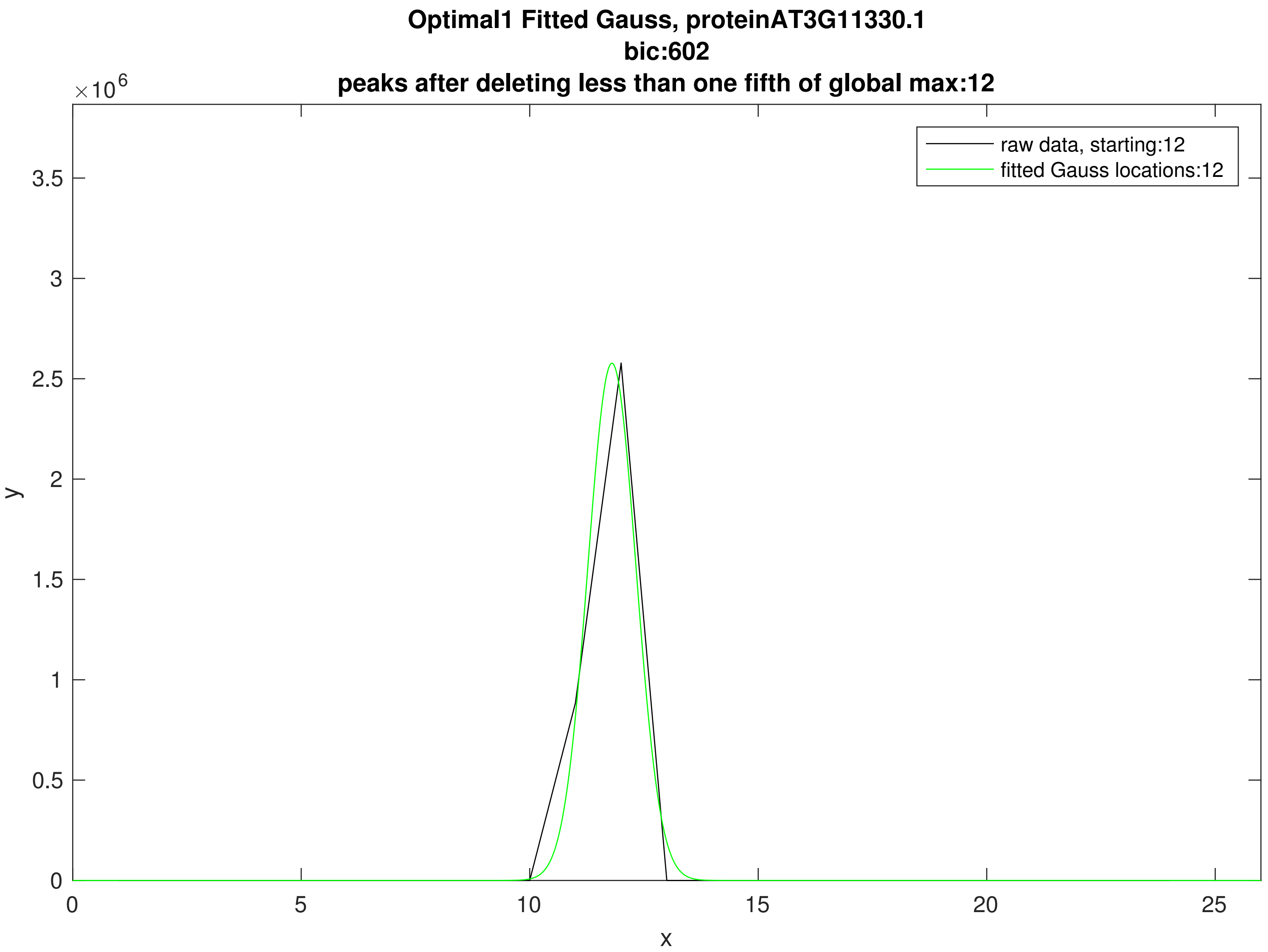

Optimal1 Fitted Gauss, proteinAT3G11330.1
bic:602
peaks after deleting less than one fifth of global max:12
×10^6
raw data, starting:12
fitted Gauss locations:12
3.5
3
2.5
2
1.5
1
0.5
0
y
0
5
10
15
20
25
x

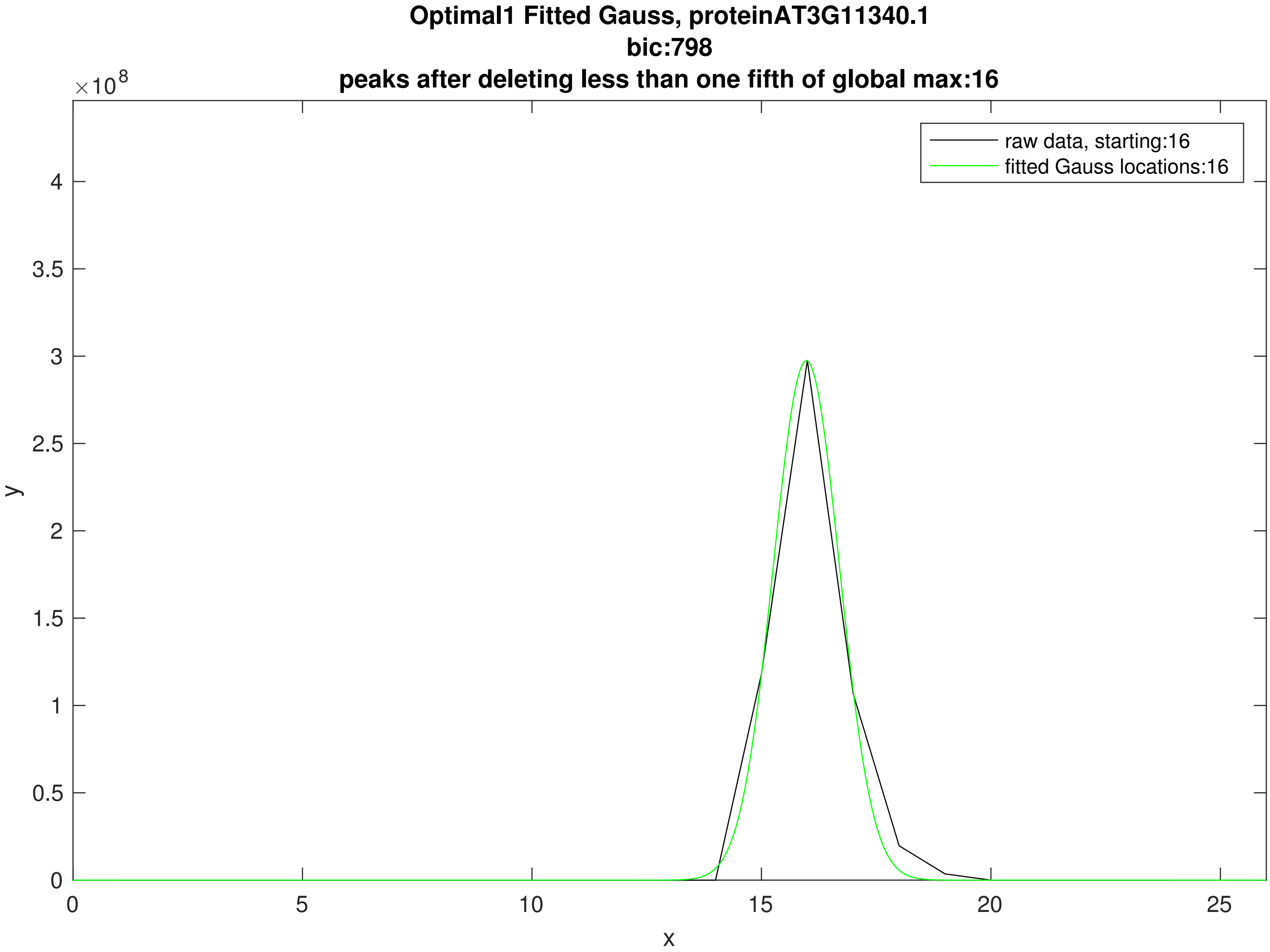
Optimal1 Fitted Gauss, proteinAT3G11340.1
bic:798
peaks after deleting less than one fifth of global max:16
×10^8
raw data, starting:16
fitted Gauss locations:16
4
3.5
3
2.5
2
1.5
1
0.5
0
y
0
5
10
15
20
25
x

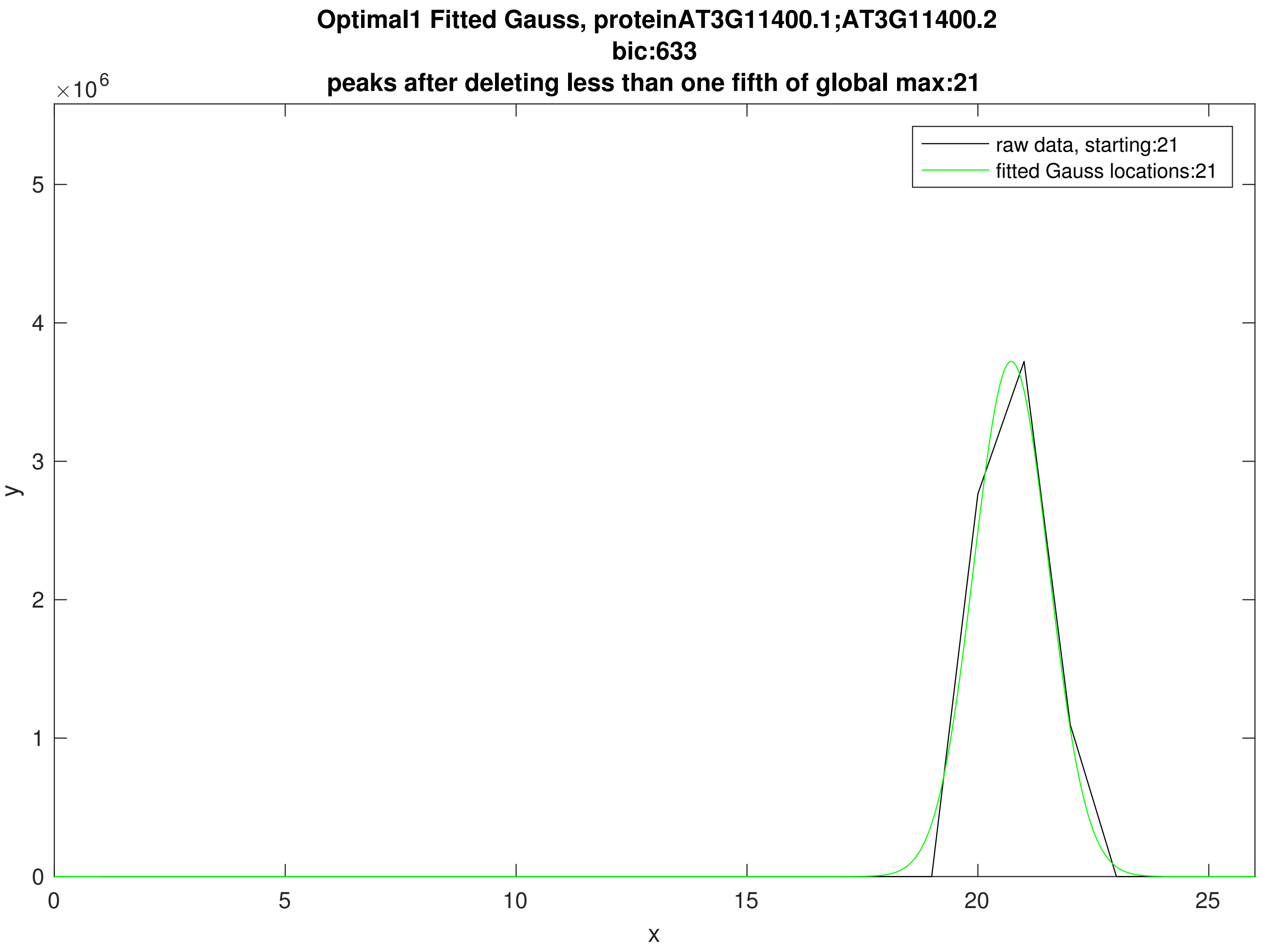

Optimal1 Fitted Gauss, proteinAT3G11400.1;AT3G11400.2
bic:633
peaks after deleting less than one fifth of global max:21
×10^6
raw data, starting:21
fitted Gauss locations:21
y
x
0
1
2
3
4
5
0
5
10
15
20
25

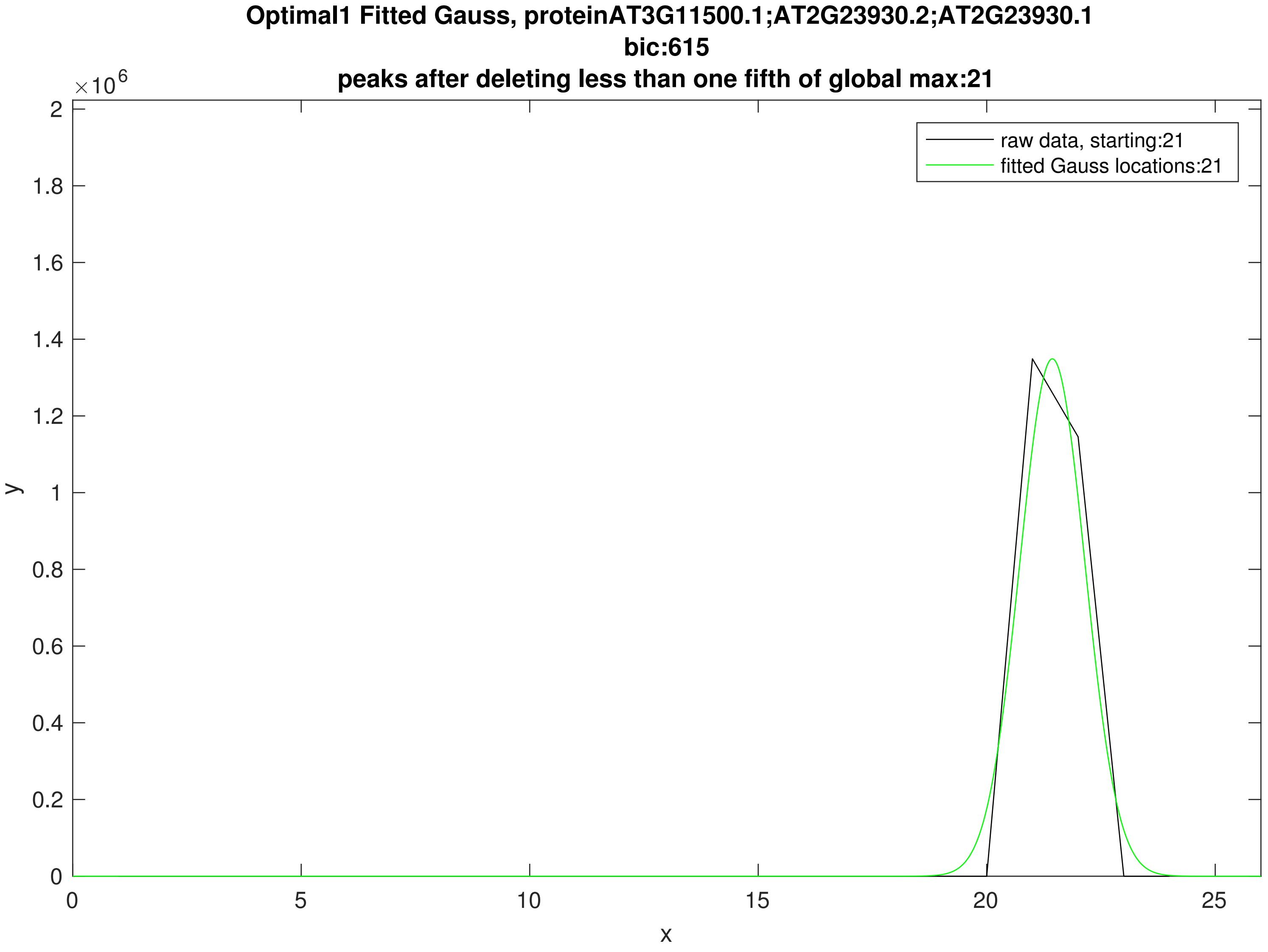
Optimal1 Fitted Gauss, proteinAT3G11500.1;AT2G23930.2;AT2G23930.1
bic:615
peaks after deleting less than one fifth of global max:21
×10^6
raw data, starting:21
fitted Gauss locations:21
y
x
0
0.2
0.4
0.6
0.8
1
1.2
1.4
1.6
1.8
2
0
5
10
15
20
25

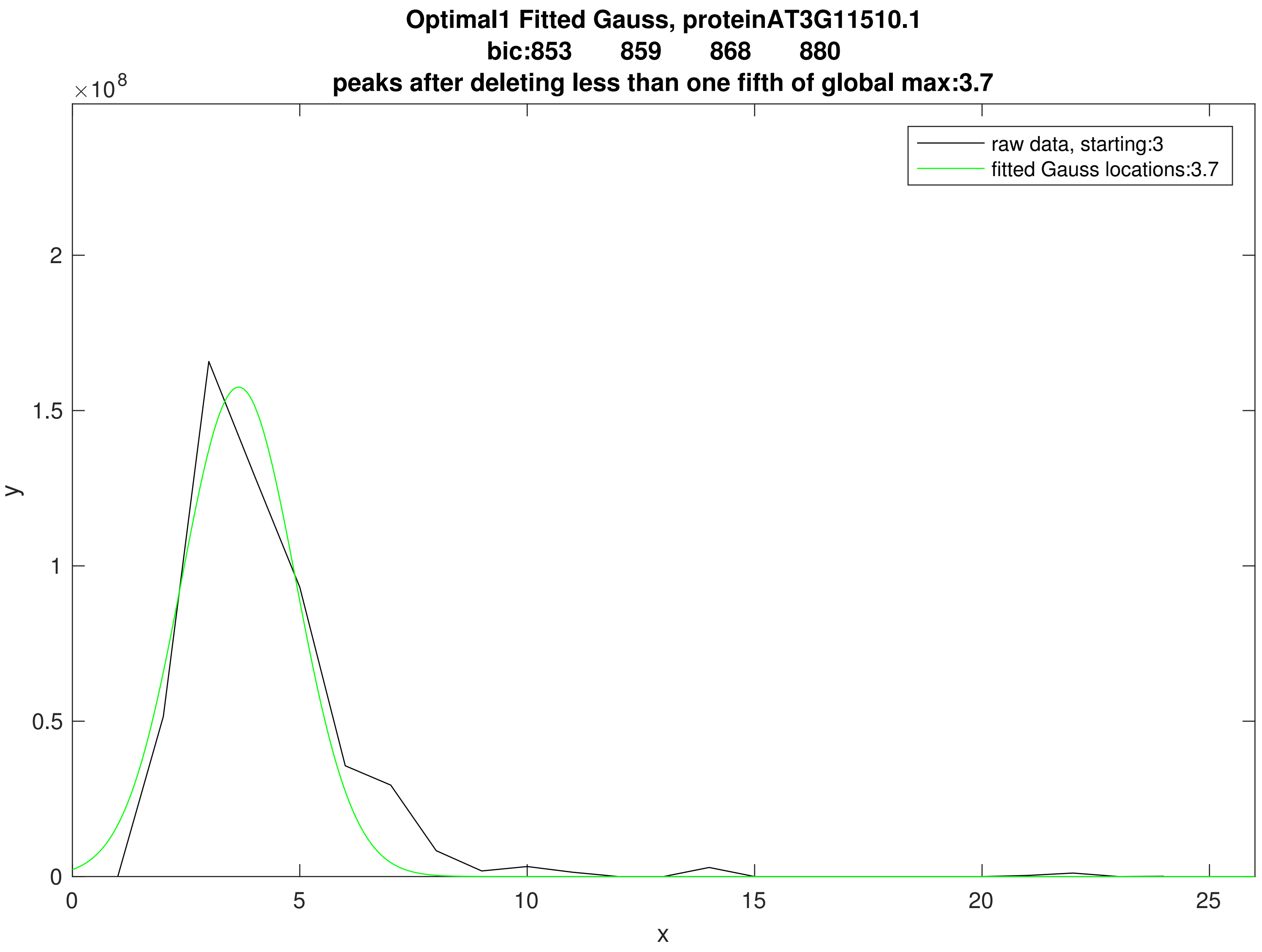

Optimal1 Fitted Gauss, proteinAT3G11510.1
bic:853 859 868 880
peaks after deleting less than one fifth of global max:3.7
×10^8
raw data, starting:3
fitted Gauss locations:3.7
y
x
0
0.5
1
1.5
2
5
10
15
20
25

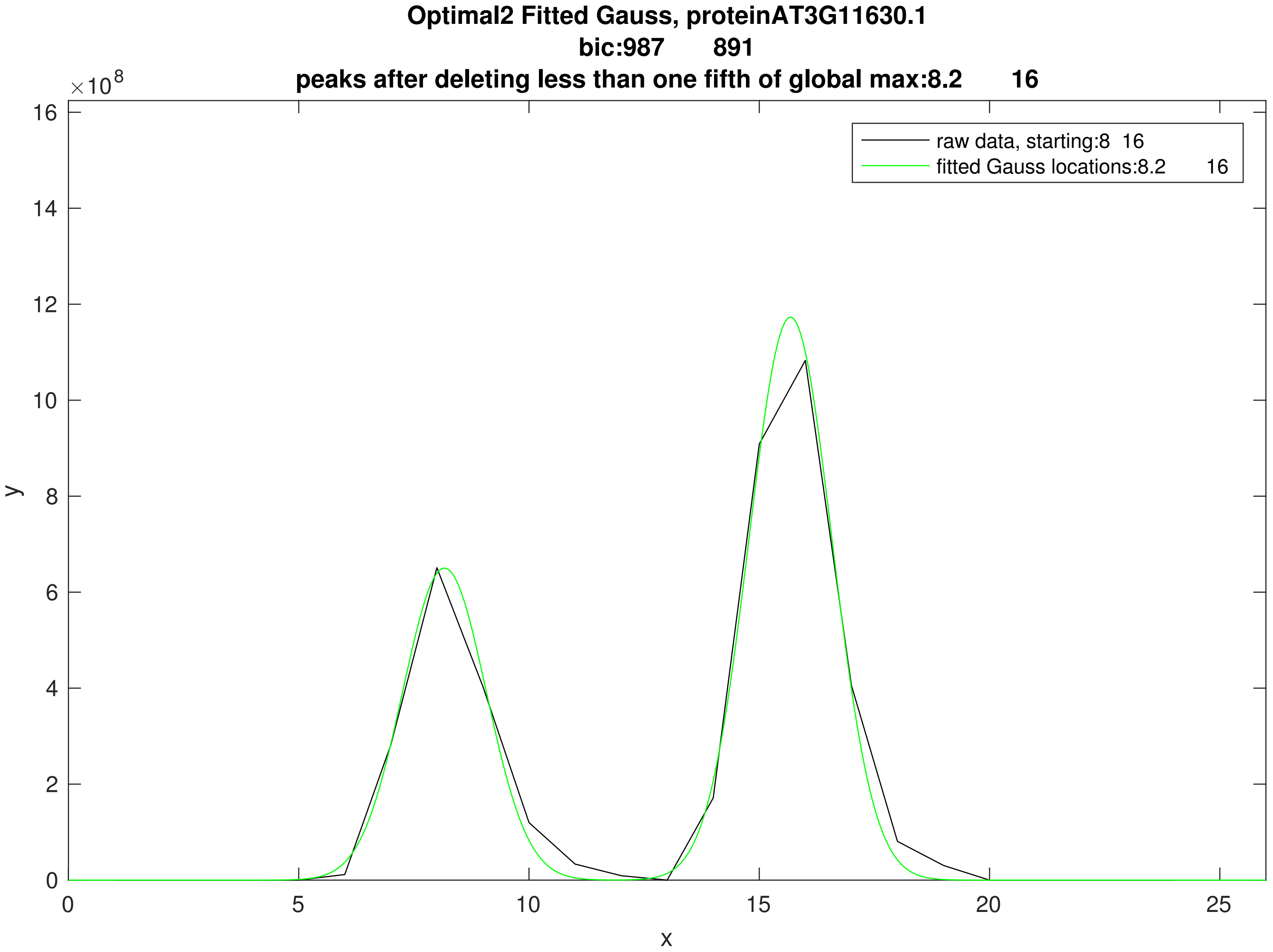
Optimal2 Fitted Gauss, proteinAT3G11630.1
bic:987 891
peaks after deleting less than one fifth of global max:8.2 16
×10^8
raw data, starting:8 16
fitted Gauss locations:8.2 16
y
x
0
2
4
6
8
10
12
14
16
0
5
10
15
20
25

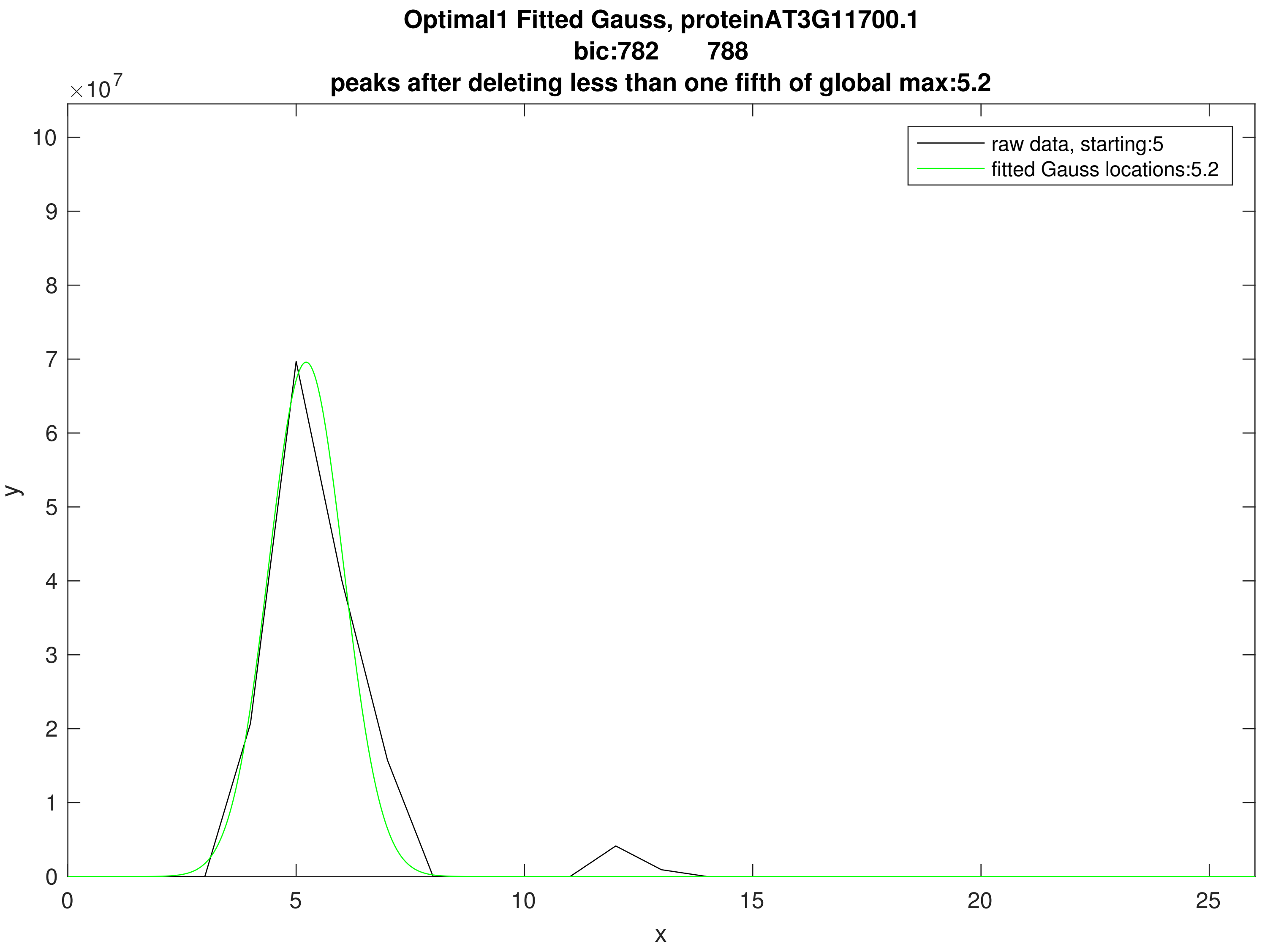

Optimal1 Fitted Gauss, proteinAT3G11700.1
bic:782 788
peaks after deleting less than one fifth of global max:5.2
×10^7
raw data, starting:5
fitted Gauss locations:5.2
y
x

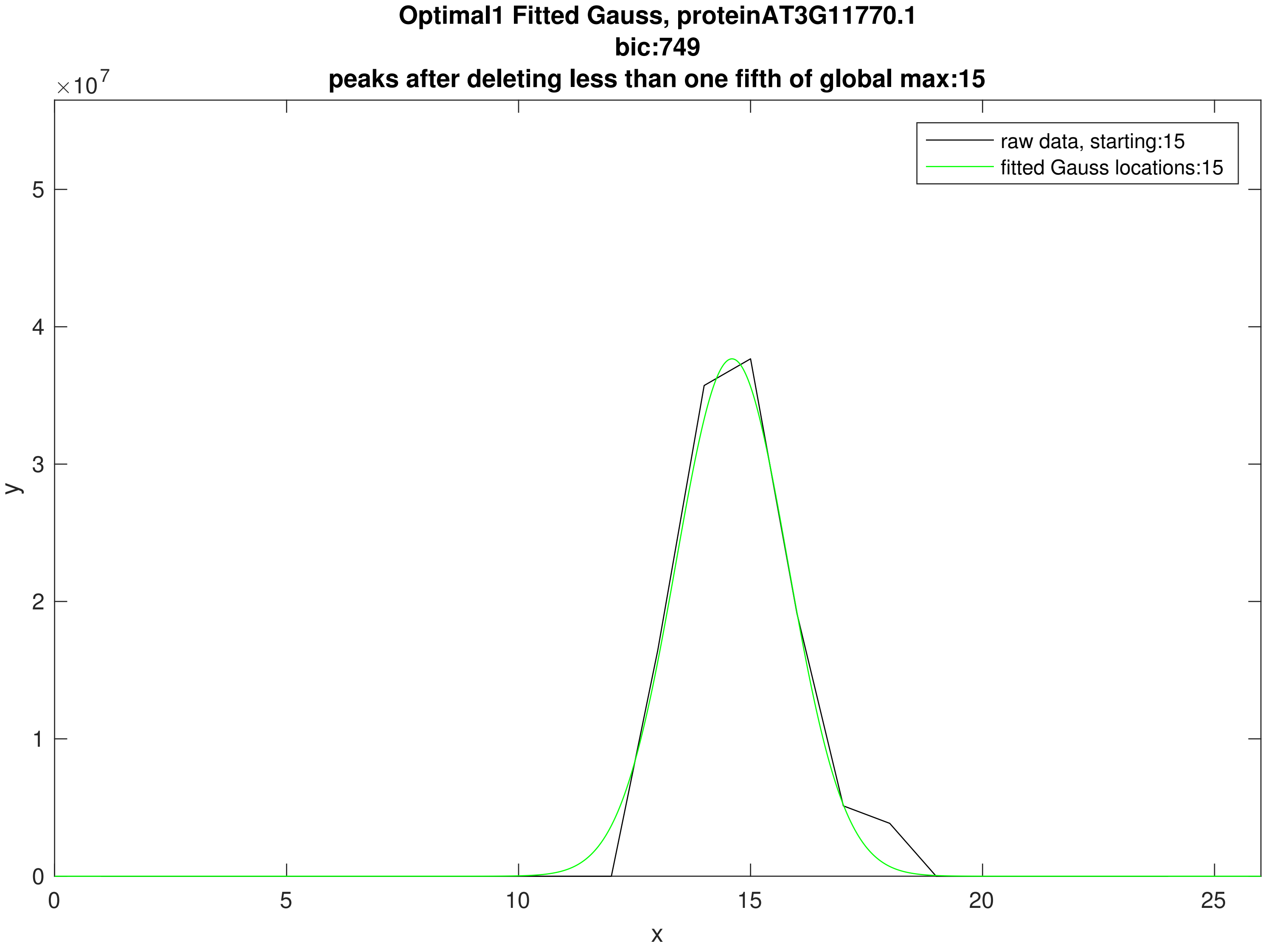
Optimal1 Fitted Gauss, proteinAT3G11770.1
bic:749
peaks after deleting less than one fifth of global max:15
×10^7
raw data, starting:15
fitted Gauss locations:15
y
x
0
1
2
3
4
5
0
5
10
15
20
25

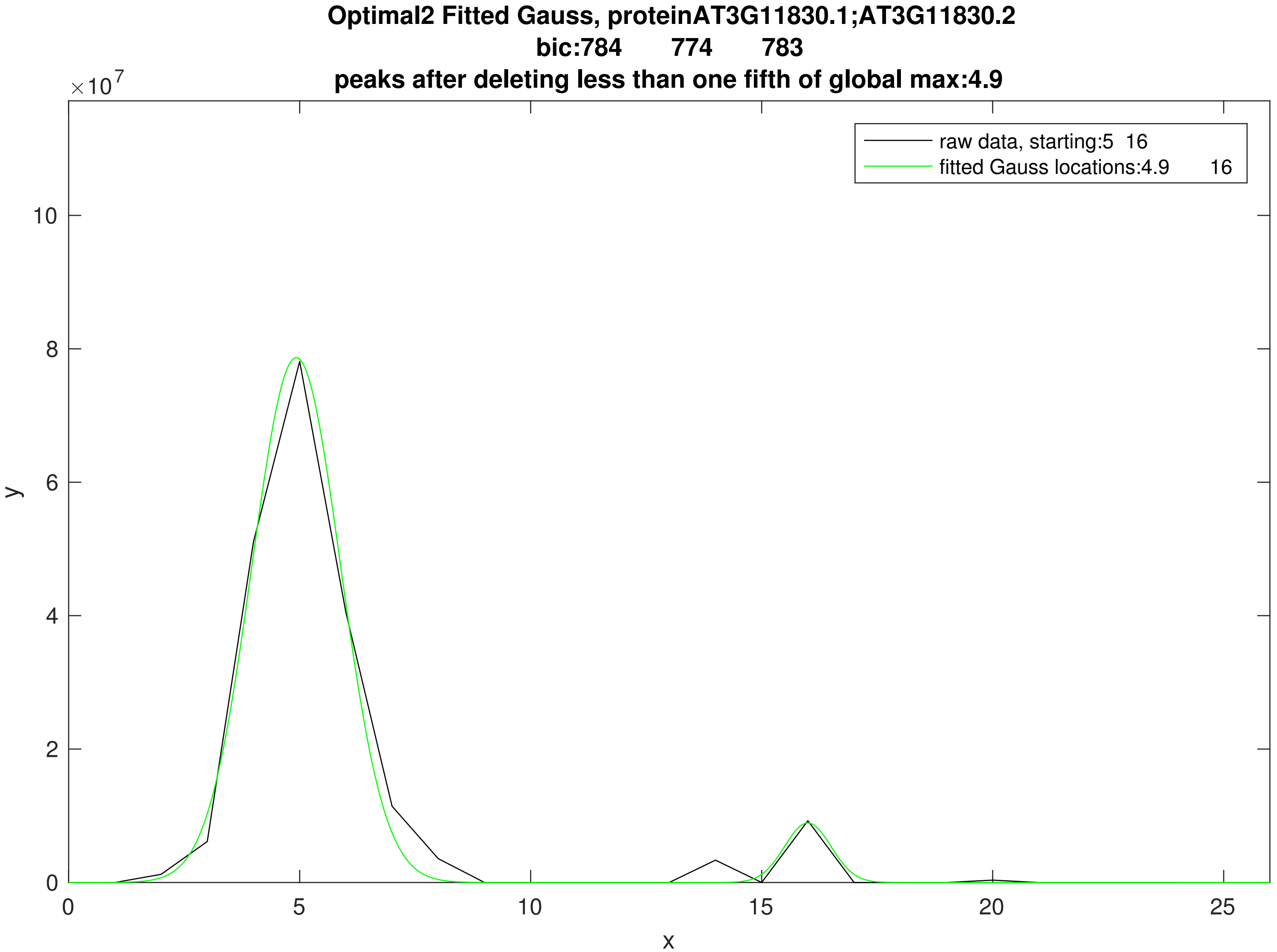

Optimal2 Fitted Gauss, proteinAT3G11830.1;AT3G11830.2
bic:784 774 783
peaks after deleting less than one fifth of global max:4.9
×10^7
raw data, starting:5 16
fitted Gauss locations:4.9 16
y
x
0
2
4
6
8
10
0
5
10
15
20
25

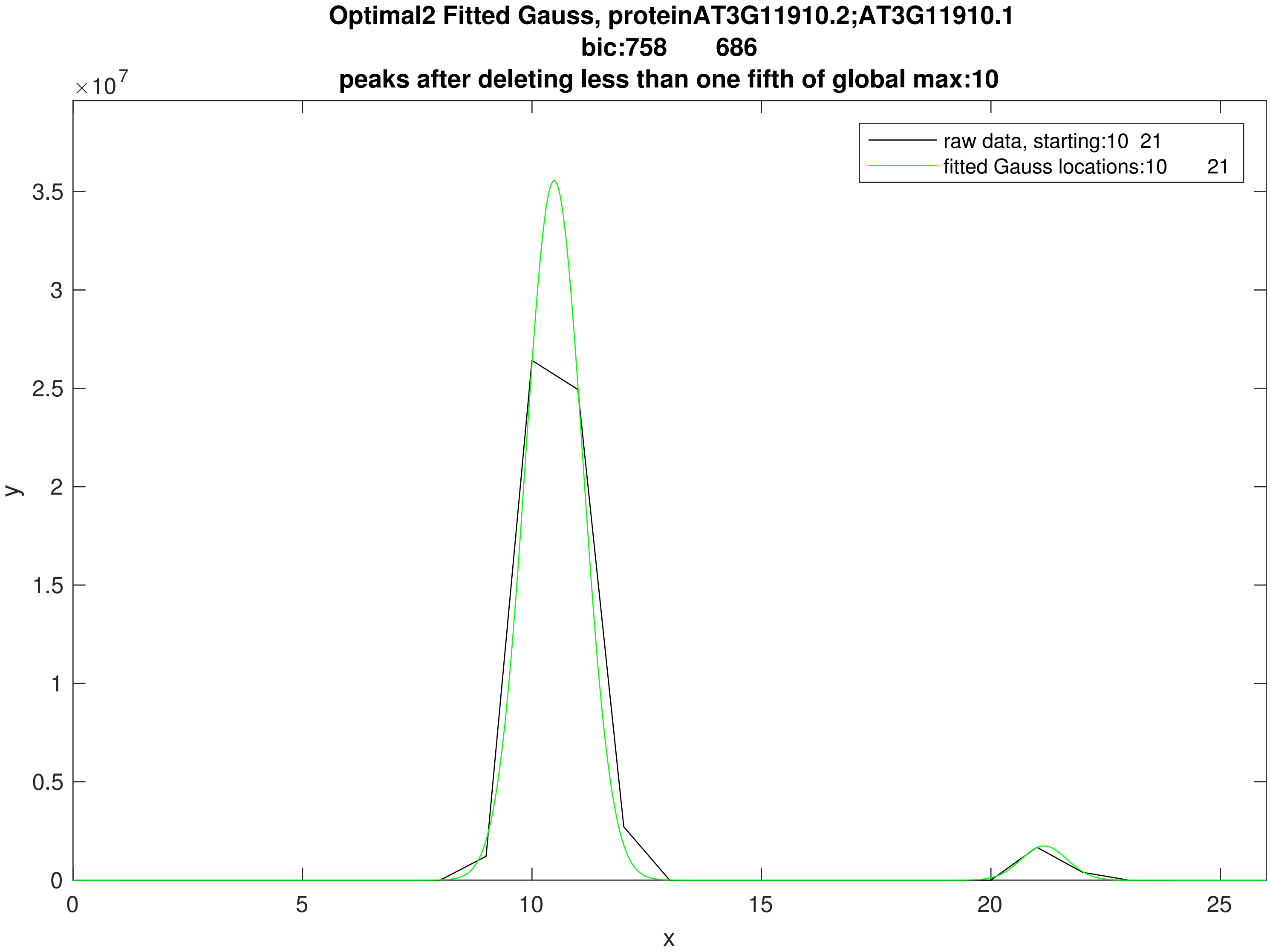

Optimal2 Fitted Gauss, proteinAT3G11910.2;AT3G11910.1
bic:758 686
peaks after deleting less than one fifth of global max:10
×10^7
raw data, starting:10 21
fitted Gauss locations:10 21
3.5
3
2.5
2
1.5
1
0.5
0
y
0
5
10
15
20
25
x

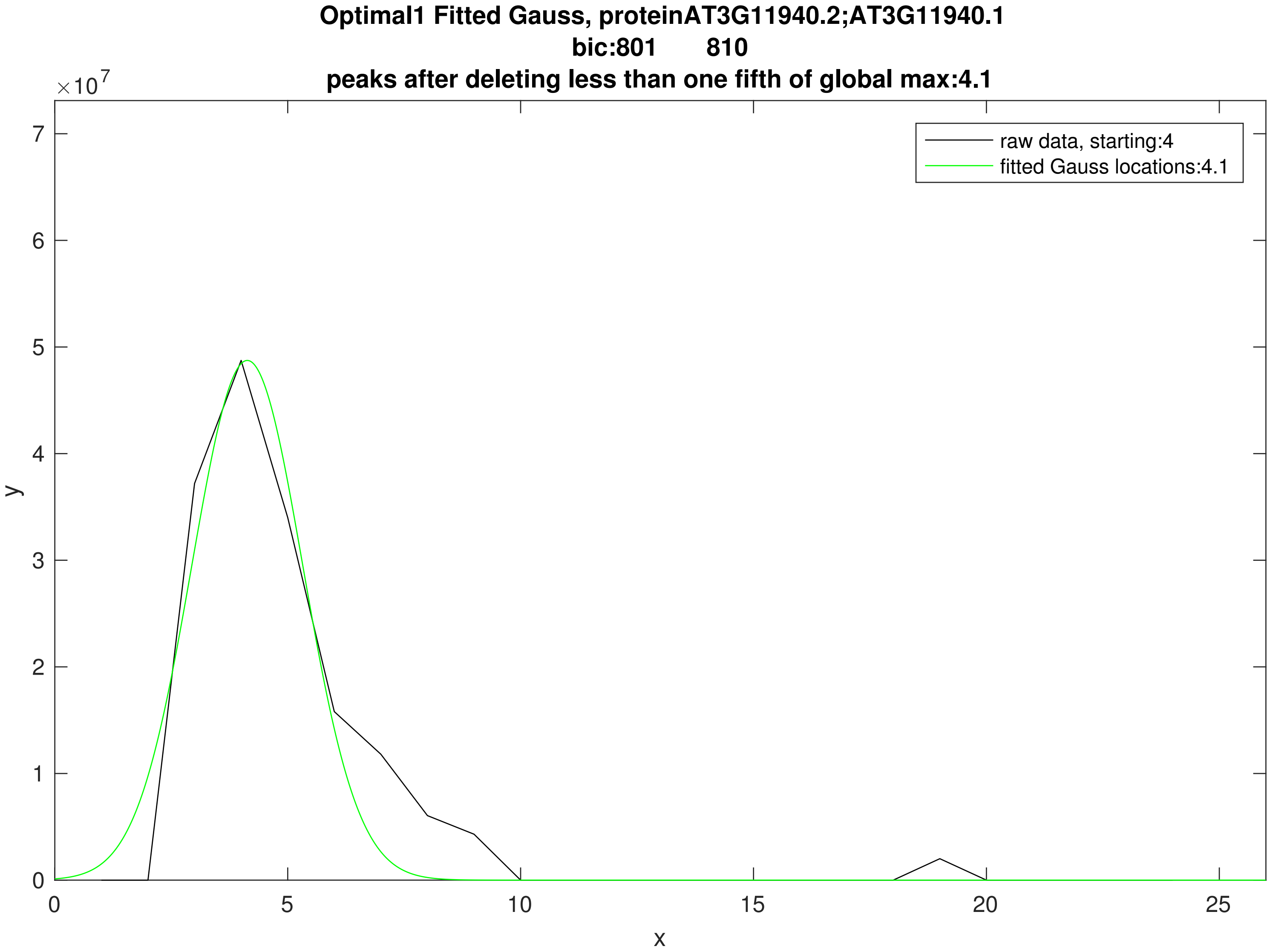
Optimal1 Fitted Gauss, proteinAT3G11940.2;AT3G11940.1
bic:801 810
peaks after deleting less than one fifth of global max:4.1
×10^7
raw data, starting:4
fitted Gauss locations:4.1
y
x
0
1
2
3
4
5
6
7
0
5
10
15
20
25

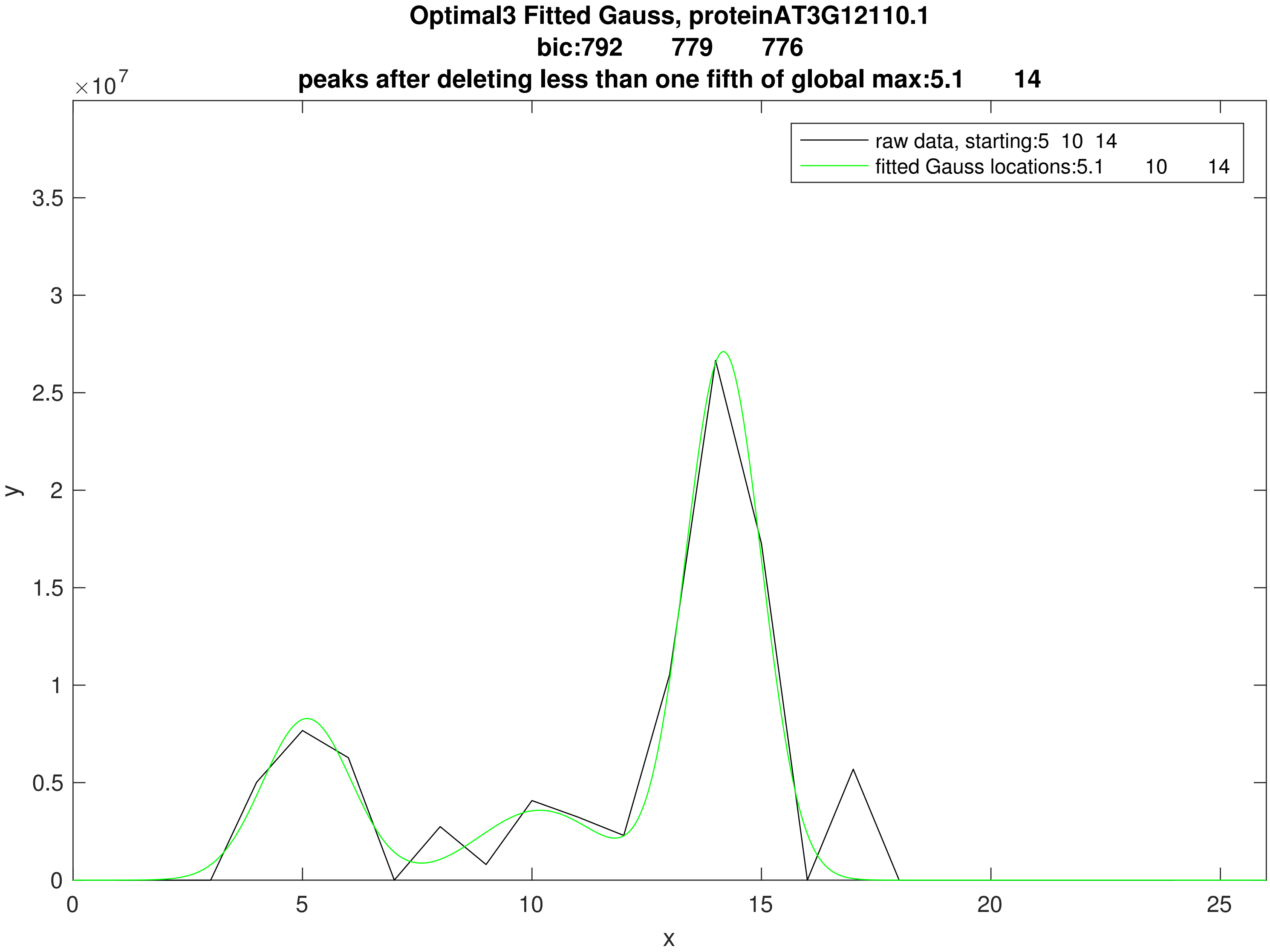

Optimal3 Fitted Gauss, proteinAT3G12110.1
bic:792 779 776
peaks after deleting less than one fifth of global max:5.1 14
×10^7
raw data, starting:5 10 14
fitted Gauss locations:5.1 10 14
3.5
3
2.5
2
1.5
1
0.5
0
y
0
5
10
15
20
25
x

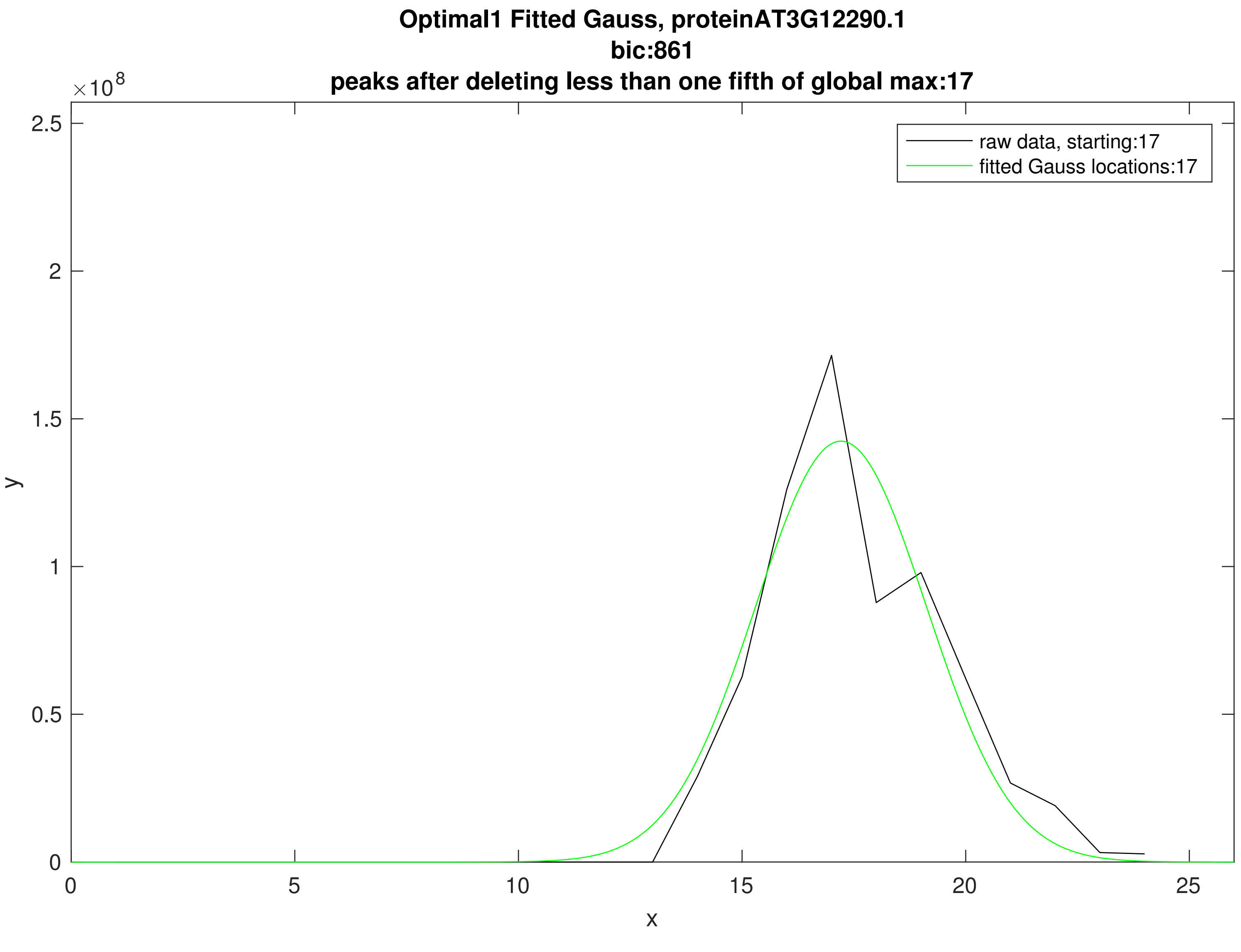

Optimal1 Fitted Gauss, proteinAT3G12290.1
bic:861
peaks after deleting less than one fifth of global max:17
$\times 10^8$
2.5
2
1.5
1
0.5
0
y
0
5
10
15
20
25
x
raw data, starting:17
fitted Gauss locations:17

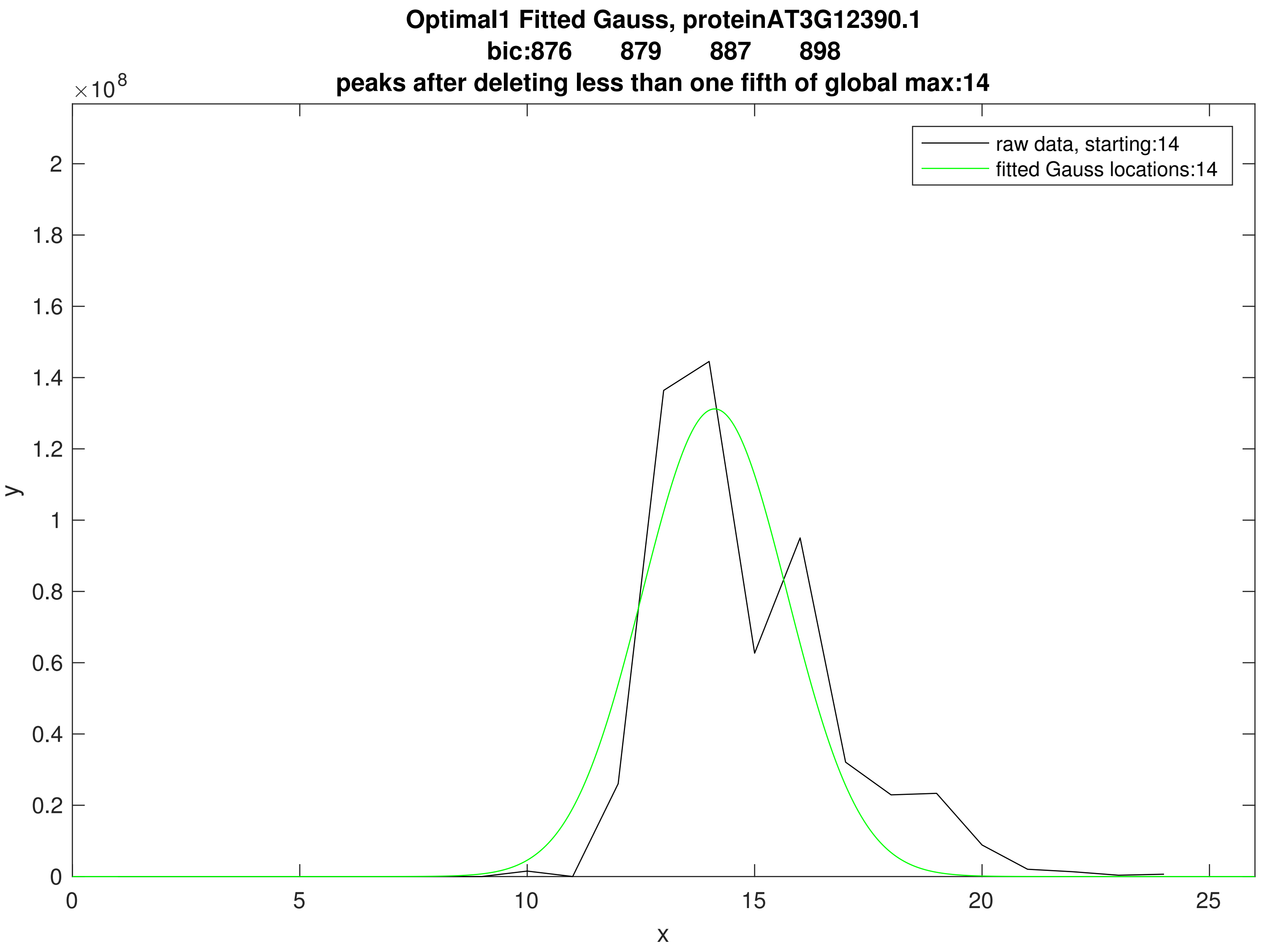
Optimal1 Fitted Gauss, proteinAT3G12390.1
bic:876 879 887 898
peaks after deleting less than one fifth of global max:14
×10^8
raw data, starting:14
fitted Gauss locations:14
y
x
0
0.2
0.4
0.6
0.8
1
1.2
1.4
1.6
1.8
2
0
5
10
15
20
25

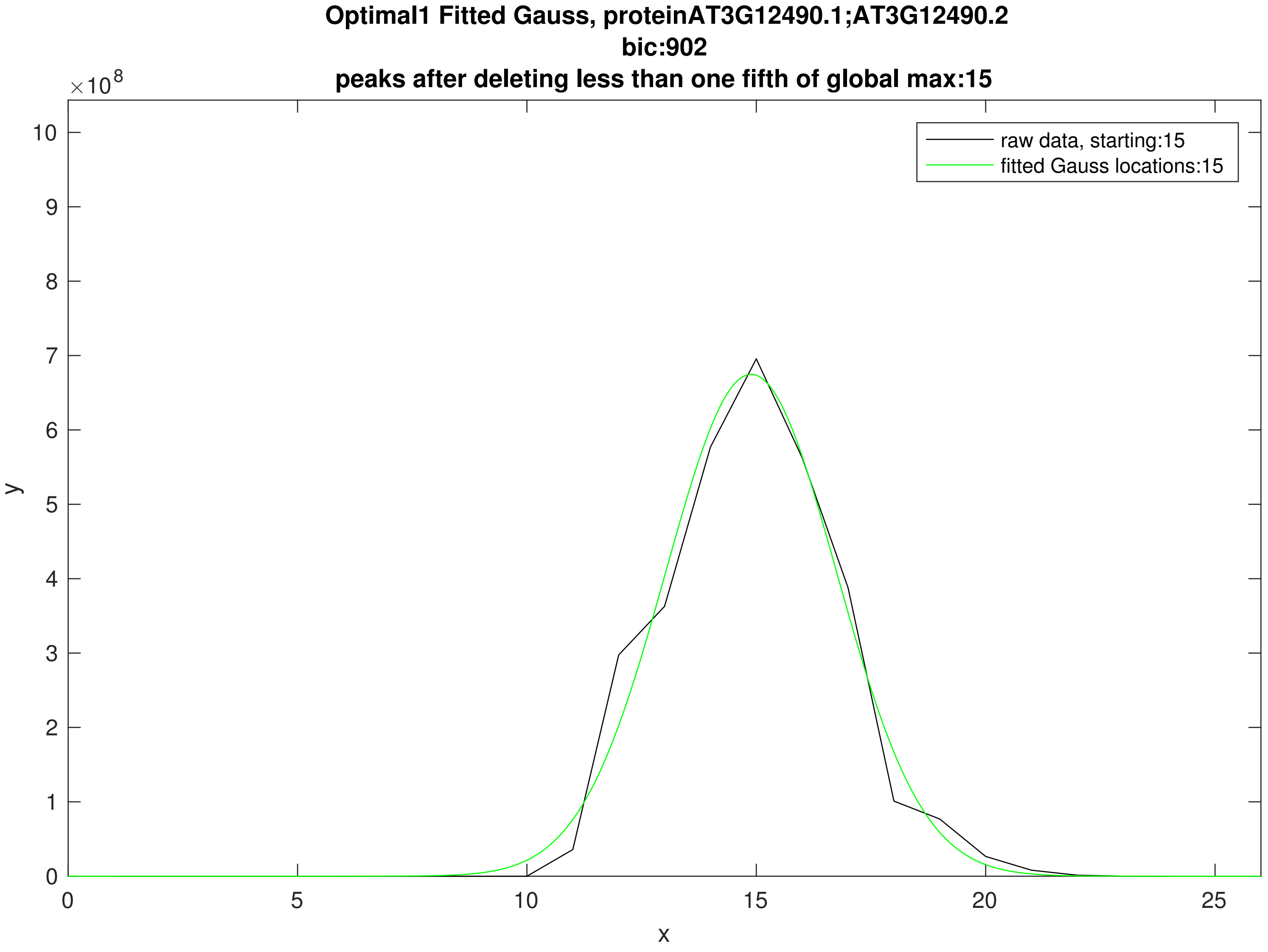

Optimal1 Fitted Gauss, proteinAT3G12490.1;AT3G12490.2
bic:902
peaks after deleting less than one fifth of global max:15
×10^8
10
9
8
7
6
5
4
3
2
1
0
y
0
5
10
15
20
25
x
raw data, starting:15
fitted Gauss locations:15

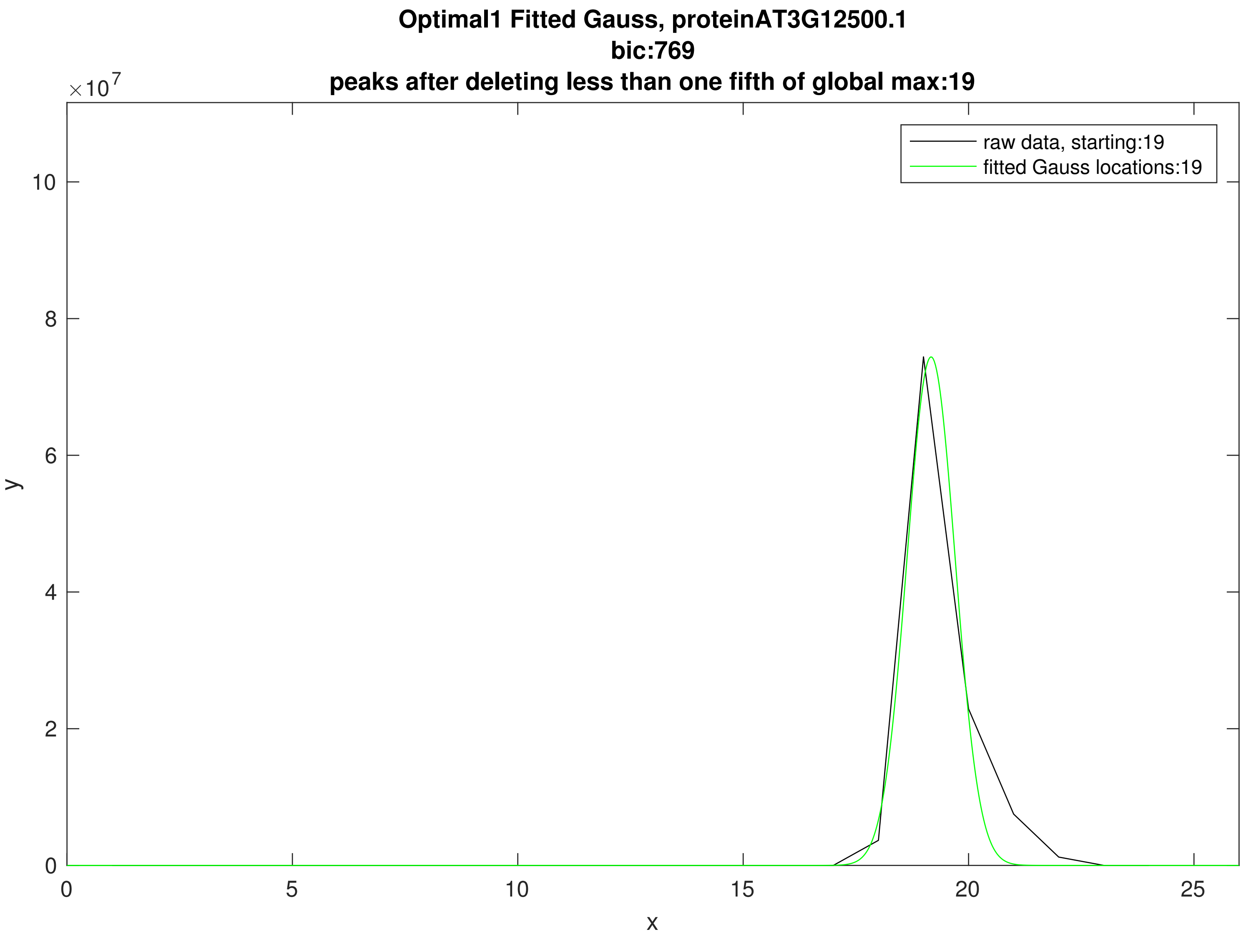
Optimal1 Fitted Gauss, proteinAT3G12500.1
bic:769
peaks after deleting less than one fifth of global max:19
×10^7
raw data, starting:19
fitted Gauss locations:19
10
8
6
4
2
0
y
0
5
10
15
20
25
x

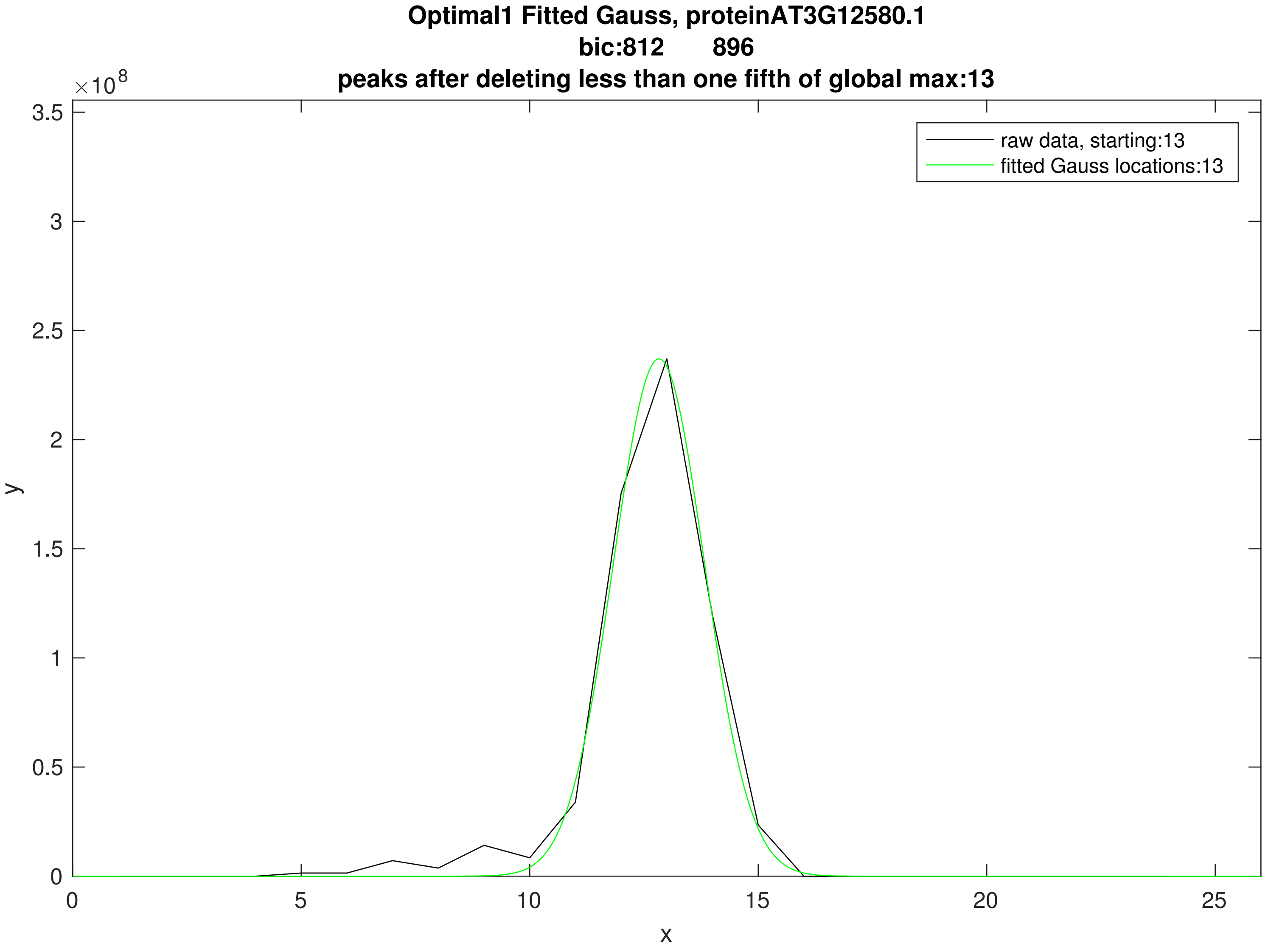
Optimal1 Fitted Gauss, proteinAT3G12580.1
bic:812 896
peaks after deleting less than one fifth of global max:13
×10^8
3.5
3
2.5
2
1.5
1
0.5
0
y
0
5
10
15
20
25
x
raw data, starting:13
fitted Gauss locations:13

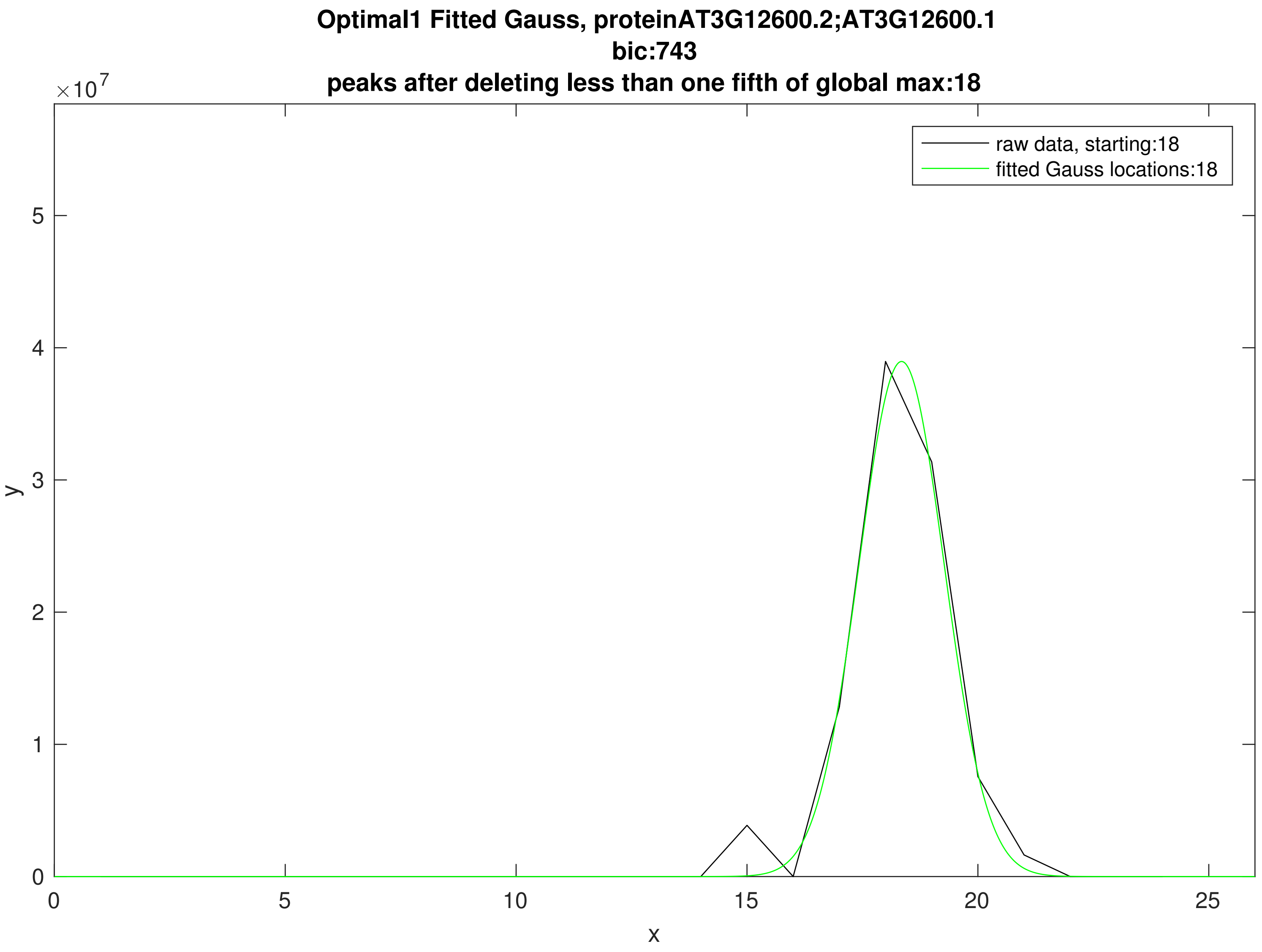
Optimal1 Fitted Gauss, proteinAT3G12600.2;AT3G12600.1
bic:743
peaks after deleting less than one fifth of global max:18
×10^7
raw data, starting:18
fitted Gauss locations:18
y
x
0
1
2
3
4
5
0
5
10
15
20
25

Optimal1 Fitted Gauss, proteinAT3G12610.1
bic:808
peaks after deleting less than one fifth of global max:16

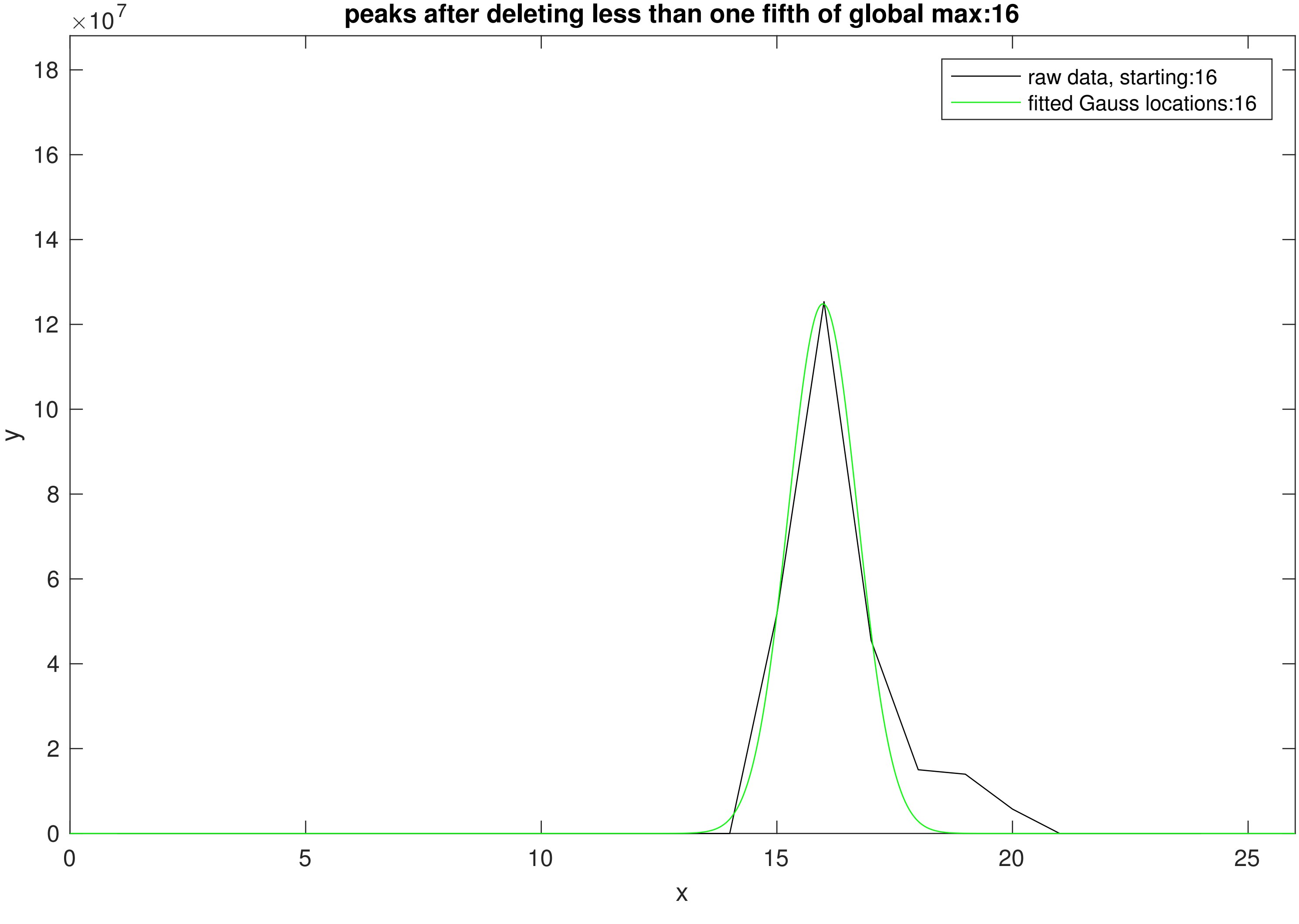

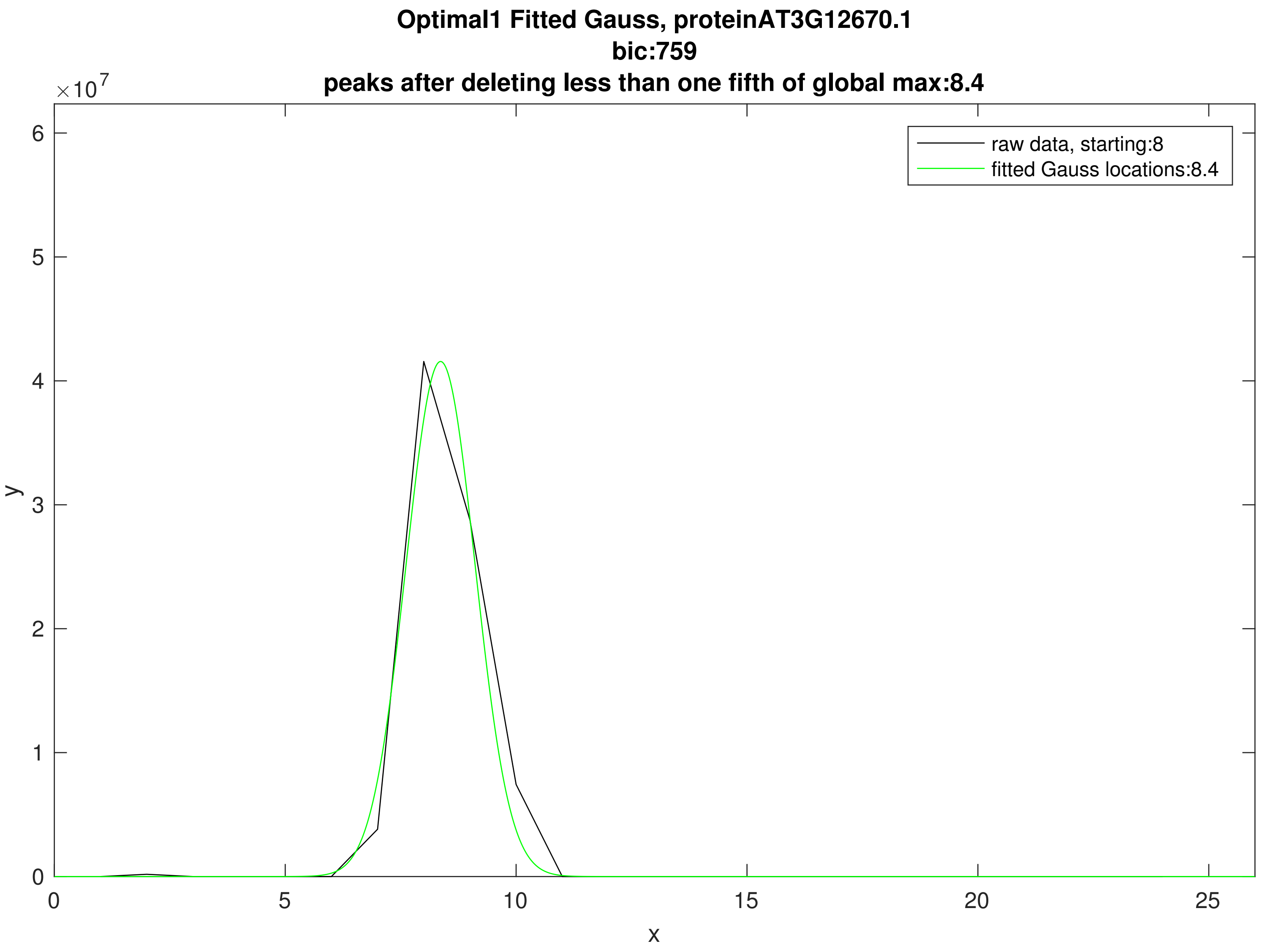
Optimal1 Fitted Gauss, proteinAT3G12670.1
bic:759
peaks after deleting less than one fifth of global max:8.4
×10^7
raw data, starting:8
fitted Gauss locations:8.4
y
x
0
1
2
3
4
5
6
0
5
10
15
20
25

**Optimal1 Fitted Gauss, proteinAT3G12780.1**
**bic:967 968 977**
**peaks after deleting less than one fifth of global max:16**

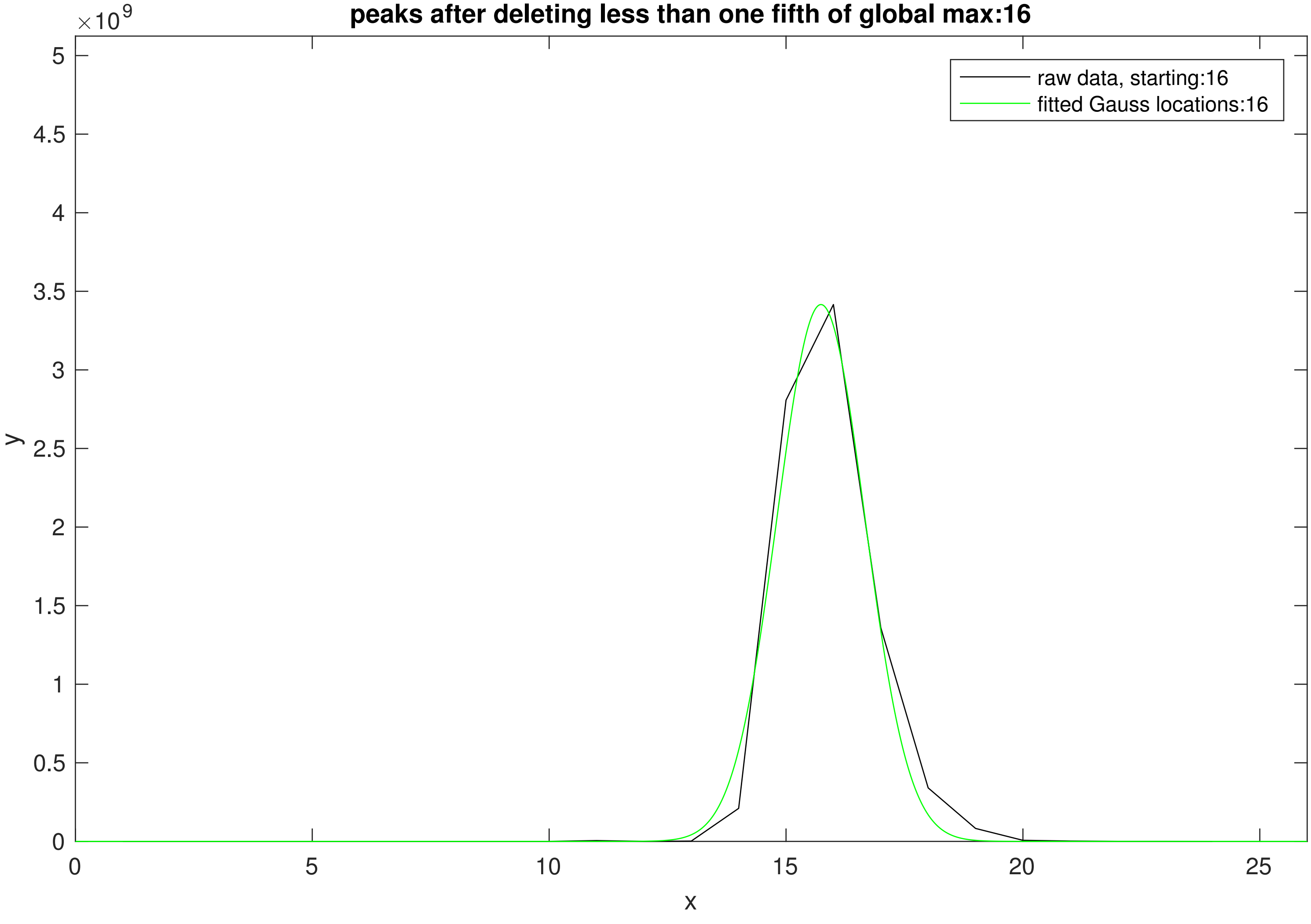

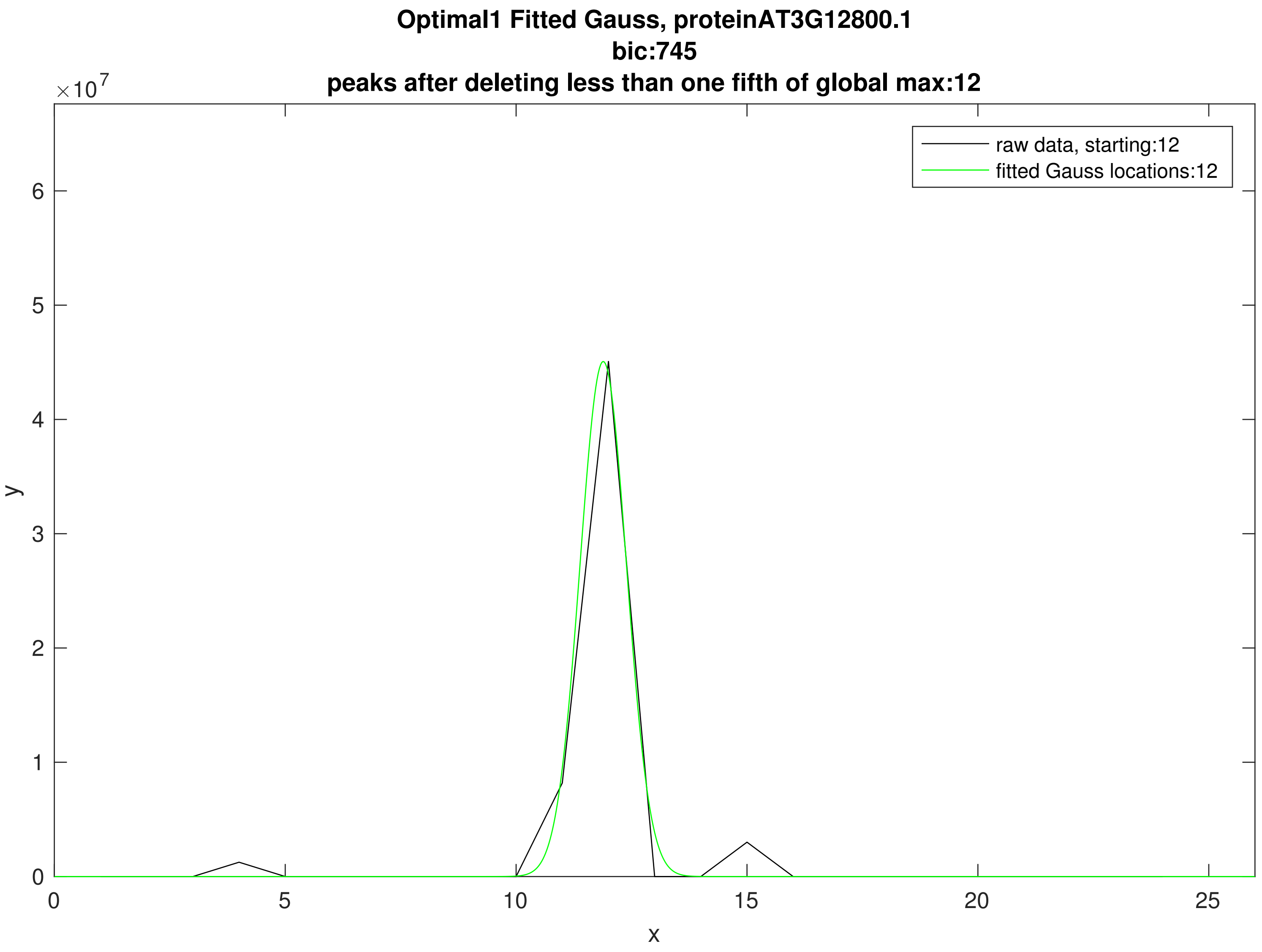
Optimal1 Fitted Gauss, proteinAT3G12800.1
bic:745
peaks after deleting less than one fifth of global max:12
×10^7
raw data, starting:12
fitted Gauss locations:12
y
x
0
1
2
3
4
5
6
0
5
10
15
20
25

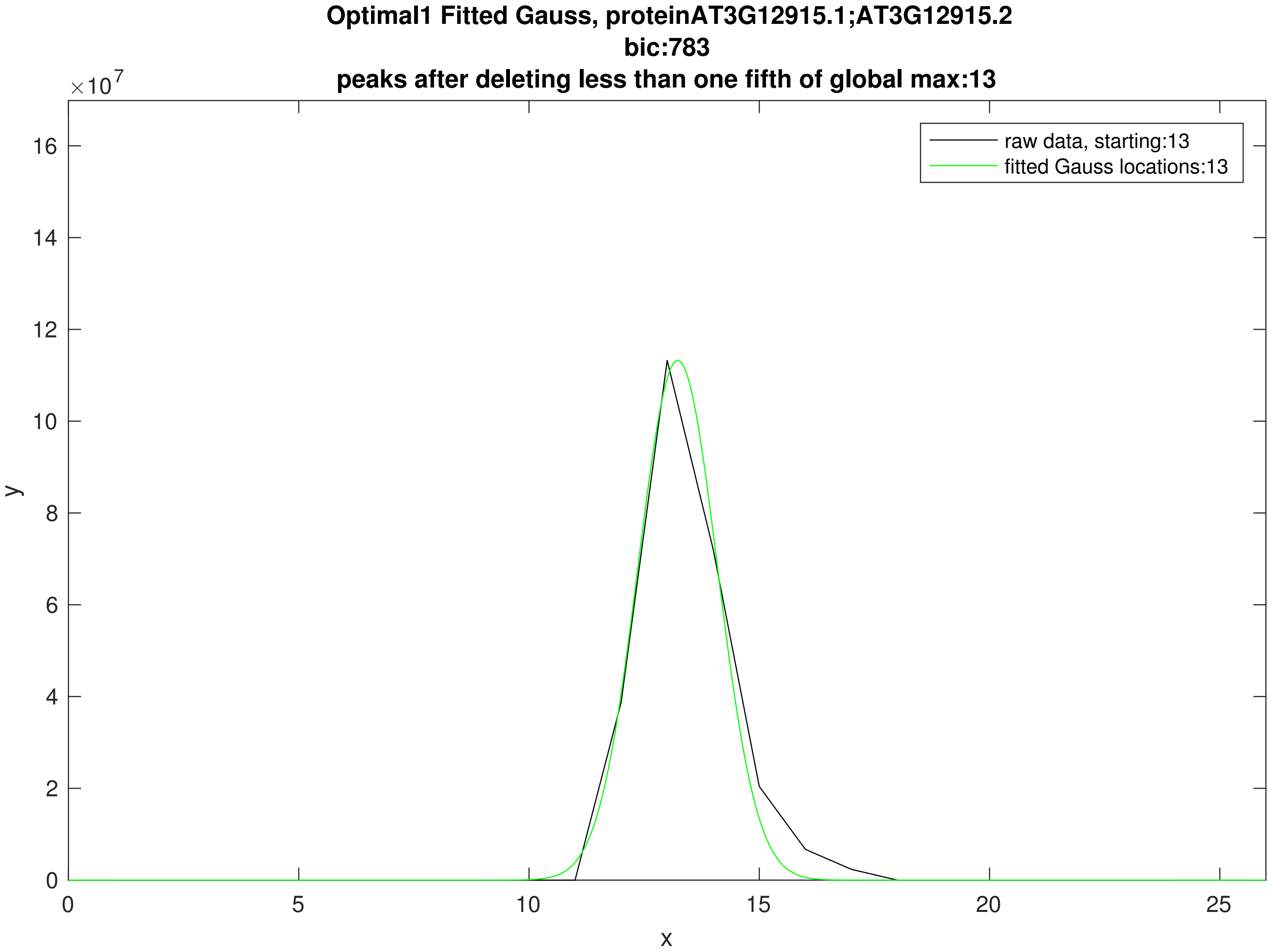

Optimal1 Fitted Gauss, proteinAT3G12915.1;AT3G12915.2
bic:783
peaks after deleting less than one fifth of global max:13
×10^7
raw data, starting:13
fitted Gauss locations:13
y
x
0
2
4
6
8
10
12
14
16
0
5
10
15
20
25

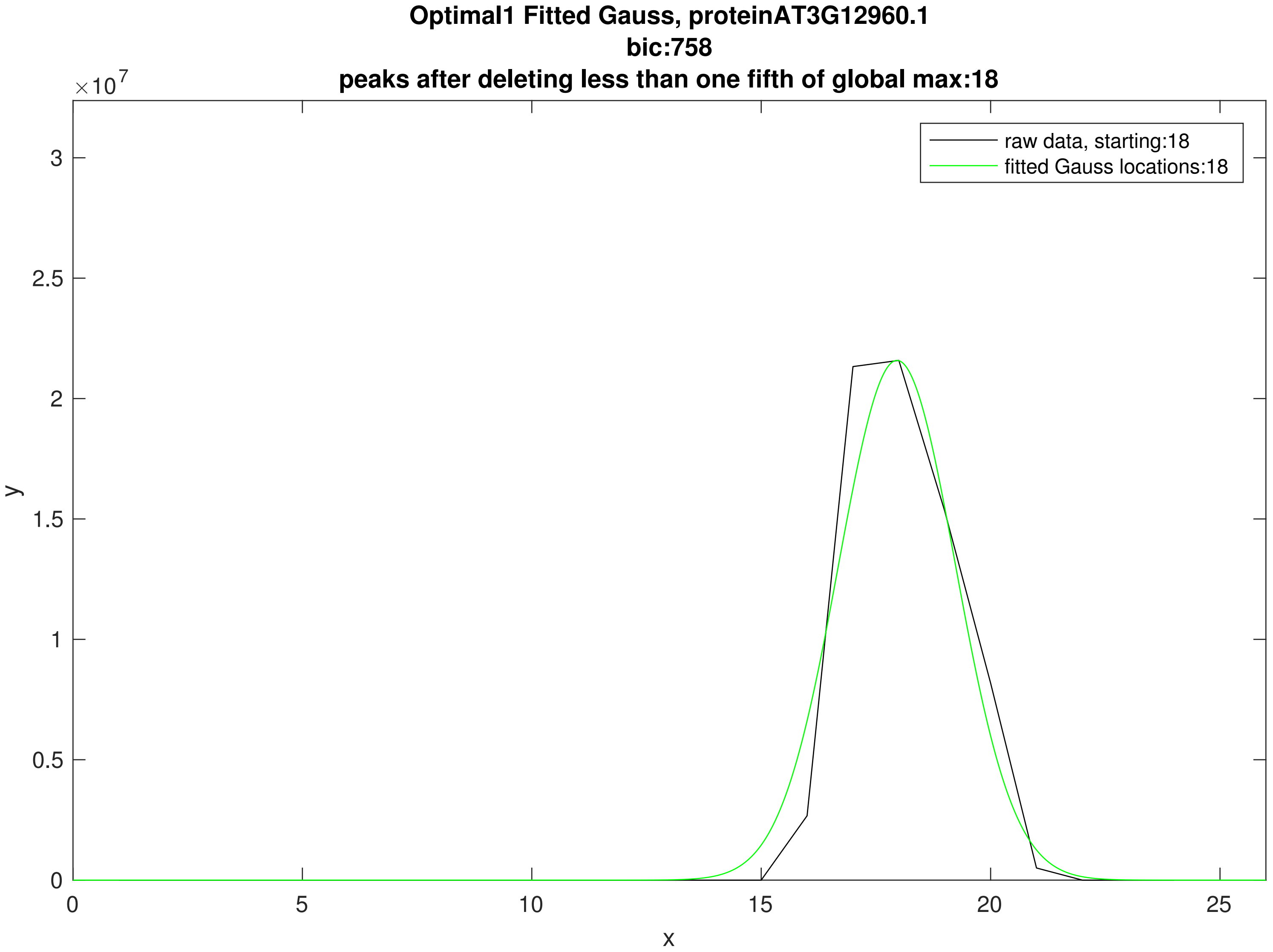

Optimal1 Fitted Gauss, proteinAT3G12960.1
bic:758
peaks after deleting less than one fifth of global max:18
×10^7
raw data, starting:18
fitted Gauss locations:18
3
2.5
2
1.5
1
0.5
0
y
0
5
10
15
20
25
x

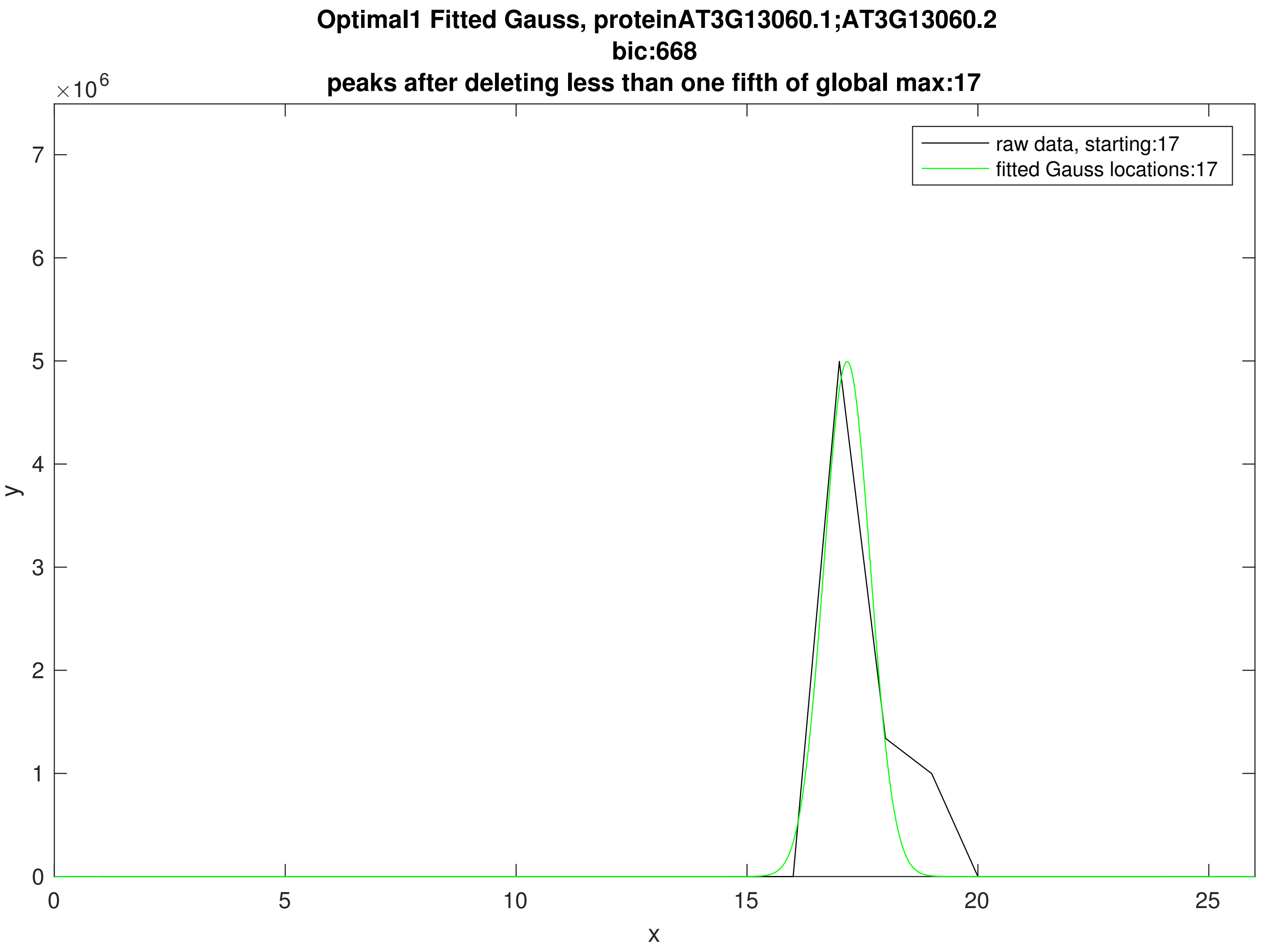

Optimal1 Fitted Gauss, proteinAT3G13060.1;AT3G13060.2
bic:668
peaks after deleting less than one fifth of global max:17
$\times 10^6$
raw data, starting:17
fitted Gauss locations:17
y
x
0
1
2
3
4
5
6
7
0
5
10
15
20
25

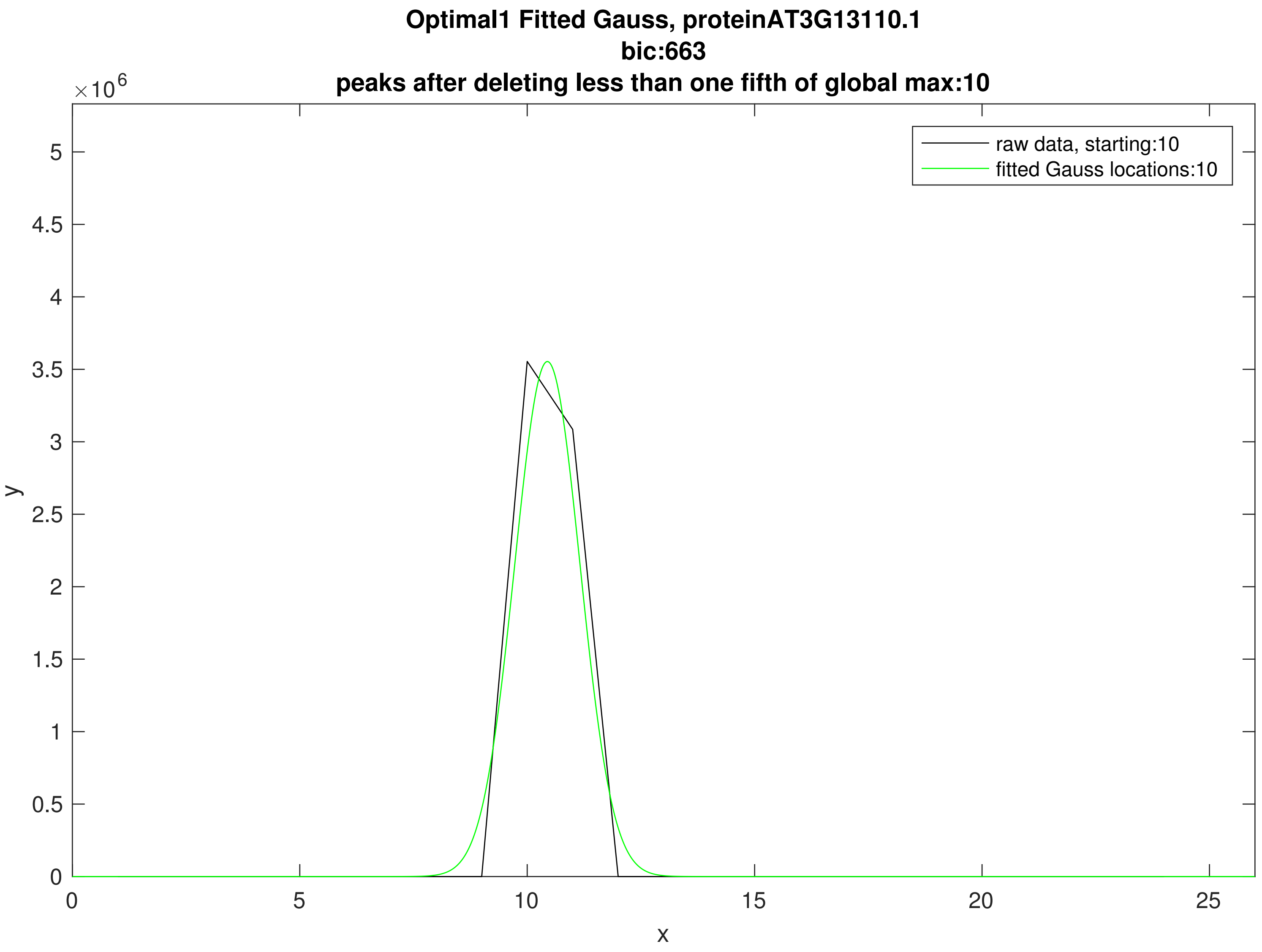
Optimal1 Fitted Gauss, proteinAT3G13110.1
bic:663
peaks after deleting less than one fifth of global max:10
×10^6
raw data, starting:10
fitted Gauss locations:10
y
x
0
0.5
1
1.5
2
2.5
3
3.5
4
4.5
5
0
5
10
15
20
25

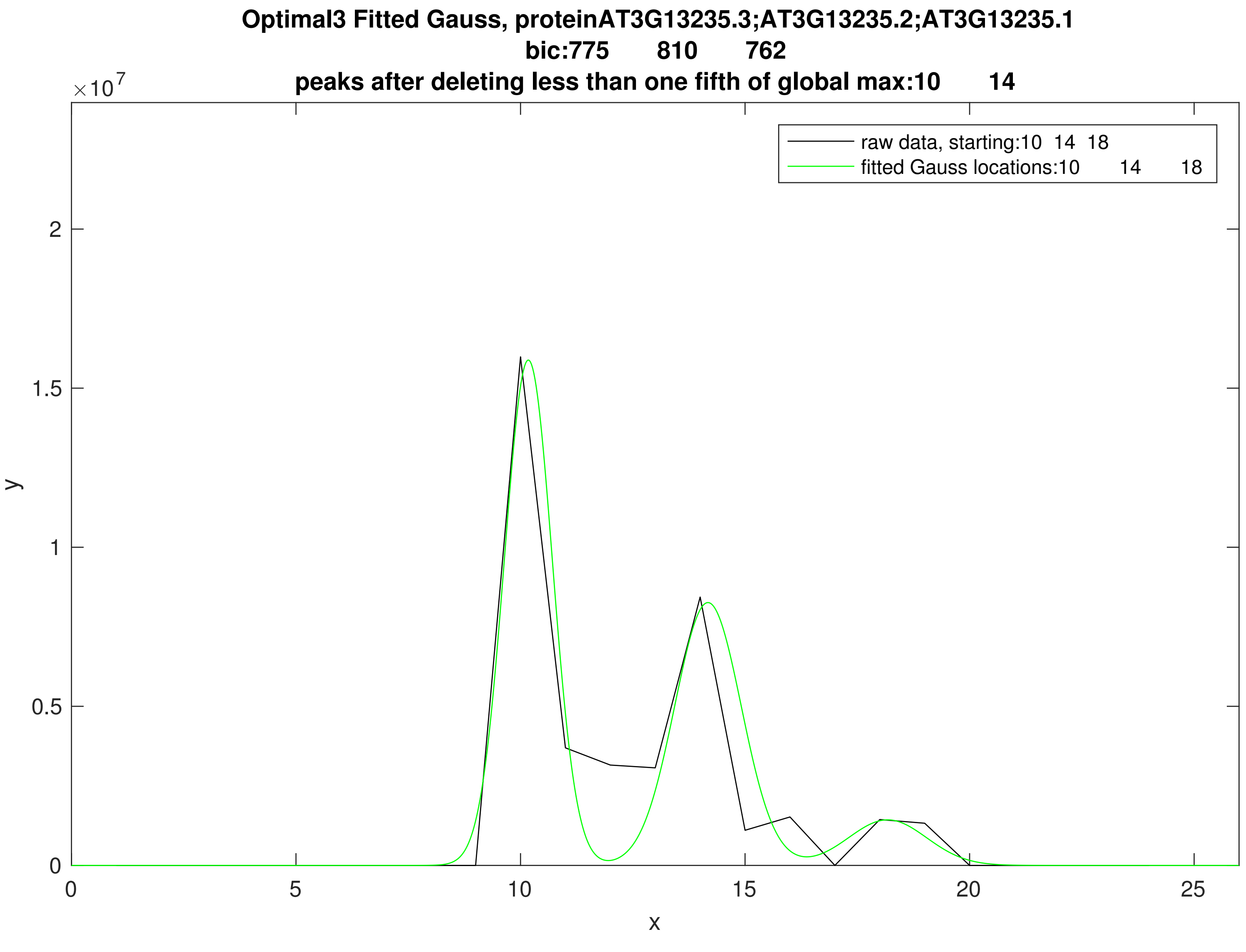
Optimal3 Fitted Gauss, proteinAT3G13235.3;AT3G13235.2;AT3G13235.1
bic:775 810 762
peaks after deleting less than one fifth of global max:10 14
×10^7
raw data, starting:10 14 18
fitted Gauss locations:10 14 18
y
x
0
0.5
1
1.5
2
0
5
10
15
20
25

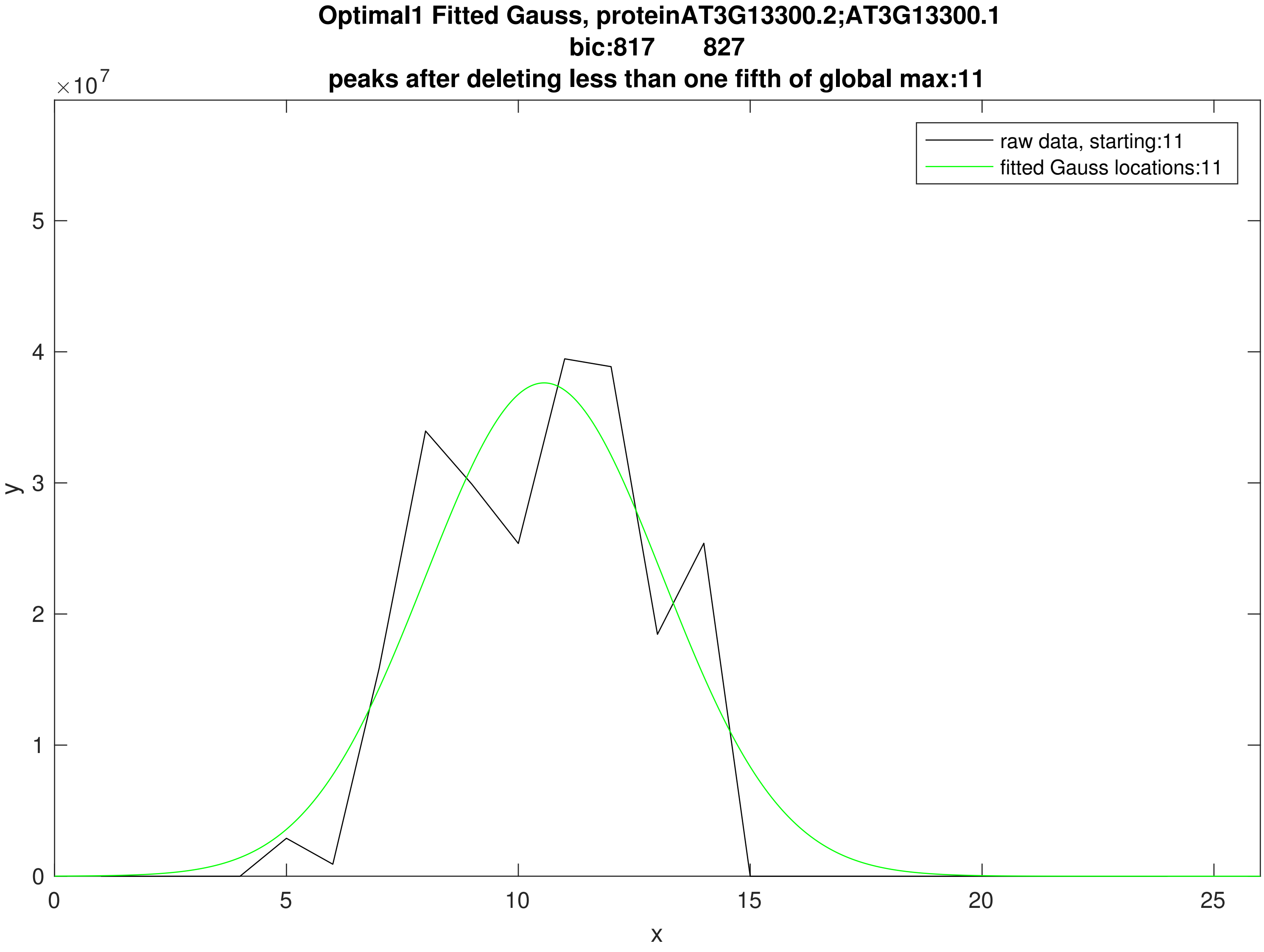

Optimal1 Fitted Gauss, proteinAT3G13300.2;AT3G13300.1
bic:817 827
peaks after deleting less than one fifth of global max:11
×10^7
raw data, starting:11
fitted Gauss locations:11
y
x
0
1
2
3
4
5
0
5
10
15
20
25

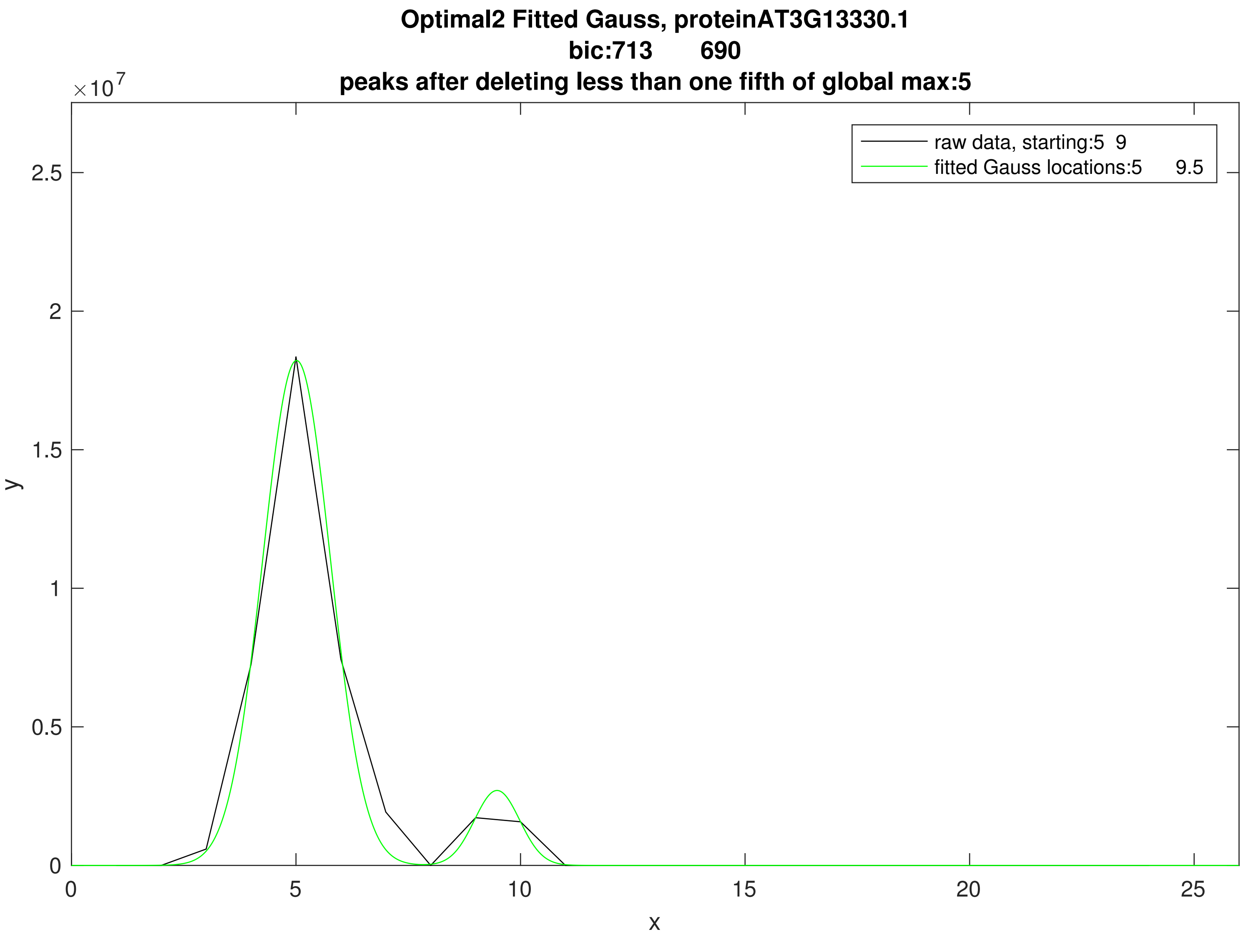

Optimal2 Fitted Gauss, proteinAT3G13330.1
bic:713 690
peaks after deleting less than one fifth of global max:5
×10^7
raw data, starting:5 9
fitted Gauss locations:5 9.5
2.5
2
1.5
1
0.5
0
y
0
5
10
15
20
25
x

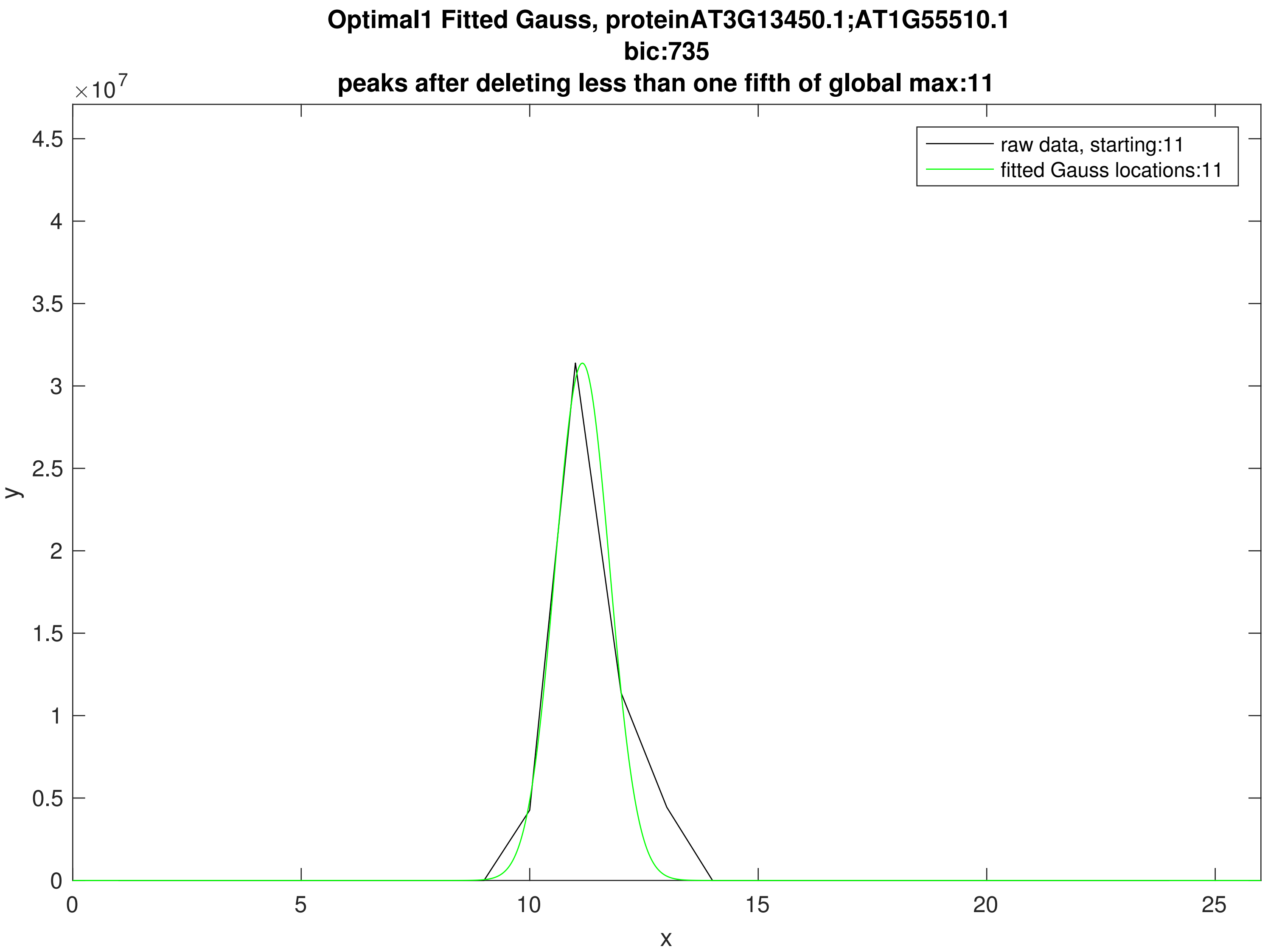
Optimal1 Fitted Gauss, proteinAT3G13450.1;AT1G55510.1
bic:735
peaks after deleting less than one fifth of global max:11
×10^7
4.5
4
3.5
3
2.5
2
1.5
1
0.5
0
y
0
5
10
15
20
25
x
raw data, starting:11
fitted Gauss locations:11

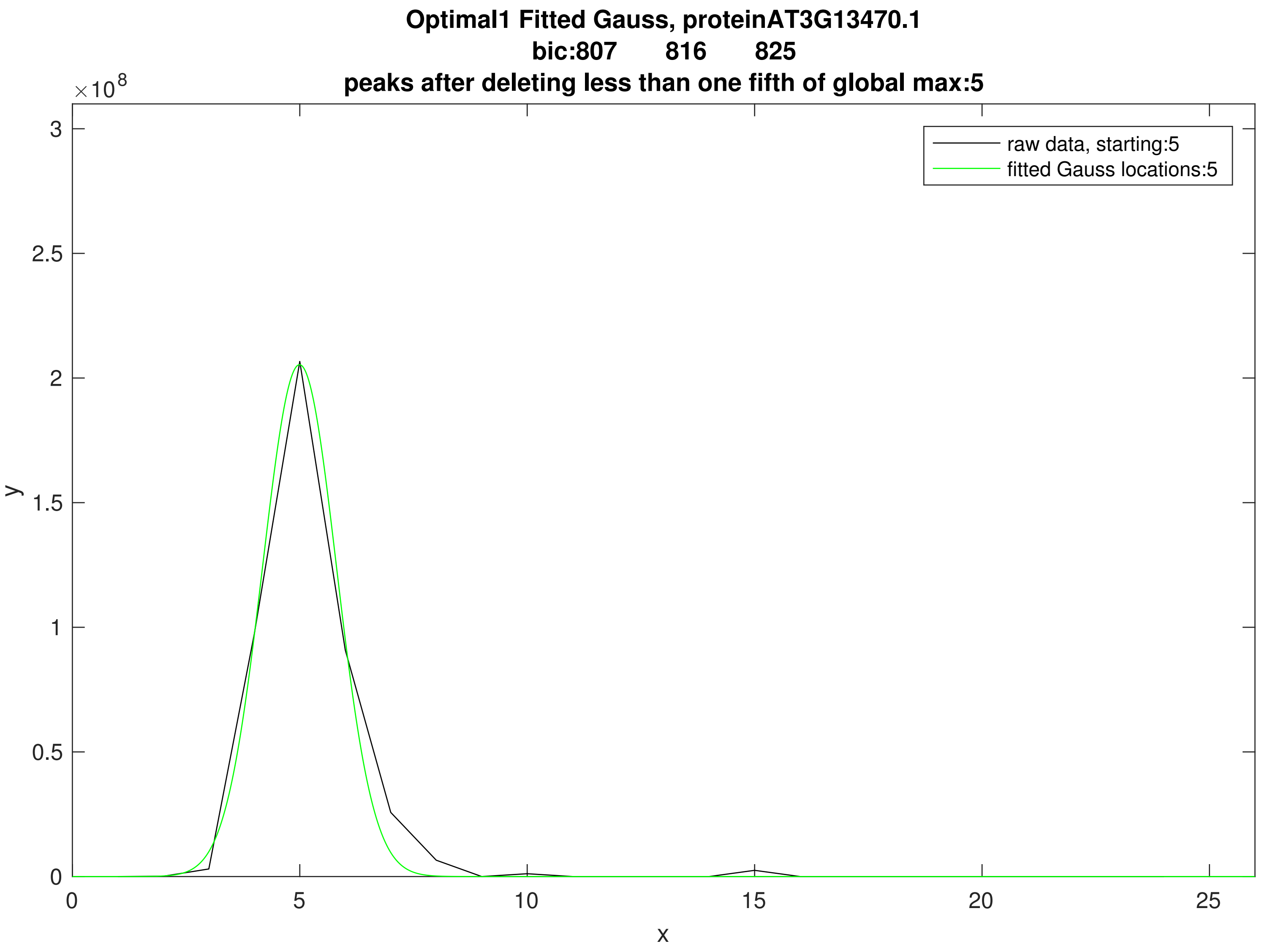
Optimal1 Fitted Gauss, proteinAT3G13470.1
bic:807 816 825
peaks after deleting less than one fifth of global max:5
×10^8
raw data, starting:5
fitted Gauss locations:5
y
x
0
0.5
1
1.5
2
2.5
3
5
10
15
20
25

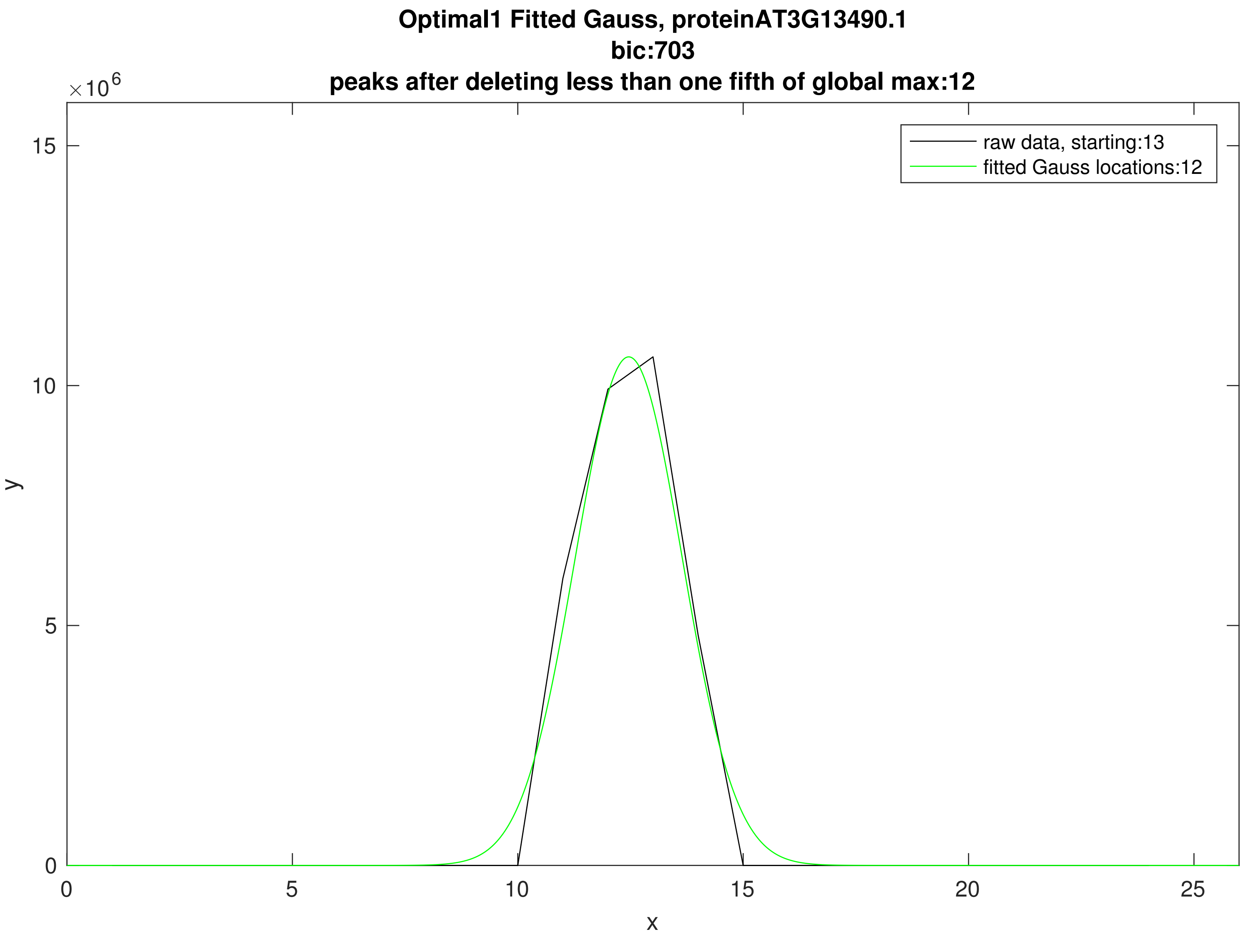
Optimal1 Fitted Gauss, proteinAT3G13490.1
bic:703
peaks after deleting less than one fifth of global max:12
×10^6
15
10
5
0
y
0
5
10
15
20
25
x
raw data, starting:13
fitted Gauss locations:12

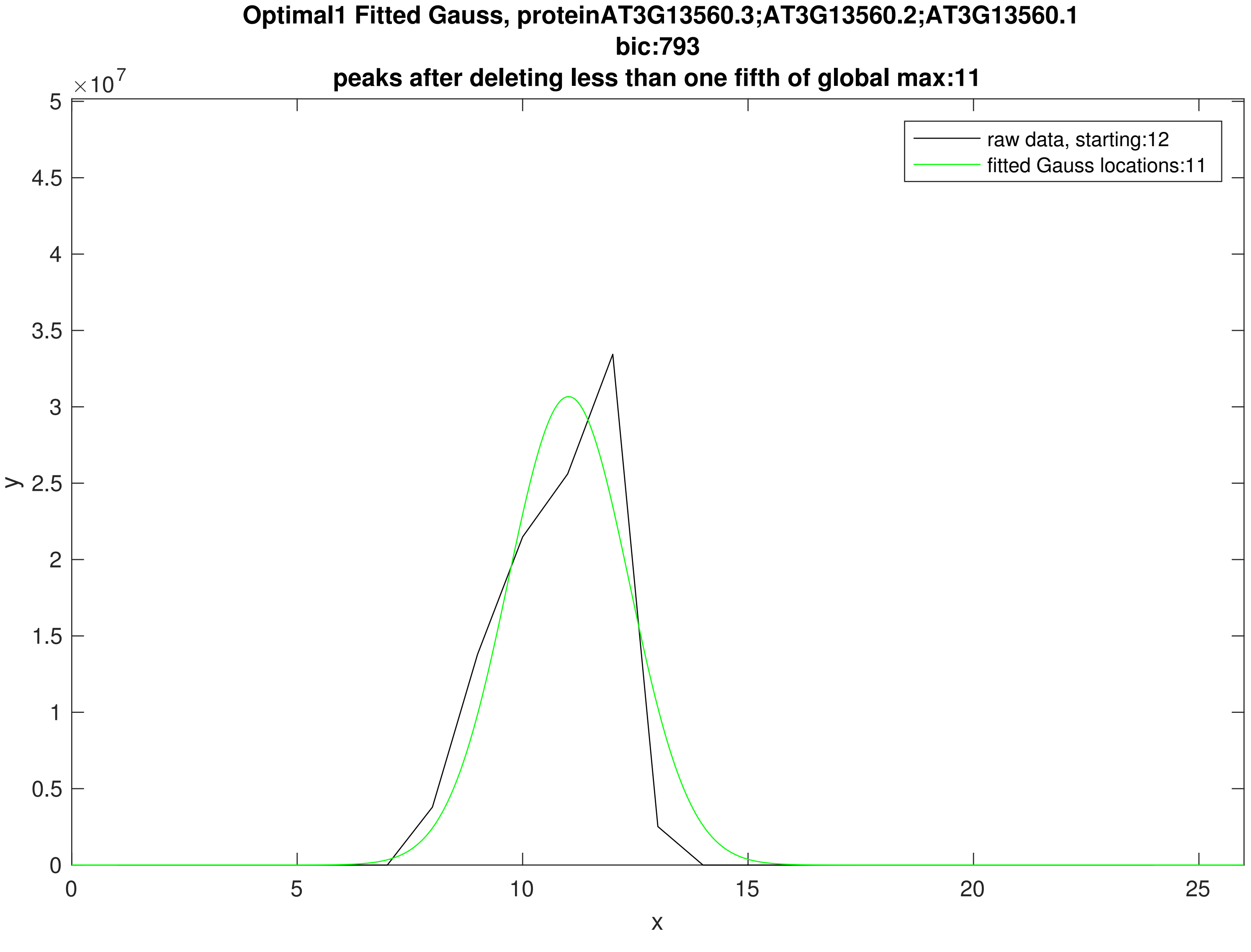

Optimal1 Fitted Gauss, proteinAT3G13560.3;AT3G13560.2;AT3G13560.1
bic:793
peaks after deleting less than one fifth of global max:11
×10^7
raw data, starting:12
fitted Gauss locations:11
y
x
0
0.5
1
1.5
2
2.5
3
3.5
4
4.5
5
0
5
10
15
20
25

**Optimal1 Fitted Gauss, proteinAT3G13750.1**
**bic:753 853**
**peaks after deleting less than one fifth of global max:17**

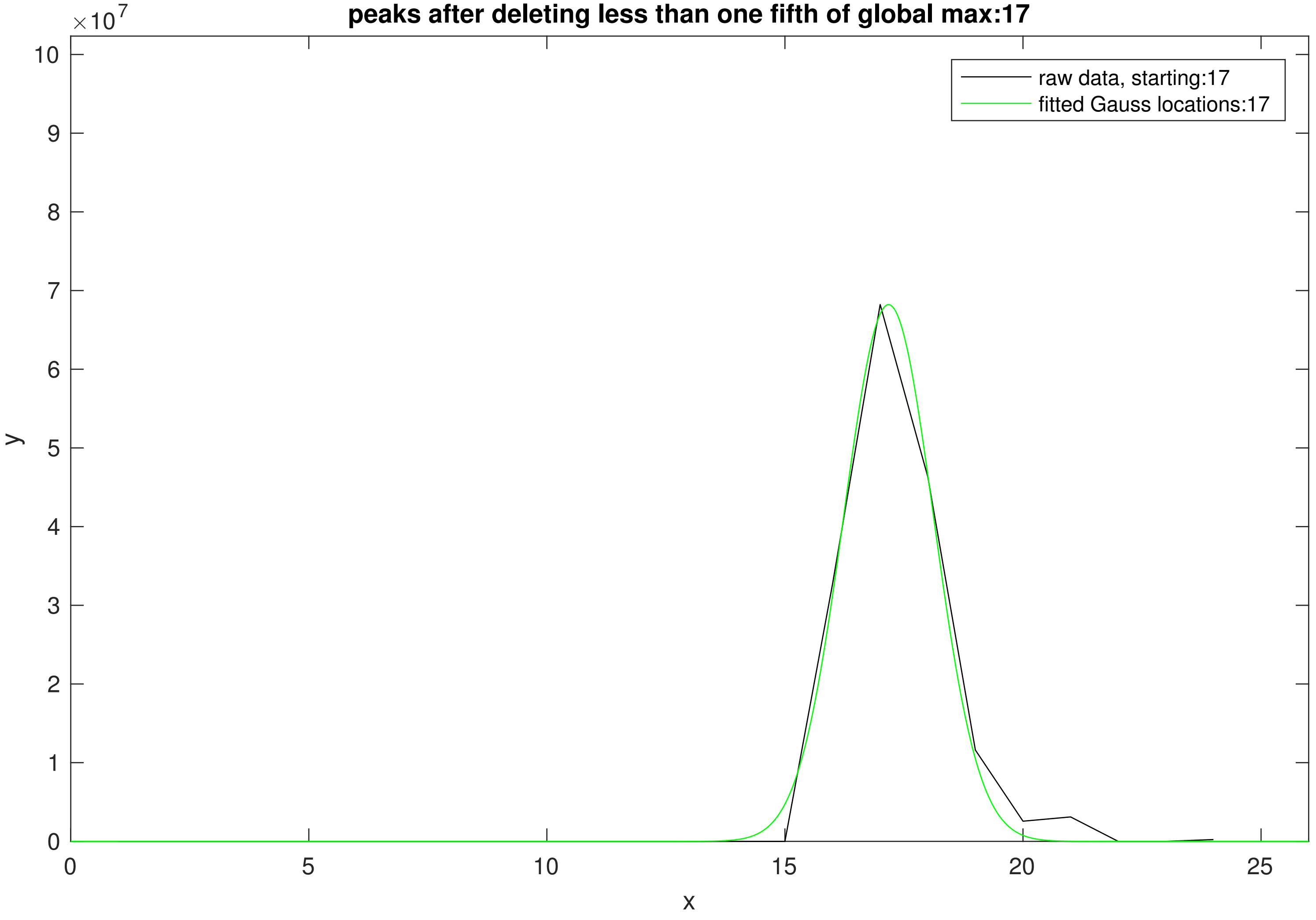

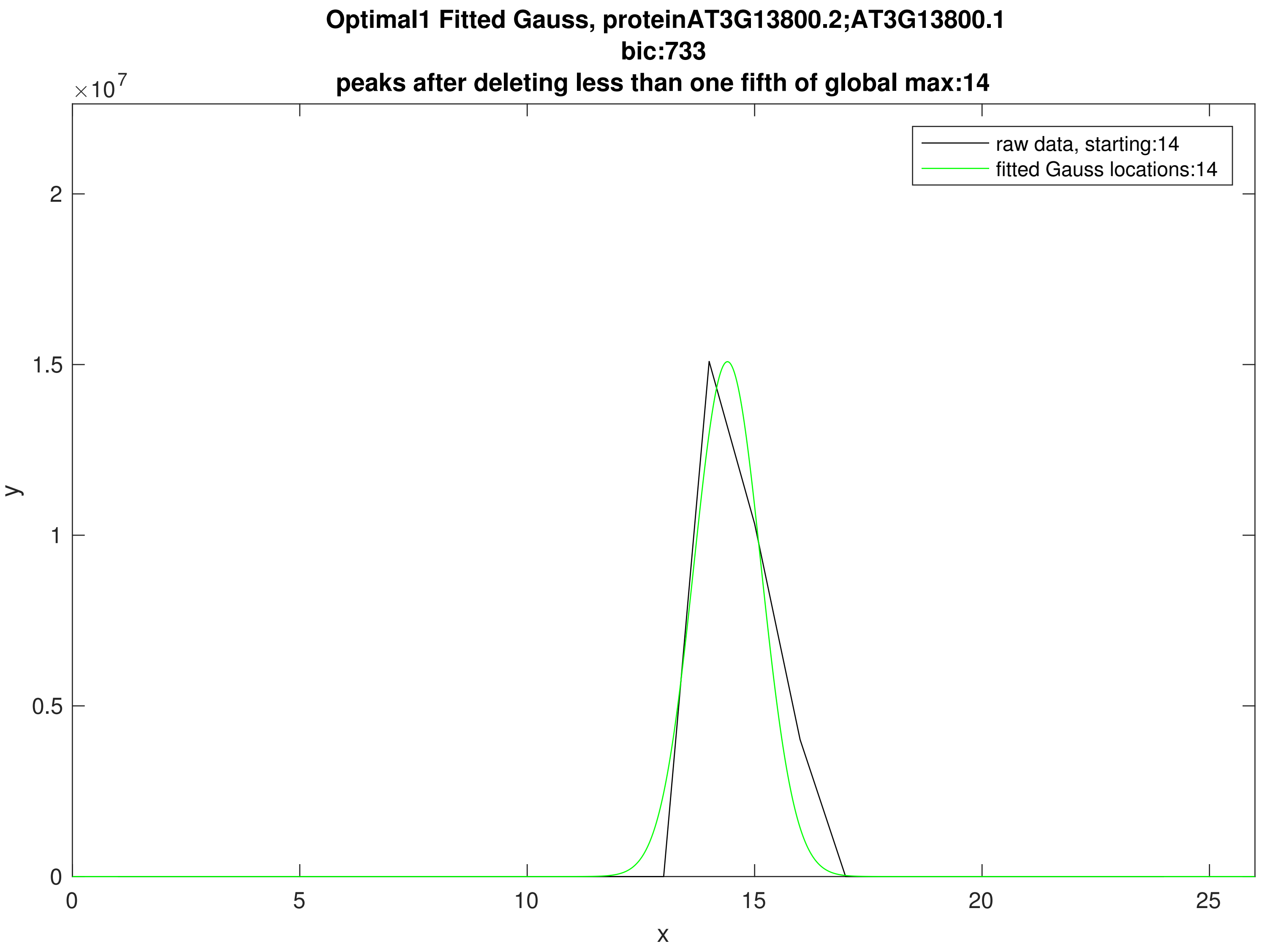
Optimal1 Fitted Gauss, proteinAT3G13800.2;AT3G13800.1
bic:733
peaks after deleting less than one fifth of global max:14
×10^7
raw data, starting:14
fitted Gauss locations:14
2
1.5
1
0.5
0
y
0
5
10
15
20
25
x

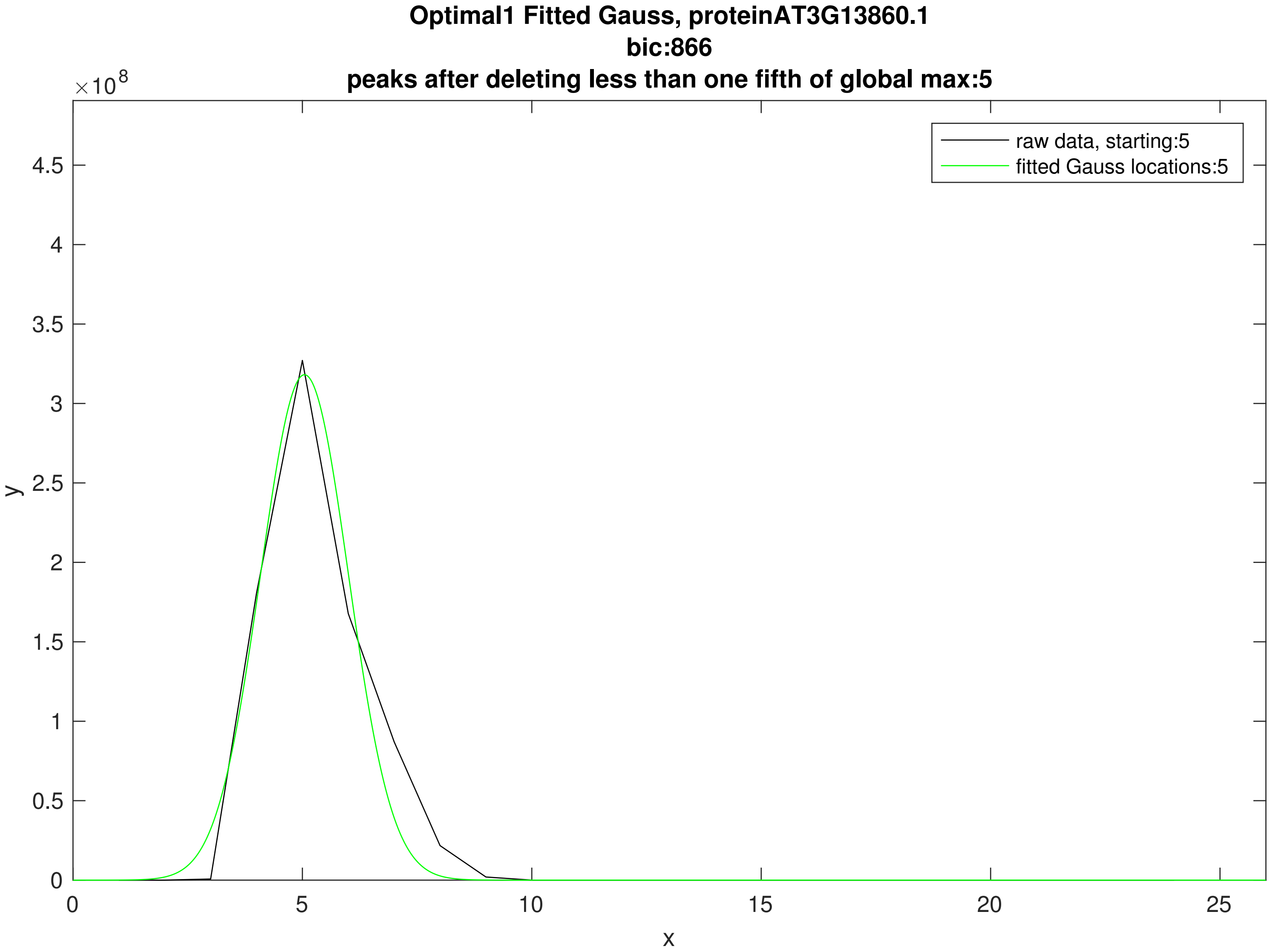
Optimal1 Fitted Gauss, proteinAT3G13860.1
bic:866
peaks after deleting less than one fifth of global max:5
×10^8
raw data, starting:5
fitted Gauss locations:5
4.5
4
3.5
3
2.5
2
1.5
1
0.5
0
y
0
5
10
15
20
25
x

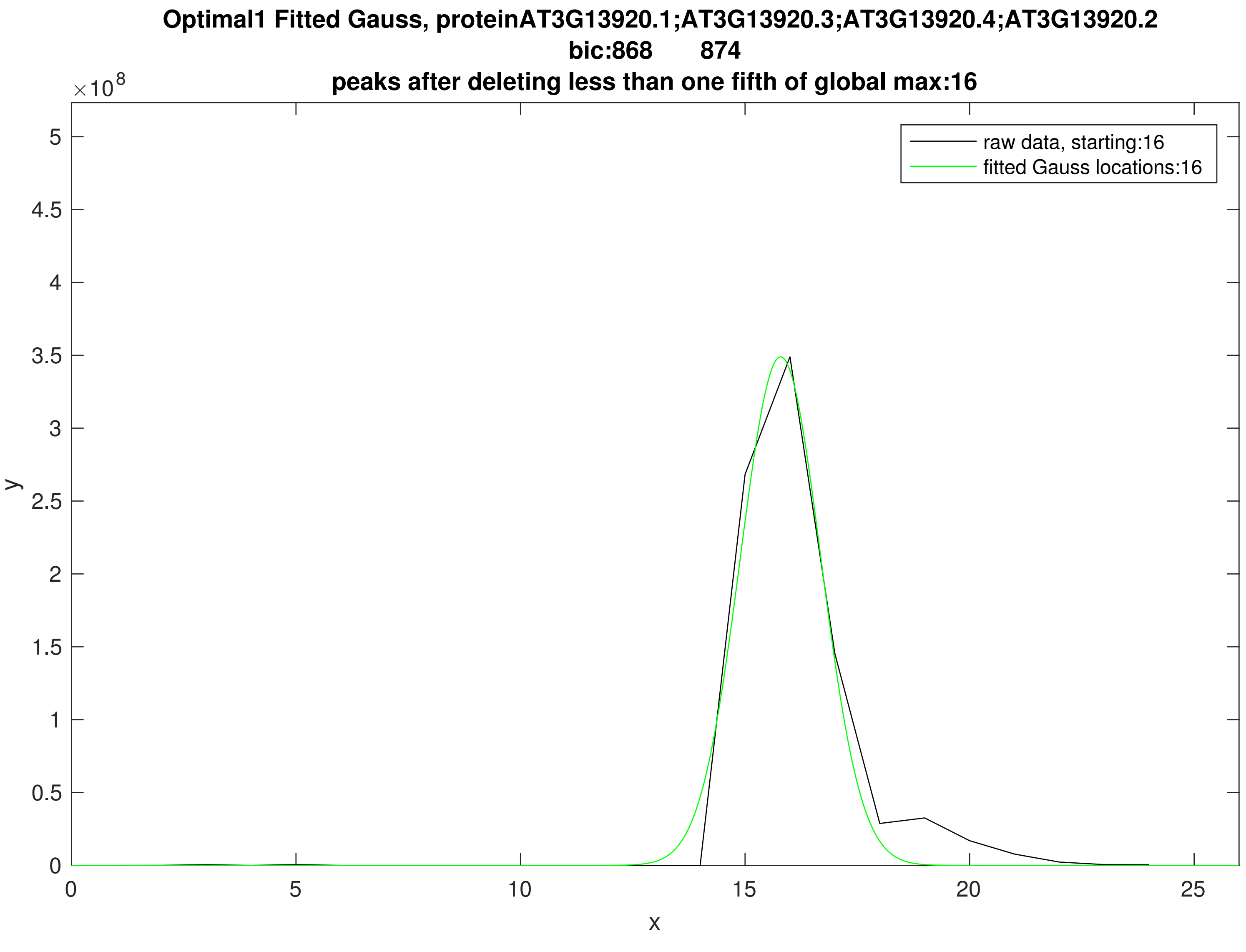

Optimal1 Fitted Gauss, proteinAT3G13920.1;AT3G13920.3;AT3G13920.4;AT3G13920.2
bic:868 874
peaks after deleting less than one fifth of global max:16
×10^8
raw data, starting:16
fitted Gauss locations:16
y
x
0
0.5
1
1.5
2
2.5
3
3.5
4
4.5
5
0
5
10
15
20
25

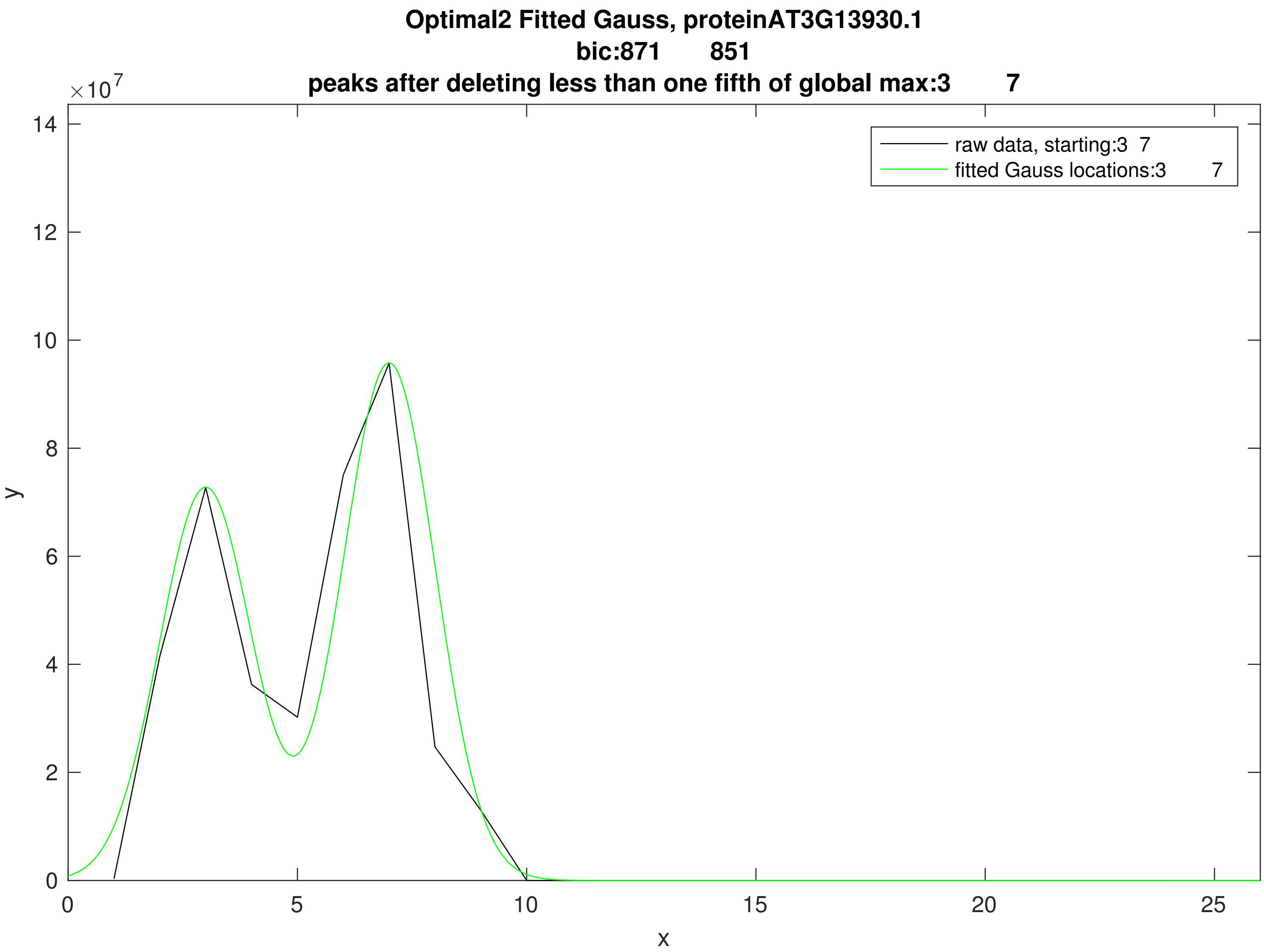

Optimal2 Fitted Gauss, proteinAT3G13930.1
bic:871 851
peaks after deleting less than one fifth of global max:3 7
×10^7
14
12
10
8
6
4
2
0
y
0
5
10
15
20
25
x
raw data, starting:3 7
fitted Gauss locations:3 7

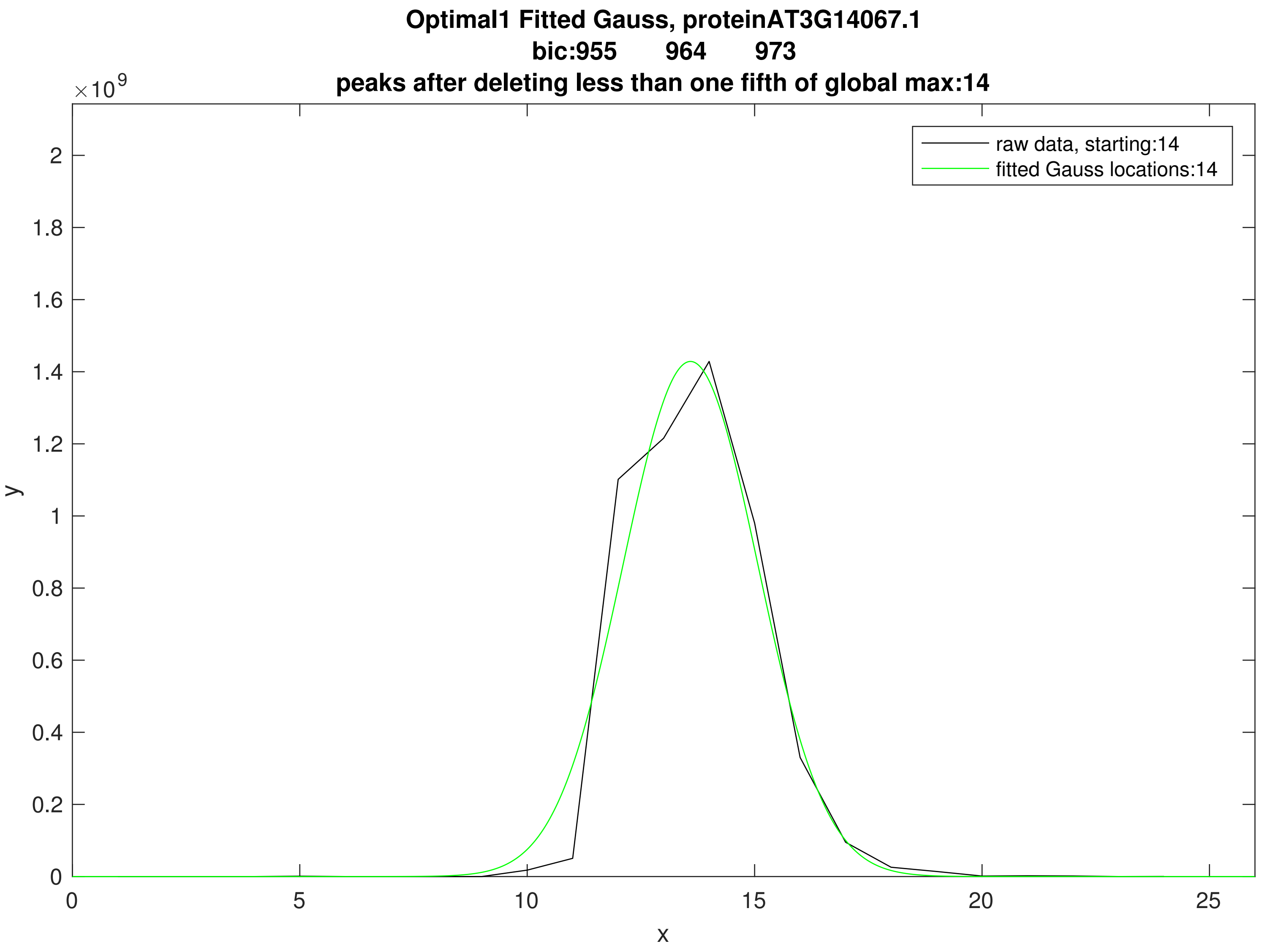
Optimal1 Fitted Gauss, proteinAT3G14067.1
bic:955 964 973
peaks after deleting less than one fifth of global max:14
×10^9
raw data, starting:14
fitted Gauss locations:14
y
x
2
1.8
1.6
1.4
1.2
1
0.8
0.6
0.4
0.2
0
0
5
10
15
20
25

**Optimal1 Fitted Gauss, proteinAT3G14290.1**
**bic:835 840**
**peaks after deleting less than one fifth of global max:5.6**

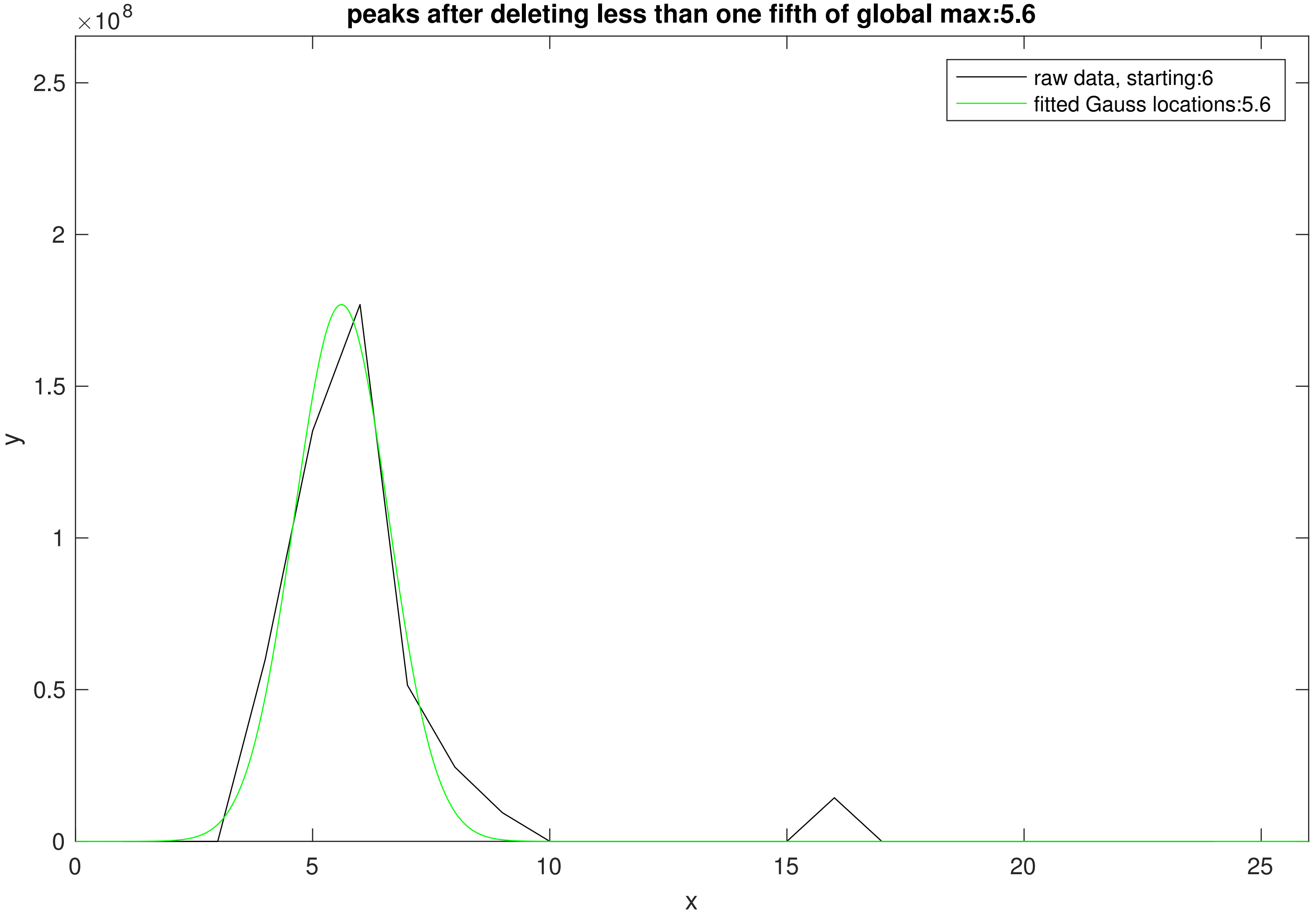

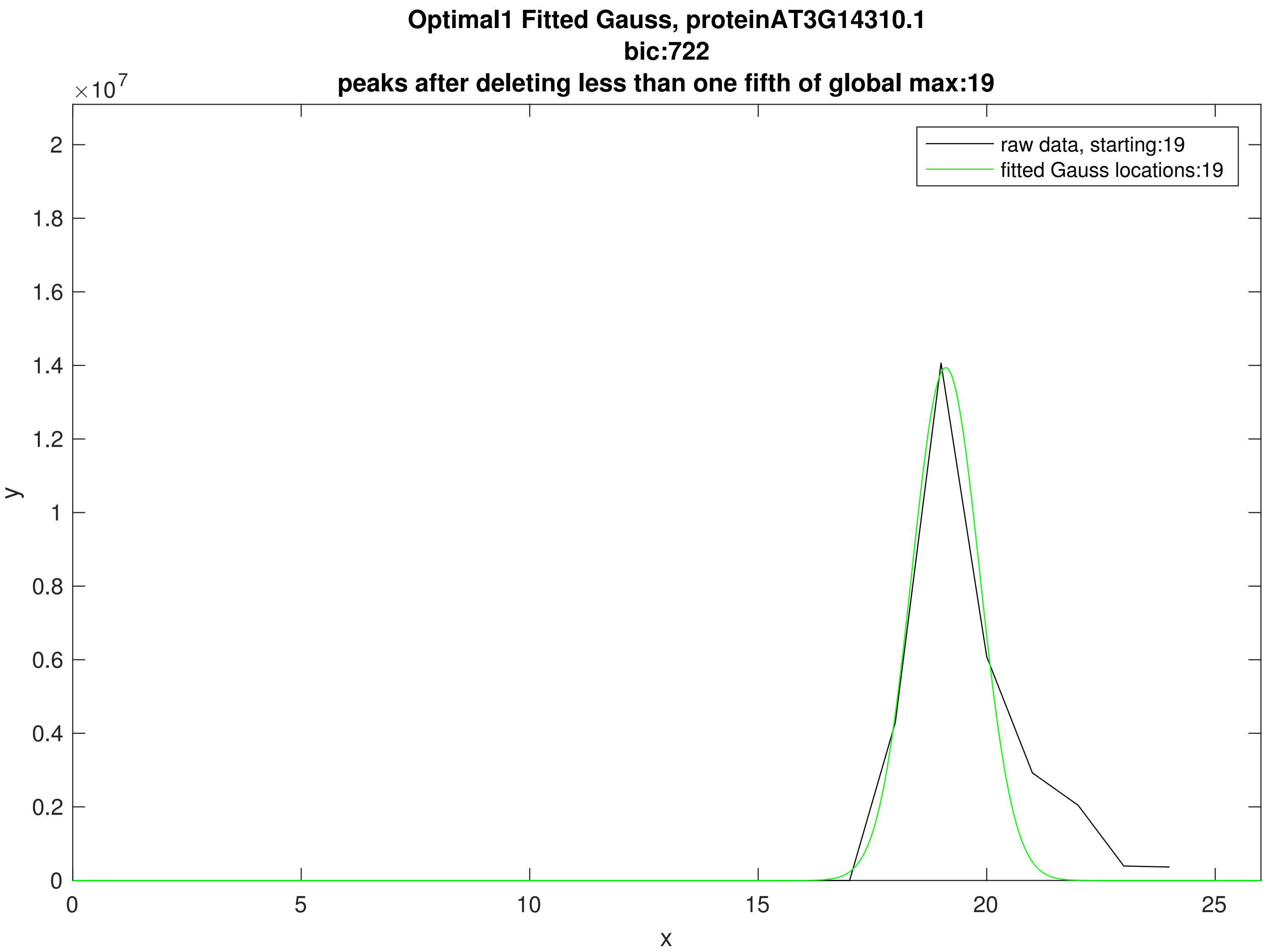
Optimal1 Fitted Gauss, proteinAT3G14310.1
bic:722
peaks after deleting less than one fifth of global max:19
×10^7
raw data, starting:19
fitted Gauss locations:19
y
x
0
0.2
0.4
0.6
0.8
1
1.2
1.4
1.6
1.8
2
0
5
10
15
20
25

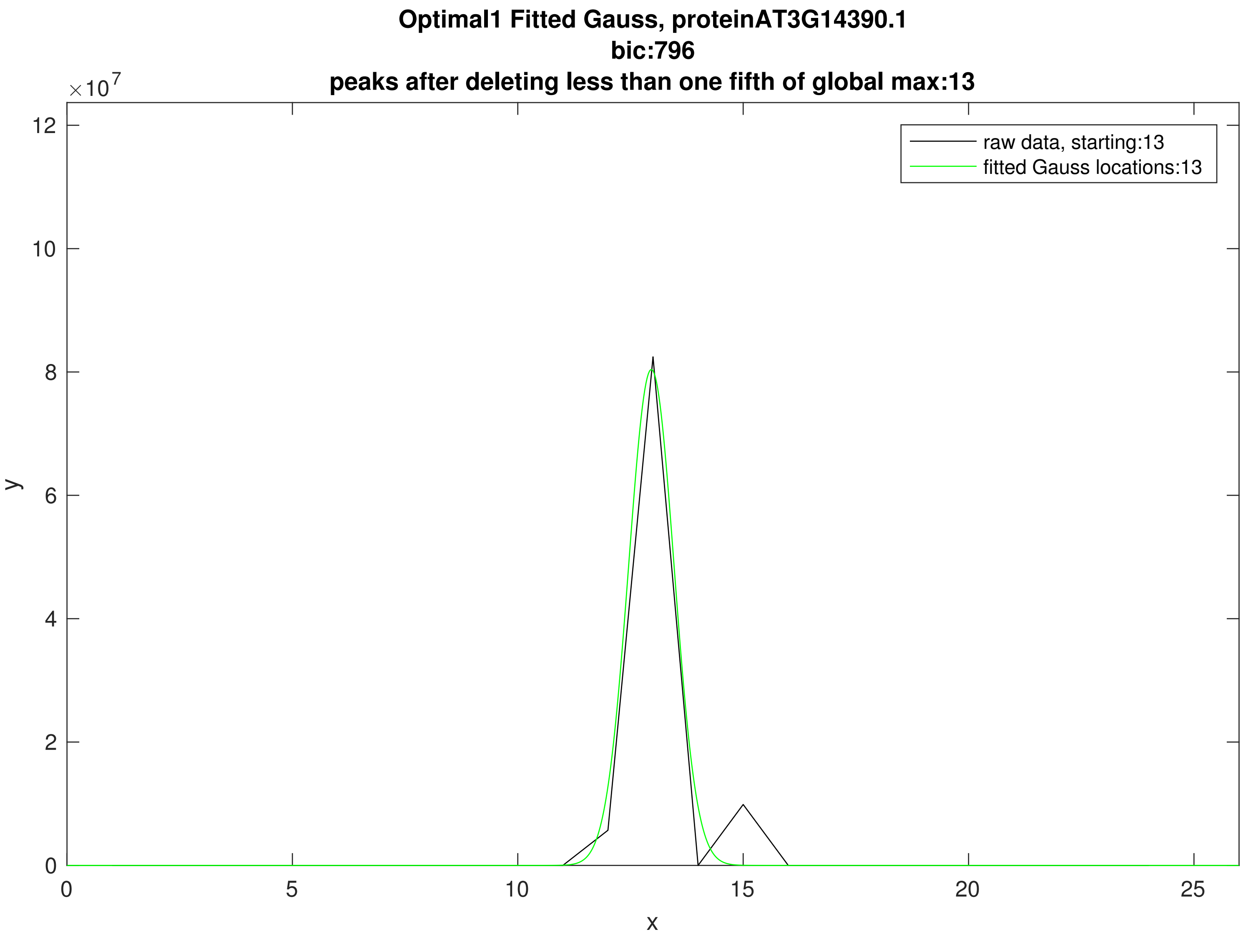

Optimal1 Fitted Gauss, proteinAT3G14390.1
bic:796
peaks after deleting less than one fifth of global max:13
×10^7
12
10
8
6
4
2
0
y
0
5
10
15
20
25
x
raw data, starting:13
fitted Gauss locations:13

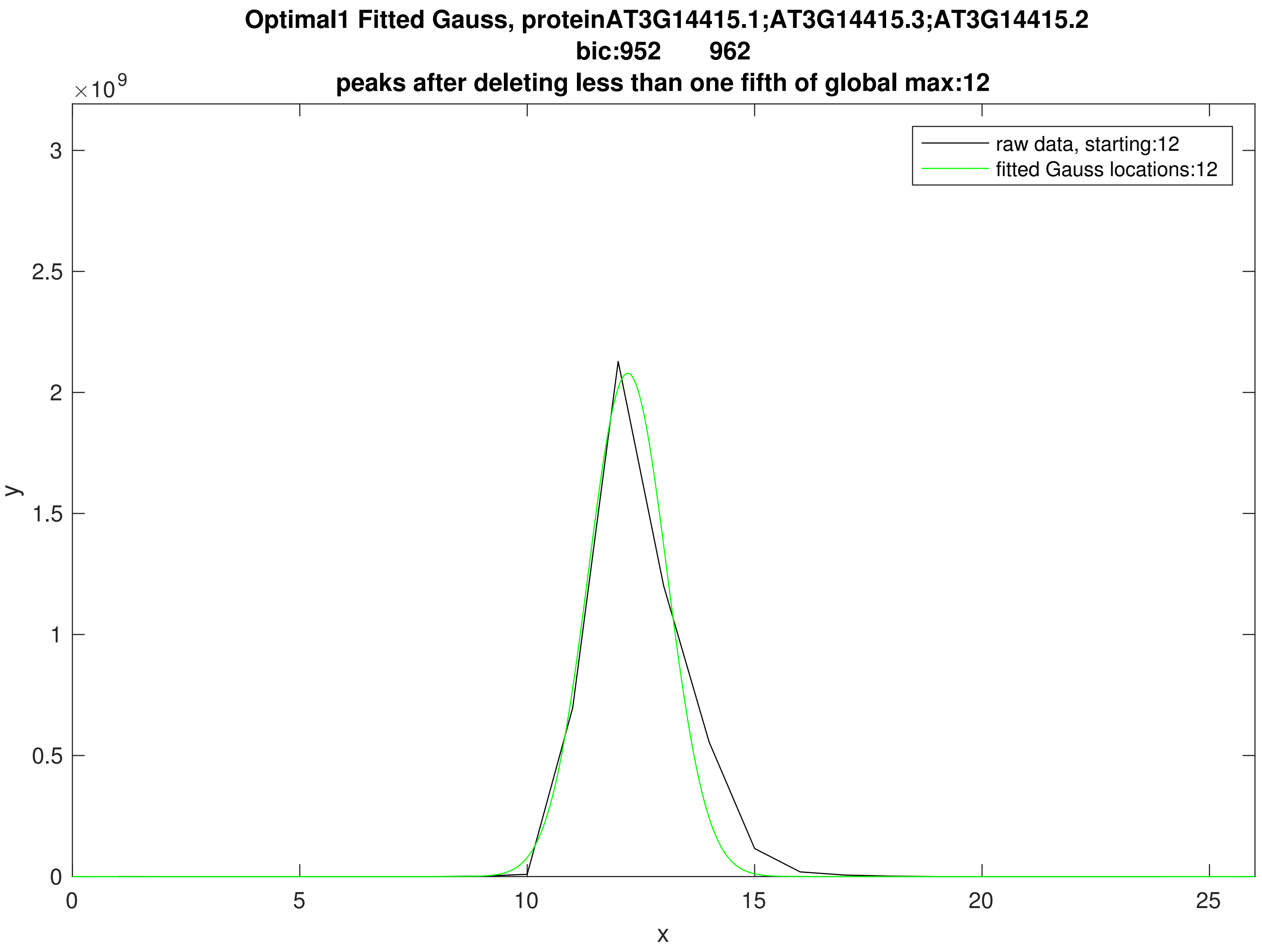

Optimal1 Fitted Gauss, proteinAT3G14415.1;AT3G14415.3;AT3G14415.2
bic:952 962
peaks after deleting less than one fifth of global max:12
×10^9
3
2.5
2
1.5
1
0.5
0
y
0
5
10
15
20
25
x
raw data, starting:12
fitted Gauss locations:12

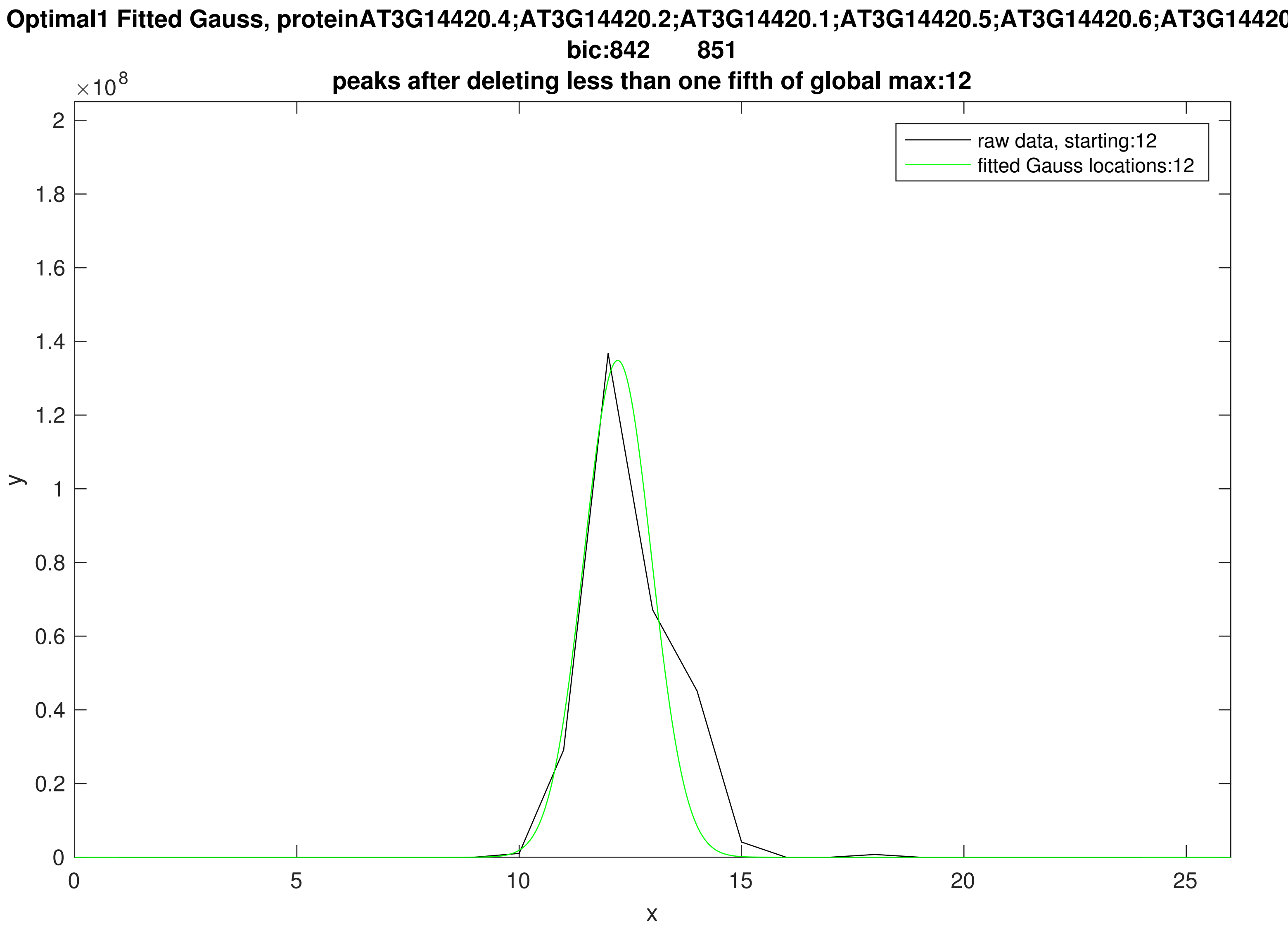
Optimal1 Fitted Gauss, proteinAT3G14420.4;AT3G14420.2;AT3G14420.1;AT3G14420.5;AT3G14420.6;AT3G1442
bic:842 851
peaks after deleting less than one fifth of global max:12
×10^8
2
1.8
1.6
1.4
1.2
1
0.8
0.6
0.4
0.2
0
y
0
5
10
15
20
25
x
raw data, starting:12
fitted Gauss locations:12

Optimal1 Fitted Gauss, proteinAT3G14790.1
bic:719 764
peaks after deleting less than one fifth of global max:6.7

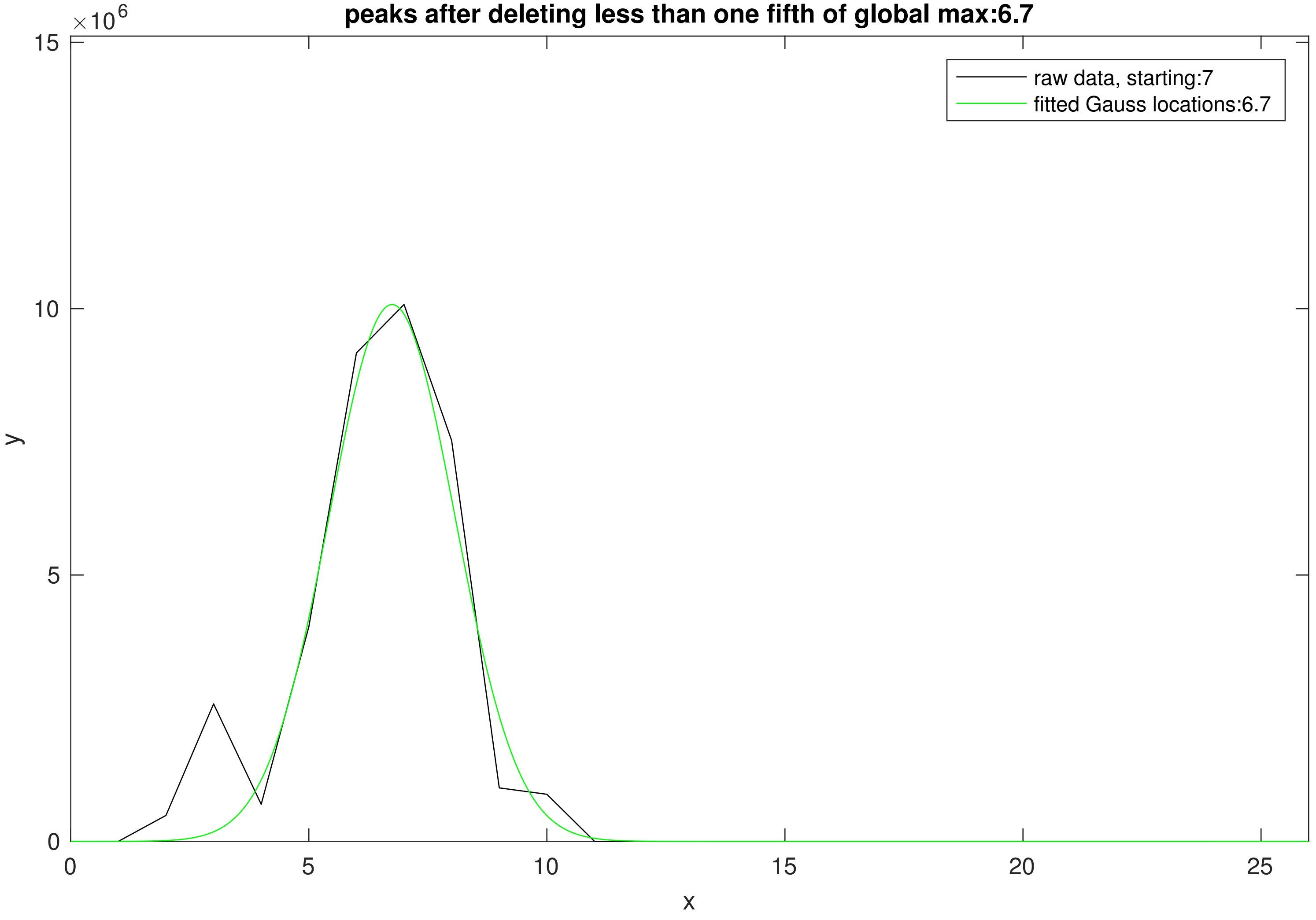

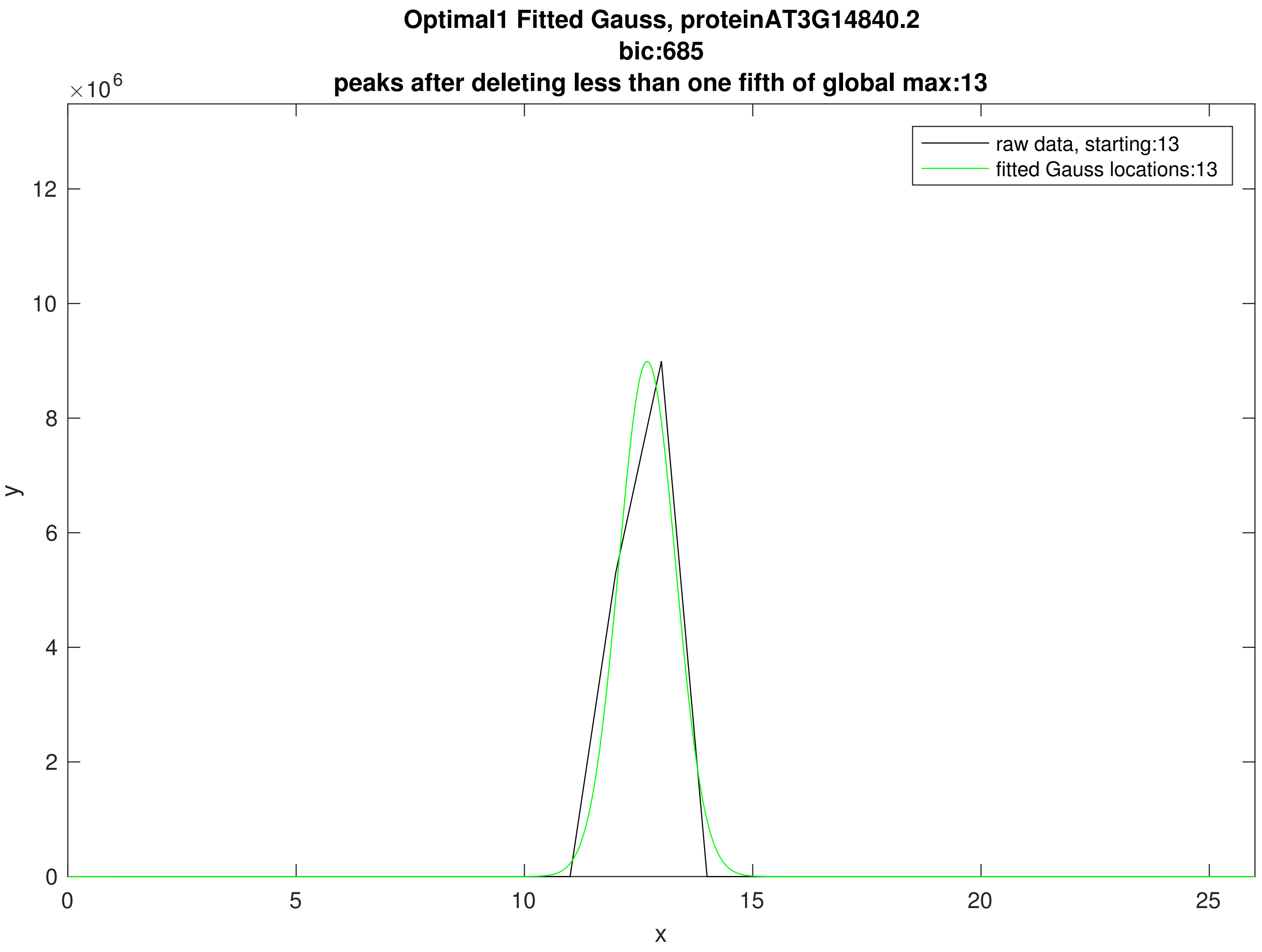
Optimal1 Fitted Gauss, proteinAT3G14840.2
bic:685
peaks after deleting less than one fifth of global max:13
×10^6
raw data, starting:13
fitted Gauss locations:13
y
x
0
2
4
6
8
10
12
0
5
10
15
20
25

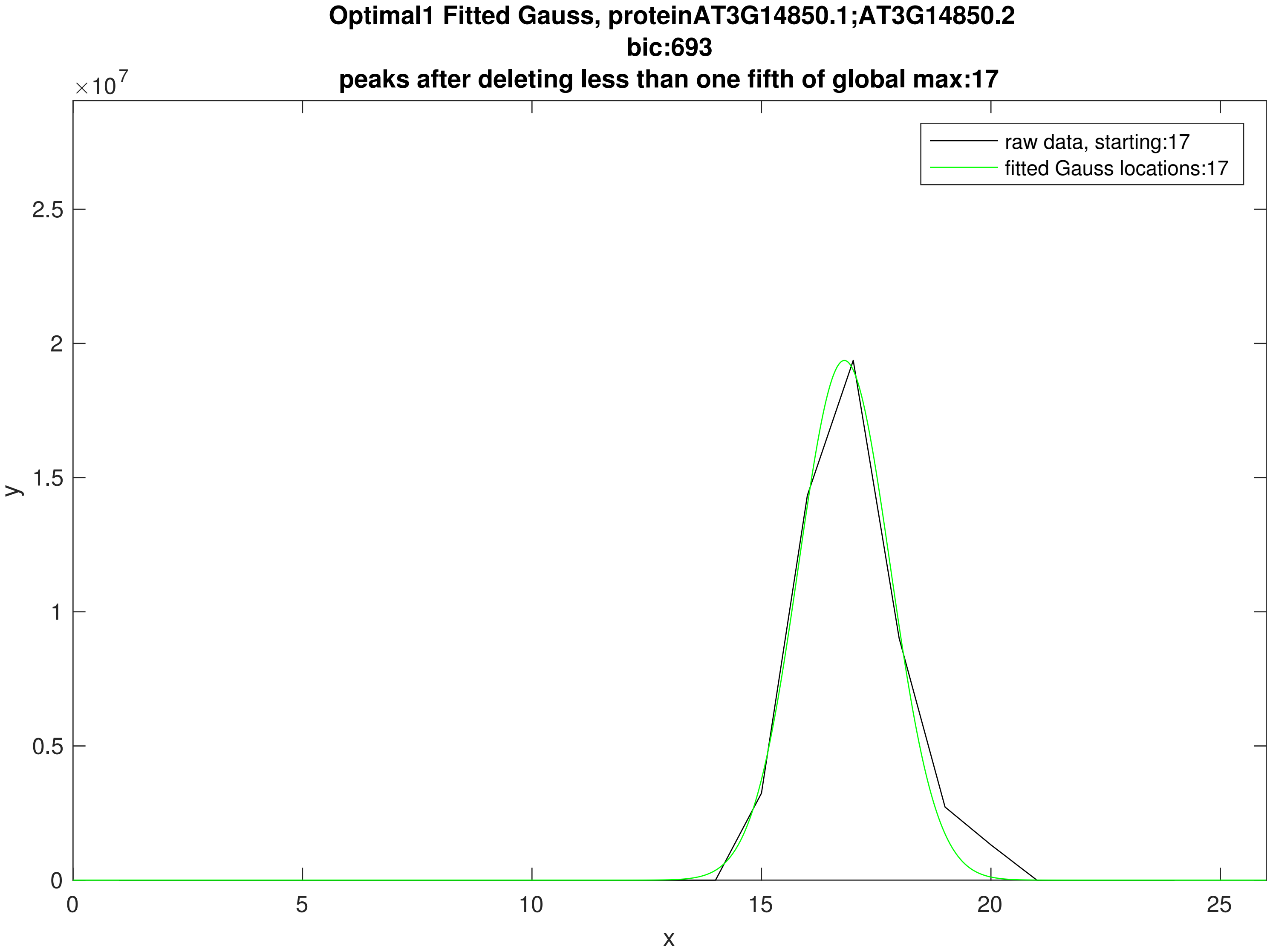
Optimal1 Fitted Gauss, proteinAT3G14850.1;AT3G14850.2
bic:693
peaks after deleting less than one fifth of global max:17
×10^7
raw data, starting:17
fitted Gauss locations:17
2.5
2
1.5
1
0.5
0
y
0
5
10
15
20
25
x

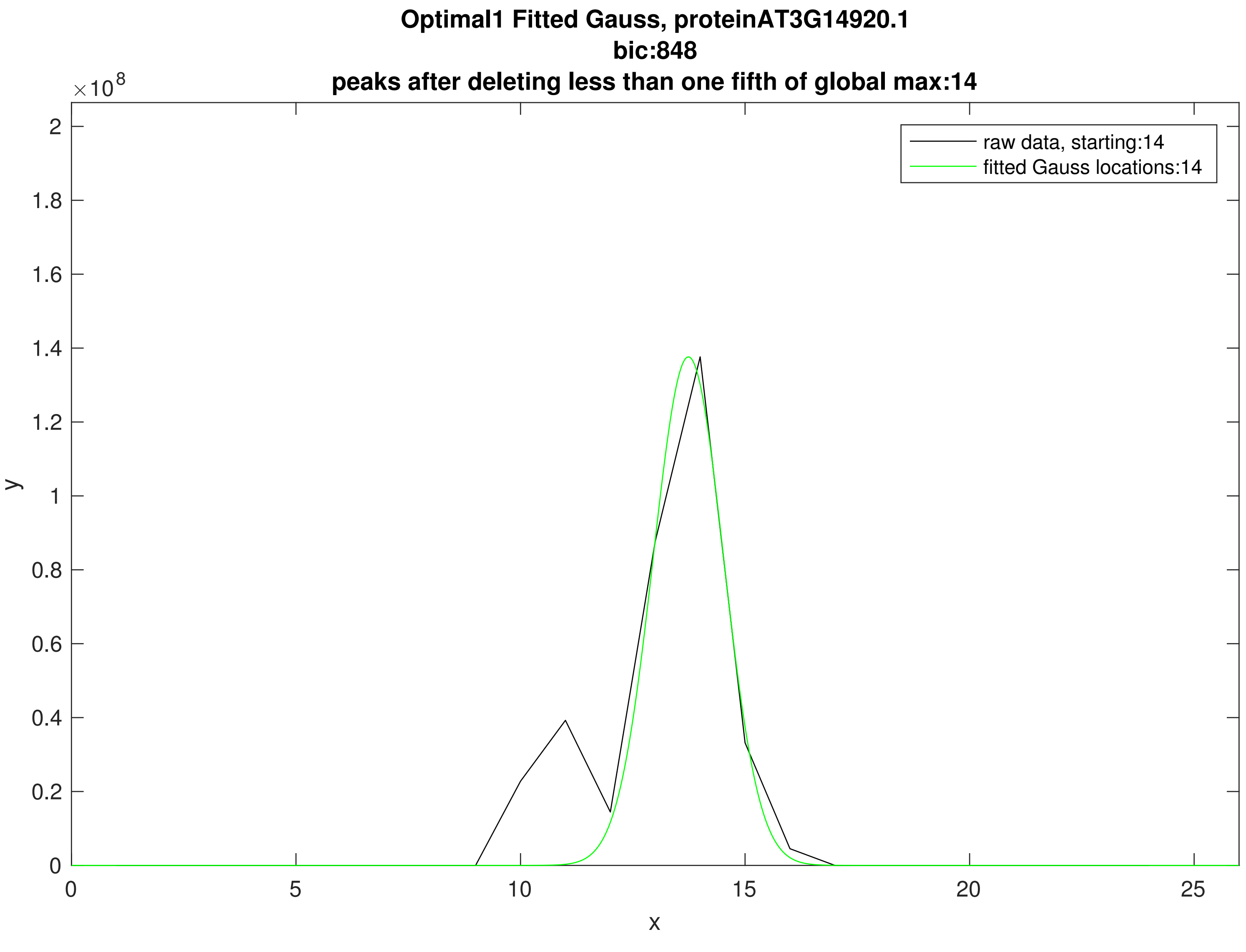
Optimal1 Fitted Gauss, proteinAT3G14920.1
bic:848
peaks after deleting less than one fifth of global max:14
×10^8
2
1.8
1.6
1.4
1.2
1
0.8
0.6
0.4
0.2
0
y
0
5
10
15
20
25
x
raw data, starting:14
fitted Gauss locations:14

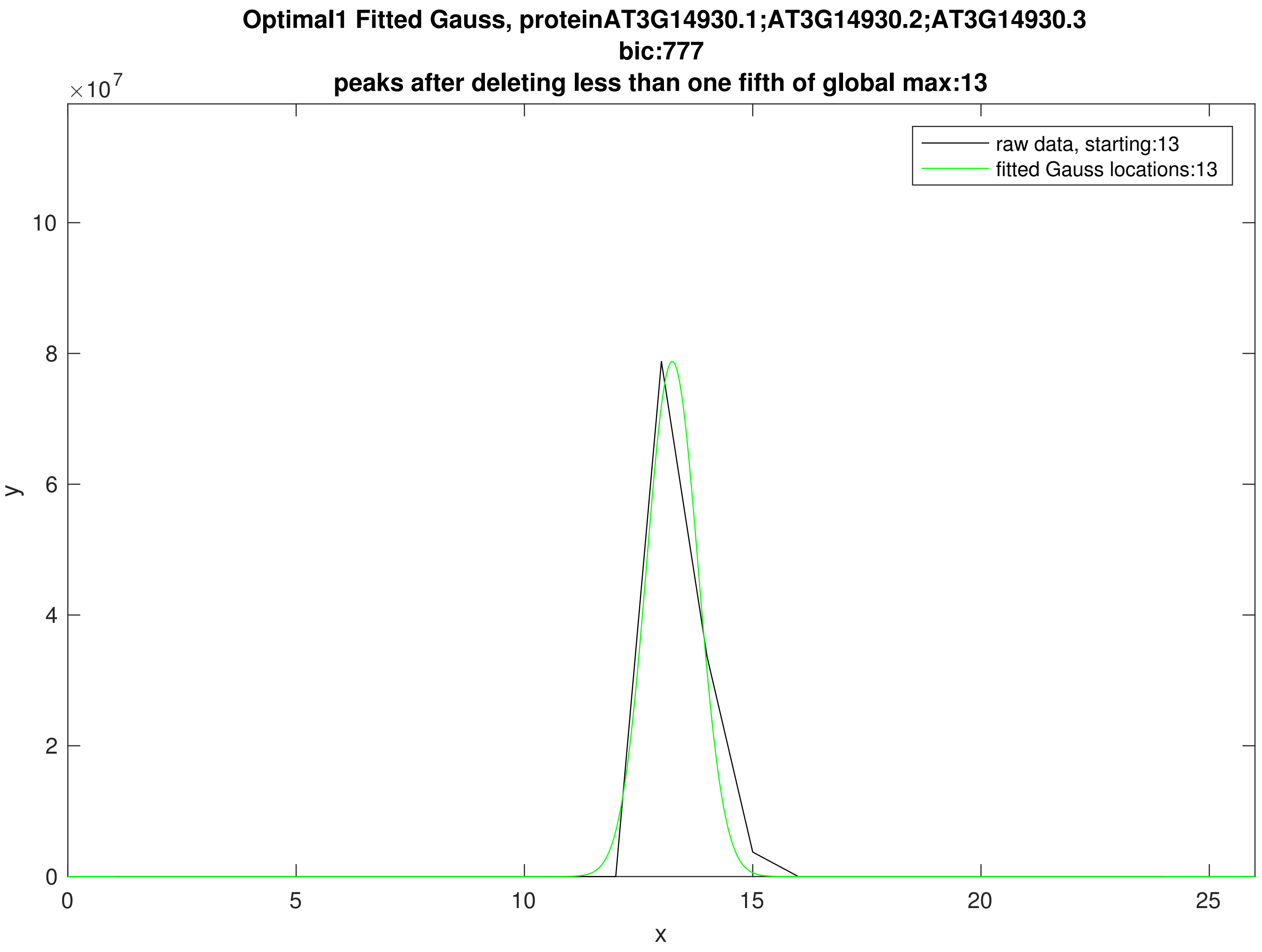
Optimal1 Fitted Gauss, proteinAT3G14930.1;AT3G14930.2;AT3G14930.3
bic:777
peaks after deleting less than one fifth of global max:13
×10^7
raw data, starting:13
fitted Gauss locations:13
y
x
0
2
4
6
8
10
0
5
10
15
20
25

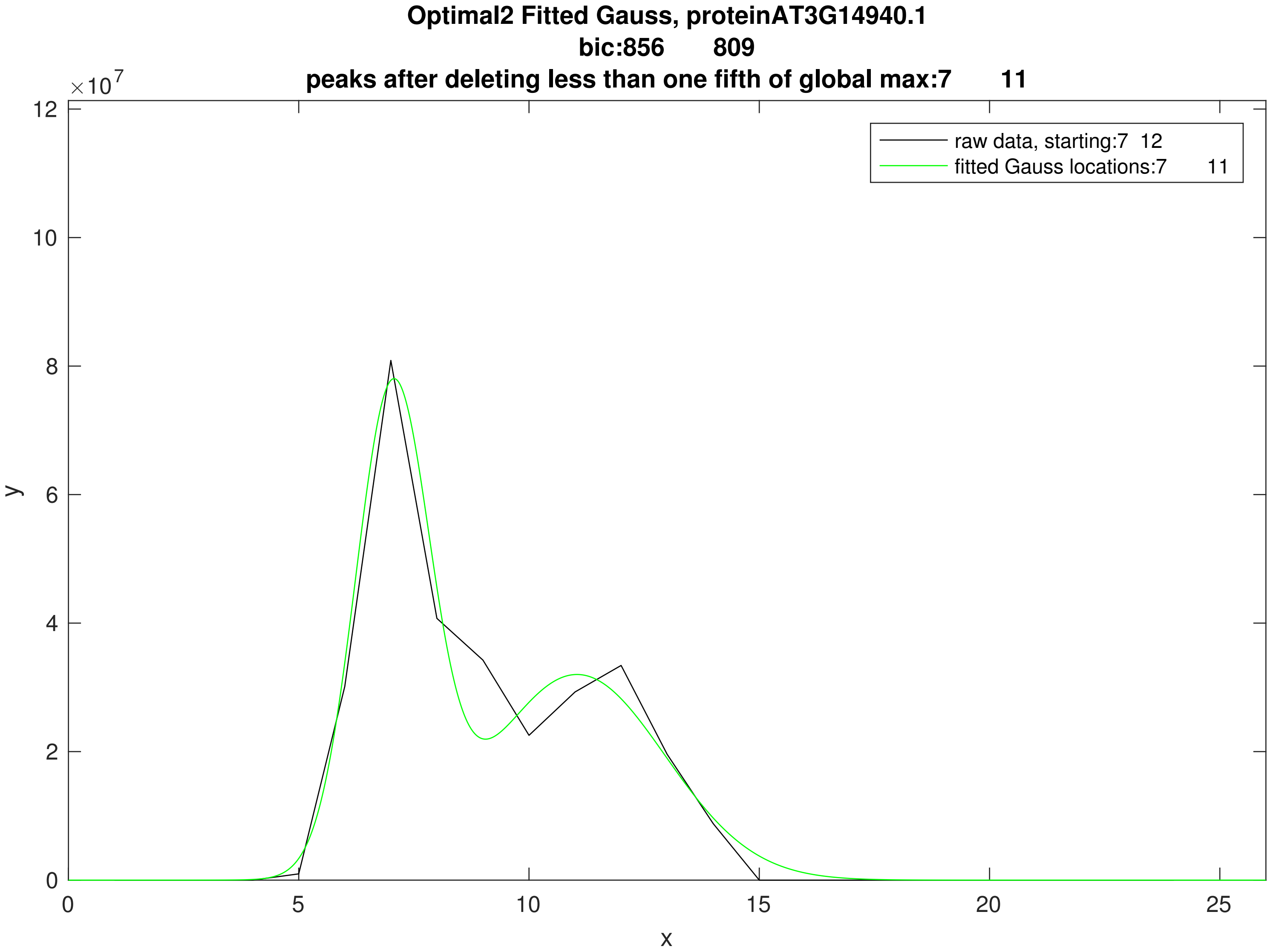
Optimal2 Fitted Gauss, proteinAT3G14940.1
bic:856 809
peaks after deleting less than one fifth of global max:7 11
×10^7
raw data, starting:7 12
fitted Gauss locations:7 11
y
x
0
2
4
6
8
10
12
5
15
20
25

**Optimal1 Fitted Gauss, proteinAT3G14990.1;AT3G14990.3;AT3G14990.2**
**bic:917 993 925**
**peaks after deleting less than one fifth of global max:17**

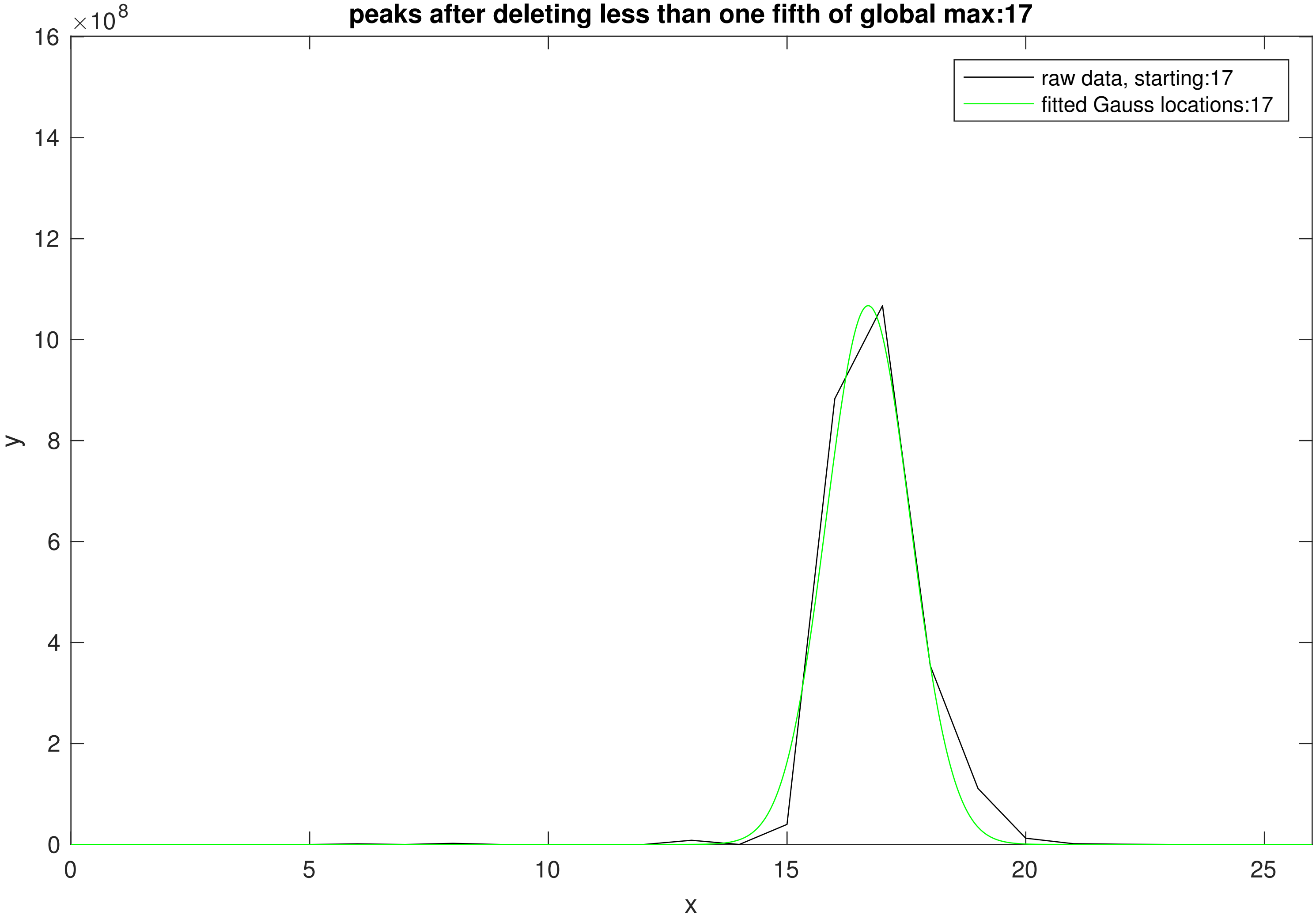

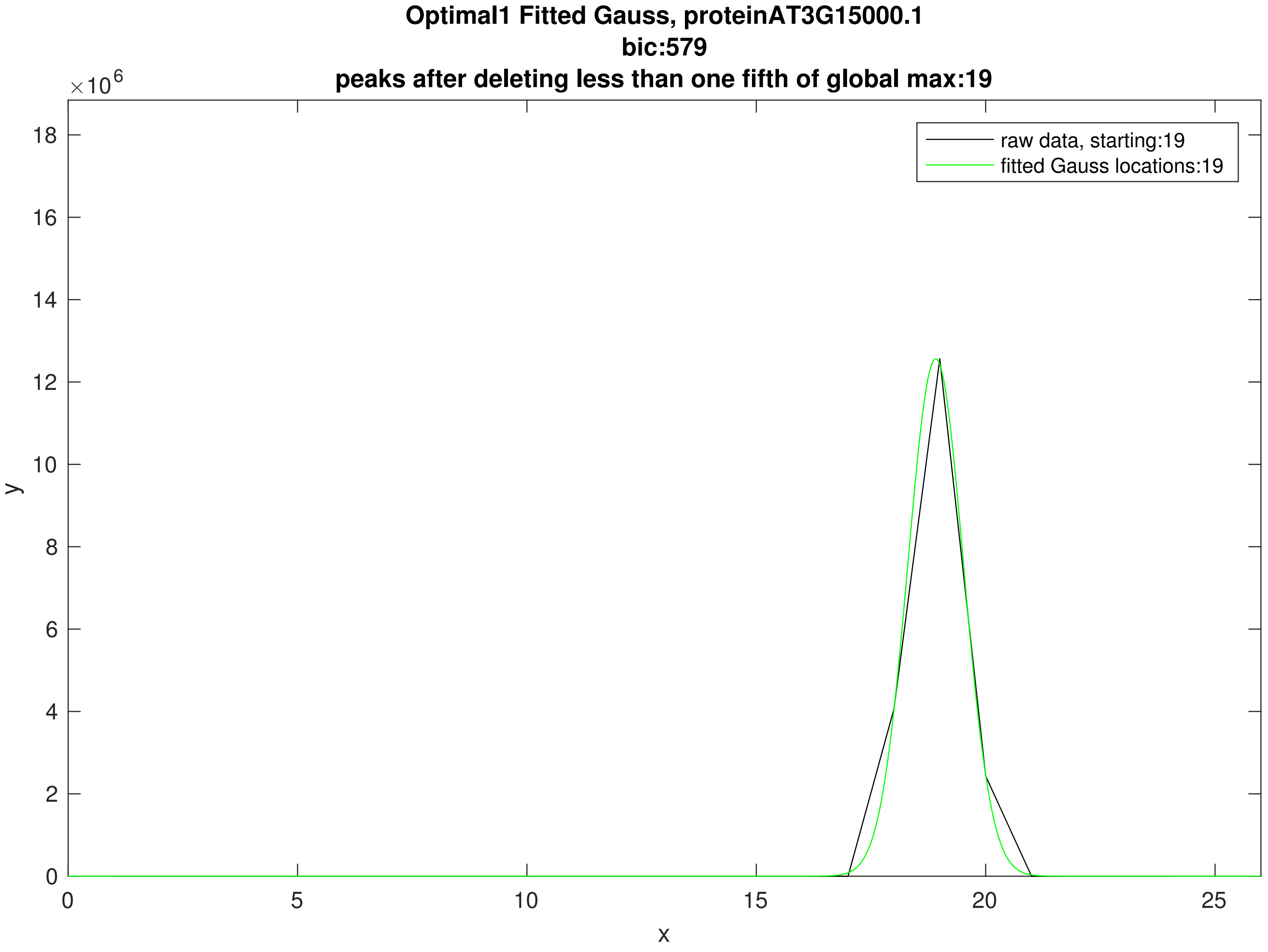

Optimal1 Fitted Gauss, proteinAT3G15000.1
bic:579
peaks after deleting less than one fifth of global max:19
×10^6
raw data, starting:19
fitted Gauss locations:19
y
x
0
2
4
6
8
10
12
14
16
18
0
5
10
15
20
25

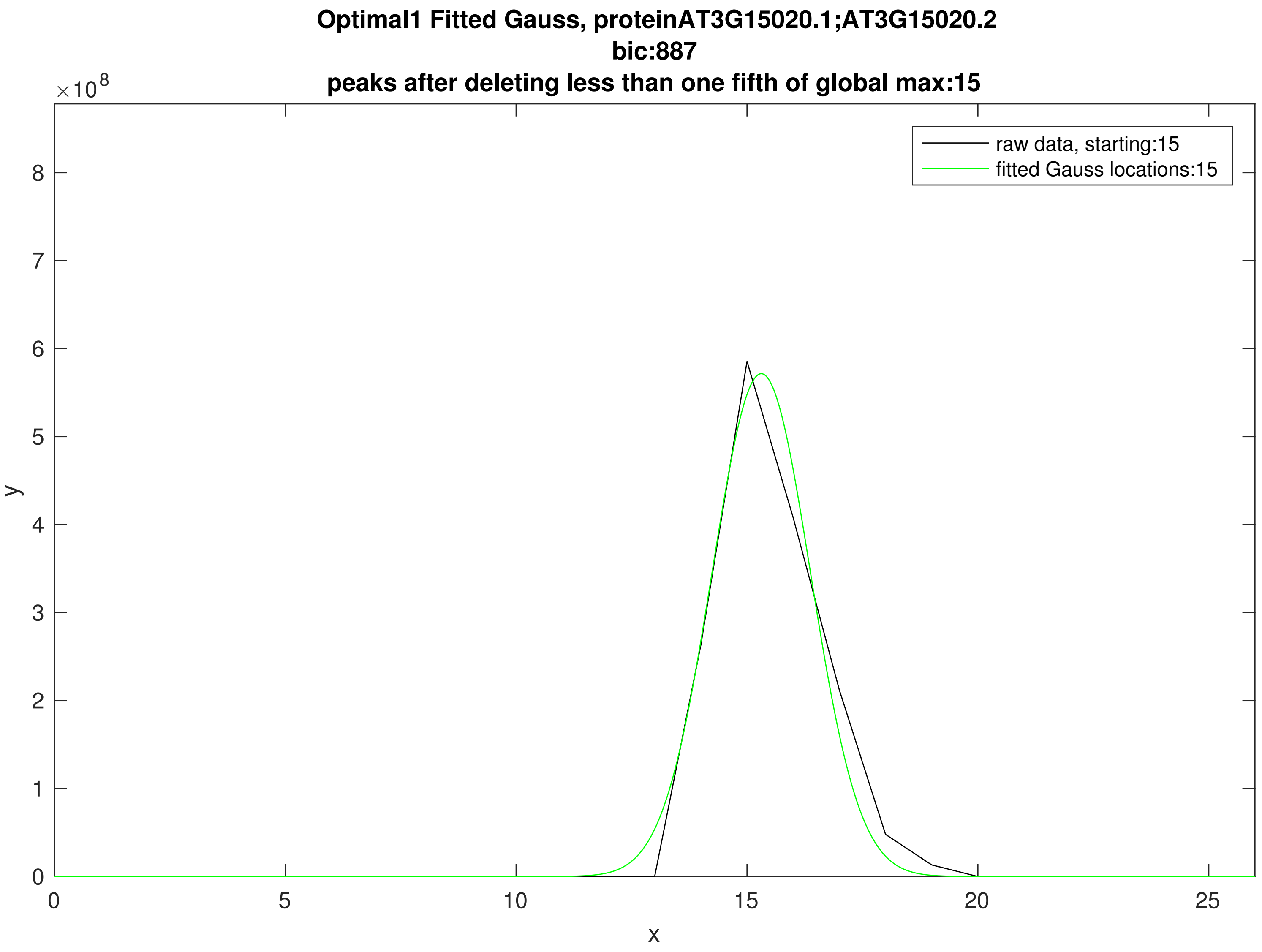
Optimal1 Fitted Gauss, proteinAT3G15020.1;AT3G15020.2
bic:887
peaks after deleting less than one fifth of global max:15
×10^8
raw data, starting:15
fitted Gauss locations:15
y
x
0
1
2
3
4
5
6
7
8
0
5
10
15
20
25

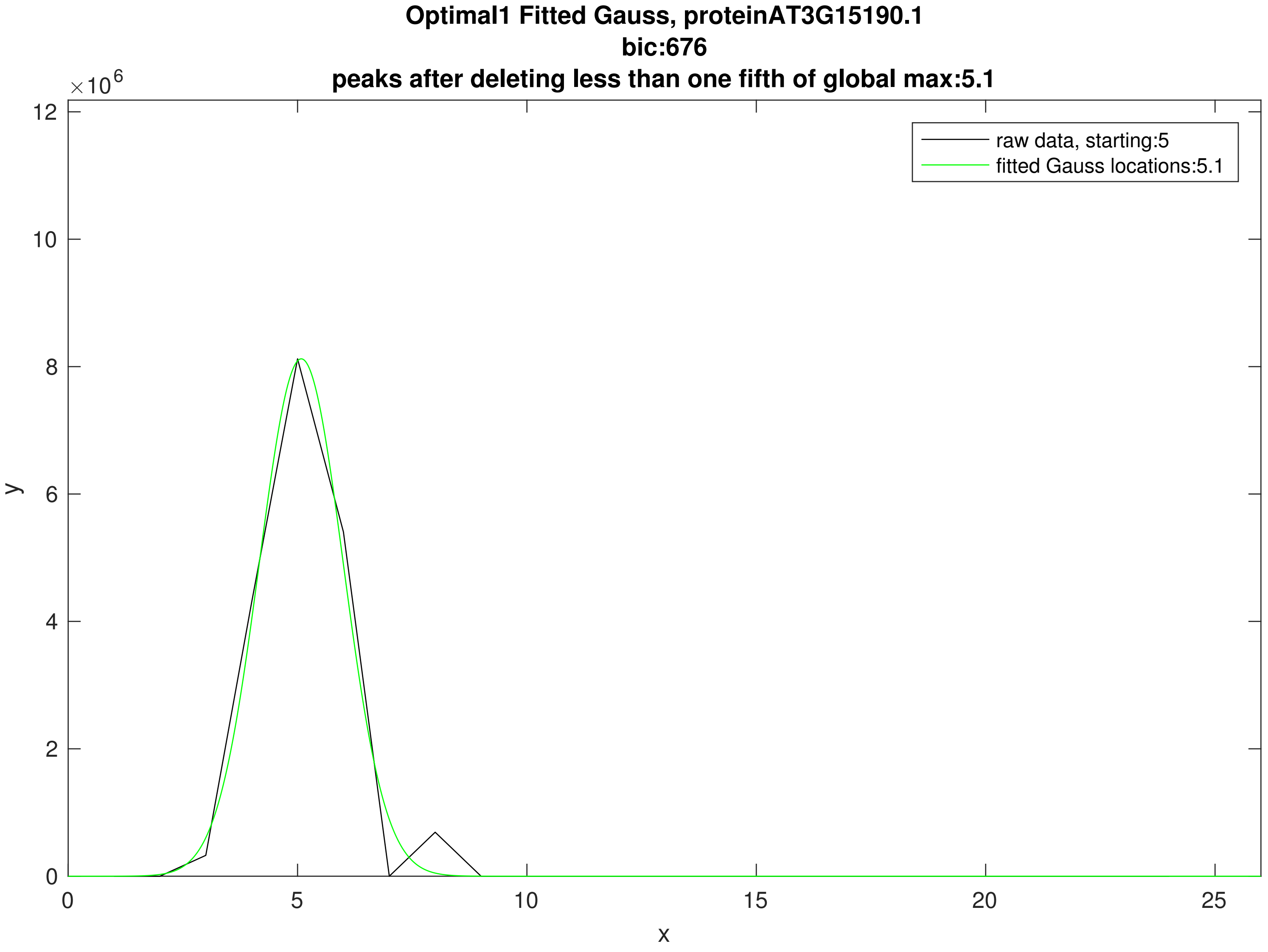

Optimal1 Fitted Gauss, proteinAT3G15190.1
bic:676
peaks after deleting less than one fifth of global max:5.1
×10^6
12
10
8
6
4
2
0
y
0
5
10
15
20
25
x
raw data, starting:5
fitted Gauss locations:5.1

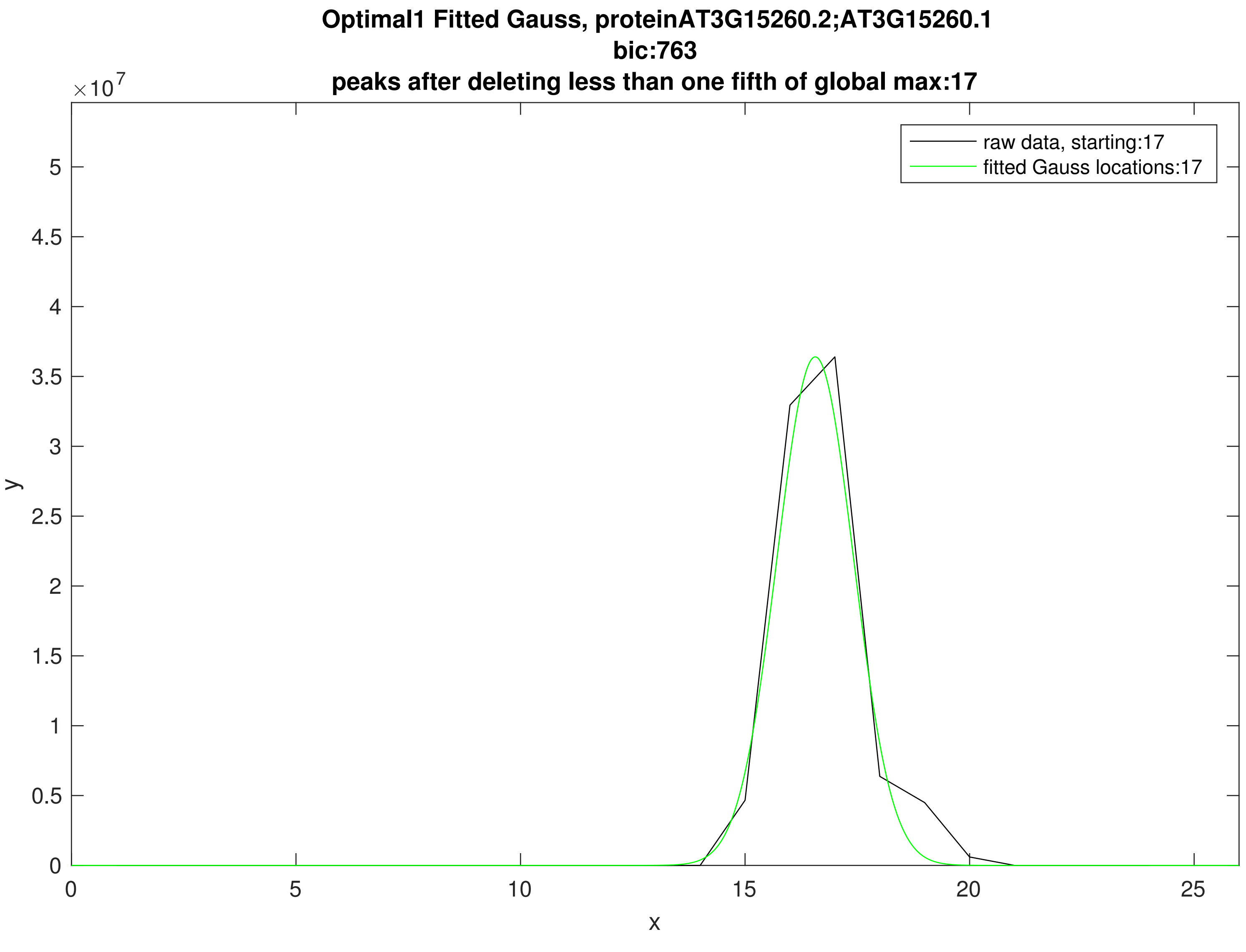

Optimal1 Fitted Gauss, proteinAT3G15260.2;AT3G15260.1
bic:763
peaks after deleting less than one fifth of global max:17
×10^7
raw data, starting:17
fitted Gauss locations:17
y
x
0
0.5
1
1.5
2
2.5
3
3.5
4
4.5
5
0
5
10
15
20
25

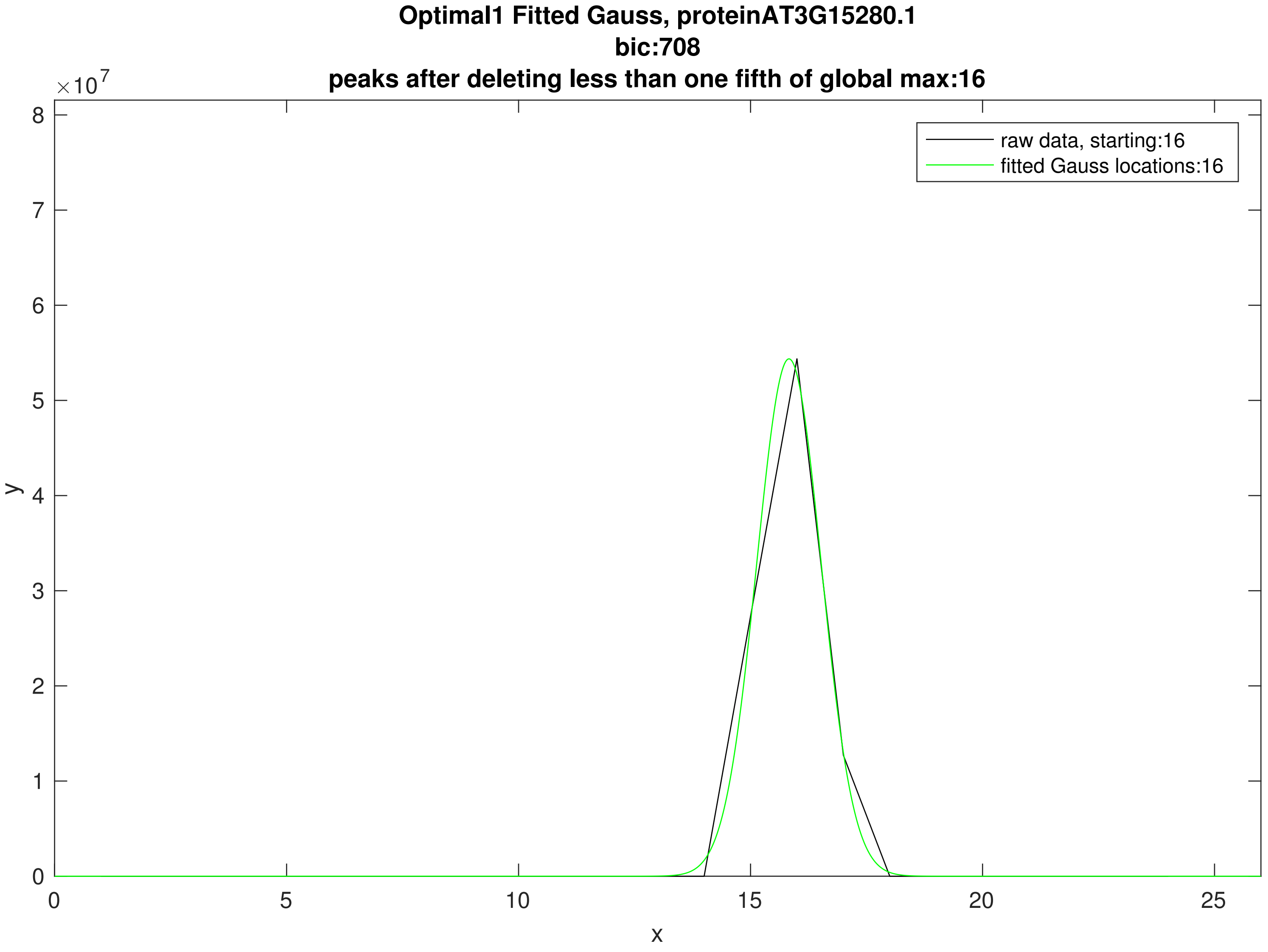
Optimal1 Fitted Gauss, proteinAT3G15280.1
bic:708
peaks after deleting less than one fifth of global max:16
×10^7
raw data, starting:16
fitted Gauss locations:16
y
x
0
1
2
3
4
5
6
7
8
0
5
10
15
20
25

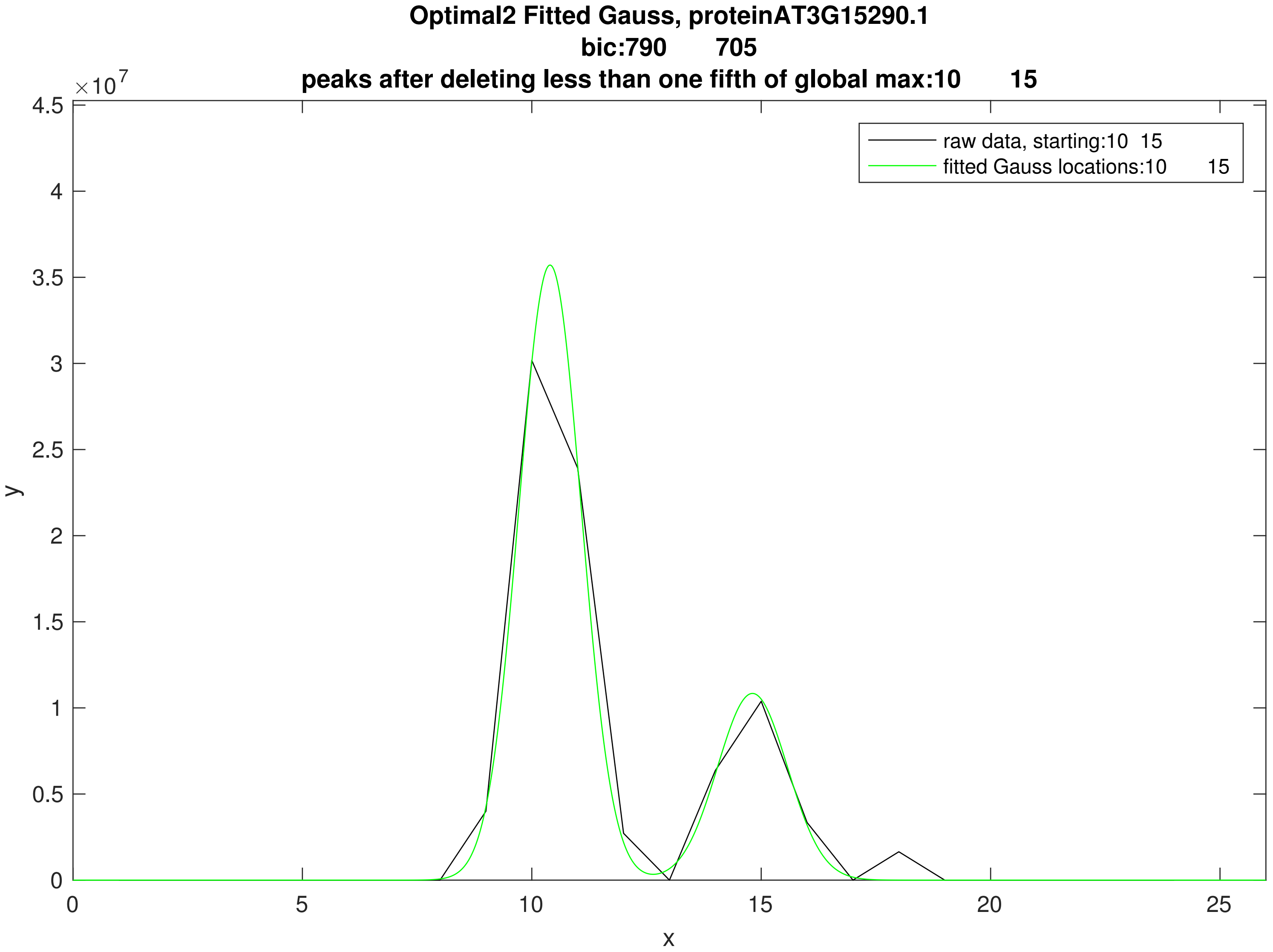

Optimal2 Fitted Gauss, proteinAT3G15290.1
bic:790 705
peaks after deleting less than one fifth of global max:10 15
×10^7
4.5
4
3.5
3
2.5
2
1.5
1
0.5
0
y
0
5
10
15
20
25
x
raw data, starting:10 15
fitted Gauss locations:10 15

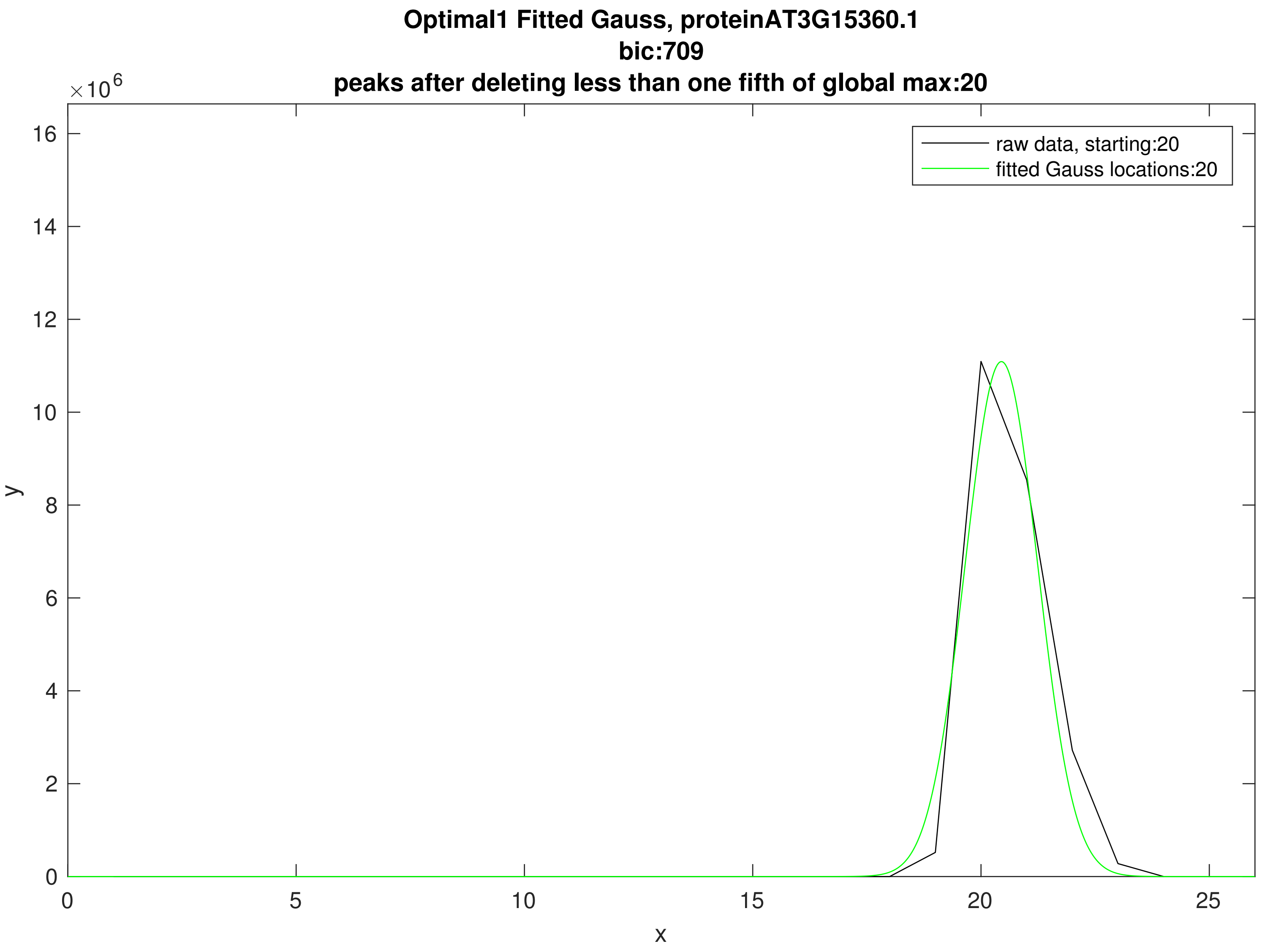
Optimal1 Fitted Gauss, proteinAT3G15360.1
bic:709
peaks after deleting less than one fifth of global max:20
×10^6
16
14
12
10
8
6
4
2
0
y
0
5
10
15
20
25
x
raw data, starting:20
fitted Gauss locations:20

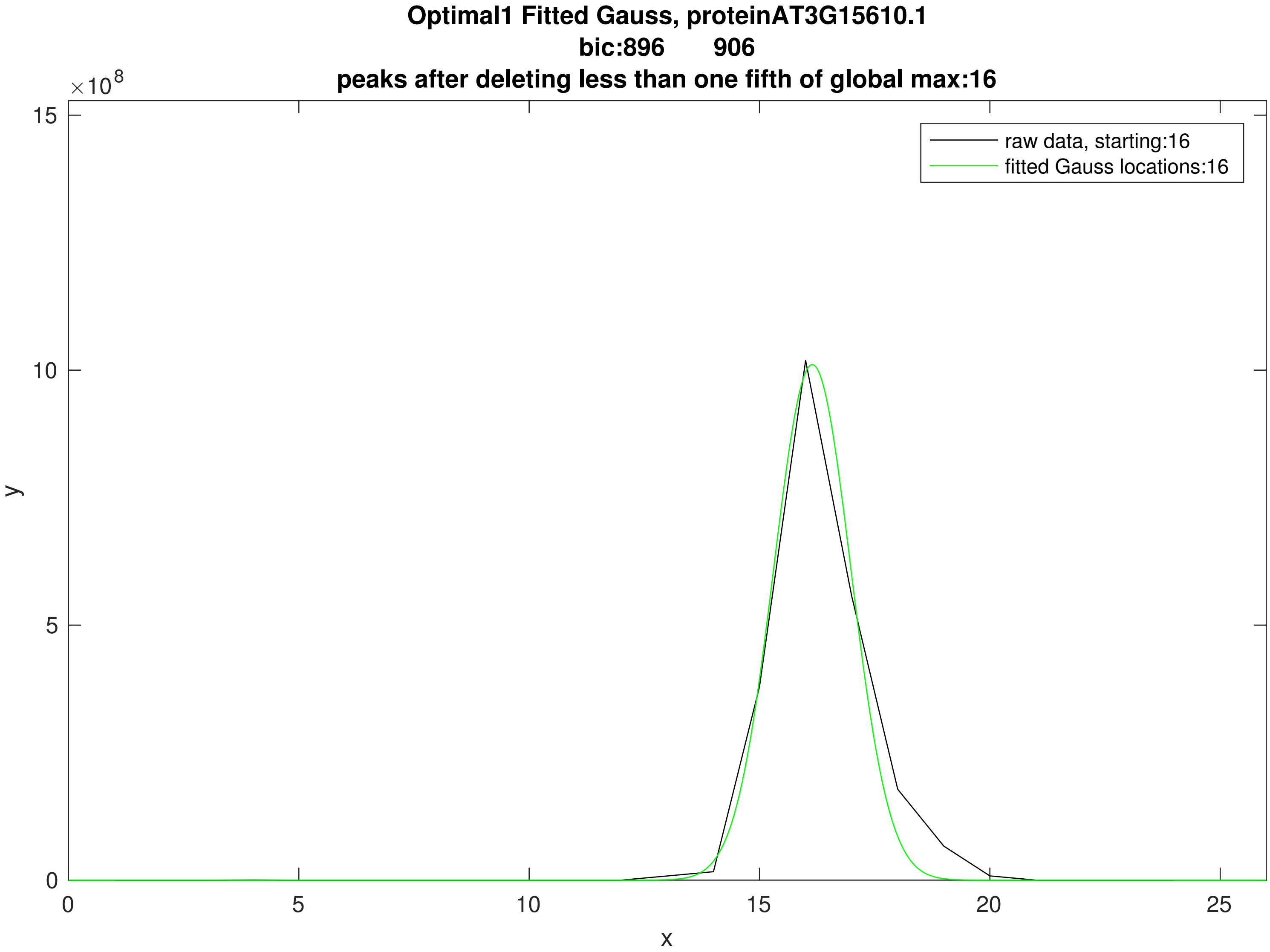

Optimal1 Fitted Gauss, proteinAT3G15610.1
bic:896 906
peaks after deleting less than one fifth of global max:16
×10^8
15
10
5
0
y
0
5
10
15
20
25
x
raw data, starting:16
fitted Gauss locations:16

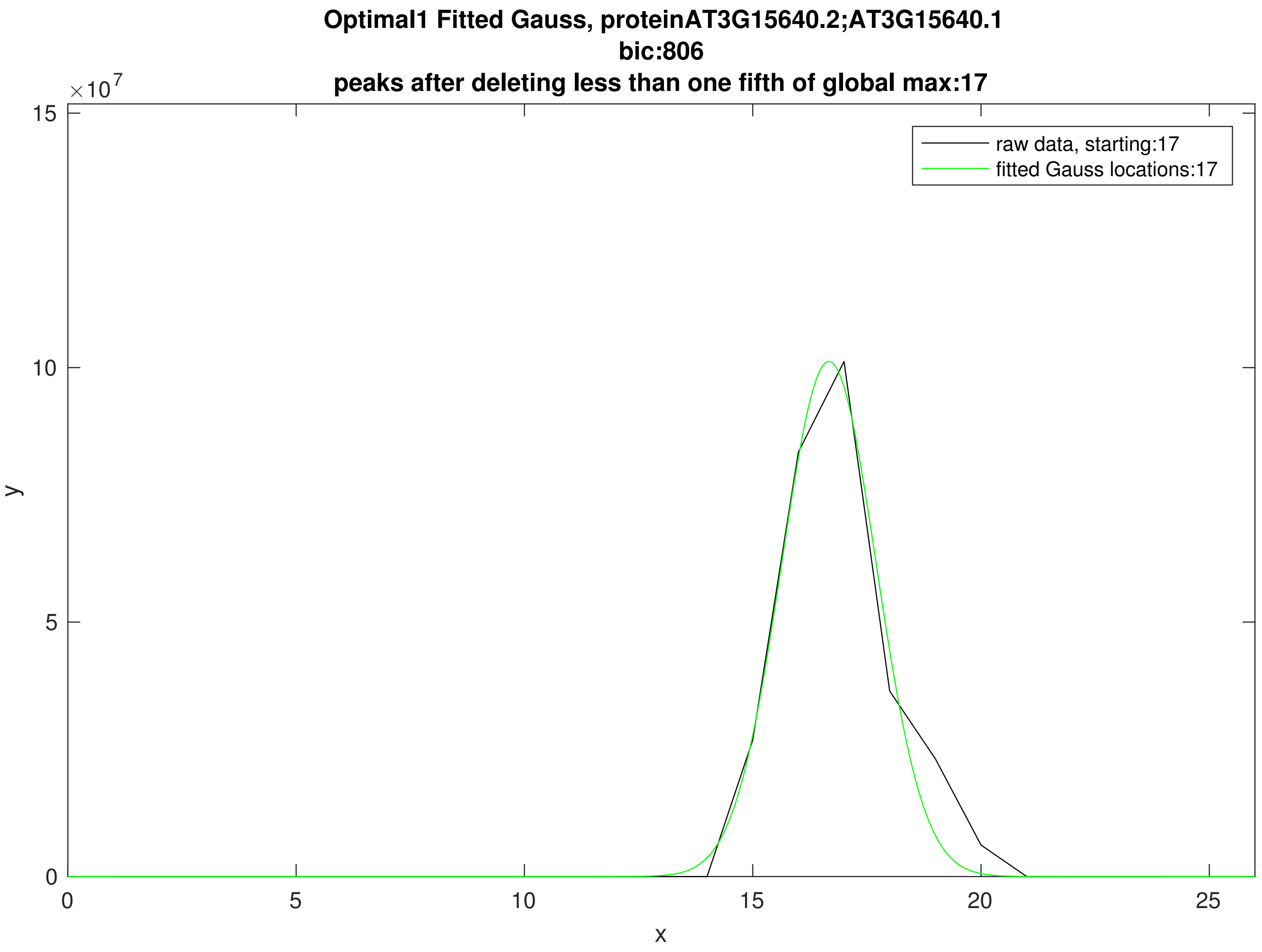

Optimal1 Fitted Gauss, proteinAT3G15640.2;AT3G15640.1
bic:806
peaks after deleting less than one fifth of global max:17
×10^7
15
10
5
0
y
0
5
10
15
20
25
x
raw data, starting:17
fitted Gauss locations:17

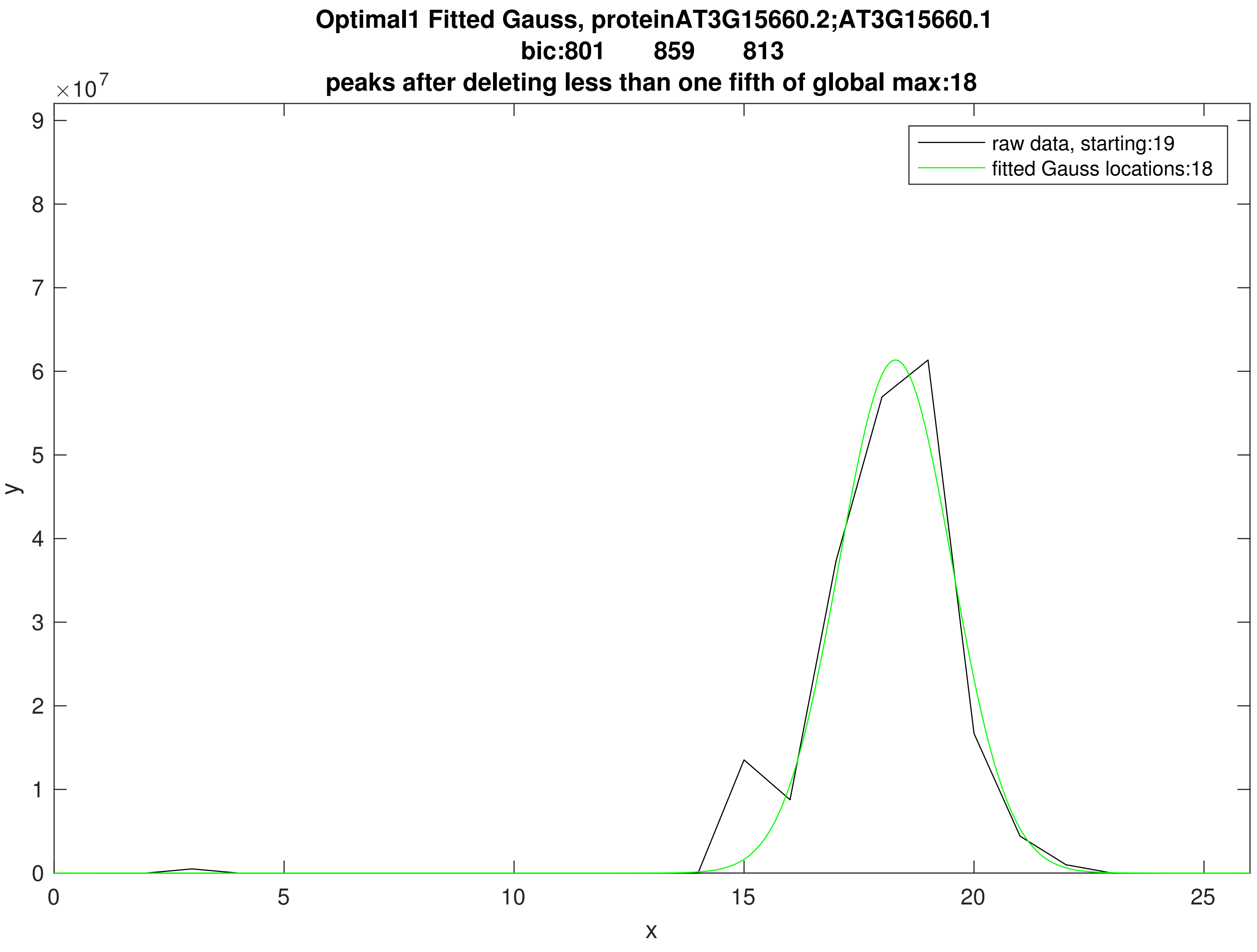
Optimal1 Fitted Gauss, proteinAT3G15660.2;AT3G15660.1
bic:801 859 813
peaks after deleting less than one fifth of global max:18
×10^7
9
8
7
6
5
4
3
2
1
0
y
0
5
10
15
20
25
x
raw data, starting:19
fitted Gauss locations:18

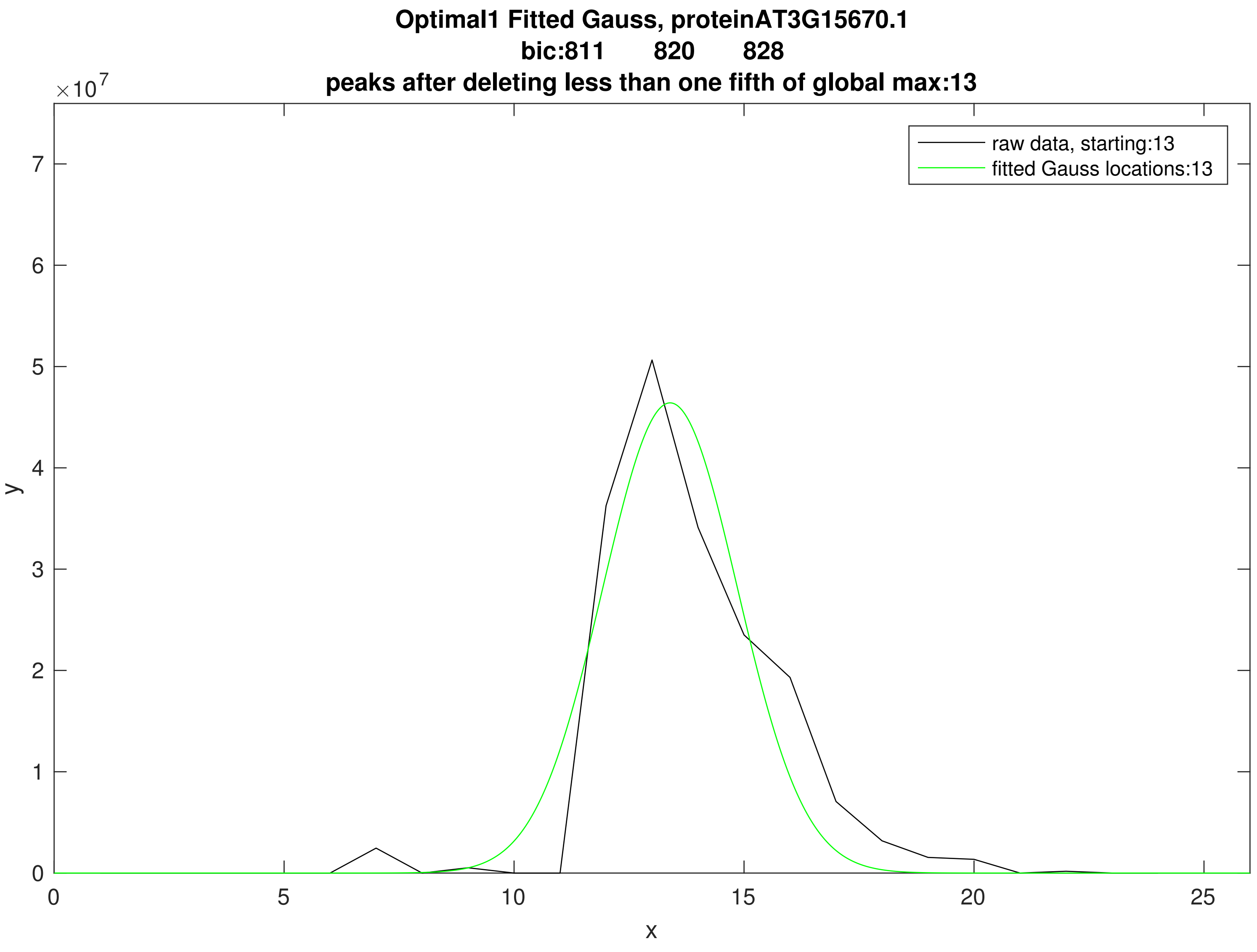

Optimal1 Fitted Gauss, proteinAT3G15670.1
bic:811 820 828
peaks after deleting less than one fifth of global max:13
×10^7
raw data, starting:13
fitted Gauss locations:13
y
x
0
1
2
3
4
5
6
7
0
5
10
15
20
25

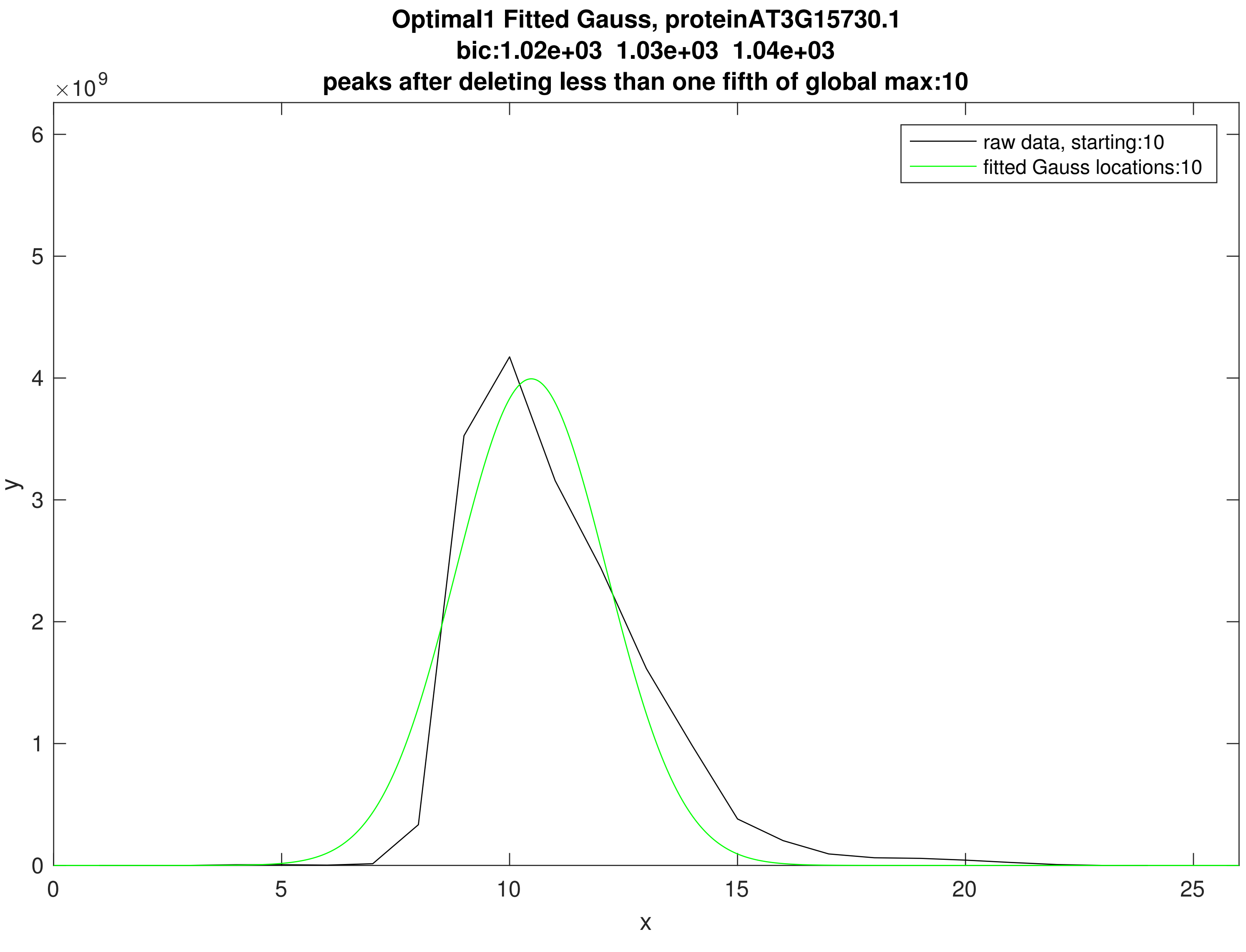
Optimal1 Fitted Gauss, proteinAT3G15730.1
bic:1.02e+03 1.03e+03 1.04e+03
peaks after deleting less than one fifth of global max:10
×10^9
raw data, starting:10
fitted Gauss locations:10
y
x
0
1
2
3
4
5
6
0
5
10
15
20
25

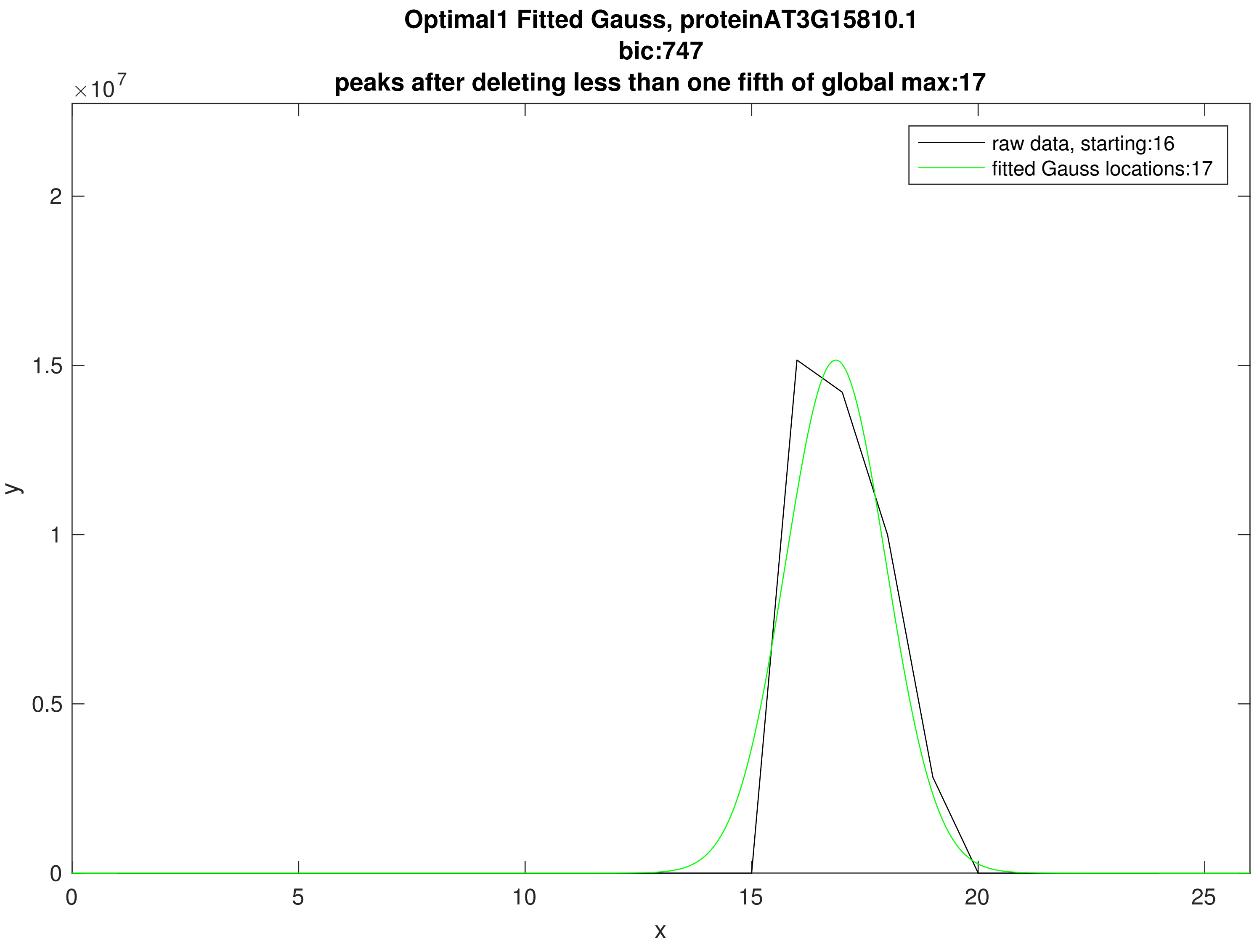

Optimal1 Fitted Gauss, proteinAT3G15810.1
bic:747
peaks after deleting less than one fifth of global max:17
×10^7
raw data, starting:16
fitted Gauss locations:17
2
1.5
1
0.5
0
y
0
5
10
15
20
25
x

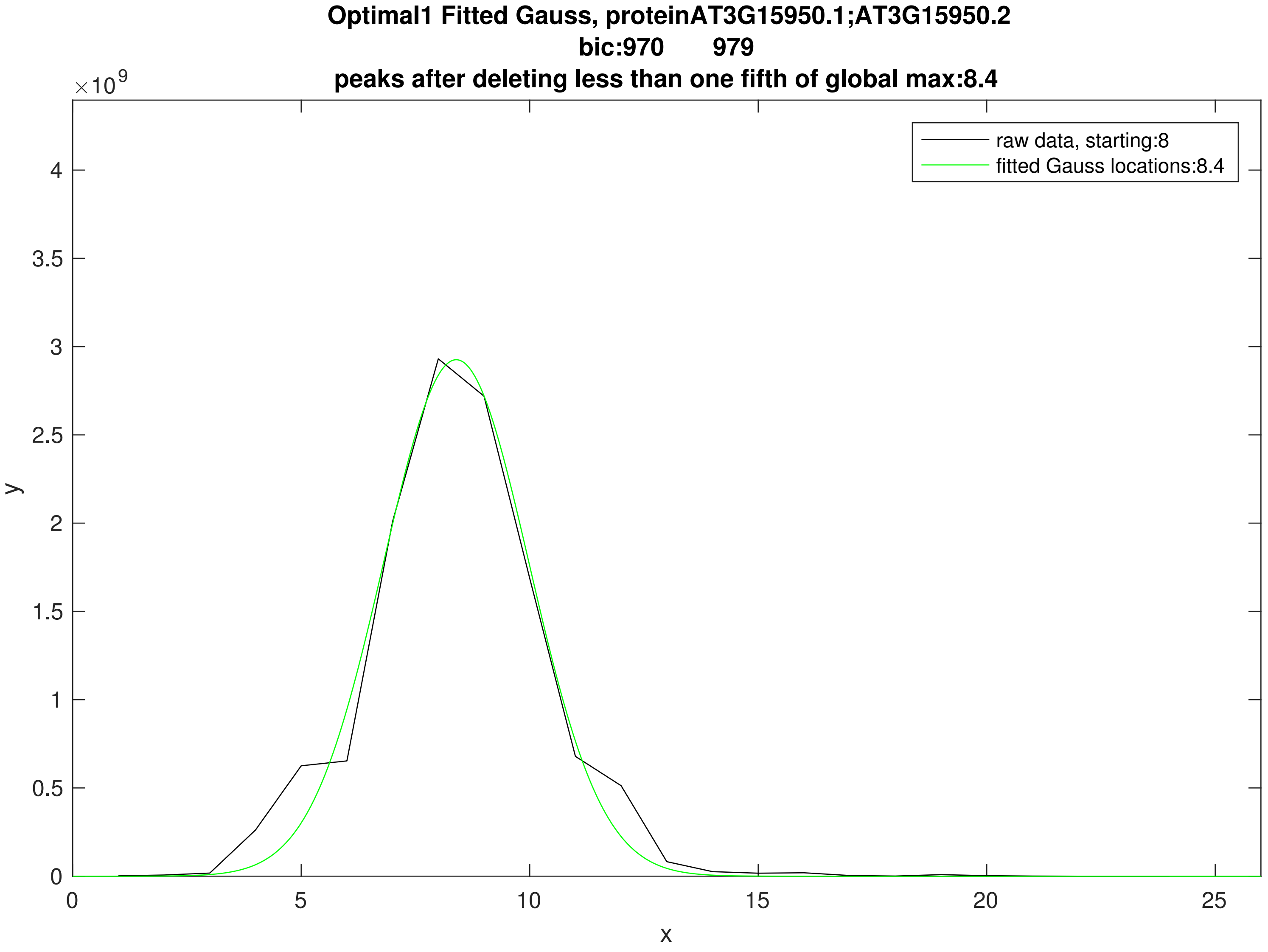

Optimal1 Fitted Gauss, proteinAT3G15950.1;AT3G15950.2
bic:970 979
peaks after deleting less than one fifth of global max:8.4
×10^9
raw data, starting:8
fitted Gauss locations:8.4
y
x
0
0.5
1
1.5
2
2.5
3
3.5
4
5
10
15
20
25

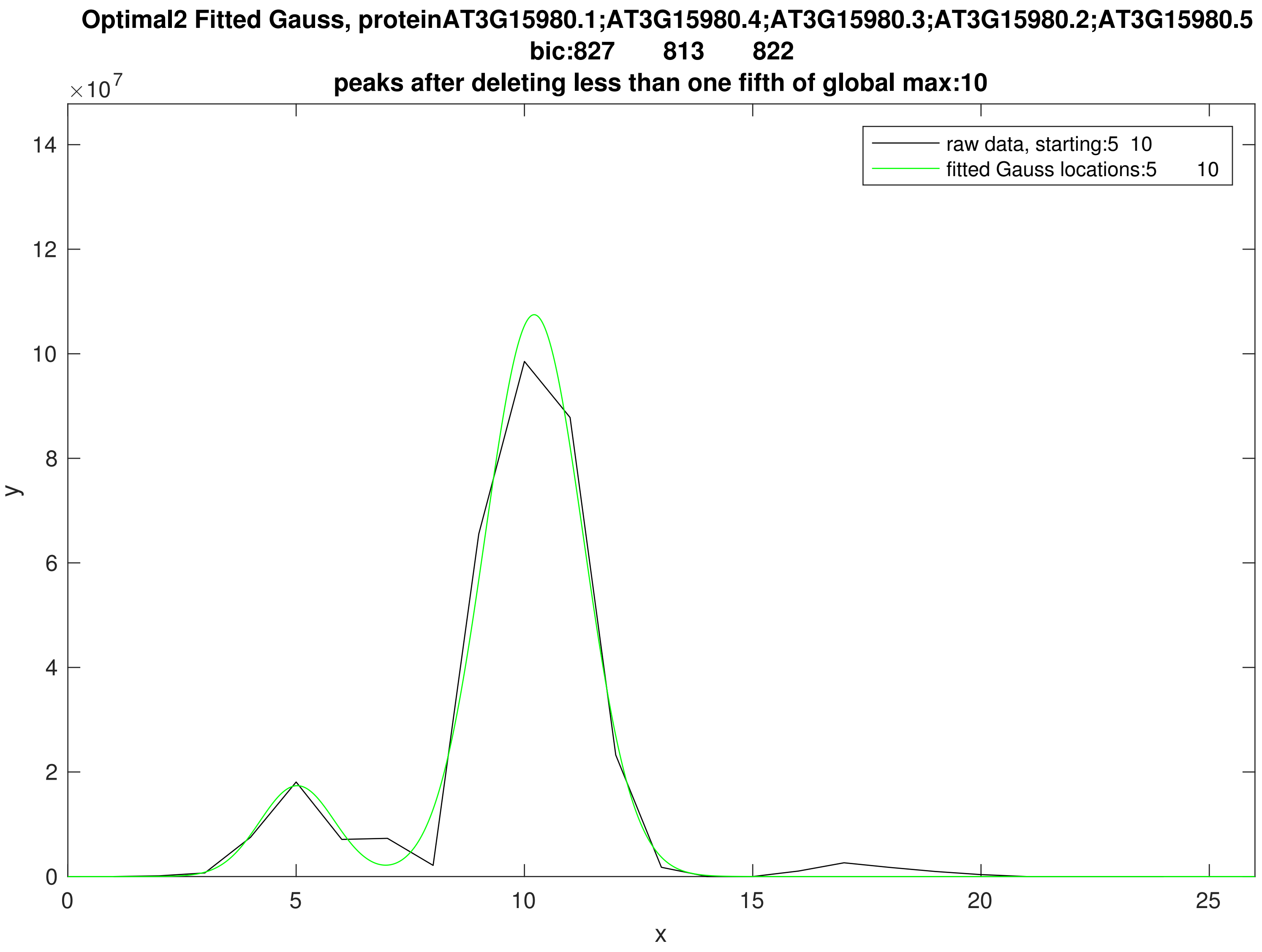
Optimal2 Fitted Gauss, proteinAT3G15980.1;AT3G15980.4;AT3G15980.3;AT3G15980.2;AT3G15980.5
bic:827 813 822
peaks after deleting less than one fifth of global max:10
raw data, starting:5 10
fitted Gauss locations:5 10
×10^7
y
x
0
2
4
6
8
10
12
14
0
5
10
15
20
25

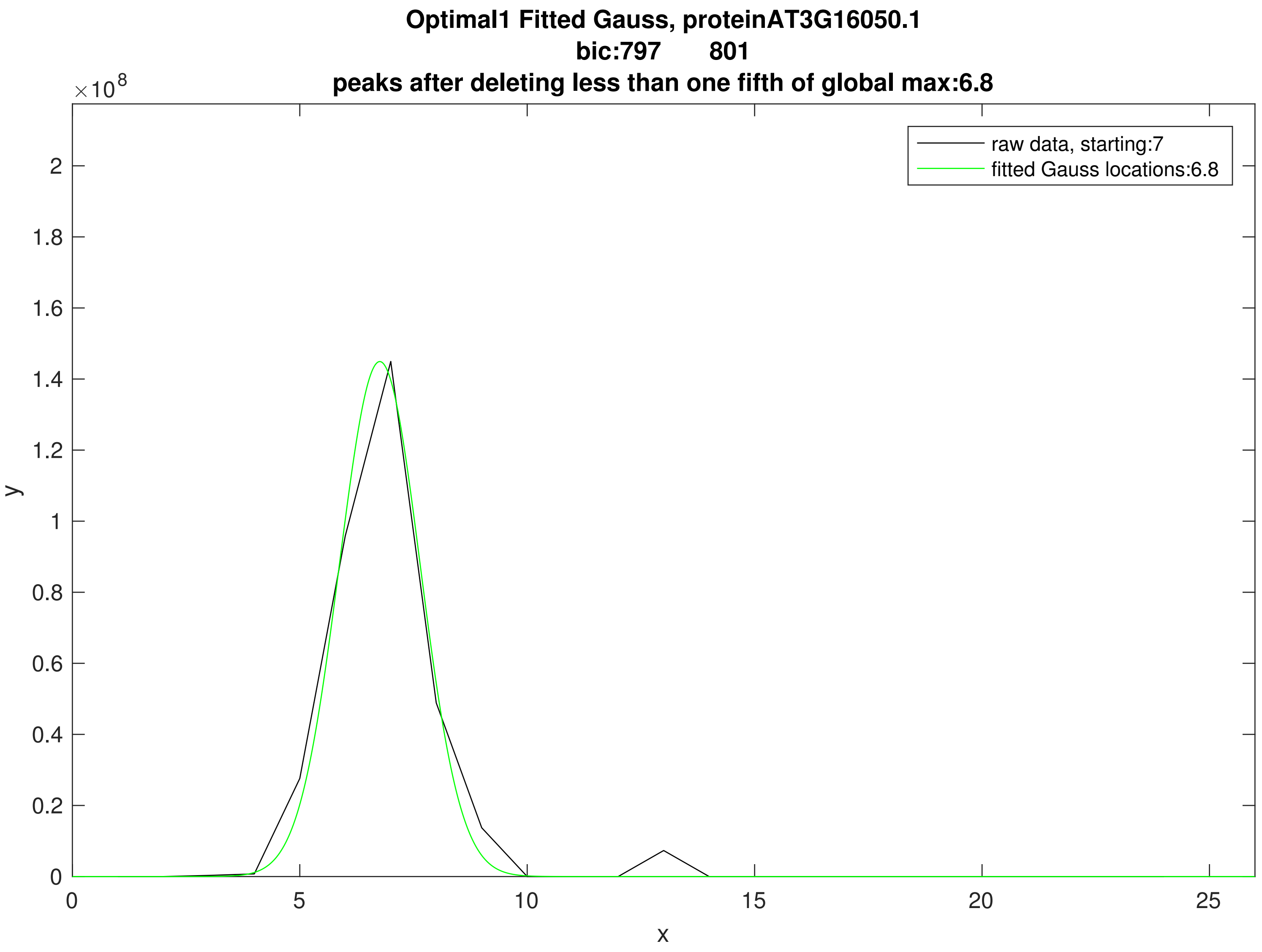

Optimal1 Fitted Gauss, proteinAT3G16050.1
bic:797 801
peaks after deleting less than one fifth of global max:6.8
×10^8
raw data, starting:7
fitted Gauss locations:6.8
2
1.8
1.6
1.4
1.2
1
0.8
0.6
0.4
0.2
0
y
0
5
10
15
20
25
x

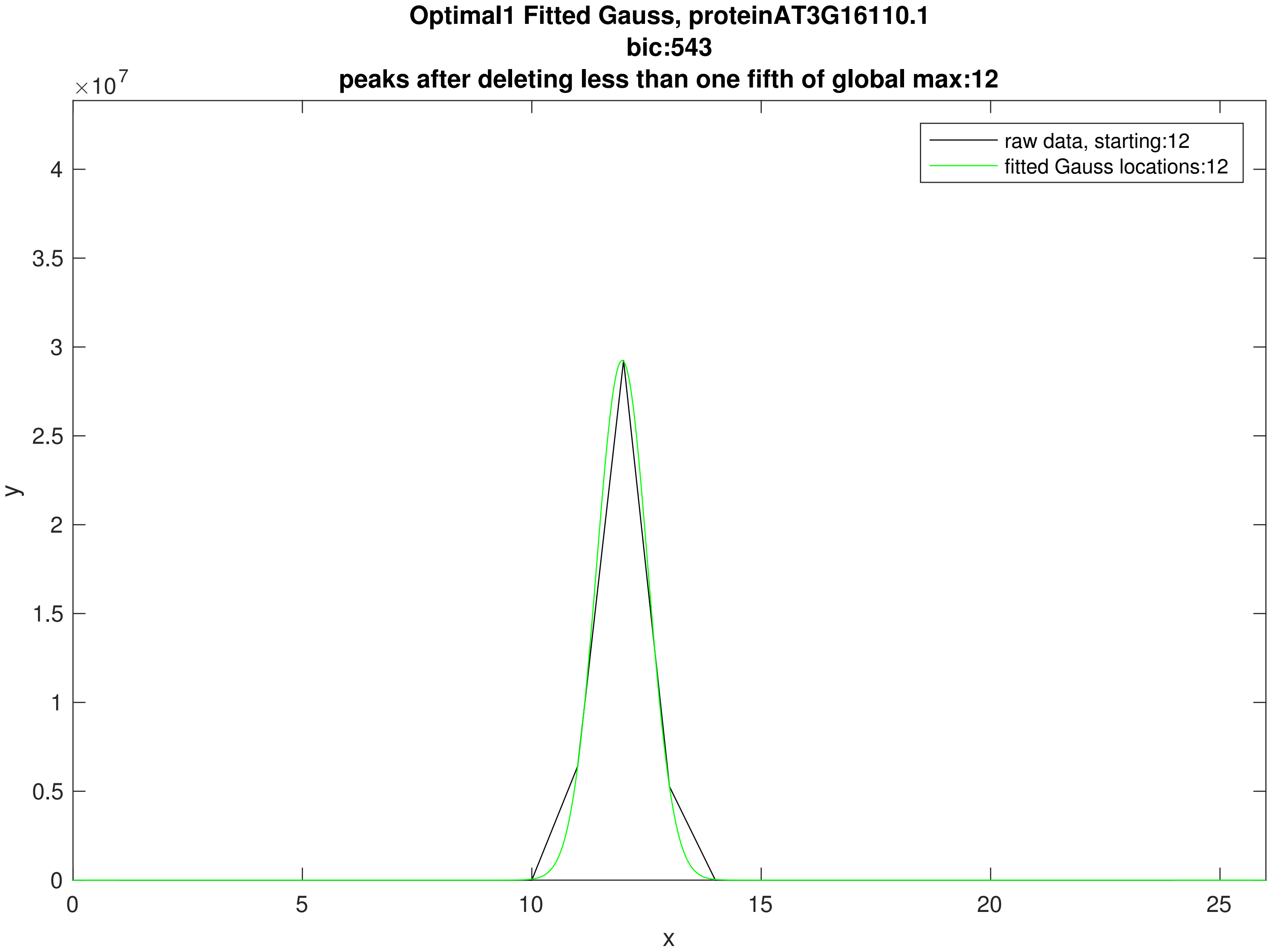

Optimal1 Fitted Gauss, proteinAT3G16110.1
bic:543
peaks after deleting less than one fifth of global max:12
×10^7
4
3.5
3
2.5
2
1.5
1
0.5
0
y
0
5
10
15
20
25
x
raw data, starting:12
fitted Gauss locations:12

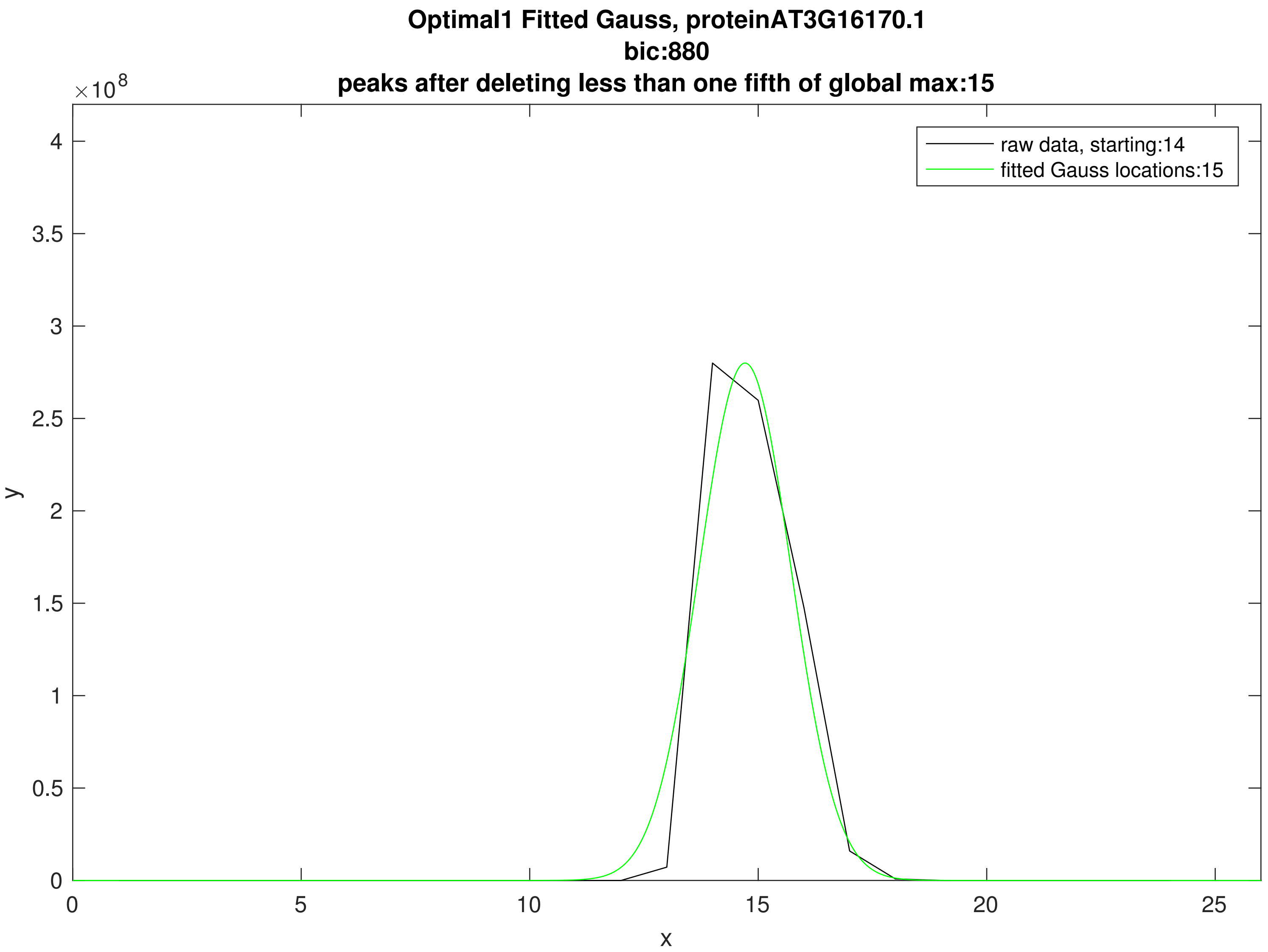

Optimal1 Fitted Gauss, proteinAT3G16170.1
bic:880
peaks after deleting less than one fifth of global max:15
×10^8
raw data, starting:14
fitted Gauss locations:15
y
x
0
0.5
1
1.5
2
2.5
3
3.5
4
0
5
10
15
20
25

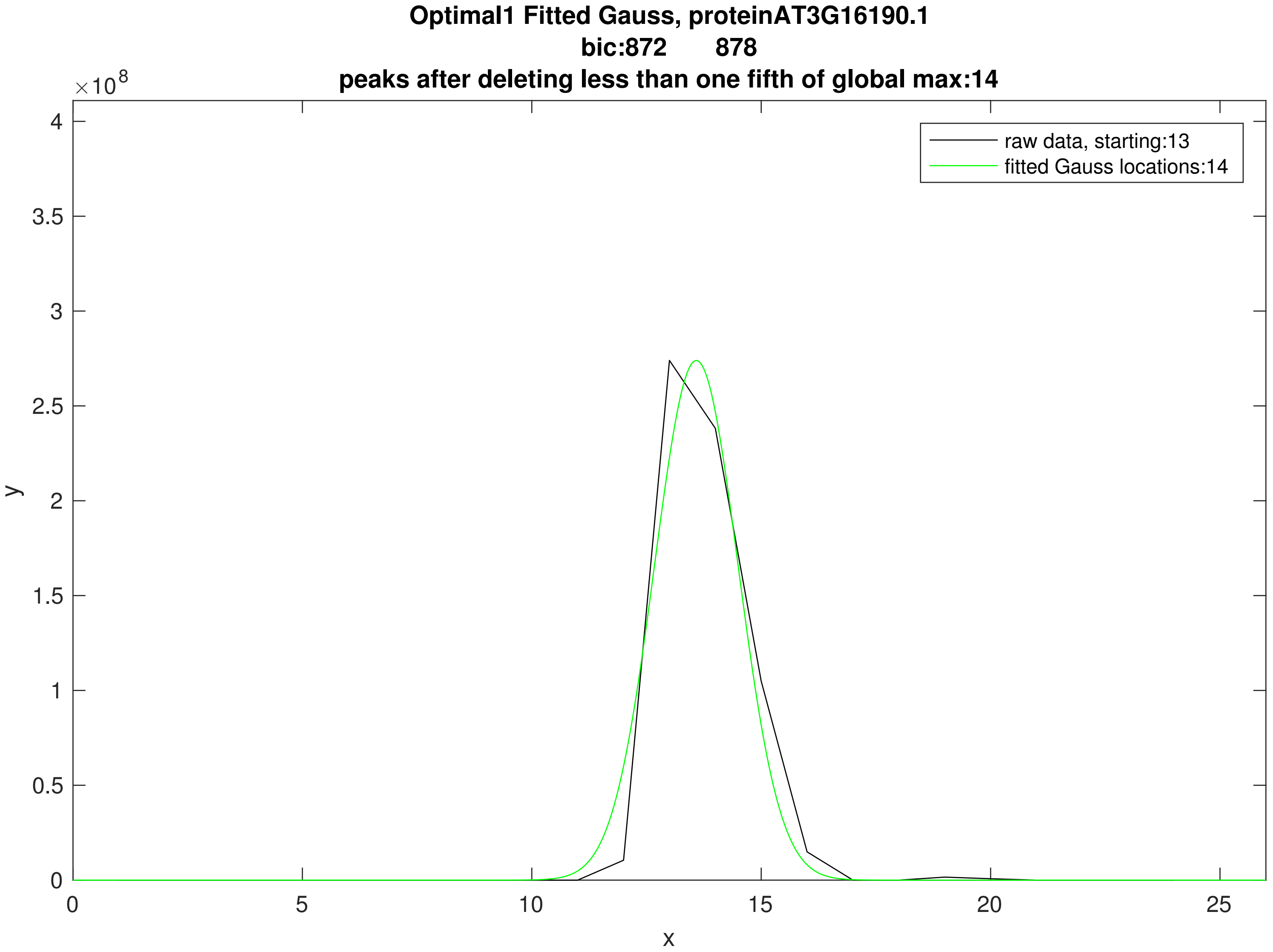

Optimal1 Fitted Gauss, proteinAT3G16190.1
bic:872 878
peaks after deleting less than one fifth of global max:14
×10^8
raw data, starting:13
fitted Gauss locations:14
y
x
0
0.5
1
1.5
2
2.5
3
3.5
4
0
5
10
15
20
25

**Optimal1 Fitted Gauss, proteinAT3G16270.1**
**bic:600**
**peaks after deleting less than one fifth of global max:19**

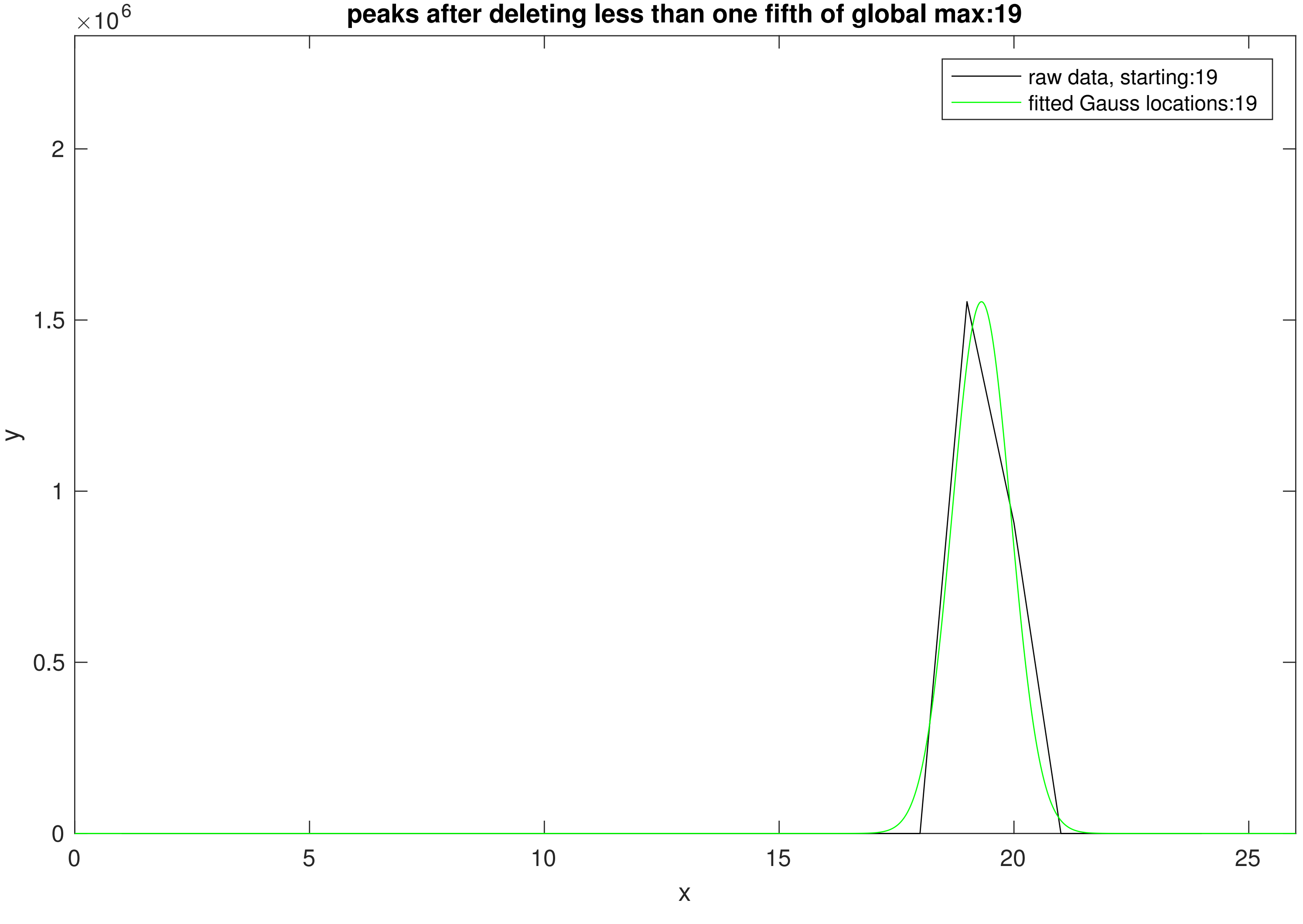

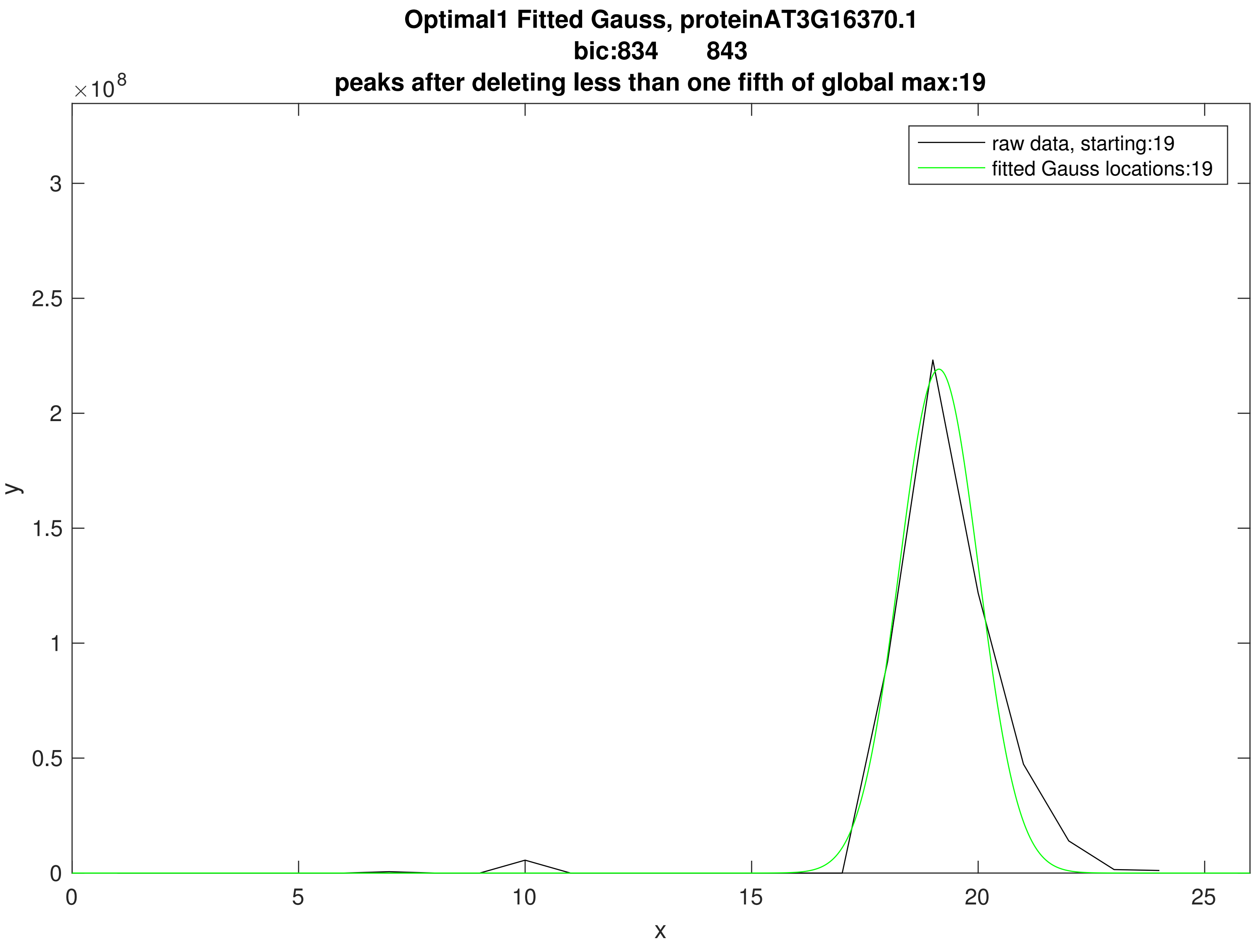

Optimal1 Fitted Gauss, proteinAT3G16370.1
bic:834 843
peaks after deleting less than one fifth of global max:19
×10^8
raw data, starting:19
fitted Gauss locations:19
y
x
0
0.5
1
1.5
2
2.5
3
0
5
10
15
20
25

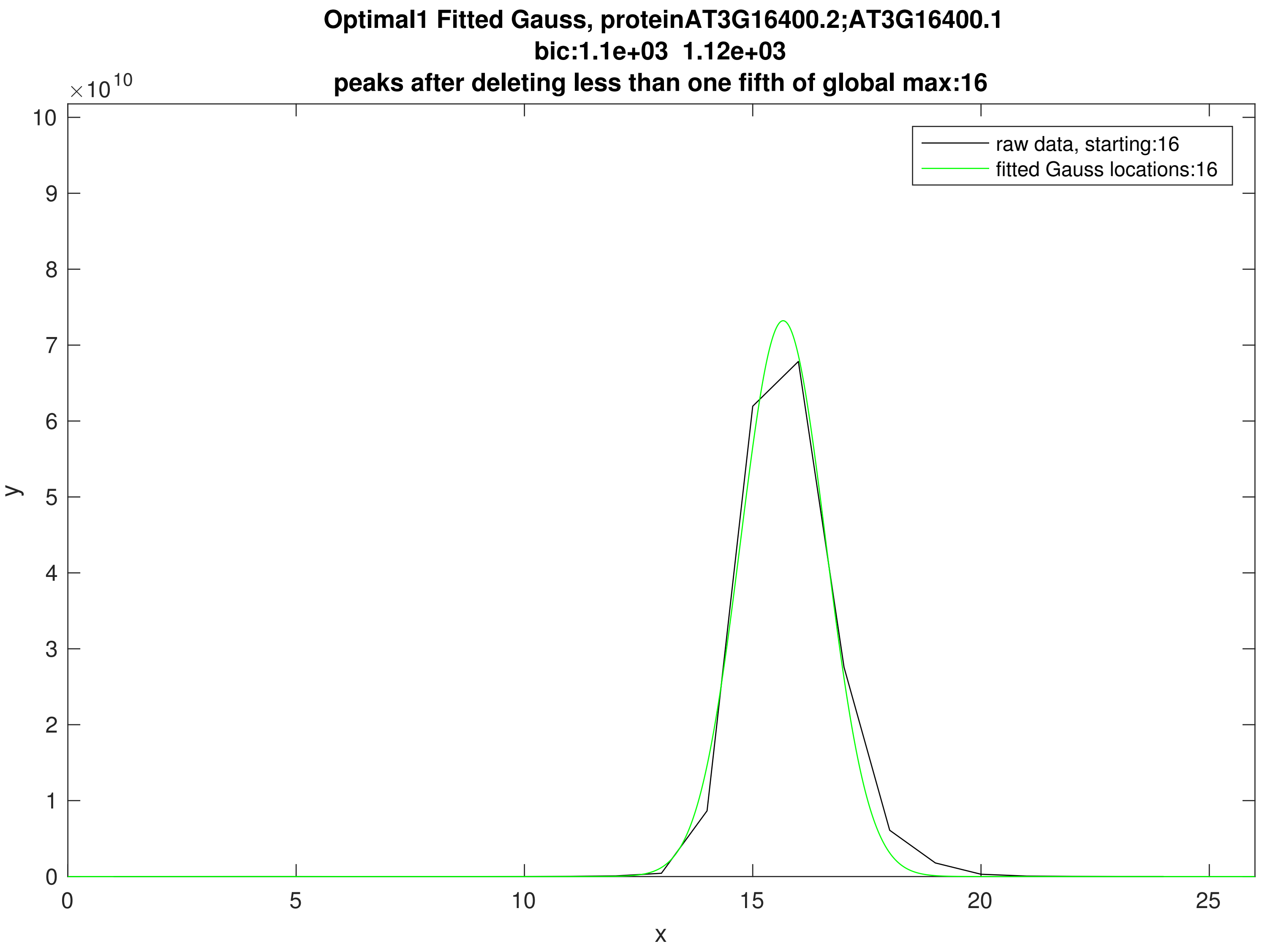

Optimal1 Fitted Gauss, proteinAT3G16400.2;AT3G16400.1
bic:1.1e+03  1.12e+03
peaks after deleting less than one fifth of global max:16
×10^10
raw data, starting:16
fitted Gauss locations:16
y
x

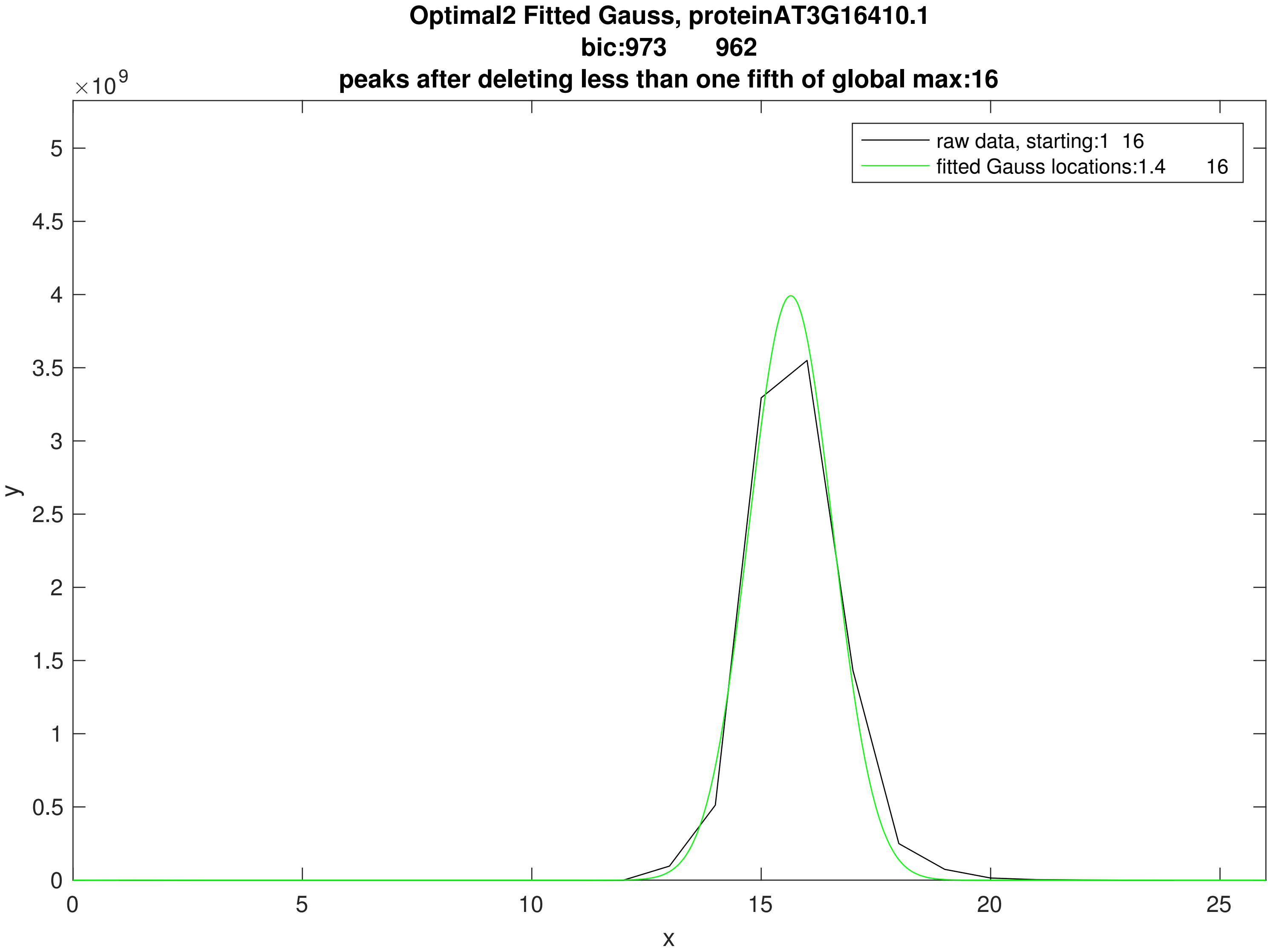
Optimal2 Fitted Gauss, proteinAT3G16410.1
bic:973 962
peaks after deleting less than one fifth of global max:16
×10^9
raw data, starting:1 16
fitted Gauss locations:1.4 16
y
x
0
5
10
15
20
25
0.5
1
1.5
2
2.5
3
3.5
4
4.5
5

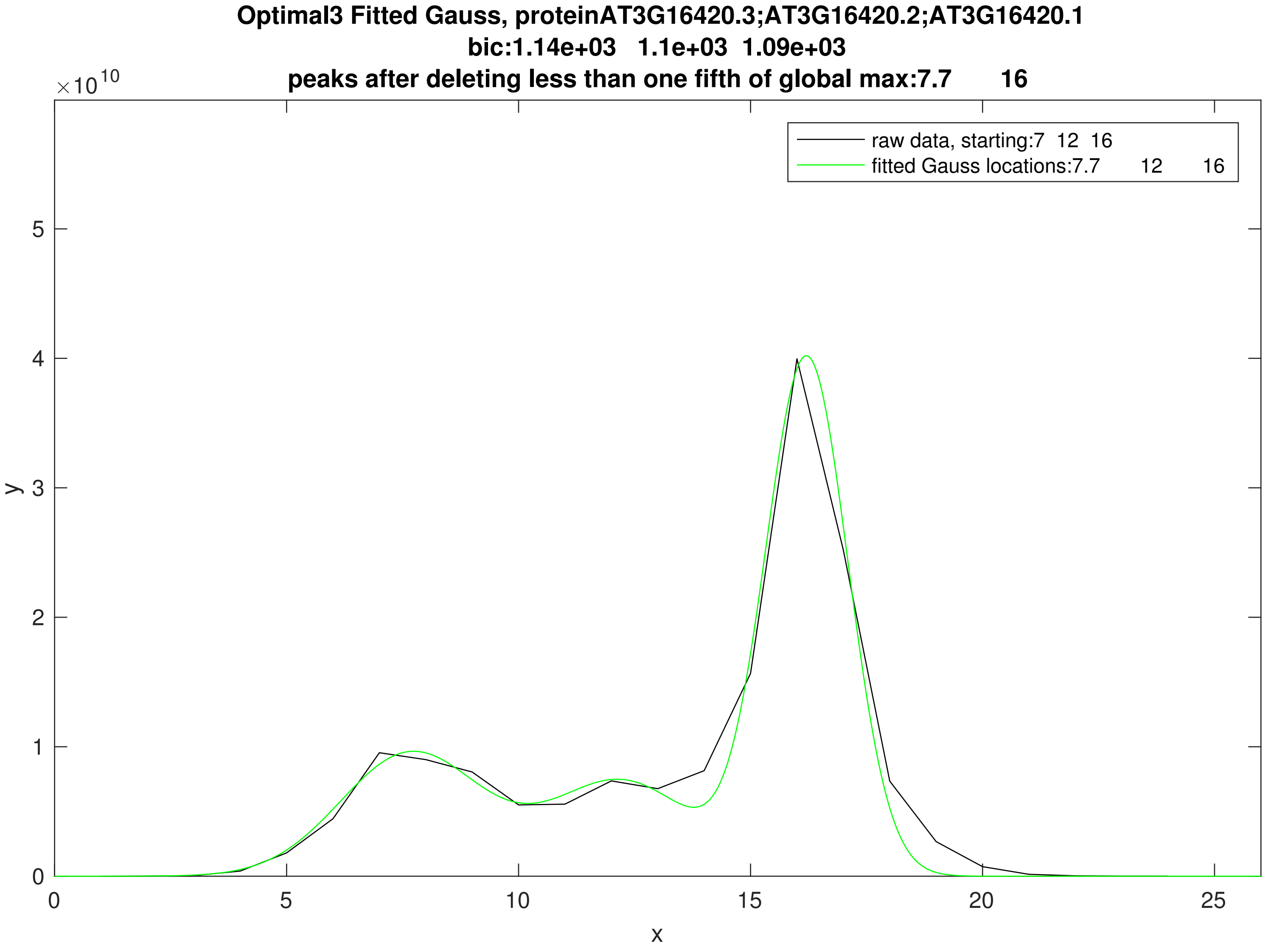
Optimal3 Fitted Gauss, proteinAT3G16420.3;AT3G16420.2;AT3G16420.1
bic:1.14e+03 1.1e+03 1.09e+03
peaks after deleting less than one fifth of global max:7.7 16
×10^10
raw data, starting:7 12 16
fitted Gauss locations:7.7 12 16
y
x
0
1
2
3
4
5
10
15
20
25

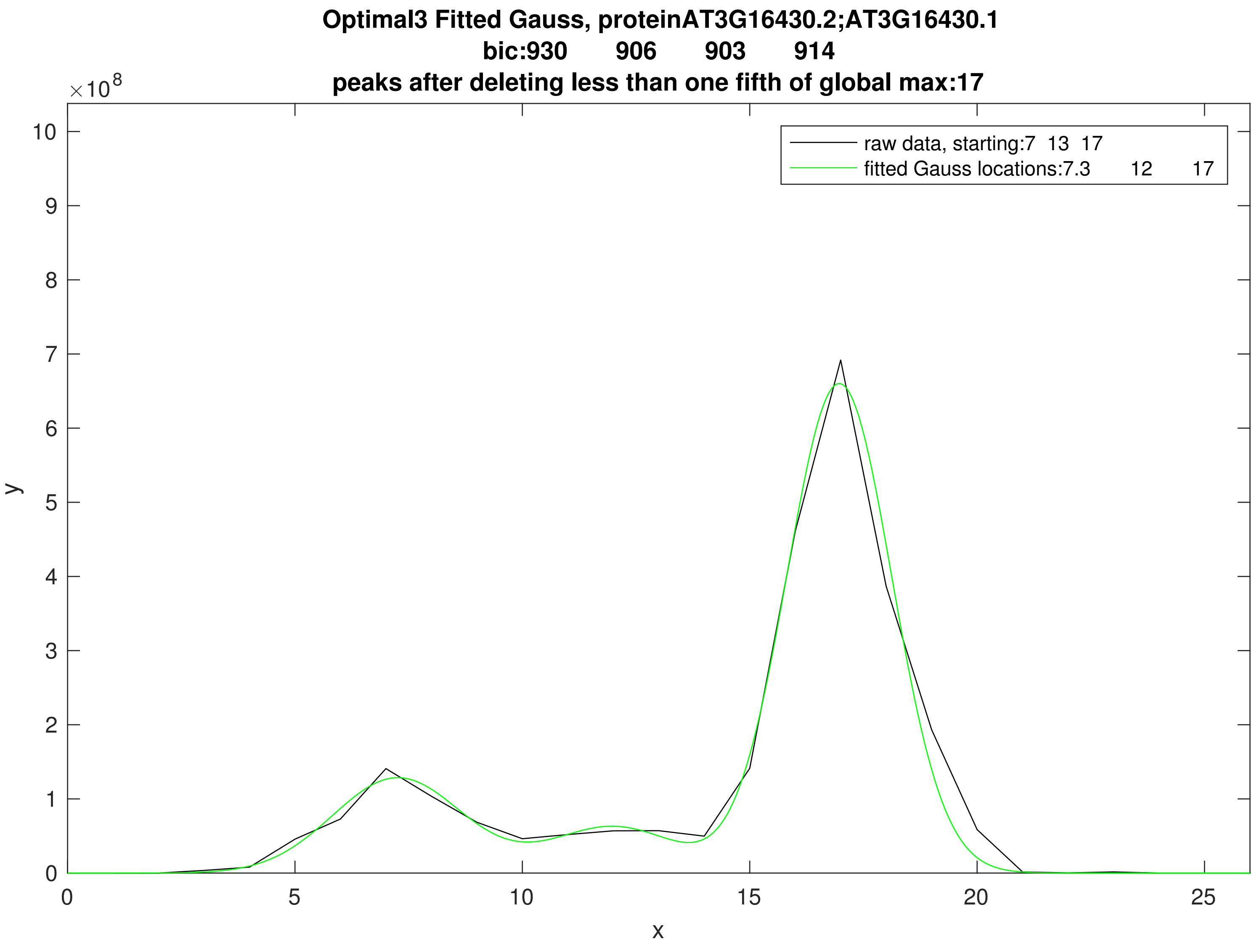

Optimal3 Fitted Gauss, proteinAT3G16430.2;AT3G16430.1
bic:930 906 903 914
peaks after deleting less than one fifth of global max:17
raw data, starting:7 13 17
fitted Gauss locations:7.3 12 17
×10^8
y
x

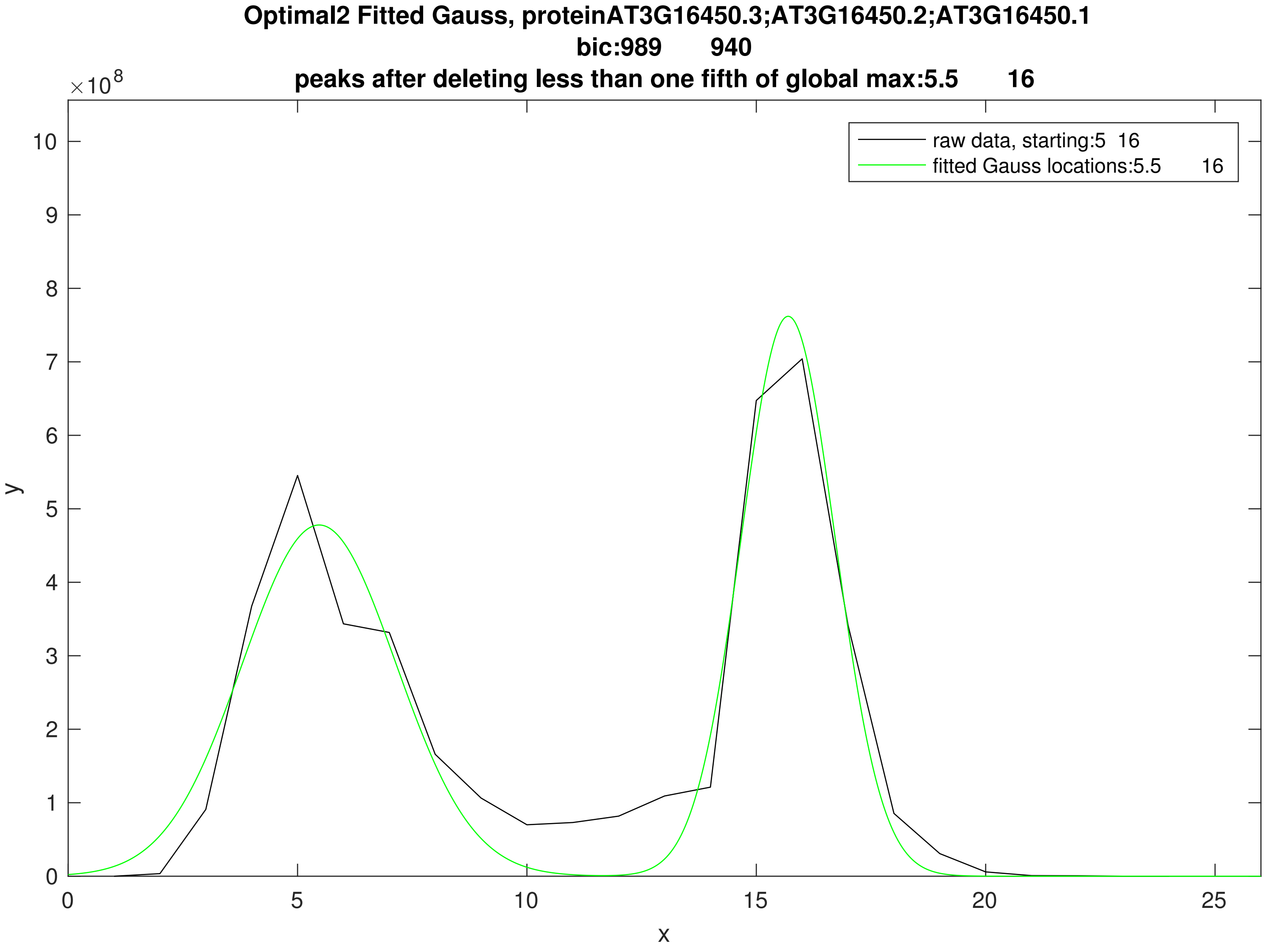

Optimal2 Fitted Gauss, proteinAT3G16450.3;AT3G16450.2;AT3G16450.1
bic:989 940
peaks after deleting less than one fifth of global max:5.5 16
×10^8
raw data, starting:5 16
fitted Gauss locations:5.5 16
y
x

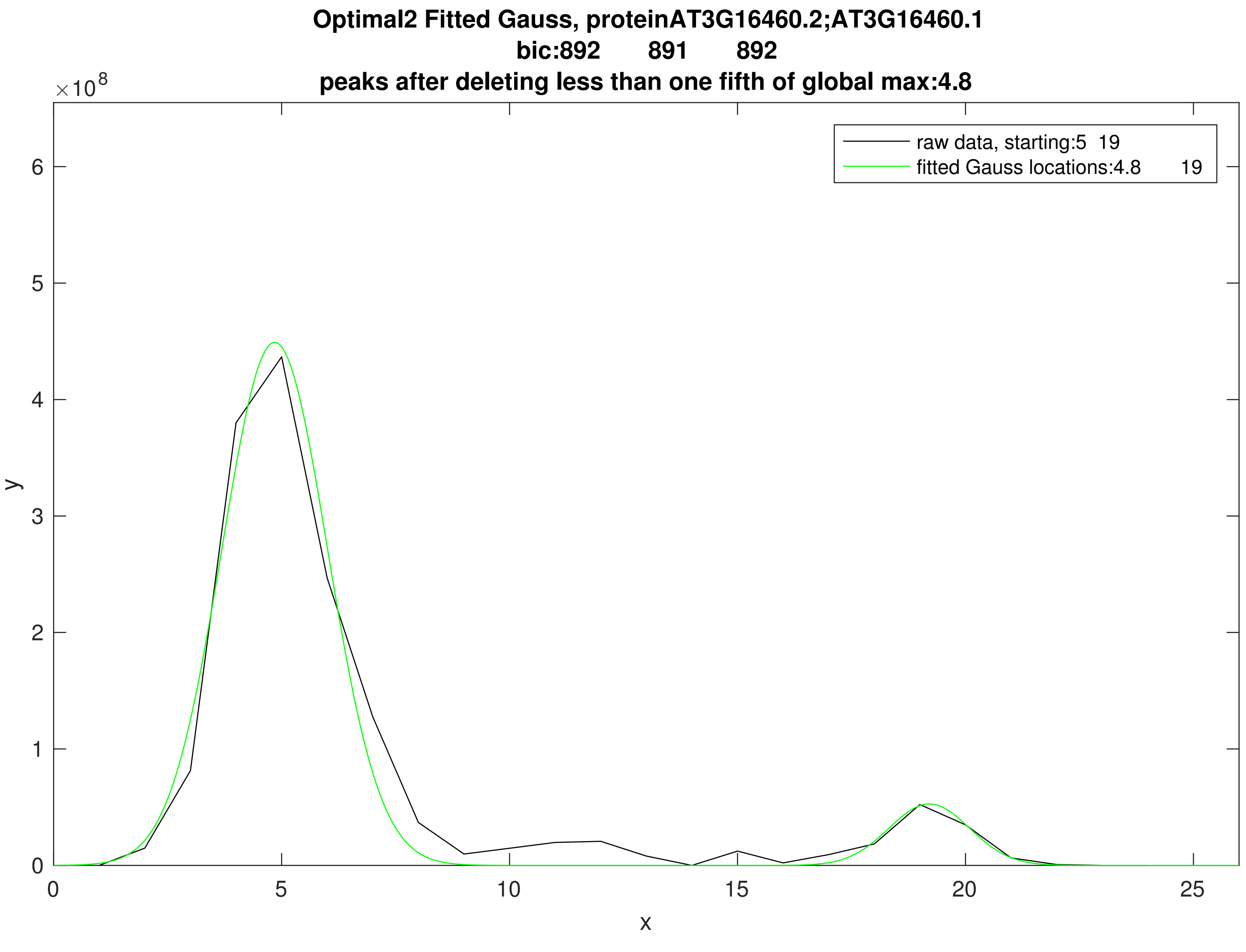
Optimal2 Fitted Gauss, proteinAT3G16460.2;AT3G16460.1
bic:892 891 892
peaks after deleting less than one fifth of global max:4.8
×10^8
raw data, starting:5 19
fitted Gauss locations:4.8 19
y
x
0
1
2
3
4
5
6
10
15
20
25

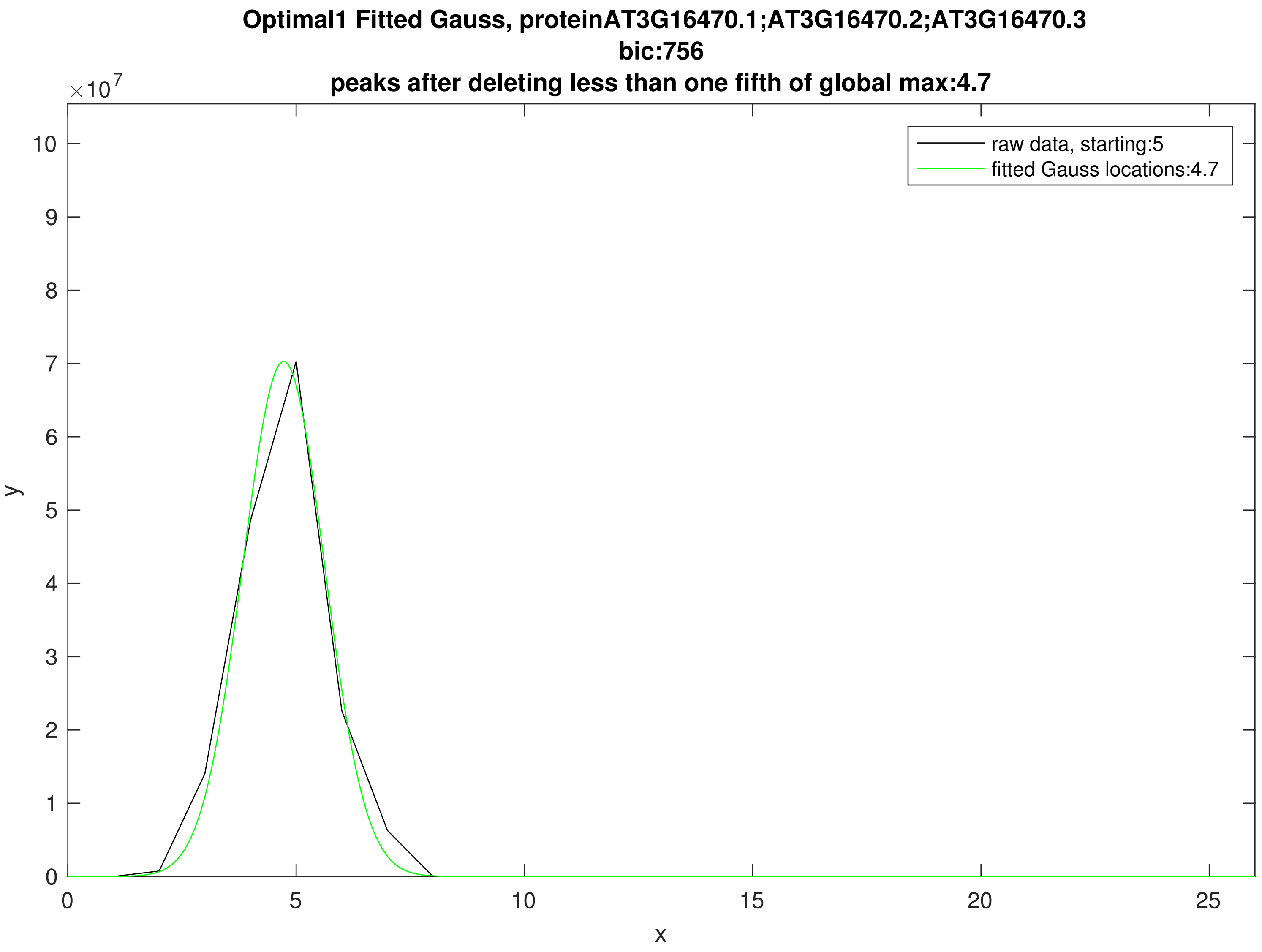
Optimal1 Fitted Gauss, proteinAT3G16470.1;AT3G16470.2;AT3G16470.3
bic:756
peaks after deleting less than one fifth of global max:4.7
×10^7
raw data, starting:5
fitted Gauss locations:4.7
y
x
0
1
2
3
4
5
6
7
8
9
10
0
5
10
15
20
25

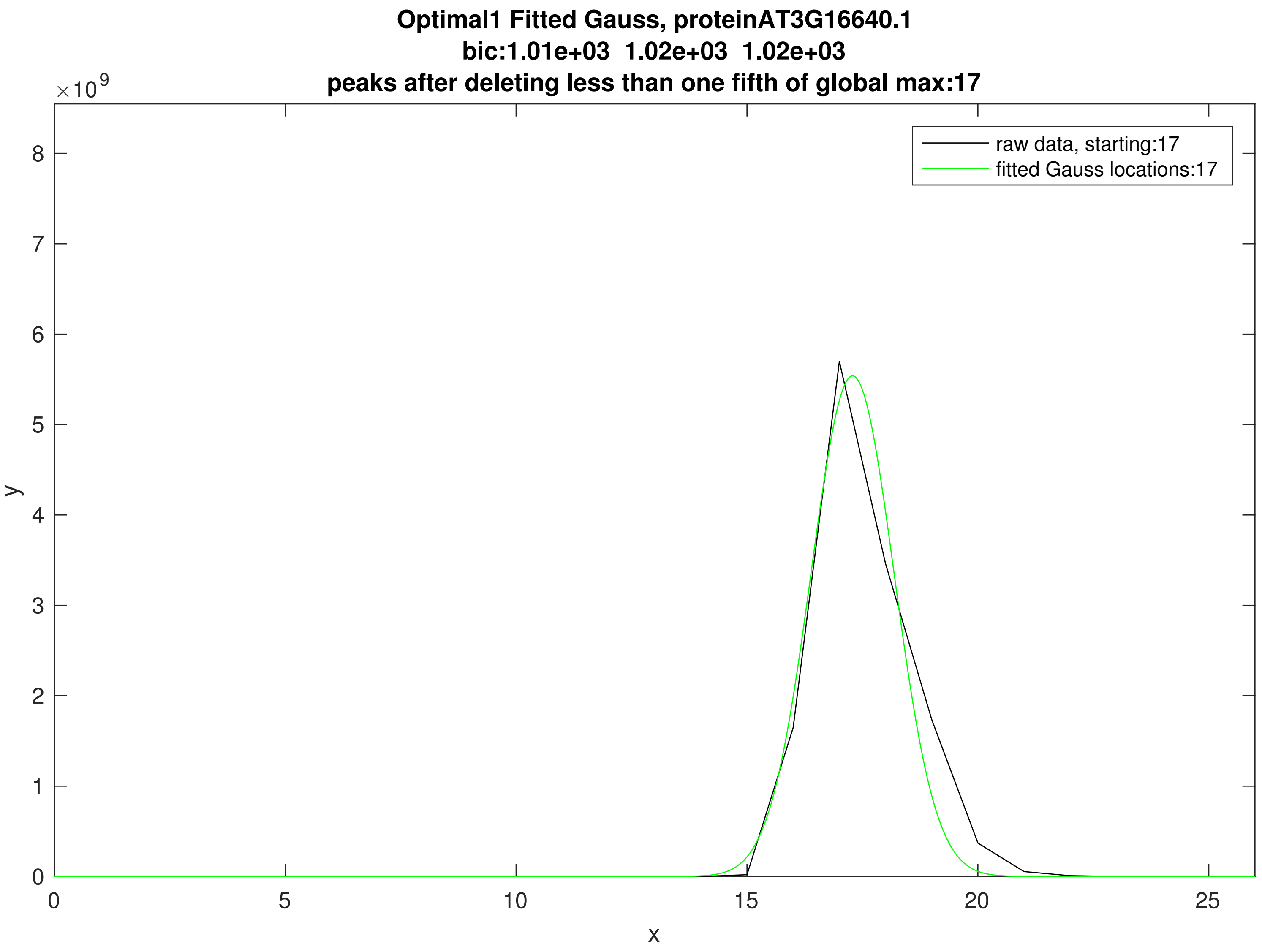

Optimal1 Fitted Gauss, proteinAT3G16640.1
bic:1.01e+03  1.02e+03  1.02e+03
peaks after deleting less than one fifth of global max:17
×10^9
raw data, starting:17
fitted Gauss locations:17
y
x
0
1
2
3
4
5
6
7
8
0
5
10
15
20
25

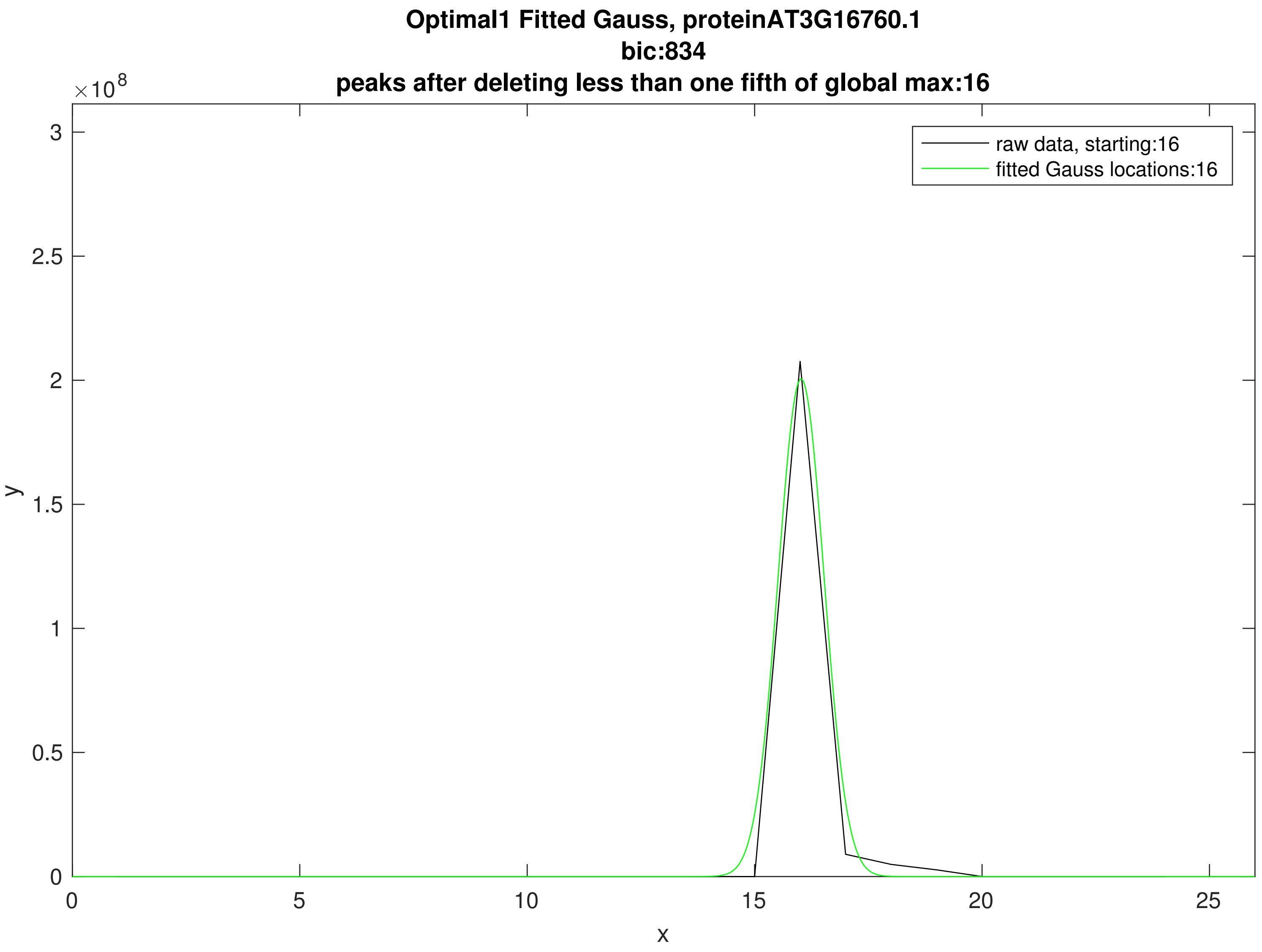
Optimal1 Fitted Gauss, proteinAT3G16760.1
bic:834
peaks after deleting less than one fifth of global max:16
×10^8
3
2.5
2
1.5
1
0.5
0
y
0
5
10
15
20
25
x
raw data, starting:16
fitted Gauss locations:16

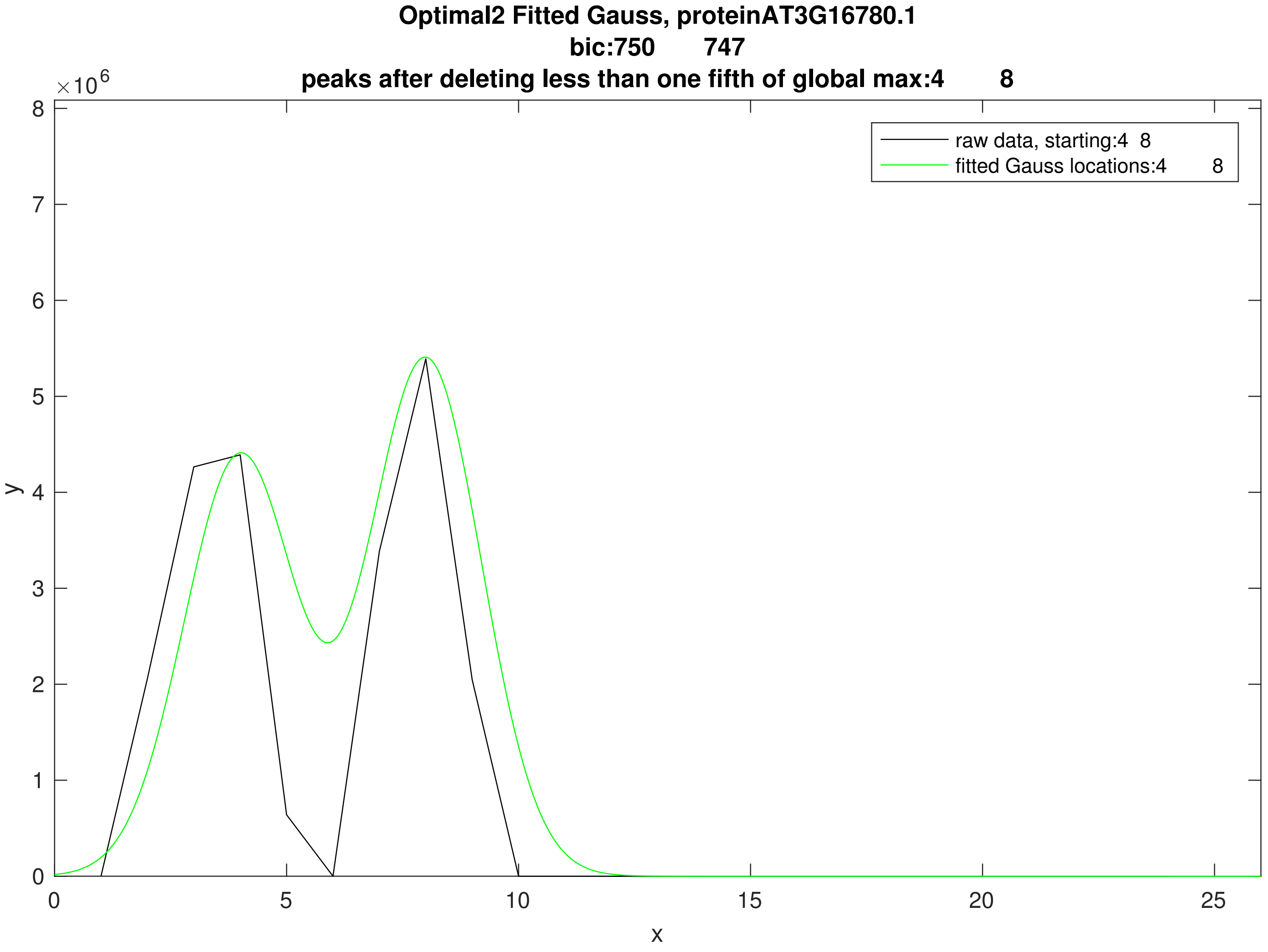

Optimal2 Fitted Gauss, proteinAT3G16780.1
bic:750 747
peaks after deleting less than one fifth of global max:4 8
×10^6
raw data, starting:4 8
fitted Gauss locations:4 8
y
x
0
1
2
3
4
5
6
7
8
0
5
10
15
20
25

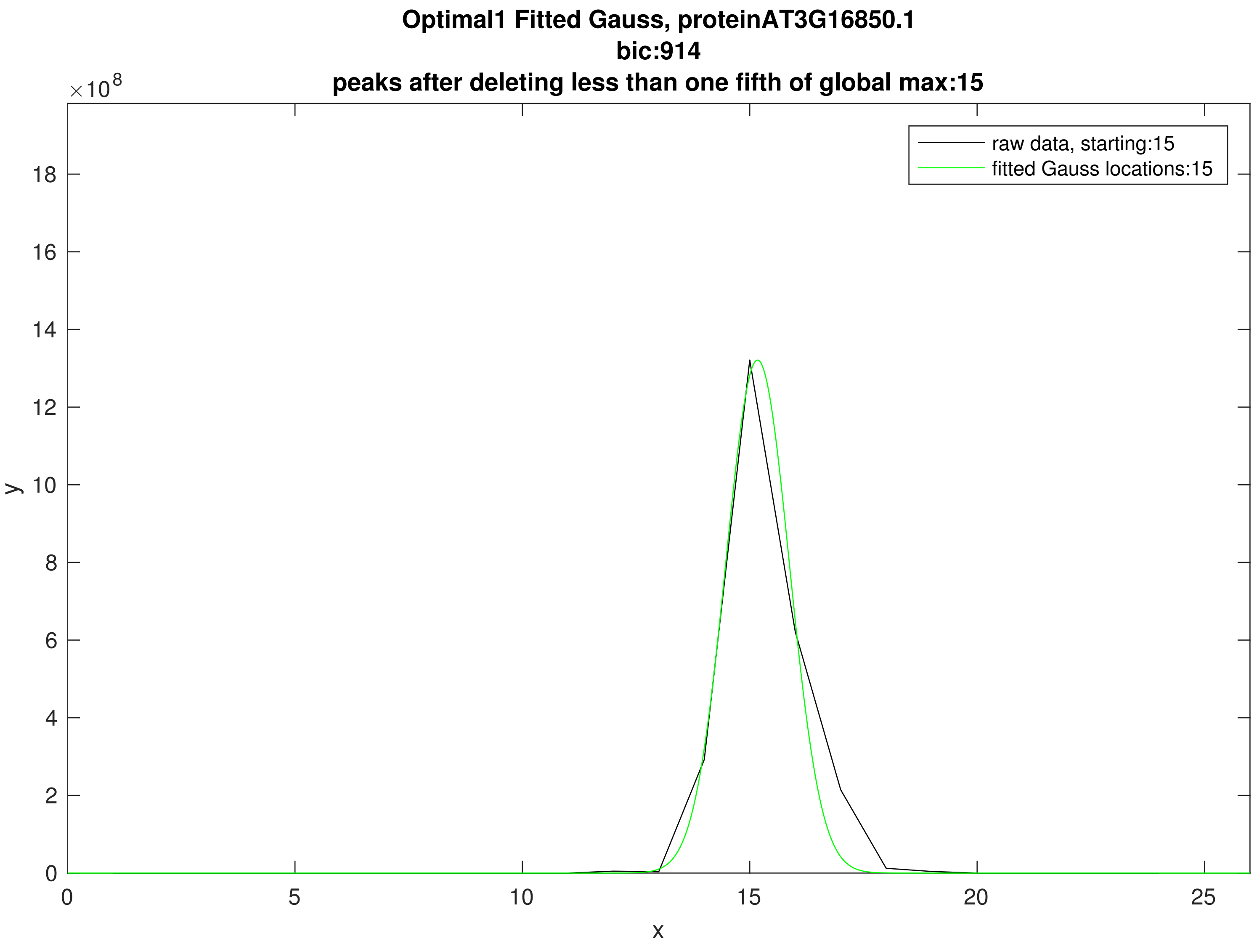

Optimal1 Fitted Gauss, proteinAT3G16850.1
bic:914
peaks after deleting less than one fifth of global max:15
×10^8
raw data, starting:15
fitted Gauss locations:15
y
x
0
2
4
6
8
10
12
14
16
18
0
5
10
15
20
25

Optimal1 Fitted Gauss, proteinAT3G16910.1
bic:749
peaks after deleting less than one fifth of global max:12

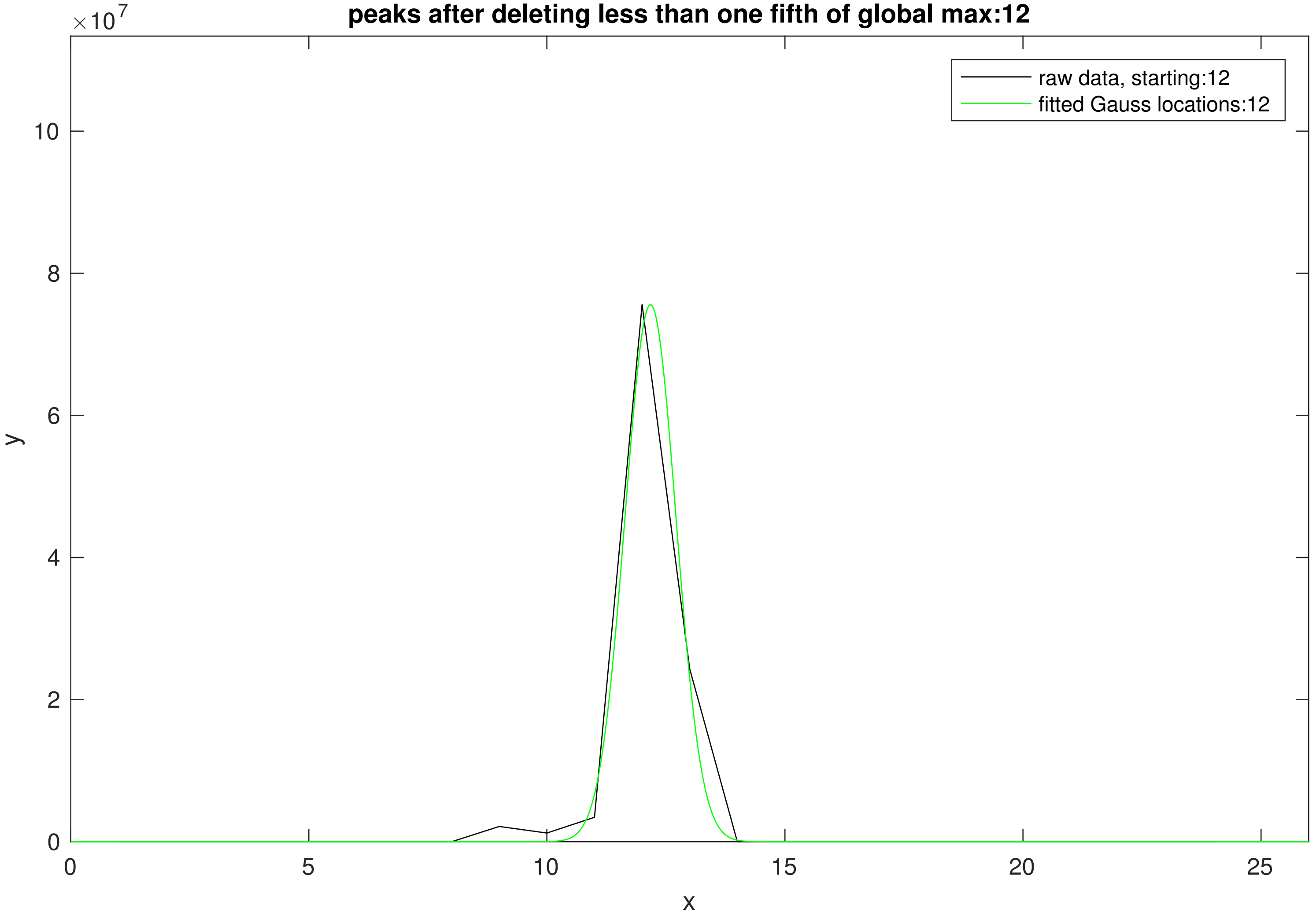

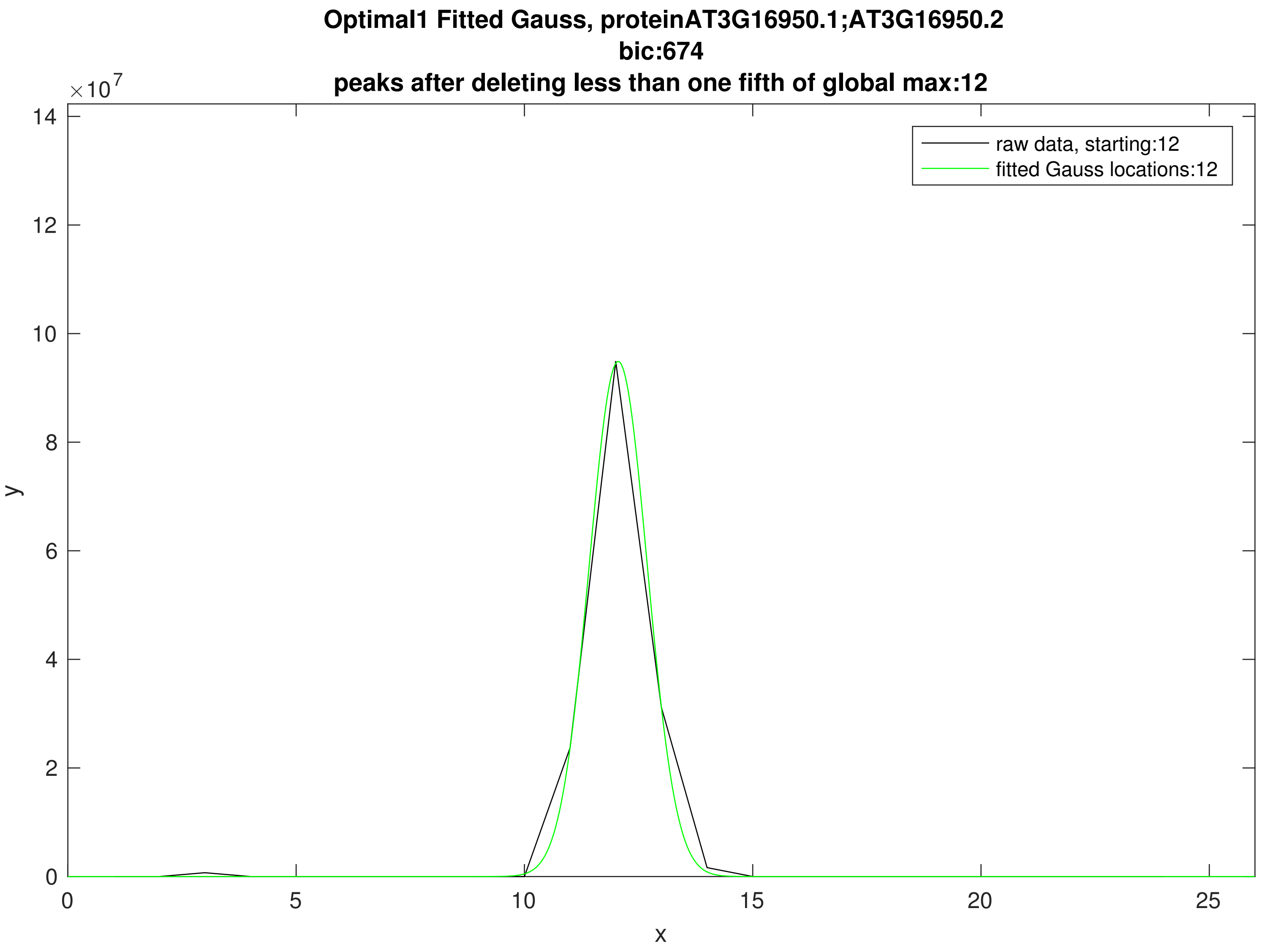

Optimal1 Fitted Gauss, proteinAT3G16950.1;AT3G16950.2
bic:674
peaks after deleting less than one fifth of global max:12
×10^7
raw data, starting:12
fitted Gauss locations:12
y
x
0
2
4
6
8
10
12
14
0
5
10
15
20
25

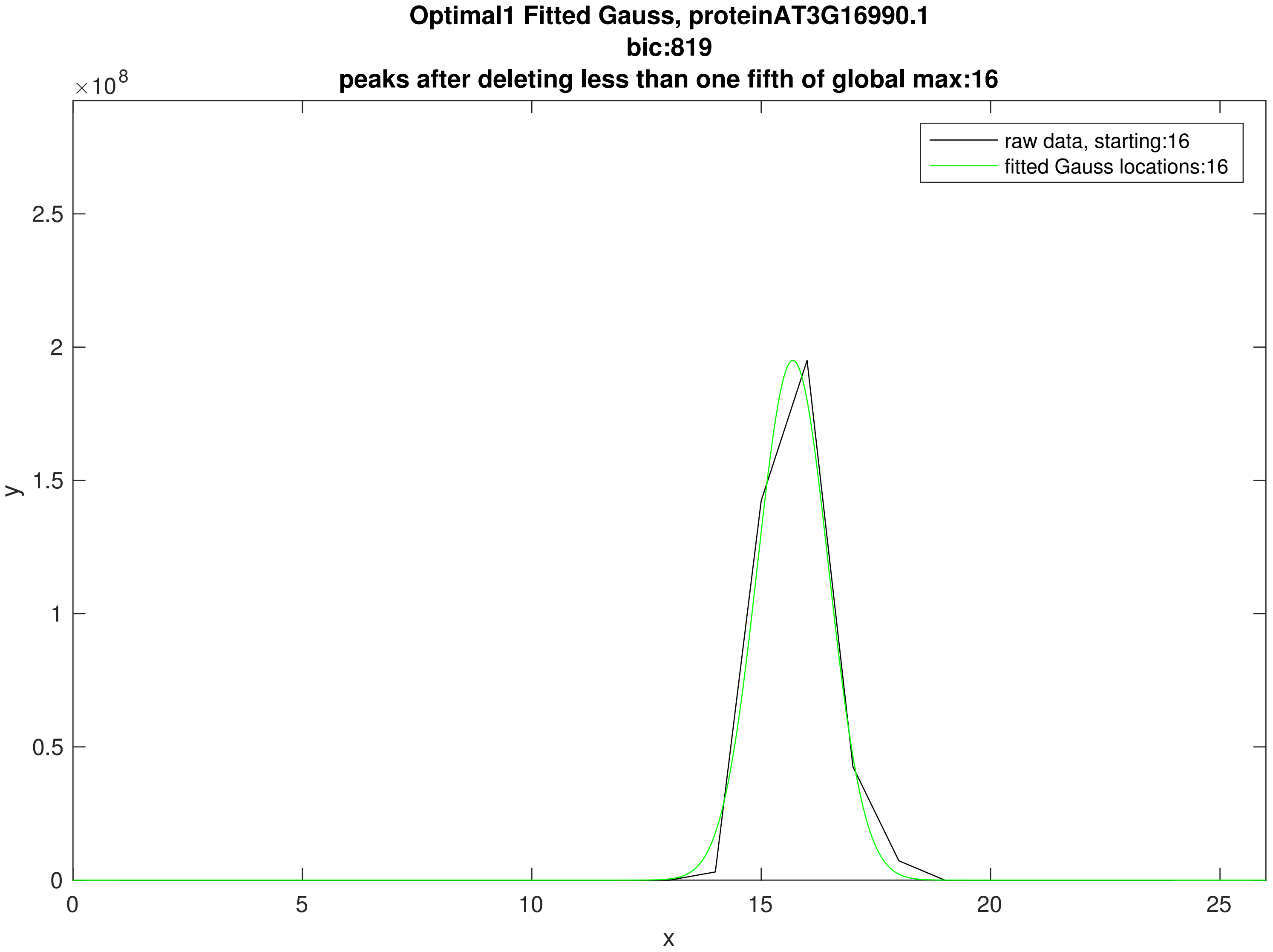

Optimal1 Fitted Gauss, proteinAT3G16990.1
bic:819
peaks after deleting less than one fifth of global max:16
×10^8
raw data, starting:16
fitted Gauss locations:16
y
x
0
0.5
1
1.5
2
2.5
0
5
10
15
20
25

**Optimal1 Fitted Gauss, proteinAT3G17020.1**
**bic:684**
**peaks after deleting less than one fifth of global max:17**

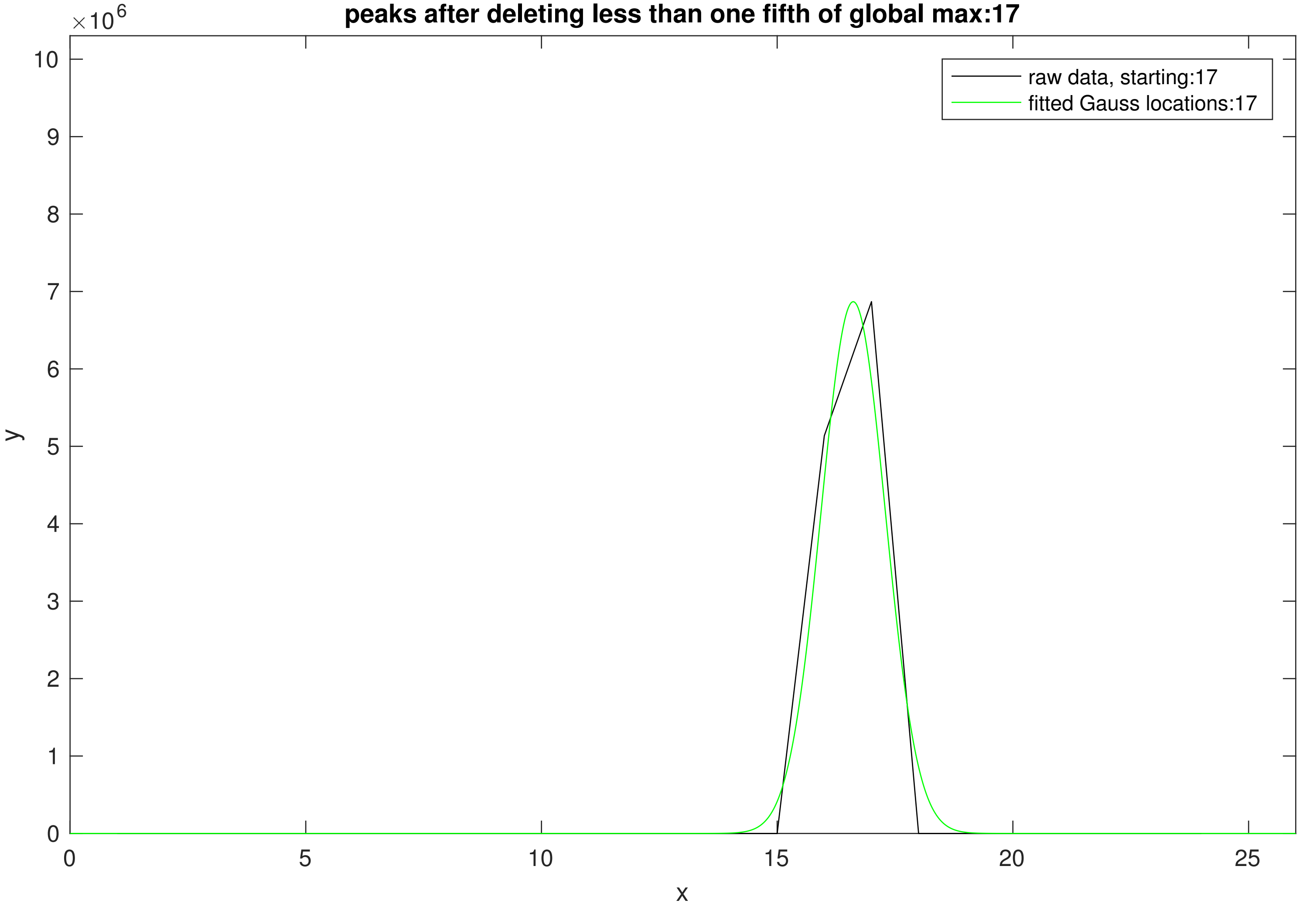

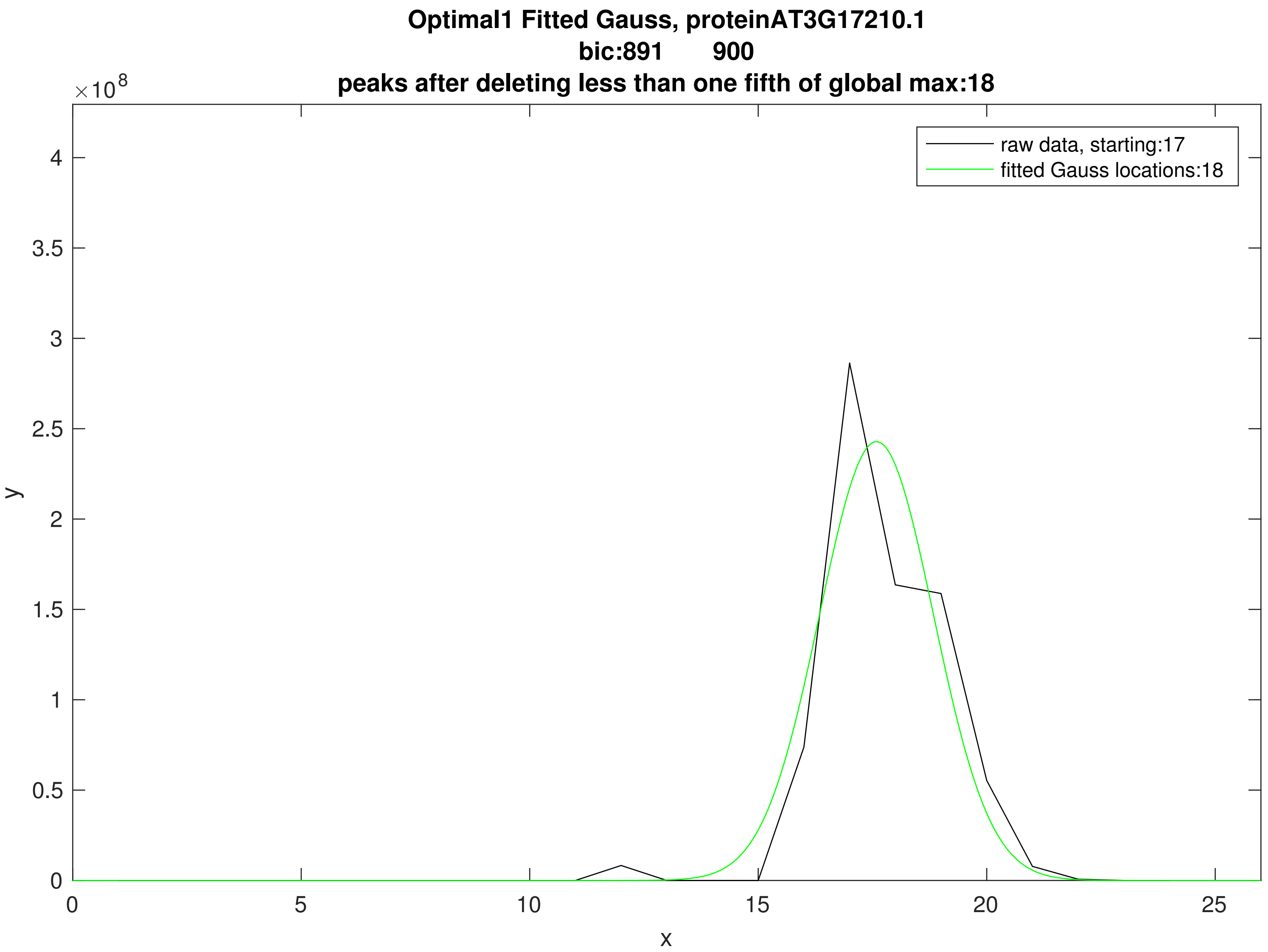
Optimal1 Fitted Gauss, proteinAT3G17210.1
bic:891 900
peaks after deleting less than one fifth of global max:18
×10^8
raw data, starting:17
fitted Gauss locations:18
y
x
0
0.5
1
1.5
2
2.5
3
3.5
4
0
5
10
15
20
25

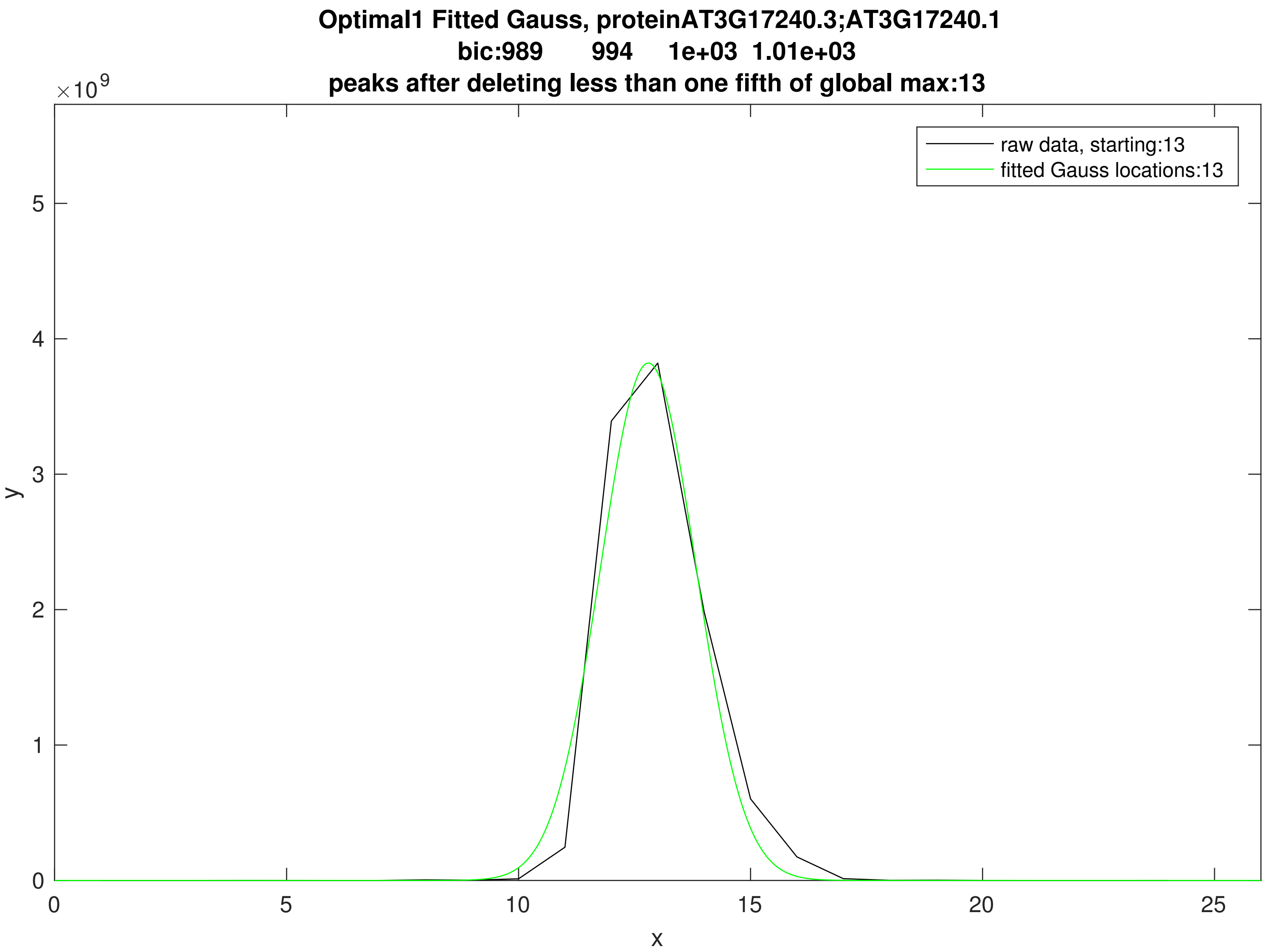
Optimal1 Fitted Gauss, proteinAT3G17240.3;AT3G17240.1
bic:989 994 1e+03 1.01e+03
peaks after deleting less than one fifth of global max:13
×10^9
raw data, starting:13
fitted Gauss locations:13
y
x
0
1
2
3
4
5
0
5
10
15
20
25

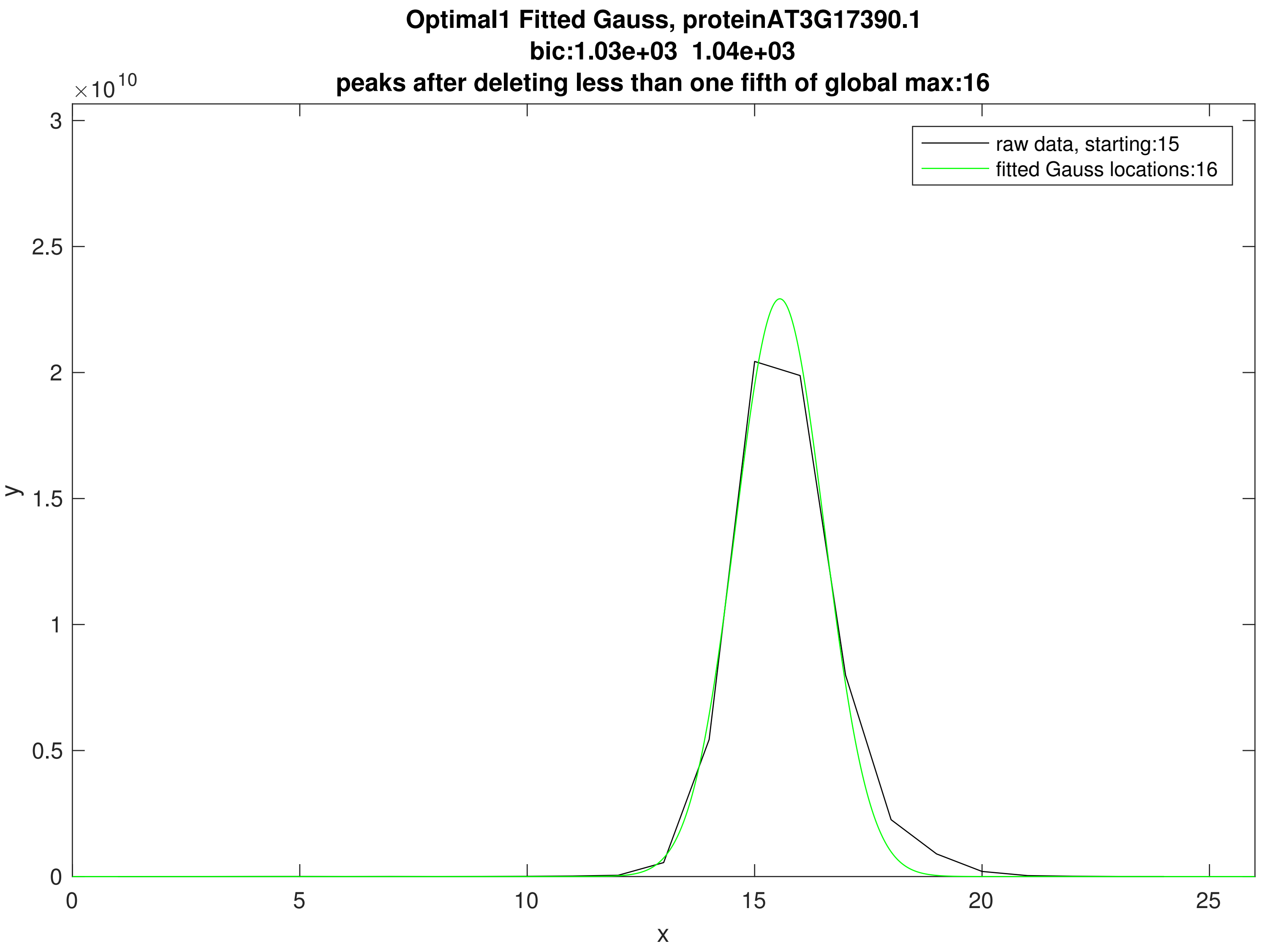
Optimal1 Fitted Gauss, proteinAT3G17390.1
bic:1.03e+03  1.04e+03
peaks after deleting less than one fifth of global max:16
×10^10
raw data, starting:15
fitted Gauss locations:16
y
x
0
0.5
1
1.5
2
2.5
3
5
10
15
20
25

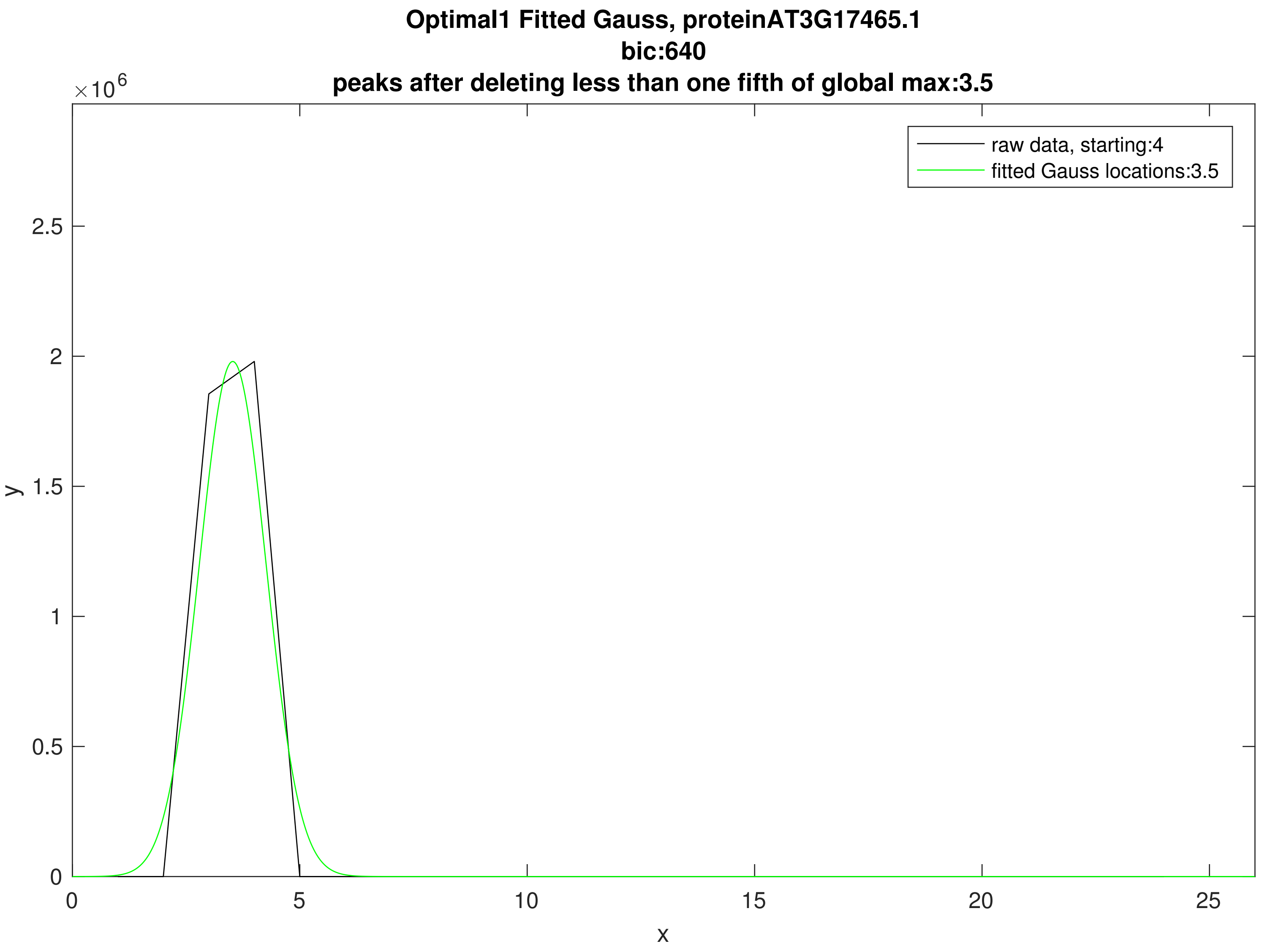
Optimal1 Fitted Gauss, proteinAT3G17465.1
bic:640
peaks after deleting less than one fifth of global max:3.5
×10^6
raw data, starting:4
fitted Gauss locations:3.5
2.5
2
1.5
1
0.5
0
y
0
5
10
15
20
25
x

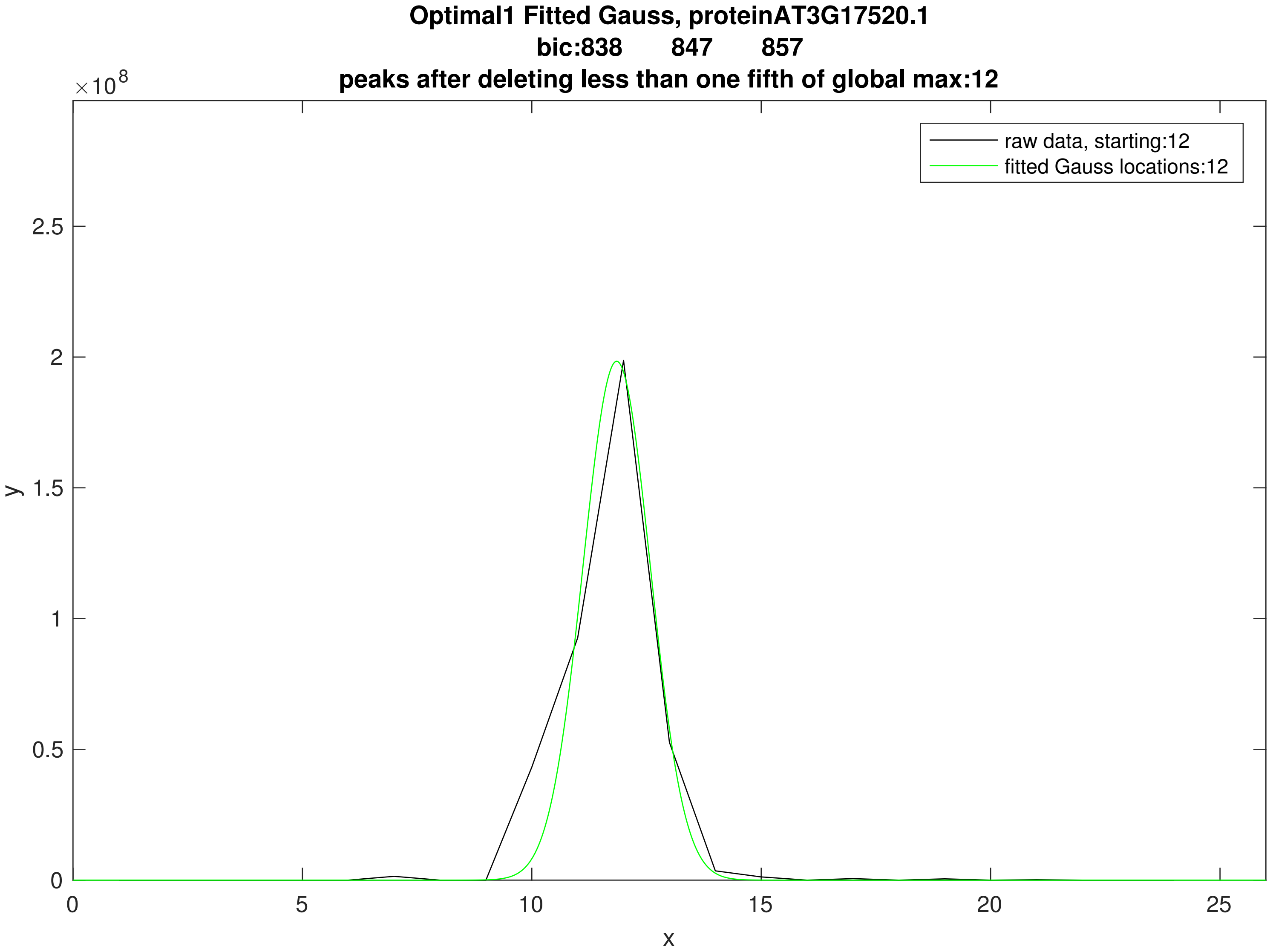
Optimal1 Fitted Gauss, proteinAT3G17520.1
bic:838 847 857
peaks after deleting less than one fifth of global max:12
×10^8
raw data, starting:12
fitted Gauss locations:12
y
x
0
0.5
1
1.5
2
2.5
5
10
15
20
25

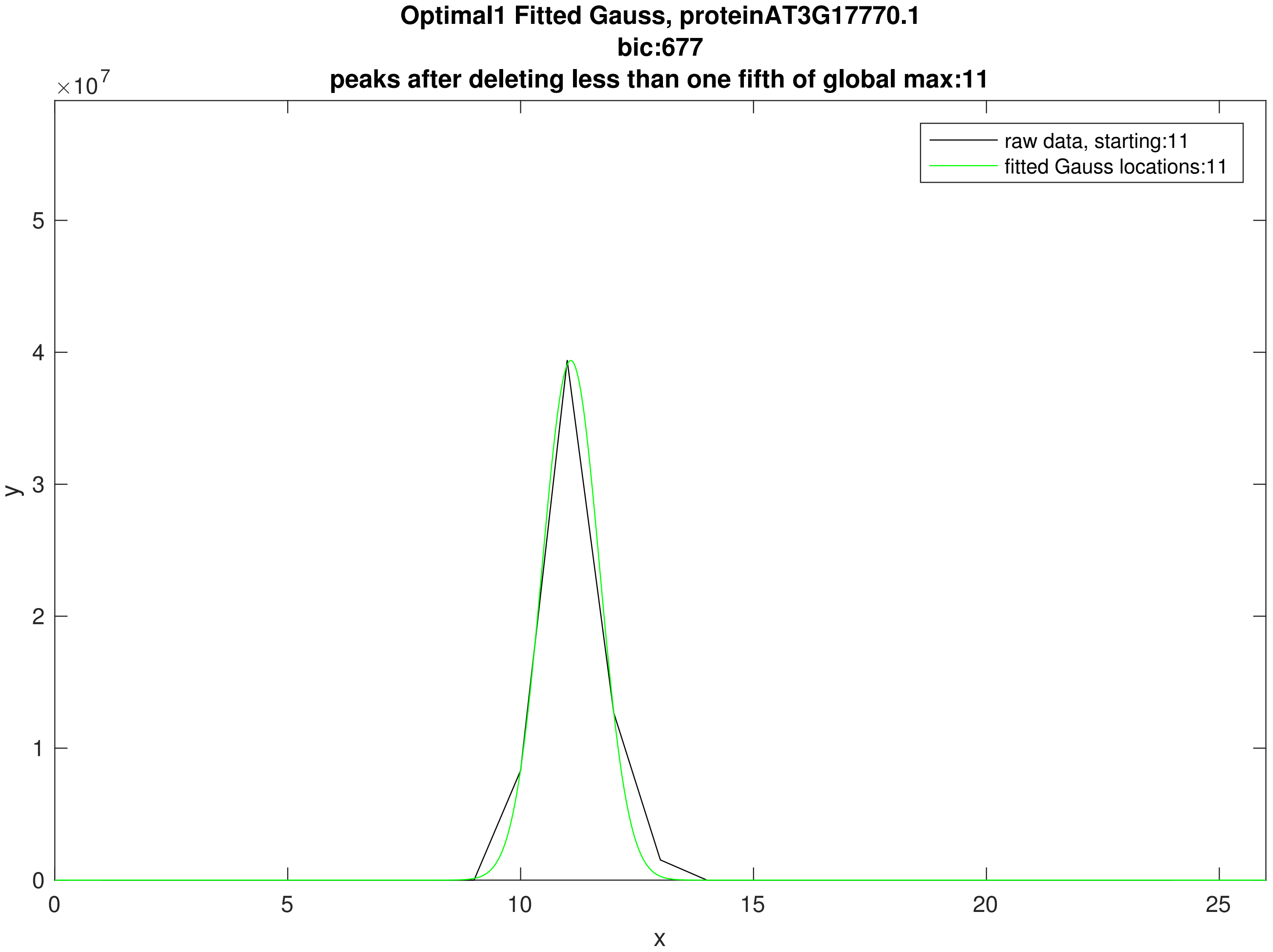
Optimal1 Fitted Gauss, proteinAT3G17770.1
bic:677
peaks after deleting less than one fifth of global max:11
×10^7
raw data, starting:11
fitted Gauss locations:11
y
x
0
1
2
3
4
5
0
5
10
15
20
25

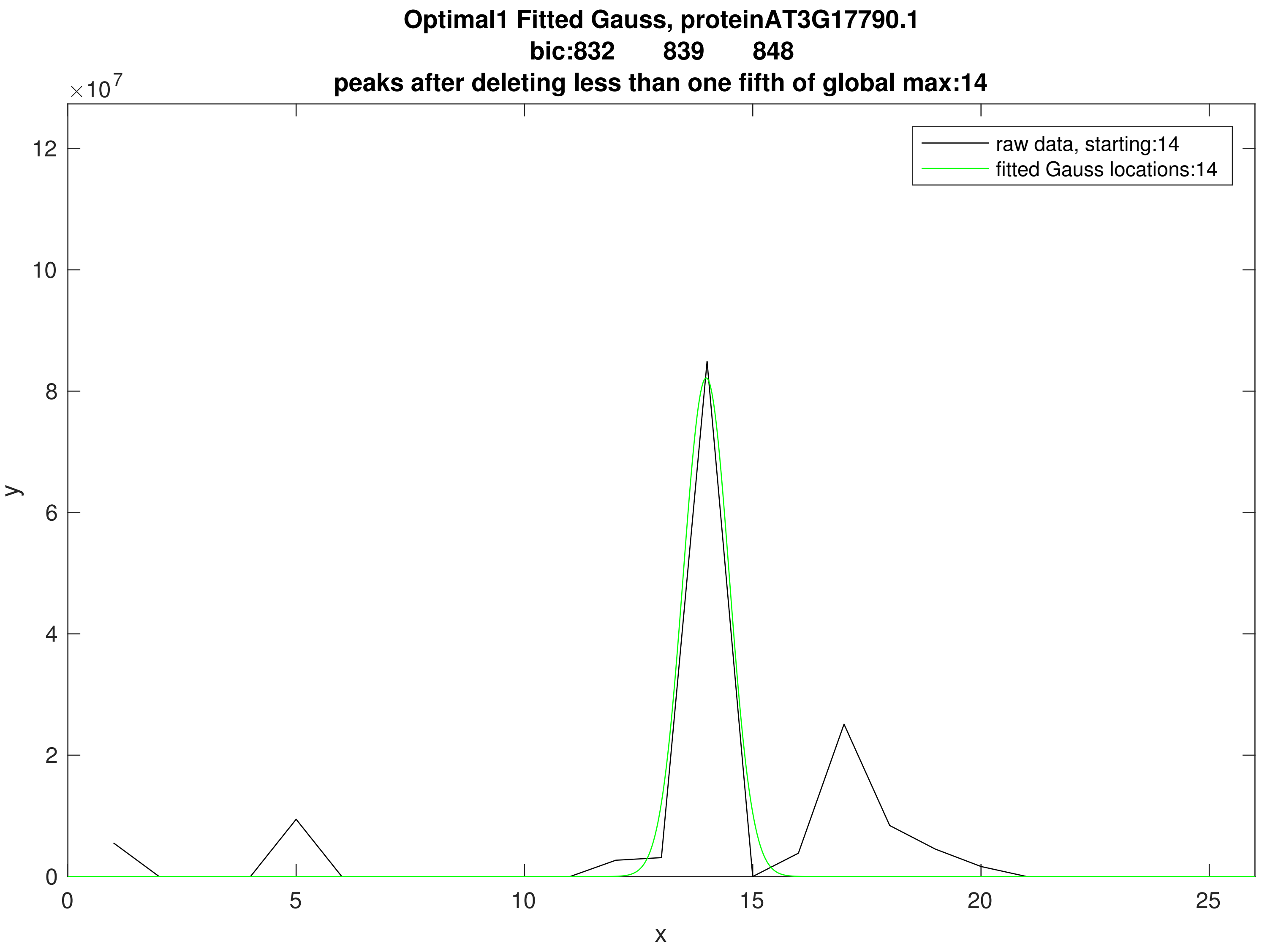
Optimal1 Fitted Gauss, proteinAT3G17790.1
bic:832 839 848
peaks after deleting less than one fifth of global max:14
×10^7
raw data, starting:14
fitted Gauss locations:14
y
x
0
2
4
6
8
10
12
0
5
10
15
20
25

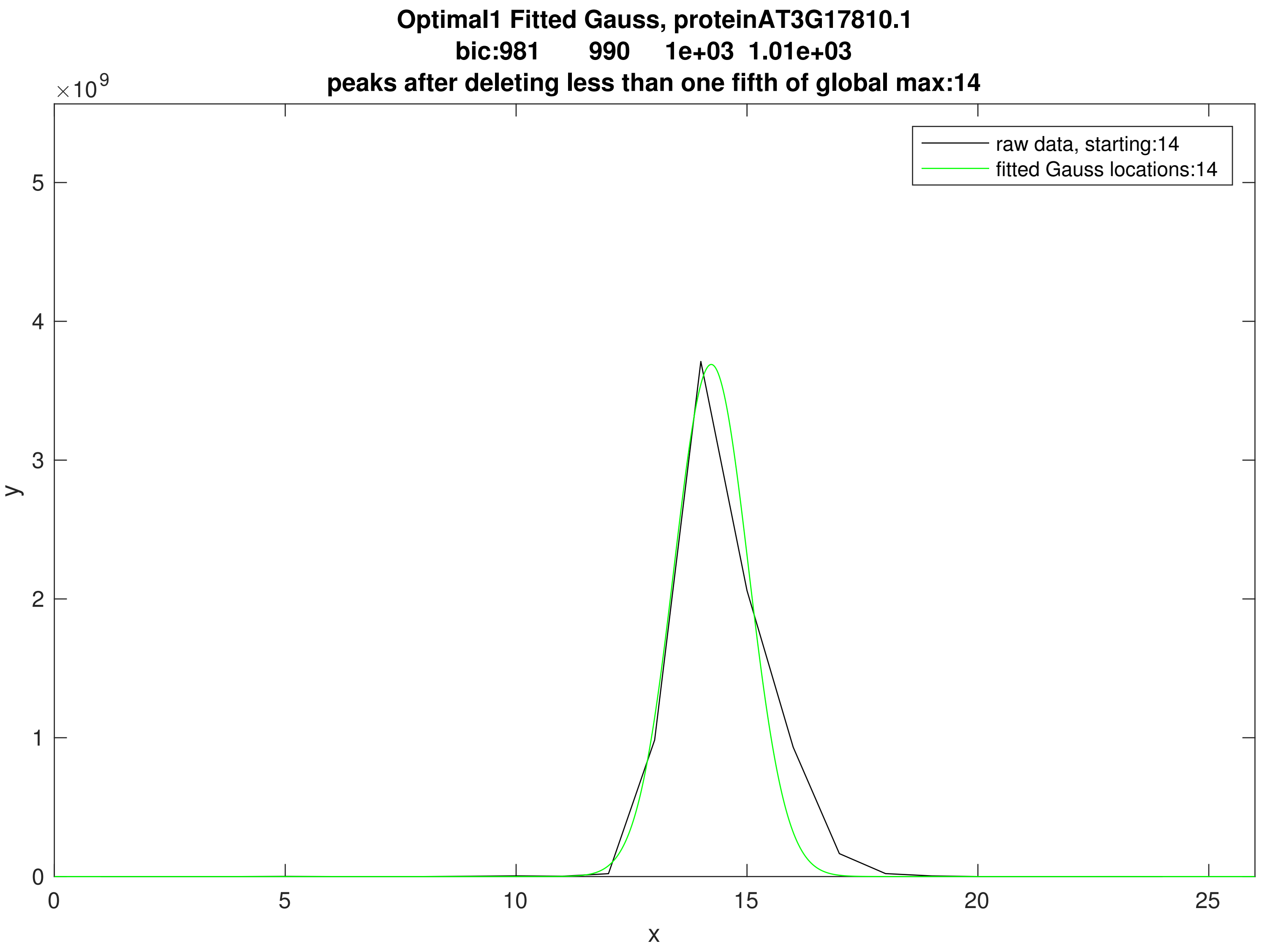
Optimal1 Fitted Gauss, proteinAT3G17810.1
bic:981 990 1e+03 1.01e+03
peaks after deleting less than one fifth of global max:14
×10^9
raw data, starting:14
fitted Gauss locations:14
y
x
0
1
2
3
4
5
0
5
10
15
20
25

Optimal1 Fitted Gauss, proteinAT3G17820.1
bic:1.01e+03  1.01e+03  1.02e+03
peaks after deleting less than one fifth of global max:7.8

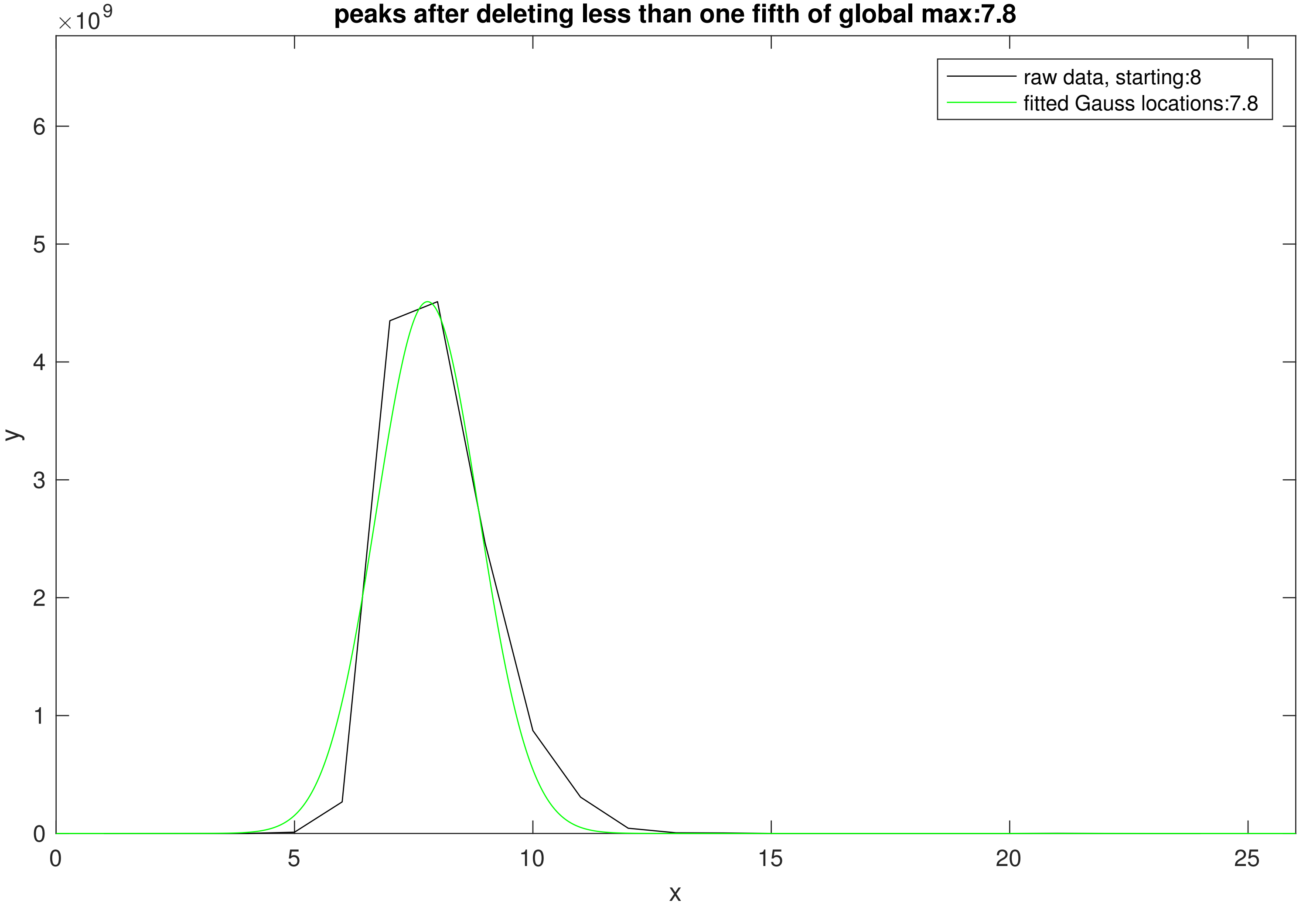

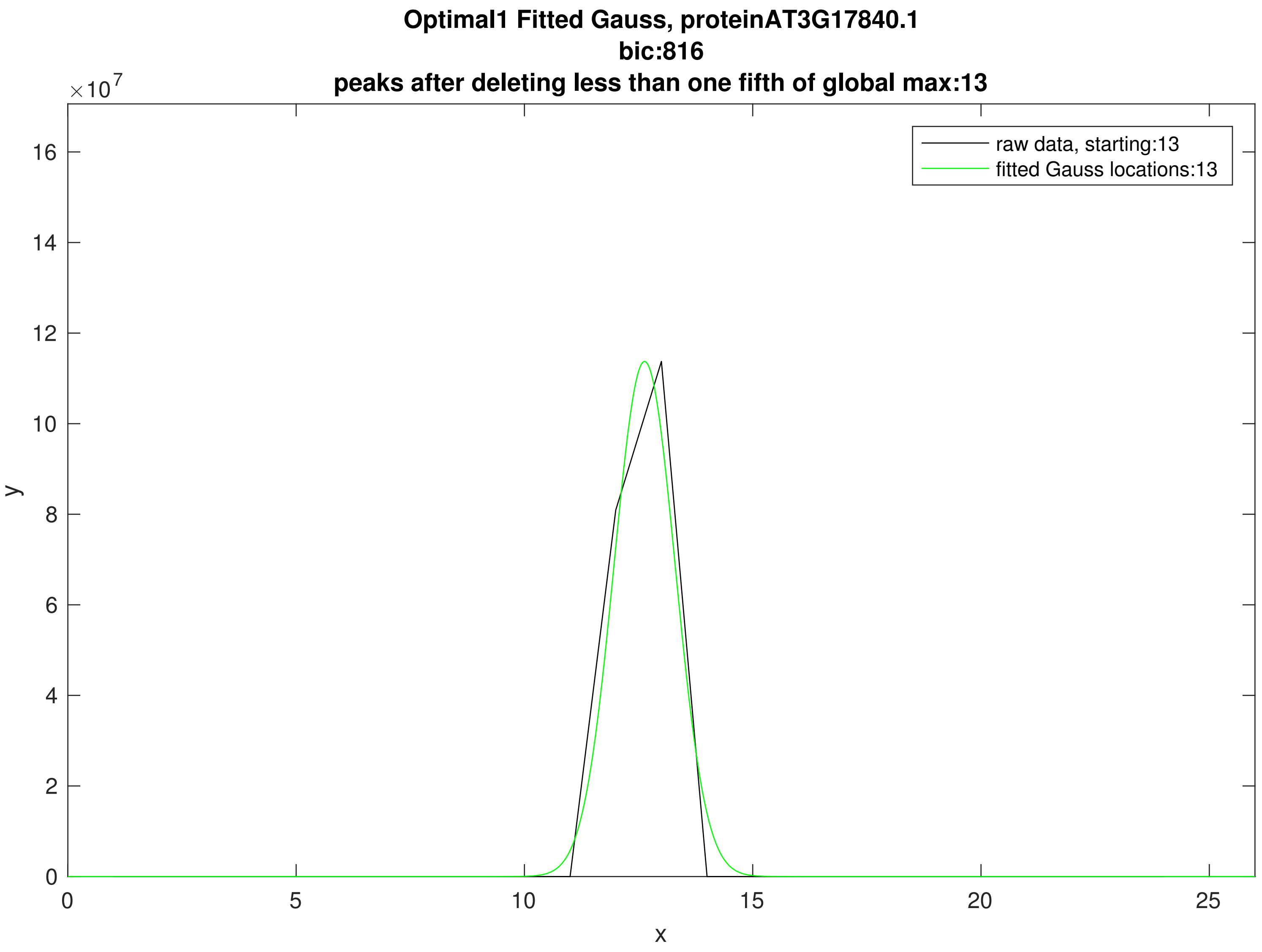
Optimal1 Fitted Gauss, proteinAT3G17840.1
bic:816
peaks after deleting less than one fifth of global max:13
×10^7
raw data, starting:13
fitted Gauss locations:13
y
x
0
2
4
6
8
10
12
14
16
0
5
10
15
20
25

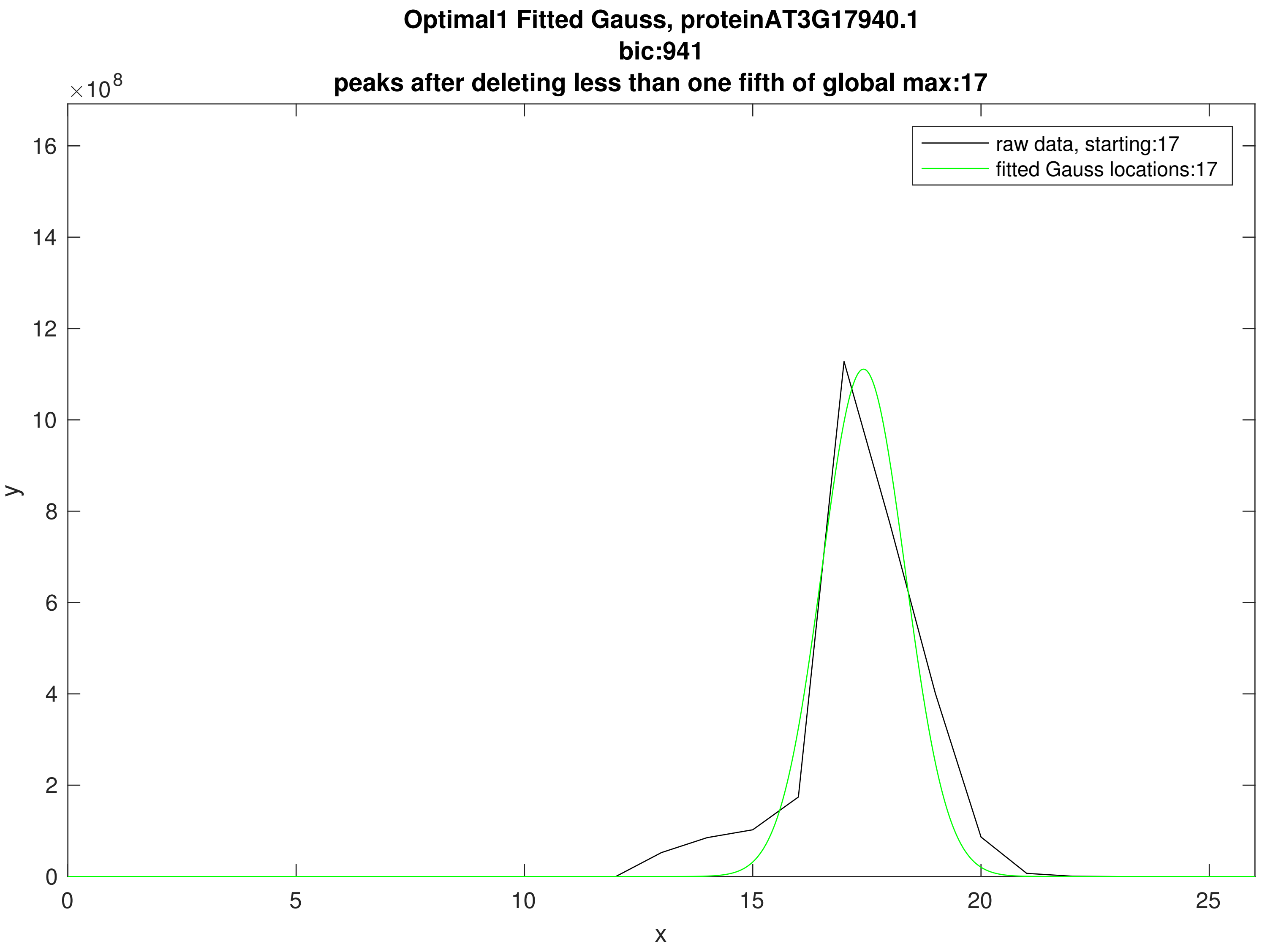
Optimal1 Fitted Gauss, proteinAT3G17940.1
bic:941
peaks after deleting less than one fifth of global max:17
×10^8
16
14
12
10
8
6
4
2
0
y
0
5
10
15
20
25
x
raw data, starting:17
fitted Gauss locations:17

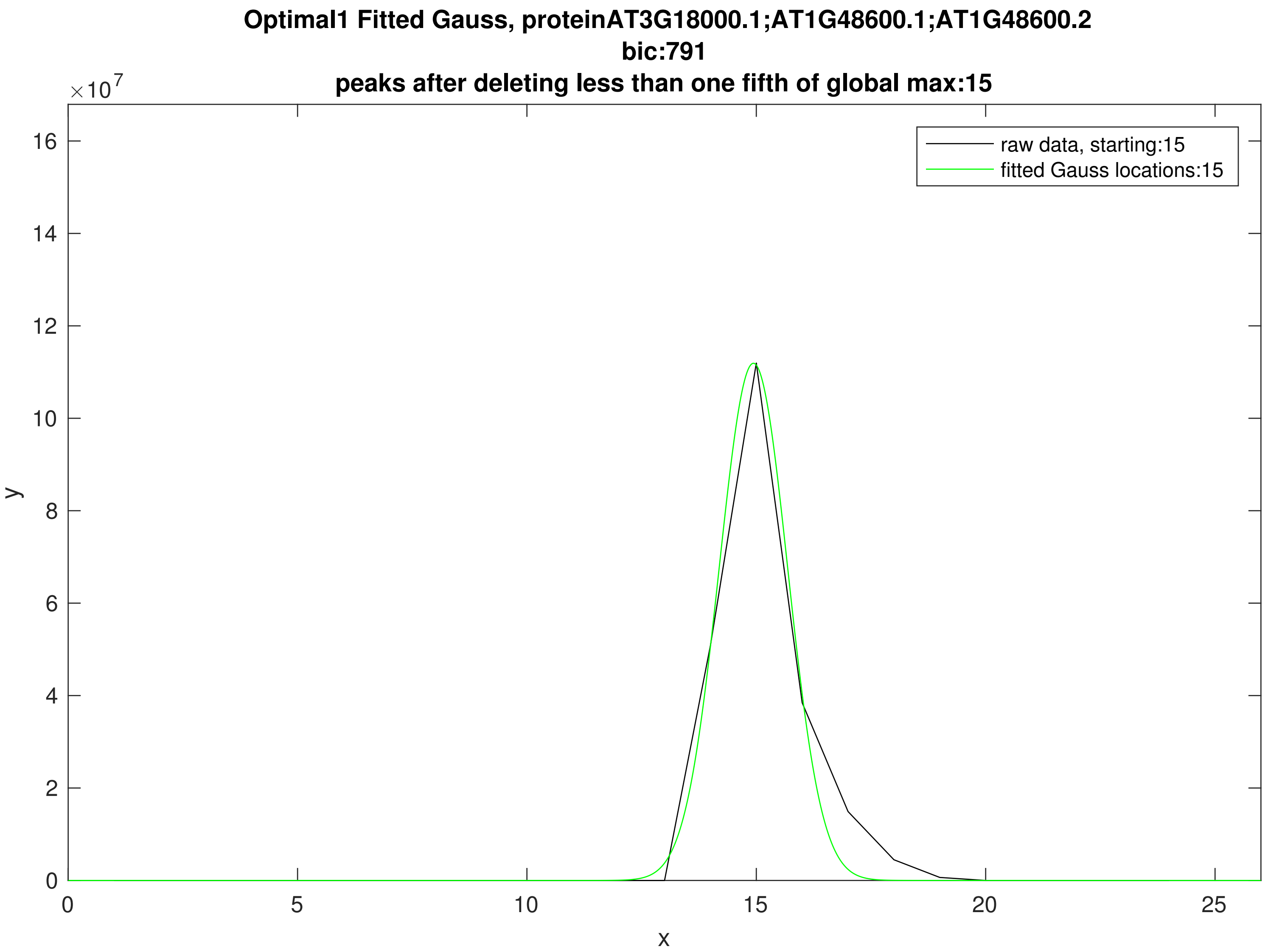
Optimal1 Fitted Gauss, proteinAT3G18000.1;AT1G48600.1;AT1G48600.2
bic:791
peaks after deleting less than one fifth of global max:15
×10^7
16
14
12
10
8
6
4
2
0
y
0
5
10
15
20
25
x
raw data, starting:15
fitted Gauss locations:15

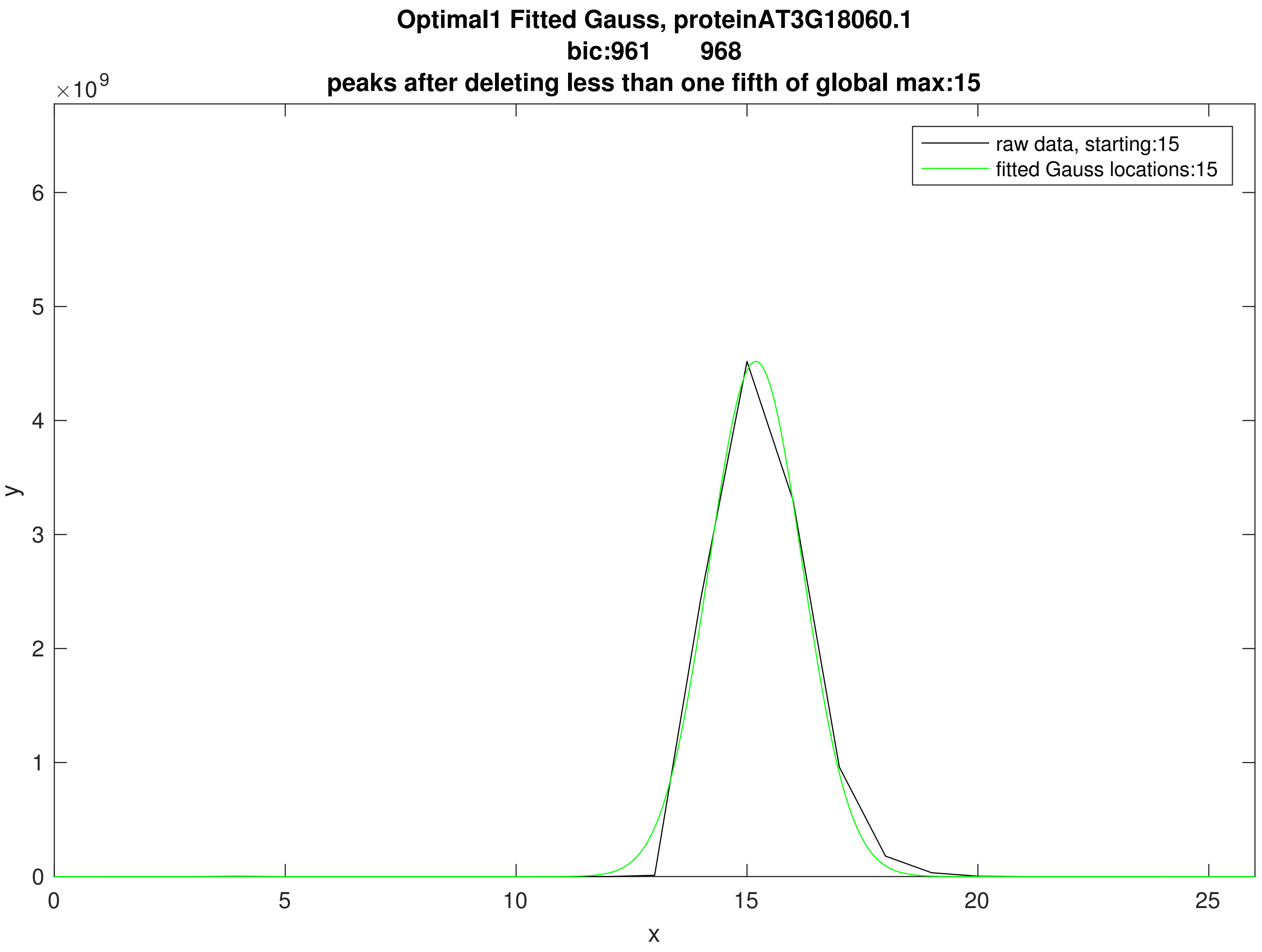

Optimal1 Fitted Gauss, proteinAT3G18060.1
bic:961 968
peaks after deleting less than one fifth of global max:15
×10^9
raw data, starting:15
fitted Gauss locations:15
y
x
0
1
2
3
4
5
6
0
5
10
15
20
25

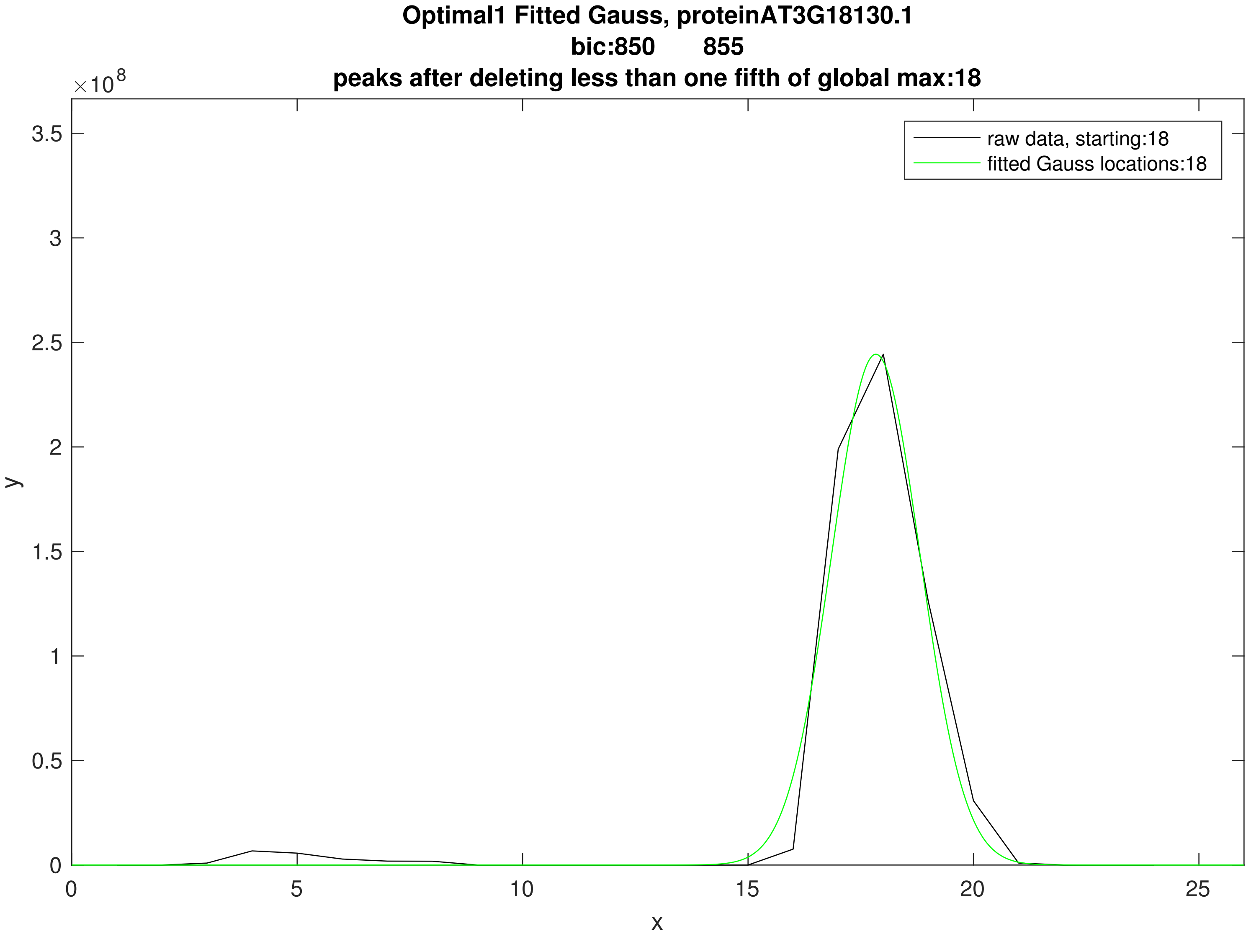

Optimal1 Fitted Gauss, proteinAT3G18130.1
bic:850 855
peaks after deleting less than one fifth of global max:18
×10^8
3.5
3
2.5
2
1.5
1
0.5
0
y
0
5
10
15
20
25
x
raw data, starting:18
fitted Gauss locations:18

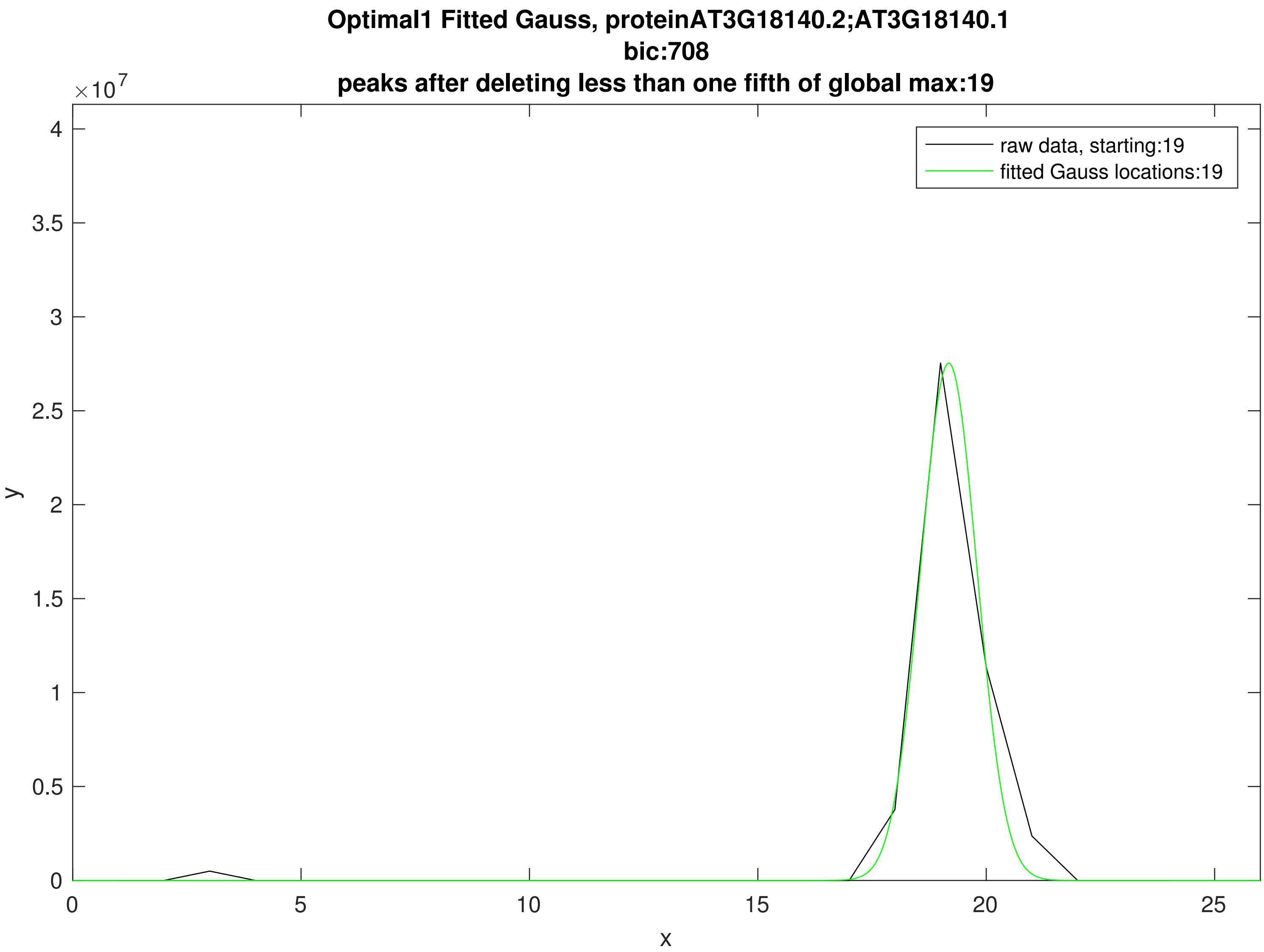

Optimal1 Fitted Gauss, proteinAT3G18140.2;AT3G18140.1
bic:708
peaks after deleting less than one fifth of global max:19
×10^7
4
3.5
3
2.5
2
1.5
1
0.5
0
y
0
5
10
15
20
25
x
raw data, starting:19
fitted Gauss locations:19

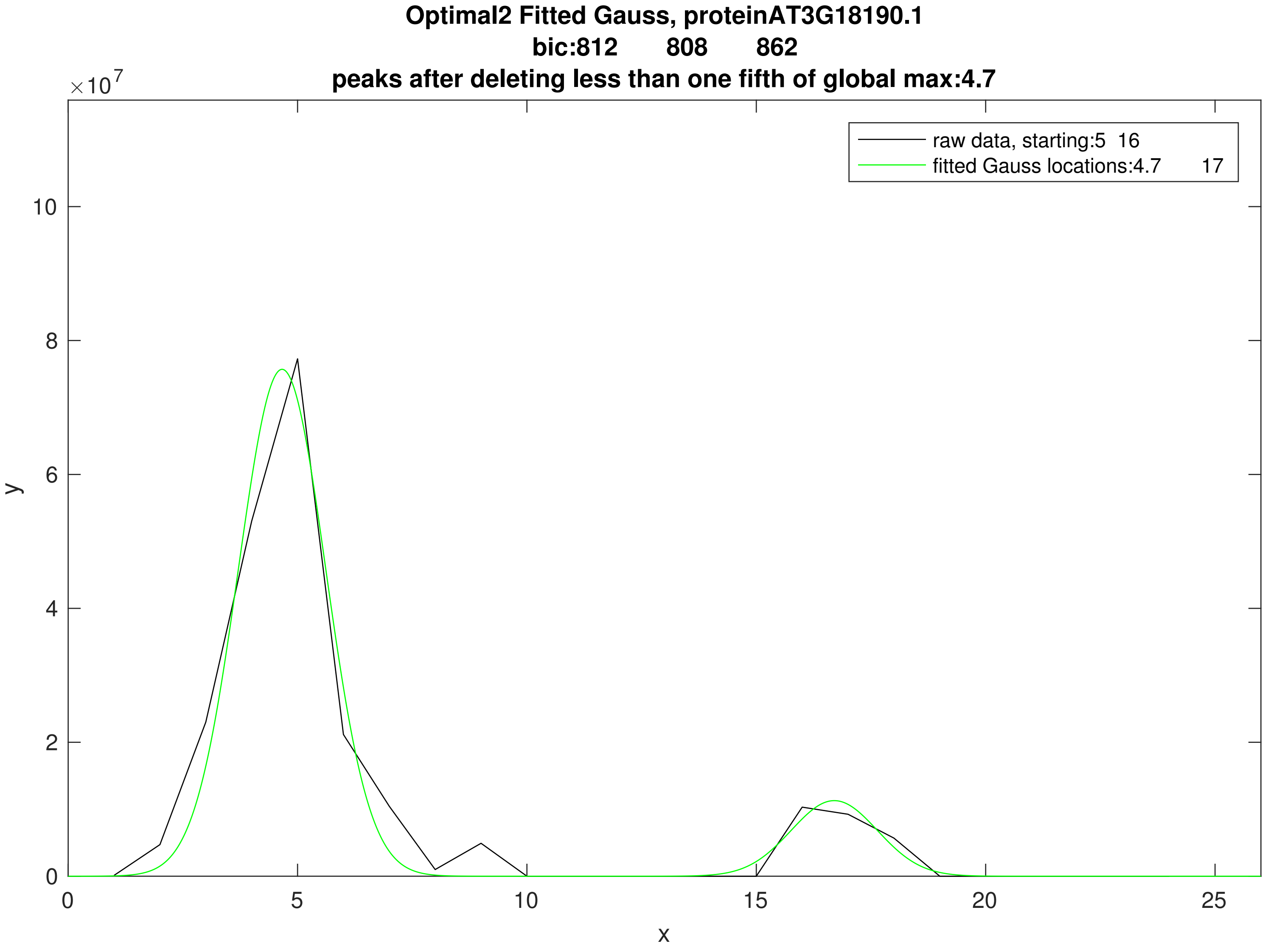
Optimal2 Fitted Gauss, proteinAT3G18190.1
bic:812 808 862
peaks after deleting less than one fifth of global max:4.7
×10^7
raw data, starting:5 16
fitted Gauss locations:4.7 17
y
x
0
2
4
6
8
10
0
5
10
15
20
25

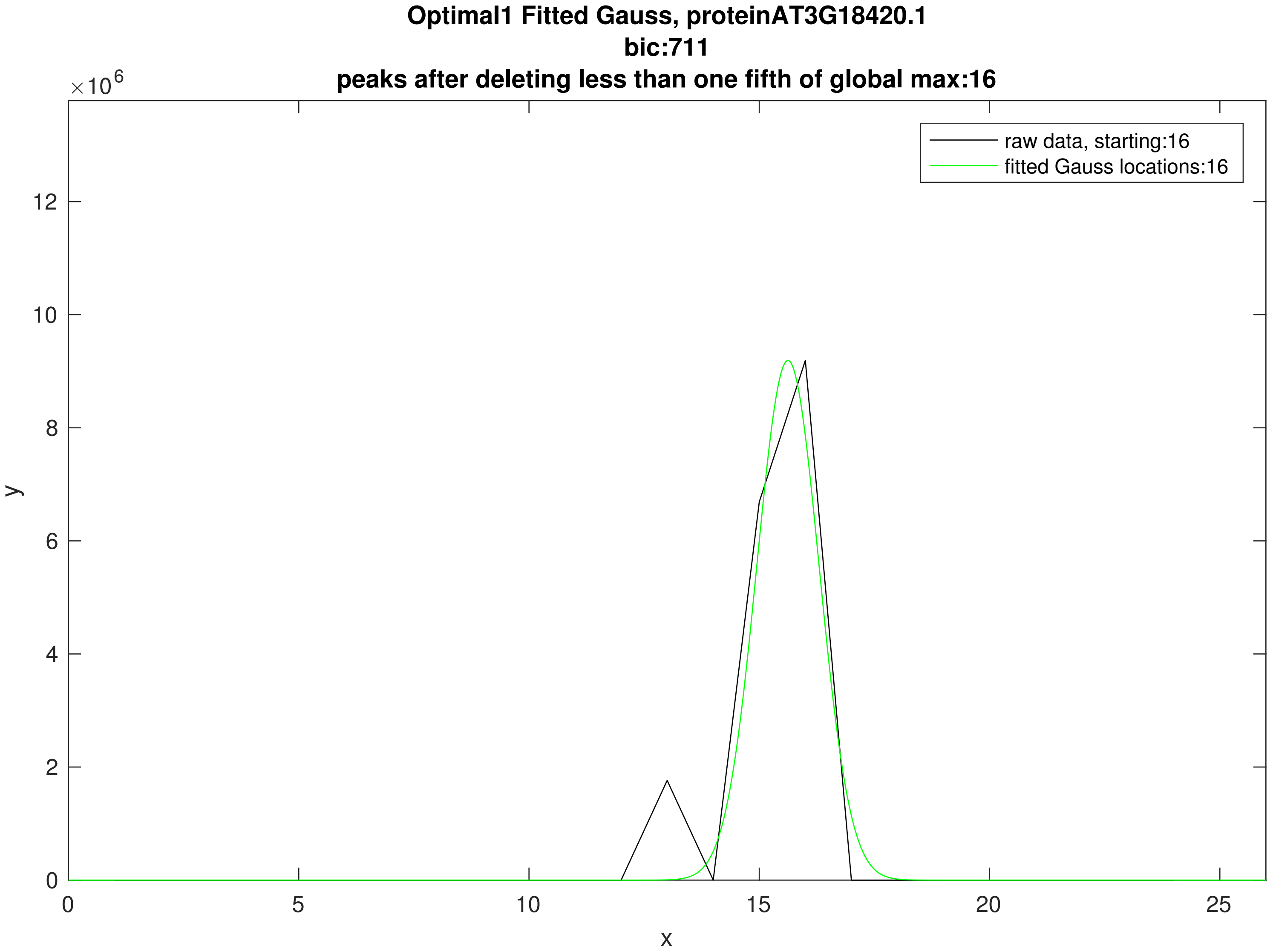

Optimal1 Fitted Gauss, proteinAT3G18420.1
bic:711
peaks after deleting less than one fifth of global max:16
×10^6
raw data, starting:16
fitted Gauss locations:16
12
10
8
6
4
2
0
y
0
5
10
15
20
25
x

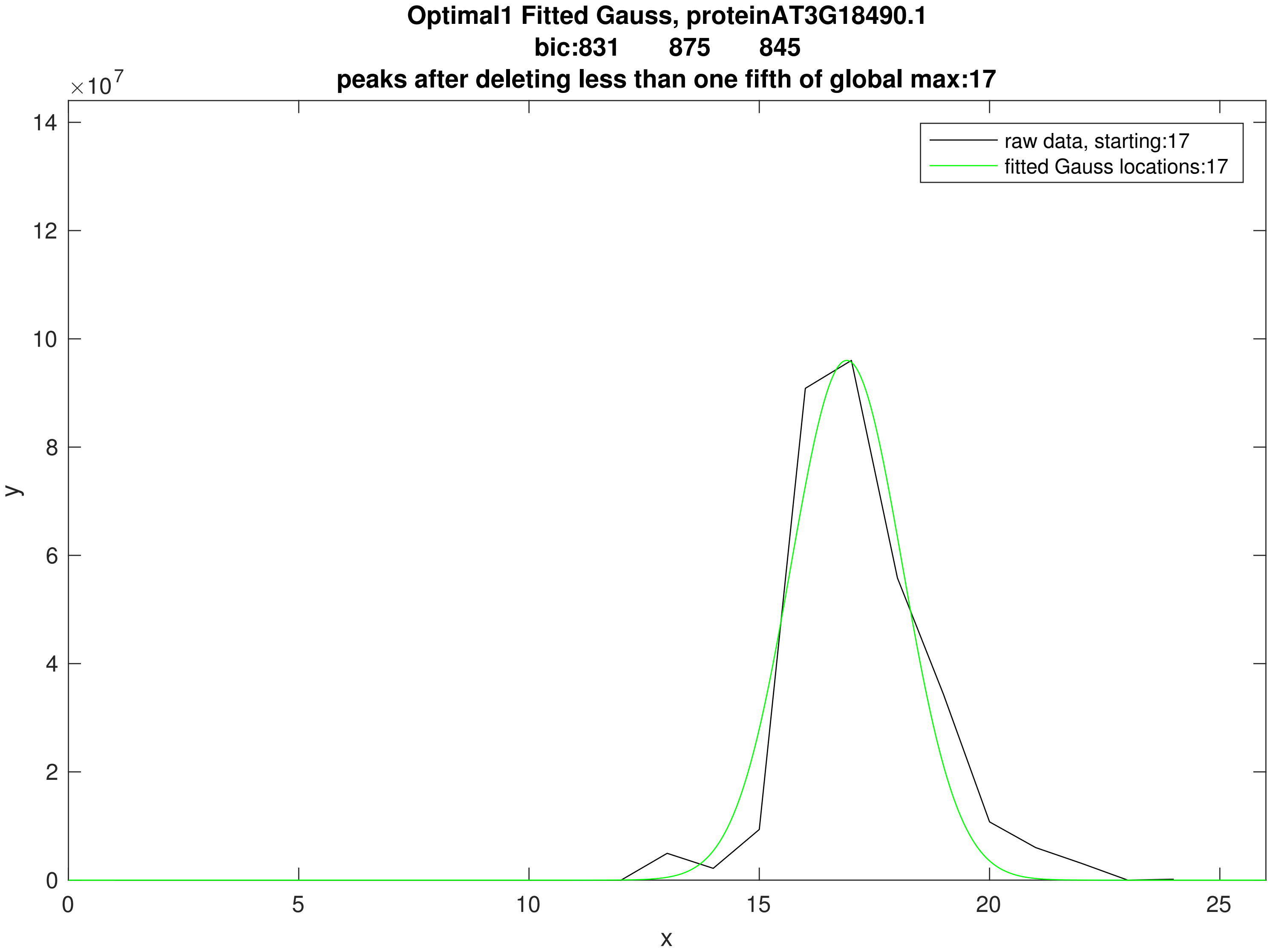
Optimal1 Fitted Gauss, proteinAT3G18490.1
bic:831 875 845
peaks after deleting less than one fifth of global max:17
×10^7
14
12
10
8
6
4
2
0
y
0
5
10
15
20
25
x
raw data, starting:17
fitted Gauss locations:17

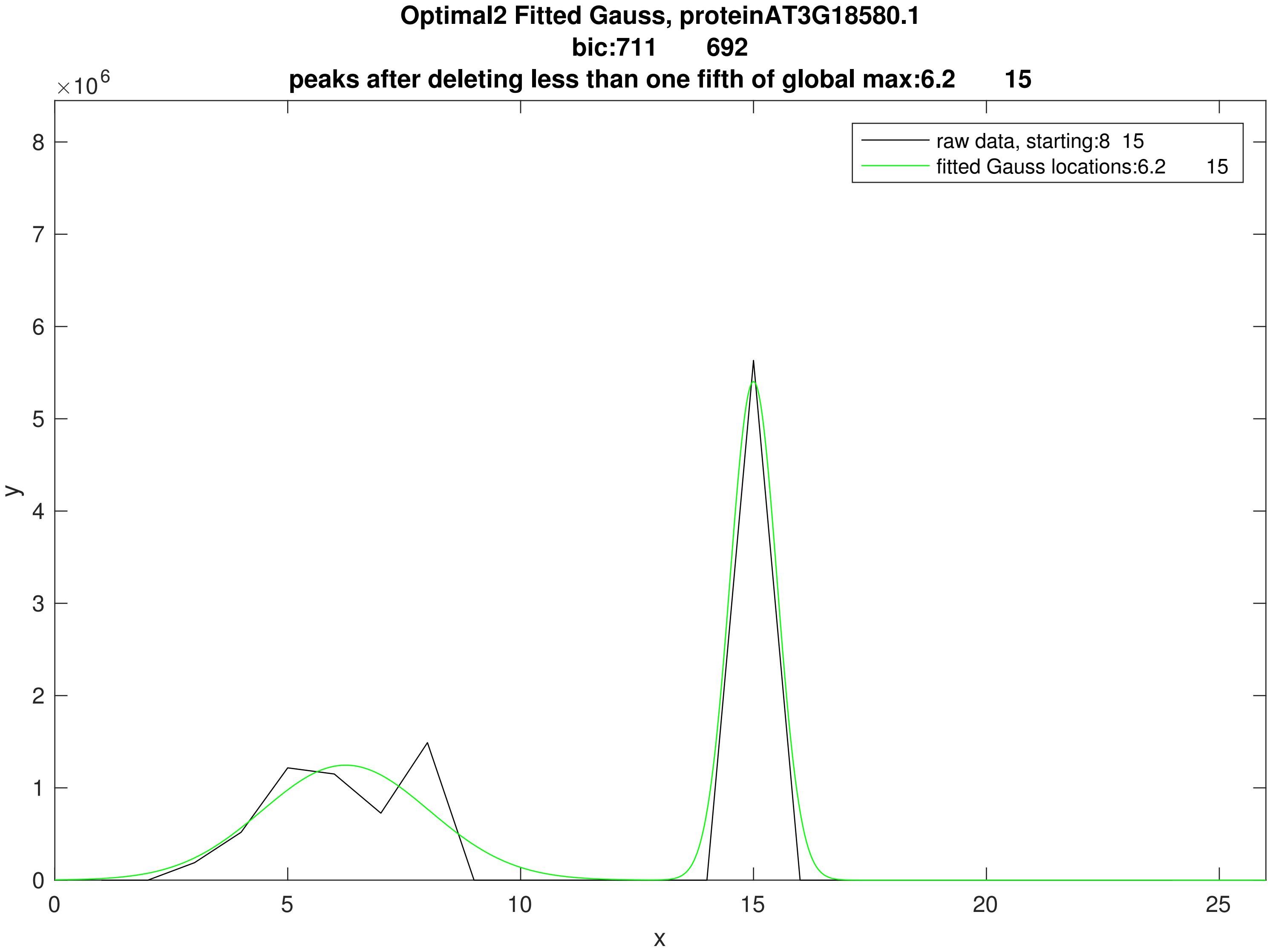
Optimal2 Fitted Gauss, proteinAT3G18580.1
bic:711 692
peaks after deleting less than one fifth of global max:6.2 15
×10^6
raw data, starting:8 15
fitted Gauss locations:6.2 15
y
x
0
1
2
3
4
5
6
7
8
0
5
10
15
20
25

Optimal1 Fitted Gauss, proteinAT3G18740.1;AT1G36240.1
bic:765
peaks after deleting less than one fifth of global max:3.9

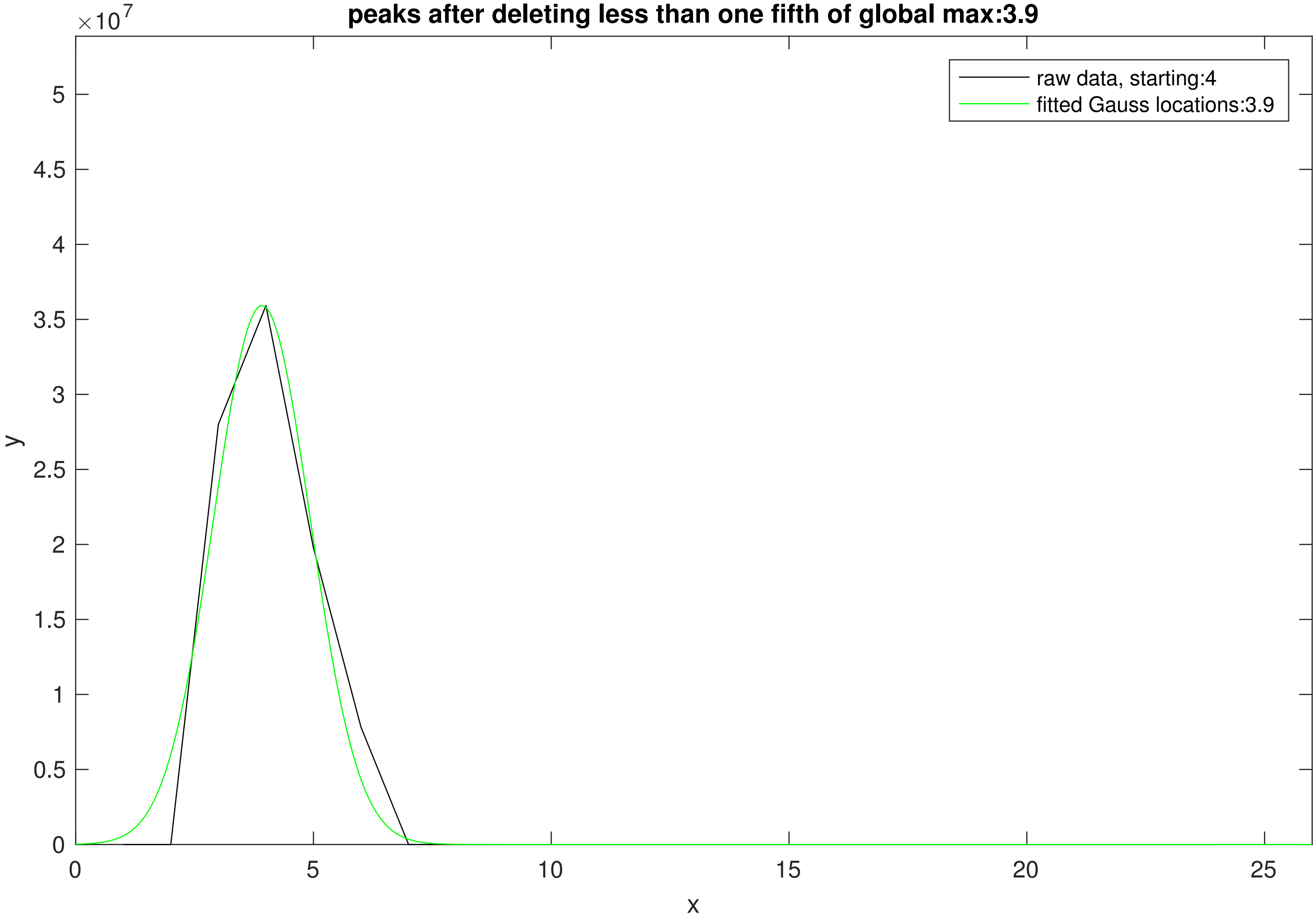

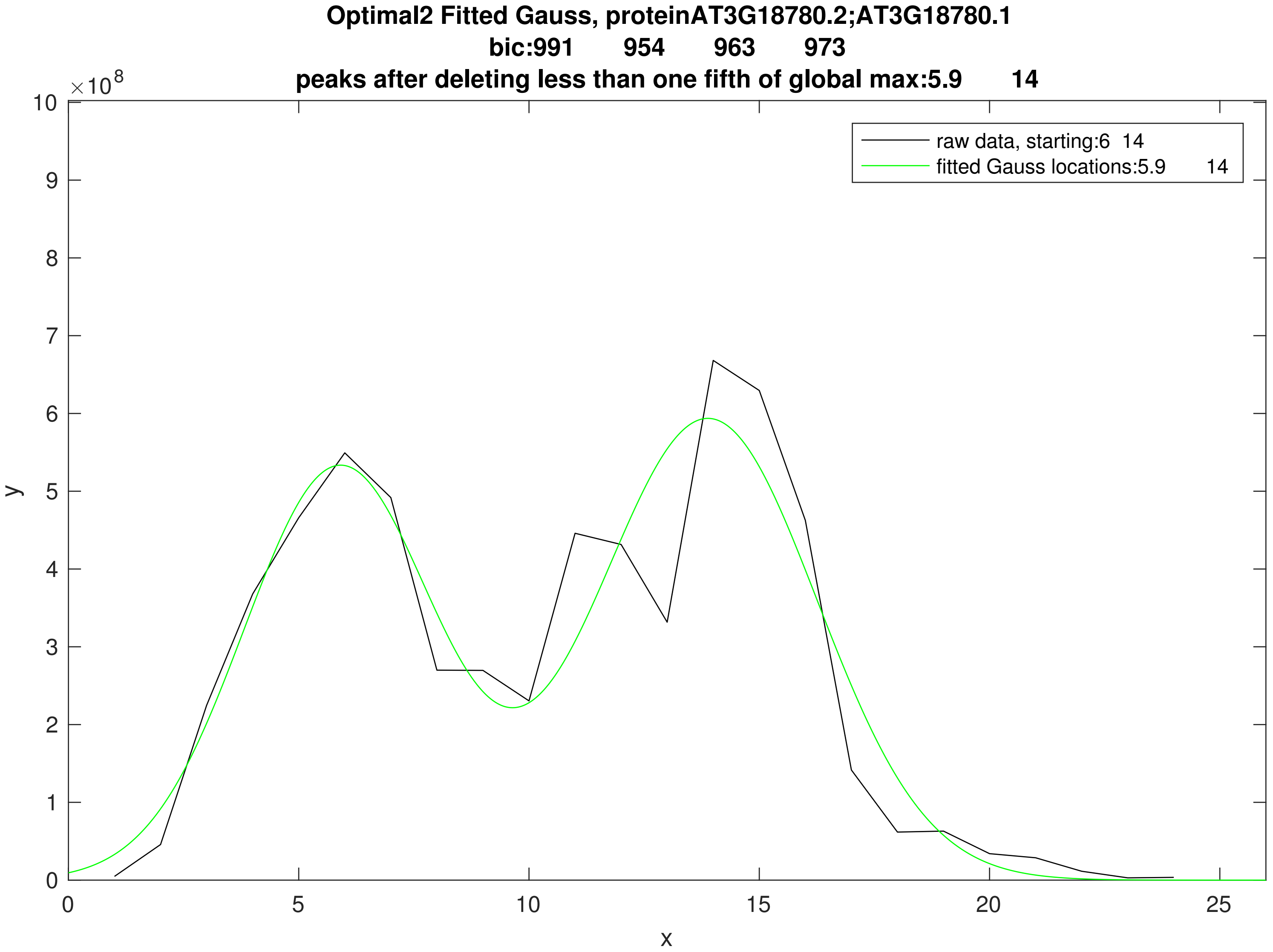
Optimal2 Fitted Gauss, proteinAT3G18780.2;AT3G18780.1
bic:991 954 963 973
peaks after deleting less than one fifth of global max:5.9 14
×10^8
raw data, starting:6 14
fitted Gauss locations:5.9 14
y
x
0
1
2
3
4
5
6
7
8
9
10
15
20
25

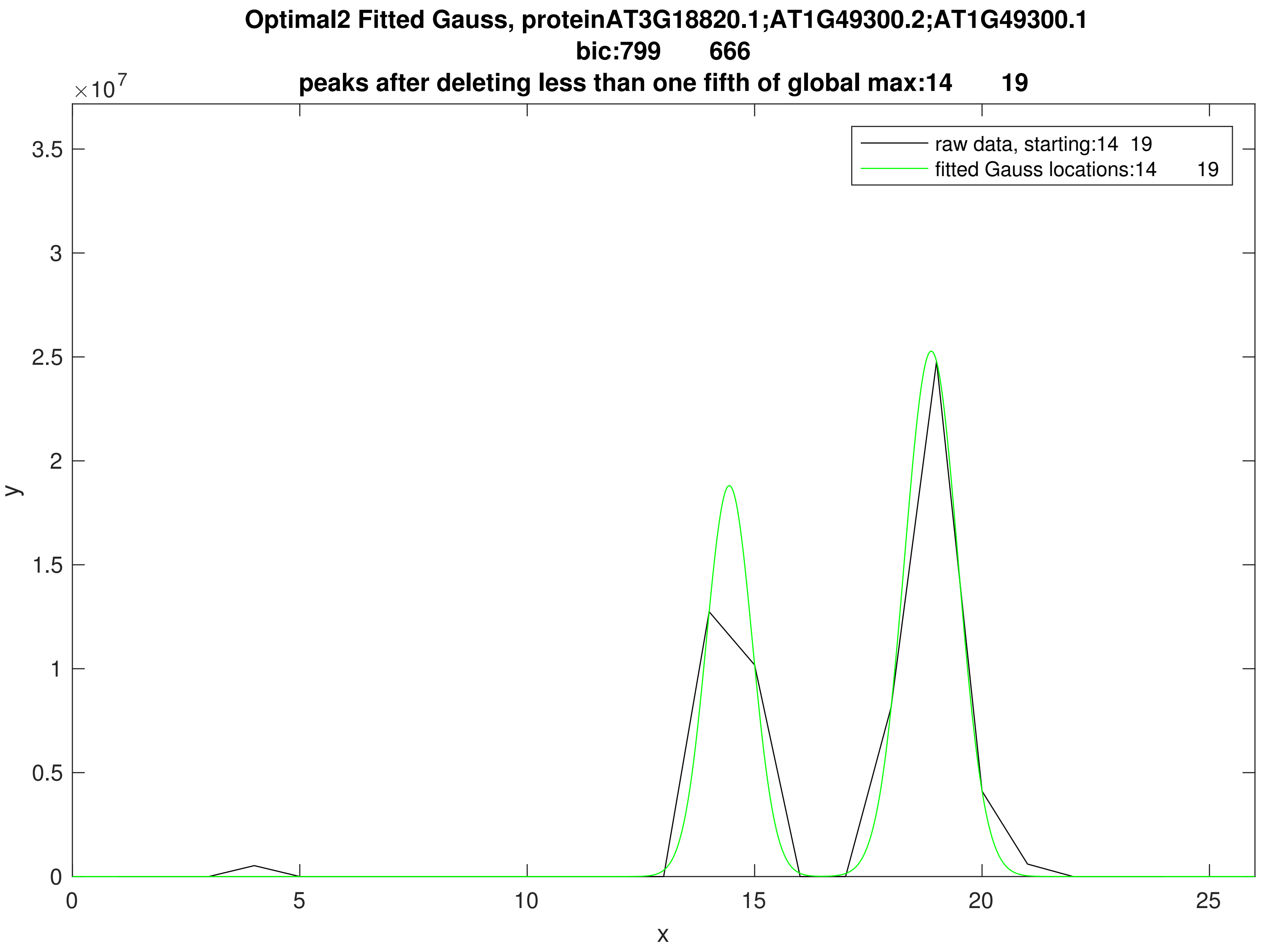

Optimal2 Fitted Gauss, proteinAT3G18820.1;AT1G49300.2;AT1G49300.1
bic:799 666
peaks after deleting less than one fifth of global max:14 19
×10^7
3.5
3
2.5
2
1.5
1
0.5
0
y
0
5
10
15
20
25
x
raw data, starting:14 19
fitted Gauss locations:14 19

Optimal1 Fitted Gauss, proteinAT3G18860.2;AT3G18860.1
bic:764
peaks after deleting less than one fifth of global max:6.7

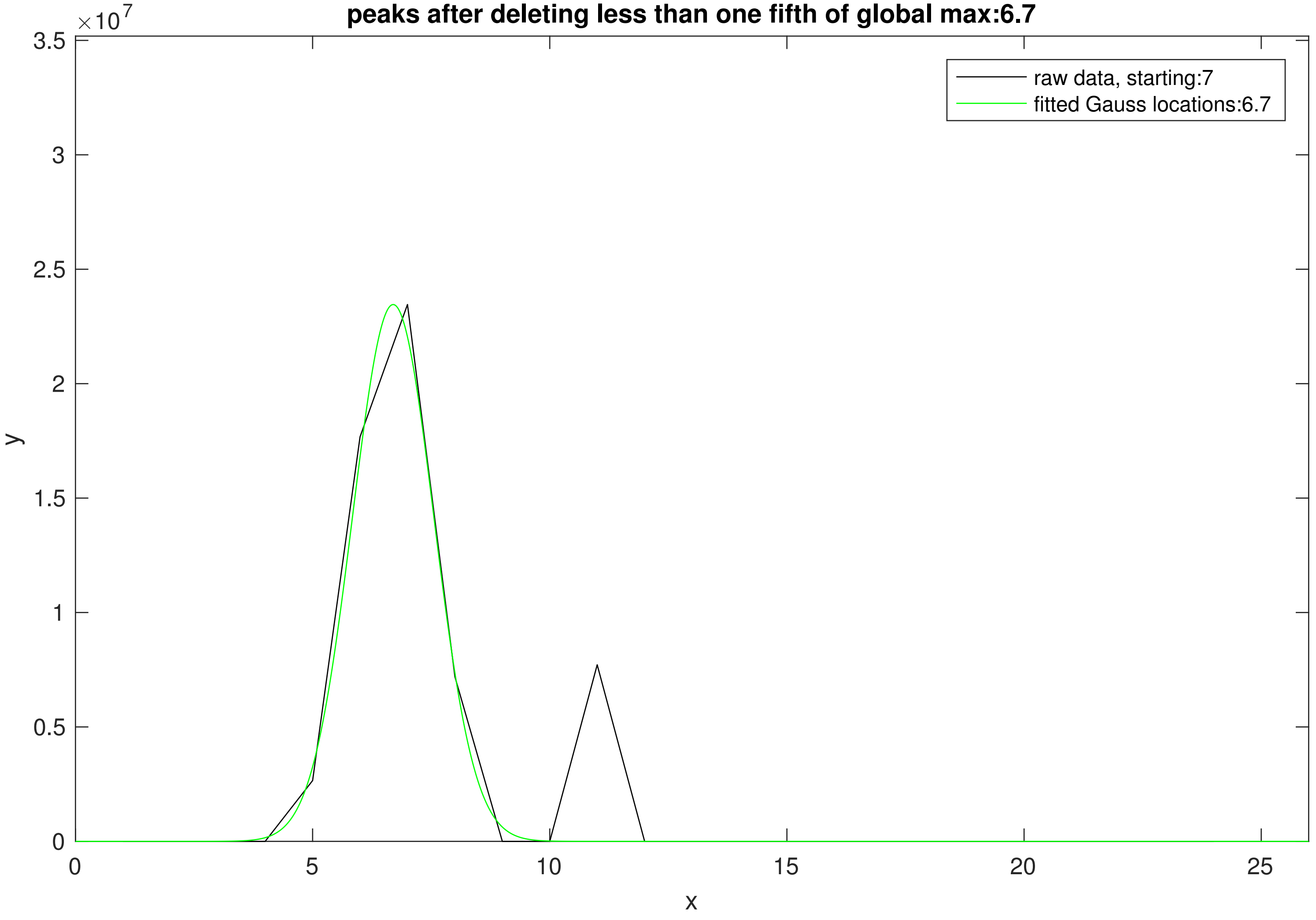

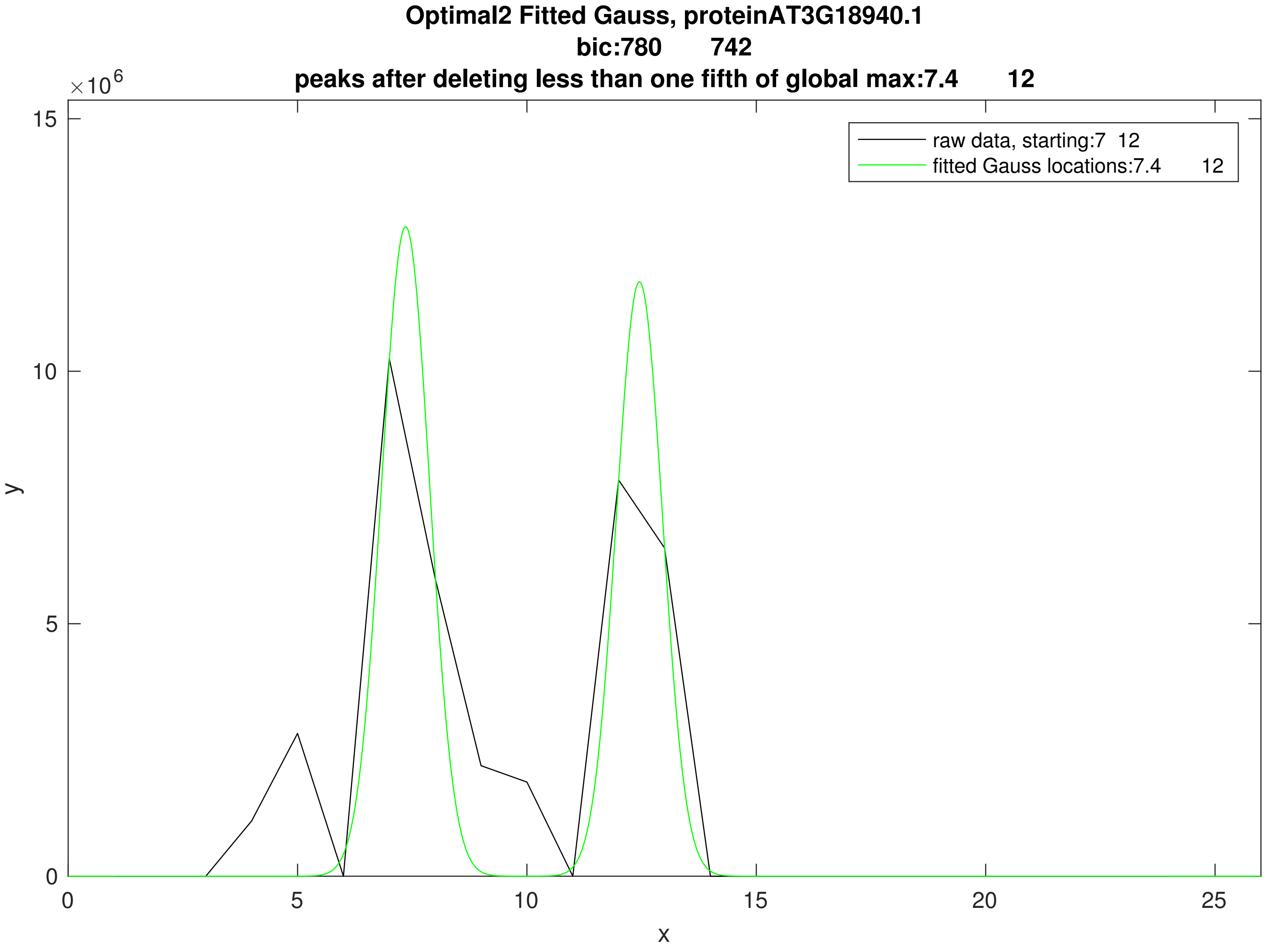
Optimal2 Fitted Gauss, proteinAT3G18940.1
bic:780 742
peaks after deleting less than one fifth of global max:7.4 12
×10^6
raw data, starting:7 12
fitted Gauss locations:7.4 12
y
x
0
5
10
15
20
25

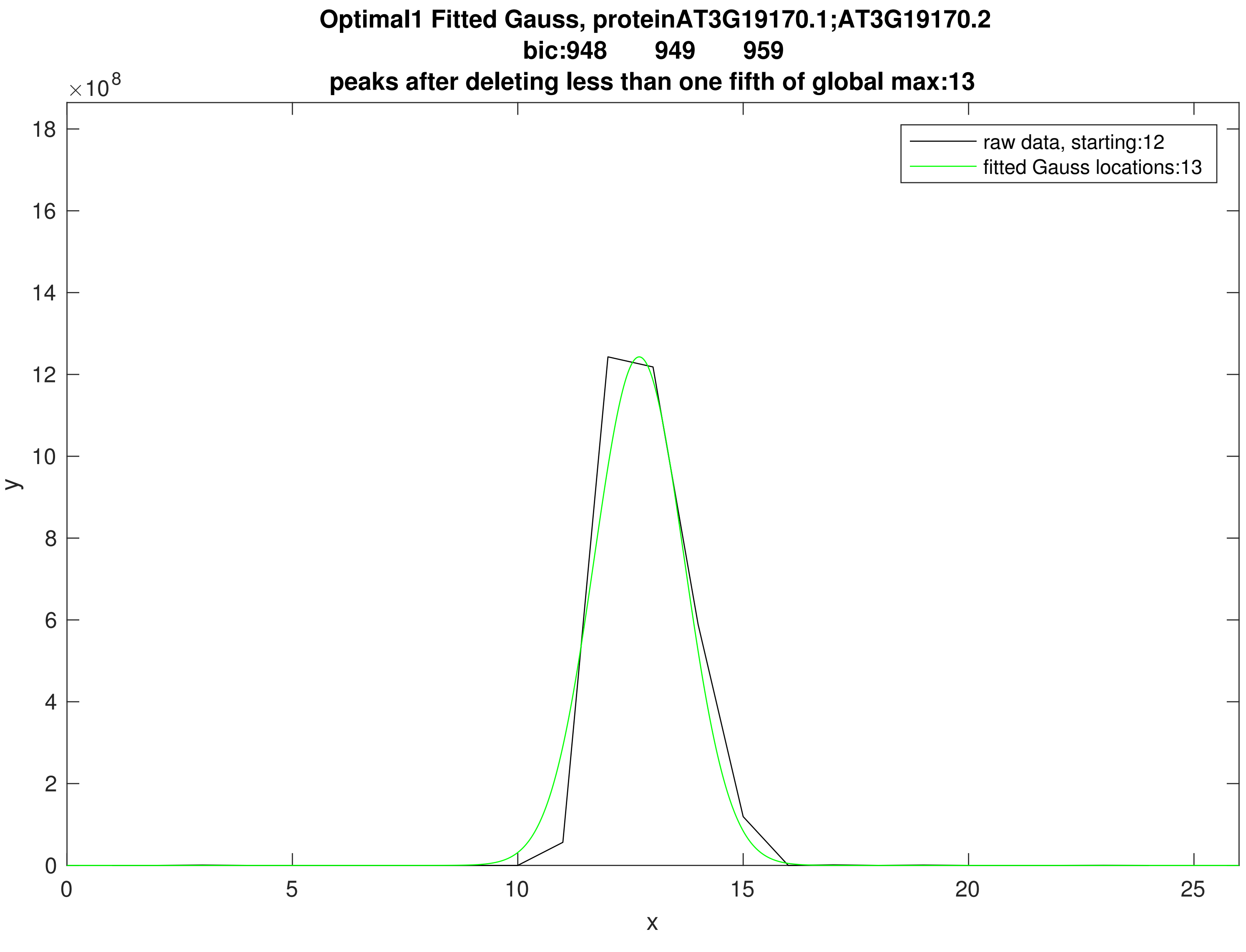
Optimal1 Fitted Gauss, proteinAT3G19170.1;AT3G19170.2
bic:948 949 959
peaks after deleting less than one fifth of global max:13
×10^8
raw data, starting:12
fitted Gauss locations:13
y
x
0
2
4
6
8
10
12
14
16
18
0
5
10
15
20
25

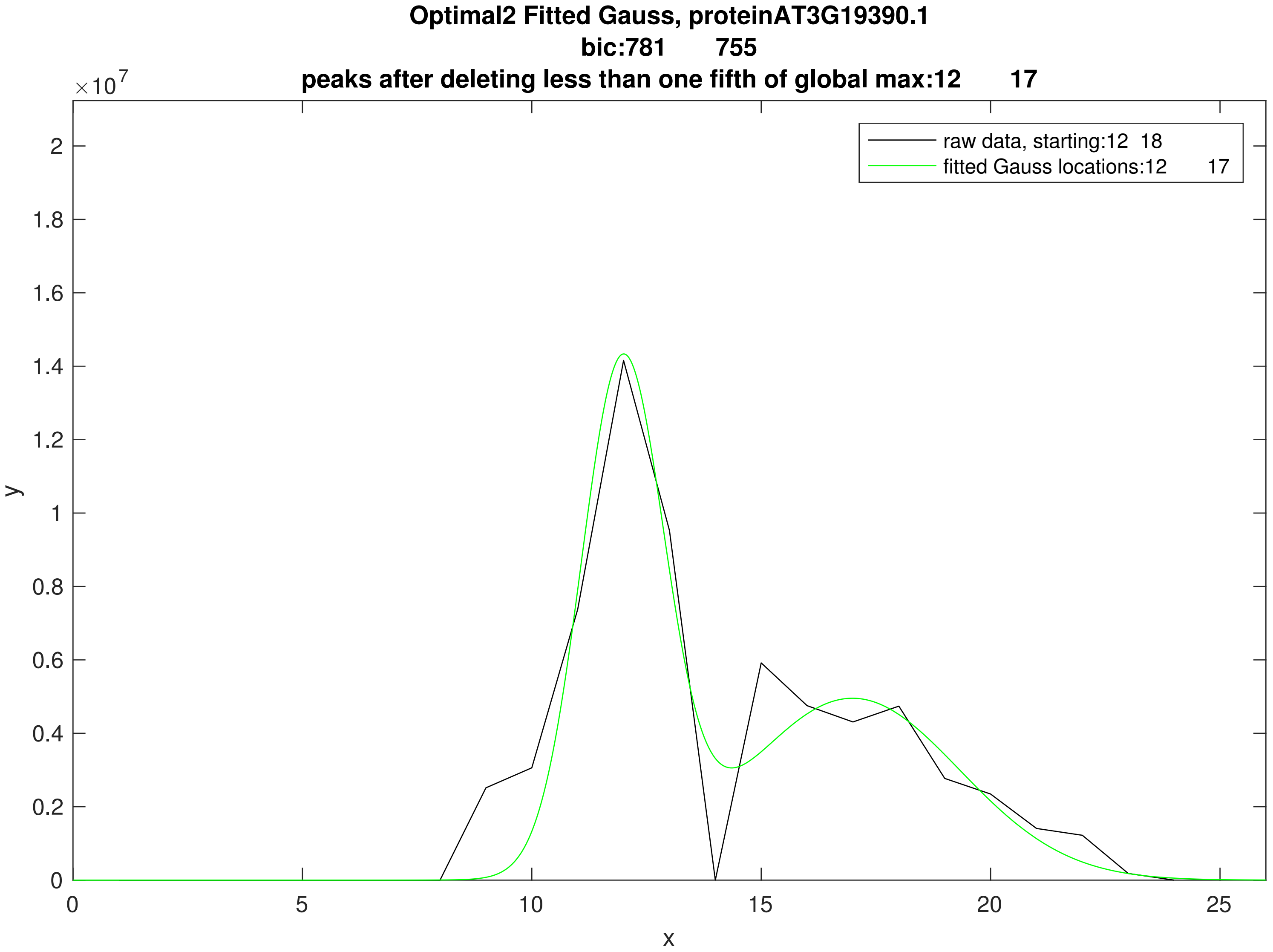
Optimal2 Fitted Gauss, proteinAT3G19390.1
bic:781 755
peaks after deleting less than one fifth of global max:12 17
×10^7
raw data, starting:12 18
fitted Gauss locations:12 17
y
x
0
0.2
0.4
0.6
0.8
1
1.2
1.4
1.6
1.8
2
0
5
10
15
20
25

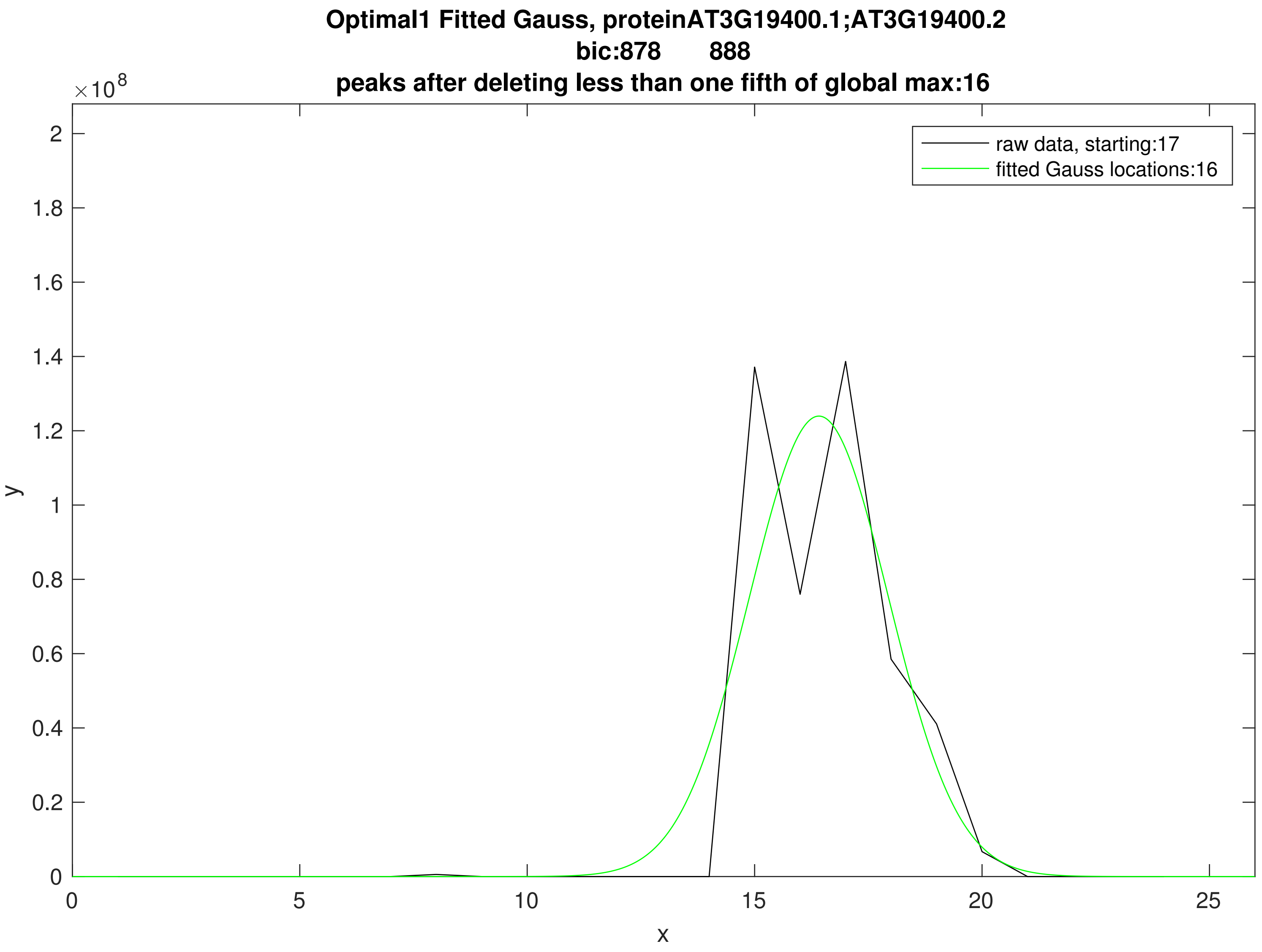

Optimal1 Fitted Gauss, proteinAT3G19400.1;AT3G19400.2
bic:878 888
peaks after deleting less than one fifth of global max:16
×10^8
raw data, starting:17
fitted Gauss locations:16
2
1.8
1.6
1.4
1.2
1
0.8
0.6
0.4
0.2
0
y
0
5
10
15
20
25
x

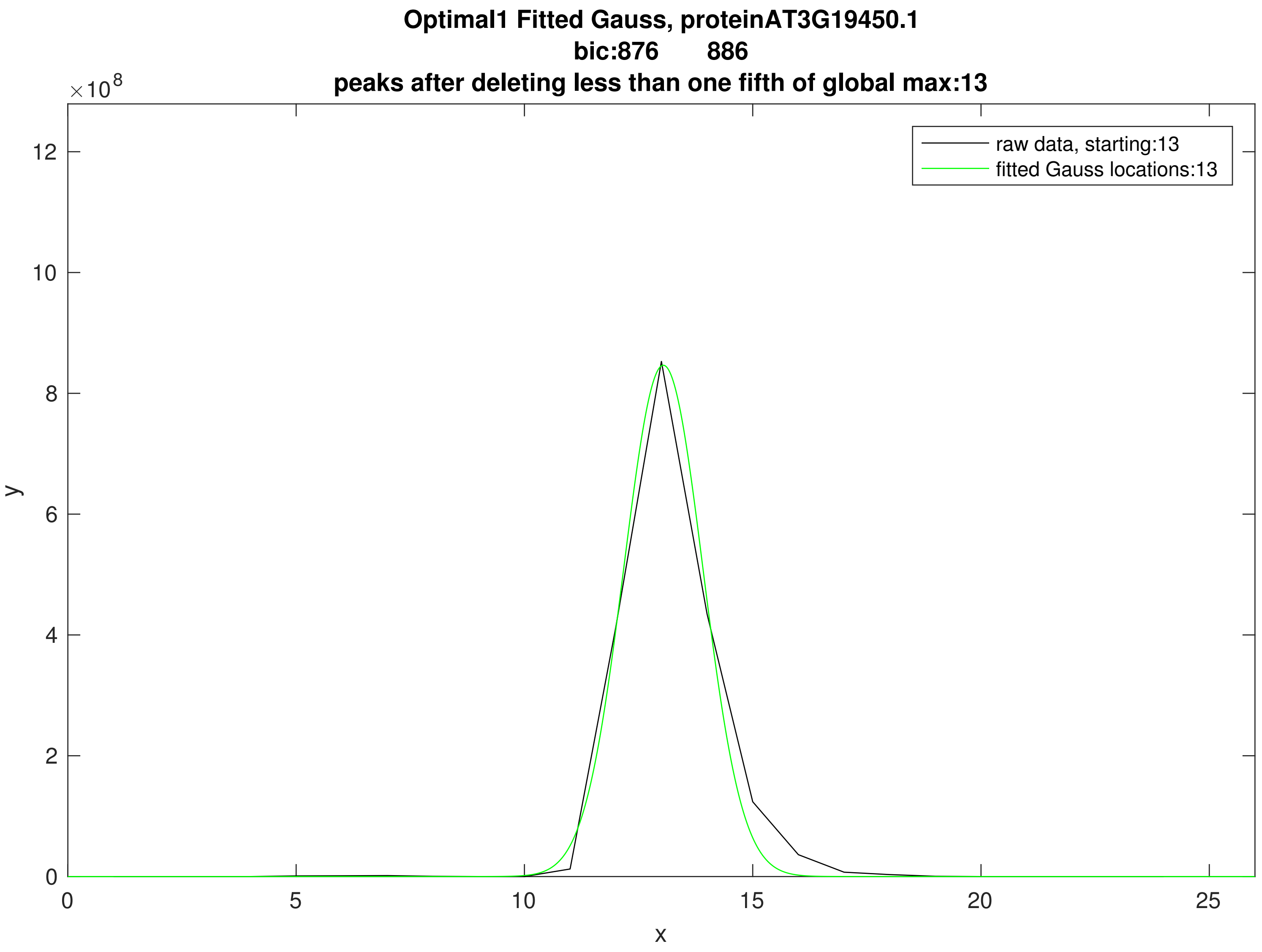
Optimal1 Fitted Gauss, proteinAT3G19450.1
bic:876 886
peaks after deleting less than one fifth of global max:13
×10^8
12
10
8
6
4
2
0
y
0
5
10
15
20
25
x
raw data, starting:13
fitted Gauss locations:13

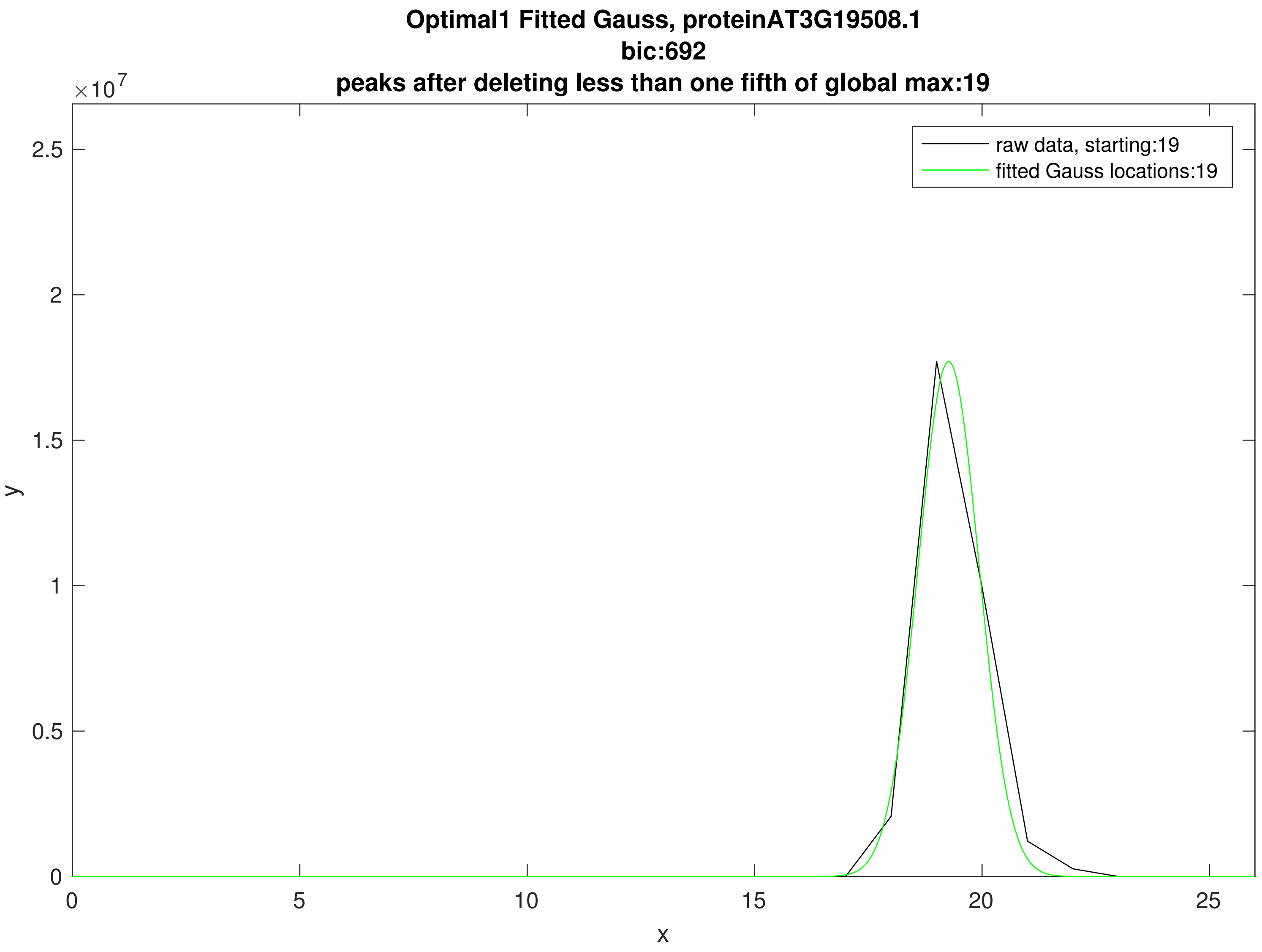
Optimal1 Fitted Gauss, proteinAT3G19508.1
bic:692
peaks after deleting less than one fifth of global max:19
×10^7
raw data, starting:19
fitted Gauss locations:19
2.5
2
1.5
1
0.5
0
y
0
5
10
15
20
25
x

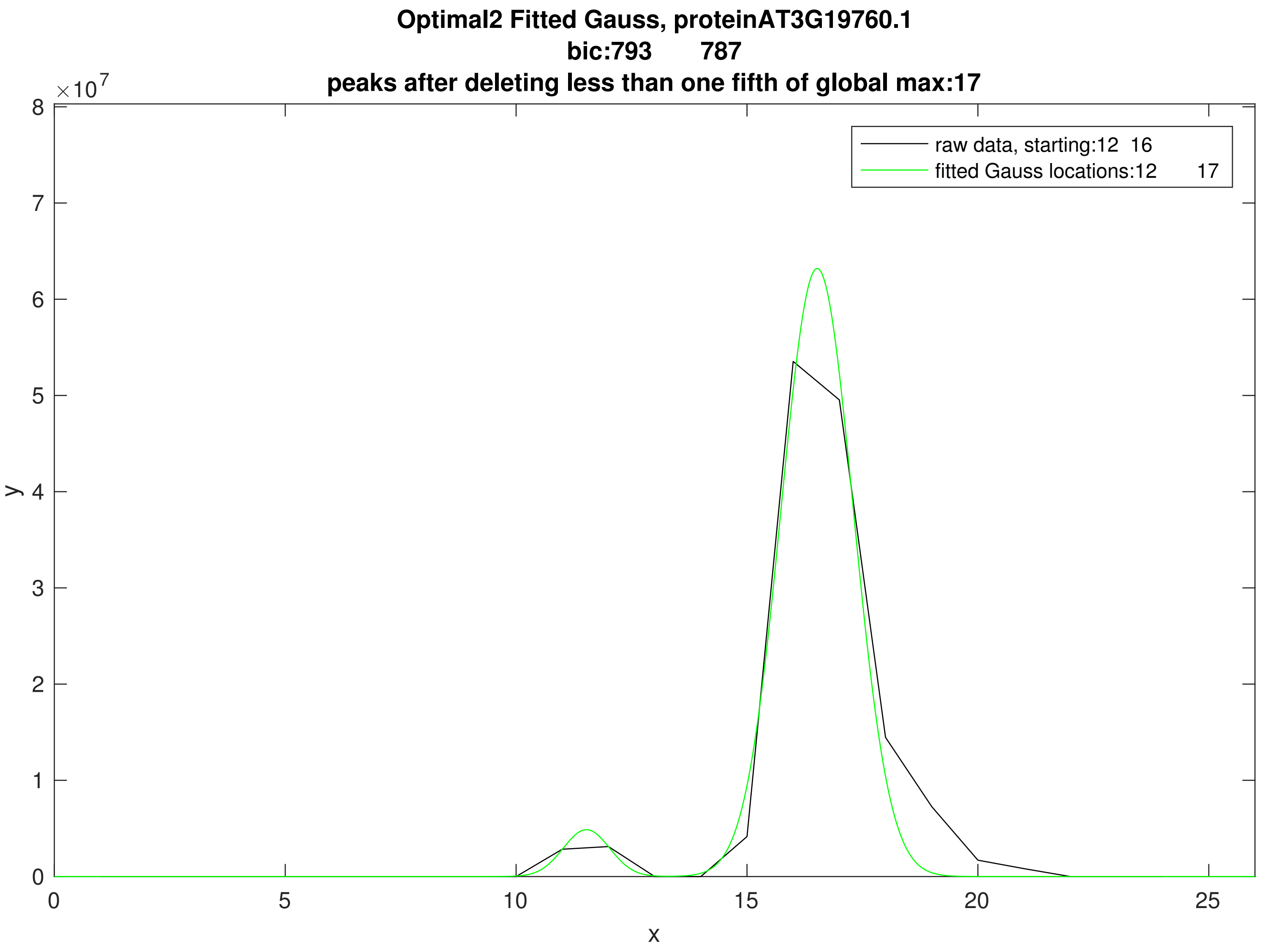
Optimal2 Fitted Gauss, proteinAT3G19760.1
bic:793 787
peaks after deleting less than one fifth of global max:17
×10^7
raw data, starting:12 16
fitted Gauss locations:12 17
y
x
0
1
2
3
4
5
6
7
8
0
5
10
15
20
25

**Optimal2 Fitted Gauss, proteinAT3G19820.3;AT3G19820.2;AT3G19820.1**
**bic:741 711**
**peaks after deleting less than one fifth of global max:3.6 14**

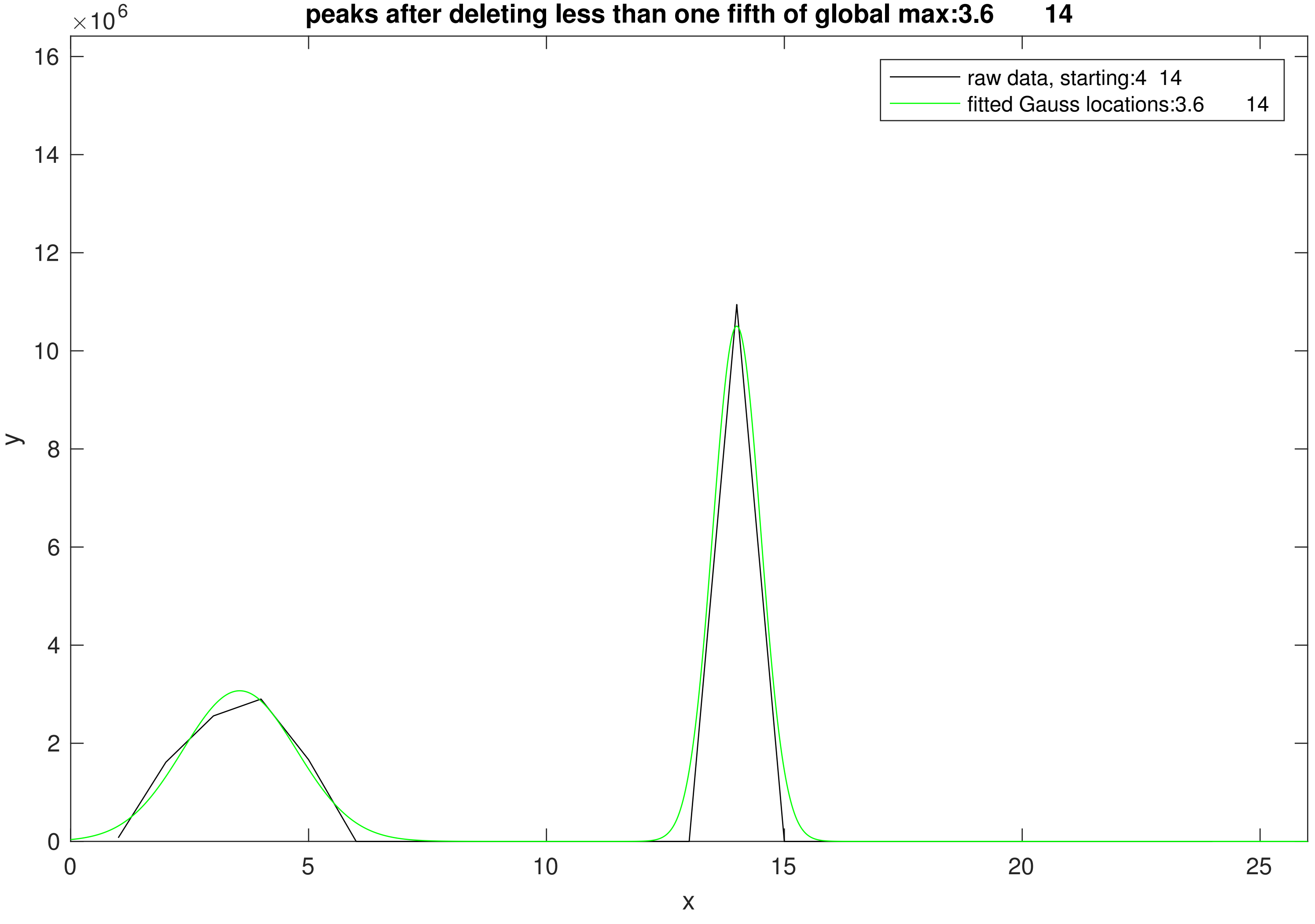

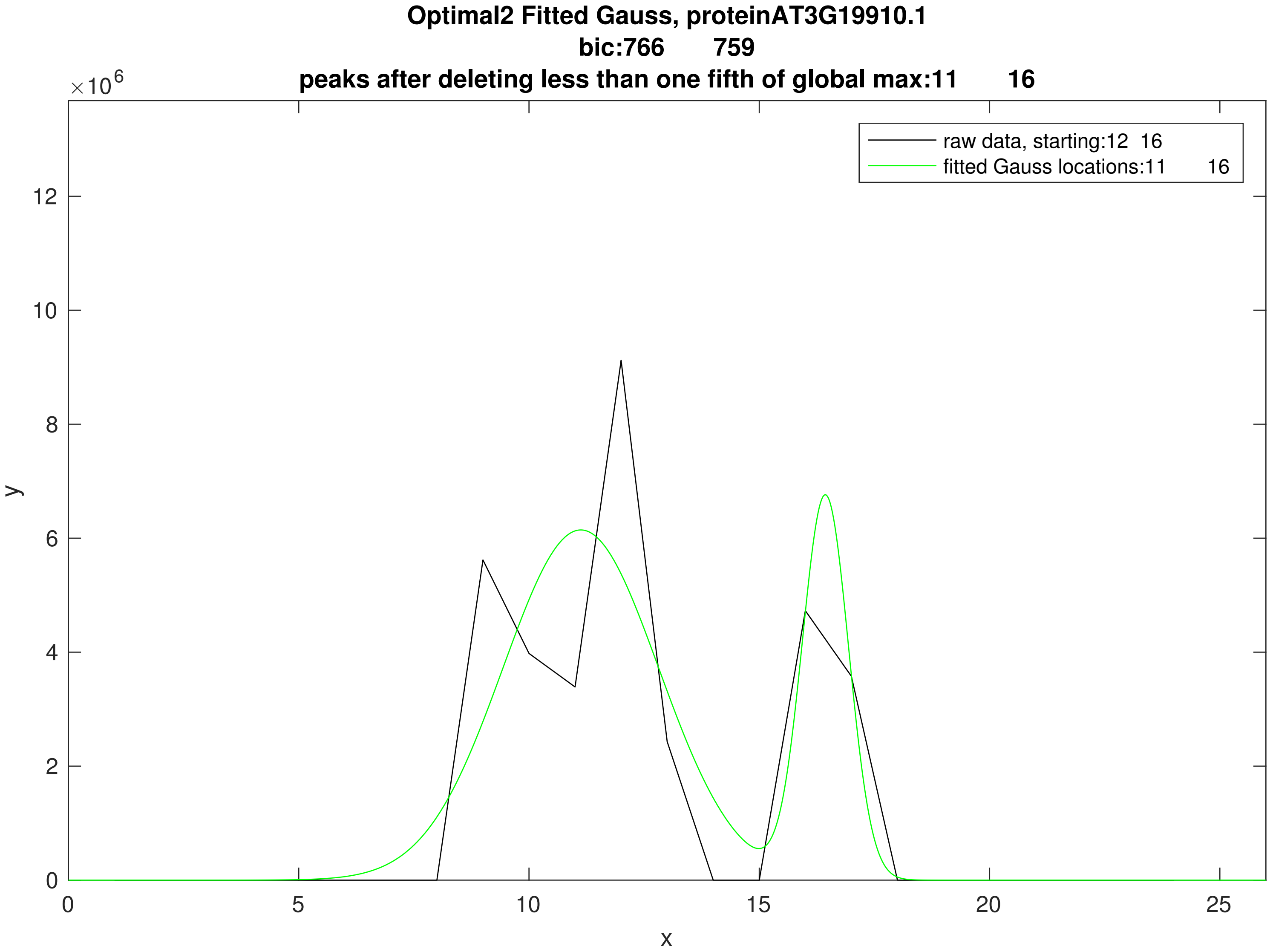
Optimal2 Fitted Gauss, proteinAT3G19910.1
bic:766 759
peaks after deleting less than one fifth of global max:11 16
×10^6
raw data, starting:12 16
fitted Gauss locations:11 16
y
x
0
2
4
6
8
10
12
0
5
10
15
20
25

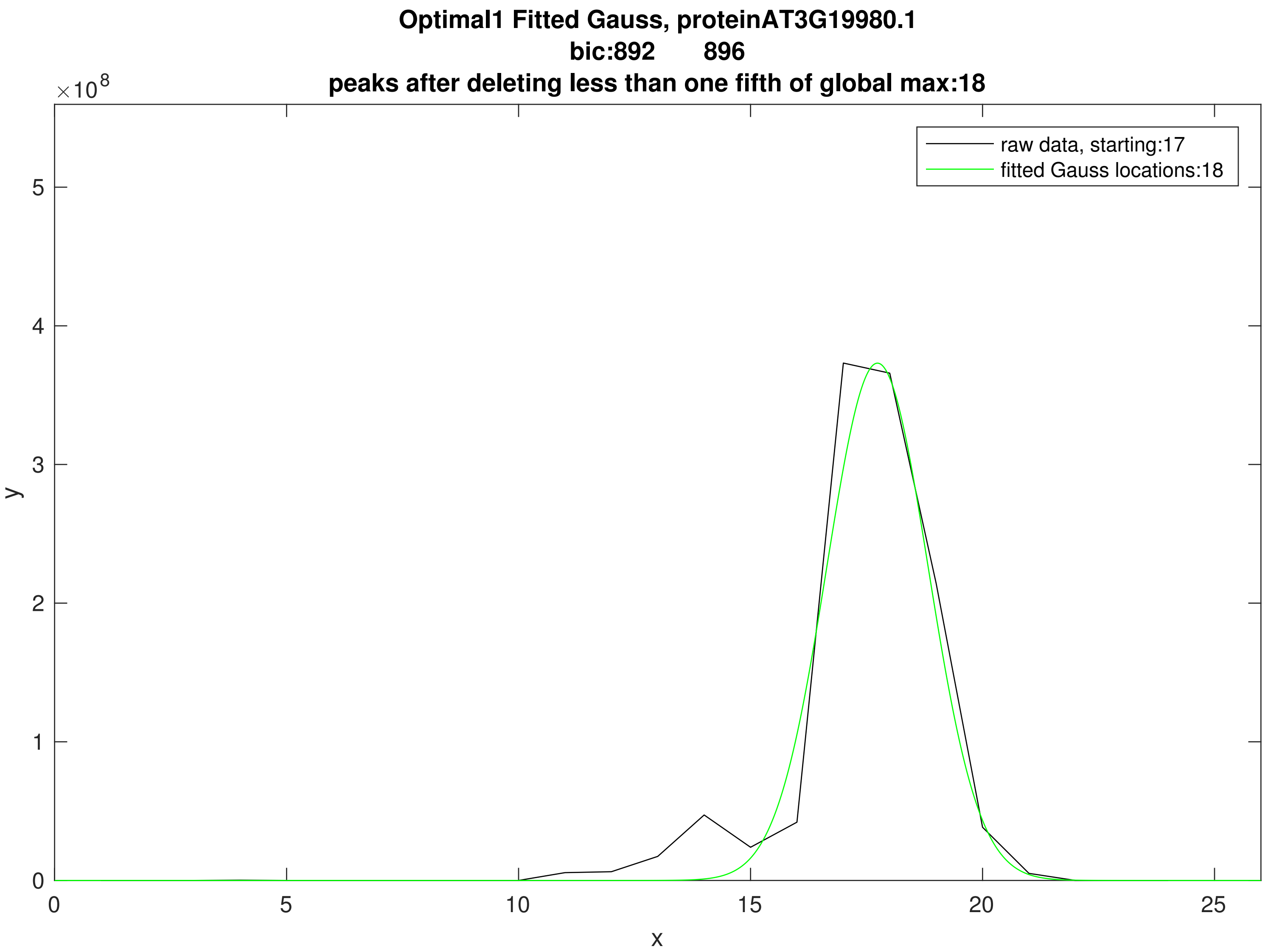

Optimal1 Fitted Gauss, proteinAT3G19980.1
bic:892 896
peaks after deleting less than one fifth of global max:18
×10^8
raw data, starting:17
fitted Gauss locations:18
5
4
3
2
1
0
y
0
5
10
15
20
25
x

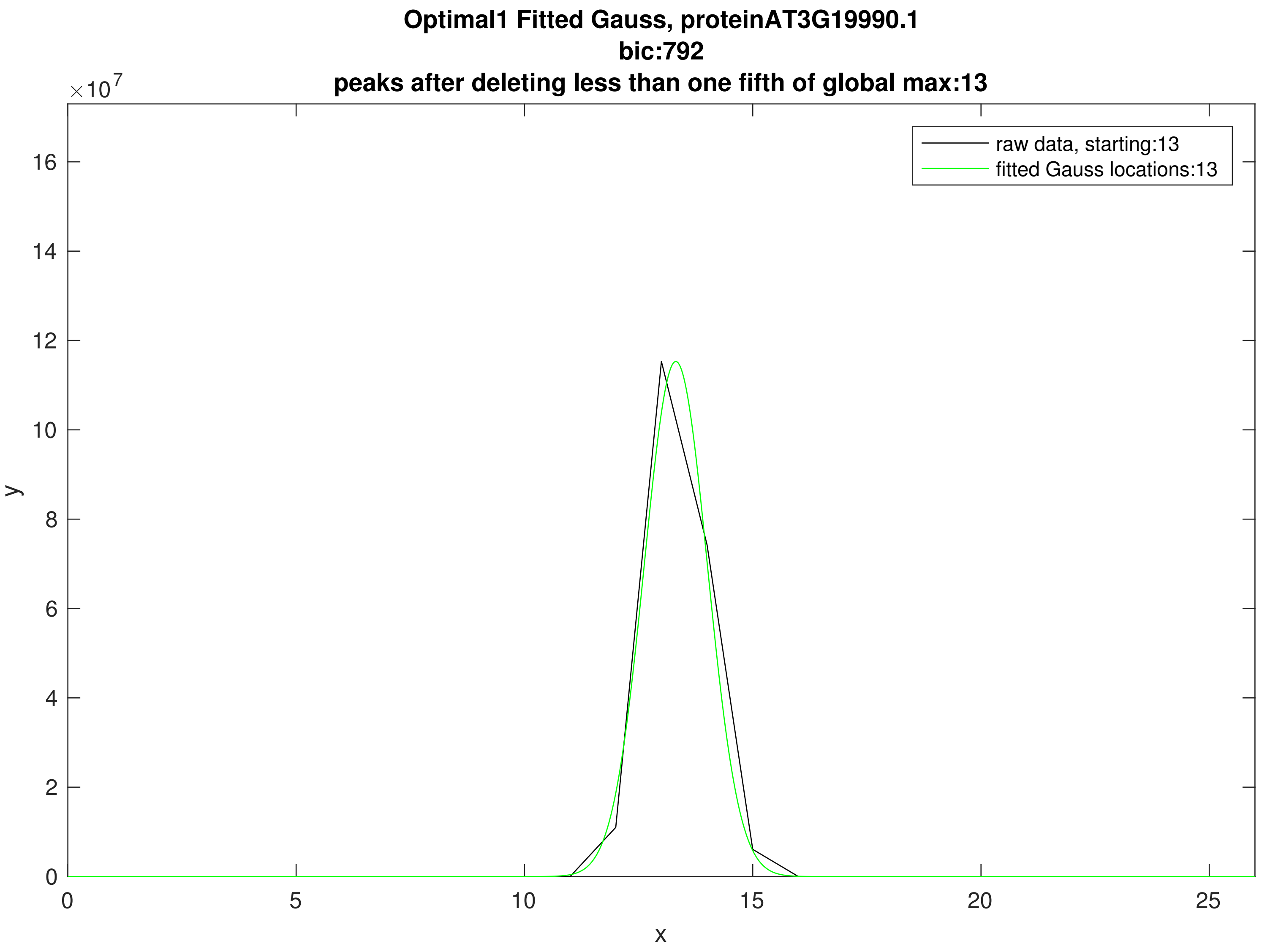

Optimal1 Fitted Gauss, proteinAT3G19990.1
bic:792
peaks after deleting less than one fifth of global max:13
×10^7
16
14
12
10
8
6
4
2
0
y
0
5
10
15
20
25
x
raw data, starting:13
fitted Gauss locations:13

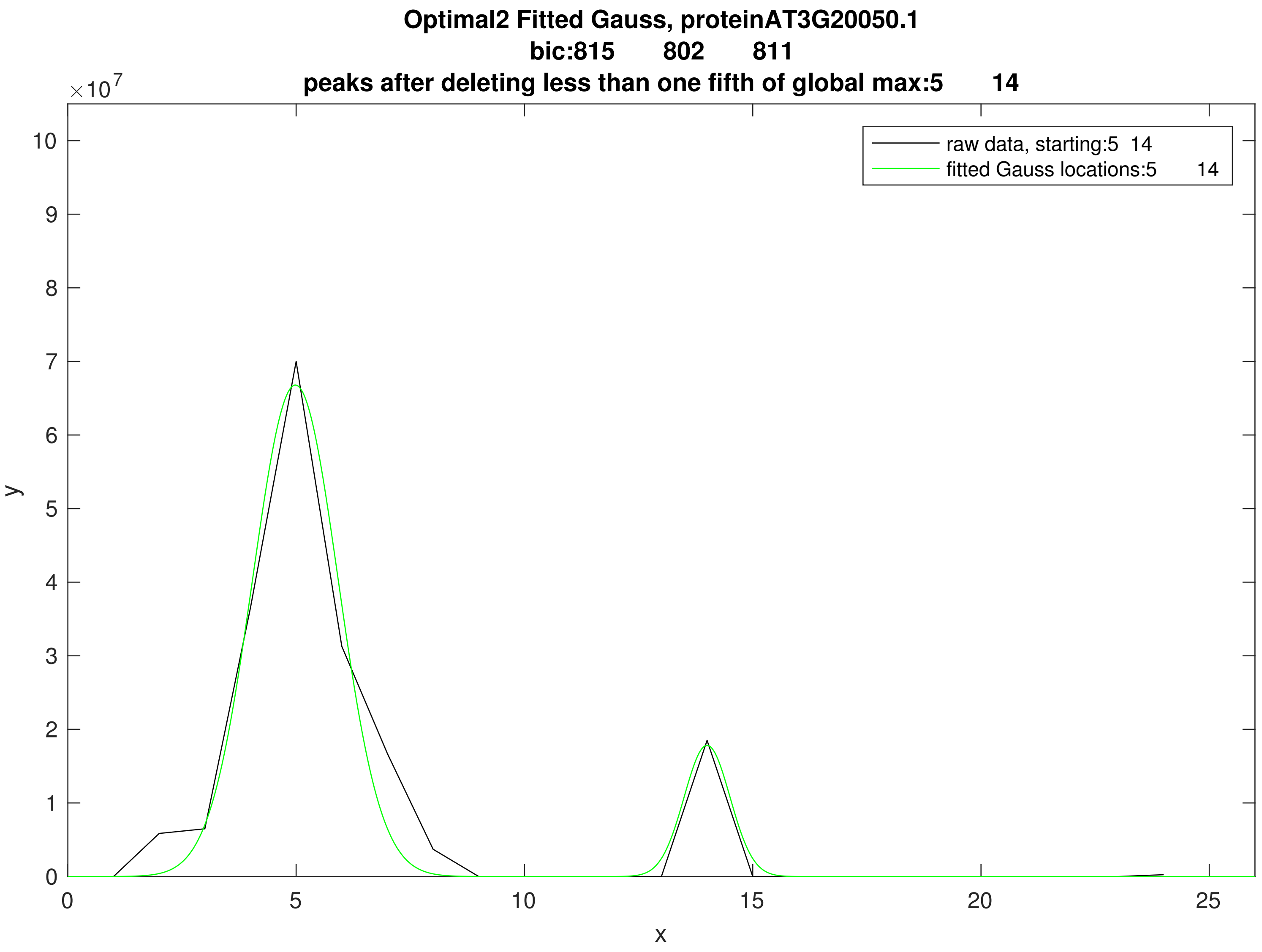

Optimal2 Fitted Gauss, proteinAT3G20050.1
bic:815 802 811
peaks after deleting less than one fifth of global max:5 14
×10^7
raw data, starting:5 14
fitted Gauss locations:5 14
y
x
0
1
2
3
4
5
6
7
8
9
10
0
5
10
15
20
25

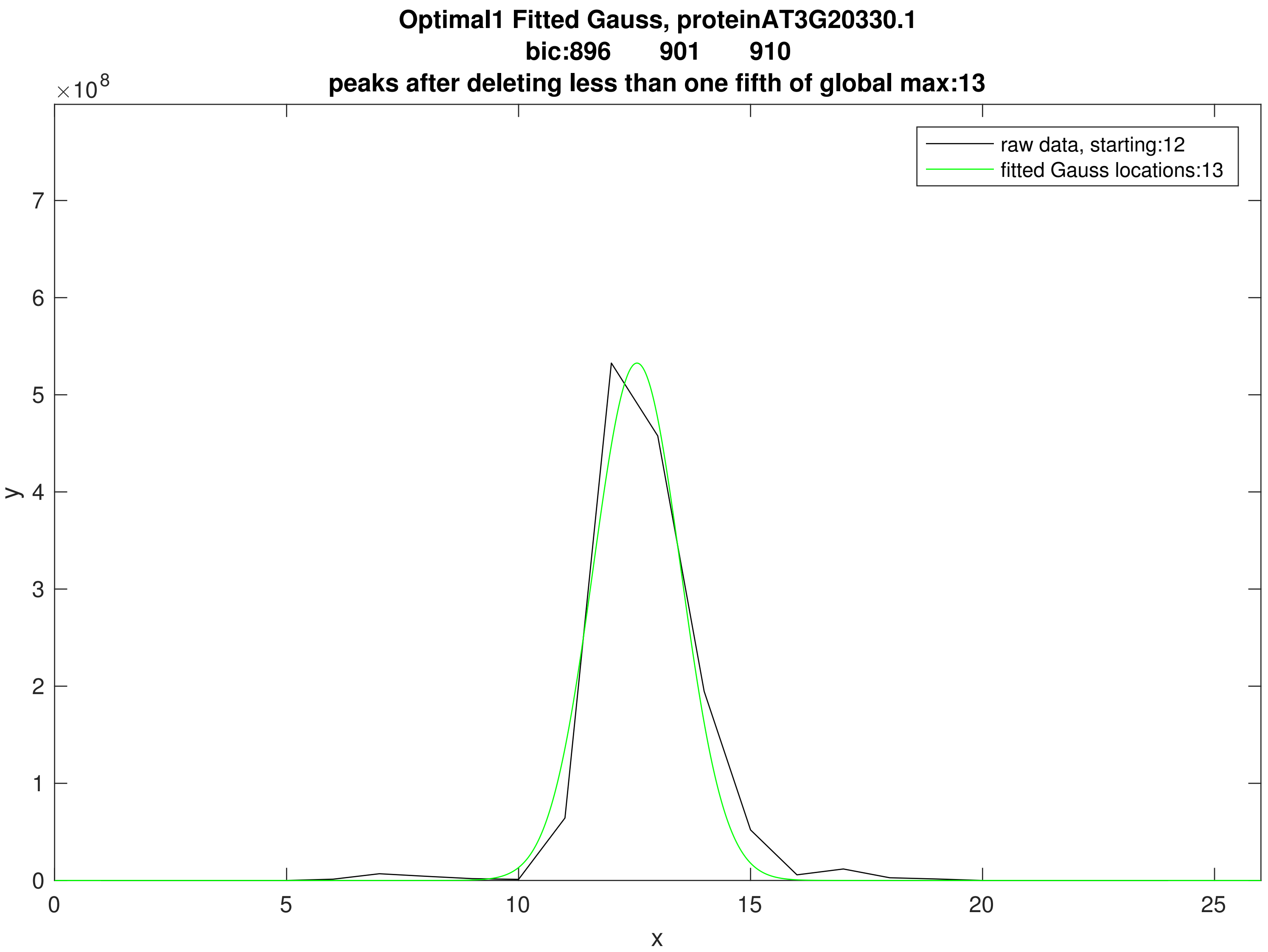
Optimal1 Fitted Gauss, proteinAT3G20330.1
bic:896 901 910
peaks after deleting less than one fifth of global max:13
×10^8
raw data, starting:12
fitted Gauss locations:13
y
x
0
1
2
3
4
5
6
7
0
5
10
15
20
25

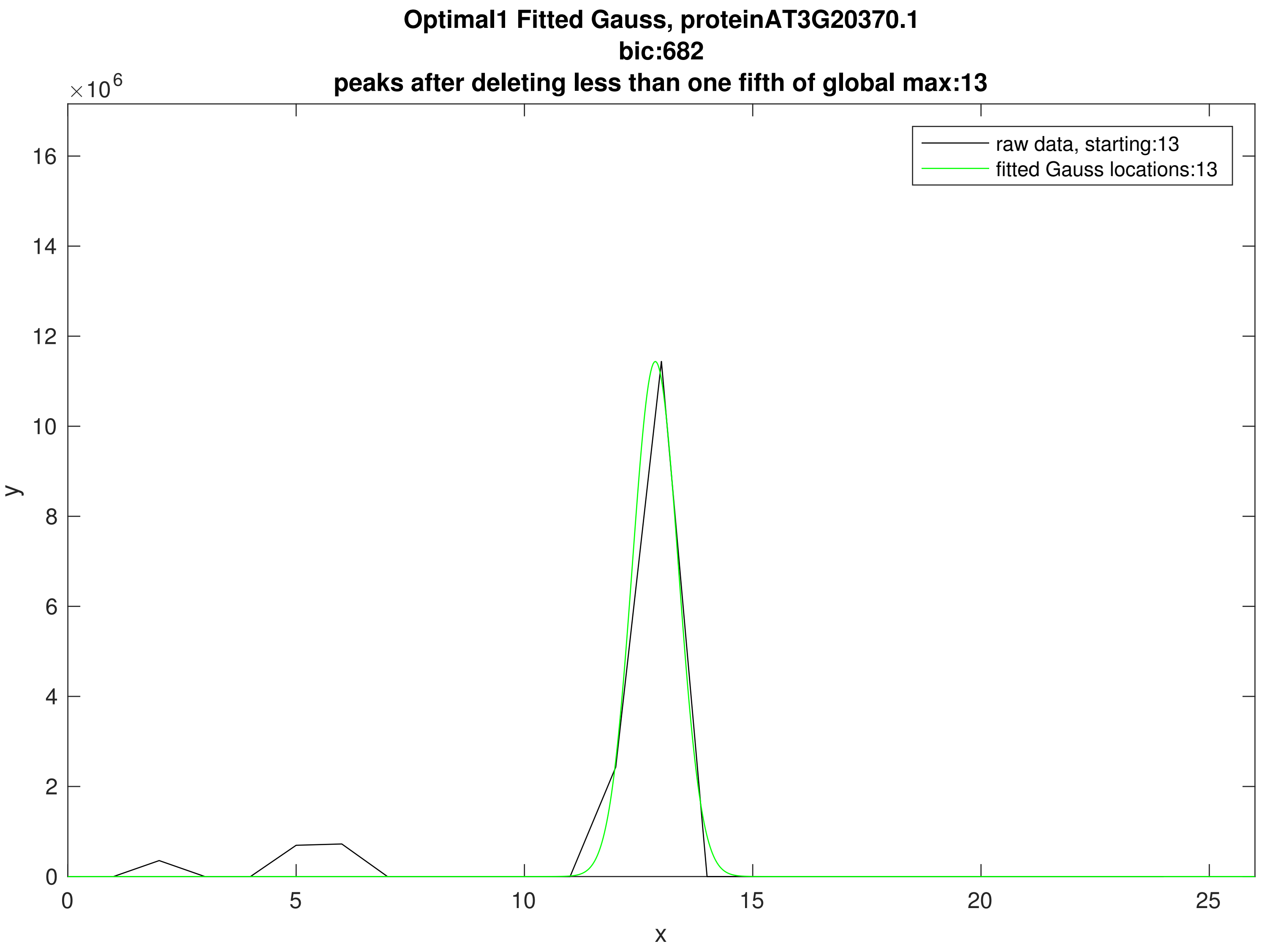

Optimal1 Fitted Gauss, proteinAT3G20370.1
bic:682
peaks after deleting less than one fifth of global max:13
×10^6
raw data, starting:13
fitted Gauss locations:13
16
14
12
10
8
6
4
2
0
y
0
5
10
15
20
25
x

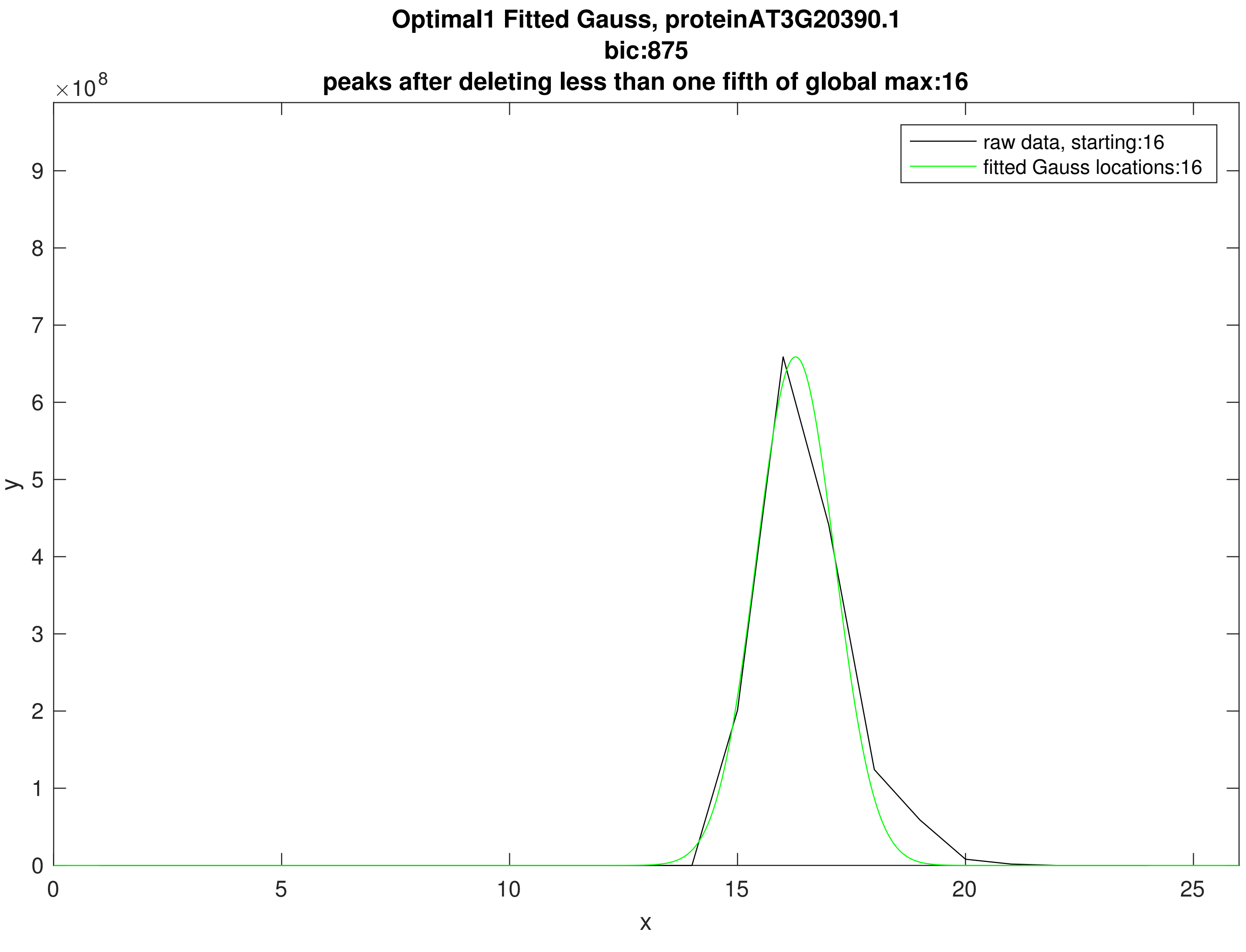
Optimal1 Fitted Gauss, proteinAT3G20390.1
bic:875
peaks after deleting less than one fifth of global max:16
×10^8
raw data, starting:16
fitted Gauss locations:16
y
x
0
1
2
3
4
5
6
7
8
9
0
5
10
15
20
25

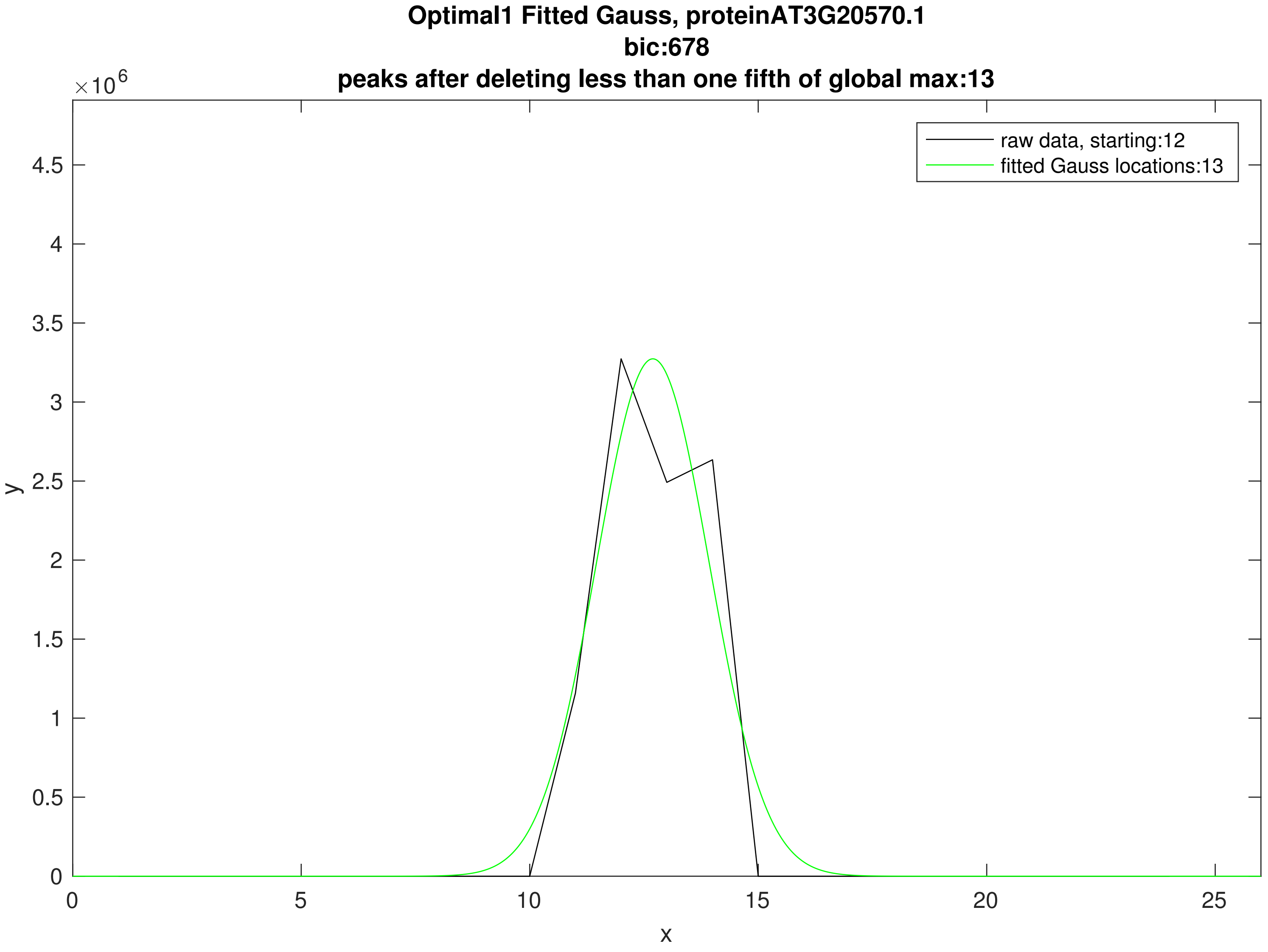

Optimal1 Fitted Gauss, proteinAT3G20570.1
bic:678
peaks after deleting less than one fifth of global max:13
×10^6
raw data, starting:12
fitted Gauss locations:13
4.5
4
3.5
3
2.5
2
1.5
1
0.5
0
y
0
5
10
15
20
25
x

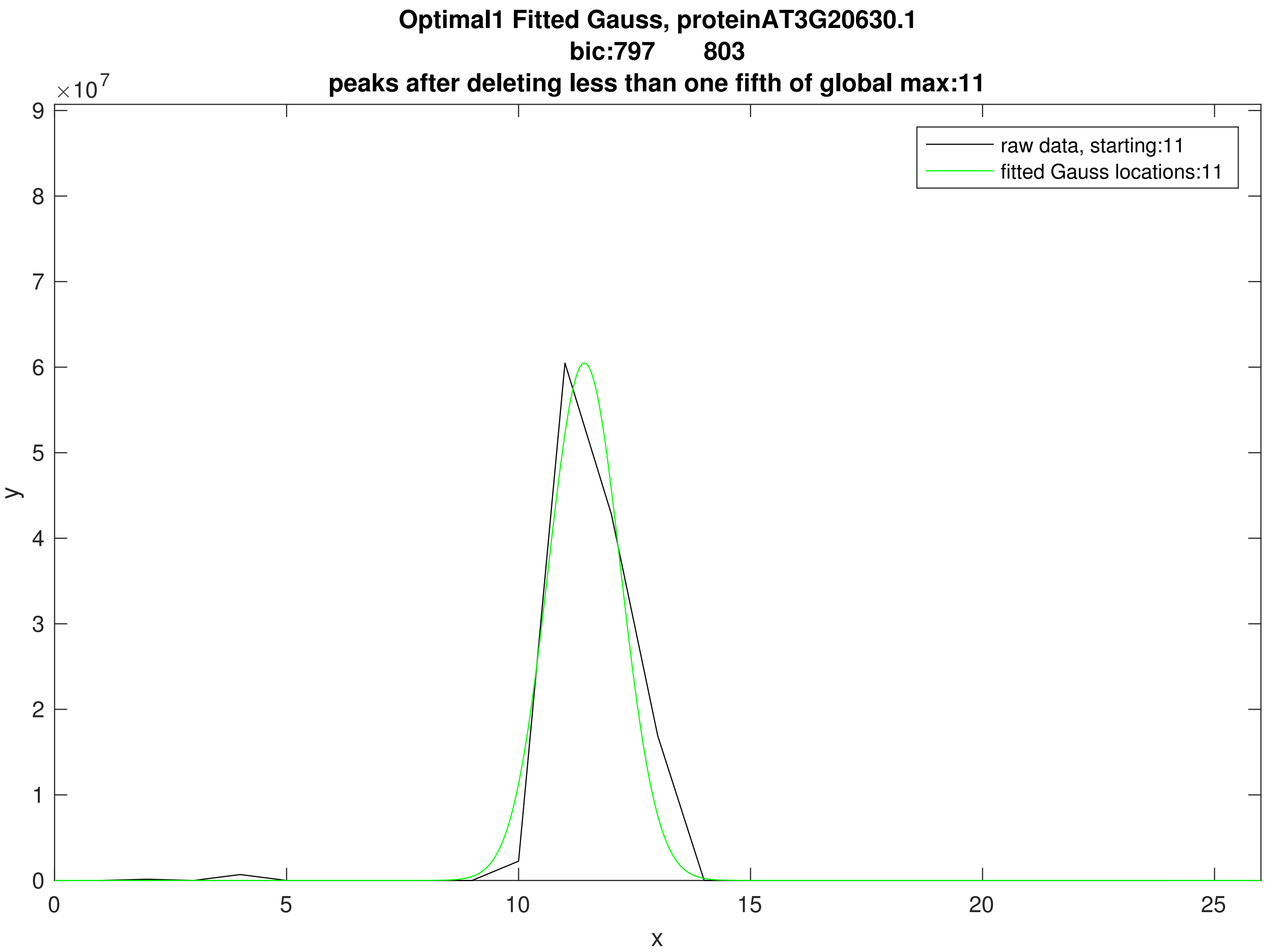
Optimal1 Fitted Gauss, proteinAT3G20630.1
bic:797 803
peaks after deleting less than one fifth of global max:11
×10^7
raw data, starting:11
fitted Gauss locations:11
y
x
0
1
2
3
4
5
6
7
8
9
0
5
10
15
20
25

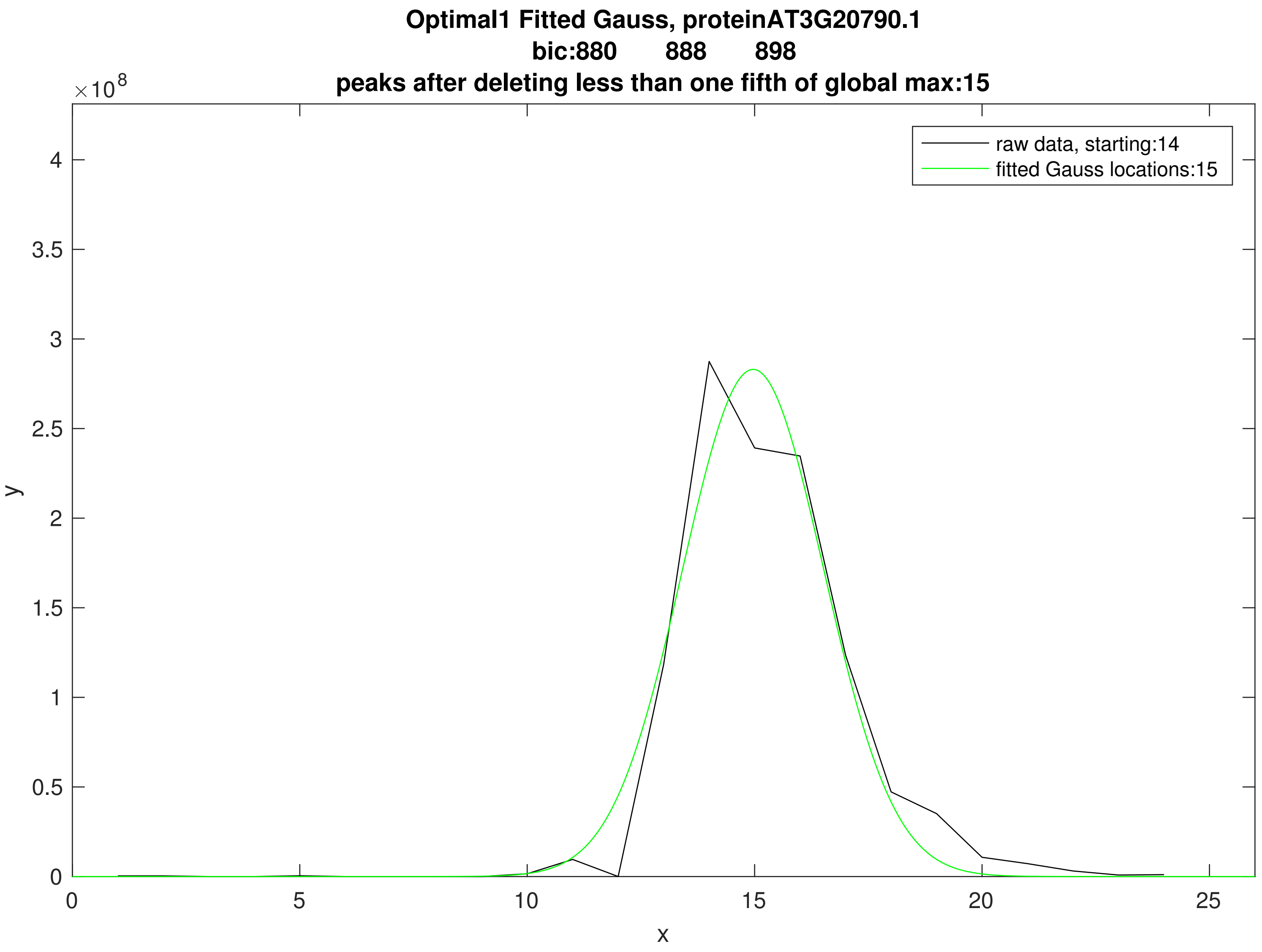
Optimal1 Fitted Gauss, proteinAT3G20790.1
bic:880 888 898
peaks after deleting less than one fifth of global max:15
×10^8
raw data, starting:14
fitted Gauss locations:15
y
x
0
0.5
1
1.5
2
2.5
3
3.5
4
0
5
10
15
20
25

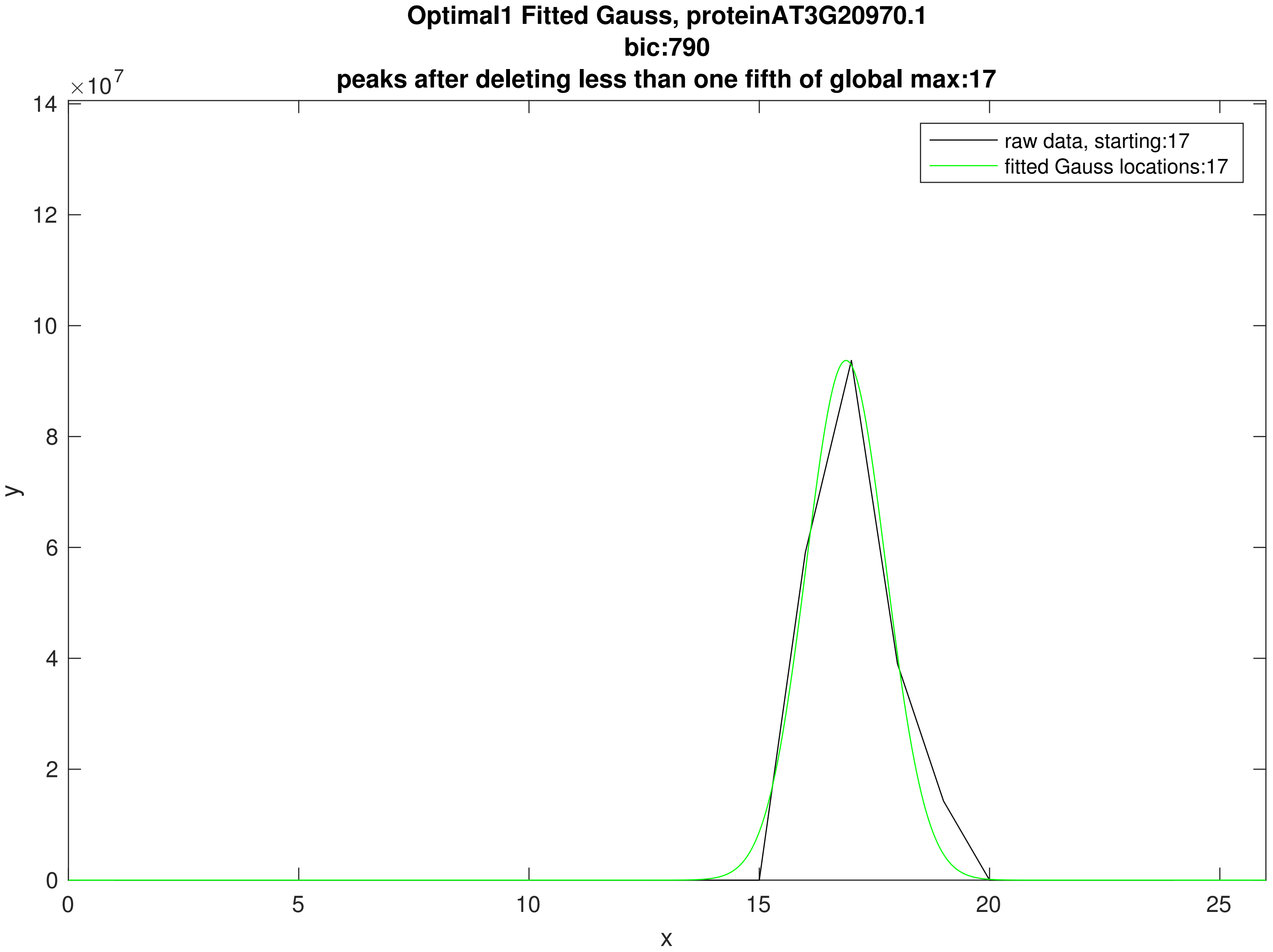

Optimal1 Fitted Gauss, proteinAT3G20970.1
bic:790
peaks after deleting less than one fifth of global max:17
×10^7
raw data, starting:17
fitted Gauss locations:17
y
x
0
2
4
6
8
10
12
14
5
15
20
25

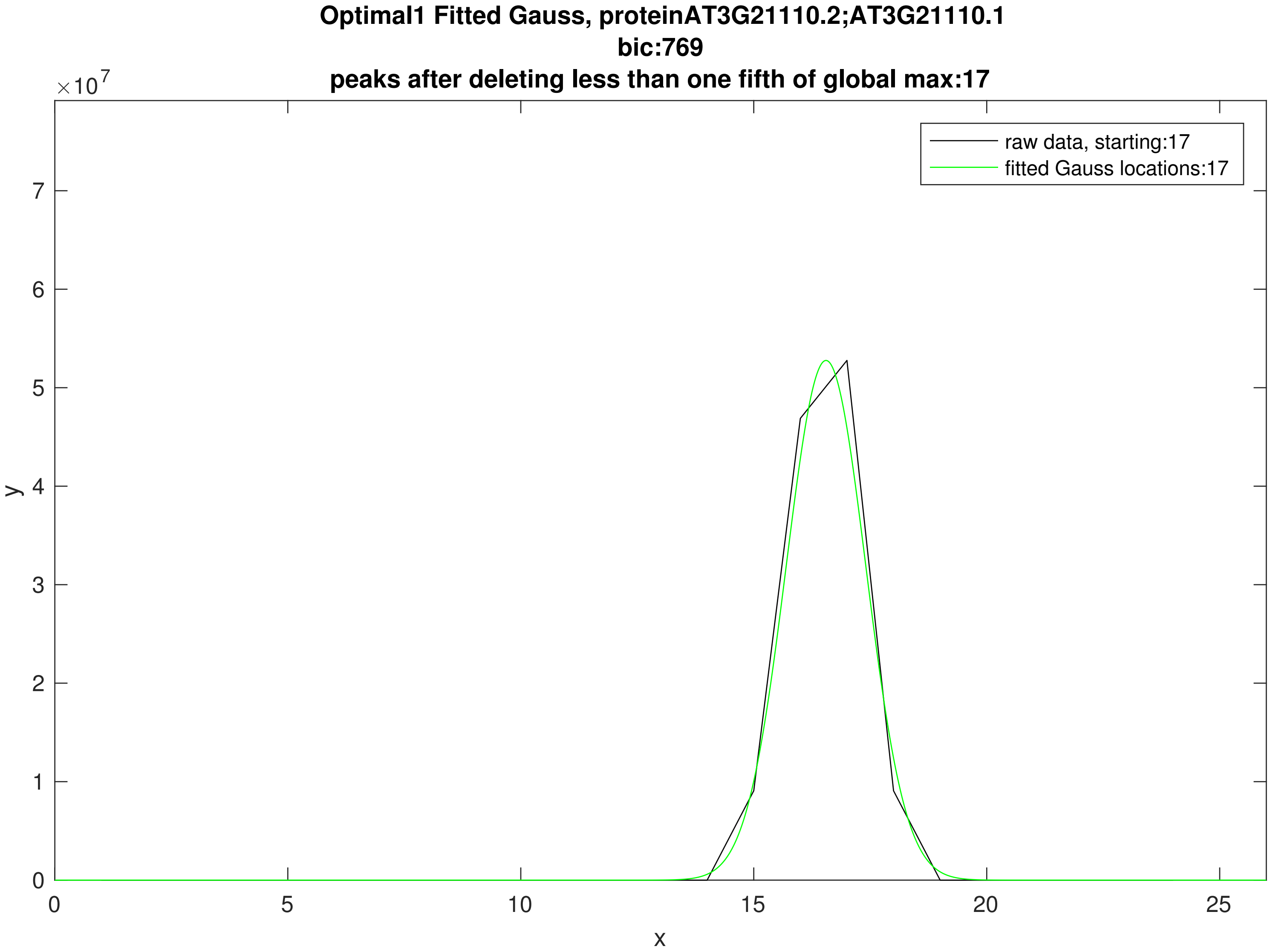
Optimal1 Fitted Gauss, proteinAT3G21110.2;AT3G21110.1
bic:769
peaks after deleting less than one fifth of global max:17
×10^7
raw data, starting:17
fitted Gauss locations:17
y
x
0
1
2
3
4
5
6
7
0
5
10
15
20
25

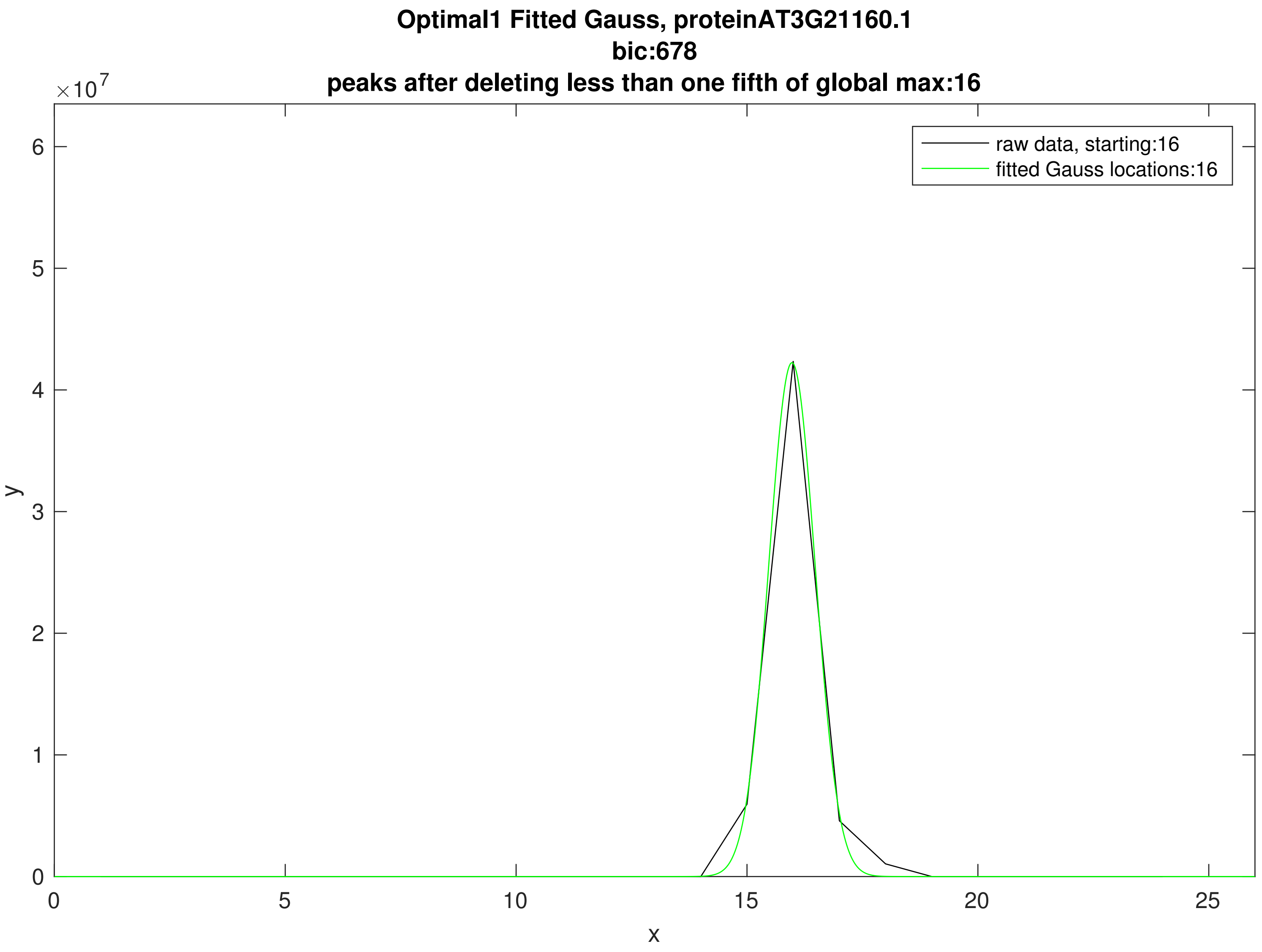

Optimal1 Fitted Gauss, proteinAT3G21160.1
bic:678
peaks after deleting less than one fifth of global max:16
×10^7
6
5
4
3
2
1
0
y
0
5
10
15
20
25
x
raw data, starting:16
fitted Gauss locations:16

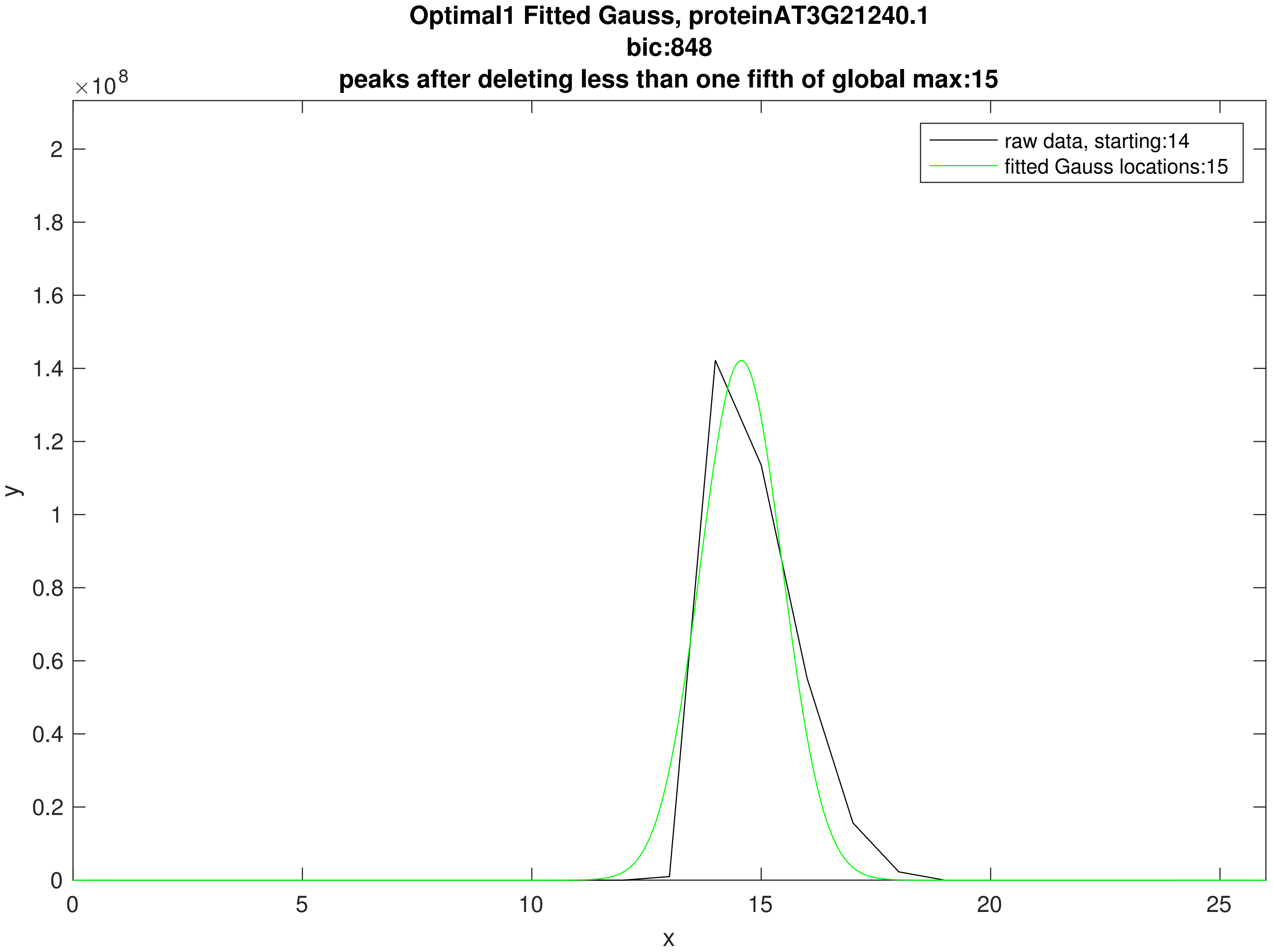
Optimal1 Fitted Gauss, proteinAT3G21240.1
bic:848
peaks after deleting less than one fifth of global max:15
×10^8
raw data, starting:14
fitted Gauss locations:15
y
x
0
0.2
0.4
0.6
0.8
1
1.2
1.4
1.6
1.8
2
0
5
10
15
20
25

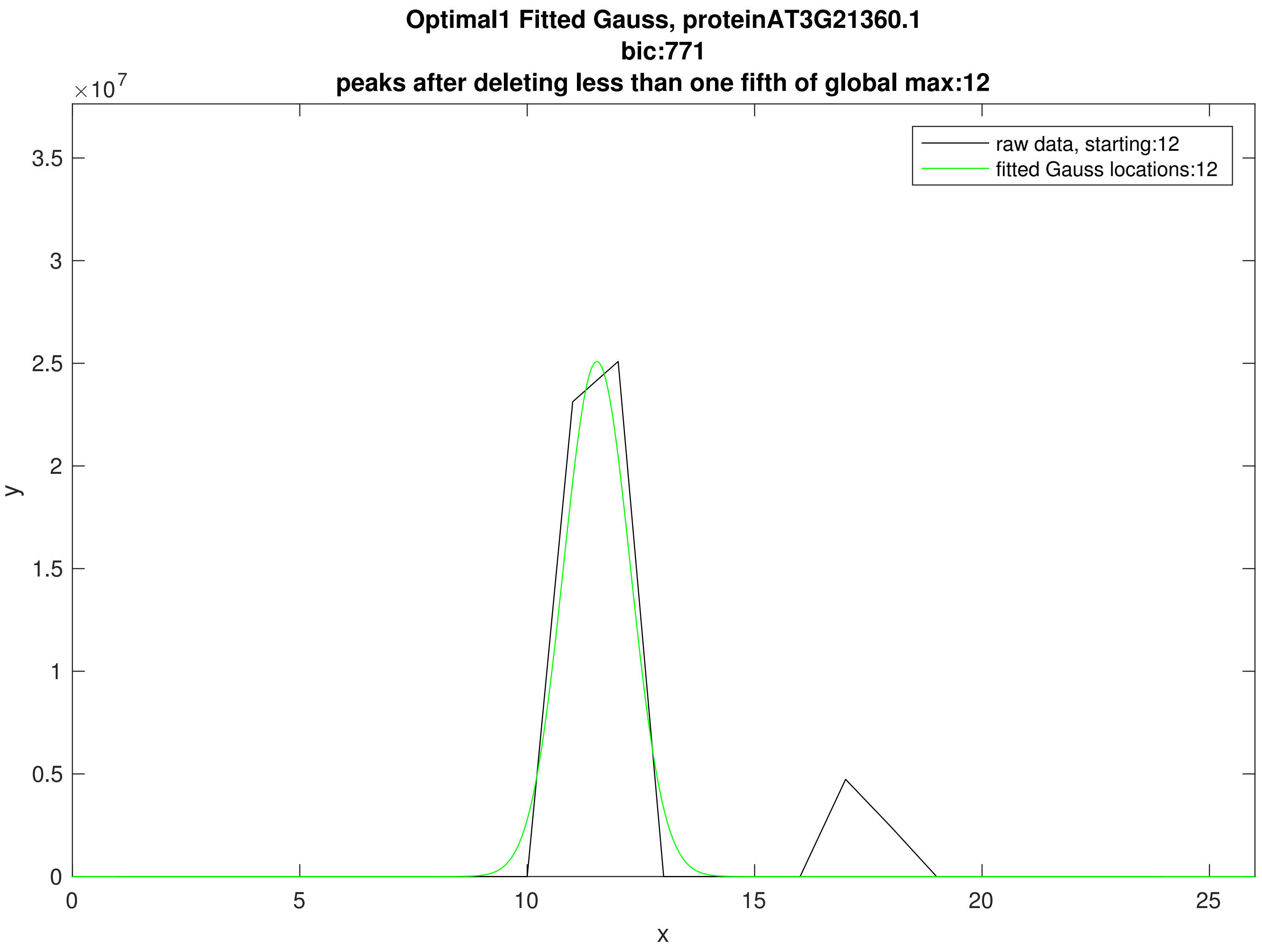

Optimal1 Fitted Gauss, proteinAT3G21360.1
bic:771
peaks after deleting less than one fifth of global max:12
×10^7
3.5
3
2.5
2
1.5
1
0.5
0
y
0
5
10
15
20
25
x
raw data, starting:12
fitted Gauss locations:12

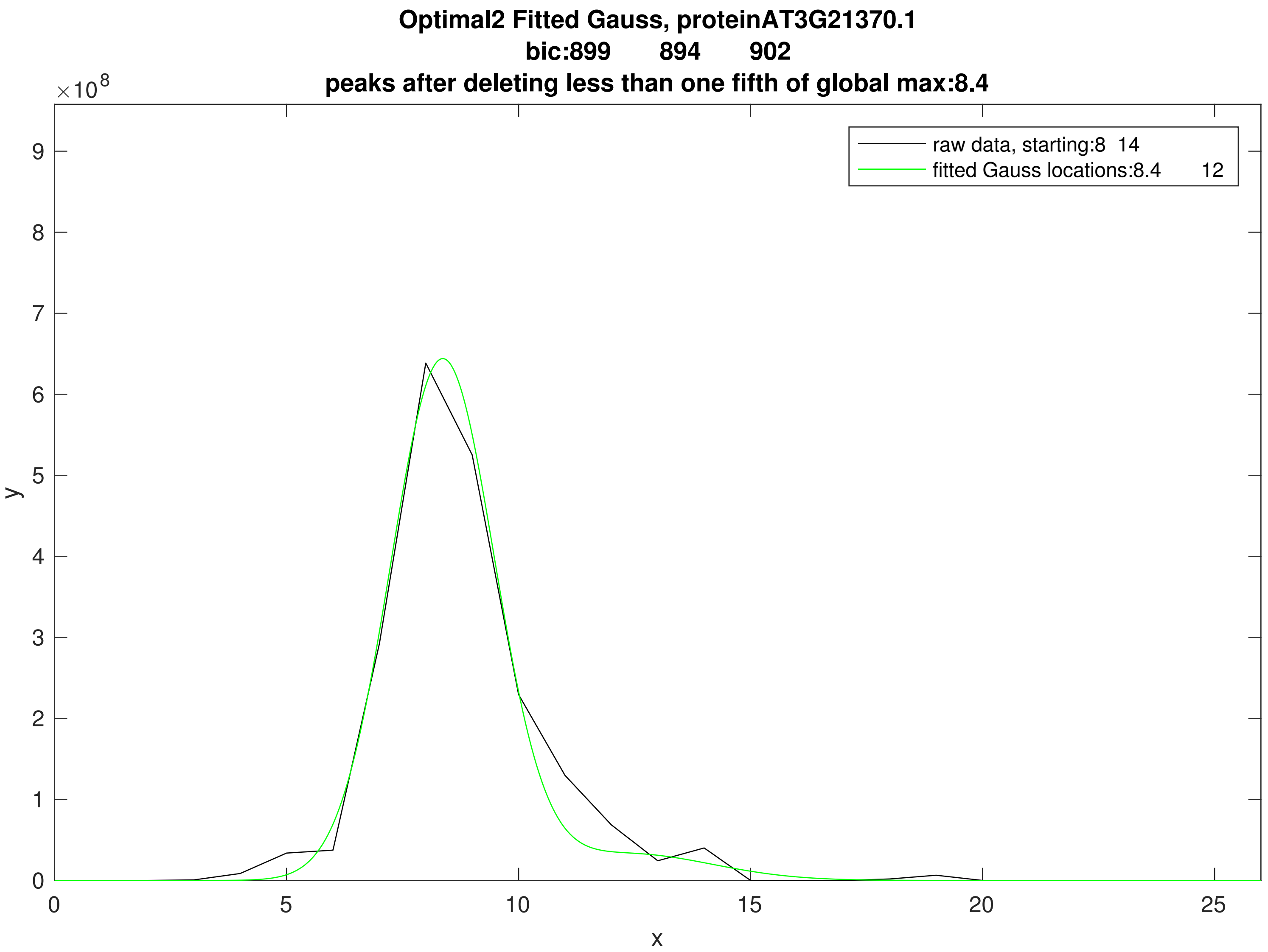
Optimal2 Fitted Gauss, proteinAT3G21370.1
bic:899 894 902
peaks after deleting less than one fifth of global max:8.4
×10^8
raw data, starting:8 14
fitted Gauss locations:8.4 12
y
x
0
1
2
3
4
5
6
7
8
9
0
5
10
15
20
25

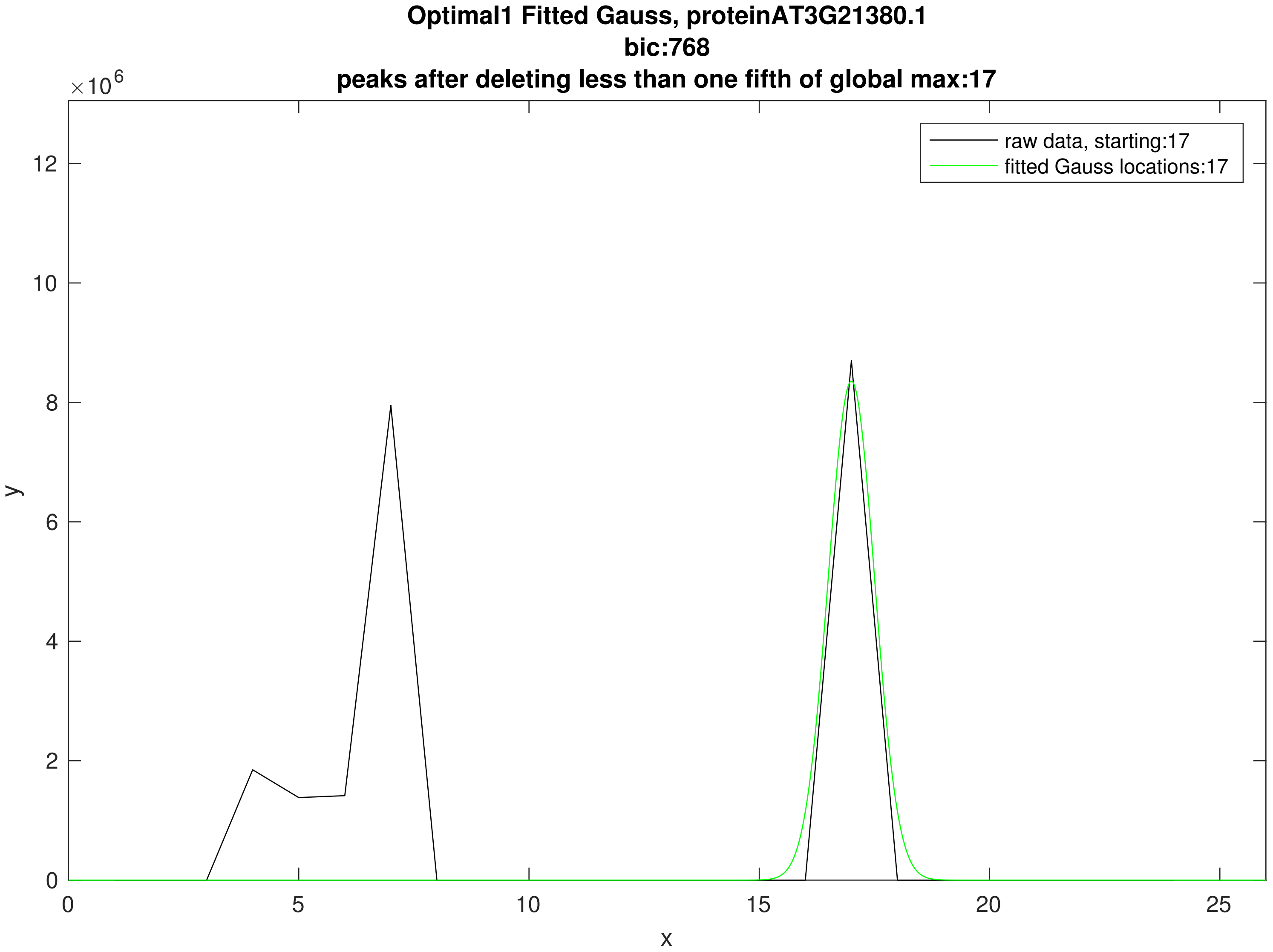
Optimal1 Fitted Gauss, proteinAT3G21380.1
bic:768
peaks after deleting less than one fifth of global max:17
$\times 10^6$
raw data, starting:17
fitted Gauss locations:17
y
x
0
2
4
6
8
10
12
0
5
10
15
20
25

#### Optimal1 Fitted Gauss, proteinAT3G21630.1
#### bic:722
#### peaks after deleting less than one fifth of global max:17

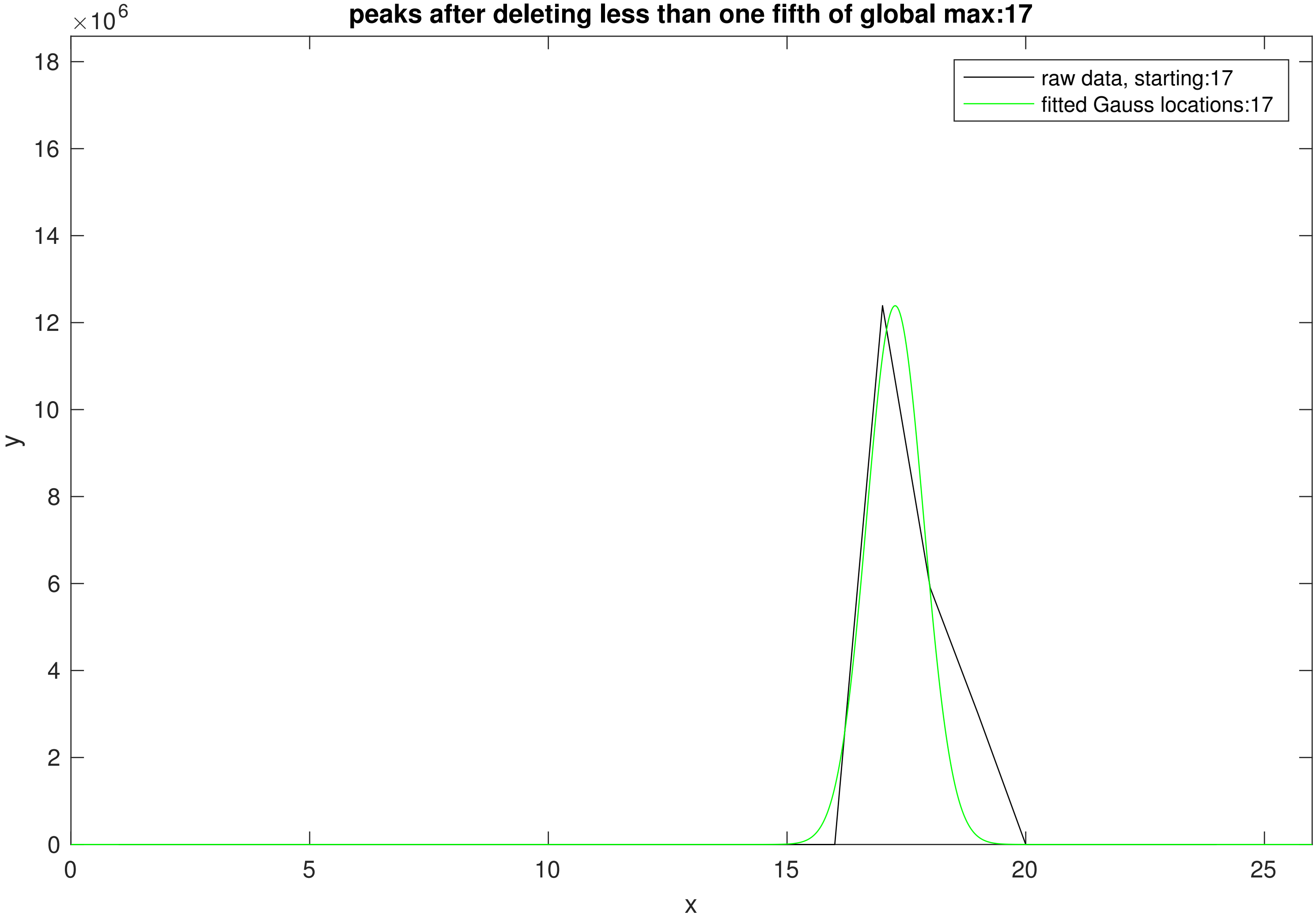

Optimal1 Fitted Gauss, proteinAT3G21720.1
bic:1.07e+03 1.08e+03 1.09e+03
peaks after deleting less than one fifth of global max:9.3

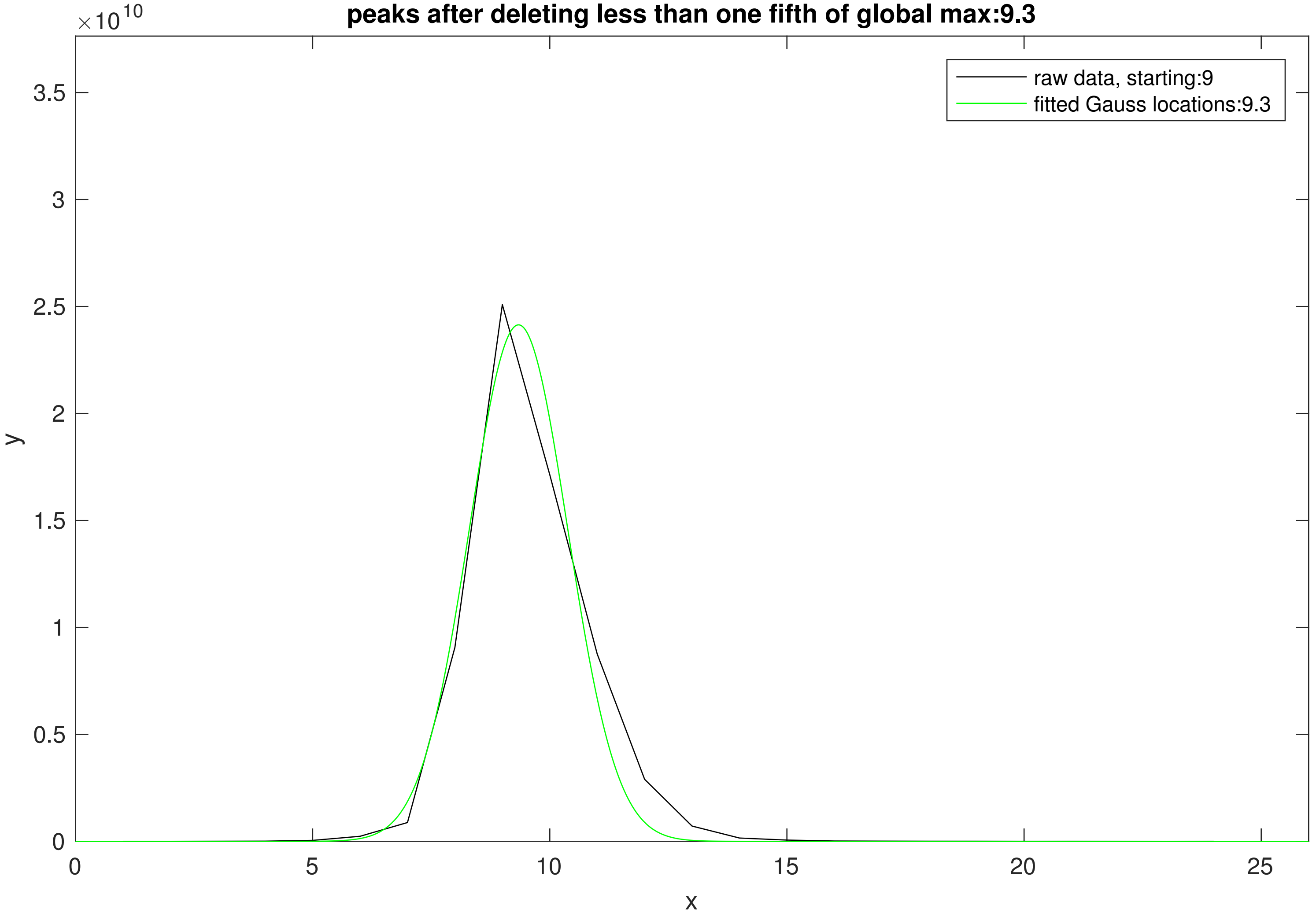

#### Optimal1 Fitted Gauss, proteinAT3G21750.1
#### bic:638
#### peaks after deleting less than one fifth of global max:16

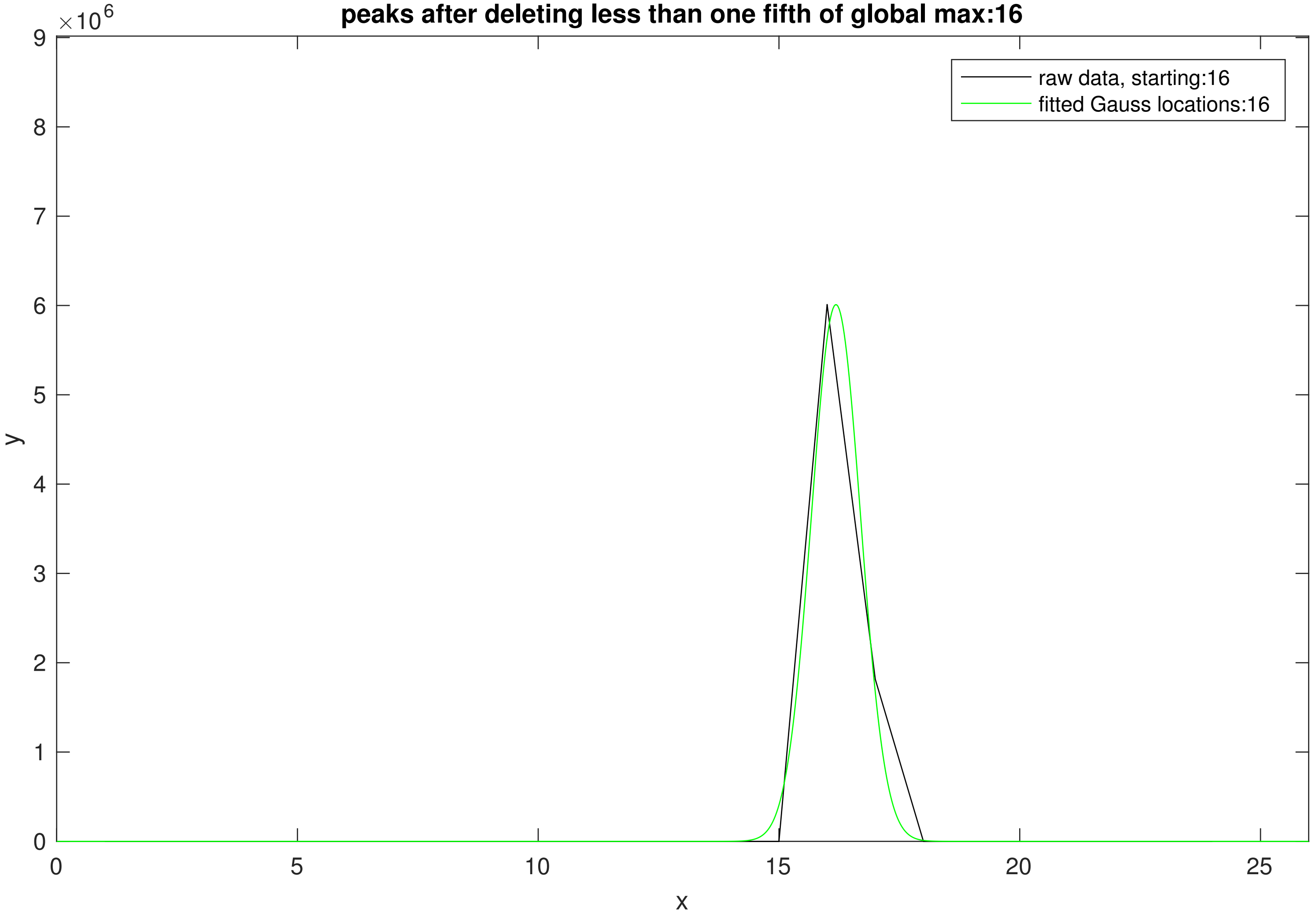

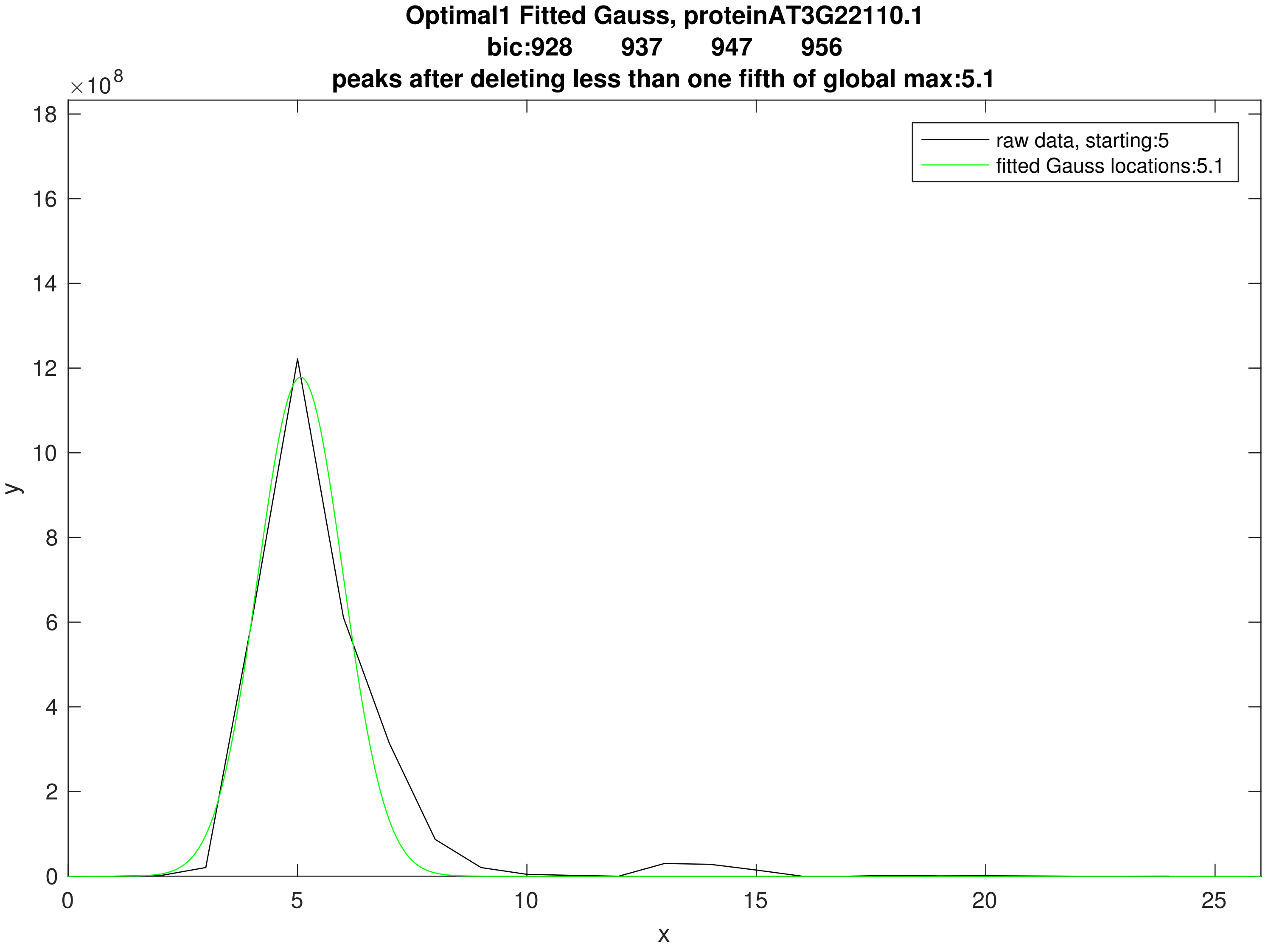

Optimal1 Fitted Gauss, proteinAT3G22110.1
bic:928 937 947 956
peaks after deleting less than one fifth of global max:5.1
×10^8
raw data, starting:5
fitted Gauss locations:5.1
y
x
0
2
4
6
8
10
12
14
16
18
0
5
10
15
20
25

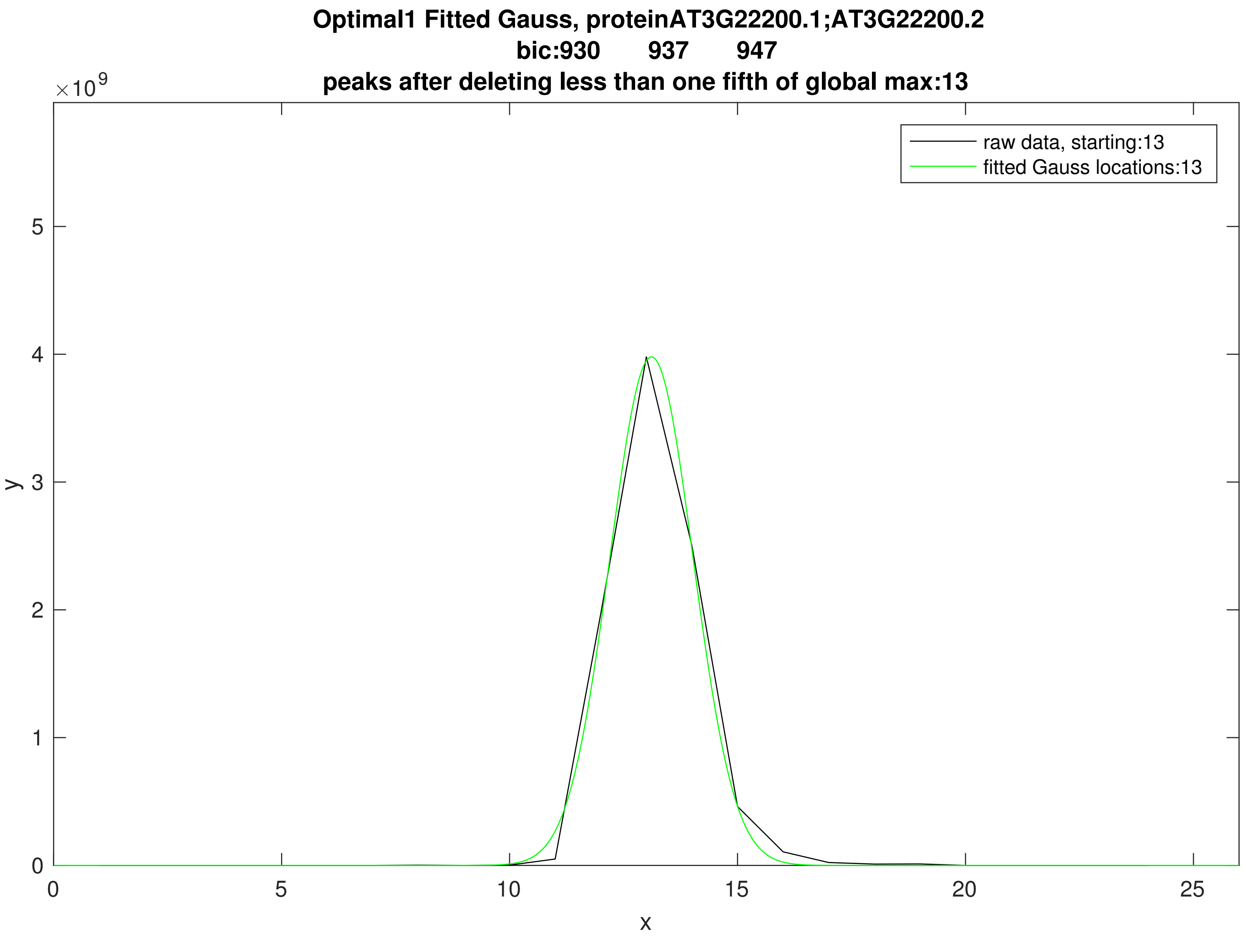

Optimal1 Fitted Gauss, proteinAT3G22200.1;AT3G22200.2
bic:930 937 947
peaks after deleting less than one fifth of global max:13
×10^9
raw data, starting:13
fitted Gauss locations:13
y
x
0
1
2
3
4
5
10
15
20
25

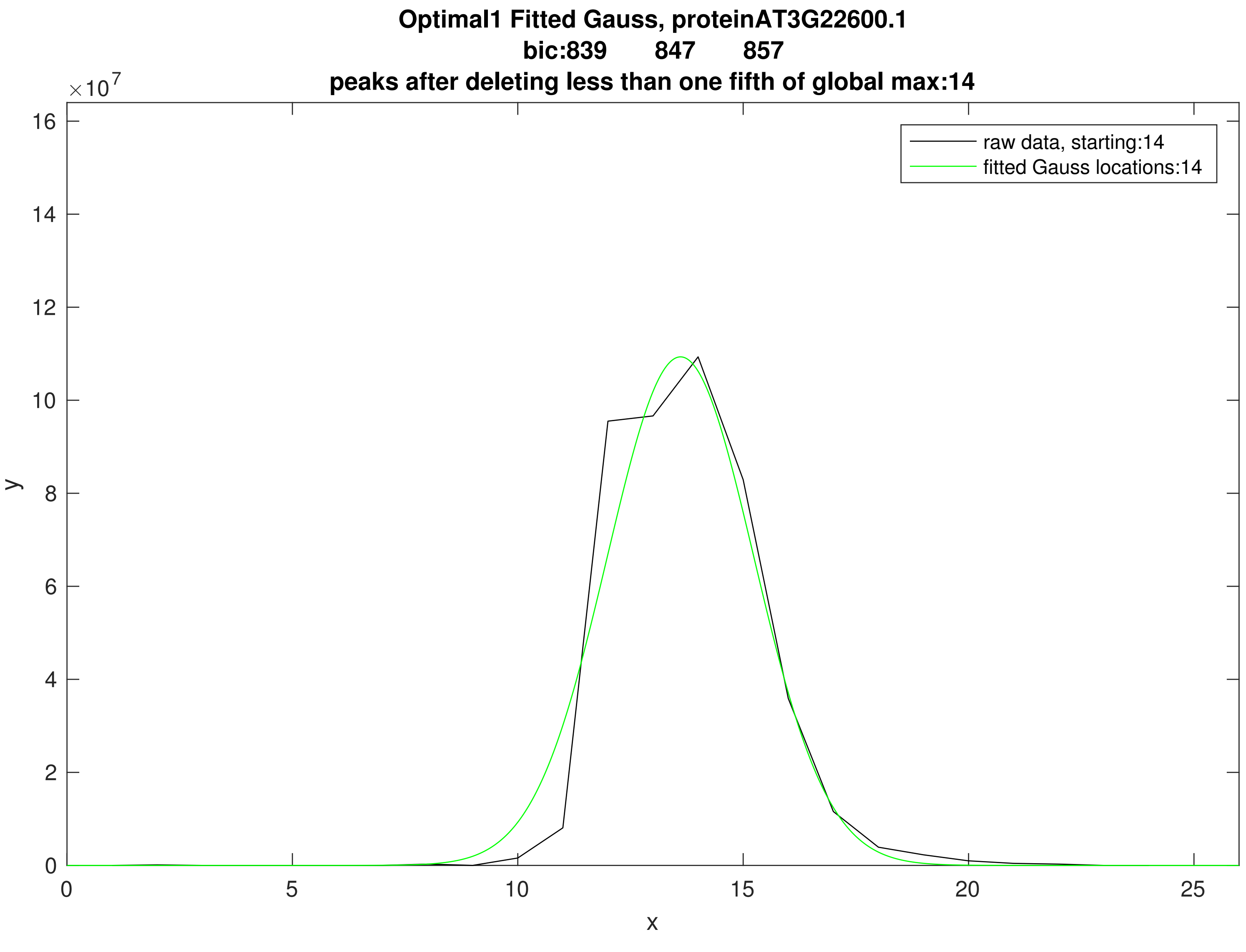
Optimal1 Fitted Gauss, proteinAT3G22600.1
bic:839 847 857
peaks after deleting less than one fifth of global max:14
×10^7
raw data, starting:14
fitted Gauss locations:14
y
x
0
2
4
6
8
10
12
14
16
0
5
10
15
20
25

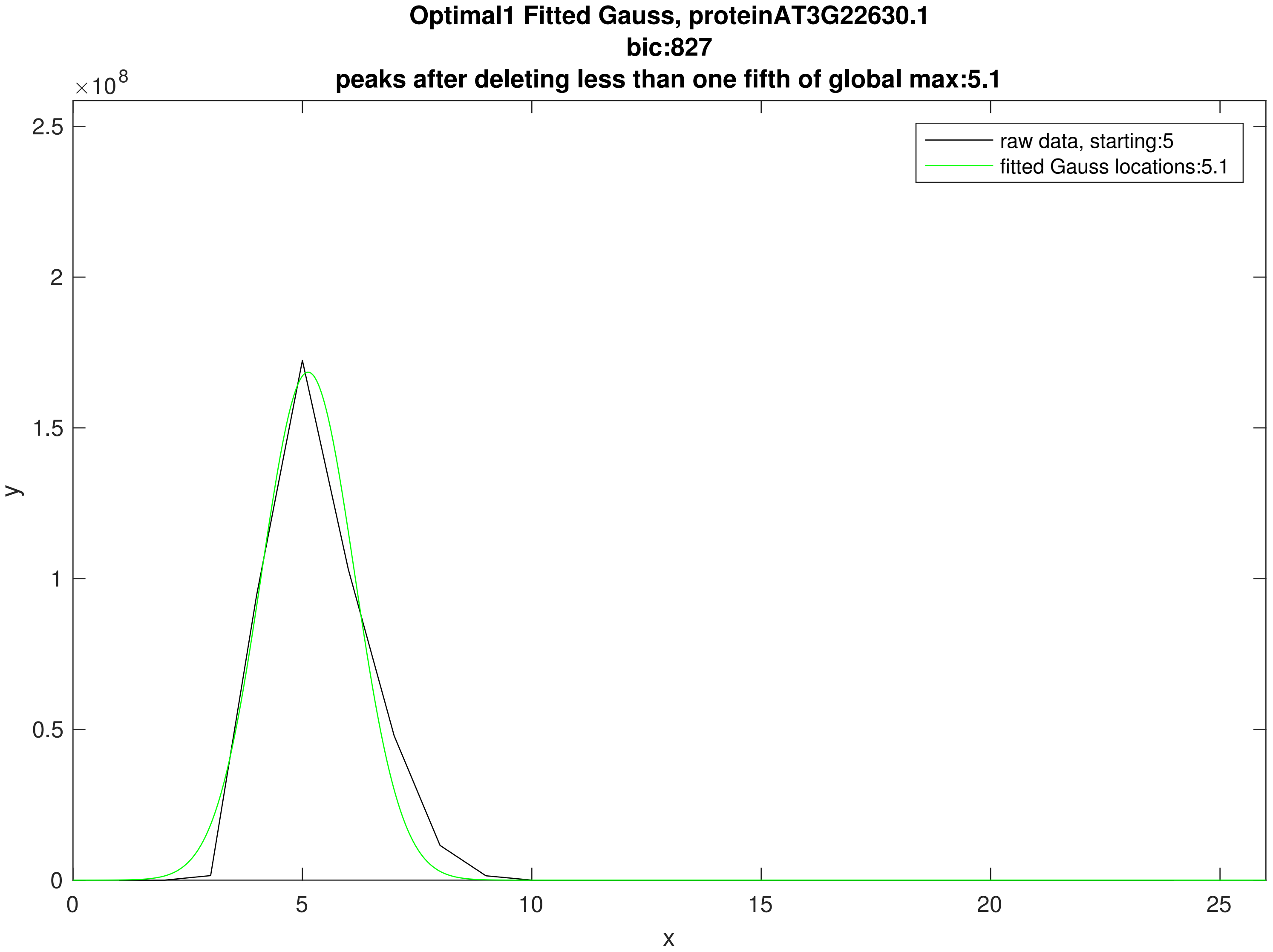
Optimal1 Fitted Gauss, proteinAT3G22630.1
bic:827
peaks after deleting less than one fifth of global max:5.1
×10^8
2.5
2
1.5
1
0.5
0
y
0
5
10
15
20
25
x
raw data, starting:5
fitted Gauss locations:5.1

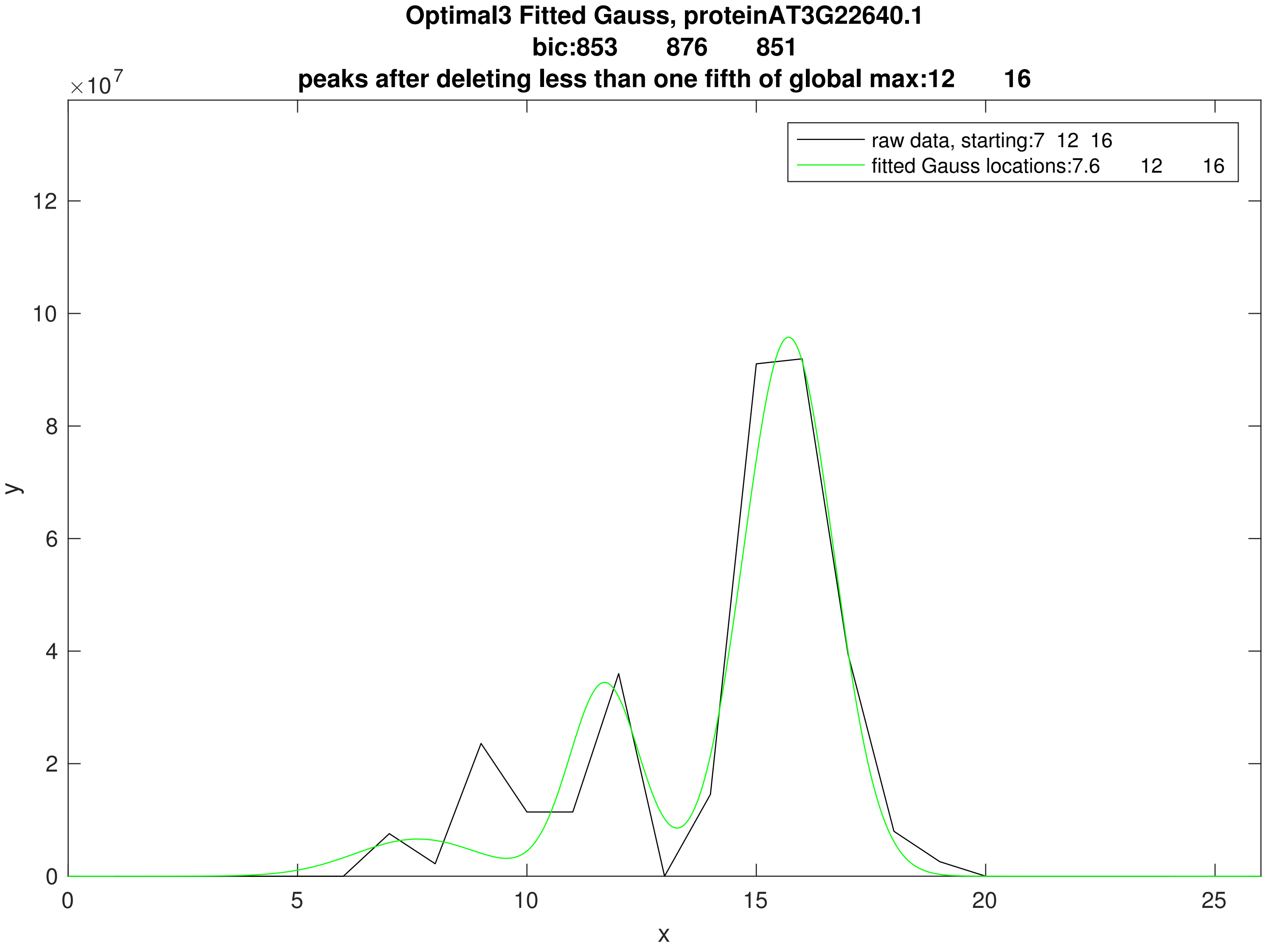
Optimal3 Fitted Gauss, proteinAT3G22640.1
bic:853 876 851
peaks after deleting less than one fifth of global max:12 16
×10^7
raw data, starting:7 12 16
fitted Gauss locations:7.6 12 16
y
x
0
2
4
6
8
10
12
5
15
20
25

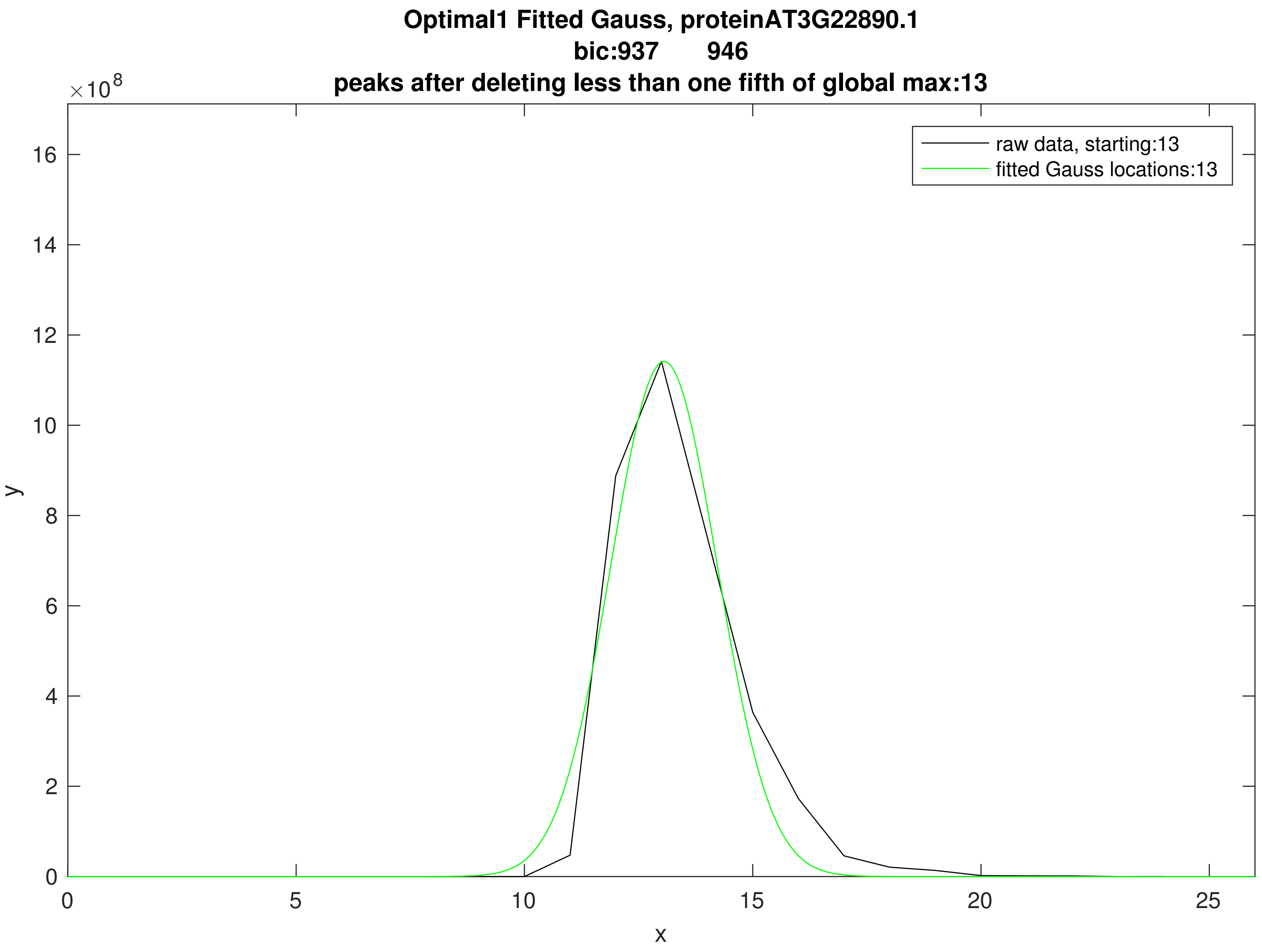

Optimal1 Fitted Gauss, proteinAT3G22890.1
bic:937 946
peaks after deleting less than one fifth of global max:13
×10^8
raw data, starting:13
fitted Gauss locations:13
16
14
12
10
8
6
4
2
0
y
0
5
10
15
20
25
x

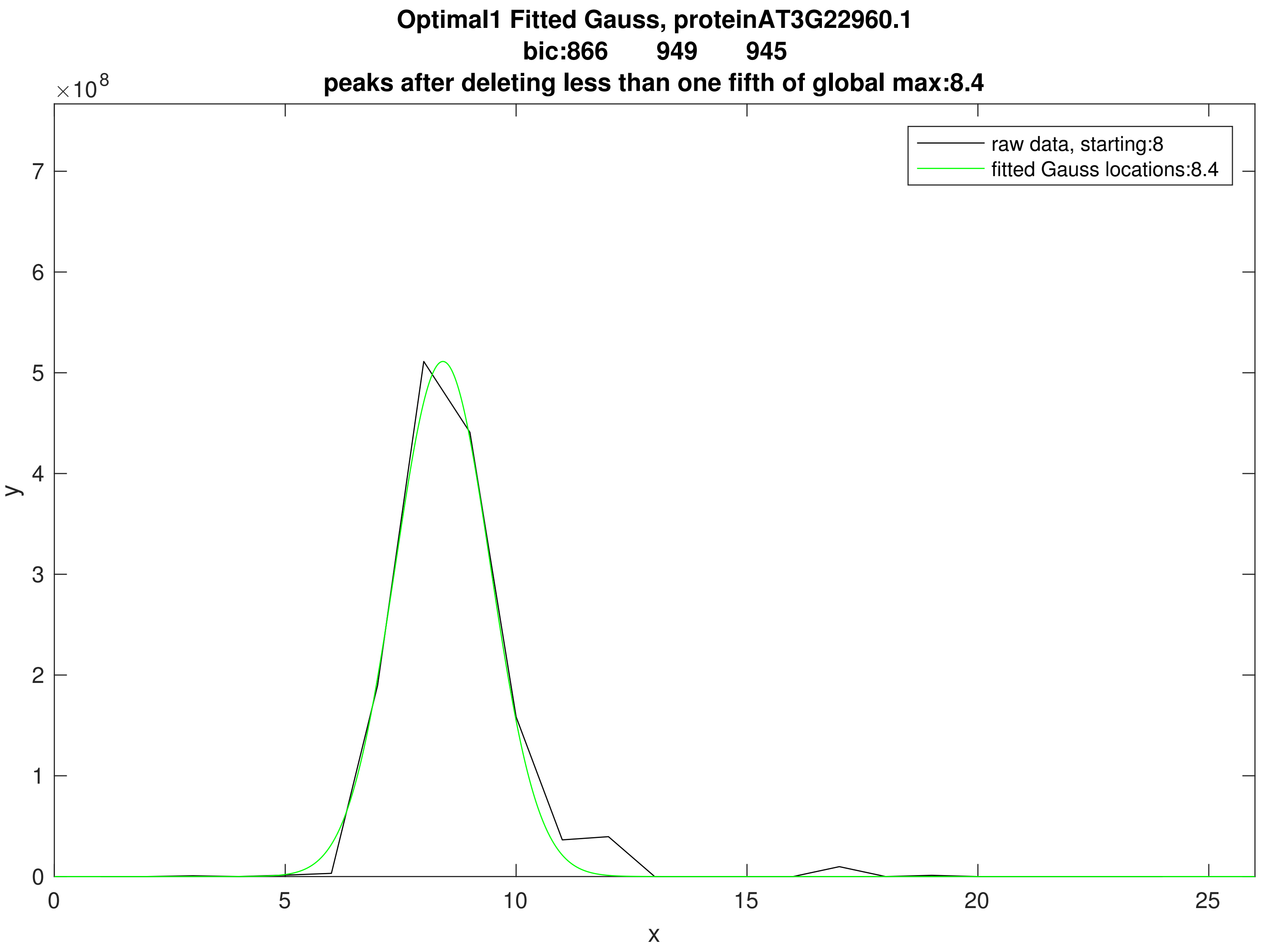
Optimal1 Fitted Gauss, proteinAT3G22960.1
bic:866 949 945
peaks after deleting less than one fifth of global max:8.4
×10^8
raw data, starting:8
fitted Gauss locations:8.4
y
x
0
1
2
3
4
5
6
7
0
5
10
15
20
25

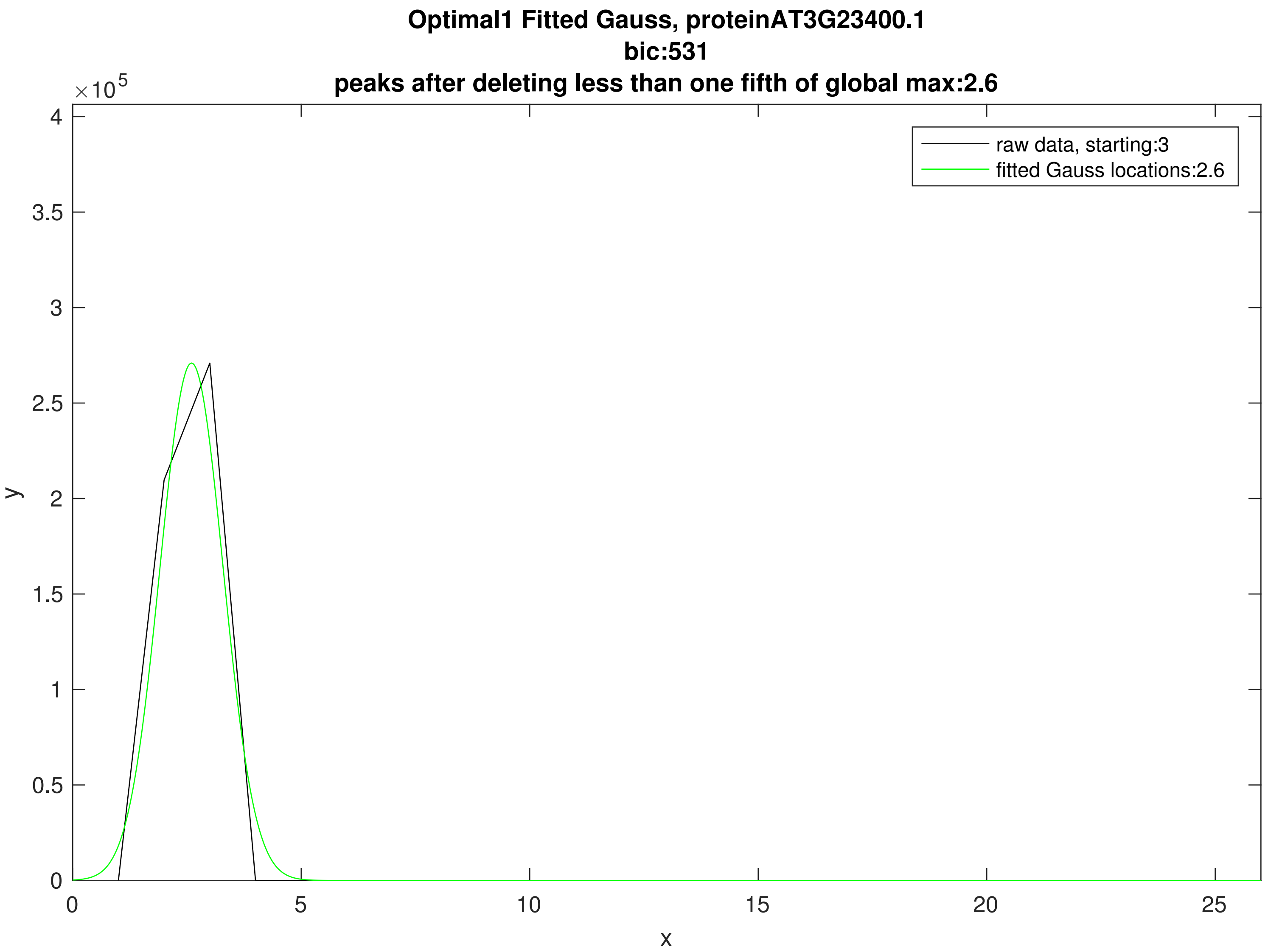
Optimal1 Fitted Gauss, proteinAT3G23400.1
bic:531
peaks after deleting less than one fifth of global max:2.6
raw data, starting:3
fitted Gauss locations:2.6
×10^5
y
x
0
0.5
1
1.5
2
2.5
3
3.5
4
0
5
10
15
20
25

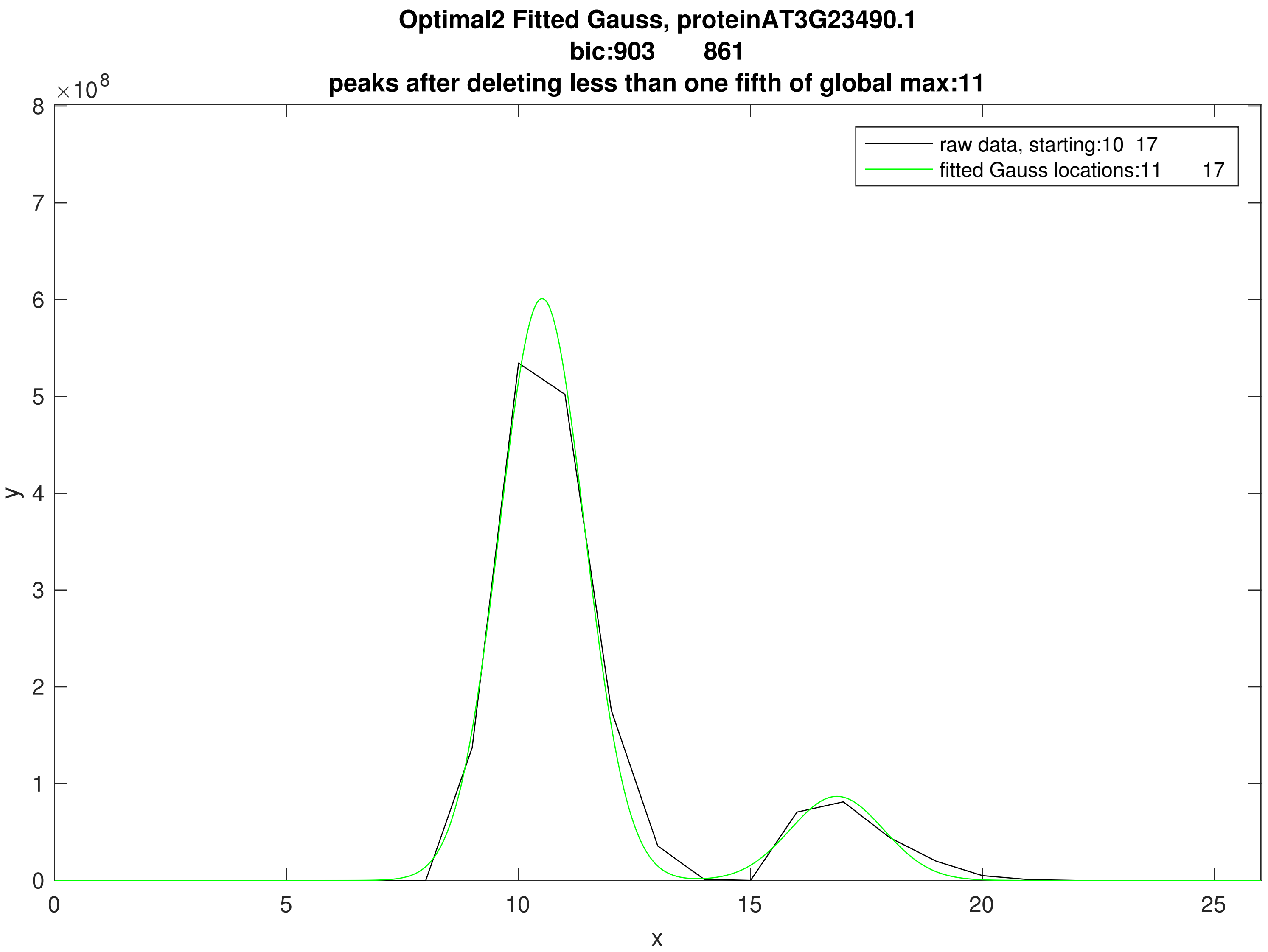

Optimal2 Fitted Gauss, proteinAT3G23490.1
bic:903 861
peaks after deleting less than one fifth of global max:11
×10^8
raw data, starting:10 17
fitted Gauss locations:11 17
y
x

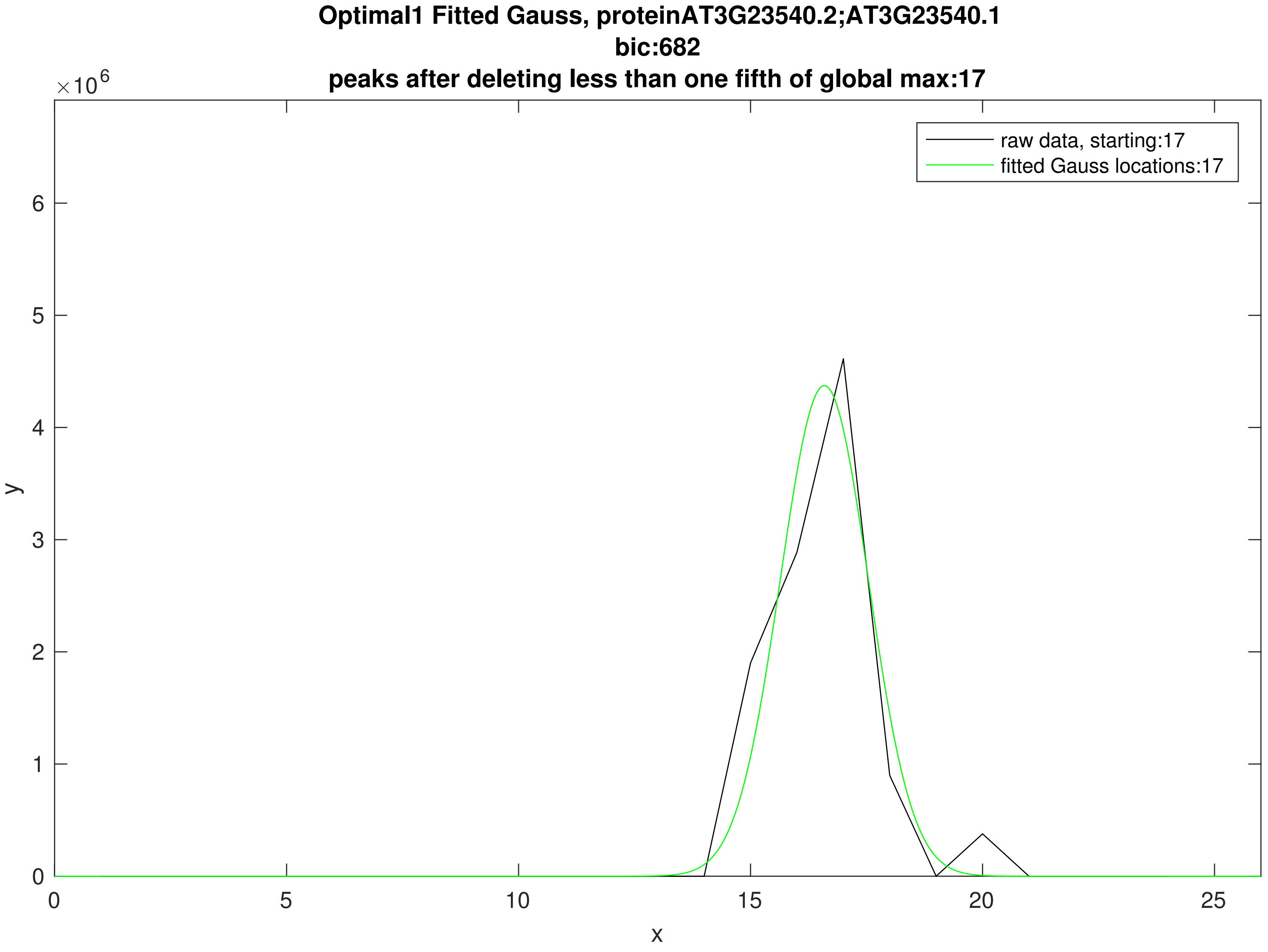

Optimal1 Fitted Gauss, proteinAT3G23540.2;AT3G23540.1
bic:682
peaks after deleting less than one fifth of global max:17
×10^6
raw data, starting:17
fitted Gauss locations:17
y
x
0
1
2
3
4
5
6
0
5
10
15
20
25

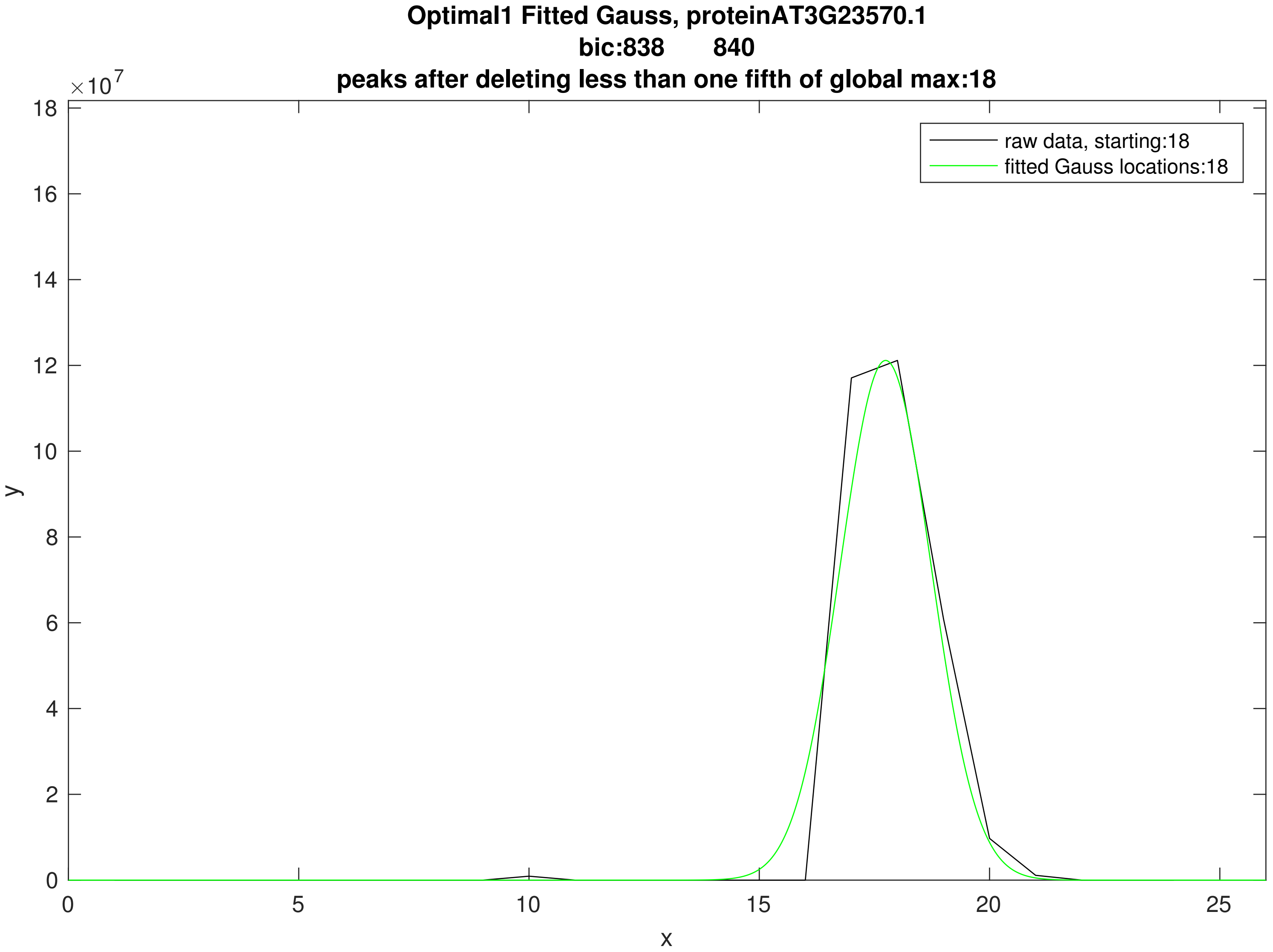
Optimal1 Fitted Gauss, proteinAT3G23570.1
bic:838 840
peaks after deleting less than one fifth of global max:18
×10^7
raw data, starting:18
fitted Gauss locations:18
y
x
0
2
4
6
8
10
12
14
16
18
0
5
10
15
20
25

**Optimal1 Fitted Gauss, proteinAT3G23600.1;AT3G23600.2**
**bic:872**
**peaks after deleting less than one fifth of global max:18**

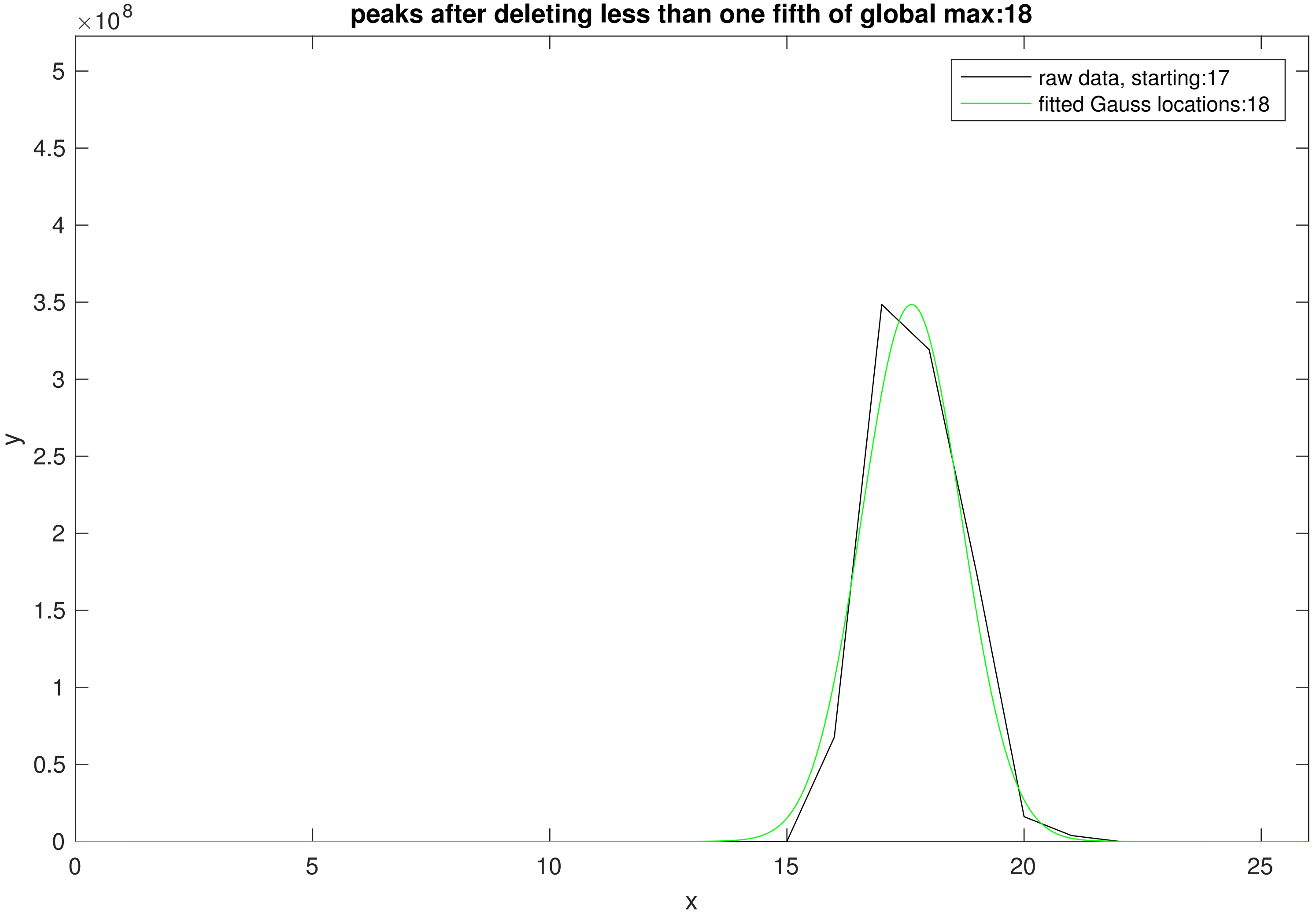

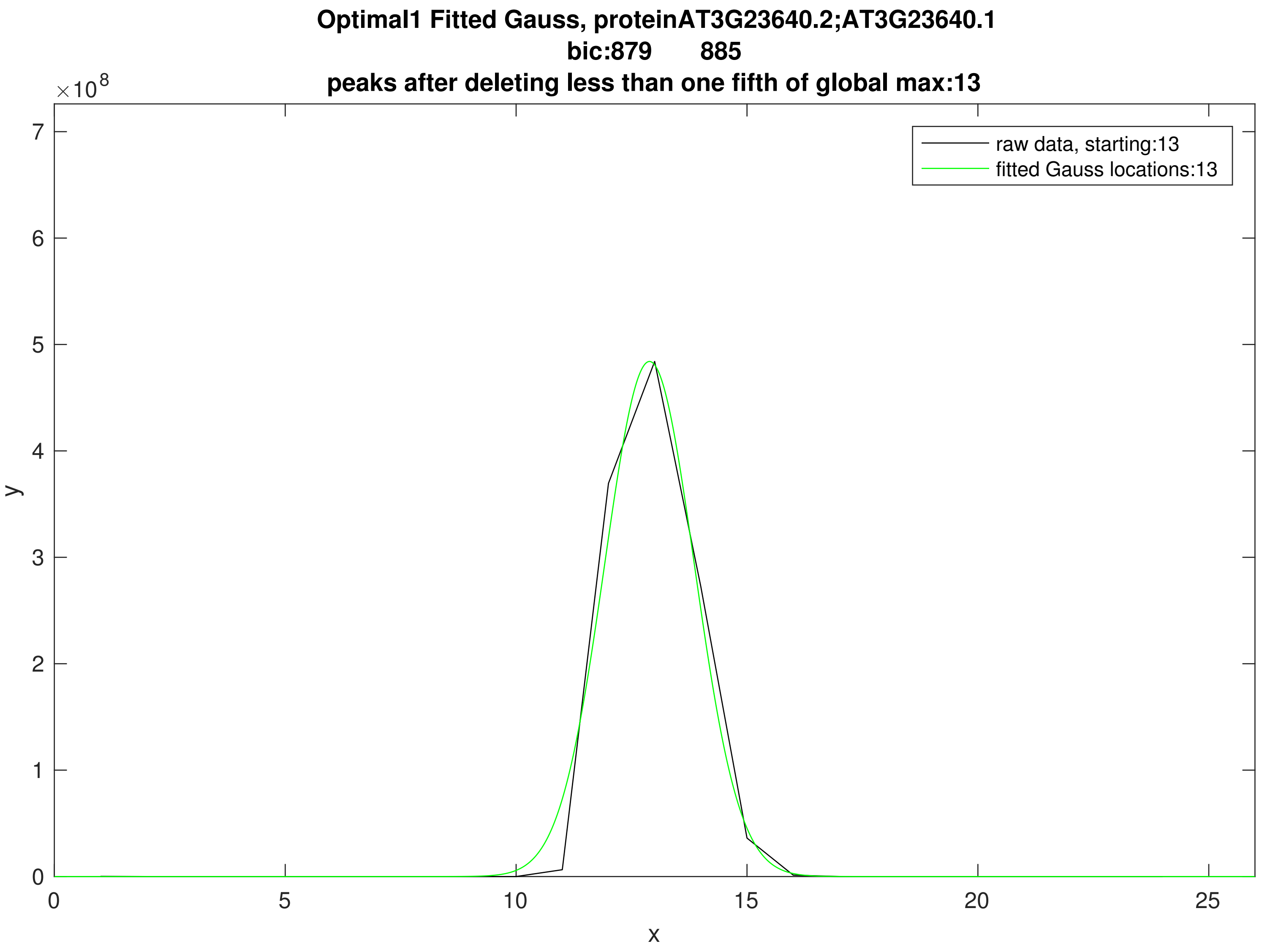

Optimal1 Fitted Gauss, proteinAT3G23640.2;AT3G23640.1
bic:879 885
peaks after deleting less than one fifth of global max:13
×10^8
raw data, starting:13
fitted Gauss locations:13
y
x
0
1
2
3
4
5
6
7
0
5
10
15
20
25

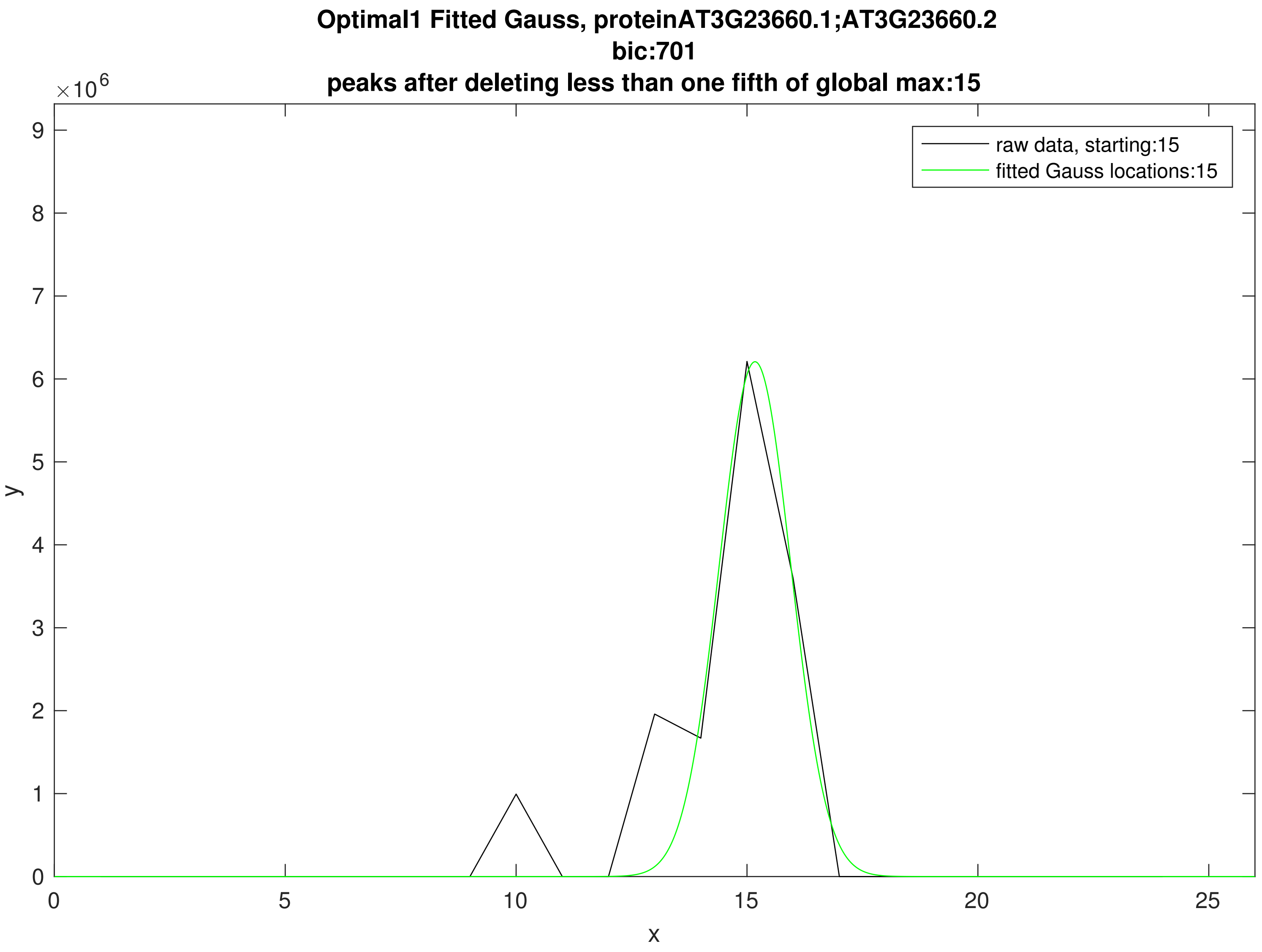

Optimal1 Fitted Gauss, proteinAT3G23660.1;AT3G23660.2
bic:701
peaks after deleting less than one fifth of global max:15
×10^6
raw data, starting:15
fitted Gauss locations:15
y
x
0
1
2
3
4
5
6
7
8
9
0
5
10
15
20
25

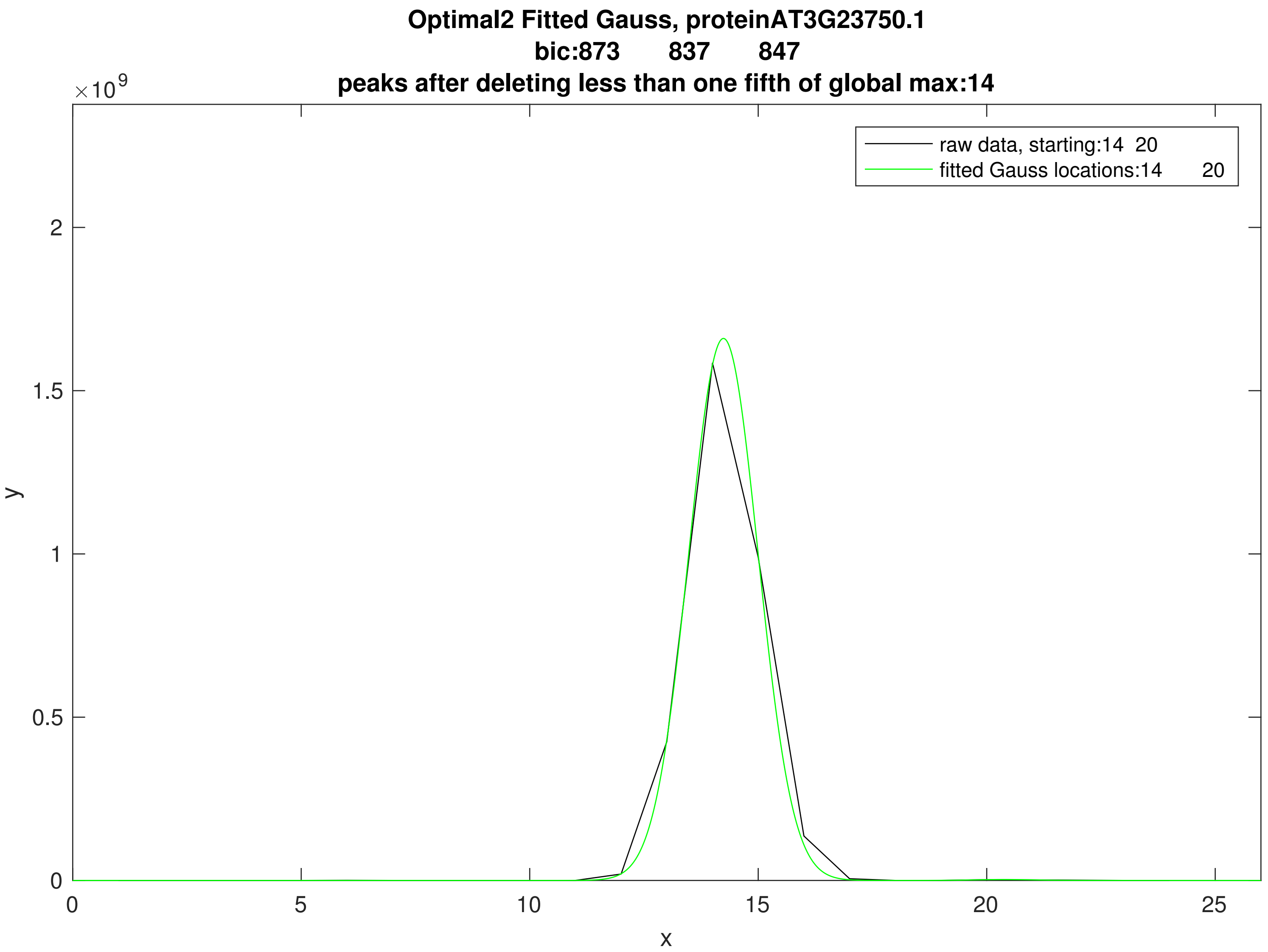

Optimal2 Fitted Gauss, proteinAT3G23750.1
bic:873 837 847
peaks after deleting less than one fifth of global max:14
×10^9
raw data, starting:14 20
fitted Gauss locations:14 20
y
x
0
0.5
1
1.5
2
5
10
15
20
25

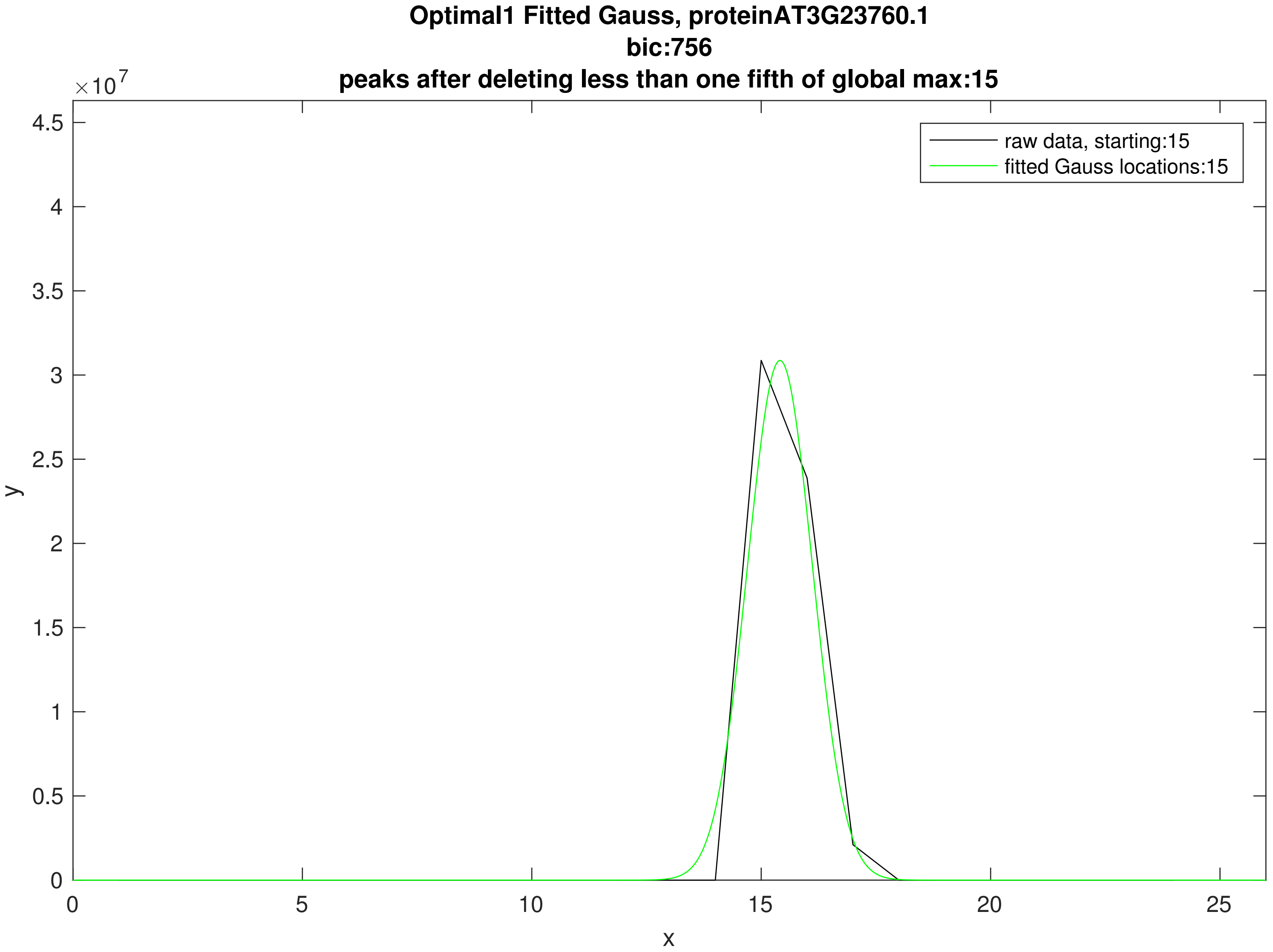
Optimal1 Fitted Gauss, proteinAT3G23760.1
bic:756
peaks after deleting less than one fifth of global max:15
×10^7
4.5
4
3.5
3
2.5
2
1.5
1
0.5
0
y
0
5
10
15
20
25
x
raw data, starting:15
fitted Gauss locations:15

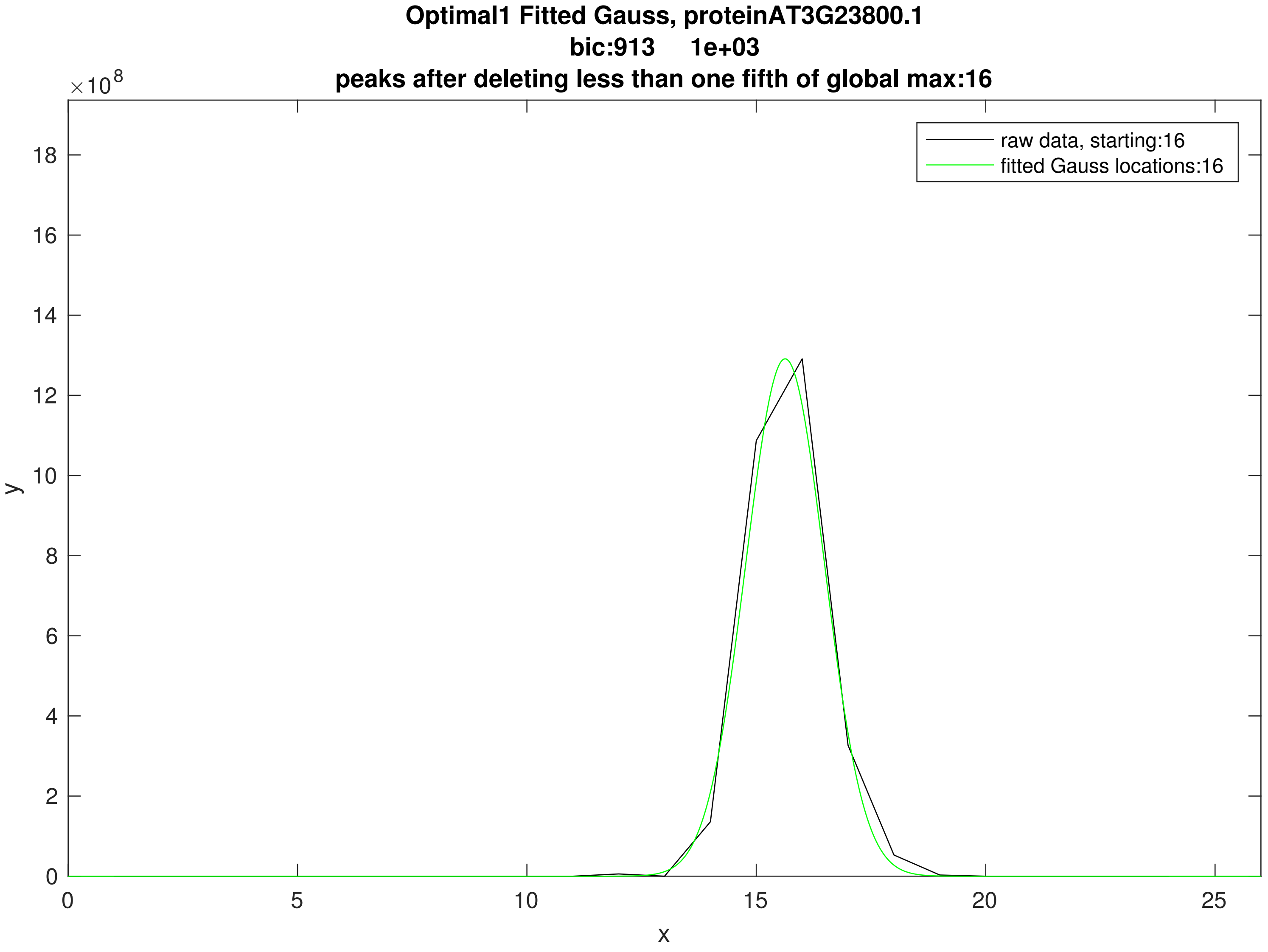
Optimal1 Fitted Gauss, proteinAT3G23800.1
bic:913 1e+03
peaks after deleting less than one fifth of global max:16
×10^8
18
16
14
12
10
8
6
4
2
0
y
0
5
10
15
20
25
x
raw data, starting:16
fitted Gauss locations:16

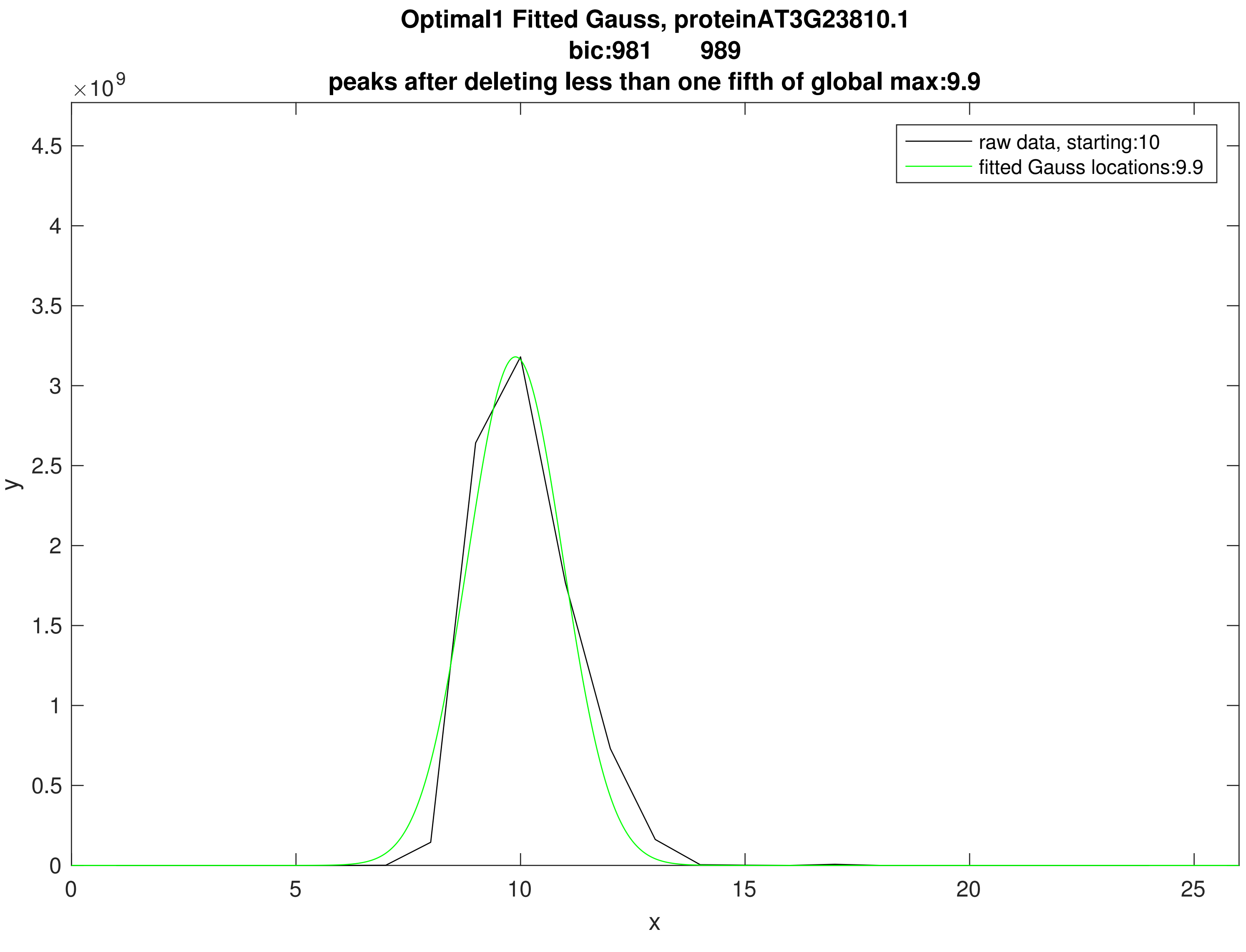
Optimal1 Fitted Gauss, proteinAT3G23810.1
bic:981 989
peaks after deleting less than one fifth of global max:9.9
×10^9
raw data, starting:10
fitted Gauss locations:9.9
y
x
0
0.5
1
1.5
2
2.5
3
3.5
4
4.5
0
5
10
15
20
25

**Optimal1 Fitted Gauss, proteinAT3G23830.2;AT3G23830.1**
**bic:659**
**peaks after deleting less than one fifth of global max:22**

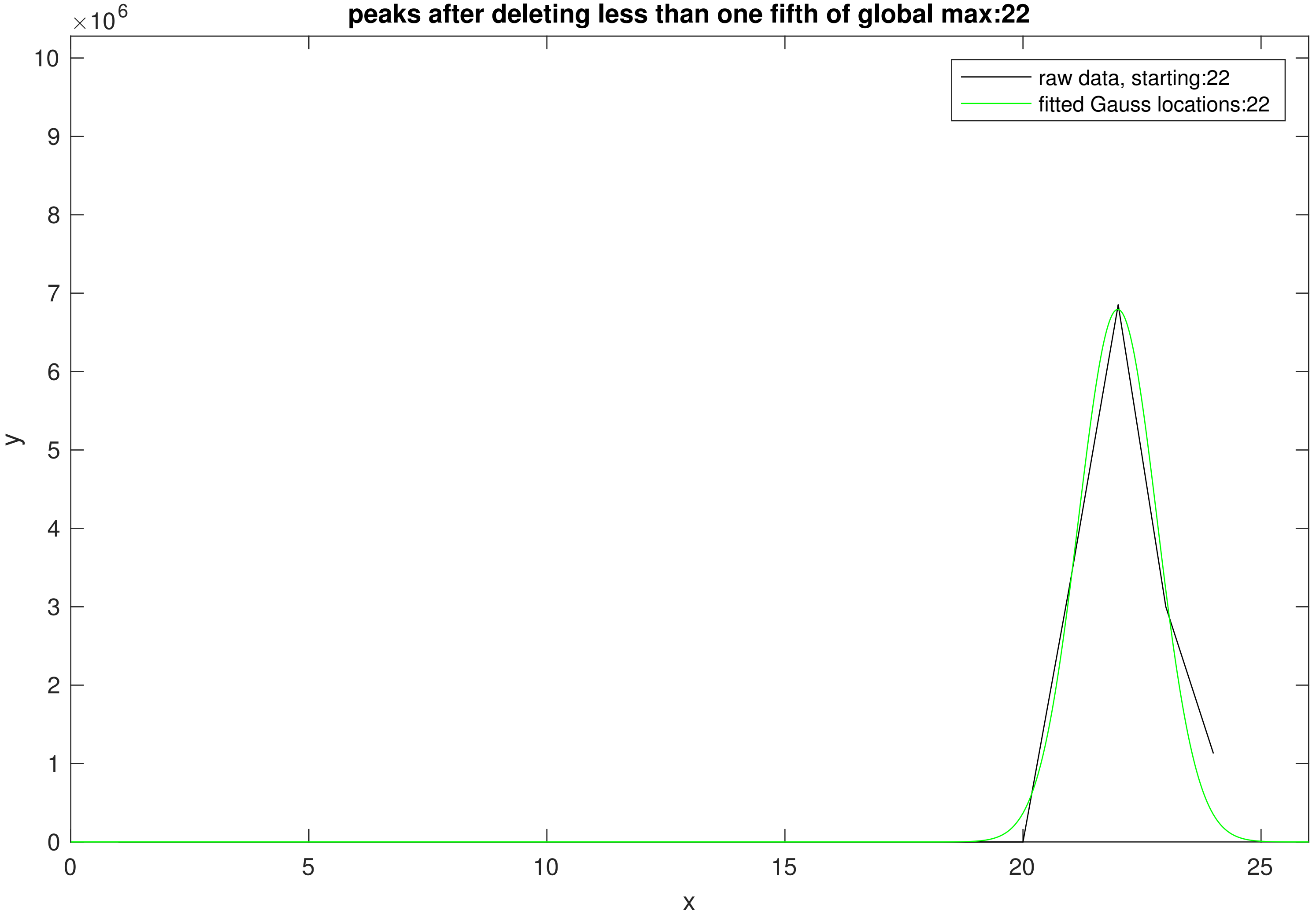

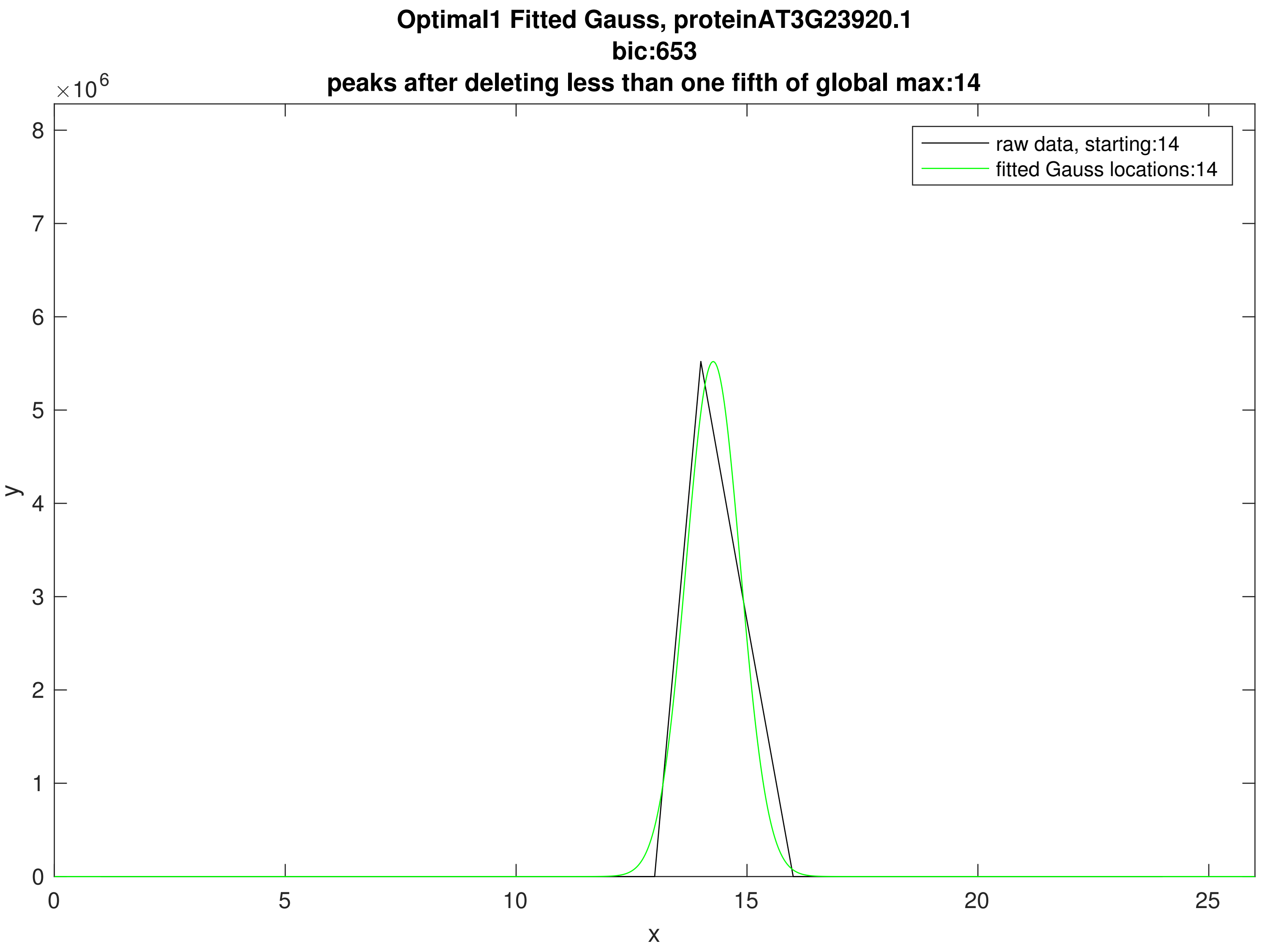
Optimal1 Fitted Gauss, proteinAT3G23920.1
bic:653
peaks after deleting less than one fifth of global max:14
×10^6
raw data, starting:14
fitted Gauss locations:14
y
x
0
1
2
3
4
5
6
7
8
0
5
10
15
20
25

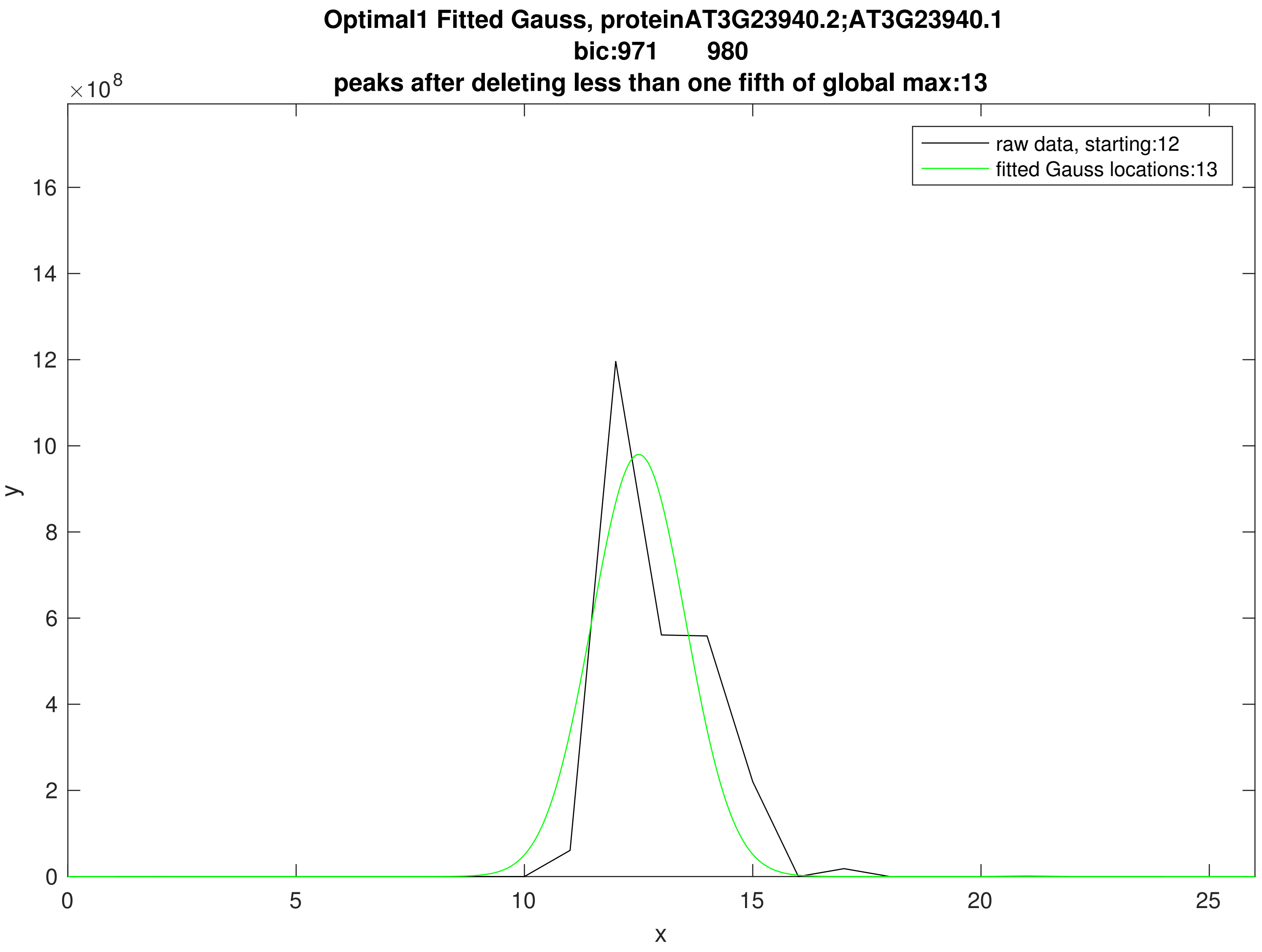
Optimal1 Fitted Gauss, proteinAT3G23940.2;AT3G23940.1
bic:971 980
peaks after deleting less than one fifth of global max:13
×10^8
raw data, starting:12
fitted Gauss locations:13
y
x
0
2
4
6
8
10
12
14
16
0
5
10
15
20
25

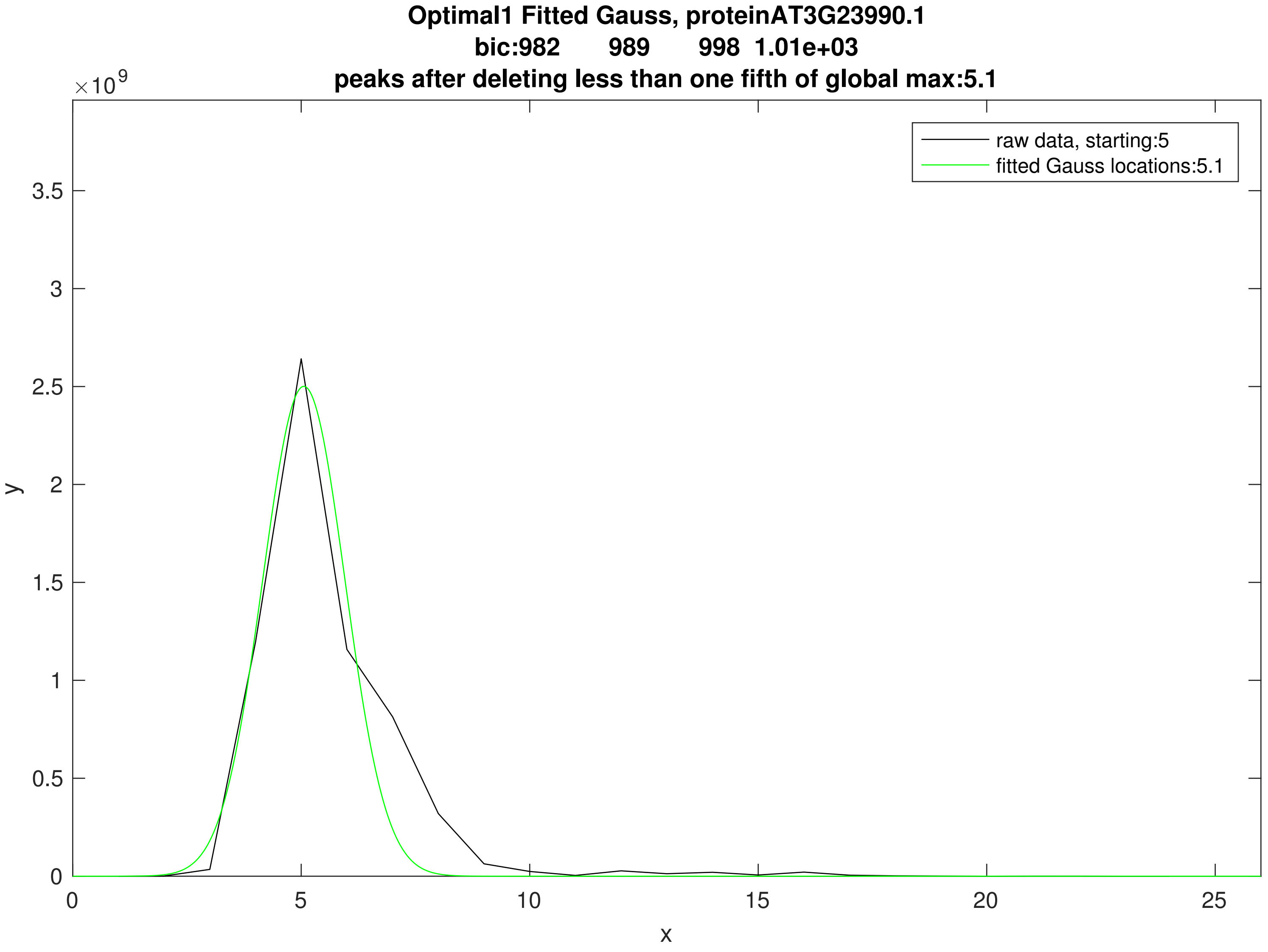
Optimal1 Fitted Gauss, proteinAT3G23990.1
bic:982 989 998 1.01e+03
peaks after deleting less than one fifth of global max:5.1
×10^9
raw data, starting:5
fitted Gauss locations:5.1
y
x
0
0.5
1
1.5
2
2.5
3
3.5
5
10
15
20
25

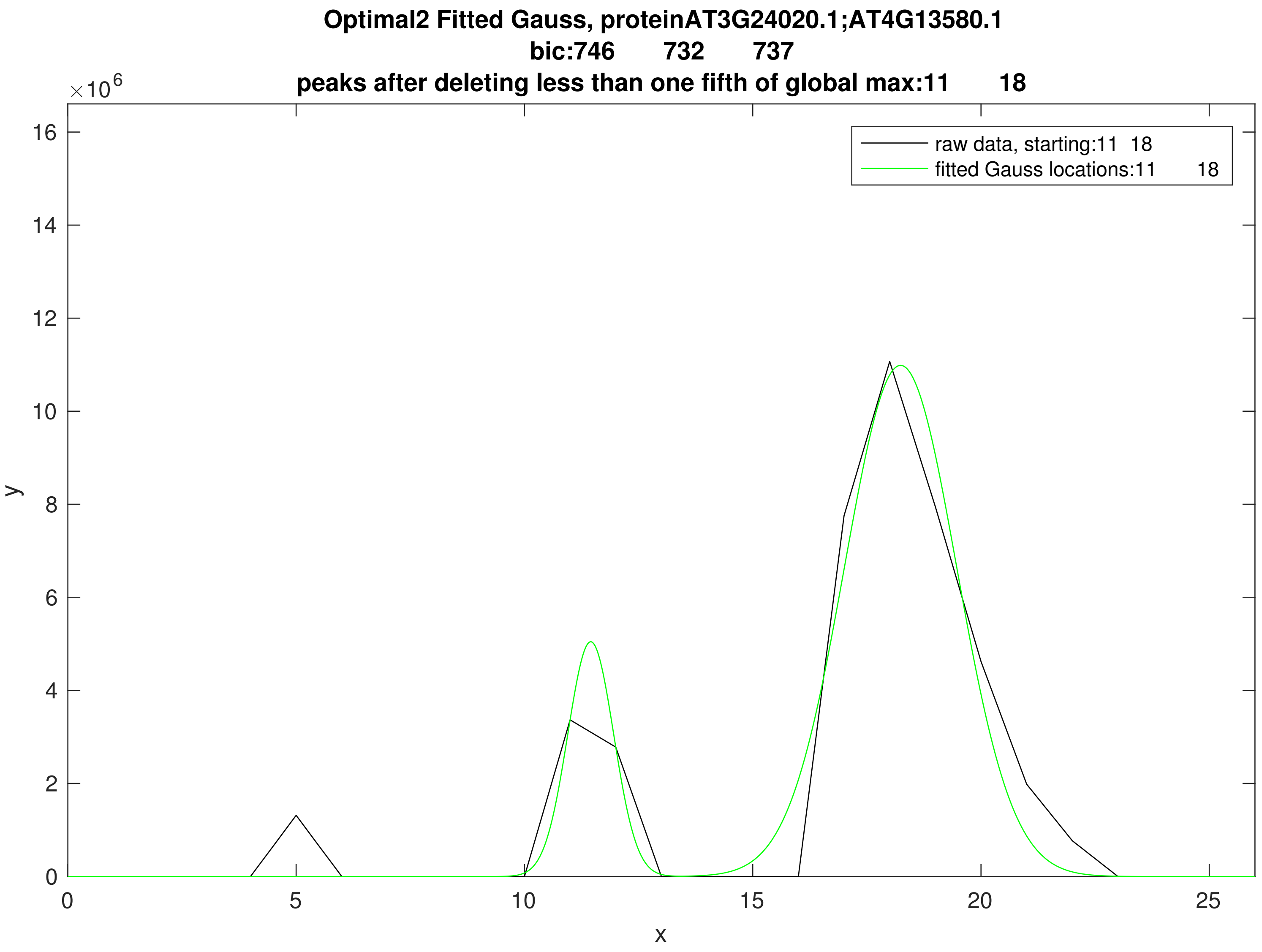
Optimal2 Fitted Gauss, proteinAT3G24020.1;AT4G13580.1
bic:746 732 737
peaks after deleting less than one fifth of global max:11 18
×10^6
raw data, starting:11 18
fitted Gauss locations:11 18
y
x
0
2
4
6
8
10
12
14
16
0
5
10
15
20
25

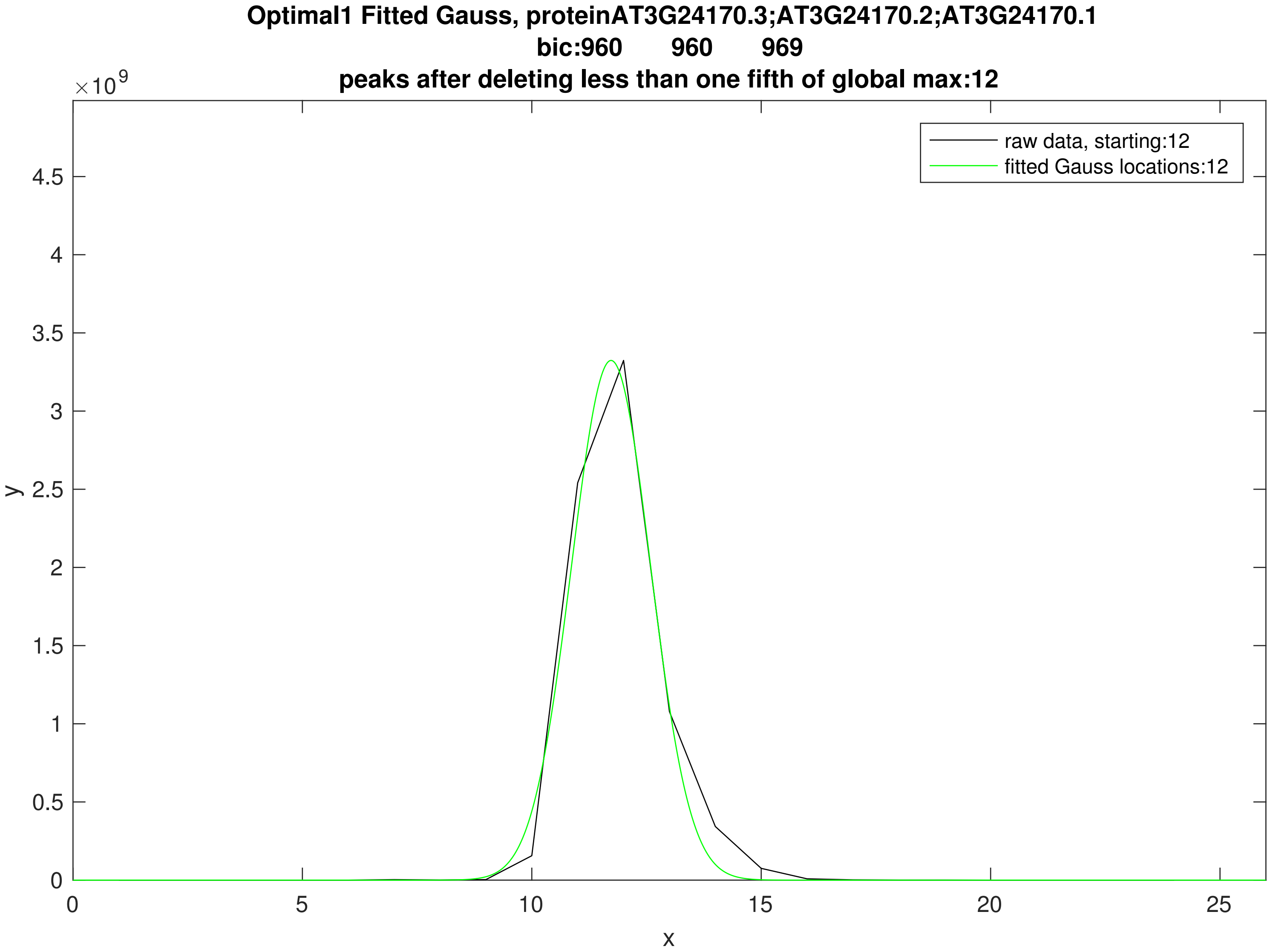

Optimal1 Fitted Gauss, proteinAT3G24170.3;AT3G24170.2;AT3G24170.1
bic:960 960 969
peaks after deleting less than one fifth of global max:12
×10^9
raw data, starting:12
fitted Gauss locations:12
y
x
0
0.5
1
1.5
2
2.5
3
3.5
4
4.5
5
10
15
20
25

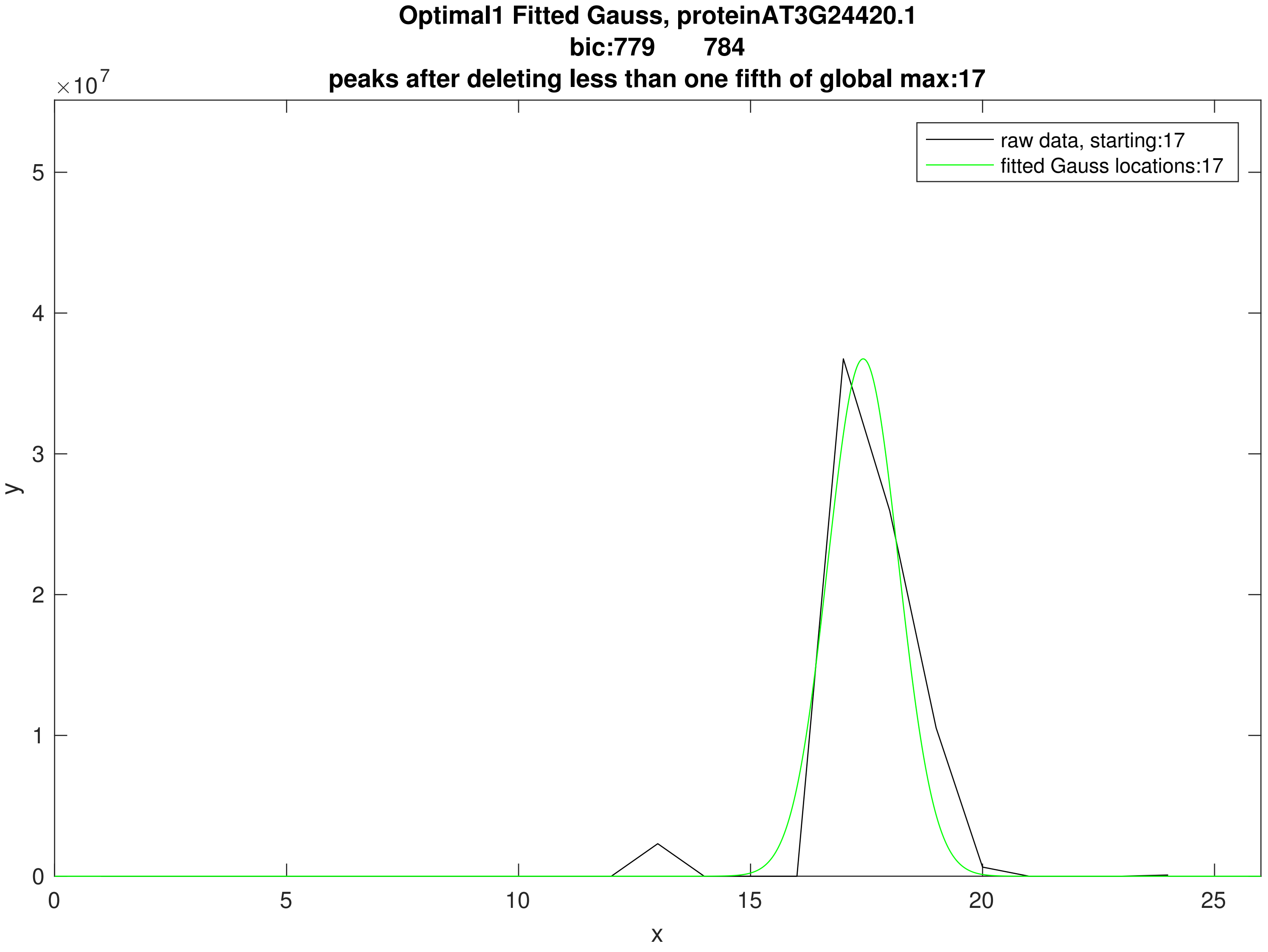

Optimal1 Fitted Gauss, proteinAT3G24420.1
bic:779 784
peaks after deleting less than one fifth of global max:17
×10^7
raw data, starting:17
fitted Gauss locations:17
y
x
0
1
2
3
4
5
0
5
10
15
20
25

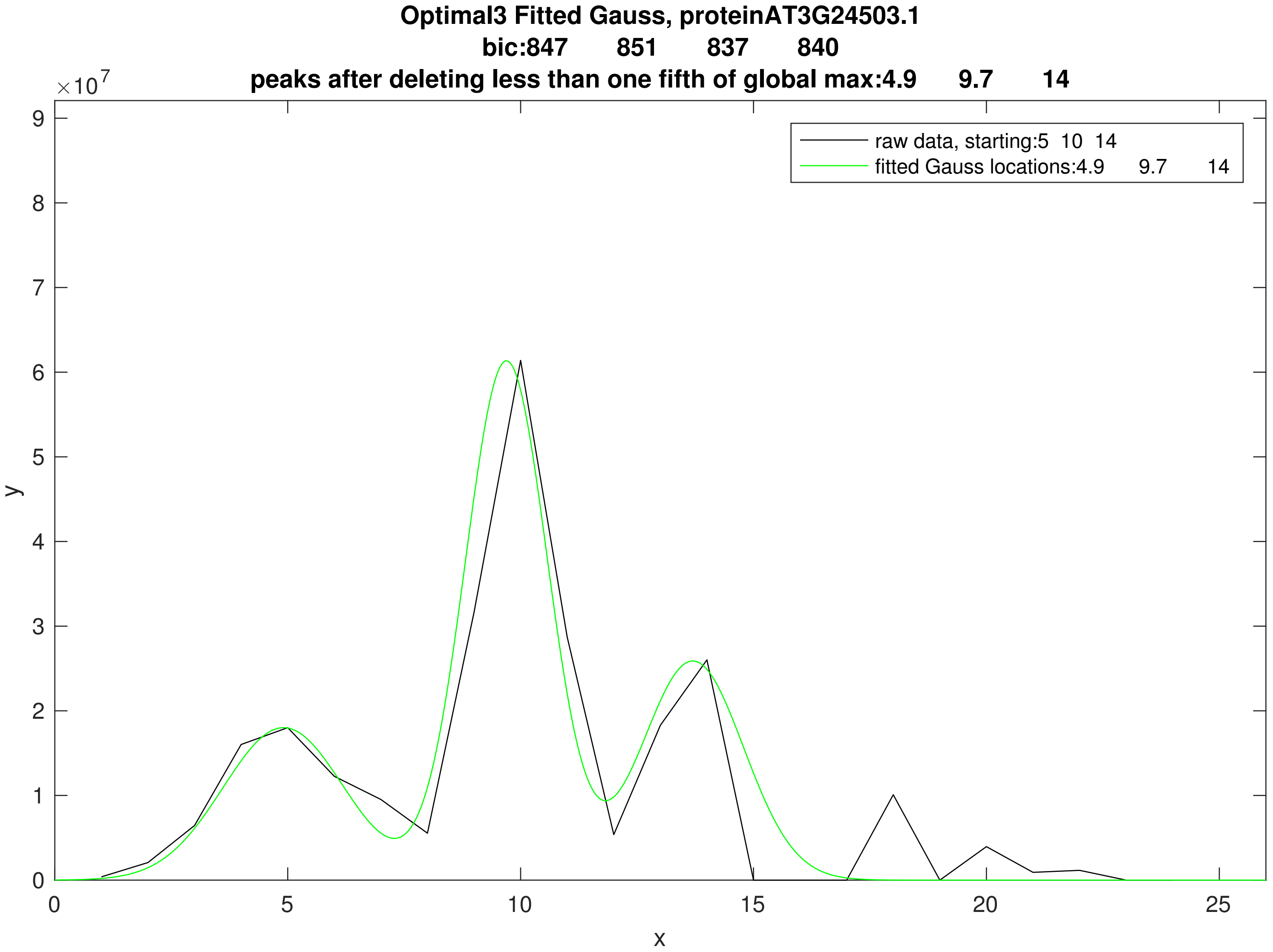
Optimal3 Fitted Gauss, proteinAT3G24503.1
bic:847 851 837 840
peaks after deleting less than one fifth of global max:4.9 9.7 14
×10^7
y
x
raw data, starting:5 10 14
fitted Gauss locations:4.9 9.7 14
0
1
2
3
4
5
6
7
8
9
0
5
10
15
20
25

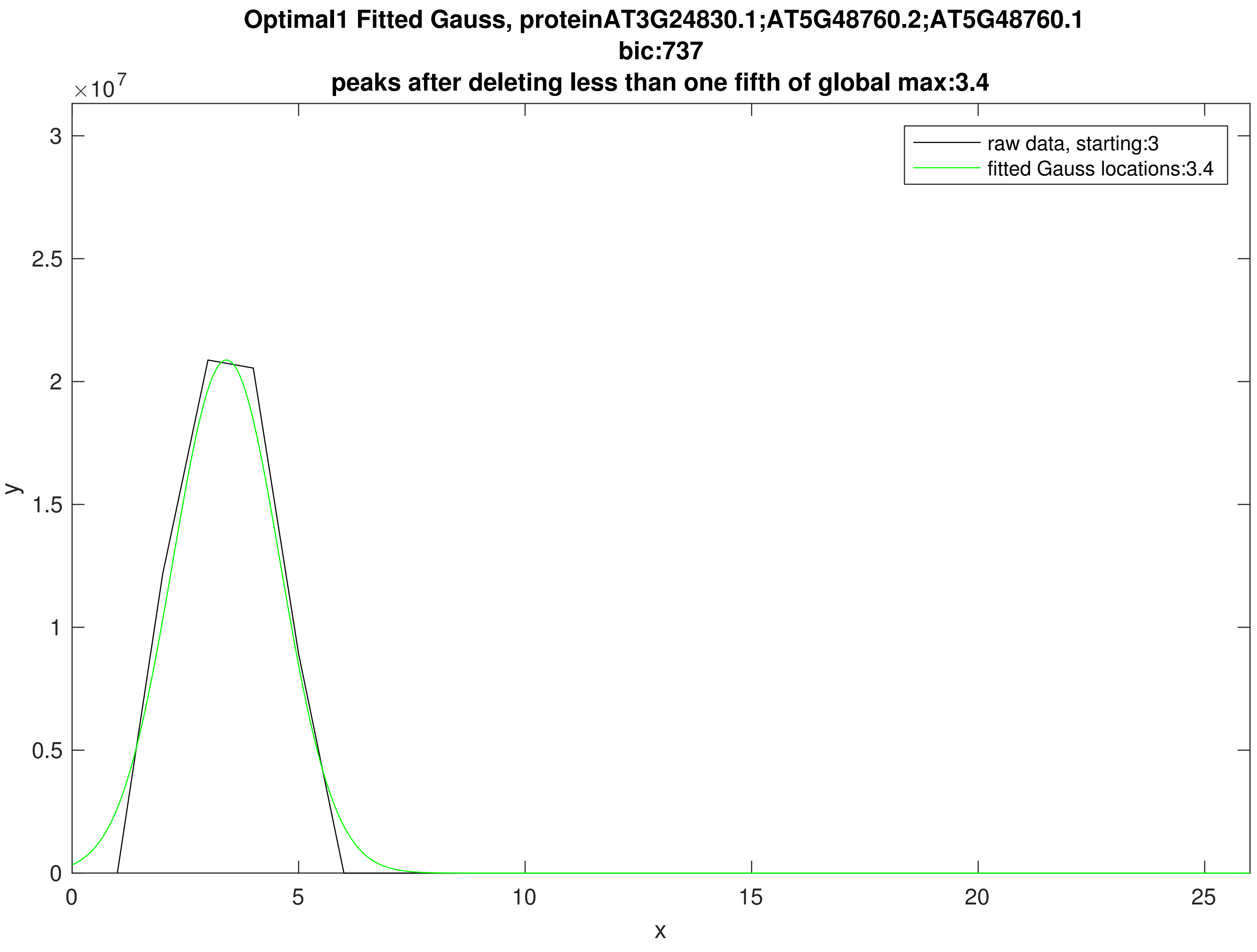
Optimal1 Fitted Gauss, proteinAT3G24830.1;AT5G48760.2;AT5G48760.1
bic:737
peaks after deleting less than one fifth of global max:3.4
×10^7
3
2.5
2
1.5
1
0.5
0
y
0
5
10
15
20
25
x
raw data, starting:3
fitted Gauss locations:3.4

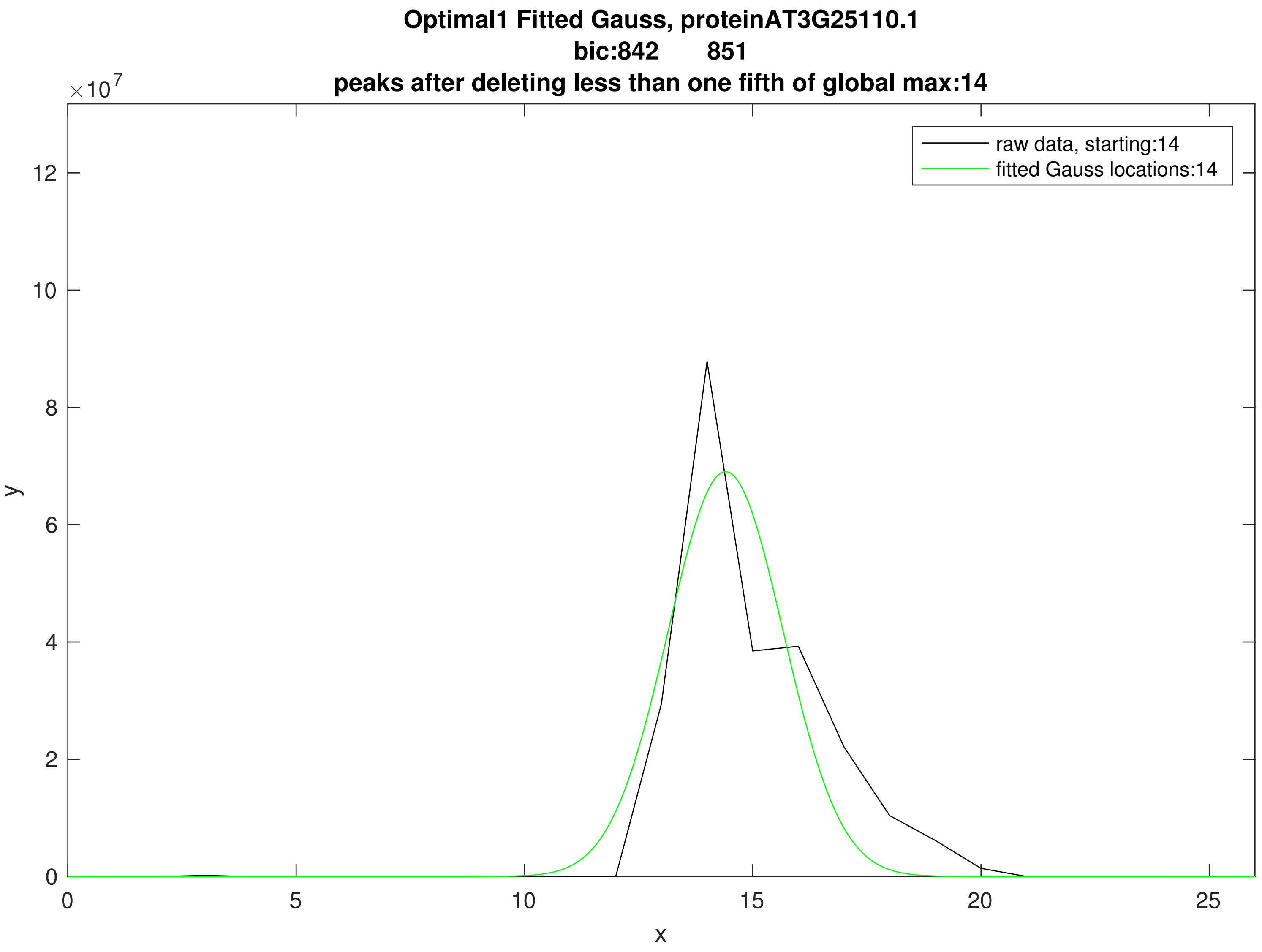
Optimal1 Fitted Gauss, proteinAT3G25110.1
bic:842 851
peaks after deleting less than one fifth of global max:14
×10^7
12
10
8
6
4
2
0
y
0
5
10
15
20
25
x
raw data, starting:14
fitted Gauss locations:14

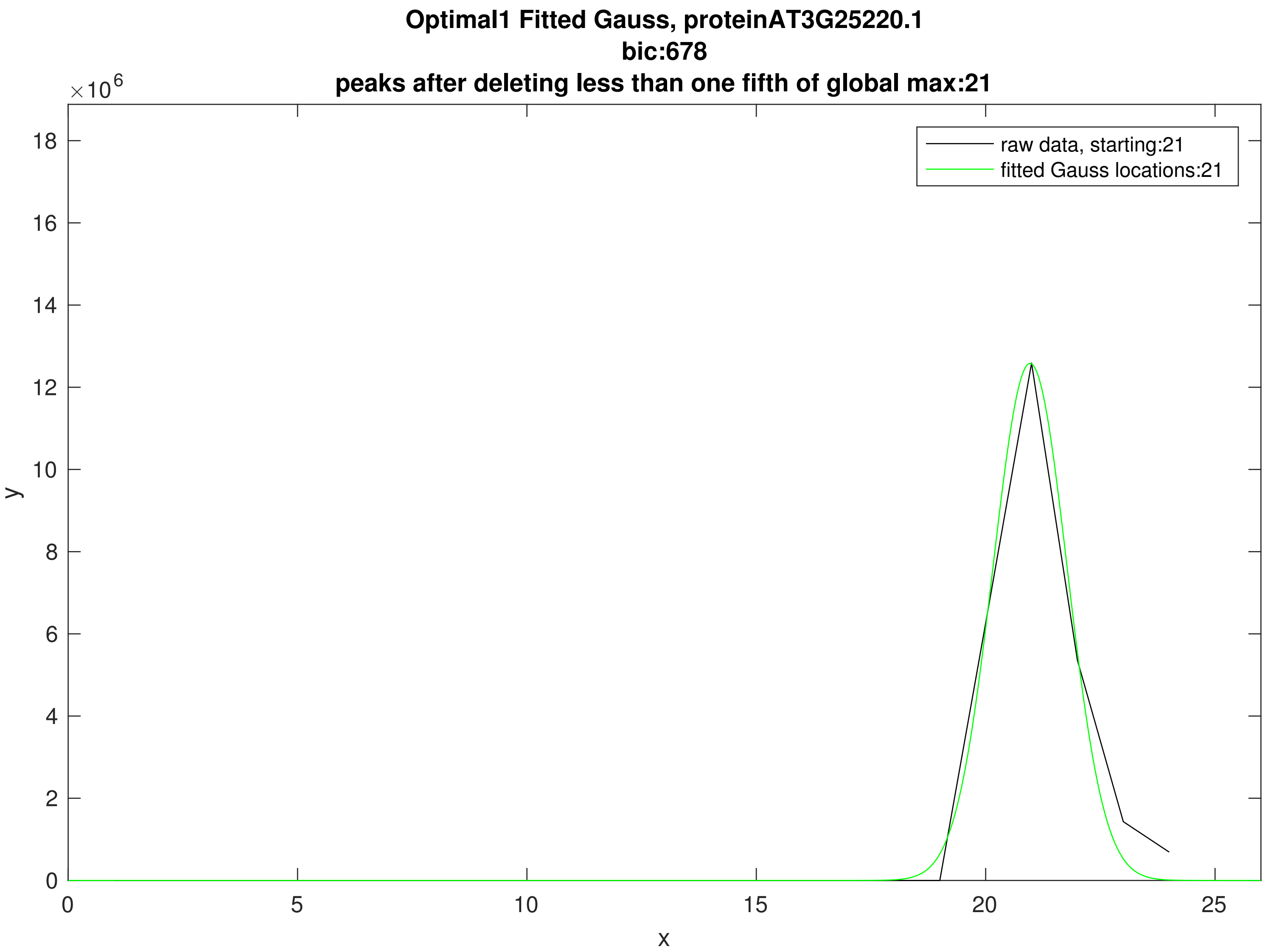
Optimal1 Fitted Gauss, proteinAT3G25220.1
bic:678
peaks after deleting less than one fifth of global max:21
×10^6
raw data, starting:21
fitted Gauss locations:21
y
x
0
2
4
6
8
10
12
14
16
18
0
5
10
15
20
25

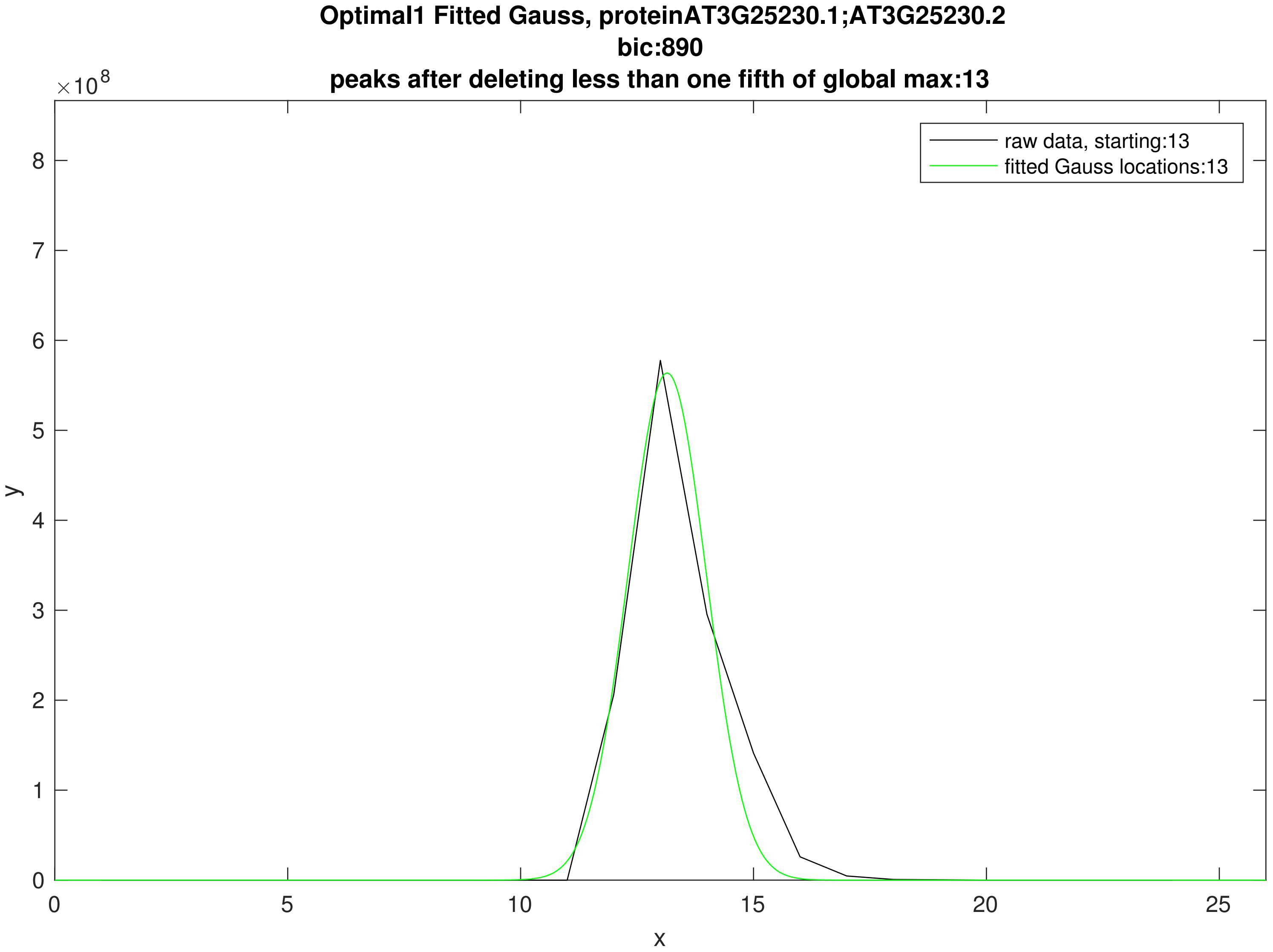
Optimal1 Fitted Gauss, proteinAT3G25230.1;AT3G25230.2
bic:890
peaks after deleting less than one fifth of global max:13
×10^8
raw data, starting:13
fitted Gauss locations:13
y
x
0
1
2
3
4
5
6
7
8
0
5
10
15
20
25

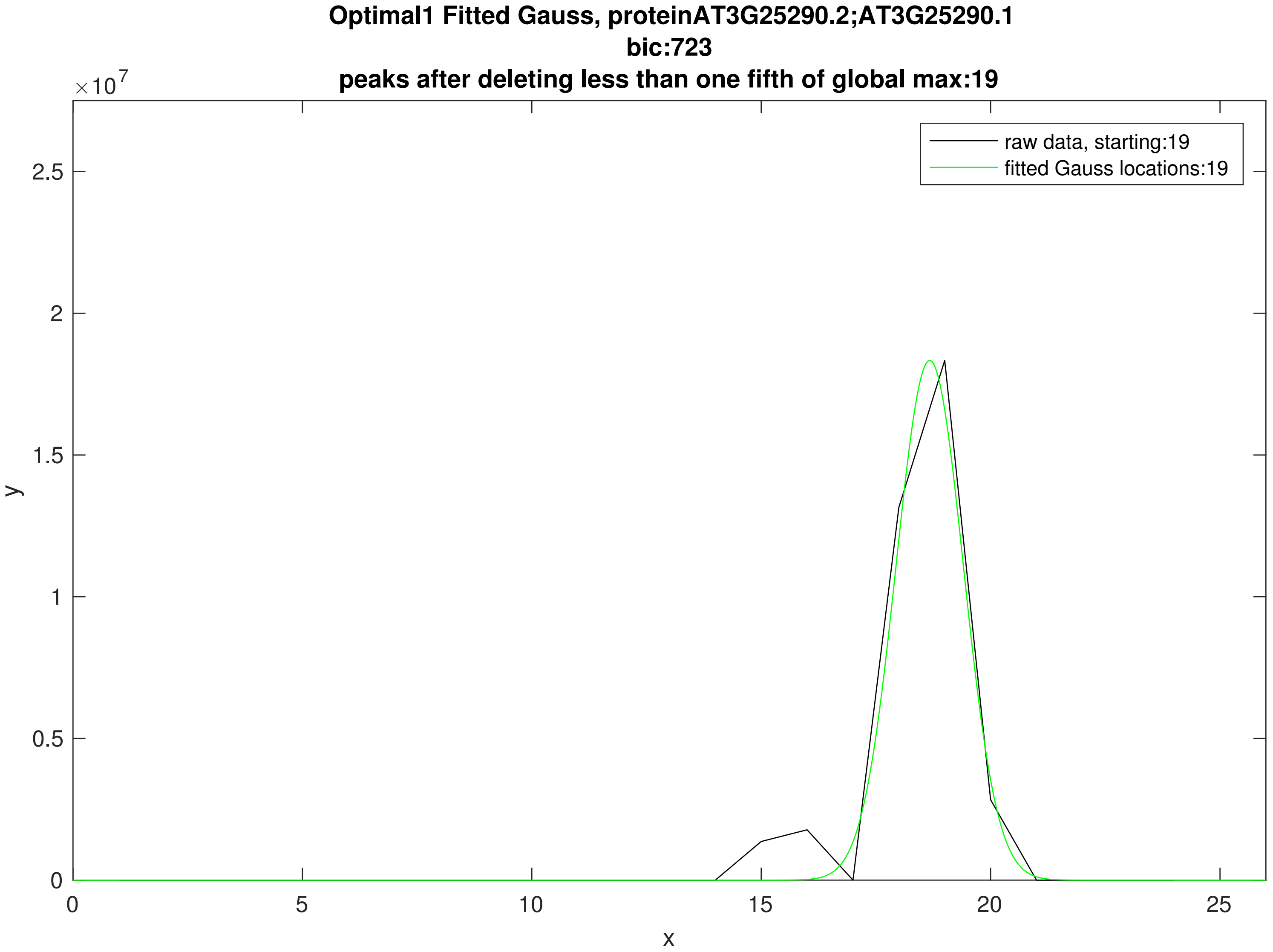

Optimal1 Fitted Gauss, proteinAT3G25290.2;AT3G25290.1
bic:723
peaks after deleting less than one fifth of global max:19
×10^7
raw data, starting:19
fitted Gauss locations:19
2.5
2
1.5
1
0.5
0
y
0
5
10
15
20
25
x

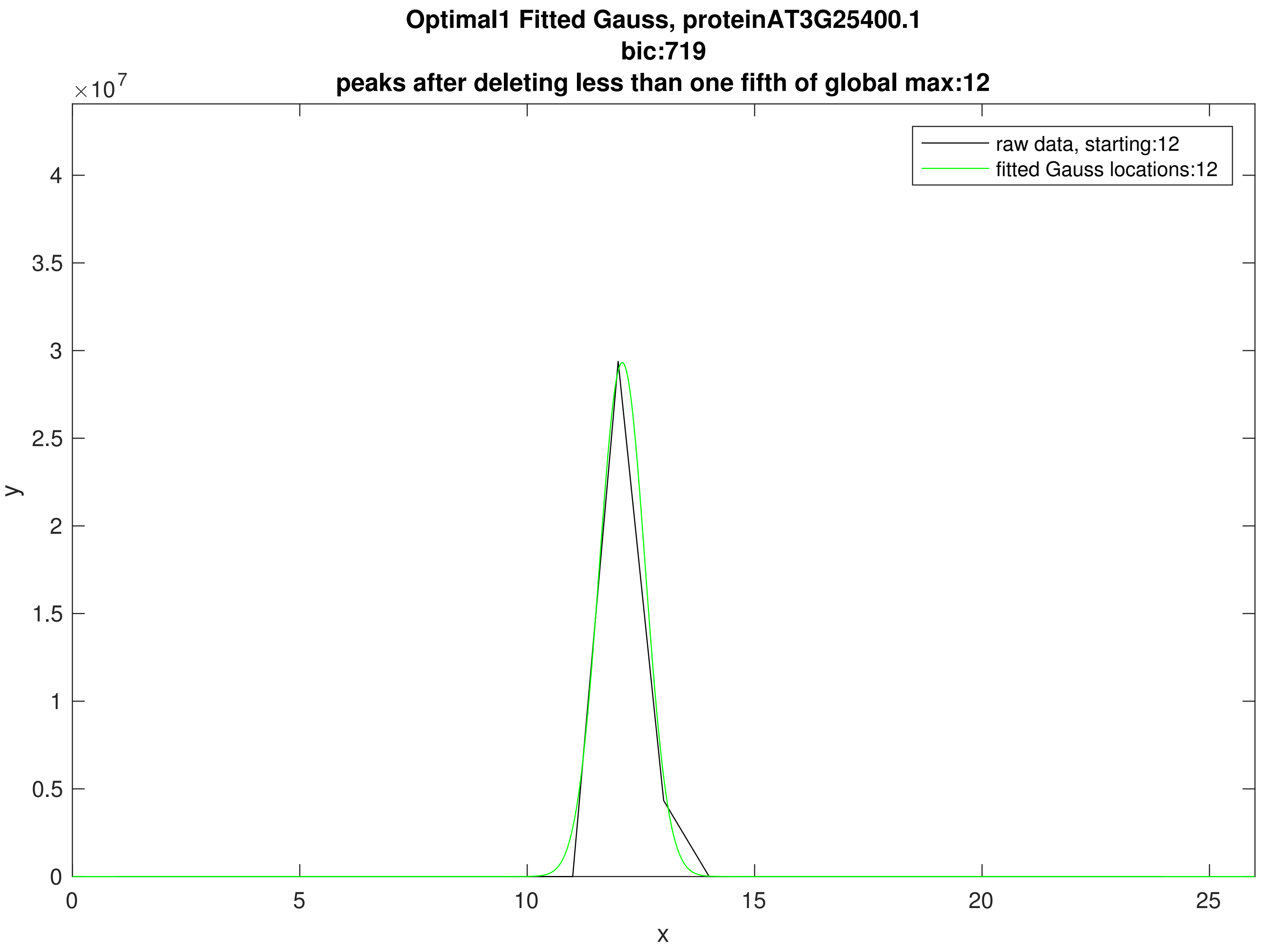

Optimal1 Fitted Gauss, proteinAT3G25400.1
bic:719
peaks after deleting less than one fifth of global max:12
×10^7
raw data, starting:12
fitted Gauss locations:12
4
3.5
3
2.5
2
1.5
1
0.5
0
y
0
5
10
15
20
25
x

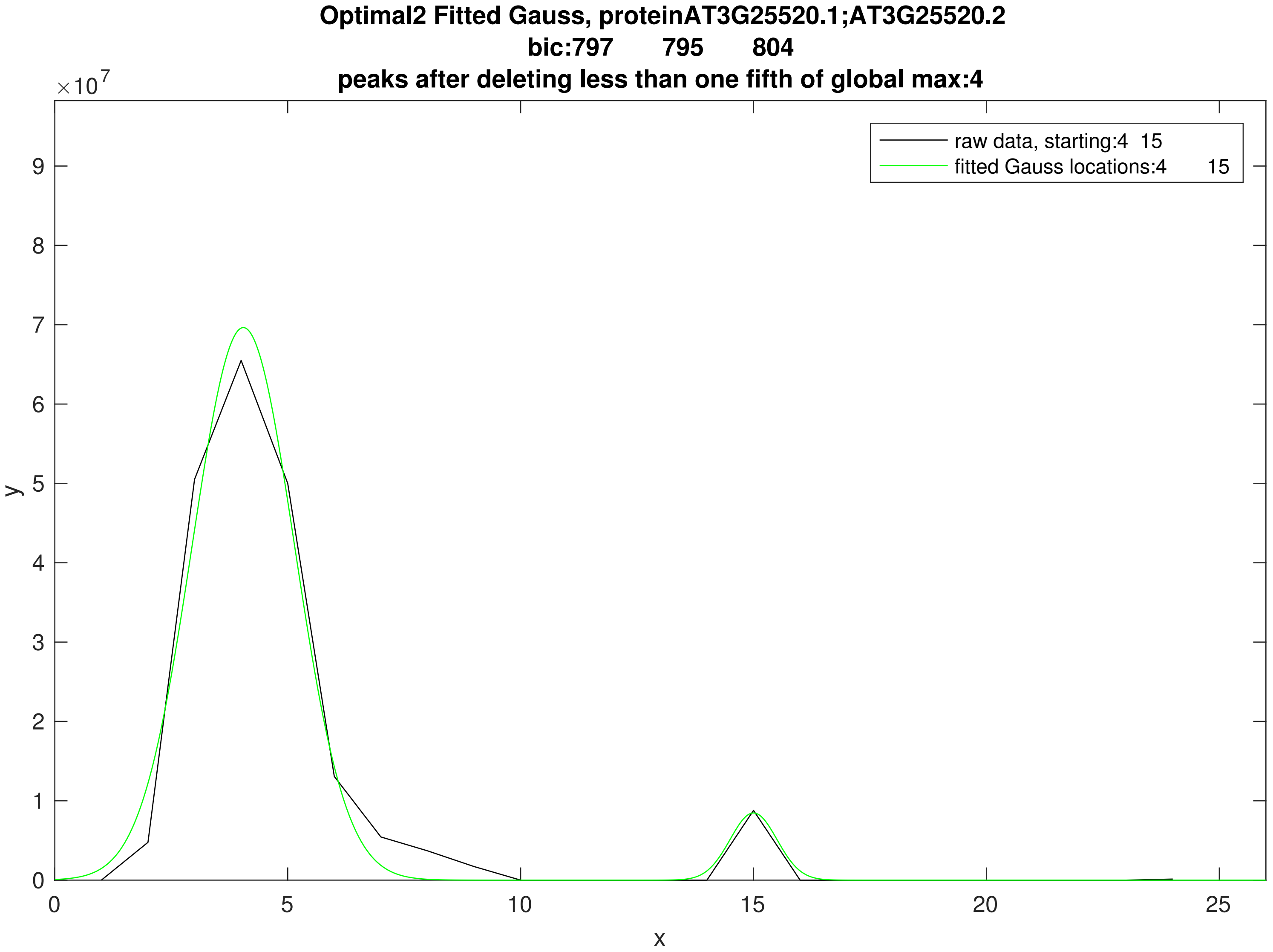
Optimal2 Fitted Gauss, proteinAT3G25520.1;AT3G25520.2
bic:797 795 804
peaks after deleting less than one fifth of global max:4
×10^7
raw data, starting:4 15
fitted Gauss locations:4 15
y
x
0
1
2
3
4
5
6
7
8
9
0
5
10
15
20
25

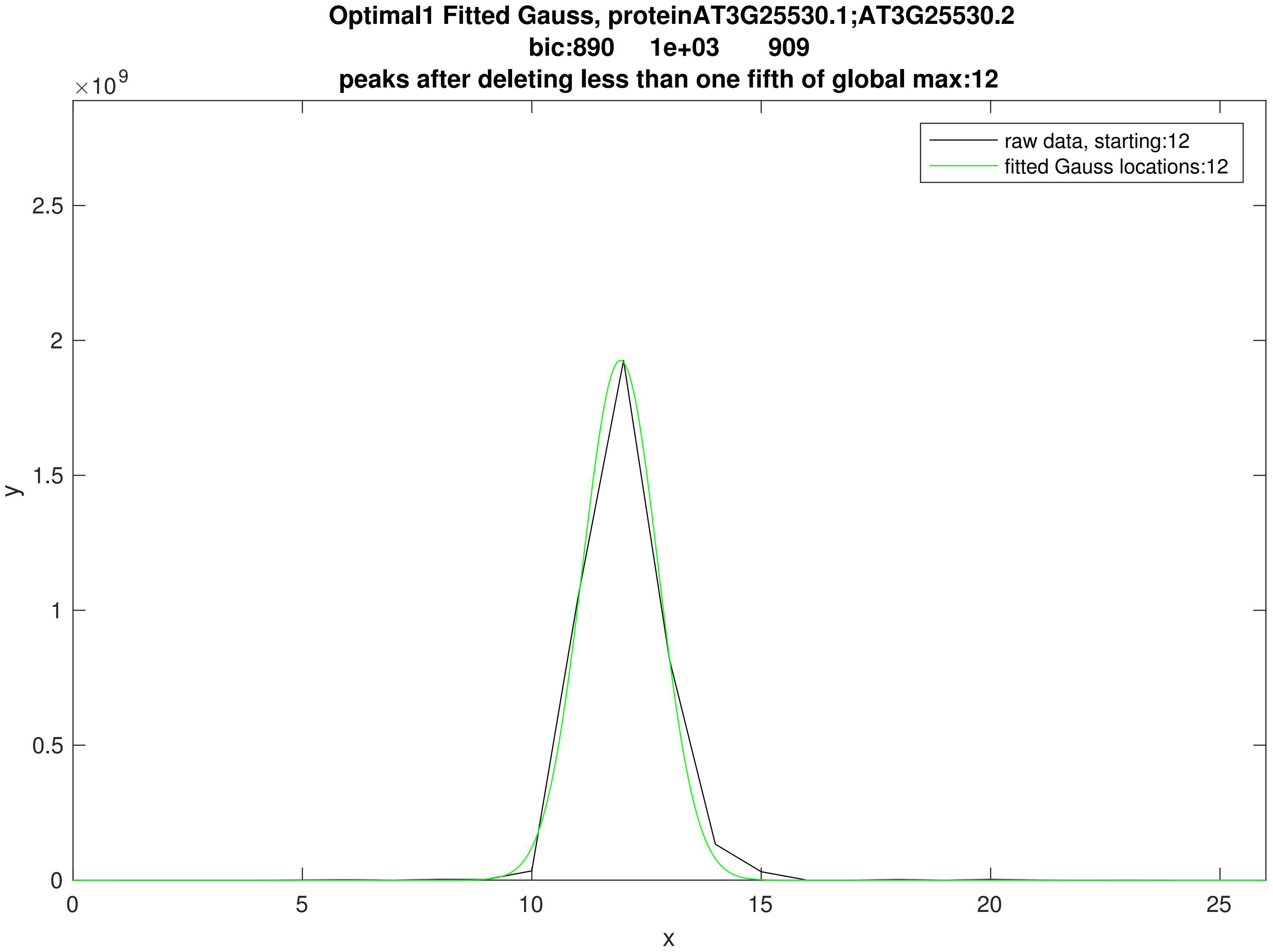

Optimal1 Fitted Gauss, proteinAT3G25530.1;AT3G25530.2
bic:890 1e+03 909
peaks after deleting less than one fifth of global max:12
×10^9
raw data, starting:12
fitted Gauss locations:12
y
x
0
0.5
1
1.5
2
2.5
5
10
15
20
25

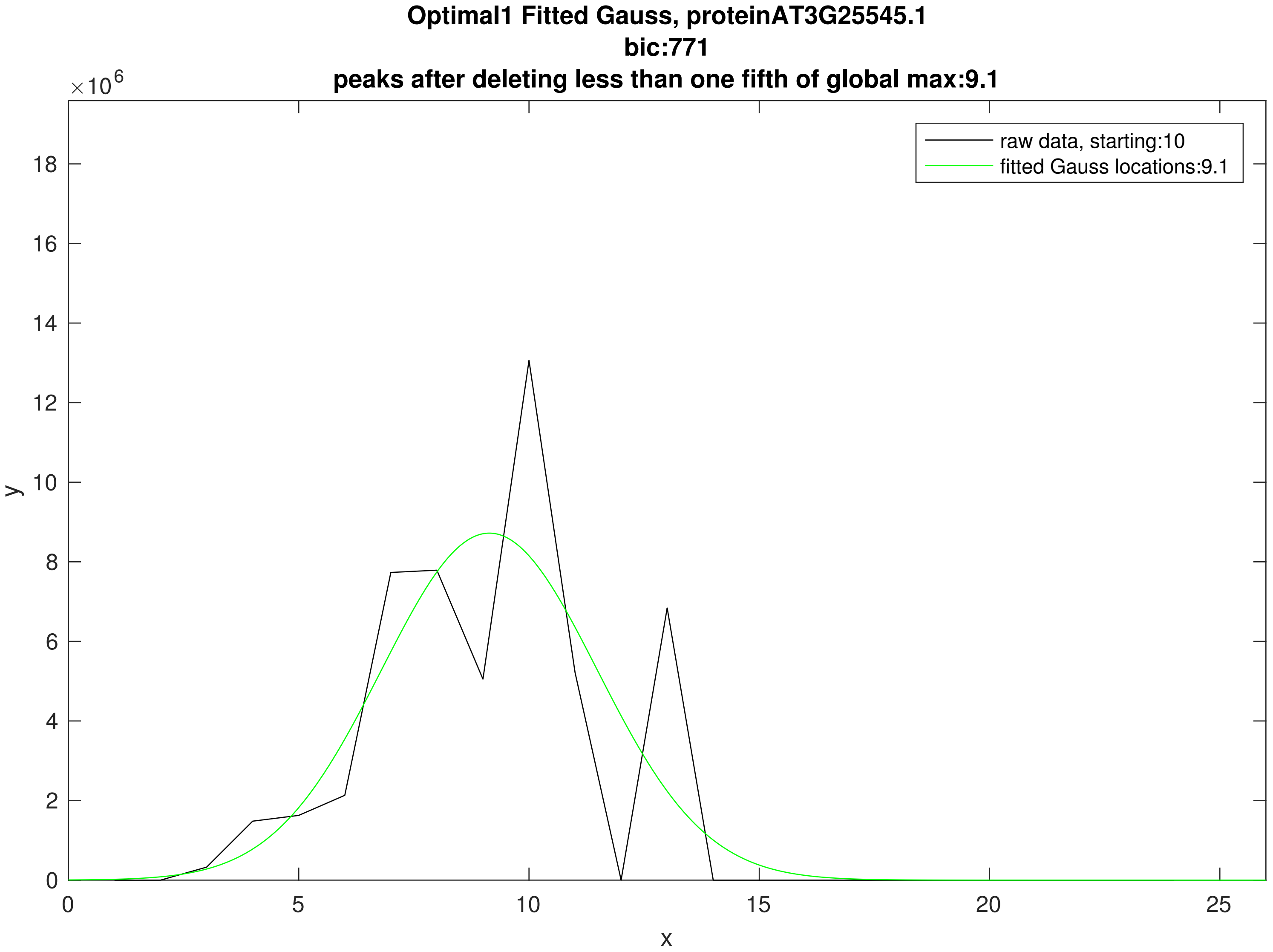

Optimal1 Fitted Gauss, proteinAT3G25545.1
bic:771
peaks after deleting less than one fifth of global max:9.1
×10^6
raw data, starting:10
fitted Gauss locations:9.1
y
x
0
2
4
6
8
10
12
14
16
18
0
5
10
15
20
25

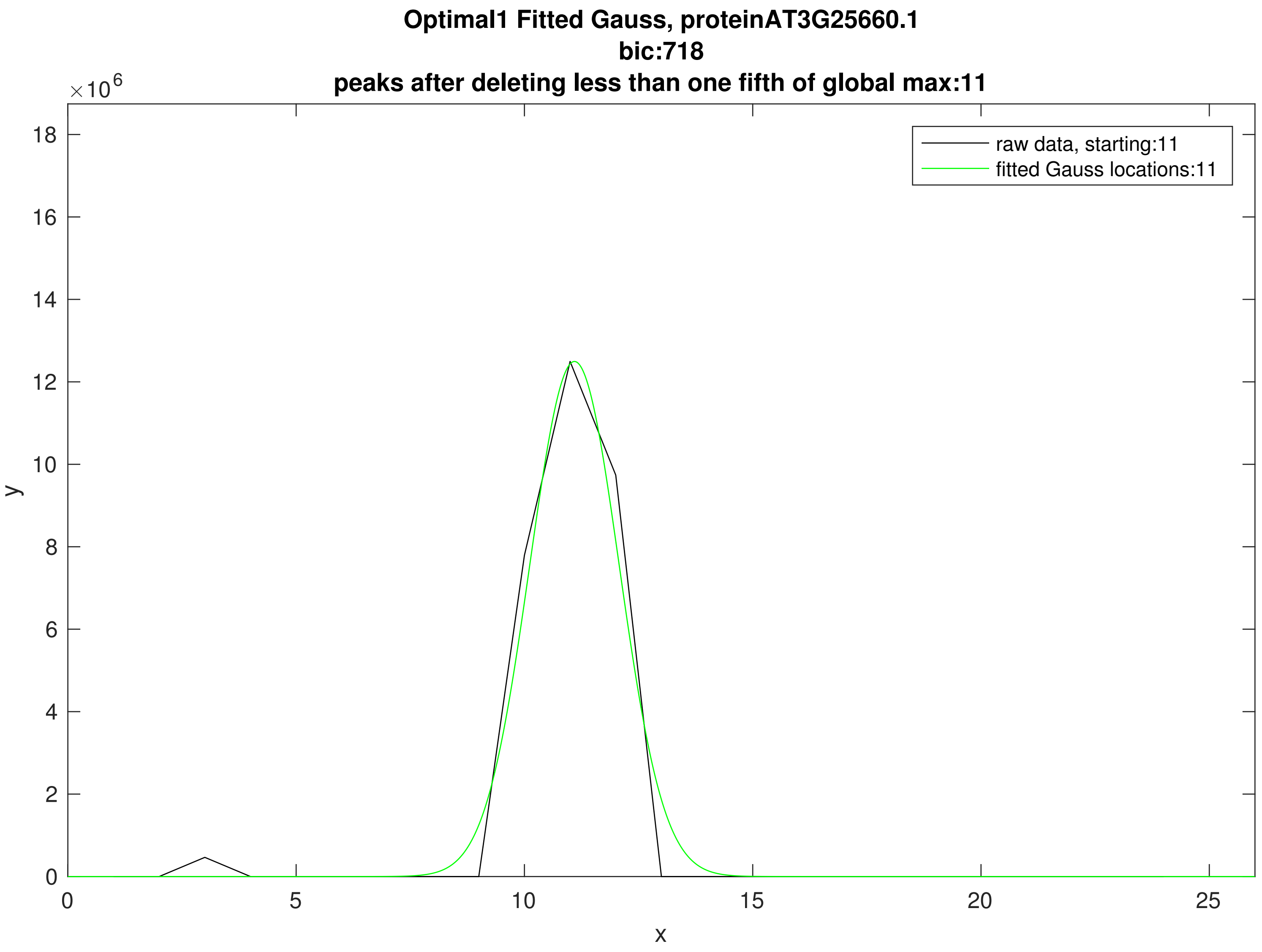

Optimal1 Fitted Gauss, proteinAT3G25660.1
bic:718
peaks after deleting less than one fifth of global max:11
×10^6
raw data, starting:11
fitted Gauss locations:11
18
16
14
12
10
8
6
4
2
0
y
0
5
10
15
20
25
x

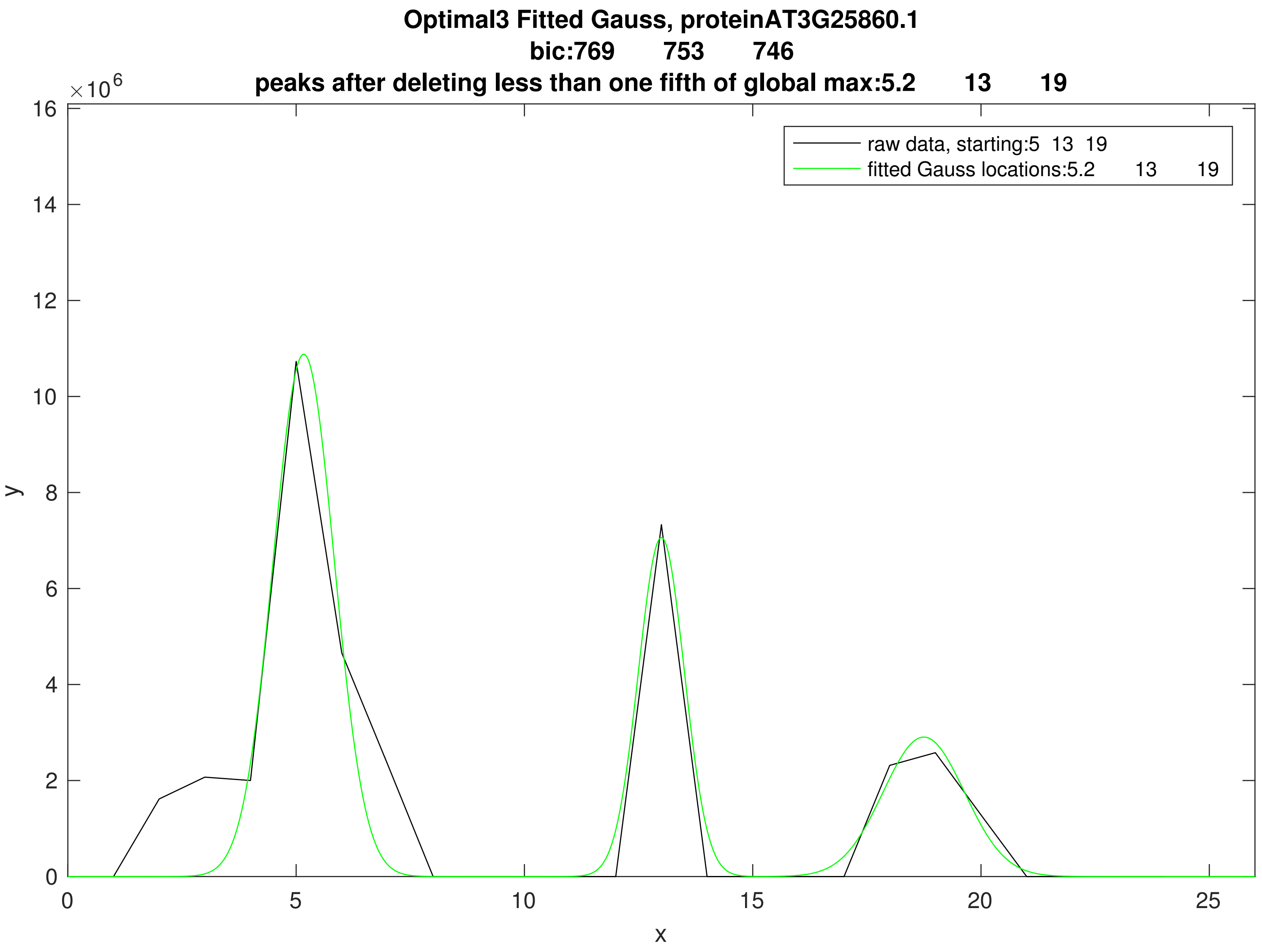
Optimal3 Fitted Gauss, proteinAT3G25860.1
bic:769 753 746
peaks after deleting less than one fifth of global max:5.2 13 19
×10^6
raw data, starting:5 13 19
fitted Gauss locations:5.2 13 19
y
x
0
2
4
6
8
10
12
14
16
0
5
10
15
20
25

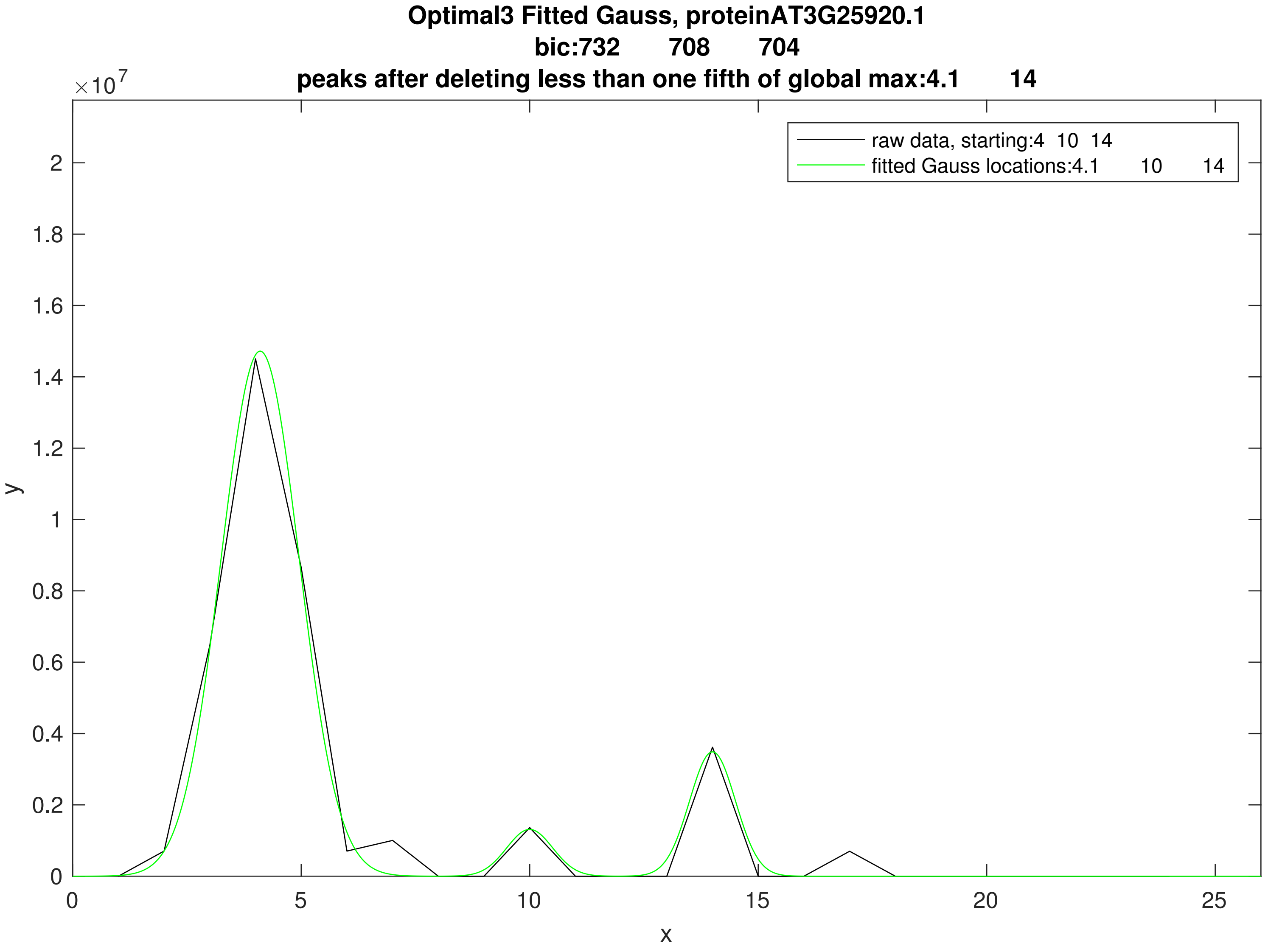

Optimal3 Fitted Gauss, proteinAT3G25920.1
bic:732 708 704
peaks after deleting less than one fifth of global max:4.1 14
×10^7
raw data, starting:4 10 14
fitted Gauss locations:4.1 10 14
y
x

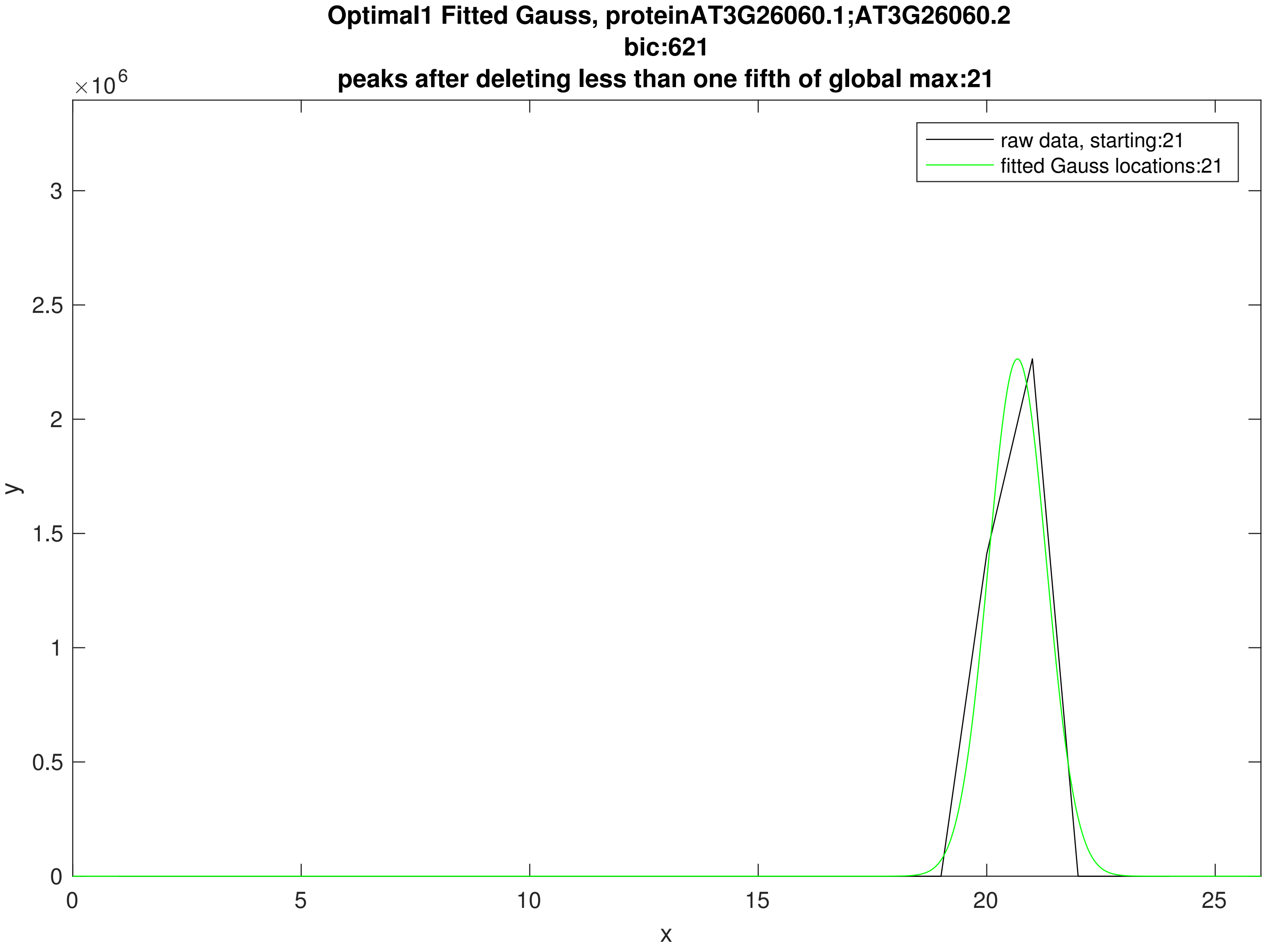
Optimal1 Fitted Gauss, proteinAT3G26060.1;AT3G26060.2
bic:621
peaks after deleting less than one fifth of global max:21
×10^6
raw data, starting:21
fitted Gauss locations:21
y
x
0
0.5
1
1.5
2
2.5
3
5
10
15
20
25

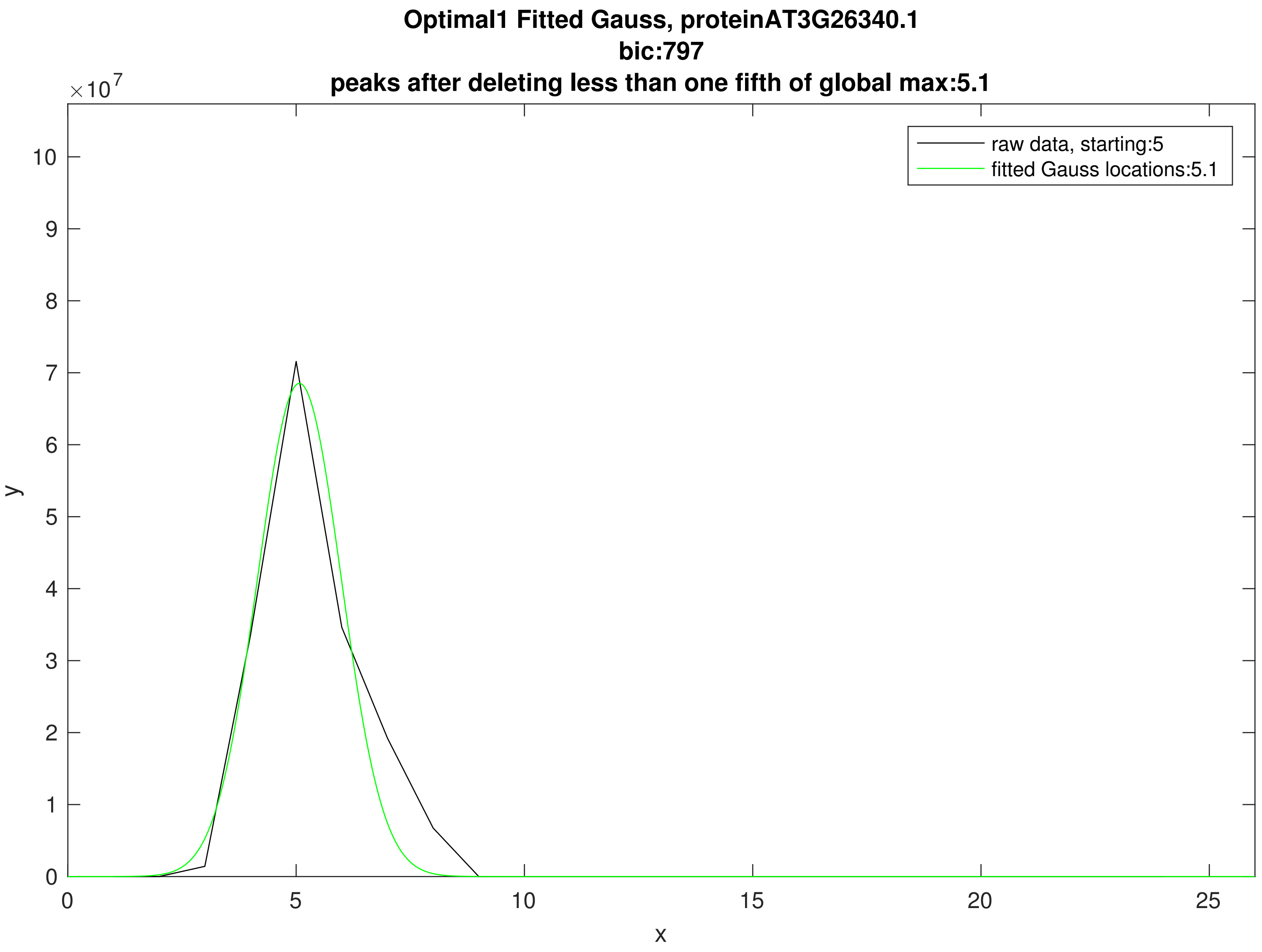
Optimal1 Fitted Gauss, proteinAT3G26340.1
bic:797
peaks after deleting less than one fifth of global max:5.1
×10^7
raw data, starting:5
fitted Gauss locations:5.1
y
x
0
1
2
3
4
5
6
7
8
9
10
0
5
10
15
20
25

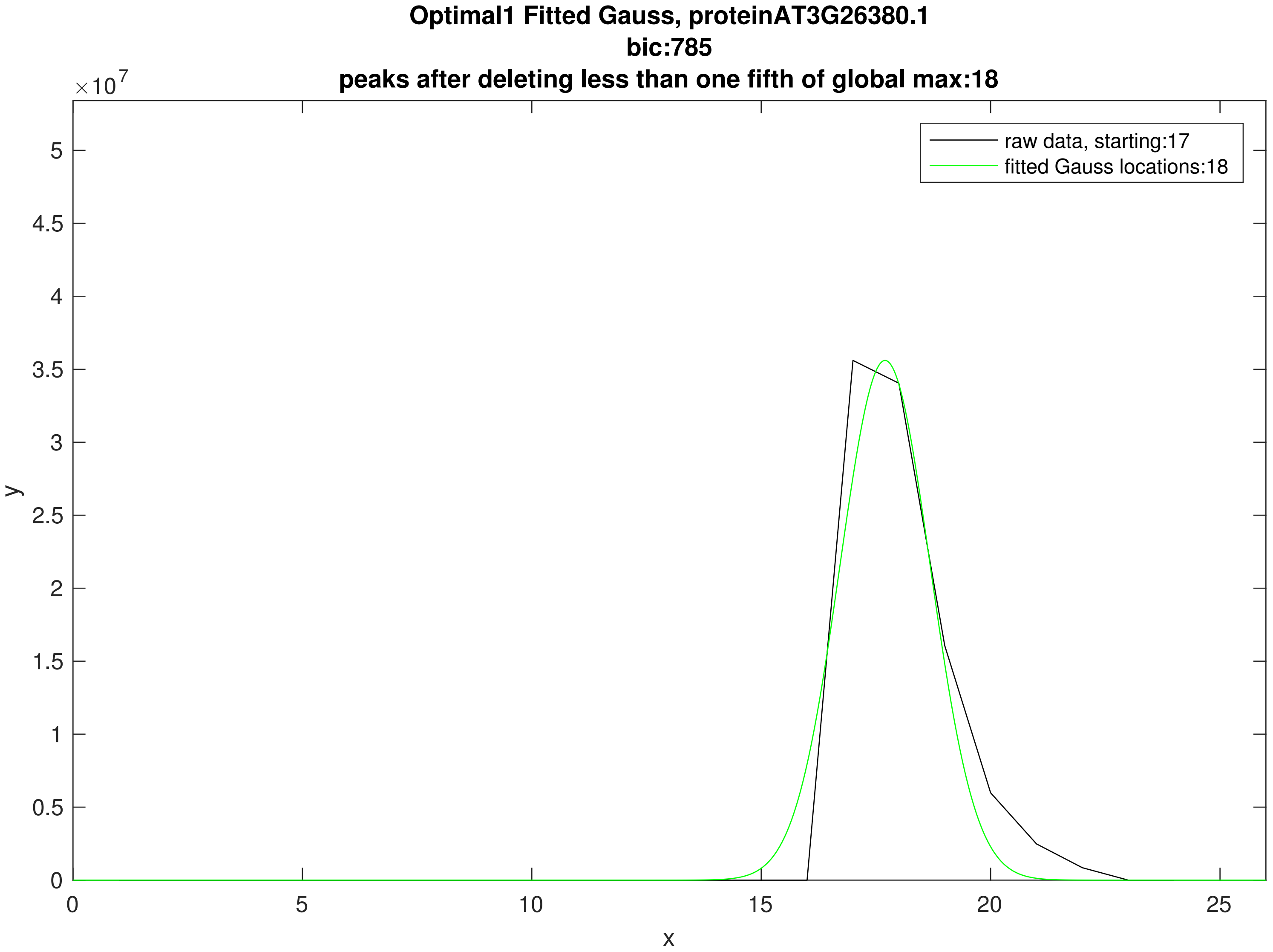
Optimal1 Fitted Gauss, proteinAT3G26380.1
bic:785
peaks after deleting less than one fifth of global max:18
×10^7
raw data, starting:17
fitted Gauss locations:18
y
x
0
0.5
1
1.5
2
2.5
3
3.5
4
4.5
5
0
5
10
15
20
25

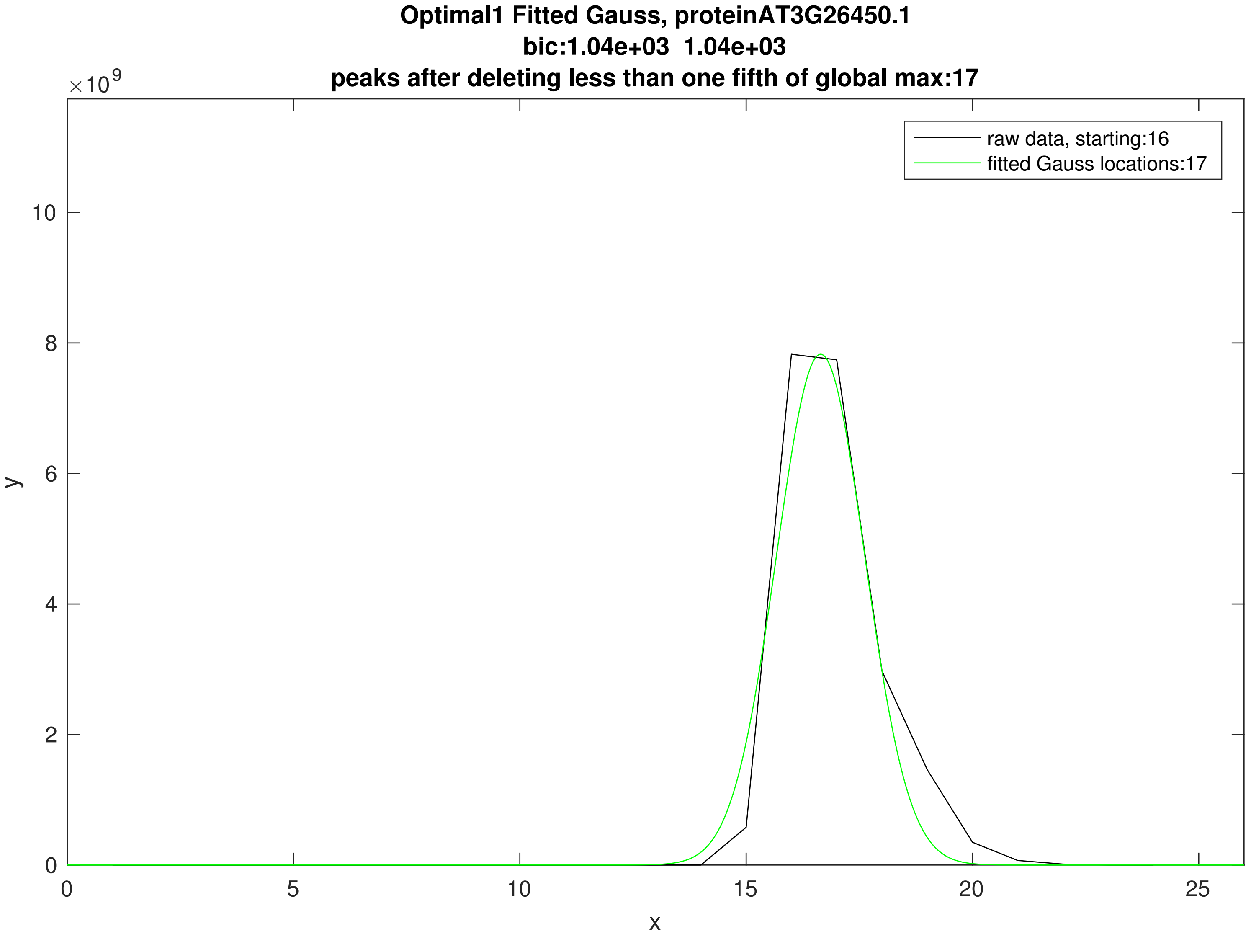
Optimal1 Fitted Gauss, proteinAT3G26450.1
bic:1.04e+03  1.04e+03
peaks after deleting less than one fifth of global max:17
×10^9
raw data, starting:16
fitted Gauss locations:17
y
x
0
2
4
6
8
10
0
5
10
15
20
25

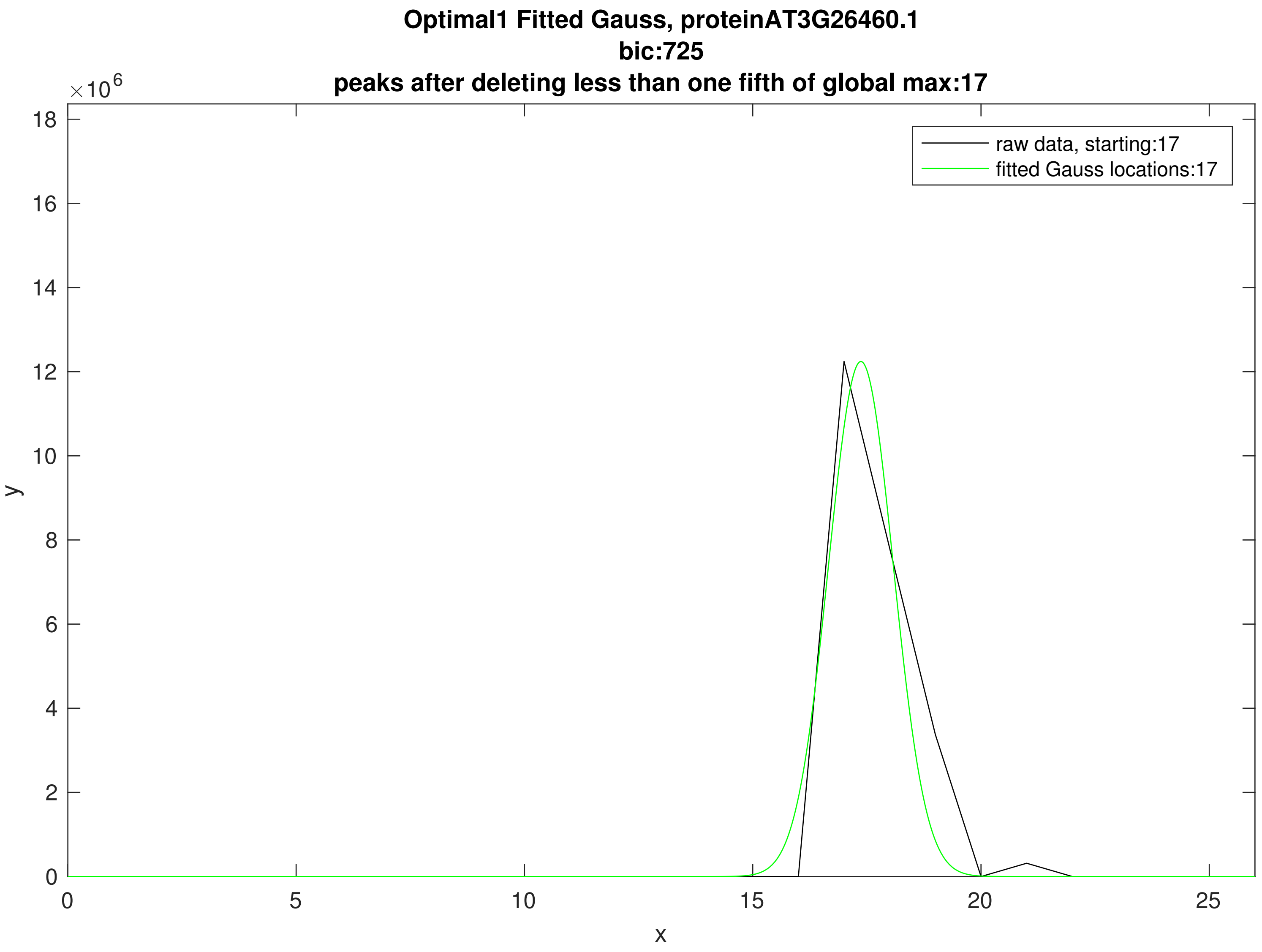
Optimal1 Fitted Gauss, proteinAT3G26460.1
bic:725
peaks after deleting less than one fifth of global max:17
×10^6
18
16
14
12
10
8
6
4
2
0
y
0
5
10
15
20
25
x
raw data, starting:17
fitted Gauss locations:17

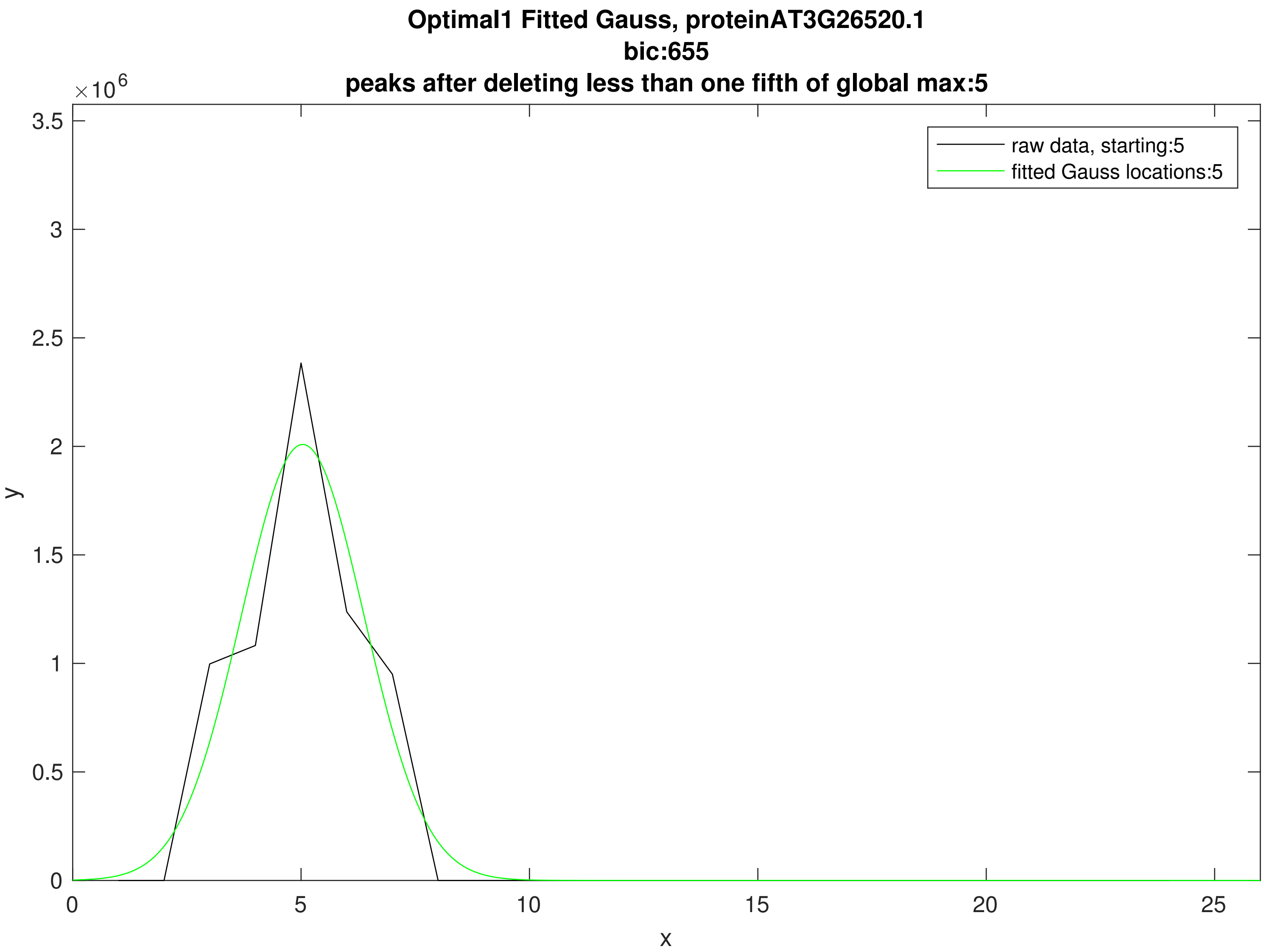
Optimal1 Fitted Gauss, proteinAT3G26520.1
bic:655
peaks after deleting less than one fifth of global max:5
×10^6
3.5
3
2.5
2
1.5
1
0.5
0
y
0
5
10
15
20
25
x
raw data, starting:5
fitted Gauss locations:5

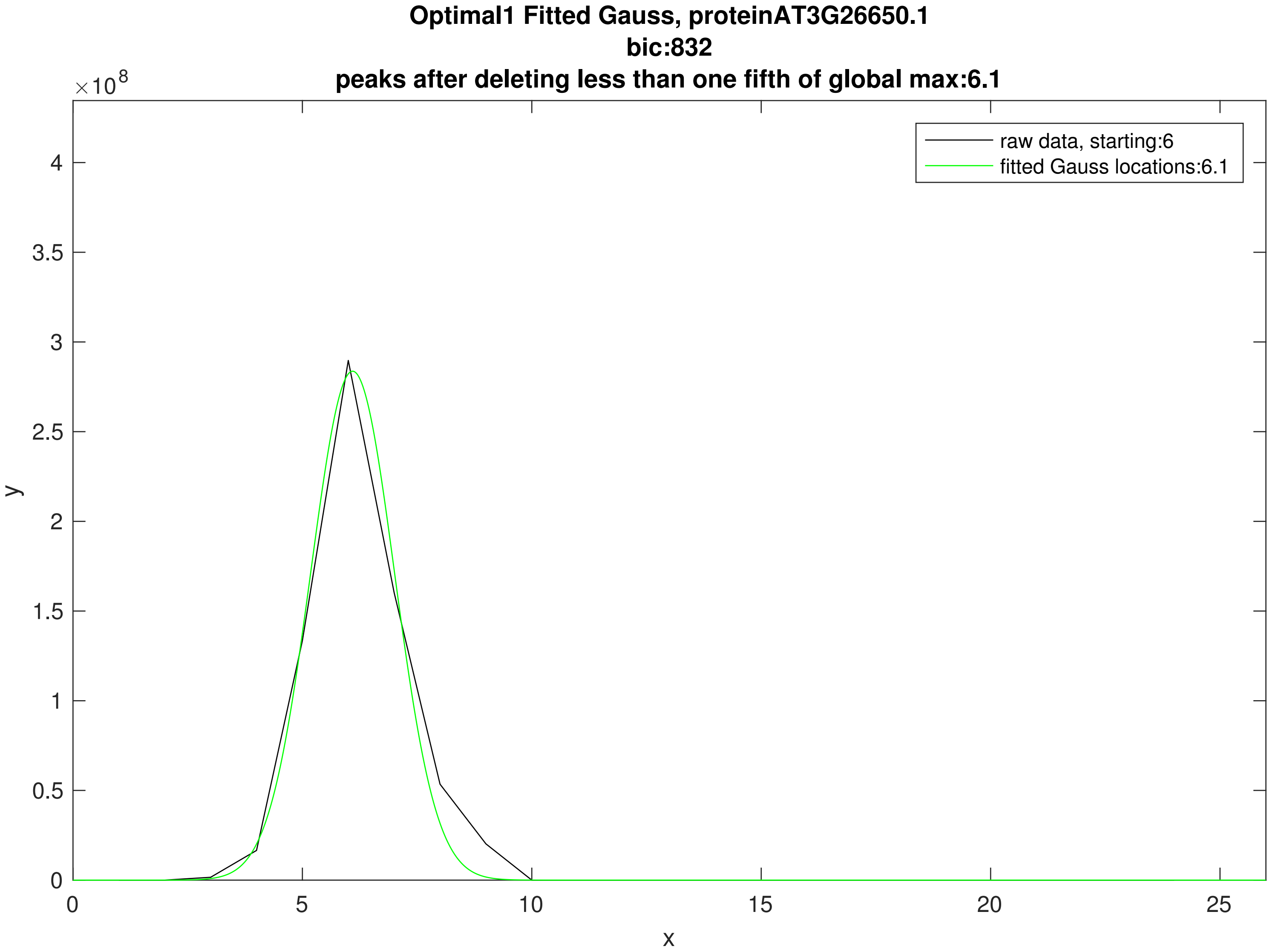
Optimal1 Fitted Gauss, proteinAT3G26650.1
bic:832
peaks after deleting less than one fifth of global max:6.1
×10^8
raw data, starting:6
fitted Gauss locations:6.1
y
x
0
0.5
1
1.5
2
2.5
3
3.5
4
0
5
10
15
20
25

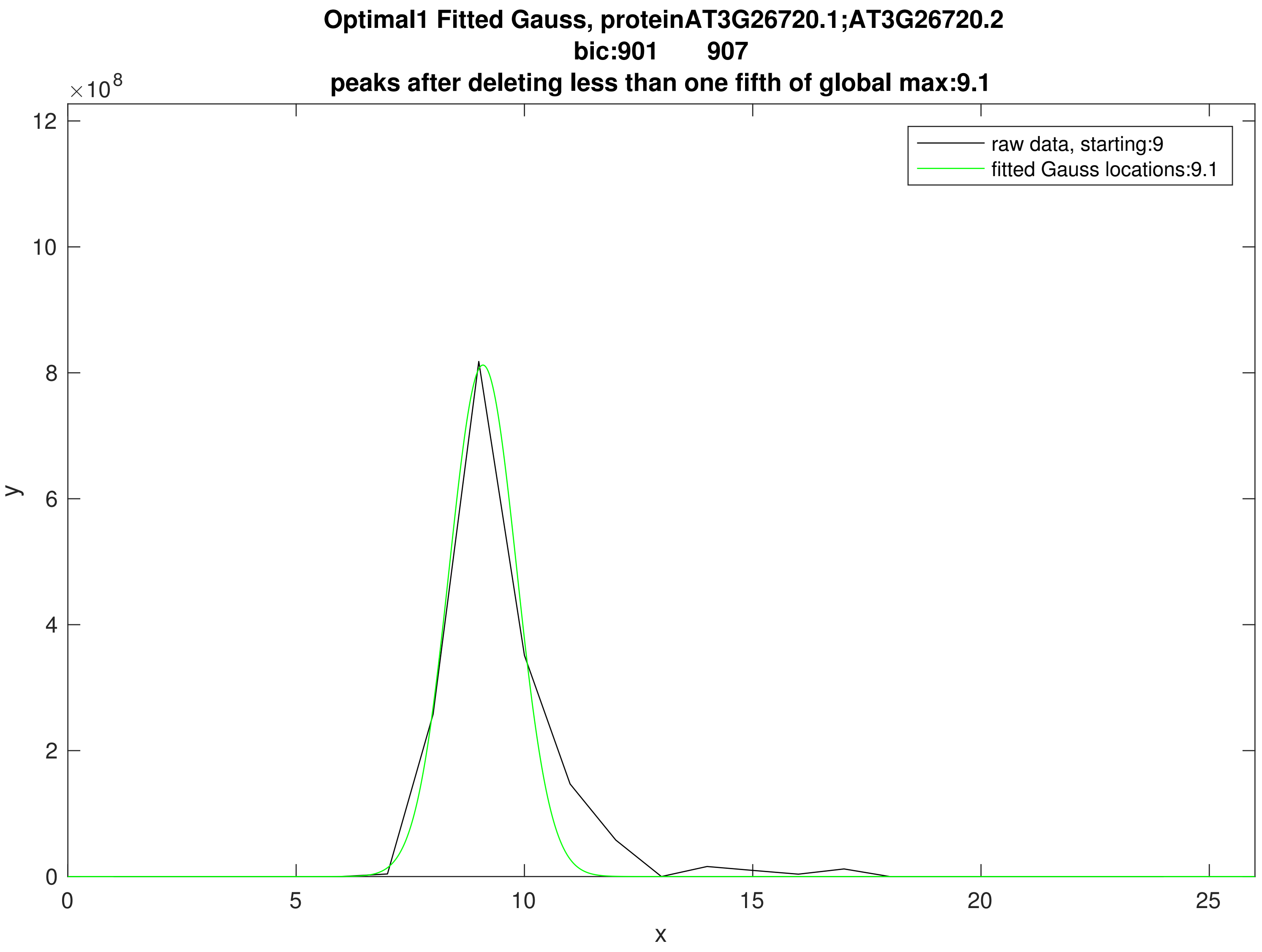

Optimal1 Fitted Gauss, proteinAT3G26720.1;AT3G26720.2
bic:901 907
peaks after deleting less than one fifth of global max:9.1
×10^8
12
10
8
6
4
2
0
y
0
5
10
15
20
25
x
raw data, starting:9
fitted Gauss locations:9.1

**Optimal1 Fitted Gauss, proteinAT3G26760.1**
**bic:751 760**
**peaks after deleting less than one fifth of global max:13**

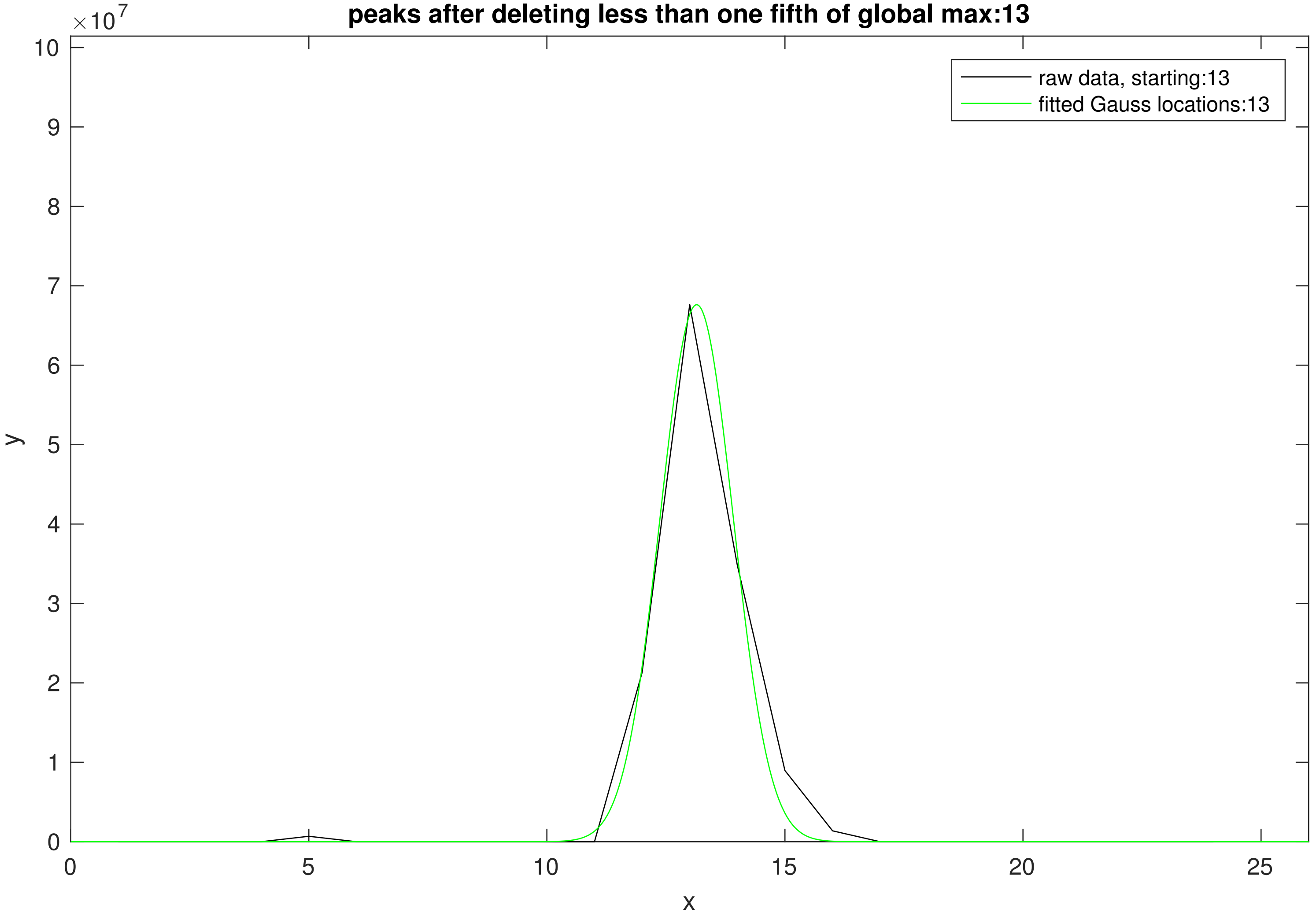

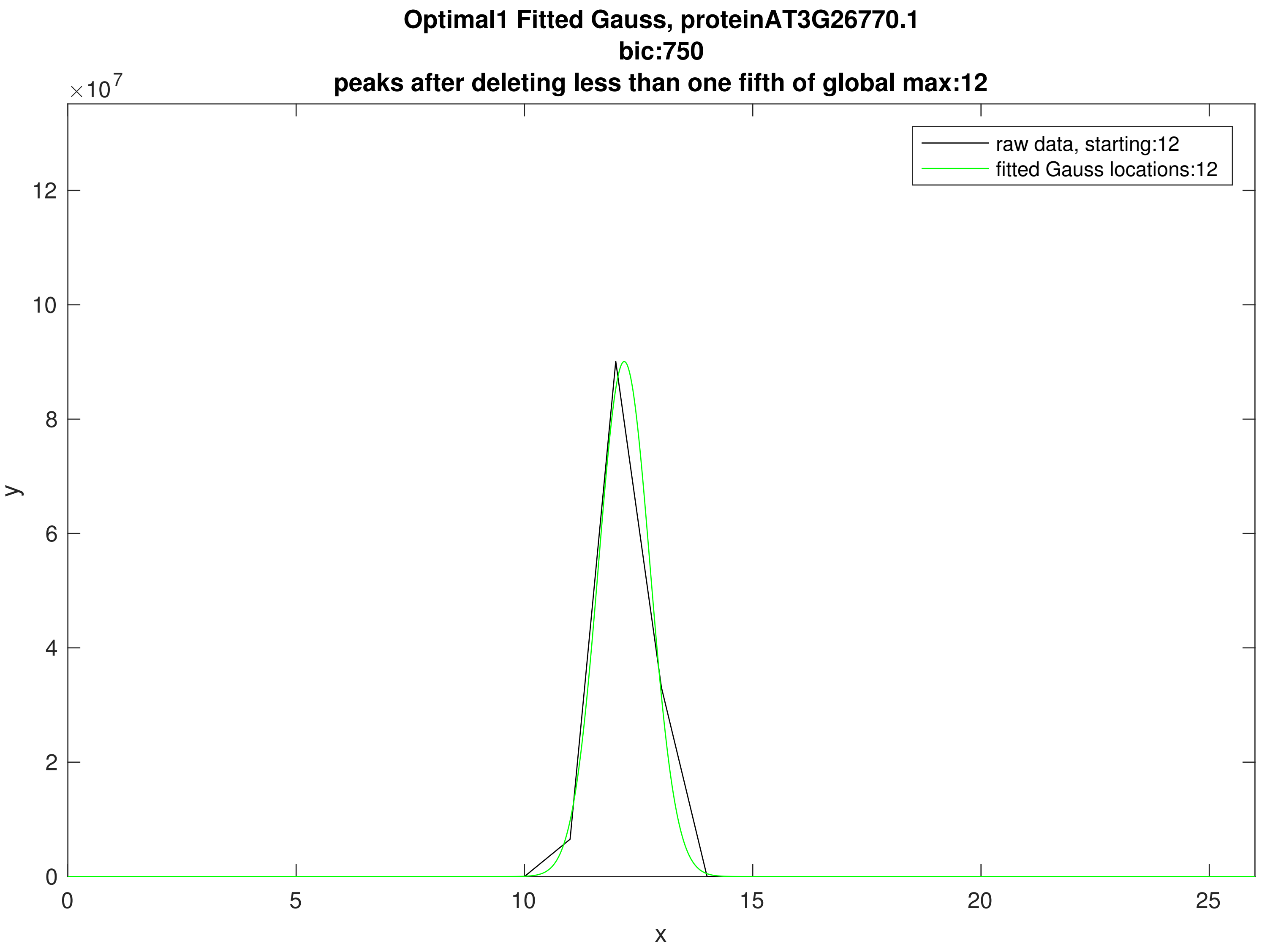
Optimal1 Fitted Gauss, proteinAT3G26770.1
bic:750
peaks after deleting less than one fifth of global max:12
×10^7
raw data, starting:12
fitted Gauss locations:12
y
x
0
2
4
6
8
10
12
0
5
10
15
20
25

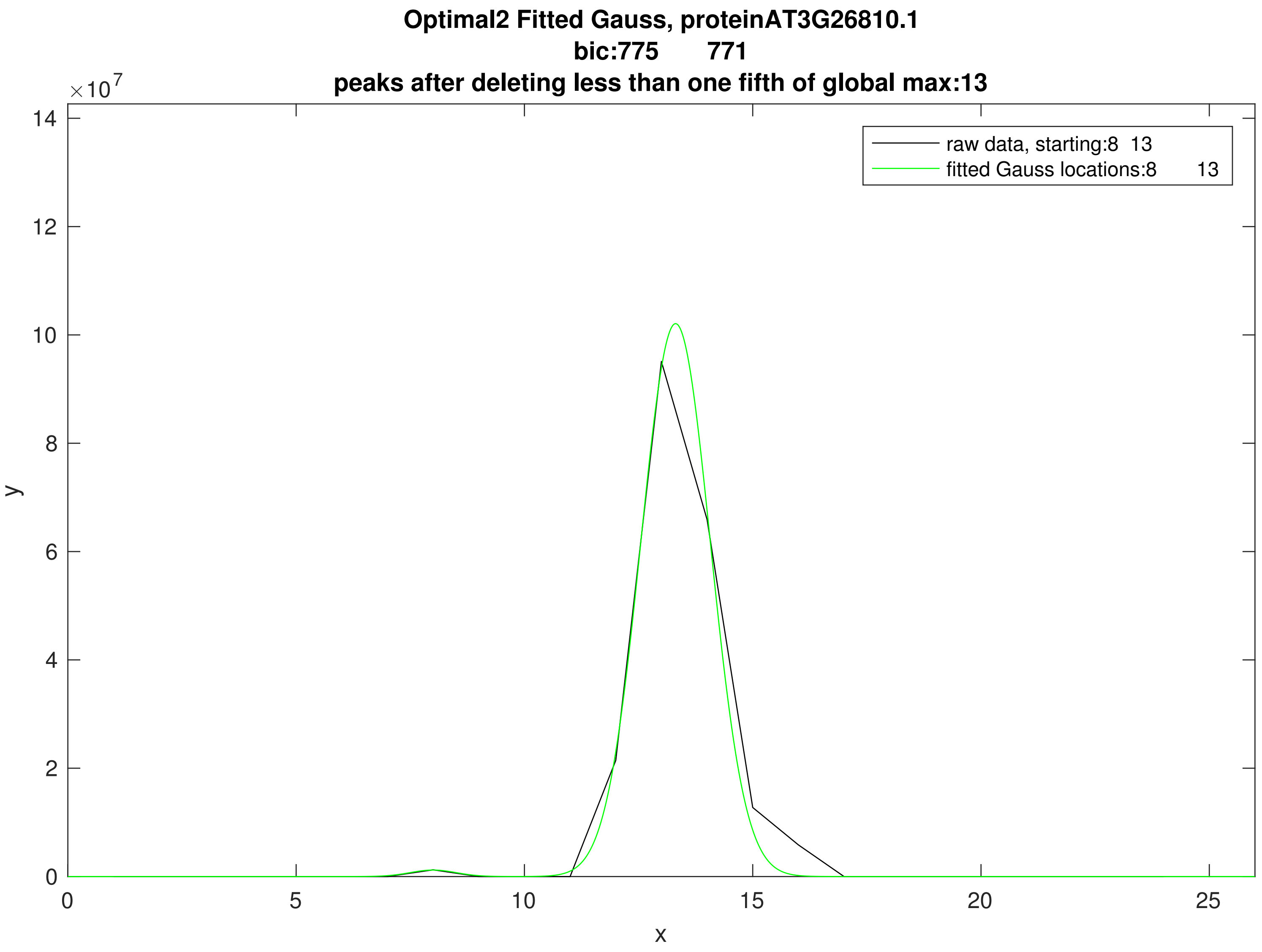

Optimal2 Fitted Gauss, proteinAT3G26810.1
bic:775 771
peaks after deleting less than one fifth of global max:13
×10^7
raw data, starting:8 13
fitted Gauss locations:8 13
14
12
10
8
6
4
2
0
y
0
5
10
15
20
25
x

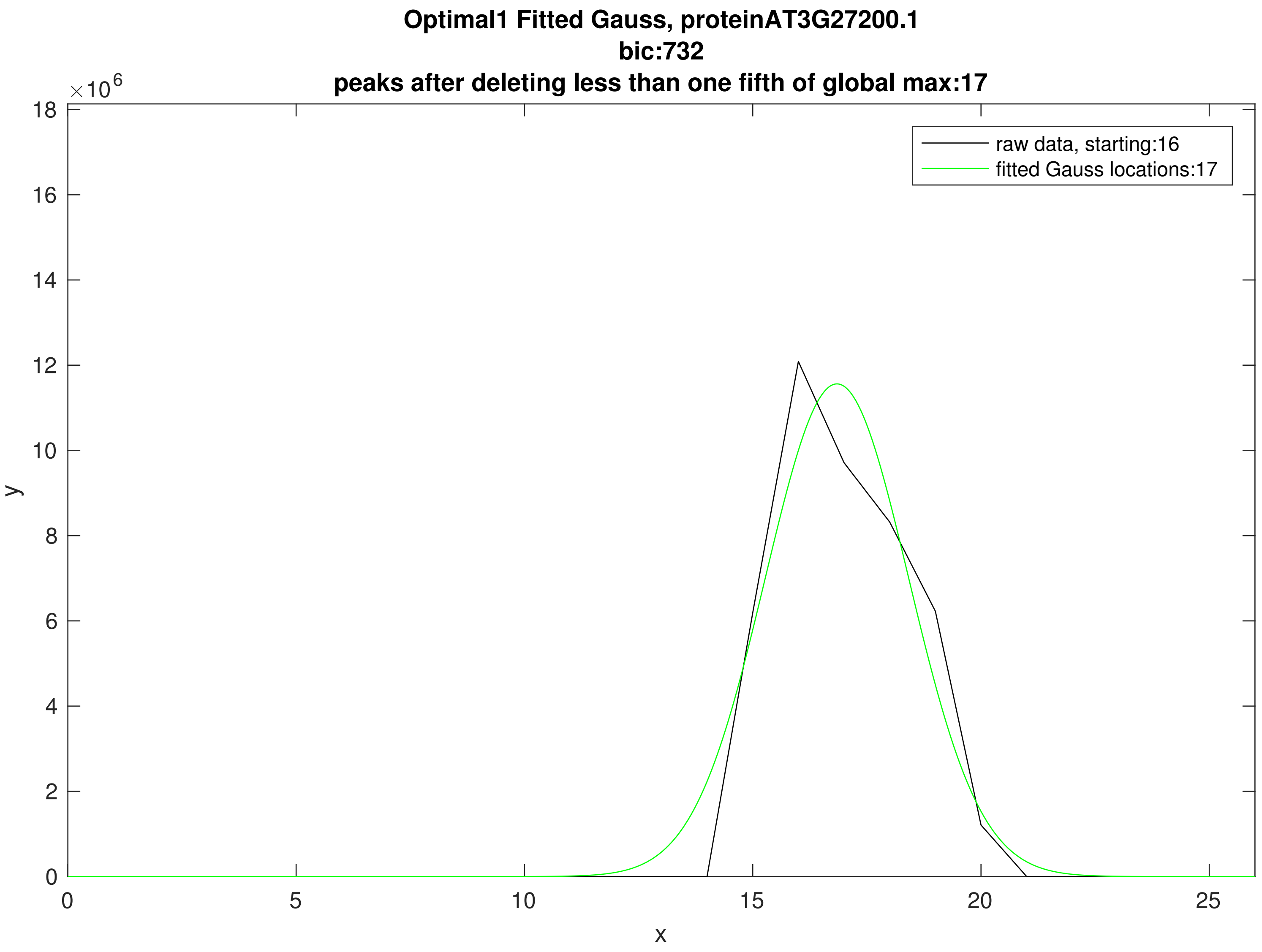

Optimal1 Fitted Gauss, proteinAT3G27200.1
bic:732
peaks after deleting less than one fifth of global max:17
×10^6
raw data, starting:16
fitted Gauss locations:17
18
16
14
12
10
8
6
4
2
0
y
0
5
10
15
20
25
x

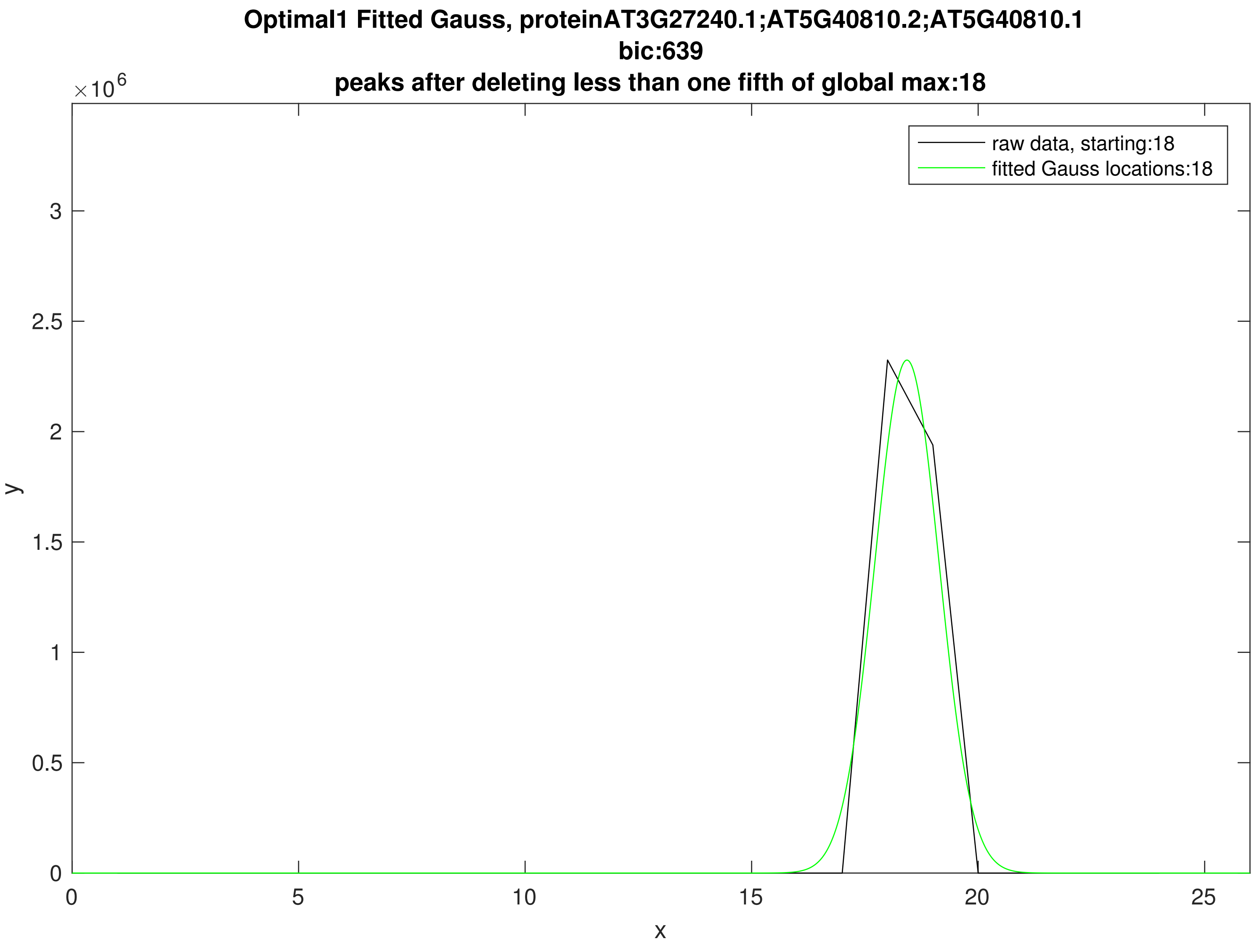
Optimal1 Fitted Gauss, proteinAT3G27240.1;AT5G40810.2;AT5G40810.1
bic:639
peaks after deleting less than one fifth of global max:18
×10^6
raw data, starting:18
fitted Gauss locations:18
y
x
0
0.5
1
1.5
2
2.5
3
0
5
10
15
20
25

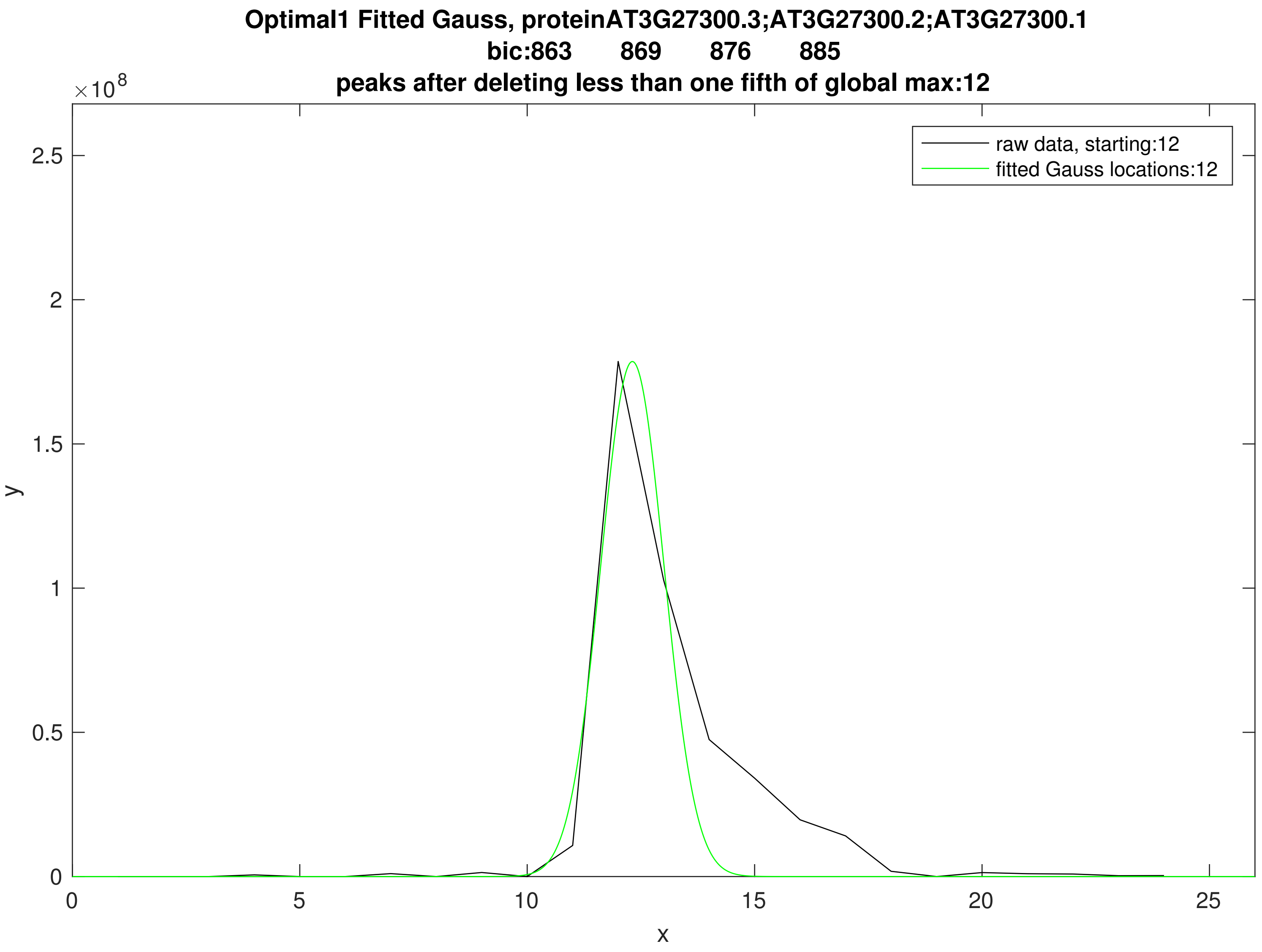

Optimal1 Fitted Gauss, proteinAT3G27300.3;AT3G27300.2;AT3G27300.1
bic:863 869 876 885
peaks after deleting less than one fifth of global max:12
×10^8
2.5
2
1.5
1
0.5
0
y
0
5
10
15
20
25
x
raw data, starting:12
fitted Gauss locations:12

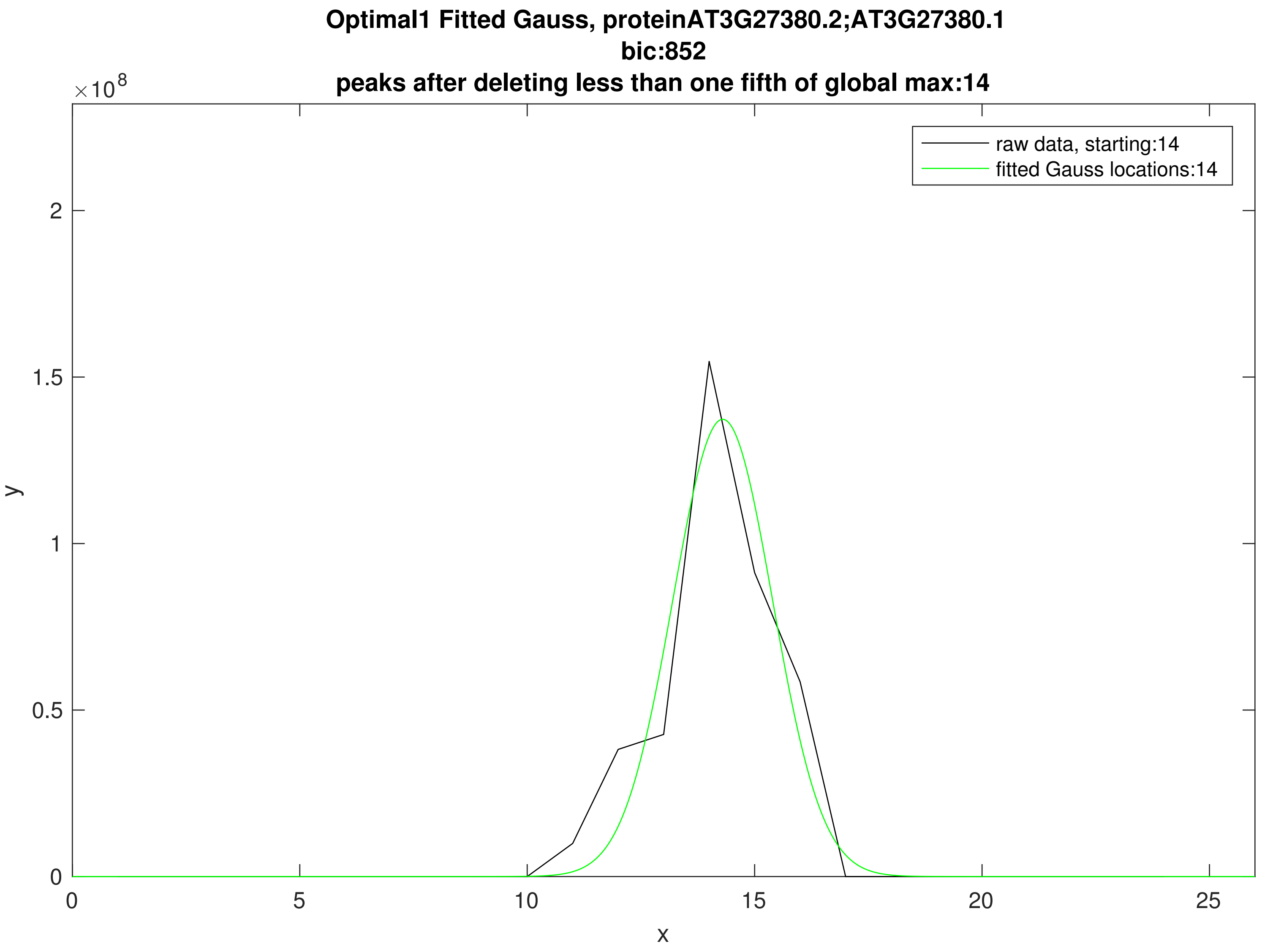
Optimal1 Fitted Gauss, proteinAT3G27380.2;AT3G27380.1
bic:852
peaks after deleting less than one fifth of global max:14
×10^8
raw data, starting:14
fitted Gauss locations:14
y
x
0
0.5
1
1.5
2
0
5
10
15
20
25

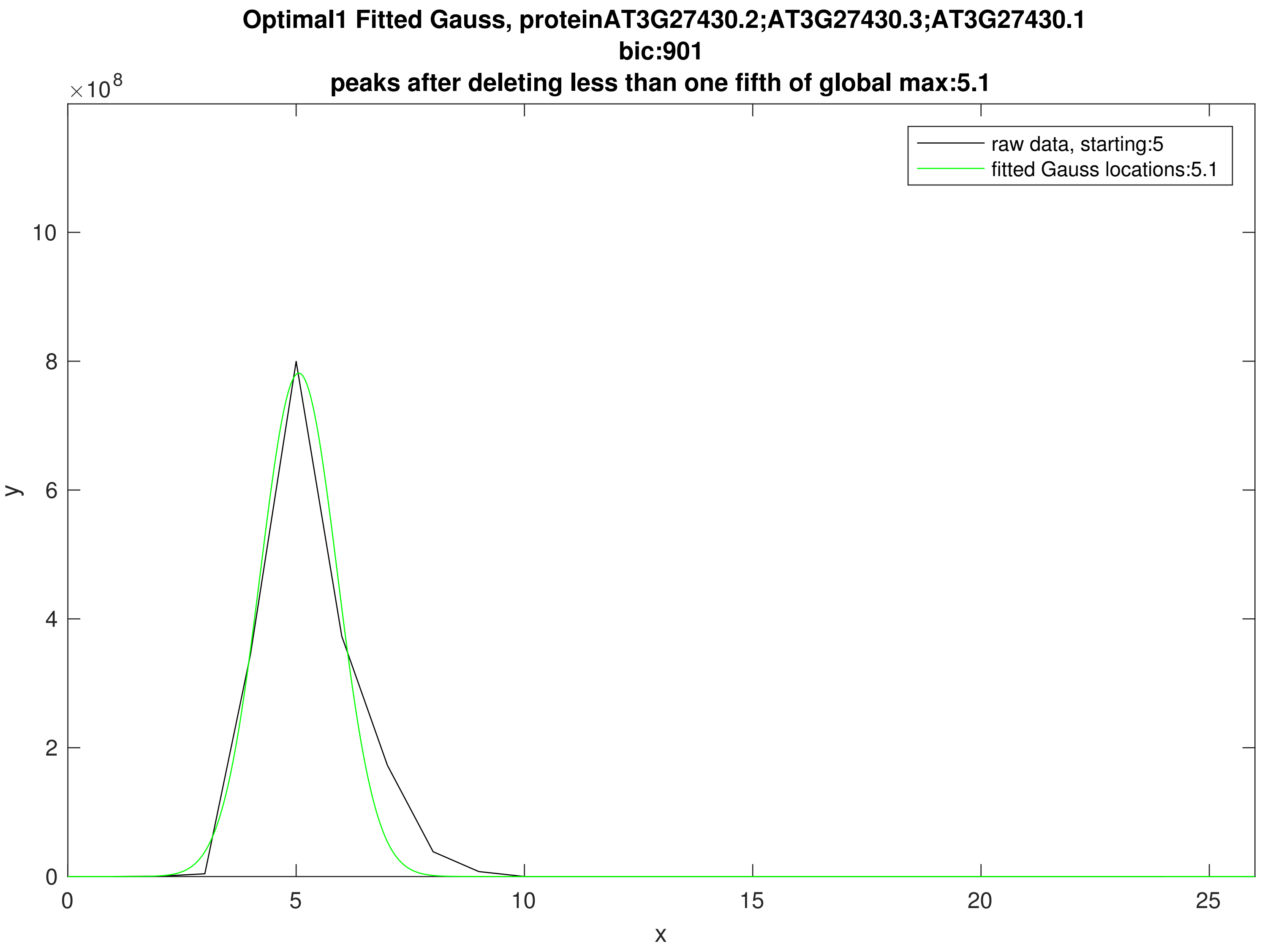
Optimal1 Fitted Gauss, proteinAT3G27430.2;AT3G27430.3;AT3G27430.1
bic:901
peaks after deleting less than one fifth of global max:5.1
×10^8
raw data, starting:5
fitted Gauss locations:5.1
y
x
0
2
4
6
8
10
0
5
10
15
20
25

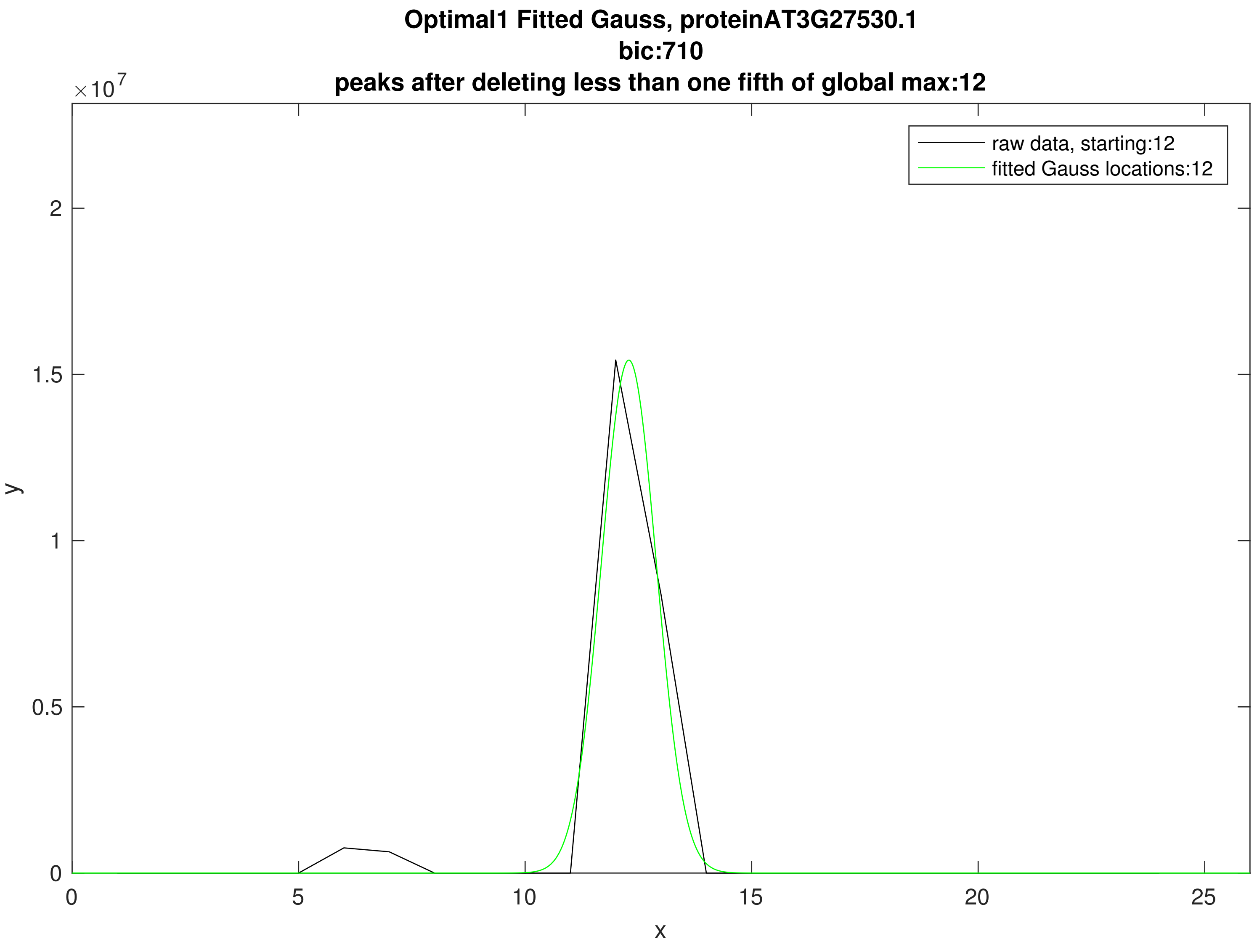

Optimal1 Fitted Gauss, proteinAT3G27530.1
bic:710
peaks after deleting less than one fifth of global max:12
×10^7
raw data, starting:12
fitted Gauss locations:12
2
1.5
1
0.5
0
y
0
5
10
15
20
25
x

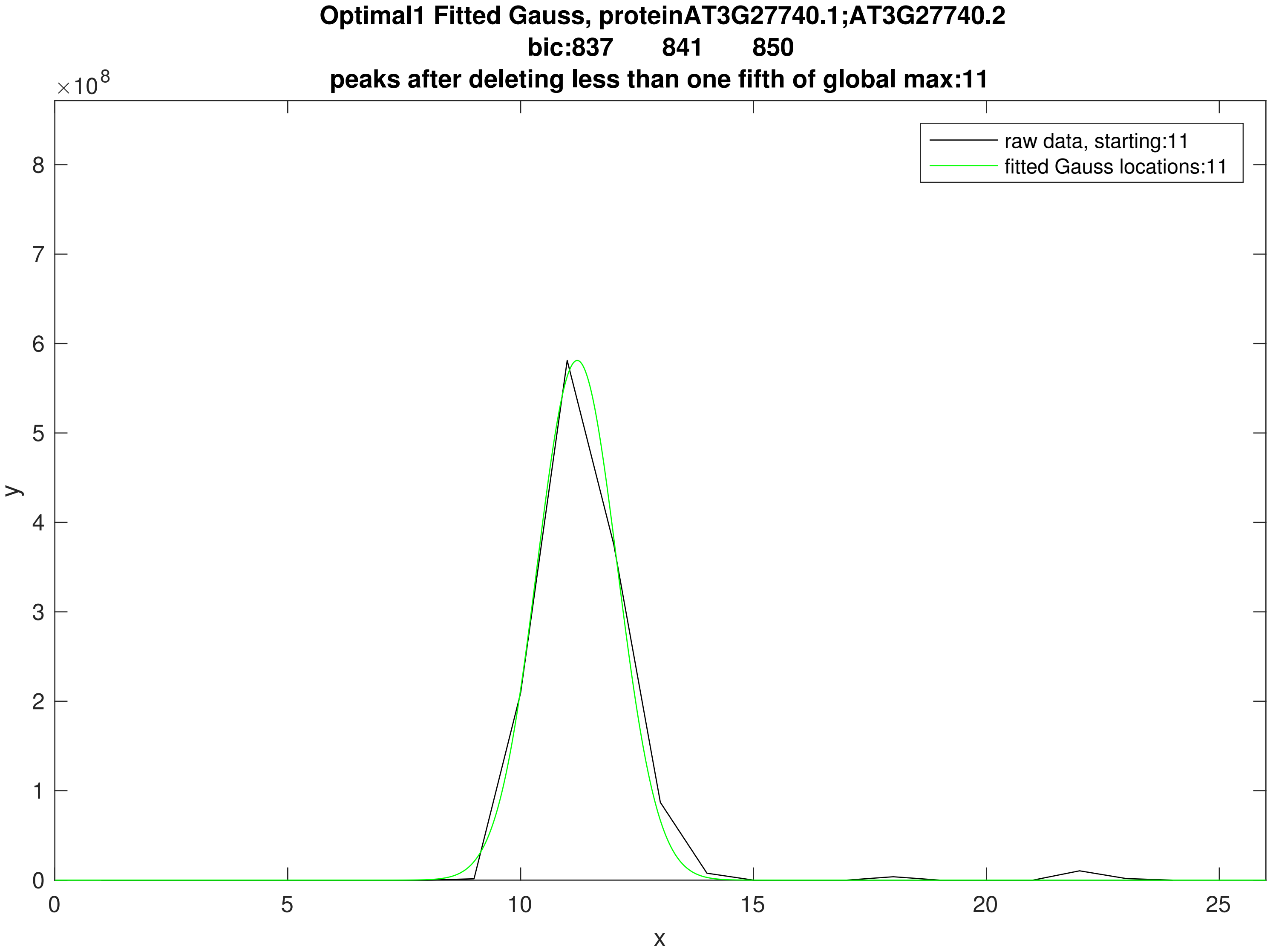

Optimal1 Fitted Gauss, proteinAT3G27740.1;AT3G27740.2
bic:837 841 850
peaks after deleting less than one fifth of global max:11
×10^8
y
x
raw data, starting:11
fitted Gauss locations:11
0
1
2
3
4
5
6
7
8
0
5
10
15
20
25

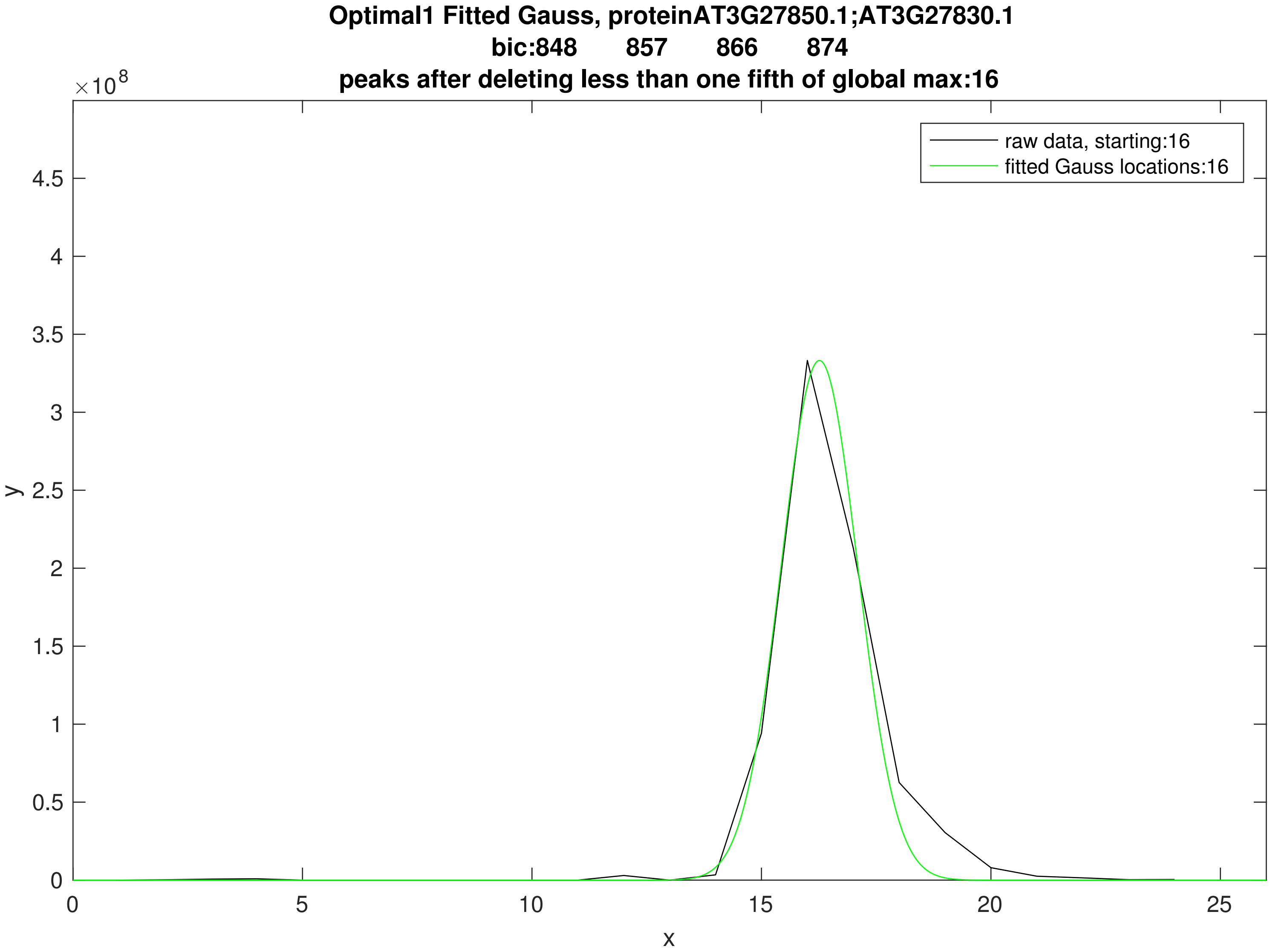
Optimal1 Fitted Gauss, proteinAT3G27850.1;AT3G27830.1
bic:848 857 866 874
peaks after deleting less than one fifth of global max:16
×10^8
raw data, starting:16
fitted Gauss locations:16
4.5
4
3.5
3
2.5
2
1.5
1
0.5
0
y
0
5
10
15
20
25
x

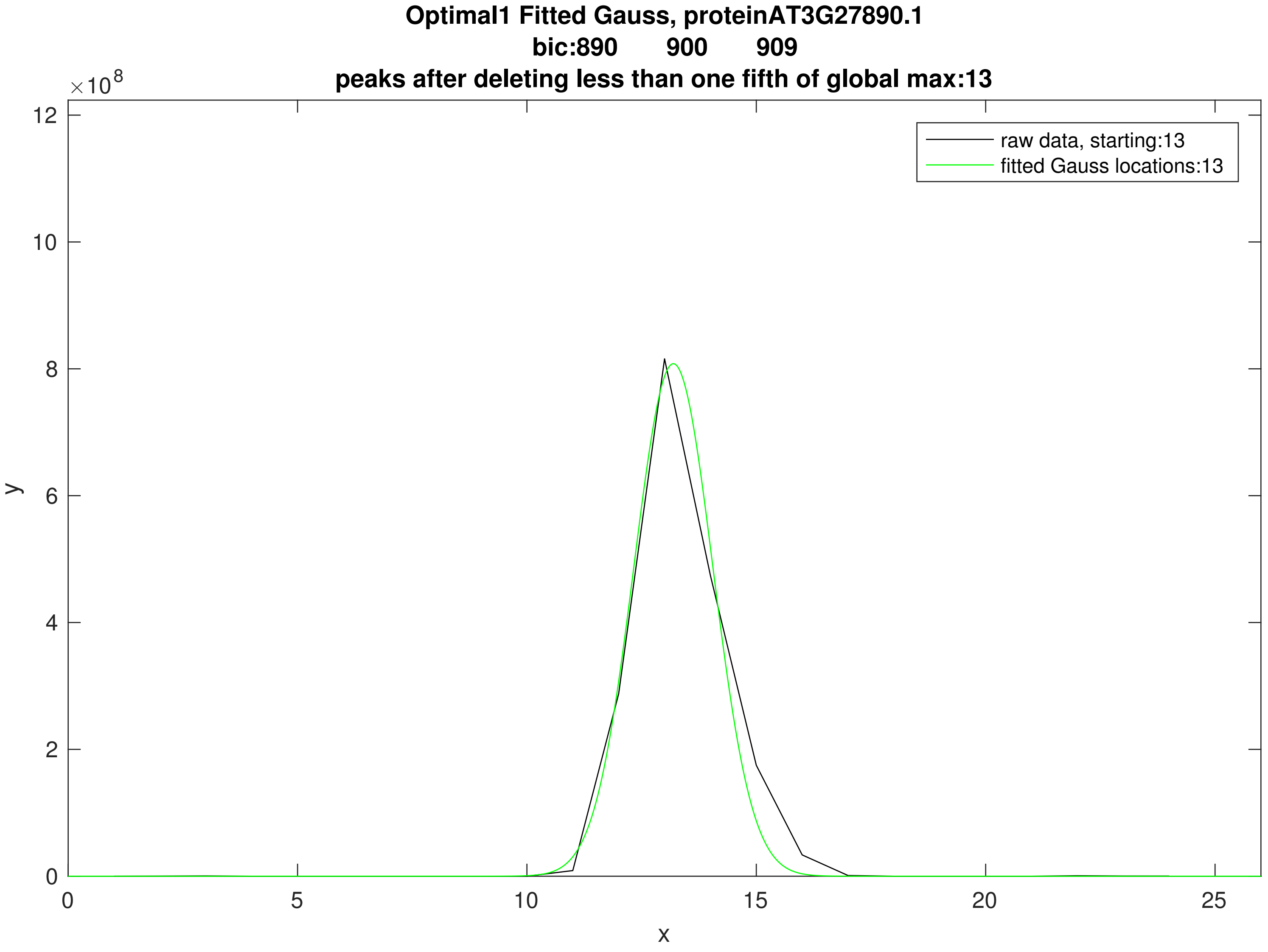

Optimal1 Fitted Gauss, proteinAT3G27890.1
bic:890 900 909
peaks after deleting less than one fifth of global max:13
×10^8
12
10
8
6
4
2
0
y
0
5
10
15
20
25
x
raw data, starting:13
fitted Gauss locations:13

#### Optimal1 Fitted Gauss, proteinAT3G28390.1
#### bic:735
#### peaks after deleting less than one fifth of global max:14

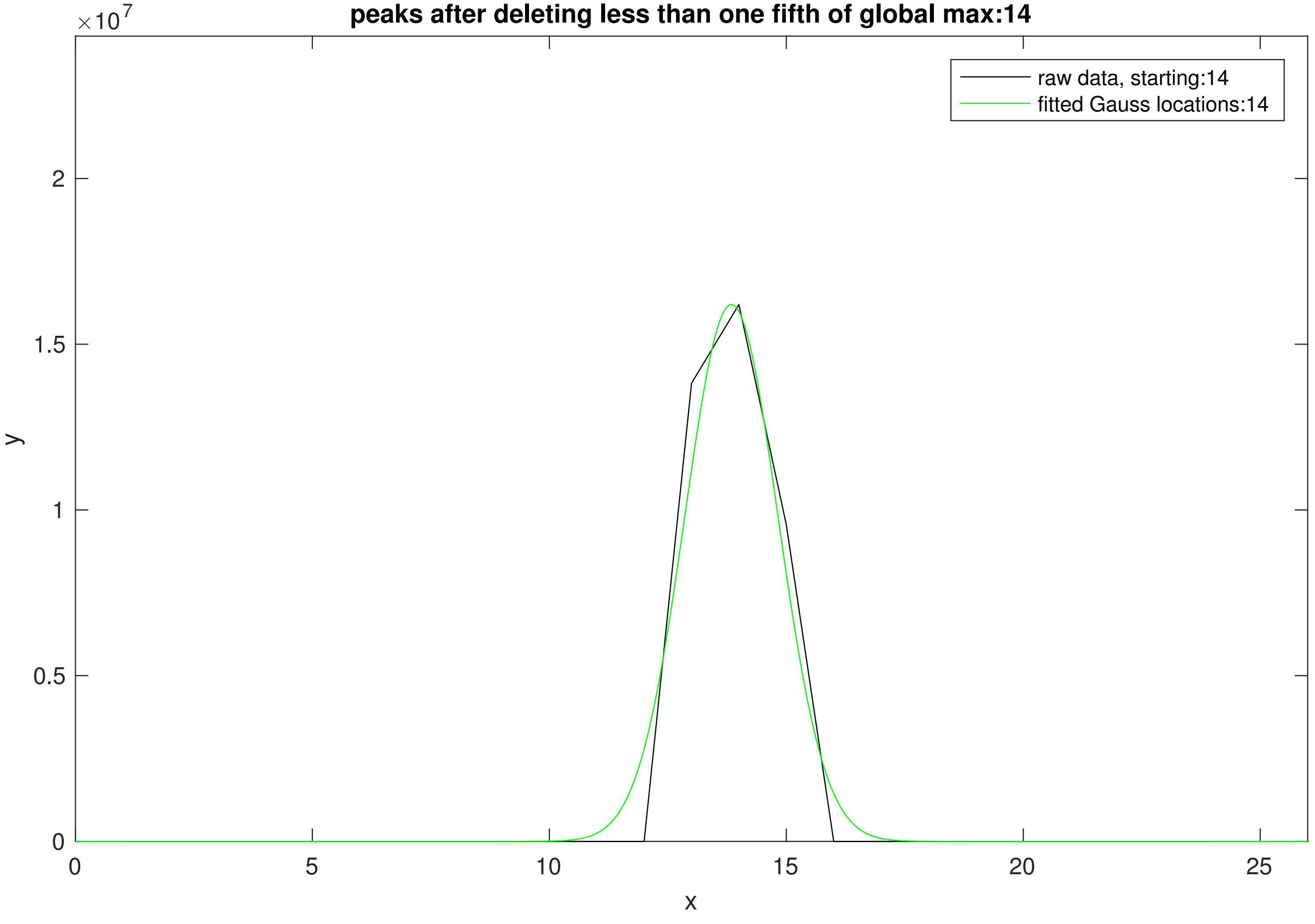

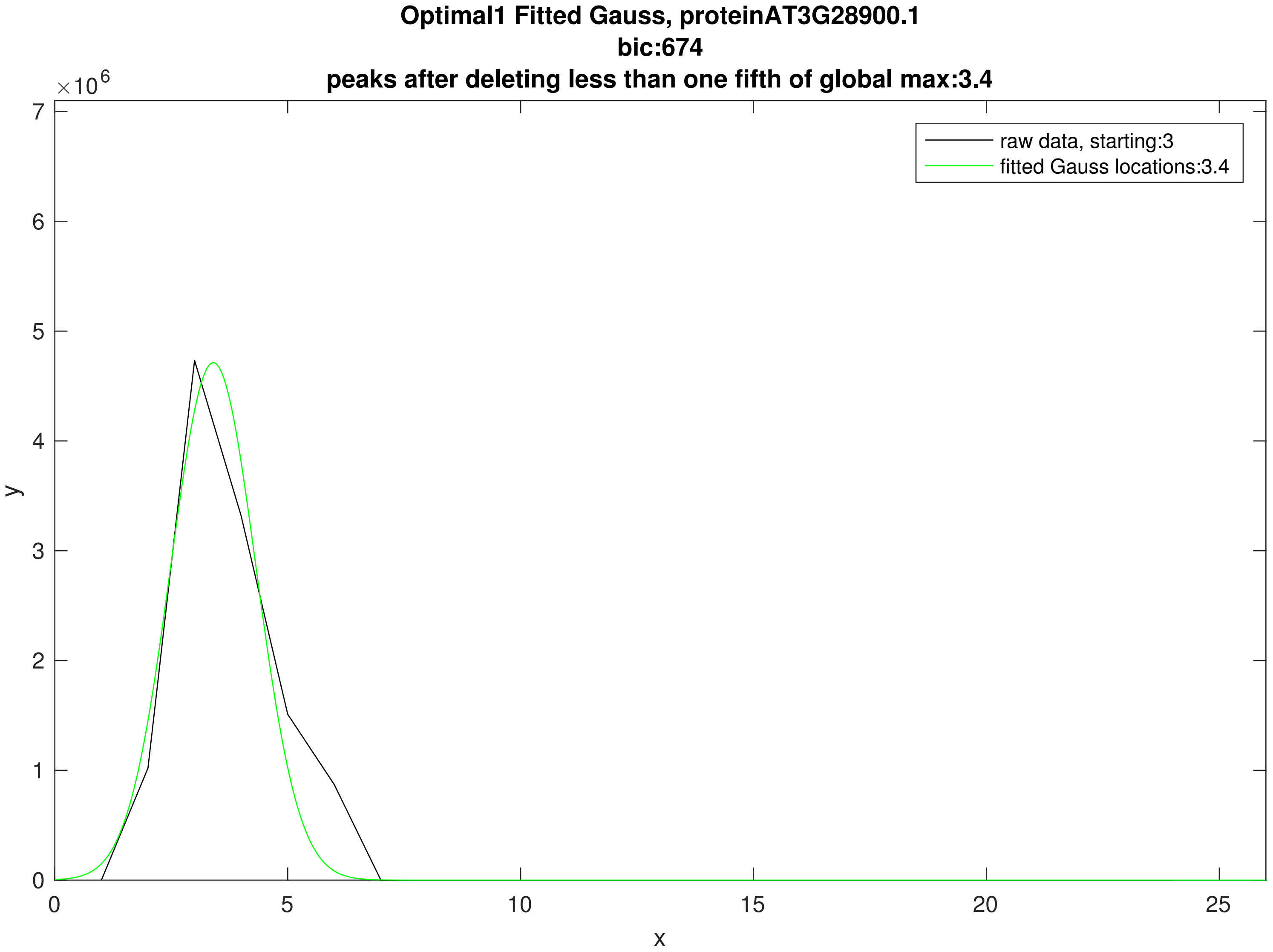
Optimal1 Fitted Gauss, proteinAT3G28900.1
bic:674
peaks after deleting less than one fifth of global max:3.4
×10^6
raw data, starting:3
fitted Gauss locations:3.4
y
x
0
1
2
3
4
5
6
7
0
5
10
15
20
25

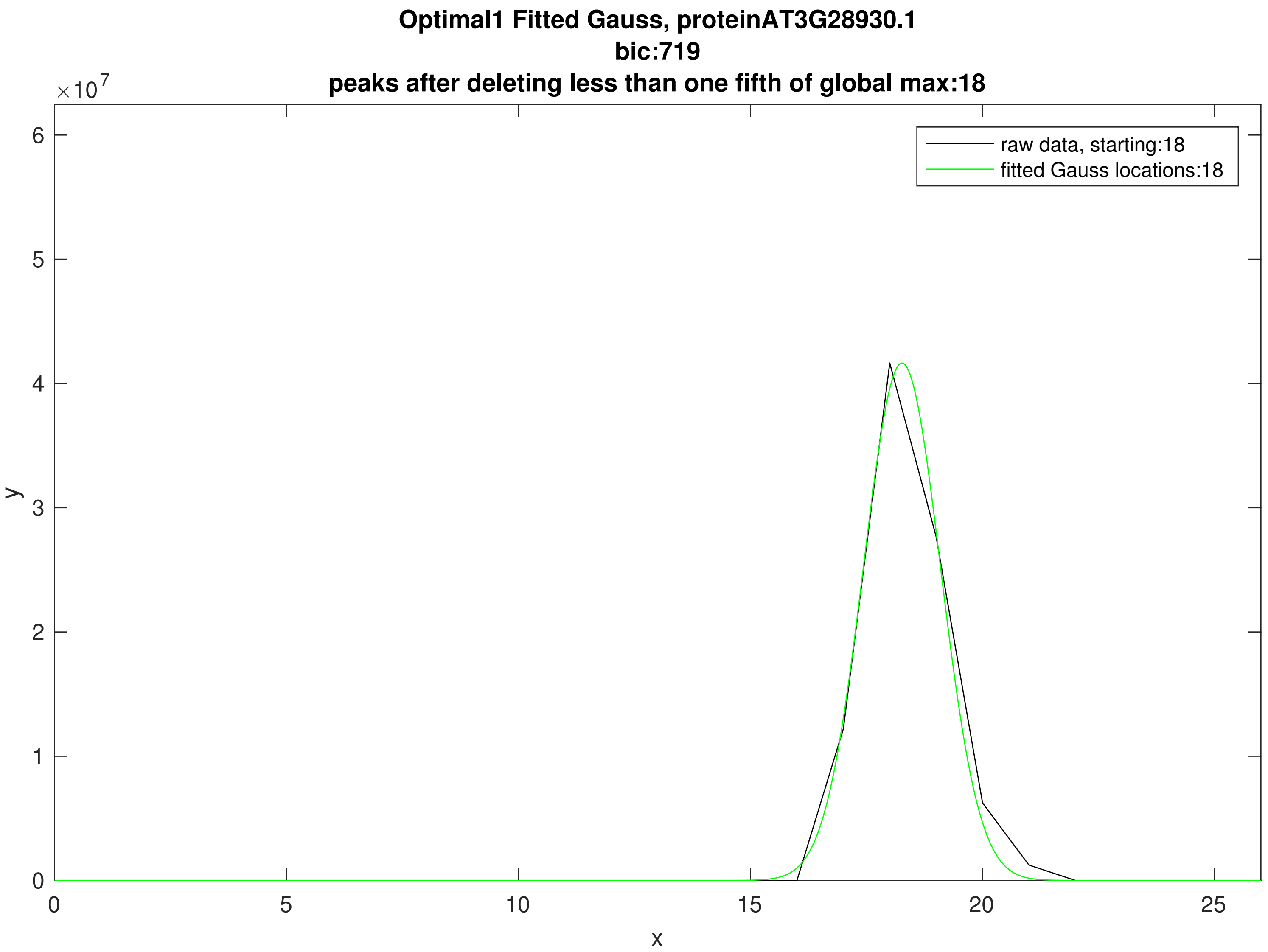

Optimal1 Fitted Gauss, proteinAT3G28930.1
bic:719
peaks after deleting less than one fifth of global max:18
×10^7
raw data, starting:18
fitted Gauss locations:18
y
x
0
1
2
3
4
5
6
0
5
10
15
20
25

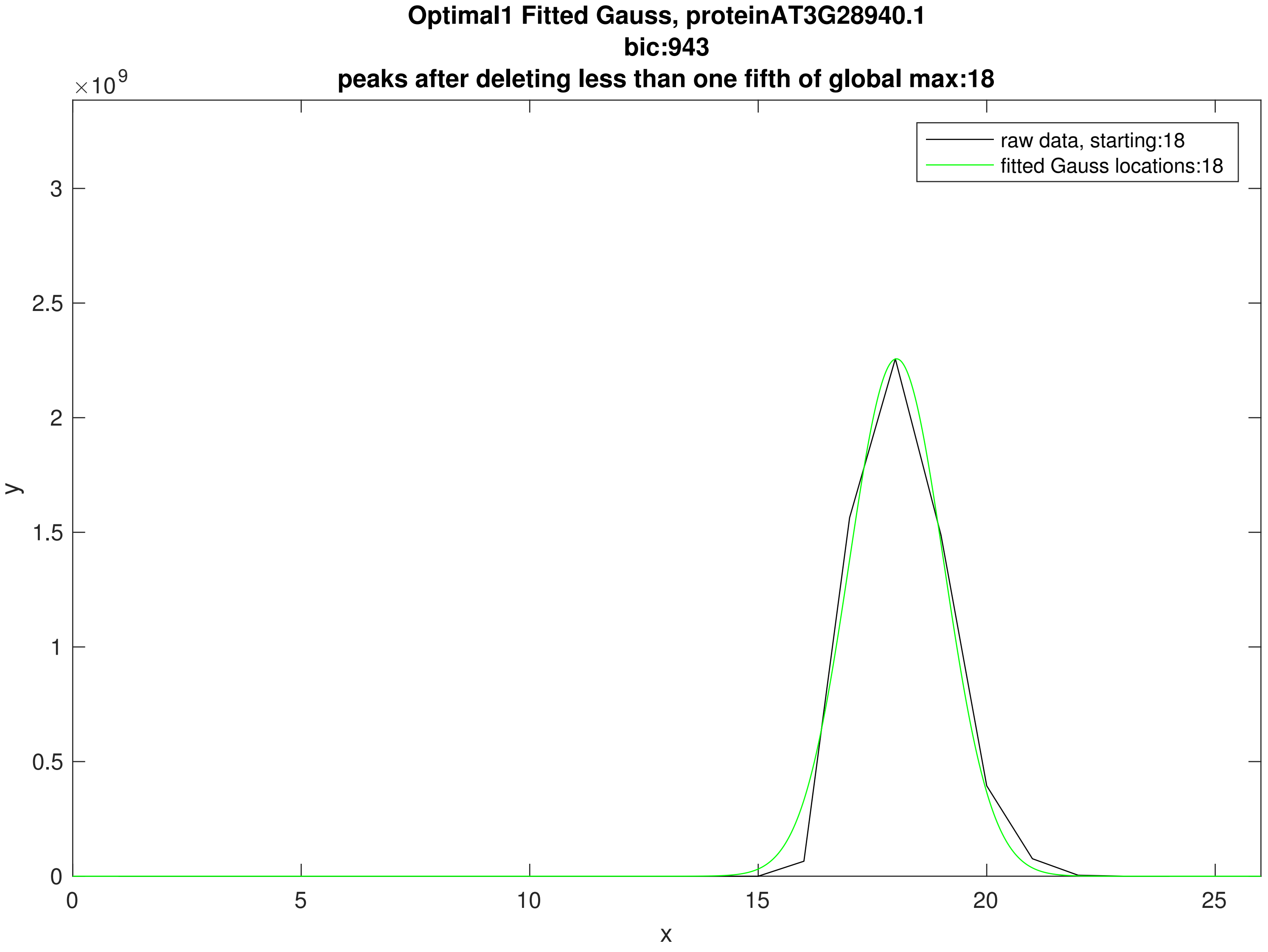
Optimal1 Fitted Gauss, proteinAT3G28940.1
bic:943
peaks after deleting less than one fifth of global max:18
×10^9
raw data, starting:18
fitted Gauss locations:18
y
x
0
0.5
1
1.5
2
2.5
3
0
5
10
15
20
25

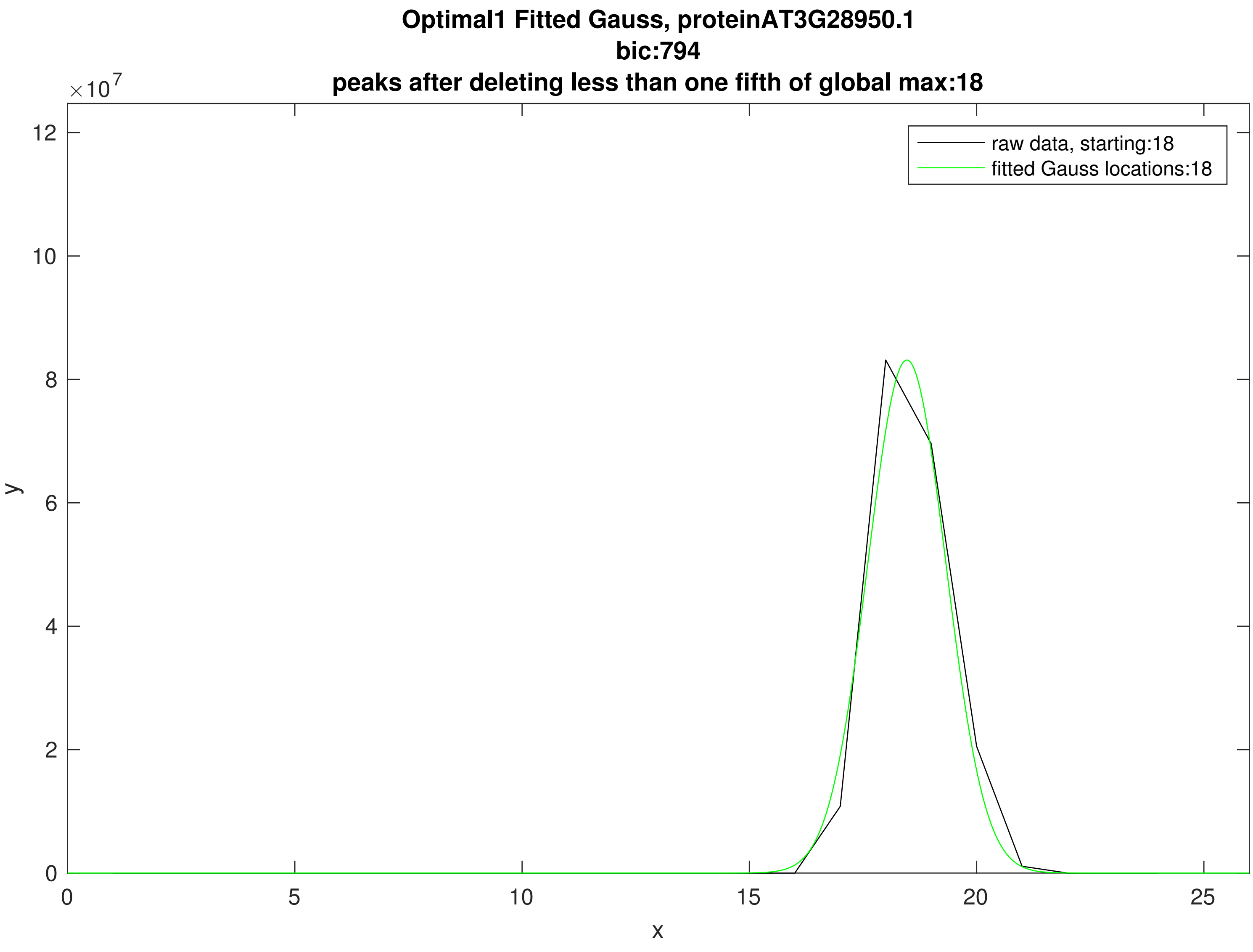

Optimal1 Fitted Gauss, proteinAT3G28950.1
bic:794
peaks after deleting less than one fifth of global max:18
×10^7
12
10
8
6
4
2
0
y
0
5
10
15
20
25
x
raw data, starting:18
fitted Gauss locations:18

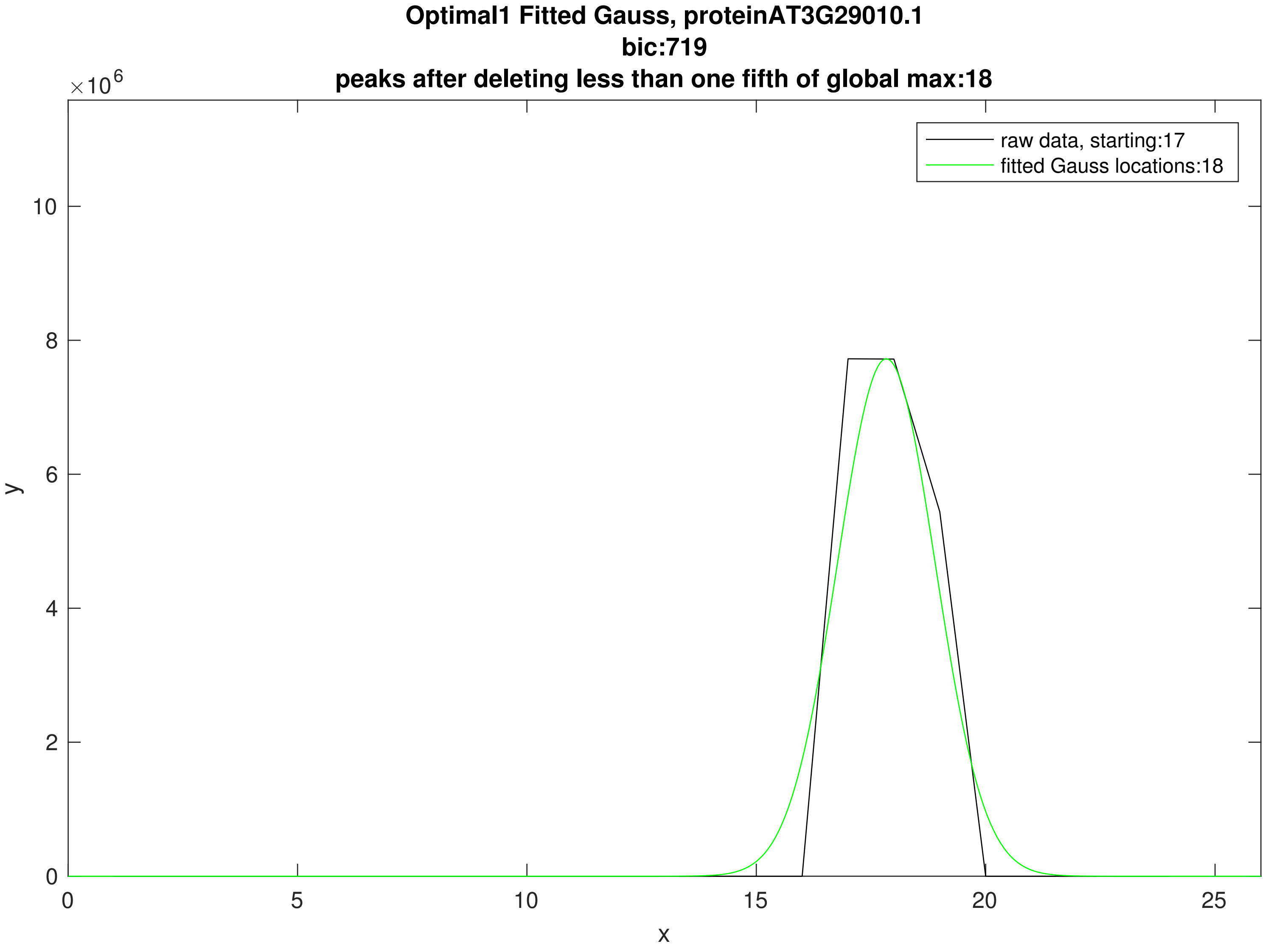

Optimal1 Fitted Gauss, proteinAT3G29010.1
bic:719
peaks after deleting less than one fifth of global max:18
×10^6
raw data, starting:17
fitted Gauss locations:18
10
8
6
4
2
0
y
0
5
10
15
20
25
x

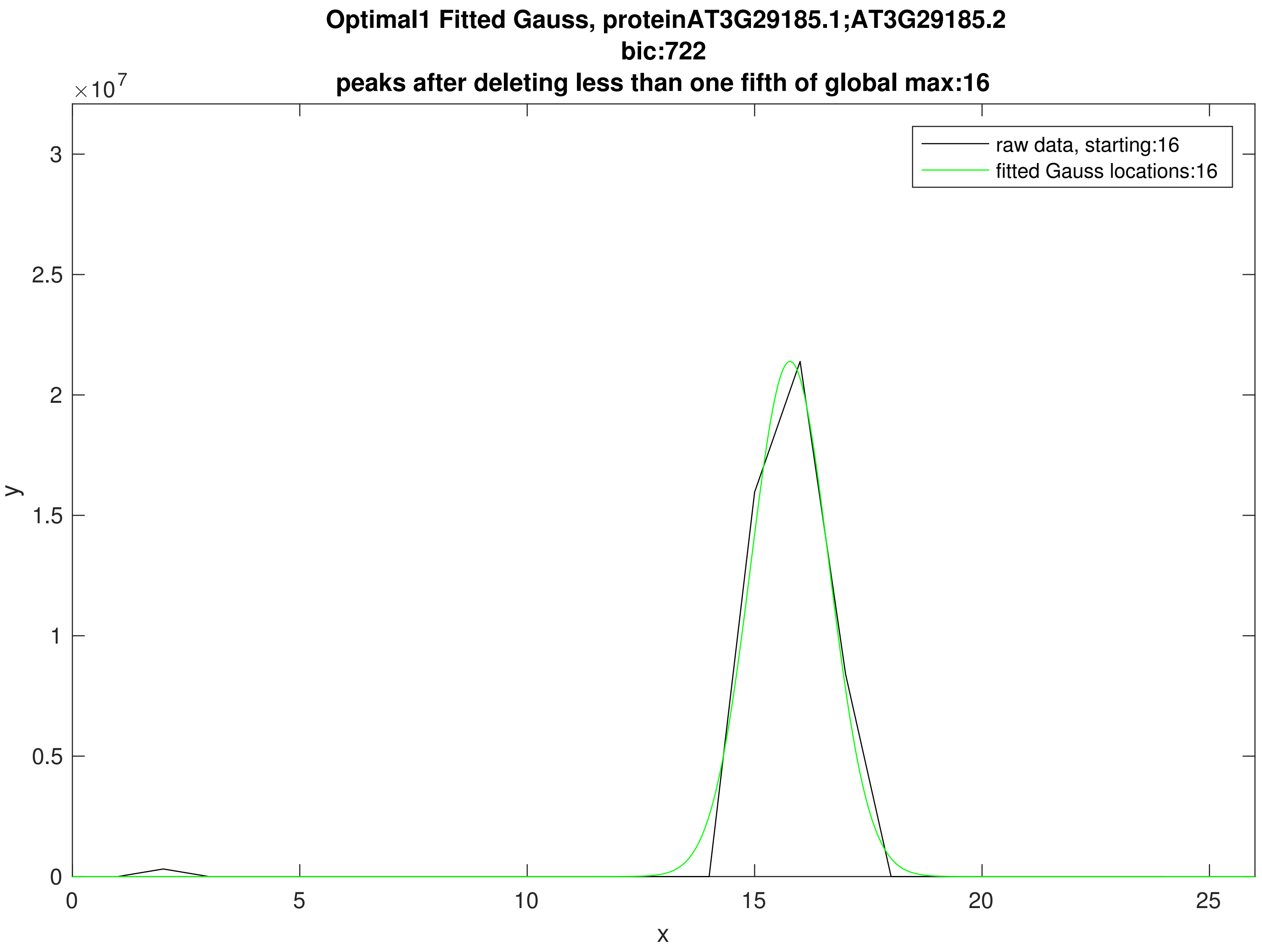
Optimal1 Fitted Gauss, proteinAT3G29185.1;AT3G29185.2
bic:722
peaks after deleting less than one fifth of global max:16
×10^7
3
2.5
2
1.5
1
0.5
0
y
0
5
10
15
20
25
x
raw data, starting:16
fitted Gauss locations:16

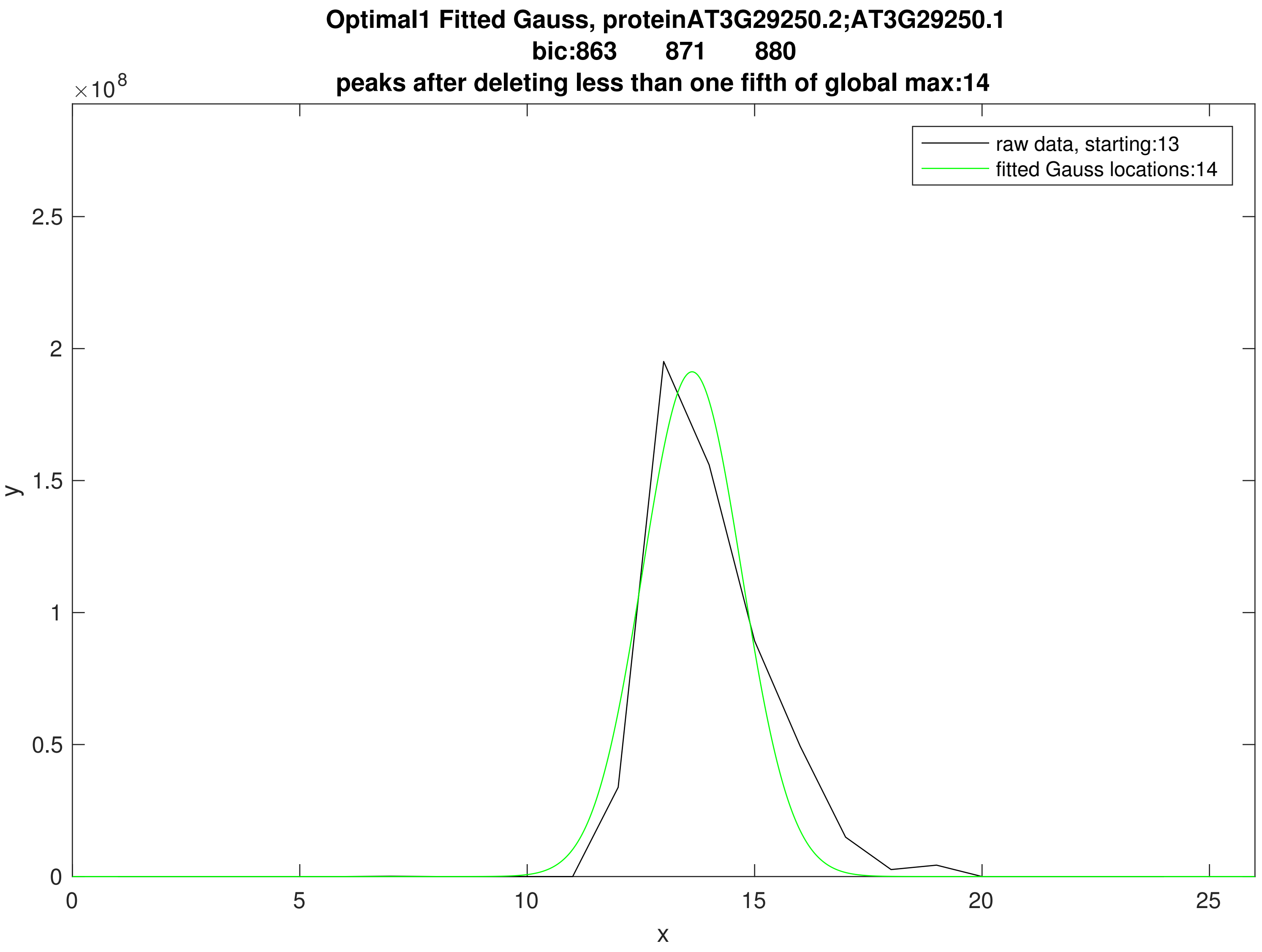
Optimal1 Fitted Gauss, proteinAT3G29250.2;AT3G29250.1
bic:863 871 880
peaks after deleting less than one fifth of global max:14
×10^8
raw data, starting:13
fitted Gauss locations:14
y
x
0
0.5
1
1.5
2
2.5
0
5
10
15
20
25

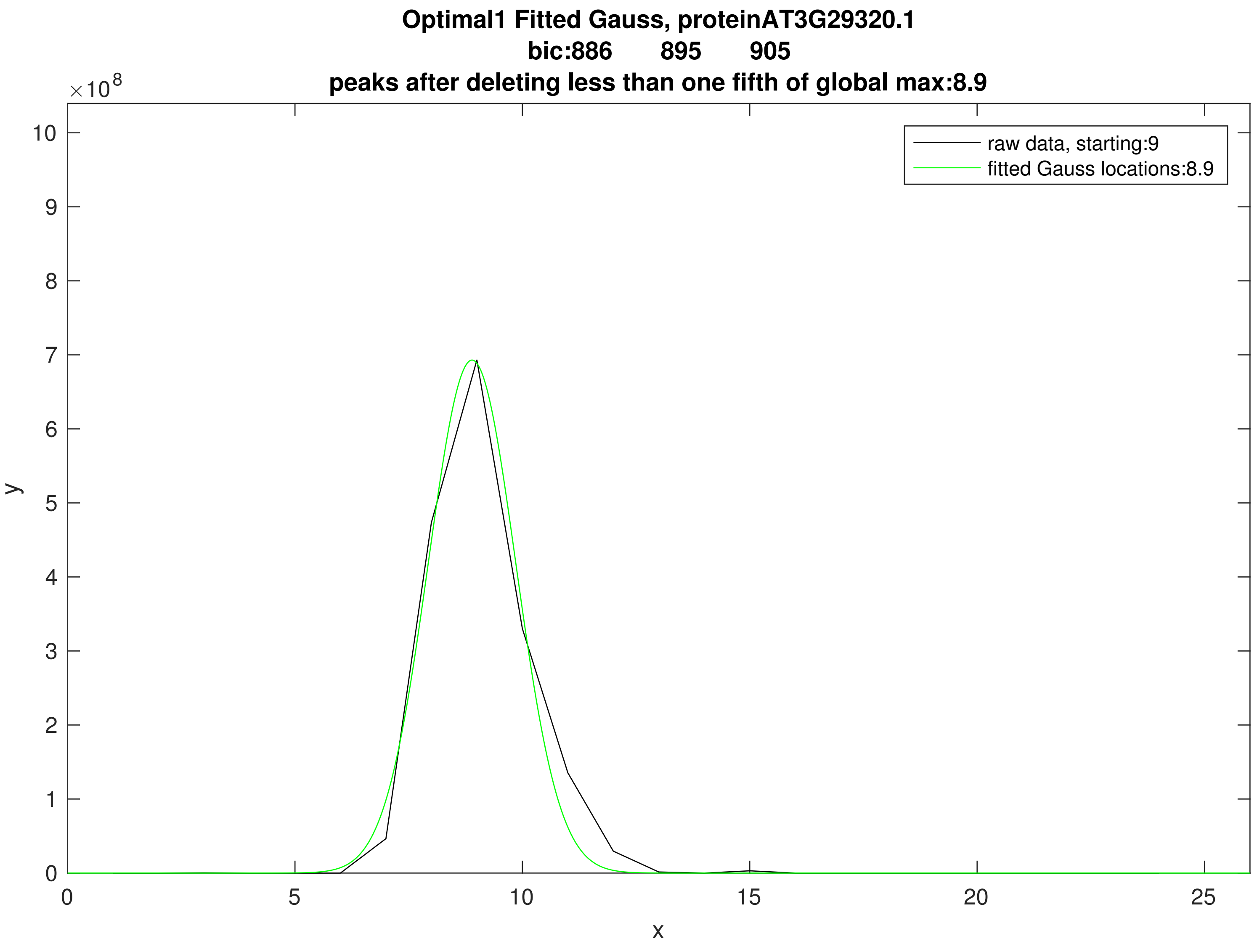
Optimal1 Fitted Gauss, proteinAT3G29320.1
bic:886 895 905
peaks after deleting less than one fifth of global max:8.9
×10^8
raw data, starting:9
fitted Gauss locations:8.9
y
x
0
1
2
3
4
5
6
7
8
9
10
15
20
25

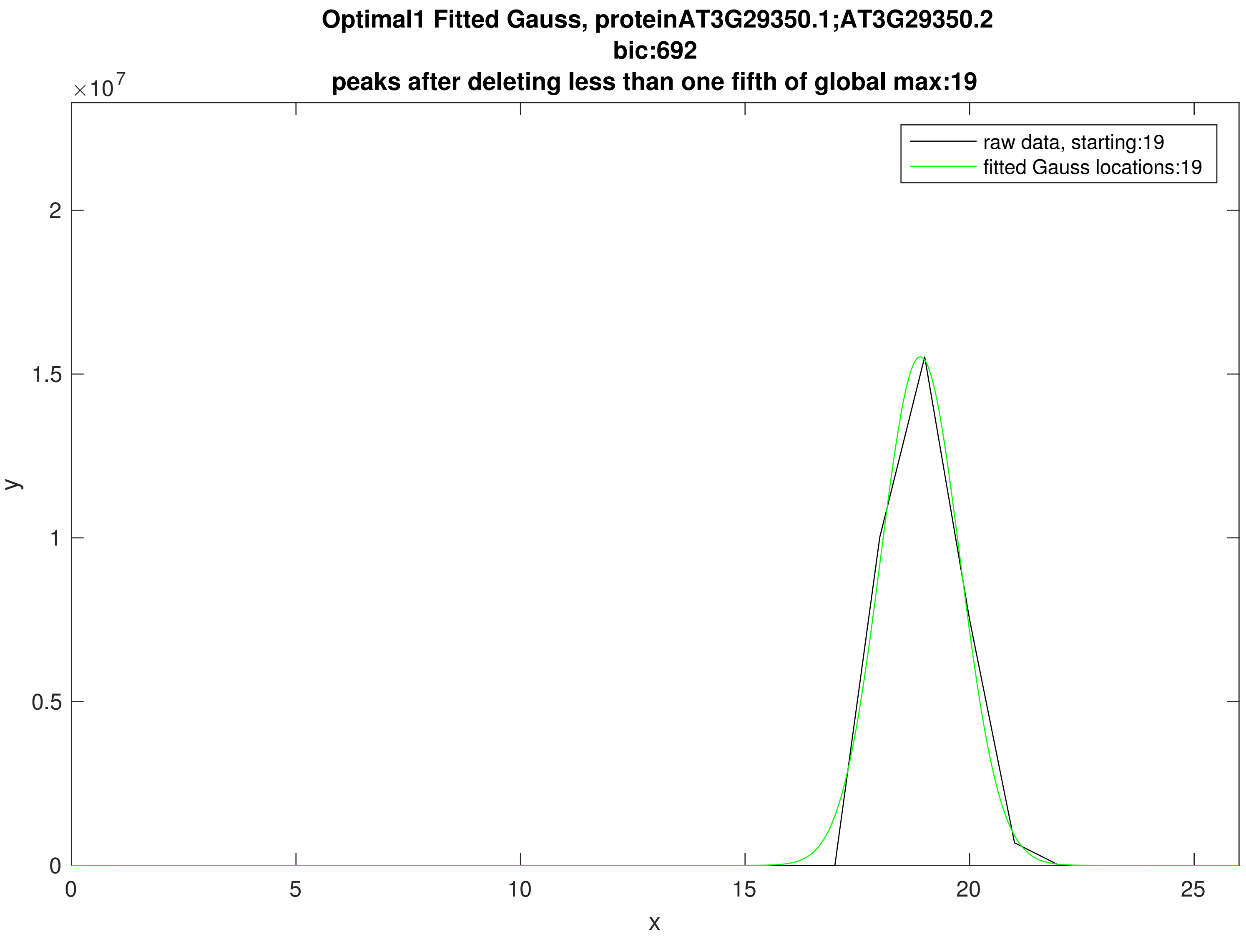
Optimal1 Fitted Gauss, proteinAT3G29350.1;AT3G29350.2
bic:692
peaks after deleting less than one fifth of global max:19
×10^7
raw data, starting:19
fitted Gauss locations:19
y
x
0
0.5
1
1.5
2
0
5
10
15
20
25

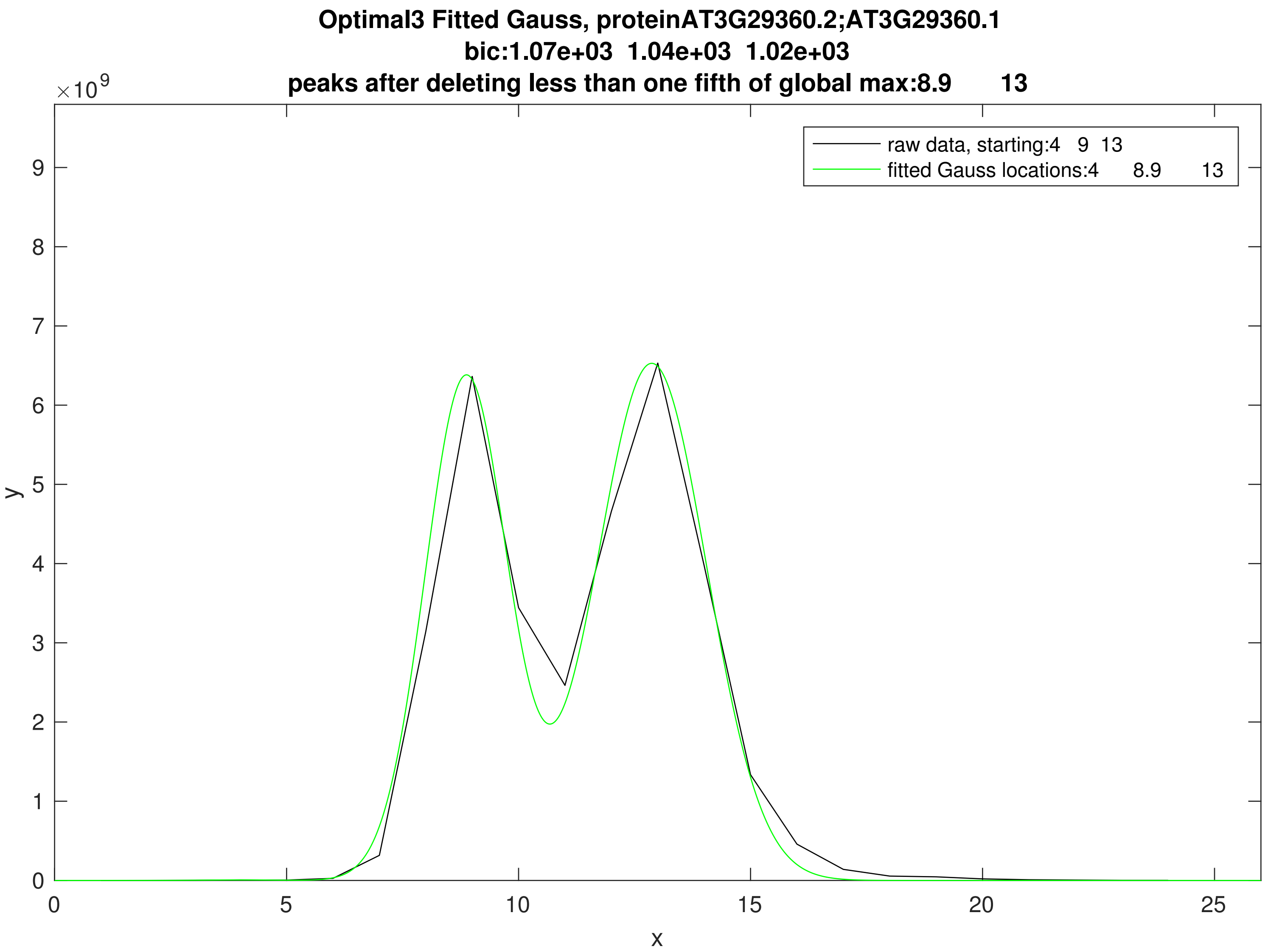
Optimal3 Fitted Gauss, proteinAT3G29360.2;AT3G29360.1
bic:1.07e+03 1.04e+03 1.02e+03
peaks after deleting less than one fifth of global max:8.9 13
×10^9
raw data, starting:4 9 13
fitted Gauss locations:4 8.9 13
0
1
2
3
4
5
6
7
8
9
y
0
5
10
15
20
25
x

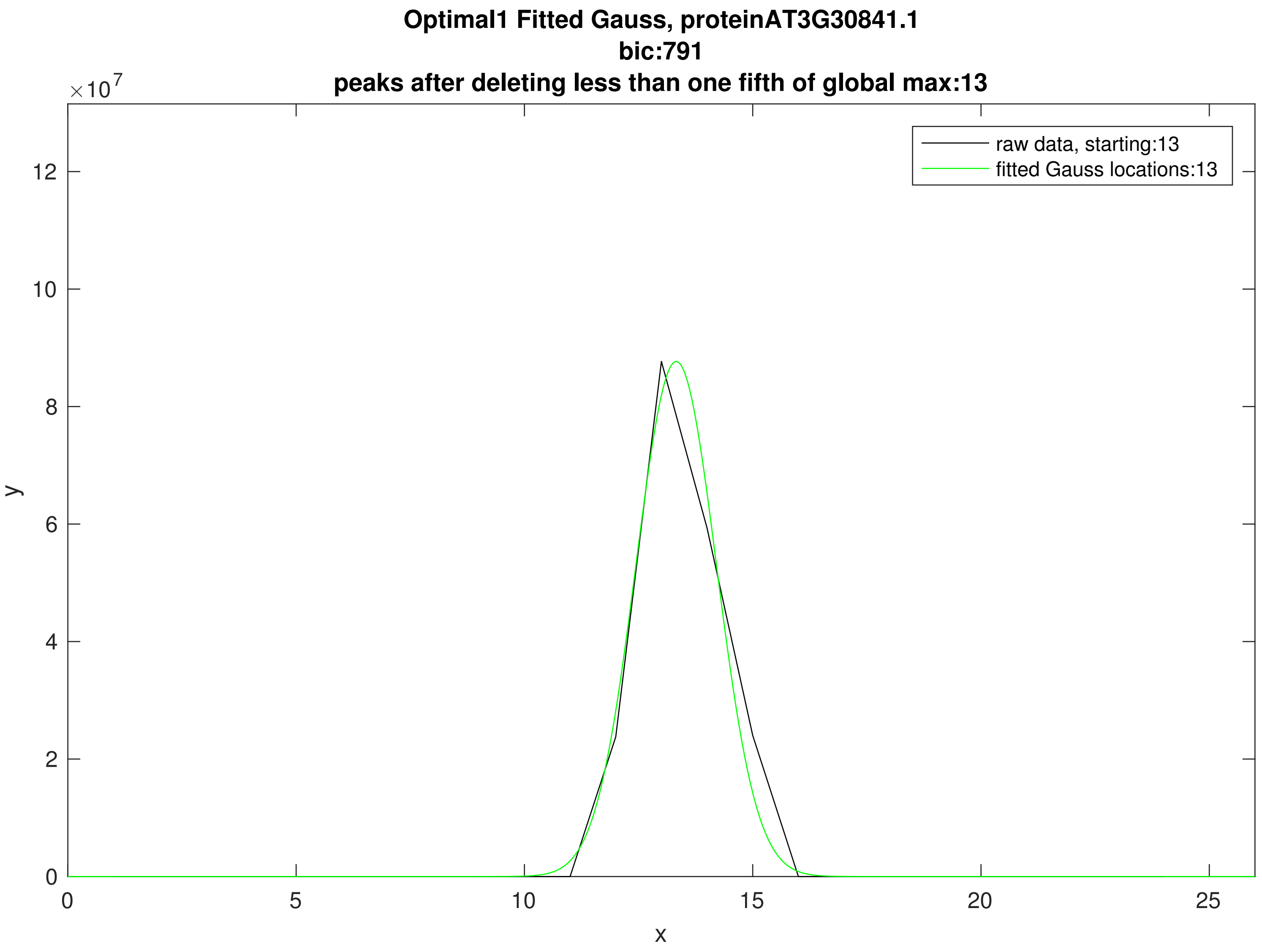

Optimal1 Fitted Gauss, proteinAT3G30841.1
bic:791
peaks after deleting less than one fifth of global max:13
×10^7
12
10
8
6
4
2
0
y
0
5
10
15
20
25
x
raw data, starting:13
fitted Gauss locations:13

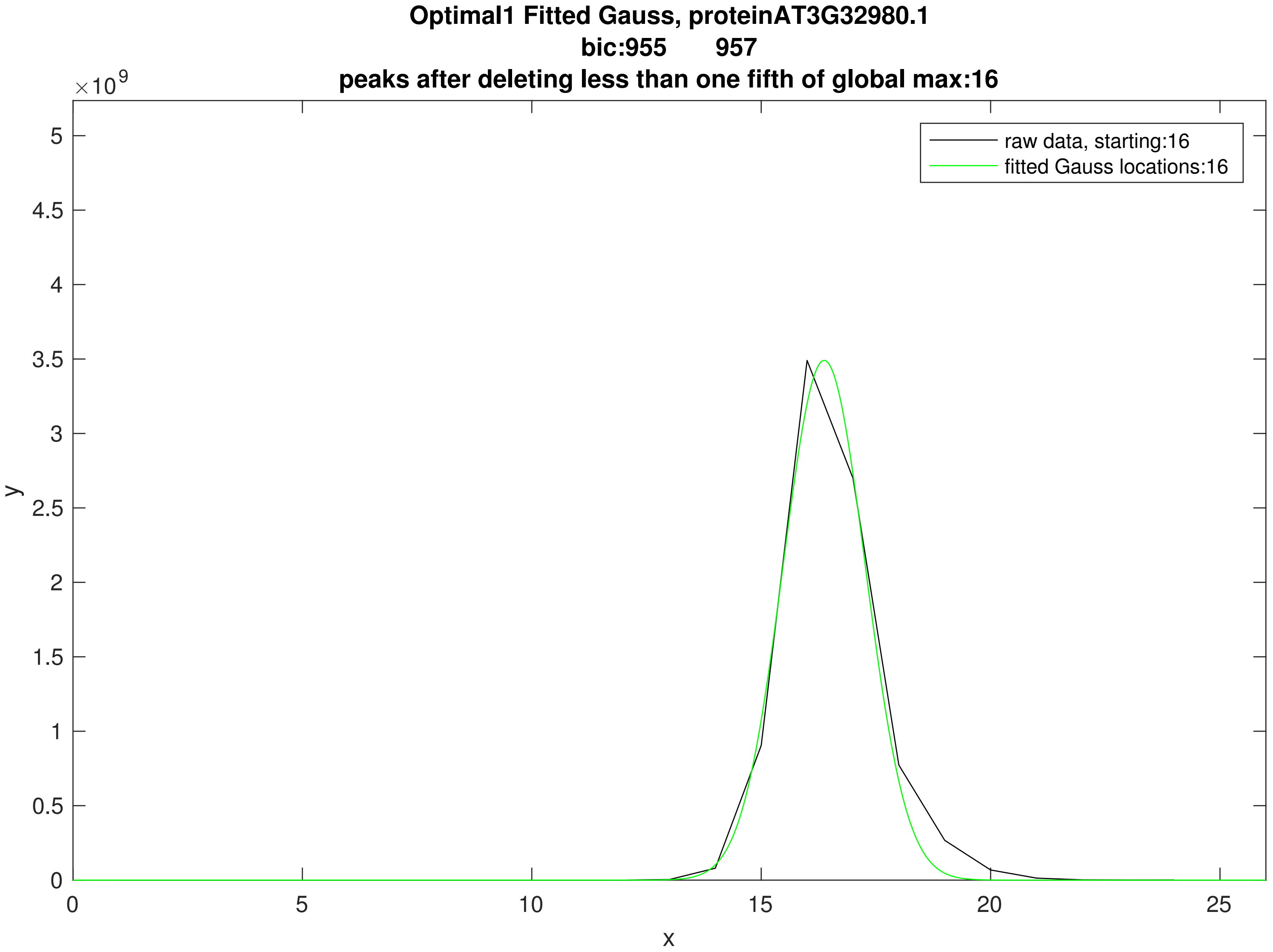
Optimal1 Fitted Gauss, proteinAT3G32980.1
bic:955 957
peaks after deleting less than one fifth of global max:16
×10^9
raw data, starting:16
fitted Gauss locations:16
y
x
0
0.5
1
1.5
2
2.5
3
3.5
4
4.5
5
0
5
10
15
20
25

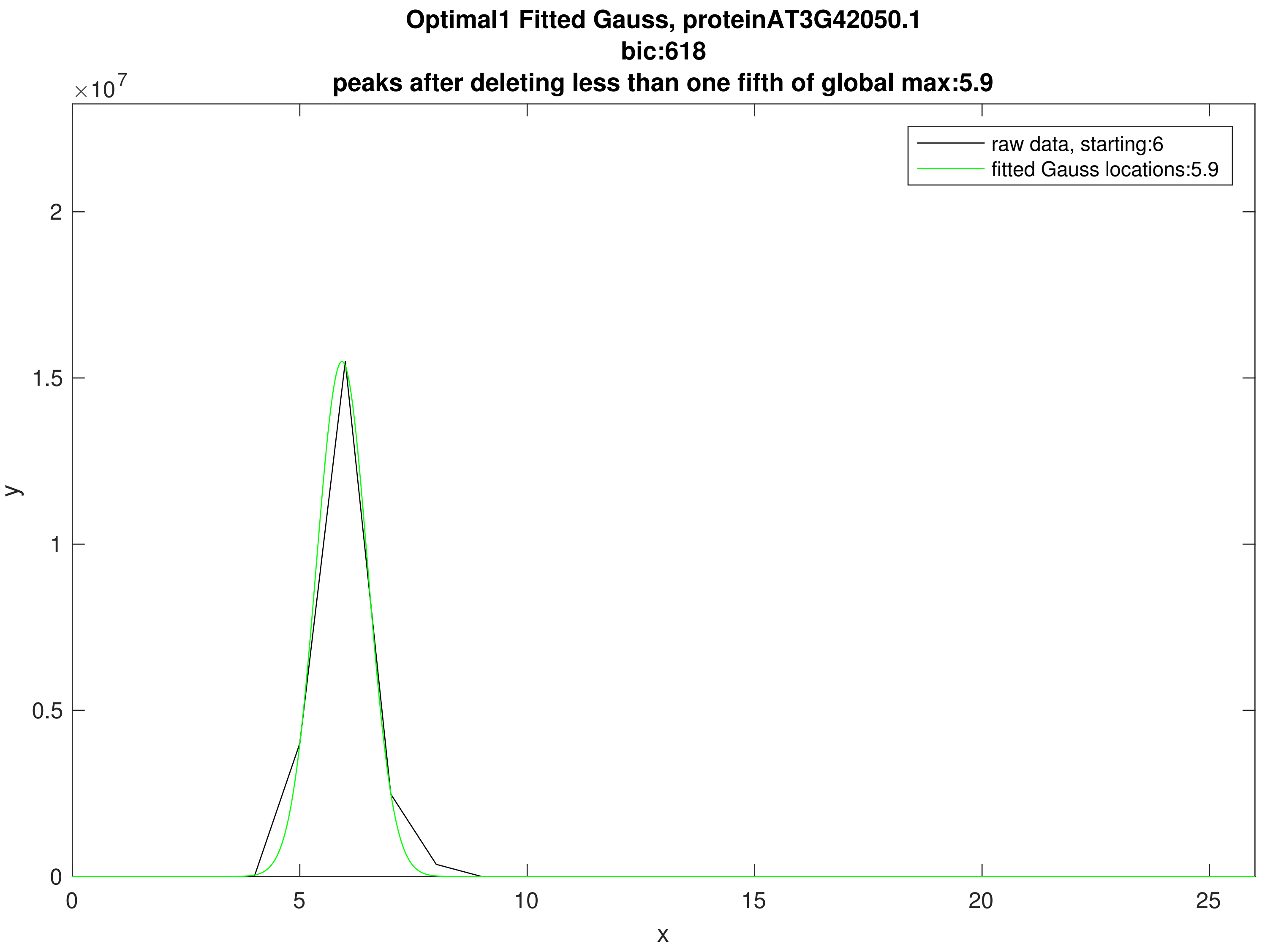
Optimal1 Fitted Gauss, proteinAT3G42050.1
bic:618
peaks after deleting less than one fifth of global max:5.9
×10^7
raw data, starting:6
fitted Gauss locations:5.9
2
1.5
1
0.5
0
y
0
5
10
15
20
25
x

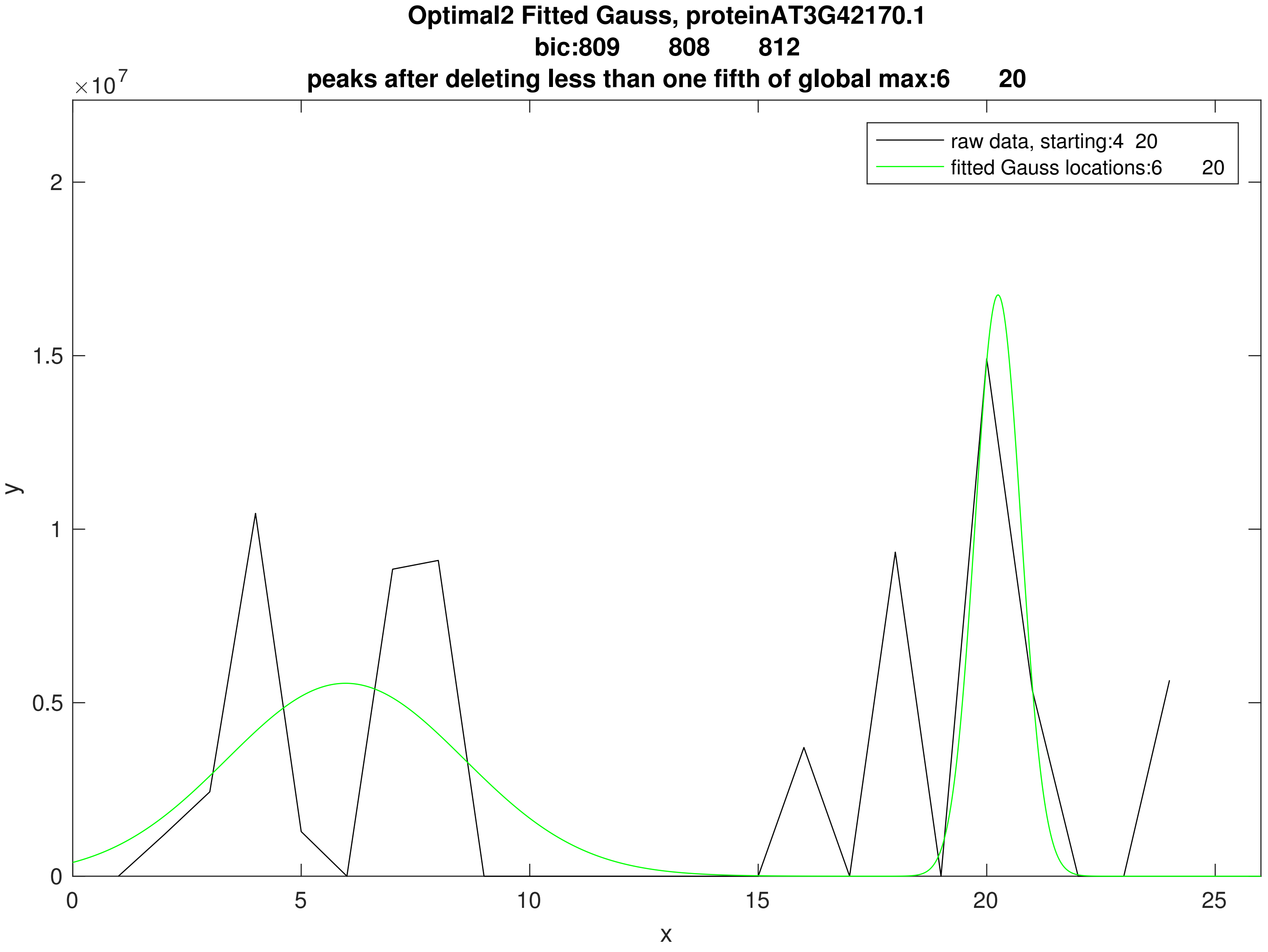
Optimal2 Fitted Gauss, proteinAT3G42170.1
bic:809 808 812
peaks after deleting less than one fifth of global max:6 20
×10^7
raw data, starting:4 20
fitted Gauss locations:6 20
y
x
0
0.5
1
1.5
2
0
5
10
15
20
25

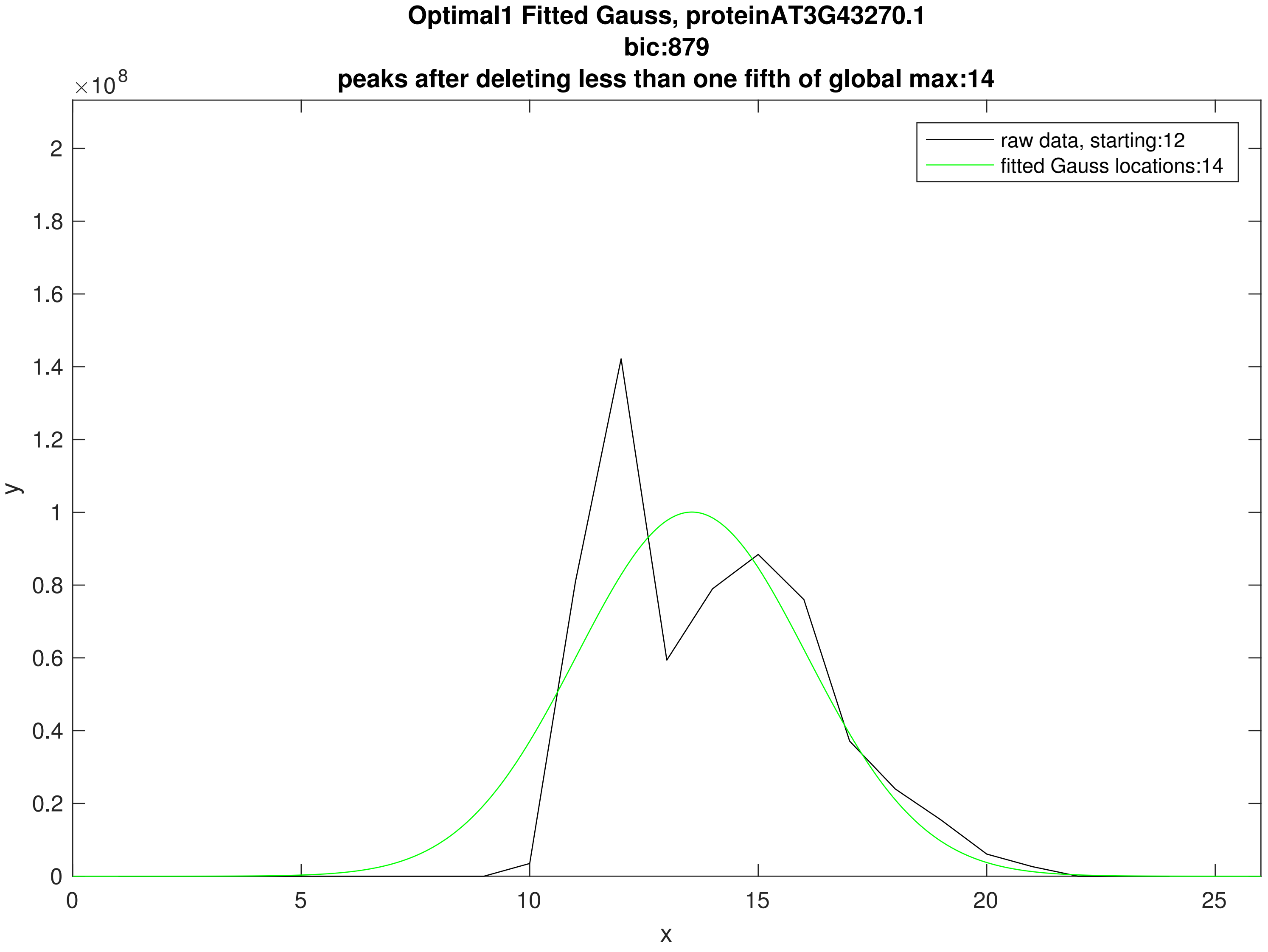
Optimal1 Fitted Gauss, proteinAT3G43270.1
bic:879
peaks after deleting less than one fifth of global max:14
×10^8
2
1.8
1.6
1.4
1.2
1
0.8
0.6
0.4
0.2
0
y
0
5
10
15
20
25
x
raw data, starting:12
fitted Gauss locations:14

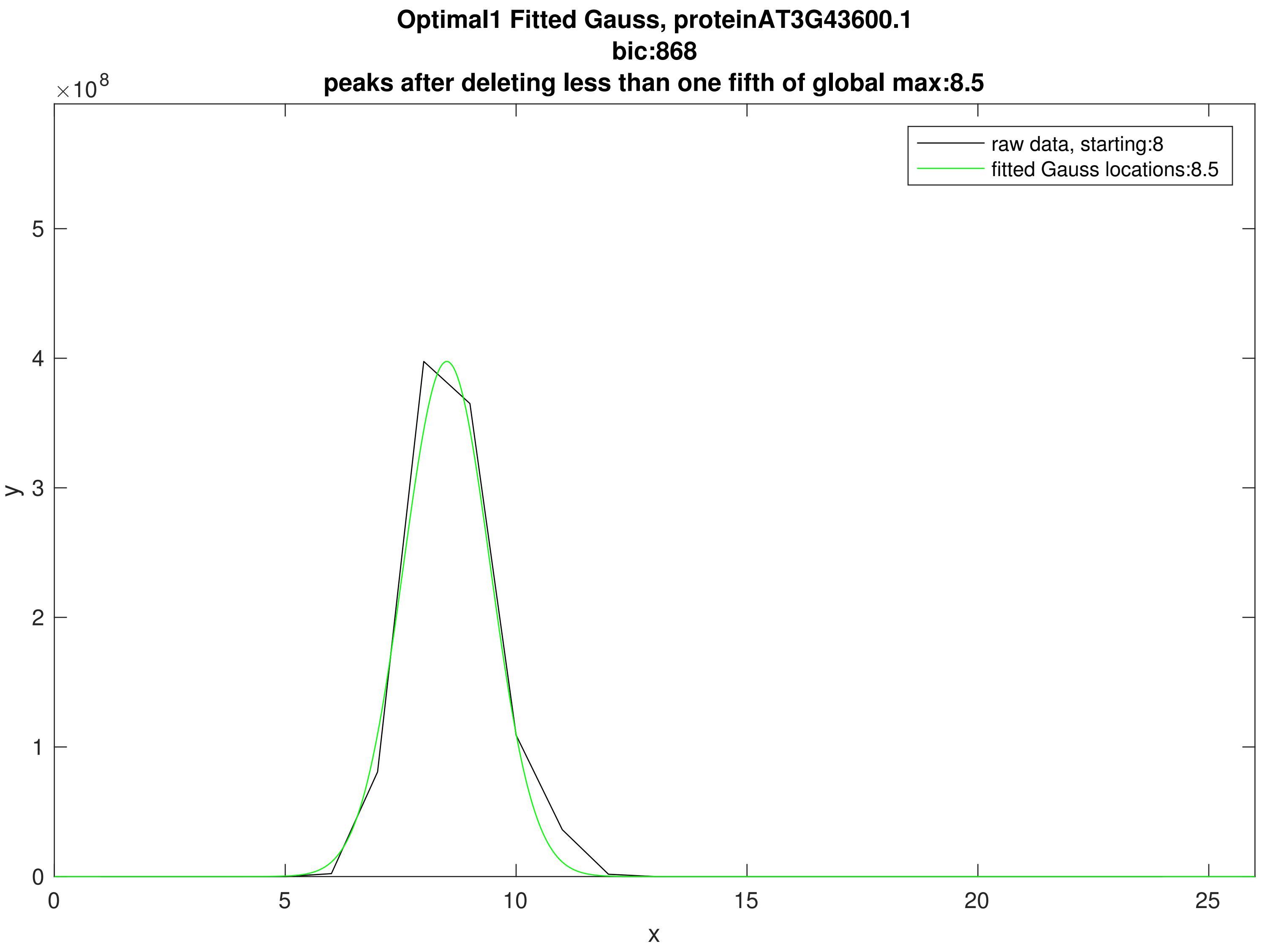

Optimal1 Fitted Gauss, proteinAT3G43600.1
bic:868
peaks after deleting less than one fifth of global max:8.5
×10^8
raw data, starting:8
fitted Gauss locations:8.5
y
x
0
1
2
3
4
5
0
5
10
15
20
25

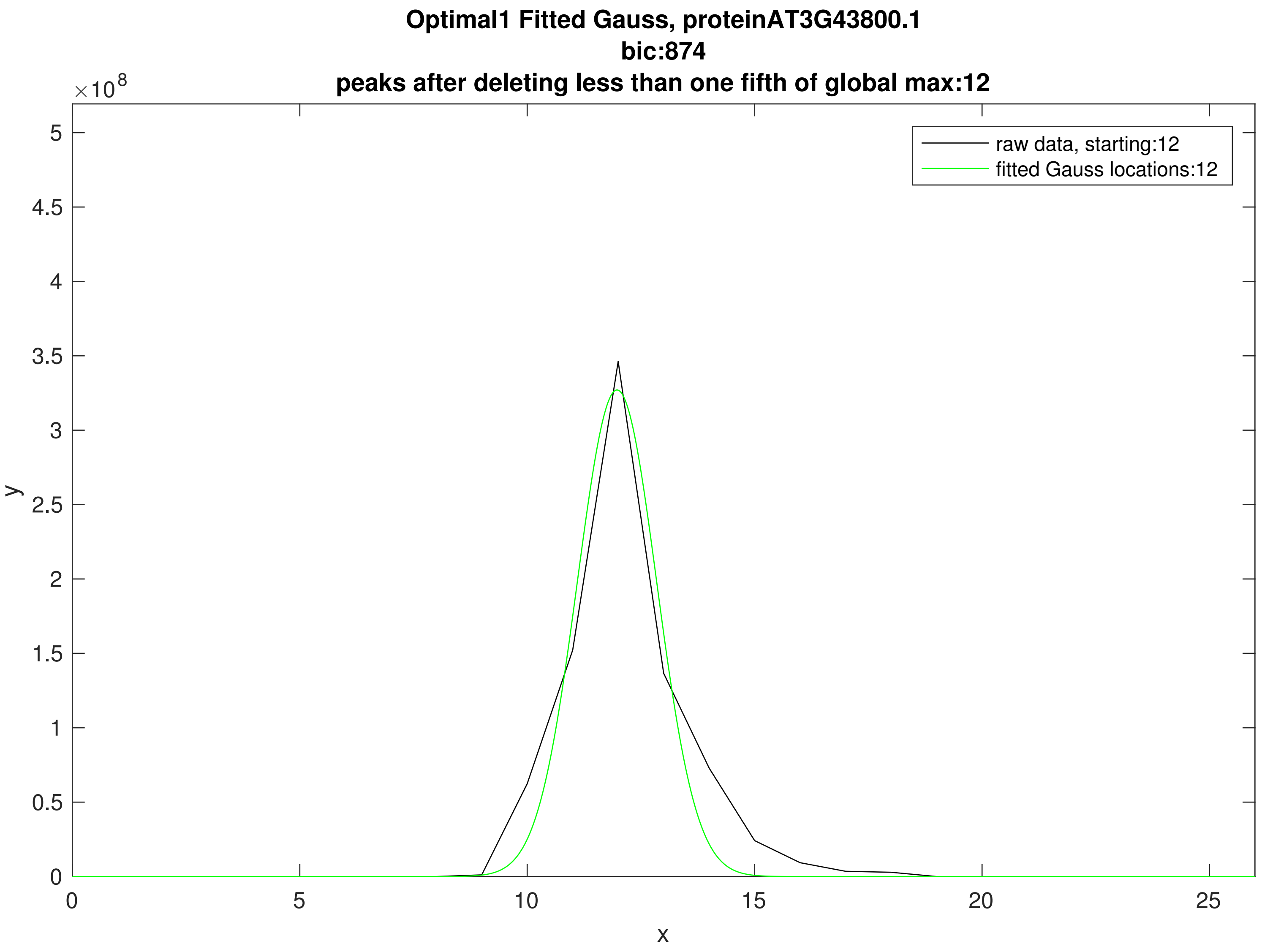
Optimal1 Fitted Gauss, proteinAT3G43800.1
bic:874
peaks after deleting less than one fifth of global max:12
×10^8
raw data, starting:12
fitted Gauss locations:12
y
x
0
0.5
1
1.5
2
2.5
3
3.5
4
4.5
5
0
5
10
15
20
25

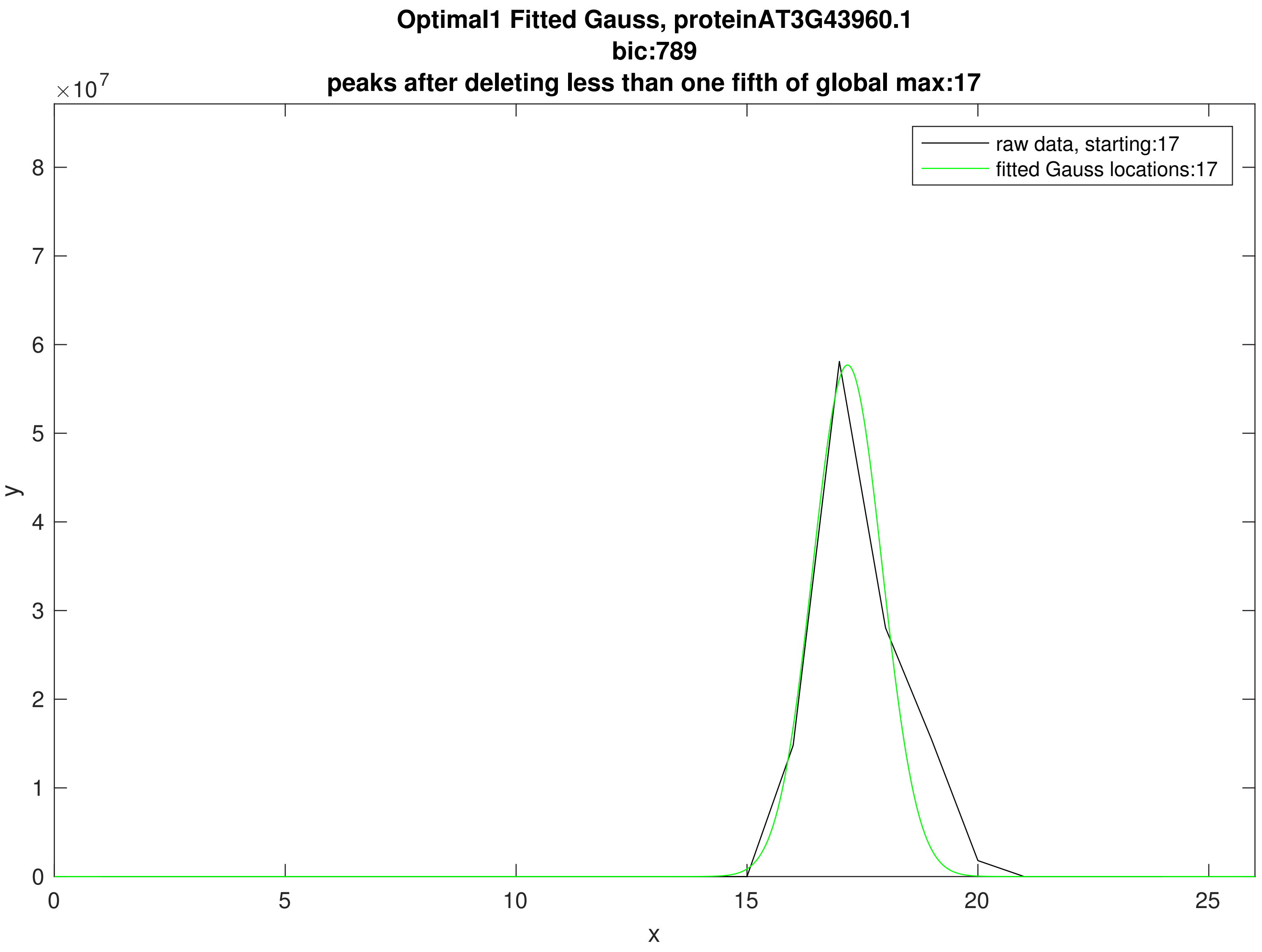
Optimal1 Fitted Gauss, proteinAT3G43960.1
bic:789
peaks after deleting less than one fifth of global max:17
×10^7
raw data, starting:17
fitted Gauss locations:17
y
x
0
1
2
3
4
5
6
7
8
0
5
10
15
20
25

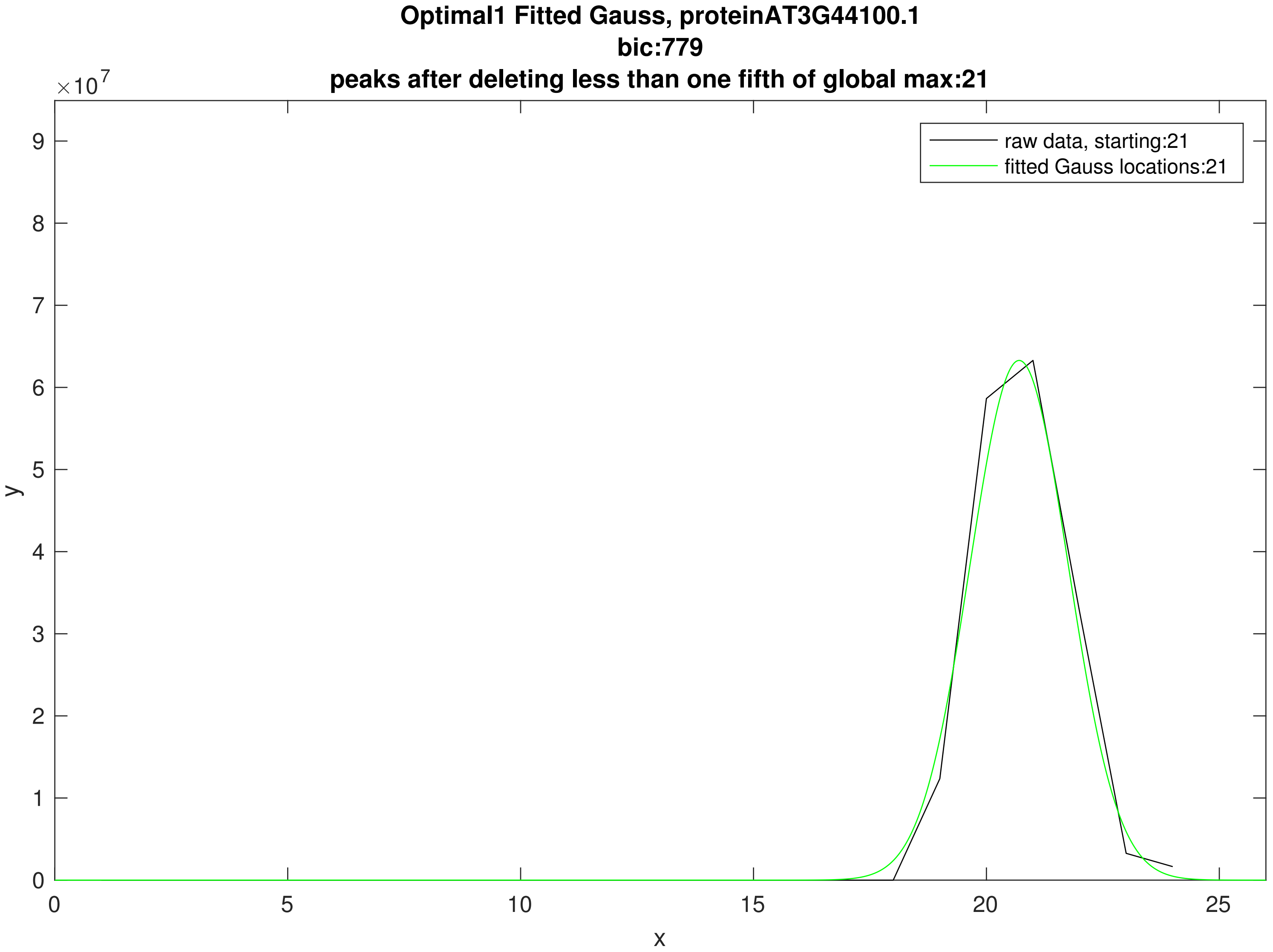
Optimal1 Fitted Gauss, proteinAT3G44100.1
bic:779
peaks after deleting less than one fifth of global max:21
×10^7
raw data, starting:21
fitted Gauss locations:21
y
x
0
1
2
3
4
5
6
7
8
9
0
5
10
15
20
25

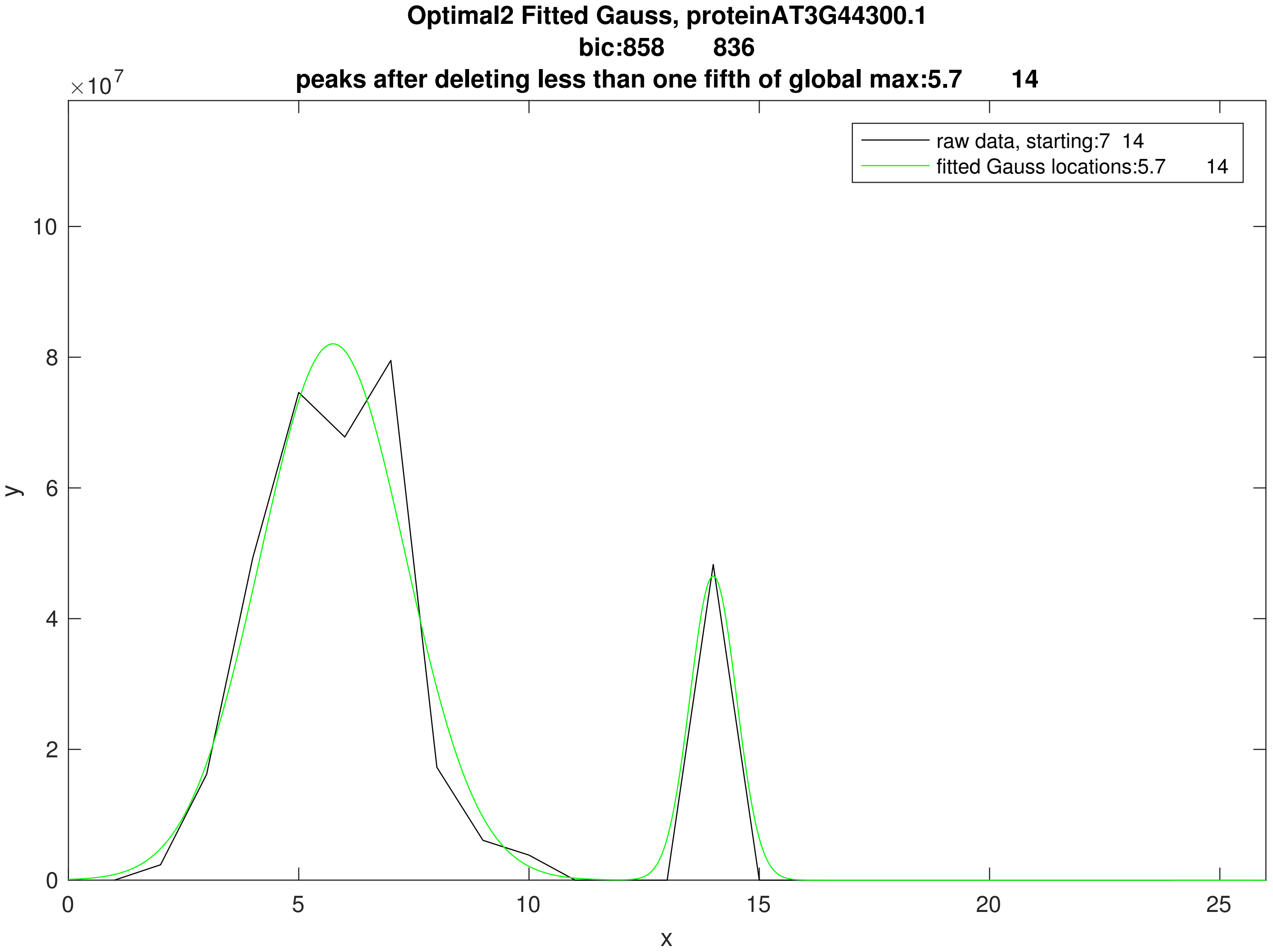
Optimal2 Fitted Gauss, proteinAT3G44300.1
bic:858 836
peaks after deleting less than one fifth of global max:5.7 14
×10^7
raw data, starting:7 14
fitted Gauss locations:5.7 14
y
x
0
2
4
6
8
10
0
5
10
15
20
25

**Optimal1 Fitted Gauss, proteinAT3G44310.1;AT3G44310.3;AT3G44310.2**
**bic:949 958 962**
**peaks after deleting less than one fifth of global max:6.3**

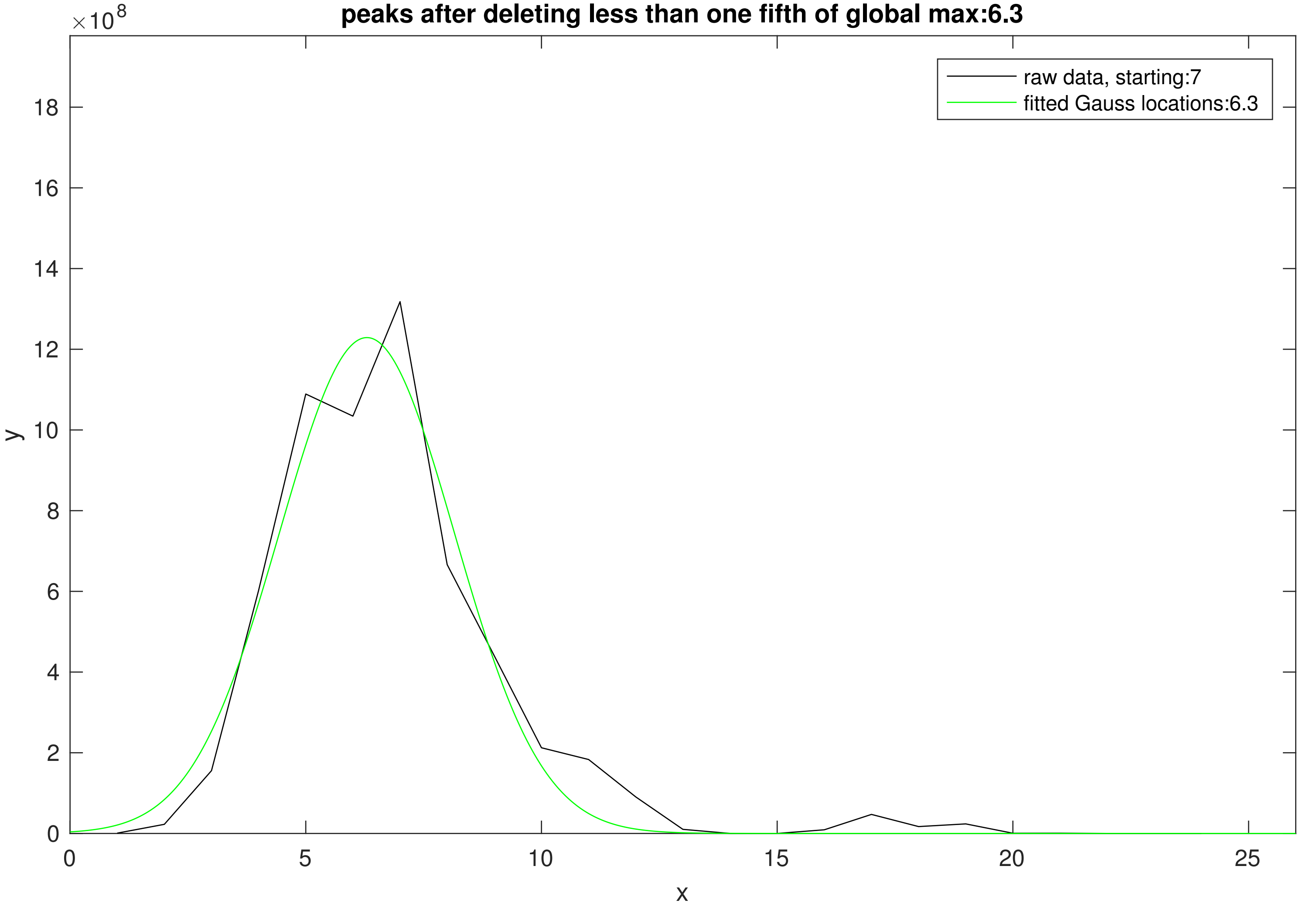

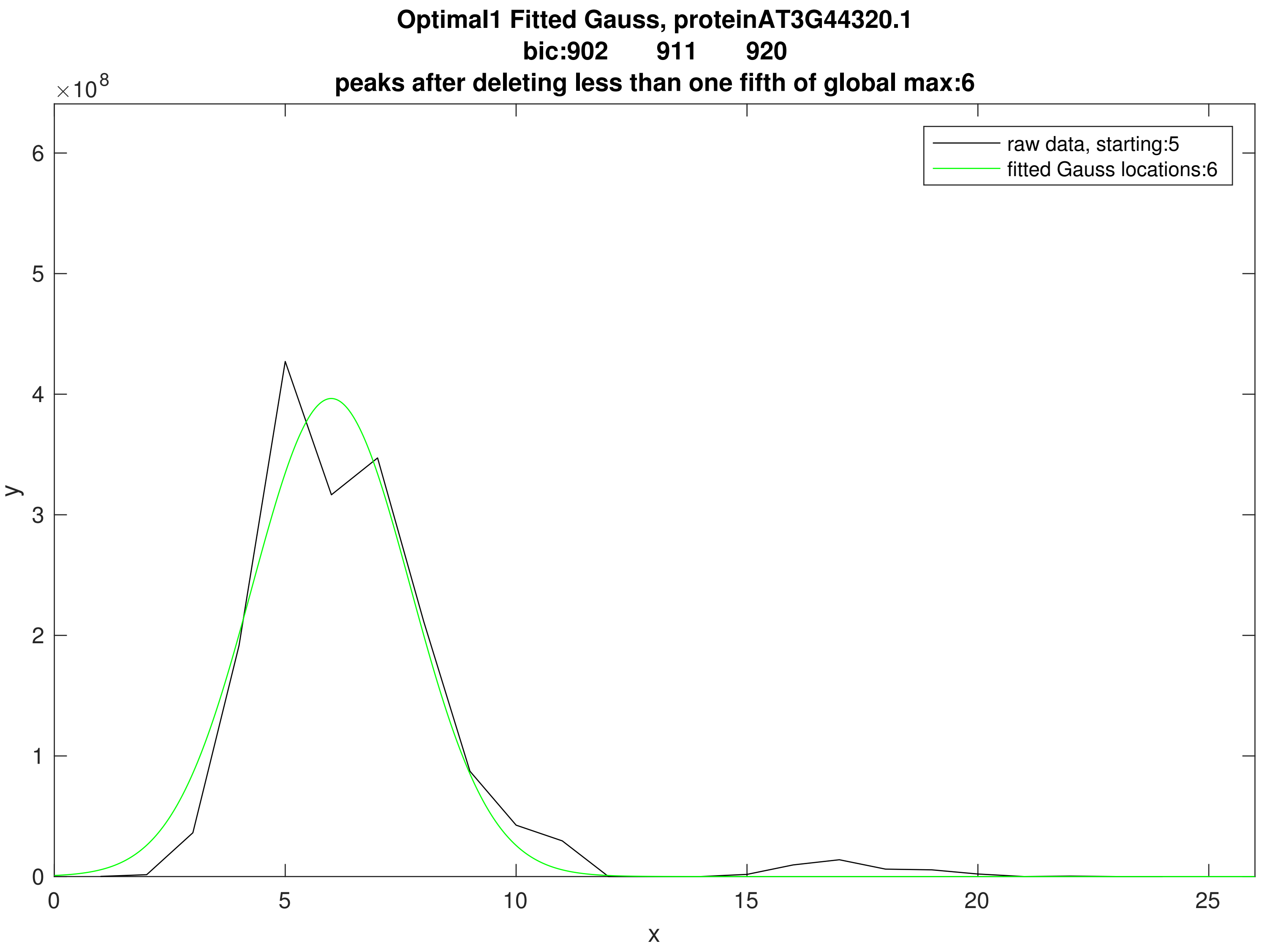

Optimal1 Fitted Gauss, proteinAT3G44320.1
bic:902 911 920
peaks after deleting less than one fifth of global max:6
×10^8
raw data, starting:5
fitted Gauss locations:6
y
x
0
1
2
3
4
5
6
10
15
20
25

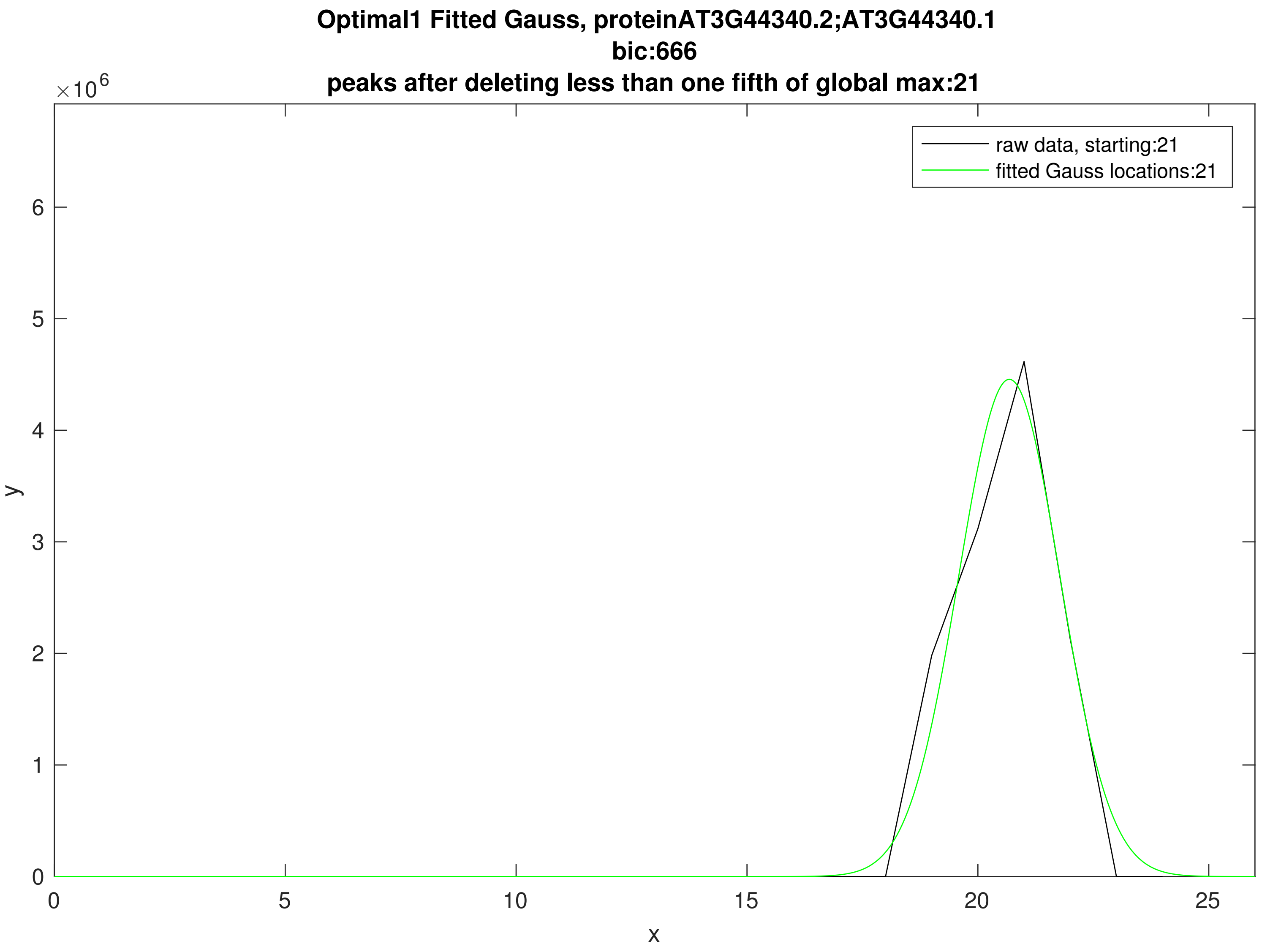

Optimal1 Fitted Gauss, proteinAT3G44340.2;AT3G44340.1
bic:666
peaks after deleting less than one fifth of global max:21
×10^6
raw data, starting:21
fitted Gauss locations:21
y
x
0
1
2
3
4
5
6
0
5
10
15
20
25

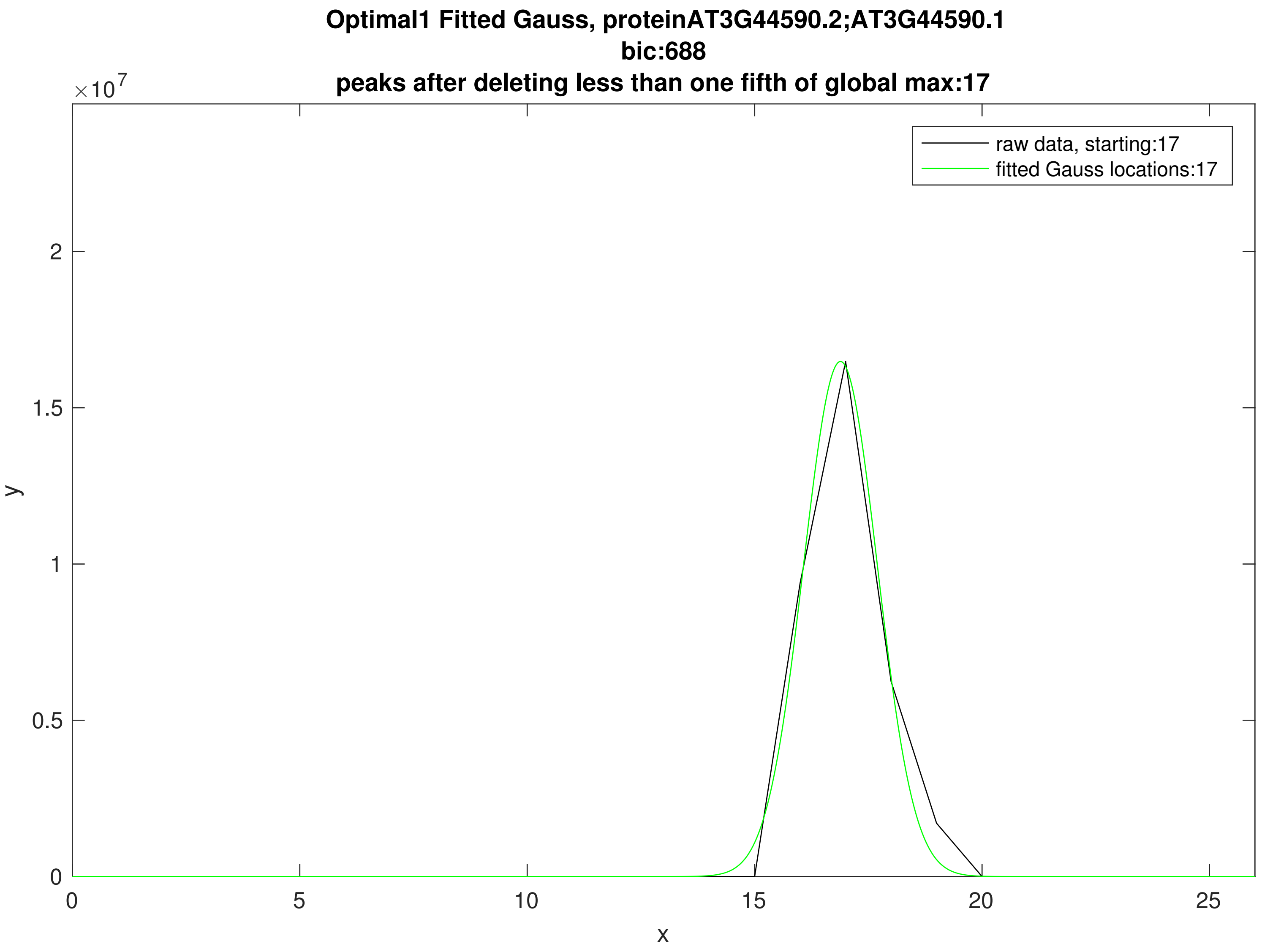
Optimal1 Fitted Gauss, proteinAT3G44590.2;AT3G44590.1
bic:688
peaks after deleting less than one fifth of global max:17
×10^7
raw data, starting:17
fitted Gauss locations:17
2
1.5
1
0.5
0
y
0
5
10
15
20
25
x

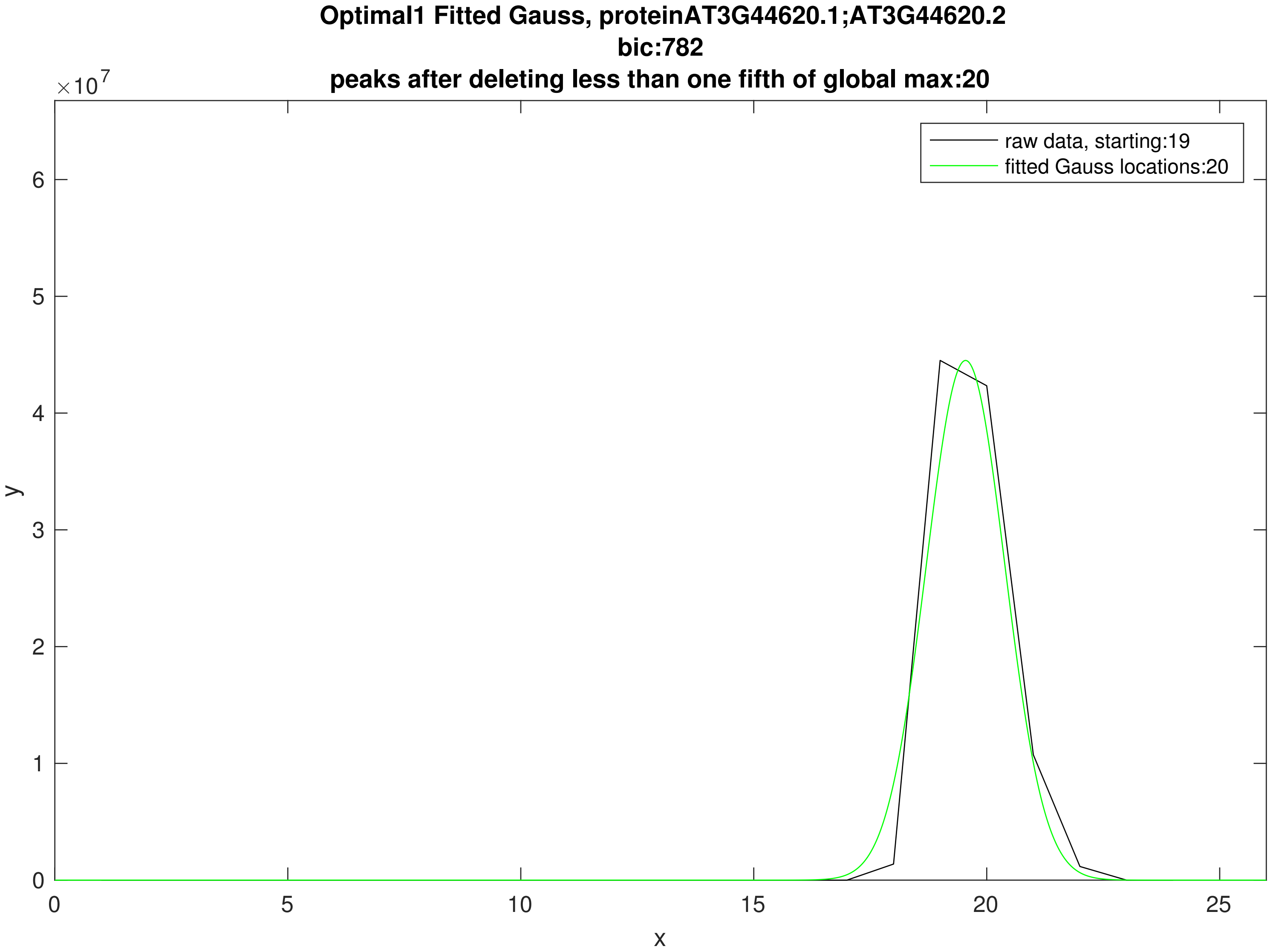
Optimal1 Fitted Gauss, proteinAT3G44620.1;AT3G44620.2
bic:782
peaks after deleting less than one fifth of global max:20
×10^7
raw data, starting:19
fitted Gauss locations:20
y
x
0
1
2
3
4
5
6
0
5
10
15
20
25

**Optimal1 Fitted Gauss, proteinAT3G44713.1**
**bic:702**
**peaks after deleting less than one fifth of global max:9.5**

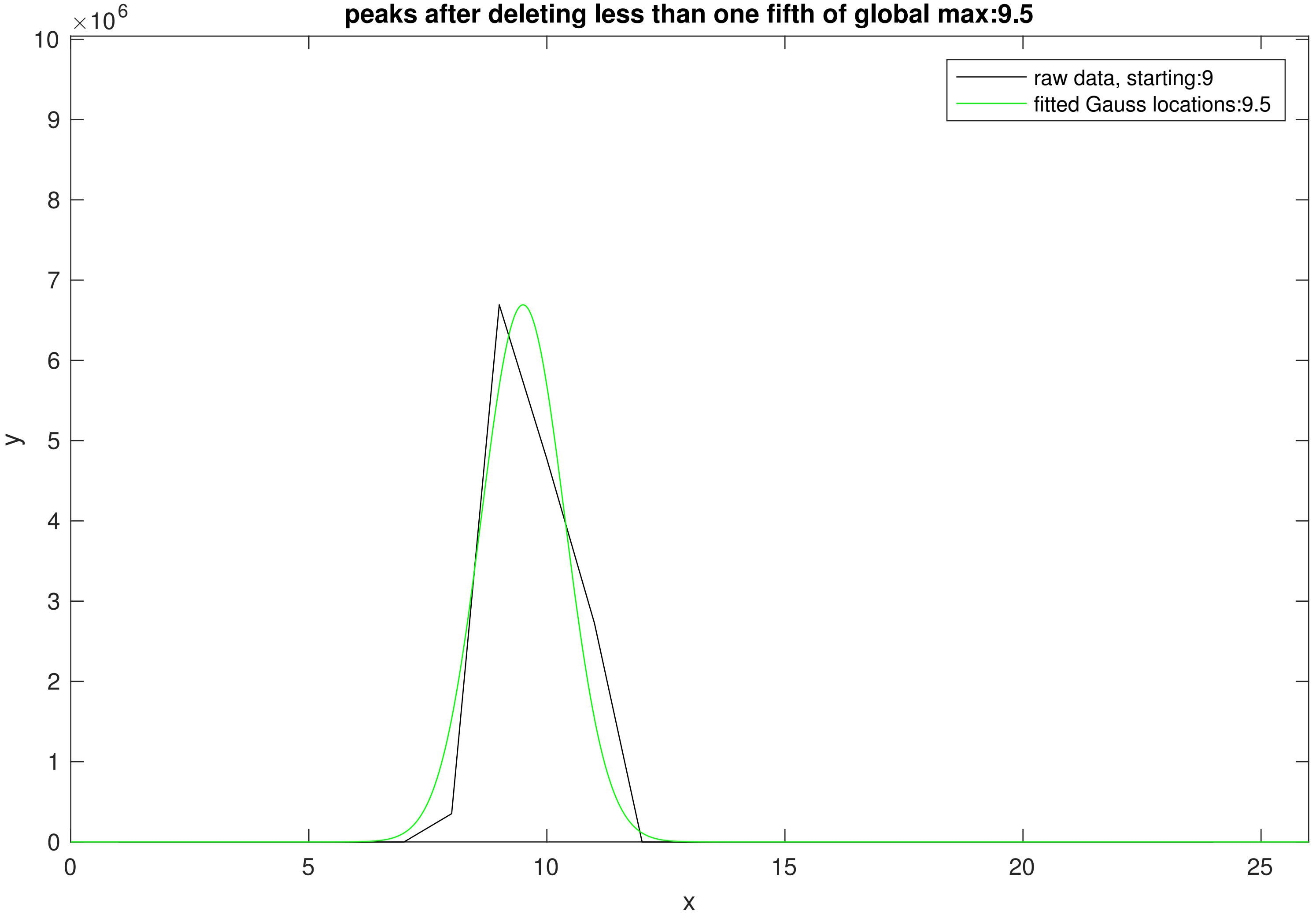

**Optimal1 Fitted Gauss, proteinAT3G45010.1**
**bic:769**
**peaks after deleting less than one fifth of global max:16**

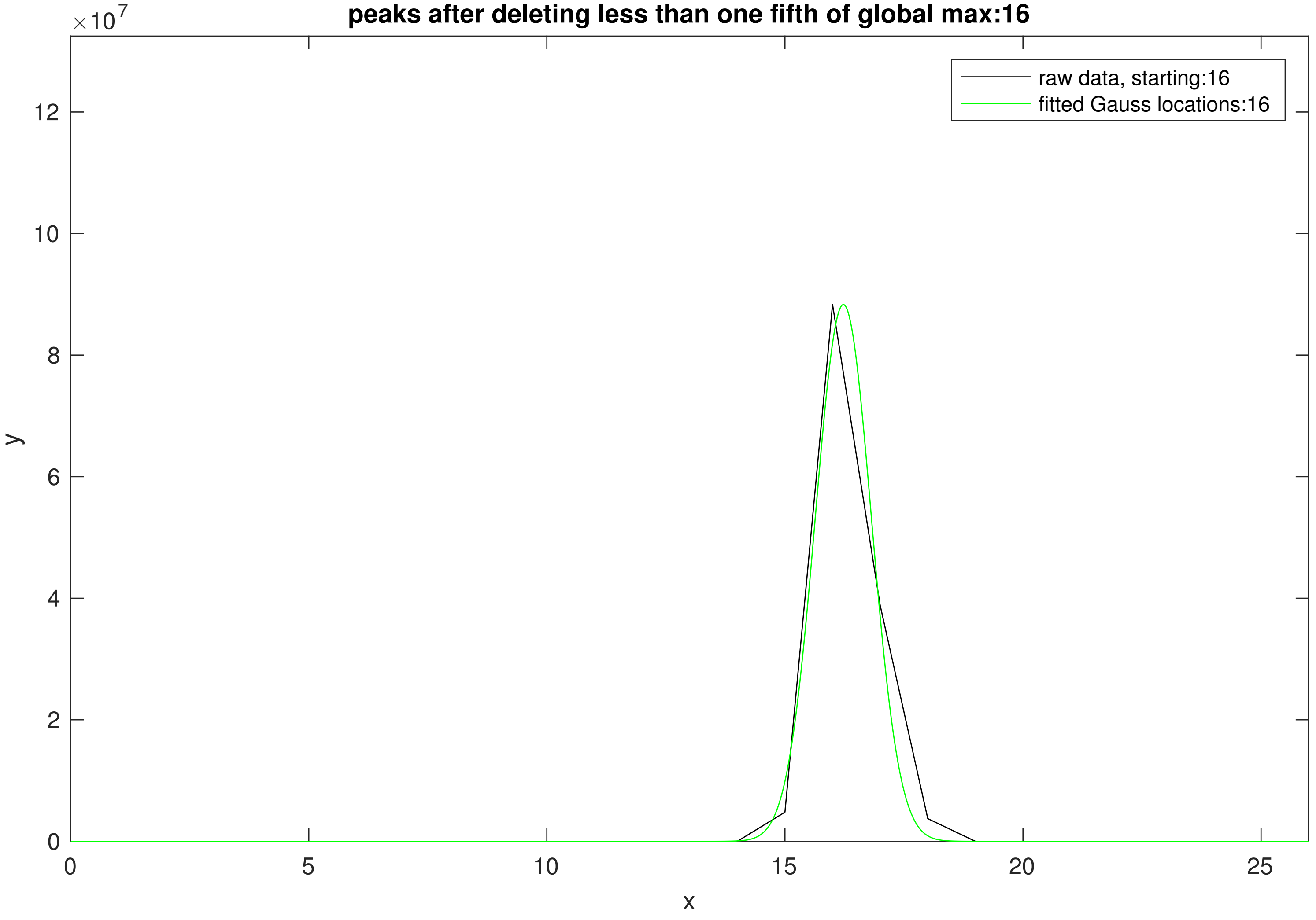

**Optimal1 Fitted Gauss, proteinAT3G45190.1**
**bic:696**
**peaks after deleting less than one fifth of global max:12**

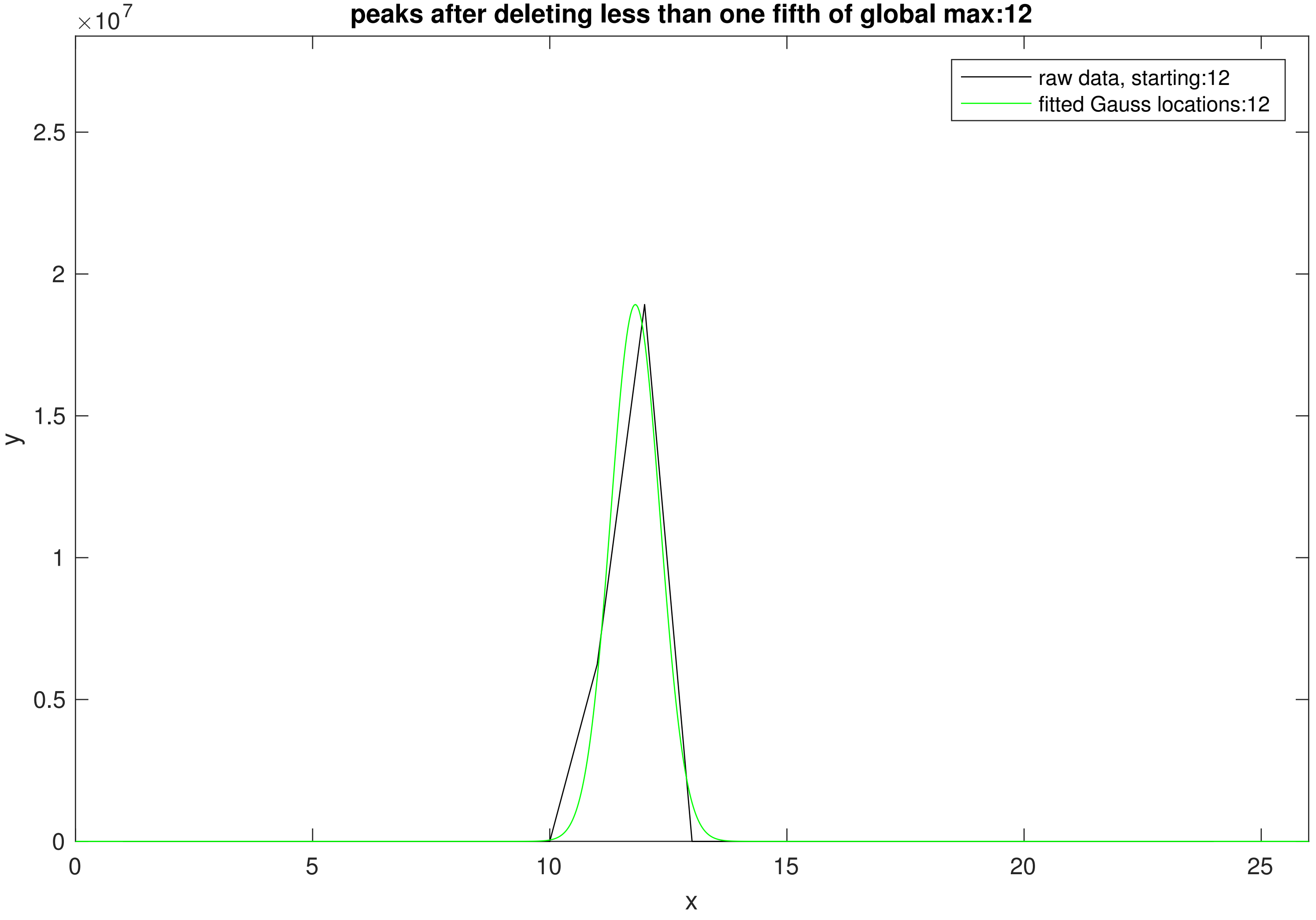

**Optimal1 Fitted Gauss, proteinAT3G45300.1**
**bic:853**
**peaks after deleting less than one fifth of global max:11**

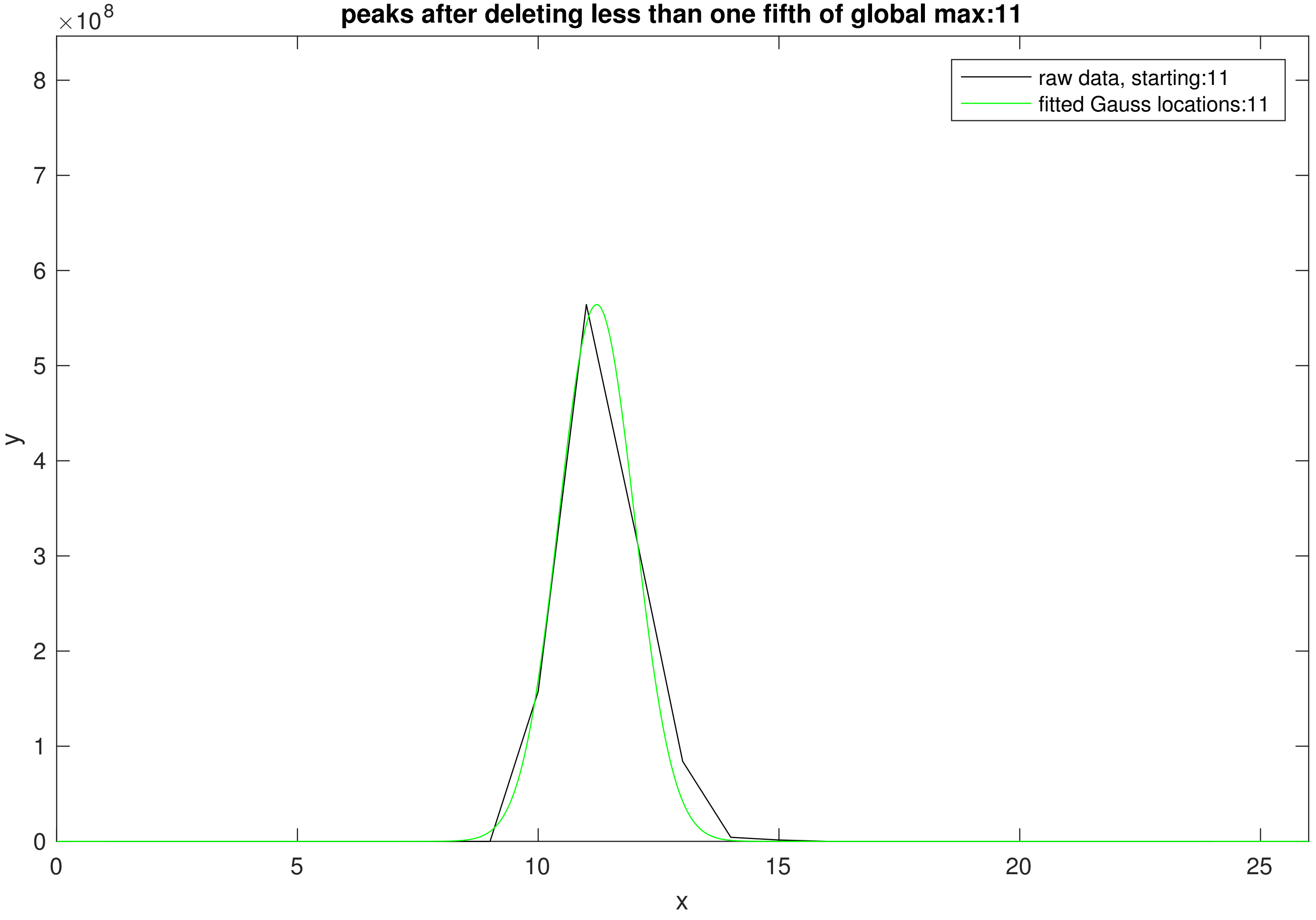

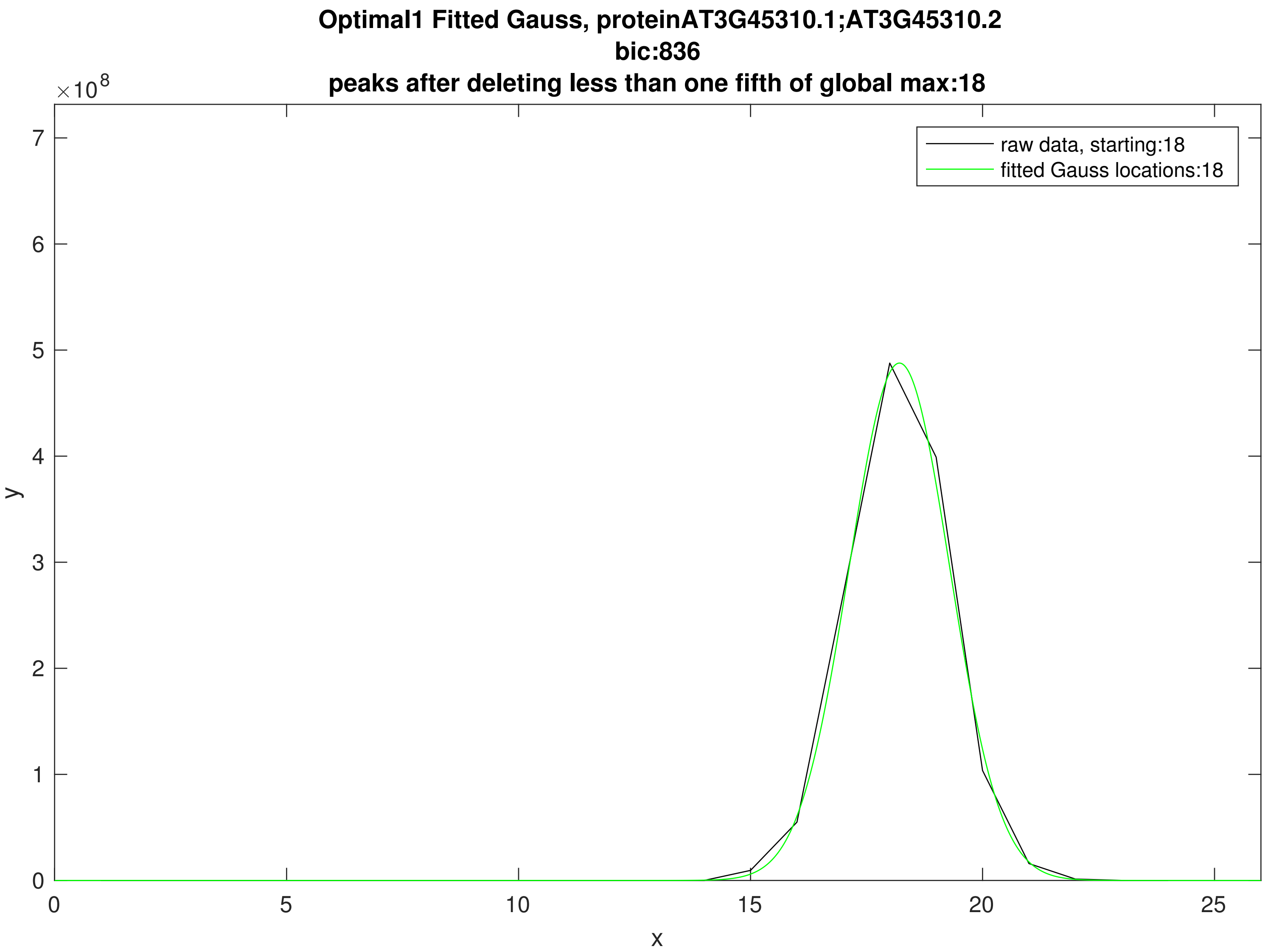

Optimal1 Fitted Gauss, proteinAT3G45310.1;AT3G45310.2
bic:836
peaks after deleting less than one fifth of global max:18
×10^8
raw data, starting:18
fitted Gauss locations:18
y
x
0
1
2
3
4
5
6
7
0
5
10
15
20
25

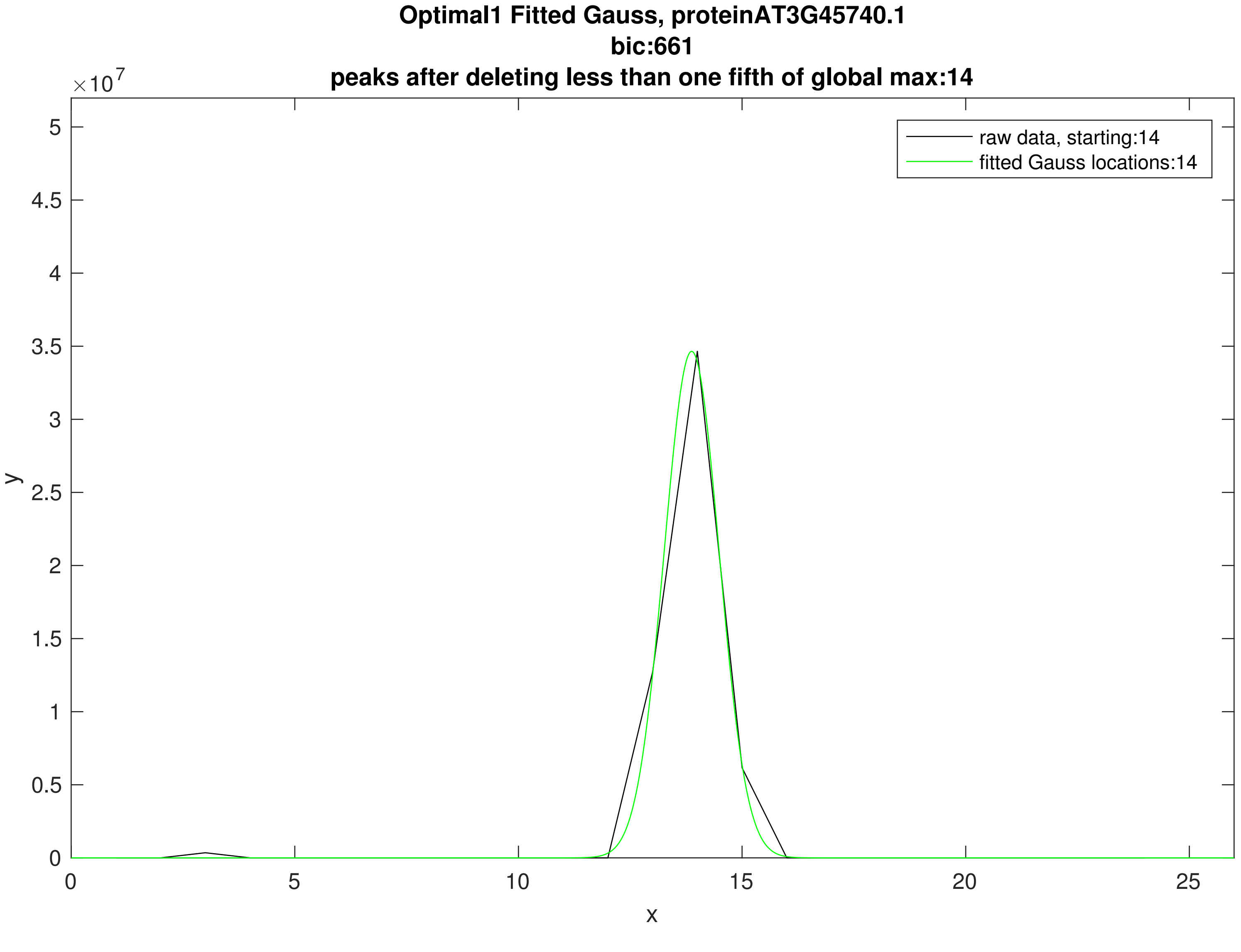
Optimal1 Fitted Gauss, proteinAT3G45740.1
bic:661
peaks after deleting less than one fifth of global max:14
×10^7
5
4.5
4
3.5
3
2.5
2
1.5
1
0.5
0
y
0
5
10
15
20
25
x
raw data, starting:14
fitted Gauss locations:14

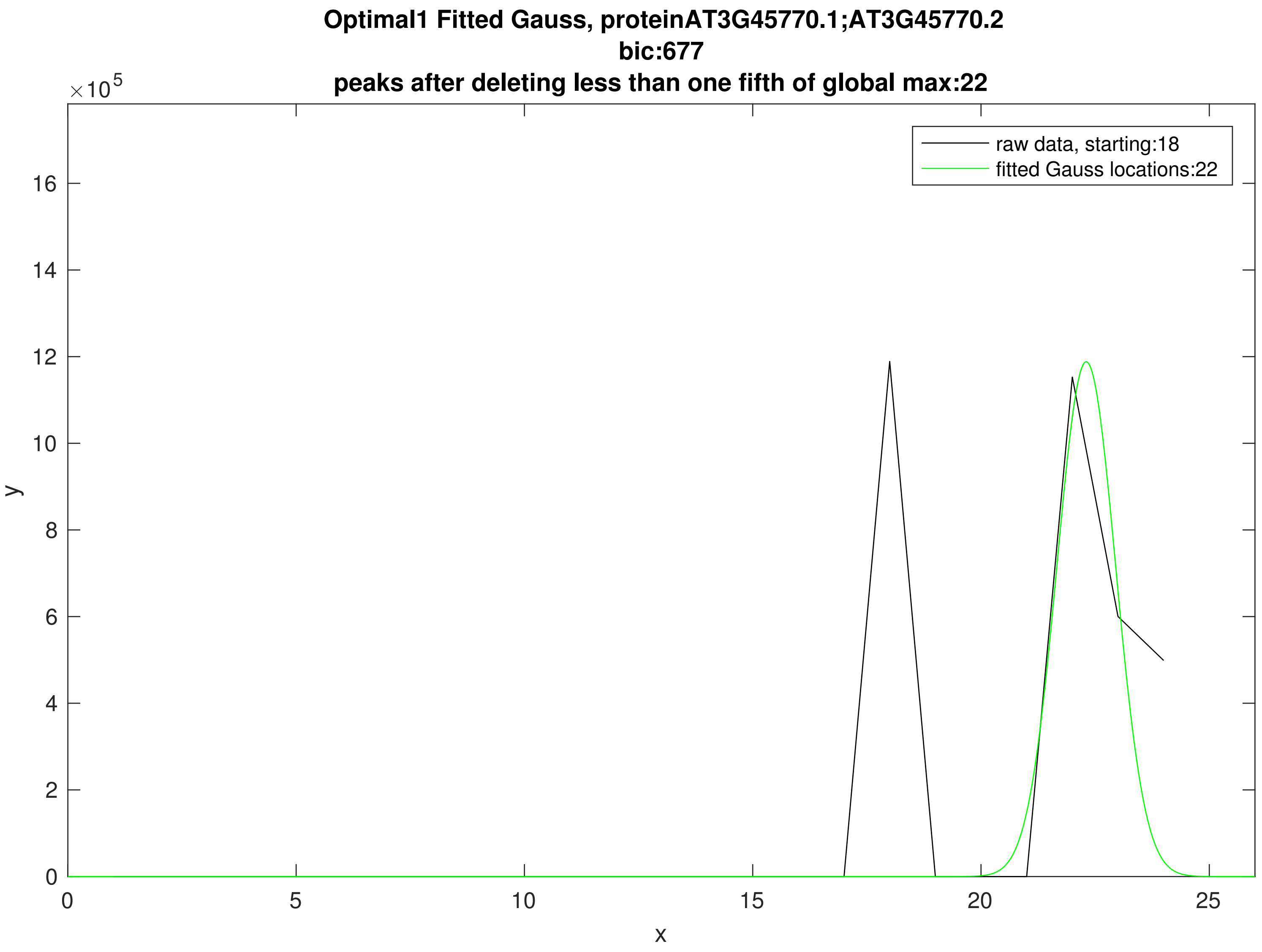

Optimal1 Fitted Gauss, proteinAT3G45770.1;AT3G45770.2
bic:677
peaks after deleting less than one fifth of global max:22
×10^5
raw data, starting:18
fitted Gauss locations:22
16
14
12
10
8
6
4
2
0
y
0
5
10
15
20
25
x

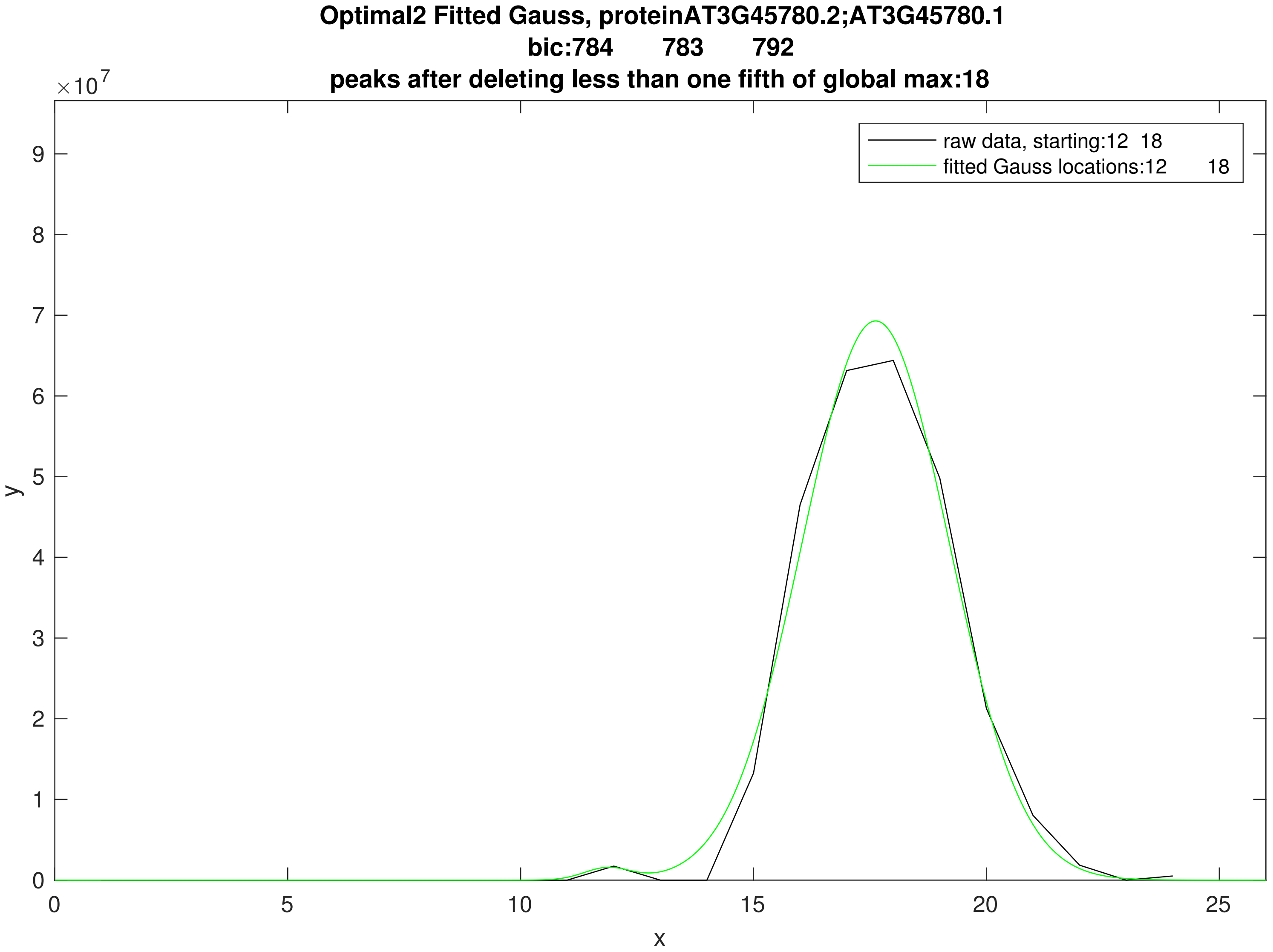
Optimal2 Fitted Gauss, proteinAT3G45780.2;AT3G45780.1
bic:784 783 792
peaks after deleting less than one fifth of global max:18
×10^7
raw data, starting:12 18
fitted Gauss locations:12 18
y
x
0
1
2
3
4
5
6
7
8
9
0
5
10
15
20
25

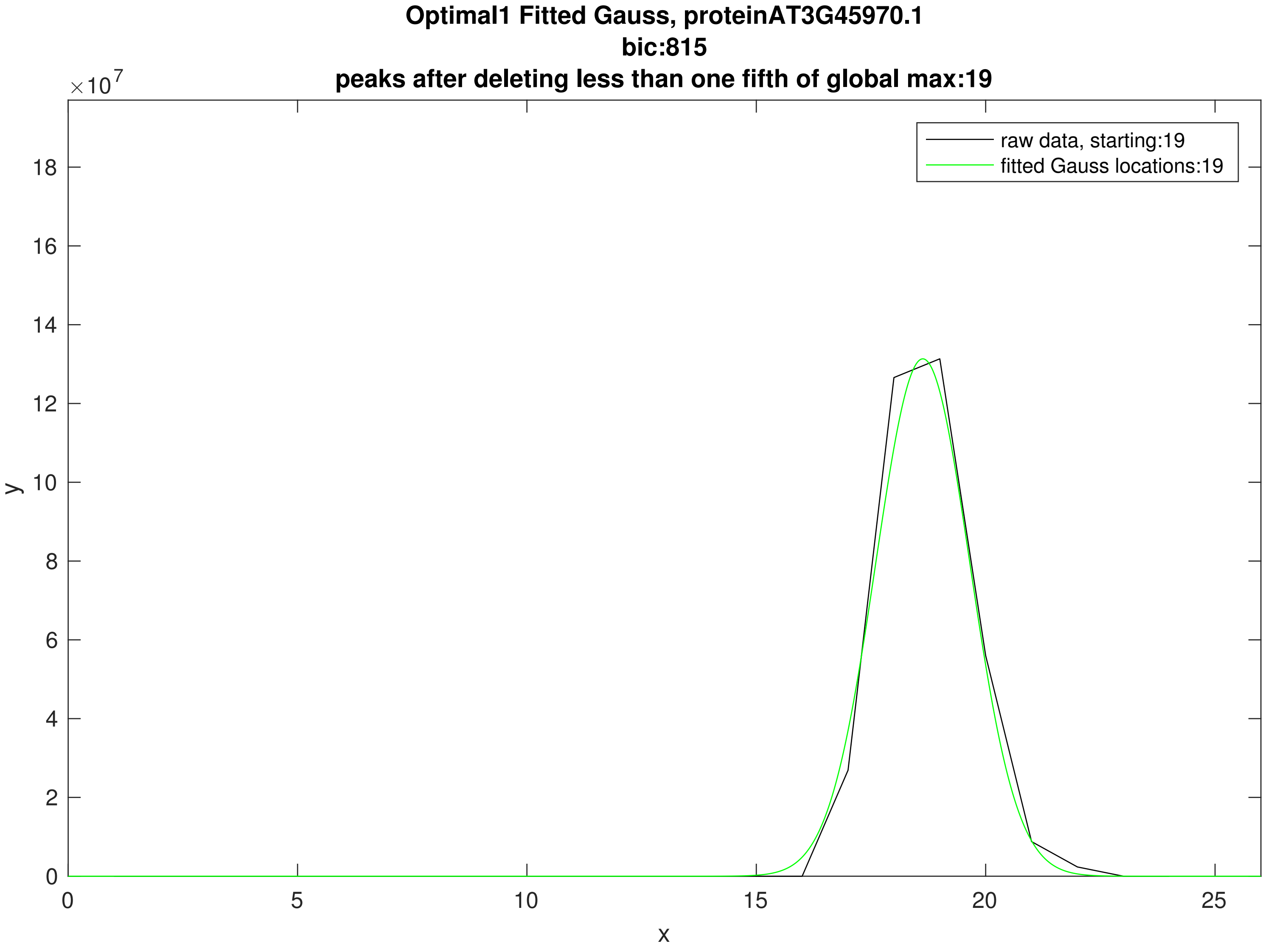
Optimal1 Fitted Gauss, proteinAT3G45970.1
bic:815
peaks after deleting less than one fifth of global max:19
×10^7
raw data, starting:19
fitted Gauss locations:19
y
x
0
2
4
6
8
10
12
14
16
18
0
5
10
15
20
25

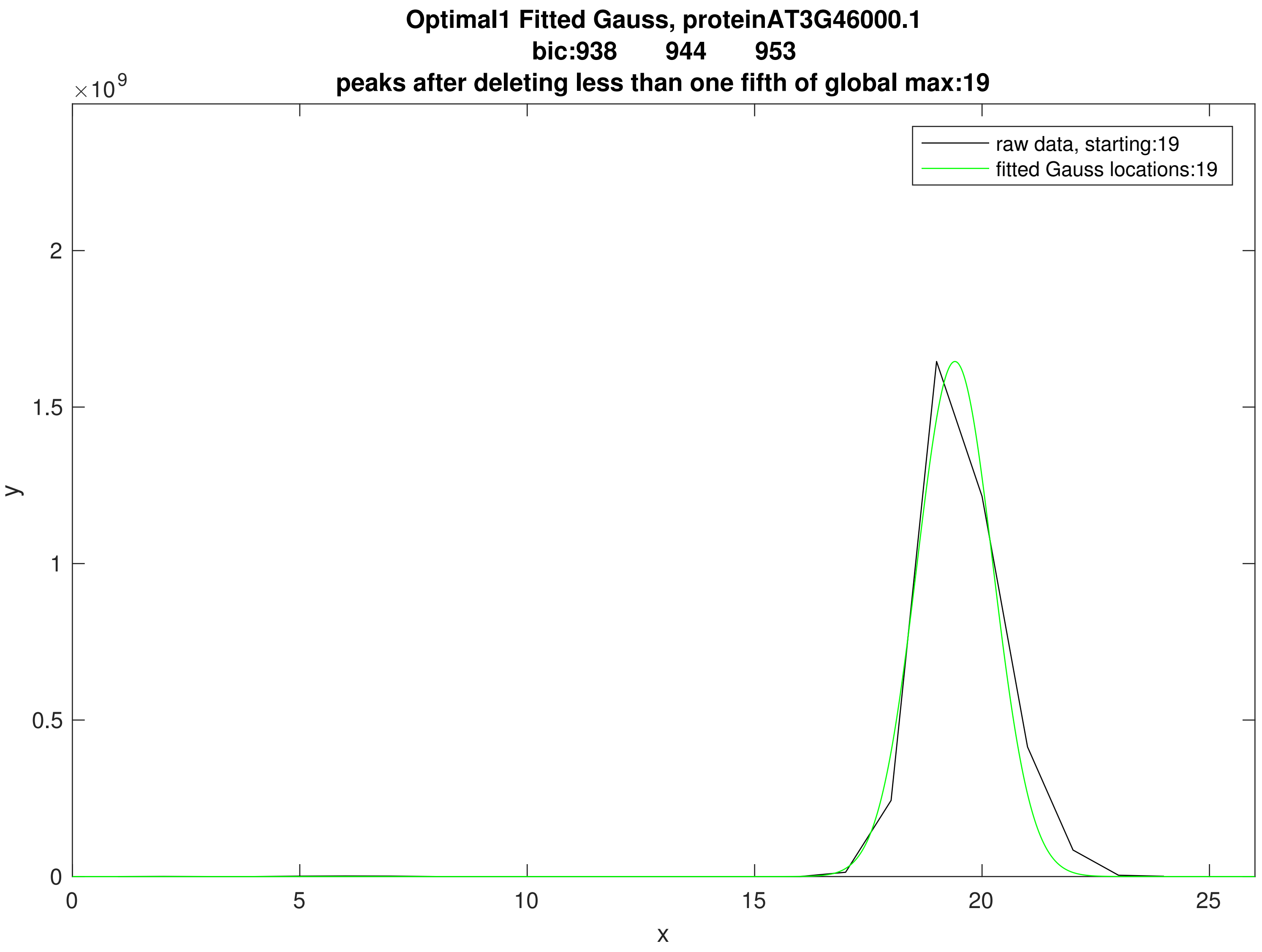

Optimal1 Fitted Gauss, proteinAT3G46000.1
bic:938 944 953
peaks after deleting less than one fifth of global max:19
×10^9
raw data, starting:19
fitted Gauss locations:19
2
1.5
1
0.5
0
y
0
5
10
15
20
25
x

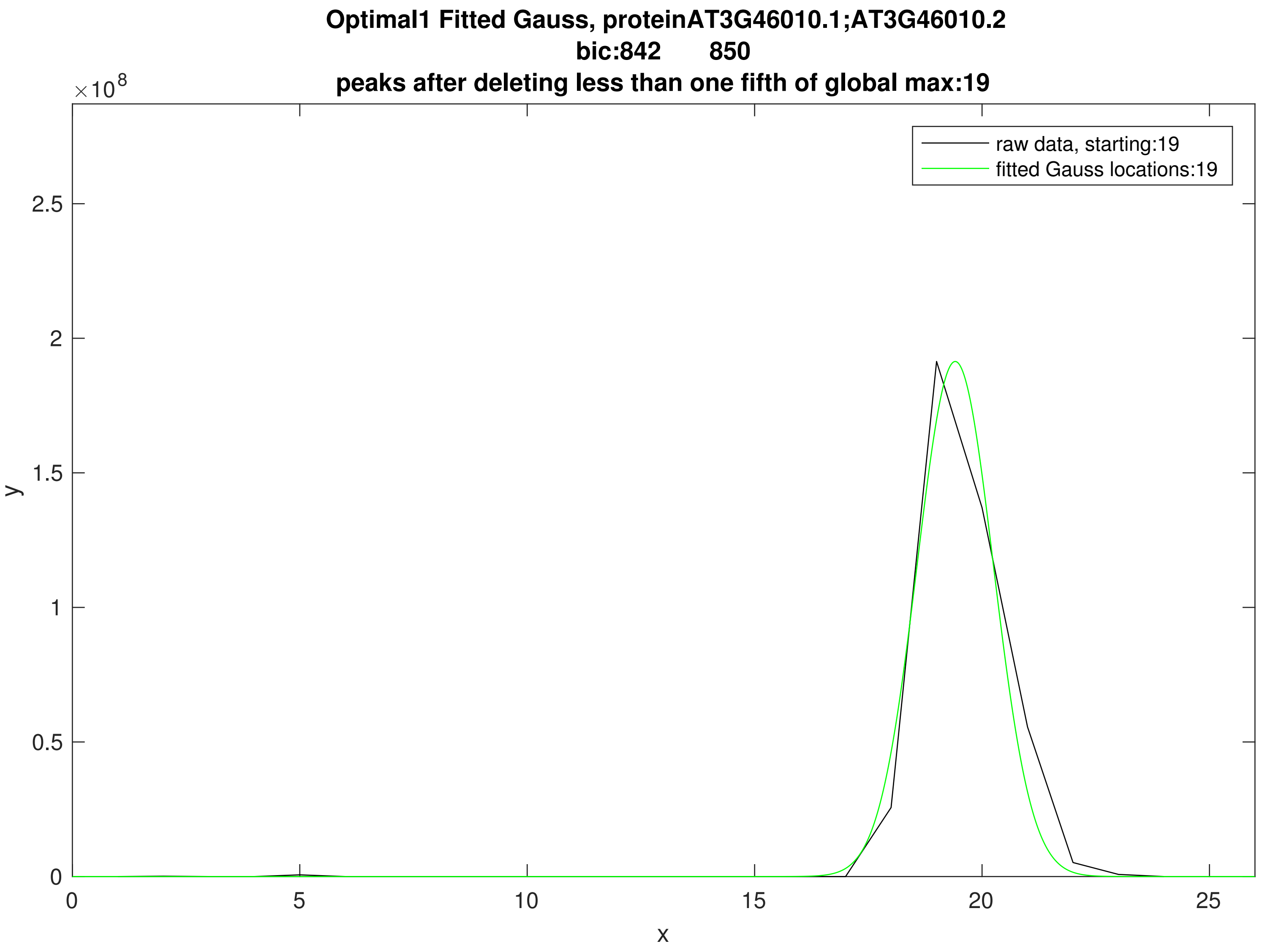
Optimal1 Fitted Gauss, proteinAT3G46010.1;AT3G46010.2
bic:842 850
peaks after deleting less than one fifth of global max:19
×10^8
2.5
2
1.5
1
0.5
0
y
0
5
10
15
20
25
x
raw data, starting:19
fitted Gauss locations:19

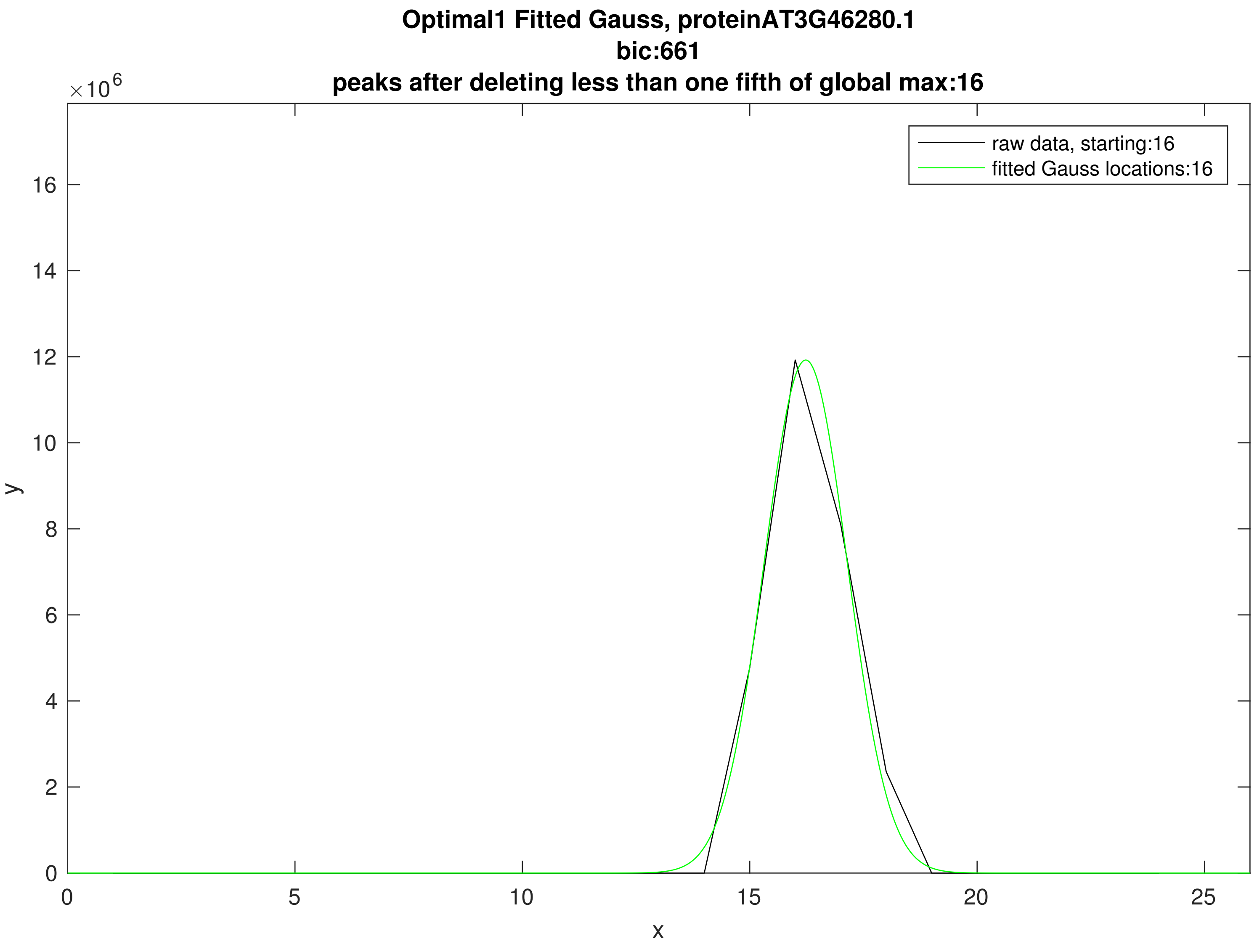
Optimal1 Fitted Gauss, proteinAT3G46280.1
bic:661
peaks after deleting less than one fifth of global max:16
×10^6
raw data, starting:16
fitted Gauss locations:16
y
x
0
2
4
6
8
10
12
14
16
0
5
10
15
20
25

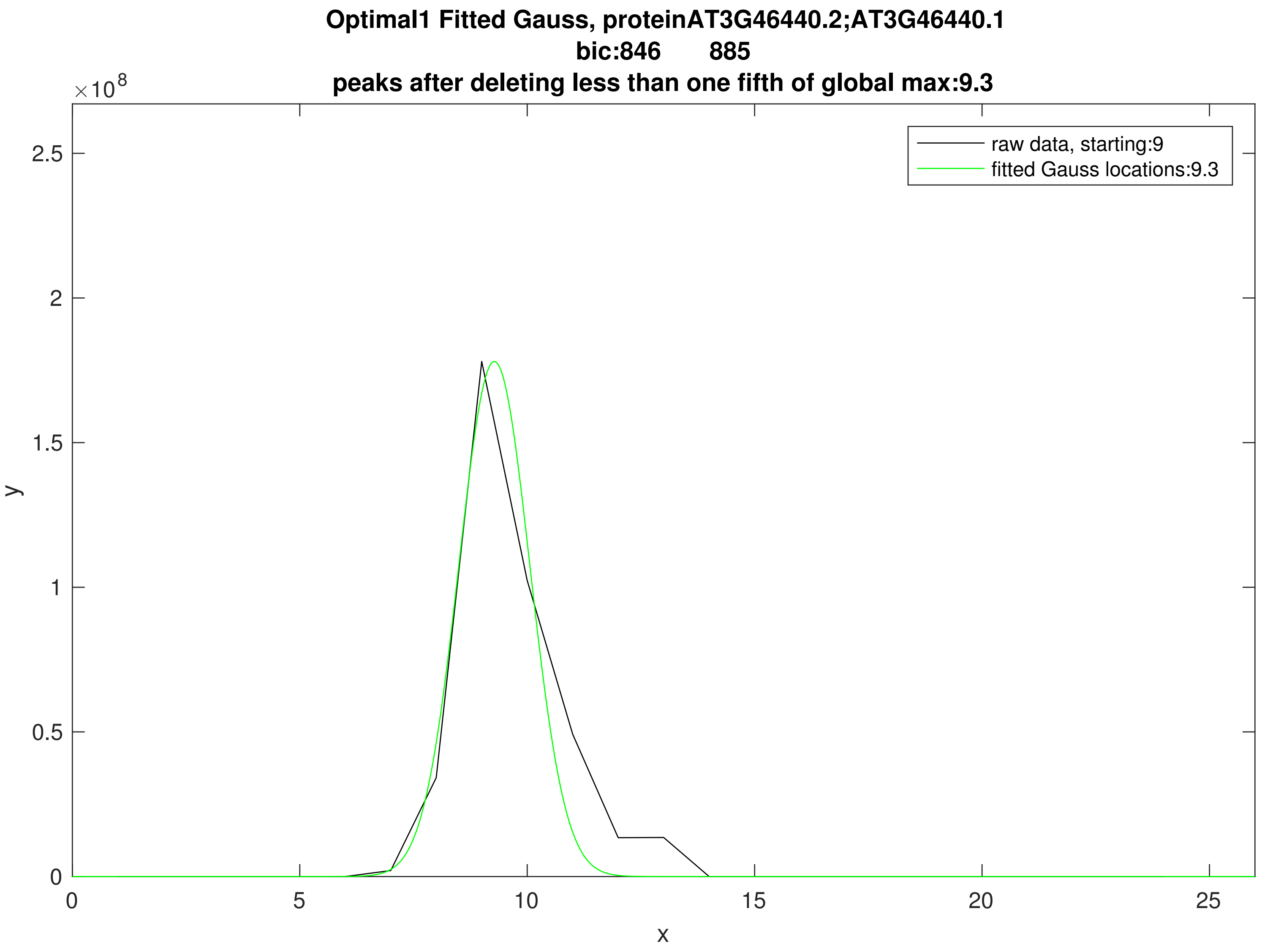

Optimal1 Fitted Gauss, proteinAT3G46440.2;AT3G46440.1
bic:846 885
peaks after deleting less than one fifth of global max:9.3
×10^8
2.5
2
1.5
1
0.5
0
y
0
5
10
15
20
25
x
raw data, starting:9
fitted Gauss locations:9.3

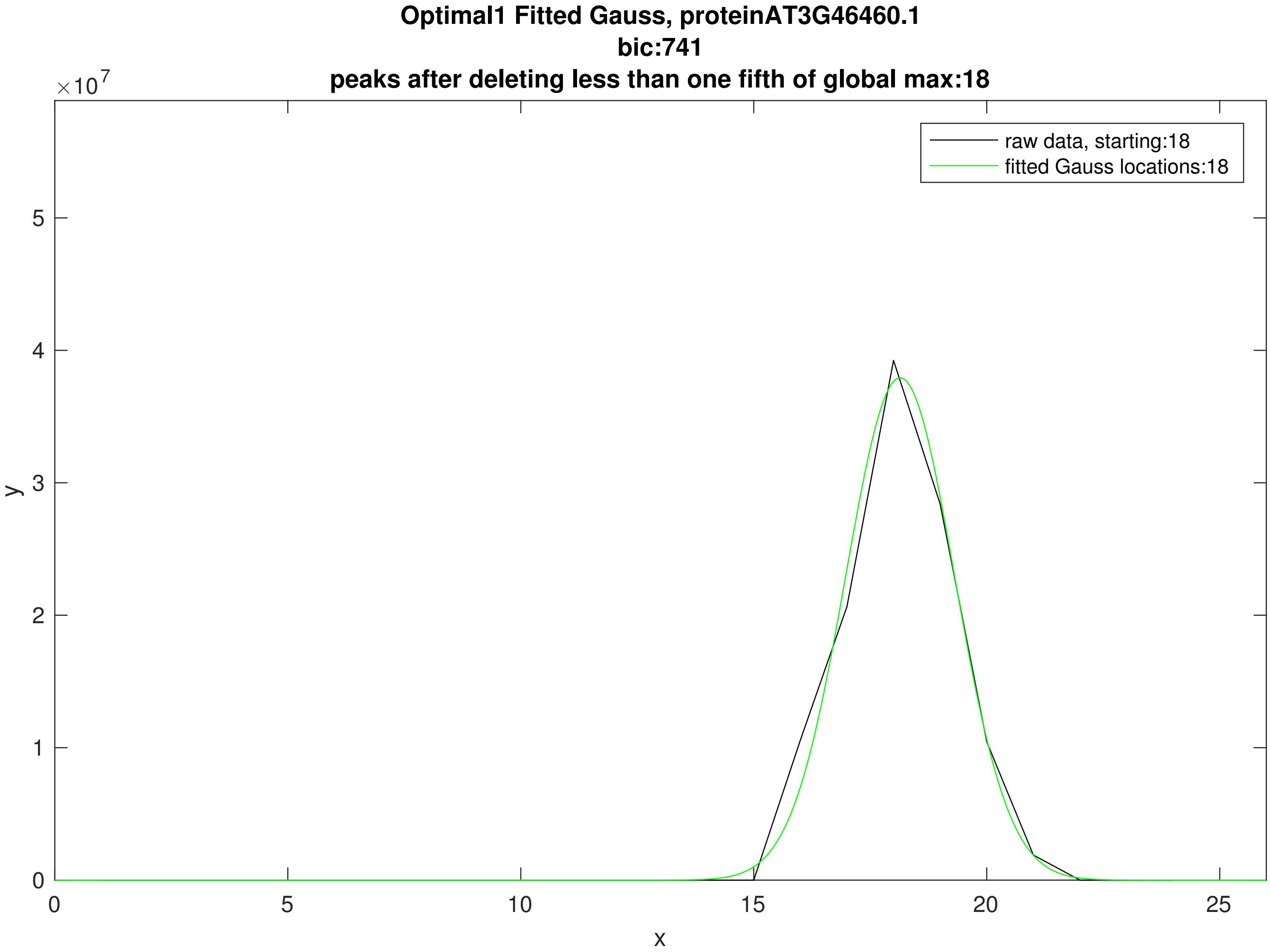
Optimal1 Fitted Gauss, proteinAT3G46460.1
bic:741
peaks after deleting less than one fifth of global max:18
×10^7
raw data, starting:18
fitted Gauss locations:18
y
x
0
1
2
3
4
5
0
5
10
15
20
25

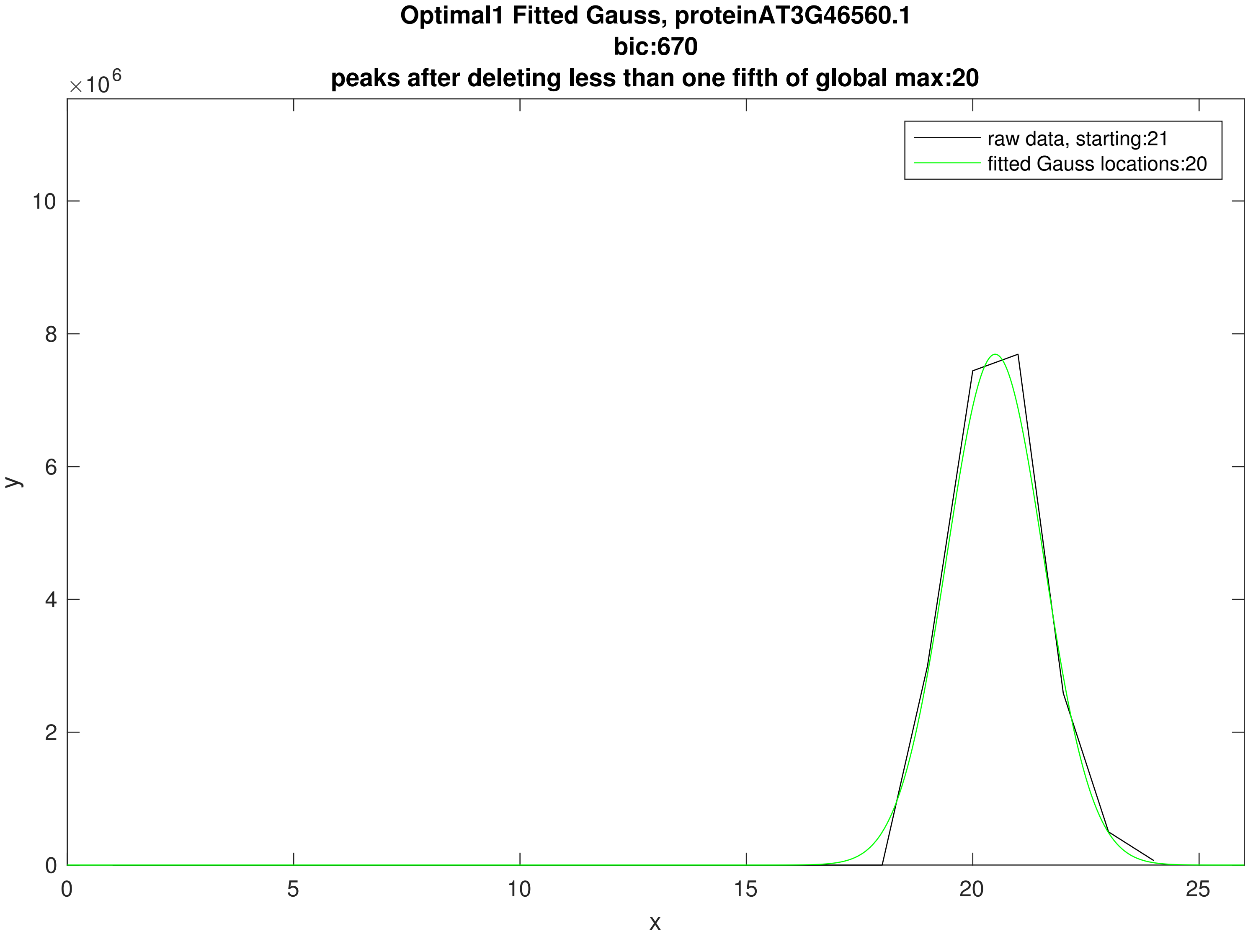
Optimal1 Fitted Gauss, proteinAT3G46560.1
bic:670
peaks after deleting less than one fifth of global max:20
×10^6
raw data, starting:21
fitted Gauss locations:20
10
8
6
4
2
0
y
0
5
10
15
20
25
x

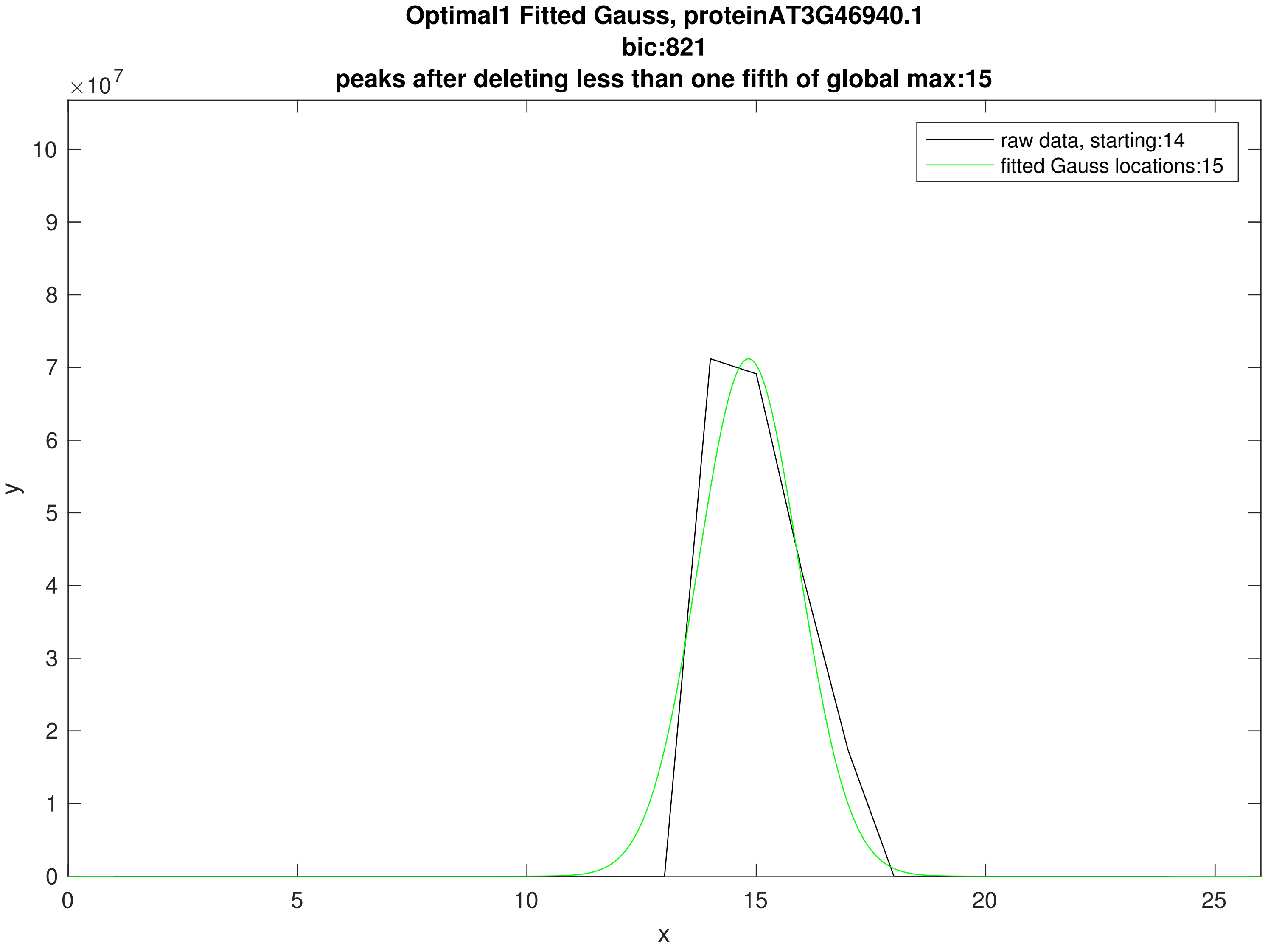
Optimal1 Fitted Gauss, proteinAT3G46940.1
bic:821
peaks after deleting less than one fifth of global max:15
×10^7
raw data, starting:14
fitted Gauss locations:15
y
x
0
1
2
3
4
5
6
7
8
9
10
0
5
10
15
20
25

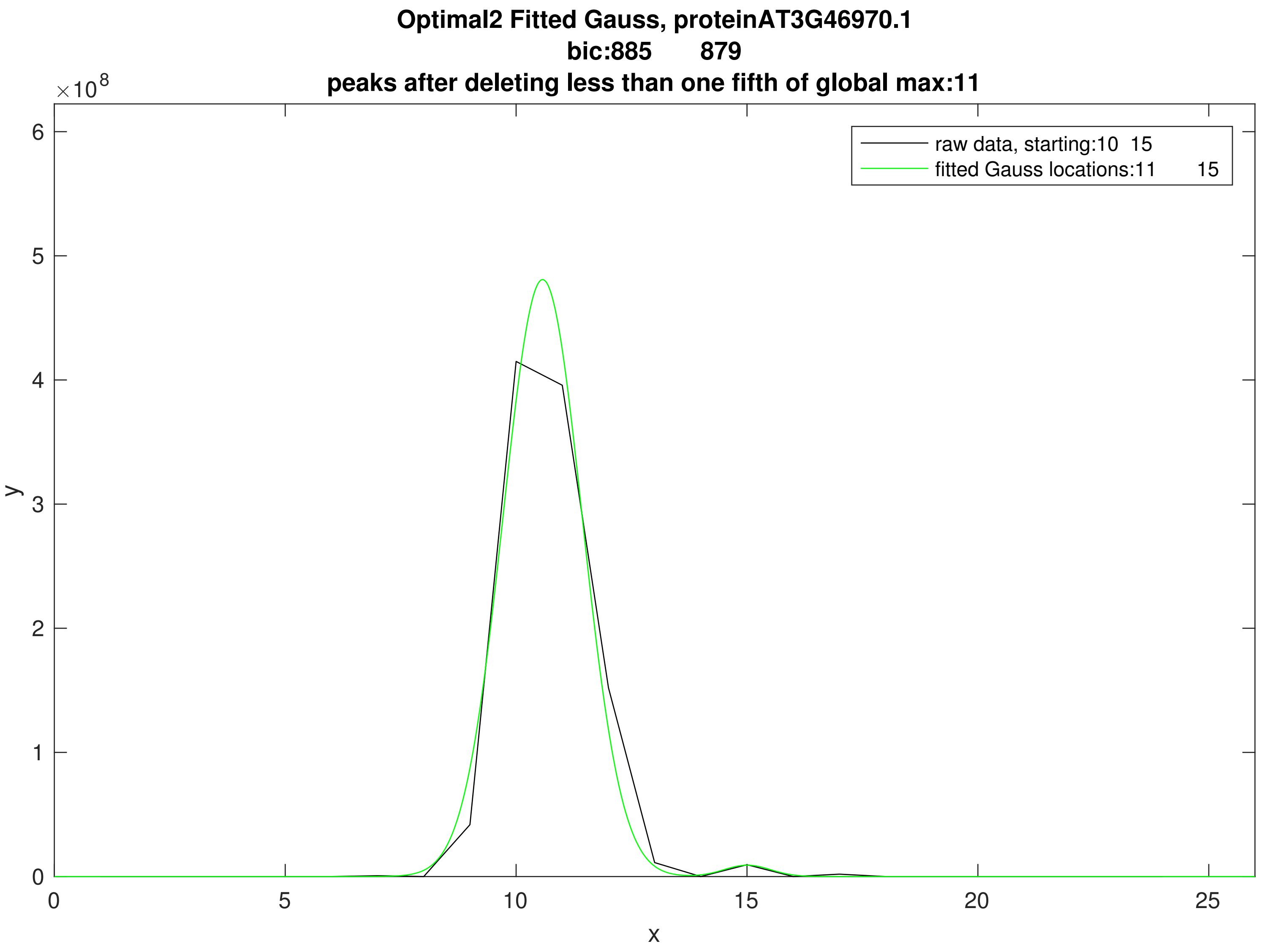

Optimal2 Fitted Gauss, proteinAT3G46970.1
bic:885 879
peaks after deleting less than one fifth of global max:11
×10^8
raw data, starting:10 15
fitted Gauss locations:11 15
y
x
0
1
2
3
4
5
6
0
5
10
15
20
25

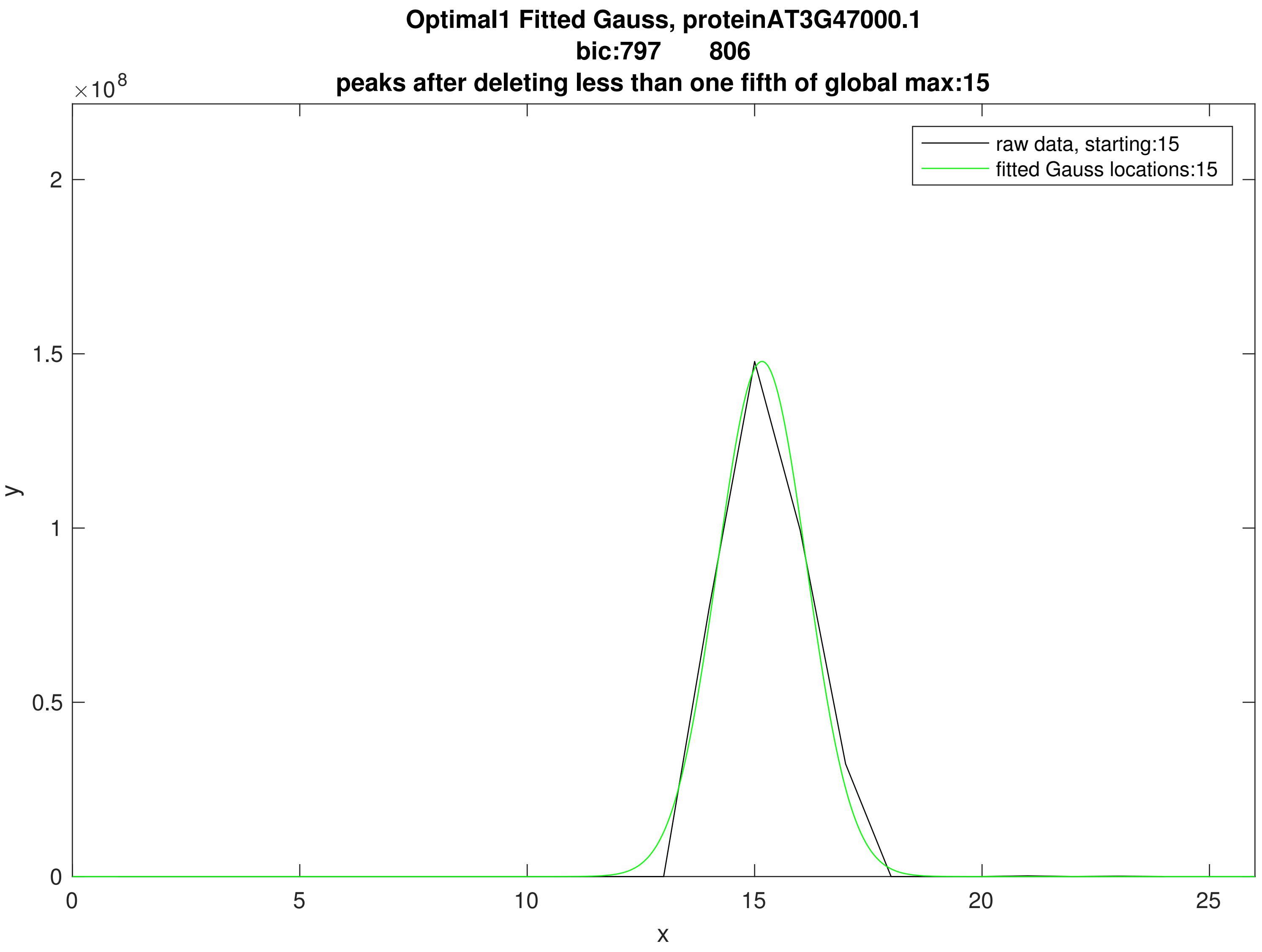

Optimal1 Fitted Gauss, proteinAT3G47000.1
bic:797 806
peaks after deleting less than one fifth of global max:15
×10^8
2
1.5
1
0.5
0
y
0
5
10
15
20
25
x
raw data, starting:15
fitted Gauss locations:15

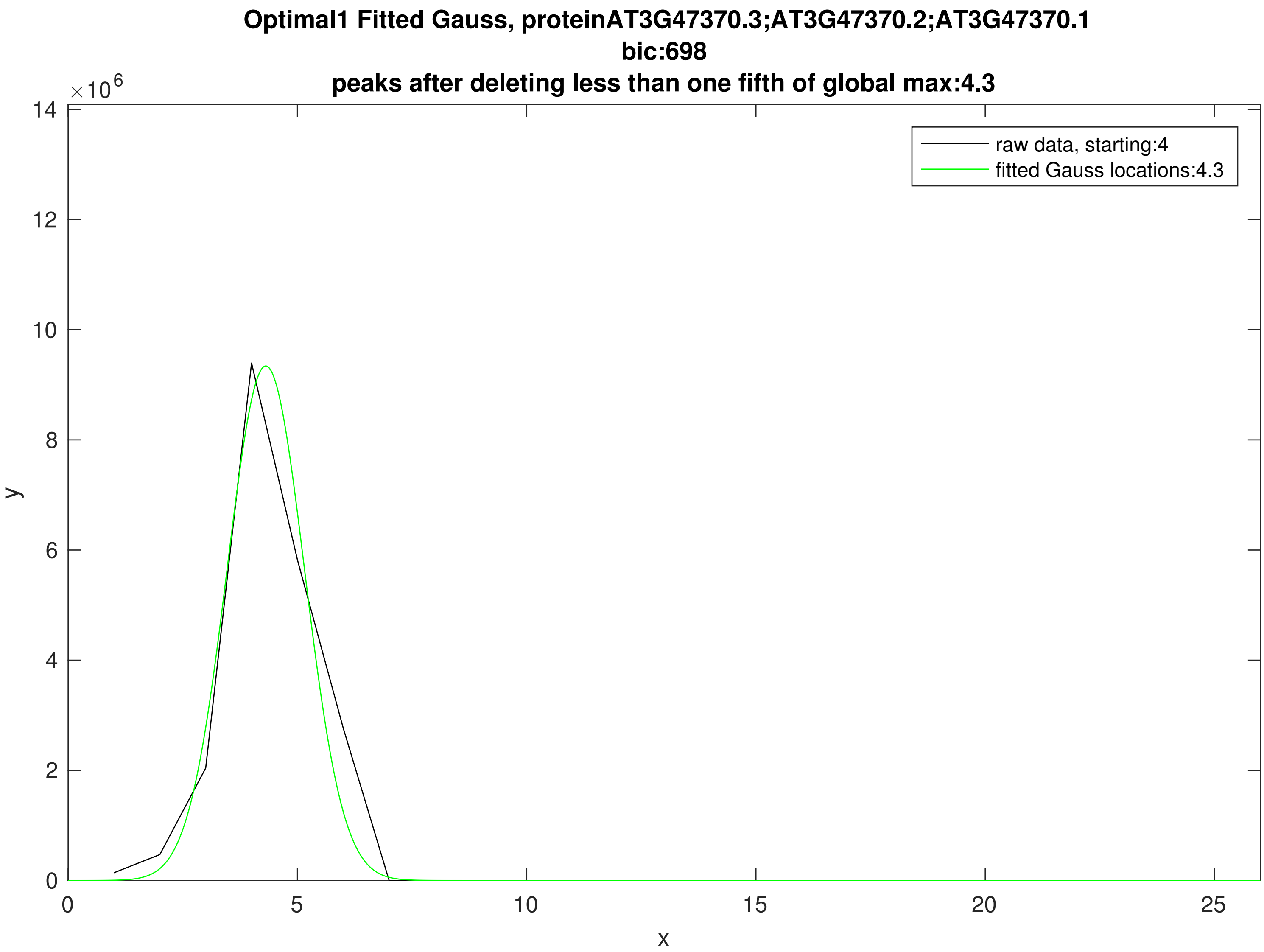
Optimal1 Fitted Gauss, proteinAT3G47370.3;AT3G47370.2;AT3G47370.1
bic:698
peaks after deleting less than one fifth of global max:4.3
×10^6
14
12
10
8
6
4
2
0
y
0
5
10
15
20
25
x
raw data, starting:4
fitted Gauss locations:4.3

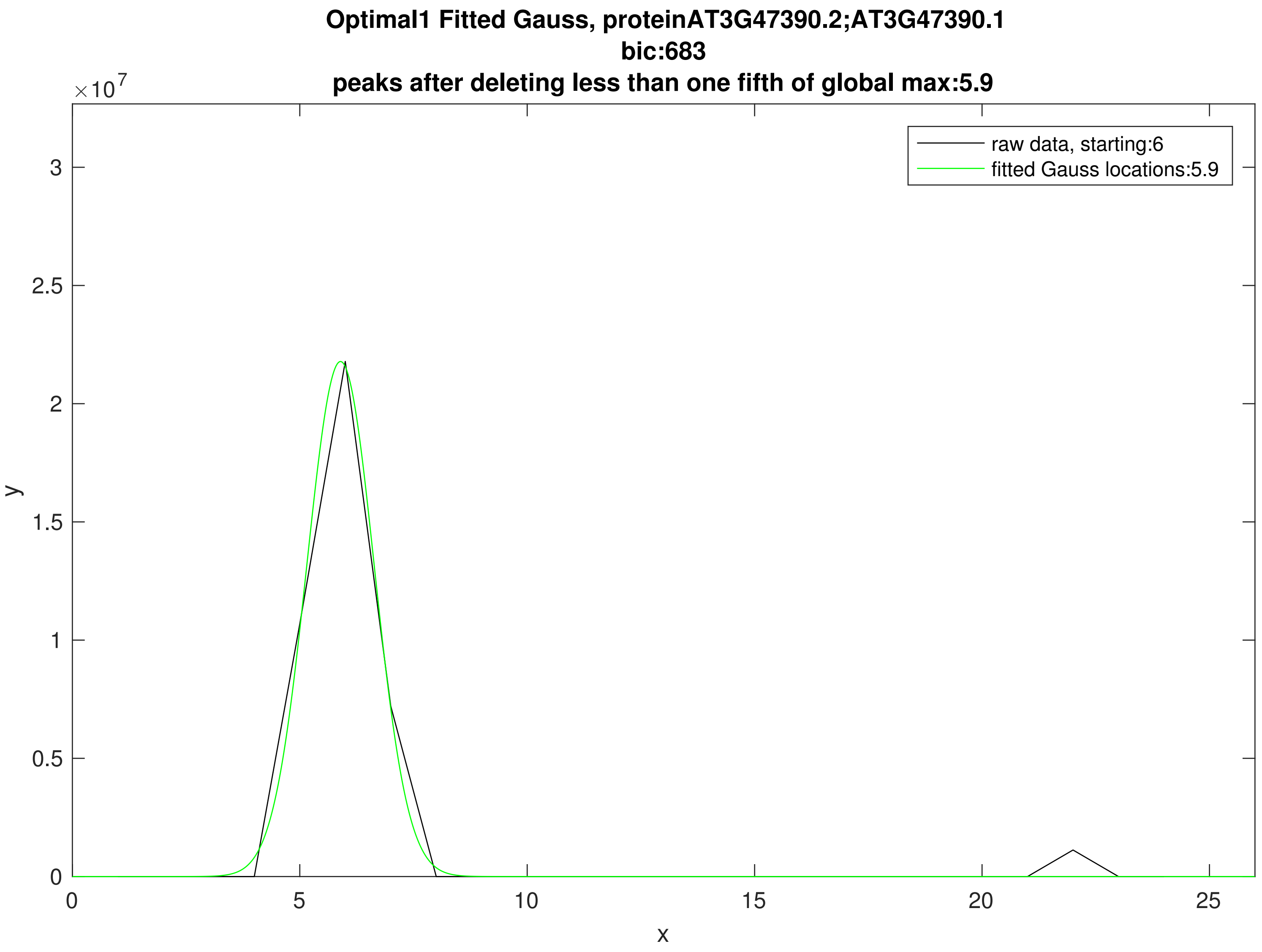
Optimal1 Fitted Gauss, proteinAT3G47390.2;AT3G47390.1
bic:683
peaks after deleting less than one fifth of global max:5.9
×10^7
raw data, starting:6
fitted Gauss locations:5.9
y
x
0
0.5
1
1.5
2
2.5
3
5
10
15
20
25

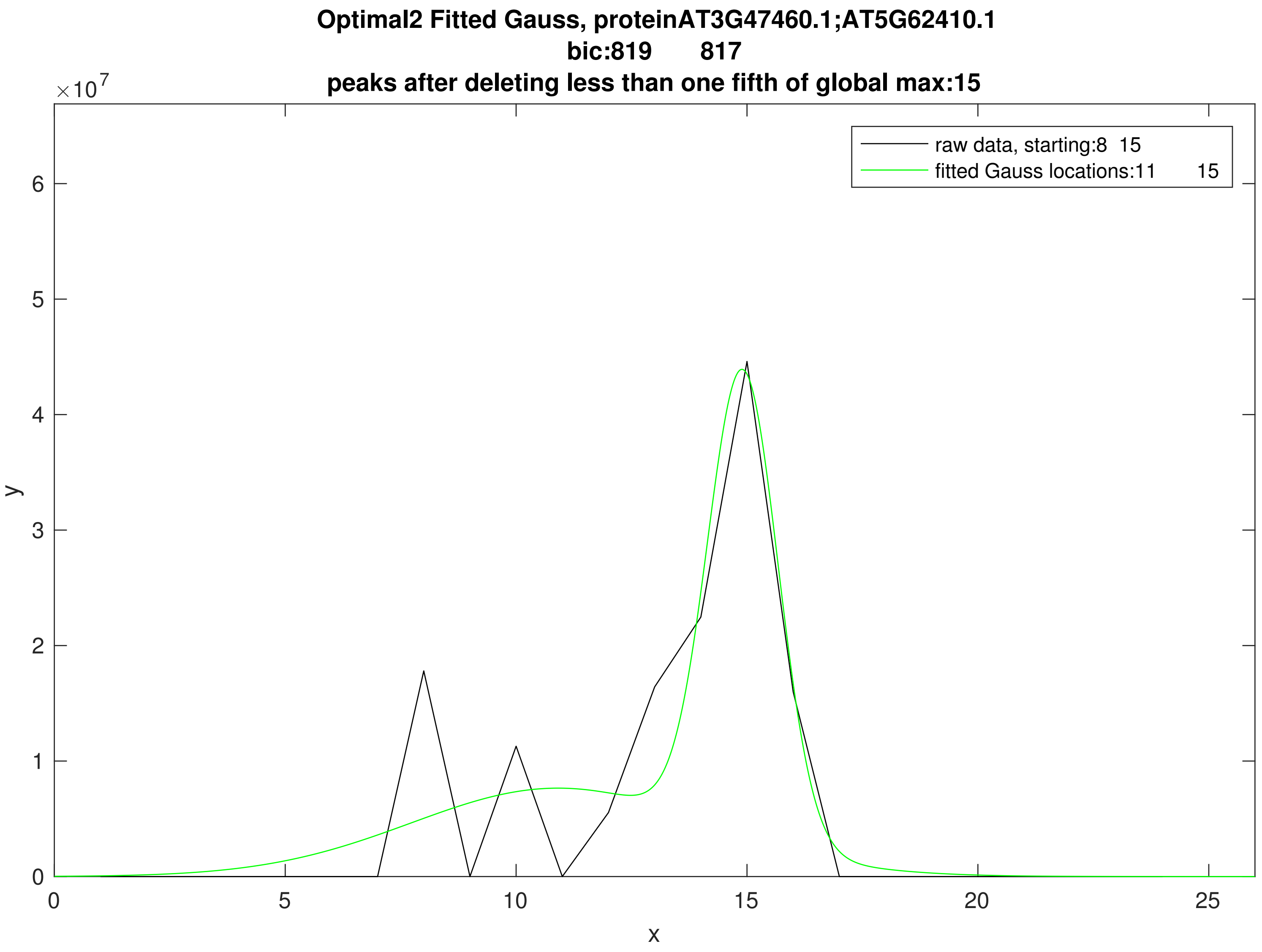
Optimal2 Fitted Gauss, proteinAT3G47460.1;AT5G62410.1
bic:819 817
peaks after deleting less than one fifth of global max:15
×10^7
raw data, starting:8 15
fitted Gauss locations:11 15
y
x
0
1
2
3
4
5
6
0
5
10
15
20
25

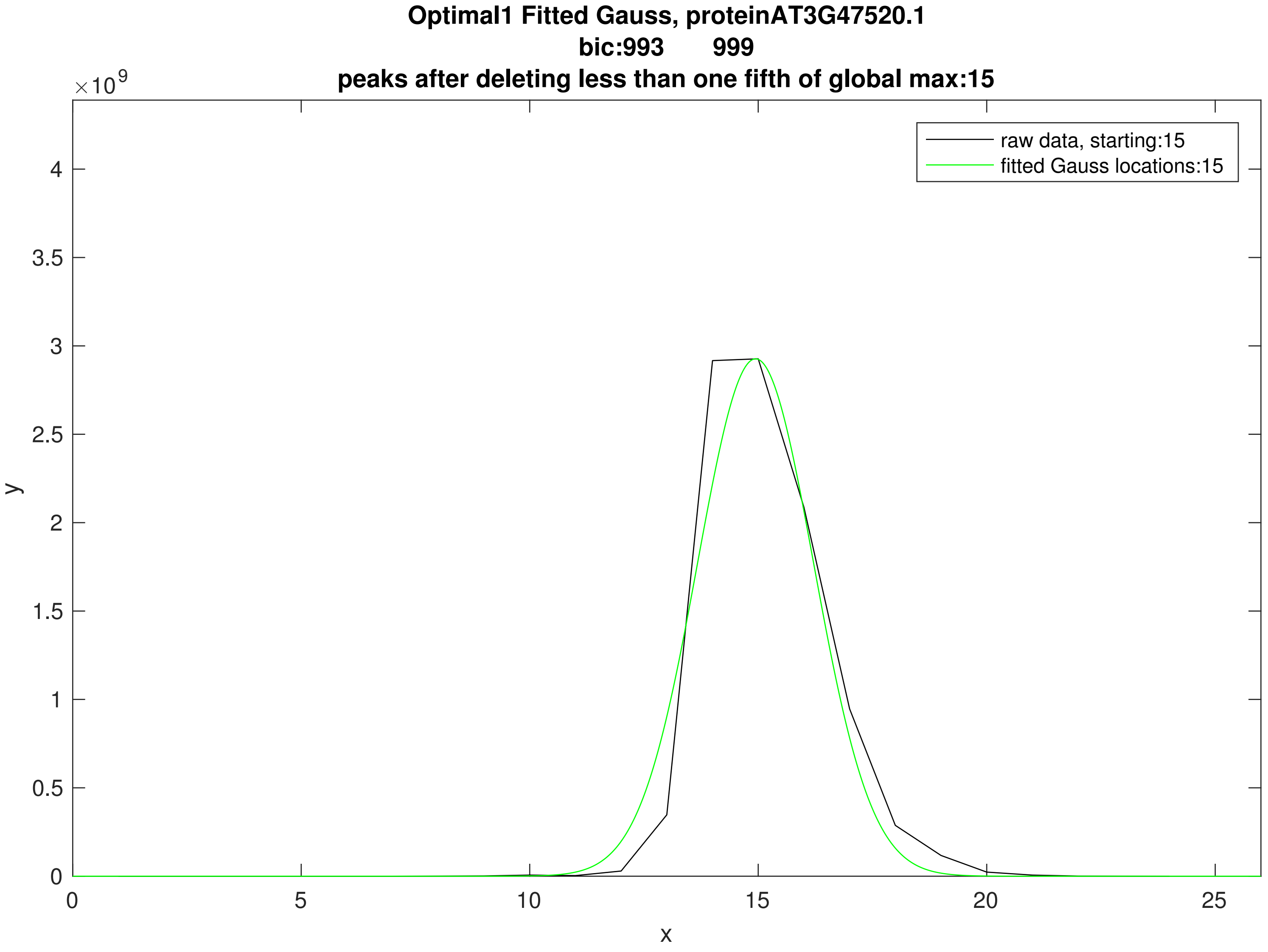

Optimal1 Fitted Gauss, proteinAT3G47520.1
bic:993 999
peaks after deleting less than one fifth of global max:15
×10^9
4
3.5
3
2.5
2
1.5
1
0.5
0
y
0
5
10
15
20
25
x
raw data, starting:15
fitted Gauss locations:15

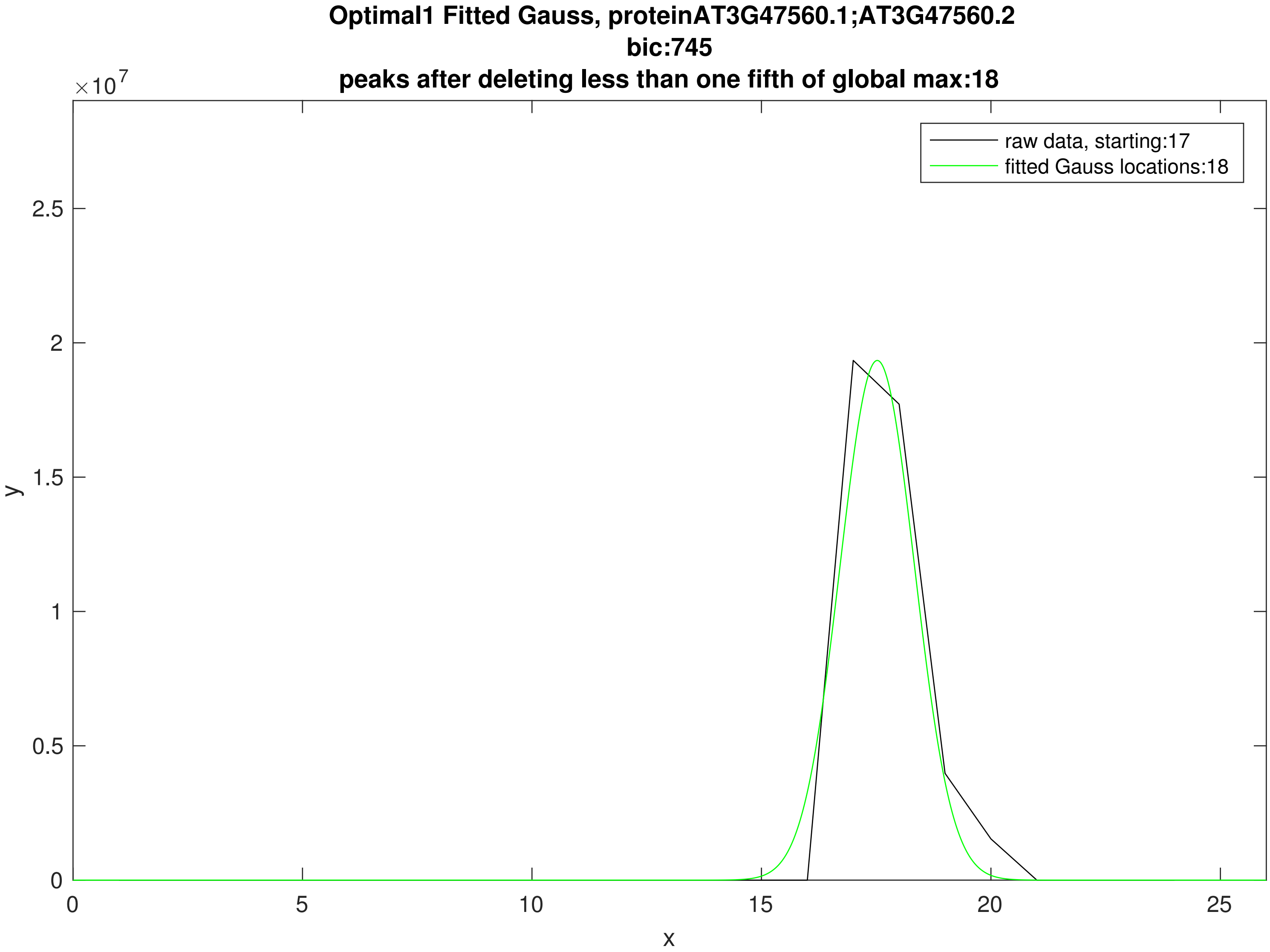
Optimal1 Fitted Gauss, proteinAT3G47560.1;AT3G47560.2
bic:745
peaks after deleting less than one fifth of global max:18
×10^7
raw data, starting:17
fitted Gauss locations:18
2.5
2
1.5
1
0.5
0
y
0
5
10
15
20
25
x

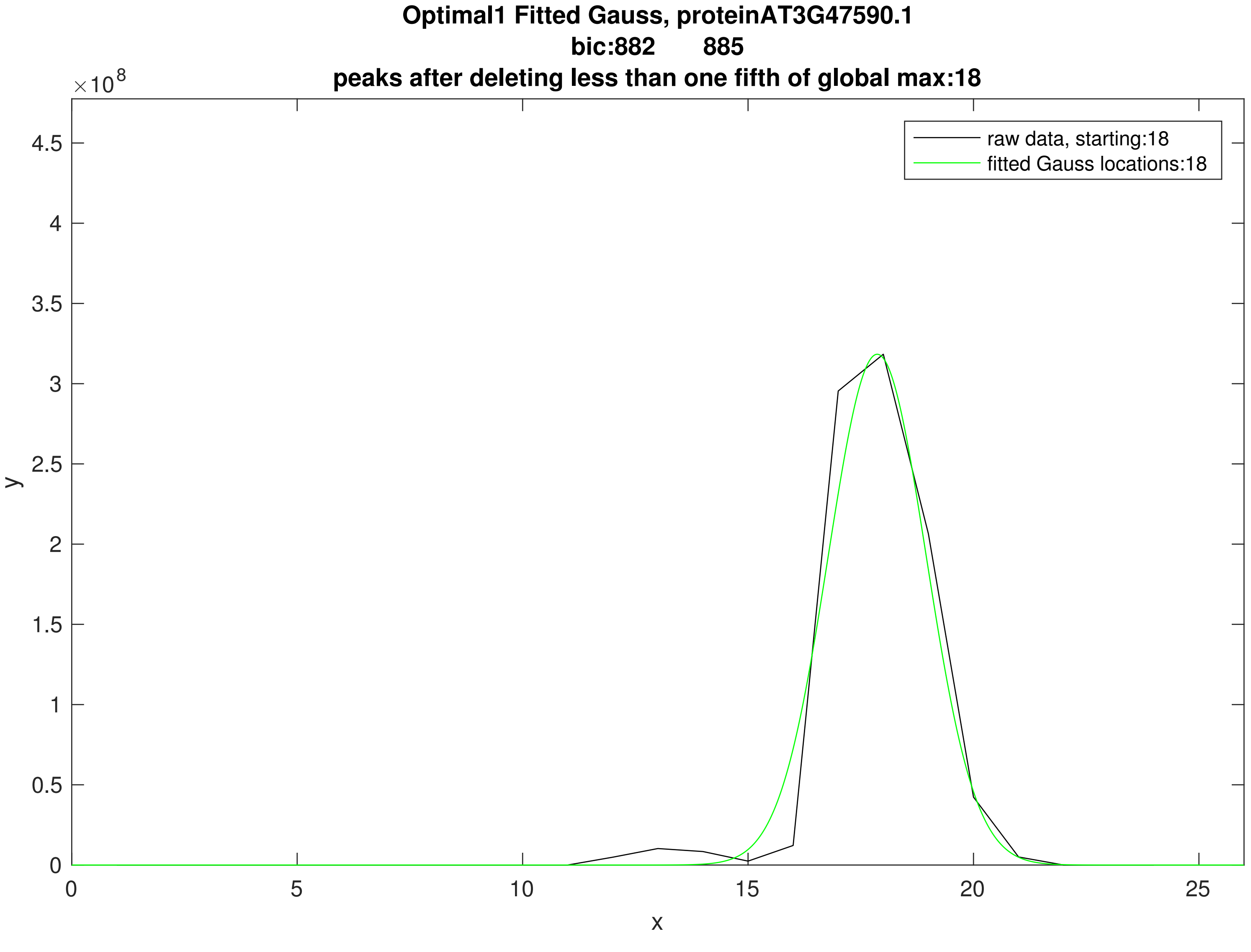

Optimal1 Fitted Gauss, proteinAT3G47590.1
bic:882 885
peaks after deleting less than one fifth of global max:18
×10^8
raw data, starting:18
fitted Gauss locations:18
y
x
0
0.5
1
1.5
2
2.5
3
3.5
4
4.5
5
10
15
20
25

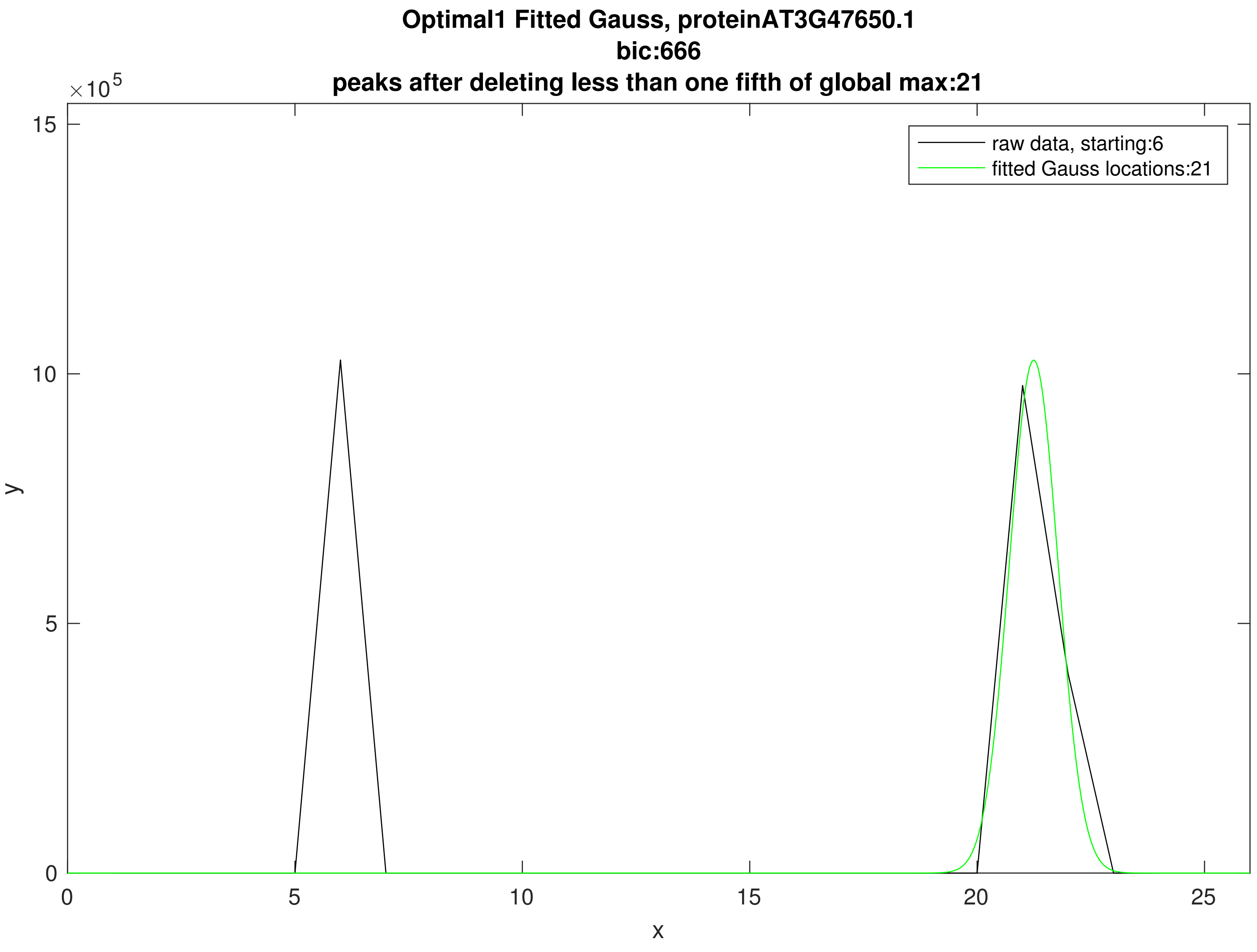

Optimal1 Fitted Gauss, proteinAT3G47650.1
bic:666
peaks after deleting less than one fifth of global max:21
×10^5
15
10
5
0
y
0
5
10
15
20
25
x
raw data, starting:6
fitted Gauss locations:21

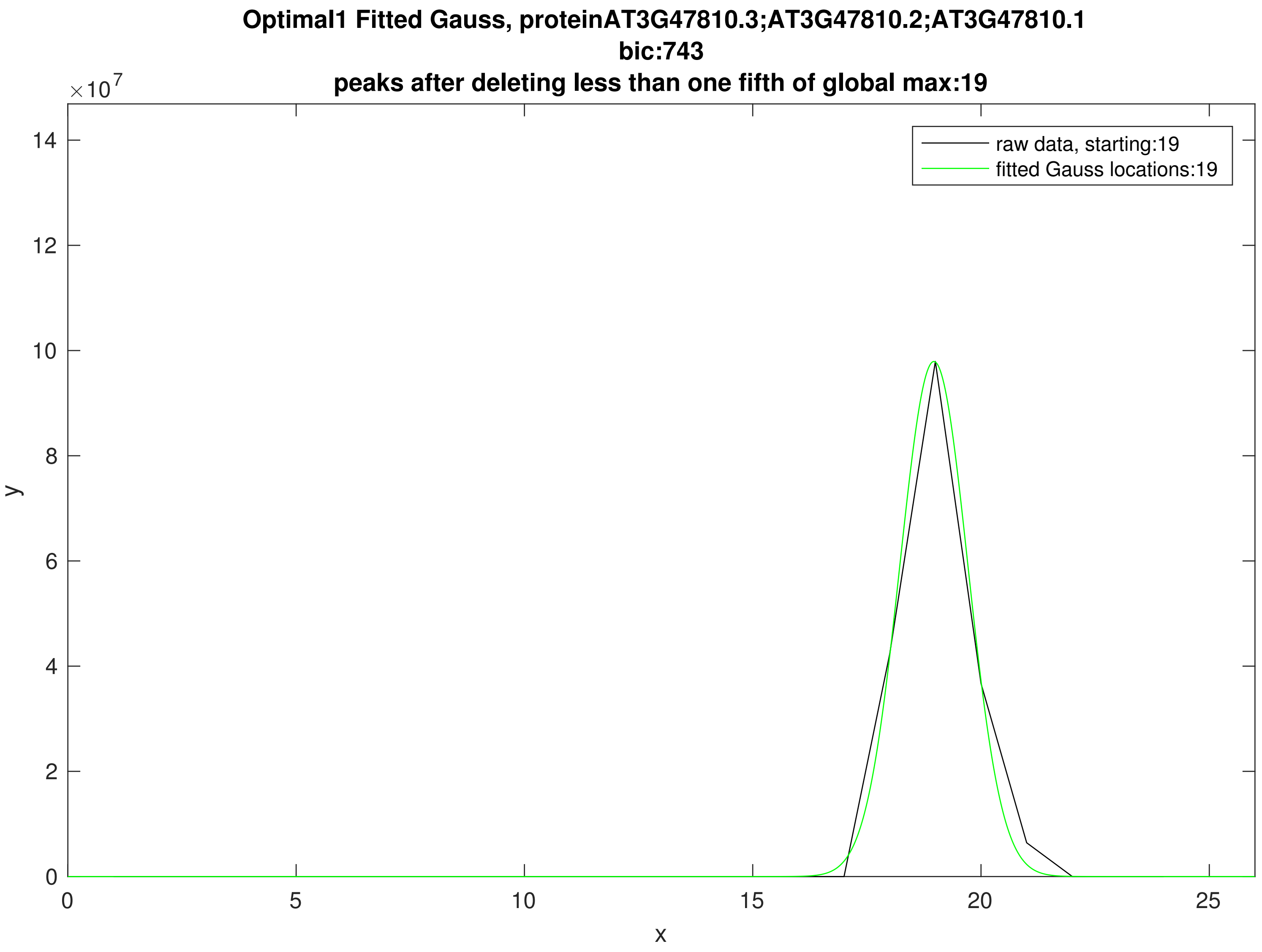

Optimal1 Fitted Gauss, proteinAT3G47810.3;AT3G47810.2;AT3G47810.1
bic:743
peaks after deleting less than one fifth of global max:19
×10^7
raw data, starting:19
fitted Gauss locations:19
14
12
10
8
6
4
2
0
y
0
5
10
15
20
25
x

**Optimal1 Fitted Gauss, proteinAT3G48000.1**
**bic:919 927**
**peaks after deleting less than one fifth of global max:11**

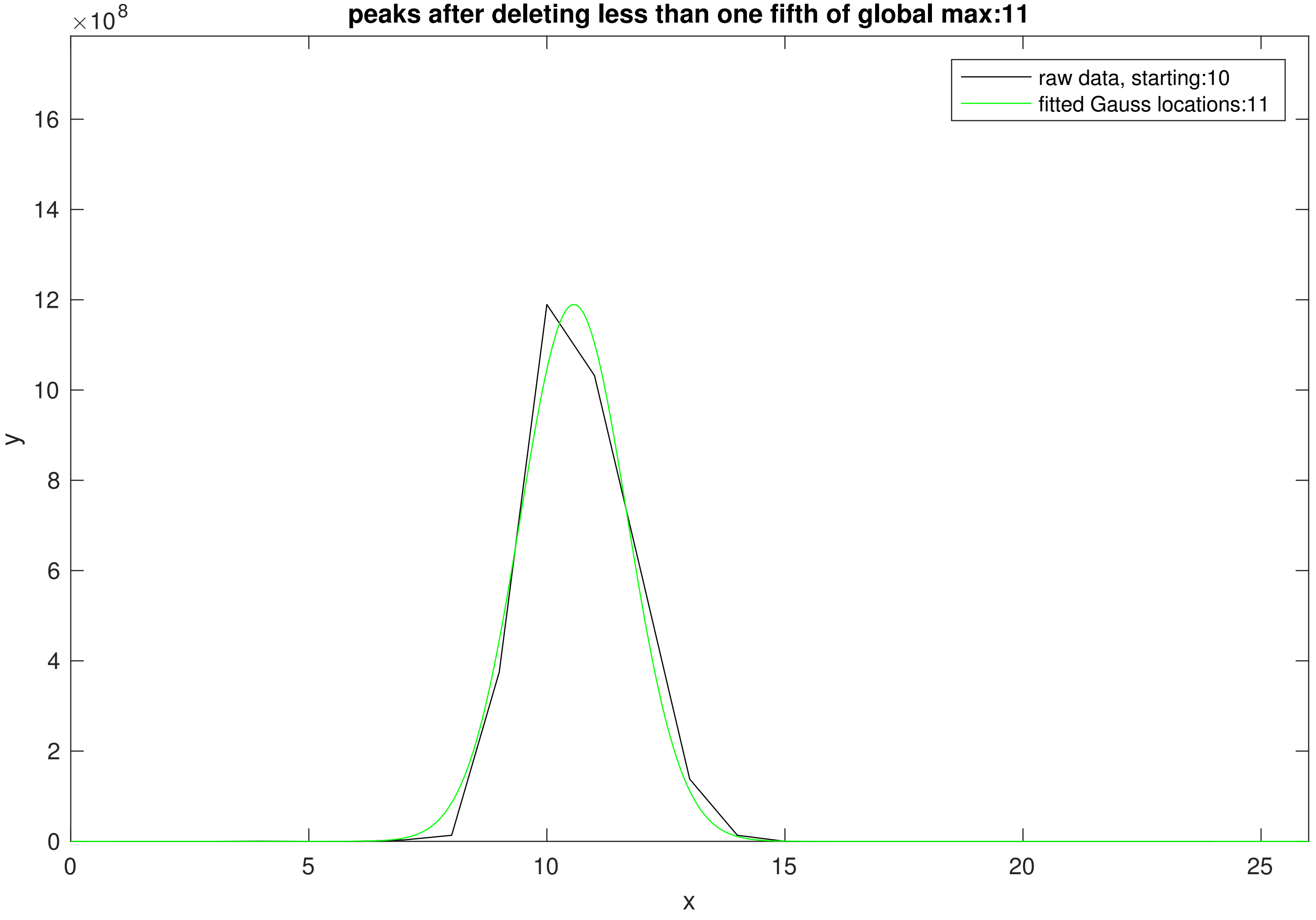

**Optimal1 Fitted Gauss, proteinAT3G48110.1**
**bic:859 860**
**peaks after deleting less than one fifth of global max:7.7**

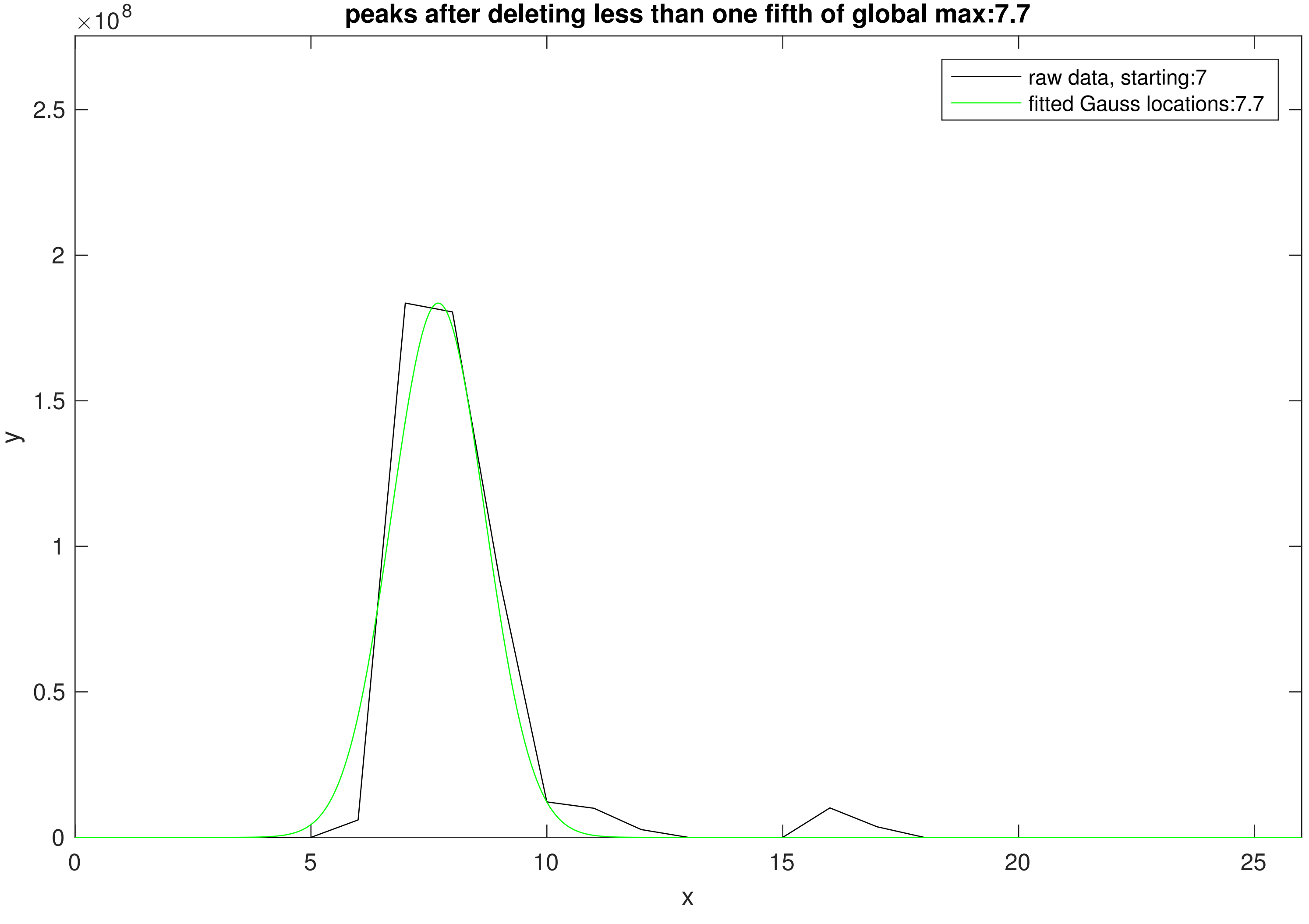

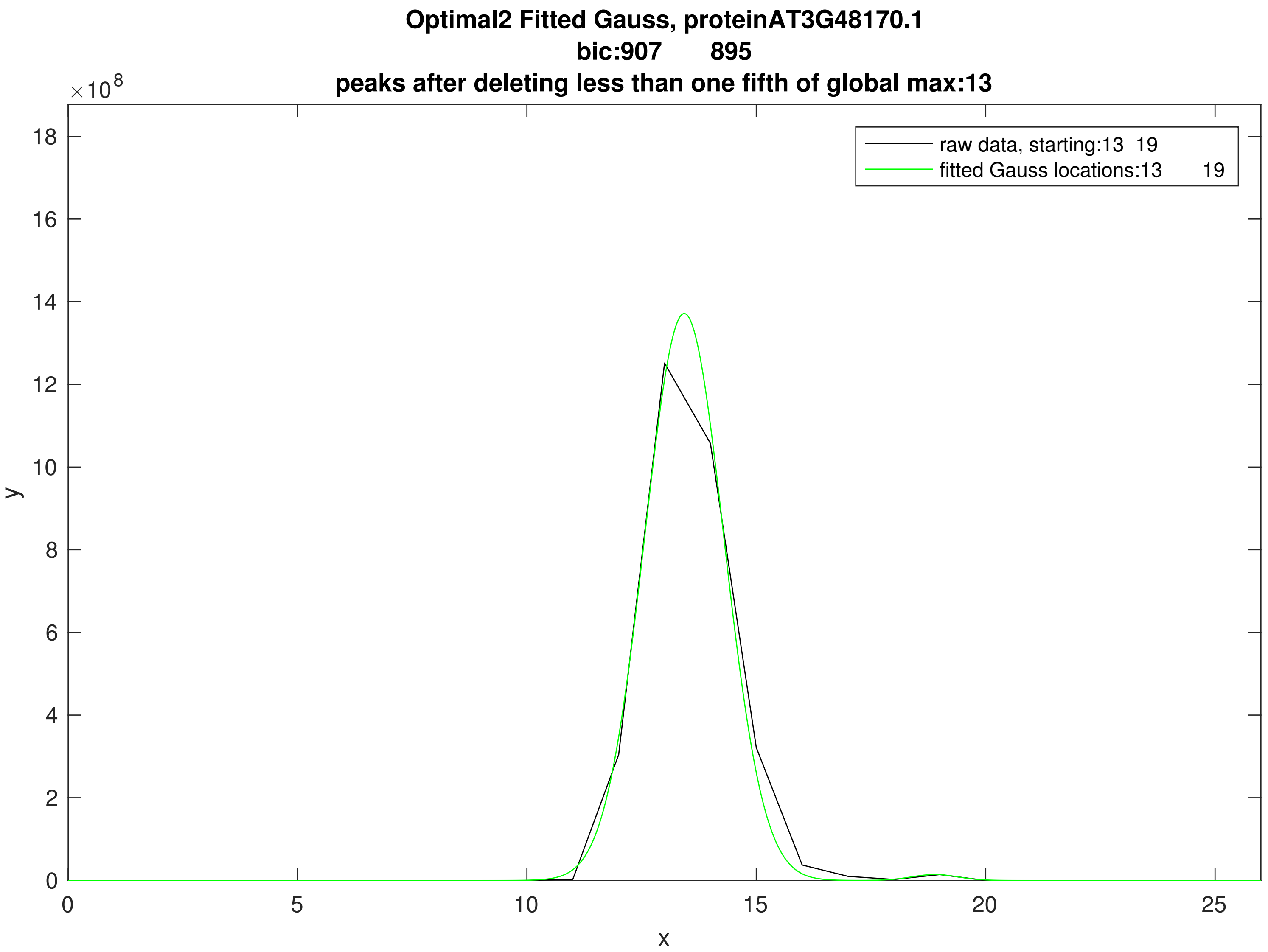

Optimal2 Fitted Gauss, proteinAT3G48170.1
bic:907 895
peaks after deleting less than one fifth of global max:13
×10^8
raw data, starting:13 19
fitted Gauss locations:13 19
y
x
0
2
4
6
8
10
12
14
16
18
0
5
10
15
20
25

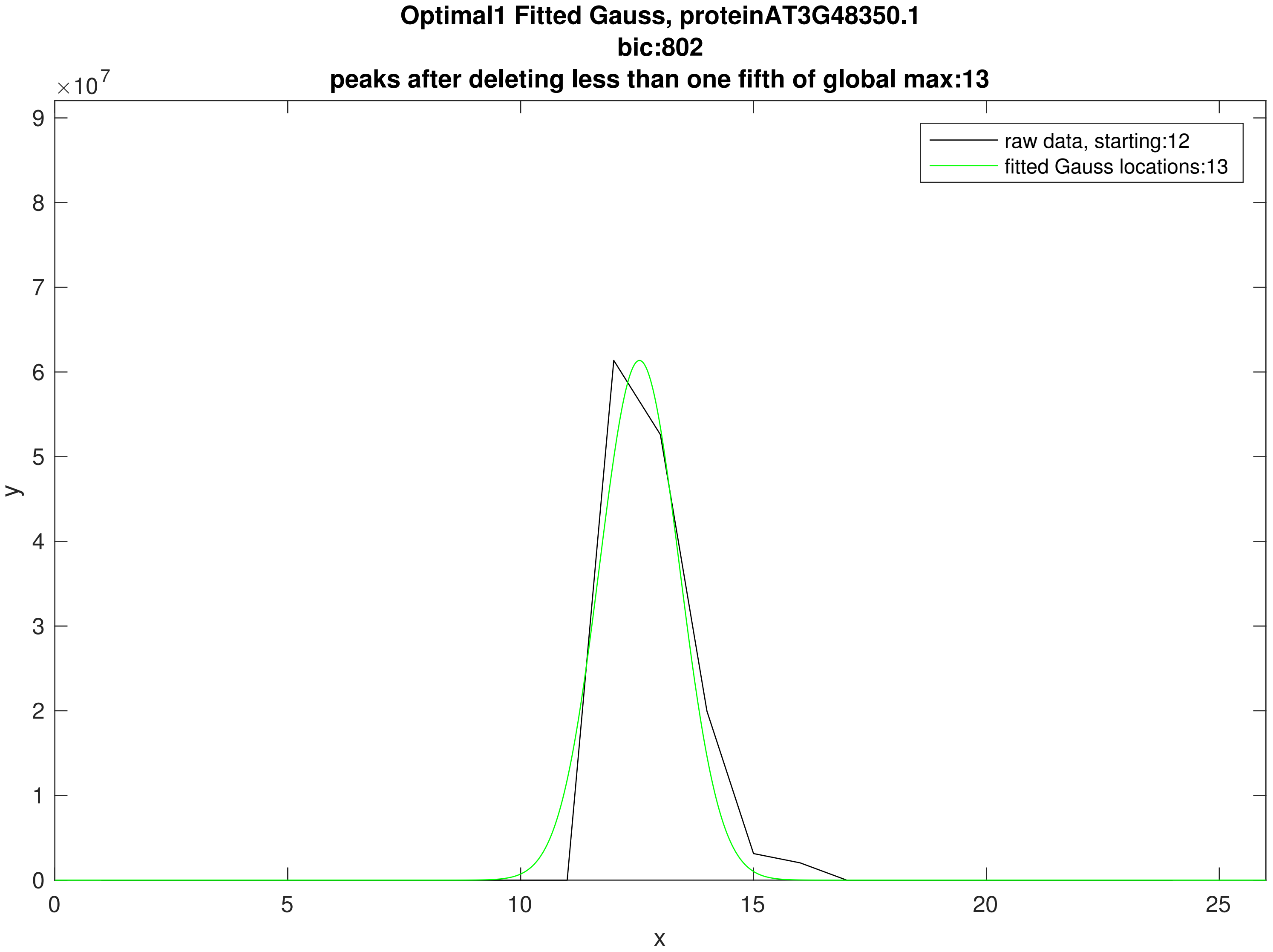

Optimal1 Fitted Gauss, proteinAT3G48350.1
bic:802
peaks after deleting less than one fifth of global max:13
×10^7
raw data, starting:12
fitted Gauss locations:13
y
x
0
1
2
3
4
5
6
7
8
9
0
5
10
15
20
25

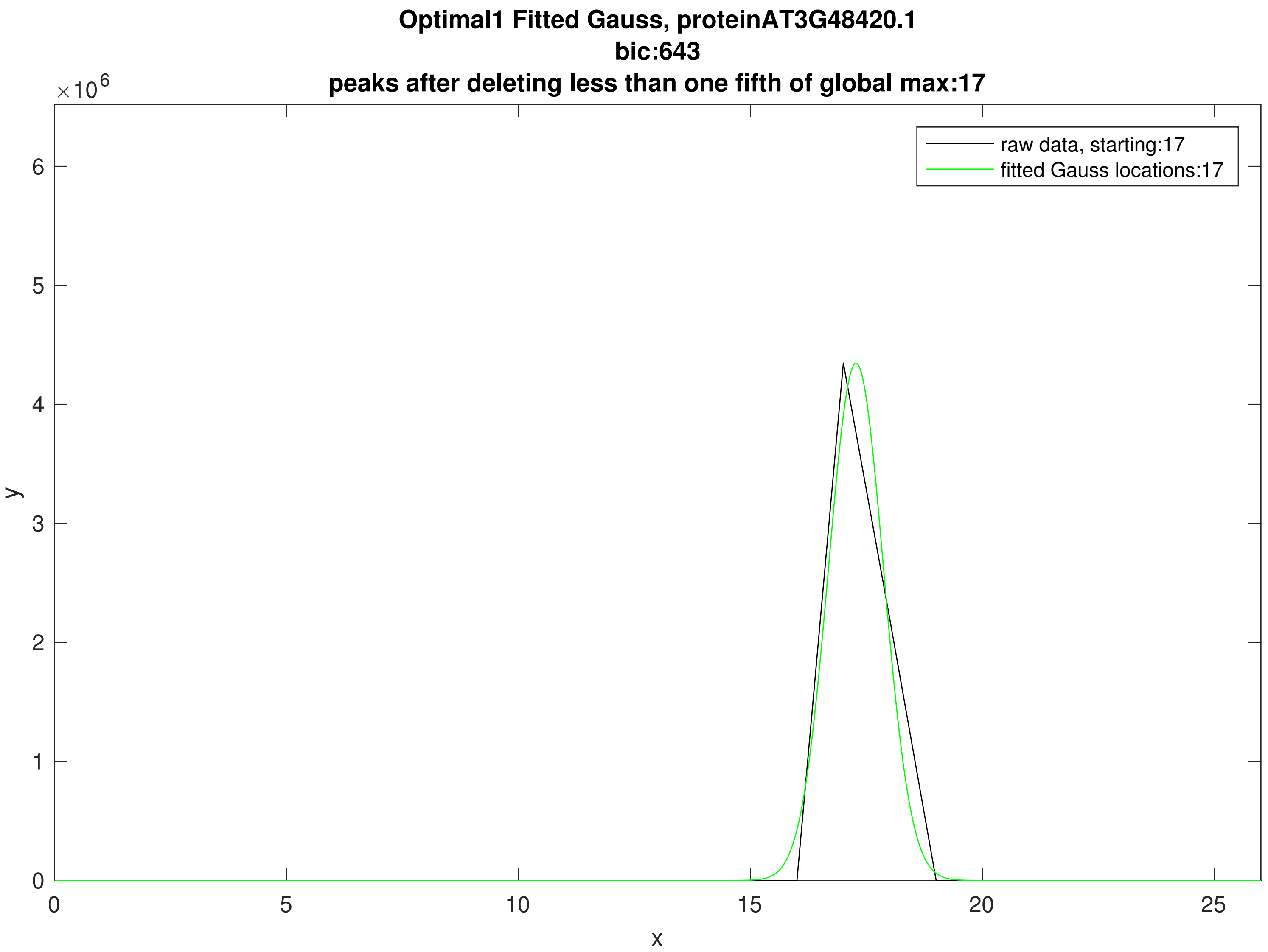
Optimal1 Fitted Gauss, proteinAT3G48420.1
bic:643
peaks after deleting less than one fifth of global max:17
×10^6
raw data, starting:17
fitted Gauss locations:17
y
x
0
1
2
3
4
5
6
0
5
10
15
20
25

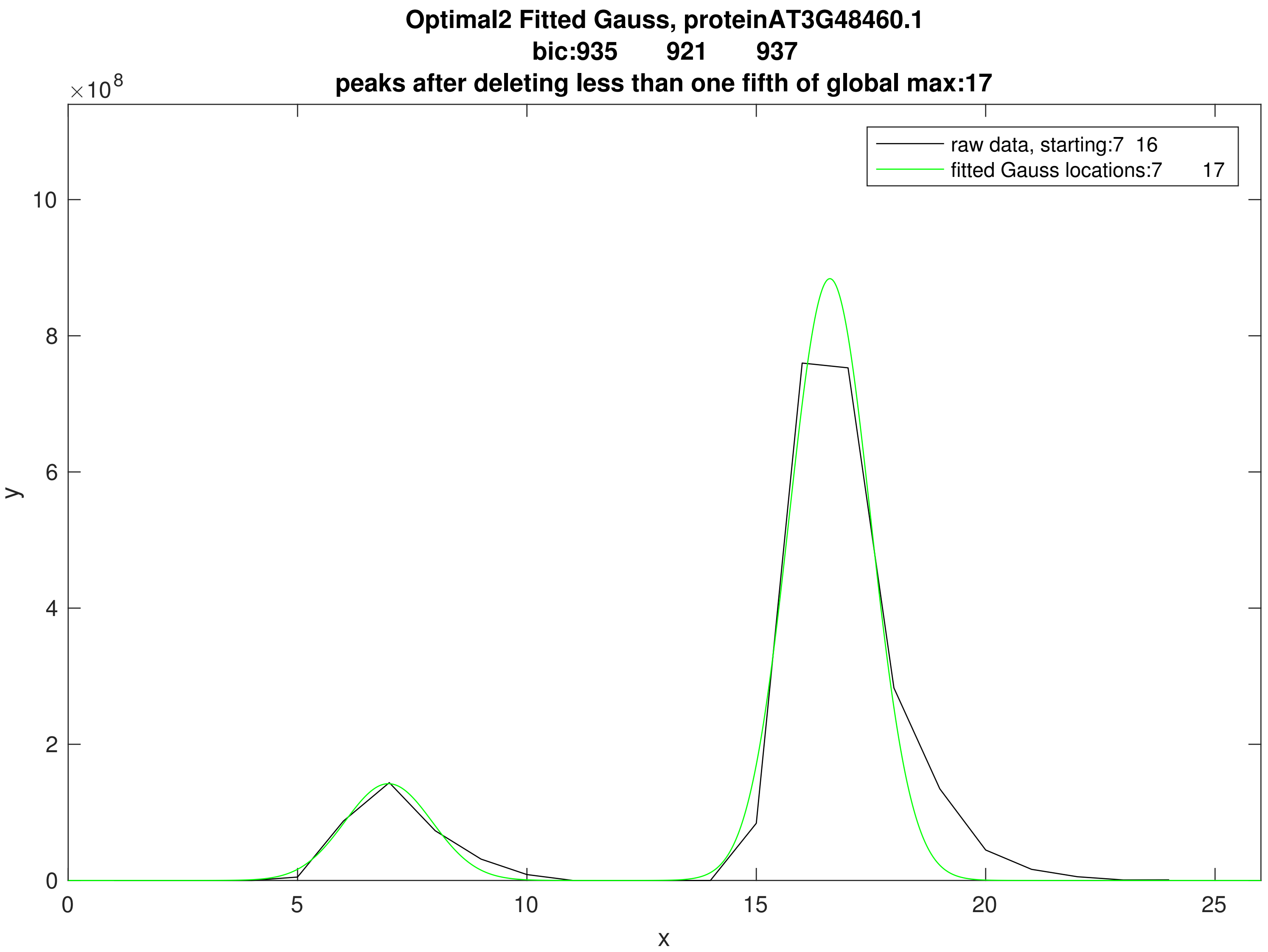
Optimal2 Fitted Gauss, proteinAT3G48460.1
bic:935 921 937
peaks after deleting less than one fifth of global max:17
×10^8
raw data, starting:7 16
fitted Gauss locations:7 17
y
x
0
2
4
6
8
10
0
5
10
15
20
25

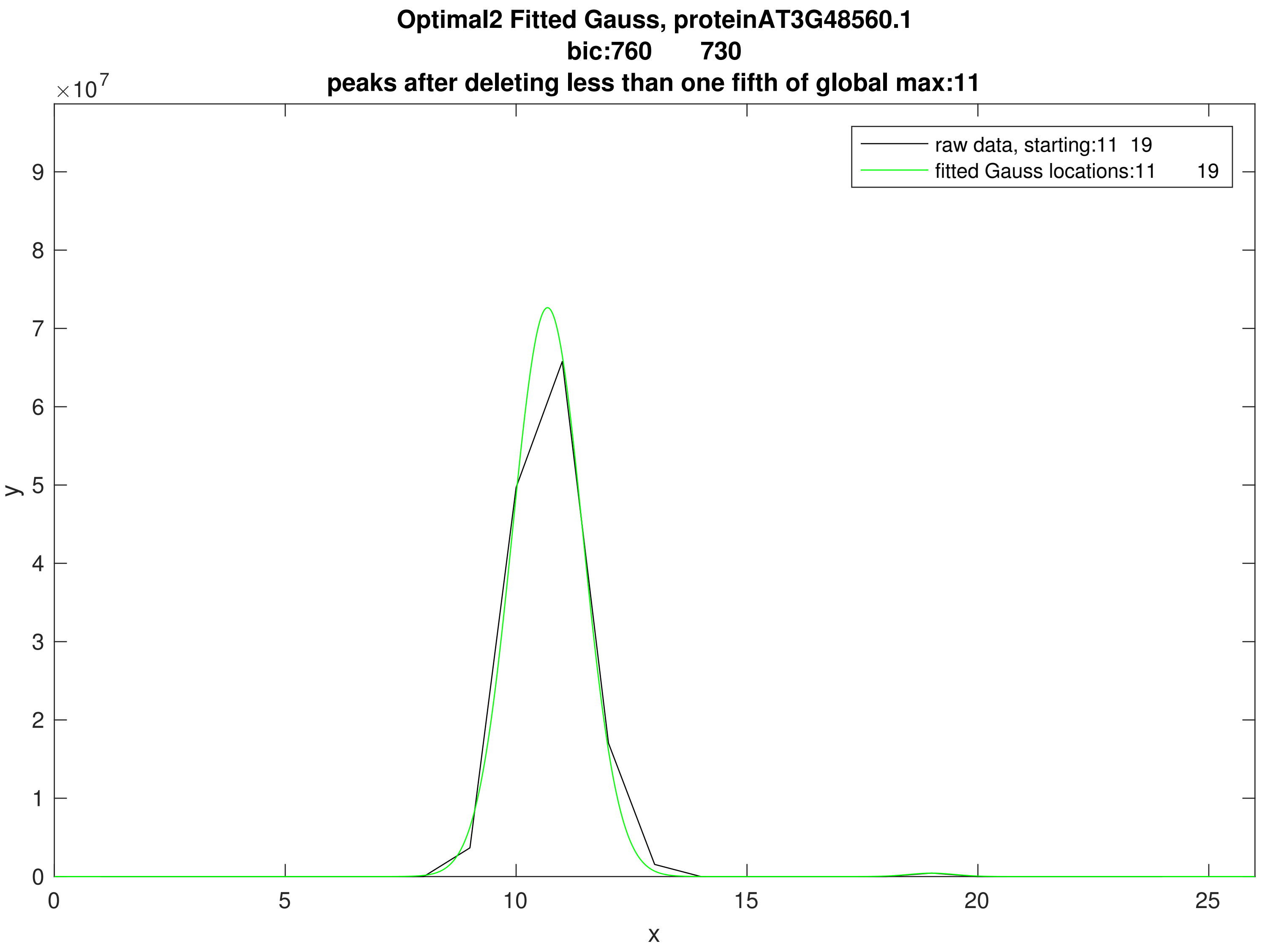

Optimal2 Fitted Gauss, proteinAT3G48560.1
bic:760 730
peaks after deleting less than one fifth of global max:11
$\times 10^7$
raw data, starting:11 19
fitted Gauss locations:11 19
y
x
0
1
2
3
4
5
6
7
8
9
0
5
10
15
20
25

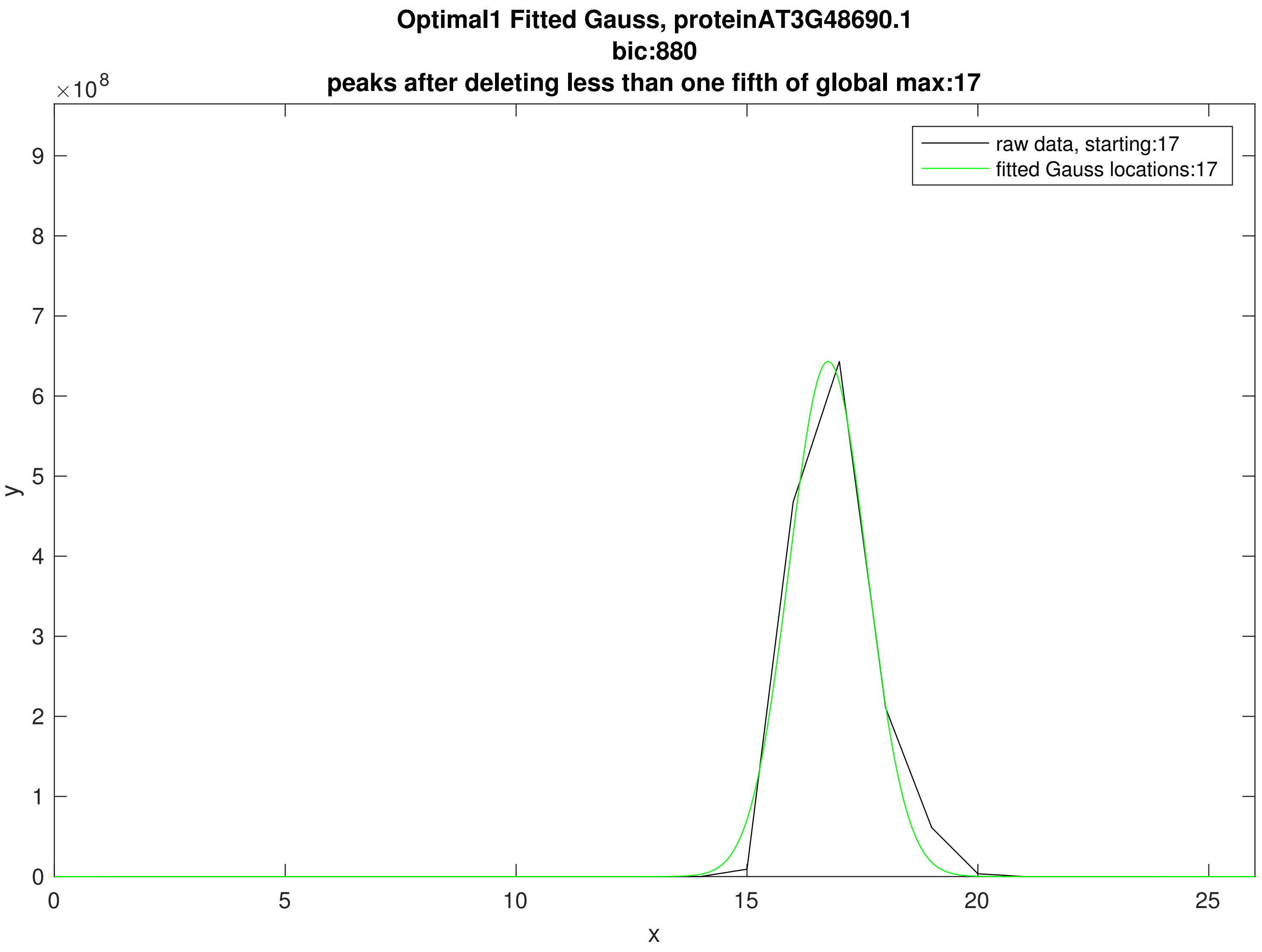

Optimal1 Fitted Gauss, proteinAT3G48690.1
bic:880
peaks after deleting less than one fifth of global max:17
×10^8
raw data, starting:17
fitted Gauss locations:17
y
x
0
1
2
3
4
5
6
7
8
9
0
5
10
15
20
25

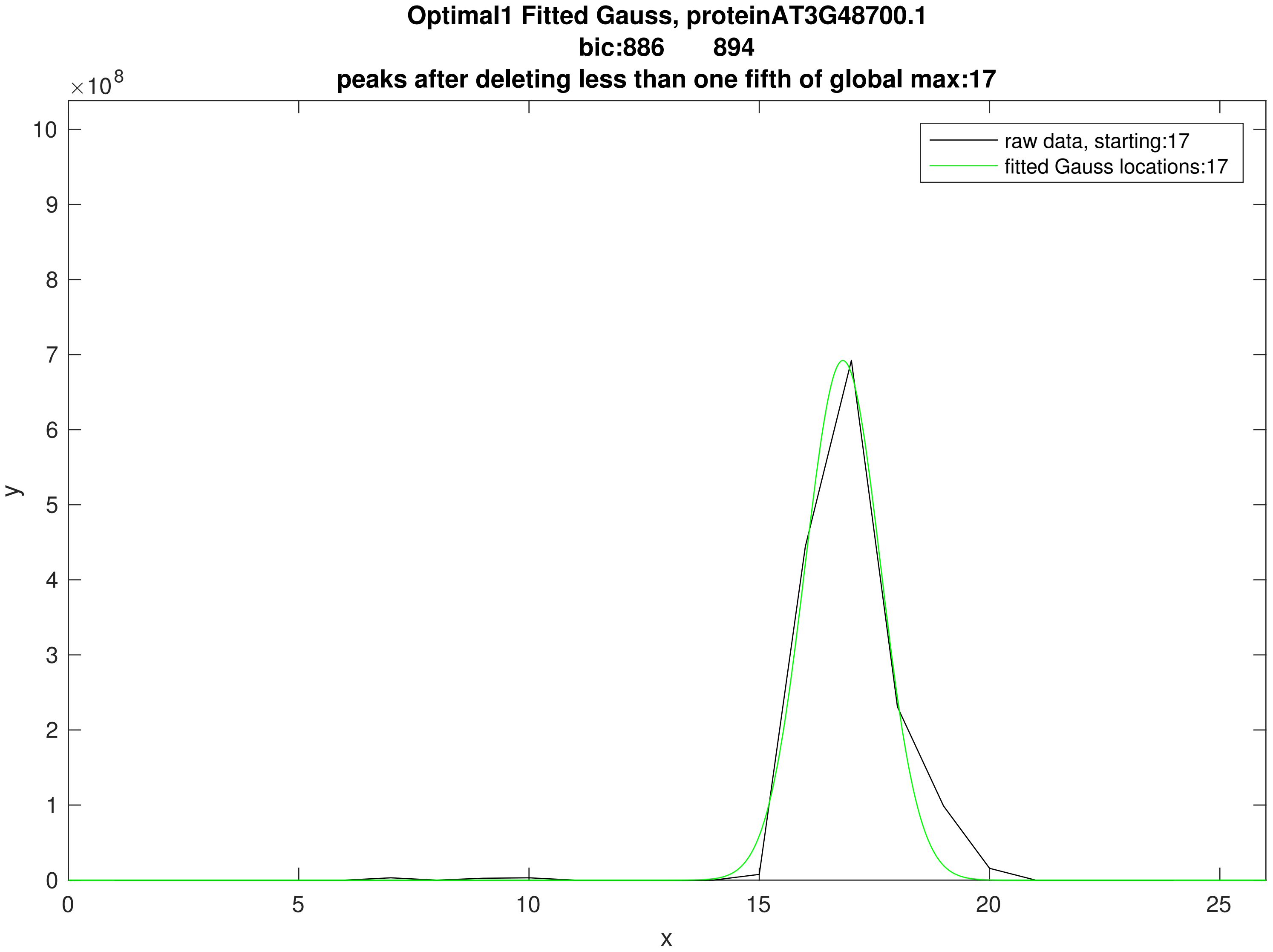

Optimal1 Fitted Gauss, proteinAT3G48700.1
bic:886 894
peaks after deleting less than one fifth of global max:17
×10^8
10
9
8
7
6
5
4
3
2
1
0
y
0
5
10
15
20
25
x
raw data, starting:17
fitted Gauss locations:17

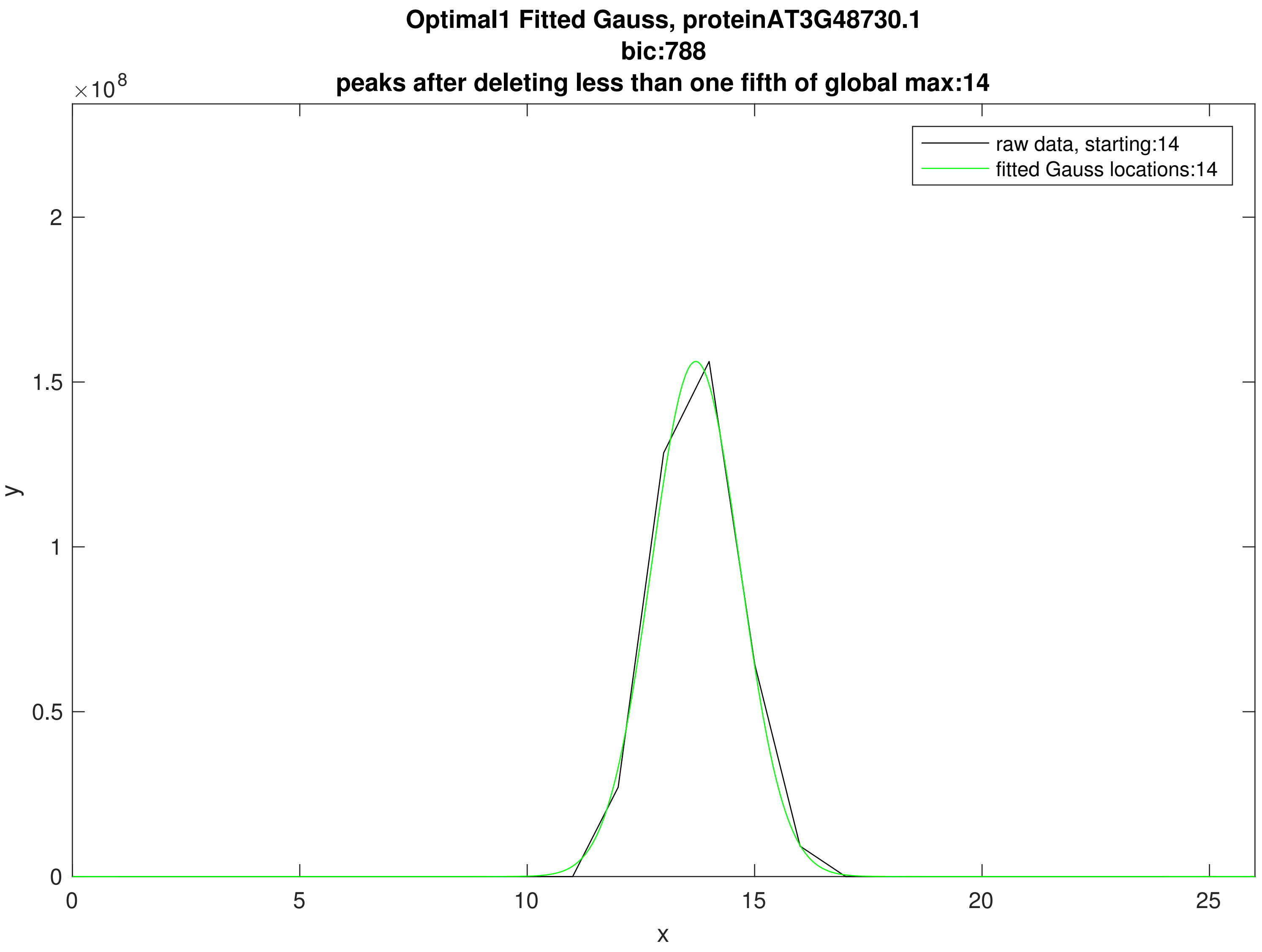

Optimal1 Fitted Gauss, proteinAT3G48730.1
bic:788
peaks after deleting less than one fifth of global max:14
×10^8
raw data, starting:14
fitted Gauss locations:14
2
1.5
1
0.5
0
y
0
5
10
15
20
25
x

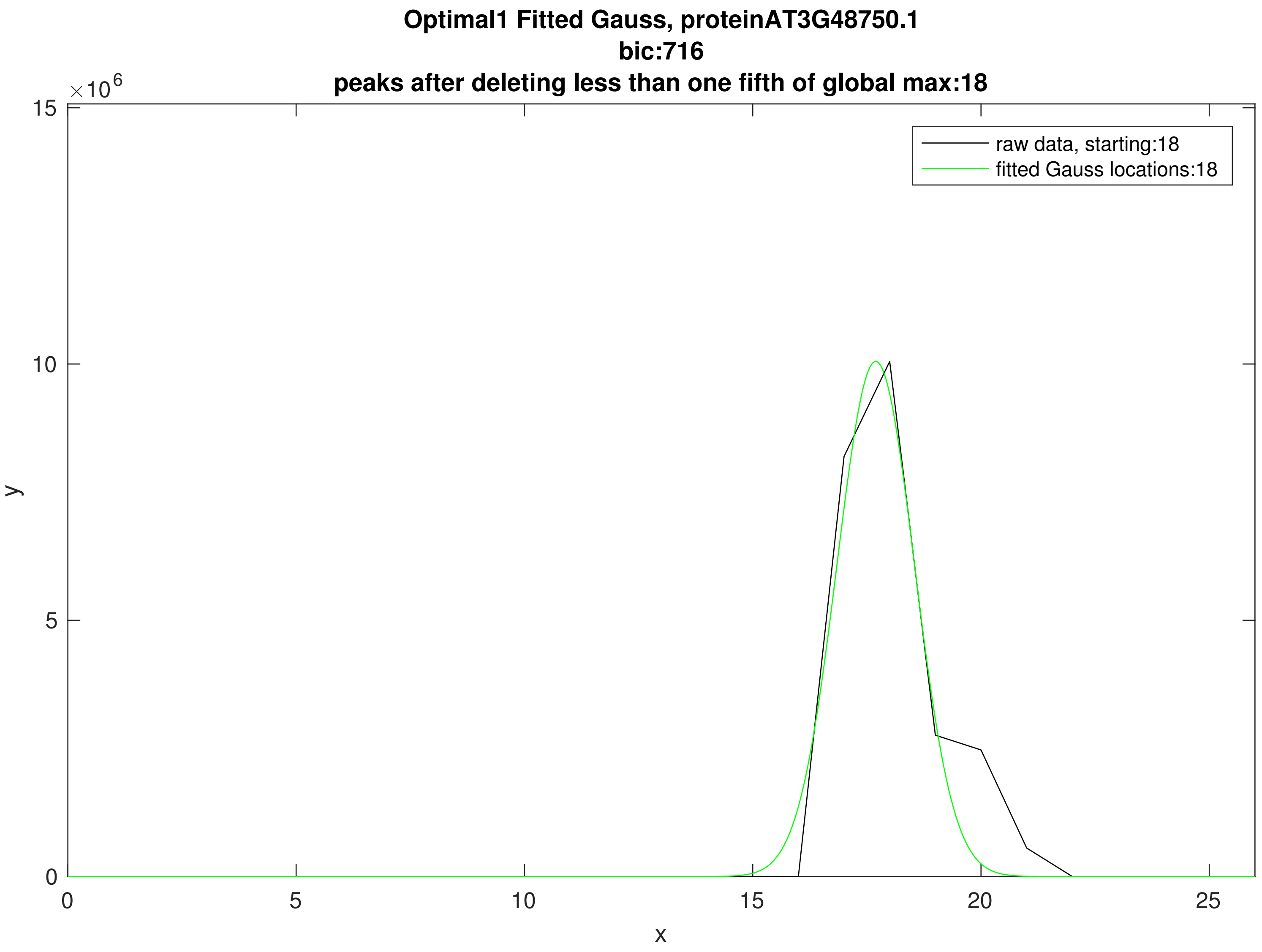

Optimal1 Fitted Gauss, proteinAT3G48750.1
bic:716
peaks after deleting less than one fifth of global max:18
×10^6
15
10
5
0
y
0
5
10
15
20
25
x
raw data, starting:18
fitted Gauss locations:18

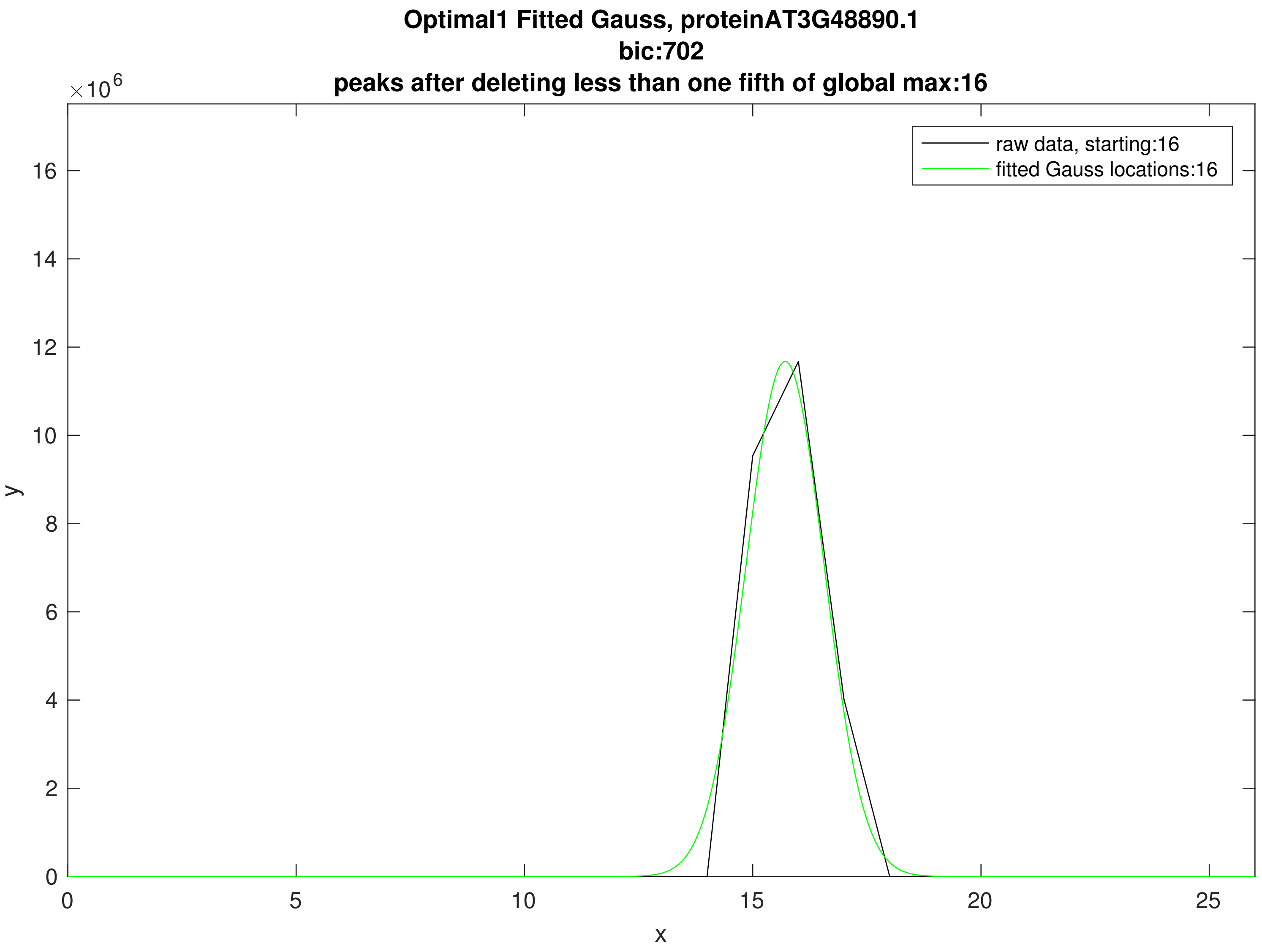
Optimal1 Fitted Gauss, proteinAT3G48890.1
bic:702
peaks after deleting less than one fifth of global max:16
×10^6
16
14
12
10
8
6
4
2
0
y
0
5
10
15
20
25
x
raw data, starting:16
fitted Gauss locations:16

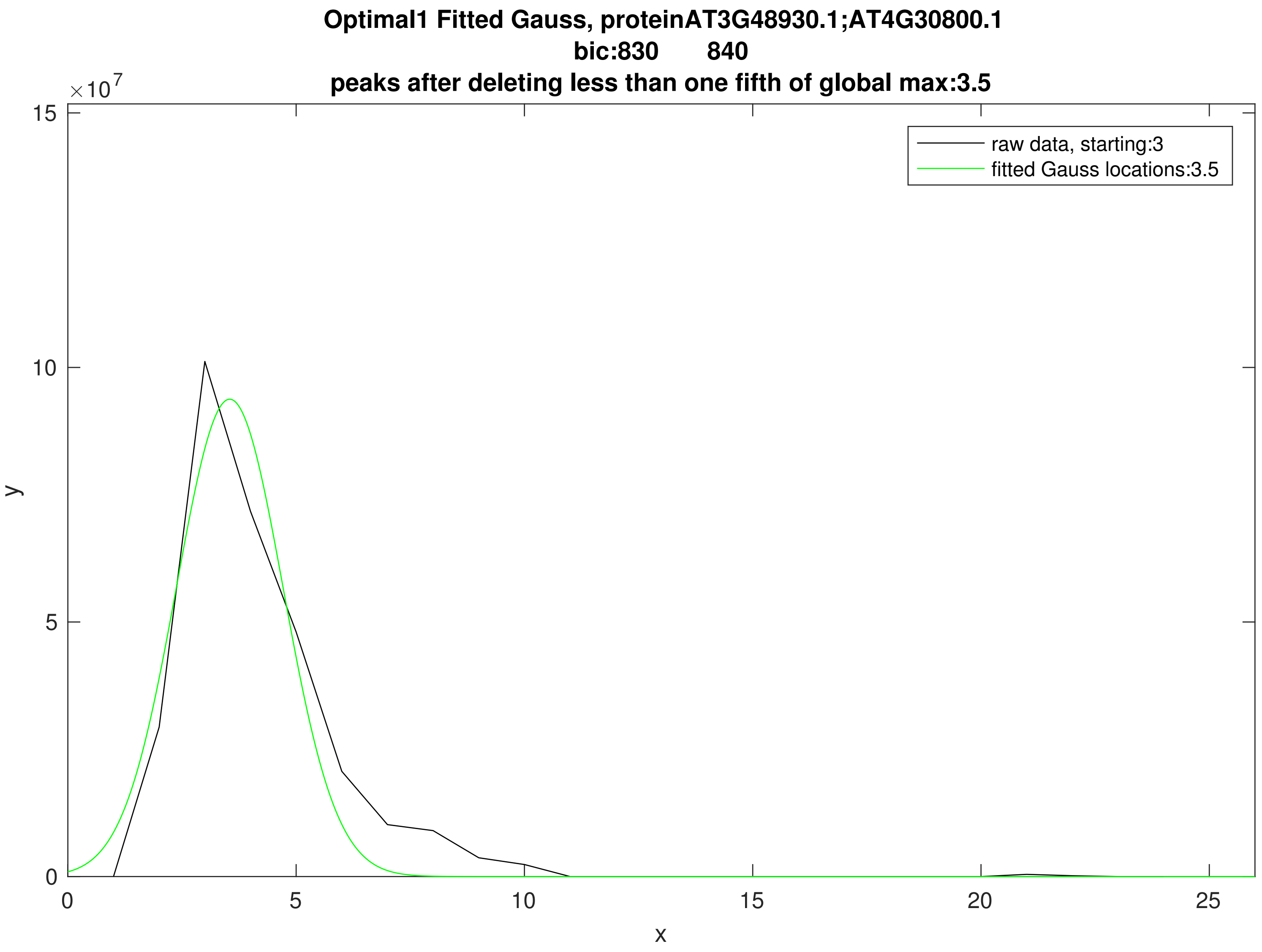
Optimal1 Fitted Gauss, proteinAT3G48930.1;AT4G30800.1
bic:830 840
peaks after deleting less than one fifth of global max:3.5
×10^7
15
10
5
0
y
0
5
10
15
20
25
x
raw data, starting:3
fitted Gauss locations:3.5

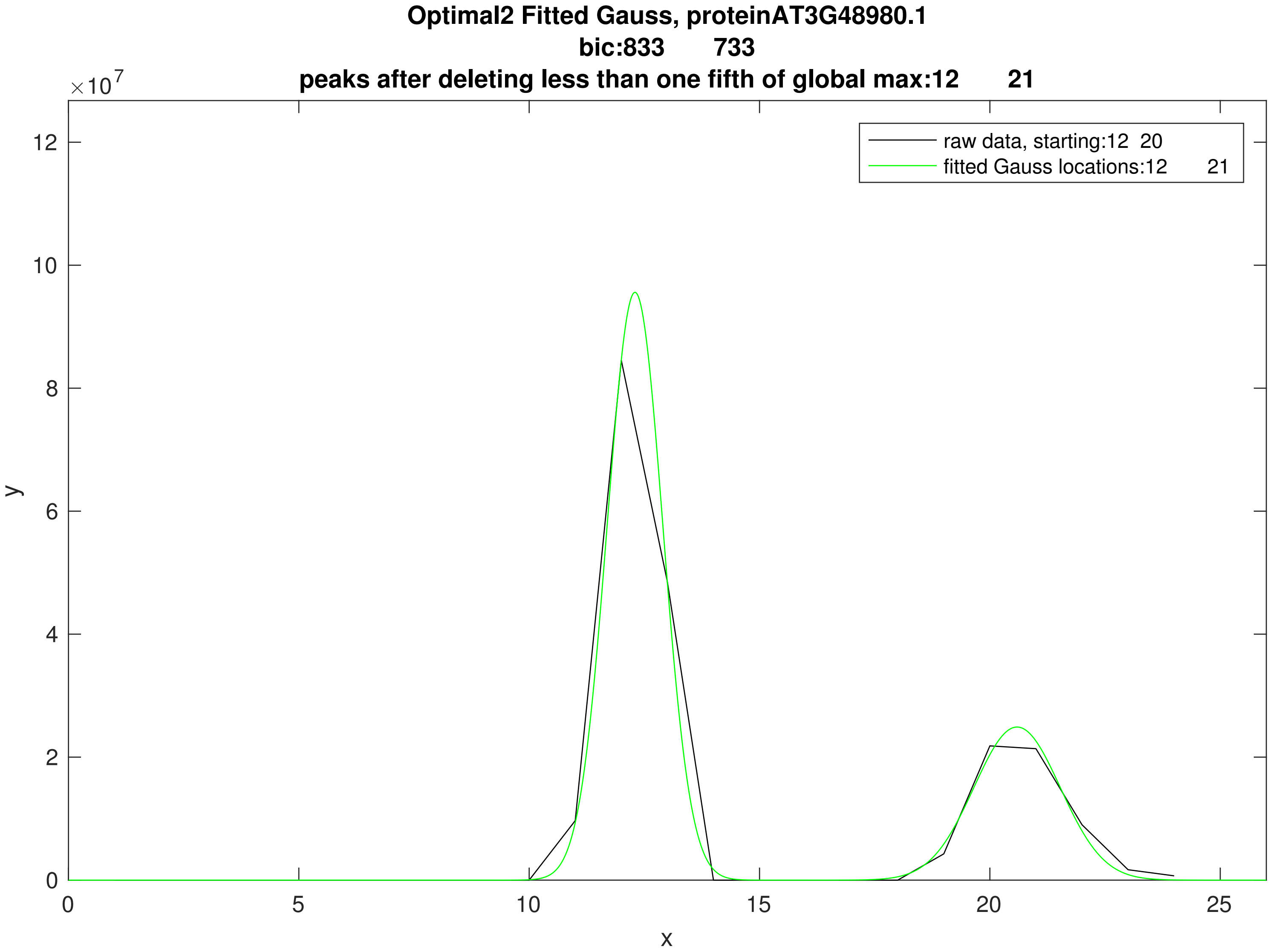

Optimal2 Fitted Gauss, proteinAT3G48980.1
bic:833 733
peaks after deleting less than one fifth of global max:12 21
×10^7
raw data, starting:12 20
fitted Gauss locations:12 21
y
x
0
2
4
6
8
10
12
0
5
10
15
20
25

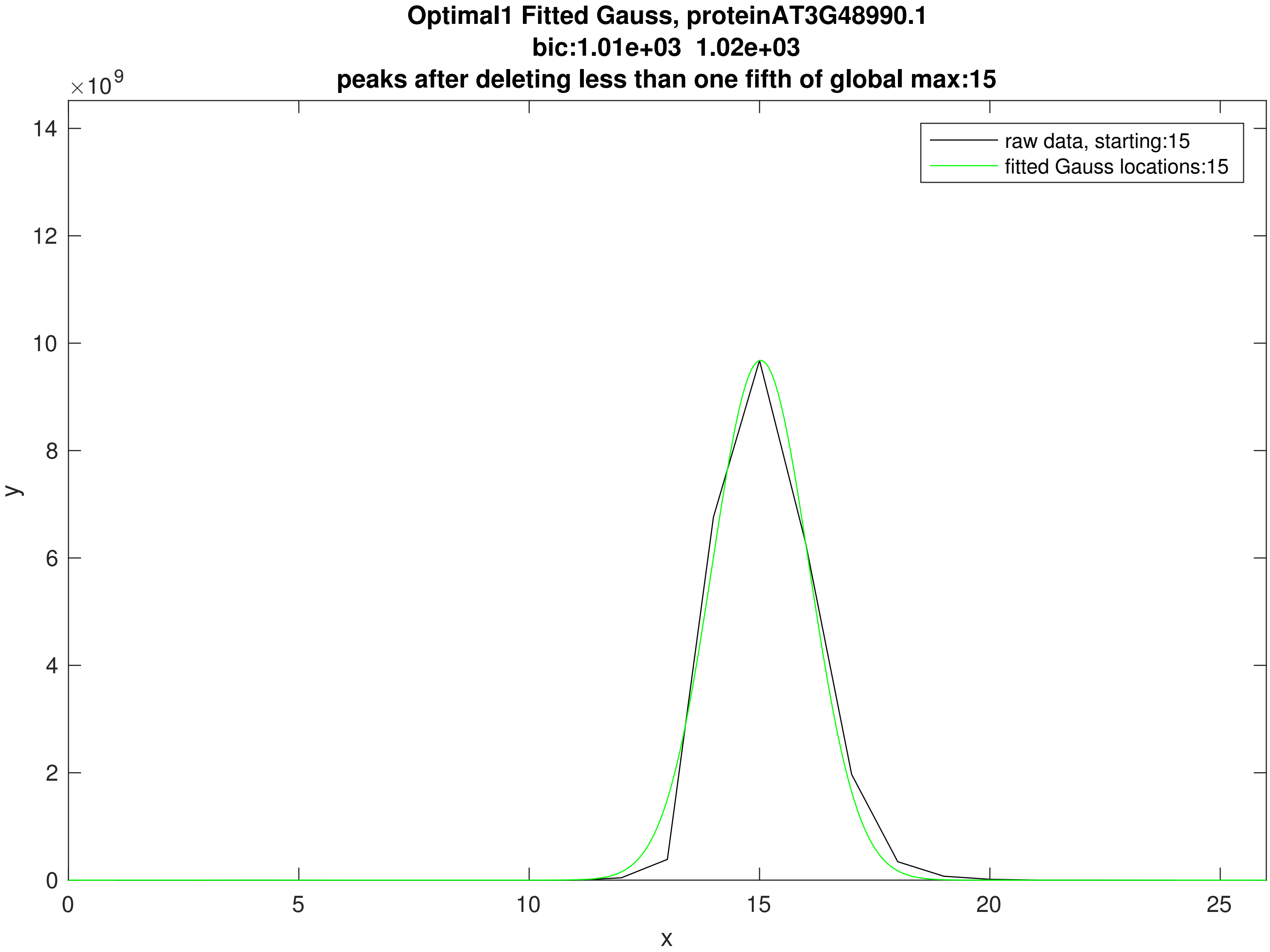

Optimal1 Fitted Gauss, proteinAT3G48990.1
bic:1.01e+03  1.02e+03
peaks after deleting less than one fifth of global max:15
×10^9
raw data, starting:15
fitted Gauss locations:15
y
x
0
2
4
6
8
10
12
14
0
5
10
15
20
25

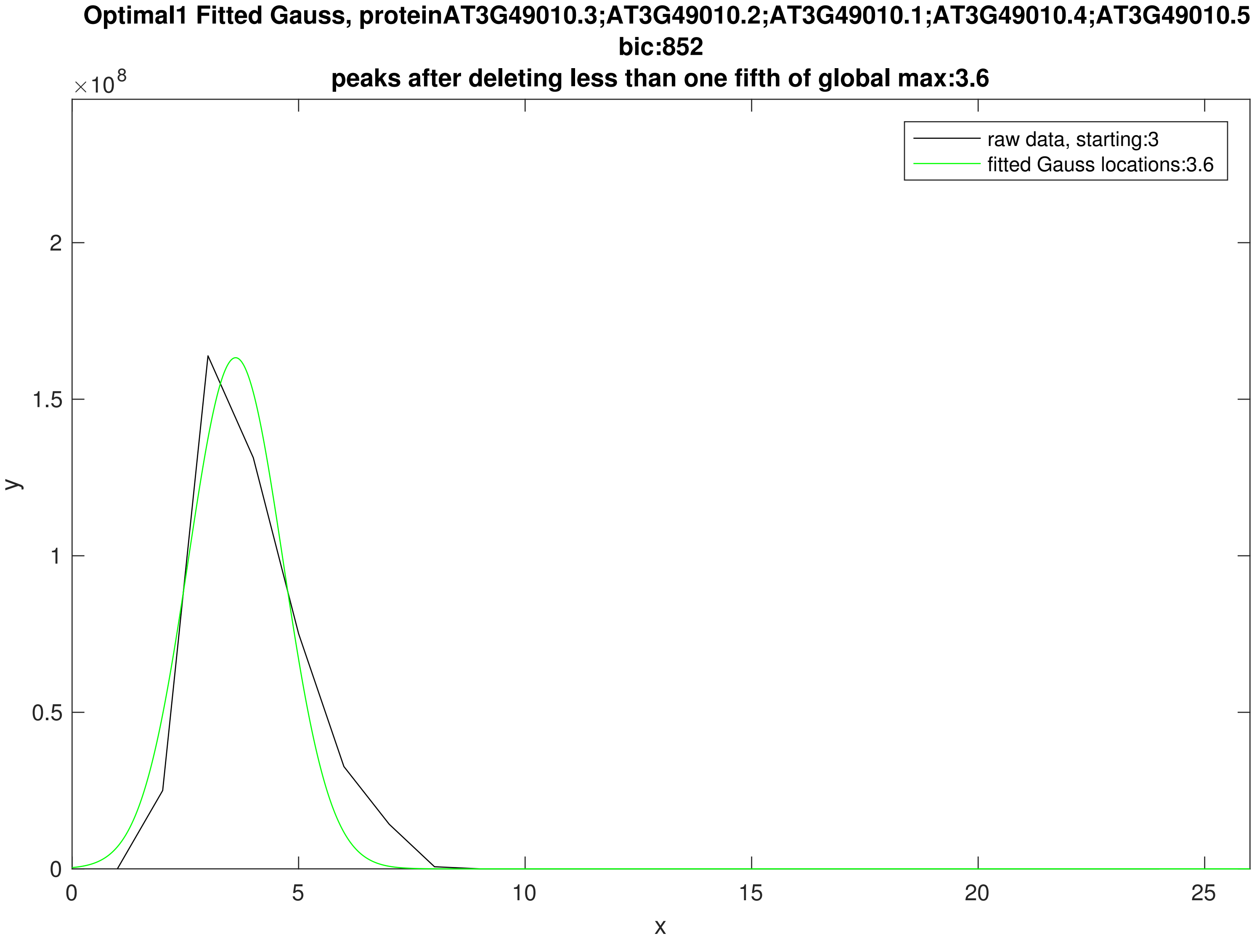
Optimal1 Fitted Gauss, proteinAT3G49010.3;AT3G49010.2;AT3G49010.1;AT3G49010.4;AT3G49010.5
bic:852
peaks after deleting less than one fifth of global max:3.6
×10^8
raw data, starting:3
fitted Gauss locations:3.6
y
x
0
0.5
1
1.5
2
0
5
10
15
20
25

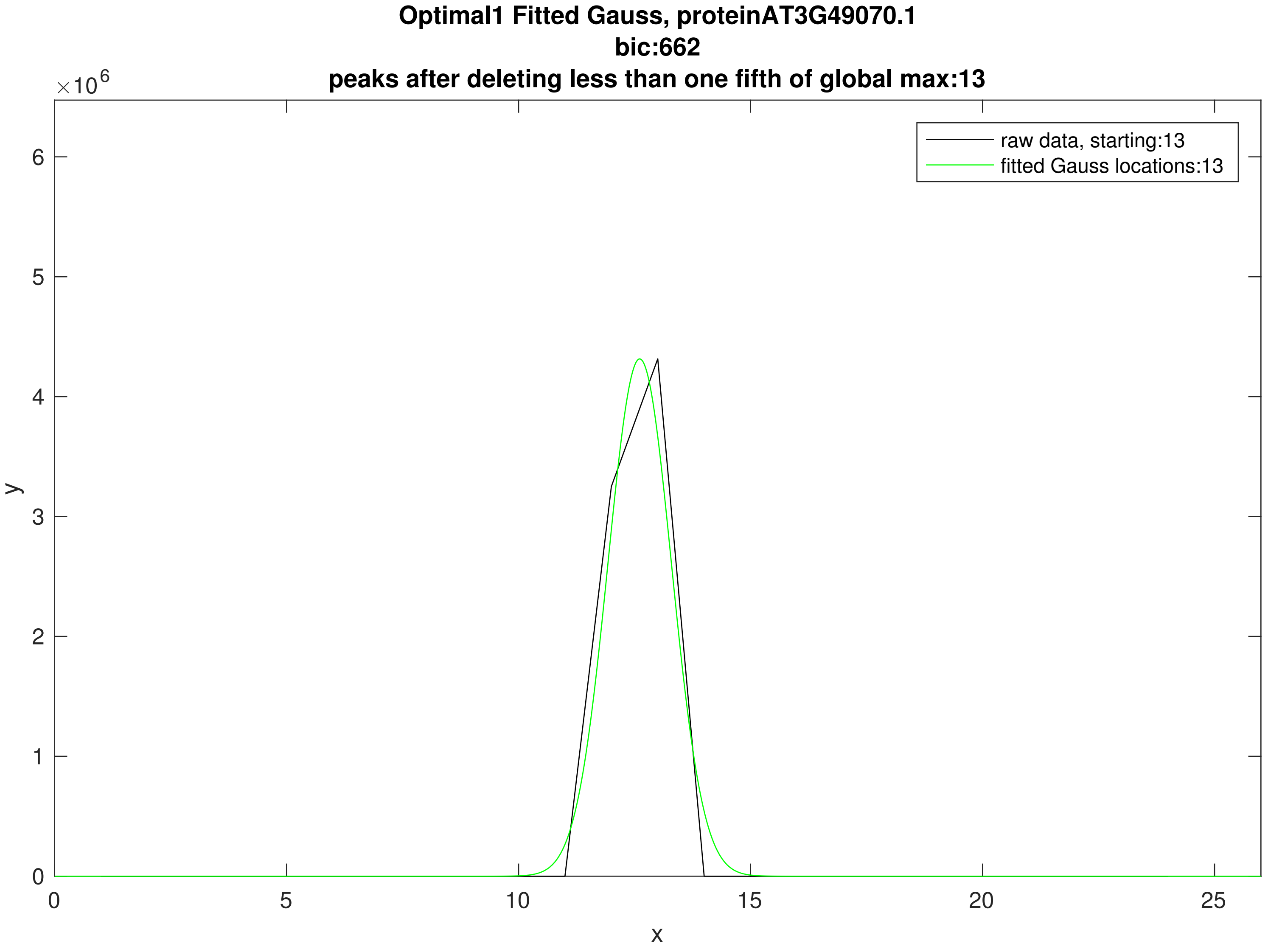
Optimal1 Fitted Gauss, proteinAT3G49070.1
bic:662
peaks after deleting less than one fifth of global max:13
×10^6
raw data, starting:13
fitted Gauss locations:13
y
x
0
1
2
3
4
5
6
0
5
10
15
20
25

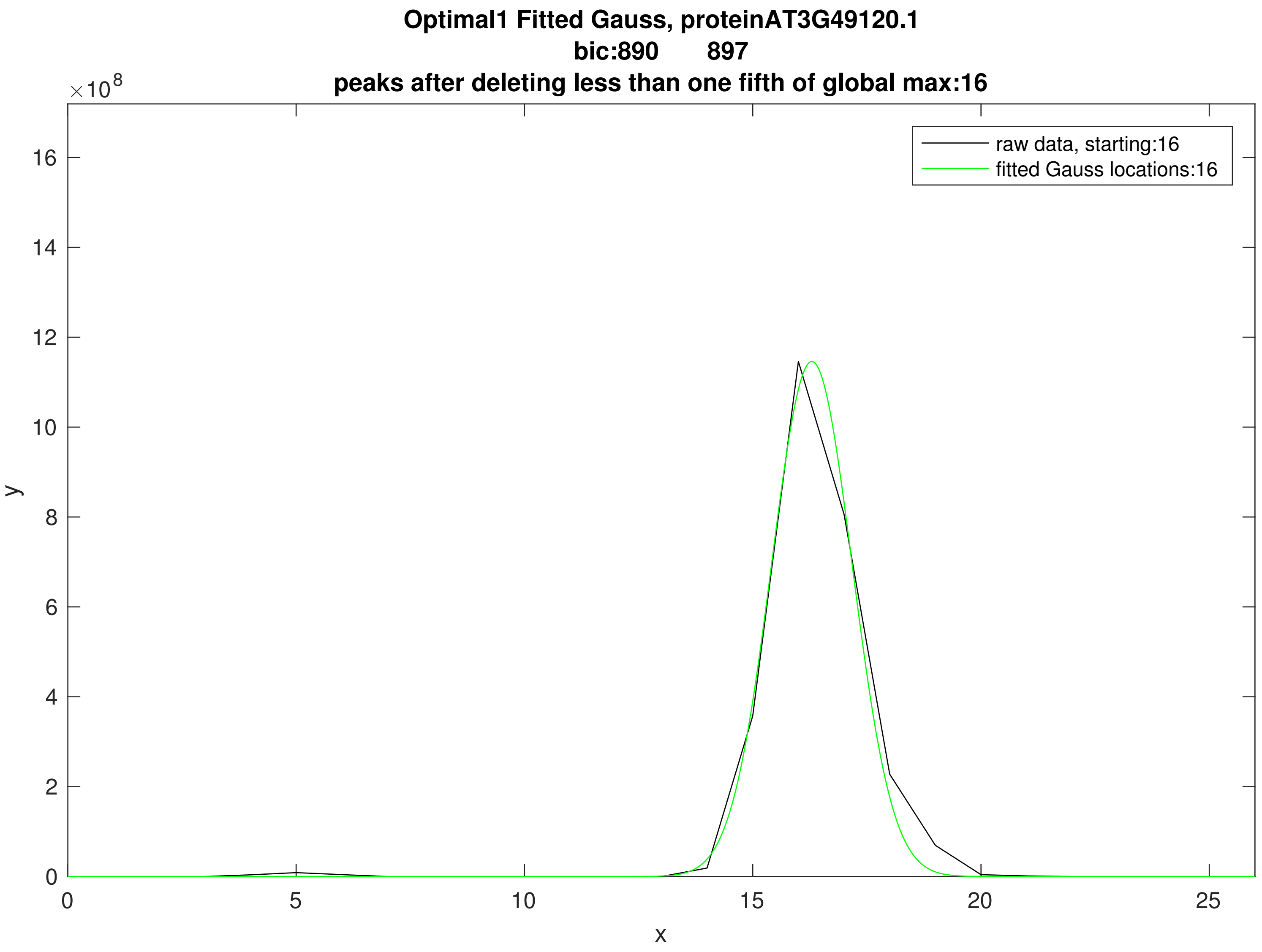

Optimal1 Fitted Gauss, proteinAT3G49120.1
bic:890 897
peaks after deleting less than one fifth of global max:16
×10^8
raw data, starting:16
fitted Gauss locations:16
y
x
0
2
4
6
8
10
12
14
16
0
5
10
15
20
25

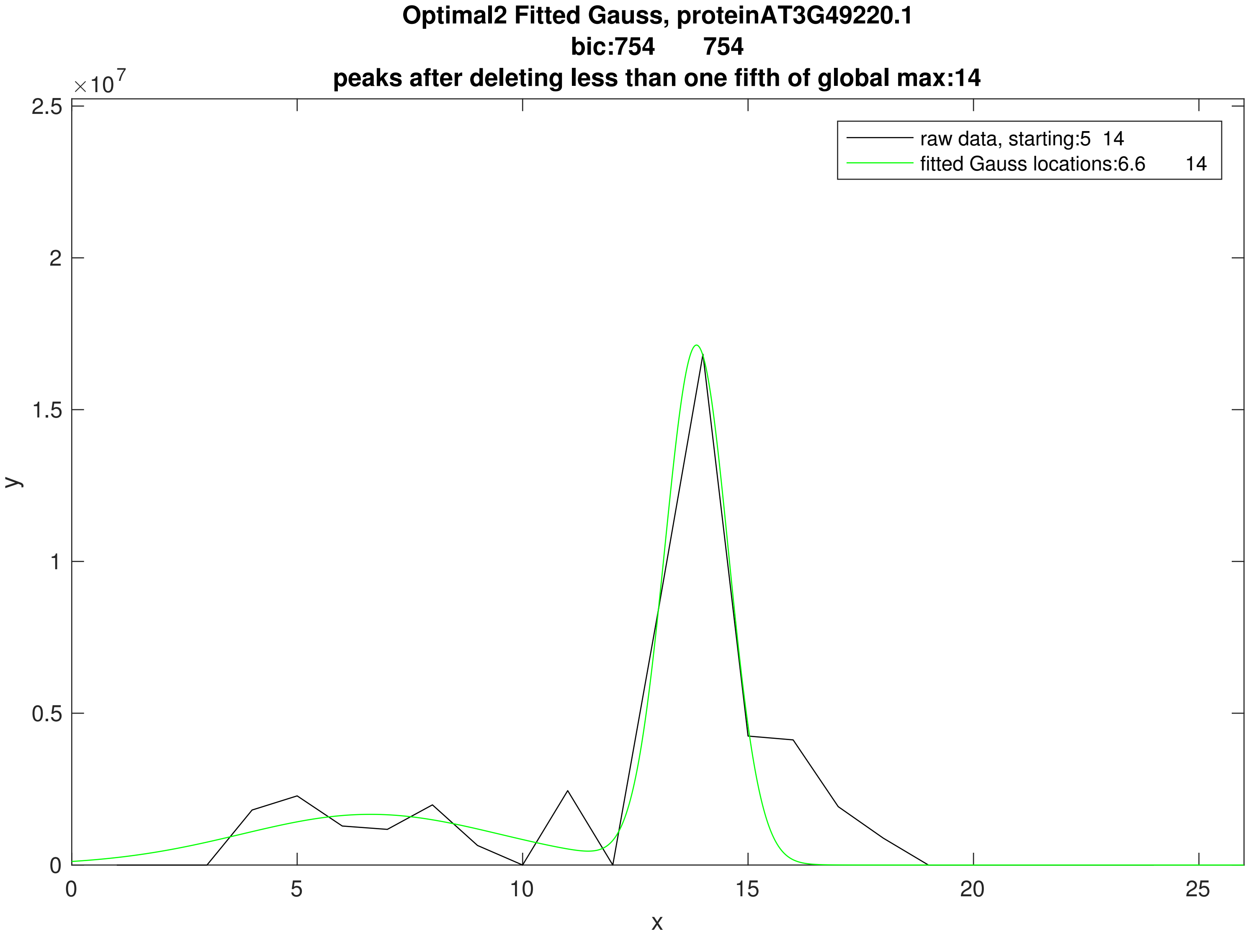

Optimal2 Fitted Gauss, proteinAT3G49220.1
bic:754 754
peaks after deleting less than one fifth of global max:14
×10^7
2.5
2
1.5
1
0.5
0
y
0
5
10
15
20
25
x
raw data, starting:5 14
fitted Gauss locations:6.6 14

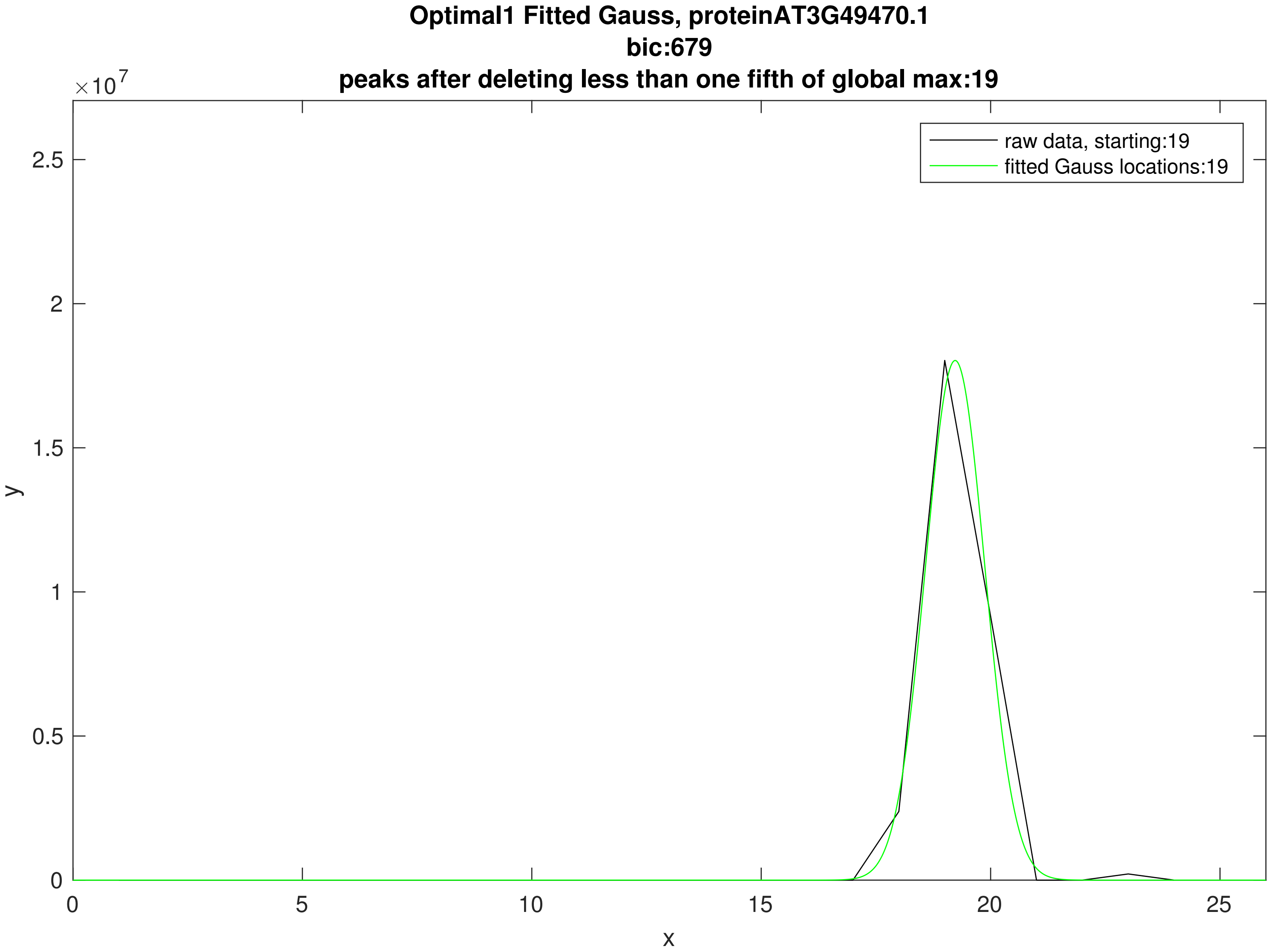

Optimal1 Fitted Gauss, proteinAT3G49470.1
bic:679
peaks after deleting less than one fifth of global max:19
×10^7
raw data, starting:19
fitted Gauss locations:19
2.5
2
1.5
1
0.5
0
y
0
5
10
15
20
25
x

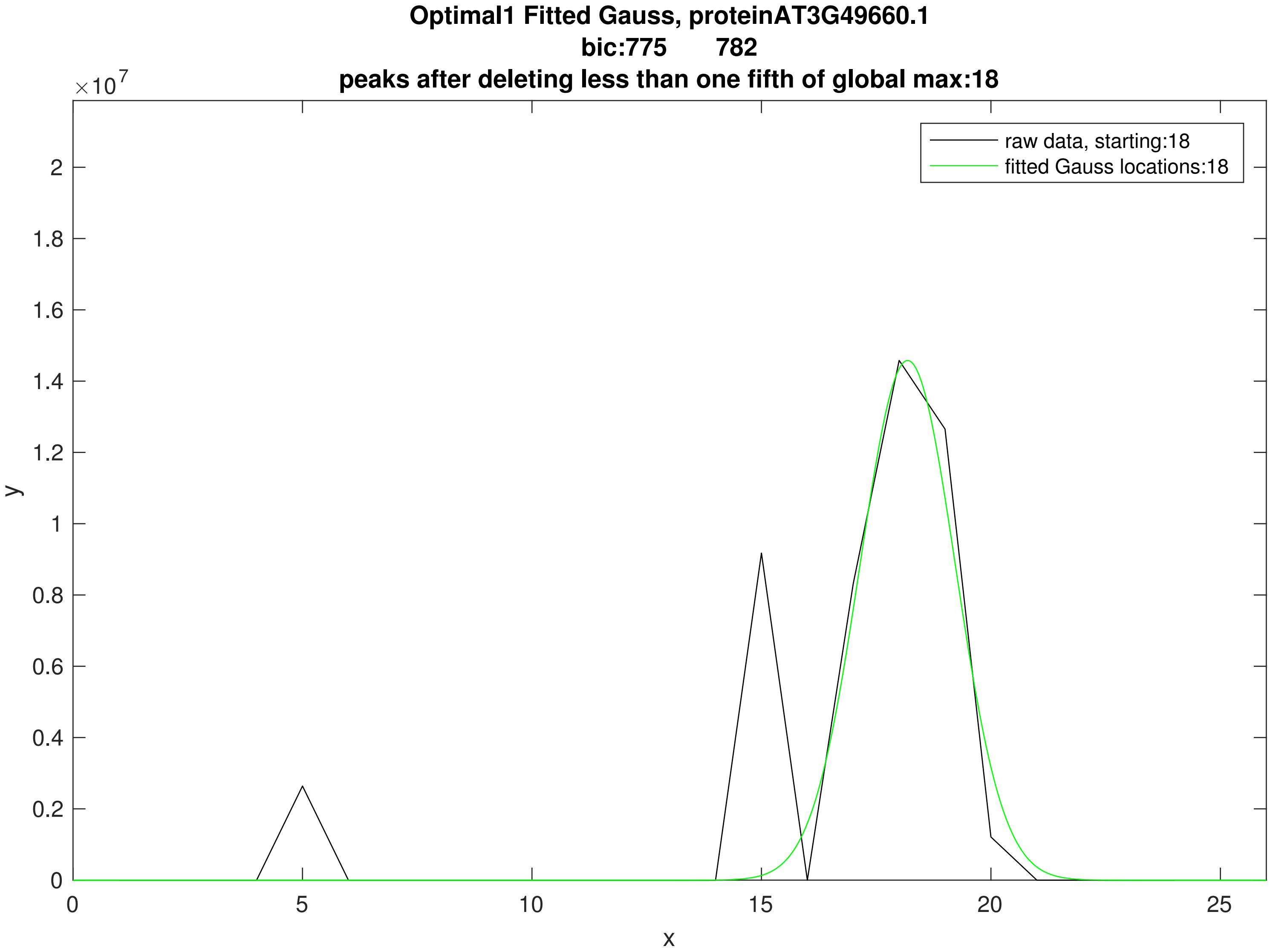

Optimal1 Fitted Gauss, proteinAT3G49660.1
bic:775 782
peaks after deleting less than one fifth of global max:18
×10^7
raw data, starting:18
fitted Gauss locations:18
y
x
0
0.2
0.4
0.6
0.8
1
1.2
1.4
1.6
1.8
2
5
10
15
20
25

#### Optimal1 Fitted Gauss, proteinAT3G49680.2;AT3G49680.1
#### bic:803 812
#### peaks after deleting less than one fifth of global max:14

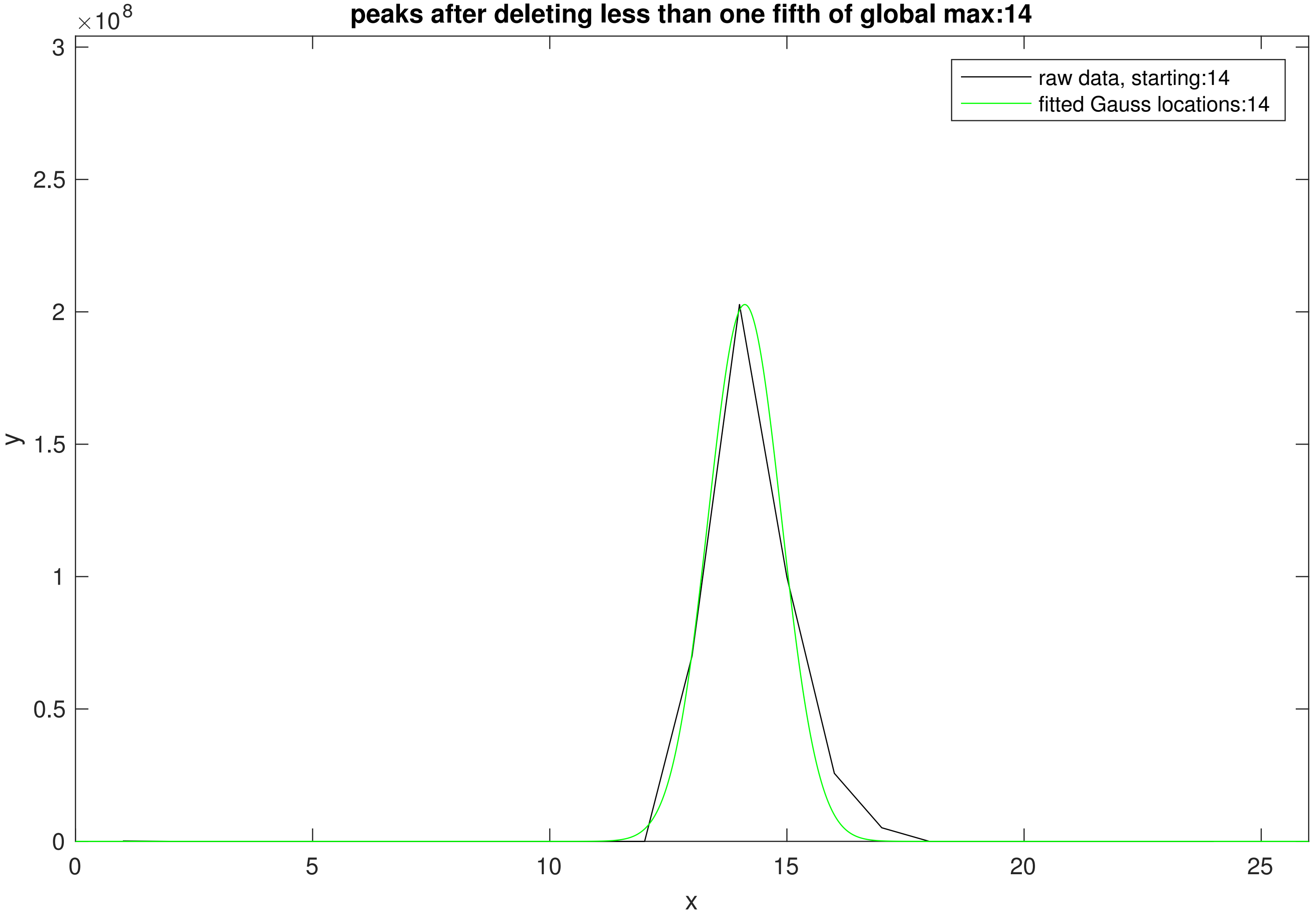

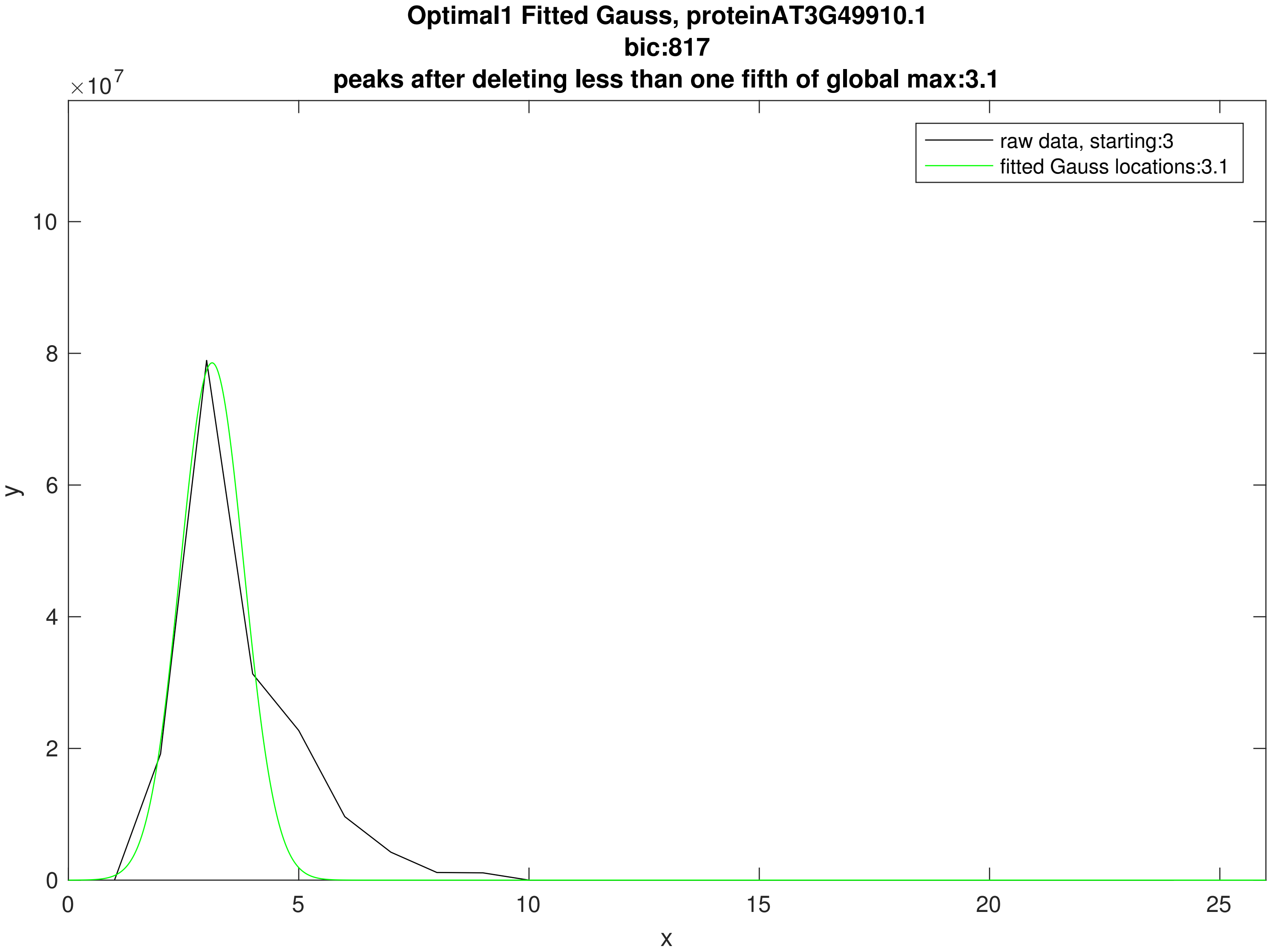
Optimal1 Fitted Gauss, proteinAT3G49910.1
bic:817
peaks after deleting less than one fifth of global max:3.1
×10^7
raw data, starting:3
fitted Gauss locations:3.1
y
x
0
2
4
6
8
10
0
5
10
15
20
25

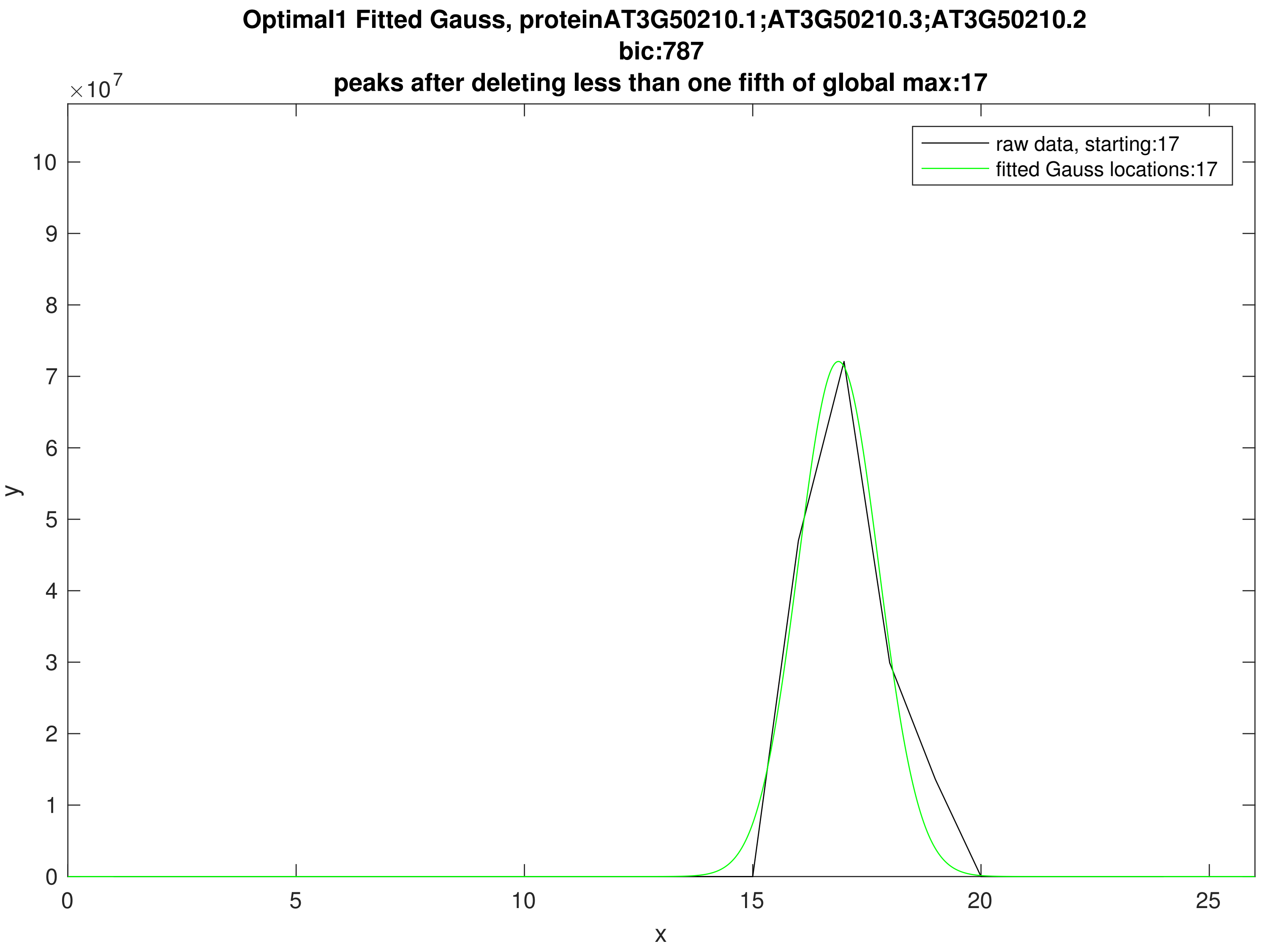

Optimal1 Fitted Gauss, proteinAT3G50210.1;AT3G50210.3;AT3G50210.2
bic:787
peaks after deleting less than one fifth of global max:17
×10^7
raw data, starting:17
fitted Gauss locations:17
y
x
0
1
2
3
4
5
6
7
8
9
10
0
5
10
15
20
25

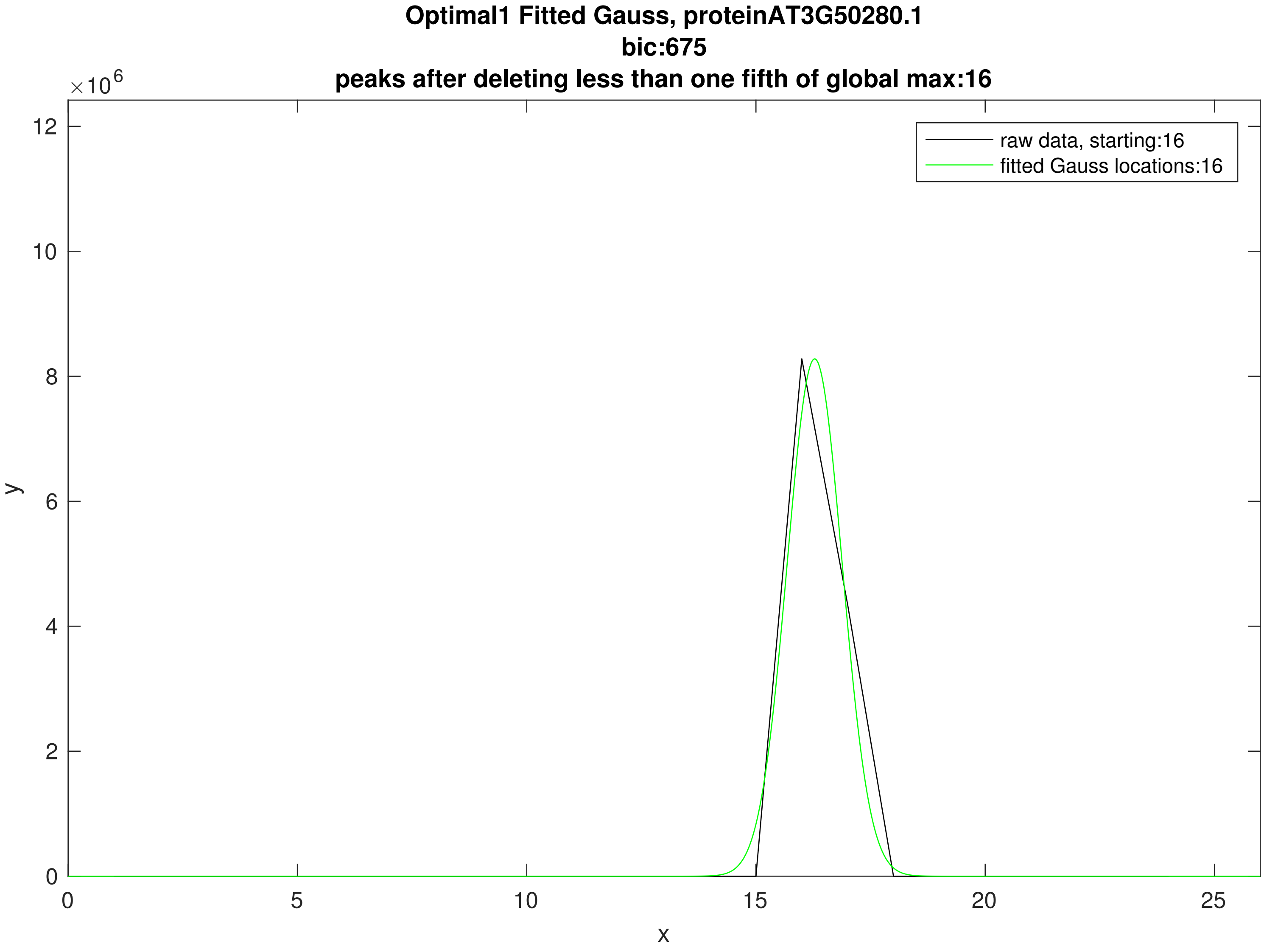

Optimal1 Fitted Gauss, proteinAT3G50280.1
bic:675
peaks after deleting less than one fifth of global max:16
×10^6
12
10
8
6
4
2
0
y
0
5
10
15
20
25
x
raw data, starting:16
fitted Gauss locations:16

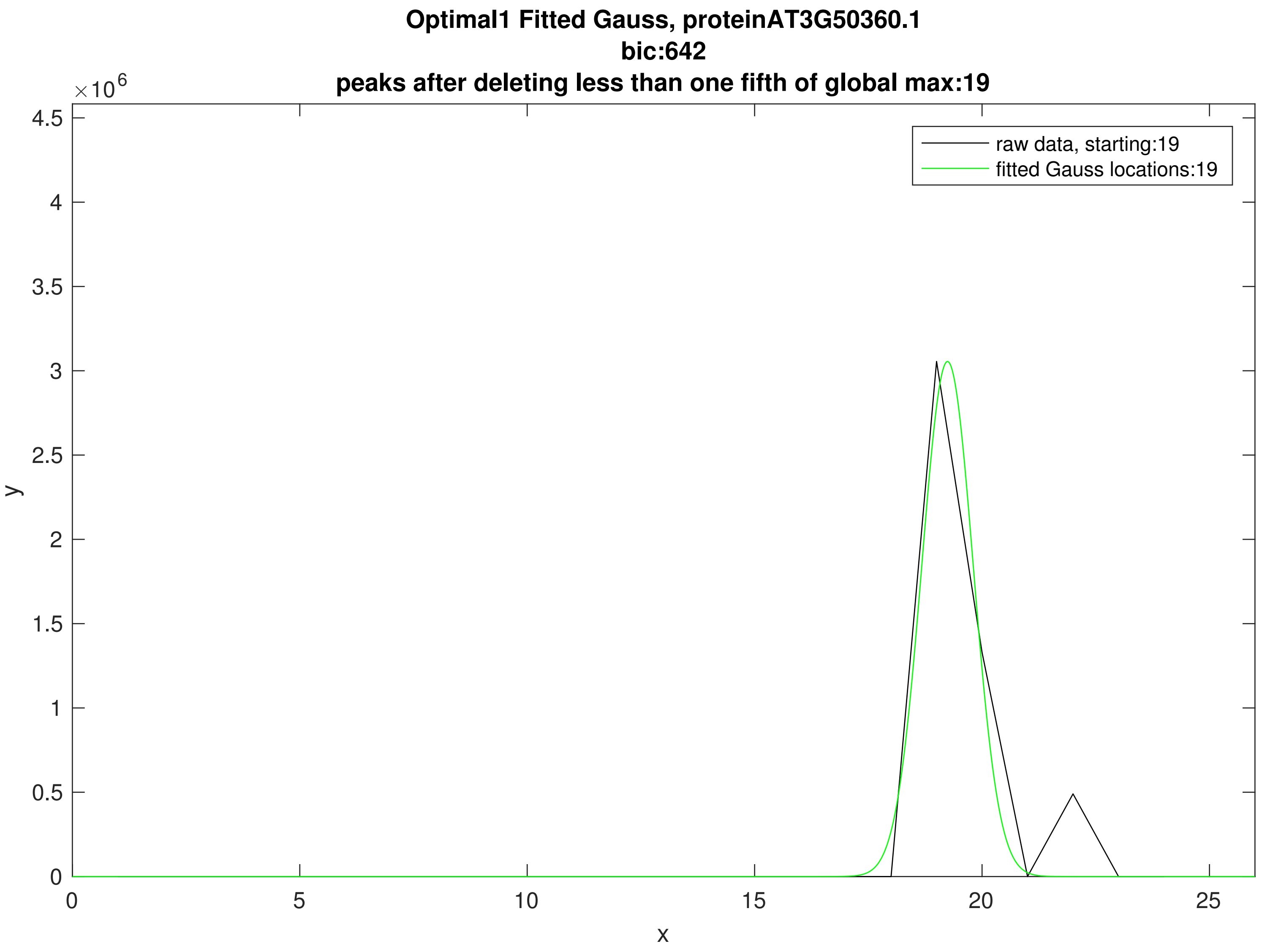
Optimal1 Fitted Gauss, proteinAT3G50360.1
bic:642
peaks after deleting less than one fifth of global max:19
×10^6
4.5
4
3.5
3
2.5
2
1.5
1
0.5
0
y
0
5
10
15
20
25
x
raw data, starting:19
fitted Gauss locations:19

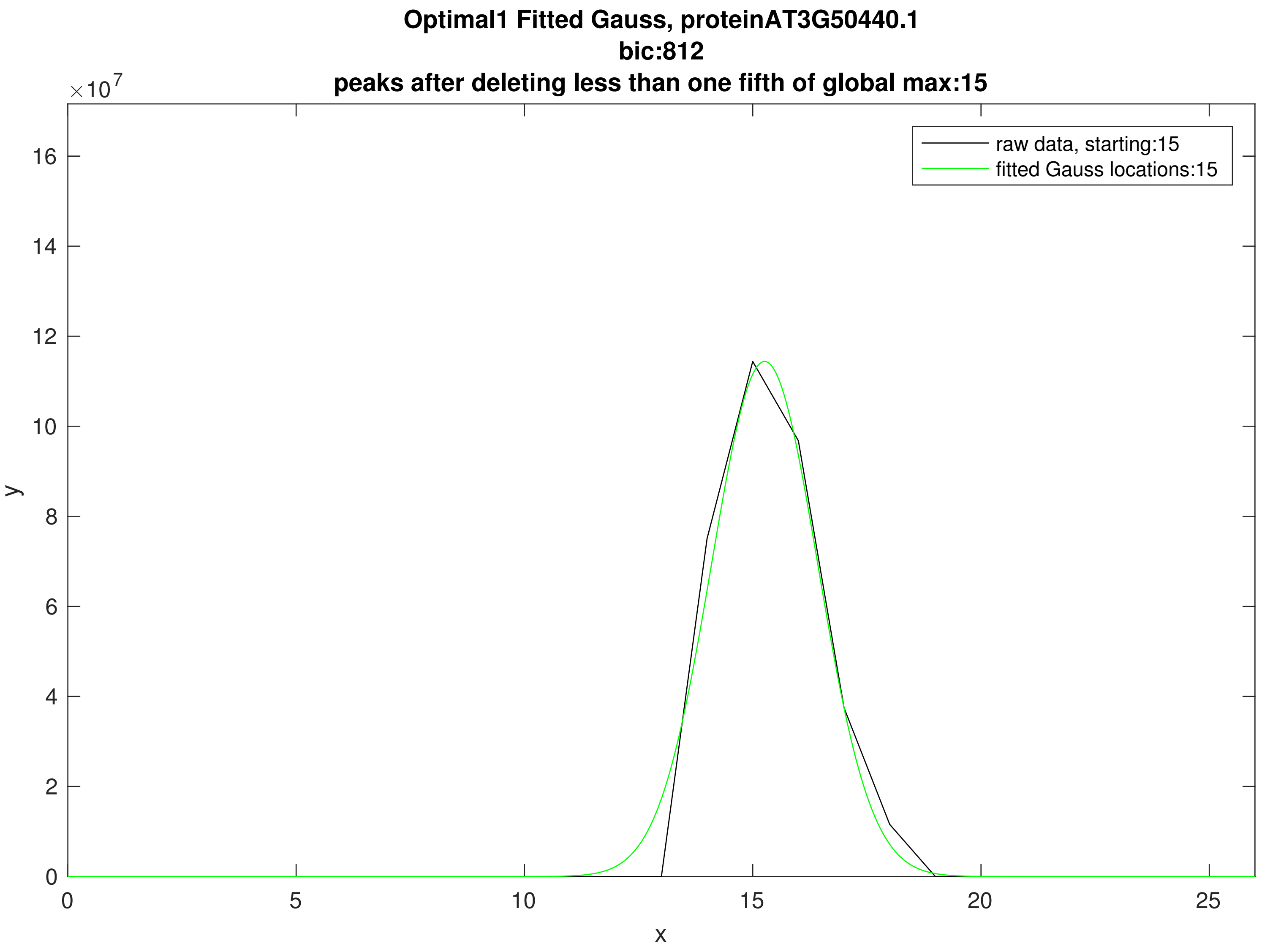
Optimal1 Fitted Gauss, proteinAT3G50440.1
bic:812
peaks after deleting less than one fifth of global max:15
×10^7
raw data, starting:15
fitted Gauss locations:15
y
x
0
2
4
6
8
10
12
14
16
0
5
10
15
20
25

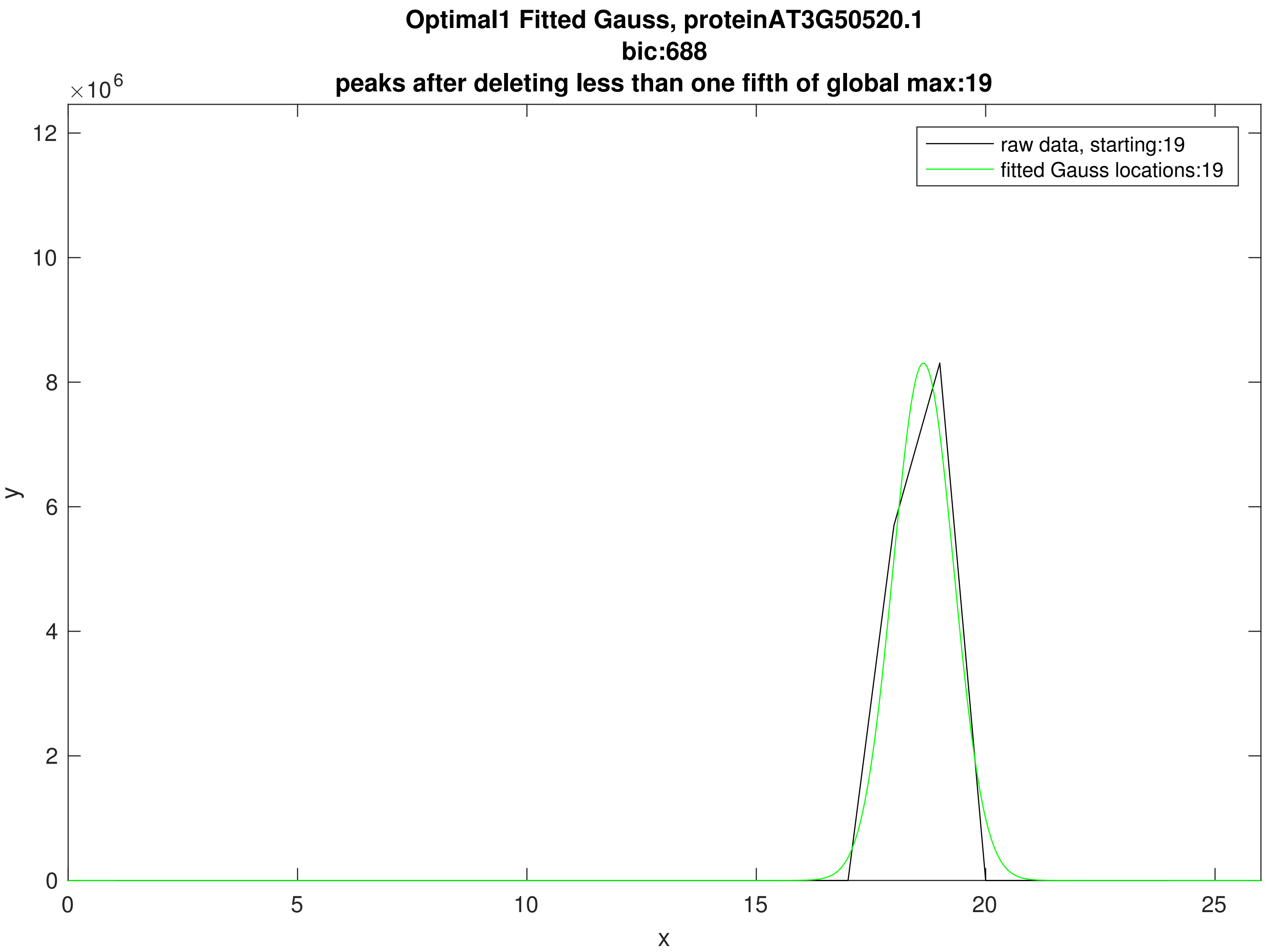

Optimal1 Fitted Gauss, proteinAT3G50520.1
bic:688
peaks after deleting less than one fifth of global max:19
×10^6
12
10
8
6
4
2
0
y
0
5
10
15
20
25
x
raw data, starting:19
fitted Gauss locations:19

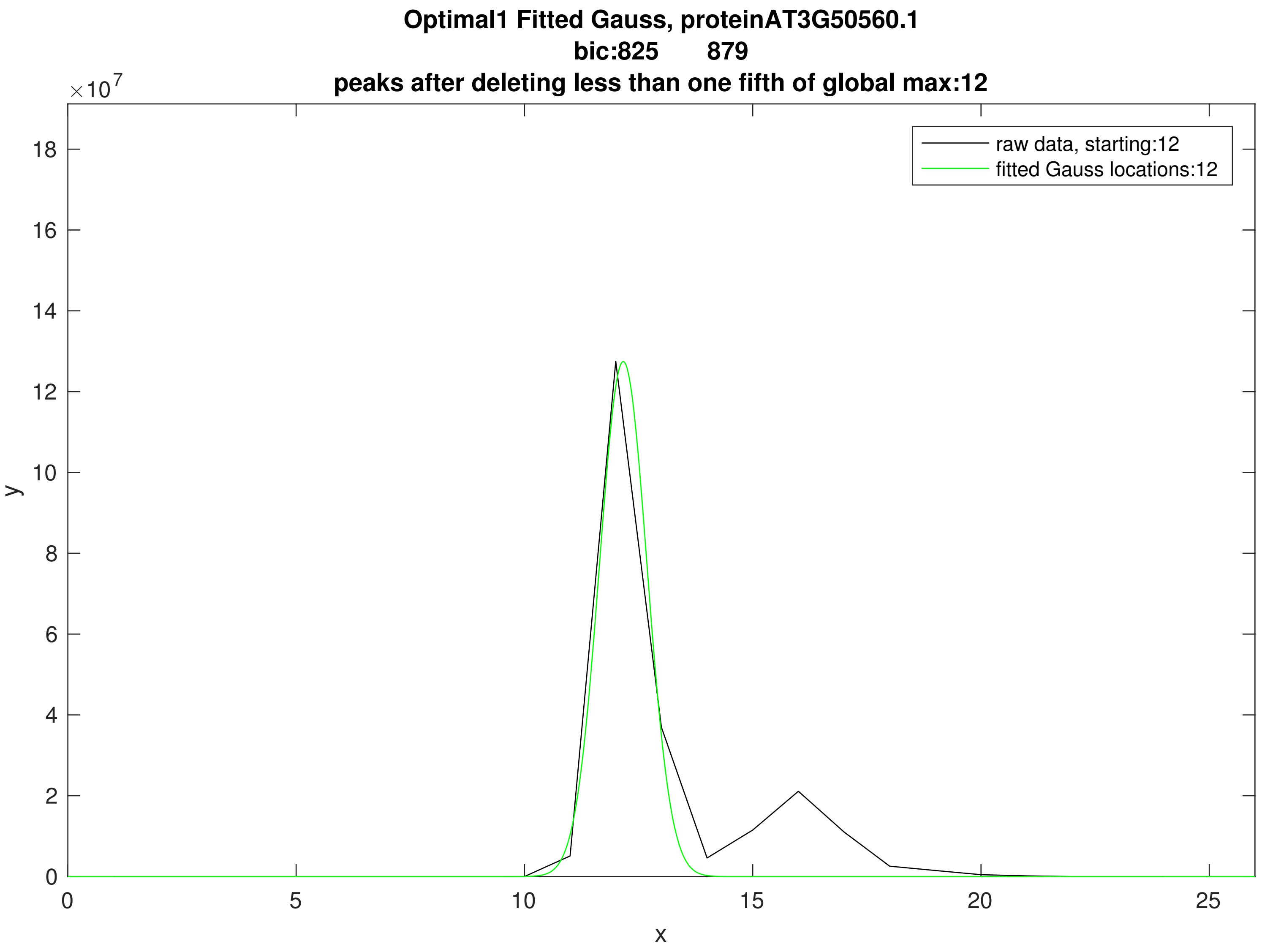
Optimal1 Fitted Gauss, proteinAT3G50560.1
bic:825 879
peaks after deleting less than one fifth of global max:12
×10^7
raw data, starting:12
fitted Gauss locations:12
y
x
0
2
4
6
8
10
12
14
16
18
0
5
10
15
20
25

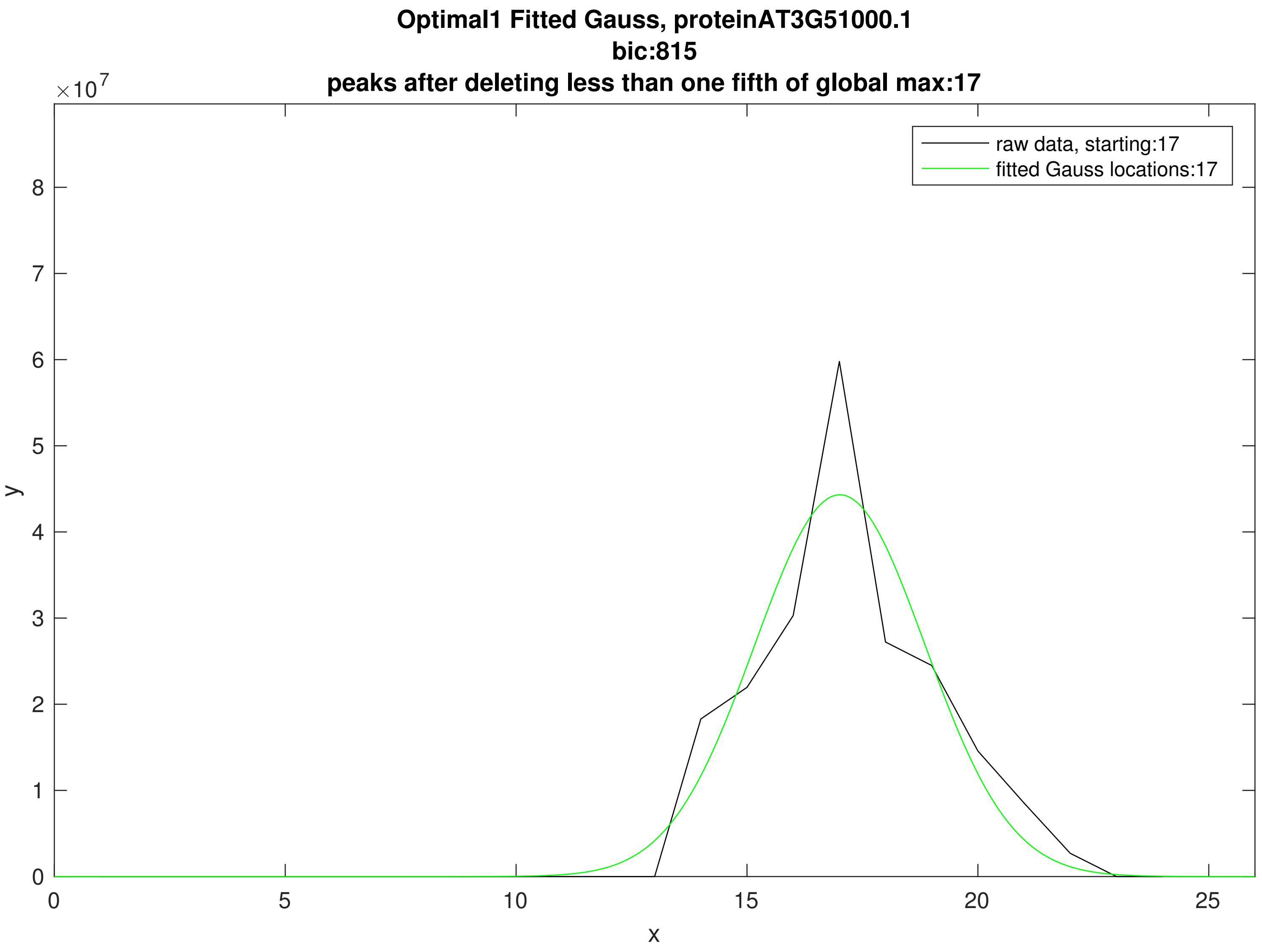

Optimal1 Fitted Gauss, proteinAT3G51000.1
bic:815
peaks after deleting less than one fifth of global max:17
×10^7
raw data, starting:17
fitted Gauss locations:17
8
7
6
5
4
3
2
1
0
y
0
5
10
15
20
25
x

**Optimal2 Fitted Gauss, proteinAT3G51160.1**
**bic:962 933 942 957**
**peaks after deleting less than one fifth of global max:11**

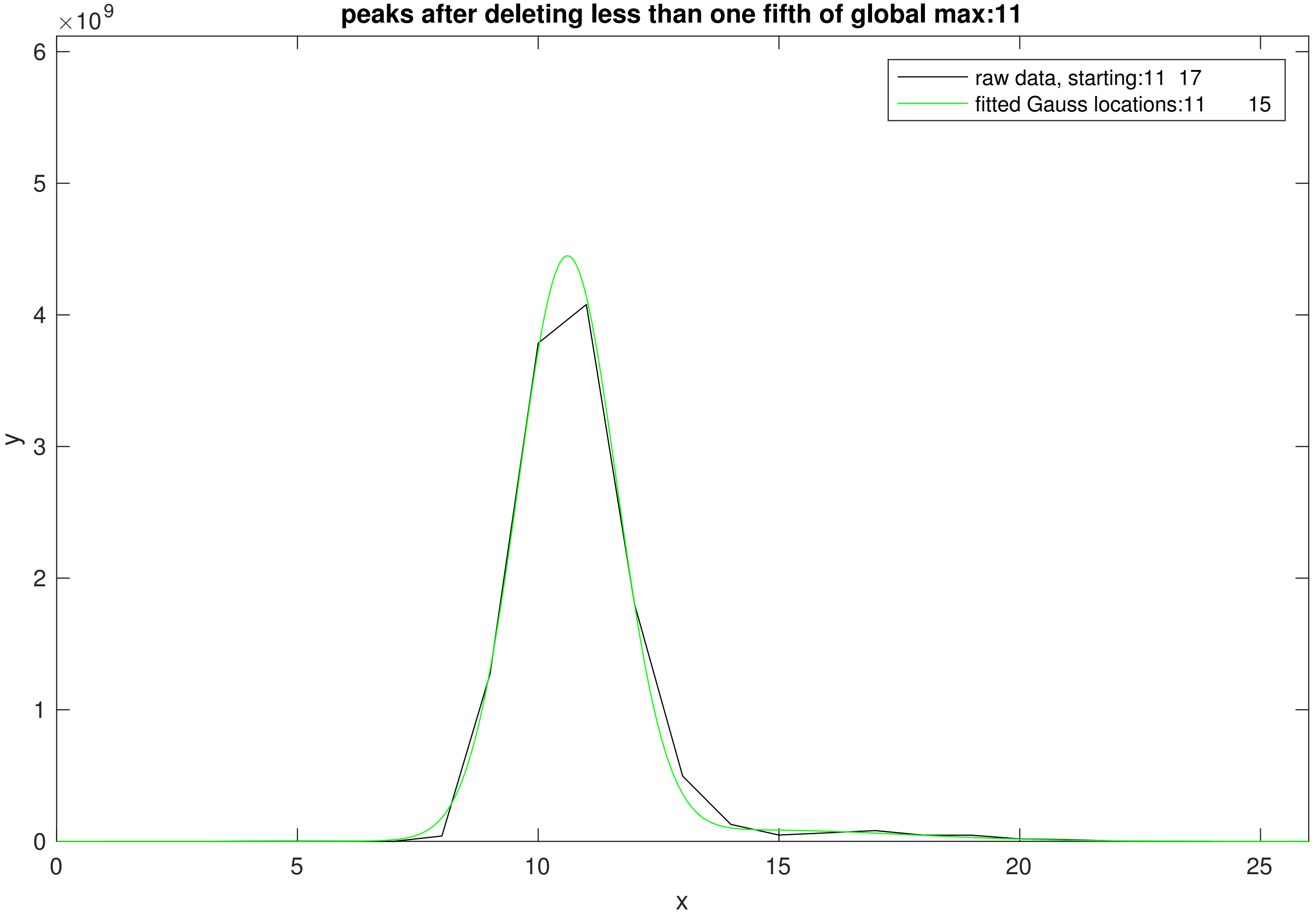

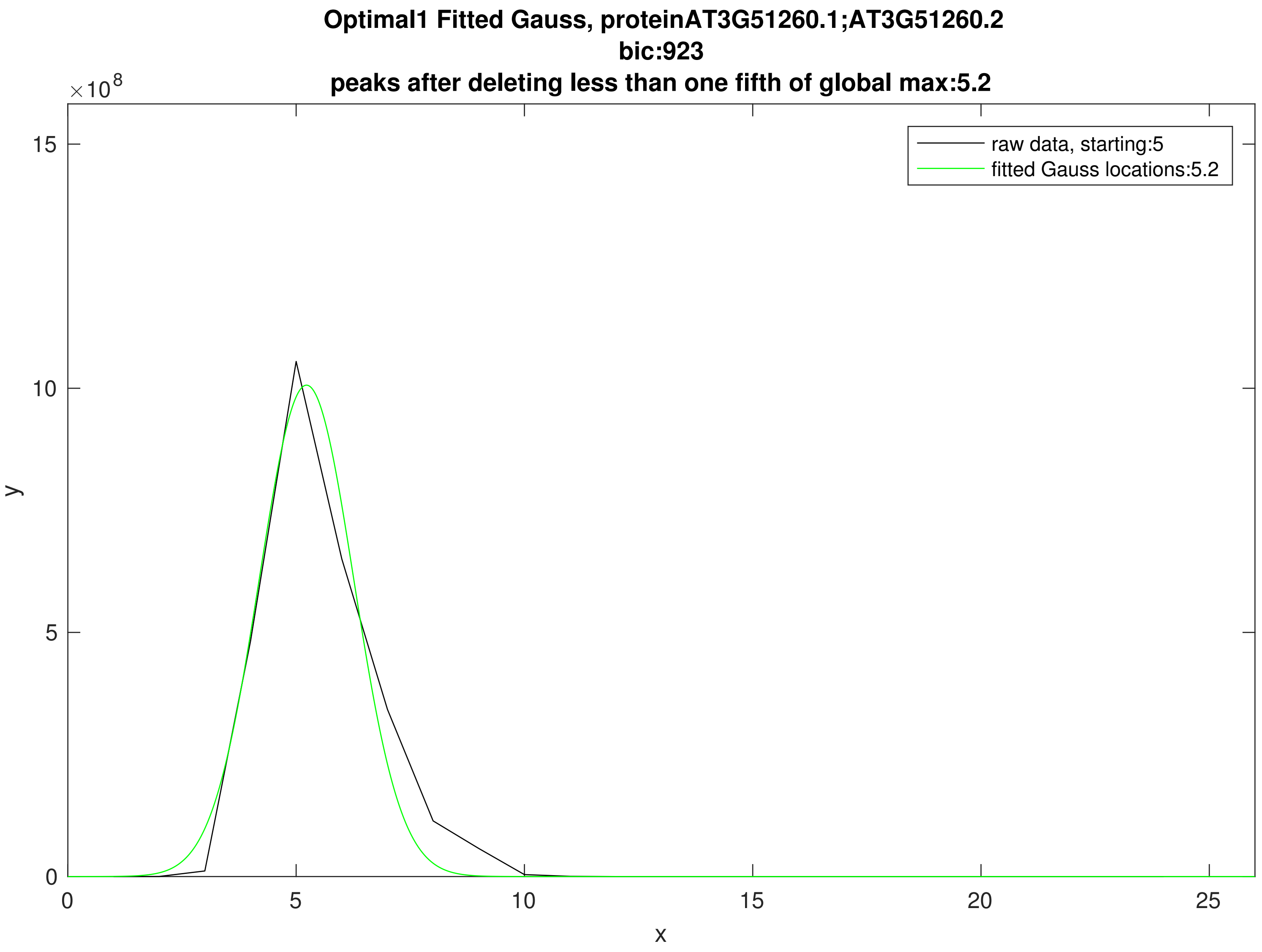

Optimal1 Fitted Gauss, proteinAT3G51260.1;AT3G51260.2
bic:923
peaks after deleting less than one fifth of global max:5.2
×10^8
raw data, starting:5
fitted Gauss locations:5.2
y
x
0
5
10
15
20
25

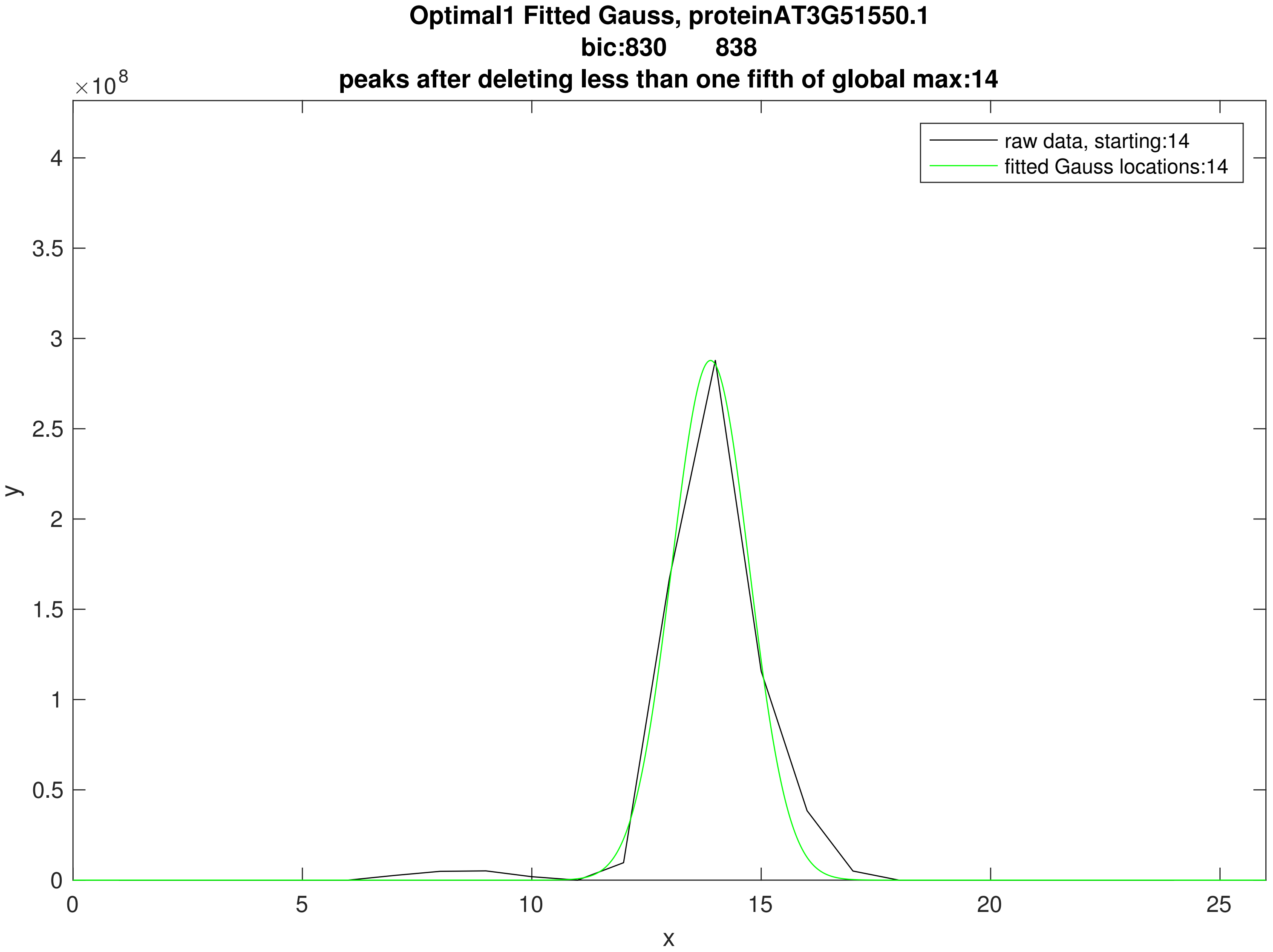

Optimal1 Fitted Gauss, proteinAT3G51550.1
bic:830 838
peaks after deleting less than one fifth of global max:14
×10^8
raw data, starting:14
fitted Gauss locations:14
4
3.5
3
2.5
2
1.5
1
0.5
0
y
0
5
10
15
20
25
x

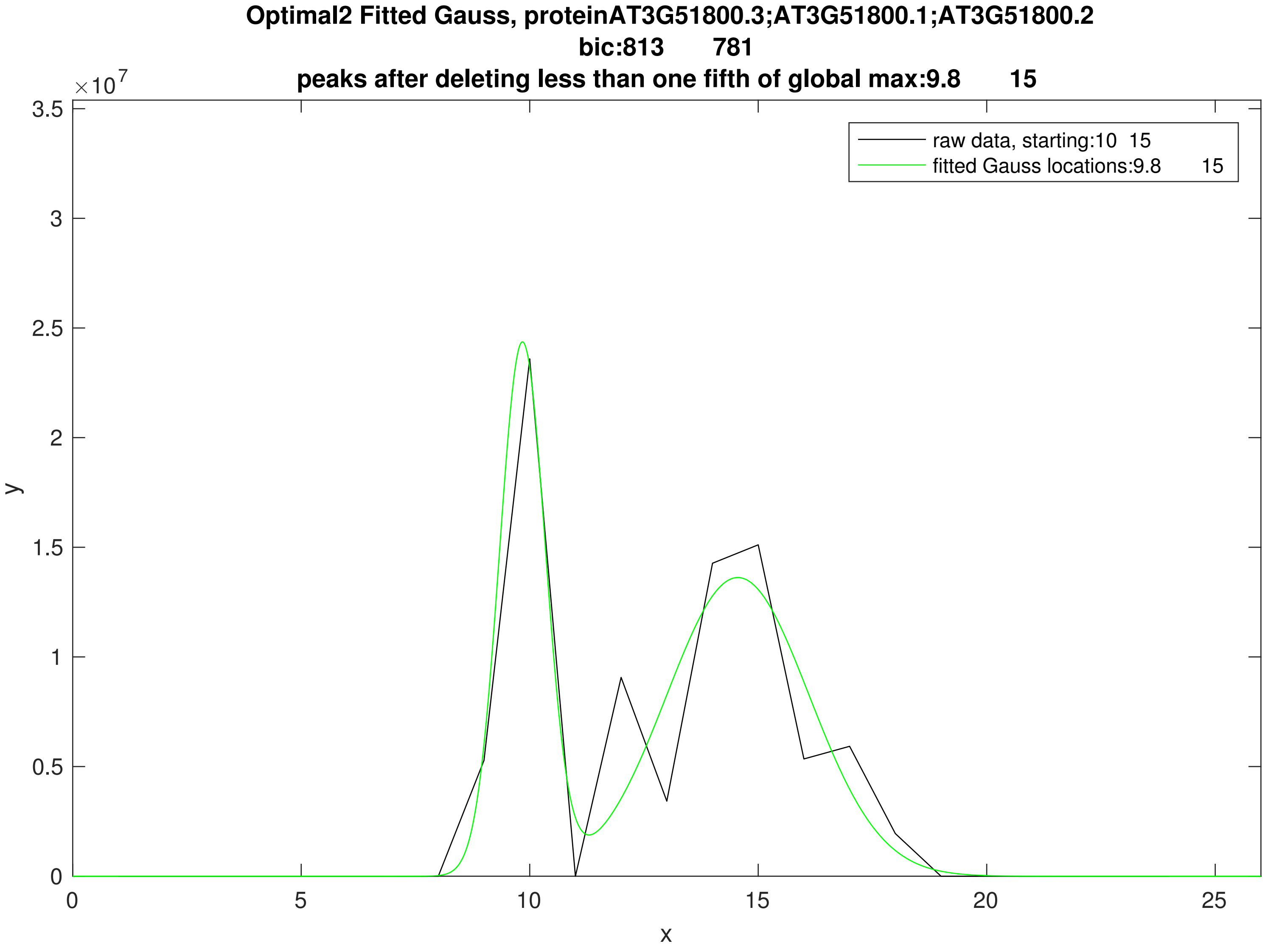
Optimal2 Fitted Gauss, proteinAT3G51800.3;AT3G51800.1;AT3G51800.2
bic:813 781
peaks after deleting less than one fifth of global max:9.8 15
×10^7
raw data, starting:10 15
fitted Gauss locations:9.8 15
y
x
0
0.5
1
1.5
2
2.5
3
3.5
5
10
15
20
25

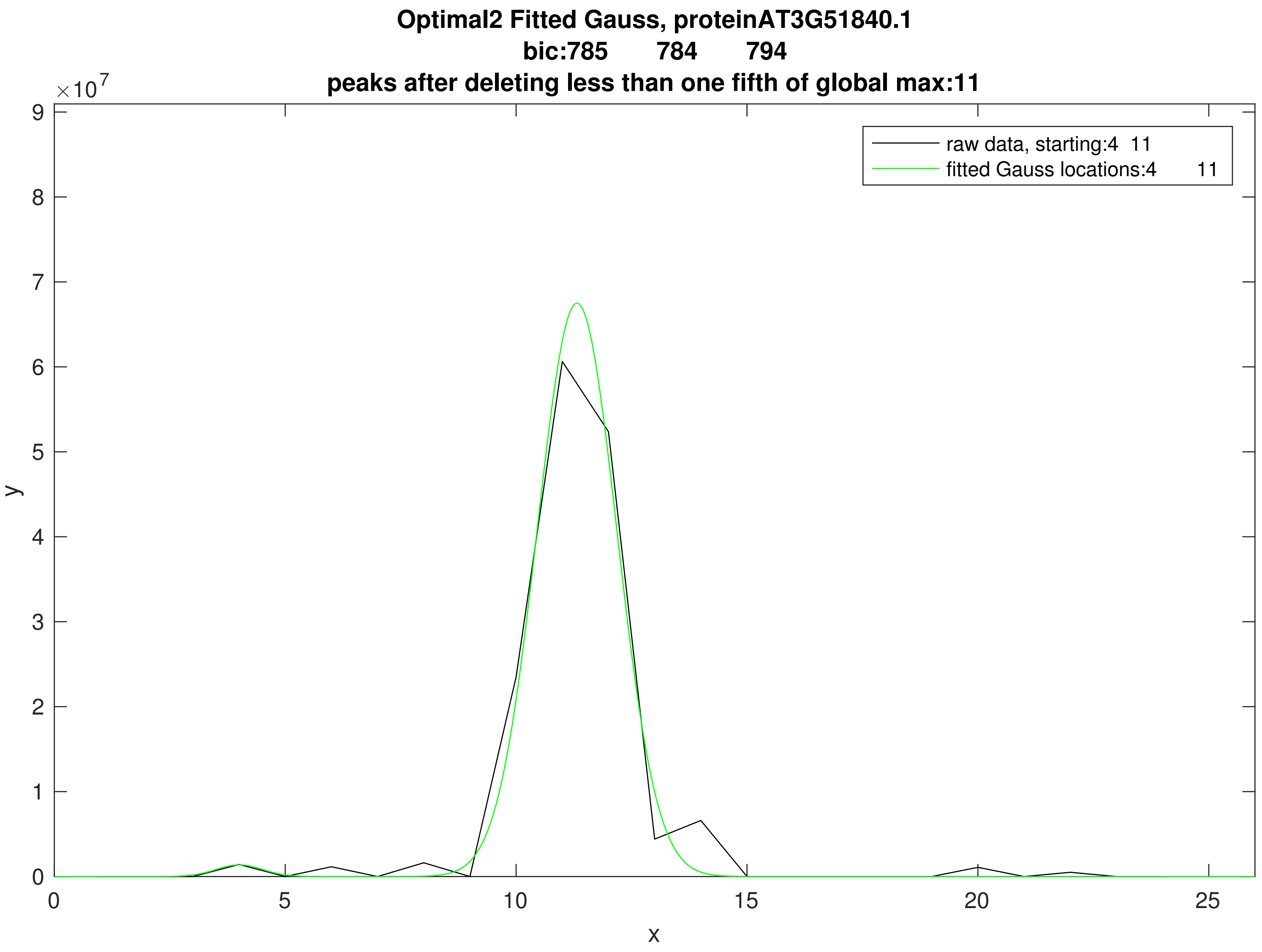

Optimal2 Fitted Gauss, proteinAT3G51840.1
bic:785 784 794
peaks after deleting less than one fifth of global max:11
×10^7
raw data, starting:4 11
fitted Gauss locations:4 11
y
x
0
1
2
3
4
5
6
7
8
9
0
5
10
15
20
25

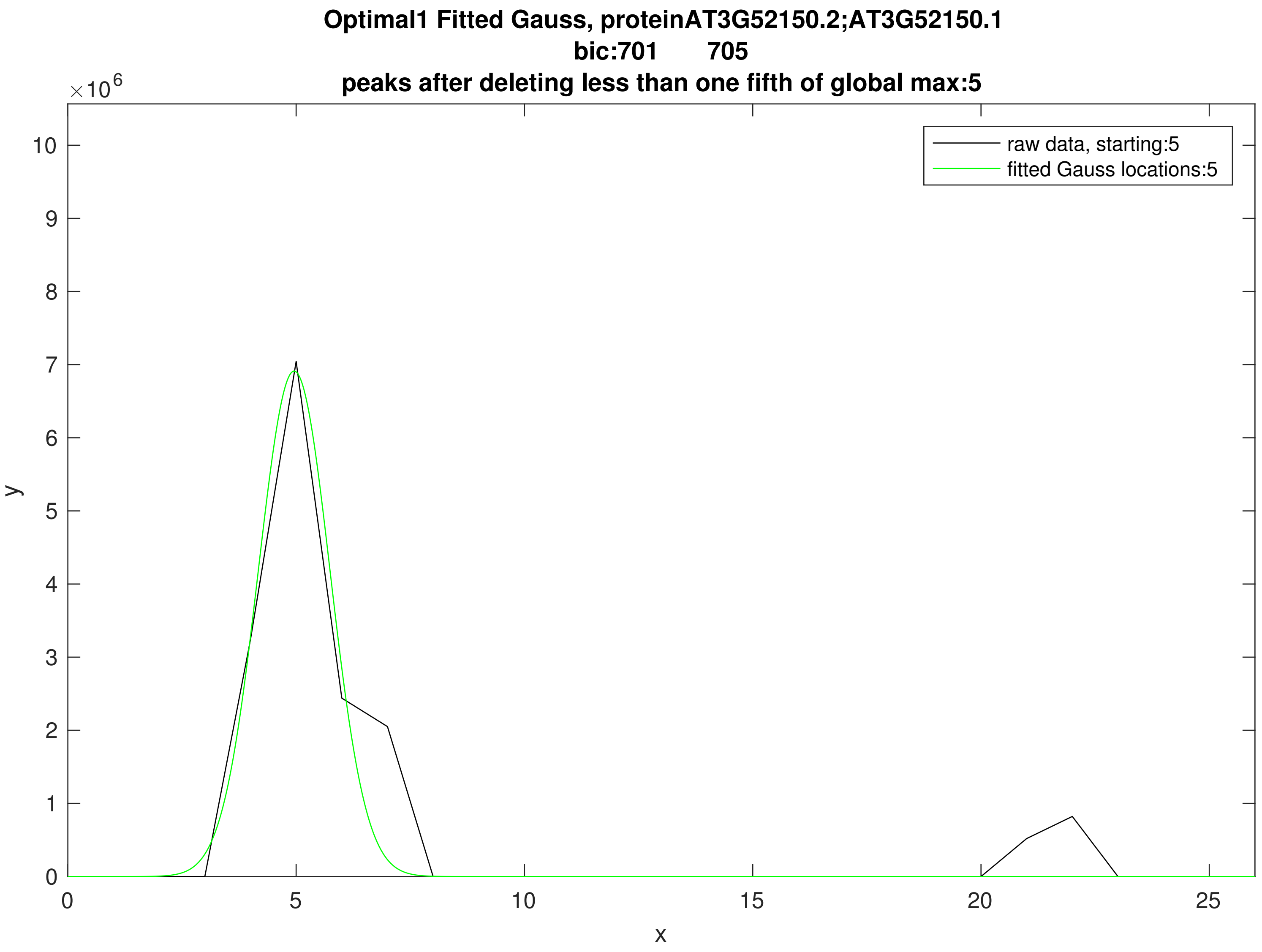
Optimal1 Fitted Gauss, proteinAT3G52150.2;AT3G52150.1
bic:701 705
peaks after deleting less than one fifth of global max:5
×10^6
raw data, starting:5
fitted Gauss locations:5
y
x
0
1
2
3
4
5
6
7
8
9
10
0
5
10
15
20
25

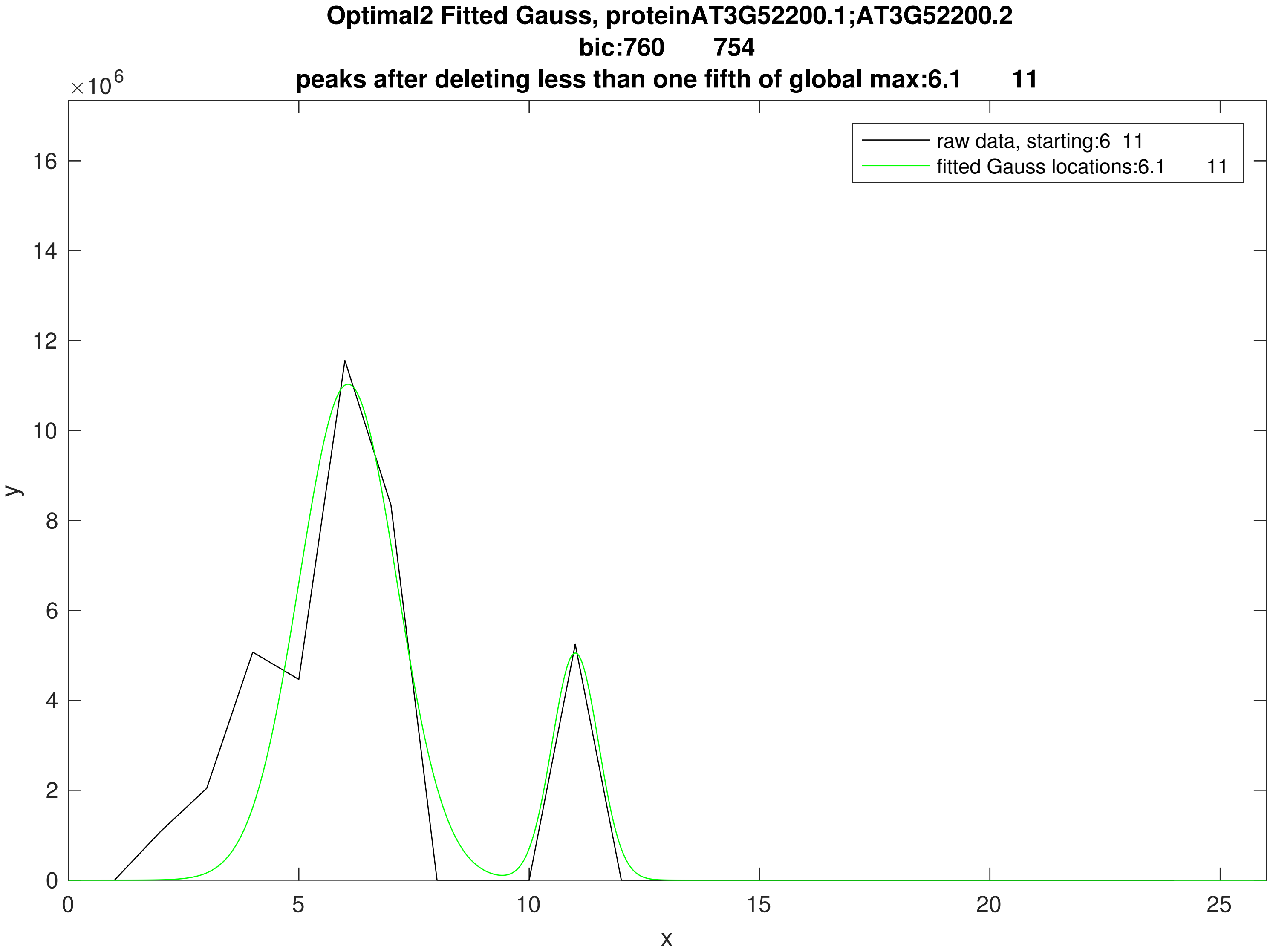

Optimal2 Fitted Gauss, proteinAT3G52200.1;AT3G52200.2
bic:760 754
peaks after deleting less than one fifth of global max:6.1 11
×10^6
raw data, starting:6 11
fitted Gauss locations:6.1 11
y
x
0
2
4
6
8
10
12
14
16
0
5
10
15
20
25

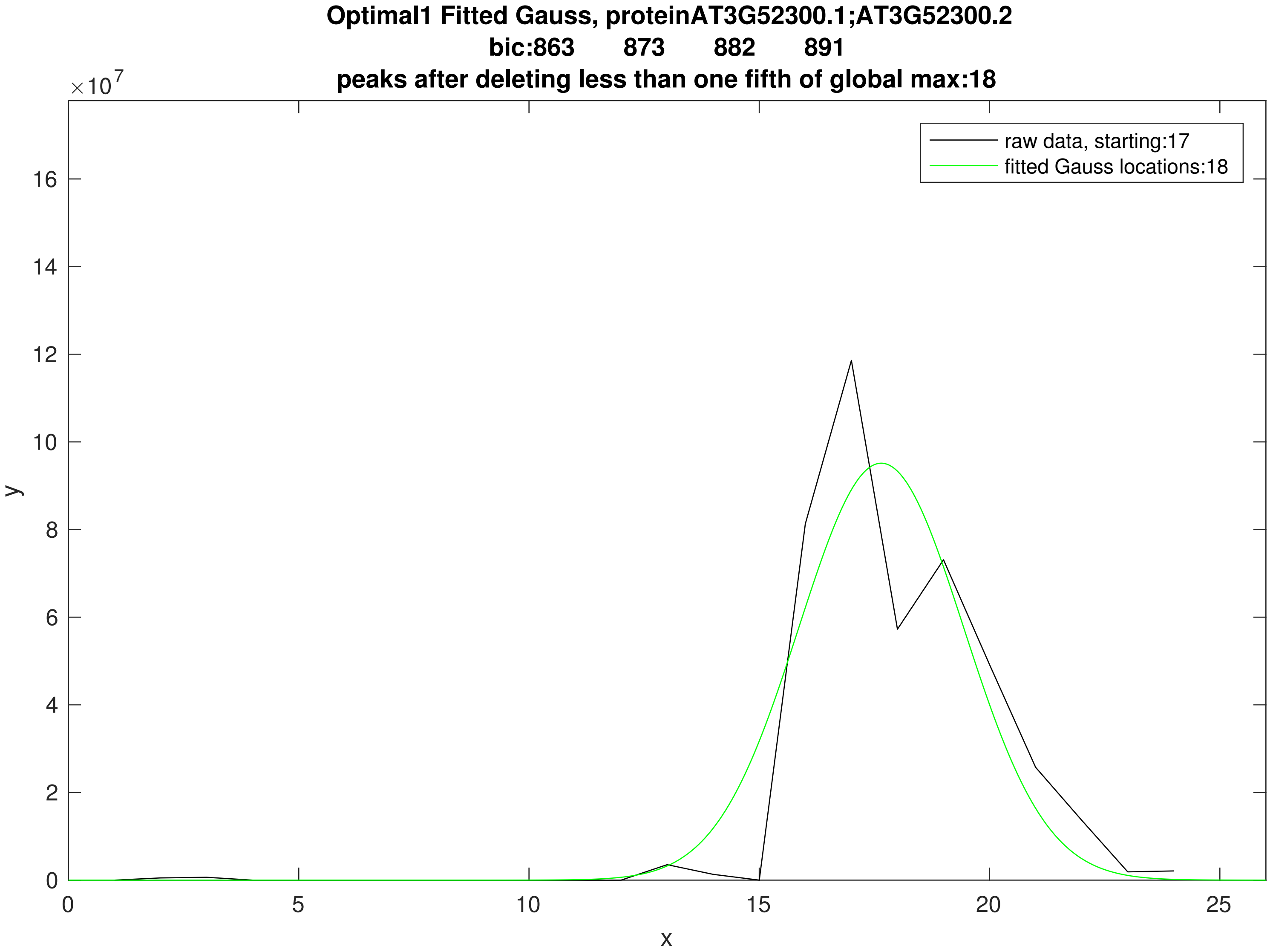

Optimal1 Fitted Gauss, proteinAT3G52300.1;AT3G52300.2
bic:863 873 882 891
peaks after deleting less than one fifth of global max:18
×10^7
raw data, starting:17
fitted Gauss locations:18
y
x
0
2
4
6
8
10
12
14
16
0
5
10
15
20
25

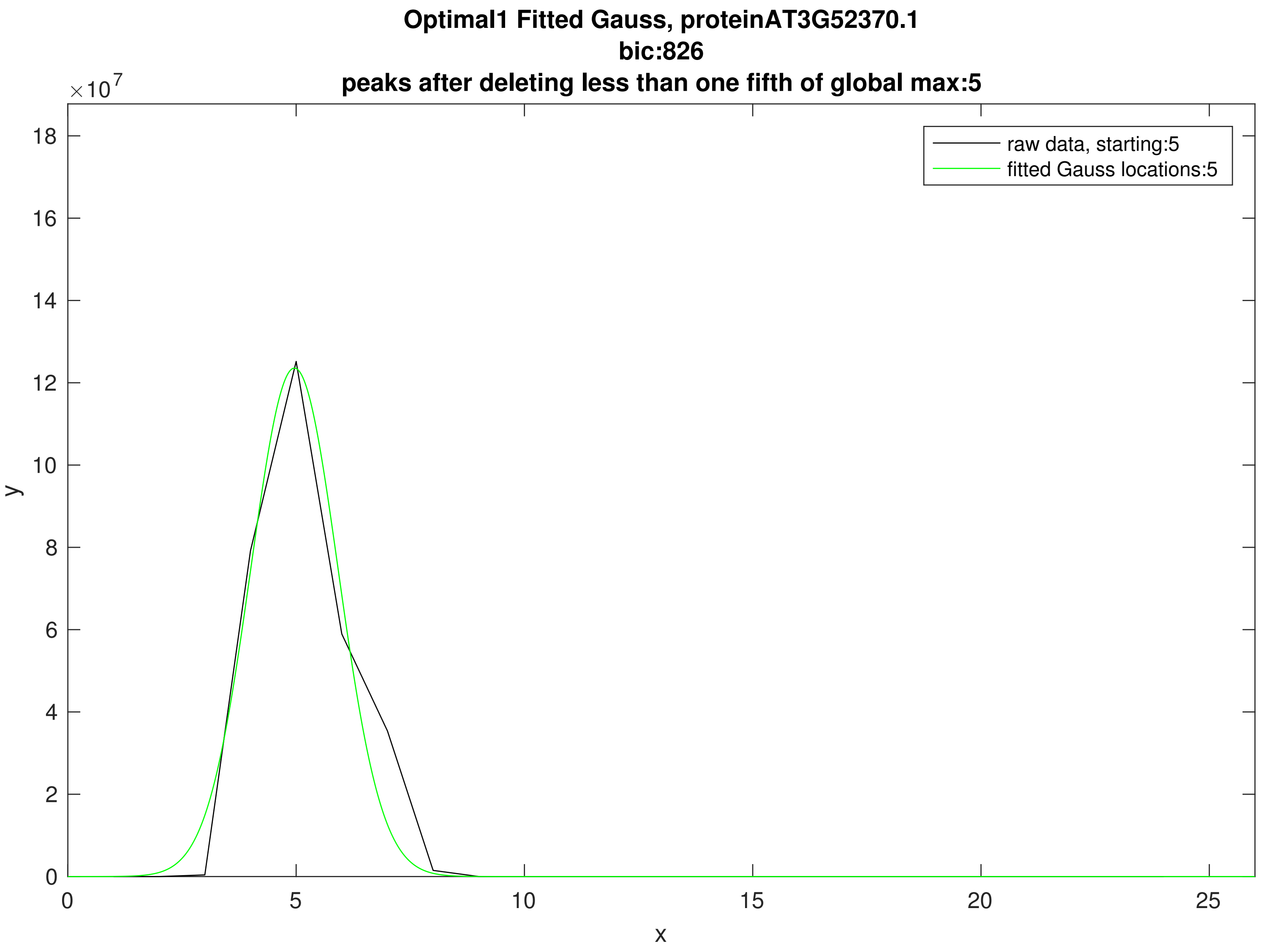
Optimal1 Fitted Gauss, proteinAT3G52370.1
bic:826
peaks after deleting less than one fifth of global max:5
×10^7
18
16
14
12
10
8
6
4
2
0
y
0
5
10
15
20
25
x
raw data, starting:5
fitted Gauss locations:5

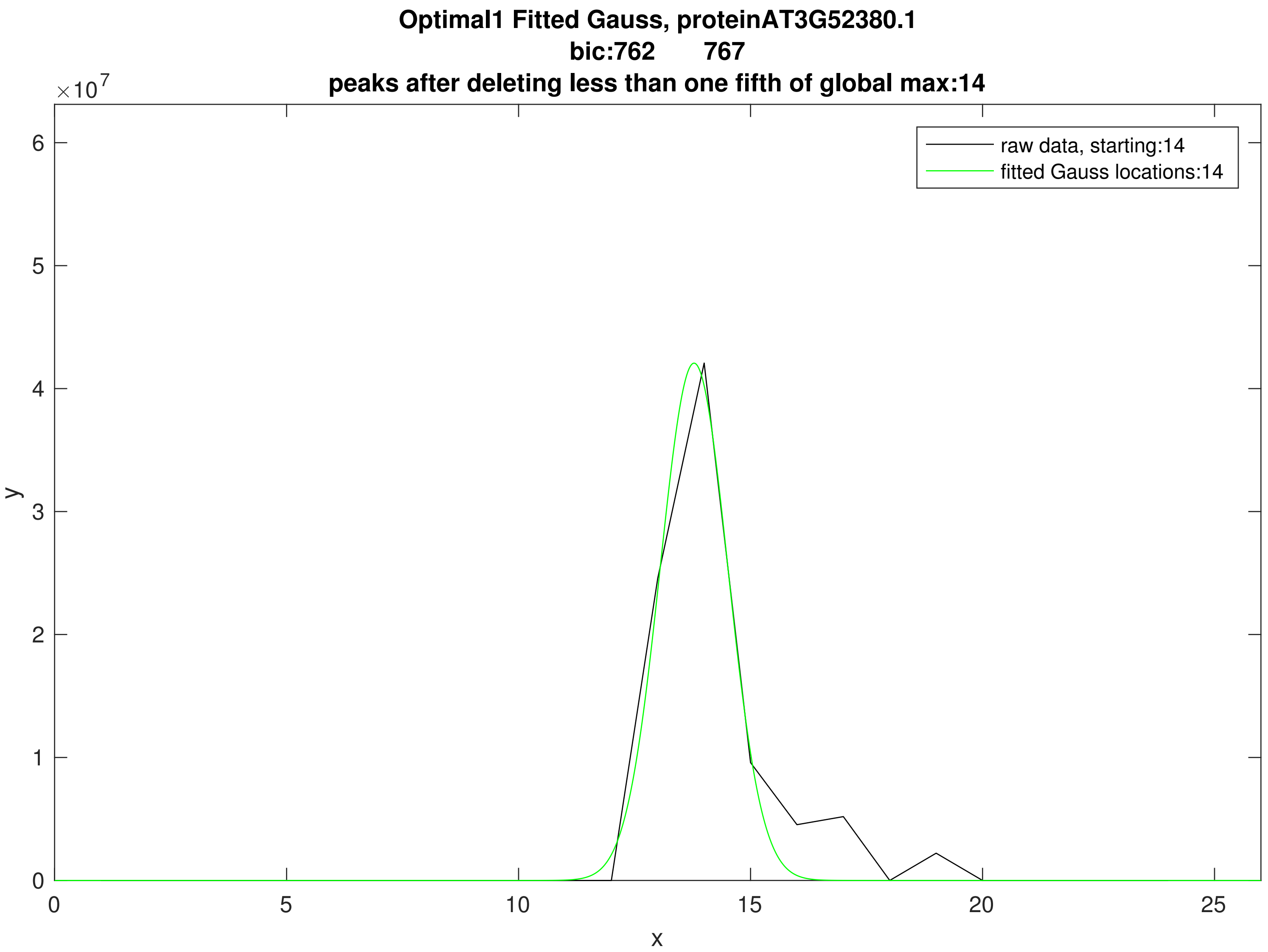
Optimal1 Fitted Gauss, proteinAT3G52380.1
bic:762 767
peaks after deleting less than one fifth of global max:14
×10^7
6
5
4
3
2
1
0
y
0
5
10
15
20
25
x
raw data, starting:14
fitted Gauss locations:14

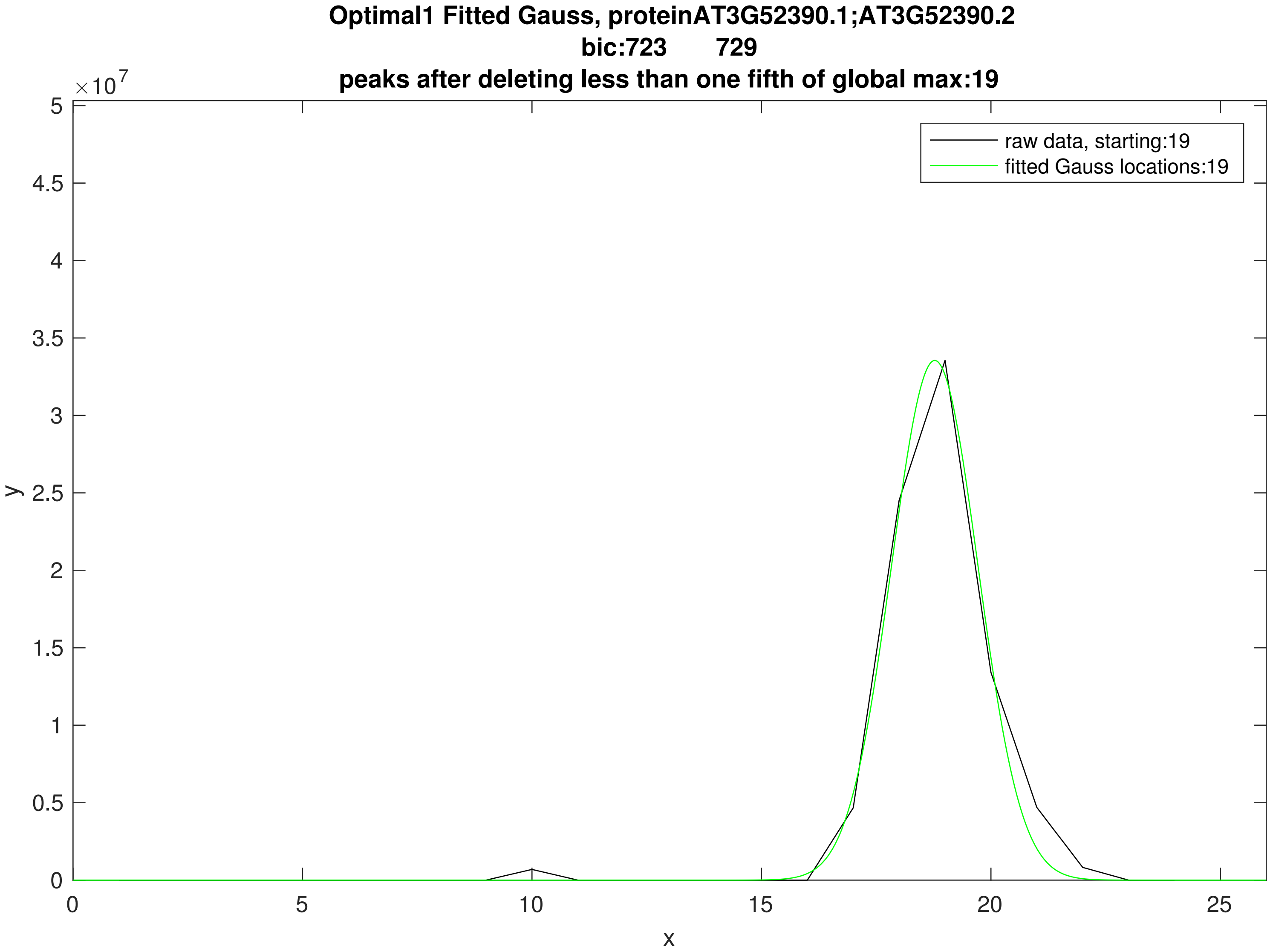
Optimal1 Fitted Gauss, proteinAT3G52390.1;AT3G52390.2
bic:723 729
peaks after deleting less than one fifth of global max:19
×10^7
5
4.5
4
3.5
3
2.5
2
1.5
1
0.5
0
y
0
5
10
15
20
25
x
raw data, starting:19
fitted Gauss locations:19

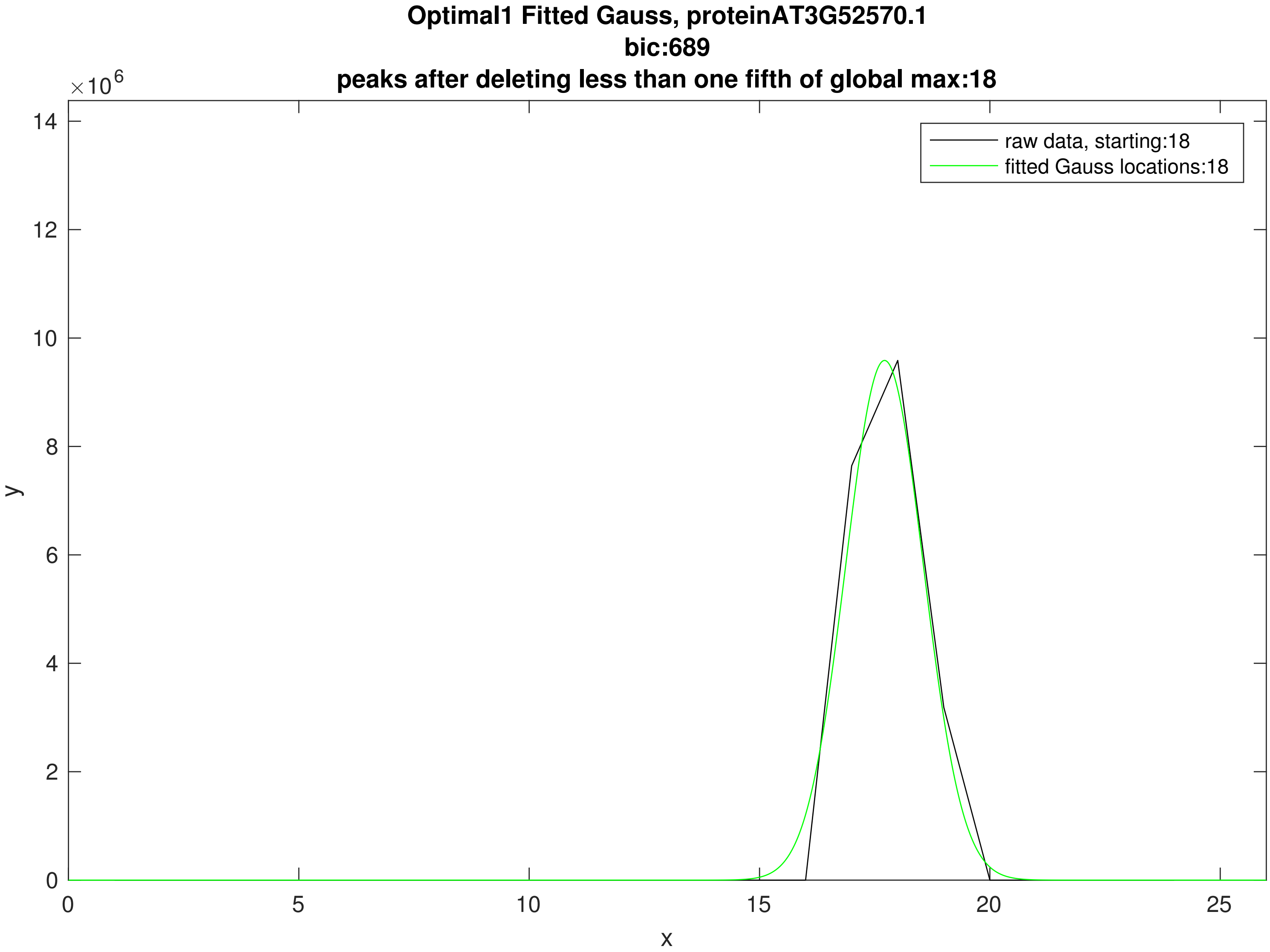

Optimal1 Fitted Gauss, proteinAT3G52570.1
bic:689
peaks after deleting less than one fifth of global max:18
×10^6
raw data, starting:18
fitted Gauss locations:18
y
x
0
2
4
6
8
10
12
14
0
5
10
15
20
25

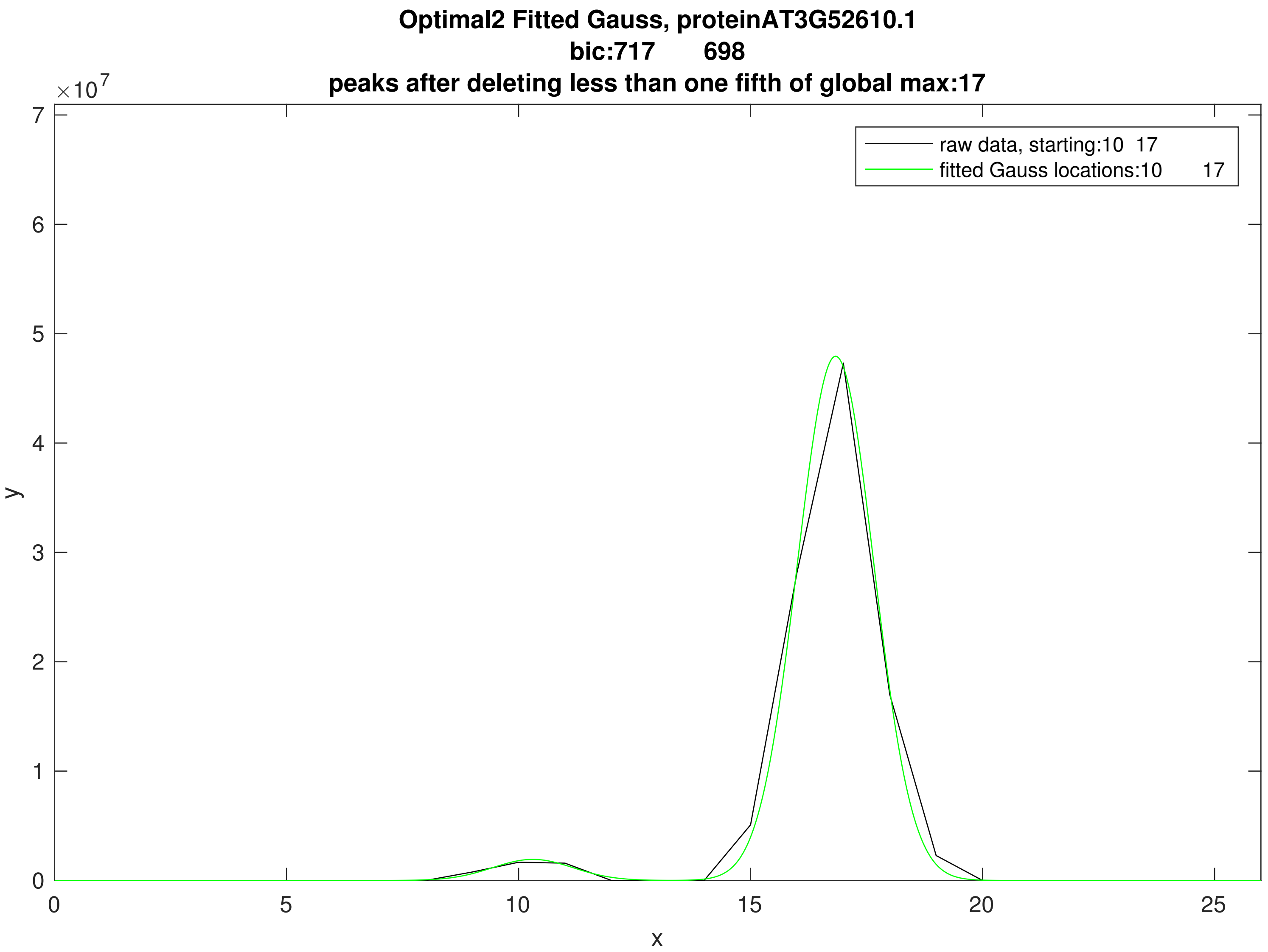

Optimal2 Fitted Gauss, proteinAT3G52610.1
bic:717 698
peaks after deleting less than one fifth of global max:17
×10^7
raw data, starting:10 17
fitted Gauss locations:10 17
y
x
0
1
2
3
4
5
6
7
0
5
10
15
20
25

Optimal1 Fitted Gauss, proteinAT3G52630.2;AT3G52630.1
bic:735
peaks after deleting less than one fifth of global max:21

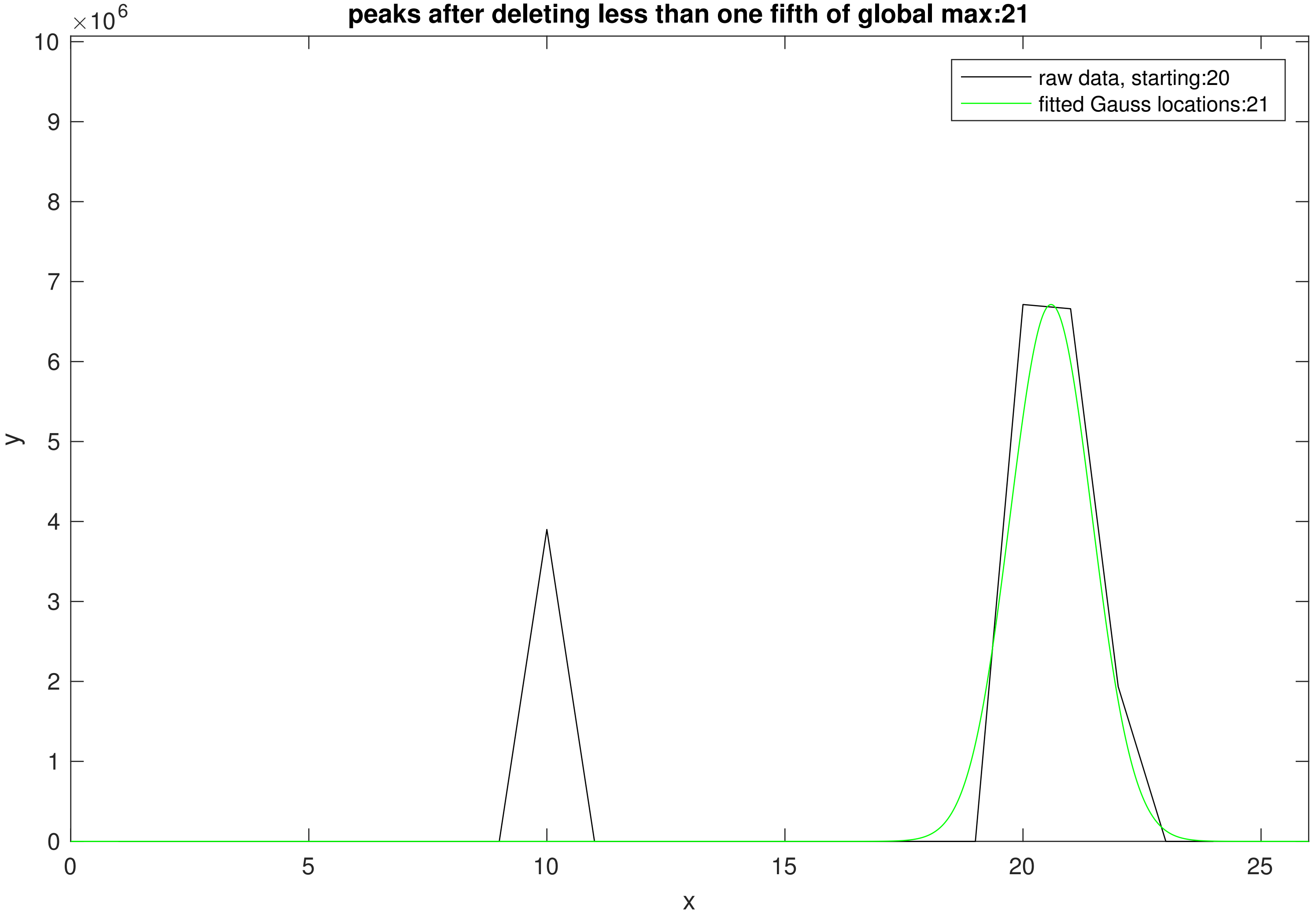

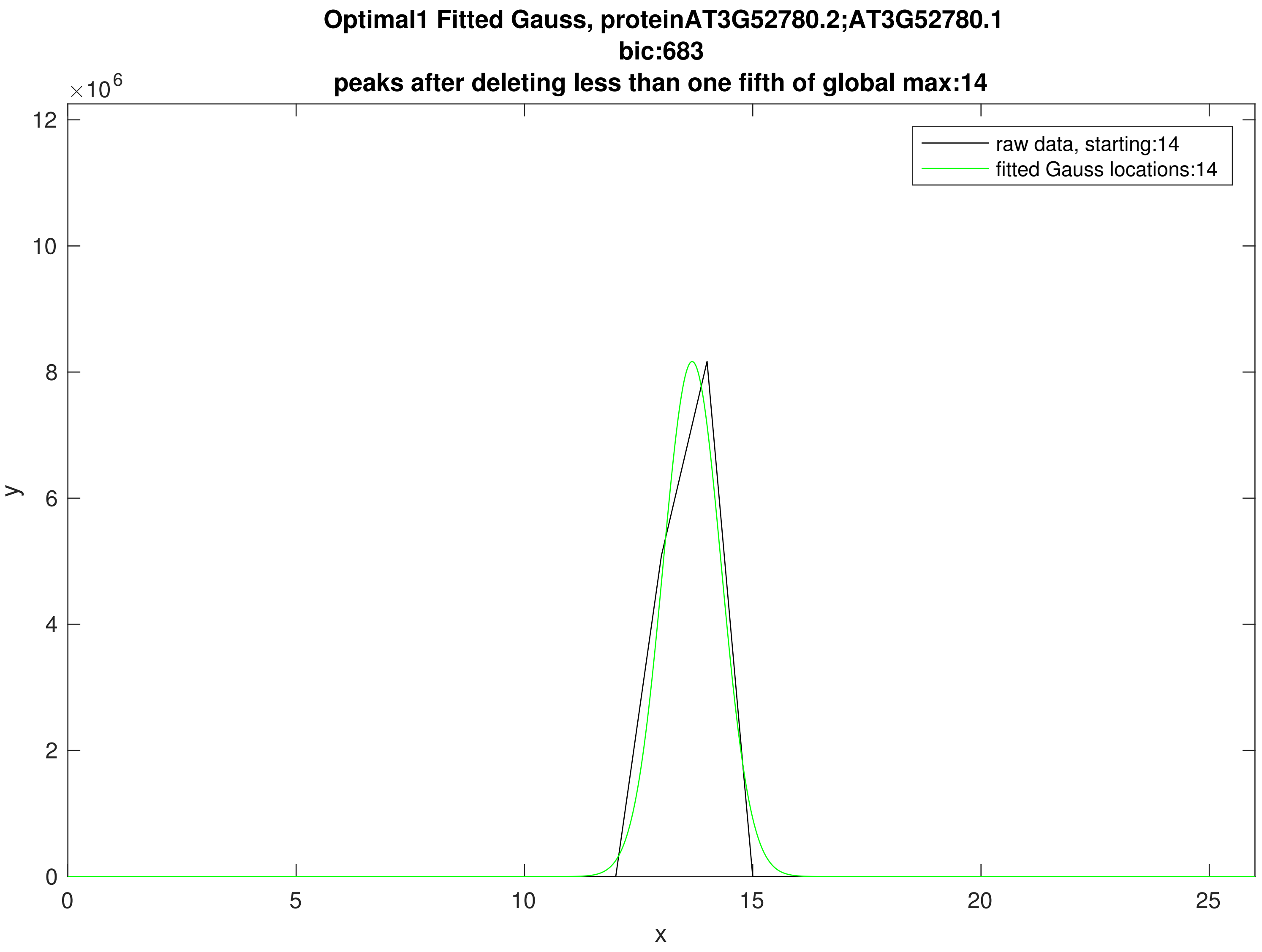
Optimal1 Fitted Gauss, proteinAT3G52780.2;AT3G52780.1
bic:683
peaks after deleting less than one fifth of global max:14
$\times 10^6$
raw data, starting:14
fitted Gauss locations:14
y
x
0
2
4
6
8
10
12
0
5
10
15
20
25

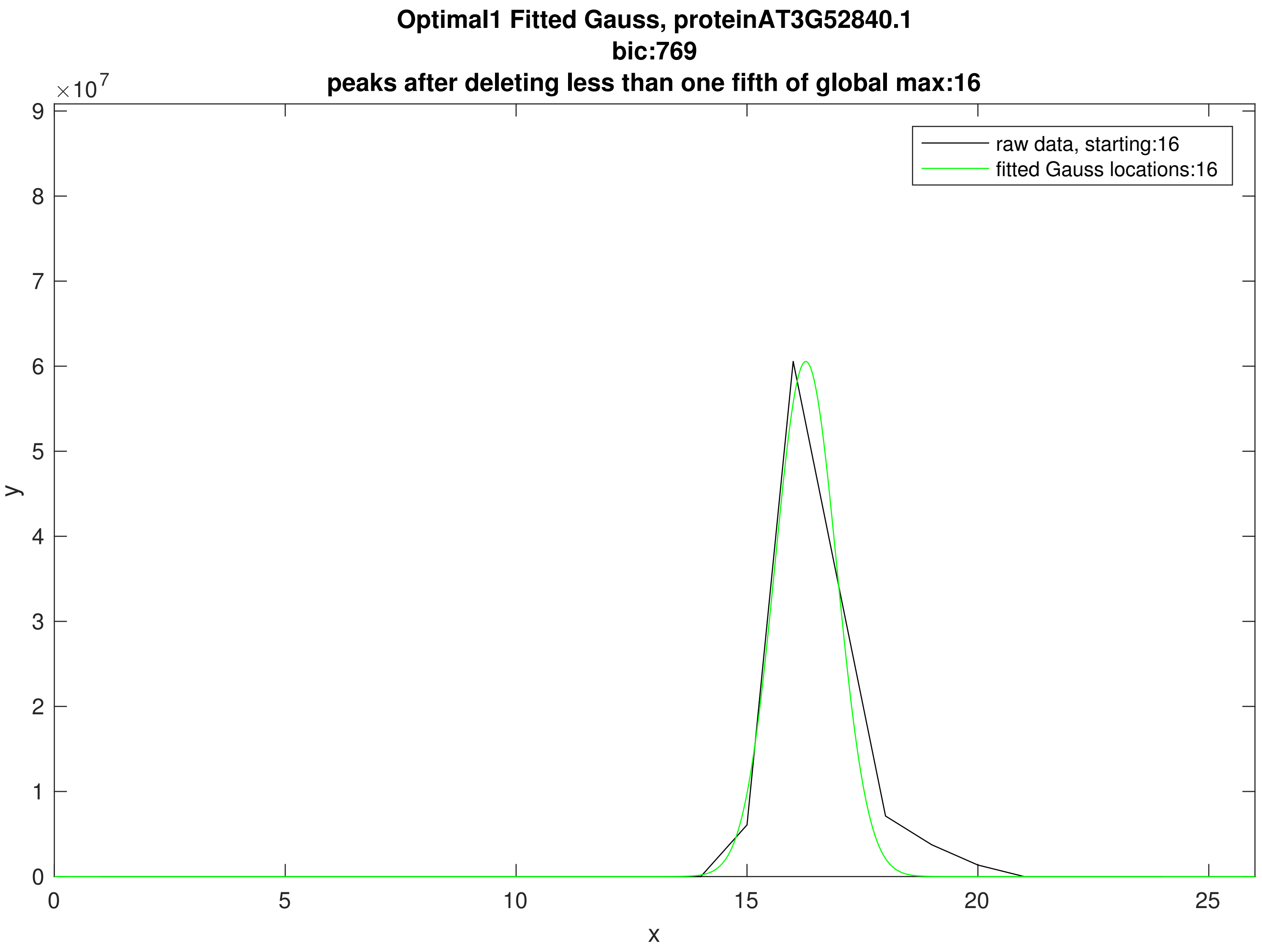

Optimal1 Fitted Gauss, proteinAT3G52840.1
bic:769
peaks after deleting less than one fifth of global max:16
×10^7
9
8
7
6
5
4
3
2
1
0
y
0
5
10
15
20
25
x
raw data, starting:16
fitted Gauss locations:16

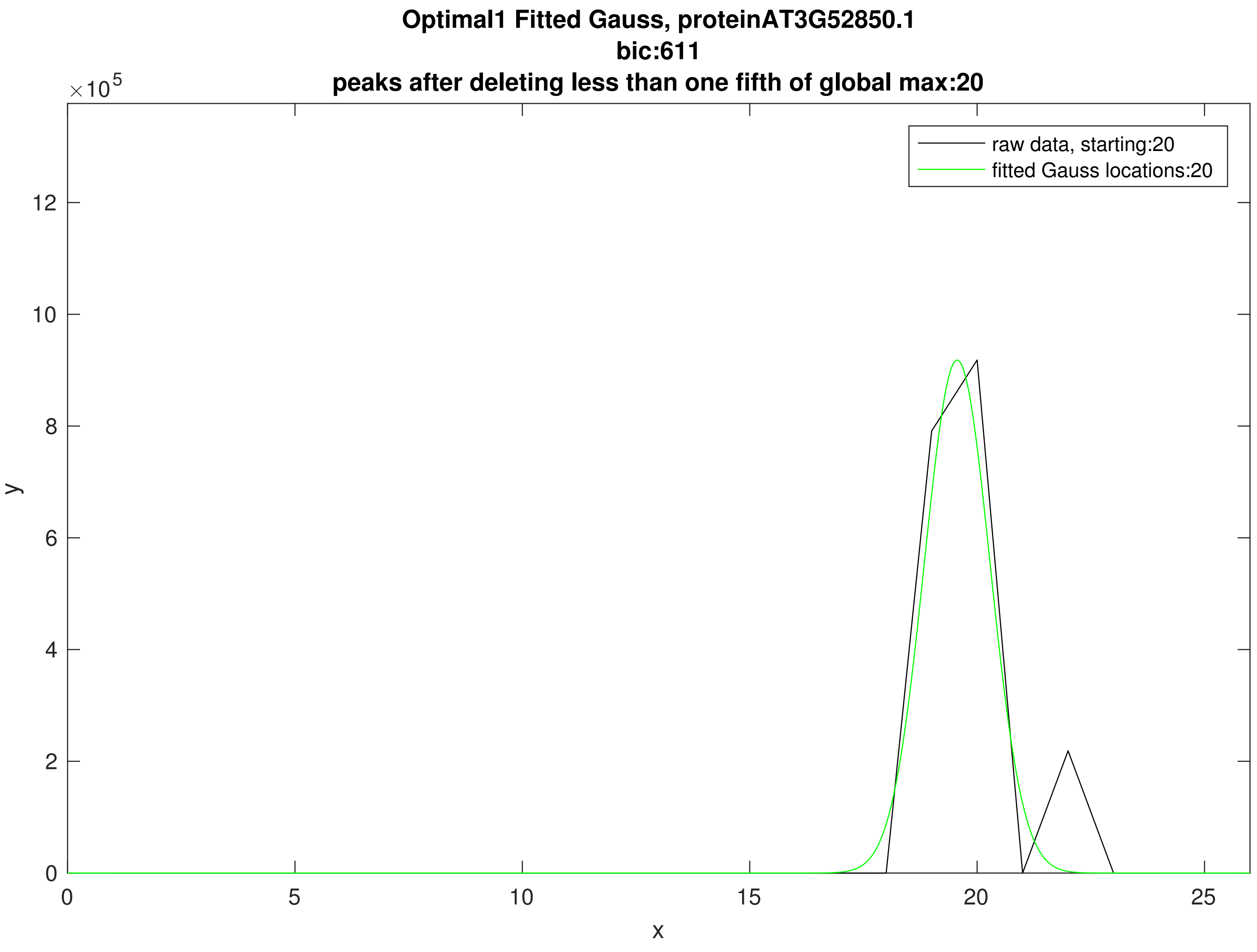
Optimal1 Fitted Gauss, proteinAT3G52850.1
bic:611
peaks after deleting less than one fifth of global max:20
×10^5
raw data, starting:20
fitted Gauss locations:20
y
x
0
2
4
6
8
10
12
0
5
10
15
20
25

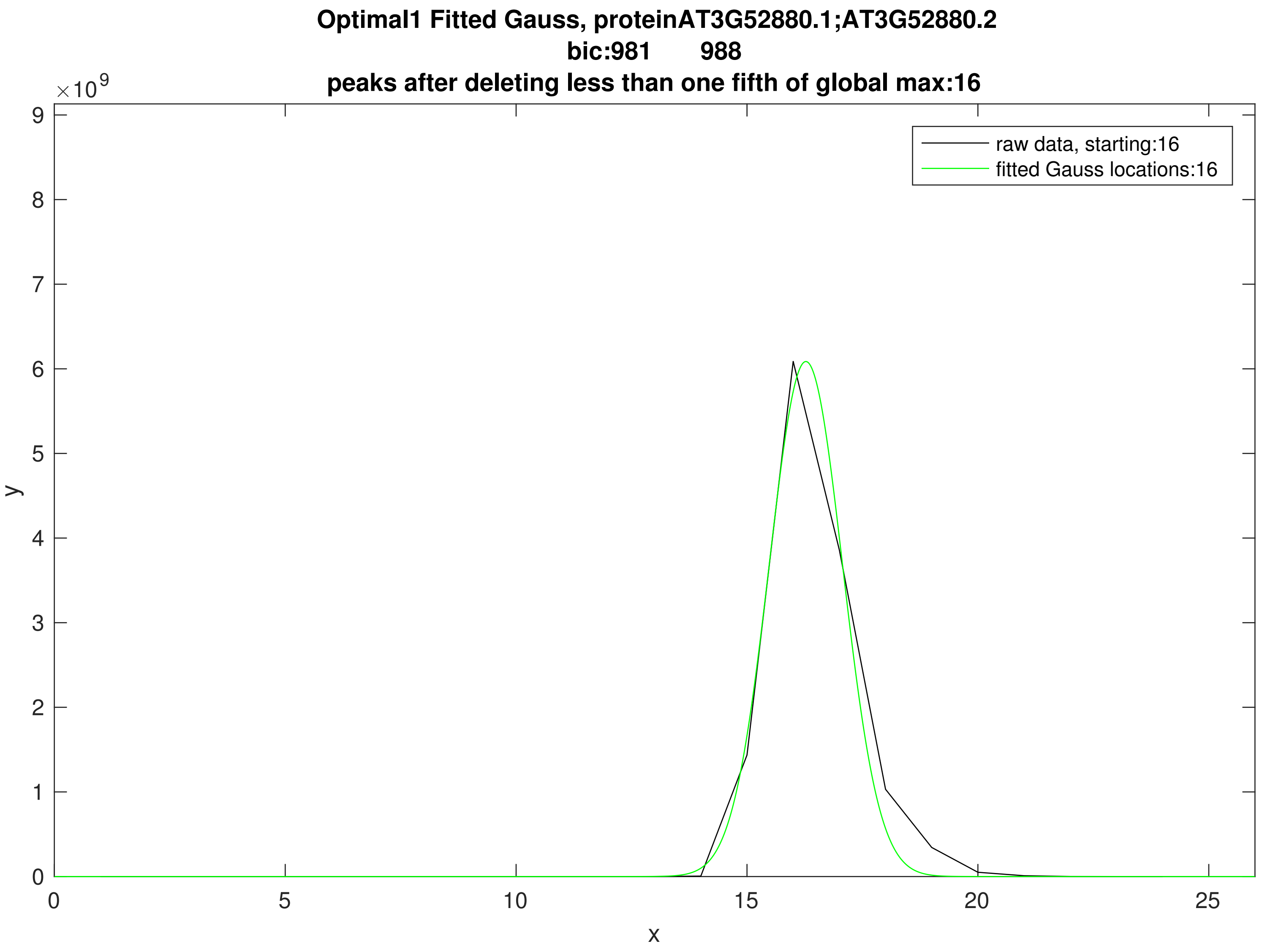
Optimal1 Fitted Gauss, proteinAT3G52880.1;AT3G52880.2
bic:981 988
peaks after deleting less than one fifth of global max:16
×10^9
raw data, starting:16
fitted Gauss locations:16
y
x
0
5
10
15
20
25
0
1
2
3
4
5
6
7
8
9

**Optimal3 Fitted Gauss, proteinAT3G52930.1**
**bic:1.1e+03  1.07e+03  1.06e+03  1.09e+03**
**peaks after deleting less than one fifth of global max:11**

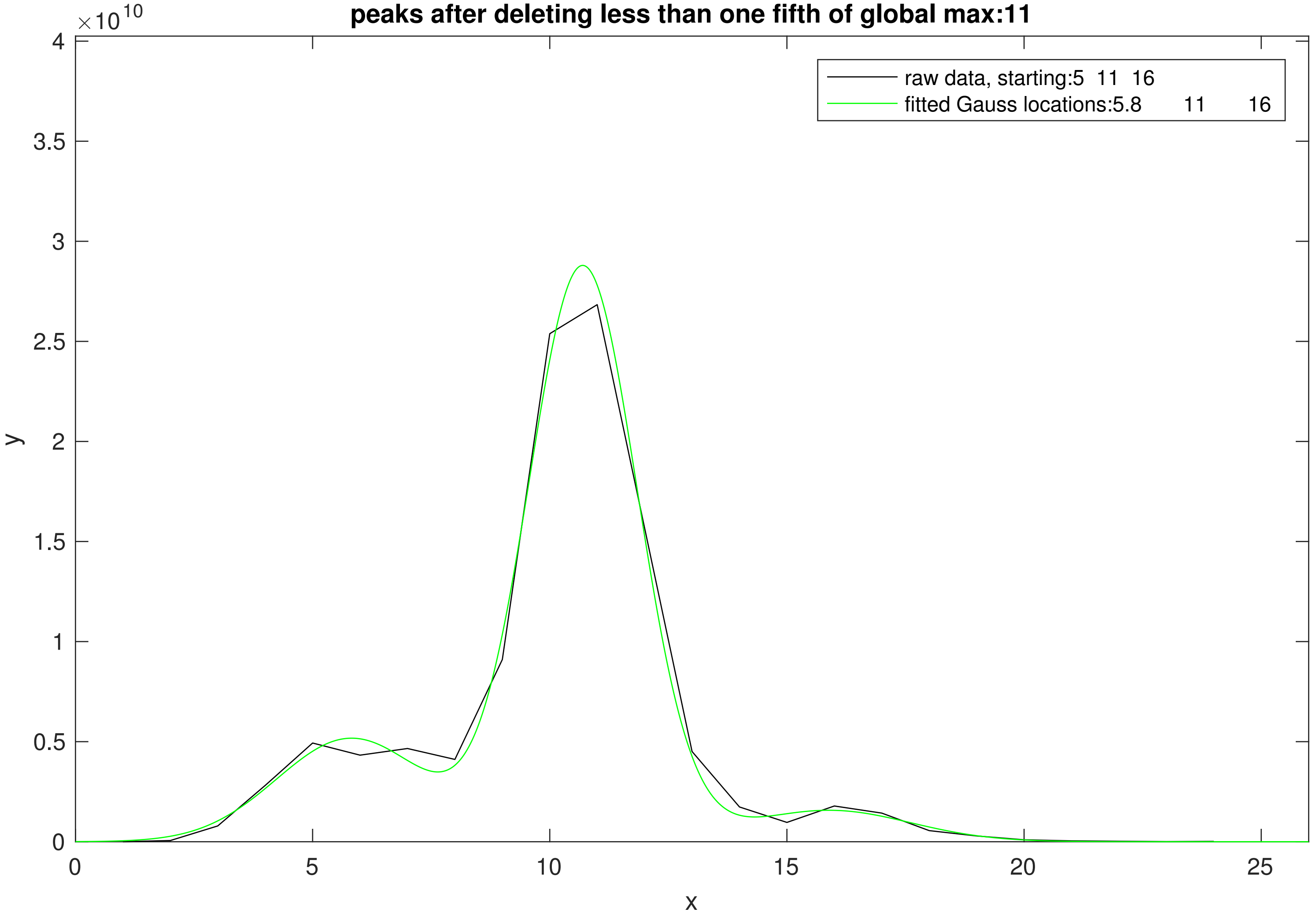

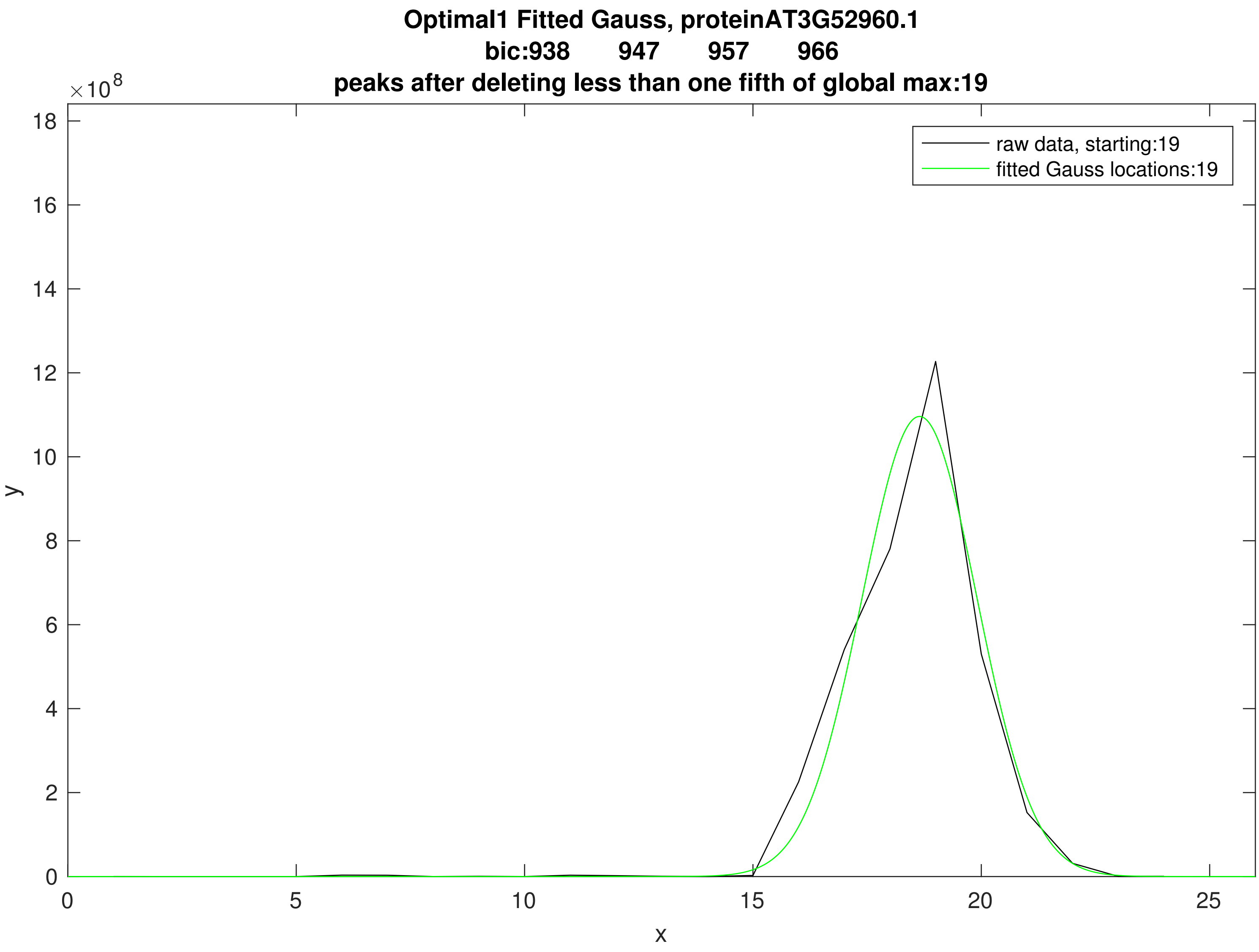

Optimal1 Fitted Gauss, proteinAT3G52960.1
bic:938 947 957 966
peaks after deleting less than one fifth of global max:19
×10^8
18
16
14
12
10
8
6
4
2
0
y
0
5
10
15
20
25
x
raw data, starting:19
fitted Gauss locations:19

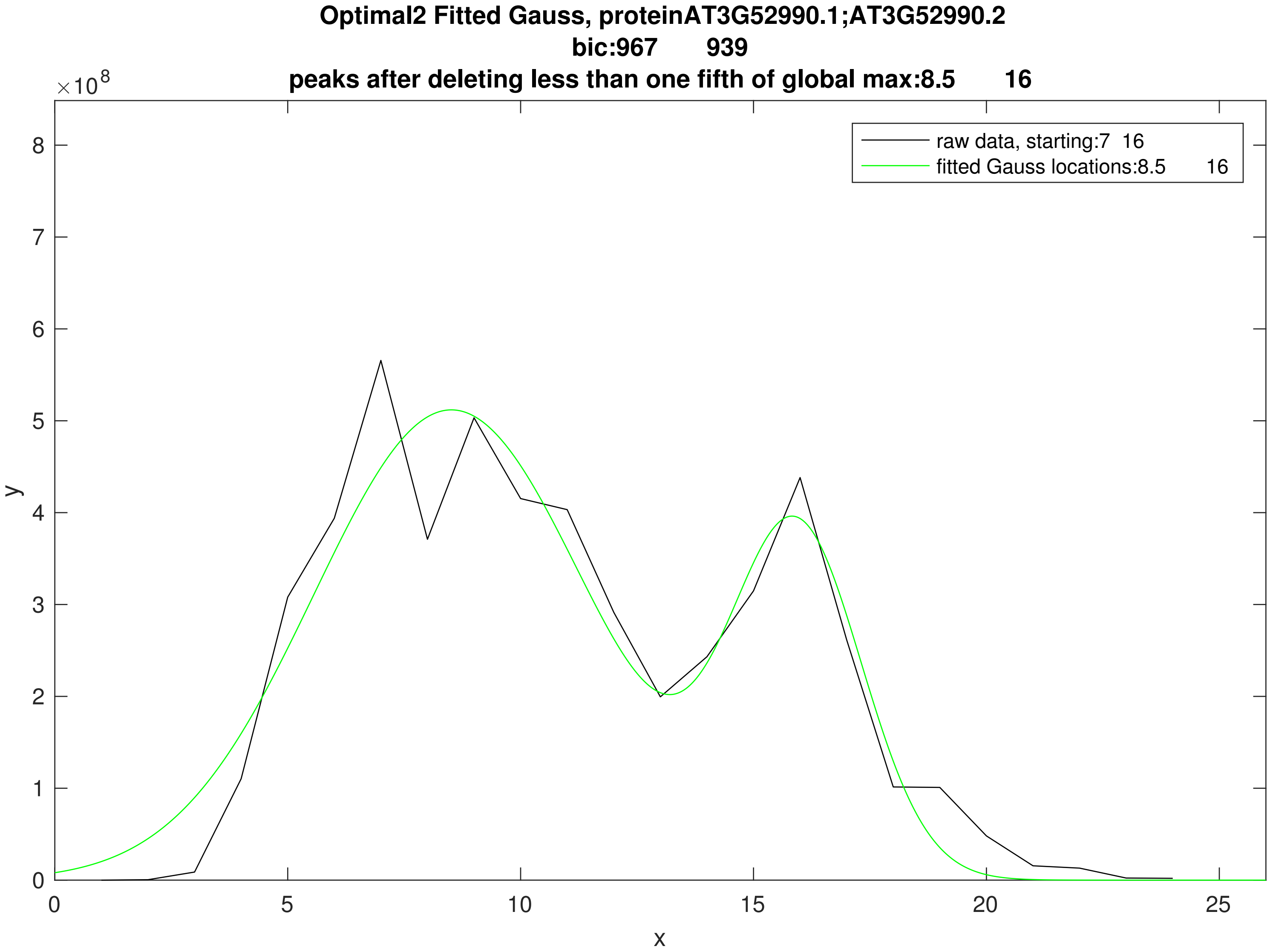
Optimal2 Fitted Gauss, proteinAT3G52990.1;AT3G52990.2
bic:967 939
peaks after deleting less than one fifth of global max:8.5 16
×10^8
raw data, starting:7 16
fitted Gauss locations:8.5 16
y
x

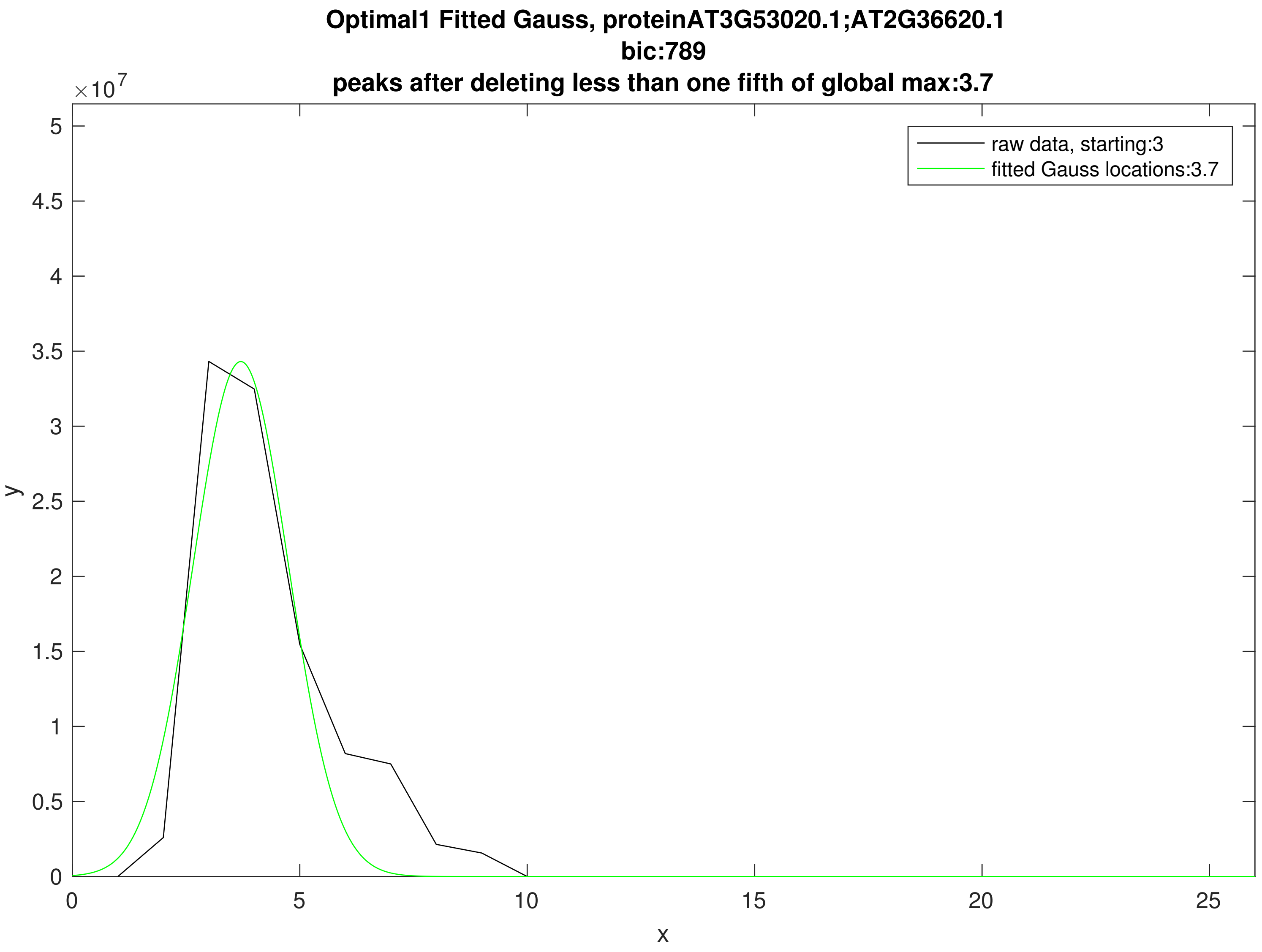
Optimal1 Fitted Gauss, proteinAT3G53020.1;AT2G36620.1
bic:789
peaks after deleting less than one fifth of global max:3.7
×10^7
raw data, starting:3
fitted Gauss locations:3.7
y
x
0
0.5
1
1.5
2
2.5
3
3.5
4
4.5
5
0
5
10
15
20
25

**Optimal1 Fitted Gauss, proteinAT3G53040.1**
**bic:897 899 907 917**
**peaks after deleting less than one fifth of global max:7.7**

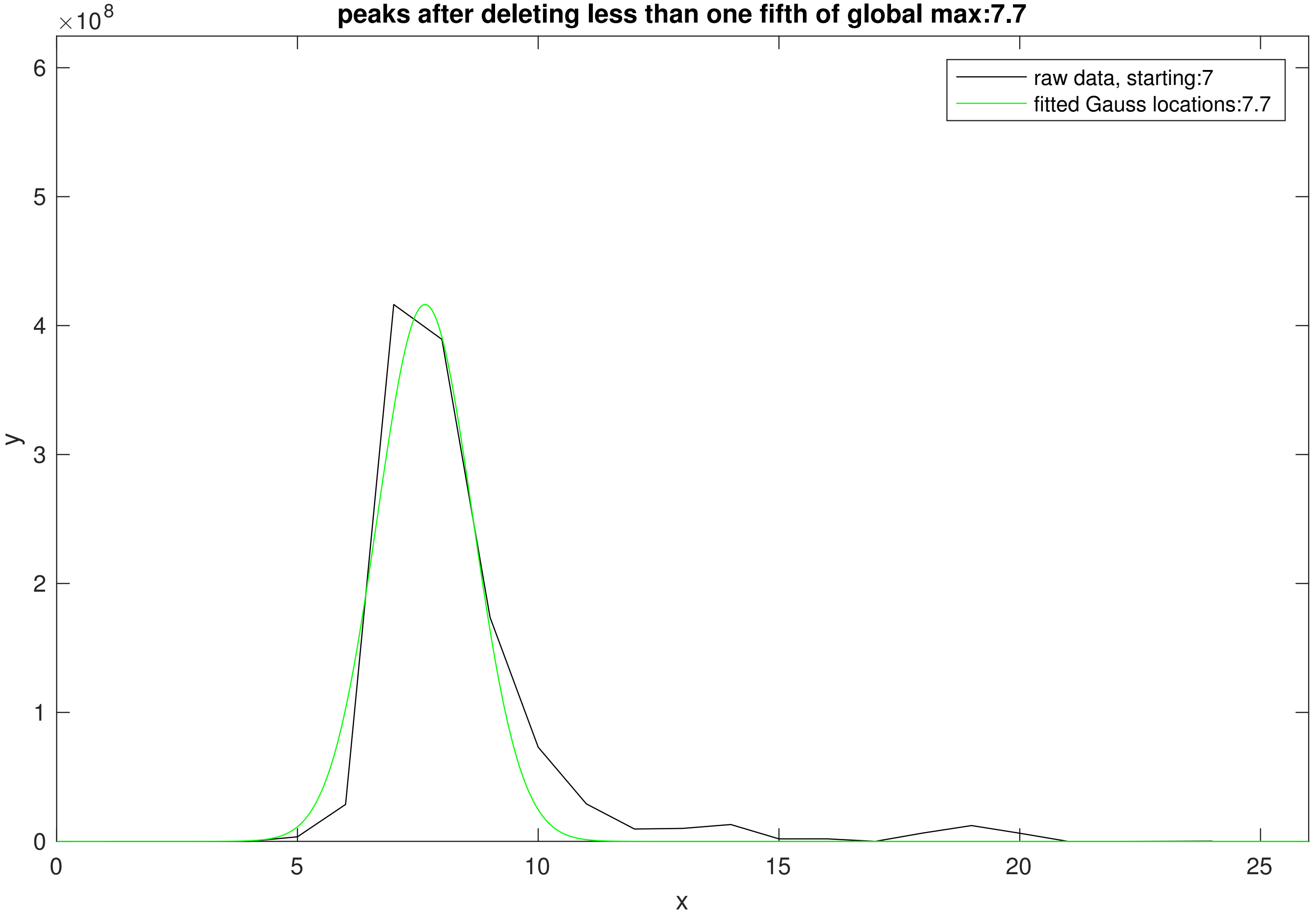

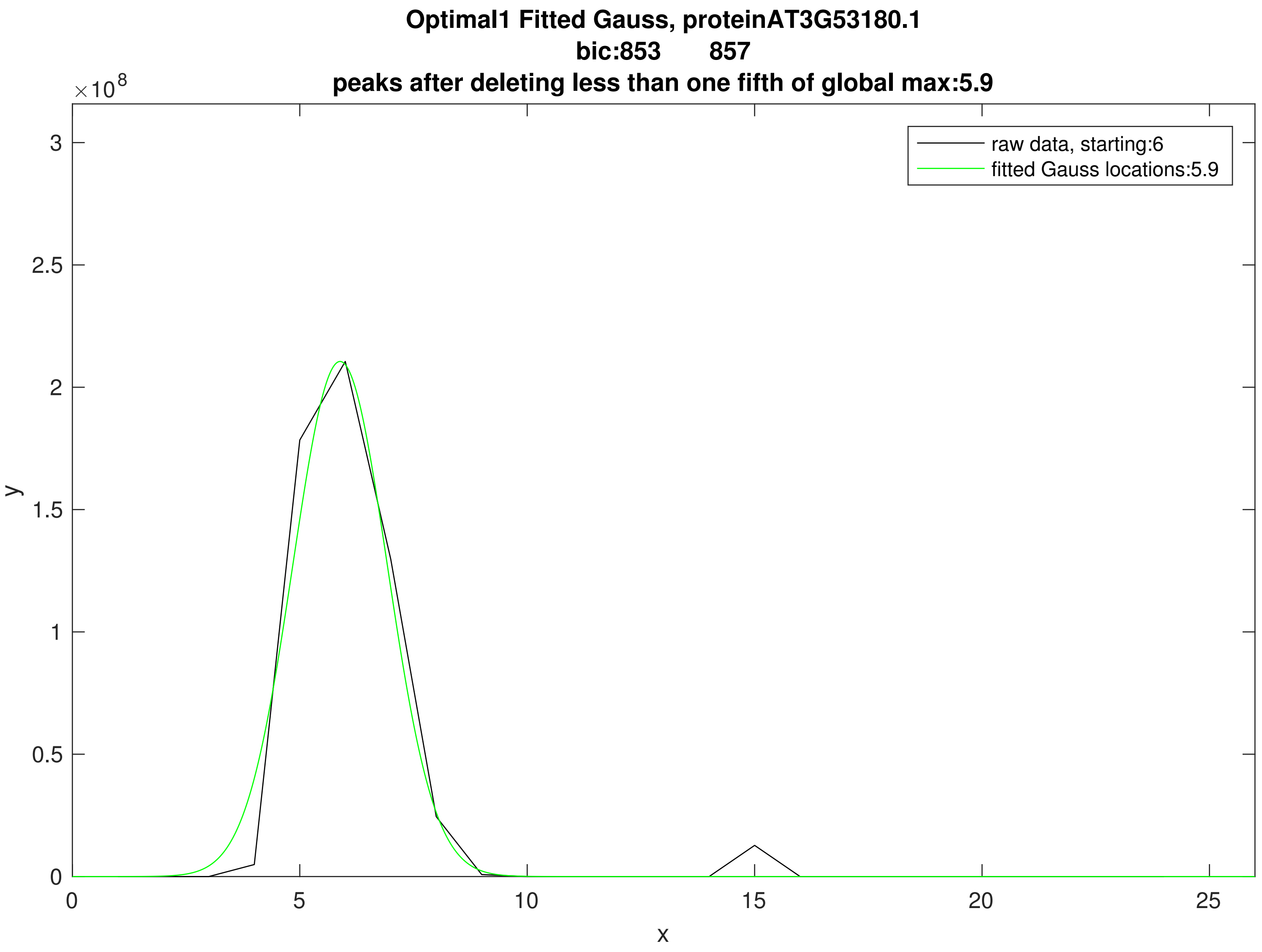

Optimal1 Fitted Gauss, proteinAT3G53180.1
bic:853 857
peaks after deleting less than one fifth of global max:5.9
×10^8
3
2.5
2
1.5
1
0.5
0
y
0
5
10
15
20
25
x
raw data, starting:6
fitted Gauss locations:5.9

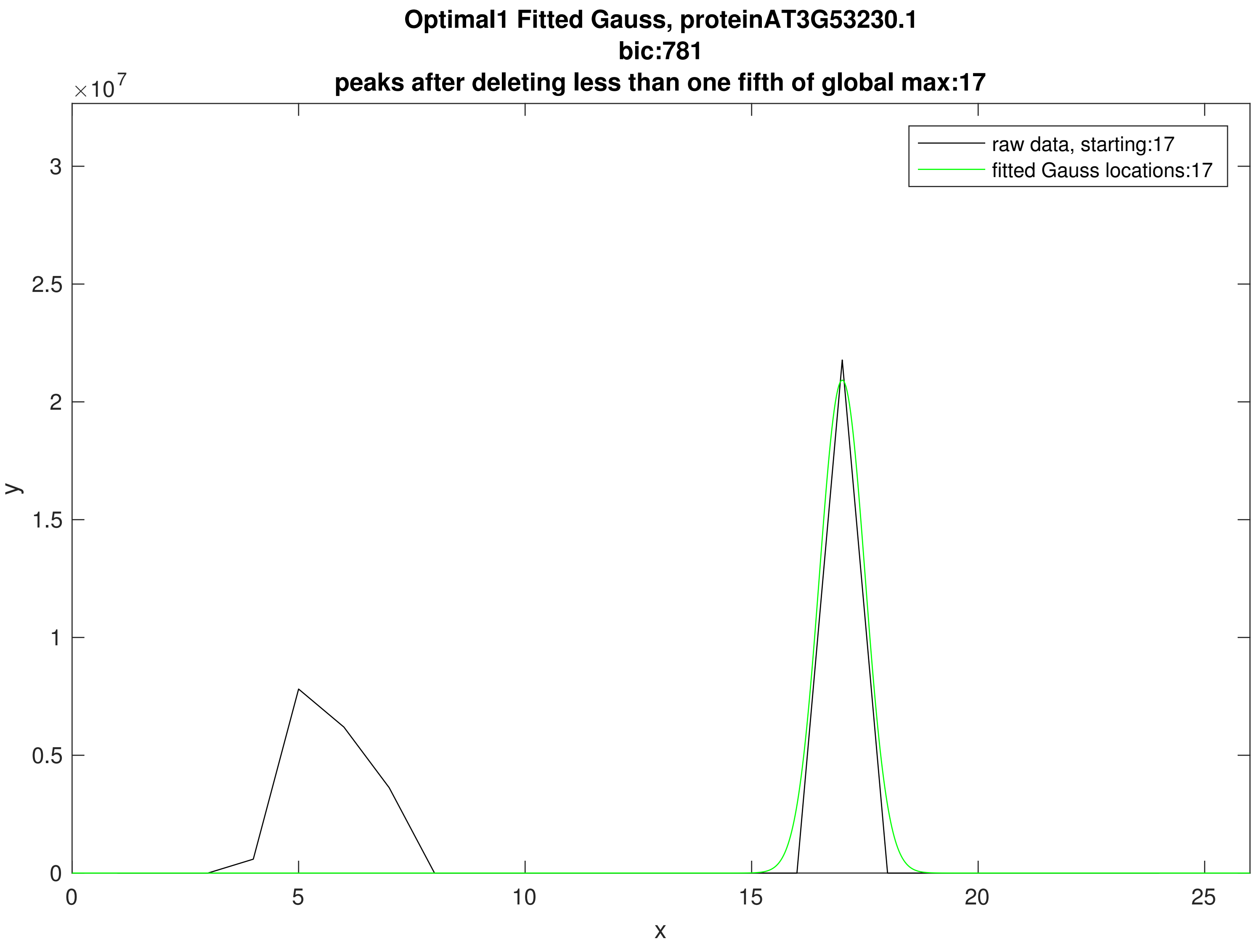
Optimal1 Fitted Gauss, proteinAT3G53230.1
bic:781
peaks after deleting less than one fifth of global max:17
×10^7
3
2.5
2
1.5
1
0.5
0
y
0
5
10
15
20
25
x
raw data, starting:17
fitted Gauss locations:17

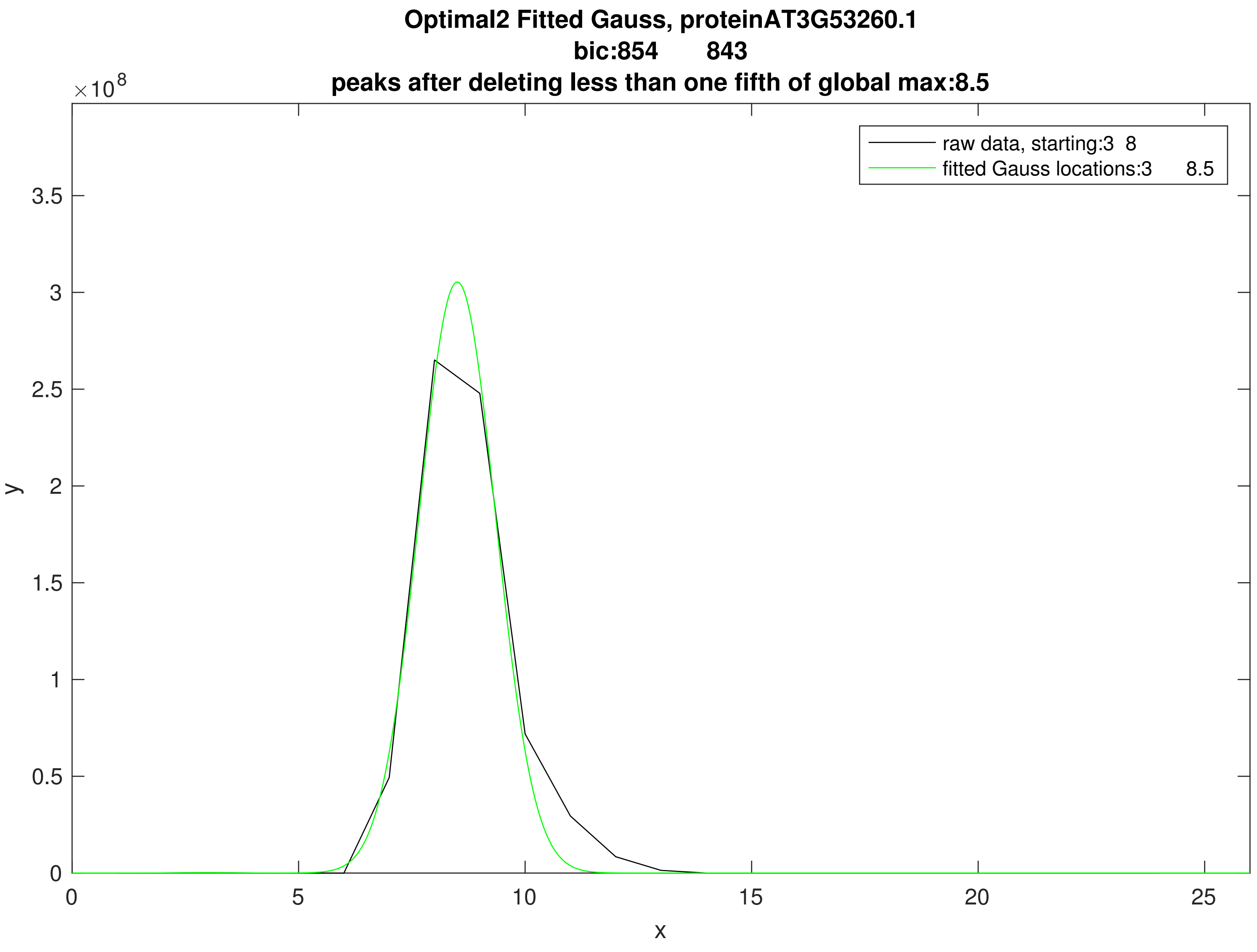

Optimal2 Fitted Gauss, proteinAT3G53260.1
bic:854 843
peaks after deleting less than one fifth of global max:8.5
×10^8
raw data, starting:3 8
fitted Gauss locations:3 8.5
y
x
0
0.5
1
1.5
2
2.5
3
3.5
5
10
15
20
25

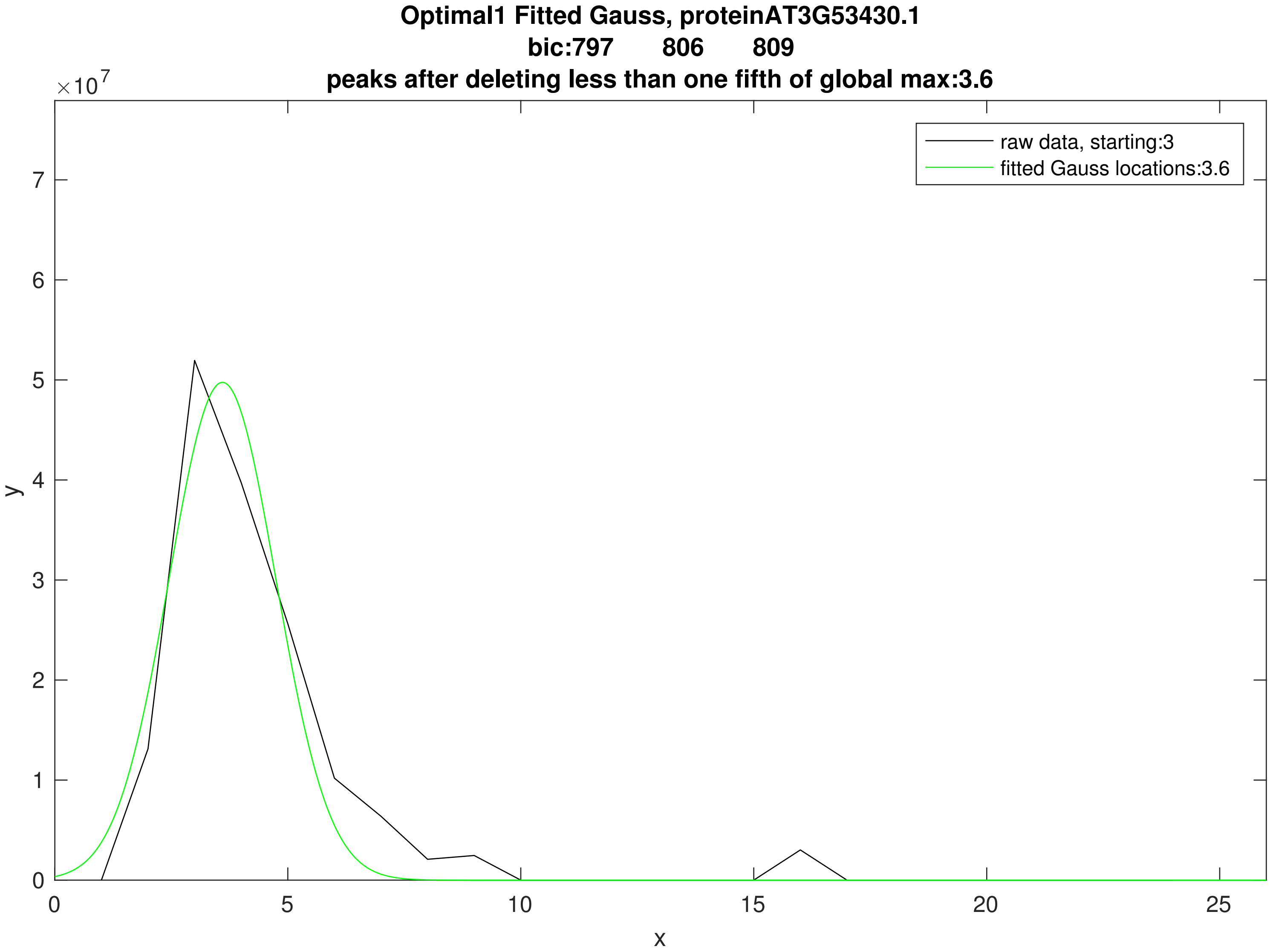

Optimal1 Fitted Gauss, proteinAT3G53430.1
bic:797 806 809
peaks after deleting less than one fifth of global max:3.6
×10^7
raw data, starting:3
fitted Gauss locations:3.6
y
x
0
1
2
3
4
5
6
7
0
5
10
15
20
25

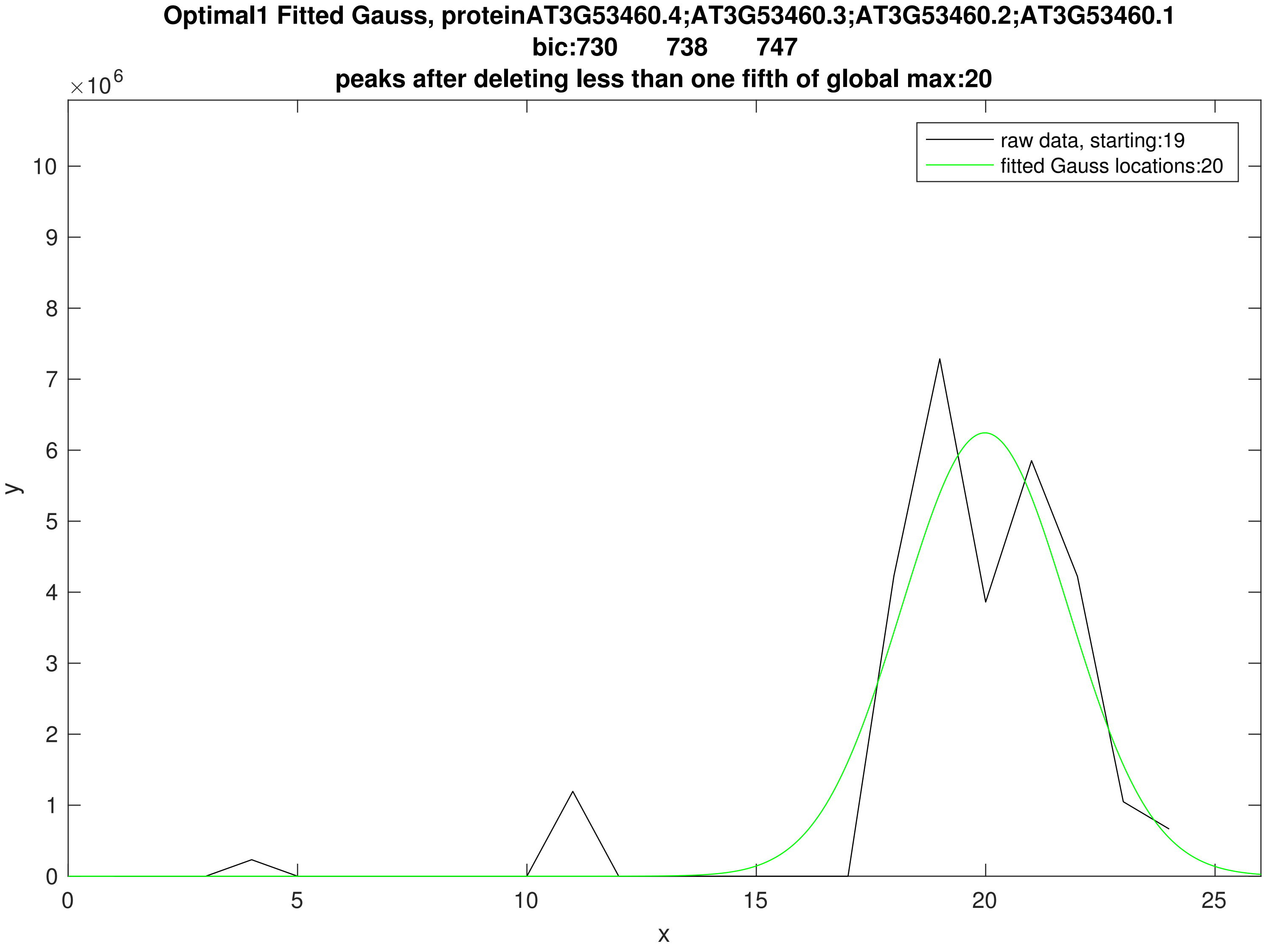
Optimal1 Fitted Gauss, proteinAT3G53460.4;AT3G53460.3;AT3G53460.2;AT3G53460.1
bic:730 738 747
peaks after deleting less than one fifth of global max:20
×10^6
raw data, starting:19
fitted Gauss locations:20
y
x
0
1
2
3
4
5
6
7
8
9
10
0
5
10
15
20
25

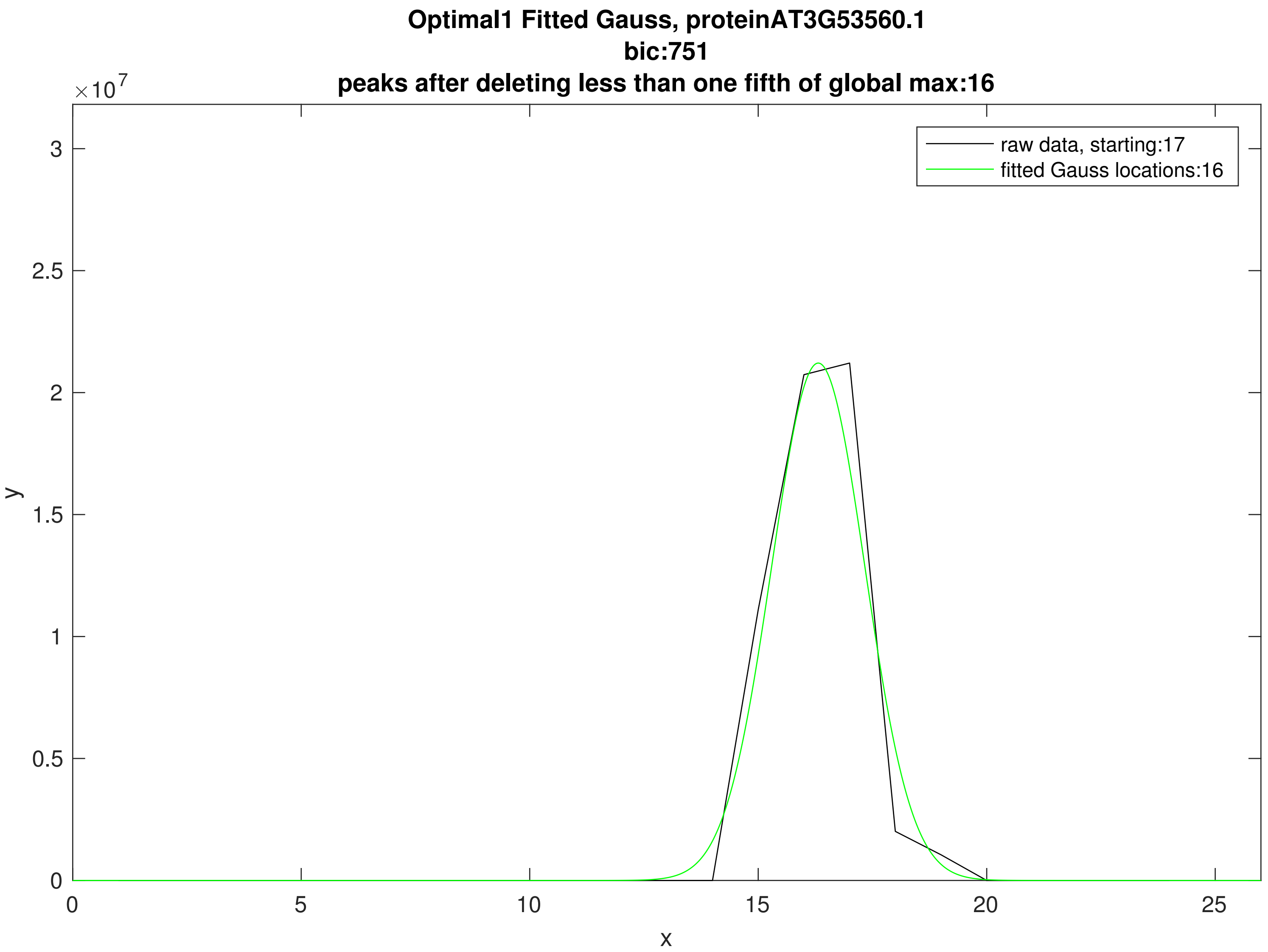
Optimal1 Fitted Gauss, proteinAT3G53560.1
bic:751
peaks after deleting less than one fifth of global max:16
×10^7
3
2.5
2
1.5
1
0.5
0
y
0
5
10
15
20
25
x
raw data, starting:17
fitted Gauss locations:16

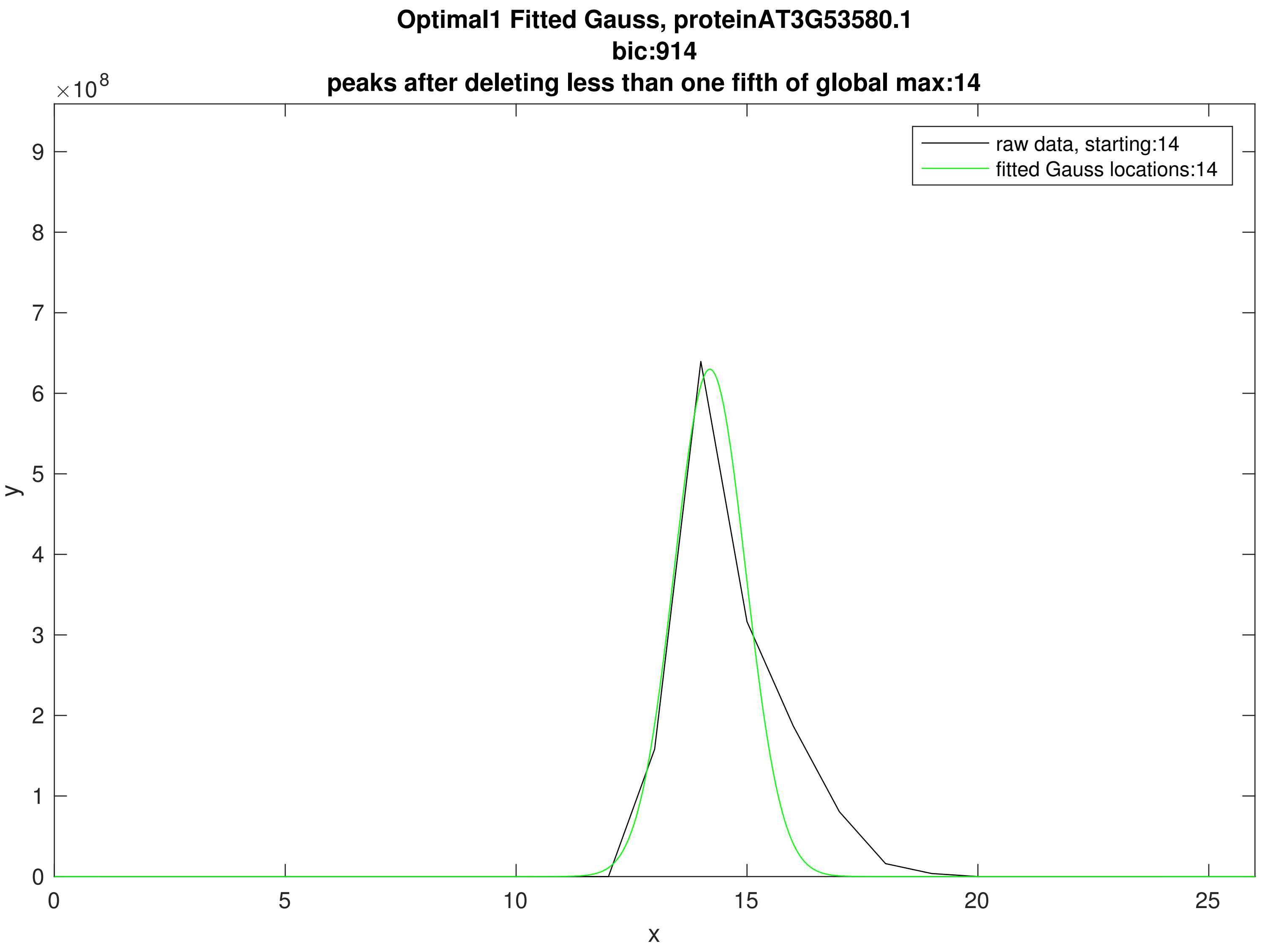
Optimal1 Fitted Gauss, proteinAT3G53580.1
bic:914
peaks after deleting less than one fifth of global max:14
×10^8
raw data, starting:14
fitted Gauss locations:14
y
x
0
1
2
3
4
5
6
7
8
9
0
5
10
15
20
25

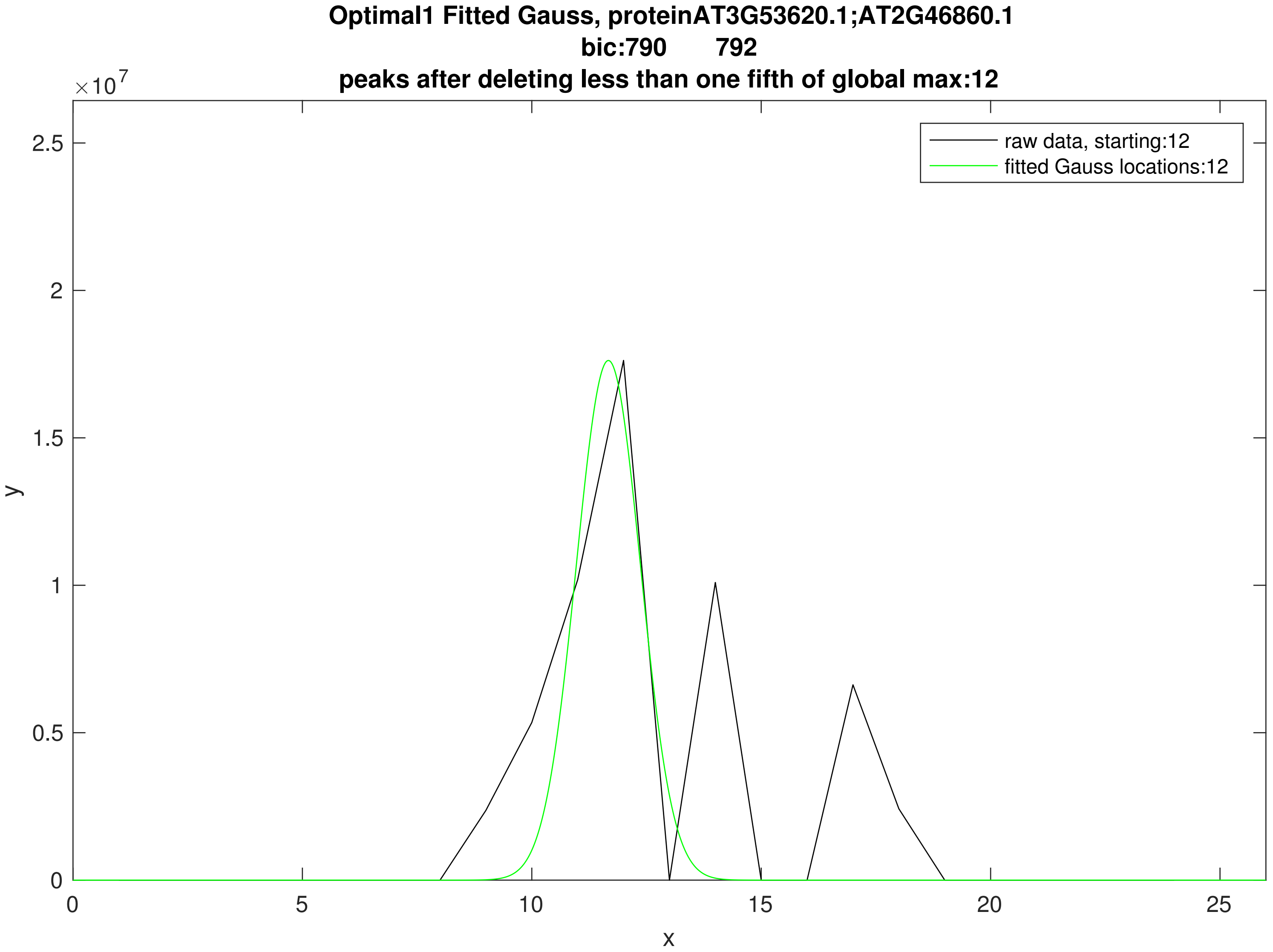

Optimal1 Fitted Gauss, proteinAT3G53620.1;AT2G46860.1
bic:790 792
peaks after deleting less than one fifth of global max:12
×10^7
2.5
2
1.5
1
0.5
0
y
0
5
10
15
20
25
x
raw data, starting:12
fitted Gauss locations:12

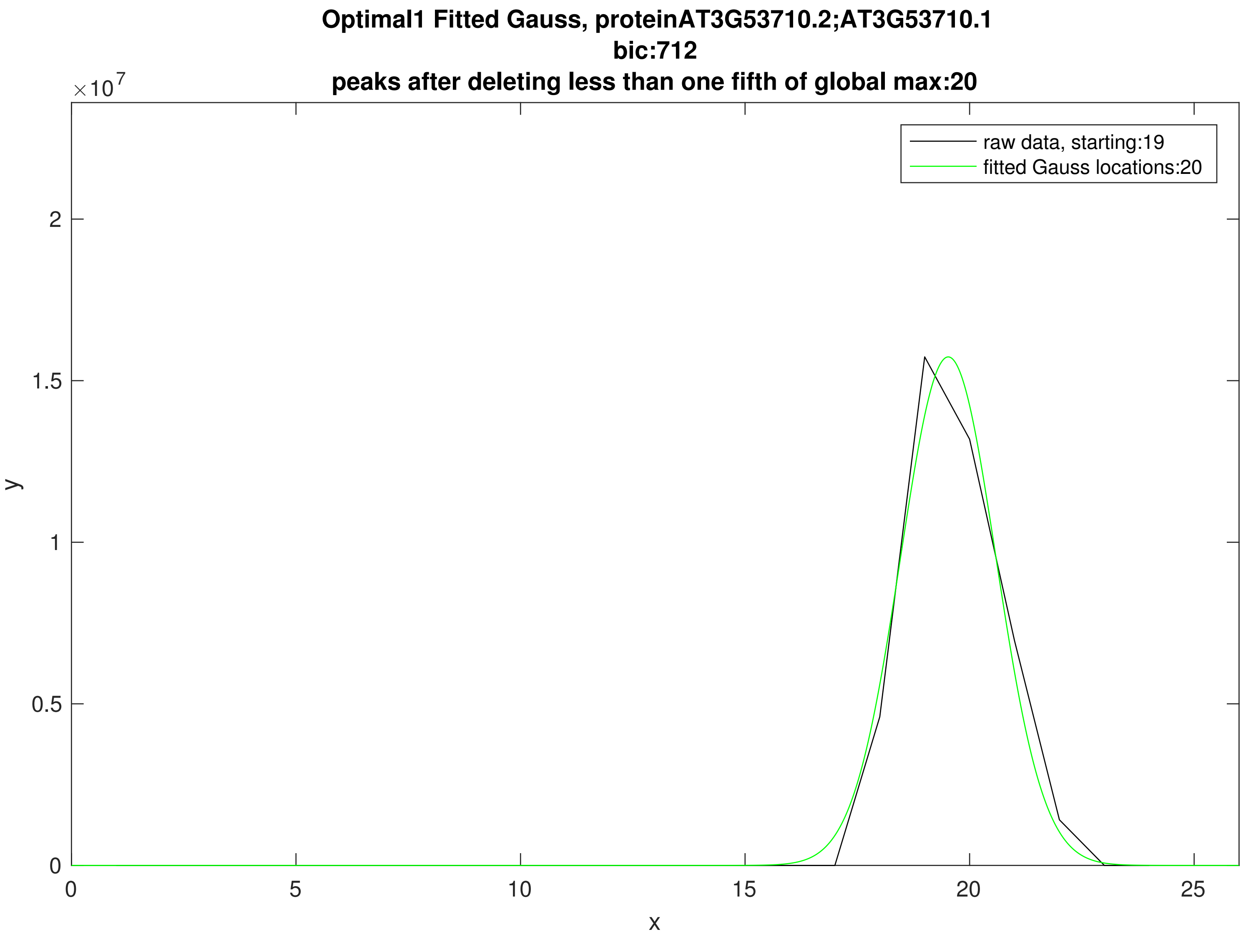
Optimal1 Fitted Gauss, proteinAT3G53710.2;AT3G53710.1
bic:712
peaks after deleting less than one fifth of global max:20
×10^7
raw data, starting:19
fitted Gauss locations:20
y
x
0
0.5
1
1.5
2
5
10
15
20
25

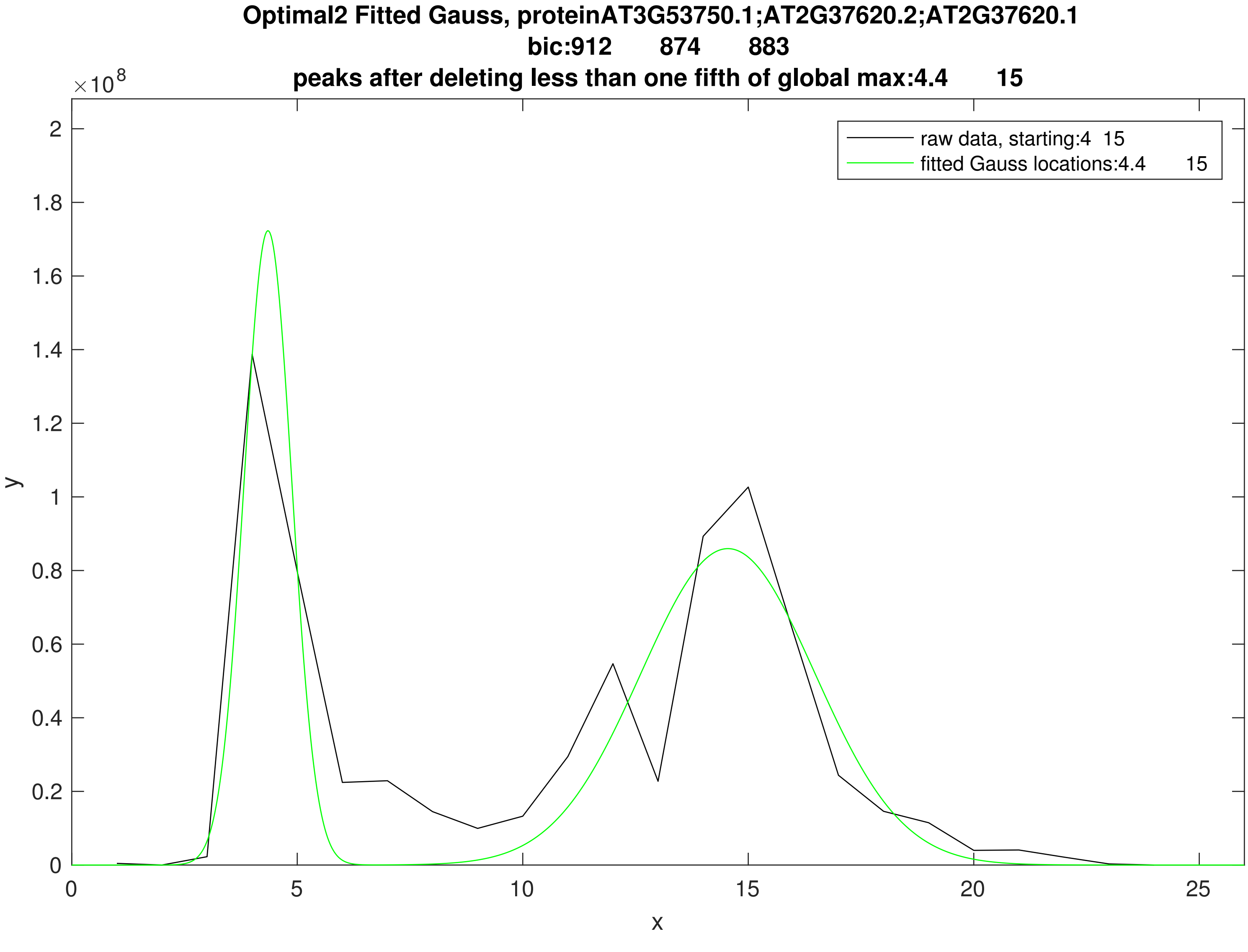

Optimal2 Fitted Gauss, proteinAT3G53750.1;AT2G37620.2;AT2G37620.1
bic:912 874 883
peaks after deleting less than one fifth of global max:4.4 15
×10^8
raw data, starting:4 15
fitted Gauss locations:4.4 15
y
x
0
0.2
0.4
0.6
0.8
1
1.2
1.4
1.6
1.8
2
5
10
15
20
25

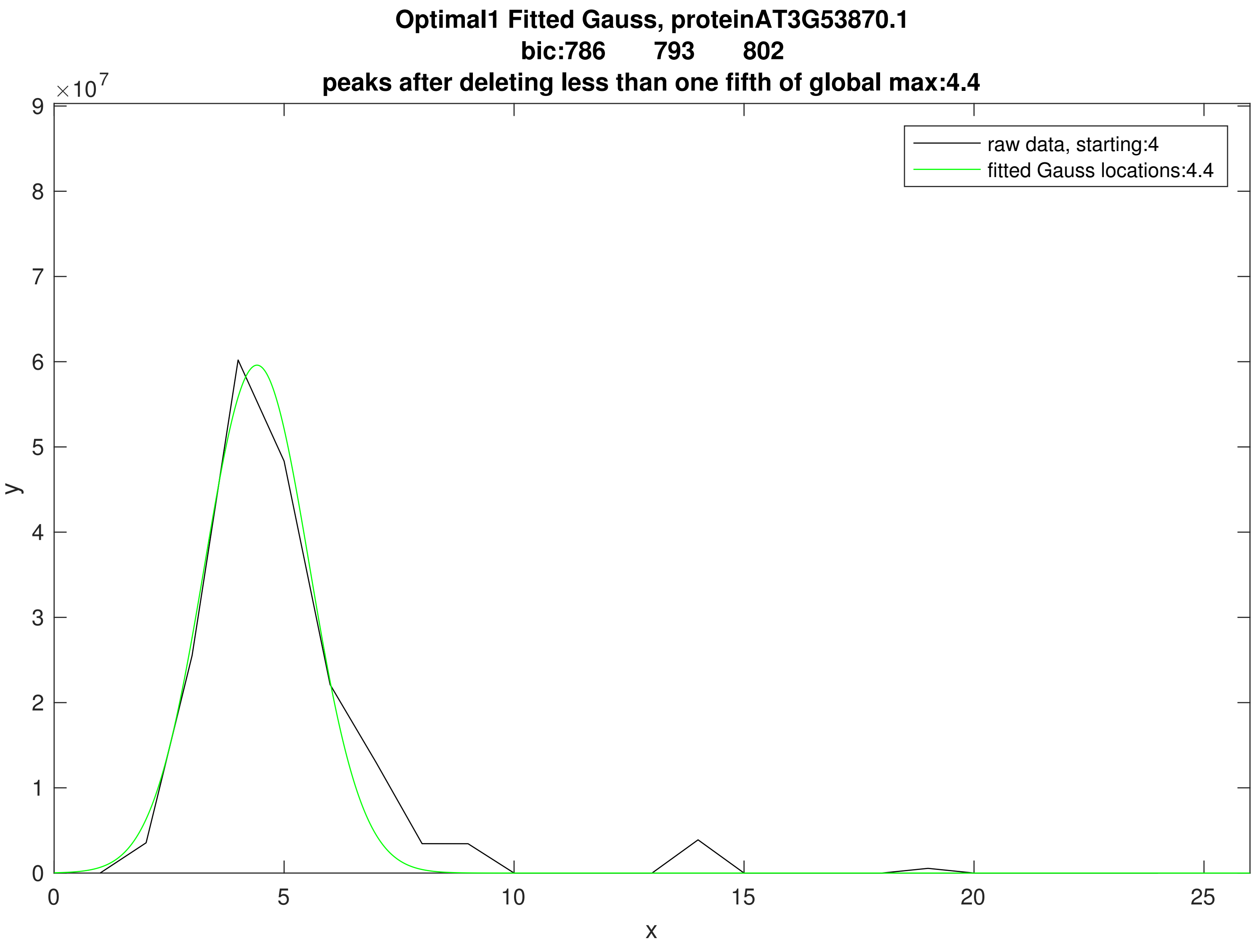
Optimal1 Fitted Gauss, proteinAT3G53870.1
bic:786 793 802
peaks after deleting less than one fifth of global max:4.4
×10^7
9
8
7
6
5
4
3
2
1
0
y
0
5
10
15
20
25
x
raw data, starting:4
fitted Gauss locations:4.4

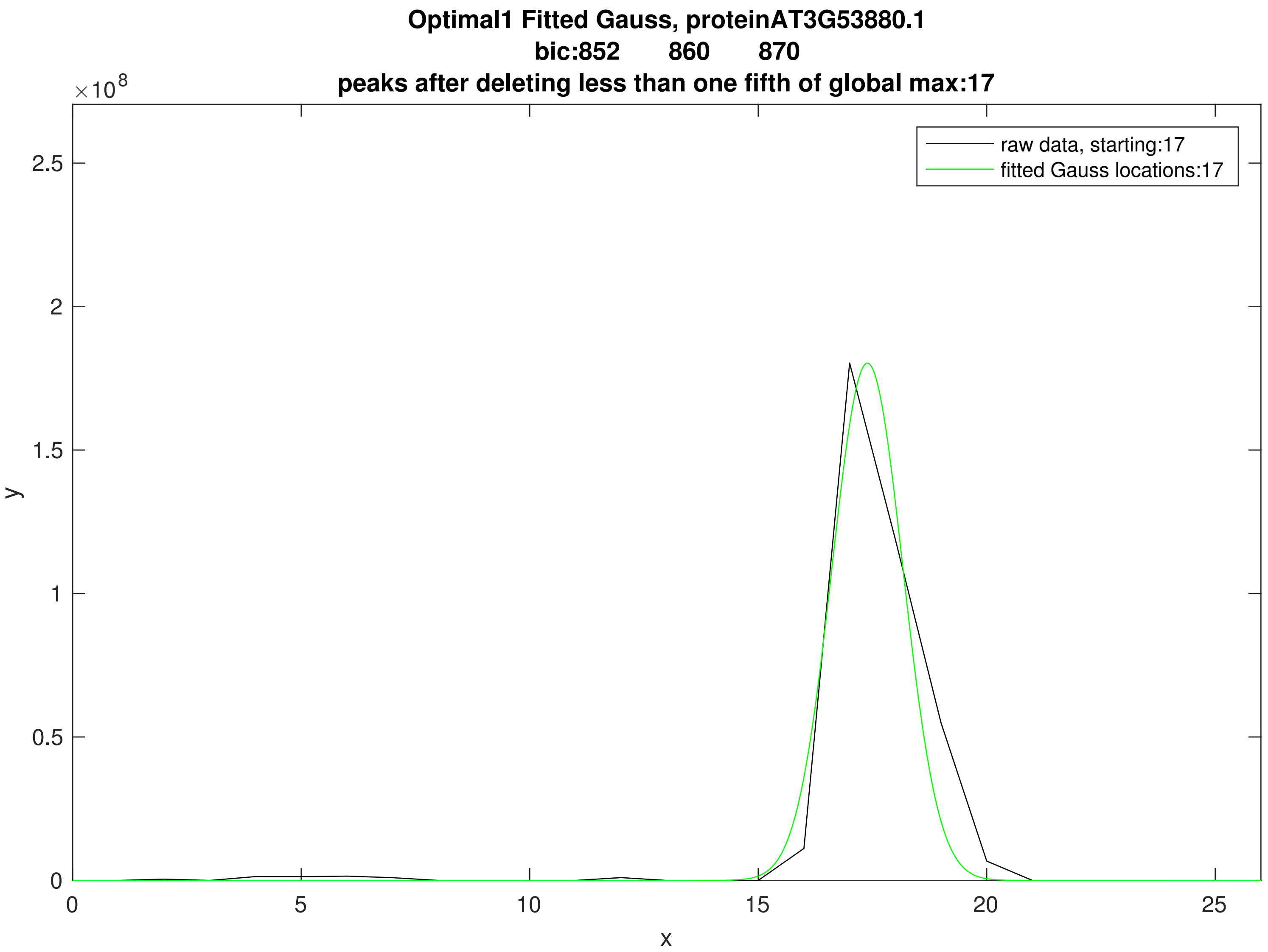
Optimal1 Fitted Gauss, proteinAT3G53880.1
bic:852 860 870
peaks after deleting less than one fifth of global max:17
×10^8
raw data, starting:17
fitted Gauss locations:17
y
x
0
0.5
1
1.5
2
2.5
5
10
15
20
25

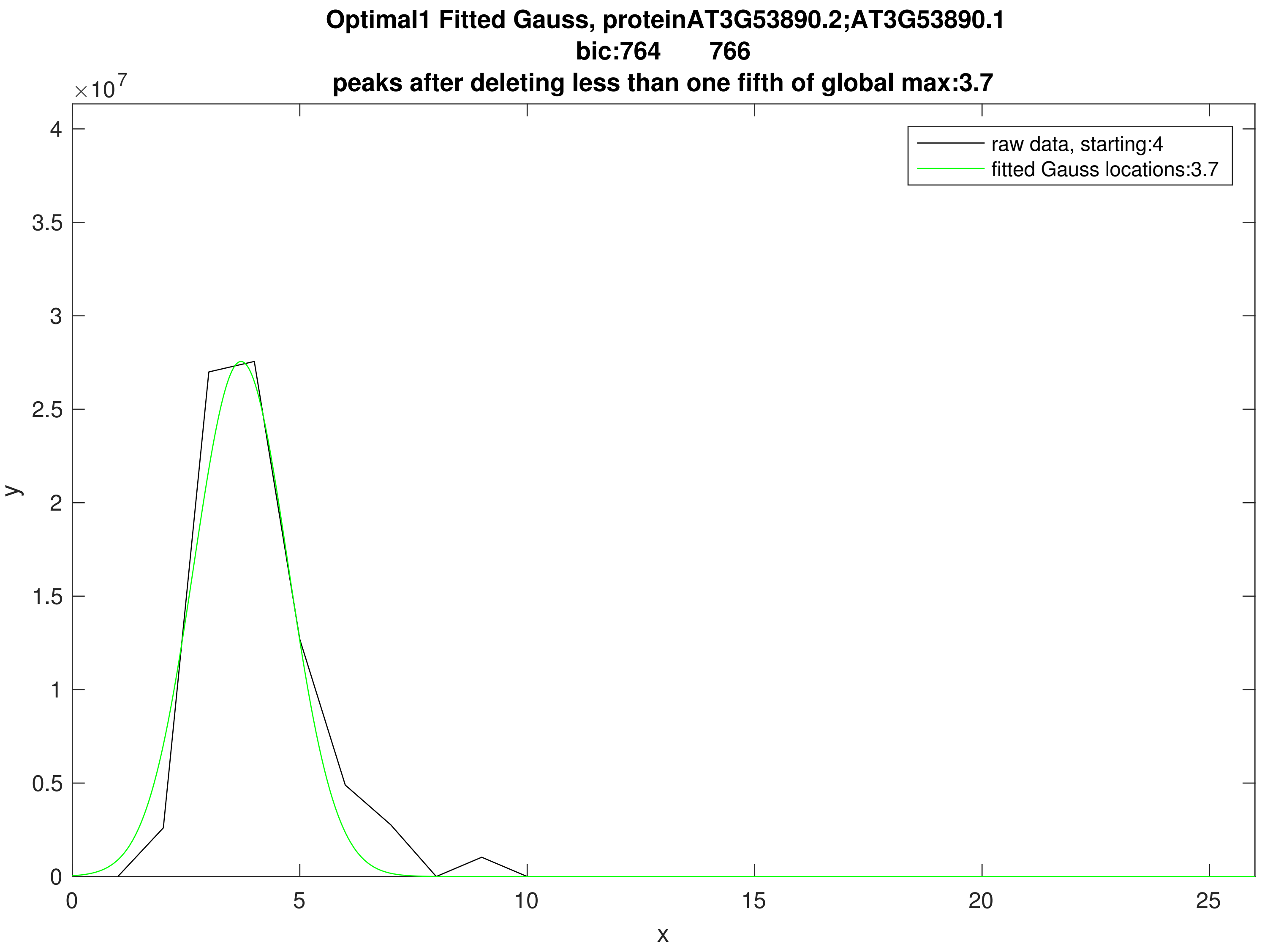

Optimal1 Fitted Gauss, proteinAT3G53890.2;AT3G53890.1
bic:764 766
peaks after deleting less than one fifth of global max:3.7
×10^7
raw data, starting:4
fitted Gauss locations:3.7
y
x
0
0.5
1
1.5
2
2.5
3
3.5
4
0
5
10
15
20
25

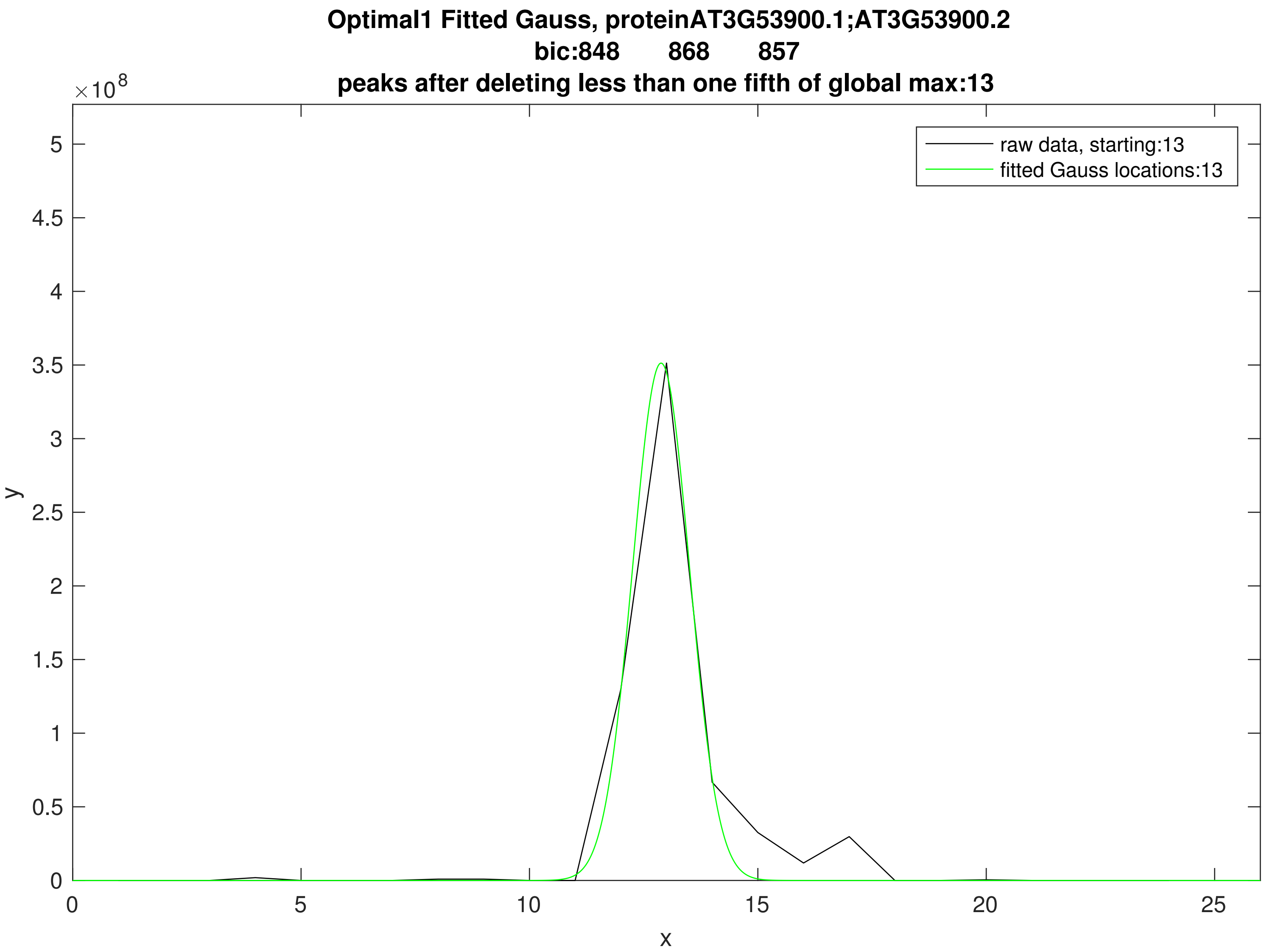

Optimal1 Fitted Gauss, proteinAT3G53900.1;AT3G53900.2
bic:848 868 857
peaks after deleting less than one fifth of global max:13
×10^8
raw data, starting:13
fitted Gauss locations:13
y
x
0
0.5
1
1.5
2
2.5
3
3.5
4
4.5
5
0
5
10
15
20
25

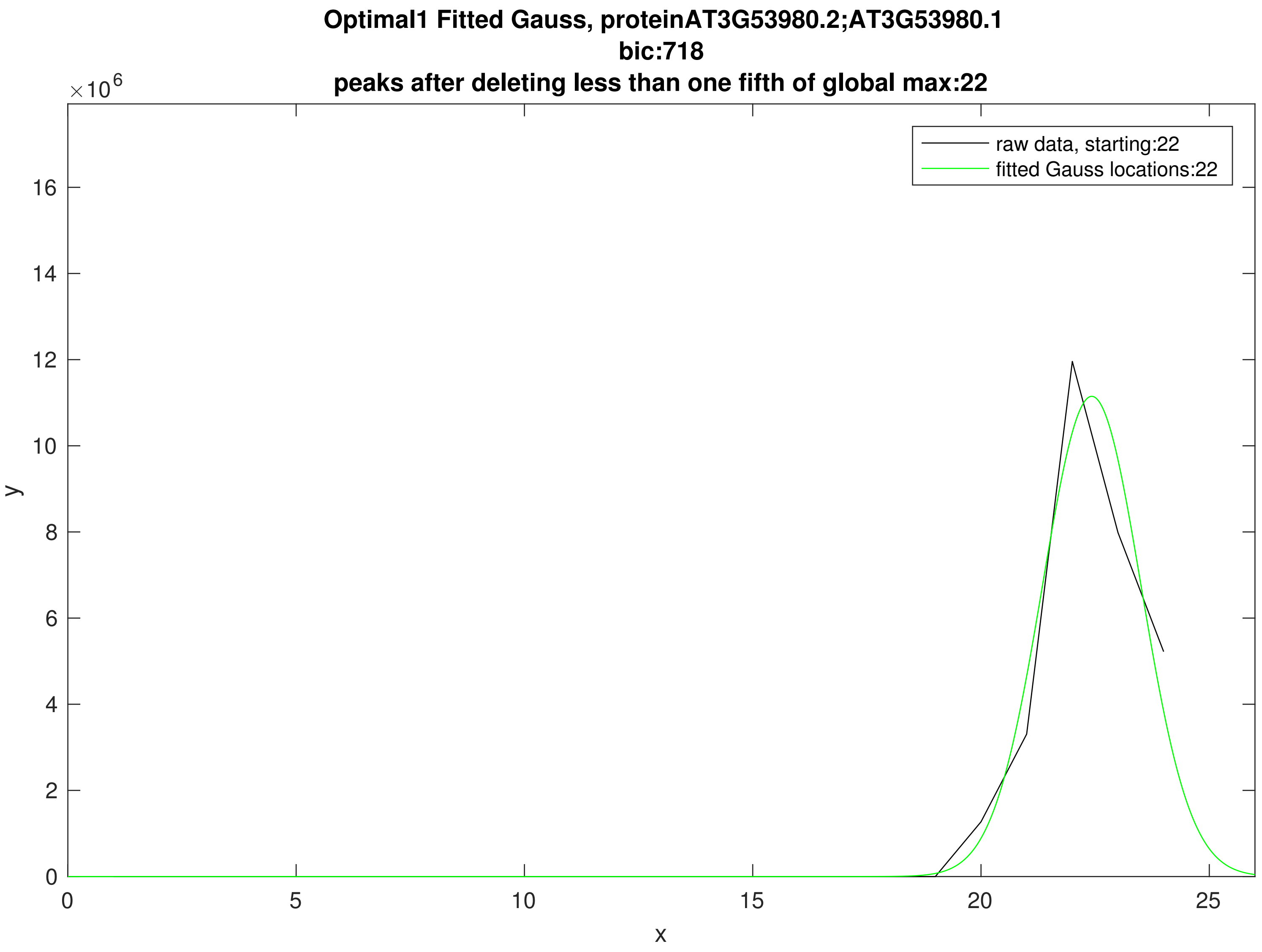

Optimal1 Fitted Gauss, proteinAT3G53980.2;AT3G53980.1
bic:718
peaks after deleting less than one fifth of global max:22
×10^6
raw data, starting:22
fitted Gauss locations:22
y
x
0
2
4
6
8
10
12
14
16
0
5
10
15
20
25

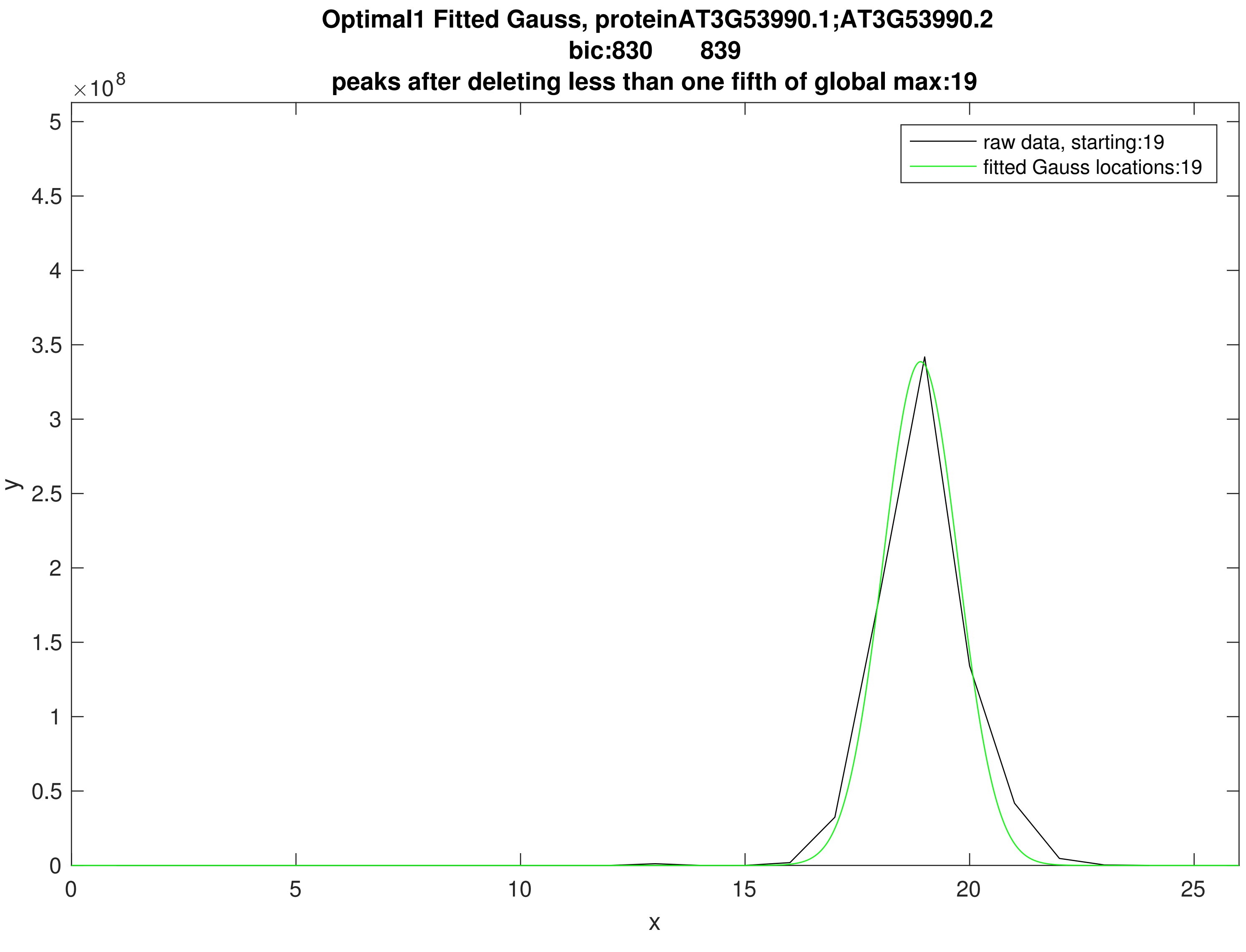
Optimal1 Fitted Gauss, proteinAT3G53990.1;AT3G53990.2
bic:830 839
peaks after deleting less than one fifth of global max:19
×10^8
5
4.5
4
3.5
3
2.5
2
1.5
1
0.5
0
y
0
5
10
15
20
25
x
raw data, starting:19
fitted Gauss locations:19

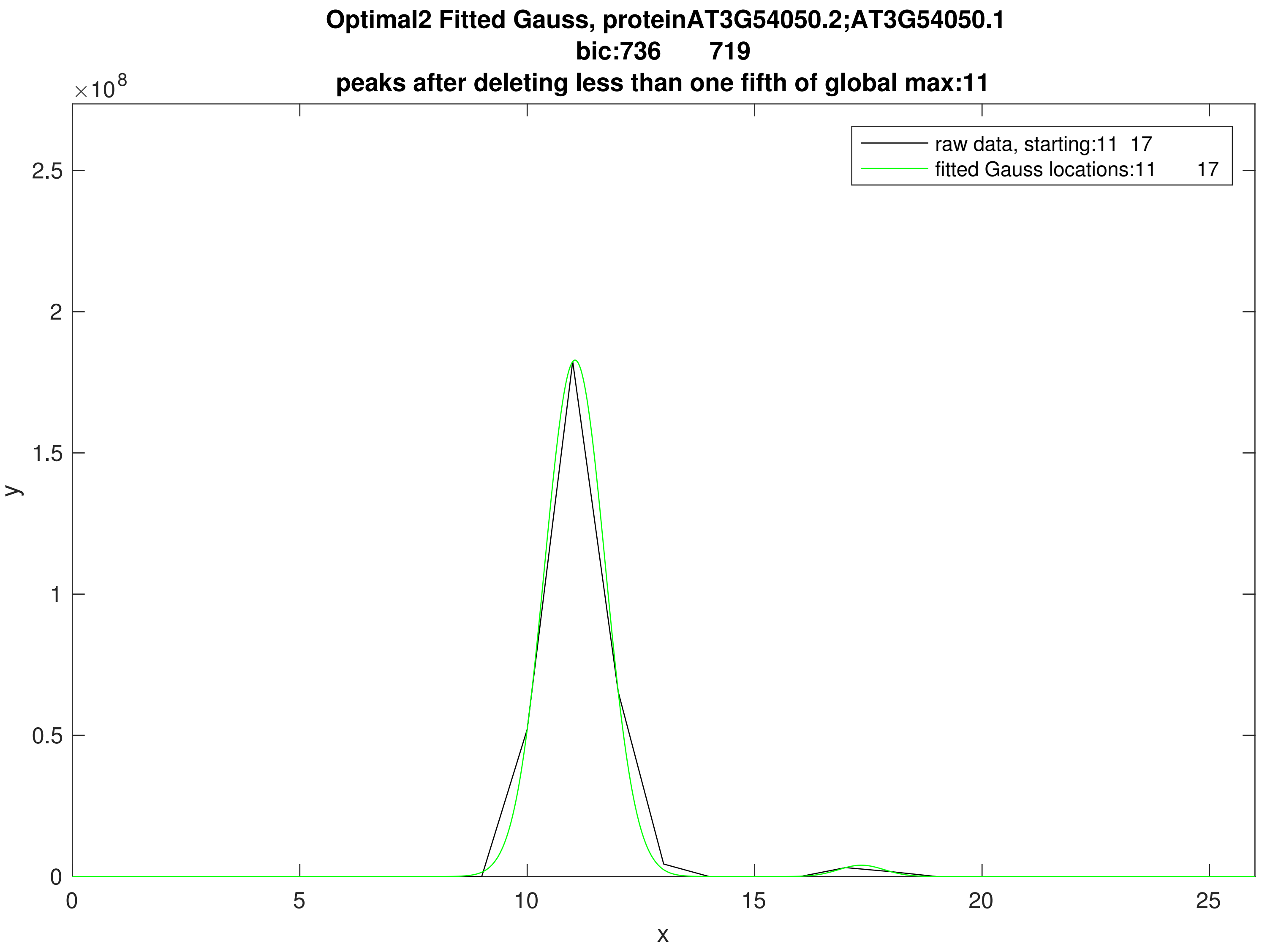
Optimal2 Fitted Gauss, proteinAT3G54050.2;AT3G54050.1
bic:736 719
peaks after deleting less than one fifth of global max:11
×10^8
2.5
2
1.5
1
0.5
0
y
0
5
10
15
20
25
x
raw data, starting:11 17
fitted Gauss locations:11 17

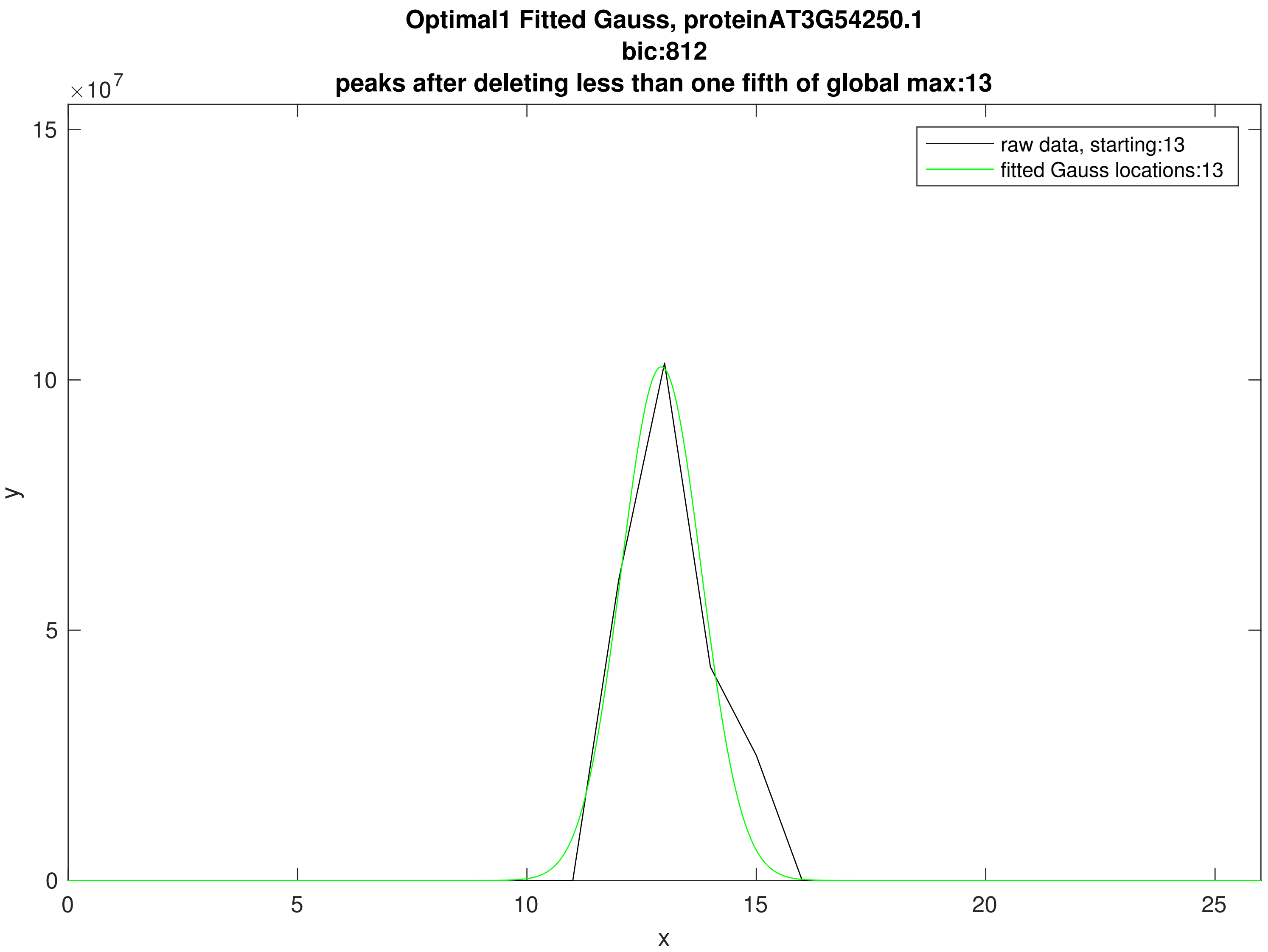
Optimal1 Fitted Gauss, proteinAT3G54250.1
bic:812
peaks after deleting less than one fifth of global max:13
×10^7
15
10
5
0
y
0
5
10
15
20
25
x
raw data, starting:13
fitted Gauss locations:13

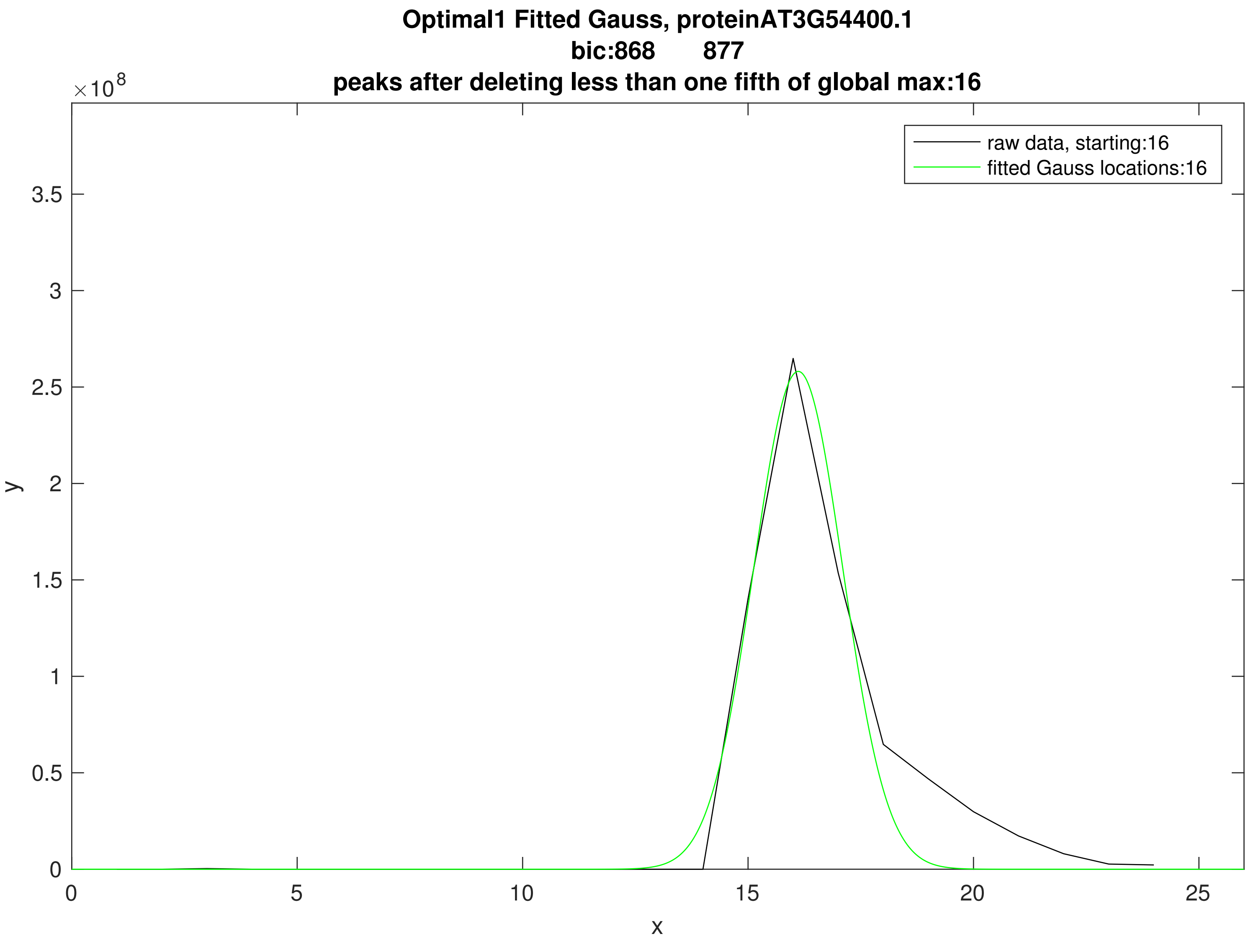

Optimal1 Fitted Gauss, proteinAT3G54400.1
bic:868 877
peaks after deleting less than one fifth of global max:16
×10^8
3.5
3
2.5
2
1.5
1
0.5
0
y
0
5
10
15
20
25
x
raw data, starting:16
fitted Gauss locations:16

**Optimal1 Fitted Gauss, proteinAT3G54440.1;AT3G54440.2;AT3G54440.3**
**bic:680**
**peaks after deleting less than one fifth of global max:8.9**

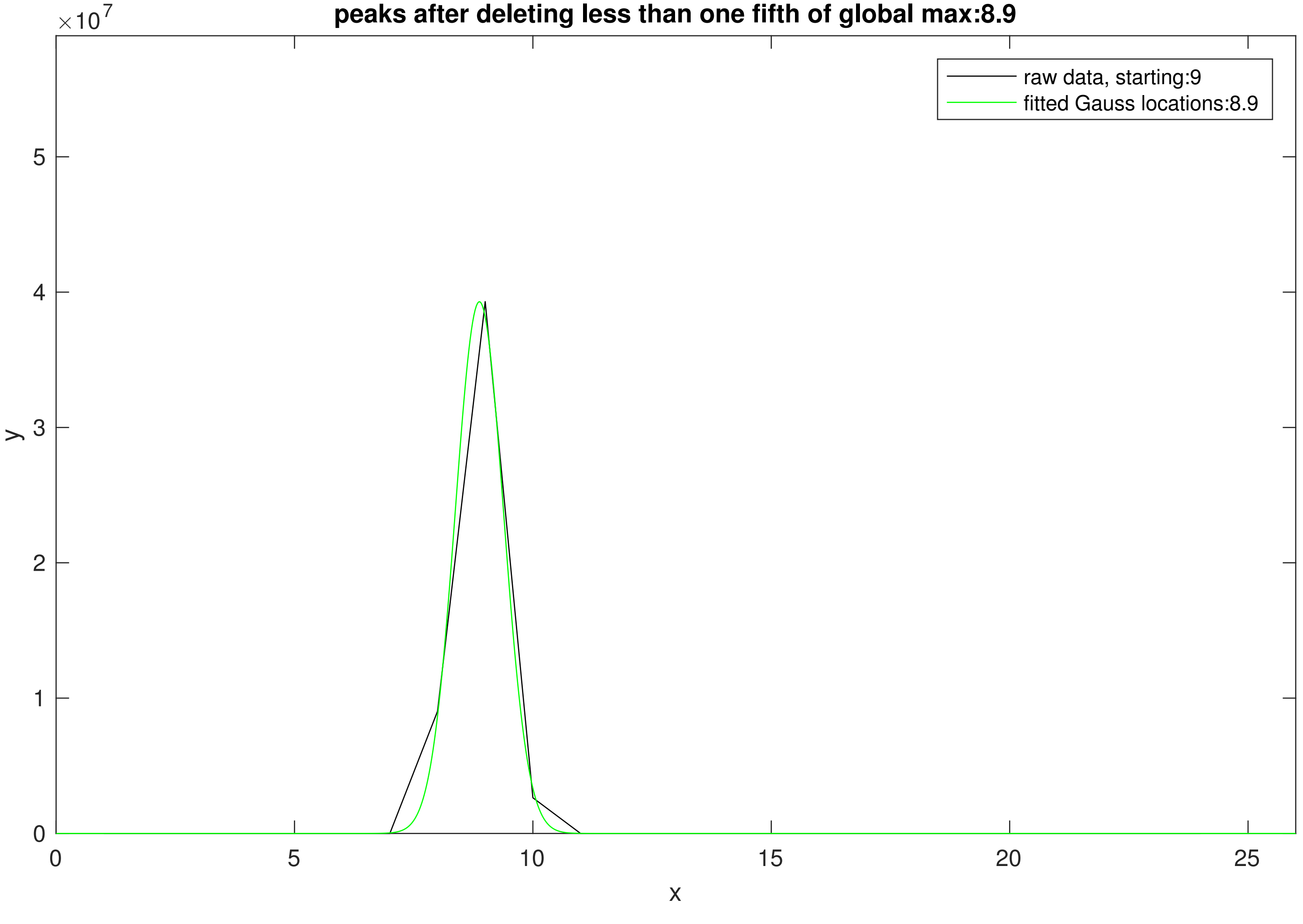

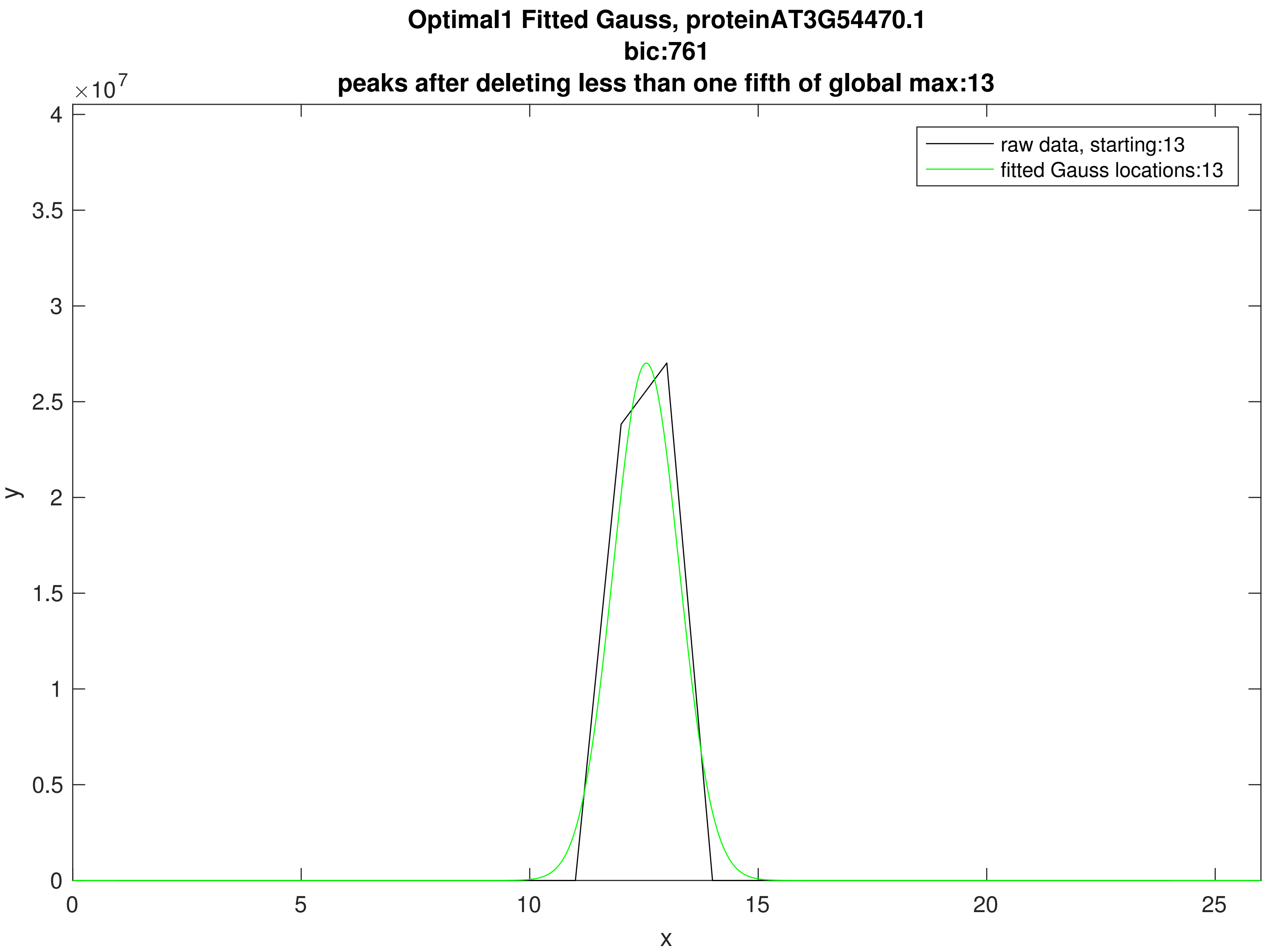

Optimal1 Fitted Gauss, proteinAT3G54470.1
bic:761
peaks after deleting less than one fifth of global max:13
×10^7
raw data, starting:13
fitted Gauss locations:13
y
x
0
0.5
1
1.5
2
2.5
3
3.5
4
0
5
10
15
20
25

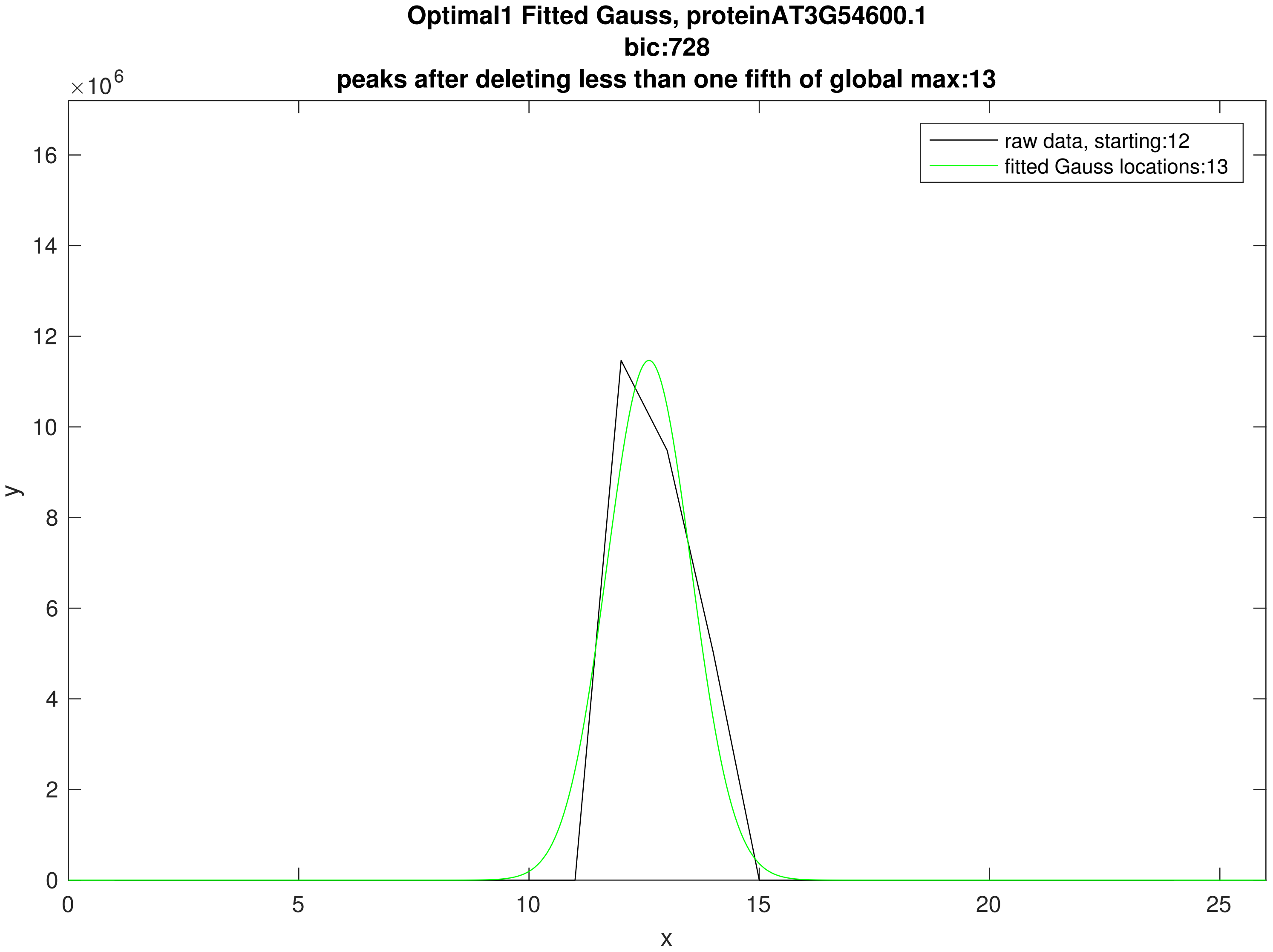
Optimal1 Fitted Gauss, proteinAT3G54600.1
bic:728
peaks after deleting less than one fifth of global max:13
$\times 10^6$
raw data, starting:12
fitted Gauss locations:13
y
x
0
2
4
6
8
10
12
14
16
0
5
10
15
20
25

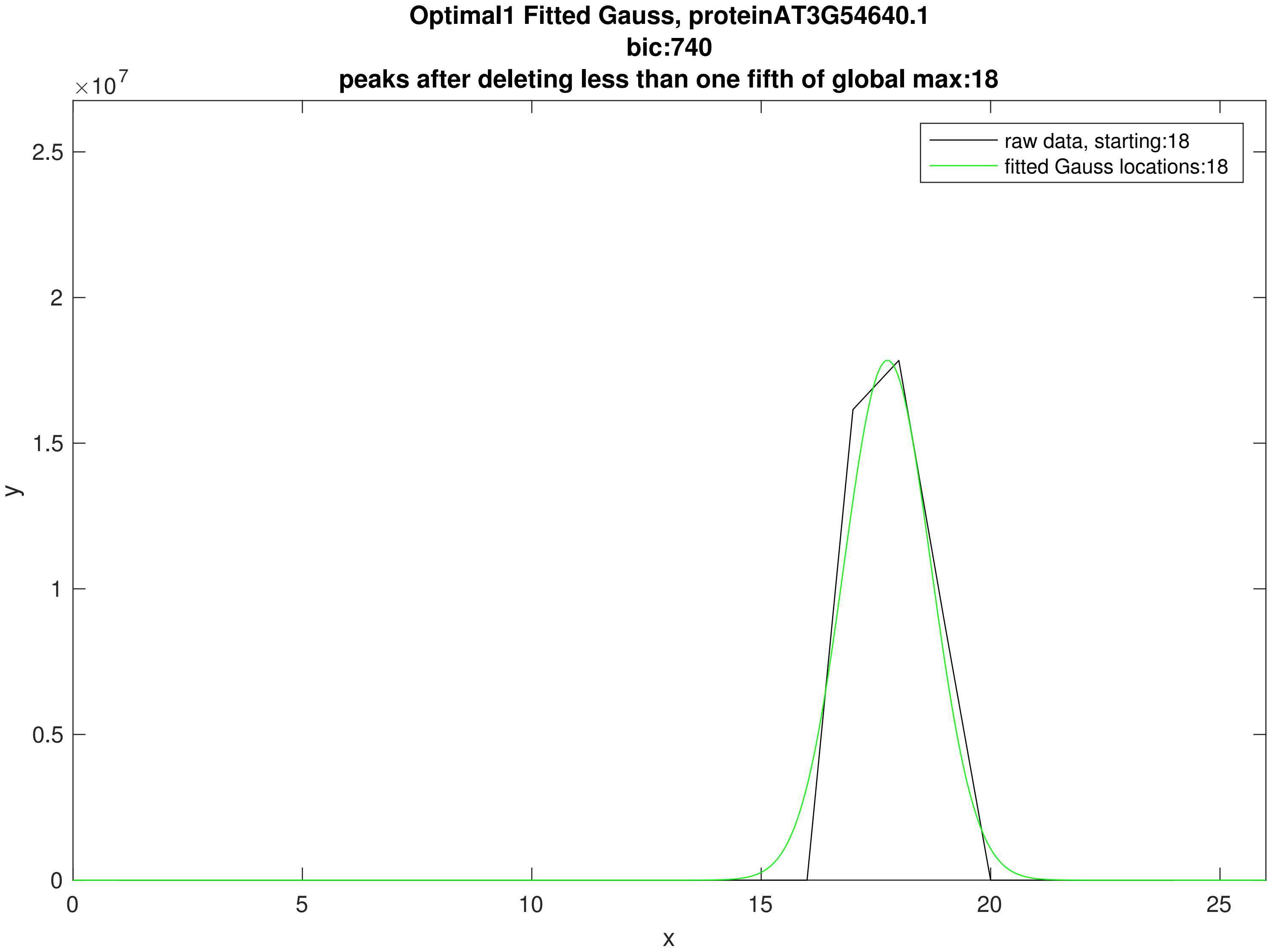
Optimal1 Fitted Gauss, proteinAT3G54640.1
bic:740
peaks after deleting less than one fifth of global max:18
×10^7
2.5
2
1.5
1
0.5
0
y
0
5
10
15
20
25
x
raw data, starting:18
fitted Gauss locations:18

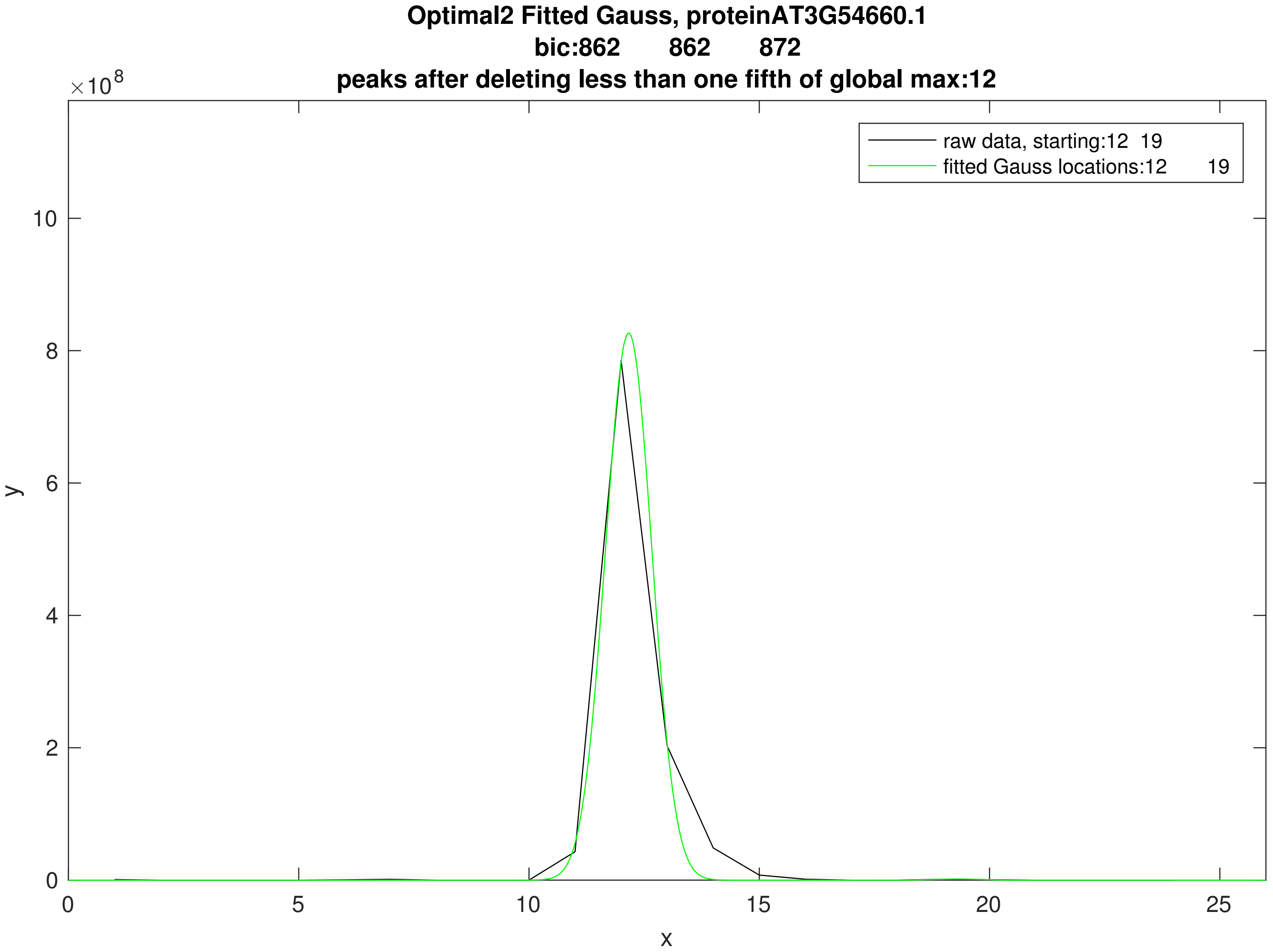
Optimal2 Fitted Gauss, proteinAT3G54660.1
bic:862 862 872
peaks after deleting less than one fifth of global max:12
×10^8
raw data, starting:12 19
fitted Gauss locations:12 19
10
8
6
4
2
0
y
0
5
10
15
20
25
x

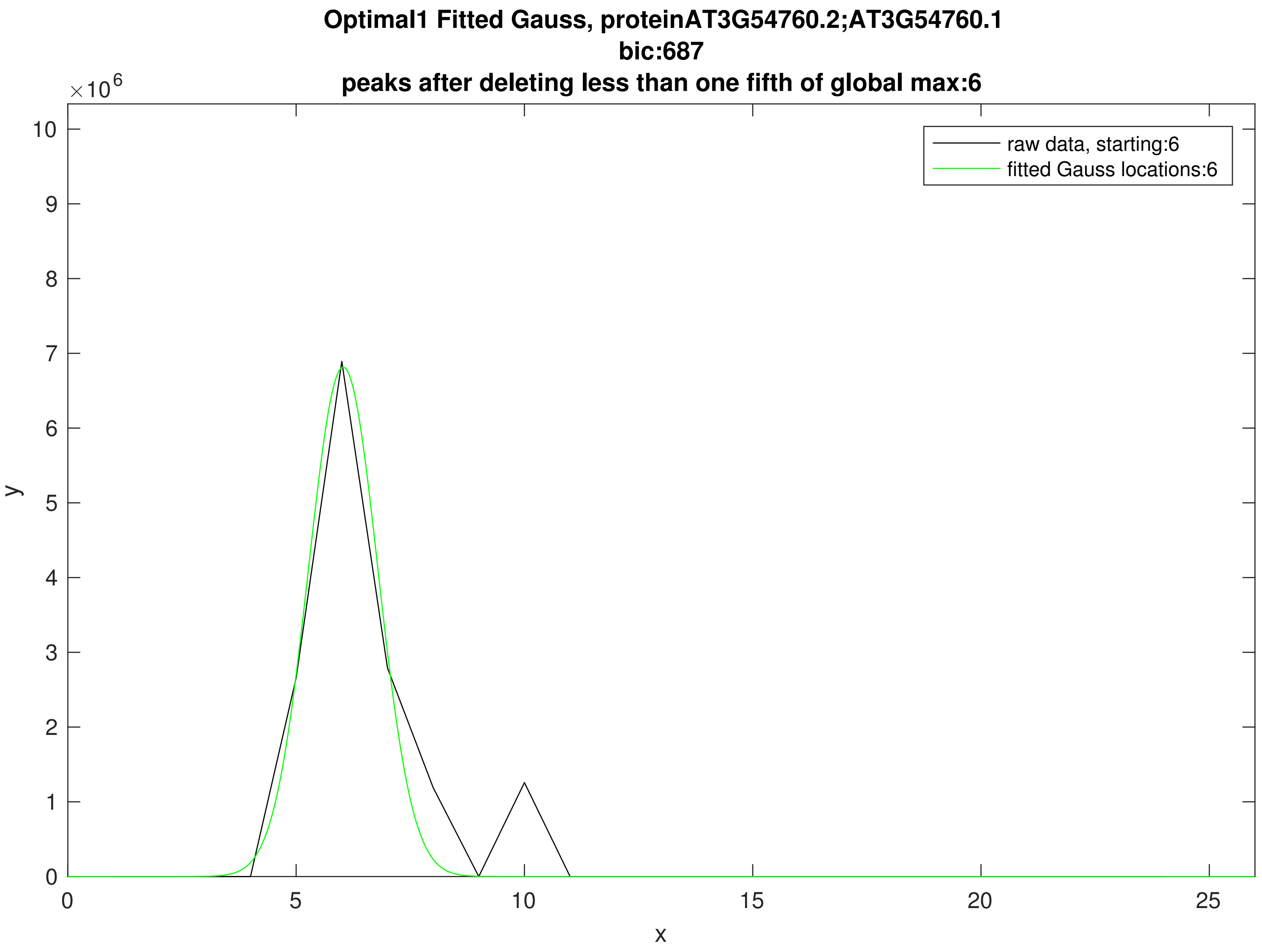
Optimal1 Fitted Gauss, proteinAT3G54760.2;AT3G54760.1
bic:687
peaks after deleting less than one fifth of global max:6
×10^6
raw data, starting:6
fitted Gauss locations:6
y
x
0
1
2
3
4
5
6
7
8
9
10
0
5
10
15
20
25

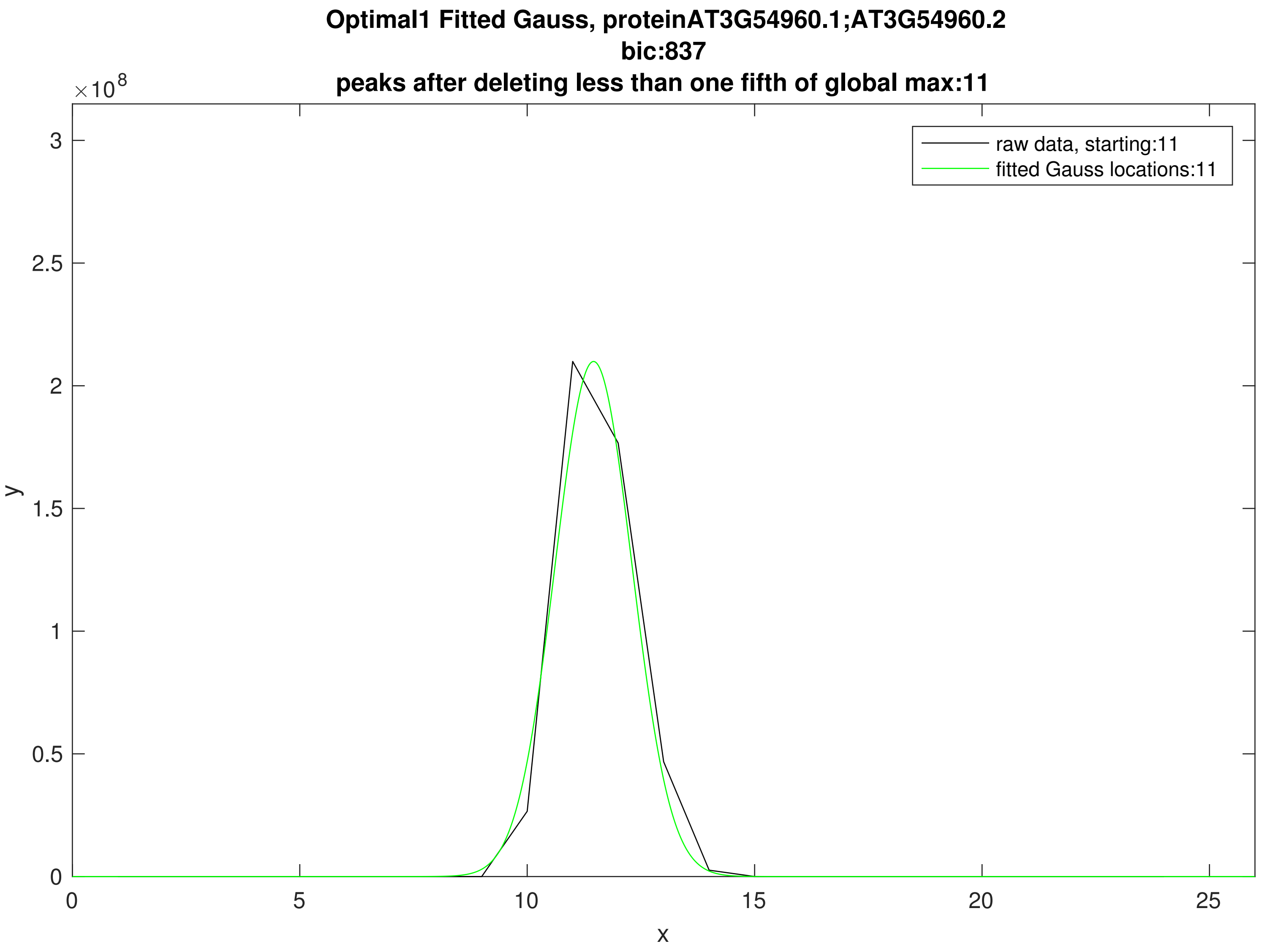
Optimal1 Fitted Gauss, proteinAT3G54960.1;AT3G54960.2
bic:837
peaks after deleting less than one fifth of global max:11
×10^8
raw data, starting:11
fitted Gauss locations:11
y
x
0
0.5
1
1.5
2
2.5
3
0
5
10
15
20
25

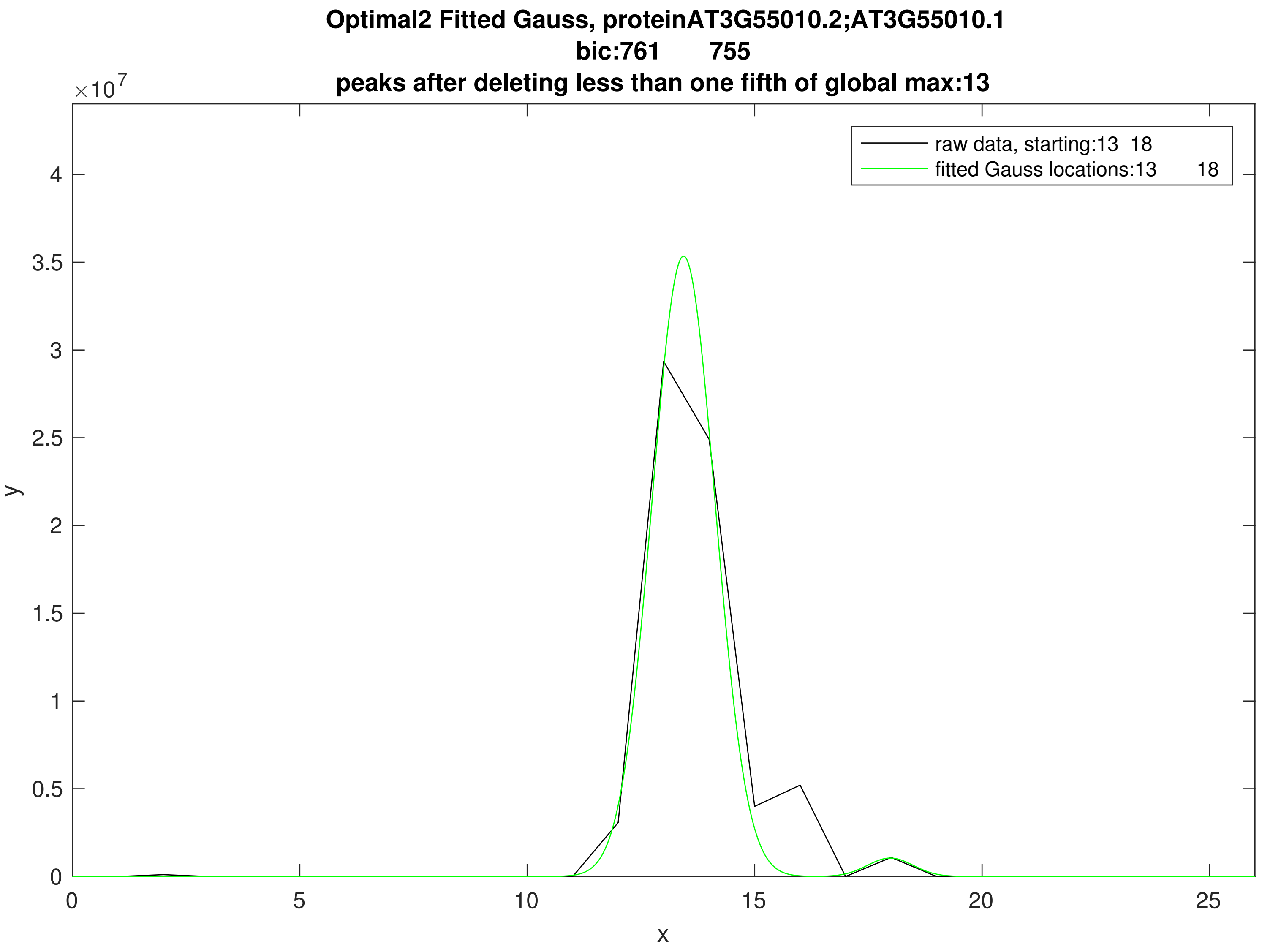
Optimal2 Fitted Gauss, proteinAT3G55010.2;AT3G55010.1
bic:761 755
peaks after deleting less than one fifth of global max:13
×10^7
raw data, starting:13 18
fitted Gauss locations:13 18
y
x
0
0.5
1
1.5
2
2.5
3
3.5
4
0
5
10
15
20
25

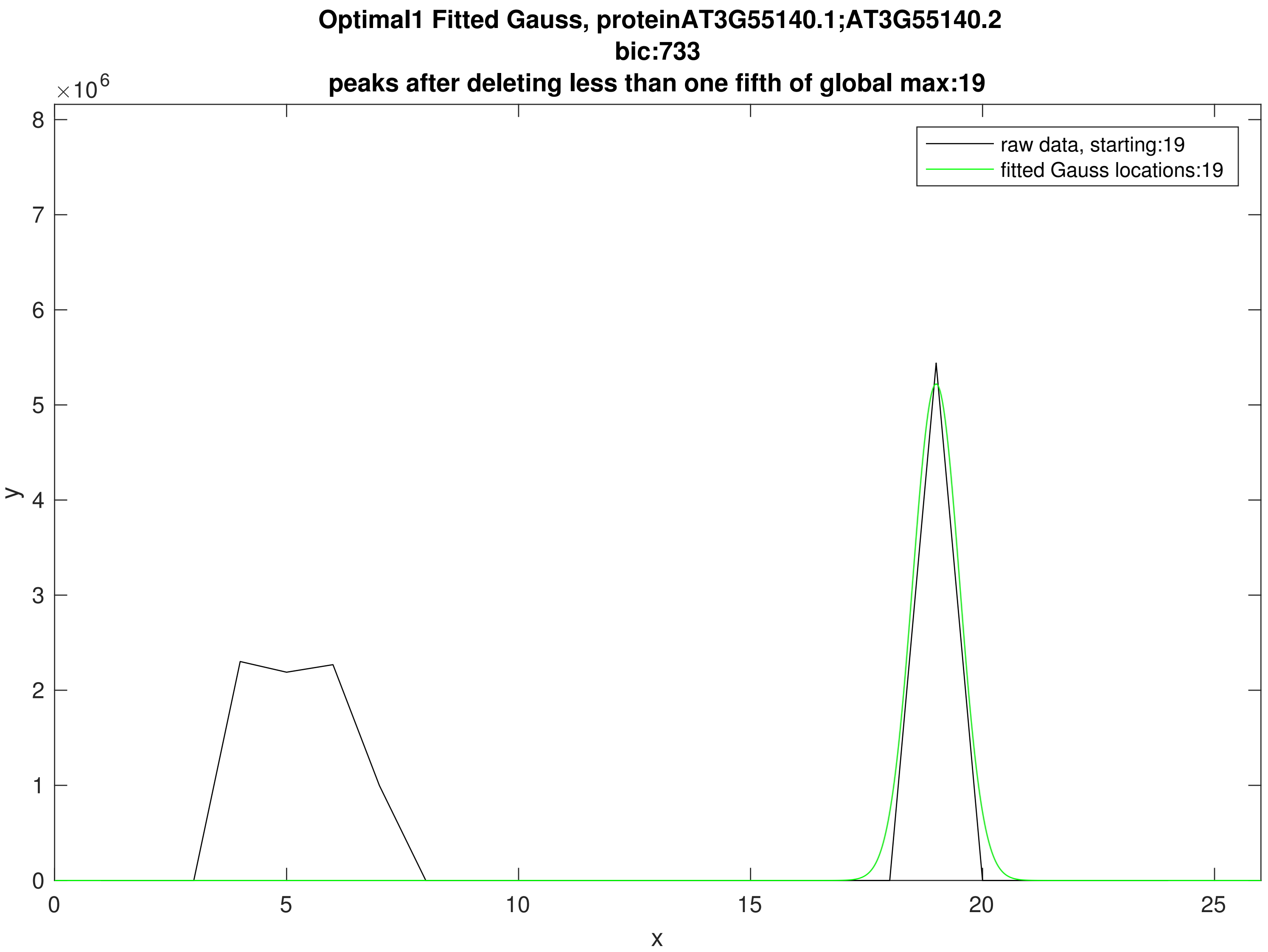
Optimal1 Fitted Gauss, proteinAT3G55140.1;AT3G55140.2
bic:733
peaks after deleting less than one fifth of global max:19
×10^6
raw data, starting:19
fitted Gauss locations:19
y
x
0
1
2
3
4
5
6
7
8
0
5
10
15
20
25

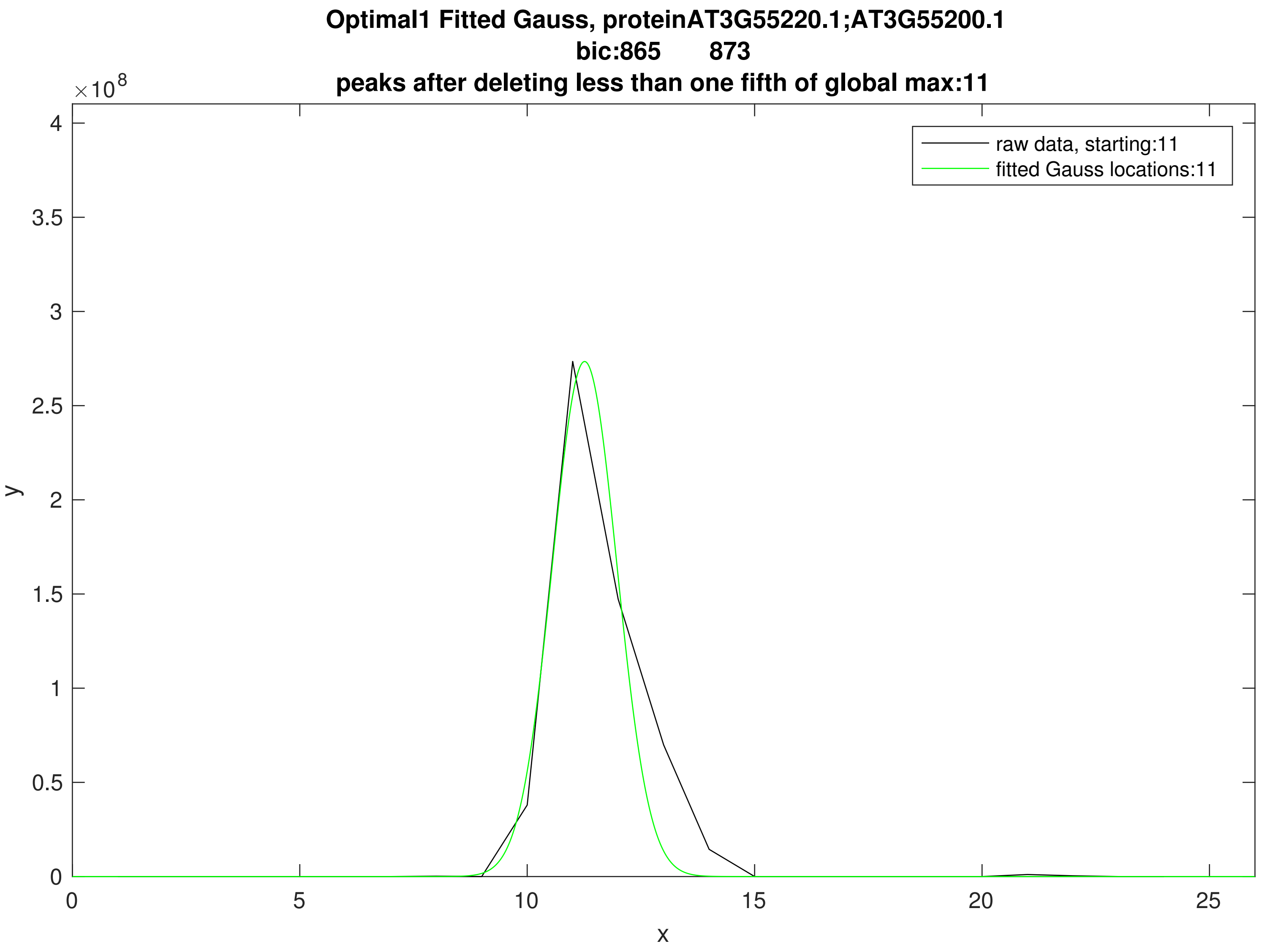
Optimal1 Fitted Gauss, proteinAT3G55220.1;AT3G55200.1
bic:865 873
peaks after deleting less than one fifth of global max:11
×10^8
4
3.5
3
2.5
2
1.5
1
0.5
0
y
0
5
10
15
20
25
x
raw data, starting:11
fitted Gauss locations:11

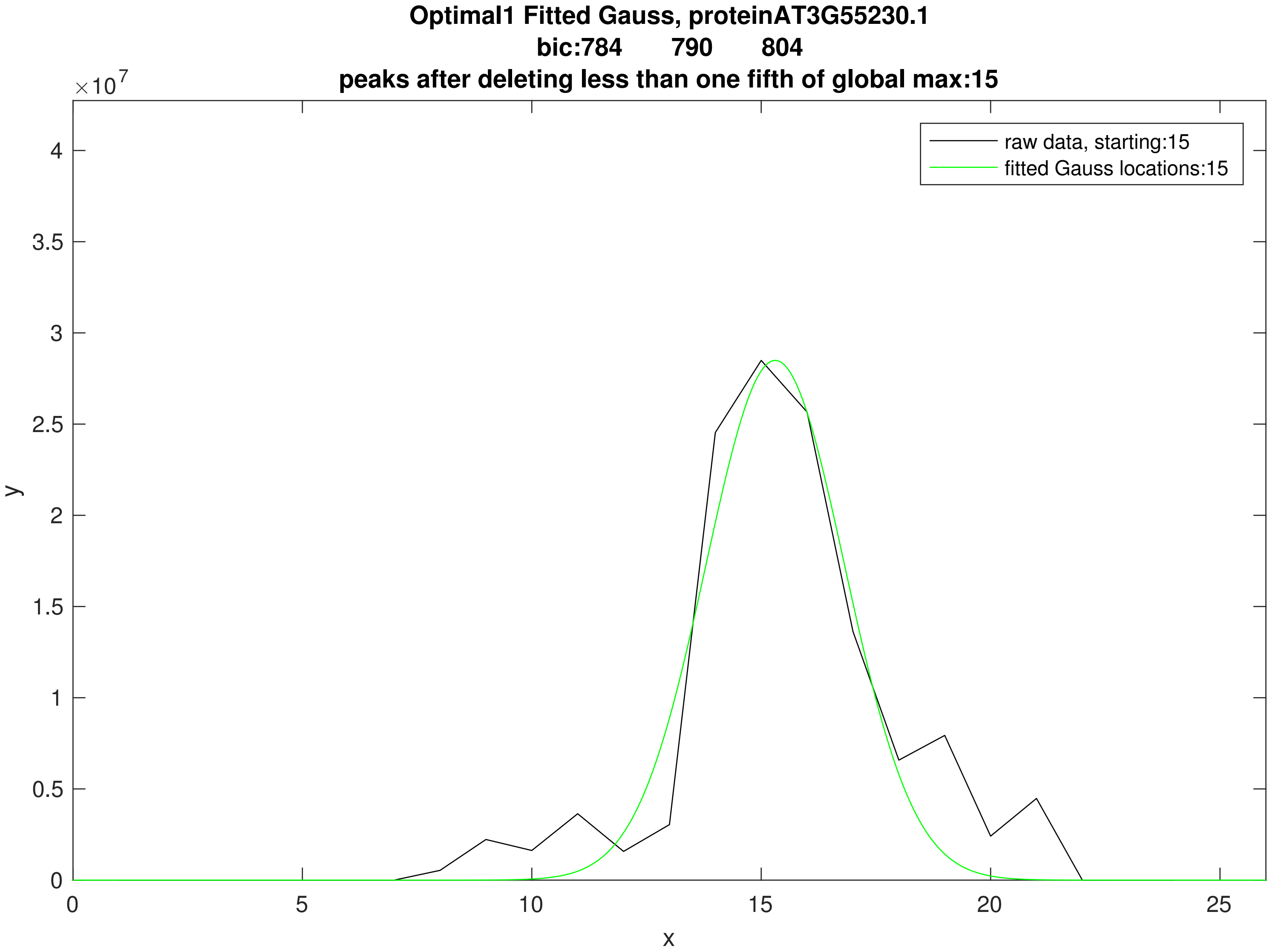
Optimal1 Fitted Gauss, proteinAT3G55230.1
bic:784 790 804
peaks after deleting less than one fifth of global max:15
×10^7
raw data, starting:15
fitted Gauss locations:15
y
x
0
0.5
1
1.5
2
2.5
3
3.5
4
0
5
10
15
20
25

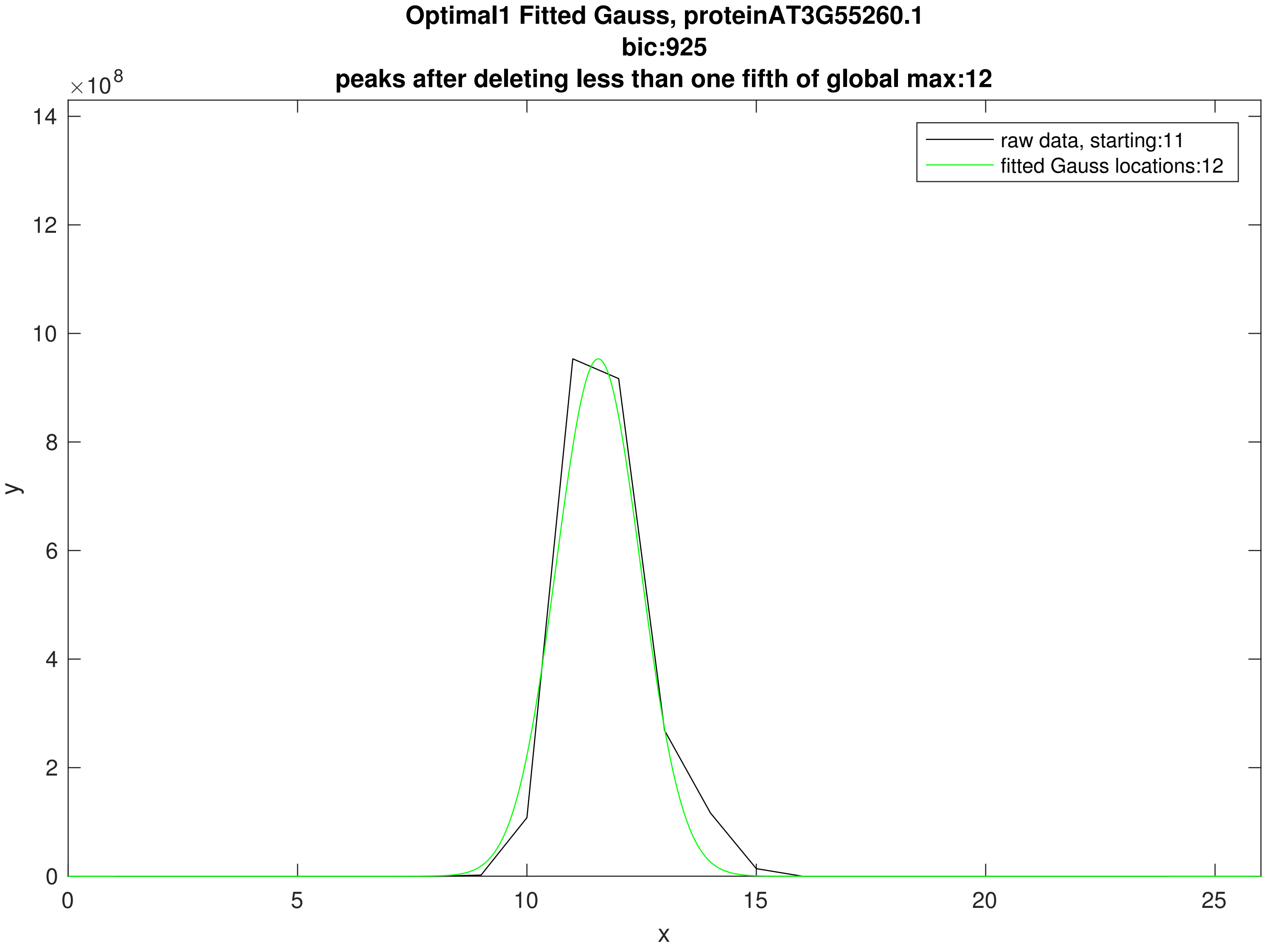
Optimal1 Fitted Gauss, proteinAT3G55260.1
bic:925
peaks after deleting less than one fifth of global max:12
×10^8
raw data, starting:11
fitted Gauss locations:12
y
x
0
2
4
6
8
10
12
14
0
5
10
15
20
25

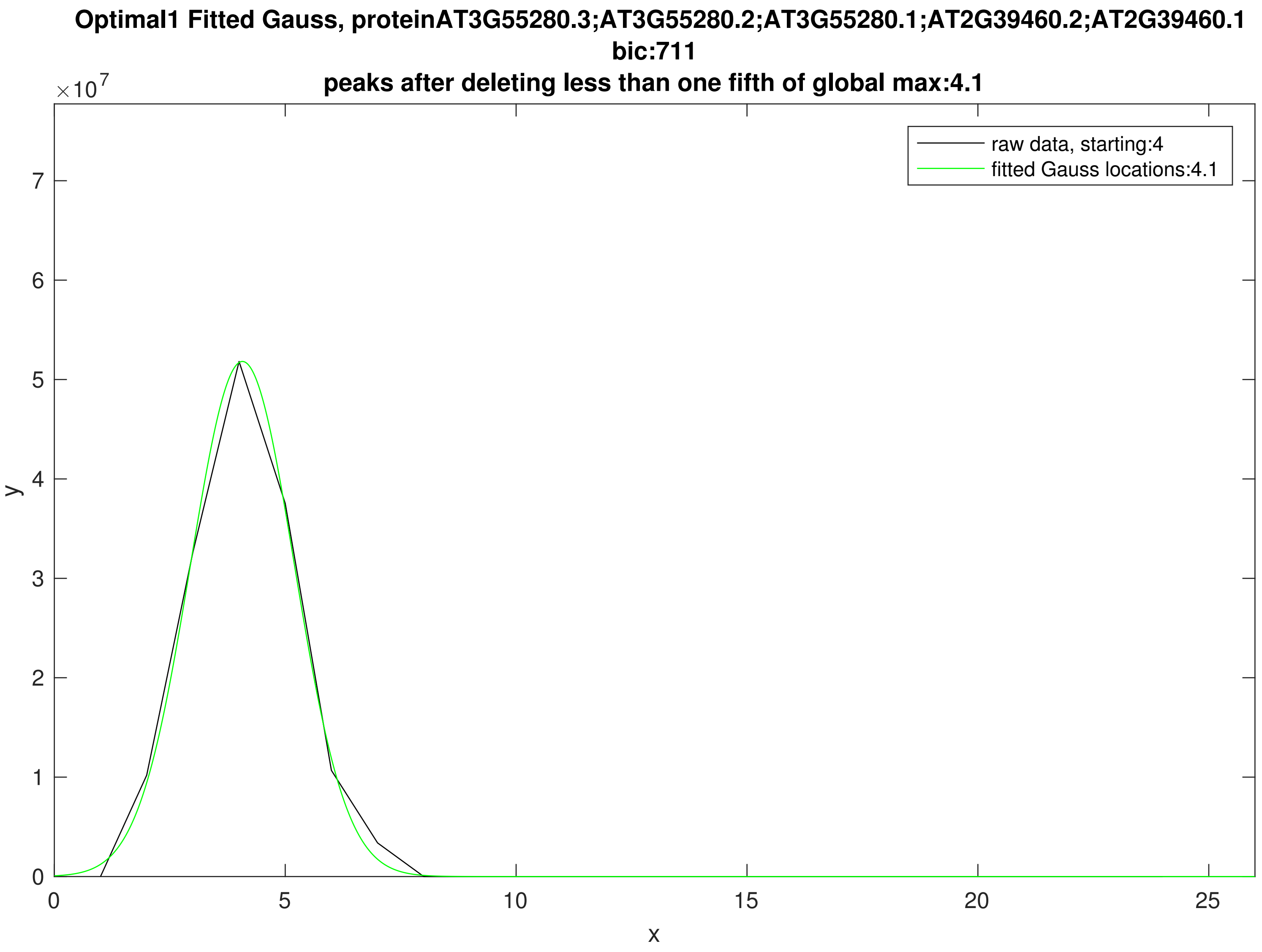

Optimal1 Fitted Gauss, proteinAT3G55280.3;AT3G55280.2;AT3G55280.1;AT2G39460.2;AT2G39460.1
bic:711
peaks after deleting less than one fifth of global max:4.1
×10^7
raw data, starting:4
fitted Gauss locations:4.1
y
x
0
1
2
3
4
5
6
7
0
5
10
15
20
25

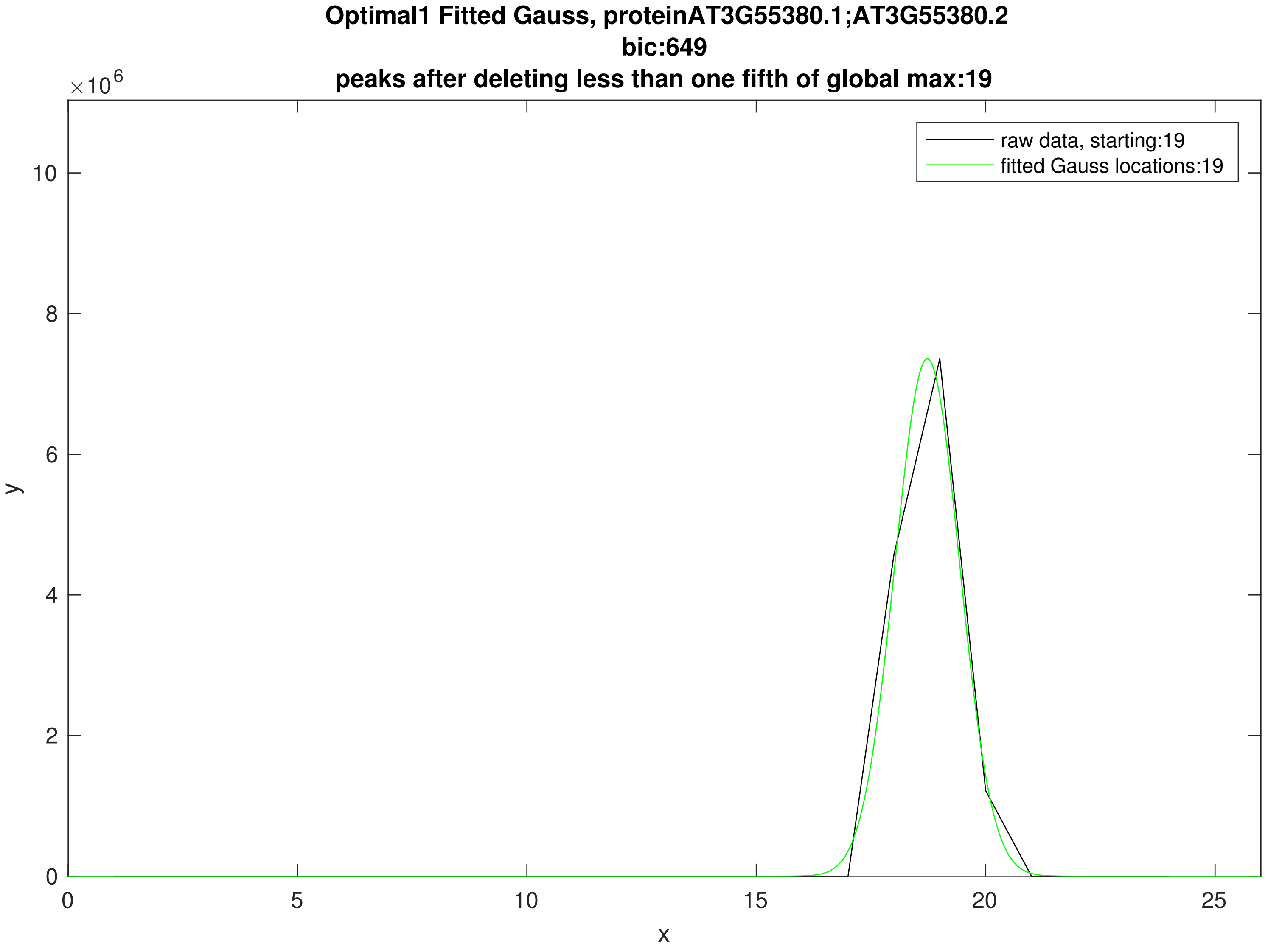
Optimal1 Fitted Gauss, proteinAT3G55380.1;AT3G55380.2
bic:649
peaks after deleting less than one fifth of global max:19
×10^6
raw data, starting:19
fitted Gauss locations:19
y
x
0
2
4
6
8
10
0
5
10
15
20
25

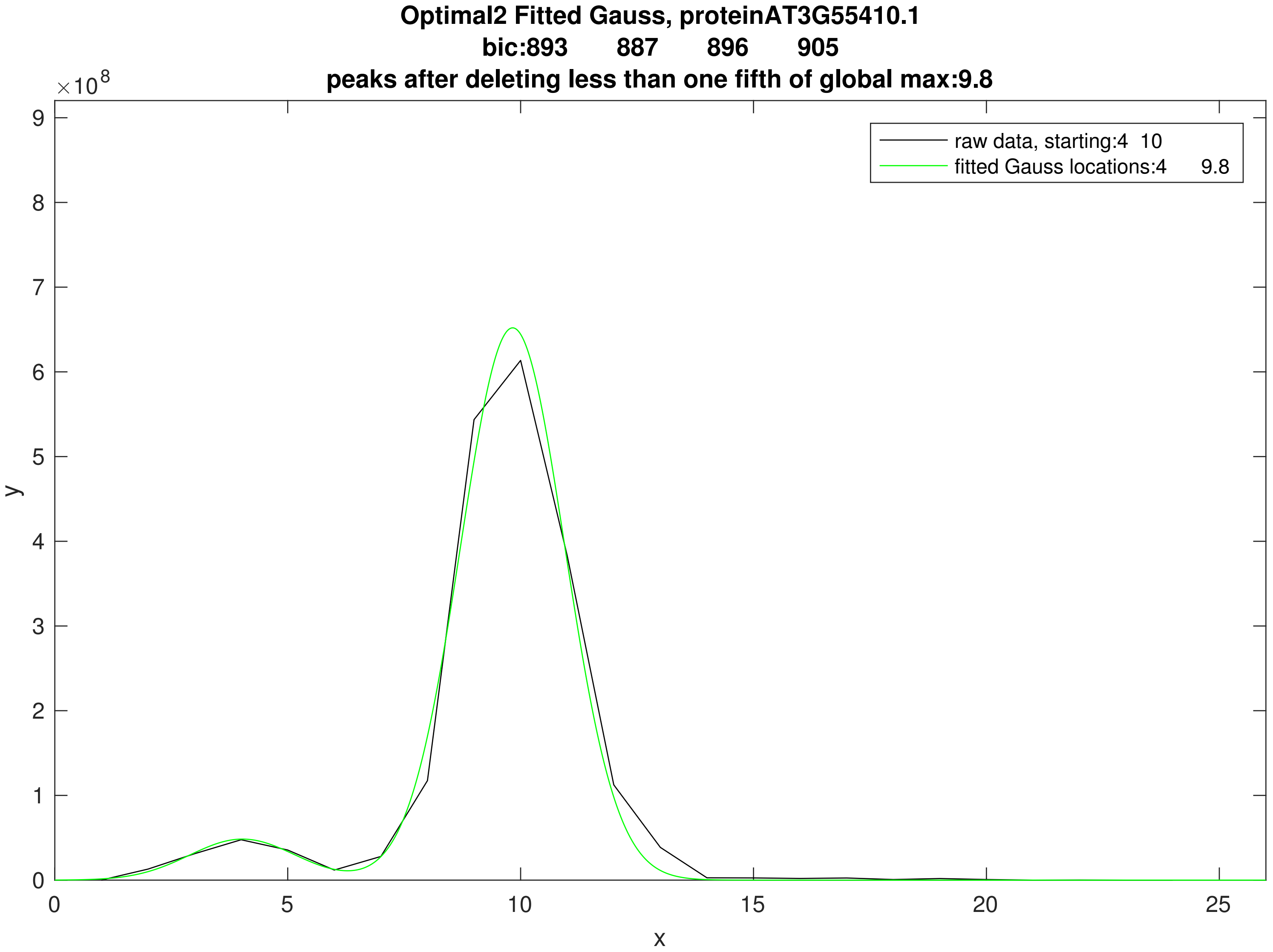
Optimal2 Fitted Gauss, proteinAT3G55410.1
bic:893 887 896 905
peaks after deleting less than one fifth of global max:9.8
×10^8
raw data, starting:4 10
fitted Gauss locations:4 9.8
y
x
0
1
2
3
4
5
6
7
8
9
0
5
10
15
20
25

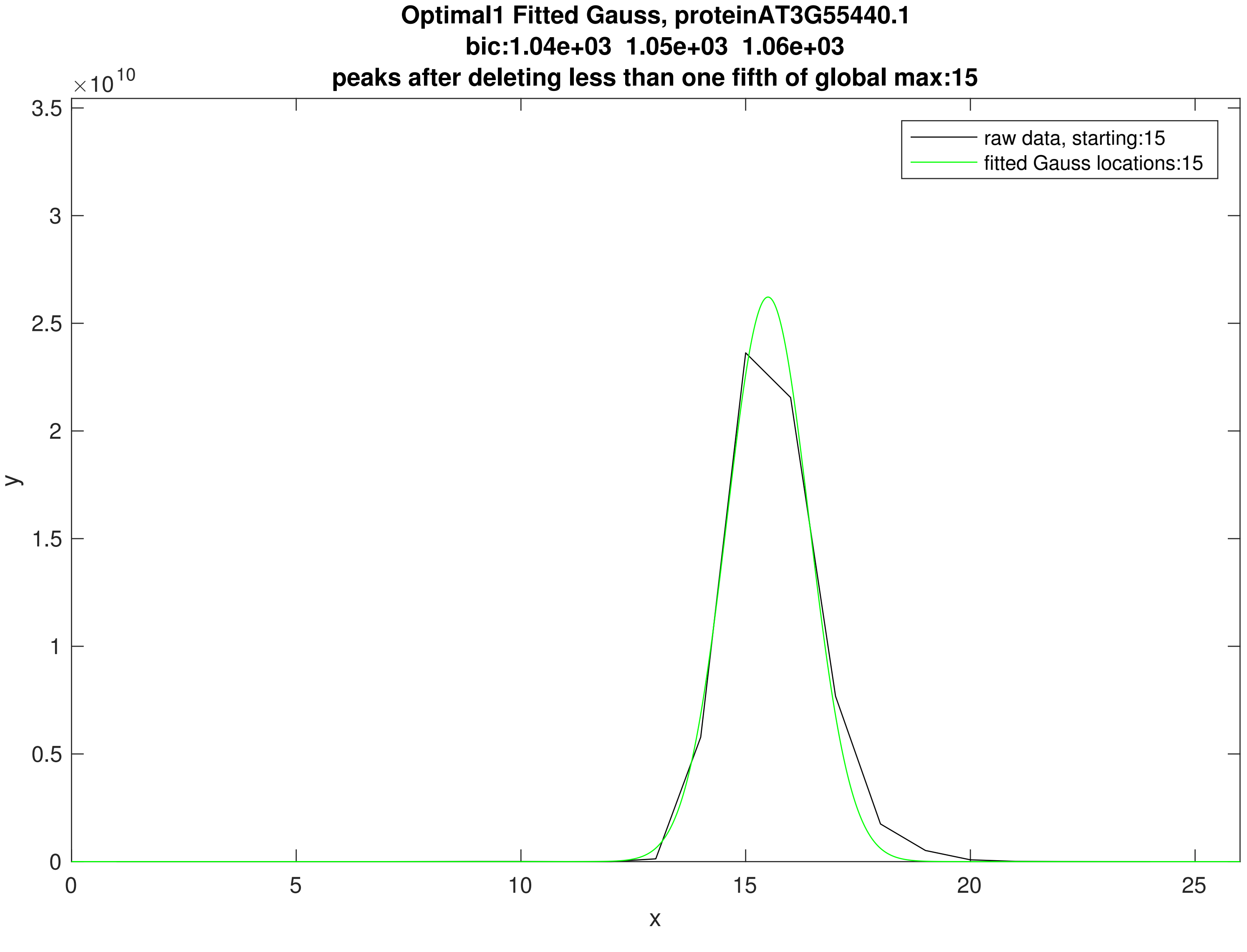
Optimal1 Fitted Gauss, proteinAT3G55440.1
bic:1.04e+03  1.05e+03  1.06e+03
peaks after deleting less than one fifth of global max:15
×10^10
3.5
3
2.5
2
1.5
1
0.5
0
y
0
5
10
15
20
25
x
raw data, starting:15
fitted Gauss locations:15

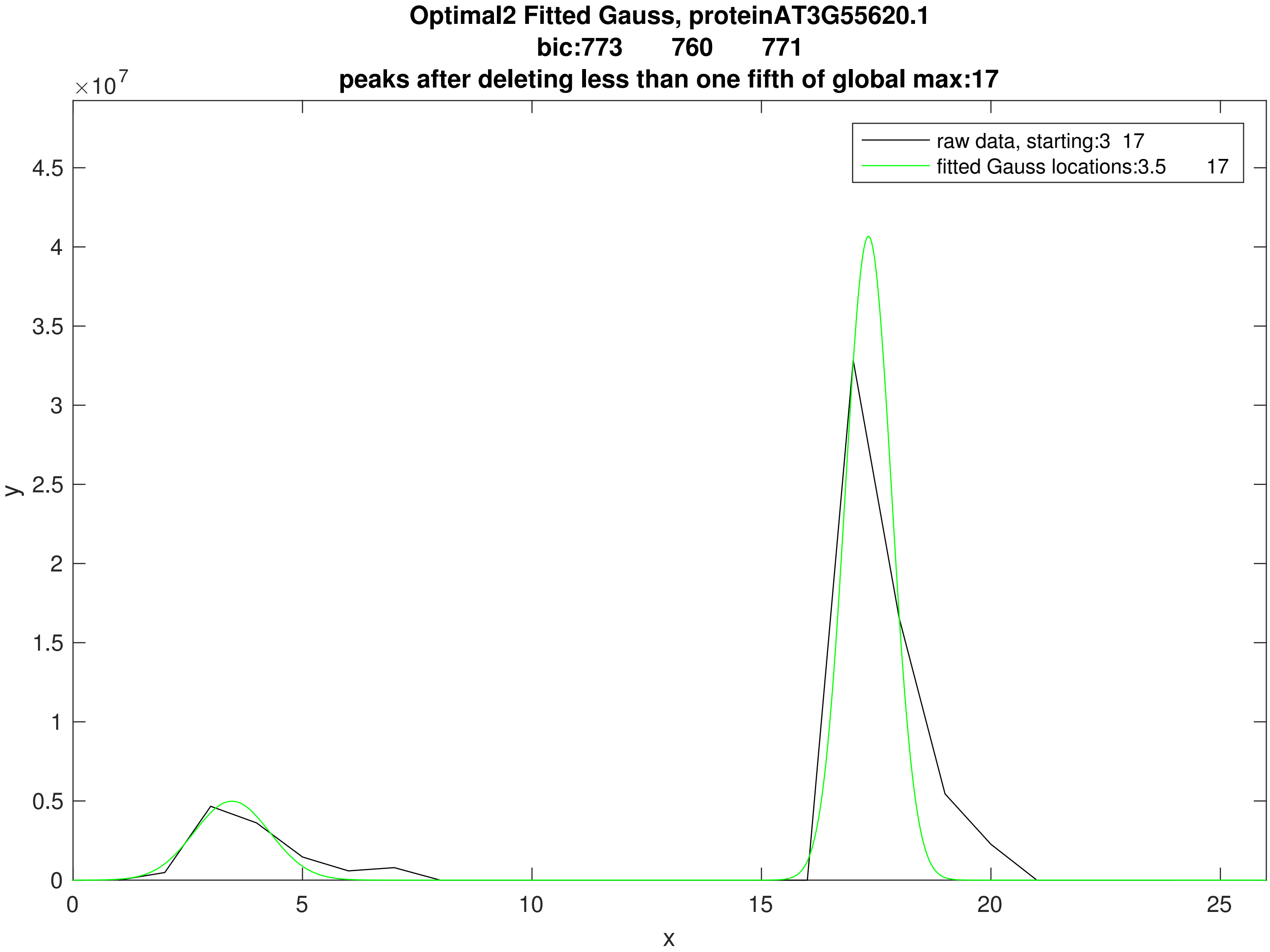

Optimal2 Fitted Gauss, proteinAT3G55620.1
bic:773 760 771
peaks after deleting less than one fifth of global max:17
×10^7
raw data, starting:3 17
fitted Gauss locations:3.5 17
y
x
0
0.5
1
1.5
2
2.5
3
3.5
4
4.5
5
10
15
20
25

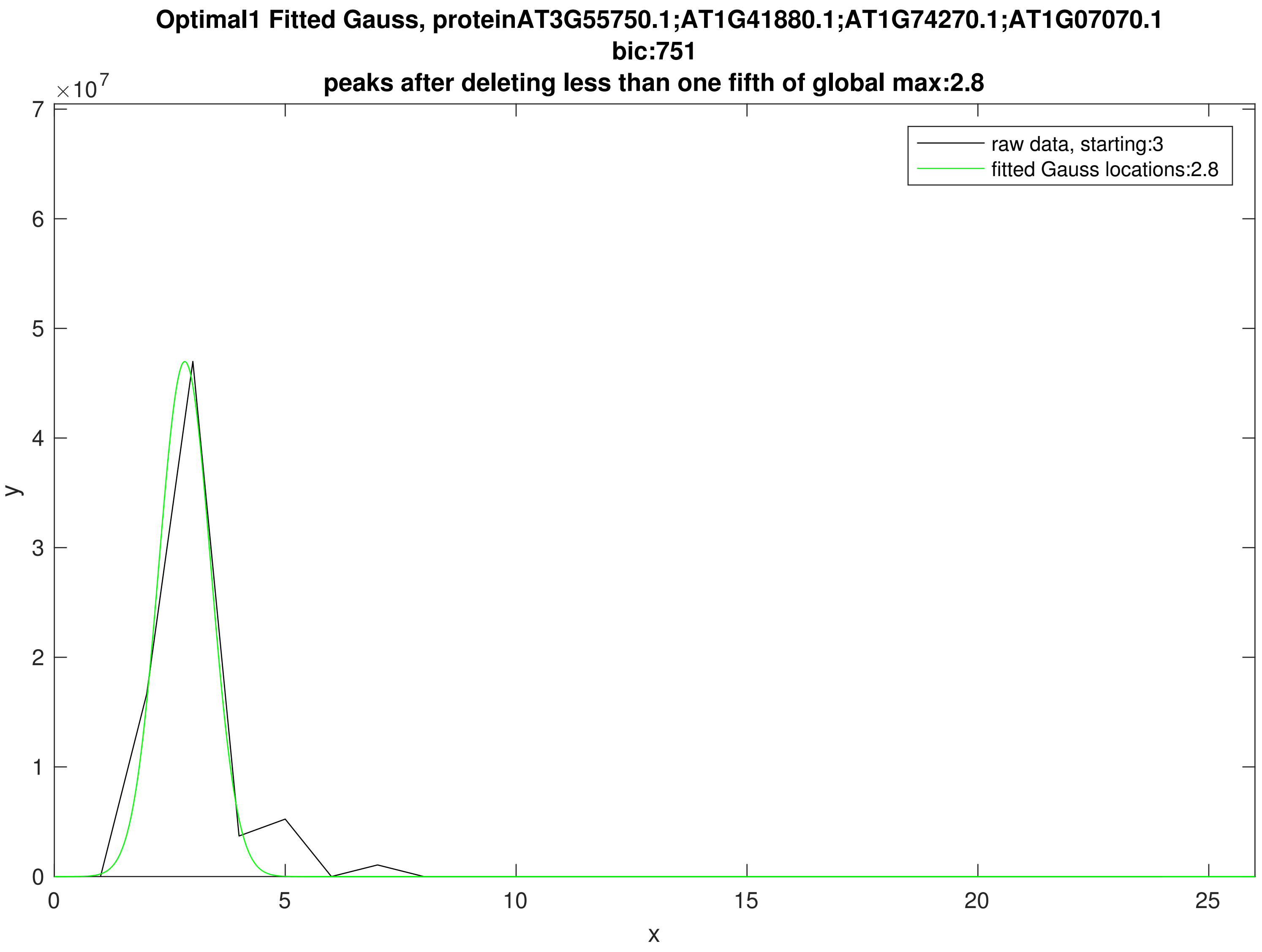
Optimal1 Fitted Gauss, proteinAT3G55750.1;AT1G41880.1;AT1G74270.1;AT1G07070.1
bic:751
peaks after deleting less than one fifth of global max:2.8
×10^7
raw data, starting:3
fitted Gauss locations:2.8
y
x
0
1
2
3
4
5
6
7
0
5
10
15
20
25

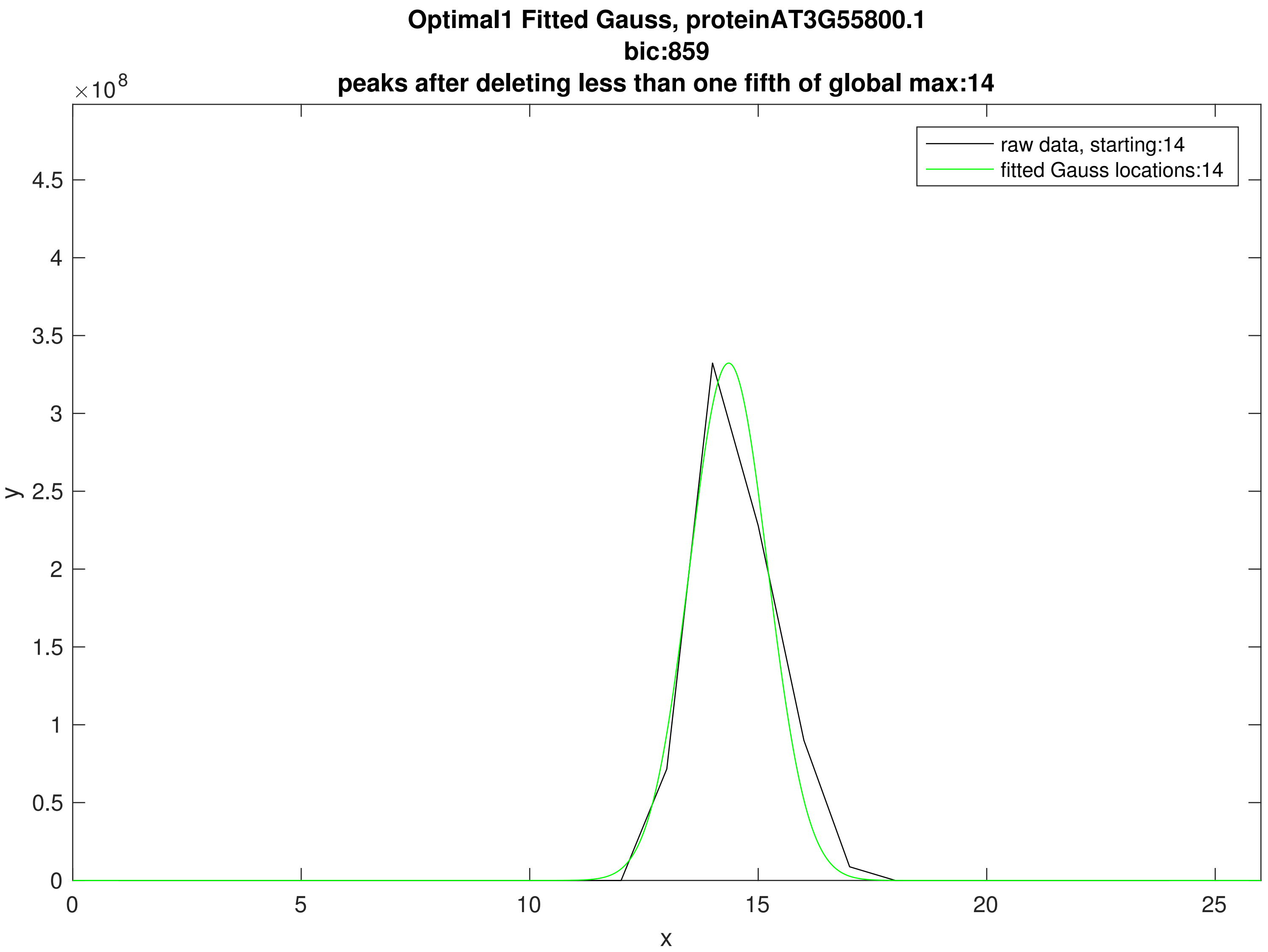
Optimal1 Fitted Gauss, proteinAT3G55800.1
bic:859
peaks after deleting less than one fifth of global max:14
×10^8
raw data, starting:14
fitted Gauss locations:14
4.5
4
3.5
3
2.5
2
1.5
1
0.5
0
y
0
5
10
15
20
25
x

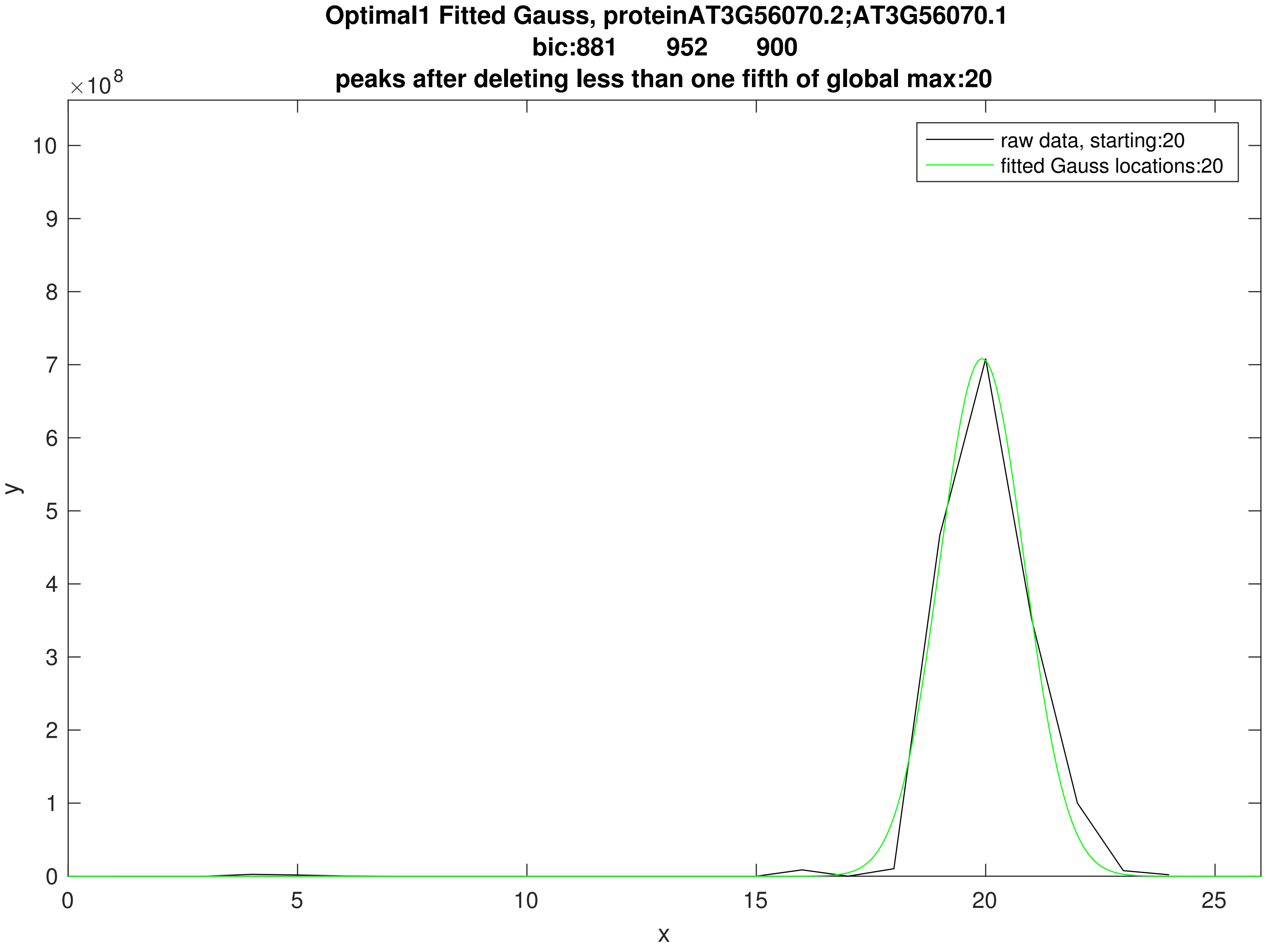
Optimal1 Fitted Gauss, proteinAT3G56070.2;AT3G56070.1
bic:881 952 900
peaks after deleting less than one fifth of global max:20
×10^8
raw data, starting:20
fitted Gauss locations:20
y
x
0
1
2
3
4
5
6
7
8
9
10
0
5
10
15
20
25

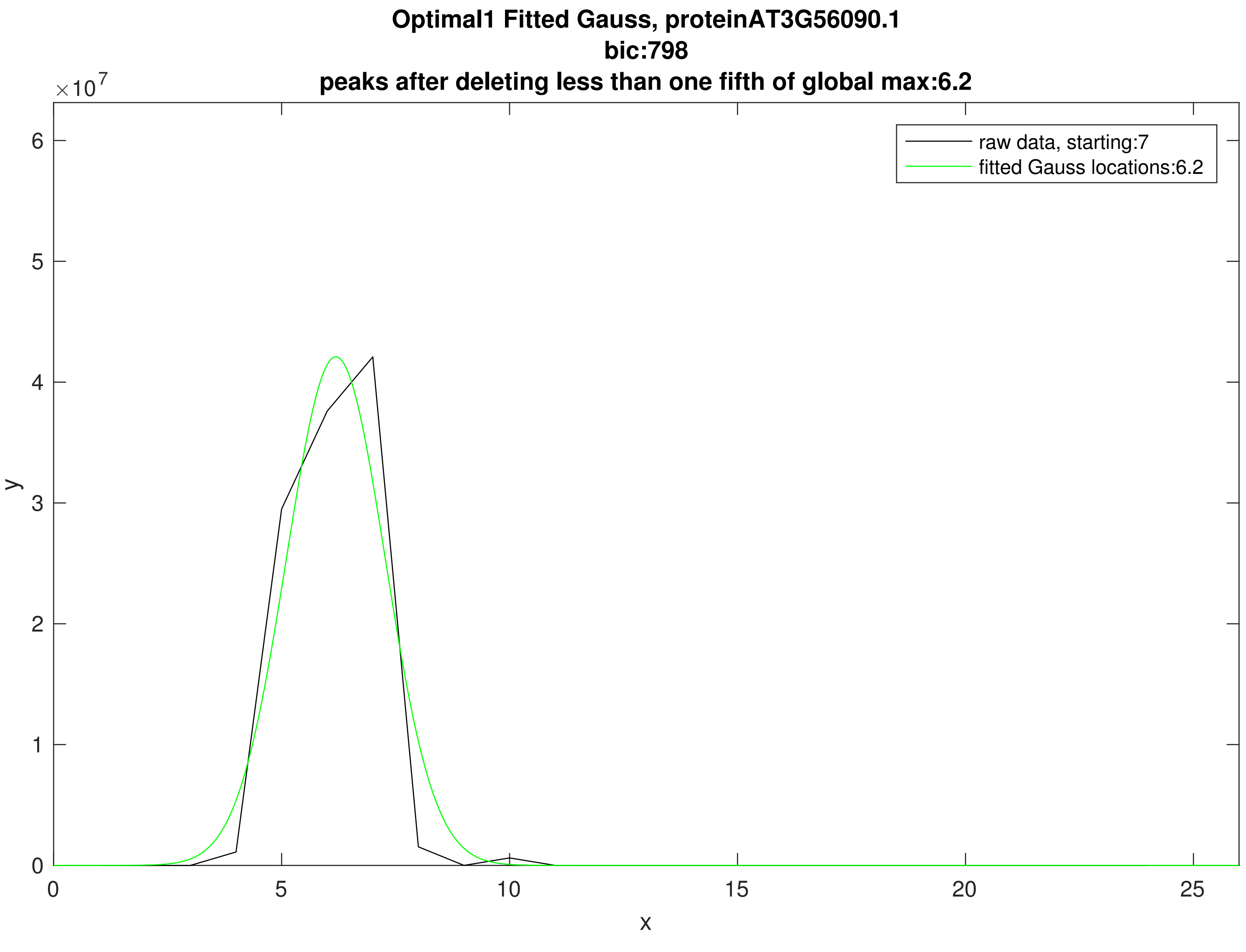

Optimal1 Fitted Gauss, proteinAT3G56090.1
bic:798
peaks after deleting less than one fifth of global max:6.2
×10^7
raw data, starting:7
fitted Gauss locations:6.2
y
x
0
1
2
3
4
5
6
0
5
10
15
20
25

**Optimal2 Fitted Gauss, proteinAT3G56120.1**
**bic:894 886**
**peaks after deleting less than one fifth of global max:14**

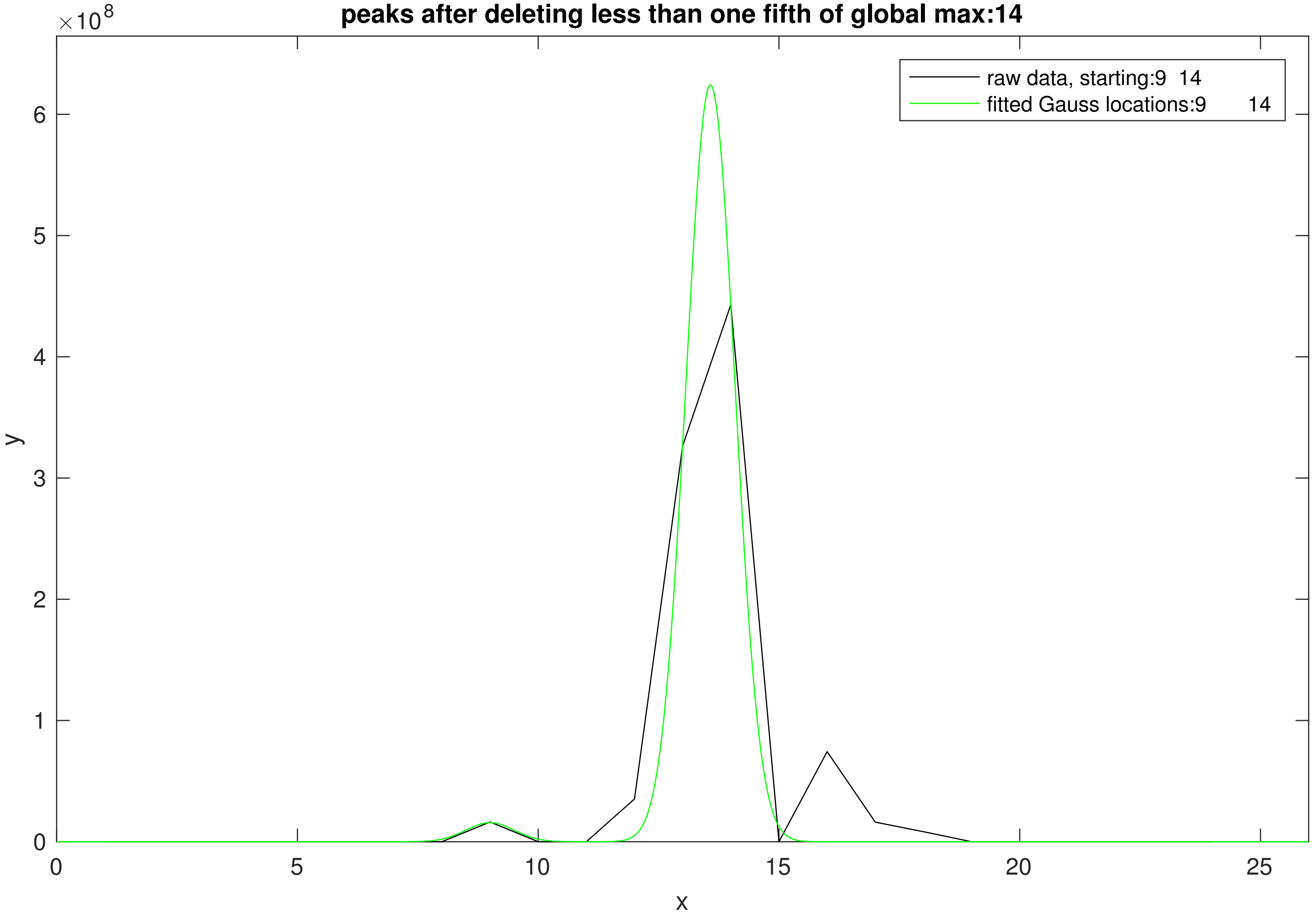

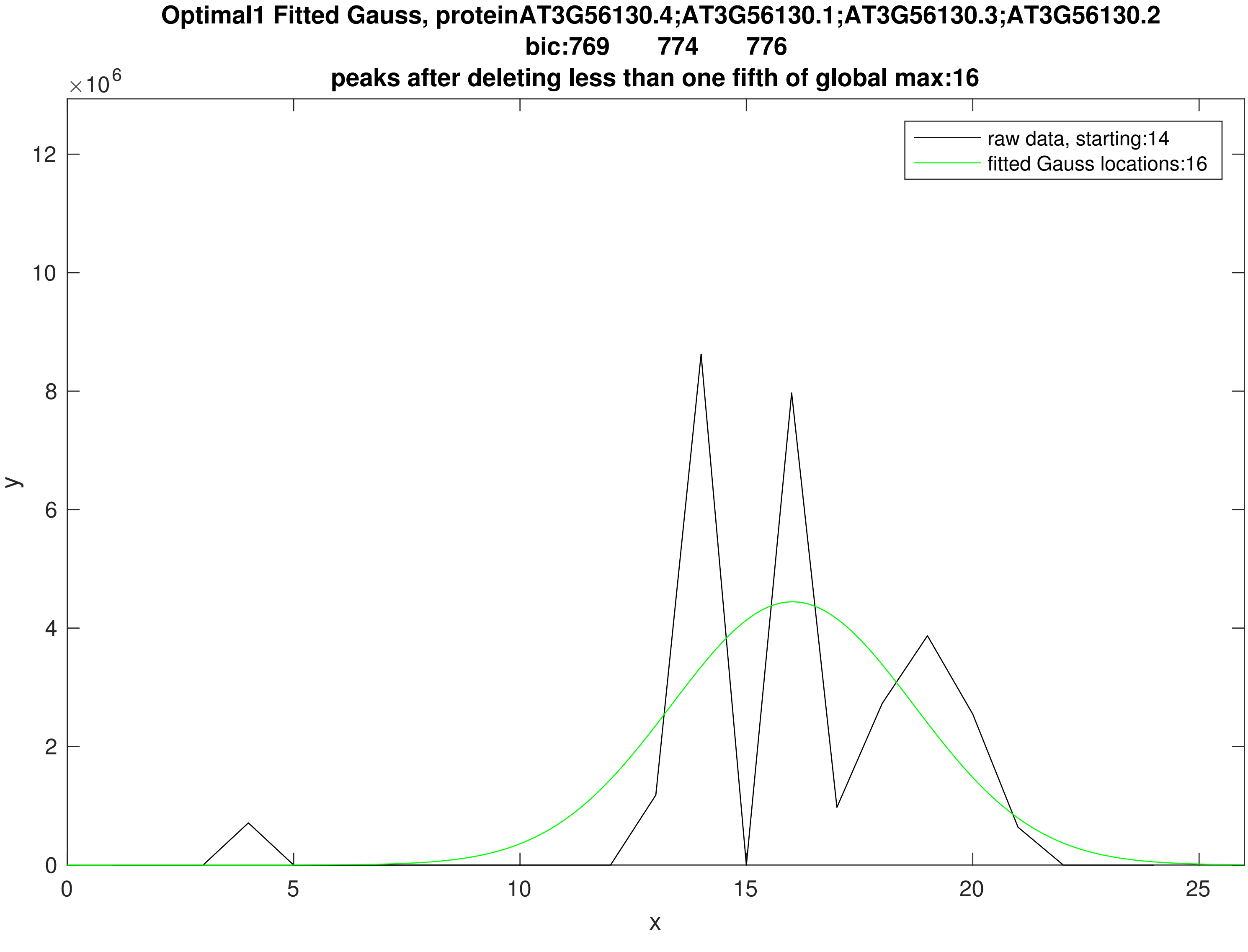
Optimal1 Fitted Gauss, proteinAT3G56130.4;AT3G56130.1;AT3G56130.3;AT3G56130.2
bic:769 774 776
peaks after deleting less than one fifth of global max:16
×10^6
raw data, starting:14
fitted Gauss locations:16
y
x
0
2
4
6
8
10
12
0
5
10
15
20
25

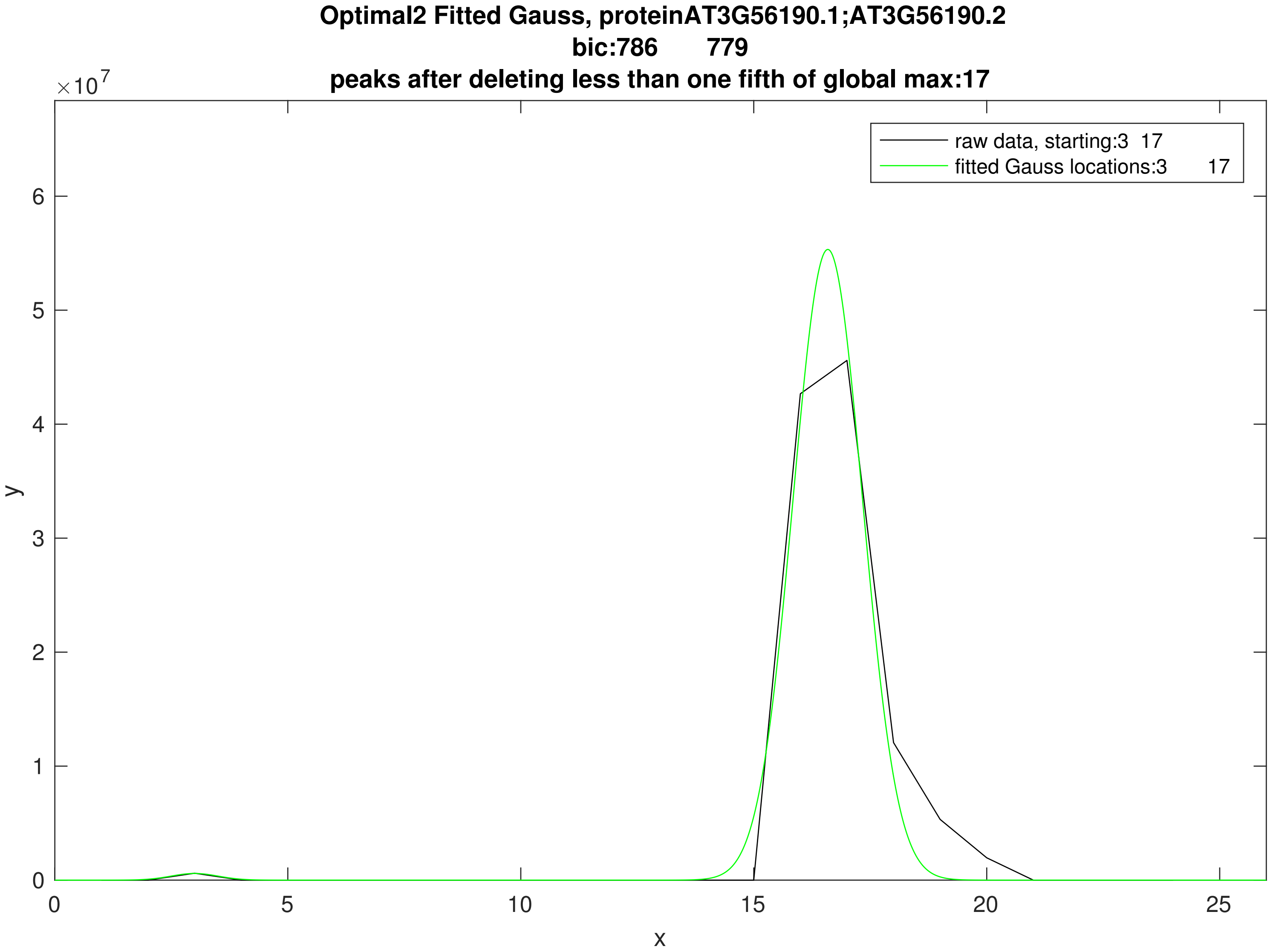
Optimal2 Fitted Gauss, proteinAT3G56190.1;AT3G56190.2
bic:786 779
peaks after deleting less than one fifth of global max:17
×10^7
raw data, starting:3 17
fitted Gauss locations:3 17
y
x
0
1
2
3
4
5
6
0
5
10
15
20
25

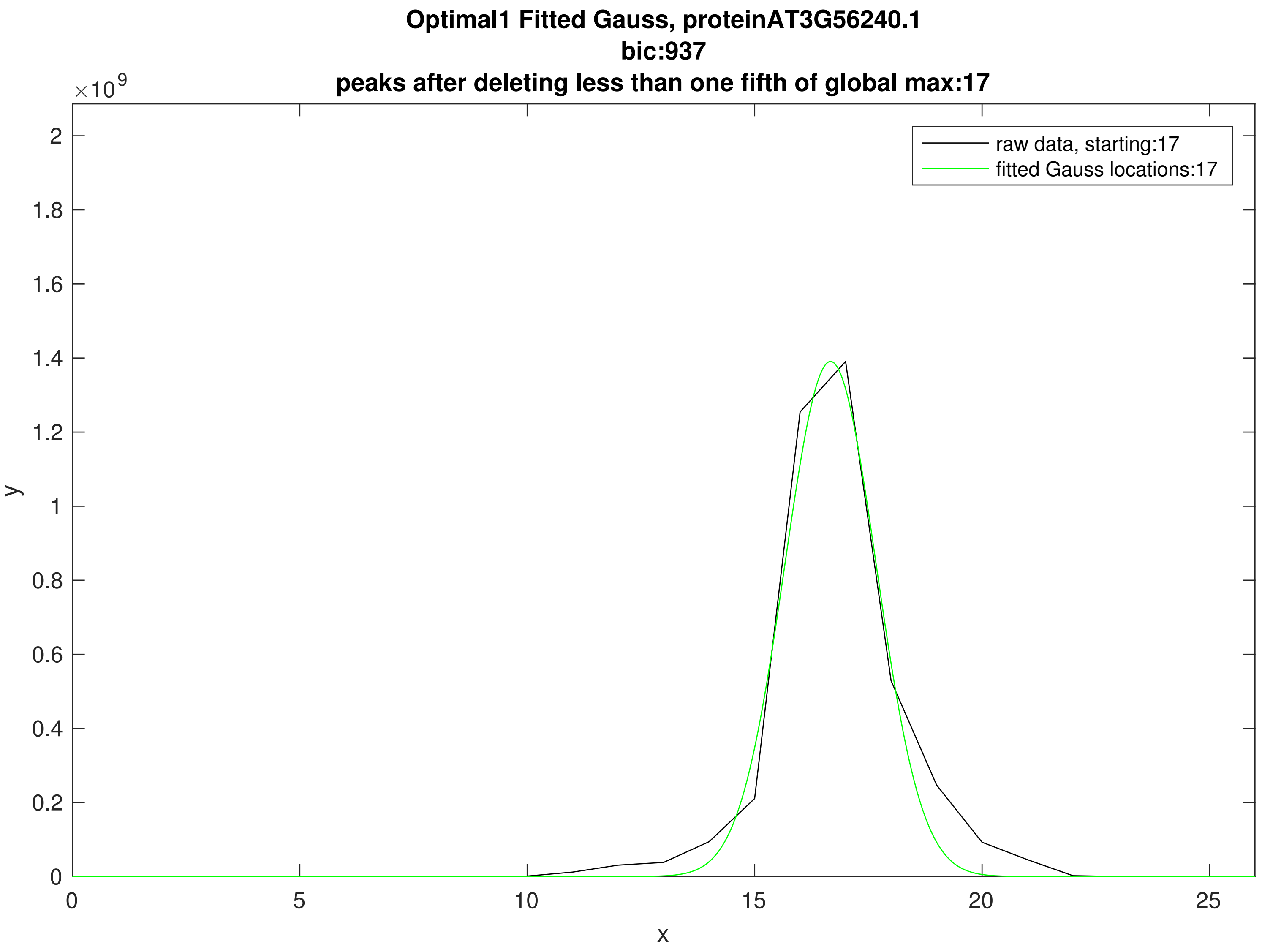

Optimal1 Fitted Gauss, proteinAT3G56240.1
bic:937
peaks after deleting less than one fifth of global max:17
×10^9
raw data, starting:17
fitted Gauss locations:17
y
x
0
0.2
0.4
0.6
0.8
1
1.2
1.4
1.6
1.8
2
0
5
10
15
20
25

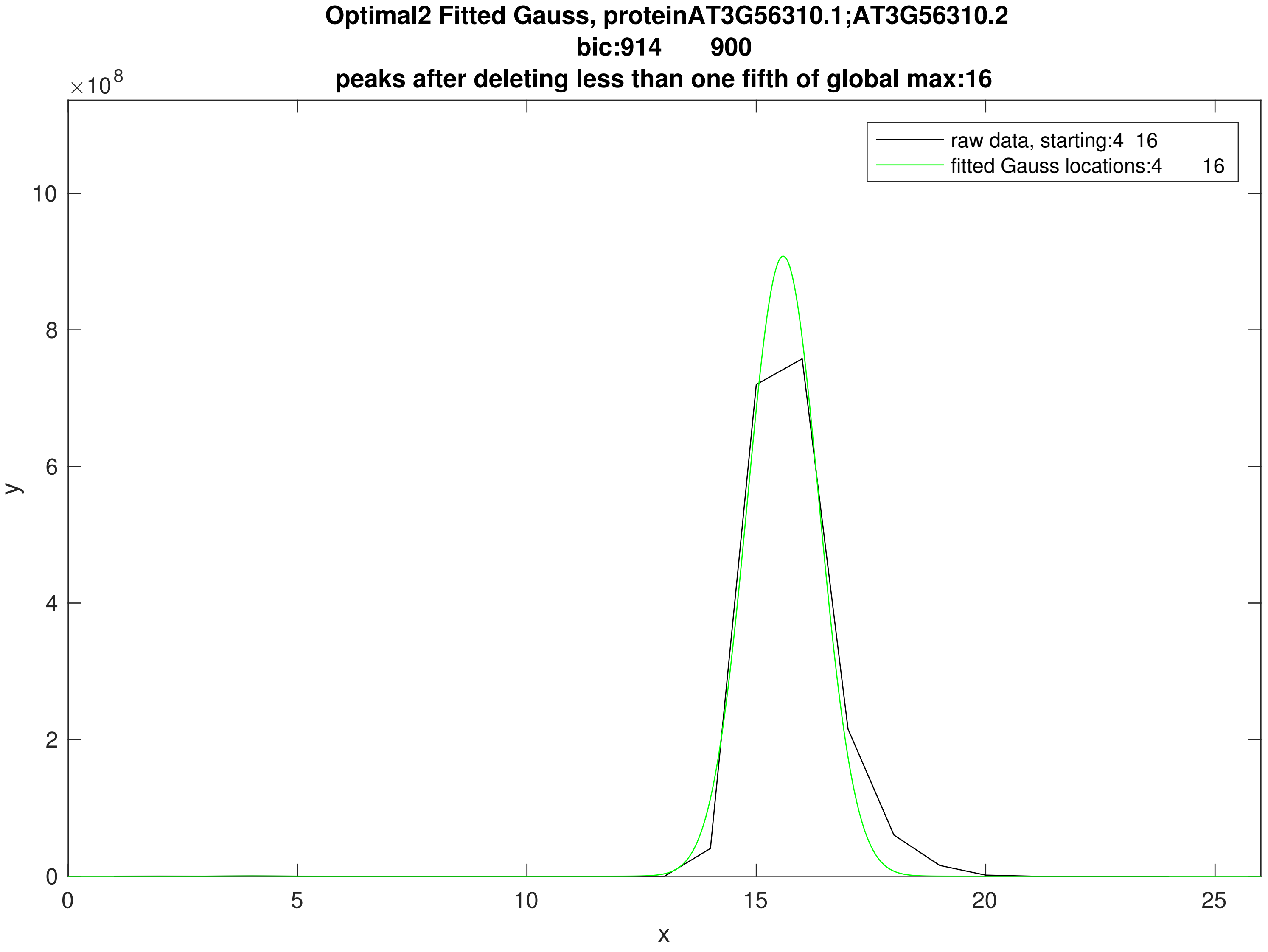
Optimal2 Fitted Gauss, proteinAT3G56310.1;AT3G56310.2
bic:914 900
peaks after deleting less than one fifth of global max:16
×10^8
raw data, starting:4 16
fitted Gauss locations:4 16
y
x
0
2
4
6
8
10
0
5
10
15
20
25

**Optimal1 Fitted Gauss, proteinAT3G56350.1**
**bic:915 924 934**
**peaks after deleting less than one fifth of global max:13**

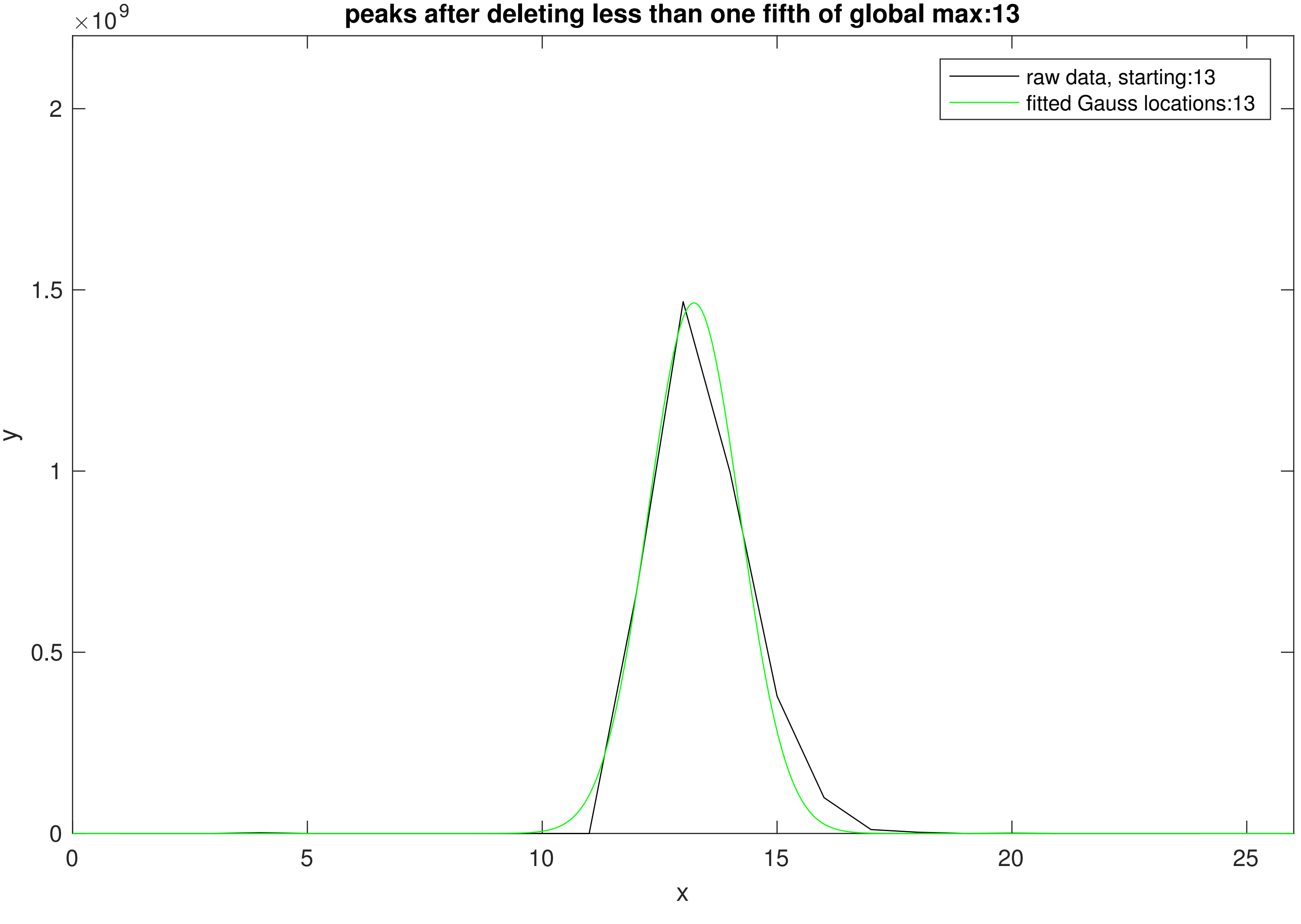

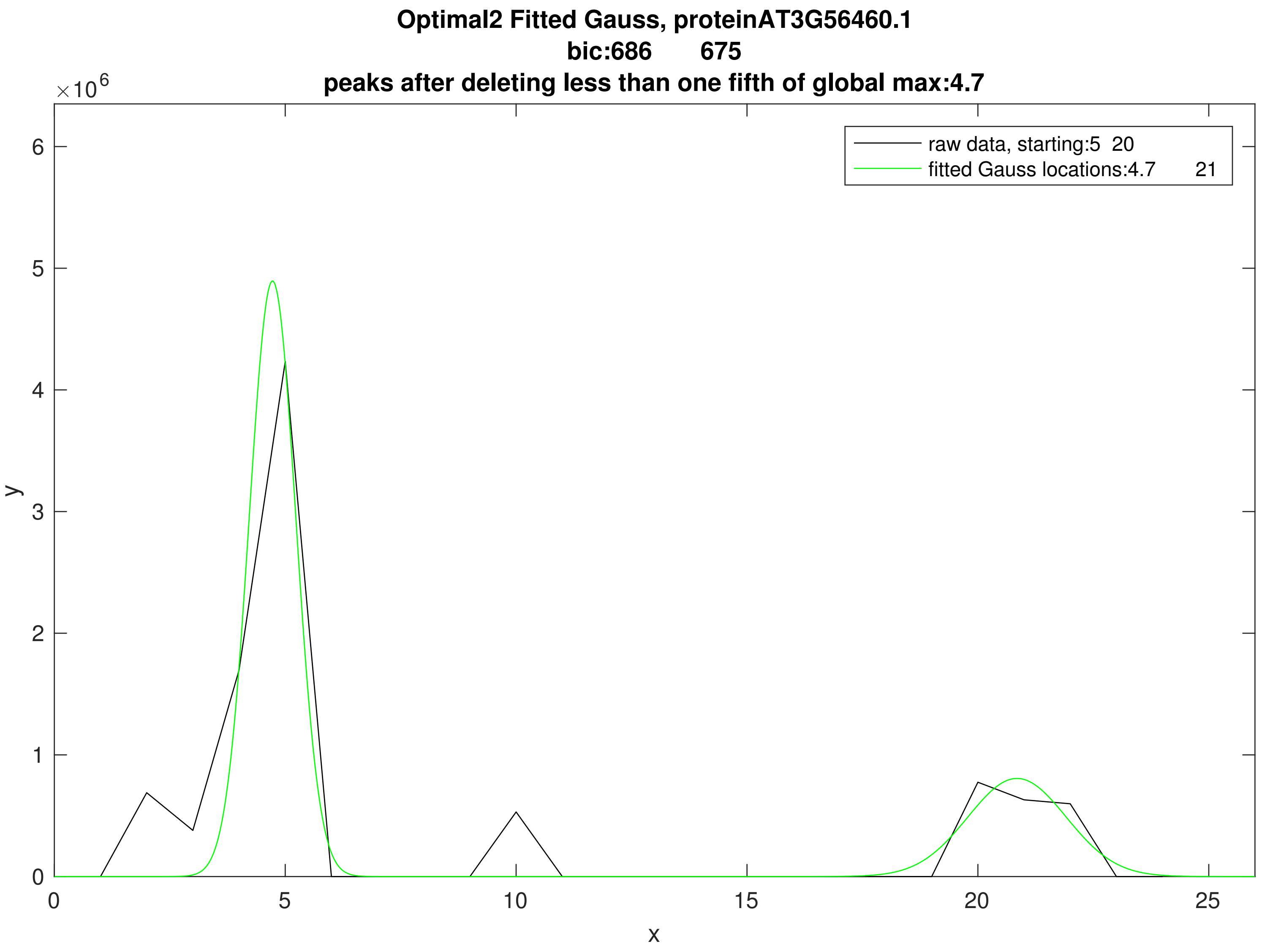
Optimal2 Fitted Gauss, proteinAT3G56460.1
bic:686 675
peaks after deleting less than one fifth of global max:4.7
×10^6
raw data, starting:5 20
fitted Gauss locations:4.7 21
y
x
0
1
2
3
4
5
6
0
5
10
15
20
25

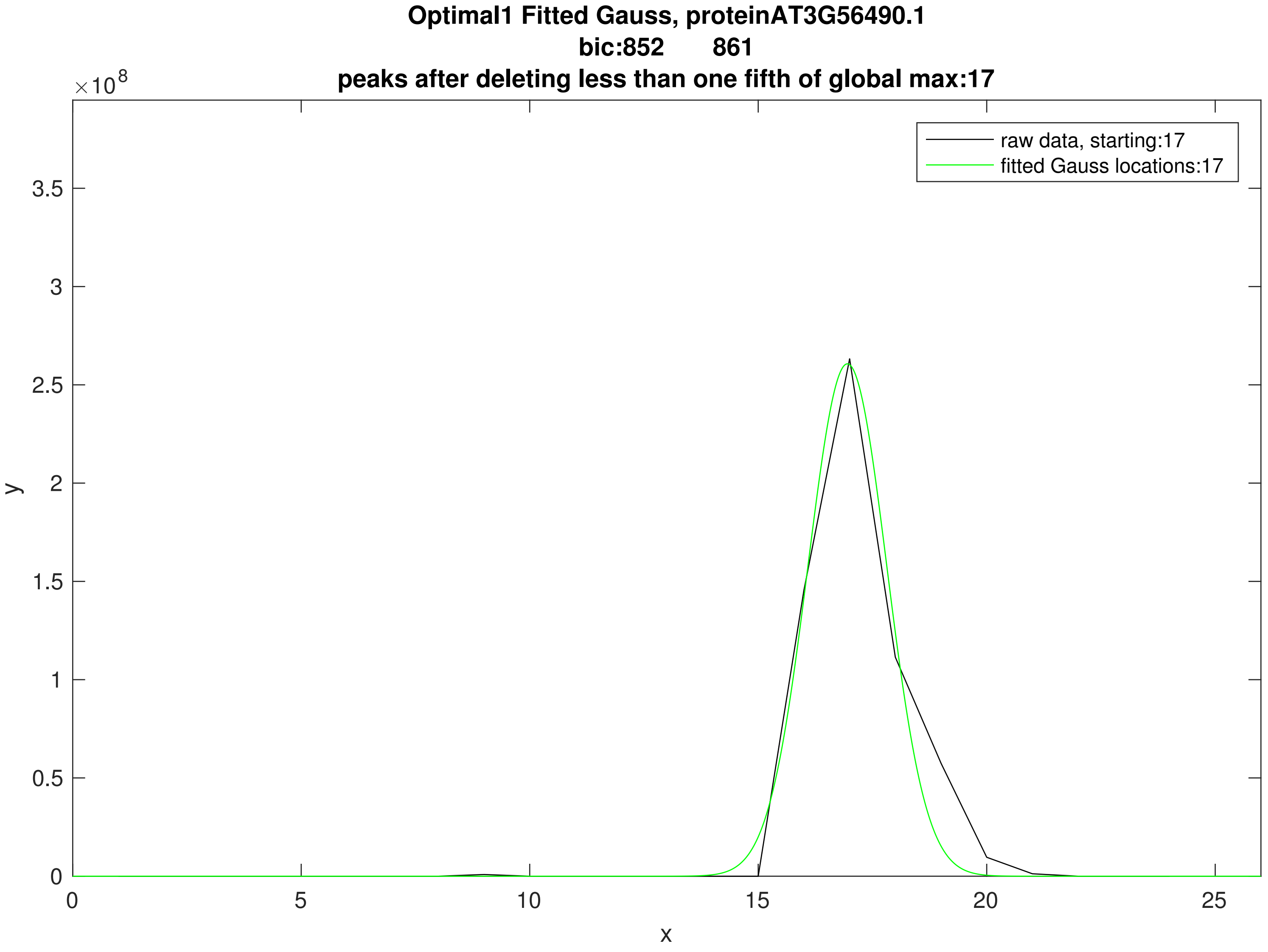

Optimal1 Fitted Gauss, proteinAT3G56490.1
bic:852 861
peaks after deleting less than one fifth of global max:17
×10^8
raw data, starting:17
fitted Gauss locations:17
y
x
0
0.5
1
1.5
2
2.5
3
3.5
5
10
15
20
25

**Fitted Gauss, proteinAT3G56800.1;AT2G41110.1;AT2G27030.1;AT2G27030.3;AT5G21274.1;AT3G43810.1;AT2G41110.**
**bic:1.02e+03  1.03e+03  1.04e+03  1.05e+03**
**peaks after deleting less than one fifth of global max:18**

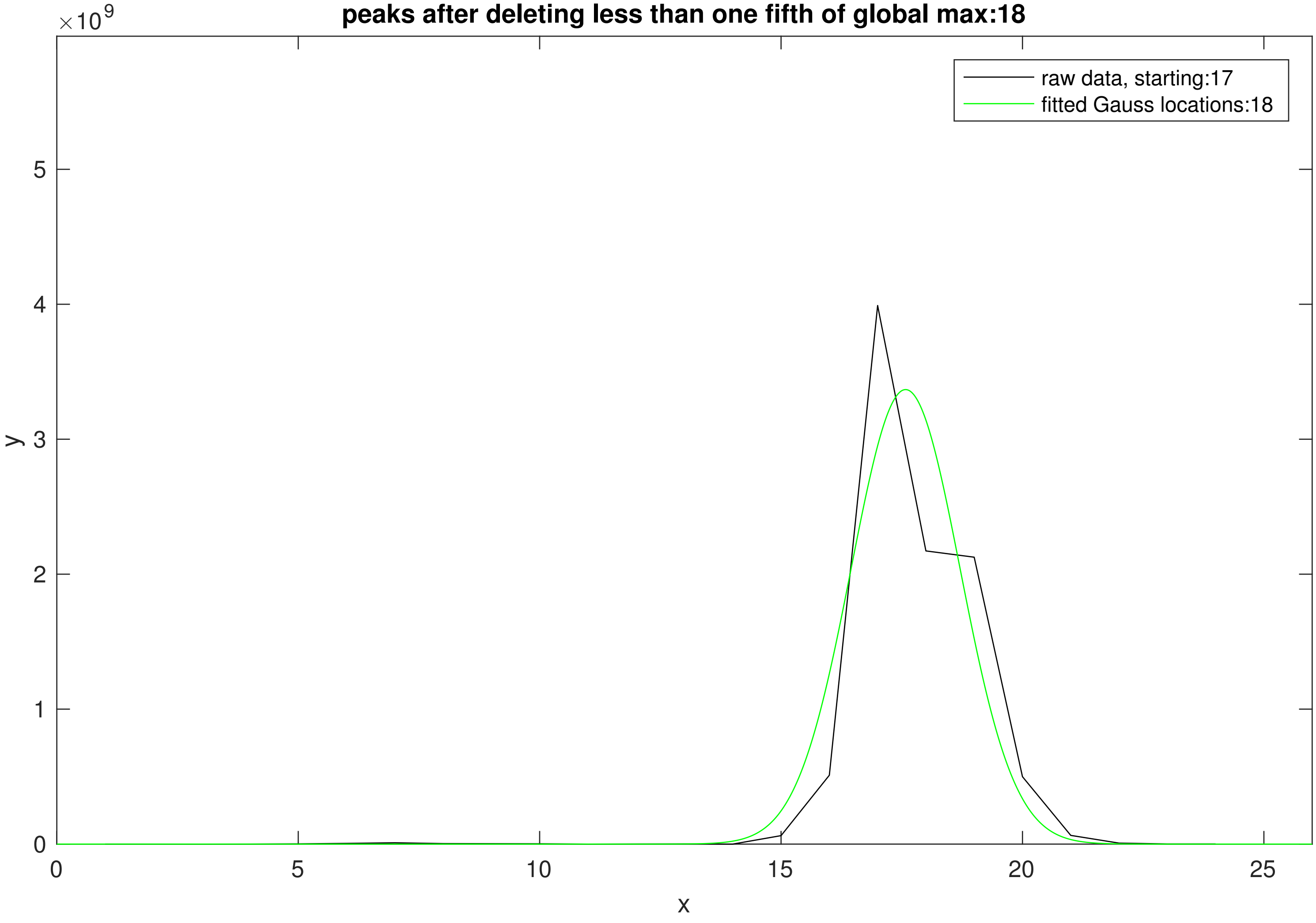

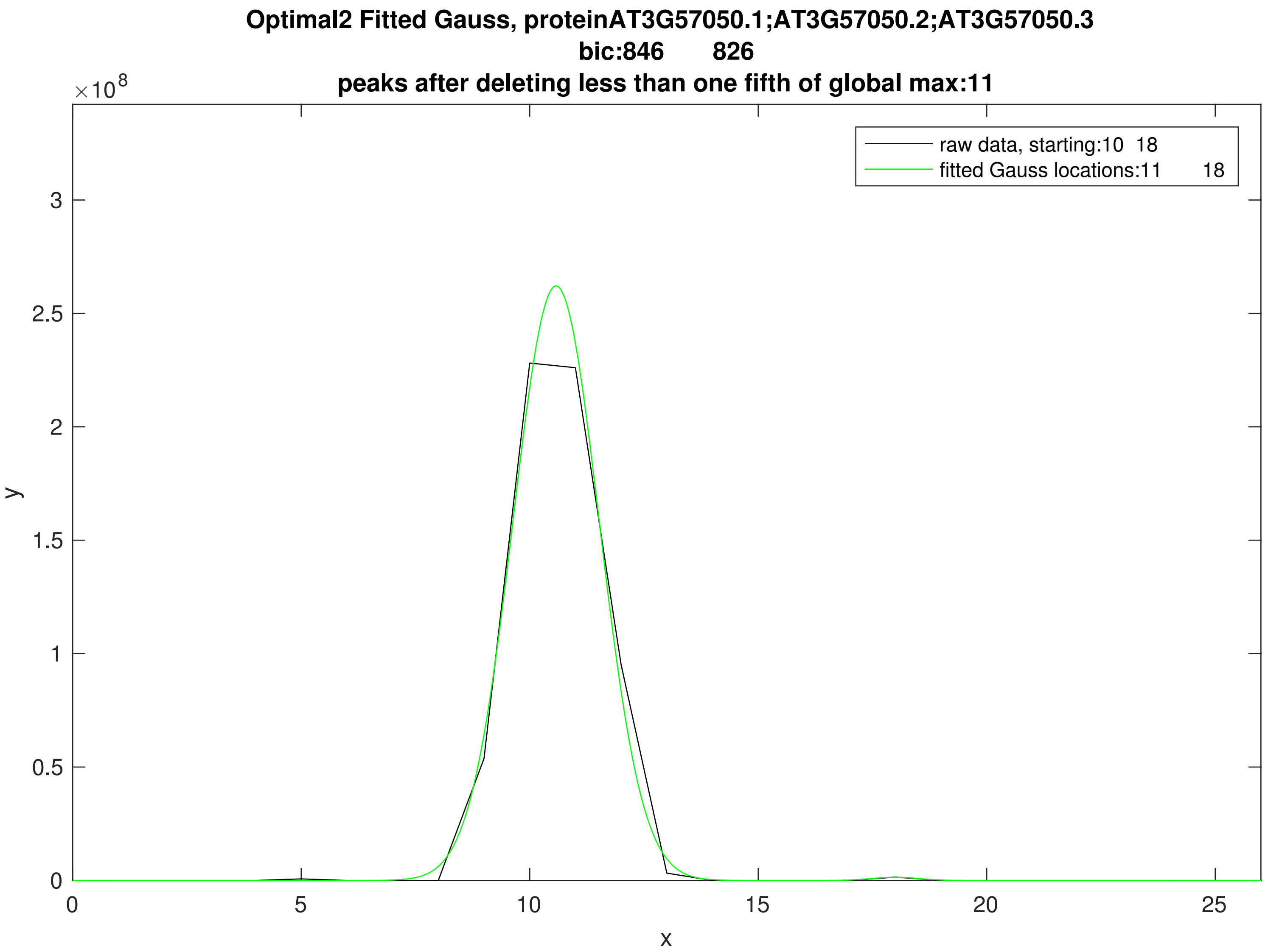

Optimal2 Fitted Gauss, proteinAT3G57050.1;AT3G57050.2;AT3G57050.3
bic:846 826
peaks after deleting less than one fifth of global max:11
×10^8
raw data, starting:10 18
fitted Gauss locations:11 18
y
x
0
0.5
1
1.5
2
2.5
3
0
5
10
15
20
25

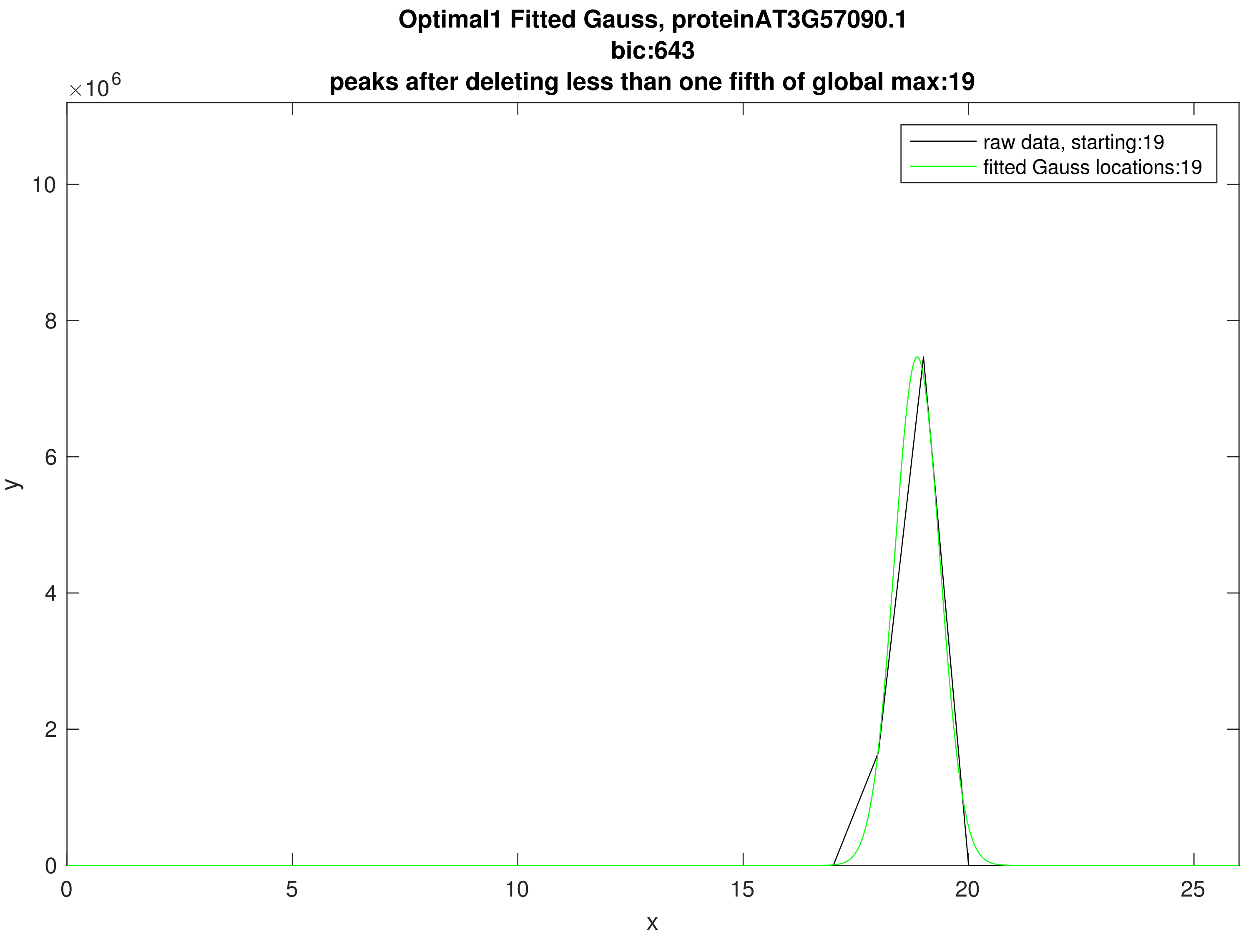

Optimal1 Fitted Gauss, proteinAT3G57090.1
bic:643
peaks after deleting less than one fifth of global max:19
×10^6
raw data, starting:19
fitted Gauss locations:19
10
8
6
4
2
0
y
0
5
10
15
20
25
x

#### Optimal1 Fitted Gauss, proteinAT3G57410.1
#### bic:768
#### peaks after deleting less than one fifth of global max:16

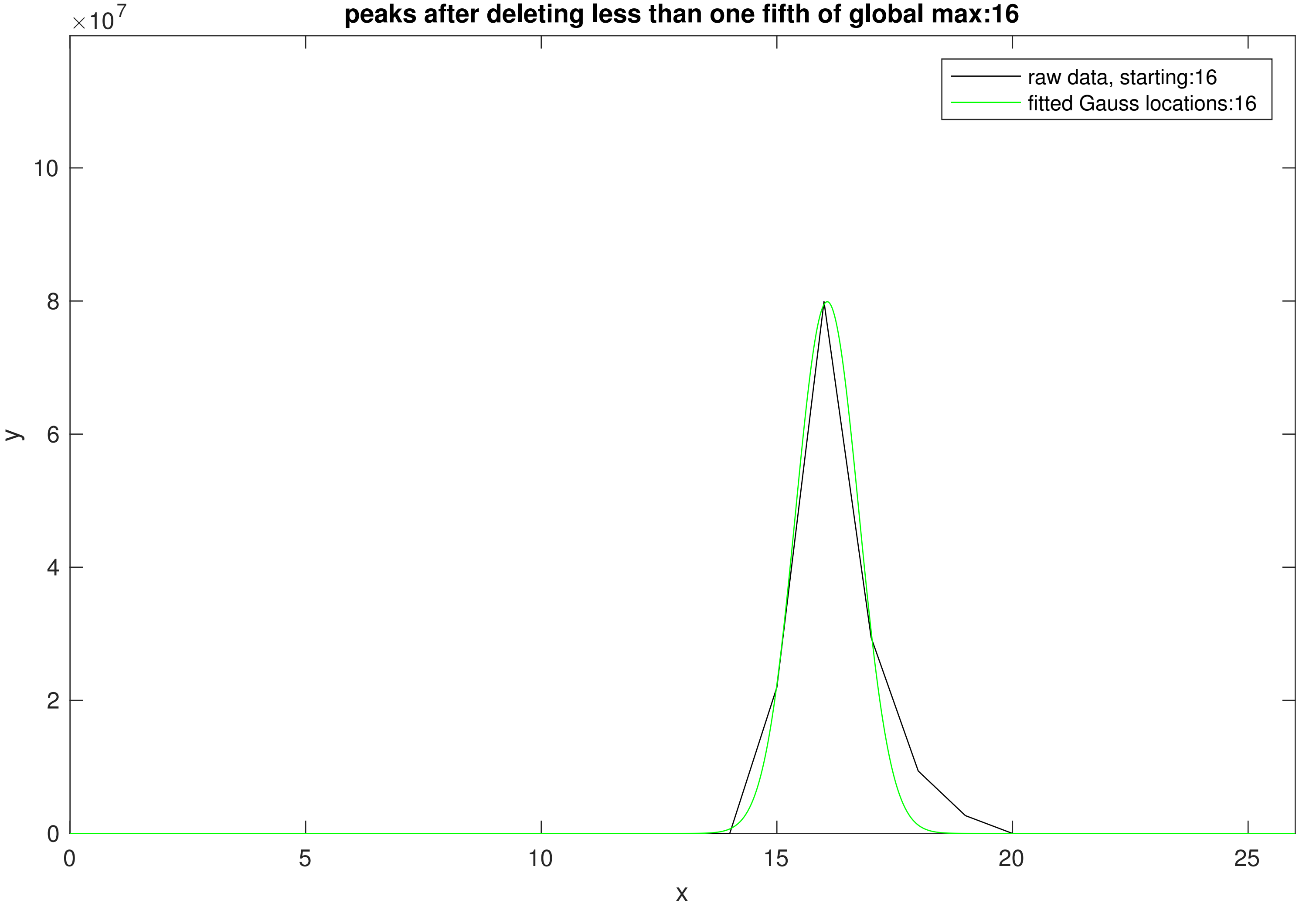

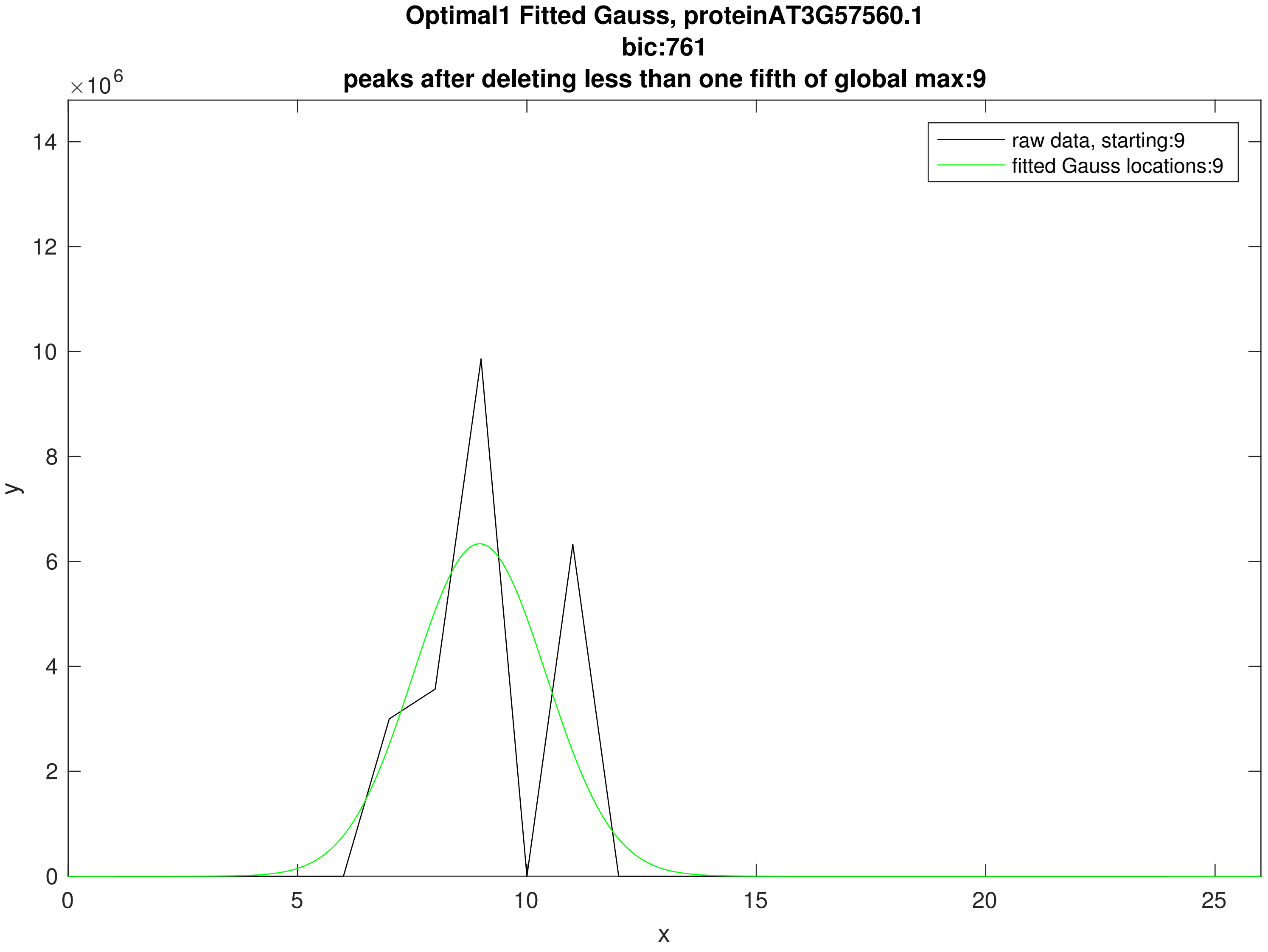

Optimal1 Fitted Gauss, proteinAT3G57560.1
bic:761
peaks after deleting less than one fifth of global max:9
×10^6
14
12
10
8
6
4
2
0
y
0
5
10
15
20
25
x
raw data, starting:9
fitted Gauss locations:9

**Optimal1 Fitted Gauss, proteinAT3G57610.1**
**bic:865 870**
**peaks after deleting less than one fifth of global max:13**

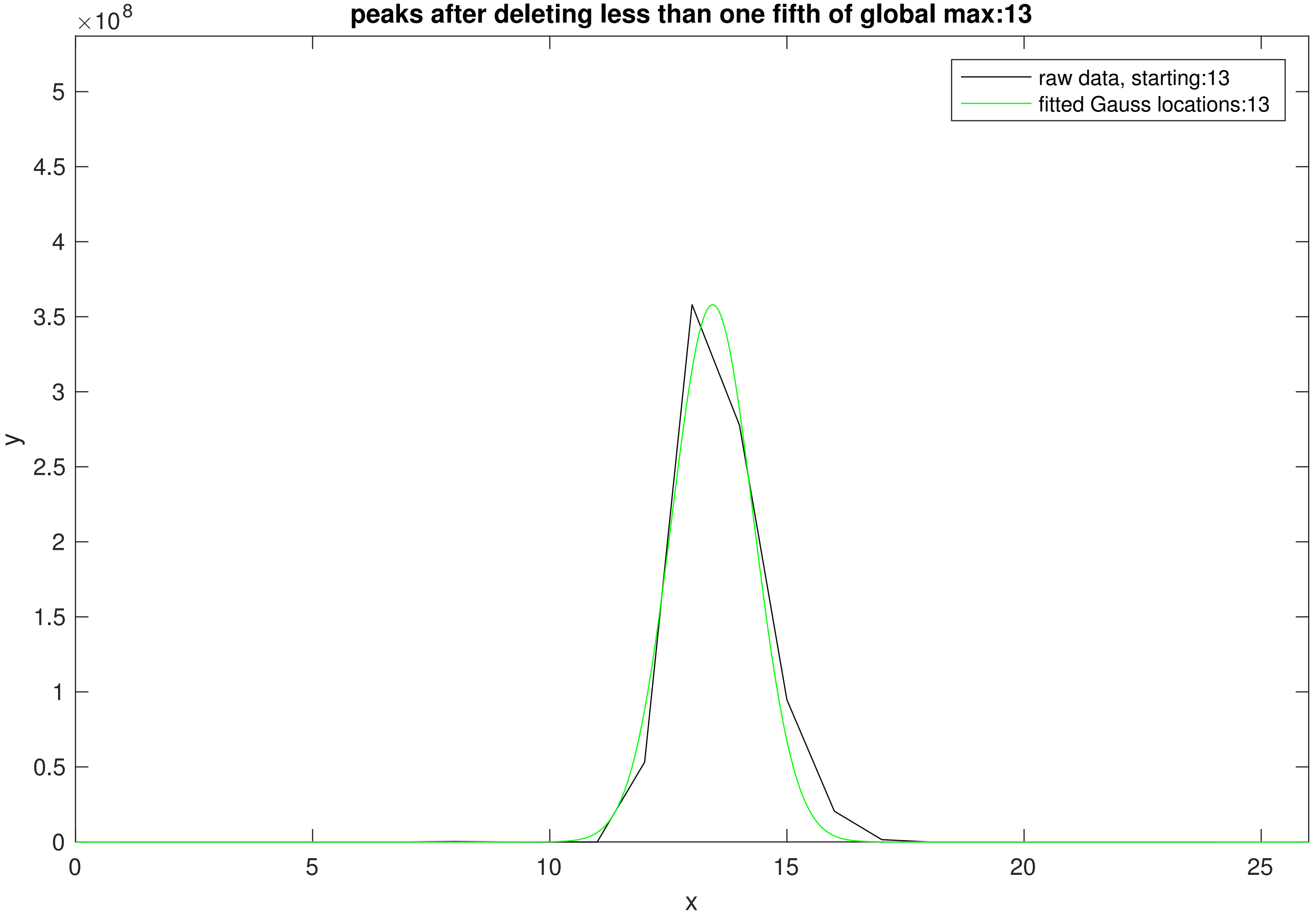

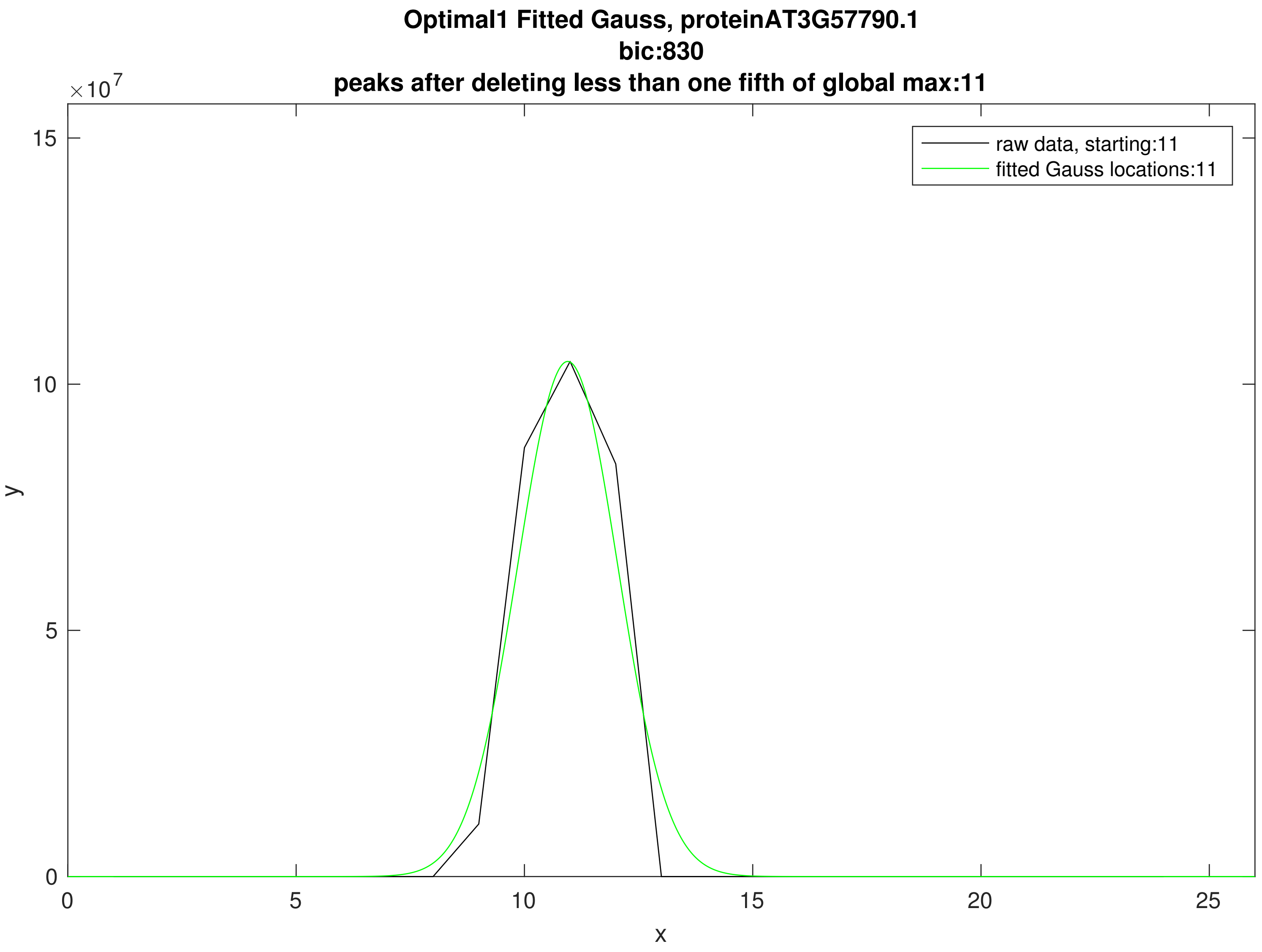
Optimal1 Fitted Gauss, proteinAT3G57790.1
bic:830
peaks after deleting less than one fifth of global max:11
×10^7
15
10
5
0
y
0
5
10
15
20
25
x
raw data, starting:11
fitted Gauss locations:11

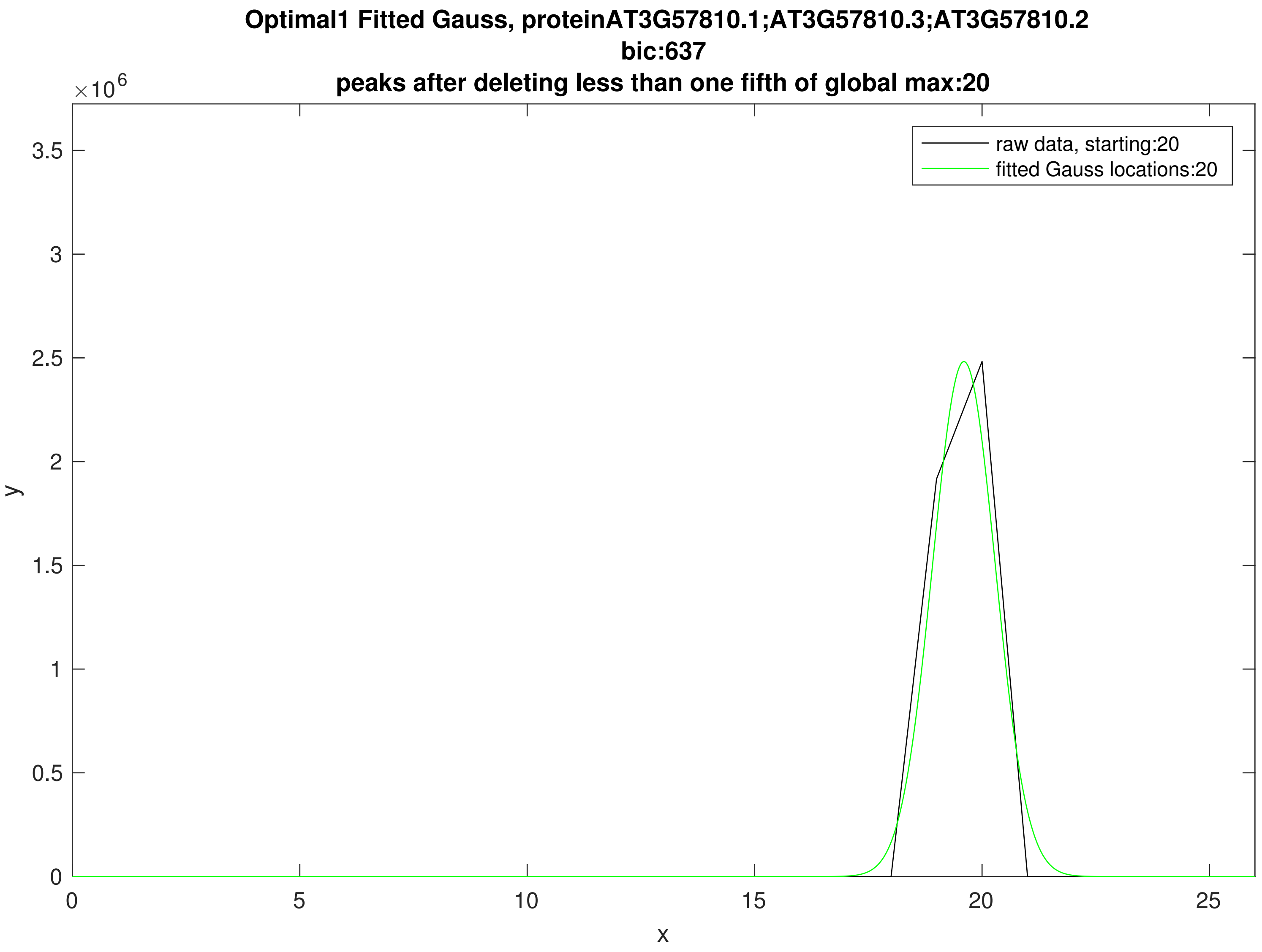
Optimal1 Fitted Gauss, proteinAT3G57810.1;AT3G57810.3;AT3G57810.2
bic:637
peaks after deleting less than one fifth of global max:20
×10^6
raw data, starting:20
fitted Gauss locations:20
3.5
3
2.5
2
1.5
1
0.5
0
y
0
5
10
15
20
25
x

Optimal1 Fitted Gauss, proteinAT3G57890.1;AT3G57890.2
bic:676
peaks after deleting less than one fifth of global max:9.7

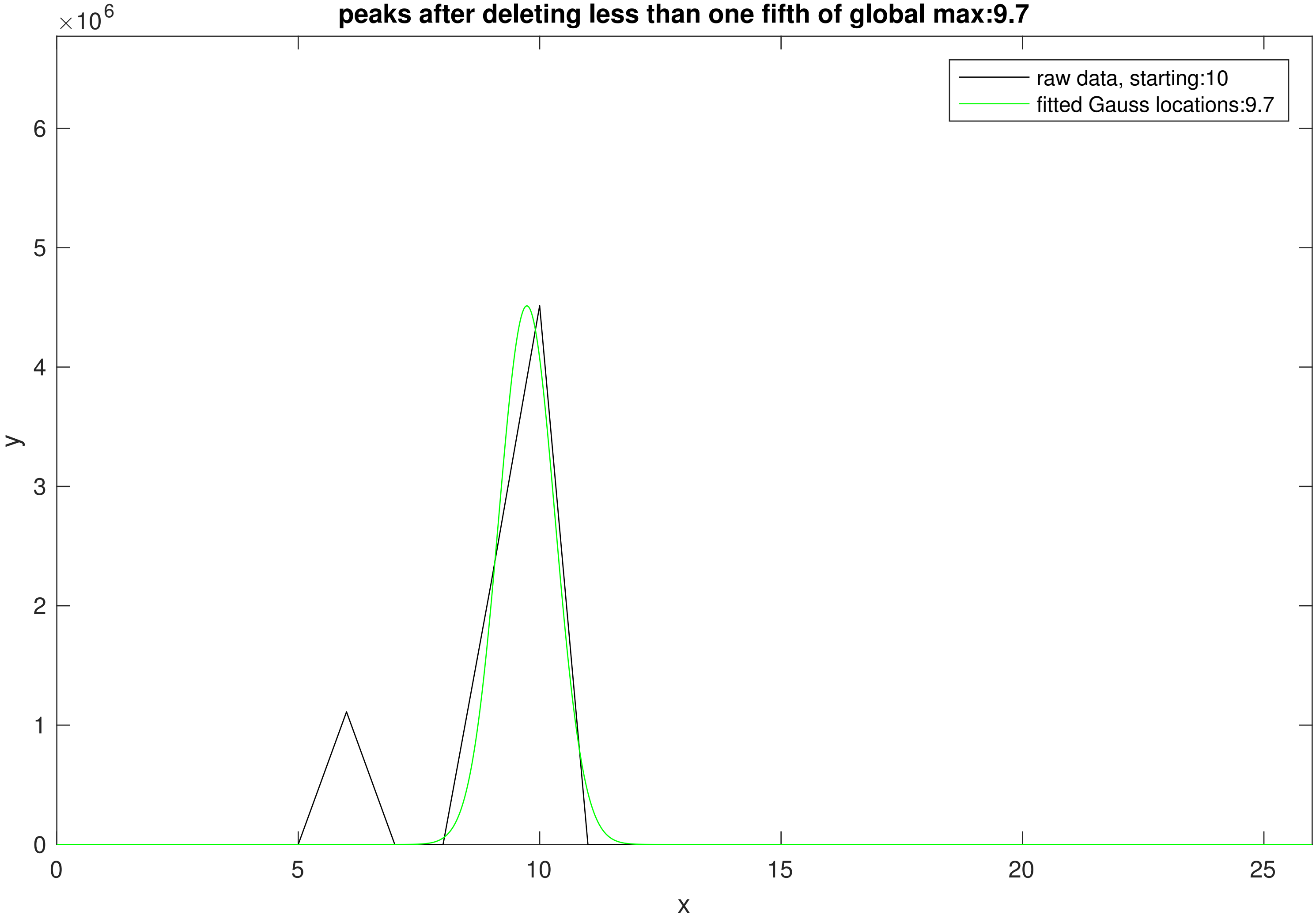

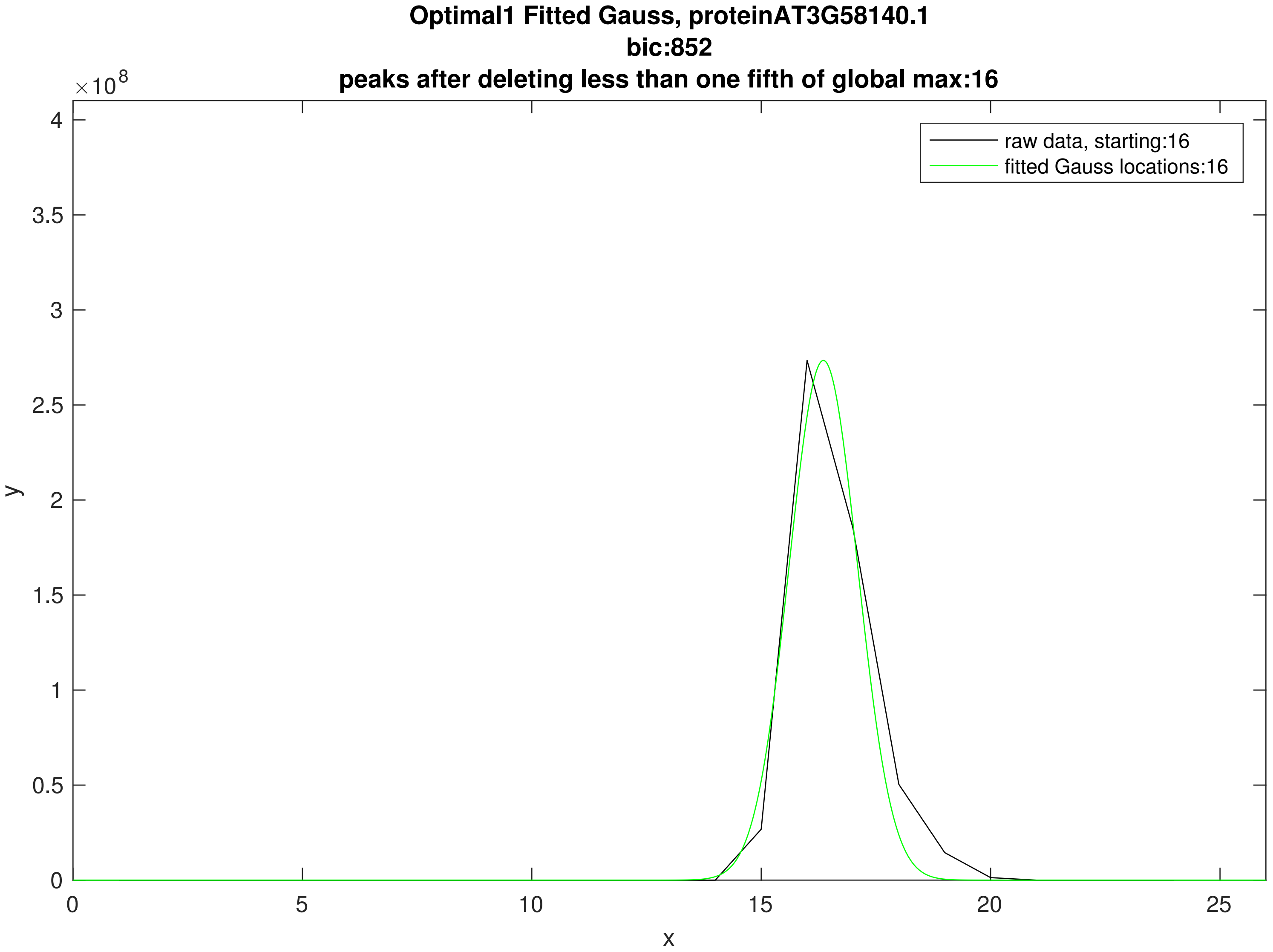
Optimal1 Fitted Gauss, proteinAT3G58140.1
bic:852
peaks after deleting less than one fifth of global max:16
×10^8
raw data, starting:16
fitted Gauss locations:16
y
x
0
0.5
1
1.5
2
2.5
3
3.5
4
0
5
10
15
20
25

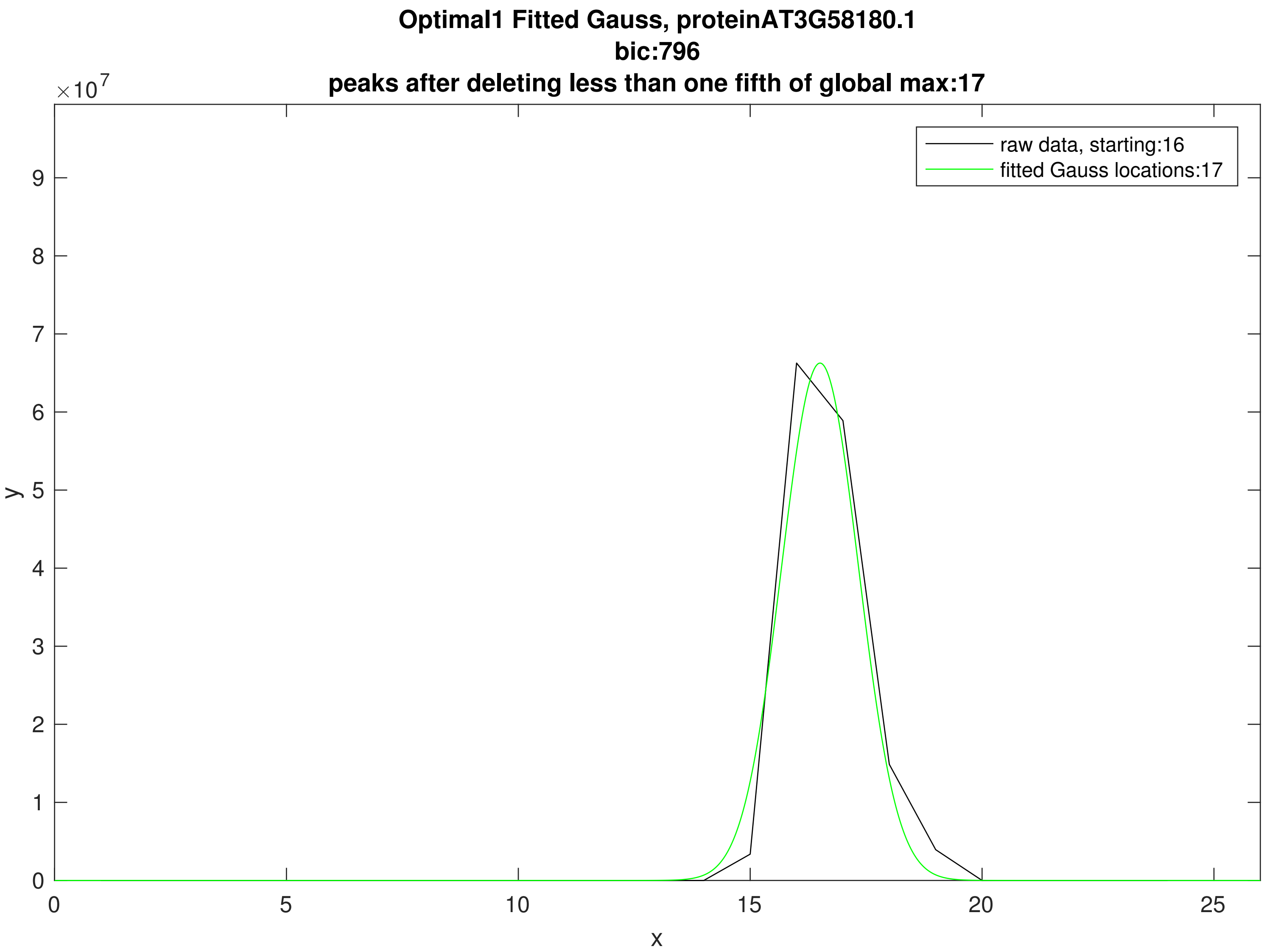
Optimal1 Fitted Gauss, proteinAT3G58180.1
bic:796
peaks after deleting less than one fifth of global max:17
×10^7
raw data, starting:16
fitted Gauss locations:17
y
x
0
1
2
3
4
5
6
7
8
9
0
5
10
15
20
25

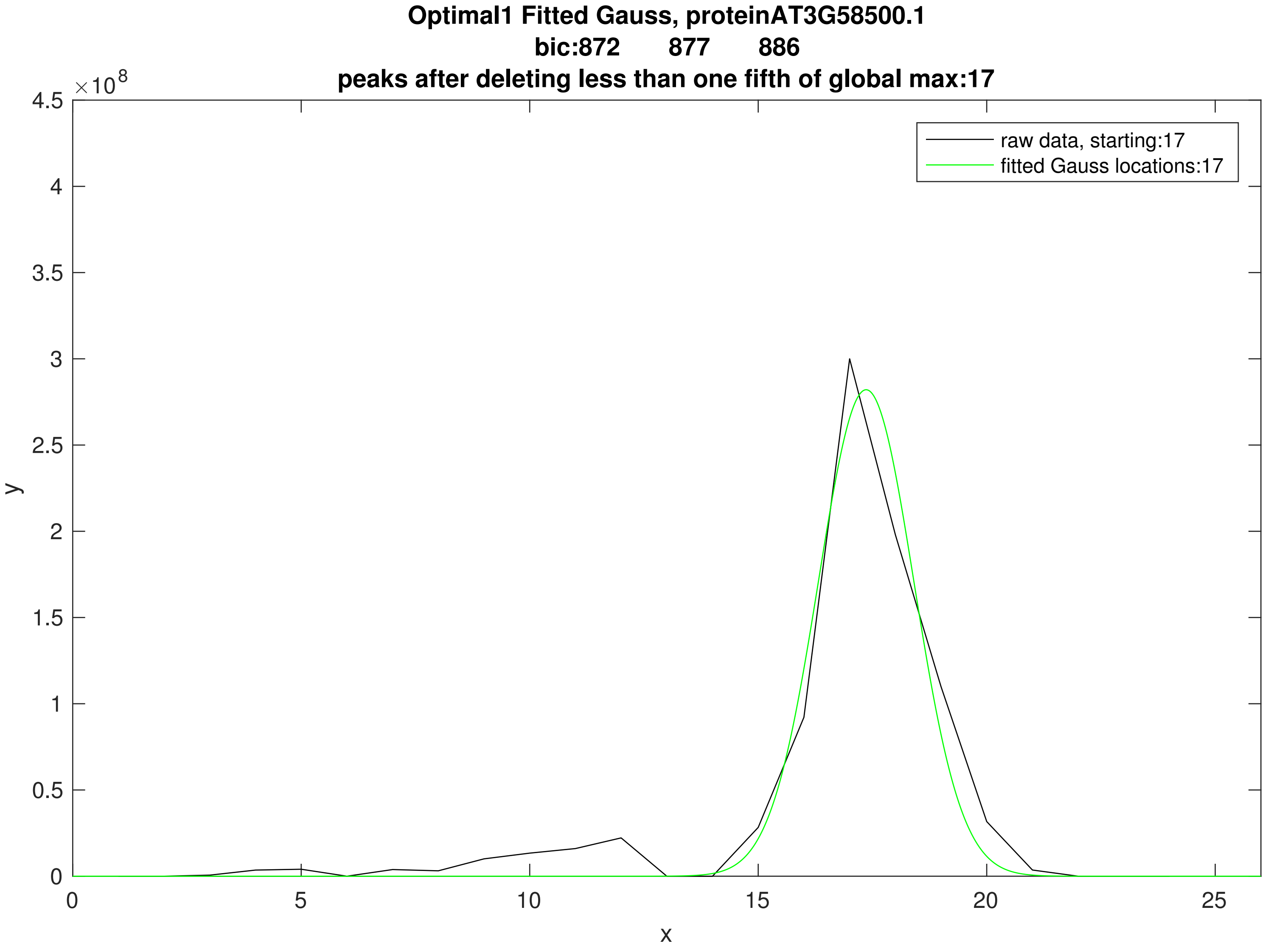
Optimal1 Fitted Gauss, proteinAT3G58500.1
bic:872 877 886
peaks after deleting less than one fifth of global max:17
×10^8
4.5
4
3.5
3
2.5
2
1.5
1
0.5
0
y
0
5
10
15
20
25
x
raw data, starting:17
fitted Gauss locations:17

Optimal1 Fitted Gauss, proteinAT3G58510.3;AT3G58510.2;AT3G58510.1;AT2G42520.1;AT3G58570.1
bic:902 906 917 928
peaks after deleting less than one fifth of global max:14

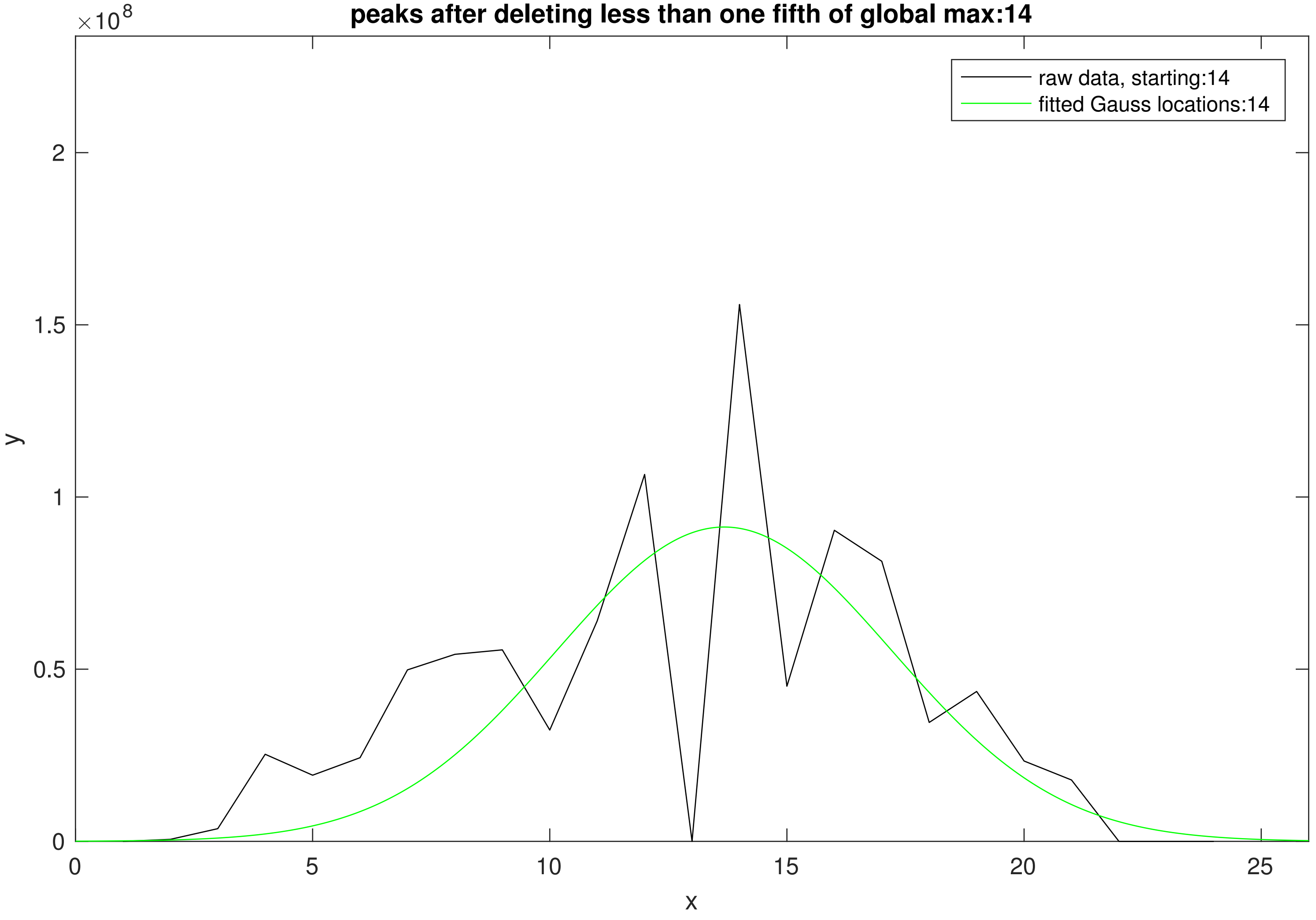

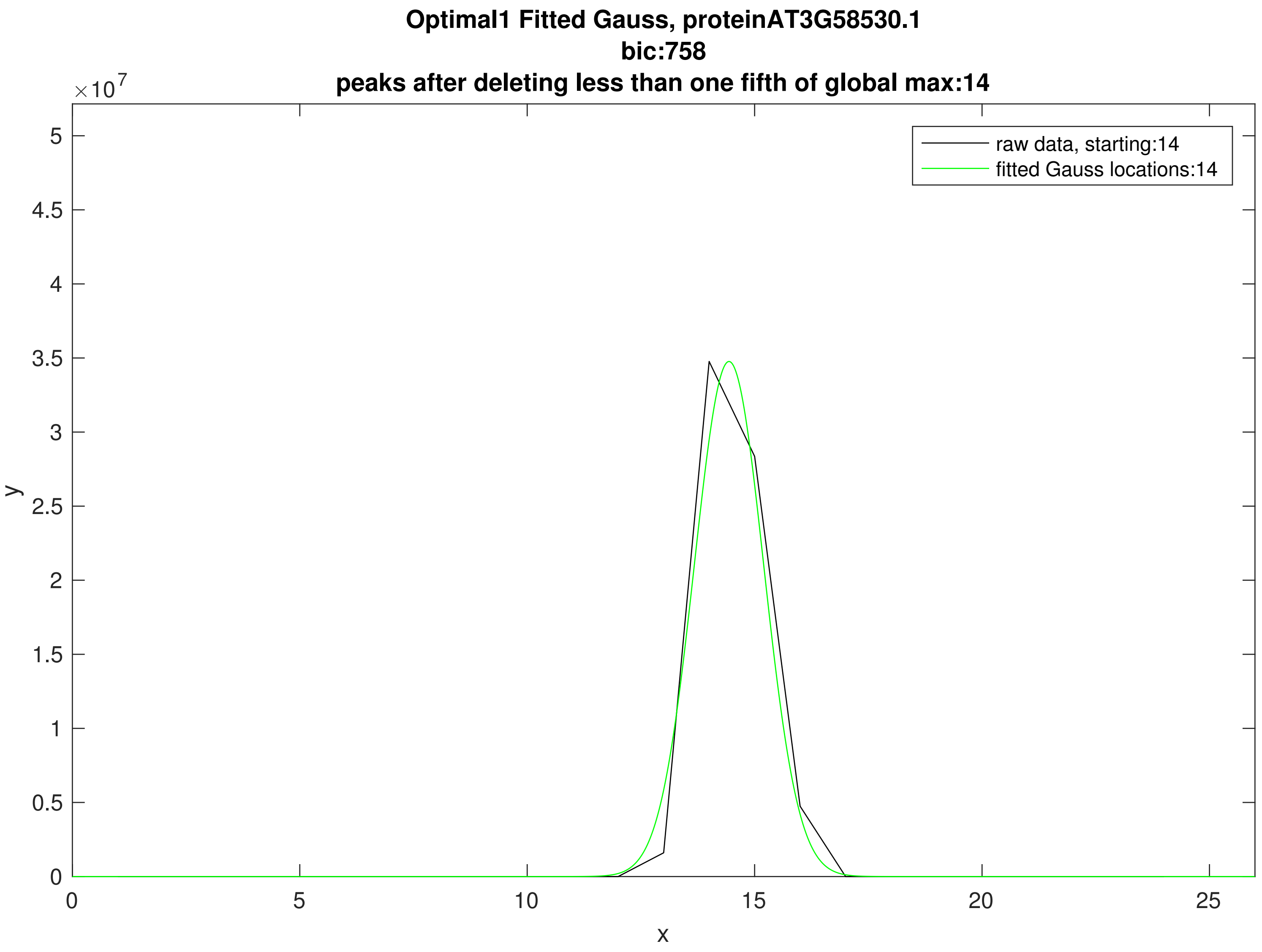
Optimal1 Fitted Gauss, proteinAT3G58530.1
bic:758
peaks after deleting less than one fifth of global max:14
×10^7
5
4.5
4
3.5
3
2.5
2
1.5
1
0.5
0
y
0
5
10
15
20
25
x
raw data, starting:14
fitted Gauss locations:14

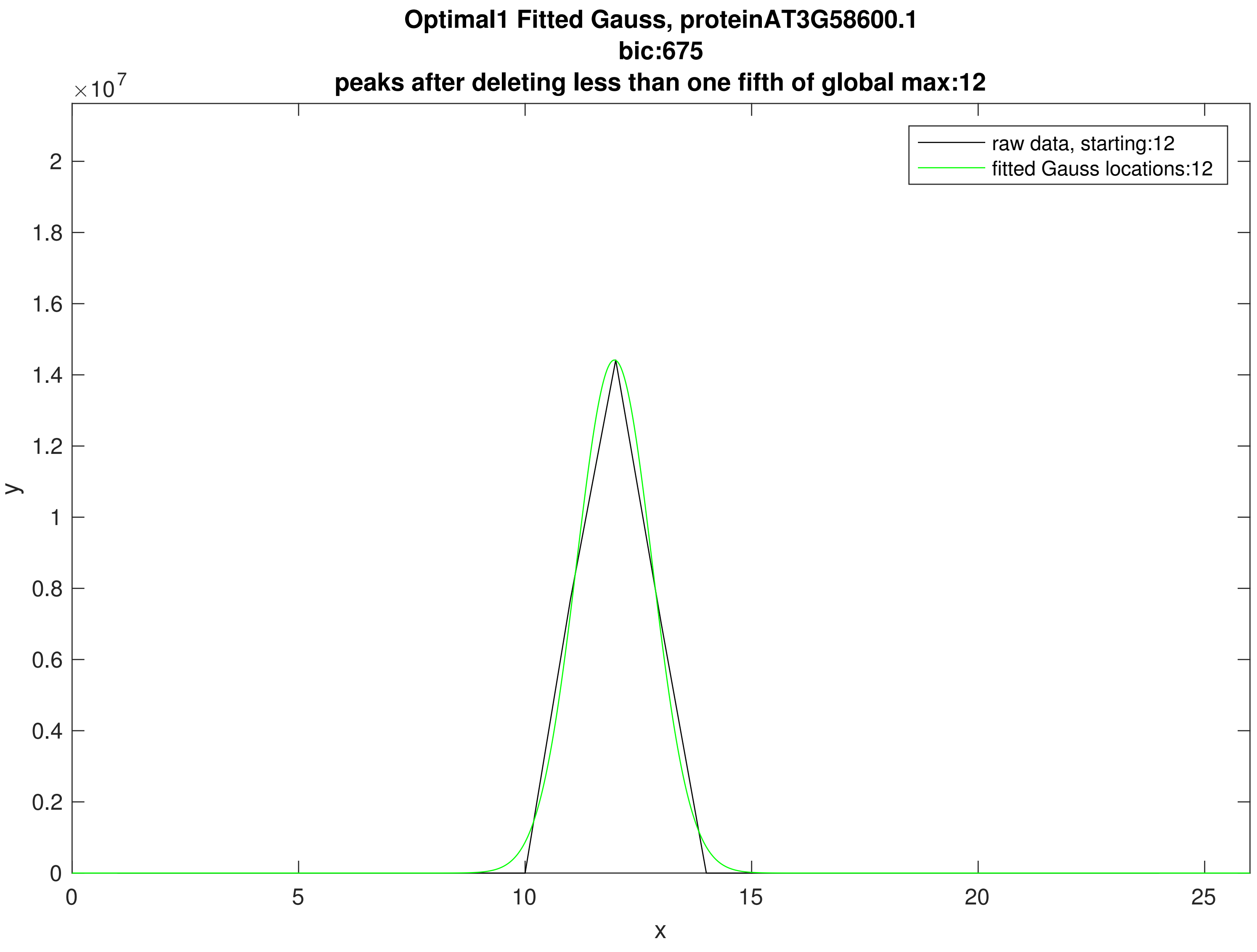

Optimal1 Fitted Gauss, proteinAT3G58600.1
bic:675
peaks after deleting less than one fifth of global max:12
×10^7
raw data, starting:12
fitted Gauss locations:12
y
x
0
0.2
0.4
0.6
0.8
1
1.2
1.4
1.6
1.8
2
0
5
10
15
20
25

Optimal1 Fitted Gauss, proteinAT3G58610.3;AT3G58610.2;AT3G58610.1
bic:977  1.05e+03
peaks after deleting less than one fifth of global max:13

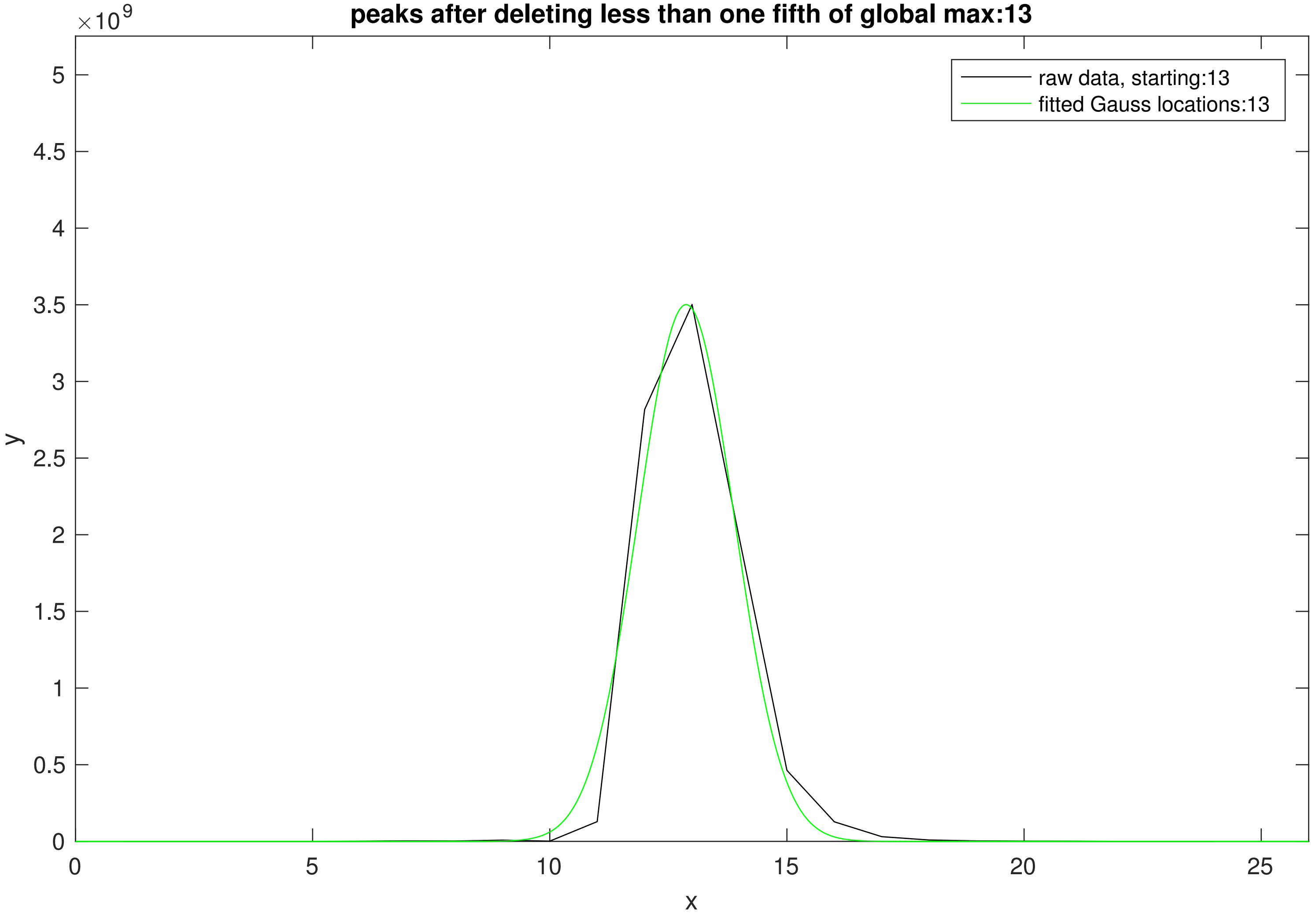

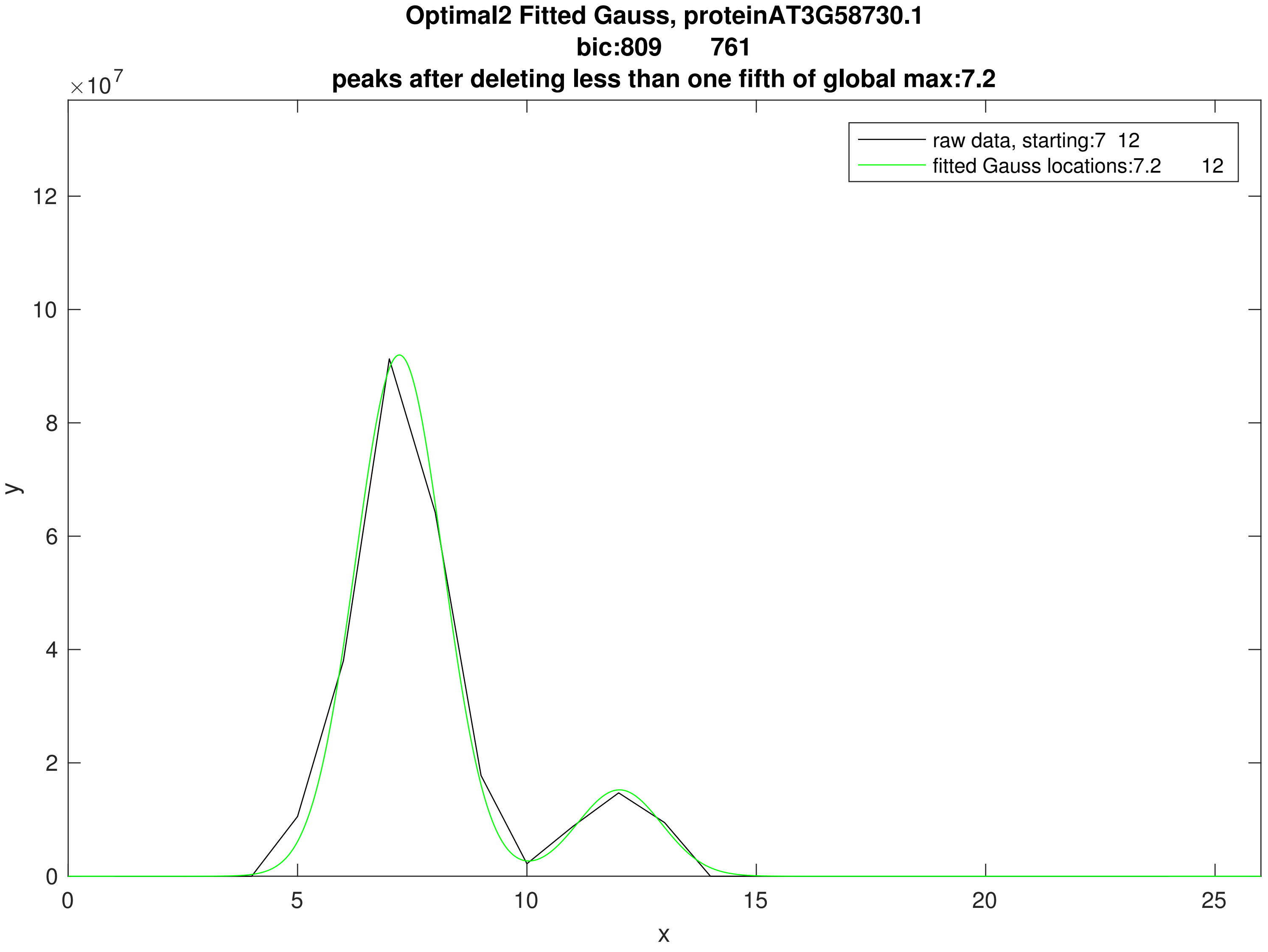
Optimal2 Fitted Gauss, proteinAT3G58730.1
bic:809 761
peaks after deleting less than one fifth of global max:7.2
raw data, starting:7 12
fitted Gauss locations:7.2 12
×10^7
y
x
0
2
4
6
8
10
12
0
5
10
15
20
25

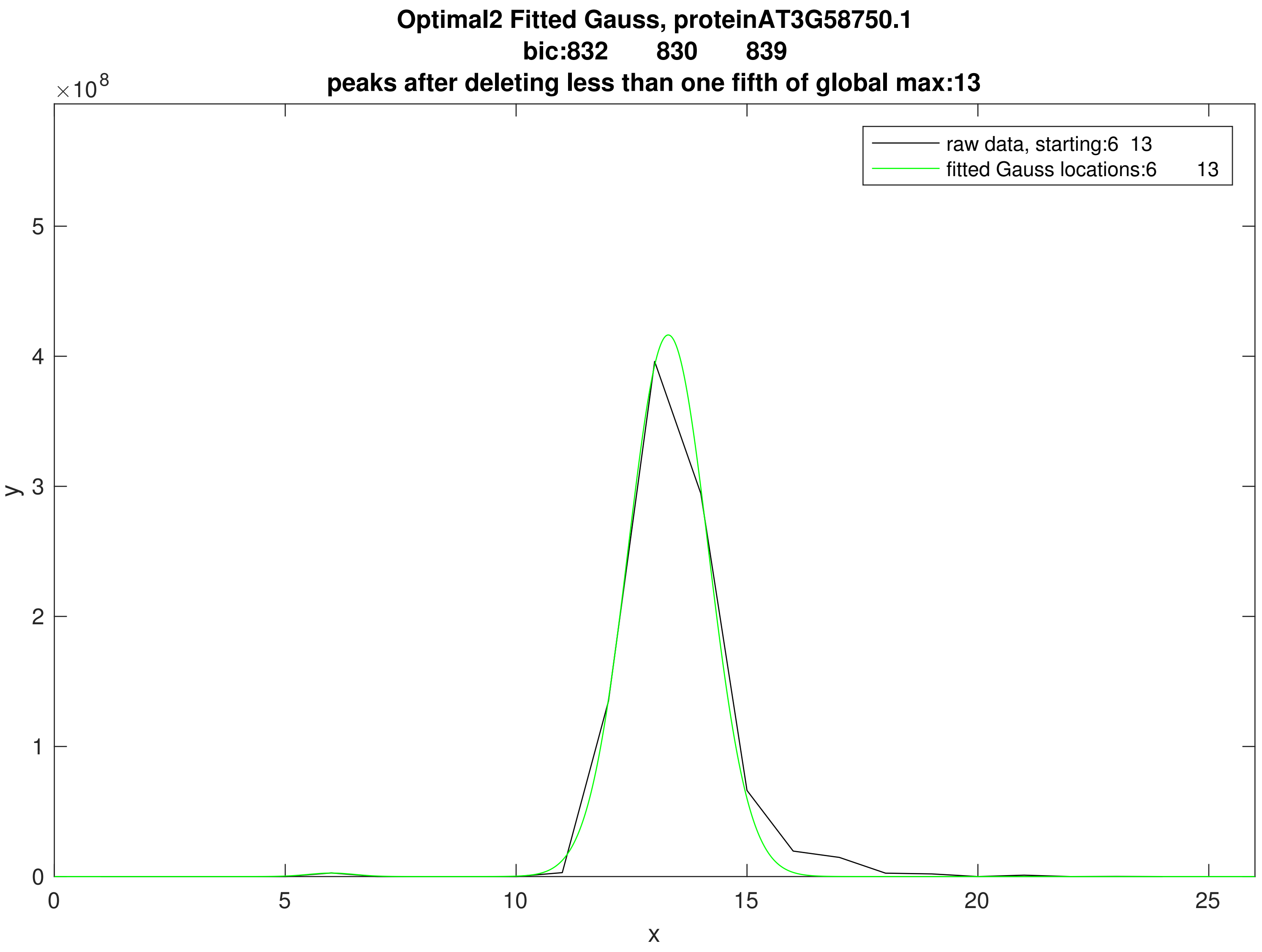
Optimal2 Fitted Gauss, proteinAT3G58750.1
bic:832 830 839
peaks after deleting less than one fifth of global max:13
raw data, starting:6 13
fitted Gauss locations:6 13
×10^8
y
x
0
1
2
3
4
5
0
5
10
15
20
25

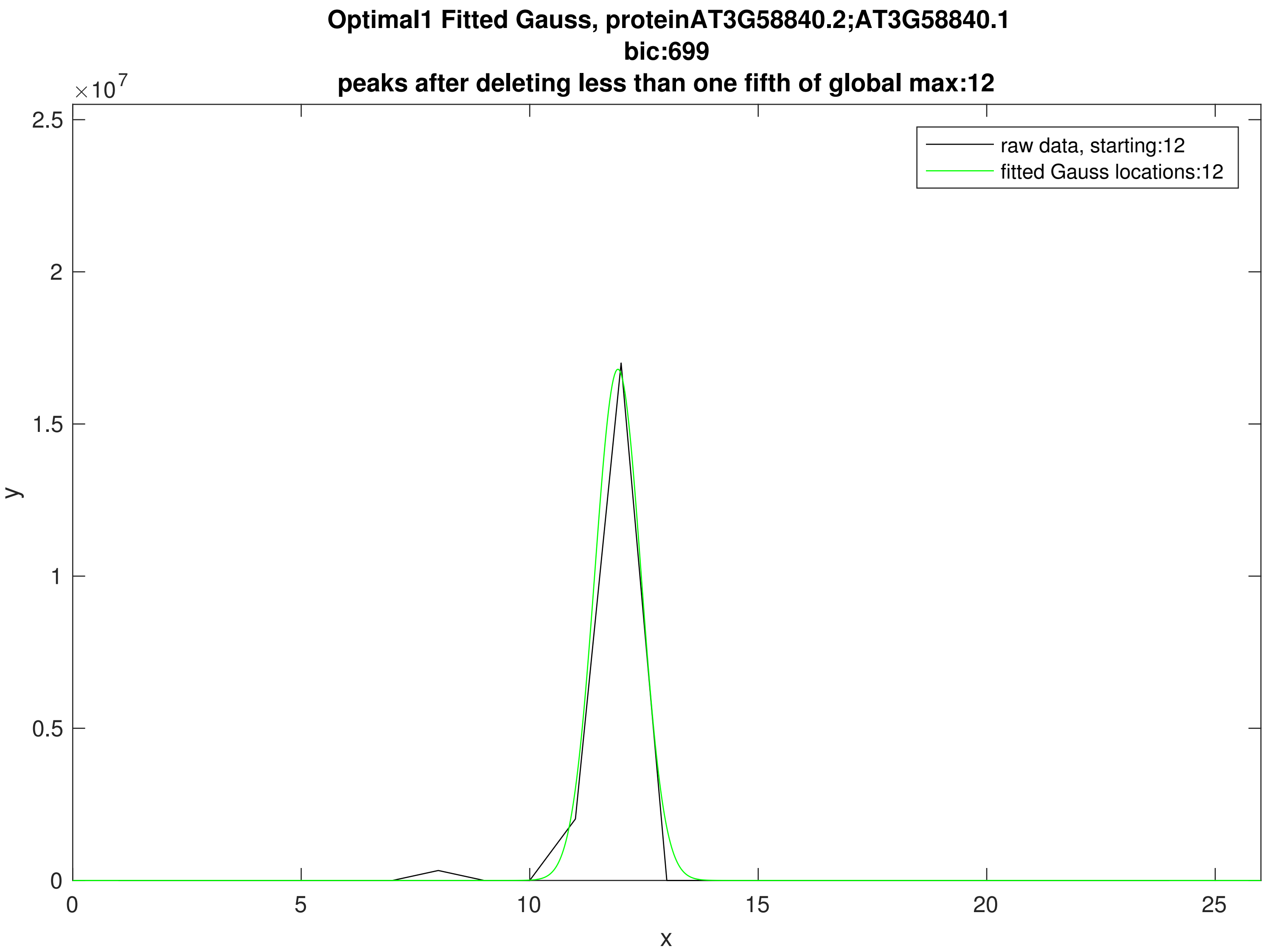
Optimal1 Fitted Gauss, proteinAT3G58840.2;AT3G58840.1
bic:699
peaks after deleting less than one fifth of global max:12
×10^7
2.5
2
1.5
1
0.5
0
y
0
5
10
15
20
25
x
raw data, starting:12
fitted Gauss locations:12

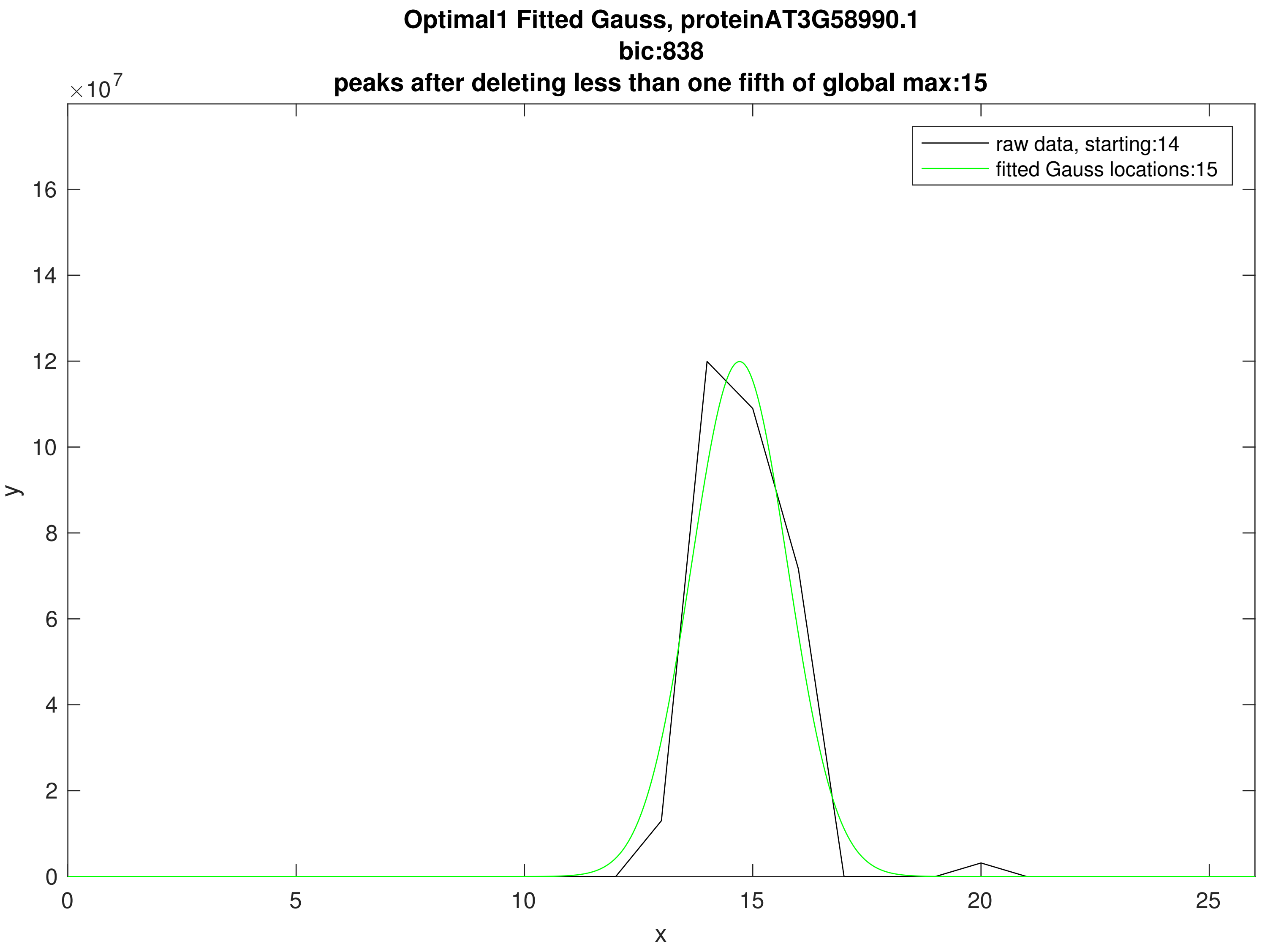

Optimal1 Fitted Gauss, proteinAT3G58990.1
bic:838
peaks after deleting less than one fifth of global max:15
×10^7
raw data, starting:14
fitted Gauss locations:15
16
14
12
10
8
6
4
2
0
y
0
5
10
15
20
25
x

**Optimal1 Fitted Gauss, proteinAT3G59050.1**
**bic:759**
**peaks after deleting less than one fifth of global max:15**

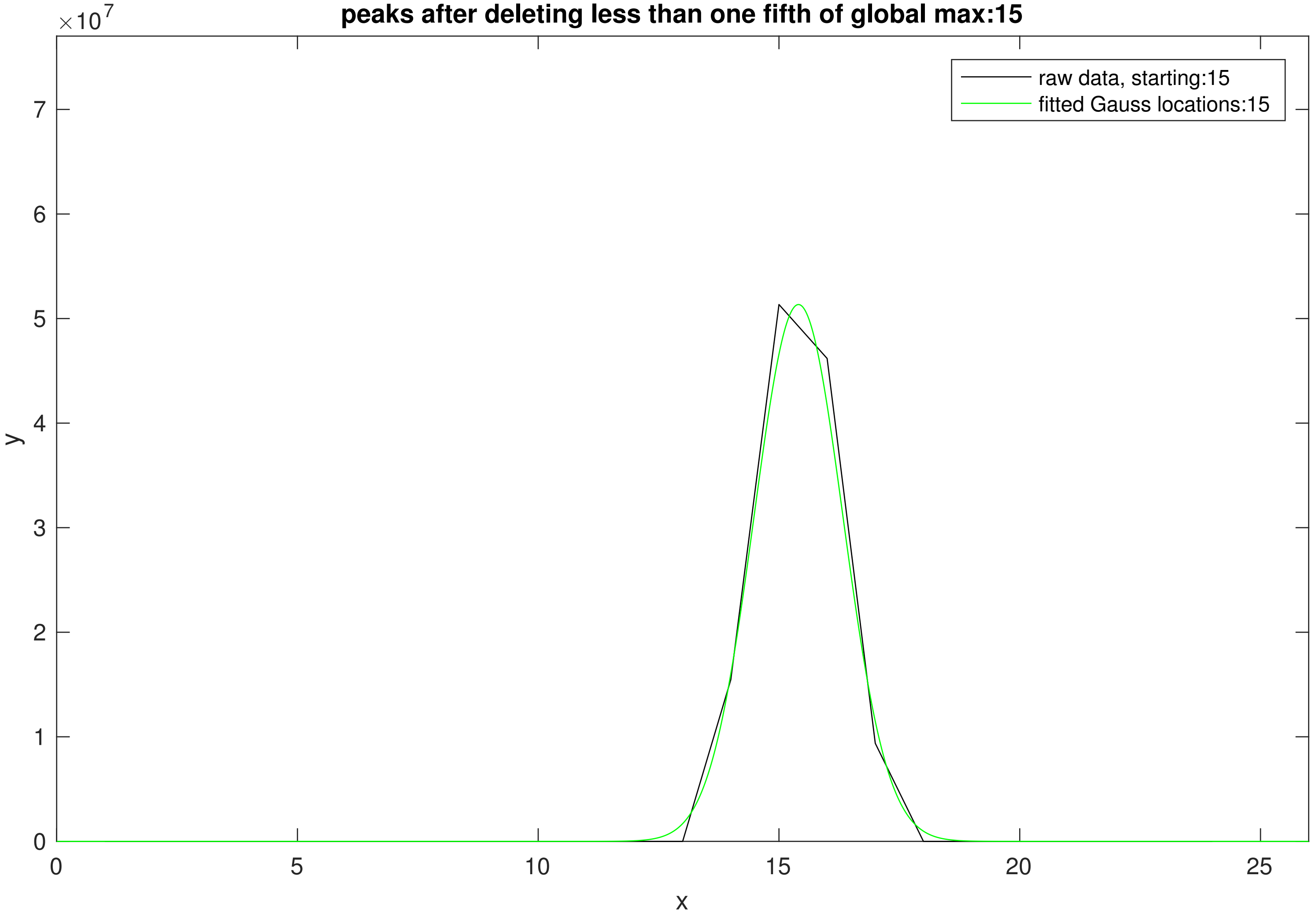

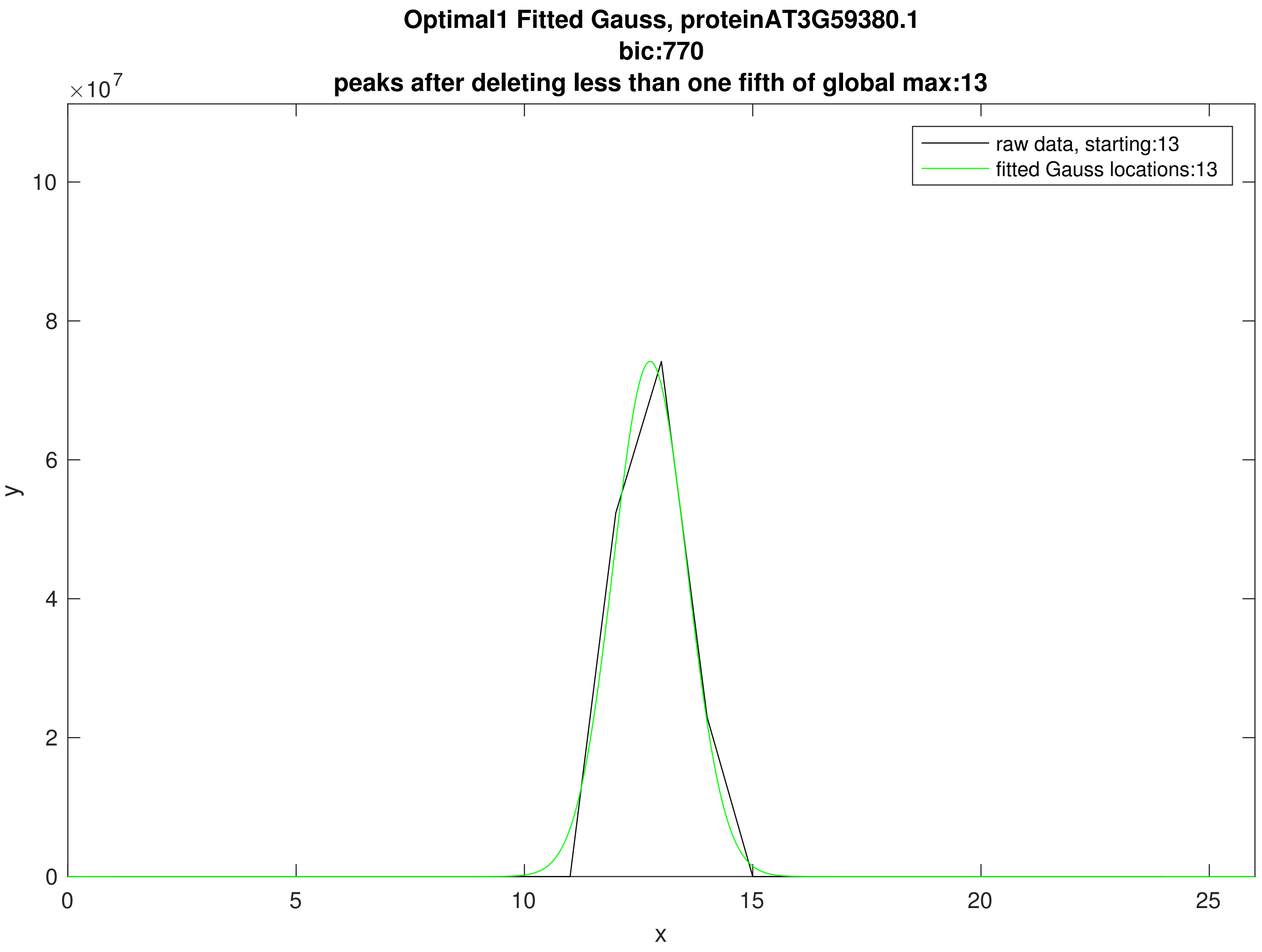
Optimal1 Fitted Gauss, proteinAT3G59380.1
bic:770
peaks after deleting less than one fifth of global max:13
×10^7
raw data, starting:13
fitted Gauss locations:13
y
x
0
2
4
6
8
10
5
15
20
25

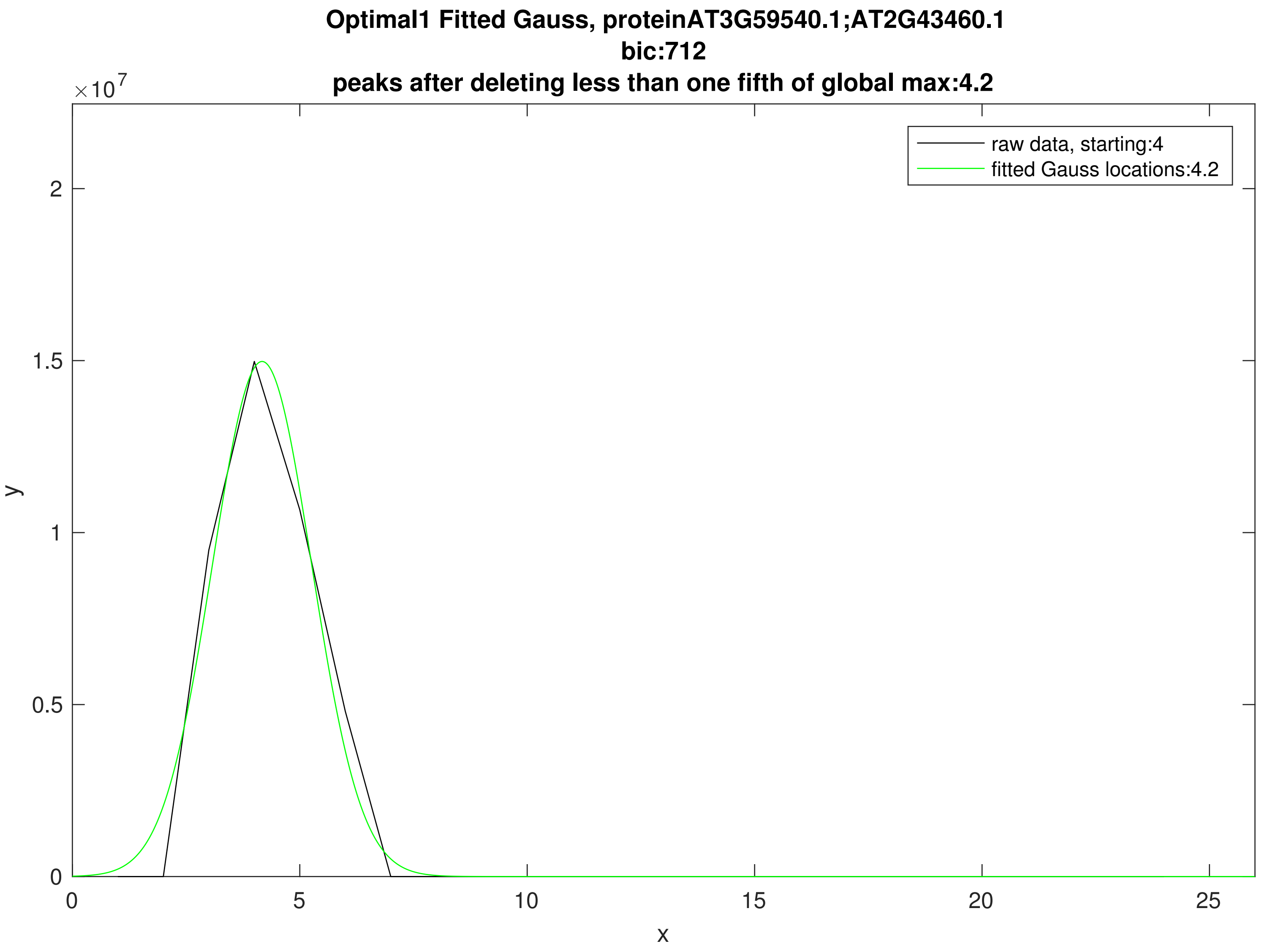

Optimal1 Fitted Gauss, proteinAT3G59540.1;AT2G43460.1
bic:712
peaks after deleting less than one fifth of global max:4.2
×10^7
raw data, starting:4
fitted Gauss locations:4.2
y
x
0
0.5
1
1.5
2
0
5
10
15
20
25

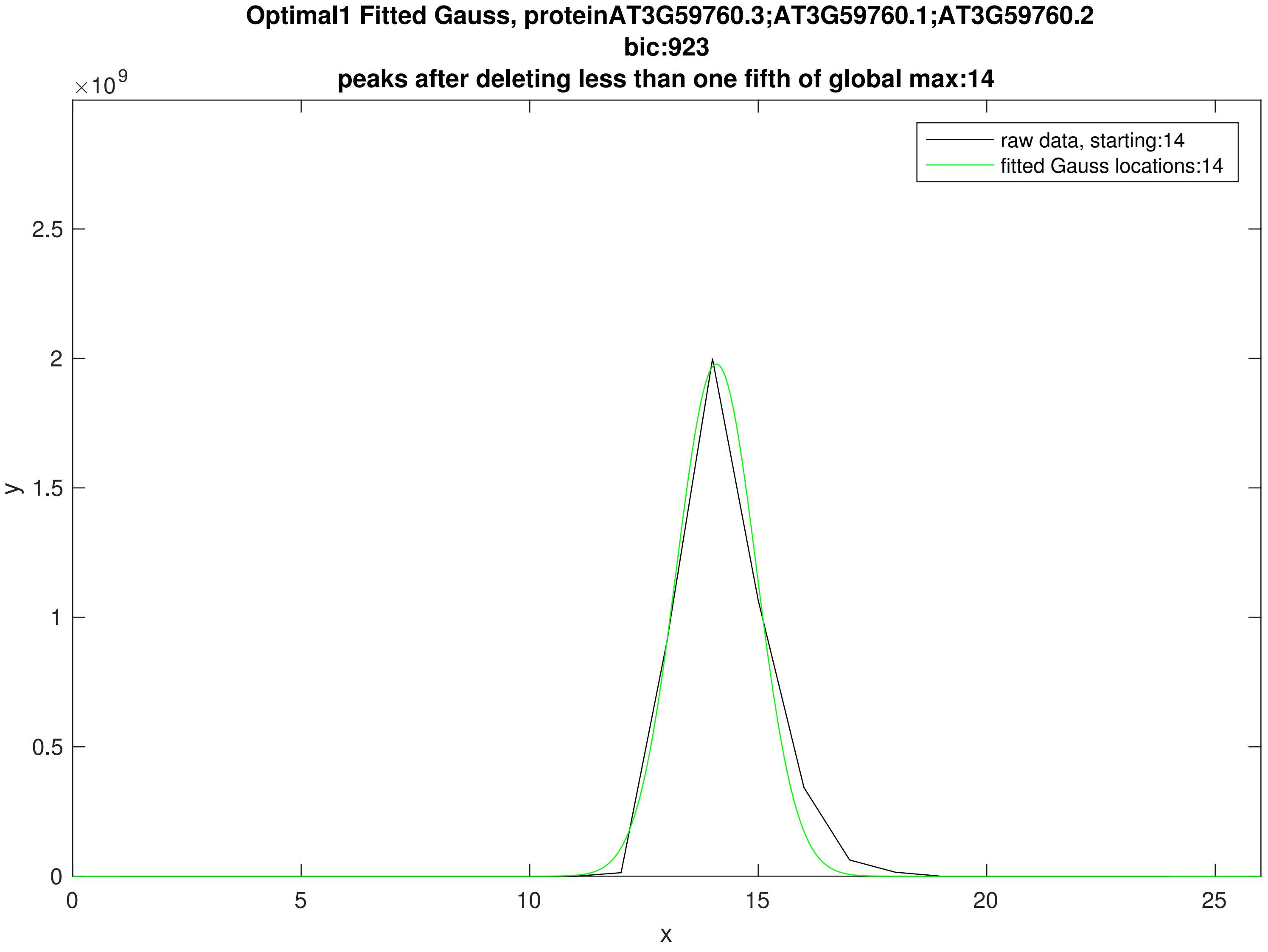

Optimal1 Fitted Gauss, proteinAT3G59760.3;AT3G59760.1;AT3G59760.2
bic:923
peaks after deleting less than one fifth of global max:14
×10^9
raw data, starting:14
fitted Gauss locations:14
2.5
2
1.5
1
0.5
0
y
0
5
10
15
20
25
x

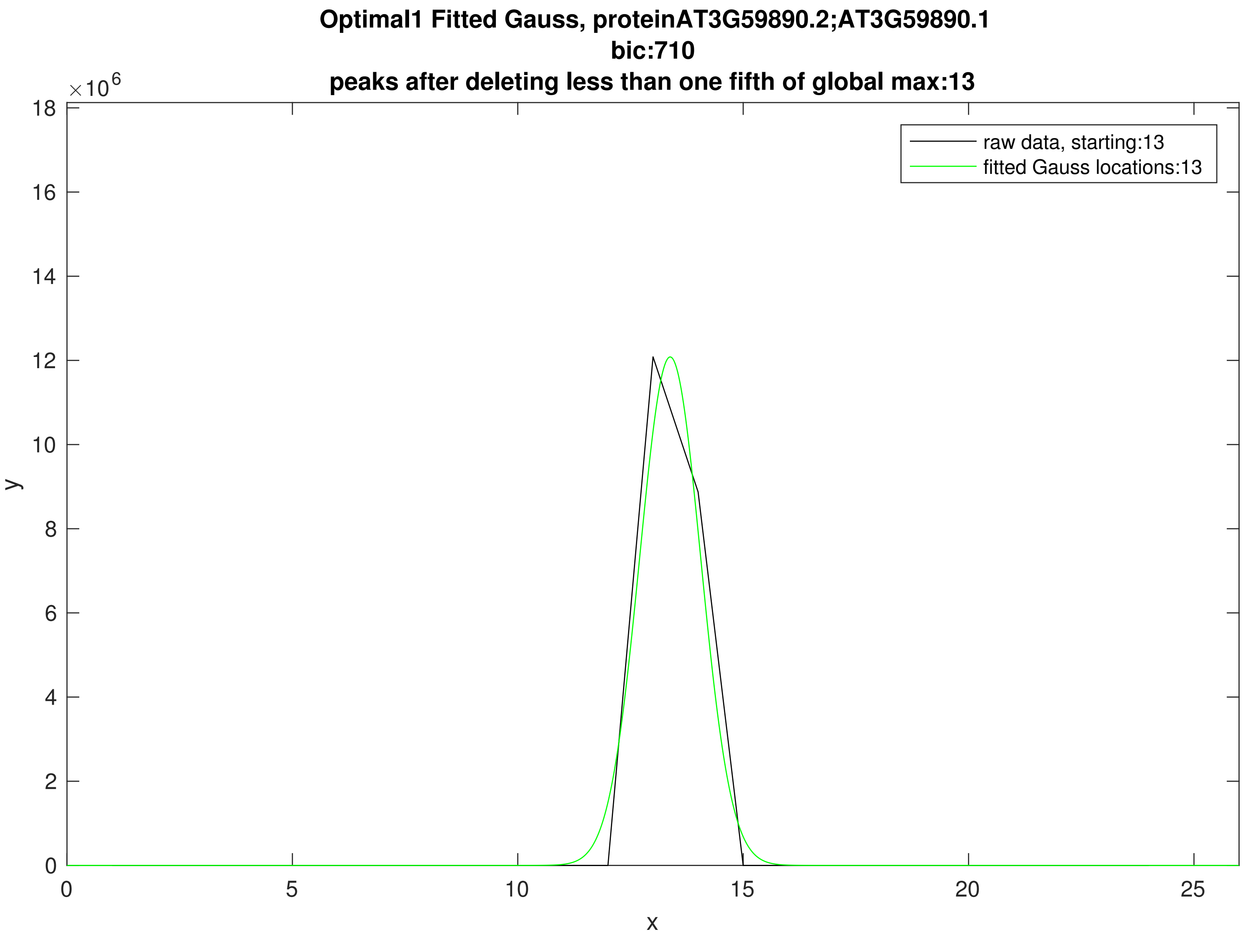
Optimal1 Fitted Gauss, proteinAT3G59890.2;AT3G59890.1
bic:710
peaks after deleting less than one fifth of global max:13
×10^6
raw data, starting:13
fitted Gauss locations:13
y
x
0
2
4
6
8
10
12
14
16
18
0
5
10
15
20
25

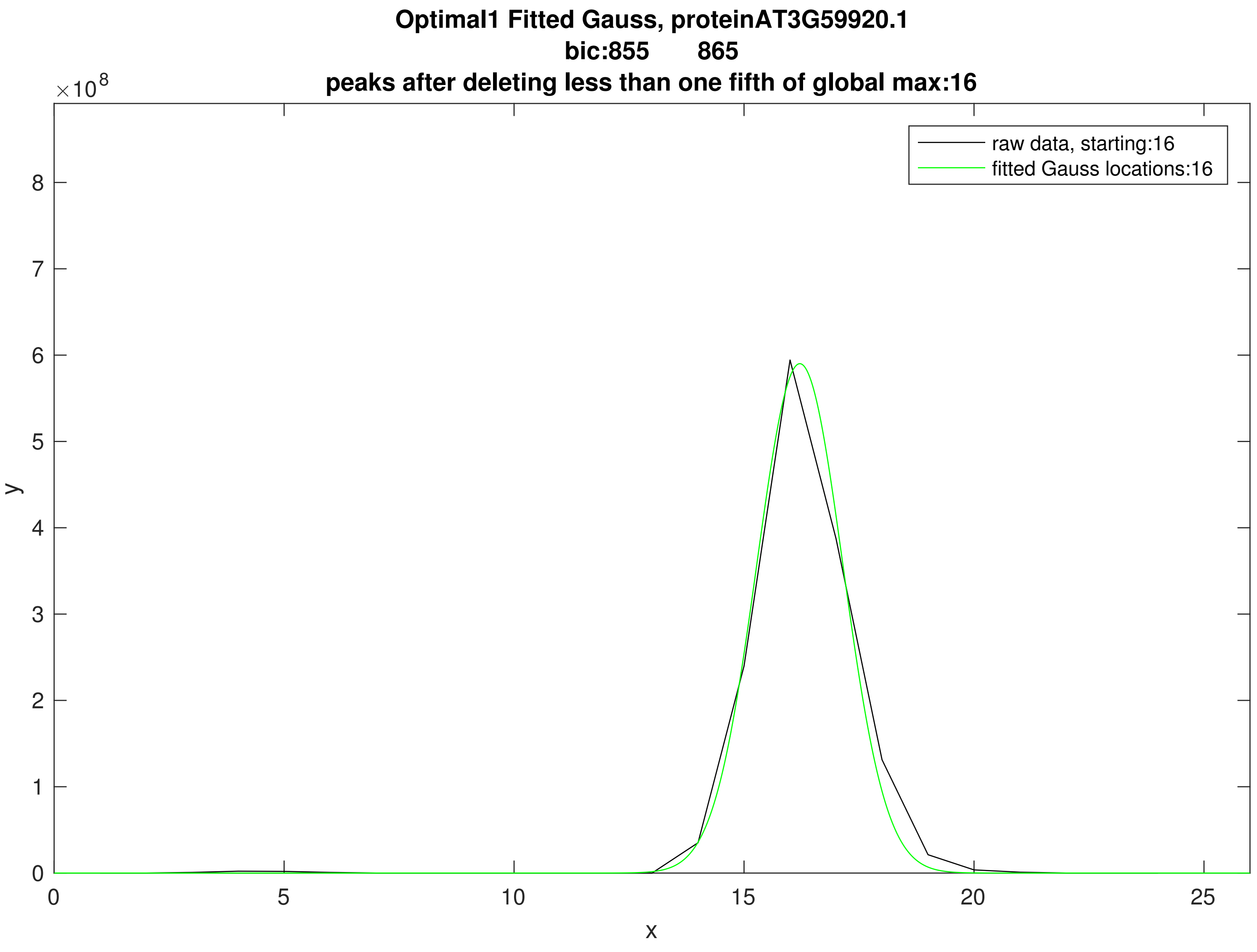
Optimal1 Fitted Gauss, proteinAT3G59920.1
bic:855 865
peaks after deleting less than one fifth of global max:16
×10^8
raw data, starting:16
fitted Gauss locations:16
y
x
0
1
2
3
4
5
6
7
8
0
5
10
15
20
25

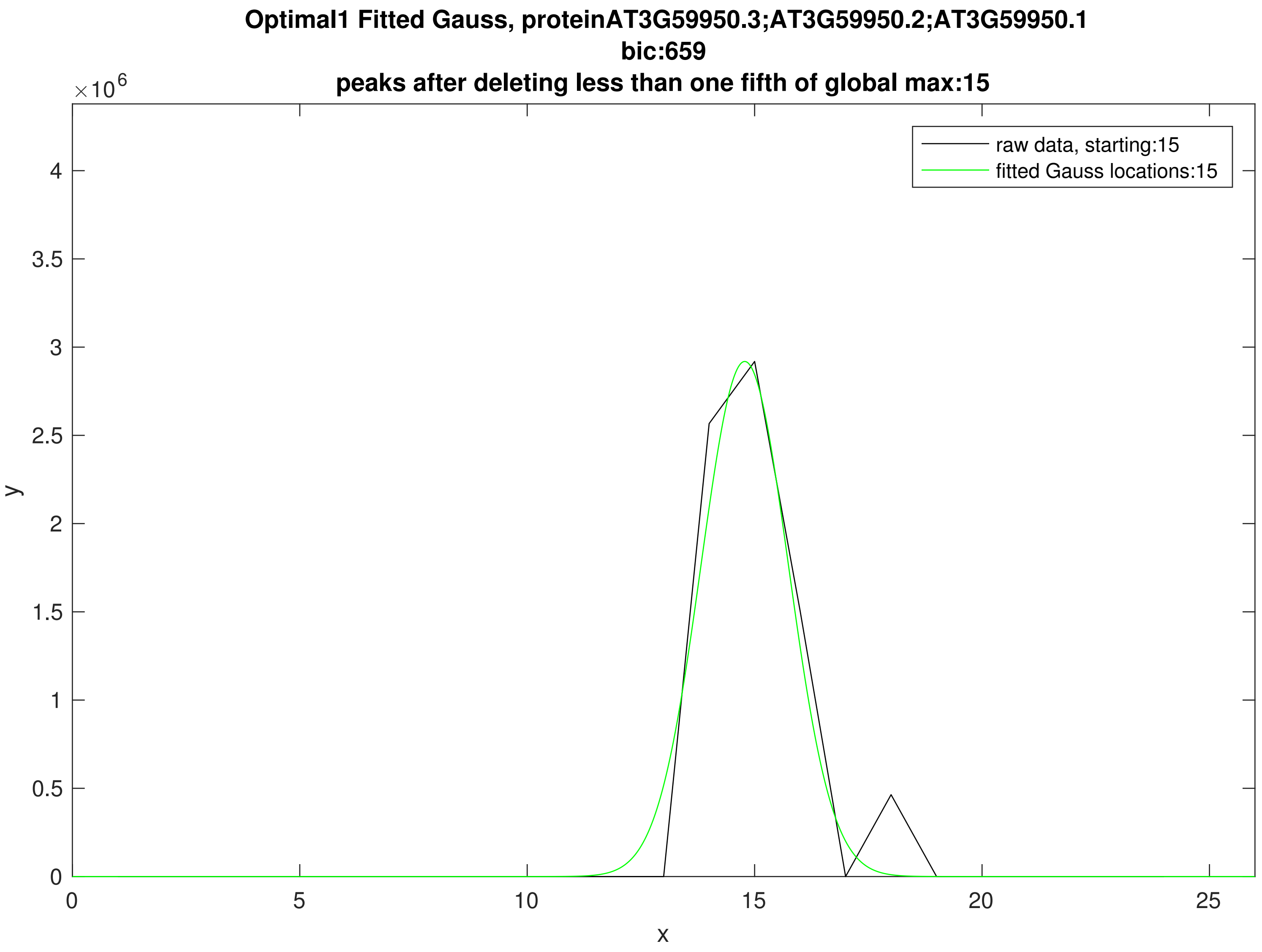

Optimal1 Fitted Gauss, proteinAT3G59950.3;AT3G59950.2;AT3G59950.1
bic:659
peaks after deleting less than one fifth of global max:15
×10^6
raw data, starting:15
fitted Gauss locations:15
y
x
0
0.5
1
1.5
2
2.5
3
3.5
4
0
5
10
15
20
25

**Optimal1 Fitted Gauss, proteinAT3G59970.3;AT3G59970.2;AT3G59970.1**
**bic:1.02e+03**
**peaks after deleting less than one fifth of global max:12**

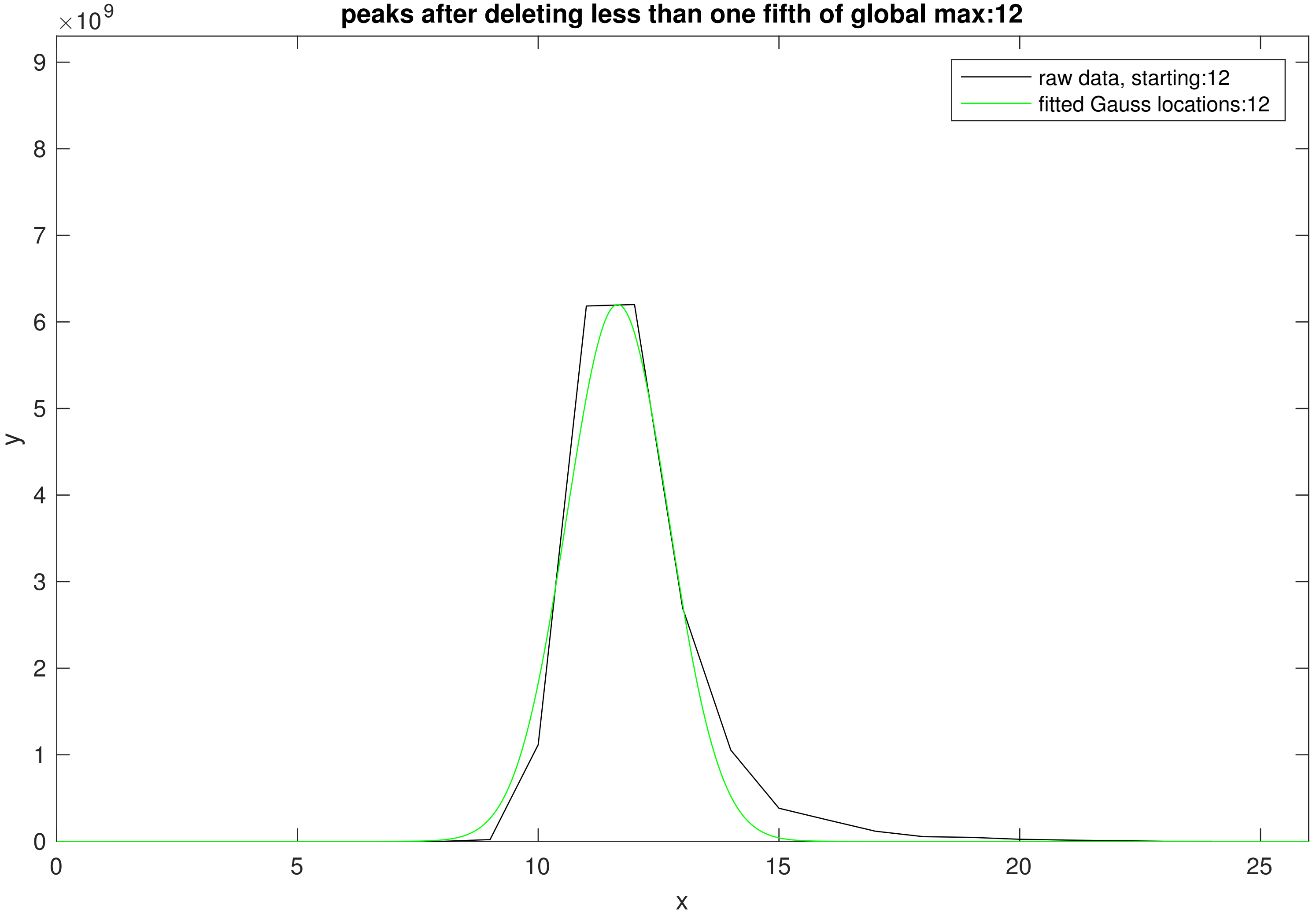

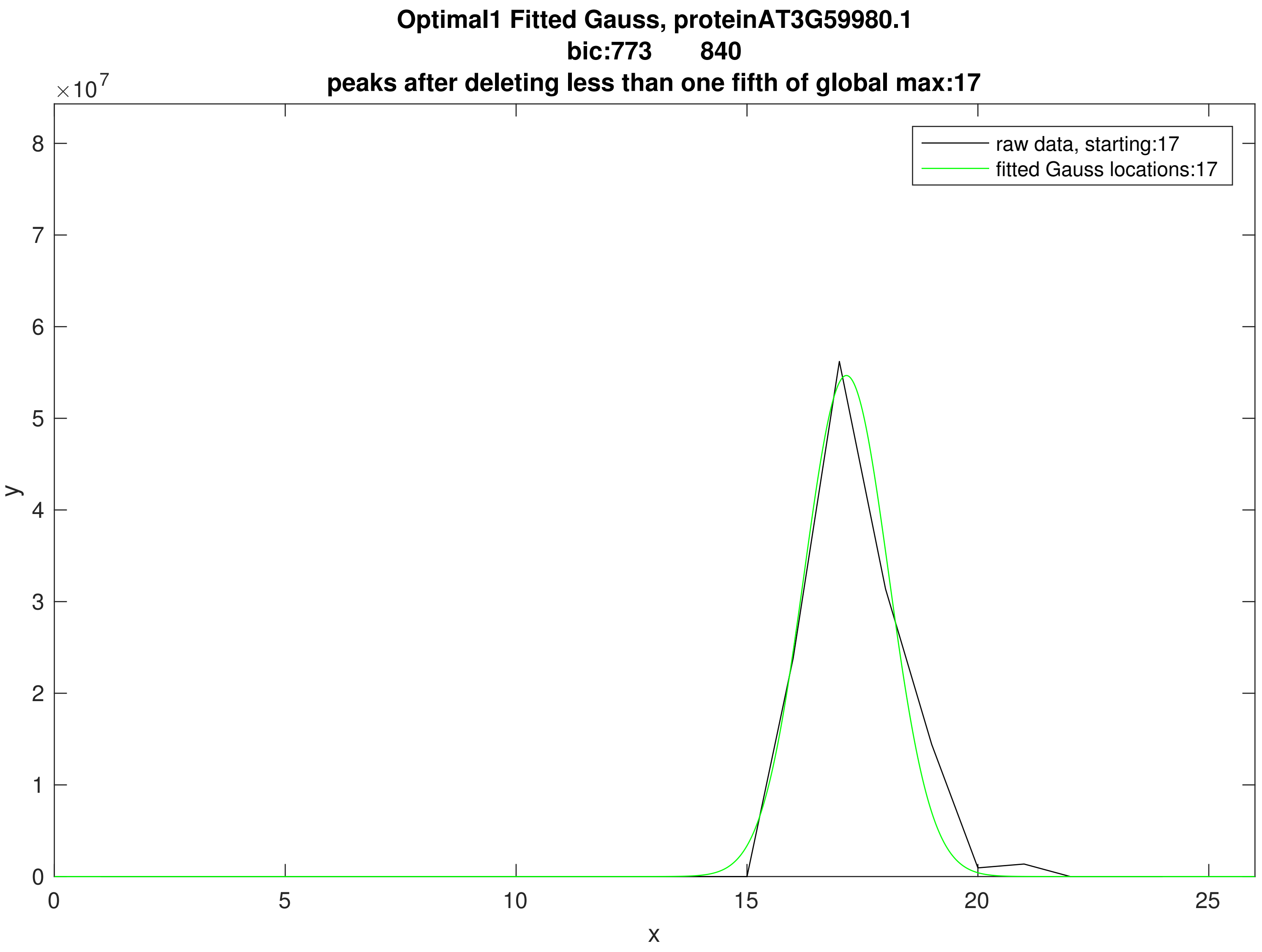
Optimal1 Fitted Gauss, proteinAT3G59980.1
bic:773 840
peaks after deleting less than one fifth of global max:17
×10^7
raw data, starting:17
fitted Gauss locations:17
y
x
0
1
2
3
4
5
6
7
8
0
5
10
15
20
25

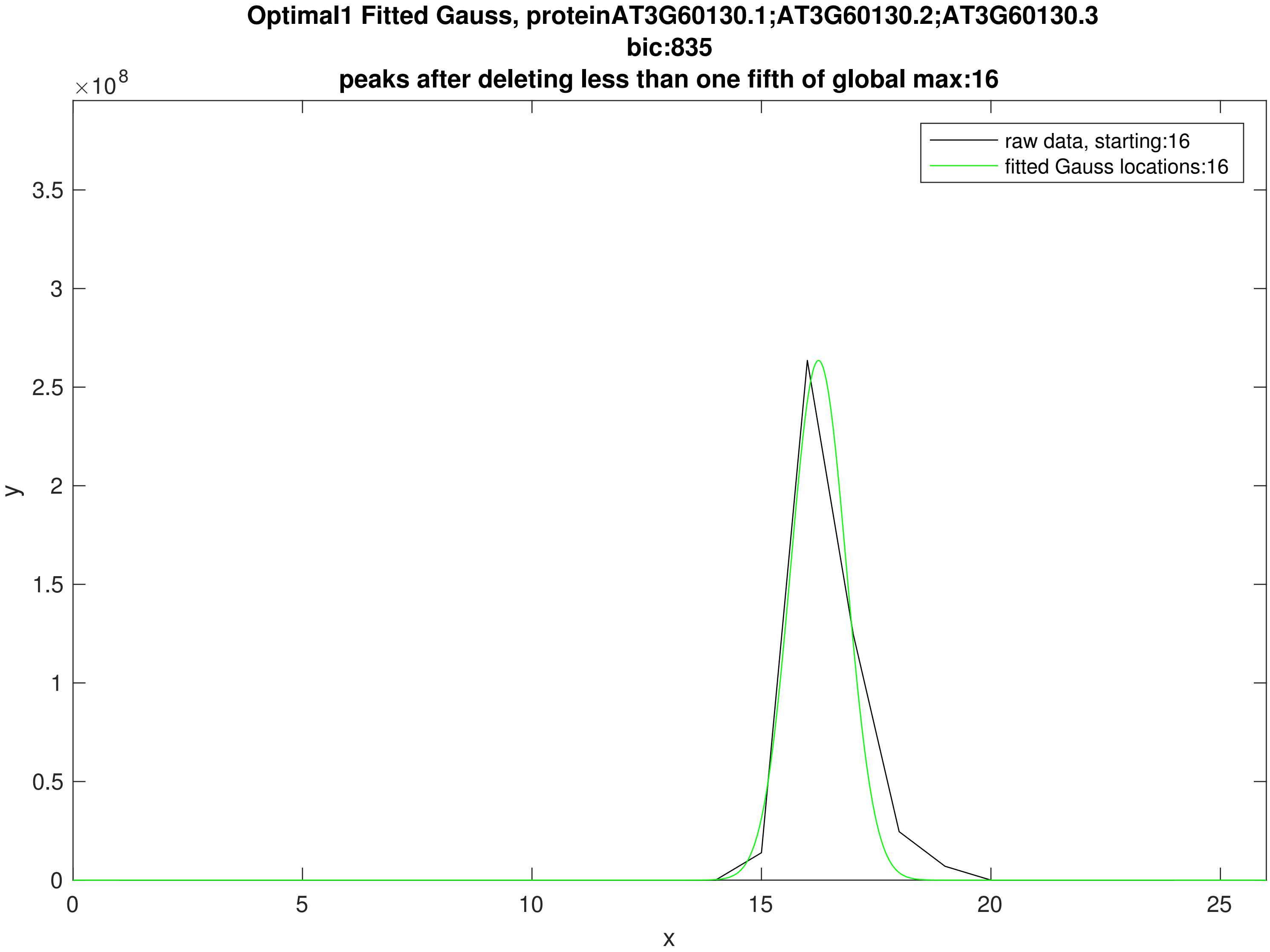
Optimal1 Fitted Gauss, proteinAT3G60130.1;AT3G60130.2;AT3G60130.3
bic:835
peaks after deleting less than one fifth of global max:16
×10^8
raw data, starting:16
fitted Gauss locations:16
y
x
0
0.5
1
1.5
2
2.5
3
3.5
0
5
10
15
20
25

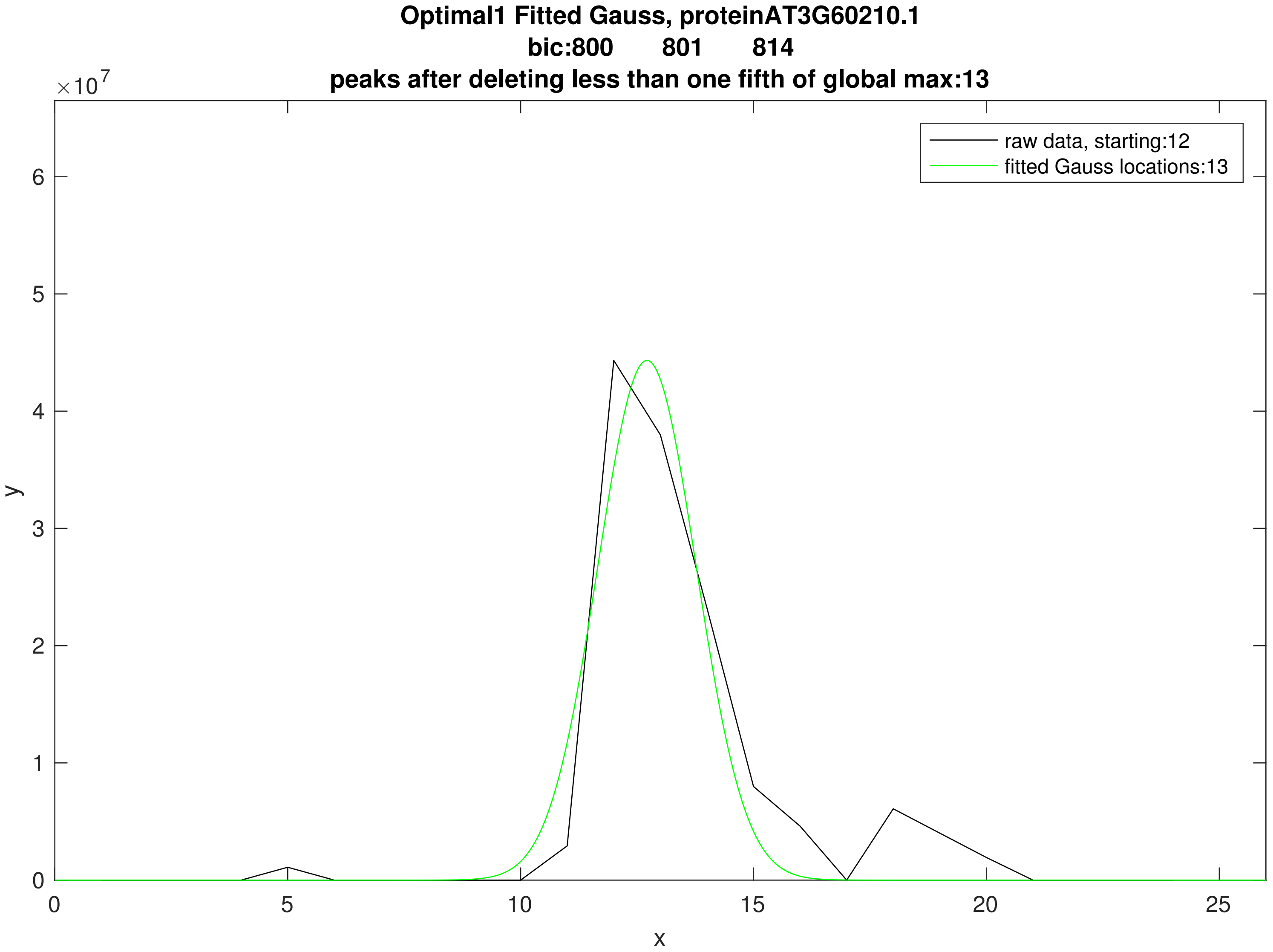
Optimal1 Fitted Gauss, proteinAT3G60210.1
bic:800 801 814
peaks after deleting less than one fifth of global max:13
×10^7
raw data, starting:12
fitted Gauss locations:13
6
5
4
3
2
1
0
y
0
5
10
15
20
25
x

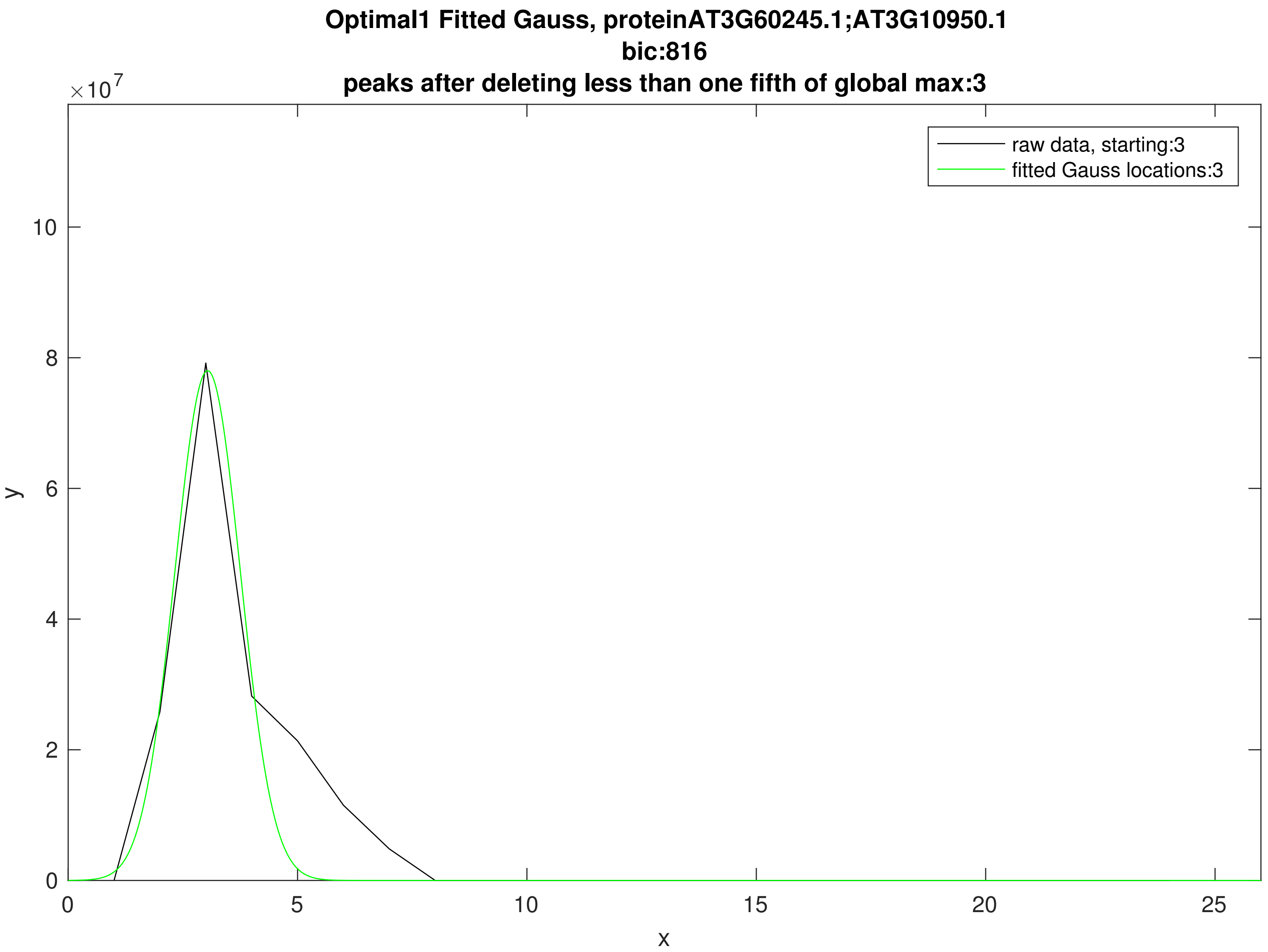
Optimal1 Fitted Gauss, proteinAT3G60245.1;AT3G10950.1
bic:816
peaks after deleting less than one fifth of global max:3
×10^7
raw data, starting:3
fitted Gauss locations:3
y
x
0
2
4
6
8
10
0
5
10
15
20
25

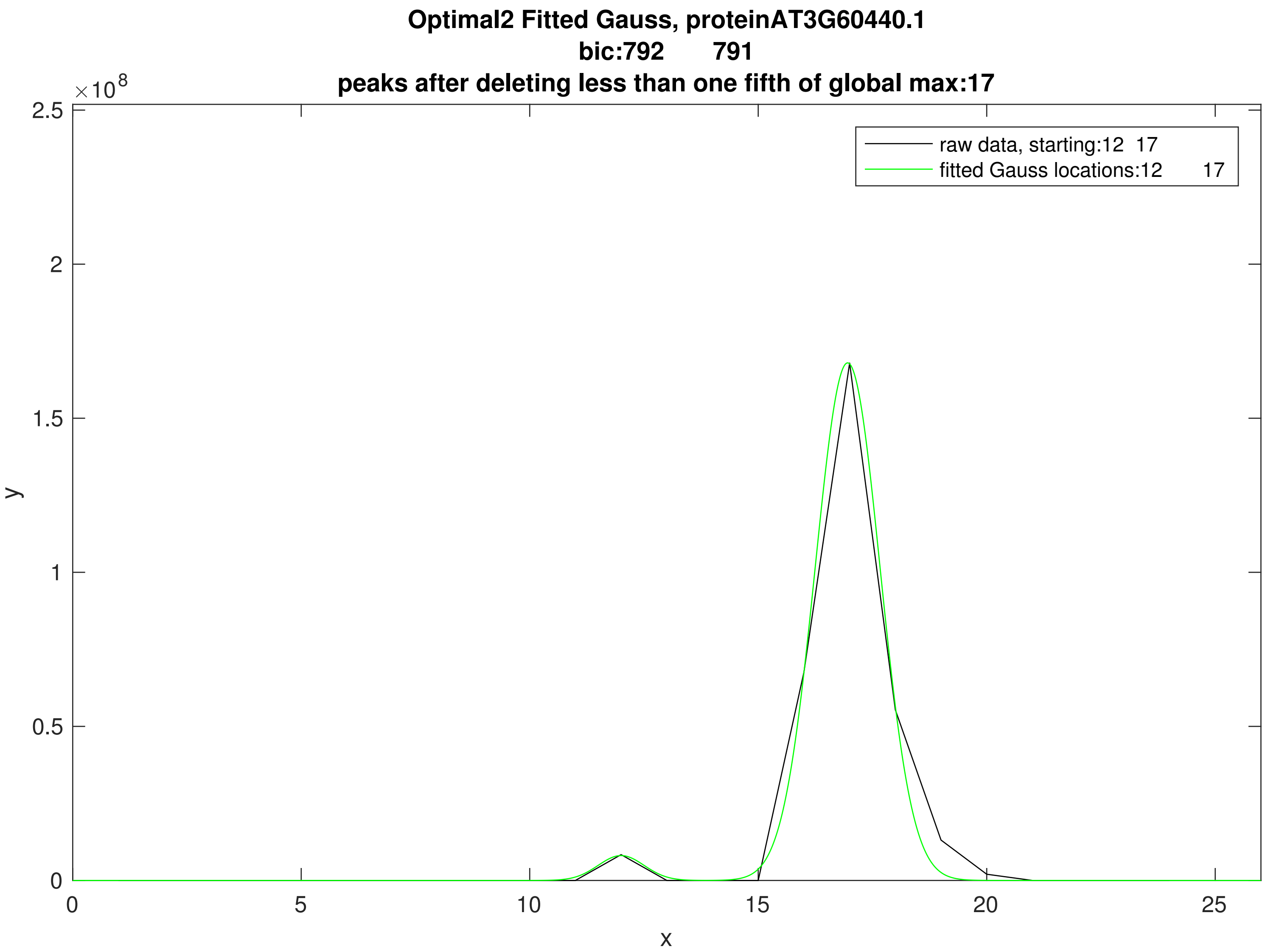
Optimal2 Fitted Gauss, proteinAT3G60440.1
bic:792 791
peaks after deleting less than one fifth of global max:17
×10^8
2.5
2
1.5
1
0.5
0
y
0
5
10
15
20
25
x
raw data, starting:12 17
fitted Gauss locations:12 17

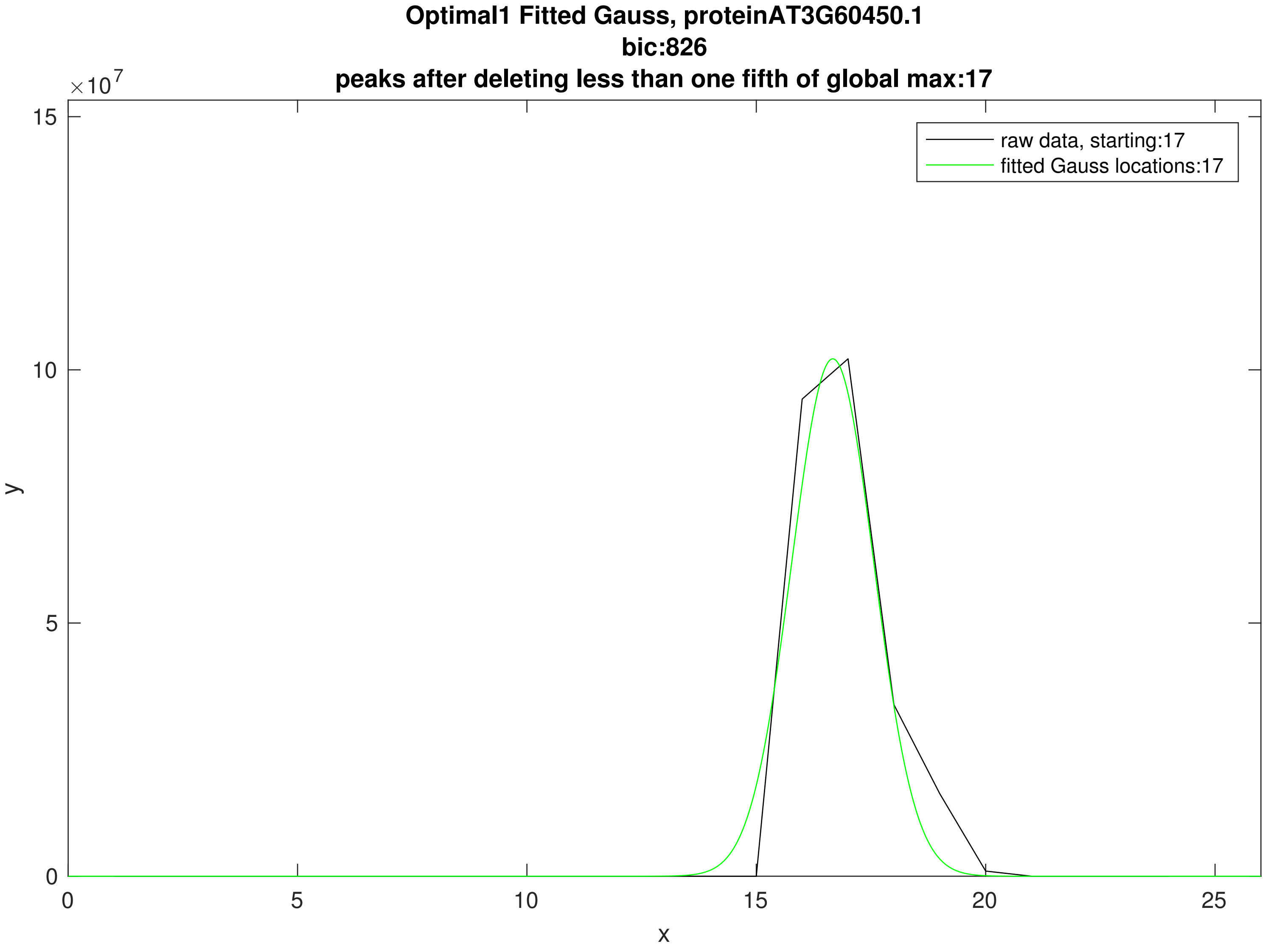
Optimal1 Fitted Gauss, proteinAT3G60450.1
bic:826
peaks after deleting less than one fifth of global max:17
×10^7
raw data, starting:17
fitted Gauss locations:17
15
10
5
0
y
0
5
10
15
20
25
x

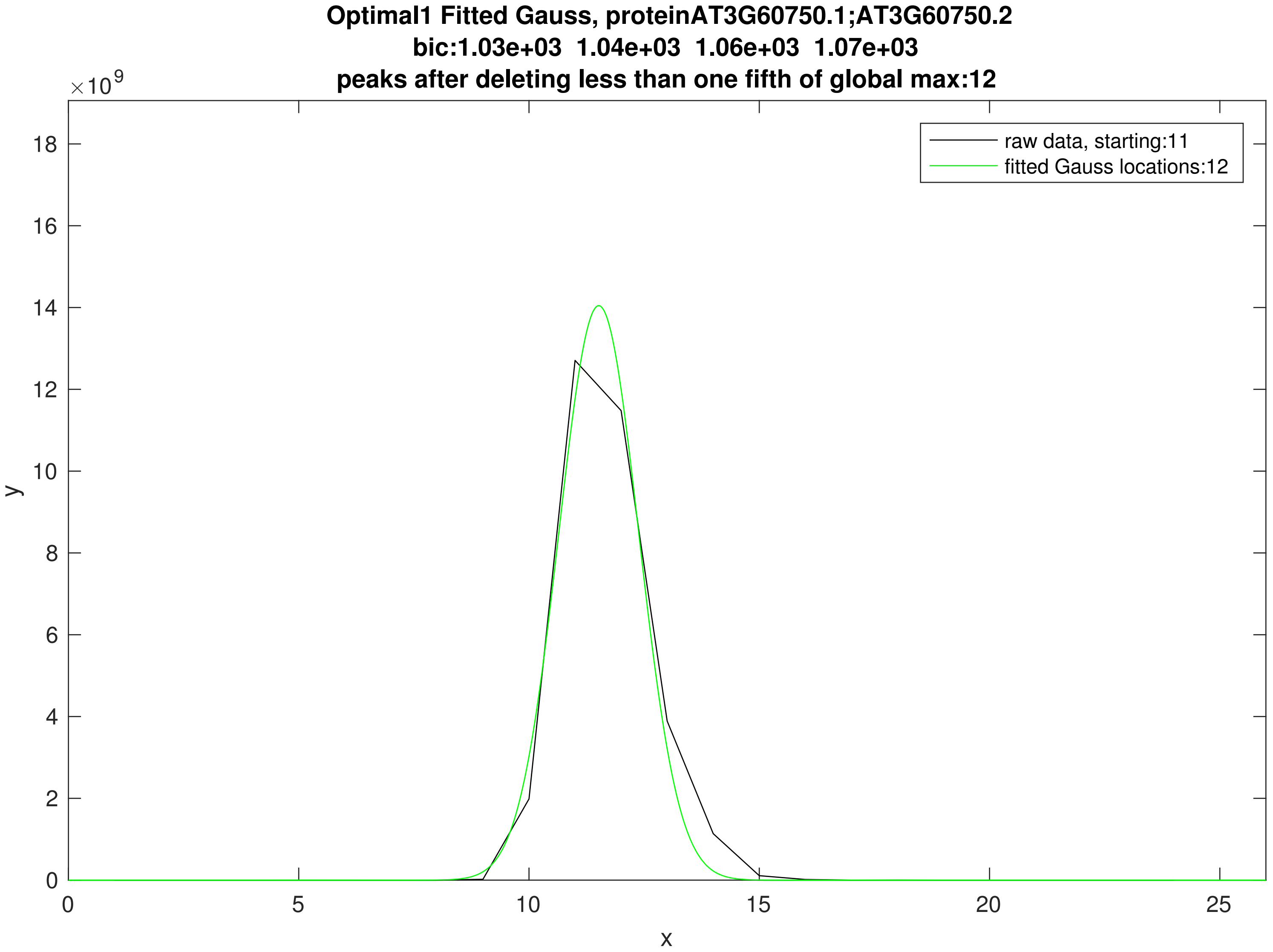
Optimal1 Fitted Gauss, proteinAT3G60750.1;AT3G60750.2
bic:1.03e+03 1.04e+03 1.06e+03 1.07e+03
peaks after deleting less than one fifth of global max:12
×10^9
raw data, starting:11
fitted Gauss locations:12
y
x
0
2
4
6
8
10
12
14
16
18
0
5
10
15
20
25

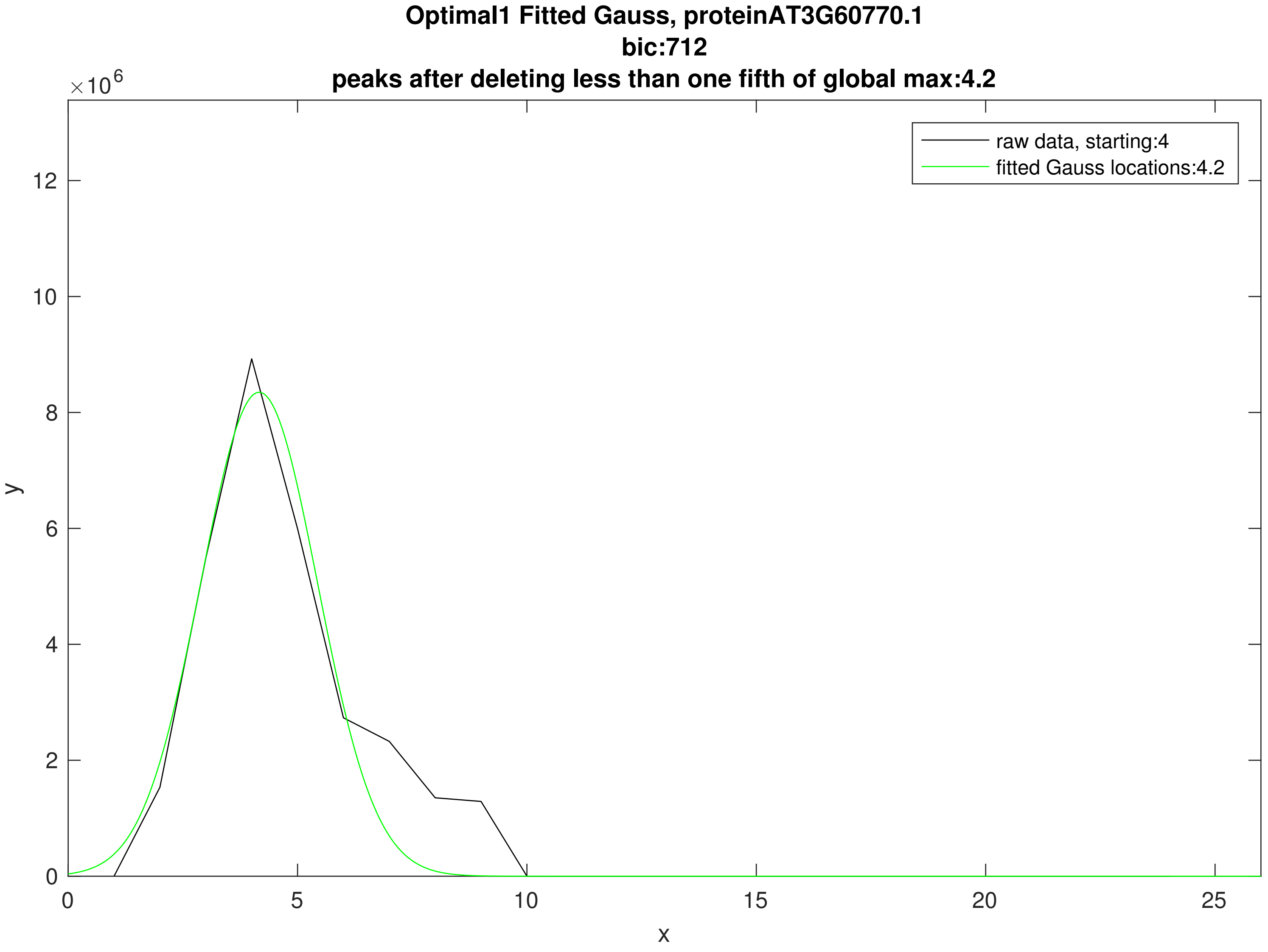
Optimal1 Fitted Gauss, proteinAT3G60770.1
bic:712
peaks after deleting less than one fifth of global max:4.2
×10^6
raw data, starting:4
fitted Gauss locations:4.2
y
x
0
2
4
6
8
10
12
0
5
10
15
20
25

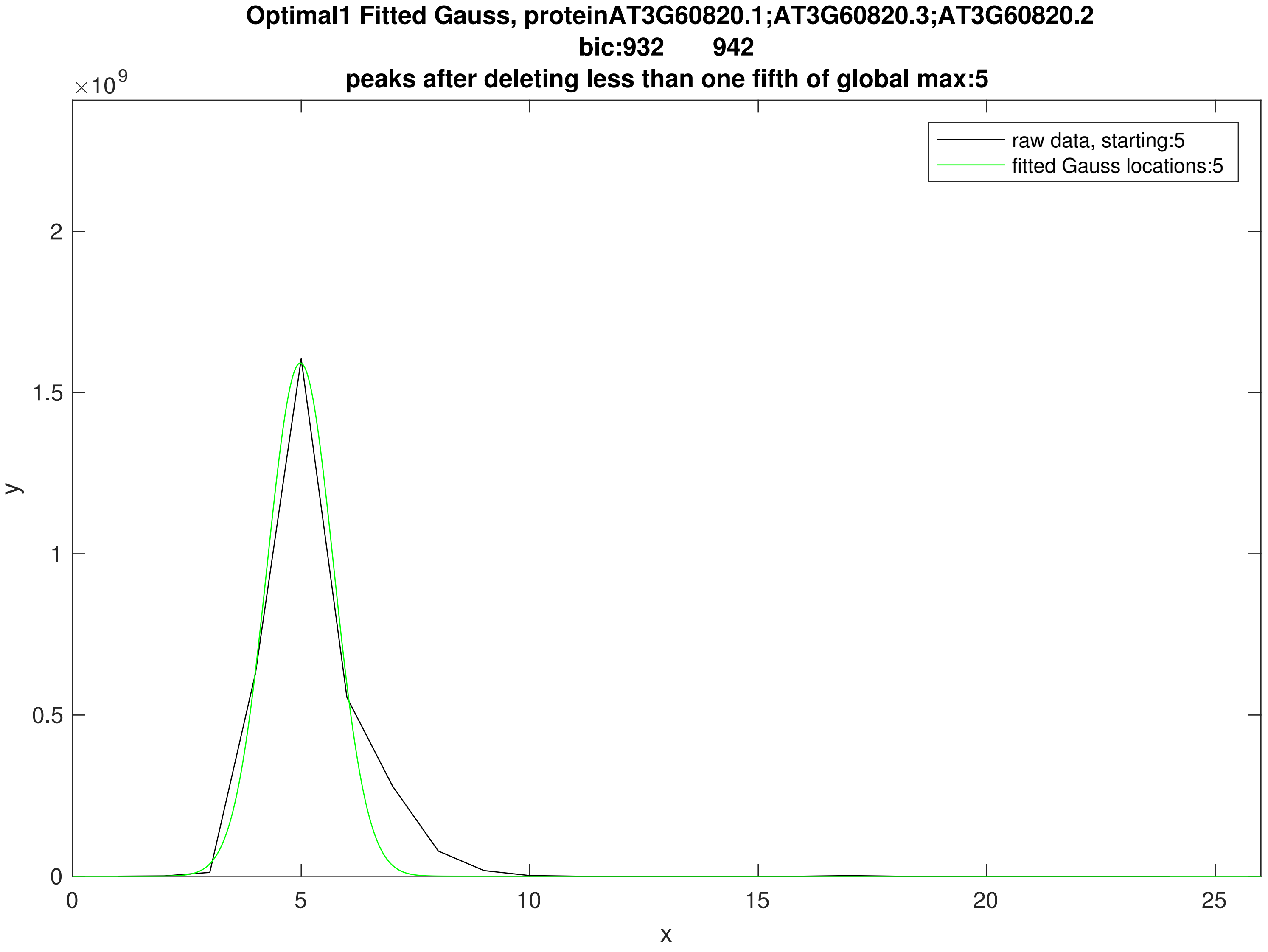

Optimal1 Fitted Gauss, proteinAT3G60820.1;AT3G60820.3;AT3G60820.2
bic:932 942
peaks after deleting less than one fifth of global max:5
×10^9
raw data, starting:5
fitted Gauss locations:5
y
x
0
0.5
1
1.5
2
0
5
10
15
20
25

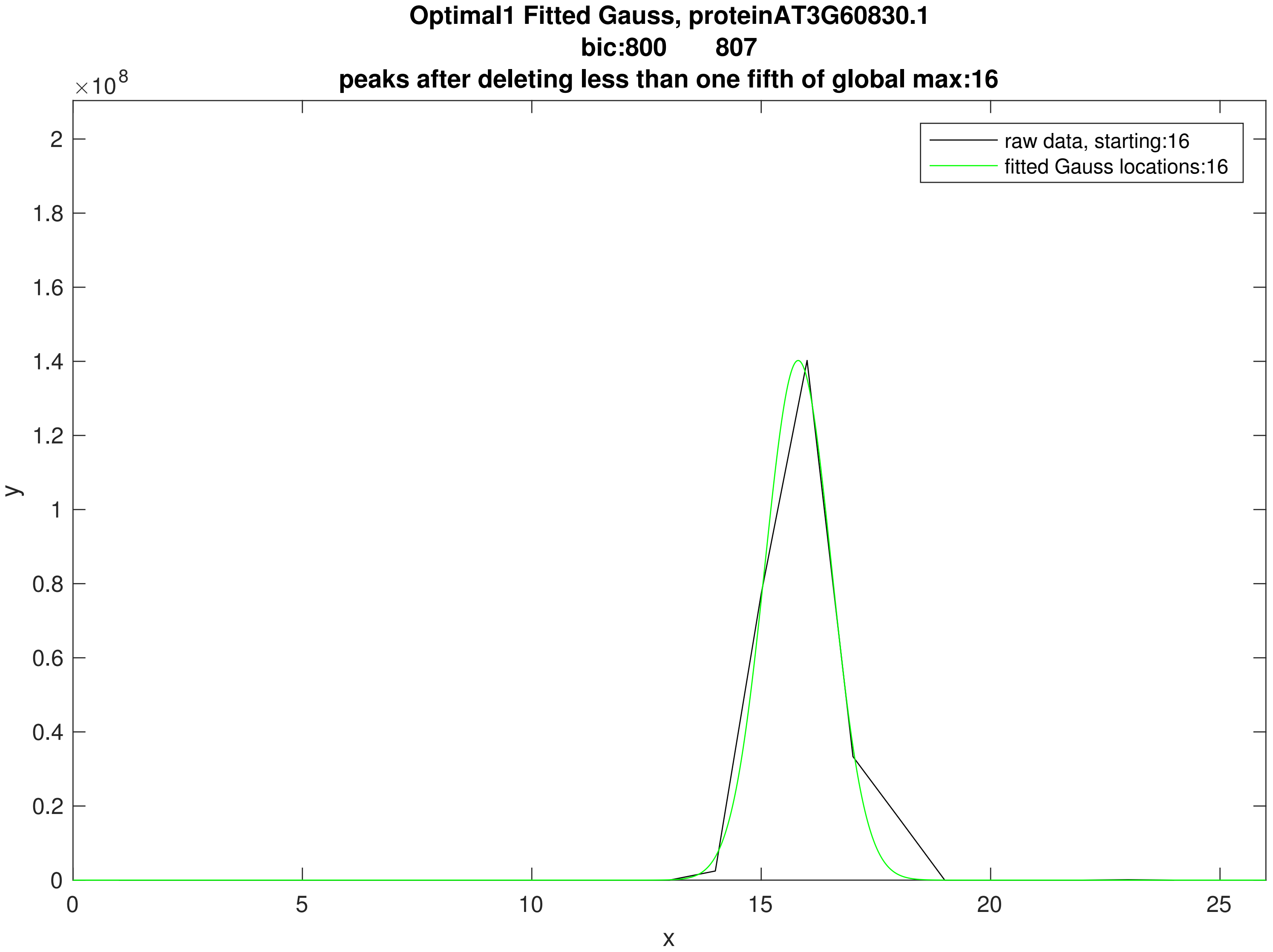
Optimal1 Fitted Gauss, proteinAT3G60830.1
bic:800 807
peaks after deleting less than one fifth of global max:16
×10^8
raw data, starting:16
fitted Gauss locations:16
y
x
0
0.2
0.4
0.6
0.8
1
1.2
1.4
1.6
1.8
2
5
10
15
20
25

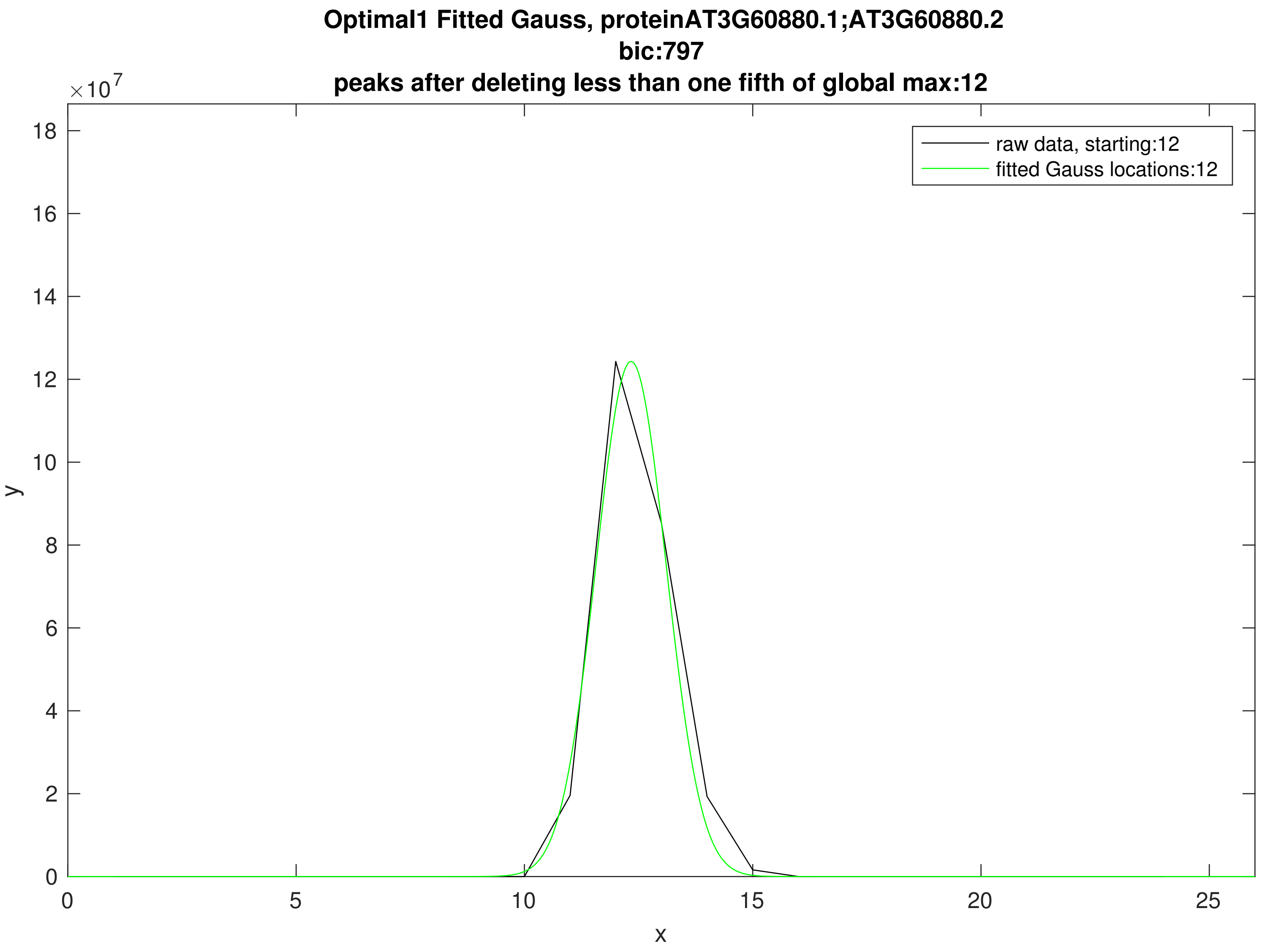
Optimal1 Fitted Gauss, proteinAT3G60880.1;AT3G60880.2
bic:797
peaks after deleting less than one fifth of global max:12
×10^7
raw data, starting:12
fitted Gauss locations:12
y
x
0
2
4
6
8
10
12
14
16
18
0
5
10
15
20
25

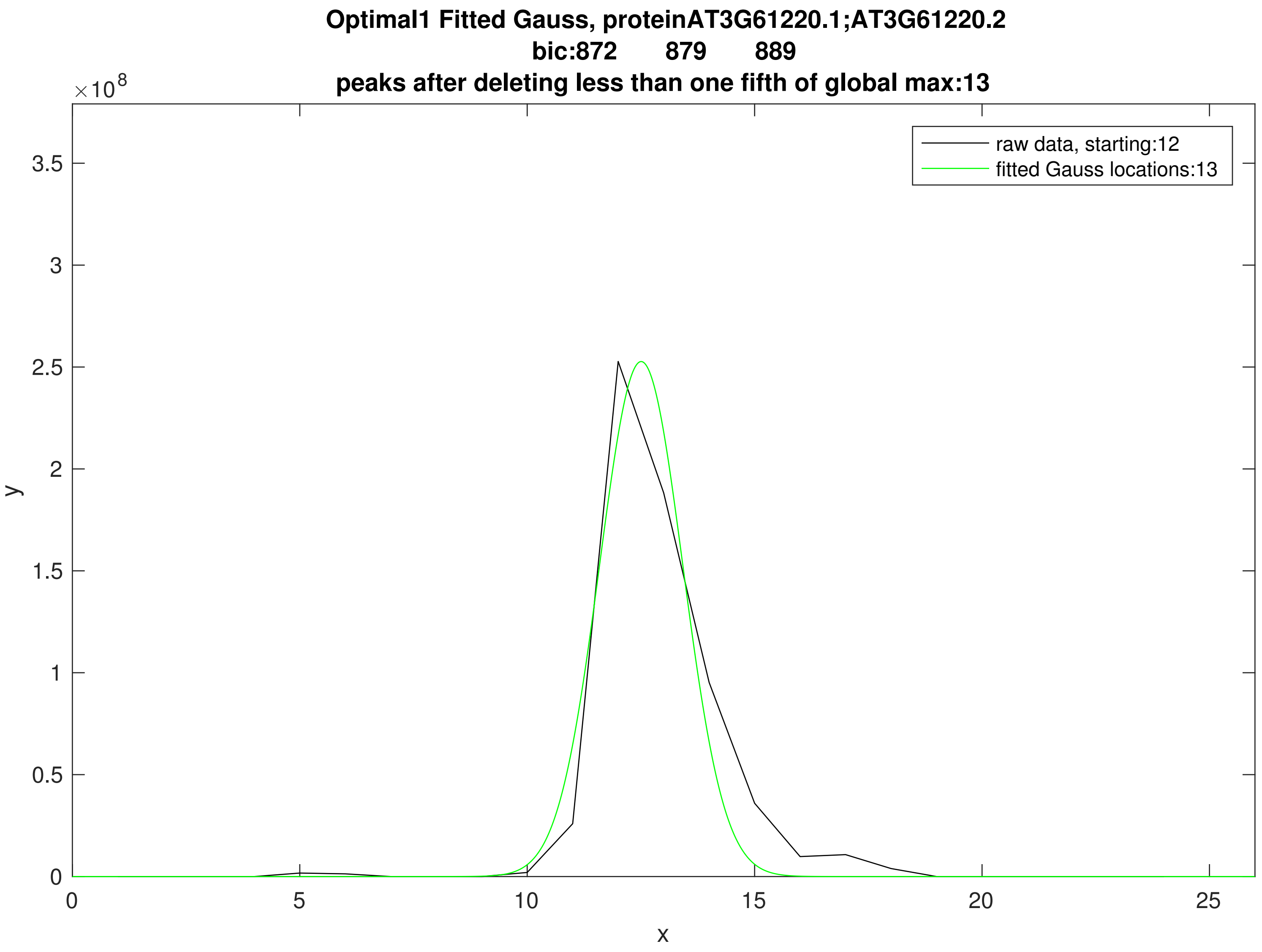
Optimal1 Fitted Gauss, proteinAT3G61220.1;AT3G61220.2
bic:872 879 889
peaks after deleting less than one fifth of global max:13
×10^8
3.5
3
2.5
2
1.5
1
0.5
0
y
0
5
10
15
20
25
x
raw data, starting:12
fitted Gauss locations:13

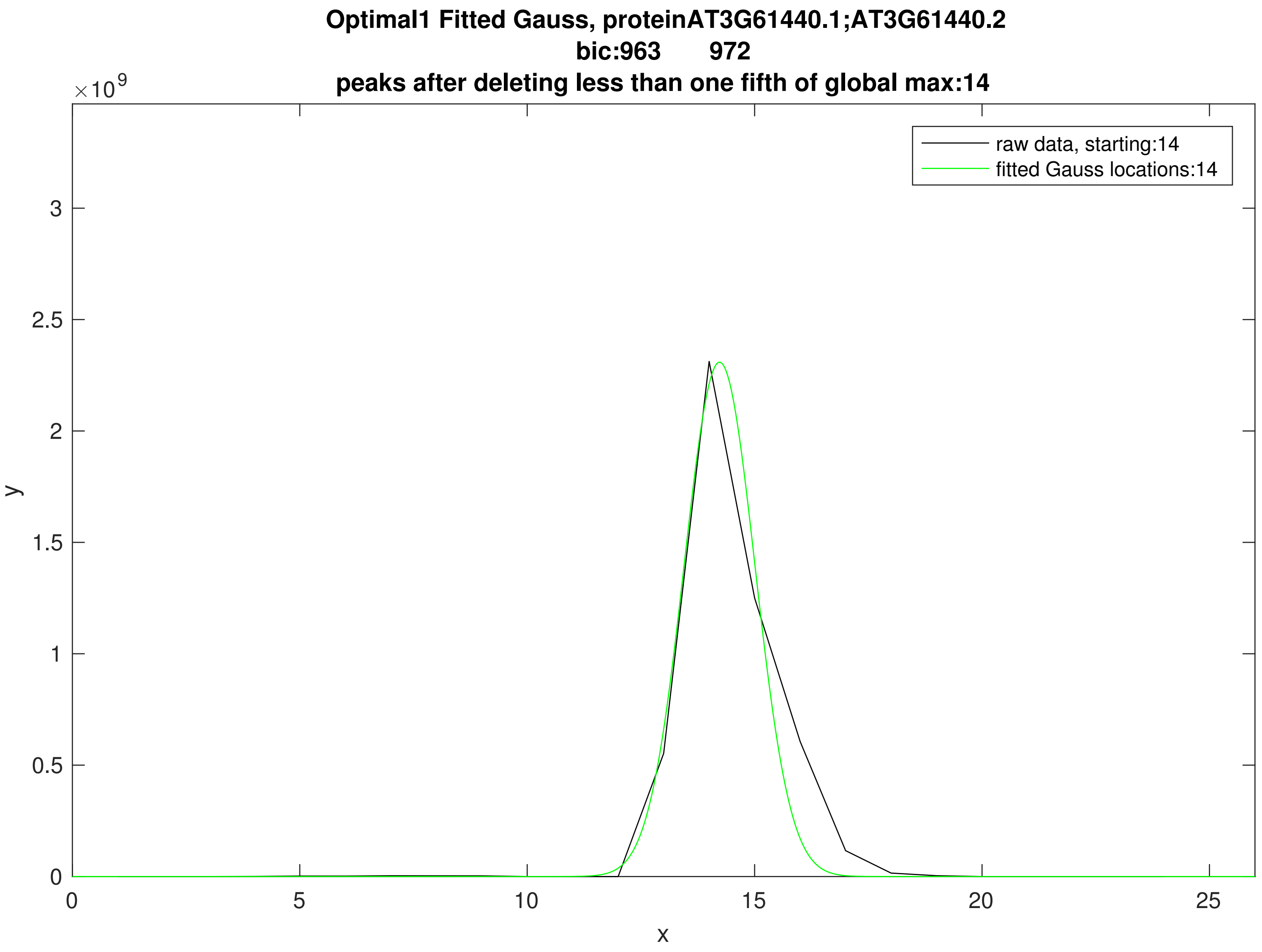
Optimal1 Fitted Gauss, proteinAT3G61440.1;AT3G61440.2
bic:963 972
peaks after deleting less than one fifth of global max:14
×10^9
raw data, starting:14
fitted Gauss locations:14
y
x
0
0.5
1
1.5
2
2.5
3
0
5
10
15
20
25

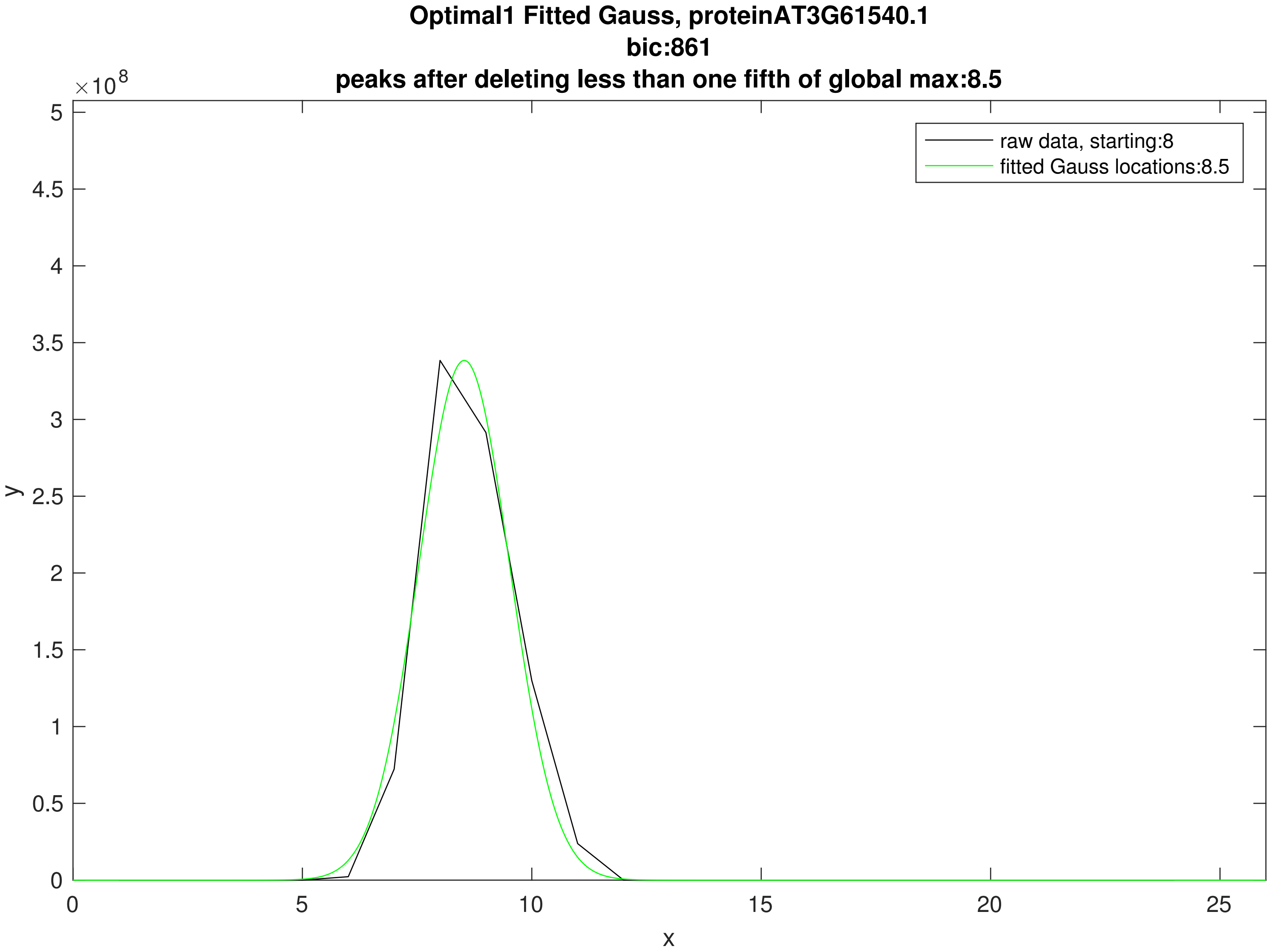
Optimal1 Fitted Gauss, proteinAT3G61540.1
bic:861
peaks after deleting less than one fifth of global max:8.5
×10^8
0
0.5
1
1.5
2
2.5
3
3.5
4
4.5
5
y
0
5
10
15
20
25
x
raw data, starting:8
fitted Gauss locations:8.5

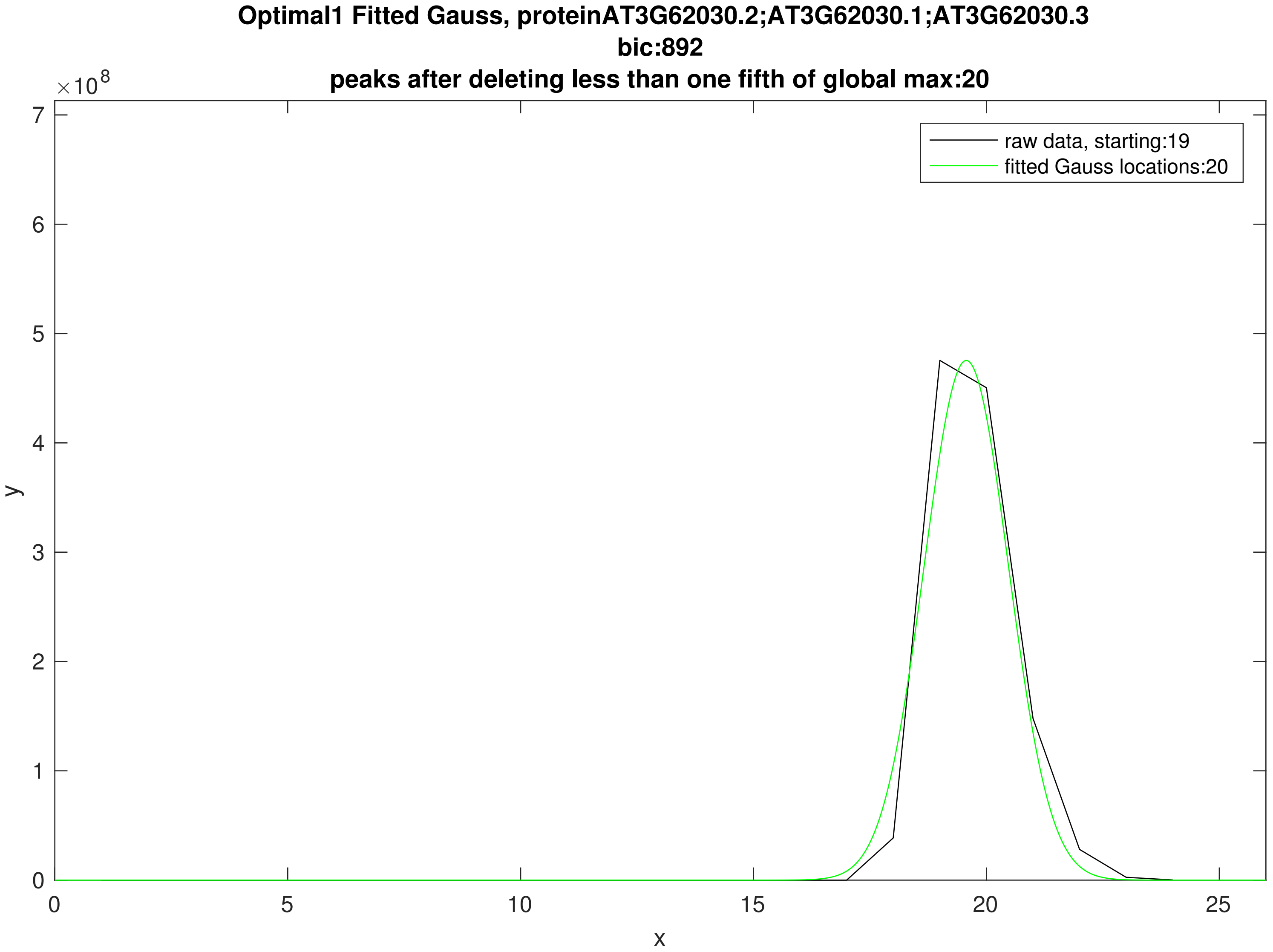

Optimal1 Fitted Gauss, proteinAT3G62030.2;AT3G62030.1;AT3G62030.3
bic:892
peaks after deleting less than one fifth of global max:20
×10^8
raw data, starting:19
fitted Gauss locations:20
y
x
0
1
2
3
4
5
6
7
0
5
10
15
20
25

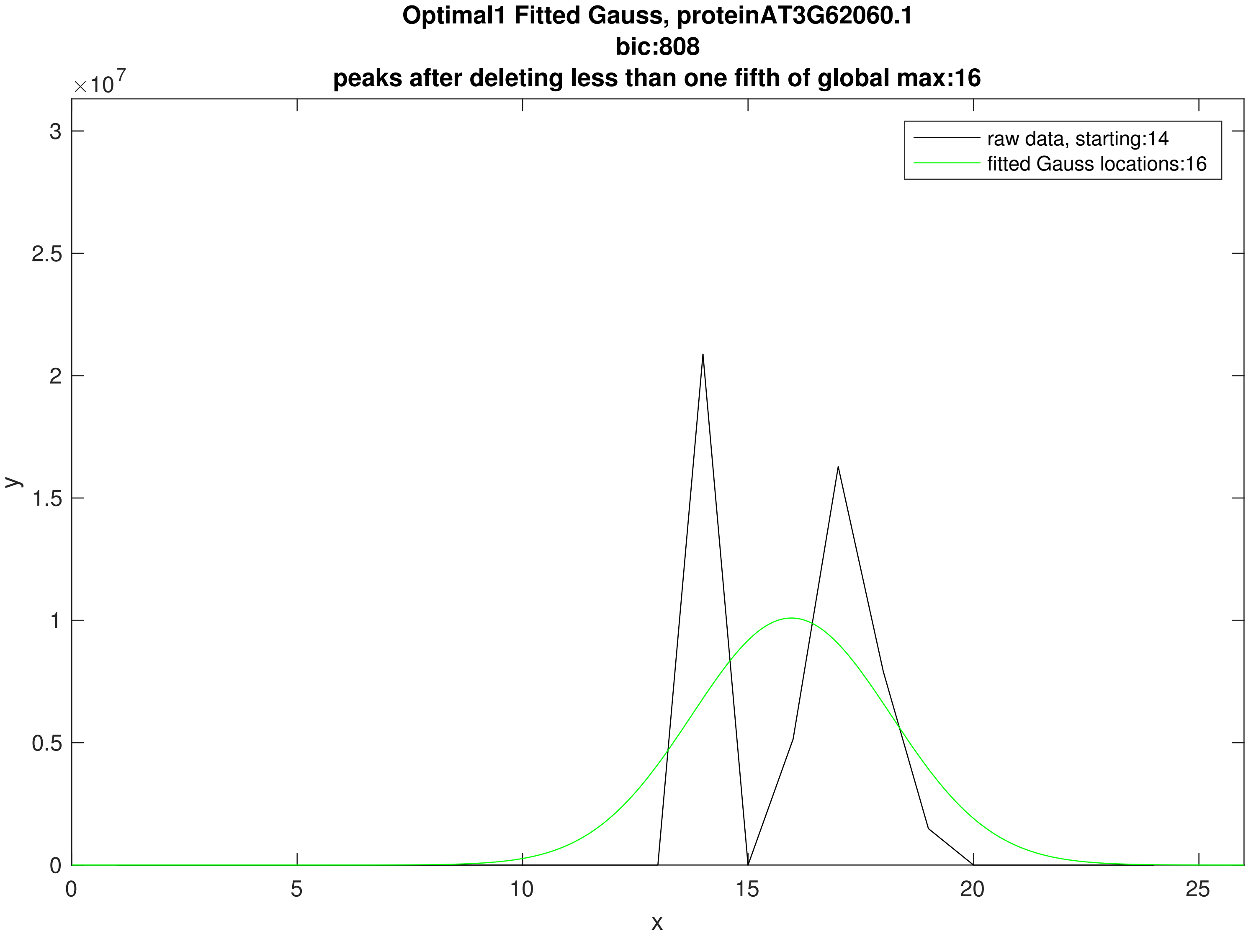
Optimal1 Fitted Gauss, proteinAT3G62060.1
bic:808
peaks after deleting less than one fifth of global max:16
×10^7
3
2.5
2
1.5
1
0.5
0
y
0
5
10
15
20
25
x
raw data, starting:14
fitted Gauss locations:16

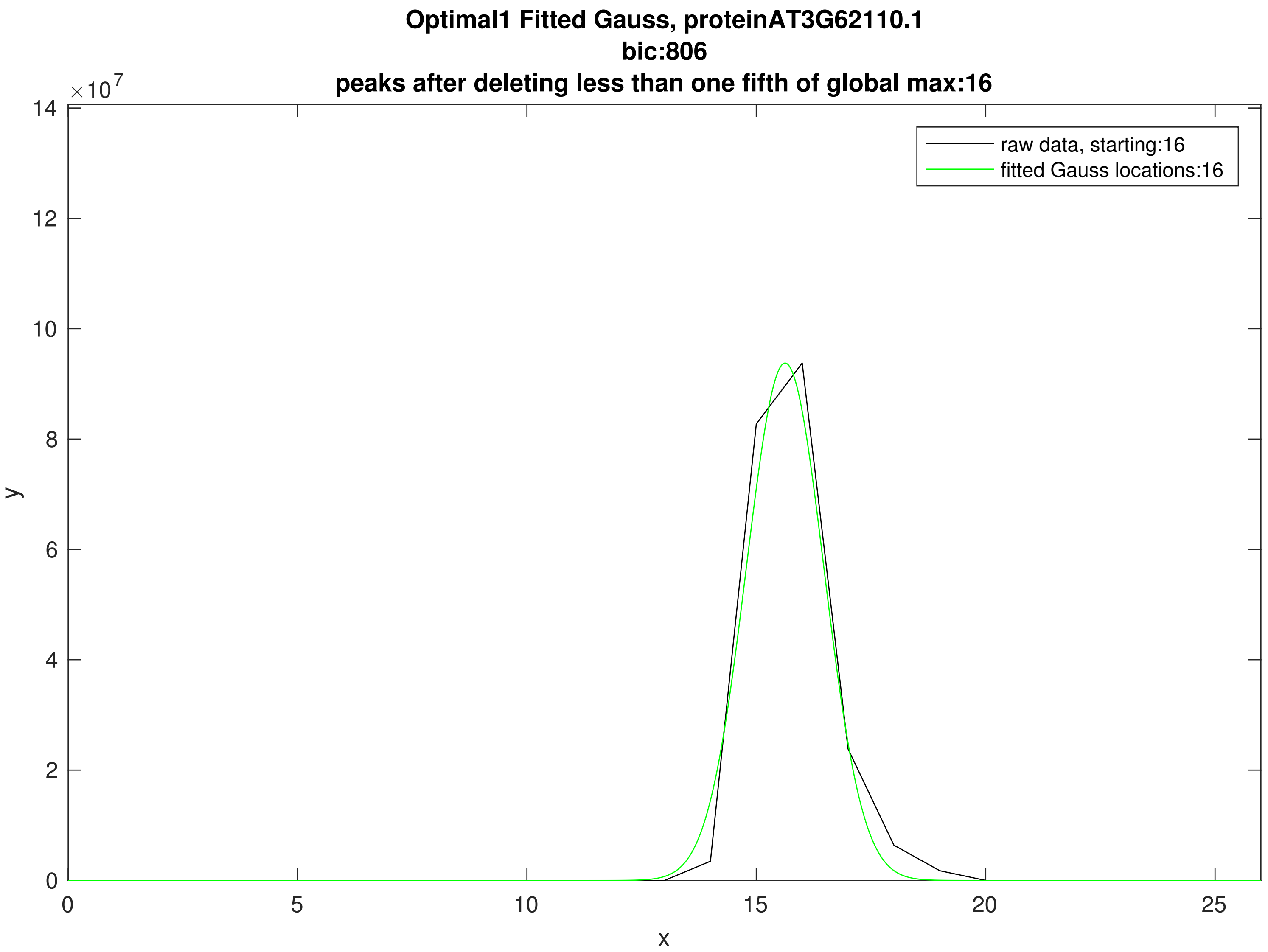
Optimal1 Fitted Gauss, proteinAT3G62110.1
bic:806
peaks after deleting less than one fifth of global max:16
×10^7
14
12
10
8
6
4
2
0
y
0
5
10
15
20
25
x
raw data, starting:16
fitted Gauss locations:16

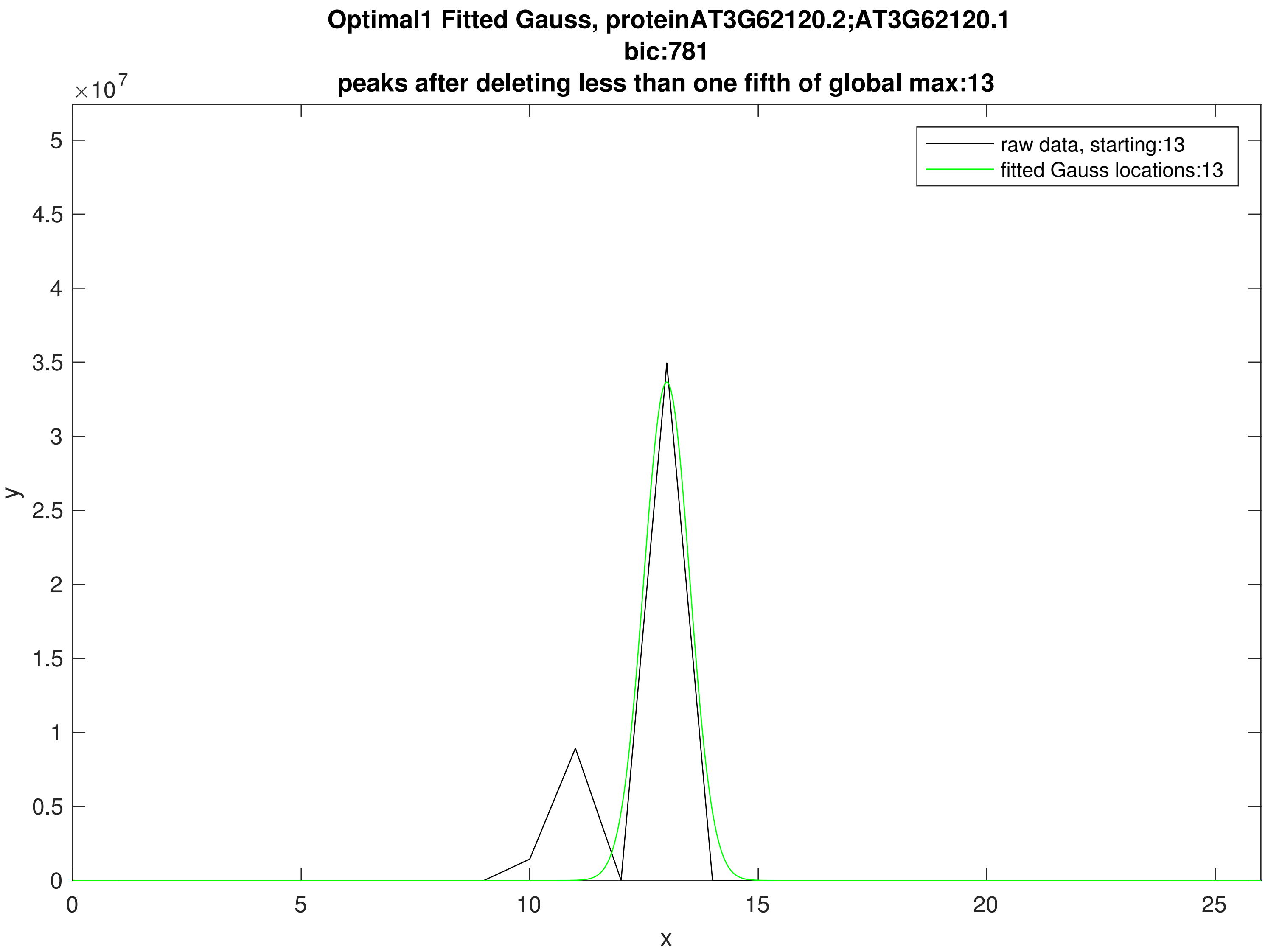
Optimal1 Fitted Gauss, proteinAT3G62120.2;AT3G62120.1
bic:781
peaks after deleting less than one fifth of global max:13
×10^7
raw data, starting:13
fitted Gauss locations:13
y
x
0
0.5
1
1.5
2
2.5
3
3.5
4
4.5
5
0
5
10
15
20
25

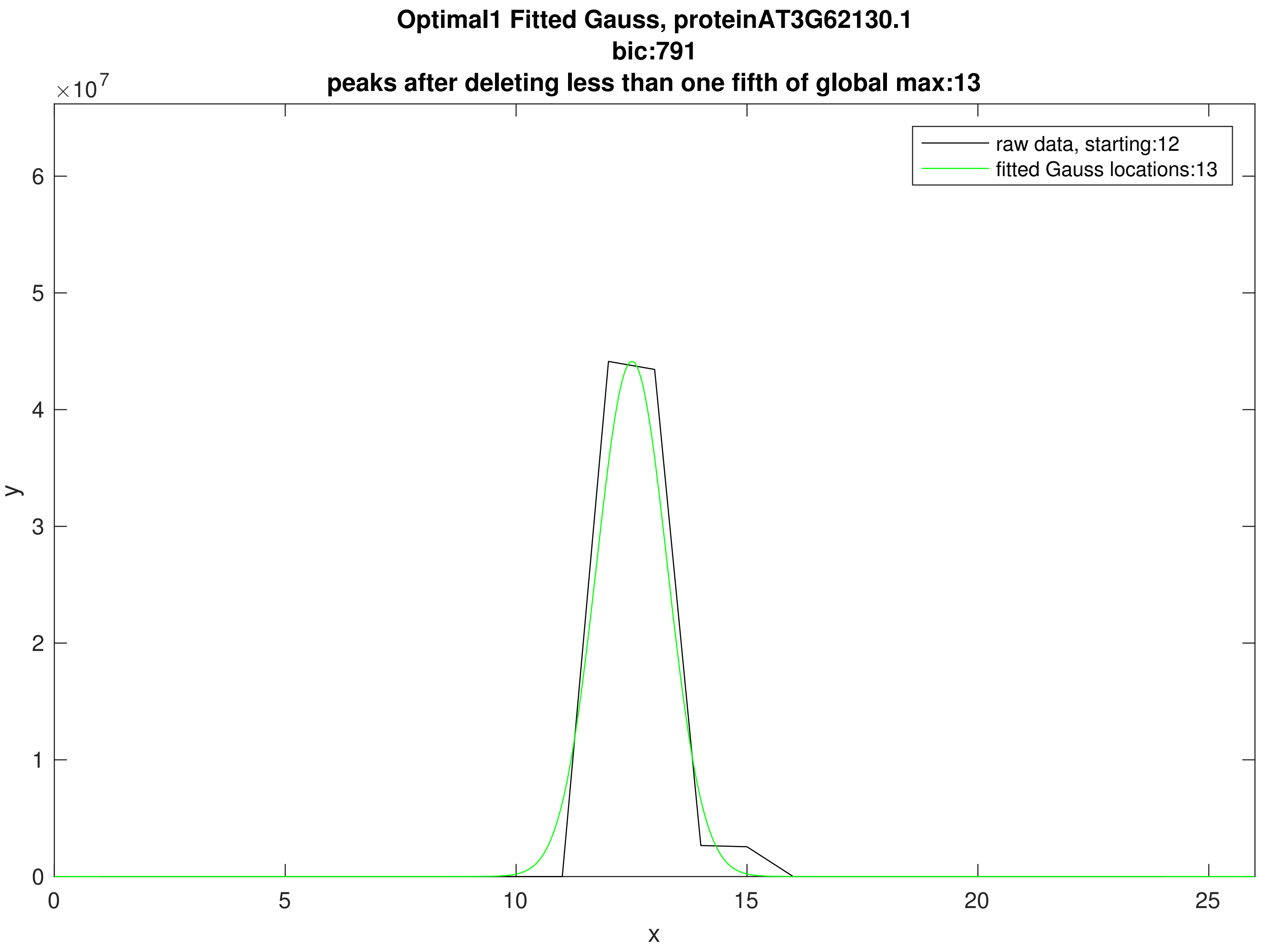
Optimal1 Fitted Gauss, proteinAT3G62130.1
bic:791
peaks after deleting less than one fifth of global max:13
×10^7
raw data, starting:12
fitted Gauss locations:13
y
x
0
1
2
3
4
5
6
0
5
10
15
20
25

s, proteinAT3G62290.3;AT3G62290.2;AT3G62290.1;AT2G47170.1;AT1G70490.3;AT1G70490.2;AT1G70490.1;AT1G2349

**bic:825 820 829**

**peaks after deleting less than one fifth of global max:20**

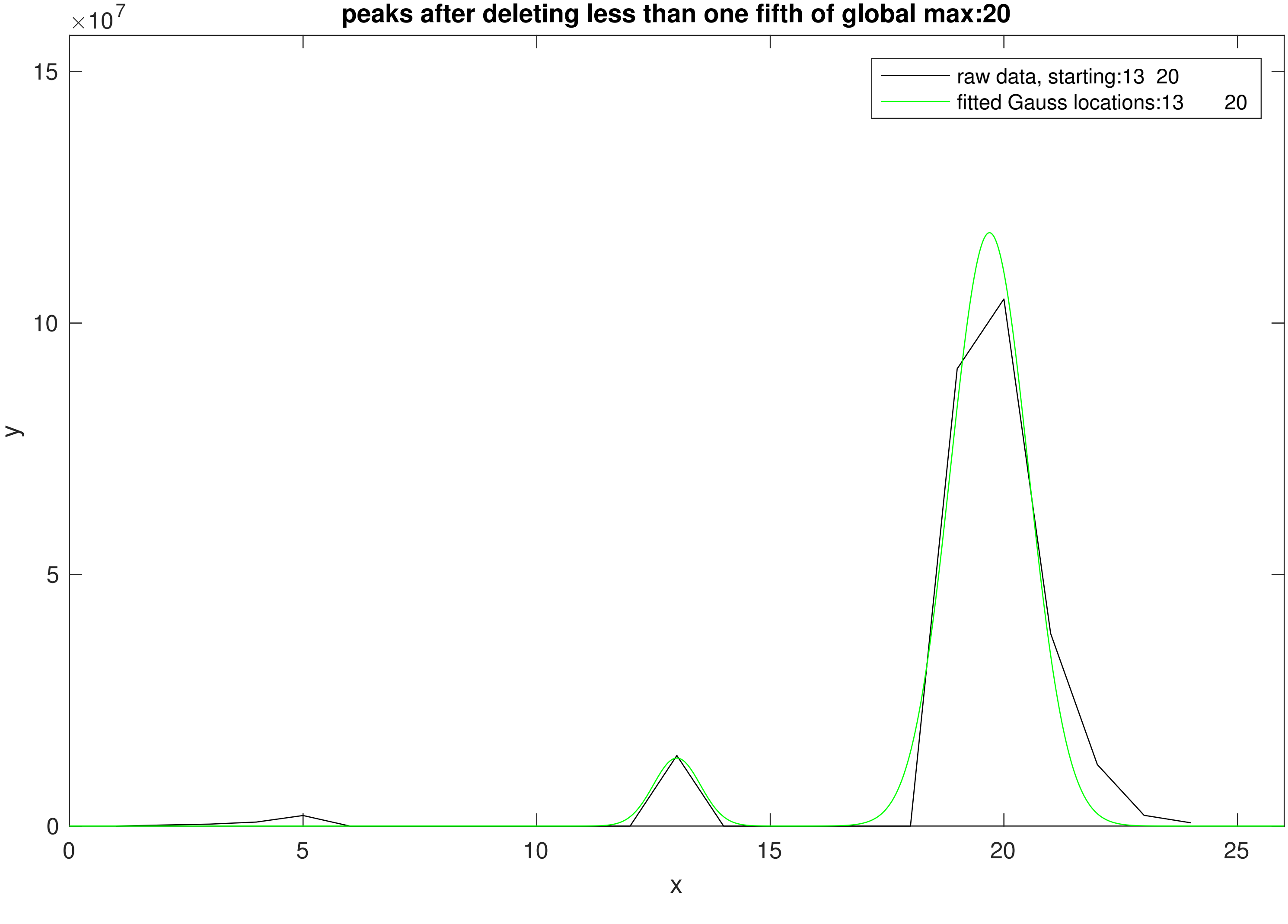

**Optimal1 Fitted Gauss, proteinAT3G62360.1**
**bic:670**
**peaks after deleting less than one fifth of global max:19**

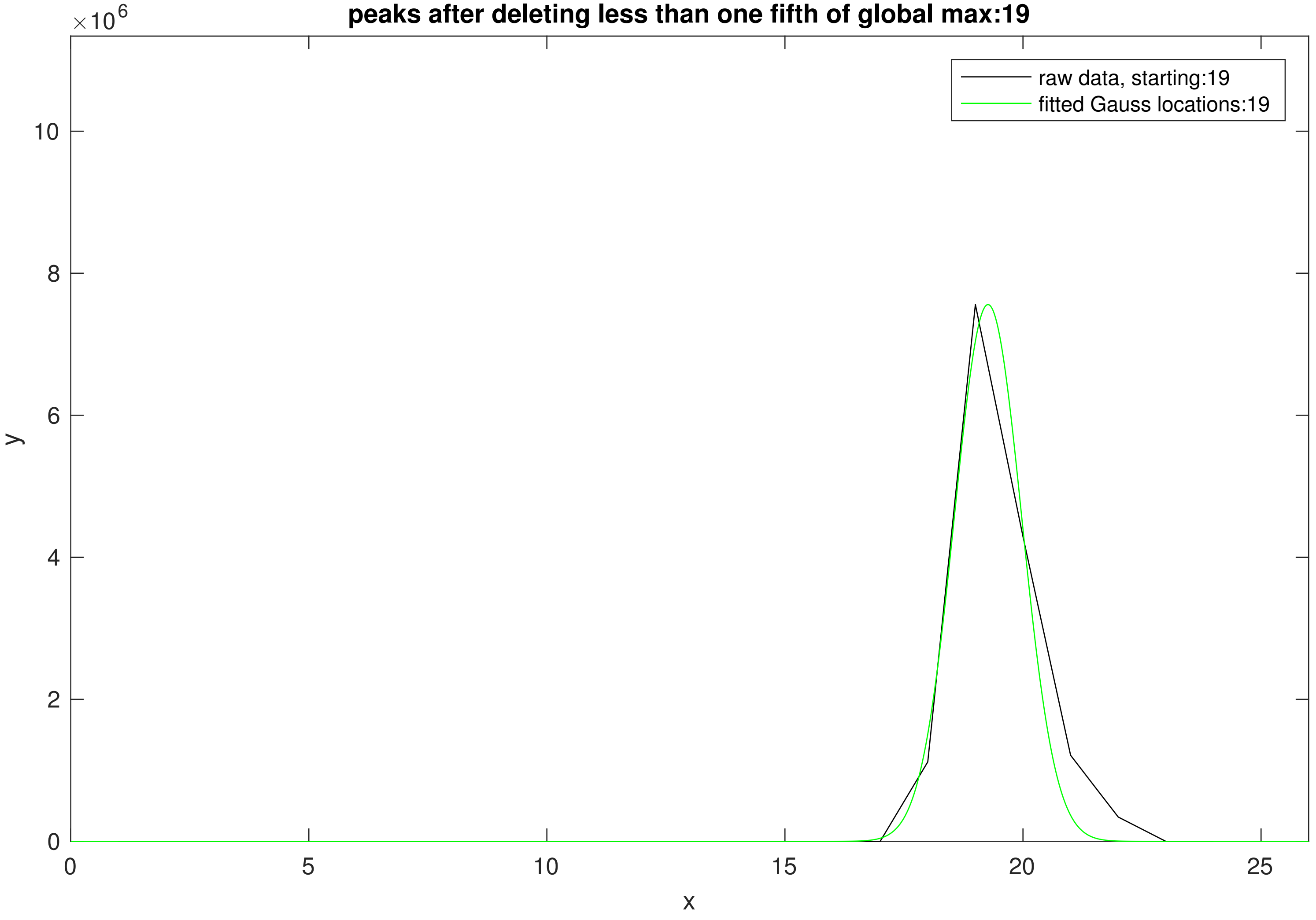

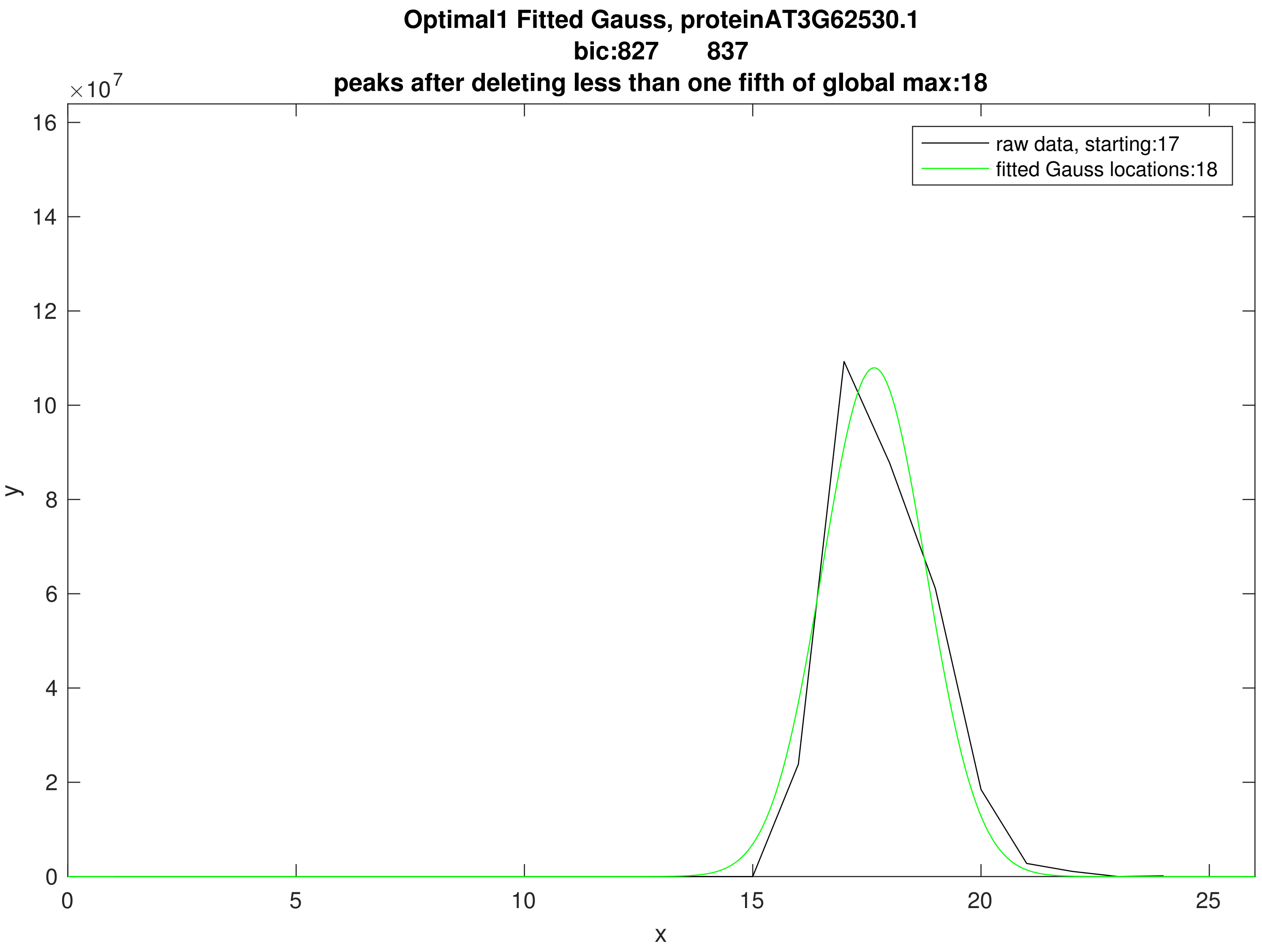

Optimal1 Fitted Gauss, proteinAT3G62530.1
bic:827 837
peaks after deleting less than one fifth of global max:18
×10^7
16
14
12
10
8
6
4
2
0
y
0
5
10
15
20
25
x
raw data, starting:17
fitted Gauss locations:18

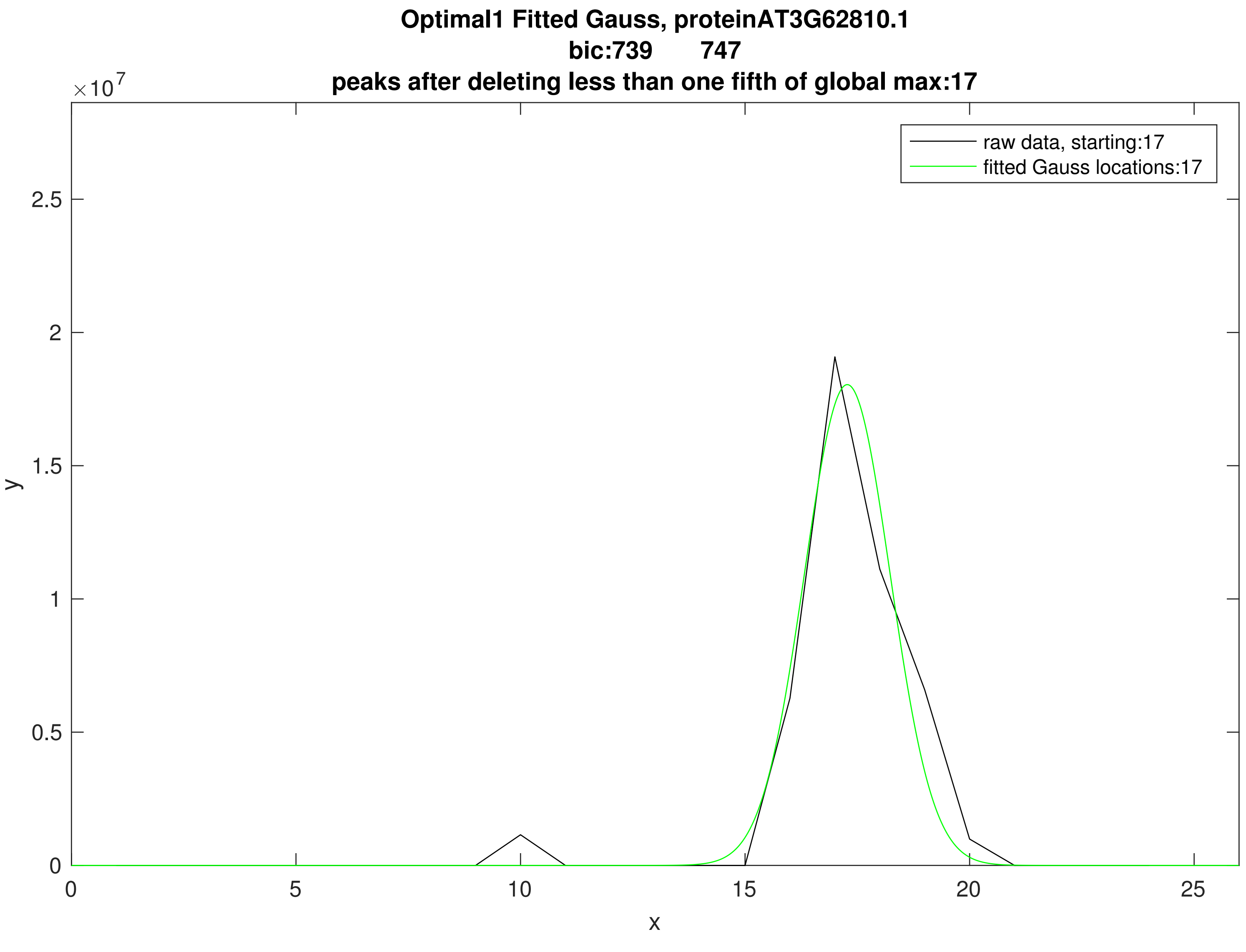

Optimal1 Fitted Gauss, proteinAT3G62810.1
bic:739 747
peaks after deleting less than one fifth of global max:17
×10^7
raw data, starting:17
fitted Gauss locations:17
2.5
2
1.5
1
0.5
0
y
0
5
10
15
20
25
x

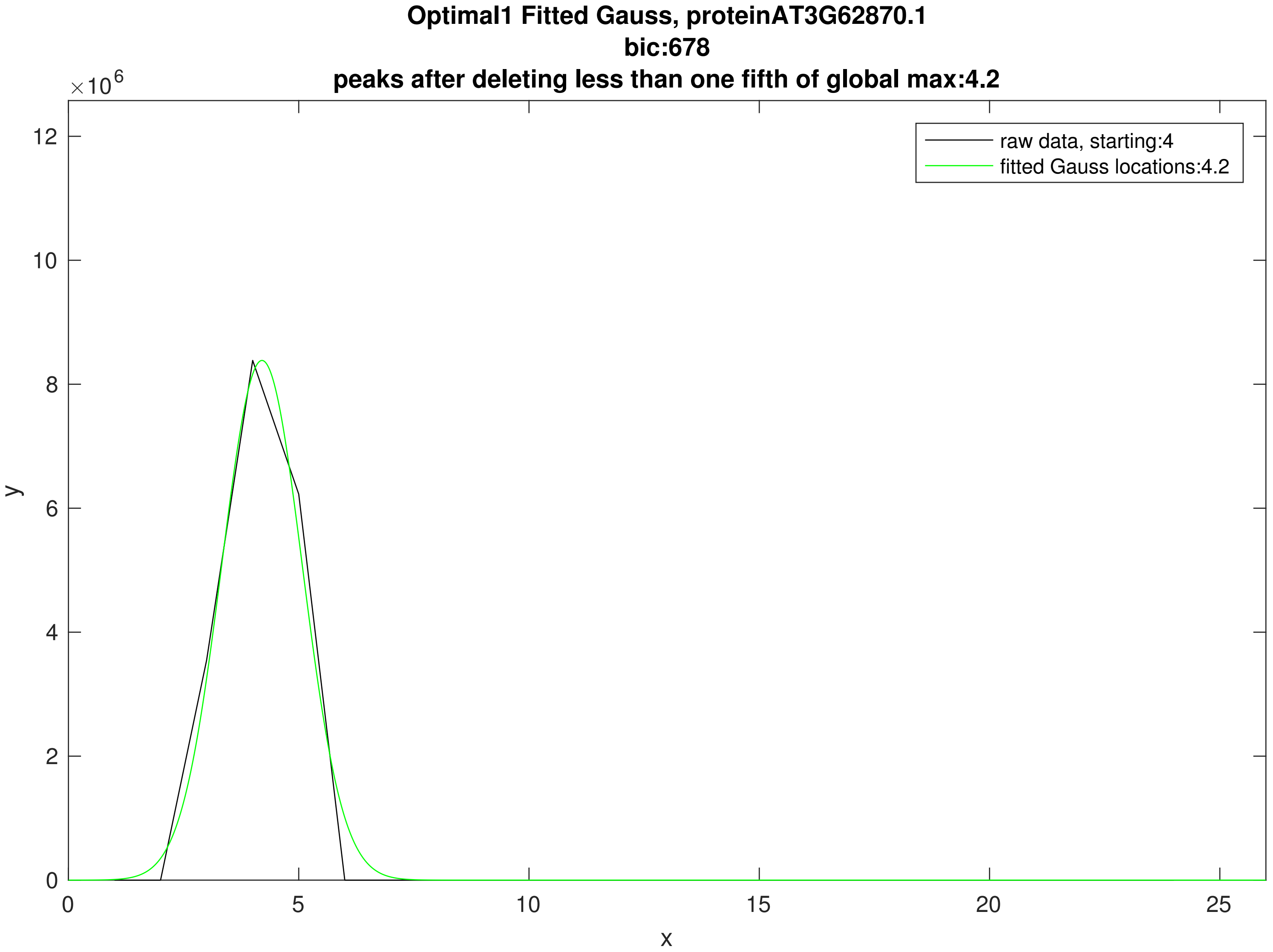
Optimal1 Fitted Gauss, proteinAT3G62870.1
bic:678
peaks after deleting less than one fifth of global max:4.2
×10^6
raw data, starting:4
fitted Gauss locations:4.2
y
x
0
2
4
6
8
10
12
0
5
10
15
20
25

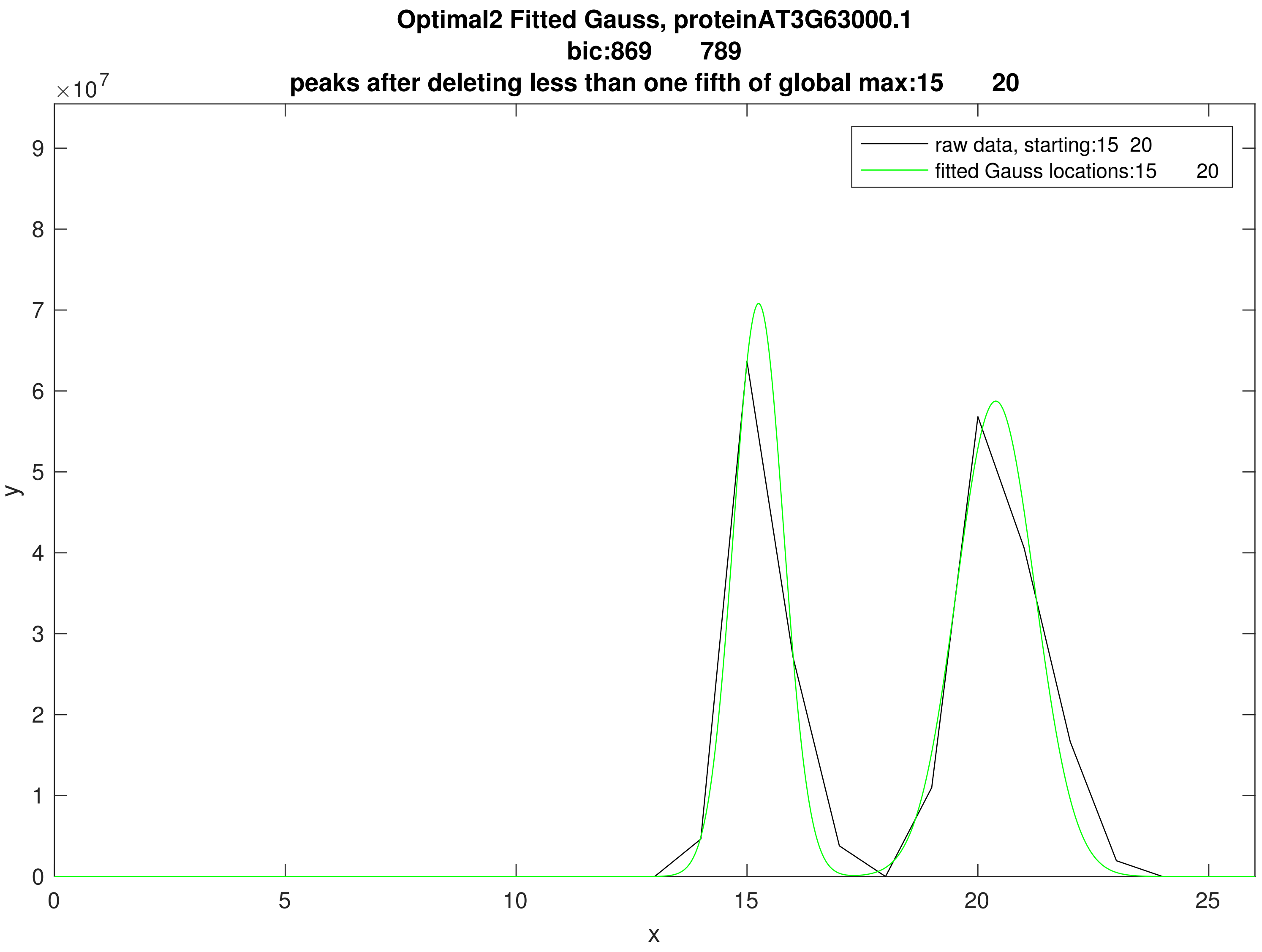

Optimal2 Fitted Gauss, proteinAT3G63000.1
bic:869 789
peaks after deleting less than one fifth of global max:15 20
×10^7
raw data, starting:15 20
fitted Gauss locations:15 20
y
x
0
1
2
3
4
5
6
7
8
9
0
5
10
15
20
25

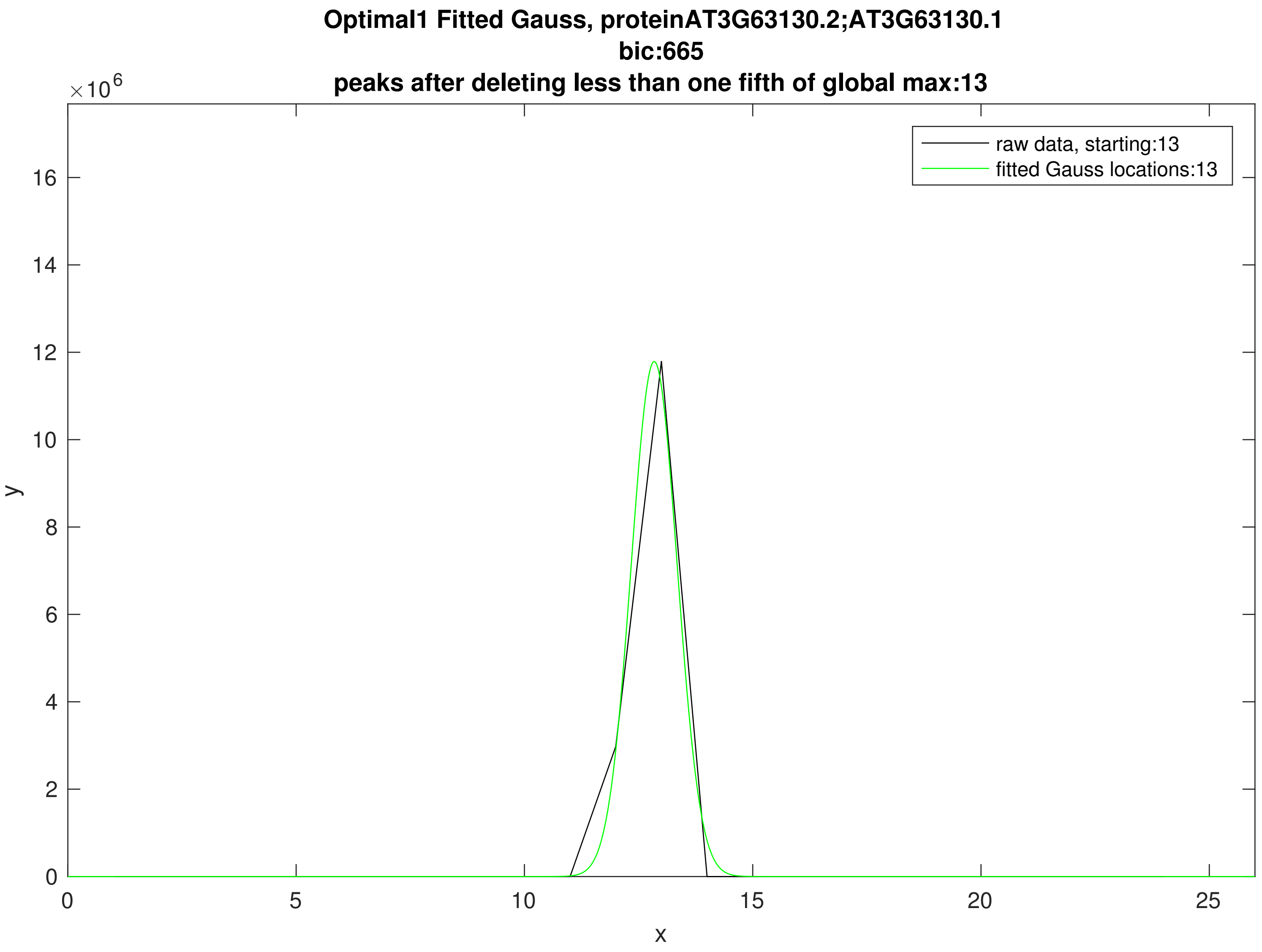

Optimal1 Fitted Gauss, proteinAT3G63130.2;AT3G63130.1
bic:665
peaks after deleting less than one fifth of global max:13
×10^6
raw data, starting:13
fitted Gauss locations:13
16
14
12
10
8
6
4
2
0
y
0
5
10
15
20
25
x

**Optimal2 Fitted Gauss, proteinAT3G63190.1**
**bic:854 804**
**peaks after deleting less than one fifth of global max:10 19**

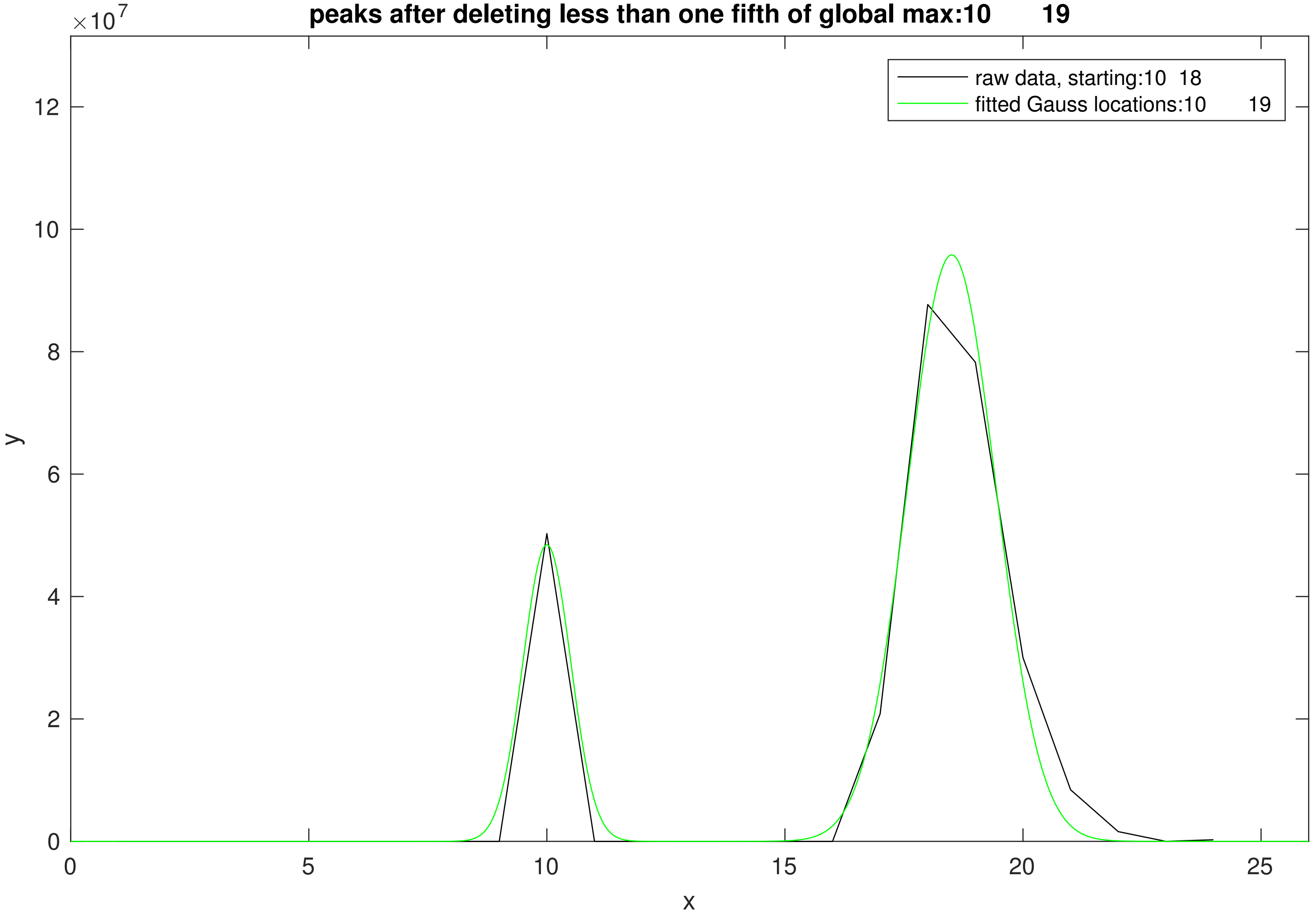

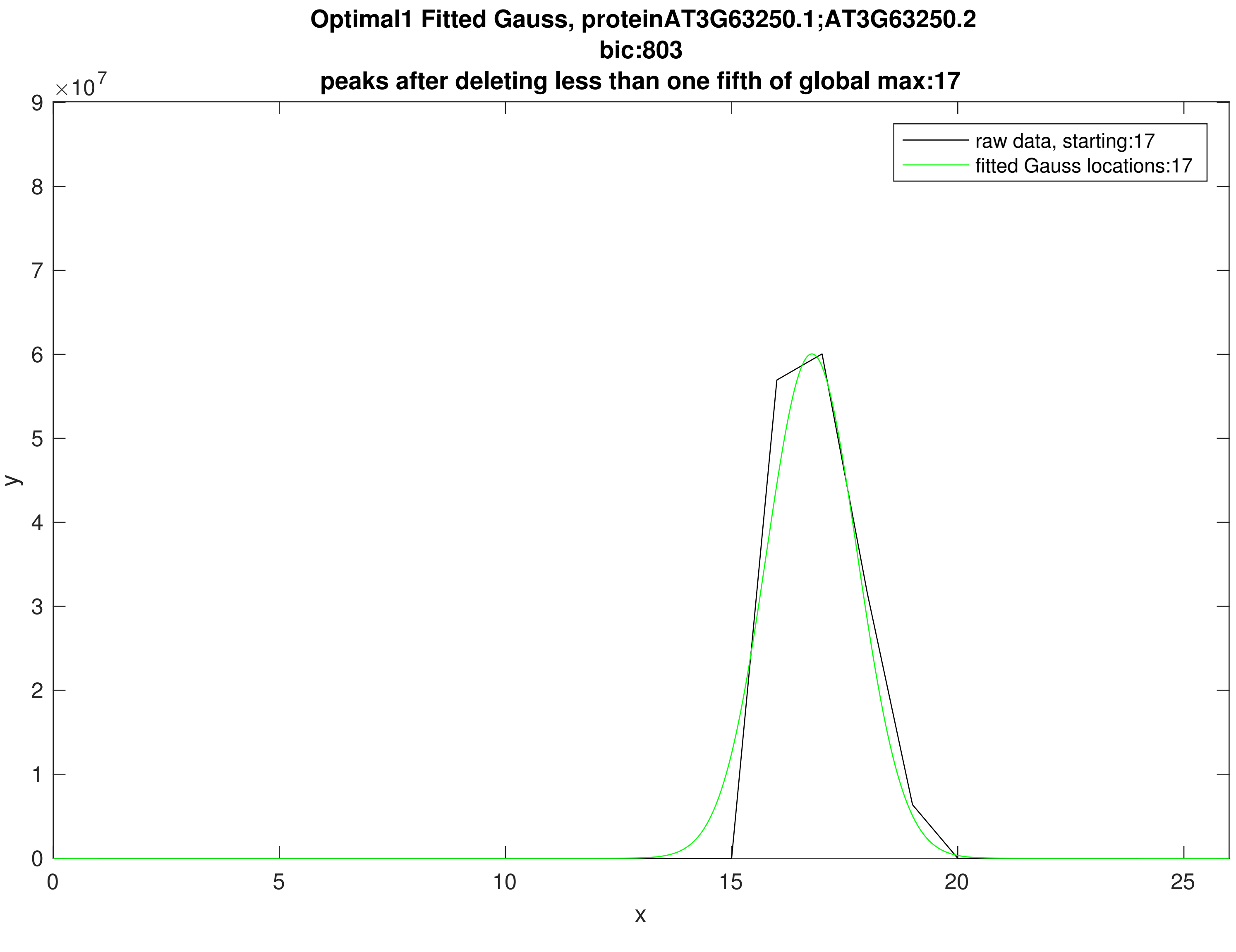
Optimal1 Fitted Gauss, proteinAT3G63250.1;AT3G63250.2
bic:803
peaks after deleting less than one fifth of global max:17
×10^7
9
8
7
6
5
4
3
2
1
0
y
0
5
10
15
20
25
x
raw data, starting:17
fitted Gauss locations:17

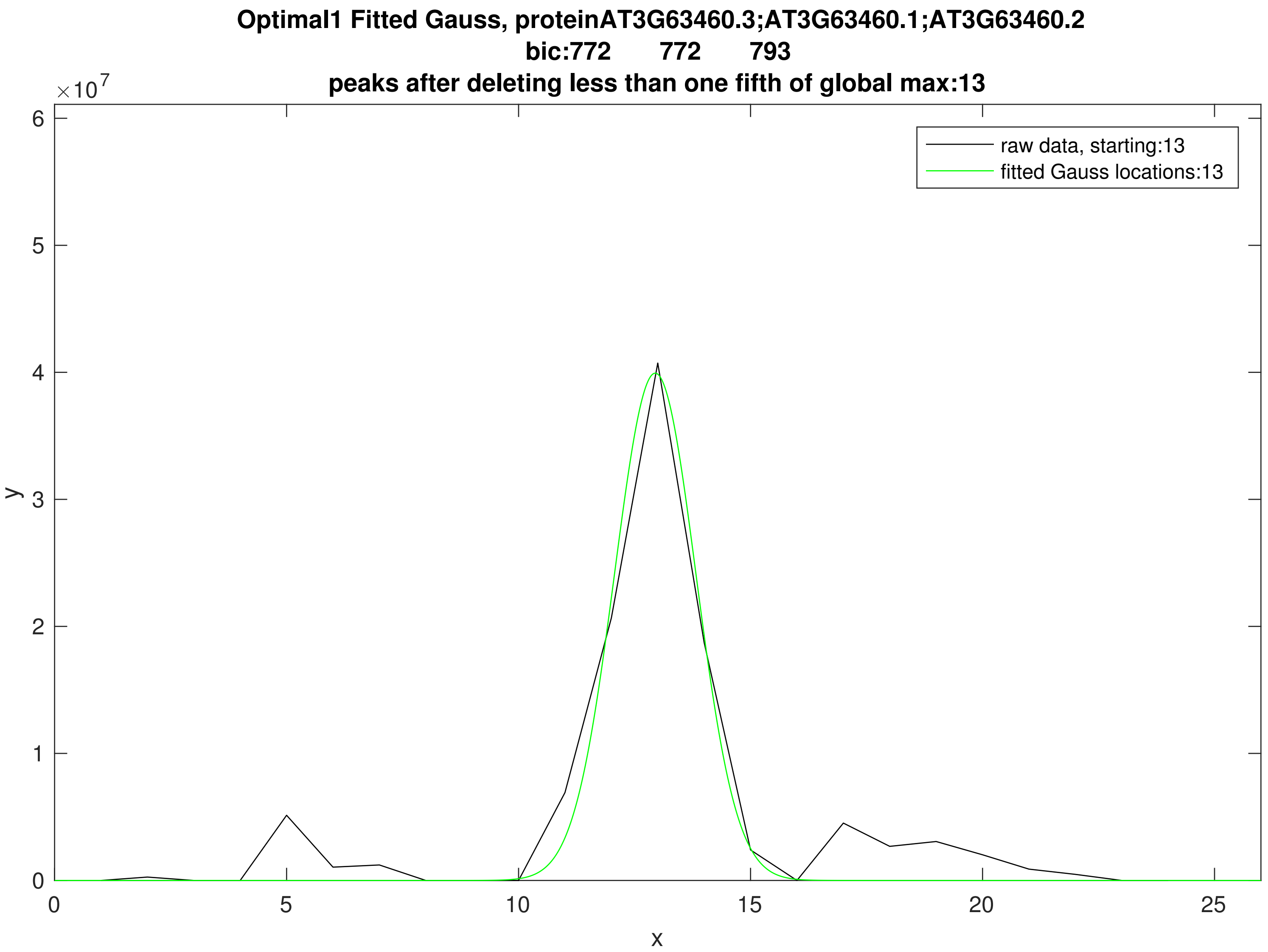
Optimal1 Fitted Gauss, proteinAT3G63460.3;AT3G63460.1;AT3G63460.2
bic:772 772 793
peaks after deleting less than one fifth of global max:13
×10^7
raw data, starting:13
fitted Gauss locations:13
y
x
0
1
2
3
4
5
6
10
15
20
25

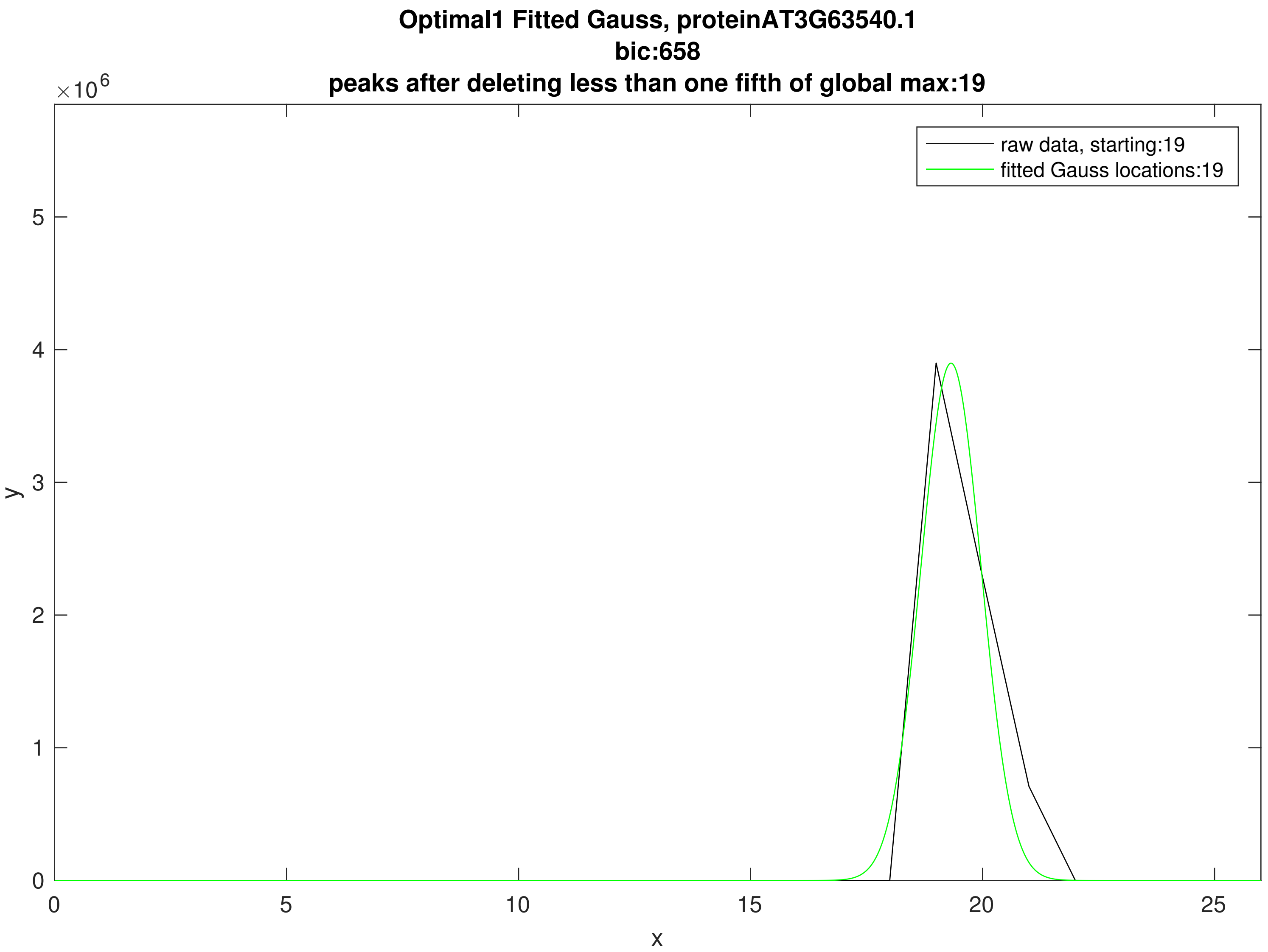
Optimal1 Fitted Gauss, proteinAT3G63540.1
bic:658
peaks after deleting less than one fifth of global max:19
×10^6
raw data, starting:19
fitted Gauss locations:19
y
x
0
1
2
3
4
5
0
5
10
15
20
25

Optimal1 Fitted Gauss, proteinAT4G00100.1
bic:818 825
peaks after deleting less than one fifth of global max:4

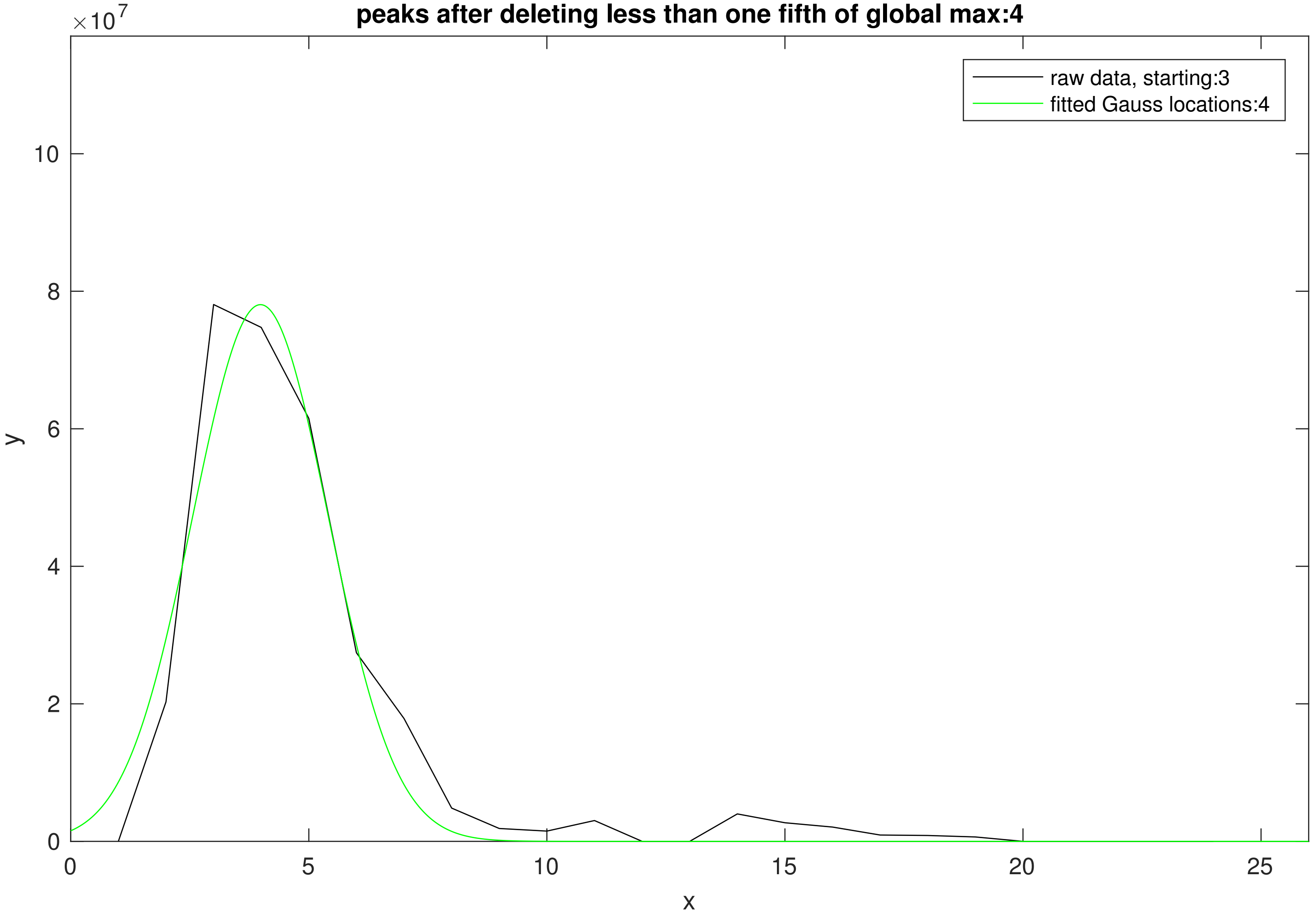

#### Optimal2 Fitted Gauss, proteinAT4G00300.2;AT4G00300.1
#### bic:773 739
#### peaks after deleting less than one fifth of global max:8.3 16

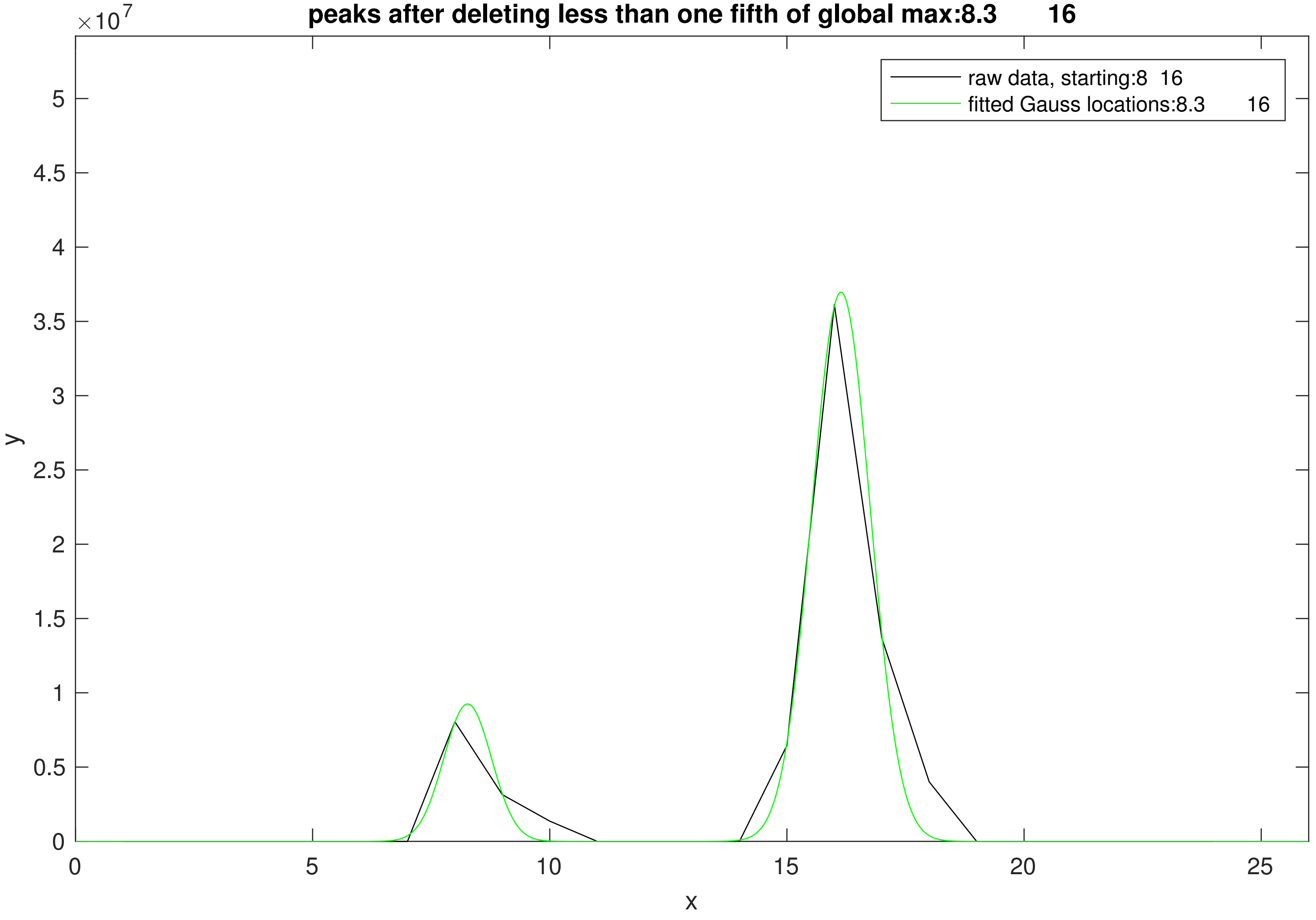

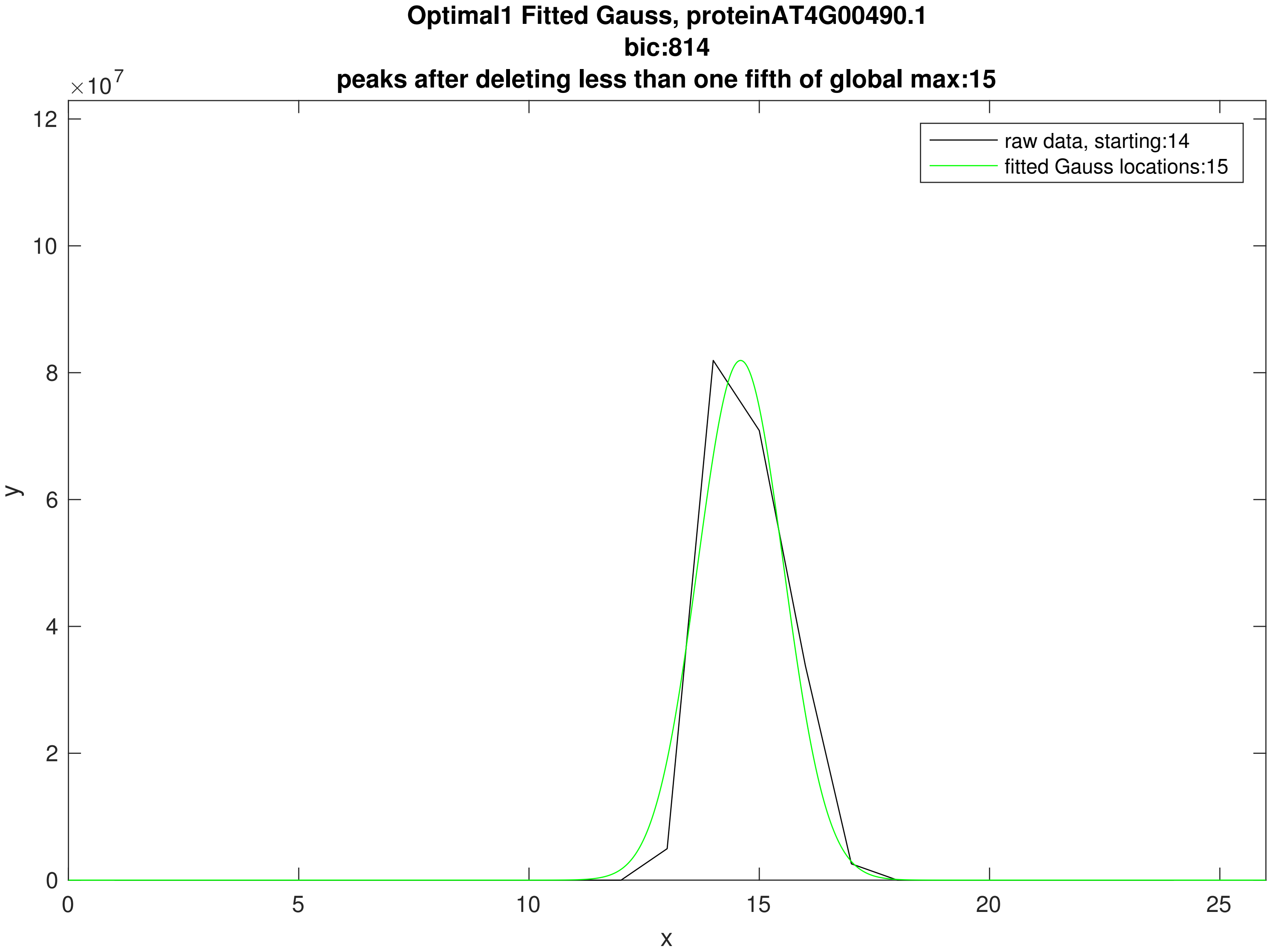
Optimal1 Fitted Gauss, proteinAT4G00490.1
bic:814
peaks after deleting less than one fifth of global max:15
×10^7
raw data, starting:14
fitted Gauss locations:15
y
x
0
2
4
6
8
10
12
0
5
10
15
20
25

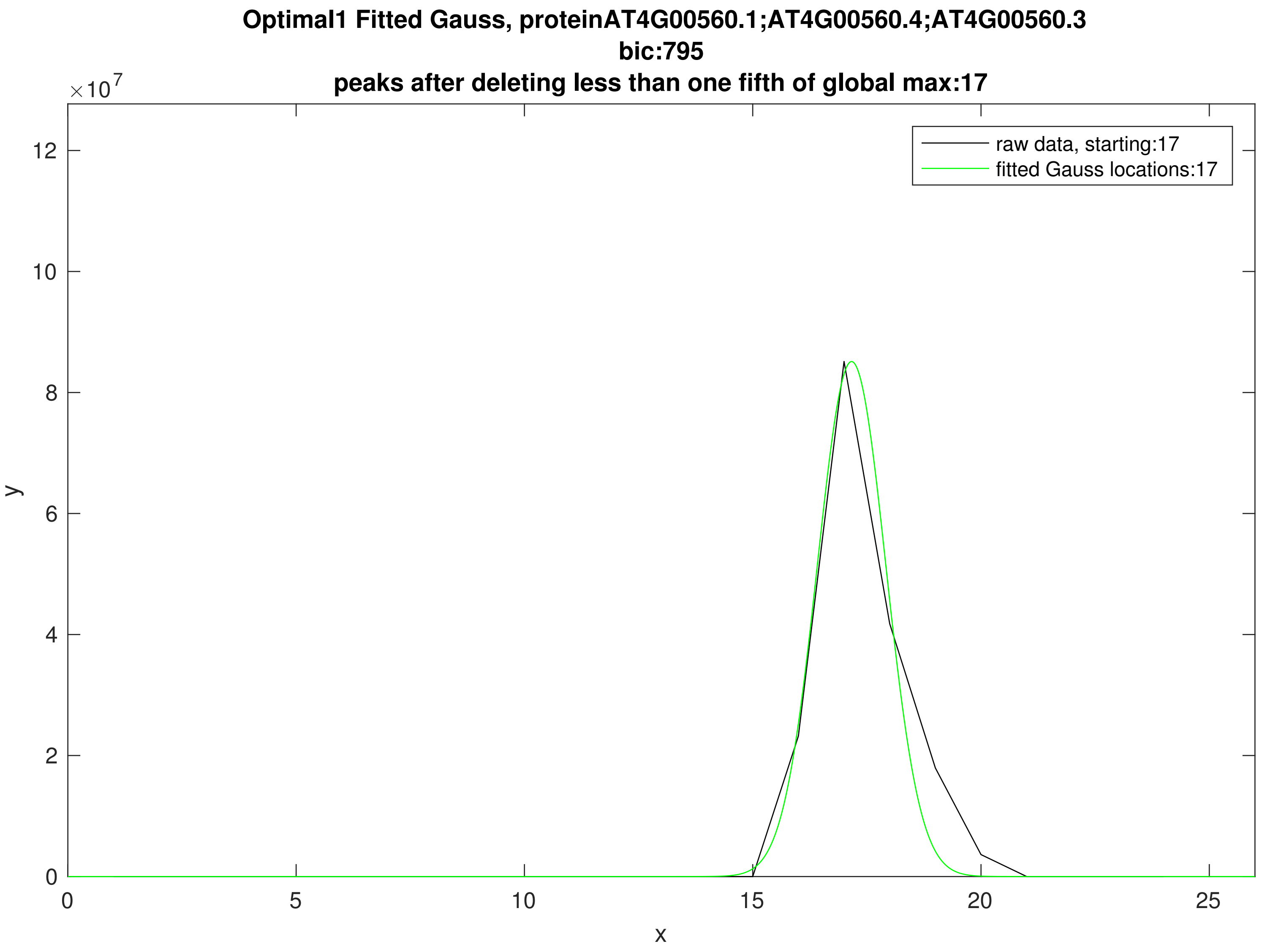
Optimal1 Fitted Gauss, proteinAT4G00560.1;AT4G00560.4;AT4G00560.3
bic:795
peaks after deleting less than one fifth of global max:17
×10^7
raw data, starting:17
fitted Gauss locations:17
y
x
0
2
4
6
8
10
12
0
5
10
15
20
25

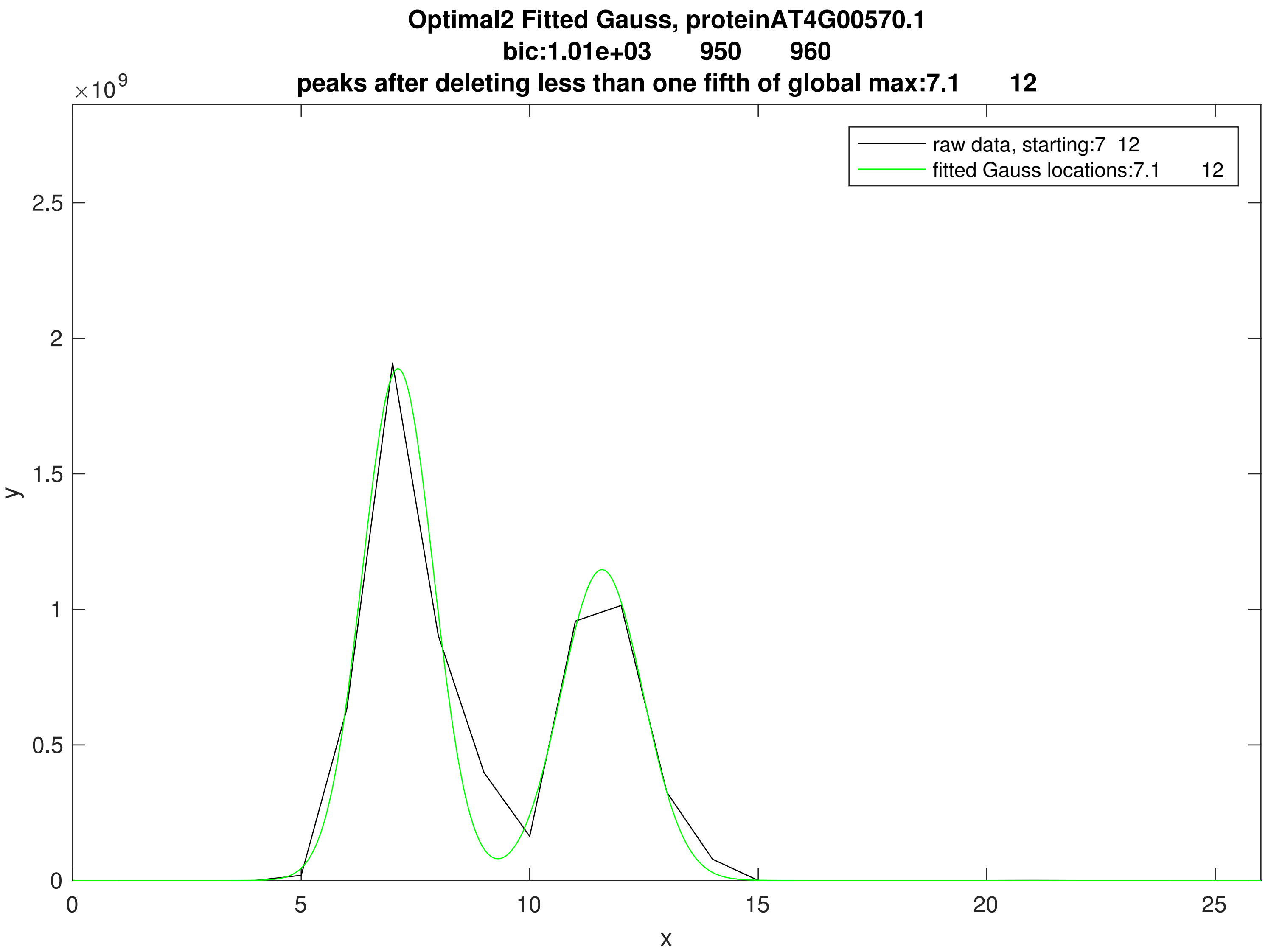
Optimal2 Fitted Gauss, proteinAT4G00570.1
bic:1.01e+03 950 960
peaks after deleting less than one fifth of global max:7.1 12
raw data, starting:7 12
fitted Gauss locations:7.1 12
×10^9
y
x

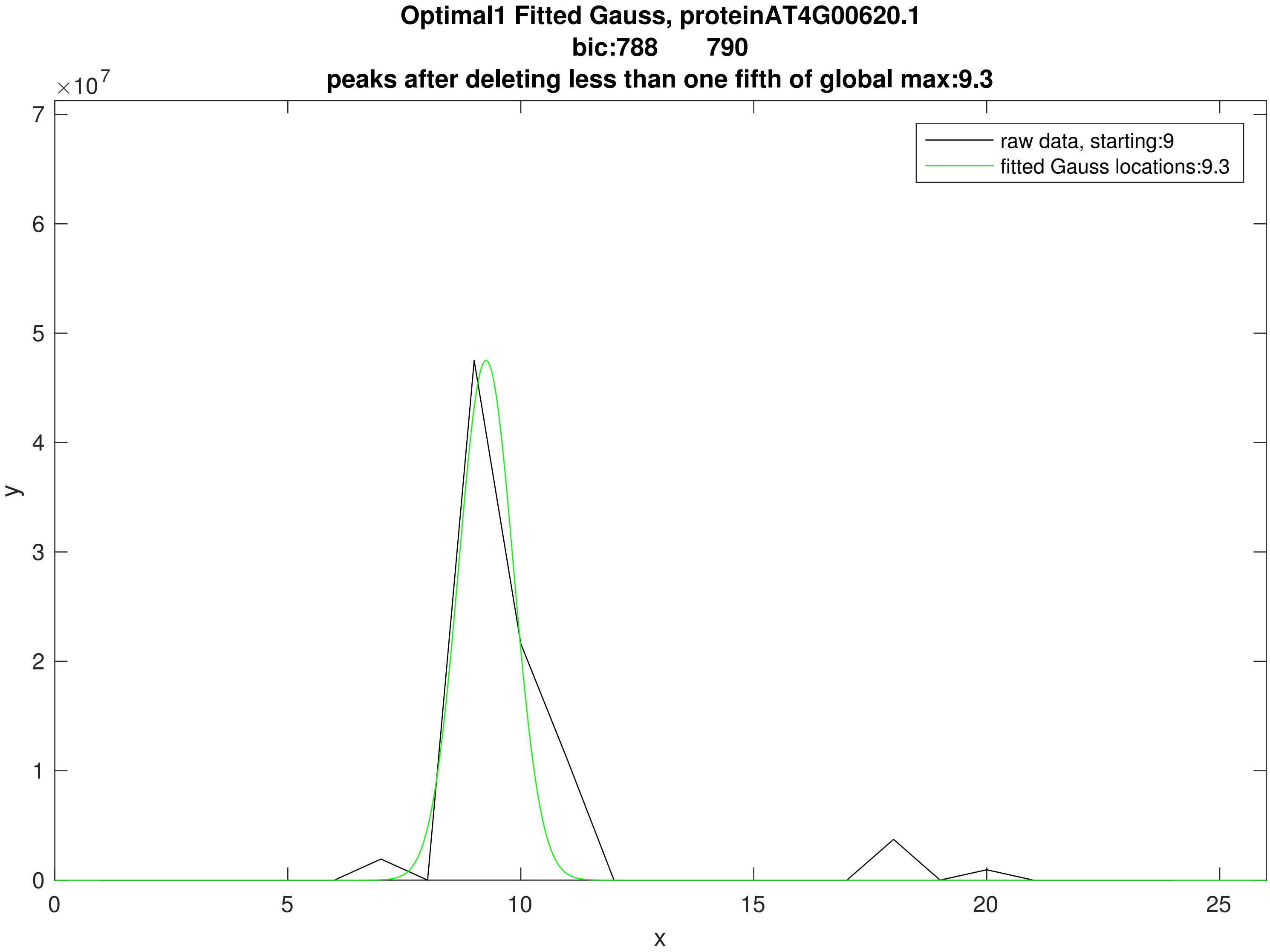

Optimal1 Fitted Gauss, proteinAT4G00620.1
bic:788 790
peaks after deleting less than one fifth of global max:9.3
×10^7
7
6
5
4
3
2
1
0
y
0
5
10
15
20
25
x
raw data, starting:9
fitted Gauss locations:9.3

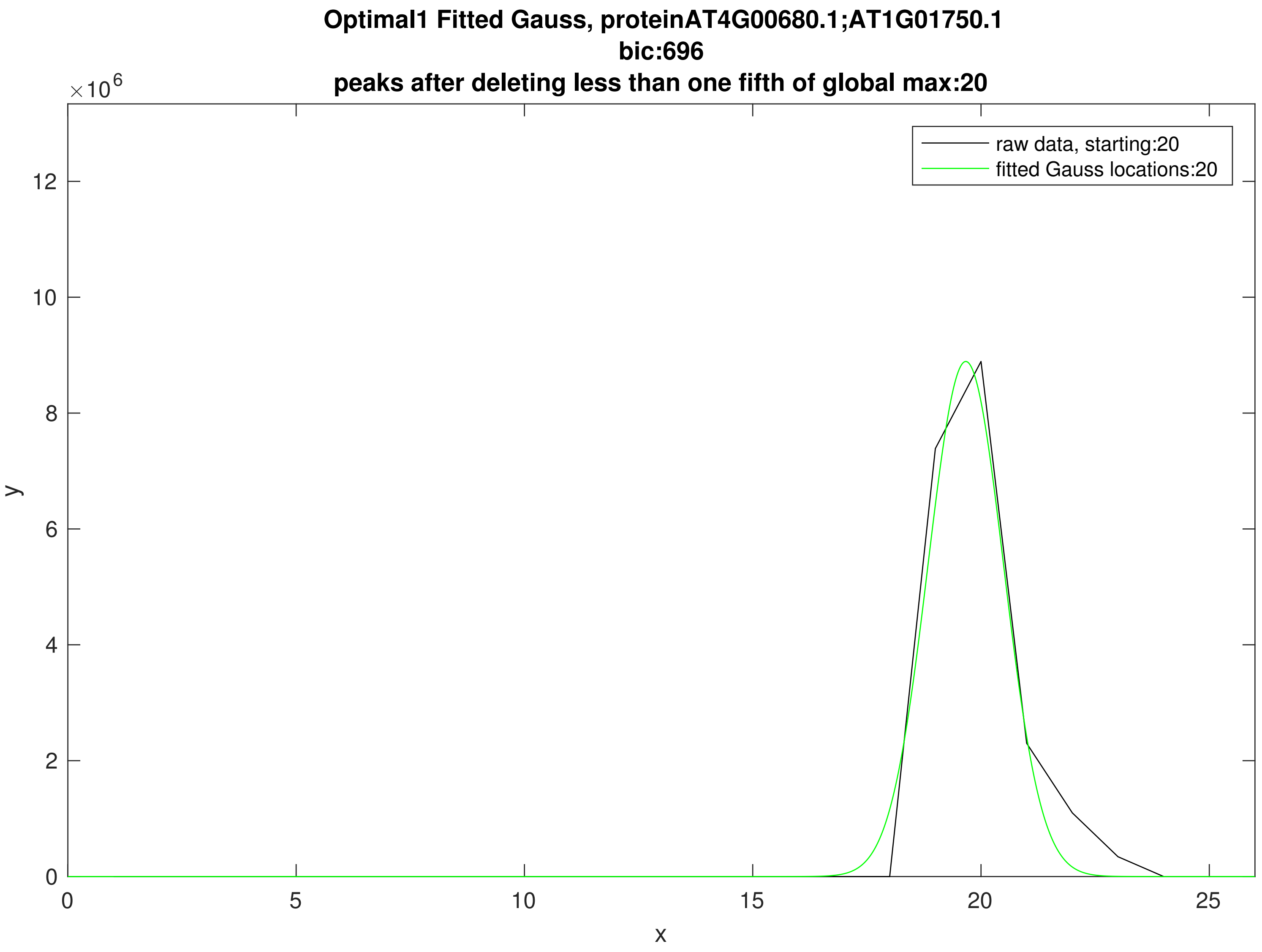

Optimal1 Fitted Gauss, proteinAT4G00680.1;AT1G01750.1
bic:696
peaks after deleting less than one fifth of global max:20
×10^6
raw data, starting:20
fitted Gauss locations:20
y
x
0
2
4
6
8
10
12
0
5
10
15
20
25

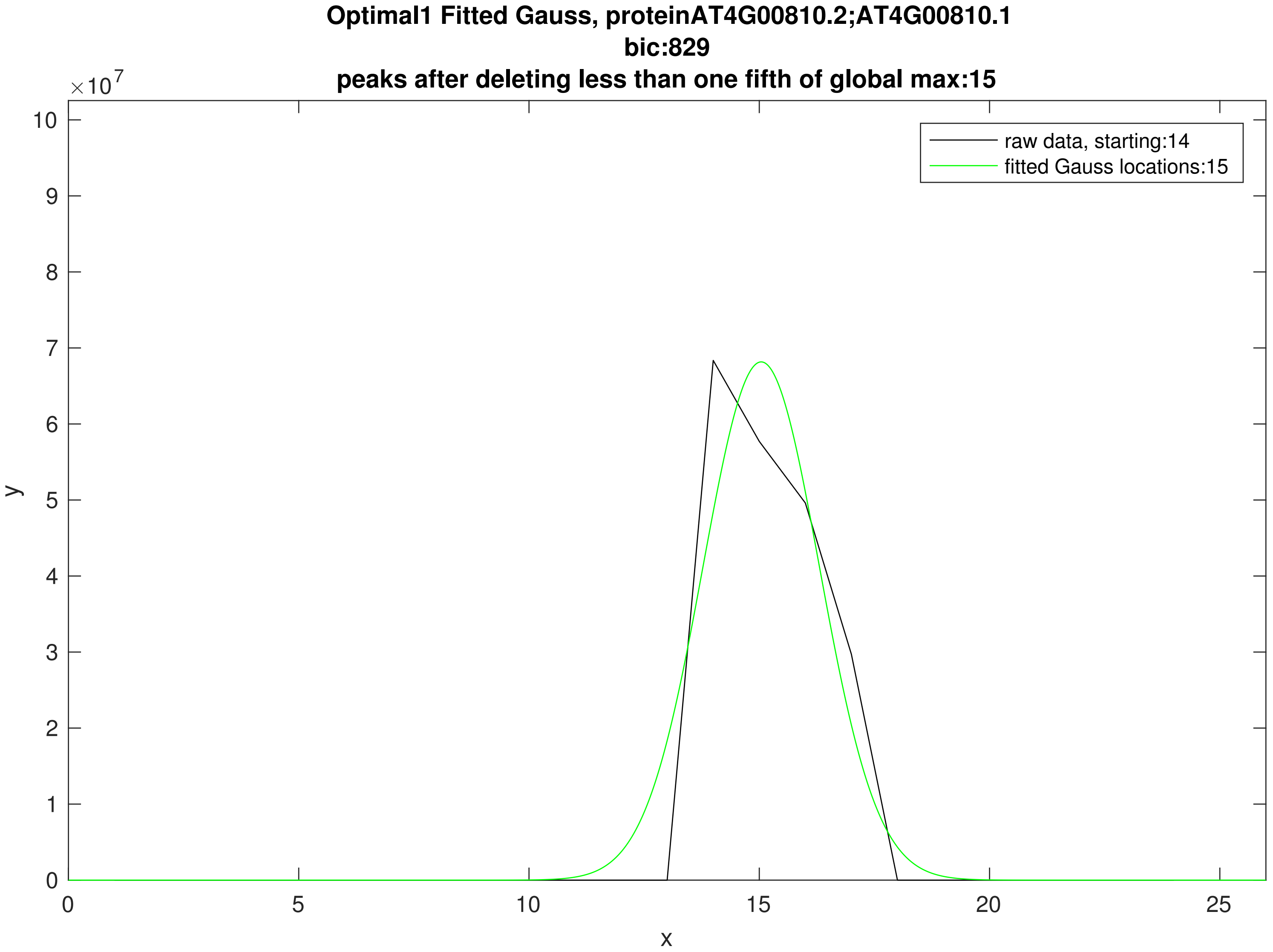

Optimal1 Fitted Gauss, proteinAT4G00810.2;AT4G00810.1
bic:829
peaks after deleting less than one fifth of global max:15
×10^7
raw data, starting:14
fitted Gauss locations:15
y
x
0
1
2
3
4
5
6
7
8
9
10
0
5
10
15
20
25

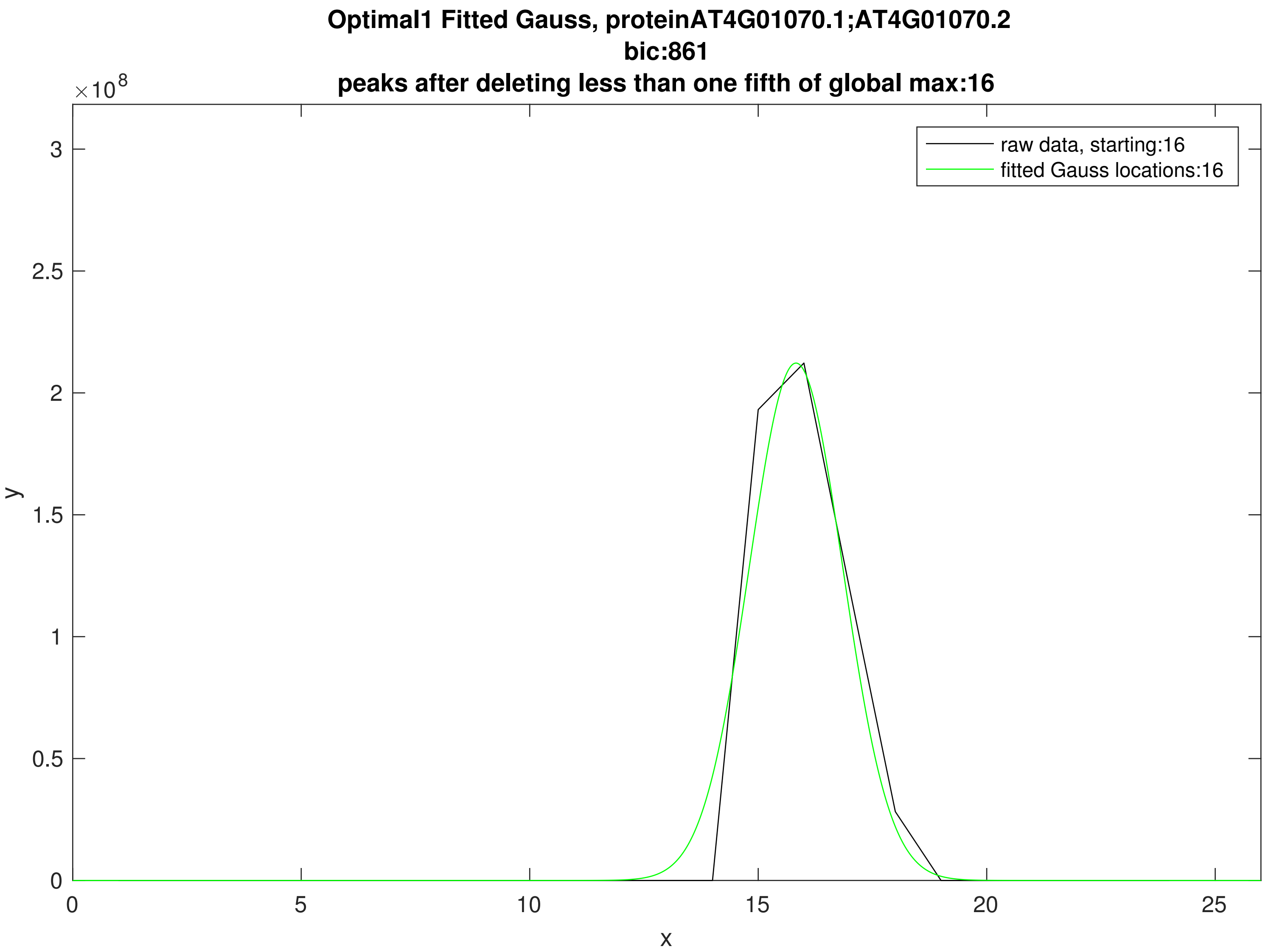
Optimal1 Fitted Gauss, proteinAT4G01070.1;AT4G01070.2
bic:861
peaks after deleting less than one fifth of global max:16
×10^8
raw data, starting:16
fitted Gauss locations:16
y
x
0
0.5
1
1.5
2
2.5
3
5
10
15
20
25

**Optimal2 Fitted Gauss, proteinAT4G01310.1**
**bic:736 709**
**peaks after deleting less than one fifth of global max:4.3 8.7**

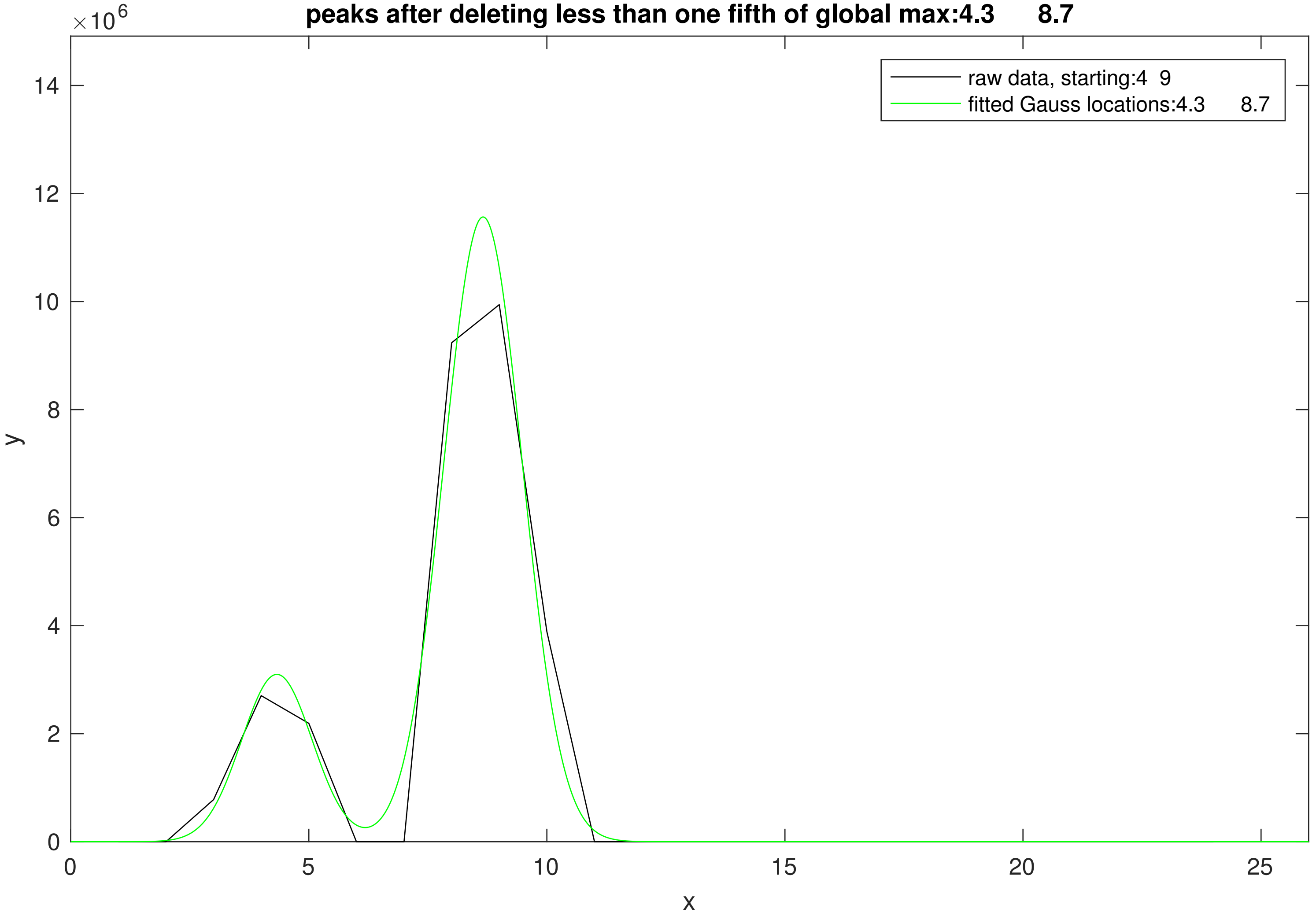

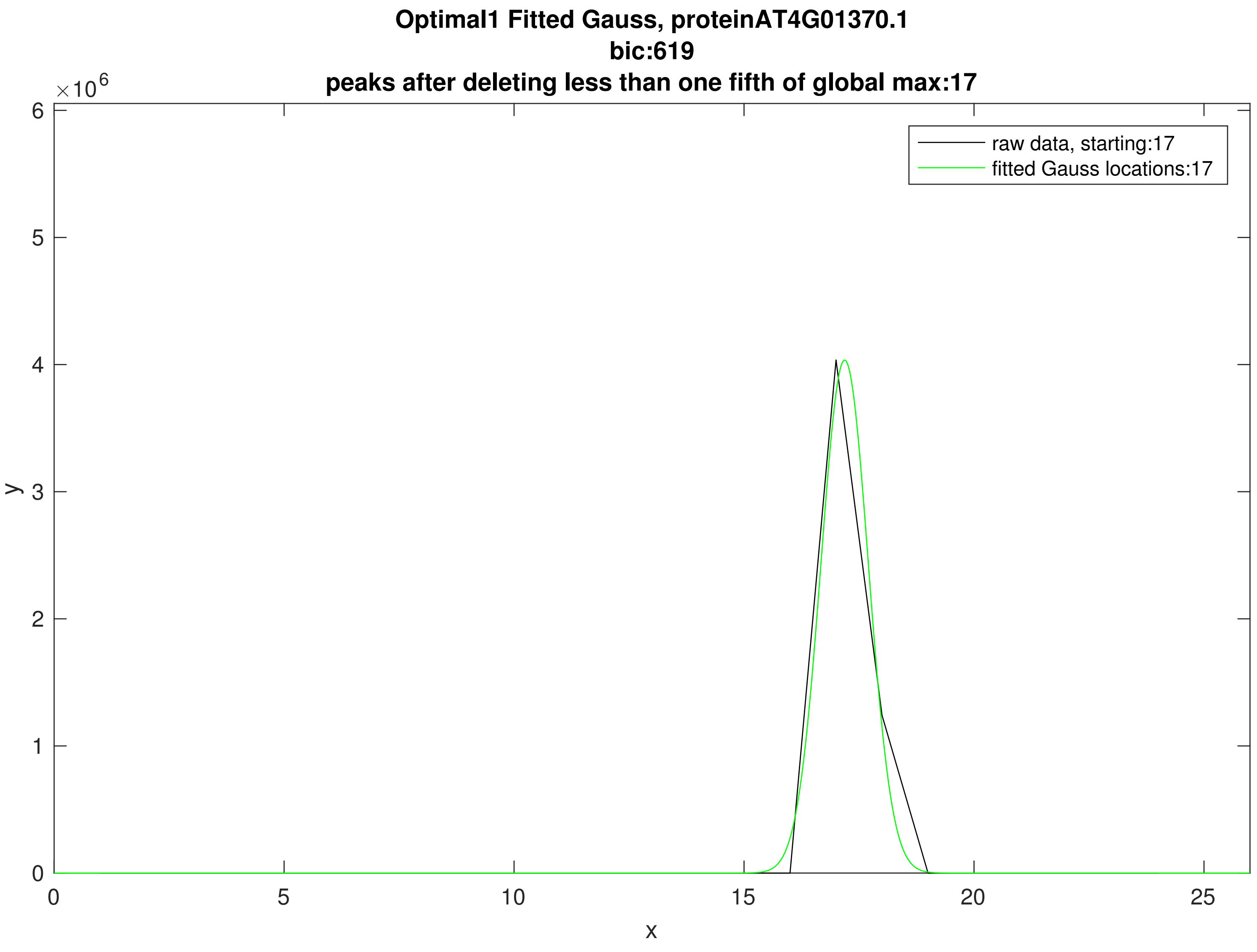
Optimal1 Fitted Gauss, proteinAT4G01370.1
bic:619
peaks after deleting less than one fifth of global max:17
$\times 10^6$
raw data, starting:17
fitted Gauss locations:17
y
x
0
1
2
3
4
5
6
0
5
10
15
20
25

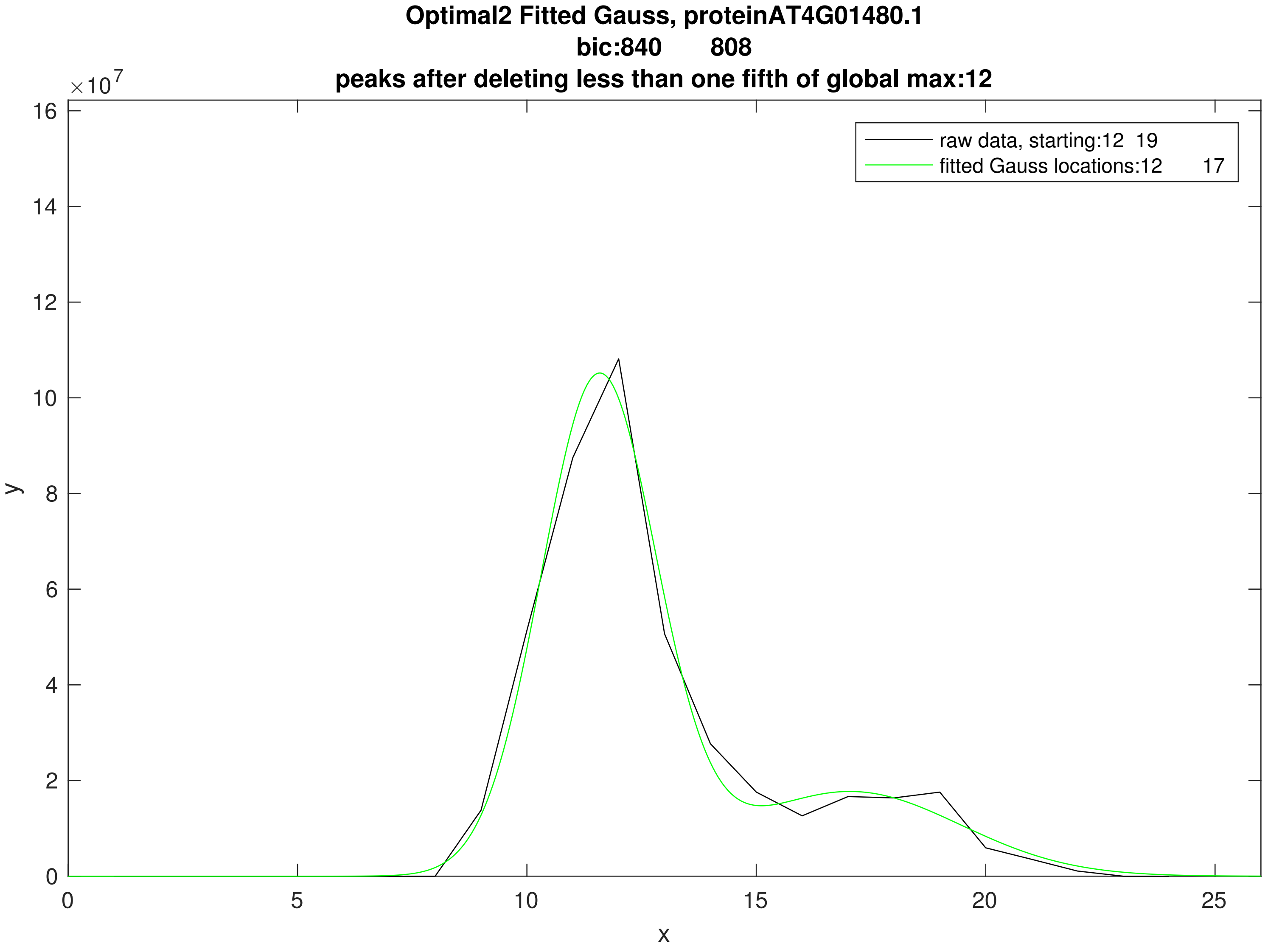

Optimal2 Fitted Gauss, proteinAT4G01480.1
bic:840 808
peaks after deleting less than one fifth of global max:12
×10^7
16
14
12
10
8
6
4
2
0
y
0
5
10
15
20
25
x
raw data, starting:12 19
fitted Gauss locations:12 17

Optimal1 Fitted Gauss, proteinAT4G01610.2;AT4G01610.1
bic:929 936
peaks after deleting less than one fifth of global max:19

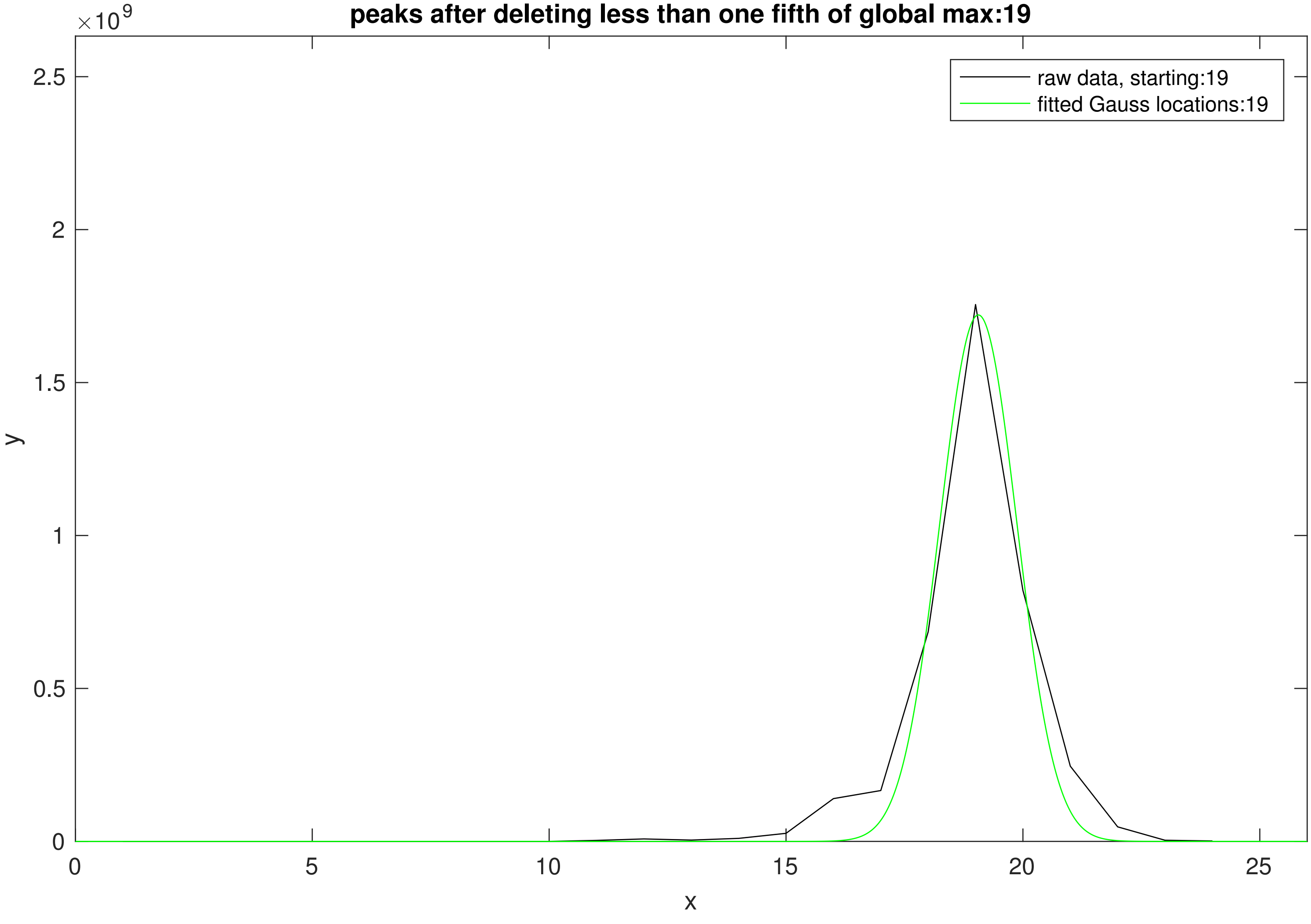

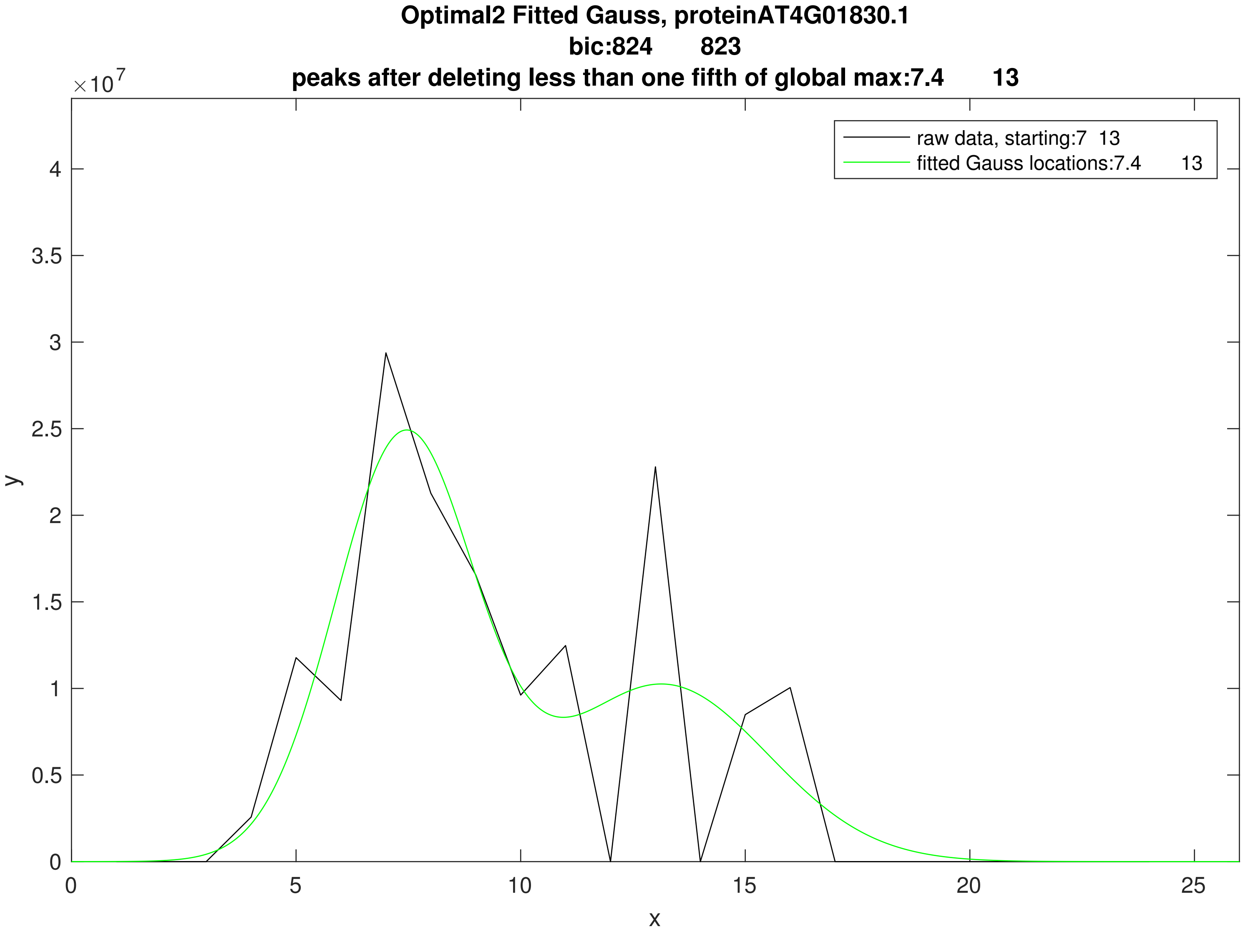
Optimal2 Fitted Gauss, proteinAT4G01830.1
bic:824 823
peaks after deleting less than one fifth of global max:7.4 13
×10^7
raw data, starting:7 13
fitted Gauss locations:7.4 13
y
x
0
0.5
1
1.5
2
2.5
3
3.5
4
0
5
10
15
20
25

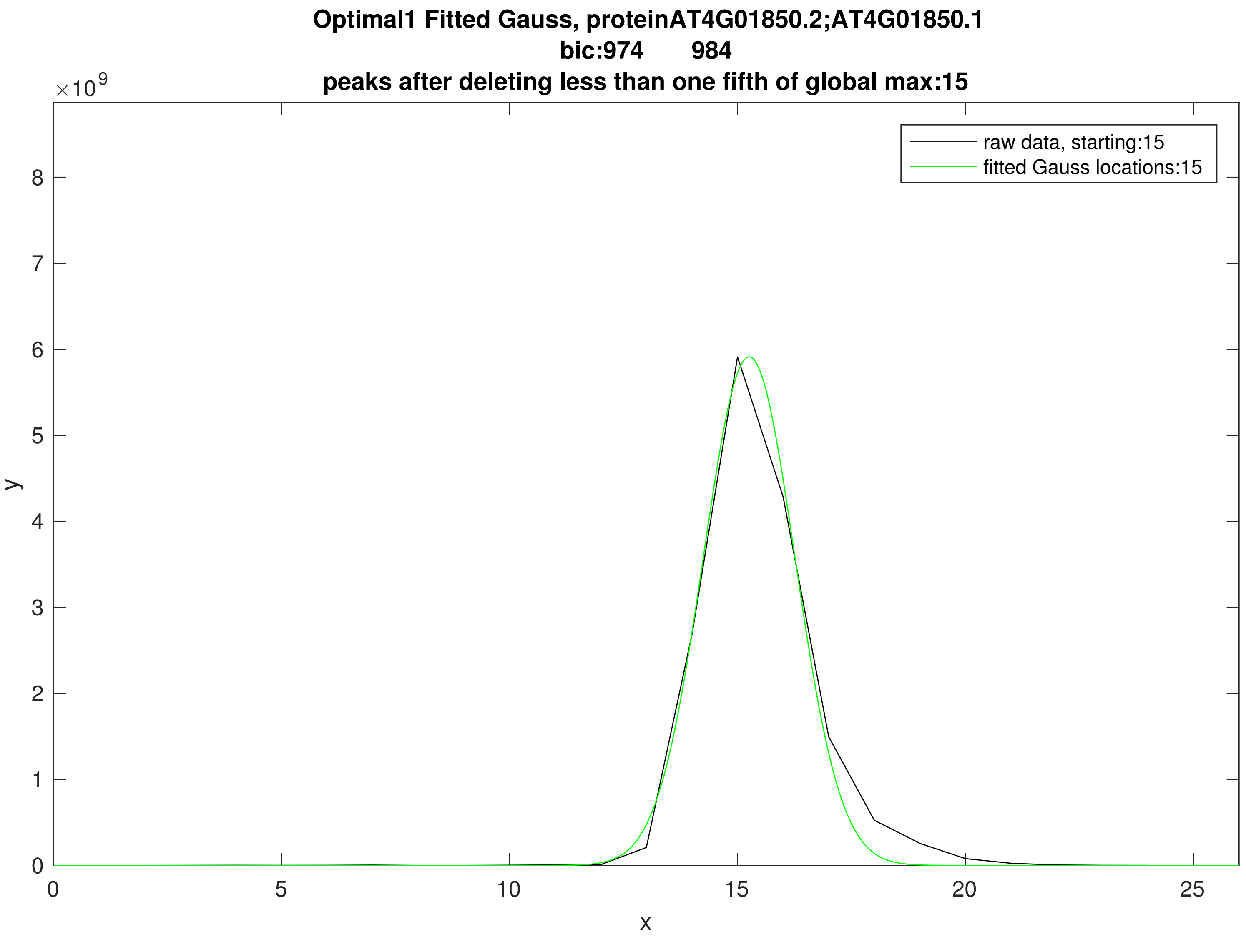
Optimal1 Fitted Gauss, proteinAT4G01850.2;AT4G01850.1
bic:974 984
peaks after deleting less than one fifth of global max:15
×10^9
raw data, starting:15
fitted Gauss locations:15
y
x
0
1
2
3
4
5
6
7
8
0
5
10
15
20
25

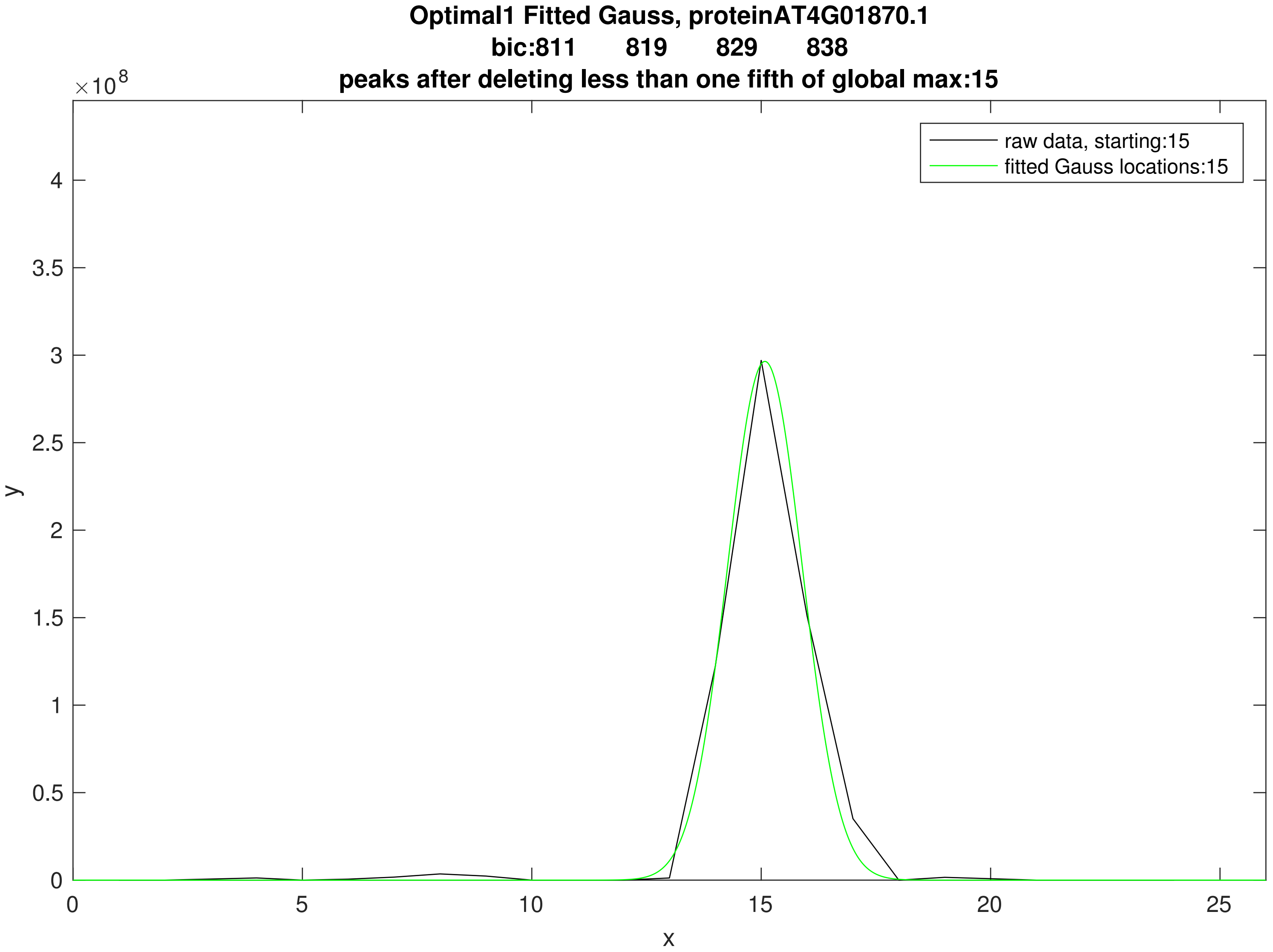
Optimal1 Fitted Gauss, proteinAT4G01870.1
bic:811 819 829 838
peaks after deleting less than one fifth of global max:15
×10^8
raw data, starting:15
fitted Gauss locations:15
y
x
0
0.5
1
1.5
2
2.5
3
3.5
4
5
10
15
20
25

**Optimal1 Fitted Gauss, proteinAT4G01897.1**
**bic:744**
**peaks after deleting less than one fifth of global max:19**

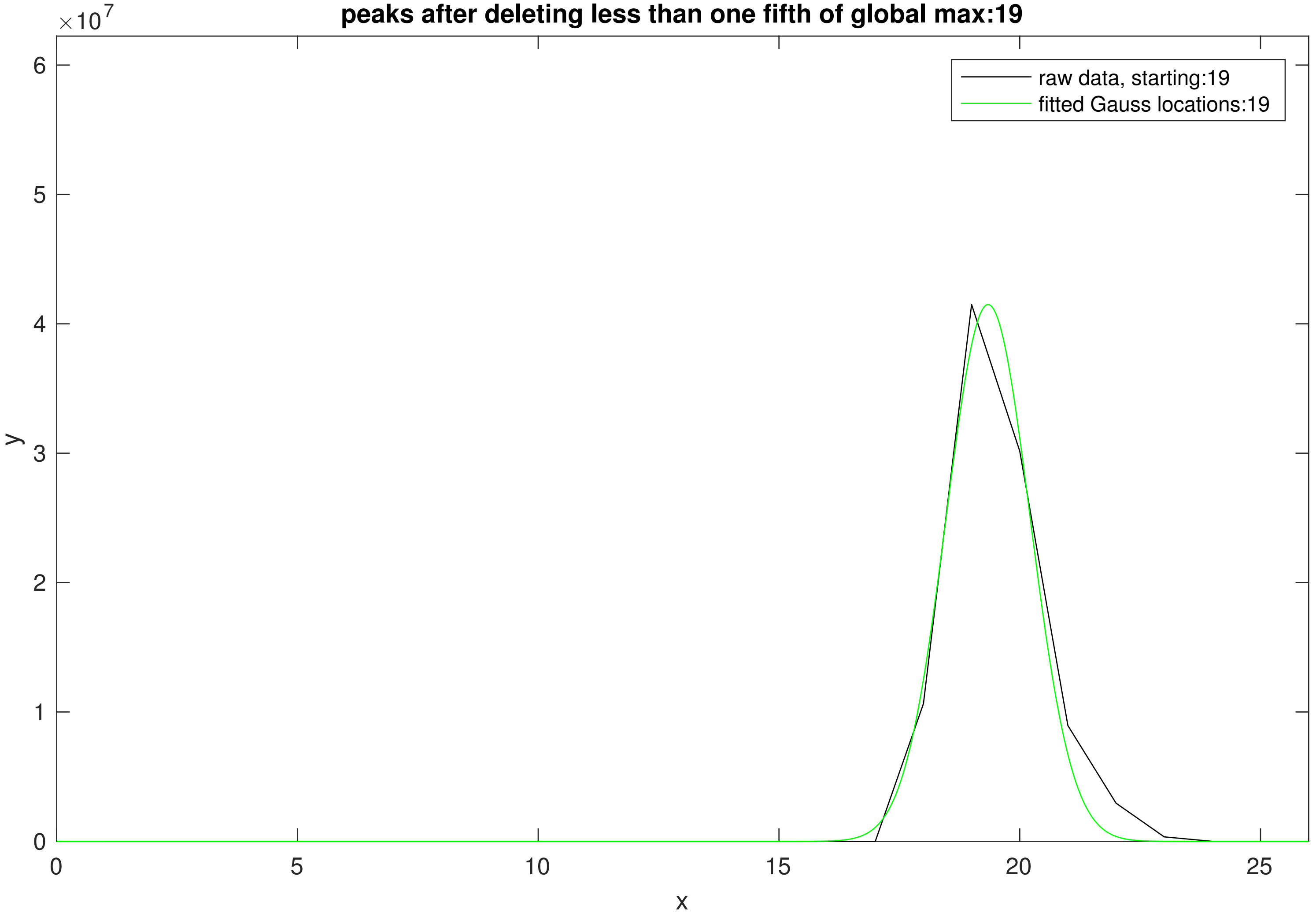

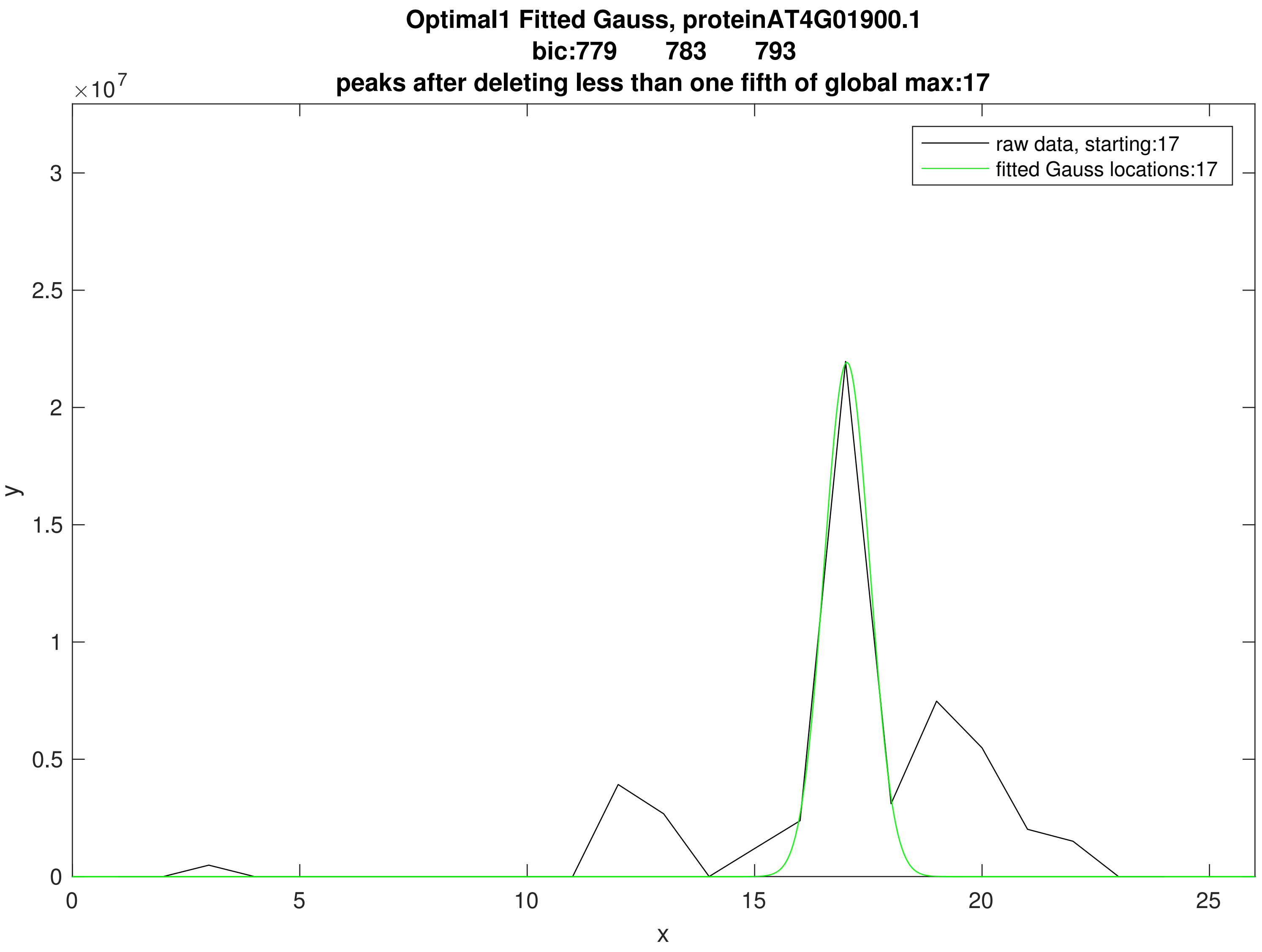
Optimal1 Fitted Gauss, proteinAT4G01900.1
bic:779 783 793
peaks after deleting less than one fifth of global max:17
×10^7
raw data, starting:17
fitted Gauss locations:17
y
x
0
0.5
1
1.5
2
2.5
3
0
5
10
15
20
25

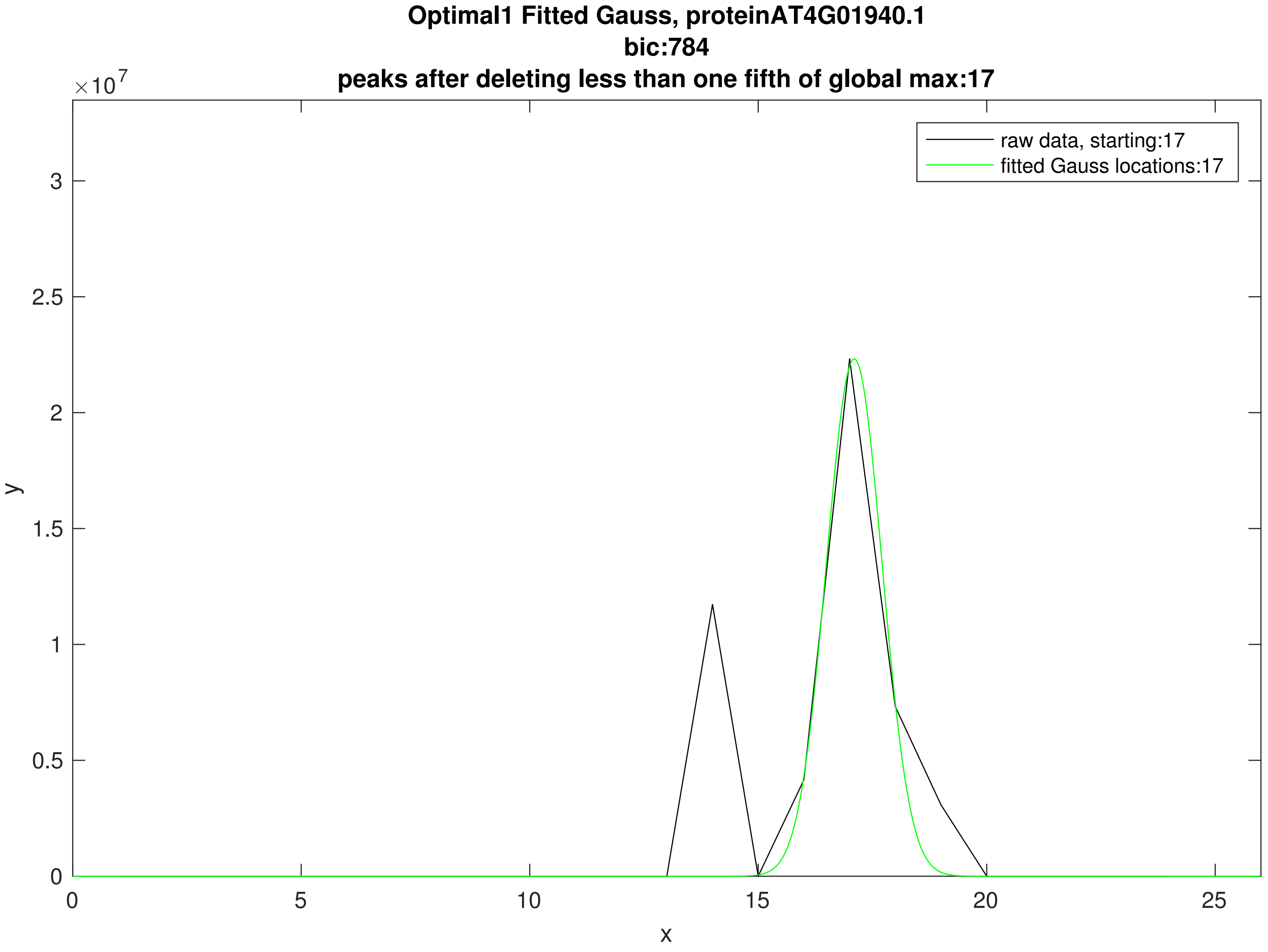
Optimal1 Fitted Gauss, proteinAT4G01940.1
bic:784
peaks after deleting less than one fifth of global max:17
×10^7
raw data, starting:17
fitted Gauss locations:17
y
x
0
0.5
1
1.5
2
2.5
3
0
5
10
15
20
25

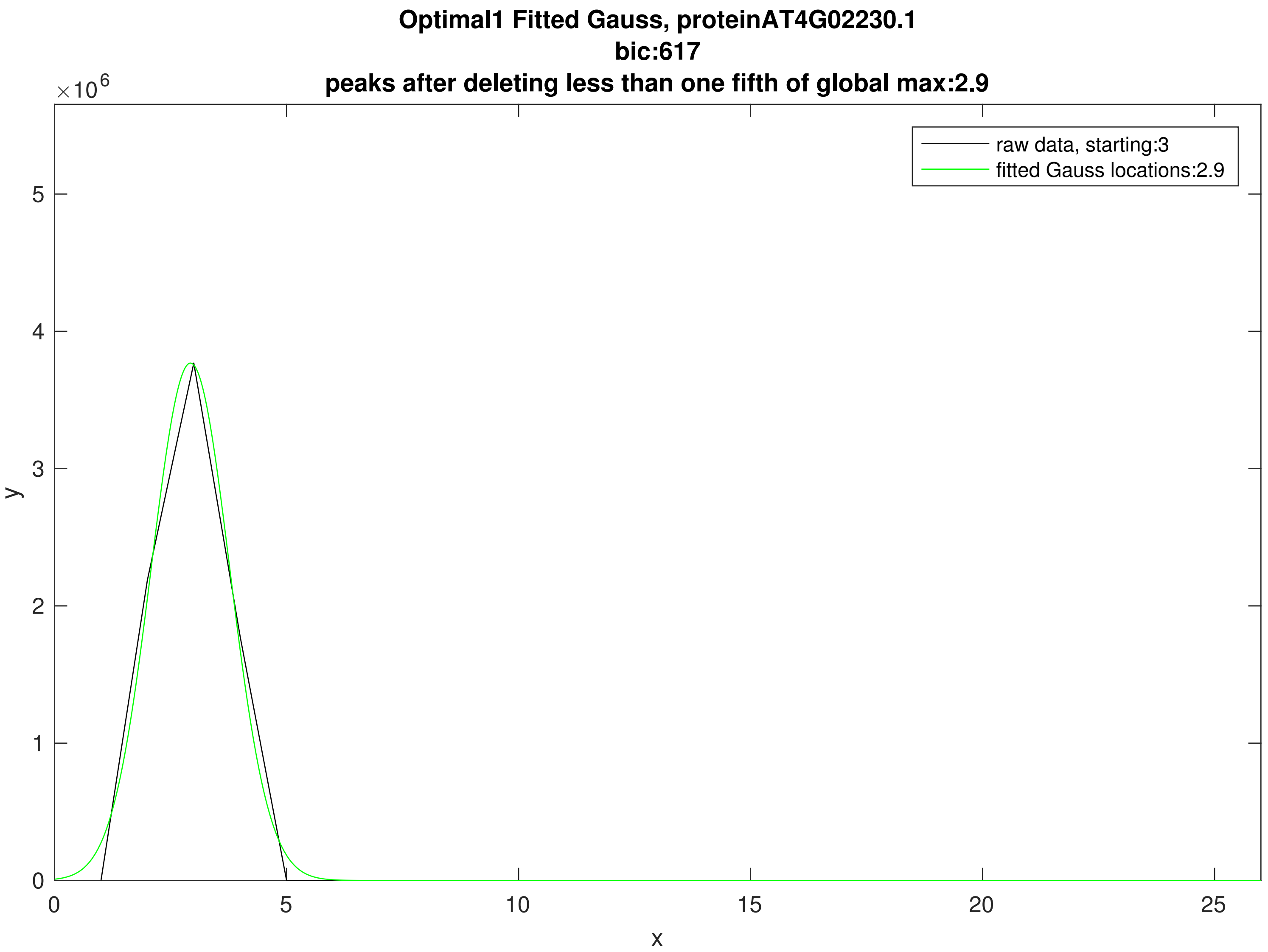
Optimal1 Fitted Gauss, proteinAT4G02230.1
bic:617
peaks after deleting less than one fifth of global max:2.9
×10^6
raw data, starting:3
fitted Gauss locations:2.9
y
x
0
1
2
3
4
5
0
5
10
15
20
25

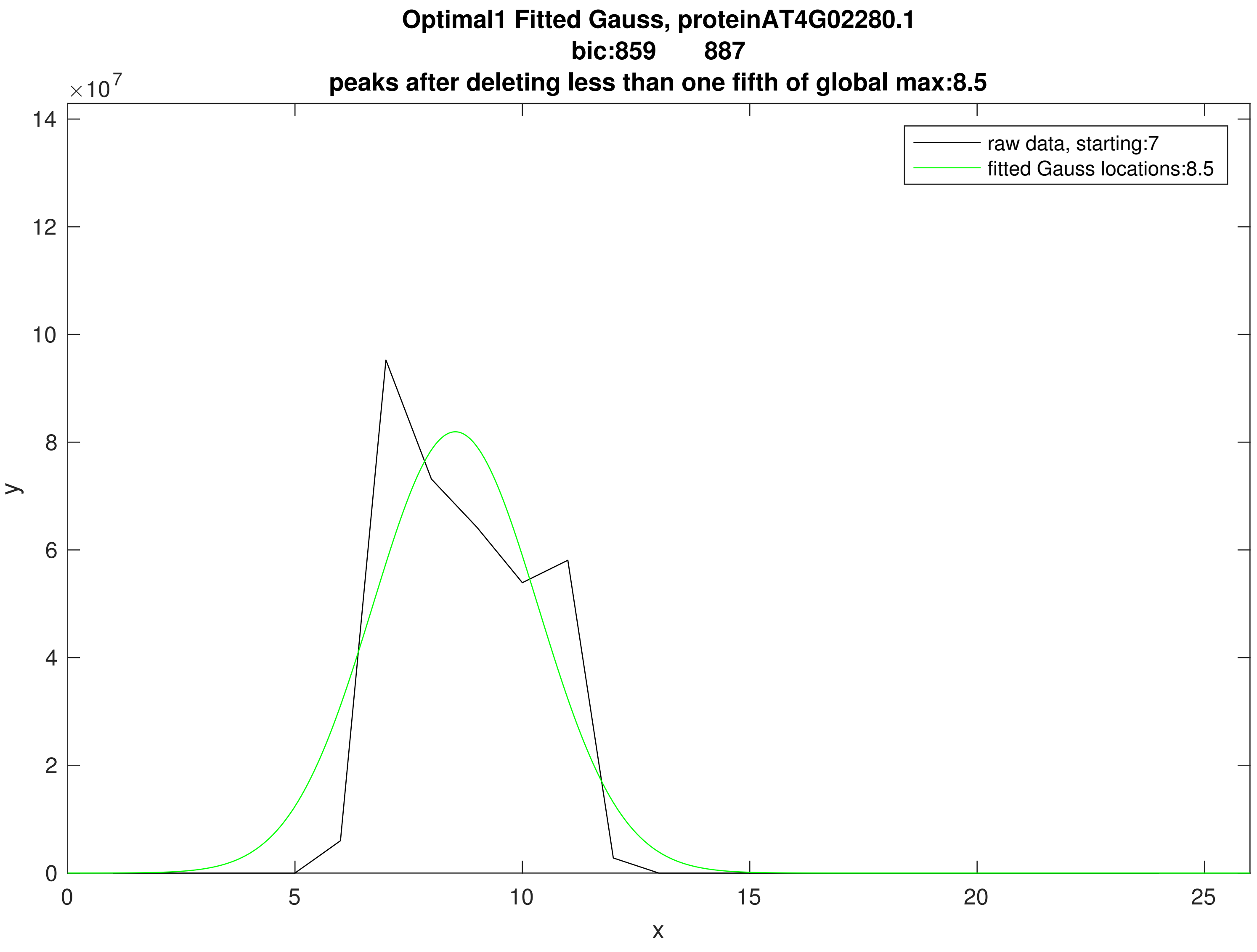

Optimal1 Fitted Gauss, proteinAT4G02280.1
bic:859 887
peaks after deleting less than one fifth of global max:8.5
×10^7
14
12
10
8
6
4
2
0
y
0
5
10
15
20
25
x
raw data, starting:7
fitted Gauss locations:8.5

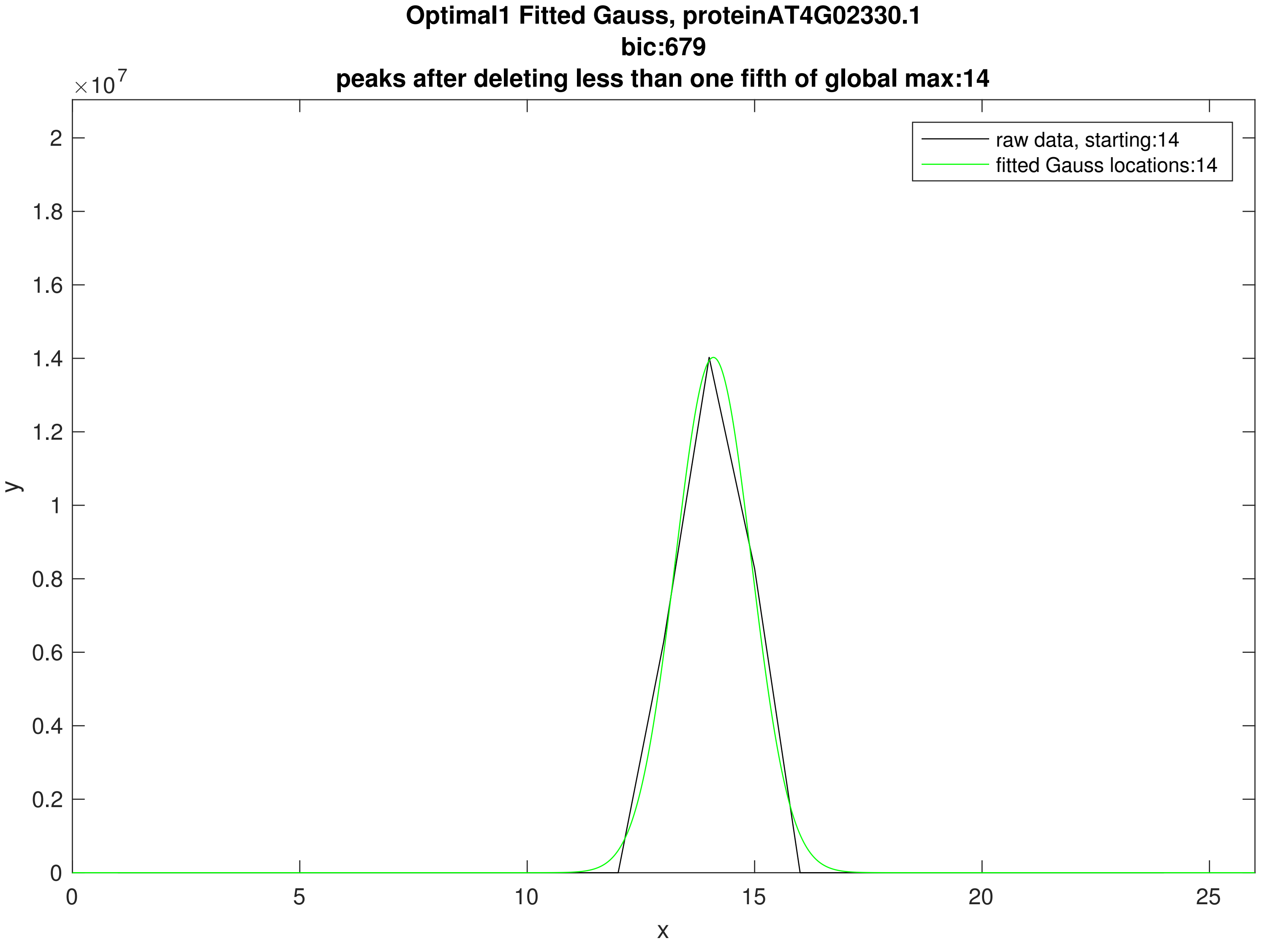
Optimal1 Fitted Gauss, proteinAT4G02330.1
bic:679
peaks after deleting less than one fifth of global max:14
×10^7
raw data, starting:14
fitted Gauss locations:14
y
x
0
0.2
0.4
0.6
0.8
1
1.2
1.4
1.6
1.8
2
5
10
15
20
25

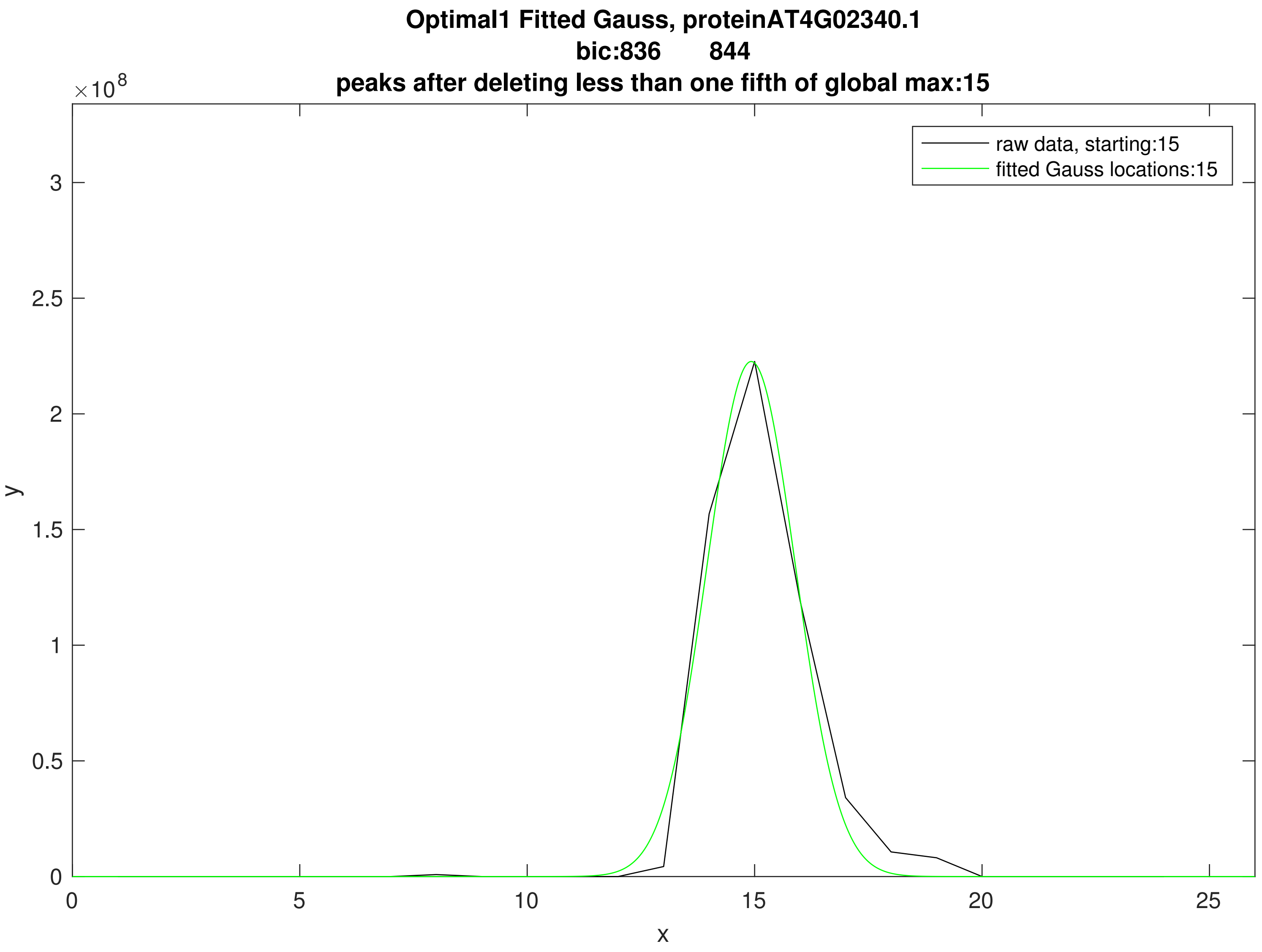
Optimal1 Fitted Gauss, proteinAT4G02340.1
bic:836 844
peaks after deleting less than one fifth of global max:15
×10^8
raw data, starting:15
fitted Gauss locations:15
y
x
0
0.5
1
1.5
2
2.5
3
0
5
10
15
20
25

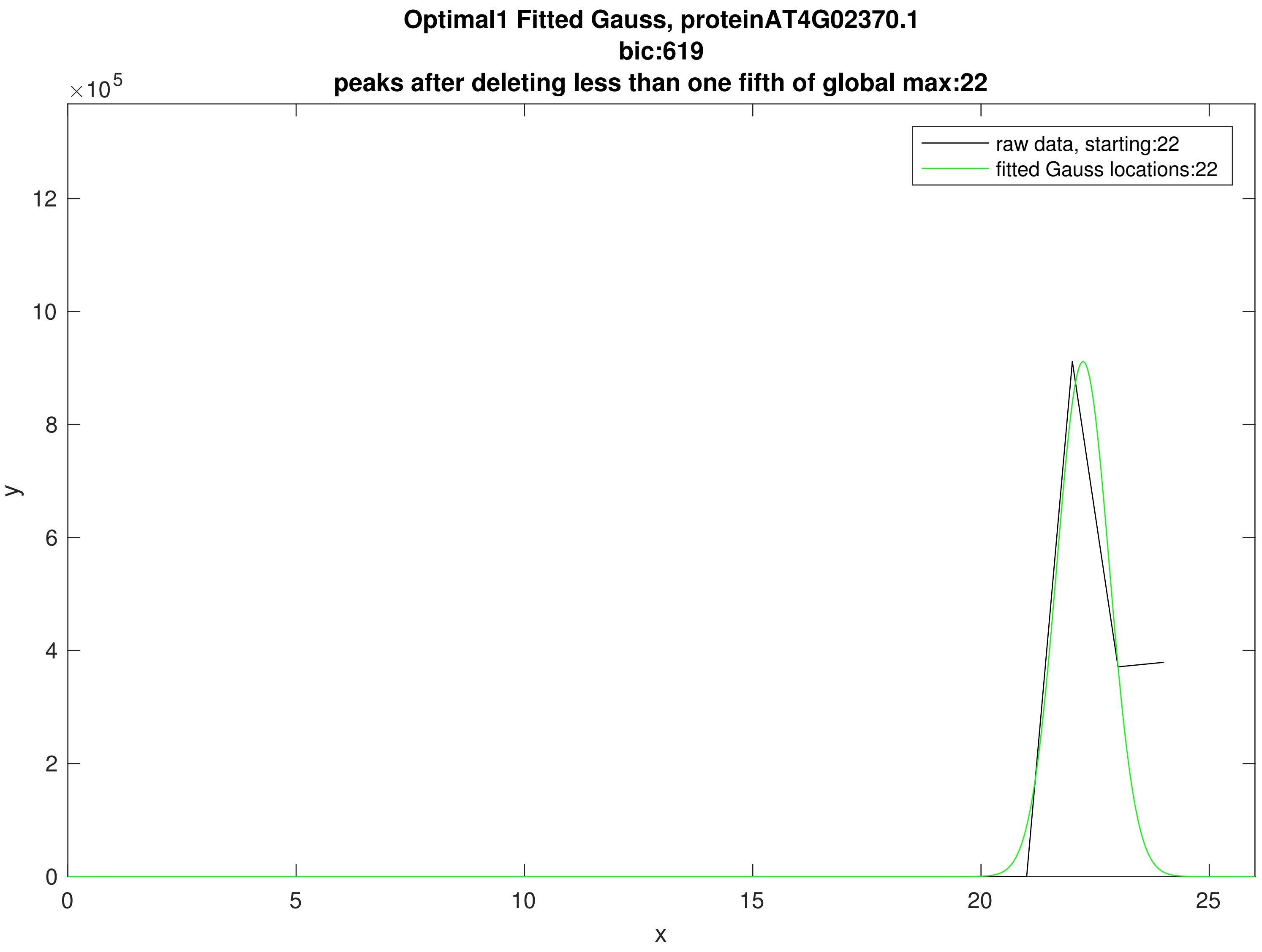
Optimal1 Fitted Gauss, proteinAT4G02370.1
bic:619
peaks after deleting less than one fifth of global max:22
×10^5
raw data, starting:22
fitted Gauss locations:22
y
x
0
2
4
6
8
10
12
0
5
10
15
20
25

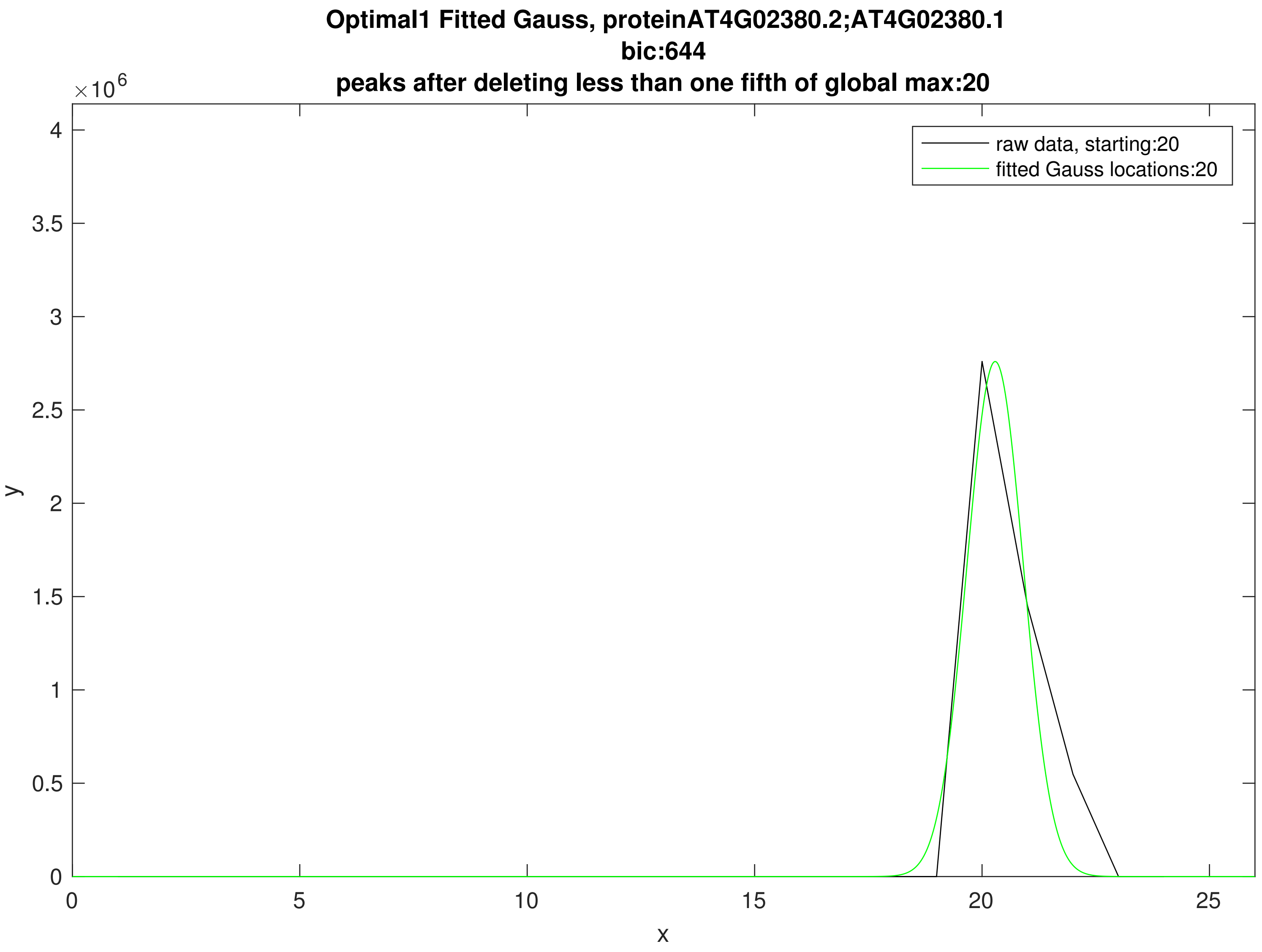

Optimal1 Fitted Gauss, proteinAT4G02380.2;AT4G02380.1
bic:644
peaks after deleting less than one fifth of global max:20
×10^6
4
3.5
3
2.5
2
1.5
1
0.5
0
y
0
5
10
15
20
25
x
raw data, starting:20
fitted Gauss locations:20

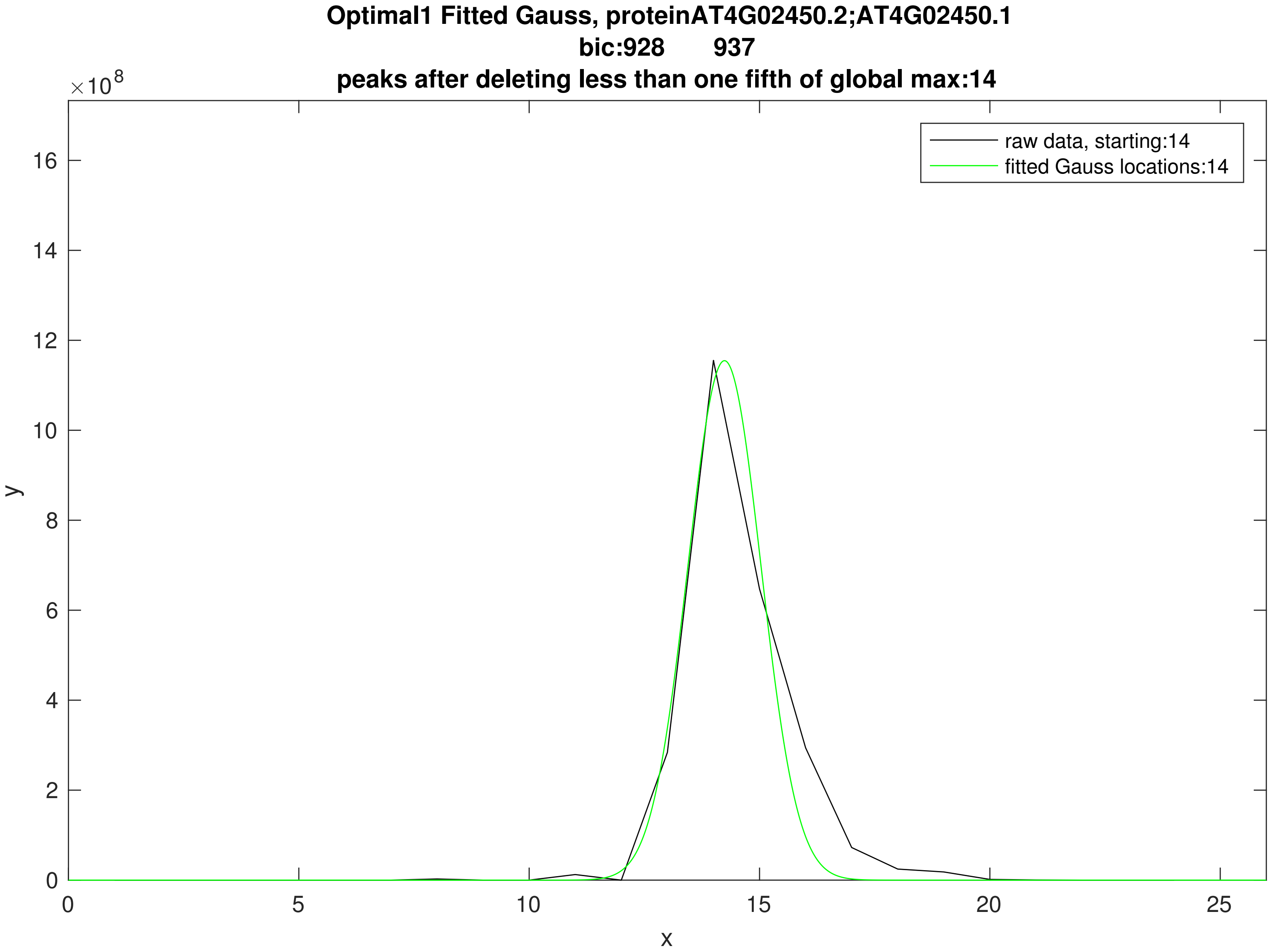

Optimal1 Fitted Gauss, proteinAT4G02450.2;AT4G02450.1
bic:928 937
peaks after deleting less than one fifth of global max:14
×10^8
raw data, starting:14
fitted Gauss locations:14
y
x
0
2
4
6
8
10
12
14
16
0
5
10
15
20
25

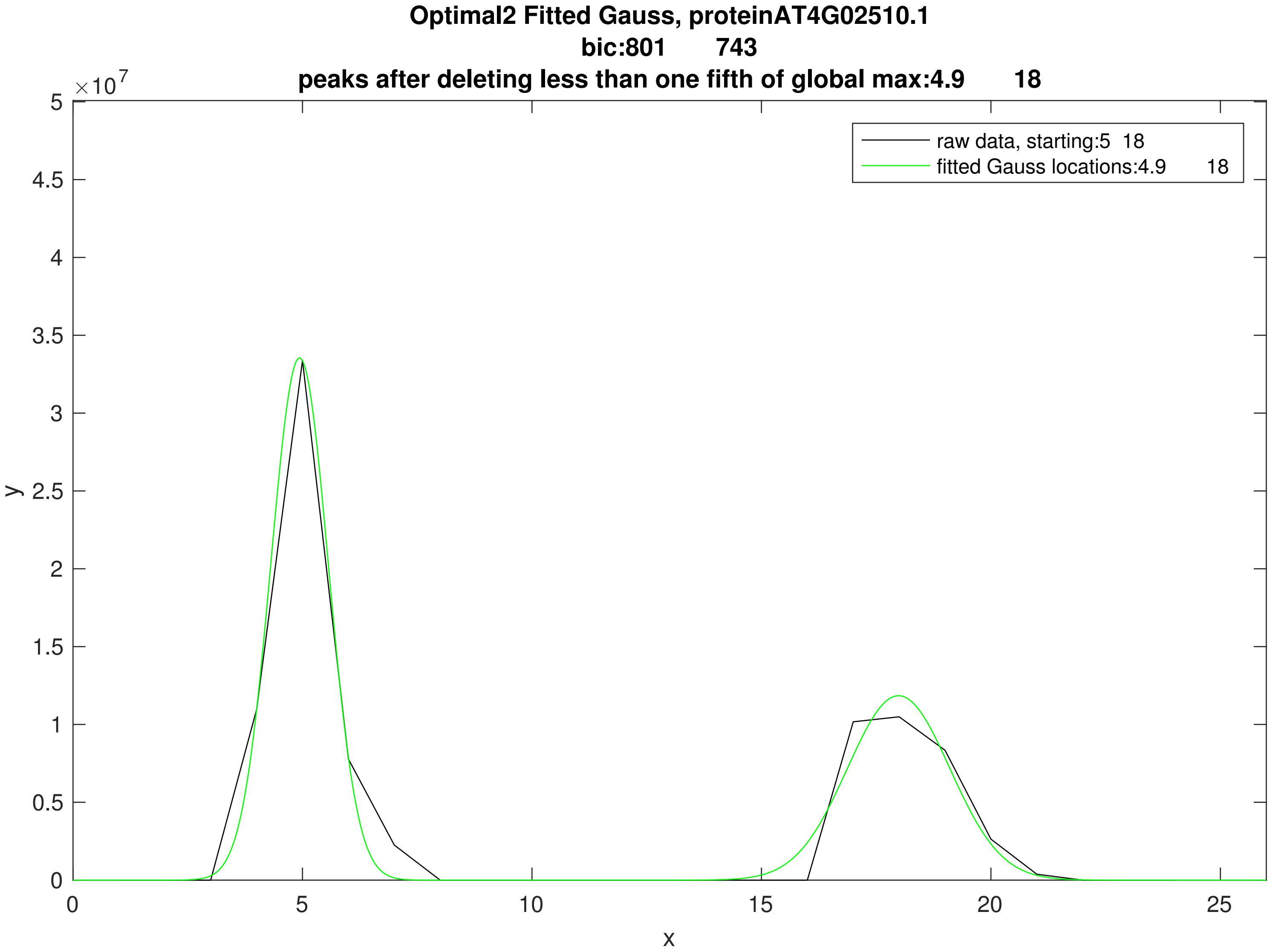

Optimal2 Fitted Gauss, proteinAT4G02510.1
bic:801 743
peaks after deleting less than one fifth of global max:4.9 18
×10^7
5
4.5
4
3.5
3
2.5
2
1.5
1
0.5
0
y
0
5
10
15
20
25
x
raw data, starting:5 18
fitted Gauss locations:4.9 18

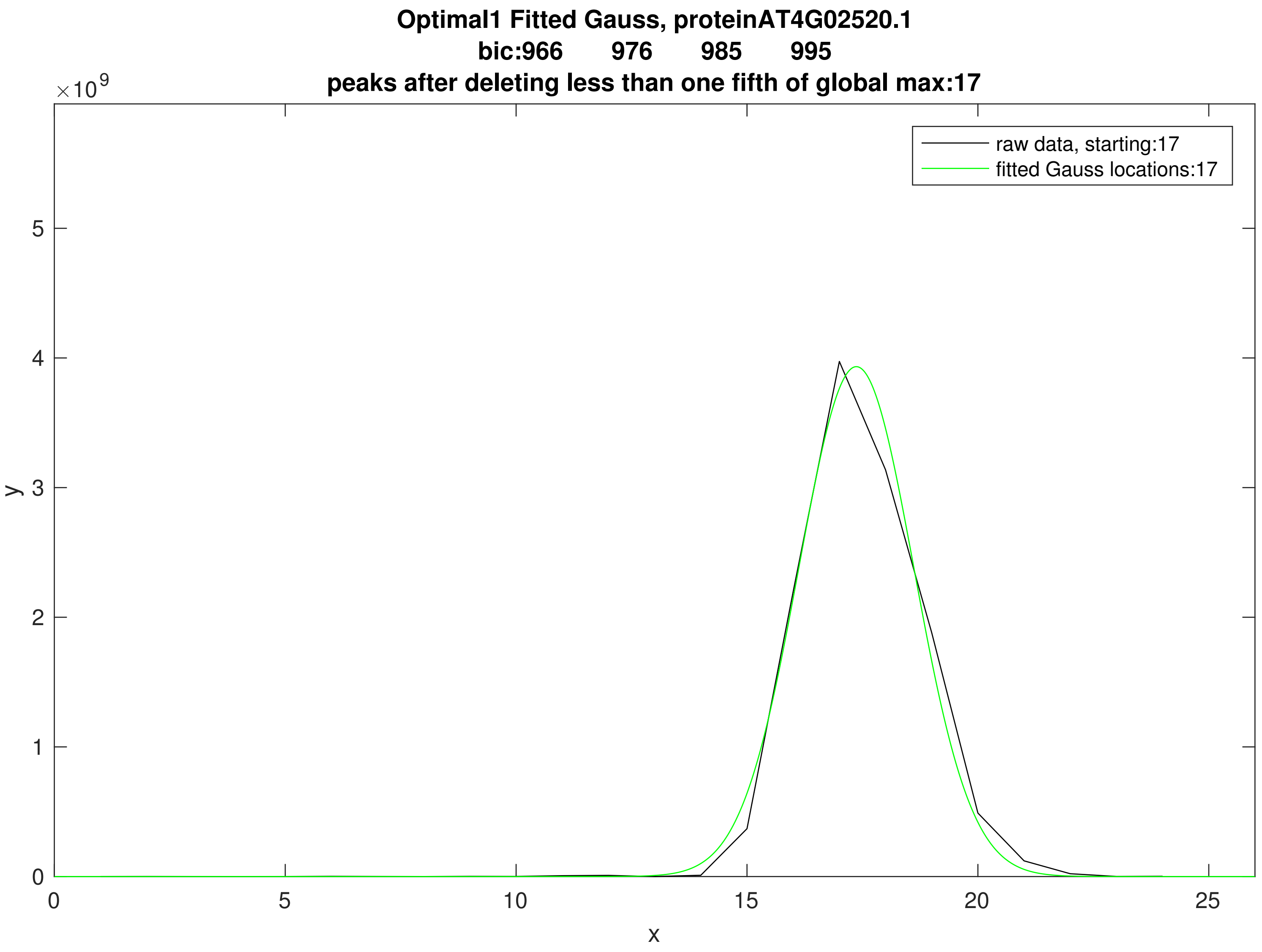

Optimal1 Fitted Gauss, proteinAT4G02520.1
bic:966 976 985 995
peaks after deleting less than one fifth of global max:17
×10^9
raw data, starting:17
fitted Gauss locations:17
y
x
0
1
2
3
4
5
0
5
10
15
20
25

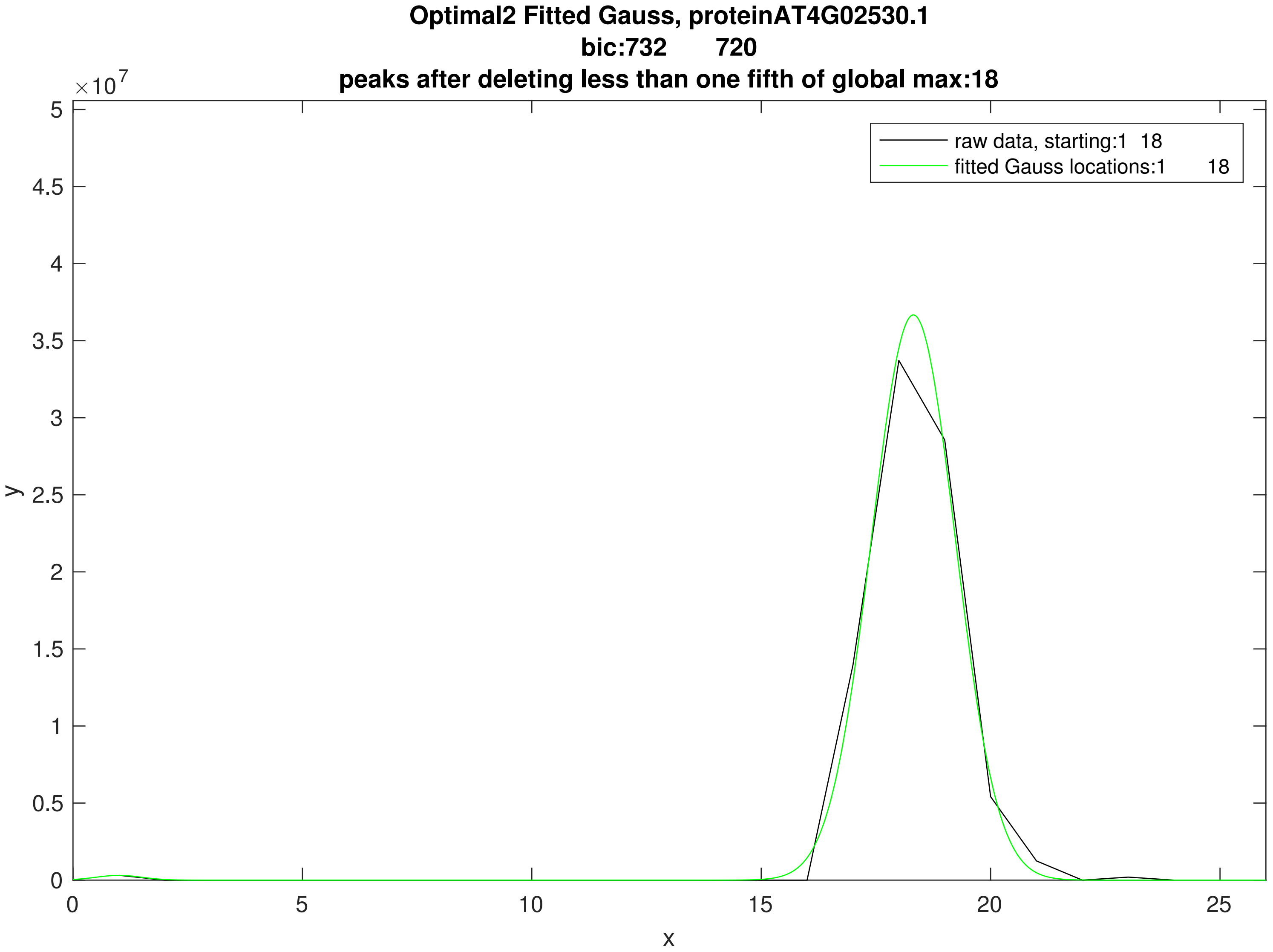

Optimal2 Fitted Gauss, proteinAT4G02530.1
bic:732 720
peaks after deleting less than one fifth of global max:18
×10^7
raw data, starting:1 18
fitted Gauss locations:1 18
y
x
0
0.5
1
1.5
2
2.5
3
3.5
4
4.5
5
0
5
10
15
20
25

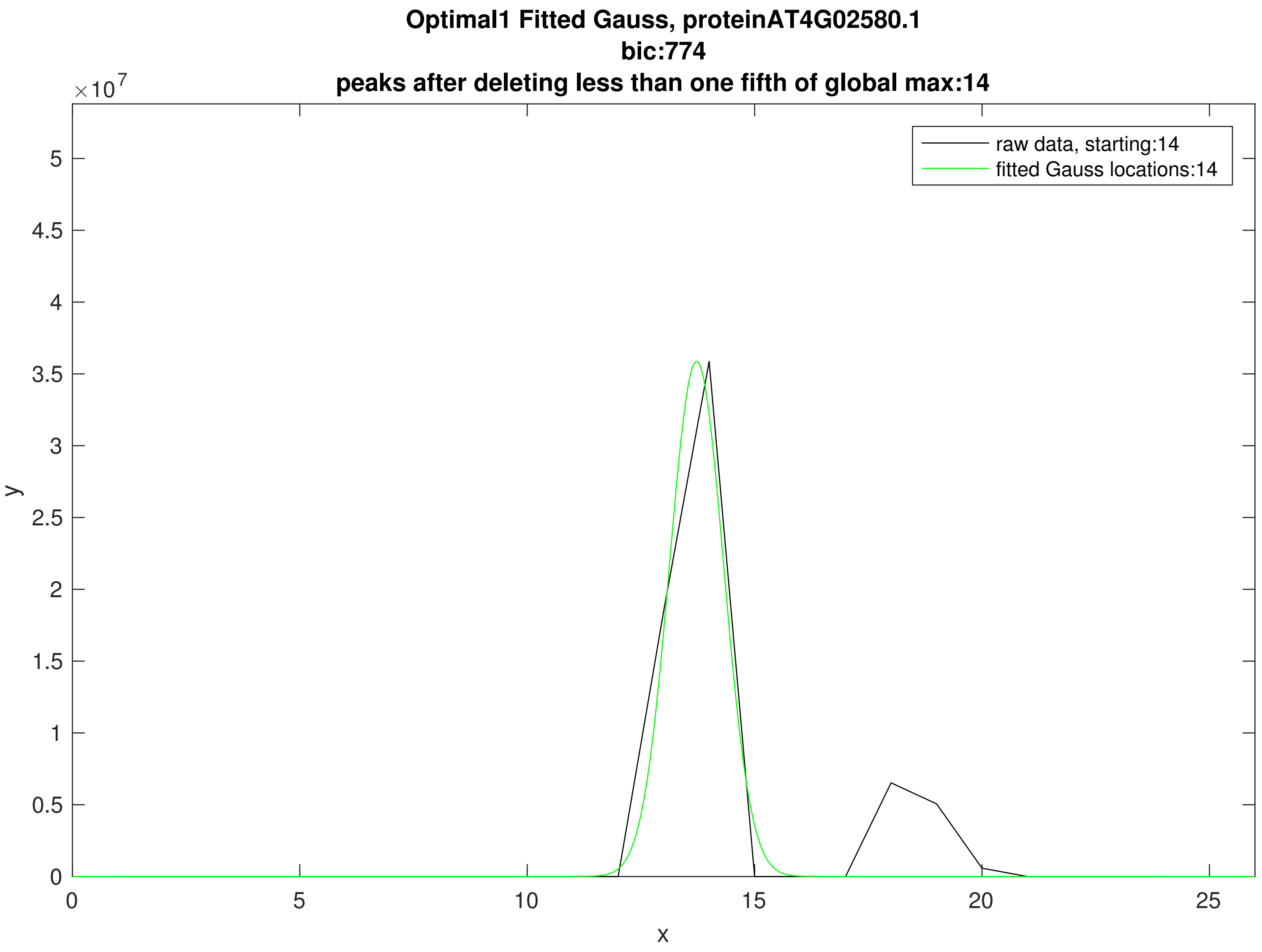
Optimal1 Fitted Gauss, proteinAT4G02580.1
bic:774
peaks after deleting less than one fifth of global max:14
×10^7
raw data, starting:14
fitted Gauss locations:14
y
x
0
0.5
1
1.5
2
2.5
3
3.5
4
4.5
5
5
10
15
20
25

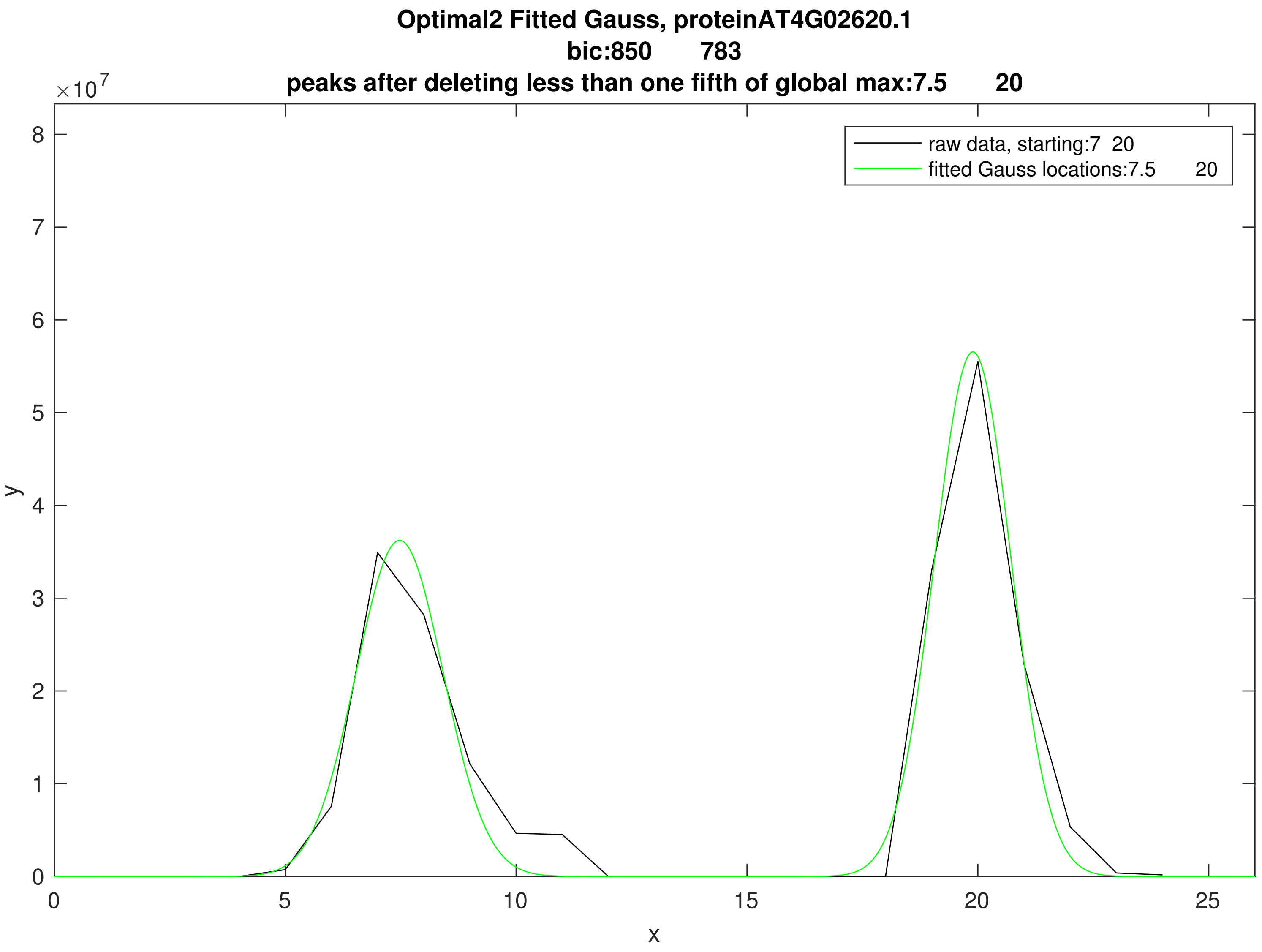

Optimal2 Fitted Gauss, proteinAT4G02620.1
bic:850 783
peaks after deleting less than one fifth of global max:7.5 20
×10^7
raw data, starting:7 20
fitted Gauss locations:7.5 20
y
x
0
1
2
3
4
5
6
7
8
0
5
10
15
20
25

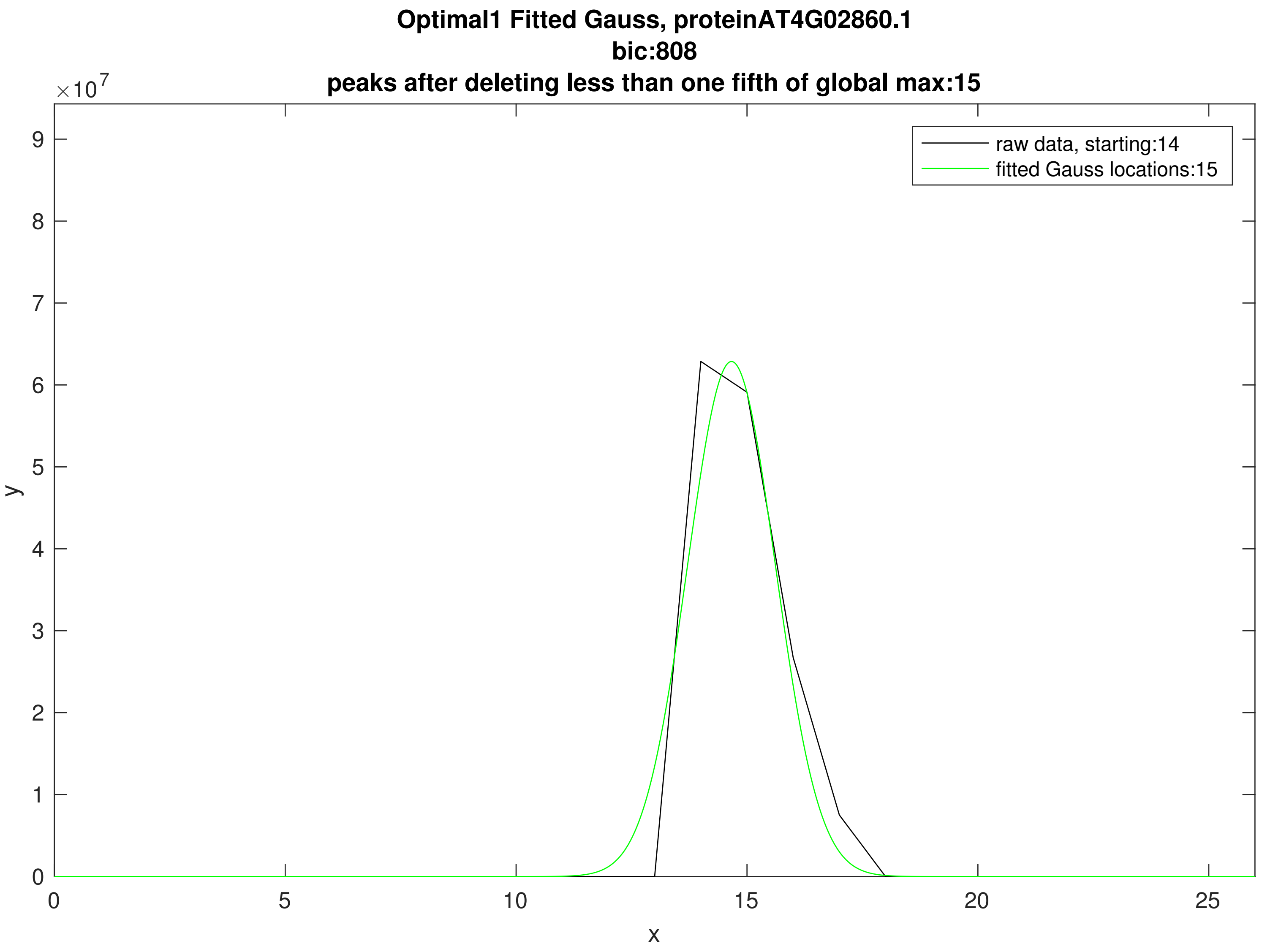

Optimal1 Fitted Gauss, proteinAT4G02860.1
bic:808
peaks after deleting less than one fifth of global max:15
×10^7
raw data, starting:14
fitted Gauss locations:15
y
x
0
1
2
3
4
5
6
7
8
9
0
5
10
15
20
25

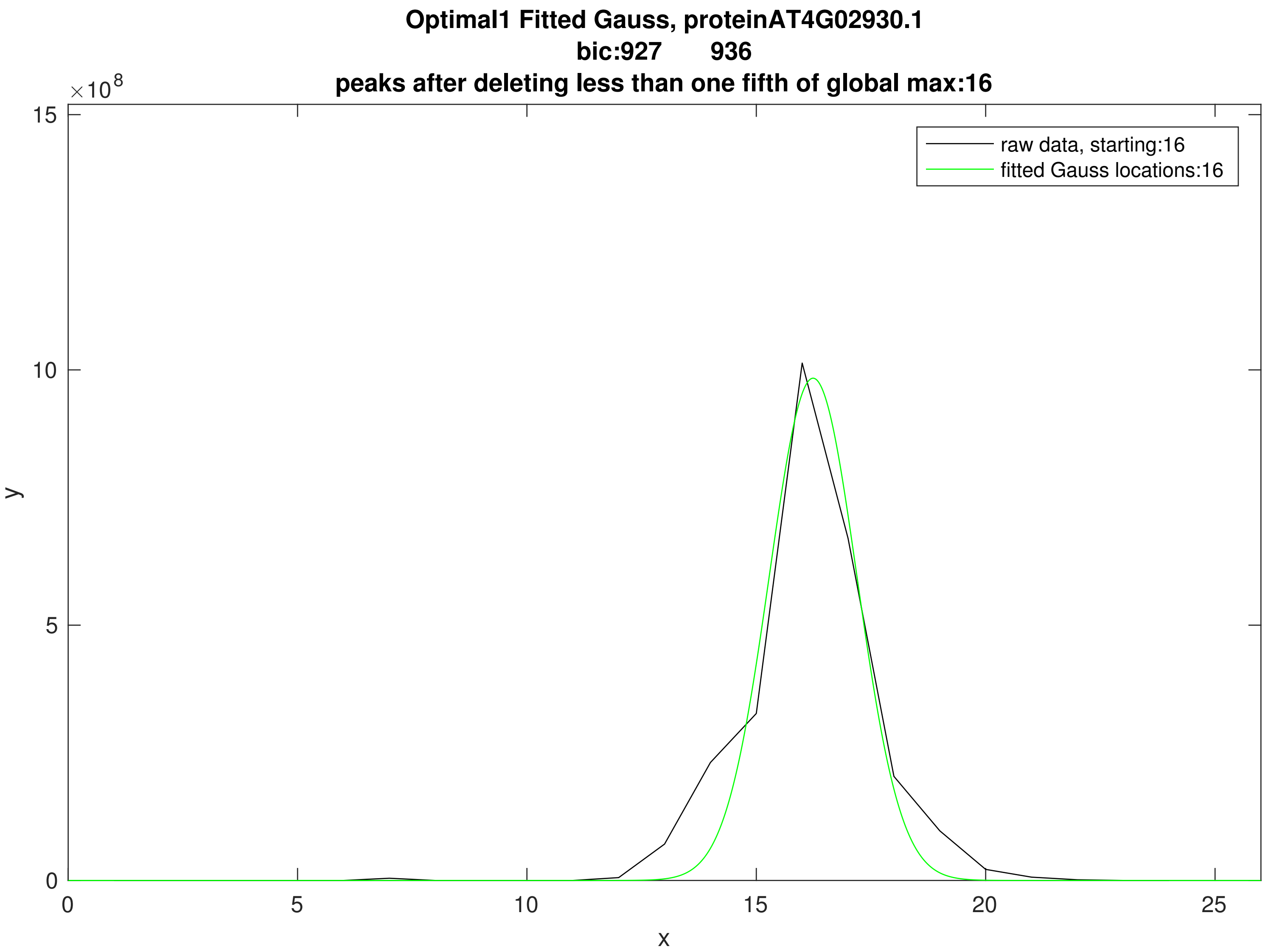

Optimal1 Fitted Gauss, proteinAT4G02930.1
bic:927 936
peaks after deleting less than one fifth of global max:16
×10^8
15
10
5
0
y
0
5
10
15
20
25
x
raw data, starting:16
fitted Gauss locations:16

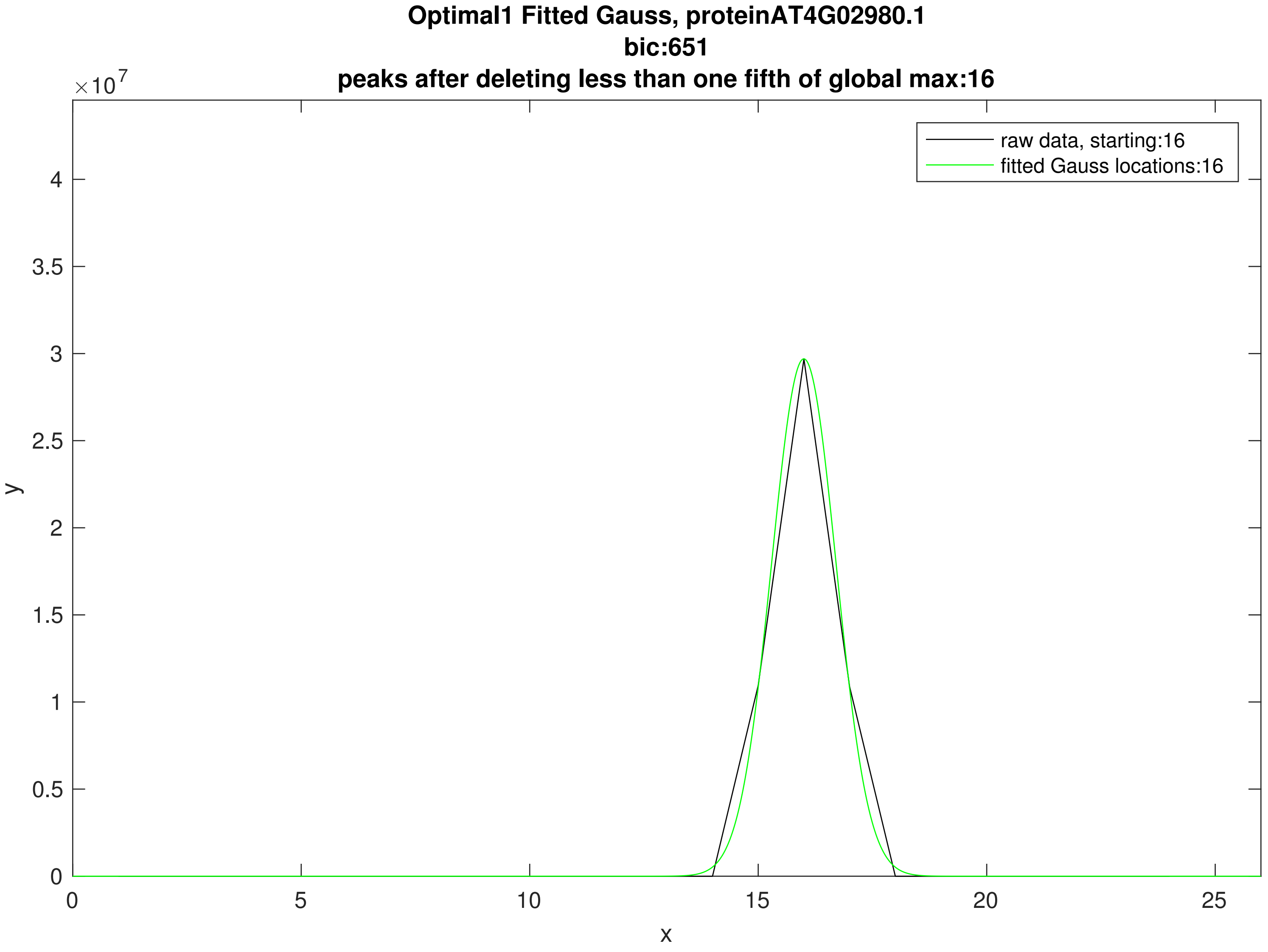

Optimal1 Fitted Gauss, proteinAT4G02980.1
bic:651
peaks after deleting less than one fifth of global max:16
×10^7
raw data, starting:16
fitted Gauss locations:16
y
x
0
0.5
1
1.5
2
2.5
3
3.5
4
0
5
10
15
20
25

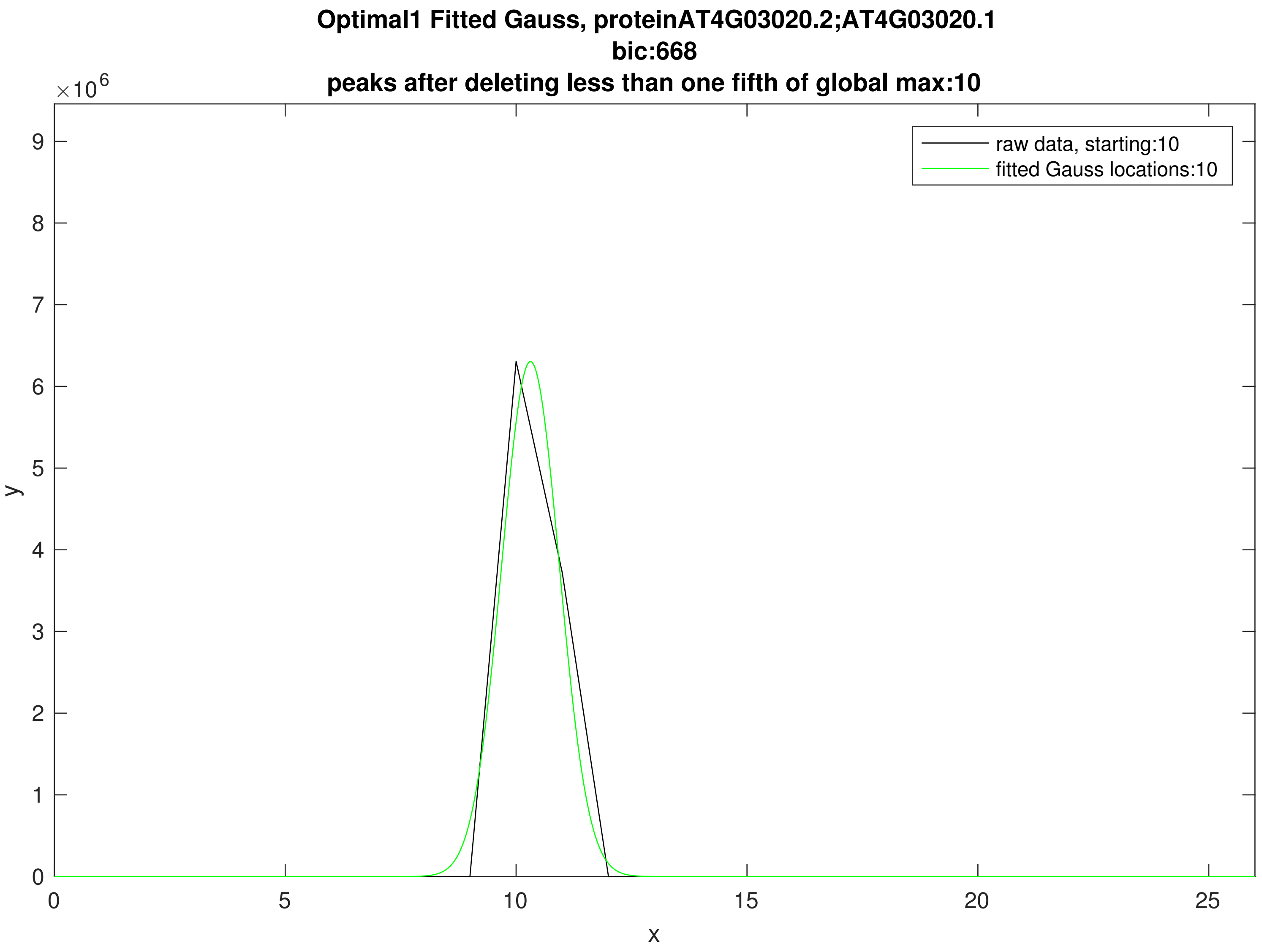
Optimal1 Fitted Gauss, proteinAT4G03020.2;AT4G03020.1
bic:668
peaks after deleting less than one fifth of global max:10
×10^6
raw data, starting:10
fitted Gauss locations:10
y
x
0
1
2
3
4
5
6
7
8
9
0
5
10
15
20
25

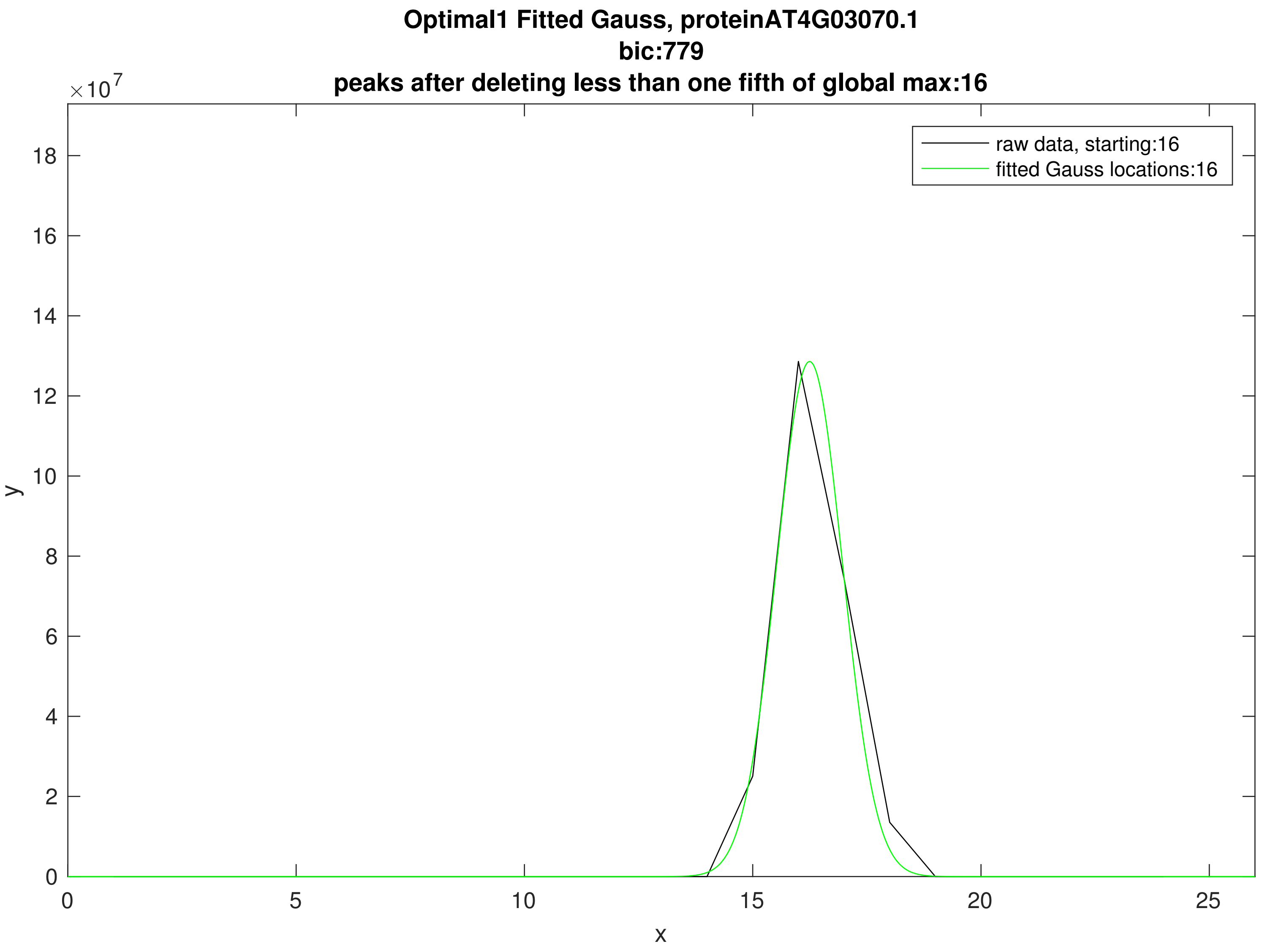
Optimal1 Fitted Gauss, proteinAT4G03070.1
bic:779
peaks after deleting less than one fifth of global max:16
×10^7
raw data, starting:16
fitted Gauss locations:16
y
x
0
2
4
6
8
10
12
14
16
18
5
15
20
25

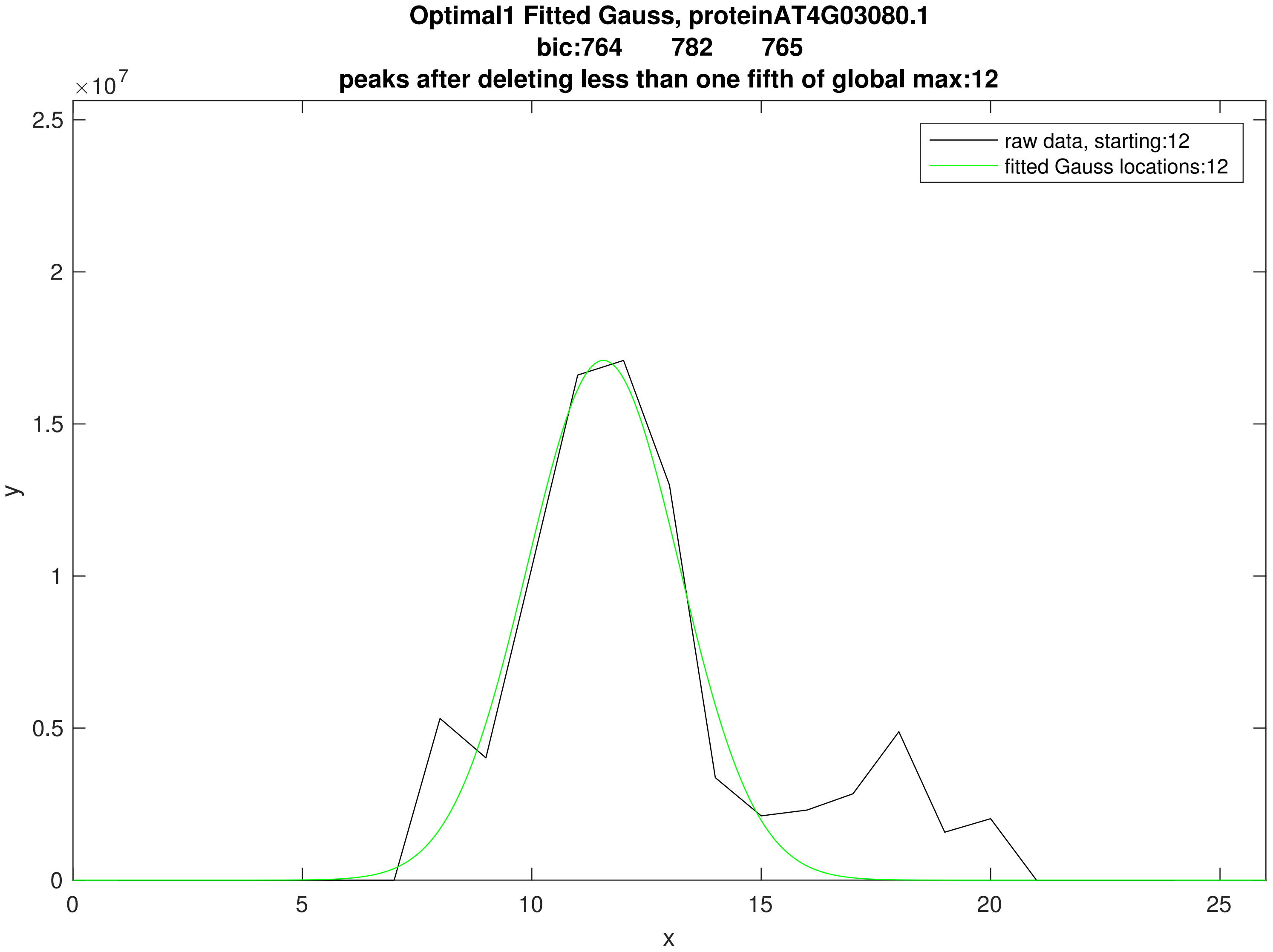
Optimal1 Fitted Gauss, proteinAT4G03080.1
bic:764 782 765
peaks after deleting less than one fifth of global max:12
×10^7
2.5
2
1.5
1
0.5
0
y
0
5
10
15
20
25
x
raw data, starting:12
fitted Gauss locations:12

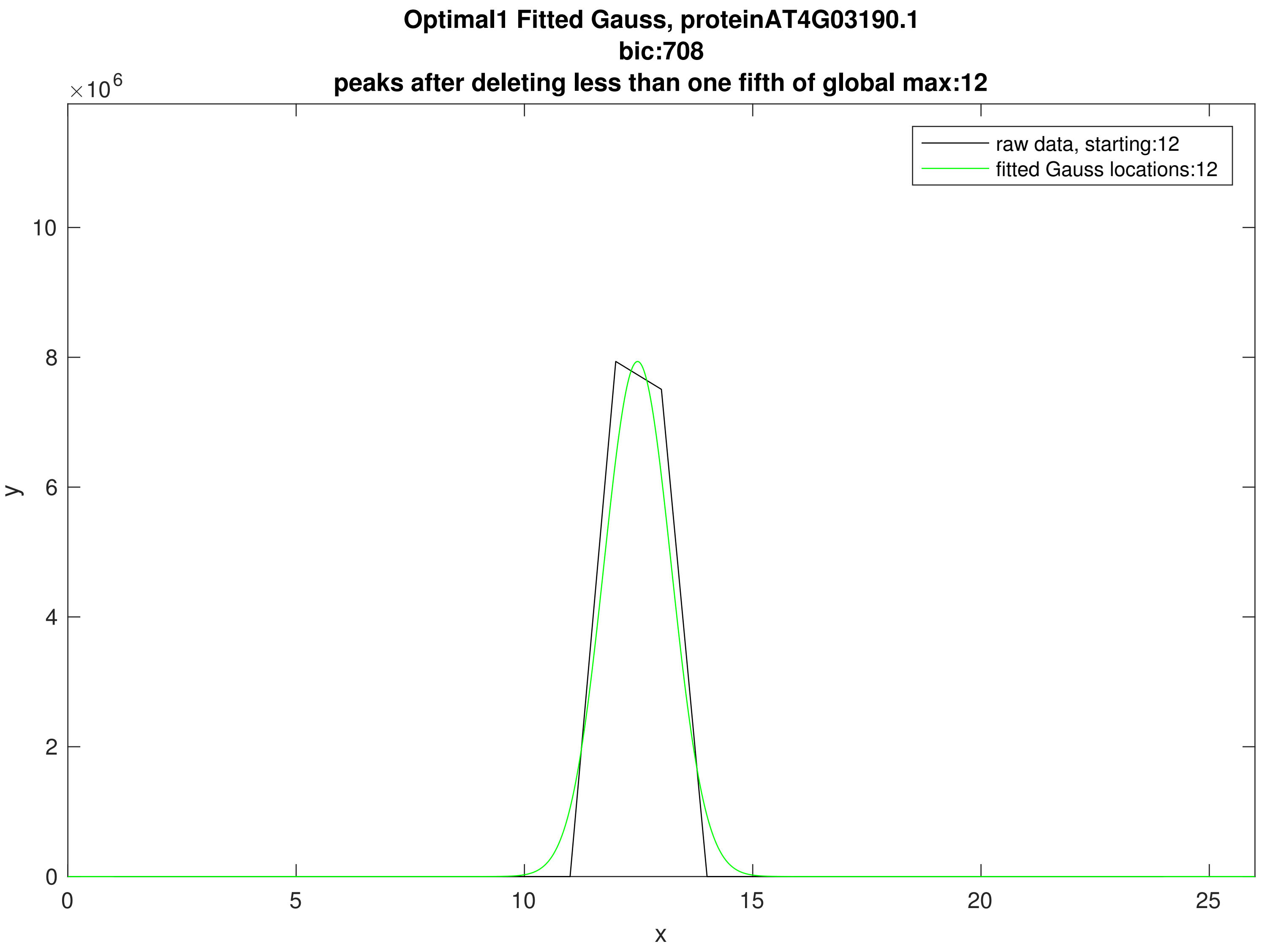

Optimal1 Fitted Gauss, proteinAT4G03190.1
bic:708
peaks after deleting less than one fifth of global max:12
×10^6
raw data, starting:12
fitted Gauss locations:12
10
8
6
4
2
0
y
0
5
10
15
20
25
x

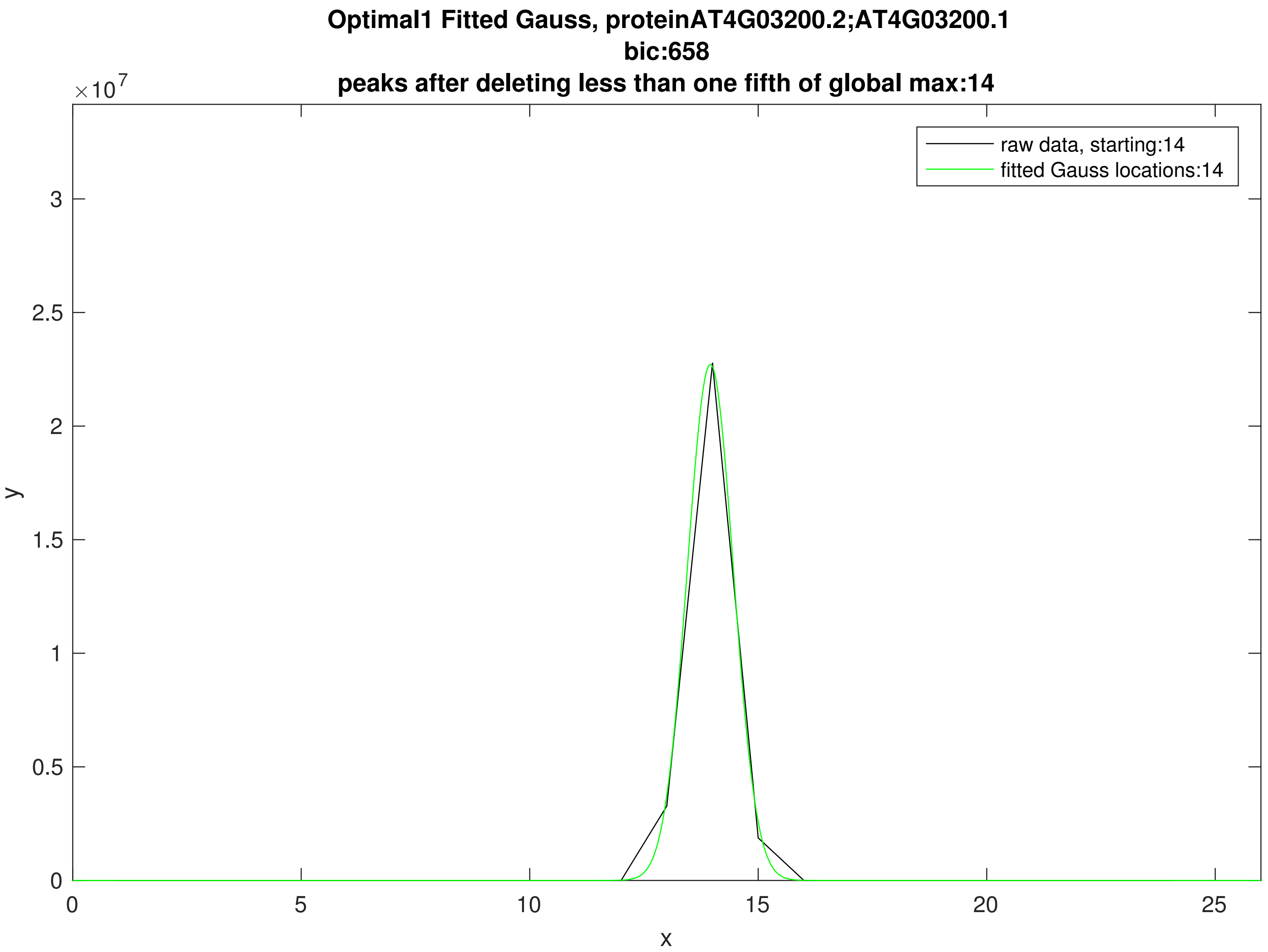
Optimal1 Fitted Gauss, proteinAT4G03200.2;AT4G03200.1
bic:658
peaks after deleting less than one fifth of global max:14
×10^7
raw data, starting:14
fitted Gauss locations:14
y
x
0
0.5
1
1.5
2
2.5
3
0
5
10
15
20
25

**Optimal1 Fitted Gauss, proteinAT4G03240.1**
**bic:689**
**peaks after deleting less than one fifth of global max:19**

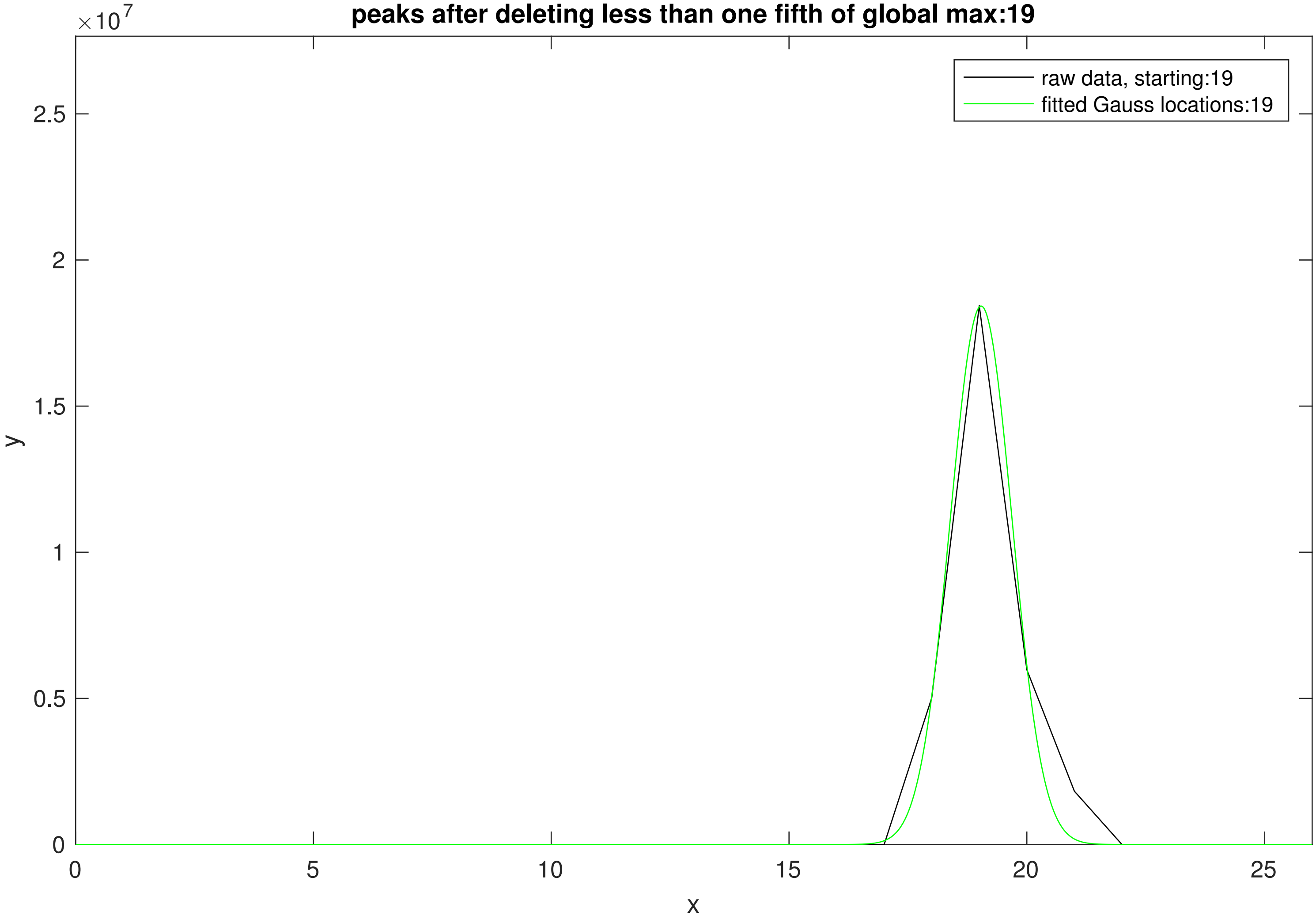

**Optimal1 Fitted Gauss, proteinAT4G03520.1;AT4G03520.2**
**bic:751**
**peaks after deleting less than one fifth of global max:21**

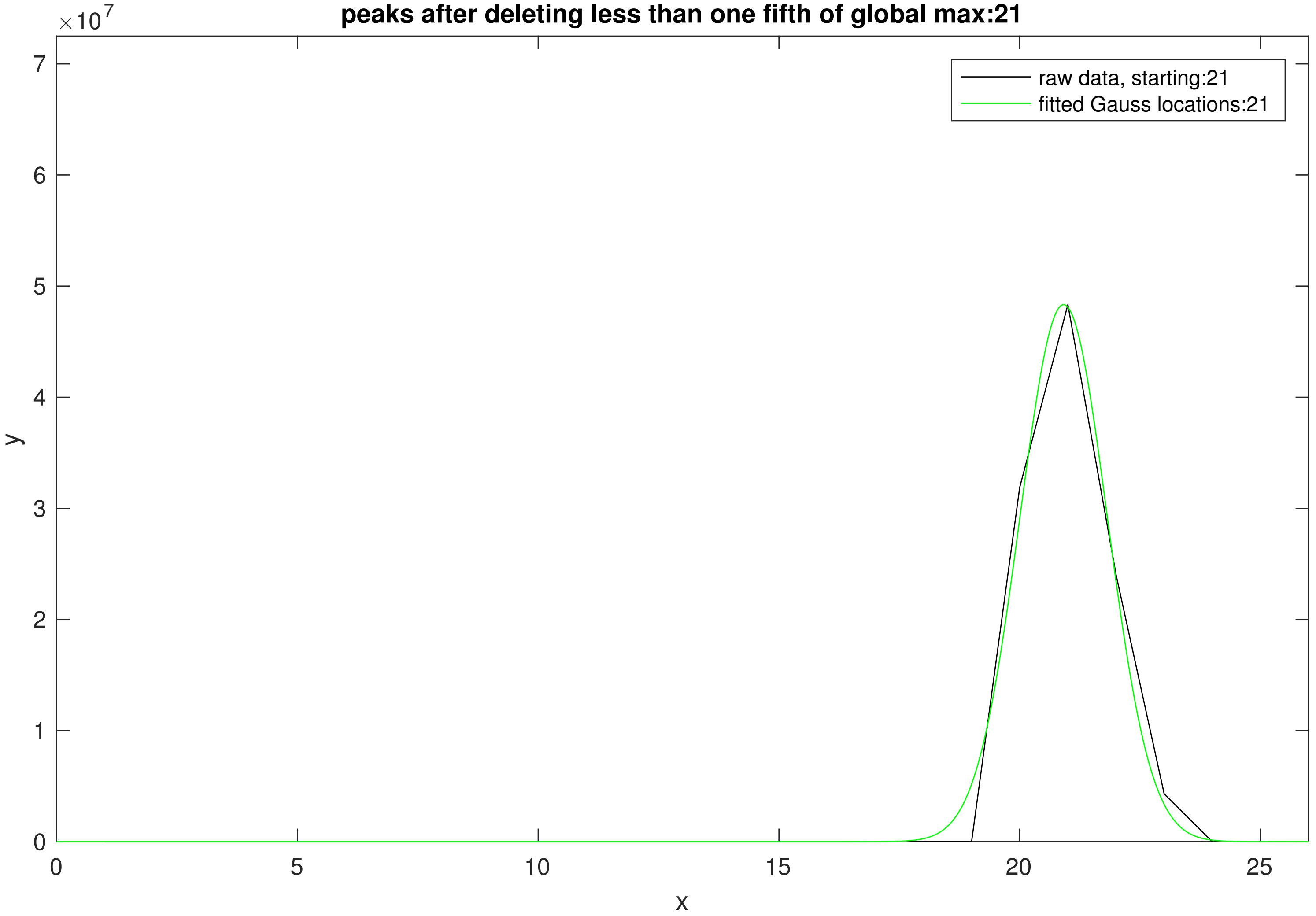

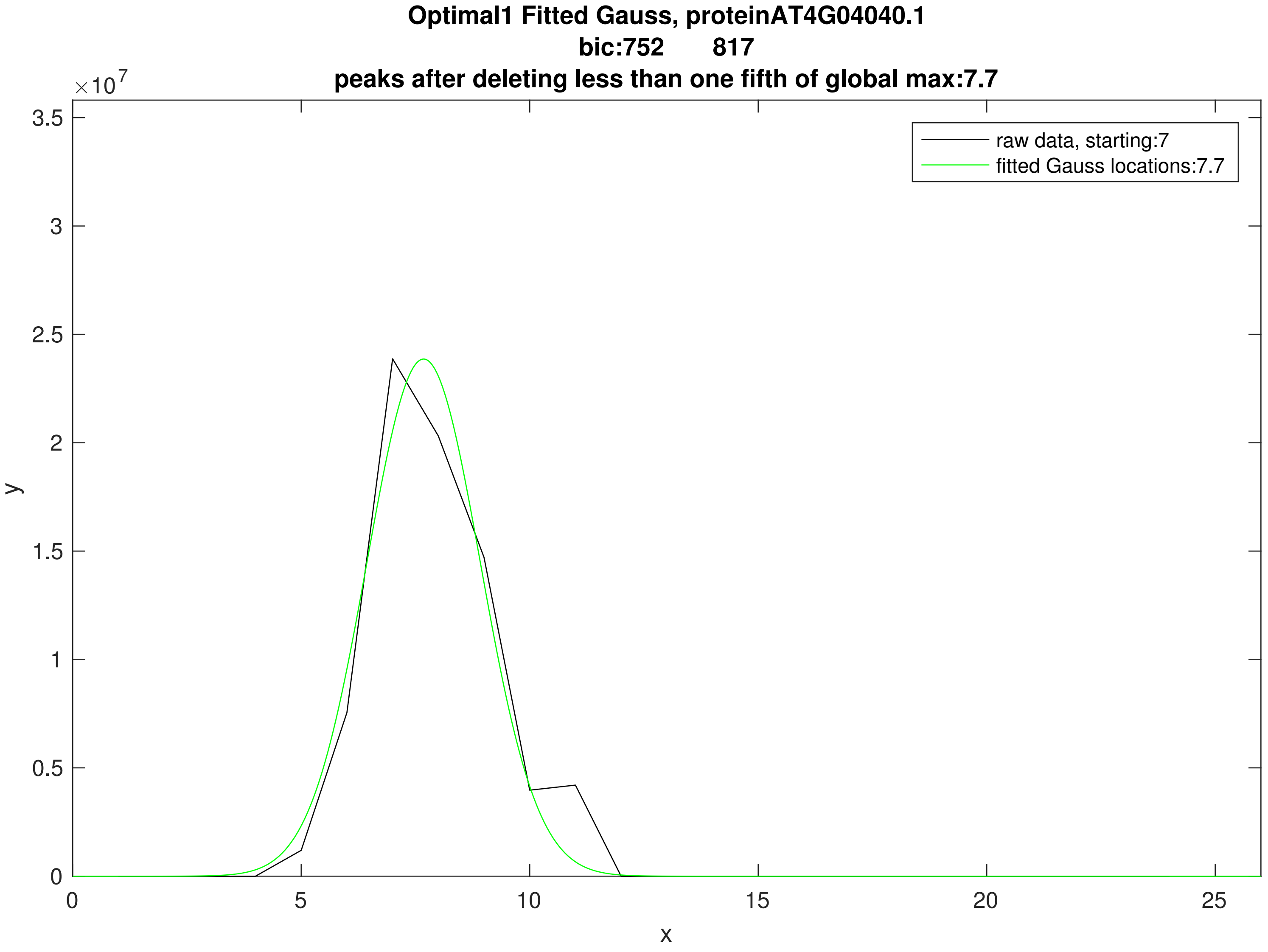

Optimal1 Fitted Gauss, proteinAT4G04040.1
bic:752 817
peaks after deleting less than one fifth of global max:7.7
×10^7
3.5
3
2.5
2
1.5
1
0.5
0
y
0
5
10
15
20
25
x
raw data, starting:7
fitted Gauss locations:7.7

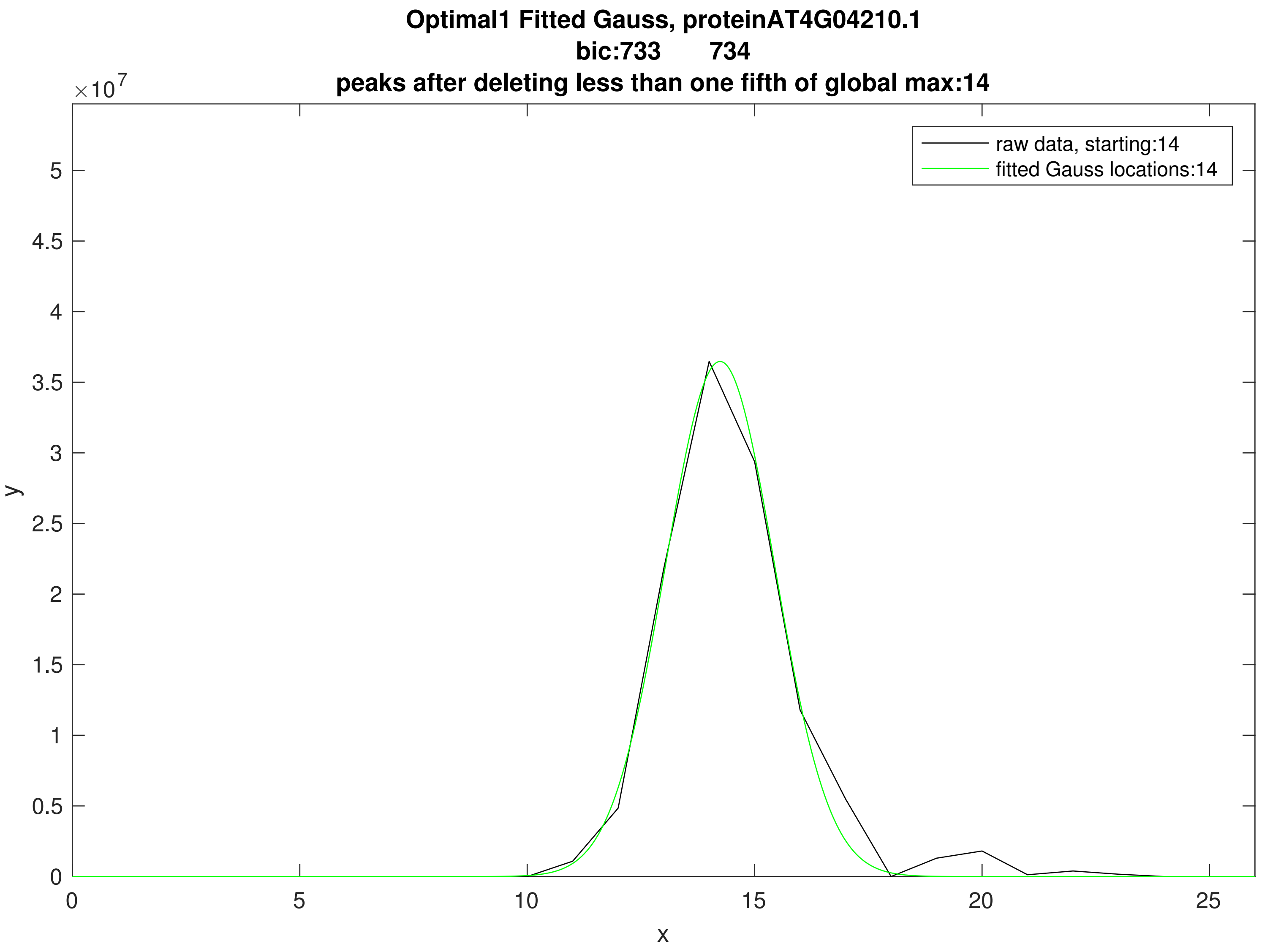
Optimal1 Fitted Gauss, proteinAT4G04210.1
bic:733 734
peaks after deleting less than one fifth of global max:14
×10^7
raw data, starting:14
fitted Gauss locations:14
5
4.5
4
3.5
3
2.5
2
1.5
1
0.5
0
y
0
5
10
15
20
25
x

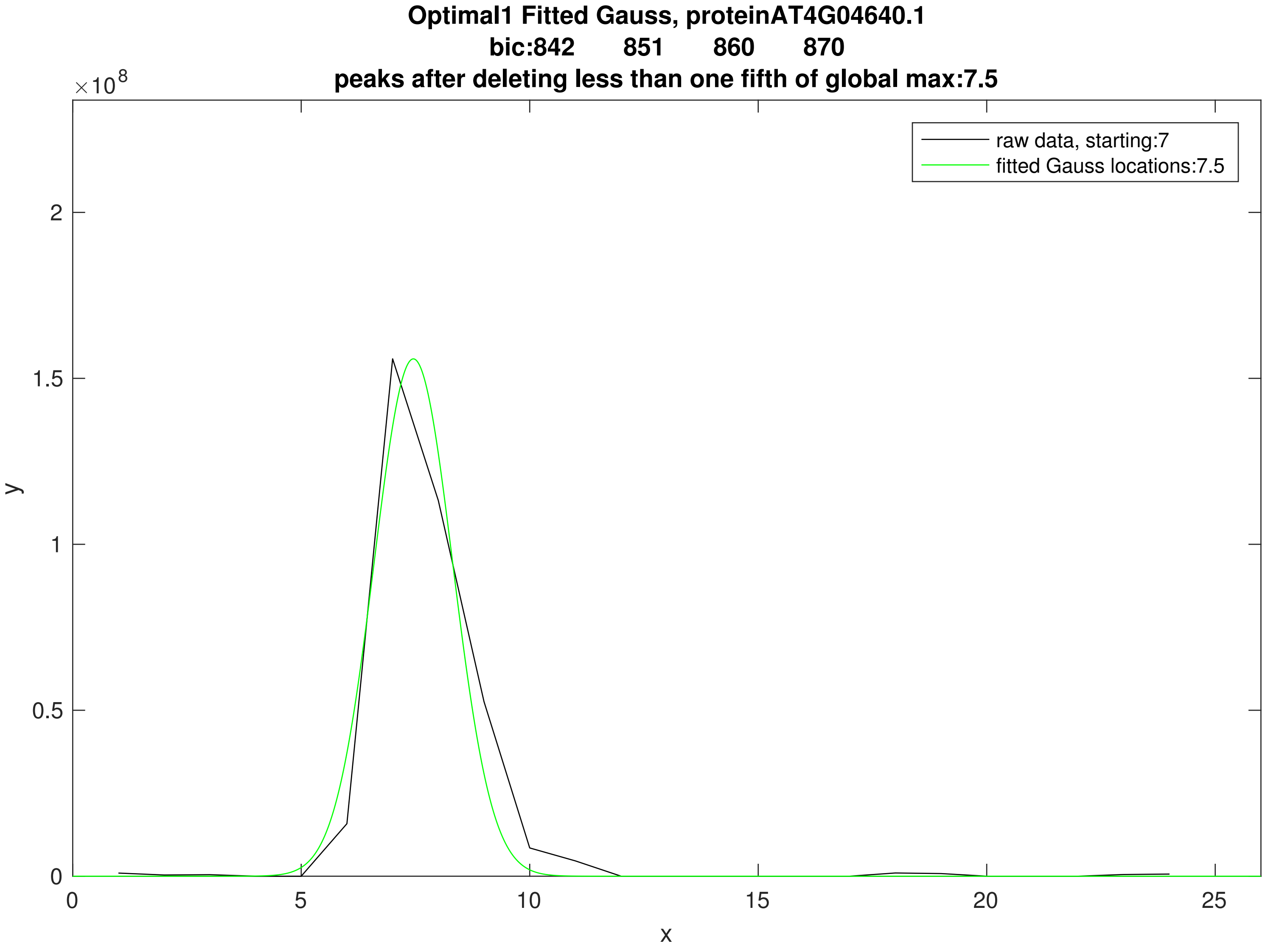
Optimal1 Fitted Gauss, proteinAT4G04640.1
bic:842 851 860 870
peaks after deleting less than one fifth of global max:7.5
×10^8
raw data, starting:7
fitted Gauss locations:7.5
y
x
0
0.5
1
1.5
2
0
5
10
15
20
25

**Optimal1 Fitted Gauss, proteinAT4G04830.1;AT4G04830.2**
**bic:723**
**peaks after deleting less than one fifth of global max:19**

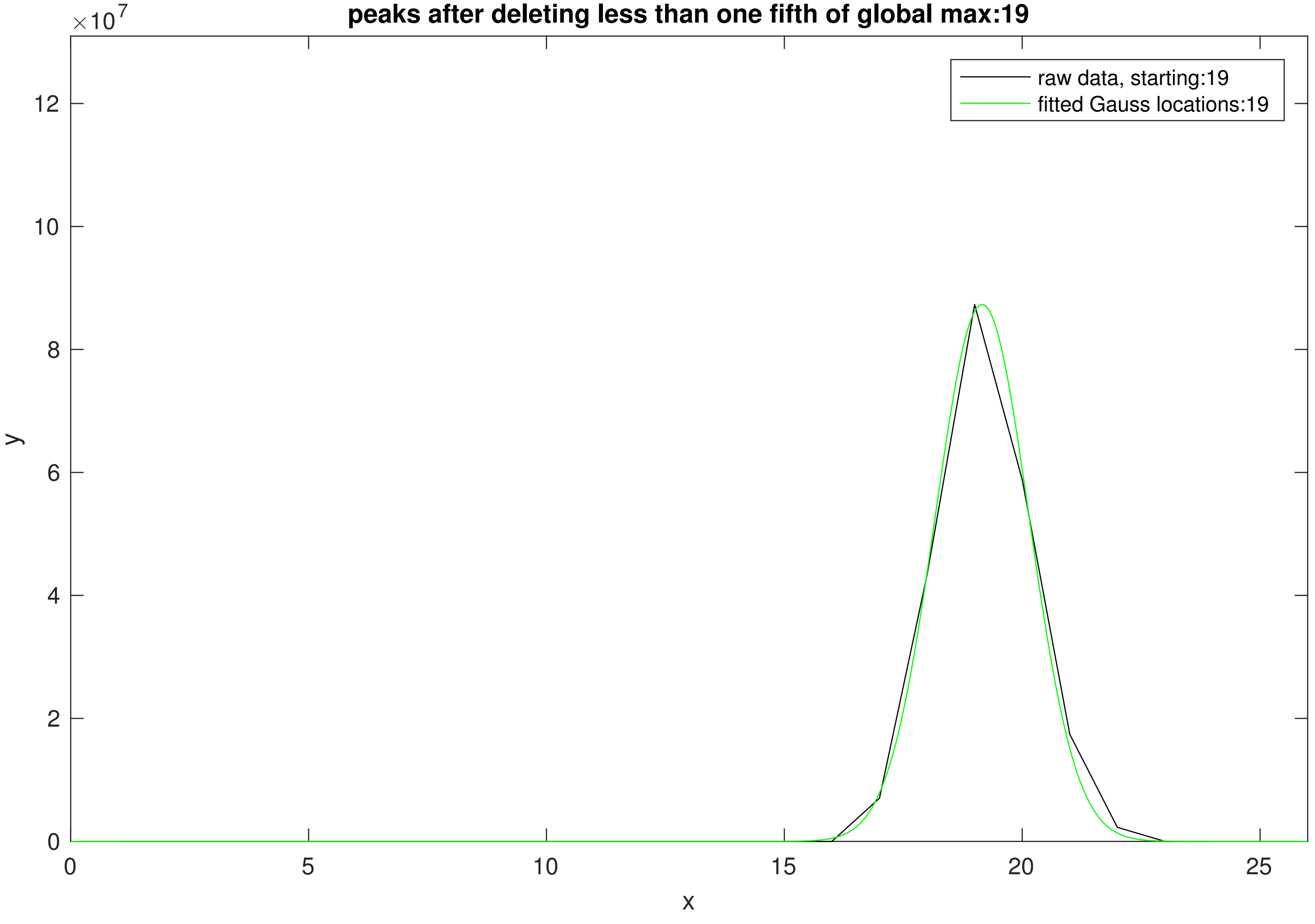

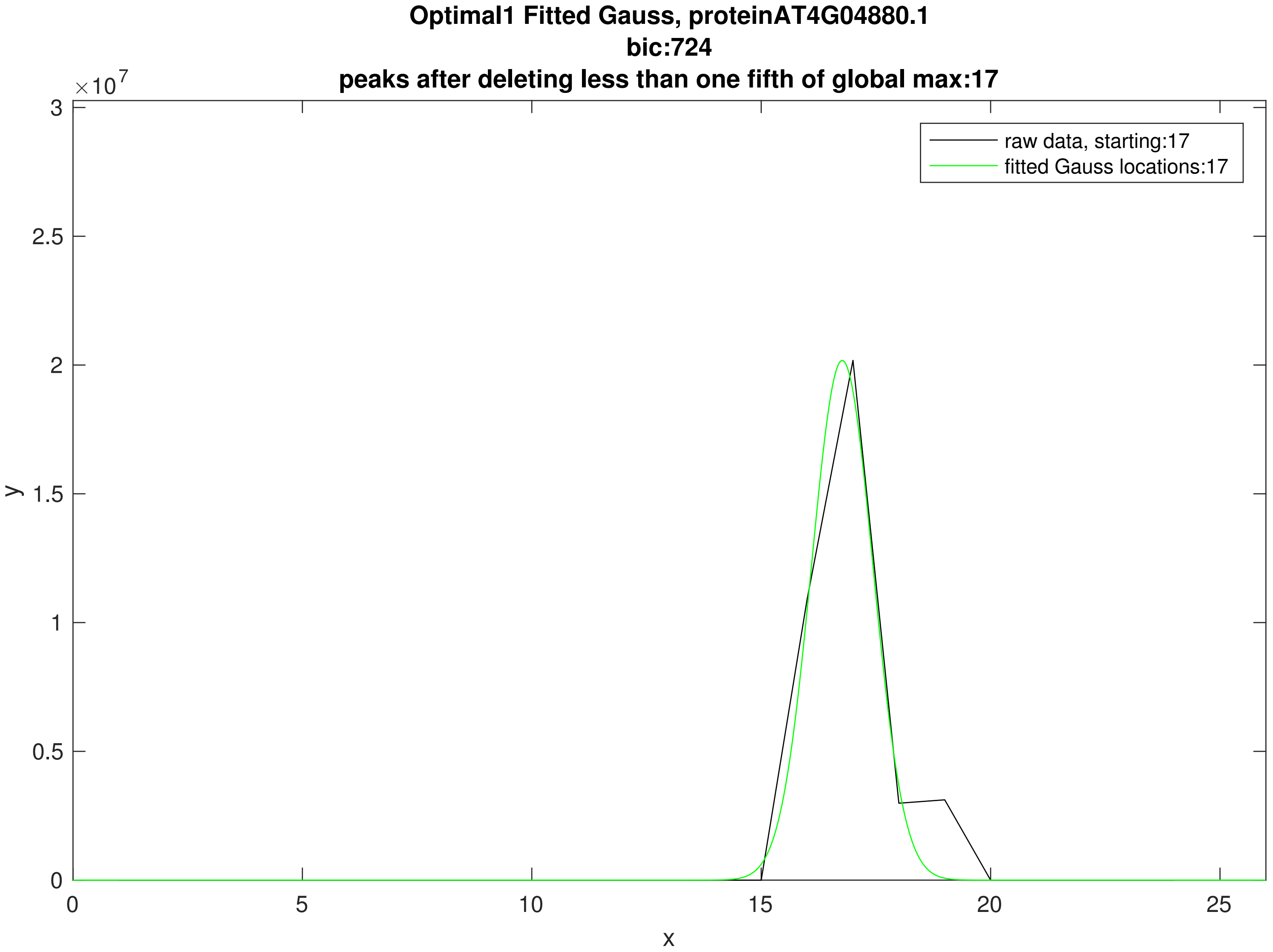
Optimal1 Fitted Gauss, proteinAT4G04880.1
bic:724
peaks after deleting less than one fifth of global max:17
×10^7
3
2.5
2
1.5
1
0.5
0
y
0
5
10
15
20
25
x
raw data, starting:17
fitted Gauss locations:17

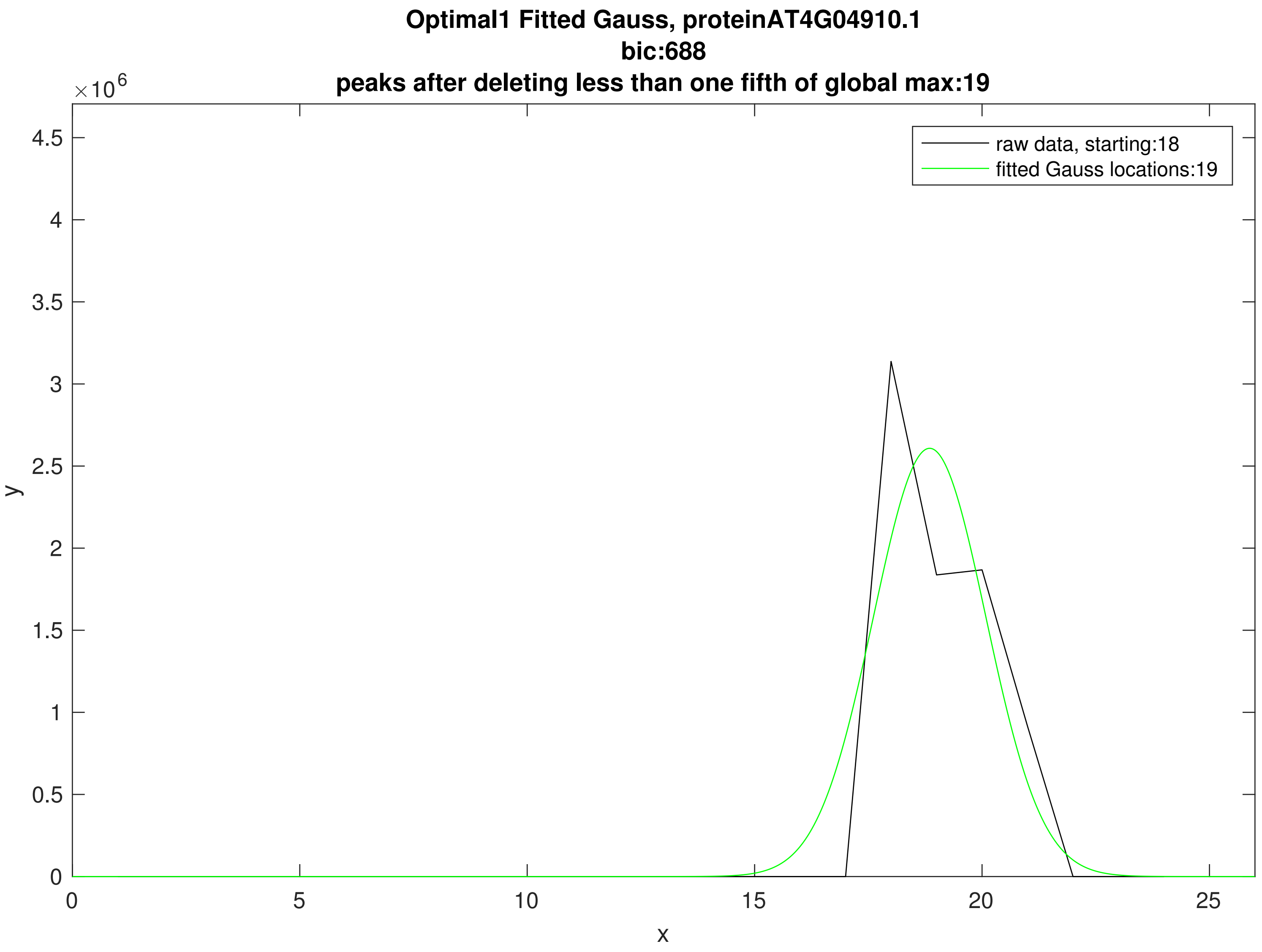
Optimal1 Fitted Gauss, proteinAT4G04910.1
bic:688
peaks after deleting less than one fifth of global max:19
×10^6
4.5
4
3.5
3
2.5
2
1.5
1
0.5
0
y
0
5
10
15
20
25
x
raw data, starting:18
fitted Gauss locations:19

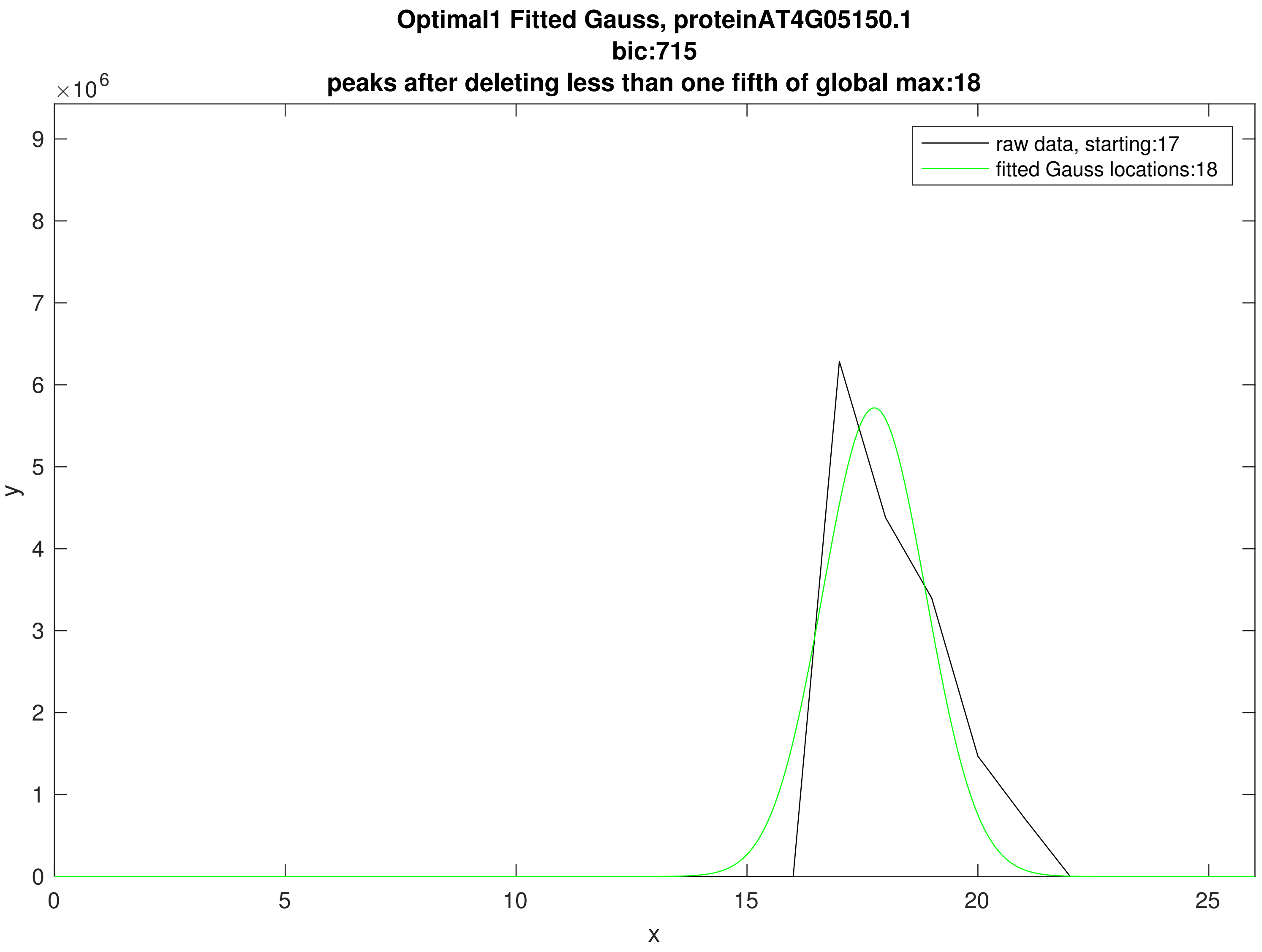
Optimal1 Fitted Gauss, proteinAT4G05150.1
bic:715
peaks after deleting less than one fifth of global max:18
×10^6
raw data, starting:17
fitted Gauss locations:18
y
x
0
1
2
3
4
5
6
7
8
9
0
5
10
15
20
25

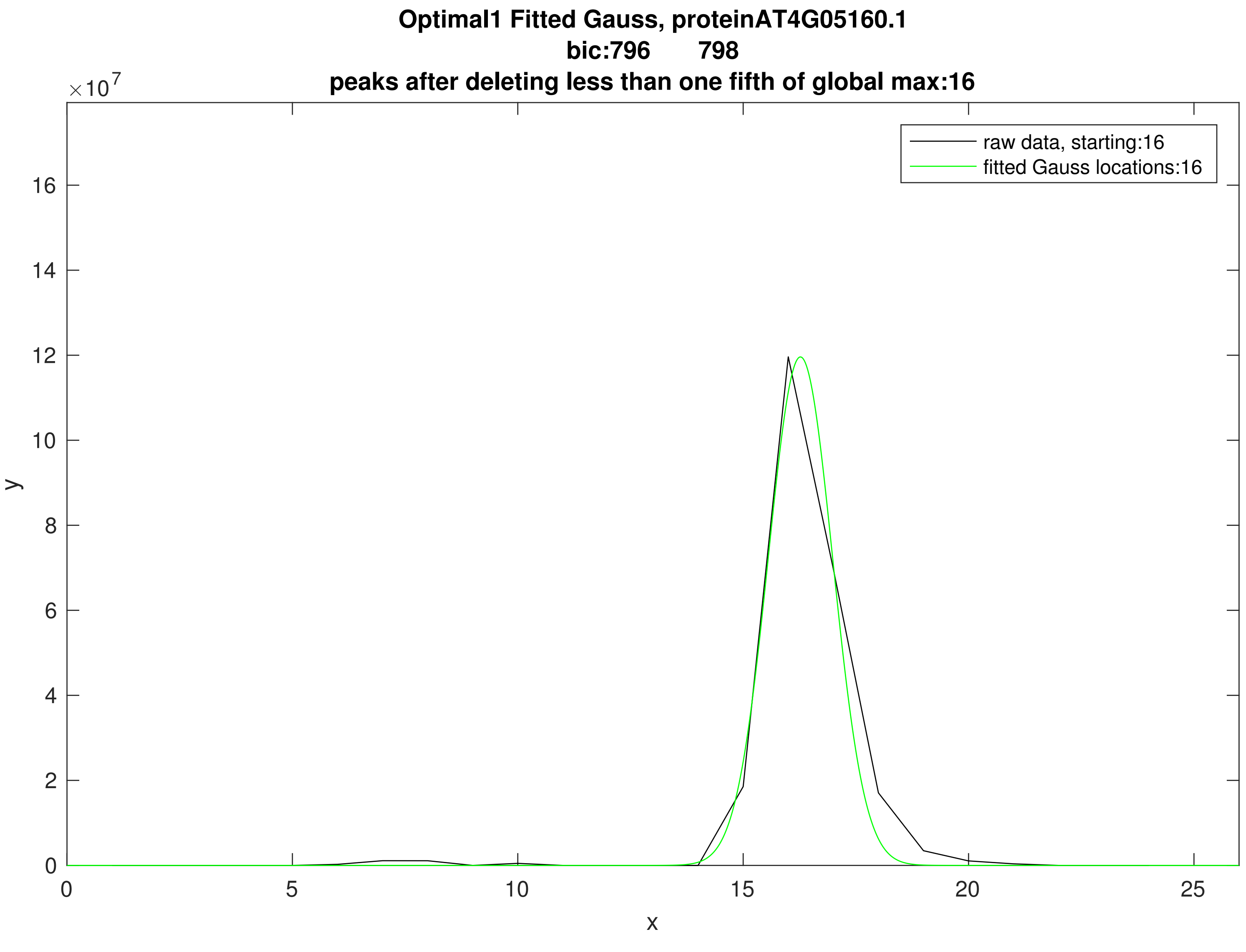
Optimal1 Fitted Gauss, proteinAT4G05160.1
bic:796 798
peaks after deleting less than one fifth of global max:16
×10^7
raw data, starting:16
fitted Gauss locations:16
16
14
12
10
8
6
4
2
0
y
0
5
10
15
20
25
x

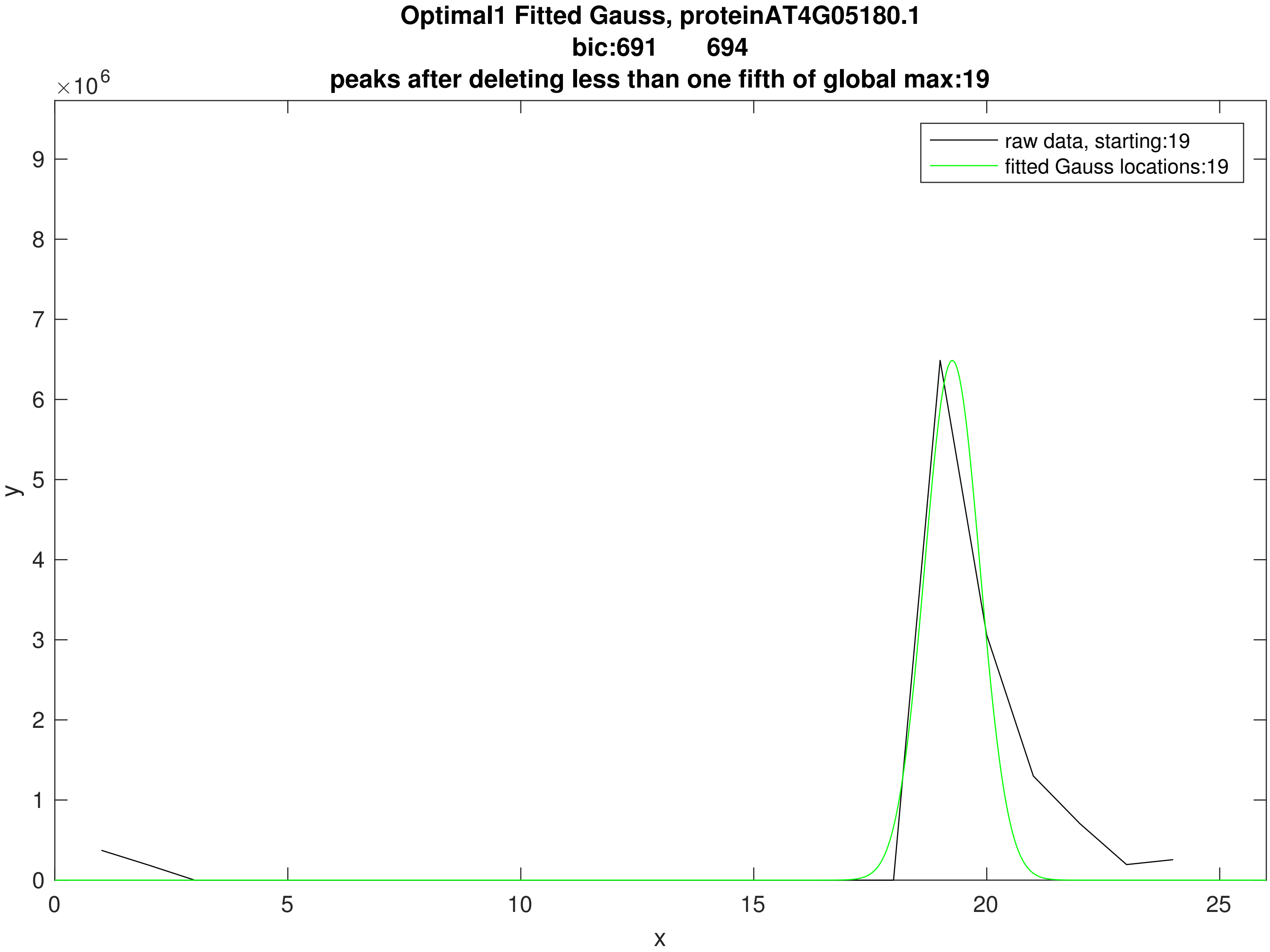
Optimal1 Fitted Gauss, proteinAT4G05180.1
bic:691 694
peaks after deleting less than one fifth of global max:19
×10^6
raw data, starting:19
fitted Gauss locations:19
y
x
0
1
2
3
4
5
6
7
8
9
0
5
10
15
20
25

#### Optimal1 Fitted Gauss, proteinAT4G05390.1;AT4G05390.2
#### bic:917 973 930
#### peaks after deleting less than one fifth of global max:17

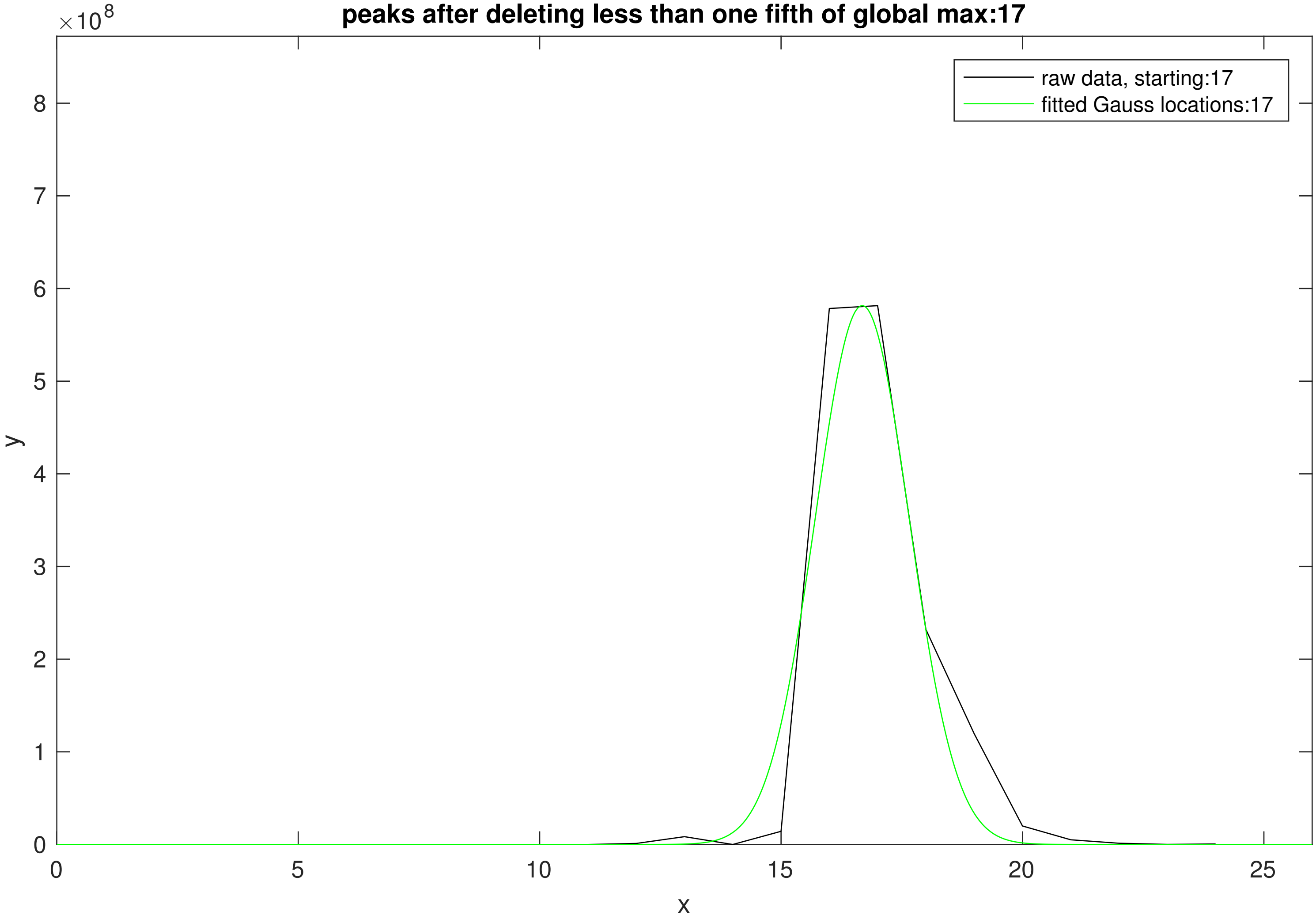

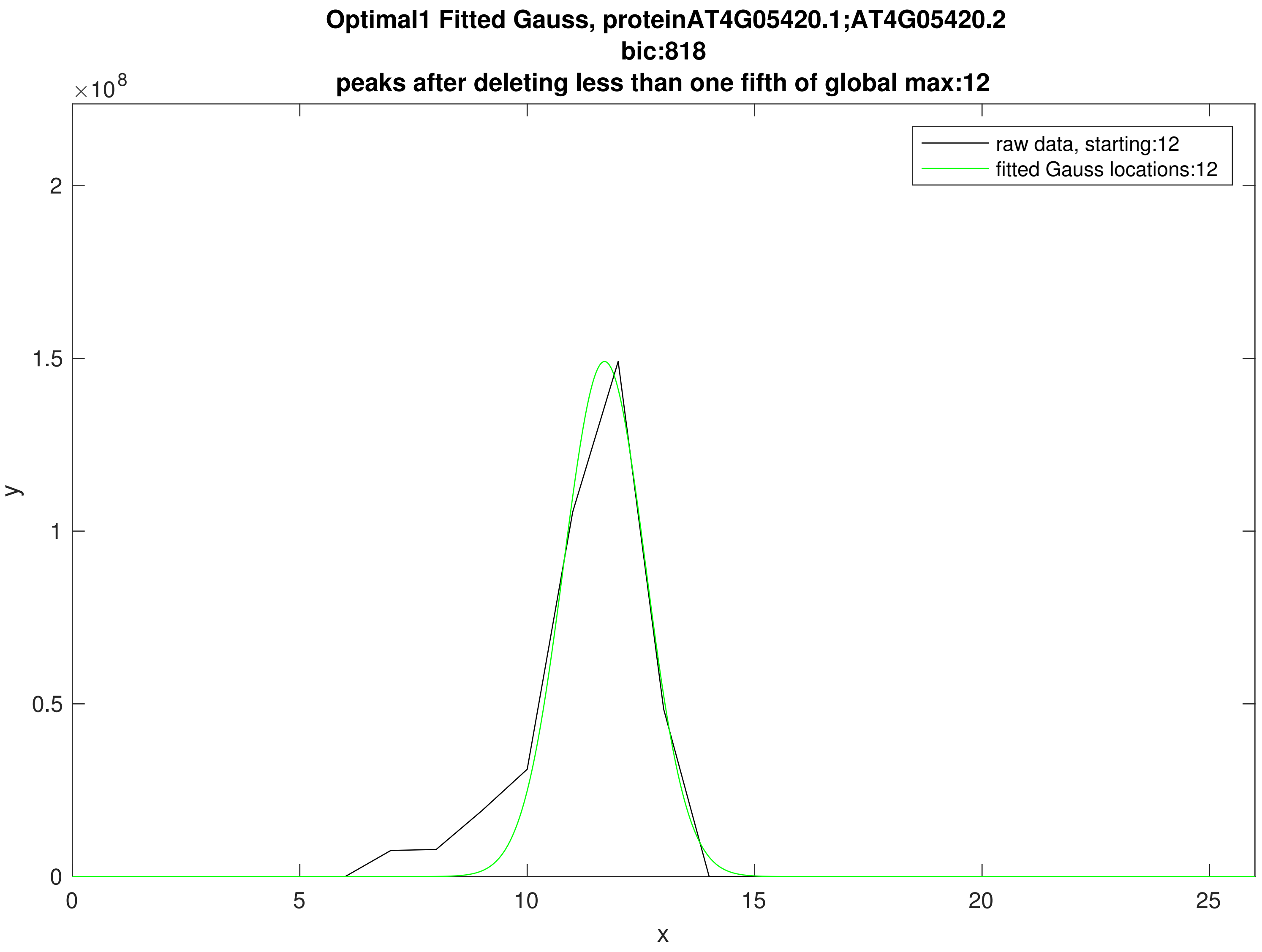
Optimal1 Fitted Gauss, proteinAT4G05420.1;AT4G05420.2
bic:818
peaks after deleting less than one fifth of global max:12
×10^8
raw data, starting:12
fitted Gauss locations:12
y
x
0
0.5
1
1.5
2
0
5
10
15
20
25

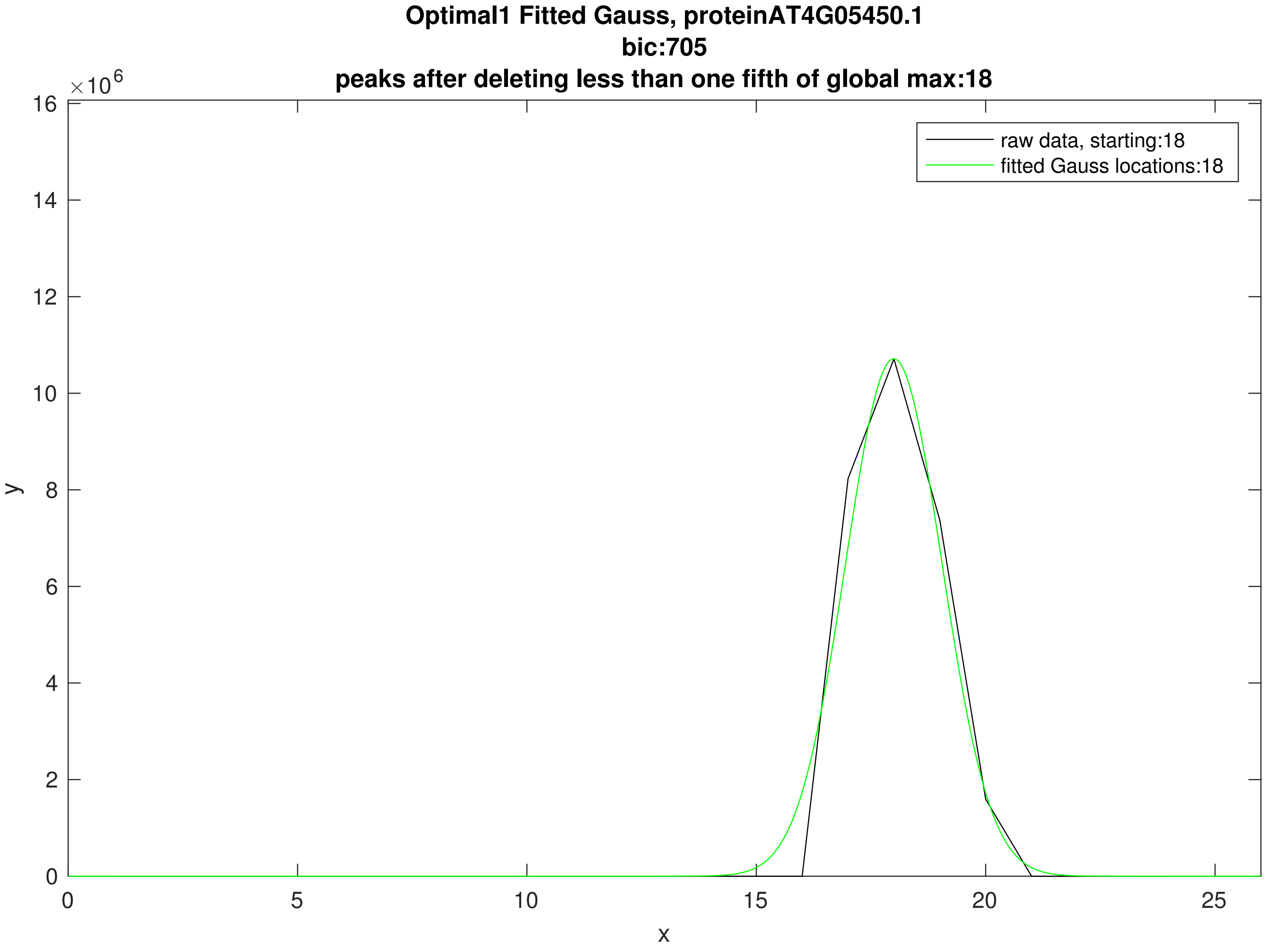
Optimal1 Fitted Gauss, proteinAT4G05450.1
bic:705
peaks after deleting less than one fifth of global max:18
×10^6
raw data, starting:18
fitted Gauss locations:18
y
x
0
2
4
6
8
10
12
14
16
5
15
20
25

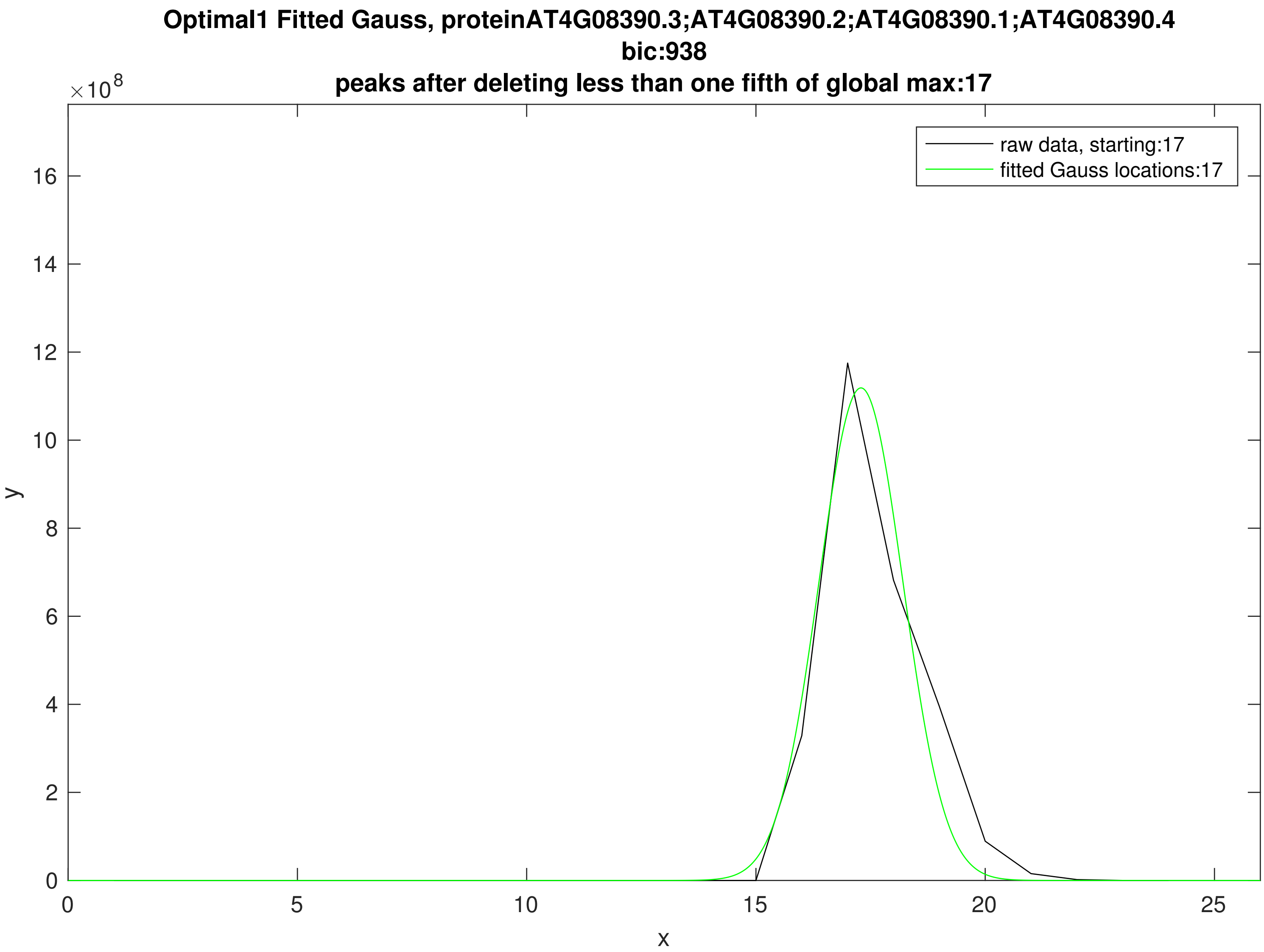
Optimal1 Fitted Gauss, proteinAT4G08390.3;AT4G08390.2;AT4G08390.1;AT4G08390.4
bic:938
peaks after deleting less than one fifth of global max:17
×10^8
raw data, starting:17
fitted Gauss locations:17
y
x
0
2
4
6
8
10
12
14
16
0
5
10
15
20
25

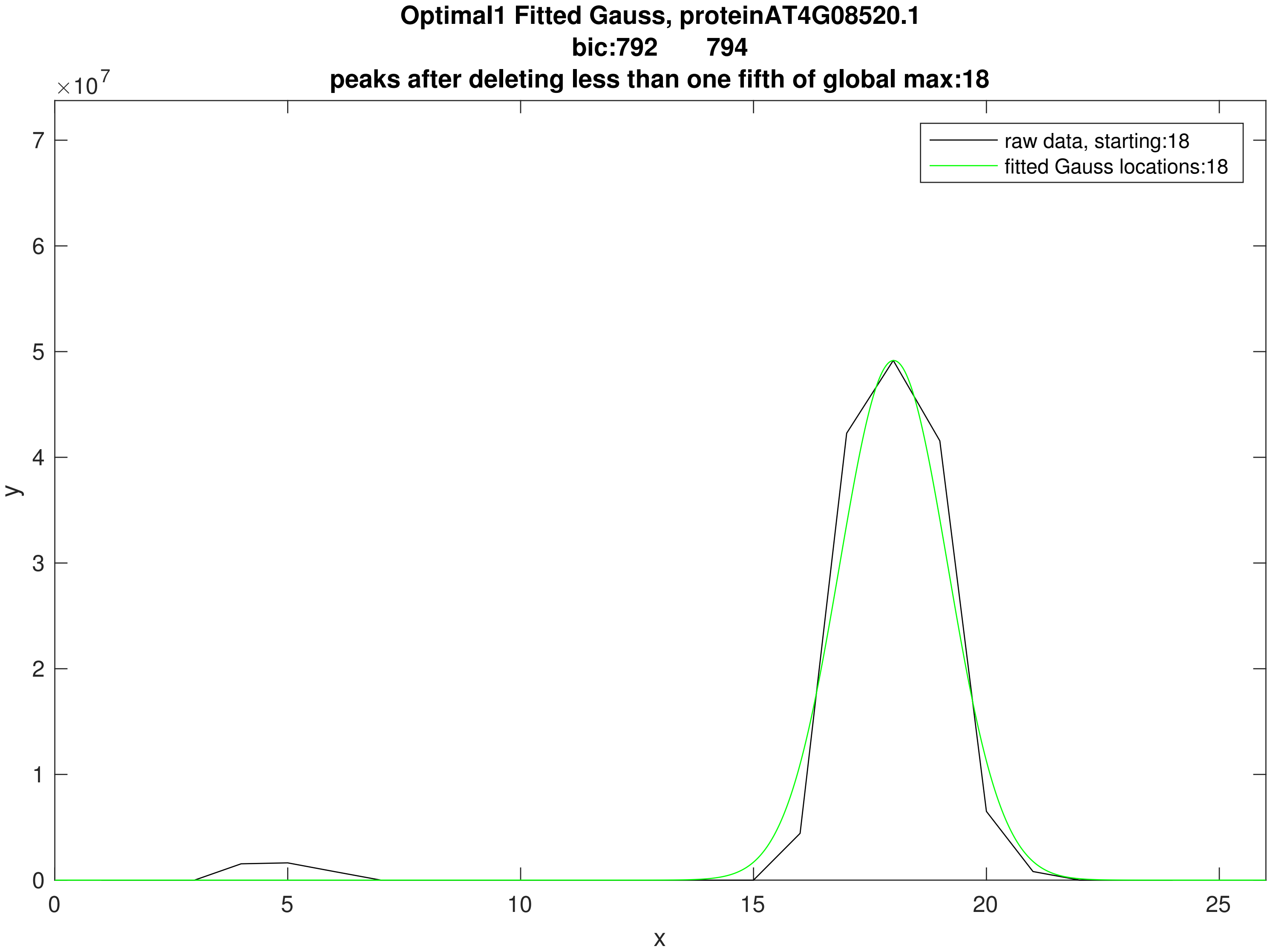

Optimal1 Fitted Gauss, proteinAT4G08520.1
bic:792 794
peaks after deleting less than one fifth of global max:18
×10^7
raw data, starting:18
fitted Gauss locations:18
y
x
0
1
2
3
4
5
6
7
0
5
10
15
20
25

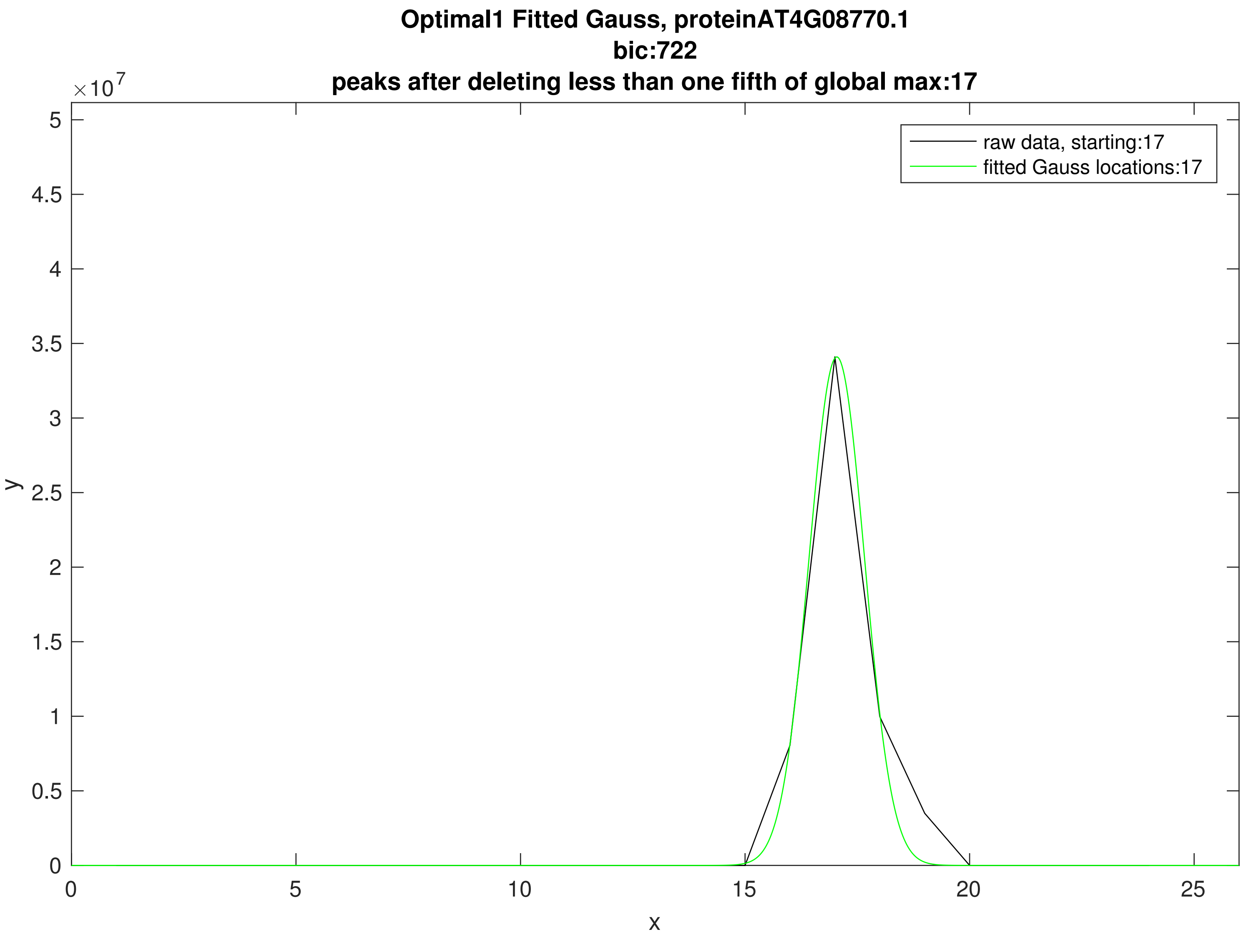
Optimal1 Fitted Gauss, proteinAT4G08770.1
bic:722
peaks after deleting less than one fifth of global max:17
×10^7
5
4.5
4
3.5
3
2.5
2
1.5
1
0.5
0
y
0
5
10
15
20
25
x
raw data, starting:17
fitted Gauss locations:17

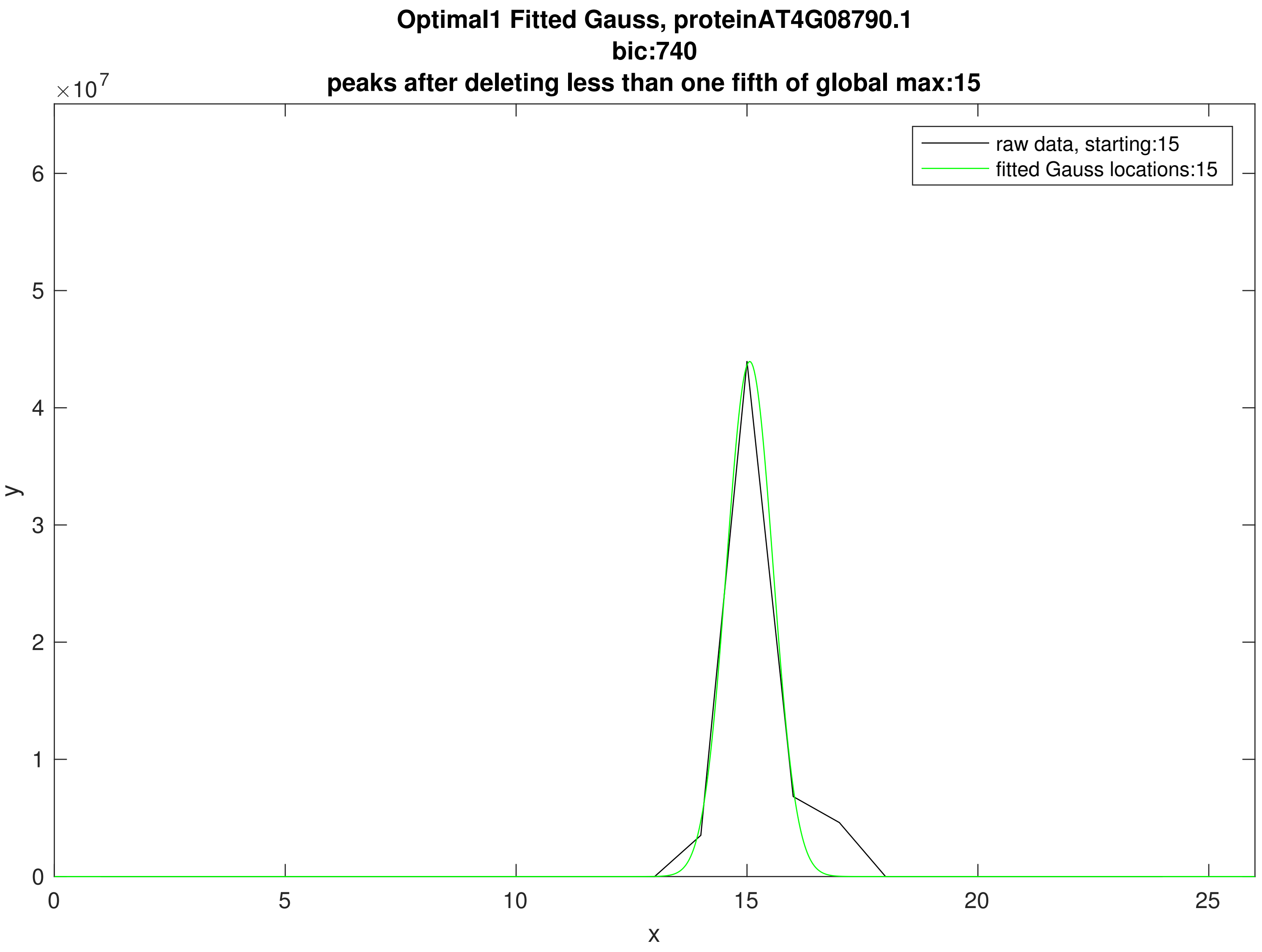

Optimal1 Fitted Gauss, proteinAT4G08790.1
bic:740
peaks after deleting less than one fifth of global max:15
×10^7
raw data, starting:15
fitted Gauss locations:15
y
x
0
1
2
3
4
5
6
0
5
10
15
20
25

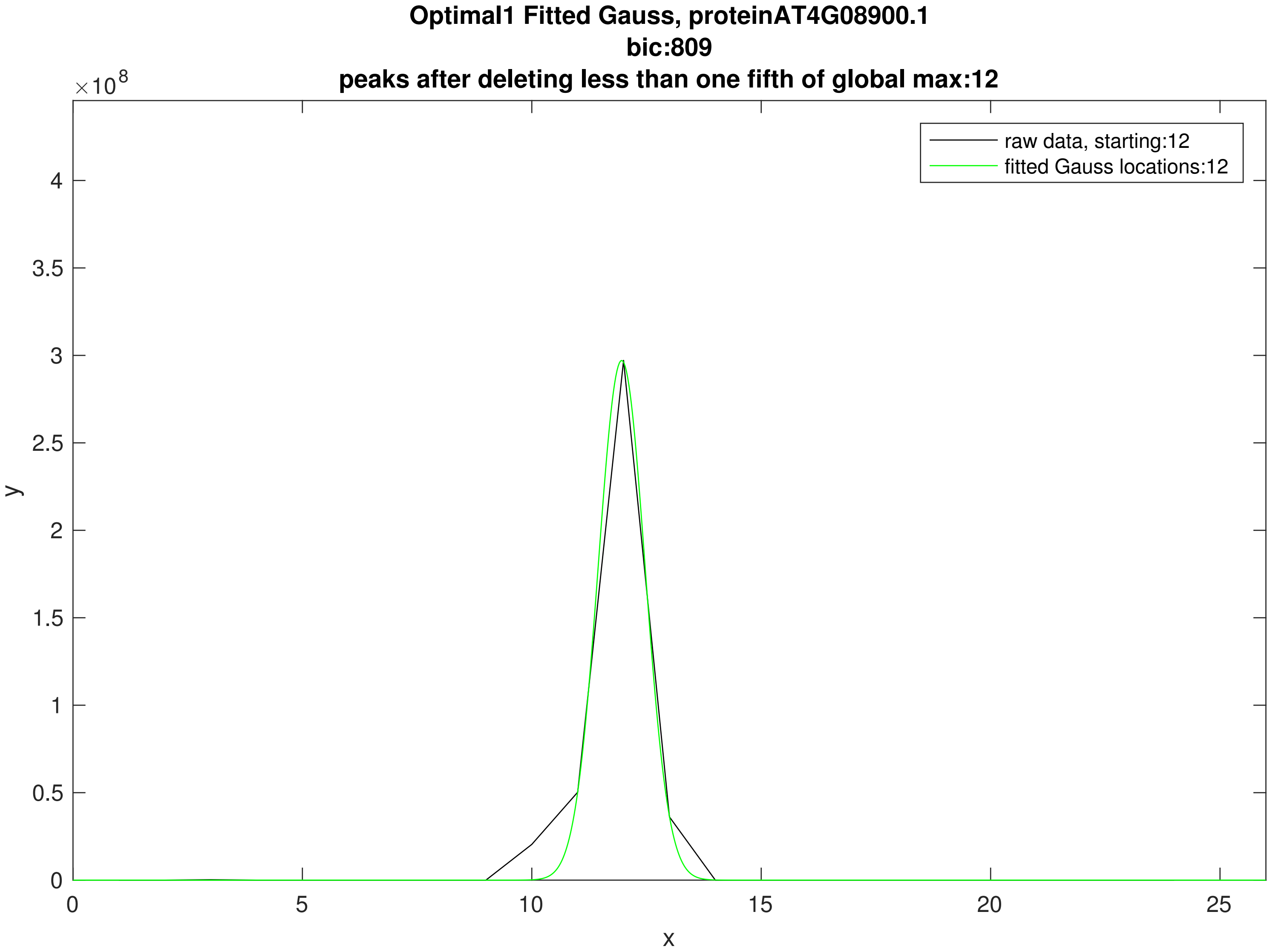
Optimal1 Fitted Gauss, proteinAT4G08900.1
bic:809
peaks after deleting less than one fifth of global max:12
×10^8
raw data, starting:12
fitted Gauss locations:12
y
x
0
0.5
1
1.5
2
2.5
3
3.5
4
0
5
10
15
20
25

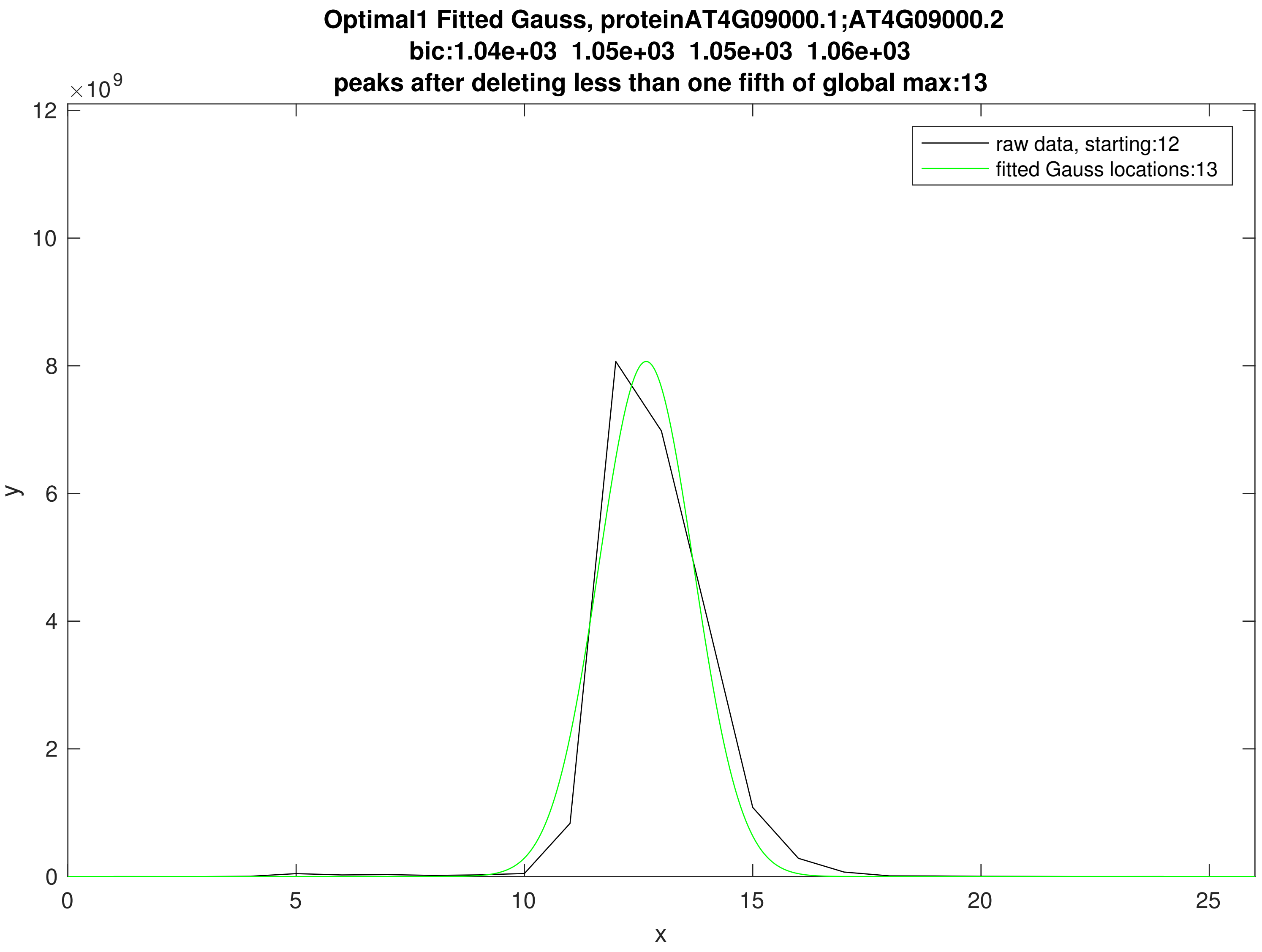
Optimal1 Fitted Gauss, proteinAT4G09000.1;AT4G09000.2
bic:1.04e+03  1.05e+03  1.05e+03  1.06e+03
peaks after deleting less than one fifth of global max:13
×10^9
raw data, starting:12
fitted Gauss locations:13
y
x
0
2
4
6
8
10
12
0
5
10
15
20
25

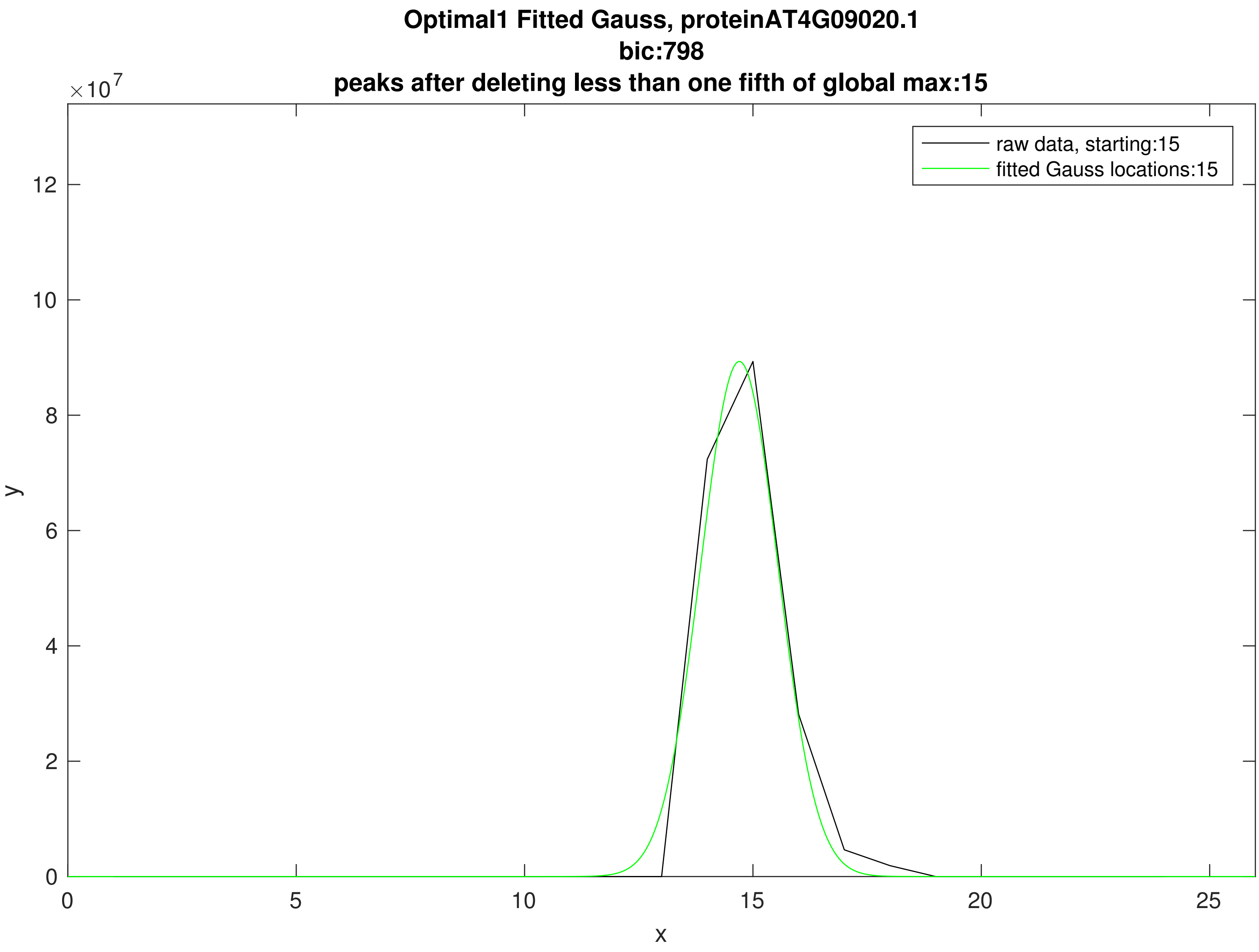
Optimal1 Fitted Gauss, proteinAT4G09020.1
bic:798
peaks after deleting less than one fifth of global max:15
×10^7
raw data, starting:15
fitted Gauss locations:15
12
10
8
6
4
2
0
y
0
5
10
15
20
25
x

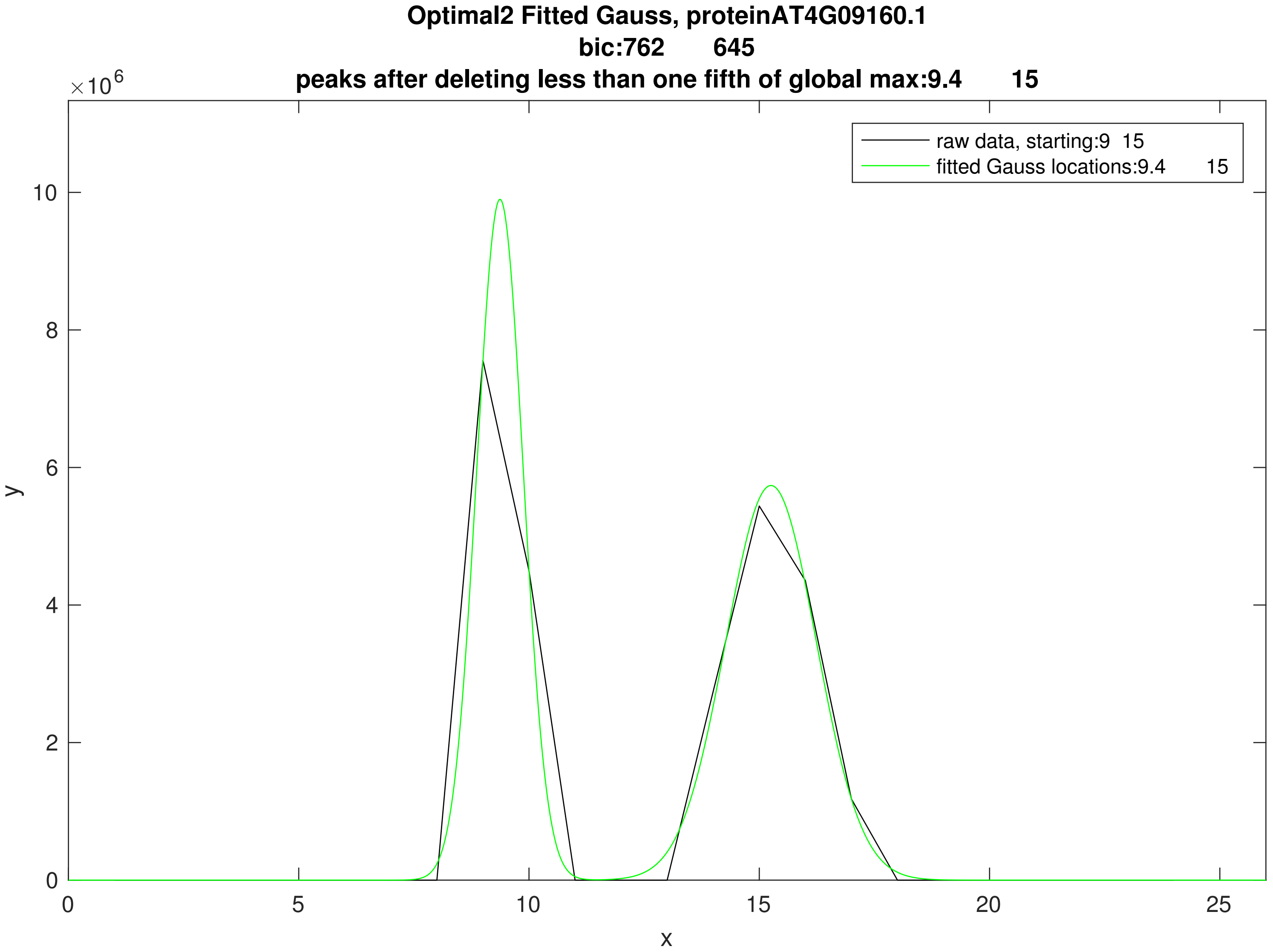
Optimal2 Fitted Gauss, proteinAT4G09160.1
bic:762 645
peaks after deleting less than one fifth of global max:9.4 15
×10^6
raw data, starting:9 15
fitted Gauss locations:9.4 15
y
x

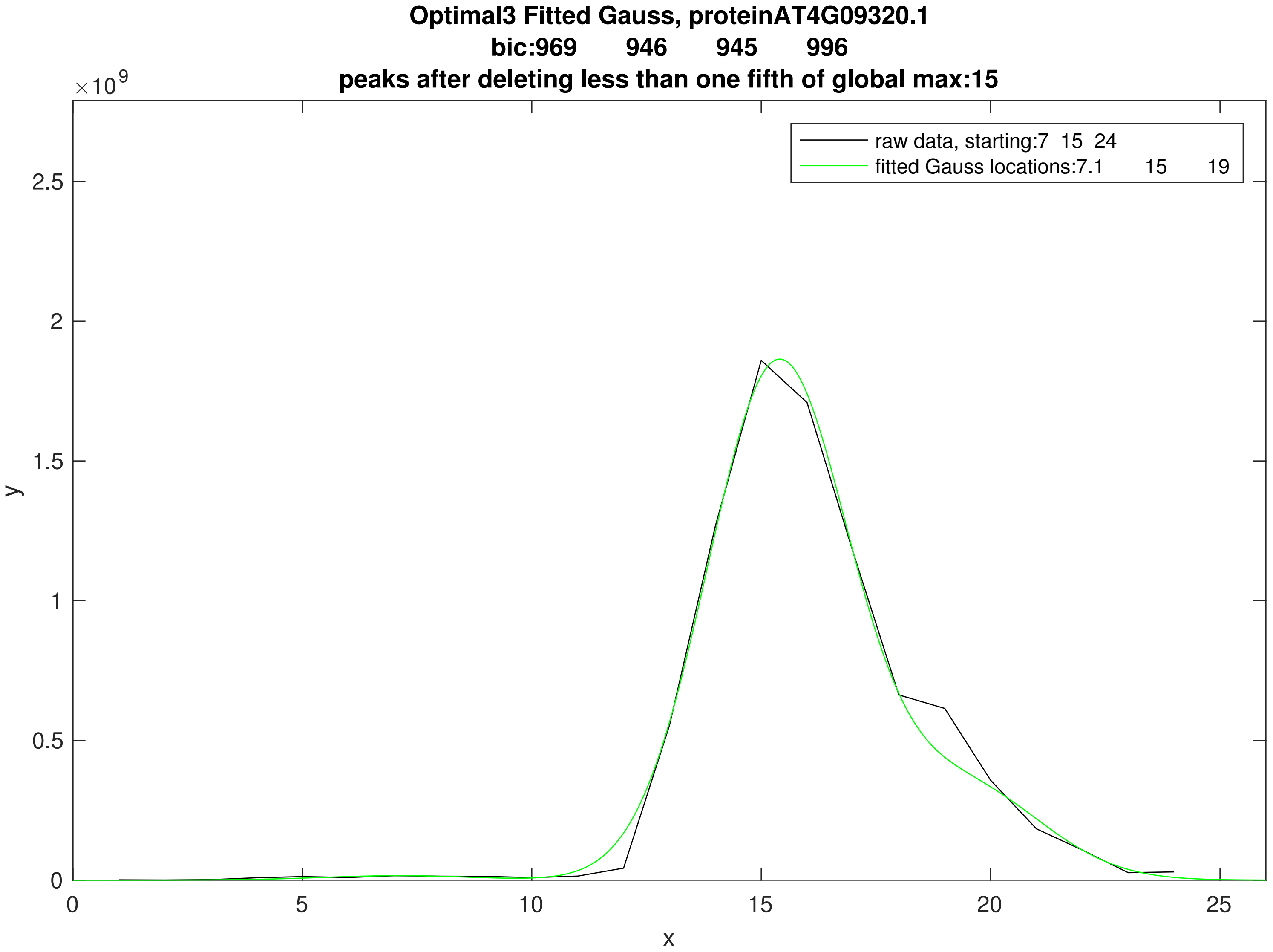

Optimal3 Fitted Gauss, proteinAT4G09320.1
bic:969 946 945 996
peaks after deleting less than one fifth of global max:15
×10^9
raw data, starting:7 15 24
fitted Gauss locations:7.1 15 19
y
x
0
0.5
1
1.5
2
2.5
0
5
10
15
20
25

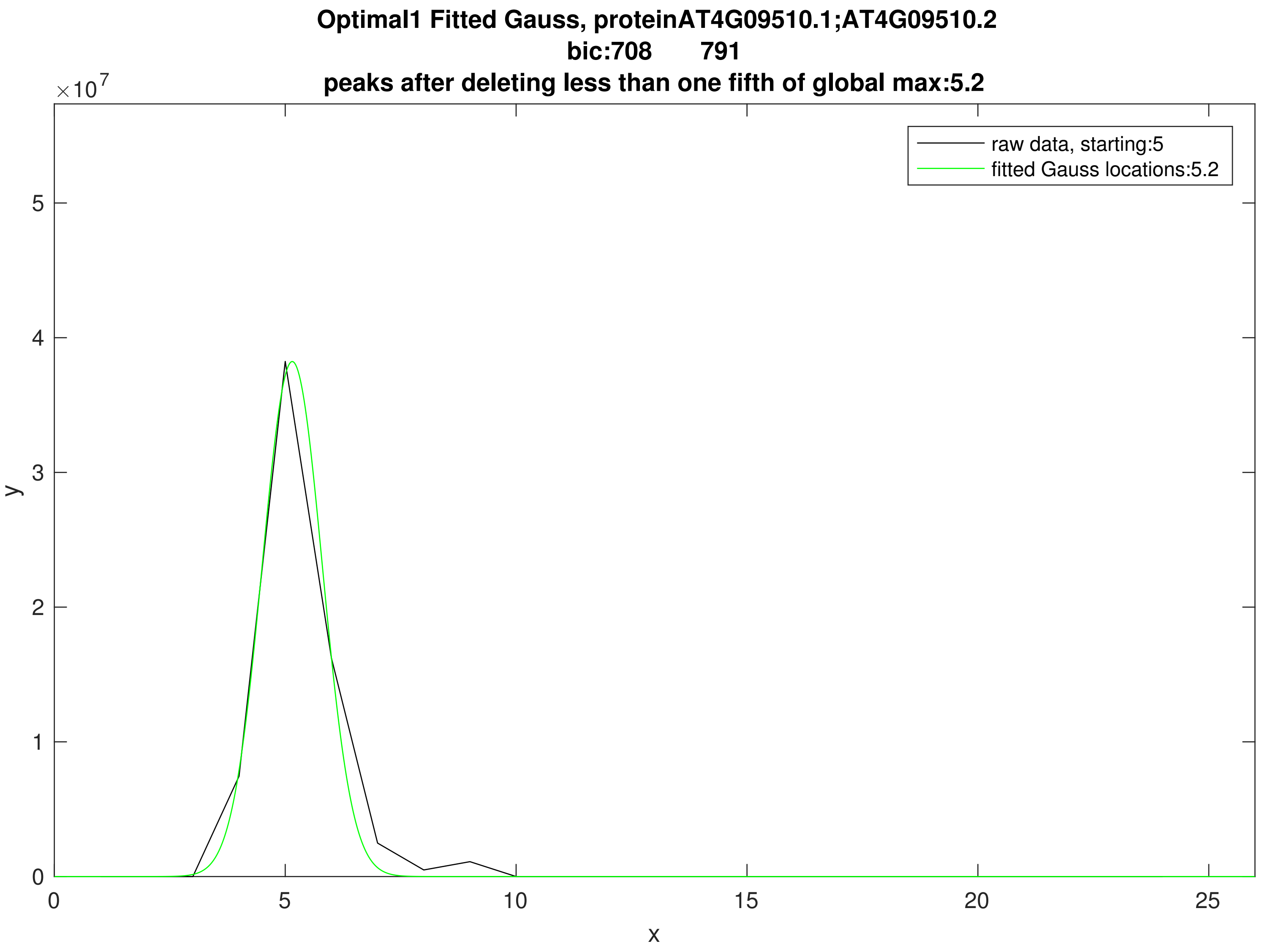
Optimal1 Fitted Gauss, proteinAT4G09510.1;AT4G09510.2
bic:708 791
peaks after deleting less than one fifth of global max:5.2
×10^7
raw data, starting:5
fitted Gauss locations:5.2
y
x
0
1
2
3
4
5
10
15
20
25

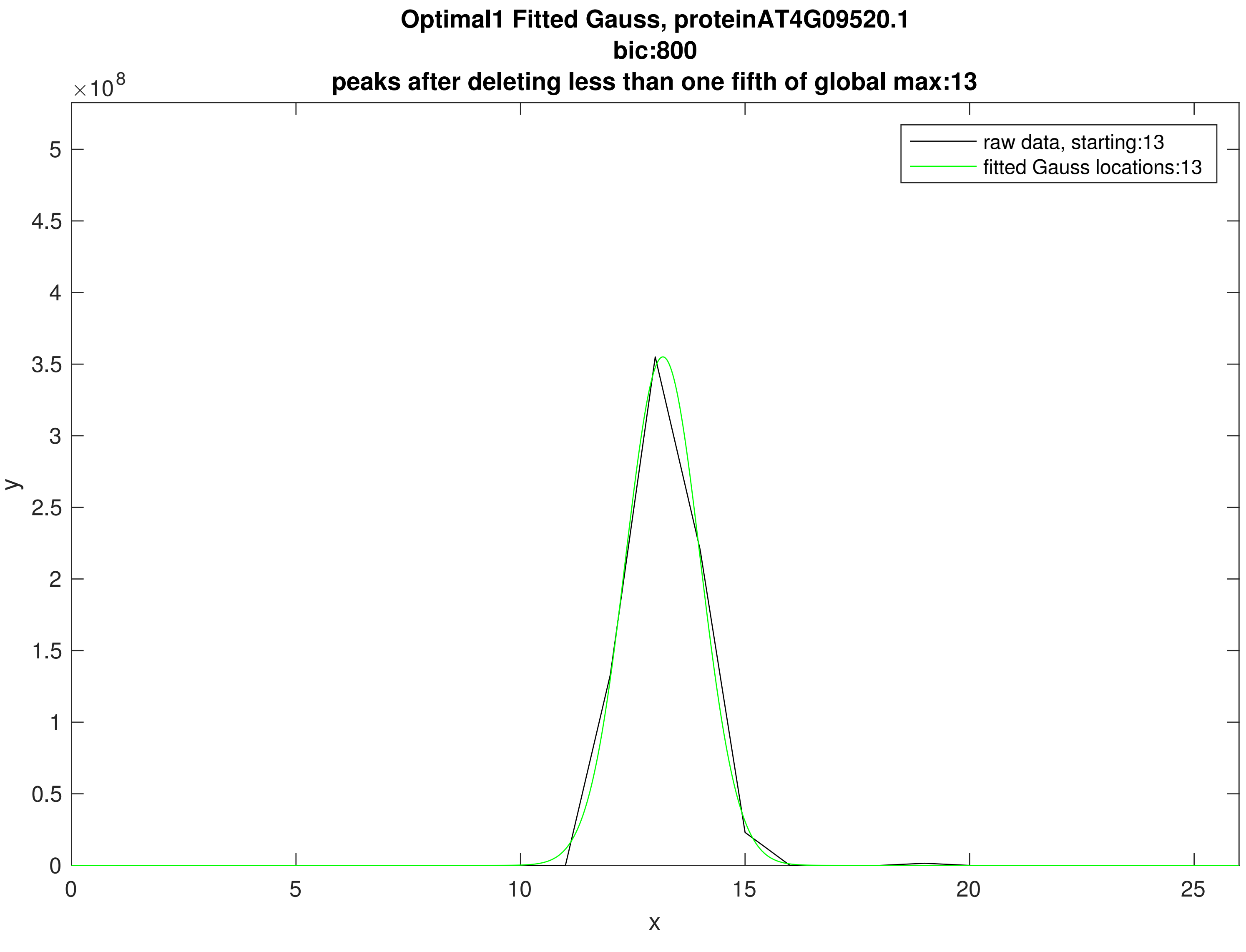

Optimal1 Fitted Gauss, proteinAT4G09520.1
bic:800
peaks after deleting less than one fifth of global max:13
×10^8
5
4.5
4
3.5
3
2.5
2
1.5
1
0.5
0
y
0
5
10
15
20
25
x
raw data, starting:13
fitted Gauss locations:13

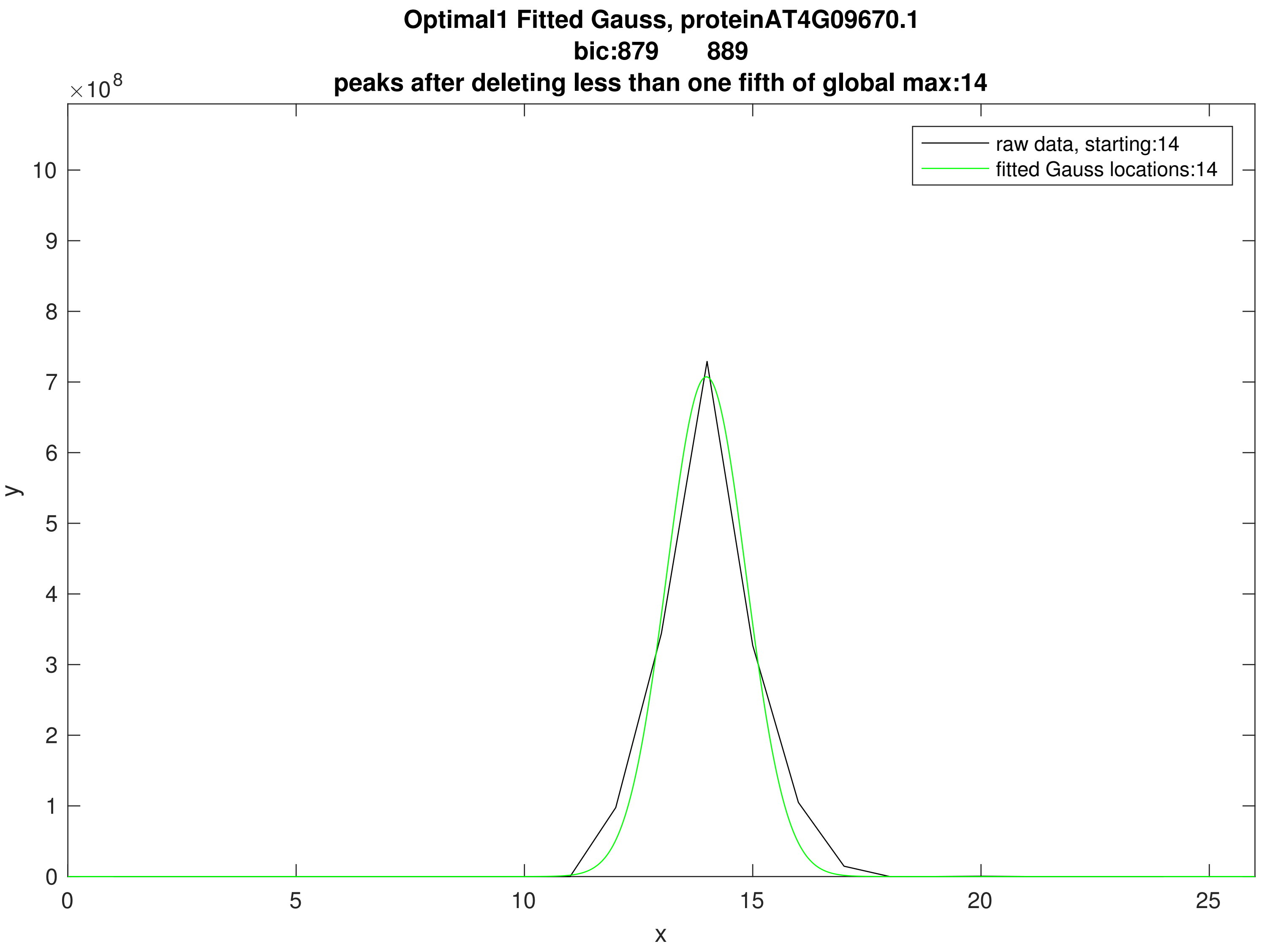
Optimal1 Fitted Gauss, proteinAT4G09670.1
bic:879 889
peaks after deleting less than one fifth of global max:14
×10^8
raw data, starting:14
fitted Gauss locations:14
y
x
0
1
2
3
4
5
6
7
8
9
10
0
5
10
15
20
25

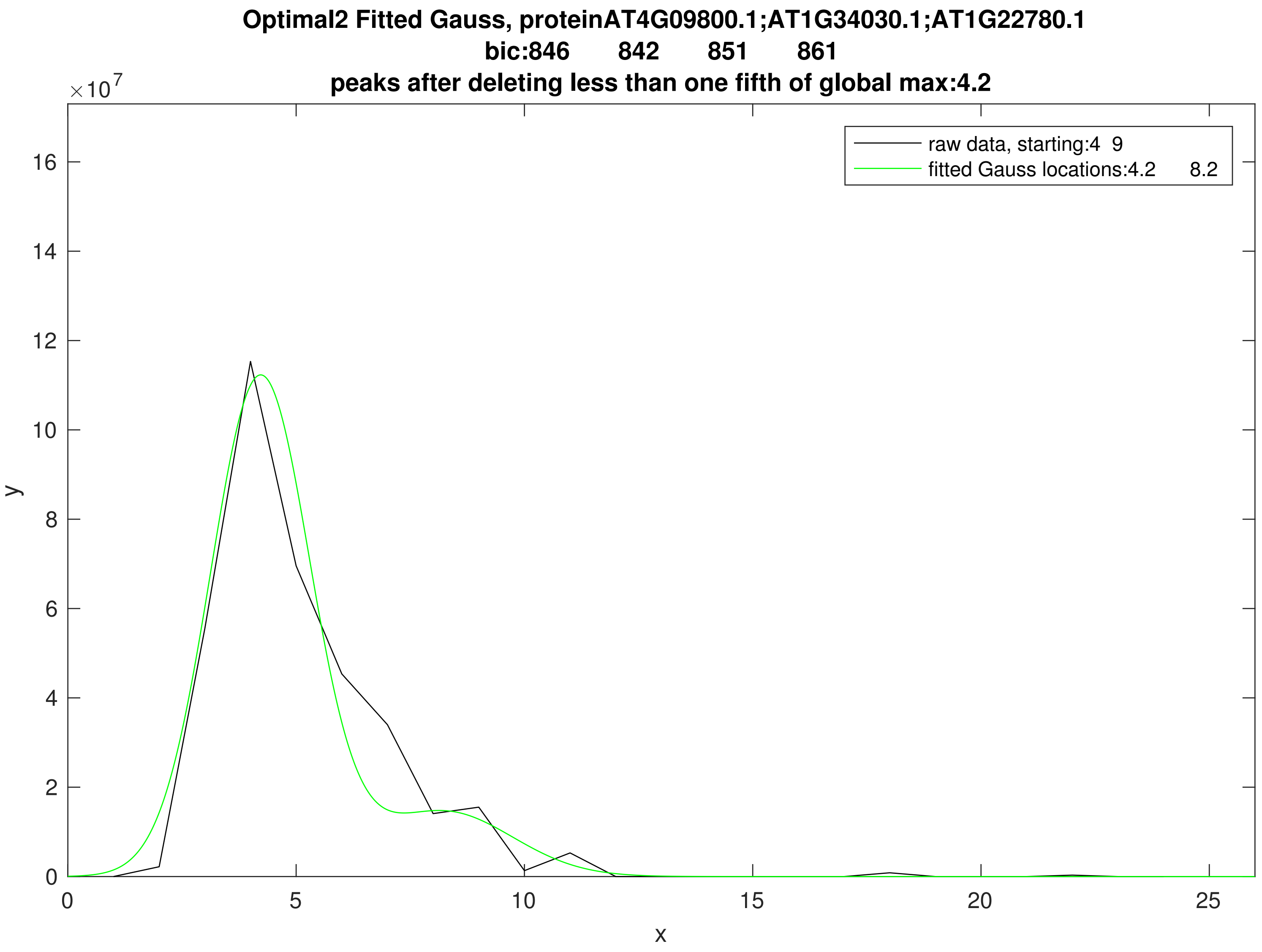
Optimal2 Fitted Gauss, proteinAT4G09800.1;AT1G34030.1;AT1G22780.1
bic:846 842 851 861
peaks after deleting less than one fifth of global max:4.2
×10^7
raw data, starting:4 9
fitted Gauss locations:4.2 8.2
y
x
0
2
4
6
8
10
12
14
16
0
5
10
15
20
25

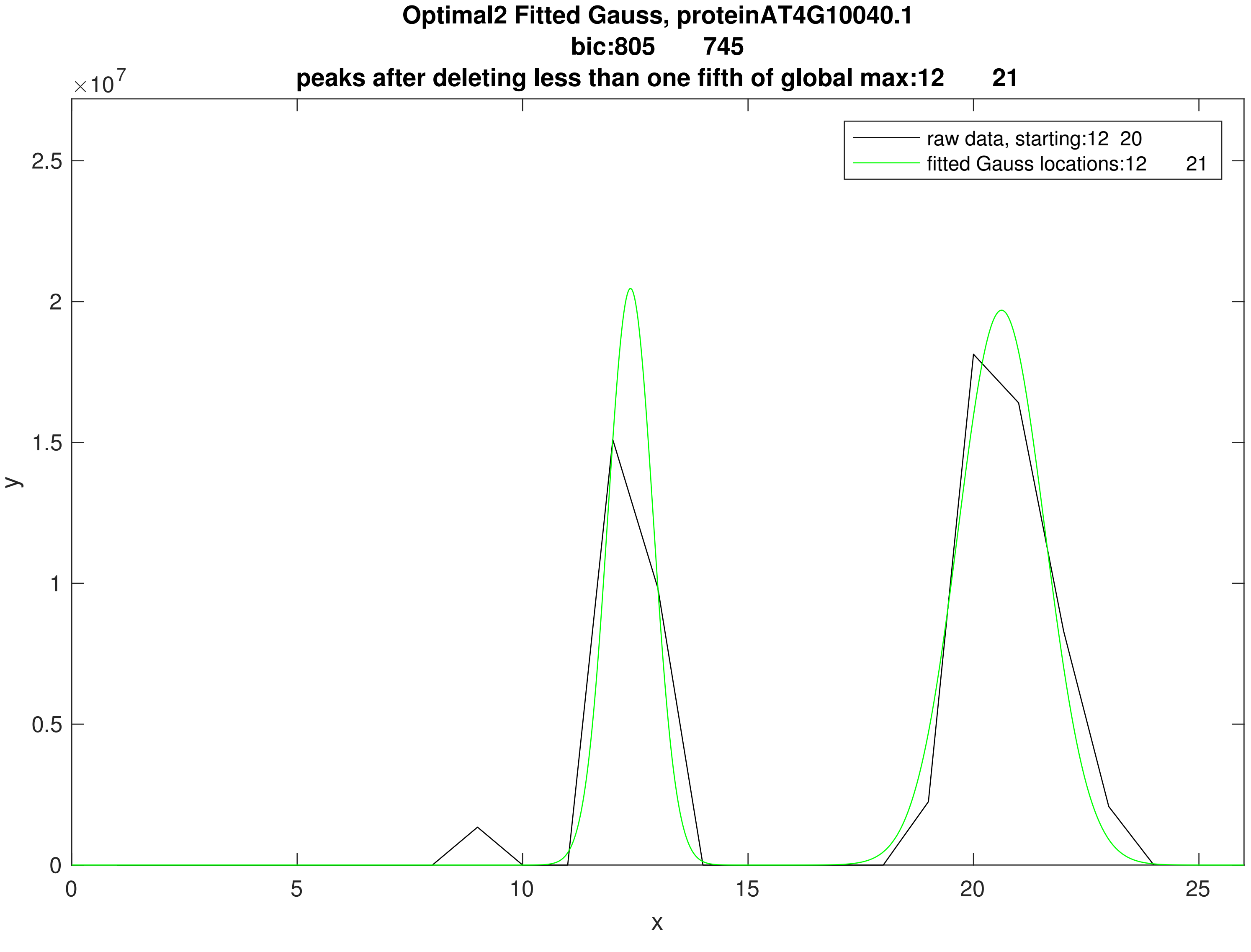
Optimal2 Fitted Gauss, proteinAT4G10040.1
bic:805 745
peaks after deleting less than one fifth of global max:12 21
×10^7
raw data, starting:12 20
fitted Gauss locations:12 21
2.5
2
1.5
1
0.5
0
y
0
5
10
15
20
25
x

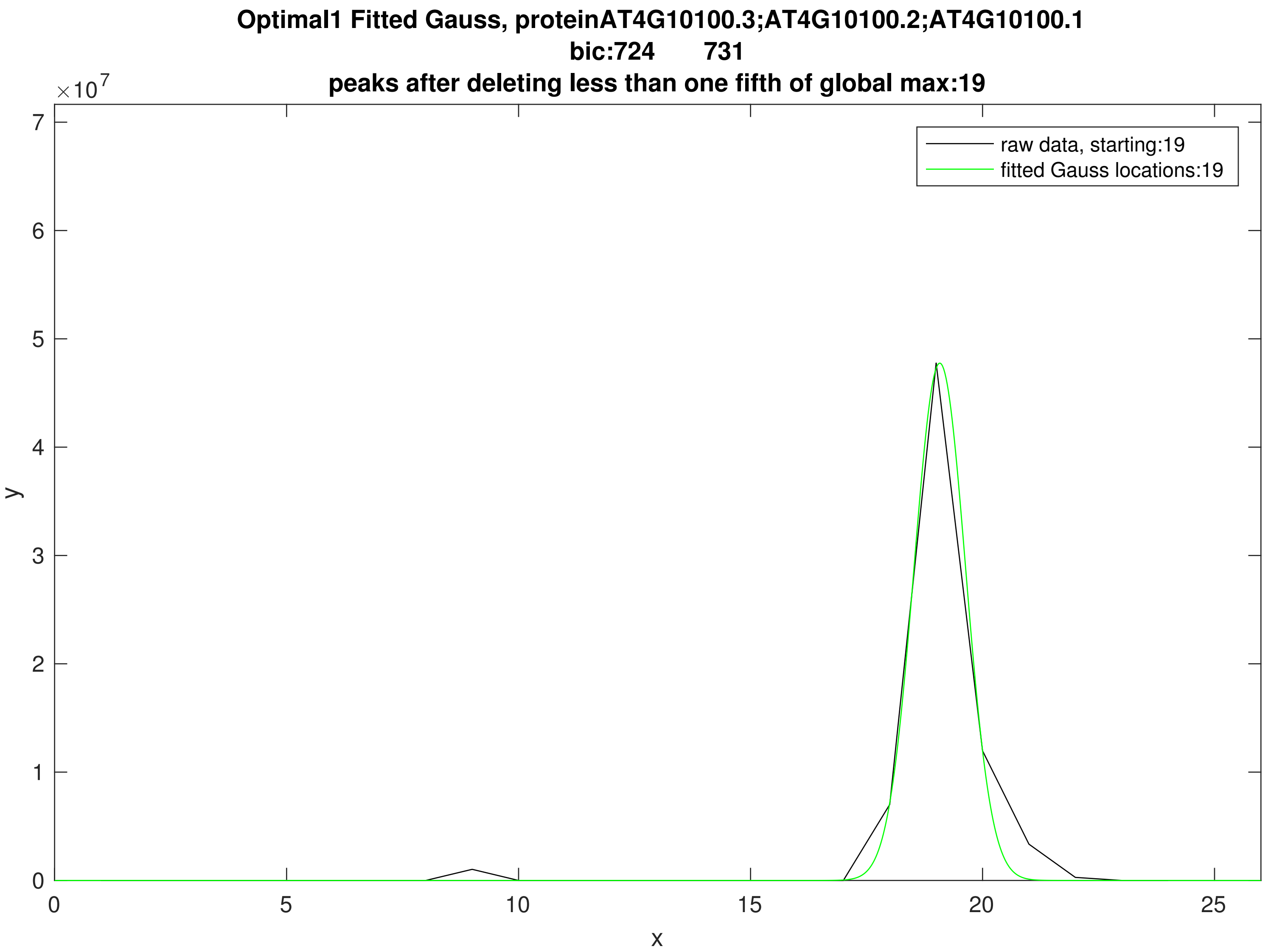
Optimal1 Fitted Gauss, proteinAT4G10100.3;AT4G10100.2;AT4G10100.1
bic:724 731
peaks after deleting less than one fifth of global max:19
×10^7
y
x
raw data, starting:19
fitted Gauss locations:19

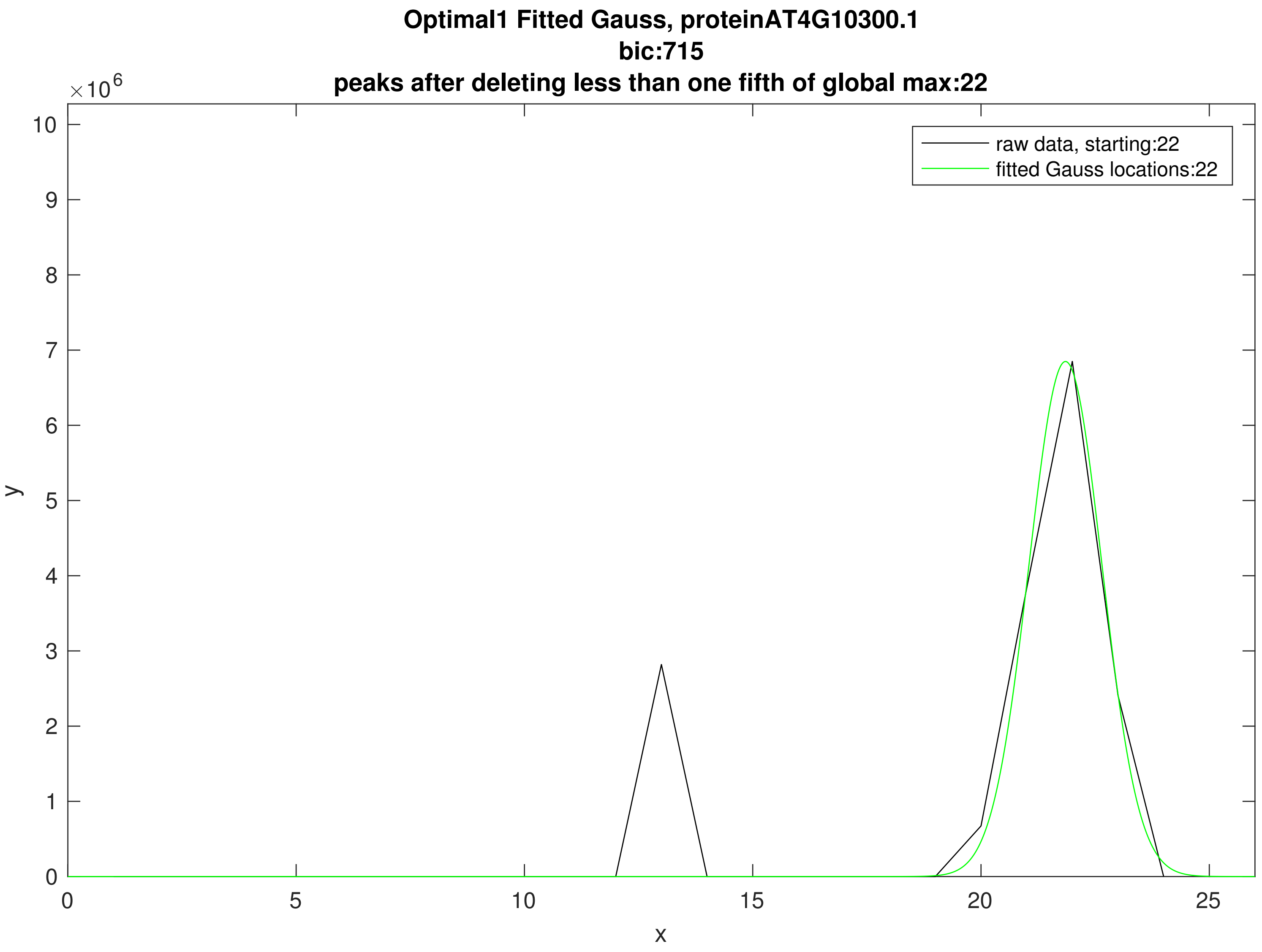

Optimal1 Fitted Gauss, proteinAT4G10300.1
bic:715
peaks after deleting less than one fifth of global max:22
×10^6
raw data, starting:22
fitted Gauss locations:22
y
x
0
1
2
3
4
5
6
7
8
9
10
0
5
10
15
20
25

**Optimal2 Fitted Gauss, proteinAT4G10320.1**
**bic:874 847**
**peaks after deleting less than one fifth of global max:5.2 9.6**

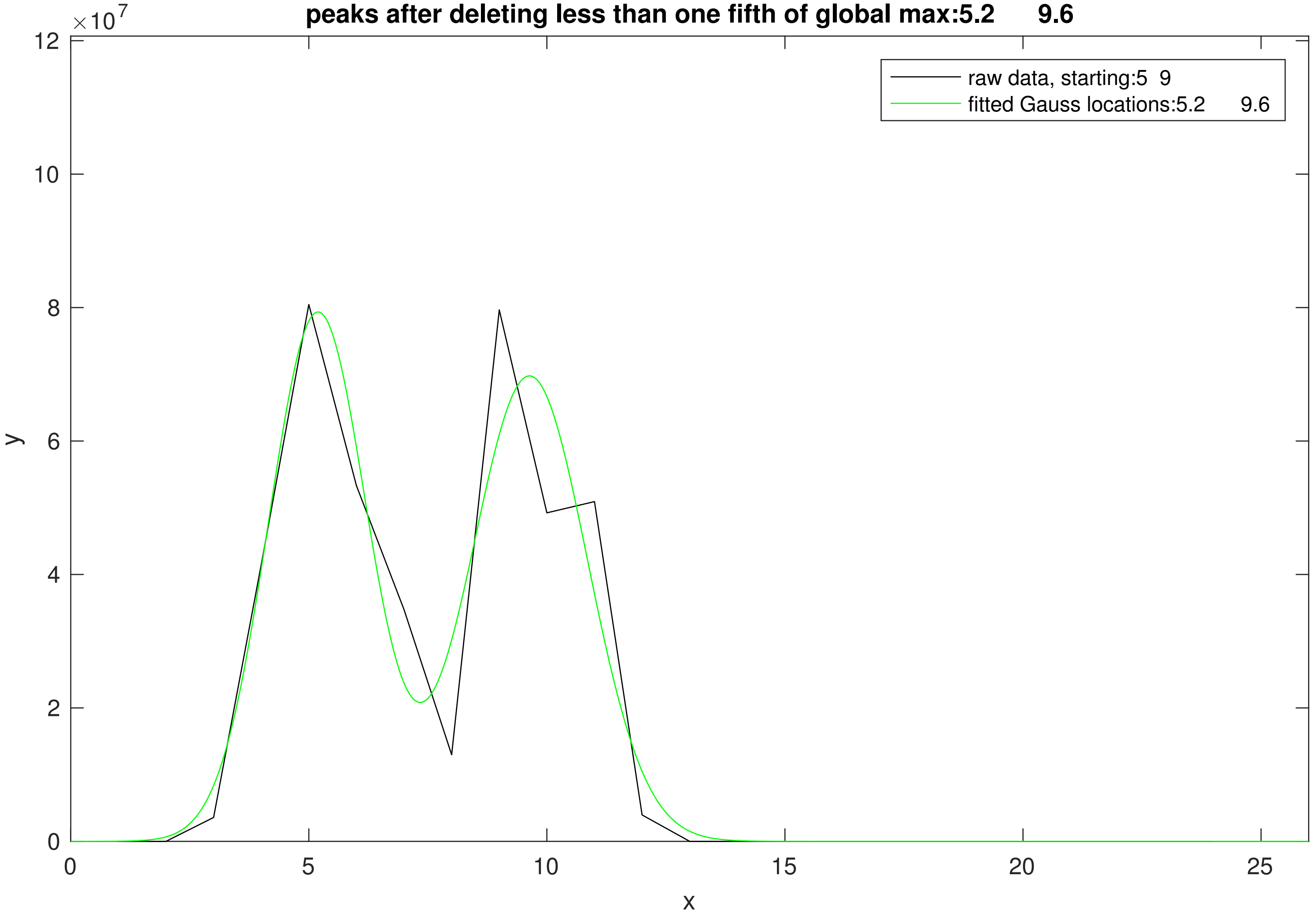

**Optimal1 Fitted Gauss, proteinAT4G10480.1;AT4G10480.2**
**bic:840 862**
**peaks after deleting less than one fifth of global max:17**

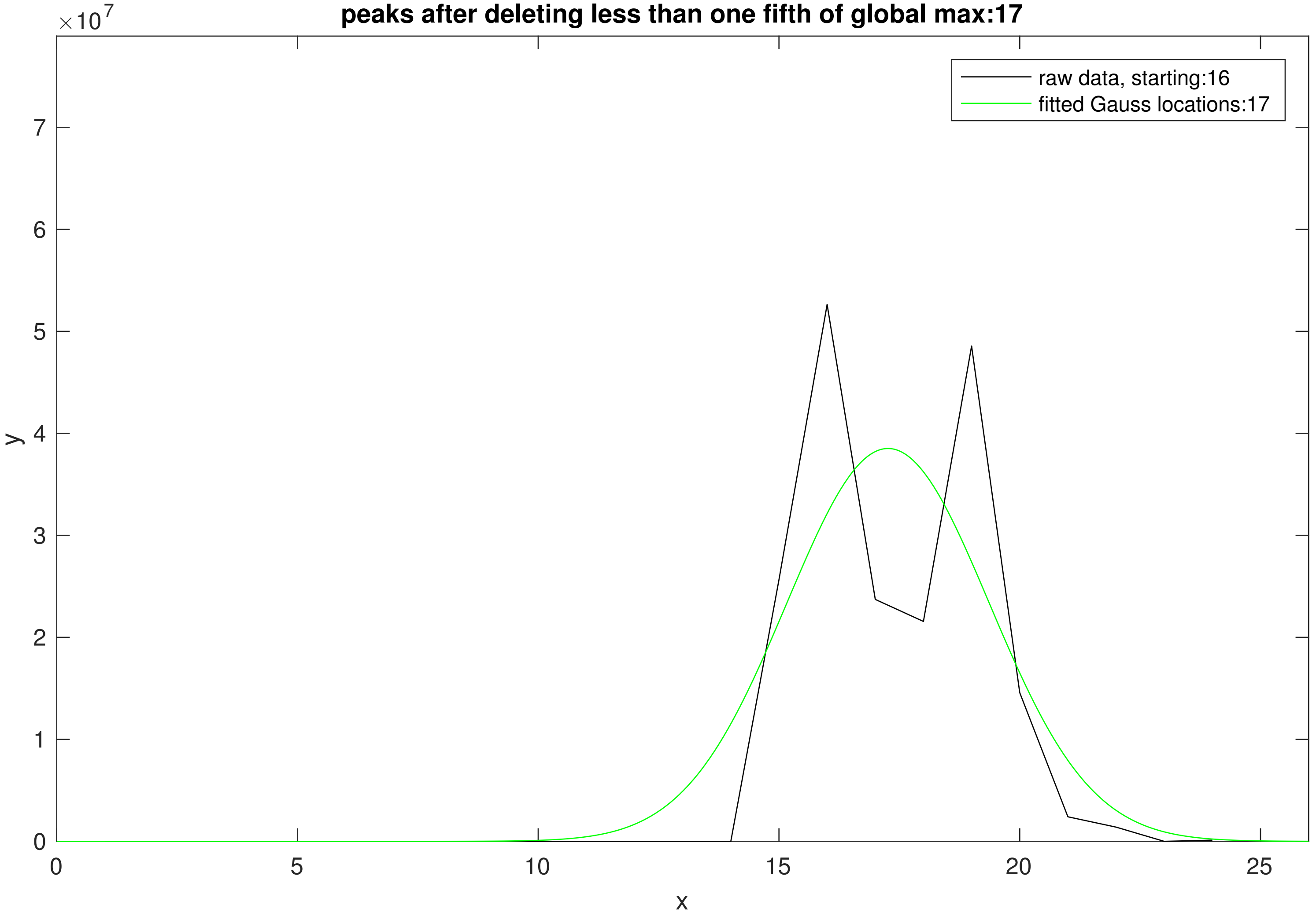

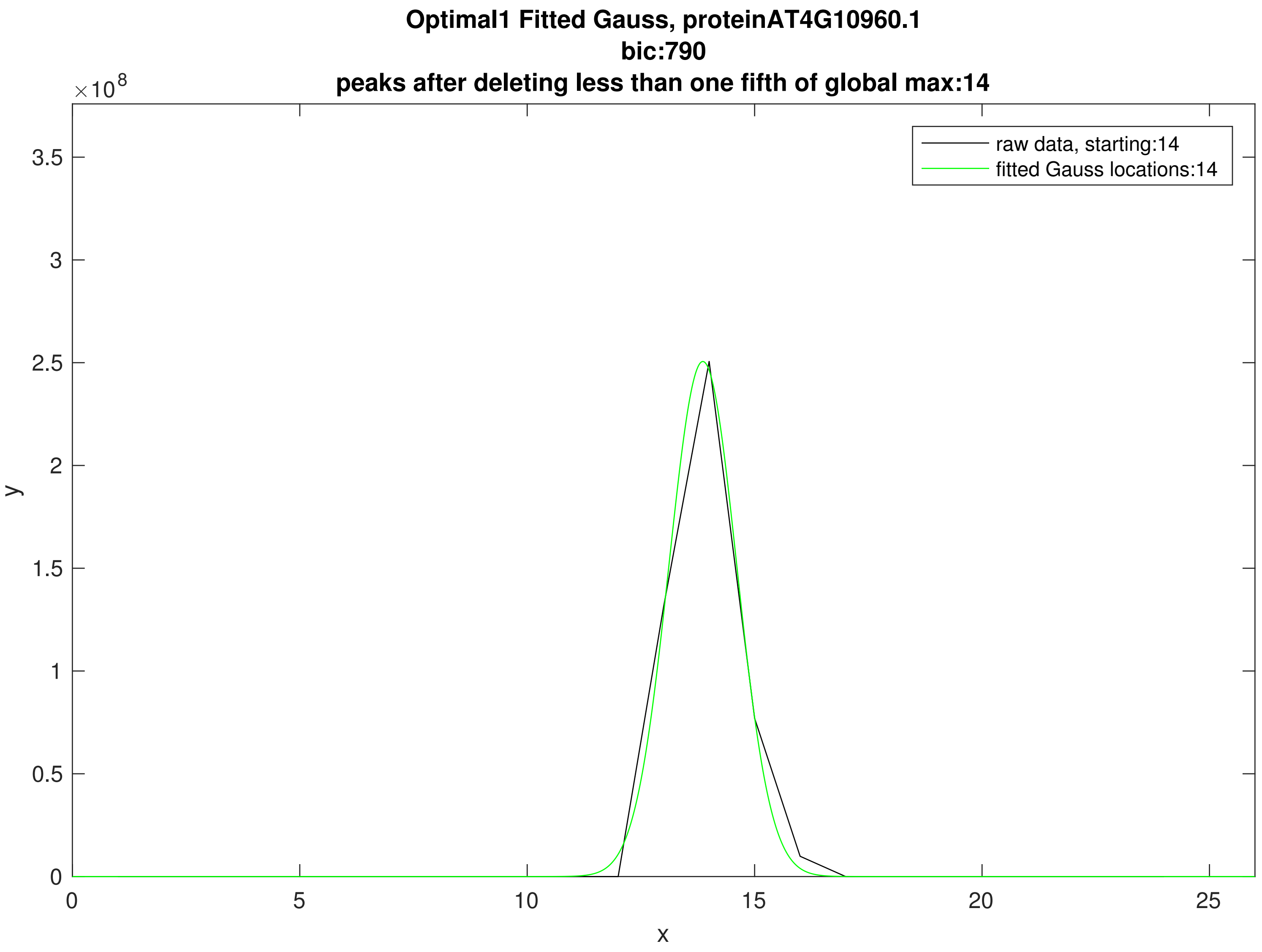
Optimal1 Fitted Gauss, proteinAT4G10960.1
bic:790
peaks after deleting less than one fifth of global max:14
×10^8
raw data, starting:14
fitted Gauss locations:14
3.5
3
2.5
2
1.5
1
0.5
0
y
0
5
10
15
20
25
x

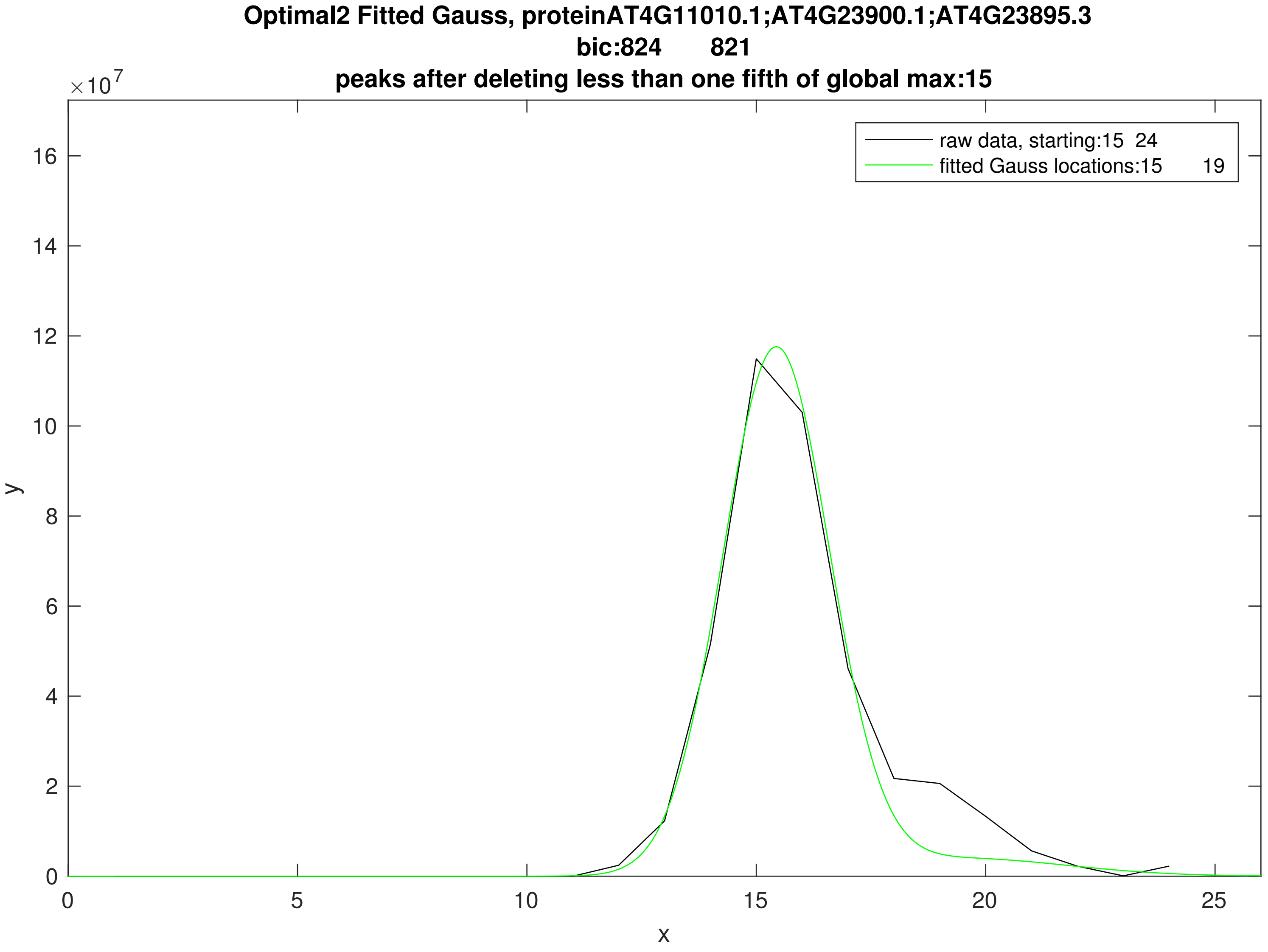
Optimal2 Fitted Gauss, proteinAT4G11010.1;AT4G23900.1;AT4G23895.3
bic:824 821
peaks after deleting less than one fifth of global max:15
×10^7
16
14
12
10
8
6
4
2
0
y
0
5
10
15
20
25
x
raw data, starting:15 24
fitted Gauss locations:15 19

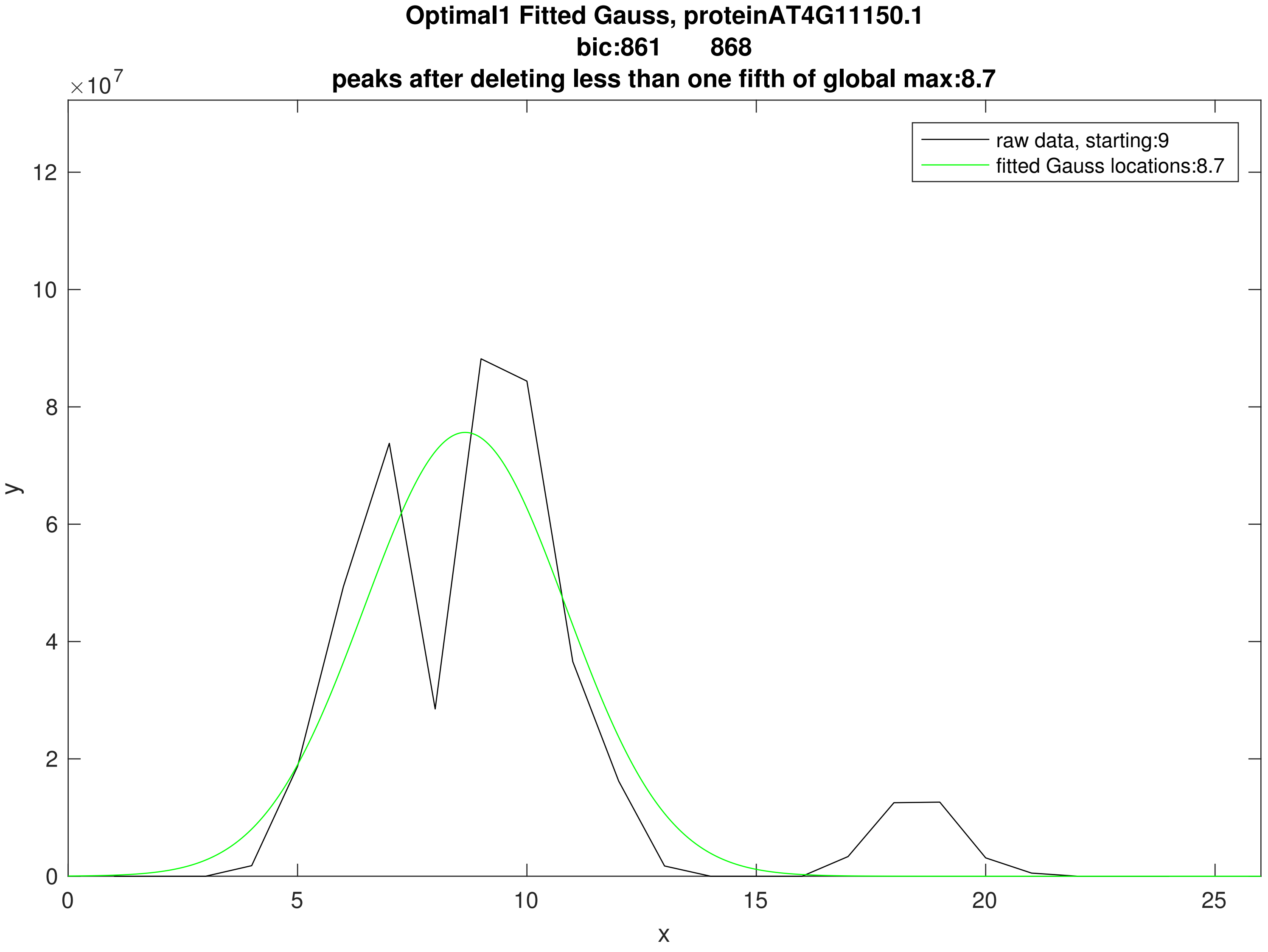

Optimal1 Fitted Gauss, proteinAT4G11150.1
bic:861 868
peaks after deleting less than one fifth of global max:8.7
×10^7
raw data, starting:9
fitted Gauss locations:8.7
y
x
0
2
4
6
8
10
12
0
5
10
15
20
25

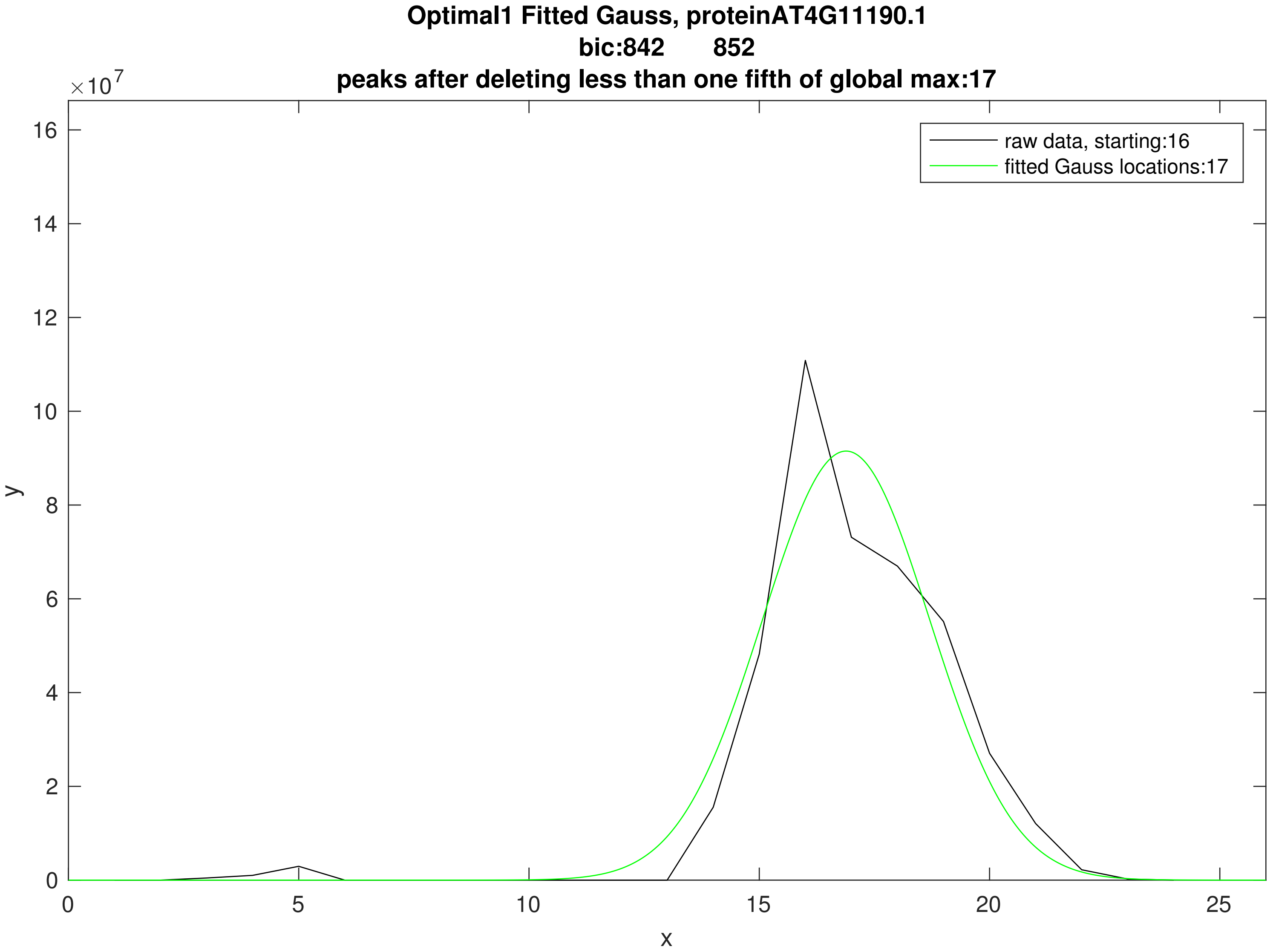

Optimal1 Fitted Gauss, proteinAT4G11190.1
bic:842 852
peaks after deleting less than one fifth of global max:17
×10^7
16
14
12
10
8
6
4
2
0
y
0
5
10
15
20
25
x
raw data, starting:16
fitted Gauss locations:17

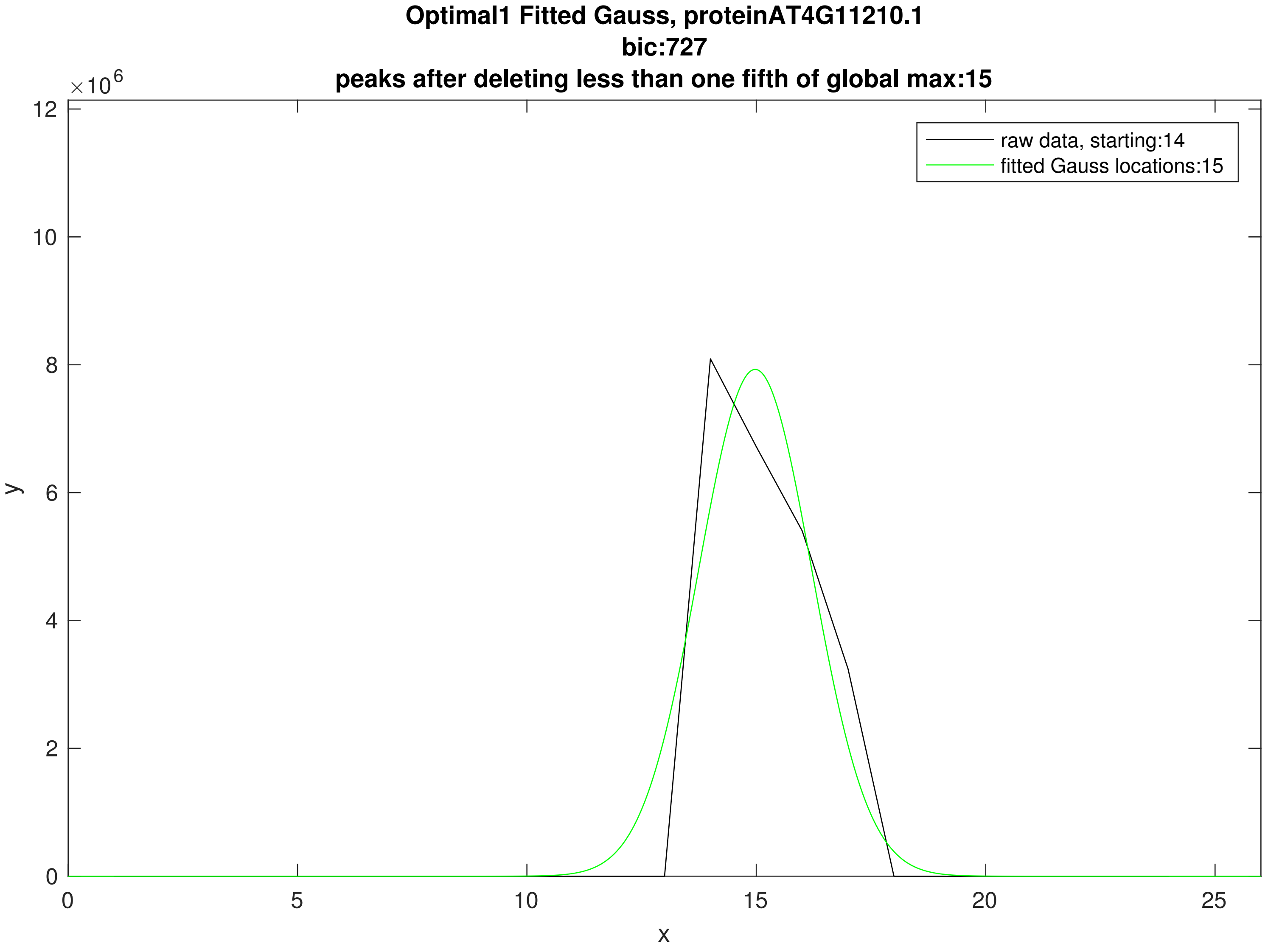
Optimal1 Fitted Gauss, proteinAT4G11210.1
bic:727
peaks after deleting less than one fifth of global max:15
×10^6
raw data, starting:14
fitted Gauss locations:15
y
x
0
2
4
6
8
10
12
0
5
10
15
20
25

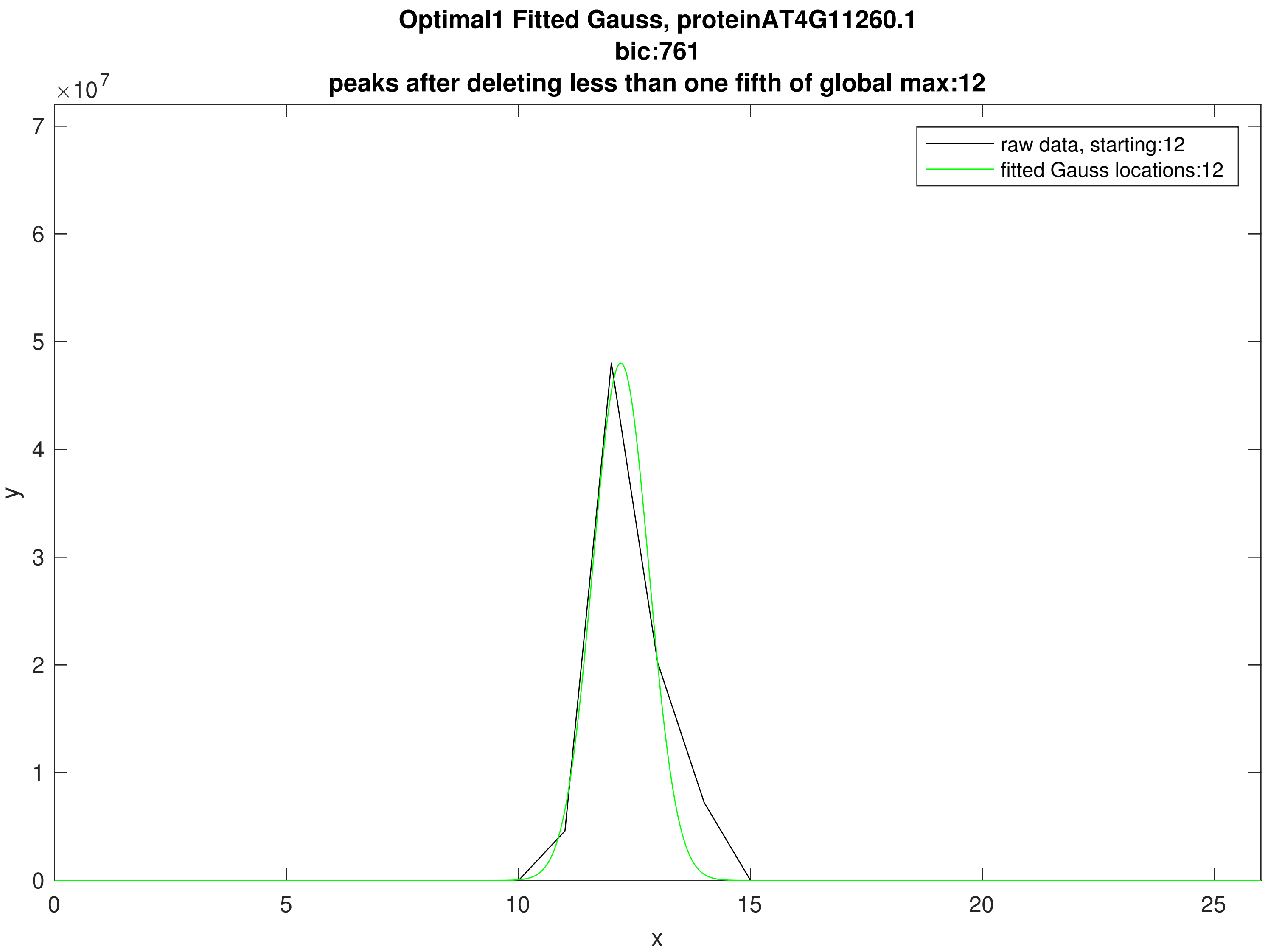
Optimal1 Fitted Gauss, proteinAT4G11260.1
bic:761
peaks after deleting less than one fifth of global max:12
×10^7
raw data, starting:12
fitted Gauss locations:12
y
x
0
1
2
3
4
5
6
7
0
5
10
15
20
25

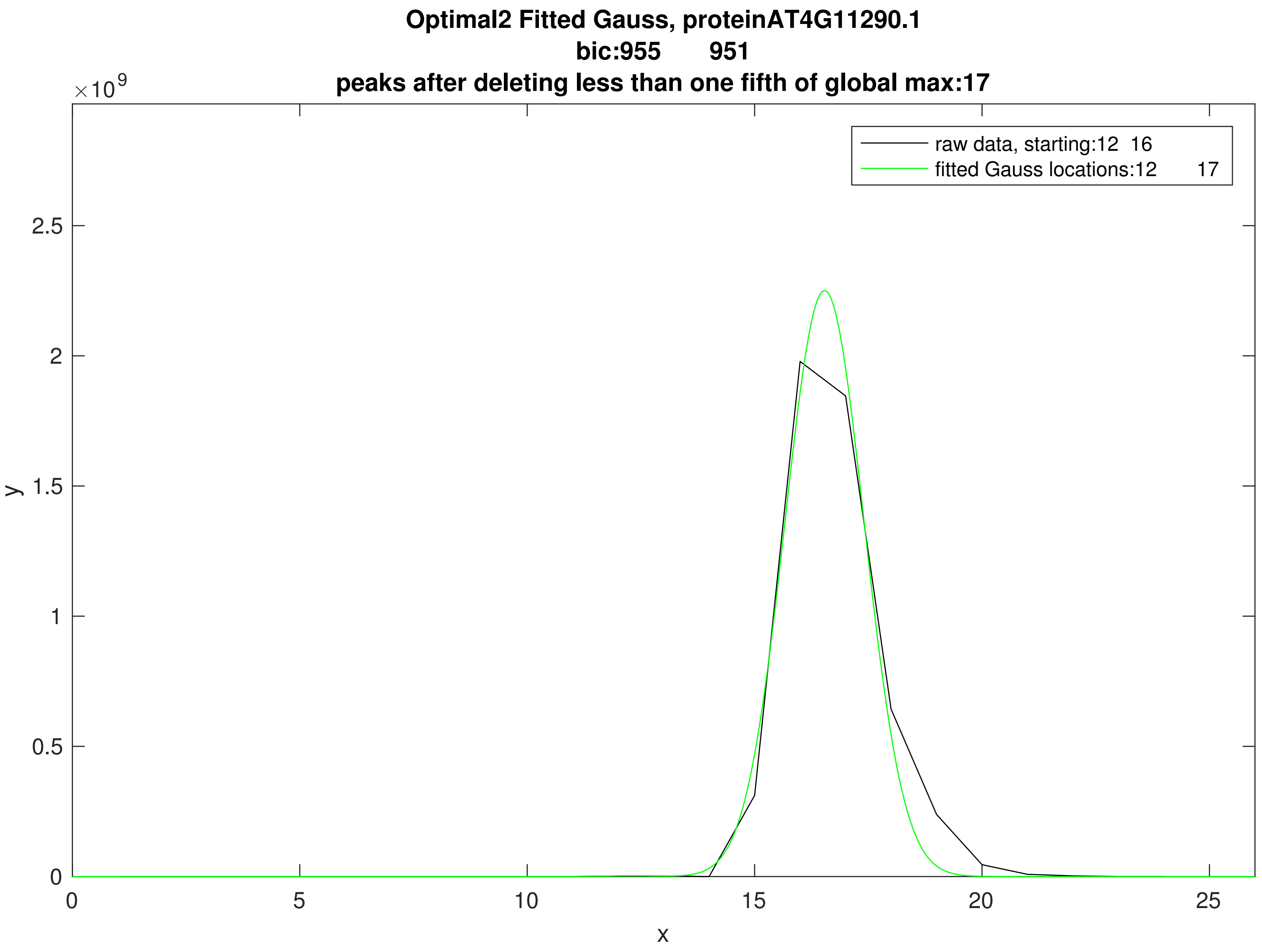

Optimal2 Fitted Gauss, proteinAT4G11290.1
bic:955 951
peaks after deleting less than one fifth of global max:17
×10^9
raw data, starting:12 16
fitted Gauss locations:12 17
2.5
2
1.5
1
0.5
0
y
0
5
10
15
20
25
x

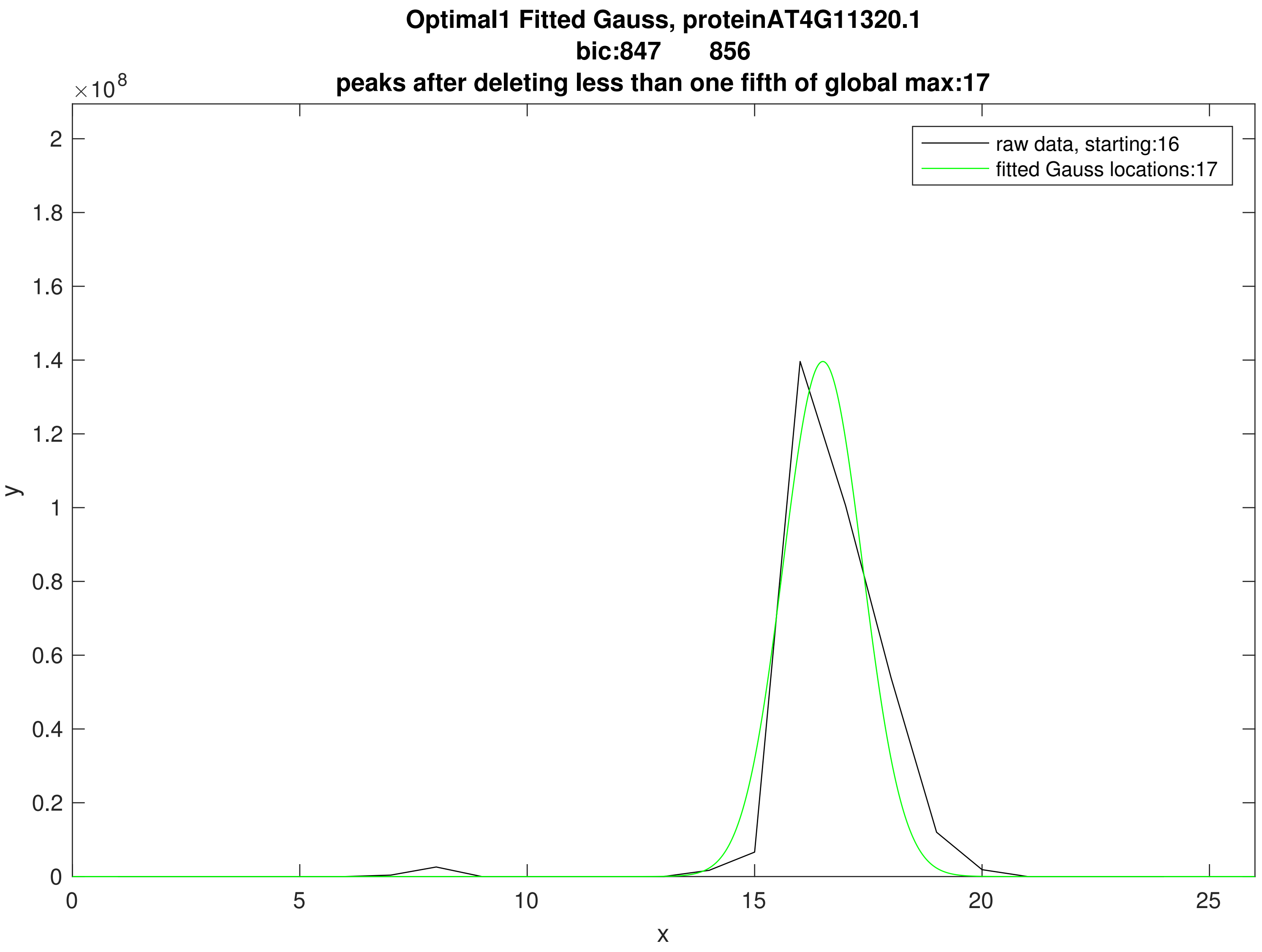

Optimal1 Fitted Gauss, proteinAT4G11320.1
bic:847 856
peaks after deleting less than one fifth of global max:17
×10^8
raw data, starting:16
fitted Gauss locations:17
y
x
0
0.2
0.4
0.6
0.8
1
1.2
1.4
1.6
1.8
2
0
5
10
15
20
25

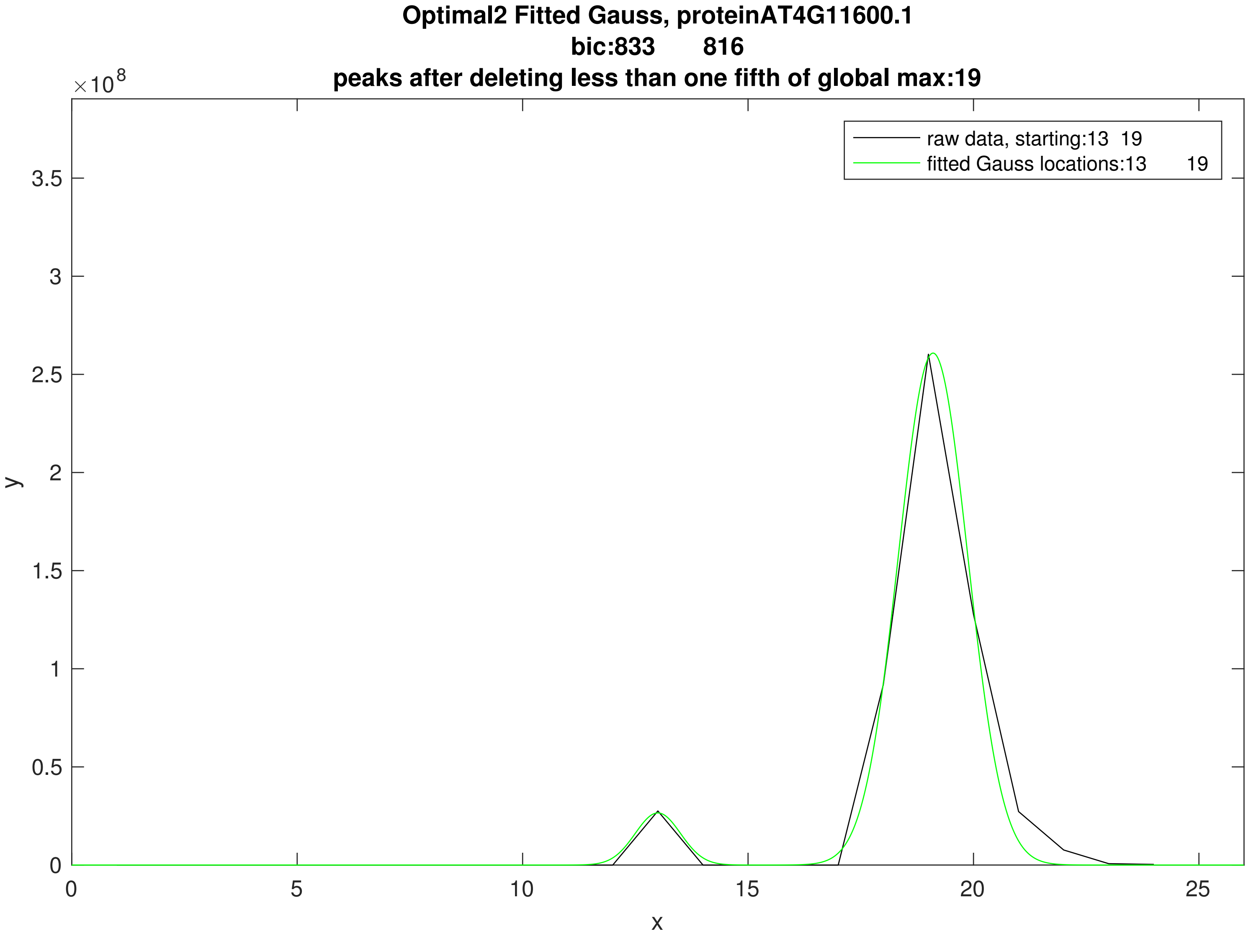
Optimal2 Fitted Gauss, proteinAT4G11600.1
bic:833 816
peaks after deleting less than one fifth of global max:19
×10^8
raw data, starting:13 19
fitted Gauss locations:13 19
y
x
0
0.5
1
1.5
2
2.5
3
3.5
5
10
15
20
25

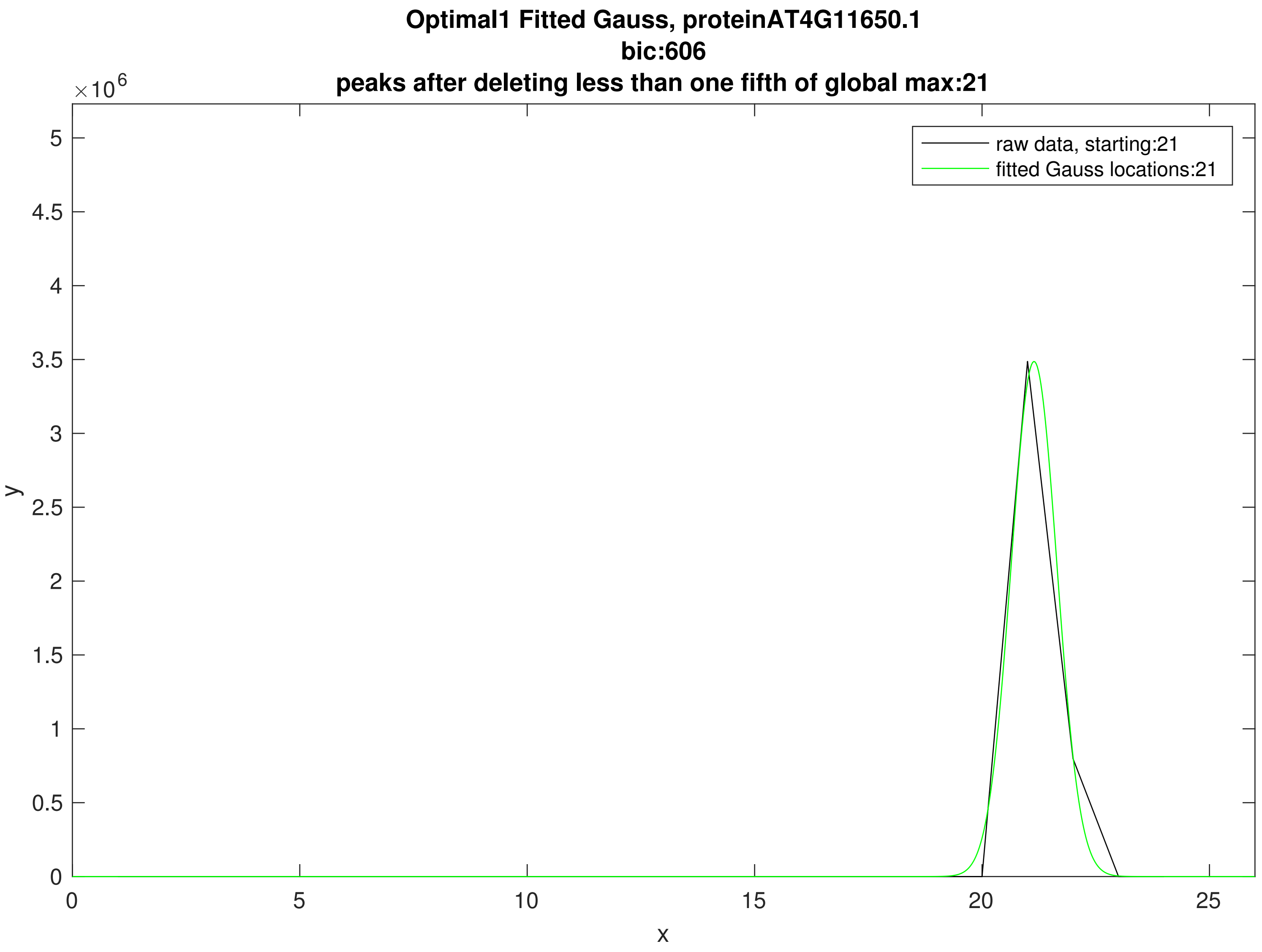
Optimal1 Fitted Gauss, proteinAT4G11650.1
bic:606
peaks after deleting less than one fifth of global max:21
×10^6
raw data, starting:21
fitted Gauss locations:21
y
x
0
0.5
1
1.5
2
2.5
3
3.5
4
4.5
5
0
5
10
15
20
25

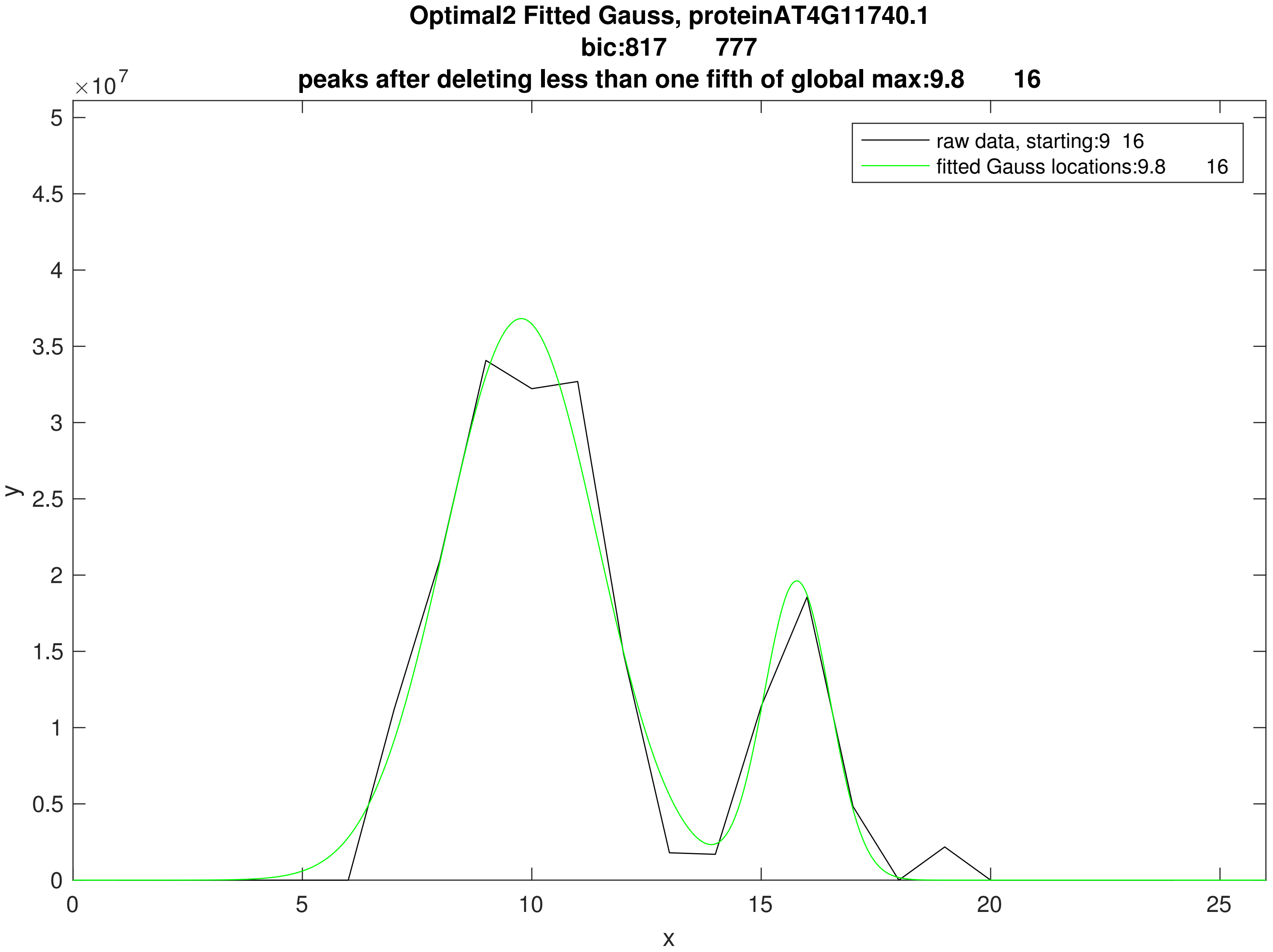

Optimal2 Fitted Gauss, proteinAT4G11740.1
bic:817 777
peaks after deleting less than one fifth of global max:9.8 16
×10^7
raw data, starting:9 16
fitted Gauss locations:9.8 16
y
x
0
0.5
1
1.5
2
2.5
3
3.5
4
4.5
5
0
5
10
15
20
25

**Optimal1 Fitted Gauss, proteinAT4G11820.2;AT4G11820.1**
**bic:973 979 989**
**peaks after deleting less than one fifth of global max:13**

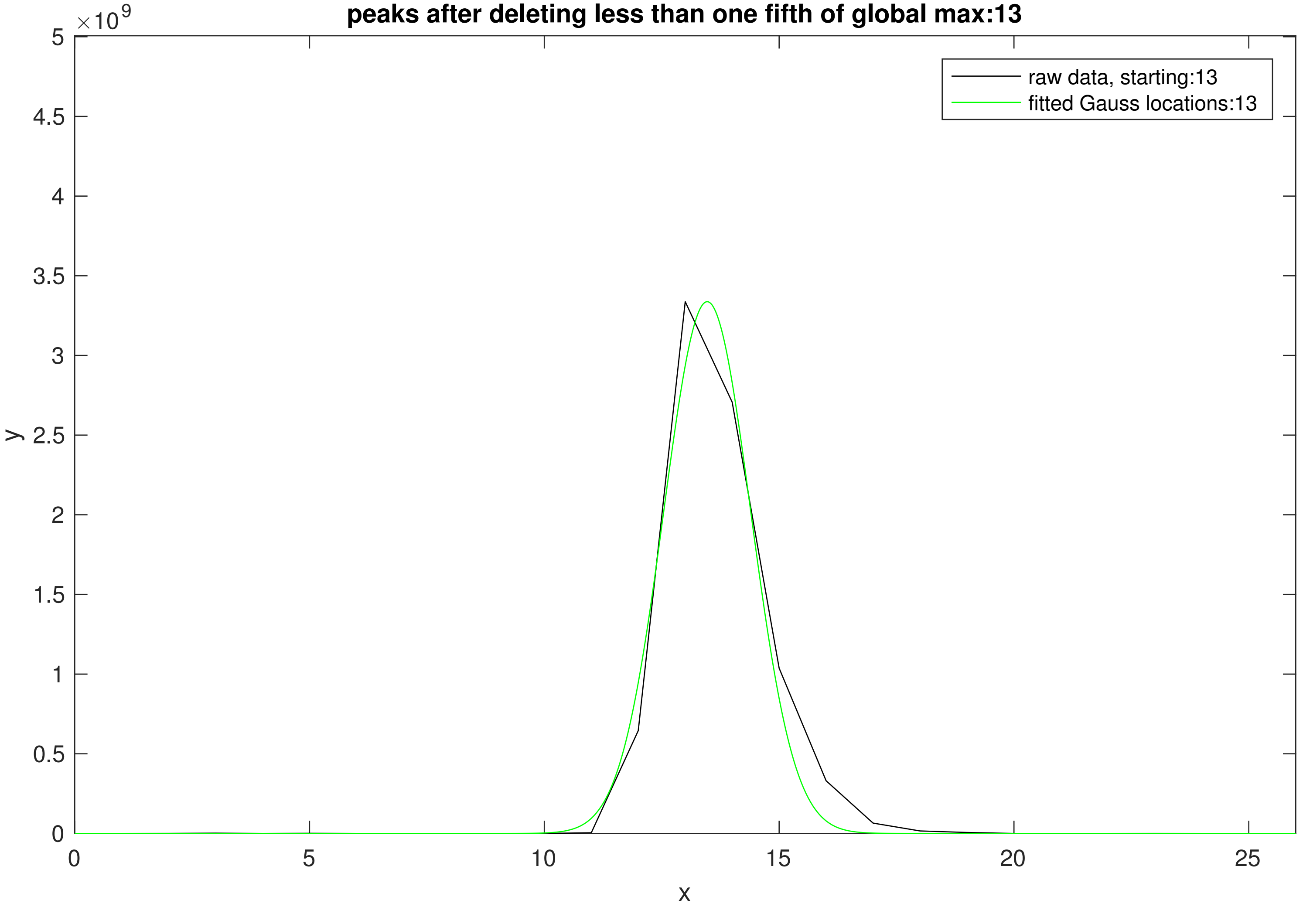

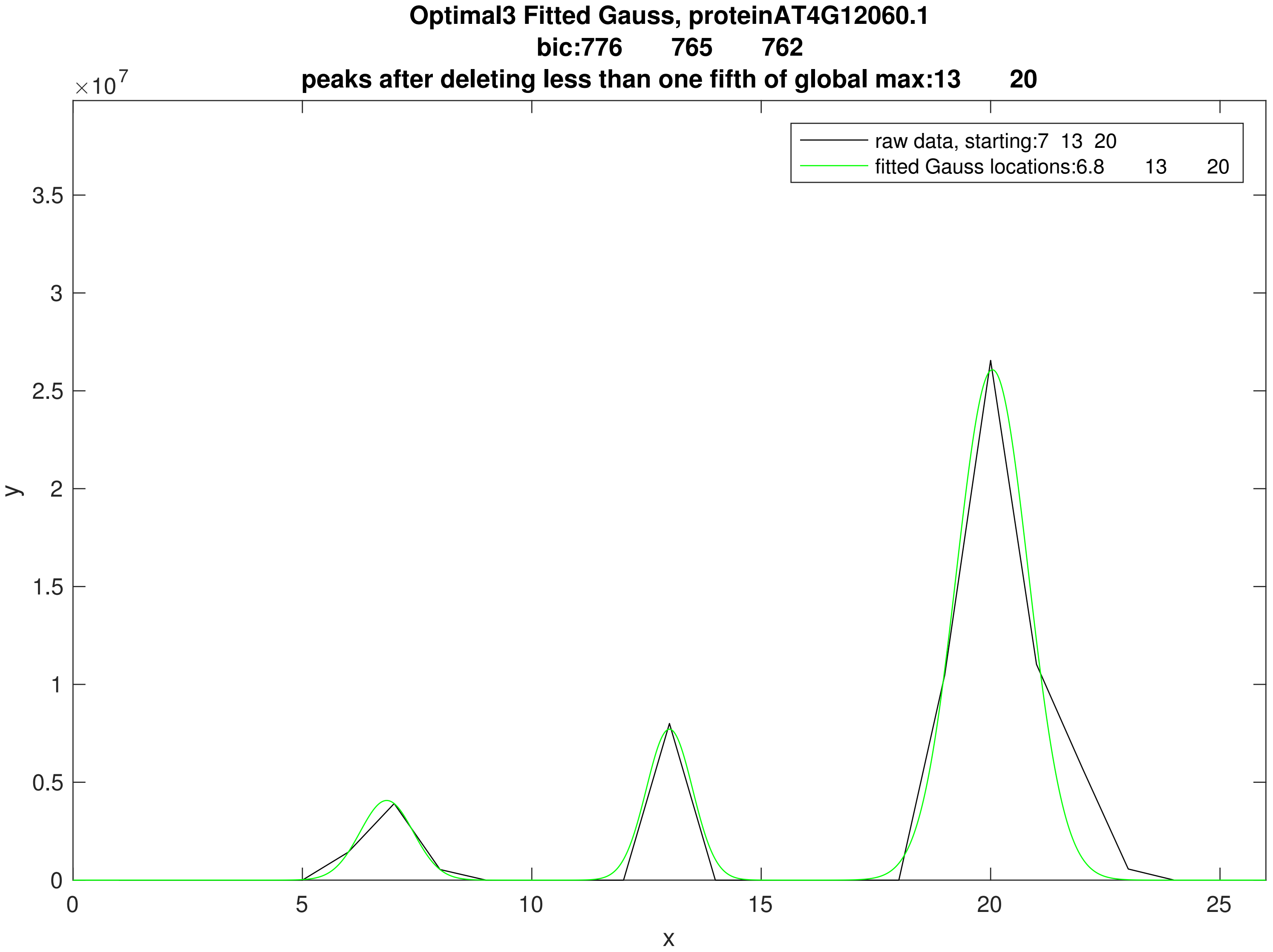
Optimal3 Fitted Gauss, proteinAT4G12060.1
bic:776 765 762
peaks after deleting less than one fifth of global max:13 20
×10^7
raw data, starting:7 13 20
fitted Gauss locations:6.8 13 20
3.5
3
2.5
2
1.5
1
0.5
0
y
0
5
10
15
20
25
x

#### Optimal1 Fitted Gauss, proteinAT4G12130.1
#### bic:741
#### peaks after deleting less than one fifth of global max:17

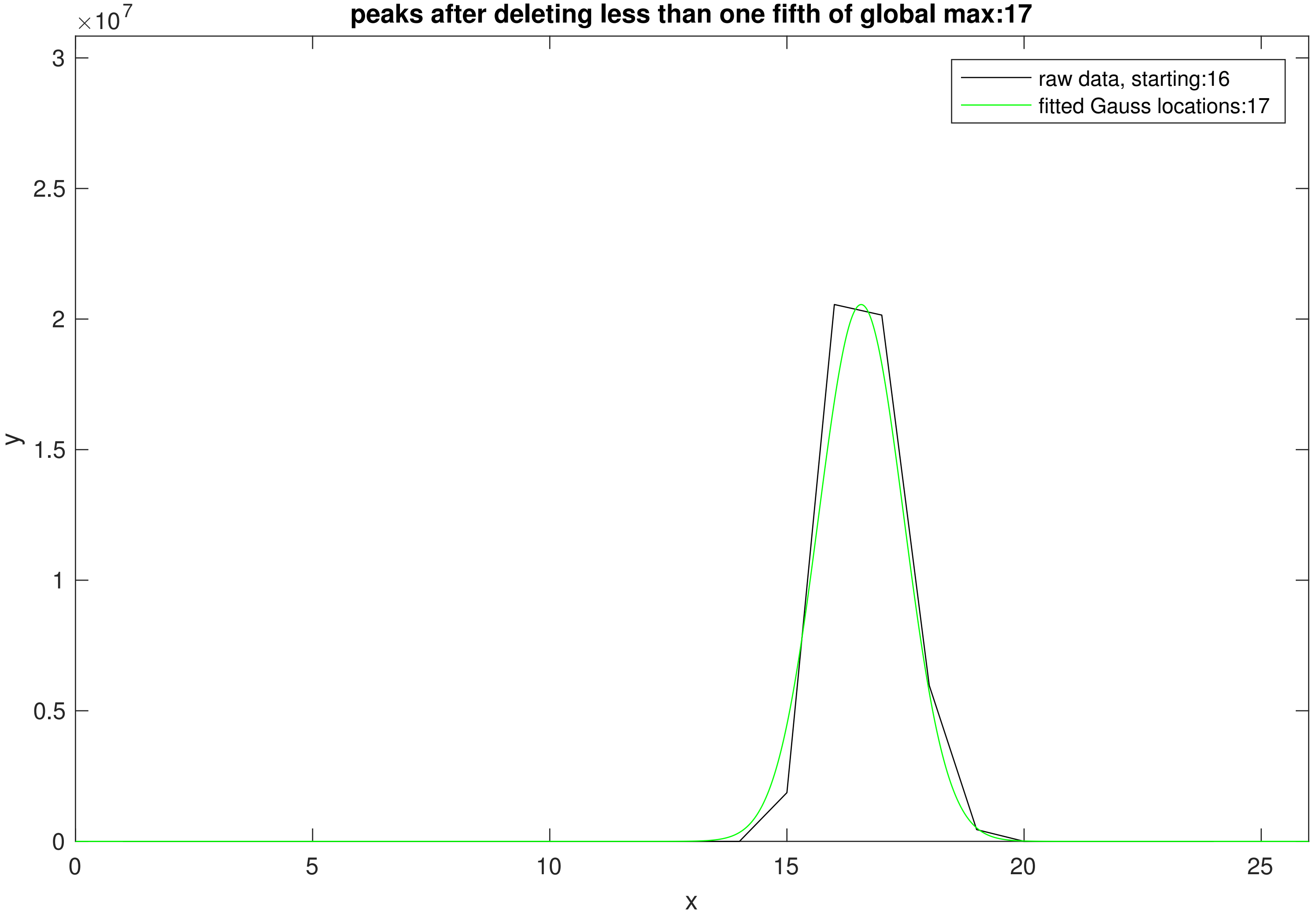

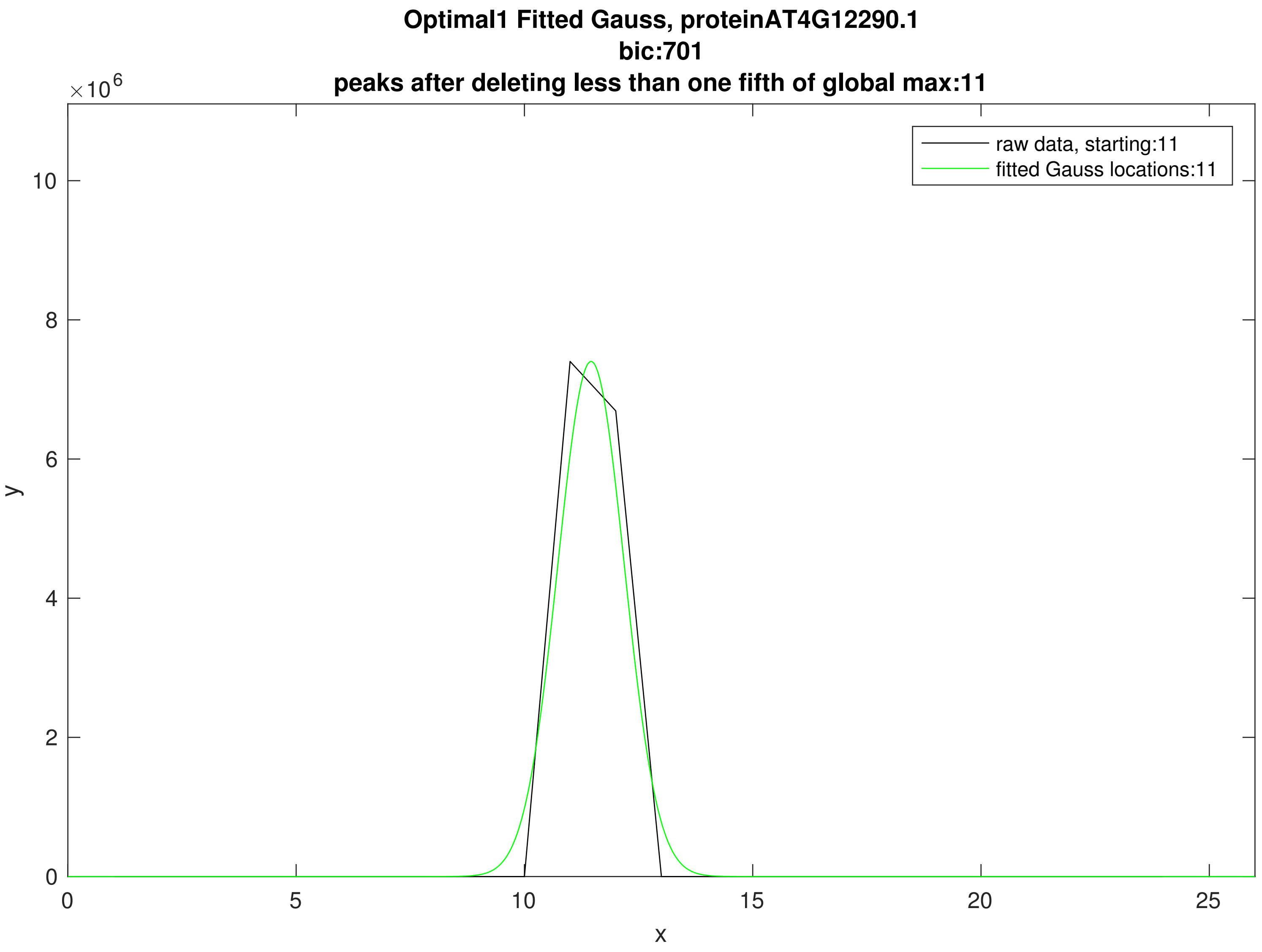

Optimal1 Fitted Gauss, proteinAT4G12290.1
bic:701
peaks after deleting less than one fifth of global max:11
×10^6
raw data, starting:11
fitted Gauss locations:11
y
x
0
2
4
6
8
10
0
5
10
15
20
25

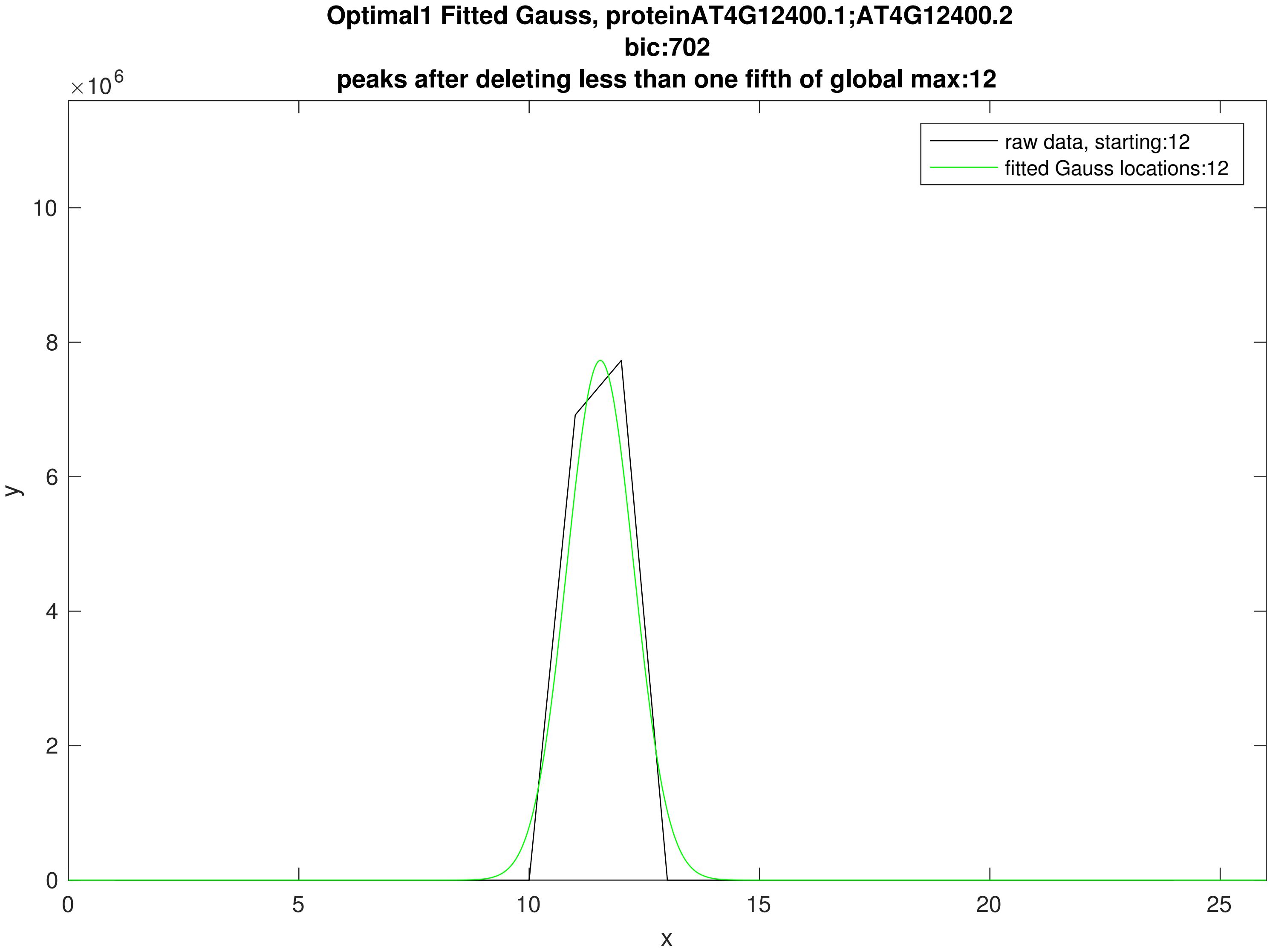
Optimal1 Fitted Gauss, proteinAT4G12400.1;AT4G12400.2
bic:702
peaks after deleting less than one fifth of global max:12
×10^6
raw data, starting:12
fitted Gauss locations:12
y
x
0
2
4
6
8
10
0
5
10
15
20
25

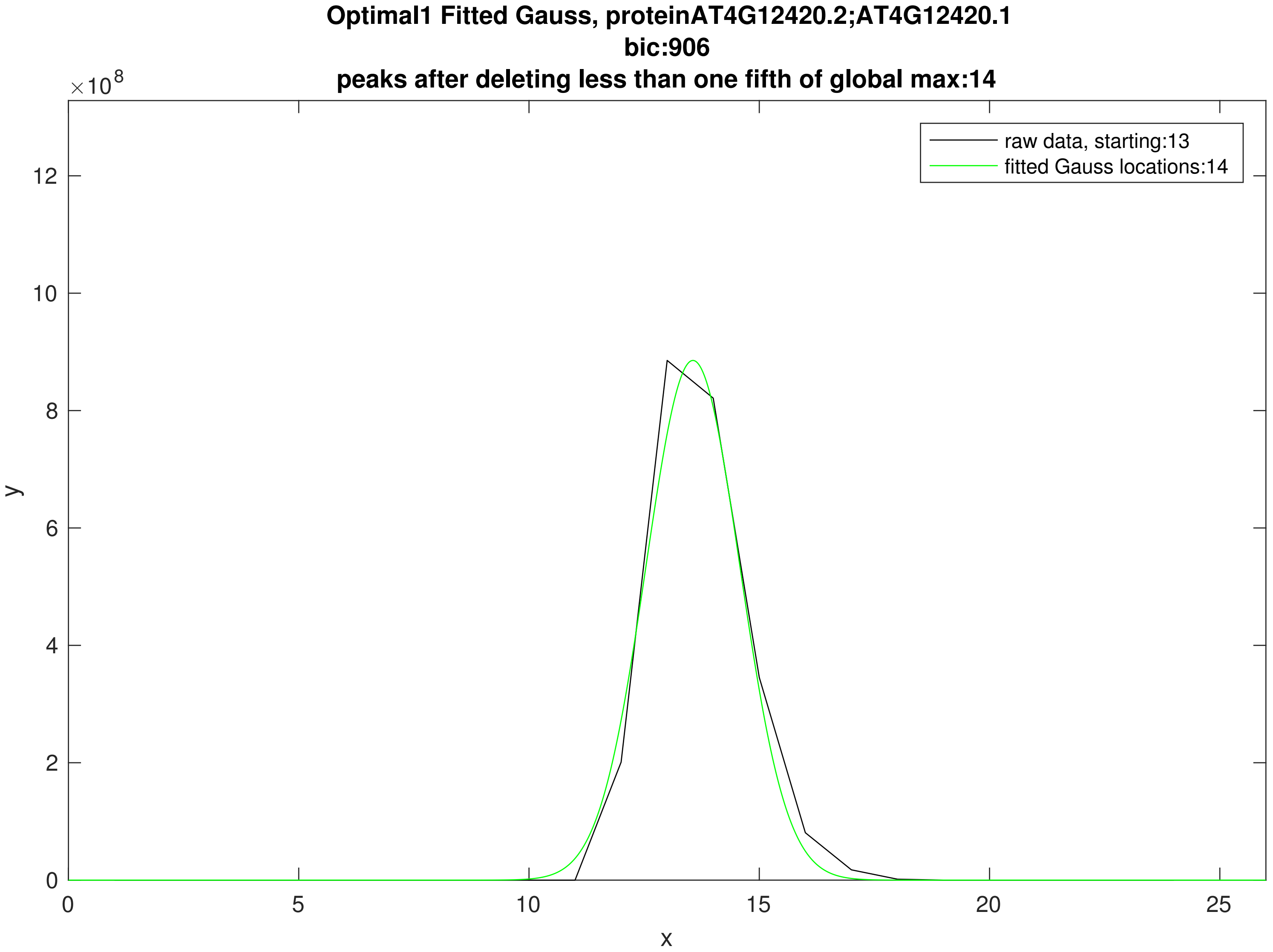
Optimal1 Fitted Gauss, proteinAT4G12420.2;AT4G12420.1
bic:906
peaks after deleting less than one fifth of global max:14
×10^8
raw data, starting:13
fitted Gauss locations:14
y
x
0
2
4
6
8
10
12
0
5
10
15
20
25

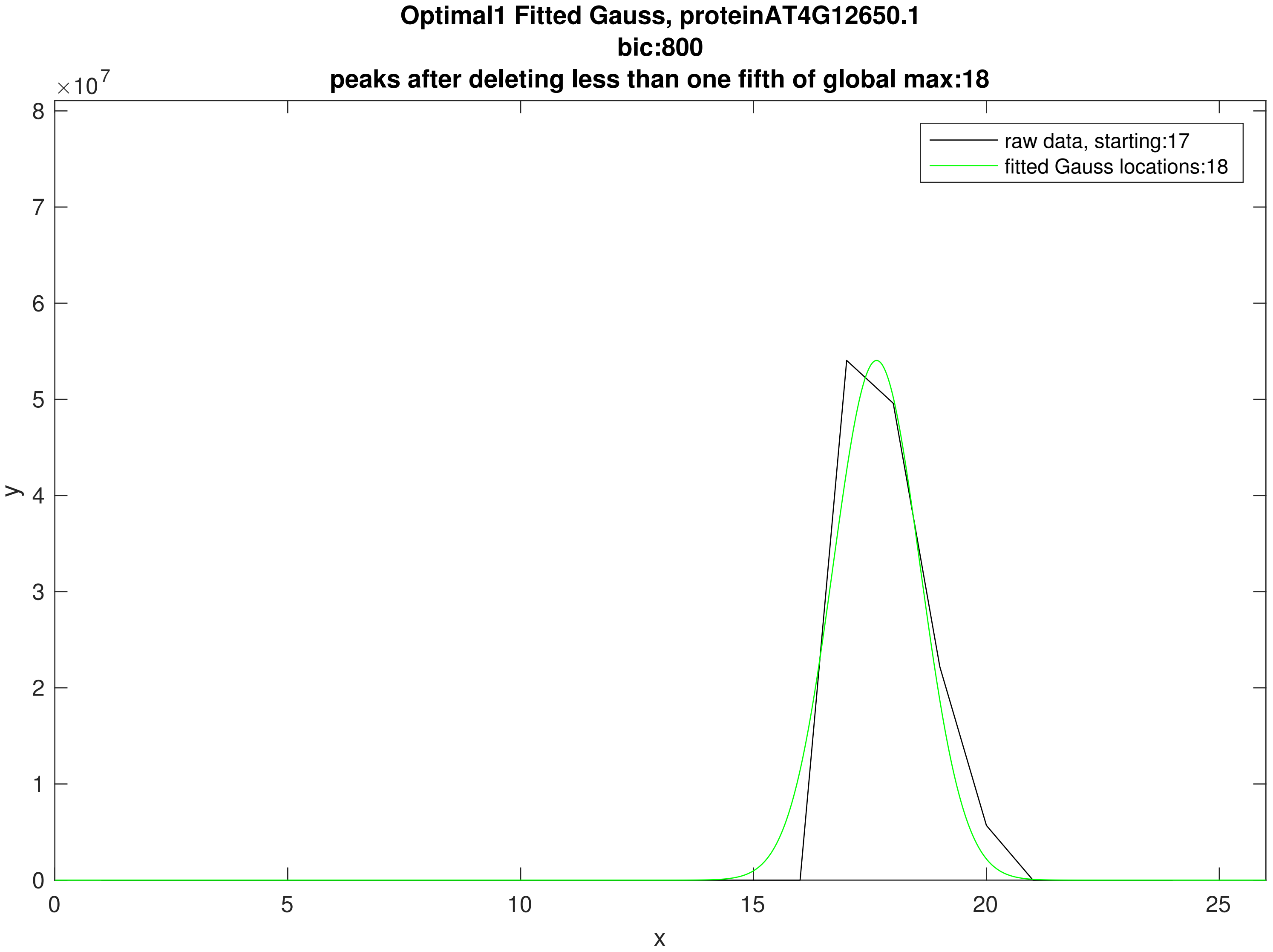

Optimal1 Fitted Gauss, proteinAT4G12650.1
bic:800
peaks after deleting less than one fifth of global max:18
×10^7
8
7
6
5
4
3
2
1
0
y
0
5
10
15
20
25
x
raw data, starting:17
fitted Gauss locations:18

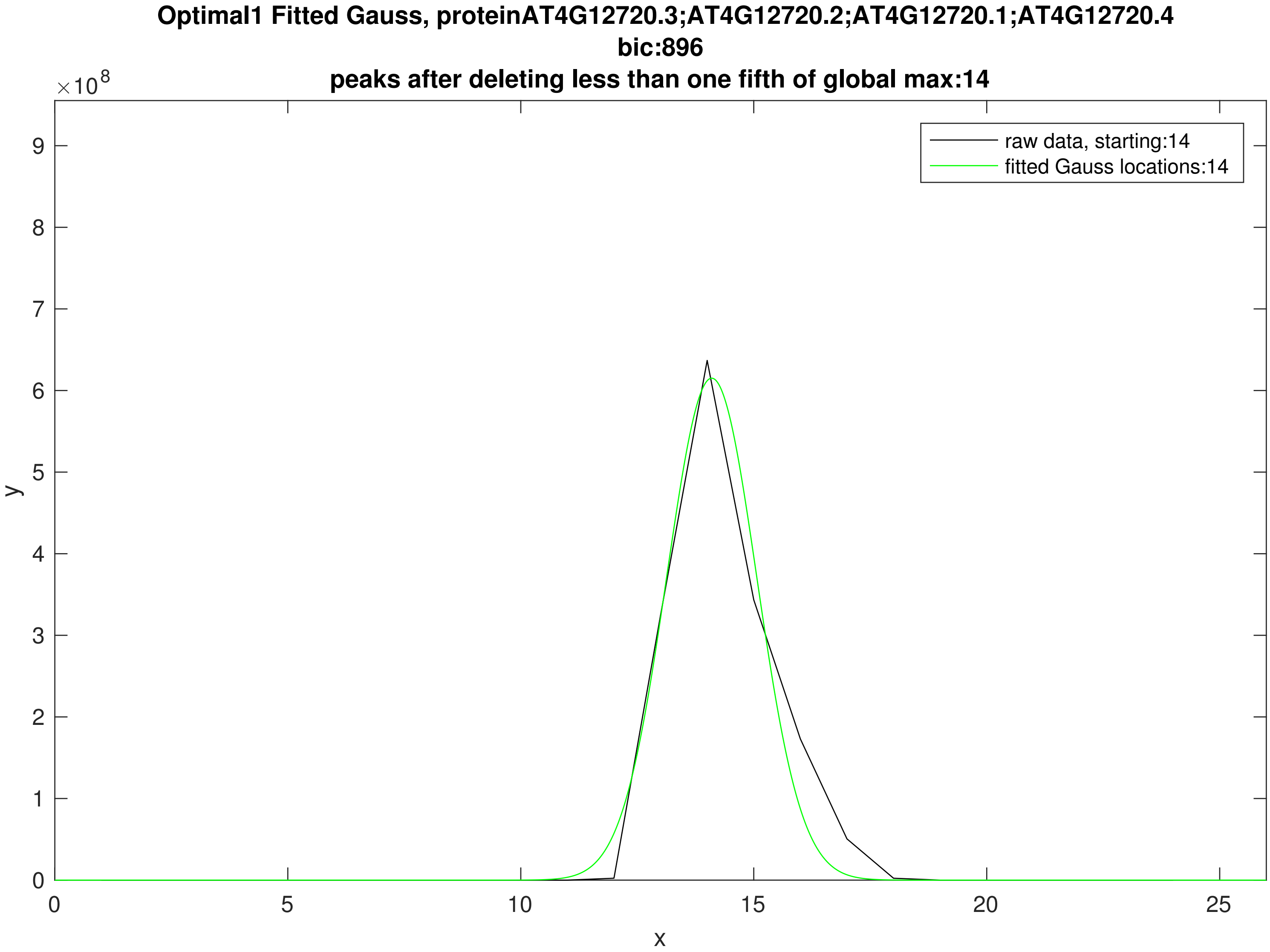
Optimal1 Fitted Gauss, proteinAT4G12720.3;AT4G12720.2;AT4G12720.1;AT4G12720.4
bic:896
peaks after deleting less than one fifth of global max:14
×10^8
raw data, starting:14
fitted Gauss locations:14
y
x
0
1
2
3
4
5
6
7
8
9
0
5
10
15
20
25

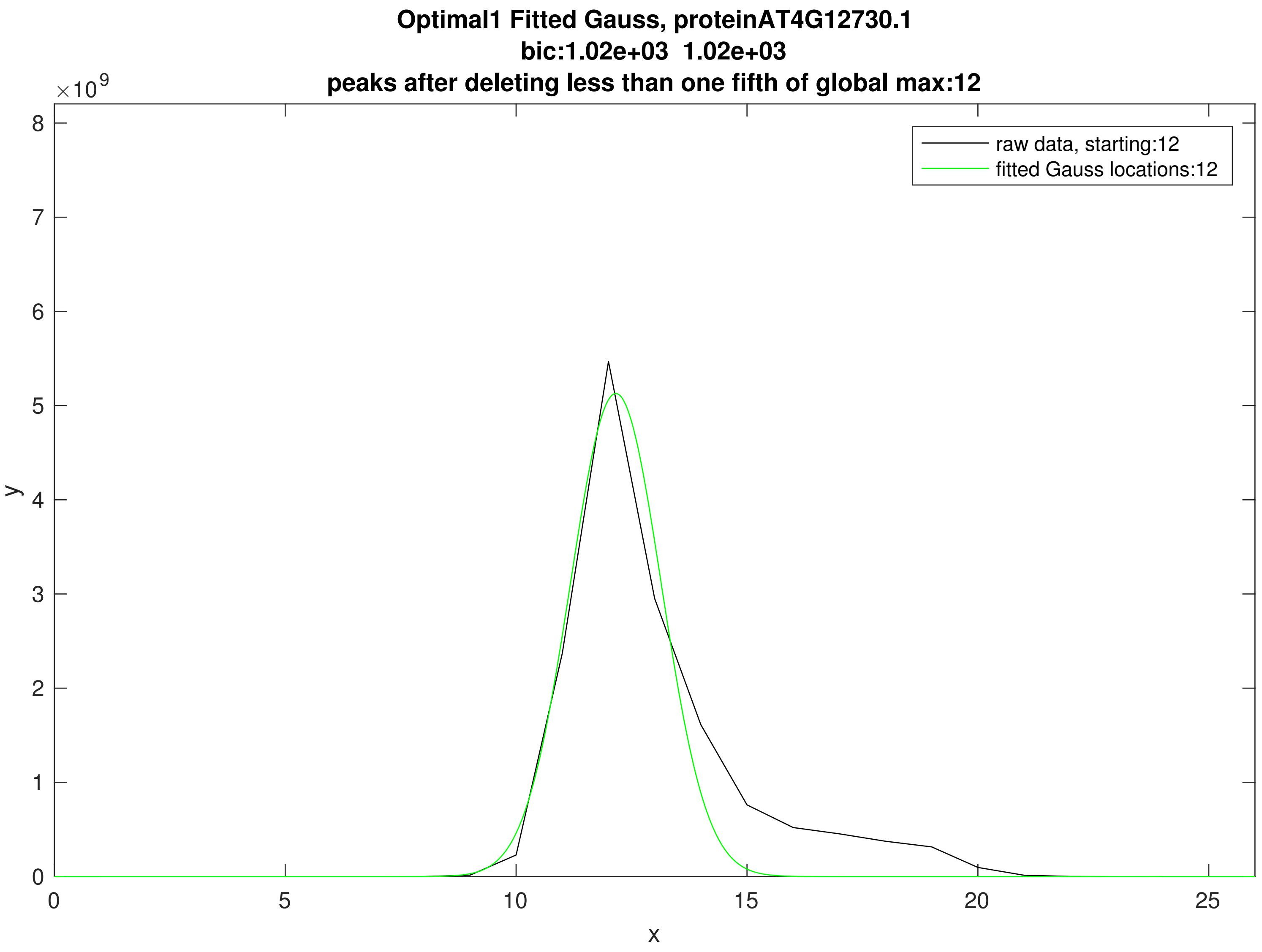
Optimal1 Fitted Gauss, proteinAT4G12730.1
bic:1.02e+03  1.02e+03
peaks after deleting less than one fifth of global max:12
×10^9
raw data, starting:12
fitted Gauss locations:12
y
x
0
1
2
3
4
5
6
7
8
0
5
10
15
20
25

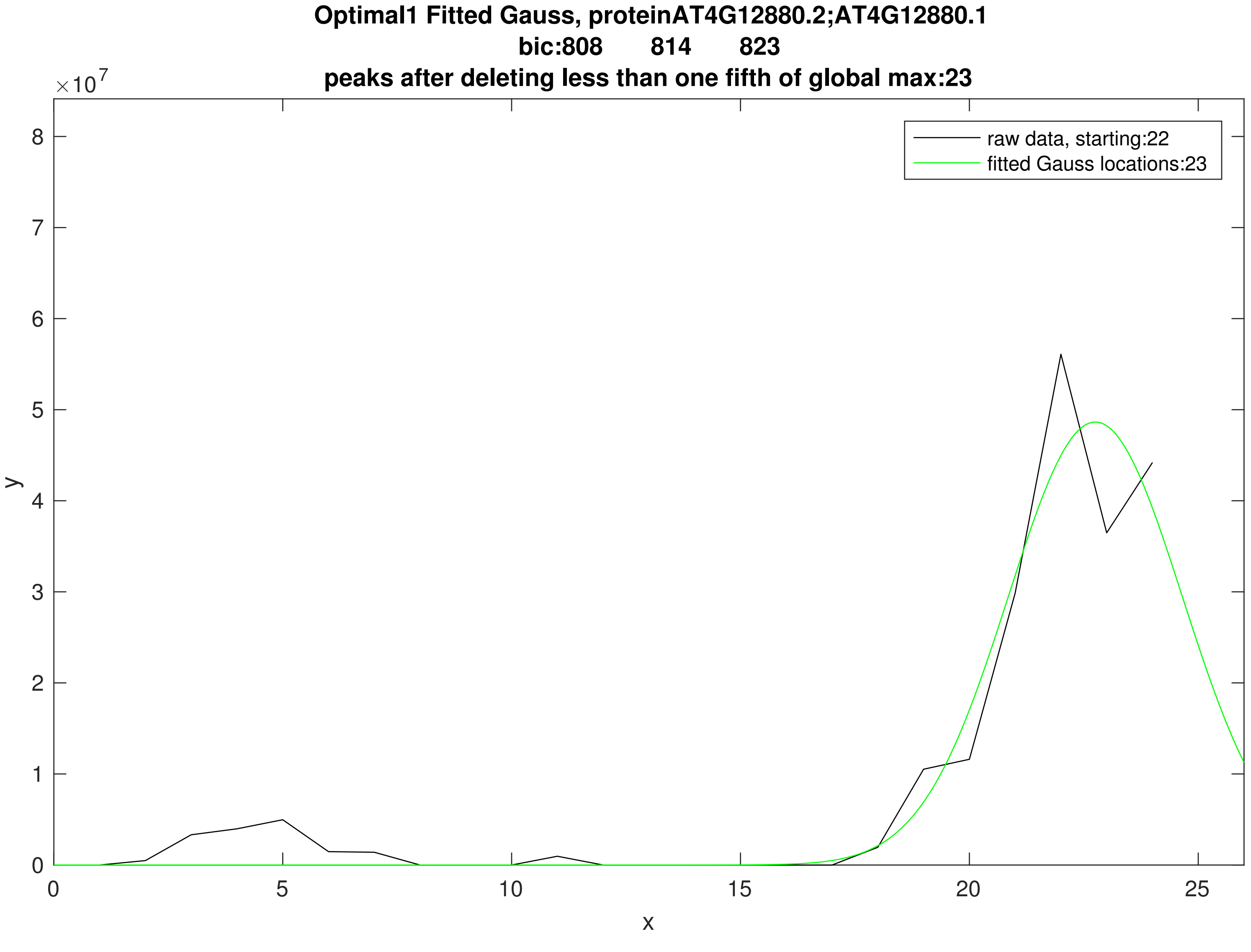

Optimal1 Fitted Gauss, proteinAT4G12880.2;AT4G12880.1
bic:808 814 823
peaks after deleting less than one fifth of global max:23
×10^7
raw data, starting:22
fitted Gauss locations:23
y
x

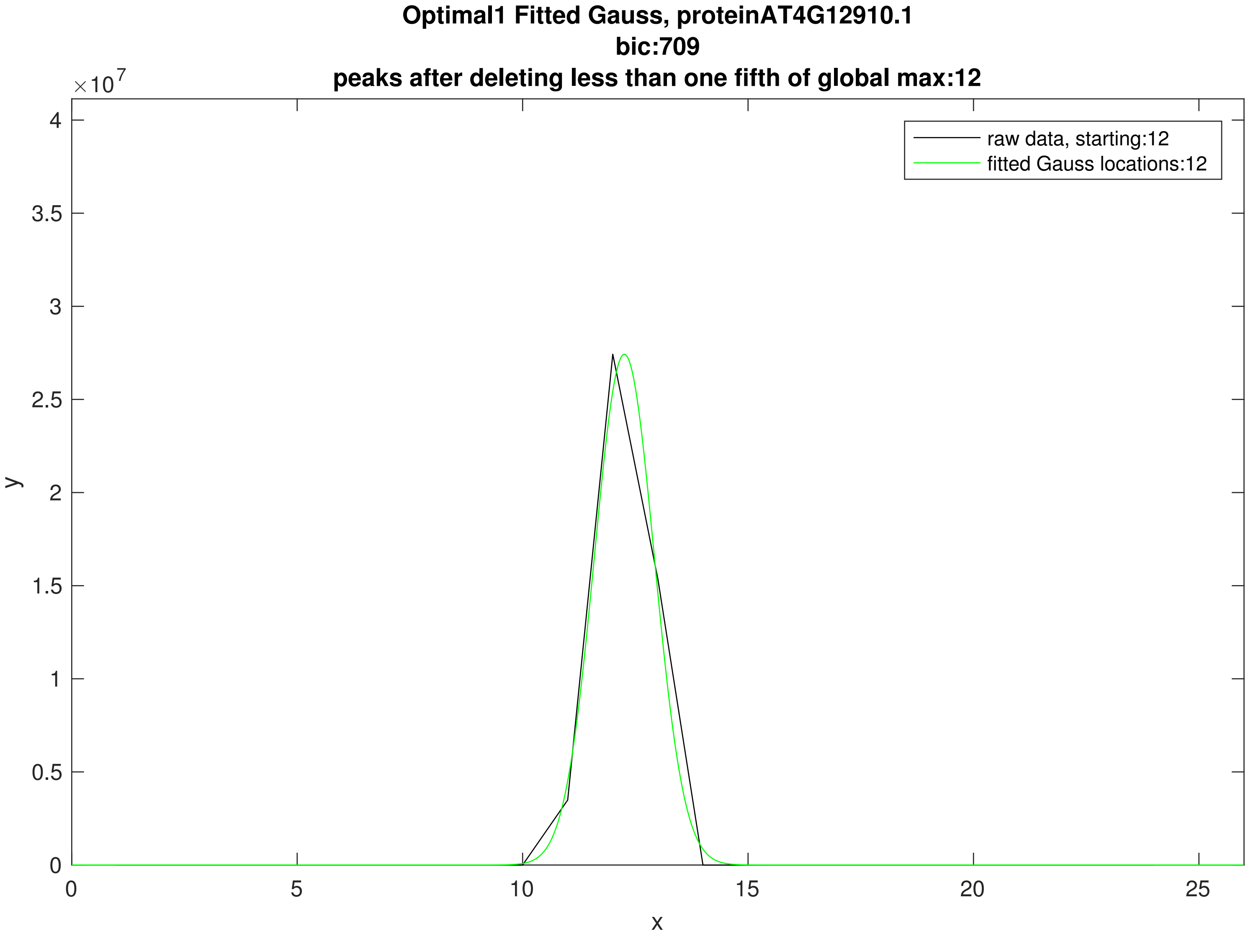
Optimal1 Fitted Gauss, proteinAT4G12910.1
bic:709
peaks after deleting less than one fifth of global max:12
×10^7
4
3.5
3
2.5
2
1.5
1
0.5
0
y
0
5
10
15
20
25
x
raw data, starting:12
fitted Gauss locations:12

**Optimal1 Fitted Gauss, proteinAT4G13050.1**
**bic:771**
**peaks after deleting less than one fifth of global max:14**

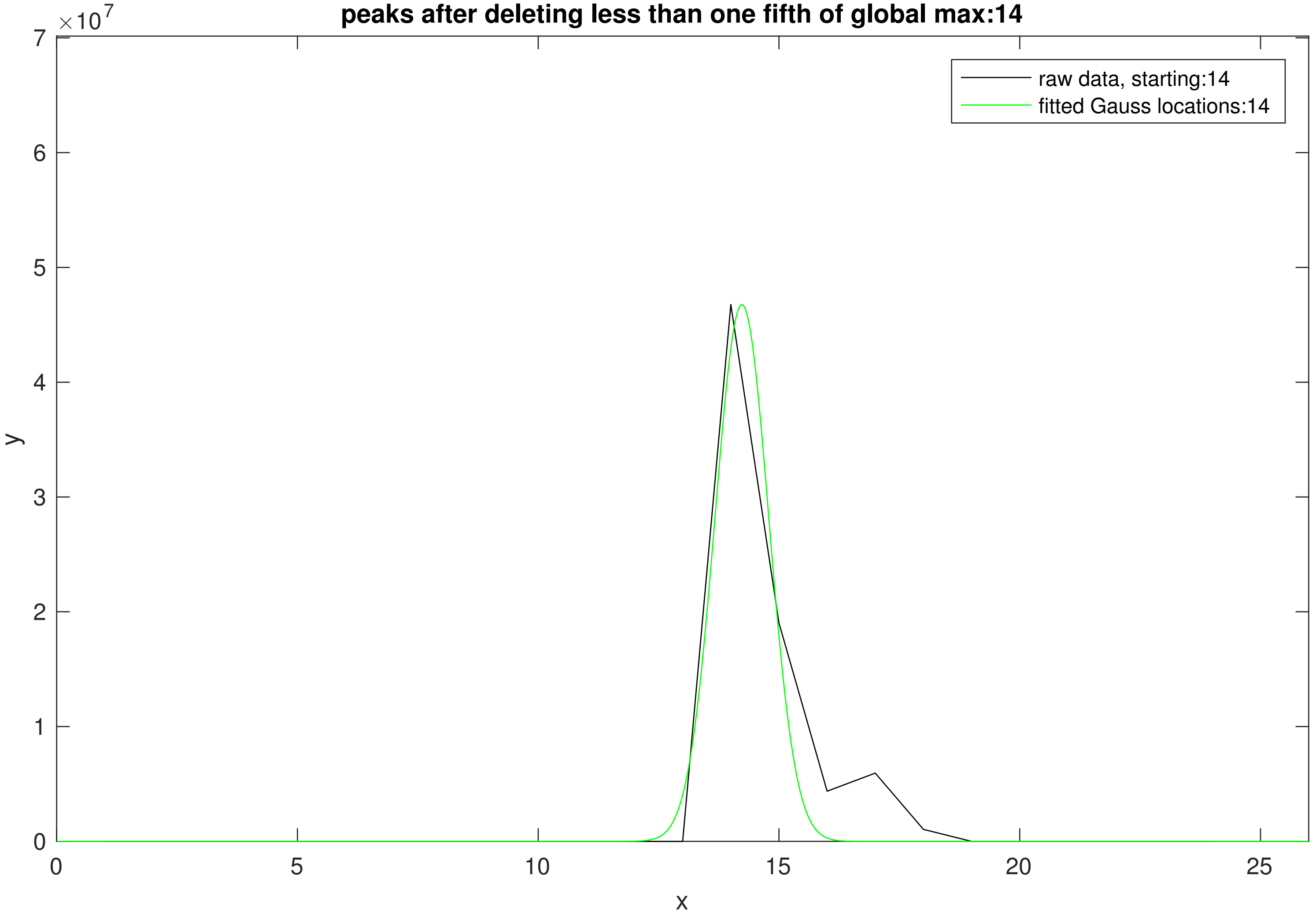

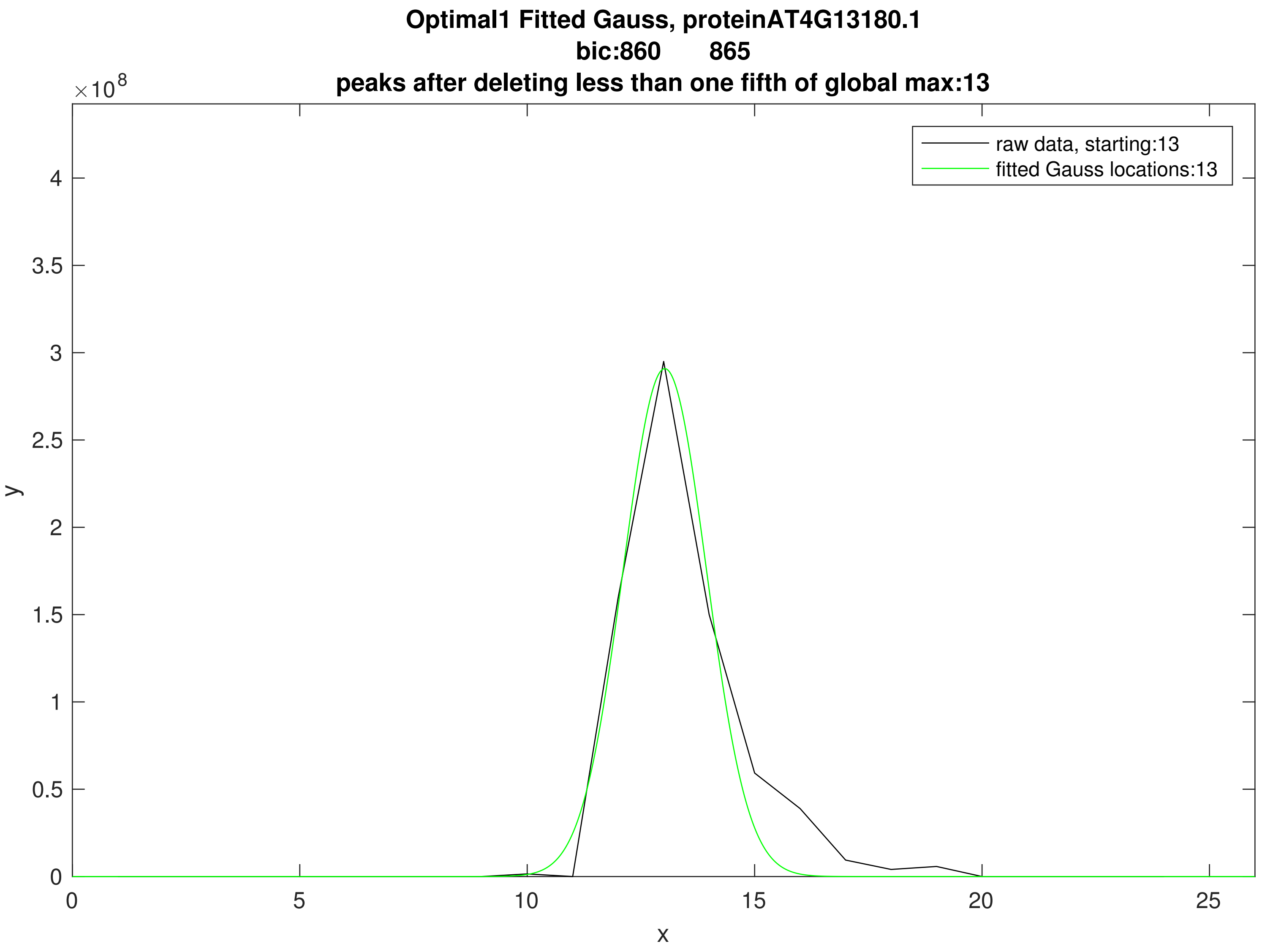

Optimal1 Fitted Gauss, proteinAT4G13180.1
bic:860 865
peaks after deleting less than one fifth of global max:13
×10^8
raw data, starting:13
fitted Gauss locations:13
y
x
0
0.5
1
1.5
2
2.5
3
3.5
4
0
5
10
15
20
25

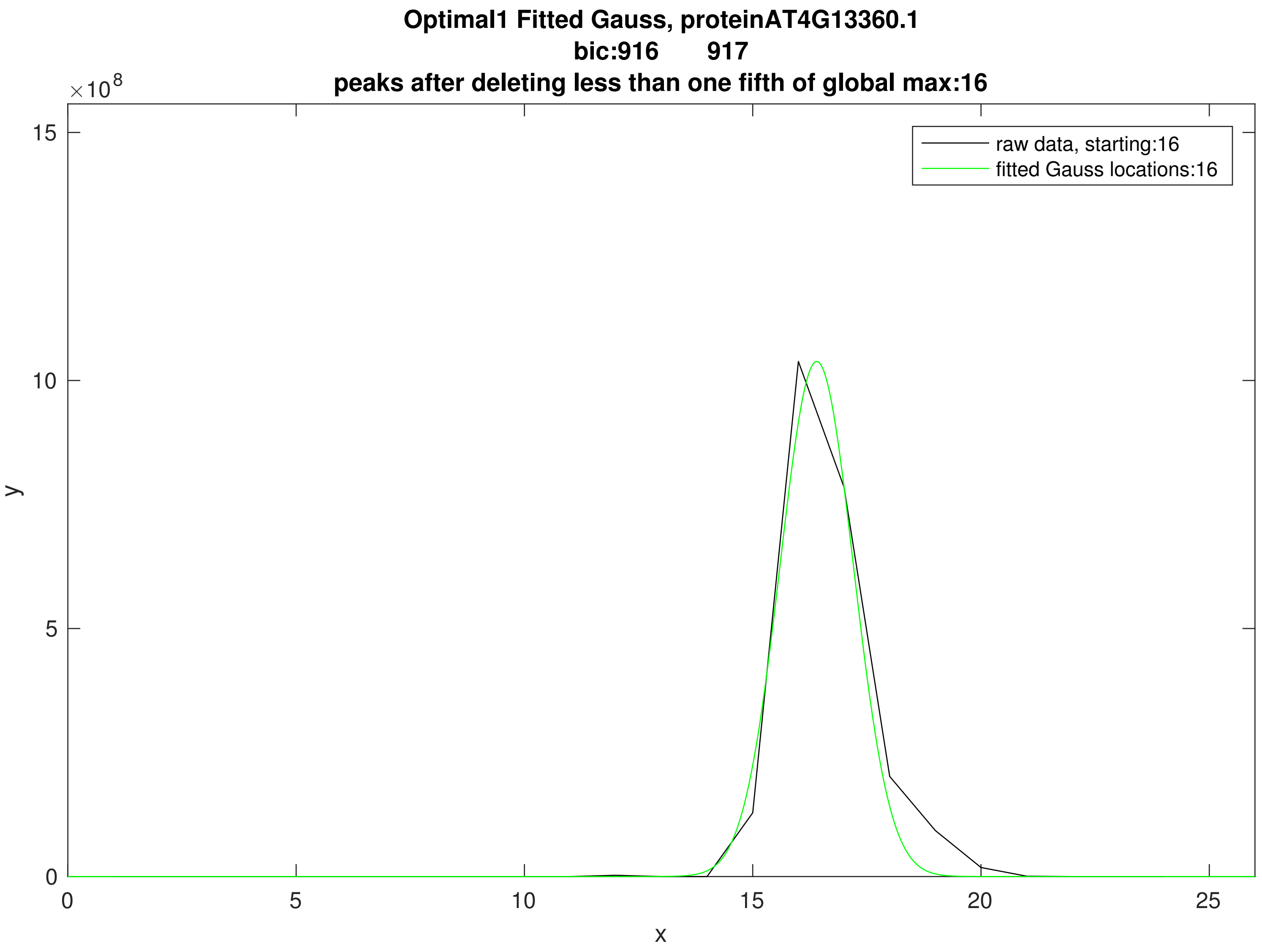
Optimal1 Fitted Gauss, proteinAT4G13360.1
bic:916 917
peaks after deleting less than one fifth of global max:16
×10^8
15
10
5
0
y
0
5
10
15
20
25
x
raw data, starting:16
fitted Gauss locations:16

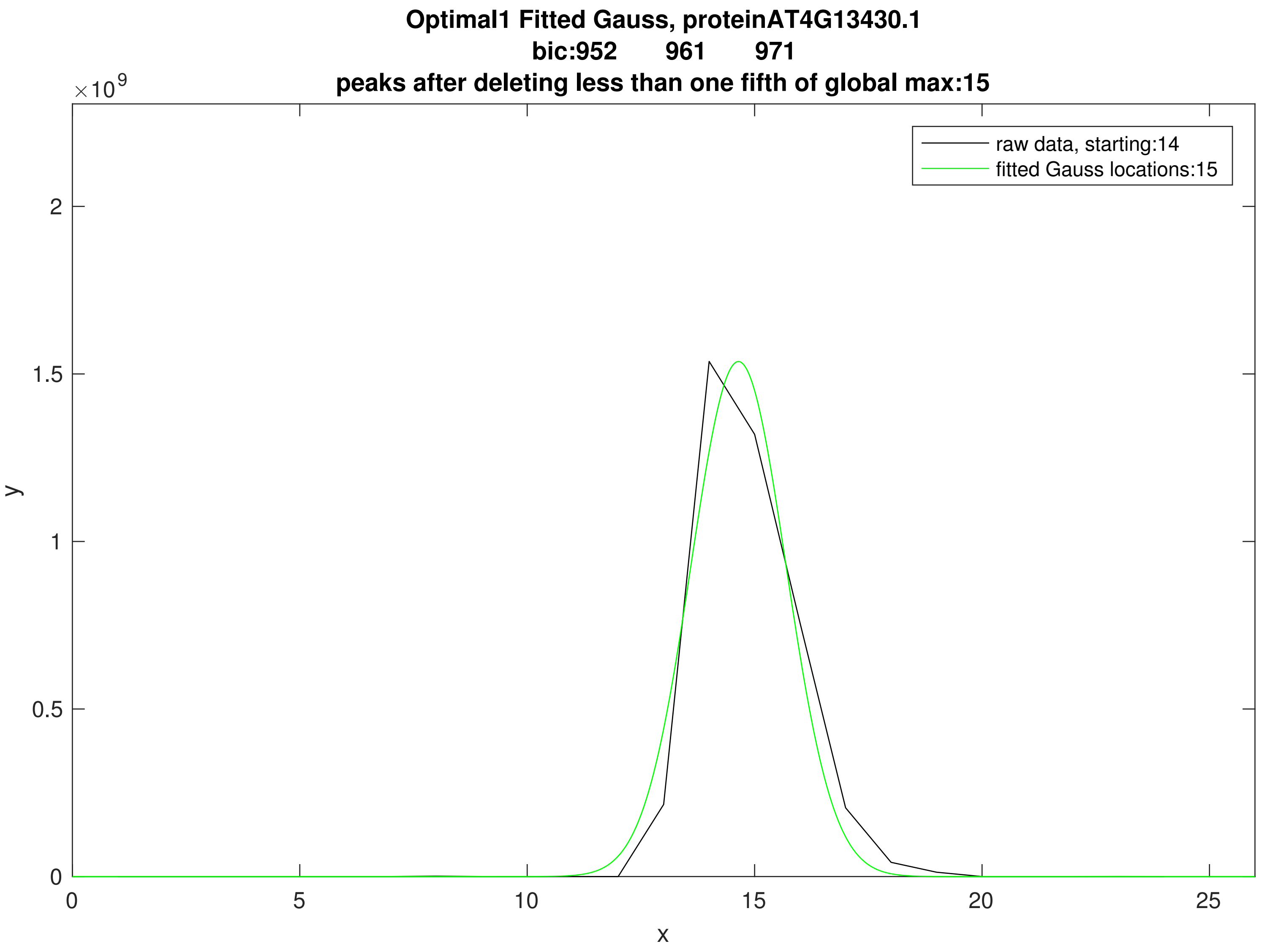
Optimal1 Fitted Gauss, proteinAT4G13430.1
bic:952 961 971
peaks after deleting less than one fifth of global max:15
×10^9
raw data, starting:14
fitted Gauss locations:15
y
x
0
0.5
1
1.5
2
5
10
15
20
25

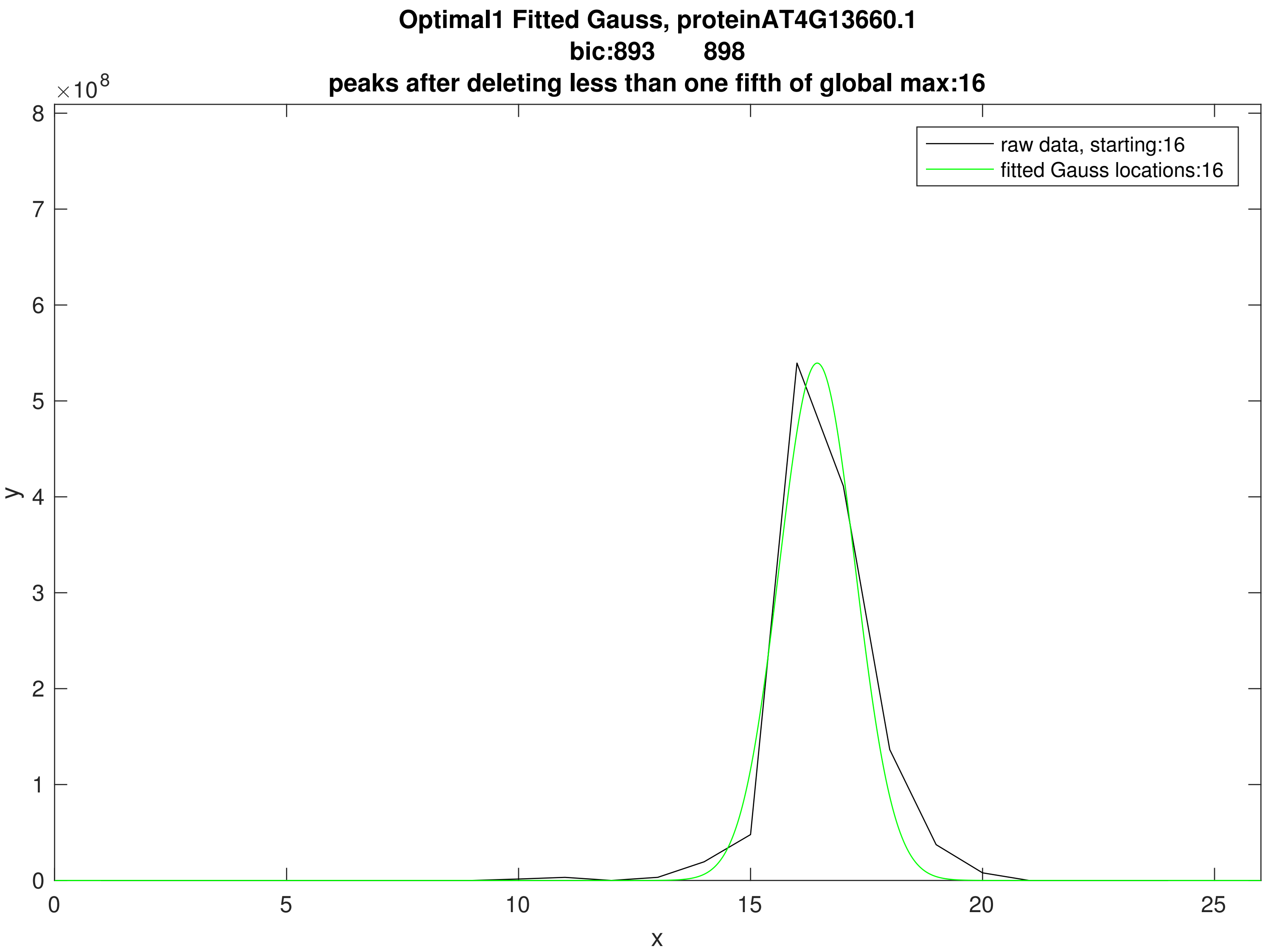
Optimal1 Fitted Gauss, proteinAT4G13660.1
bic:893 898
peaks after deleting less than one fifth of global max:16
×10^8
raw data, starting:16
fitted Gauss locations:16
y
x
0
1
2
3
4
5
6
7
8
0
5
10
15
20
25

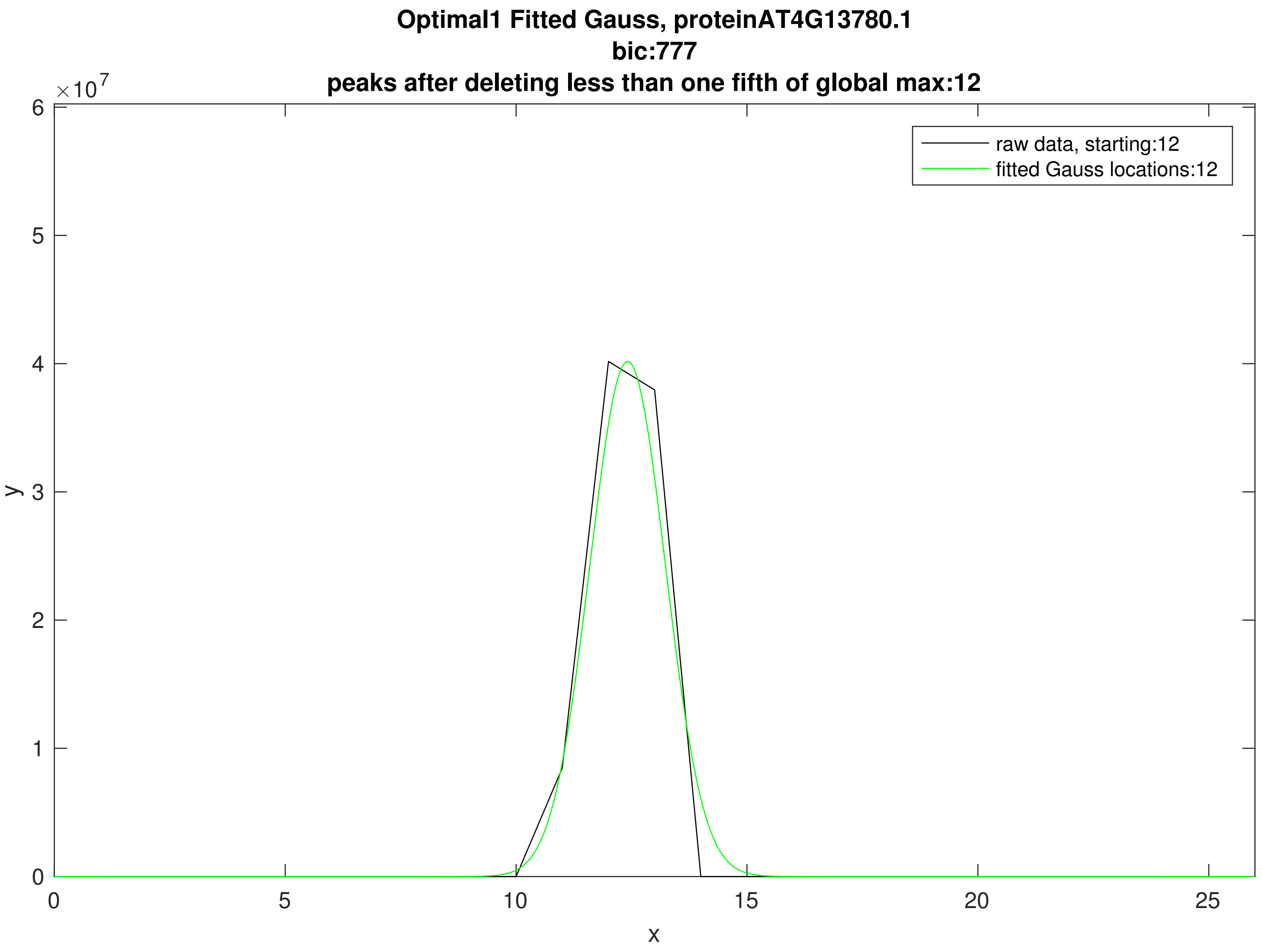

Optimal1 Fitted Gauss, proteinAT4G13780.1
bic:777
peaks after deleting less than one fifth of global max:12
×10^7
raw data, starting:12
fitted Gauss locations:12
y
x
0
1
2
3
4
5
6
0
5
10
15
20
25

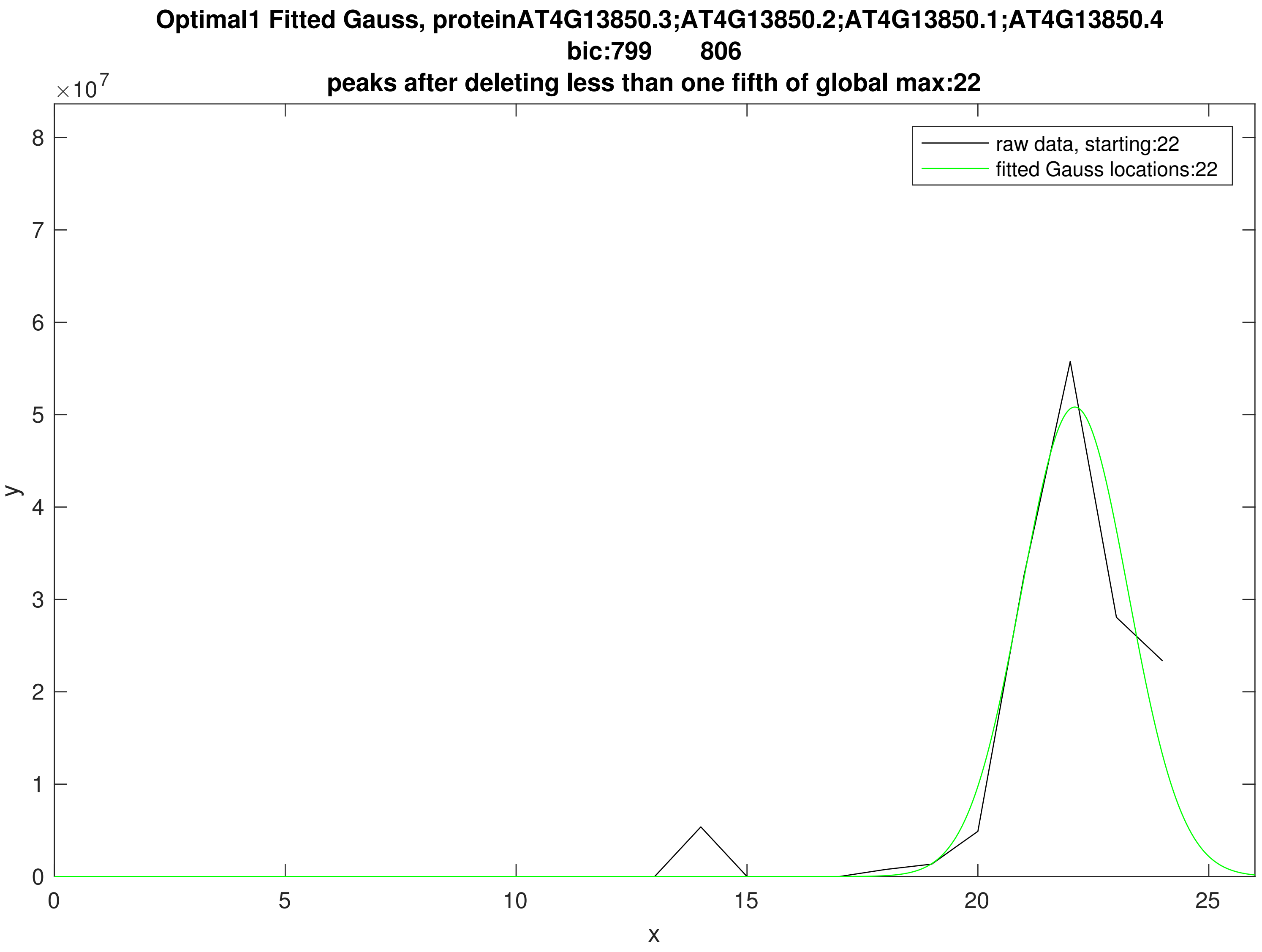
Optimal1 Fitted Gauss, proteinAT4G13850.3;AT4G13850.2;AT4G13850.1;AT4G13850.4
bic:799 806
peaks after deleting less than one fifth of global max:22
×10^7
raw data, starting:22
fitted Gauss locations:22
y
x
0
1
2
3
4
5
6
7
8
0
5
10
15
20
25

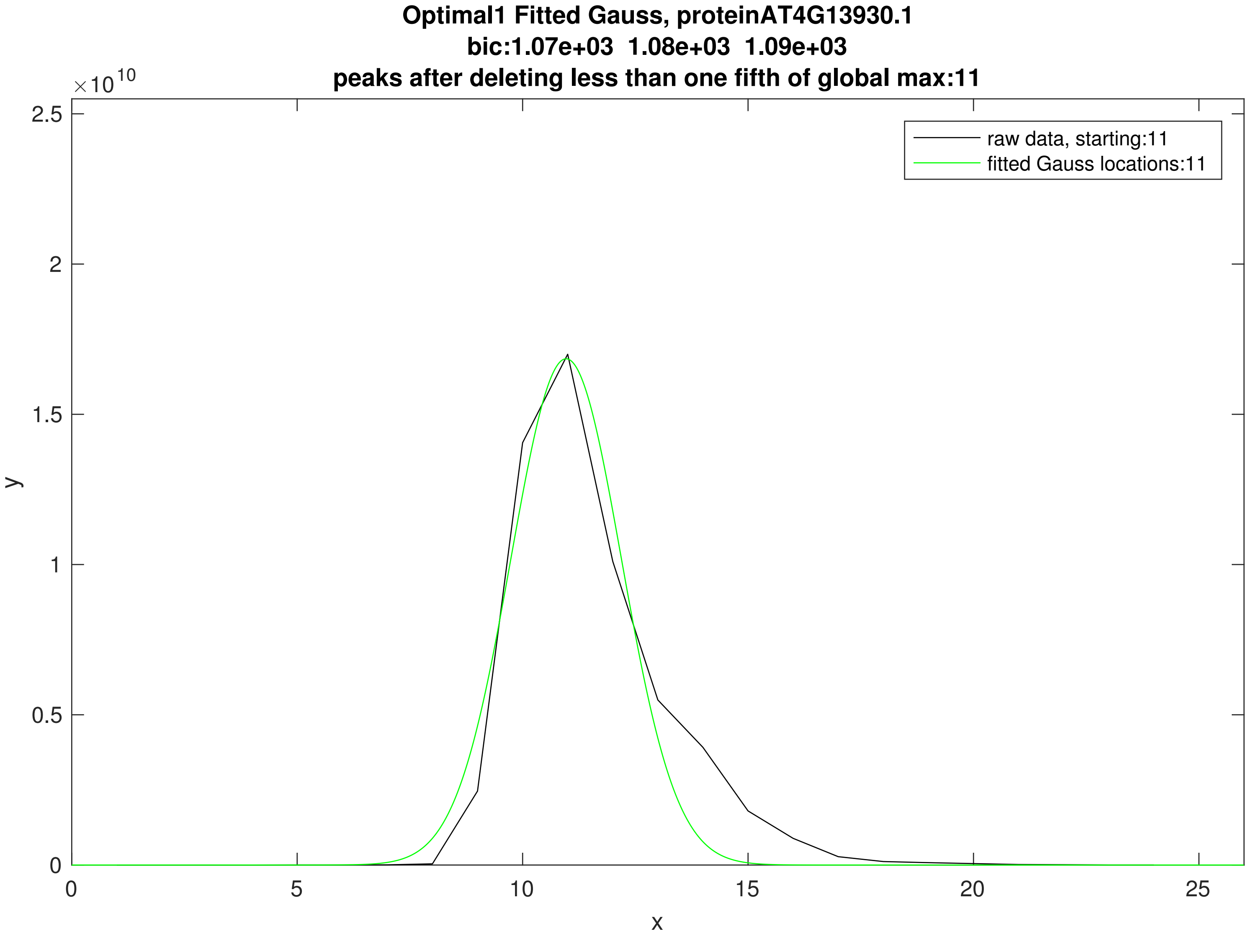
Optimal1 Fitted Gauss, proteinAT4G13930.1
bic:1.07e+03 1.08e+03 1.09e+03
peaks after deleting less than one fifth of global max:11
×10^10
2.5
2
1.5
1
0.5
0
y
0
5
10
15
20
25
x
raw data, starting:11
fitted Gauss locations:11

Optimal1 Fitted Gauss, proteinAT4G13940.1;AT4G13940.3;AT4G13940.2;AT4G13940.4
bic:1.11e+03  1.12e+03  1.13e+03
peaks after deleting less than one fifth of global max:10

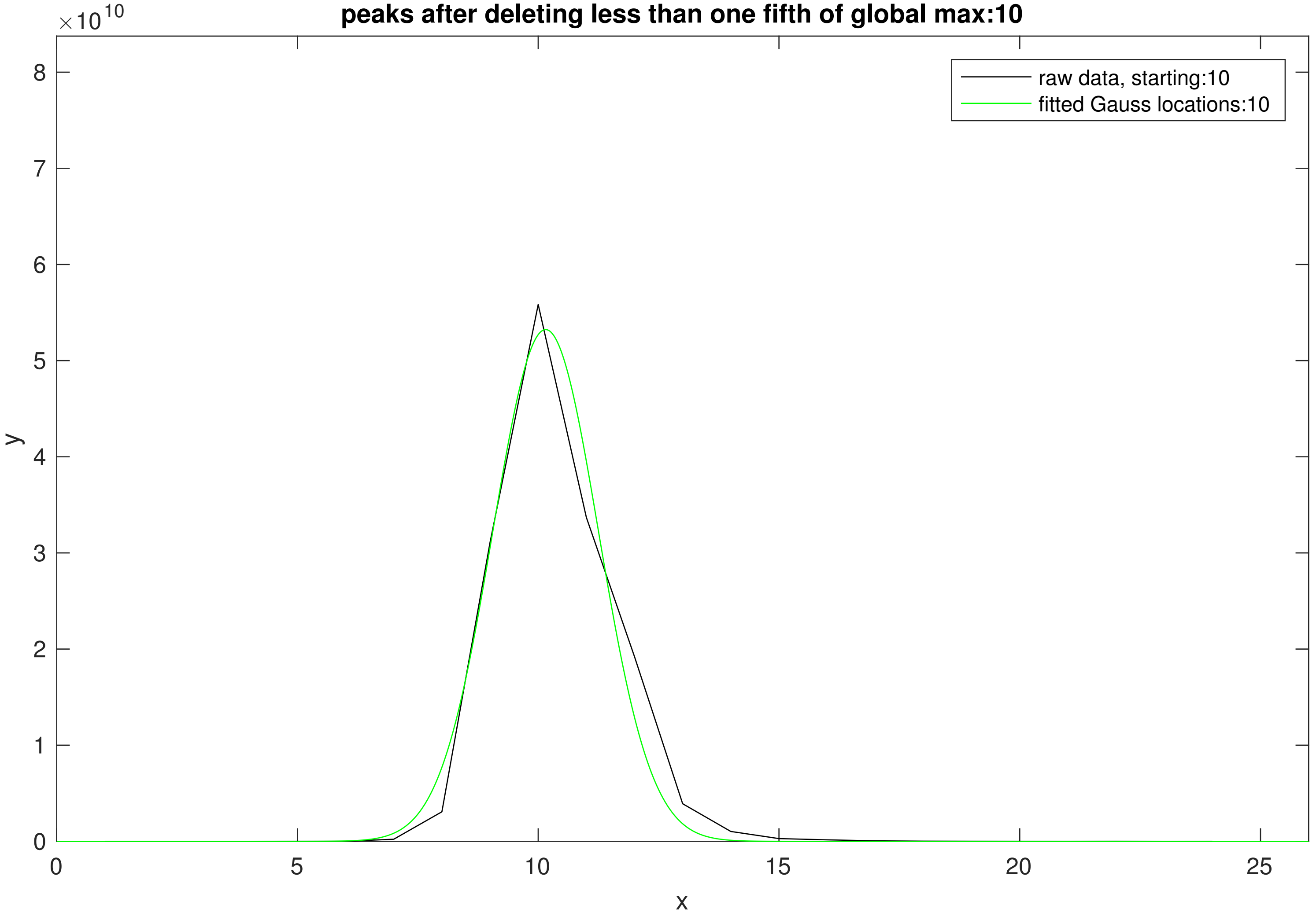

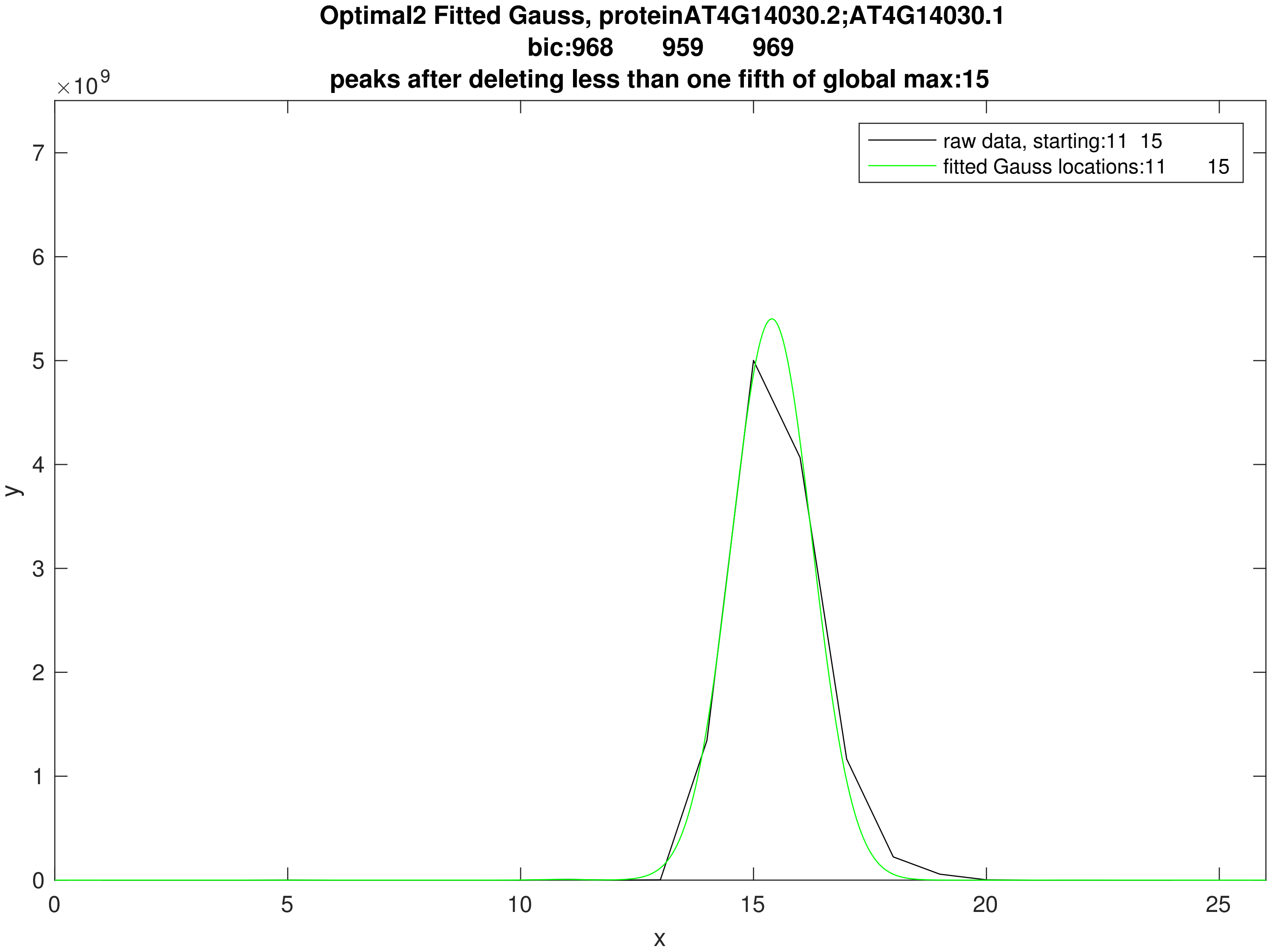
Optimal2 Fitted Gauss, proteinAT4G14030.2;AT4G14030.1
bic:968 959 969
peaks after deleting less than one fifth of global max:15
×10^9
raw data, starting:11 15
fitted Gauss locations:11 15
y
x
0
1
2
3
4
5
6
7
0
5
10
15
20
25

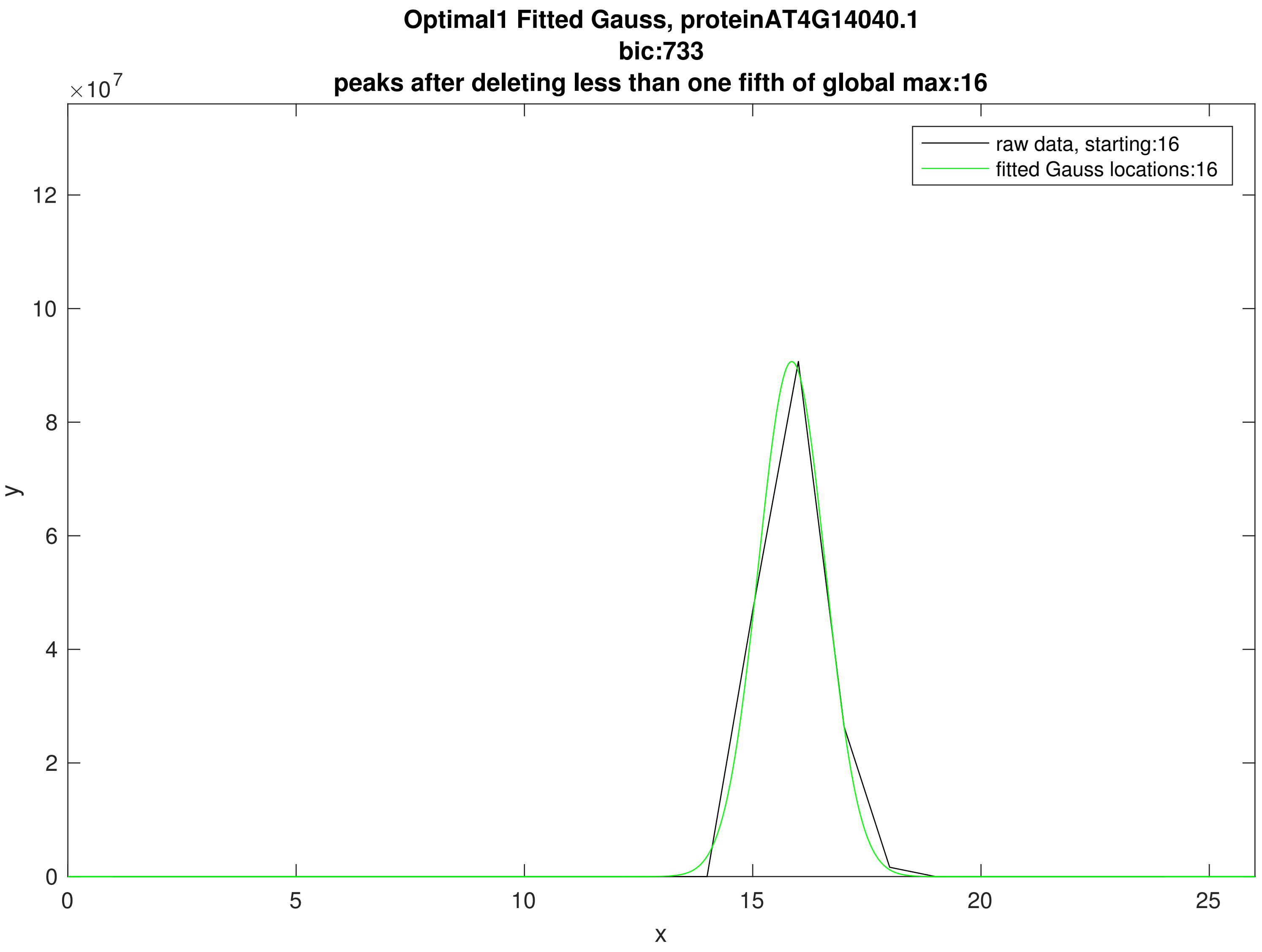

Optimal1 Fitted Gauss, proteinAT4G14040.1
bic:733
peaks after deleting less than one fifth of global max:16
×10^7
raw data, starting:16
fitted Gauss locations:16
y
x
0
2
4
6
8
10
12
0
5
10
15
20
25

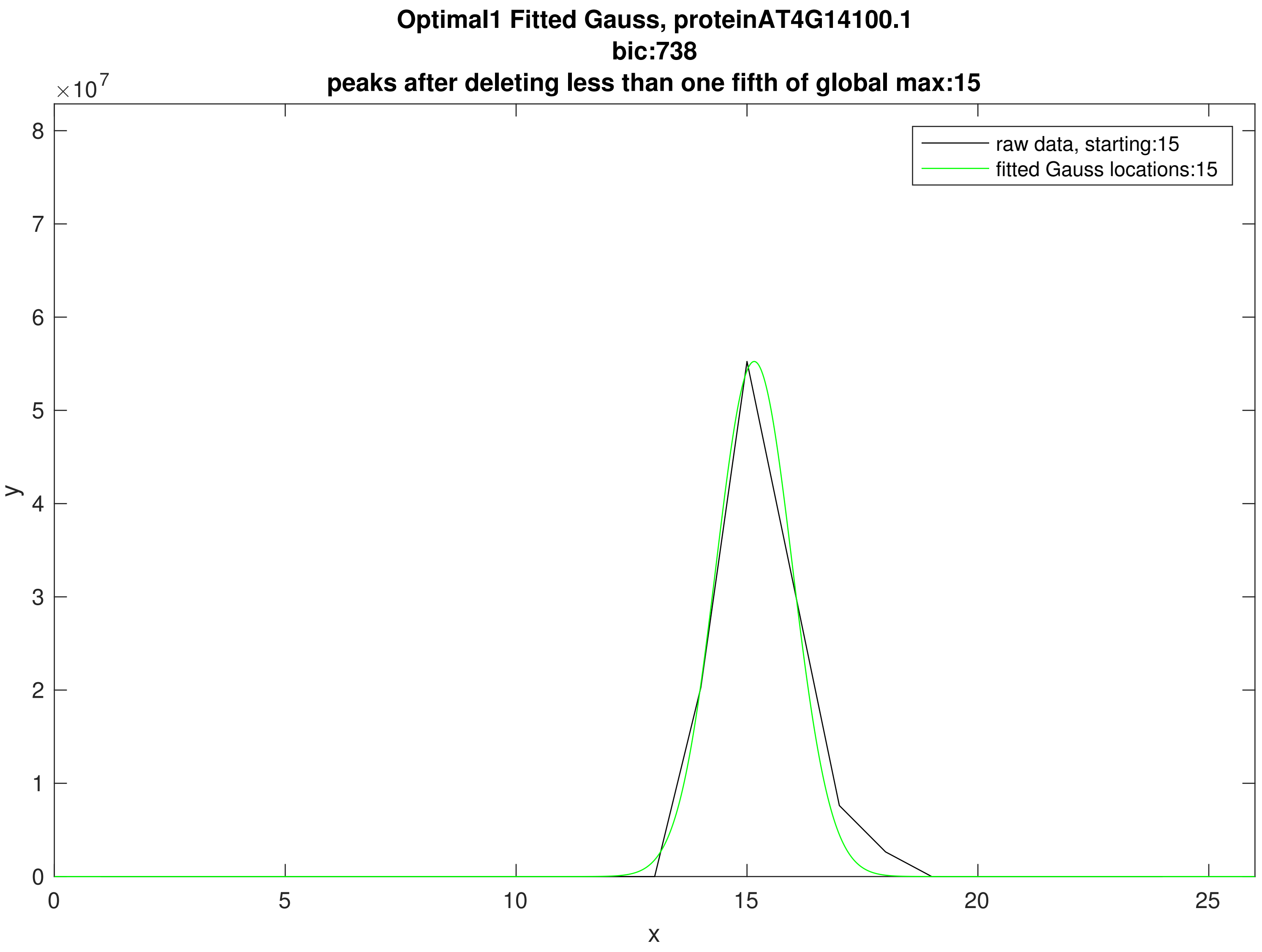
Optimal1 Fitted Gauss, proteinAT4G14100.1
bic:738
peaks after deleting less than one fifth of global max:15
×10^7
raw data, starting:15
fitted Gauss locations:15
y
x
0
1
2
3
4
5
6
7
8
0
5
10
15
20
25

**Optimal1 Fitted Gauss, proteinAT4G14300.1**
**bic:791 823**
**peaks after deleting less than one fifth of global max:18**

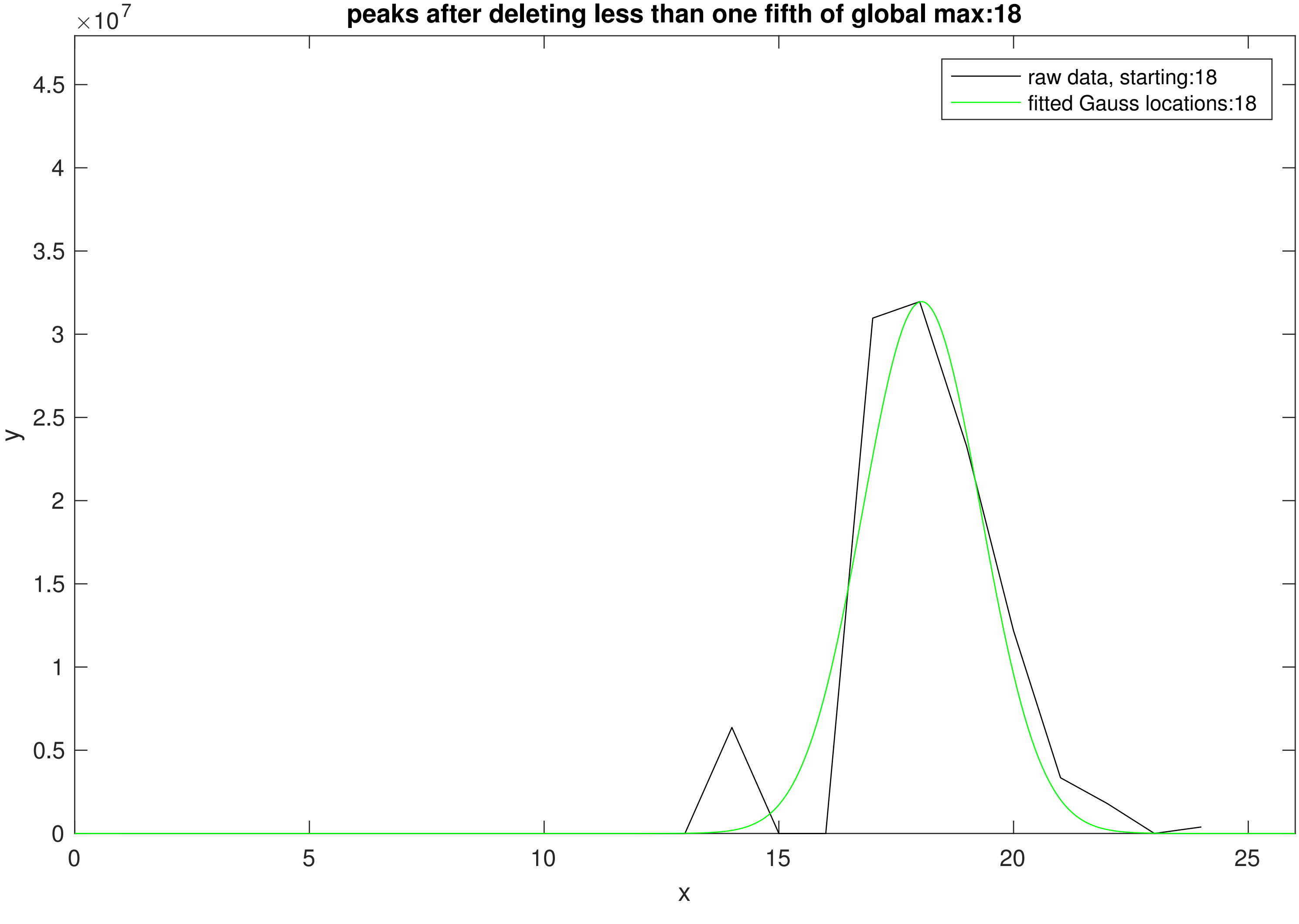

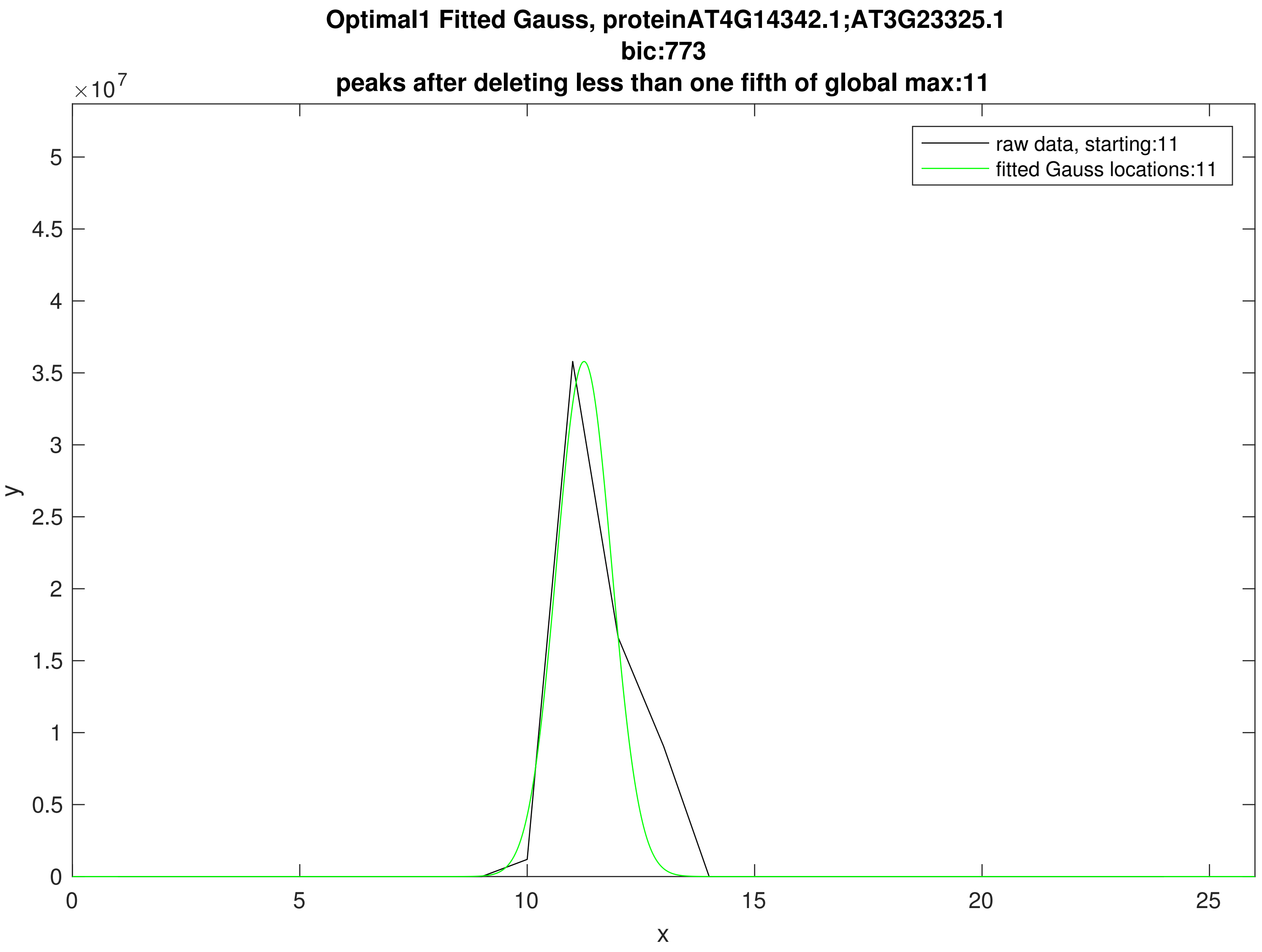
Optimal1 Fitted Gauss, proteinAT4G14342.1;AT3G23325.1
bic:773
peaks after deleting less than one fifth of global max:11
×10^7
5
4.5
4
3.5
3
2.5
2
1.5
1
0.5
0
y
0
5
10
15
20
25
x
raw data, starting:11
fitted Gauss locations:11

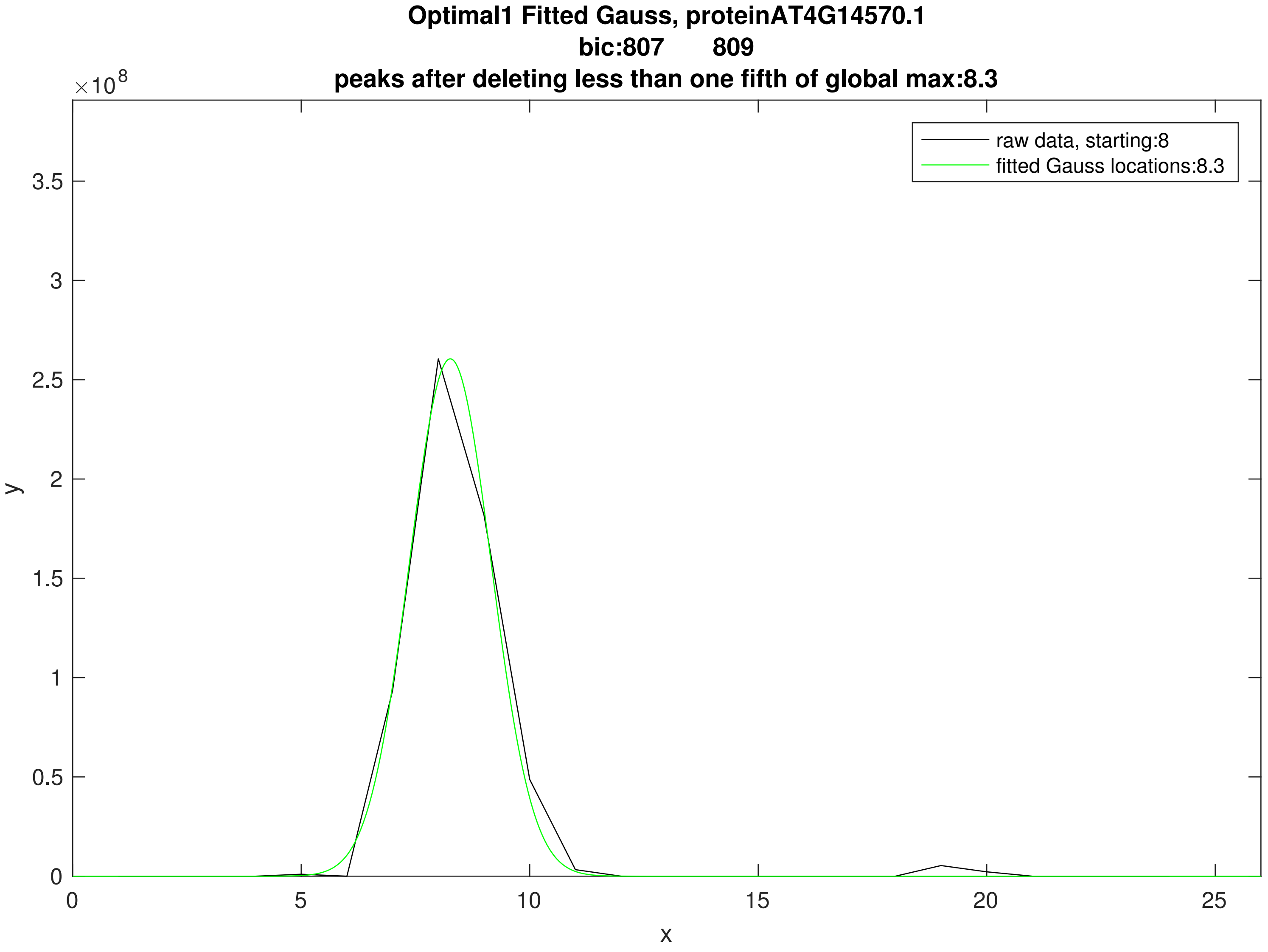

Optimal1 Fitted Gauss, proteinAT4G14570.1
bic:807 809
peaks after deleting less than one fifth of global max:8.3
×10^8
raw data, starting:8
fitted Gauss locations:8.3
3.5
3
2.5
2
1.5
1
0.5
0
y
0
5
10
15
20
25
x

Optimal1 Fitted Gauss, proteinAT4G14710.2;AT4G14710.1;AT4G14710.3;AT4G14710.5;AT4G14710.4
bic:907 911
peaks after deleting less than one fifth of global max:18

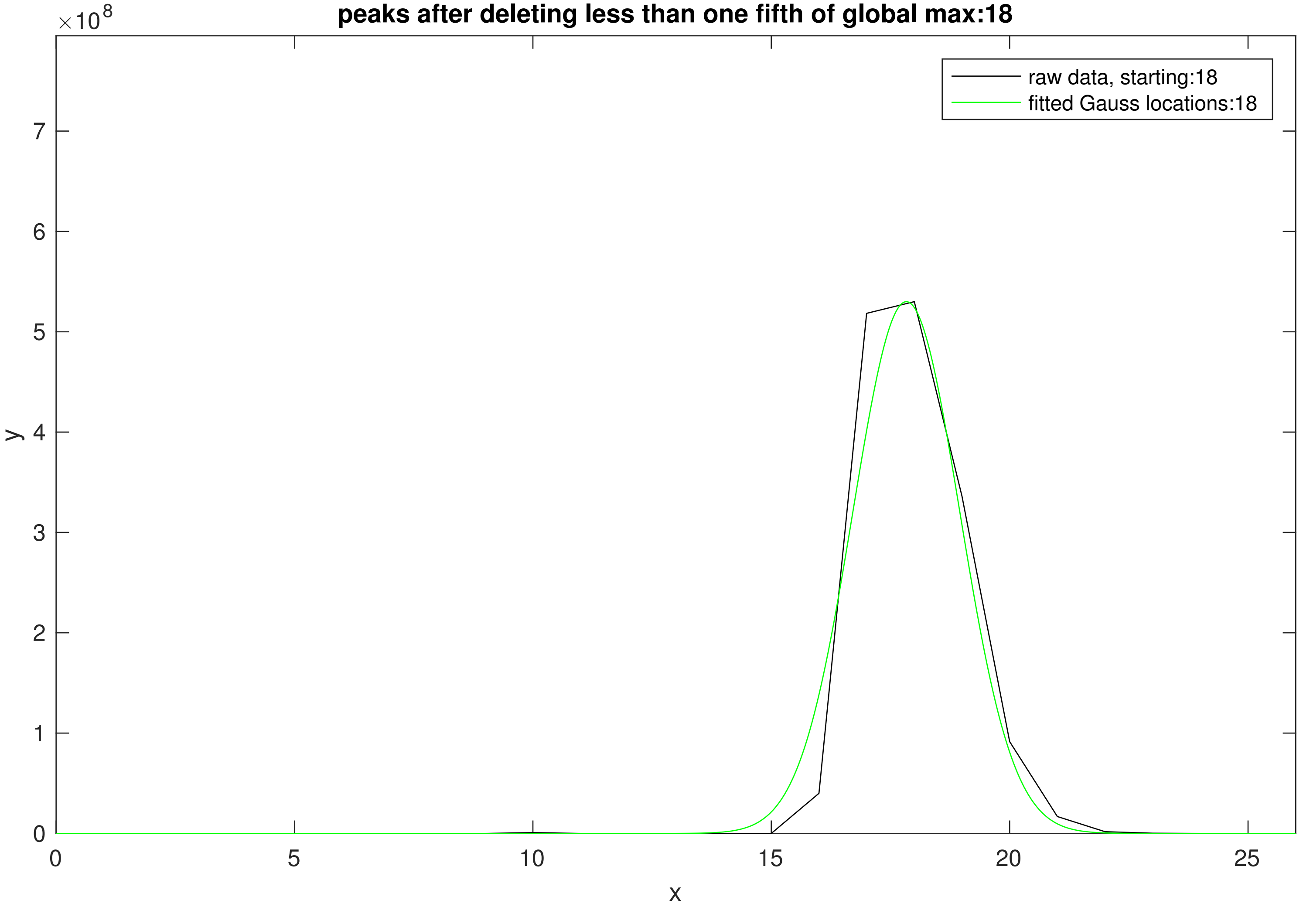

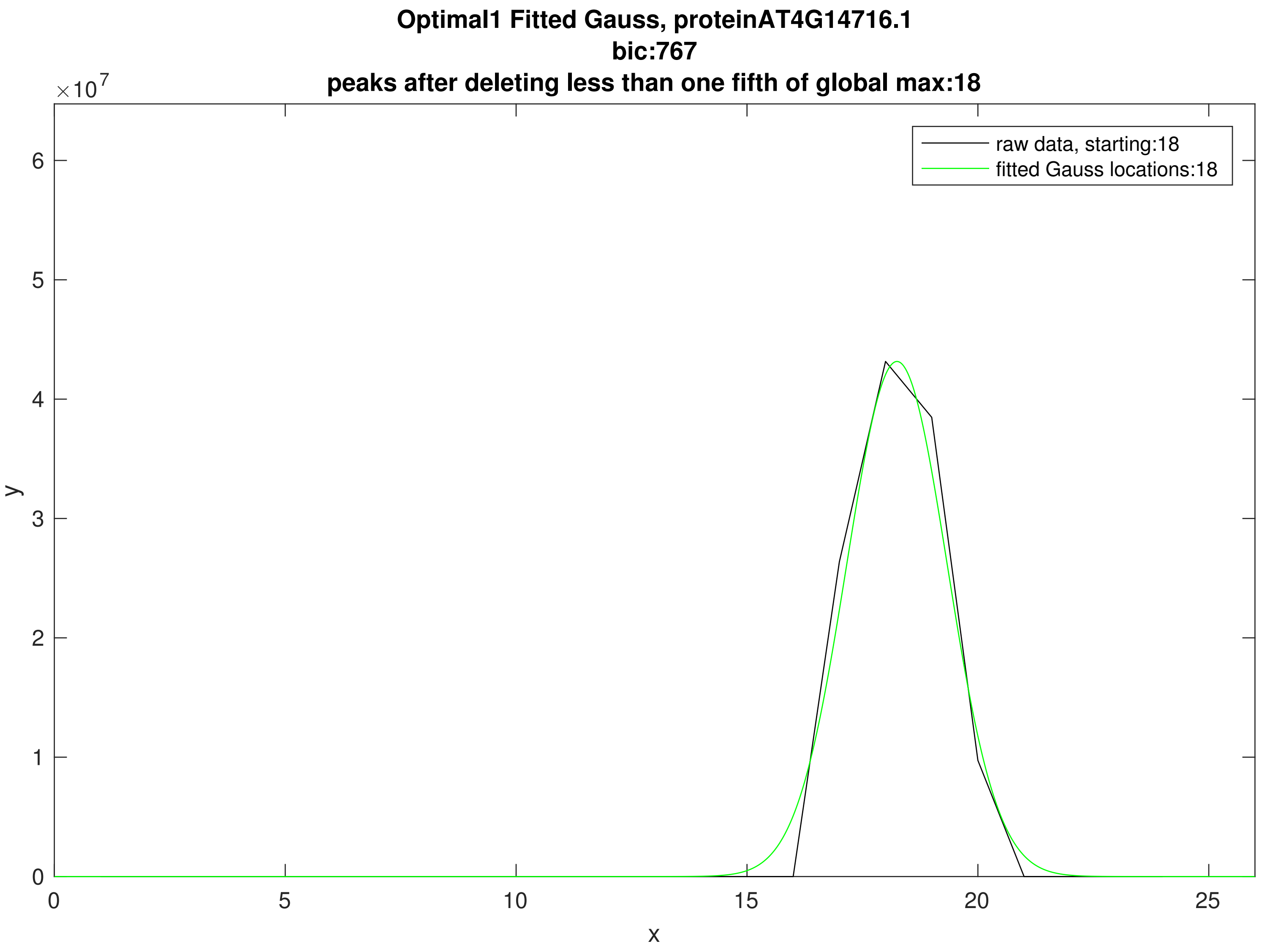
Optimal1 Fitted Gauss, proteinAT4G14716.1
bic:767
peaks after deleting less than one fifth of global max:18
×10^7
raw data, starting:18
fitted Gauss locations:18
y
x
0
1
2
3
4
5
6
0
5
10
15
20
25

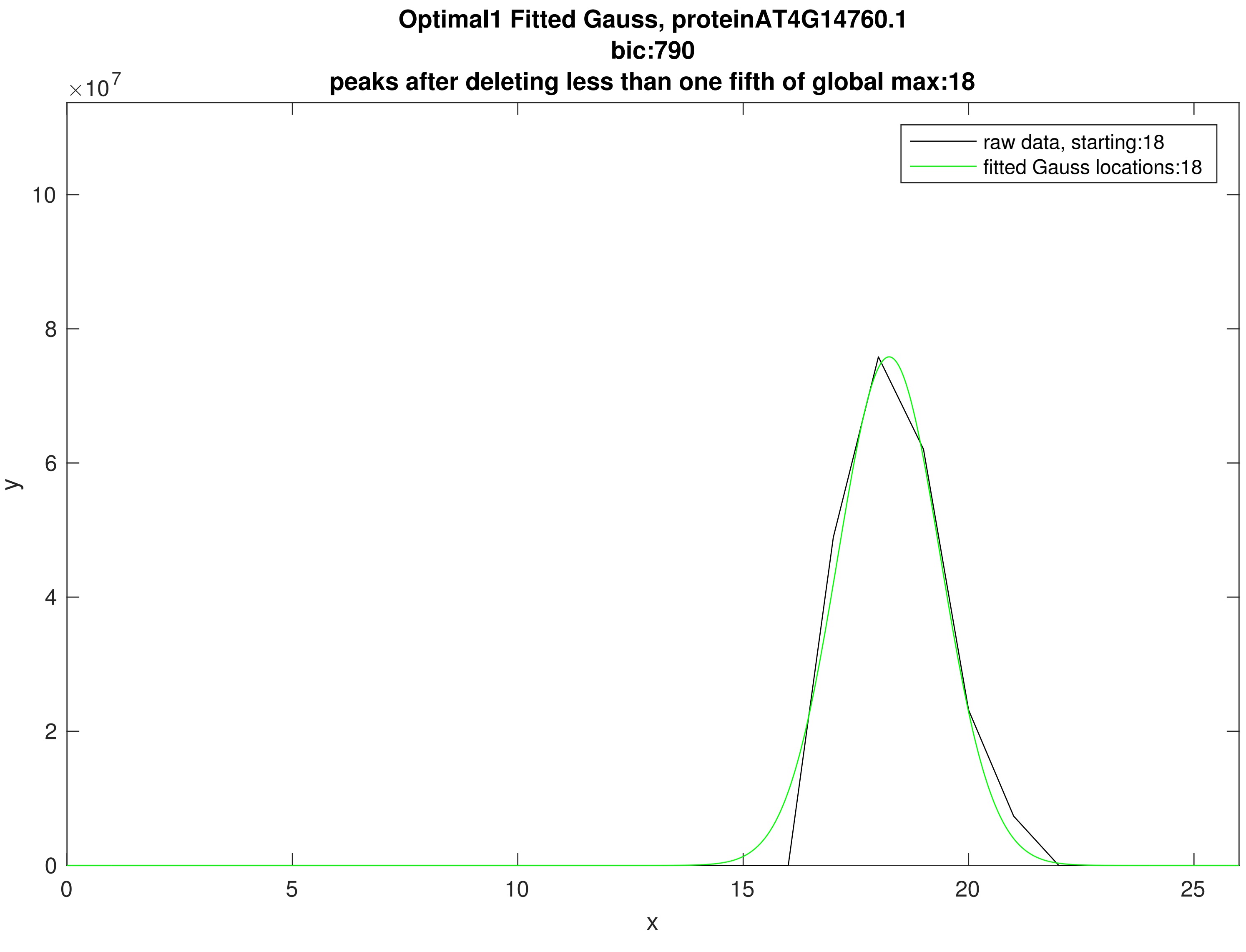
Optimal1 Fitted Gauss, proteinAT4G14760.1
bic:790
peaks after deleting less than one fifth of global max:18
×10^7
raw data, starting:18
fitted Gauss locations:18
y
x
0
2
4
6
8
10
0
5
10
15
20
25

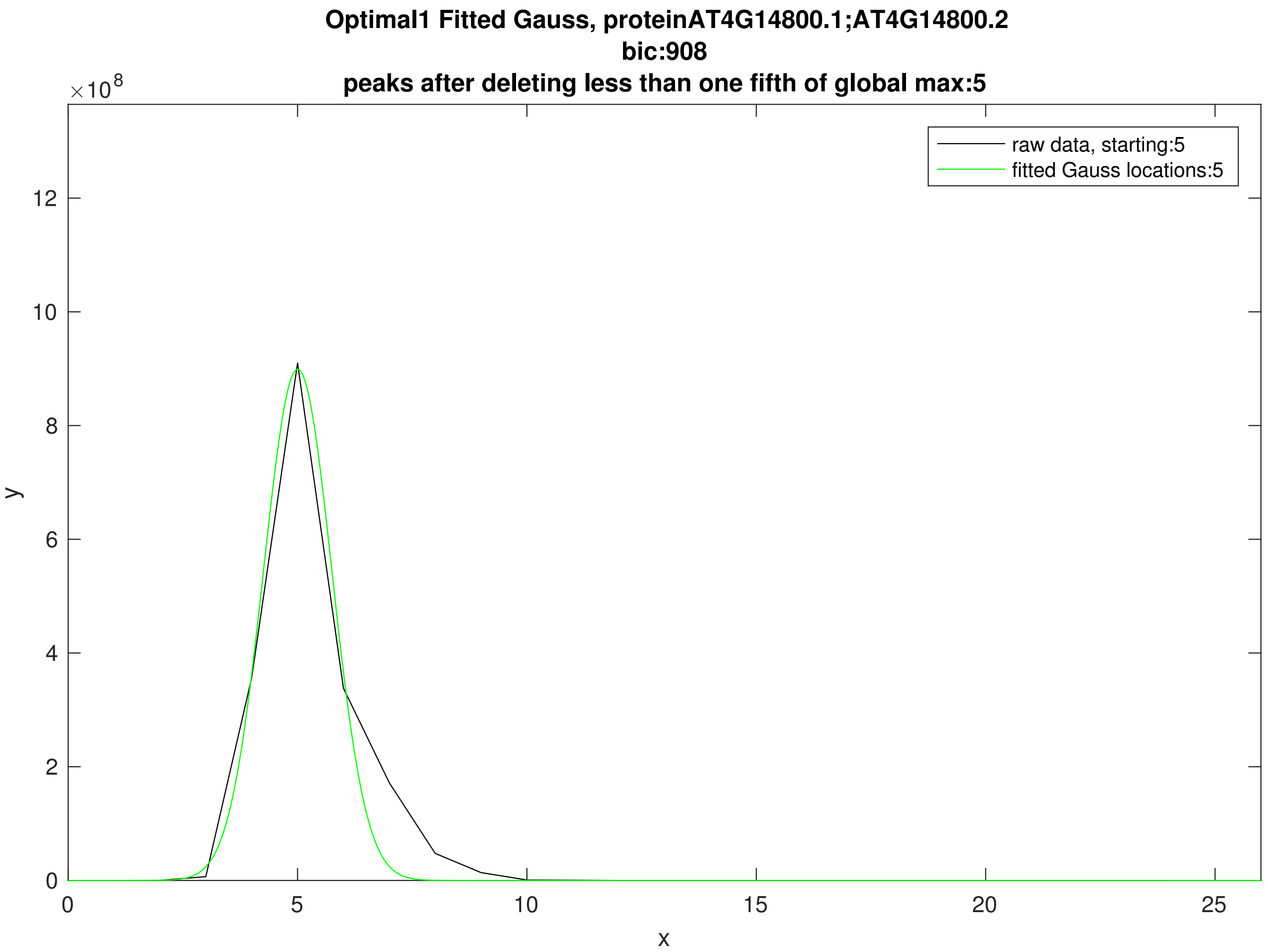

Optimal1 Fitted Gauss, proteinAT4G14800.1;AT4G14800.2
bic:908
peaks after deleting less than one fifth of global max:5
×10^8
raw data, starting:5
fitted Gauss locations:5
y
x
0
2
4
6
8
10
12
0
5
10
15
20
25

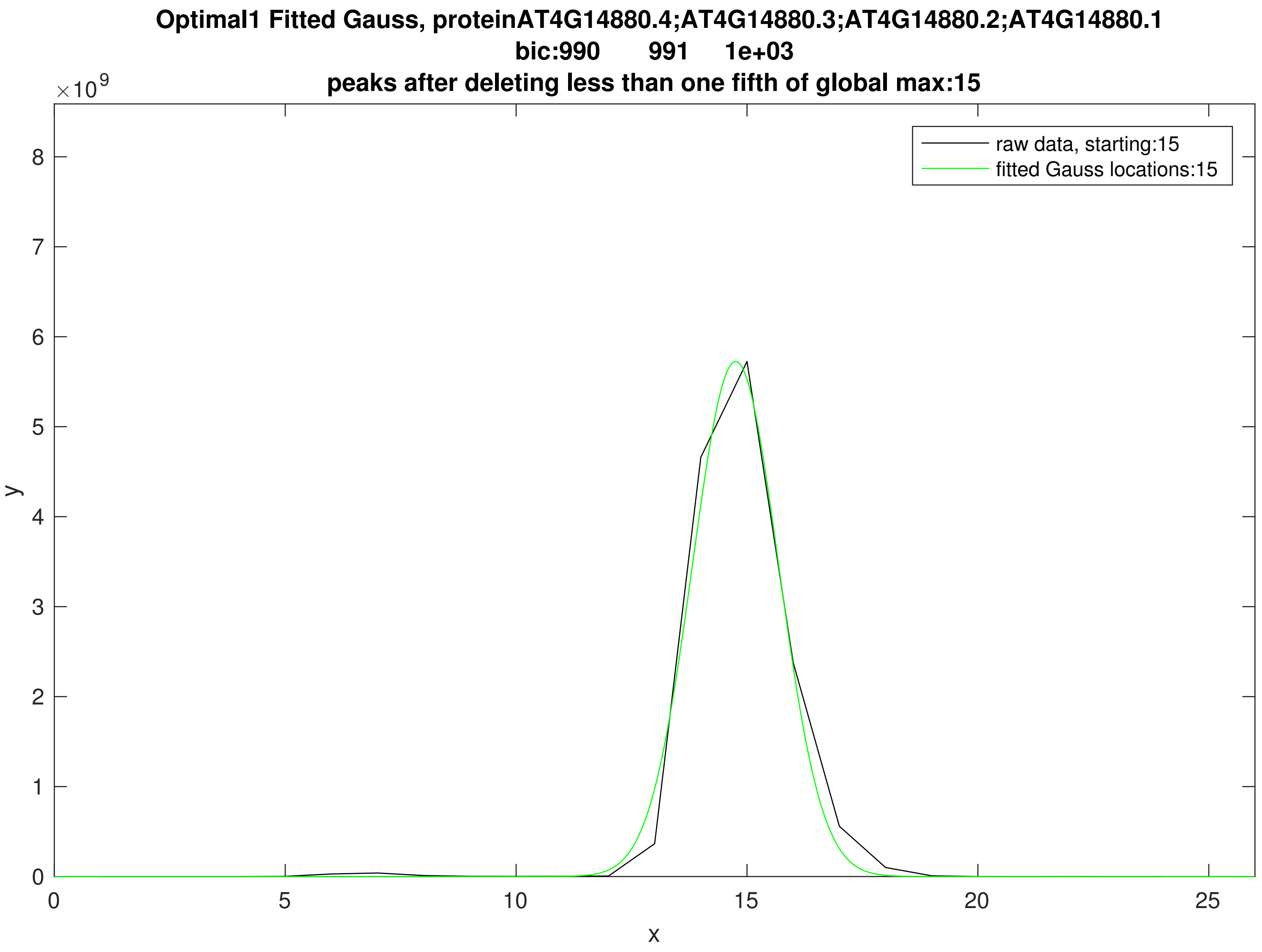

Optimal1 Fitted Gauss, proteinAT4G14880.4;AT4G14880.3;AT4G14880.2;AT4G14880.1
bic:990 991 1e+03
peaks after deleting less than one fifth of global max:15
×10^9
raw data, starting:15
fitted Gauss locations:15
y
x
0
1
2
3
4
5
6
7
8
10
15
20
25

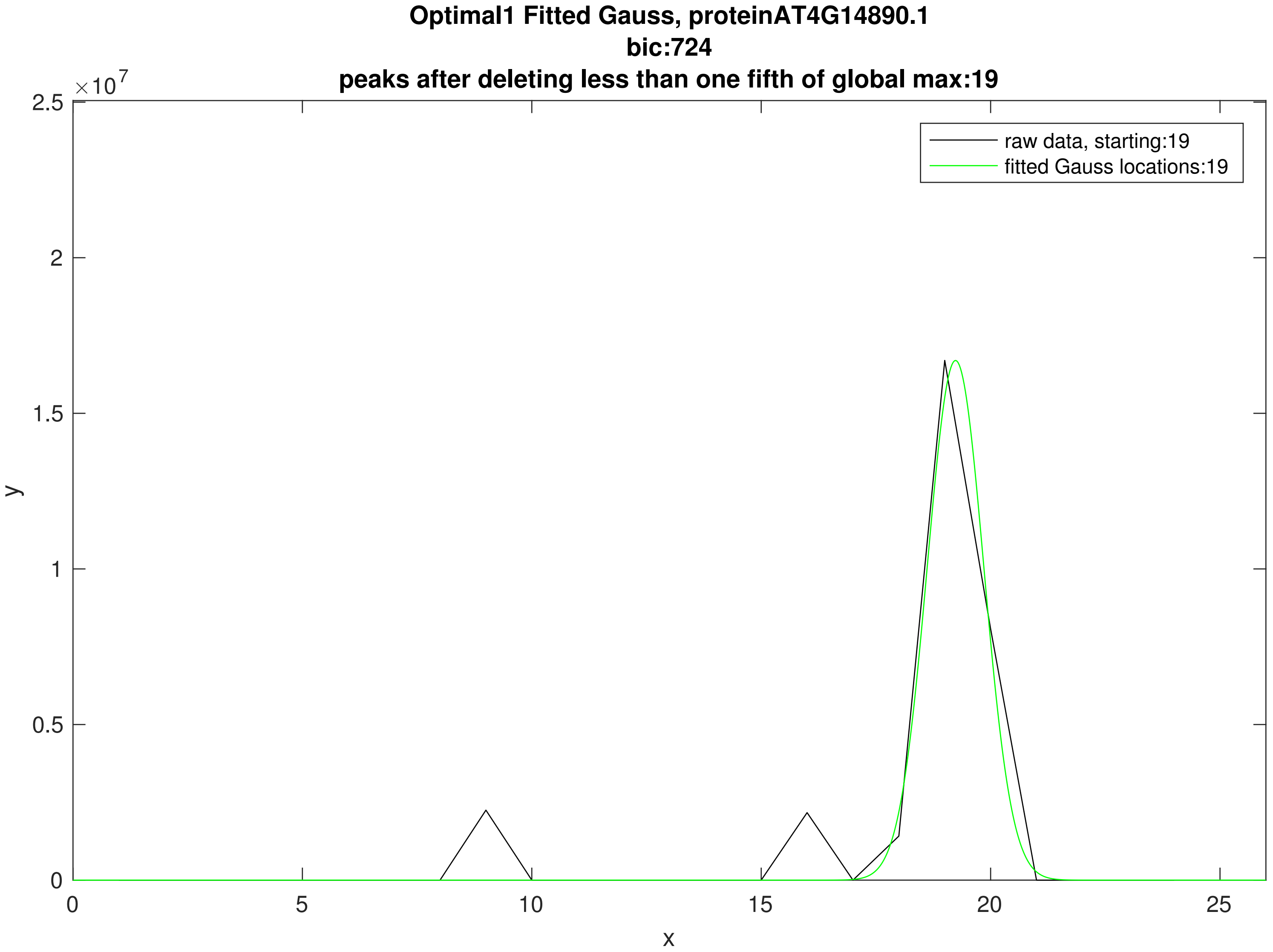
Optimal1 Fitted Gauss, proteinAT4G14890.1
bic:724
peaks after deleting less than one fifth of global max:19
×10^7
2.5
2
1.5
1
0.5
0
y
0
5
10
15
20
25
x
raw data, starting:19
fitted Gauss locations:19

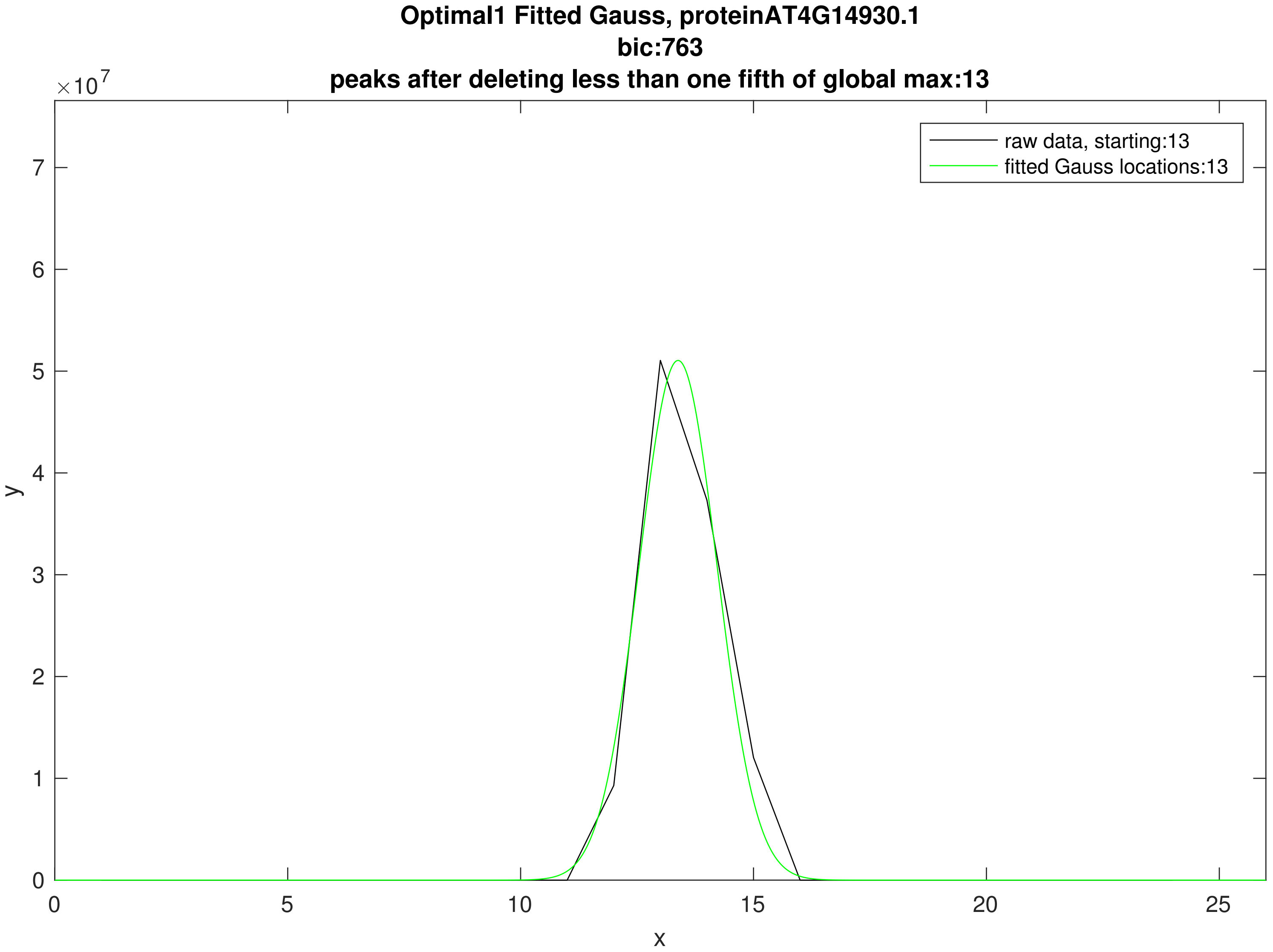
Optimal1 Fitted Gauss, proteinAT4G14930.1
bic:763
peaks after deleting less than one fifth of global max:13
×10^7
raw data, starting:13
fitted Gauss locations:13
7
6
5
4
3
2
1
0
y
0
5
10
15
20
25
x

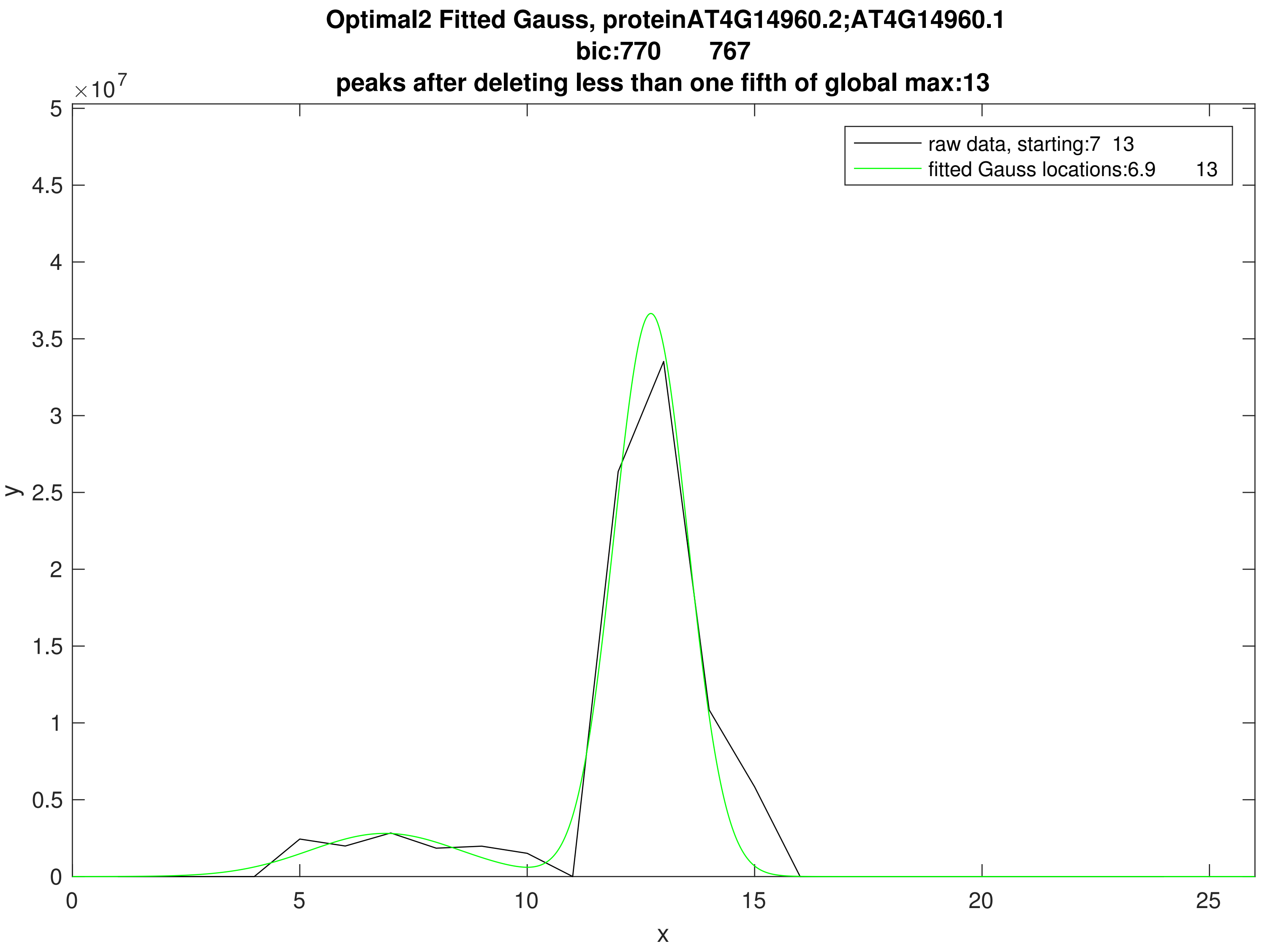
Optimal2 Fitted Gauss, proteinAT4G14960.2;AT4G14960.1
bic:770 767
peaks after deleting less than one fifth of global max:13
×10^7
raw data, starting:7 13
fitted Gauss locations:6.9 13
y
x
0
0.5
1
1.5
2
2.5
3
3.5
4
4.5
5
0
5
10
15
20
25

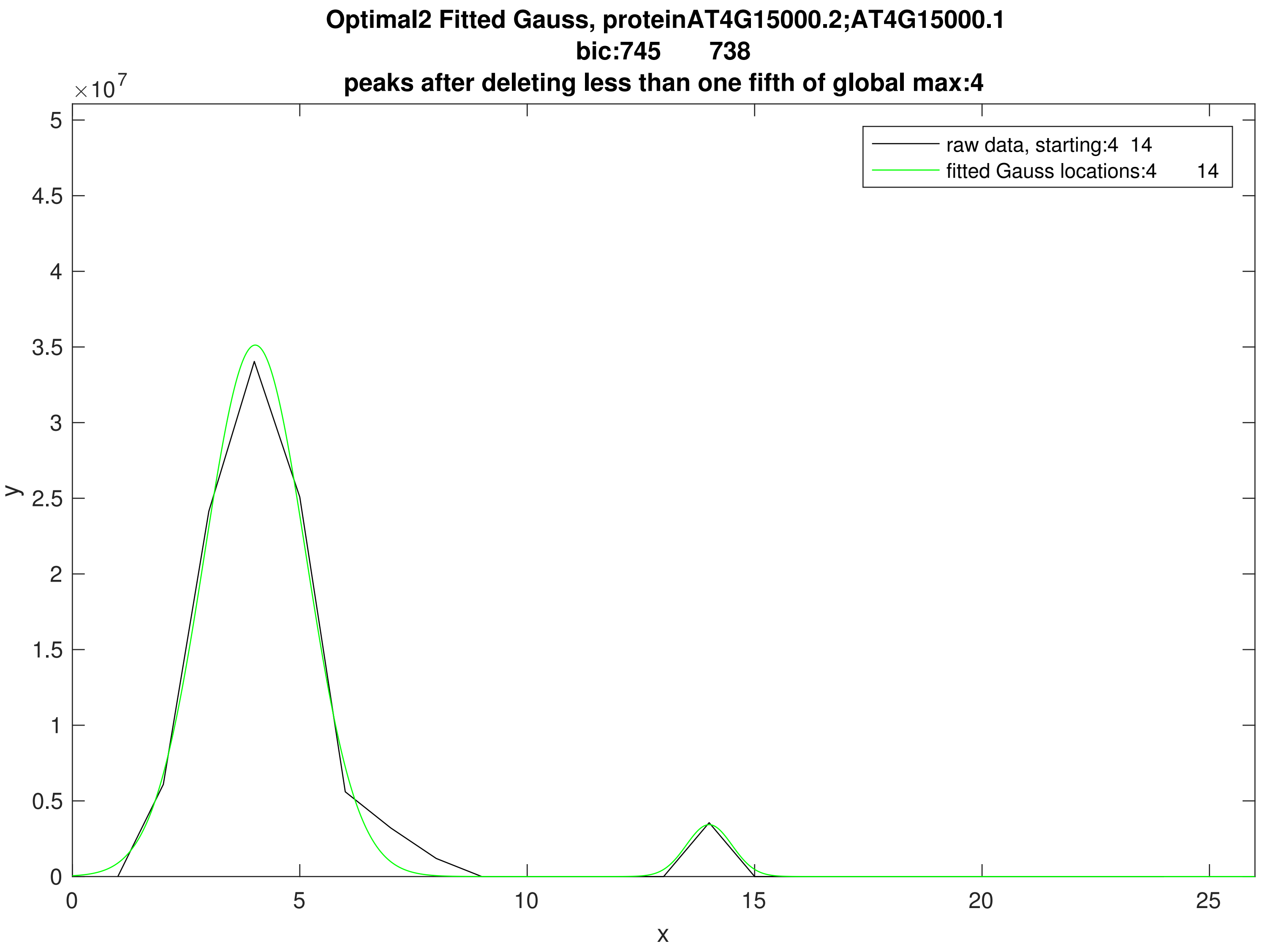
Optimal2 Fitted Gauss, proteinAT4G15000.2;AT4G15000.1
bic:745 738
peaks after deleting less than one fifth of global max:4
×10^7
raw data, starting:4 14
fitted Gauss locations:4 14
y
x
0
0.5
1
1.5
2
2.5
3
3.5
4
4.5
5
0
5
10
15
20
25

Optimal1 Fitted Gauss, proteinAT4G15093.1
bic:776
peaks after deleting less than one fifth of global max:19

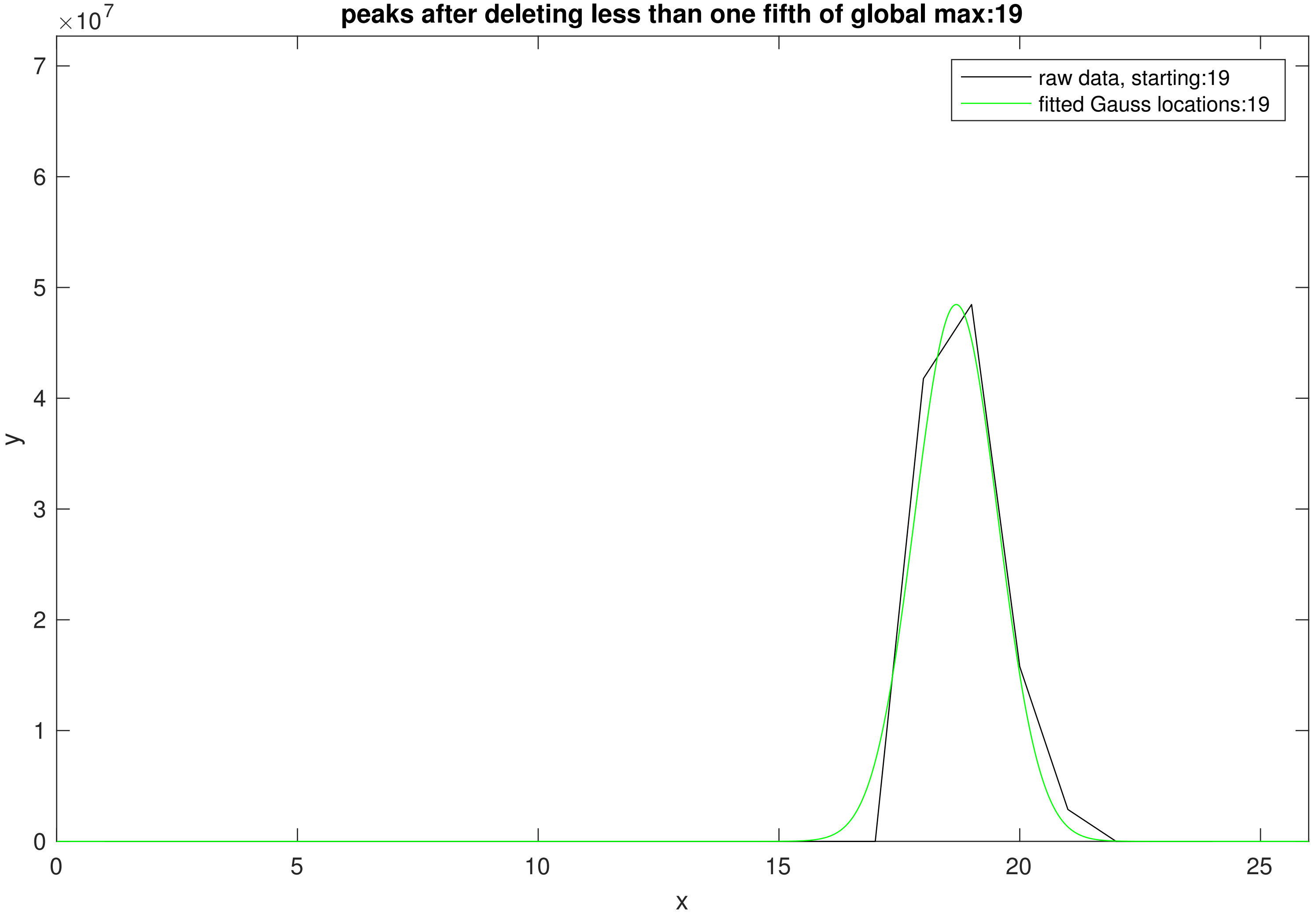

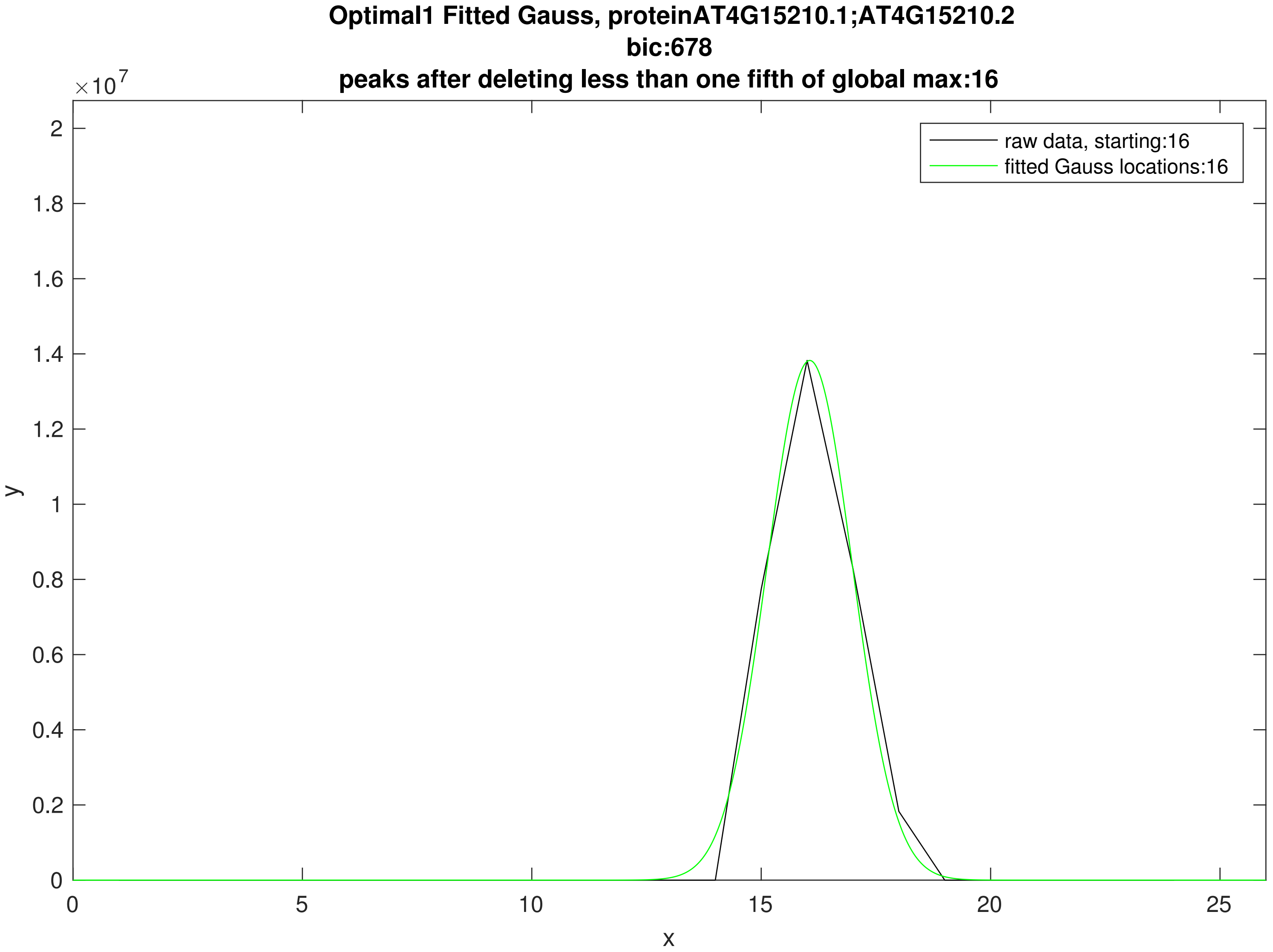
Optimal1 Fitted Gauss, proteinAT4G15210.1;AT4G15210.2
bic:678
peaks after deleting less than one fifth of global max:16
×10^7
2
1.8
1.6
1.4
1.2
1
0.8
0.6
0.4
0.2
0
y
0
5
10
15
20
25
x
raw data, starting:16
fitted Gauss locations:16

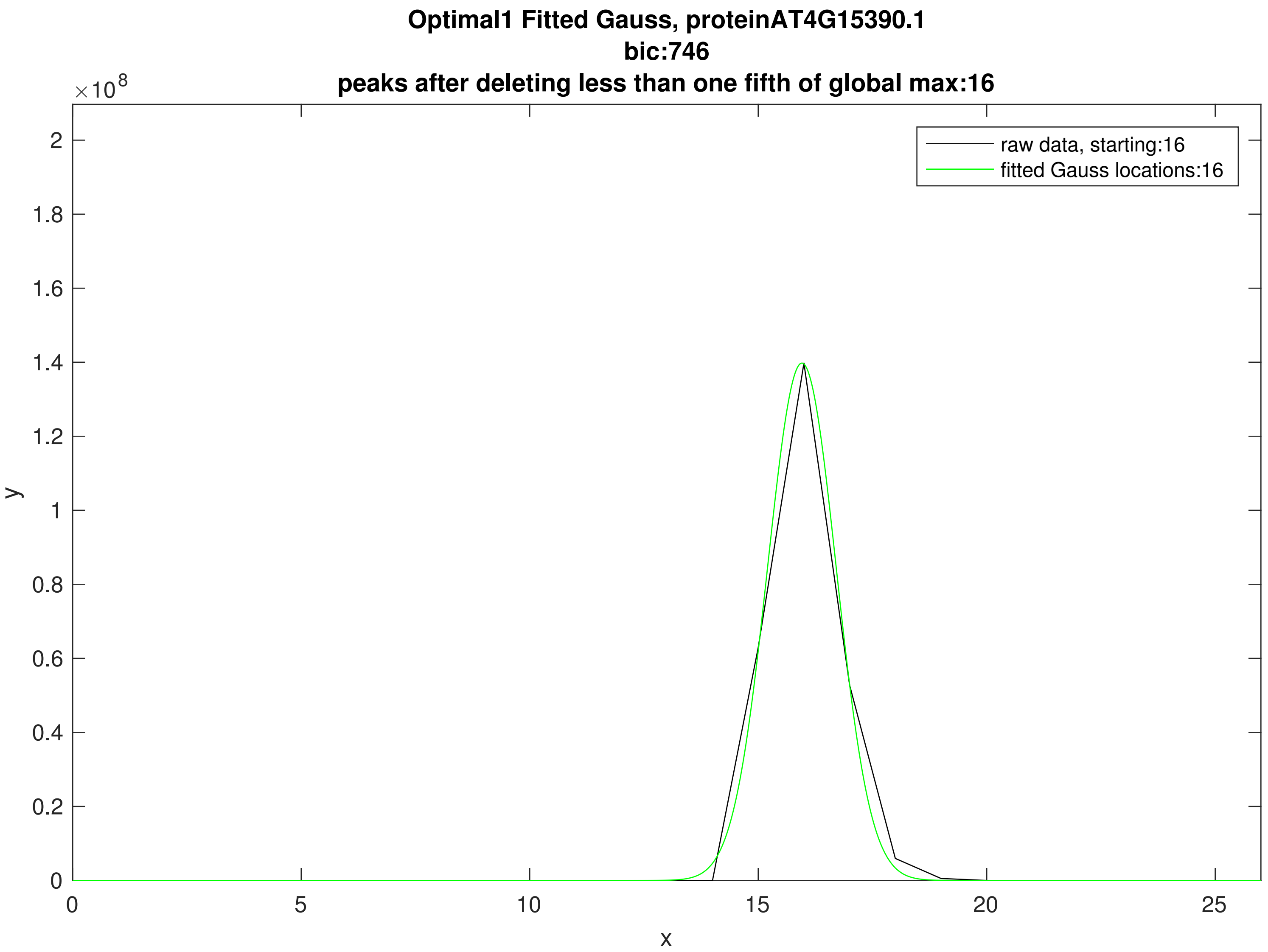
Optimal1 Fitted Gauss, proteinAT4G15390.1
bic:746
peaks after deleting less than one fifth of global max:16
×10^8
raw data, starting:16
fitted Gauss locations:16
y
x
0
0.2
0.4
0.6
0.8
1
1.2
1.4
1.6
1.8
2
0
5
10
15
20
25

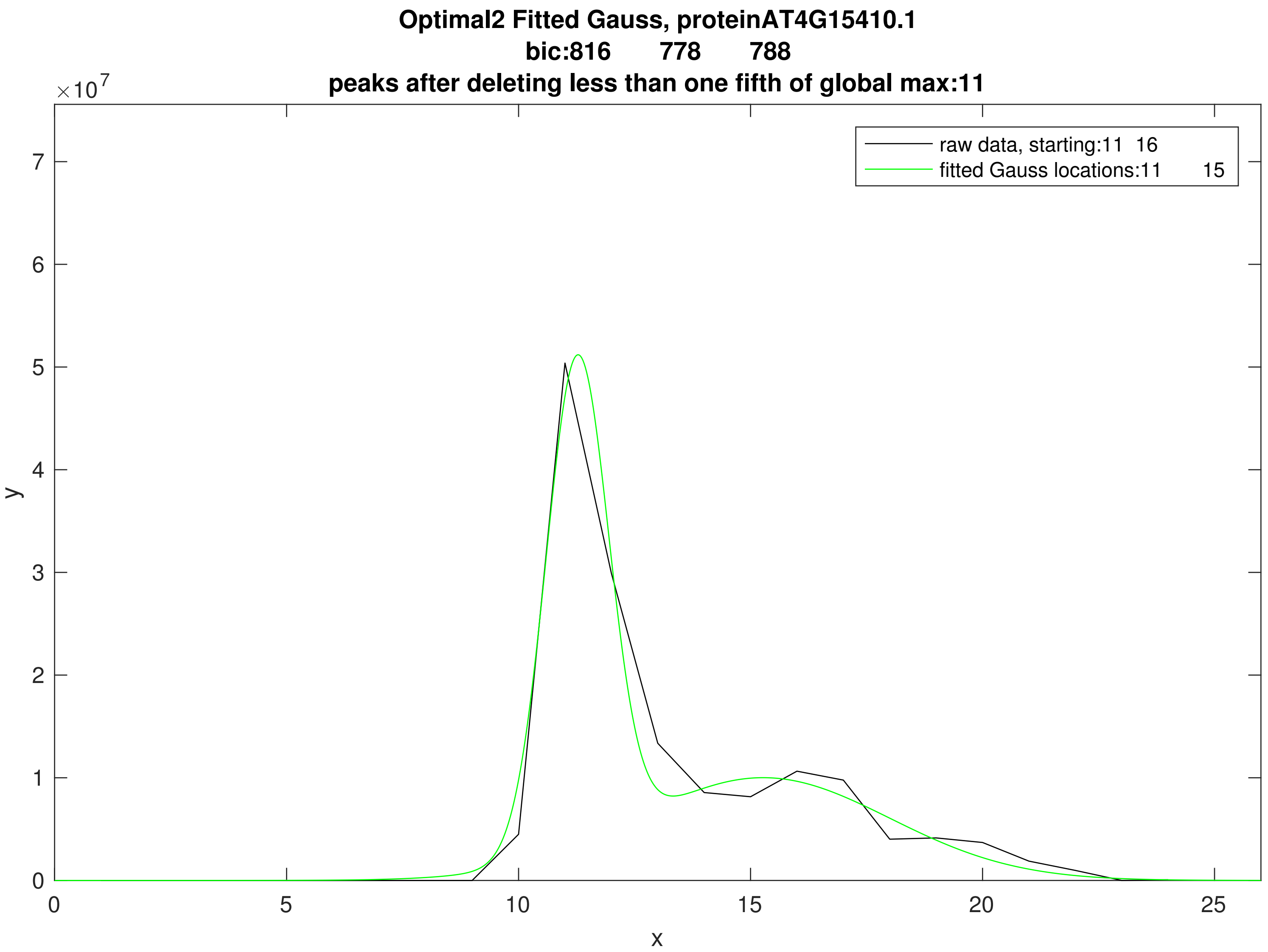
Optimal2 Fitted Gauss, proteinAT4G15410.1
bic:816 778 788
peaks after deleting less than one fifth of global max:11
×10^7
raw data, starting:11 16
fitted Gauss locations:11 15
y
x
0
1
2
3
4
5
6
7
0
5
10
15
20
25

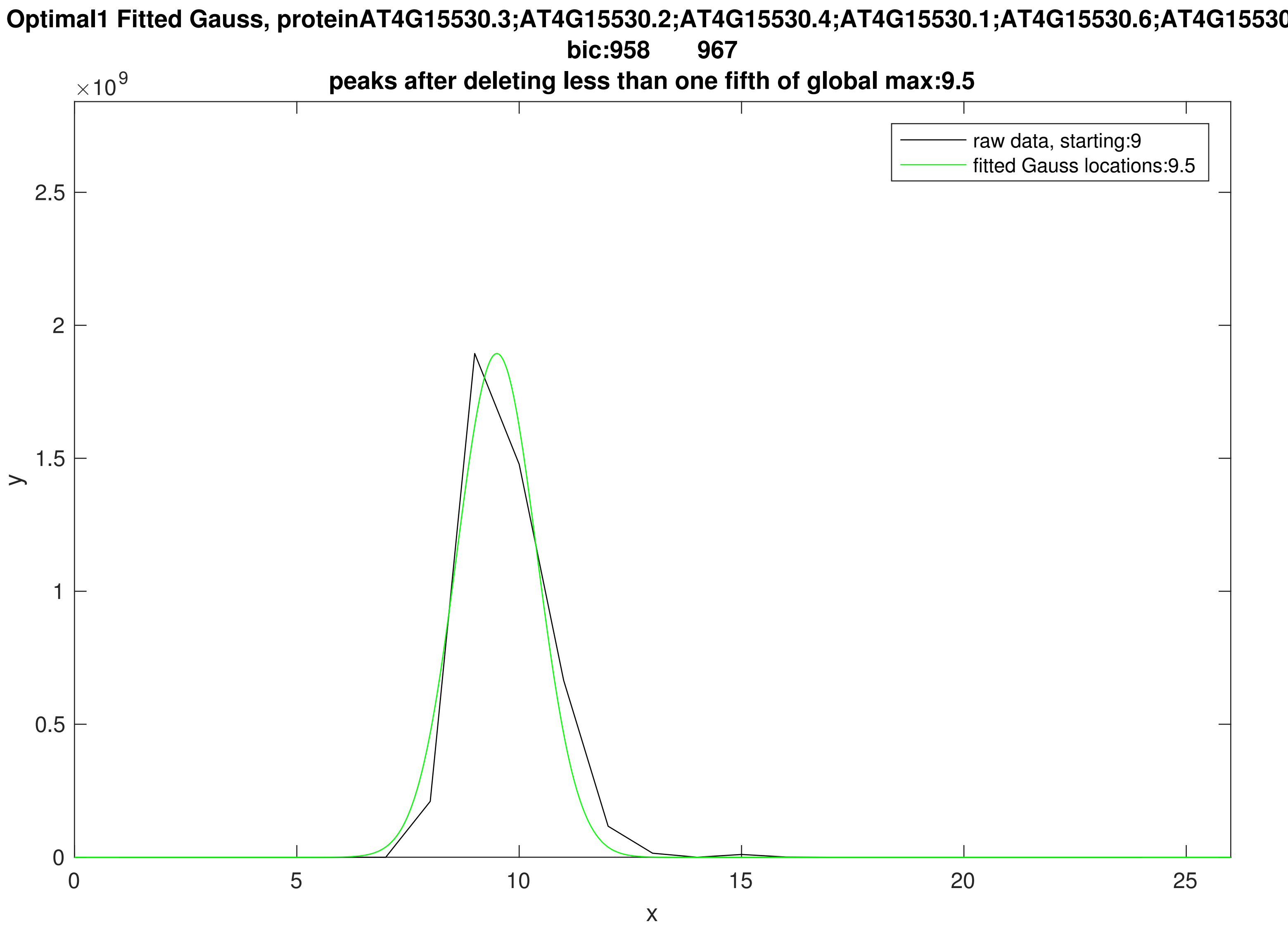

Optimal1 Fitted Gauss, proteinAT4G15530.3;AT4G15530.2;AT4G15530.4;AT4G15530.1;AT4G15530.6;AT4G15530
bic:958 967
peaks after deleting less than one fifth of global max:9.5
×10^9
raw data, starting:9
fitted Gauss locations:9.5
y
x
0
0.5
1
1.5
2
2.5
0
5
10
15
20
25

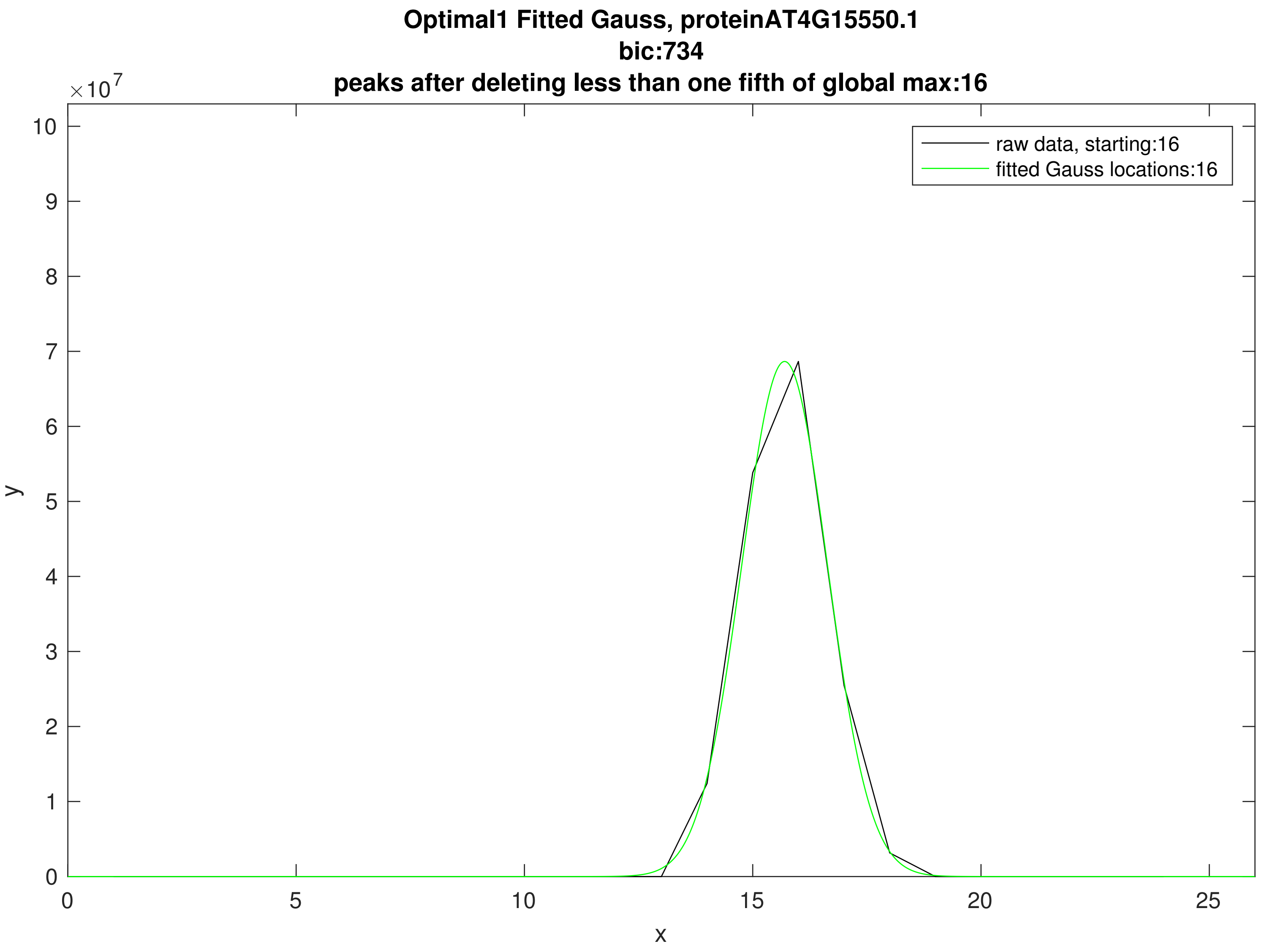

Optimal1 Fitted Gauss, proteinAT4G15550.1
bic:734
peaks after deleting less than one fifth of global max:16
×10^7
10
9
8
7
6
5
4
3
2
1
0
y
0
5
10
15
20
25
x
raw data, starting:16
fitted Gauss locations:16

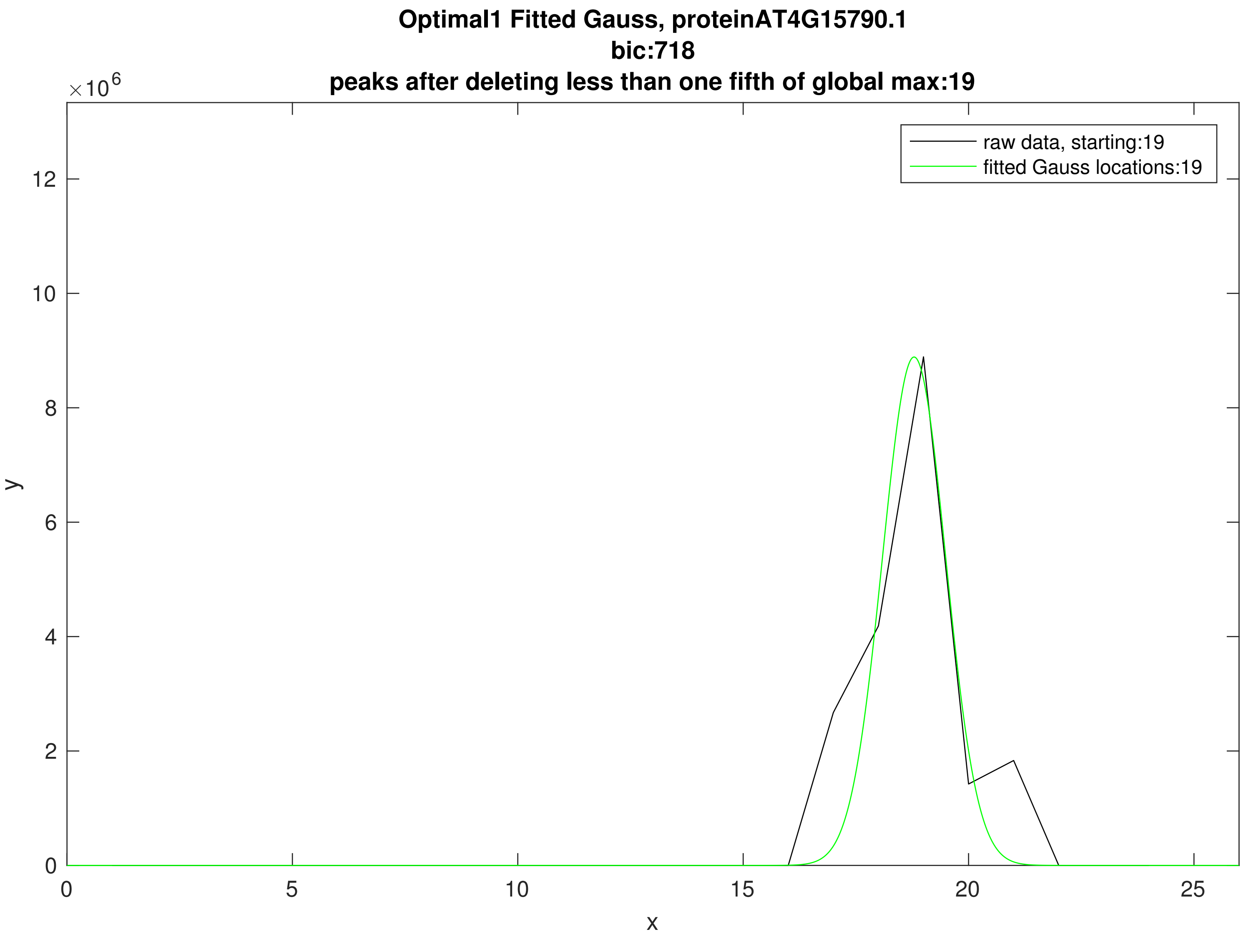

Optimal1 Fitted Gauss, proteinAT4G15790.1
bic:718
peaks after deleting less than one fifth of global max:19
×10^6
raw data, starting:19
fitted Gauss locations:19
y
x
0
2
4
6
8
10
12
0
5
10
15
20
25

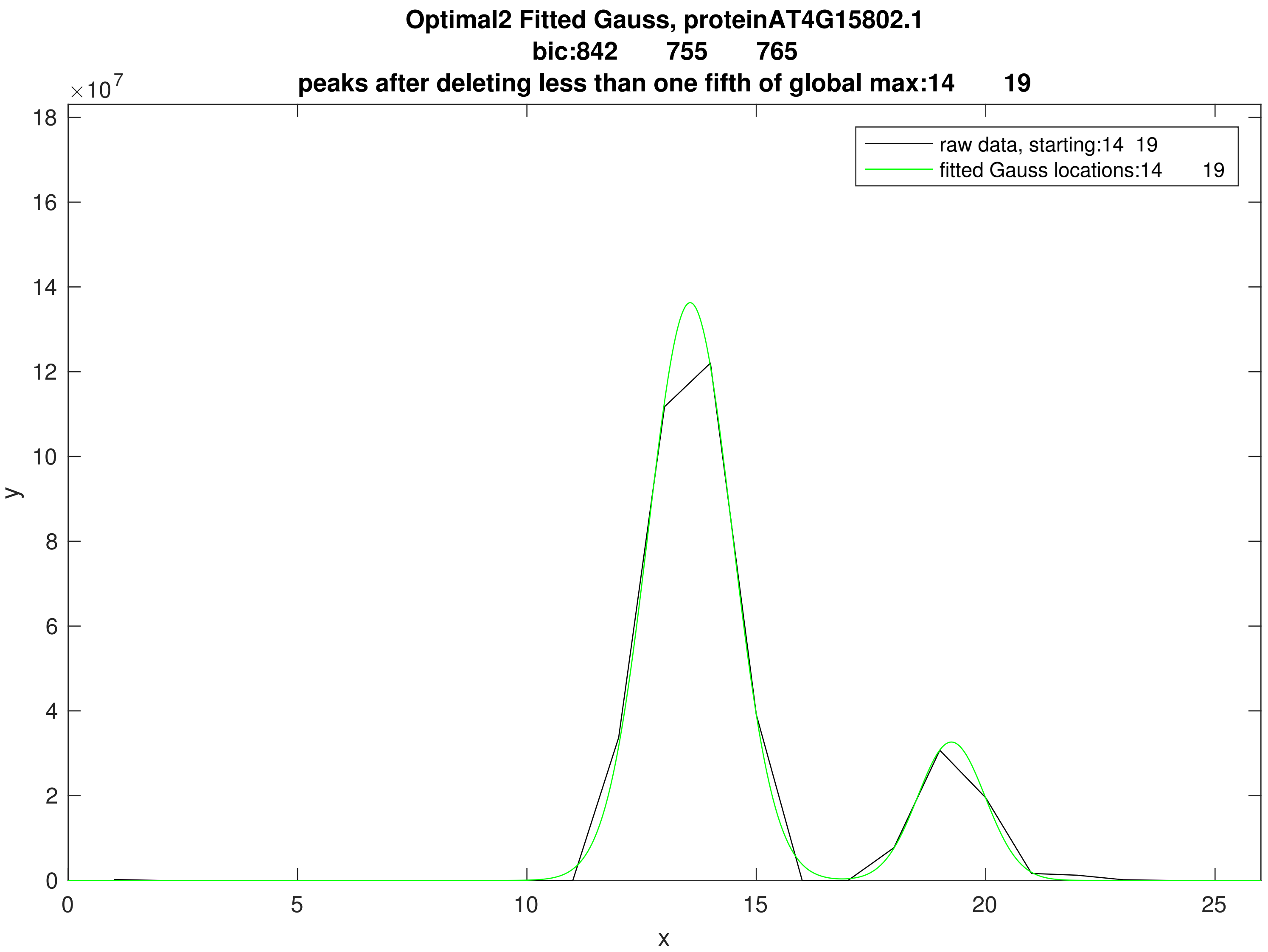

Optimal2 Fitted Gauss, proteinAT4G15802.1
bic:842 755 765
peaks after deleting less than one fifth of global max:14 19
×10^7
18
16
14
12
10
8
6
4
2
0
y
0
5
10
15
20
25
x
raw data, starting:14 19
fitted Gauss locations:14 19

**Optimal2 Fitted Gauss, proteinAT4G15940.1**
**bic:826 823**
**peaks after deleting less than one fifth of global max:16**

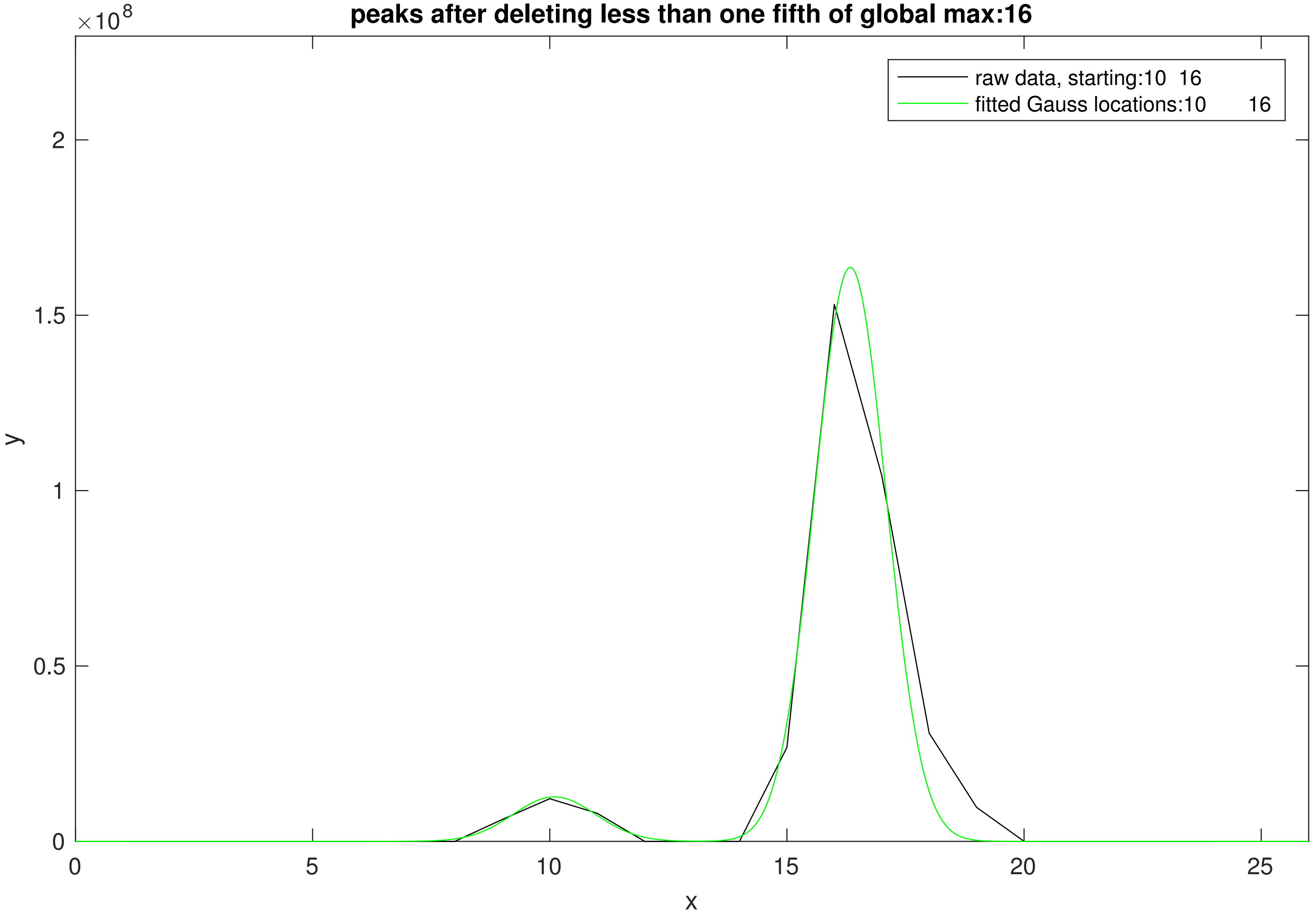

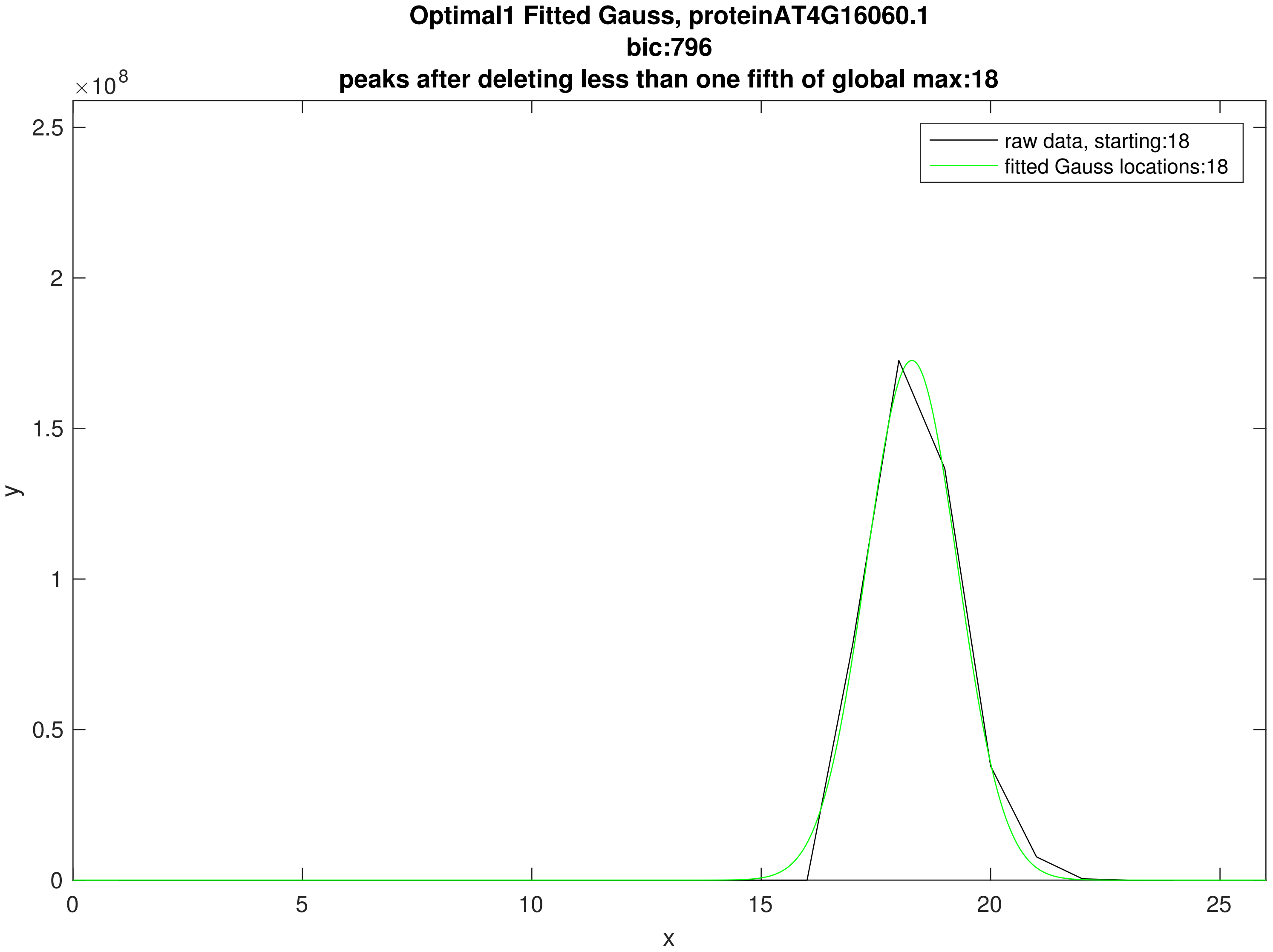
Optimal1 Fitted Gauss, proteinAT4G16060.1
bic:796
peaks after deleting less than one fifth of global max:18
×10^8
2.5
2
1.5
1
0.5
0
y
0
5
10
15
20
25
x
raw data, starting:18
fitted Gauss locations:18

**Optimal1 Fitted Gauss, proteinAT4G16130.1**
**bic:819 851**
**peaks after deleting less than one fifth of global max:9.2**

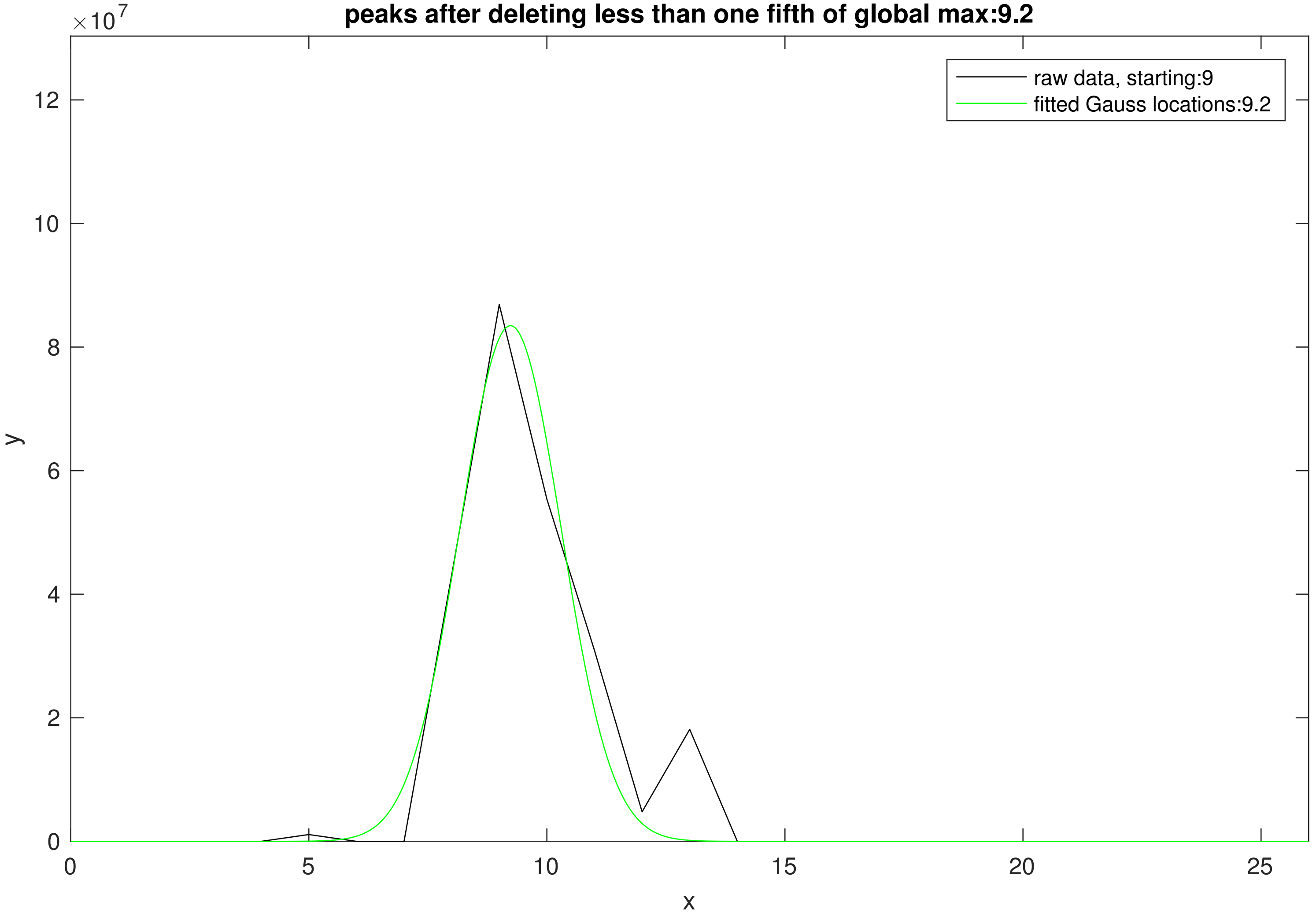

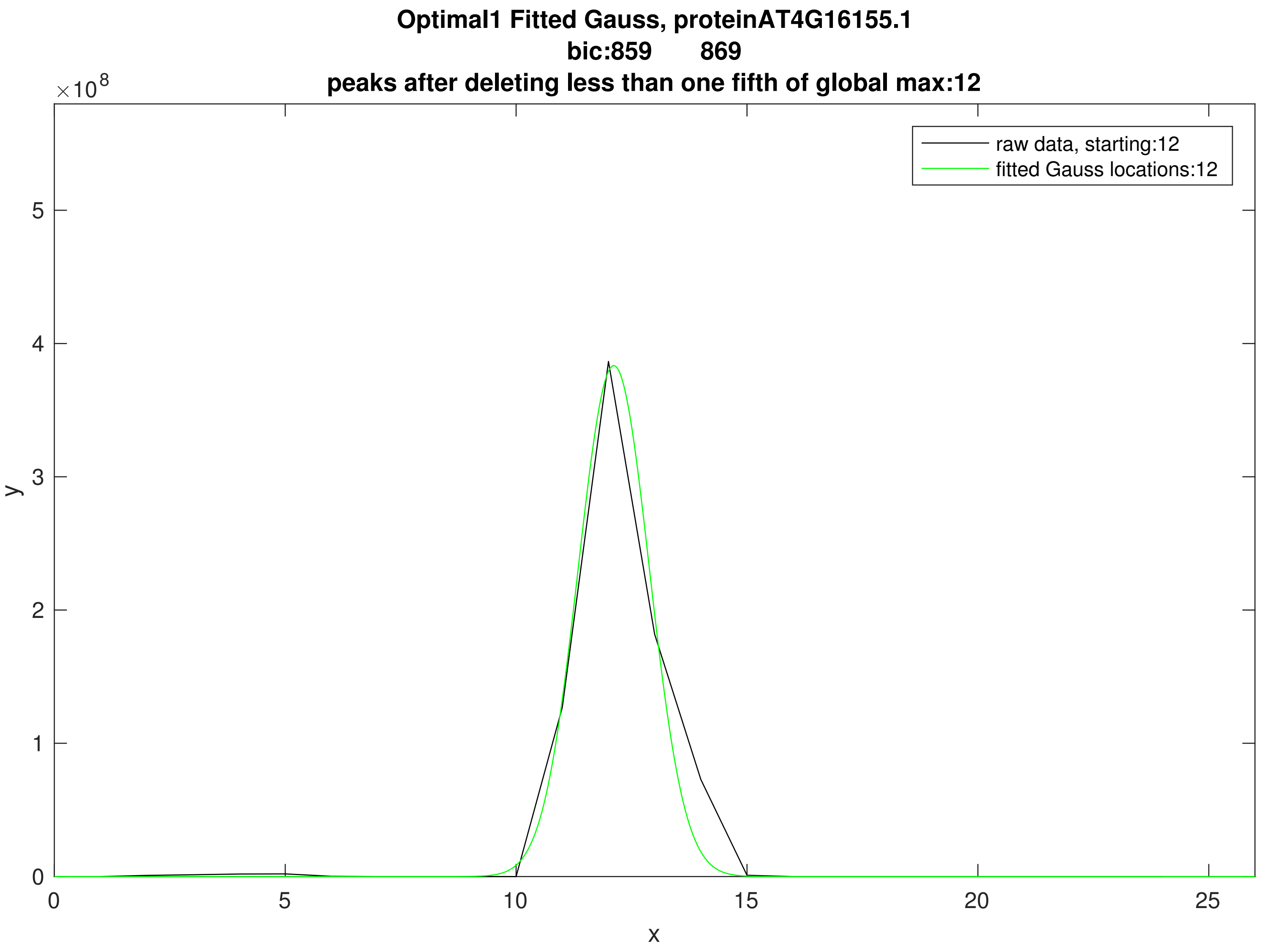
Optimal1 Fitted Gauss, proteinAT4G16155.1
bic:859 869
peaks after deleting less than one fifth of global max:12
×10^8
raw data, starting:12
fitted Gauss locations:12
5
4
3
2
1
0
y
0
5
10
15
20
25
x

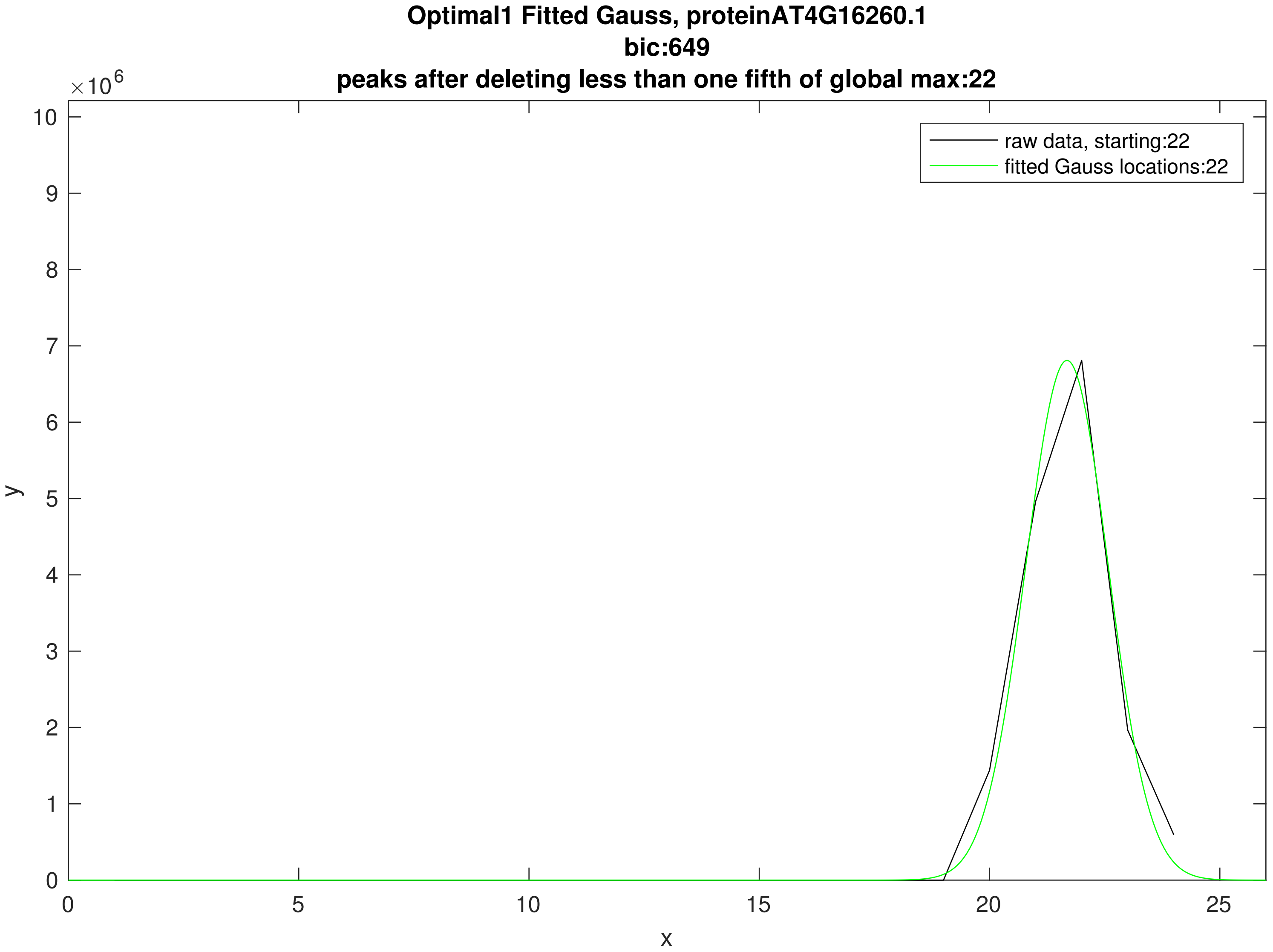

Optimal1 Fitted Gauss, proteinAT4G16260.1
bic:649
peaks after deleting less than one fifth of global max:22
×10^6
raw data, starting:22
fitted Gauss locations:22
y
x
0
1
2
3
4
5
6
7
8
9
10
0
5
10
15
20
25

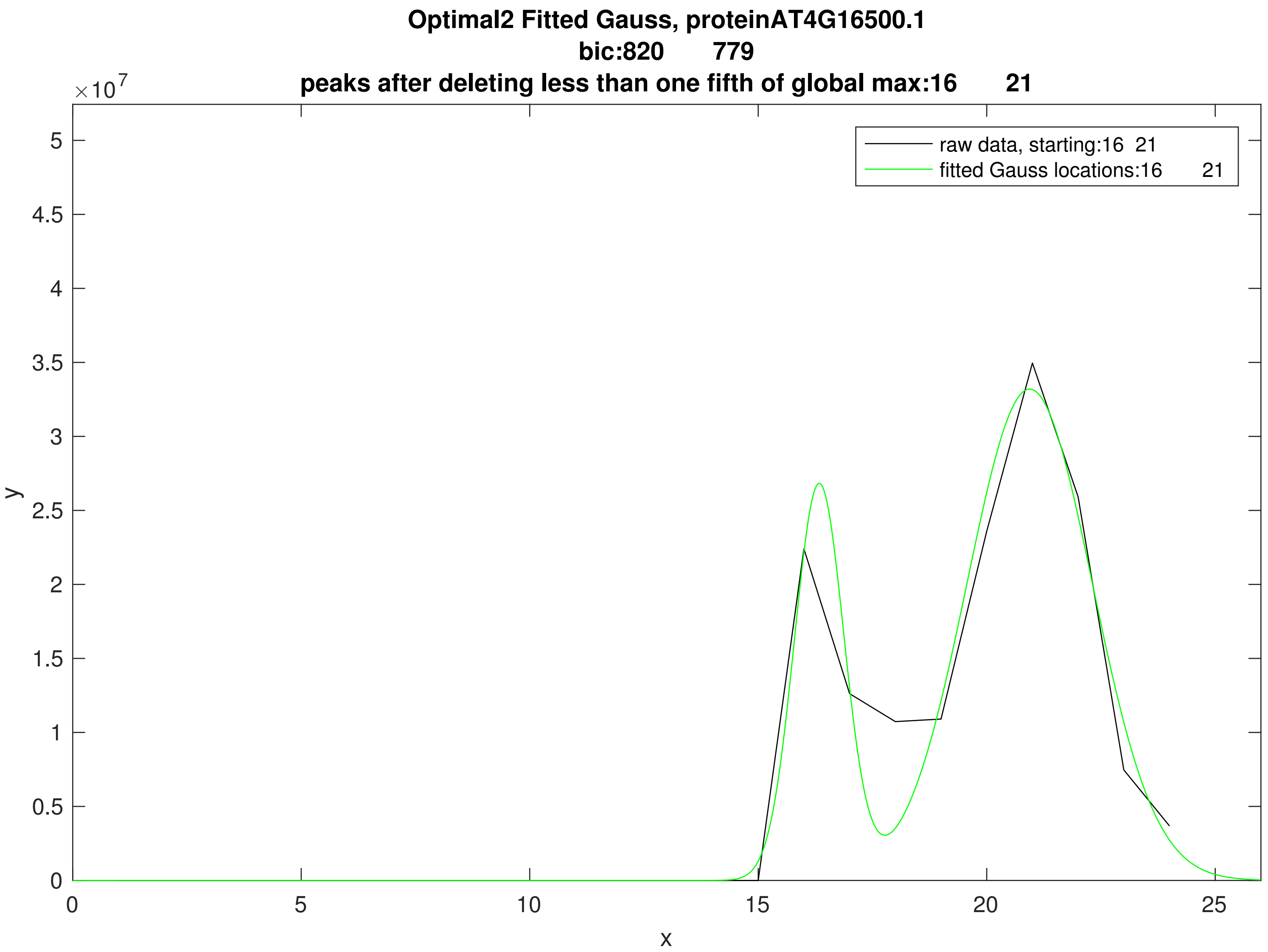

Optimal2 Fitted Gauss, proteinAT4G16500.1
bic:820 779
peaks after deleting less than one fifth of global max:16 21
×10^7
raw data, starting:16 21
fitted Gauss locations:16 21
y
x
0
0.5
1
1.5
2
2.5
3
3.5
4
4.5
5
0
5
10
15
20
25

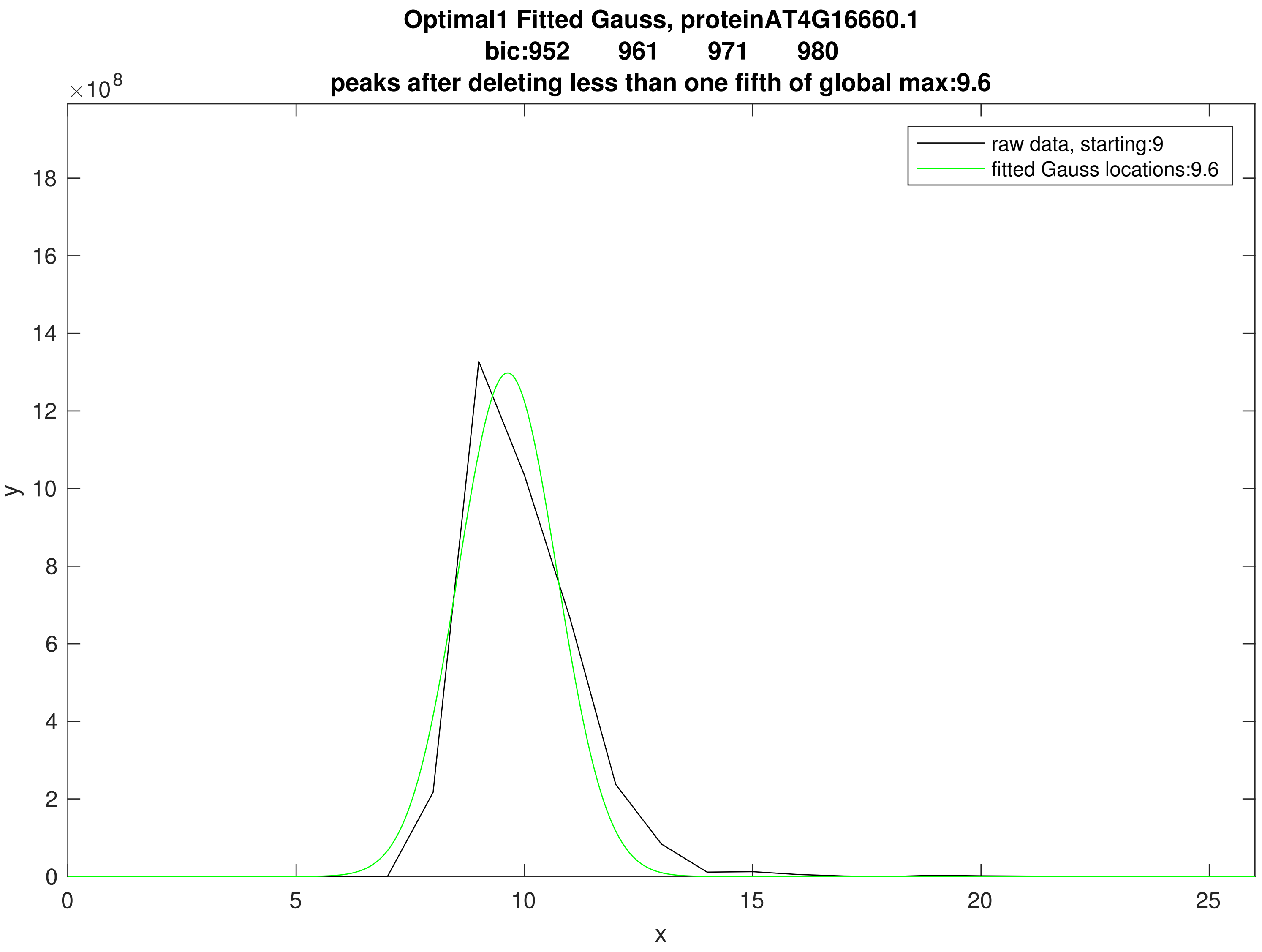
Optimal1 Fitted Gauss, proteinAT4G16660.1
bic:952 961 971 980
peaks after deleting less than one fifth of global max:9.6
×10^8
raw data, starting:9
fitted Gauss locations:9.6
y
x
0
2
4
6
8
10
12
14
16
18
0
5
10
15
20
25

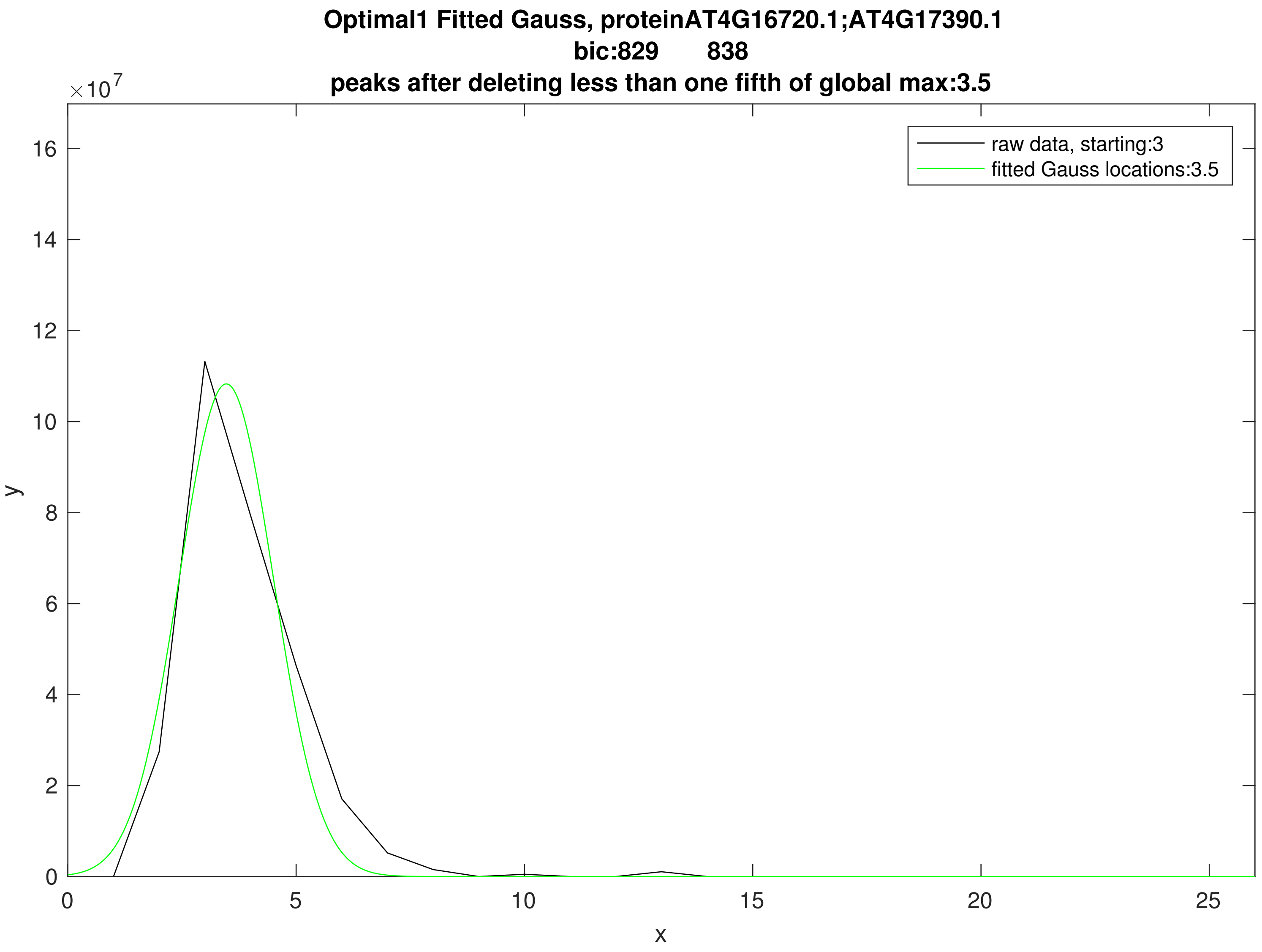
Optimal1 Fitted Gauss, proteinAT4G16720.1;AT4G17390.1
bic:829 838
peaks after deleting less than one fifth of global max:3.5
×10^7
16
14
12
10
8
6
4
2
0
y
0
5
10
15
20
25
x
raw data, starting:3
fitted Gauss locations:3.5

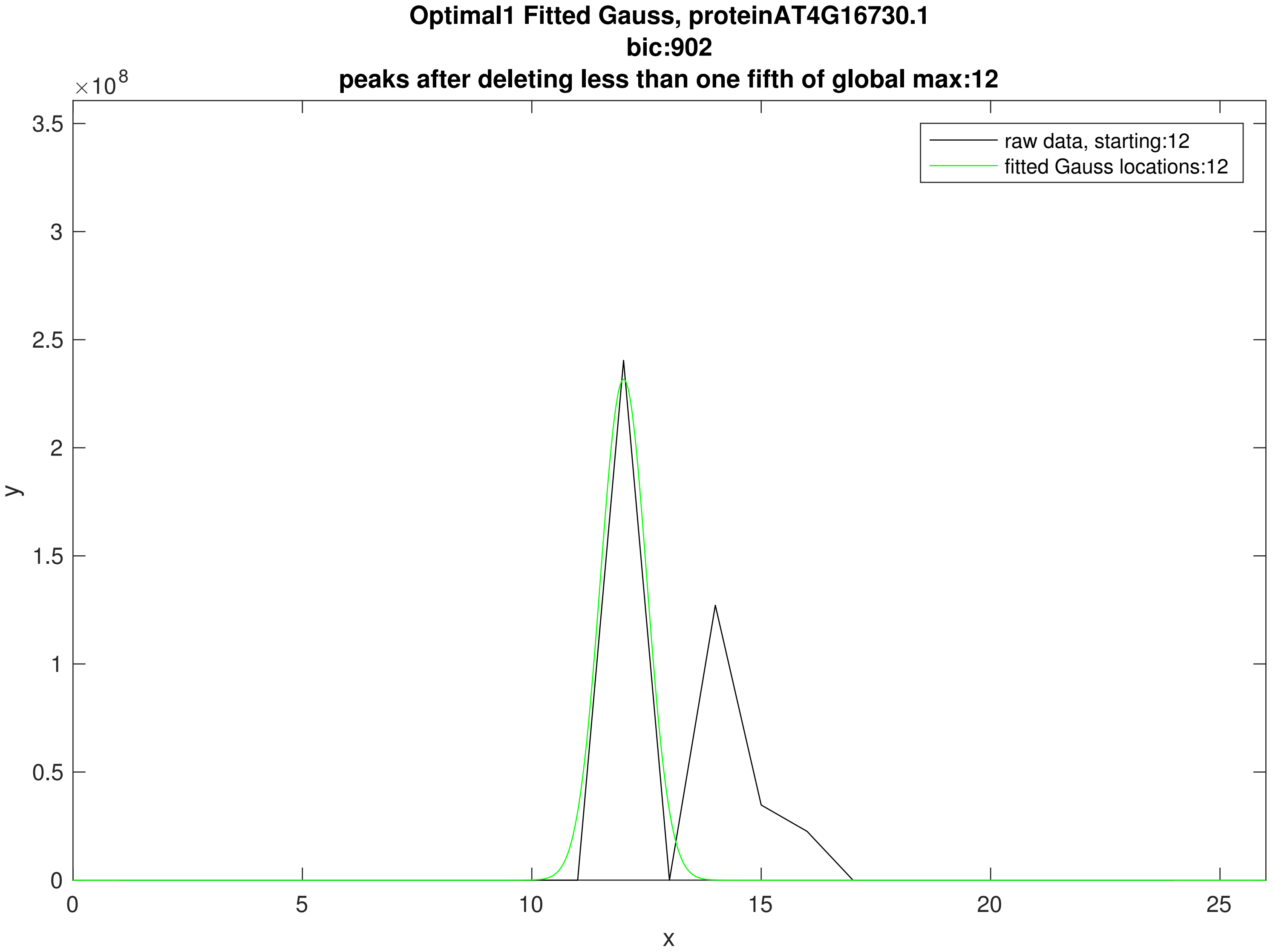
Optimal1 Fitted Gauss, proteinAT4G16730.1
bic:902
peaks after deleting less than one fifth of global max:12
×10^8
3.5
3
2.5
2
1.5
1
0.5
0
y
0
5
10
15
20
25
x
raw data, starting:12
fitted Gauss locations:12

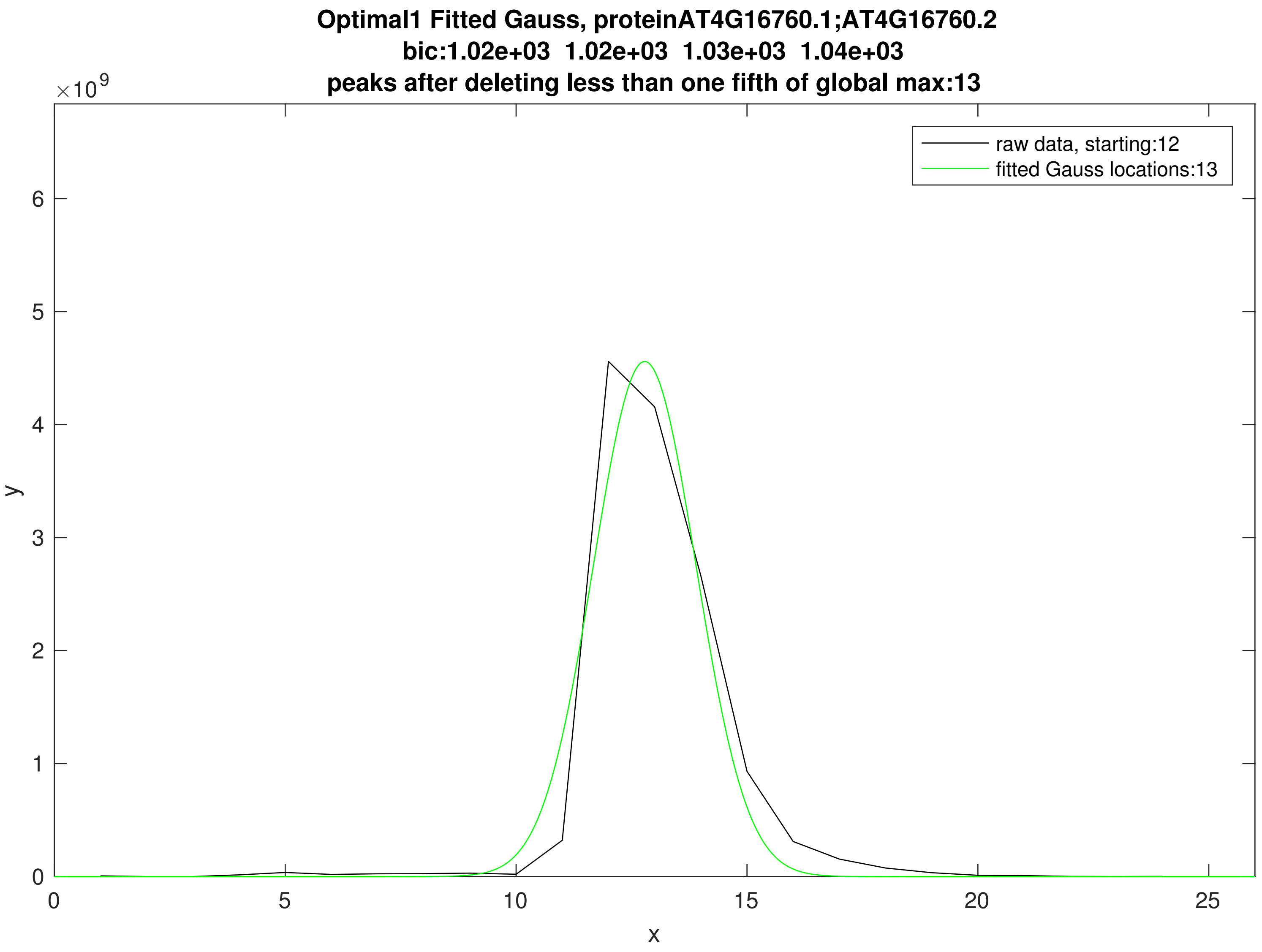
Optimal1 Fitted Gauss, proteinAT4G16760.1;AT4G16760.2
bic:1.02e+03 1.02e+03 1.03e+03 1.04e+03
peaks after deleting less than one fifth of global max:13
×10^9
raw data, starting:12
fitted Gauss locations:13
y
x
0
1
2
3
4
5
6
0
5
10
15
20
25

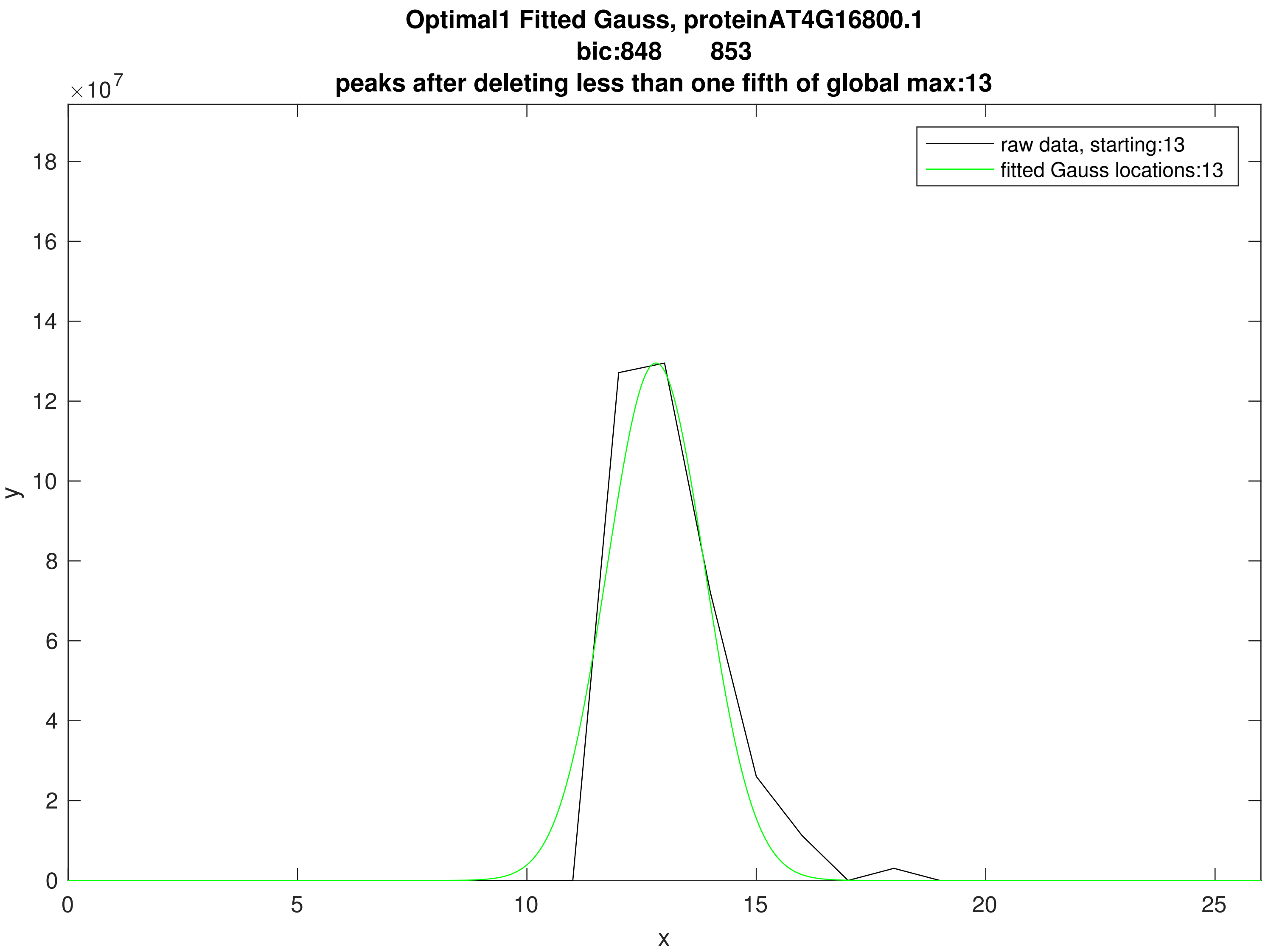
Optimal1 Fitted Gauss, proteinAT4G16800.1
bic:848 853
peaks after deleting less than one fifth of global max:13
×10^7
raw data, starting:13
fitted Gauss locations:13
y
x
0
2
4
6
8
10
12
14
16
18
0
5
10
15
20
25

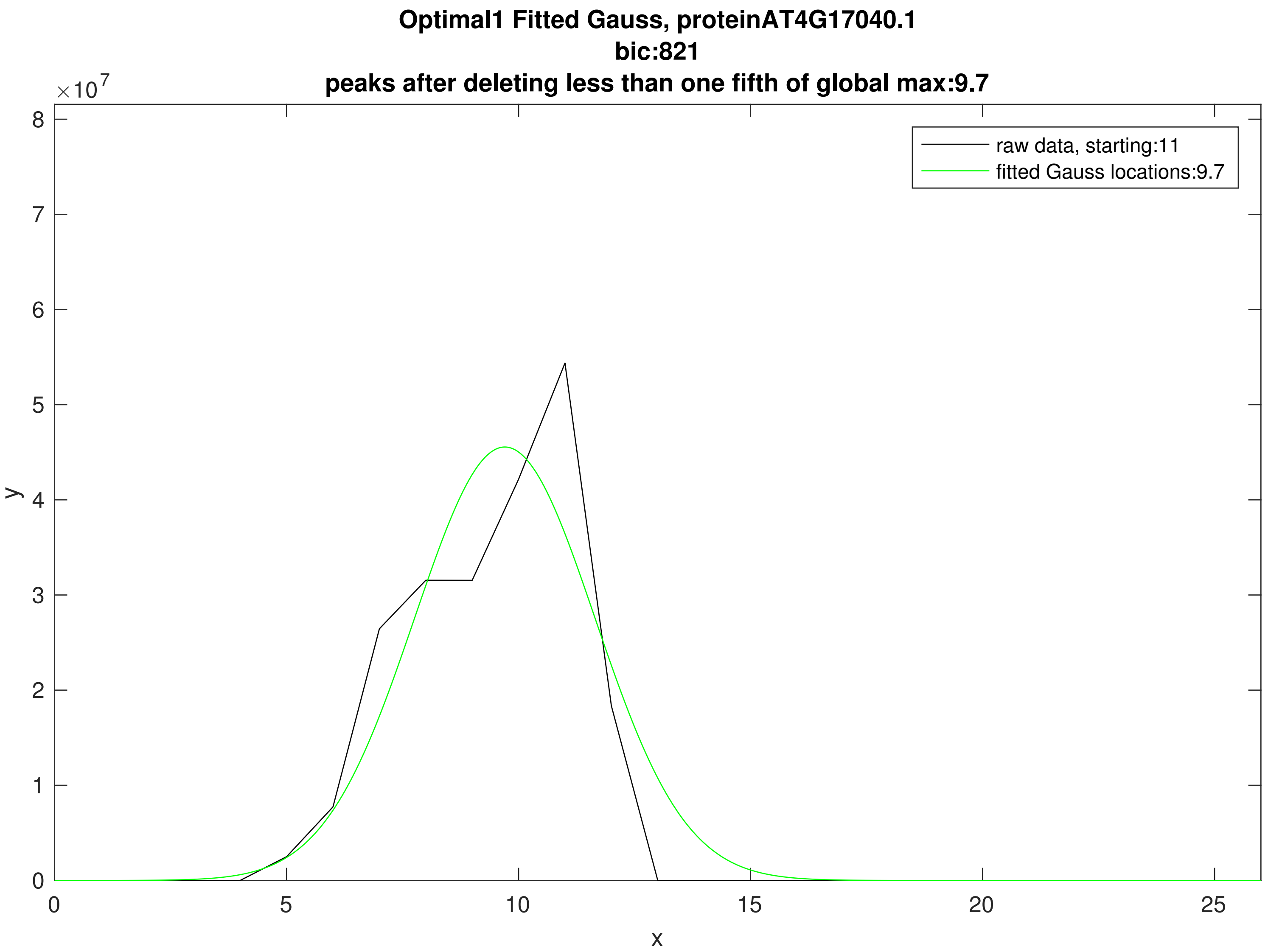

Optimal1 Fitted Gauss, proteinAT4G17040.1
bic:821
peaks after deleting less than one fifth of global max:9.7
×10^7
8
7
6
5
4
3
2
1
0
y
0
5
10
15
20
25
x
raw data, starting:11
fitted Gauss locations:9.7

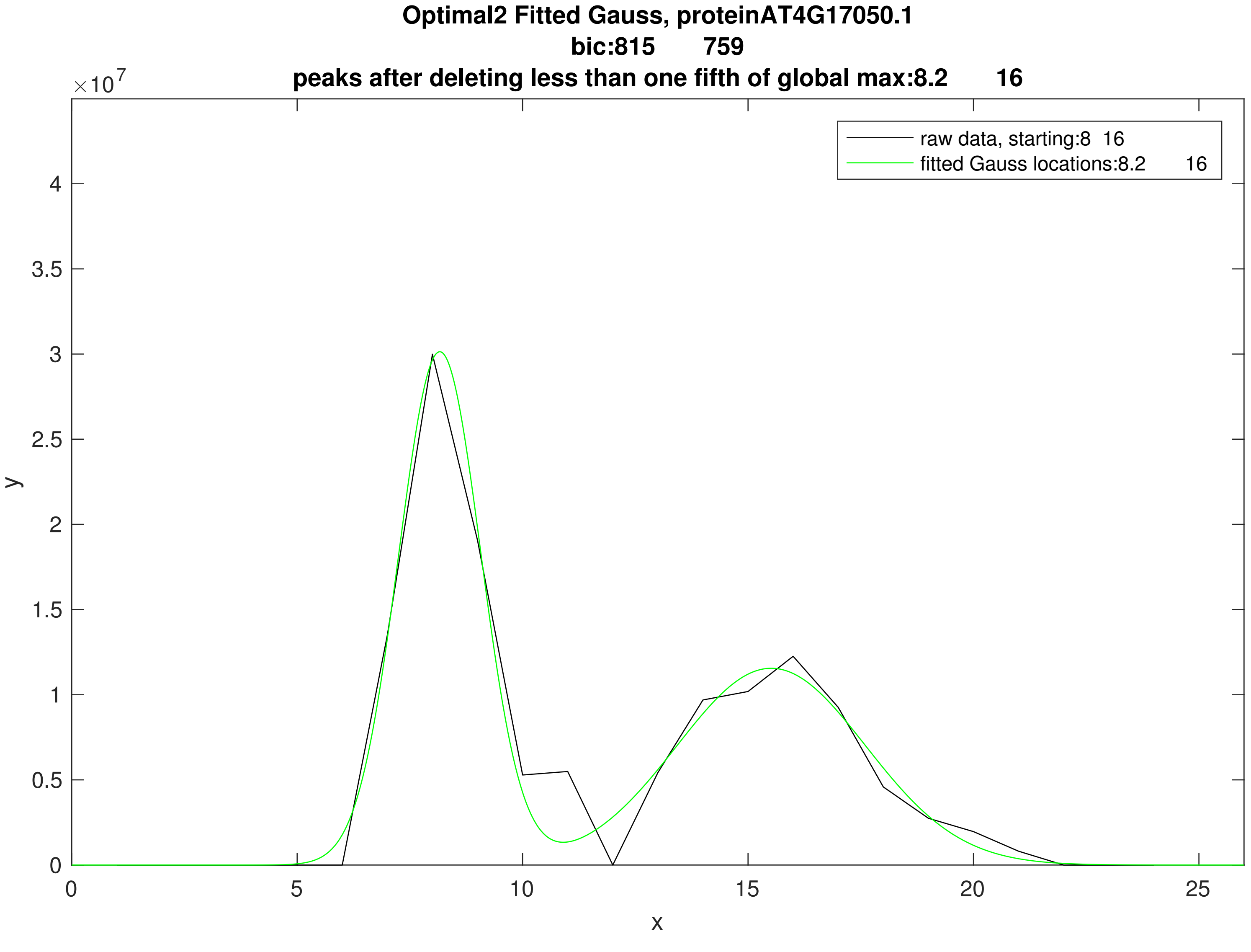
Optimal2 Fitted Gauss, proteinAT4G17050.1
bic:815 759
peaks after deleting less than one fifth of global max:8.2 16
×10^7
raw data, starting:8 16
fitted Gauss locations:8.2 16
y
x
0
0.5
1
1.5
2
2.5
3
3.5
4
5
10
15
20
25

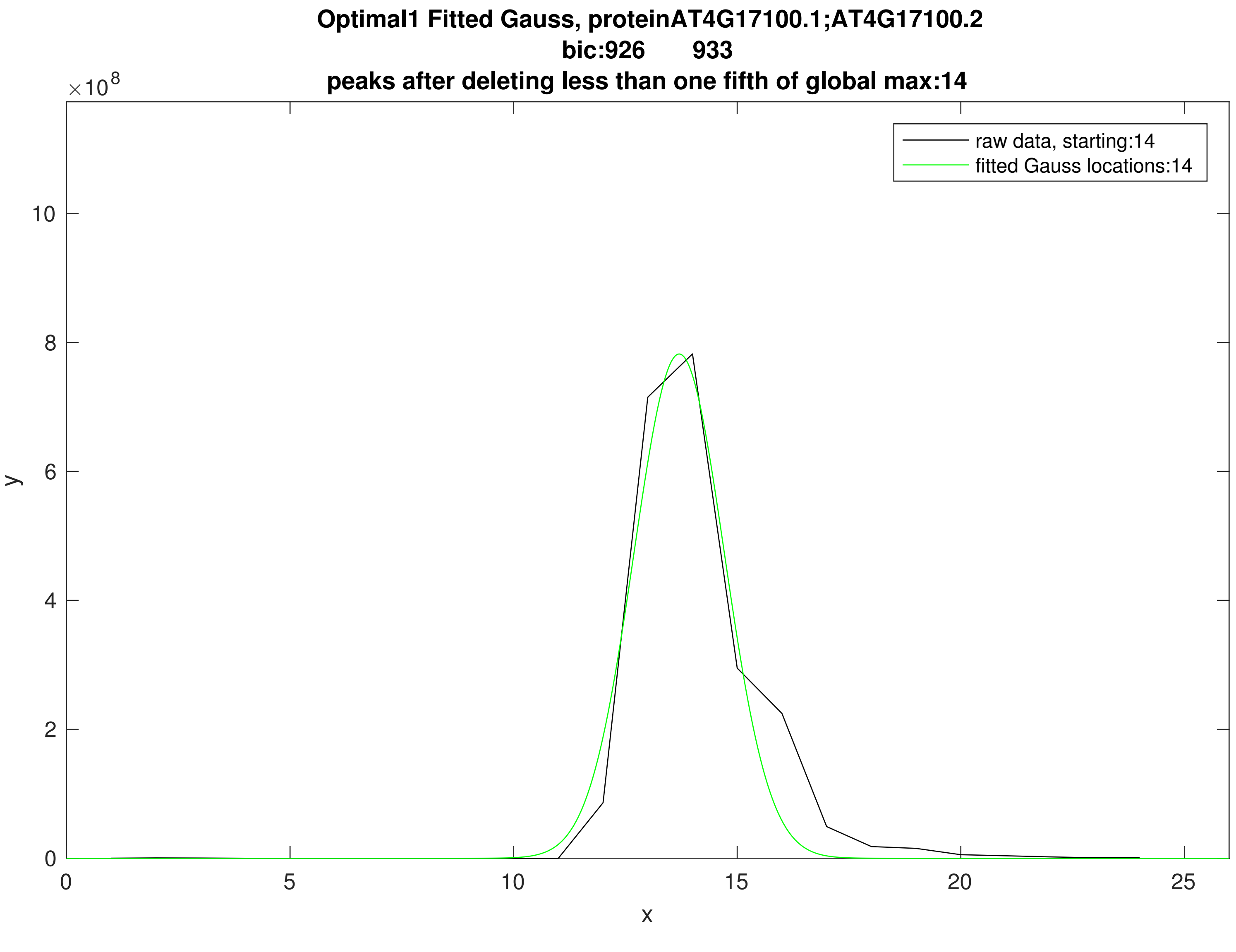

Optimal1 Fitted Gauss, proteinAT4G17100.1;AT4G17100.2
bic:926 933
peaks after deleting less than one fifth of global max:14
×10^8
raw data, starting:14
fitted Gauss locations:14
y
x
0
2
4
6
8
10
0
5
10
15
20
25

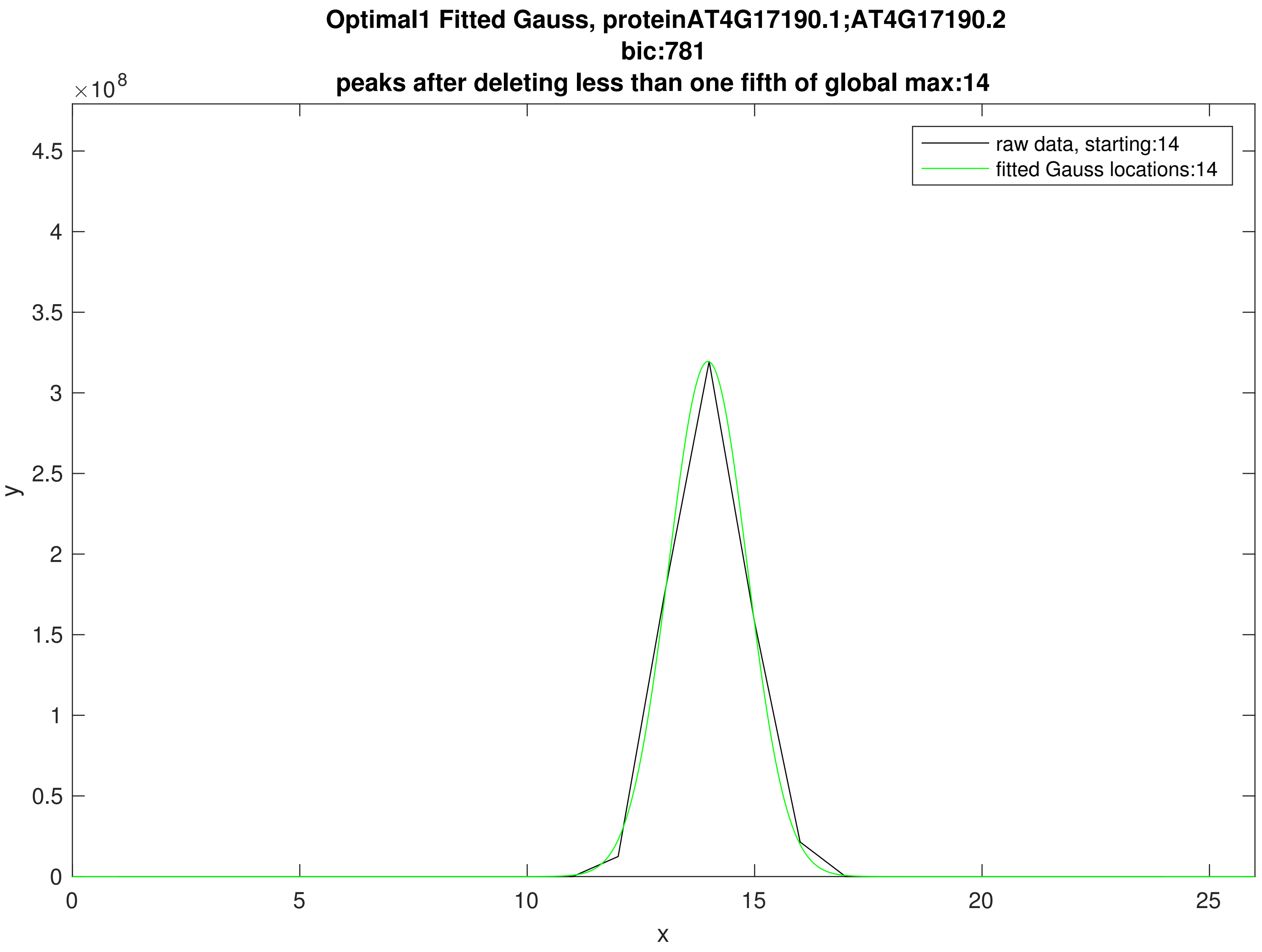
Optimal1 Fitted Gauss, proteinAT4G17190.1;AT4G17190.2
bic:781
peaks after deleting less than one fifth of global max:14
raw data, starting:14
fitted Gauss locations:14
$\times 10^8$
y
x
0
0.5
1
1.5
2
2.5
3
3.5
4
4.5
0
5
10
15
20
25

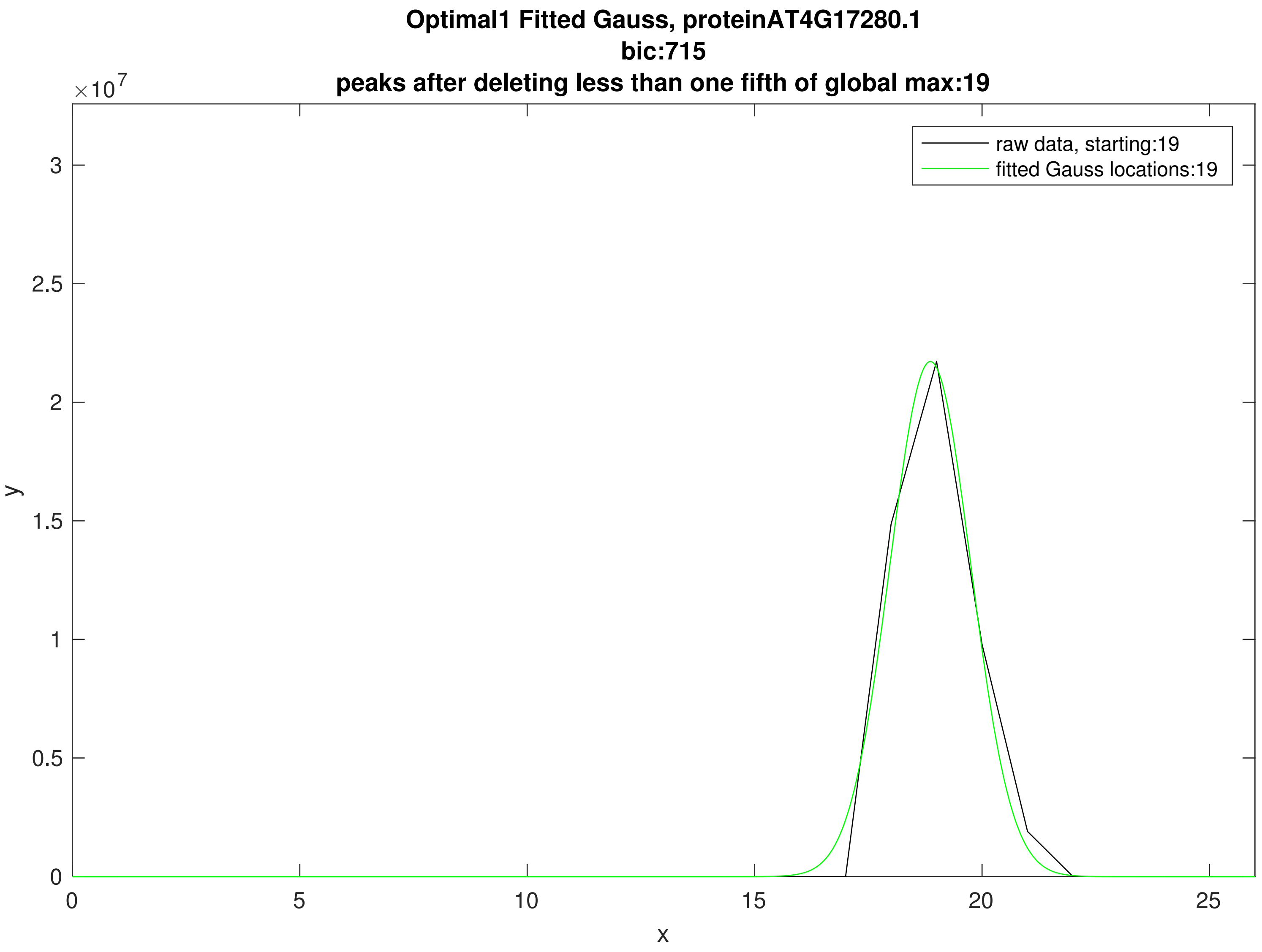

Optimal1 Fitted Gauss, proteinAT4G17280.1
bic:715
peaks after deleting less than one fifth of global max:19
×10^7
raw data, starting:19
fitted Gauss locations:19
y
x
0
0.5
1
1.5
2
2.5
3
5
10
15
20
25

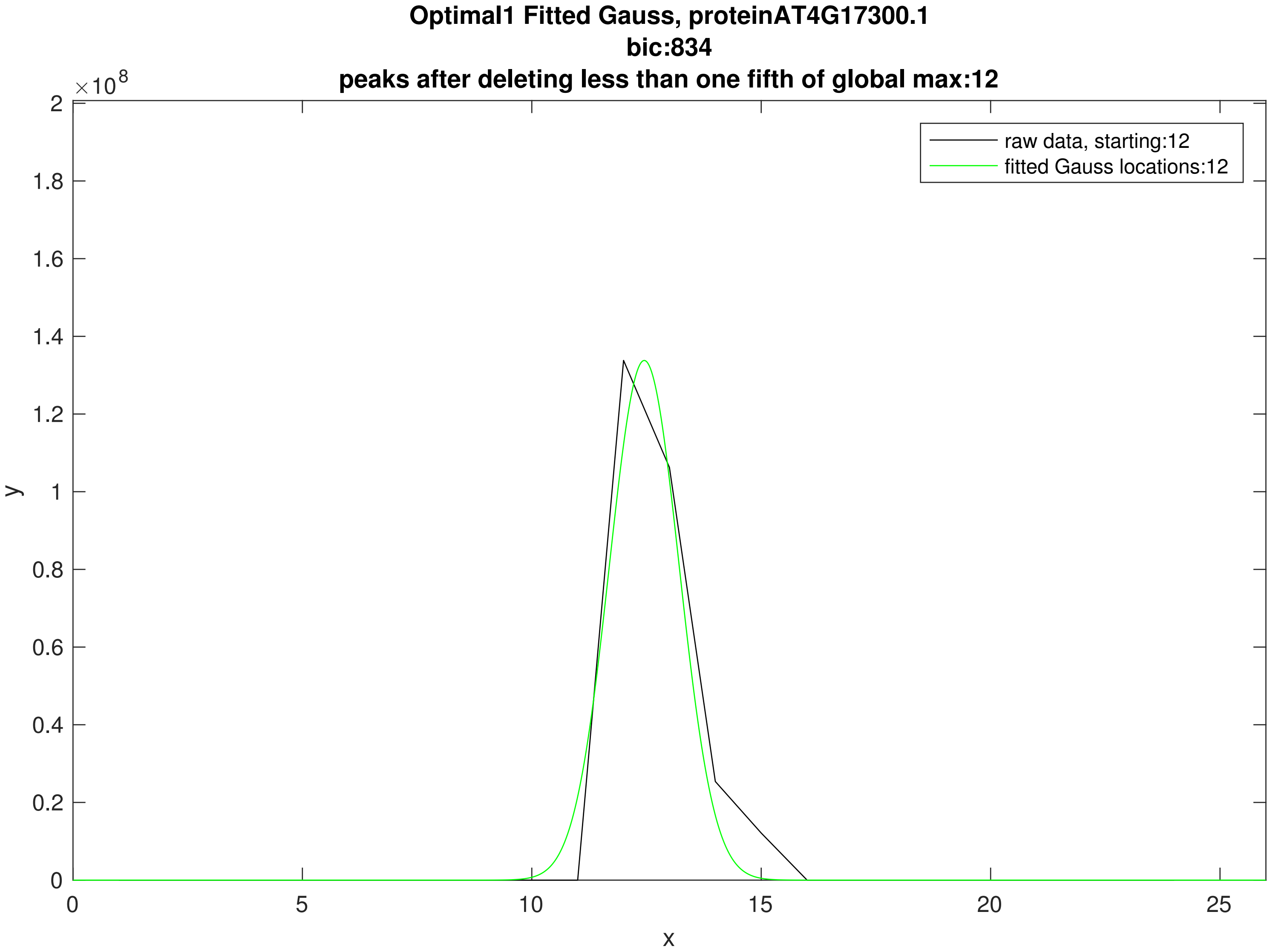
Optimal1 Fitted Gauss, proteinAT4G17300.1
bic:834
peaks after deleting less than one fifth of global max:12
×10^8
raw data, starting:12
fitted Gauss locations:12
y
x
0
0.2
0.4
0.6
0.8
1
1.2
1.4
1.6
1.8
2
0
5
10
15
20
25

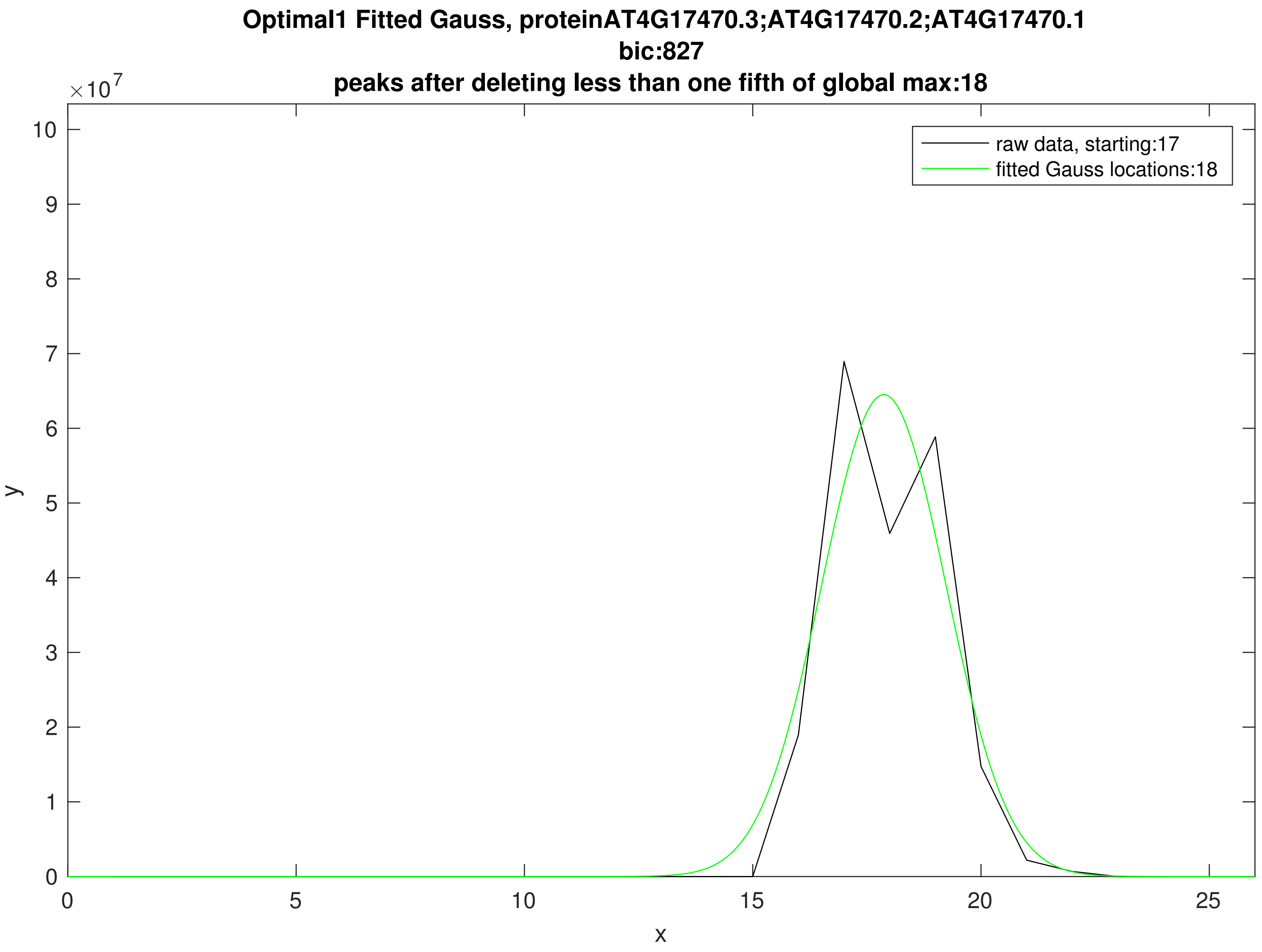
Optimal1 Fitted Gauss, proteinAT4G17470.3;AT4G17470.2;AT4G17470.1
bic:827
peaks after deleting less than one fifth of global max:18
×10^7
raw data, starting:17
fitted Gauss locations:18
y
x

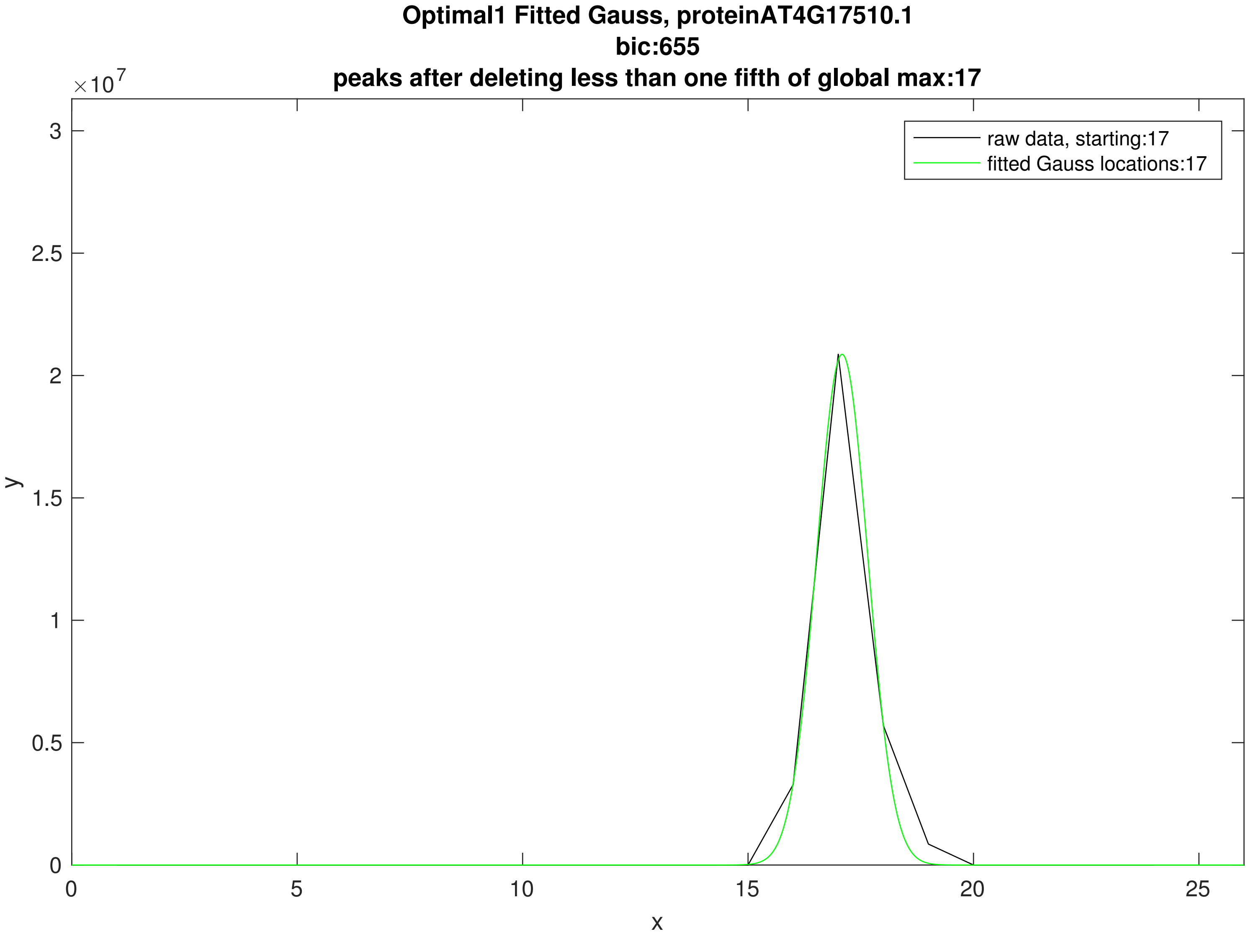

Optimal1 Fitted Gauss, proteinAT4G17510.1
bic:655
peaks after deleting less than one fifth of global max:17
×10^7
raw data, starting:17
fitted Gauss locations:17
y
x
0
0.5
1
1.5
2
2.5
3
5
10
15
20
25

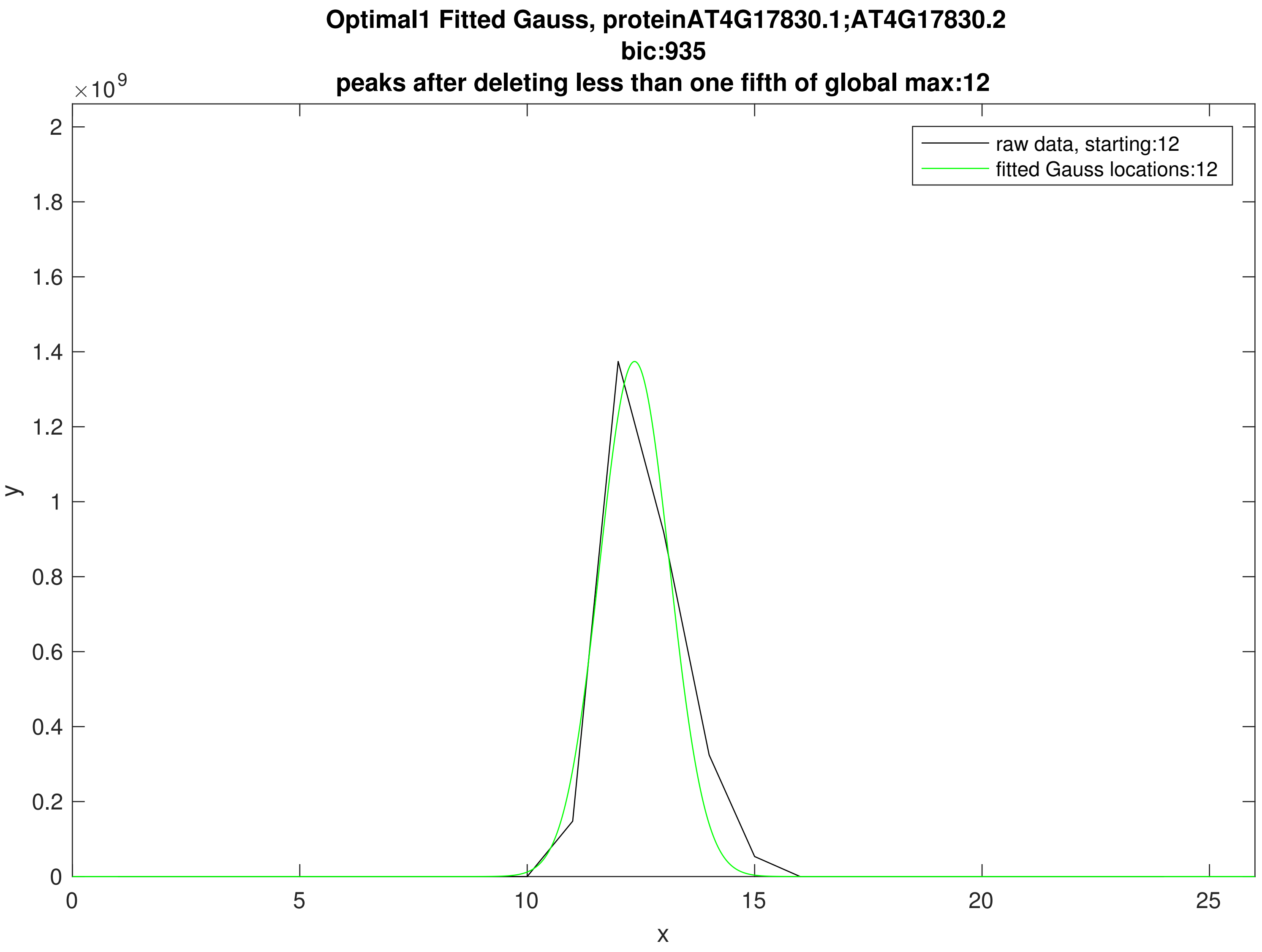

Optimal1 Fitted Gauss, proteinAT4G17830.1;AT4G17830.2
bic:935
peaks after deleting less than one fifth of global max:12
×10^9
2
1.8
1.6
1.4
1.2
1
0.8
0.6
0.4
0.2
0
y
0
5
10
15
20
25
x
raw data, starting:12
fitted Gauss locations:12

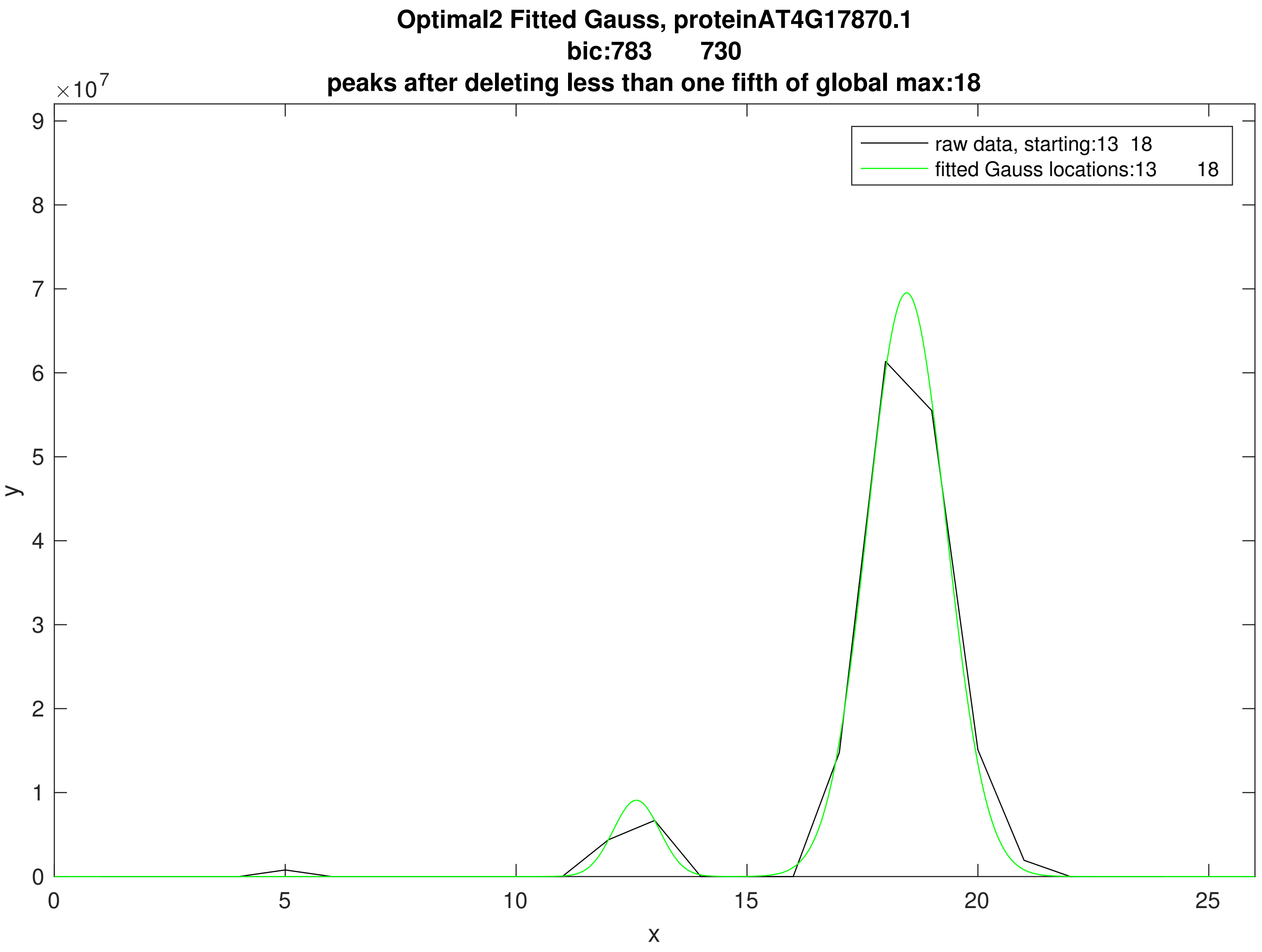
Optimal2 Fitted Gauss, proteinAT4G17870.1
bic:783 730
peaks after deleting less than one fifth of global max:18
×10^7
raw data, starting:13 18
fitted Gauss locations:13 18
y
x
0
1
2
3
4
5
6
7
8
9
0
5
10
15
20
25

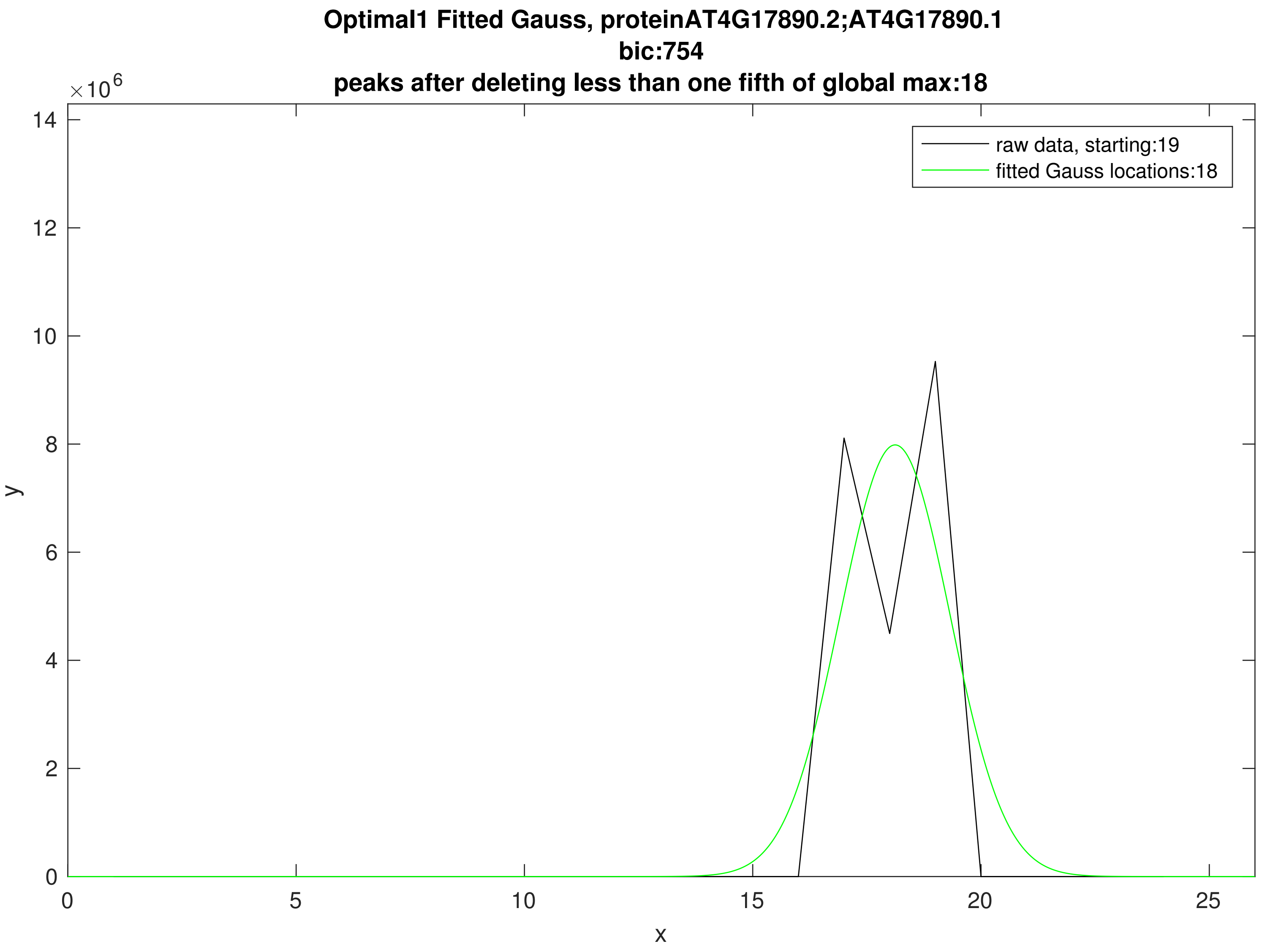

Optimal1 Fitted Gauss, proteinAT4G17890.2;AT4G17890.1
bic:754
peaks after deleting less than one fifth of global max:18
×10^6
14
12
10
8
6
4
2
0
y
0
5
10
15
20
25
x
raw data, starting:19
fitted Gauss locations:18

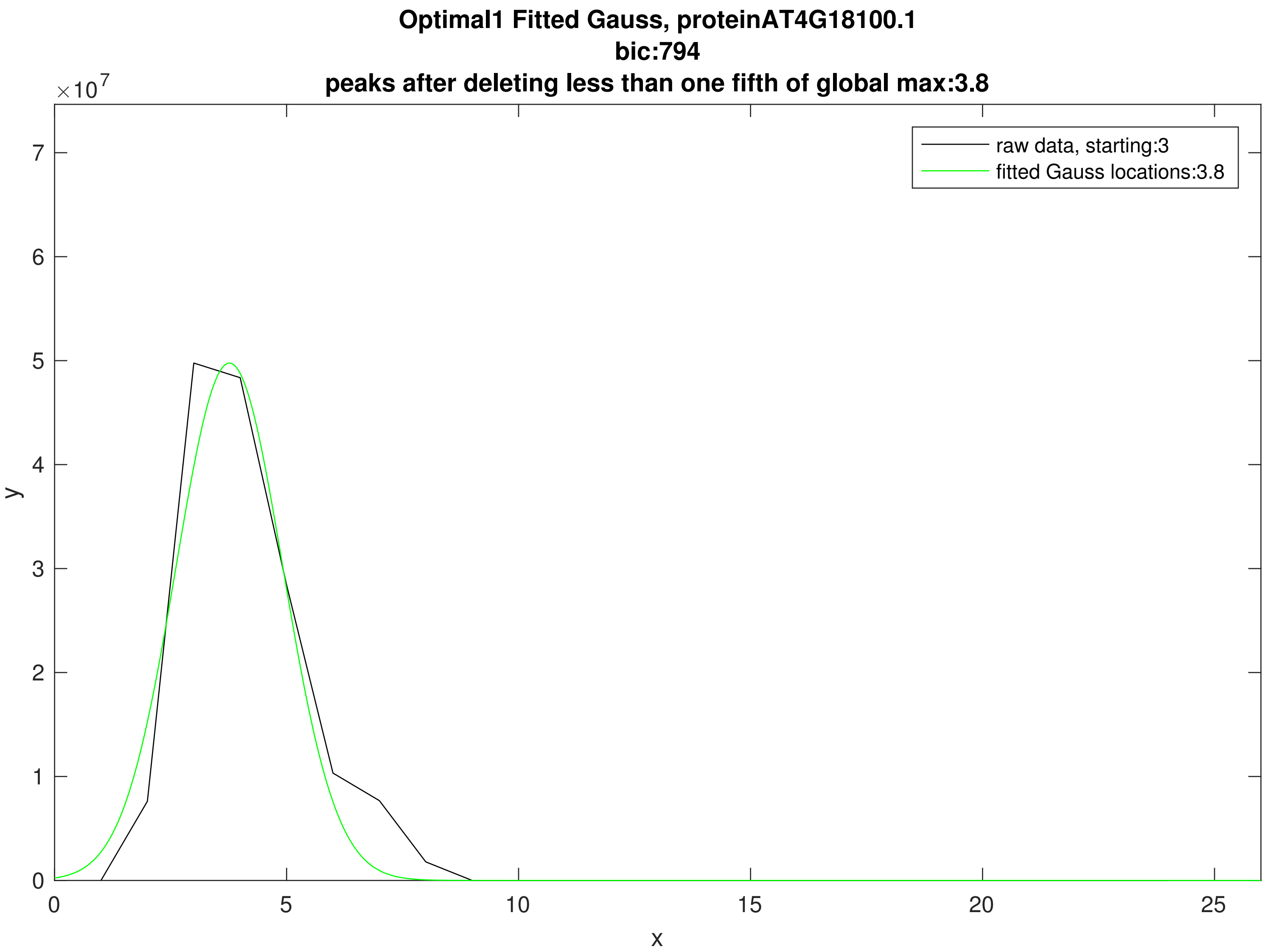
Optimal1 Fitted Gauss, proteinAT4G18100.1
bic:794
peaks after deleting less than one fifth of global max:3.8
×10^7
raw data, starting:3
fitted Gauss locations:3.8
y
x
0
1
2
3
4
5
6
7
0
5
10
15
20
25

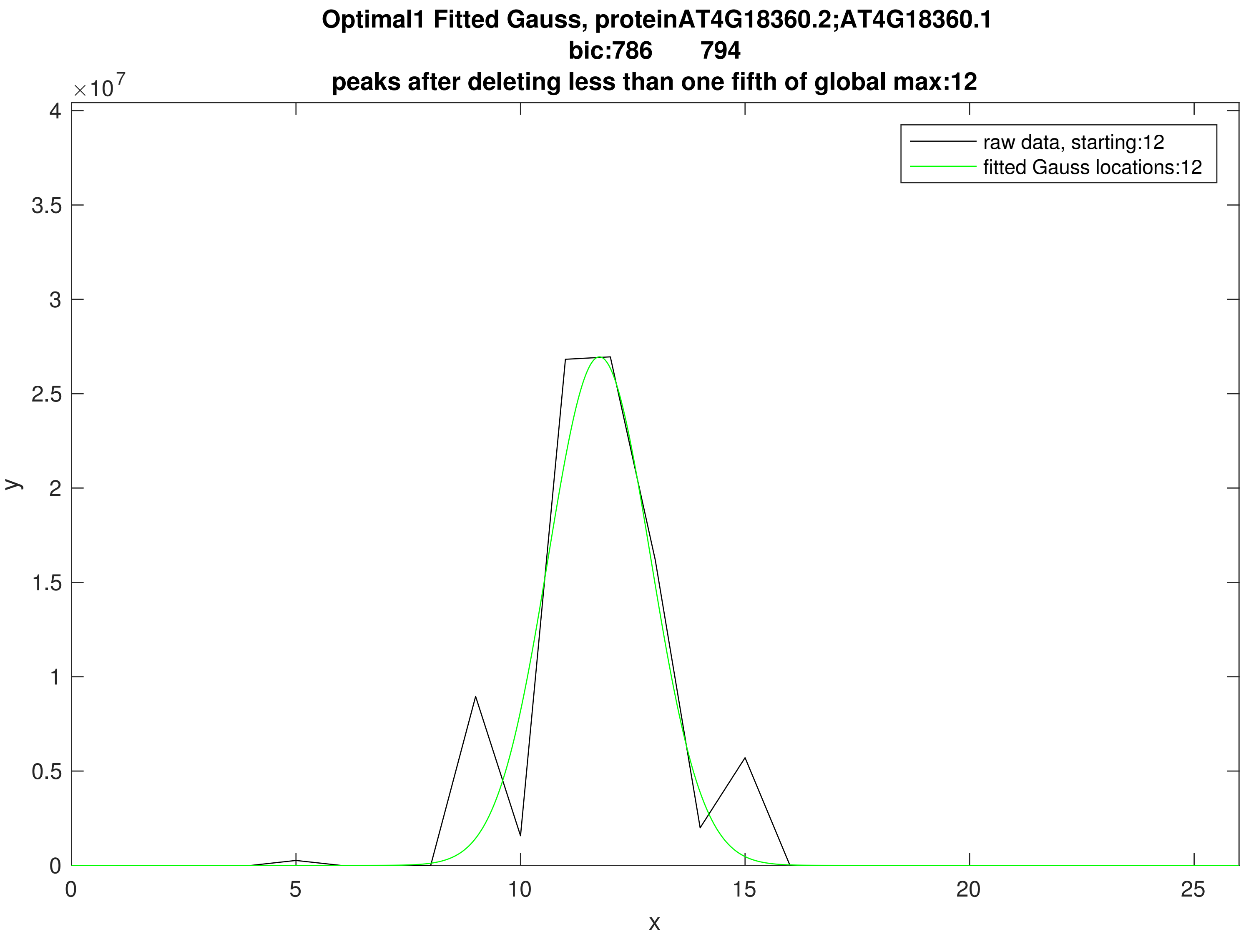

Optimal1 Fitted Gauss, proteinAT4G18360.2;AT4G18360.1
bic:786 794
peaks after deleting less than one fifth of global max:12
×10^7
raw data, starting:12
fitted Gauss locations:12
y
x
0
0.5
1
1.5
2
2.5
3
3.5
4
5
10
15
20
25

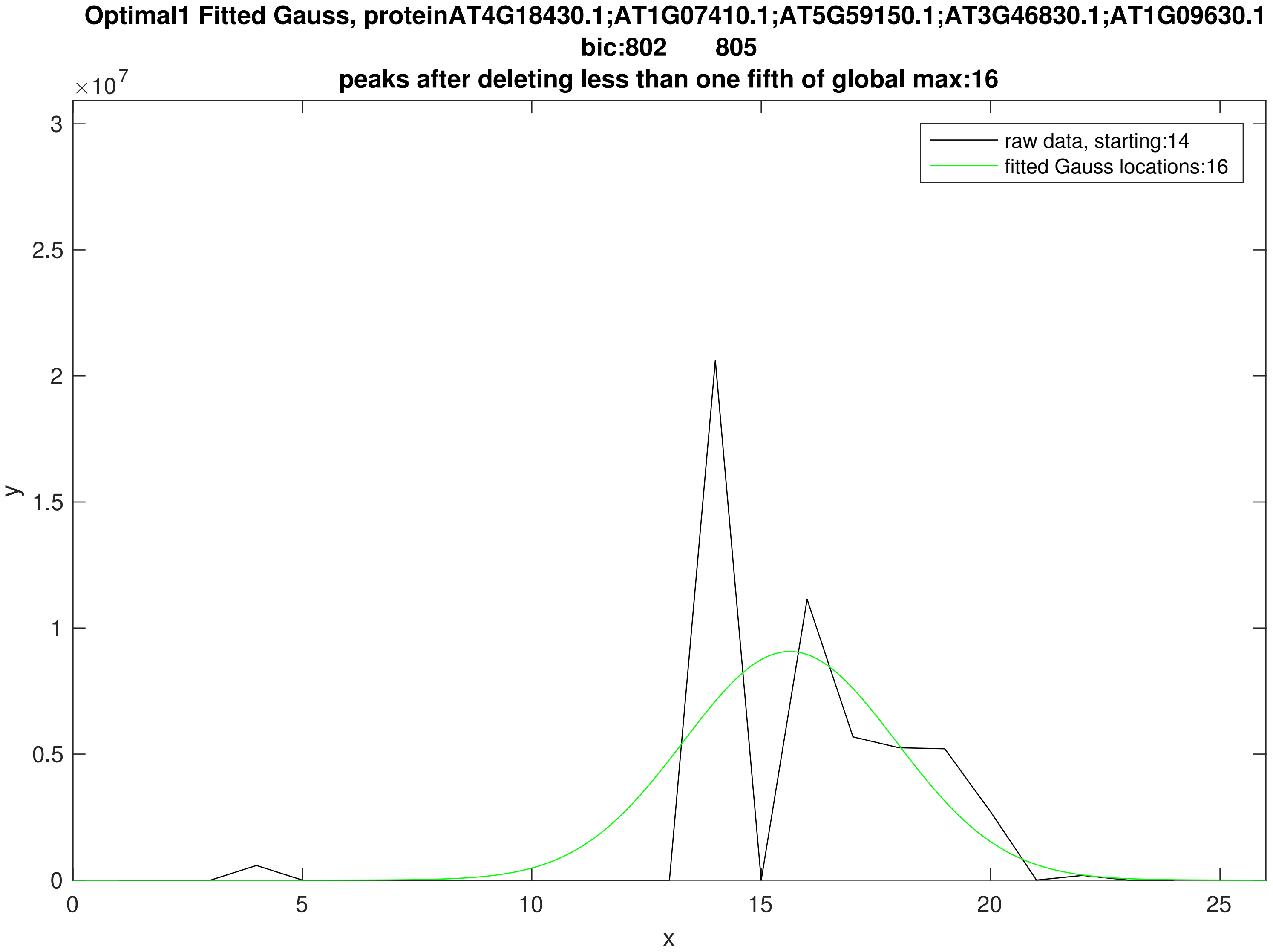
Optimal1 Fitted Gauss, proteinAT4G18430.1;AT1G07410.1;AT5G59150.1;AT3G46830.1;AT1G09630.1
bic:802 805
peaks after deleting less than one fifth of global max:16
×10^7
raw data, starting:14
fitted Gauss locations:16
y
x
0
0.5
1
1.5
2
2.5
3
5
10
15
20
25

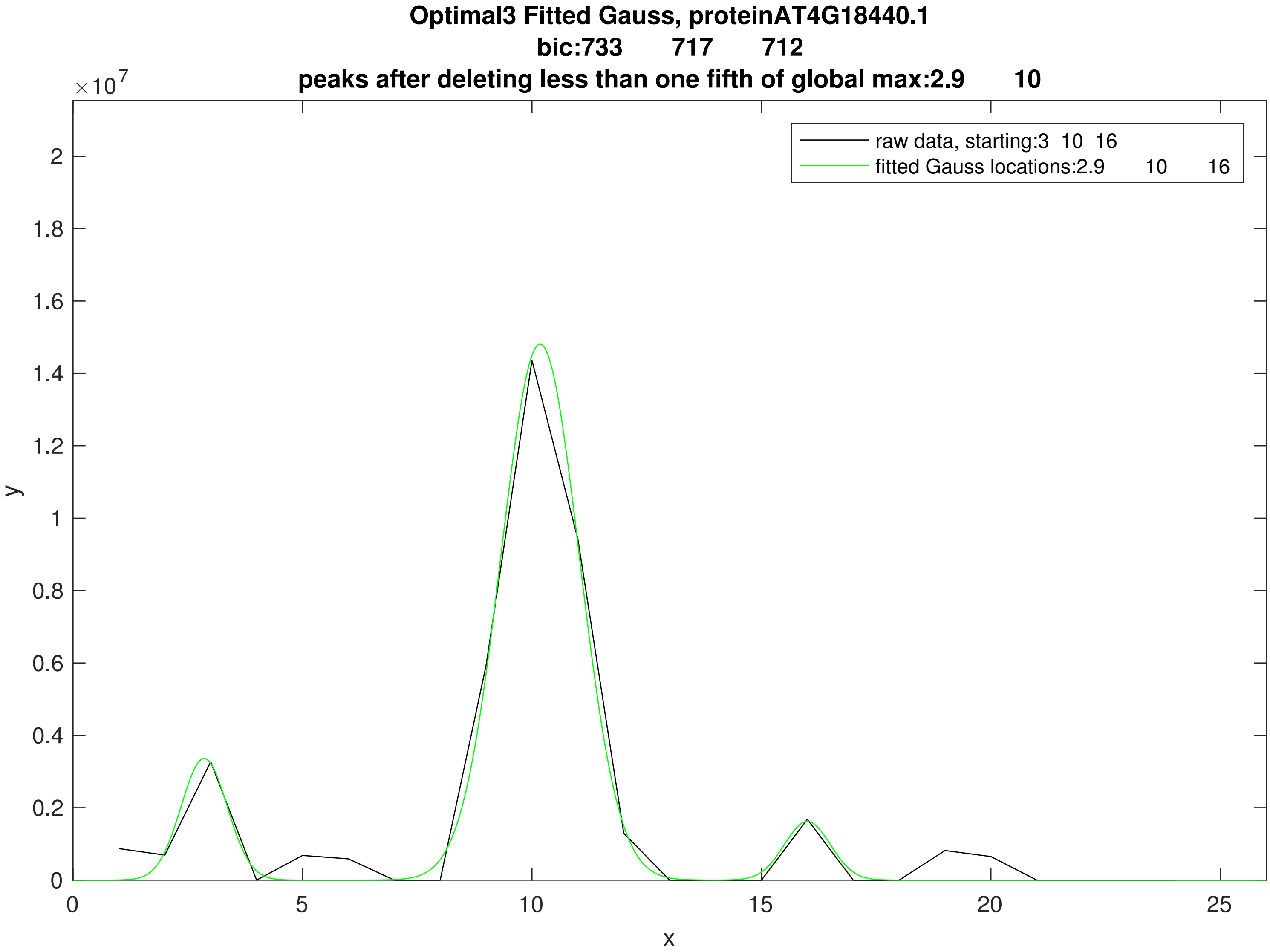
Optimal3 Fitted Gauss, proteinAT4G18440.1
bic:733 717 712
peaks after deleting less than one fifth of global max:2.9 10
×10^7
raw data, starting:3 10 16
fitted Gauss locations:2.9 10 16
y
x
0
0.2
0.4
0.6
0.8
1
1.2
1.4
1.6
1.8
2
5
10
15
20
25

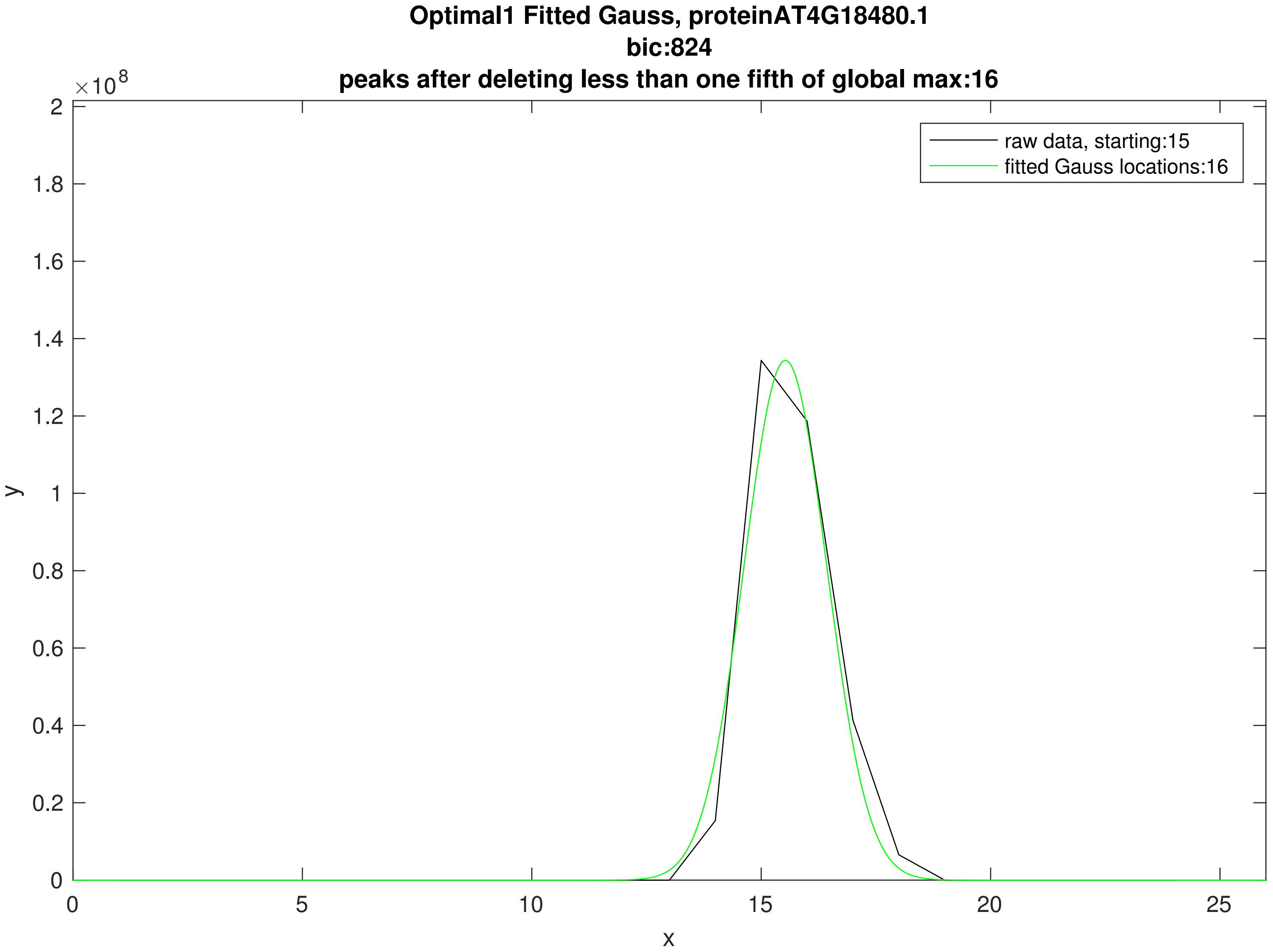

Optimal1 Fitted Gauss, proteinAT4G18480.1
bic:824
peaks after deleting less than one fifth of global max:16
×10^8
2
1.8
1.6
1.4
1.2
1
0.8
0.6
0.4
0.2
0
y
0
5
10
15
20
25
x
raw data, starting:15
fitted Gauss locations:16

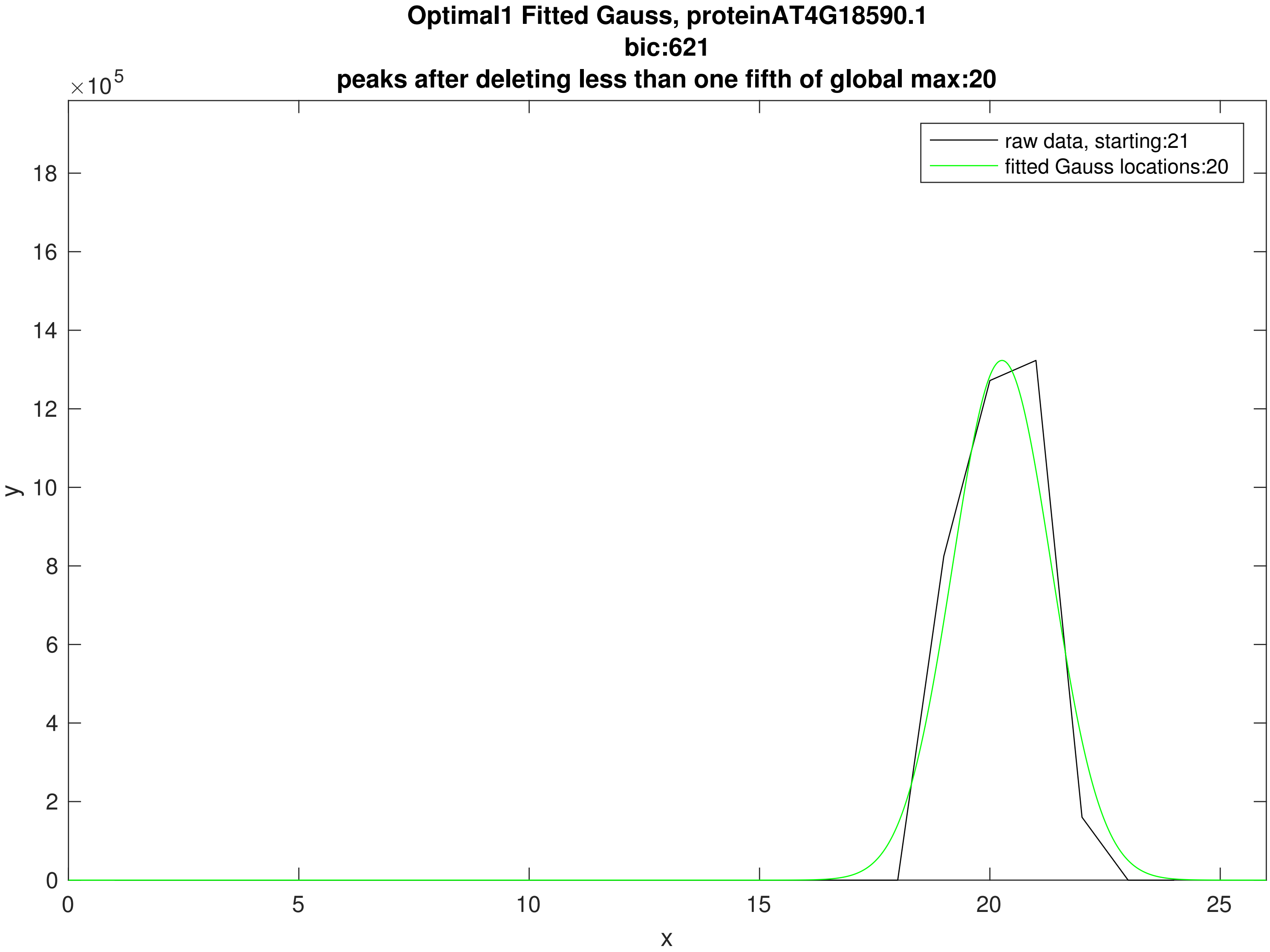

Optimal1 Fitted Gauss, proteinAT4G18590.1
bic:621
peaks after deleting less than one fifth of global max:20
×10^5
raw data, starting:21
fitted Gauss locations:20
y
x
0
2
4
6
8
10
12
14
16
18
0
5
10
15
20
25

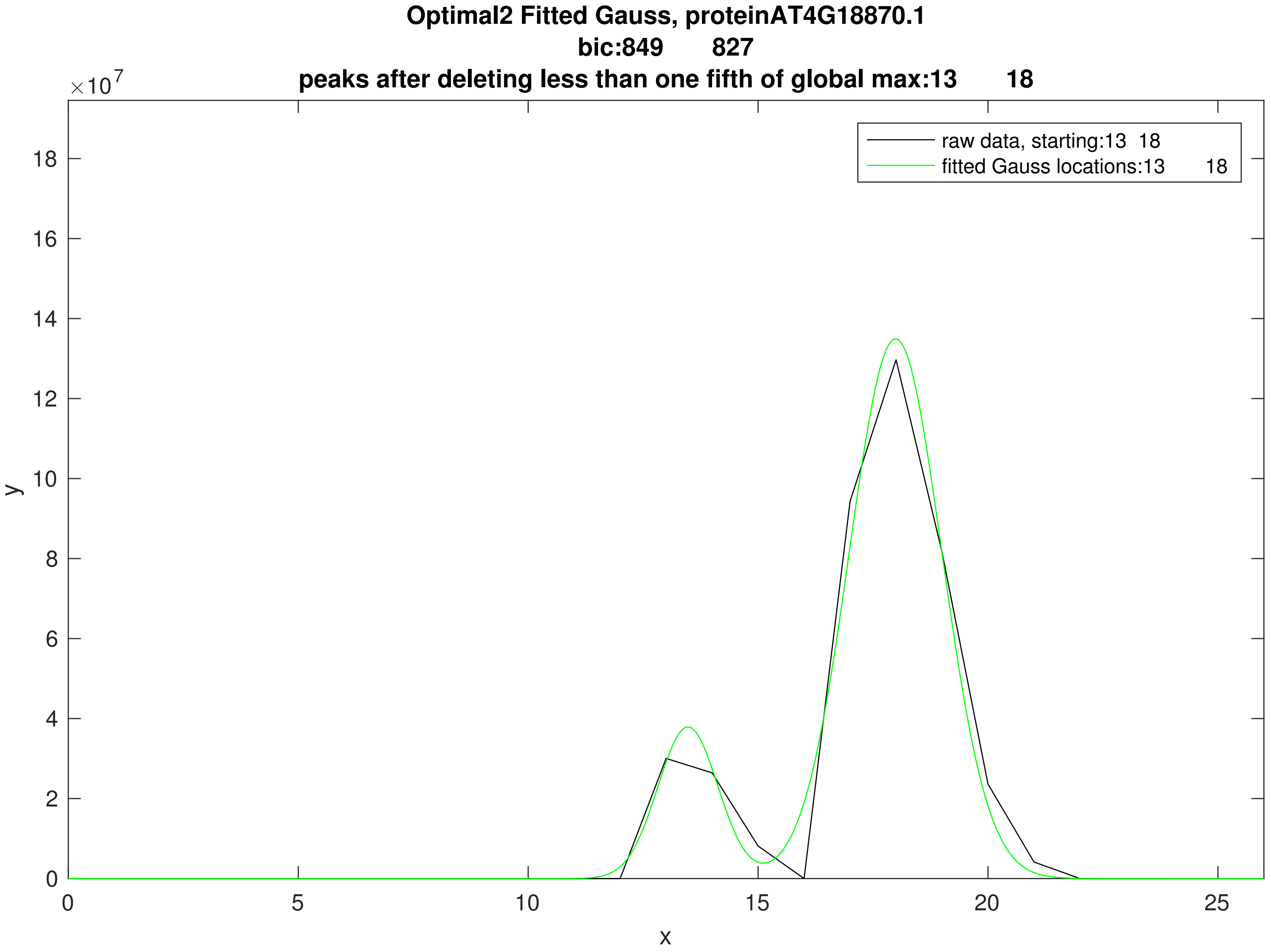
Optimal2 Fitted Gauss, proteinAT4G18870.1
bic:849 827
peaks after deleting less than one fifth of global max:13 18
×10^7
raw data, starting:13 18
fitted Gauss locations:13 18
y
x
0
2
4
6
8
10
12
14
16
18
0
5
10
15
20
25

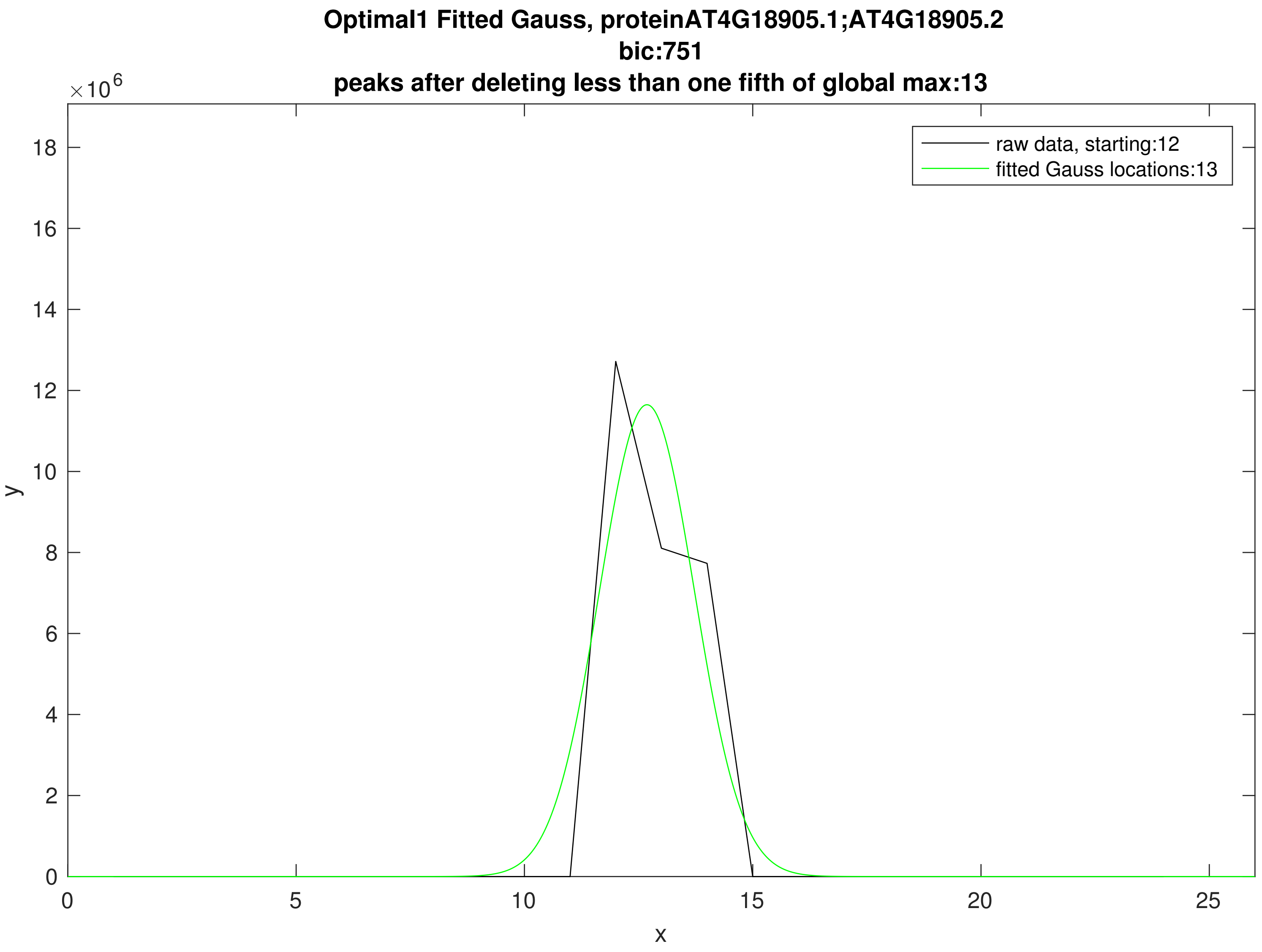
Optimal1 Fitted Gauss, proteinAT4G18905.1;AT4G18905.2
bic:751
peaks after deleting less than one fifth of global max:13
×10^6
raw data, starting:12
fitted Gauss locations:13
y
x
0
2
4
6
8
10
12
14
16
18
0
5
10
15
20
25

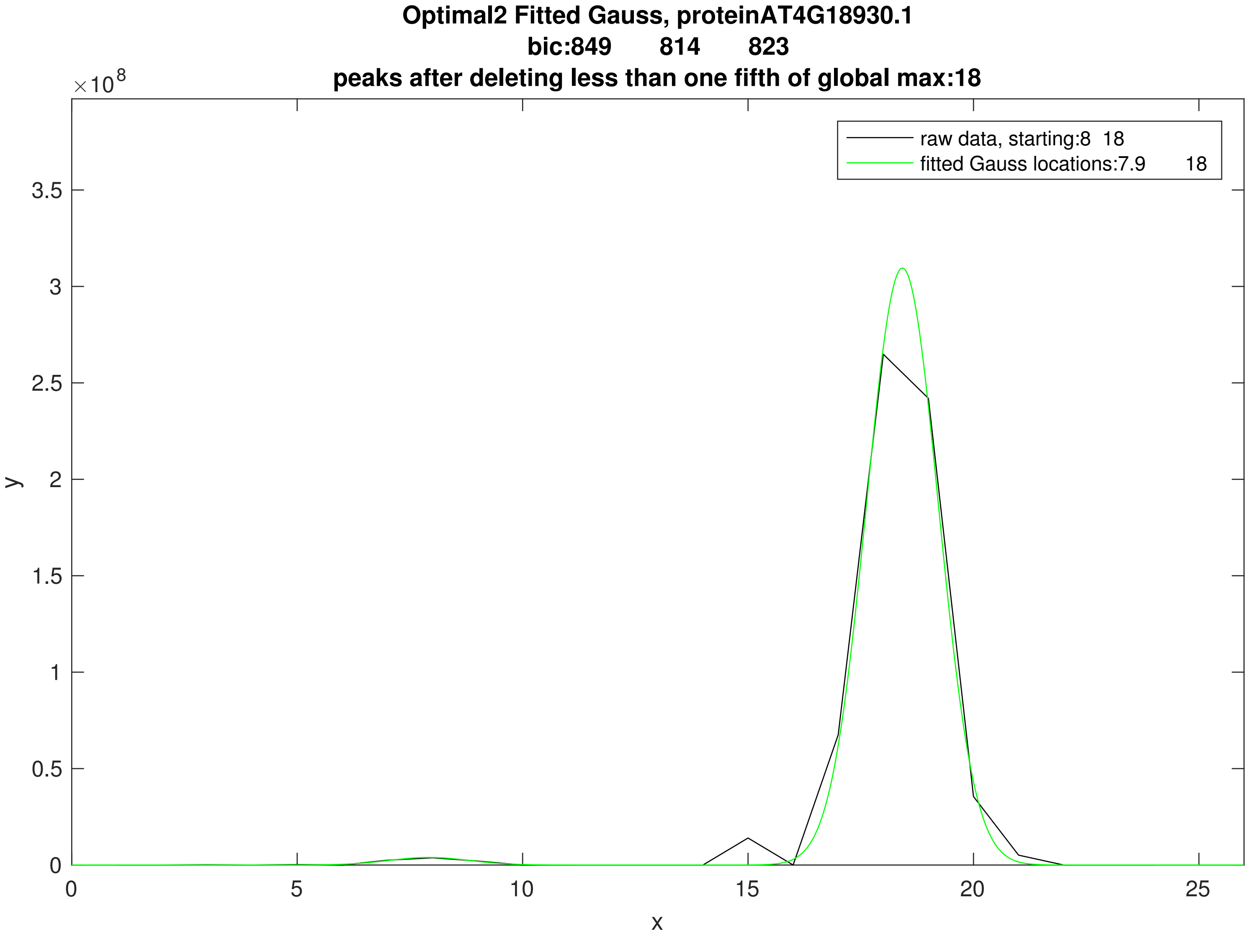
Optimal2 Fitted Gauss, proteinAT4G18930.1
bic:849 814 823
peaks after deleting less than one fifth of global max:18
raw data, starting:8 18
fitted Gauss locations:7.9 18
×10^8
3.5
3
2.5
2
1.5
1
0.5
0
y
0
5
10
15
20
25
x

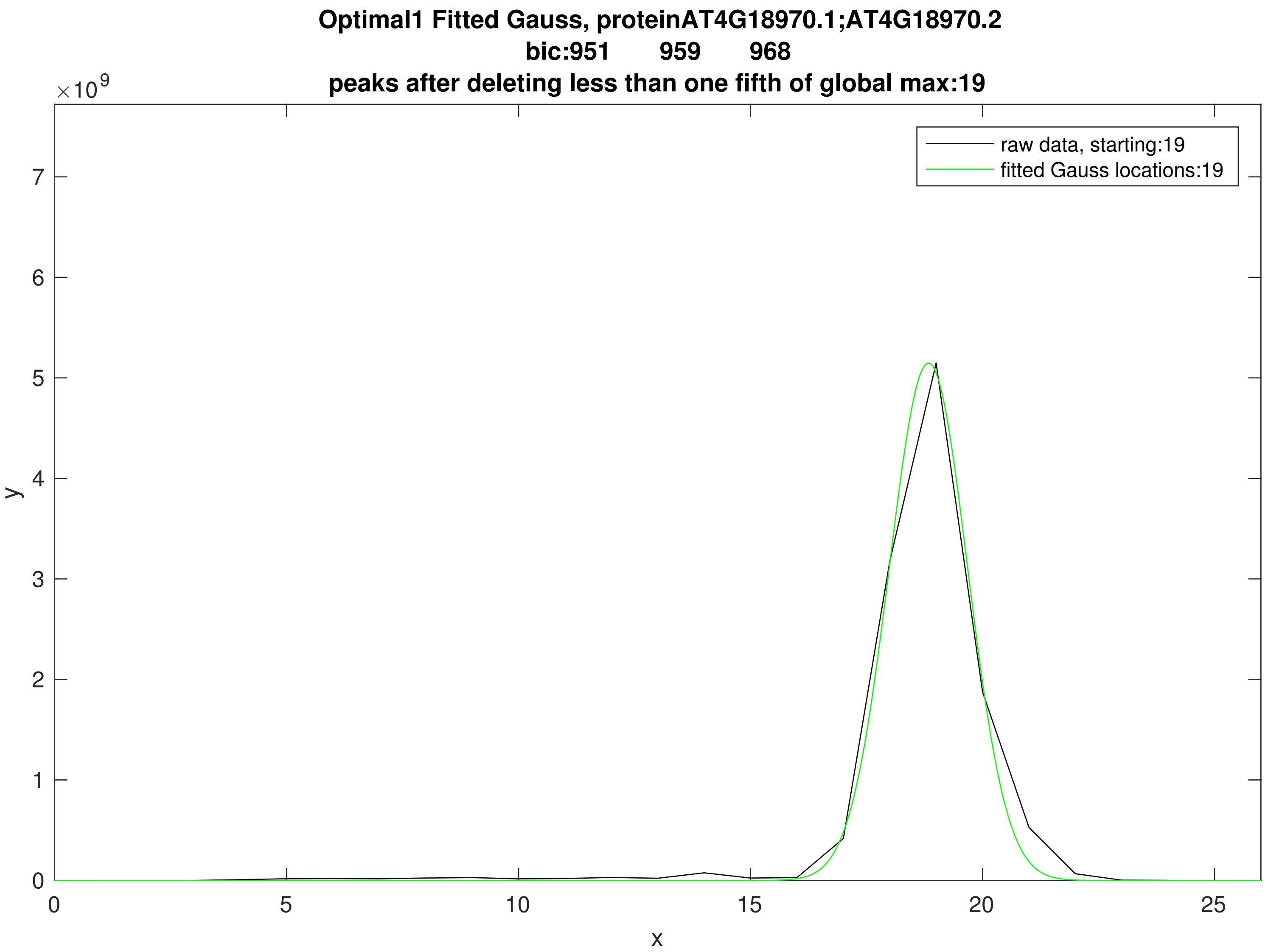

Optimal1 Fitted Gauss, proteinAT4G18970.1;AT4G18970.2
bic:951 959 968
peaks after deleting less than one fifth of global max:19
×10^9
raw data, starting:19
fitted Gauss locations:19
y
x
0
1
2
3
4
5
6
7
0
5
10
15
20
25

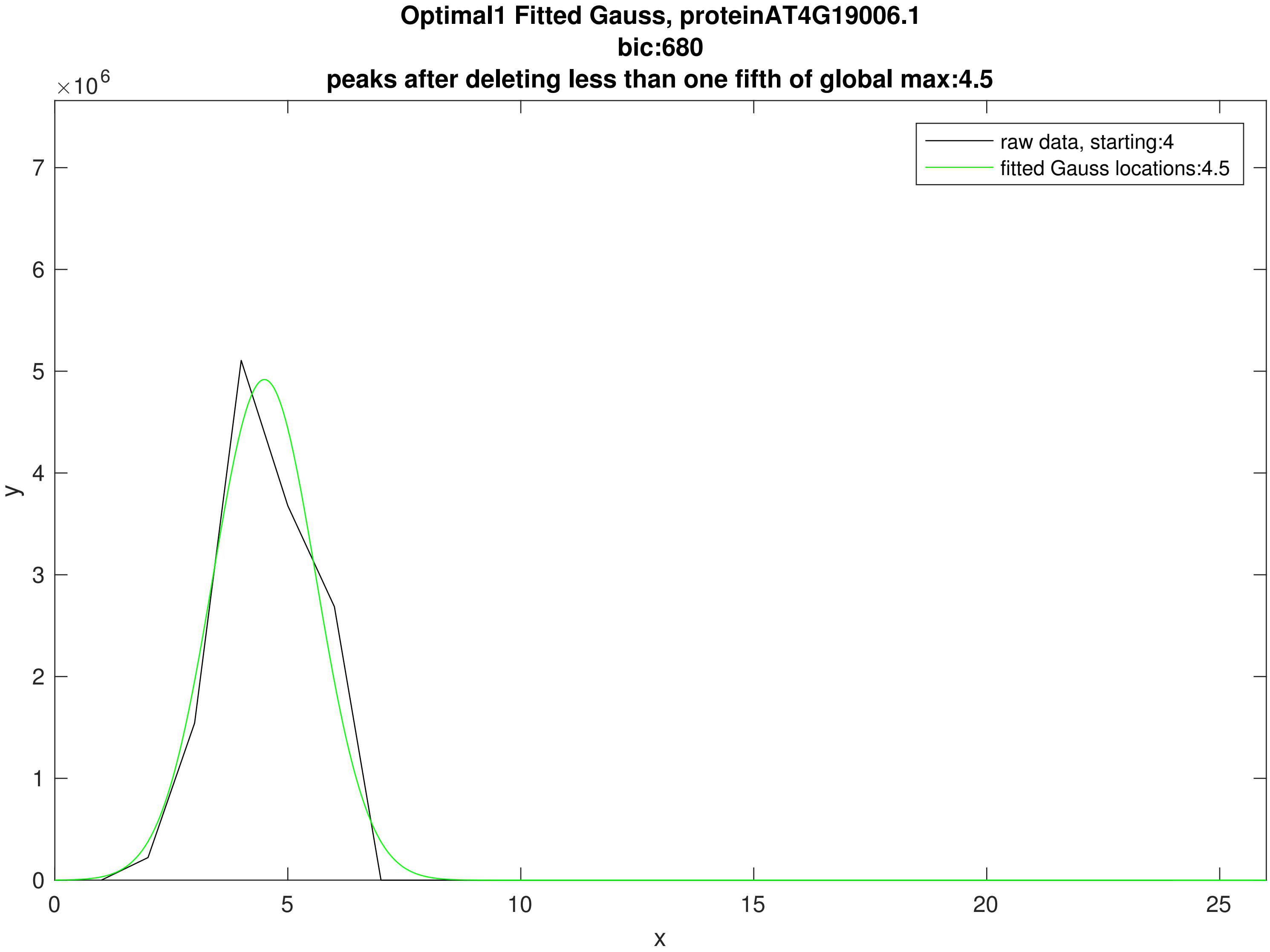

Optimal1 Fitted Gauss, proteinAT4G19006.1
bic:680
peaks after deleting less than one fifth of global max:4.5
×10^6
raw data, starting:4
fitted Gauss locations:4.5
y
x
0
1
2
3
4
5
6
7
0
5
10
15
20
25

Optimal1 Fitted Gauss, proteinAT4G19210.1
bic:787 836 828
peaks after deleting less than one fifth of global max:15

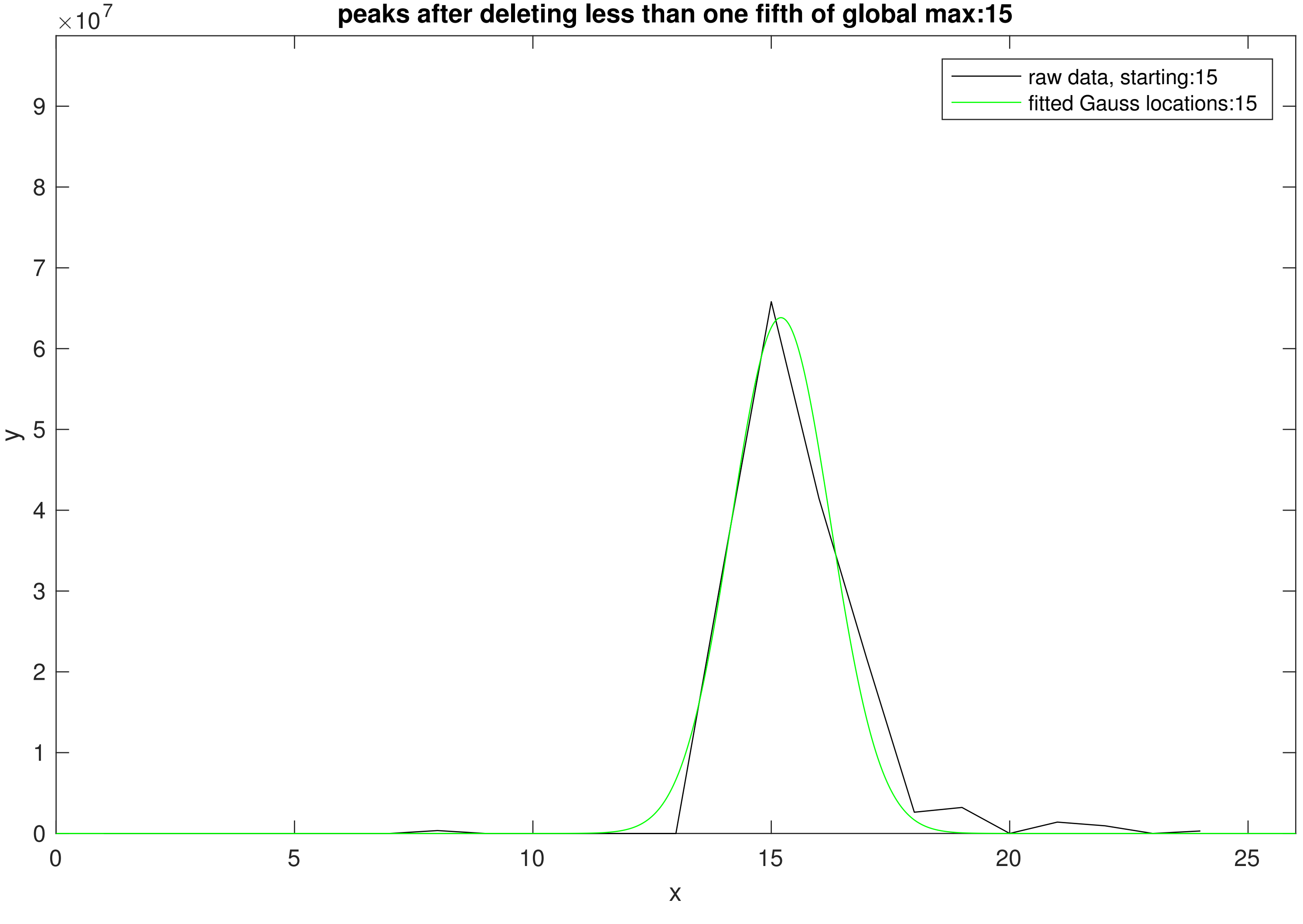

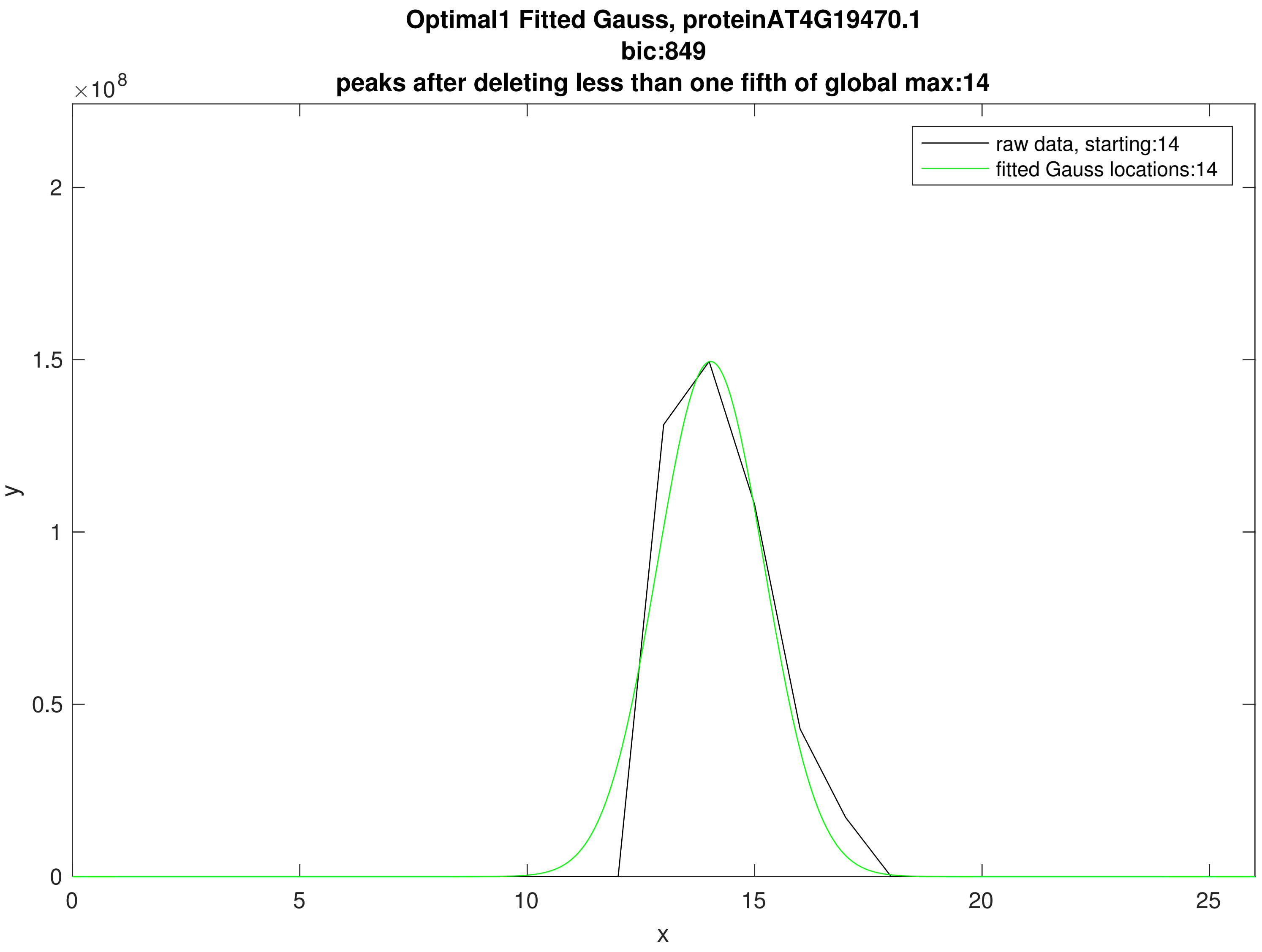

Optimal1 Fitted Gauss, proteinAT4G19470.1
bic:849
peaks after deleting less than one fifth of global max:14
×10^8
raw data, starting:14
fitted Gauss locations:14
2
1.5
1
0.5
0
y
0
5
10
15
20
25
x

**Optimal1 Fitted Gauss, proteinAT4G19710.2;AT4G19710.1**
**bic:757 762 772**
**peaks after deleting less than one fifth of global max:11**

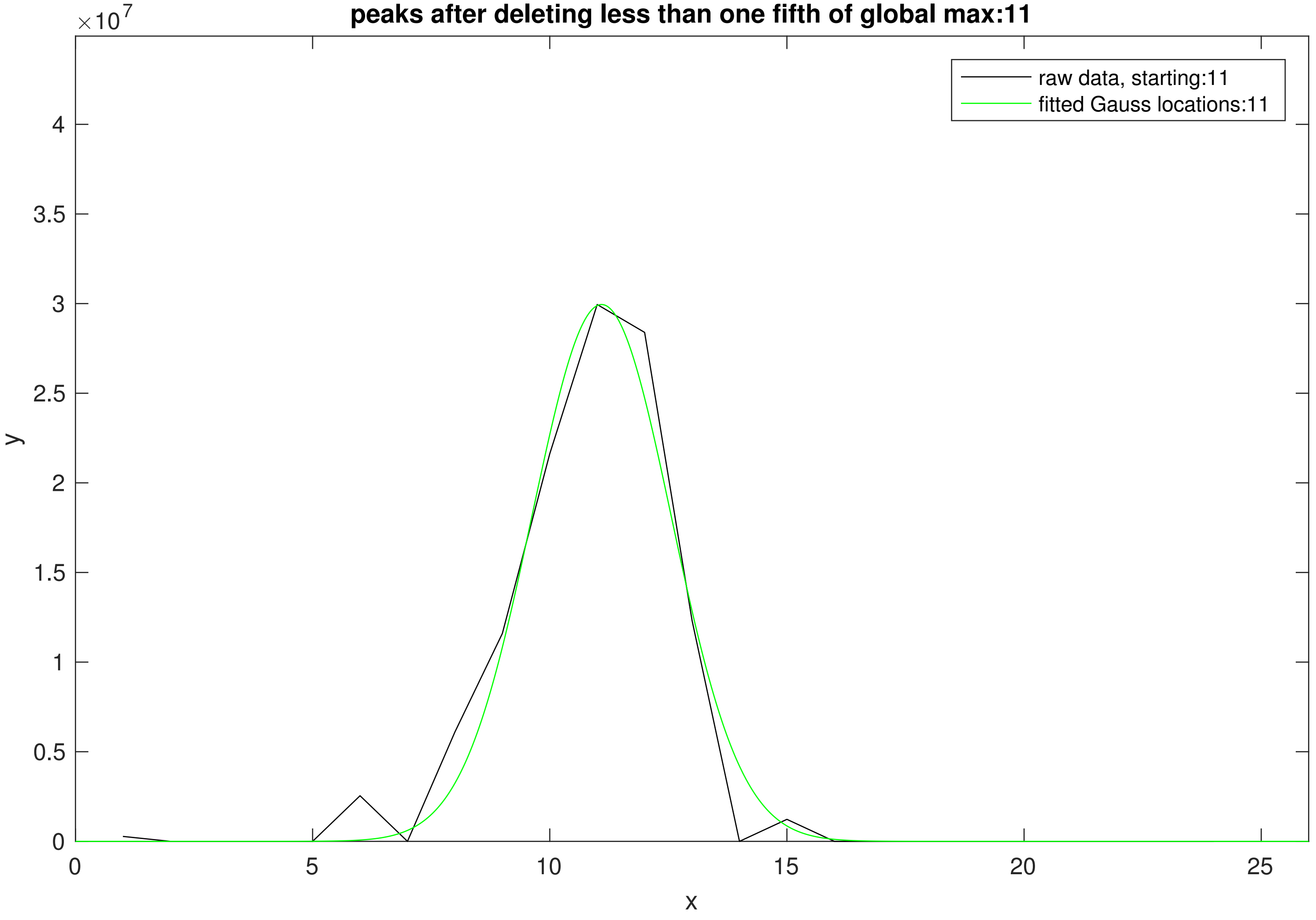

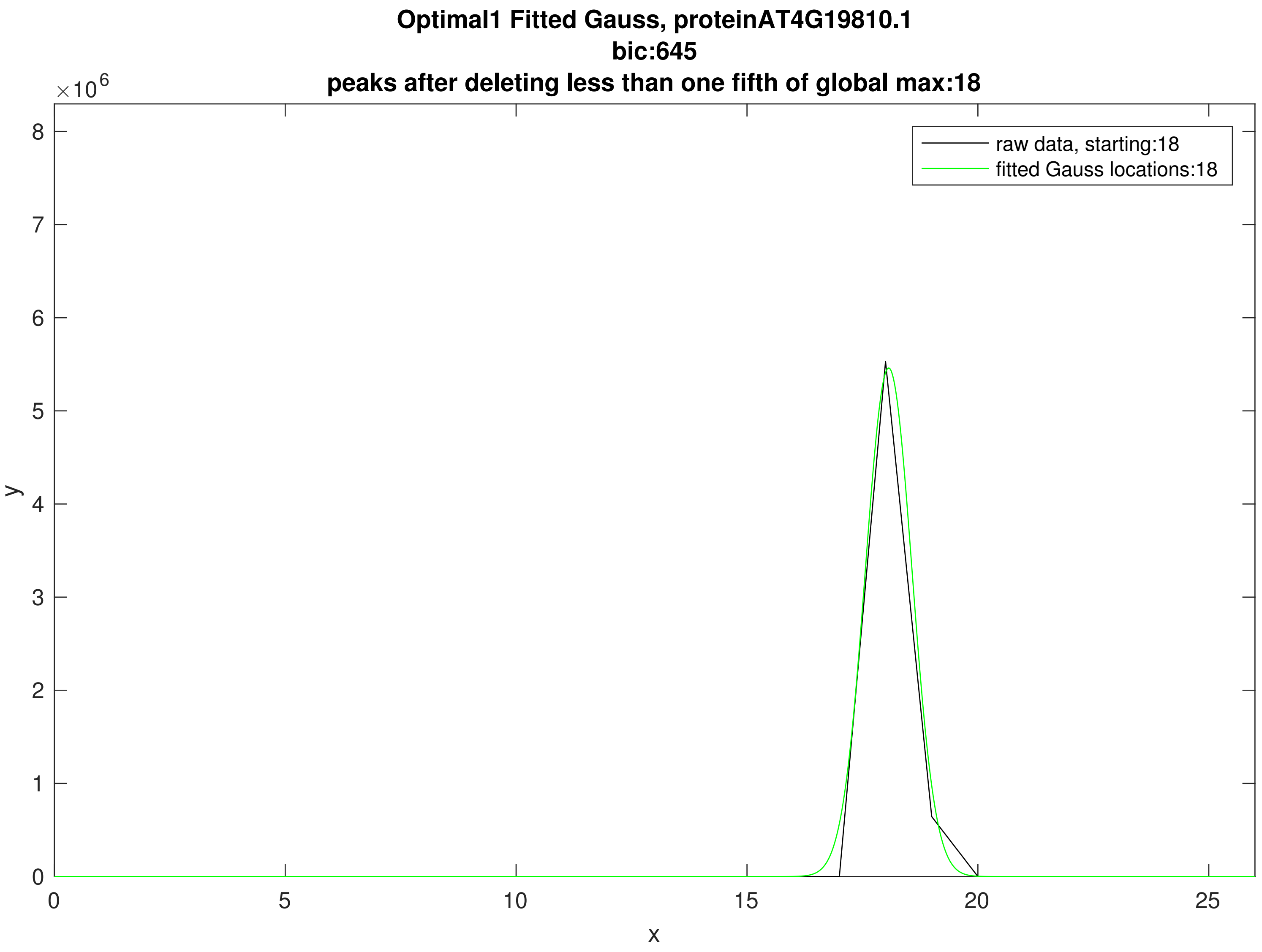

Optimal1 Fitted Gauss, proteinAT4G19810.1
bic:645
peaks after deleting less than one fifth of global max:18
×10^6
raw data, starting:18
fitted Gauss locations:18
y
x
0
1
2
3
4
5
6
7
8
0
5
10
15
20
25

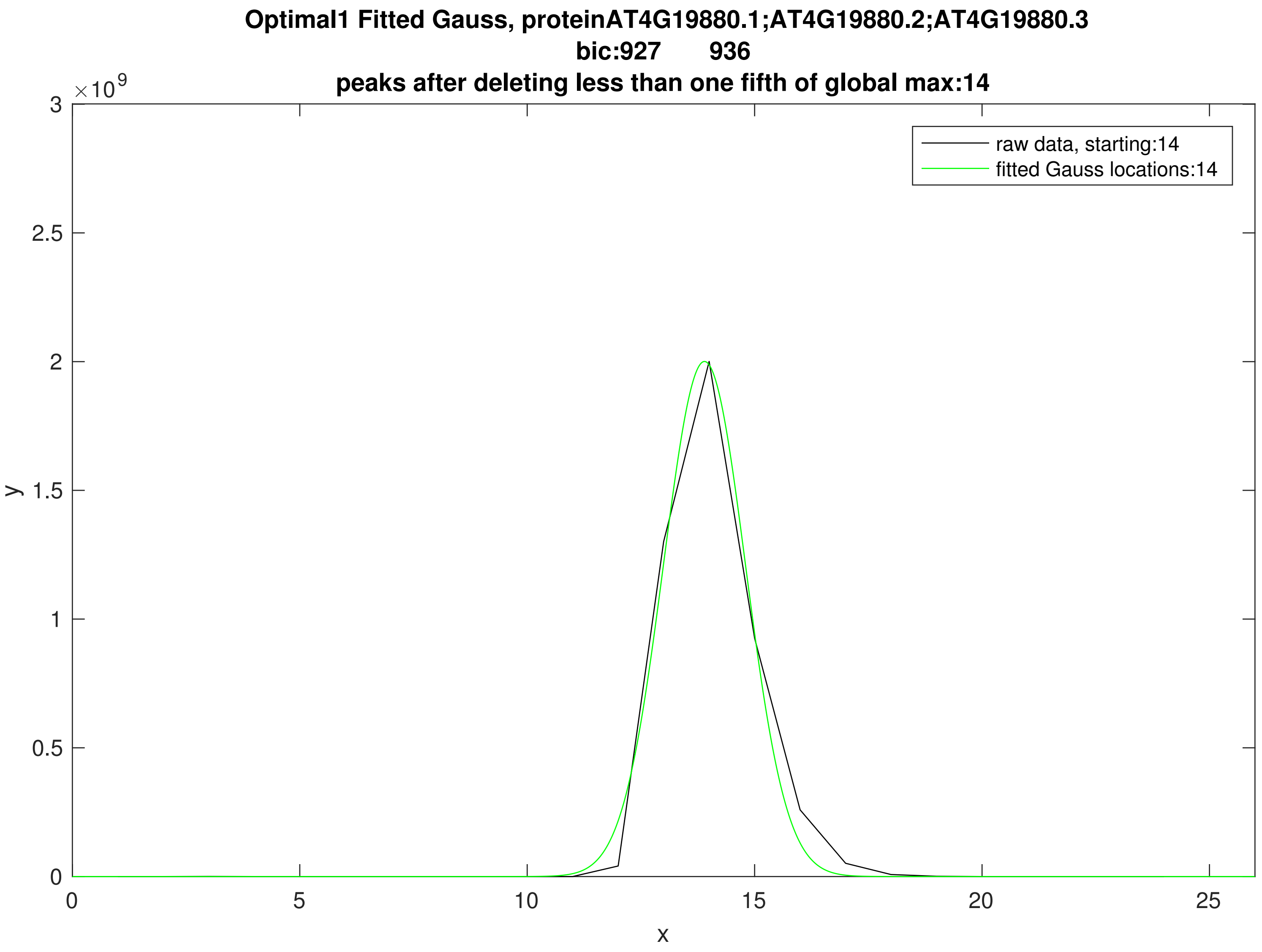
Optimal1 Fitted Gauss, proteinAT4G19880.1;AT4G19880.2;AT4G19880.3
bic:927 936
peaks after deleting less than one fifth of global max:14
×10^9
3
2.5
2
y 1.5
1
0.5
0
0
5
10
15
20
25
x
raw data, starting:14
fitted Gauss locations:14

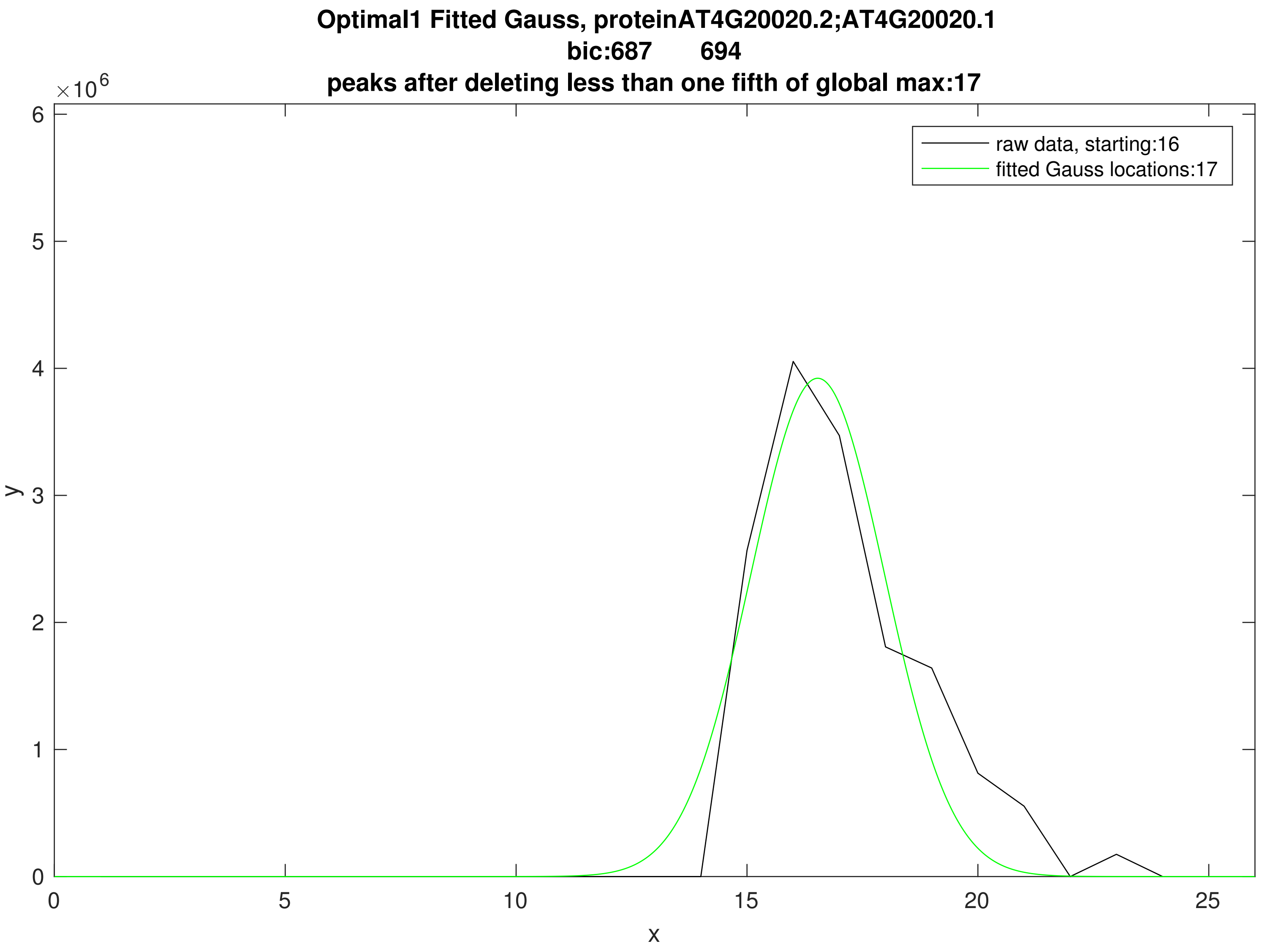
Optimal1 Fitted Gauss, proteinAT4G20020.2;AT4G20020.1
bic:687 694
peaks after deleting less than one fifth of global max:17
×10^6
raw data, starting:16
fitted Gauss locations:17
y
x
0
1
2
3
4
5
6
0
5
10
15
20
25

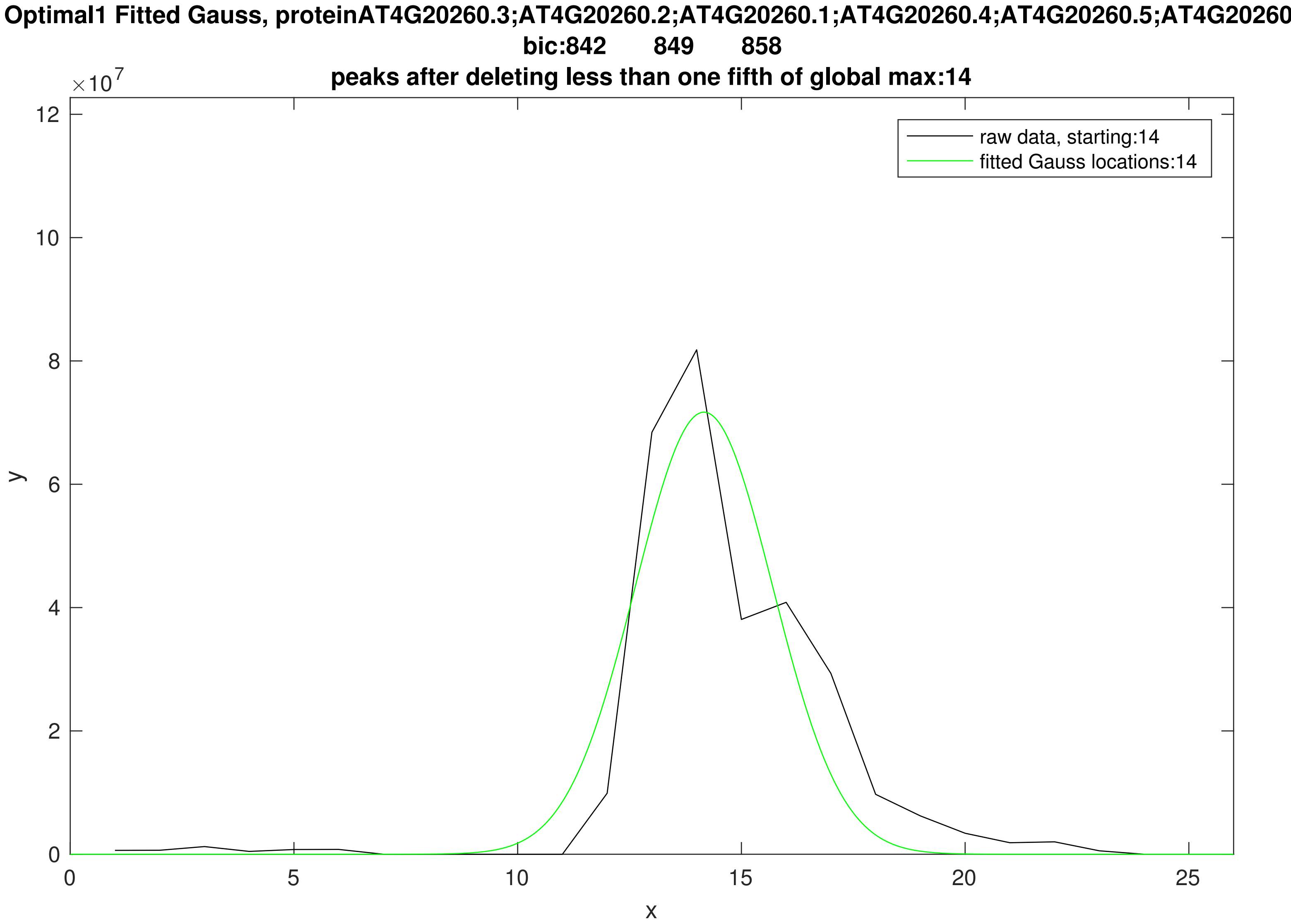
Optimal1 Fitted Gauss, proteinAT4G20260.3;AT4G20260.2;AT4G20260.1;AT4G20260.4;AT4G20260.5;AT4G20260
bic:842 849 858
peaks after deleting less than one fifth of global max:14
×10^7
raw data, starting:14
fitted Gauss locations:14
y
x
0
2
4
6
8
10
12
0
5
10
15
20
25

**Optimal1 Fitted Gauss, proteinAT4G20360.1**
**bic:910 919 929**
**peaks after deleting less than one fifth of global max:16**

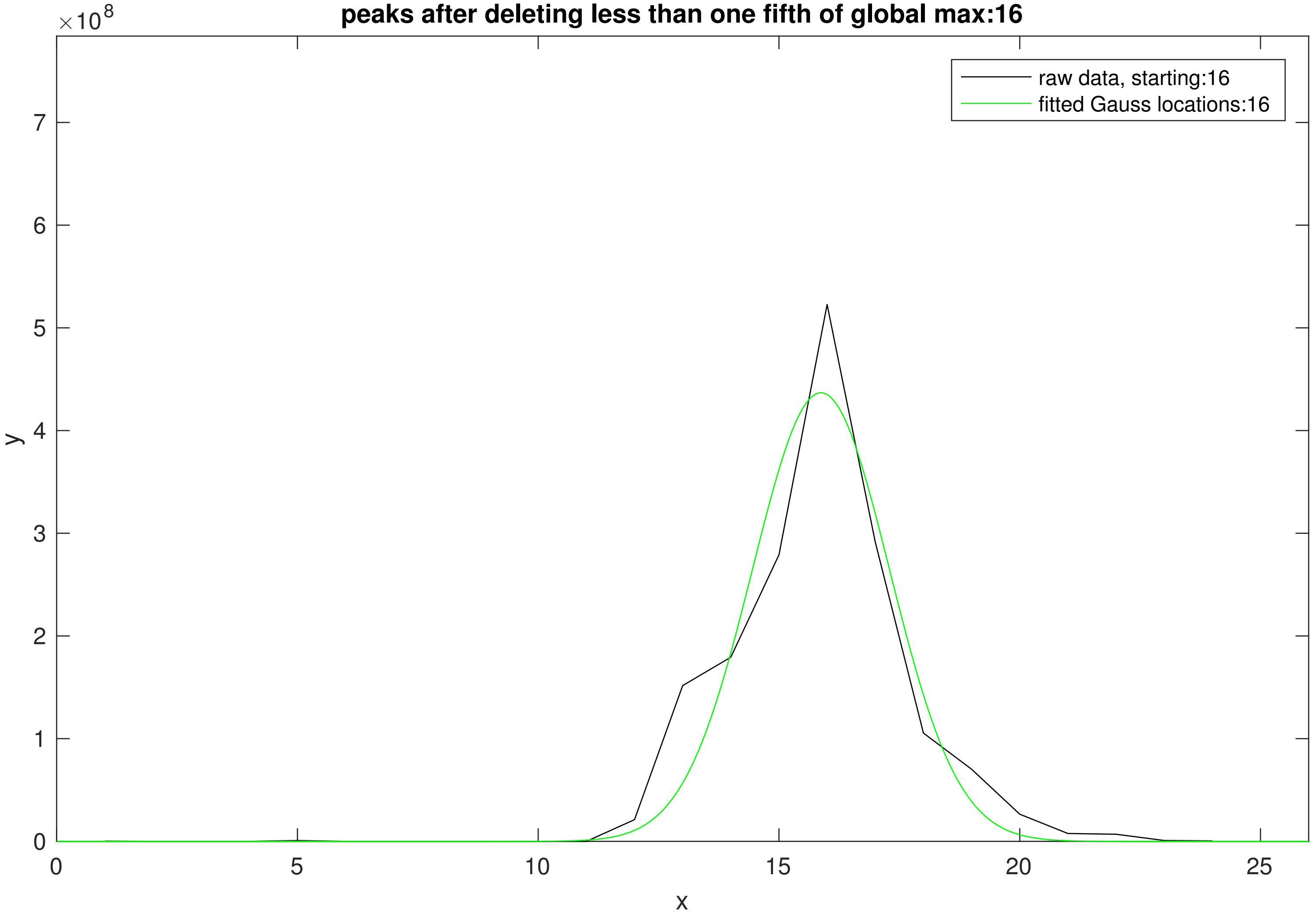

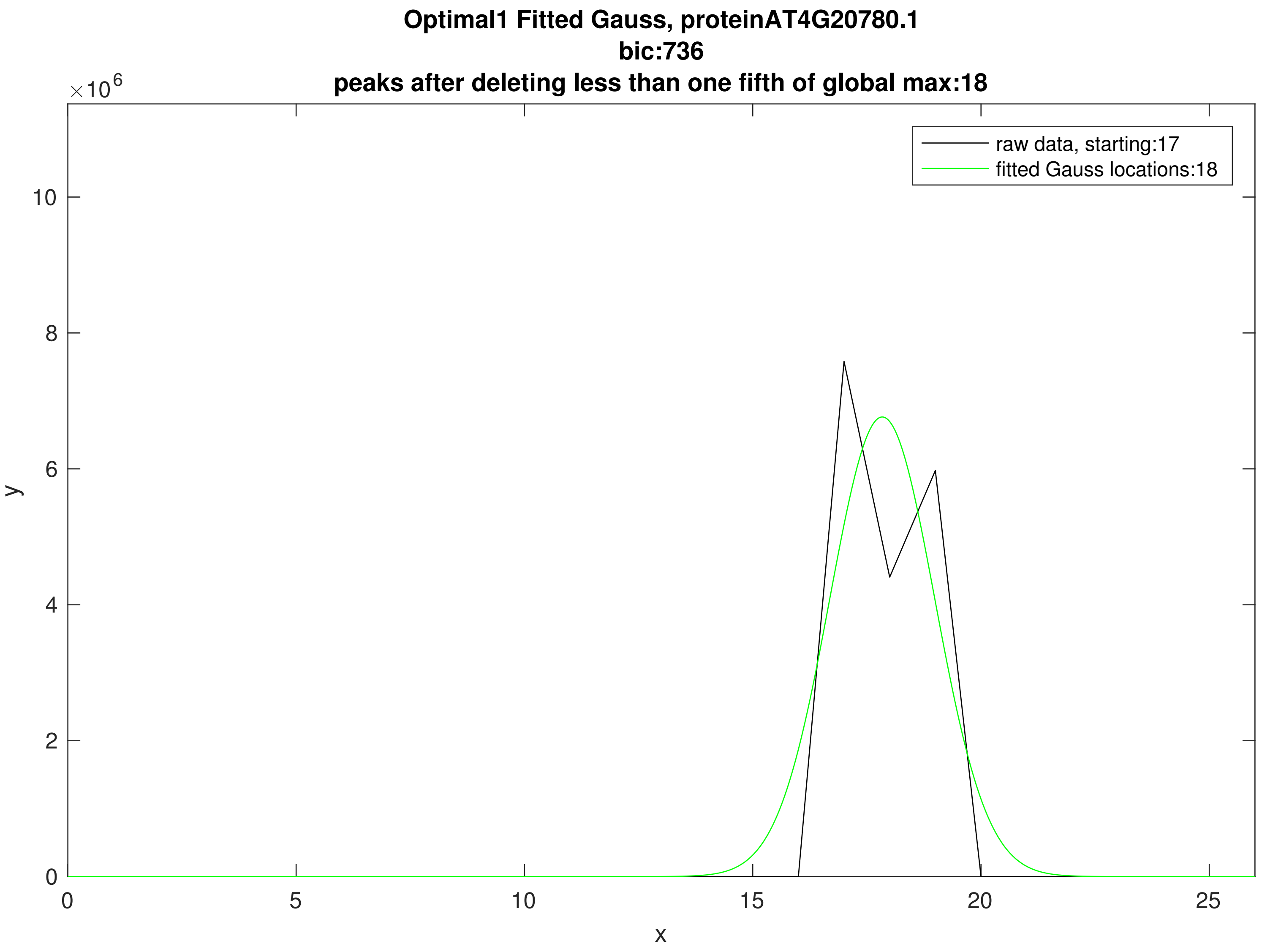

Optimal1 Fitted Gauss, proteinAT4G20780.1
bic:736
peaks after deleting less than one fifth of global max:18
×10^6
raw data, starting:17
fitted Gauss locations:18
y
x
0
2
4
6
8
10
0
5
10
15
20
25

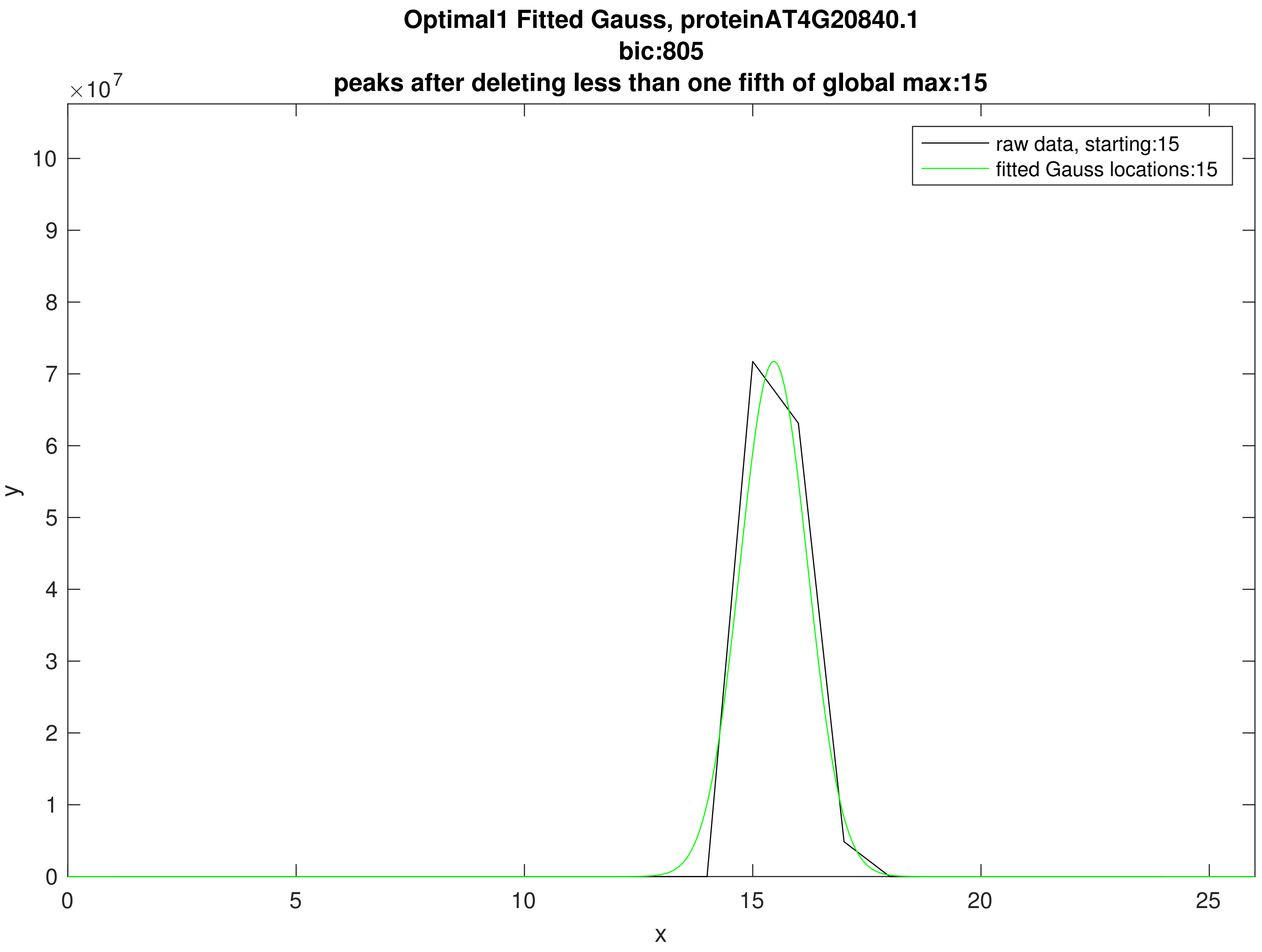
Optimal1 Fitted Gauss, proteinAT4G20840.1
bic:805
peaks after deleting less than one fifth of global max:15
×10^7
raw data, starting:15
fitted Gauss locations:15
y
x
0
1
2
3
4
5
6
7
8
9
10
0
5
10
15
20
25

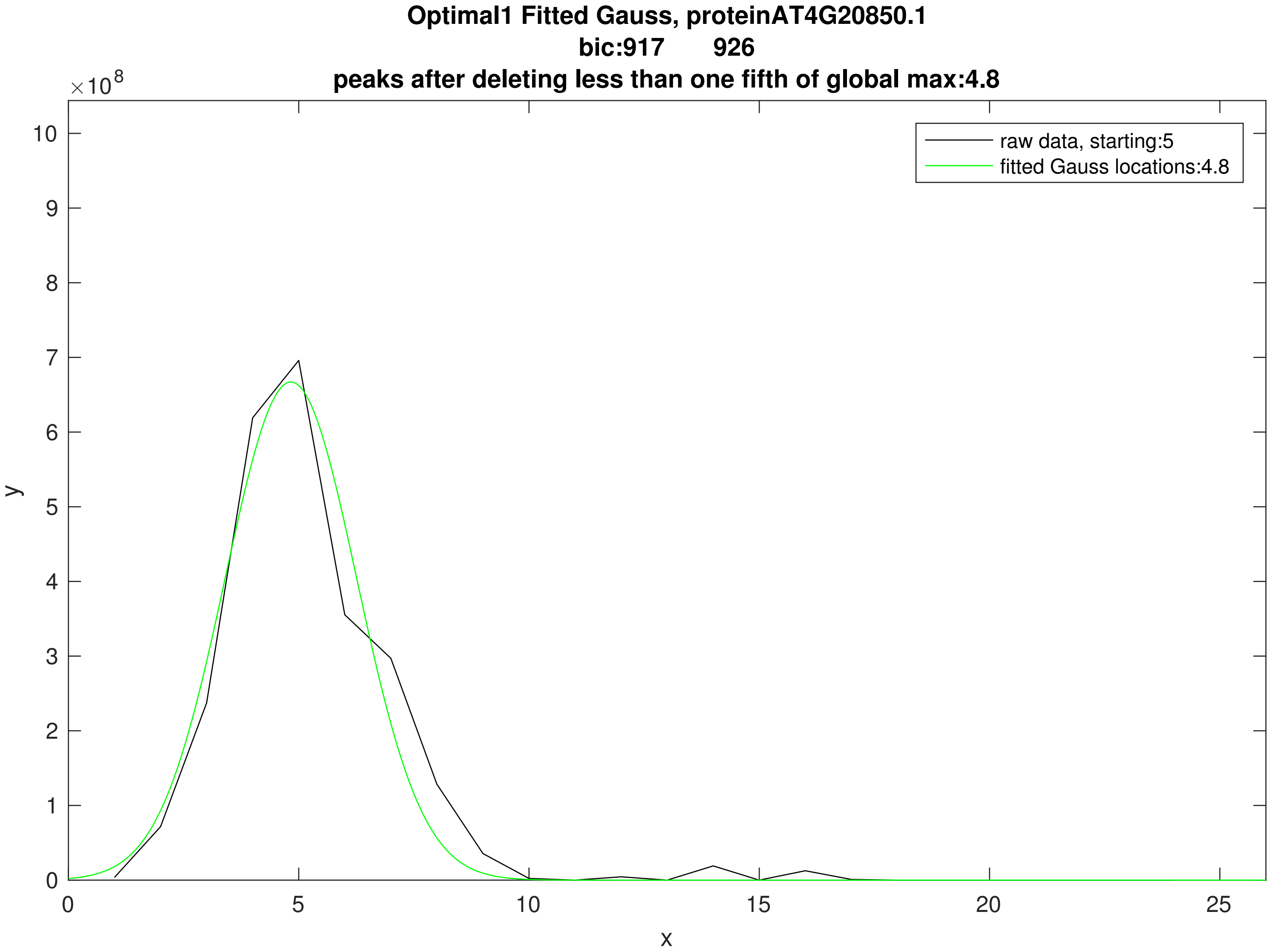
Optimal1 Fitted Gauss, proteinAT4G20850.1
bic:917 926
peaks after deleting less than one fifth of global max:4.8
×10^8
raw data, starting:5
fitted Gauss locations:4.8
y
x
0
1
2
3
4
5
6
7
8
9
10
0
5
10
15
20
25

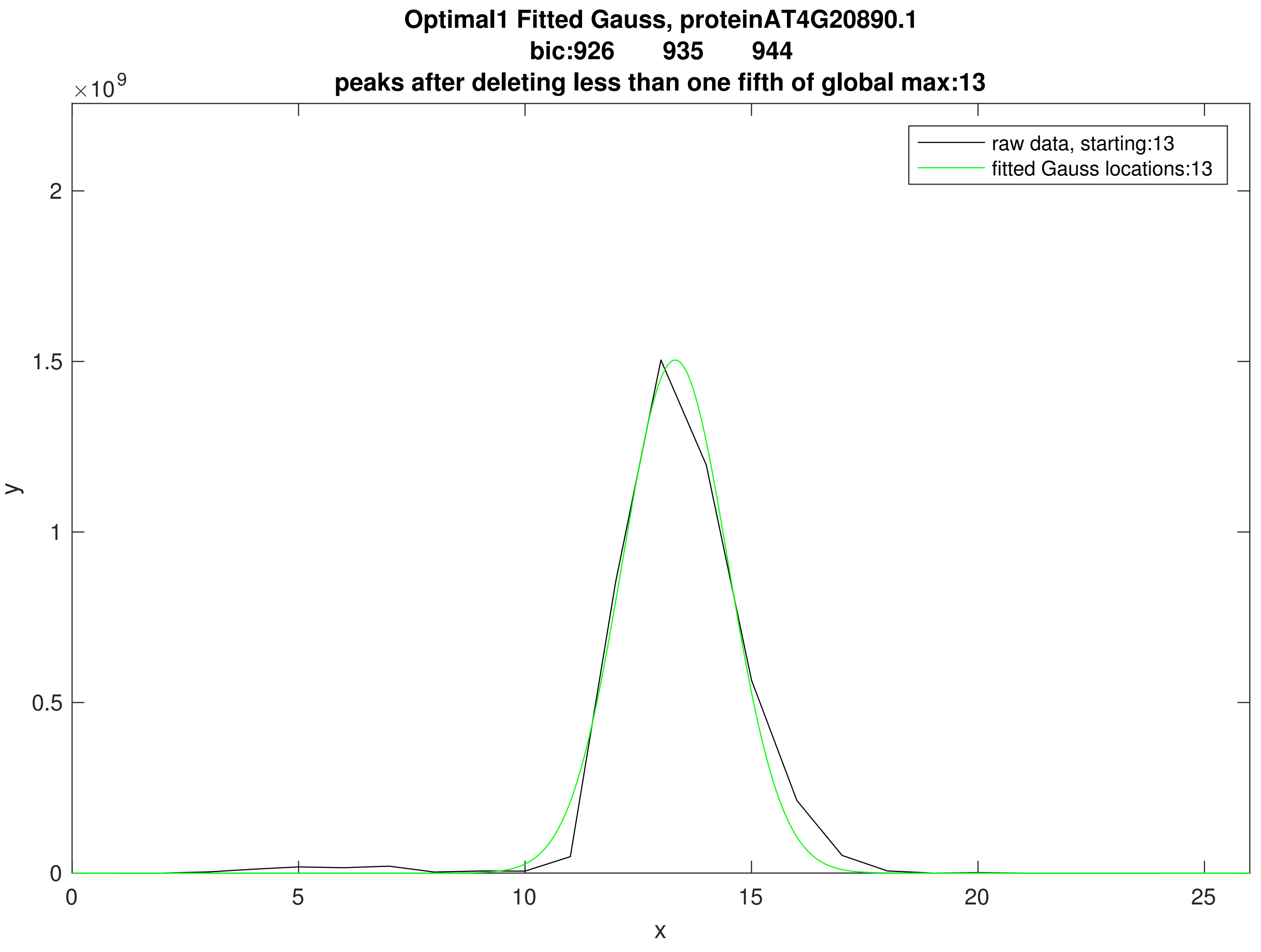
Optimal1 Fitted Gauss, proteinAT4G20890.1
bic:926 935 944
peaks after deleting less than one fifth of global max:13
×10^9
2
1.5
1
0.5
0
y
0
5
10
15
20
25
x
raw data, starting:13
fitted Gauss locations:13

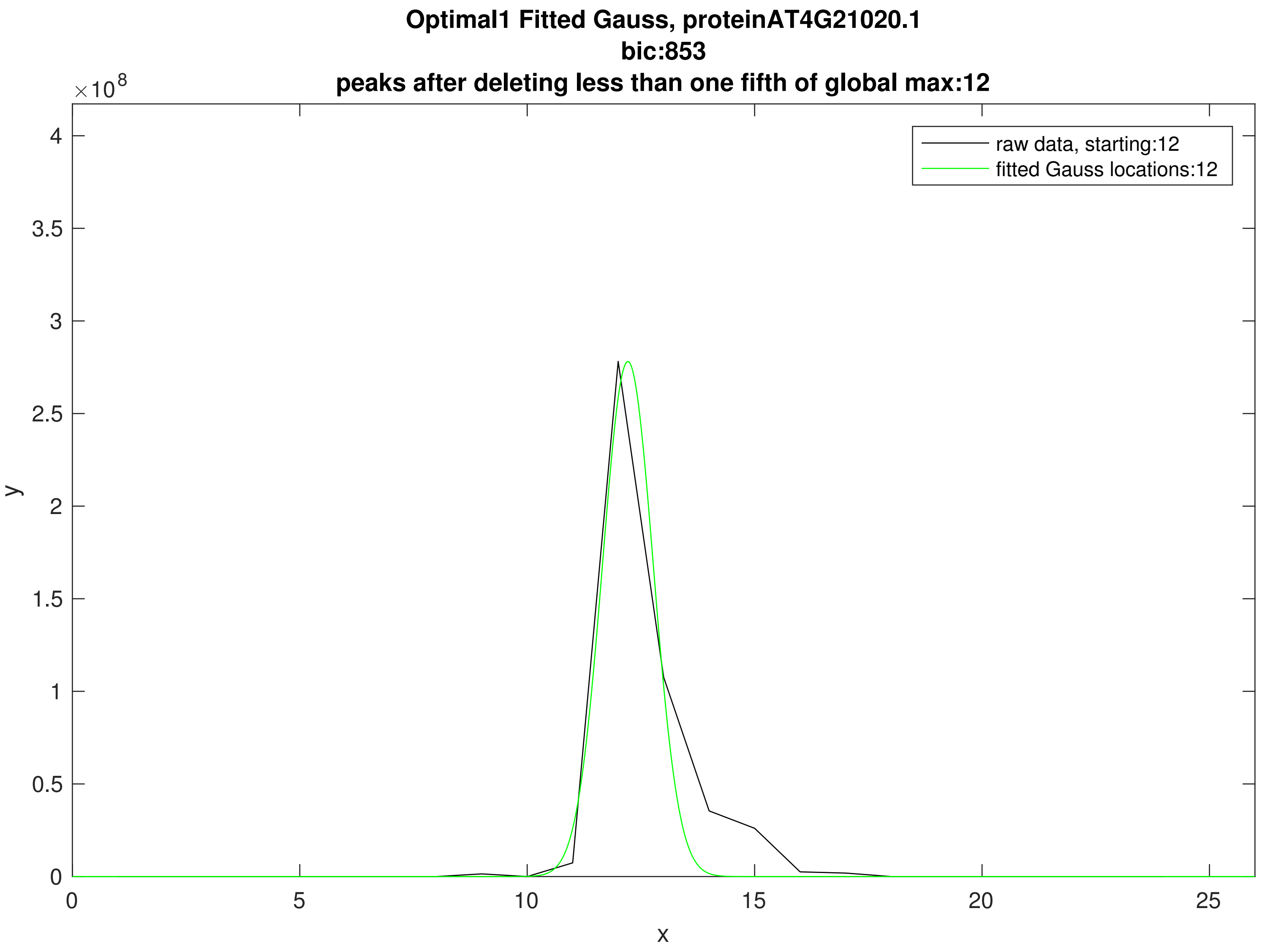
Optimal1 Fitted Gauss, proteinAT4G21020.1
bic:853
peaks after deleting less than one fifth of global max:12
×10^8
raw data, starting:12
fitted Gauss locations:12
y
x
0
0.5
1
1.5
2
2.5
3
3.5
4
5
10
15
20
25

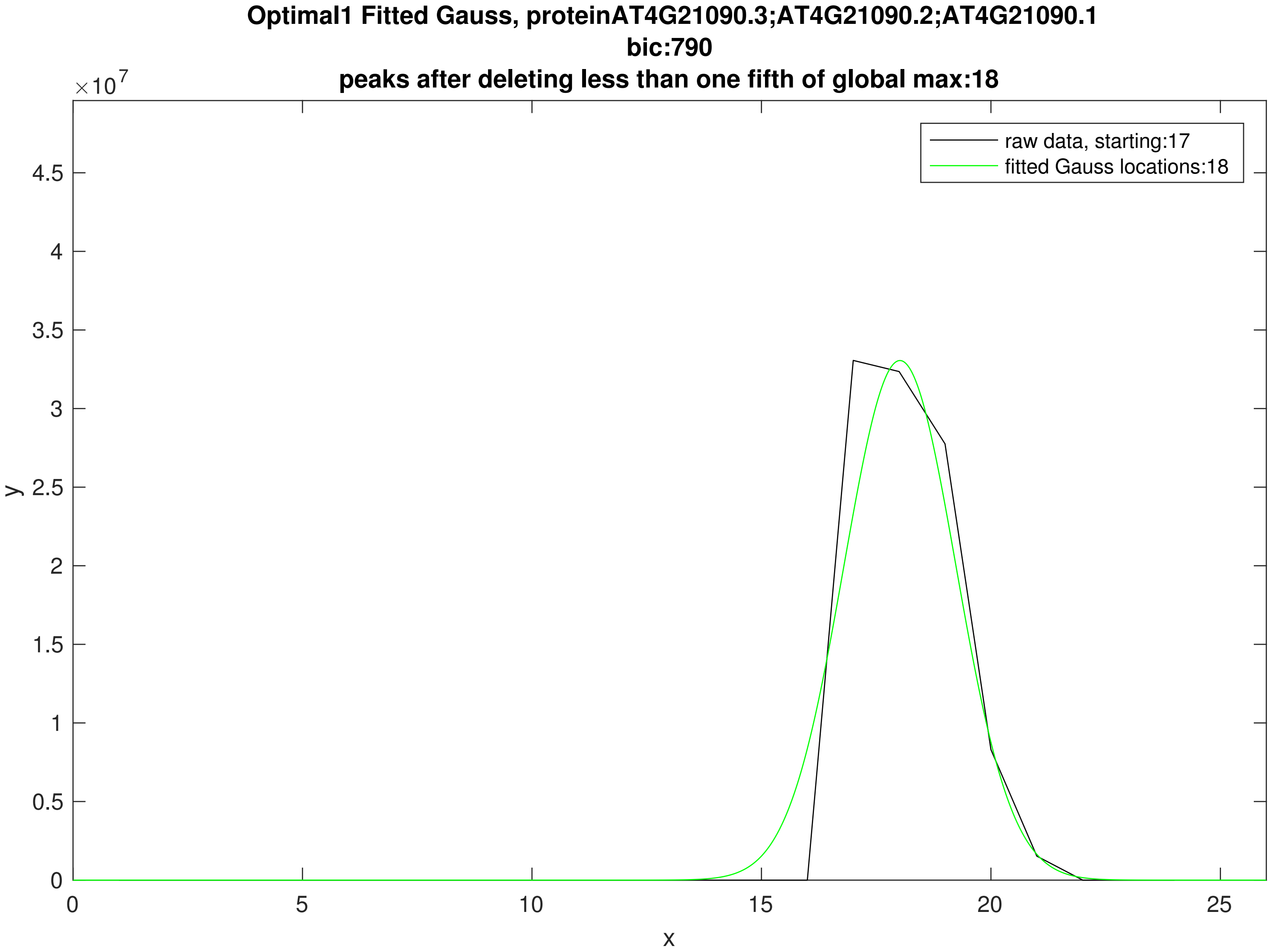

Optimal1 Fitted Gauss, proteinAT4G21090.3;AT4G21090.2;AT4G21090.1
bic:790
peaks after deleting less than one fifth of global max:18
×10^7
raw data, starting:17
fitted Gauss locations:18
y
x
0
0.5
1
1.5
2
2.5
3
3.5
4
4.5
5
10
15
20
25

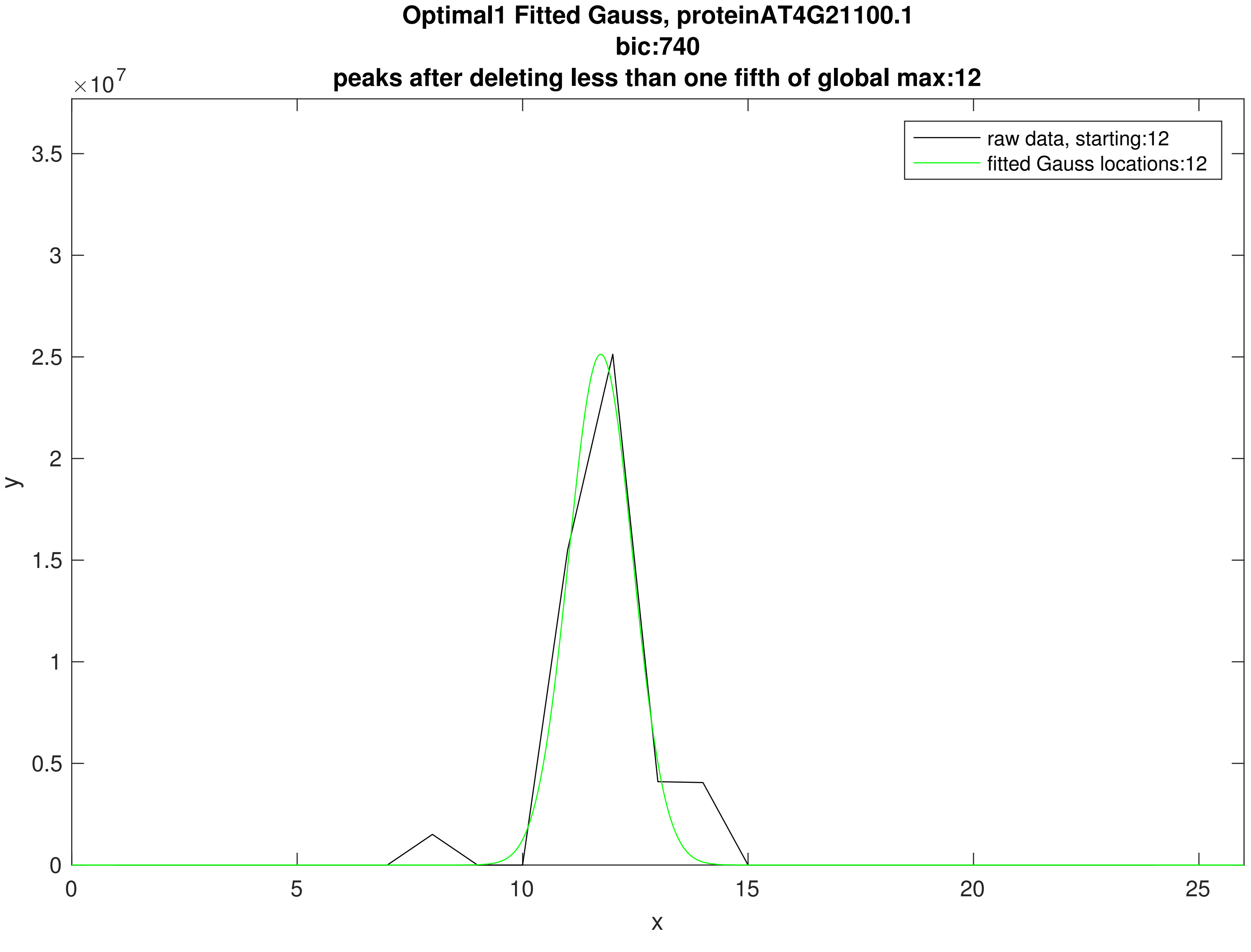
Optimal1 Fitted Gauss, proteinAT4G21100.1
bic:740
peaks after deleting less than one fifth of global max:12
×10^7
3.5
3
2.5
2
1.5
1
0.5
0
y
0
5
10
15
20
25
x
raw data, starting:12
fitted Gauss locations:12

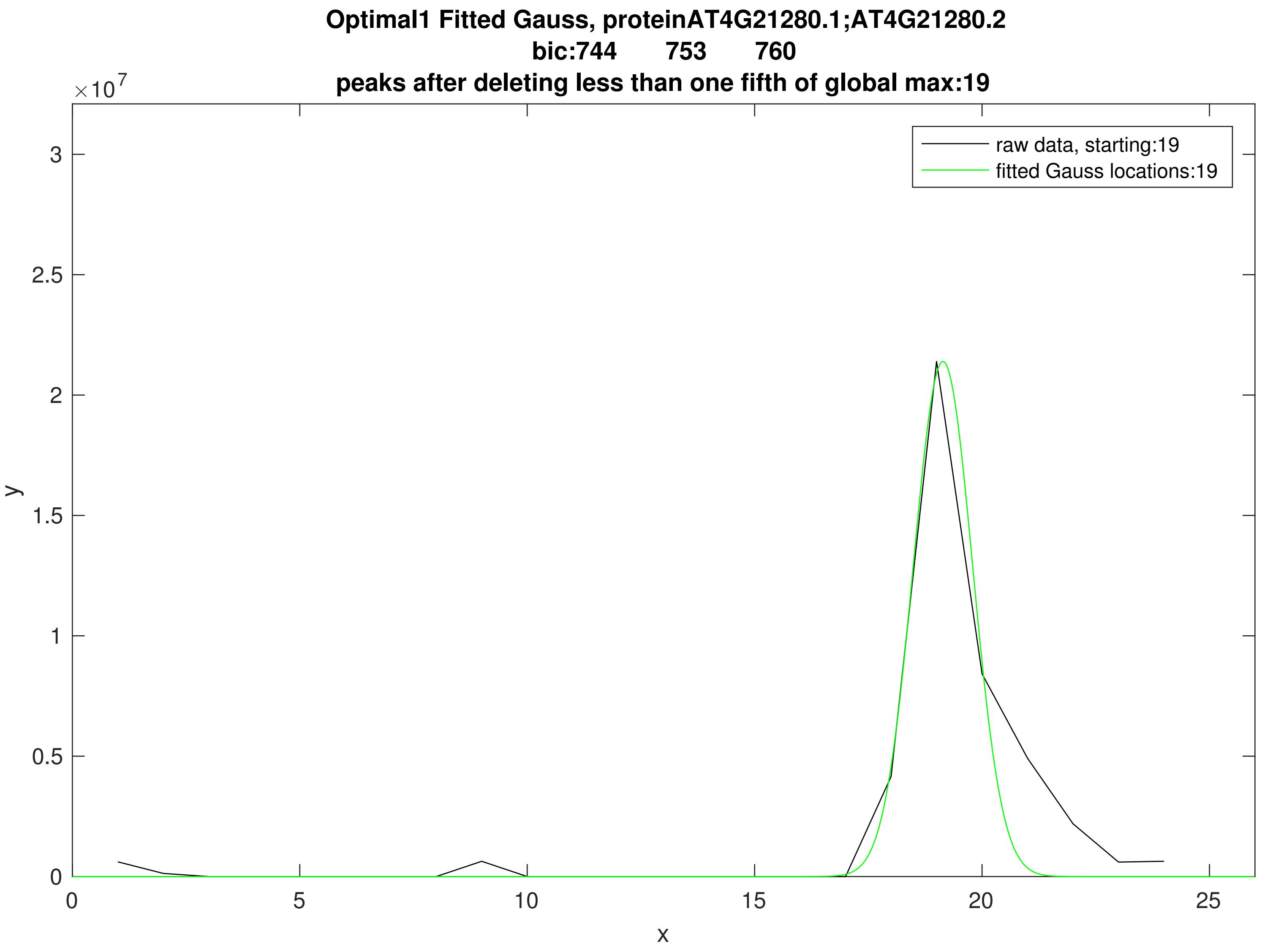
Optimal1 Fitted Gauss, proteinAT4G21280.1;AT4G21280.2
bic:744 753 760
peaks after deleting less than one fifth of global max:19
×10^7
raw data, starting:19
fitted Gauss locations:19
y
x
0
0.5
1
1.5
2
2.5
3
0
5
10
15
20
25

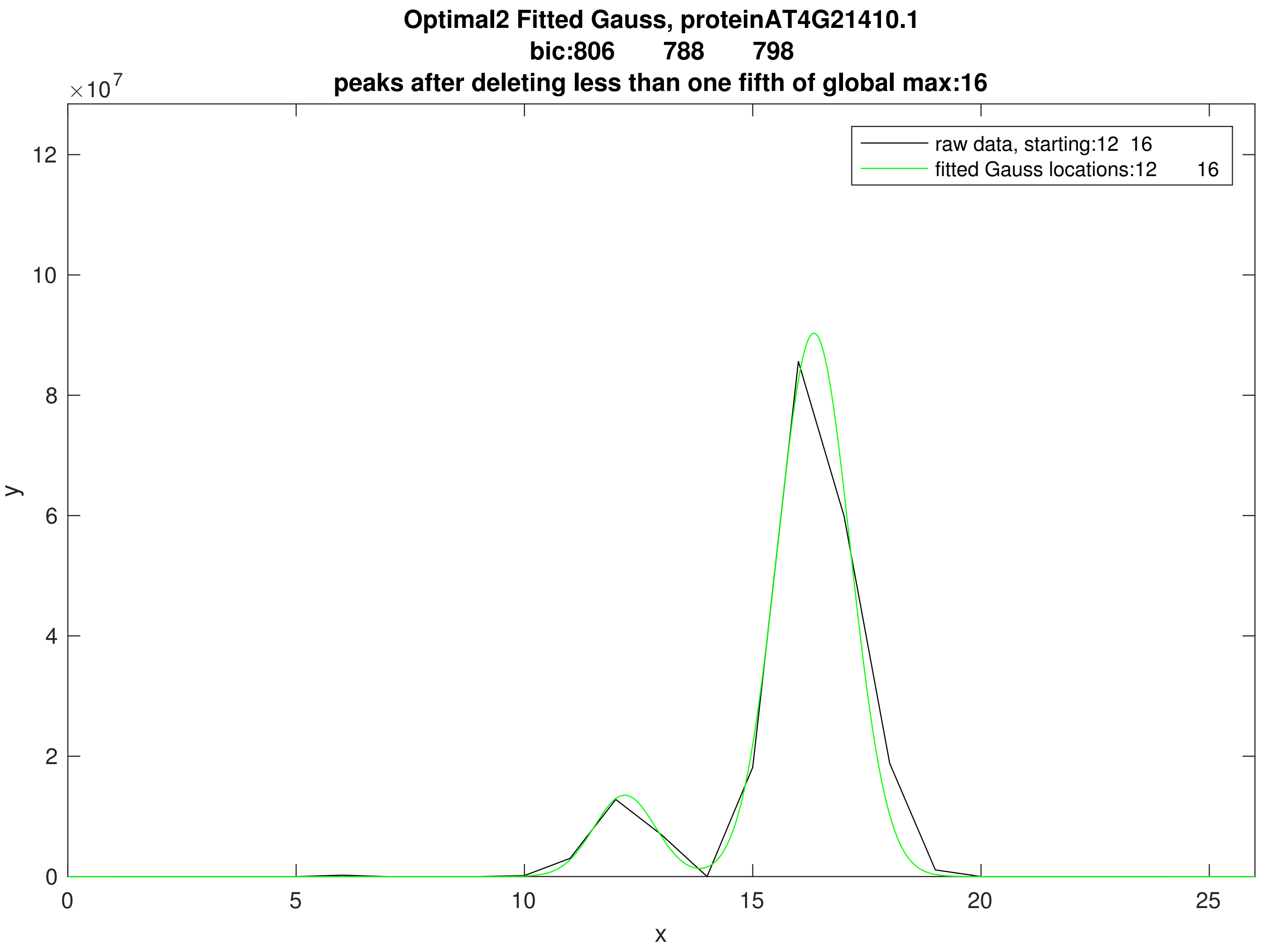
Optimal2 Fitted Gauss, proteinAT4G21410.1
bic:806 788 798
peaks after deleting less than one fifth of global max:16
×10^7
raw data, starting:12 16
fitted Gauss locations:12 16
y
x
0
2
4
6
8
10
12
0
5
10
15
20
25

Optimal1 Fitted Gauss, proteinAT4G21560.3;AT4G21560.2;AT4G21560.1
bic:703
peaks after deleting less than one fifth of global max:18

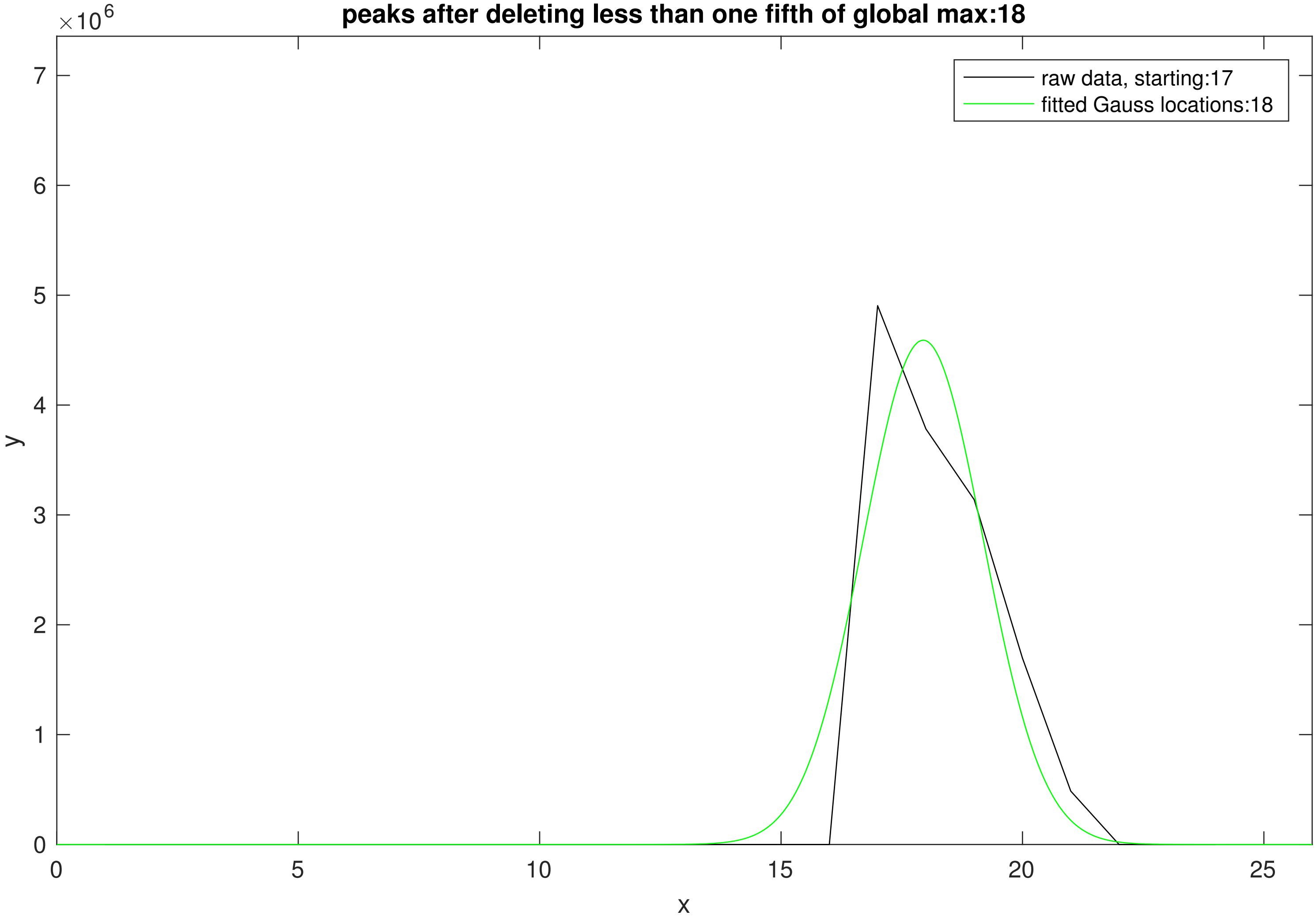

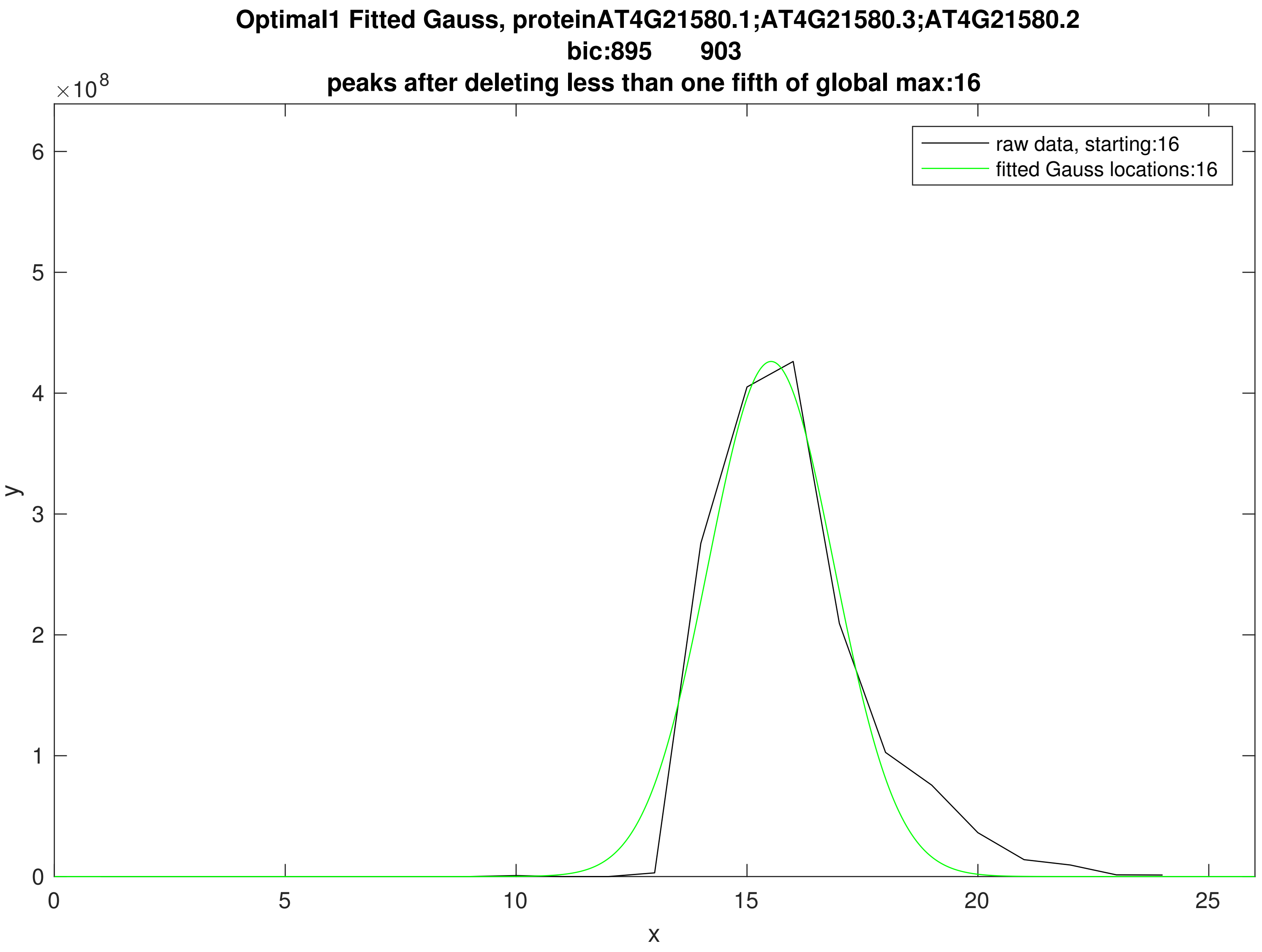
Optimal1 Fitted Gauss, proteinAT4G21580.1;AT4G21580.3;AT4G21580.2
bic:895 903
peaks after deleting less than one fifth of global max:16
×10^8
y
x
raw data, starting:16
fitted Gauss locations:16
0
1
2
3
4
5
6
0
5
10
15
20
25

**Optimal1 Fitted Gauss, proteinAT4G21590.2;AT4G21590.1**
**bic:679**
**peaks after deleting less than one fifth of global max:16**

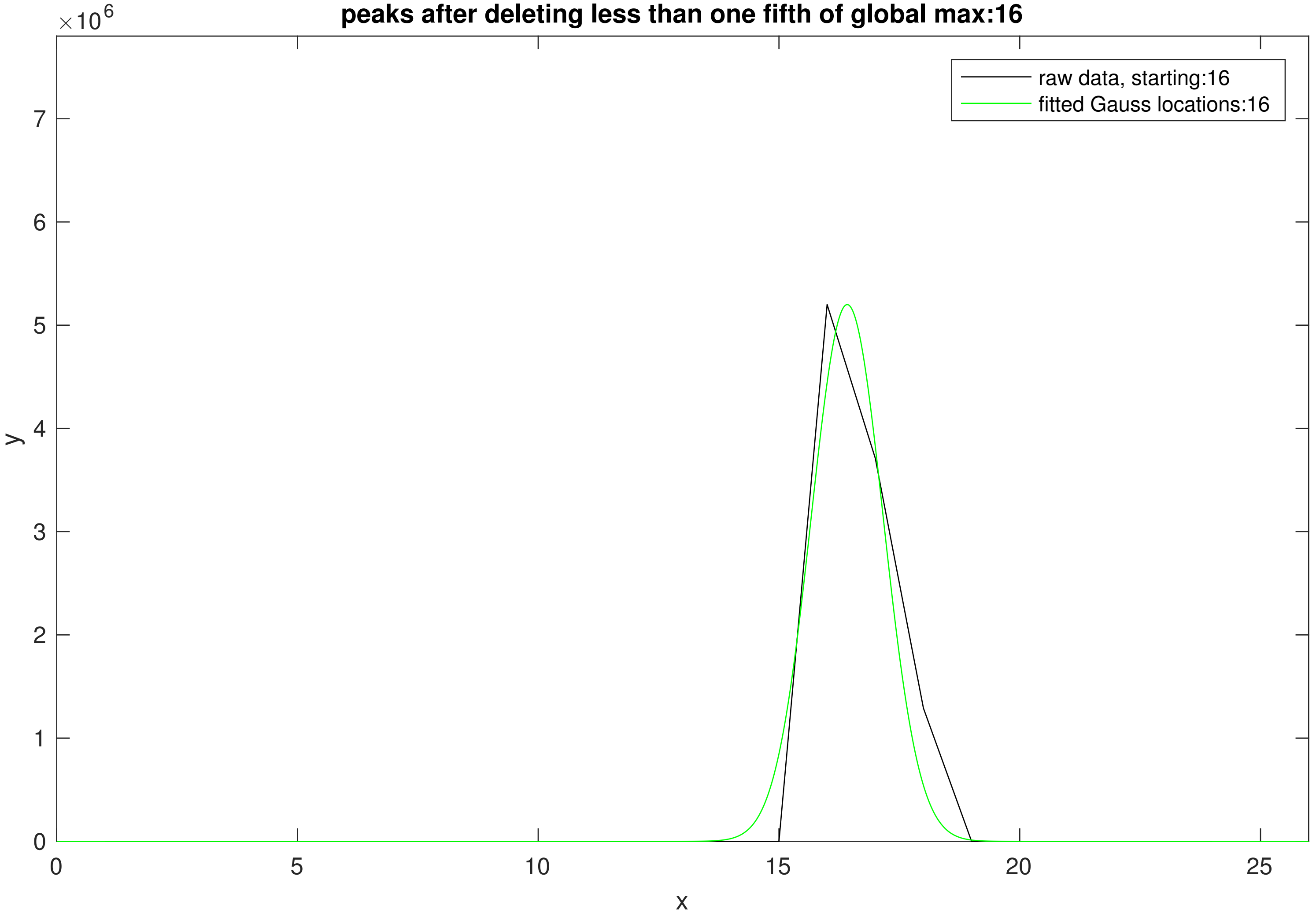

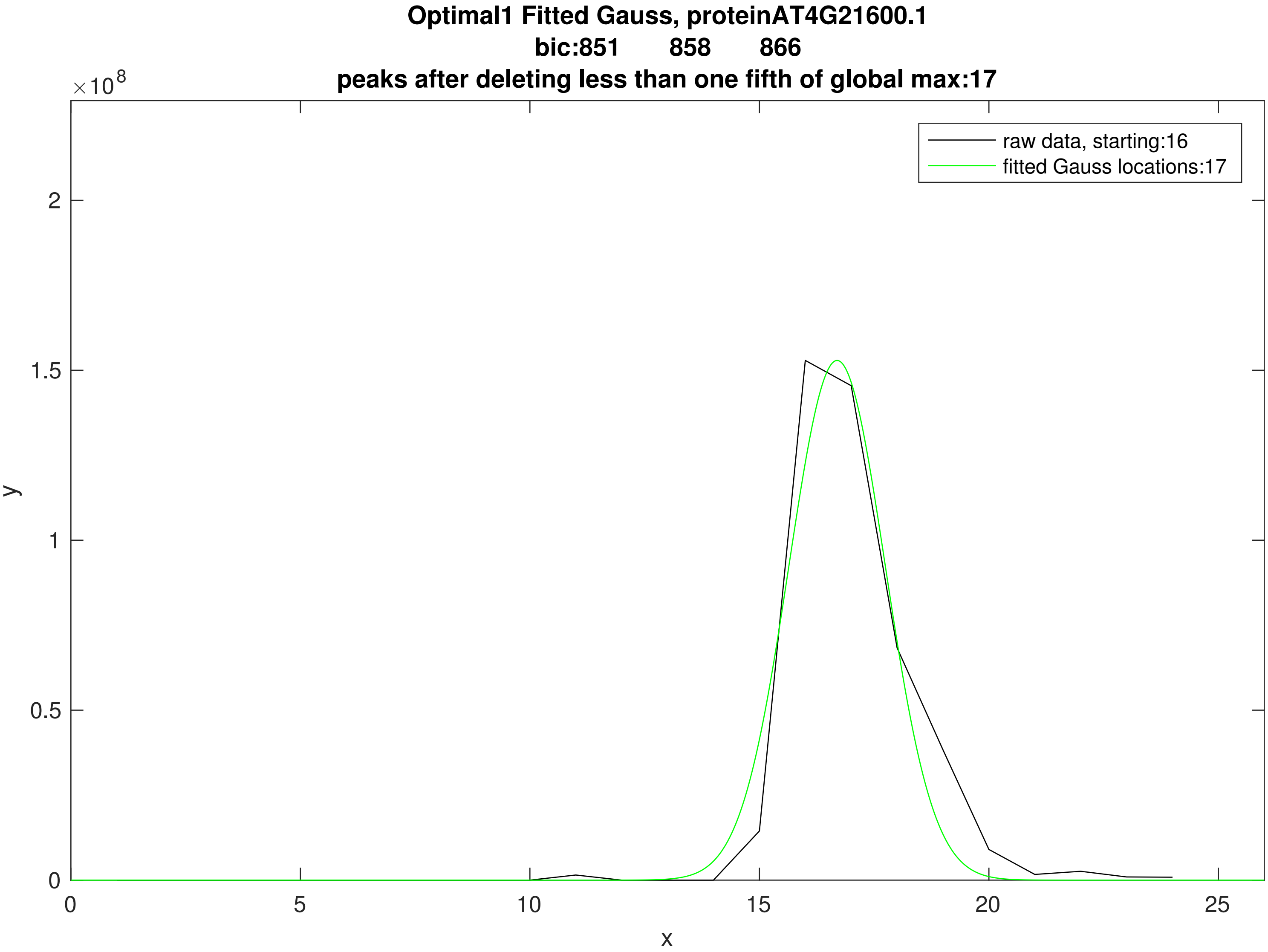
Optimal1 Fitted Gauss, proteinAT4G21600.1
bic:851 858 866
peaks after deleting less than one fifth of global max:17
×10^8
raw data, starting:16
fitted Gauss locations:17
y
x
0
0.5
1
1.5
2
0
5
10
15
20
25

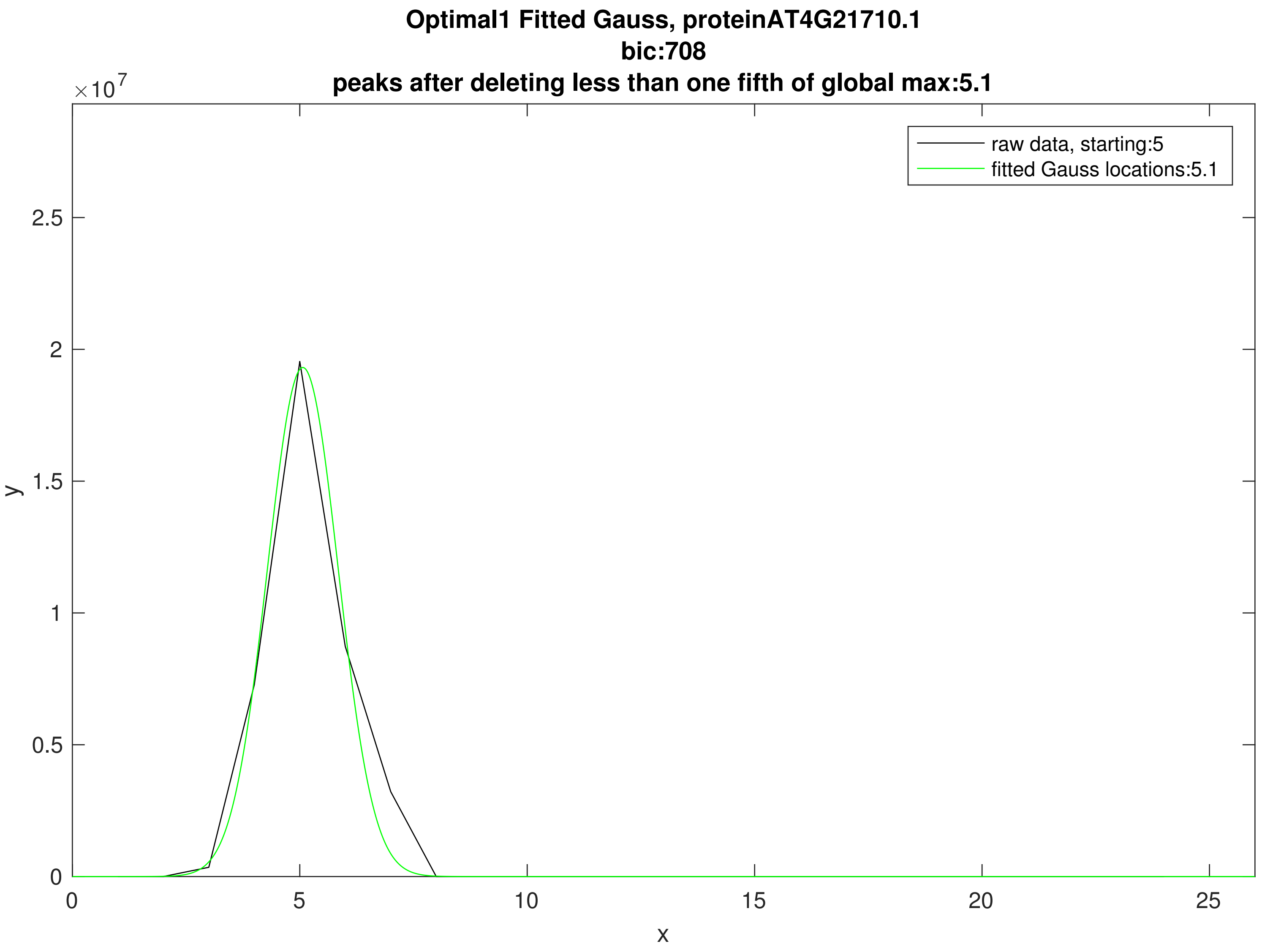

Optimal1 Fitted Gauss, proteinAT4G21710.1
bic:708
peaks after deleting less than one fifth of global max:5.1
×10^7
raw data, starting:5
fitted Gauss locations:5.1
y
2.5
2
1.5
1
0.5
0
0
5
10
15
20
25
x

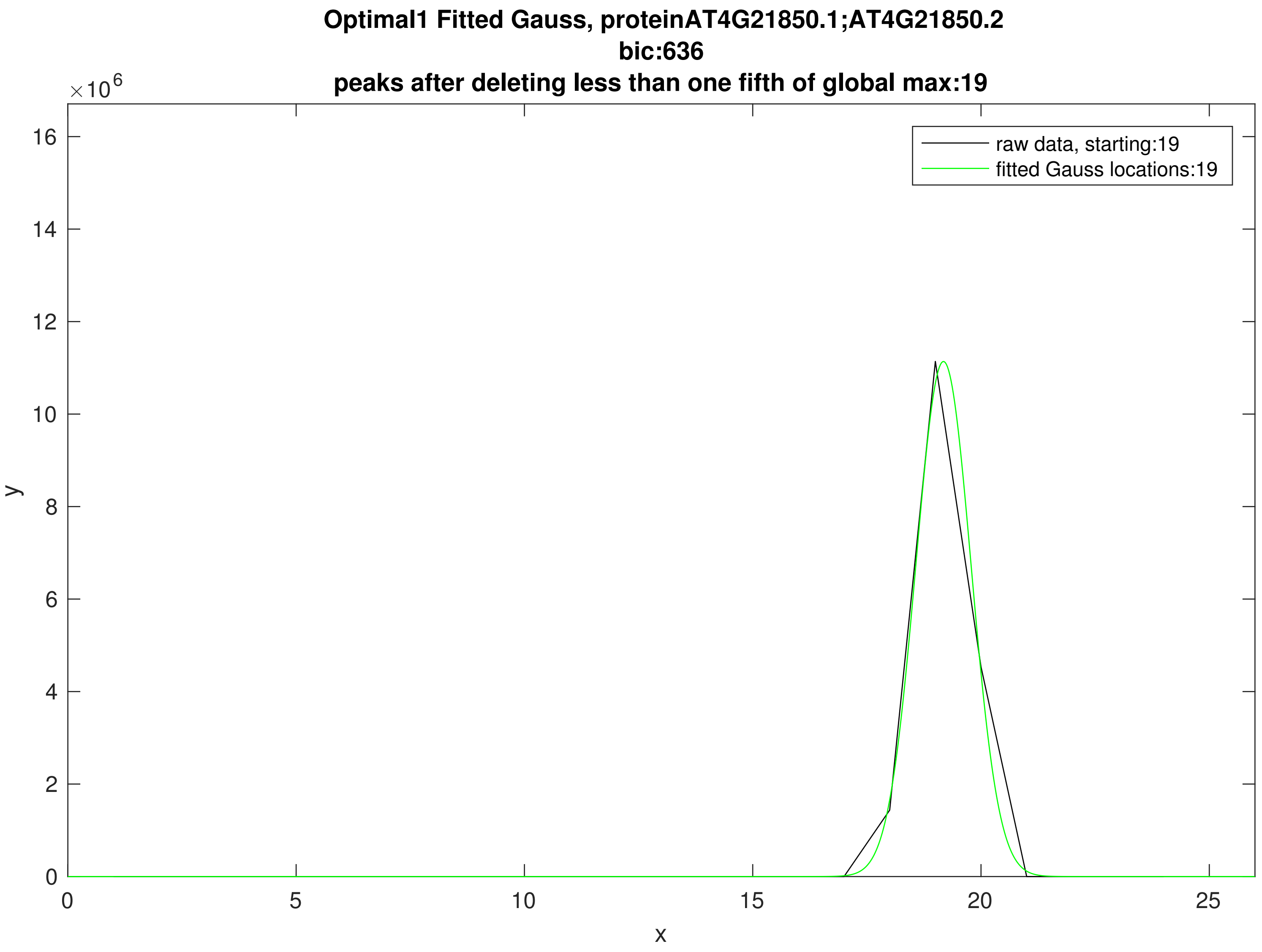
Optimal1 Fitted Gauss, proteinAT4G21850.1;AT4G21850.2
bic:636
peaks after deleting less than one fifth of global max:19
×10^6
16
14
12
10
8
6
4
2
0
y
0
5
10
15
20
25
x
raw data, starting:19
fitted Gauss locations:19

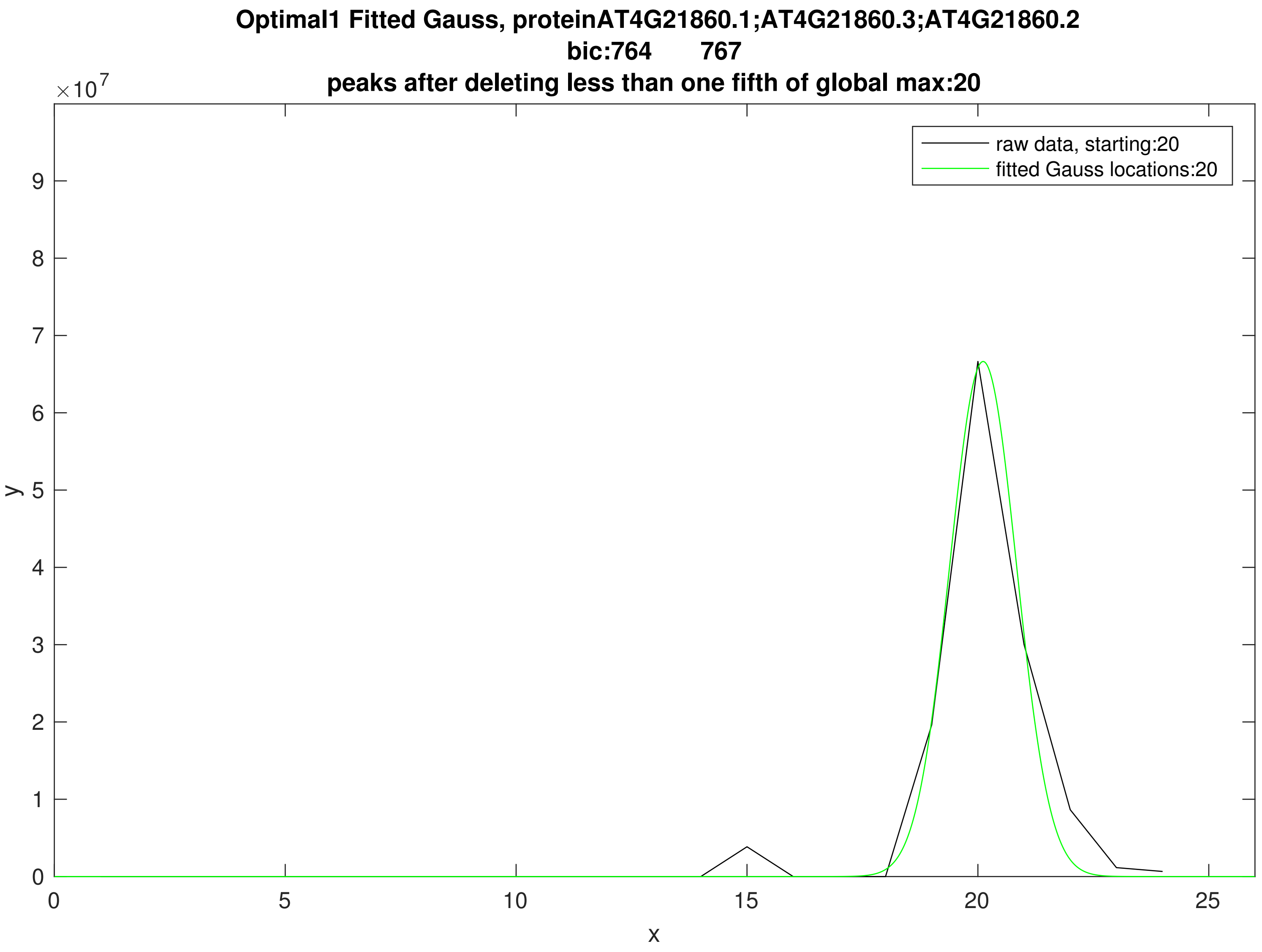

Optimal1 Fitted Gauss, proteinAT4G21860.1;AT4G21860.3;AT4G21860.2
bic:764 767
peaks after deleting less than one fifth of global max:20
×10^7
raw data, starting:20
fitted Gauss locations:20
y
x
0
1
2
3
4
5
6
7
8
9
0
5
10
15
20
25

**Optimal1 Fitted Gauss, proteinAT4G22150.1**
**bic:770 778**
**peaks after deleting less than one fifth of global max:14**

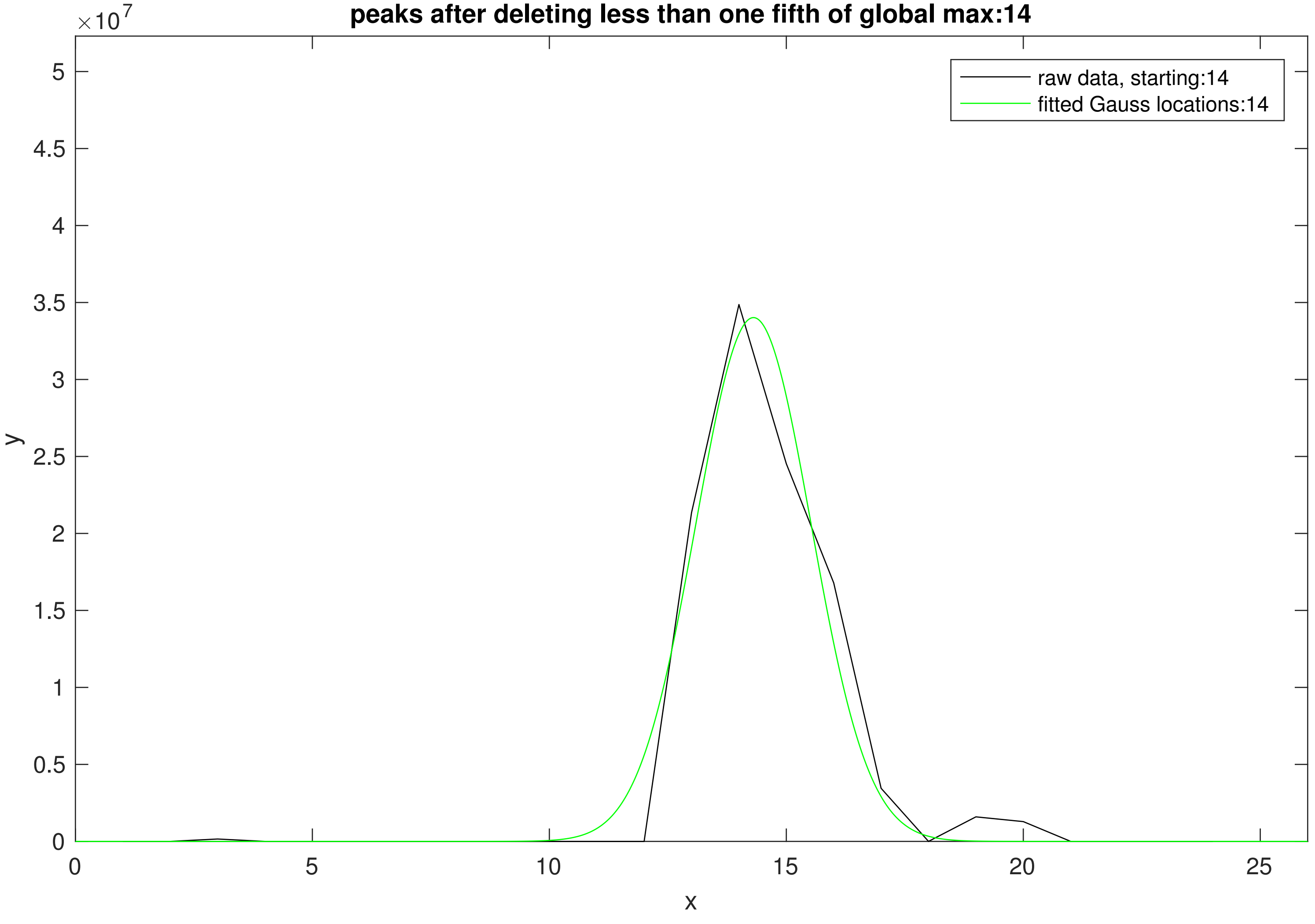

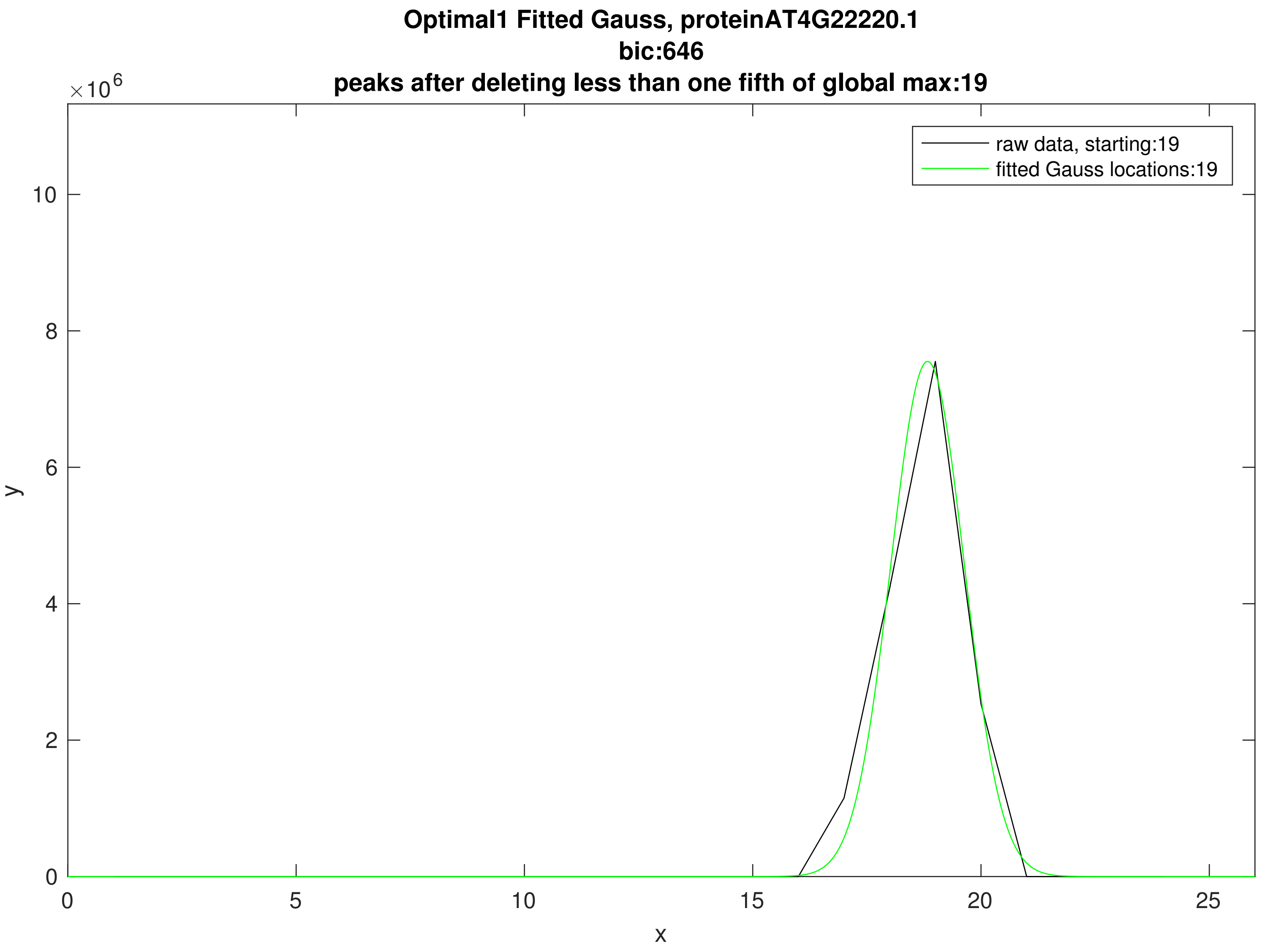
Optimal1 Fitted Gauss, proteinAT4G22220.1
bic:646
peaks after deleting less than one fifth of global max:19
×10^6
raw data, starting:19
fitted Gauss locations:19
10
8
6
4
2
0
y
0
5
10
15
20
25
x

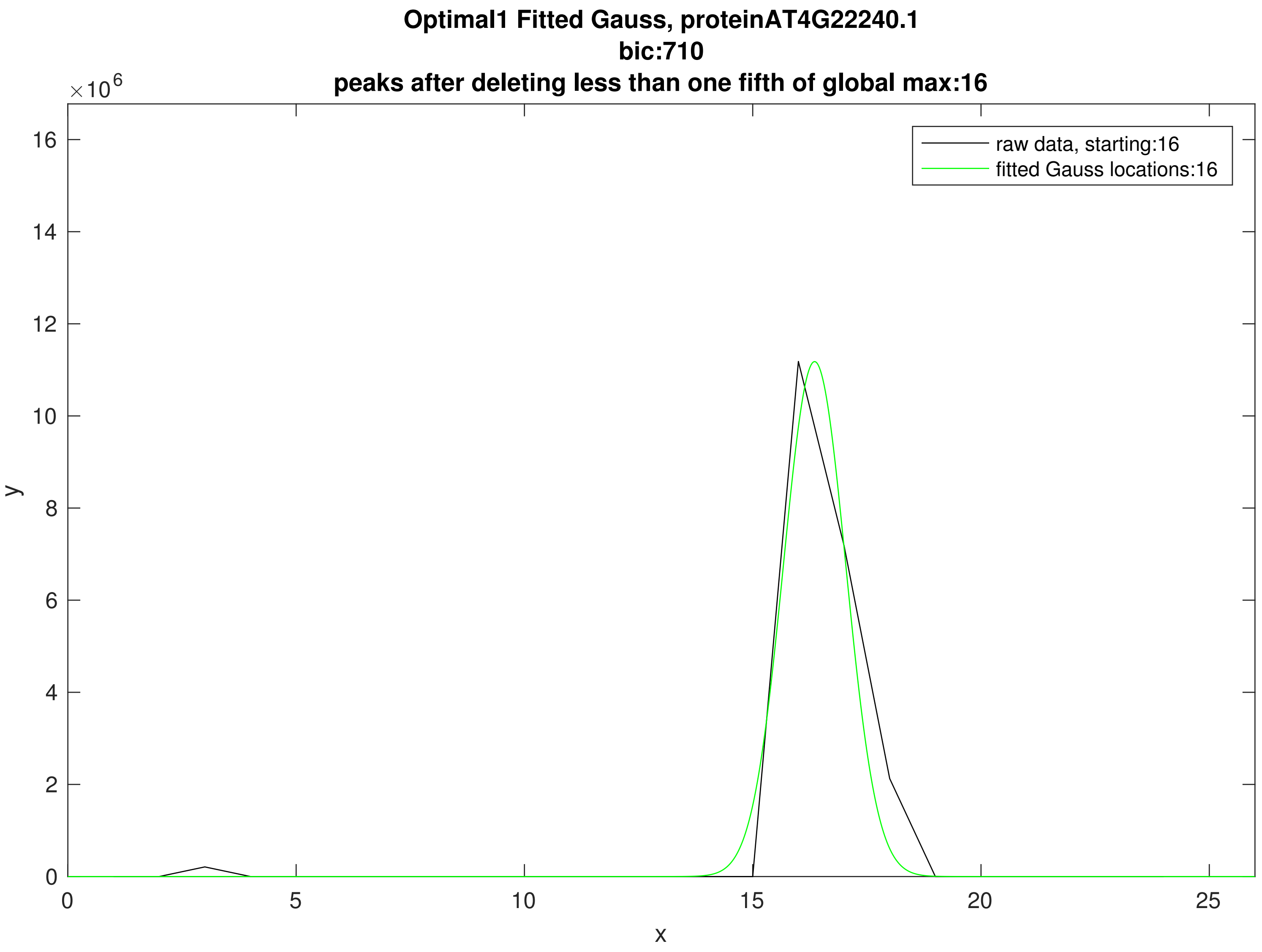

Optimal1 Fitted Gauss, proteinAT4G22240.1
bic:710
peaks after deleting less than one fifth of global max:16
×10^6
raw data, starting:16
fitted Gauss locations:16
y
x
0
2
4
6
8
10
12
14
16
0
5
10
15
20
25

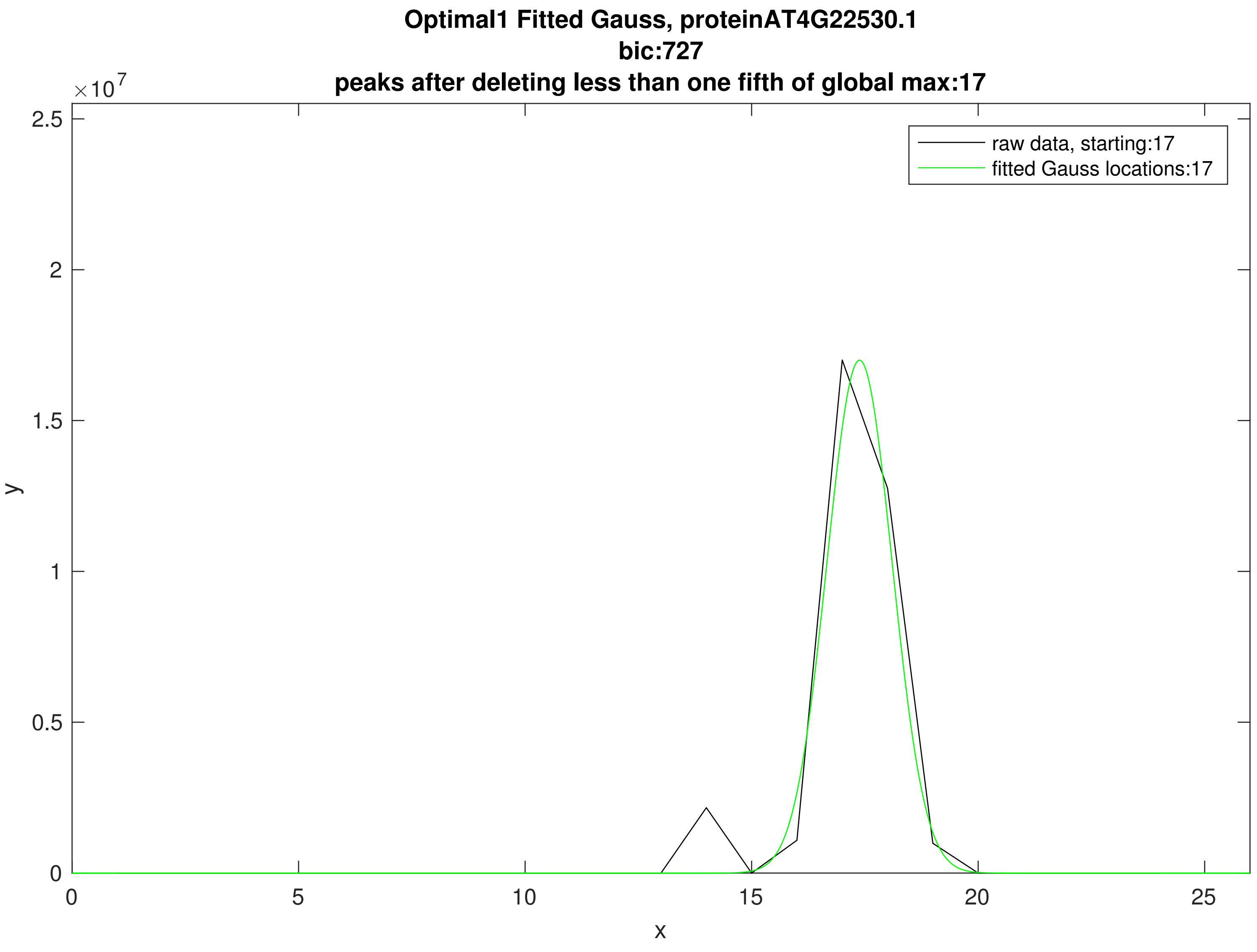
Optimal1 Fitted Gauss, proteinAT4G22530.1
bic:727
peaks after deleting less than one fifth of global max:17
×10^7
2.5
2
1.5
1
0.5
0
y
0
5
10
15
20
25
x
raw data, starting:17
fitted Gauss locations:17

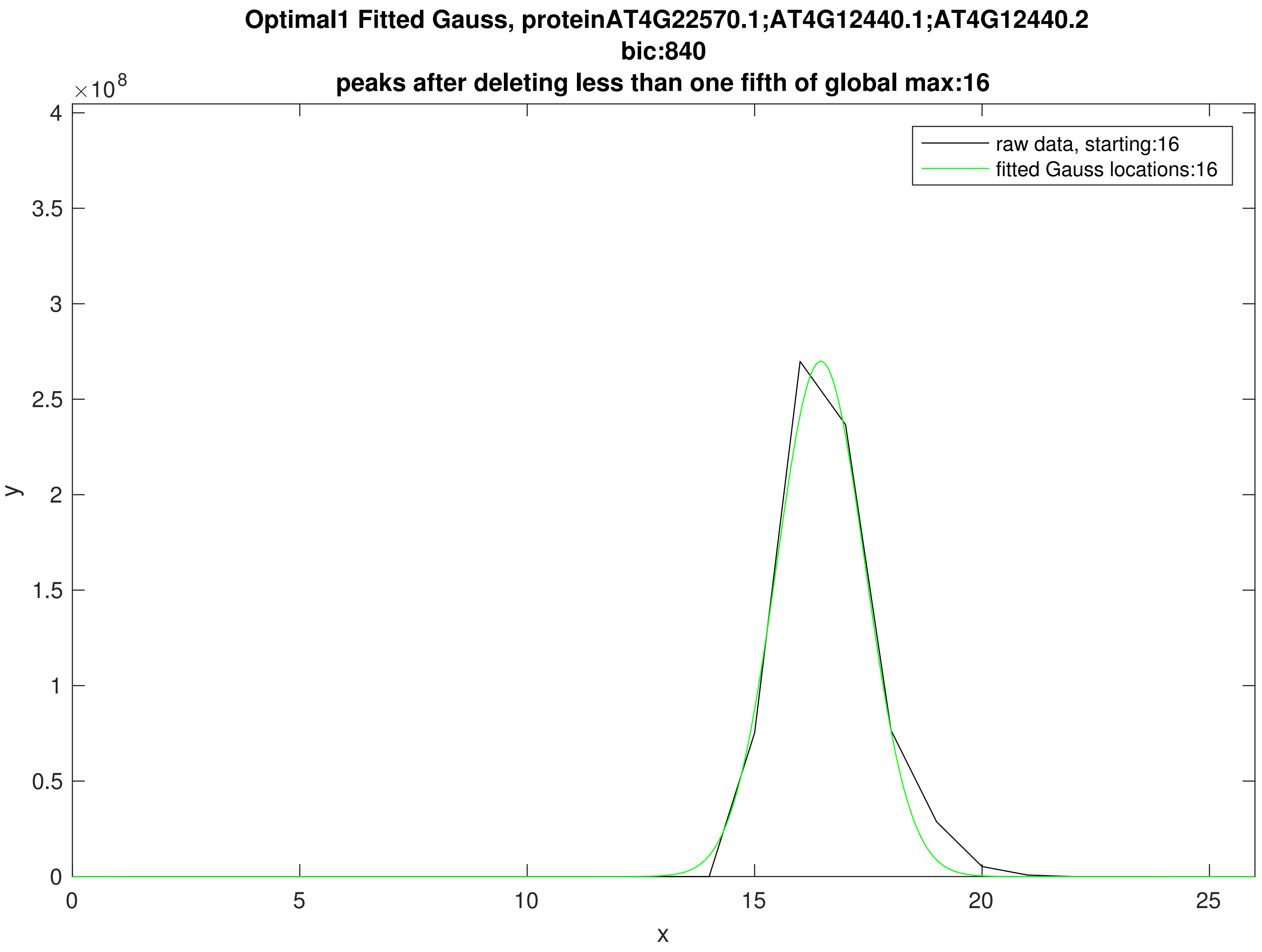

Optimal1 Fitted Gauss, proteinAT4G22570.1;AT4G12440.1;AT4G12440.2
bic:840
peaks after deleting less than one fifth of global max:16
×10^8
raw data, starting:16
fitted Gauss locations:16
y
x
0
0.5
1
1.5
2
2.5
3
3.5
4
5
10
15
20
25

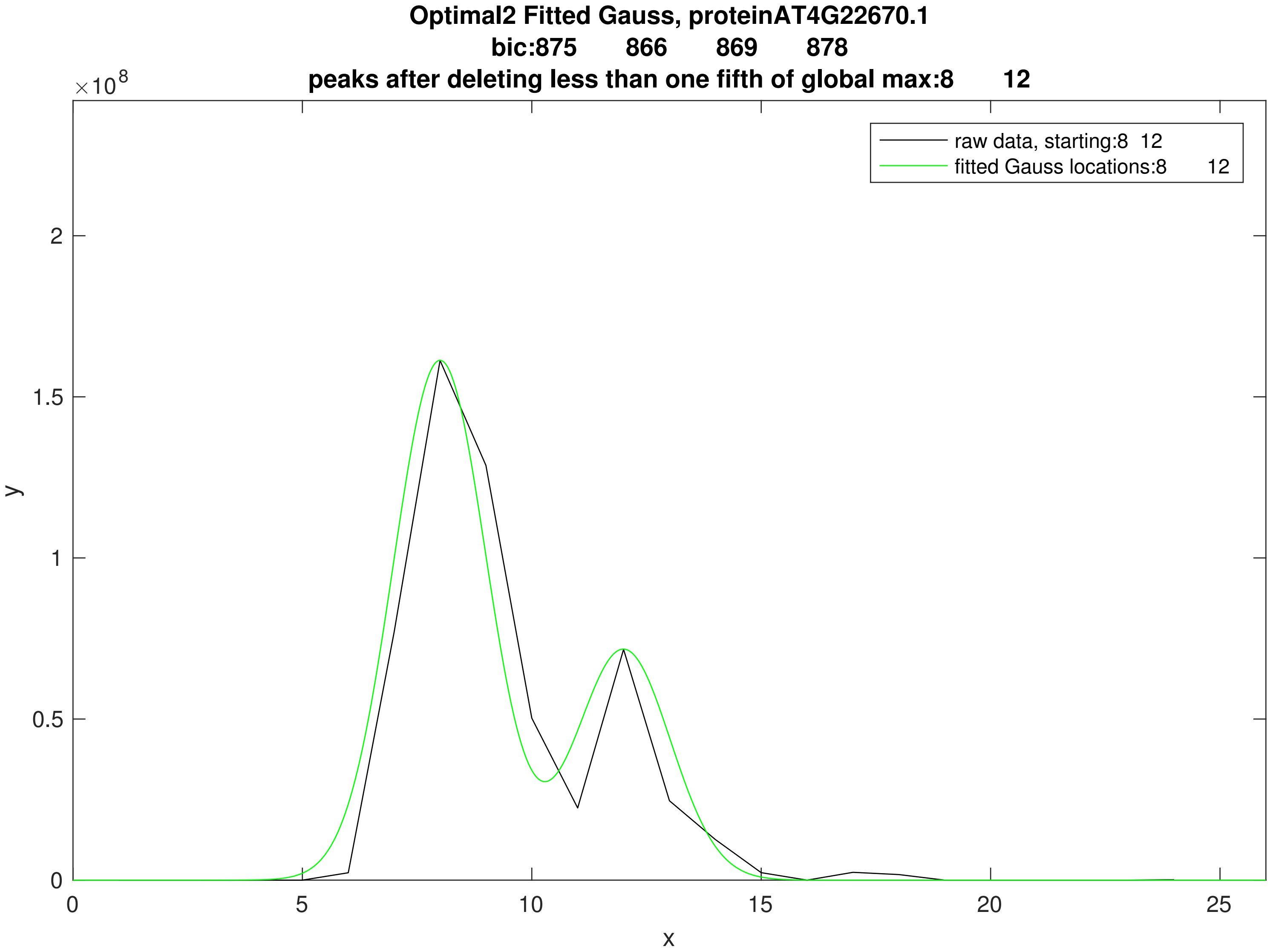
Optimal2 Fitted Gauss, proteinAT4G22670.1
bic:875 866 869 878
peaks after deleting less than one fifth of global max:8 12
×10^8
raw data, starting:8 12
fitted Gauss locations:8 12
2
1.5
1
0.5
0
y
0
5
10
15
20
25
x

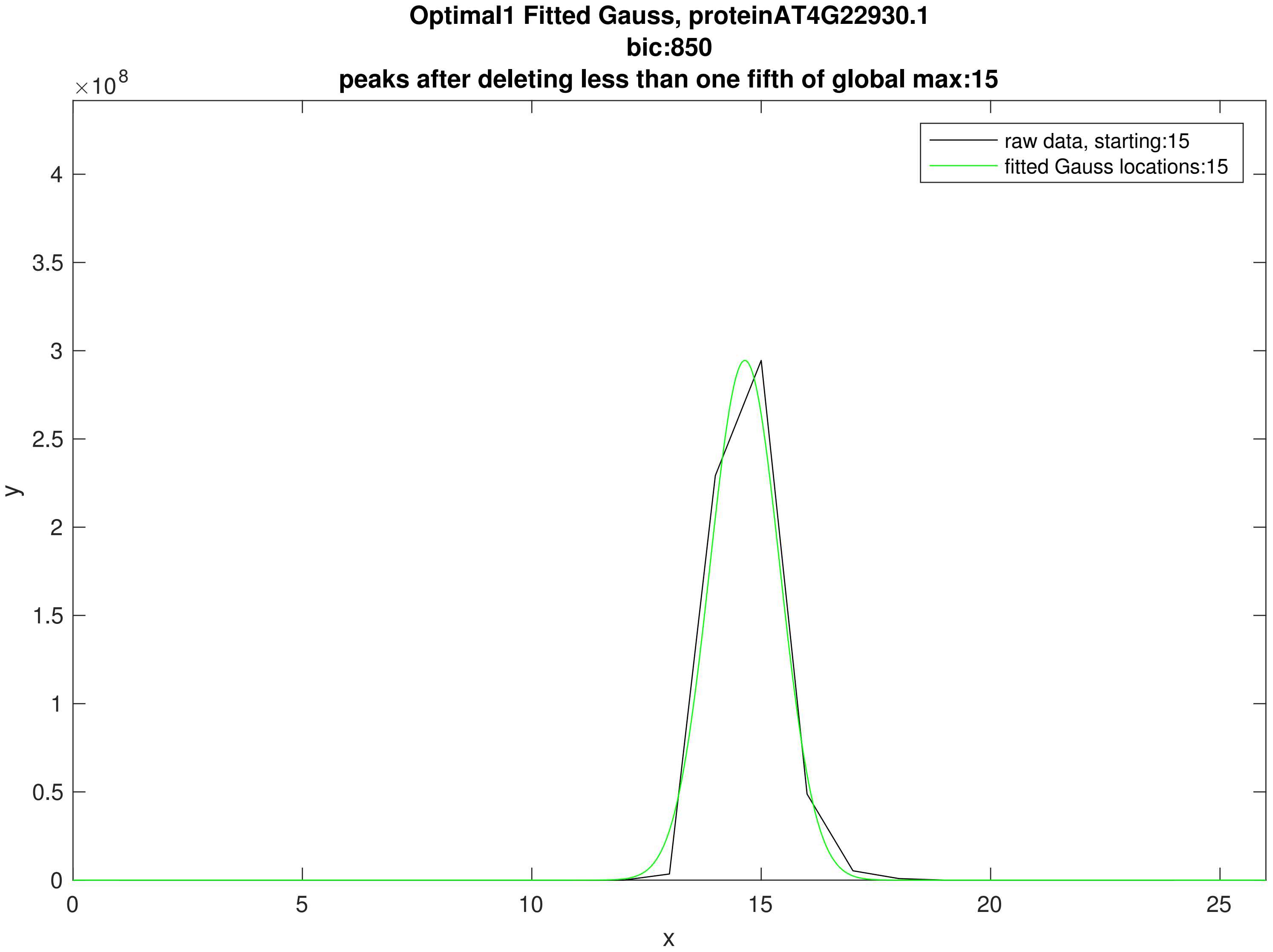
Optimal1 Fitted Gauss, proteinAT4G22930.1
bic:850
peaks after deleting less than one fifth of global max:15
×10^8
raw data, starting:15
fitted Gauss locations:15
y
x
0
0.5
1
1.5
2
2.5
3
3.5
4
0
5
10
15
20
25

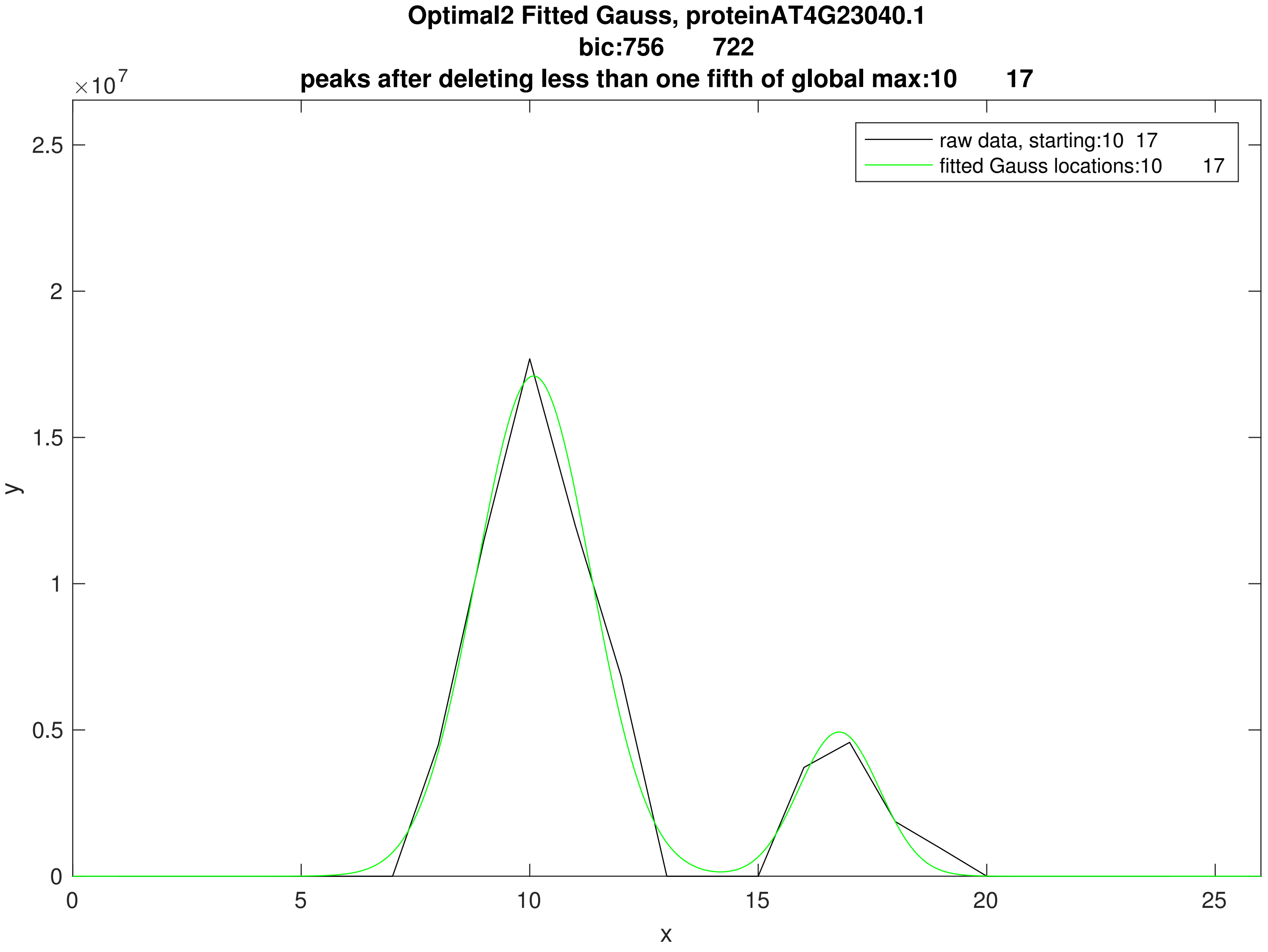

Optimal2 Fitted Gauss, proteinAT4G23040.1
bic:756 722
peaks after deleting less than one fifth of global max:10 17
×10^7
raw data, starting:10 17
fitted Gauss locations:10 17
y
x
0
0.5
1
1.5
2
2.5
5
10
15
20
25

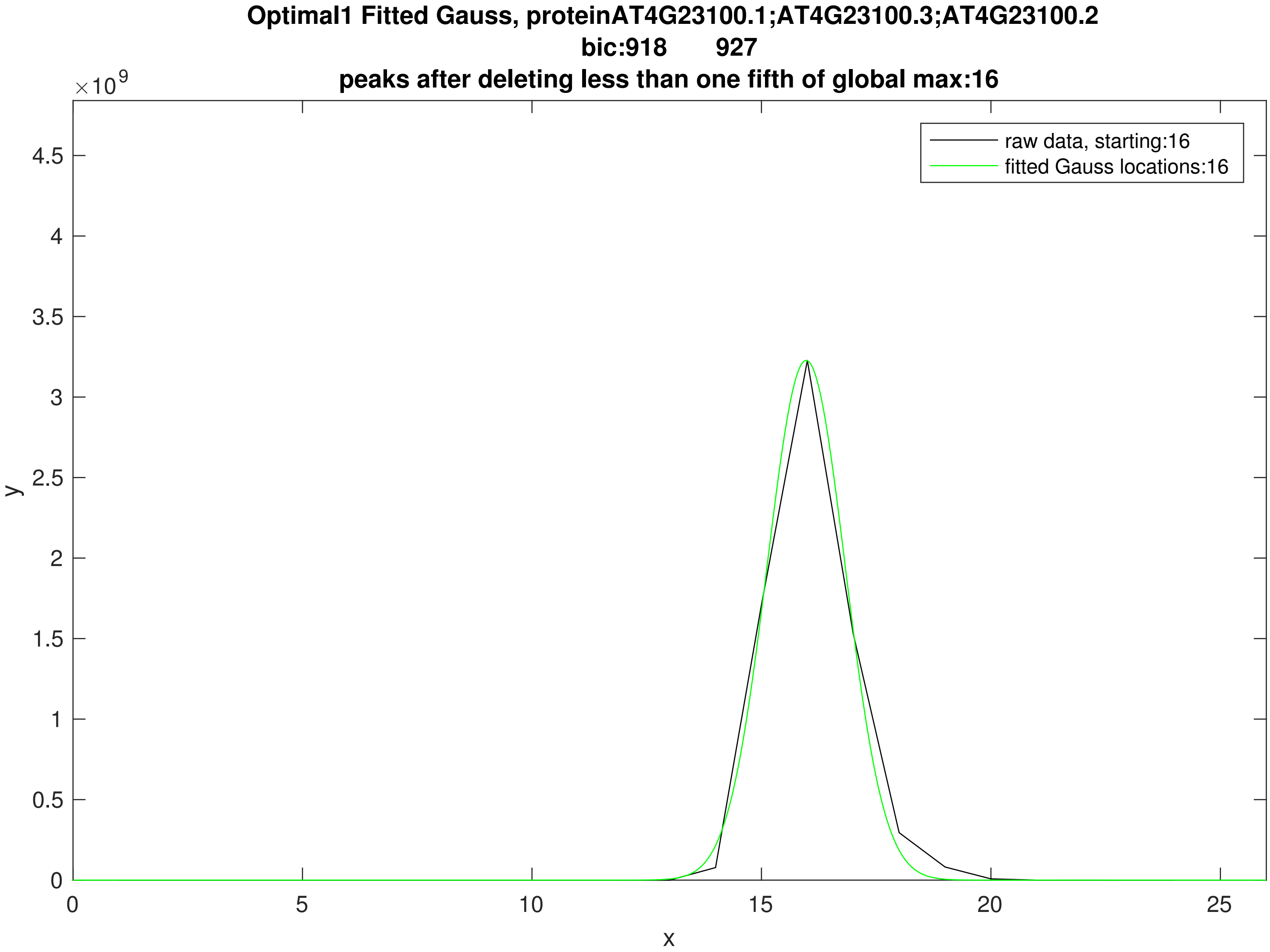

Optimal1 Fitted Gauss, proteinAT4G23100.1;AT4G23100.3;AT4G23100.2
bic:918 927
peaks after deleting less than one fifth of global max:16
×10^9
4.5
4
3.5
3
2.5
2
1.5
1
0.5
0
y
0
5
10
15
20
25
x
raw data, starting:16
fitted Gauss locations:16

**Optimal1 Fitted Gauss, proteinAT4G23270.1**
**bic:652**
**peaks after deleting less than one fifth of global max:16**

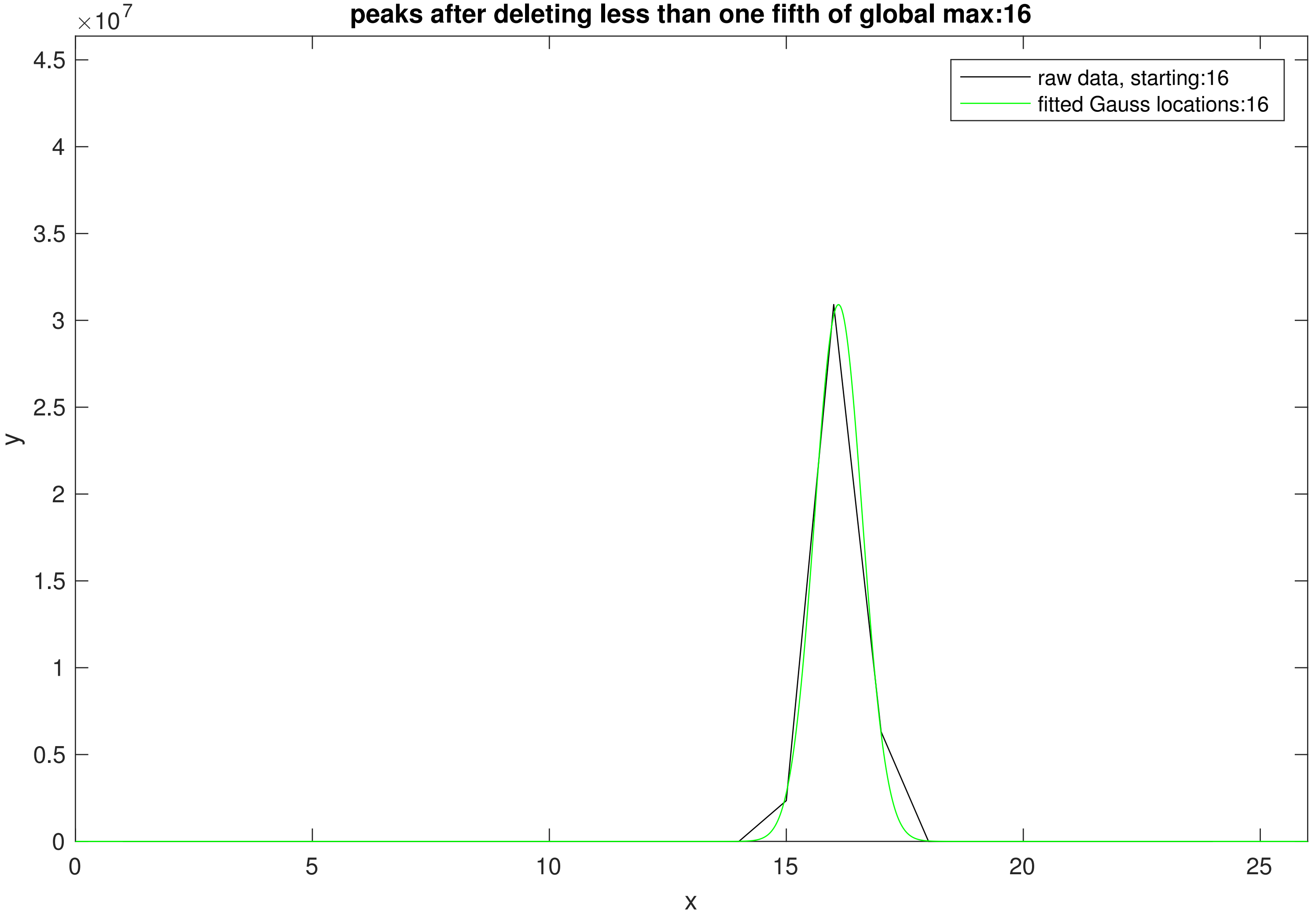

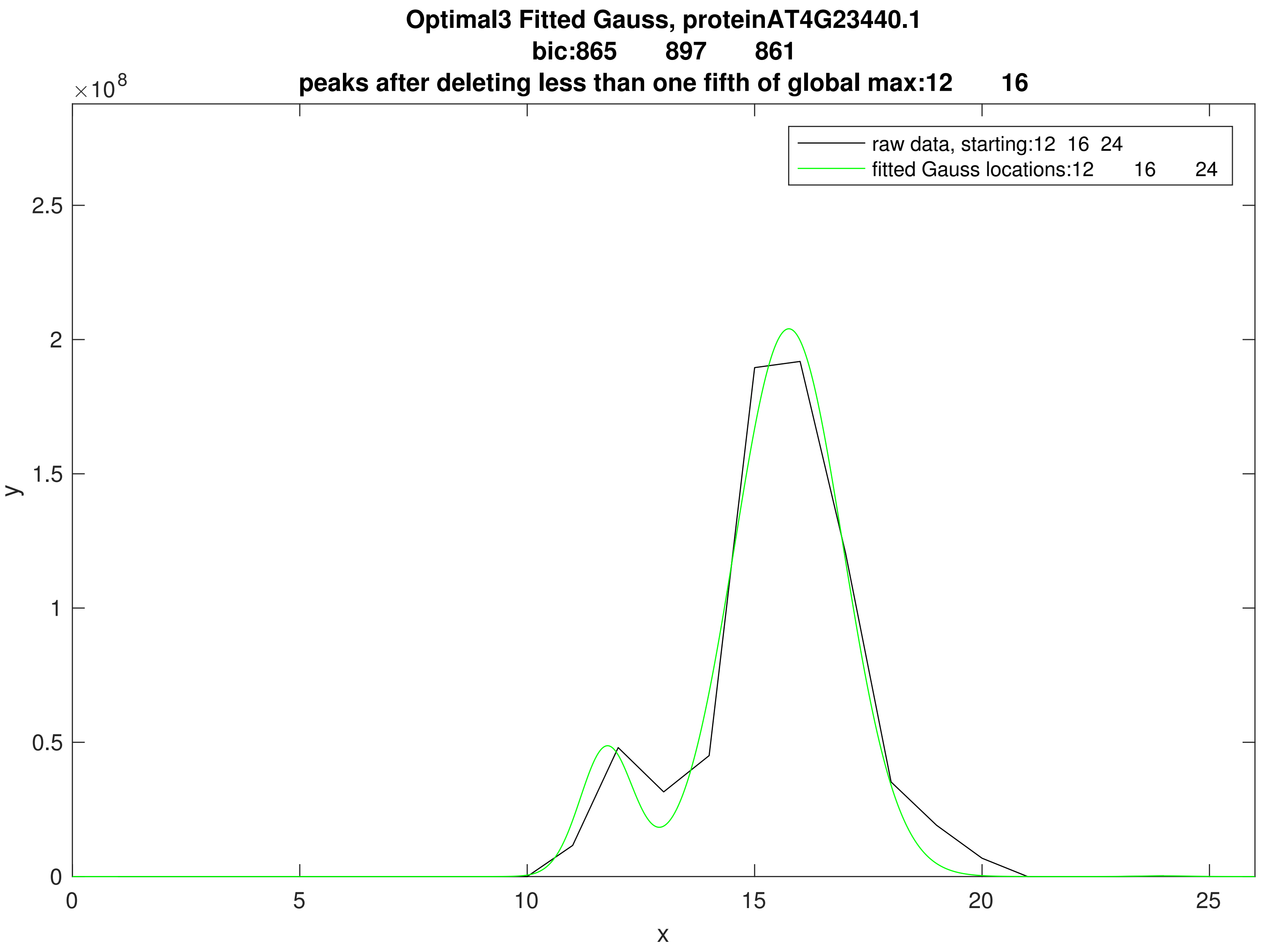

Optimal3 Fitted Gauss, proteinAT4G23440.1
bic:865 897 861
peaks after deleting less than one fifth of global max:12 16
×10^8
raw data, starting:12 16 24
fitted Gauss locations:12 16 24
y
x
0
0.5
1
1.5
2
2.5
5
10
15
20
25

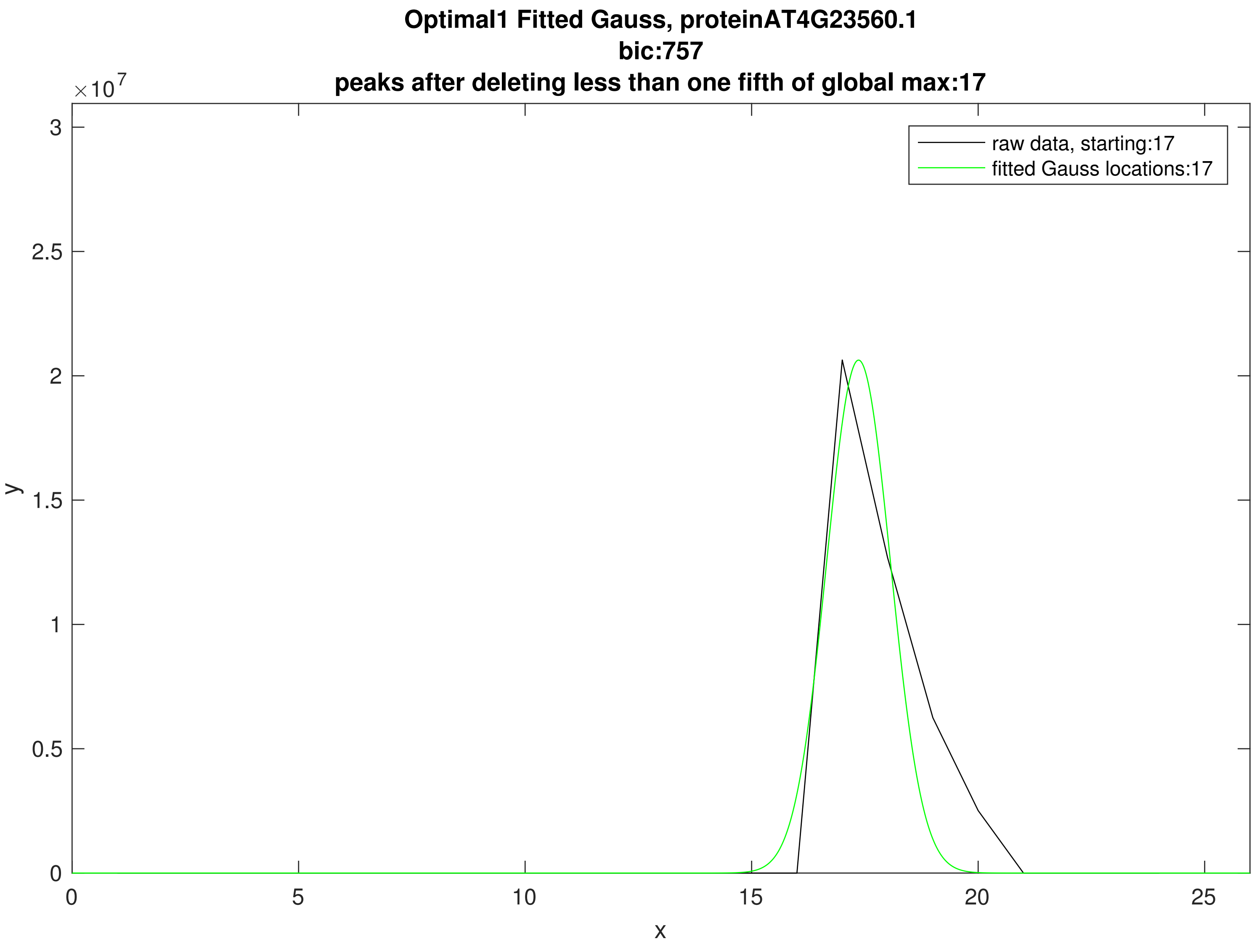

Optimal1 Fitted Gauss, proteinAT4G23560.1
bic:757
peaks after deleting less than one fifth of global max:17
×10^7
raw data, starting:17
fitted Gauss locations:17
y
x
0
0.5
1
1.5
2
2.5
3
5
10
15
20
25

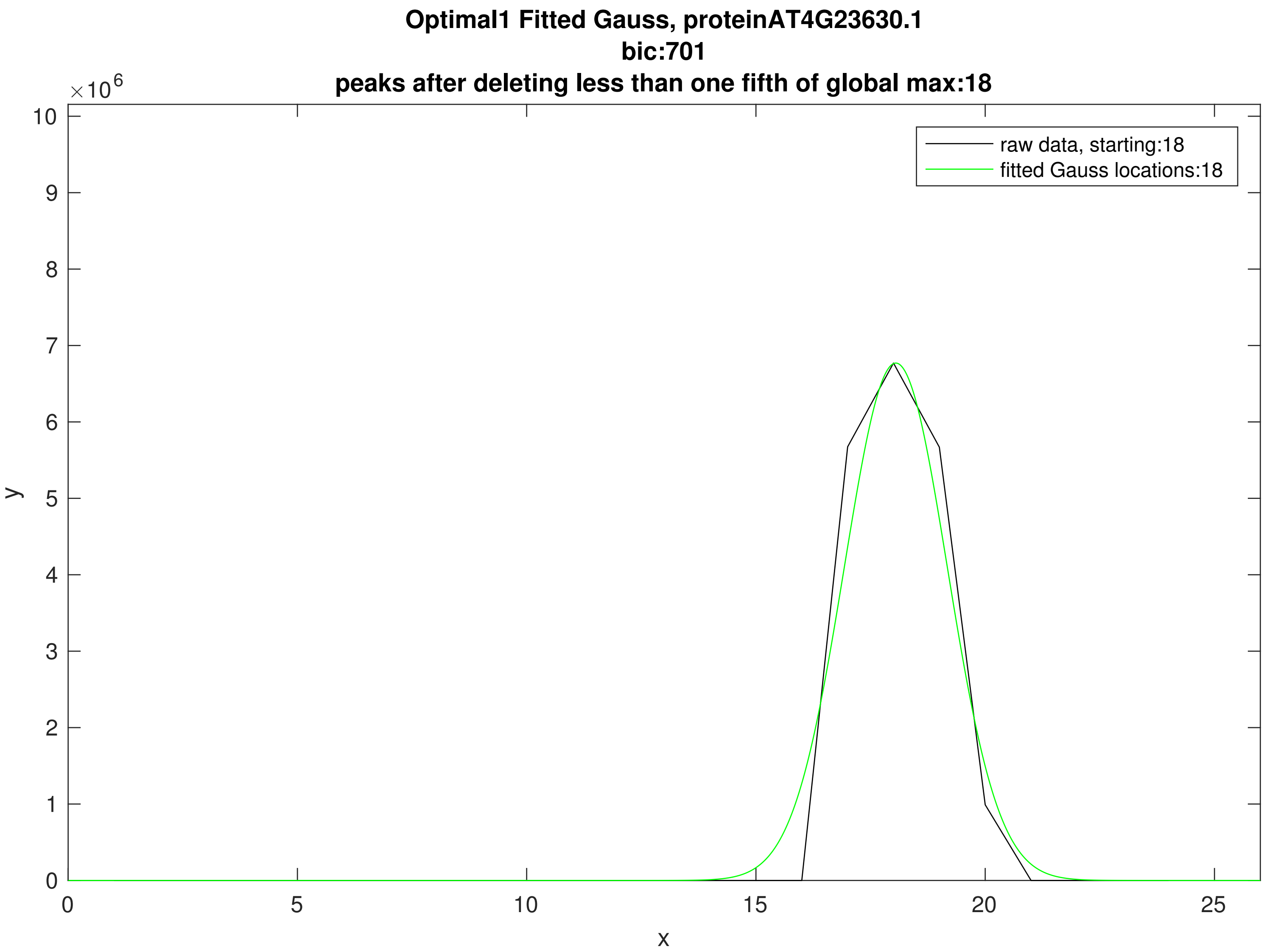
Optimal1 Fitted Gauss, proteinAT4G23630.1
bic:701
peaks after deleting less than one fifth of global max:18
×10^6
raw data, starting:18
fitted Gauss locations:18
y
x
0
1
2
3
4
5
6
7
8
9
10
0
5
10
15
20
25

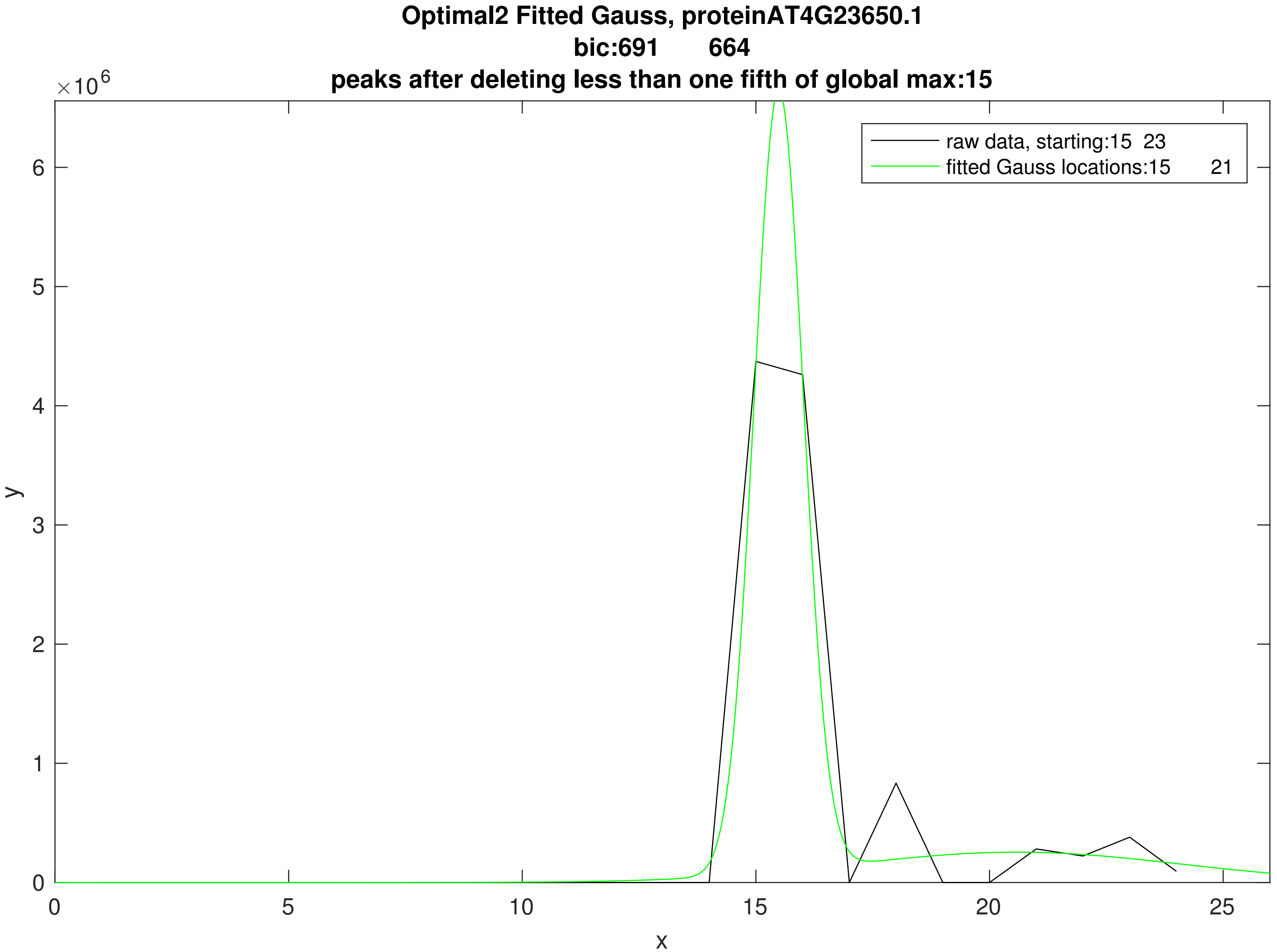
Optimal2 Fitted Gauss, proteinAT4G23650.1
bic:691 664
peaks after deleting less than one fifth of global max:15
×10^6
raw data, starting:15 23
fitted Gauss locations:15 21
y
x
0
1
2
3
4
5
6
0
5
10
15
20
25

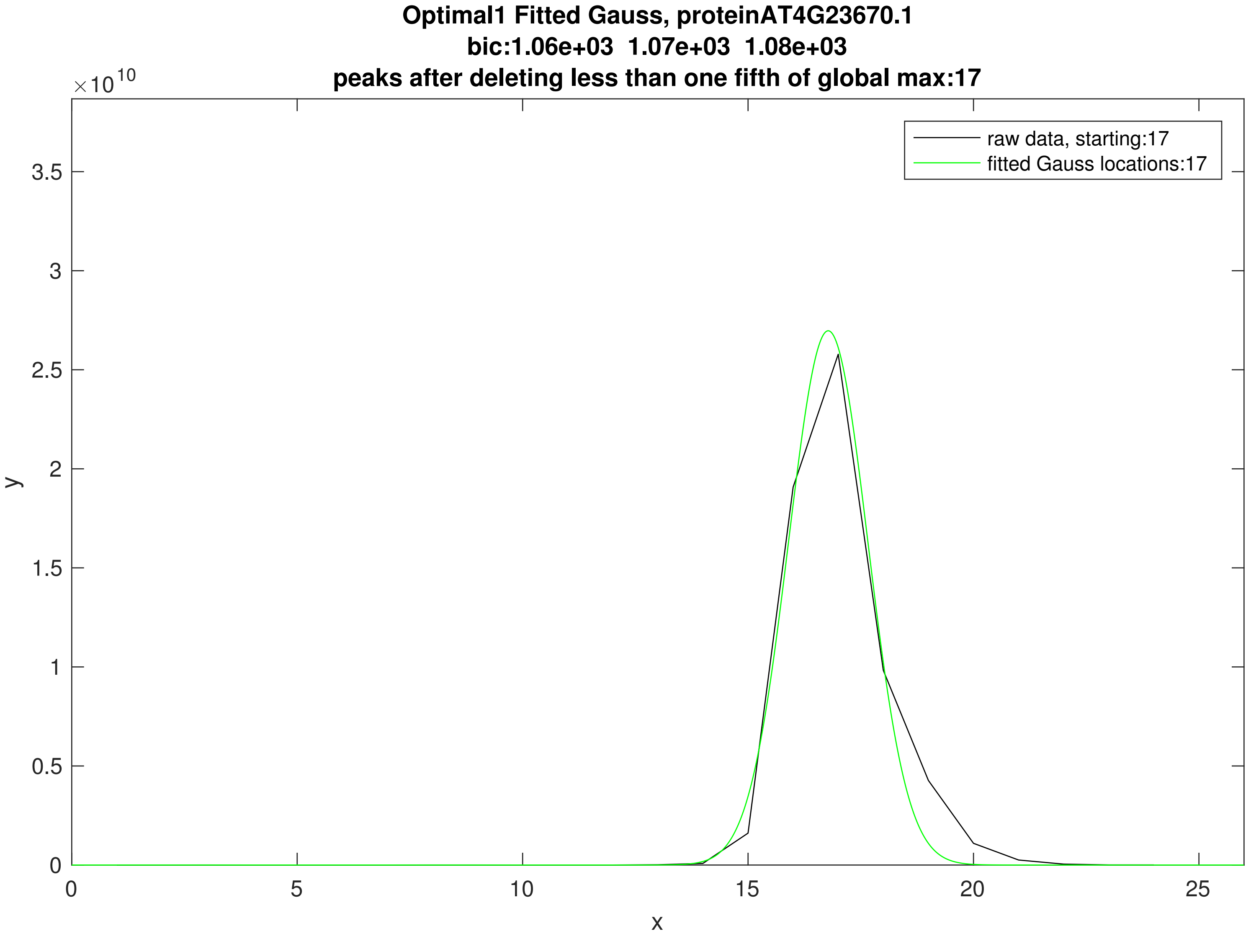
Optimal1 Fitted Gauss, proteinAT4G23670.1
bic:1.06e+03  1.07e+03  1.08e+03
peaks after deleting less than one fifth of global max:17
raw data, starting:17
fitted Gauss locations:17
×10^10
y
x
0
5
10
15
20
25
0.5
1
1.5
2
2.5
3
3.5

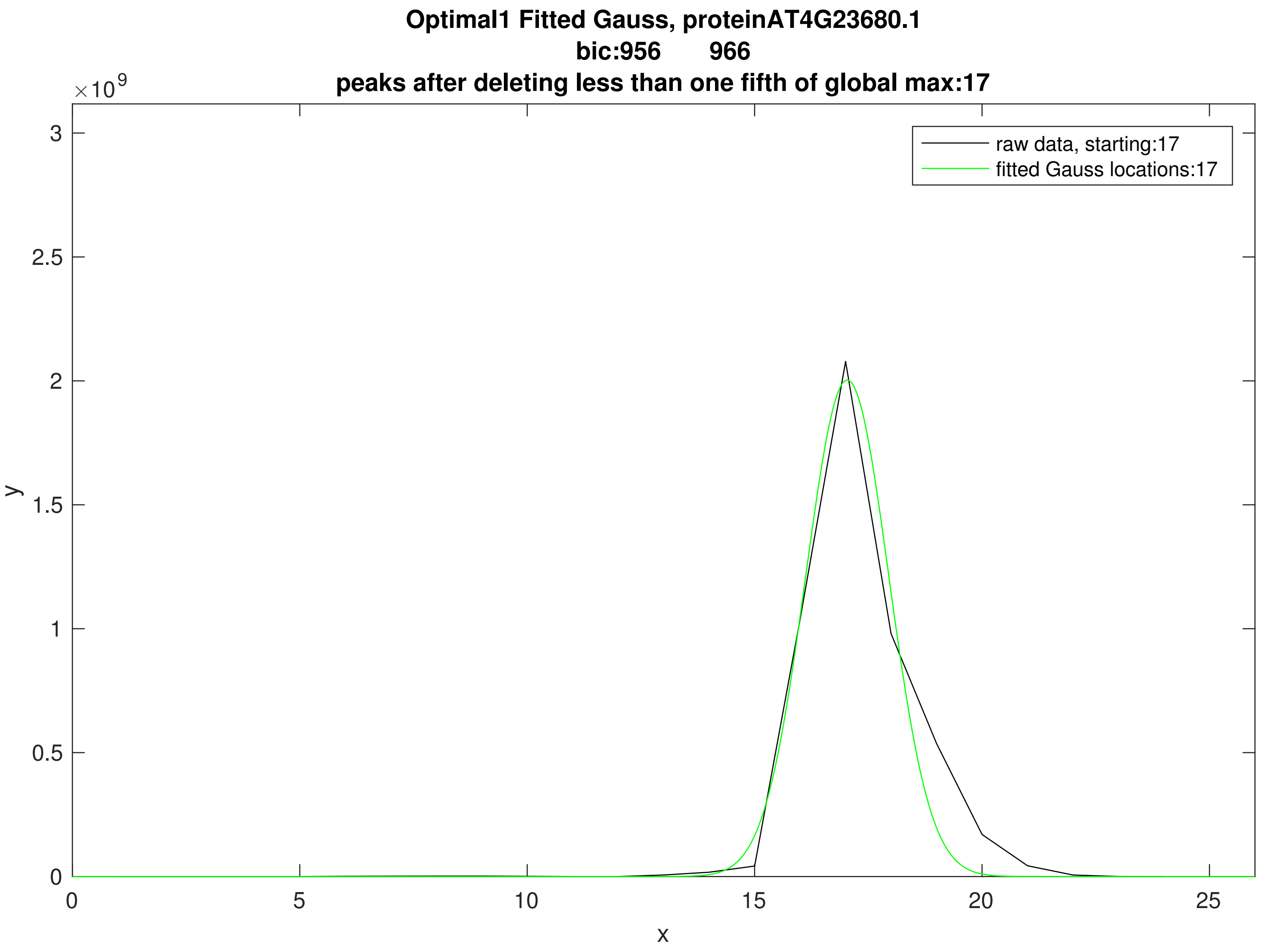
Optimal1 Fitted Gauss, proteinAT4G23680.1
bic:956 966
peaks after deleting less than one fifth of global max:17
×10^9
raw data, starting:17
fitted Gauss locations:17
y
x
0
0.5
1
1.5
2
2.5
3
0
5
10
15
20
25

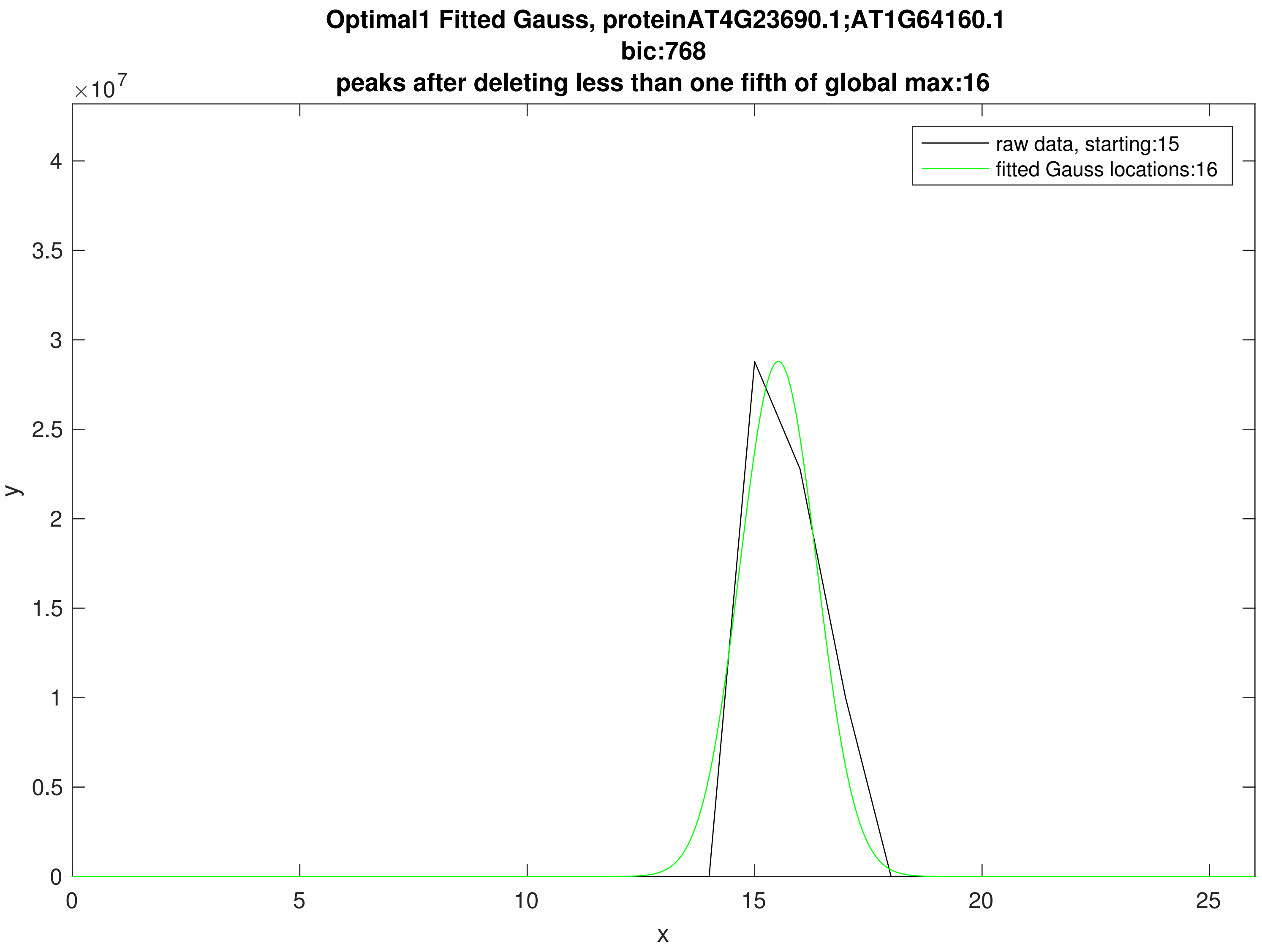
Optimal1 Fitted Gauss, proteinAT4G23690.1;AT1G64160.1
bic:768
peaks after deleting less than one fifth of global max:16
×10^7
raw data, starting:15
fitted Gauss locations:16
y
x
0
0.5
1
1.5
2
2.5
3
3.5
4
0
5
10
15
20
25

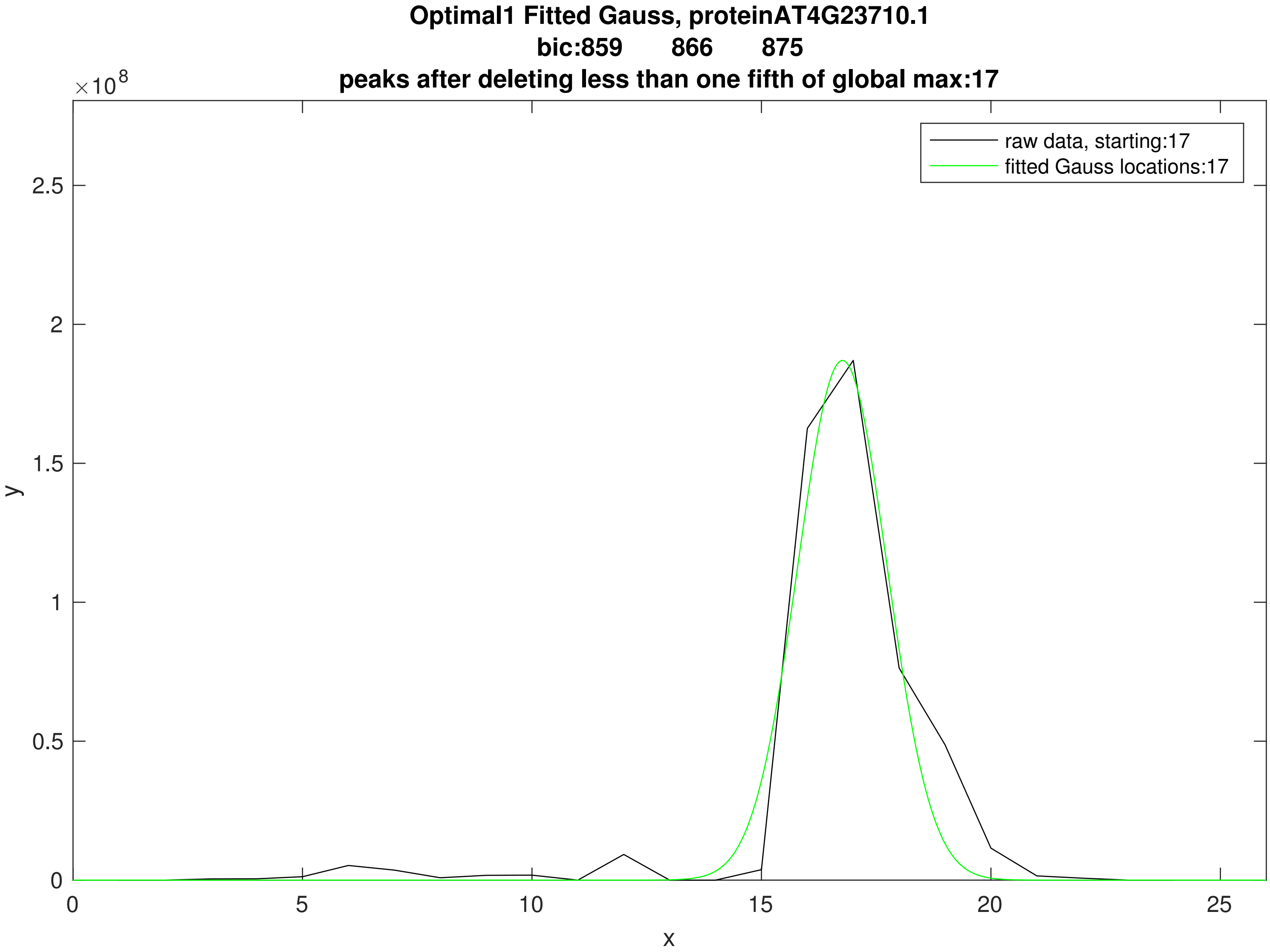
Optimal1 Fitted Gauss, proteinAT4G23710.1
bic:859 866 875
peaks after deleting less than one fifth of global max:17
×10^8
2.5
2
1.5
1
0.5
0
y
0
5
10
15
20
25
x
raw data, starting:17
fitted Gauss locations:17

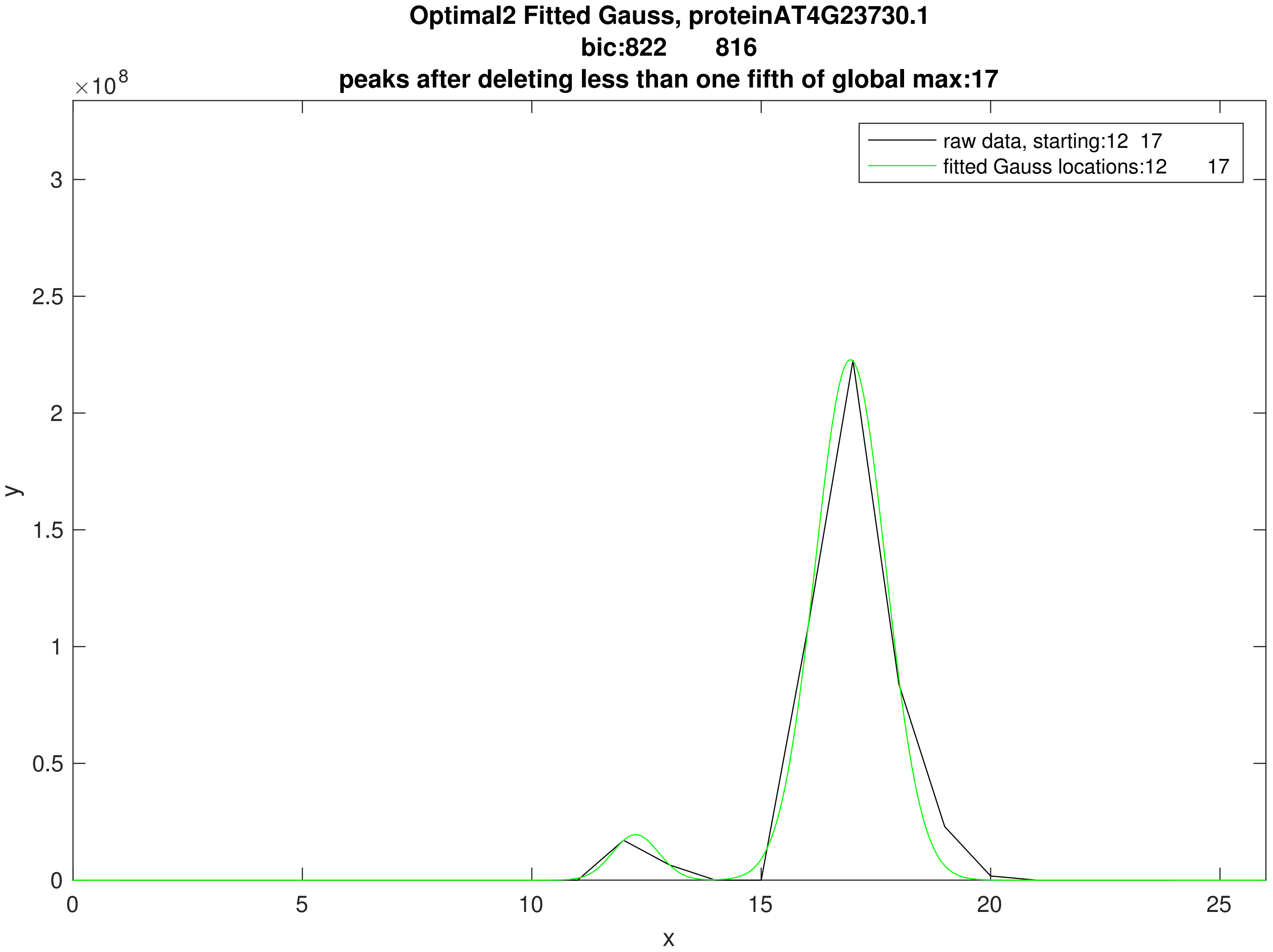
Optimal2 Fitted Gauss, proteinAT4G23730.1
bic:822 816
peaks after deleting less than one fifth of global max:17
×10^8
raw data, starting:12 17
fitted Gauss locations:12 17
y
x
0
0.5
1
1.5
2
2.5
3
5
10
15
20
25

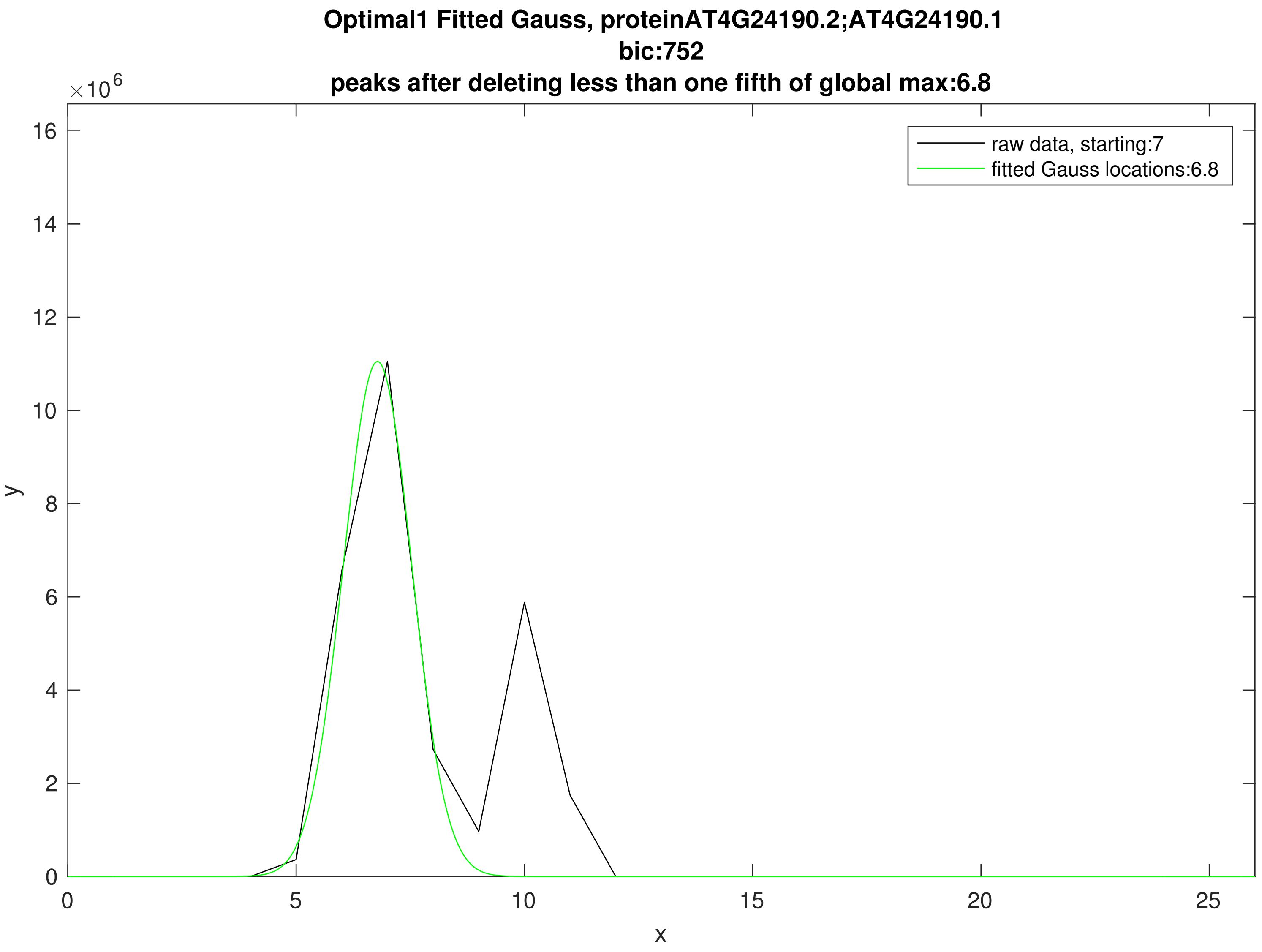
Optimal1 Fitted Gauss, proteinAT4G24190.2;AT4G24190.1
bic:752
peaks after deleting less than one fifth of global max:6.8
×10^6
raw data, starting:7
fitted Gauss locations:6.8
y
x
0
2
4
6
8
10
12
14
16
0
5
10
15
20
25

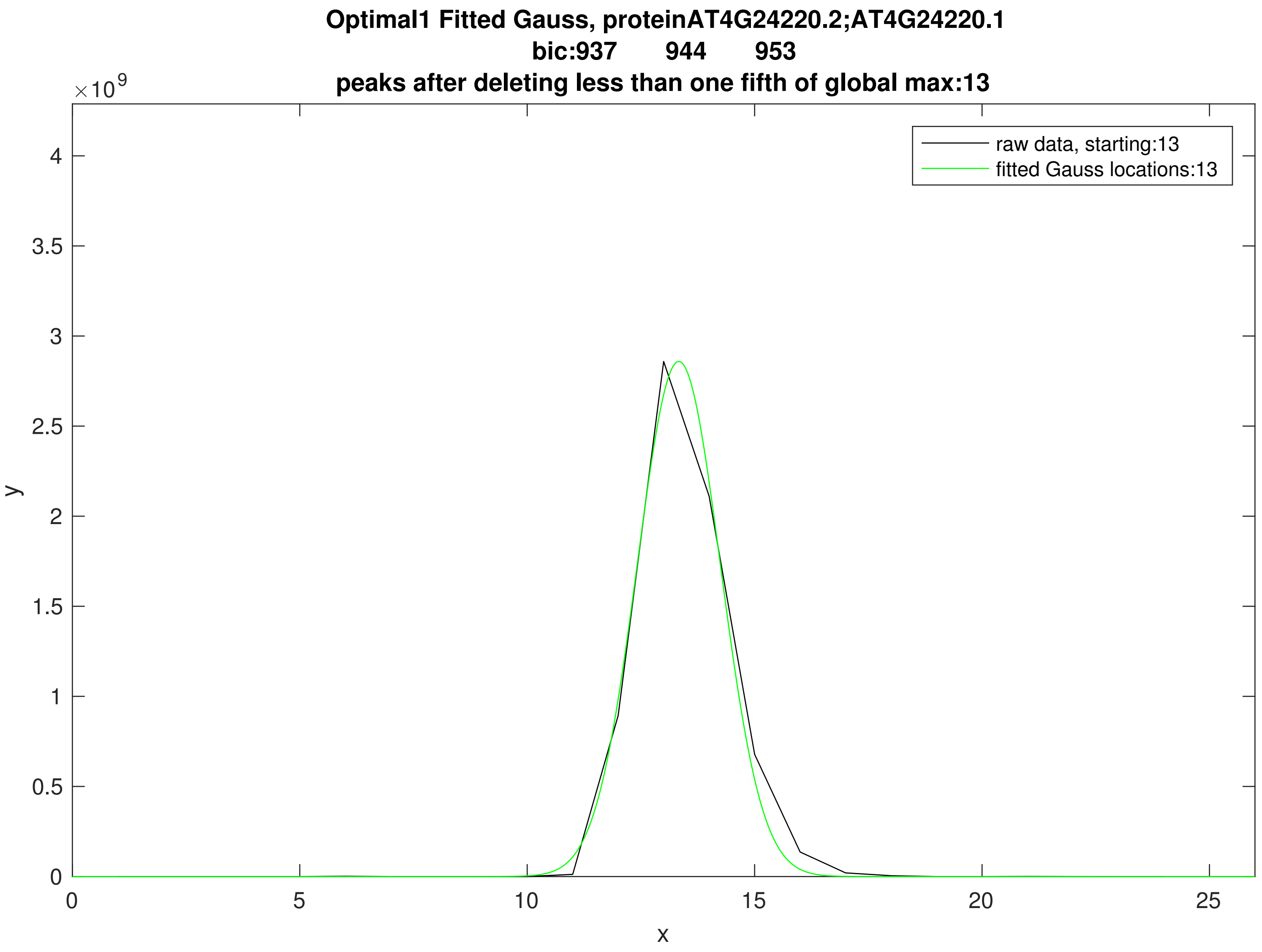
Optimal1 Fitted Gauss, proteinAT4G24220.2;AT4G24220.1
bic:937 944 953
peaks after deleting less than one fifth of global max:13
×10^9
raw data, starting:13
fitted Gauss locations:13
4
3.5
3
2.5
2
1.5
1
0.5
0
y
0
5
10
15
20
25
x

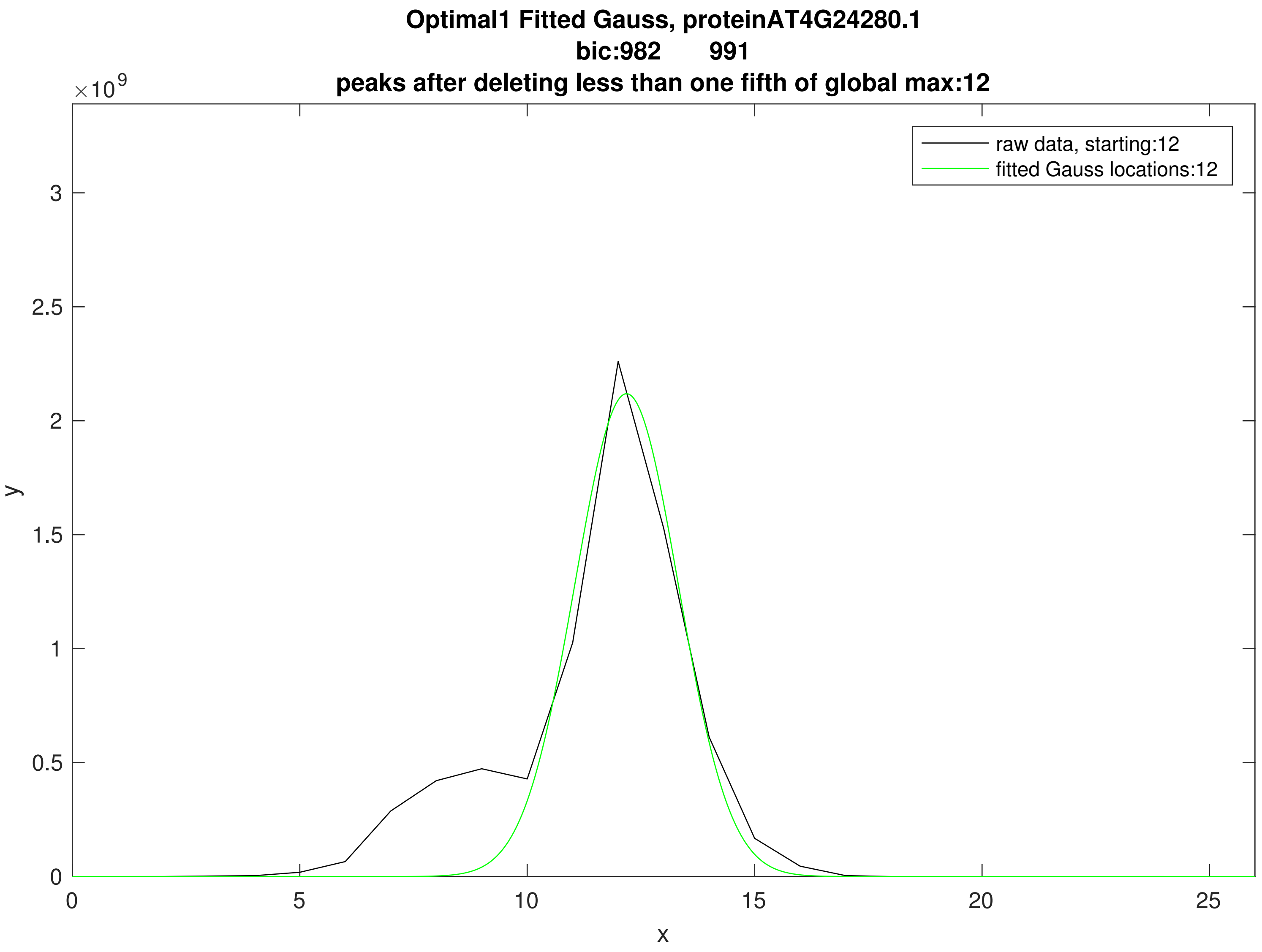
Optimal1 Fitted Gauss, proteinAT4G24280.1
bic:982 991
peaks after deleting less than one fifth of global max:12
×10^9
raw data, starting:12
fitted Gauss locations:12
y
x
0
0.5
1
1.5
2
2.5
3
0
5
10
15
20
25

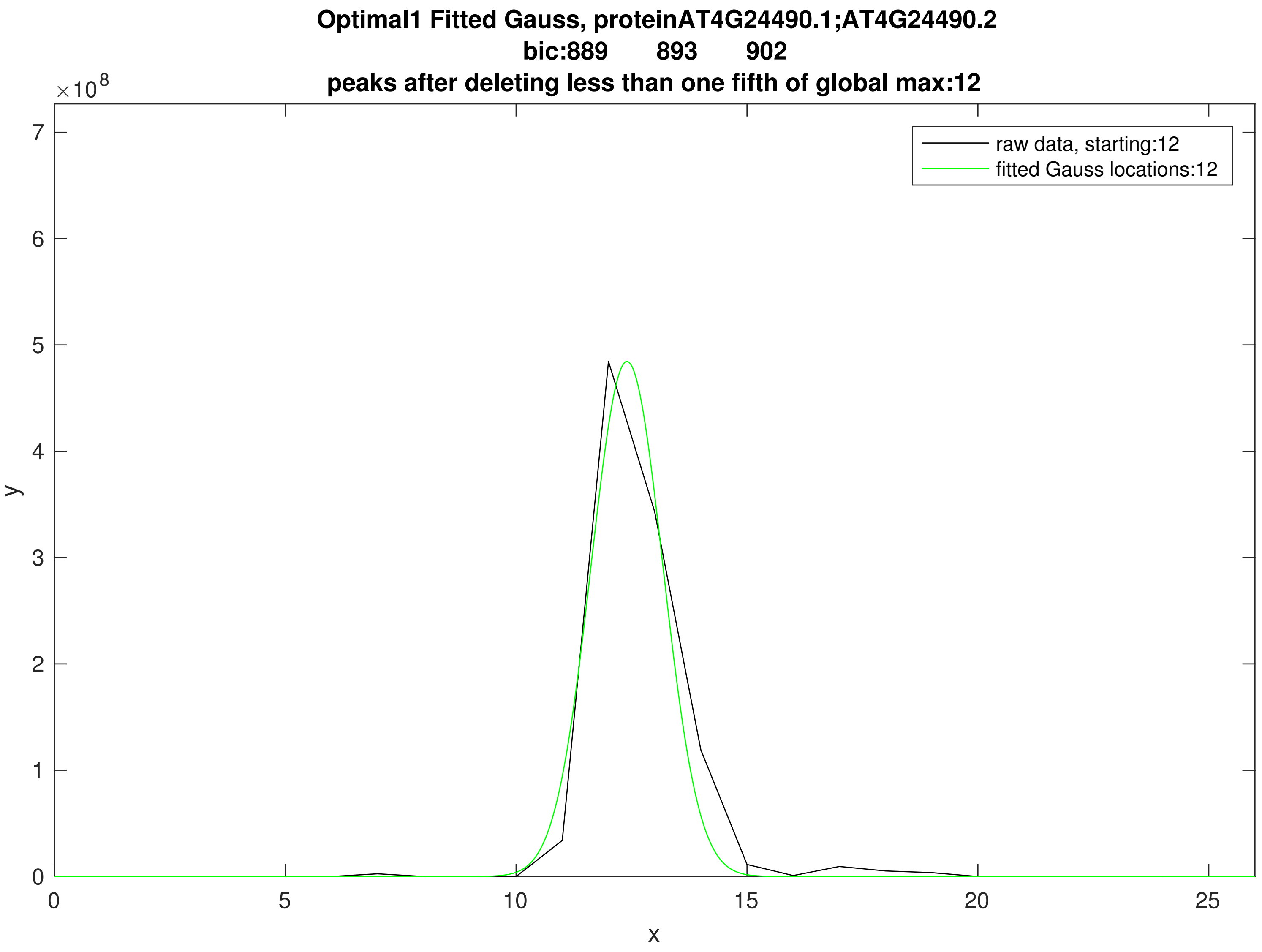
Optimal1 Fitted Gauss, proteinAT4G24490.1;AT4G24490.2
bic:889 893 902
peaks after deleting less than one fifth of global max:12
×10^8
raw data, starting:12
fitted Gauss locations:12
y
x
0
1
2
3
4
5
6
7
0
5
10
15
20
25

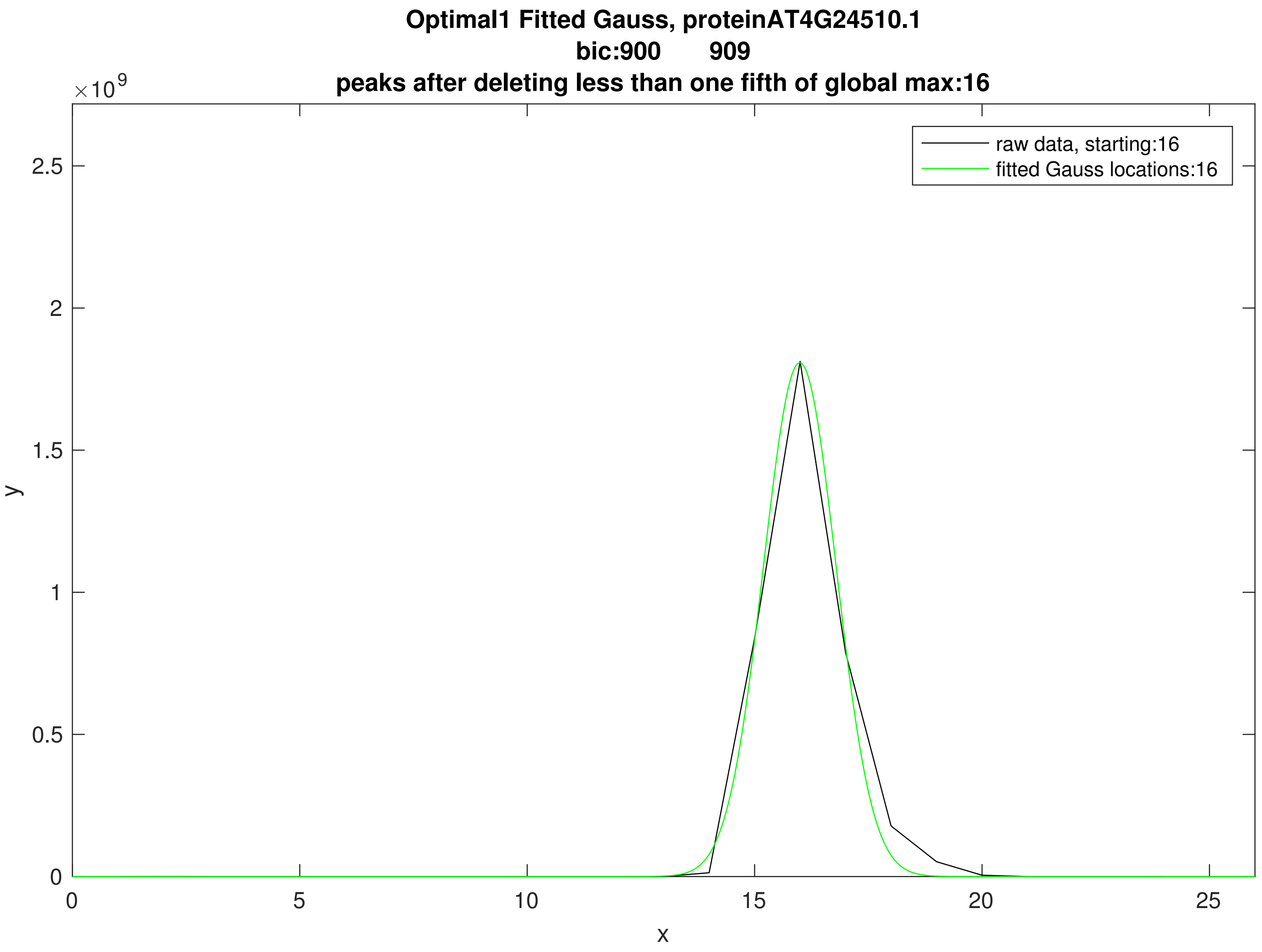
Optimal1 Fitted Gauss, proteinAT4G24510.1
bic:900 909
peaks after deleting less than one fifth of global max:16
×10^9
raw data, starting:16
fitted Gauss locations:16
y
x
2.5
2
1.5
1
0.5
0
5
10
15
20
25

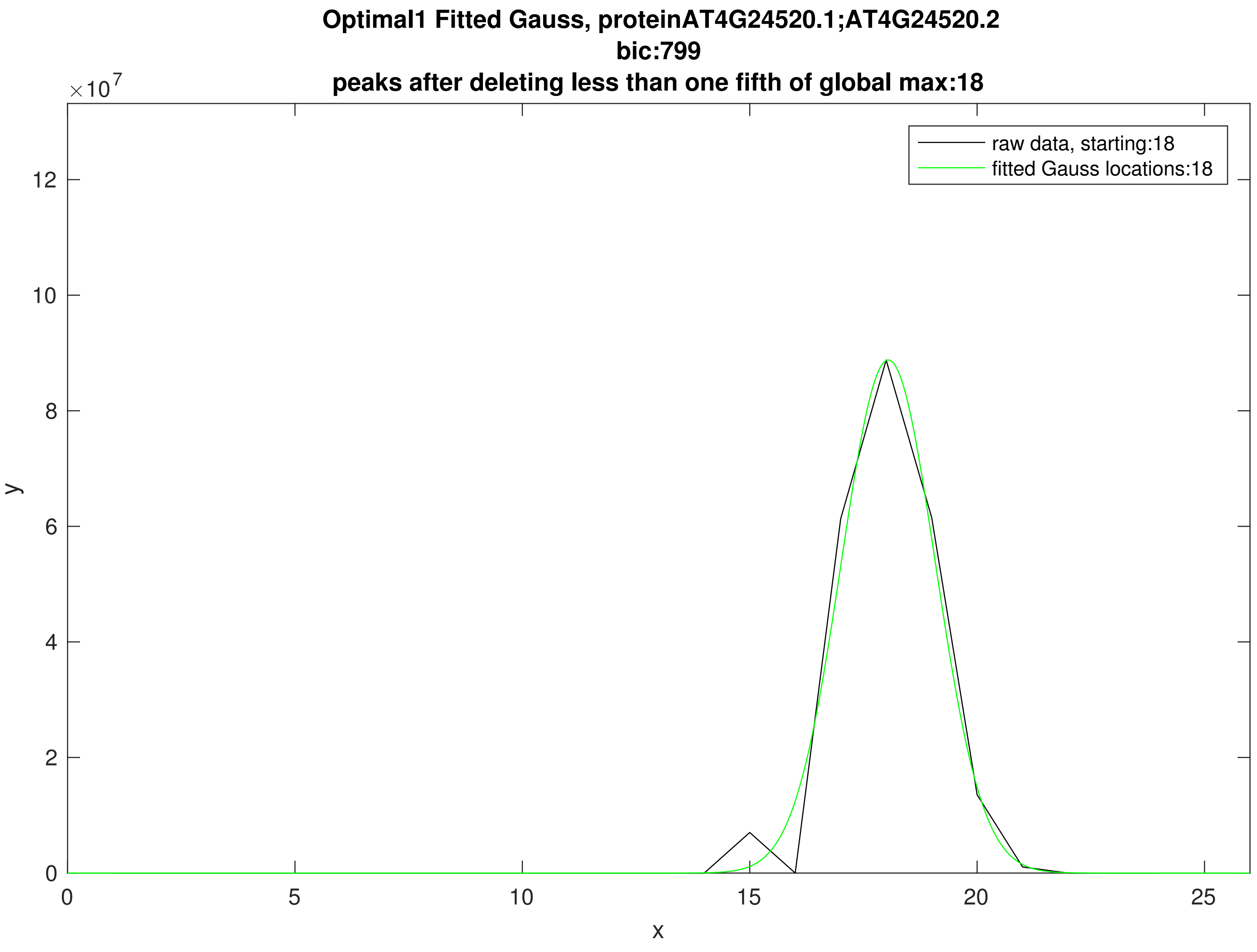
Optimal1 Fitted Gauss, proteinAT4G24520.1;AT4G24520.2
bic:799
peaks after deleting less than one fifth of global max:18
×10^7
raw data, starting:18
fitted Gauss locations:18
y
x
0
2
4
6
8
10
12
0
5
10
15
20
25

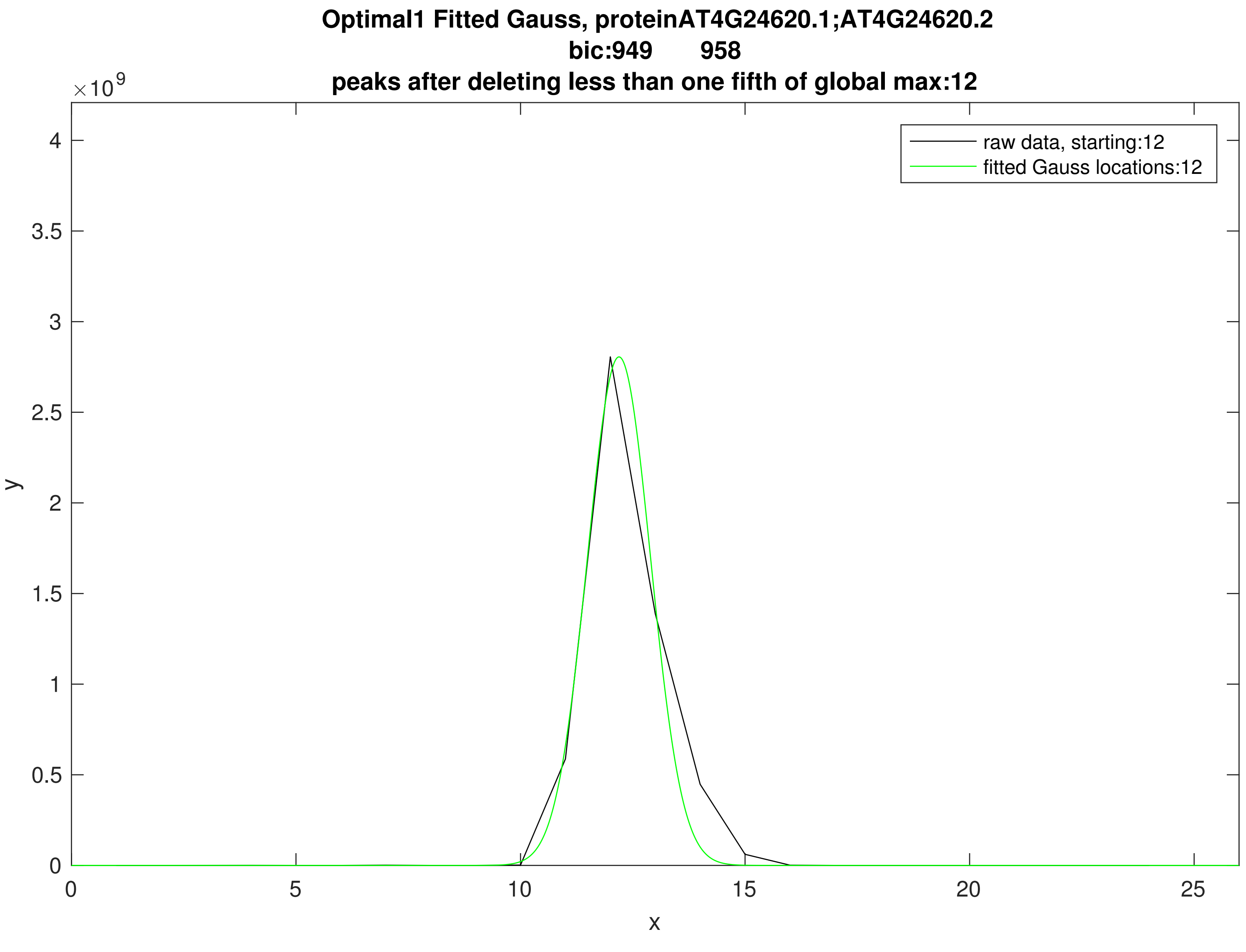

Optimal1 Fitted Gauss, proteinAT4G24620.1;AT4G24620.2
bic:949 958
peaks after deleting less than one fifth of global max:12
×10^9
raw data, starting:12
fitted Gauss locations:12
y
x
0
0.5
1
1.5
2
2.5
3
3.5
4
5
10
15
20
25

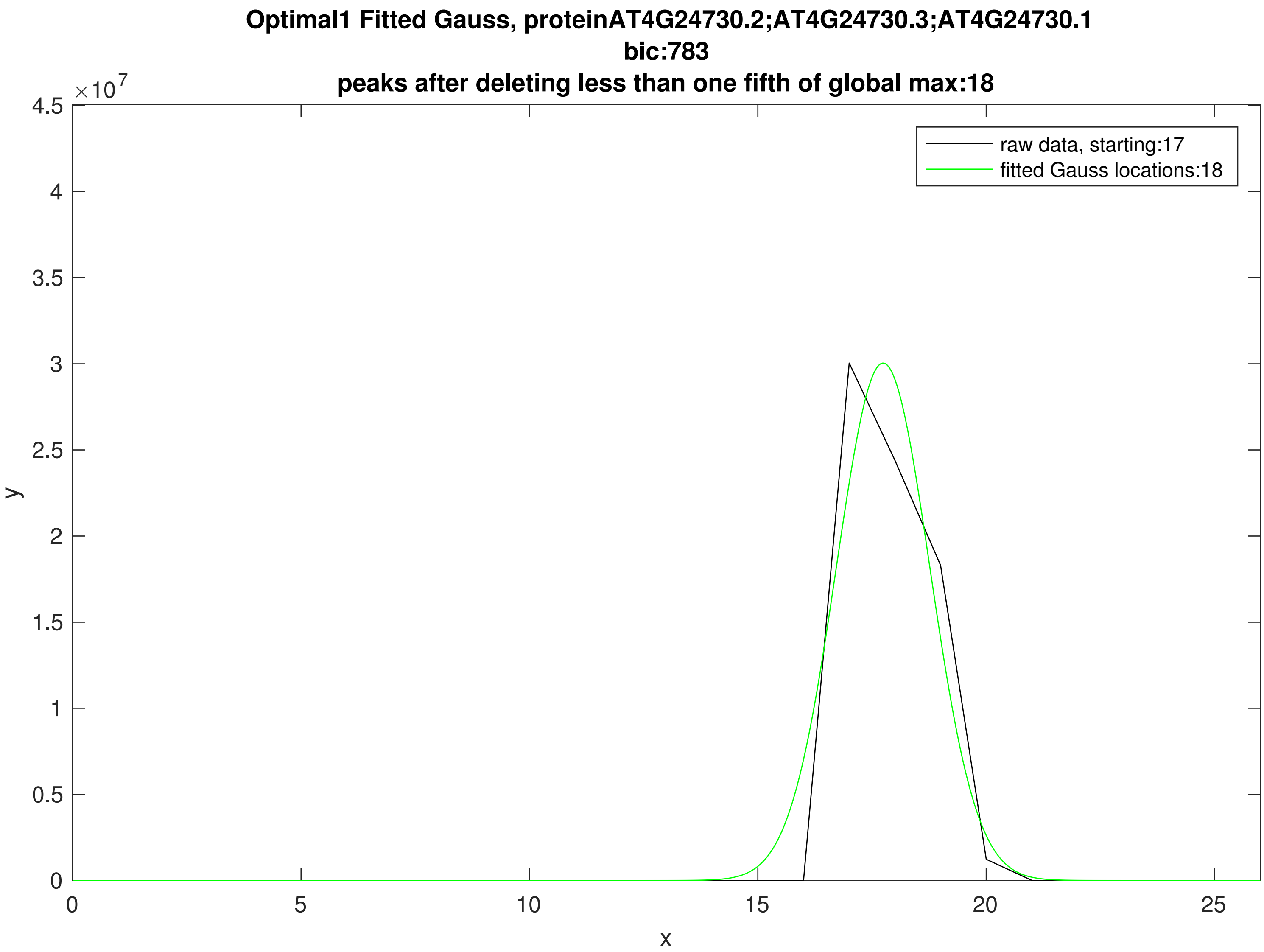
Optimal1 Fitted Gauss, proteinAT4G24730.2;AT4G24730.3;AT4G24730.1
bic:783
peaks after deleting less than one fifth of global max:18
×10^7
4.5
4
3.5
3
2.5
2
1.5
1
0.5
0
y
0
5
10
15
20
25
x
raw data, starting:17
fitted Gauss locations:18

**Optimal1 Fitted Gauss, proteinAT4G24770.1**
**bic:889 899**
**peaks after deleting less than one fifth of global max:15**

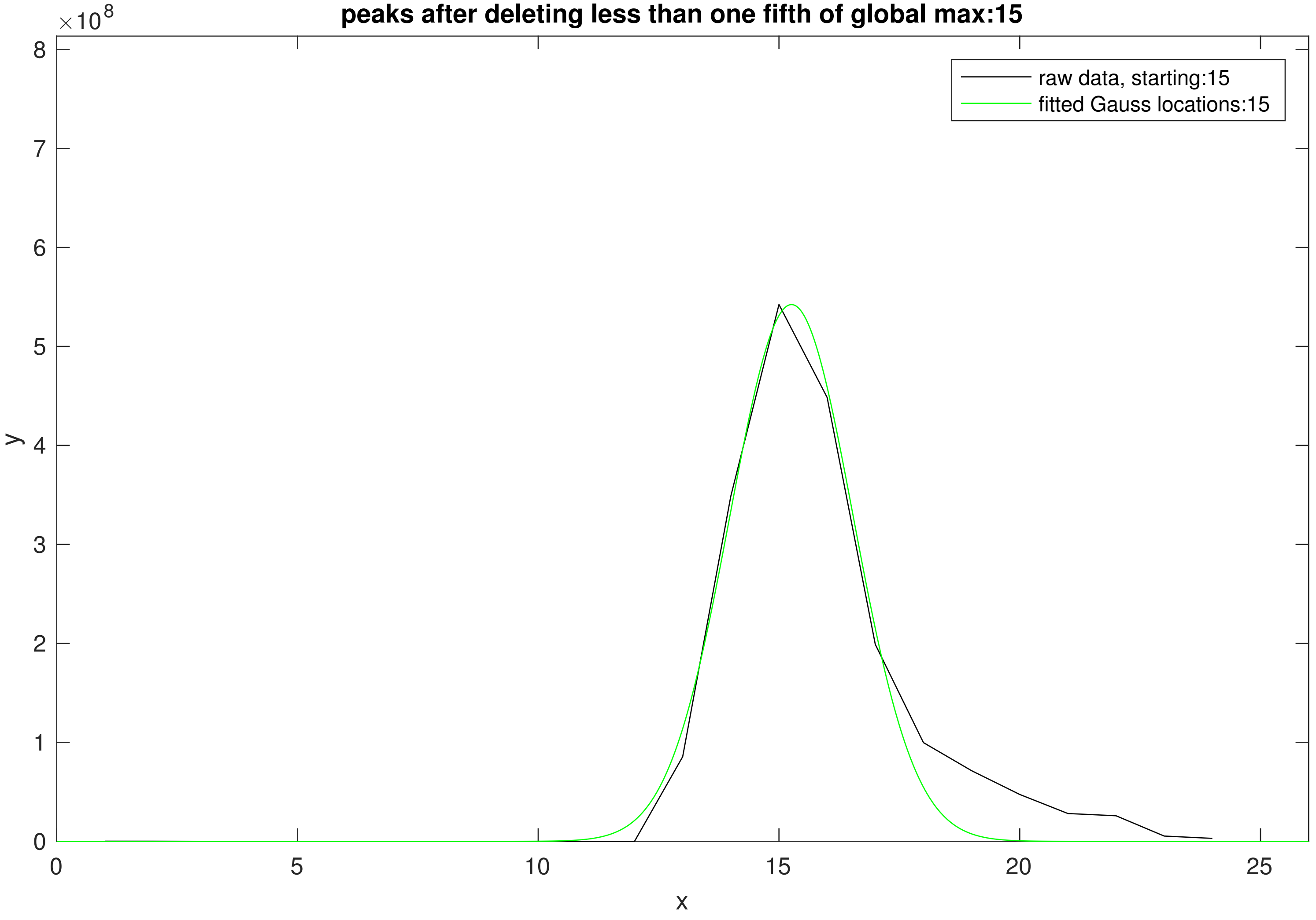

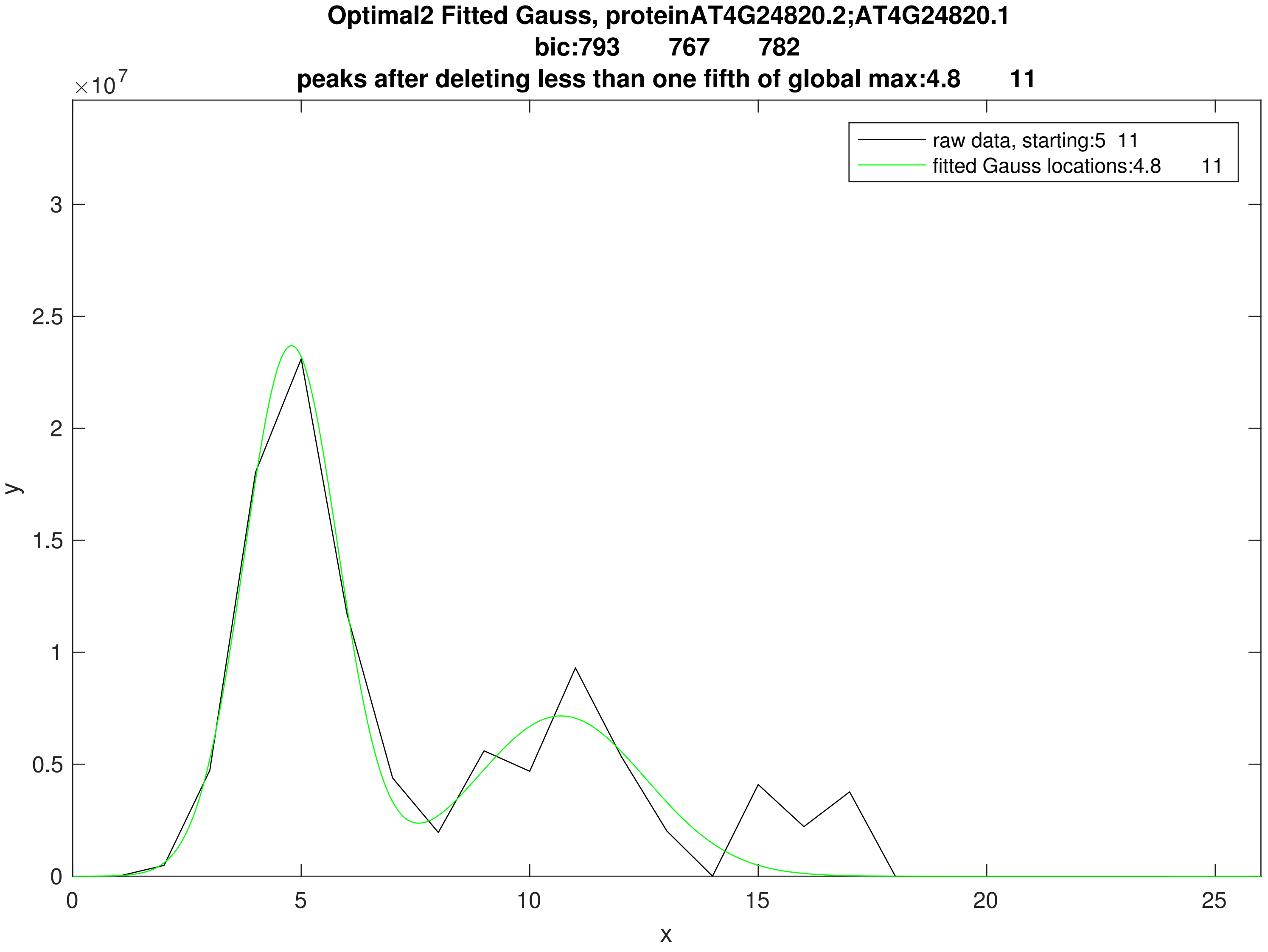
Optimal2 Fitted Gauss, proteinAT4G24820.2;AT4G24820.1
bic:793 767 782
peaks after deleting less than one fifth of global max:4.8 11
×10^7
raw data, starting:5 11
fitted Gauss locations:4.8 11
y
x
0
0.5
1
1.5
2
2.5
3
0
5
10
15
20
25

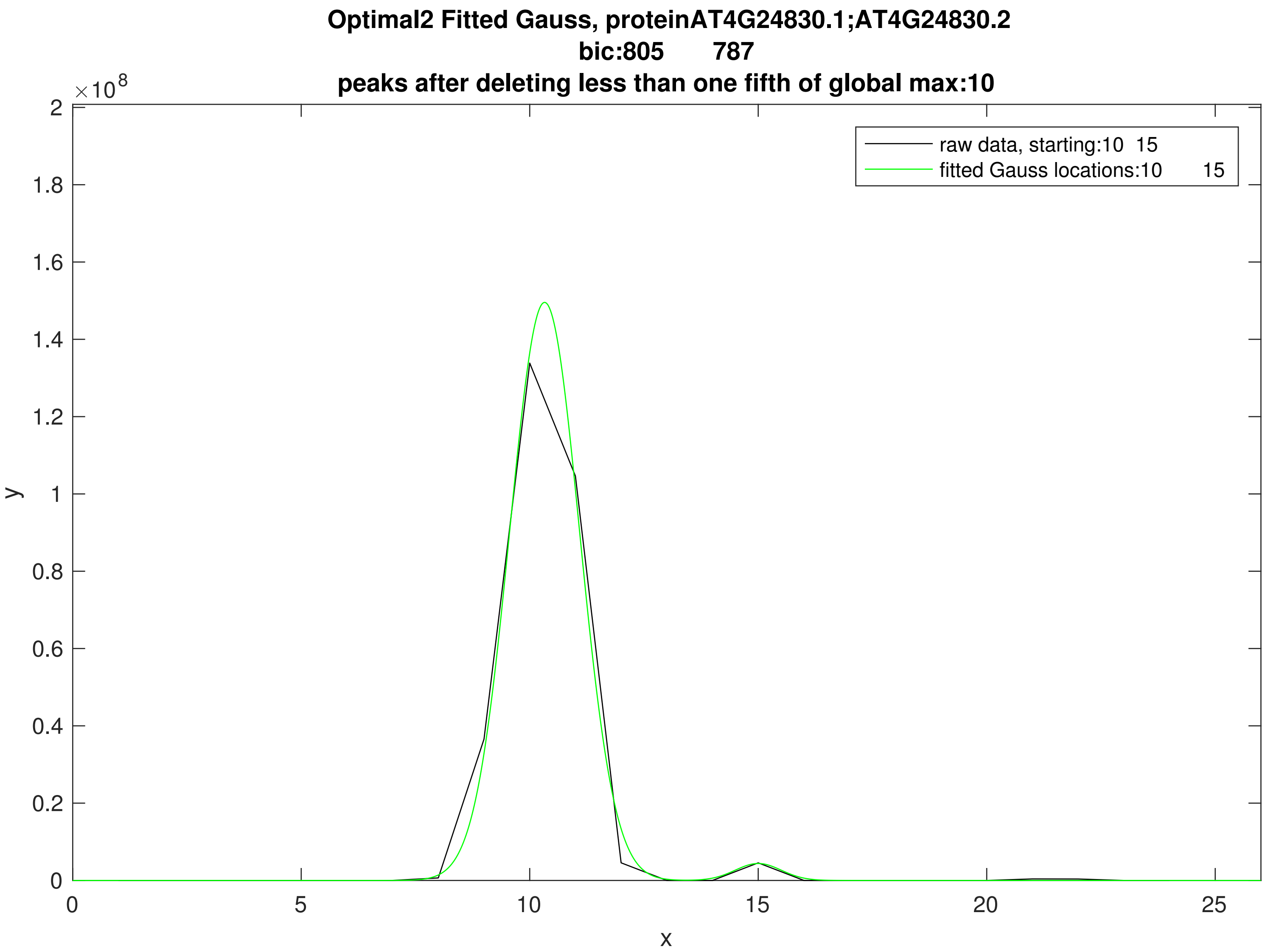
Optimal2 Fitted Gauss, proteinAT4G24830.1;AT4G24830.2
bic:805 787
peaks after deleting less than one fifth of global max:10
×10^8
raw data, starting:10 15
fitted Gauss locations:10 15
y
x
0
0.2
0.4
0.6
0.8
1
1.2
1.4
1.6
1.8
2
5
10
15
20
25

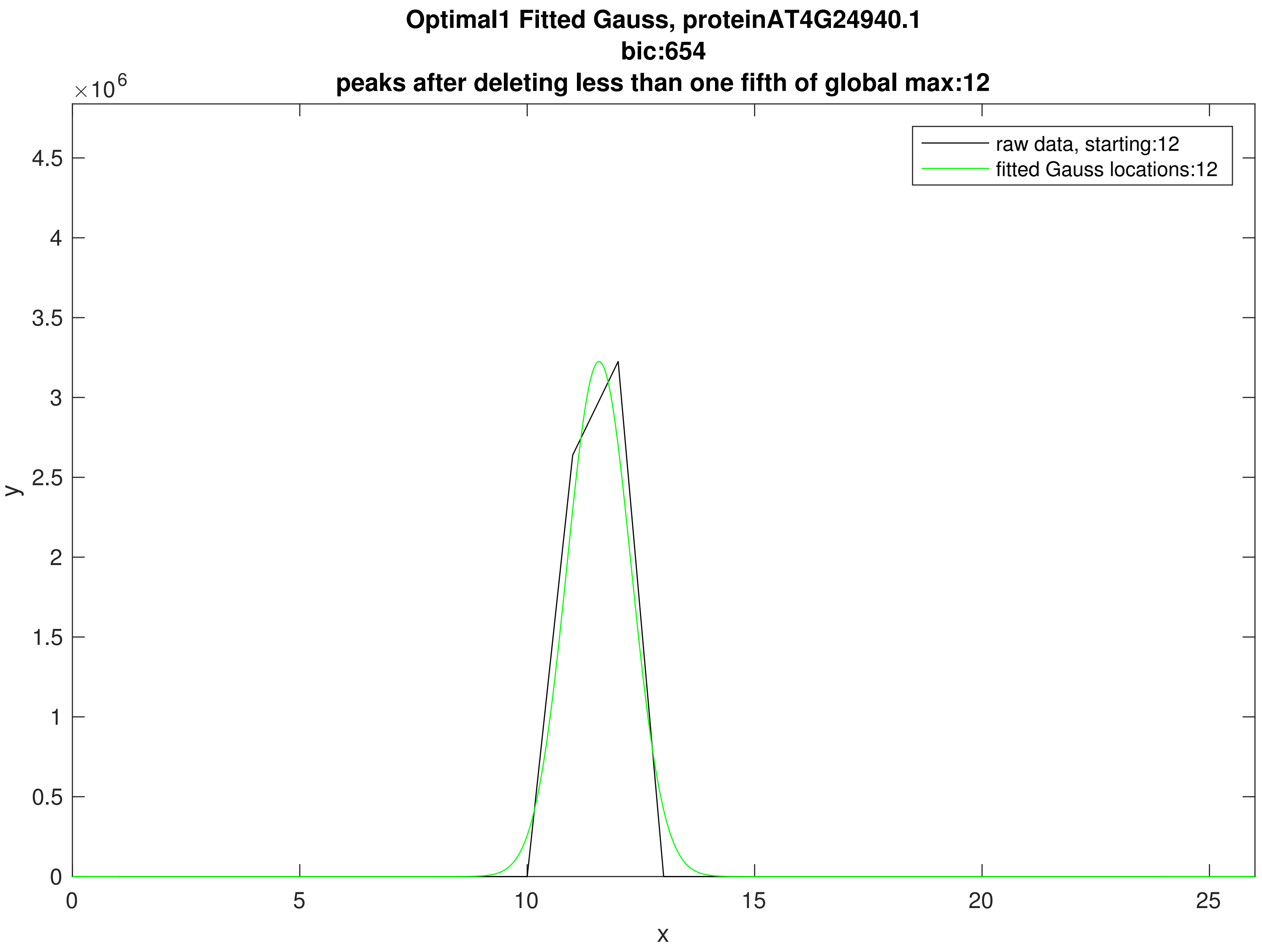

Optimal1 Fitted Gauss, proteinAT4G24940.1
bic:654
peaks after deleting less than one fifth of global max:12
×10^6
raw data, starting:12
fitted Gauss locations:12
y
x
0
0.5
1
1.5
2
2.5
3
3.5
4
4.5
0
5
10
15
20
25

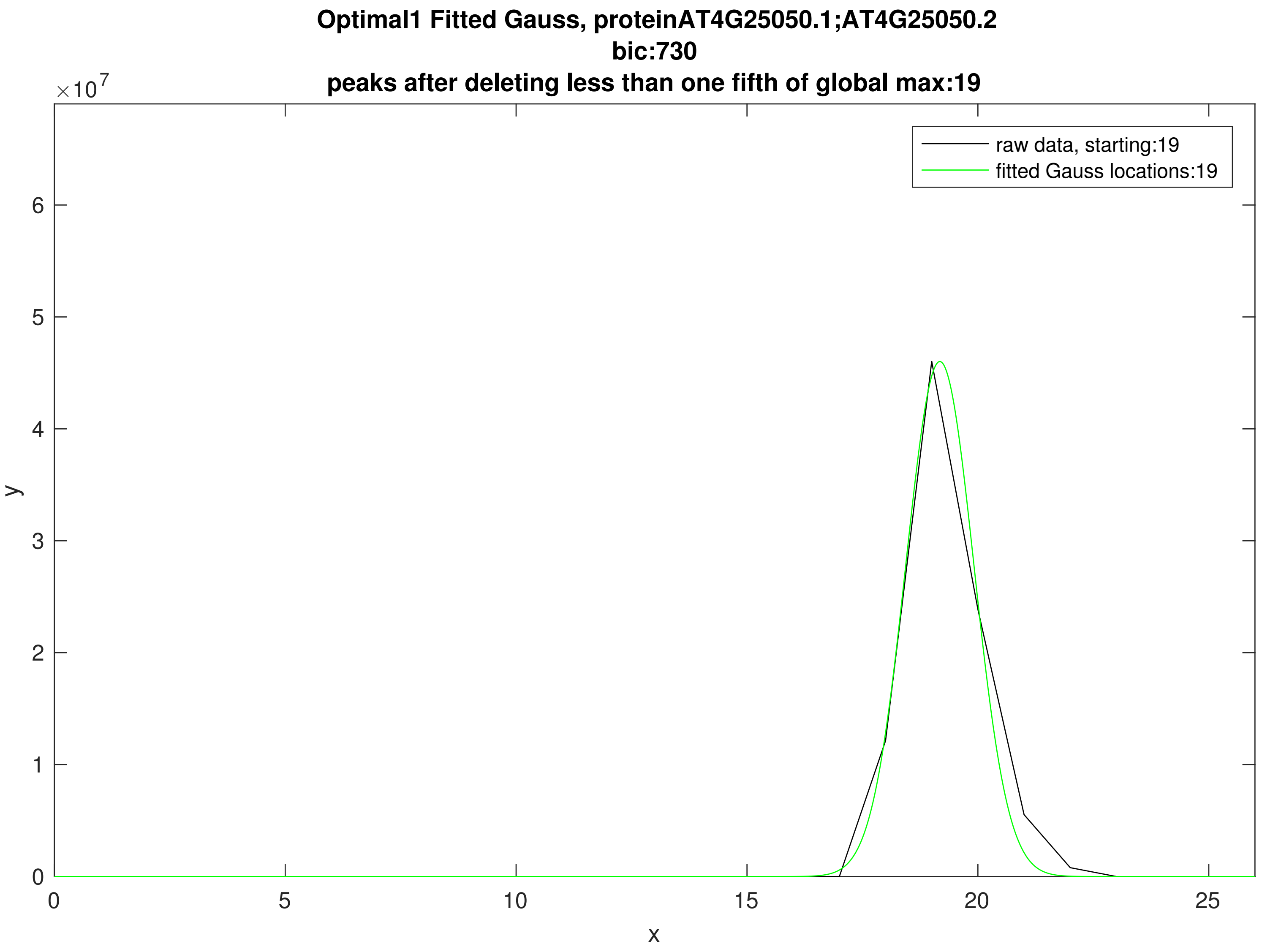
Optimal1 Fitted Gauss, proteinAT4G25050.1;AT4G25050.2
bic:730
peaks after deleting less than one fifth of global max:19
×10^7
raw data, starting:19
fitted Gauss locations:19
y
x
0
1
2
3
4
5
6
0
5
10
15
20
25

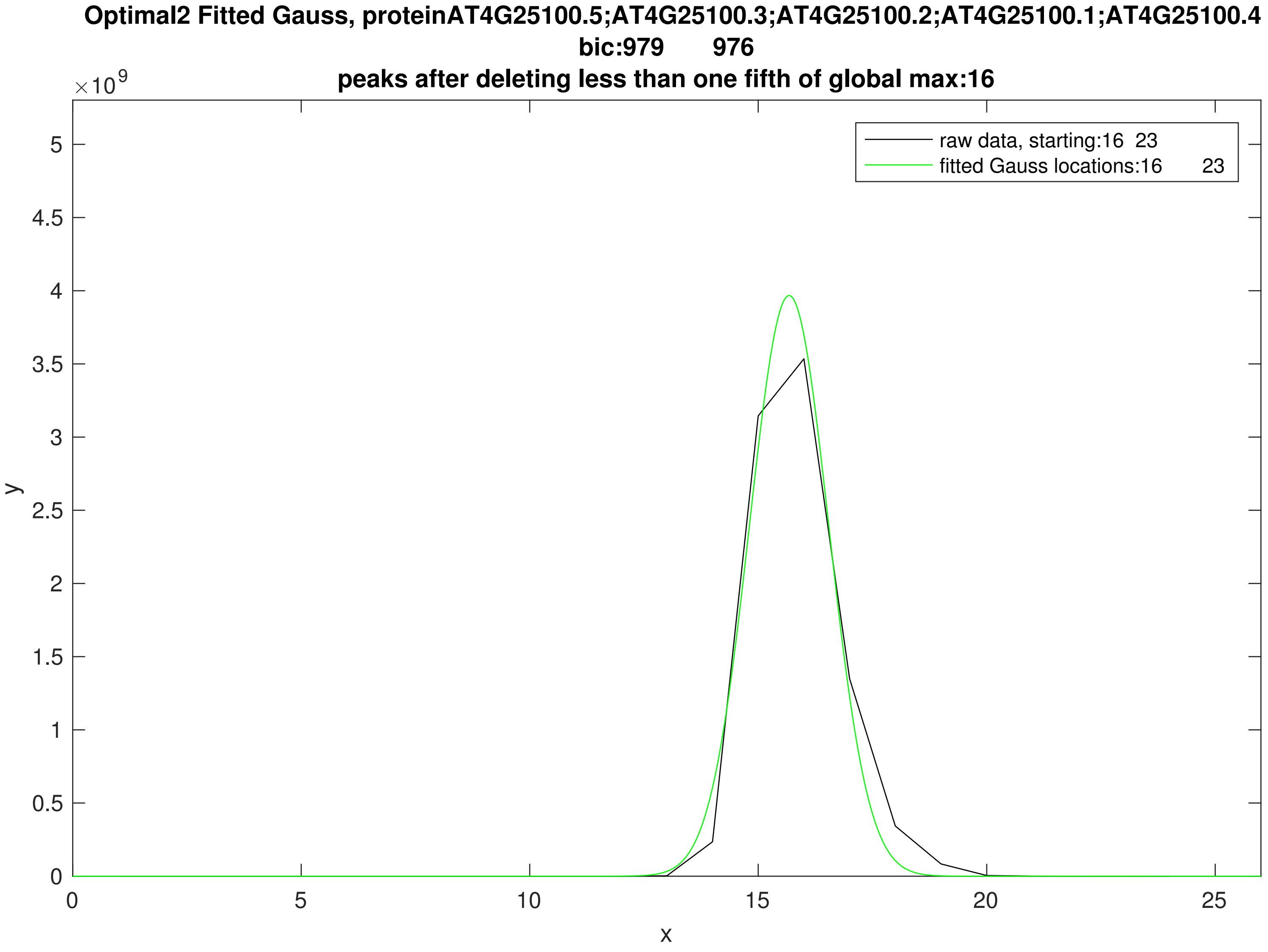
Optimal2 Fitted Gauss, proteinAT4G25100.5;AT4G25100.3;AT4G25100.2;AT4G25100.1;AT4G25100.4
bic:979 976
peaks after deleting less than one fifth of global max:16
×10^9
raw data, starting:16 23
fitted Gauss locations:16 23
y
x
0
0.5
1
1.5
2
2.5
3
3.5
4
4.5
5
0
5
10
15
20
25

**Optimal1 Fitted Gauss, proteinAT4G25130.1**
**bic:863 872**
**peaks after deleting less than one fifth of global max:20**

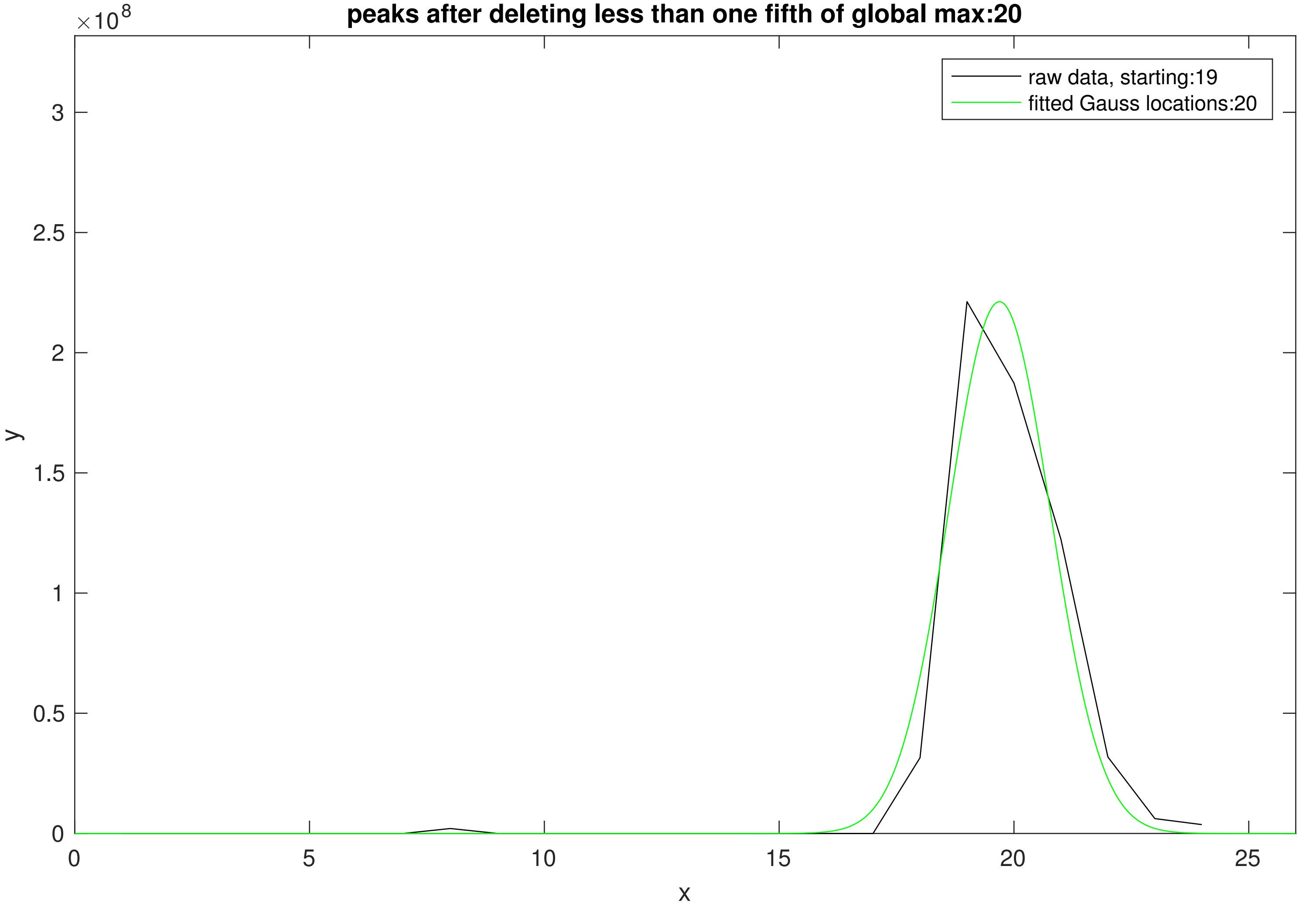

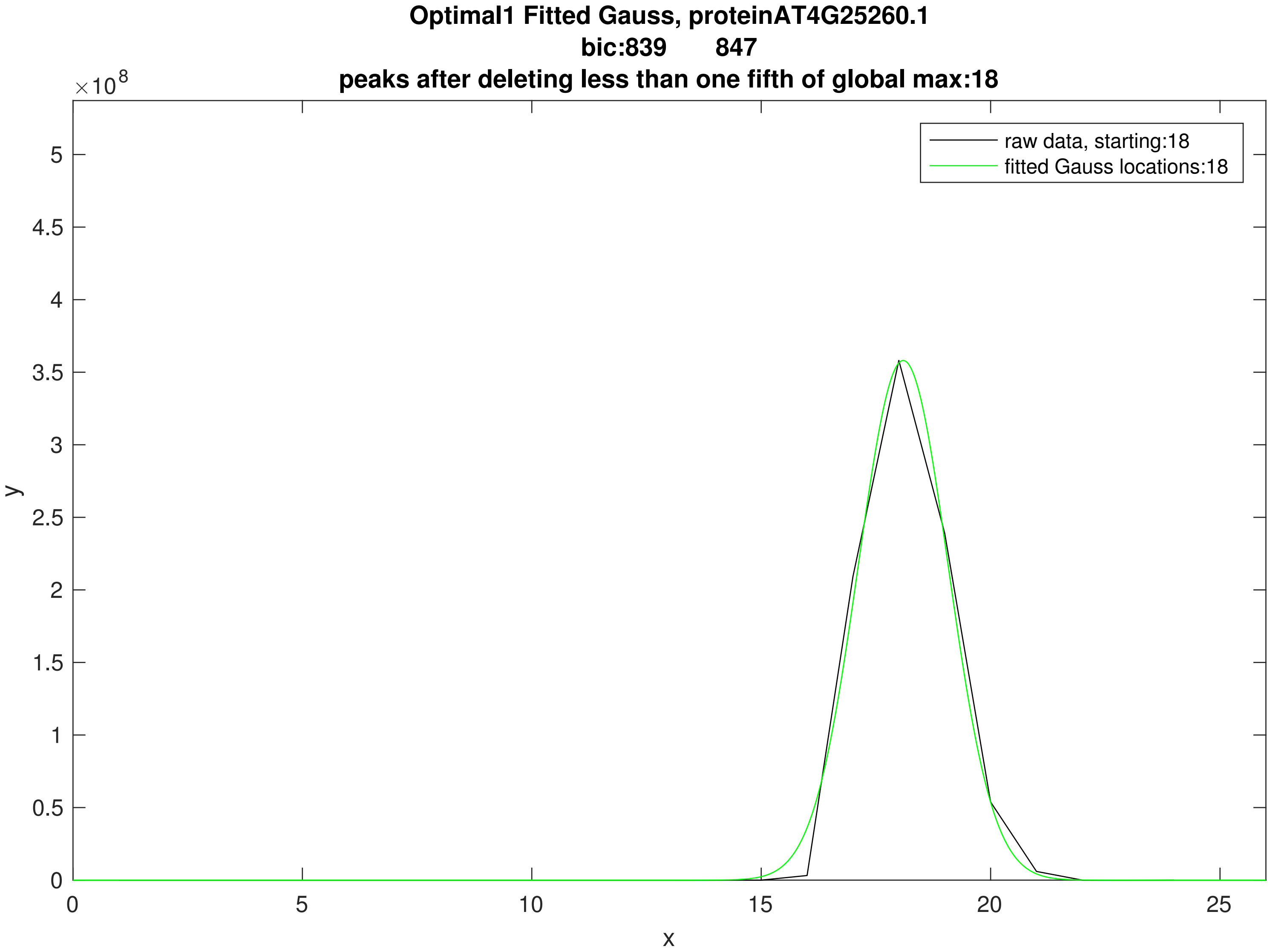
Optimal1 Fitted Gauss, proteinAT4G25260.1
bic:839 847
peaks after deleting less than one fifth of global max:18
×10^8
raw data, starting:18
fitted Gauss locations:18
y
x
0
0.5
1
1.5
2
2.5
3
3.5
4
4.5
5
0
5
10
15
20
25

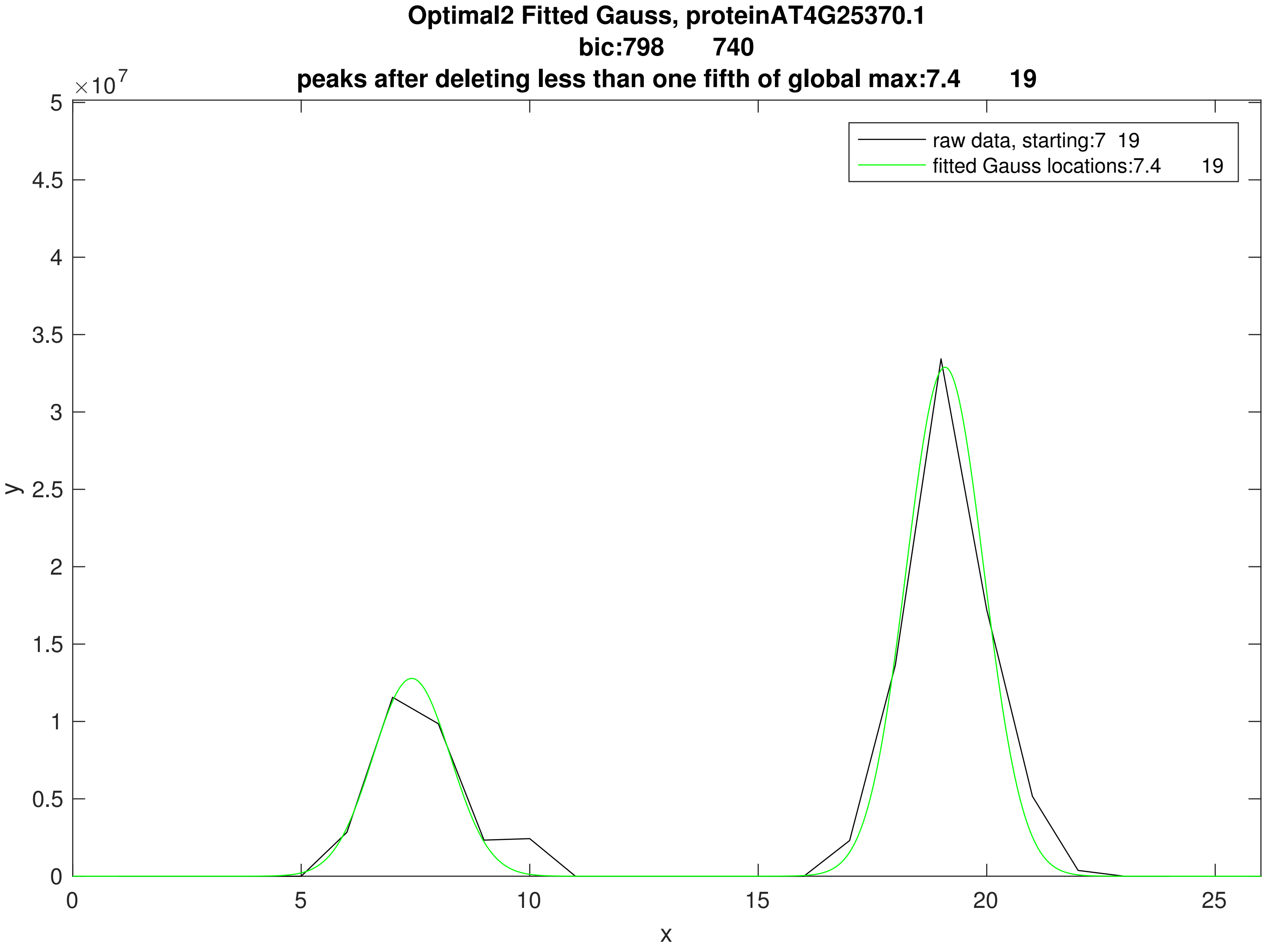
Optimal2 Fitted Gauss, proteinAT4G25370.1
bic:798 740
peaks after deleting less than one fifth of global max:7.4 19
×10^7
5
4.5
4
3.5
3
2.5
2
1.5
1
0.5
0
y
0
5
10
15
20
25
x
raw data, starting:7 19
fitted Gauss locations:7.4 19

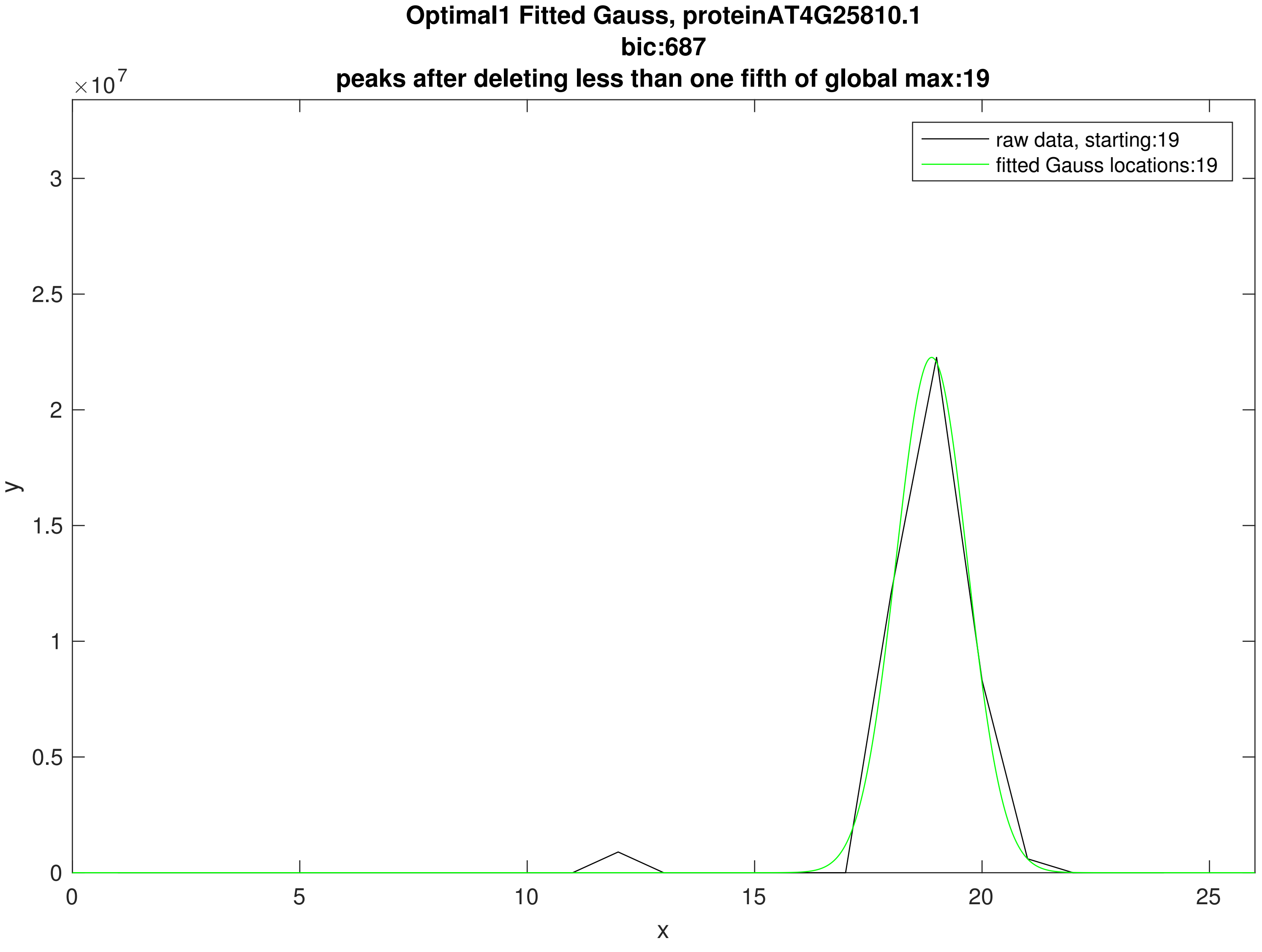
Optimal1 Fitted Gauss, proteinAT4G25810.1
bic:687
peaks after deleting less than one fifth of global max:19
×10^7
raw data, starting:19
fitted Gauss locations:19
y
x
0
0.5
1
1.5
2
2.5
3
0
5
10
15
20
25

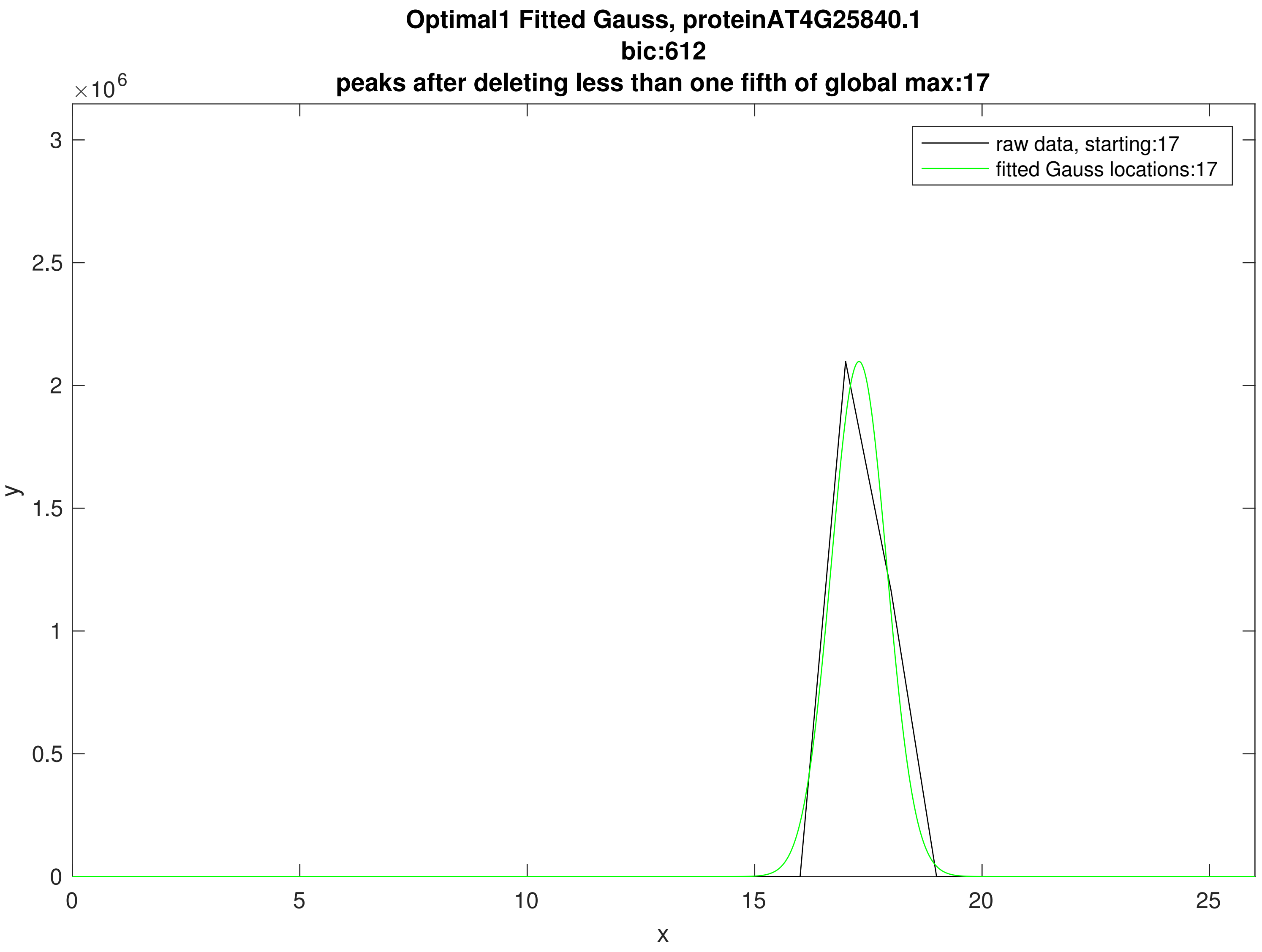

Optimal1 Fitted Gauss, proteinAT4G25840.1
bic:612
peaks after deleting less than one fifth of global max:17
×10^6
3
2.5
2
1.5
1
0.5
0
y
0
5
10
15
20
25
x
raw data, starting:17
fitted Gauss locations:17

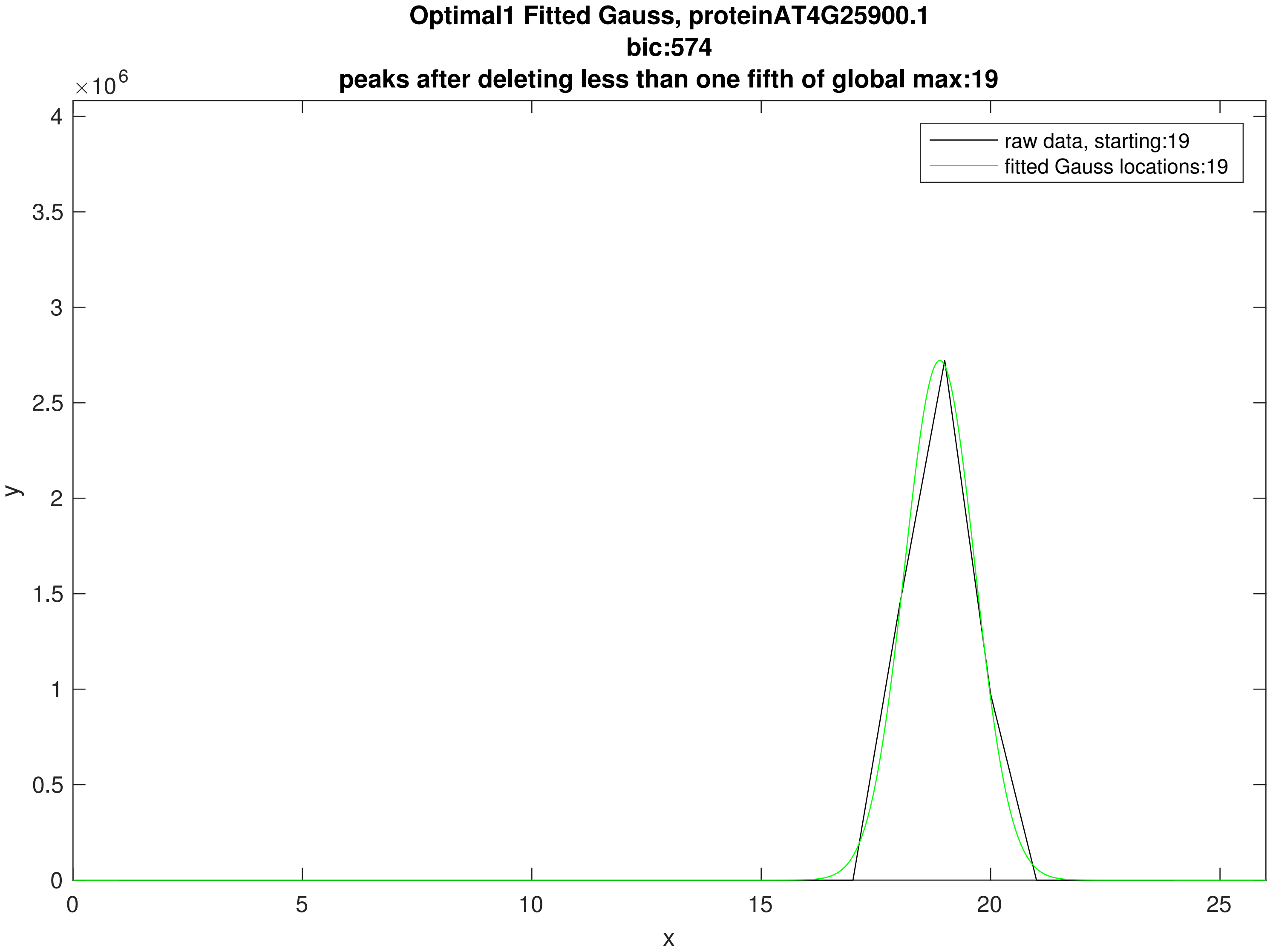
Optimal1 Fitted Gauss, proteinAT4G25900.1
bic:574
peaks after deleting less than one fifth of global max:19
×10^6
raw data, starting:19
fitted Gauss locations:19
4
3.5
3
2.5
2
1.5
1
0.5
0
y
0
5
10
15
20
25
x

Optimal1 Fitted Gauss, proteinAT4G26110.2;AT4G26110.1
bic:724
peaks after deleting less than one fifth of global max:12

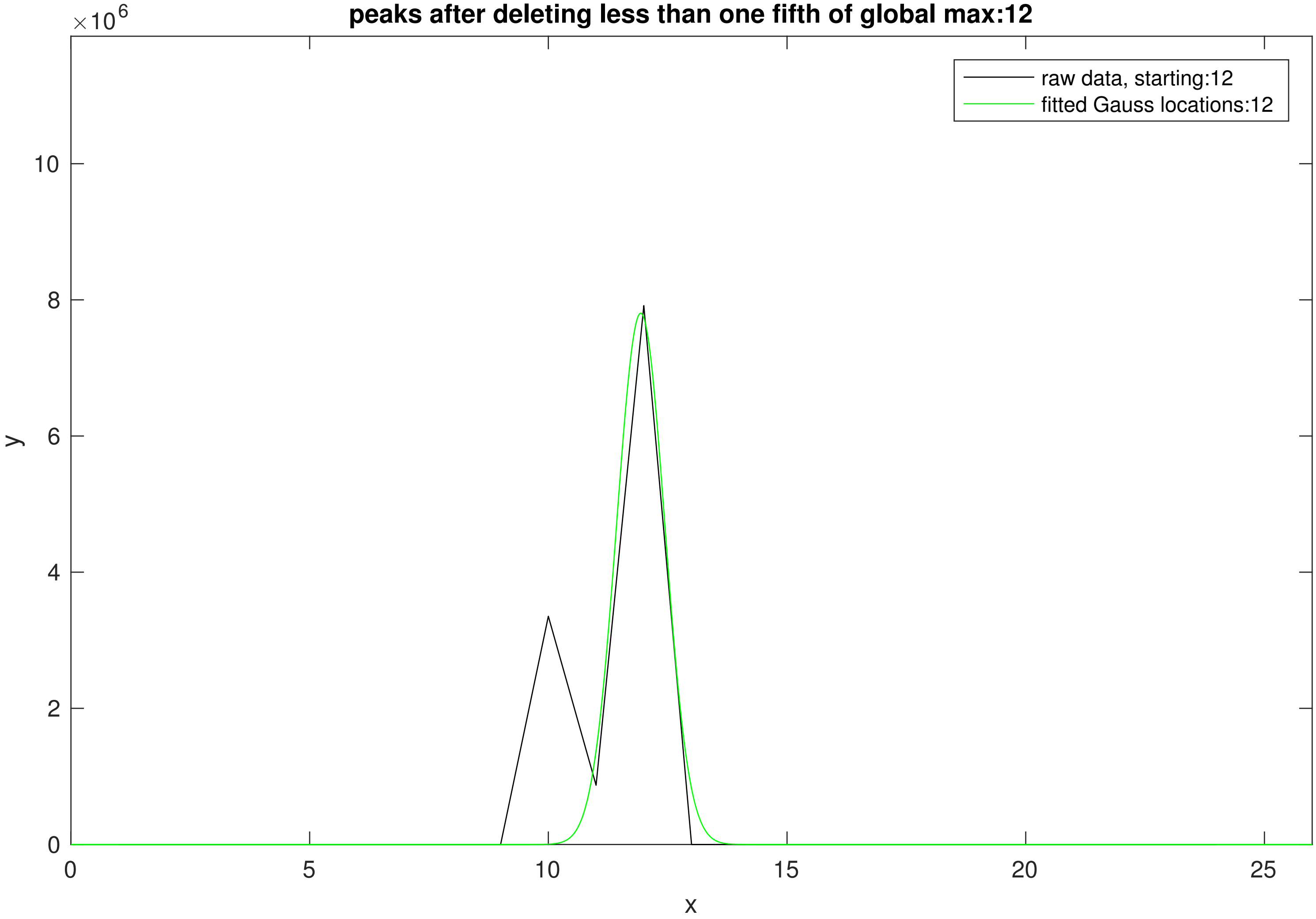

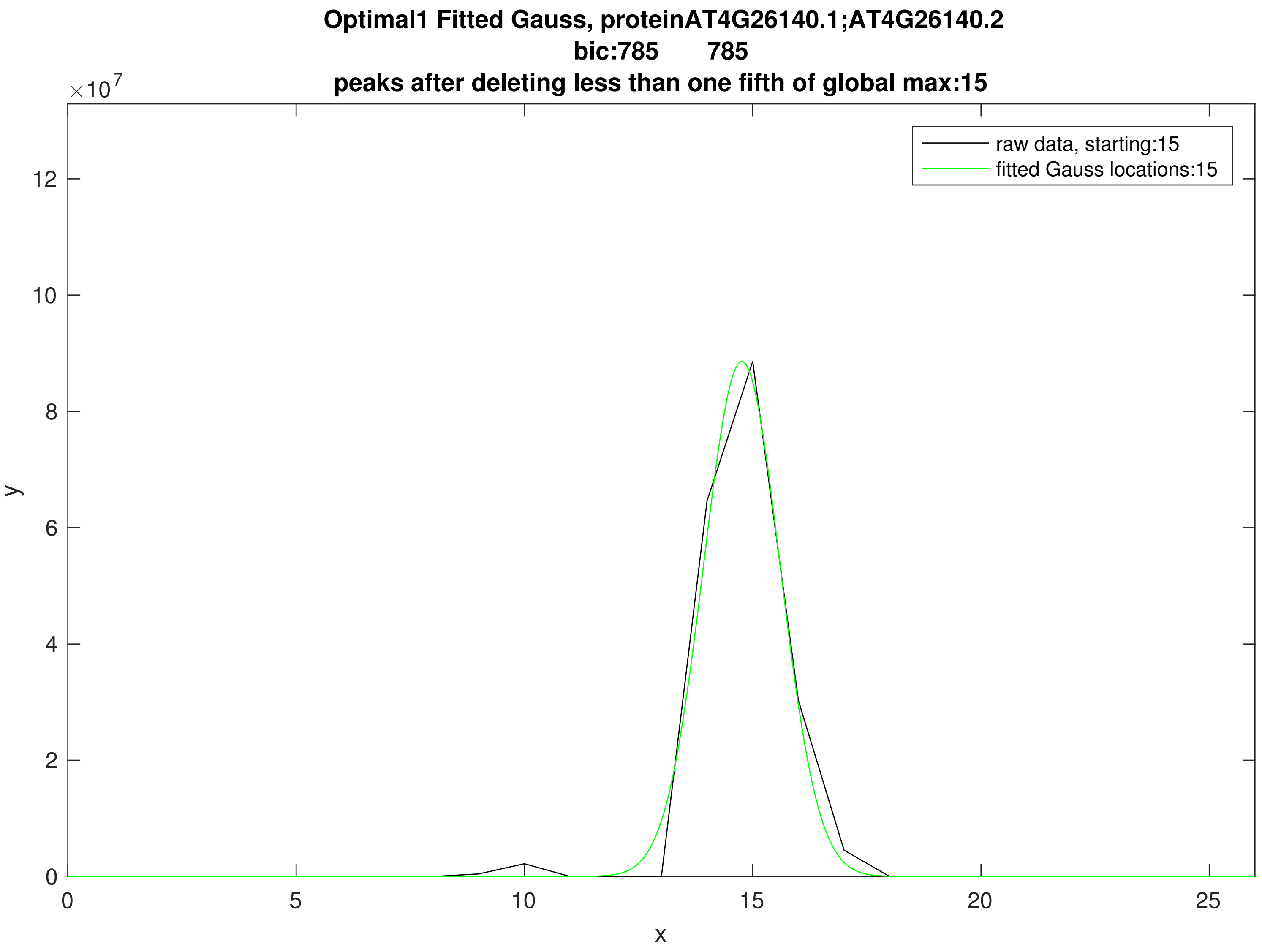

Optimal1 Fitted Gauss, proteinAT4G26140.1;AT4G26140.2
bic:785 785
peaks after deleting less than one fifth of global max:15
×10^7
12
10
8
6
4
2
0
y
0
5
10
15
20
25
x
raw data, starting:15
fitted Gauss locations:15

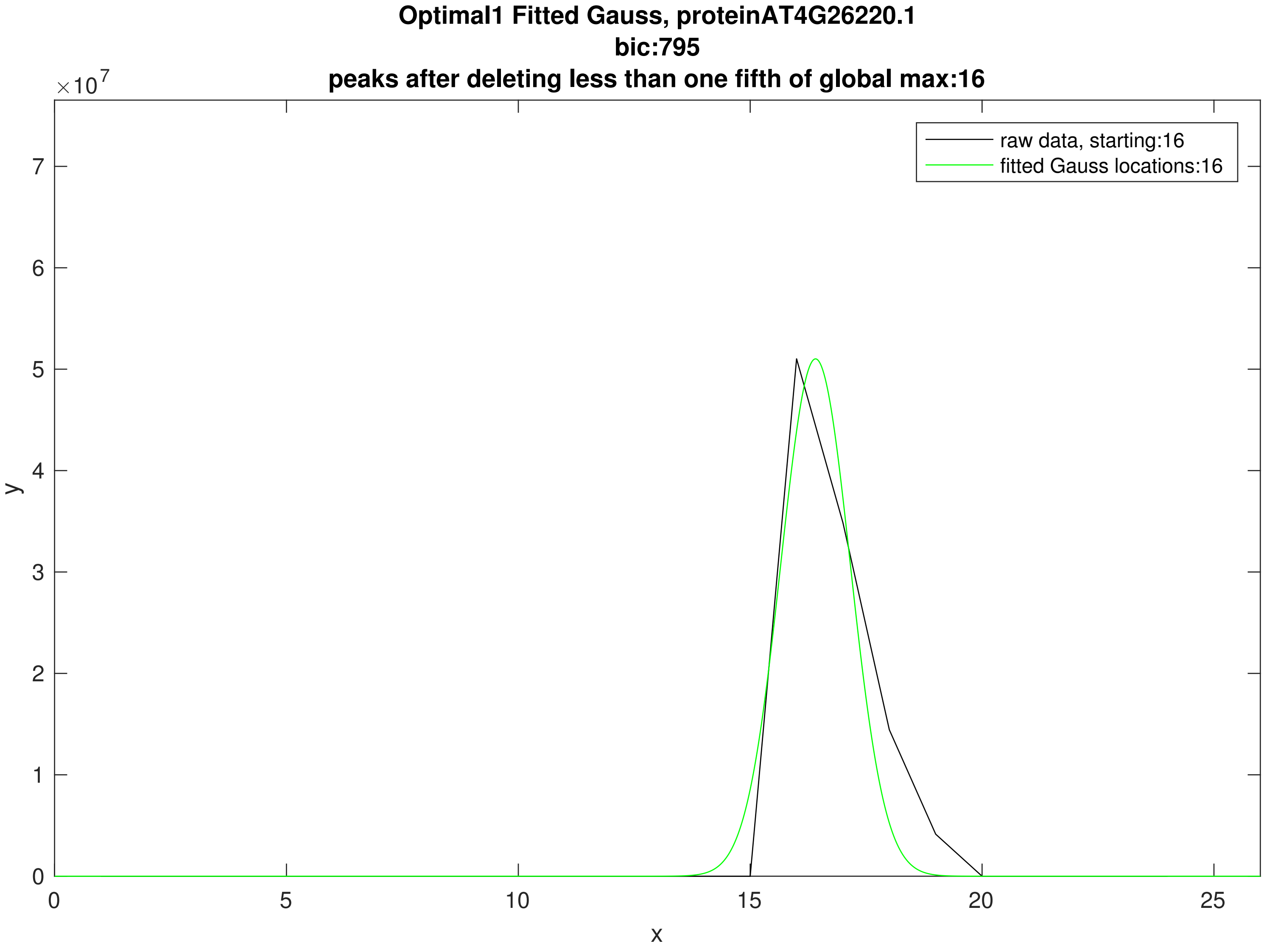

Optimal1 Fitted Gauss, proteinAT4G26220.1
bic:795
peaks after deleting less than one fifth of global max:16
×10^7
raw data, starting:16
fitted Gauss locations:16
y
x
0
1
2
3
4
5
6
7
0
5
10
15
20
25

**Optimal1 Fitted Gauss, proteinAT4G26300.1**
**bic:864 869**
**peaks after deleting less than one fifth of global max:14**

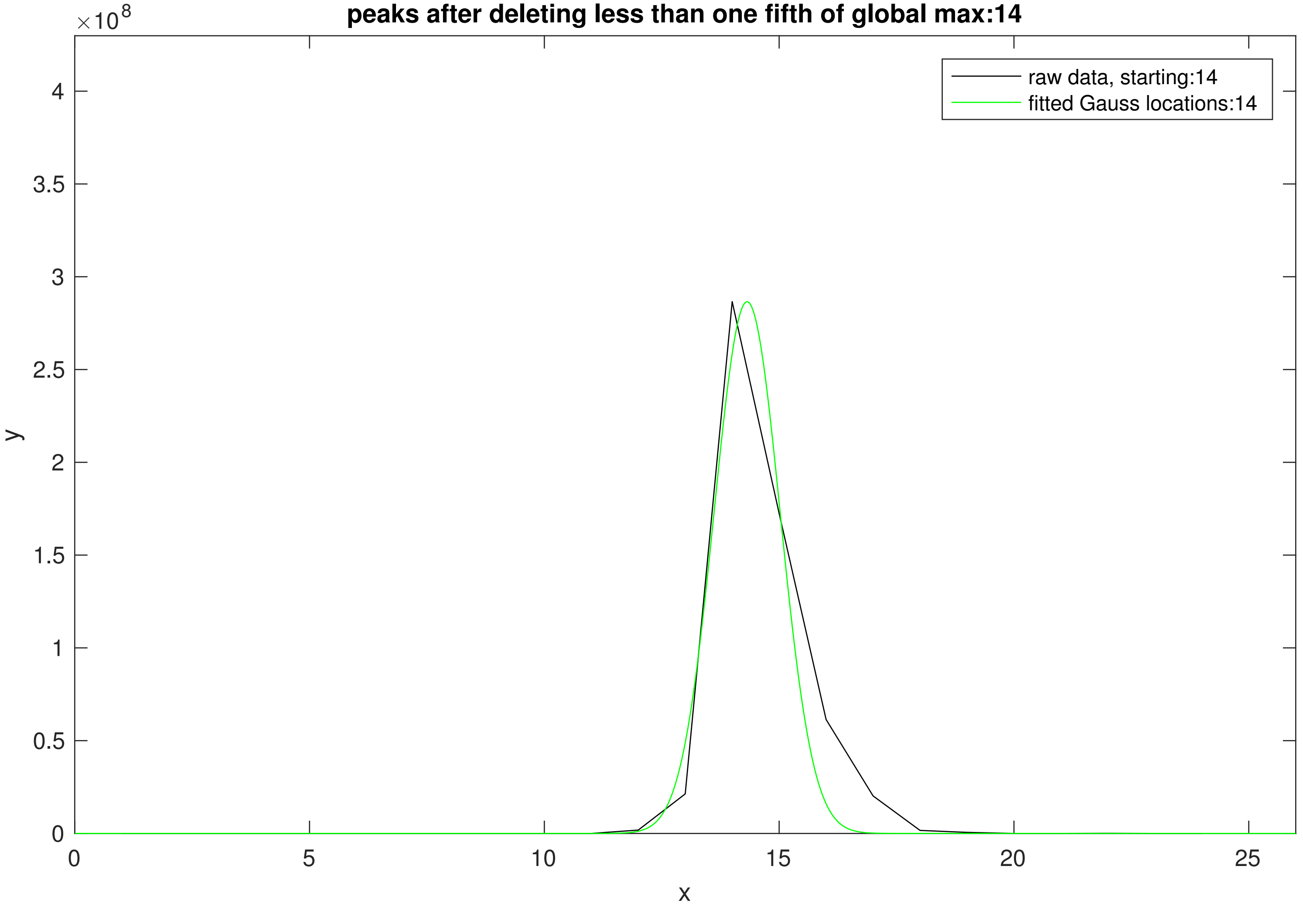

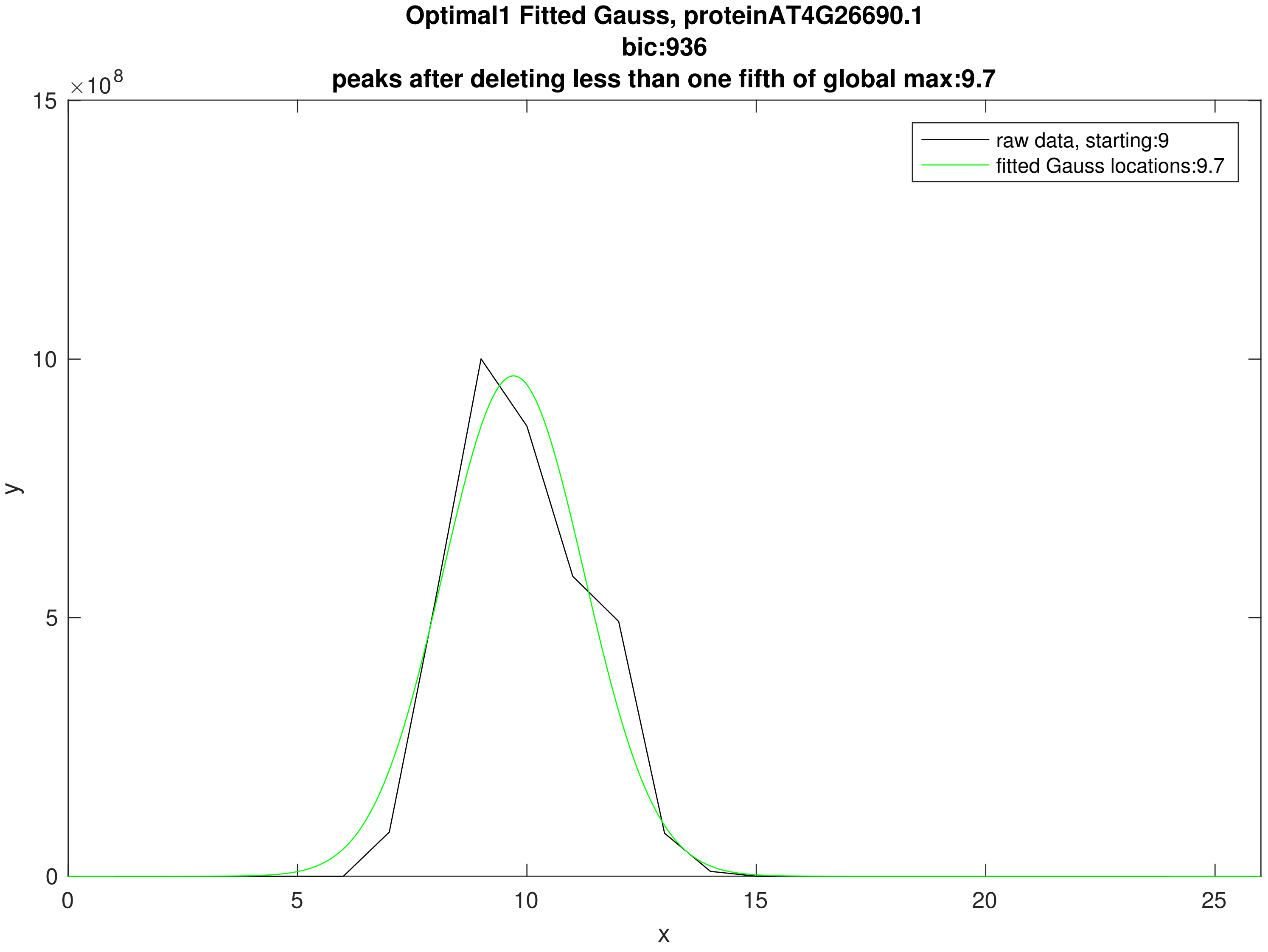

Optimal1 Fitted Gauss, proteinAT4G26690.1
bic:936
peaks after deleting less than one fifth of global max:9.7
×10^8
15
10
5
0
y
0
5
10
15
20
25
x
raw data, starting:9
fitted Gauss locations:9.7

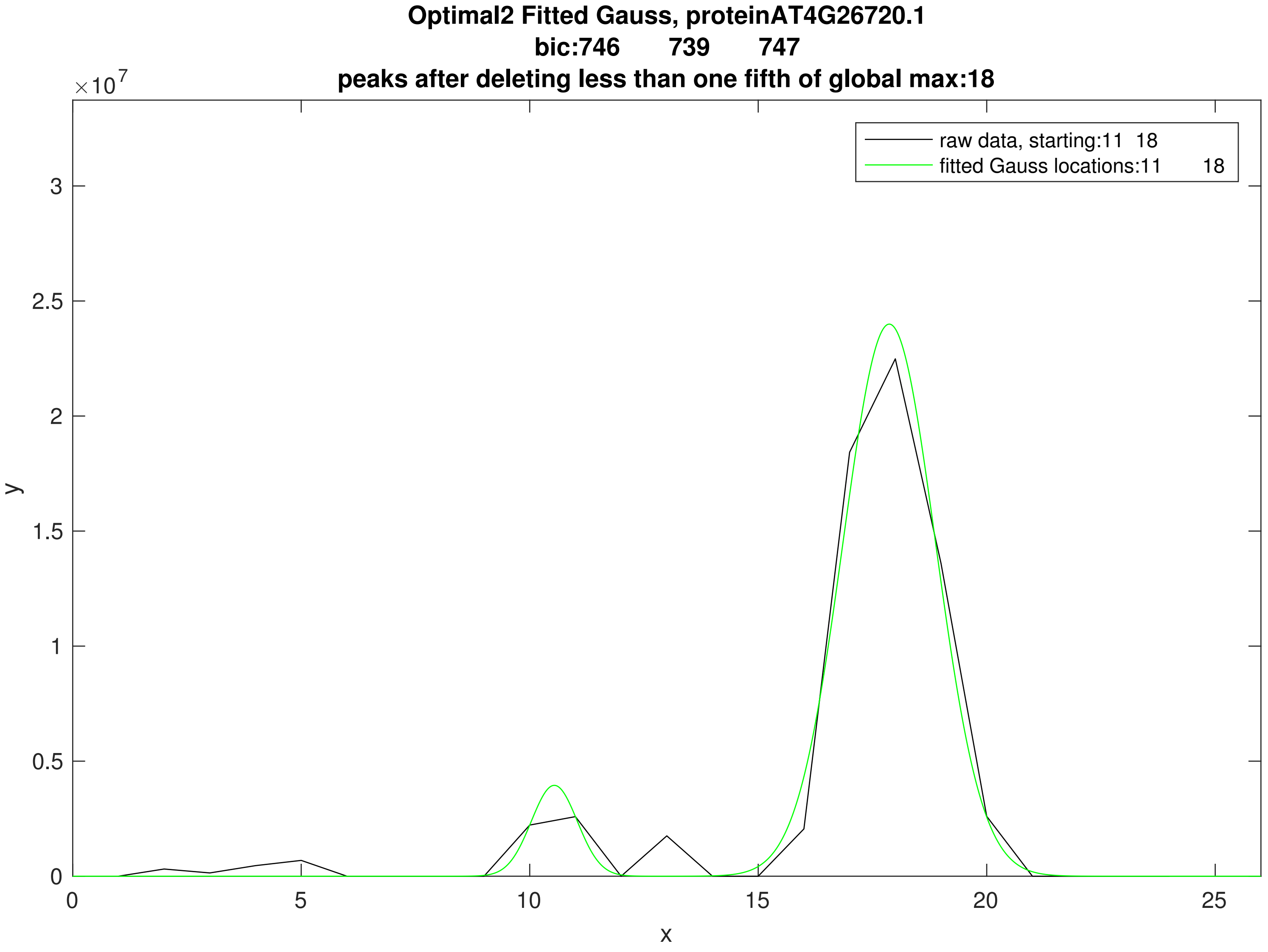
Optimal2 Fitted Gauss, proteinAT4G26720.1
bic:746 739 747
peaks after deleting less than one fifth of global max:18
×10^7
raw data, starting:11 18
fitted Gauss locations:11 18
y
x
0
0.5
1
1.5
2
2.5
3
0
5
10
15
20
25

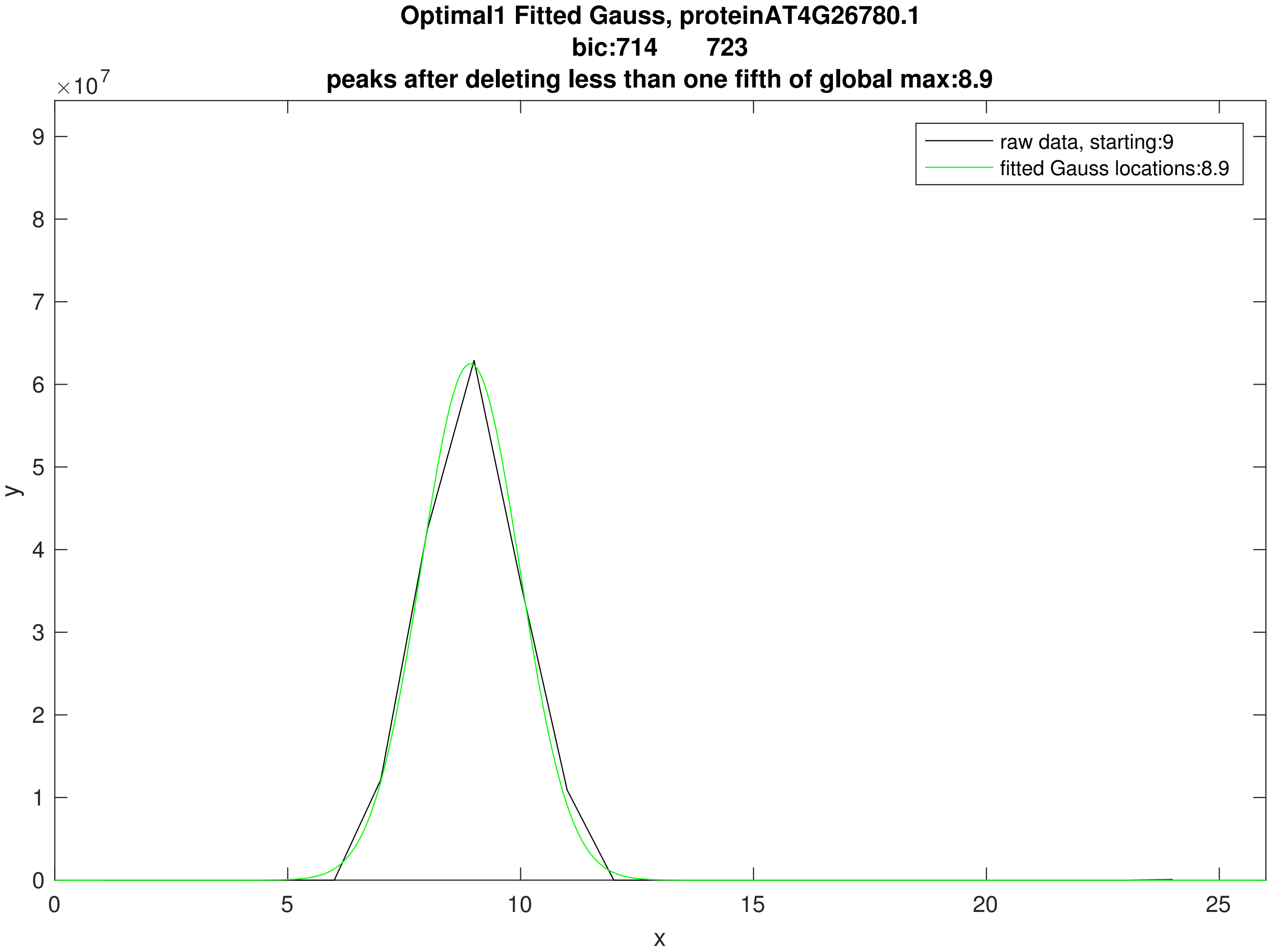

Optimal1 Fitted Gauss, proteinAT4G26780.1
bic:714 723
peaks after deleting less than one fifth of global max:8.9
×10^7
raw data, starting:9
fitted Gauss locations:8.9
y
x
0
1
2
3
4
5
6
7
8
9
0
5
10
15
20
25

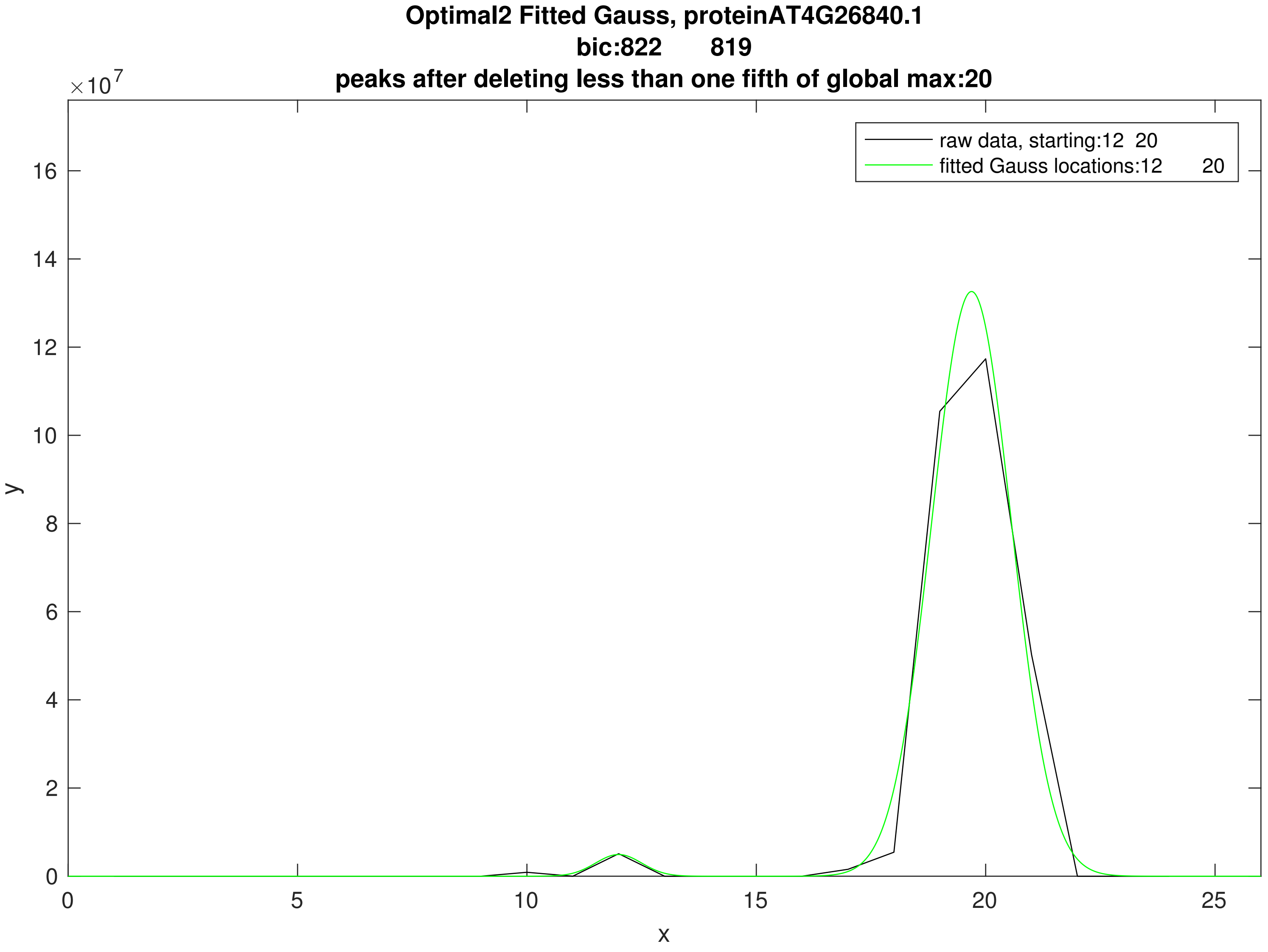
Optimal2 Fitted Gauss, proteinAT4G26840.1
bic:822 819
peaks after deleting less than one fifth of global max:20
×10^7
raw data, starting:12 20
fitted Gauss locations:12 20
y
x
0
2
4
6
8
10
12
14
16
0
5
10
15
20
25

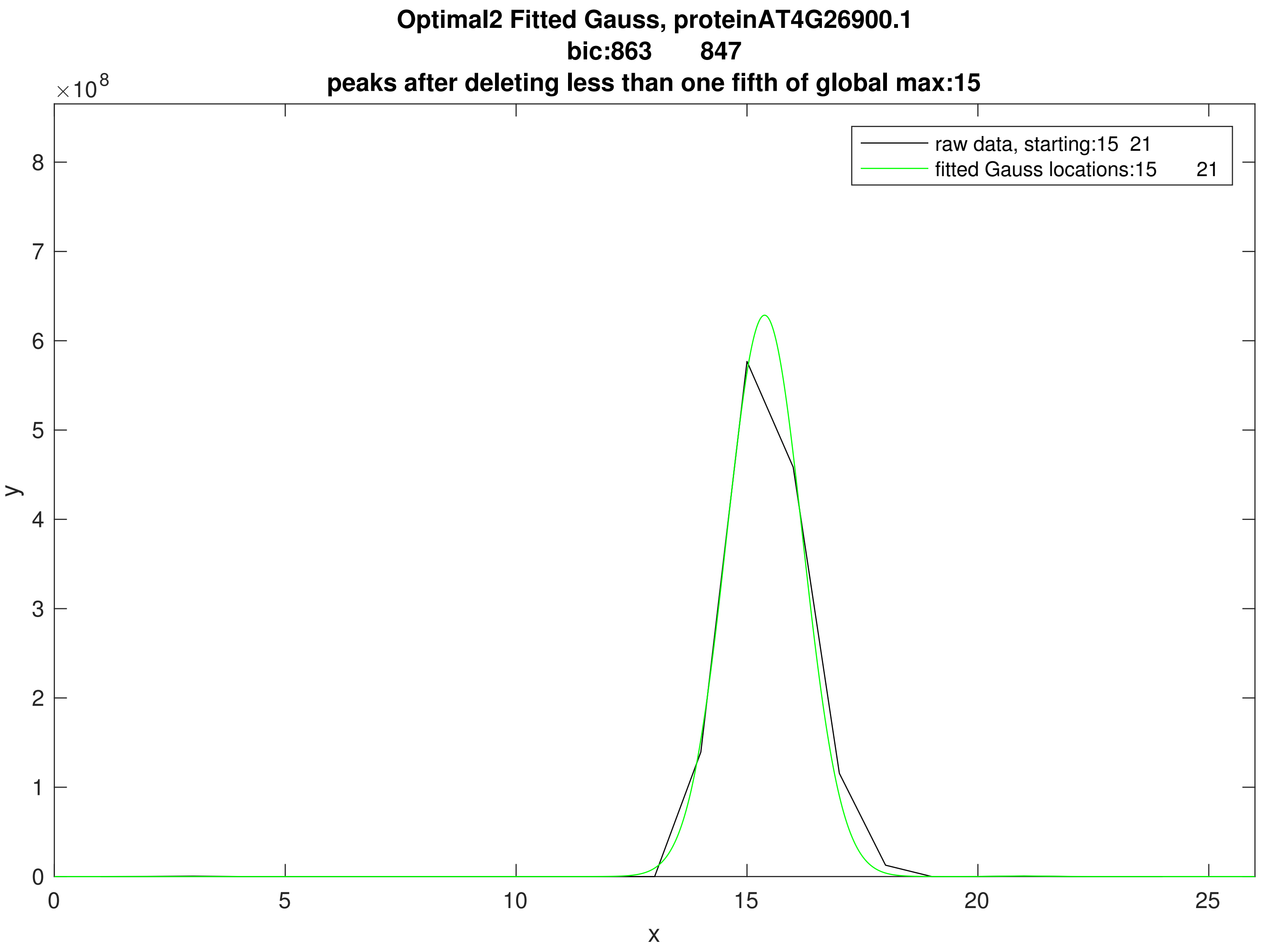

Optimal2 Fitted Gauss, proteinAT4G26900.1
bic:863 847
peaks after deleting less than one fifth of global max:15
×10^8
raw data, starting:15 21
fitted Gauss locations:15 21
y
x
0
1
2
3
4
5
6
7
8
0
5
10
15
20
25

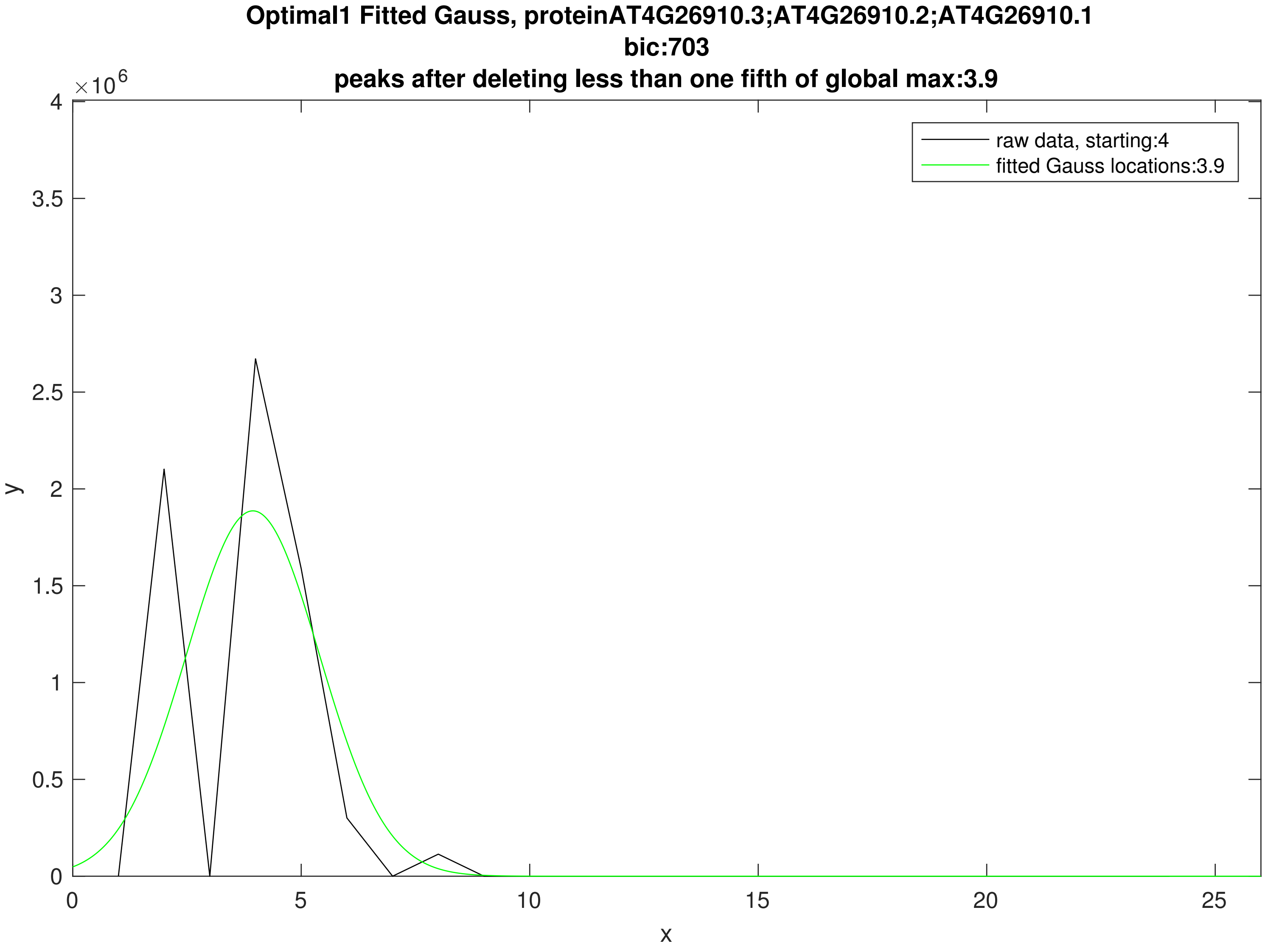
Optimal1 Fitted Gauss, proteinAT4G26910.3;AT4G26910.2;AT4G26910.1
bic:703
peaks after deleting less than one fifth of global max:3.9
×10^6
raw data, starting:4
fitted Gauss locations:3.9
y
x
0
0.5
1
1.5
2
2.5
3
3.5
4
0
5
10
15
20
25

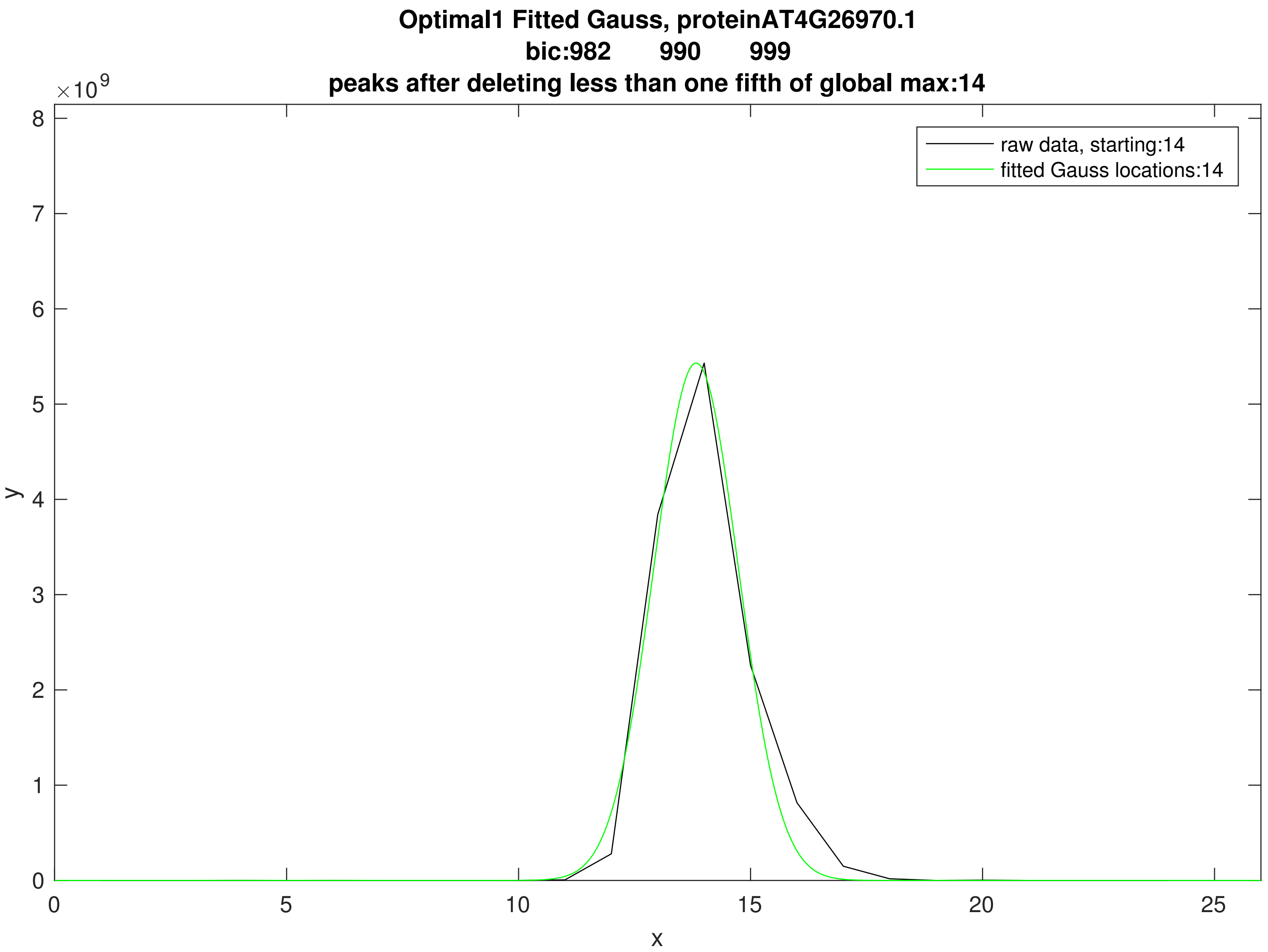

Optimal1 Fitted Gauss, proteinAT4G26970.1
bic:982 990 999
peaks after deleting less than one fifth of global max:14
×10^9
raw data, starting:14
fitted Gauss locations:14
y
x

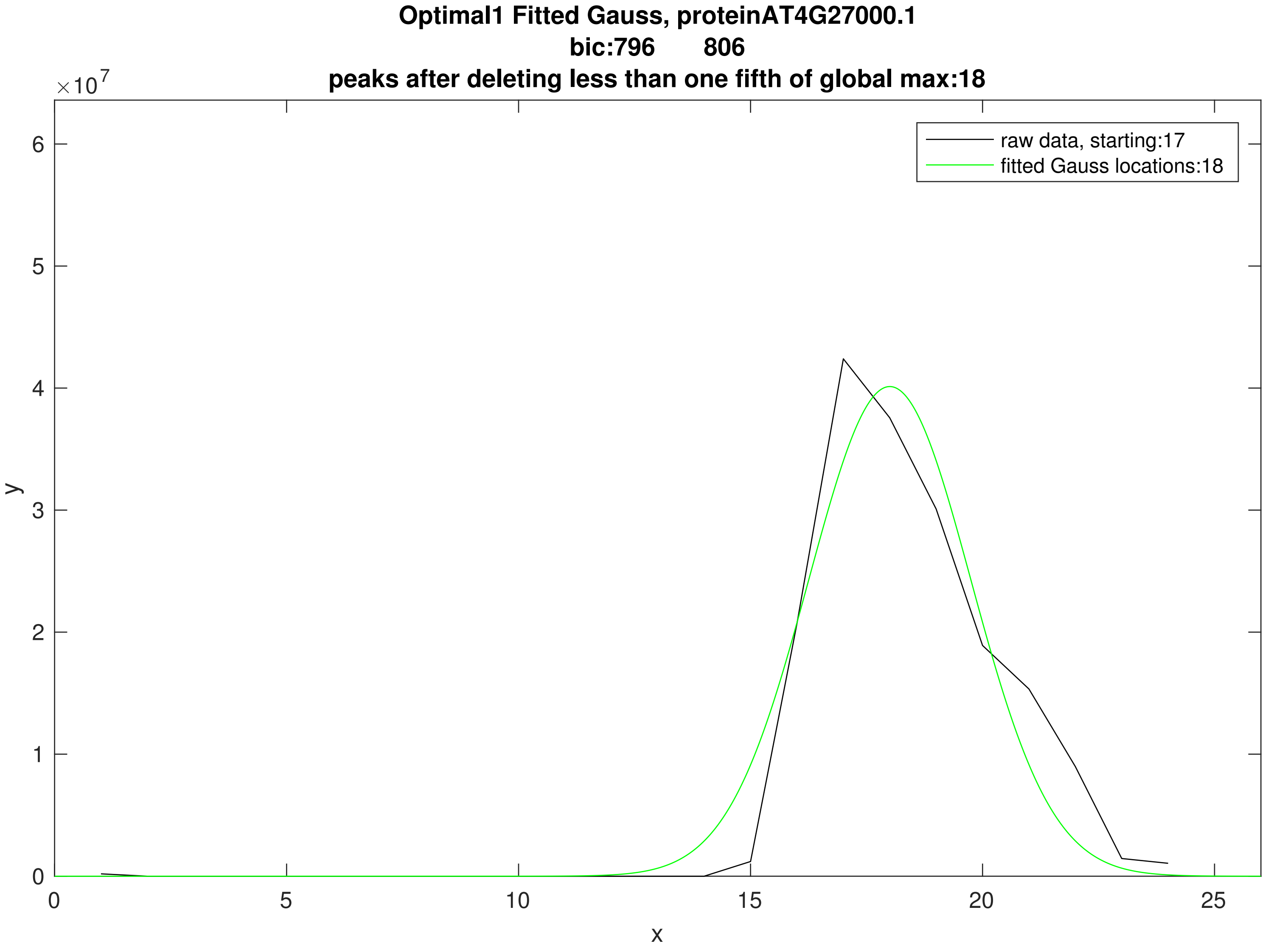
Optimal1 Fitted Gauss, proteinAT4G27000.1
bic:796 806
peaks after deleting less than one fifth of global max:18
×10^7
raw data, starting:17
fitted Gauss locations:18
y
x
0
1
2
3
4
5
6
0
5
10
15
20
25

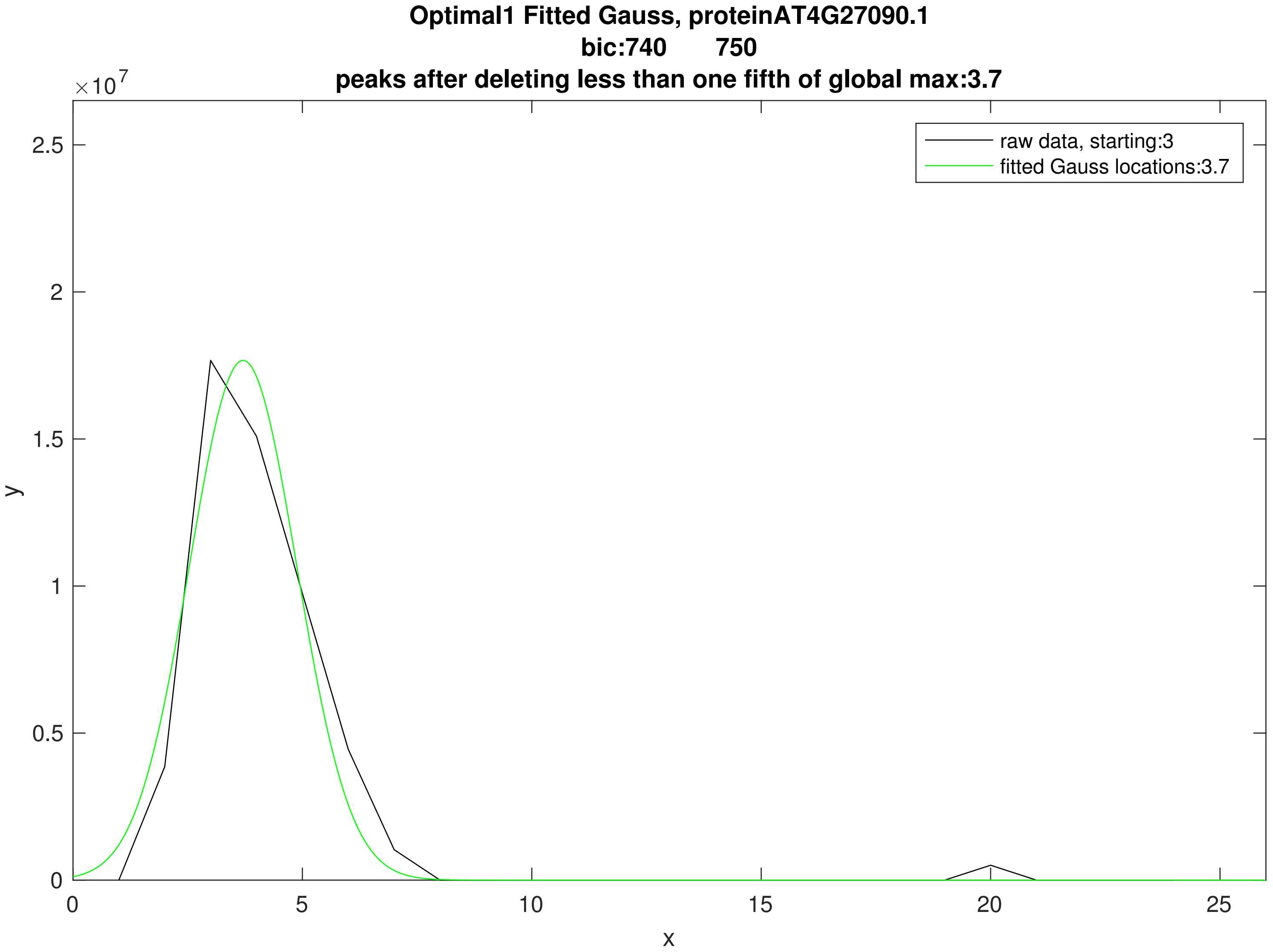

Optimal1 Fitted Gauss, proteinAT4G27090.1
bic:740 750
peaks after deleting less than one fifth of global max:3.7
×10^7
raw data, starting:3
fitted Gauss locations:3.7
2.5
2
1.5
1
0.5
0
y
0
5
10
15
20
25
x

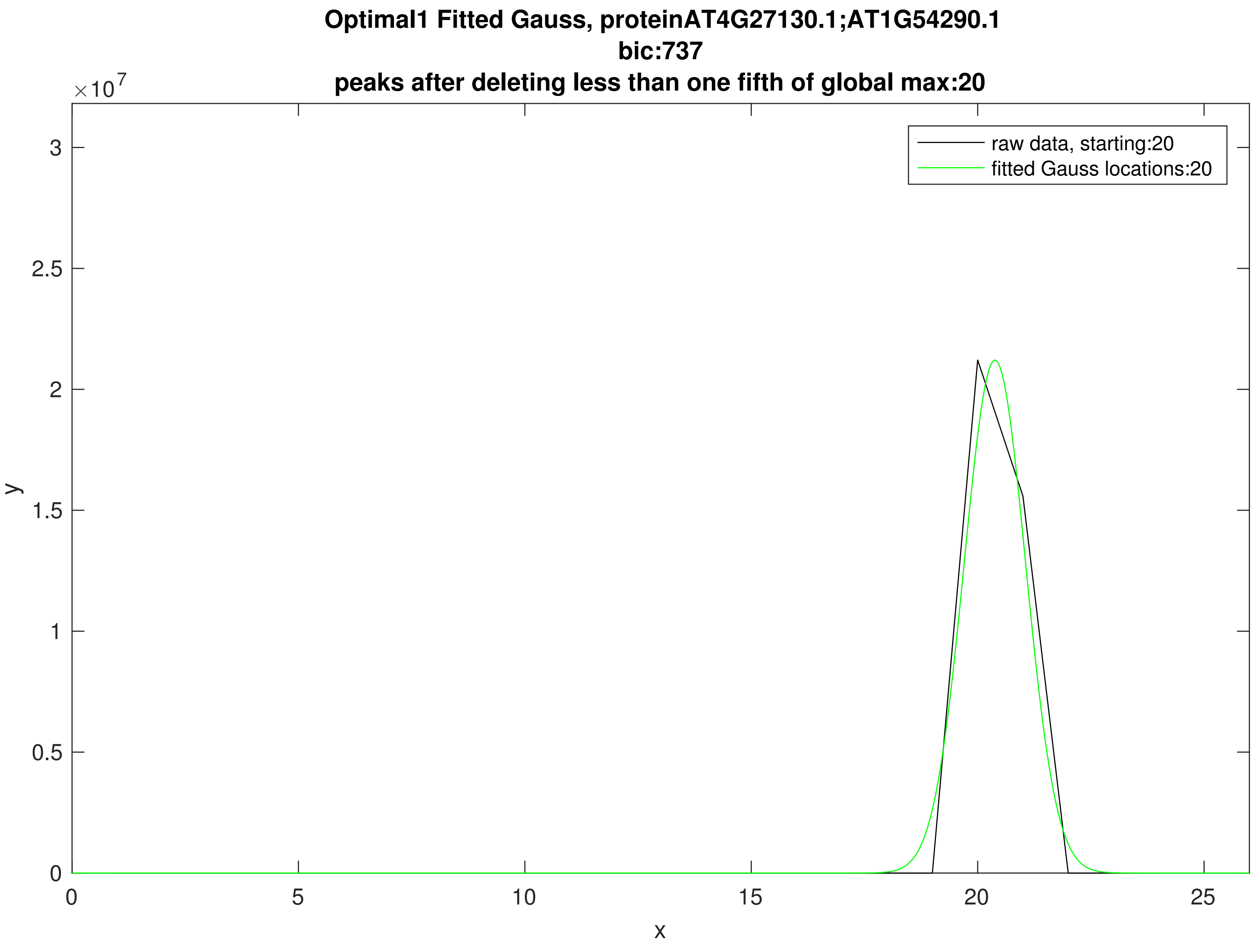

Optimal1 Fitted Gauss, proteinAT4G27130.1;AT1G54290.1
bic:737
peaks after deleting less than one fifth of global max:20
×10^7
3
2.5
2
1.5
1
0.5
0
y
0
5
10
15
20
25
x
raw data, starting:20
fitted Gauss locations:20

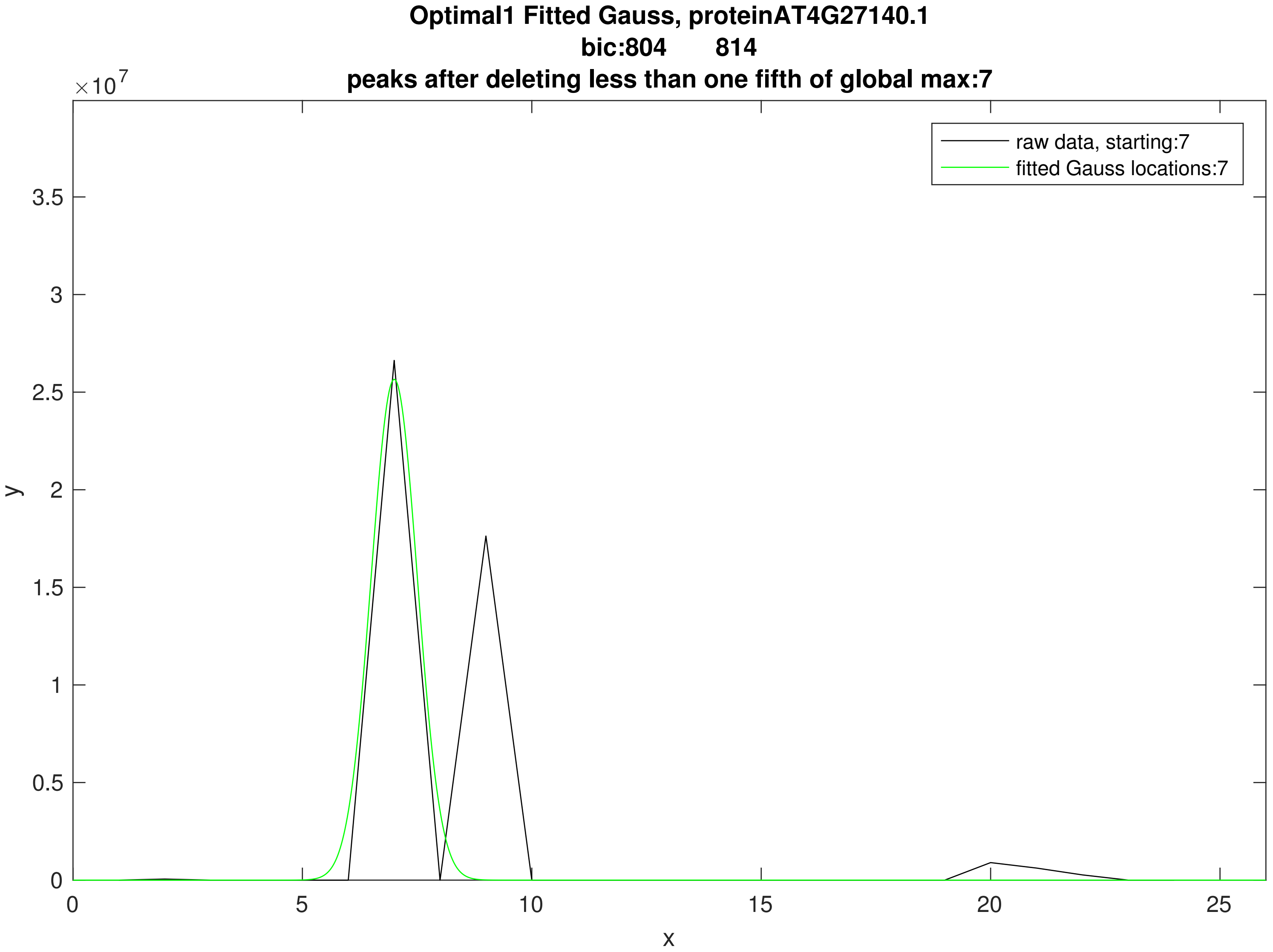
Optimal1 Fitted Gauss, proteinAT4G27140.1
bic:804 814
peaks after deleting less than one fifth of global max:7
×10^7
raw data, starting:7
fitted Gauss locations:7
y
x
0
0.5
1
1.5
2
2.5
3
3.5
0
5
10
15
20
25

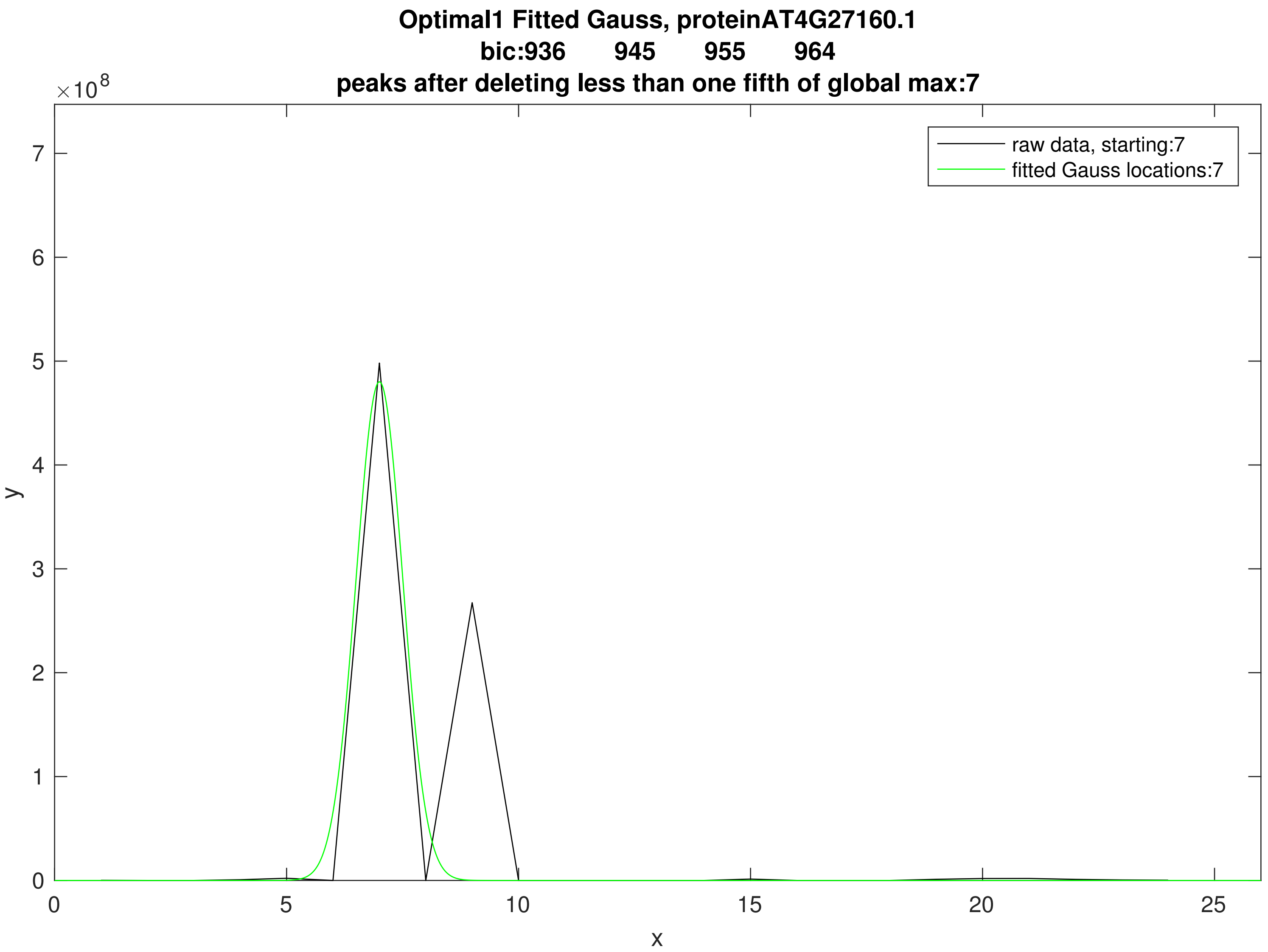
Optimal1 Fitted Gauss, proteinAT4G27160.1
bic:936 945 955 964
peaks after deleting less than one fifth of global max:7
×10^8
raw data, starting:7
fitted Gauss locations:7
y
x
0
1
2
3
4
5
6
7
10
15
20
25

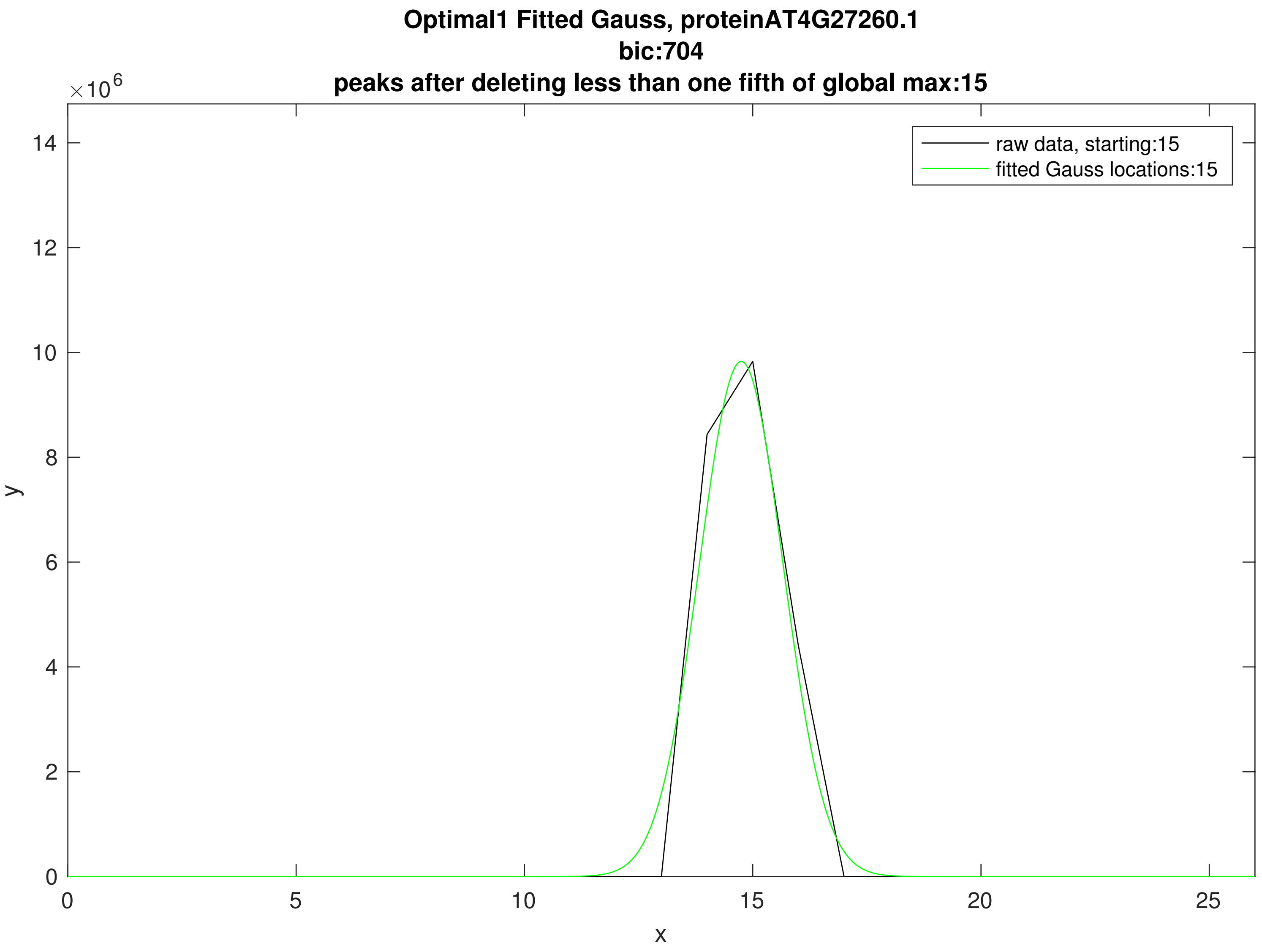

Optimal1 Fitted Gauss, proteinAT4G27260.1
bic:704
peaks after deleting less than one fifth of global max:15
×10^6
raw data, starting:15
fitted Gauss locations:15
y
x
0
2
4
6
8
10
12
14
0
5
10
15
20
25

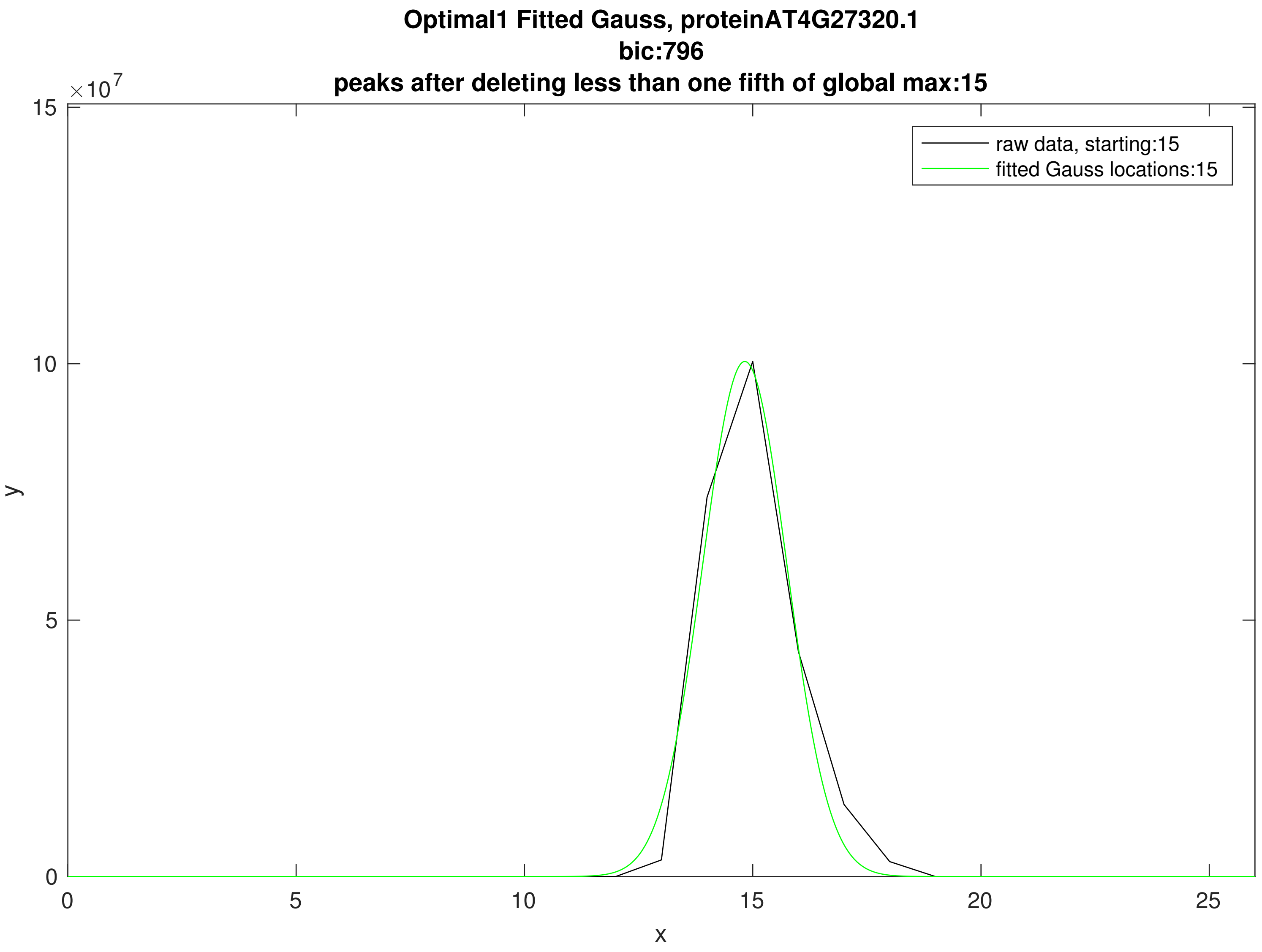

Optimal1 Fitted Gauss, proteinAT4G27320.1
bic:796
peaks after deleting less than one fifth of global max:15
×10^7
15
10
5
0
y
0
5
10
15
20
25
x
raw data, starting:15
fitted Gauss locations:15

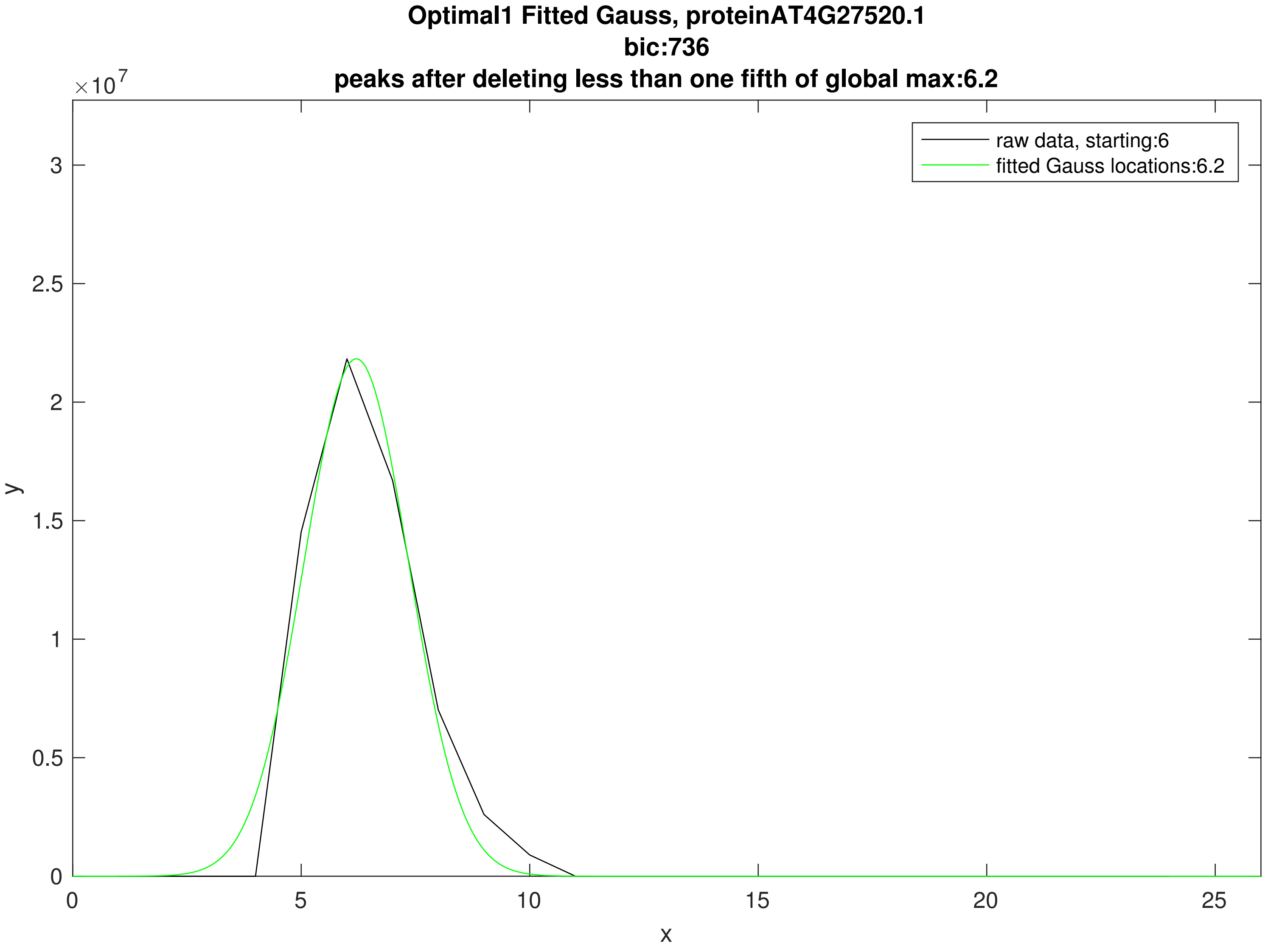

Optimal1 Fitted Gauss, proteinAT4G27520.1
bic:736
peaks after deleting less than one fifth of global max:6.2
×10^7
raw data, starting:6
fitted Gauss locations:6.2
3
2.5
2
1.5
1
0.5
0
y
0
5
10
15
20
25
x

**Optimal1 Fitted Gauss, proteinAT4G27690.2;AT4G27690.1**
**bic:832**
**peaks after deleting less than one fifth of global max:17**

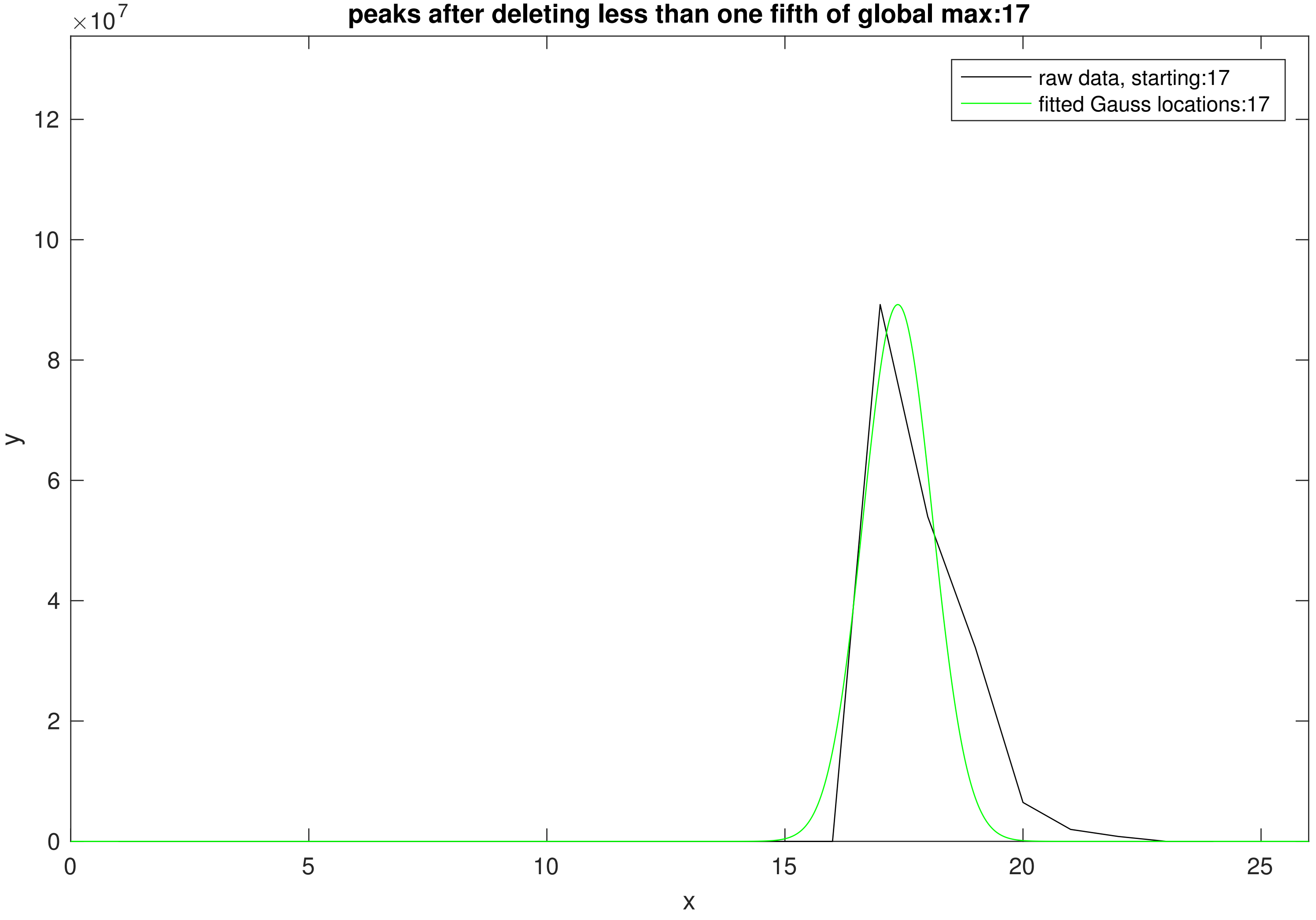

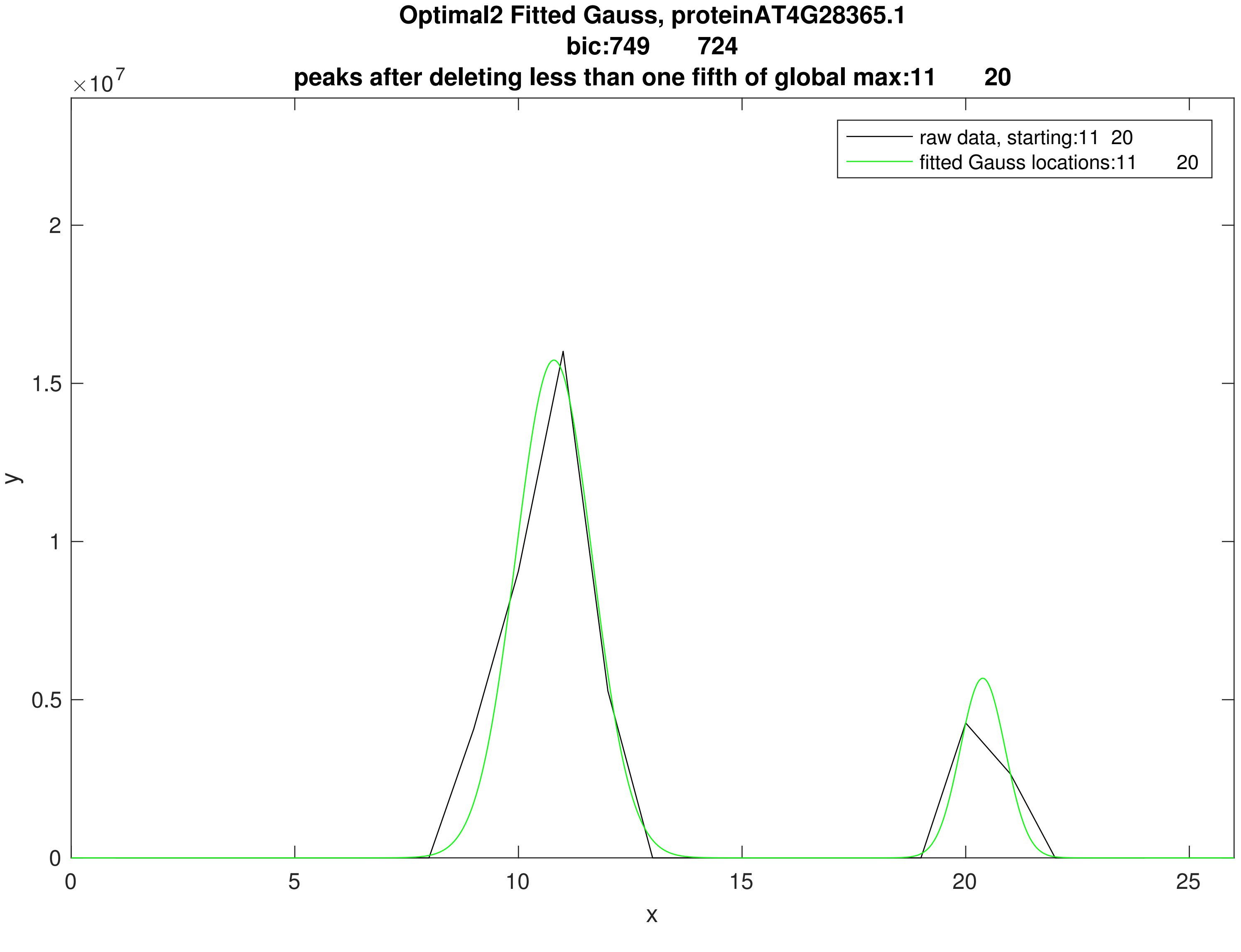
Optimal2 Fitted Gauss, proteinAT4G28365.1
bic:749 724
peaks after deleting less than one fifth of global max:11 20
×10^7
raw data, starting:11 20
fitted Gauss locations:11 20
y
x
0
0.5
1
1.5
2
5
10
15
20
25

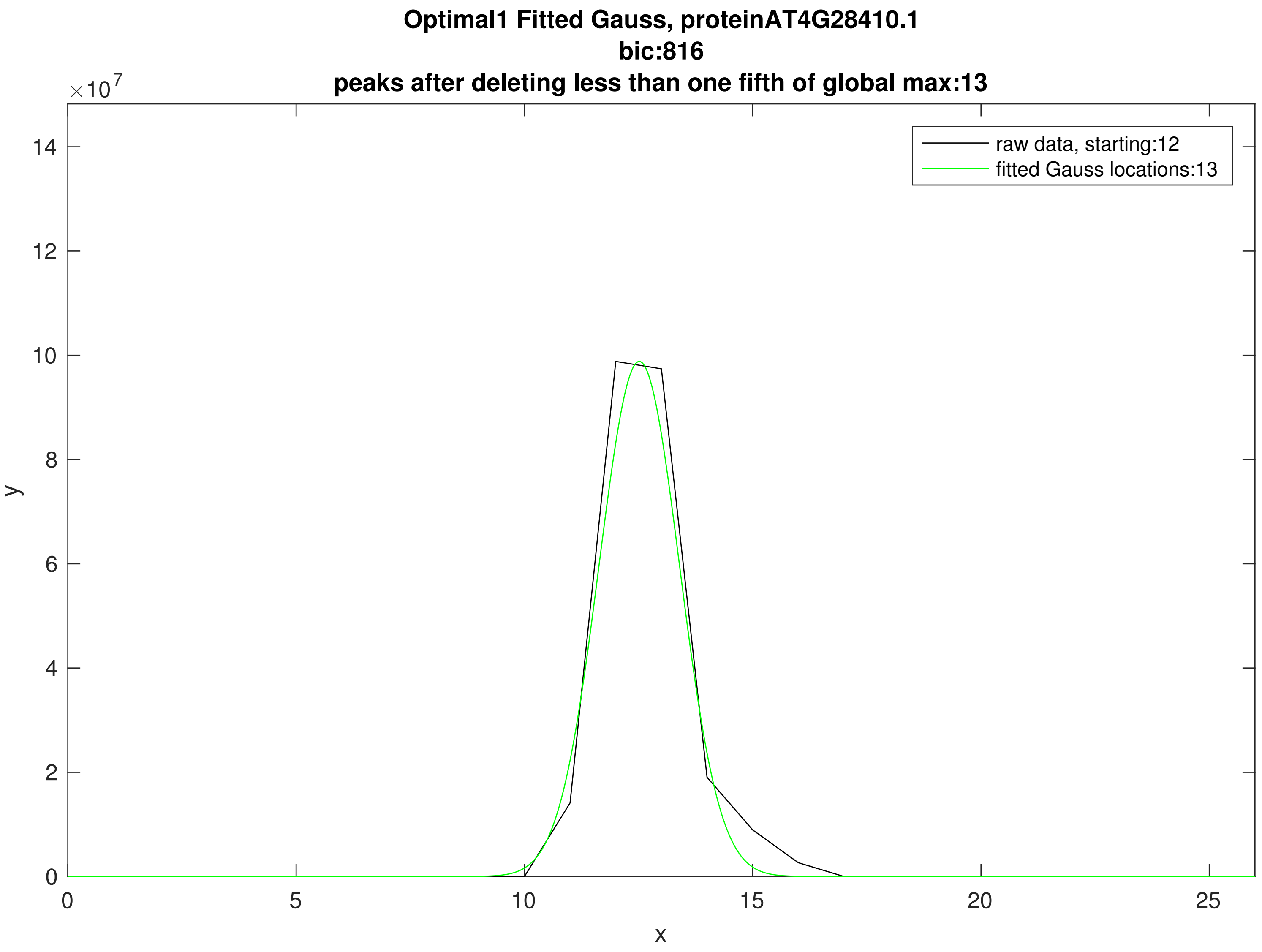

Optimal1 Fitted Gauss, proteinAT4G28410.1
bic:816
peaks after deleting less than one fifth of global max:13
×10^7
raw data, starting:12
fitted Gauss locations:13
y
x
0
2
4
6
8
10
12
14
0
5
10
15
20
25

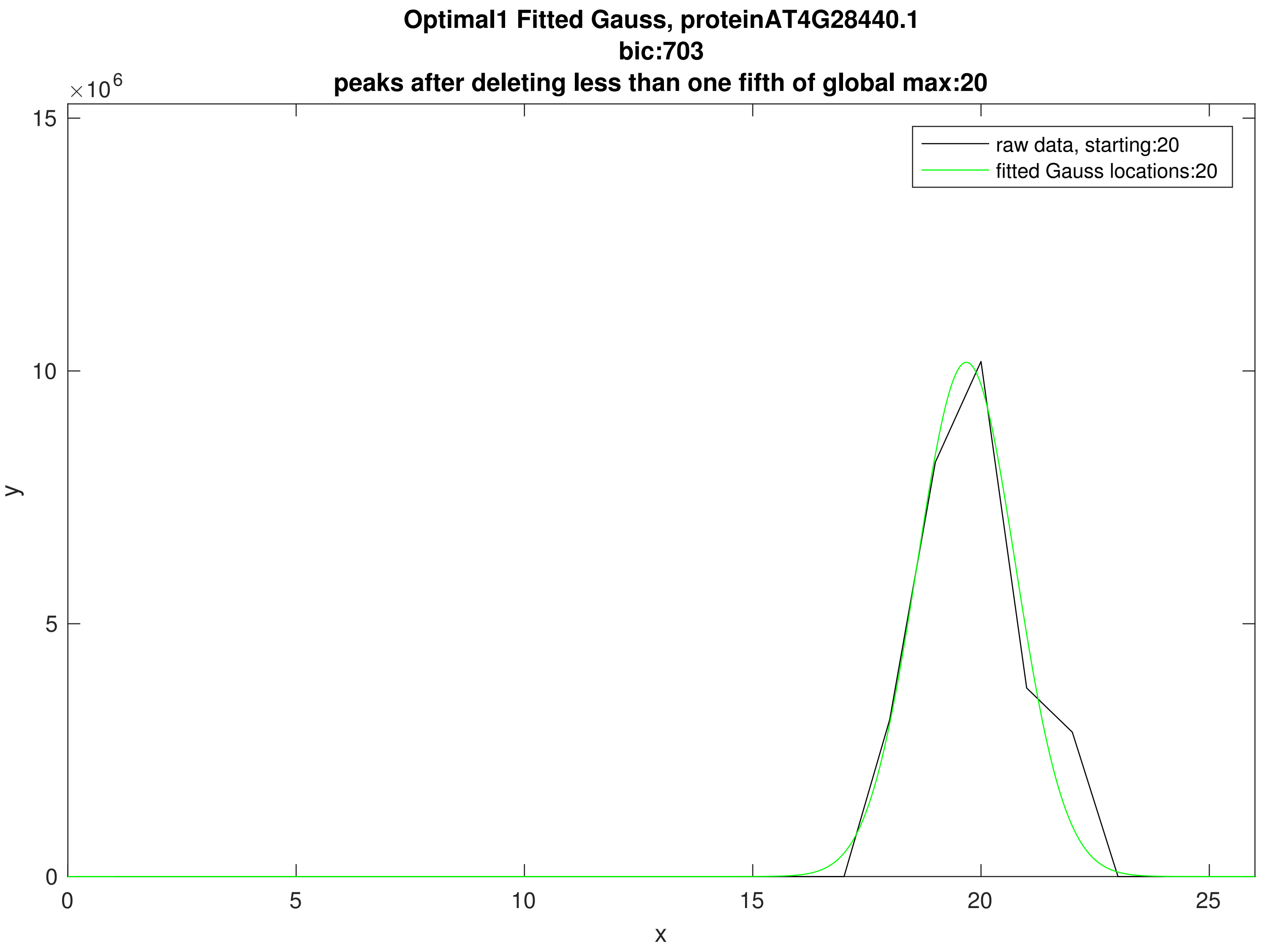
Optimal1 Fitted Gauss, proteinAT4G28440.1
bic:703
peaks after deleting less than one fifth of global max:20
×10^6
15
10
5
0
y
0
5
10
15
20
25
x
raw data, starting:20
fitted Gauss locations:20

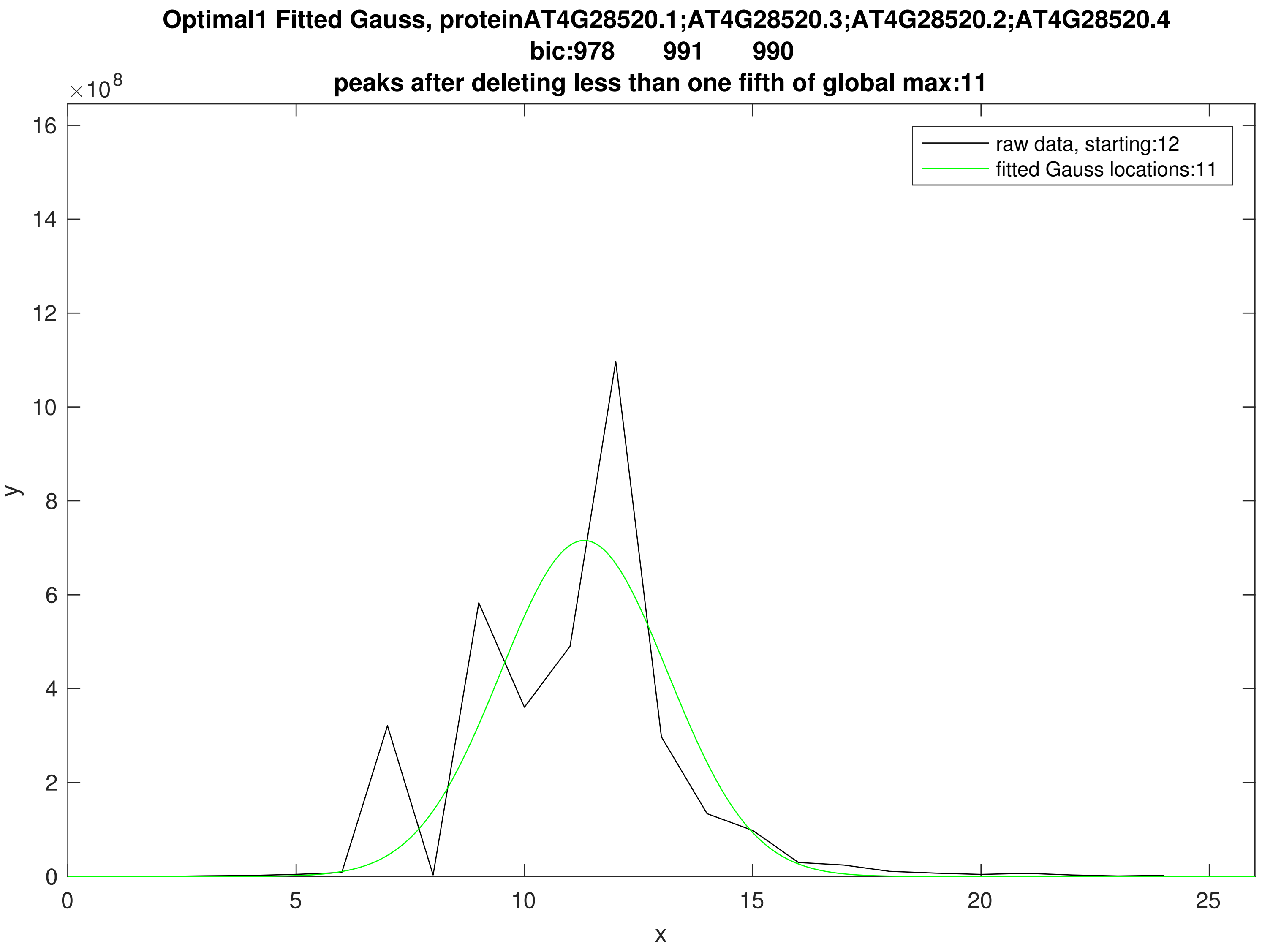

Optimal1 Fitted Gauss, proteinAT4G28520.1;AT4G28520.3;AT4G28520.2;AT4G28520.4
bic:978 991 990
peaks after deleting less than one fifth of global max:11
×10^8
raw data, starting:12
fitted Gauss locations:11
y
x

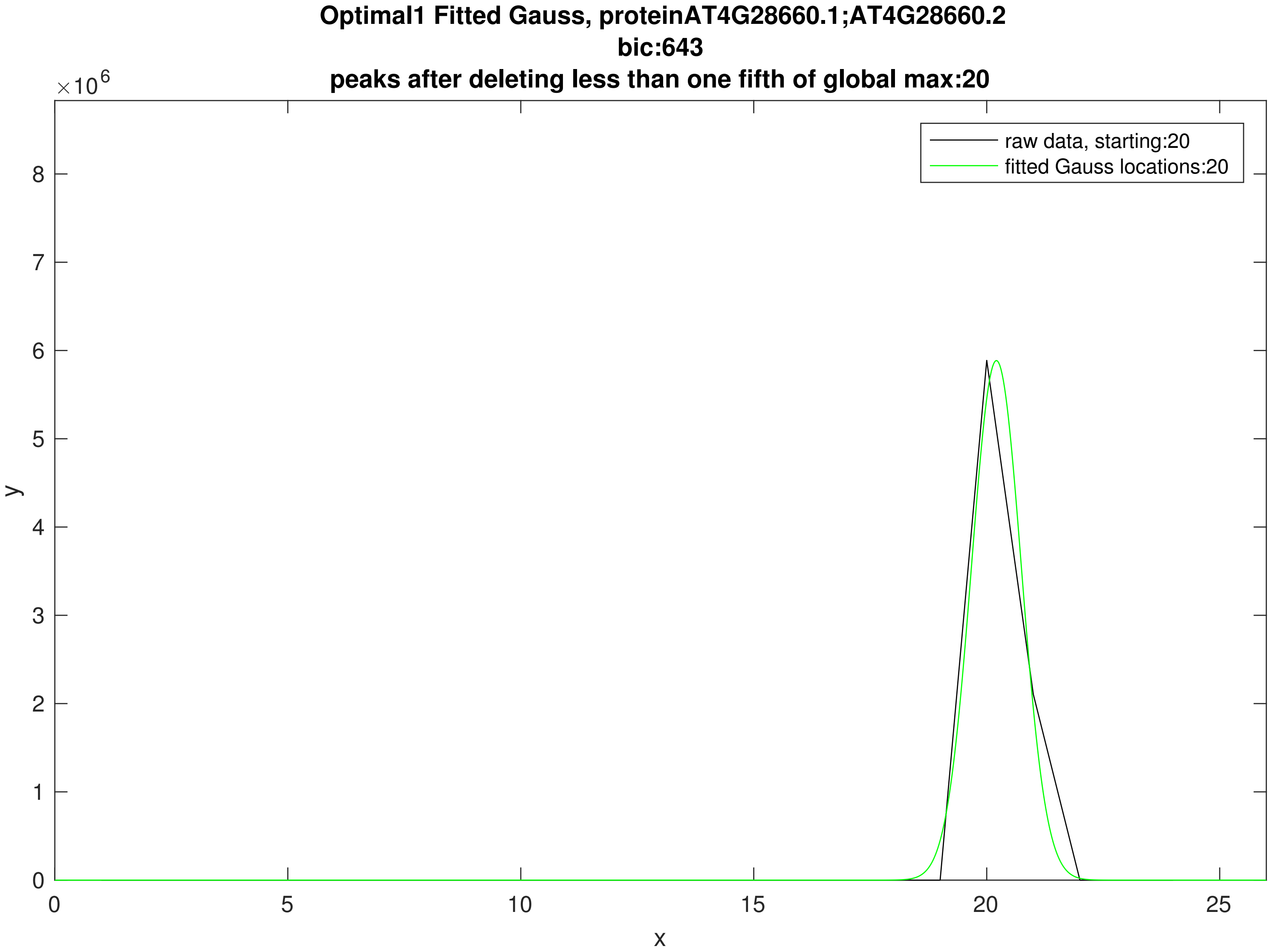
Optimal1 Fitted Gauss, proteinAT4G28660.1;AT4G28660.2
bic:643
peaks after deleting less than one fifth of global max:20
×10^6
raw data, starting:20
fitted Gauss locations:20
8
7
6
5
4
3
2
1
0
y
0
5
10
15
20
25
x

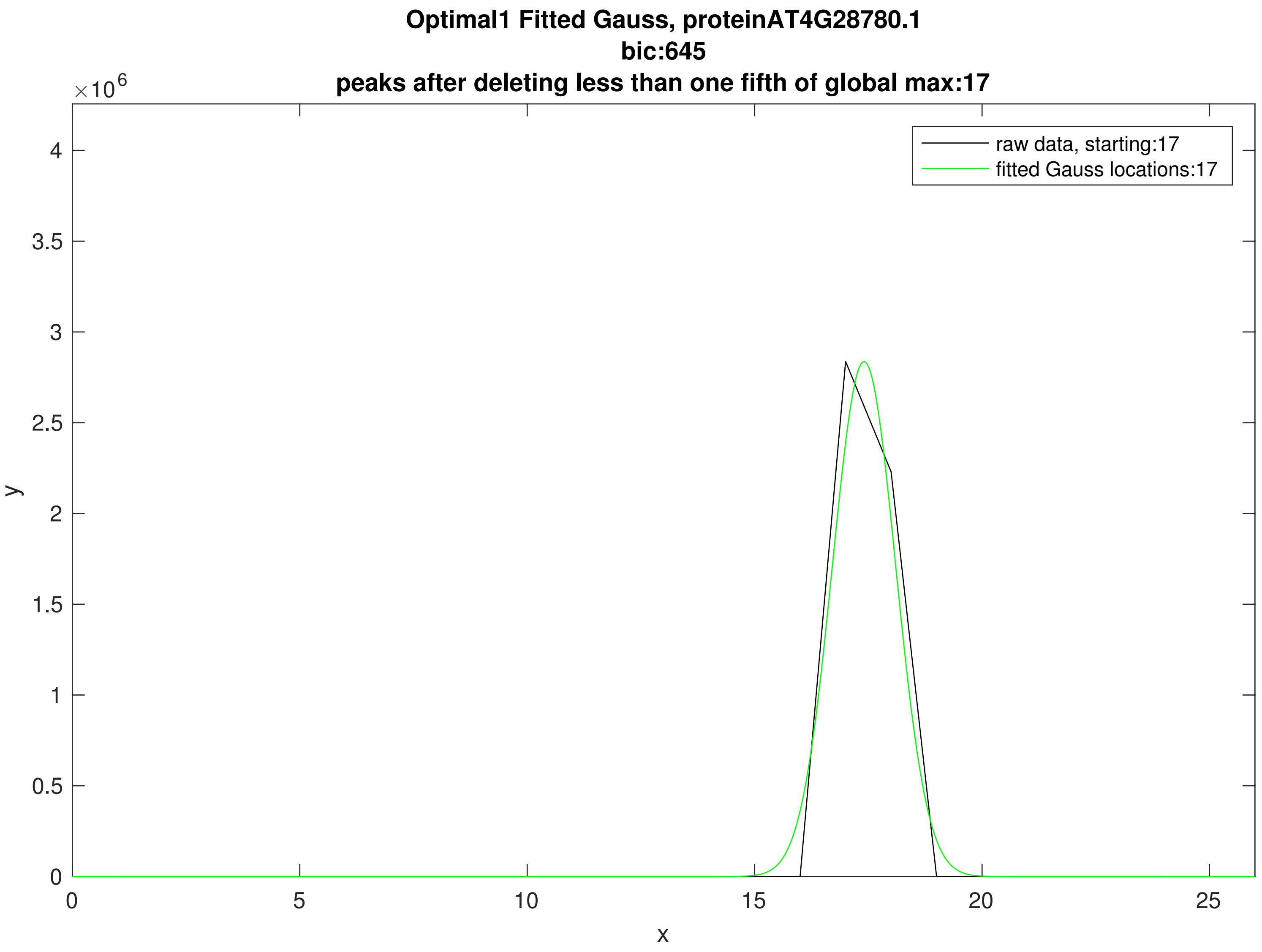

Optimal1 Fitted Gauss, proteinAT4G28780.1
bic:645
peaks after deleting less than one fifth of global max:17
×10^6
4
3.5
3
2.5
2
1.5
1
0.5
0
y
0
5
10
15
20
25
x
raw data, starting:17
fitted Gauss locations:17

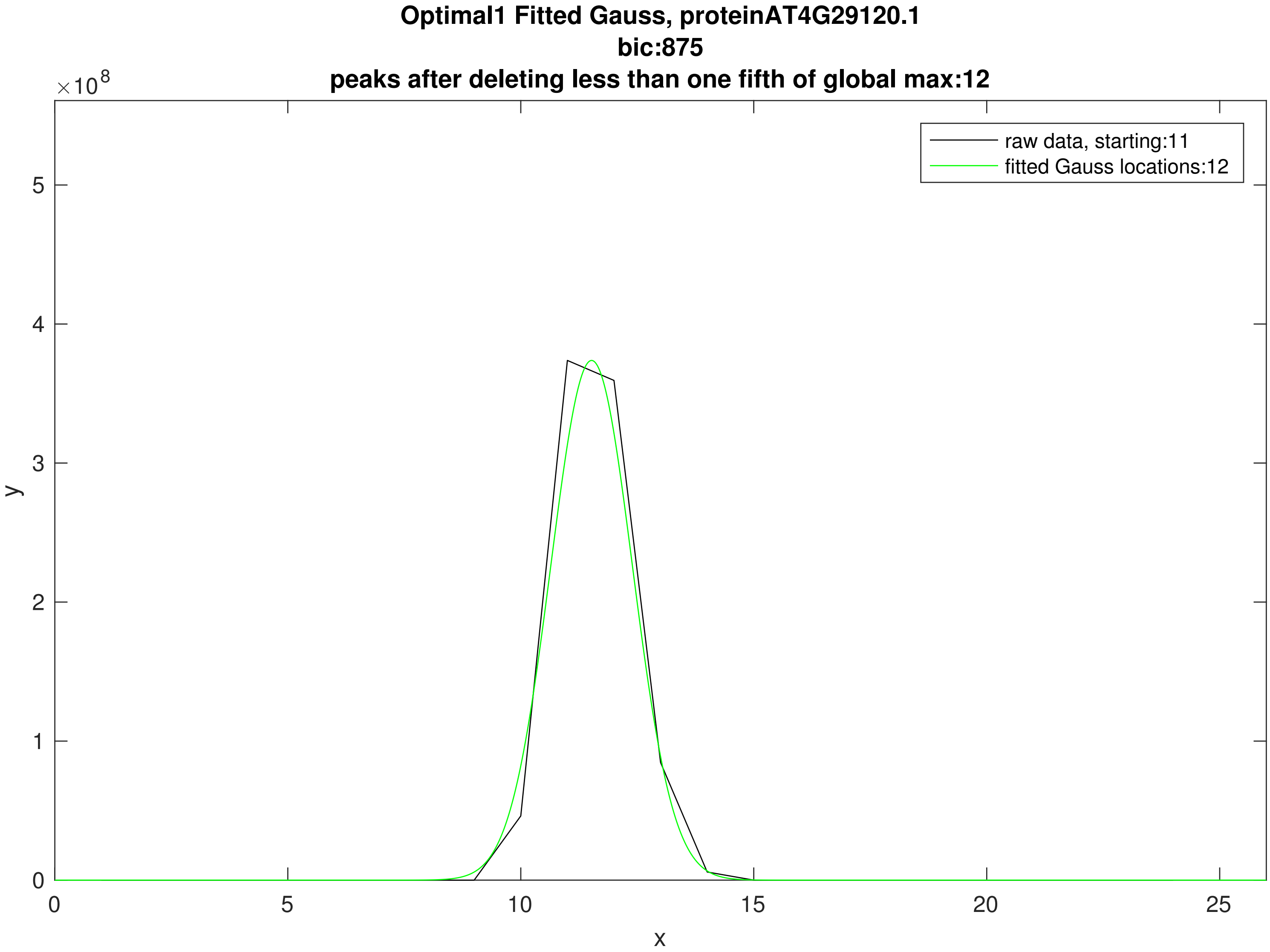
Optimal1 Fitted Gauss, proteinAT4G29120.1
bic:875
peaks after deleting less than one fifth of global max:12
×10^8
raw data, starting:11
fitted Gauss locations:12
5
4
3
2
1
0
y
0
5
10
15
20
25
x

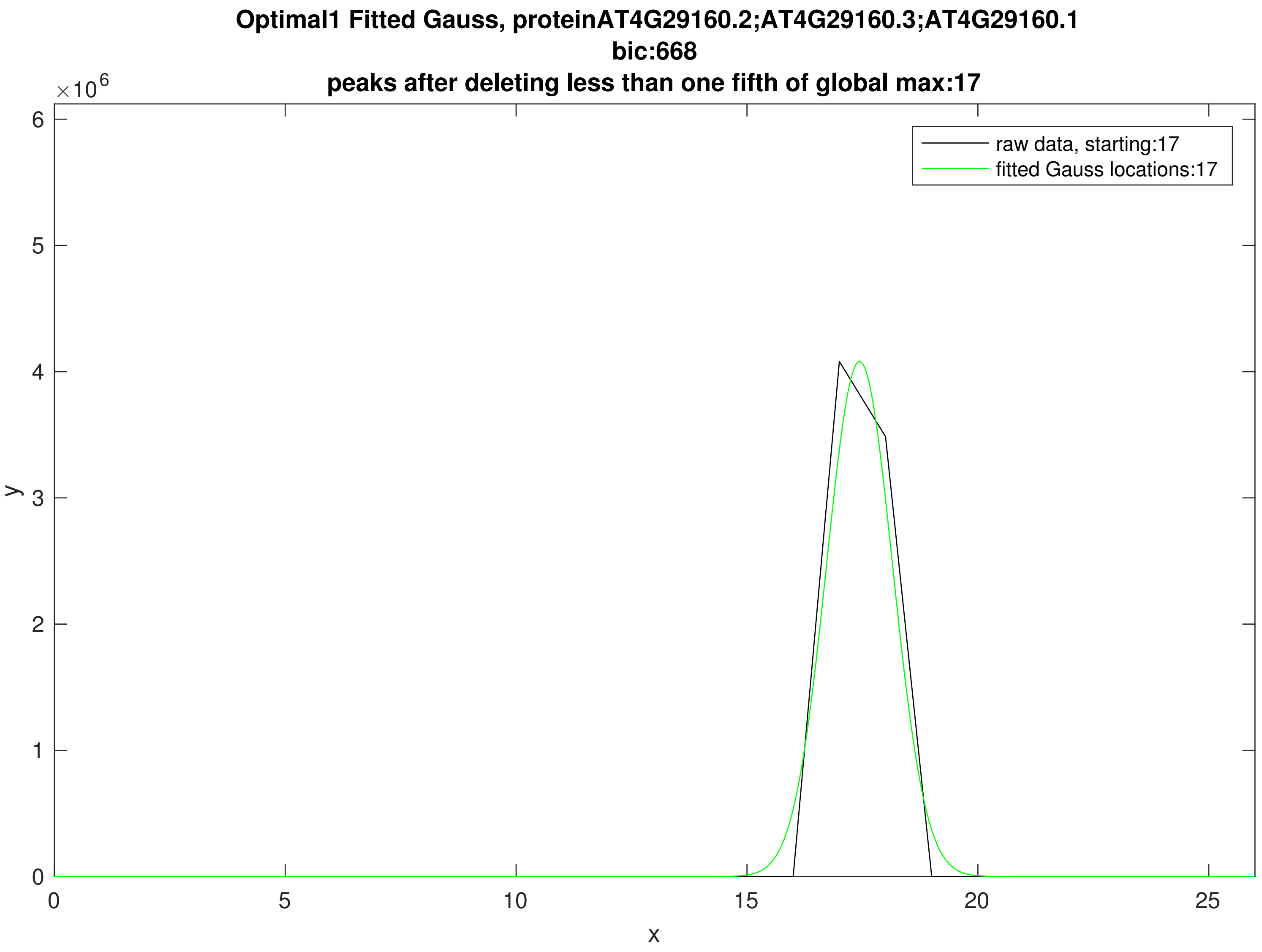

Optimal1 Fitted Gauss, proteinAT4G29160.2;AT4G29160.3;AT4G29160.1
bic:668
peaks after deleting less than one fifth of global max:17
×10^6
raw data, starting:17
fitted Gauss locations:17
y
x
0
1
2
3
4
5
6
0
5
10
15
20
25

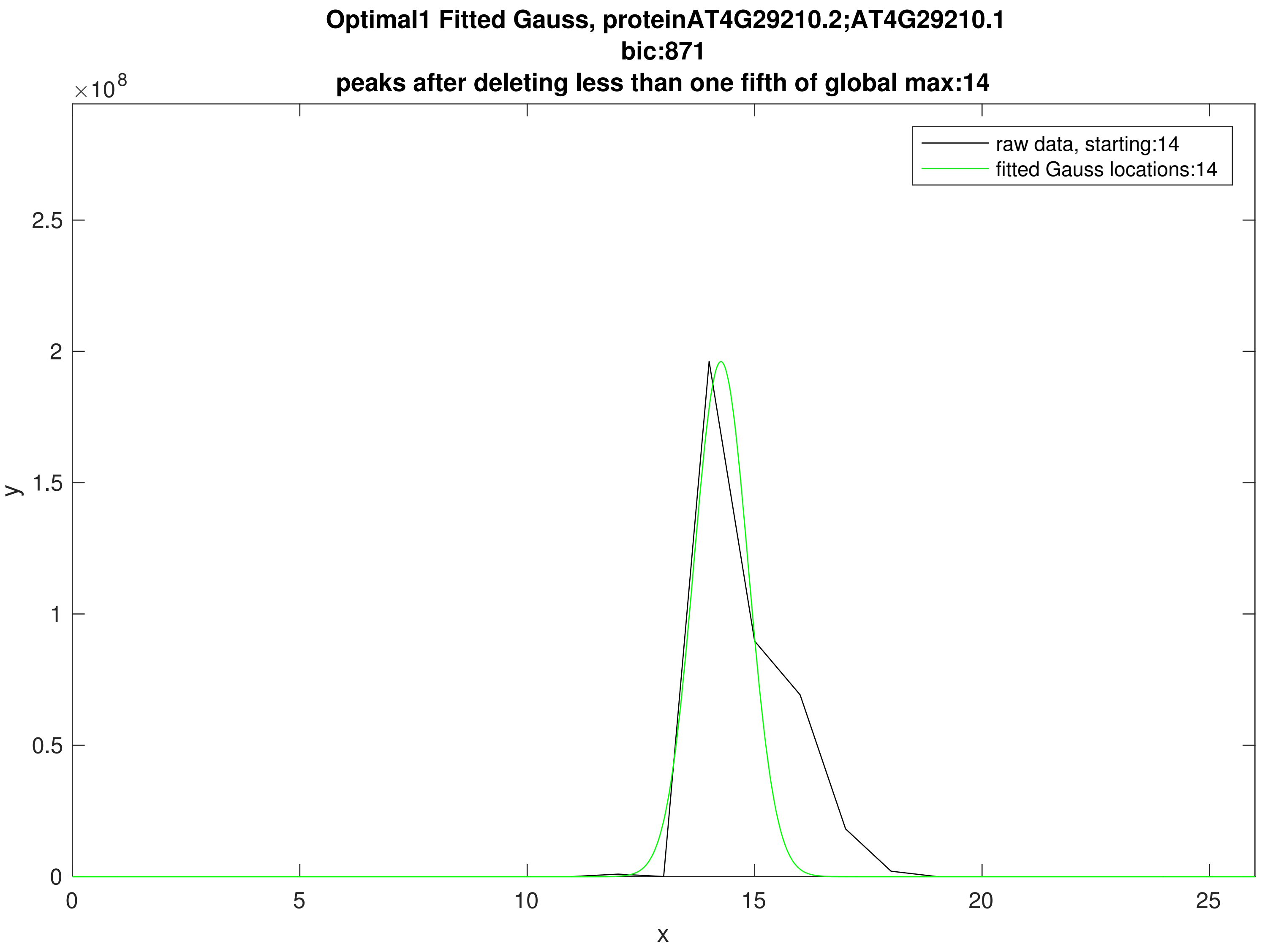
Optimal1 Fitted Gauss, proteinAT4G29210.2;AT4G29210.1
bic:871
peaks after deleting less than one fifth of global max:14
×10^8
raw data, starting:14
fitted Gauss locations:14
y
x
0
0.5
1
1.5
2
2.5
0
5
10
15
20
25

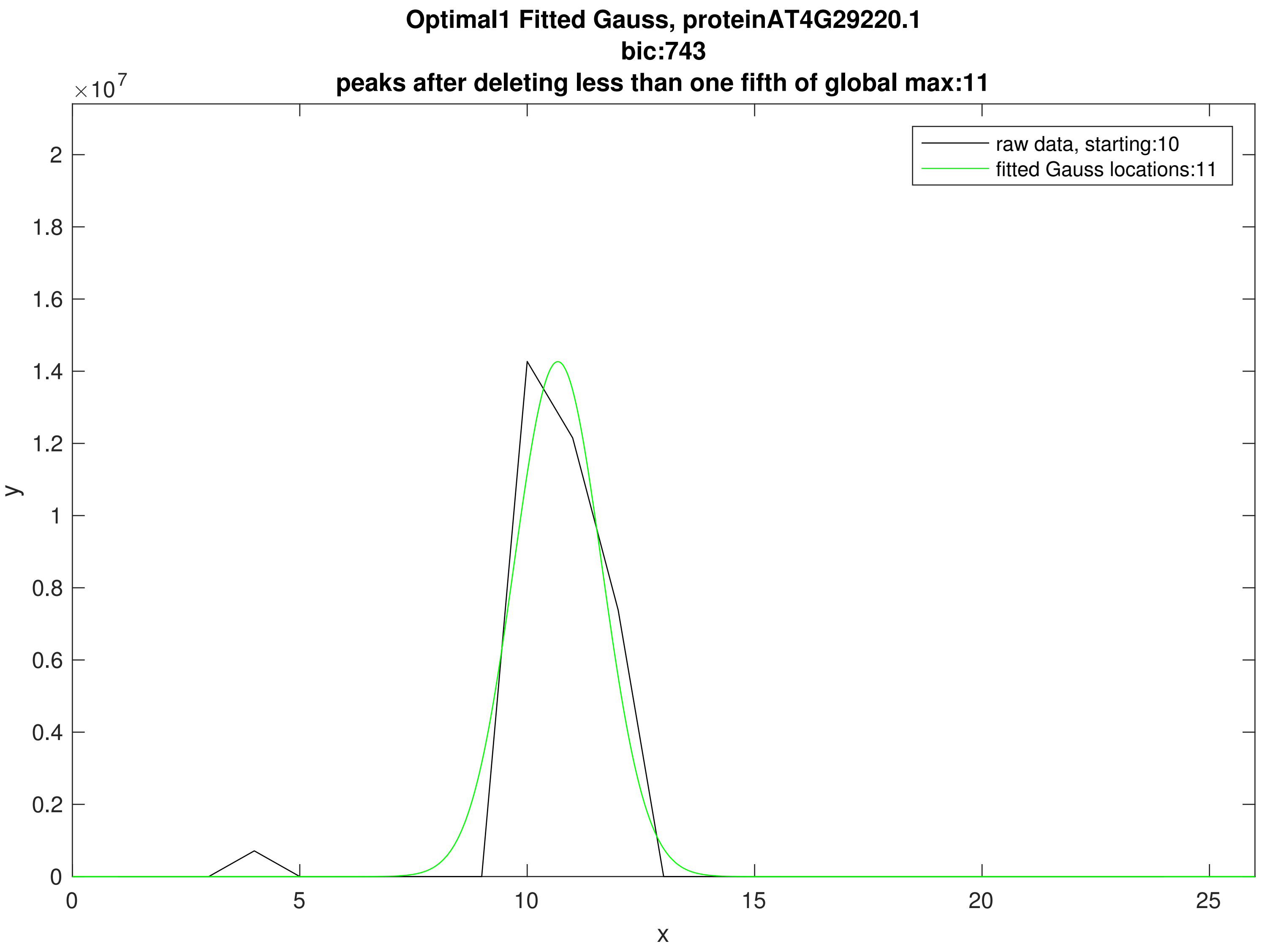
Optimal1 Fitted Gauss, proteinAT4G29220.1
bic:743
peaks after deleting less than one fifth of global max:11
×10^7
raw data, starting:10
fitted Gauss locations:11
y
x
0
0.2
0.4
0.6
0.8
1
1.2
1.4
1.6
1.8
2
0
5
10
15
20
25

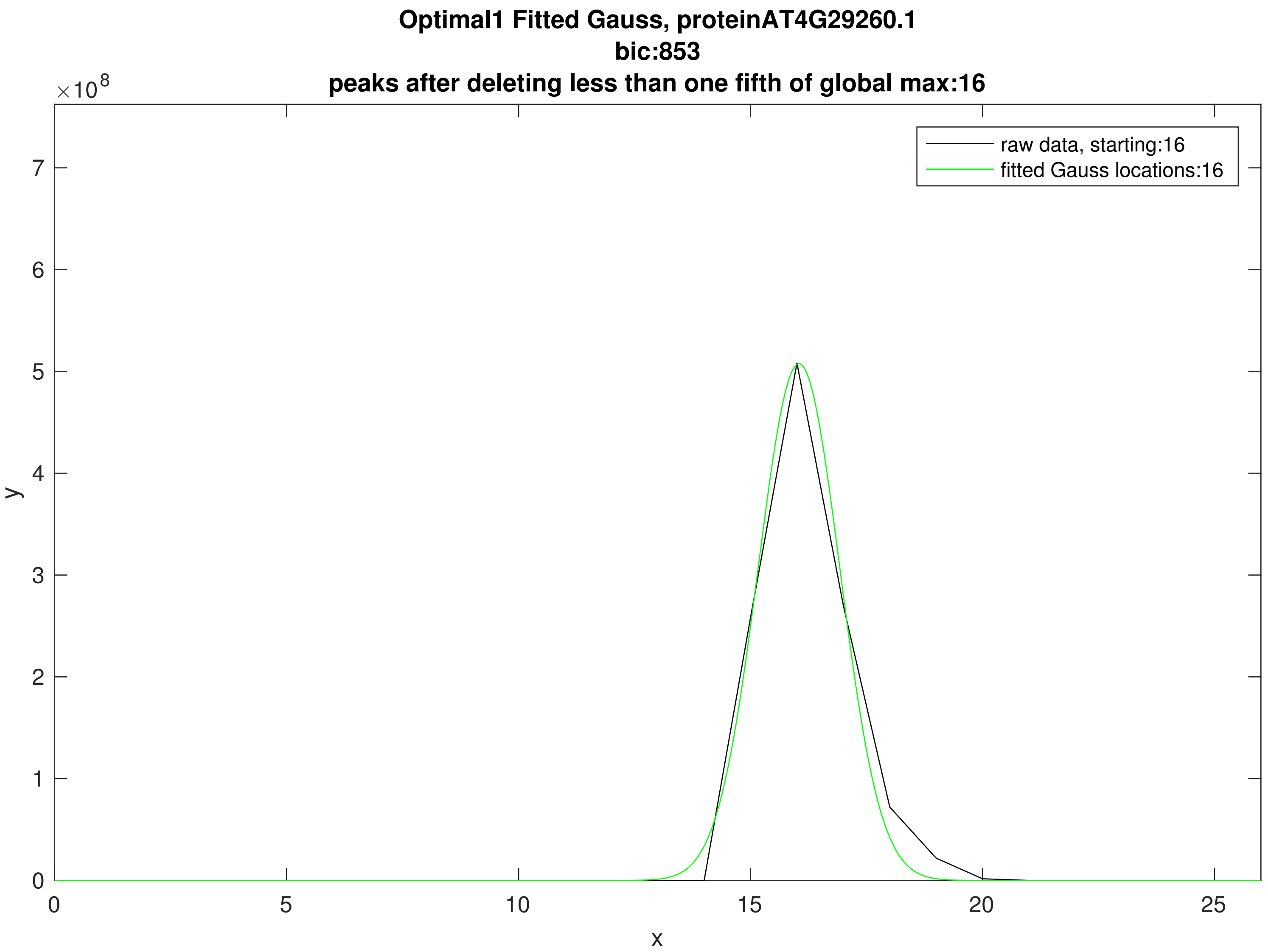
Optimal1 Fitted Gauss, proteinAT4G29260.1
bic:853
peaks after deleting less than one fifth of global max:16
×10^8
raw data, starting:16
fitted Gauss locations:16
y
x
0
1
2
3
4
5
6
7
0
5
10
15
20
25

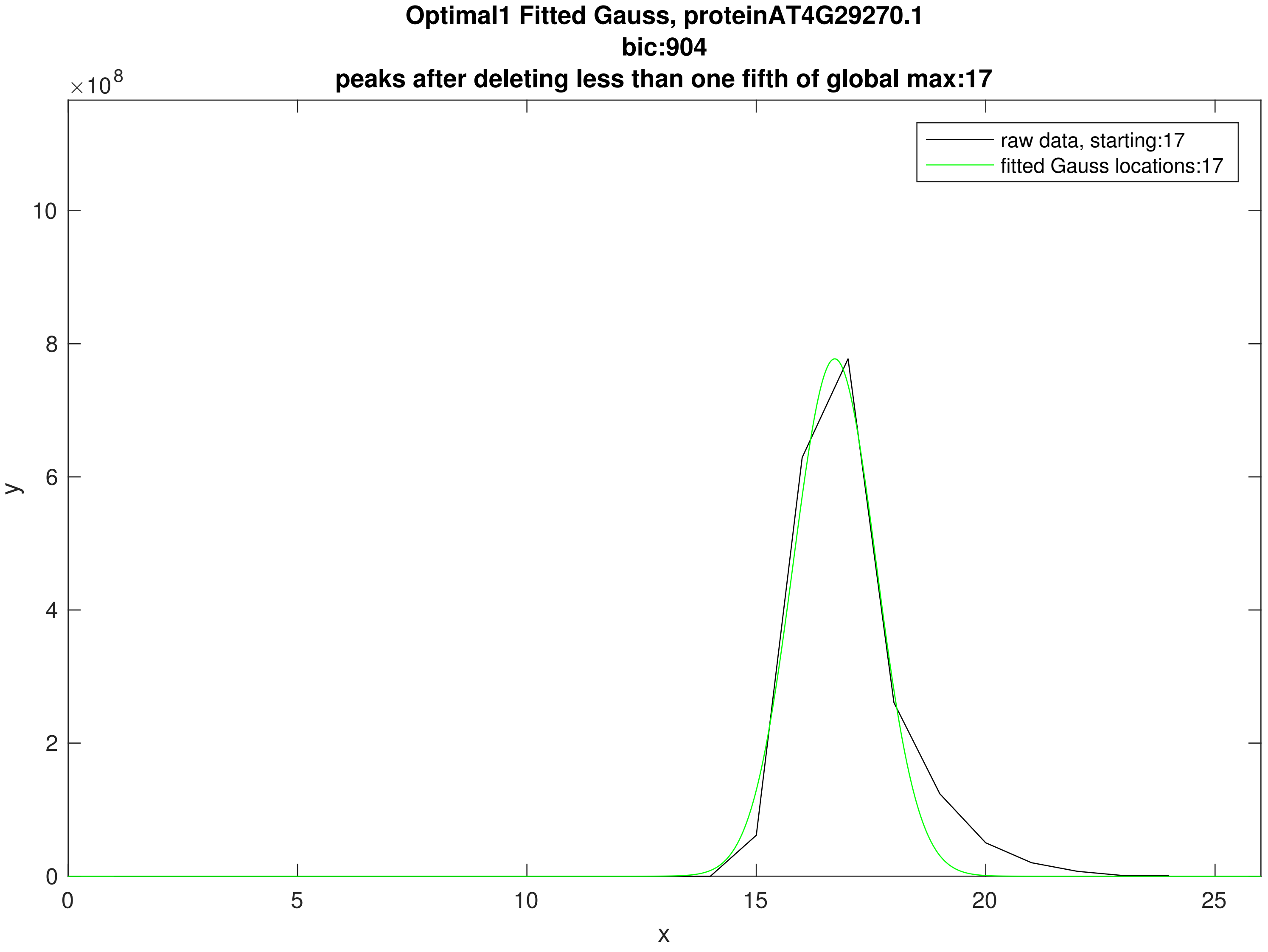
Optimal1 Fitted Gauss, proteinAT4G29270.1
bic:904
peaks after deleting less than one fifth of global max:17
×10^8
raw data, starting:17
fitted Gauss locations:17
y
x
0
2
4
6
8
10
0
5
10
15
20
25

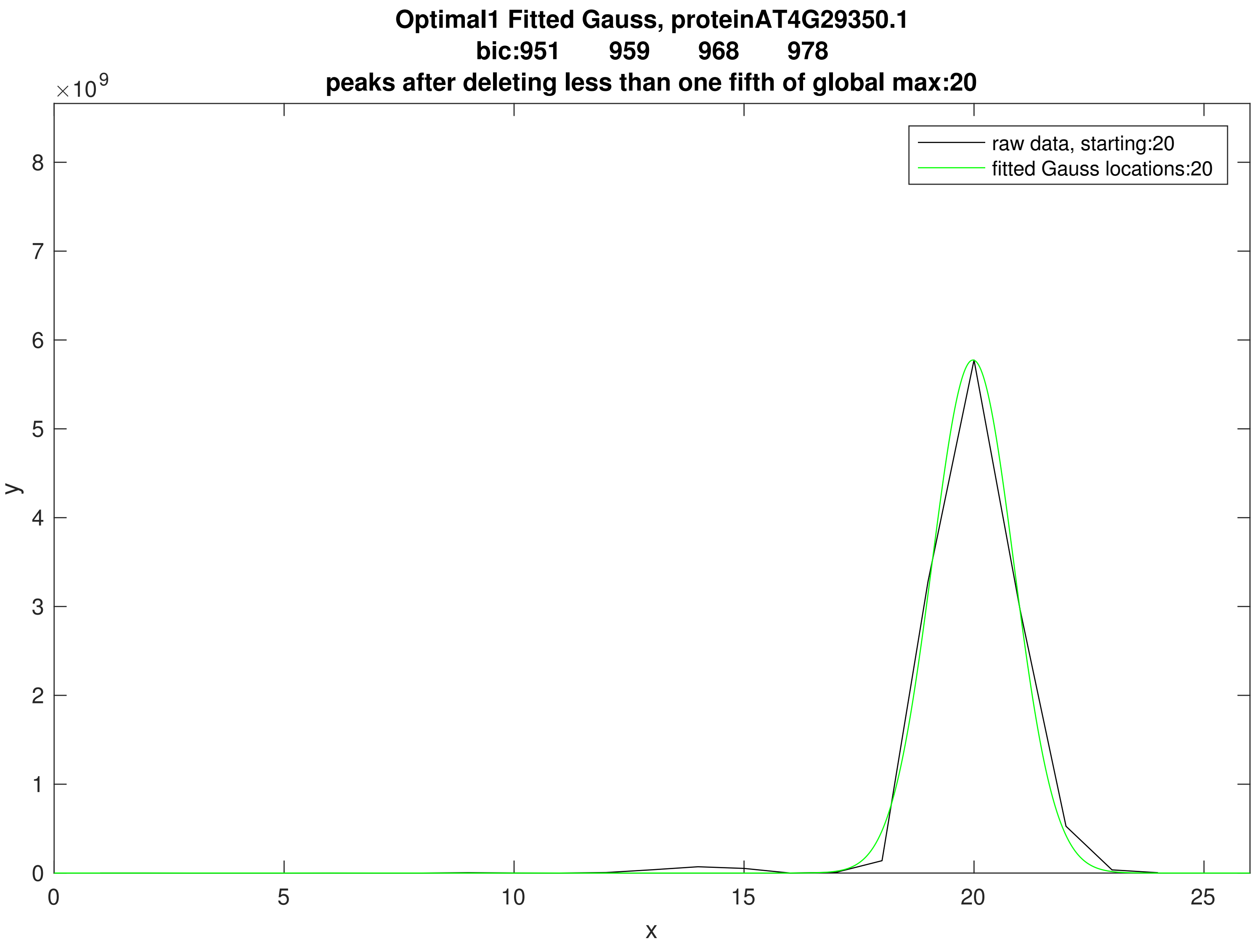
Optimal1 Fitted Gauss, proteinAT4G29350.1
bic:951 959 968 978
peaks after deleting less than one fifth of global max:20
×10^9
raw data, starting:20
fitted Gauss locations:20
y
x
0
1
2
3
4
5
6
7
8
0
5
10
15
20
25

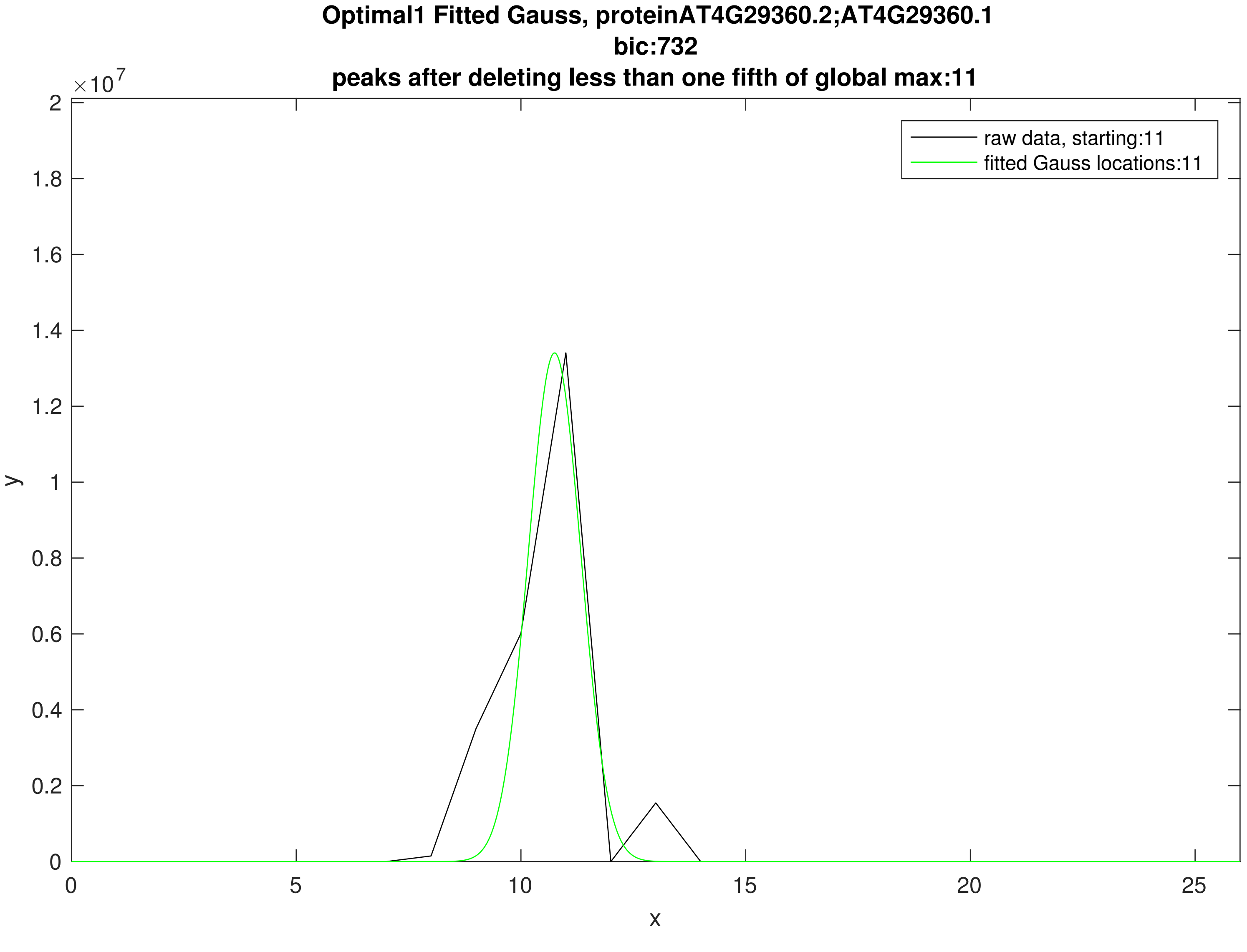
Optimal1 Fitted Gauss, proteinAT4G29360.2;AT4G29360.1
bic:732
peaks after deleting less than one fifth of global max:11
×10^7
raw data, starting:11
fitted Gauss locations:11
y
x
0
0.2
0.4
0.6
0.8
1
1.2
1.4
1.6
1.8
2
0
5
10
15
20
25

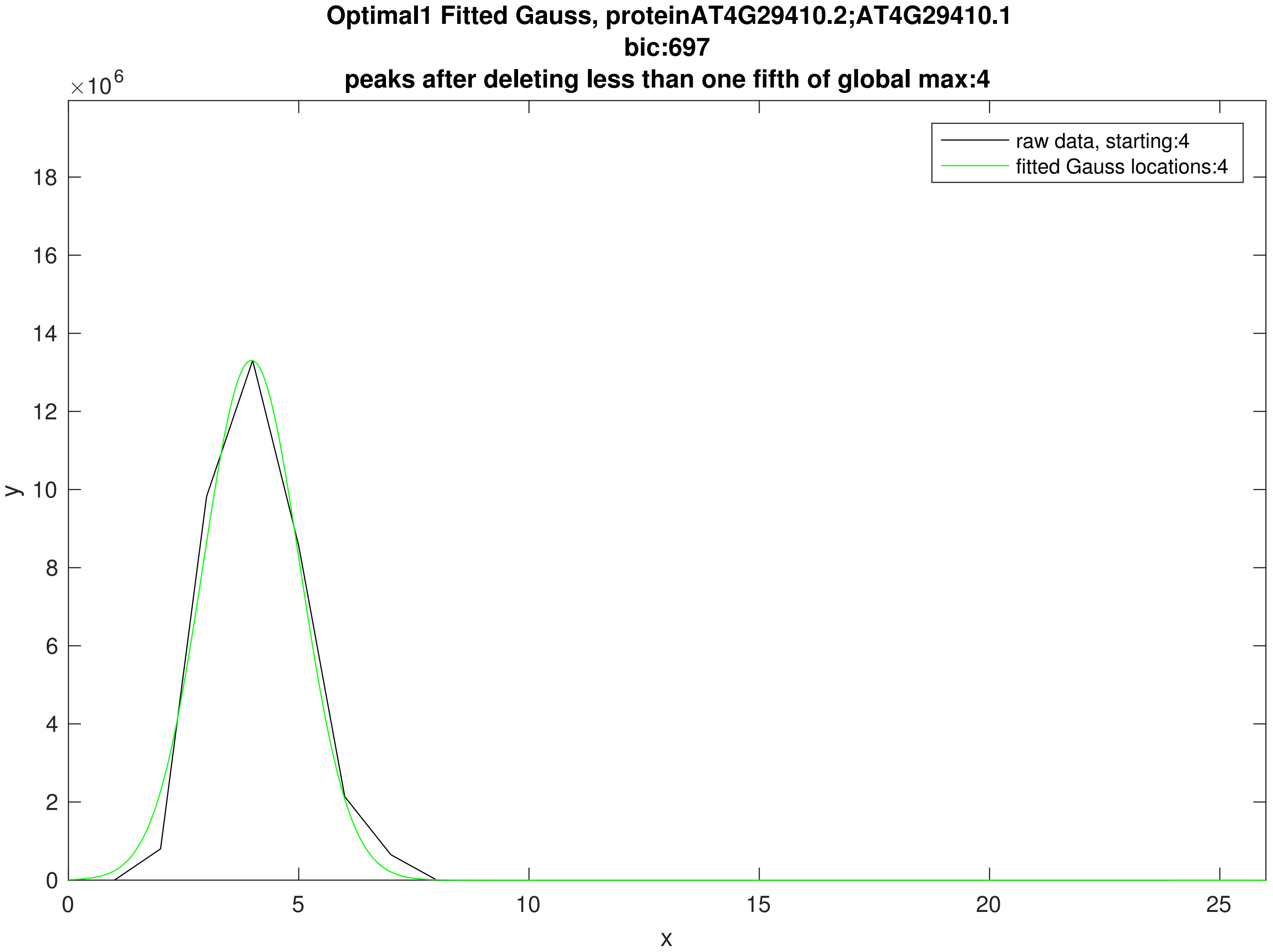
Optimal1 Fitted Gauss, proteinAT4G29410.2;AT4G29410.1
bic:697
peaks after deleting less than one fifth of global max:4
×10^6
raw data, starting:4
fitted Gauss locations:4
y
x
0
2
4
6
8
10
12
14
16
18
0
5
10
15
20
25

**Optimal1 Fitted Gauss, proteinAT4G29490.1**
**bic:762**
**peaks after deleting less than one fifth of global max:13**

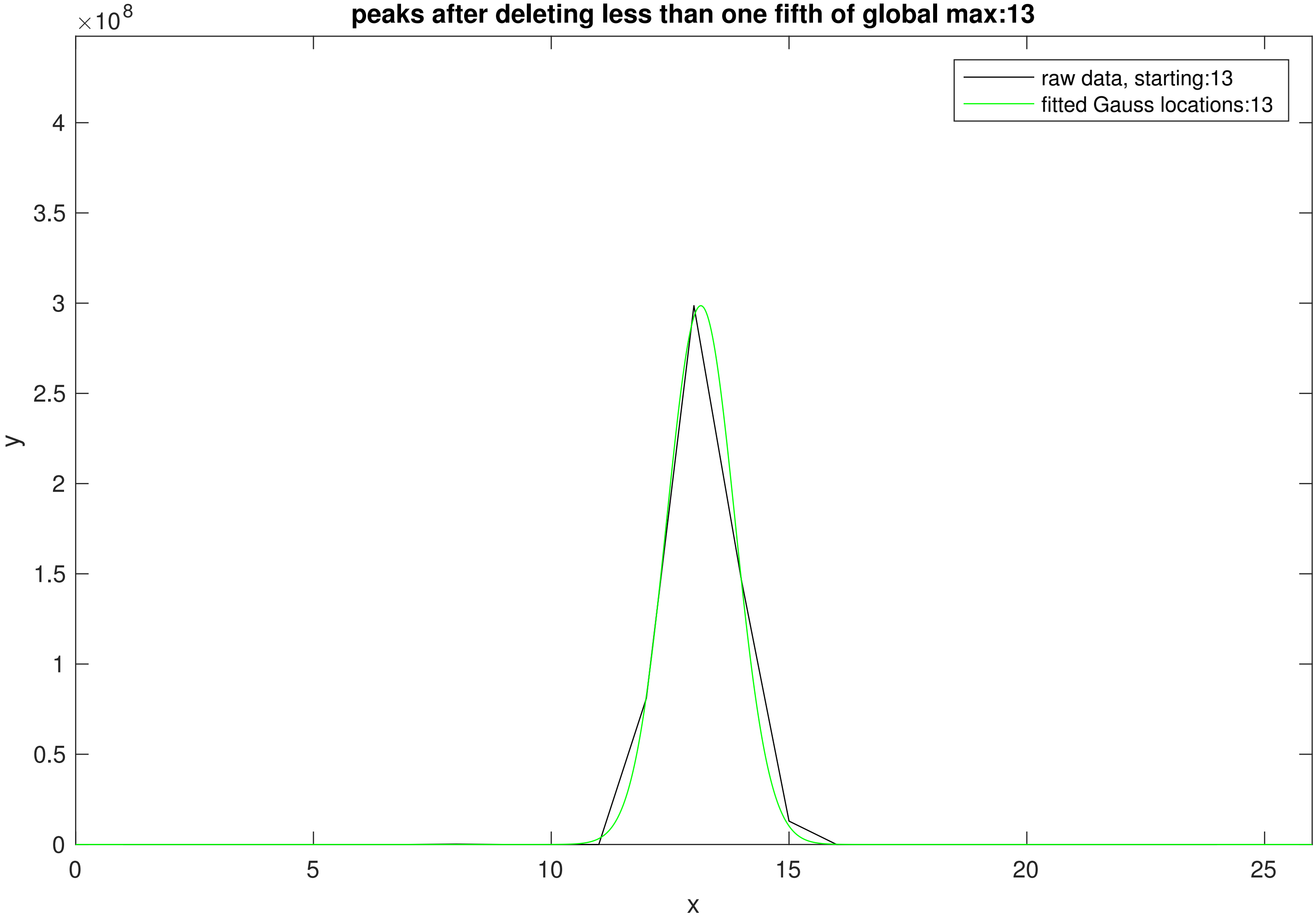

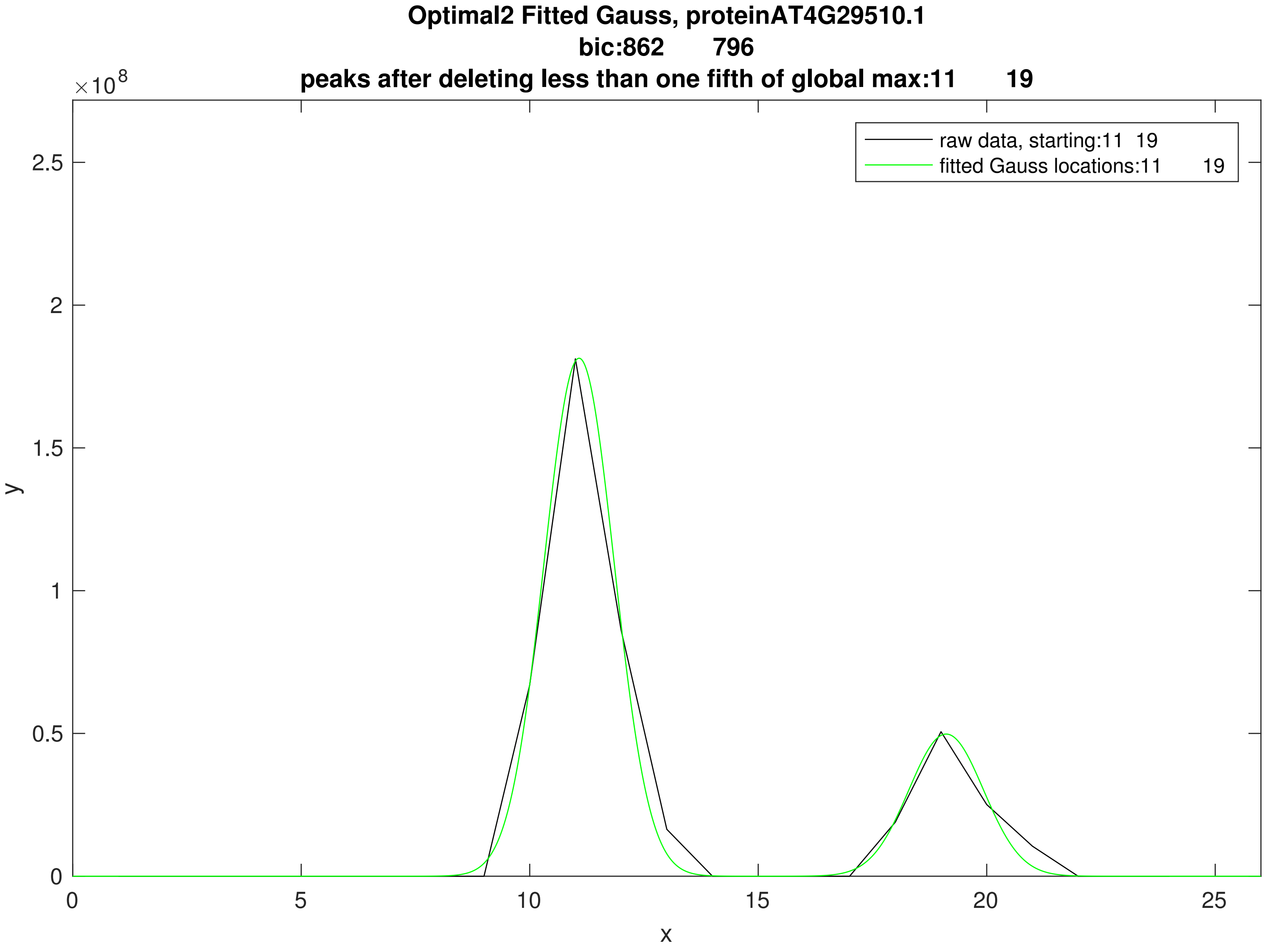

Optimal2 Fitted Gauss, proteinAT4G29510.1
bic:862 796
peaks after deleting less than one fifth of global max:11 19
×10^8
raw data, starting:11 19
fitted Gauss locations:11 19
2.5
2
1.5
1
0.5
0
y
0
5
10
15
20
25
x

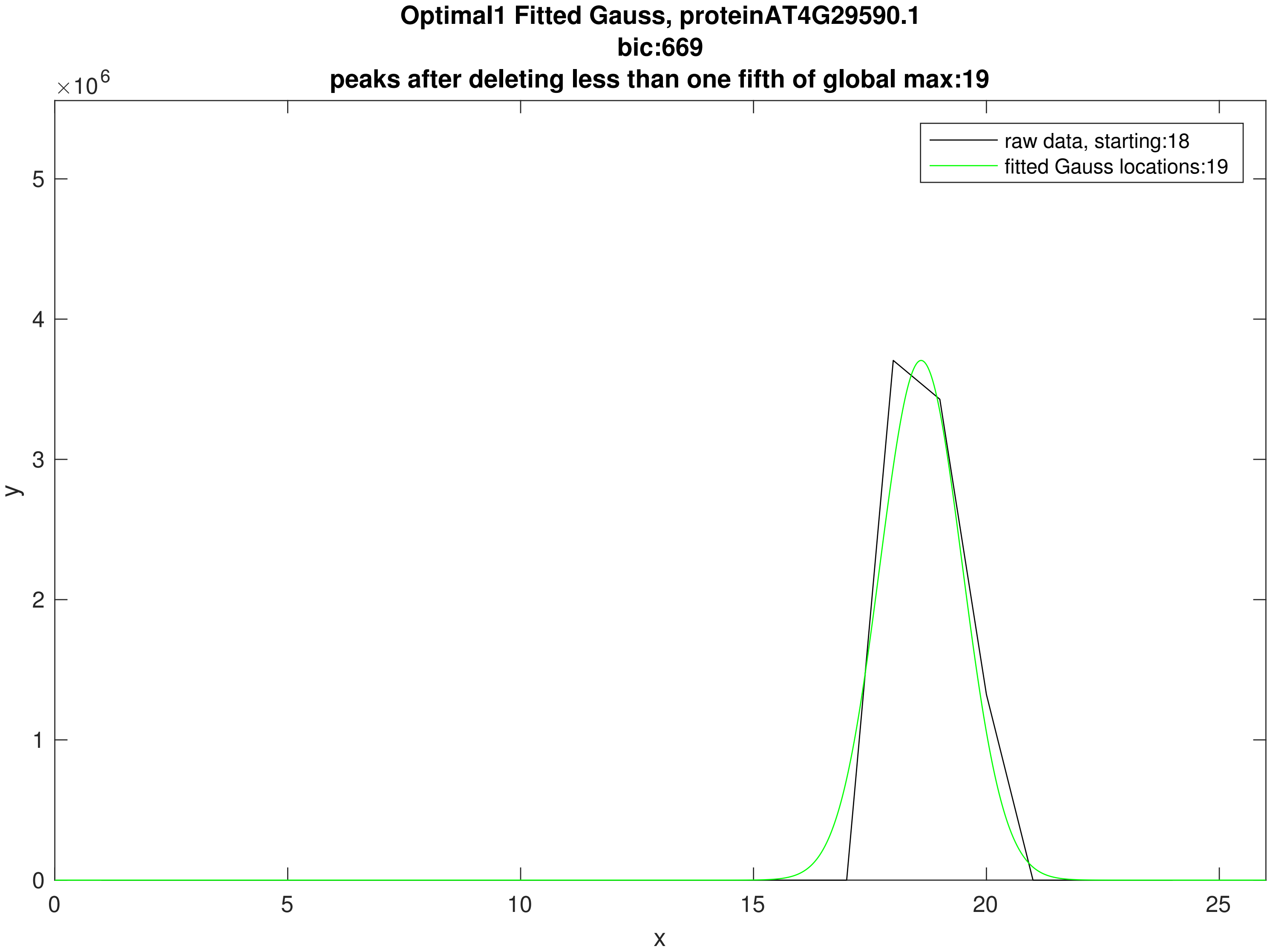
Optimal1 Fitted Gauss, proteinAT4G29590.1
bic:669
peaks after deleting less than one fifth of global max:19
×10^6
raw data, starting:18
fitted Gauss locations:19
y
x
0
1
2
3
4
5
0
5
10
15
20
25

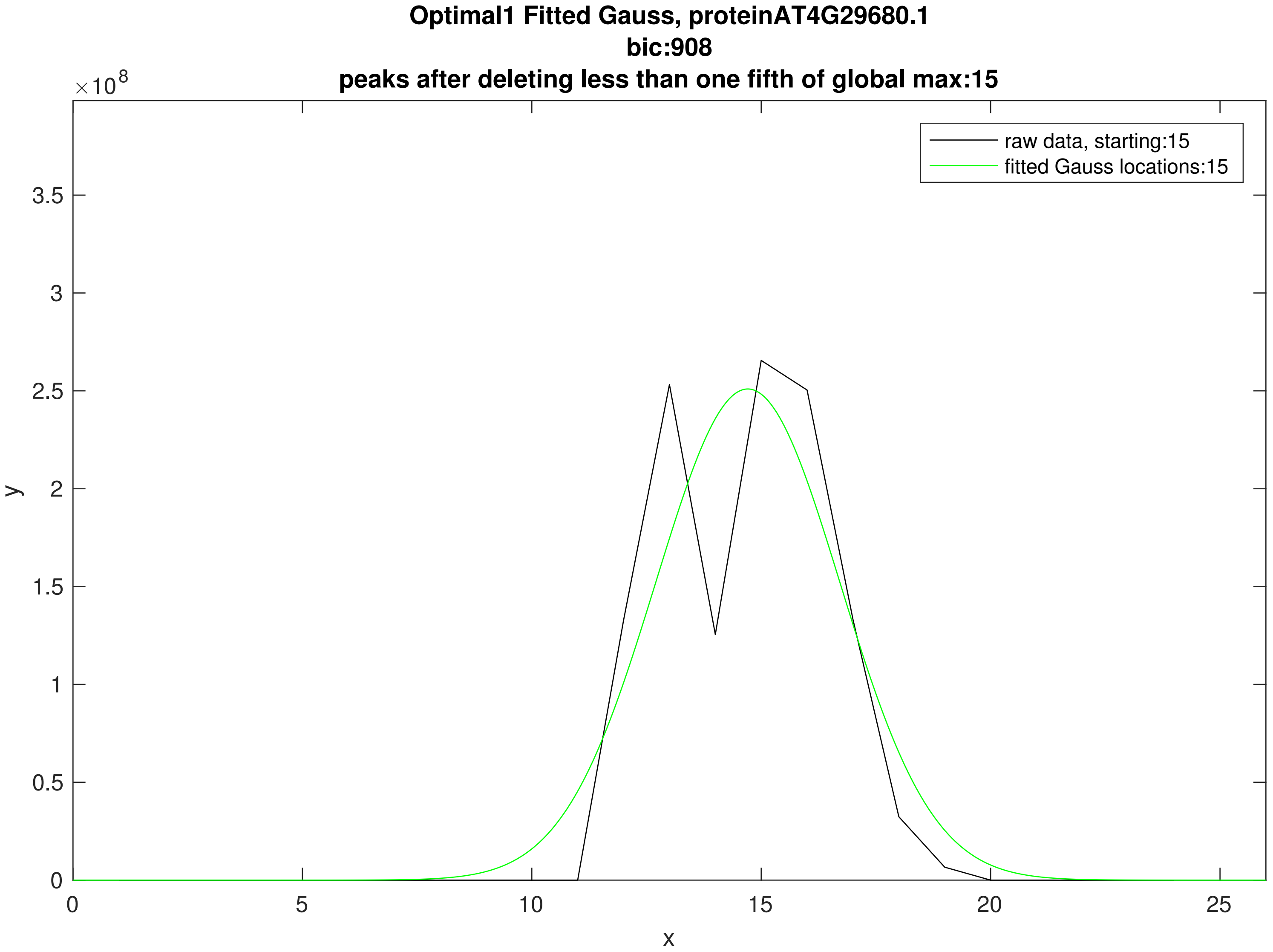
Optimal1 Fitted Gauss, proteinAT4G29680.1
bic:908
peaks after deleting less than one fifth of global max:15
×10^8
raw data, starting:15
fitted Gauss locations:15
3.5
3
2.5
2
1.5
1
0.5
0
y
0
5
10
15
20
25
x

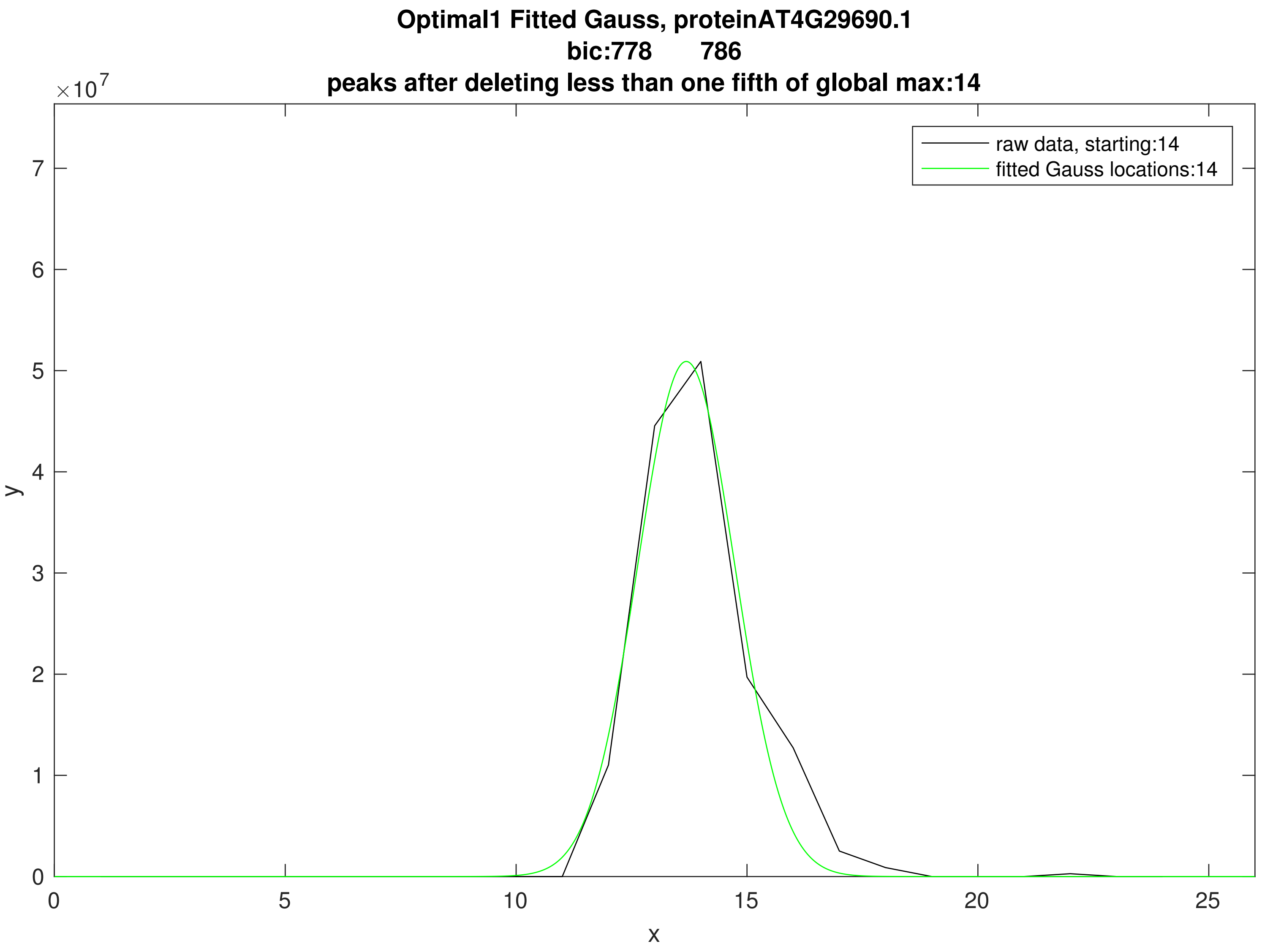

Optimal1 Fitted Gauss, proteinAT4G29690.1
bic:778 786
peaks after deleting less than one fifth of global max:14
×10^7
raw data, starting:14
fitted Gauss locations:14
y
x
0
1
2
3
4
5
6
7
0
5
10
15
20
25

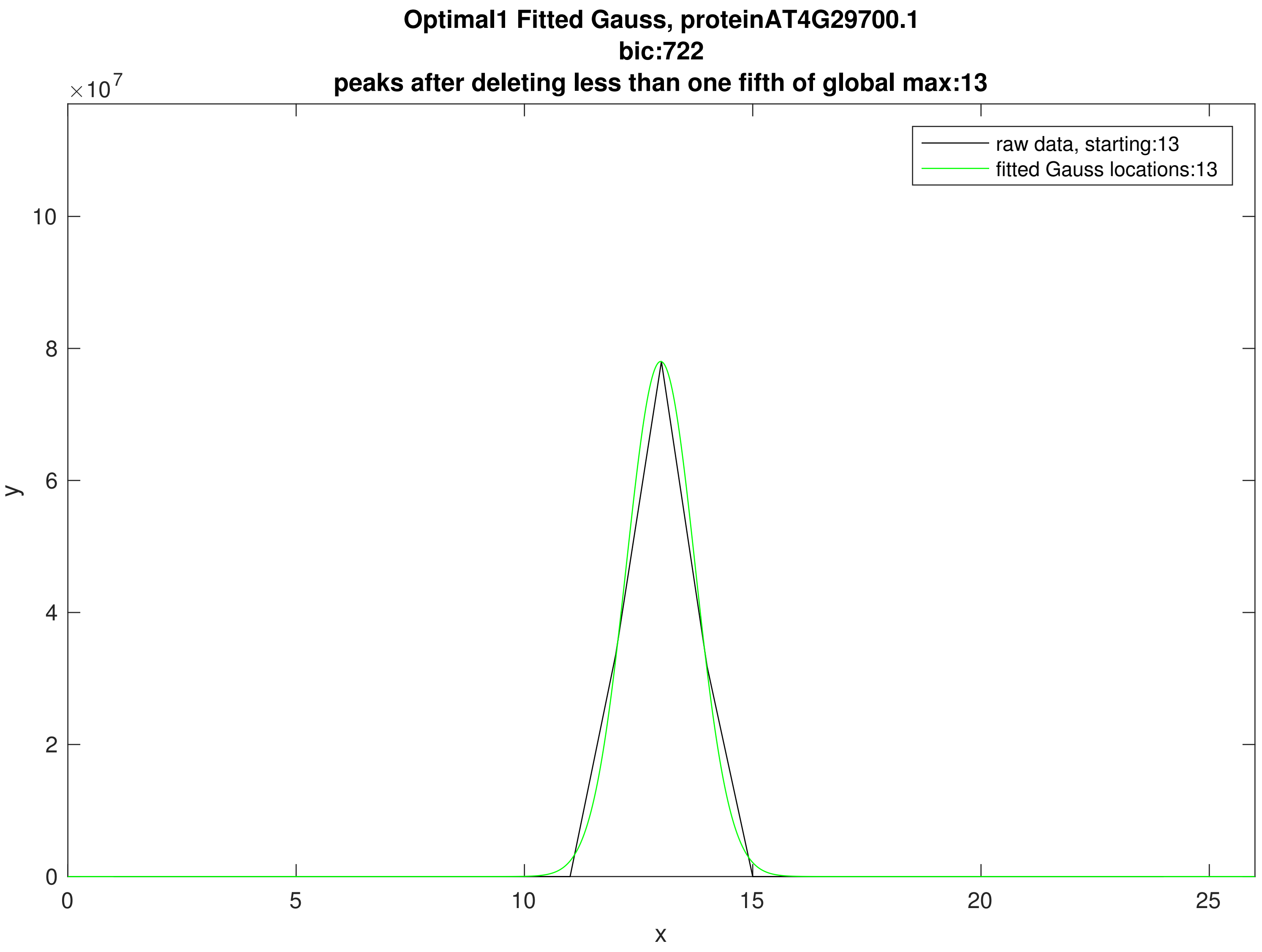
Optimal1 Fitted Gauss, proteinAT4G29700.1
bic:722
peaks after deleting less than one fifth of global max:13
×10^7
raw data, starting:13
fitted Gauss locations:13
y
x
0
2
4
6
8
10
0
5
10
15
20
25

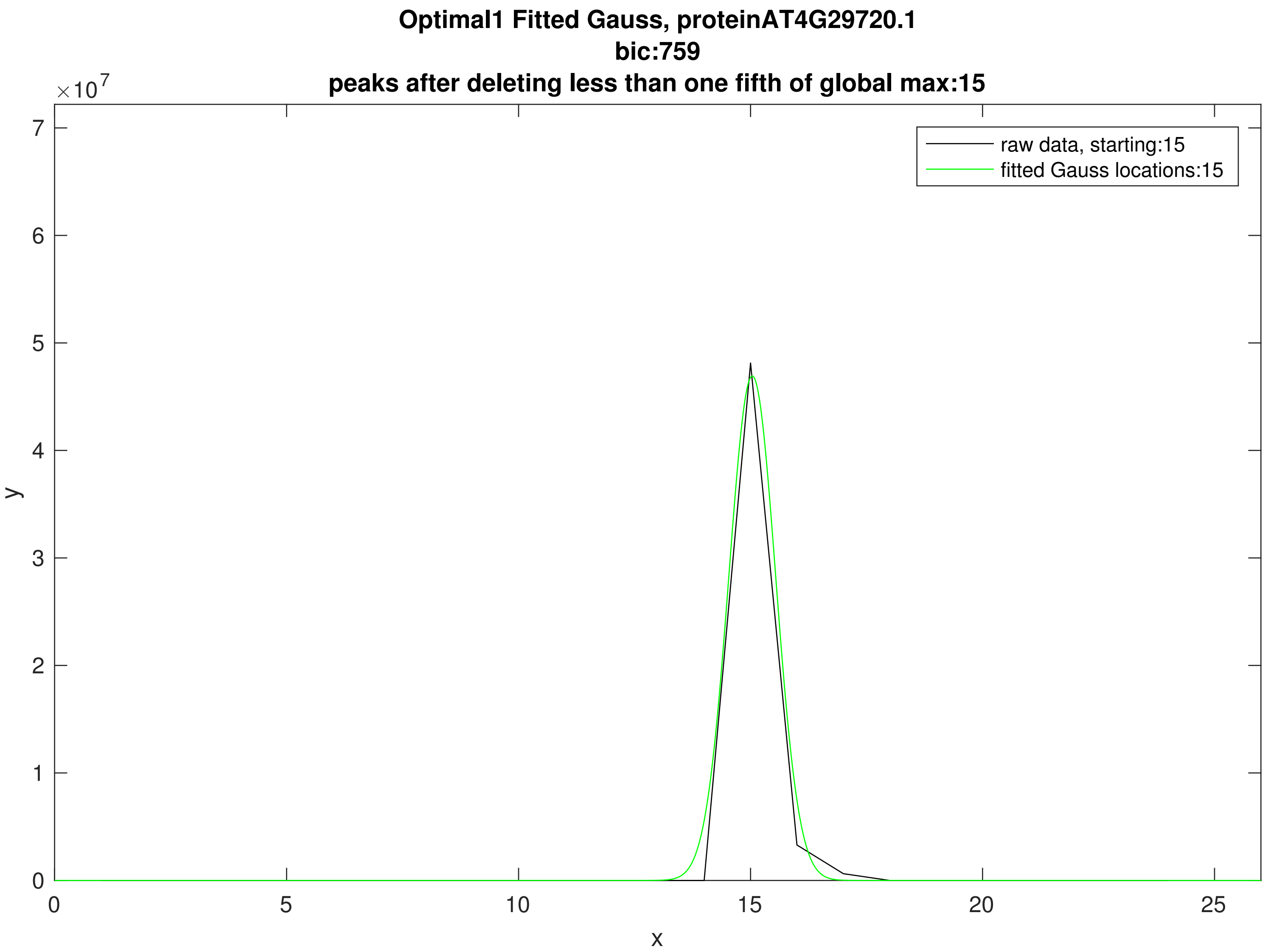
Optimal1 Fitted Gauss, proteinAT4G29720.1
bic:759
peaks after deleting less than one fifth of global max:15
raw data, starting:15
fitted Gauss locations:15
$\times 10^7$
y
x

**Optimal1 Fitted Gauss, proteinAT4G29830.1**
**bic:856**
**peaks after deleting less than one fifth of global max:18**

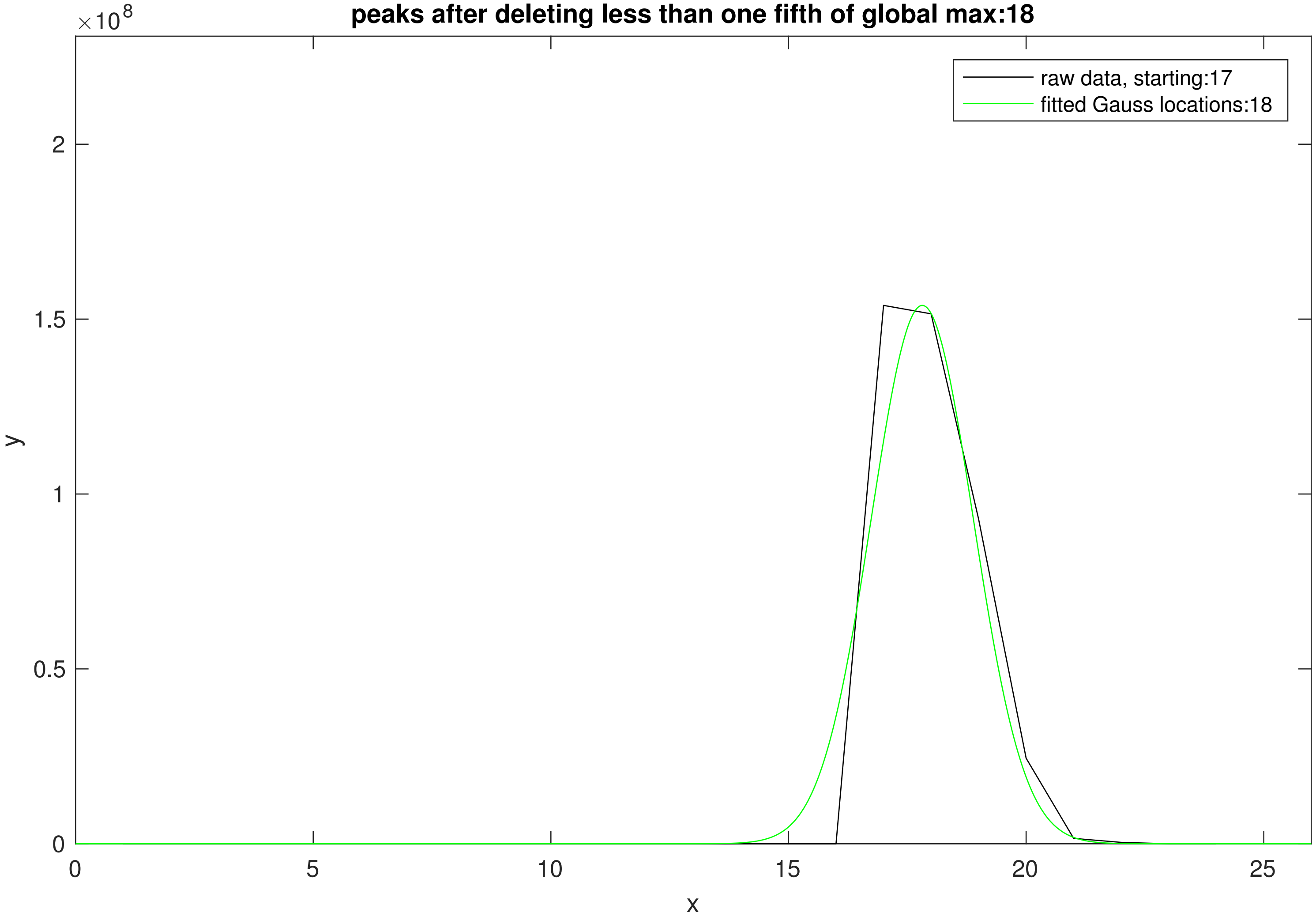

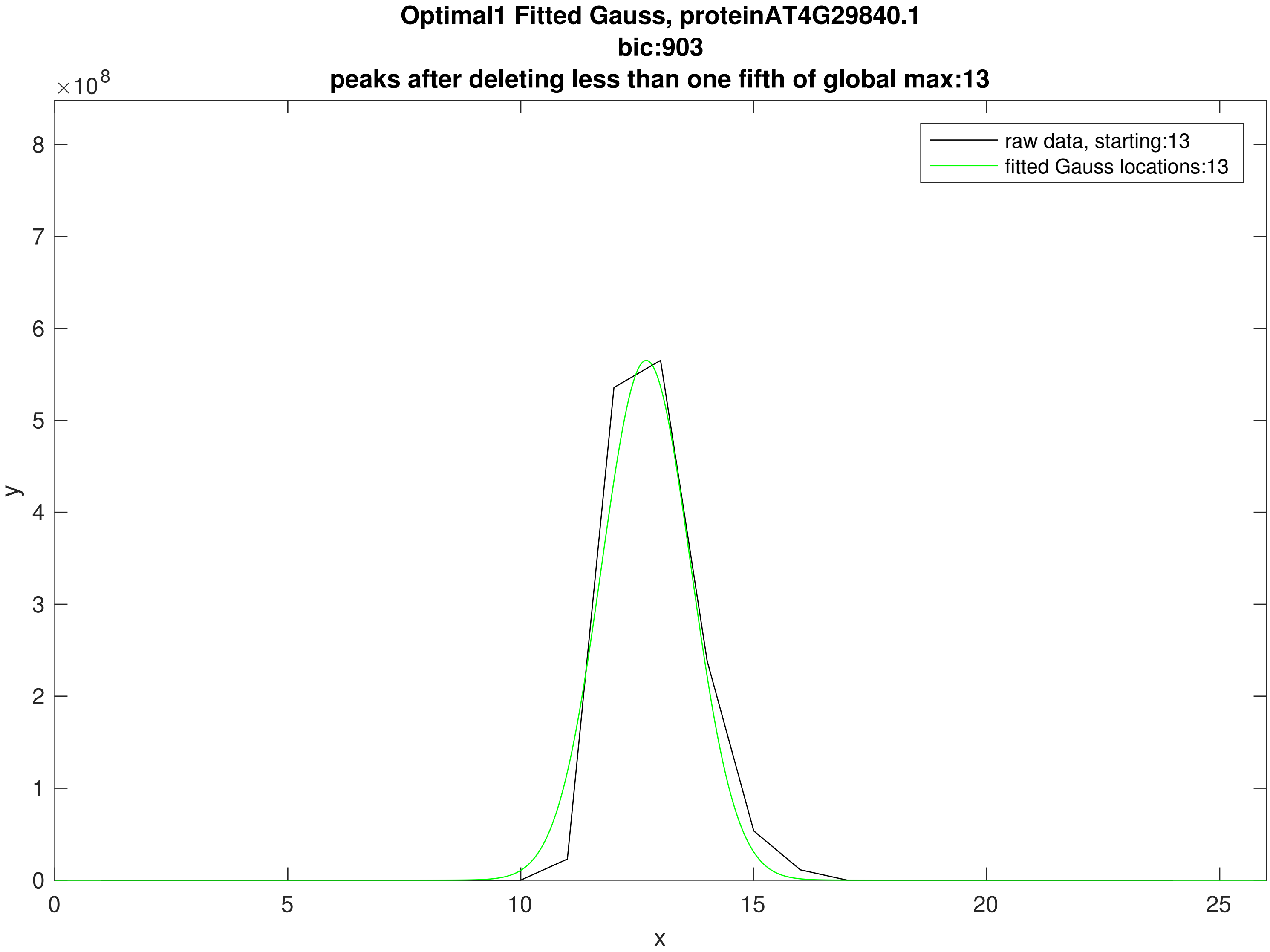

Optimal1 Fitted Gauss, proteinAT4G29840.1
bic:903
peaks after deleting less than one fifth of global max:13
×10^8
raw data, starting:13
fitted Gauss locations:13
y
x
0
1
2
3
4
5
6
7
8
0
5
10
15
20
25

Optimal1 Fitted Gauss, proteinAT4G30160.1;AT4G30160.2
bic:761 770
peaks after deleting less than one fifth of global max:16

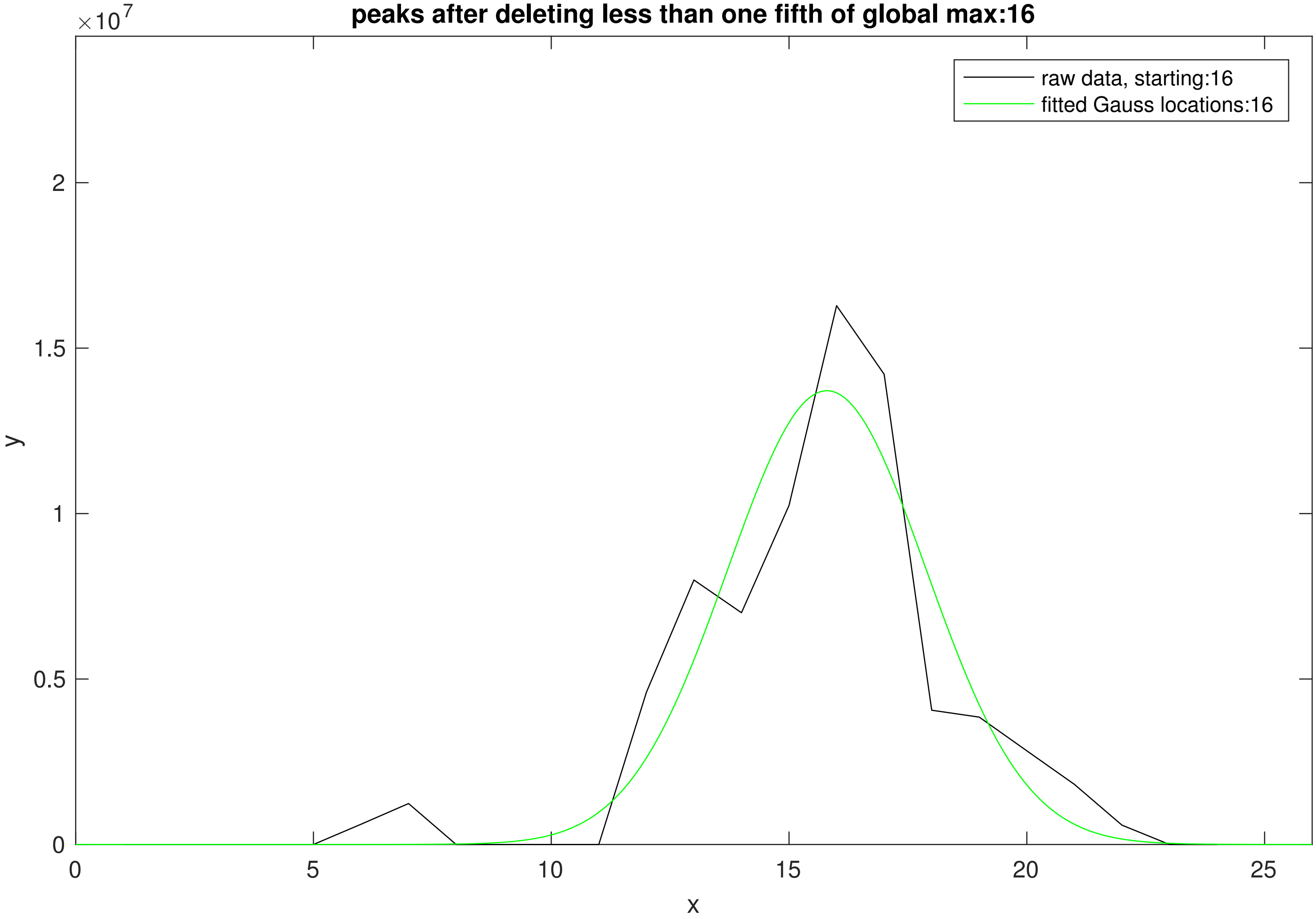

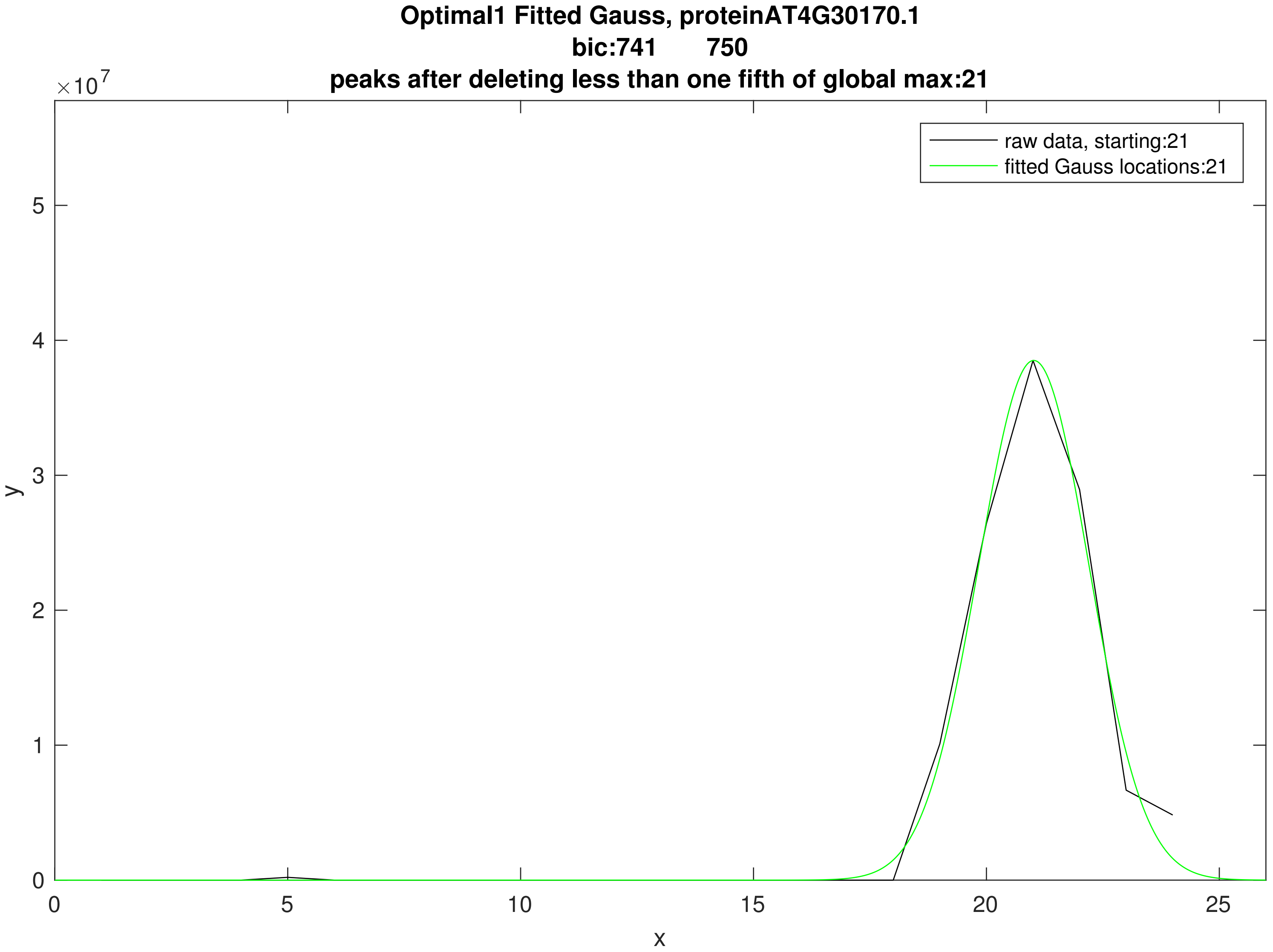
Optimal1 Fitted Gauss, proteinAT4G30170.1
bic:741 750
peaks after deleting less than one fifth of global max:21
×10^7
raw data, starting:21
fitted Gauss locations:21
y
x
0
1
2
3
4
5
0
5
10
15
20
25

Optimal1 Fitted Gauss, proteinAT4G30210.2;AT4G30210.1
bic:757 765
peaks after deleting less than one fifth of global max:18

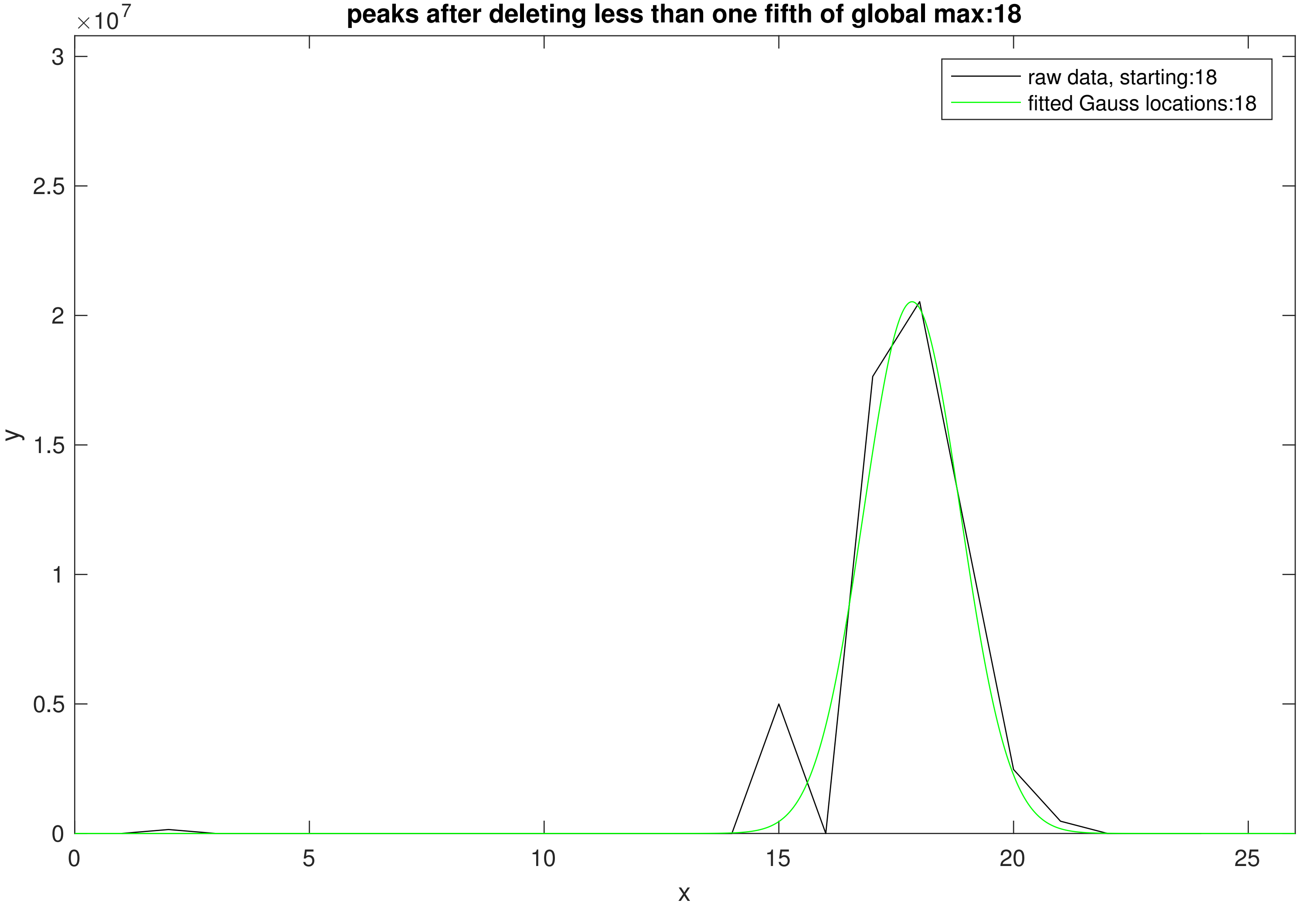

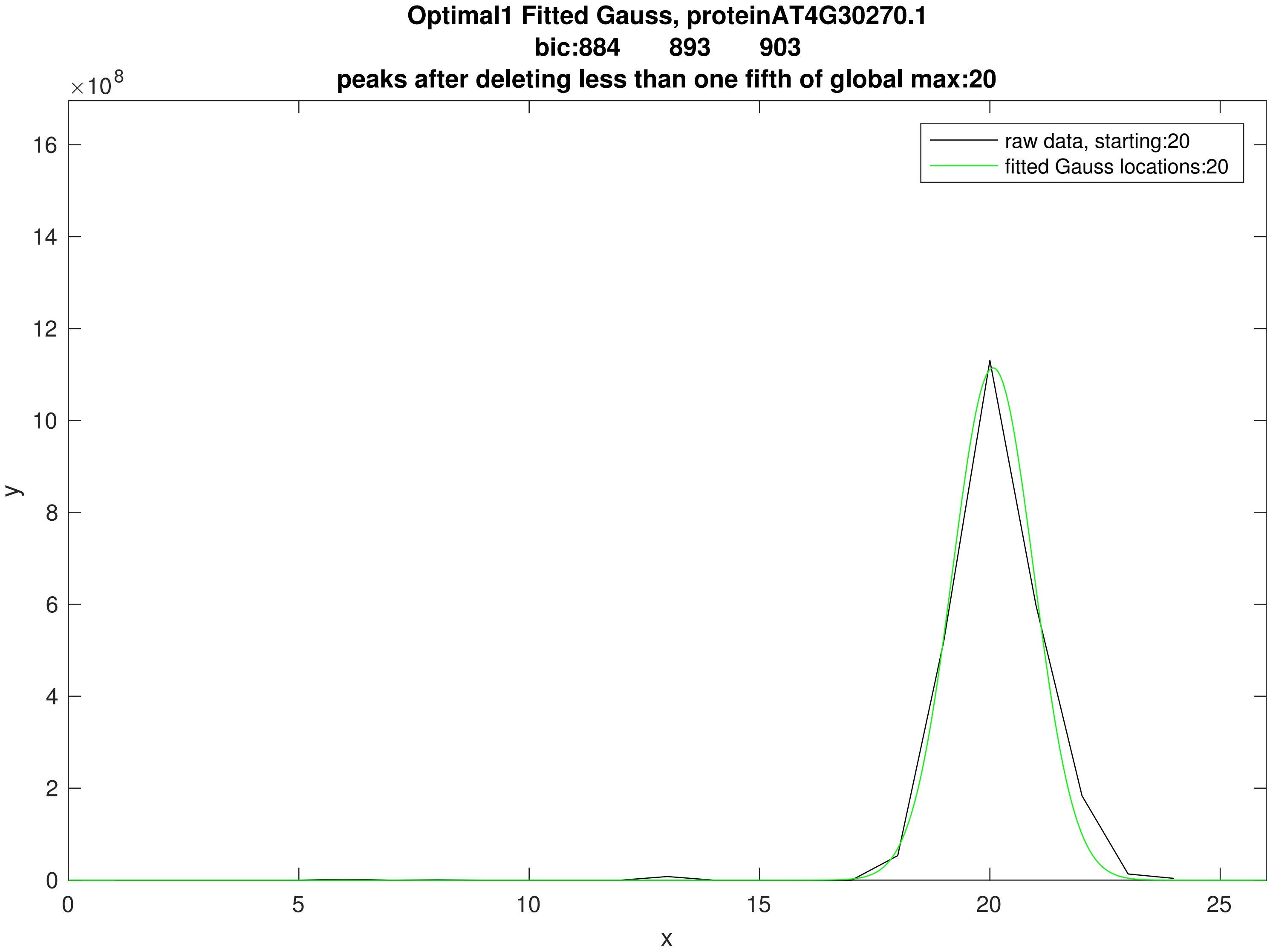

Optimal1 Fitted Gauss, proteinAT4G30270.1
bic:884 893 903
peaks after deleting less than one fifth of global max:20
×10^8
16
14
12
10
8
6
4
2
0
y
0
5
10
15
20
25
x
raw data, starting:20
fitted Gauss locations:20

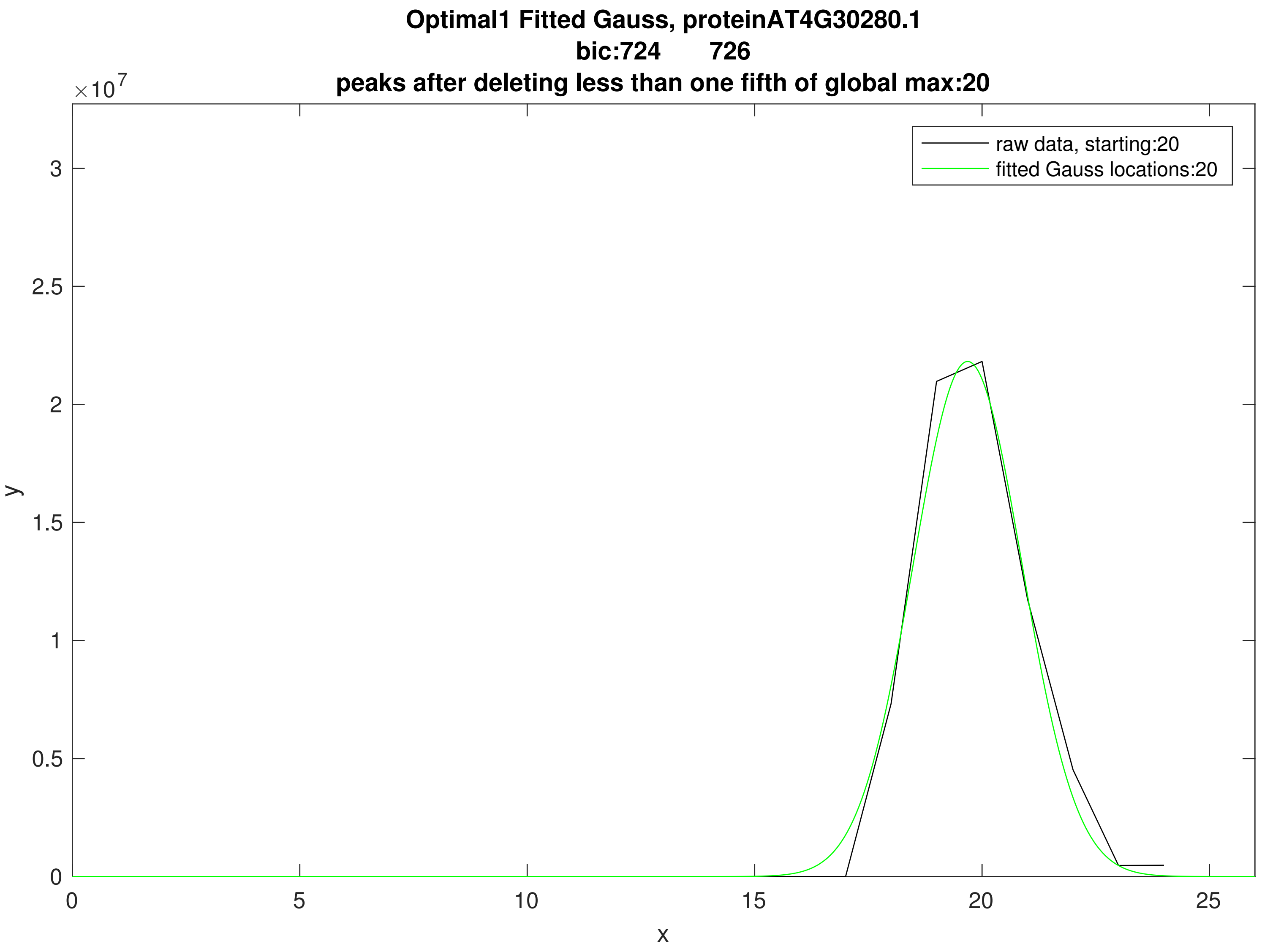
Optimal1 Fitted Gauss, proteinAT4G30280.1
bic:724 726
peaks after deleting less than one fifth of global max:20
×10^7
raw data, starting:20
fitted Gauss locations:20
y
x
0
0.5
1
1.5
2
2.5
3
0
5
10
15
20
25

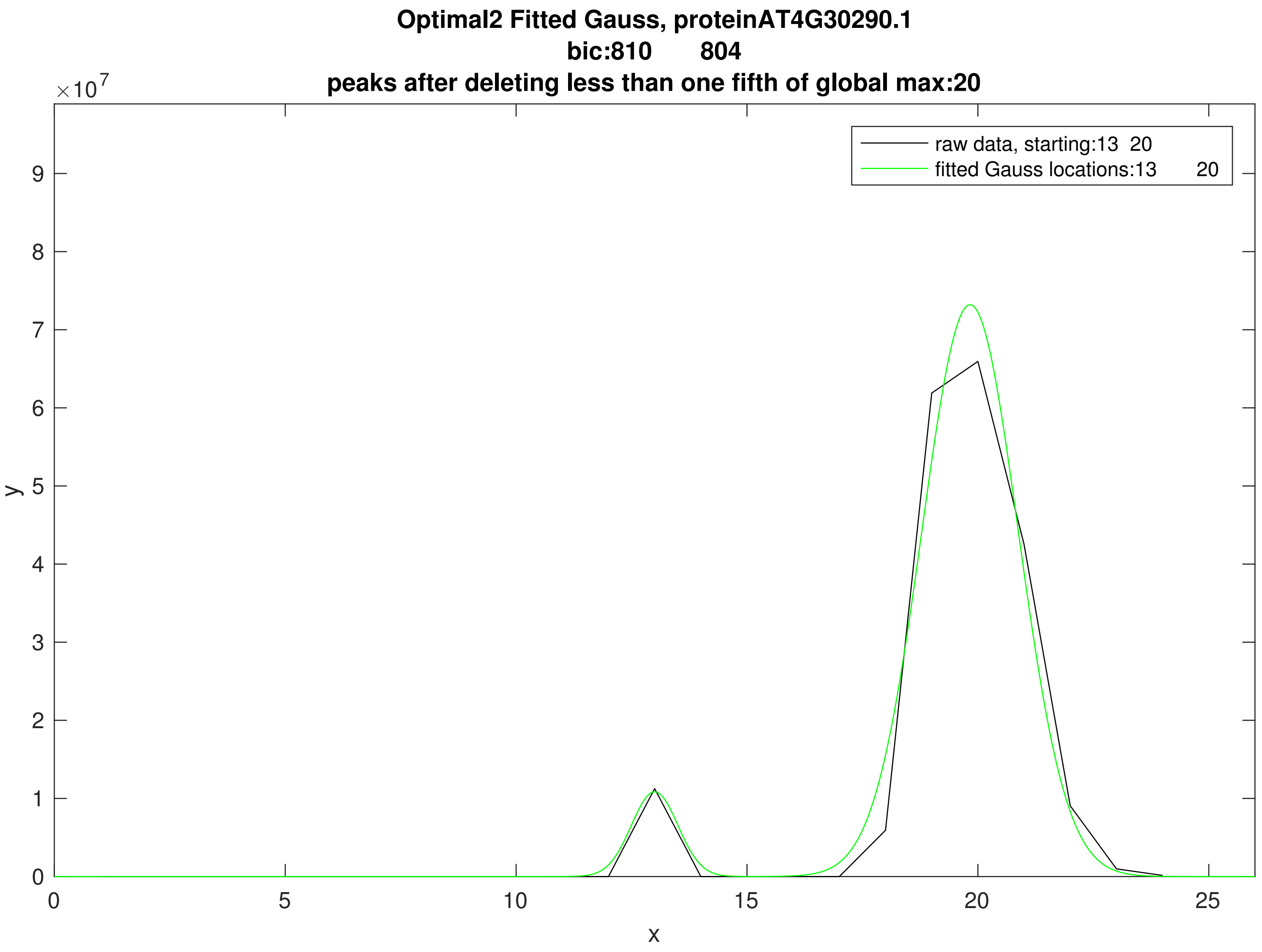
Optimal2 Fitted Gauss, proteinAT4G30290.1
bic:810 804
peaks after deleting less than one fifth of global max:20
×10^7
raw data, starting:13 20
fitted Gauss locations:13 20
y
x
0
1
2
3
4
5
6
7
8
9
0
5
10
15
20
25

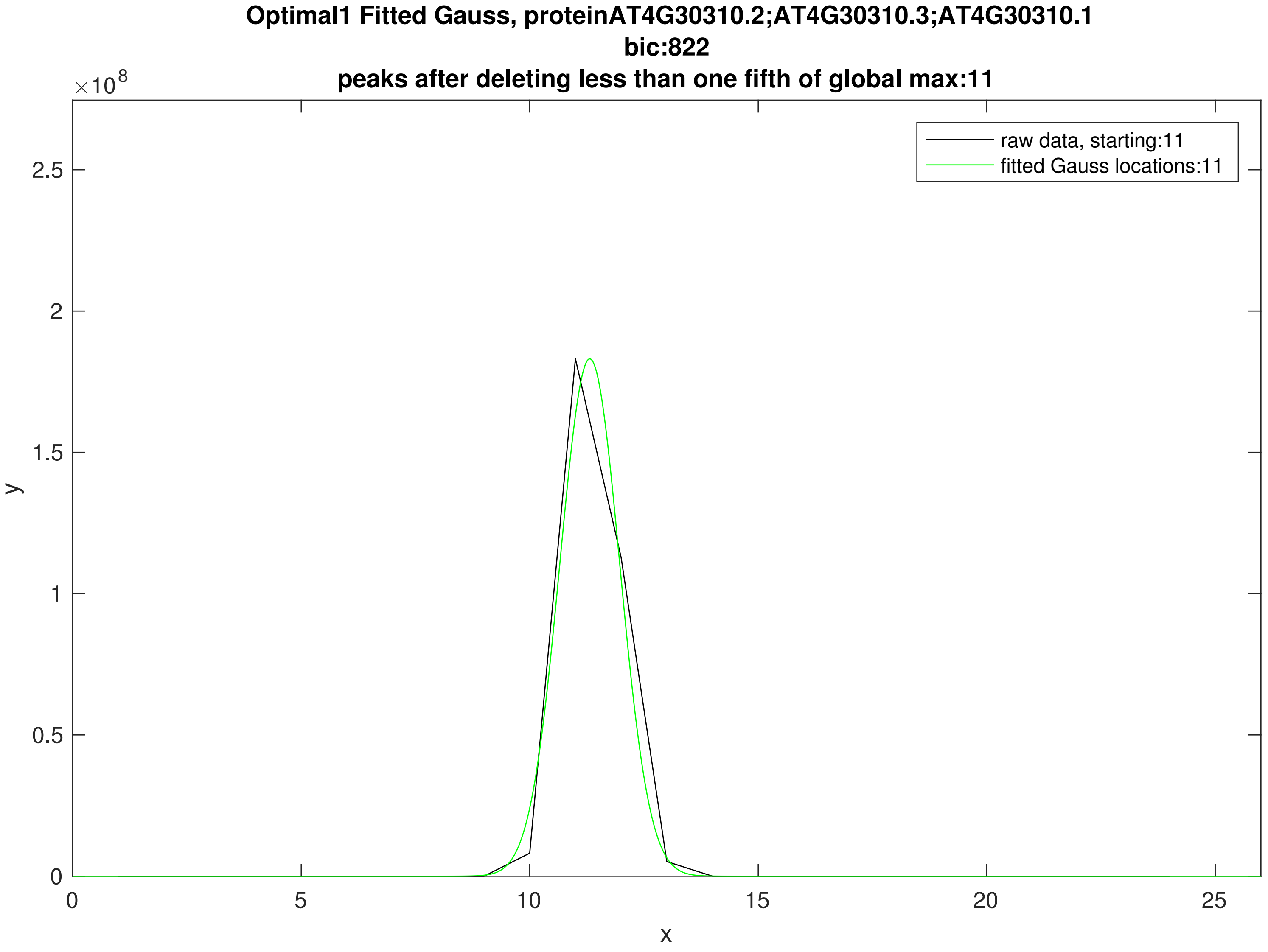

Optimal1 Fitted Gauss, proteinAT4G30310.2;AT4G30310.3;AT4G30310.1
bic:822
peaks after deleting less than one fifth of global max:11
×10^8
raw data, starting:11
fitted Gauss locations:11
2.5
2
1.5
1
0.5
0
y
0
5
10
15
20
25
x

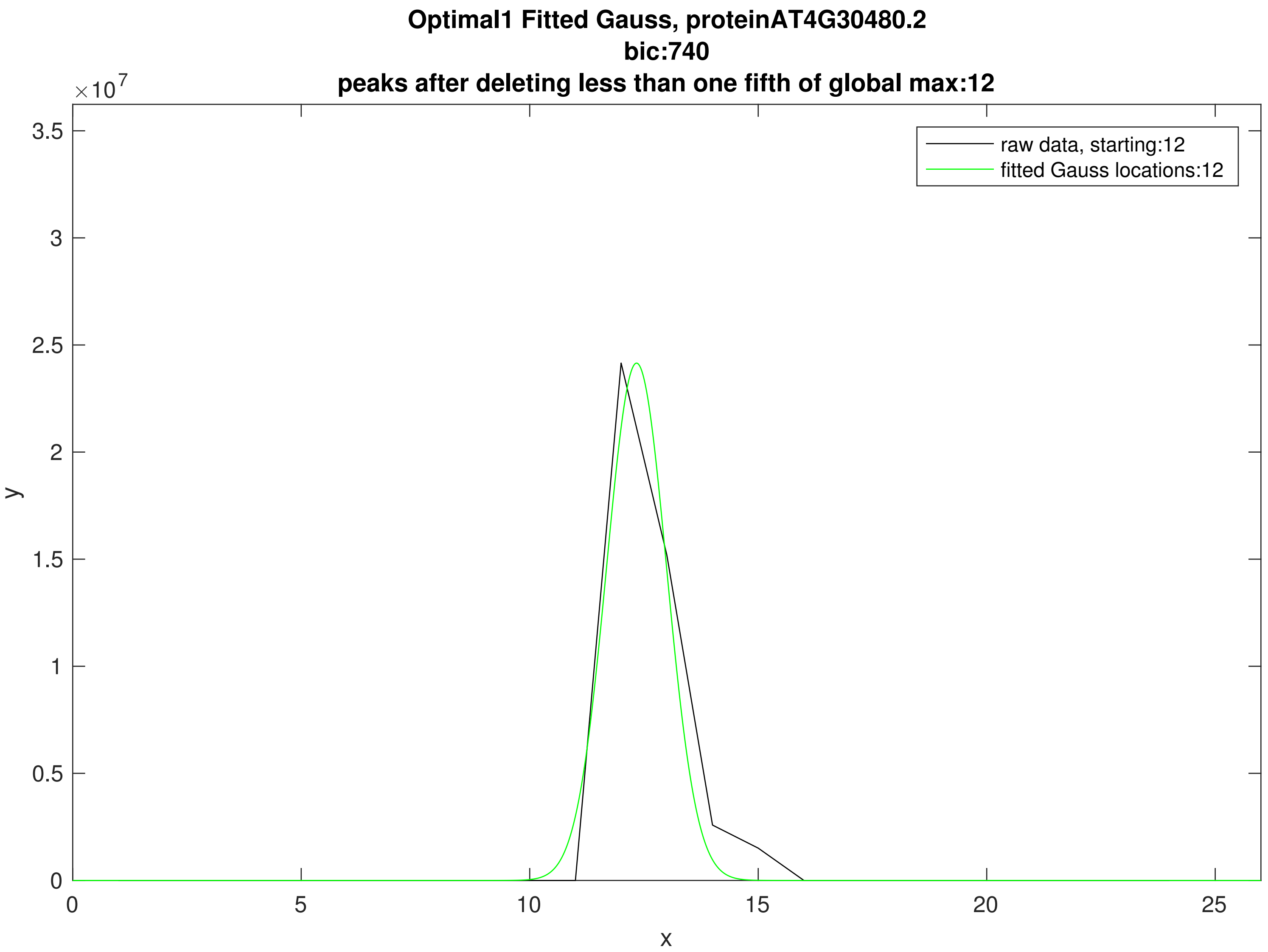
Optimal1 Fitted Gauss, proteinAT4G30480.2
bic:740
peaks after deleting less than one fifth of global max:12
×10^7
3.5
3
2.5
2
1.5
1
0.5
0
y
0
5
10
15
20
25
x
raw data, starting:12
fitted Gauss locations:12

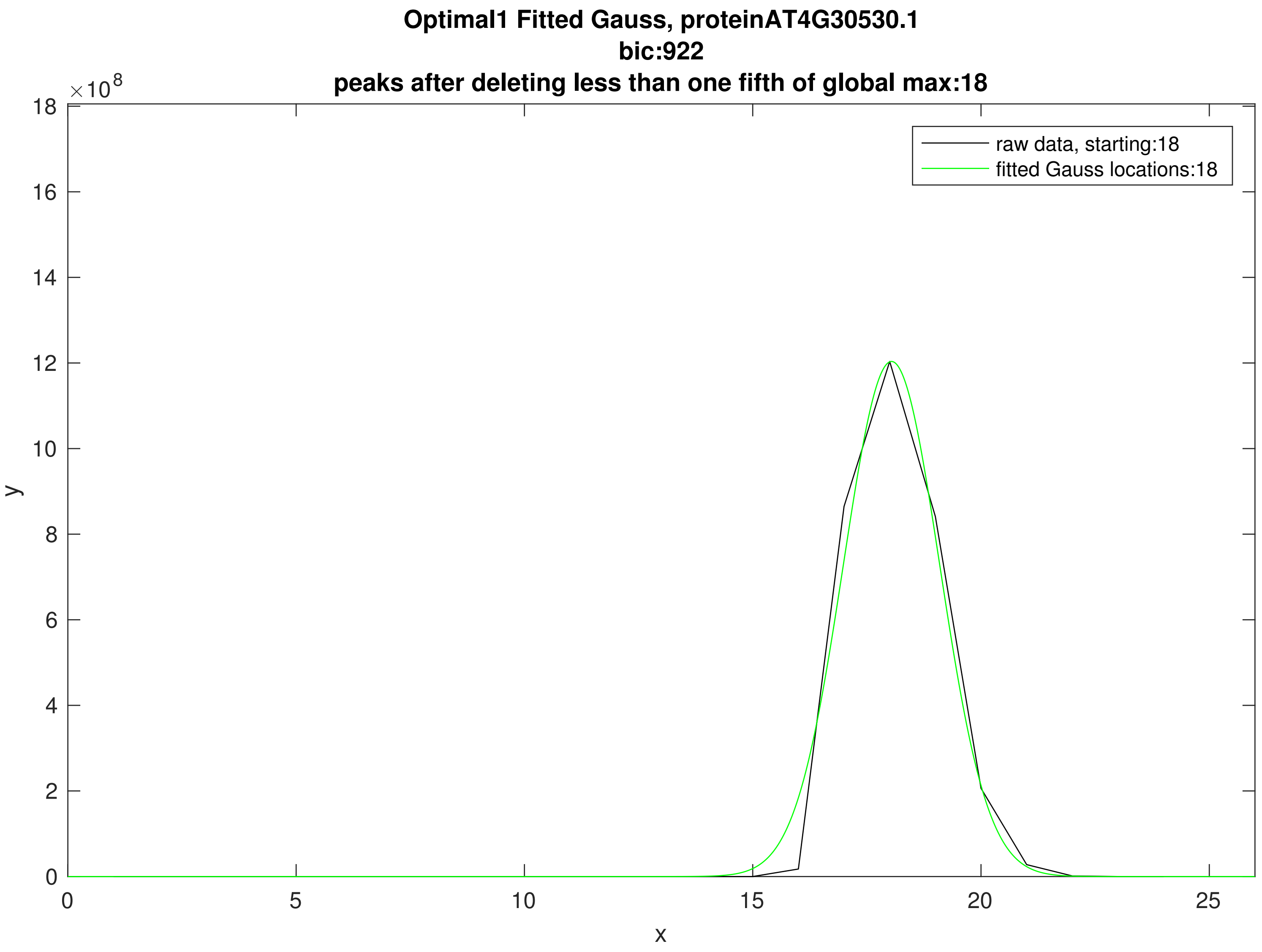

Optimal1 Fitted Gauss, proteinAT4G30530.1
bic:922
peaks after deleting less than one fifth of global max:18
×10^8
18
16
14
12
10
8
6
4
2
0
y
0
5
10
15
20
25
x
raw data, starting:18
fitted Gauss locations:18

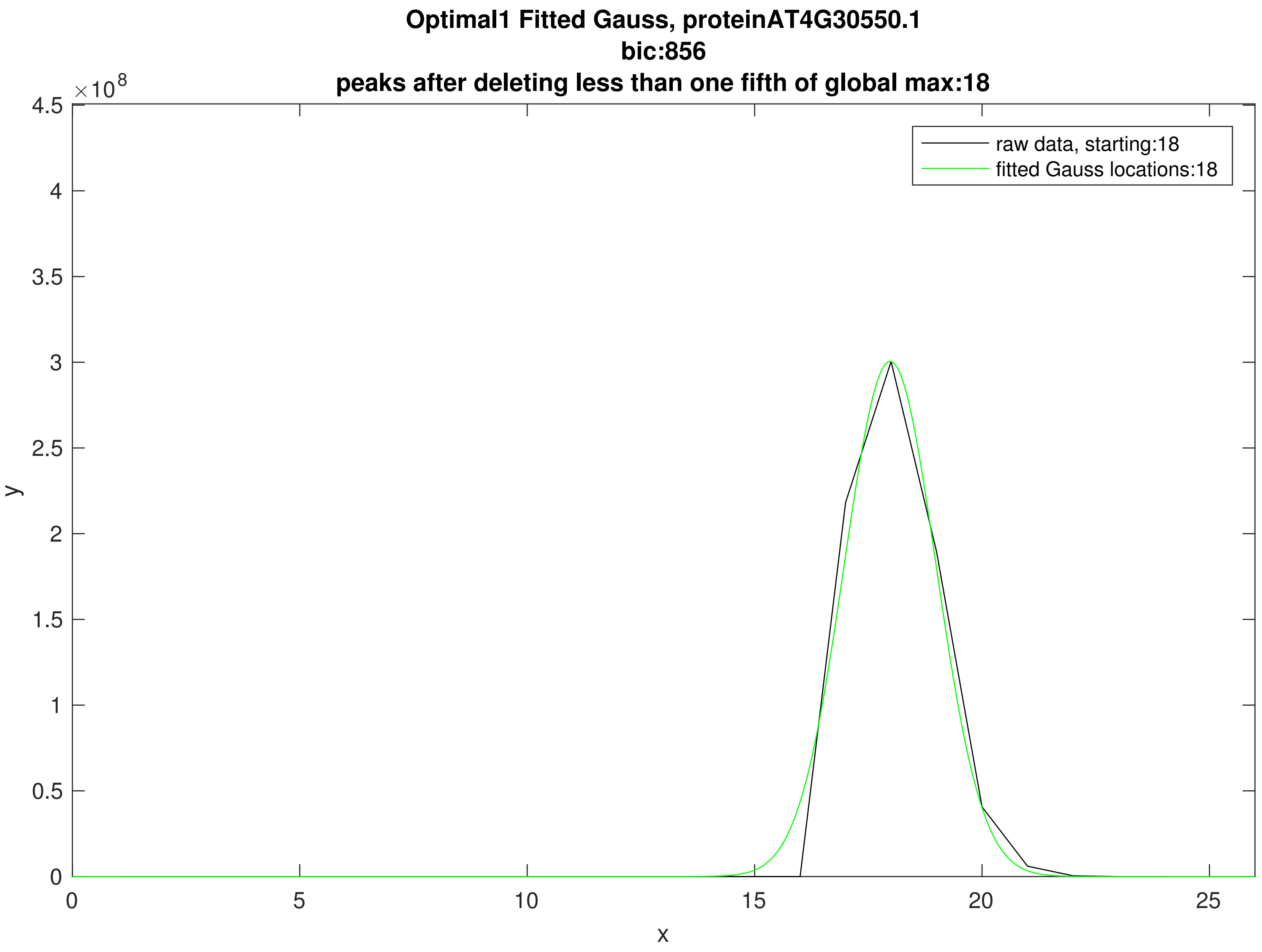

Optimal1 Fitted Gauss, proteinAT4G30550.1
bic:856
peaks after deleting less than one fifth of global max:18
×10^8
4.5
4
3.5
3
2.5
2
1.5
1
0.5
0
y
0
5
10
15
20
25
x
raw data, starting:18
fitted Gauss locations:18

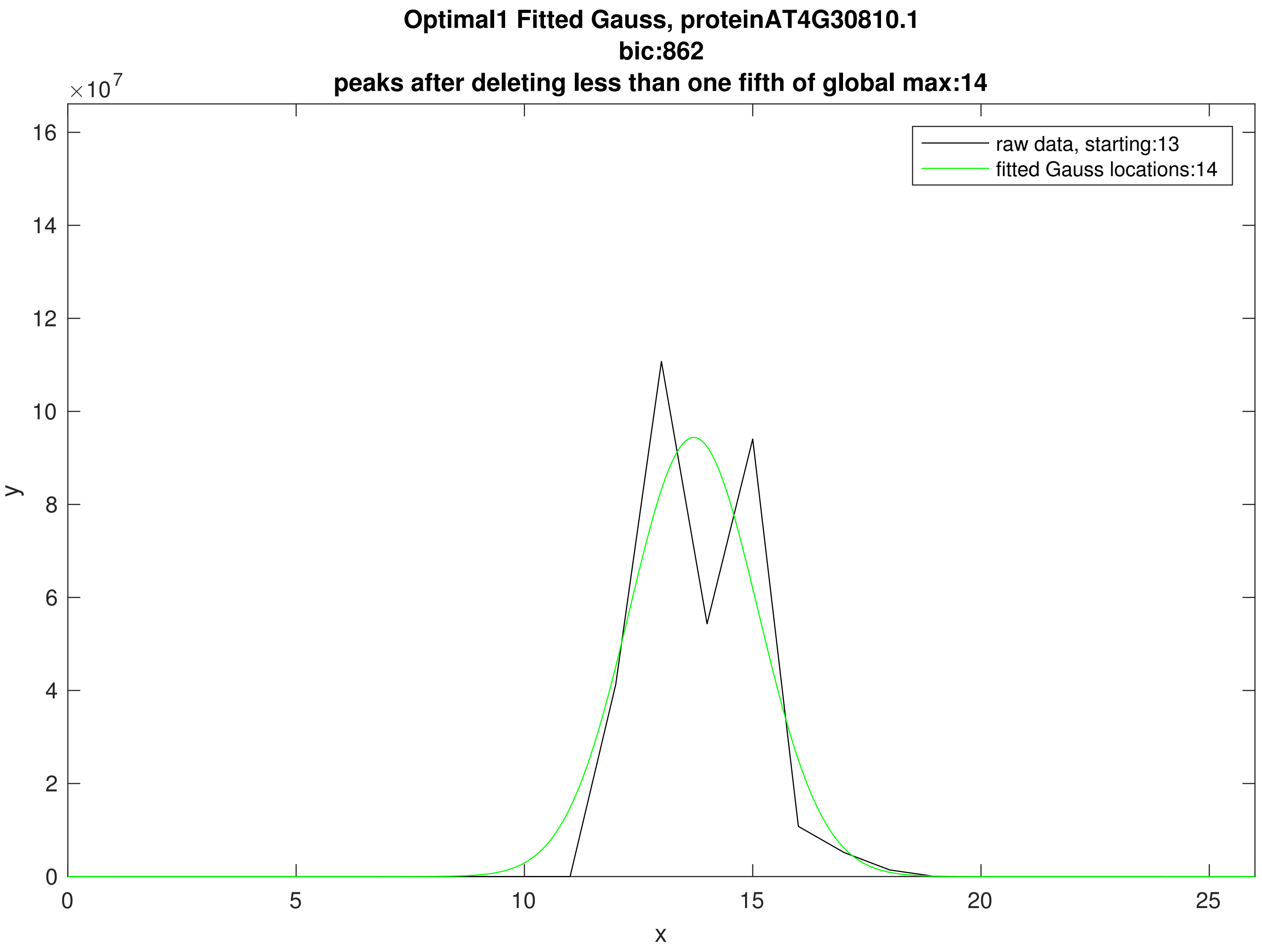

Optimal1 Fitted Gauss, proteinAT4G30810.1
bic:862
peaks after deleting less than one fifth of global max:14
×10^7
16
14
12
10
8
6
4
2
0
y
0
5
10
15
20
25
x
raw data, starting:13
fitted Gauss locations:14

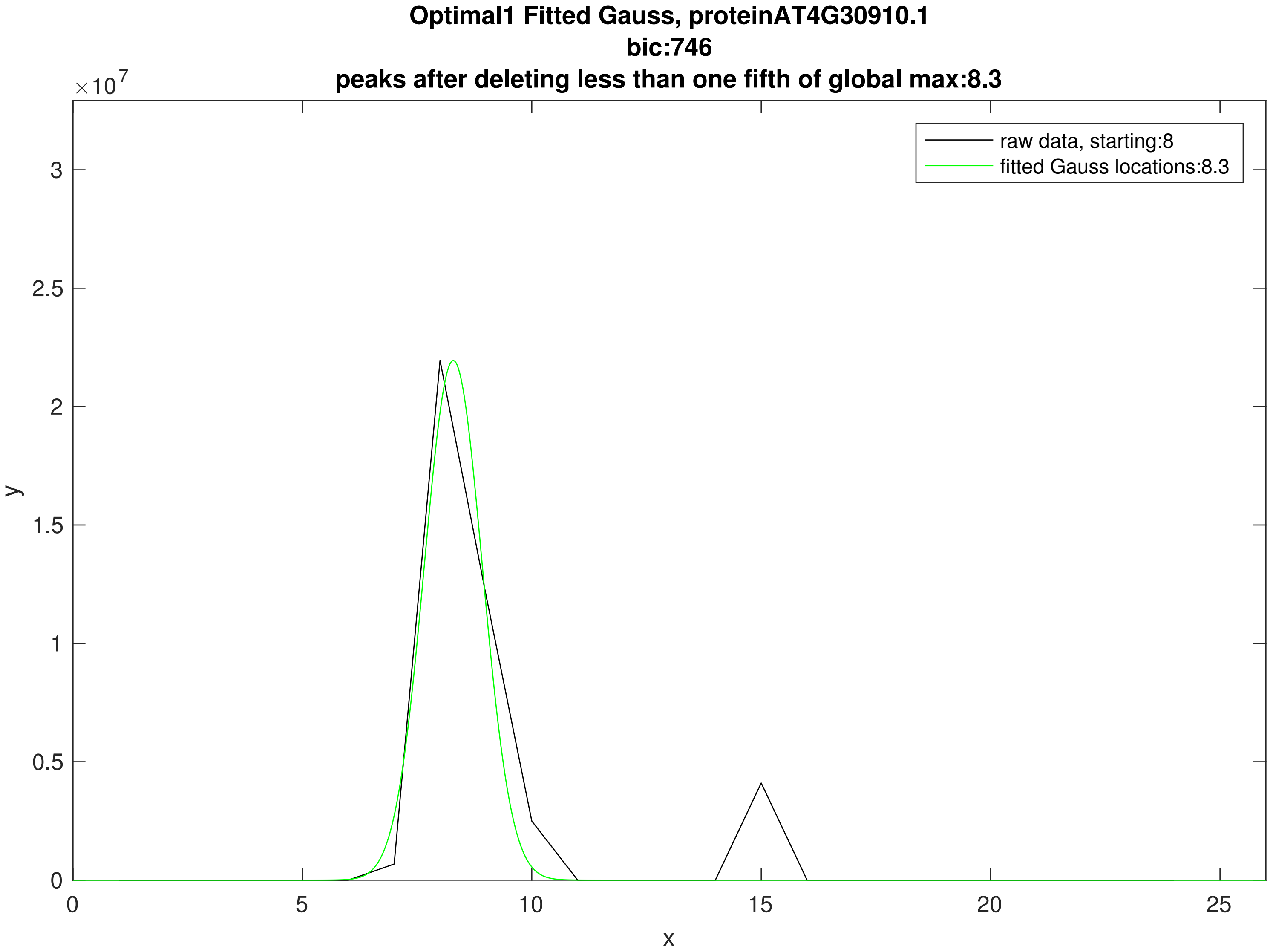

Optimal1 Fitted Gauss, proteinAT4G30910.1
bic:746
peaks after deleting less than one fifth of global max:8.3
×10^7
raw data, starting:8
fitted Gauss locations:8.3
y
x
0
0.5
1
1.5
2
2.5
3
0
5
10
15
20
25

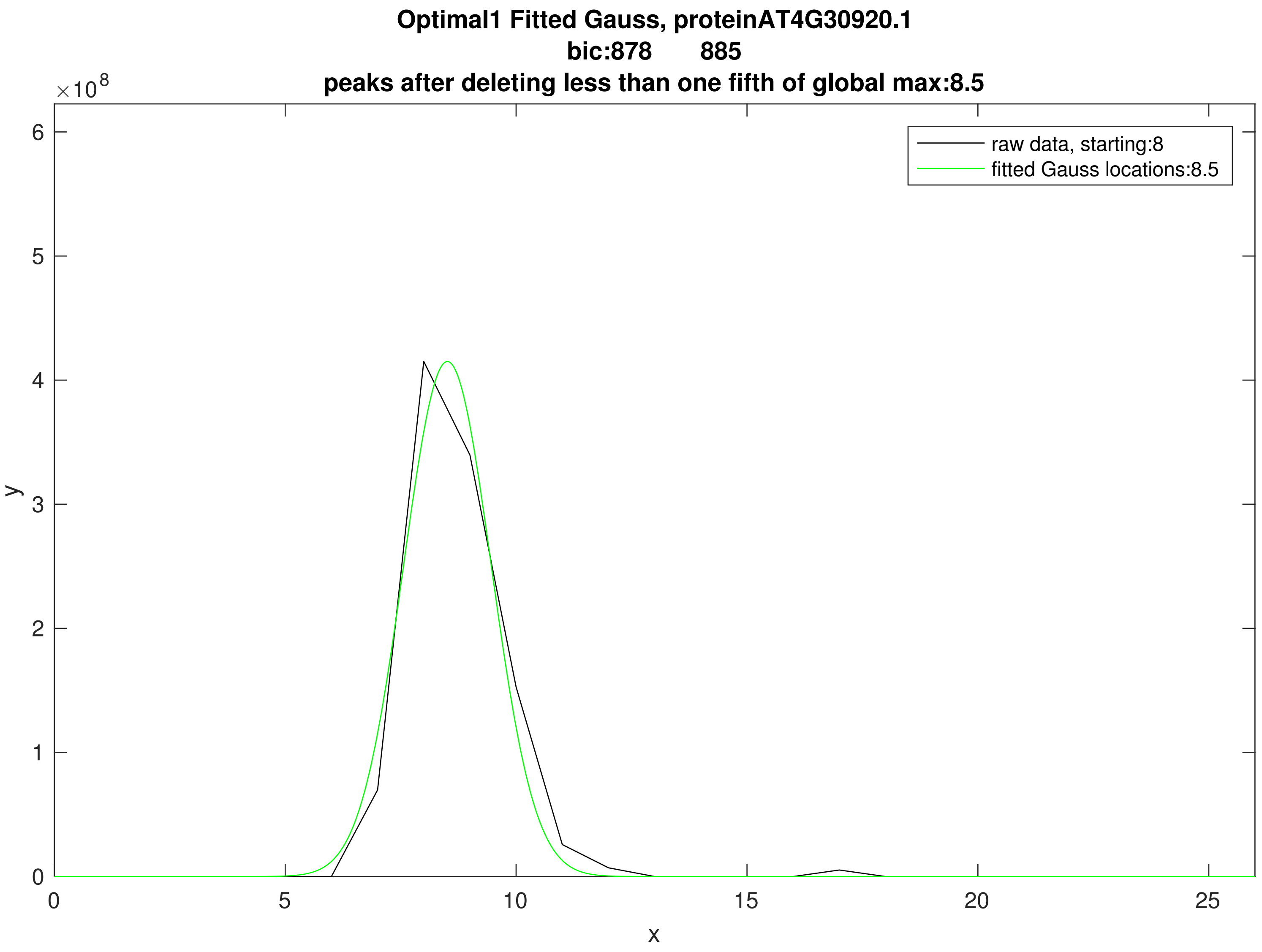

Optimal1 Fitted Gauss, proteinAT4G30920.1
bic:878 885
peaks after deleting less than one fifth of global max:8.5
×10^8
raw data, starting:8
fitted Gauss locations:8.5
y
x
0
1
2
3
4
5
6
0
5
10
15
20
25

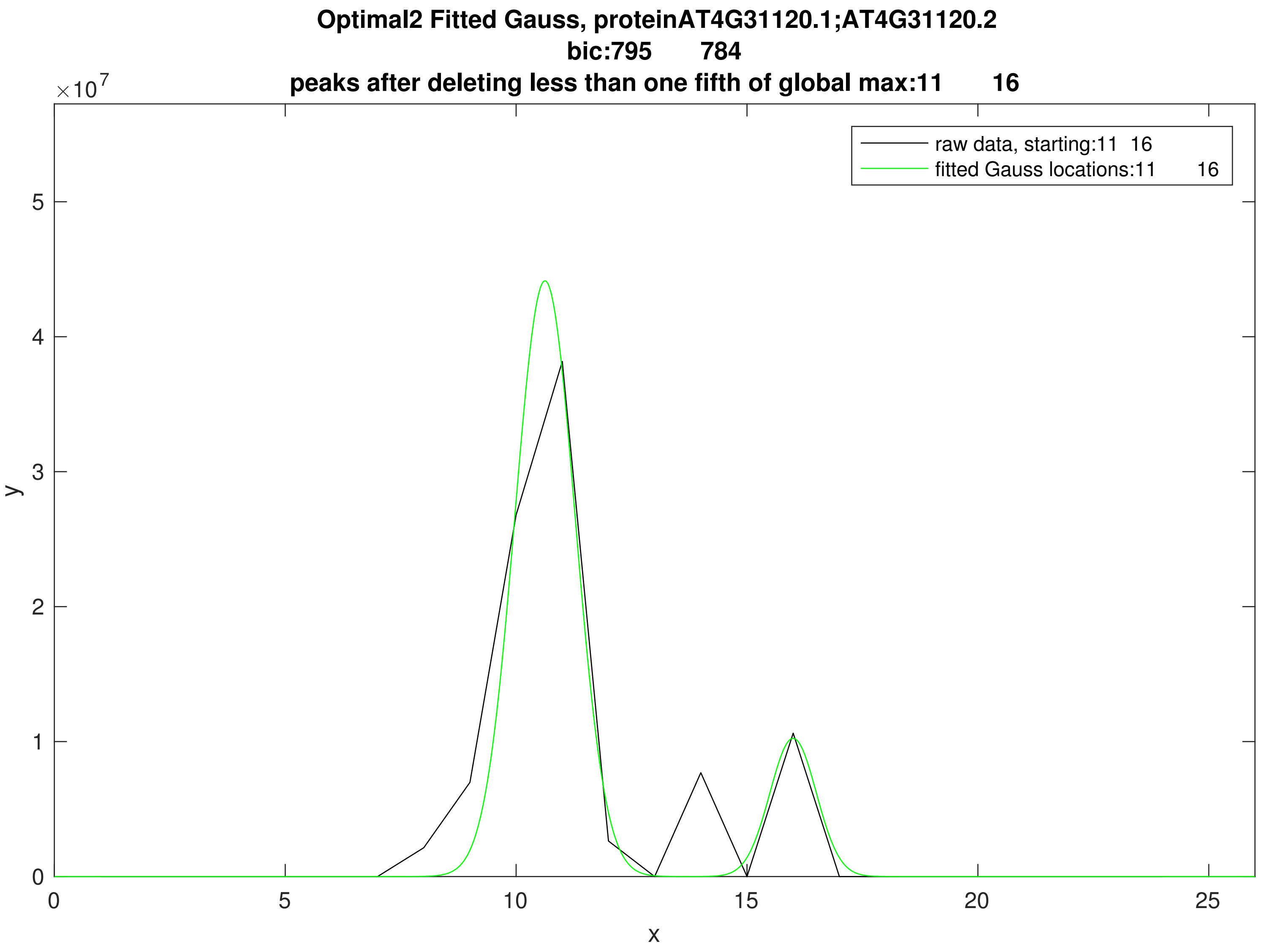
Optimal2 Fitted Gauss, proteinAT4G31120.1;AT4G31120.2
bic:795 784
peaks after deleting less than one fifth of global max:11 16
×10^7
raw data, starting:11 16
fitted Gauss locations:11 16
y
x
0
1
2
3
4
5
0
5
10
15
20
25

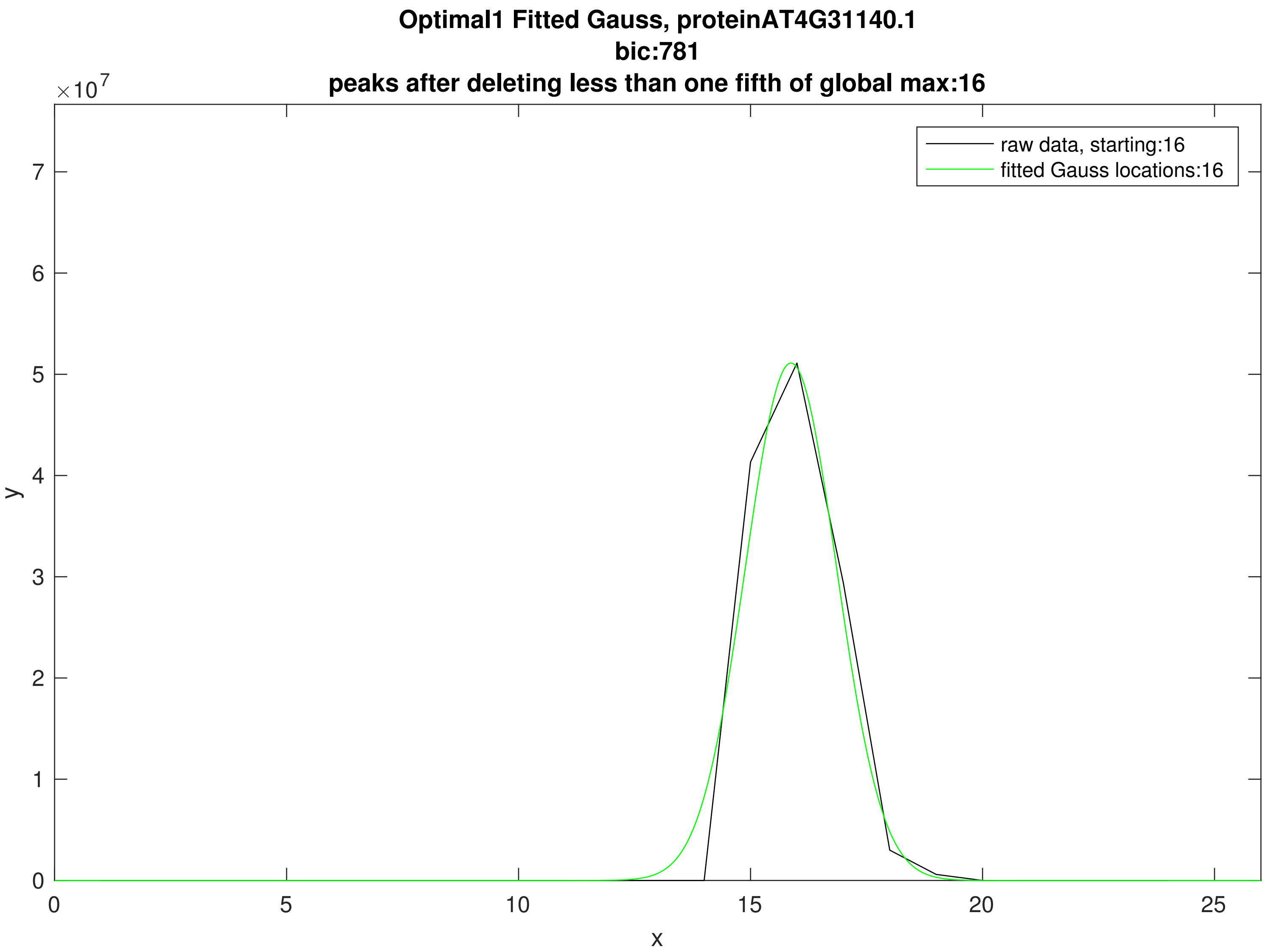
Optimal1 Fitted Gauss, proteinAT4G31140.1
bic:781
peaks after deleting less than one fifth of global max:16
×10^7
raw data, starting:16
fitted Gauss locations:16
y
x
0
1
2
3
4
5
6
7
0
5
10
15
20
25

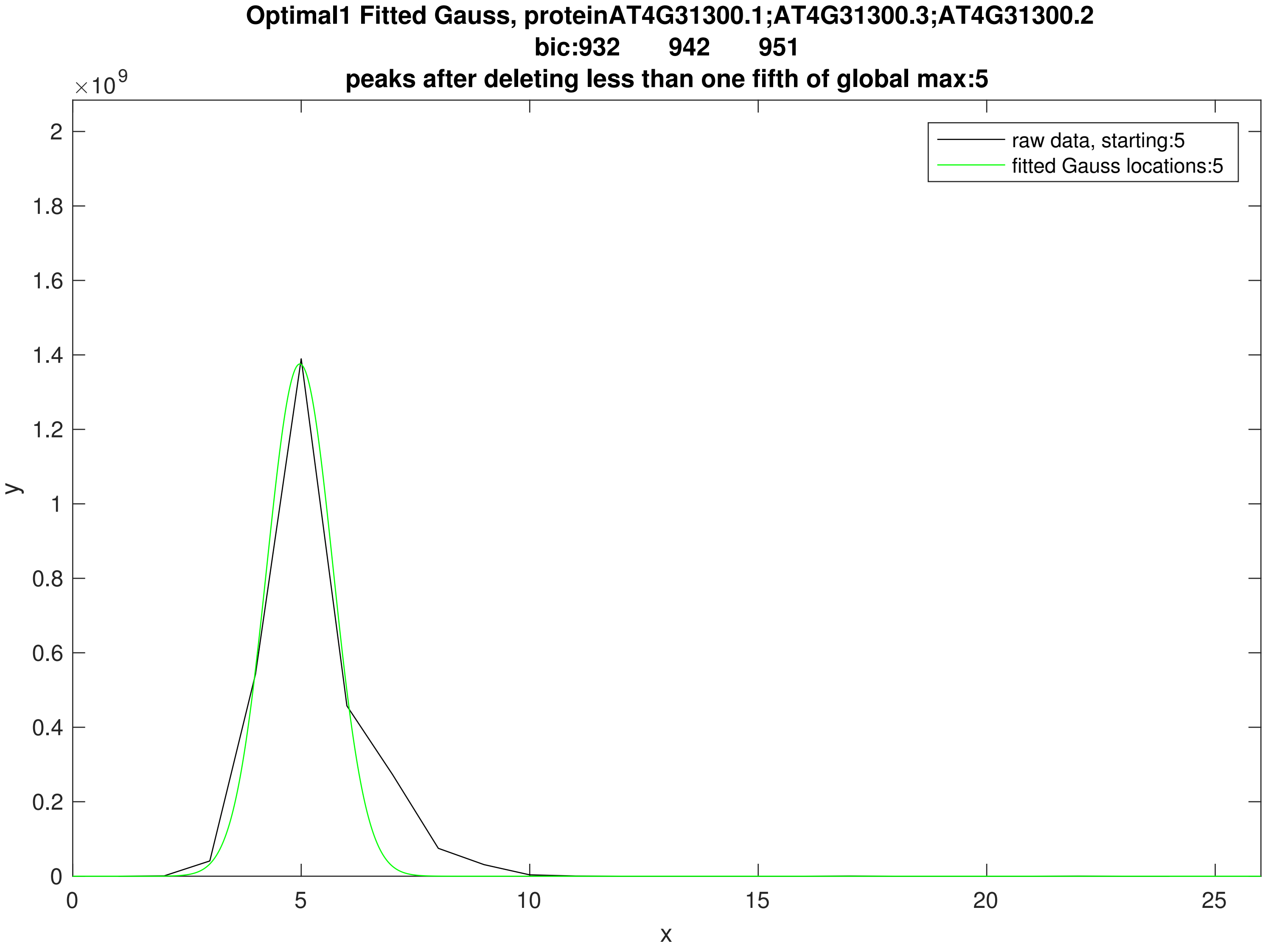

Optimal1 Fitted Gauss, proteinAT4G31300.1;AT4G31300.3;AT4G31300.2
bic:932 942 951
peaks after deleting less than one fifth of global max:5
×10^9
2
1.8
1.6
1.4
1.2
1
0.8
0.6
0.4
0.2
0
y
0
5
10
15
20
25
x
raw data, starting:5
fitted Gauss locations:5

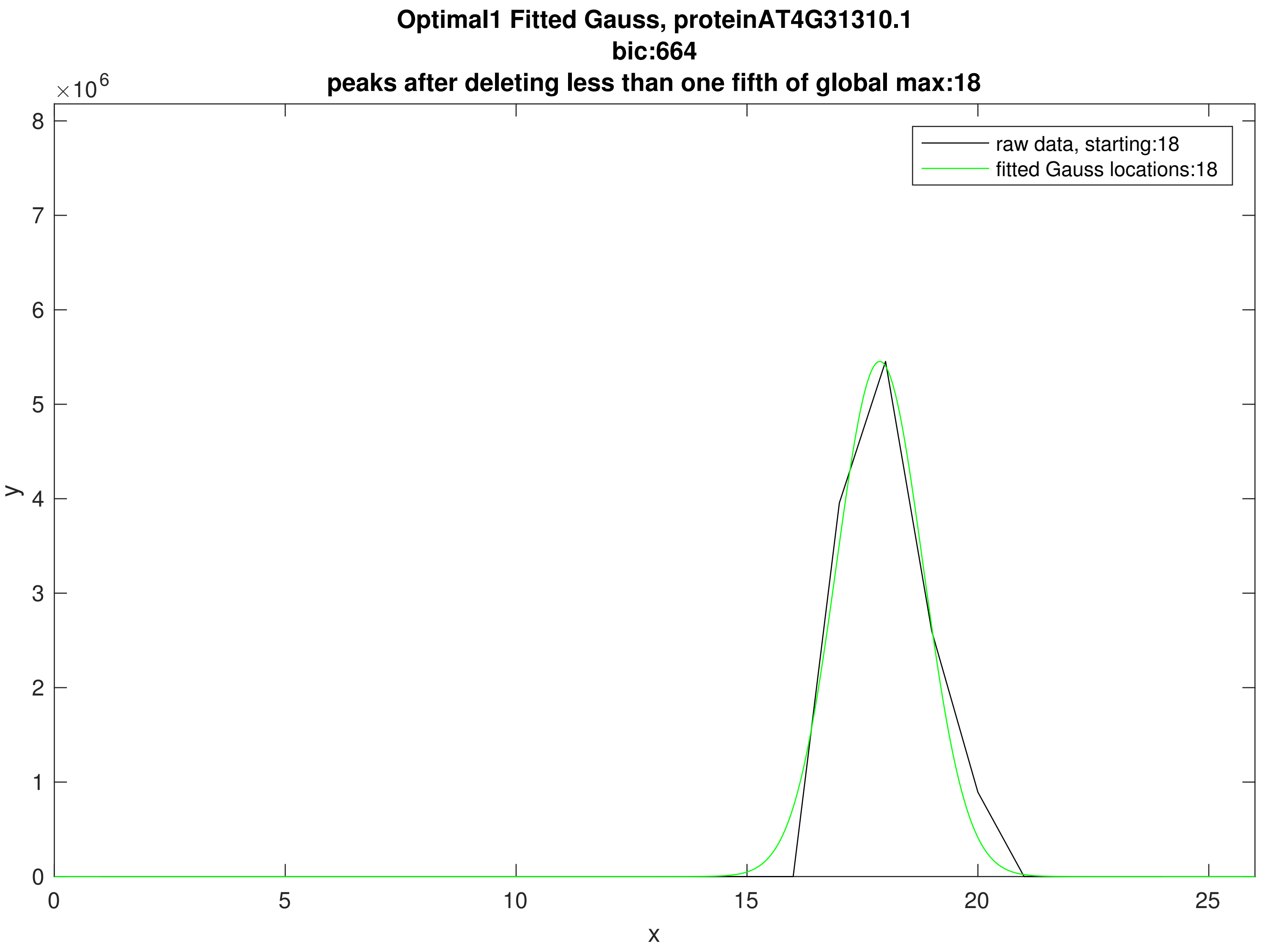
Optimal1 Fitted Gauss, proteinAT4G31310.1
bic:664
peaks after deleting less than one fifth of global max:18
×10^6
y
x
raw data, starting:18
fitted Gauss locations:18

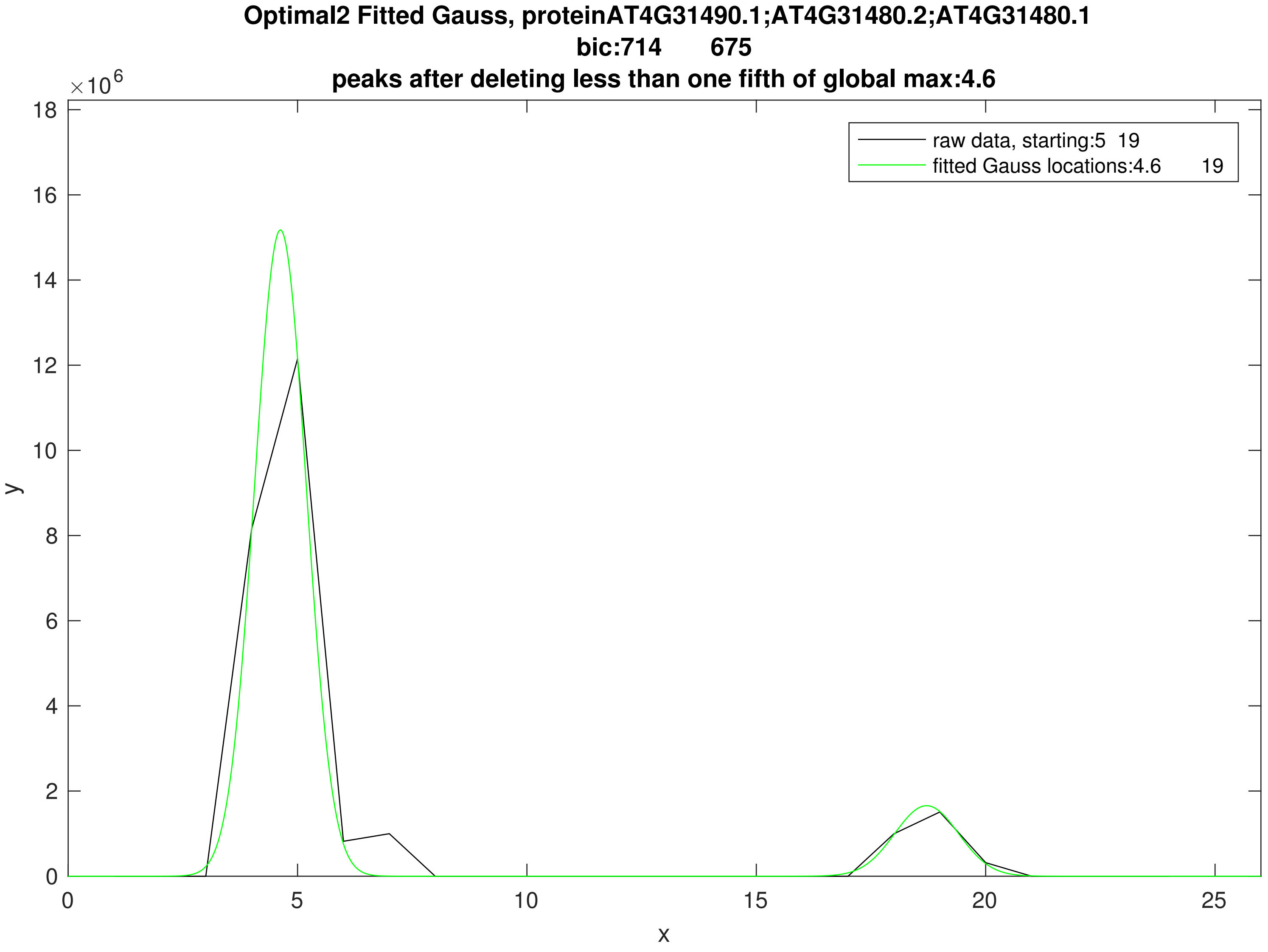
Optimal2 Fitted Gauss, proteinAT4G31490.1;AT4G31480.2;AT4G31480.1
bic:714 675
peaks after deleting less than one fifth of global max:4.6
×10^6
raw data, starting:5 19
fitted Gauss locations:4.6 19
y
x
0
2
4
6
8
10
12
14
16
18
0
5
10
15
20
25

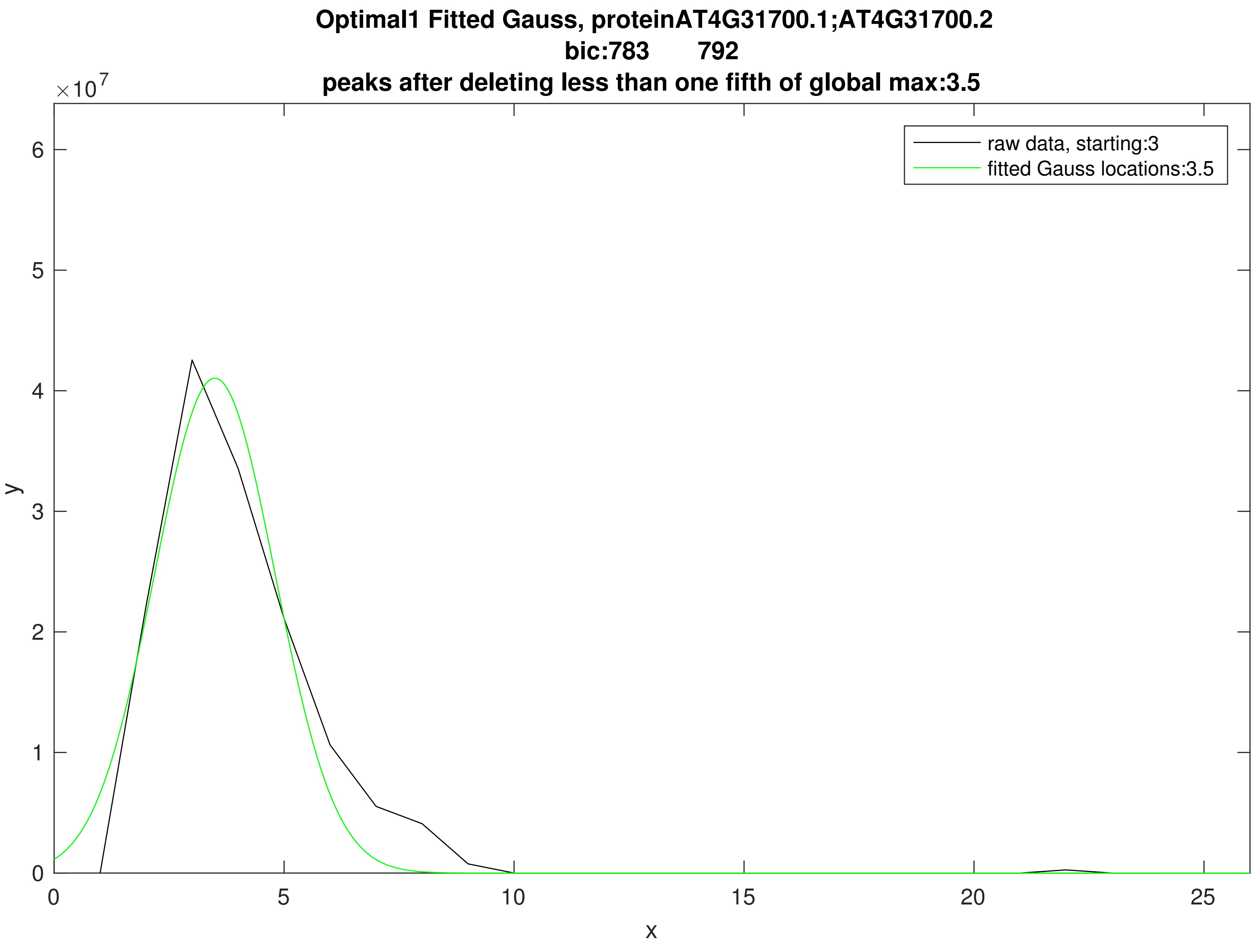
Optimal1 Fitted Gauss, proteinAT4G31700.1;AT4G31700.2
bic:783 792
peaks after deleting less than one fifth of global max:3.5
×10^7
y
x
raw data, starting:3
fitted Gauss locations:3.5

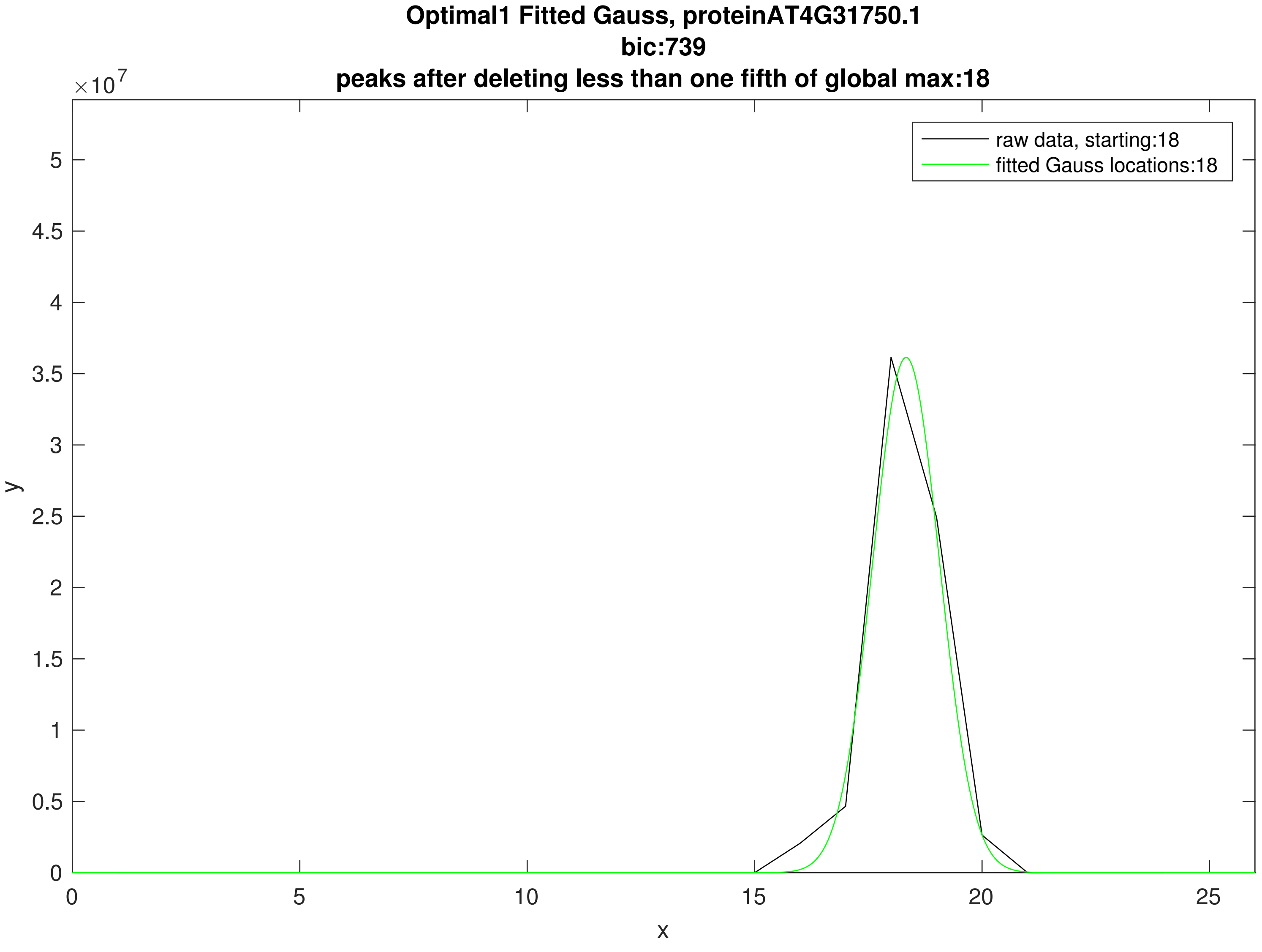
Optimal1 Fitted Gauss, proteinAT4G31750.1
bic:739
peaks after deleting less than one fifth of global max:18
×10^7
raw data, starting:18
fitted Gauss locations:18
y
x
0
0.5
1
1.5
2
2.5
3
3.5
4
4.5
5
0
5
10
15
20
25

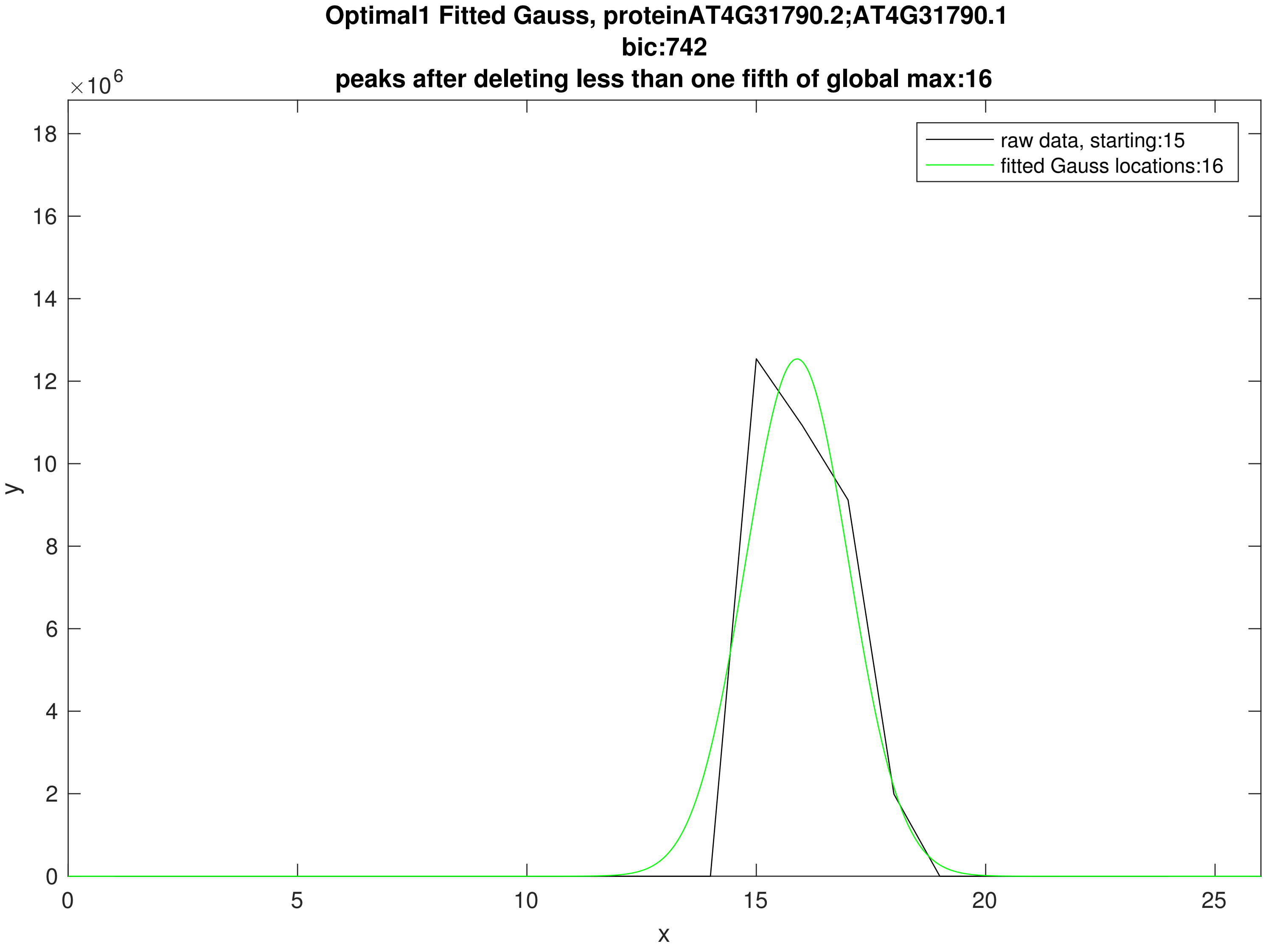
Optimal1 Fitted Gauss, proteinAT4G31790.2;AT4G31790.1
bic:742
peaks after deleting less than one fifth of global max:16
×10^6
raw data, starting:15
fitted Gauss locations:16
y
x
0
2
4
6
8
10
12
14
16
18
5
15
20
25

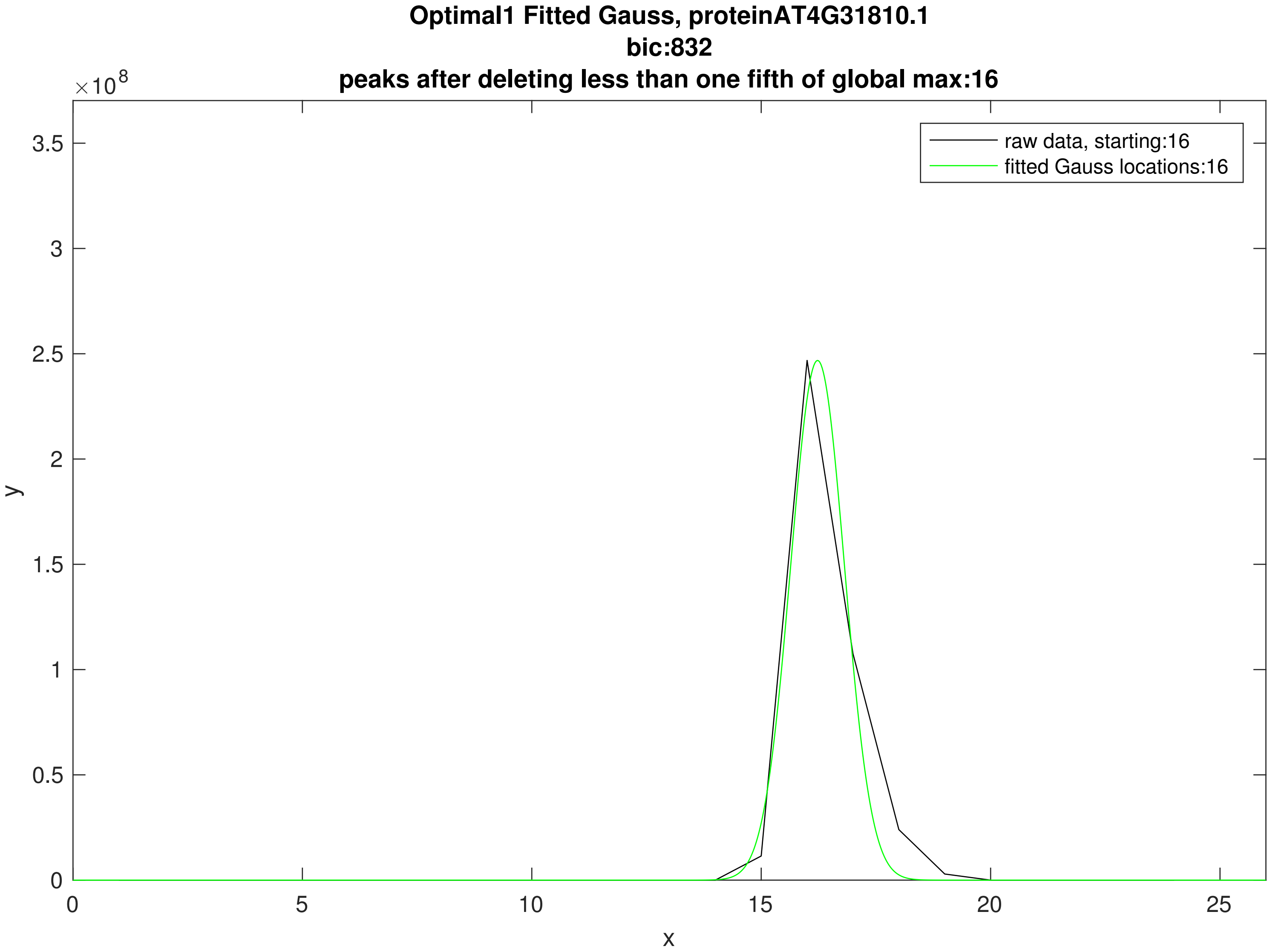
Optimal1 Fitted Gauss, proteinAT4G31810.1
bic:832
peaks after deleting less than one fifth of global max:16
×10^8
3.5
3
2.5
2
1.5
1
0.5
0
y
0
5
10
15
20
25
x
raw data, starting:16
fitted Gauss locations:16

Optimal1 Fitted Gauss, proteinAT4G31840.1
bic:732
peaks after deleting less than one fifth of global max:9

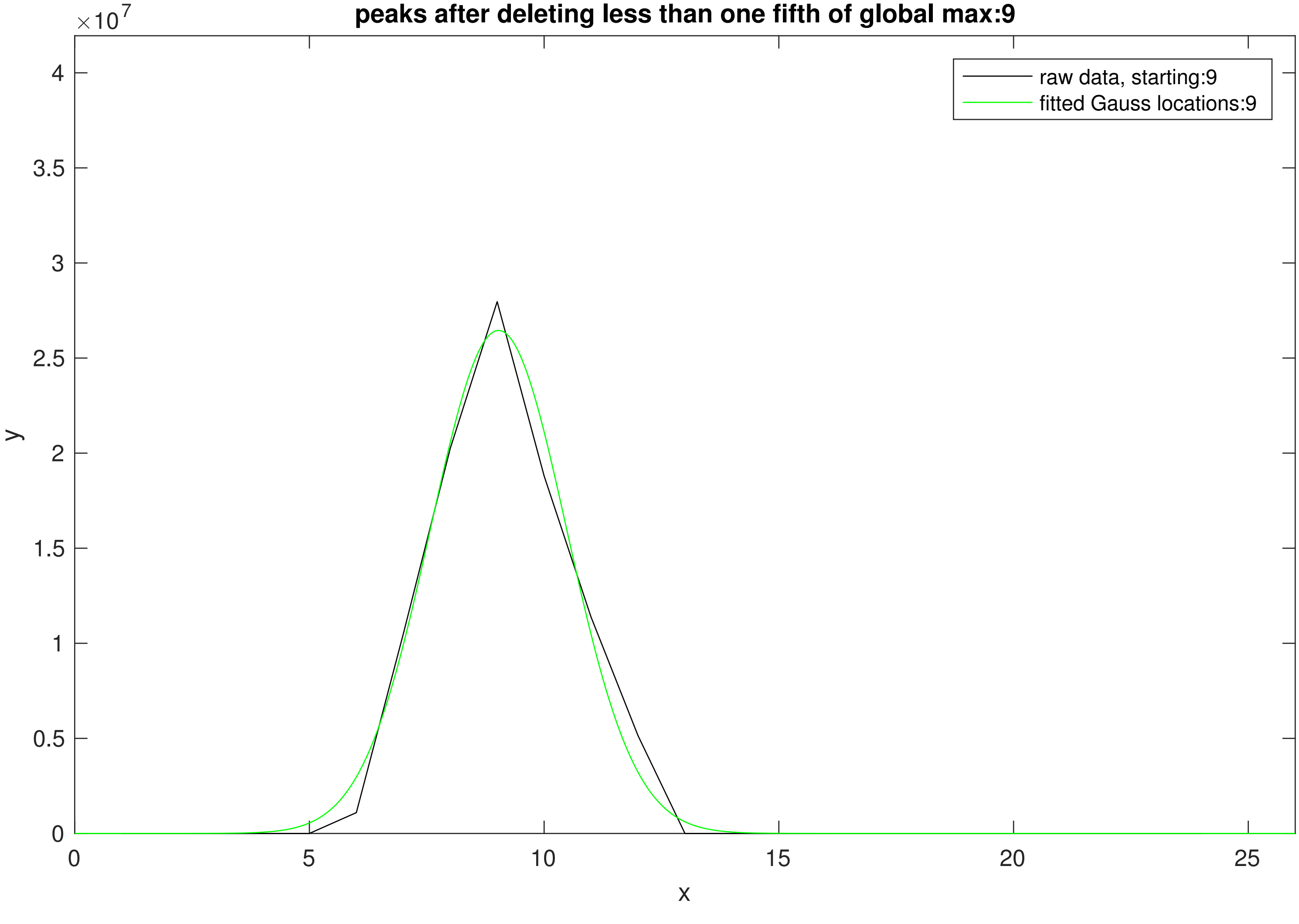

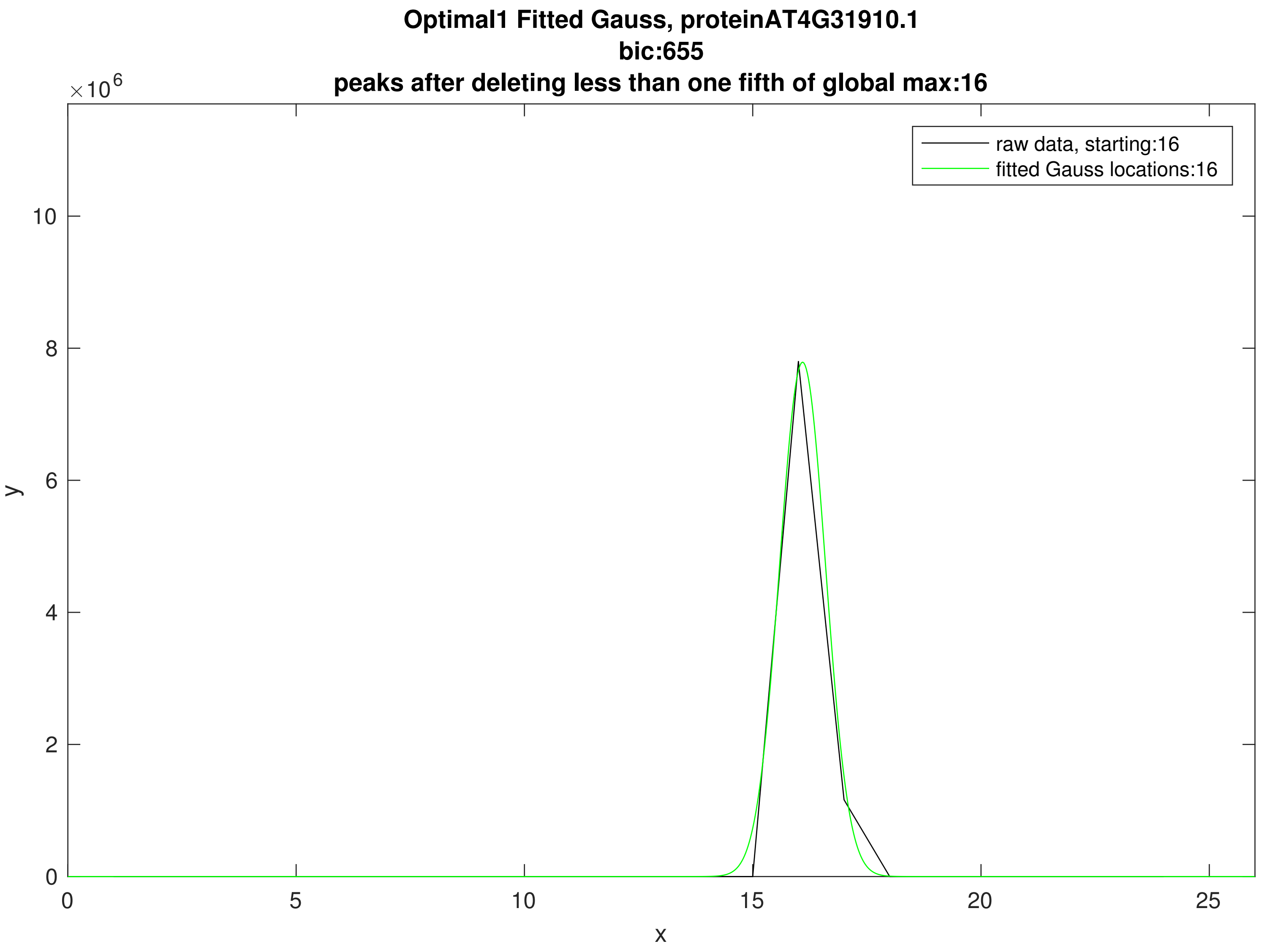
Optimal1 Fitted Gauss, proteinAT4G31910.1
bic:655
peaks after deleting less than one fifth of global max:16
×10^6
raw data, starting:16
fitted Gauss locations:16
10
8
6
4
2
0
y
0
5
10
15
20
25
x

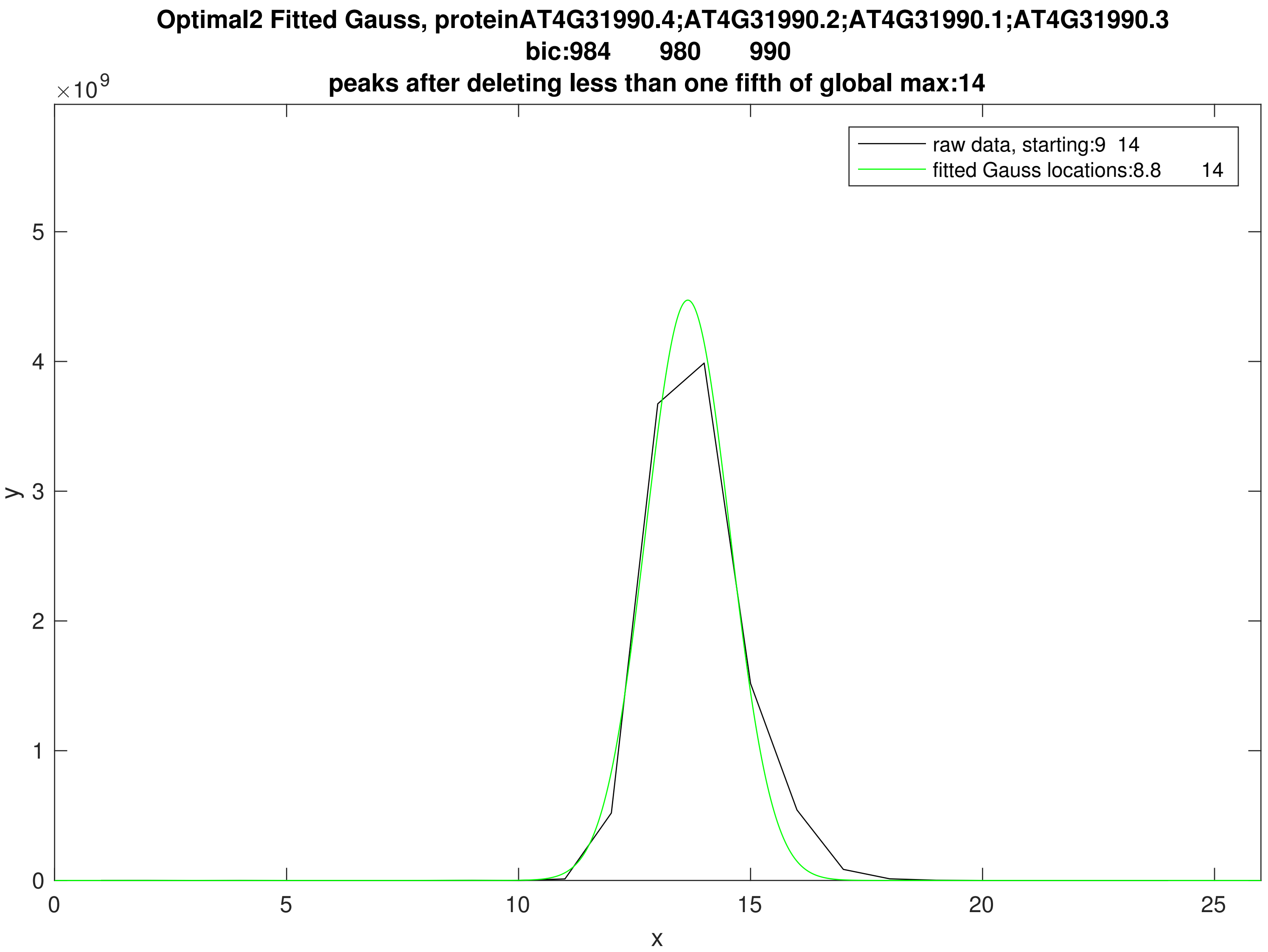

Optimal2 Fitted Gauss, proteinAT4G31990.4;AT4G31990.2;AT4G31990.1;AT4G31990.3
bic:984 980 990
peaks after deleting less than one fifth of global max:14
×10^9
raw data, starting:9 14
fitted Gauss locations:8.8 14
y
x
0
1
2
3
4
5
0
5
10
15
20
25

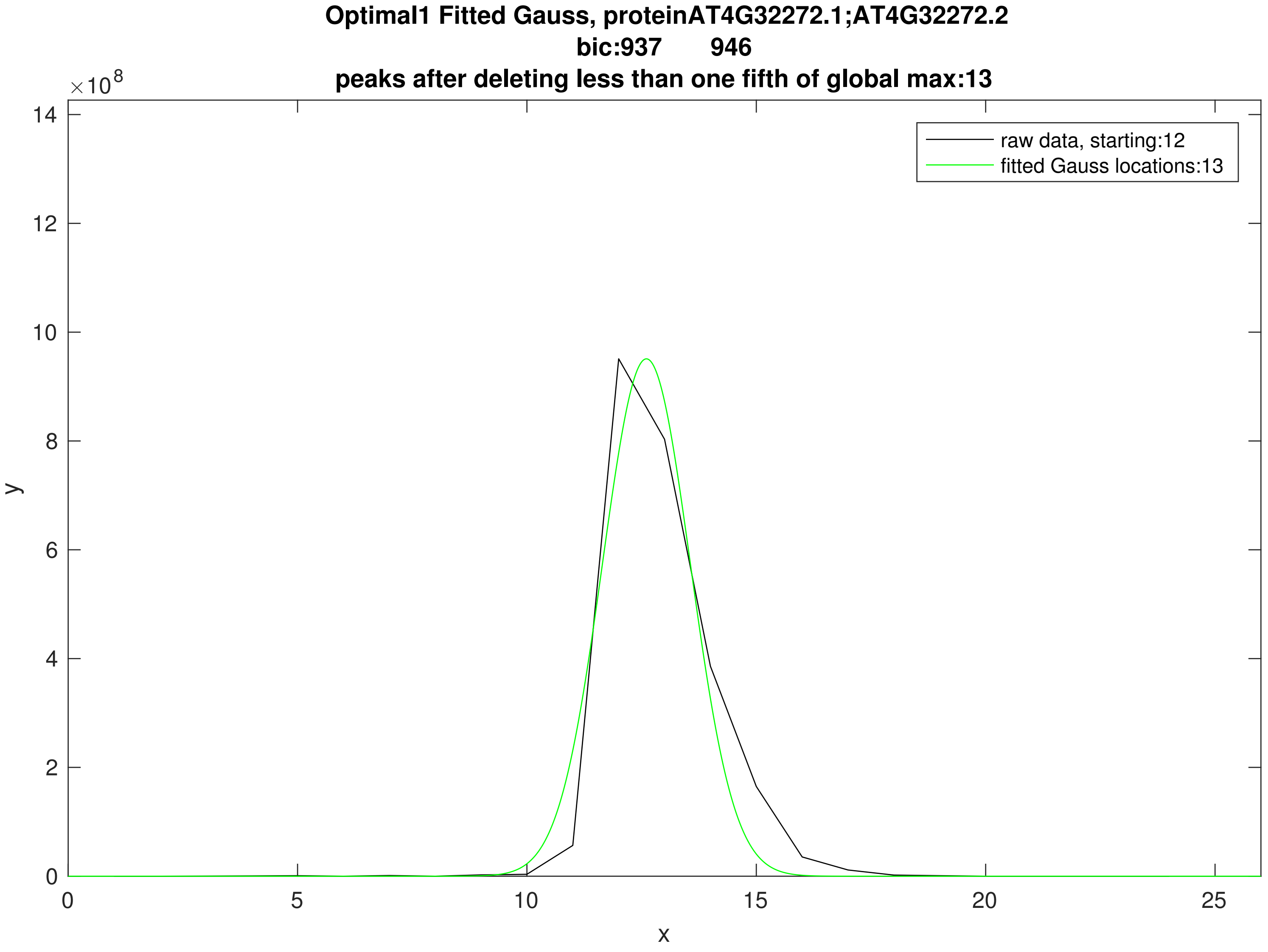

Optimal1 Fitted Gauss, proteinAT4G32272.1;AT4G32272.2
bic:937 946
peaks after deleting less than one fifth of global max:13
×10^8
raw data, starting:12
fitted Gauss locations:13
y
x
0
2
4
6
8
10
12
14
5
15
20
25

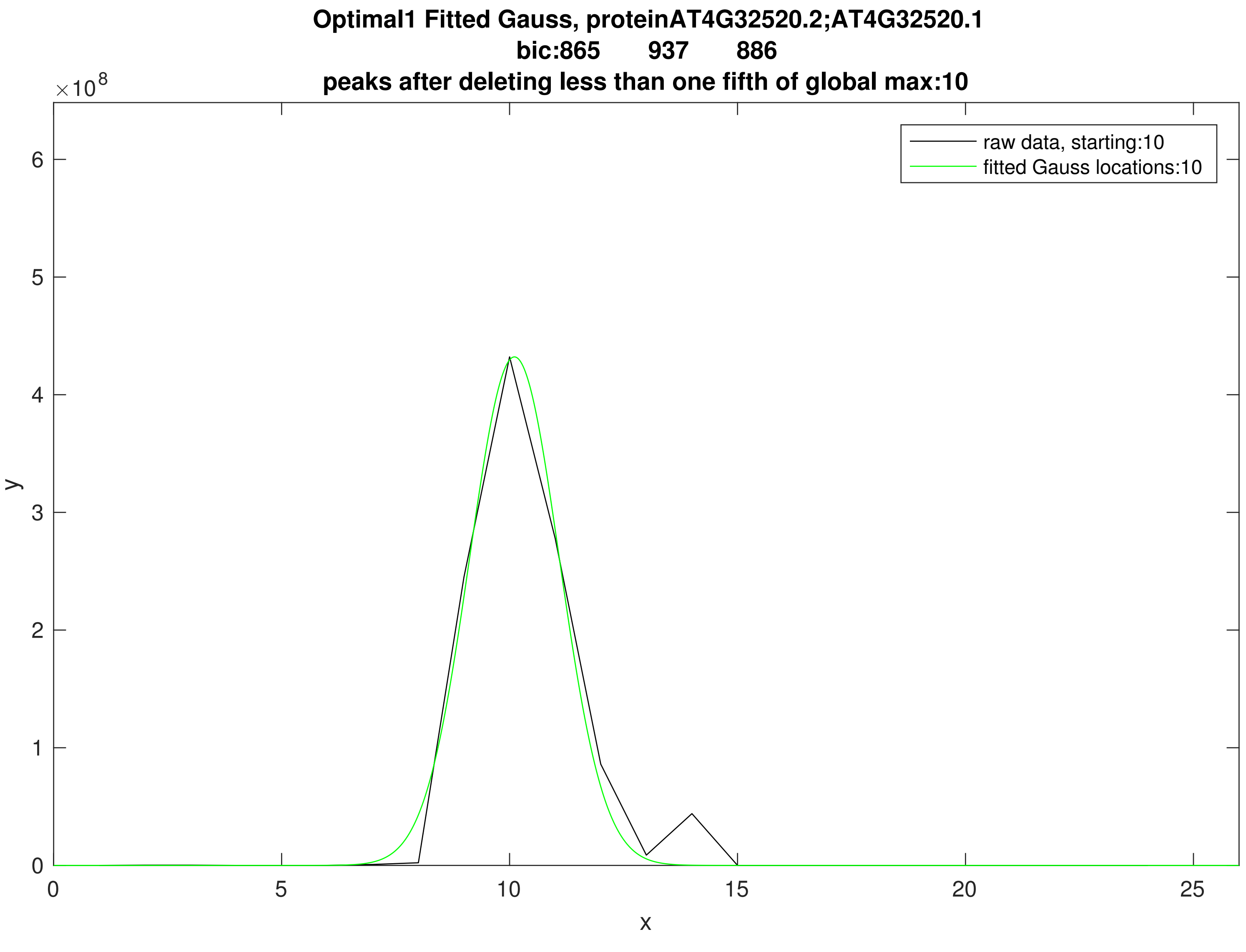

Optimal1 Fitted Gauss, proteinAT4G32520.2;AT4G32520.1
bic:865 937 886
peaks after deleting less than one fifth of global max:10
×10^8
raw data, starting:10
fitted Gauss locations:10
y
x
0
1
2
3
4
5
6
0
5
10
15
20
25

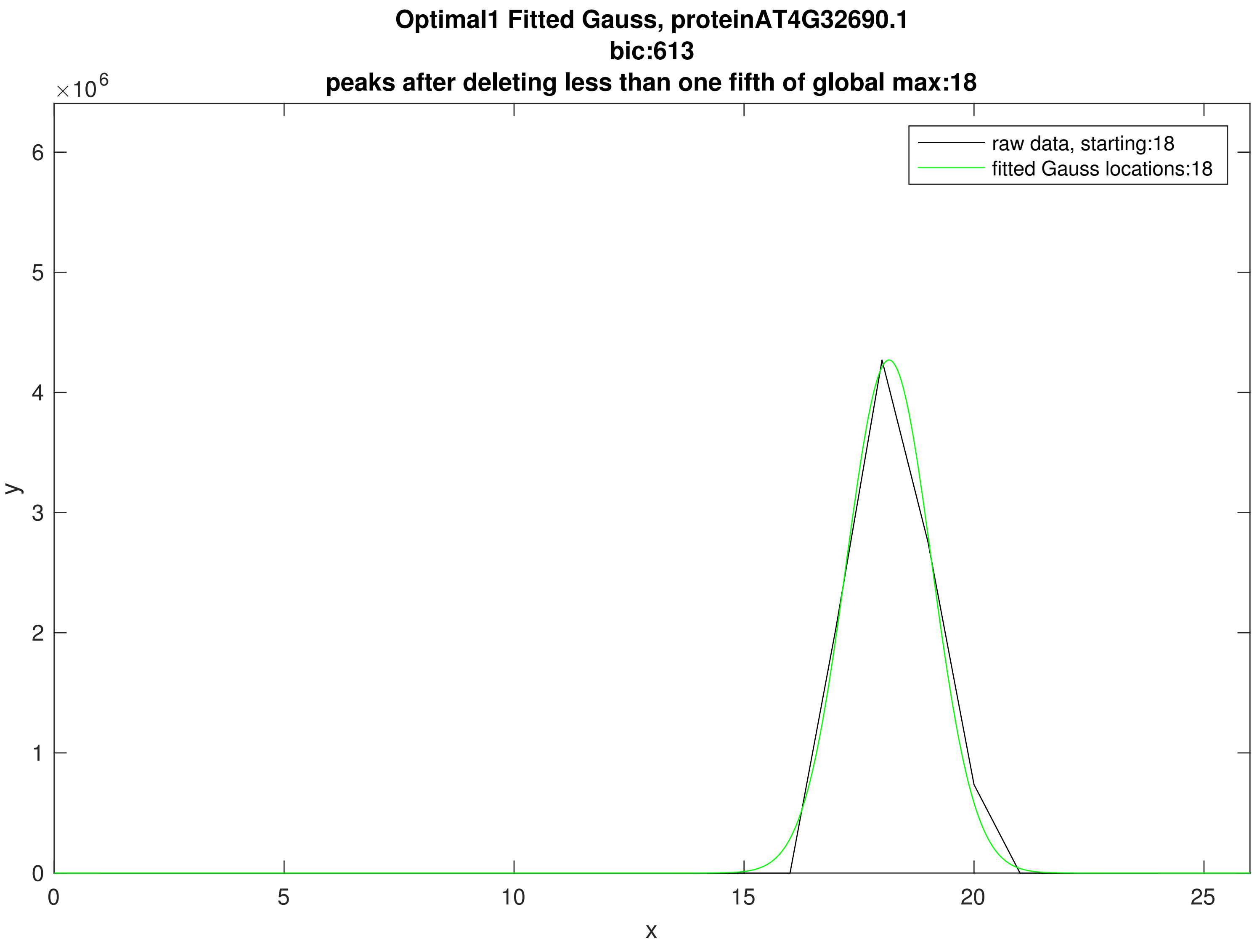
Optimal1 Fitted Gauss, proteinAT4G32690.1
bic:613
peaks after deleting less than one fifth of global max:18
×10^6
raw data, starting:18
fitted Gauss locations:18
y
x
0
1
2
3
4
5
6
0
5
10
15
20
25

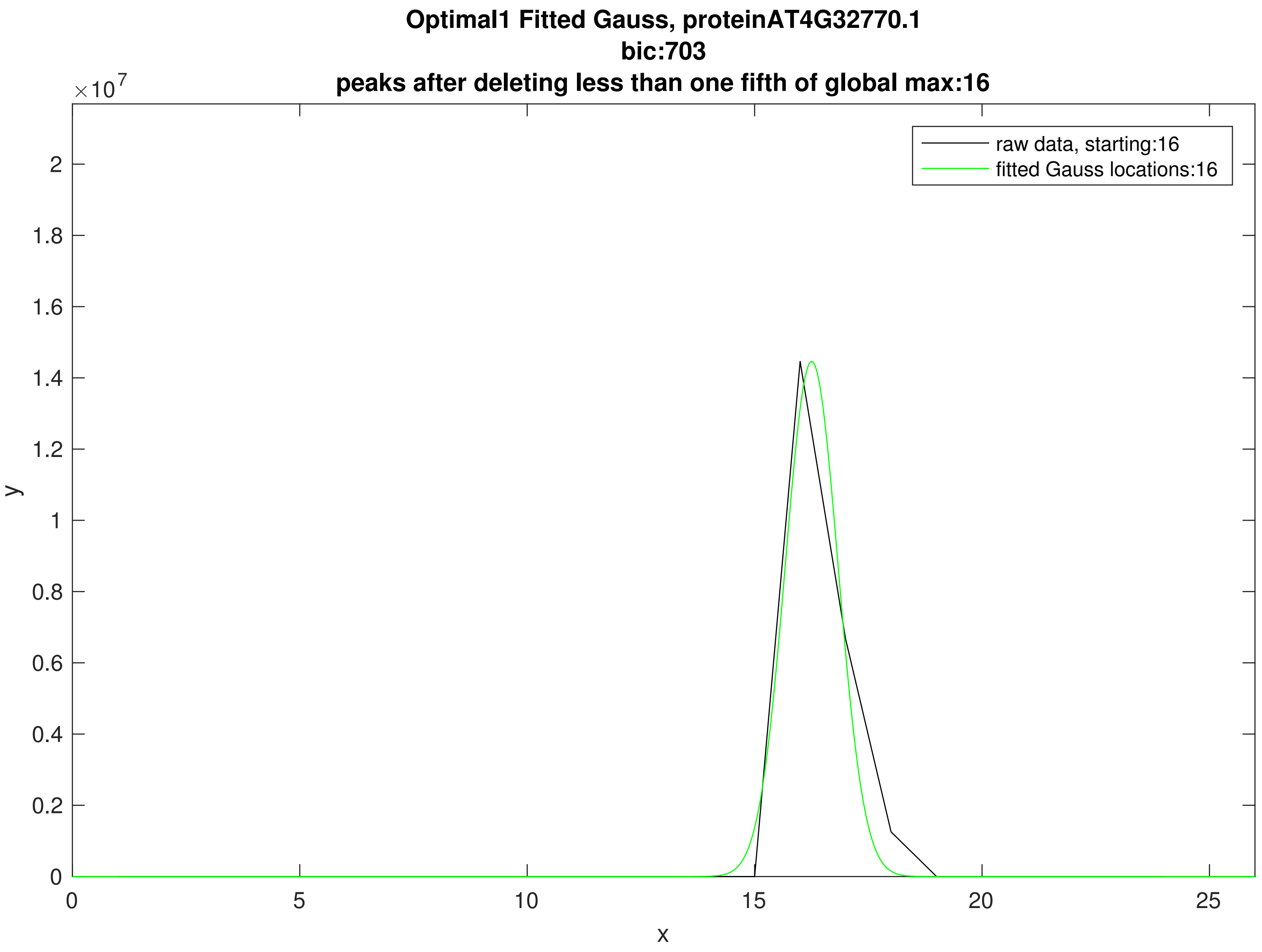
Optimal1 Fitted Gauss, proteinAT4G32770.1
bic:703
peaks after deleting less than one fifth of global max:16
×10^7
raw data, starting:16
fitted Gauss locations:16
y
x
0
0.2
0.4
0.6
0.8
1
1.2
1.4
1.6
1.8
2
0
5
10
15
20
25

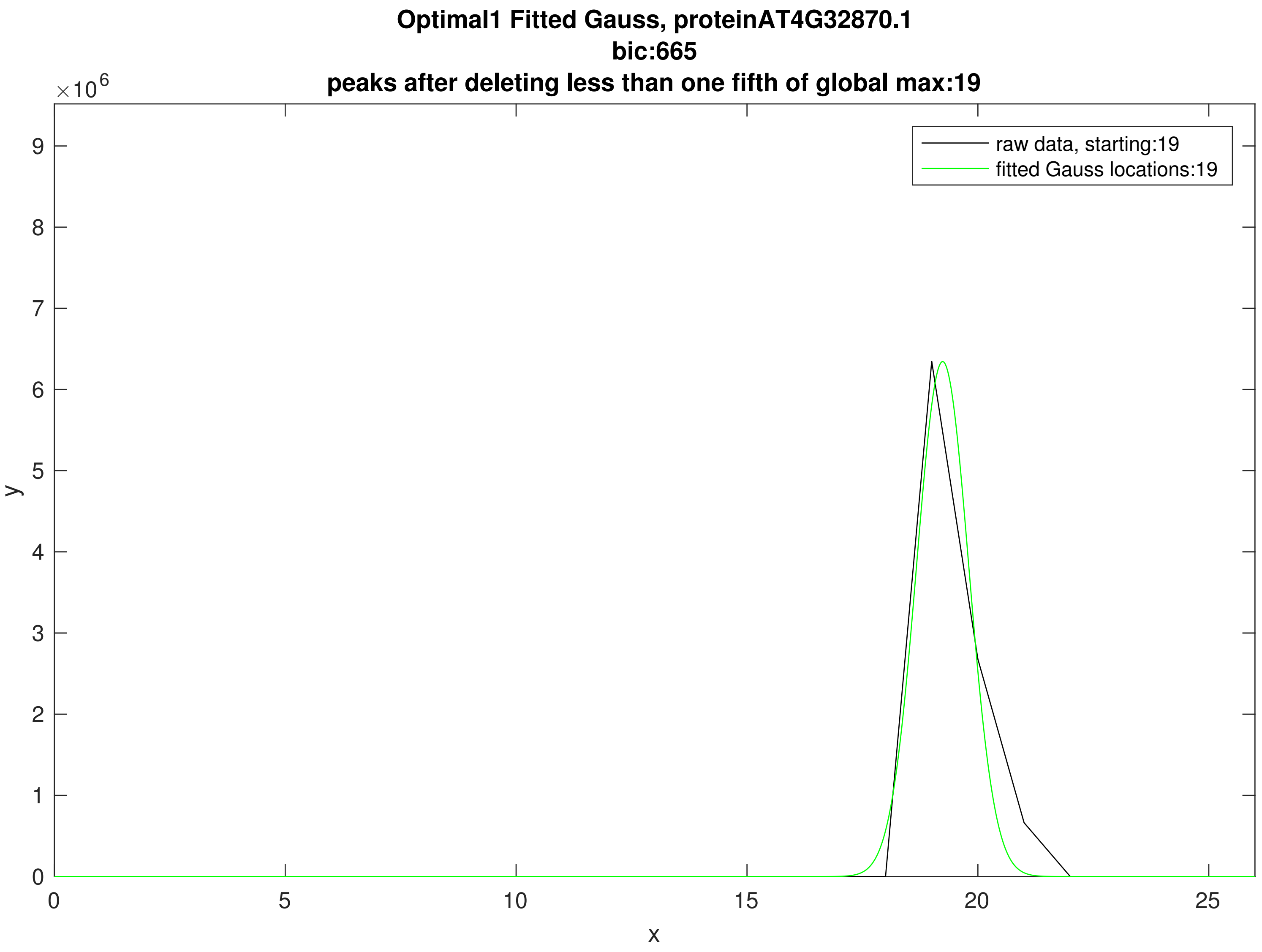
Optimal1 Fitted Gauss, proteinAT4G32870.1
bic:665
peaks after deleting less than one fifth of global max:19
×10^6
raw data, starting:19
fitted Gauss locations:19
y
x
0
1
2
3
4
5
6
7
8
9
0
5
10
15
20
25

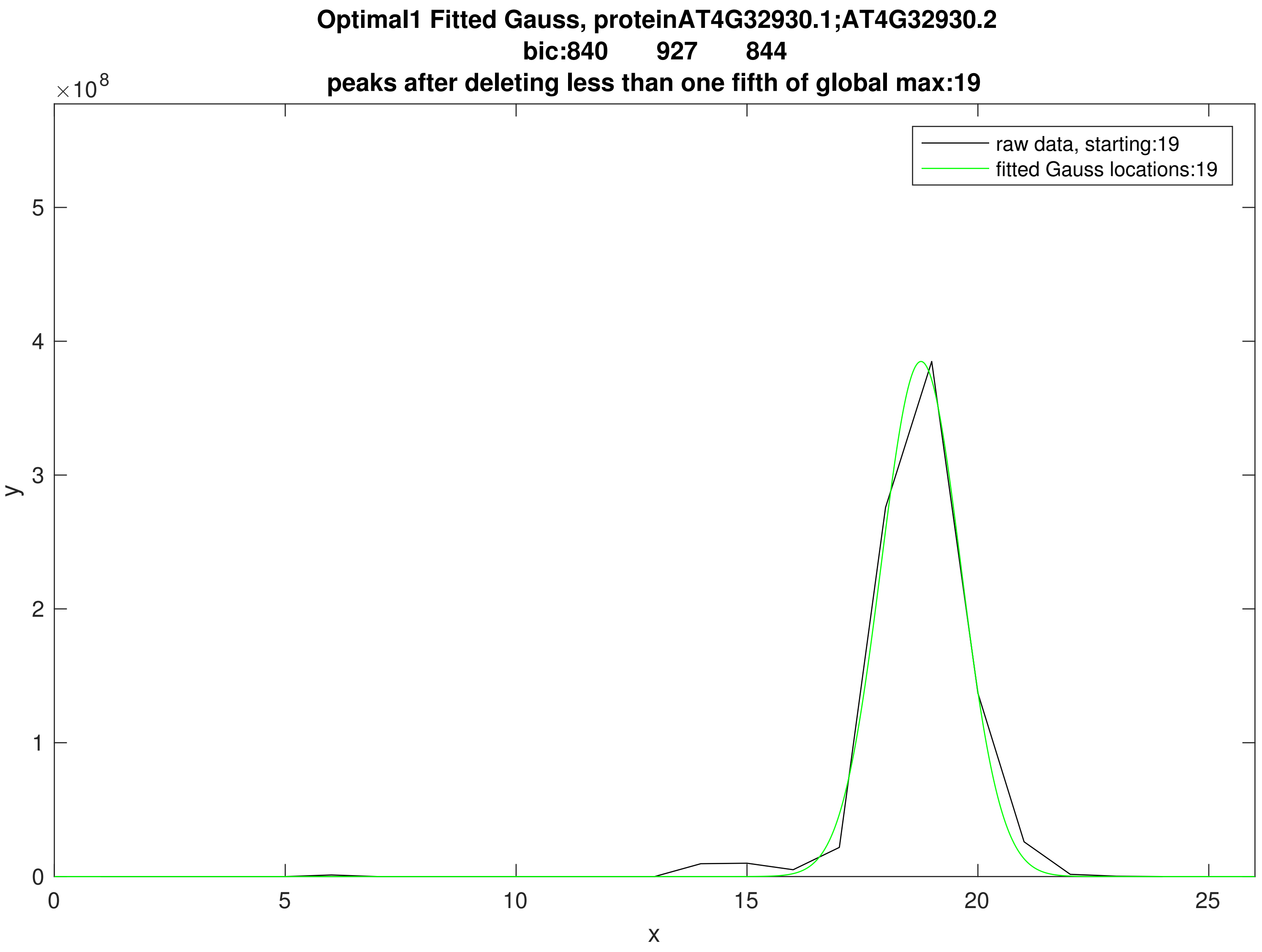
Optimal1 Fitted Gauss, proteinAT4G32930.1;AT4G32930.2
bic:840 927 844
peaks after deleting less than one fifth of global max:19
×10^8
raw data, starting:19
fitted Gauss locations:19
y
x
0
1
2
3
4
5
0
5
10
15
20
25

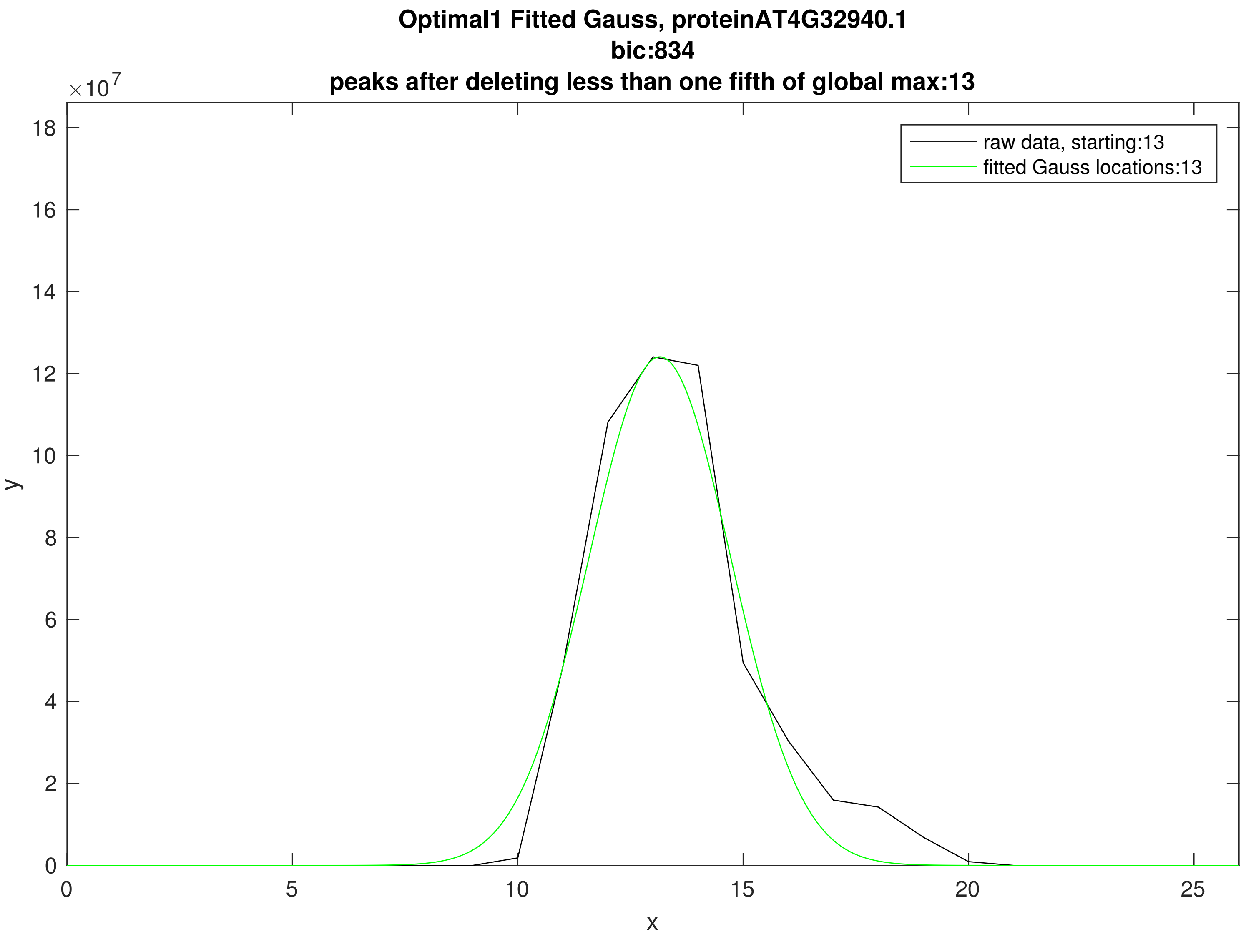

Optimal1 Fitted Gauss, proteinAT4G32940.1
bic:834
peaks after deleting less than one fifth of global max:13
×10^7
raw data, starting:13
fitted Gauss locations:13
18
16
14
12
10
8
6
4
2
0
y
0
5
10
15
20
25
x

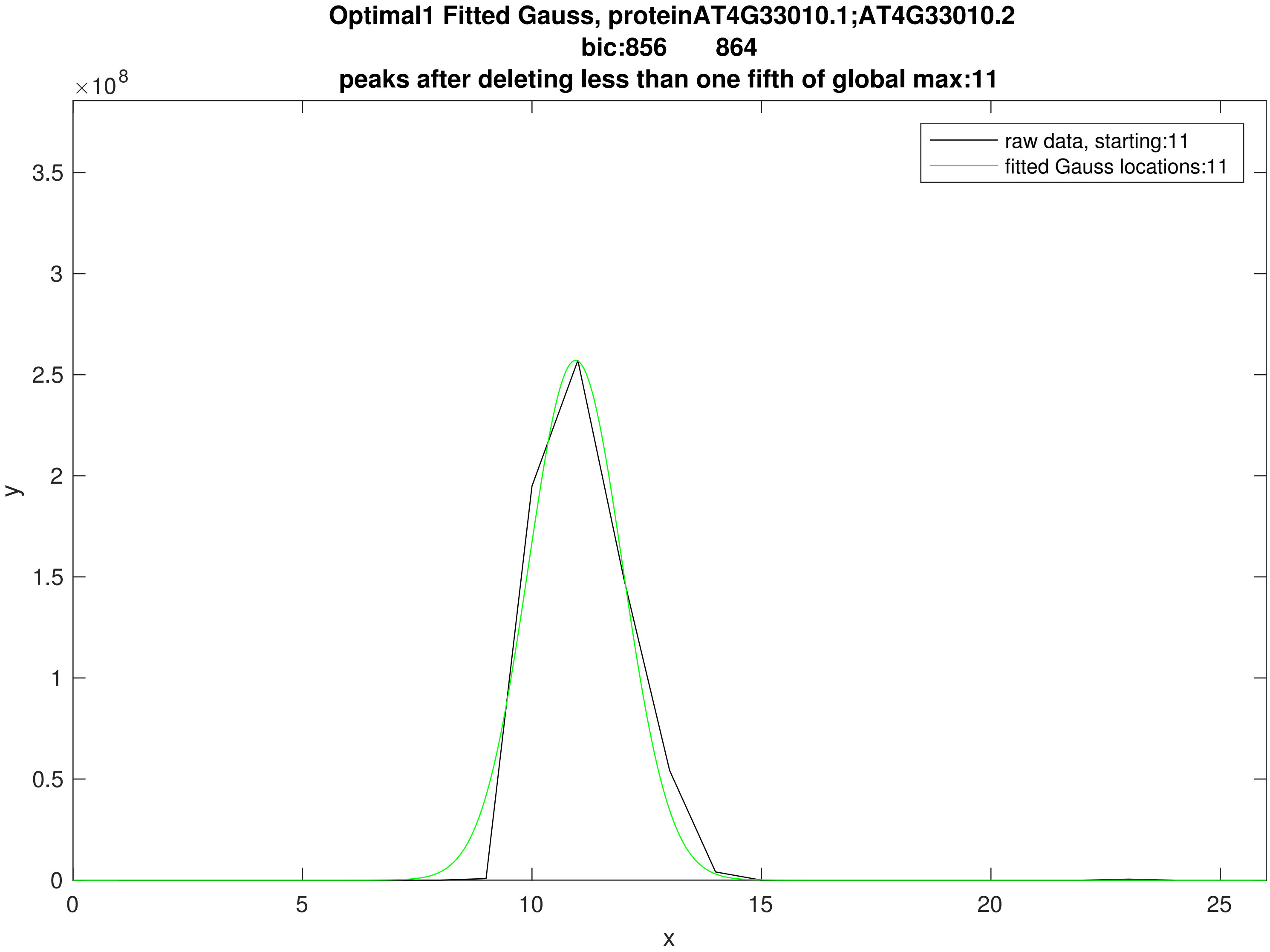
Optimal1 Fitted Gauss, proteinAT4G33010.1;AT4G33010.2
bic:856 864
peaks after deleting less than one fifth of global max:11
×10^8
3.5
3
2.5
2
1.5
1
0.5
0
y
0
5
10
15
20
25
x
raw data, starting:11
fitted Gauss locations:11

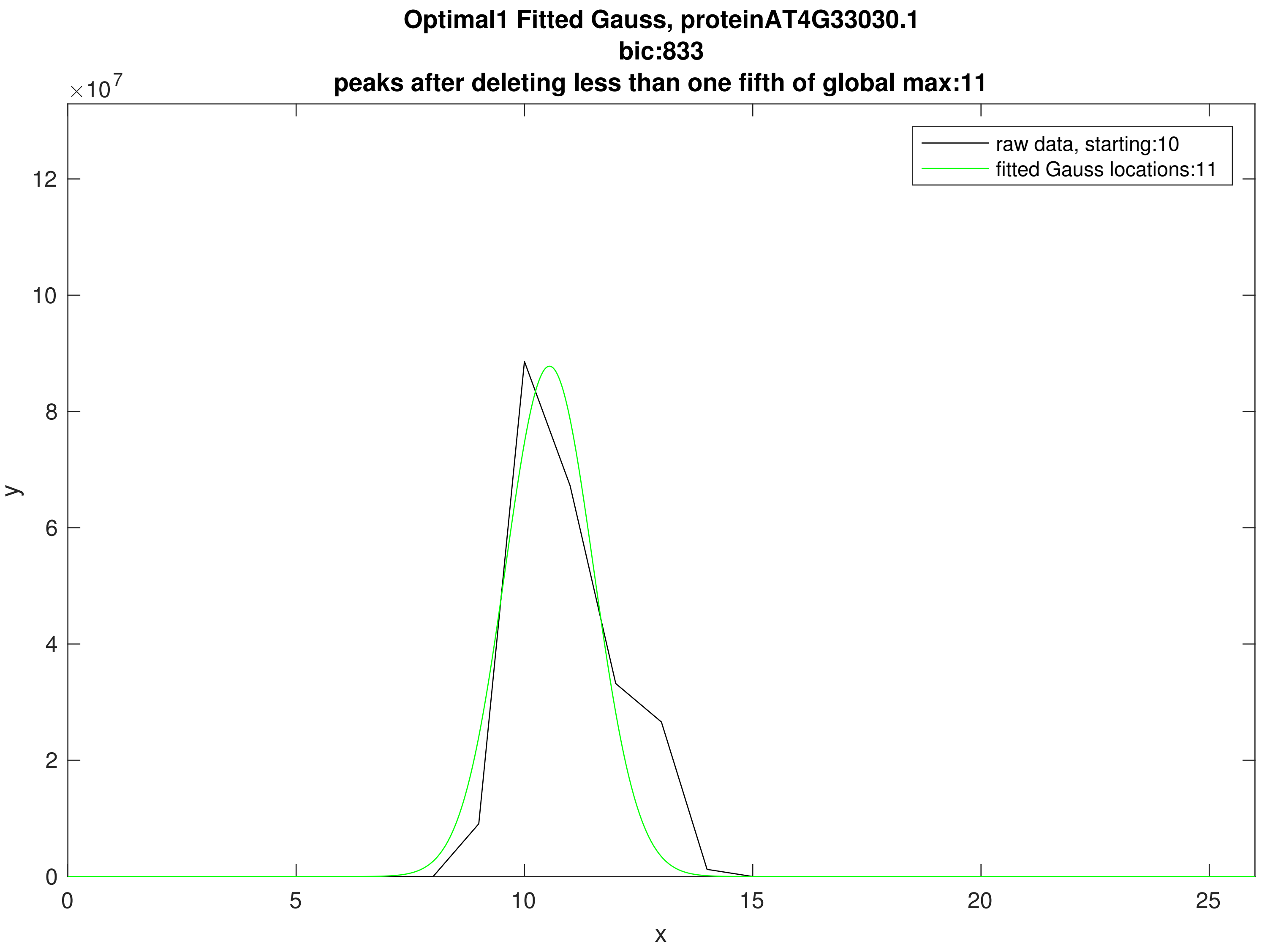
Optimal1 Fitted Gauss, proteinAT4G33030.1
bic:833
peaks after deleting less than one fifth of global max:11
×10^7
raw data, starting:10
fitted Gauss locations:11
12
10
8
6
4
2
0
y
0
5
10
15
20
25
x

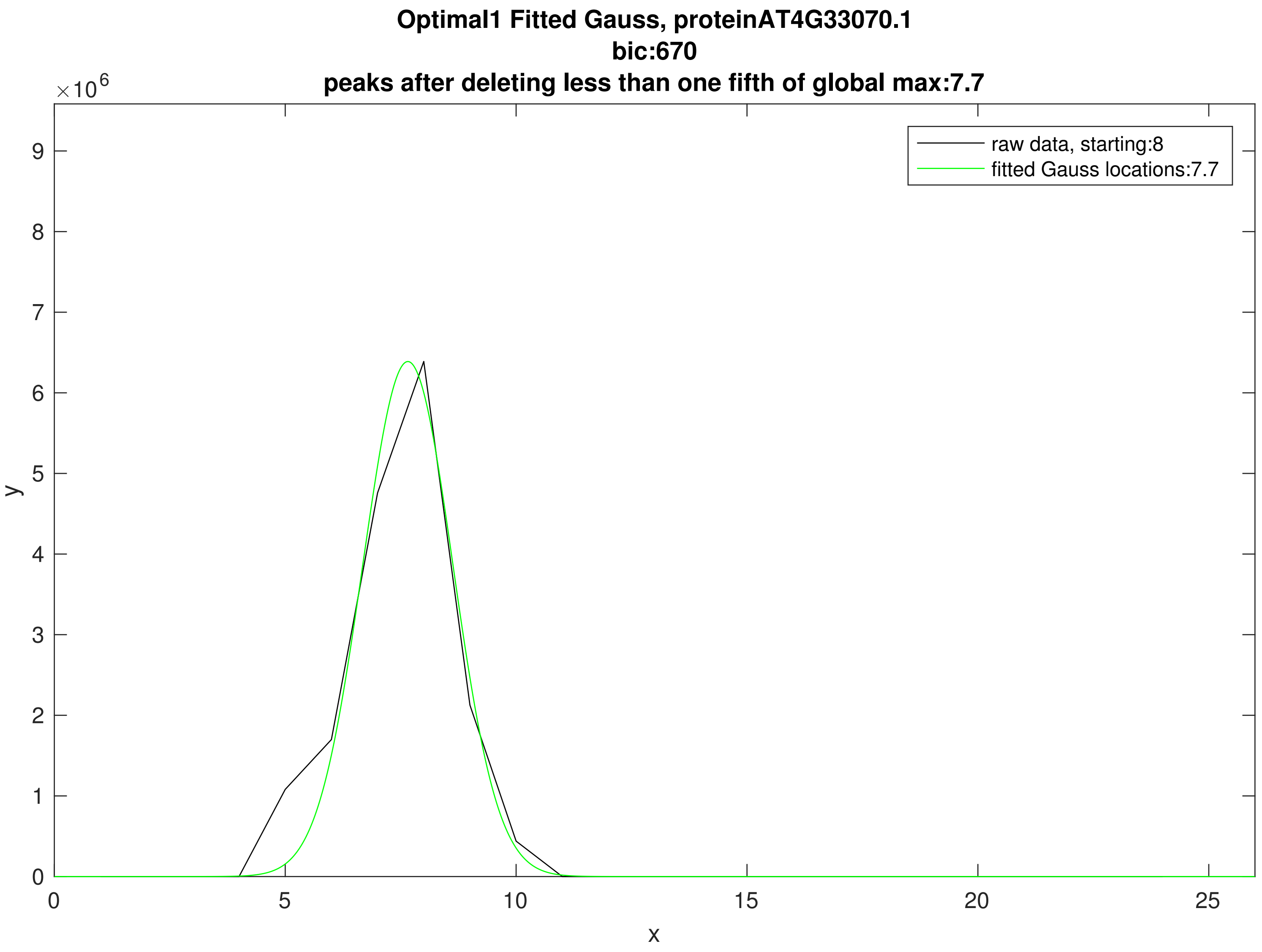
Optimal1 Fitted Gauss, proteinAT4G33070.1
bic:670
peaks after deleting less than one fifth of global max:7.7
×10^6
raw data, starting:8
fitted Gauss locations:7.7
y
x
0
1
2
3
4
5
6
7
8
9
0
5
10
15
20
25

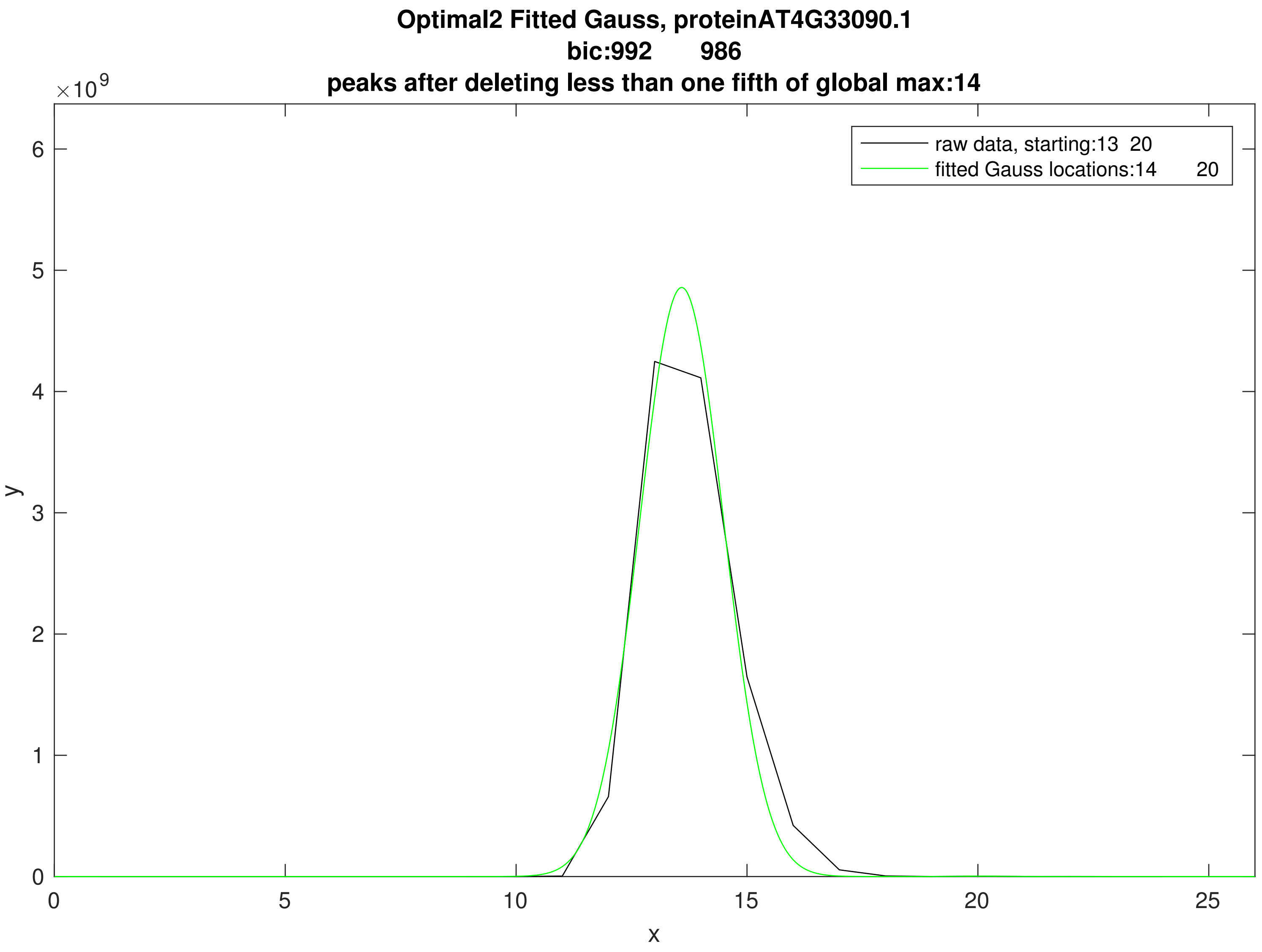

Optimal2 Fitted Gauss, proteinAT4G33090.1
bic:992 986
peaks after deleting less than one fifth of global max:14
×10^9
raw data, starting:13 20
fitted Gauss locations:14 20
y
x
0
1
2
3
4
5
6
0
5
10
15
20
25

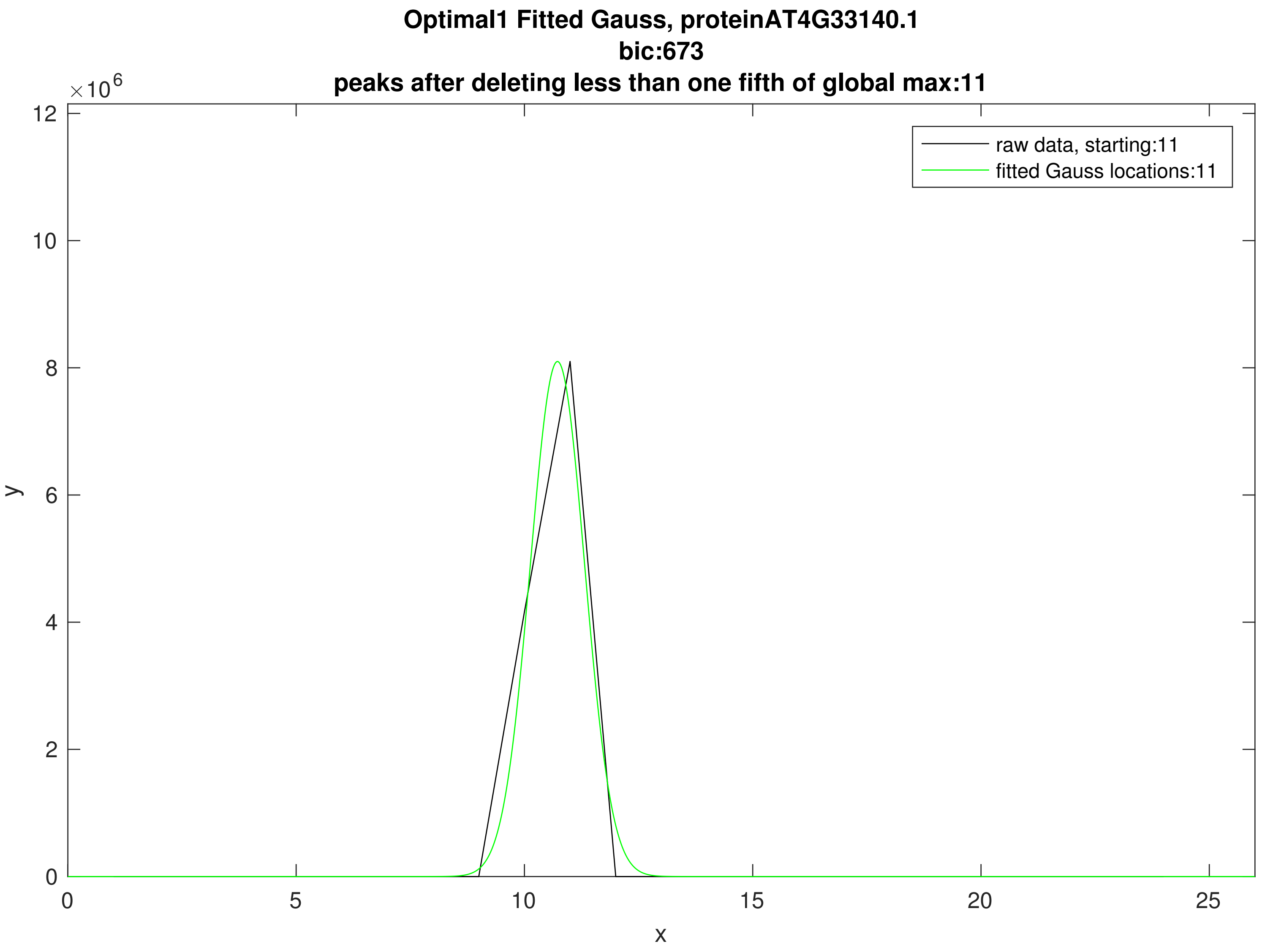

Optimal1 Fitted Gauss, proteinAT4G33140.1
bic:673
peaks after deleting less than one fifth of global max:11
×10^6
12
10
8
6
4
2
0
y
0
5
10
15
20
25
x
raw data, starting:11
fitted Gauss locations:11

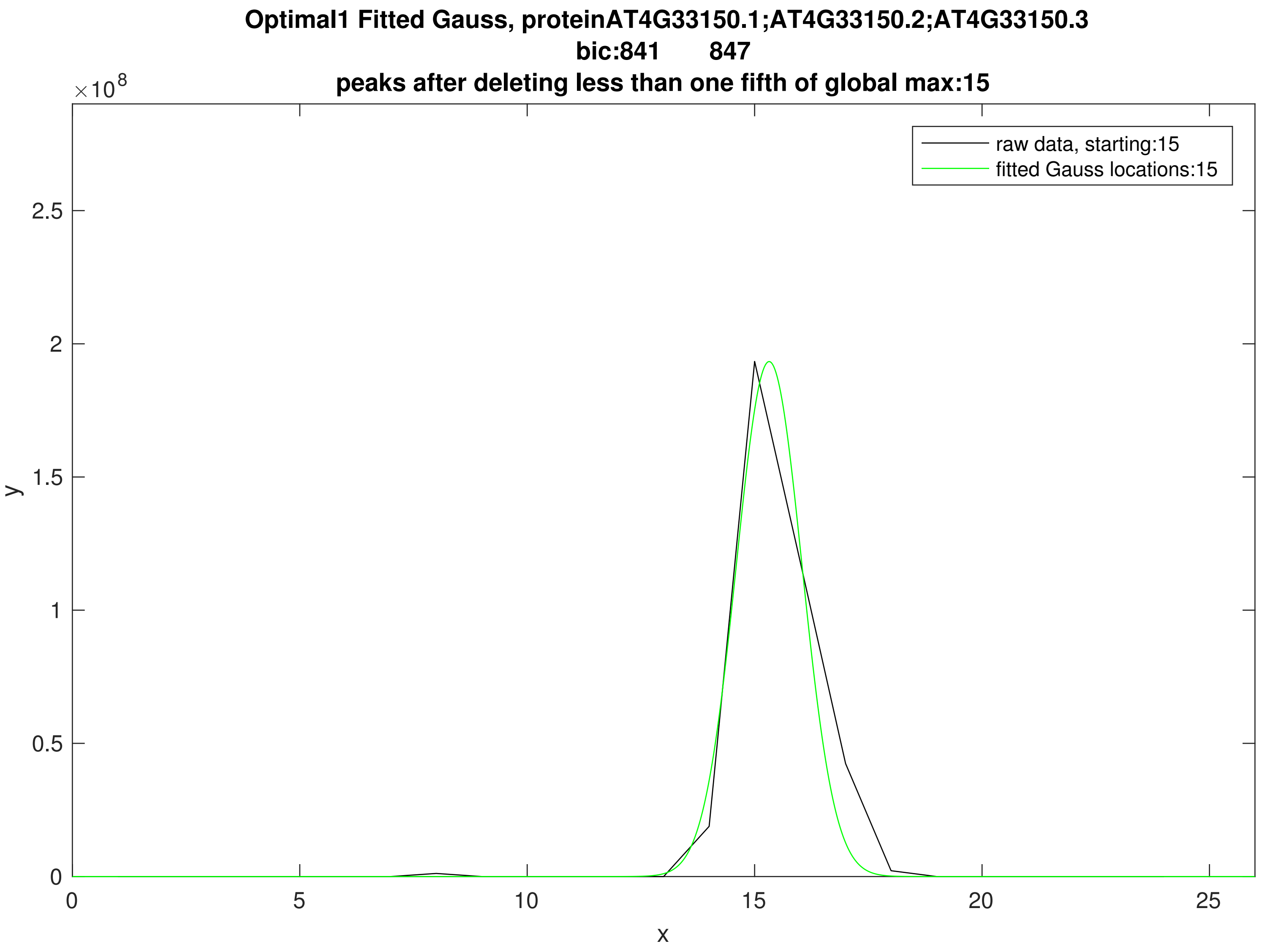
Optimal1 Fitted Gauss, proteinAT4G33150.1;AT4G33150.2;AT4G33150.3
bic:841 847
peaks after deleting less than one fifth of global max:15
×10^8
raw data, starting:15
fitted Gauss locations:15
y
x
0
0.5
1
1.5
2
2.5
5
10
15
20
25

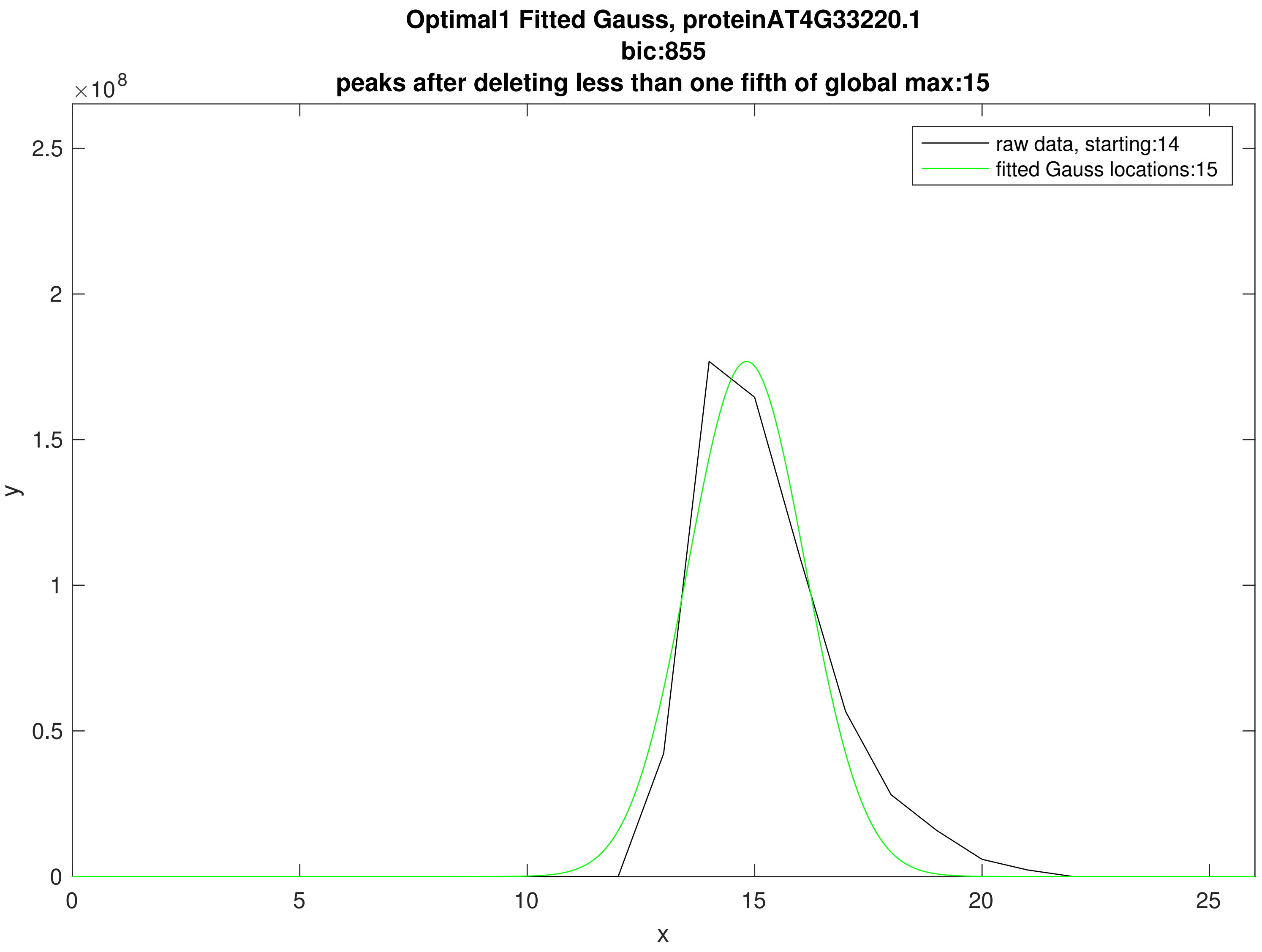

Optimal1 Fitted Gauss, proteinAT4G33220.1
bic:855
peaks after deleting less than one fifth of global max:15
×10^8
2.5
2
1.5
1
0.5
0
y
0
5
10
15
20
25
x
raw data, starting:14
fitted Gauss locations:15

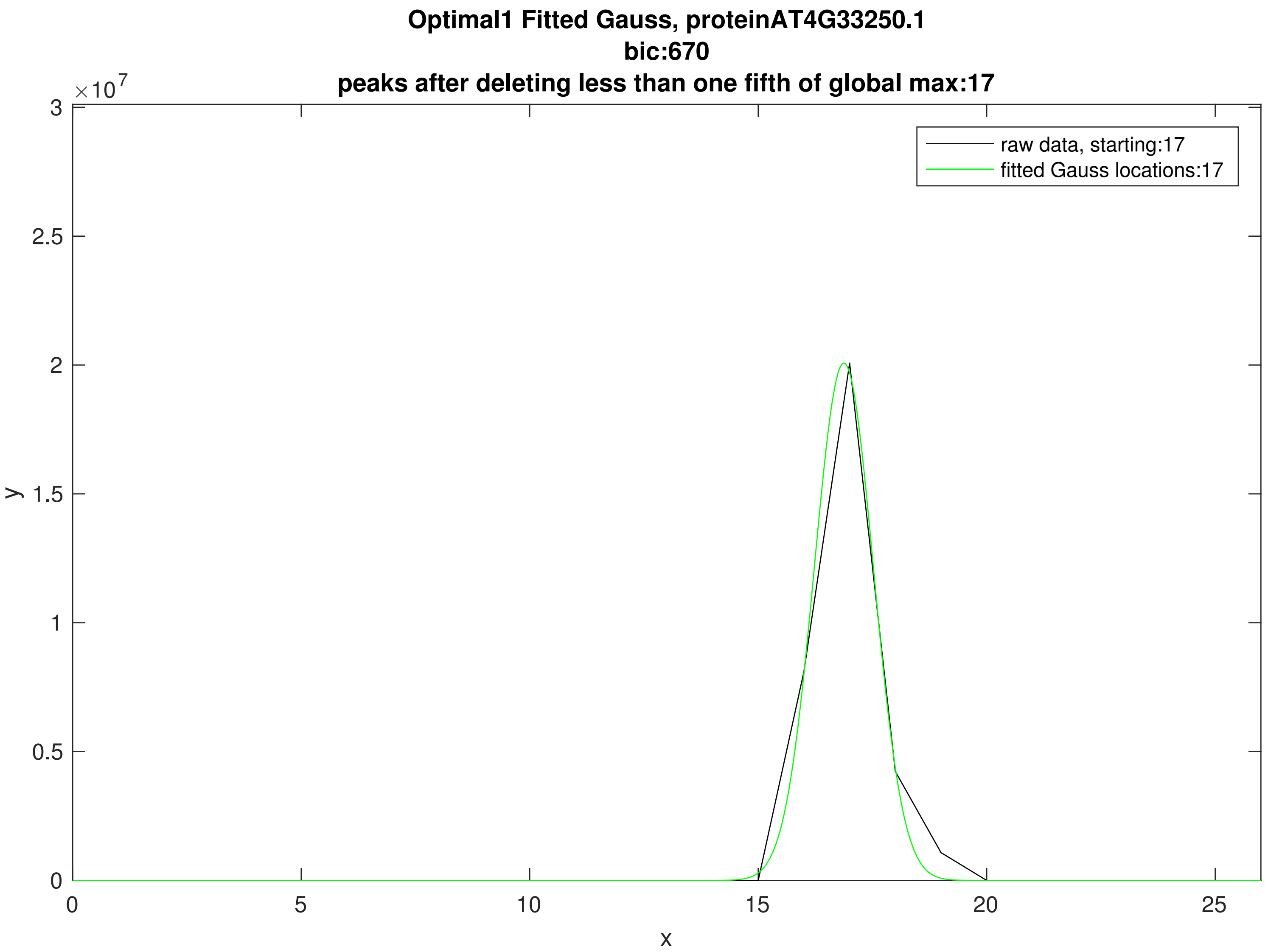
Optimal1 Fitted Gauss, proteinAT4G33250.1
bic:670
peaks after deleting less than one fifth of global max:17
raw data, starting:17
fitted Gauss locations:17
$\times 10^7$
3
2.5
2
1.5
1
0.5
0
y
0
5
10
15
20
25
x

#### Optimal1 Fitted Gauss, proteinAT4G33420.1
## bic:732       737
#### peaks after deleting less than one fifth of global max:18

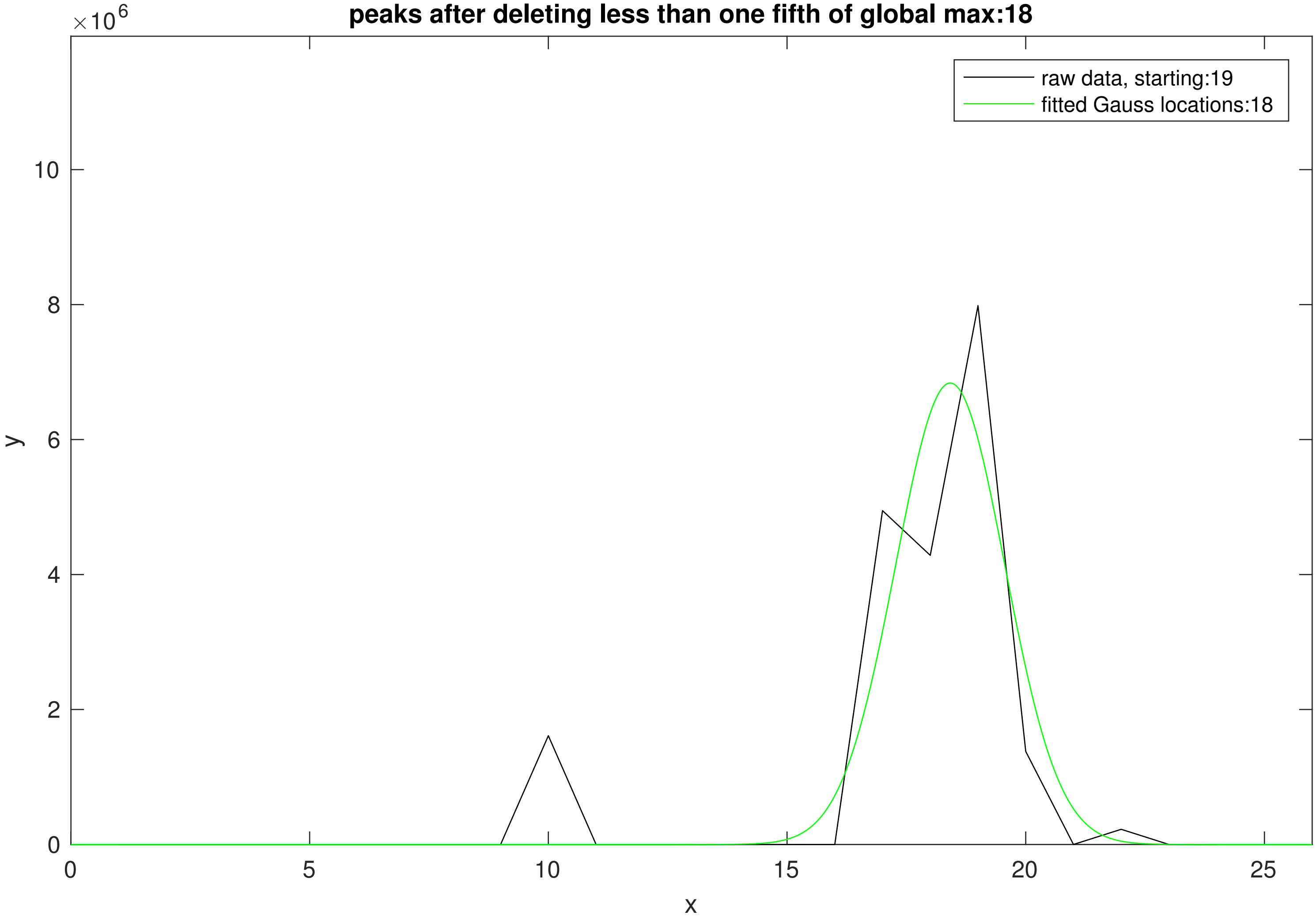

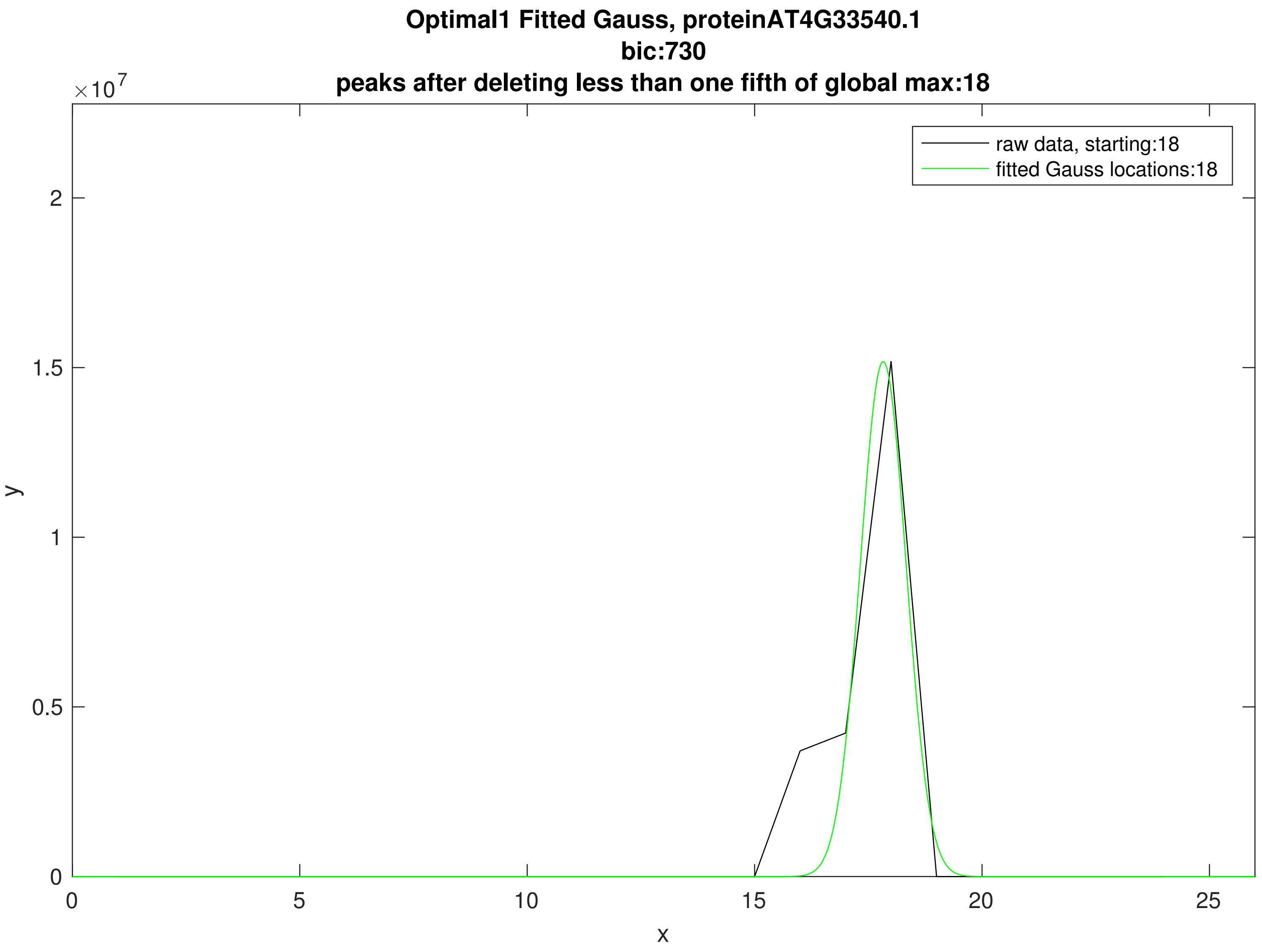
Optimal1 Fitted Gauss, proteinAT4G33540.1
bic:730
peaks after deleting less than one fifth of global max:18
×10^7
raw data, starting:18
fitted Gauss locations:18
y
x
0
0.5
1
1.5
2
0
5
10
15
20
25

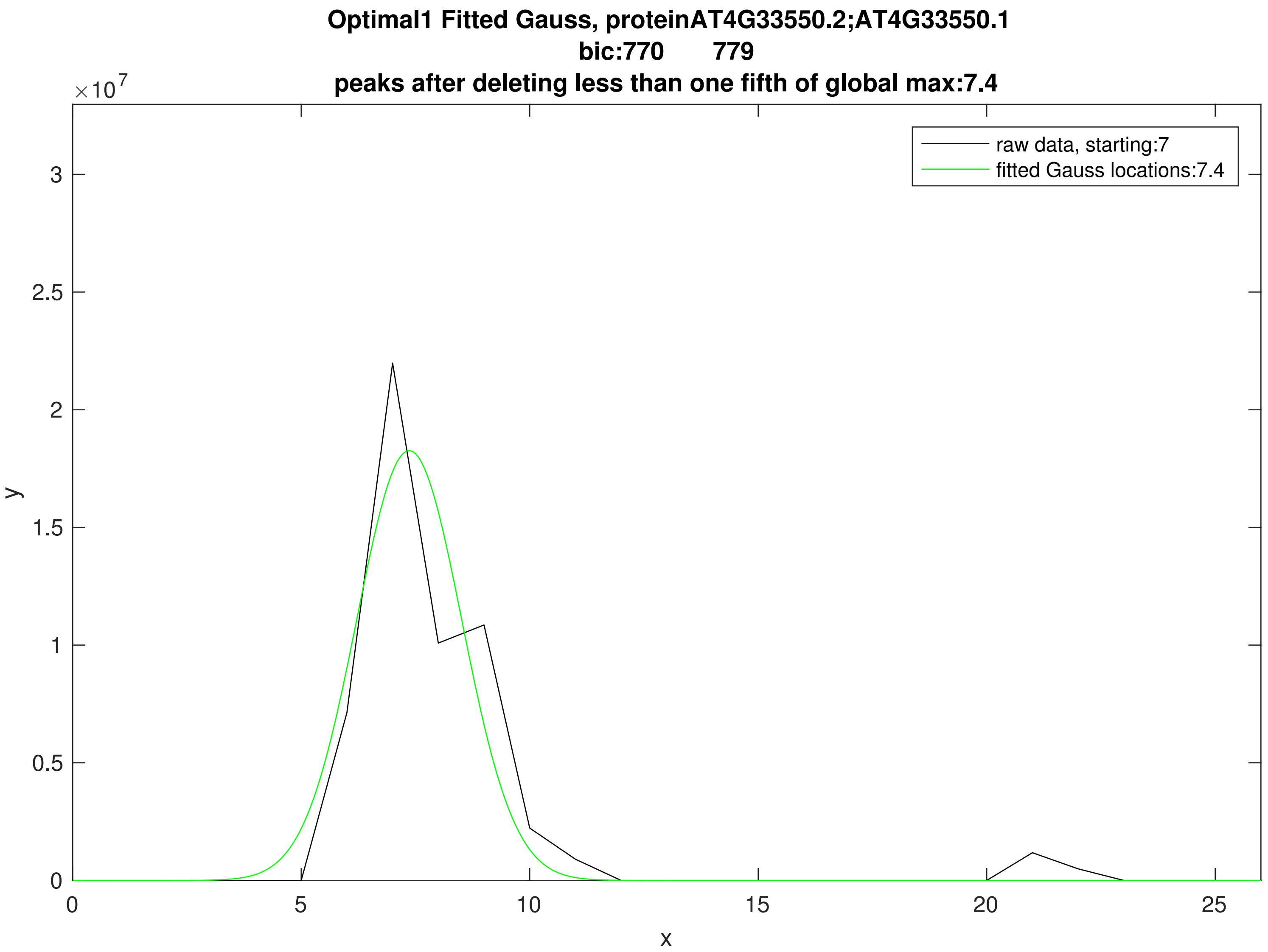

Optimal1 Fitted Gauss, proteinAT4G33550.2;AT4G33550.1
bic:770 779
peaks after deleting less than one fifth of global max:7.4
×10^7
3
2.5
2
1.5
1
0.5
0
y
0
5
10
15
20
25
x
raw data, starting:7
fitted Gauss locations:7.4

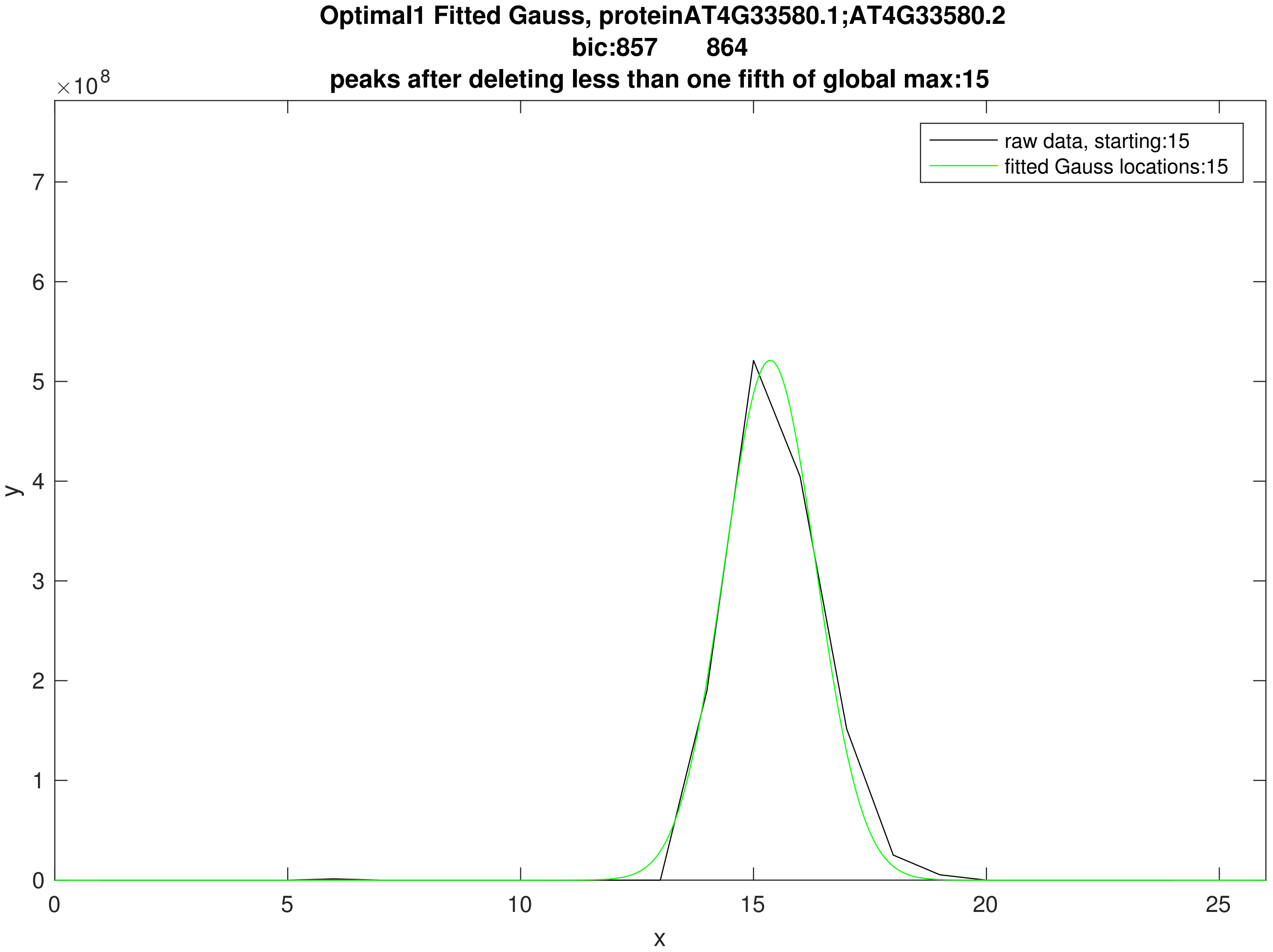
Optimal1 Fitted Gauss, proteinAT4G33580.1;AT4G33580.2
bic:857 864
peaks after deleting less than one fifth of global max:15
×10^8
raw data, starting:15
fitted Gauss locations:15
y
x
0
1
2
3
4
5
6
7
0
5
10
15
20
25

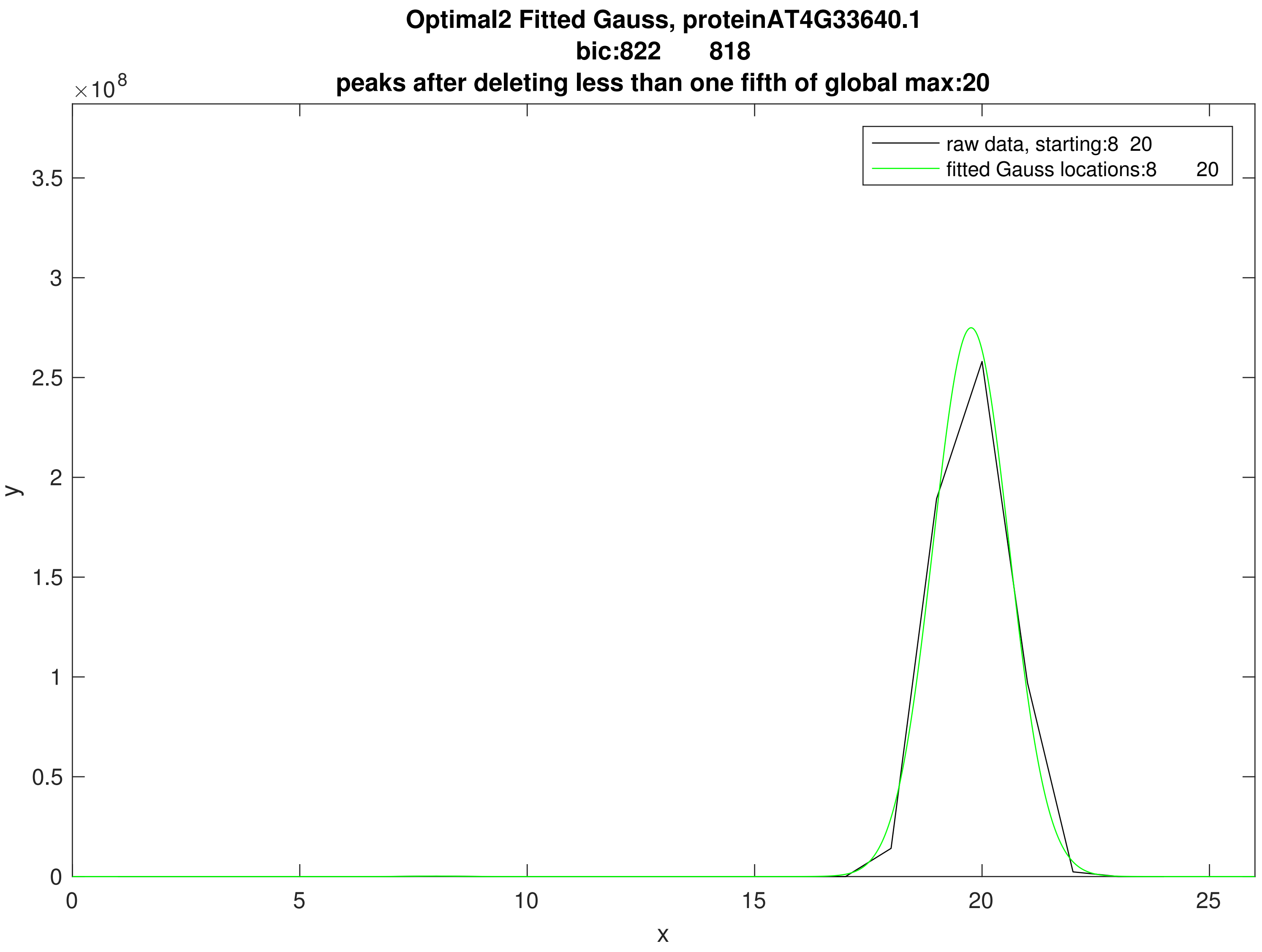
Optimal2 Fitted Gauss, proteinAT4G33640.1
bic:822 818
peaks after deleting less than one fifth of global max:20
×10^8
raw data, starting:8 20
fitted Gauss locations:8 20
y
x
0
0.5
1
1.5
2
2.5
3
3.5
5
10
15
20
25

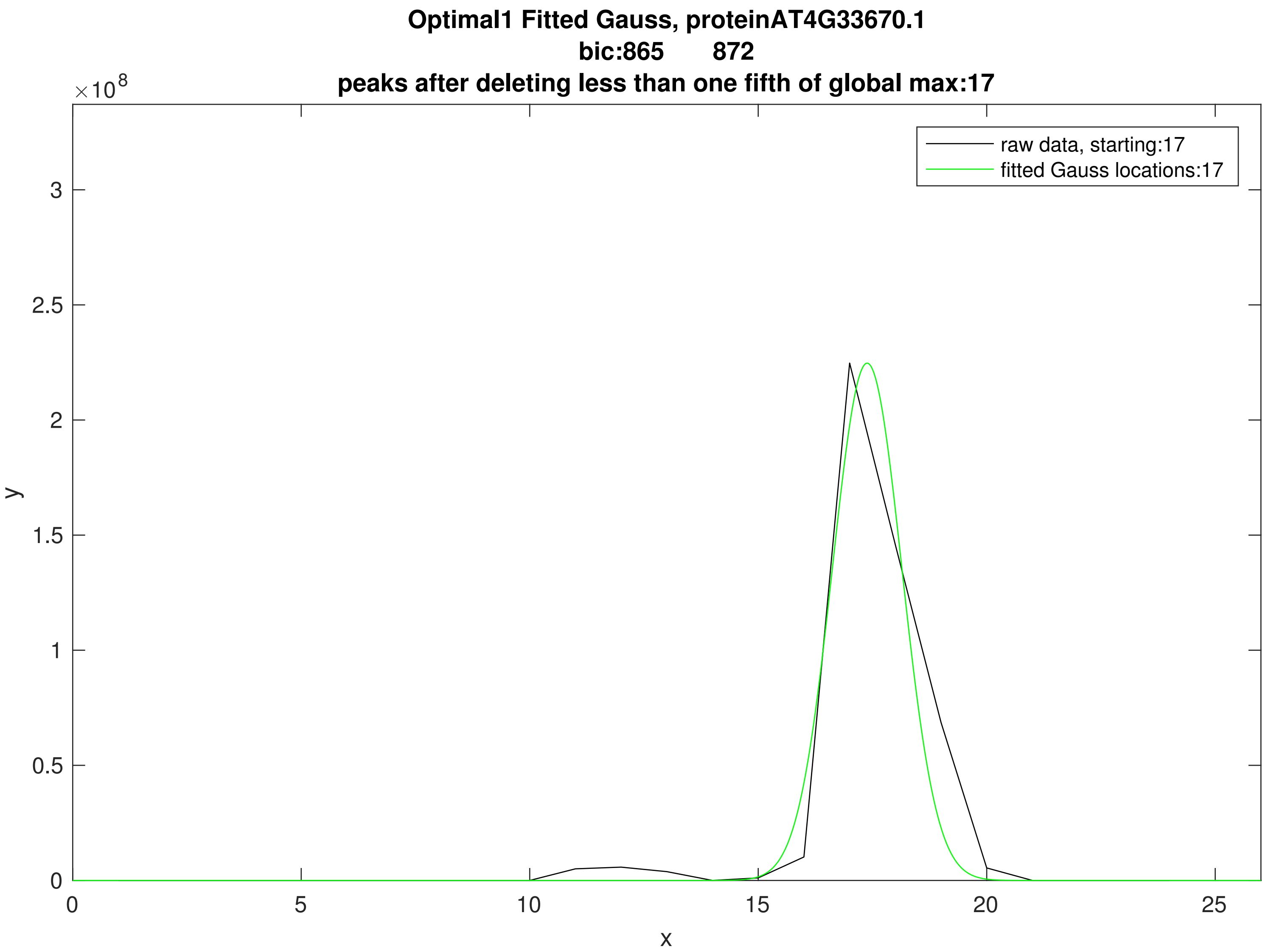
Optimal1 Fitted Gauss, proteinAT4G33670.1
bic:865 872
peaks after deleting less than one fifth of global max:17
×10^8
raw data, starting:17
fitted Gauss locations:17
3
2.5
2
1.5
1
0.5
0
y
0
5
10
15
20
25
x

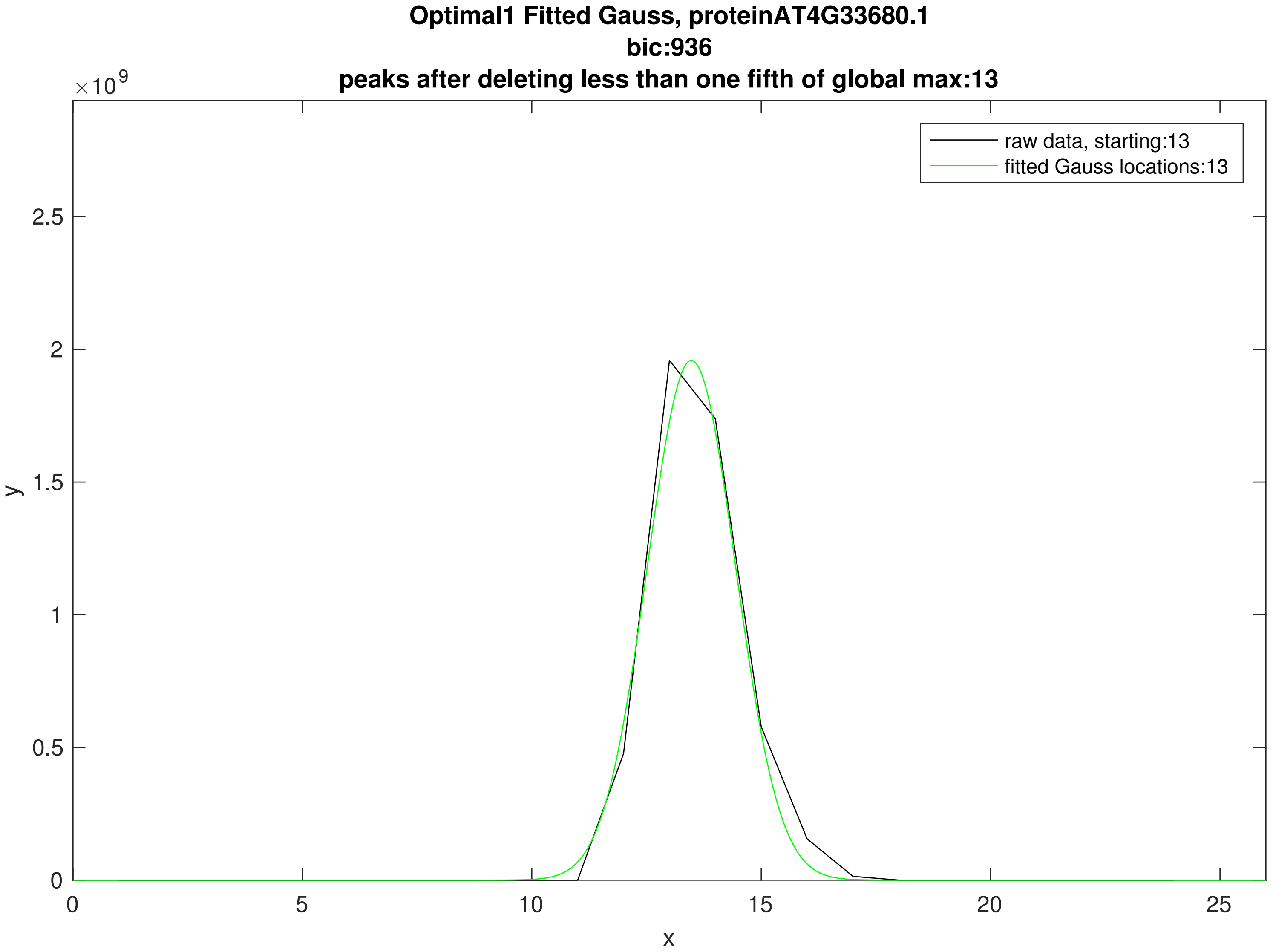

Optimal1 Fitted Gauss, proteinAT4G33680.1
bic:936
peaks after deleting less than one fifth of global max:13
×10^9
raw data, starting:13
fitted Gauss locations:13
2.5
2
1.5
1
0.5
0
y
0
5
10
15
20
25
x

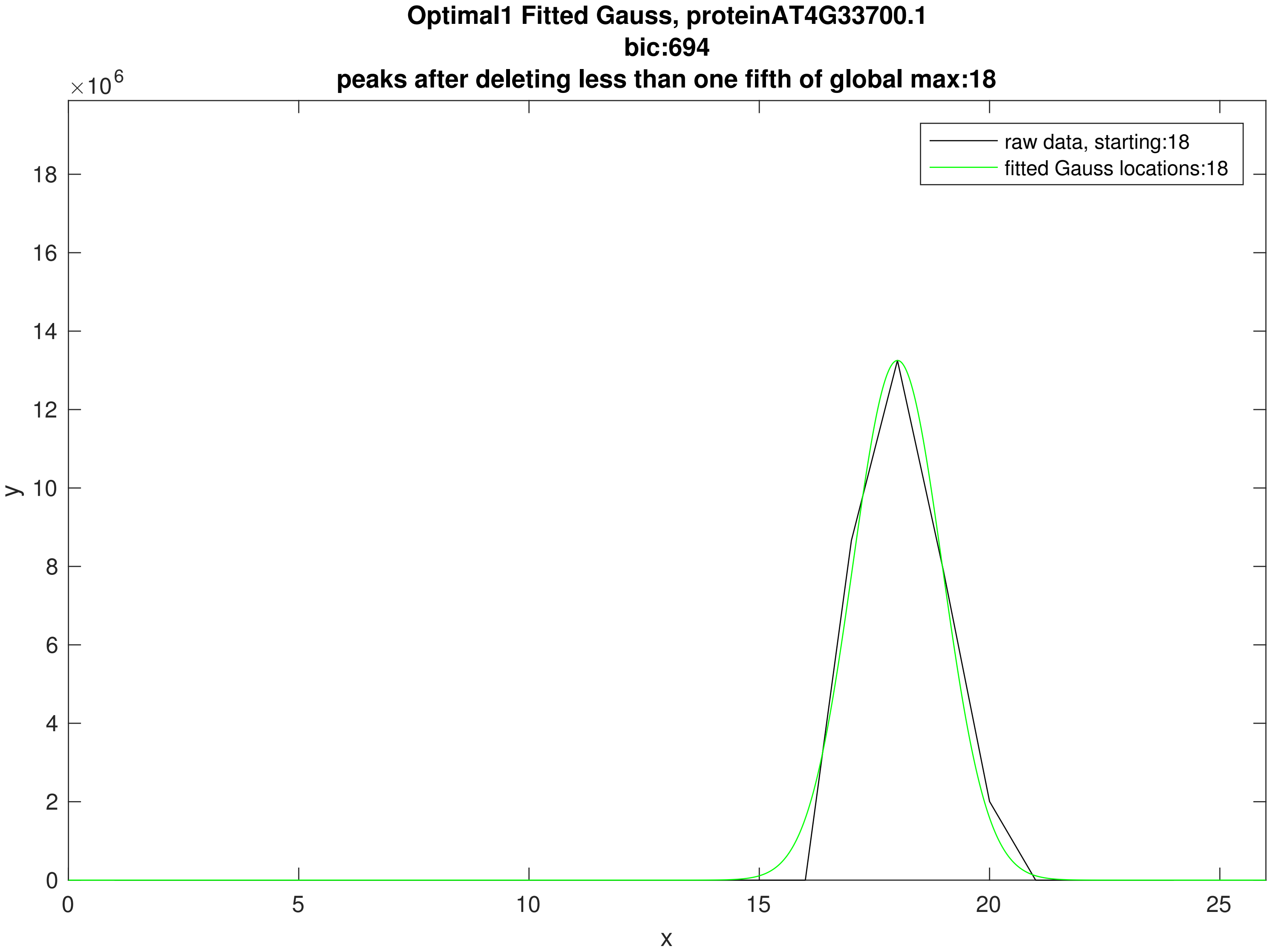
Optimal1 Fitted Gauss, proteinAT4G33700.1
bic:694
peaks after deleting less than one fifth of global max:18
×10^6
raw data, starting:18
fitted Gauss locations:18
y
x
0
2
4
6
8
10
12
14
16
18
0
5
10
15
20
25

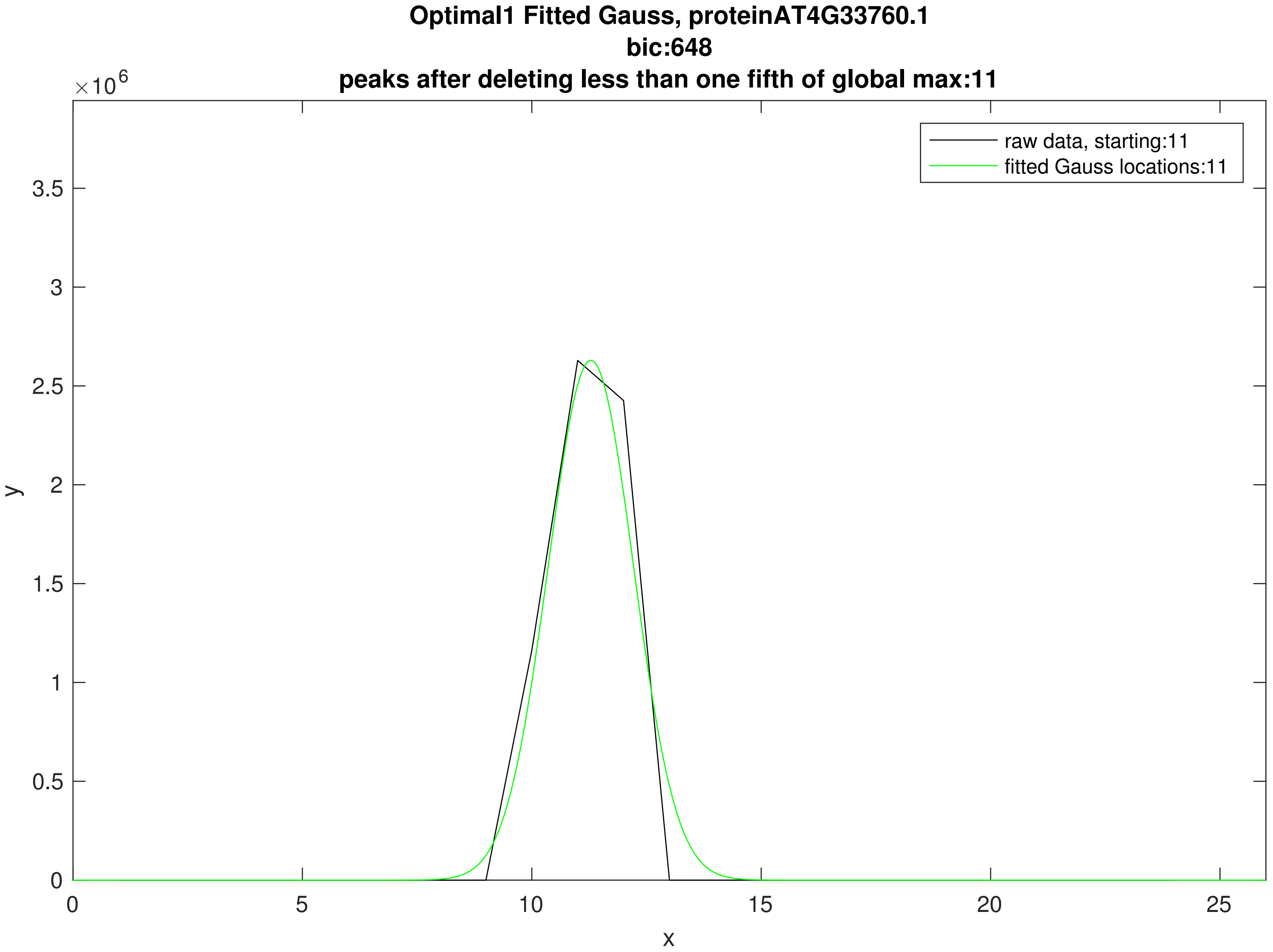
Optimal1 Fitted Gauss, proteinAT4G33760.1
bic:648
peaks after deleting less than one fifth of global max:11
×10^6
raw data, starting:11
fitted Gauss locations:11
y
x
0
0.5
1
1.5
2
2.5
3
3.5
0
5
10
15
20
25

**Optimal1 Fitted Gauss, proteinAT4G33865.1;AT3G44010.1;AT3G43980.1**
**bic:634**
**peaks after deleting less than one fifth of global max:3.1**

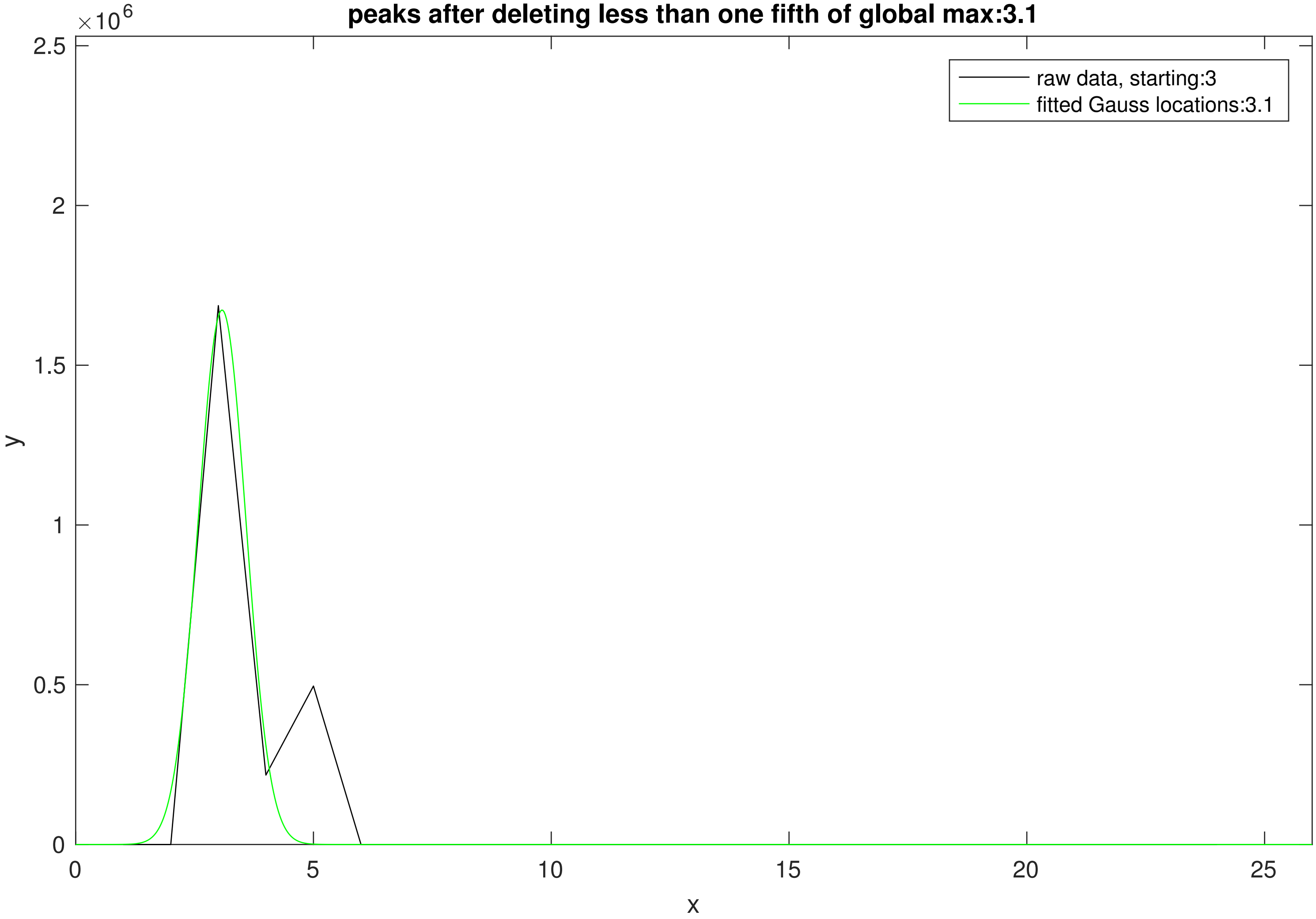

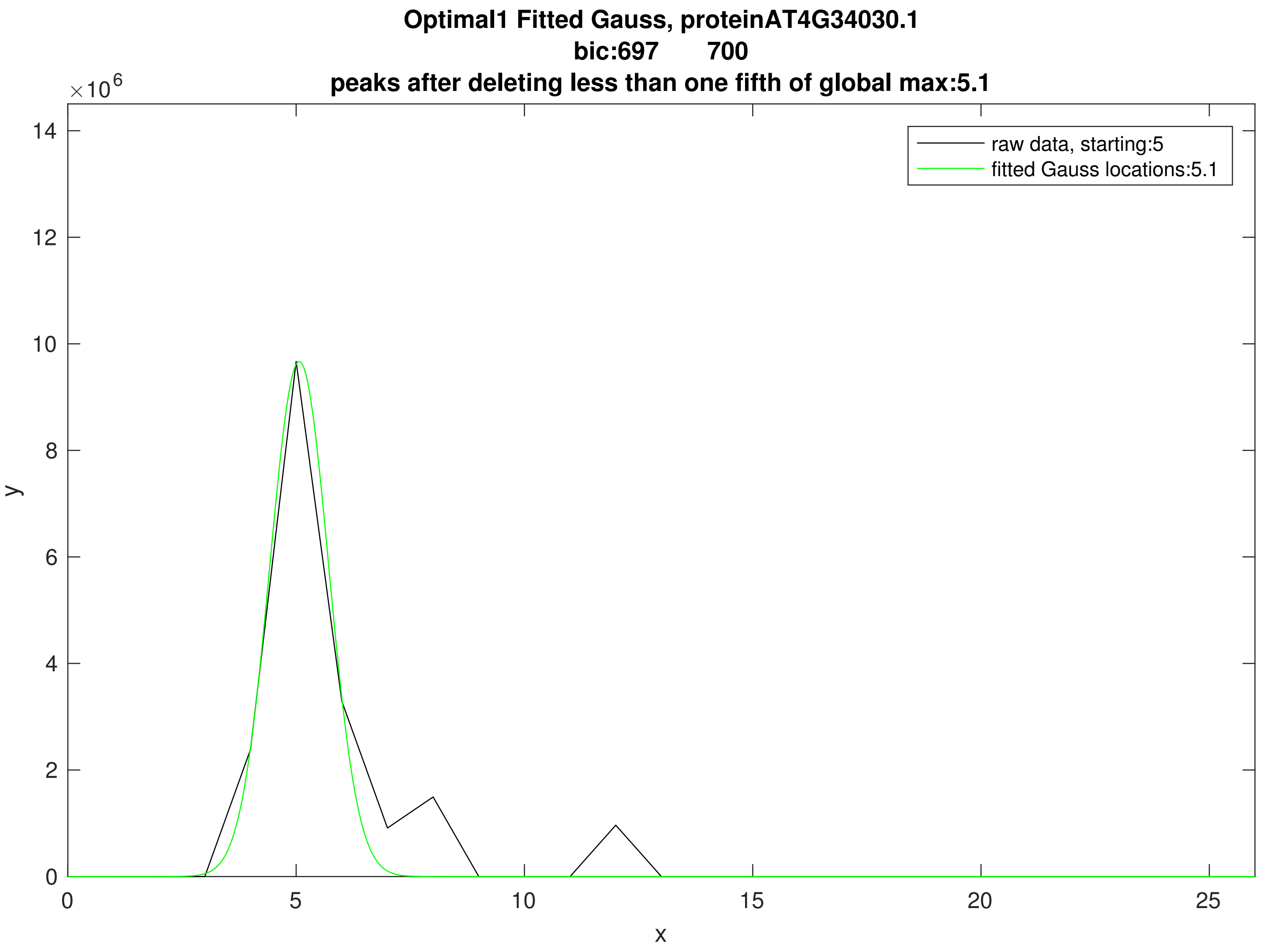
Optimal1 Fitted Gauss, proteinAT4G34030.1
bic:697 700
peaks after deleting less than one fifth of global max:5.1
×10^6
raw data, starting:5
fitted Gauss locations:5.1
y
x
0
2
4
6
8
10
12
14
0
5
10
15
20
25

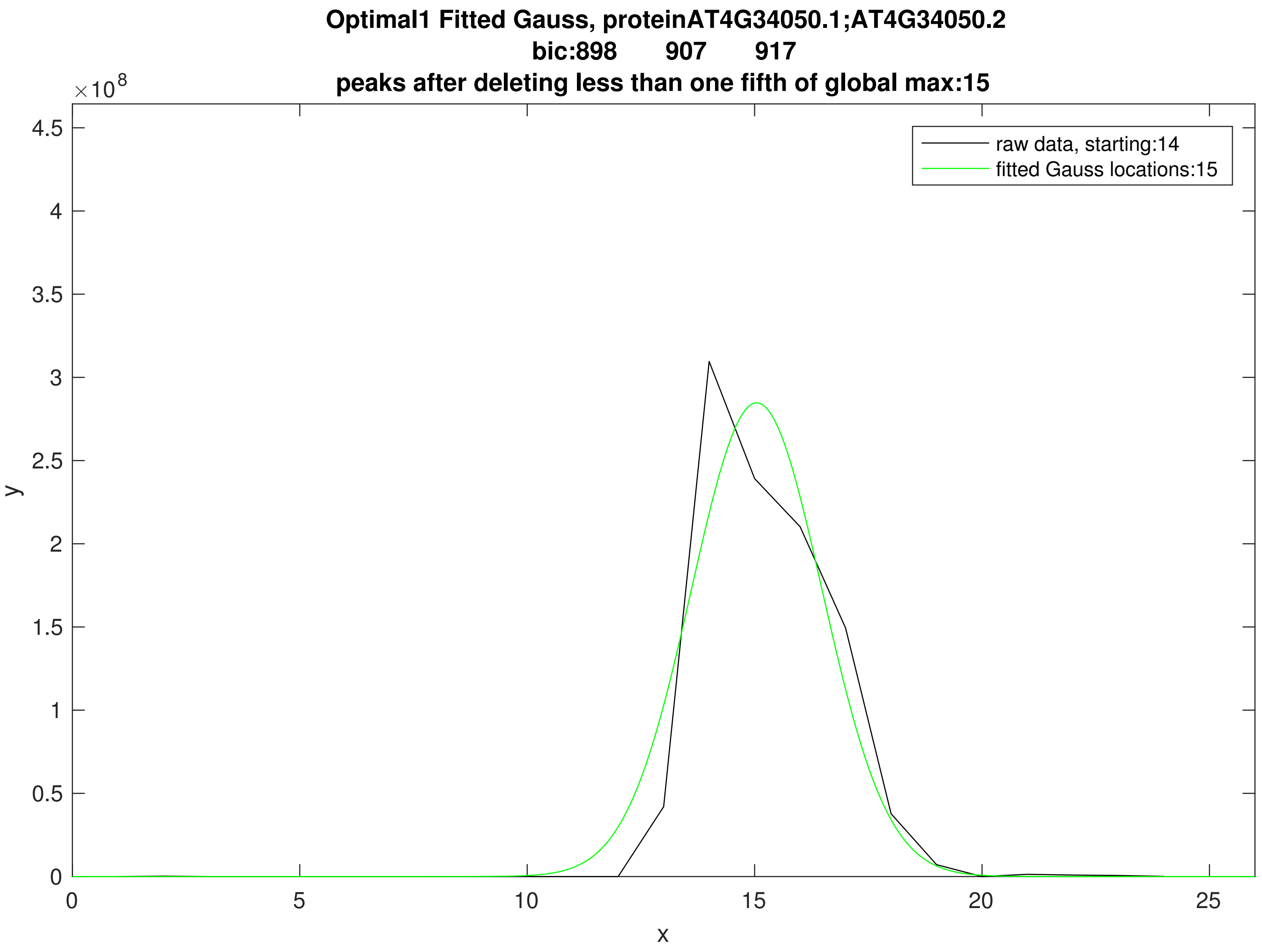
Optimal1 Fitted Gauss, proteinAT4G34050.1;AT4G34050.2
bic:898 907 917
peaks after deleting less than one fifth of global max:15
×10^8
raw data, starting:14
fitted Gauss locations:15
y
x
0
0.5
1
1.5
2
2.5
3
3.5
4
4.5
5
10
15
20
25

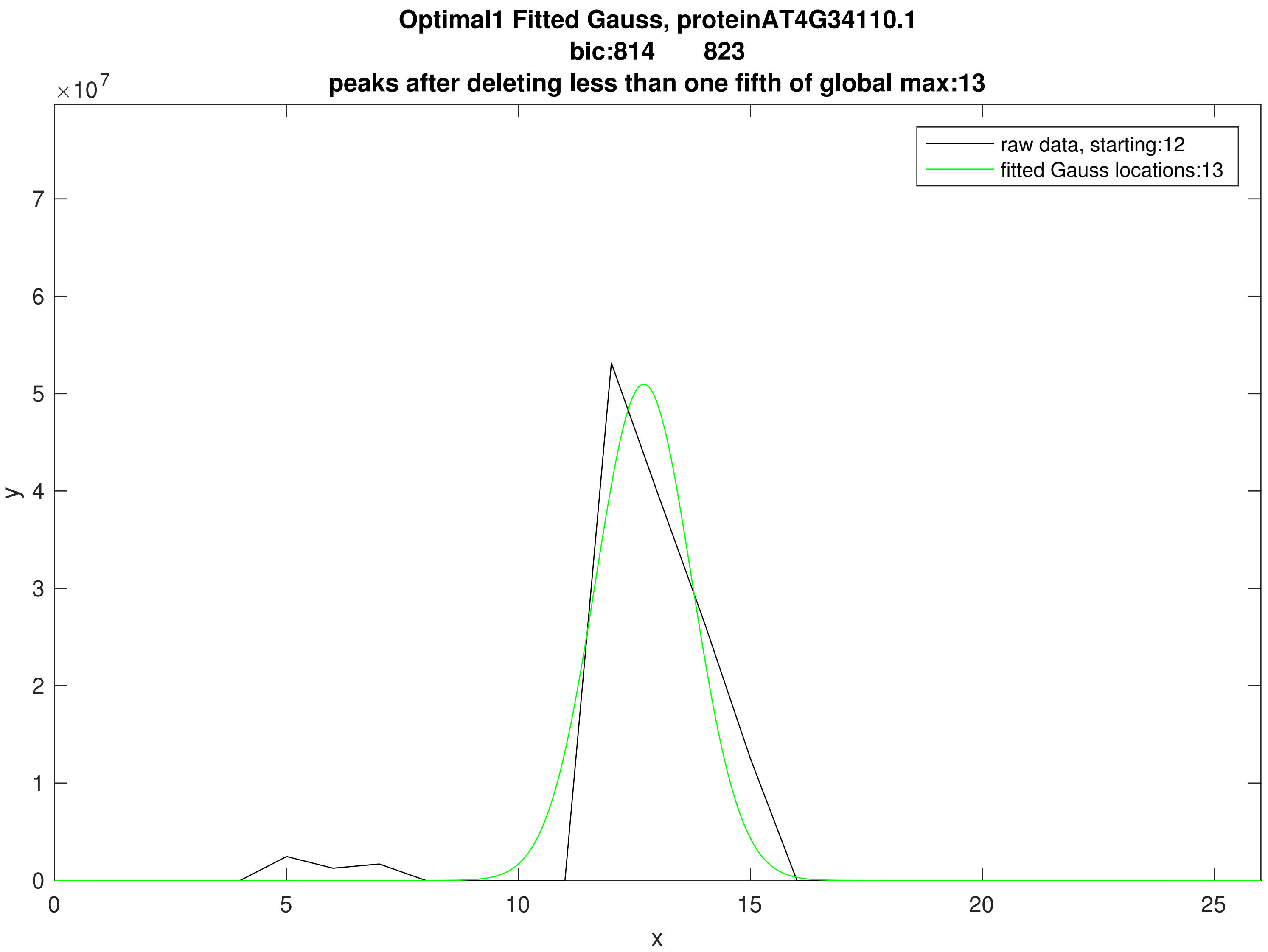

Optimal1 Fitted Gauss, proteinAT4G34110.1
bic:814 823
peaks after deleting less than one fifth of global max:13
×10^7
raw data, starting:12
fitted Gauss locations:13
y
x
0
1
2
3
4
5
6
7
0
5
10
15
20
25

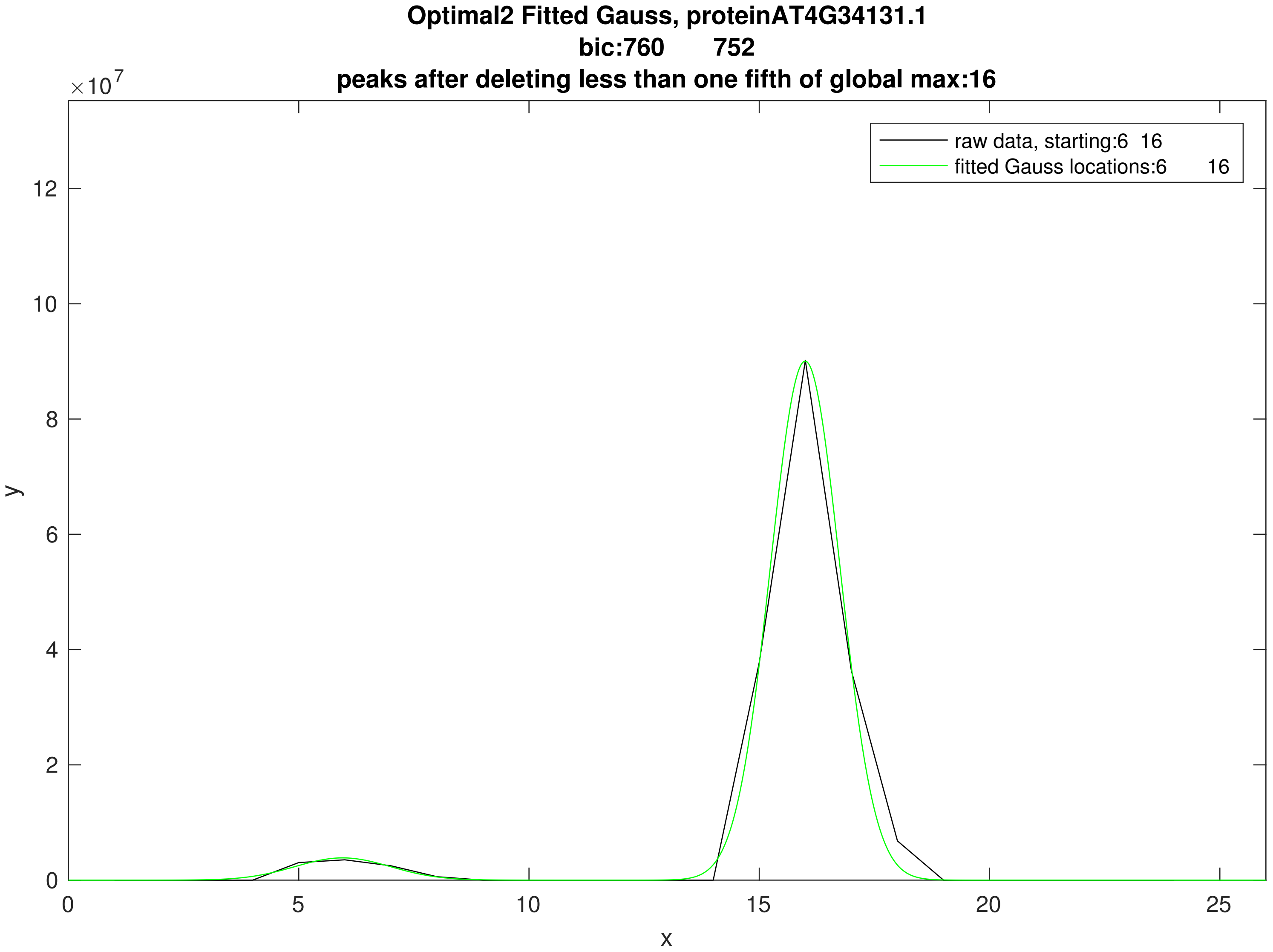

Optimal2 Fitted Gauss, proteinAT4G34131.1
bic:760 752
peaks after deleting less than one fifth of global max:16
×10^7
raw data, starting:6 16
fitted Gauss locations:6 16
y
x
0
2
4
6
8
10
12
0
5
10
15
20
25

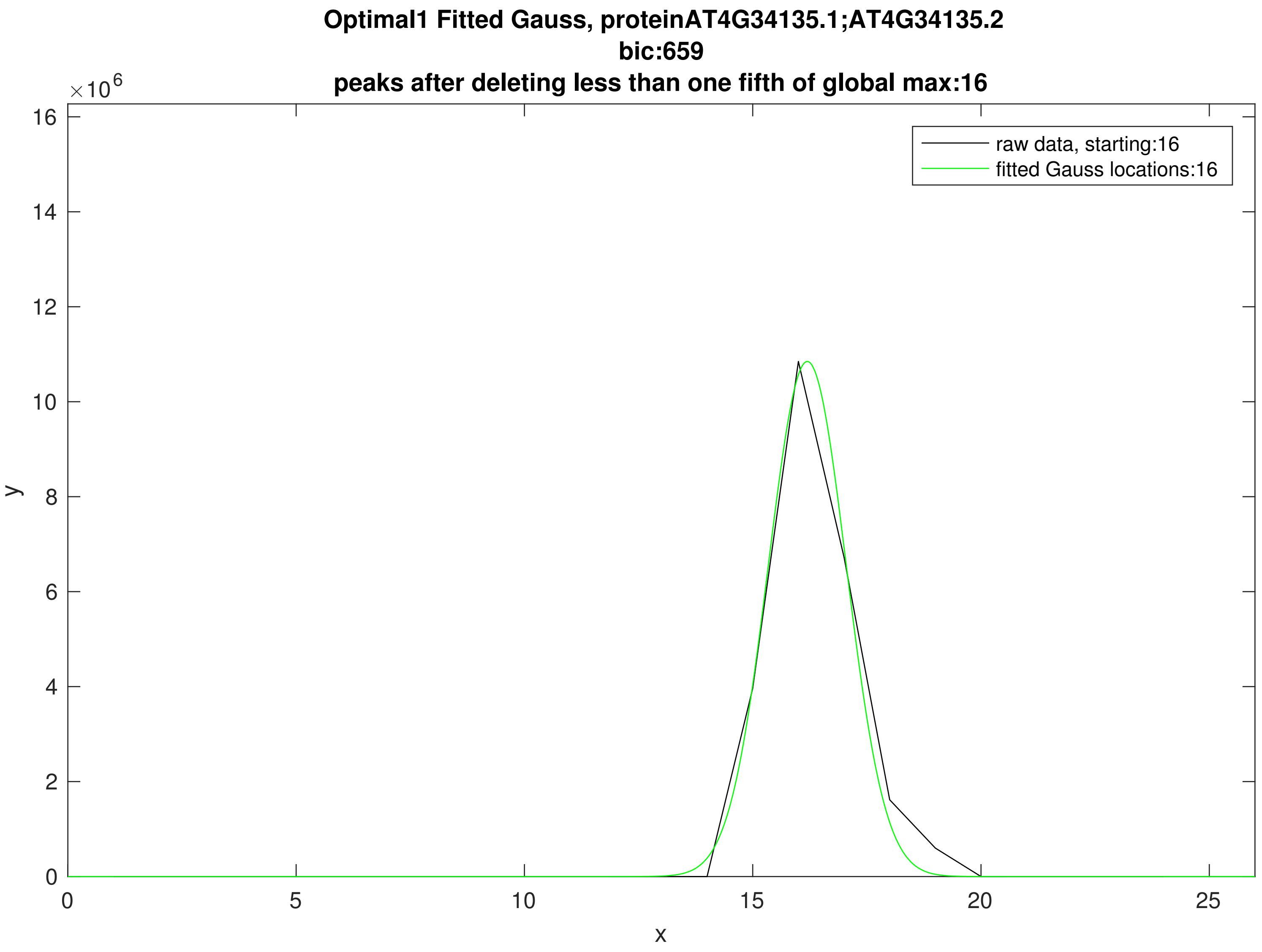
Optimal1 Fitted Gauss, proteinAT4G34135.1;AT4G34135.2
bic:659
peaks after deleting less than one fifth of global max:16
×10^6
16
14
12
10
8
6
4
2
0
y
0
5
10
15
20
25
x
raw data, starting:16
fitted Gauss locations:16

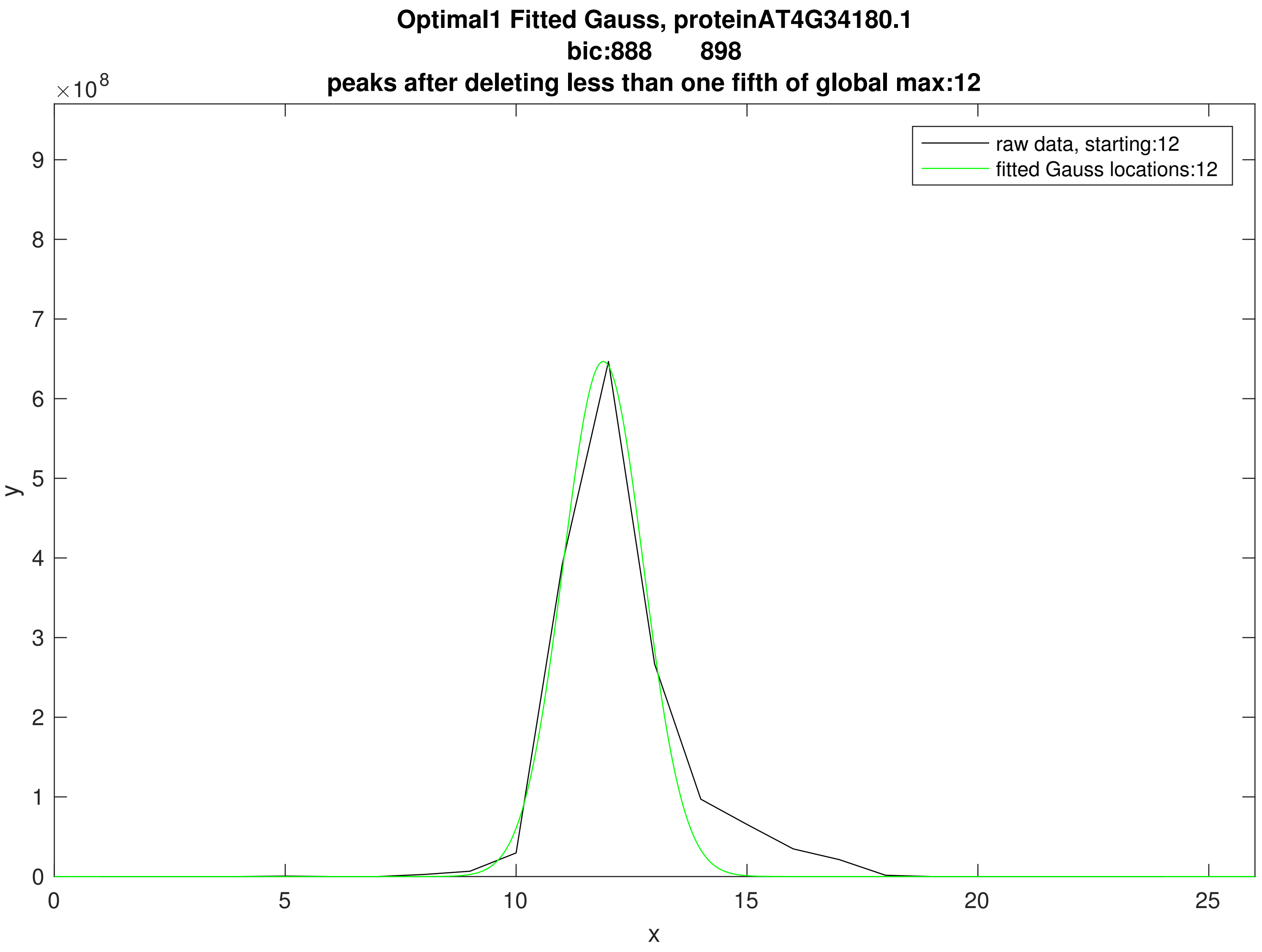
Optimal1 Fitted Gauss, proteinAT4G34180.1
bic:888 898
peaks after deleting less than one fifth of global max:12
×10^8
raw data, starting:12
fitted Gauss locations:12
y
x
0
1
2
3
4
5
6
7
8
9
0
5
10
15
20
25

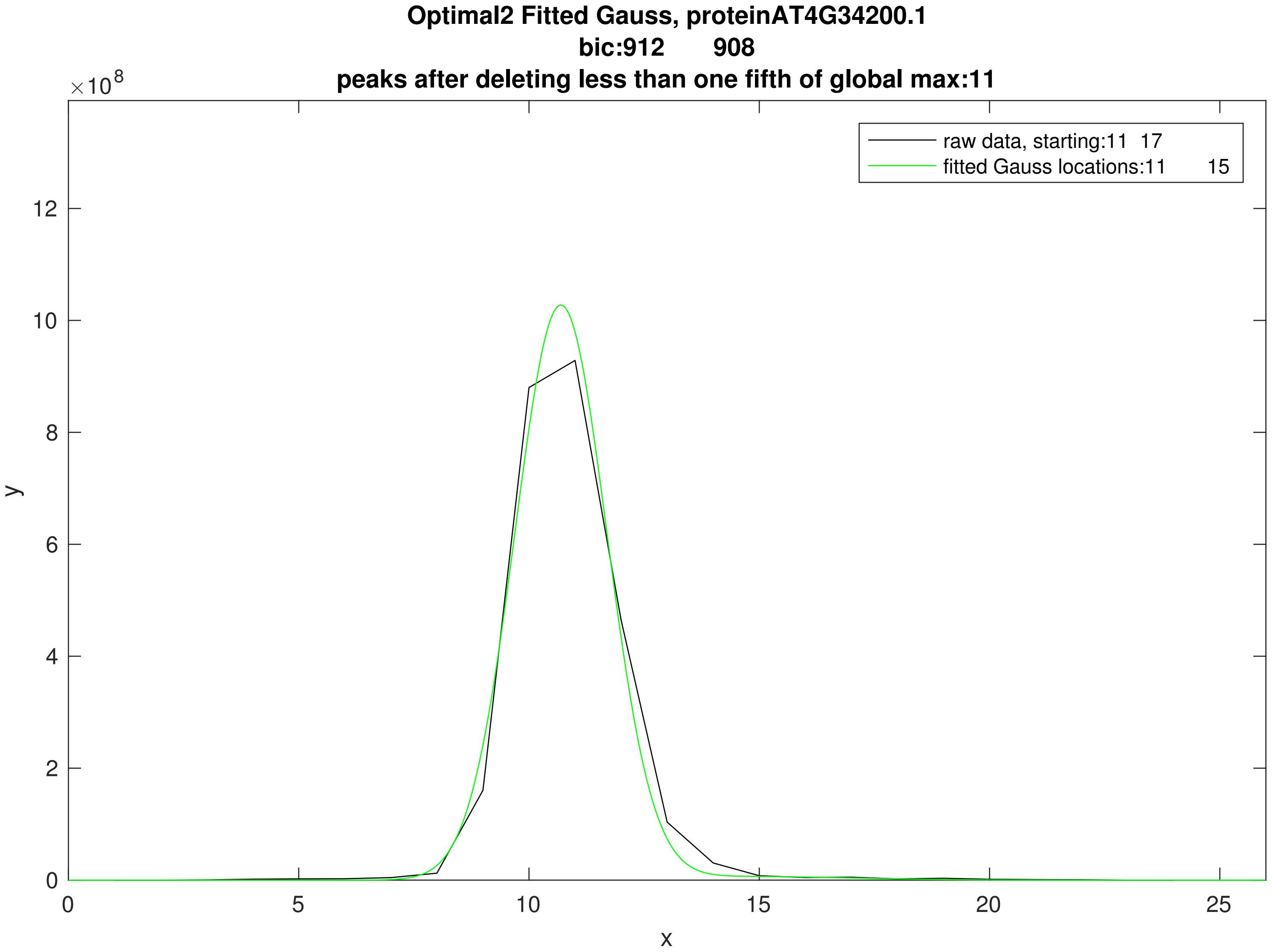

Optimal2 Fitted Gauss, proteinAT4G34200.1
bic:912 908
peaks after deleting less than one fifth of global max:11
×10^8
raw data, starting:11 17
fitted Gauss locations:11 15
12
10
8
6
4
2
0
y
0
5
10
15
20
25
x

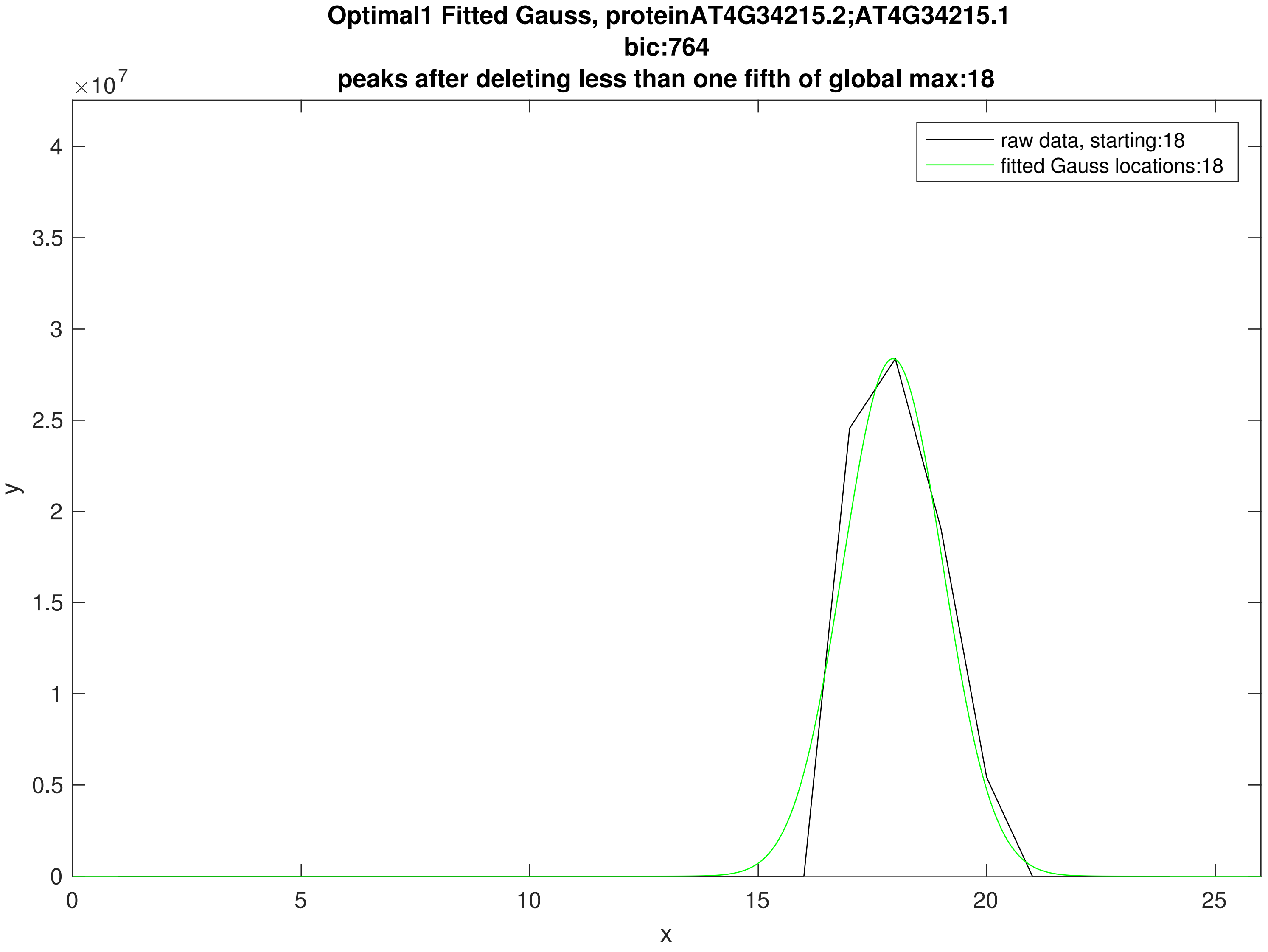
Optimal1 Fitted Gauss, proteinAT4G34215.2;AT4G34215.1
bic:764
peaks after deleting less than one fifth of global max:18
×10^7
raw data, starting:18
fitted Gauss locations:18
y
x
0
0.5
1
1.5
2
2.5
3
3.5
4
0
5
10
15
20
25

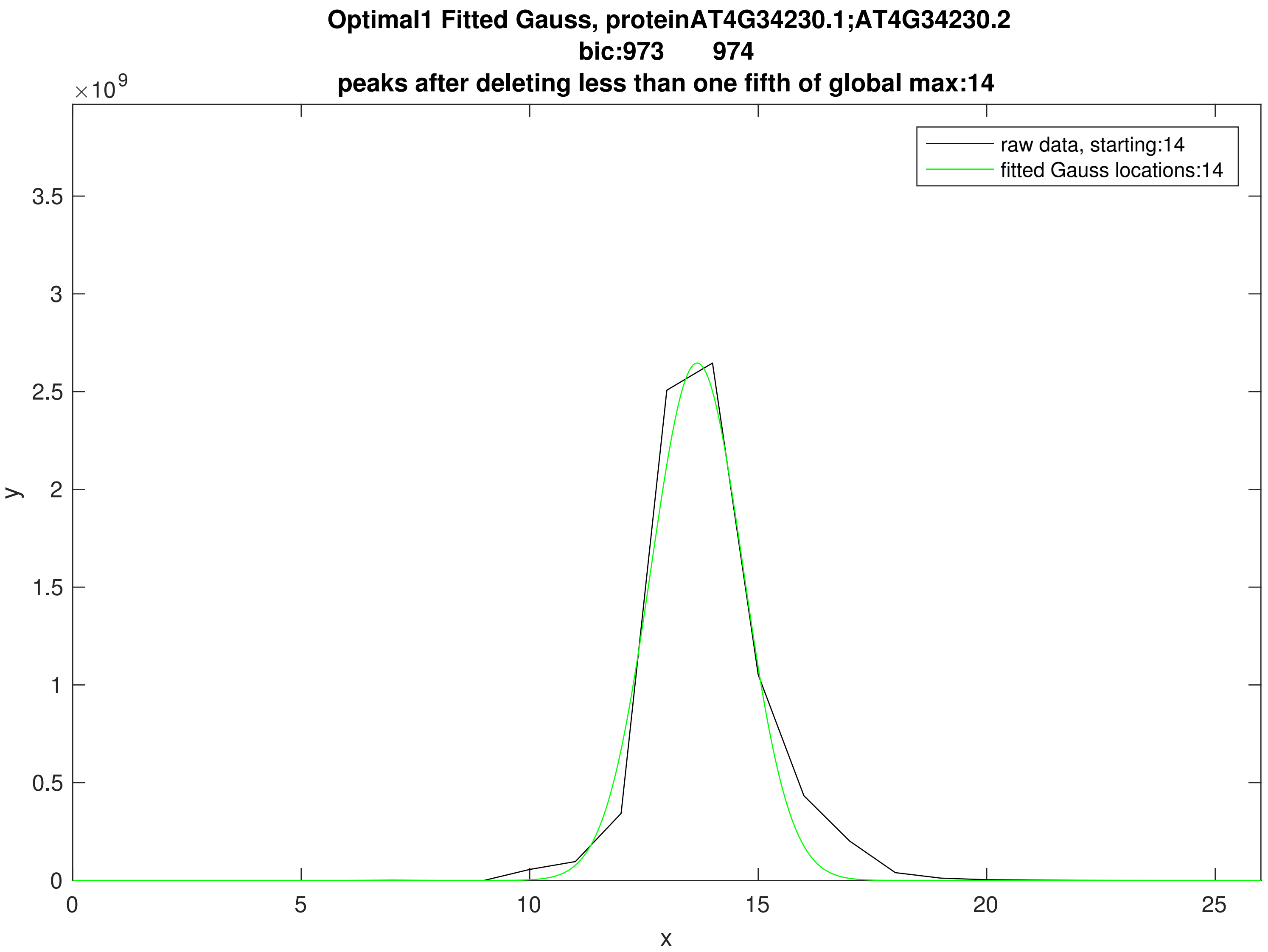

Optimal1 Fitted Gauss, proteinAT4G34230.1;AT4G34230.2
bic:973 974
peaks after deleting less than one fifth of global max:14
×10^9
3.5
3
2.5
2
1.5
1
0.5
0
y
0
5
10
15
20
25
x
raw data, starting:14
fitted Gauss locations:14

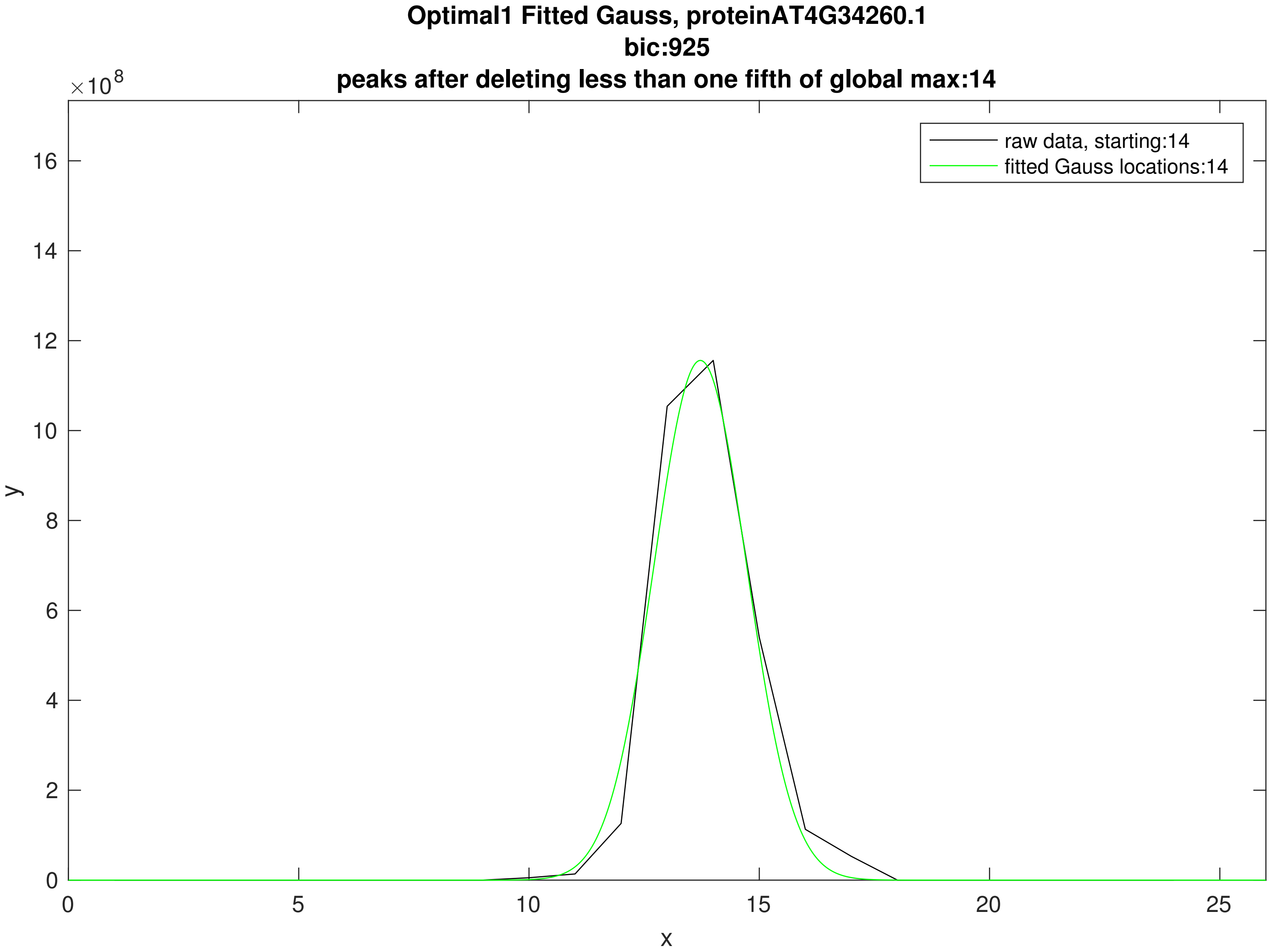
Optimal1 Fitted Gauss, proteinAT4G34260.1
bic:925
peaks after deleting less than one fifth of global max:14
×10^8
raw data, starting:14
fitted Gauss locations:14
y
x
0
2
4
6
8
10
12
14
16
0
5
10
15
20
25

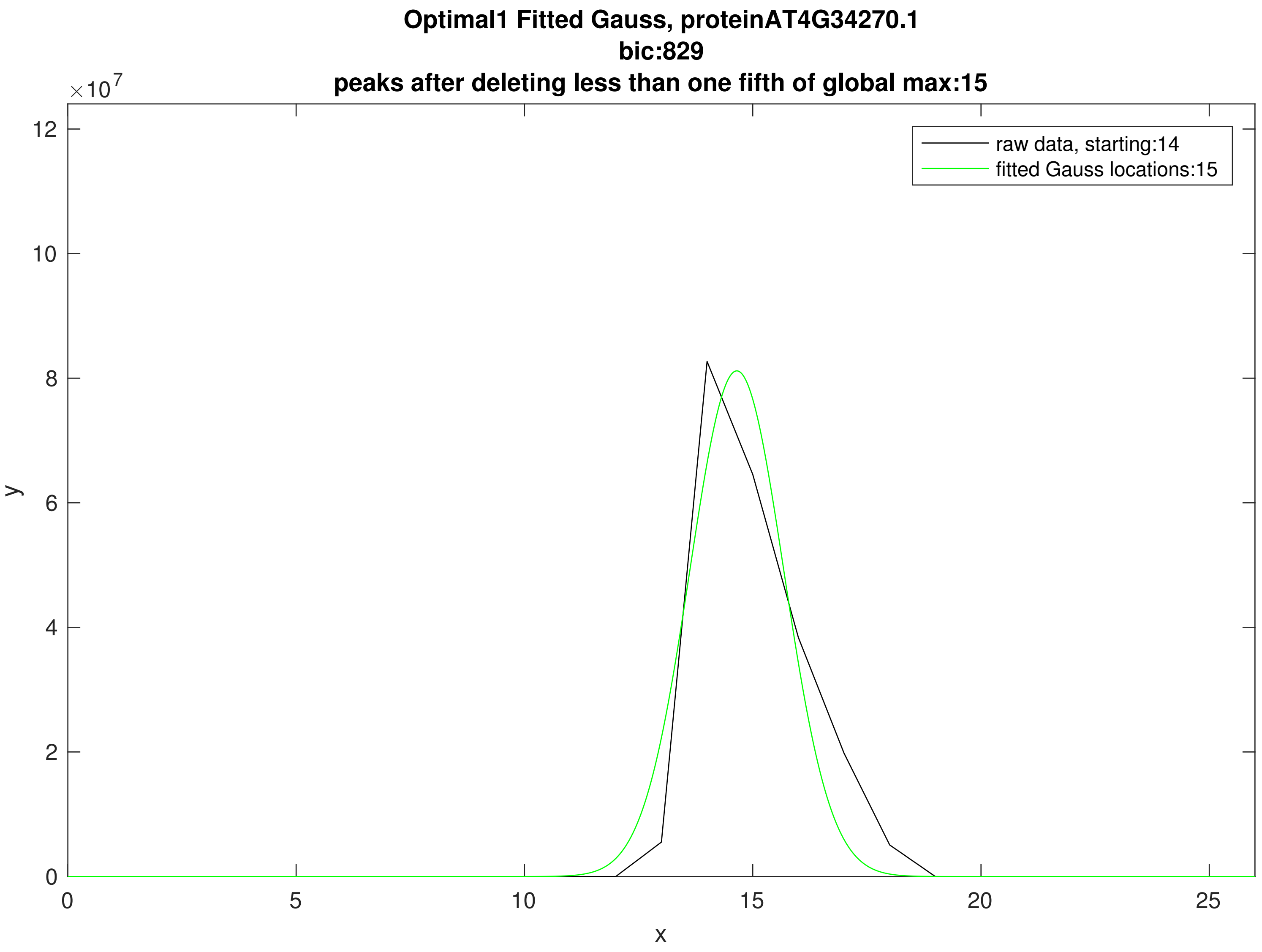

Optimal1 Fitted Gauss, proteinAT4G34270.1
bic:829
peaks after deleting less than one fifth of global max:15
×10^7
12
10
8
6
4
2
0
y
0
5
10
15
20
25
x
raw data, starting:14
fitted Gauss locations:15

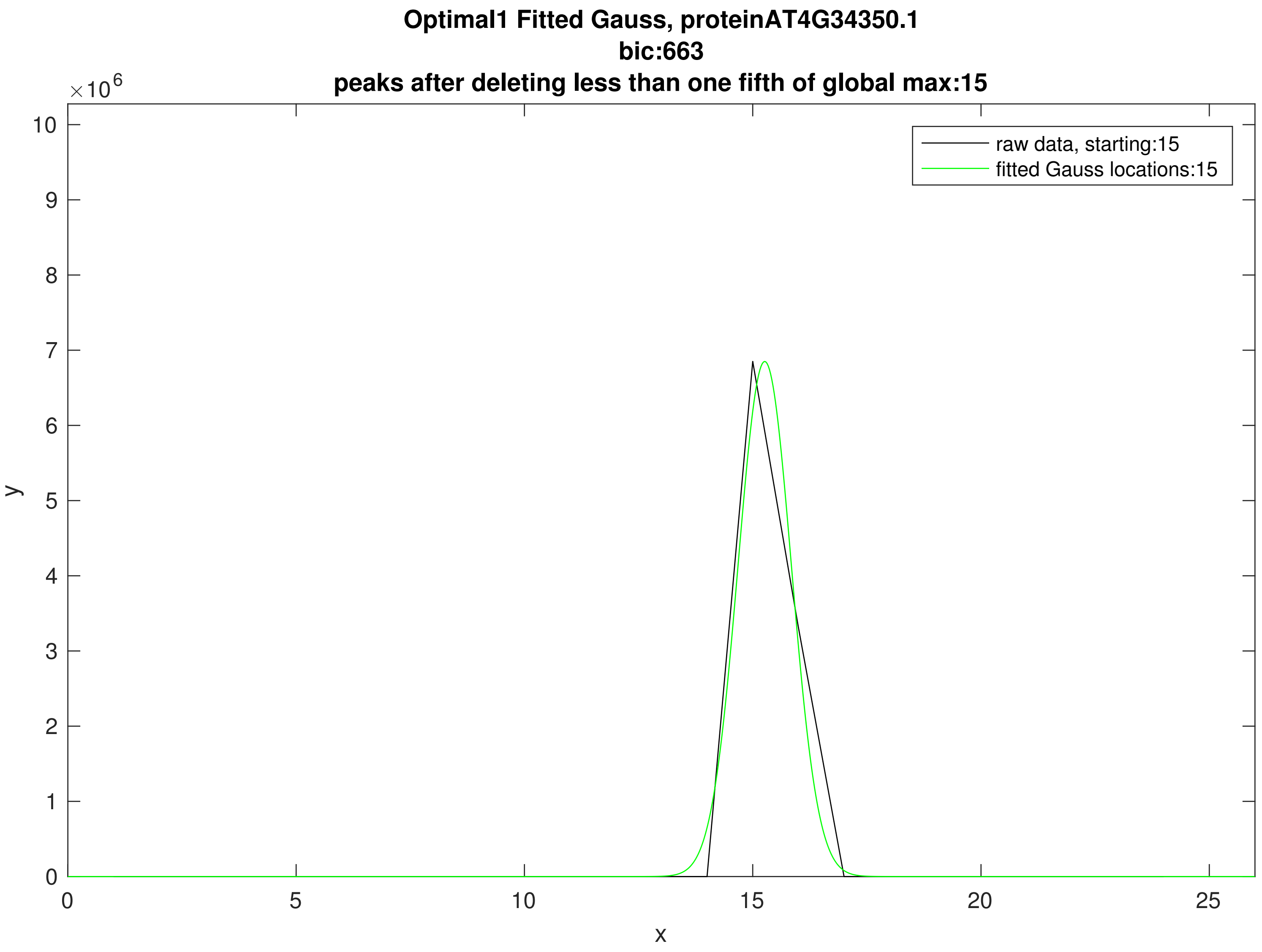
Optimal1 Fitted Gauss, proteinAT4G34350.1
bic:663
peaks after deleting less than one fifth of global max:15
×10^6
raw data, starting:15
fitted Gauss locations:15
y
x
0
1
2
3
4
5
6
7
8
9
10
0
5
10
15
20
25

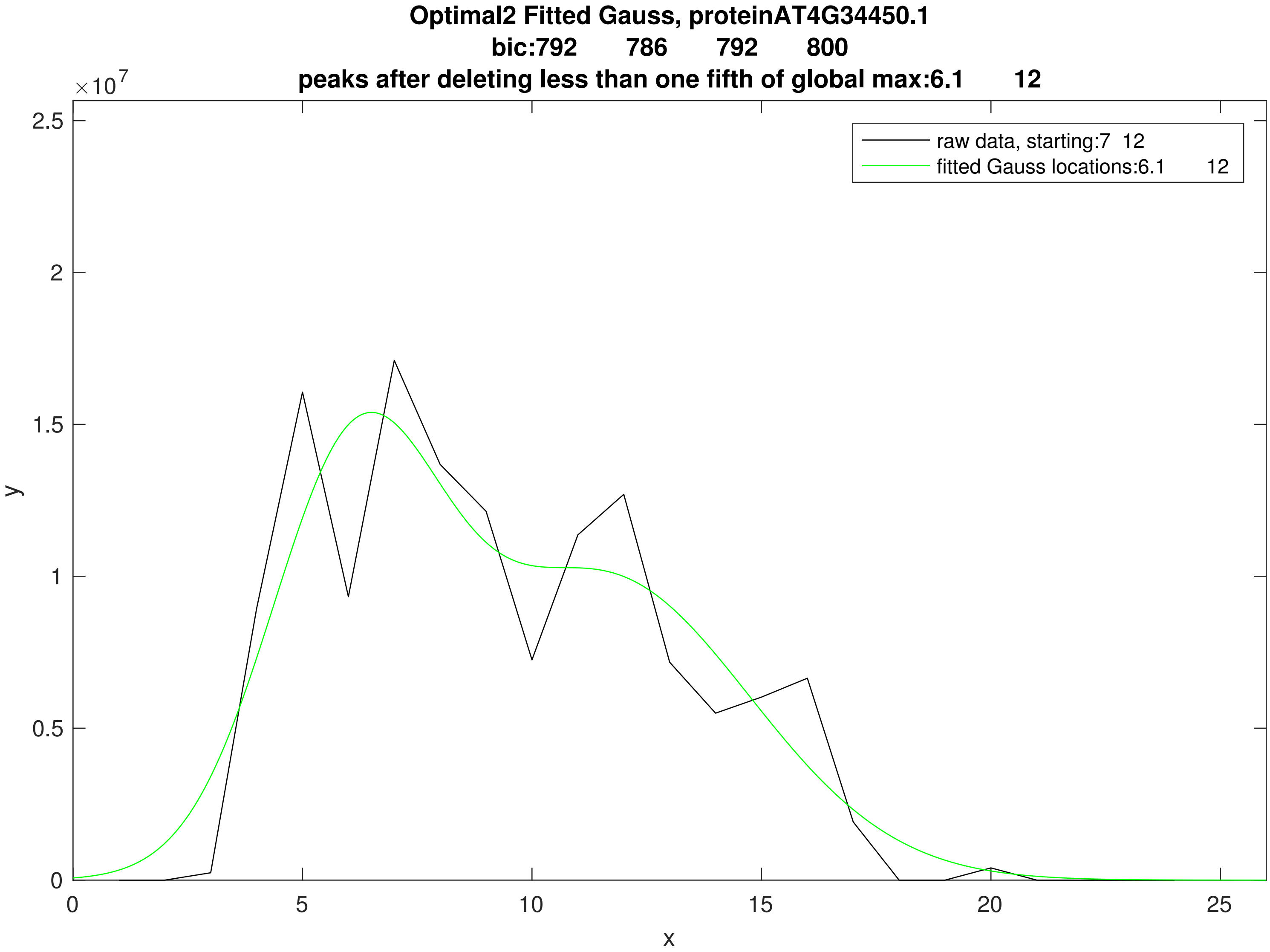
Optimal2 Fitted Gauss, proteinAT4G34450.1
bic:792 786 792 800
peaks after deleting less than one fifth of global max:6.1 12
×10^7
raw data, starting:7 12
fitted Gauss locations:6.1 12
y
x
0
0.5
1
1.5
2
2.5
5
10
15
20
25

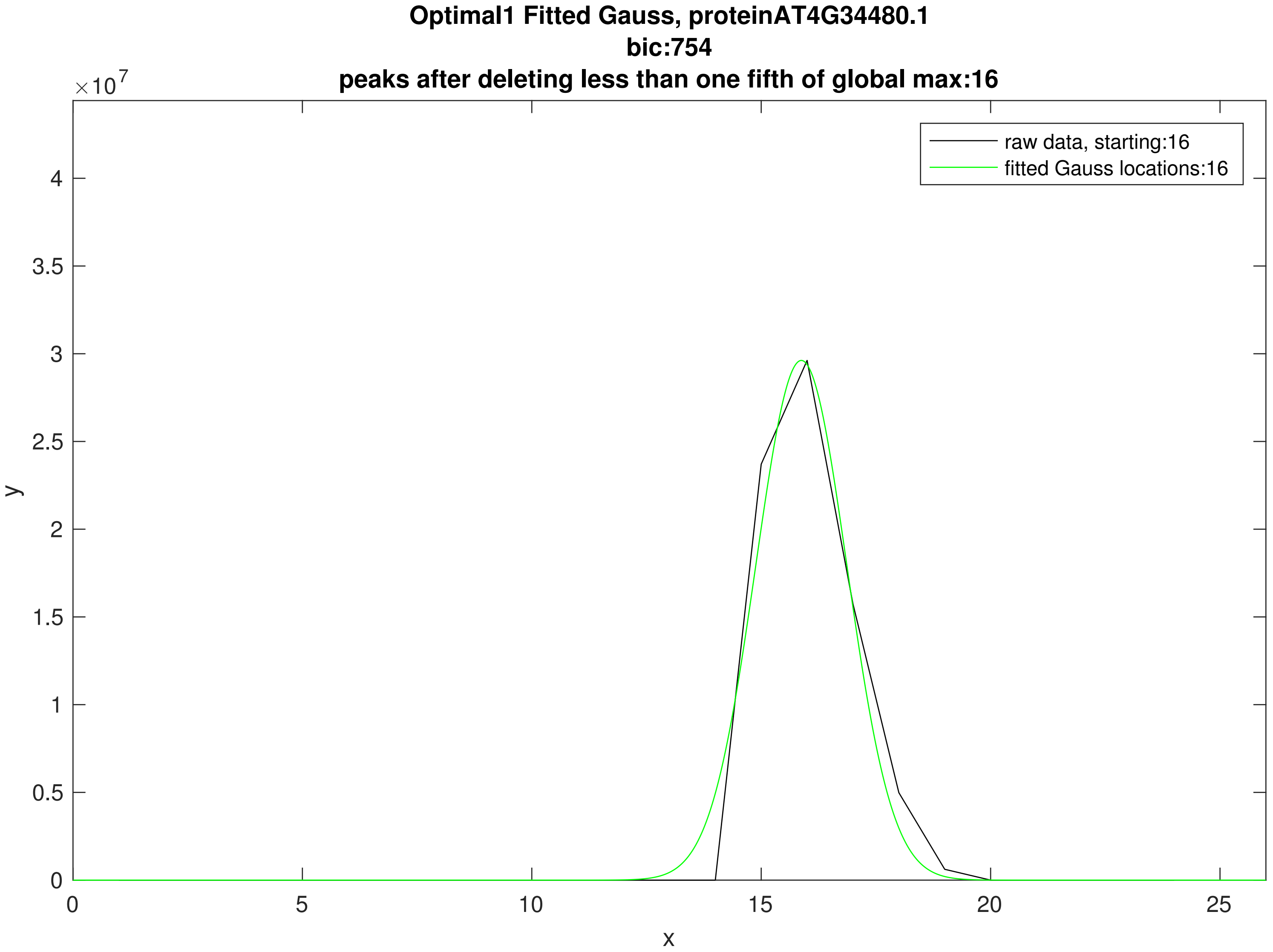

Optimal1 Fitted Gauss, proteinAT4G34480.1
bic:754
peaks after deleting less than one fifth of global max:16
×10^7
raw data, starting:16
fitted Gauss locations:16
y
x
0
0.5
1
1.5
2
2.5
3
3.5
4
0
5
10
15
20
25

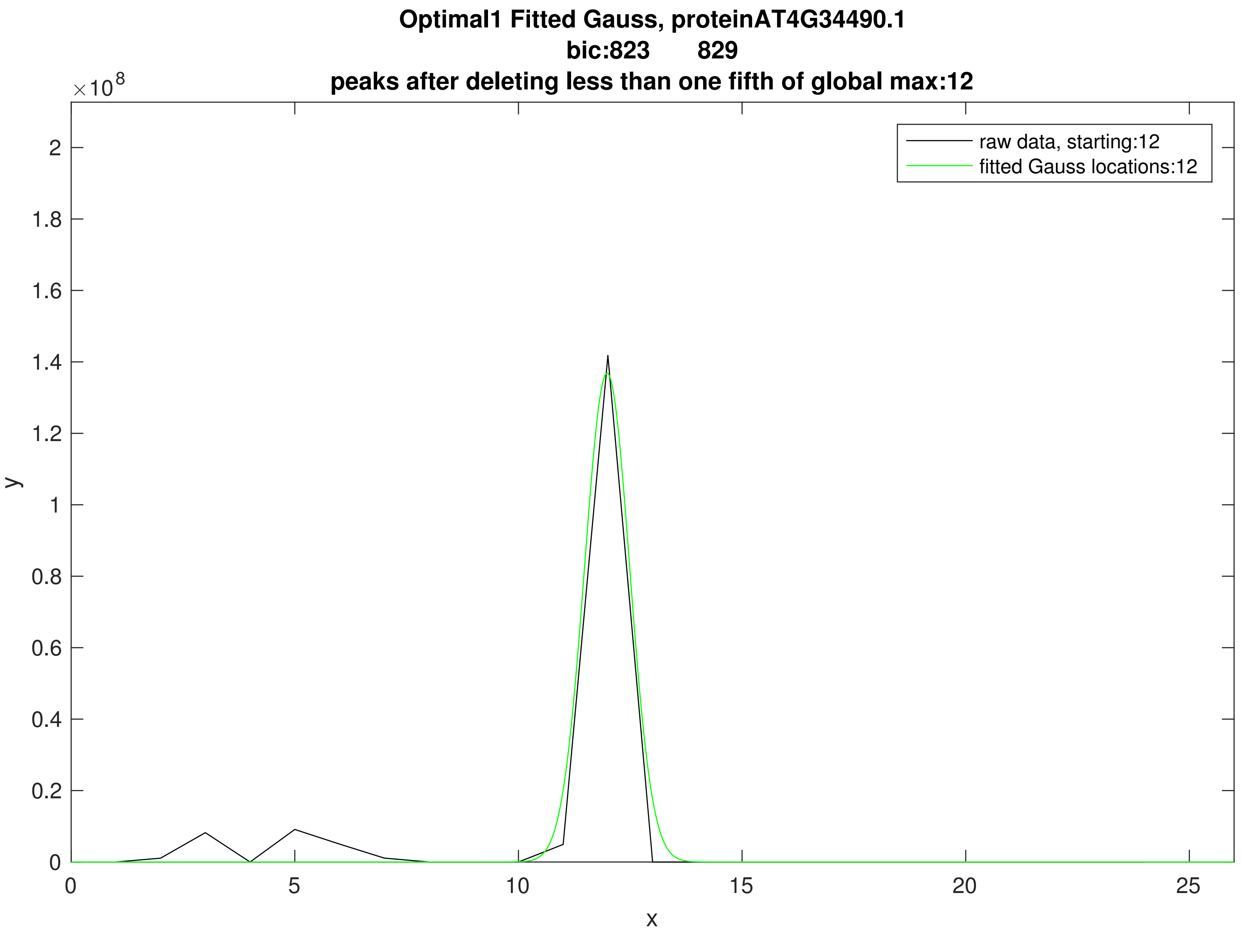

Optimal1 Fitted Gauss, proteinAT4G34490.1
bic:823 829
peaks after deleting less than one fifth of global max:12
×10^8
raw data, starting:12
fitted Gauss locations:12
y
x
0
0.2
0.4
0.6
0.8
1
1.2
1.4
1.6
1.8
2
5
10
15
20
25

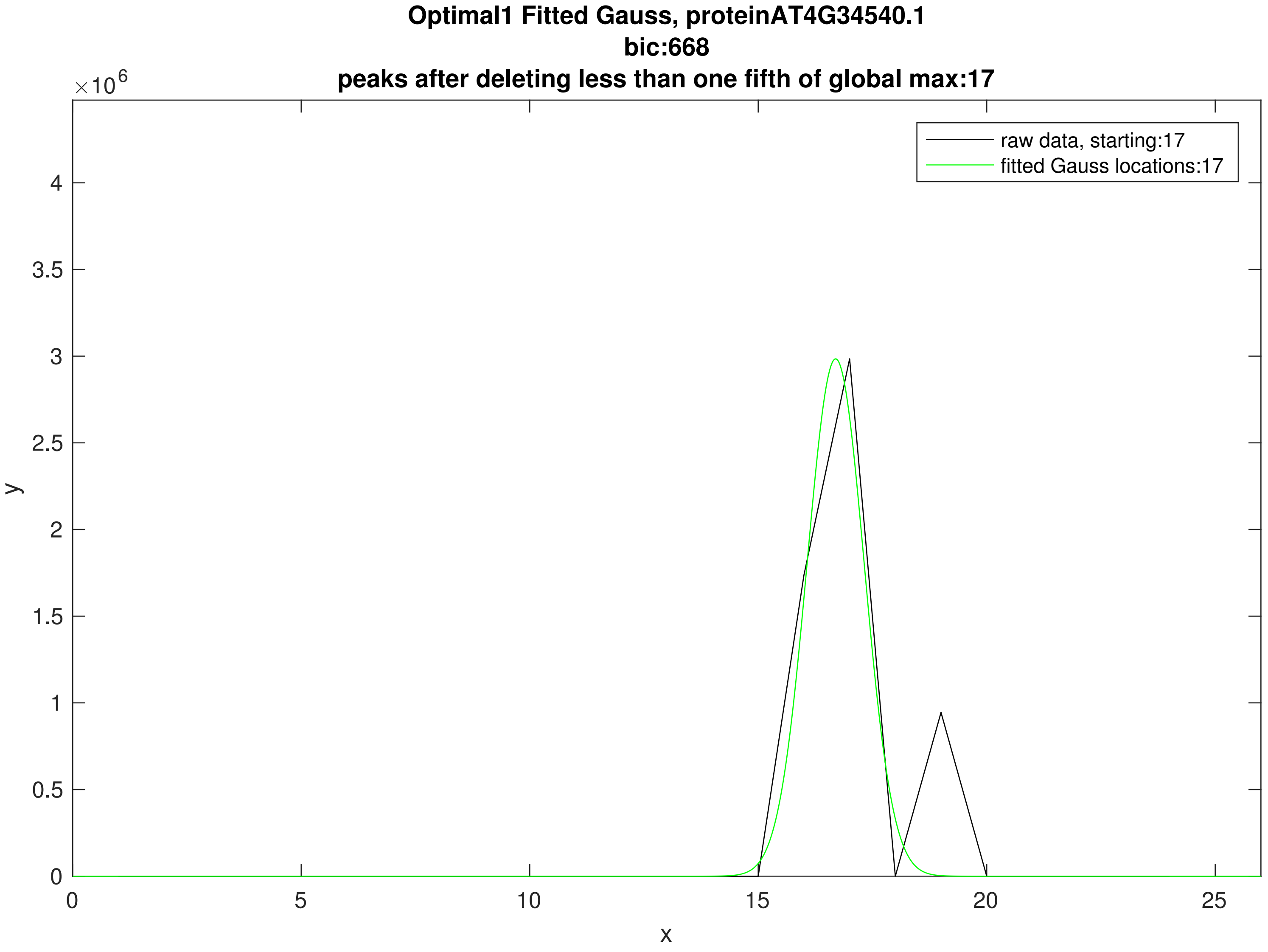
Optimal1 Fitted Gauss, proteinAT4G34540.1
bic:668
peaks after deleting less than one fifth of global max:17
×10^6
raw data, starting:17
fitted Gauss locations:17
y
x
0
0.5
1
1.5
2
2.5
3
3.5
4
0
5
10
15
20
25

**Optimal1 Fitted Gauss, proteinAT4G34620.1**
**bic:717**
**peaks after deleting less than one fifth of global max:5**

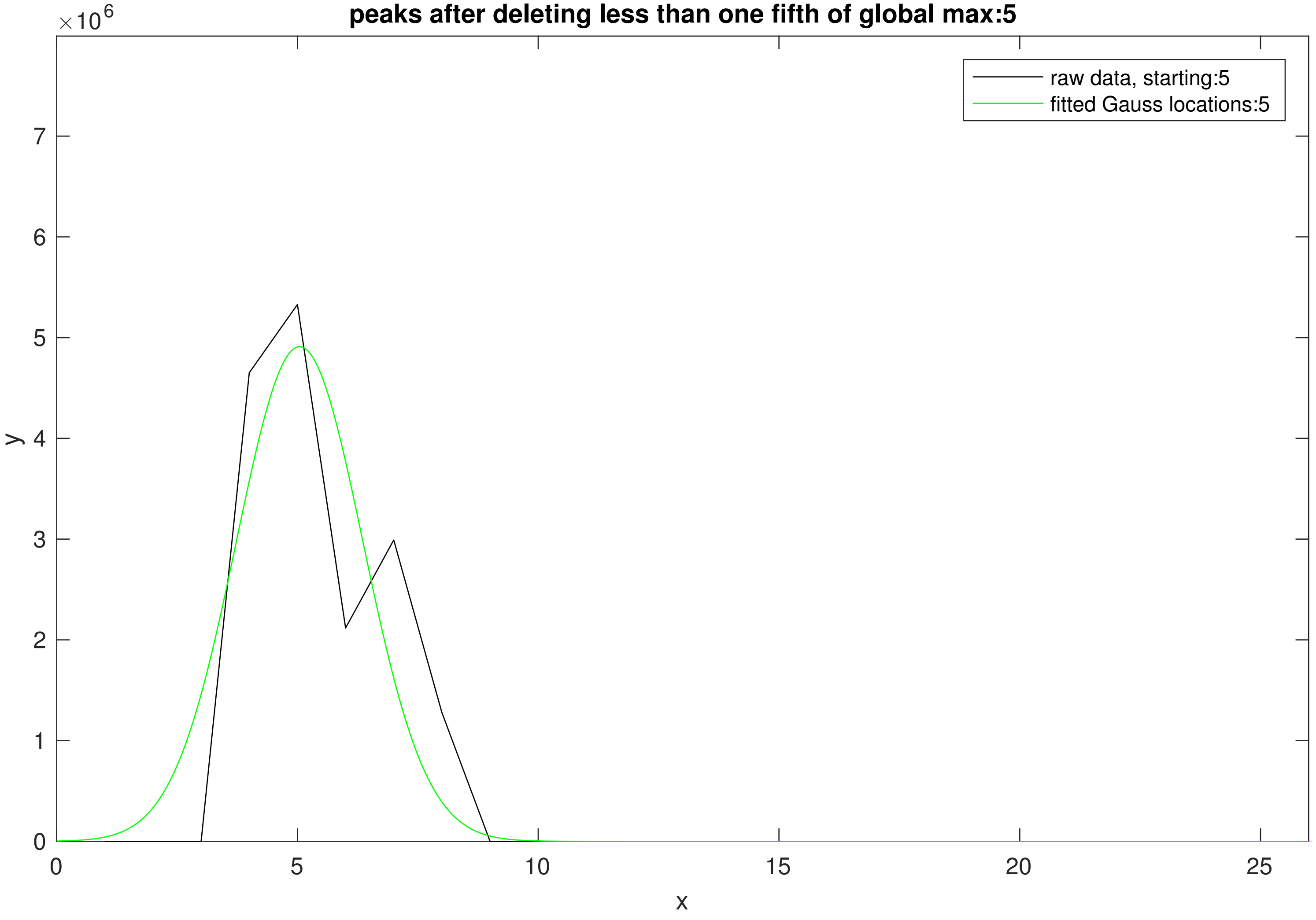

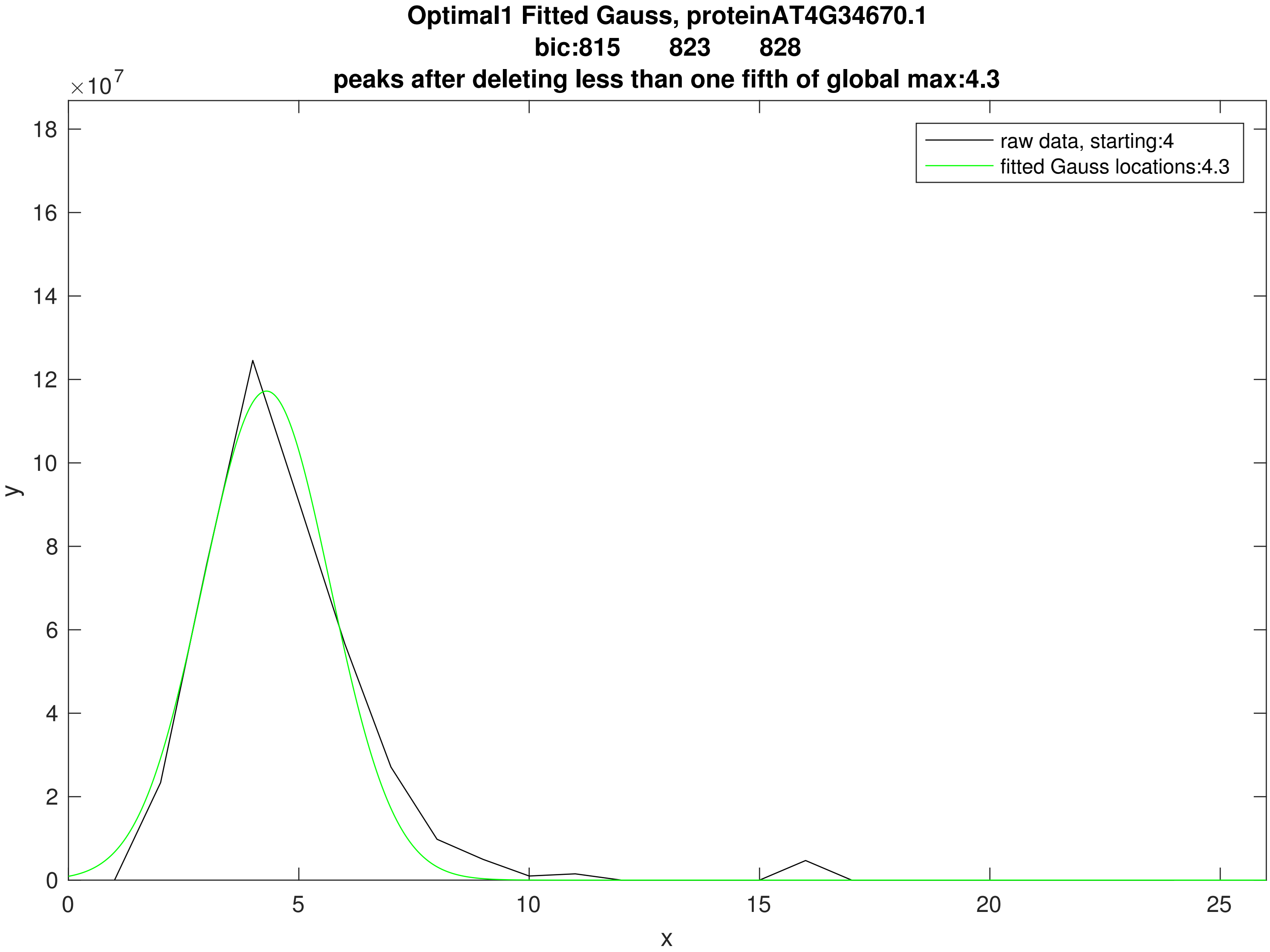

Optimal1 Fitted Gauss, proteinAT4G34670.1
bic:815 823 828
peaks after deleting less than one fifth of global max:4.3
×10^7
raw data, starting:4
fitted Gauss locations:4.3
y
x
0
2
4
6
8
10
12
14
16
18
0
5
10
15
20
25

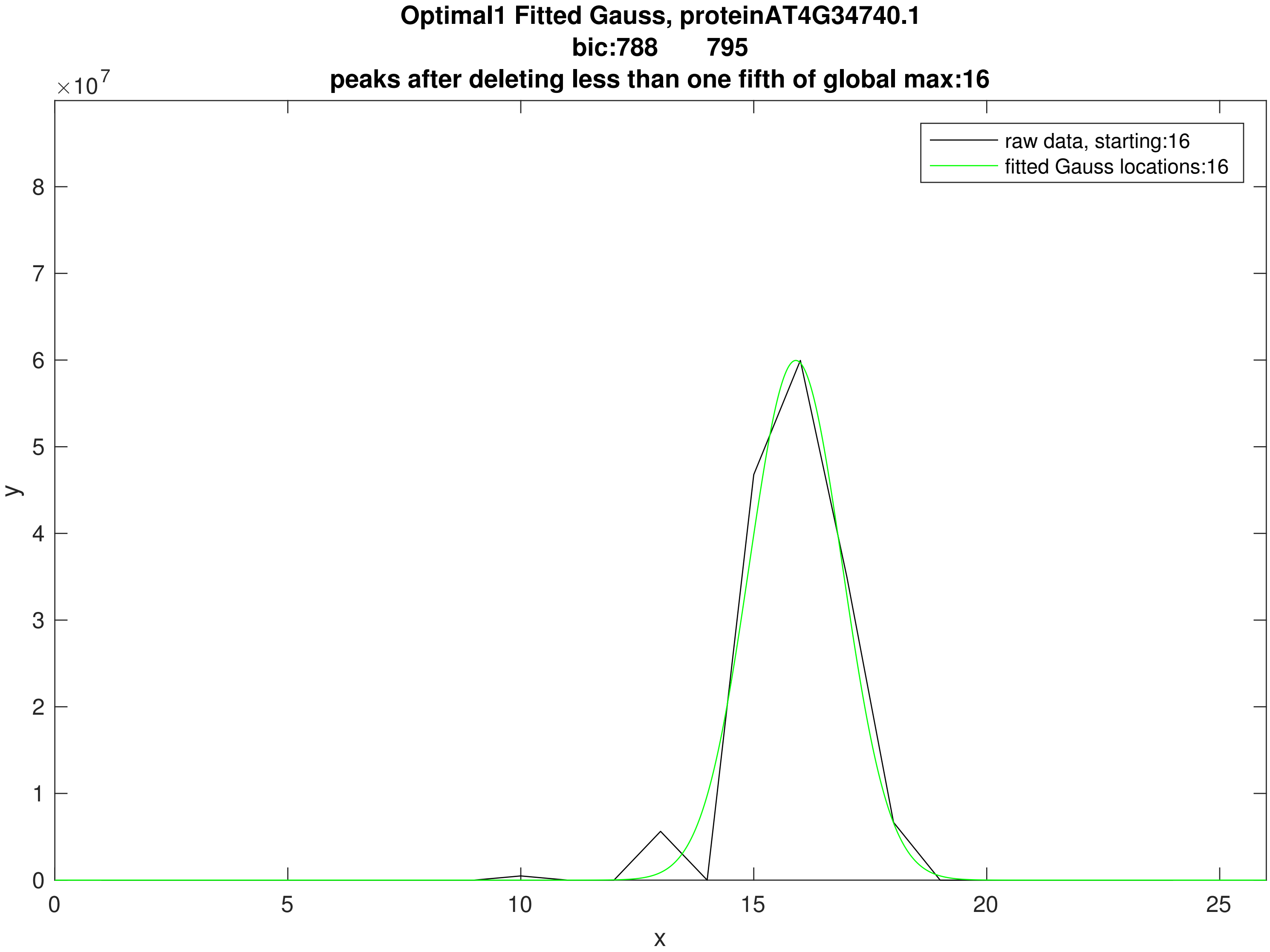

Optimal1 Fitted Gauss, proteinAT4G34740.1
bic:788 795
peaks after deleting less than one fifth of global max:16
×10^7
raw data, starting:16
fitted Gauss locations:16
y
x
0
1
2
3
4
5
6
7
8
10
15
20
25

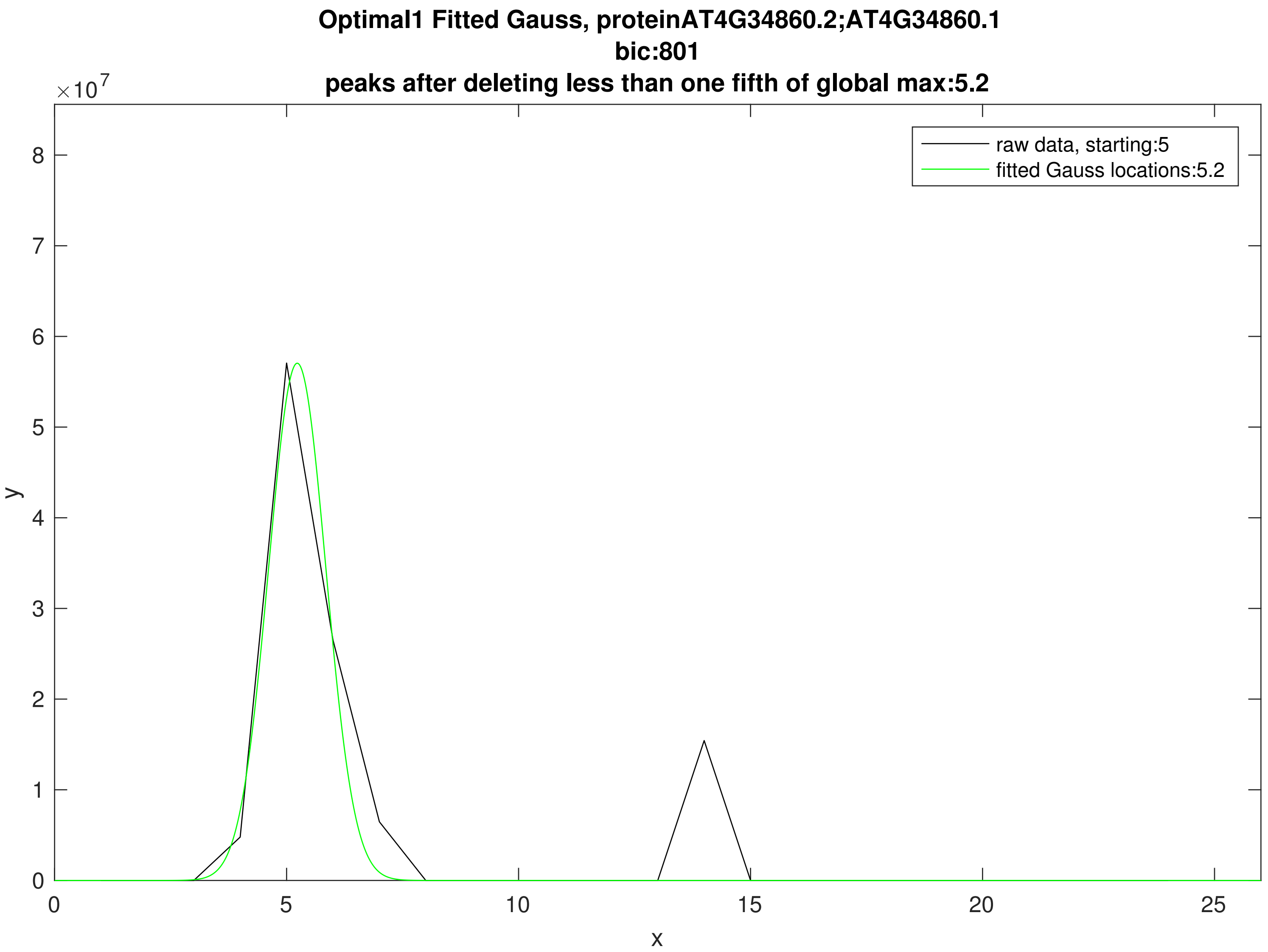

Optimal1 Fitted Gauss, proteinAT4G34860.2;AT4G34860.1
bic:801
peaks after deleting less than one fifth of global max:5.2
×10^7
raw data, starting:5
fitted Gauss locations:5.2
y
x
0
1
2
3
4
5
6
7
8
0
5
10
15
20
25

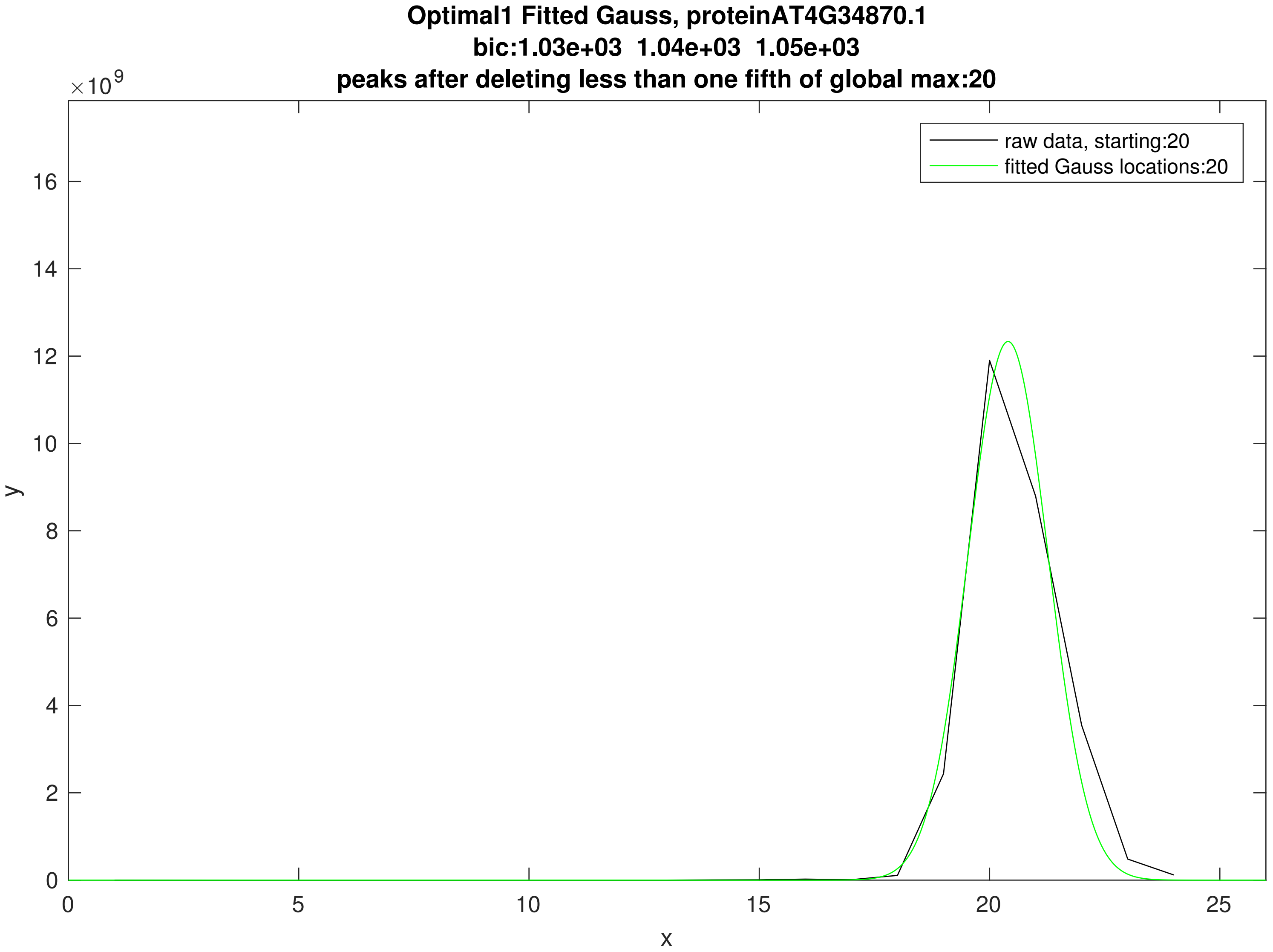

Optimal1 Fitted Gauss, proteinAT4G34870.1
bic:1.03e+03  1.04e+03  1.05e+03
peaks after deleting less than one fifth of global max:20
×10^9
raw data, starting:20
fitted Gauss locations:20
y
x
0
2
4
6
8
10
12
14
16
0
5
10
15
20
25

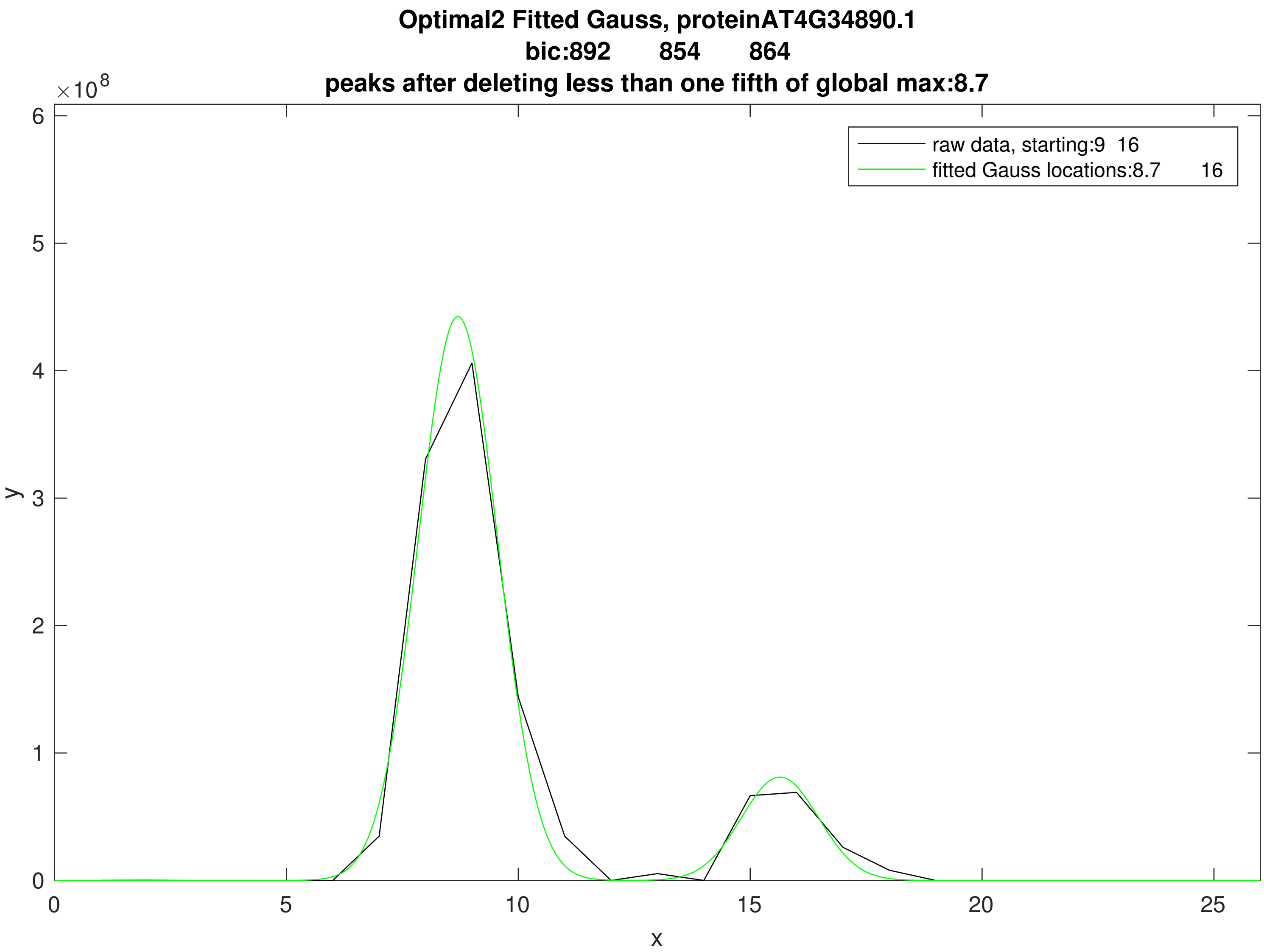
Optimal2 Fitted Gauss, proteinAT4G34890.1
bic:892 854 864
peaks after deleting less than one fifth of global max:8.7
×10^8
raw data, starting:9 16
fitted Gauss locations:8.7 16
y
x
0
1
2
3
4
5
6
10
15
20
25

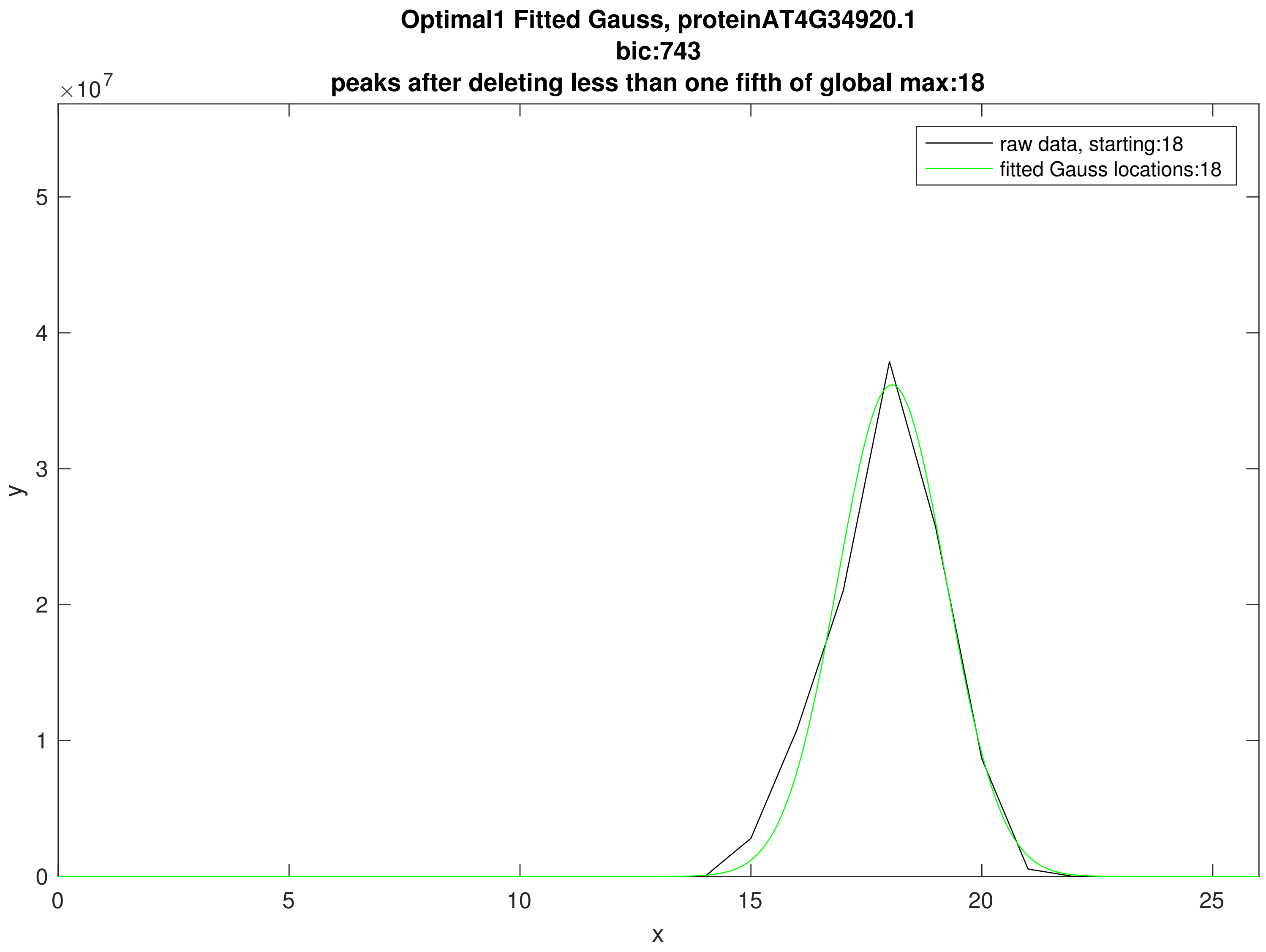
Optimal1 Fitted Gauss, proteinAT4G34920.1
bic:743
peaks after deleting less than one fifth of global max:18
×10^7
raw data, starting:18
fitted Gauss locations:18
y
x
0
1
2
3
4
5
0
5
10
15
20
25

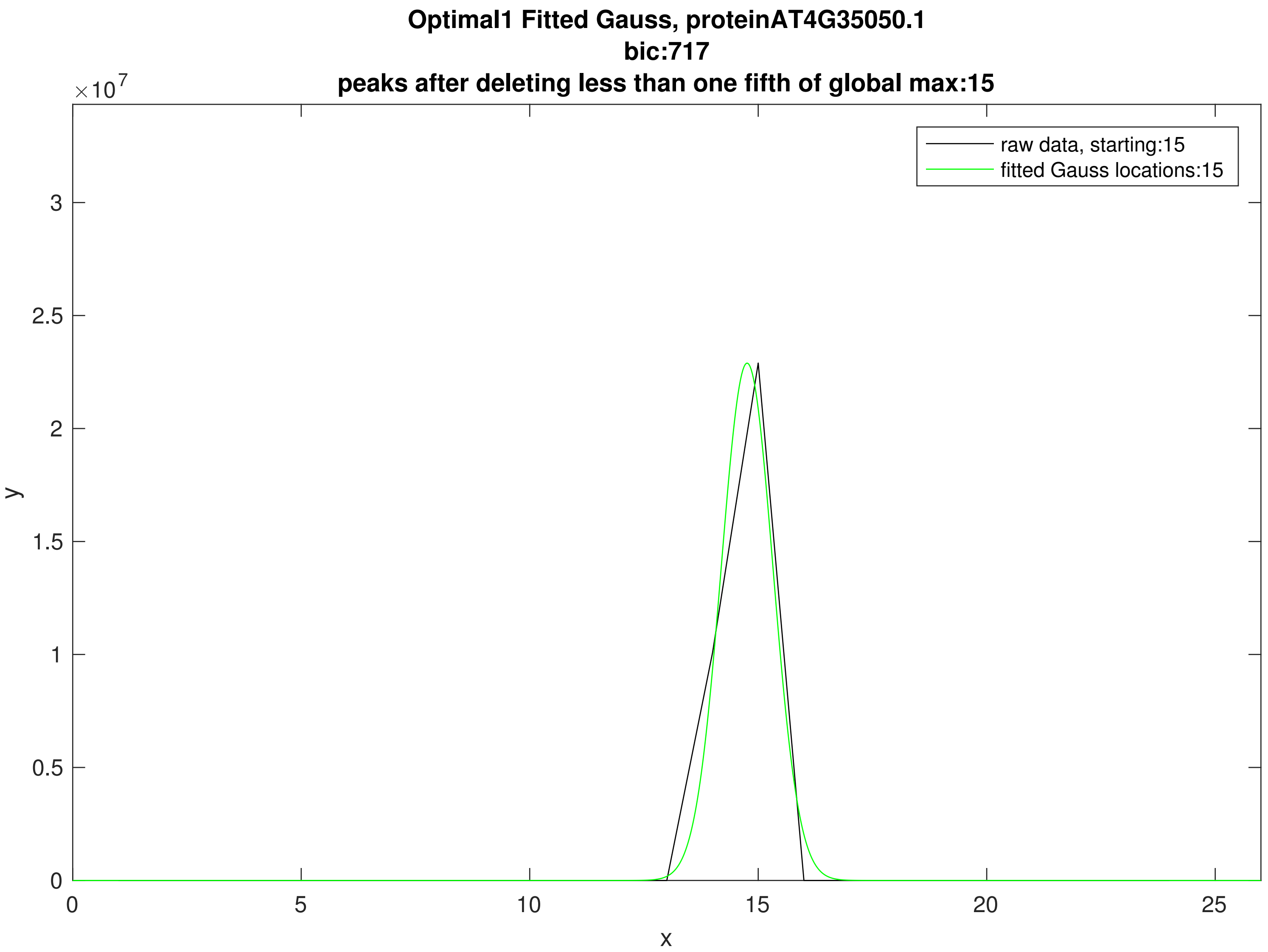
Optimal1 Fitted Gauss, proteinAT4G35050.1
bic:717
peaks after deleting less than one fifth of global max:15
×10^7
3
2.5
2
1.5
1
0.5
0
y
0
5
10
15
20
25
x
raw data, starting:15
fitted Gauss locations:15

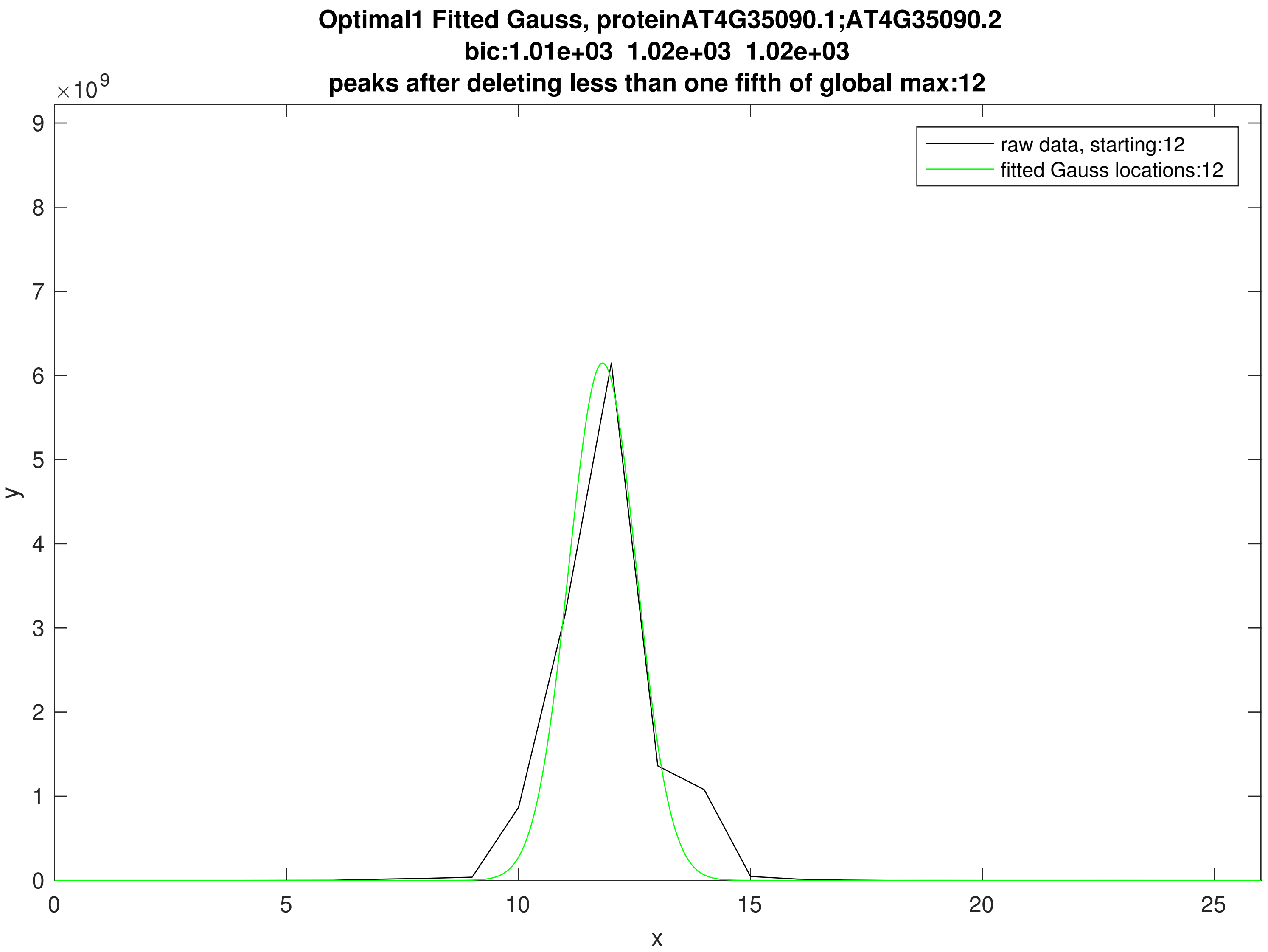
Optimal1 Fitted Gauss, proteinAT4G35090.1;AT4G35090.2
bic:1.01e+03 1.02e+03 1.02e+03
peaks after deleting less than one fifth of global max:12
×10^9
raw data, starting:12
fitted Gauss locations:12
y
x
0
1
2
3
4
5
6
7
8
9
0
5
10
15
20
25

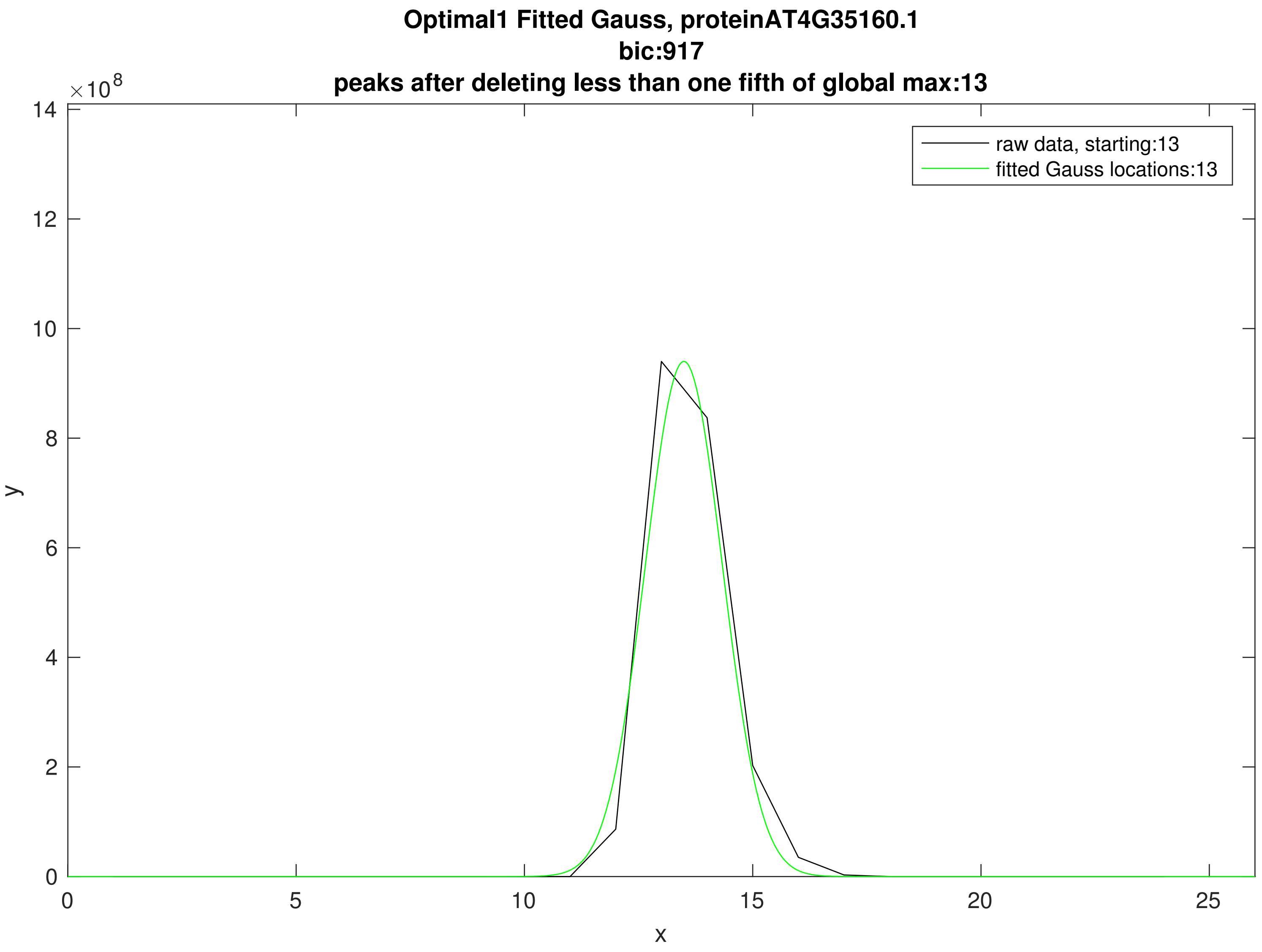
Optimal1 Fitted Gauss, proteinAT4G35160.1
bic:917
peaks after deleting less than one fifth of global max:13
×10^8
14
12
10
8
6
4
2
0
y
0
5
10
15
20
25
x
raw data, starting:13
fitted Gauss locations:13

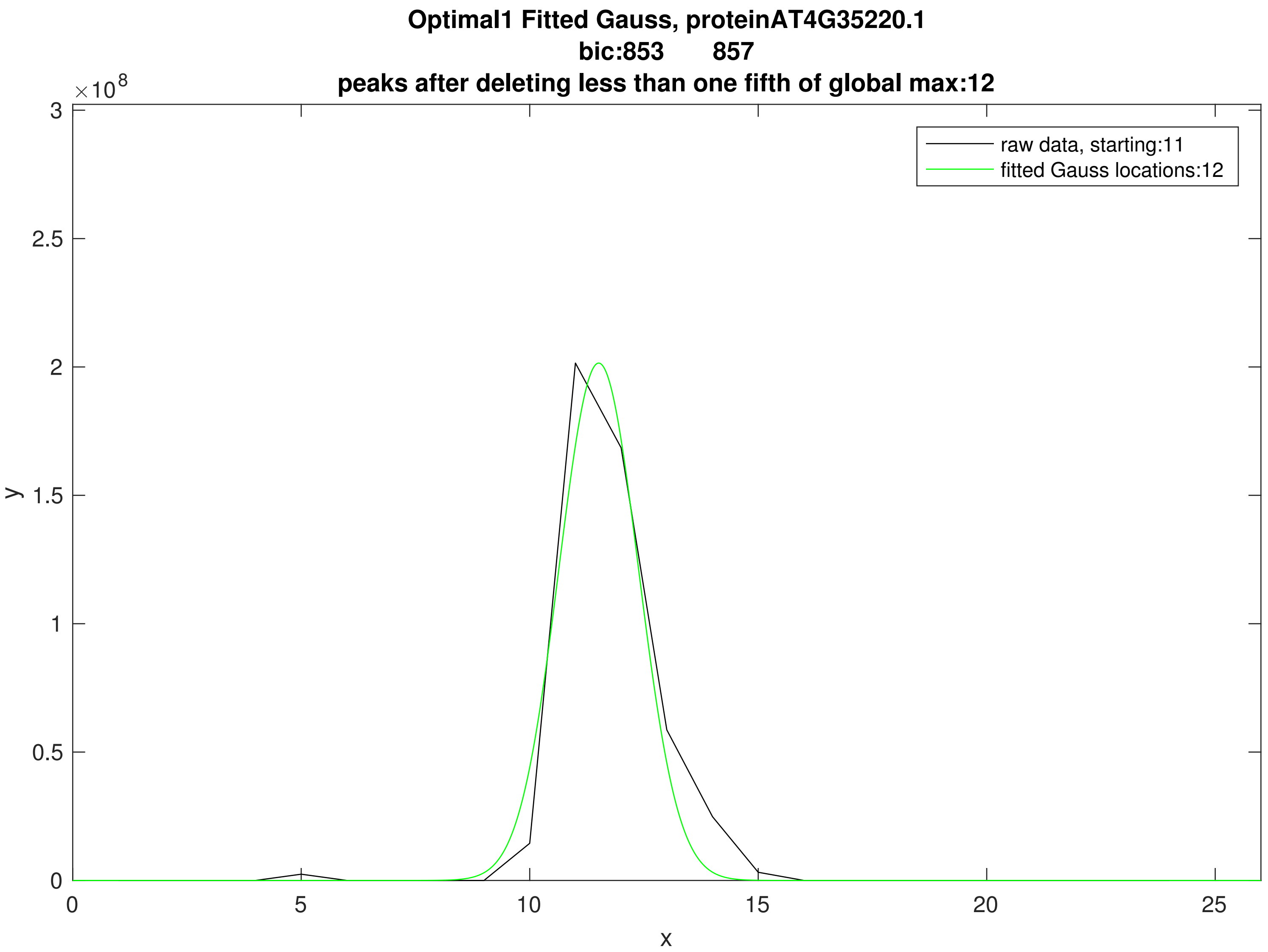

Optimal1 Fitted Gauss, proteinAT4G35220.1
bic:853 857
peaks after deleting less than one fifth of global max:12
×10^8
raw data, starting:11
fitted Gauss locations:12
y
x
0
0.5
1
1.5
2
2.5
3
0
5
10
15
20
25

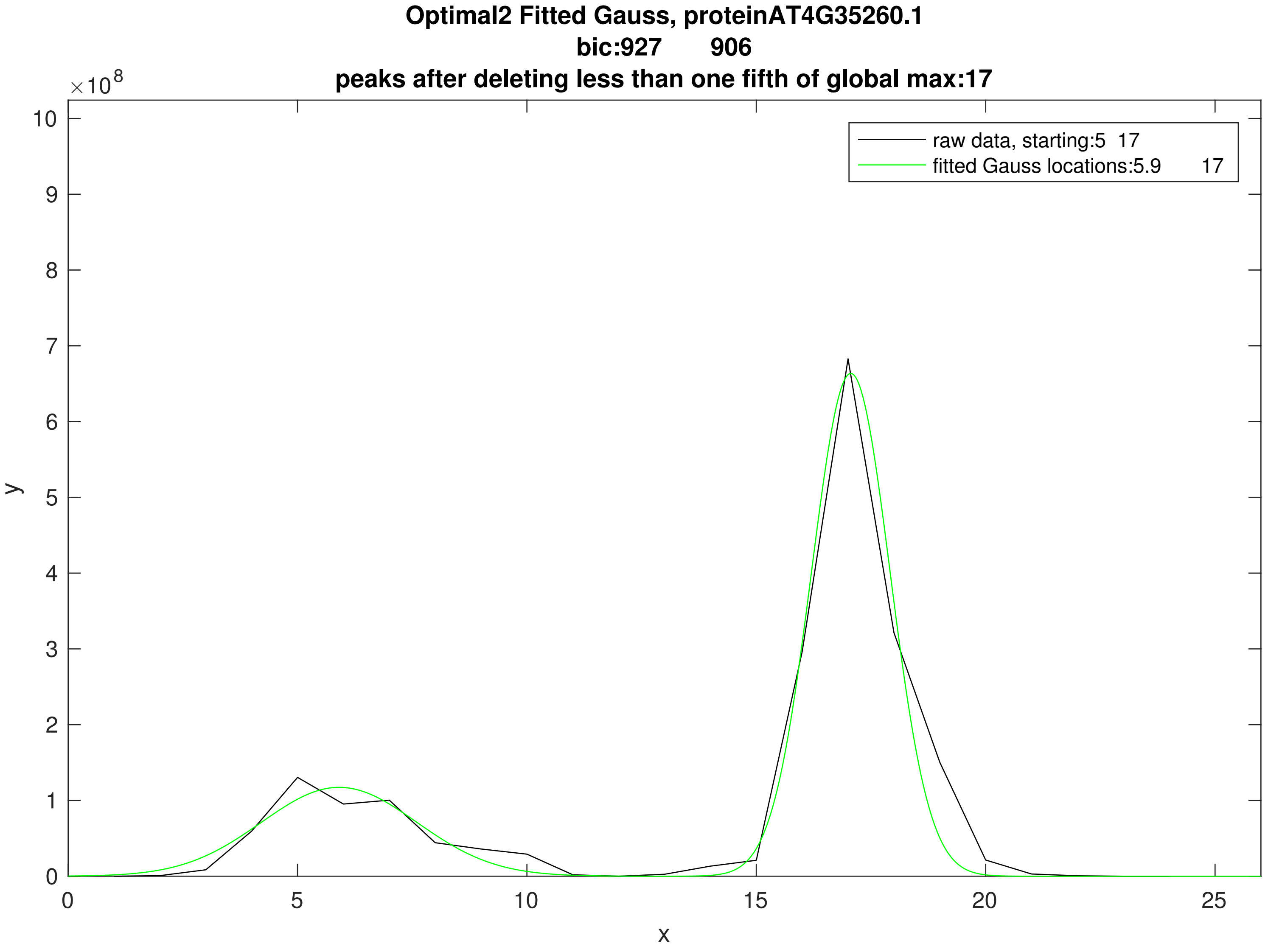

Optimal2 Fitted Gauss, proteinAT4G35260.1
bic:927 906
peaks after deleting less than one fifth of global max:17
×10^8
raw data, starting:5 17
fitted Gauss locations:5.9 17
y
x
0
1
2
3
4
5
6
7
8
9
10
0
5
10
15
20
25

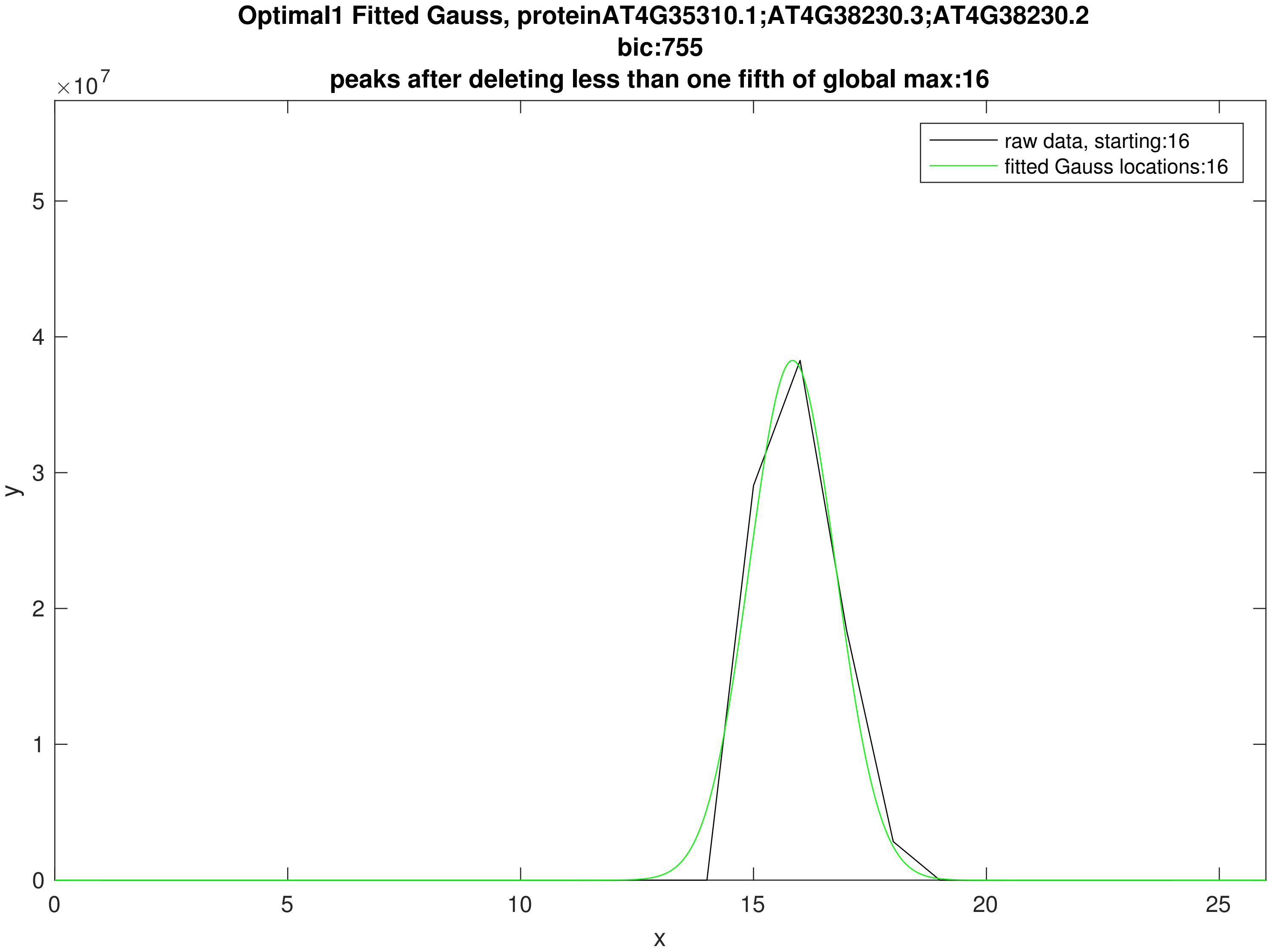

Optimal1 Fitted Gauss, proteinAT4G35310.1;AT4G38230.3;AT4G38230.2
bic:755
peaks after deleting less than one fifth of global max:16
×10^7
raw data, starting:16
fitted Gauss locations:16
y
x
0
1
2
3
4
5
0
5
10
15
20
25

Optimal1 Fitted Gauss, proteinAT4G35350.2;AT4G35350.1
bic:737
peaks after deleting less than one fifth of global max:17

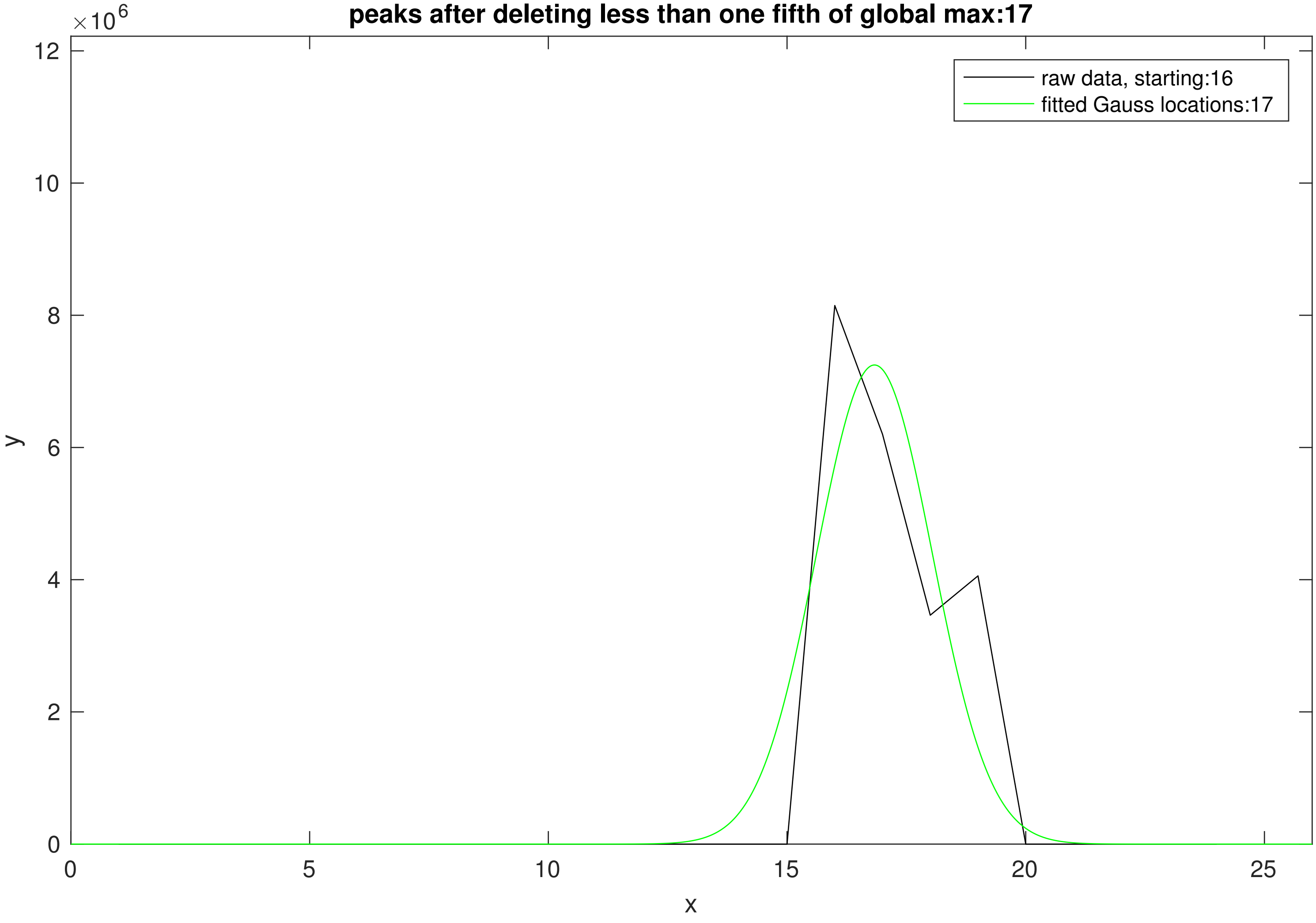

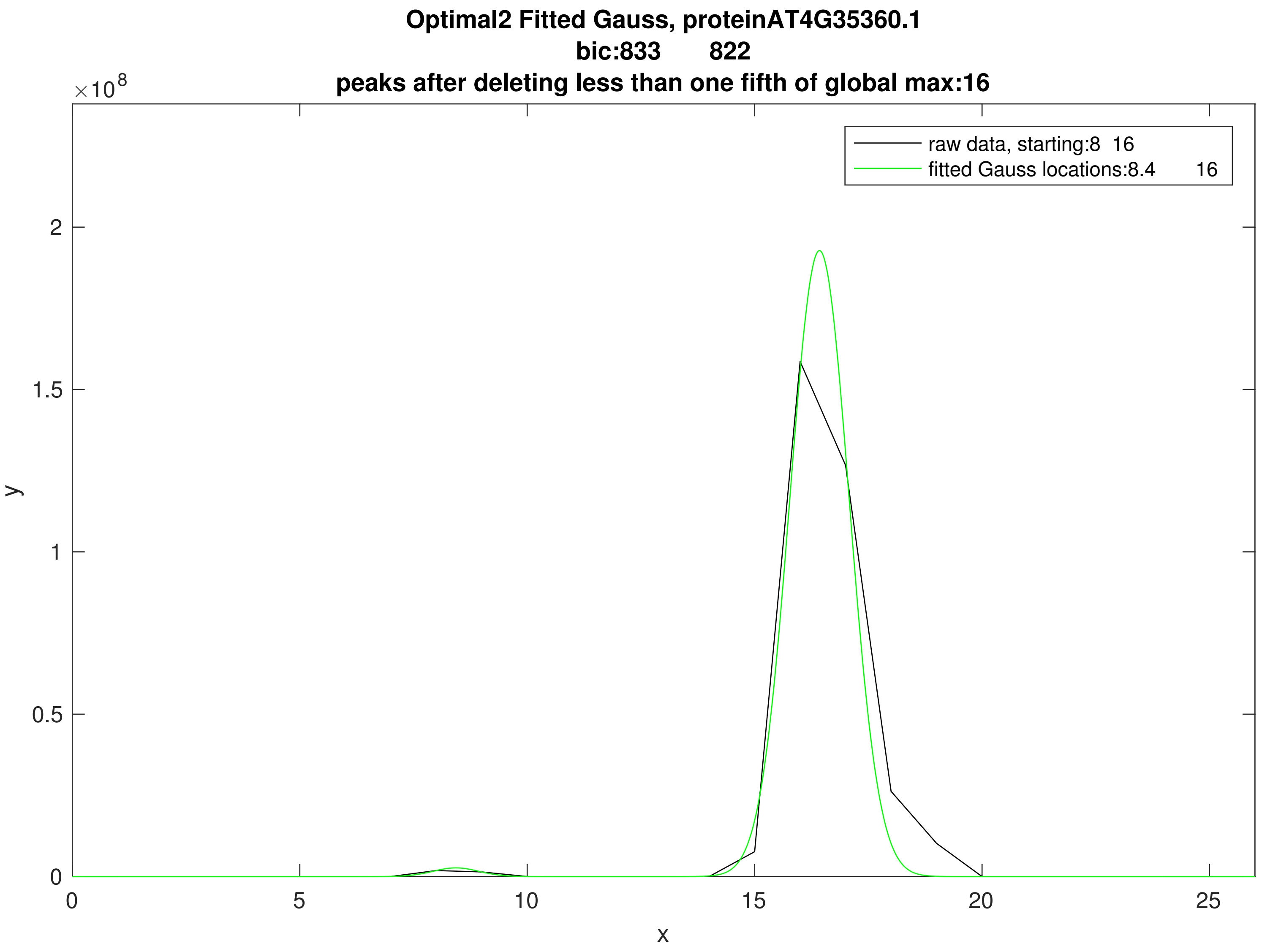

Optimal2 Fitted Gauss, proteinAT4G35360.1
bic:833 822
peaks after deleting less than one fifth of global max:16
×10^8
raw data, starting:8 16
fitted Gauss locations:8.4 16
2
1.5
1
0.5
0
y
0
5
10
15
20
25
x

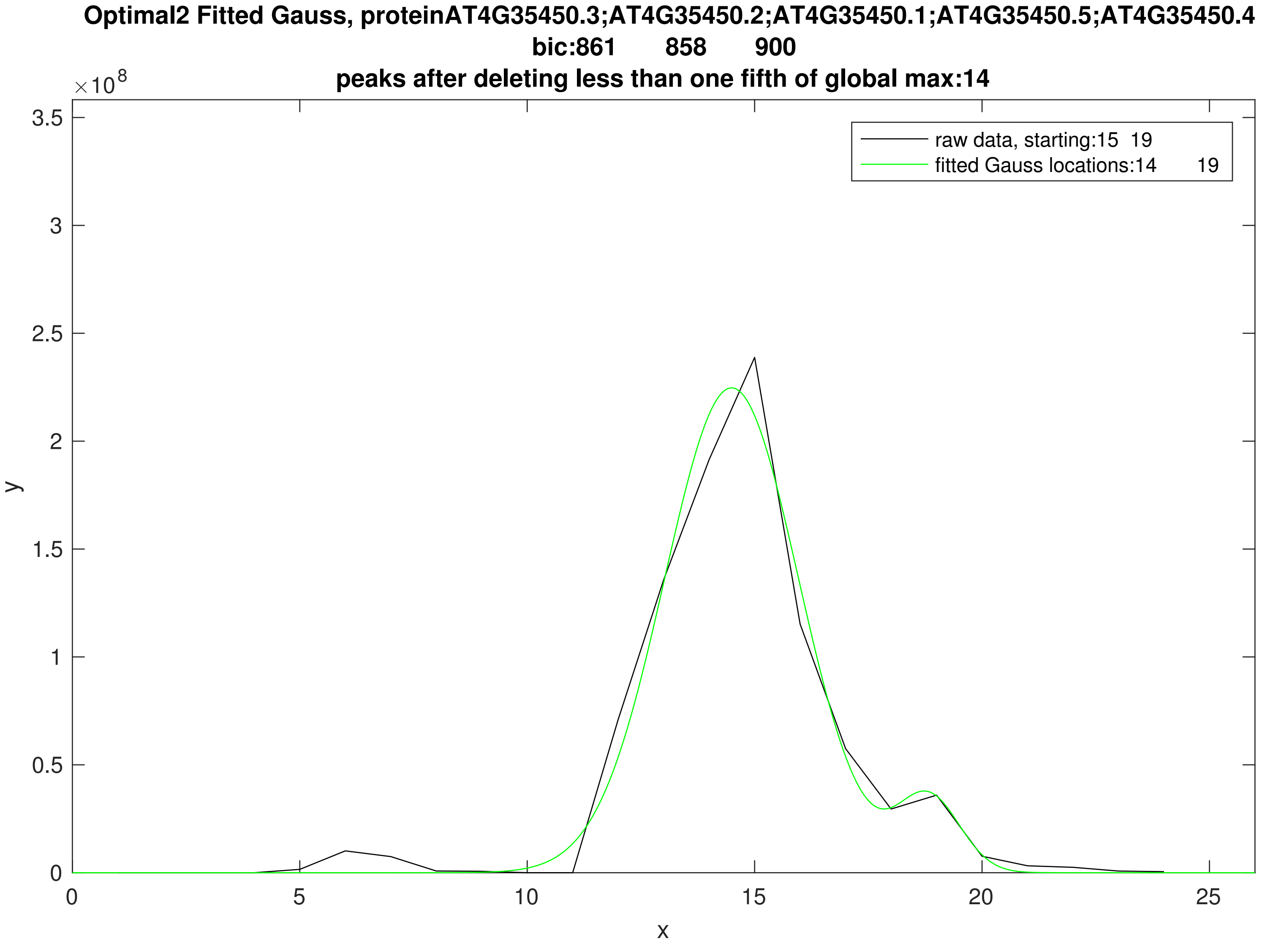
Optimal2 Fitted Gauss, proteinAT4G35450.3;AT4G35450.2;AT4G35450.1;AT4G35450.5;AT4G35450.4
bic:861 858 900
peaks after deleting less than one fifth of global max:14
×10^8
raw data, starting:15 19
fitted Gauss locations:14 19
y
x
0
0.5
1
1.5
2
2.5
3
3.5
0
5
10
15
20
25

**Optimal2 Fitted Gauss, proteinAT4G35460.1**
**bic:829 823**
**peaks after deleting less than one fifth of global max:15**

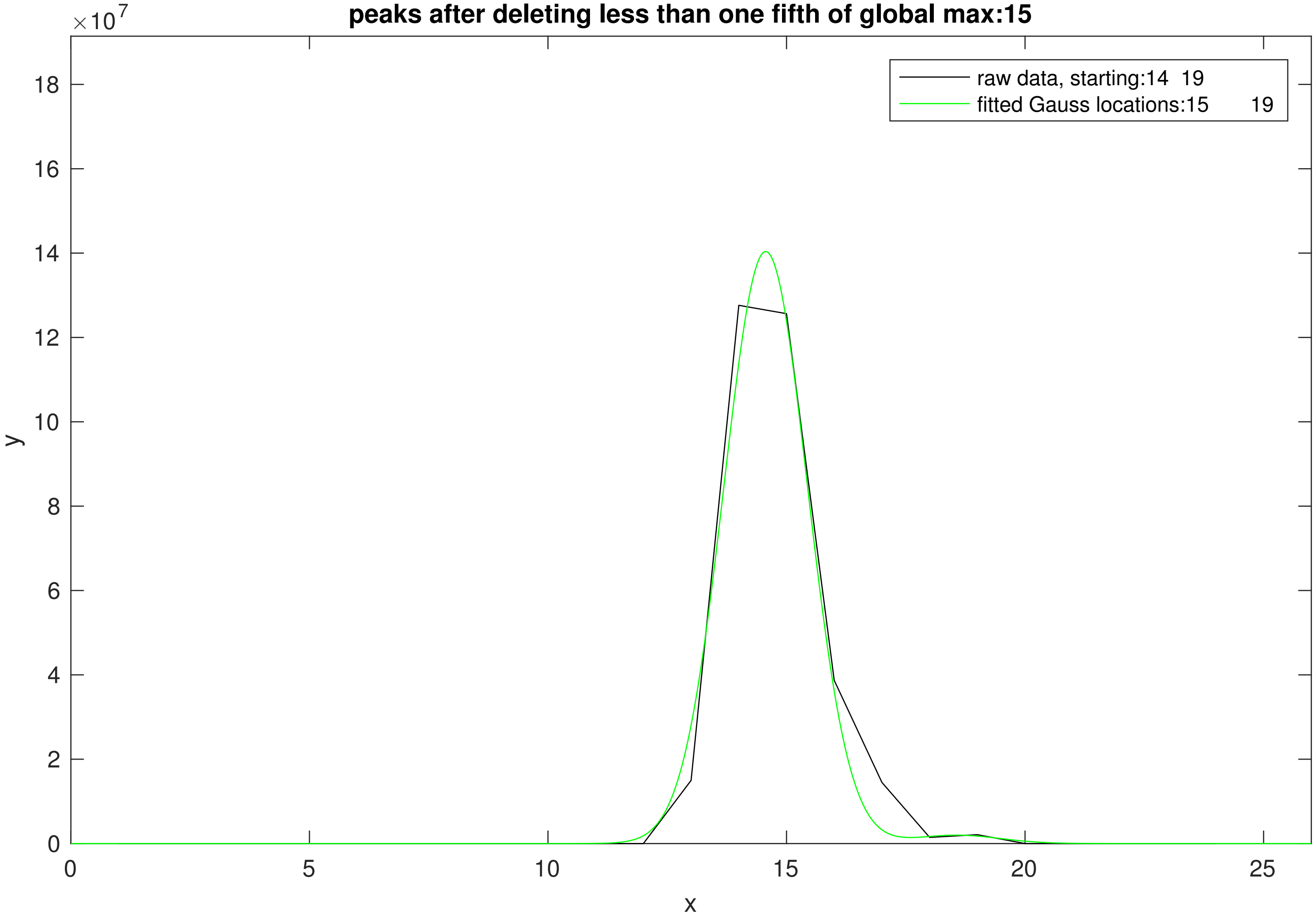

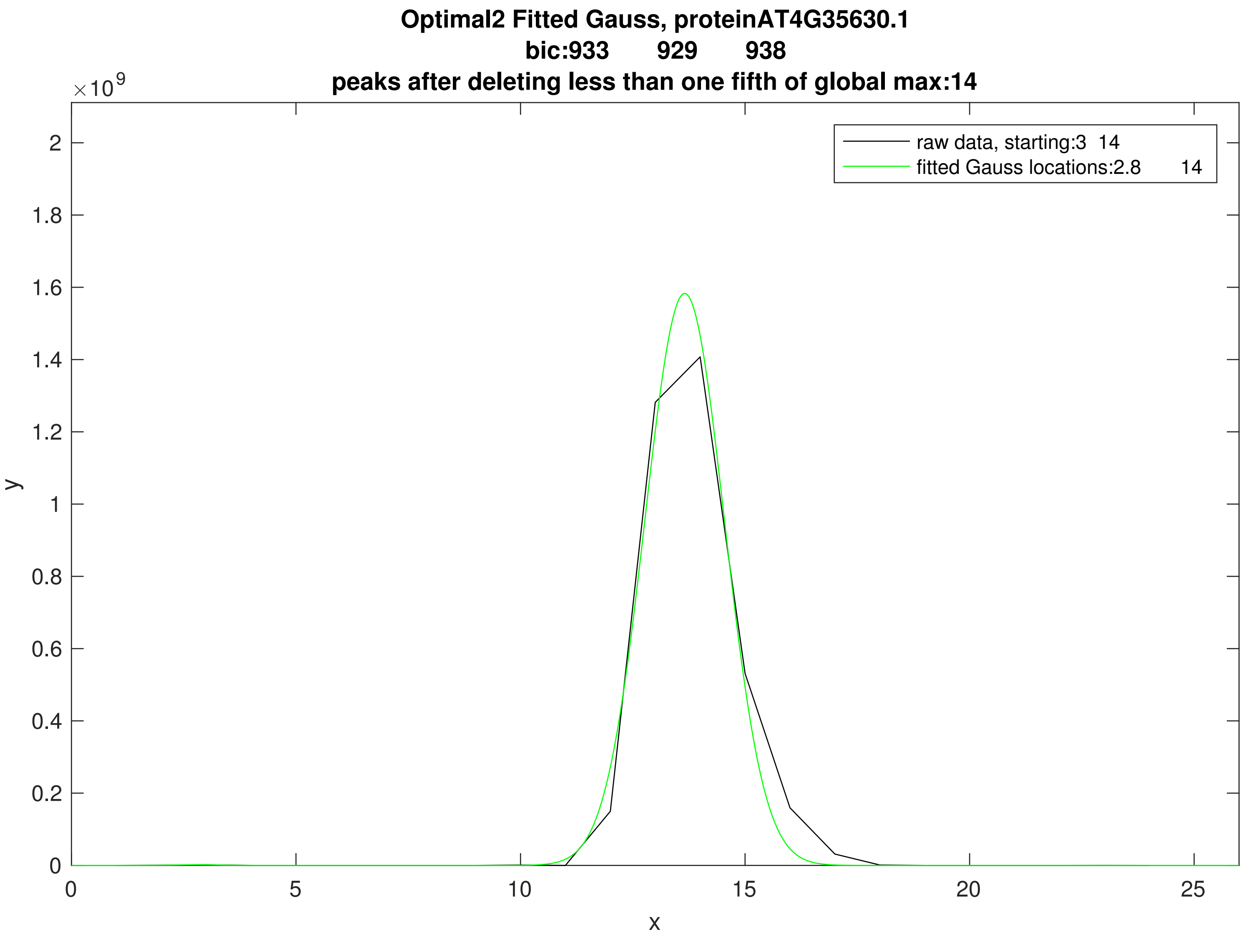

Optimal2 Fitted Gauss, proteinAT4G35630.1
bic:933 929 938
peaks after deleting less than one fifth of global max:14
×10^9
raw data, starting:3 14
fitted Gauss locations:2.8 14
y
x
0
0.2
0.4
0.6
0.8
1
1.2
1.4
1.6
1.8
2
0
5
10
15
20
25

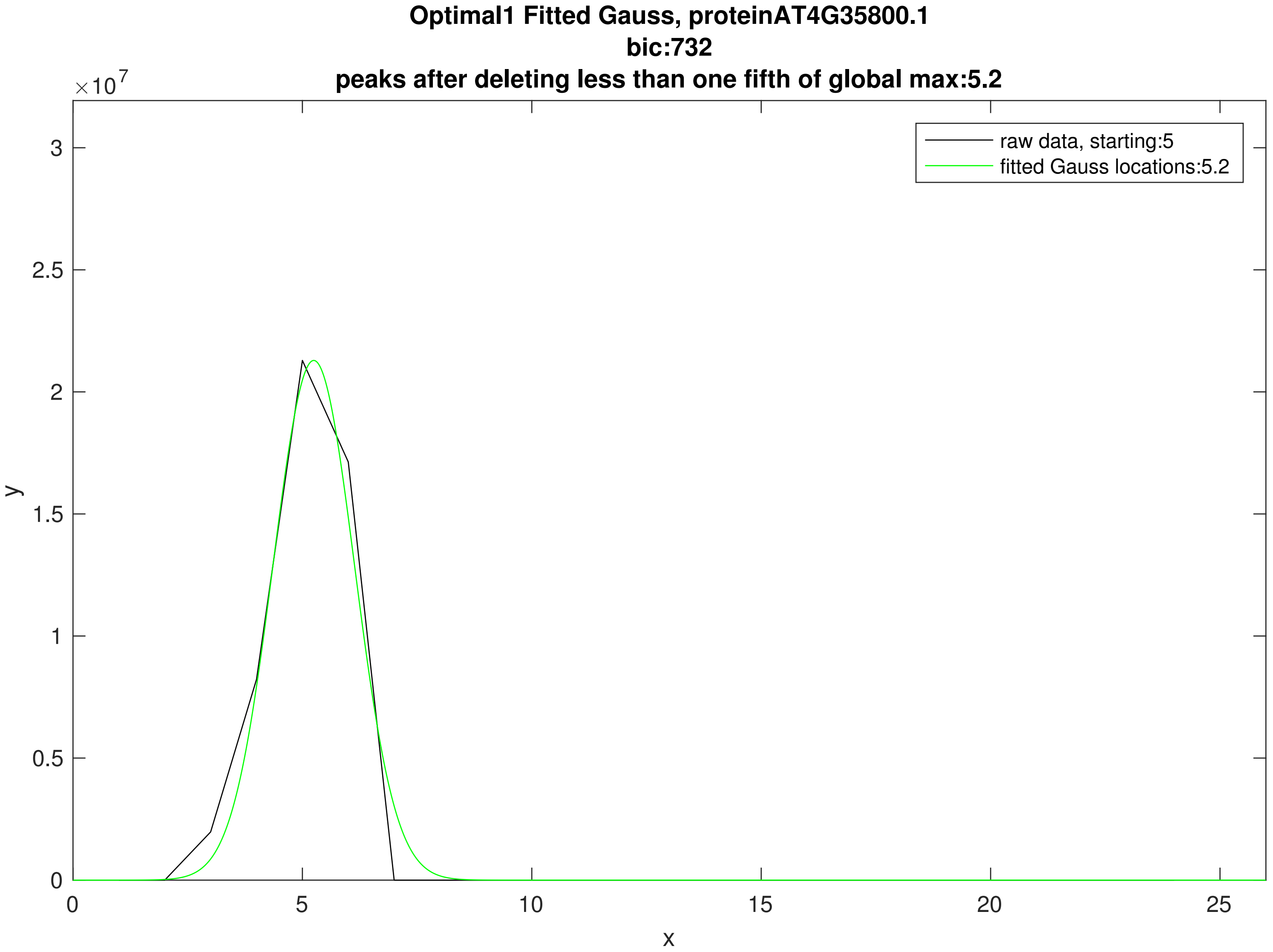
Optimal1 Fitted Gauss, proteinAT4G35800.1
bic:732
peaks after deleting less than one fifth of global max:5.2
×10^7
raw data, starting:5
fitted Gauss locations:5.2
y
x
0
0.5
1
1.5
2
2.5
3
0
5
10
15
20
25

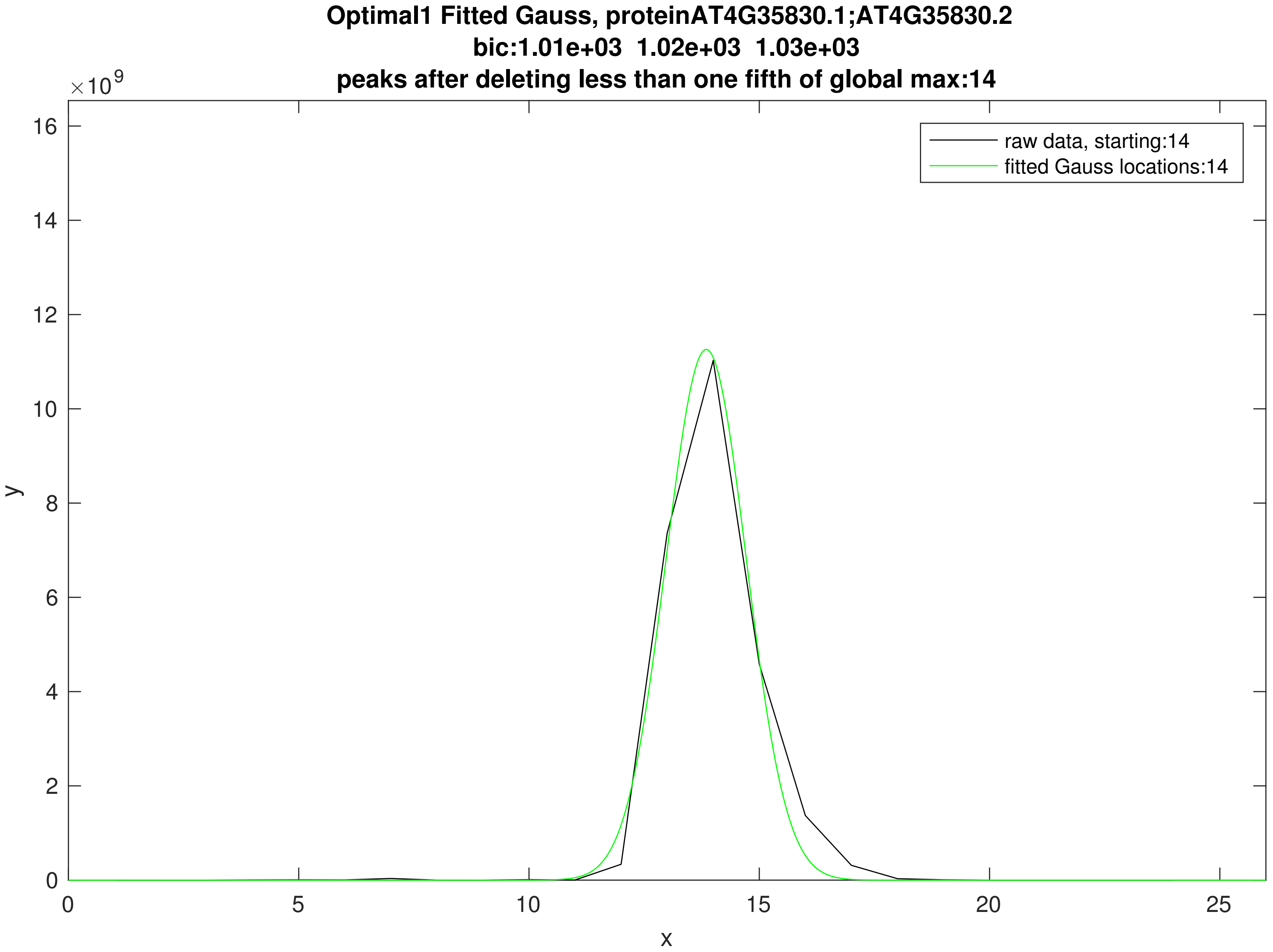
Optimal1 Fitted Gauss, proteinAT4G35830.1;AT4G35830.2
bic:1.01e+03 1.02e+03 1.03e+03
peaks after deleting less than one fifth of global max:14
×10^9
raw data, starting:14
fitted Gauss locations:14
y
x
0
2
4
6
8
10
12
14
16
5
15
20
25

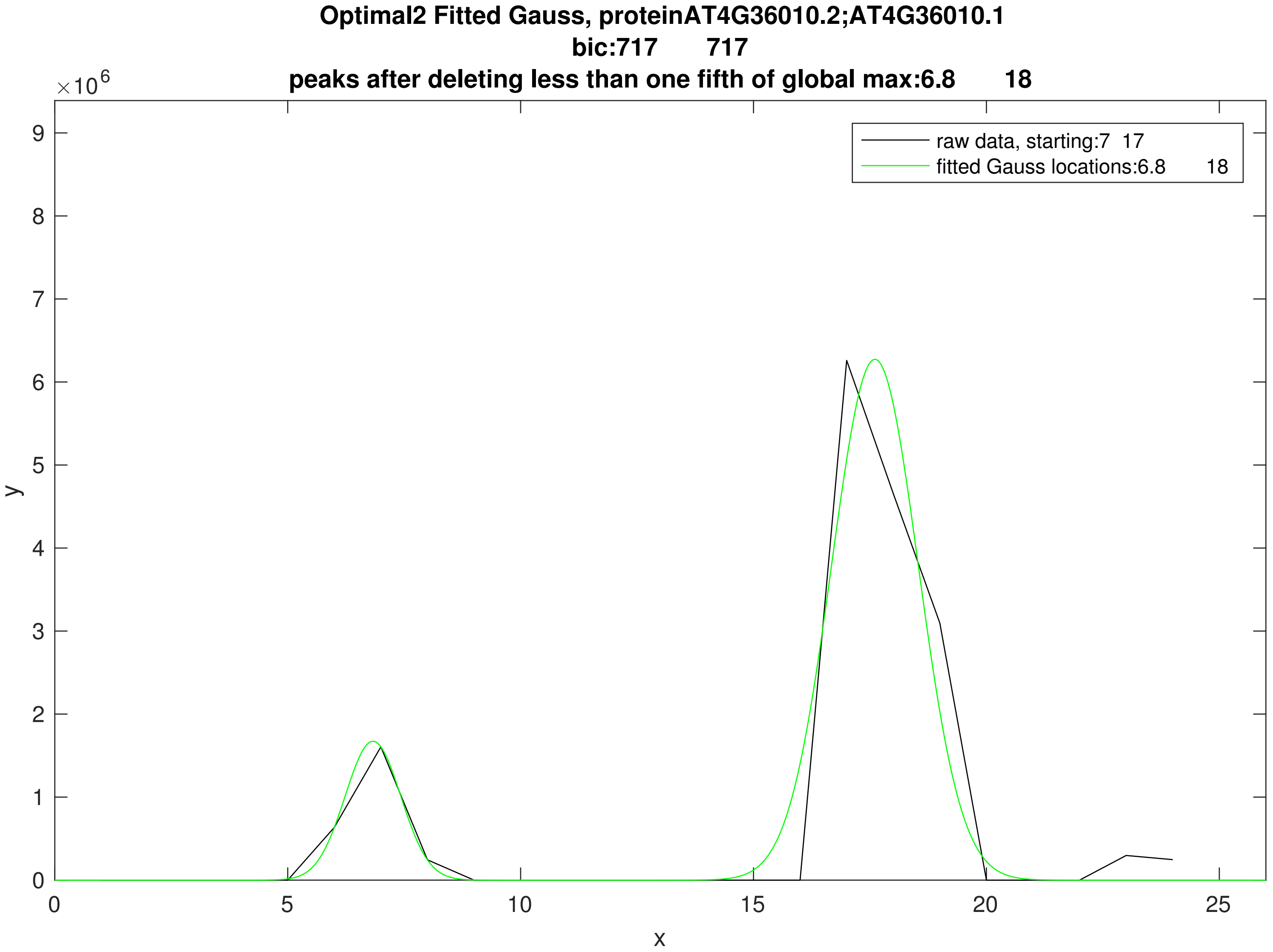

Optimal2 Fitted Gauss, proteinAT4G36010.2;AT4G36010.1
bic:717 717
peaks after deleting less than one fifth of global max:6.8 18
×10^6
raw data, starting:7 17
fitted Gauss locations:6.8 18
y
x
0
1
2
3
4
5
6
7
8
9
0
5
10
15
20
25

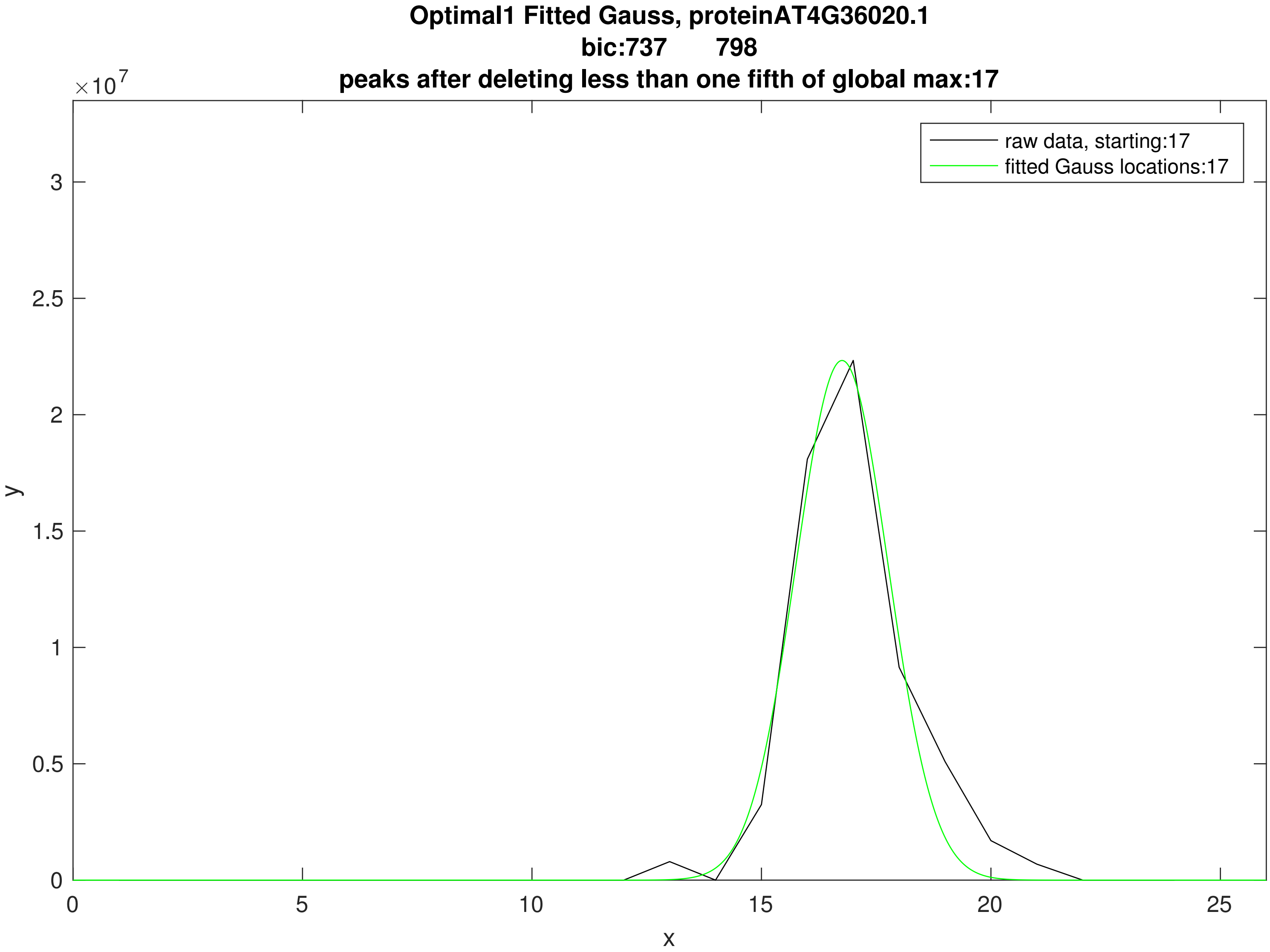
Optimal1 Fitted Gauss, proteinAT4G36020.1
bic:737 798
peaks after deleting less than one fifth of global max:17
×10^7
raw data, starting:17
fitted Gauss locations:17
y
x
0
0.5
1
1.5
2
2.5
3
0
5
10
15
20
25

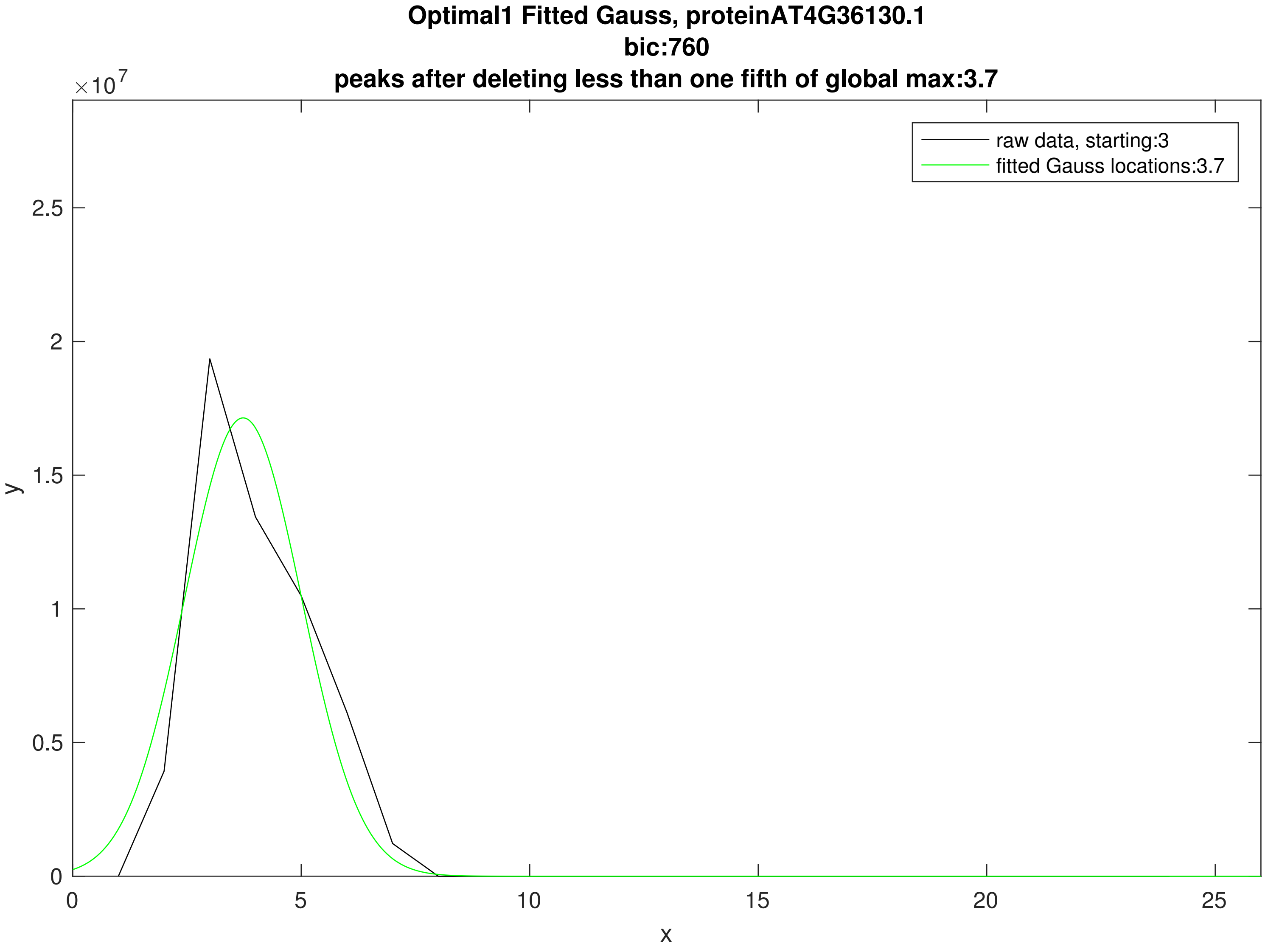
Optimal1 Fitted Gauss, proteinAT4G36130.1
bic:760
peaks after deleting less than one fifth of global max:3.7
×10^7
raw data, starting:3
fitted Gauss locations:3.7
y
x
0
0.5
1
1.5
2
2.5
0
5
10
15
20
25

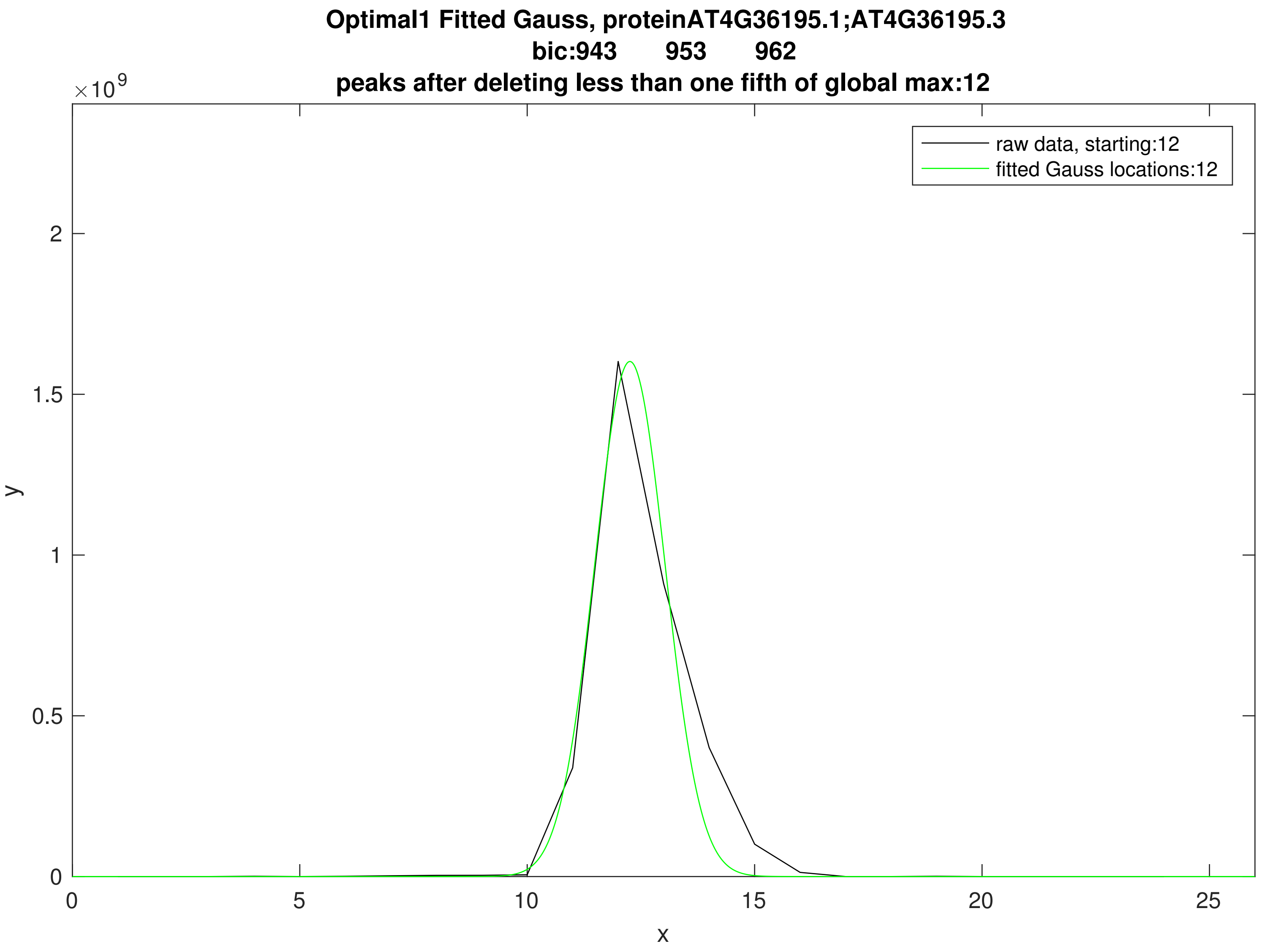
Optimal1 Fitted Gauss, proteinAT4G36195.1;AT4G36195.3
bic:943 953 962
peaks after deleting less than one fifth of global max:12
×10^9
raw data, starting:12
fitted Gauss locations:12
y
x
0
0.5
1
1.5
2
0
5
10
15
20
25

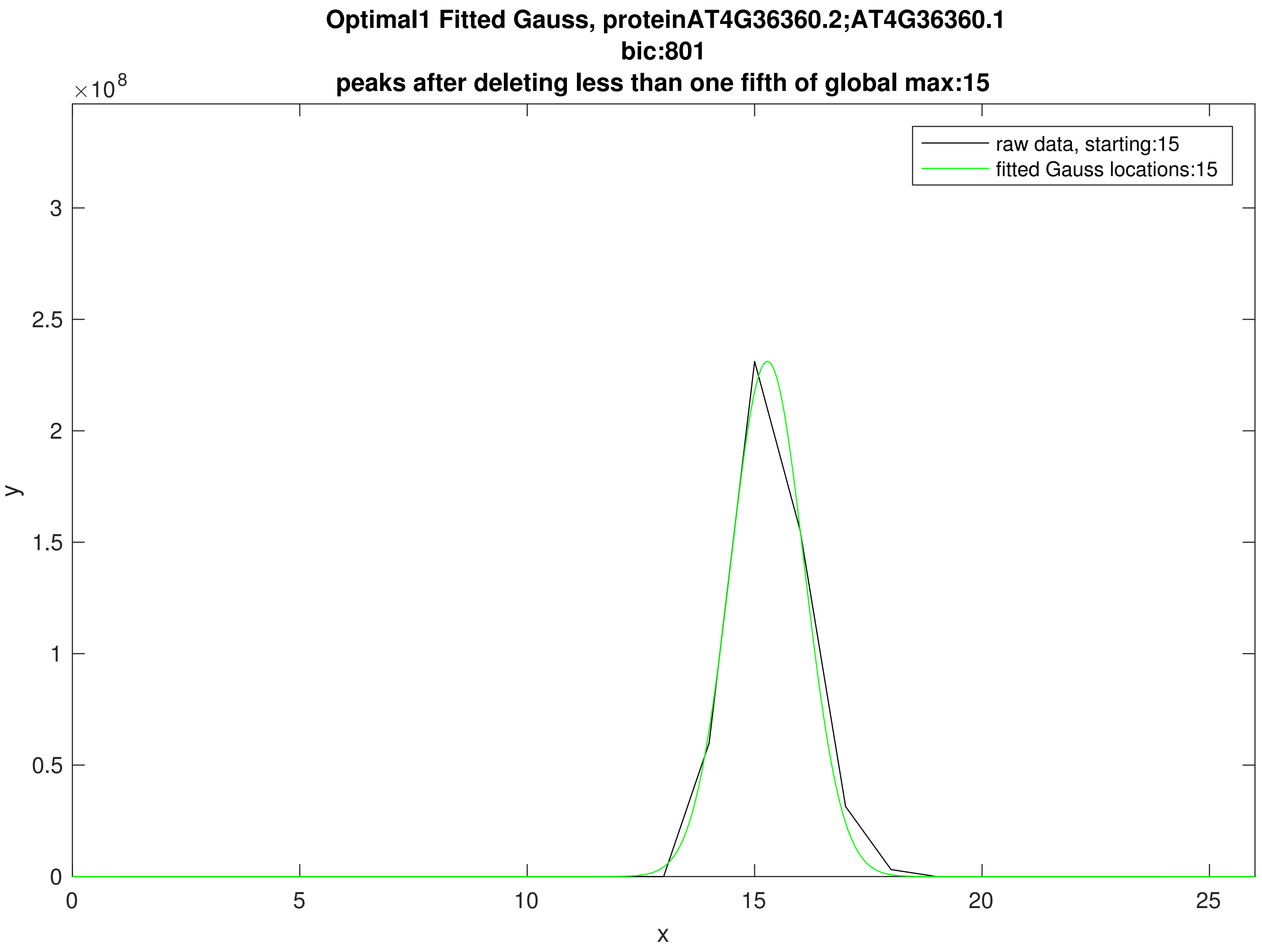
Optimal1 Fitted Gauss, proteinAT4G36360.2;AT4G36360.1
bic:801
peaks after deleting less than one fifth of global max:15
×10^8
raw data, starting:15
fitted Gauss locations:15
y
x
0
0.5
1
1.5
2
2.5
3
0
5
10
15
20
25

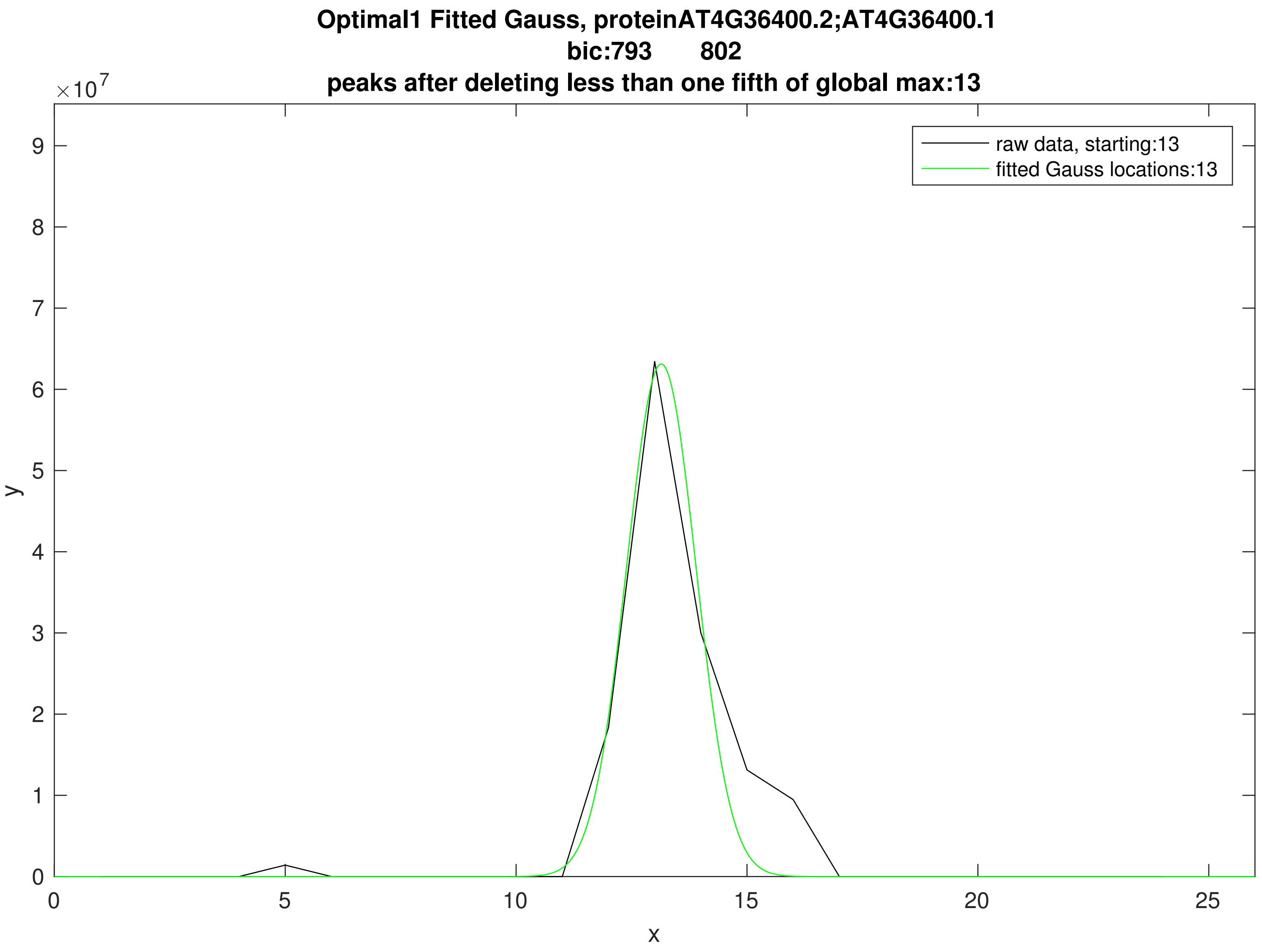

Optimal1 Fitted Gauss, proteinAT4G36400.2;AT4G36400.1
bic:793 802
peaks after deleting less than one fifth of global max:13
×10^7
9
8
7
6
5
4
3
2
1
0
y
0
5
10
15
20
25
x
raw data, starting:13
fitted Gauss locations:13

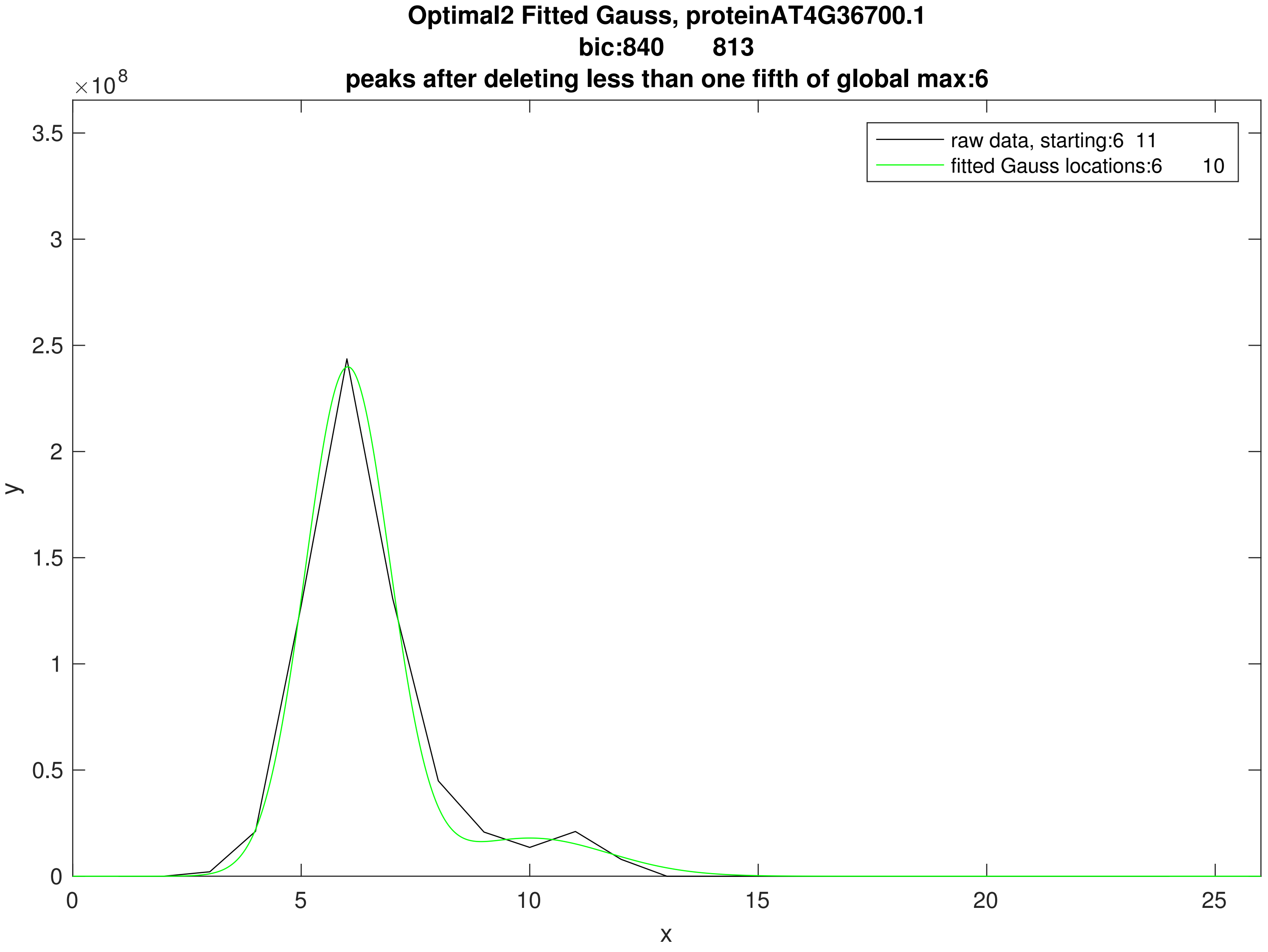

Optimal2 Fitted Gauss, proteinAT4G36700.1
bic:840 813
peaks after deleting less than one fifth of global max:6
×10^8
3.5
3
2.5
2
1.5
1
0.5
0
y
0
5
10
15
20
25
x
raw data, starting:6 11
fitted Gauss locations:6 10

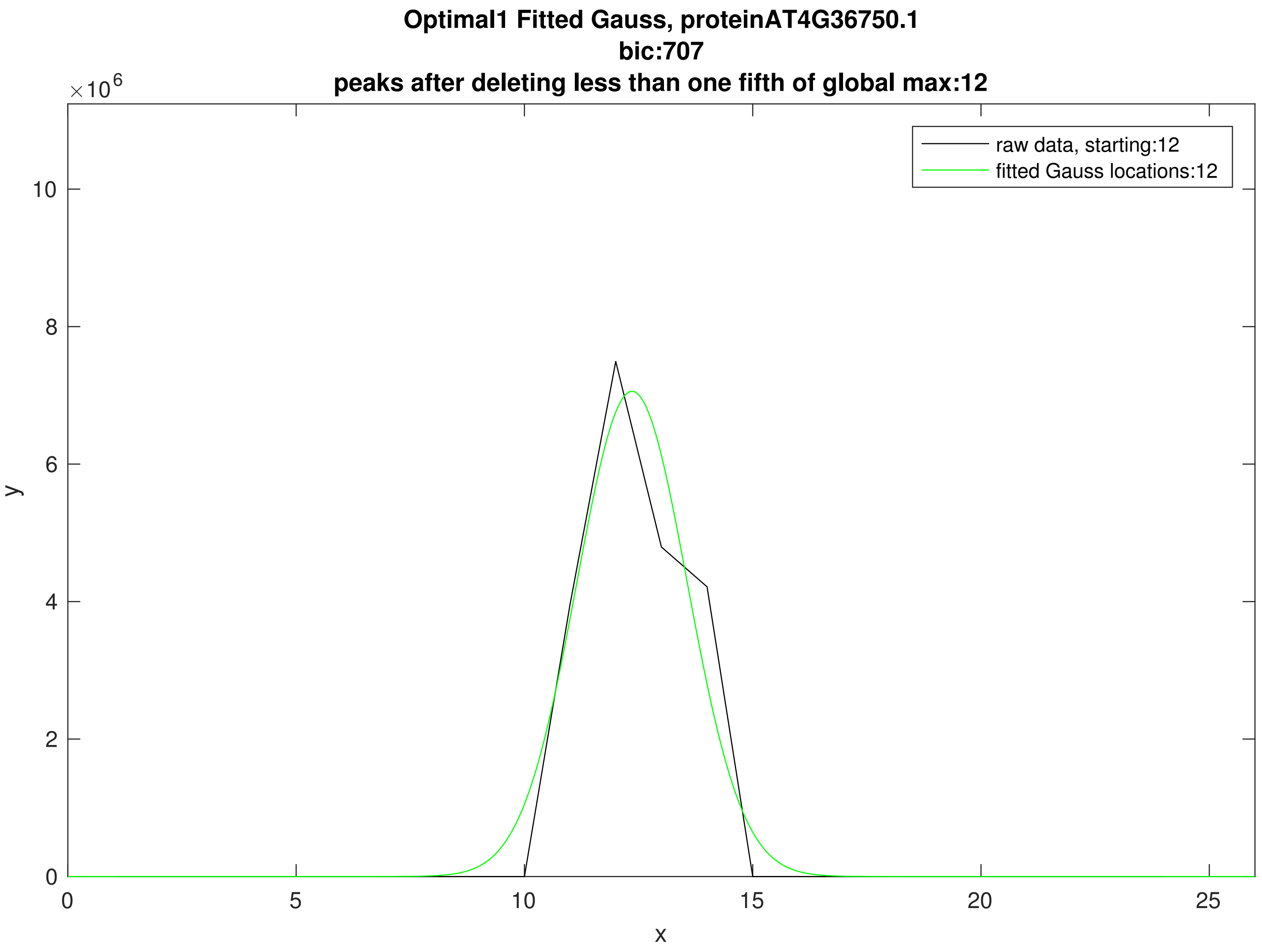
Optimal1 Fitted Gauss, proteinAT4G36750.1
bic:707
peaks after deleting less than one fifth of global max:12
×10^6
raw data, starting:12
fitted Gauss locations:12
10
8
6
4
2
0
y
0
5
10
15
20
25
x

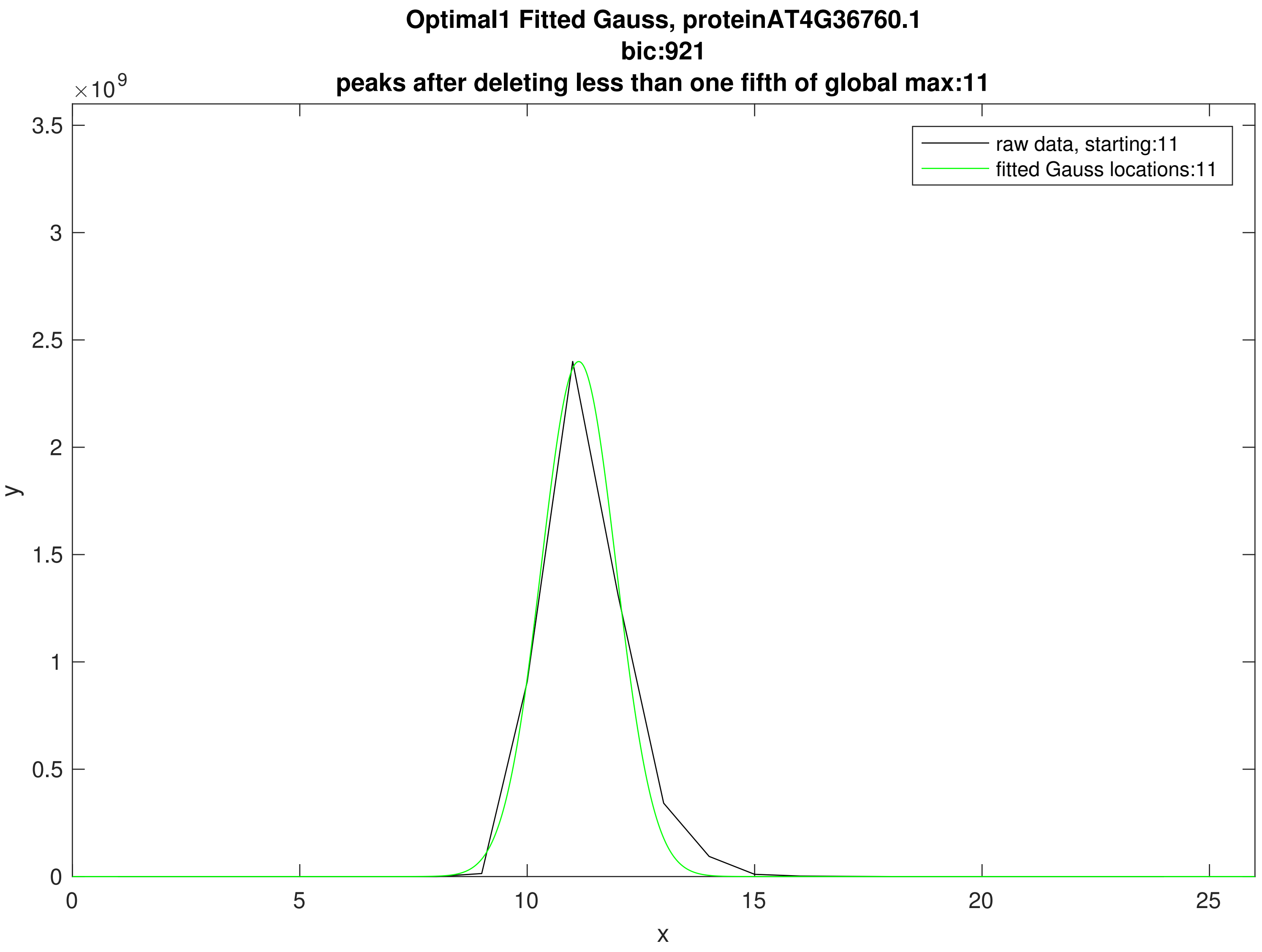
Optimal1 Fitted Gauss, proteinAT4G36760.1
bic:921
peaks after deleting less than one fifth of global max:11
×10^9
3.5
3
2.5
2
1.5
1
0.5
0
y
0
5
10
15
20
25
x
raw data, starting:11
fitted Gauss locations:11

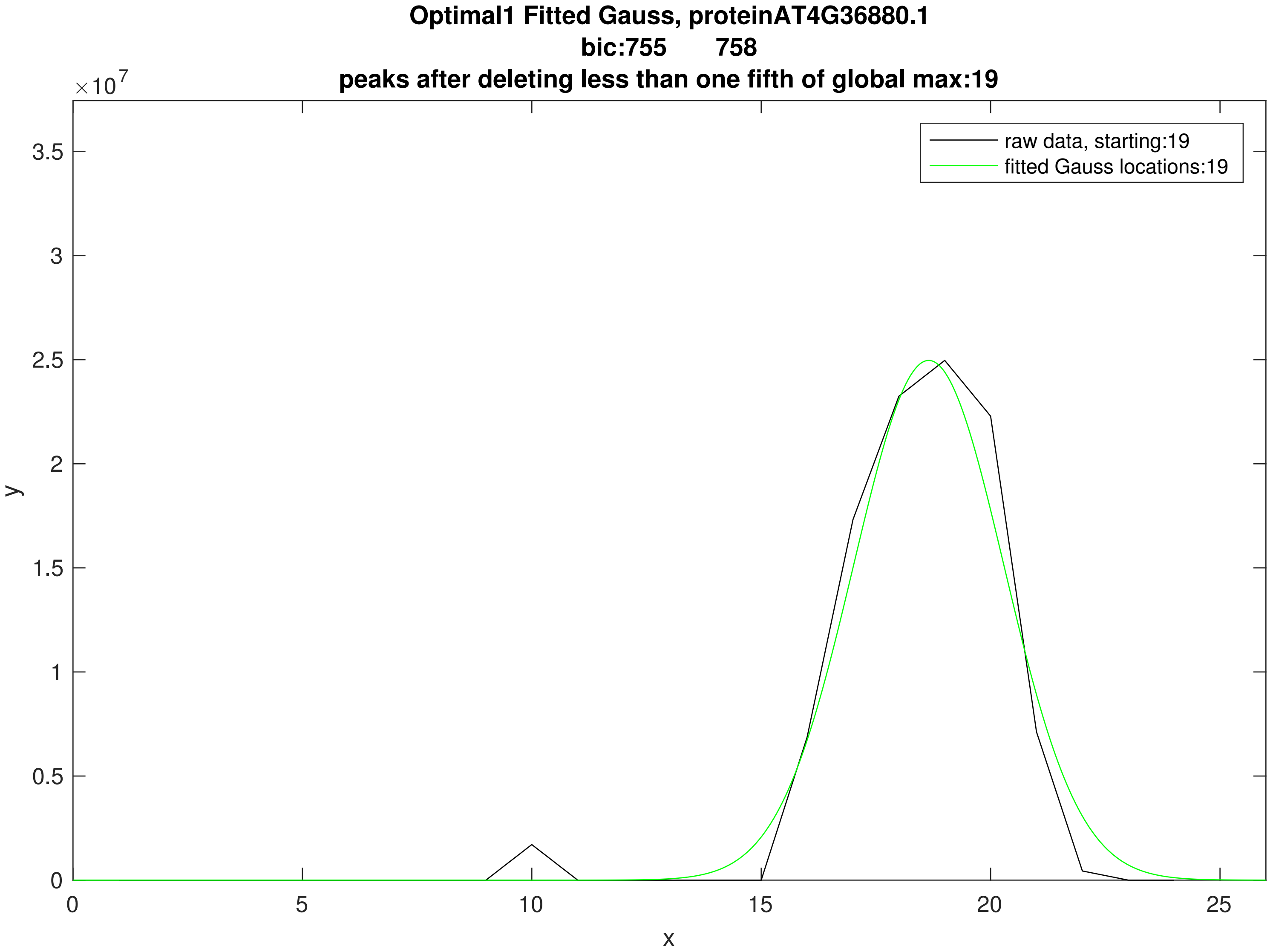

Optimal1 Fitted Gauss, proteinAT4G36880.1
bic:755 758
peaks after deleting less than one fifth of global max:19
×10^7
raw data, starting:19
fitted Gauss locations:19
y
x
0
0.5
1
1.5
2
2.5
3
3.5
0
5
10
15
20
25

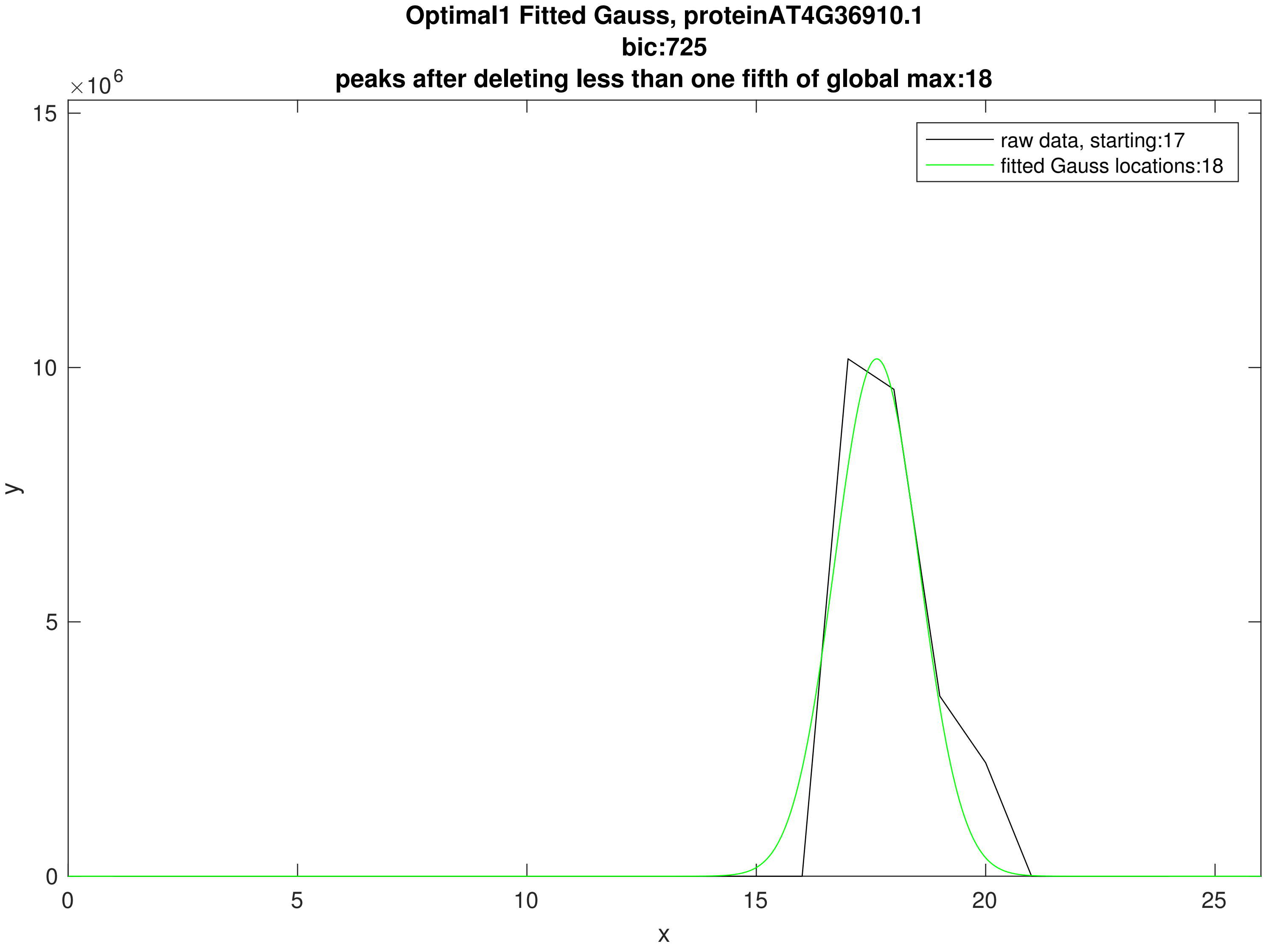
Optimal1 Fitted Gauss, proteinAT4G36910.1
bic:725
peaks after deleting less than one fifth of global max:18
×10^6
15
10
5
0
y
0
5
10
15
20
25
x
raw data, starting:17
fitted Gauss locations:18

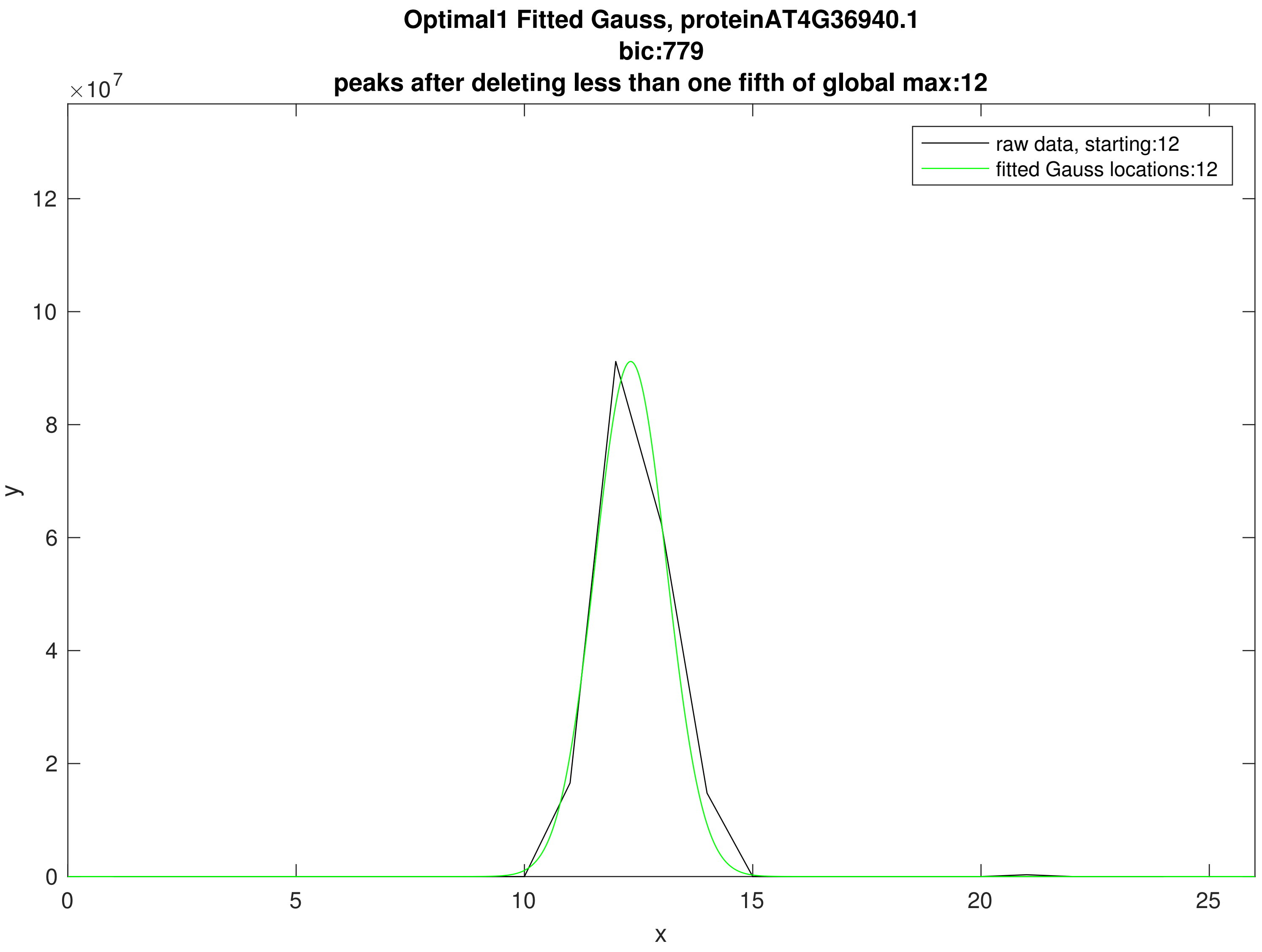

Optimal1 Fitted Gauss, proteinAT4G36940.1
bic:779
peaks after deleting less than one fifth of global max:12
×10^7
raw data, starting:12
fitted Gauss locations:12
y
x
0
2
4
6
8
10
12
0
5
10
15
20
25

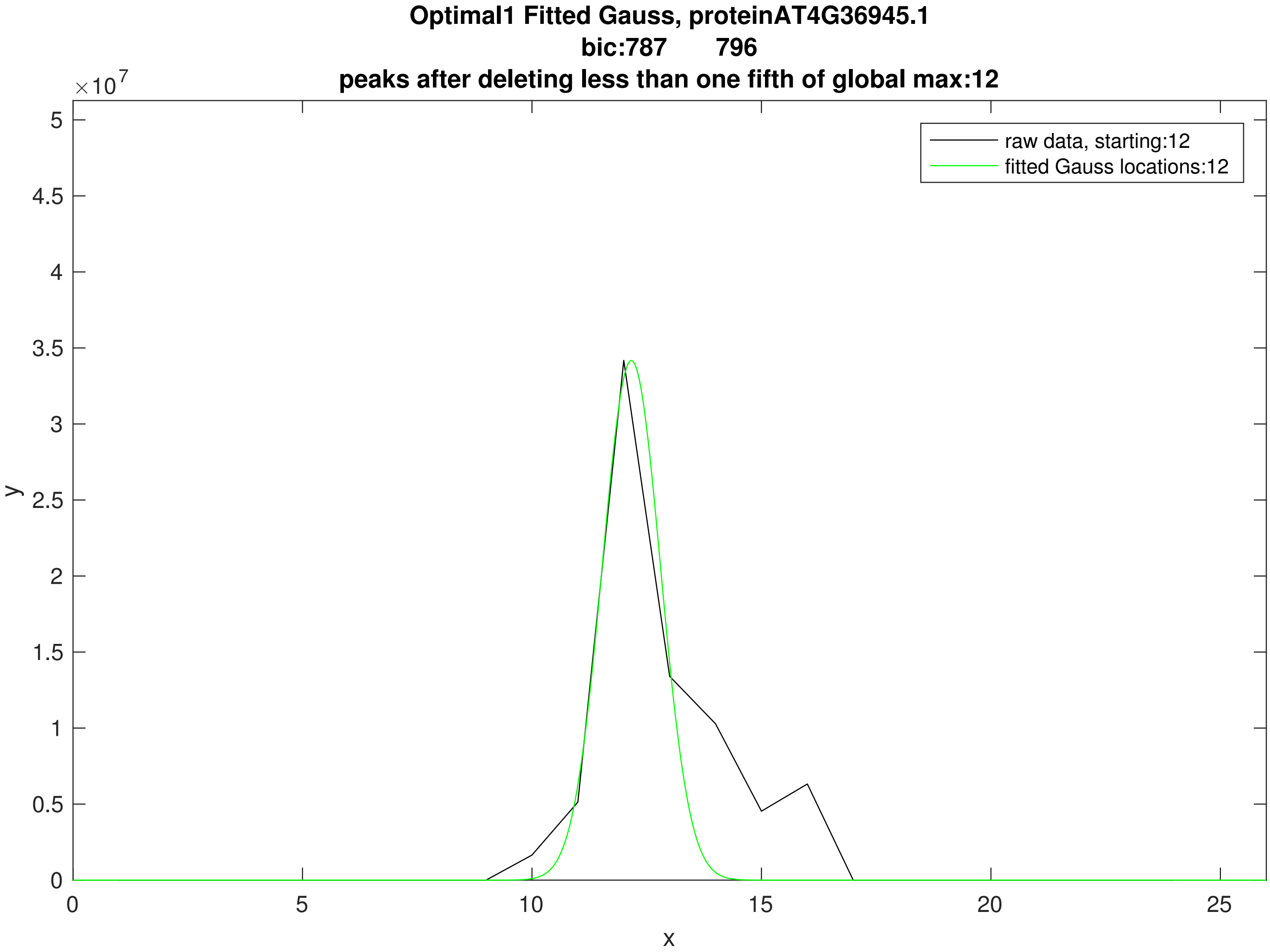
Optimal1 Fitted Gauss, proteinAT4G36945.1
bic:787 796
peaks after deleting less than one fifth of global max:12
×10^7
5
4.5
4
3.5
3
2.5
2
1.5
1
0.5
0
y
0
5
10
15
20
25
x
raw data, starting:12
fitted Gauss locations:12

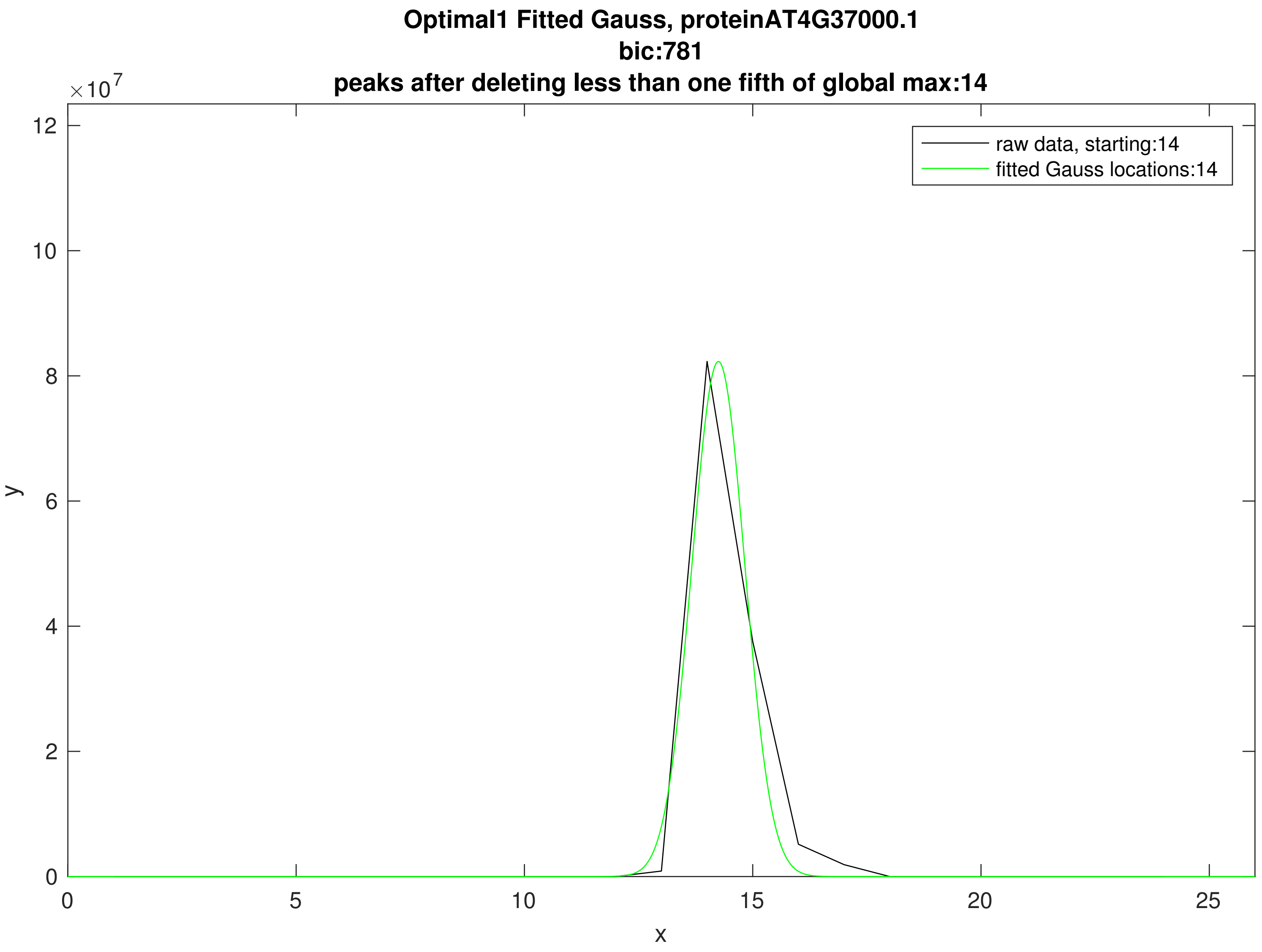

Optimal1 Fitted Gauss, proteinAT4G37000.1
bic:781
peaks after deleting less than one fifth of global max:14
×10^7
12
10
8
6
4
2
0
y
0
5
10
15
20
25
x
raw data, starting:14
fitted Gauss locations:14

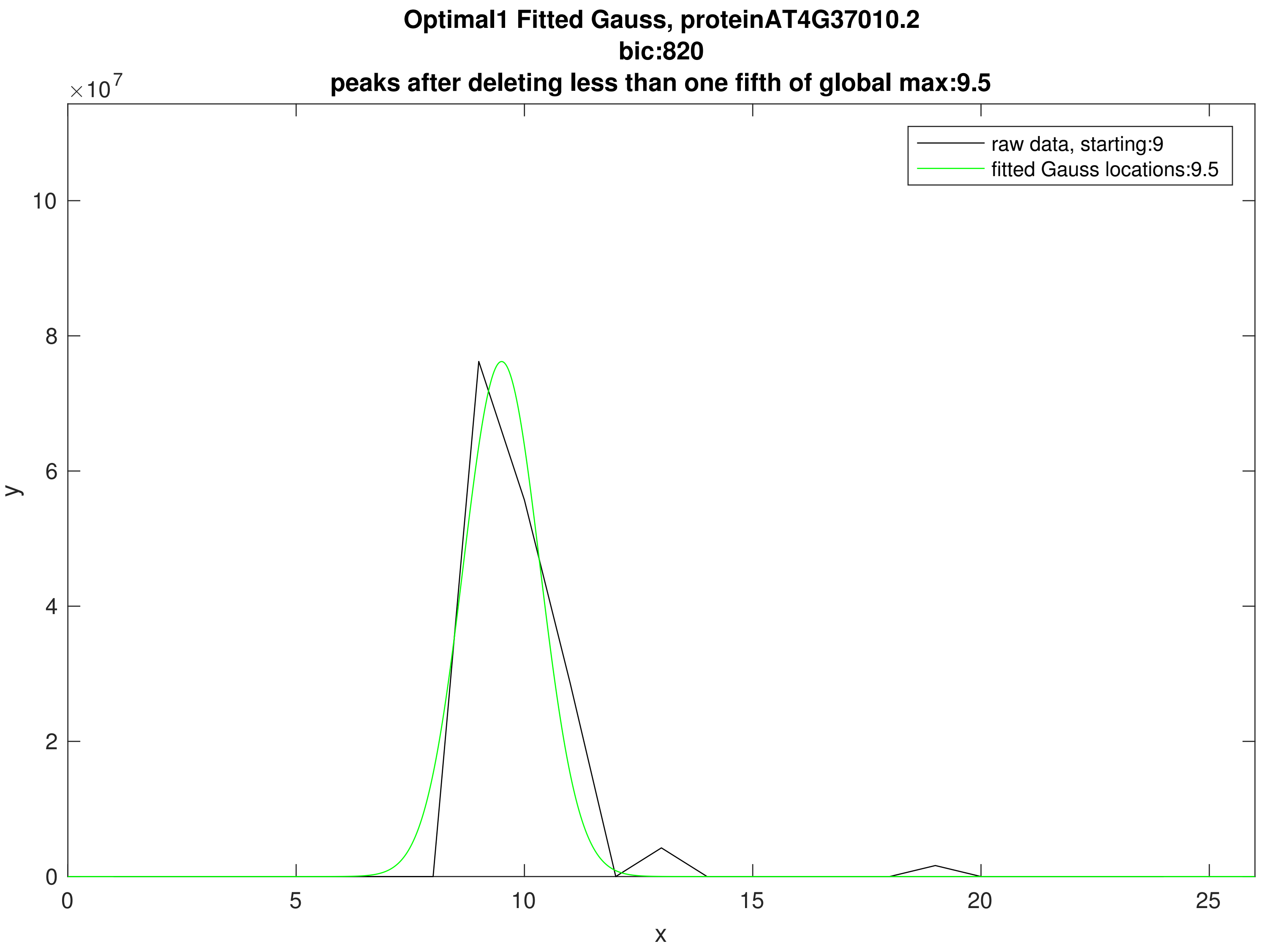

Optimal1 Fitted Gauss, proteinAT4G37010.2
bic:820
peaks after deleting less than one fifth of global max:9.5
×10^7
raw data, starting:9
fitted Gauss locations:9.5
y
x
0
2
4
6
8
10
0
5
10
15
20
25

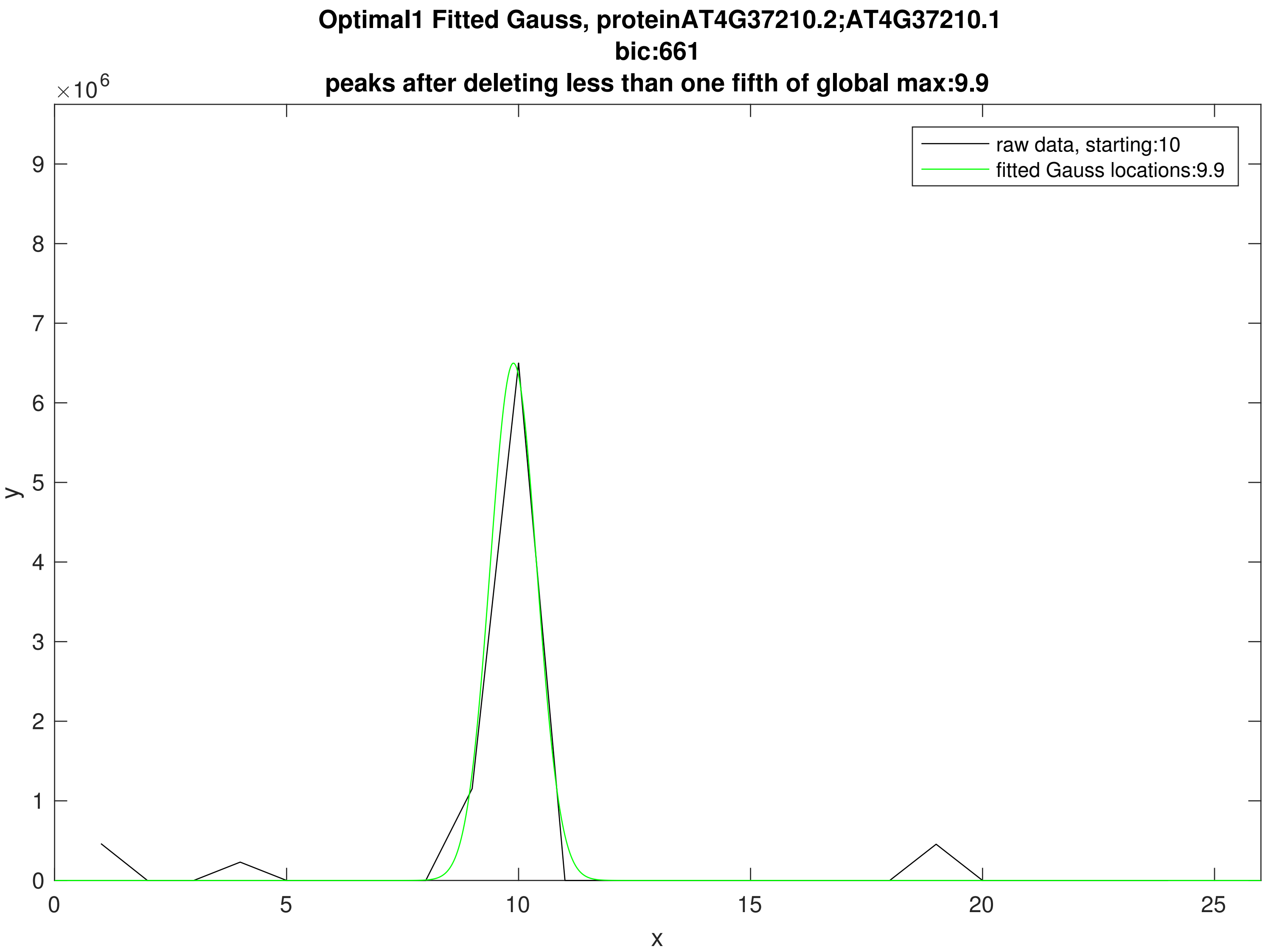

Optimal1 Fitted Gauss, proteinAT4G37210.2;AT4G37210.1
bic:661
peaks after deleting less than one fifth of global max:9.9
×10^6
raw data, starting:10
fitted Gauss locations:9.9
y
x
0
1
2
3
4
5
6
7
8
9
10
15
20
25

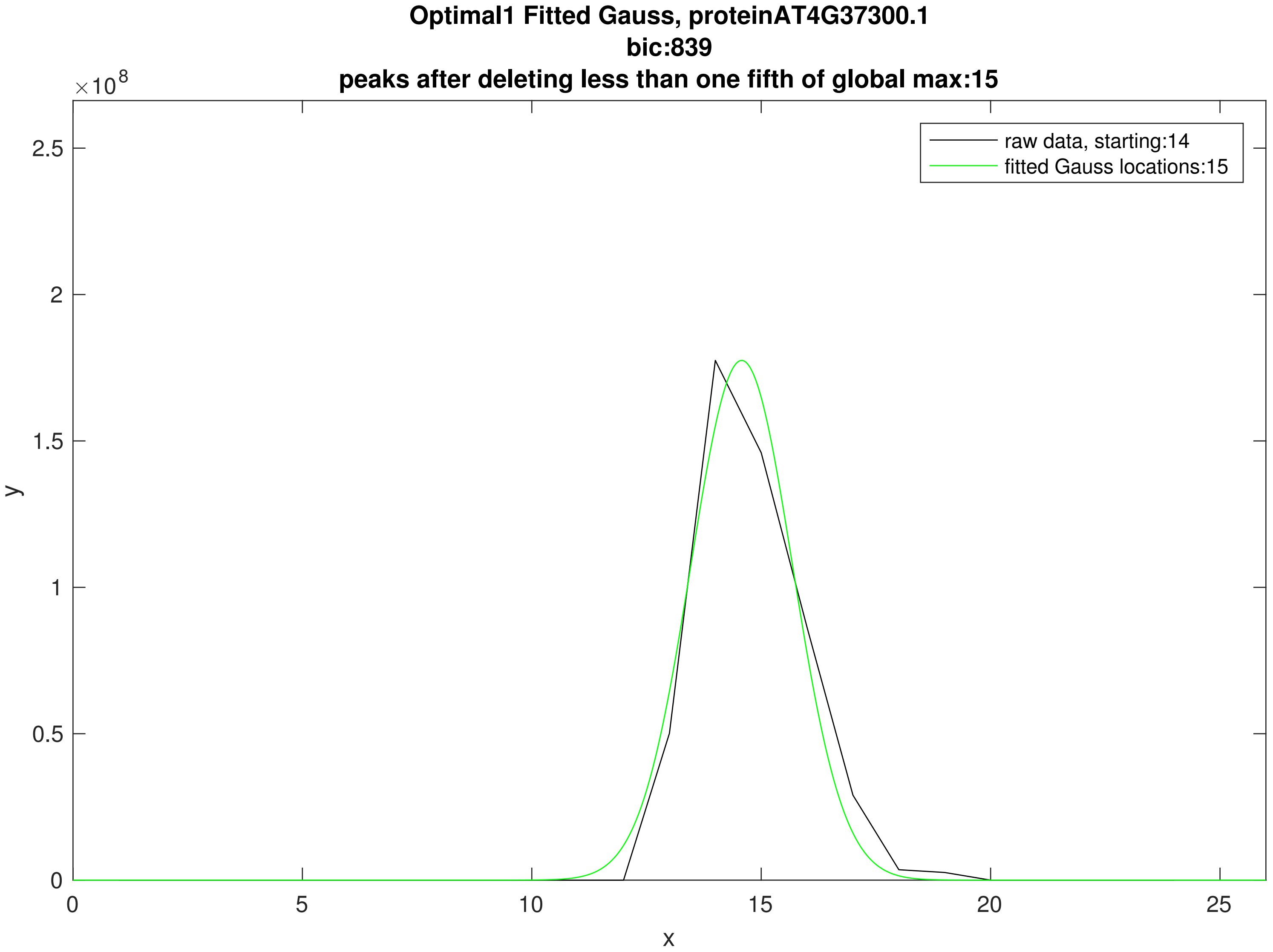
Optimal1 Fitted Gauss, proteinAT4G37300.1
bic:839
peaks after deleting less than one fifth of global max:15
×10^8
2.5
2
1.5
1
0.5
0
y
0
5
10
15
20
25
x
raw data, starting:14
fitted Gauss locations:15

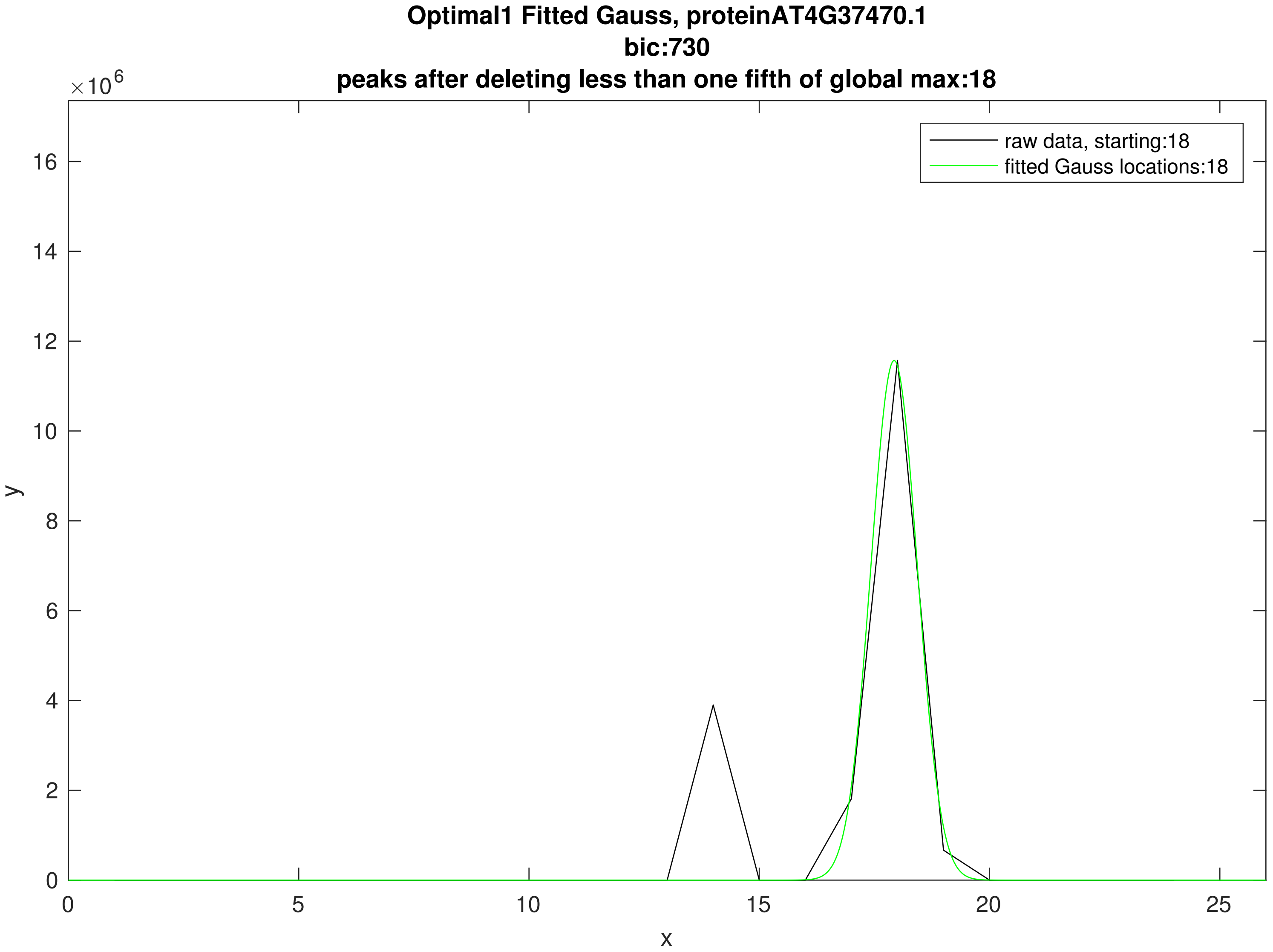
Optimal1 Fitted Gauss, proteinAT4G37470.1
bic:730
peaks after deleting less than one fifth of global max:18
×10^6
raw data, starting:18
fitted Gauss locations:18
y
x
0
2
4
6
8
10
12
14
16
0
5
10
15
20
25

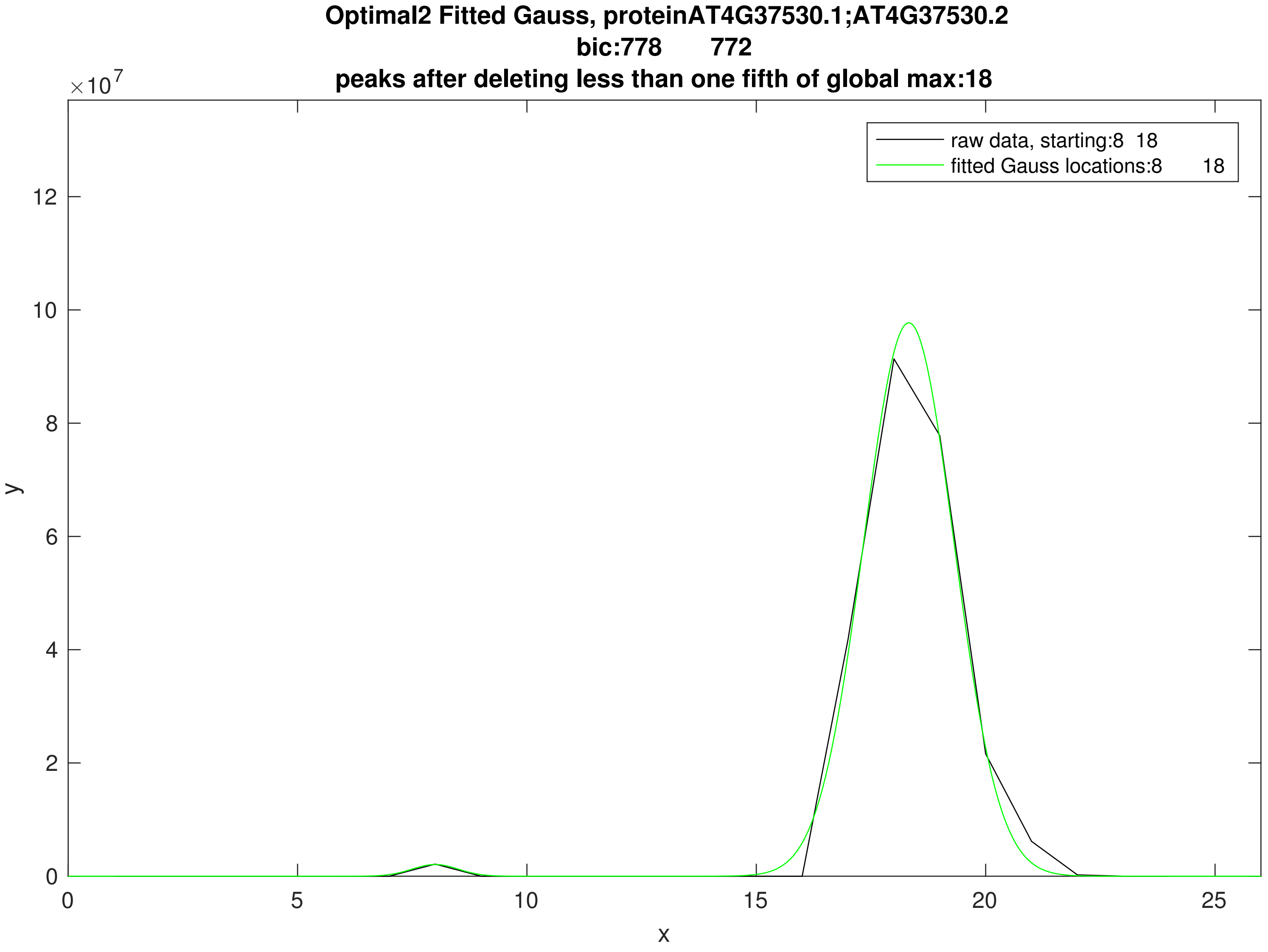
Optimal2 Fitted Gauss, proteinAT4G37530.1;AT4G37530.2
bic:778 772
peaks after deleting less than one fifth of global max:18
×10^7
raw data, starting:8 18
fitted Gauss locations:8 18
y
x
0
2
4
6
8
10
12
0
5
10
15
20
25

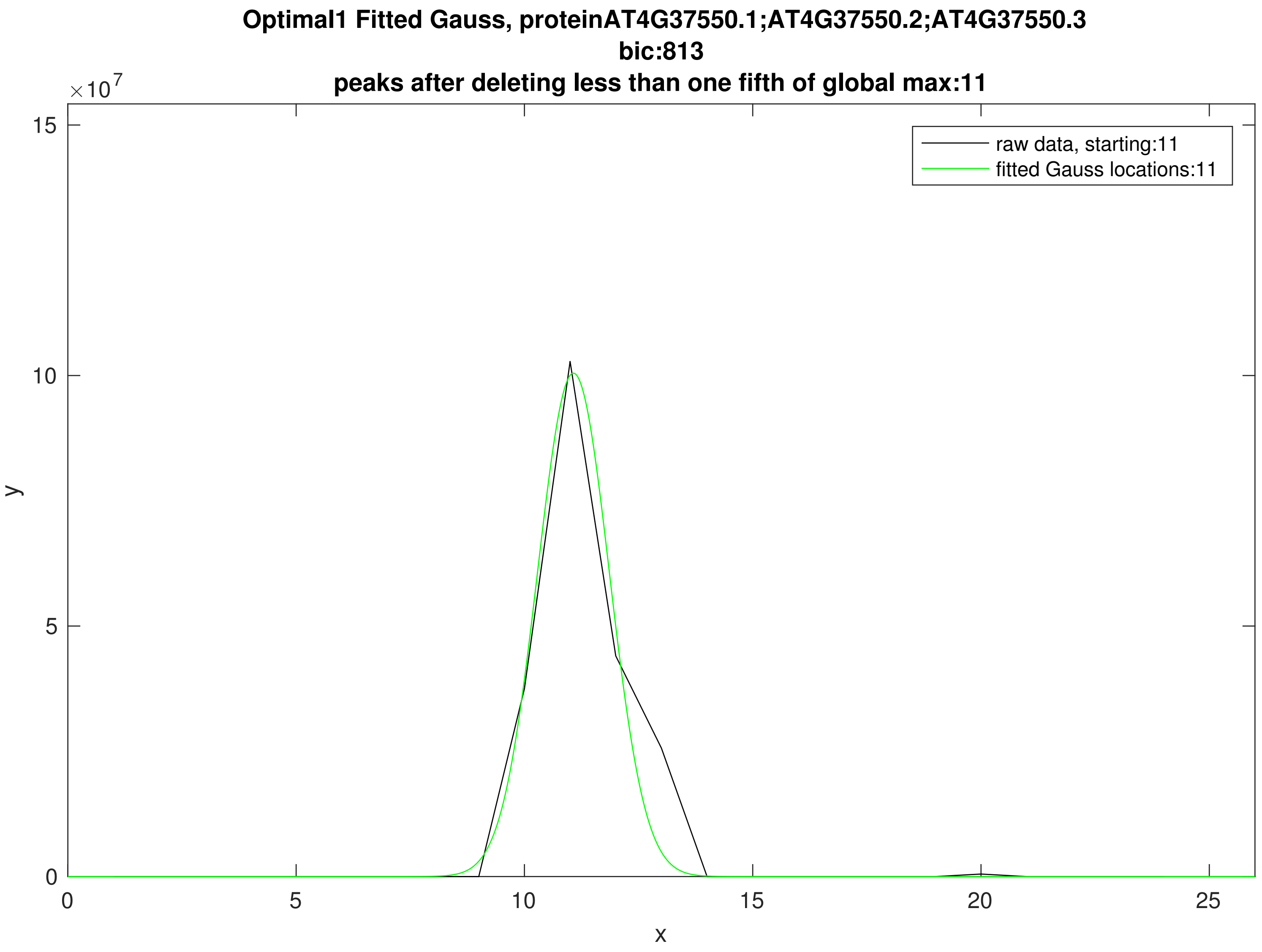
Optimal1 Fitted Gauss, proteinAT4G37550.1;AT4G37550.2;AT4G37550.3
bic:813
peaks after deleting less than one fifth of global max:11
×10^7
15
10
5
0
y
0
5
10
15
20
25
x
raw data, starting:11
fitted Gauss locations:11

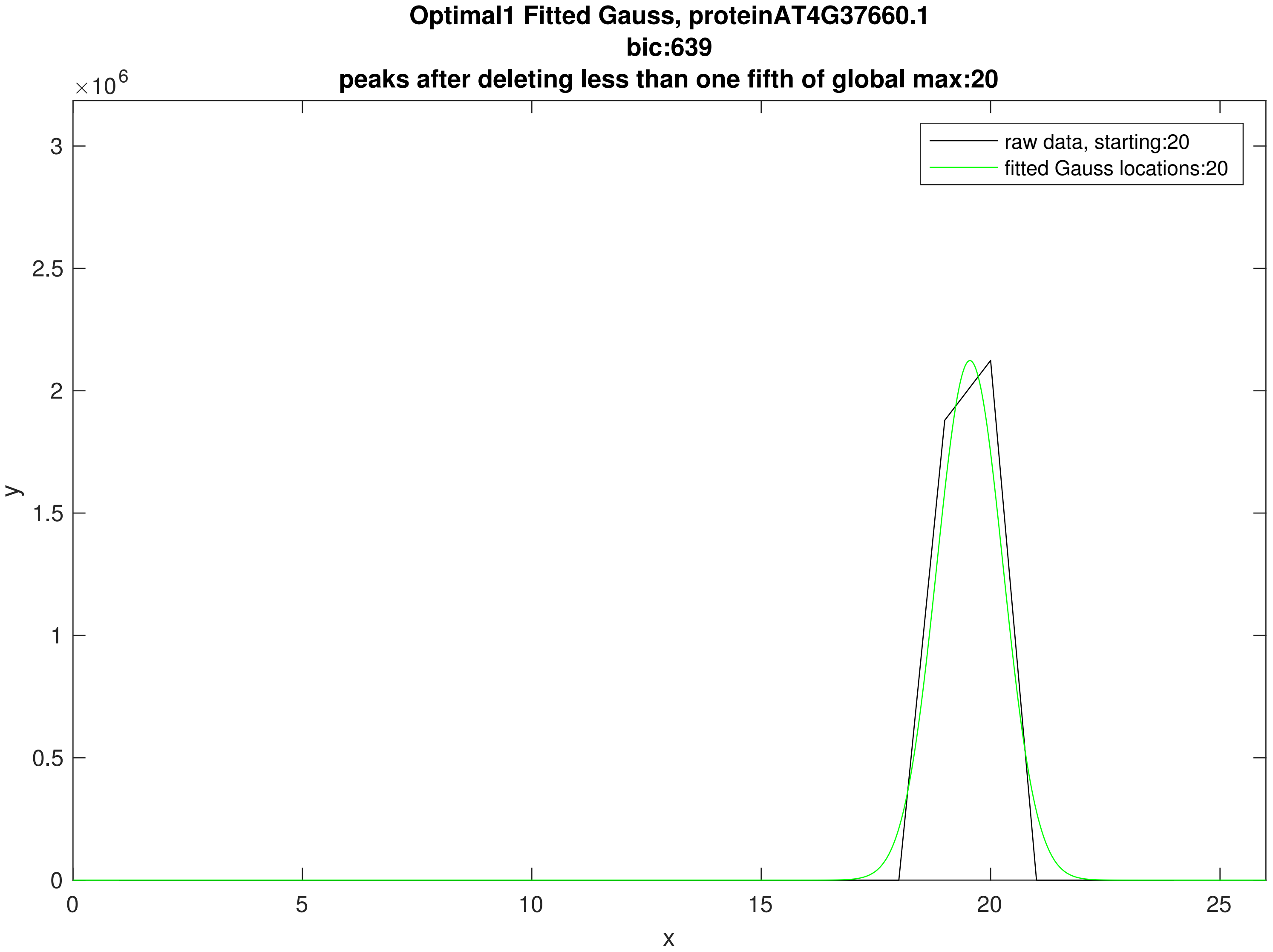
Optimal1 Fitted Gauss, proteinAT4G37660.1
bic:639
peaks after deleting less than one fifth of global max:20
×10^6
3
2.5
2
1.5
1
0.5
0
y
0
5
10
15
20
25
x
raw data, starting:20
fitted Gauss locations:20

**Optimal1 Fitted Gauss, proteinAT4G37800.1**
**bic:809**
**peaks after deleting less than one fifth of global max:20**

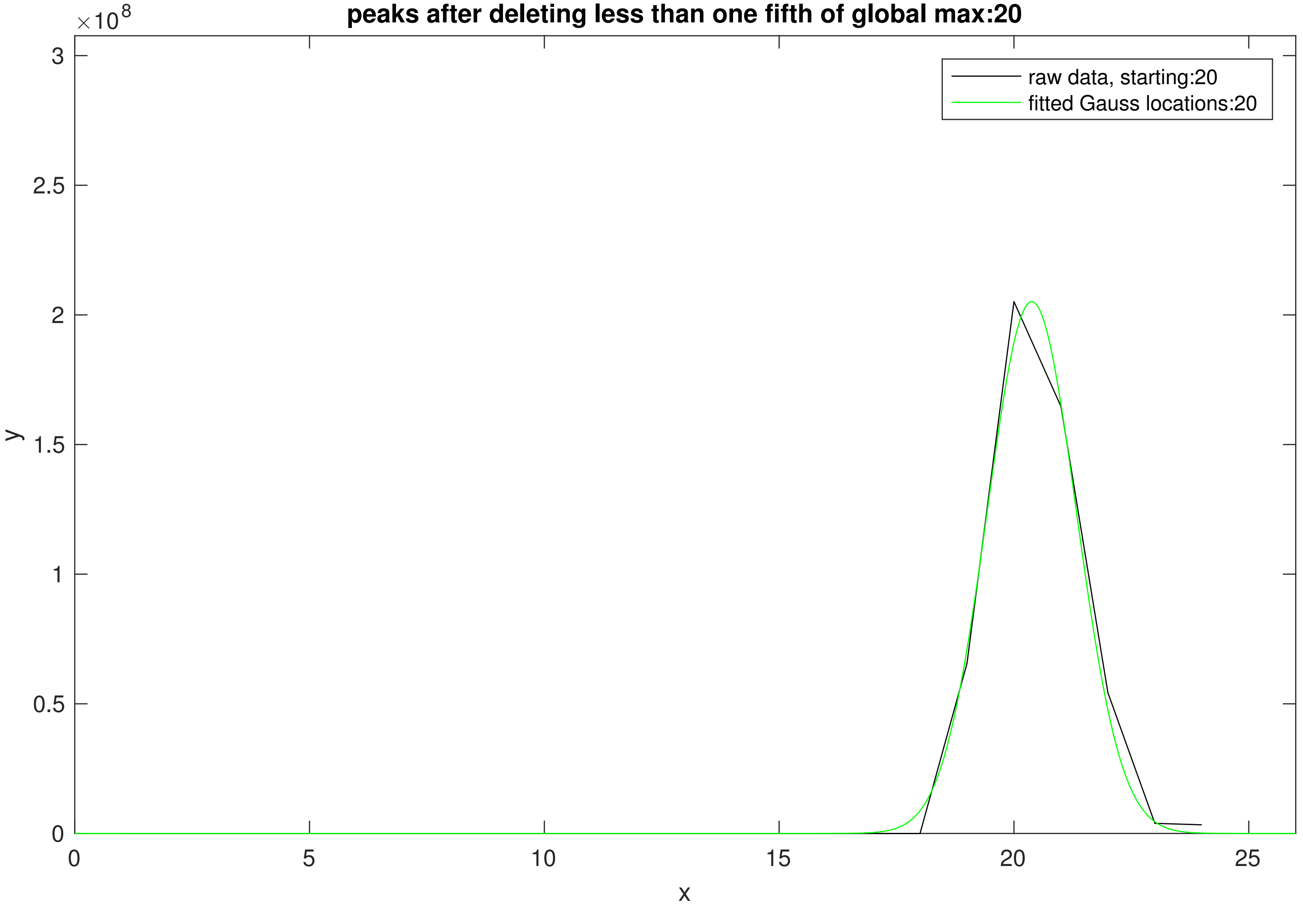

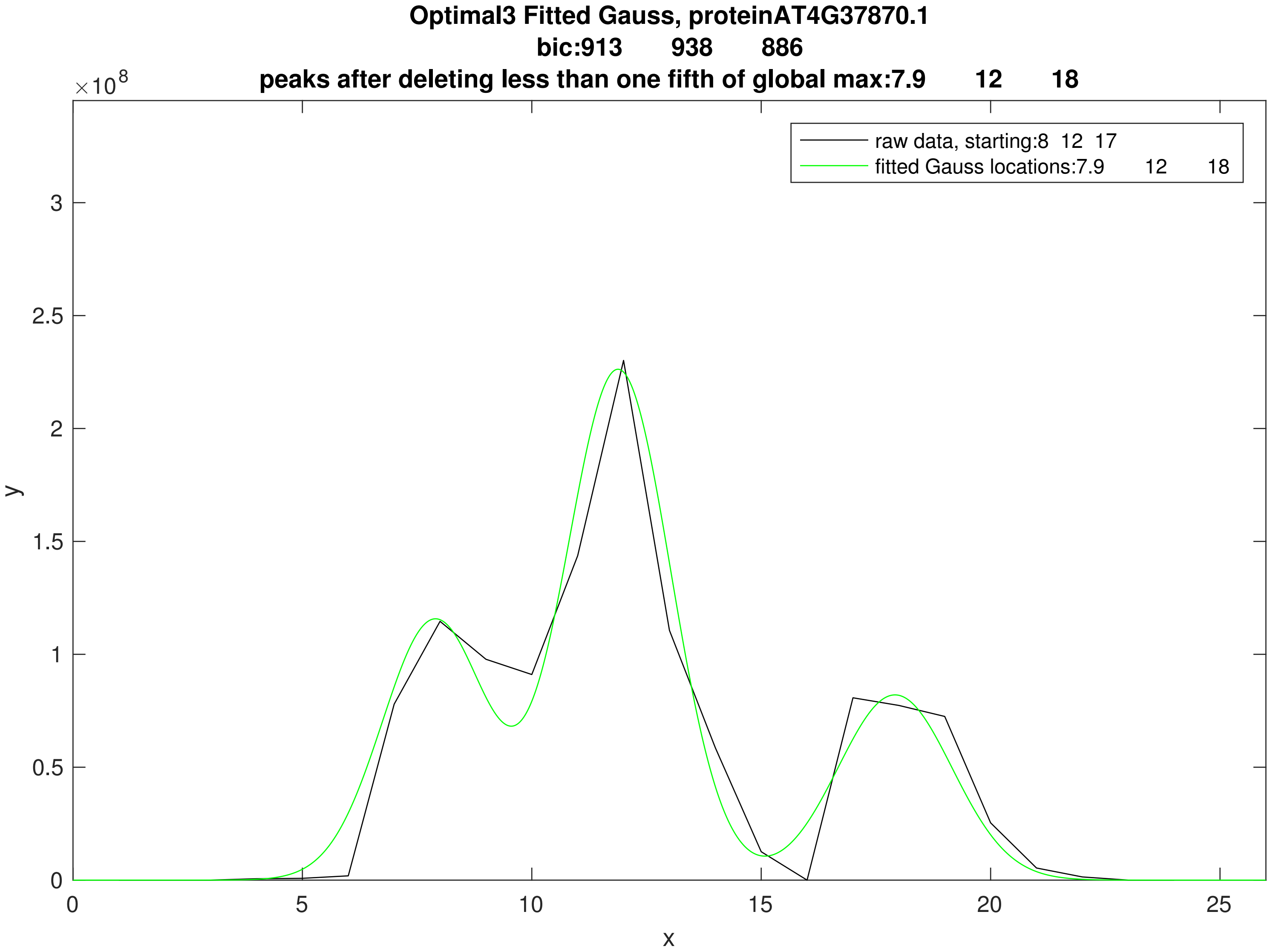

Optimal3 Fitted Gauss, proteinAT4G37870.1
bic:913 938 886
peaks after deleting less than one fifth of global max:7.9 12 18
×10^8
raw data, starting:8 12 17
fitted Gauss locations:7.9 12 18
y
x
0
0.5
1
1.5
2
2.5
3
0
5
10
15
20
25

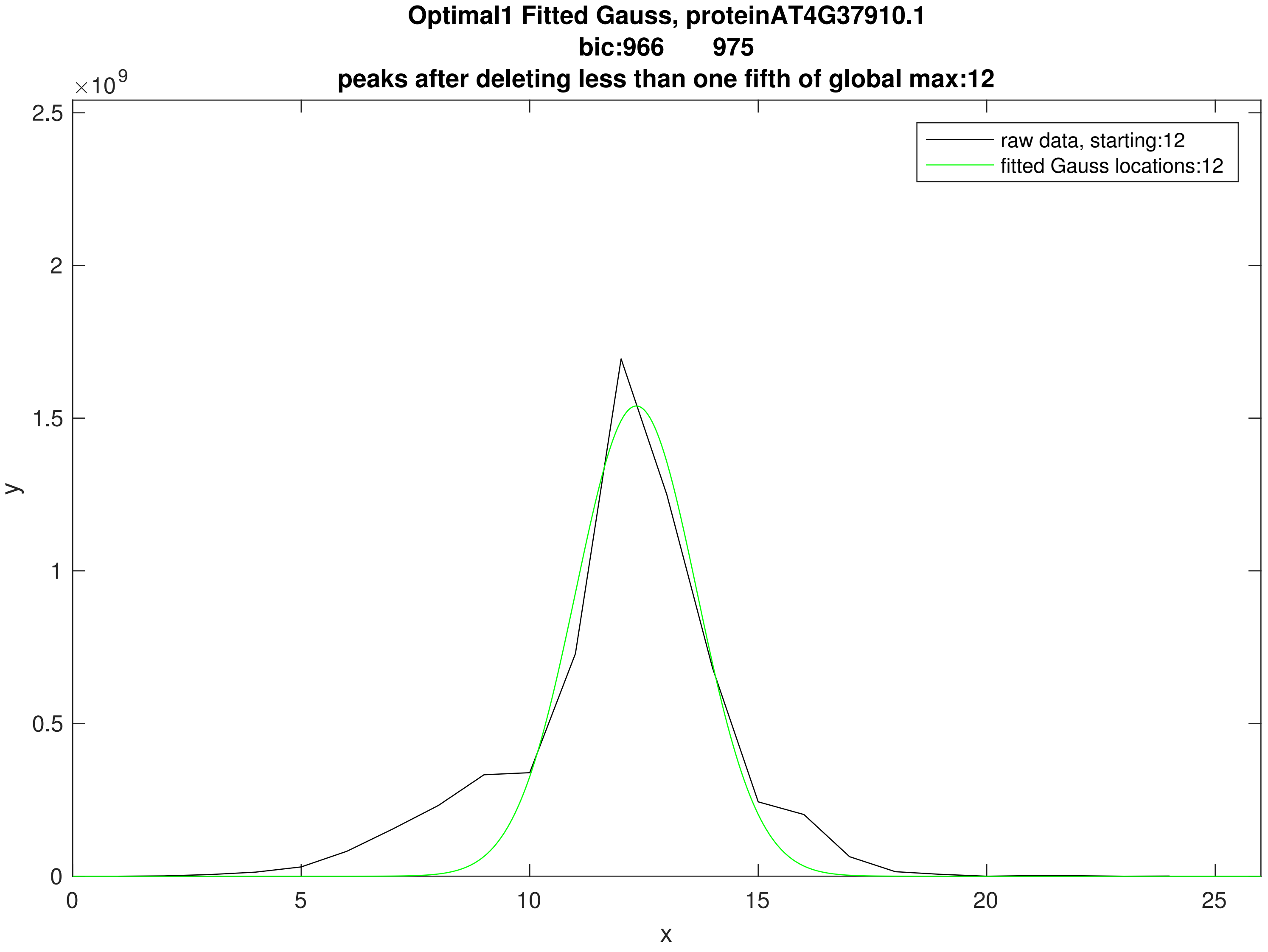
Optimal1 Fitted Gauss, proteinAT4G37910.1
bic:966 975
peaks after deleting less than one fifth of global max:12
×10^9
2.5
2
1.5
1
0.5
0
y
0
5
10
15
20
25
x
raw data, starting:12
fitted Gauss locations:12

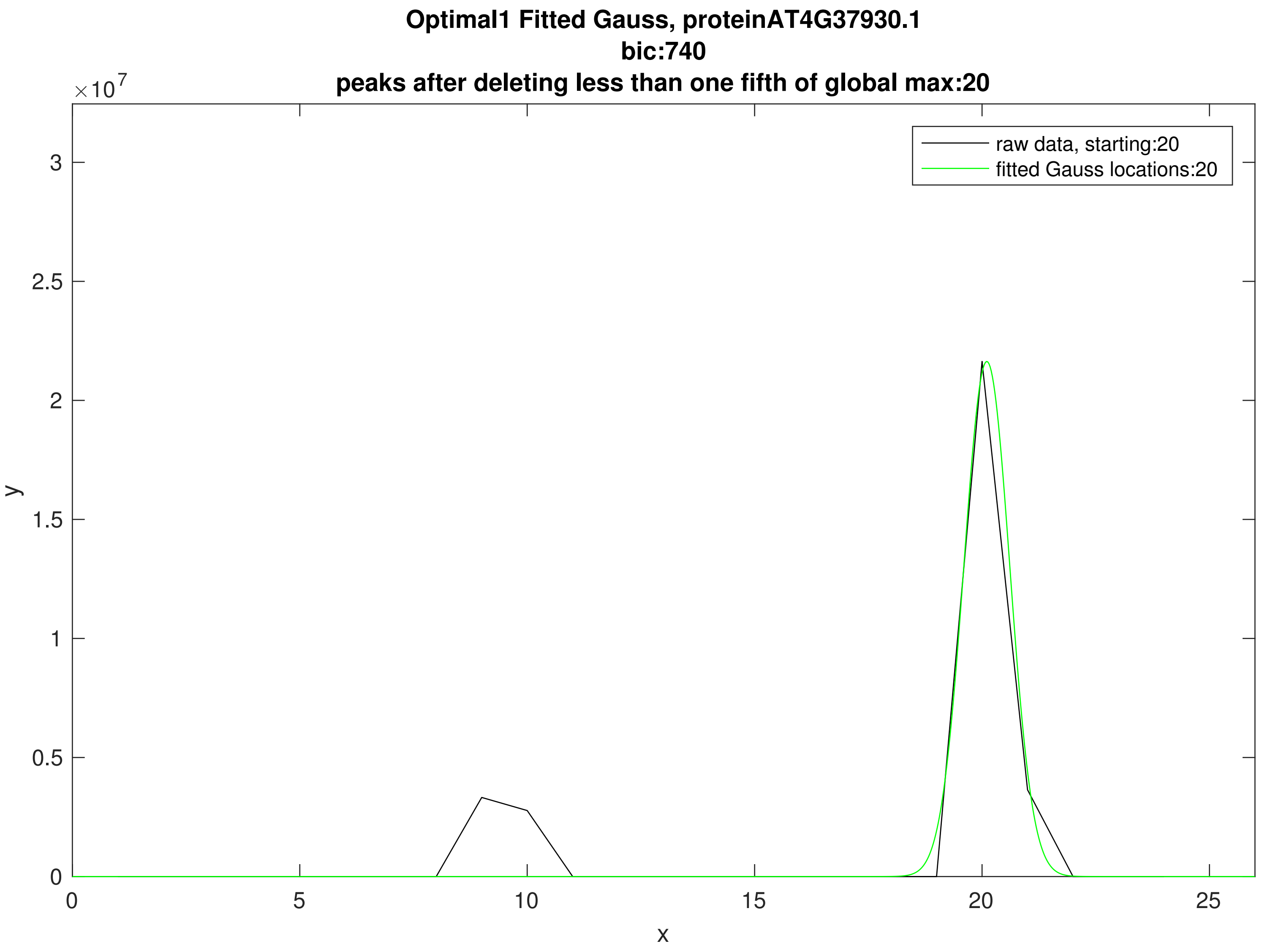

Optimal1 Fitted Gauss, proteinAT4G37930.1
bic:740
peaks after deleting less than one fifth of global max:20
×10^7
raw data, starting:20
fitted Gauss locations:20
3
2.5
2
1.5
1
0.5
0
y
0
5
10
15
20
25
x

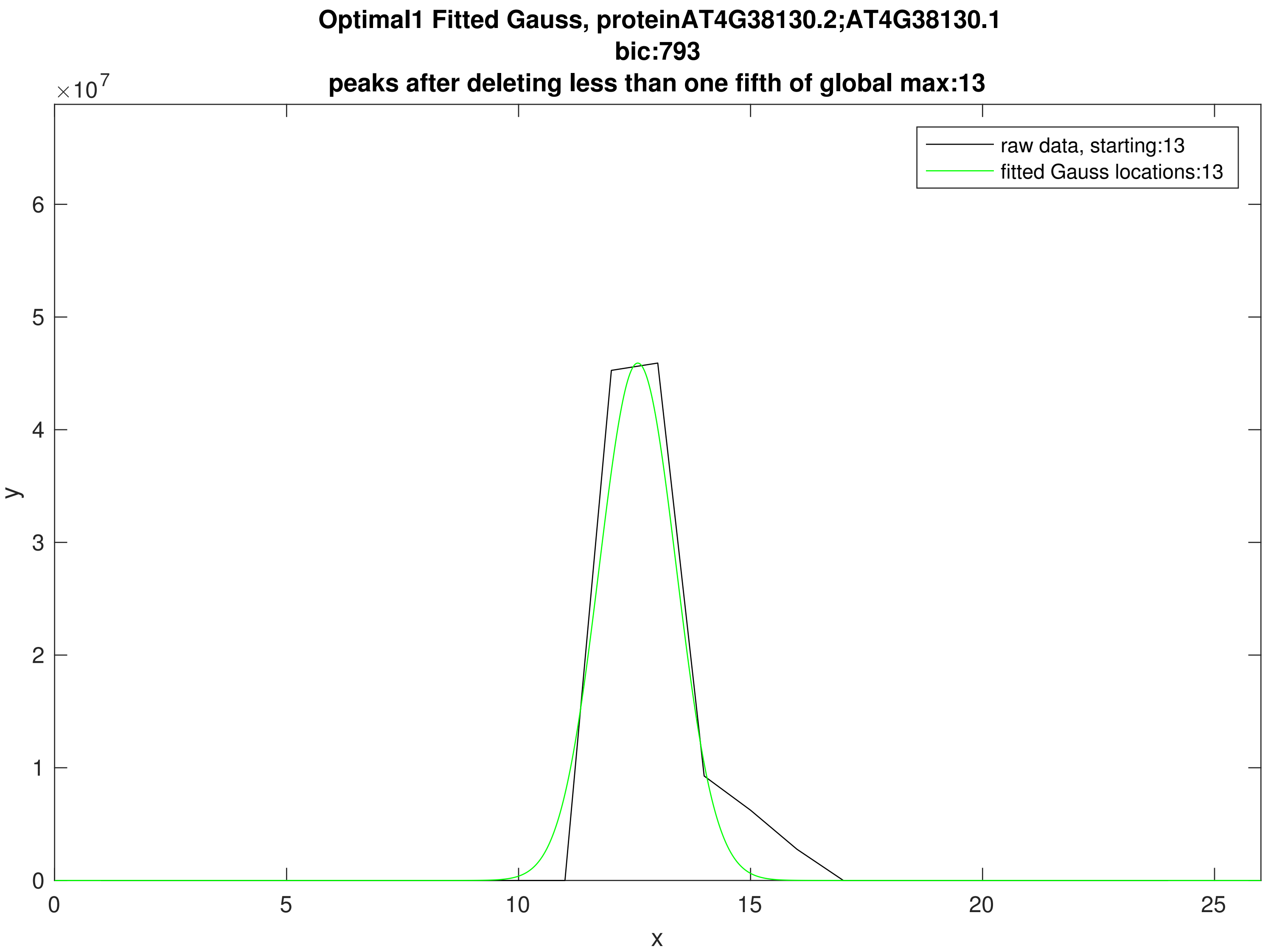
Optimal1 Fitted Gauss, proteinAT4G38130.2;AT4G38130.1
bic:793
peaks after deleting less than one fifth of global max:13
×10^7
raw data, starting:13
fitted Gauss locations:13
y
x
0
1
2
3
4
5
6
0
5
10
15
20
25

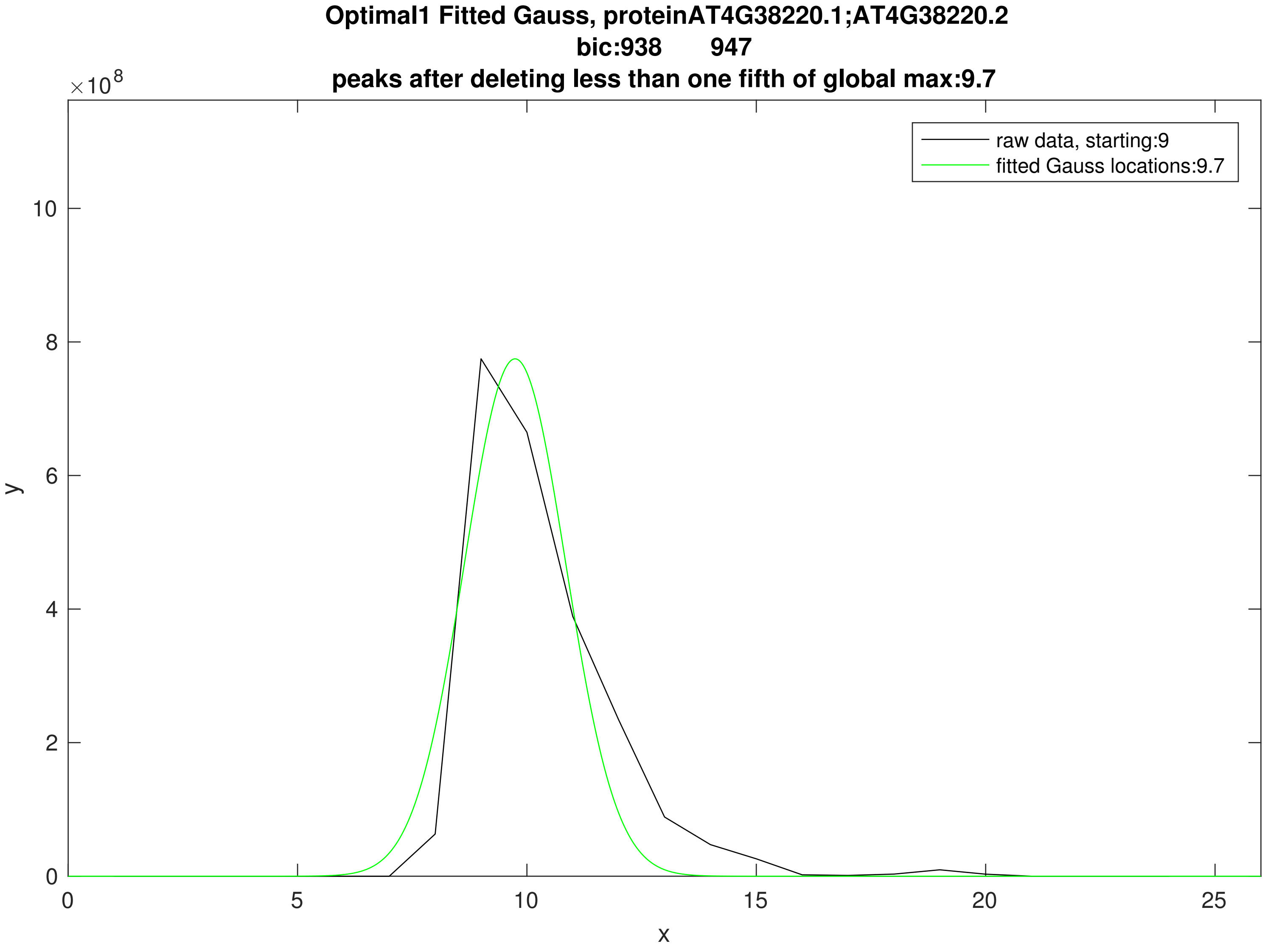

Optimal1 Fitted Gauss, proteinAT4G38220.1;AT4G38220.2
bic:938 947
peaks after deleting less than one fifth of global max:9.7
×10^8
raw data, starting:9
fitted Gauss locations:9.7
10
8
6
4
2
0
y
0
5
10
15
20
25
x

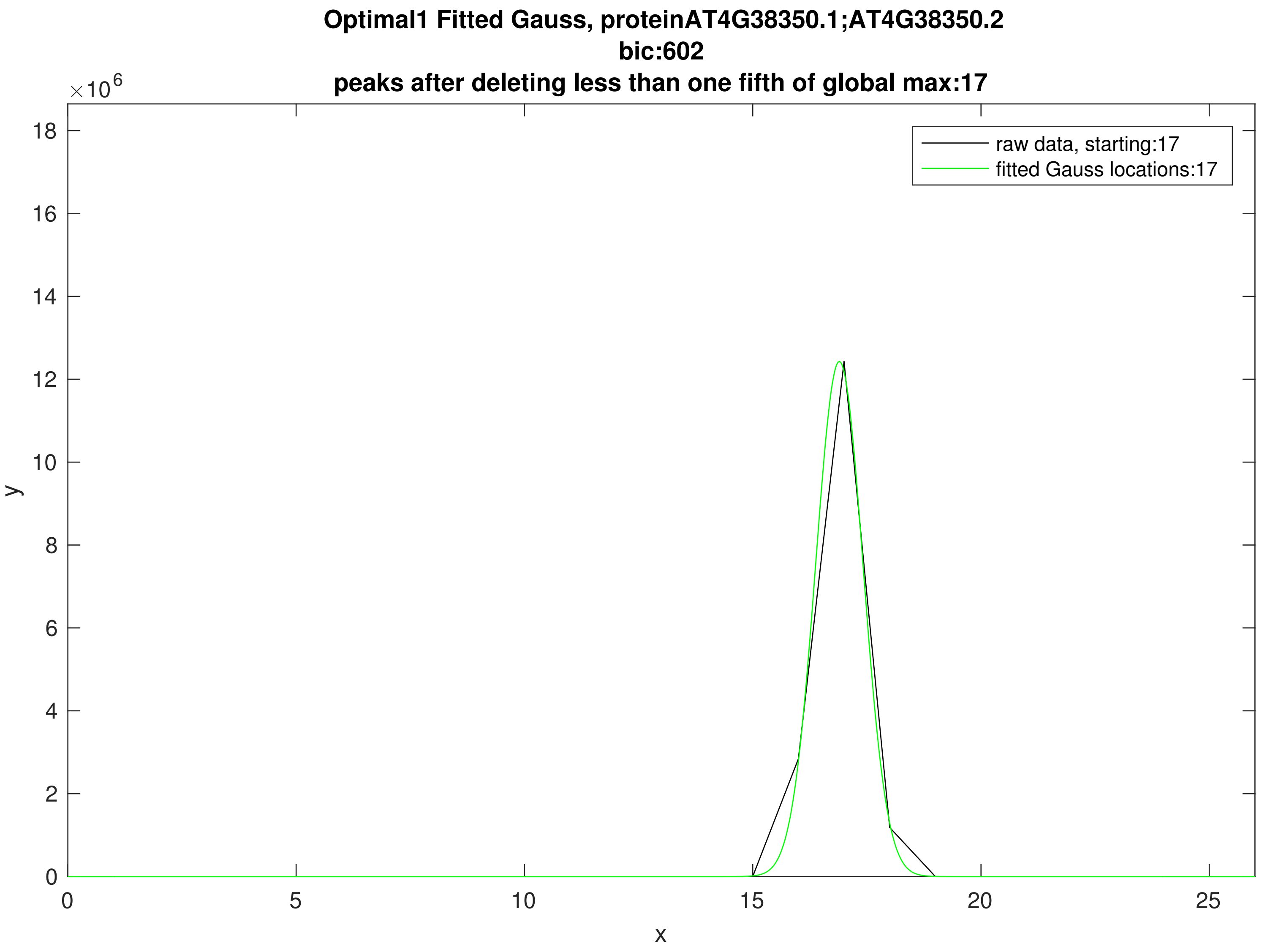
Optimal1 Fitted Gauss, proteinAT4G38350.1;AT4G38350.2
bic:602
peaks after deleting less than one fifth of global max:17
×10^6
y
x
raw data, starting:17
fitted Gauss locations:17
0
2
4
6
8
10
12
14
16
18
0
5
10
15
20
25

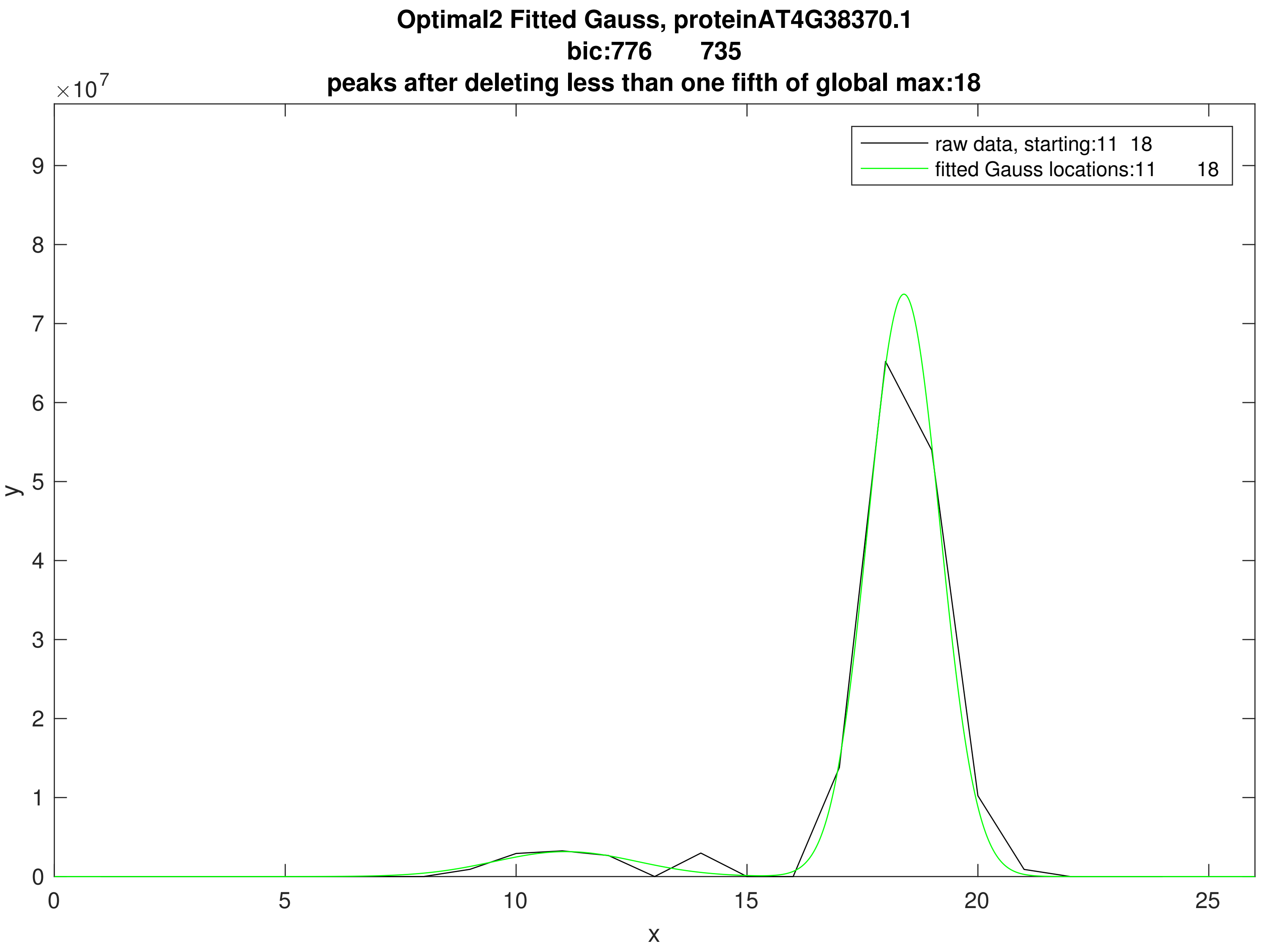
Optimal2 Fitted Gauss, proteinAT4G38370.1
bic:776 735
peaks after deleting less than one fifth of global max:18
×10^7
raw data, starting:11 18
fitted Gauss locations:11 18
y
x
0
1
2
3
4
5
6
7
8
9
0
5
10
15
20
25

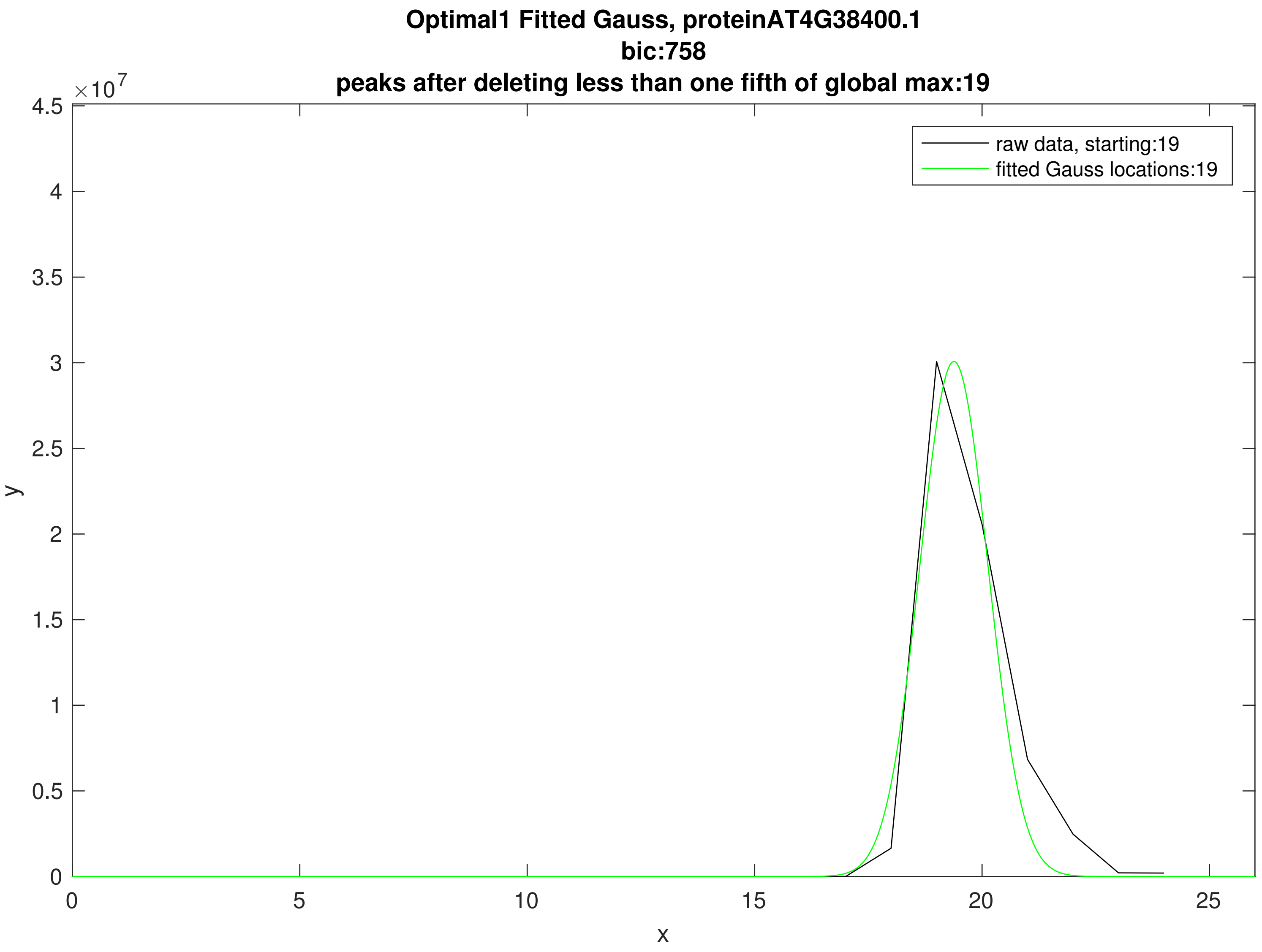

Optimal1 Fitted Gauss, proteinAT4G38400.1
bic:758
peaks after deleting less than one fifth of global max:19
×10^7
4.5
4
3.5
3
2.5
2
1.5
1
0.5
0
y
0
5
10
15
20
25
x
raw data, starting:19
fitted Gauss locations:19

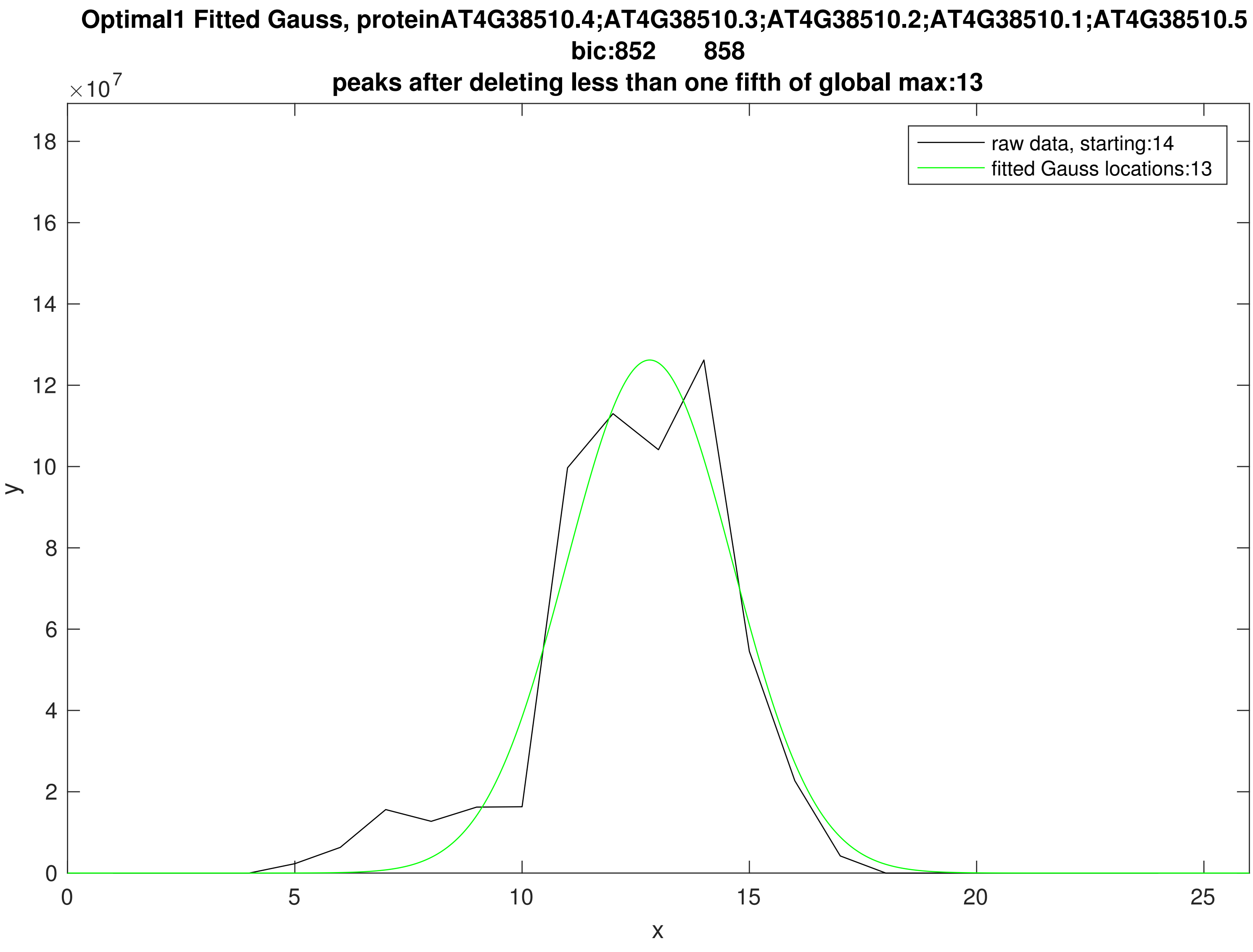
Optimal1 Fitted Gauss, proteinAT4G38510.4;AT4G38510.3;AT4G38510.2;AT4G38510.1;AT4G38510.5
bic:852 858
peaks after deleting less than one fifth of global max:13
×10^7
raw data, starting:14
fitted Gauss locations:13
y
x
0
2
4
6
8
10
12
14
16
18
0
5
10
15
20
25

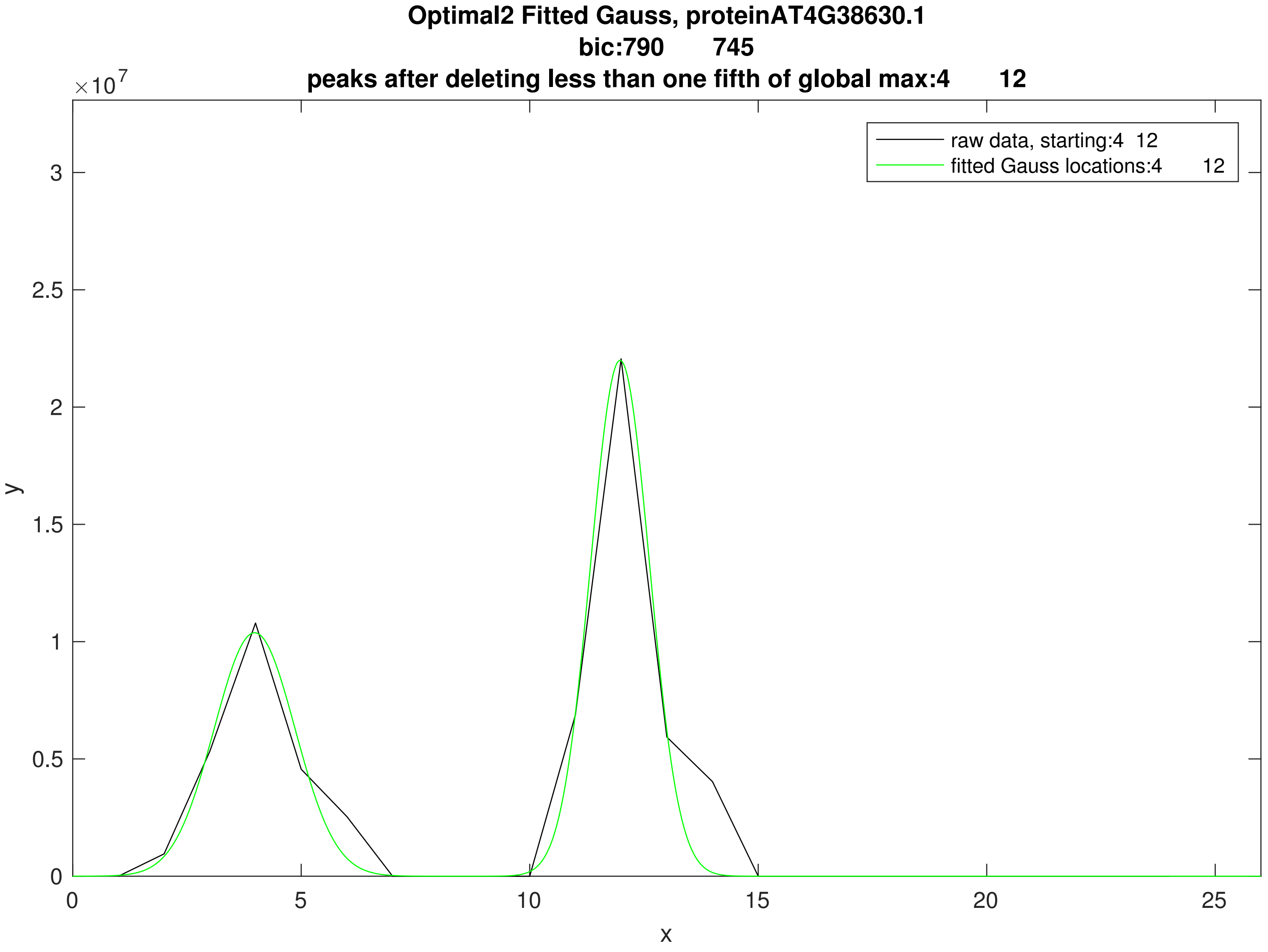
Optimal2 Fitted Gauss, proteinAT4G38630.1
bic:790 745
peaks after deleting less than one fifth of global max:4 12
×10^7
raw data, starting:4 12
fitted Gauss locations:4 12
y
x
0
0.5
1
1.5
2
2.5
3
0
5
10
15
20
25

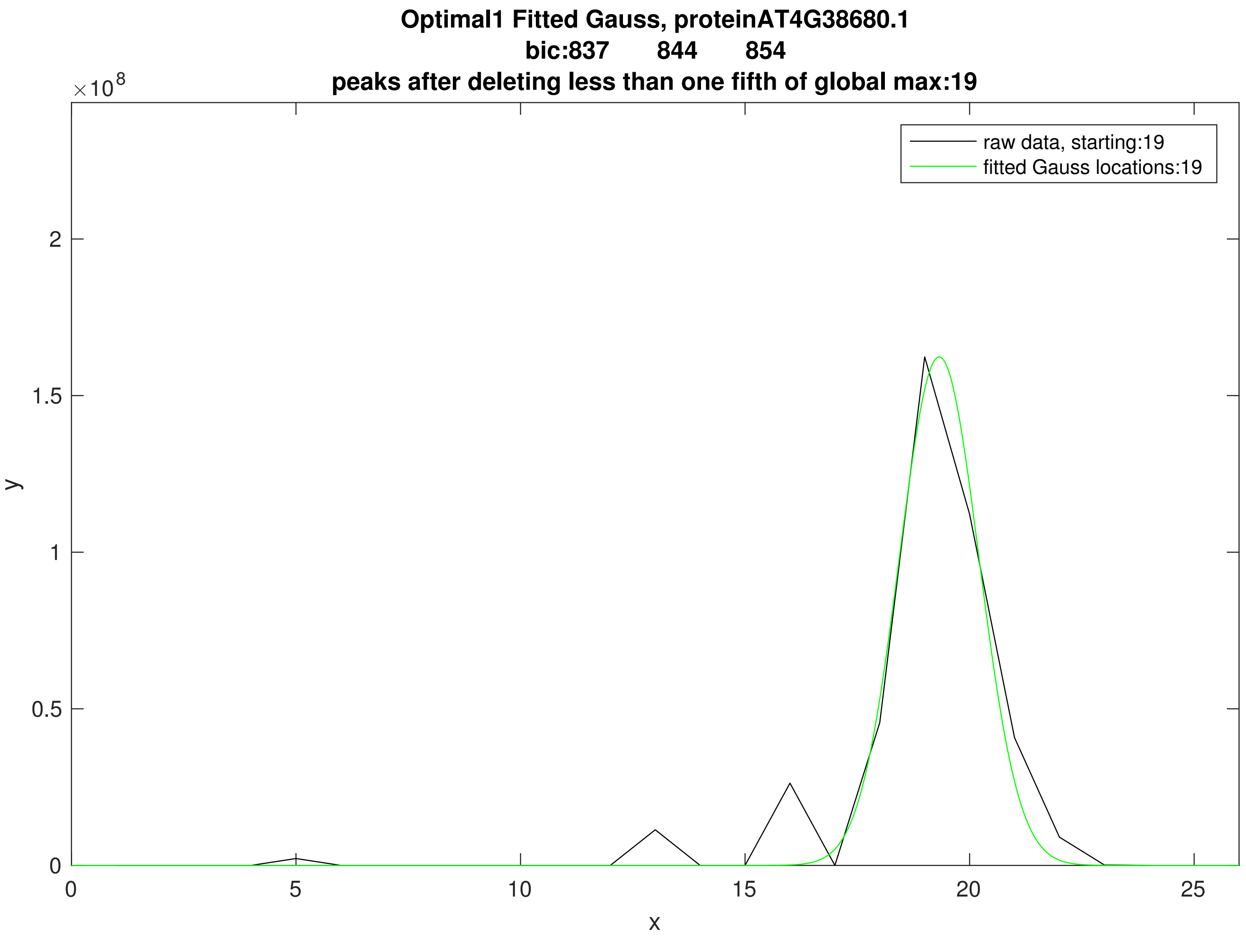

Optimal1 Fitted Gauss, proteinAT4G38680.1
bic:837 844 854
peaks after deleting less than one fifth of global max:19
×10^8
raw data, starting:19
fitted Gauss locations:19
y
x
0
0.5
1
1.5
2
0
5
10
15
20
25

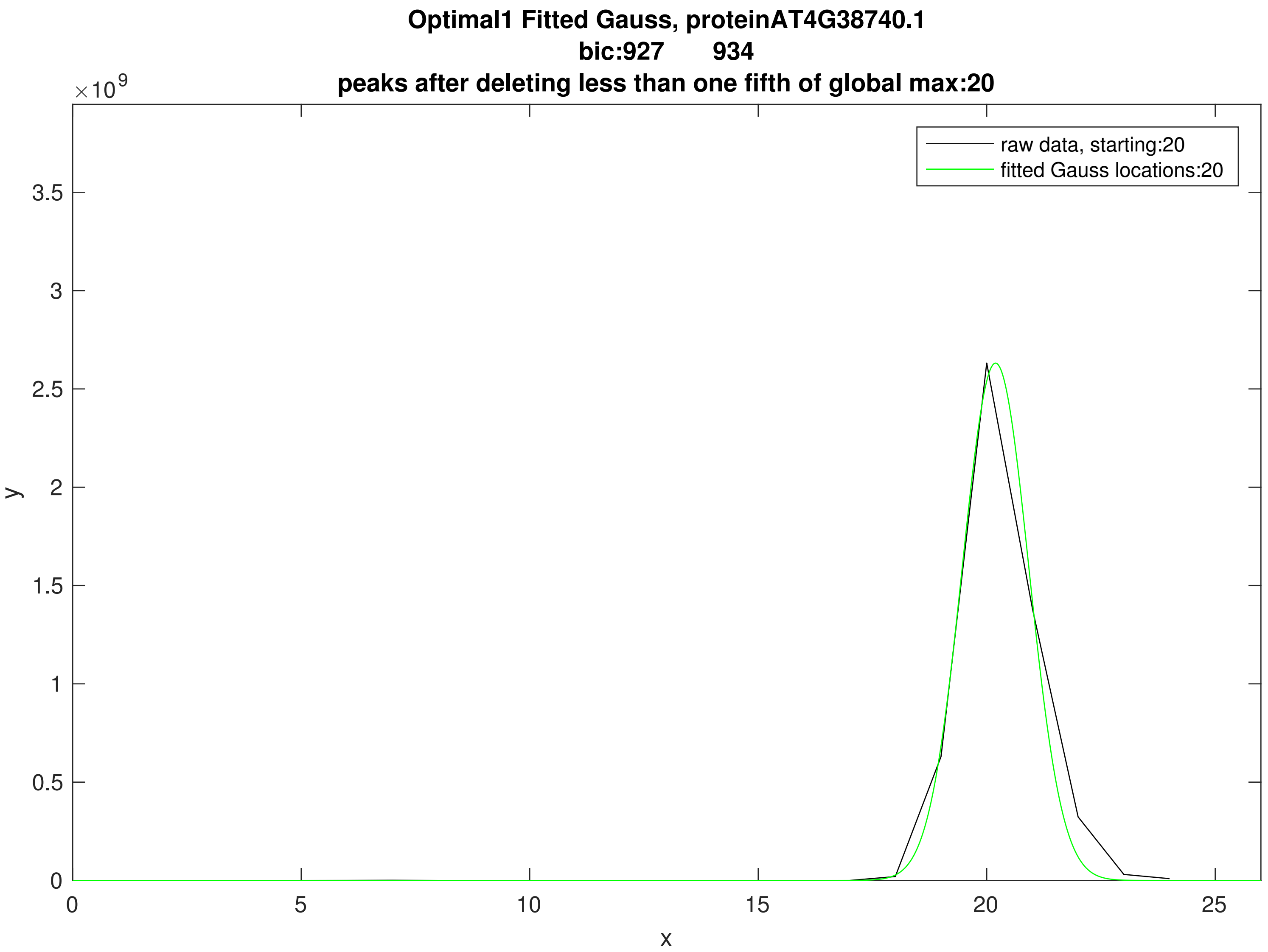

Optimal1 Fitted Gauss, proteinAT4G38740.1
bic:927 934
peaks after deleting less than one fifth of global max:20
×10^9
raw data, starting:20
fitted Gauss locations:20
3.5
3
2.5
2
1.5
1
0.5
0
y
0
5
10
15
20
25
x

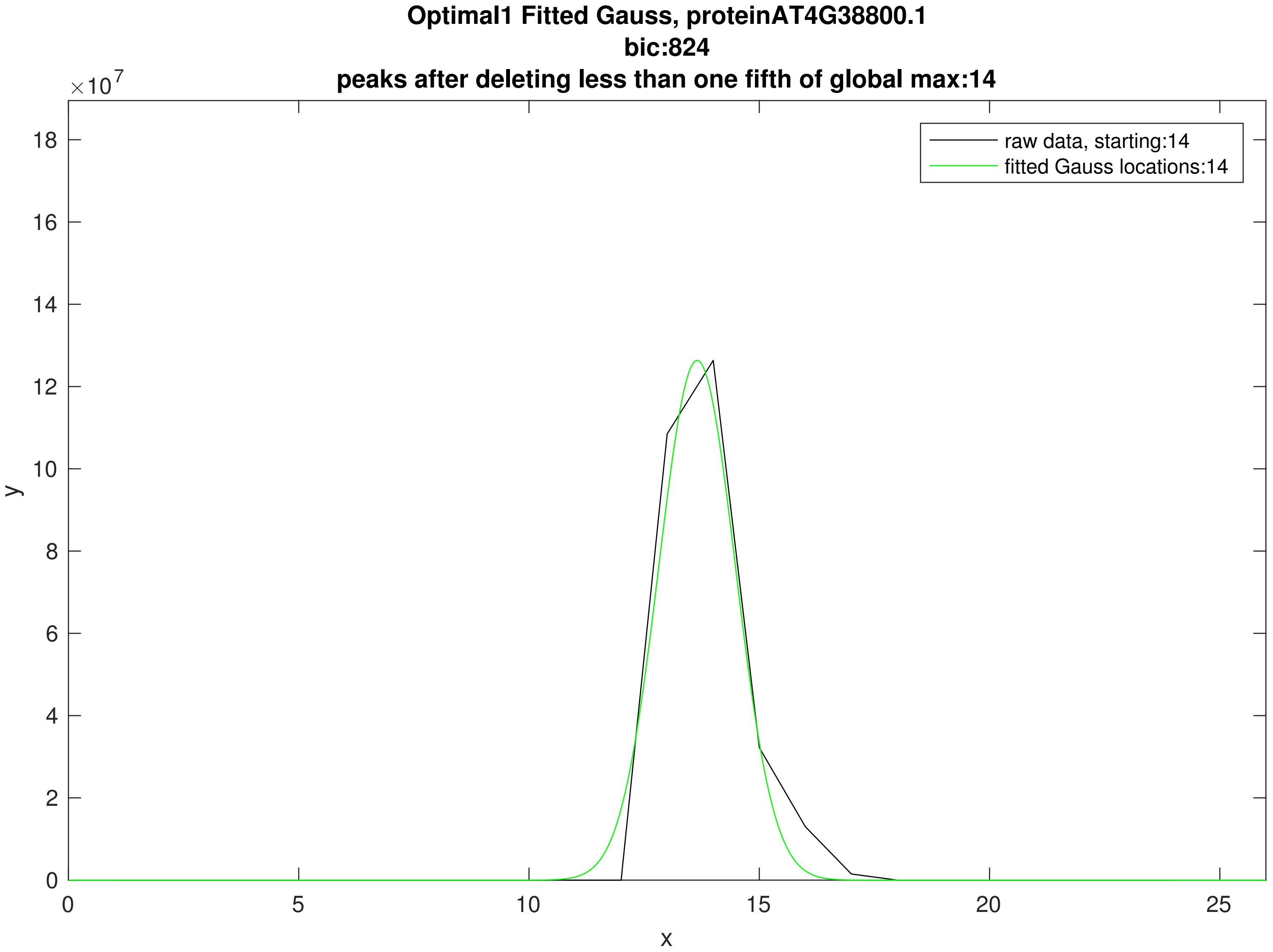
Optimal1 Fitted Gauss, proteinAT4G38800.1
bic:824
peaks after deleting less than one fifth of global max:14
×10^7
raw data, starting:14
fitted Gauss locations:14
y
x
0
2
4
6
8
10
12
14
16
18
0
5
10
15
20
25

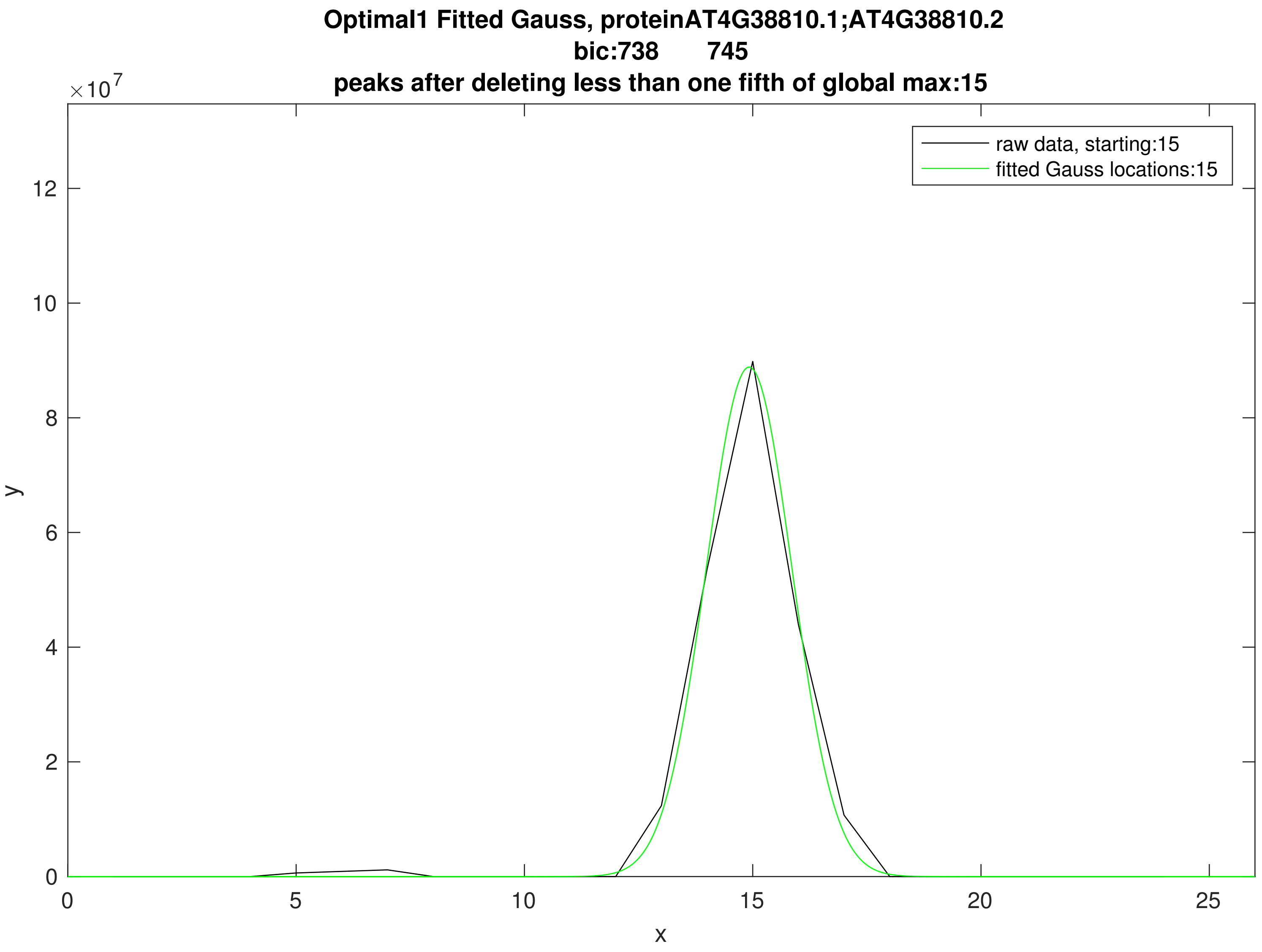

Optimal1 Fitted Gauss, proteinAT4G38810.1;AT4G38810.2
bic:738 745
peaks after deleting less than one fifth of global max:15
×10^7
raw data, starting:15
fitted Gauss locations:15
12
10
8
6
4
2
0
y
0
5
10
15
20
25
x

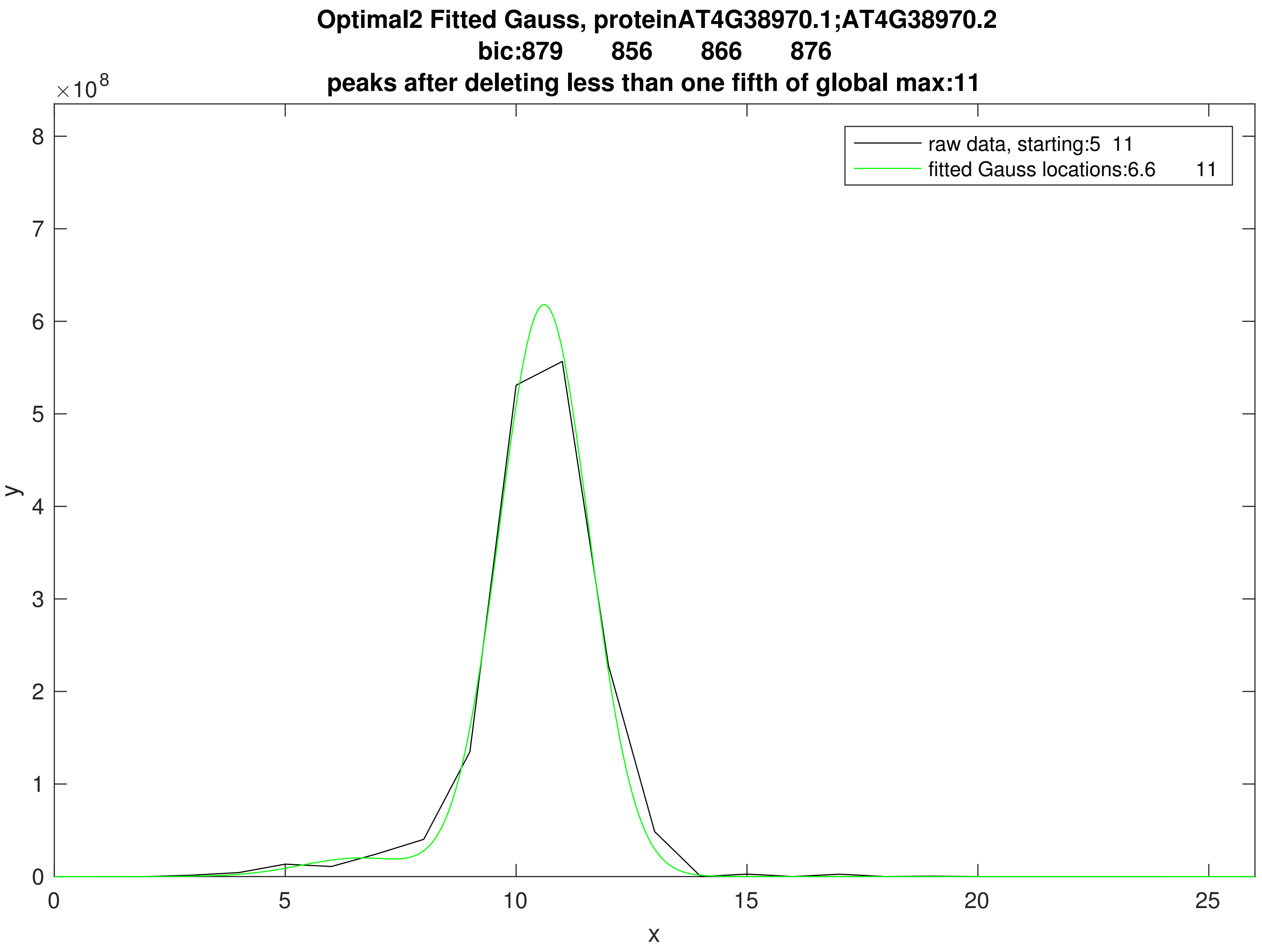
Optimal2 Fitted Gauss, proteinAT4G38970.1;AT4G38970.2
bic:879 856 866 876
peaks after deleting less than one fifth of global max:11
×10^8
raw data, starting:5 11
fitted Gauss locations:6.6 11
y
x
0
1
2
3
4
5
6
7
8
0
5
10
15
20
25

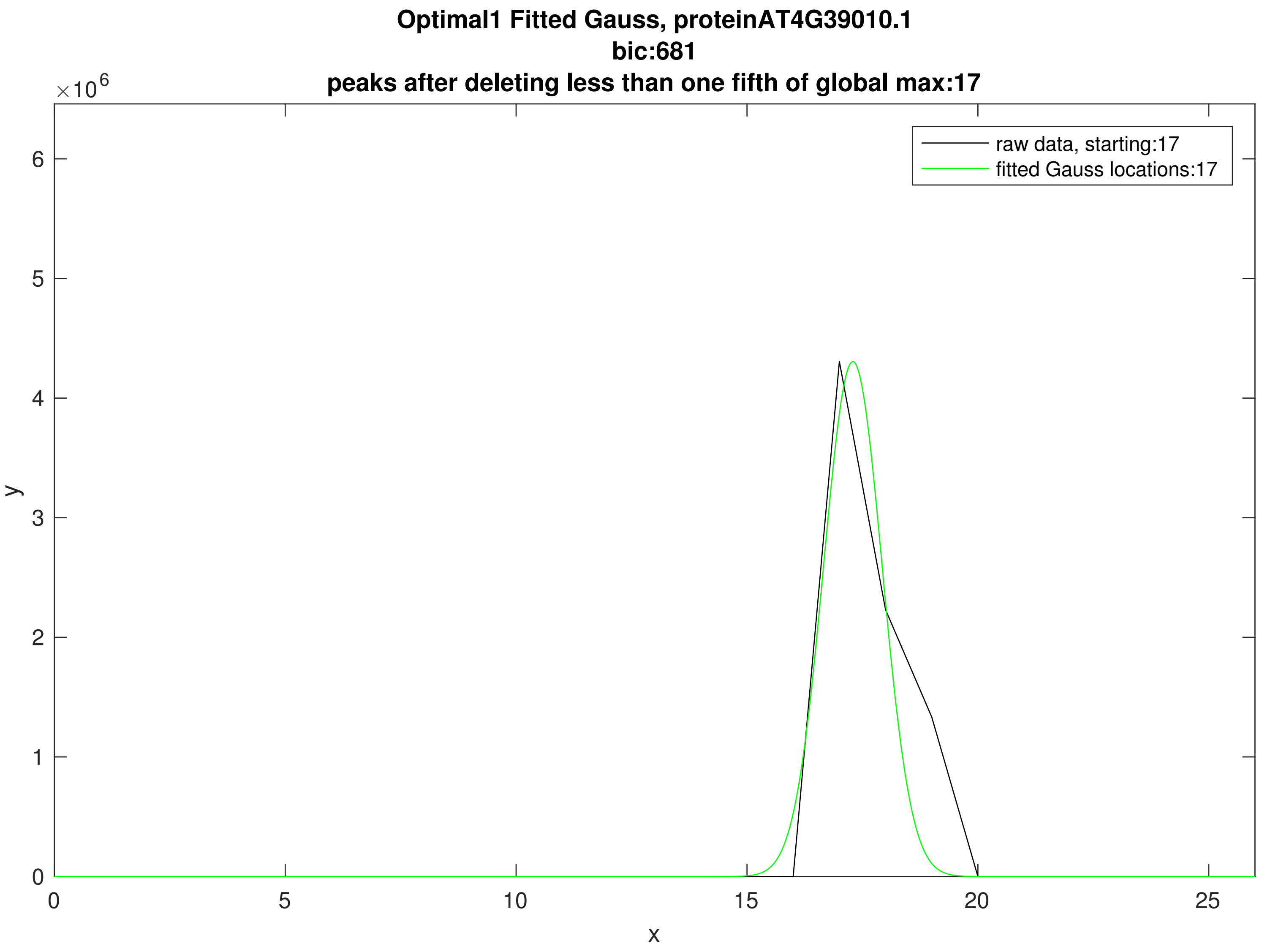
Optimal1 Fitted Gauss, proteinAT4G39010.1
bic:681
peaks after deleting less than one fifth of global max:17
×10^6
raw data, starting:17
fitted Gauss locations:17
y
x
0
1
2
3
4
5
6
0
5
10
15
20
25

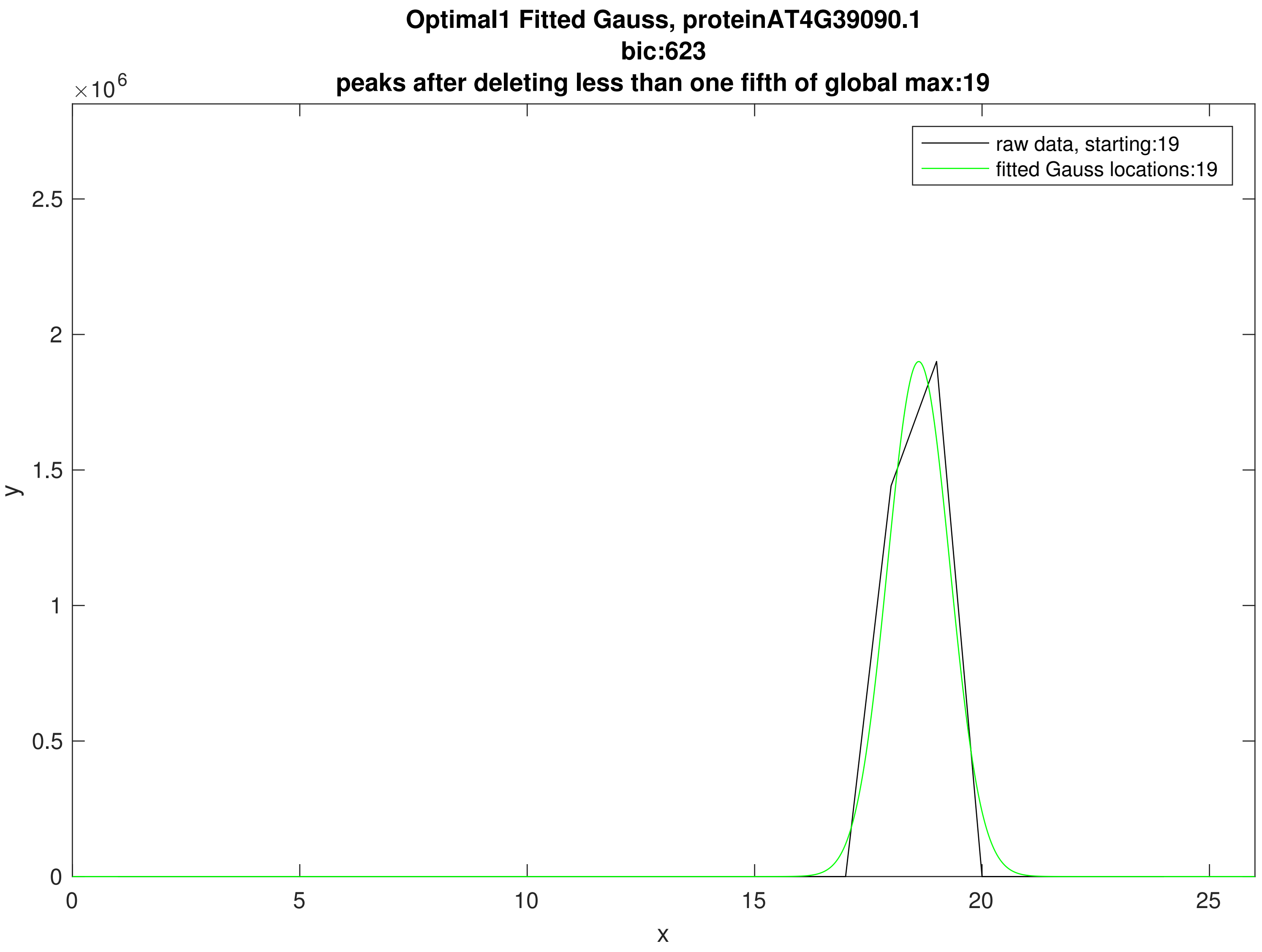

Optimal1 Fitted Gauss, proteinAT4G39090.1
bic:623
peaks after deleting less than one fifth of global max:19
×10^6
raw data, starting:19
fitted Gauss locations:19
2.5
2
1.5
1
0.5
0
y
0
5
10
15
20
25
x

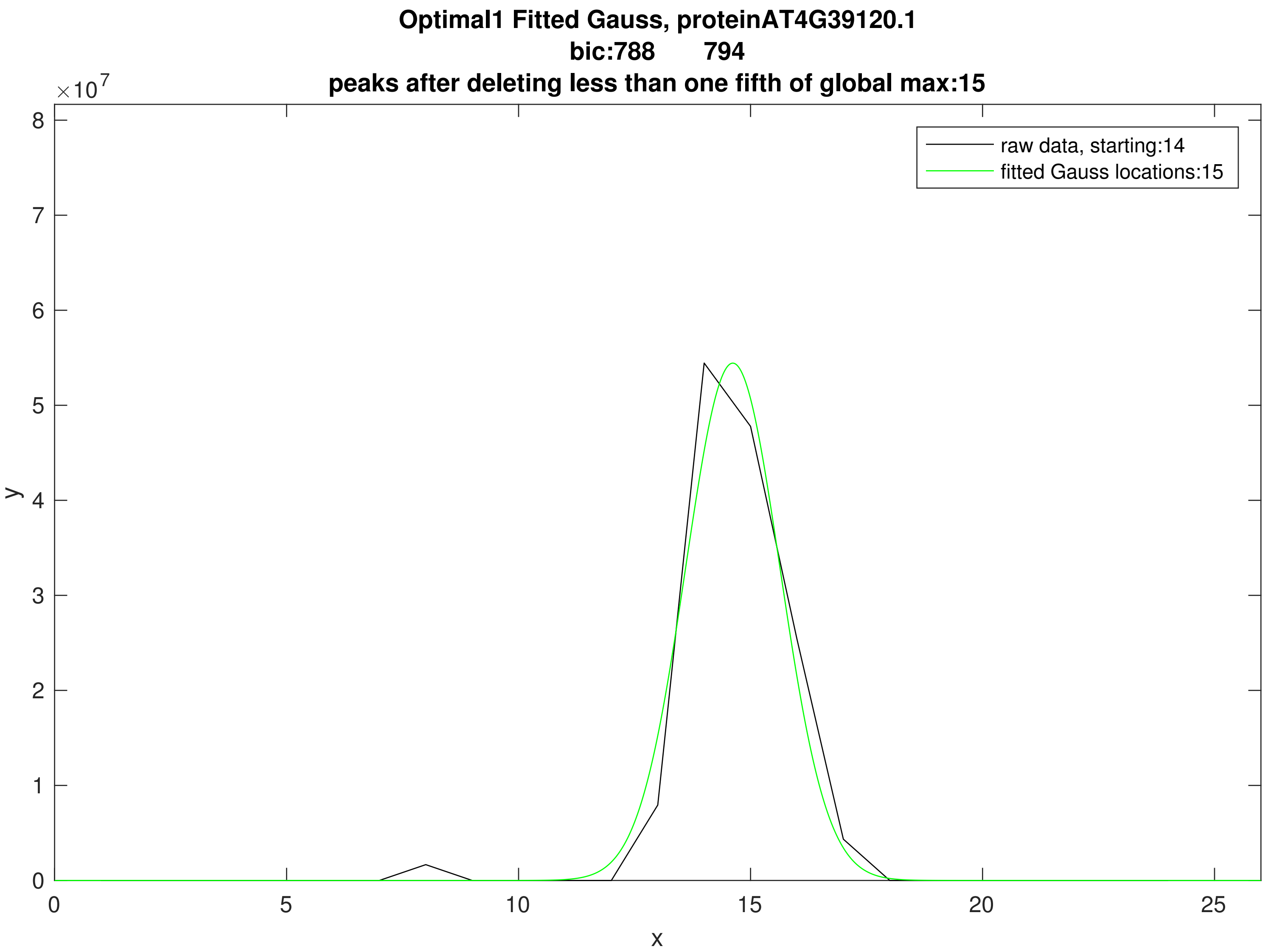

Optimal1 Fitted Gauss, proteinAT4G39120.1
bic:788 794
peaks after deleting less than one fifth of global max:15
×10^7
raw data, starting:14
fitted Gauss locations:15
y
x

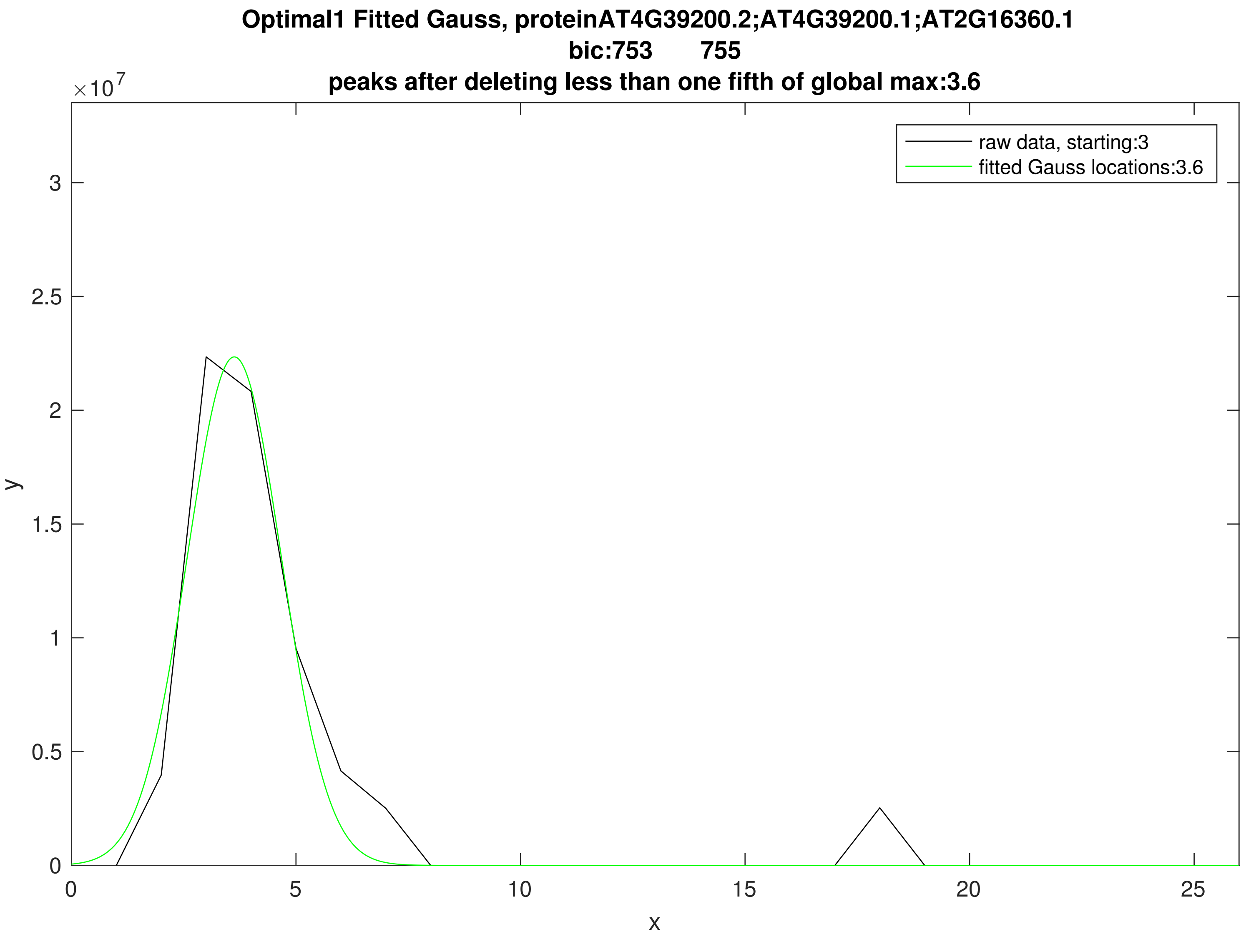

Optimal1 Fitted Gauss, proteinAT4G39200.2;AT4G39200.1;AT2G16360.1
bic:753 755
peaks after deleting less than one fifth of global max:3.6
×10^7
raw data, starting:3
fitted Gauss locations:3.6
y
x
0
0.5
1
1.5
2
2.5
3
5
10
15
20
25

Optimal1 Fitted Gauss, proteinAT4G39210.1
bic:728
peaks after deleting less than one fifth of global max:10

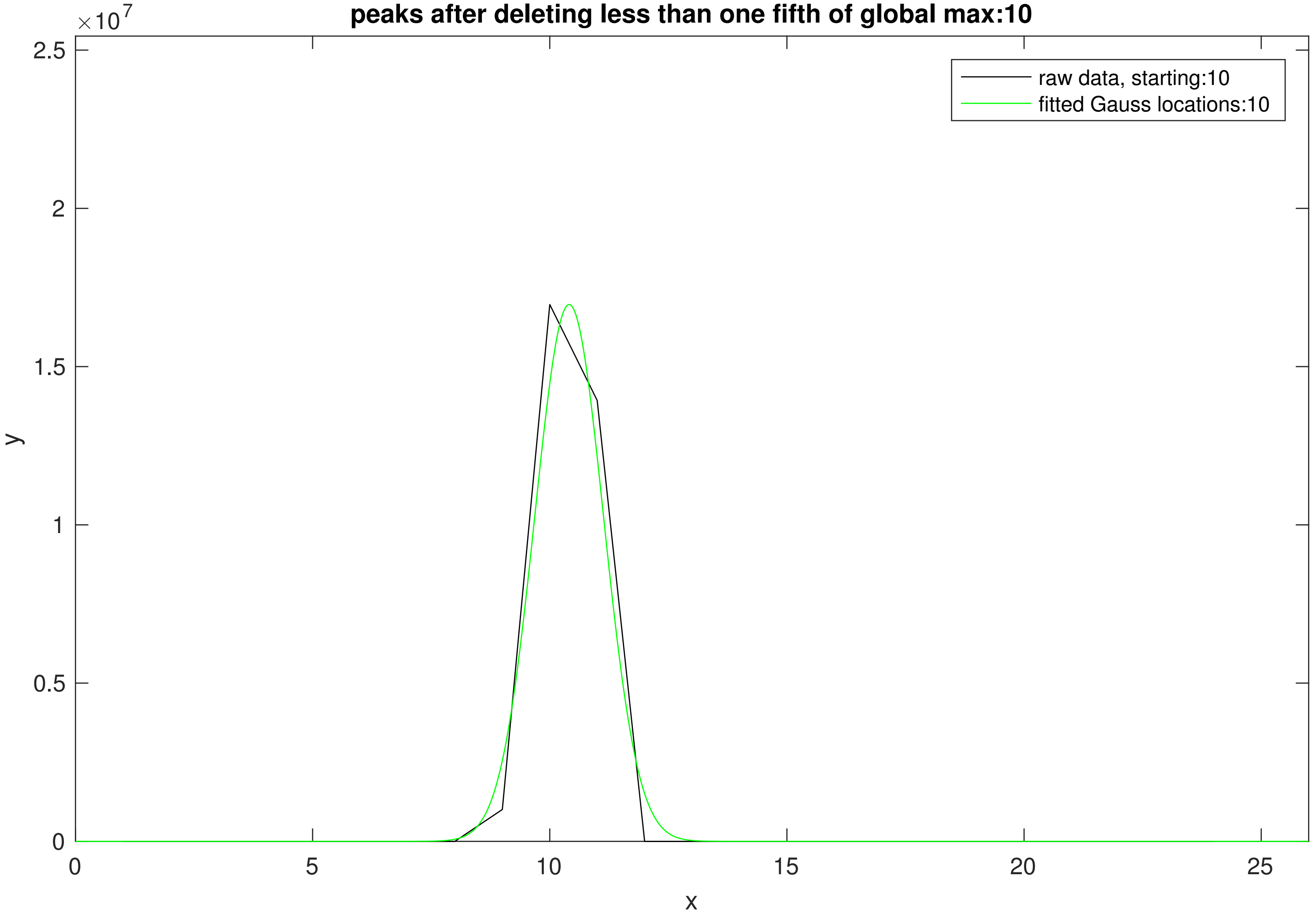

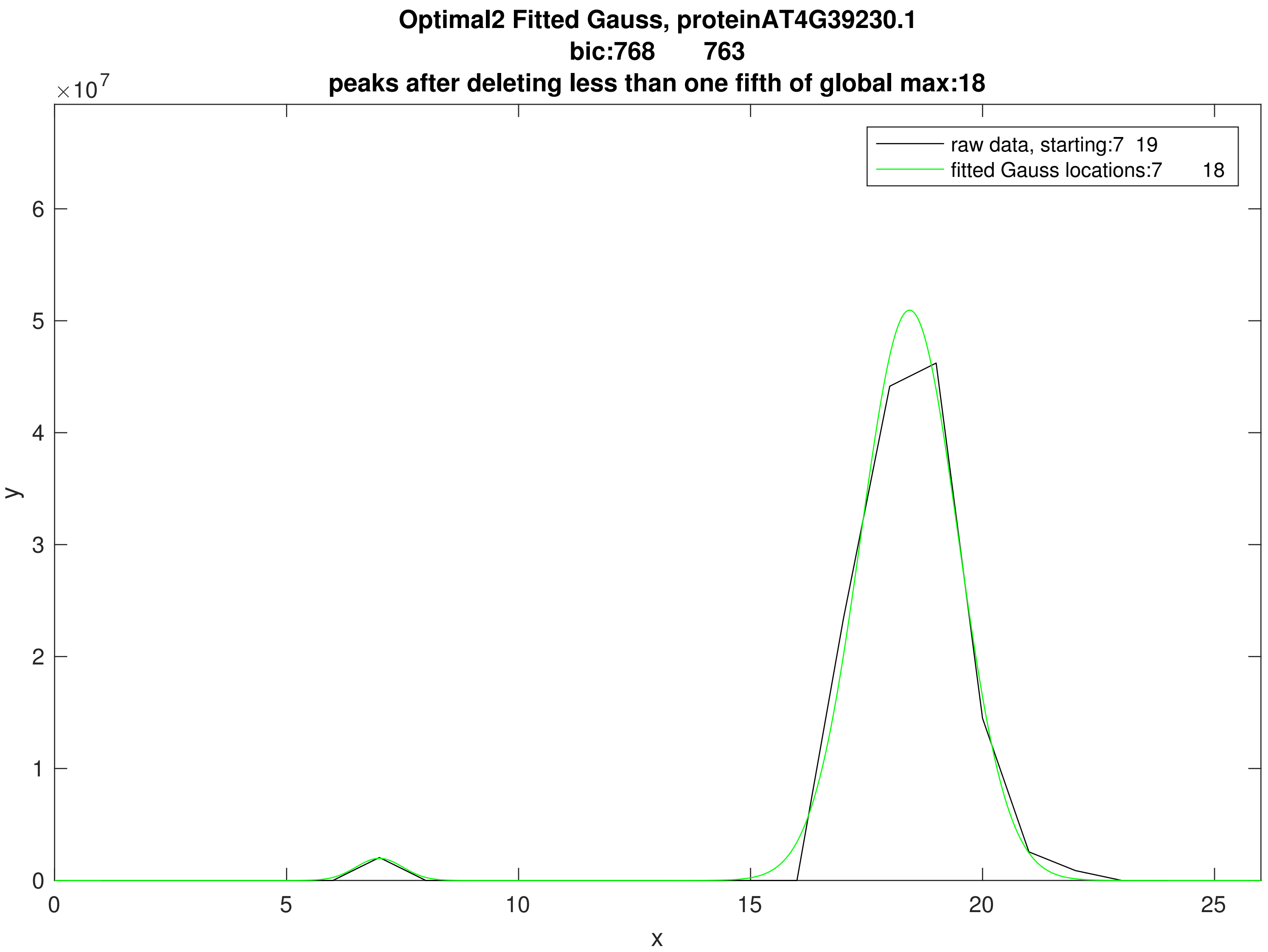

Optimal2 Fitted Gauss, proteinAT4G39230.1
bic:768 763
peaks after deleting less than one fifth of global max:18
×10^7
raw data, starting:7 19
fitted Gauss locations:7 18
y
x
0
1
2
3
4
5
6
0
5
10
15
20
25

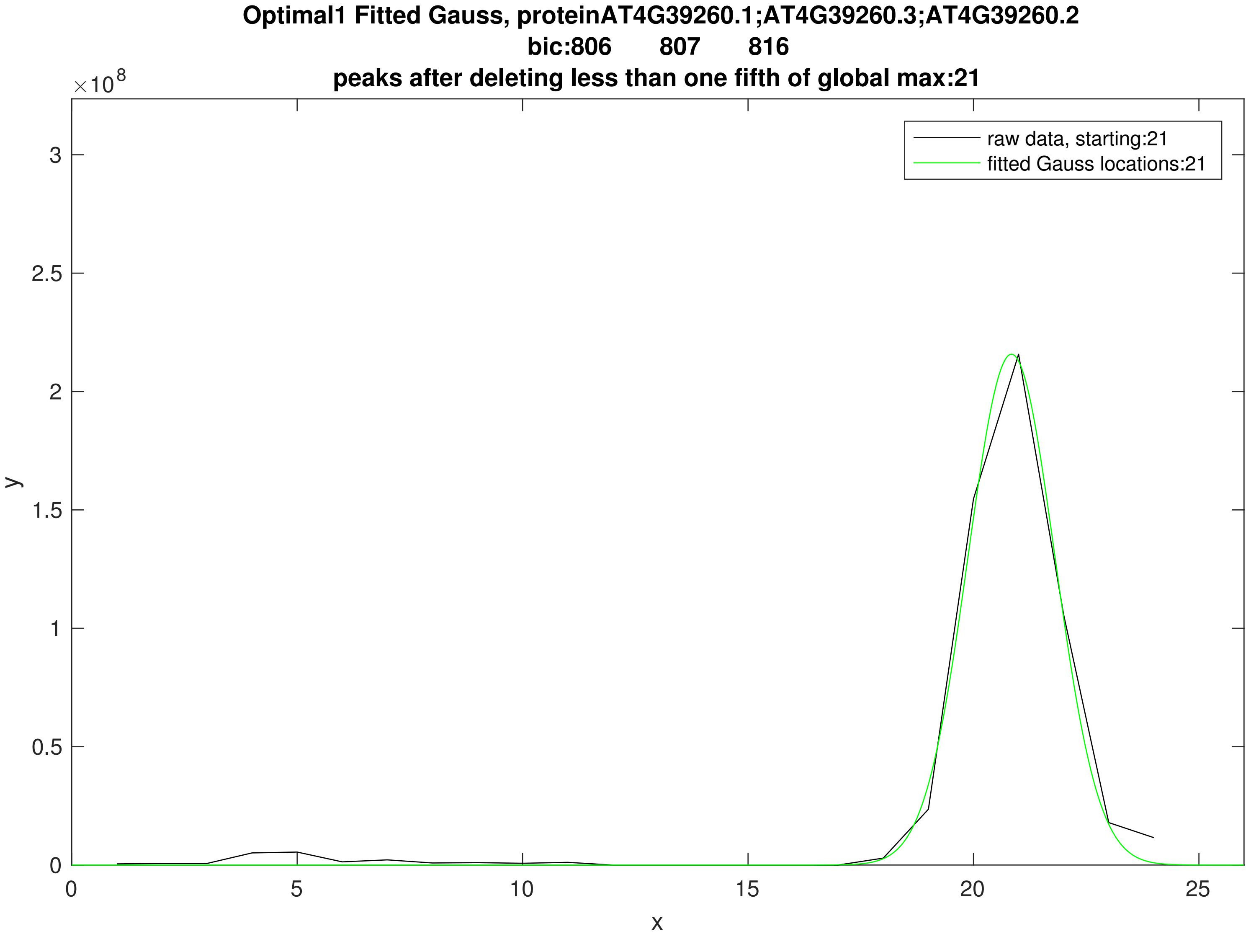
Optimal1 Fitted Gauss, proteinAT4G39260.1;AT4G39260.3;AT4G39260.2
bic:806 807 816
peaks after deleting less than one fifth of global max:21
×10^8
raw data, starting:21
fitted Gauss locations:21
y
x
0
0.5
1
1.5
2
2.5
3
5
10
15
20
25

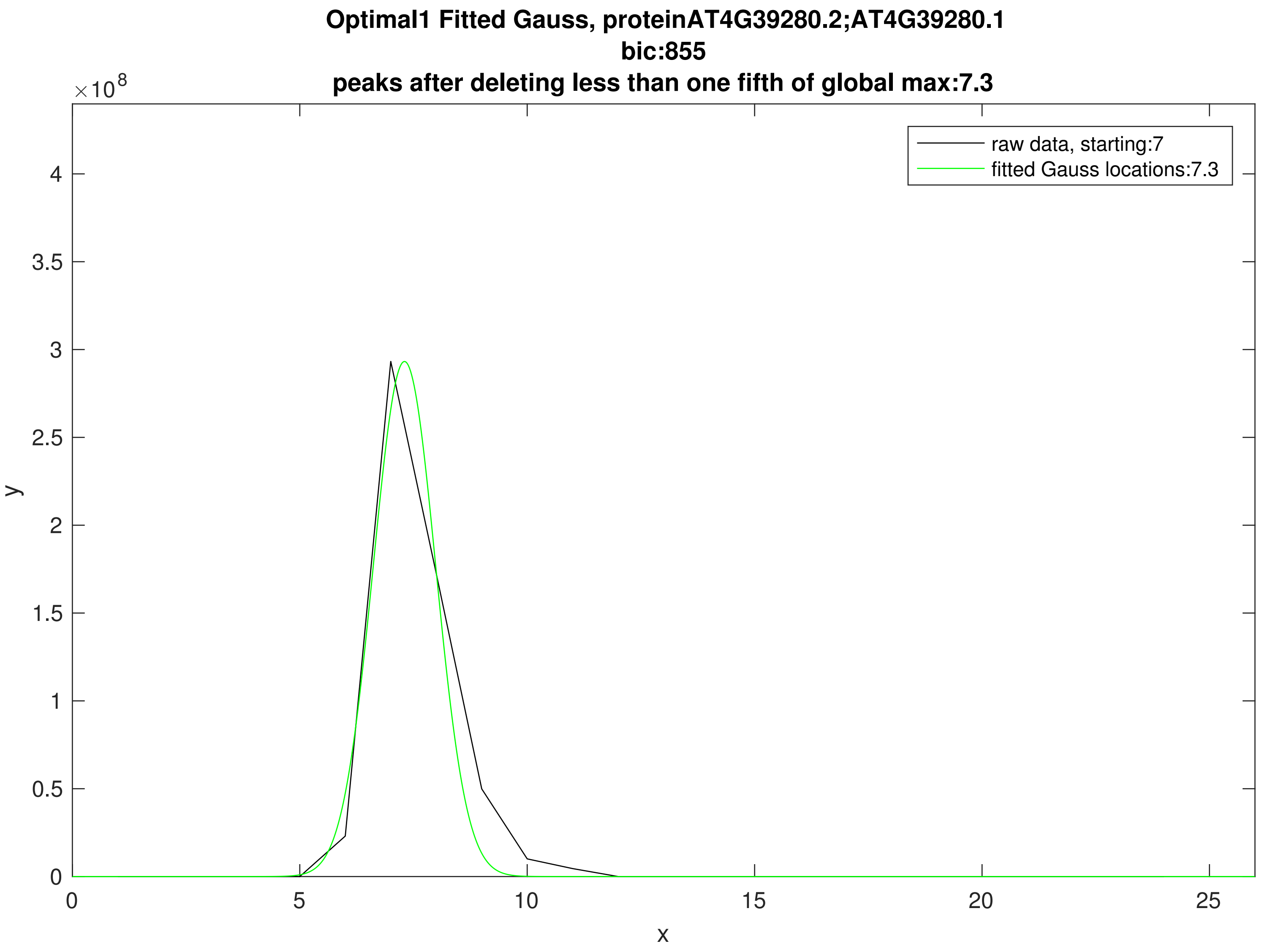
Optimal1 Fitted Gauss, proteinAT4G39280.2;AT4G39280.1
bic:855
peaks after deleting less than one fifth of global max:7.3
×10^8
raw data, starting:7
fitted Gauss locations:7.3
y
x
0
0.5
1
1.5
2
2.5
3
3.5
4
5
10
15
20
25

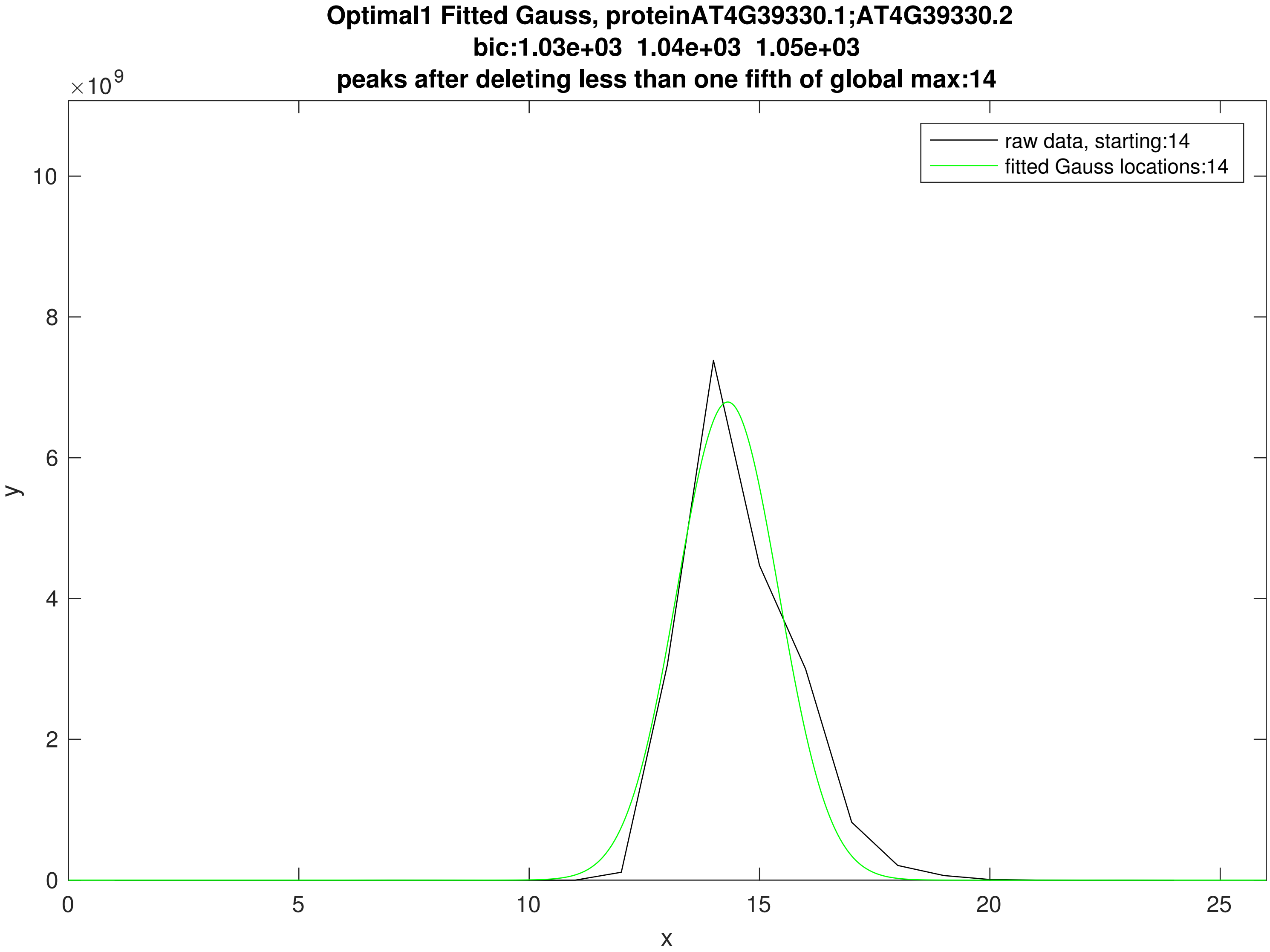
Optimal1 Fitted Gauss, proteinAT4G39330.1;AT4G39330.2
bic:1.03e+03  1.04e+03  1.05e+03
peaks after deleting less than one fifth of global max:14
×10^9
raw data, starting:14
fitted Gauss locations:14
y
x
0
2
4
6
8
10
0
5
10
15
20
25

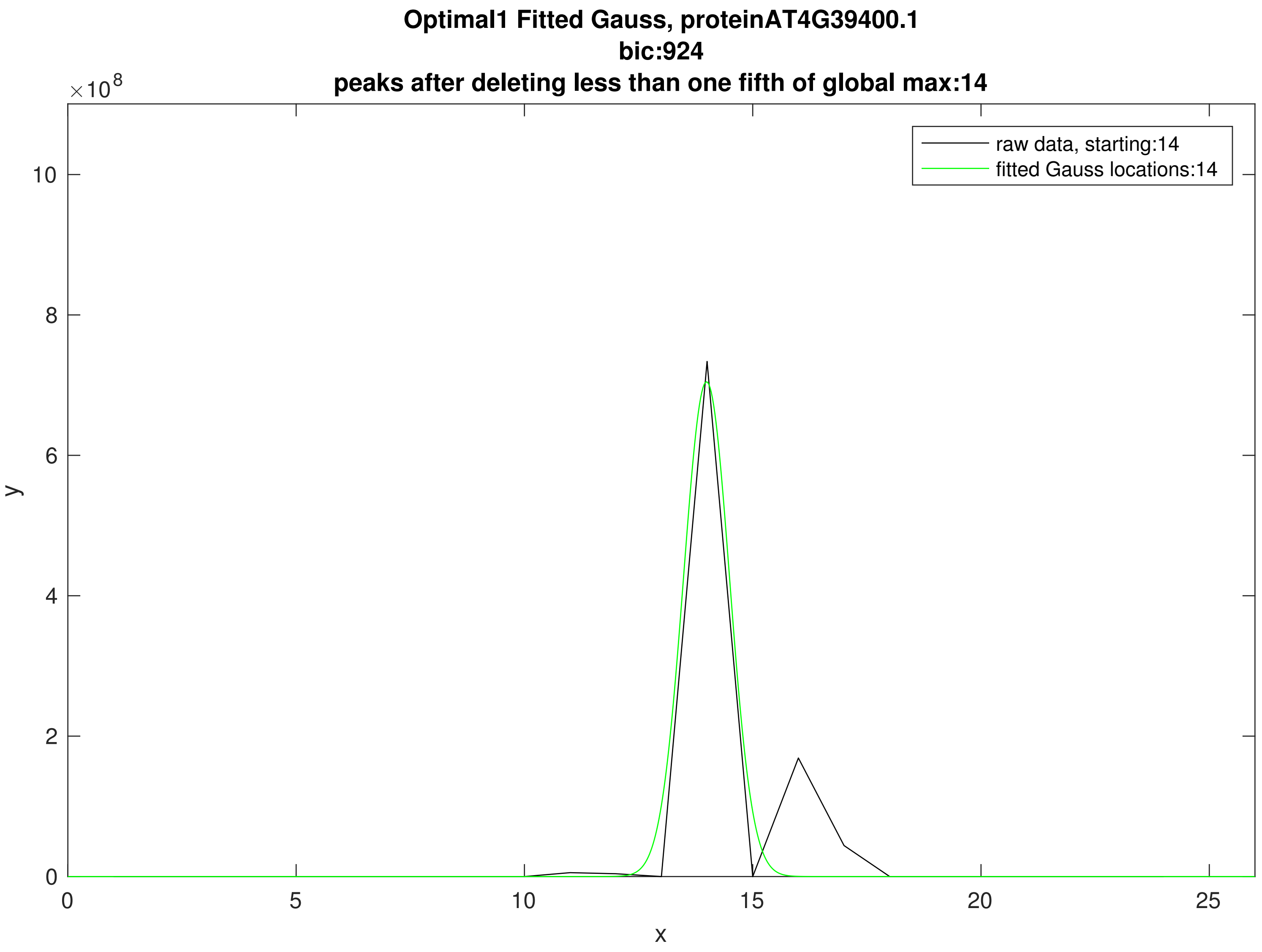
Optimal1 Fitted Gauss, proteinAT4G39400.1
bic:924
peaks after deleting less than one fifth of global max:14
×10^8
10
8
6
4
2
0
y
0
5
10
15
20
25
x
raw data, starting:14
fitted Gauss locations:14

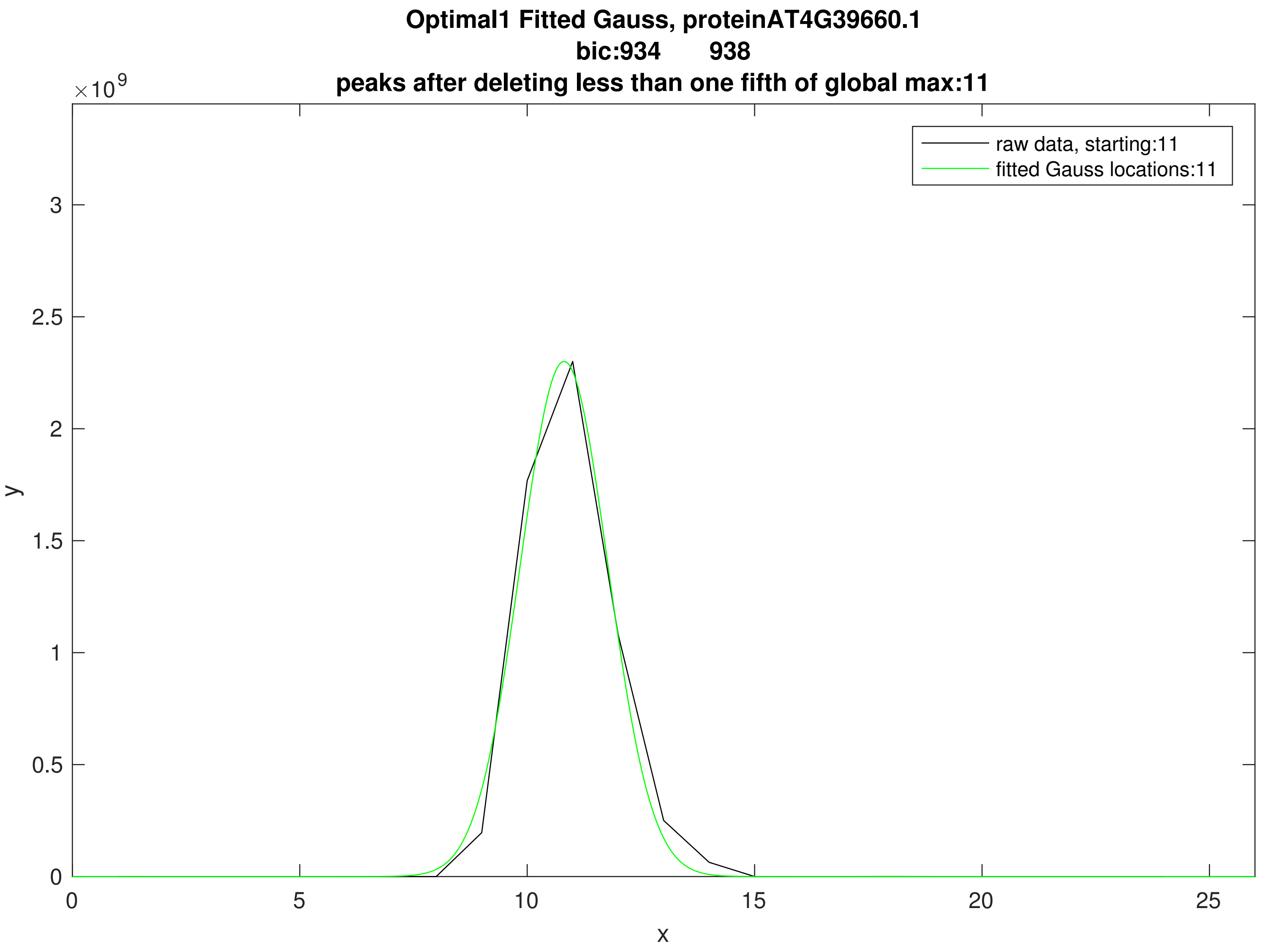

Optimal1 Fitted Gauss, proteinAT4G39660.1
bic:934 938
peaks after deleting less than one fifth of global max:11
×10^9
raw data, starting:11
fitted Gauss locations:11
y
x
0
0.5
1
1.5
2
2.5
3
5
10
15
20
25

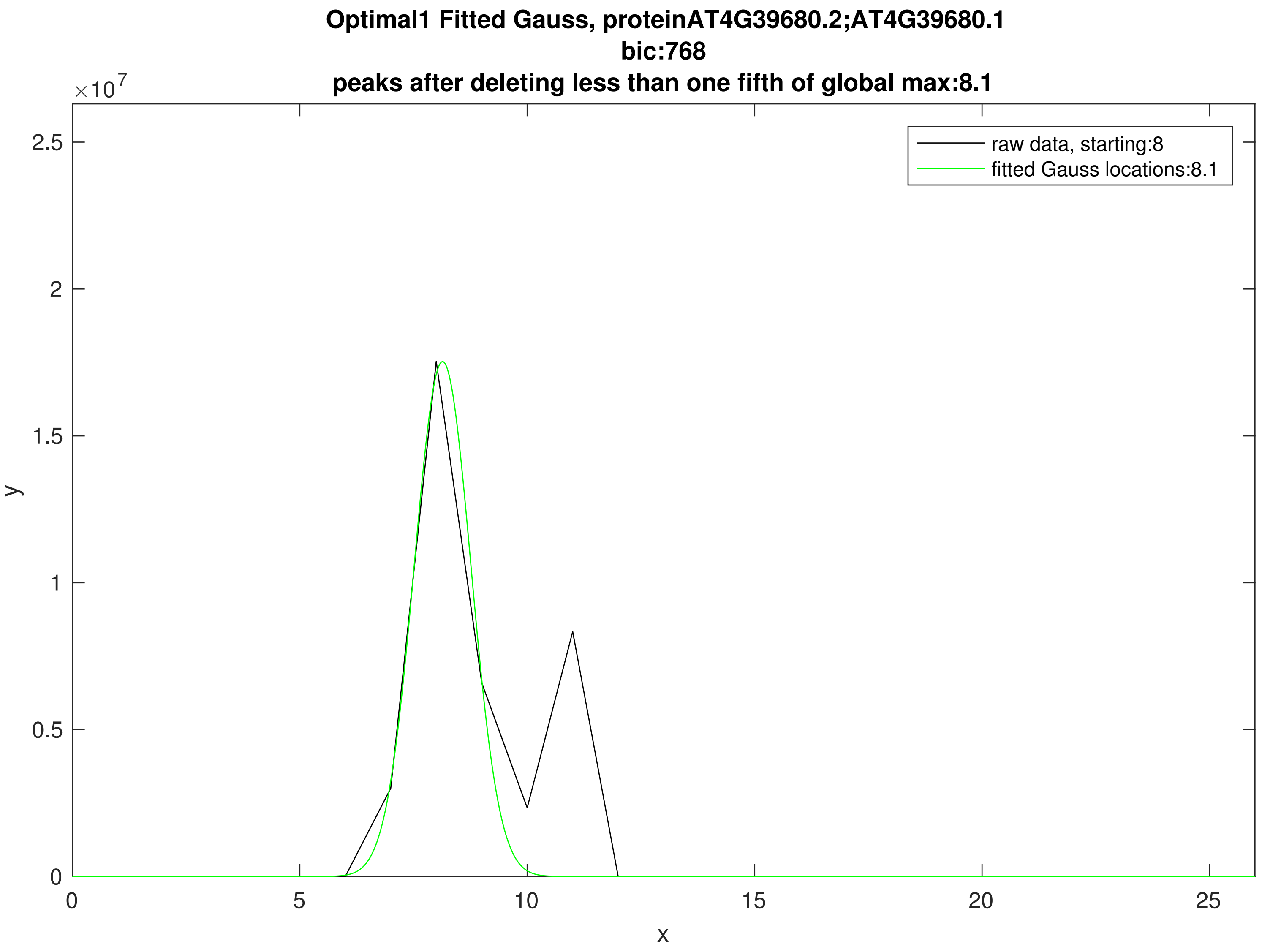
Optimal1 Fitted Gauss, proteinAT4G39680.2;AT4G39680.1
bic:768
peaks after deleting less than one fifth of global max:8.1
×10^7
2.5
2
1.5
1
0.5
0
y
0
5
10
15
20
25
x
raw data, starting:8
fitted Gauss locations:8.1

**Optimal1 Fitted Gauss, proteinAT4G39730.1**
**bic:875**
**peaks after deleting less than one fifth of global max:18**

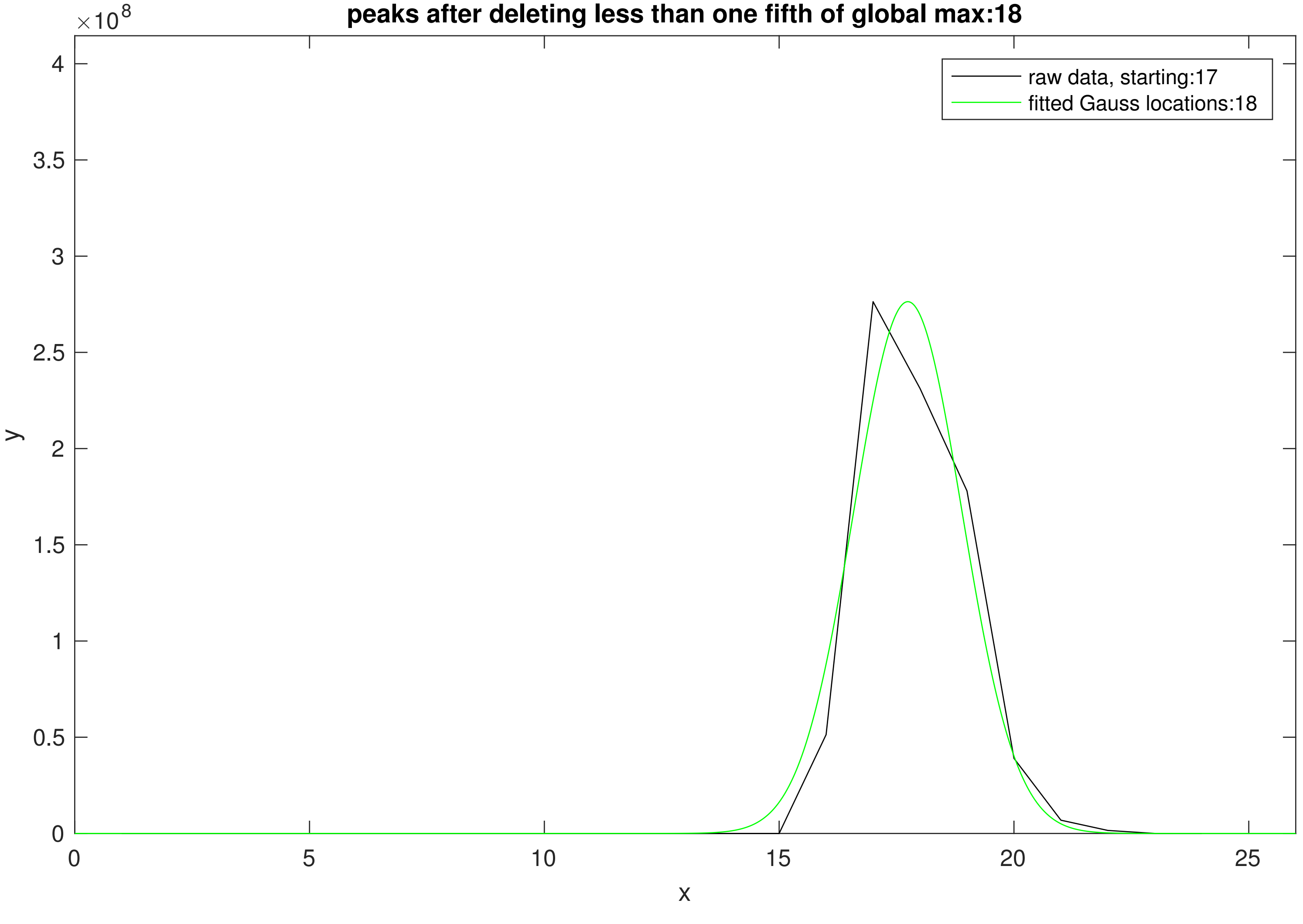

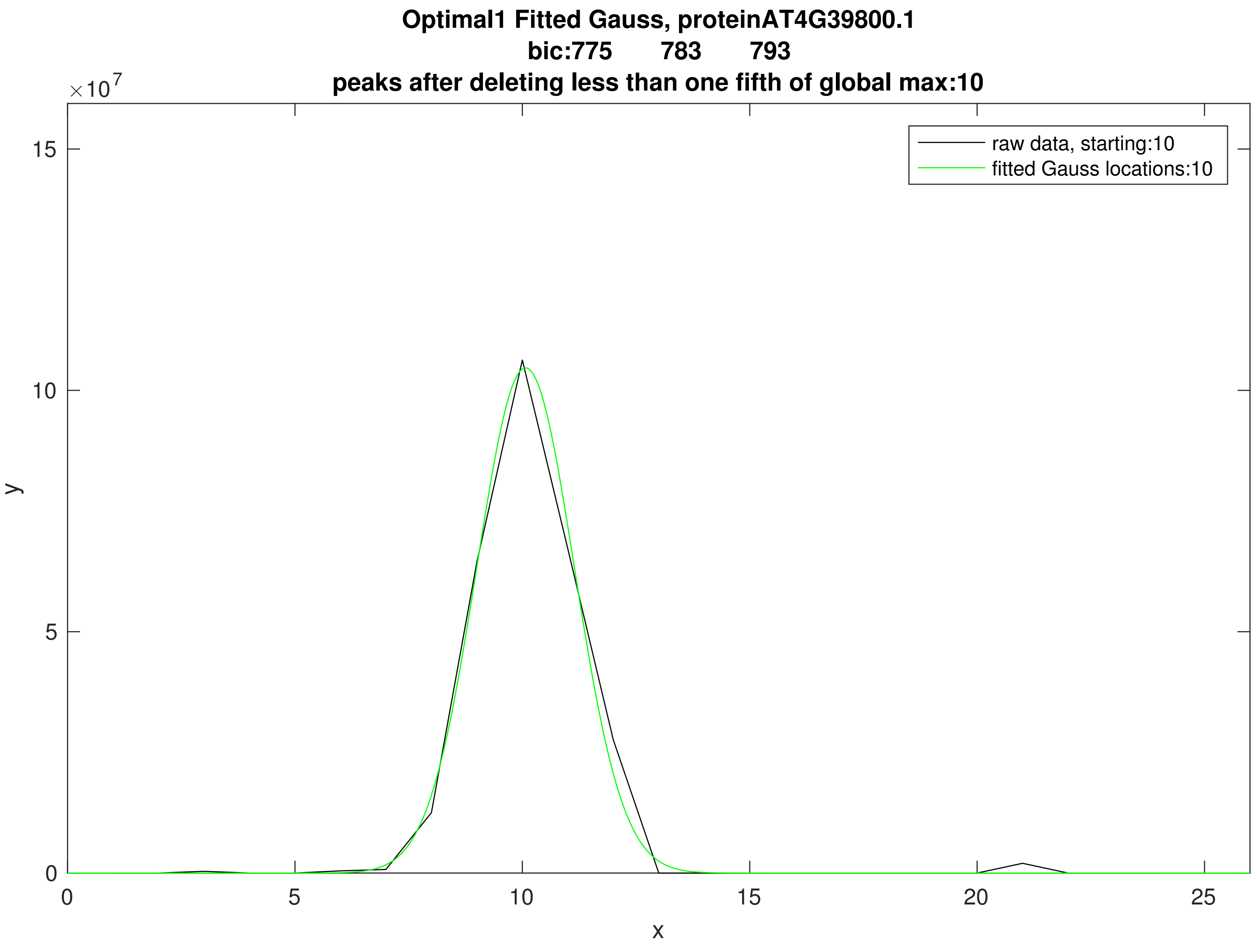
Optimal1 Fitted Gauss, proteinAT4G39800.1
bic:775 783 793
peaks after deleting less than one fifth of global max:10
×10^7
15
10
5
0
y
0
5
10
15
20
25
x
raw data, starting:10
fitted Gauss locations:10

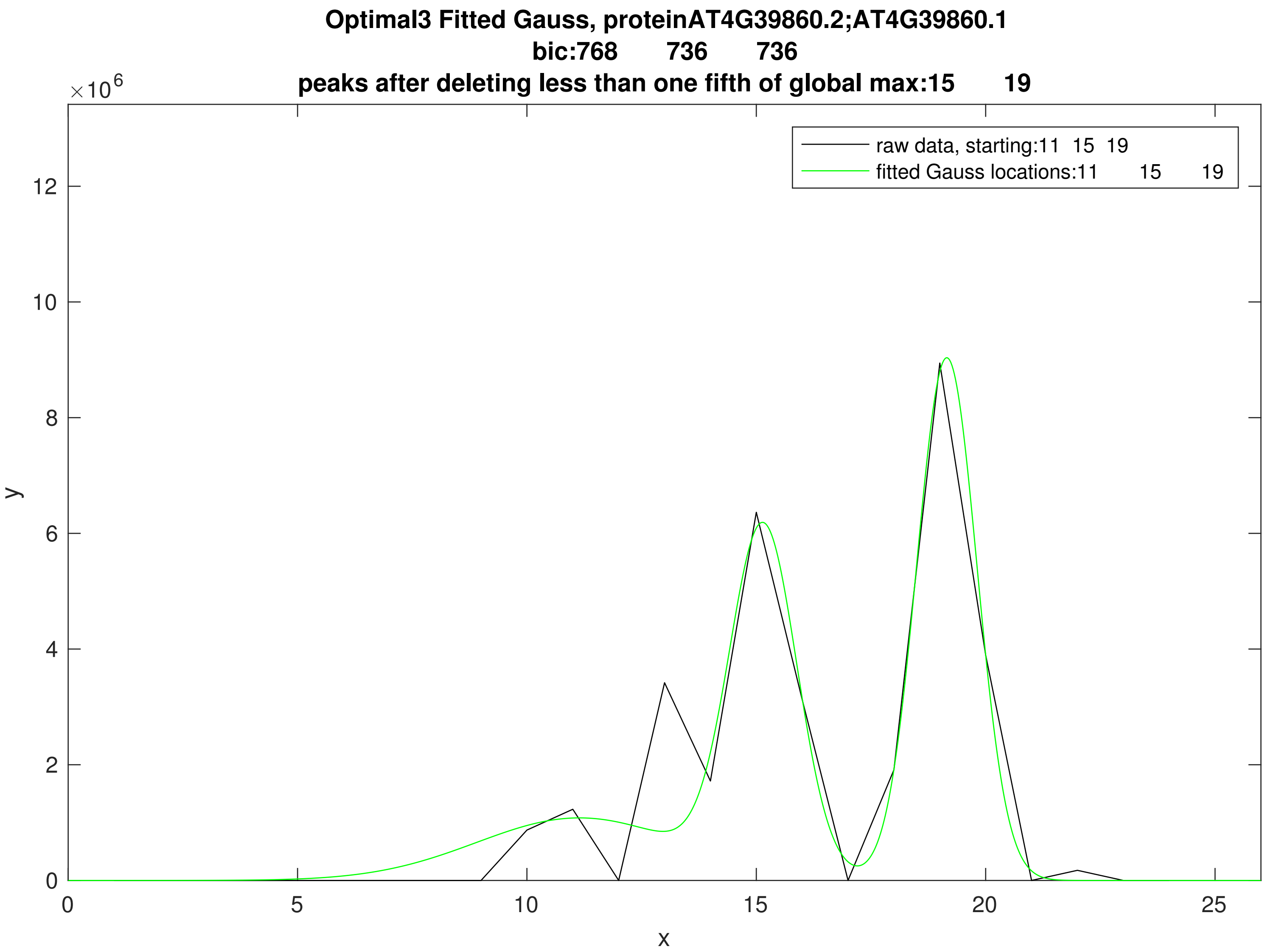

Optimal3 Fitted Gauss, proteinAT4G39860.2;AT4G39860.1
bic:768 736 736
peaks after deleting less than one fifth of global max:15 19
×10^6
raw data, starting:11 15 19
fitted Gauss locations:11 15 19
y
x
0
2
4
6
8
10
12
5
15
20
25

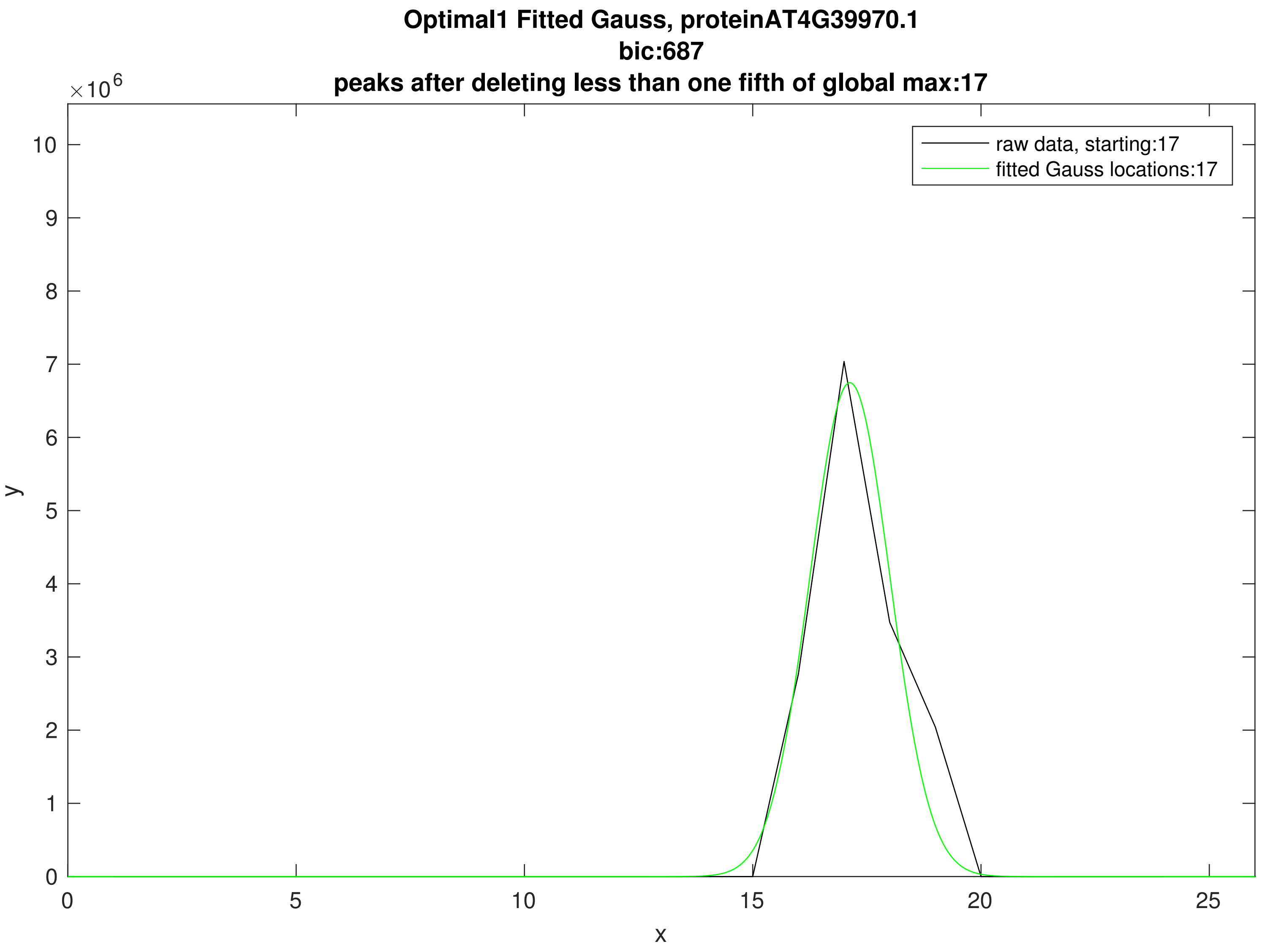
Optimal1 Fitted Gauss, proteinAT4G39970.1
bic:687
peaks after deleting less than one fifth of global max:17
×10^6
raw data, starting:17
fitted Gauss locations:17
y
x
0
1
2
3
4
5
6
7
8
9
10
0
5
10
15
20
25

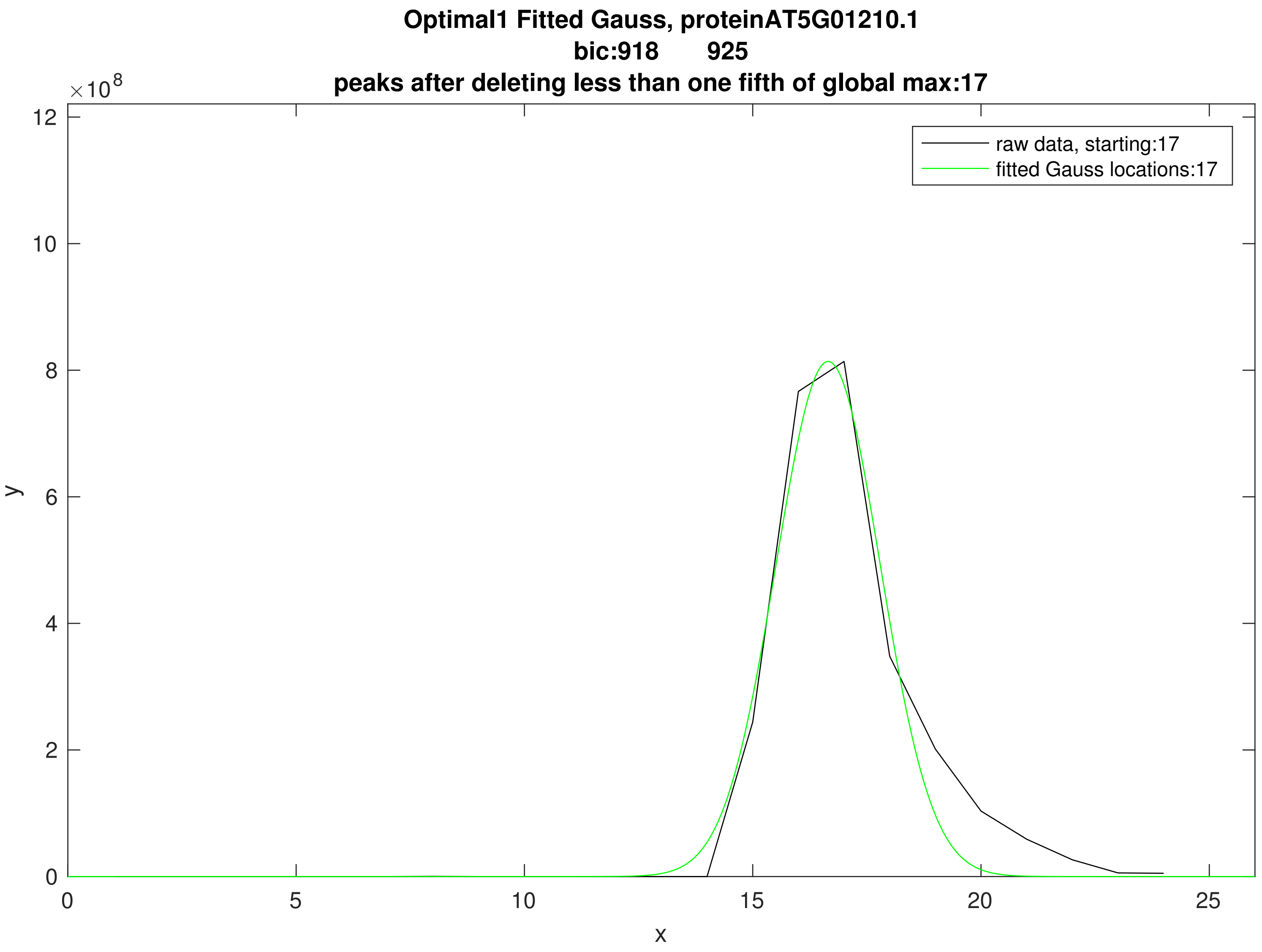
Optimal1 Fitted Gauss, proteinAT5G01210.1
bic:918 925
peaks after deleting less than one fifth of global max:17
×10^8
12
10
8
6
4
2
0
y
0
5
10
15
20
25
x
raw data, starting:17
fitted Gauss locations:17

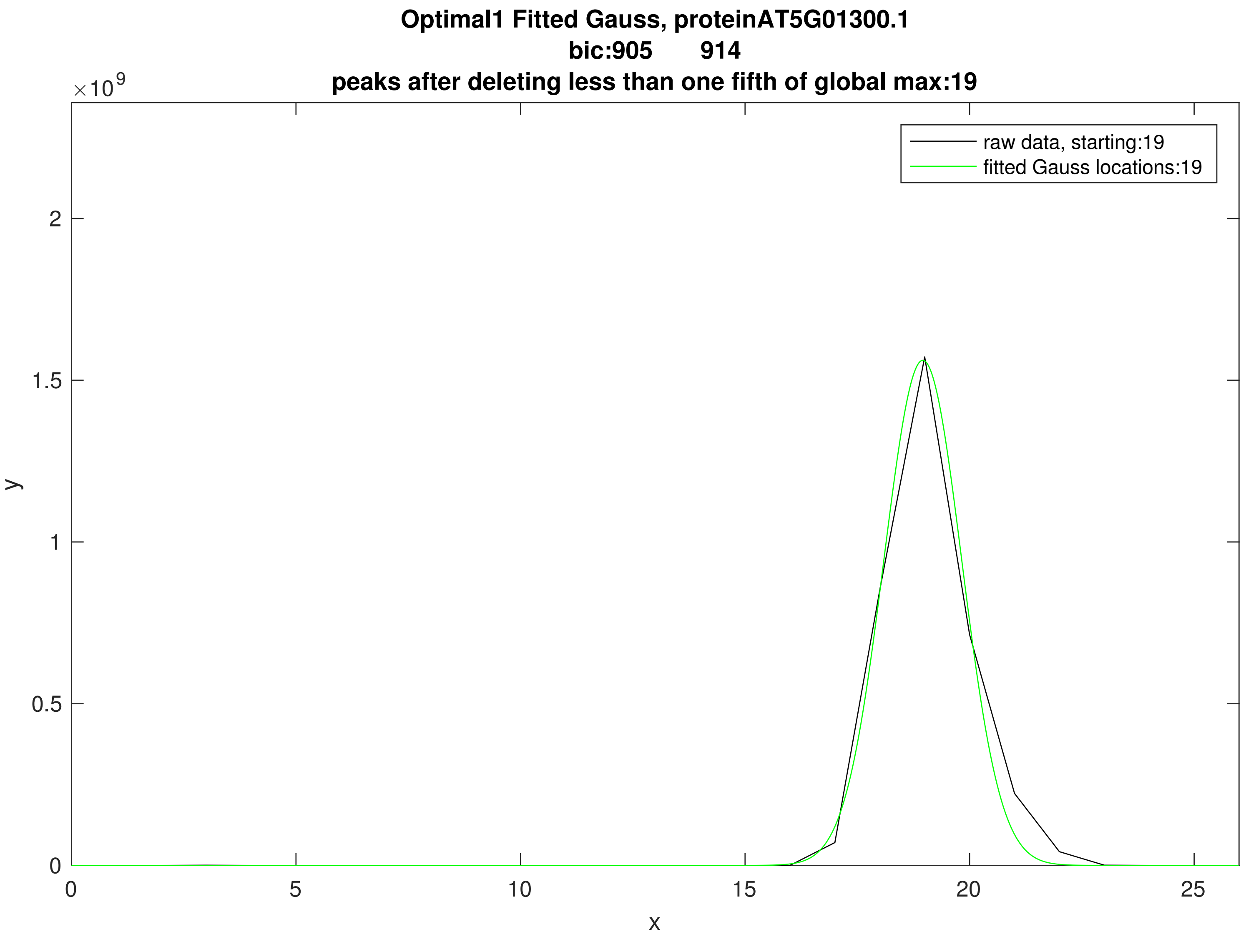

Optimal1 Fitted Gauss, proteinAT5G01300.1
bic:905 914
peaks after deleting less than one fifth of global max:19
×10^9
raw data, starting:19
fitted Gauss locations:19
2
1.5
1
0.5
0
y
0
5
10
15
20
25
x

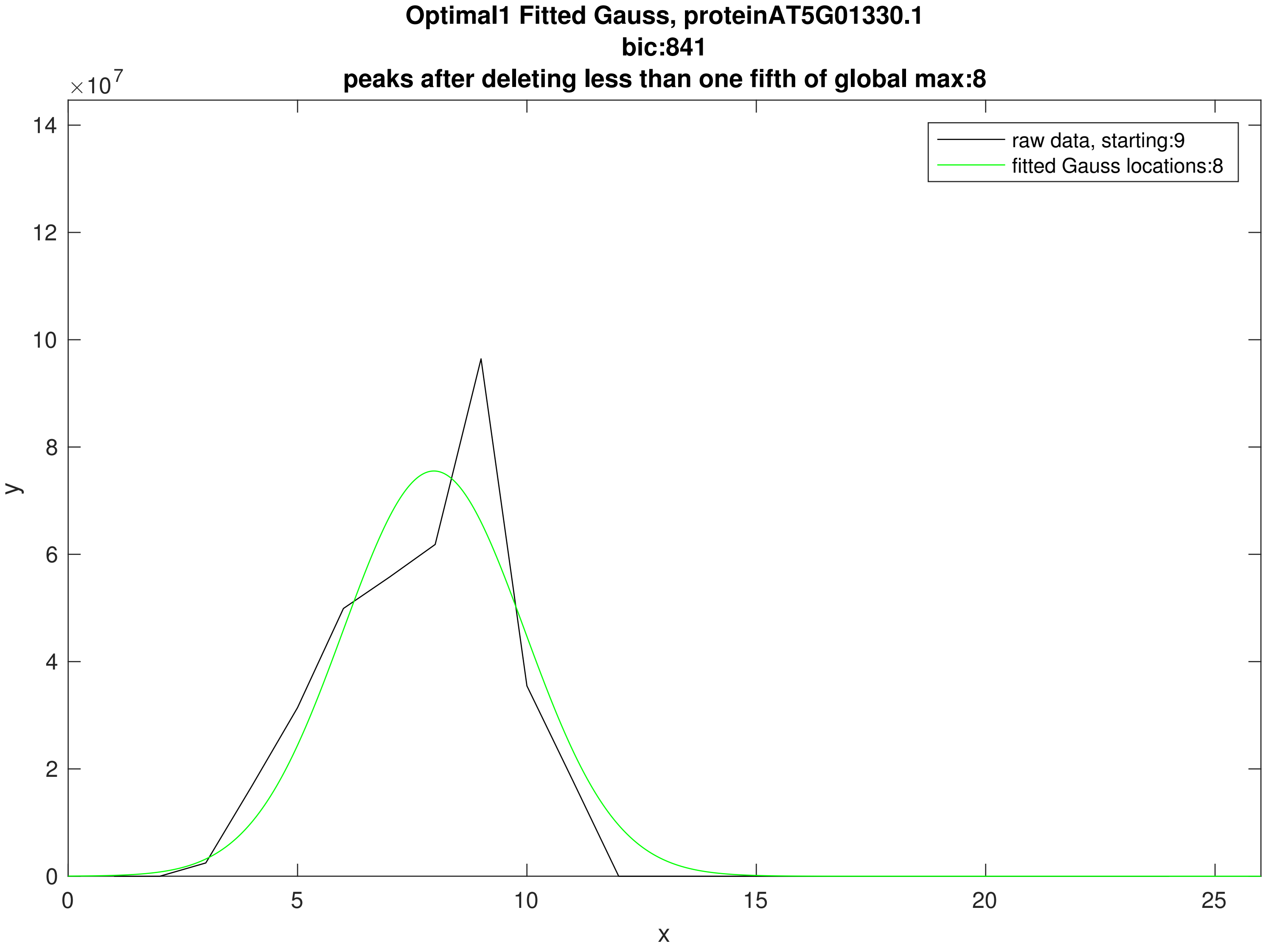
Optimal1 Fitted Gauss, proteinAT5G01330.1
bic:841
peaks after deleting less than one fifth of global max:8
×10^7
raw data, starting:9
fitted Gauss locations:8
y
x
0
2
4
6
8
10
12
14
5
15
20
25

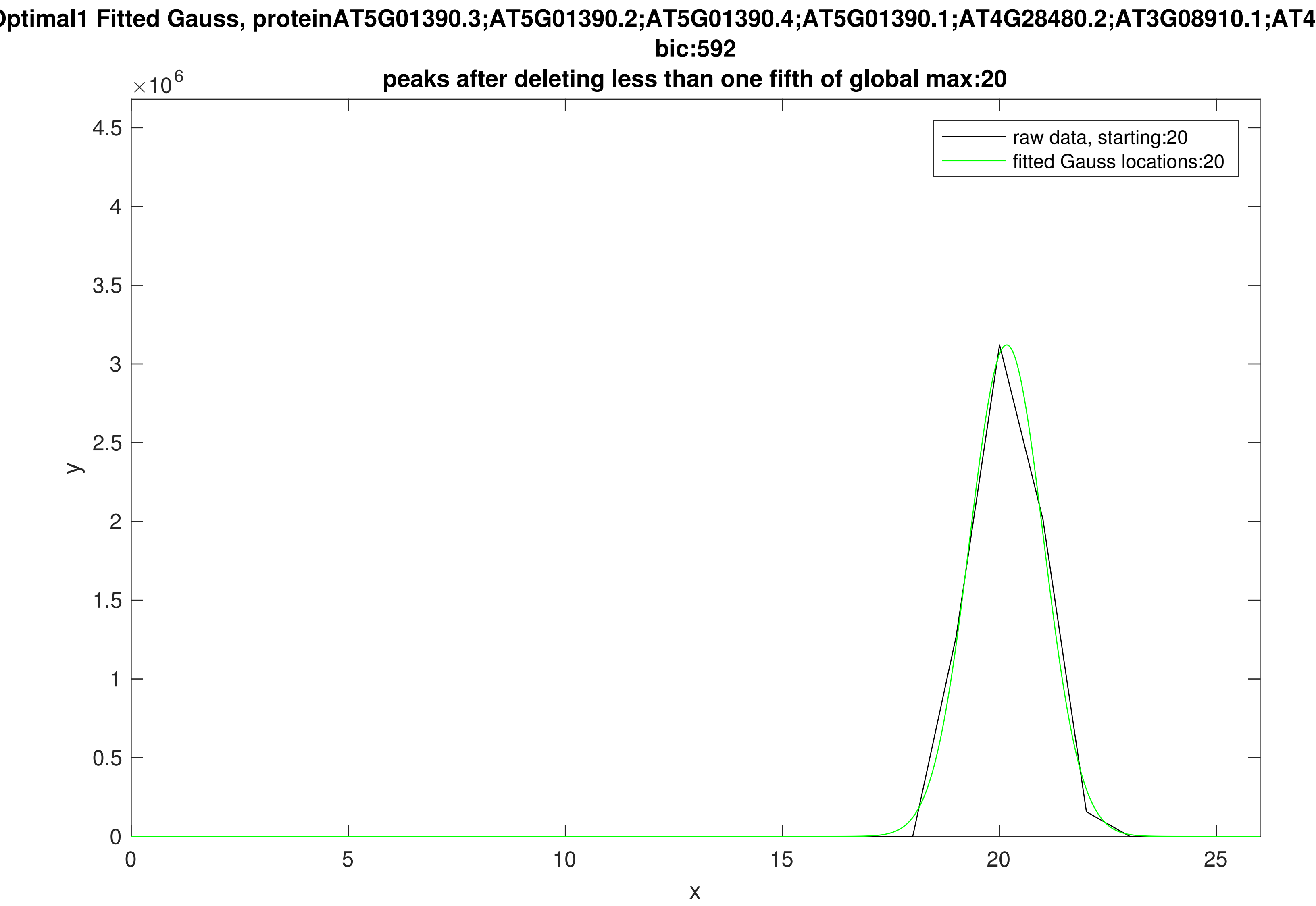
Optimal1 Fitted Gauss, proteinAT5G01390.3;AT5G01390.2;AT5G01390.4;AT5G01390.1;AT4G28480.2;AT3G08910.1;AT4
bic:592
peaks after deleting less than one fifth of global max:20
×10^6
4.5
4
3.5
3
2.5
2
1.5
1
0.5
0
y
0
5
10
15
20
25
x
raw data, starting:20
fitted Gauss locations:20

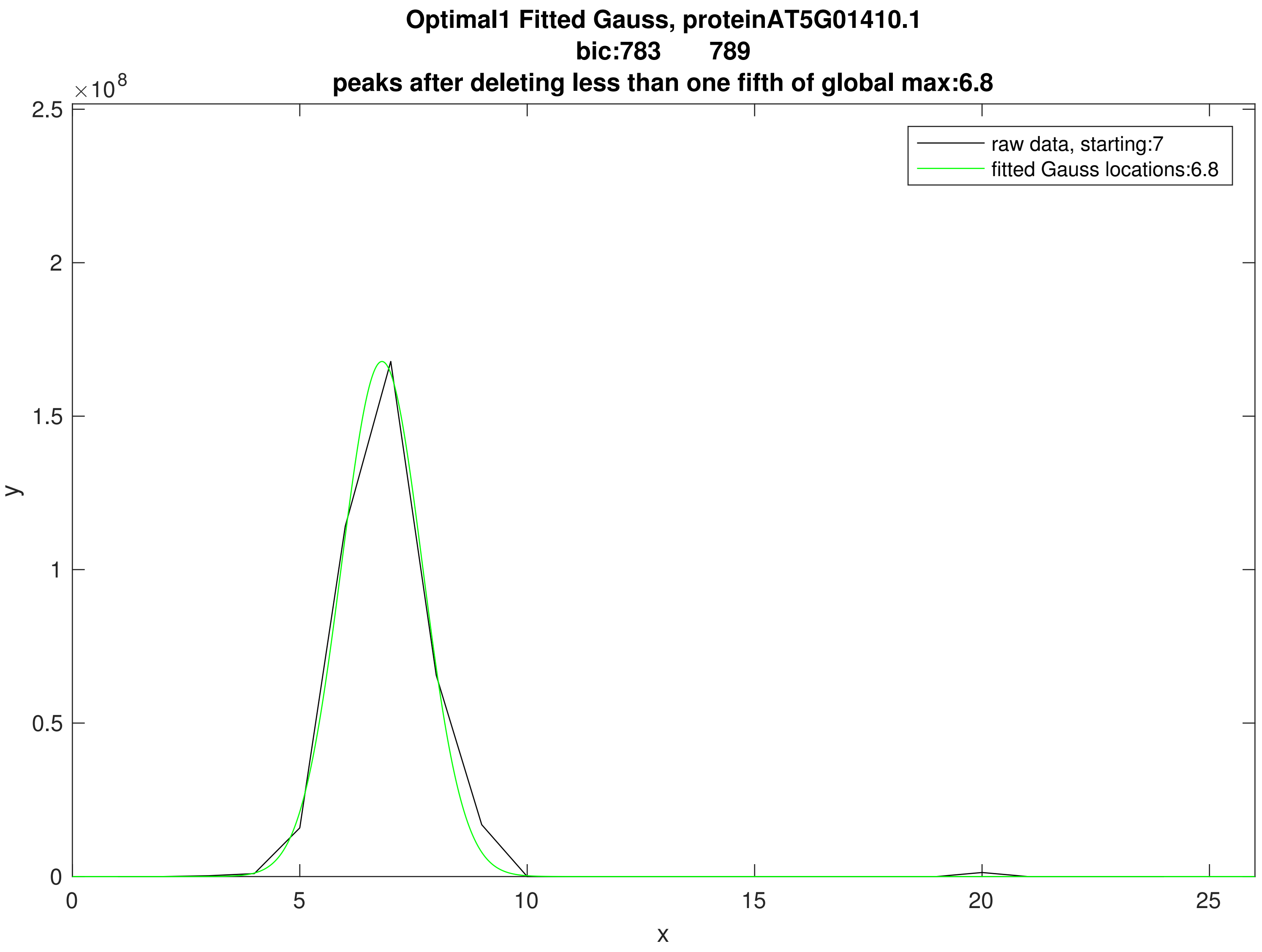
Optimal1 Fitted Gauss, proteinAT5G01410.1
bic:783 789
peaks after deleting less than one fifth of global max:6.8
×10^8
2.5
2
1.5
1
0.5
0
y
0
5
10
15
20
25
x
raw data, starting:7
fitted Gauss locations:6.8

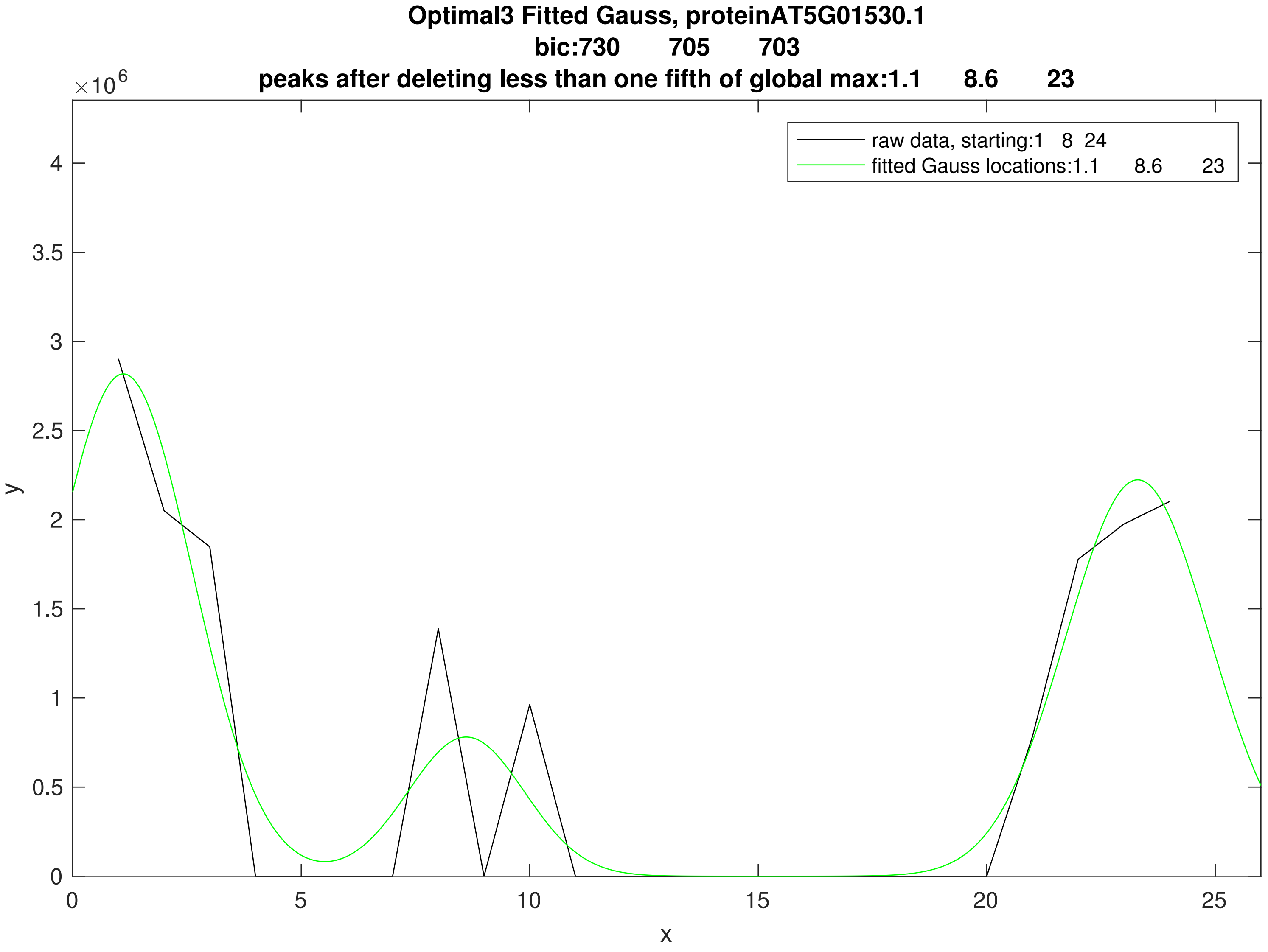

Optimal3 Fitted Gauss, proteinAT5G01530.1
bic:730 705 703
peaks after deleting less than one fifth of global max:1.1 8.6 23
$\times 10^6$
raw data, starting:1 8 24
fitted Gauss locations:1.1 8.6 23
y
x
0
0.5
1
1.5
2
2.5
3
3.5
4
0
5
10
15
20
25

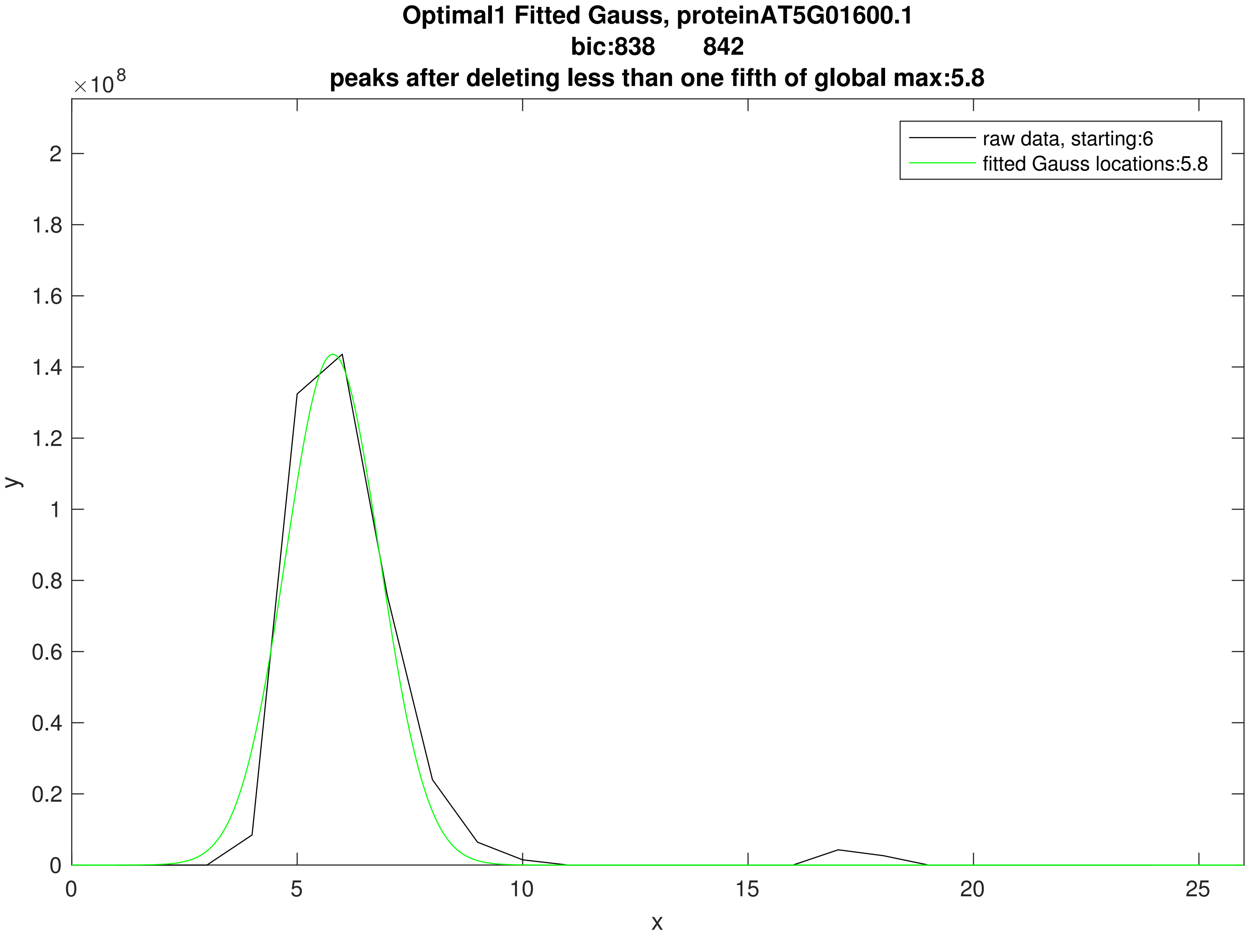

Optimal1 Fitted Gauss, proteinAT5G01600.1
bic:838 842
peaks after deleting less than one fifth of global max:5.8
×10^8
raw data, starting:6
fitted Gauss locations:5.8
y
x
0
0.2
0.4
0.6
0.8
1
1.2
1.4
1.6
1.8
2
5
10
15
20
25

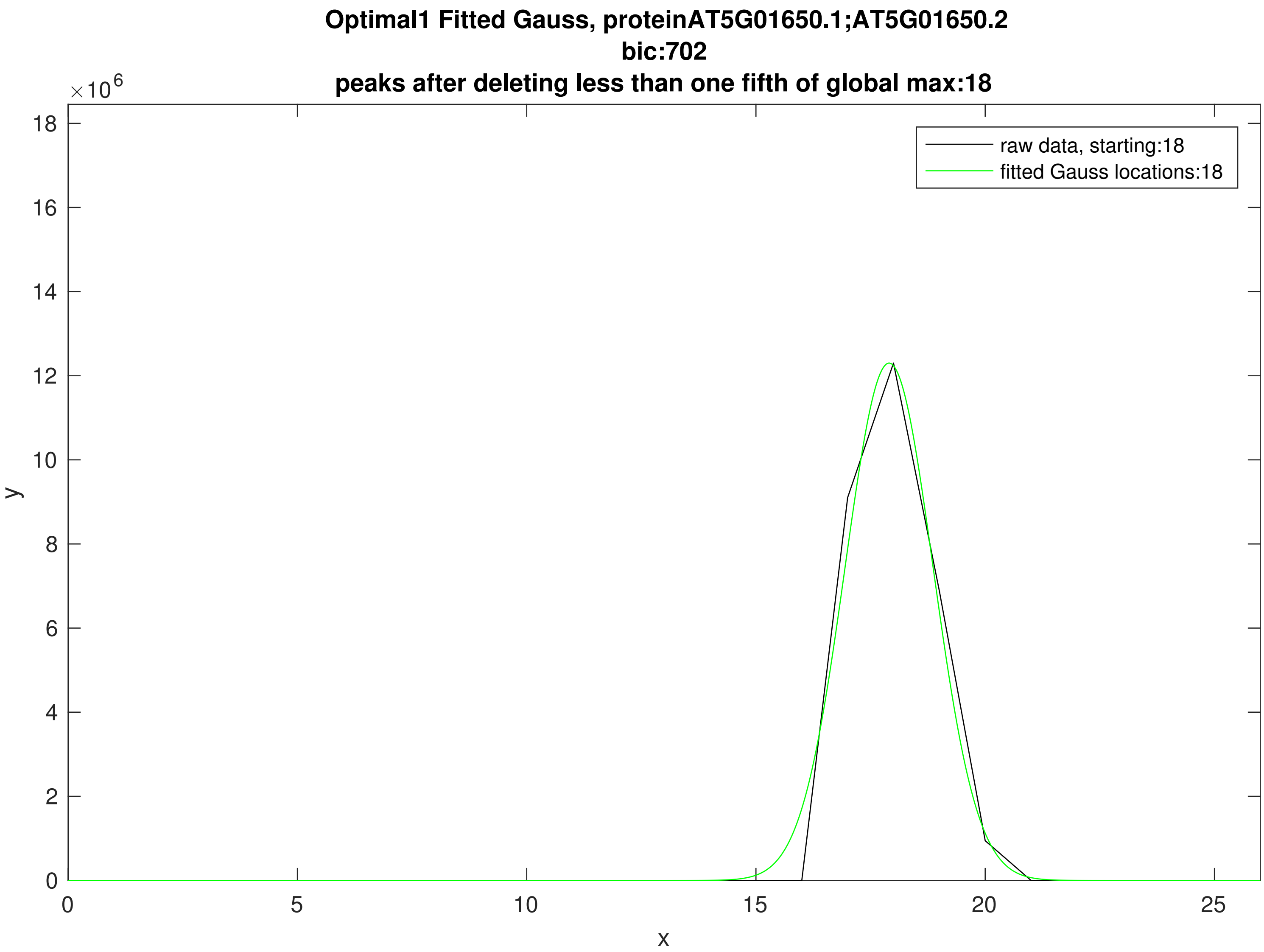

Optimal1 Fitted Gauss, proteinAT5G01650.1;AT5G01650.2
bic:702
peaks after deleting less than one fifth of global max:18
×10^6
18
16
14
12
10
8
6
4
2
0
y
0
5
10
15
20
25
x
raw data, starting:18
fitted Gauss locations:18

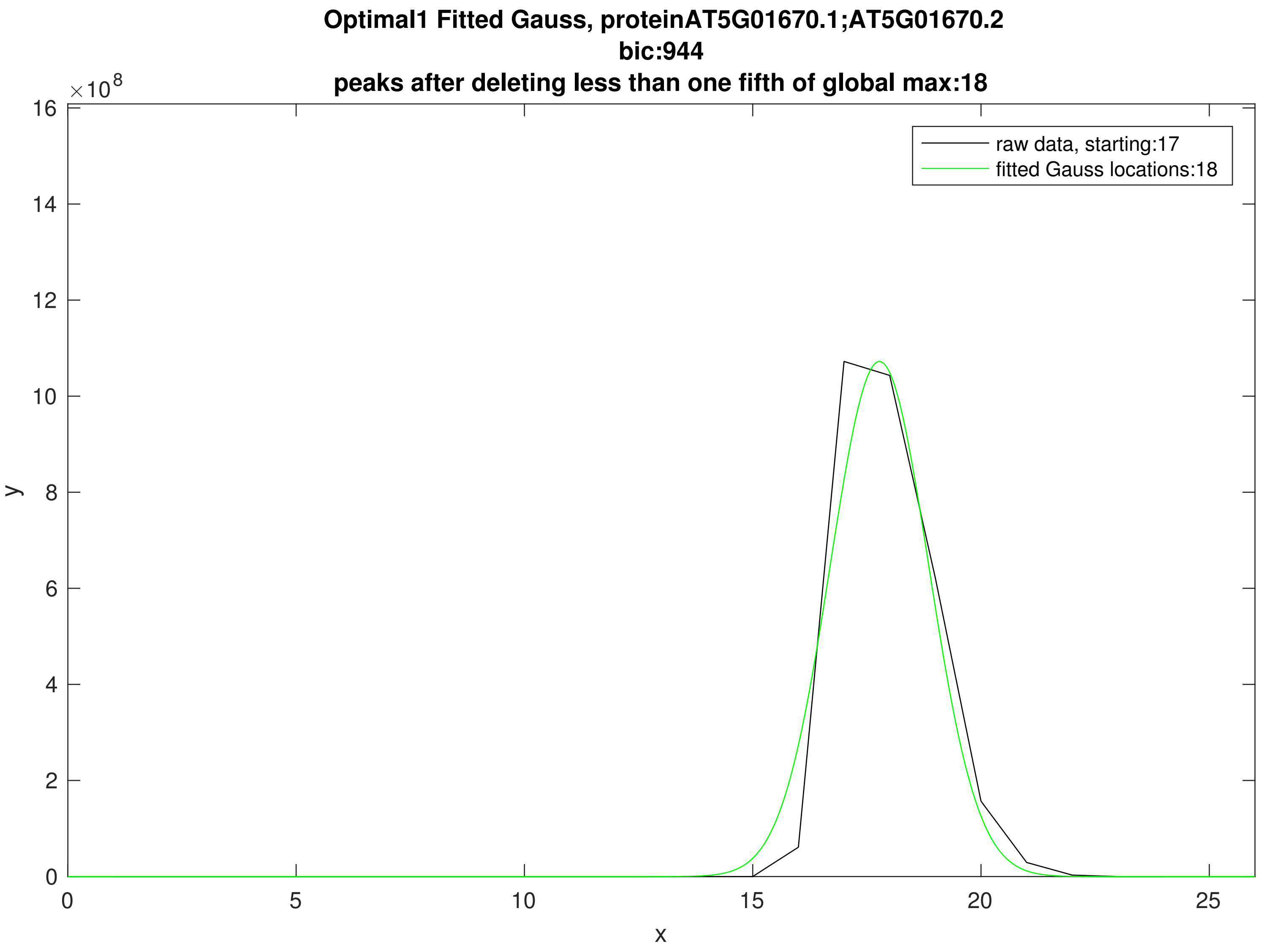
Optimal1 Fitted Gauss, proteinAT5G01670.1;AT5G01670.2
bic:944
peaks after deleting less than one fifth of global max:18
×10^8
16
14
12
10
8
6
4
2
0
y
0
5
10
15
20
25
x
raw data, starting:17
fitted Gauss locations:18

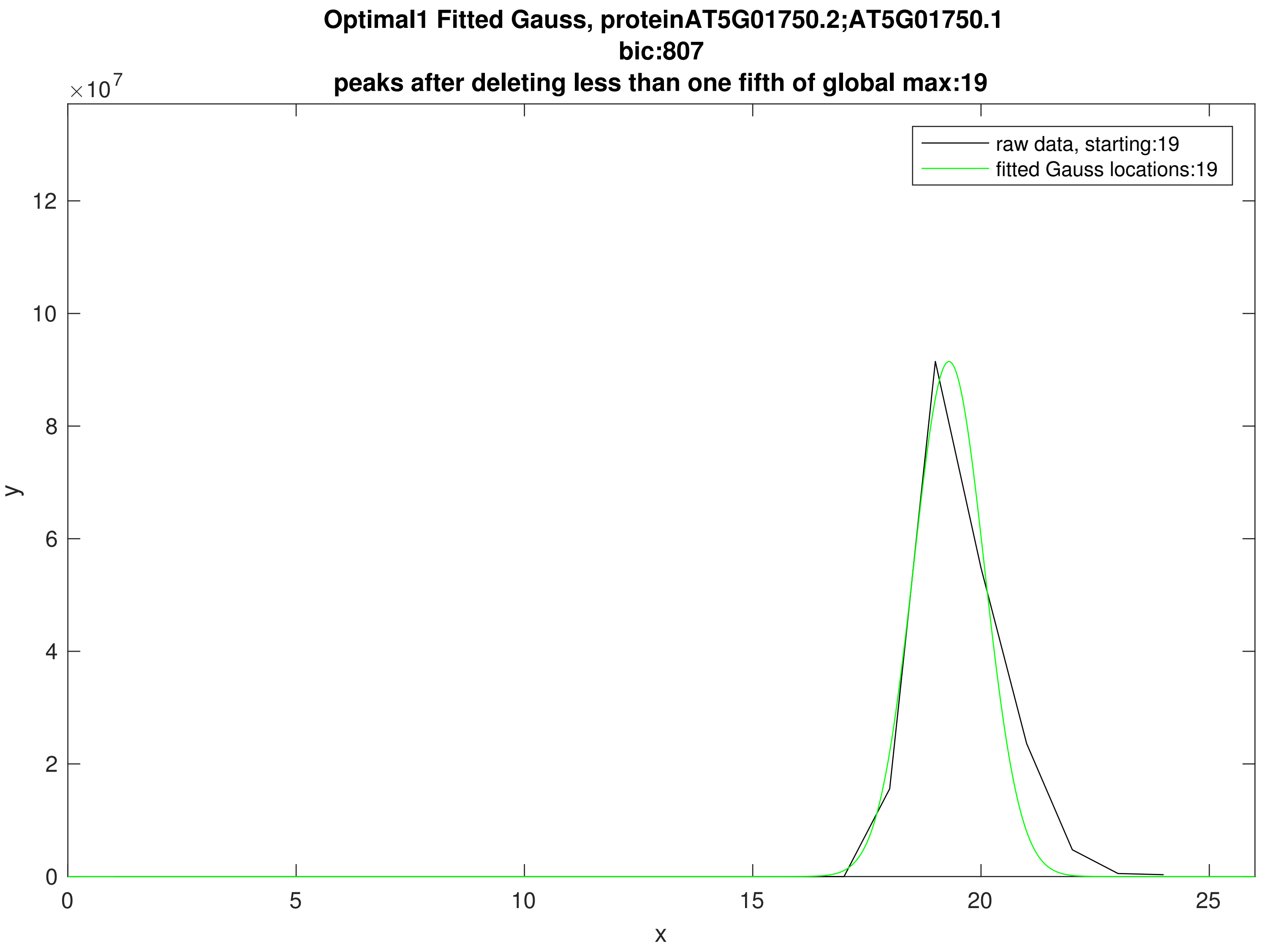

Optimal1 Fitted Gauss, proteinAT5G01750.2;AT5G01750.1
bic:807
peaks after deleting less than one fifth of global max:19
×10^7
raw data, starting:19
fitted Gauss locations:19
y
x
0
2
4
6
8
10
12
0
5
10
15
20
25

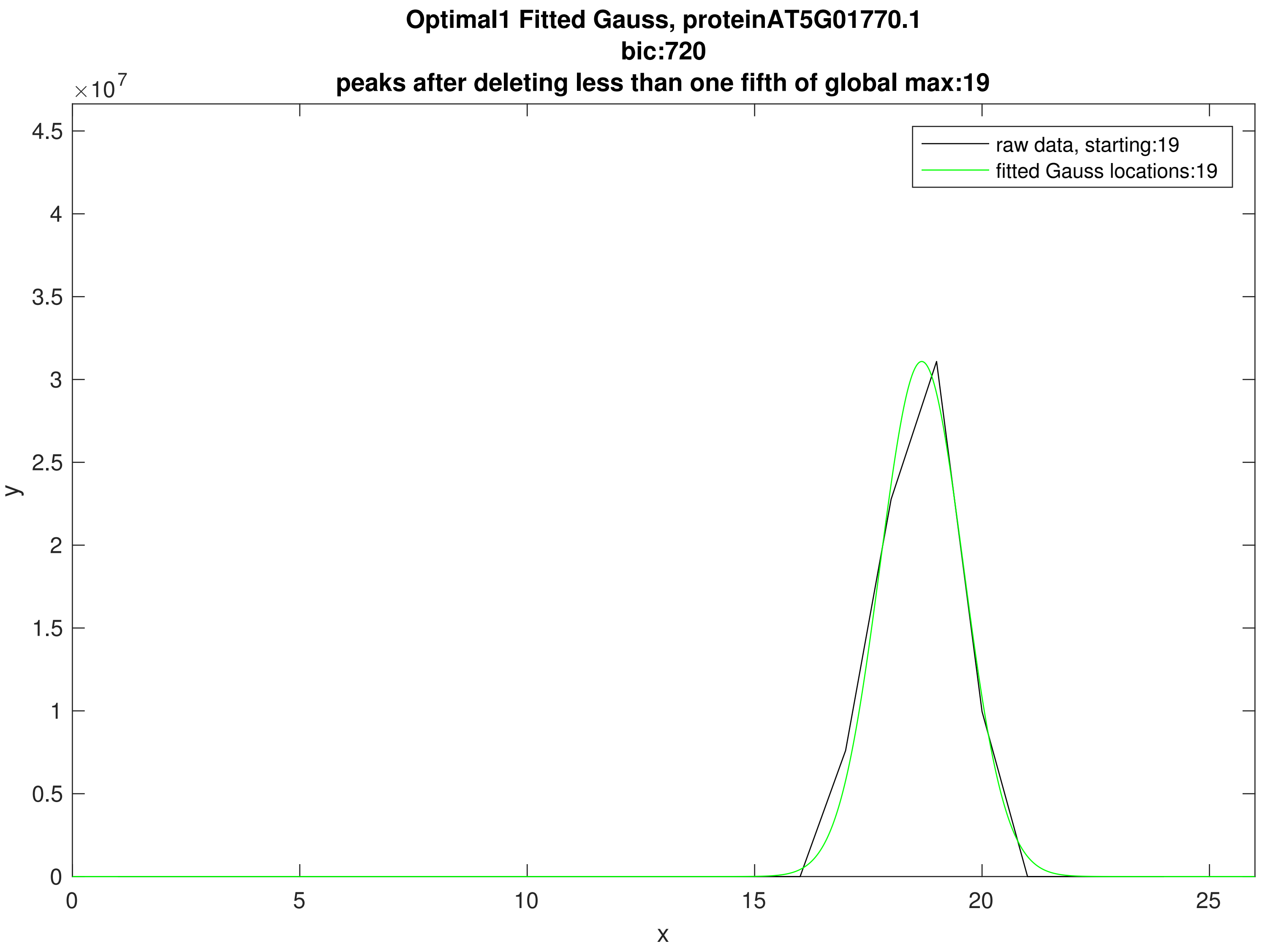
Optimal1 Fitted Gauss, proteinAT5G01770.1
bic:720
peaks after deleting less than one fifth of global max:19
×10^7
raw data, starting:19
fitted Gauss locations:19
y
x
0
0.5
1
1.5
2
2.5
3
3.5
4
4.5
0
5
10
15
20
25

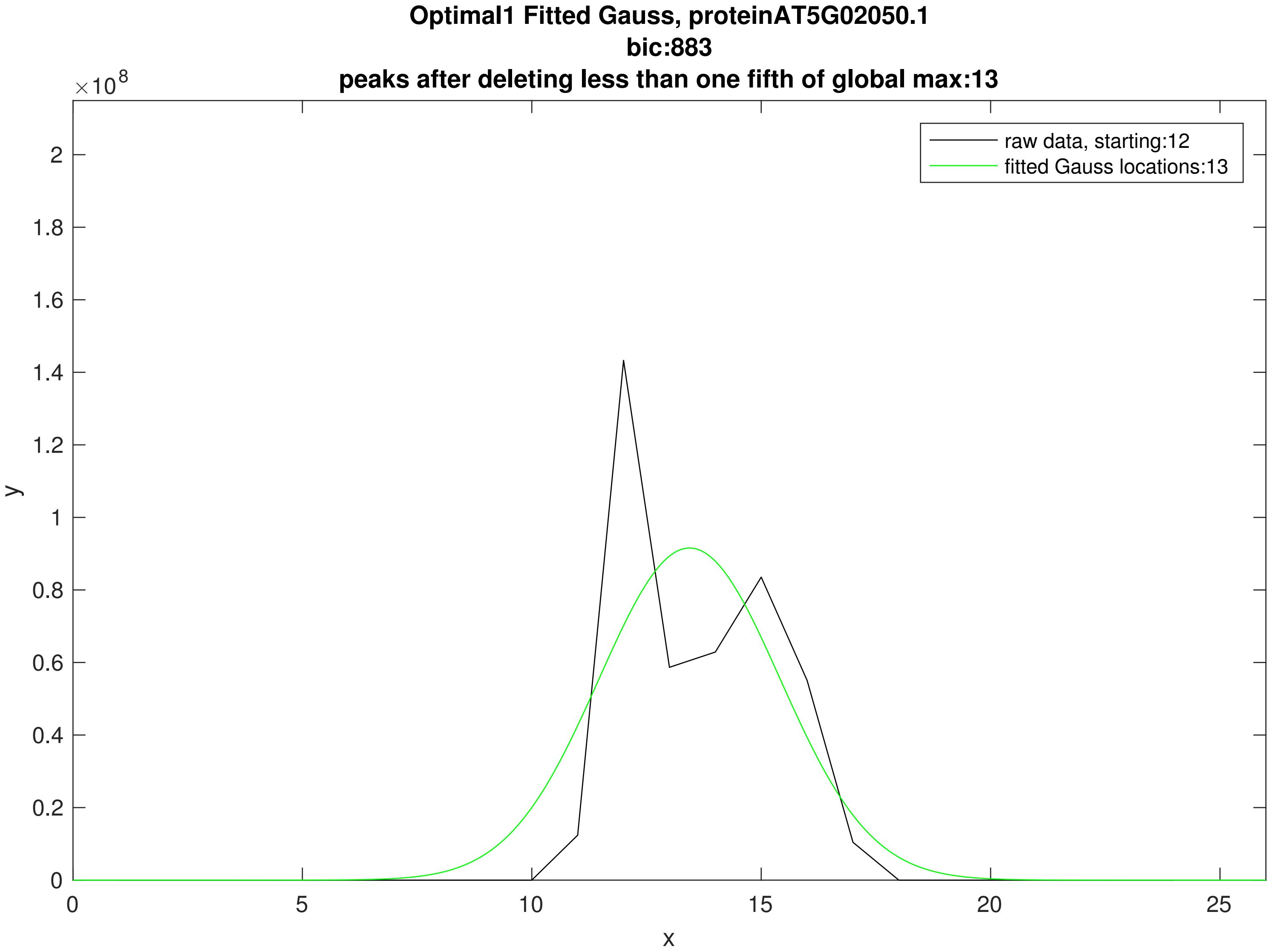
Optimal1 Fitted Gauss, proteinAT5G02050.1
bic:883
peaks after deleting less than one fifth of global max:13
×10^8
raw data, starting:12
fitted Gauss locations:13
2
1.8
1.6
1.4
1.2
1
0.8
0.6
0.4
0.2
0
y
0
5
10
15
20
25
x

**Optimal1 Fitted Gauss, proteinAT5G02130.1**
**bic:666**
**peaks after deleting less than one fifth of global max:17**

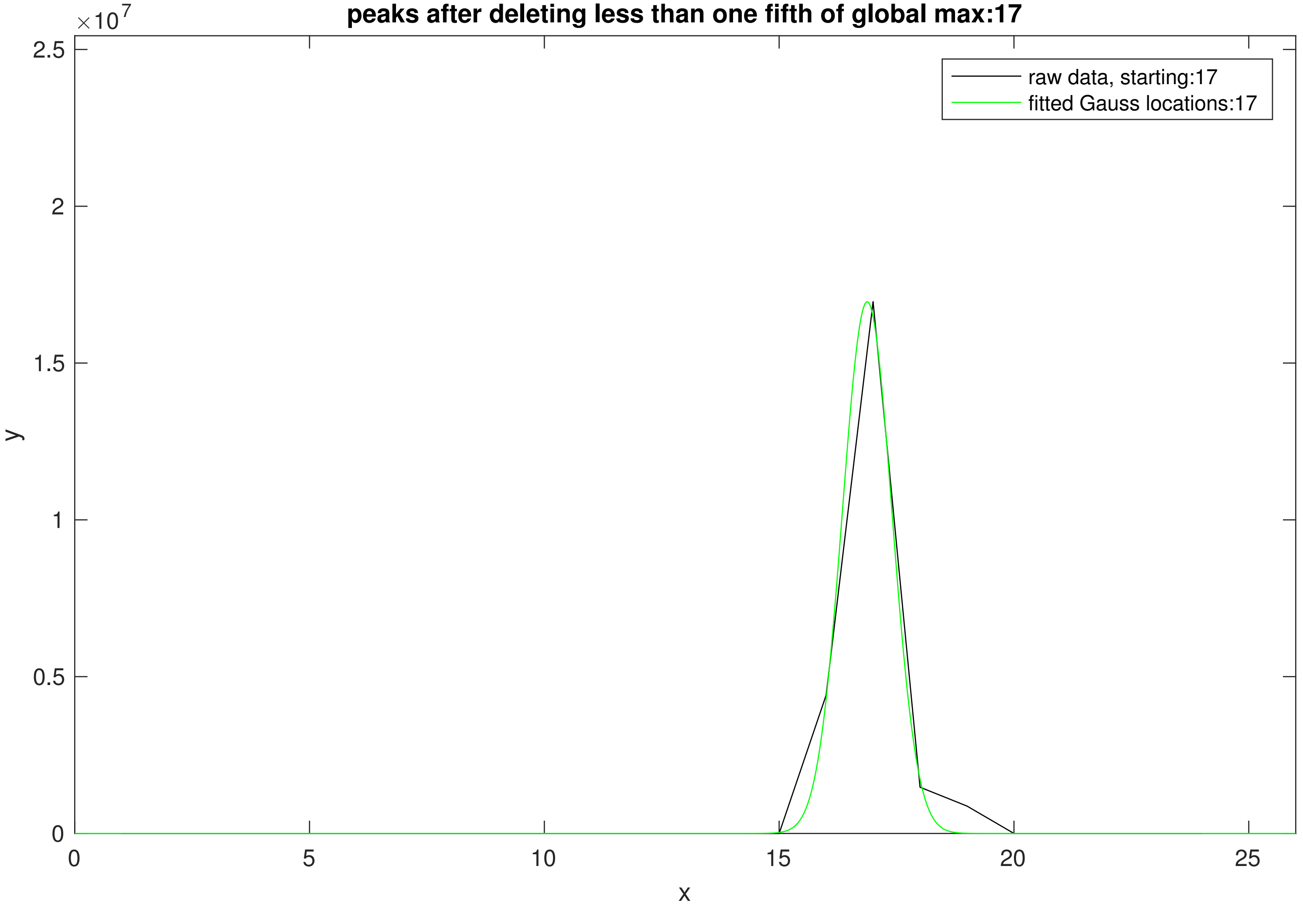

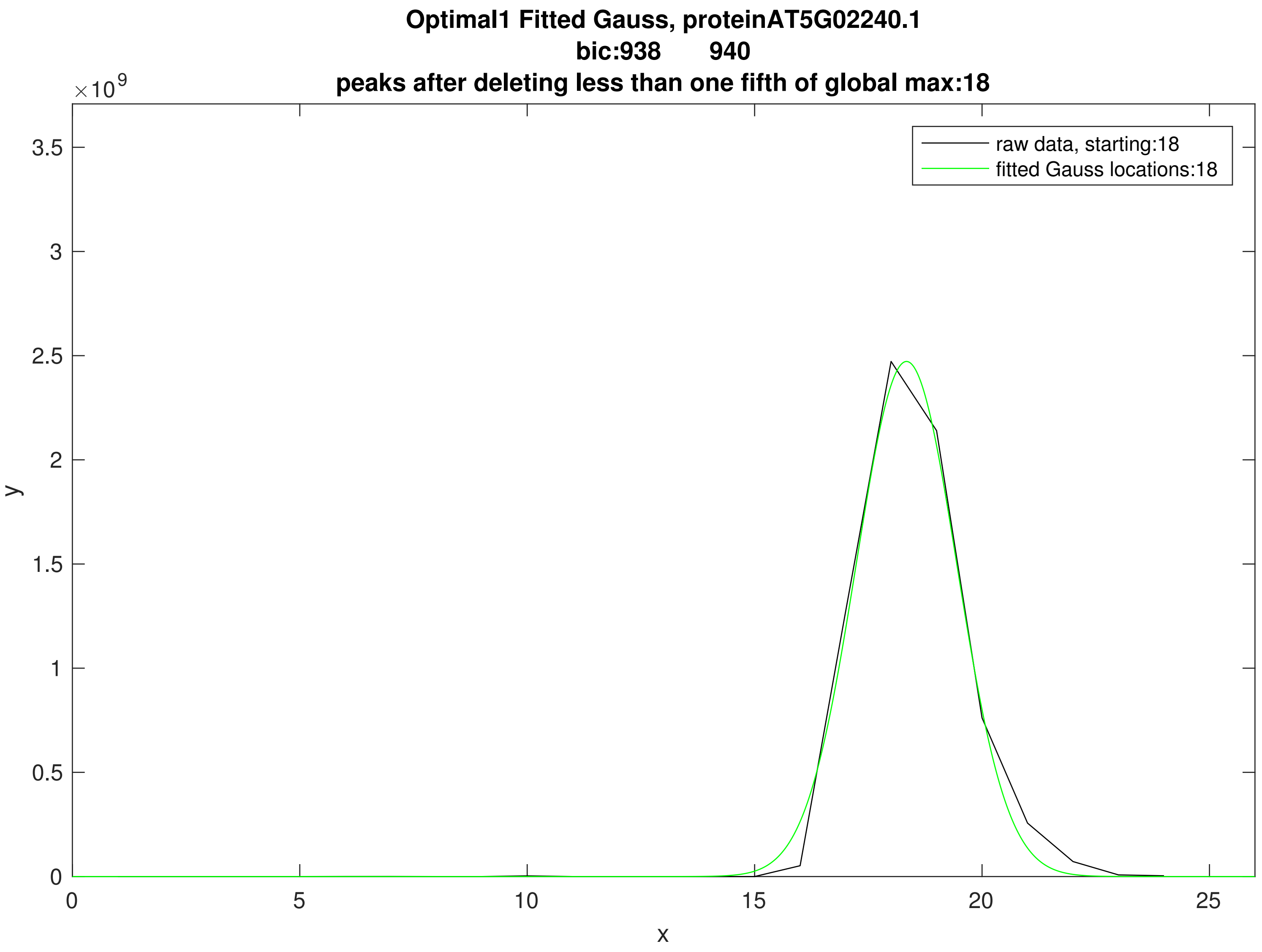

Optimal1 Fitted Gauss, proteinAT5G02240.1
bic:938 940
peaks after deleting less than one fifth of global max:18
×10^9
3.5
3
2.5
2
1.5
1
0.5
0
y
0
5
10
15
20
25
x
raw data, starting:18
fitted Gauss locations:18

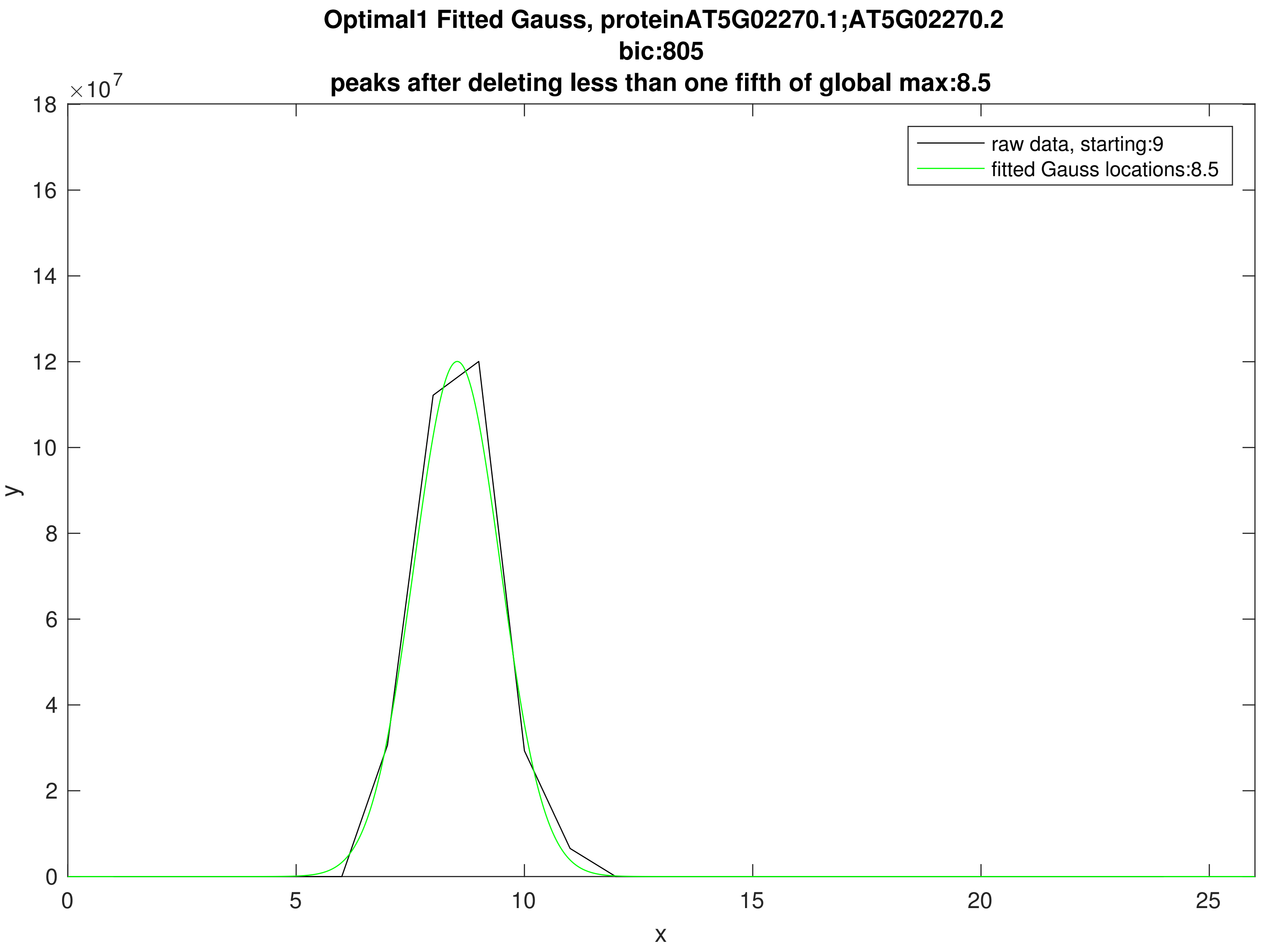
Optimal1 Fitted Gauss, proteinAT5G02270.1;AT5G02270.2
bic:805
peaks after deleting less than one fifth of global max:8.5
×10^7
18
16
14
12
10
8
6
4
2
0
y
0
5
10
15
20
25
x
raw data, starting:9
fitted Gauss locations:8.5

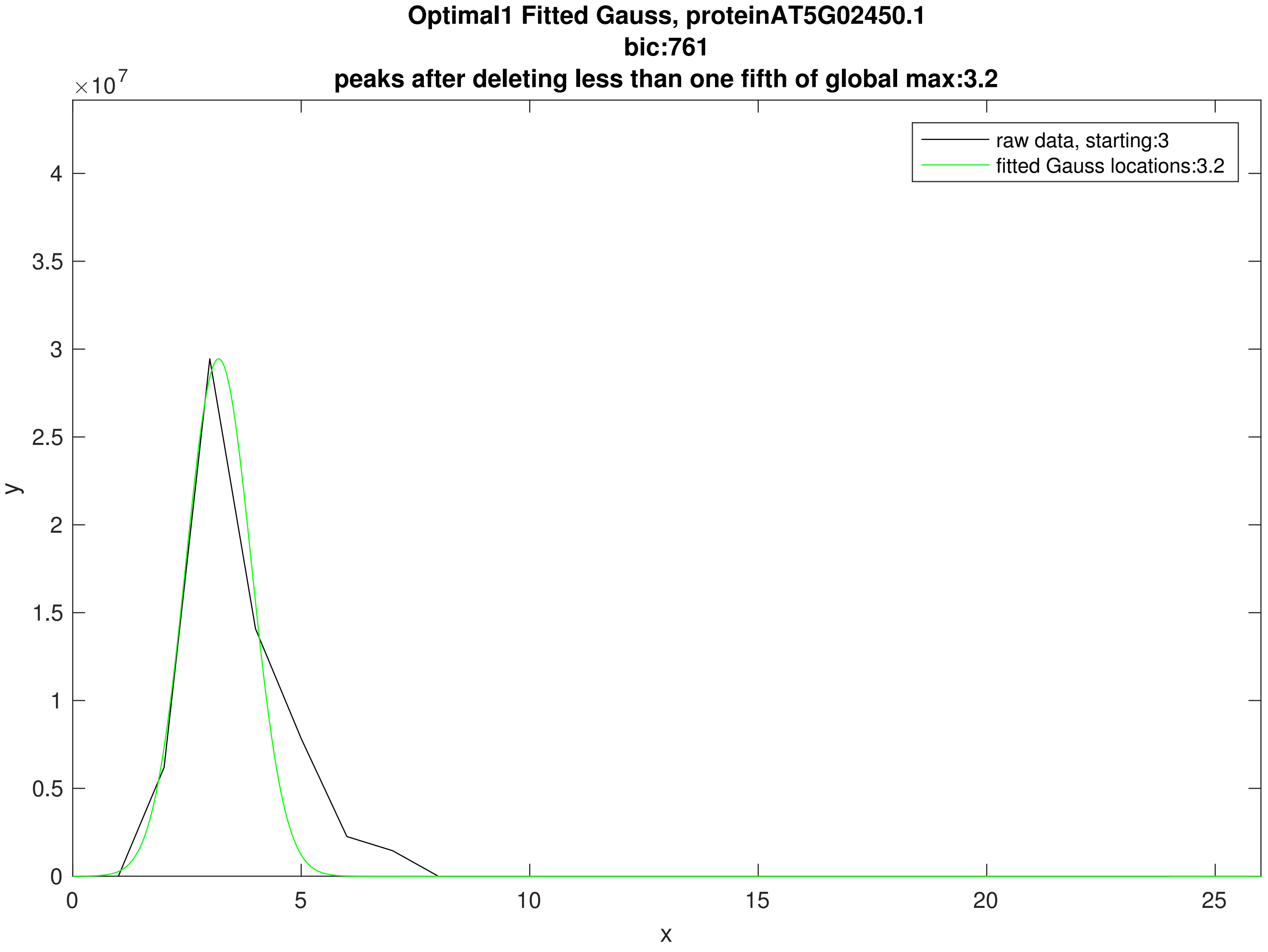

Optimal1 Fitted Gauss, proteinAT5G02450.1
bic:761
peaks after deleting less than one fifth of global max:3.2
×10^7
raw data, starting:3
fitted Gauss locations:3.2
4
3.5
3
2.5
2
1.5
1
0.5
0
y
0
5
10
15
20
25
x

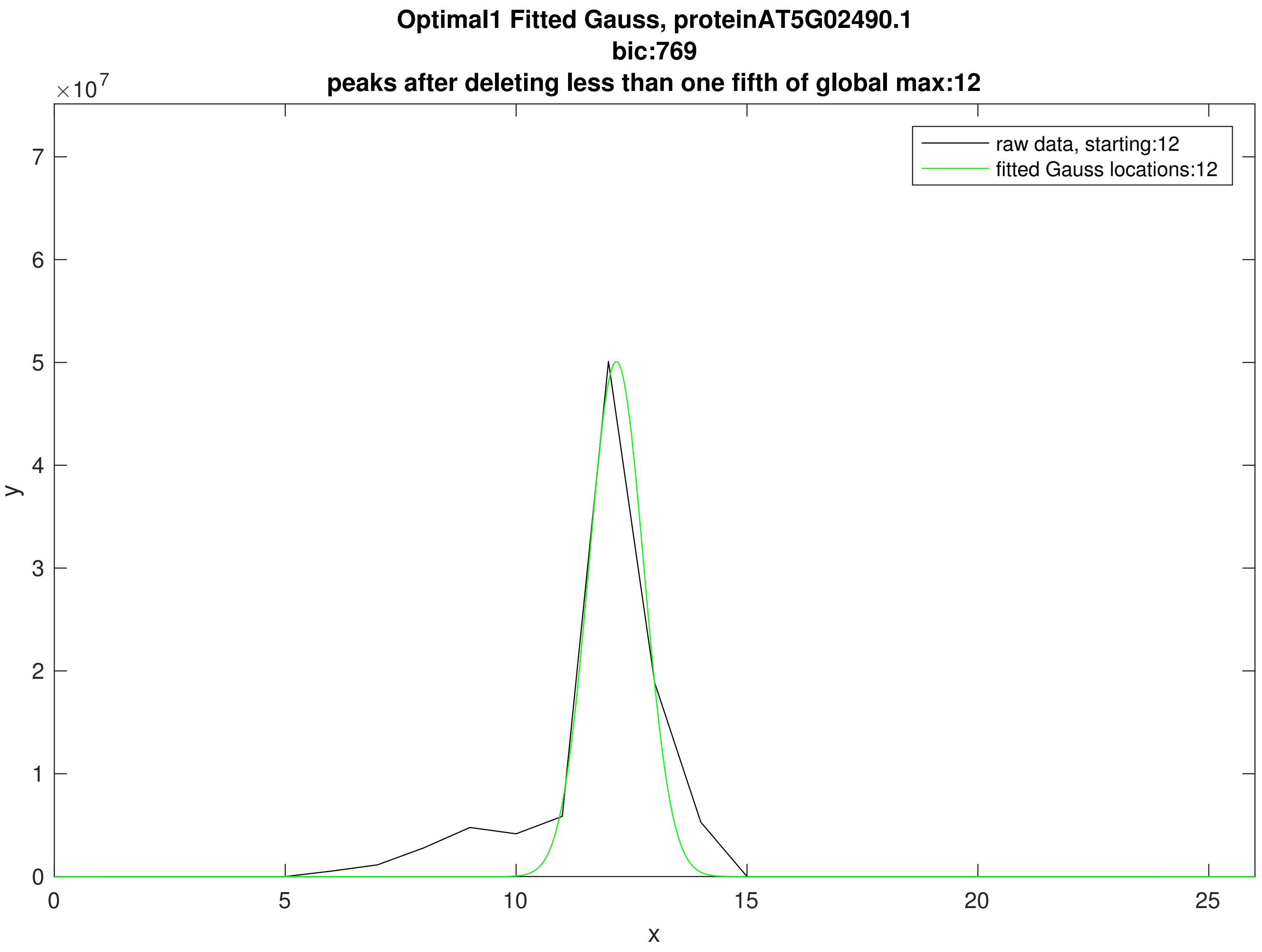
Optimal1 Fitted Gauss, proteinAT5G02490.1
bic:769
peaks after deleting less than one fifth of global max:12
×10^7
raw data, starting:12
fitted Gauss locations:12
y
x
0
1
2
3
4
5
6
7
0
5
10
15
20
25

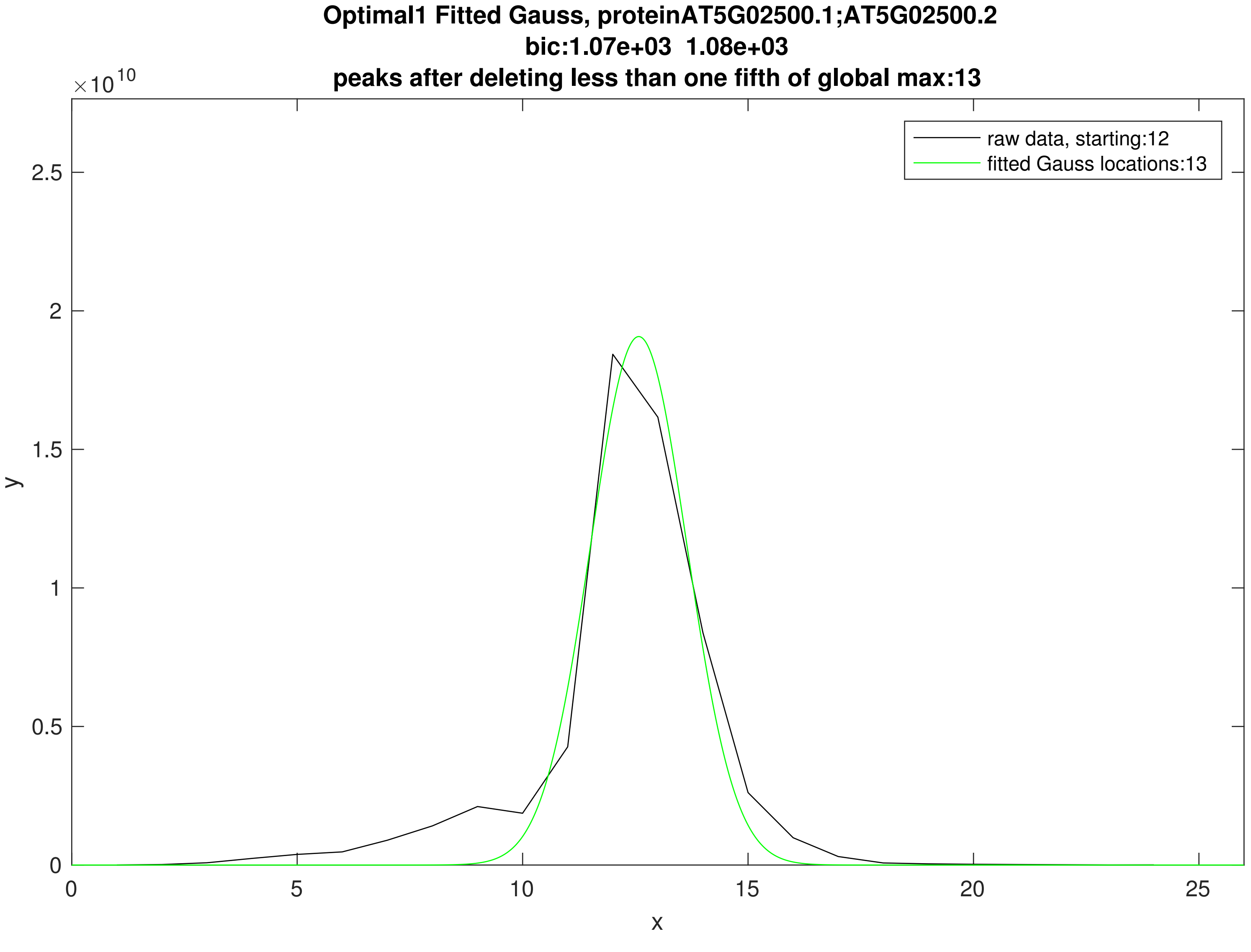

Optimal1 Fitted Gauss, proteinAT5G02500.1;AT5G02500.2
bic:1.07e+03  1.08e+03
peaks after deleting less than one fifth of global max:13
×10^10
2.5
2
1.5
1
0.5
0
y
0
5
10
15
20
25
x
raw data, starting:12
fitted Gauss locations:13

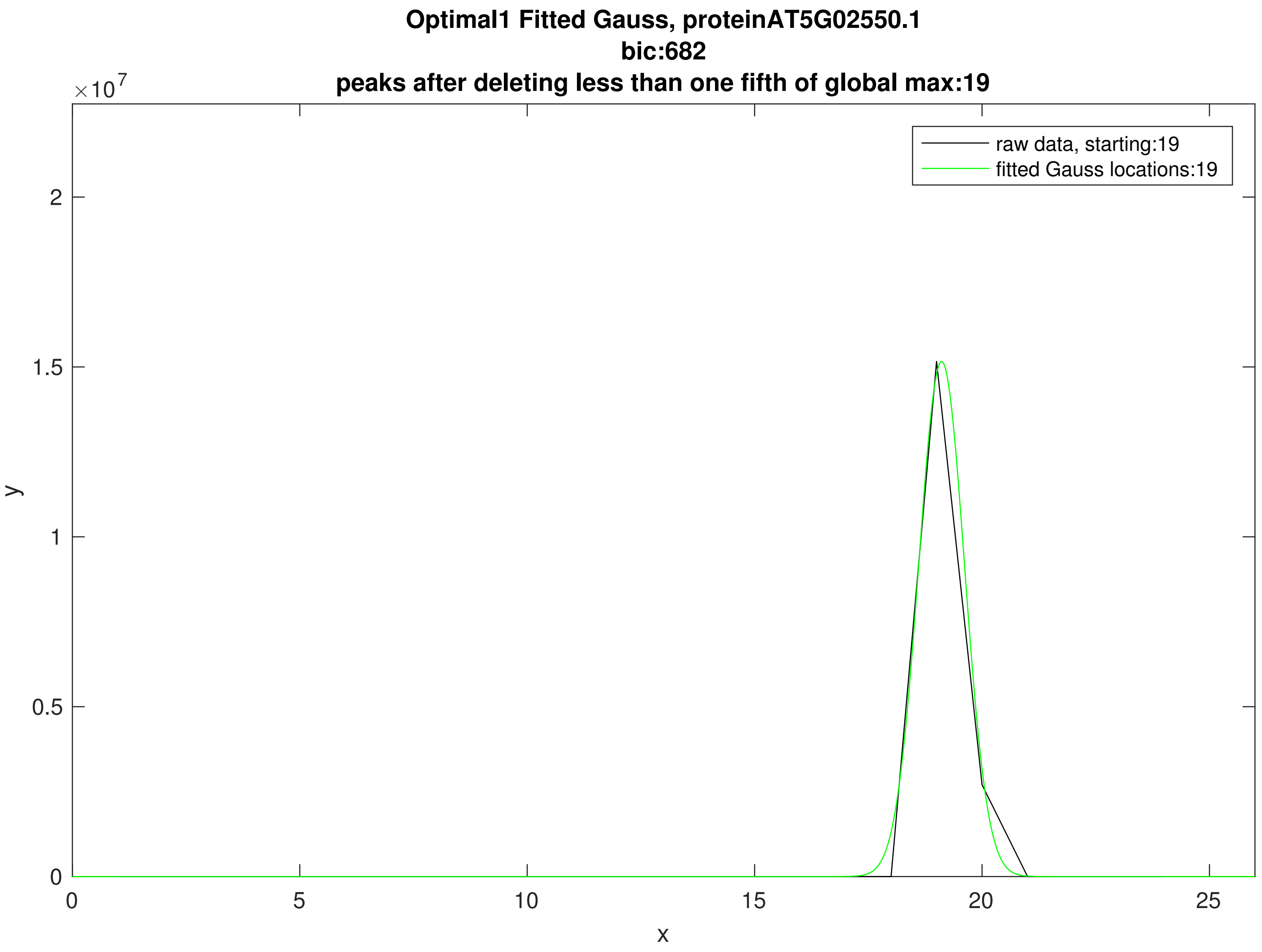
Optimal1 Fitted Gauss, proteinAT5G02550.1
bic:682
peaks after deleting less than one fifth of global max:19
×10^7
raw data, starting:19
fitted Gauss locations:19
y
x
0
0.5
1
1.5
2
0
5
10
15
20
25

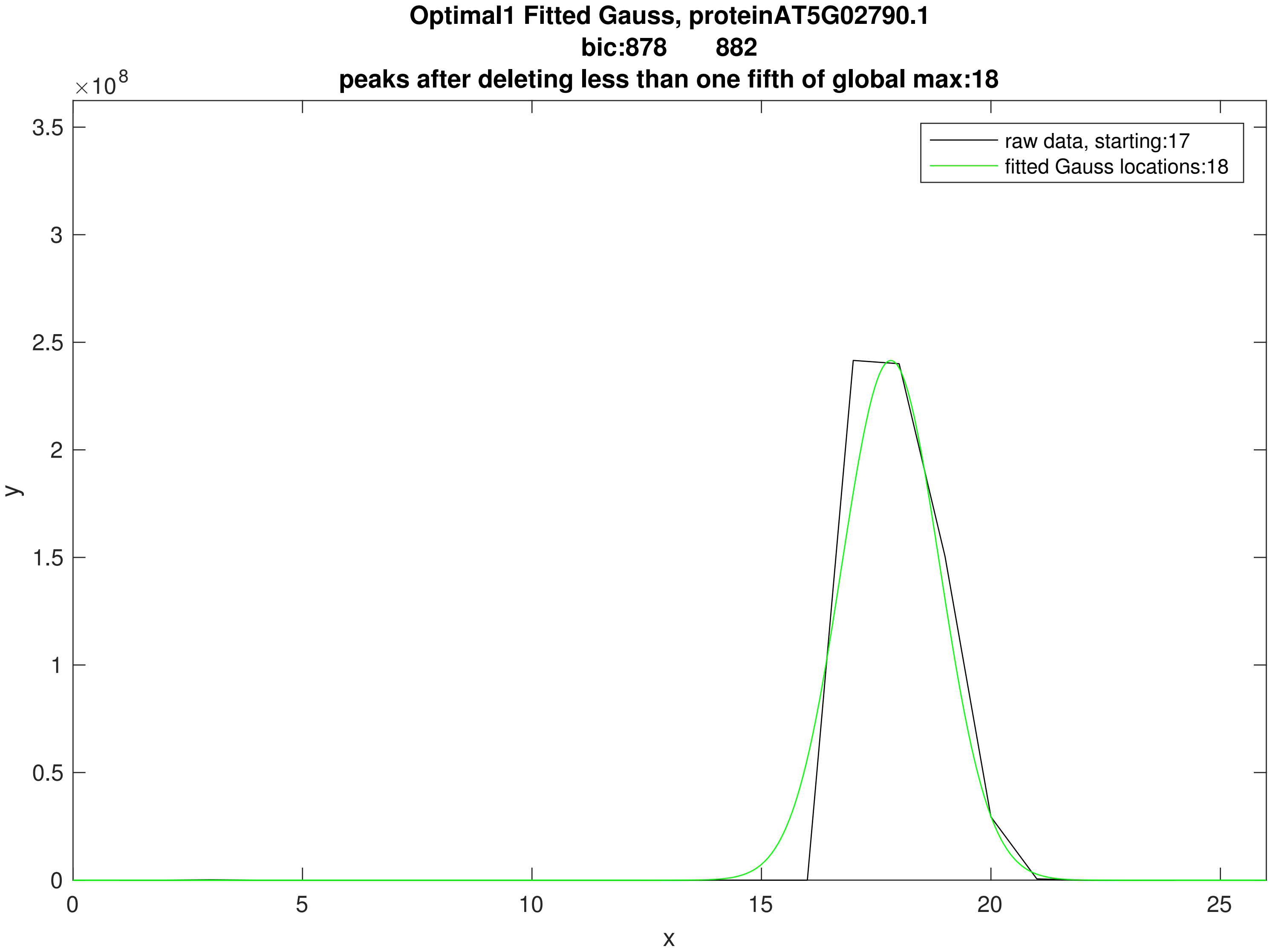

Optimal1 Fitted Gauss, proteinAT5G02790.1
bic:878 882
peaks after deleting less than one fifth of global max:18
×10^8
raw data, starting:17
fitted Gauss locations:18
y
x
0
0.5
1
1.5
2
2.5
3
3.5
5
10
15
20
25

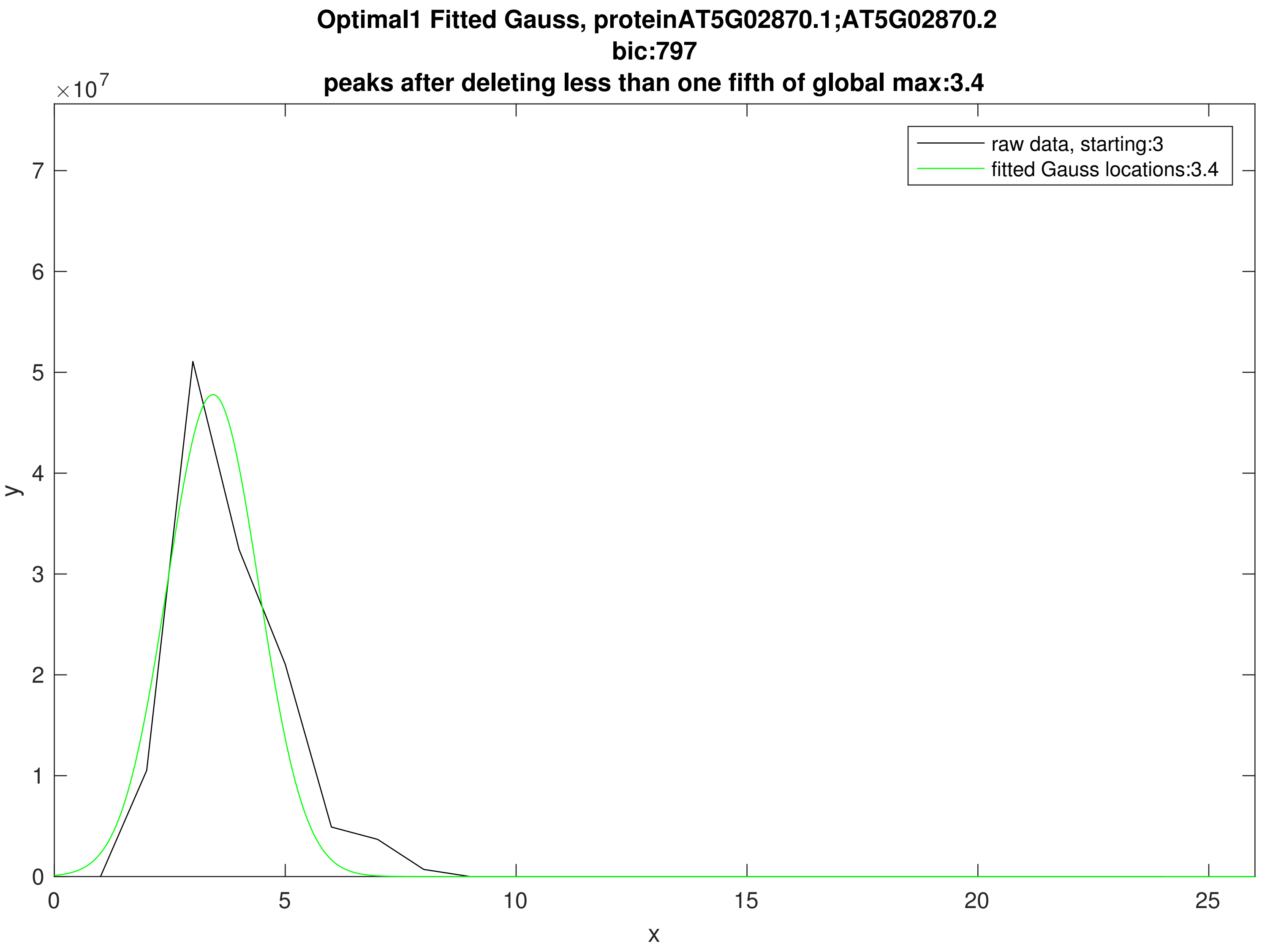
Optimal1 Fitted Gauss, proteinAT5G02870.1;AT5G02870.2
bic:797
peaks after deleting less than one fifth of global max:3.4
×10^7
raw data, starting:3
fitted Gauss locations:3.4
y
x
0
1
2
3
4
5
6
7
0
5
10
15
20
25

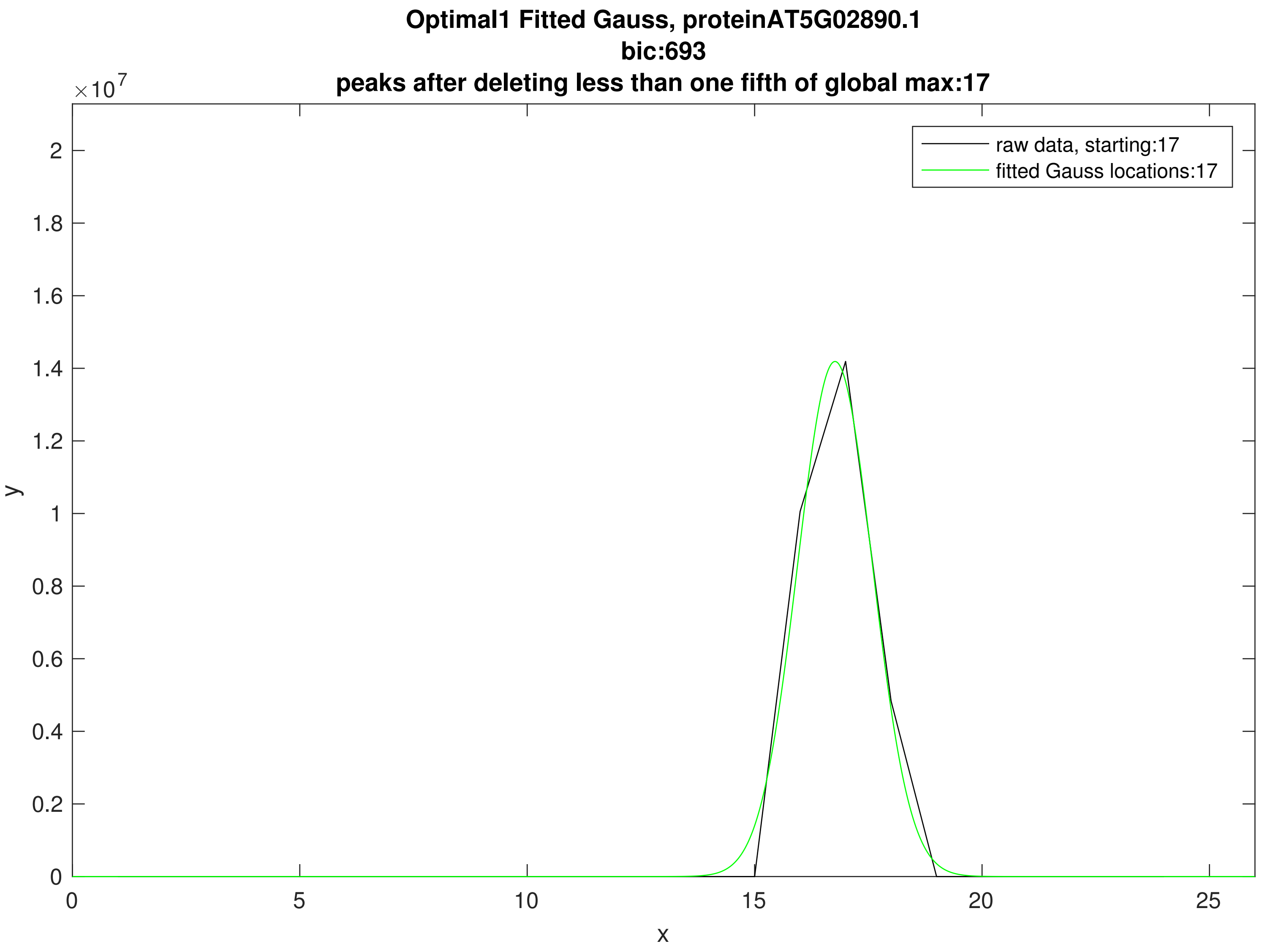

Optimal1 Fitted Gauss, proteinAT5G02890.1
bic:693
peaks after deleting less than one fifth of global max:17
×10^7
2
1.8
1.6
1.4
1.2
1
0.8
0.6
0.4
0.2
0
y
0
5
10
15
20
25
x
raw data, starting:17
fitted Gauss locations:17

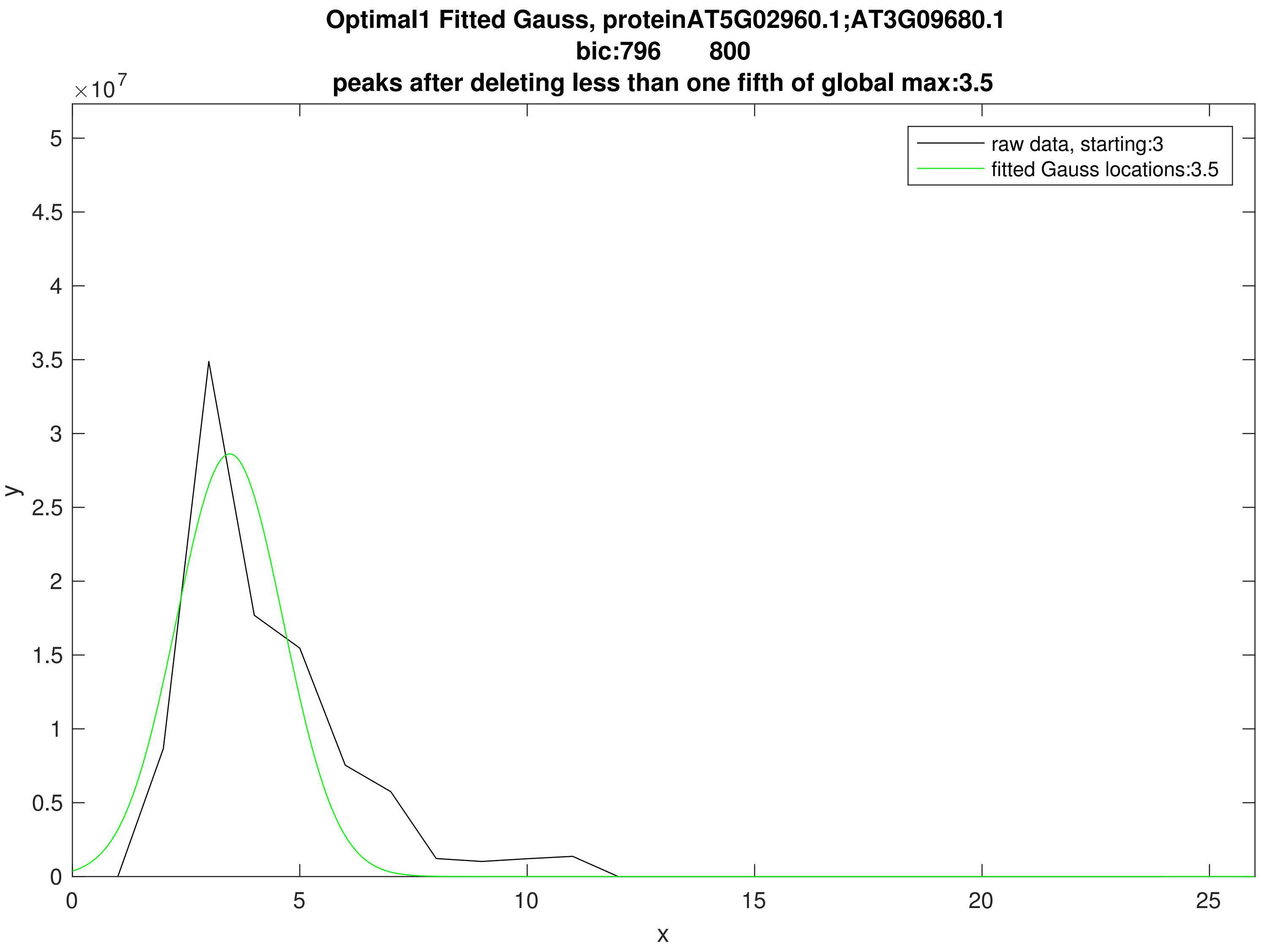
Optimal1 Fitted Gauss, proteinAT5G02960.1;AT3G09680.1
bic:796 800
peaks after deleting less than one fifth of global max:3.5
×10^7
5
4.5
4
3.5
3
2.5
2
1.5
1
0.5
0
y
0
5
10
15
20
25
x
raw data, starting:3
fitted Gauss locations:3.5

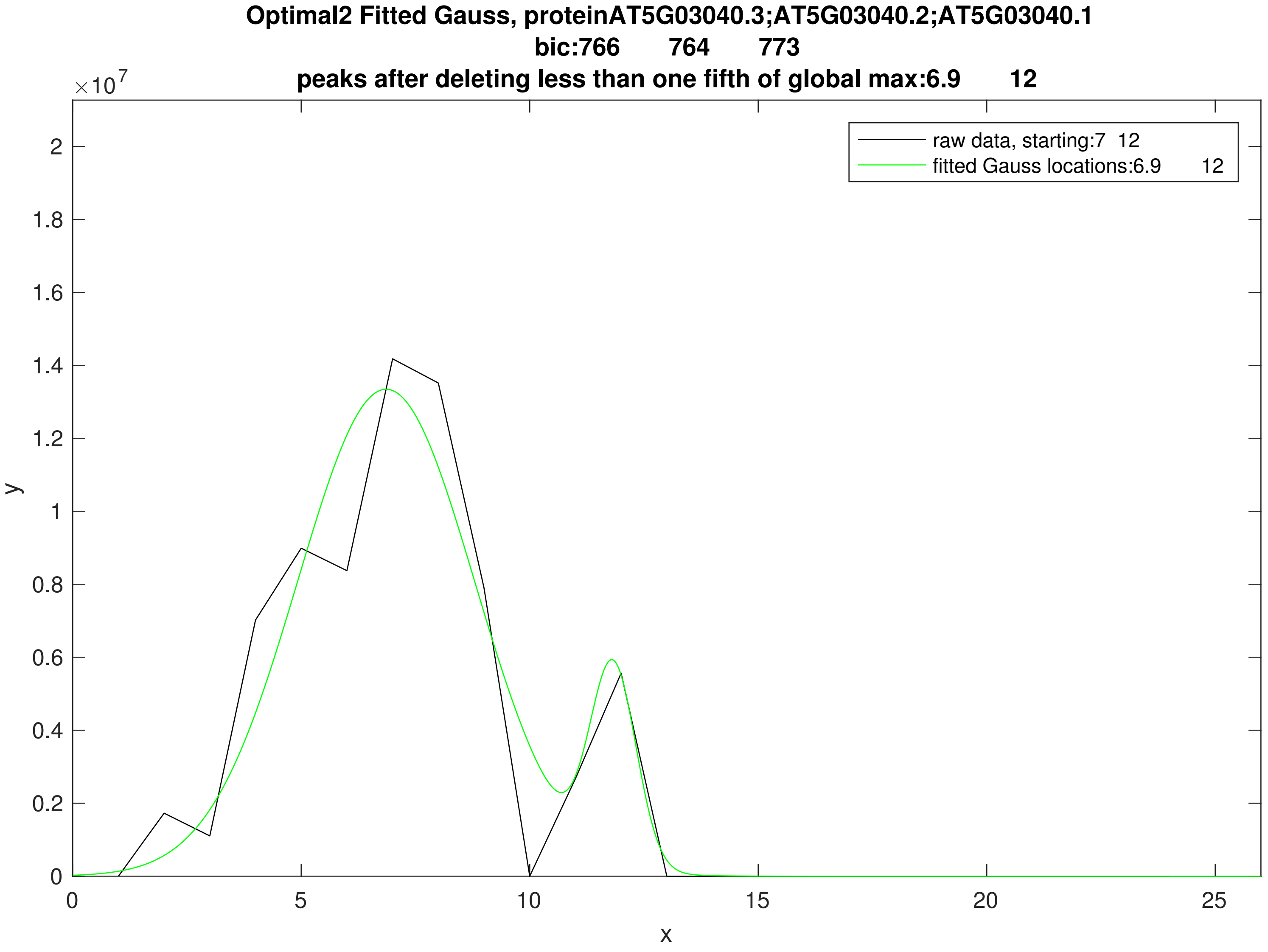
Optimal2 Fitted Gauss, proteinAT5G03040.3;AT5G03040.2;AT5G03040.1
bic:766 764 773
peaks after deleting less than one fifth of global max:6.9 12
×10^7
raw data, starting:7 12
fitted Gauss locations:6.9 12
y
x
0
0.2
0.4
0.6
0.8
1
1.2
1.4
1.6
1.8
2
0
5
10
15
20
25

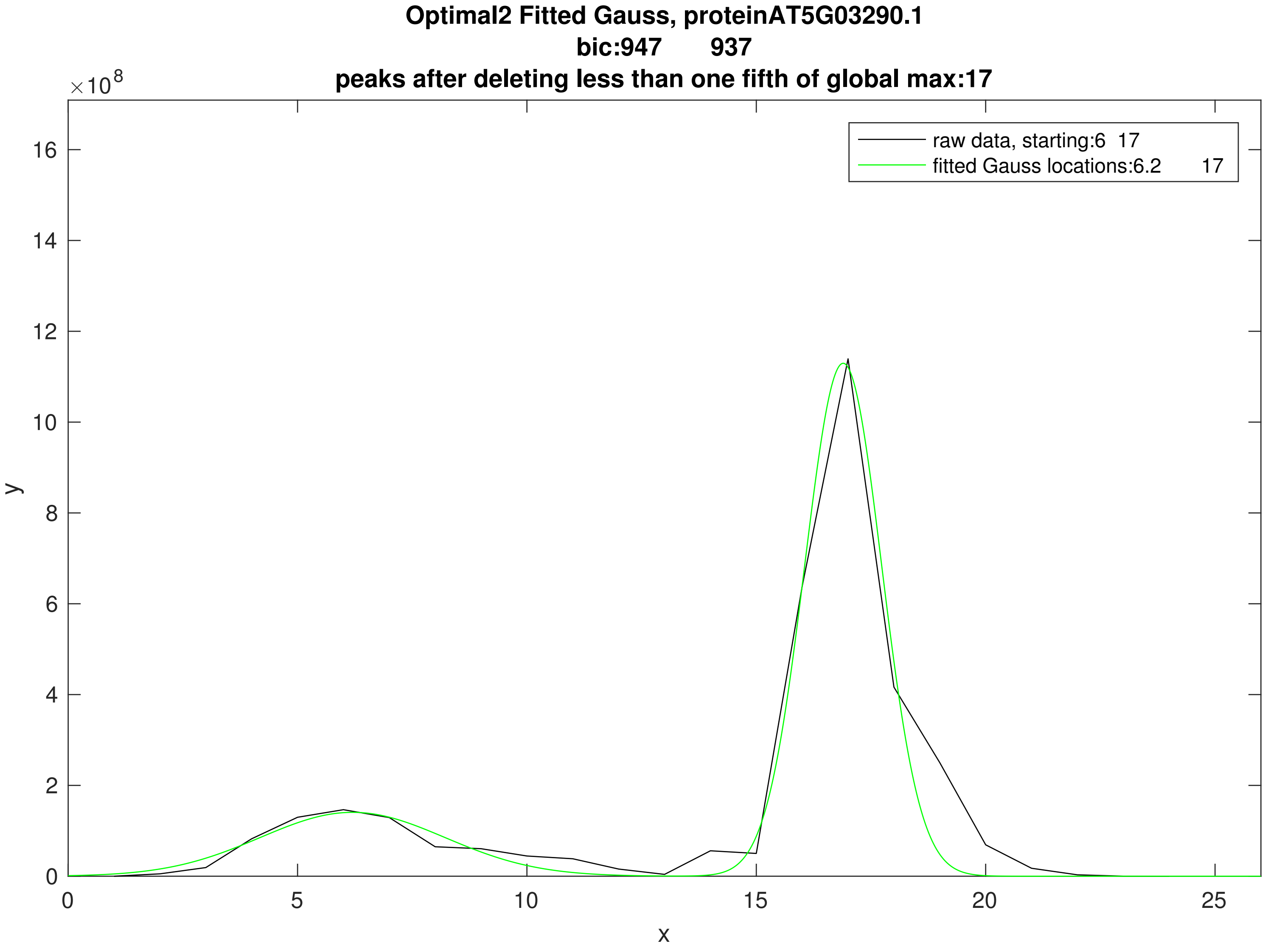
Optimal2 Fitted Gauss, proteinAT5G03290.1
bic:947 937
peaks after deleting less than one fifth of global max:17
×10^8
raw data, starting:6 17
fitted Gauss locations:6.2 17
y
x
0
2
4
6
8
10
12
14
16
0
5
10
15
20
25

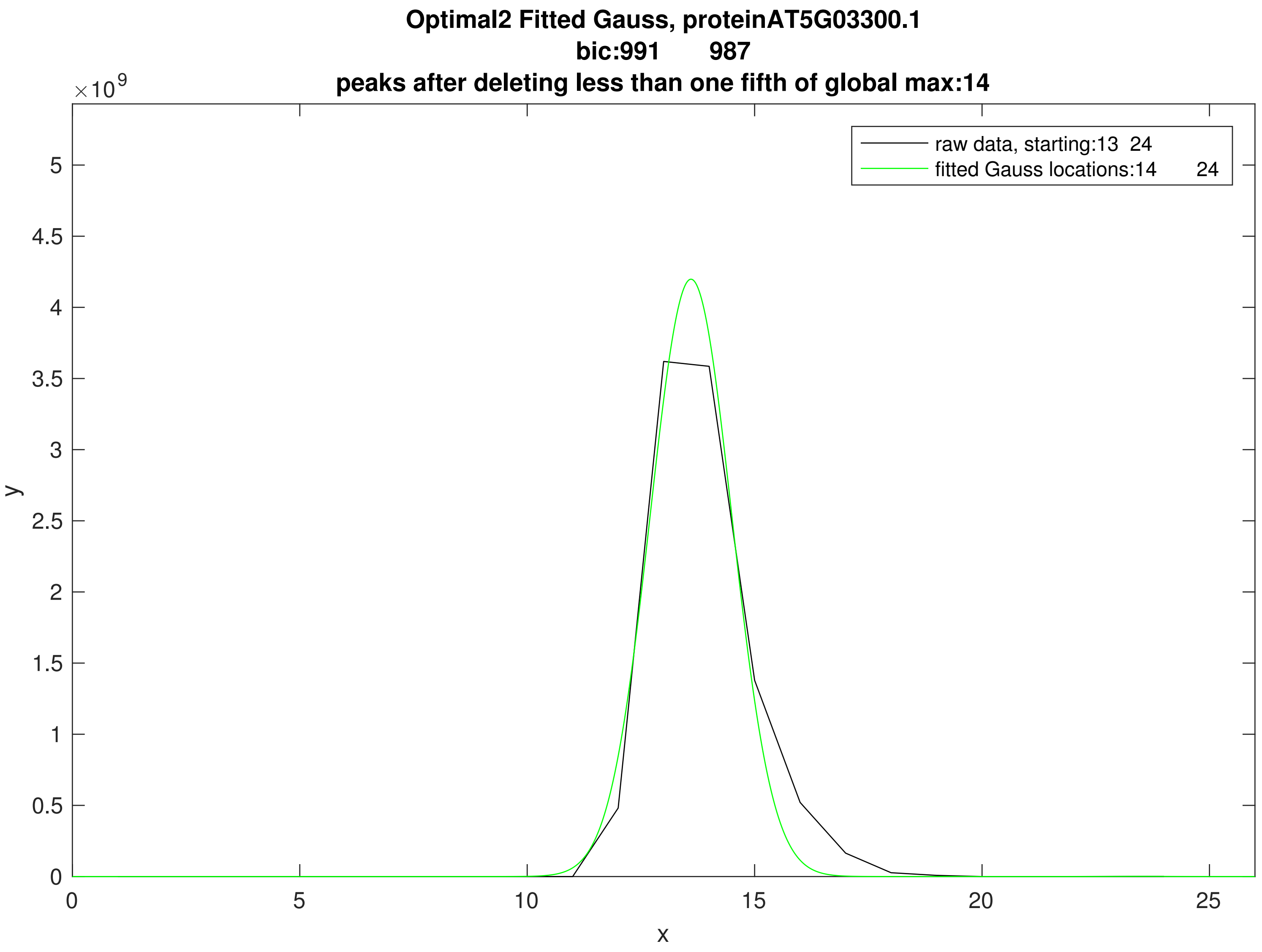
Optimal2 Fitted Gauss, proteinAT5G03300.1
bic:991 987
peaks after deleting less than one fifth of global max:14
×10^9
raw data, starting:13 24
fitted Gauss locations:14 24
y
x
0
0.5
1
1.5
2
2.5
3
3.5
4
4.5
5
0
5
10
15
20
25

**Optimal1 Fitted Gauss, proteinAT5G03340.1**
**bic:771**
**peaks after deleting less than one fifth of global max:5.3**

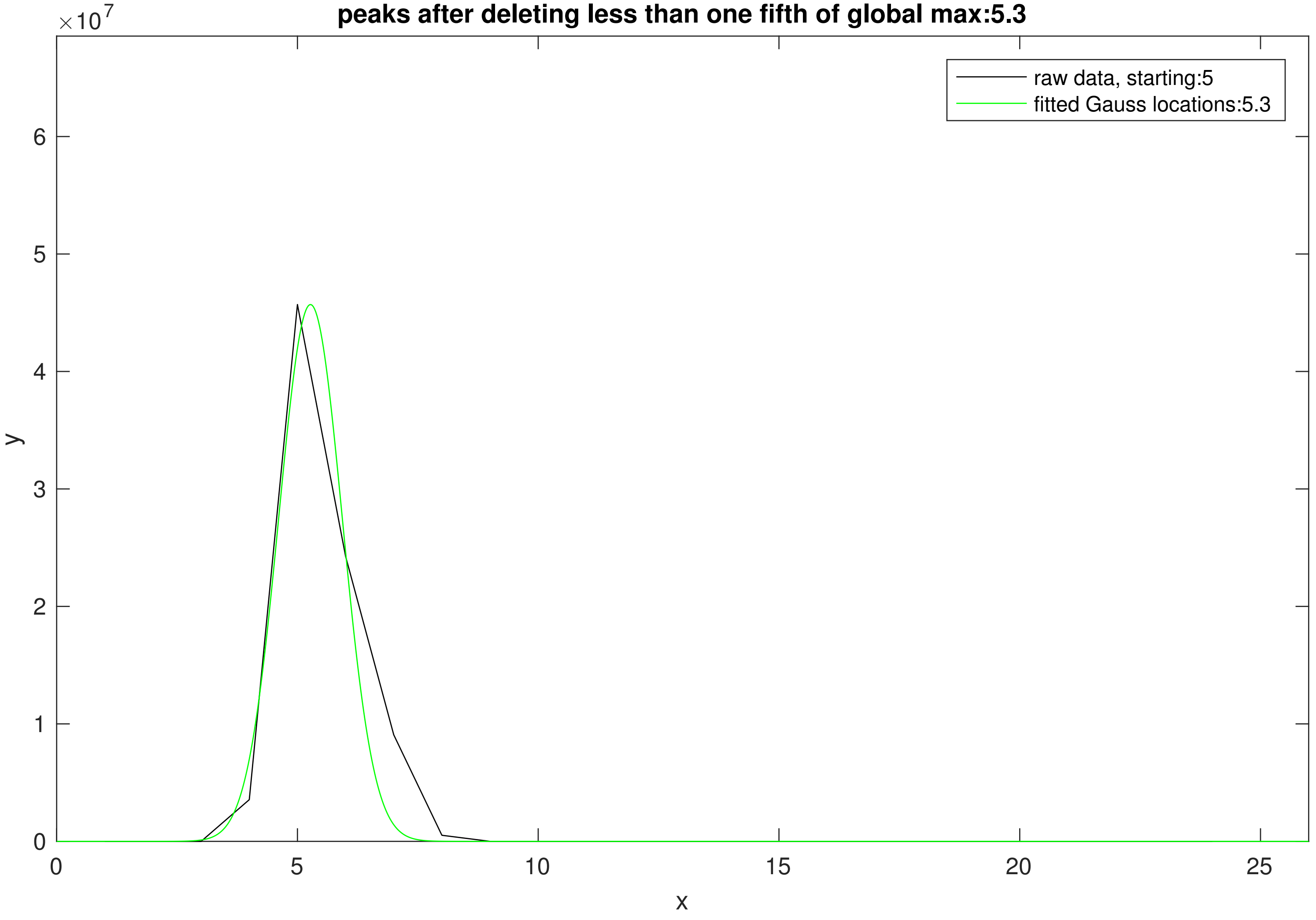

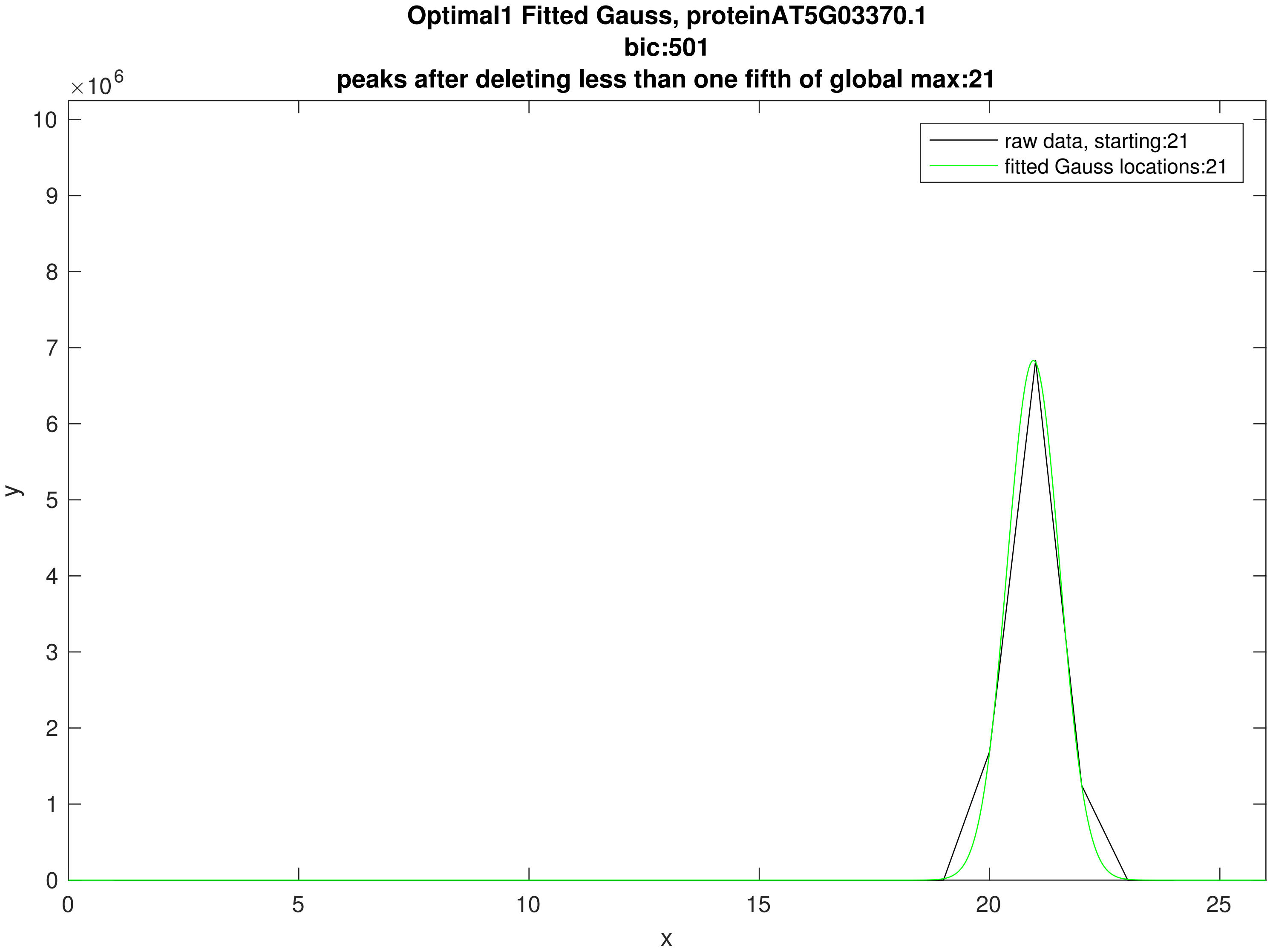

Optimal1 Fitted Gauss, proteinAT5G03370.1
bic:501
peaks after deleting less than one fifth of global max:21
×10^6
raw data, starting:21
fitted Gauss locations:21
y
x
0
1
2
3
4
5
6
7
8
9
10
0
5
10
15
20
25

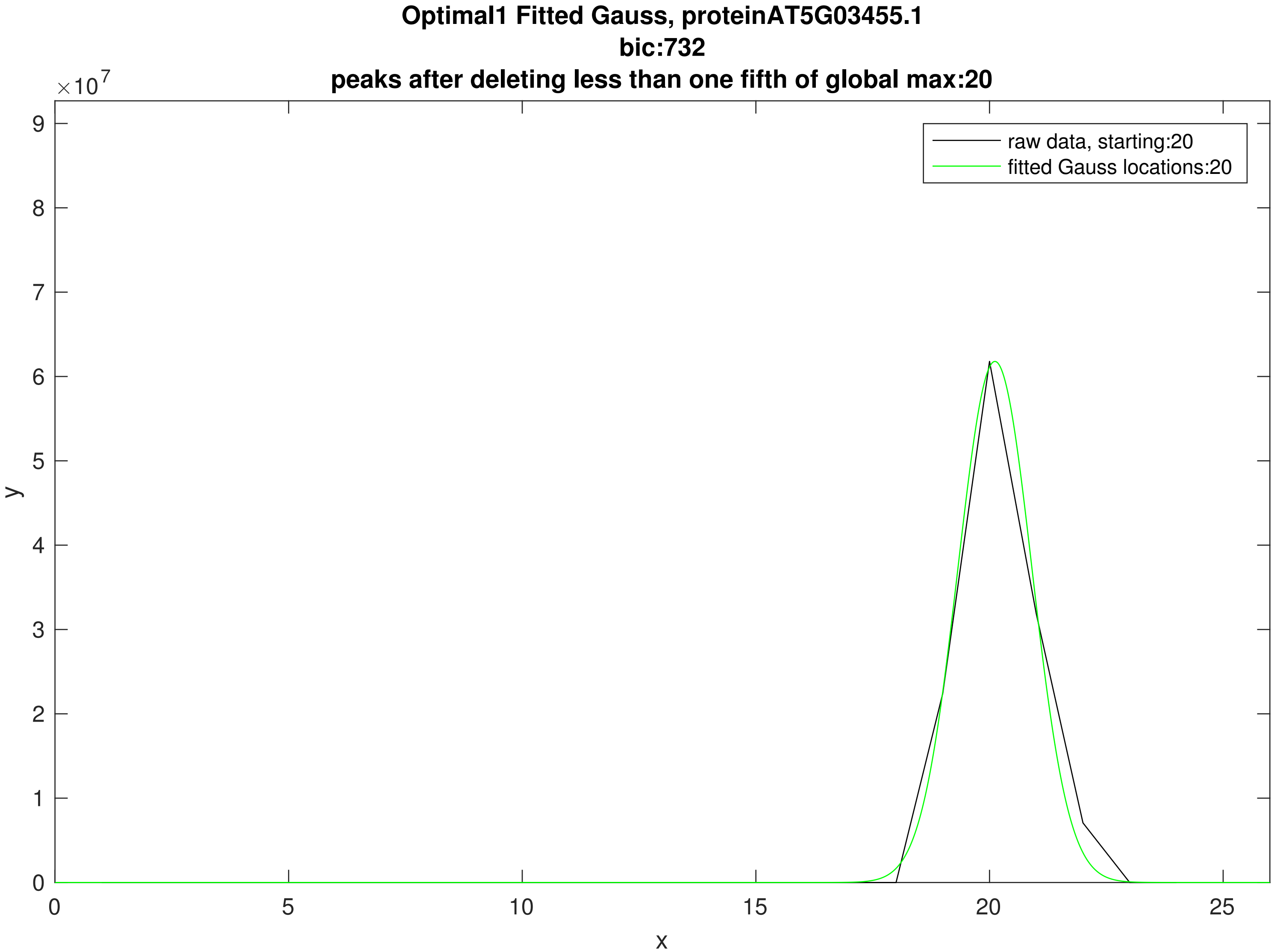
Optimal1 Fitted Gauss, proteinAT5G03455.1
bic:732
peaks after deleting less than one fifth of global max:20
$\times 10^7$
raw data, starting:20
fitted Gauss locations:20
y
x
0
1
2
3
4
5
6
7
8
9
0
5
10
15
20
25

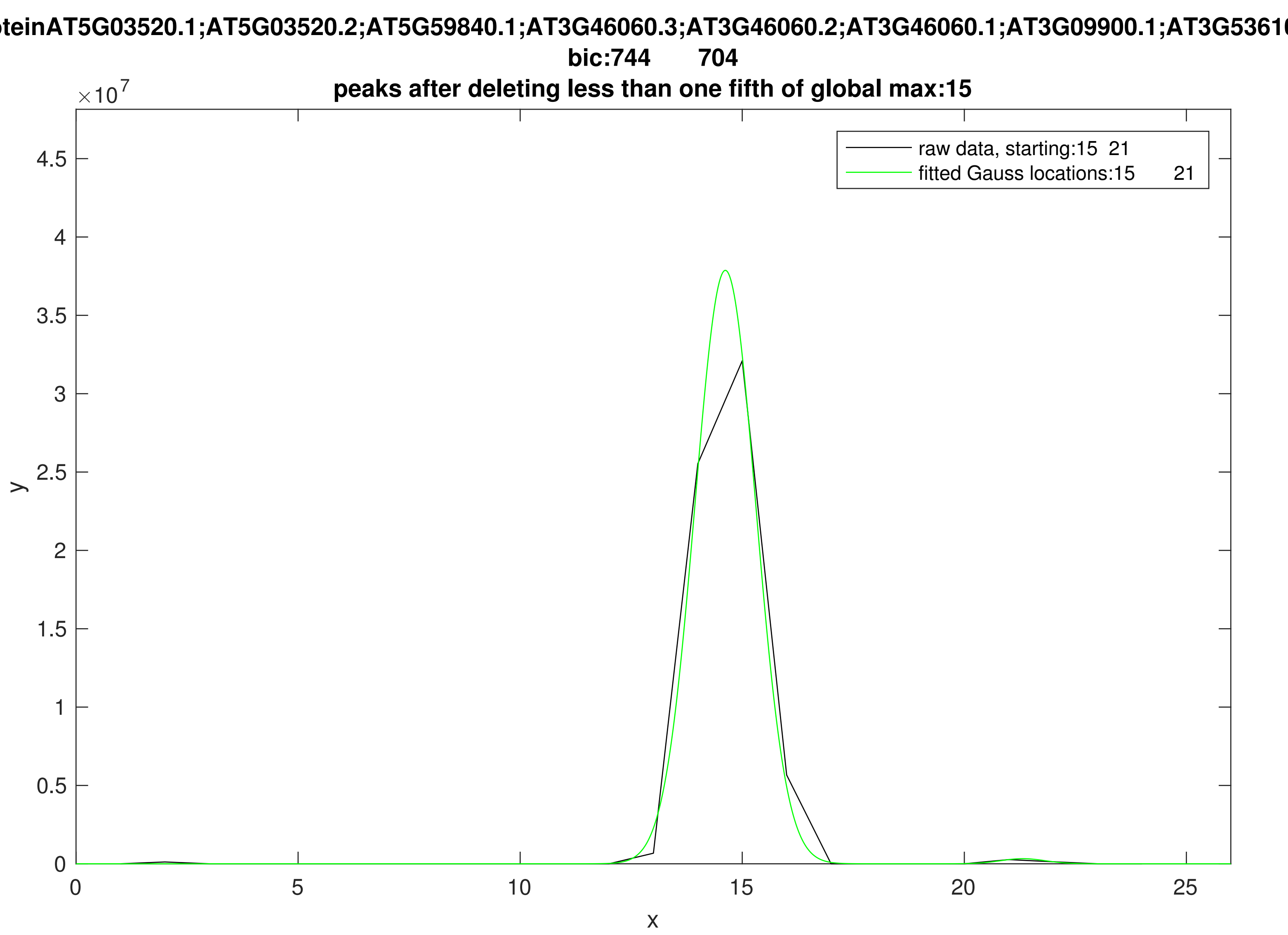
s, proteinAT5G03520.1;AT5G03520.2;AT5G59840.1;AT3G46060.3;AT3G46060.2;AT3G46060.1;AT3G09900.1;AT3G5361
bic:744 704
peaks after deleting less than one fifth of global max:15
×10^7
raw data, starting:15 21
fitted Gauss locations:15 21
y
x
0
0.5
1
1.5
2
2.5
3
3.5
4
4.5
5
10
15
20
25

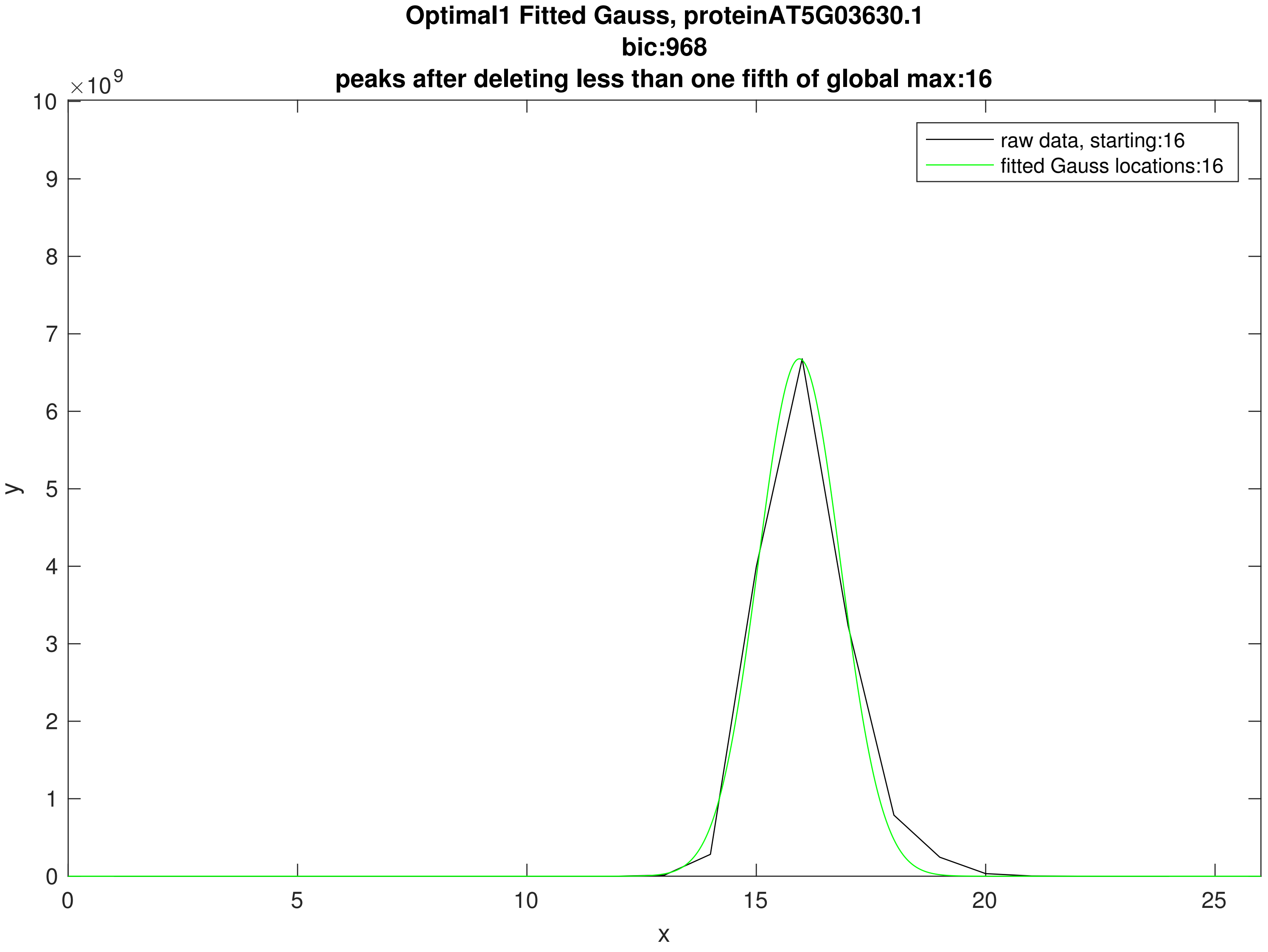
Optimal1 Fitted Gauss, proteinAT5G03630.1
bic:968
peaks after deleting less than one fifth of global max:16
×10^9
10
9
8
7
6
5
4
3
2
1
0
y
0
5
10
15
20
25
x
raw data, starting:16
fitted Gauss locations:16

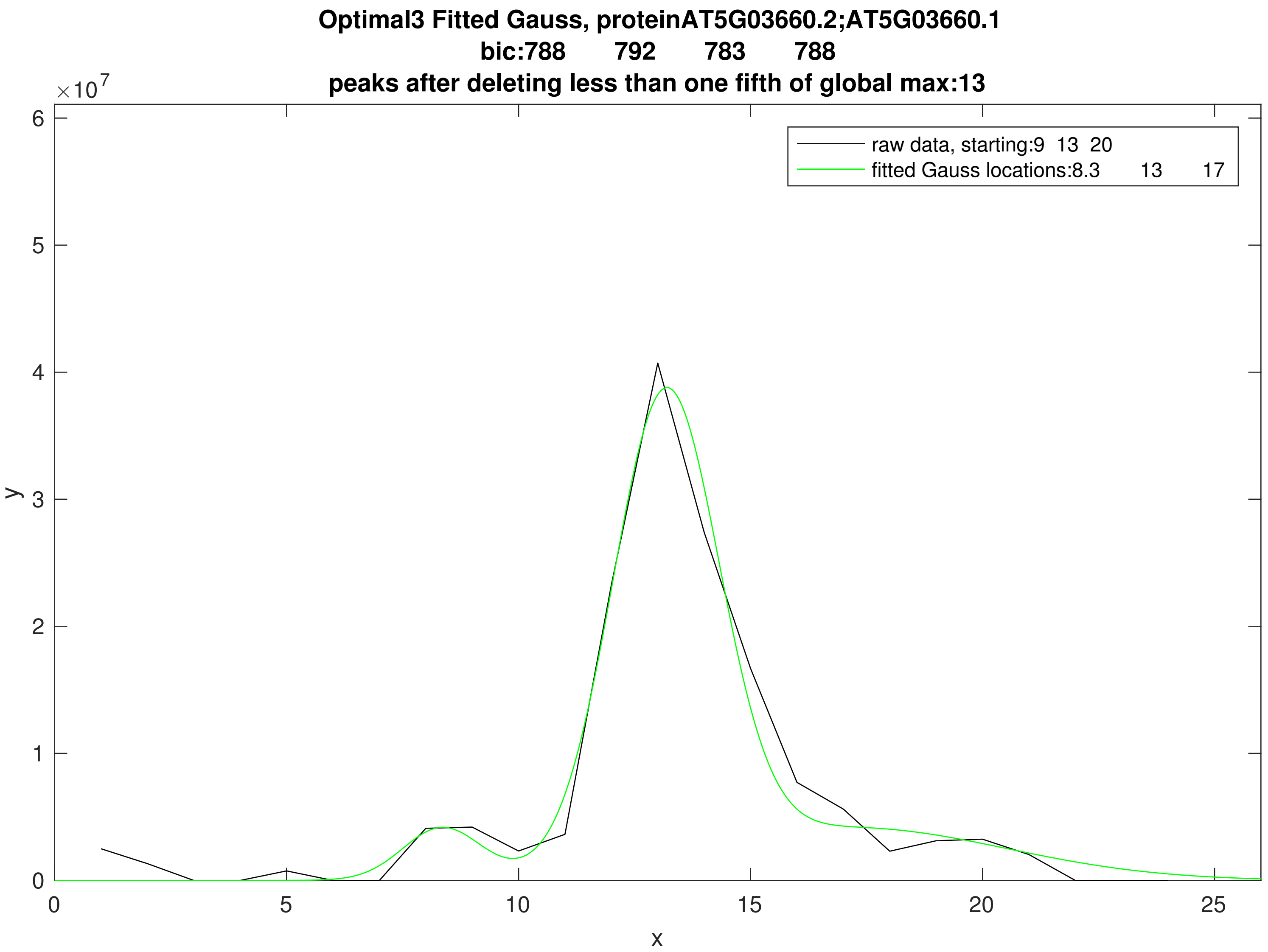
Optimal3 Fitted Gauss, proteinAT5G03660.2;AT5G03660.1
bic:788 792 783 788
peaks after deleting less than one fifth of global max:13
×10^7
raw data, starting:9 13 20
fitted Gauss locations:8.3 13 17
y
x
0
1
2
3
4
5
6
0
5
10
15
20
25

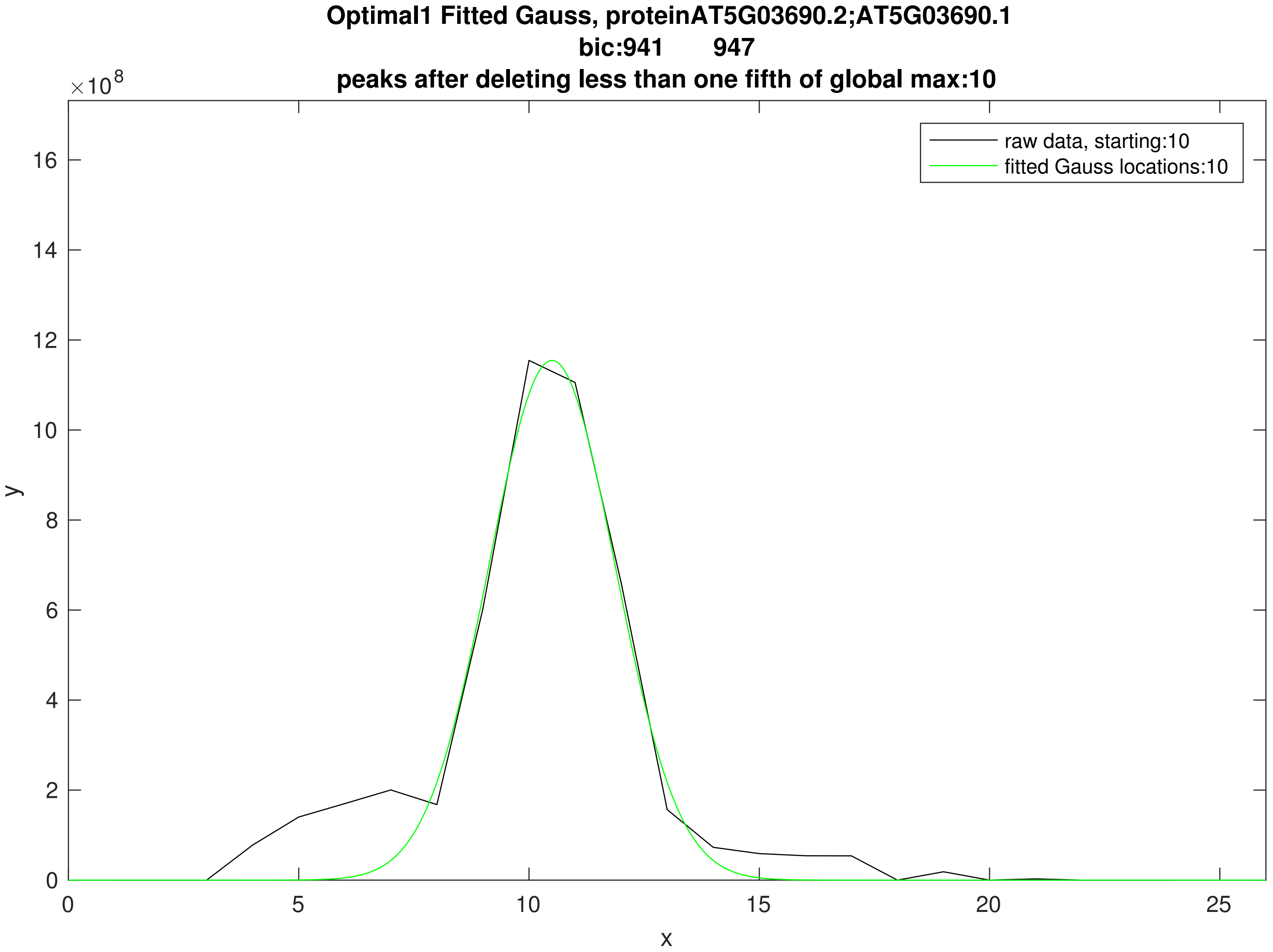
Optimal1 Fitted Gauss, proteinAT5G03690.2;AT5G03690.1
bic:941 947
peaks after deleting less than one fifth of global max:10
×10^8
raw data, starting:10
fitted Gauss locations:10
y
x
0
2
4
6
8
10
12
14
16
0
5
10
15
20
25

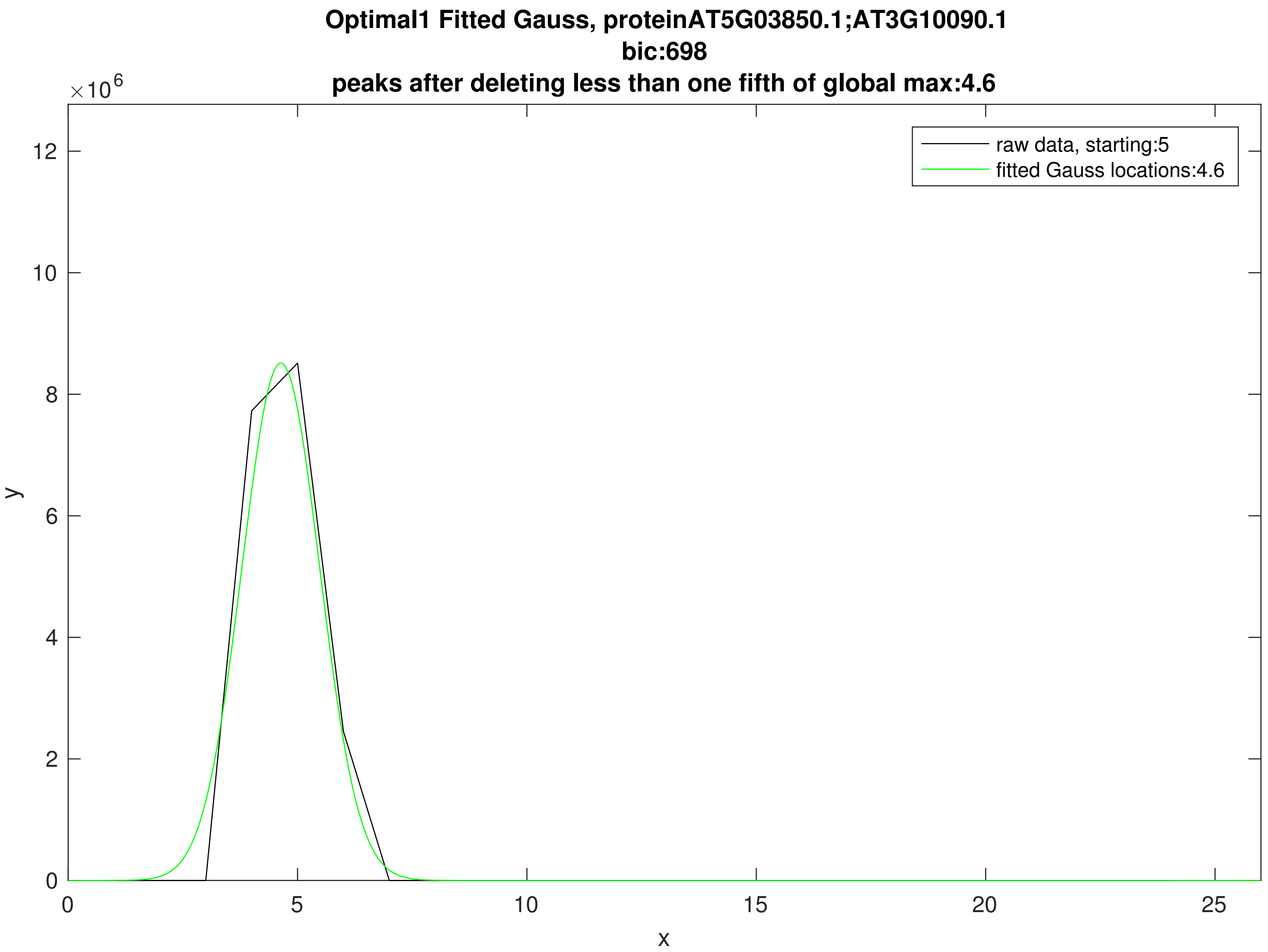
Optimal1 Fitted Gauss, proteinAT5G03850.1;AT3G10090.1
bic:698
peaks after deleting less than one fifth of global max:4.6
×10^6
raw data, starting:5
fitted Gauss locations:4.6
y
x
0
2
4
6
8
10
12
0
5
10
15
20
25

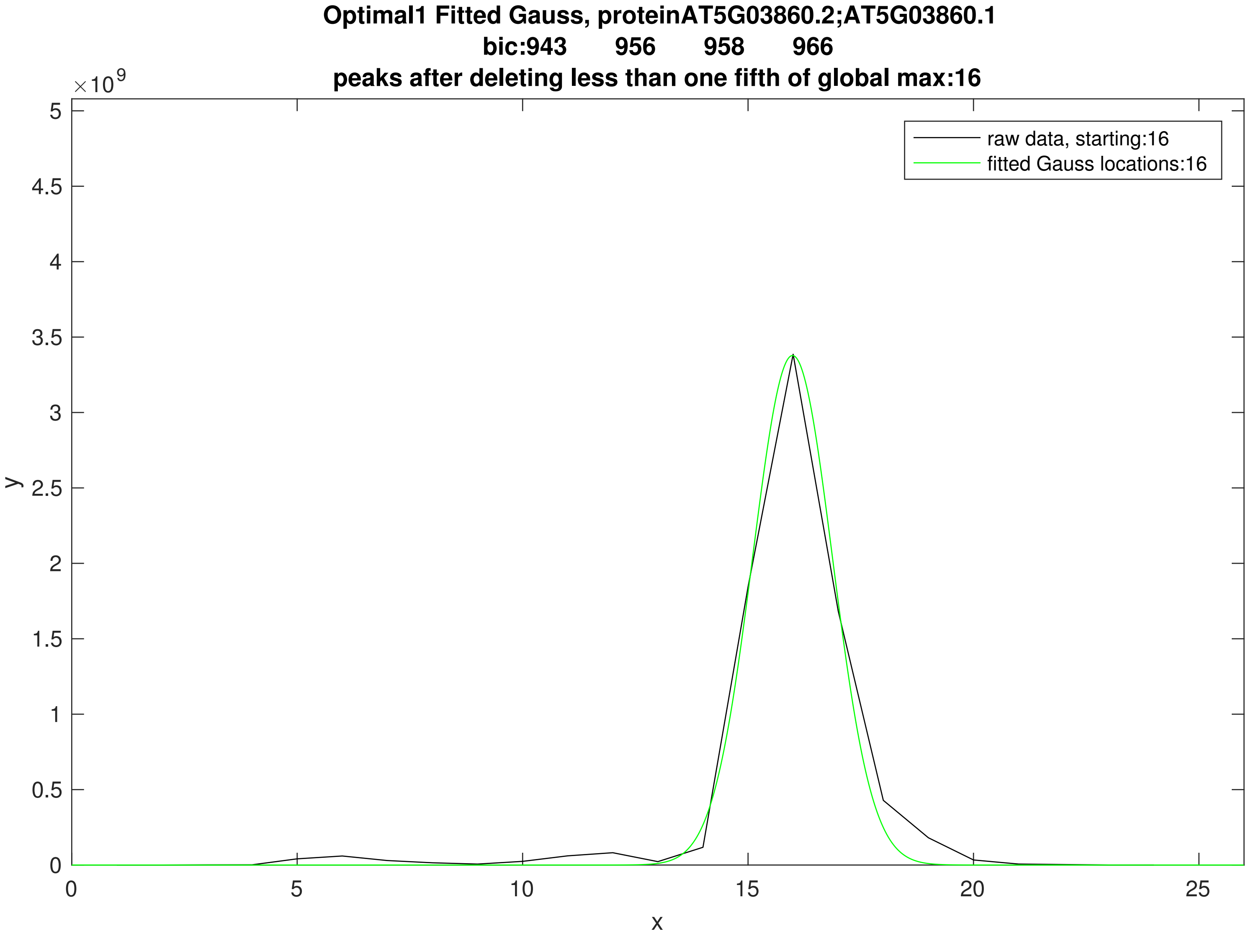

Optimal1 Fitted Gauss, proteinAT5G03860.2;AT5G03860.1
bic:943 956 958 966
peaks after deleting less than one fifth of global max:16
×10^9
5
4.5
4
3.5
3
2.5
2
1.5
1
0.5
0
y
0
5
10
15
20
25
x
raw data, starting:16
fitted Gauss locations:16

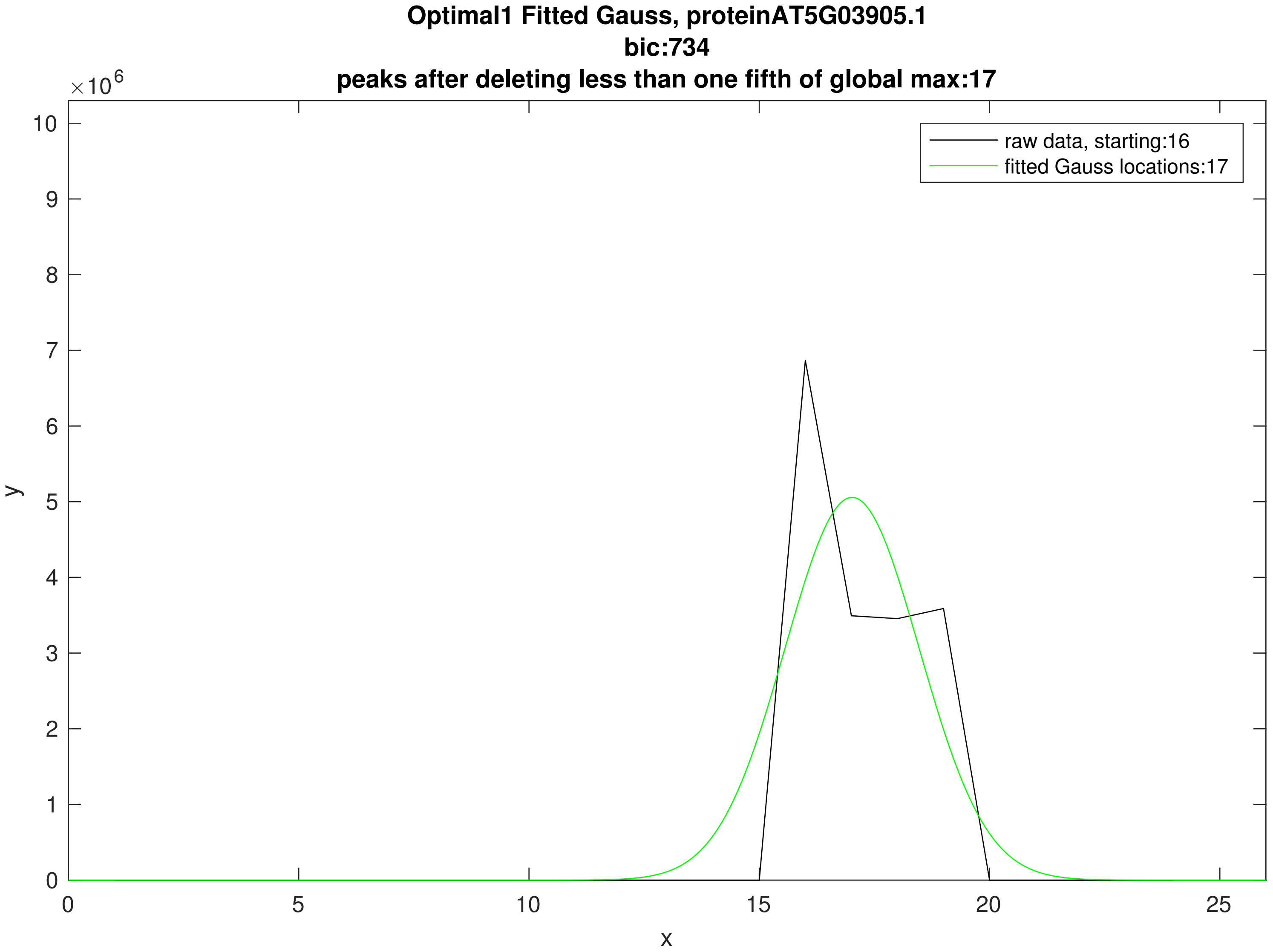
Optimal1 Fitted Gauss, proteinAT5G03905.1
bic:734
peaks after deleting less than one fifth of global max:17
×10^6
raw data, starting:16
fitted Gauss locations:17
y
x
0
1
2
3
4
5
6
7
8
9
10
0
5
10
15
20
25

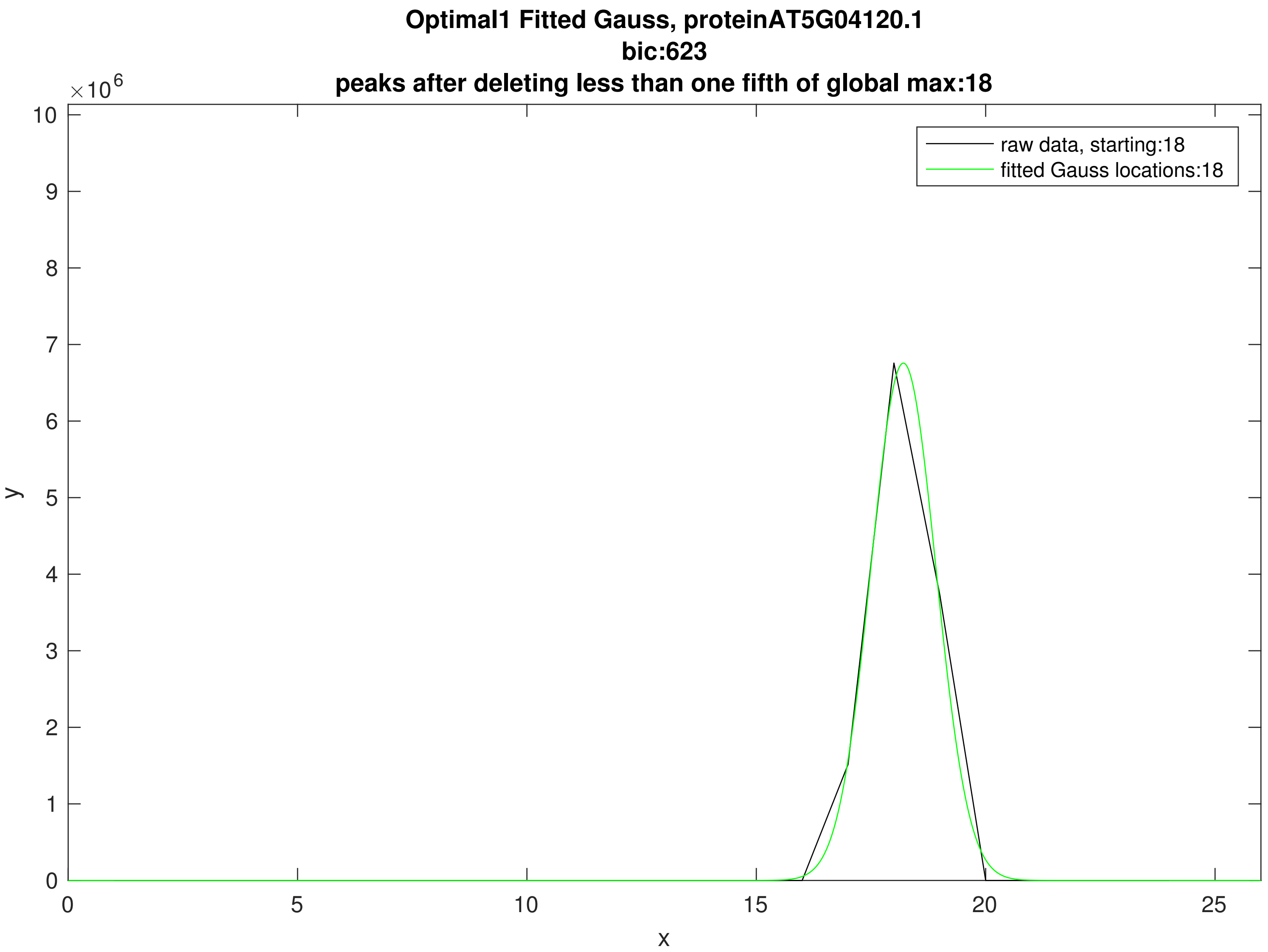
Optimal1 Fitted Gauss, proteinAT5G04120.1
bic:623
peaks after deleting less than one fifth of global max:18
×10^6
10
9
8
7
6
5
4
3
2
1
0
y
0
5
10
15
20
25
x
raw data, starting:18
fitted Gauss locations:18

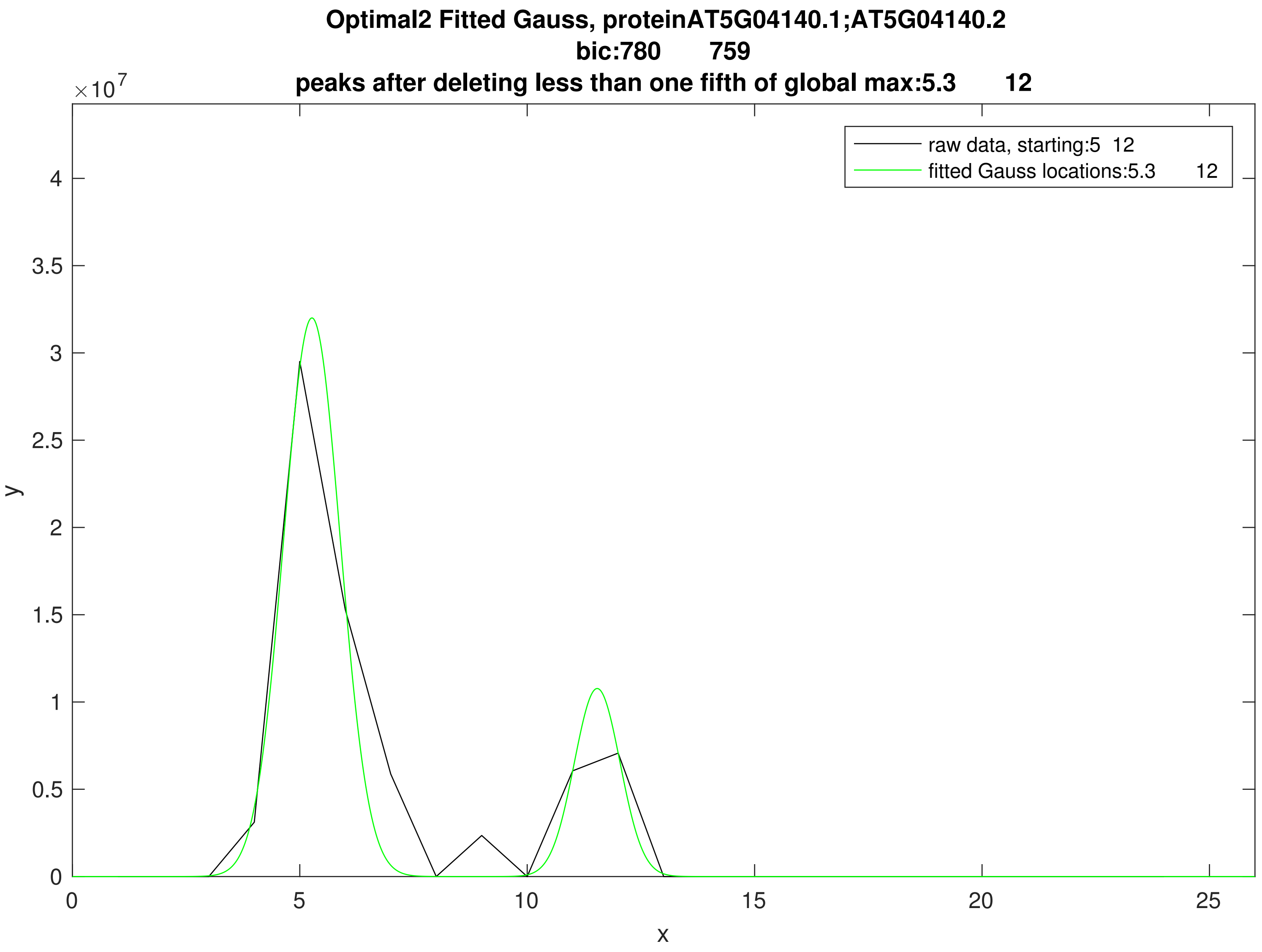
Optimal2 Fitted Gauss, proteinAT5G04140.1;AT5G04140.2
bic:780 759
peaks after deleting less than one fifth of global max:5.3 12
×10^7
raw data, starting:5 12
fitted Gauss locations:5.3 12
y
x
0
0.5
1
1.5
2
2.5
3
3.5
4
5
10
15
20
25

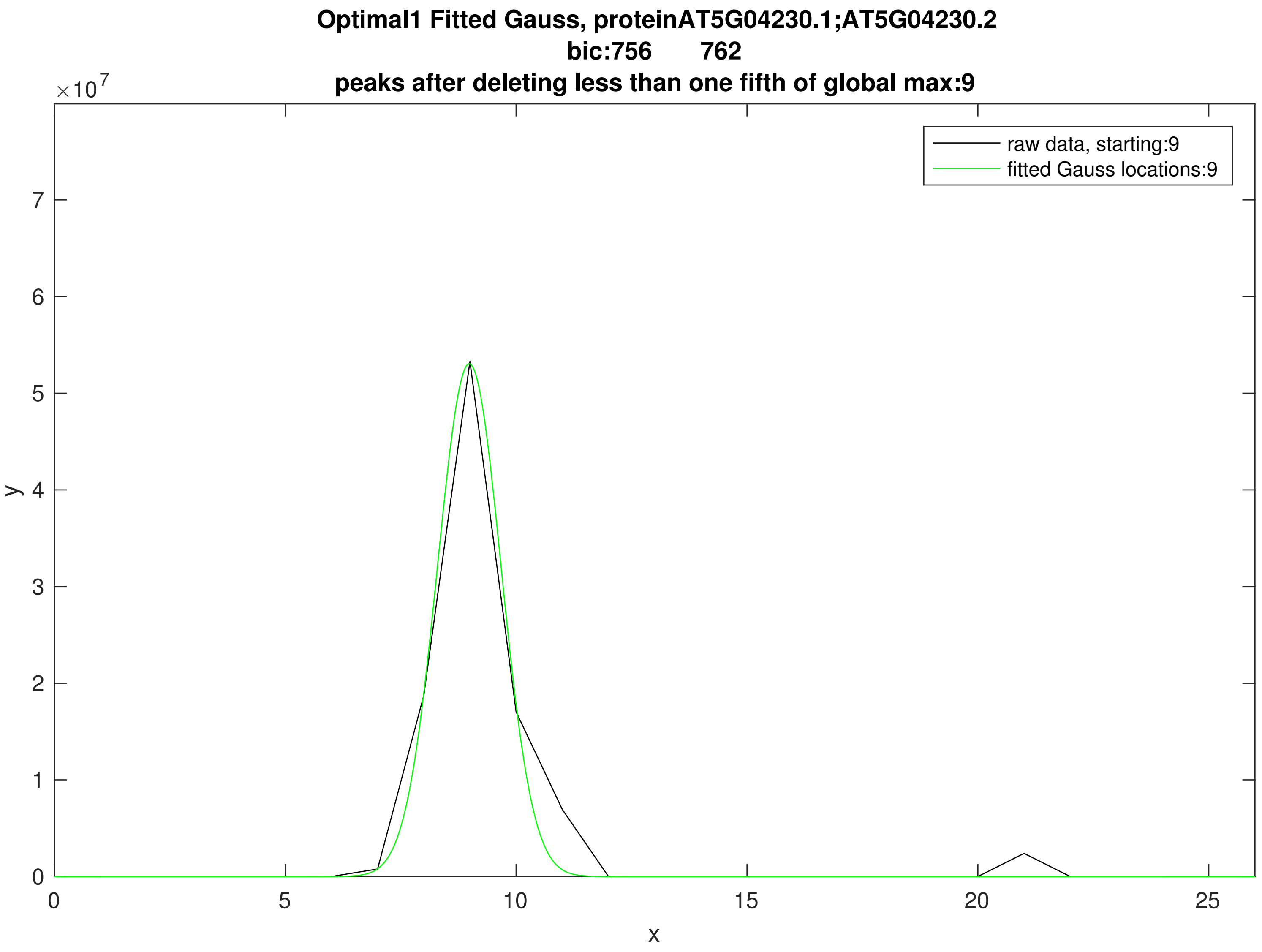
Optimal1 Fitted Gauss, proteinAT5G04230.1;AT5G04230.2
bic:756 762
peaks after deleting less than one fifth of global max:9
×10^7
raw data, starting:9
fitted Gauss locations:9
y
x
0
1
2
3
4
5
6
7
0
5
10
15
20
25

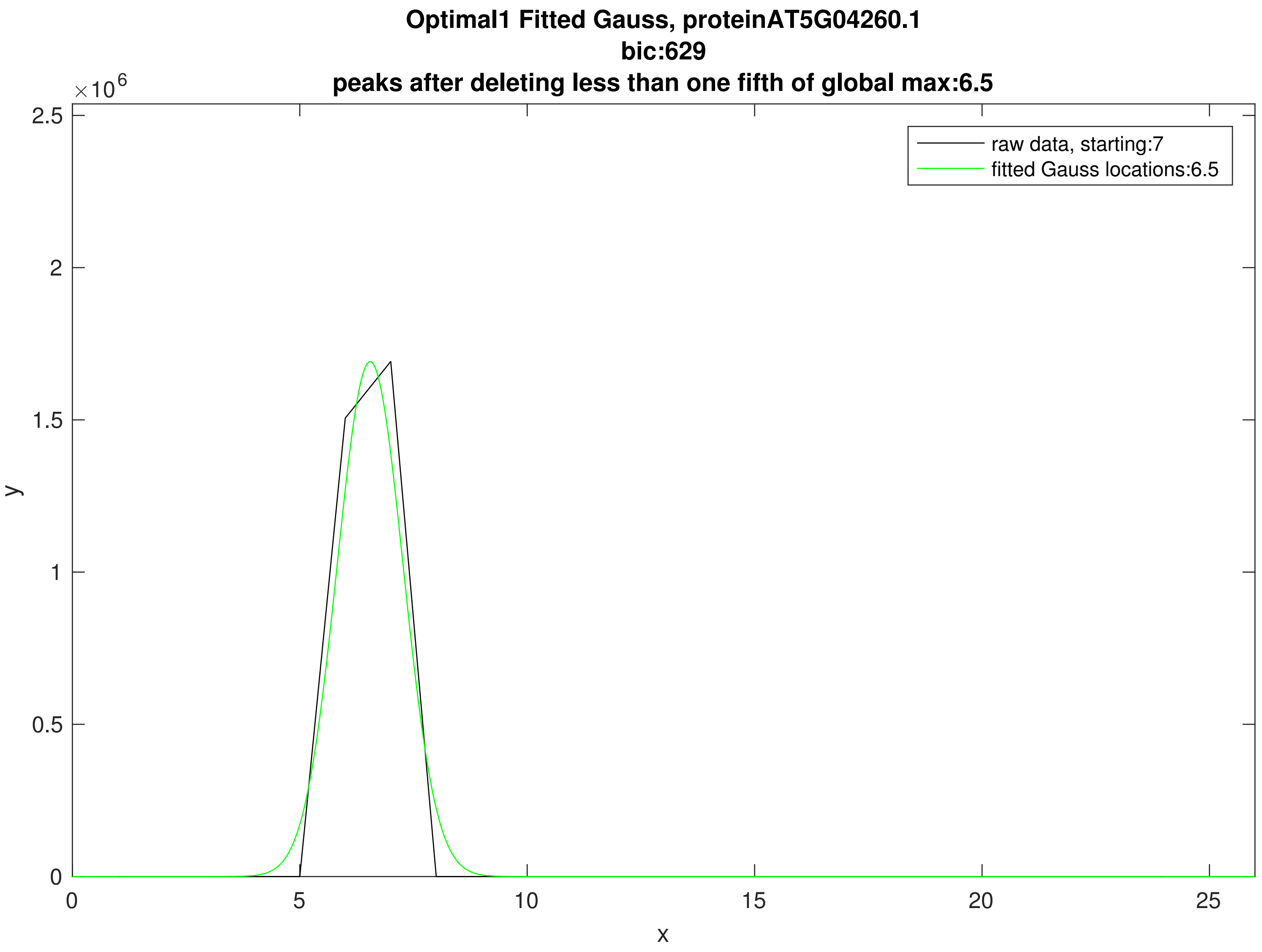
Optimal1 Fitted Gauss, proteinAT5G04260.1
bic:629
peaks after deleting less than one fifth of global max:6.5
×10^6
2.5
2
1.5
1
0.5
0
y
0
5
10
15
20
25
x
raw data, starting:7
fitted Gauss locations:6.5

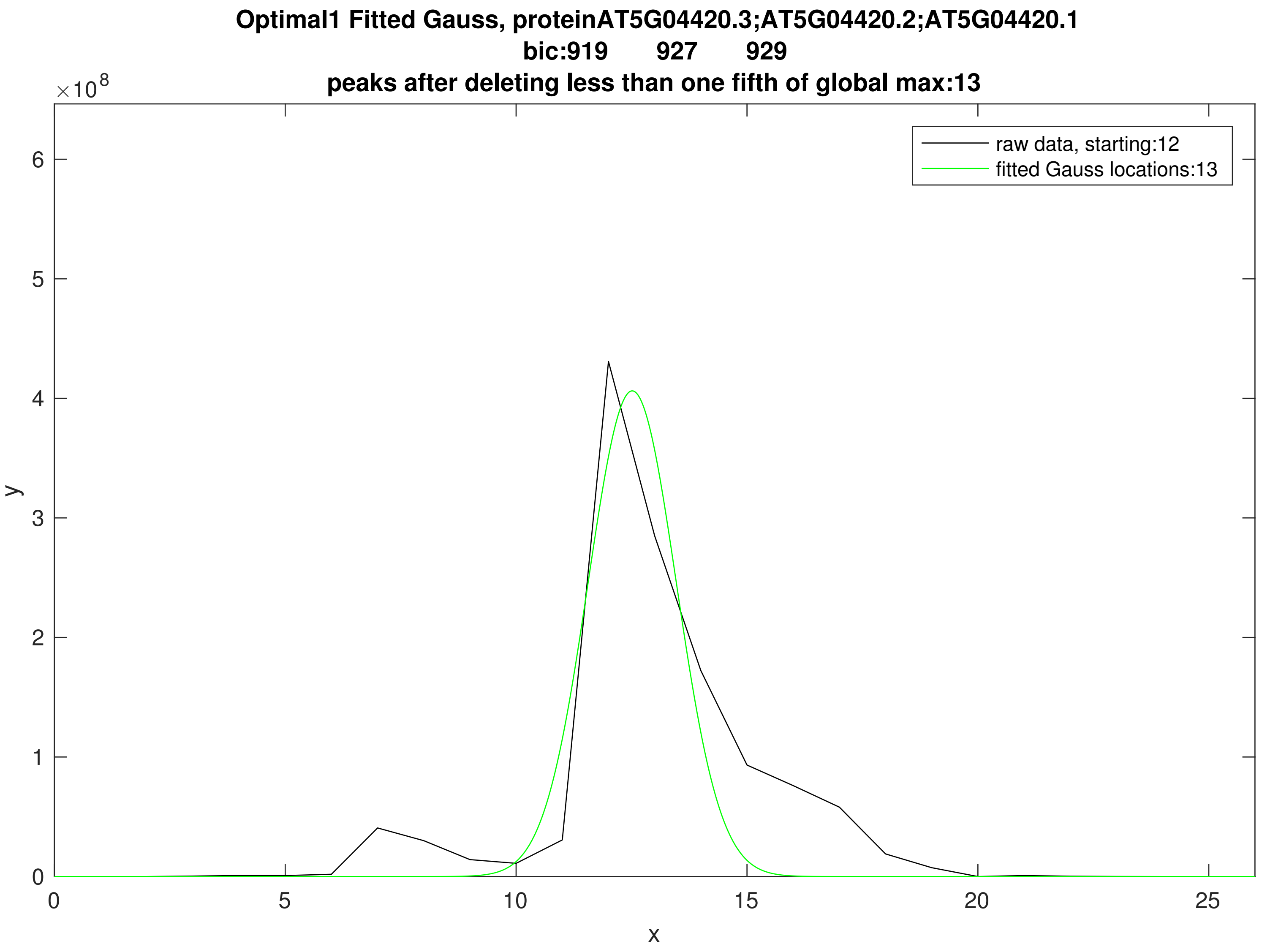

Optimal1 Fitted Gauss, proteinAT5G04420.3;AT5G04420.2;AT5G04420.1
bic:919 927 929
peaks after deleting less than one fifth of global max:13
×10^8
6
5
4
3
2
1
0
y
0
5
10
15
20
25
x
raw data, starting:12
fitted Gauss locations:13

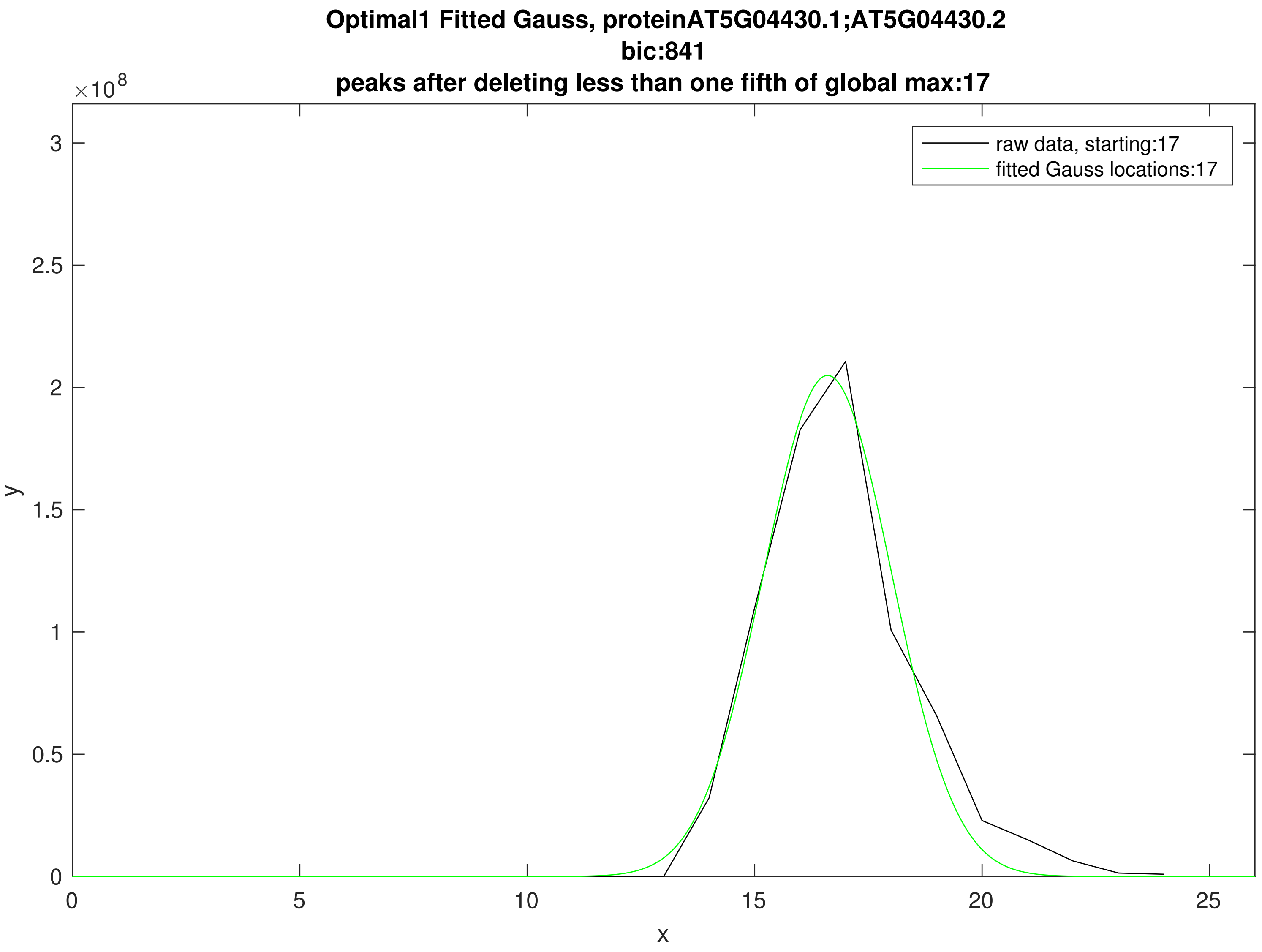
Optimal1 Fitted Gauss, proteinAT5G04430.1;AT5G04430.2
bic:841
peaks after deleting less than one fifth of global max:17
×10^8
raw data, starting:17
fitted Gauss locations:17
y
x
0
0.5
1
1.5
2
2.5
3
0
5
10
15
20
25

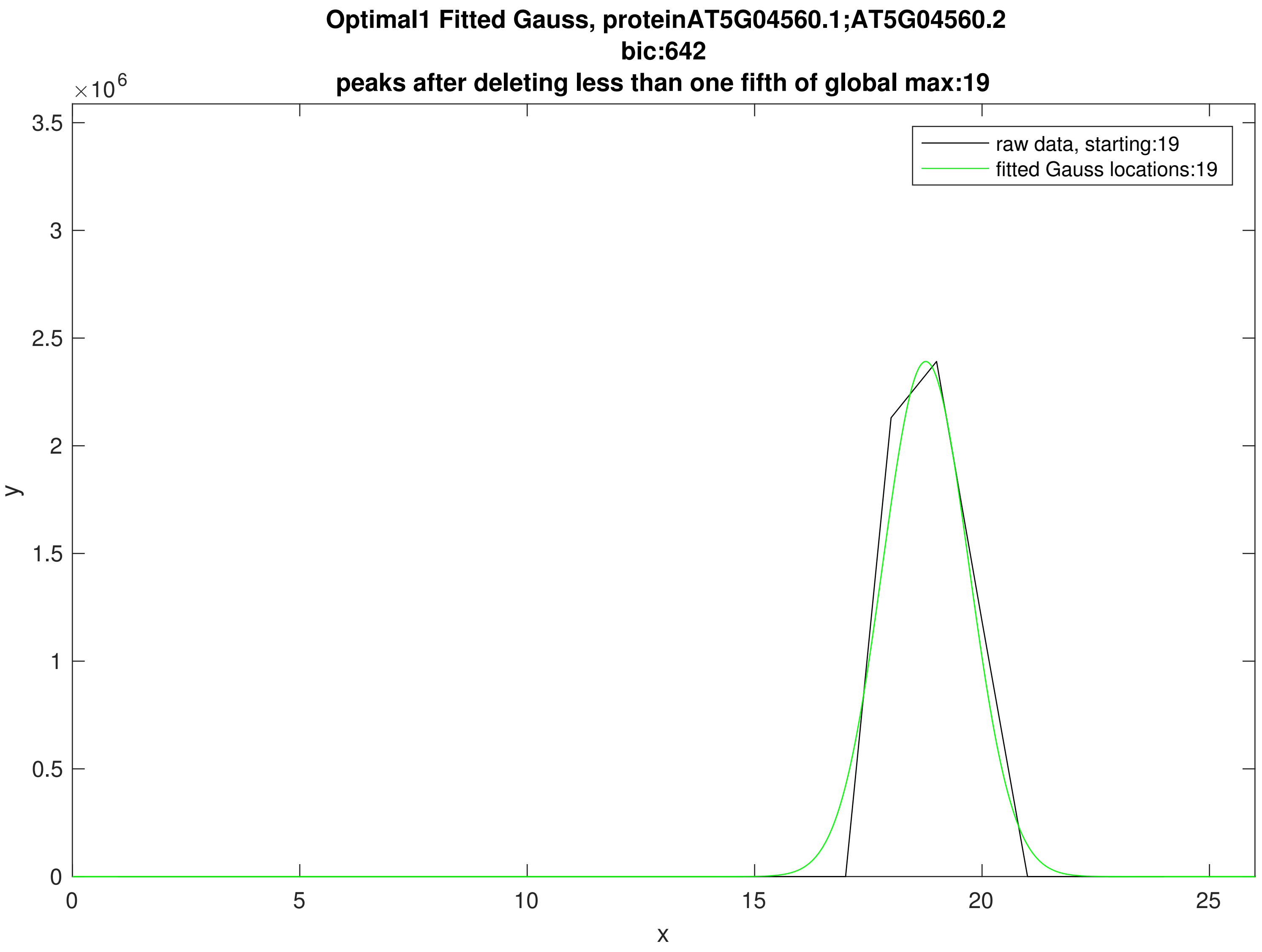
Optimal1 Fitted Gauss, proteinAT5G04560.1;AT5G04560.2
bic:642
peaks after deleting less than one fifth of global max:19
×10^6
raw data, starting:19
fitted Gauss locations:19
y
x
0
5
10
15
20
25
0.5
1
1.5
2
2.5
3
3.5

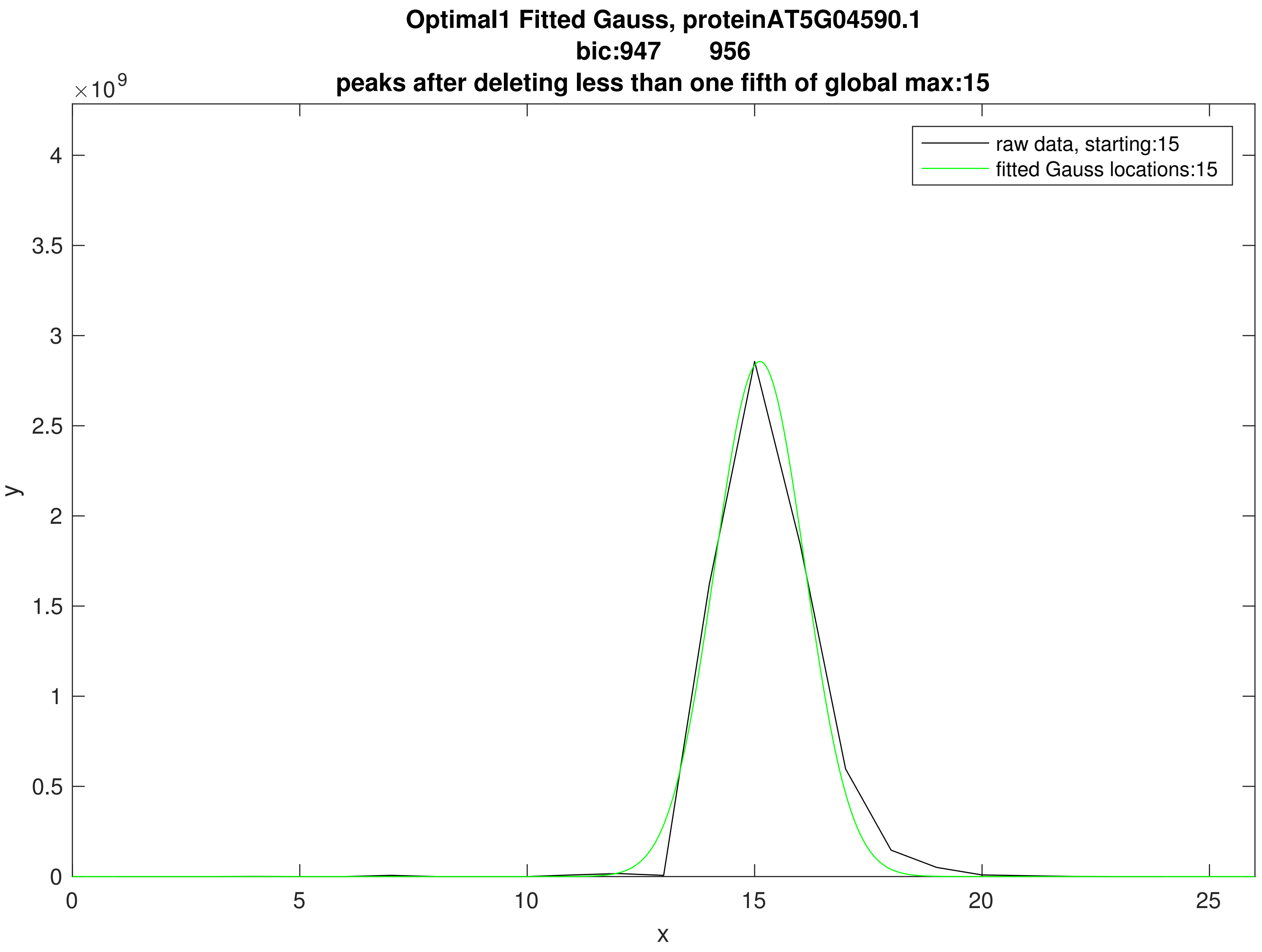

Optimal1 Fitted Gauss, proteinAT5G04590.1
bic:947 956
peaks after deleting less than one fifth of global max:15
×10^9
4
3.5
3
2.5
2
1.5
1
0.5
0
y
0
5
10
15
20
25
x
raw data, starting:15
fitted Gauss locations:15

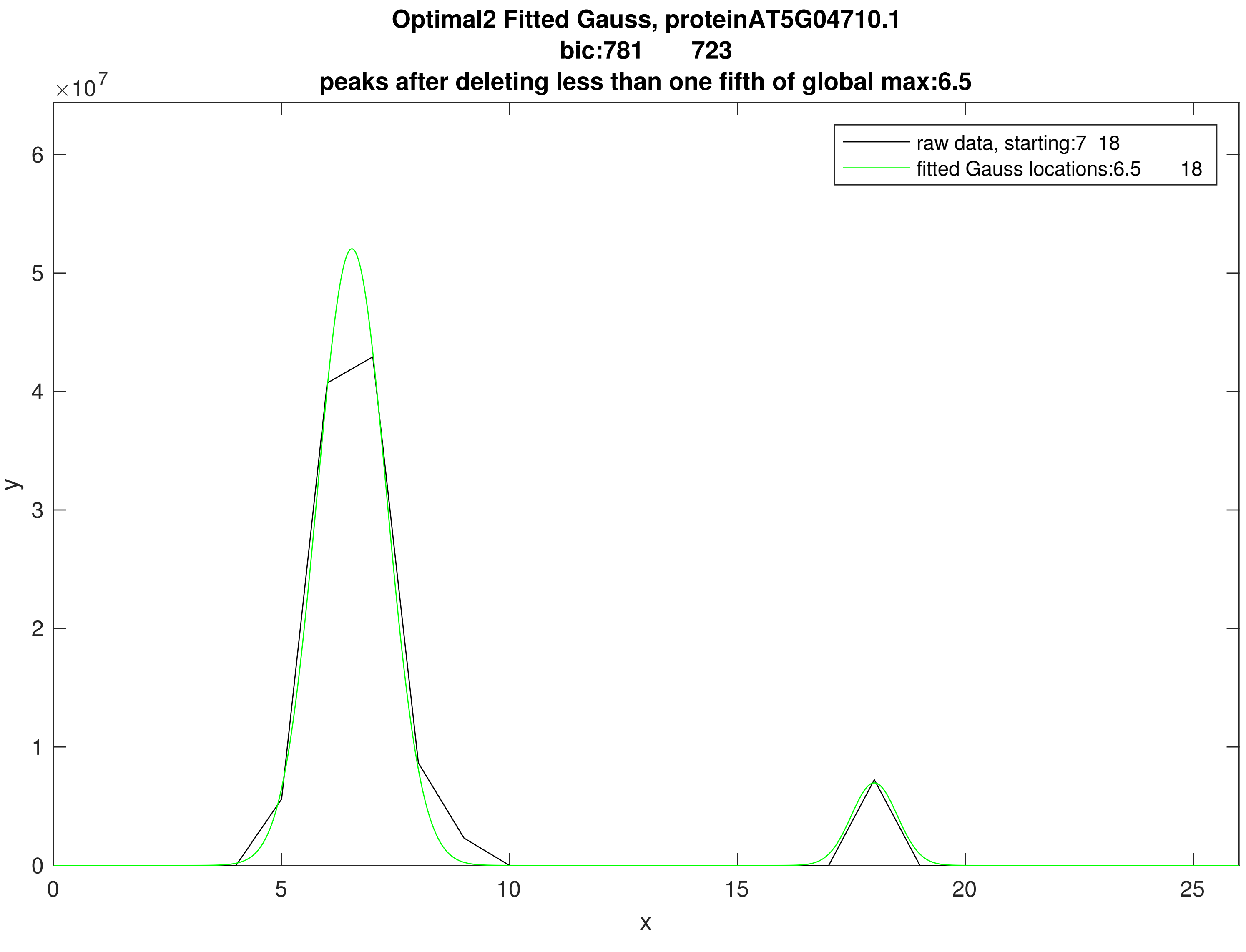
Optimal2 Fitted Gauss, proteinAT5G04710.1
bic:781 723
peaks after deleting less than one fifth of global max:6.5
×10^7
raw data, starting:7 18
fitted Gauss locations:6.5 18
y
x
0
1
2
3
4
5
6
0
5
10
15
20
25

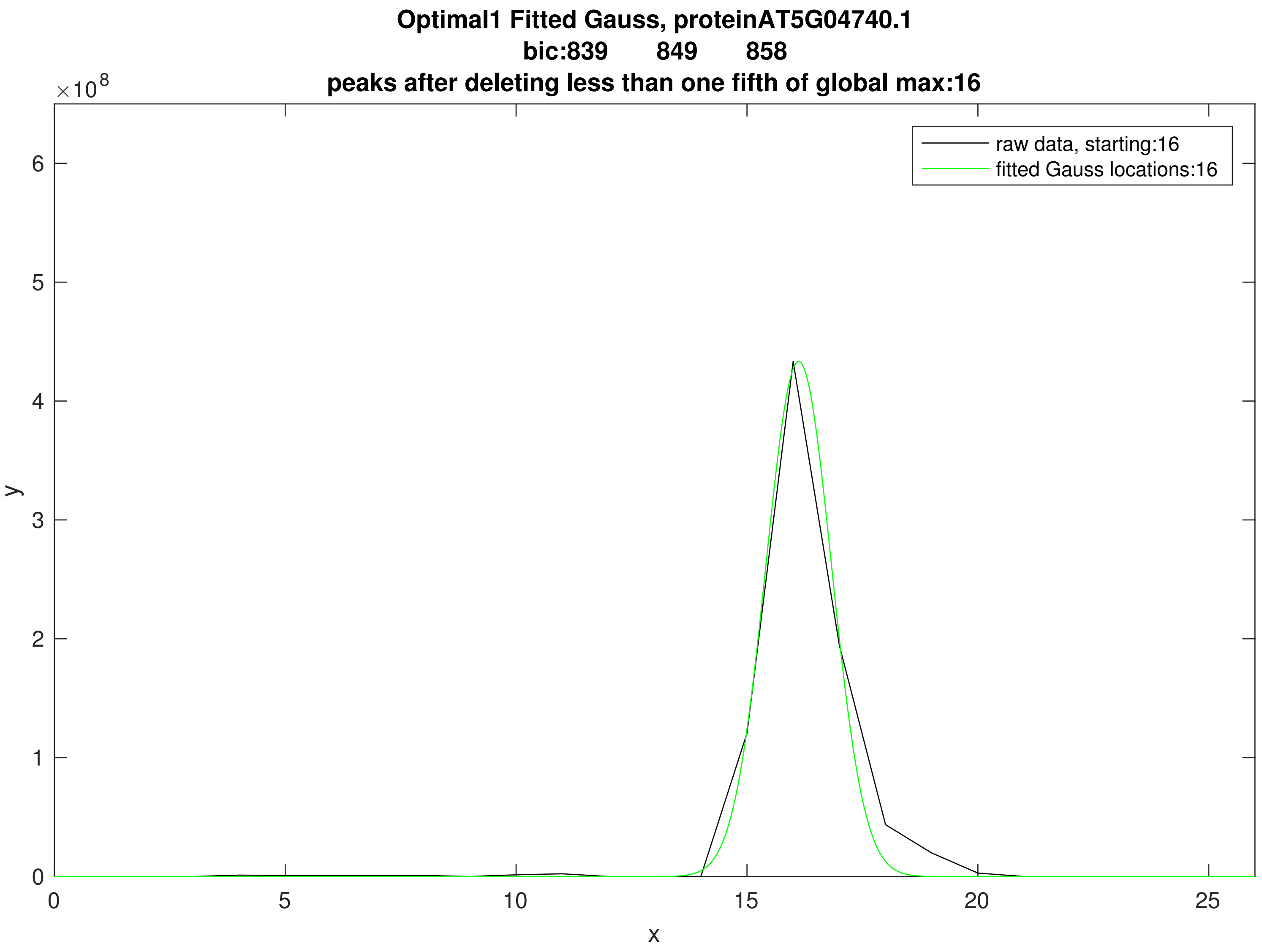
Optimal1 Fitted Gauss, proteinAT5G04740.1
bic:839 849 858
peaks after deleting less than one fifth of global max:16
×10^8
raw data, starting:16
fitted Gauss locations:16
y
x
0
1
2
3
4
5
6
0
5
10
15
20
25

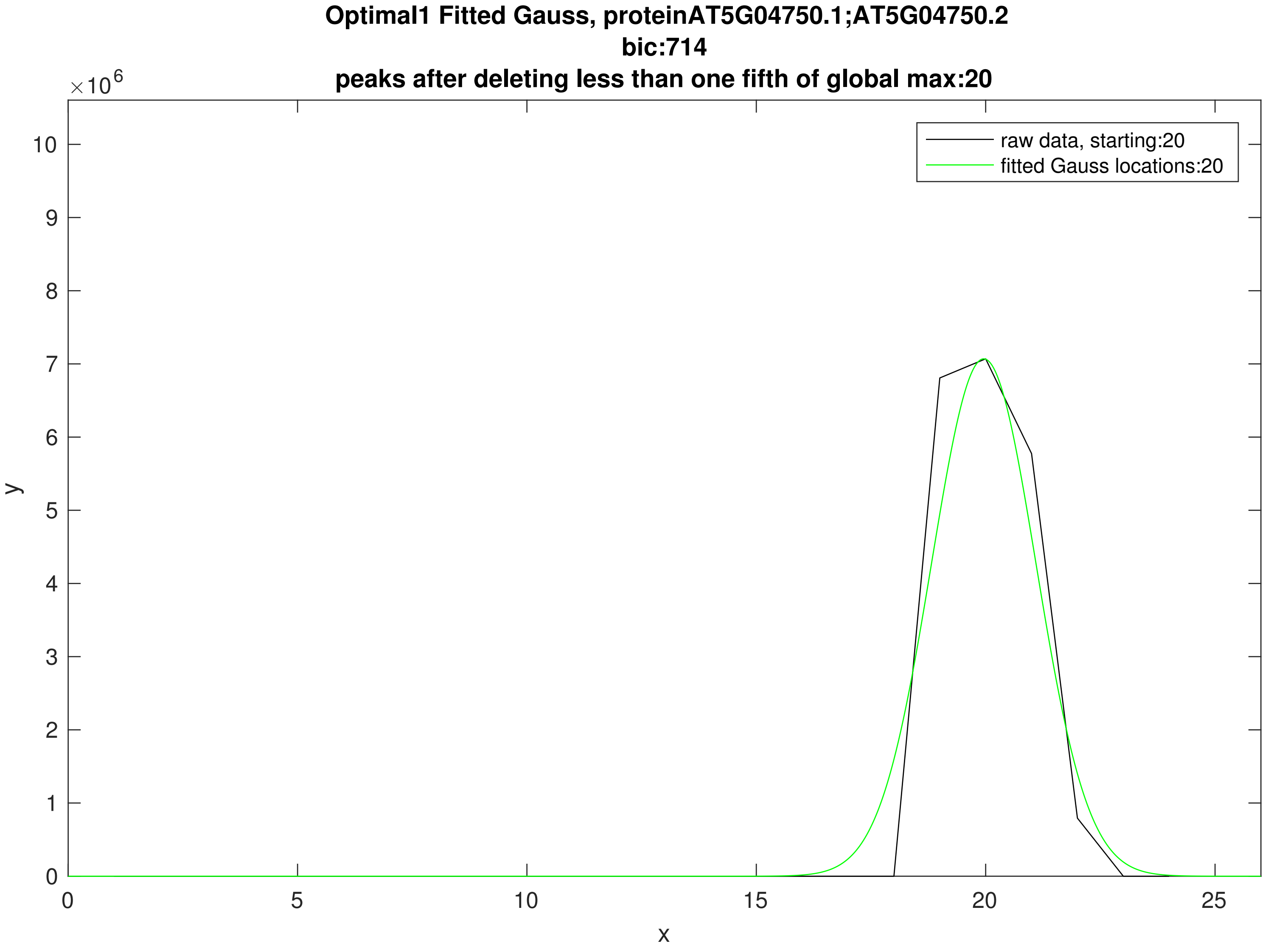
Optimal1 Fitted Gauss, proteinAT5G04750.1;AT5G04750.2
bic:714
peaks after deleting less than one fifth of global max:20
×10^6
raw data, starting:20
fitted Gauss locations:20
y
x
0
1
2
3
4
5
6
7
8
9
10
0
5
10
15
20
25

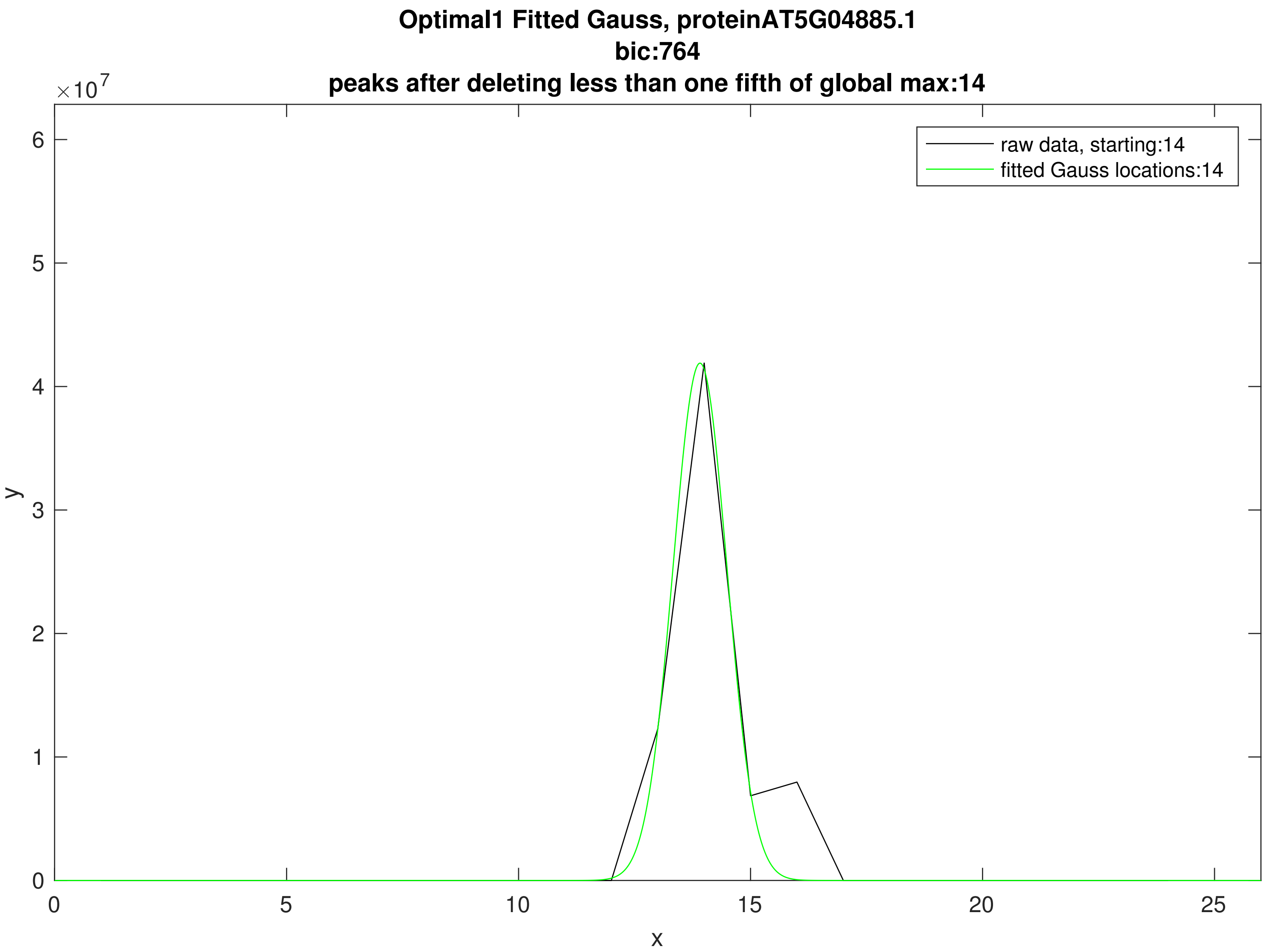
Optimal1 Fitted Gauss, proteinAT5G04885.1
bic:764
peaks after deleting less than one fifth of global max:14
×10^7
6
5
4
3
2
1
0
y
0
5
10
15
20
25
x
raw data, starting:14
fitted Gauss locations:14

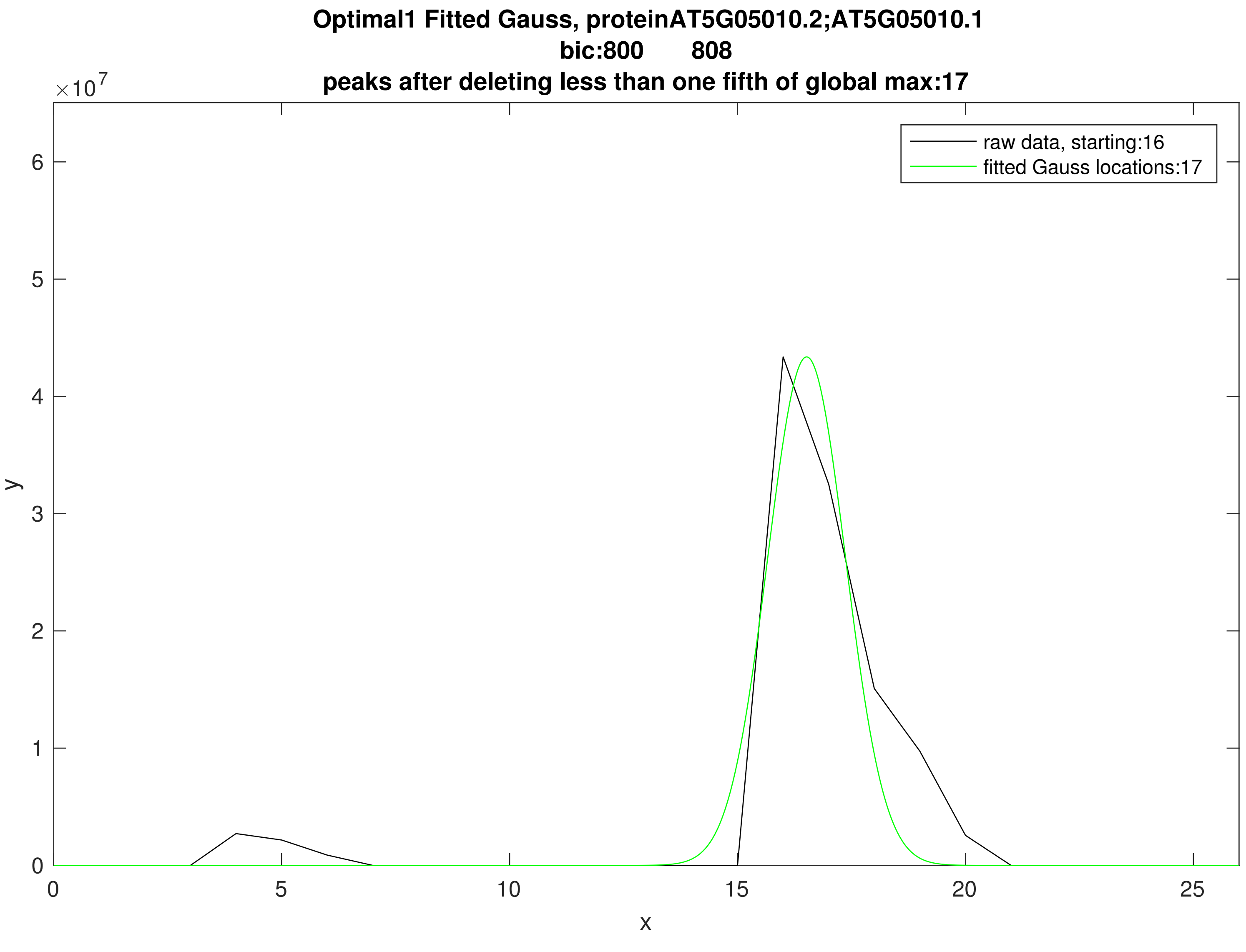

Optimal1 Fitted Gauss, proteinAT5G05010.2;AT5G05010.1
bic:800 808
peaks after deleting less than one fifth of global max:17
×10^7
raw data, starting:16
fitted Gauss locations:17
y
x
0
1
2
3
4
5
6
0
5
10
15
20
25

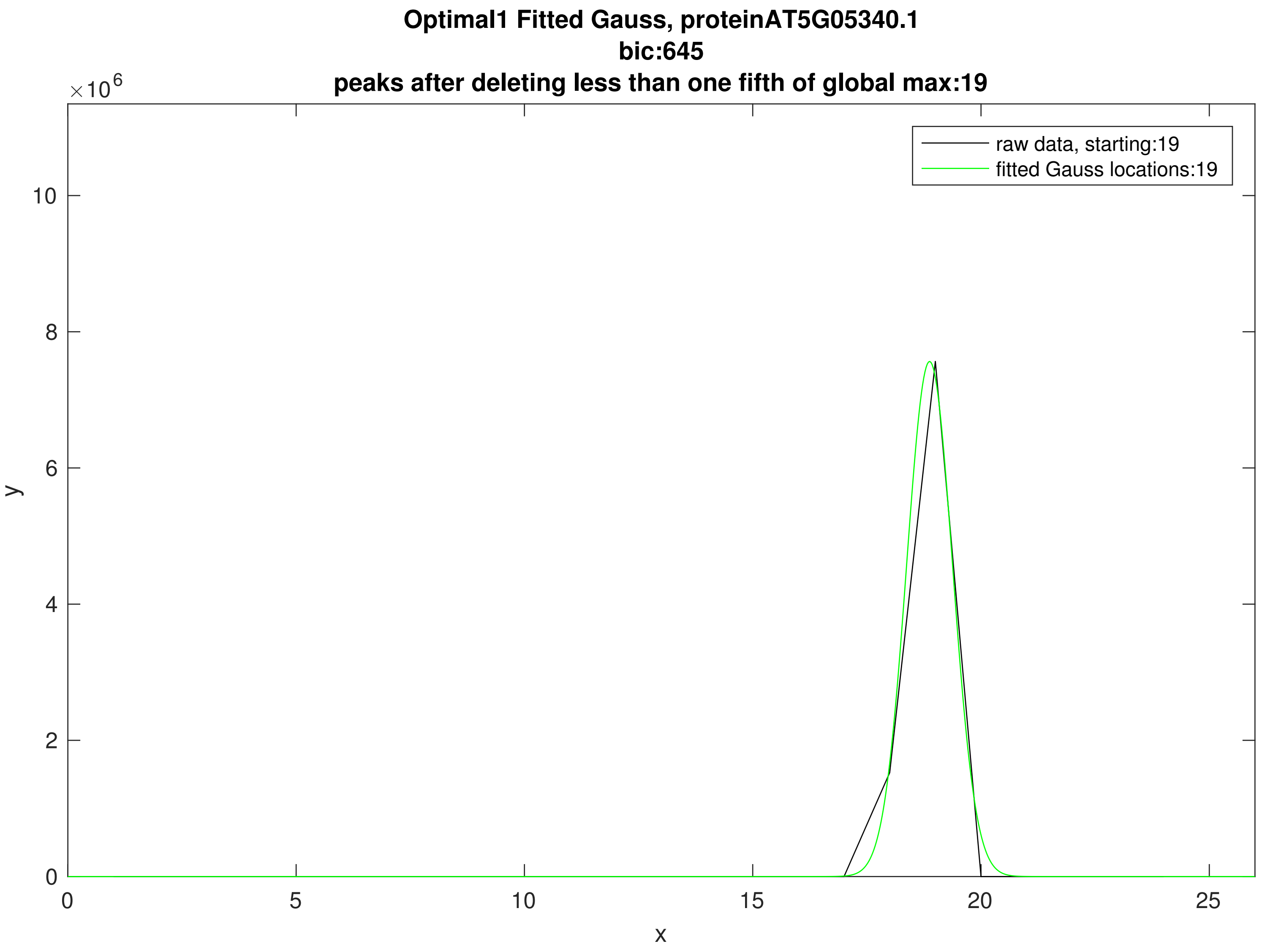
Optimal1 Fitted Gauss, proteinAT5G05340.1
bic:645
peaks after deleting less than one fifth of global max:19
×10^6
raw data, starting:19
fitted Gauss locations:19
10
8
6
4
2
0
y
0
5
10
15
20
25
x

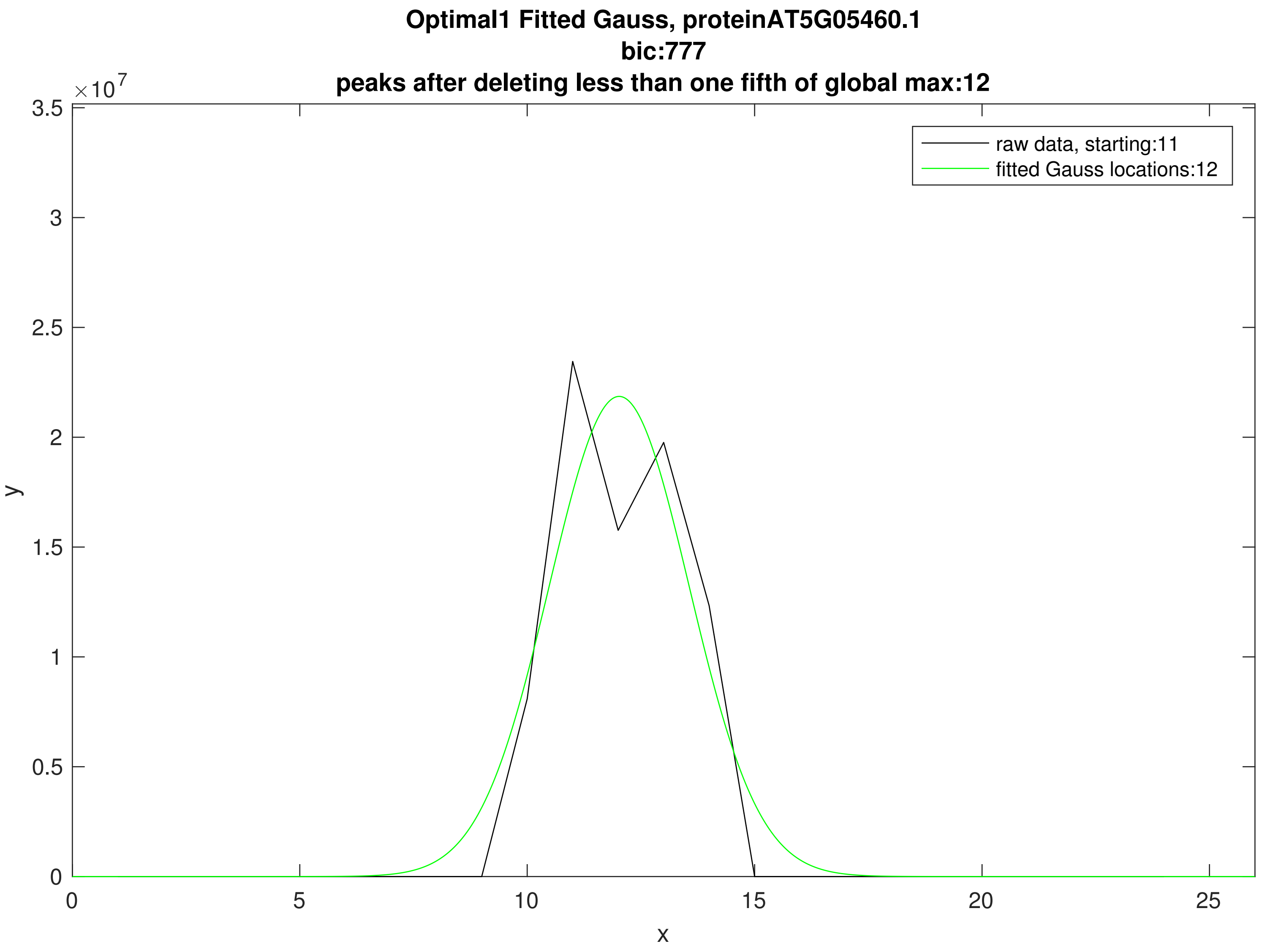
Optimal1 Fitted Gauss, proteinAT5G05460.1
bic:777
peaks after deleting less than one fifth of global max:12
×10^7
3.5
3
2.5
2
1.5
1
0.5
0
y
0
5
10
15
20
25
x
raw data, starting:11
fitted Gauss locations:12

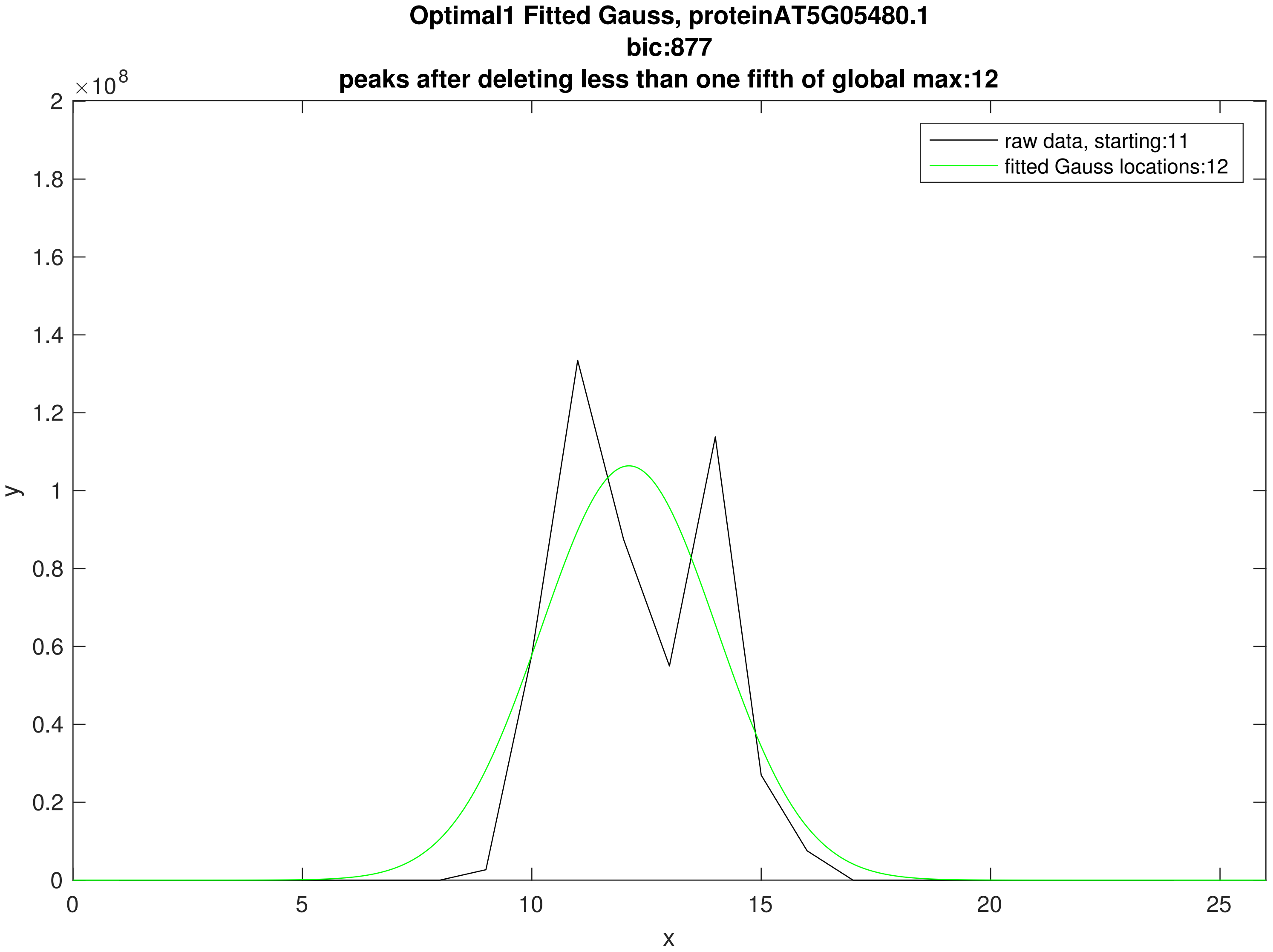
Optimal1 Fitted Gauss, proteinAT5G05480.1
bic:877
peaks after deleting less than one fifth of global max:12
×10^8
raw data, starting:11
fitted Gauss locations:12
y
x
0
0.2
0.4
0.6
0.8
1
1.2
1.4
1.6
1.8
2
5
10
15
20
25

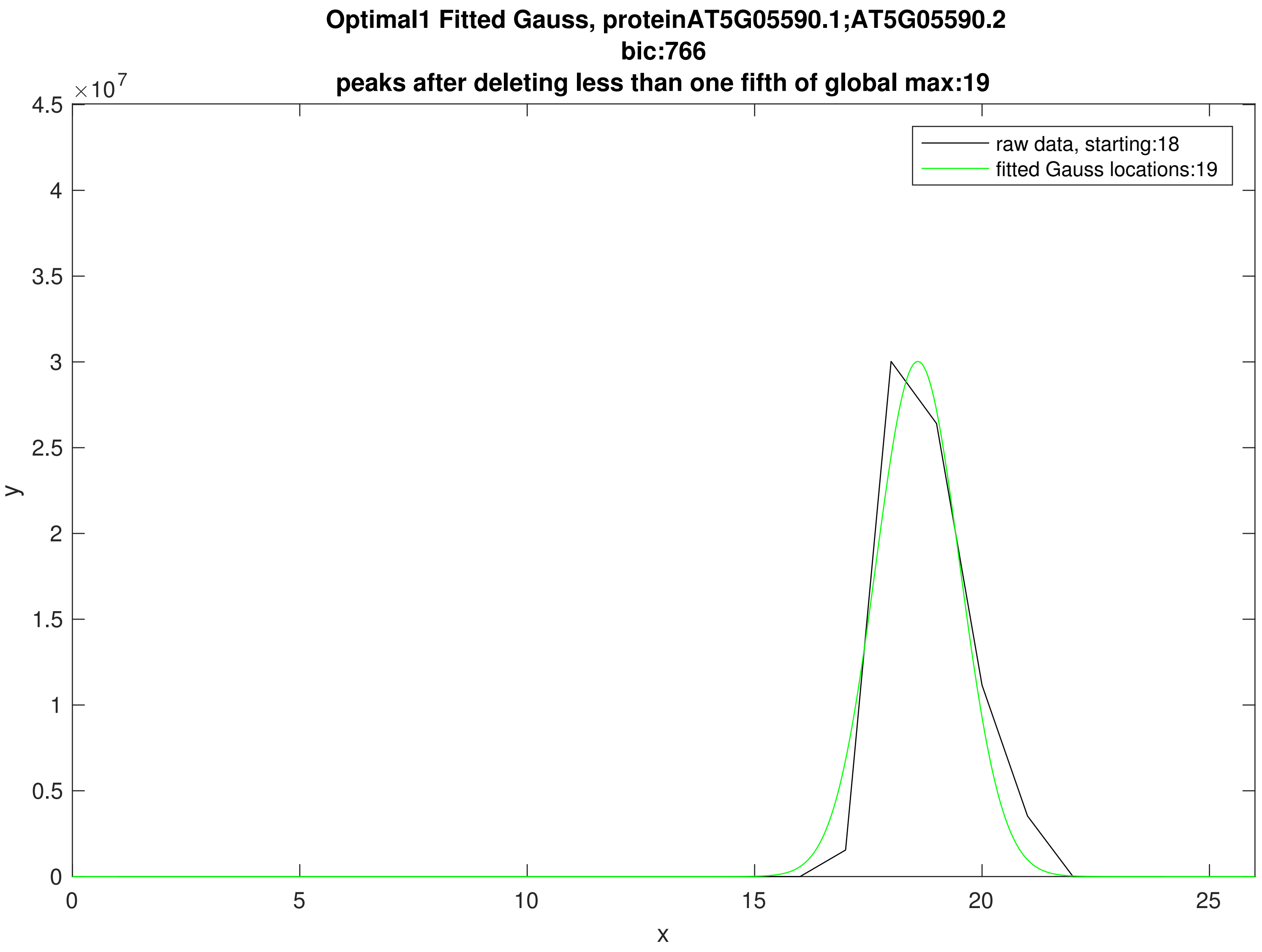

Optimal1 Fitted Gauss, proteinAT5G05590.1;AT5G05590.2
bic:766
peaks after deleting less than one fifth of global max:19
×10^7
4.5
4
3.5
3
2.5
2
1.5
1
0.5
0
y
0
5
10
15
20
25
x
raw data, starting:18
fitted Gauss locations:19

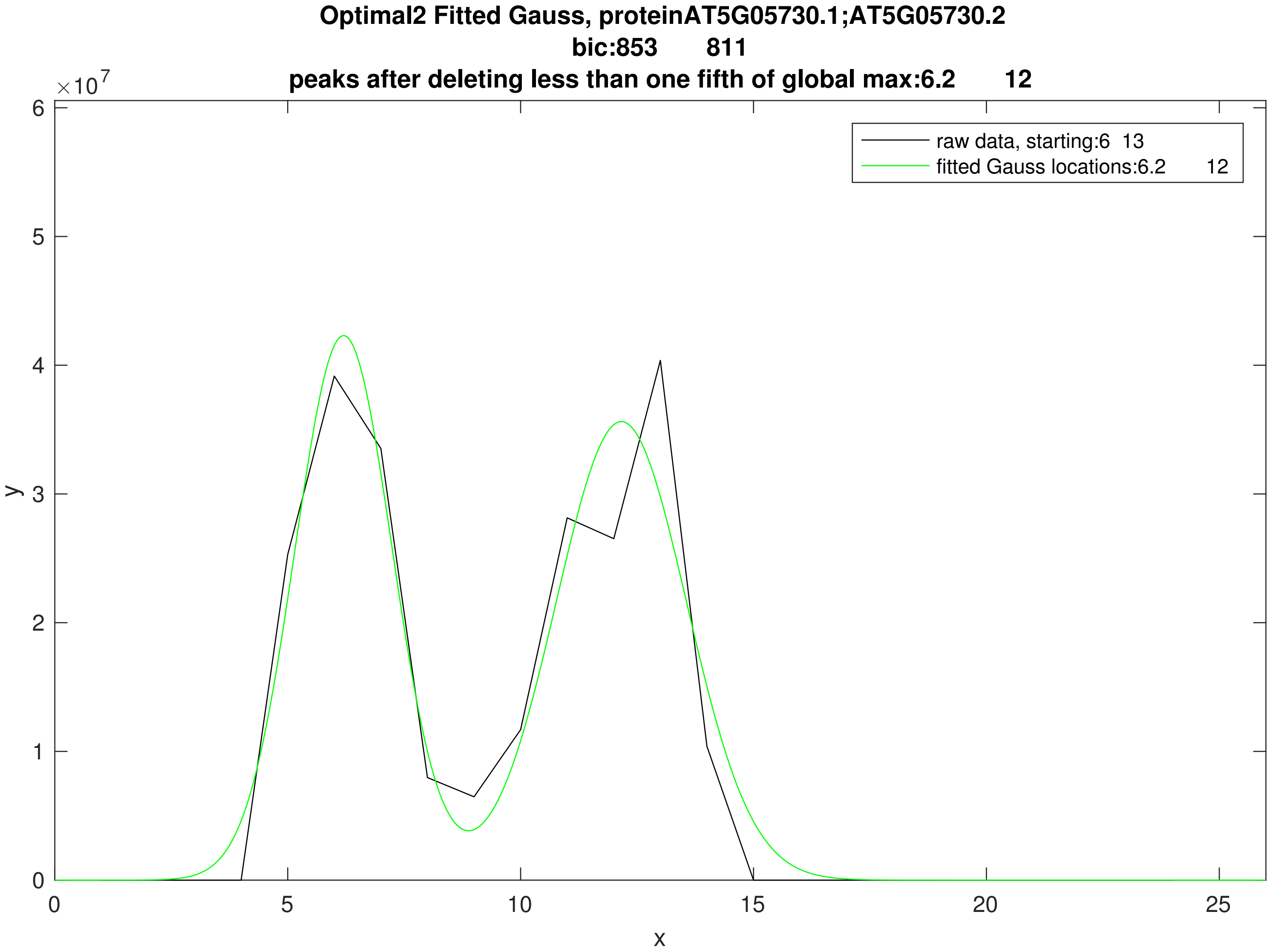
Optimal2 Fitted Gauss, proteinAT5G05730.1;AT5G05730.2
bic:853 811
peaks after deleting less than one fifth of global max:6.2 12
×10^7
raw data, starting:6 13
fitted Gauss locations:6.2 12
y
x
0
1
2
3
4
5
6
0
5
10
15
20
25

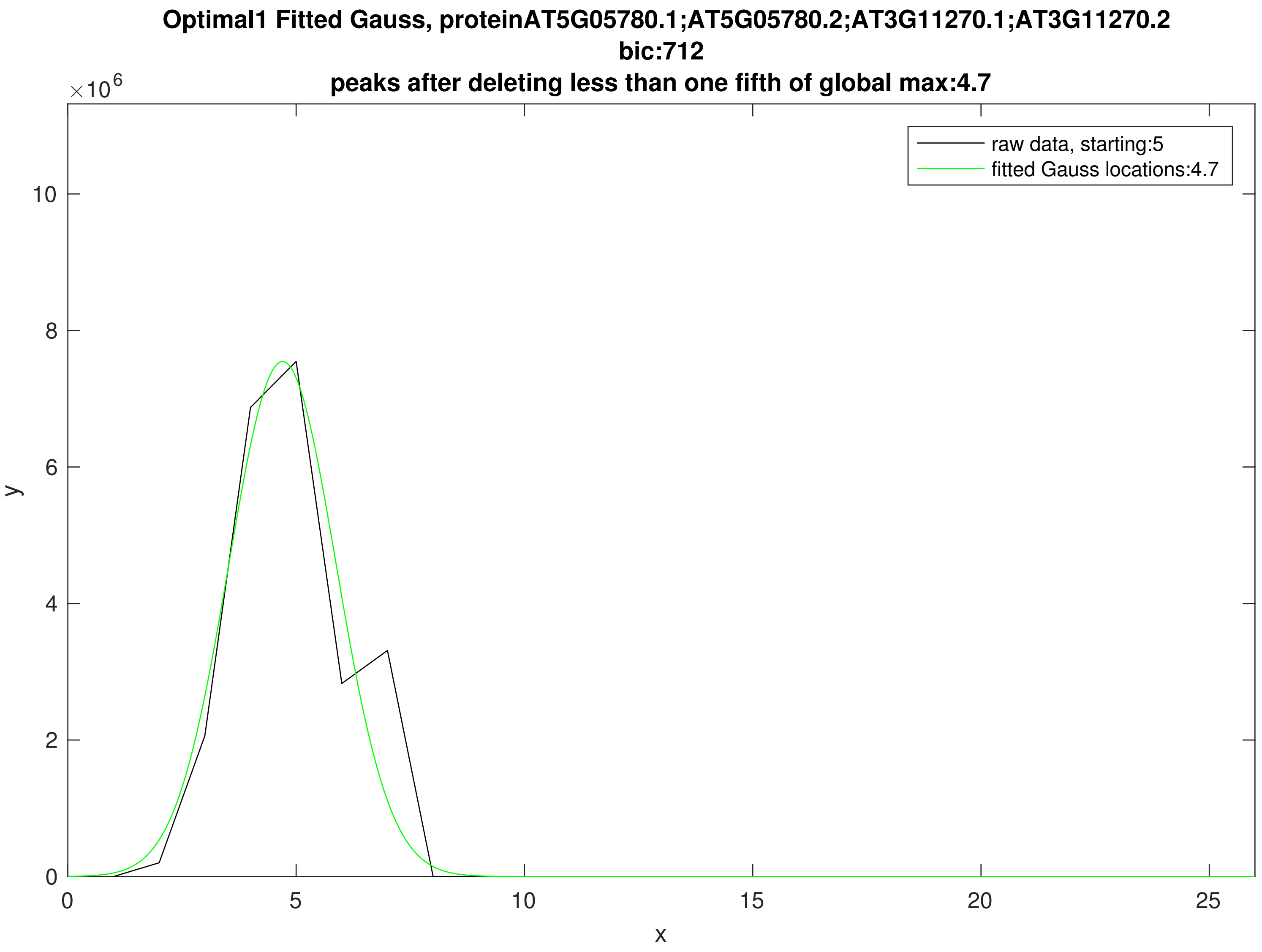
Optimal1 Fitted Gauss, proteinAT5G05780.1;AT5G05780.2;AT3G11270.1;AT3G11270.2
bic:712
peaks after deleting less than one fifth of global max:4.7
×10^6
raw data, starting:5
fitted Gauss locations:4.7
y
x
0
2
4
6
8
10
5
15
20
25

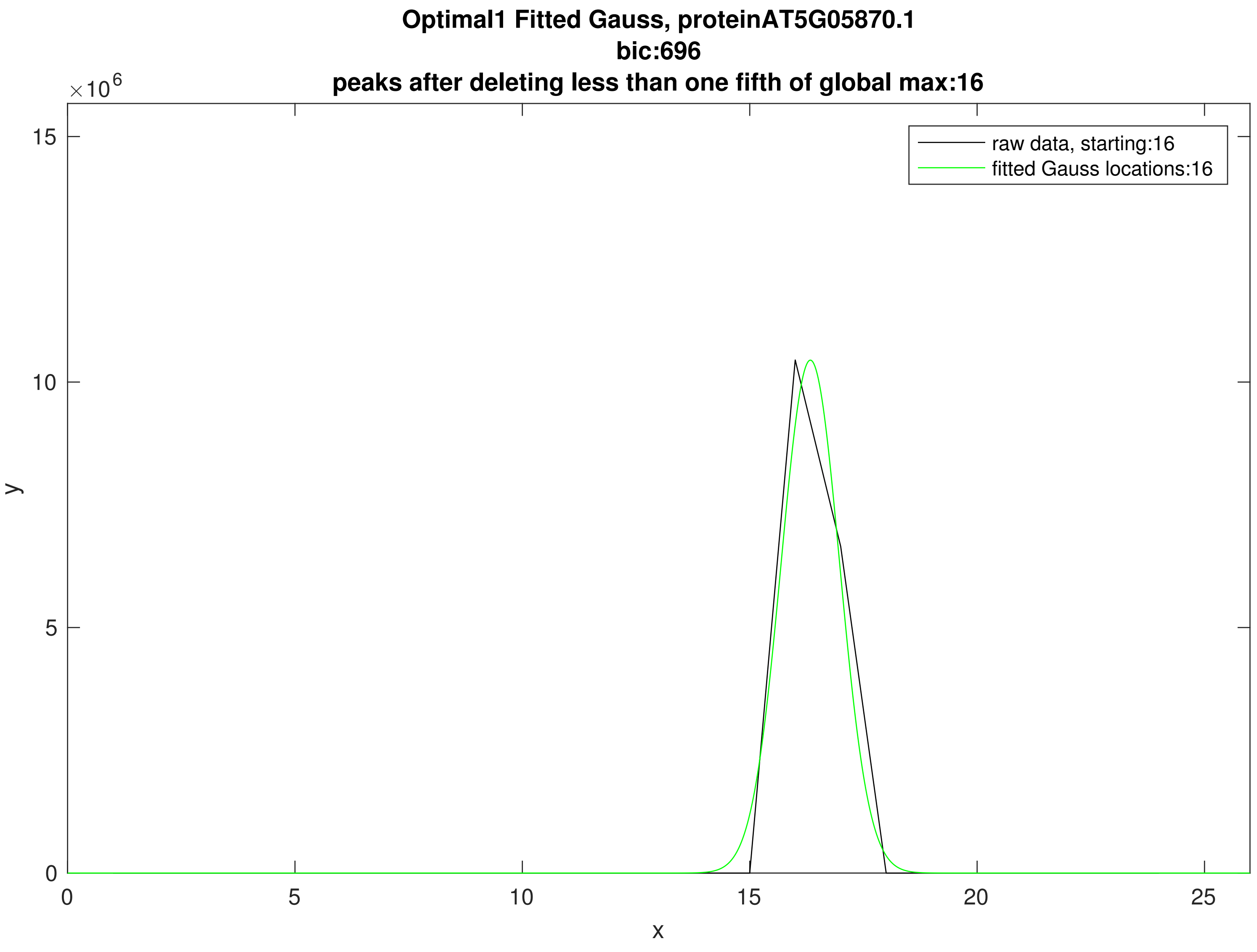
Optimal1 Fitted Gauss, proteinAT5G05870.1
bic:696
peaks after deleting less than one fifth of global max:16
×10^6
raw data, starting:16
fitted Gauss locations:16
y
x
0
5
10
15
20
25

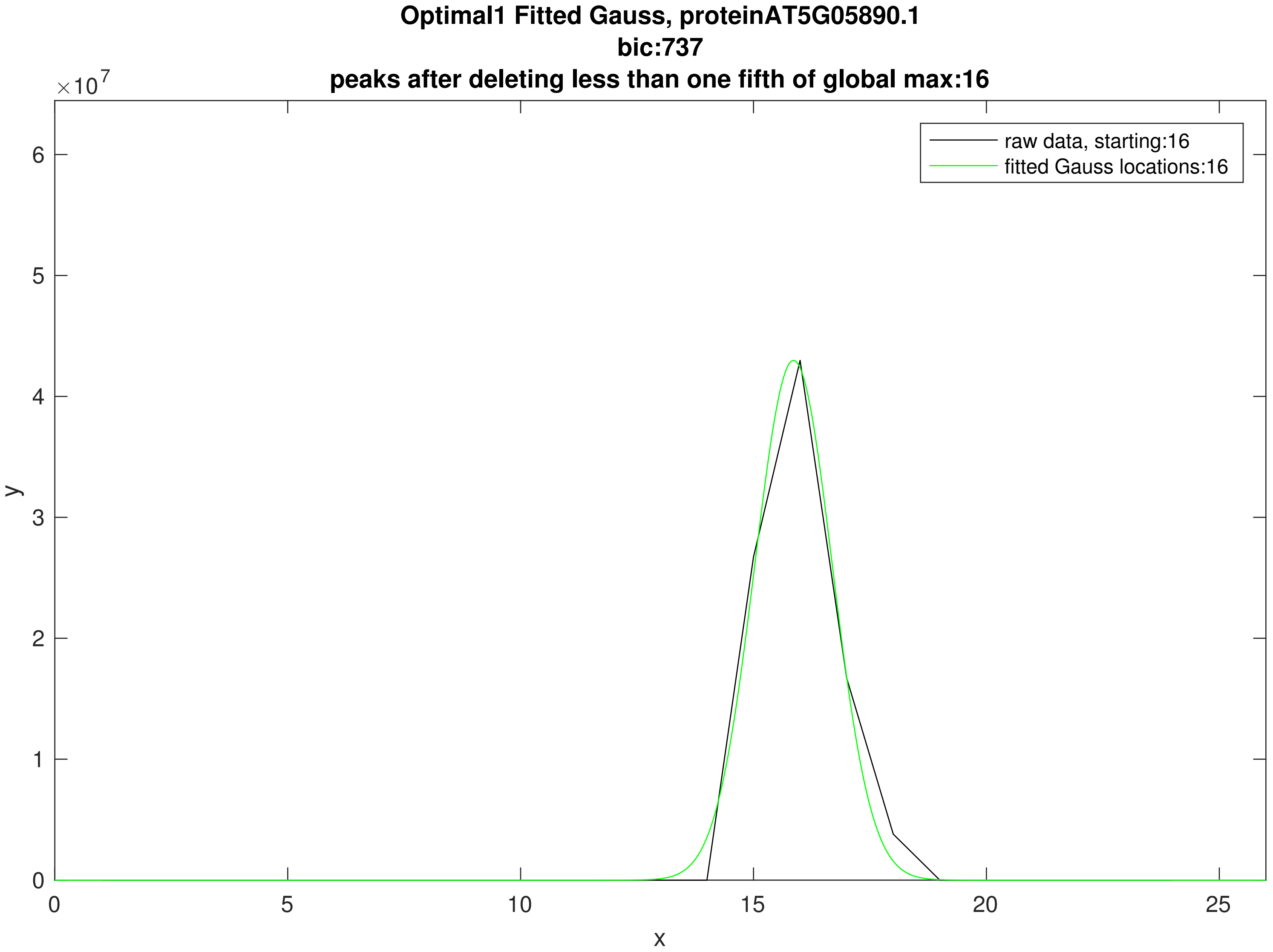

Optimal1 Fitted Gauss, proteinAT5G05890.1
bic:737
peaks after deleting less than one fifth of global max:16
×10^7
6
5
4
3
2
1
0
y
0
5
10
15
20
25
x
raw data, starting:16
fitted Gauss locations:16

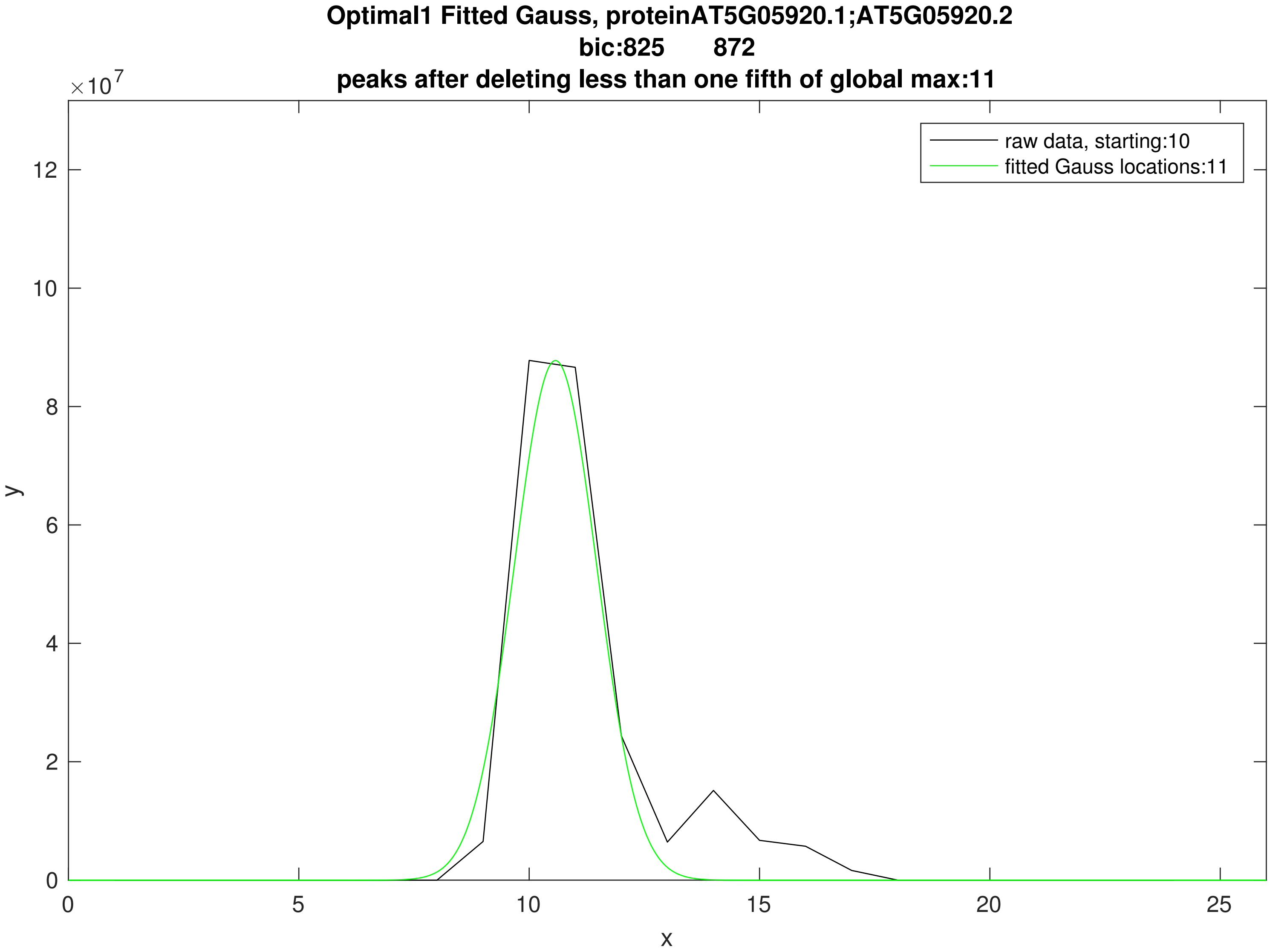

Optimal1 Fitted Gauss, proteinAT5G05920.1;AT5G05920.2
bic:825 872
peaks after deleting less than one fifth of global max:11
×10^7
raw data, starting:10
fitted Gauss locations:11
y
x
0
2
4
6
8
10
12
0
5
10
15
20
25

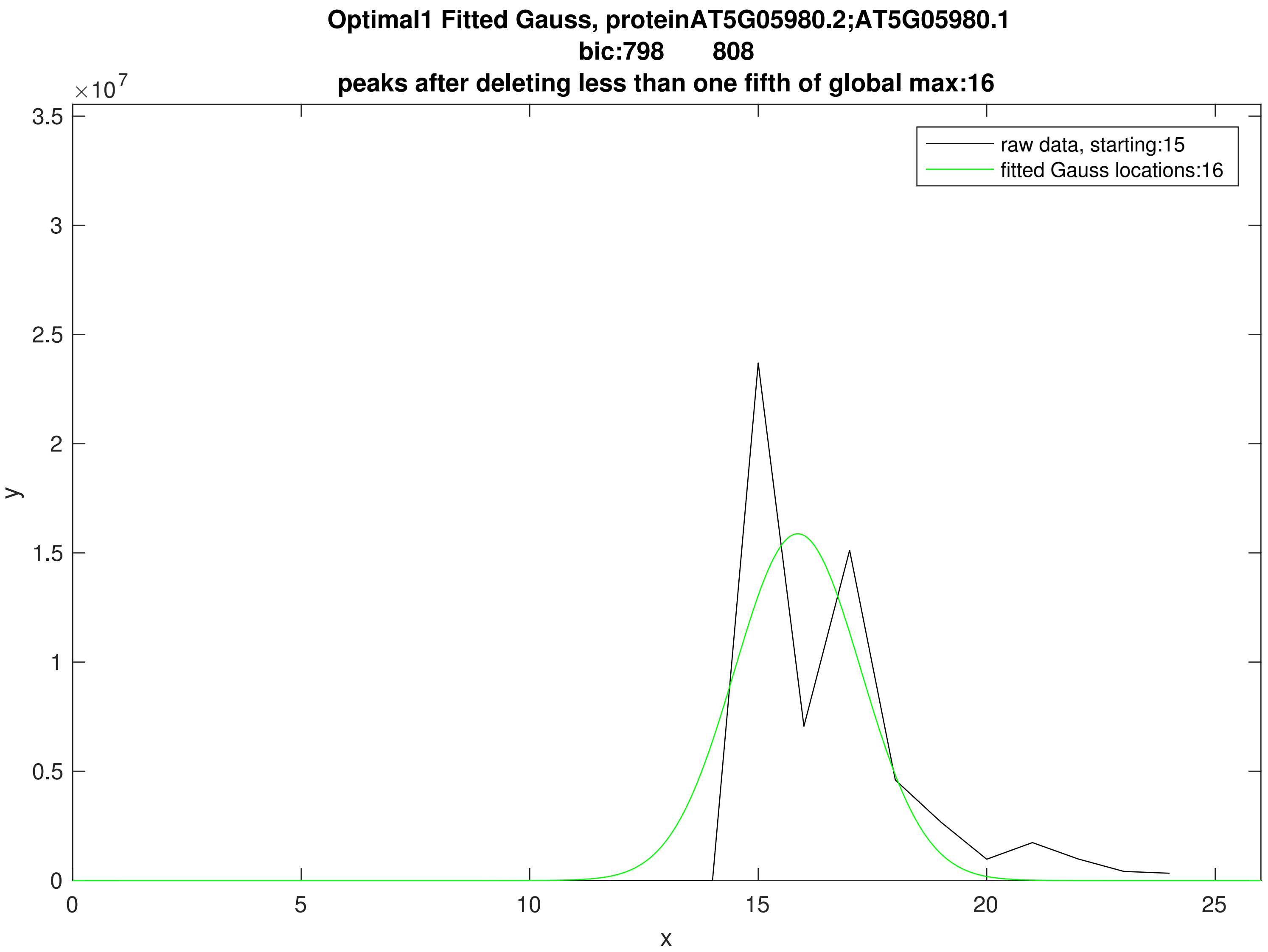
Optimal1 Fitted Gauss, proteinAT5G05980.2;AT5G05980.1
bic:798 808
peaks after deleting less than one fifth of global max:16
×10^7
3.5
3
2.5
2
1.5
1
0.5
0
y
0
5
10
15
20
25
x
raw data, starting:15
fitted Gauss locations:16

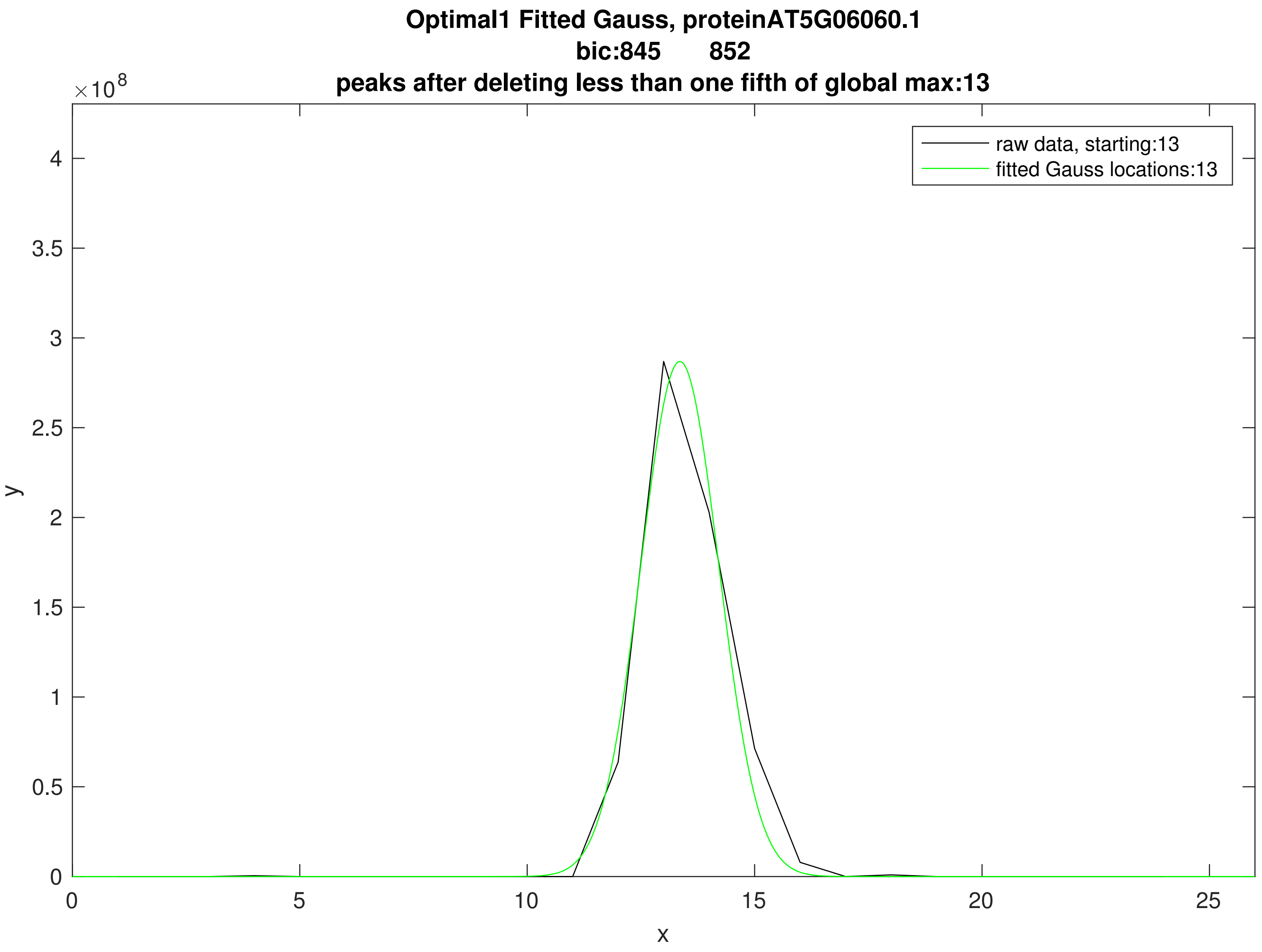

Optimal1 Fitted Gauss, proteinAT5G06060.1
bic:845 852
peaks after deleting less than one fifth of global max:13
×10^8
4
3.5
3
2.5
2
1.5
1
0.5
0
y
0
5
10
15
20
25
x
raw data, starting:13
fitted Gauss locations:13

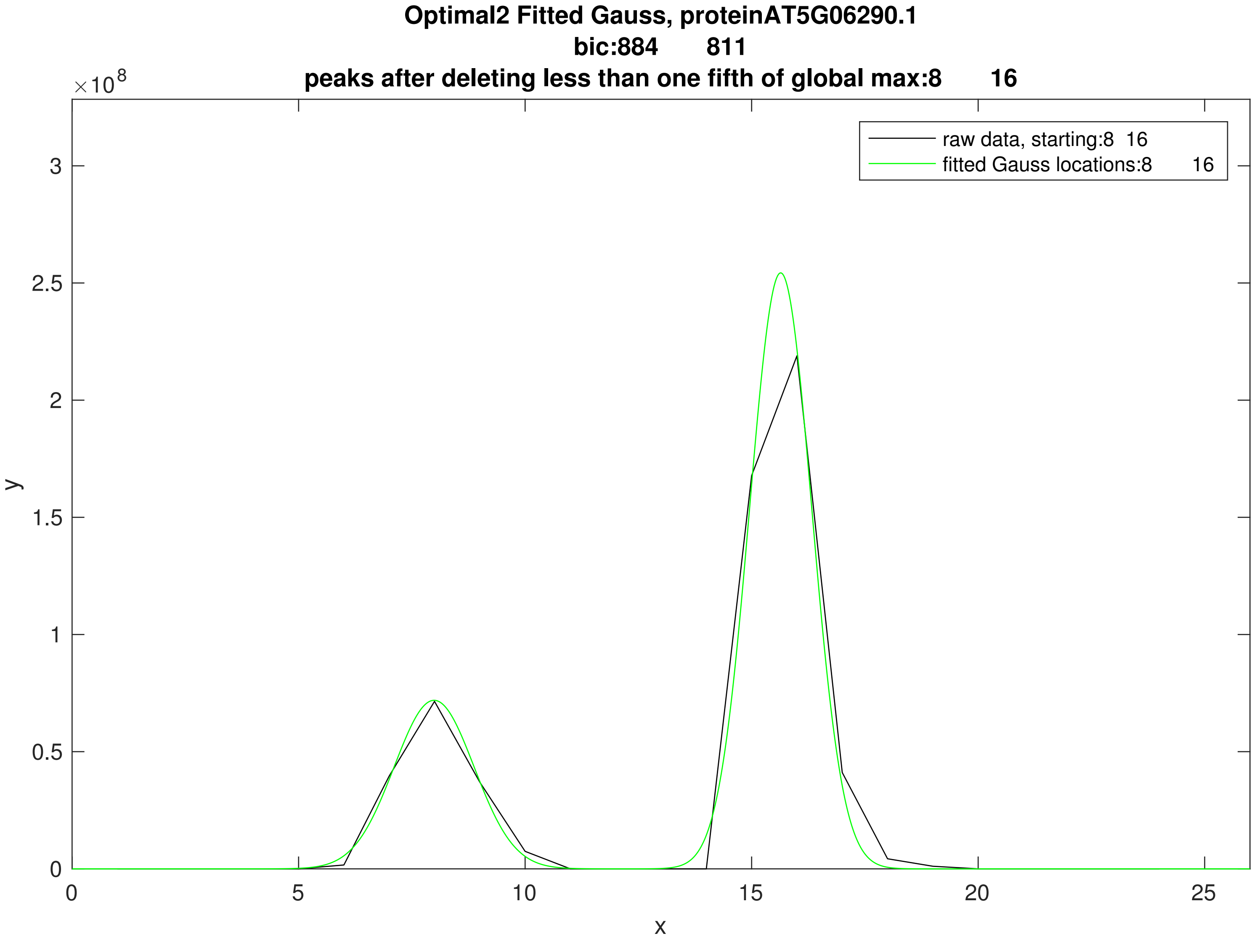
Optimal2 Fitted Gauss, proteinAT5G06290.1
bic:884 811
peaks after deleting less than one fifth of global max:8 16
×10^8
raw data, starting:8 16
fitted Gauss locations:8 16
y
x
0
0.5
1
1.5
2
2.5
3
0
5
10
15
20
25

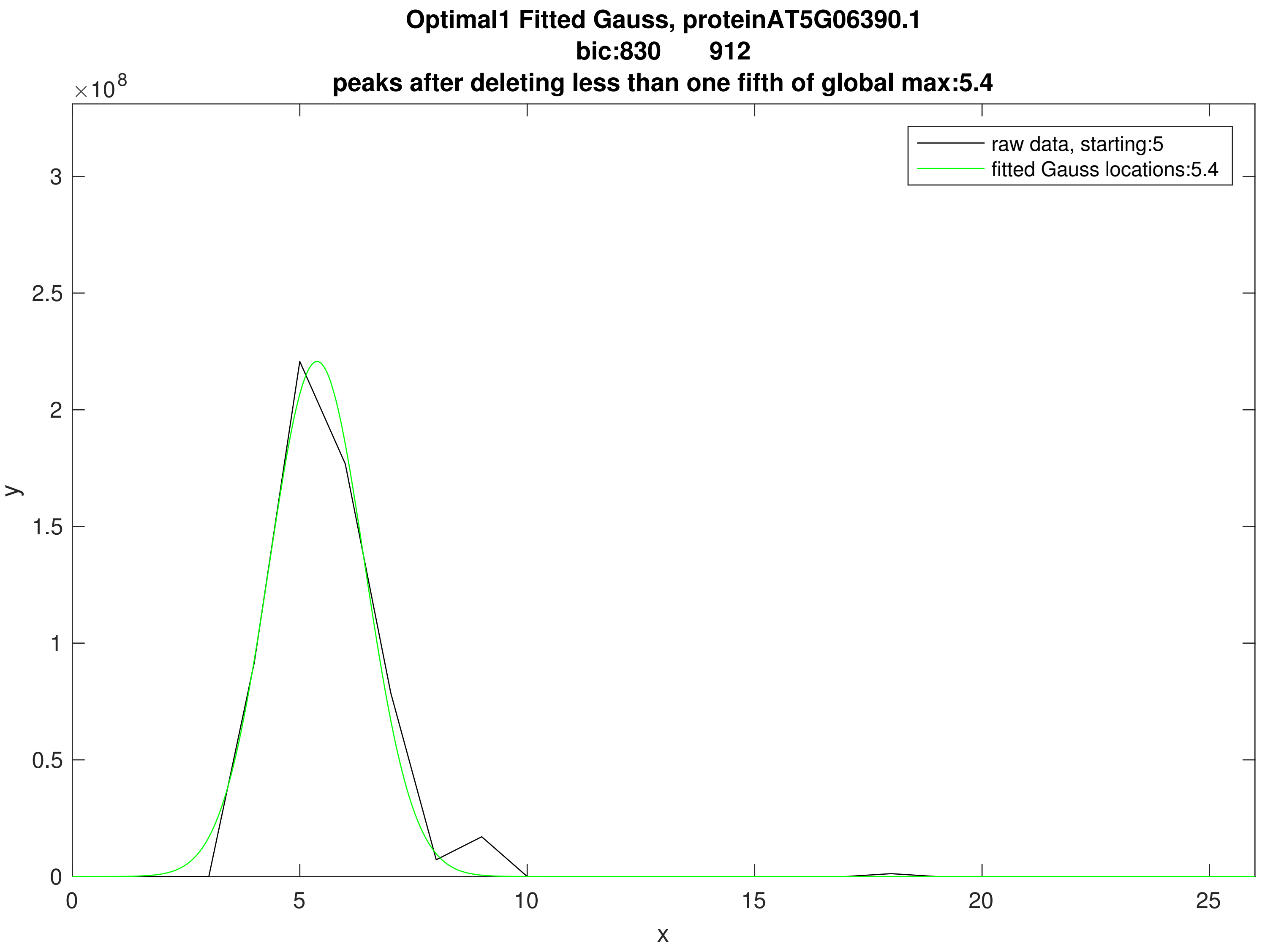

Optimal1 Fitted Gauss, proteinAT5G06390.1
bic:830 912
peaks after deleting less than one fifth of global max:5.4
×10^8
raw data, starting:5
fitted Gauss locations:5.4
y
x
0
0.5
1
1.5
2
2.5
3
0
5
10
15
20
25

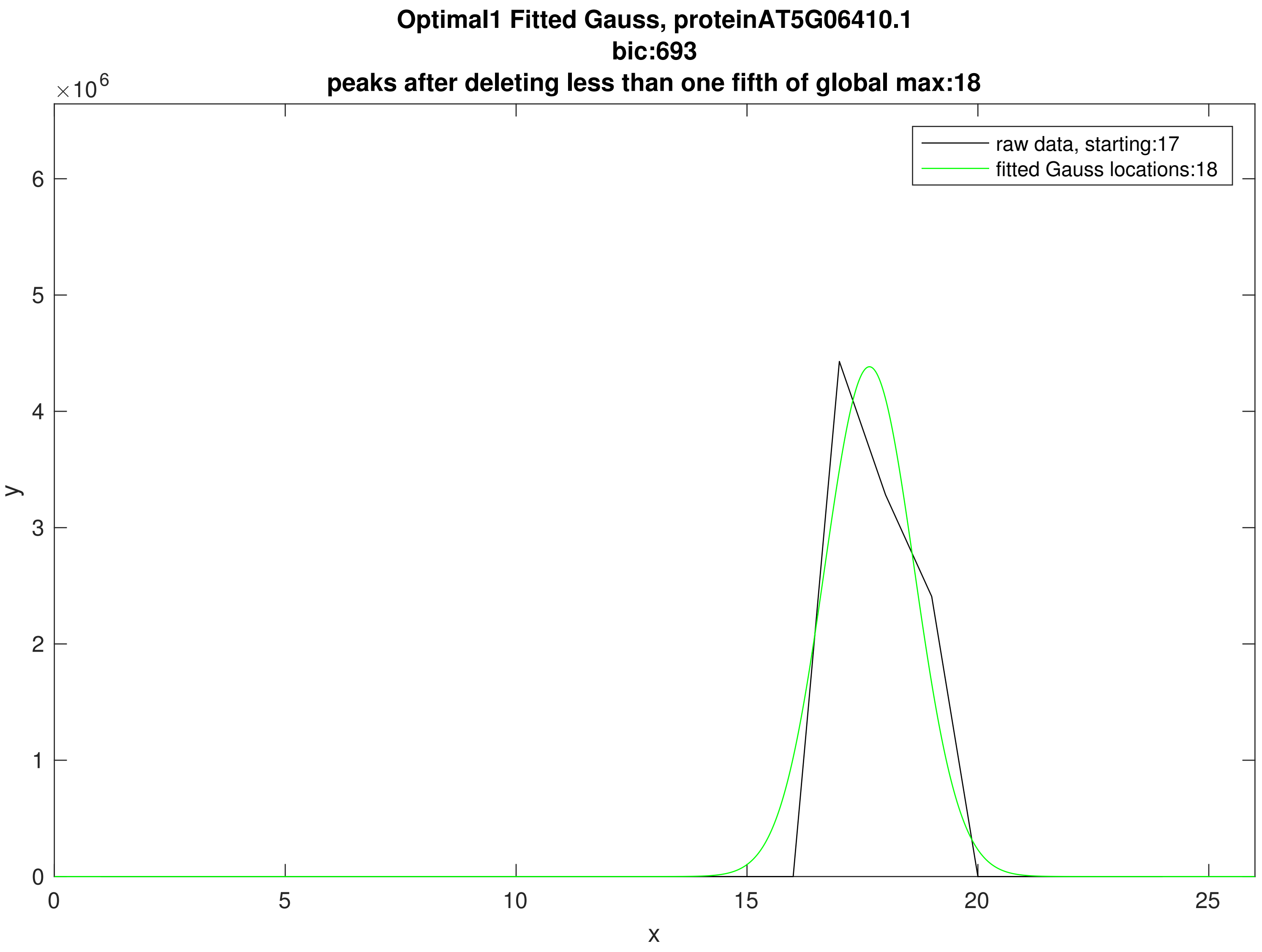

Optimal1 Fitted Gauss, proteinAT5G06410.1
bic:693
peaks after deleting less than one fifth of global max:18
×10^6
raw data, starting:17
fitted Gauss locations:18
y
x
0
1
2
3
4
5
6
0
5
10
15
20
25

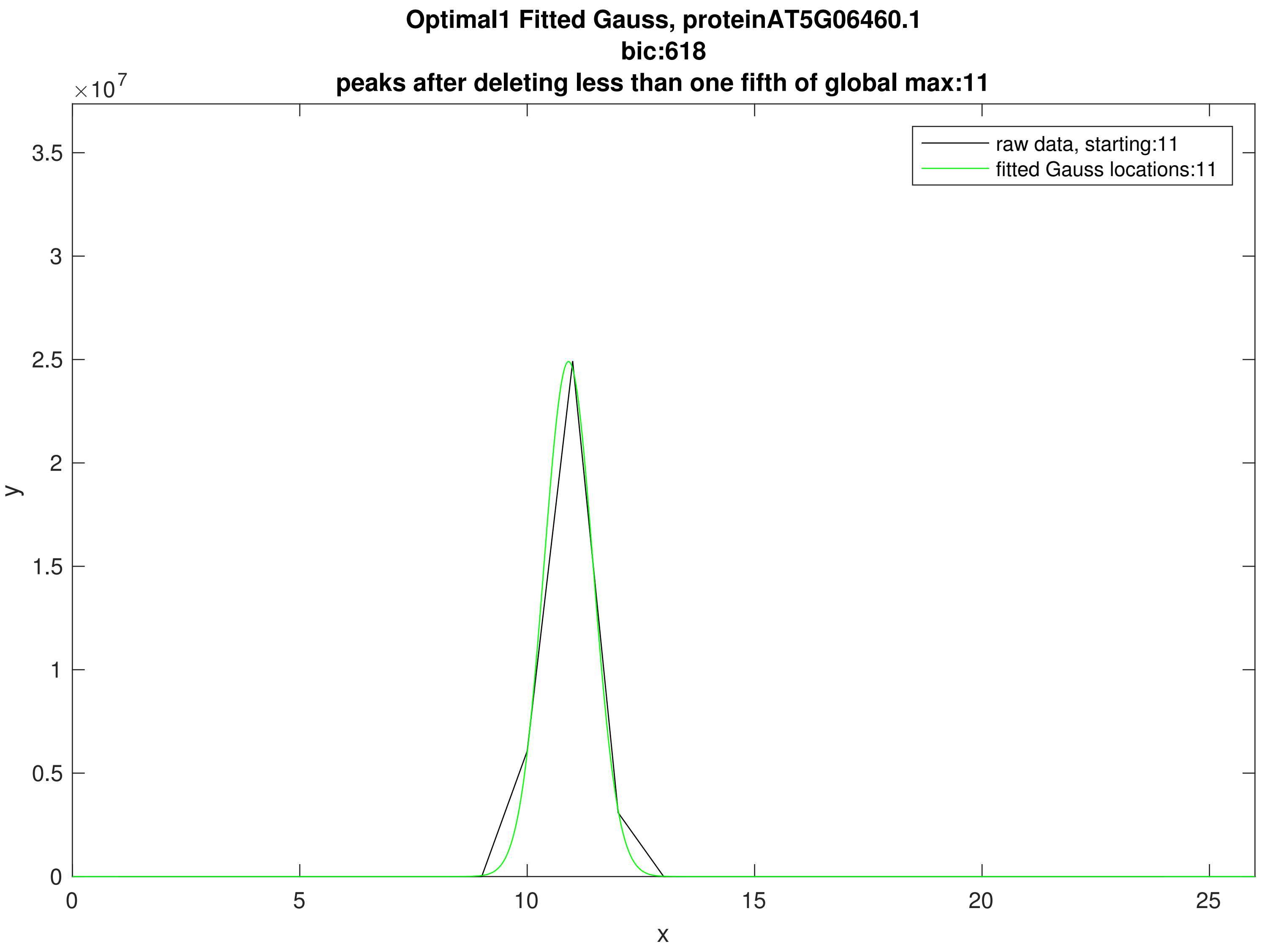
Optimal1 Fitted Gauss, proteinAT5G06460.1
bic:618
peaks after deleting less than one fifth of global max:11
×10^7
raw data, starting:11
fitted Gauss locations:11
3.5
3
2.5
2
1.5
1
0.5
0
y
0
5
10
15
20
25
x

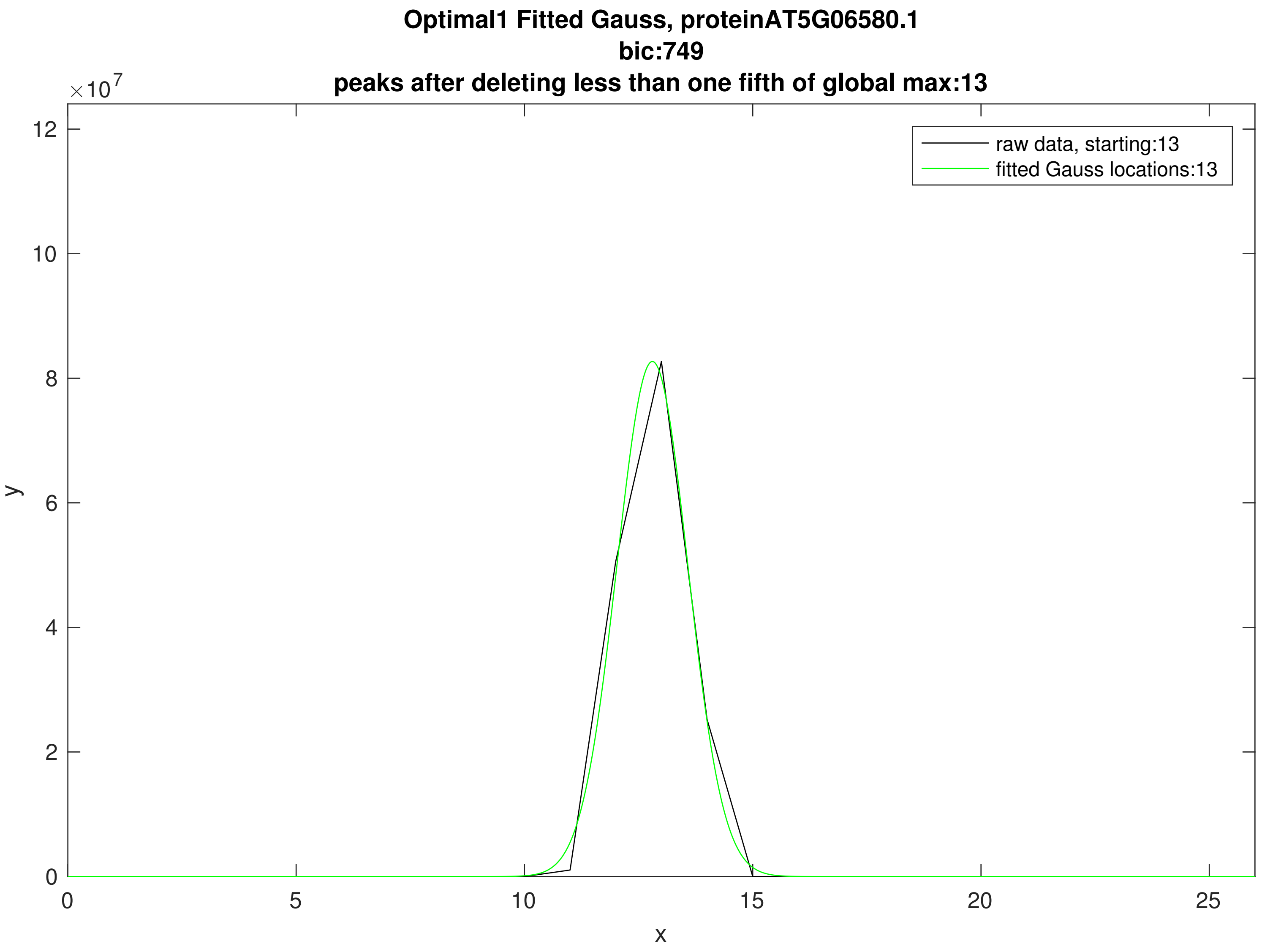
Optimal1 Fitted Gauss, proteinAT5G06580.1
bic:749
peaks after deleting less than one fifth of global max:13
×10^7
raw data, starting:13
fitted Gauss locations:13
y
x
0
2
4
6
8
10
12
0
5
10
15
20
25

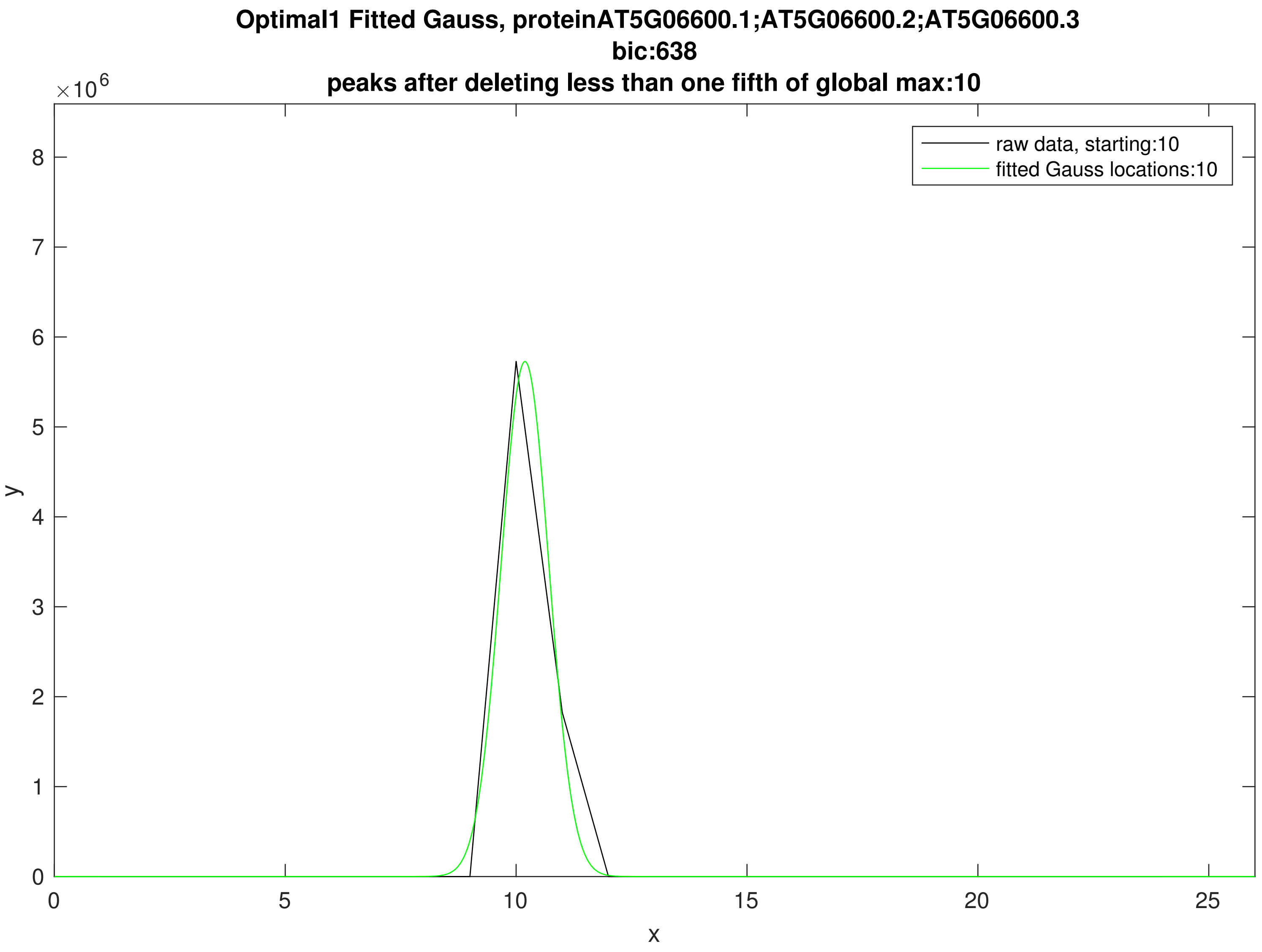
Optimal1 Fitted Gauss, proteinAT5G06600.1;AT5G06600.2;AT5G06600.3
bic:638
peaks after deleting less than one fifth of global max:10
×10^6
raw data, starting:10
fitted Gauss locations:10
y
x
0
1
2
3
4
5
6
7
8
0
5
10
15
20
25

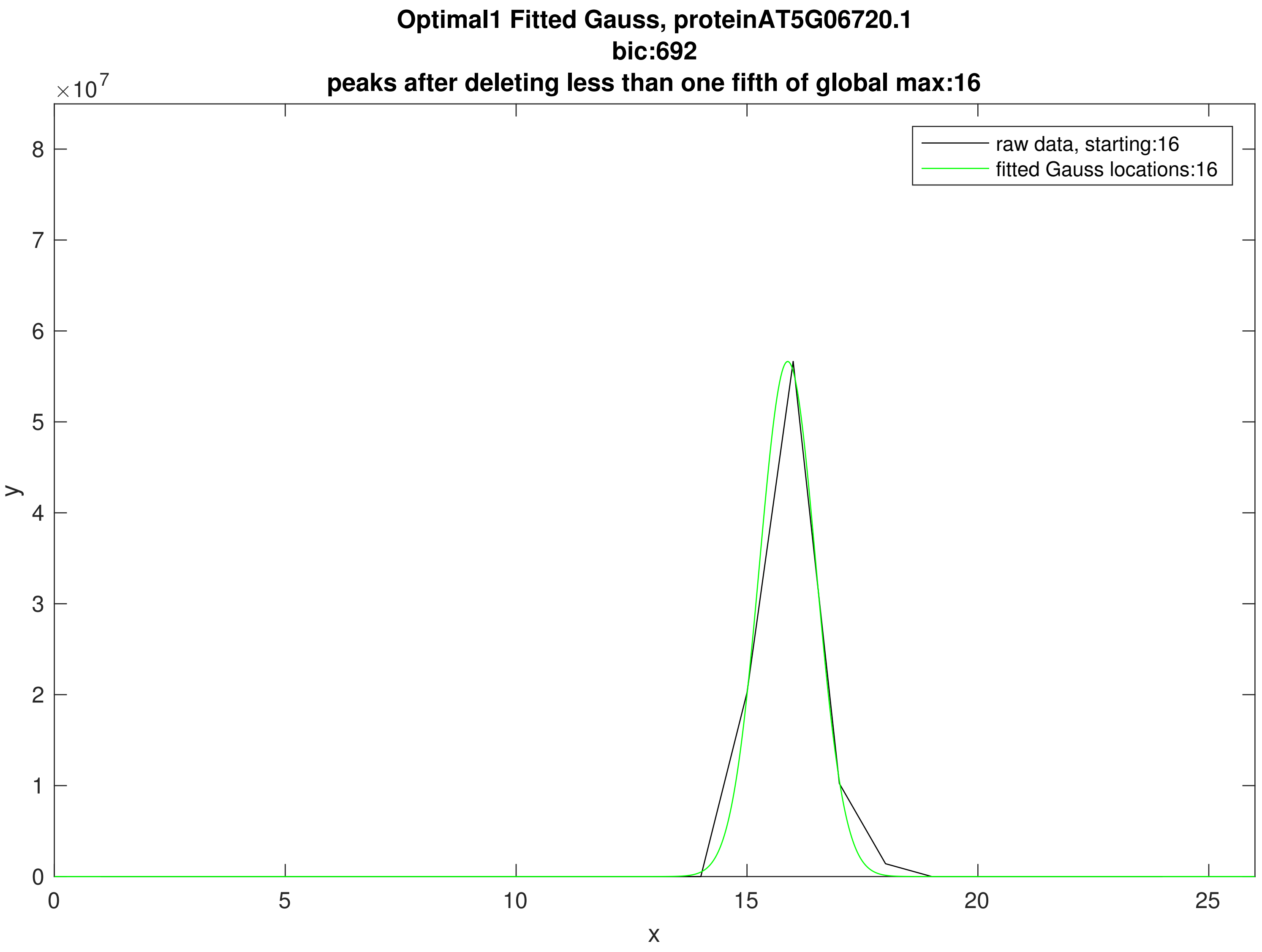
Optimal1 Fitted Gauss, proteinAT5G06720.1
bic:692
peaks after deleting less than one fifth of global max:16
×10^7
raw data, starting:16
fitted Gauss locations:16
y
x
0
1
2
3
4
5
6
7
8
10
15
20
25

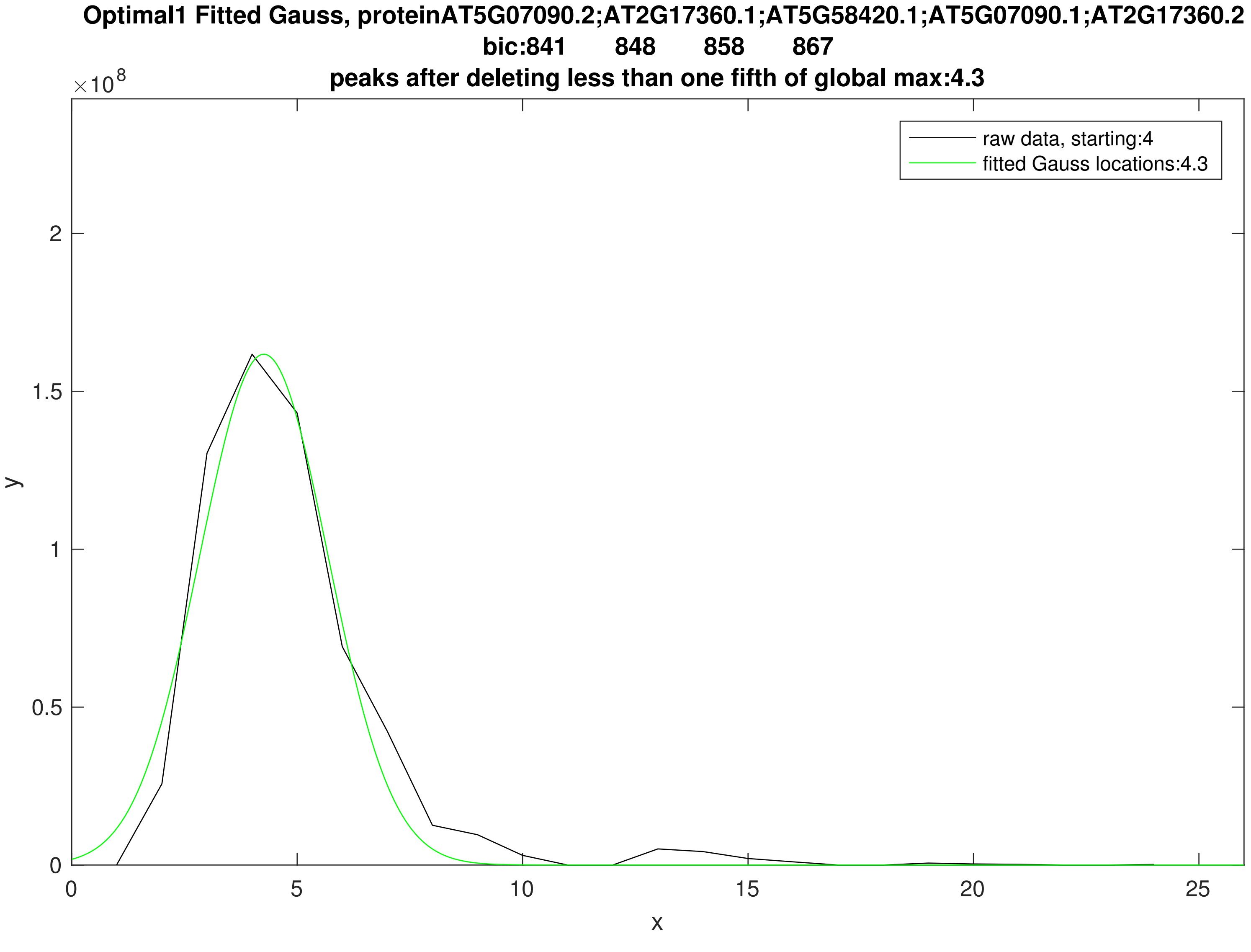
Optimal1 Fitted Gauss, proteinAT5G07090.2;AT2G17360.1;AT5G58420.1;AT5G07090.1;AT2G17360.2
bic:841 848 858 867
peaks after deleting less than one fifth of global max:4.3
×10^8
raw data, starting:4
fitted Gauss locations:4.3
2
1.5
1
0.5
0
y
0
5
10
15
20
25
x

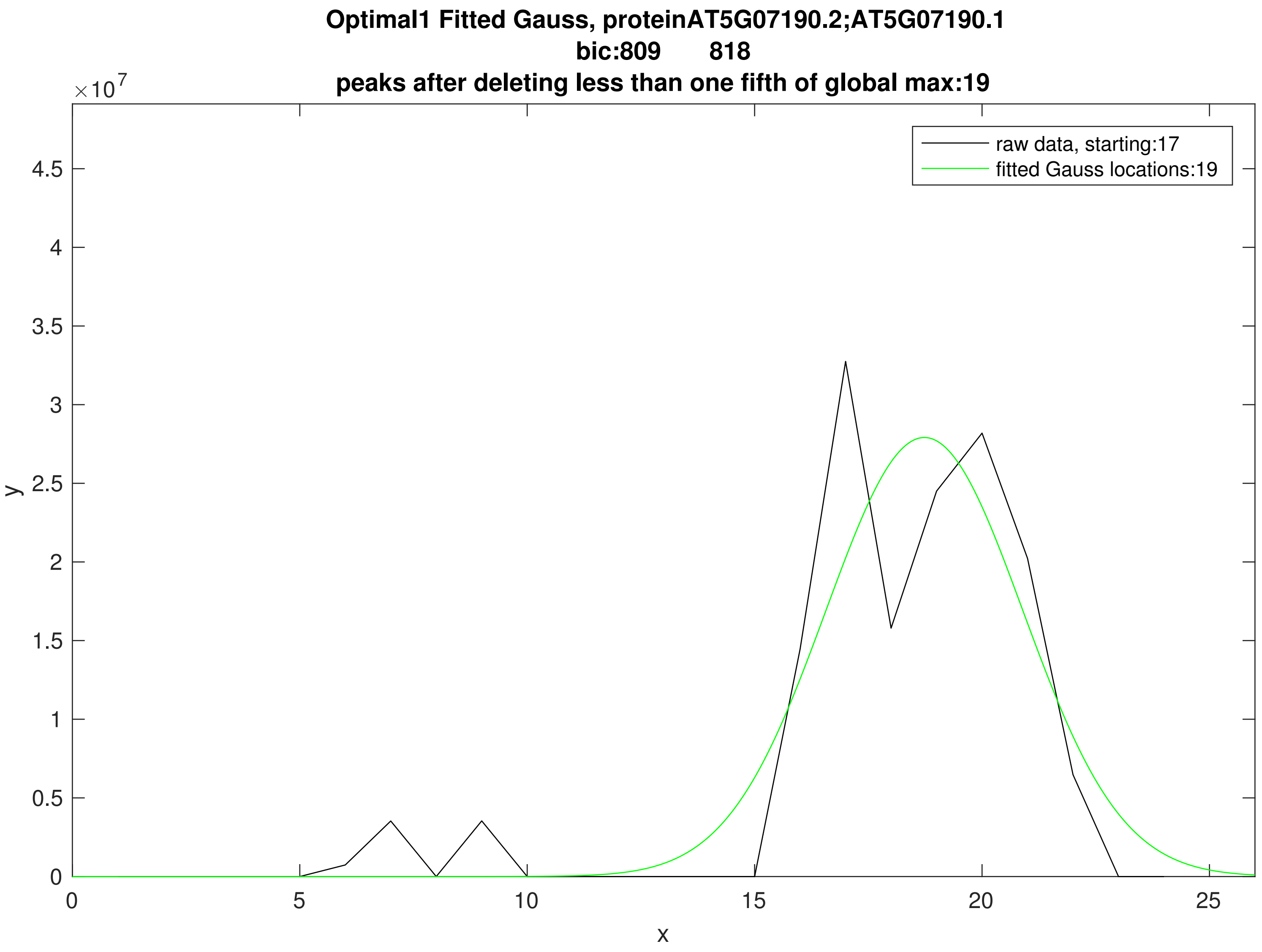

Optimal1 Fitted Gauss, proteinAT5G07190.2;AT5G07190.1
bic:809 818
peaks after deleting less than one fifth of global max:19
×10^7
raw data, starting:17
fitted Gauss locations:19
y
x
0
0.5
1
1.5
2
2.5
3
3.5
4
4.5
5
10
15
20
25

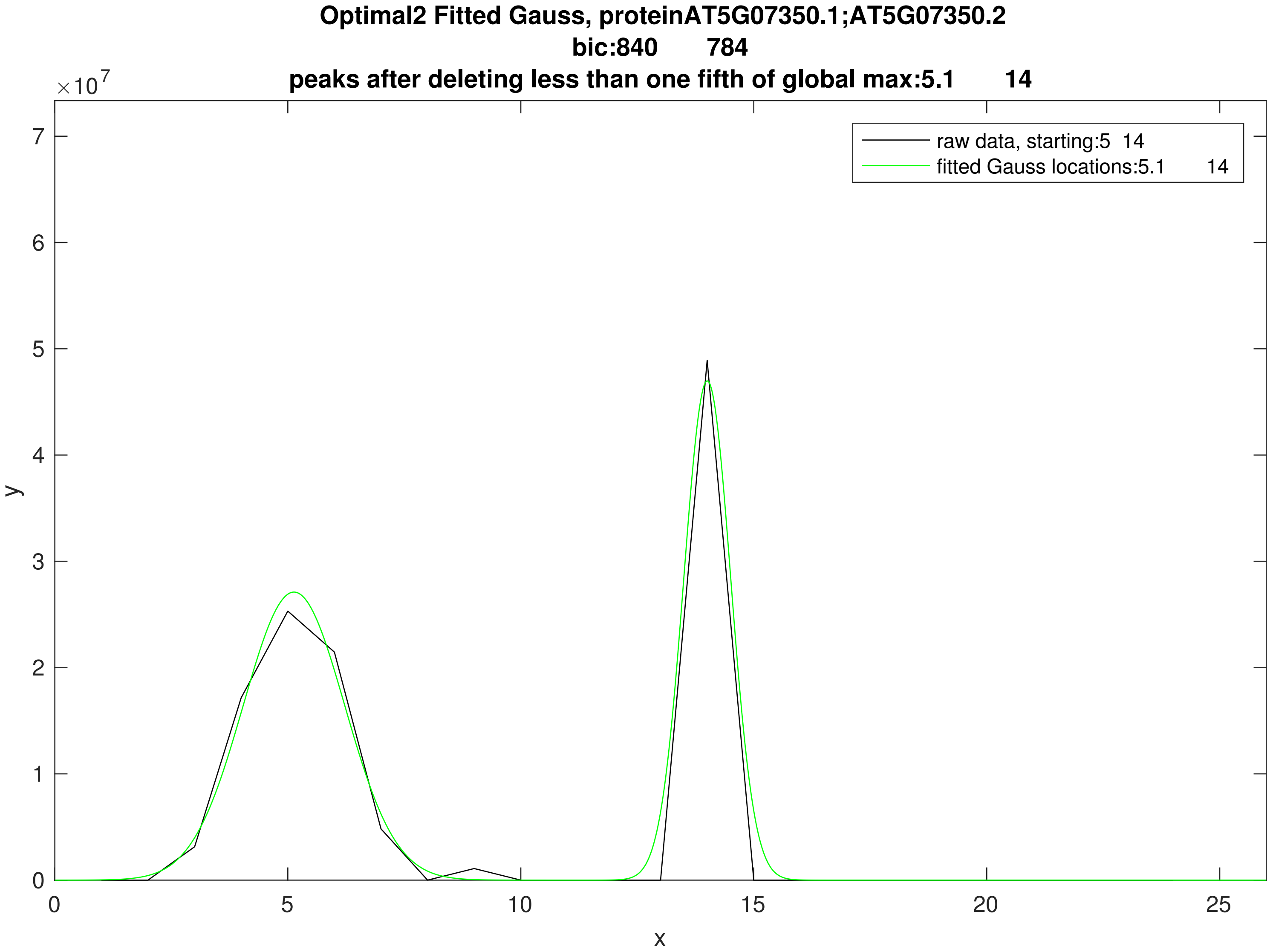

Optimal2 Fitted Gauss, proteinAT5G07350.1;AT5G07350.2
bic:840 784
peaks after deleting less than one fifth of global max:5.1 14
×10^7
raw data, starting:5 14
fitted Gauss locations:5.1 14
0
1
2
3
4
5
6
7
y
0
5
10
15
20
25
x

**Optimal2 Fitted Gauss, proteinAT5G07360.2;AT5G07360.1**
**bic:807 806**
**peaks after deleting less than one fifth of global max:12**

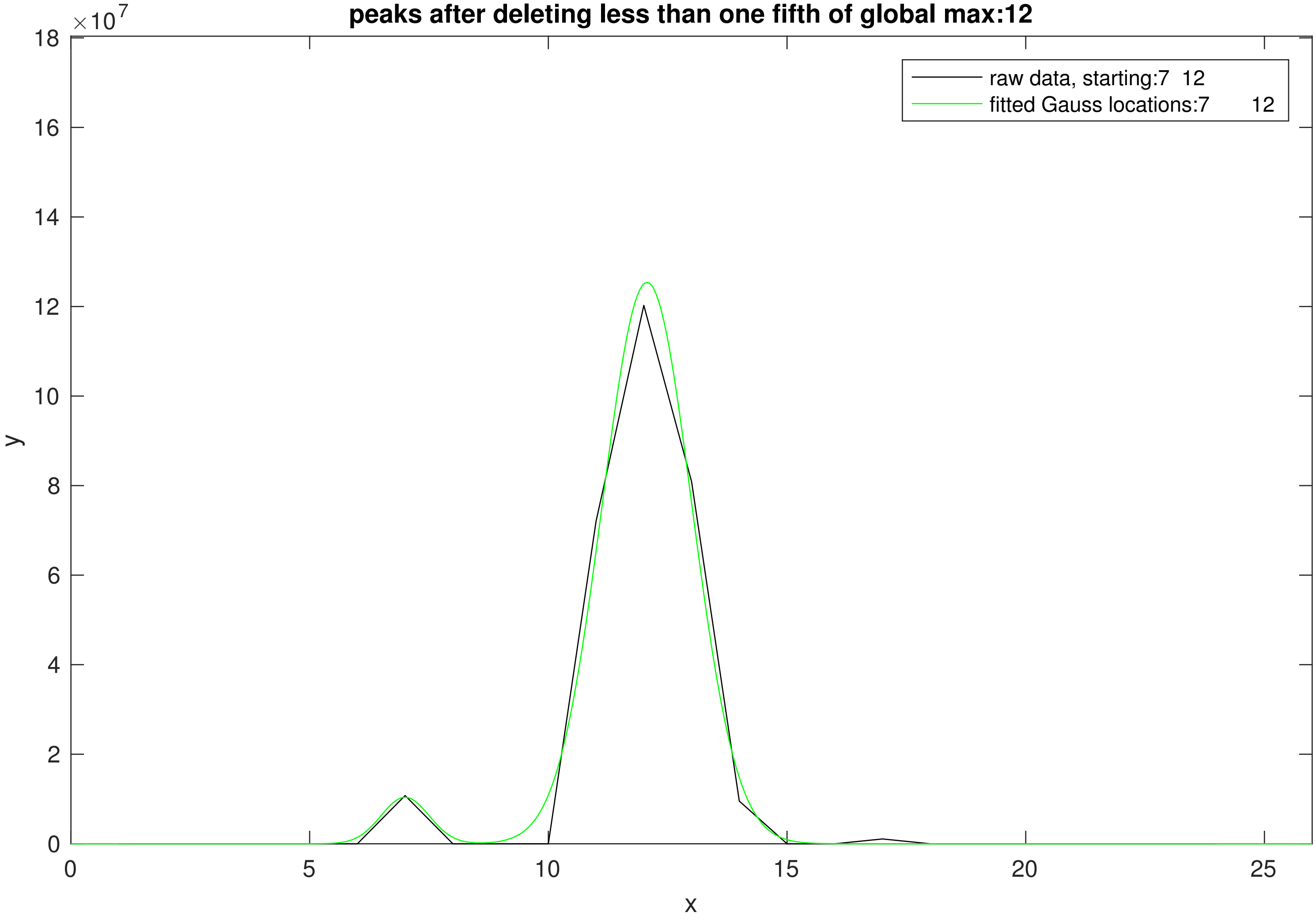

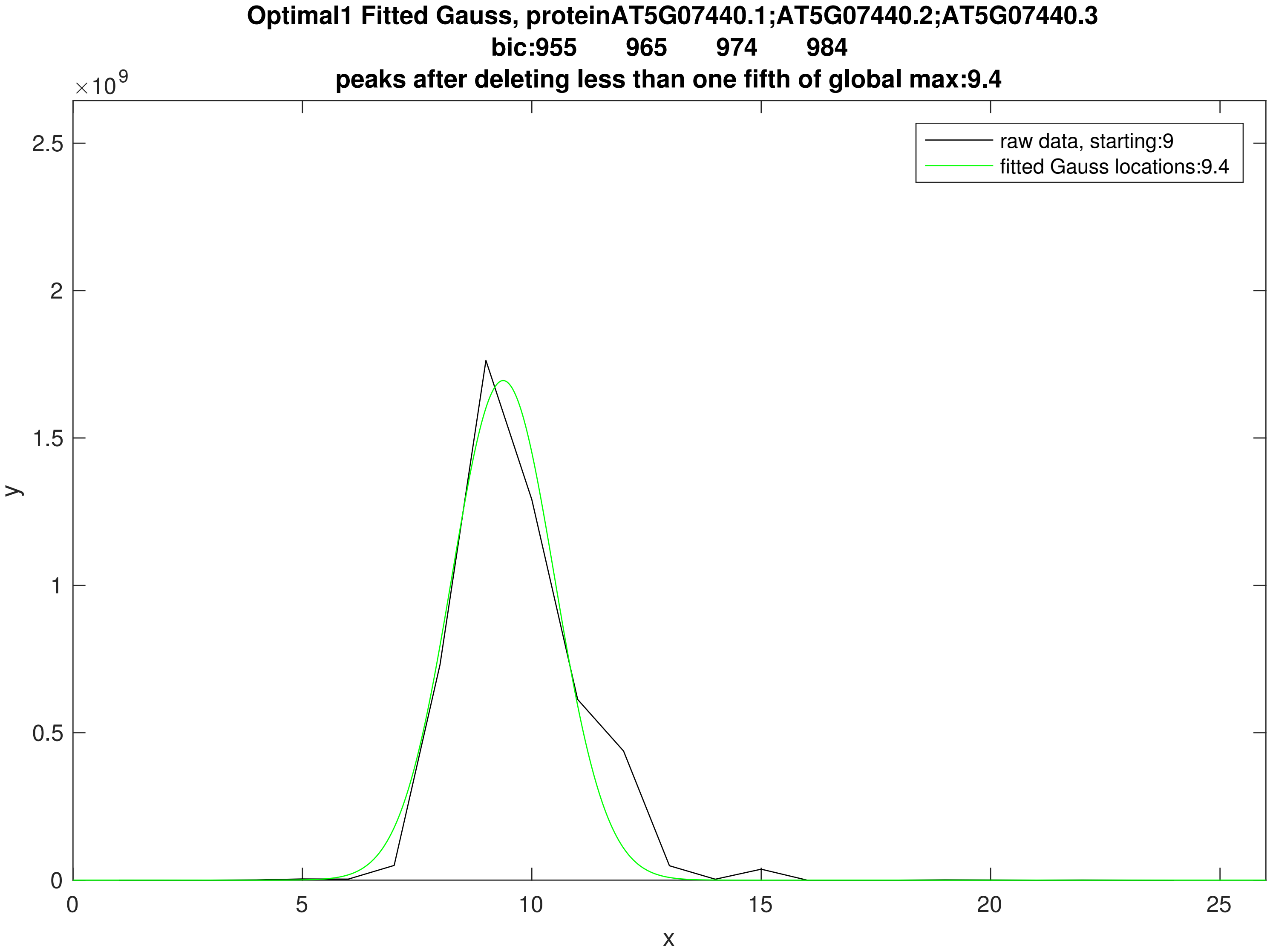
Optimal1 Fitted Gauss, proteinAT5G07440.1;AT5G07440.2;AT5G07440.3
bic:955 965 974 984
peaks after deleting less than one fifth of global max:9.4
×10^9
raw data, starting:9
fitted Gauss locations:9.4
y
x
0
0.5
1
1.5
2
2.5
5
10
15
20
25

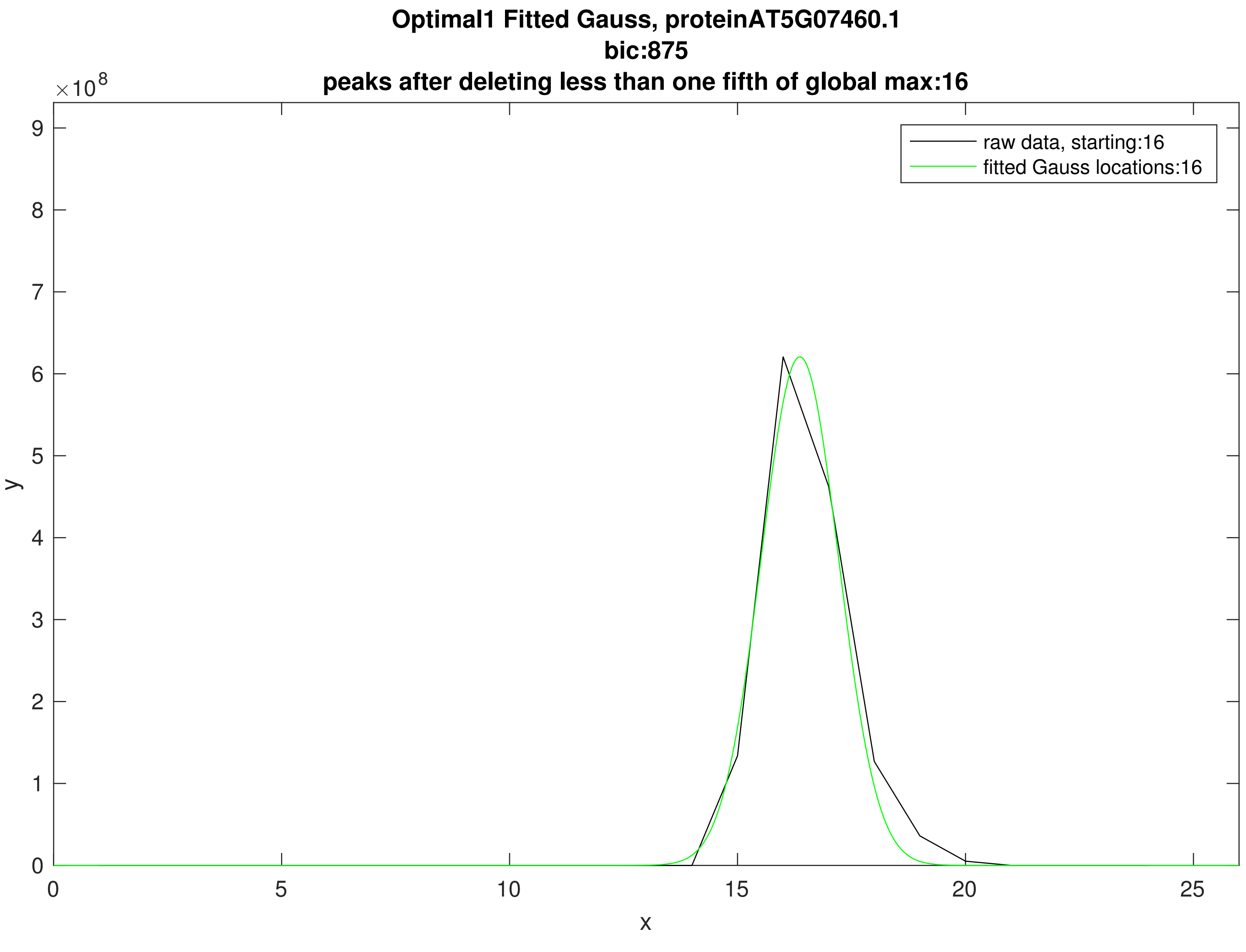
Optimal1 Fitted Gauss, proteinAT5G07460.1
bic:875
peaks after deleting less than one fifth of global max:16
×10^8
raw data, starting:16
fitted Gauss locations:16
y
x
0
1
2
3
4
5
6
7
8
9
0
5
10
15
20
25

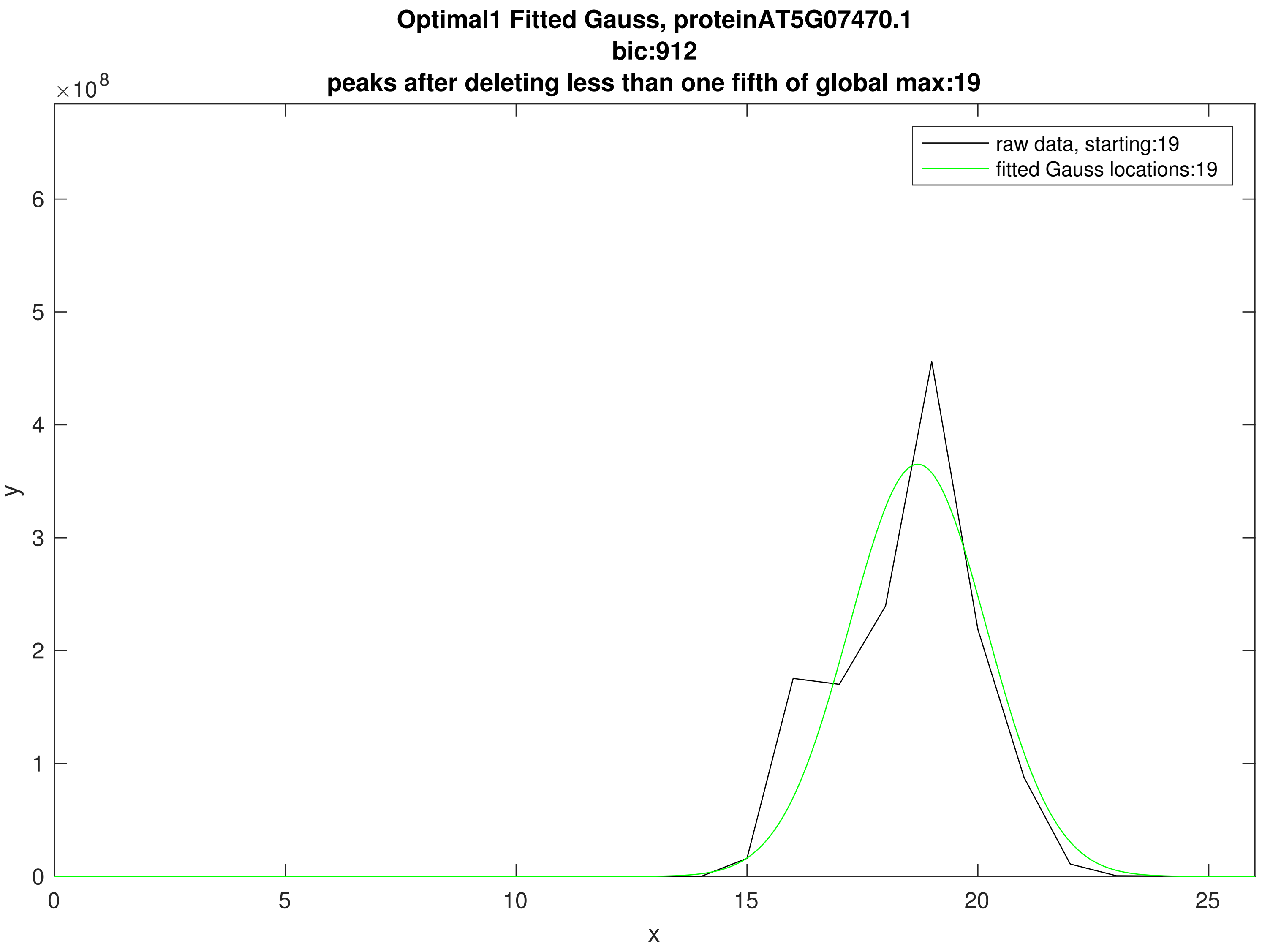
Optimal1 Fitted Gauss, proteinAT5G07470.1
bic:912
peaks after deleting less than one fifth of global max:19
×10^8
raw data, starting:19
fitted Gauss locations:19
y
x
0
1
2
3
4
5
6
0
5
10
15
20
25

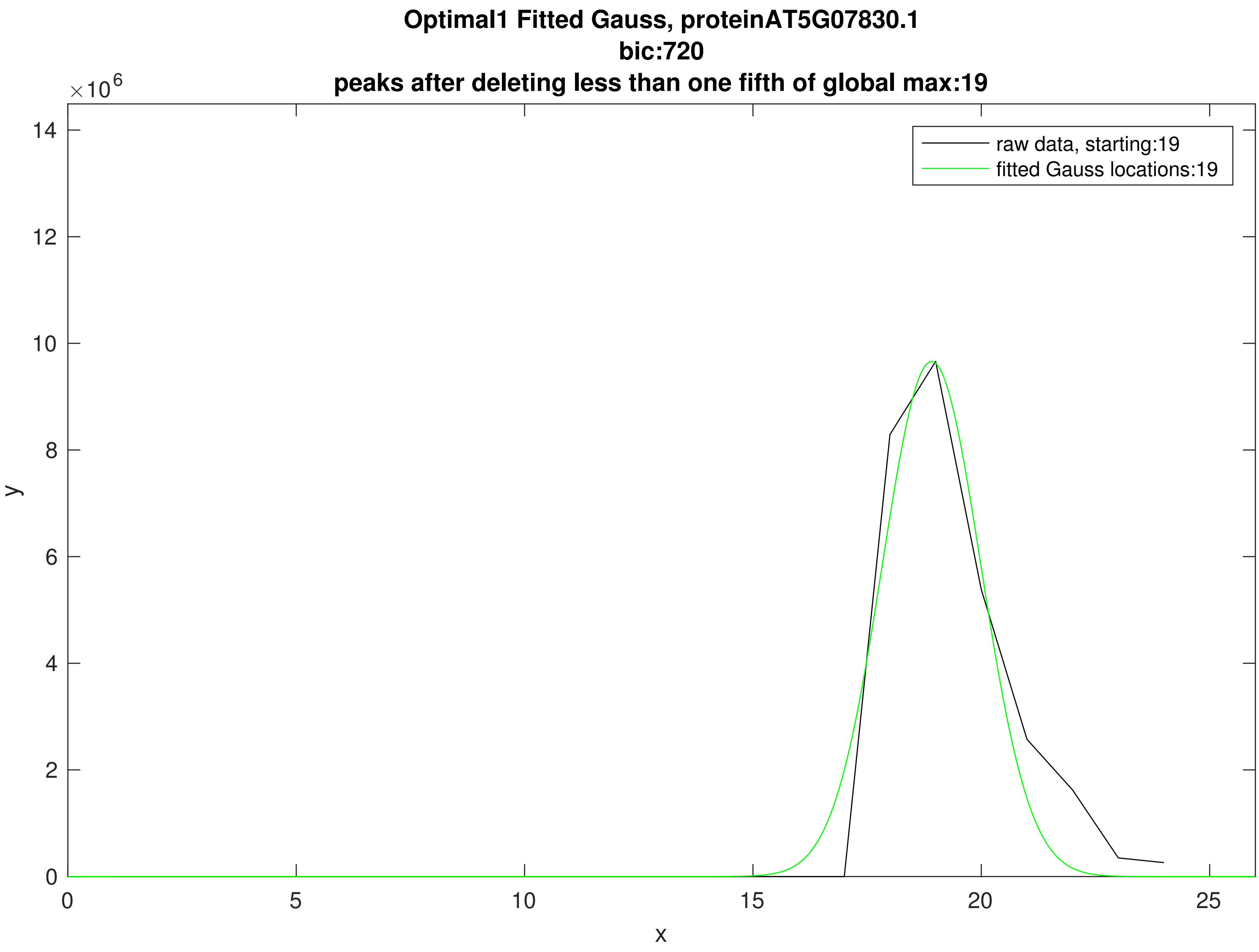

Optimal1 Fitted Gauss, proteinAT5G07830.1
bic:720
peaks after deleting less than one fifth of global max:19
×10^6
14
12
10
8
6
4
2
0
y
0
5
10
15
20
25
x
raw data, starting:19
fitted Gauss locations:19

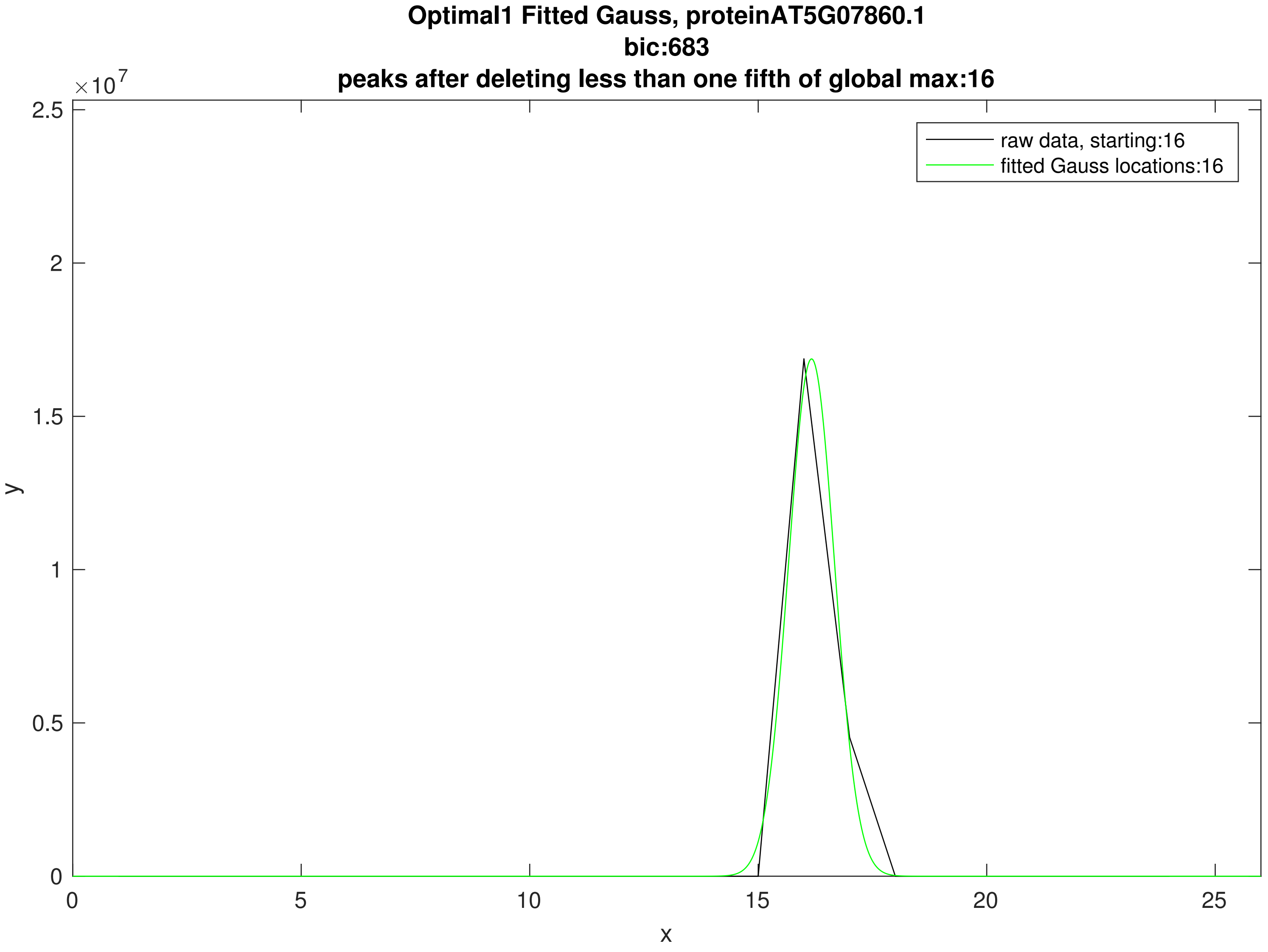

Optimal1 Fitted Gauss, proteinAT5G07860.1
bic:683
peaks after deleting less than one fifth of global max:16
×10^7
2.5
2
1.5
1
0.5
0
y
0
5
10
15
20
25
x
raw data, starting:16
fitted Gauss locations:16

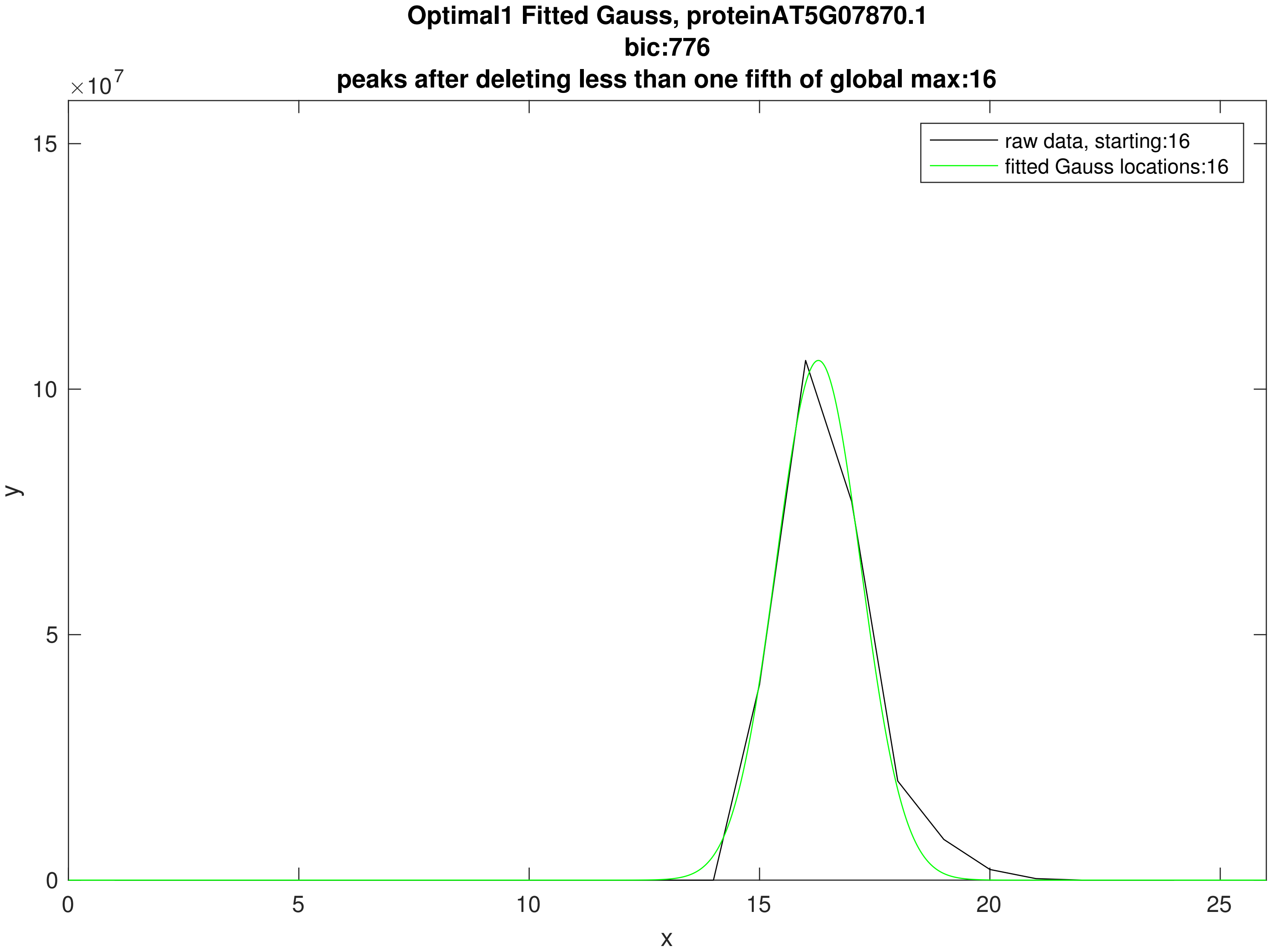

Optimal1 Fitted Gauss, proteinAT5G07870.1
bic:776
peaks after deleting less than one fifth of global max:16
×10^7
15
10
5
0
y
0
5
10
15
20
25
x
raw data, starting:16
fitted Gauss locations:16

**Optimal1 Fitted Gauss, proteinAT5G08000.1**
**bic:750**
**peaks after deleting less than one fifth of global max:17**

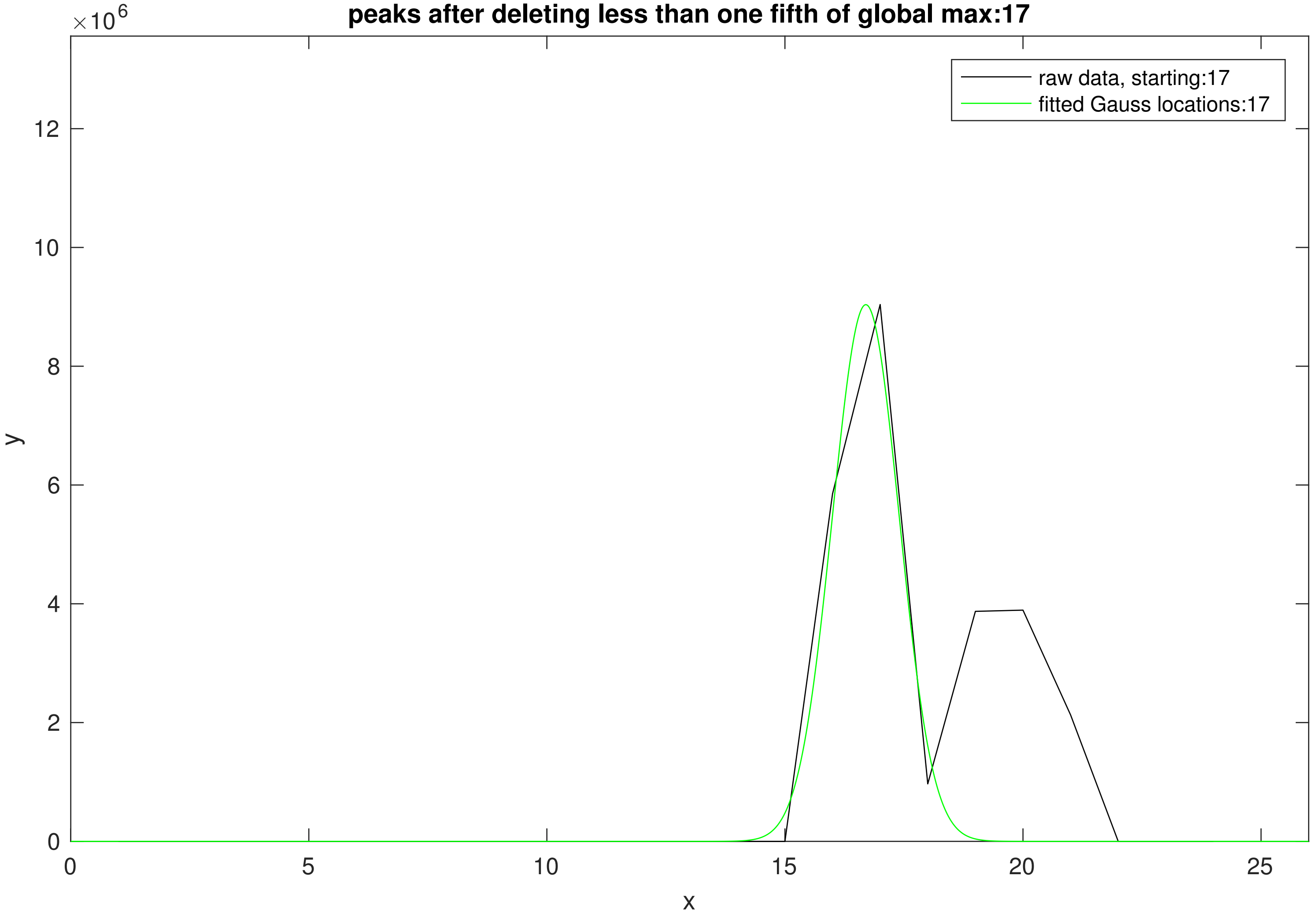

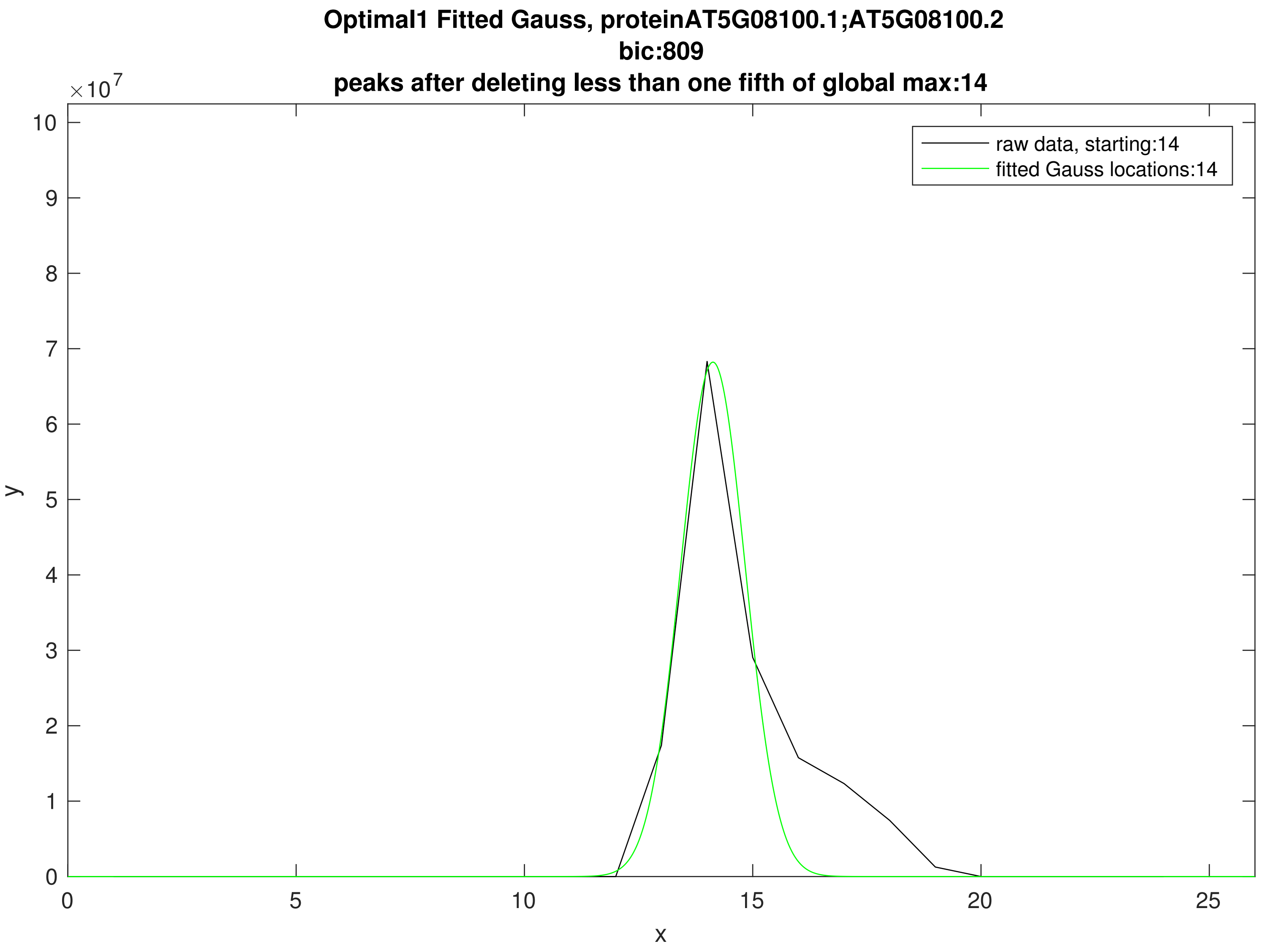
Optimal1 Fitted Gauss, proteinAT5G08100.1;AT5G08100.2
bic:809
peaks after deleting less than one fifth of global max:14
×10^7
raw data, starting:14
fitted Gauss locations:14
y
x
0
1
2
3
4
5
6
7
8
9
10
0
5
10
15
20
25

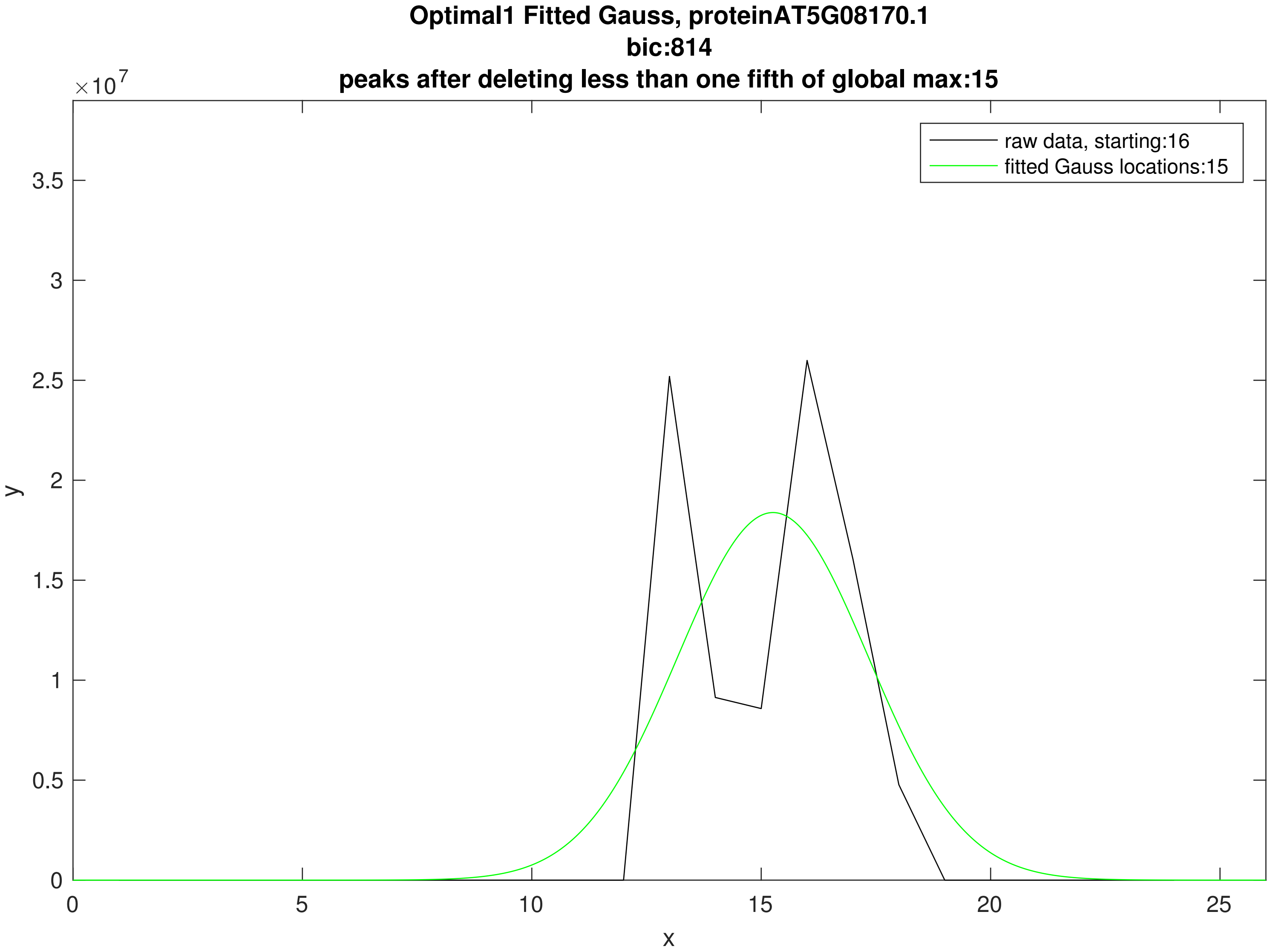

Optimal1 Fitted Gauss, proteinAT5G08170.1
bic:814
peaks after deleting less than one fifth of global max:15
×10^7
3.5
3
2.5
2
1.5
1
0.5
0
y
0
5
10
15
20
25
x
raw data, starting:16
fitted Gauss locations:15

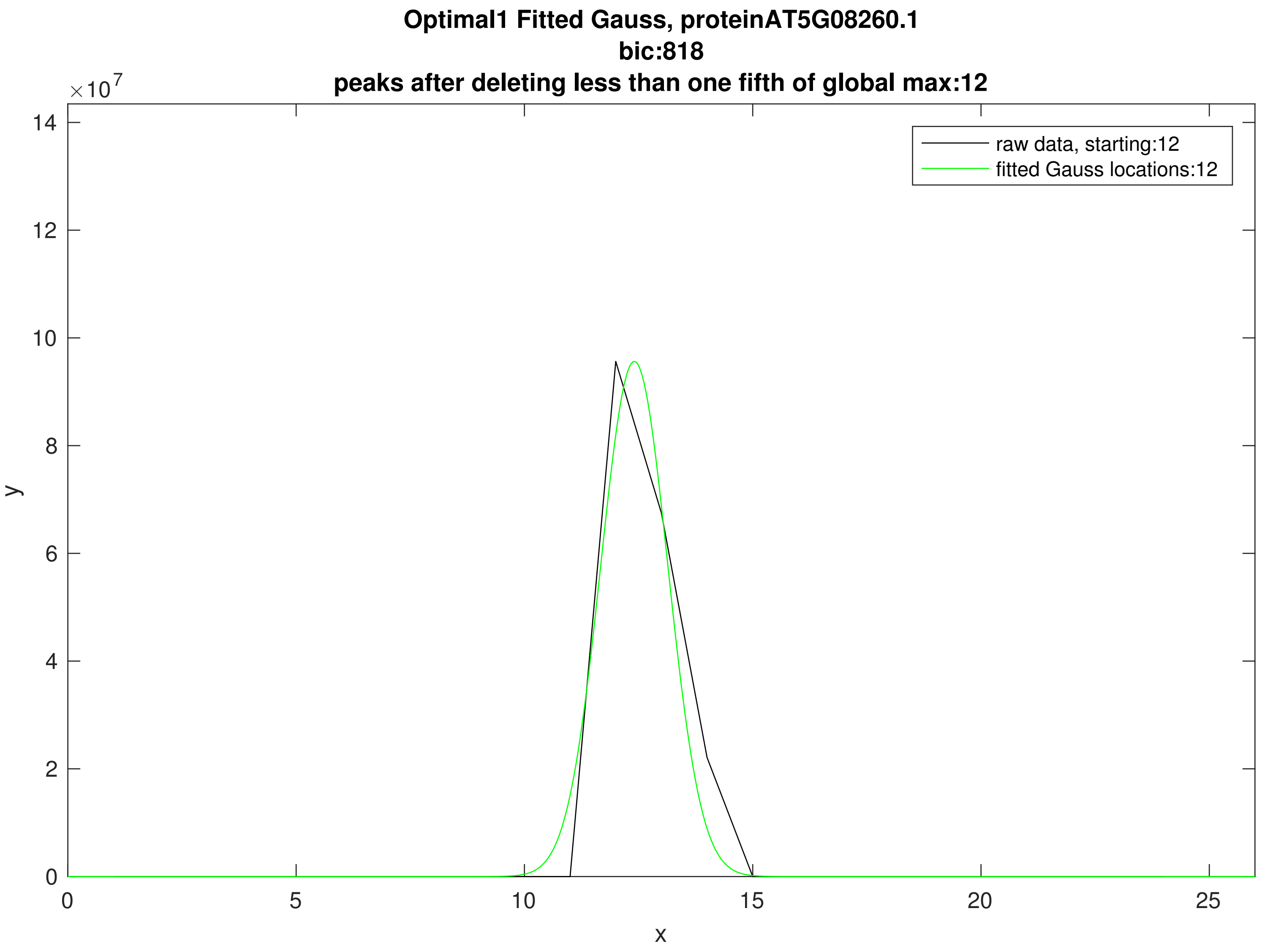
Optimal1 Fitted Gauss, proteinAT5G08260.1
bic:818
peaks after deleting less than one fifth of global max:12
×10^7
raw data, starting:12
fitted Gauss locations:12
y
x
0
2
4
6
8
10
12
14
0
5
10
15
20
25

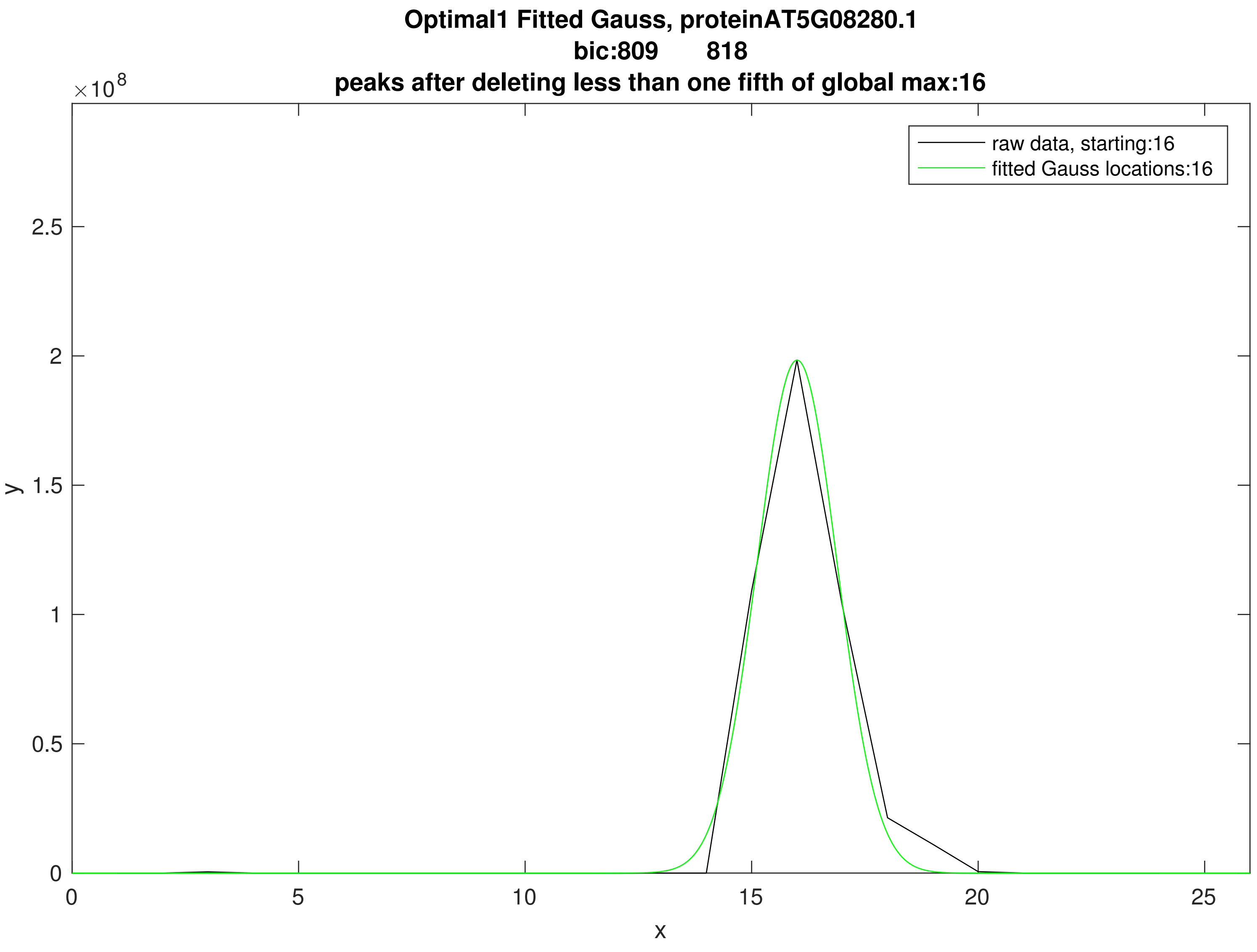
Optimal1 Fitted Gauss, proteinAT5G08280.1
bic:809 818
peaks after deleting less than one fifth of global max:16
×10^8
raw data, starting:16
fitted Gauss locations:16
2.5
2
1.5
1
0.5
0
y
0
5
10
15
20
25
x

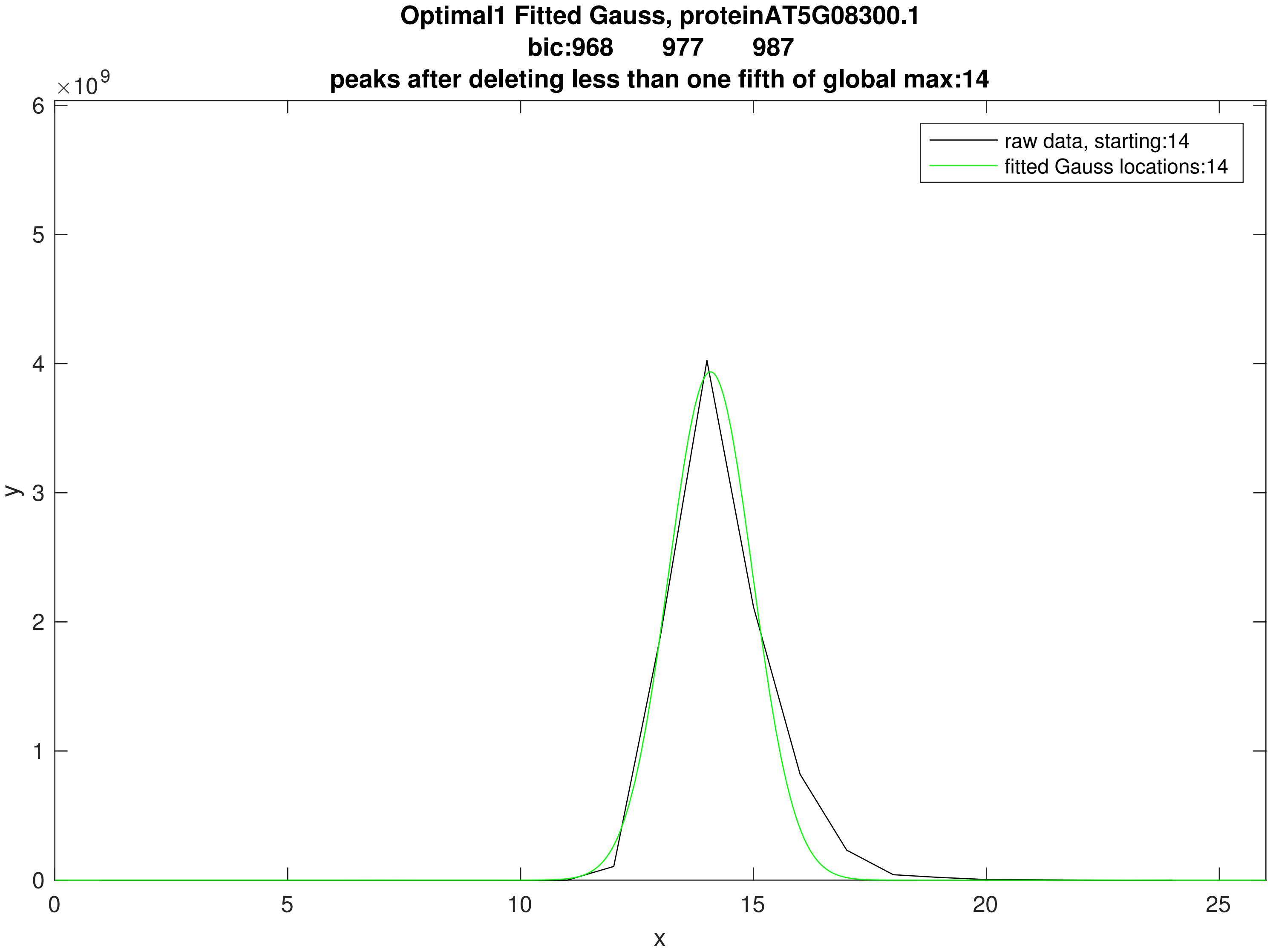

Optimal1 Fitted Gauss, proteinAT5G08300.1
bic:968 977 987
peaks after deleting less than one fifth of global max:14
×10^9
6
5
4
3
2
1
0
y
0
5
10
15
20
25
x
raw data, starting:14
fitted Gauss locations:14

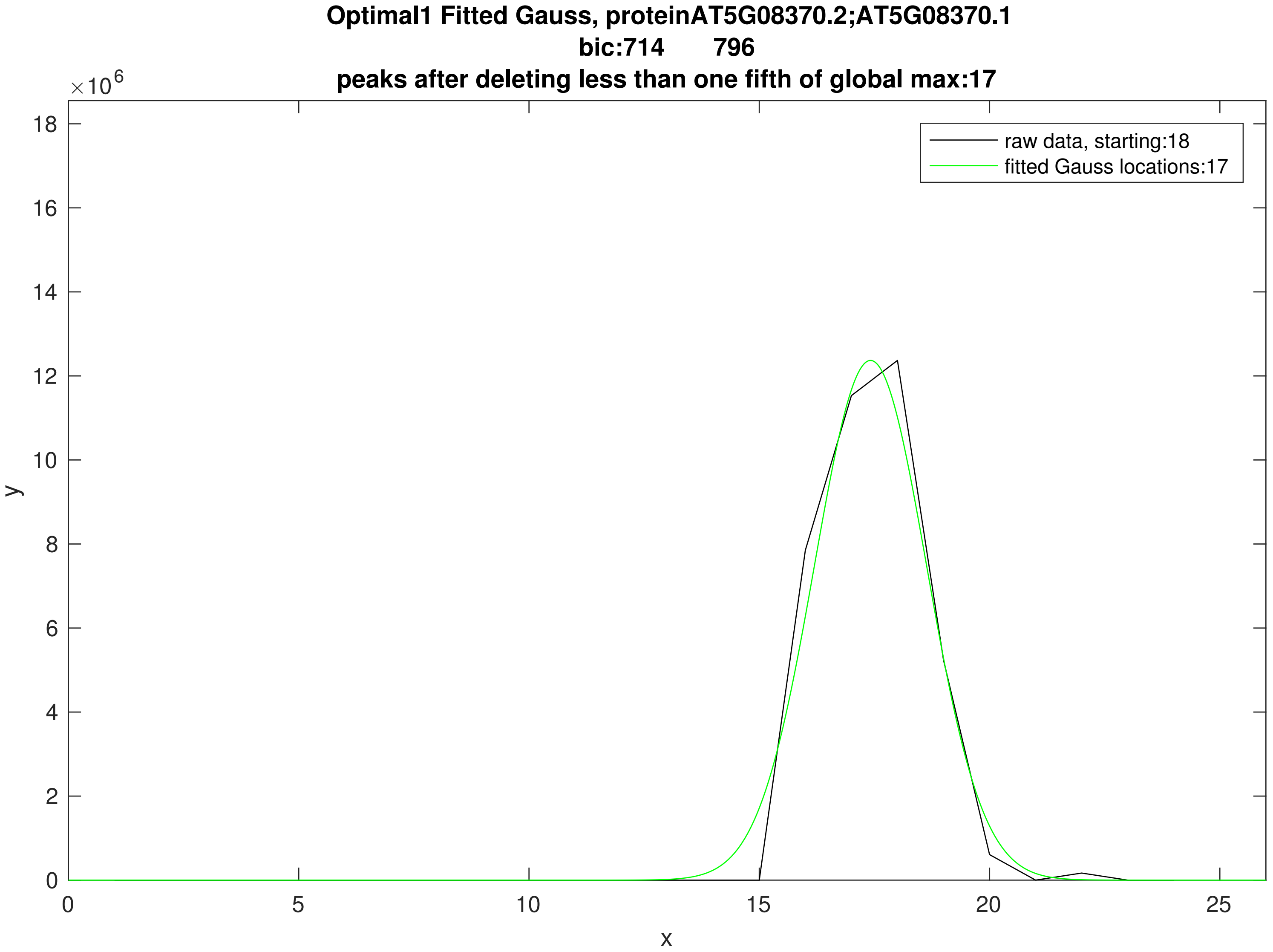
Optimal1 Fitted Gauss, proteinAT5G08370.2;AT5G08370.1
bic:714 796
peaks after deleting less than one fifth of global max:17
×10^6
raw data, starting:18
fitted Gauss locations:17
y
x
0
2
4
6
8
10
12
14
16
18
0
5
10
15
20
25

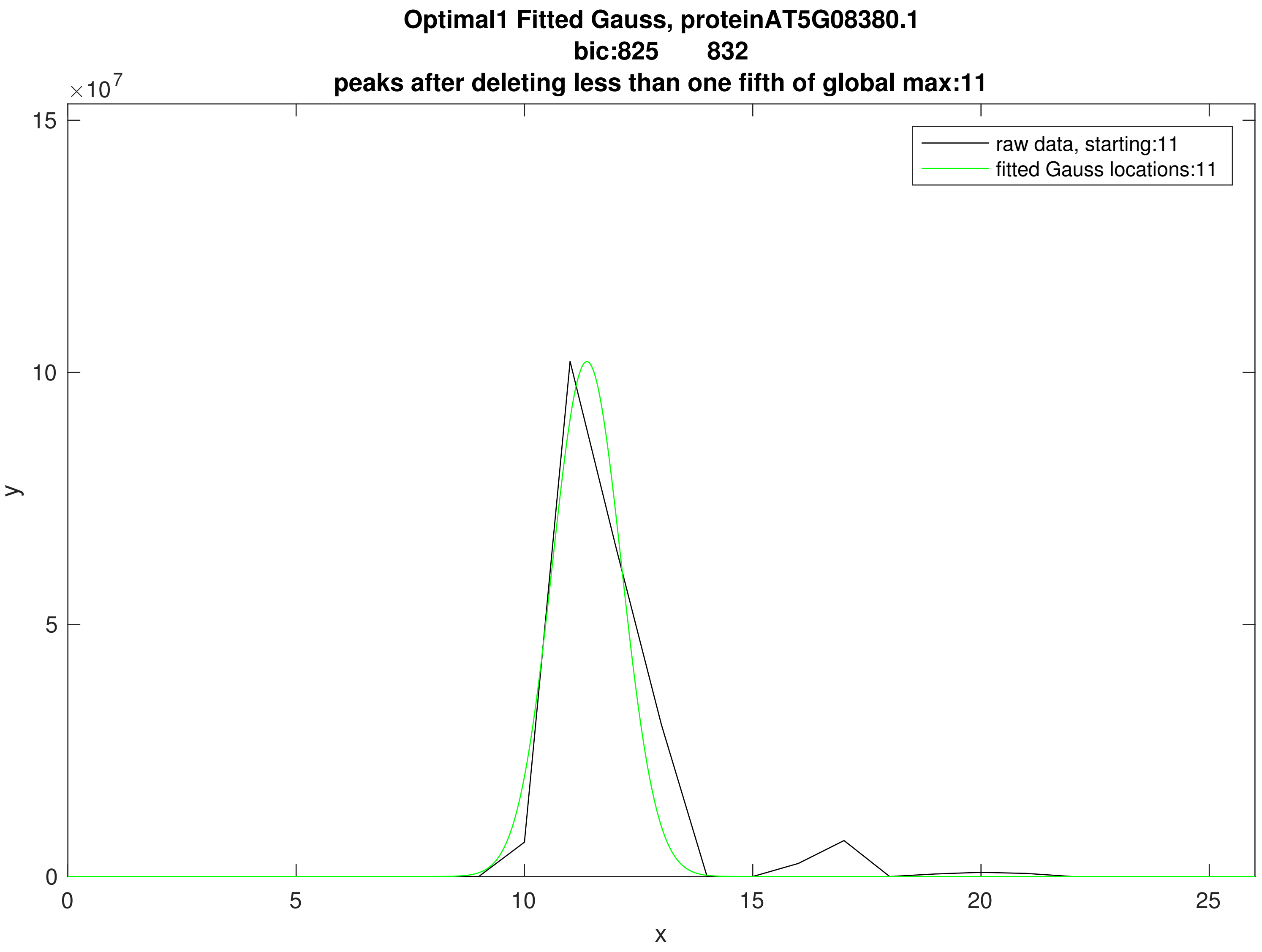

Optimal1 Fitted Gauss, proteinAT5G08380.1
bic:825 832
peaks after deleting less than one fifth of global max:11
×10^7
15
10
5
0
y
0
5
10
15
20
25
x
raw data, starting:11
fitted Gauss locations:11

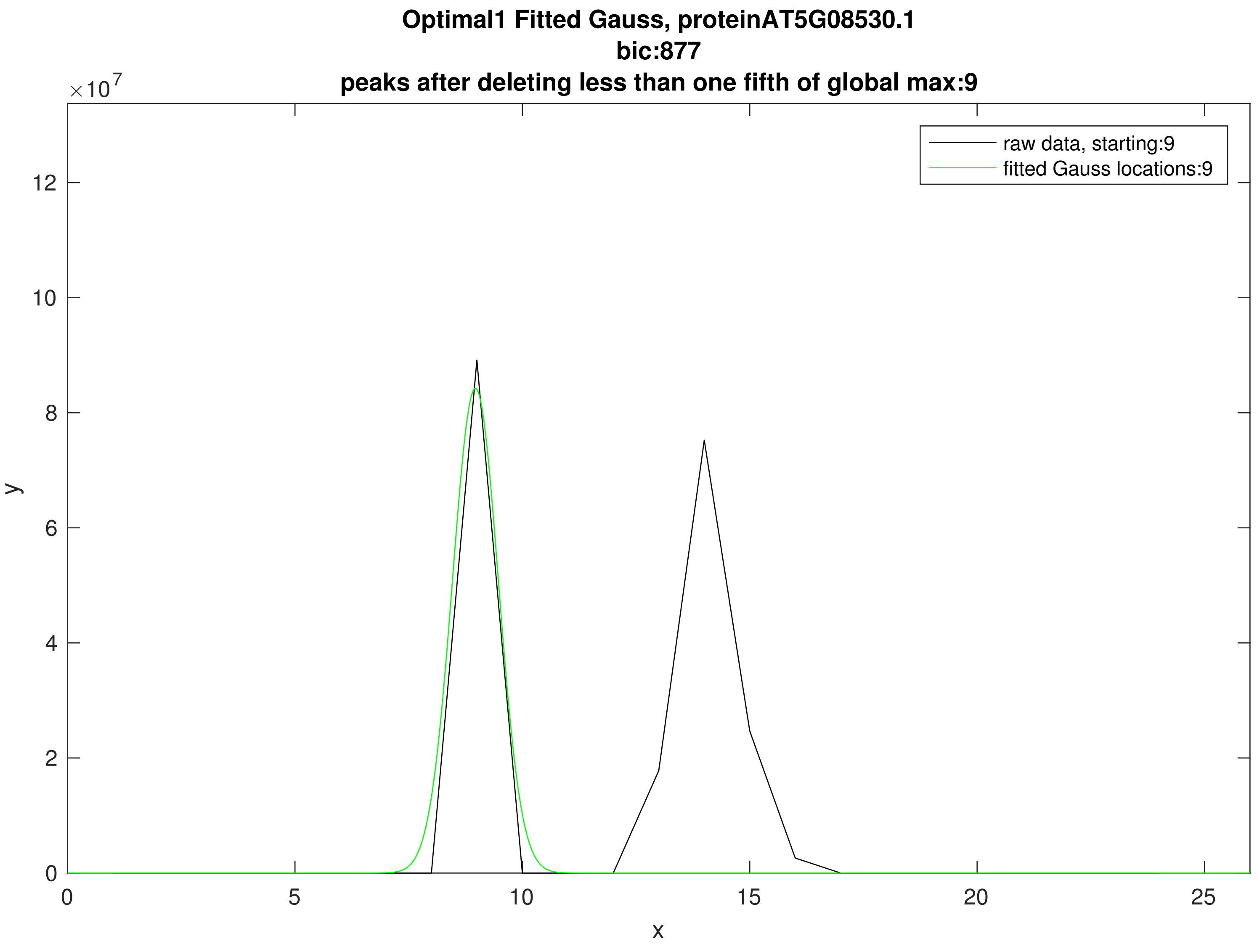

Optimal1 Fitted Gauss, proteinAT5G08530.1
bic:877
peaks after deleting less than one fifth of global max:9
×10^7
raw data, starting:9
fitted Gauss locations:9
12
10
8
6
4
2
0
y
0
5
10
15
20
25
x

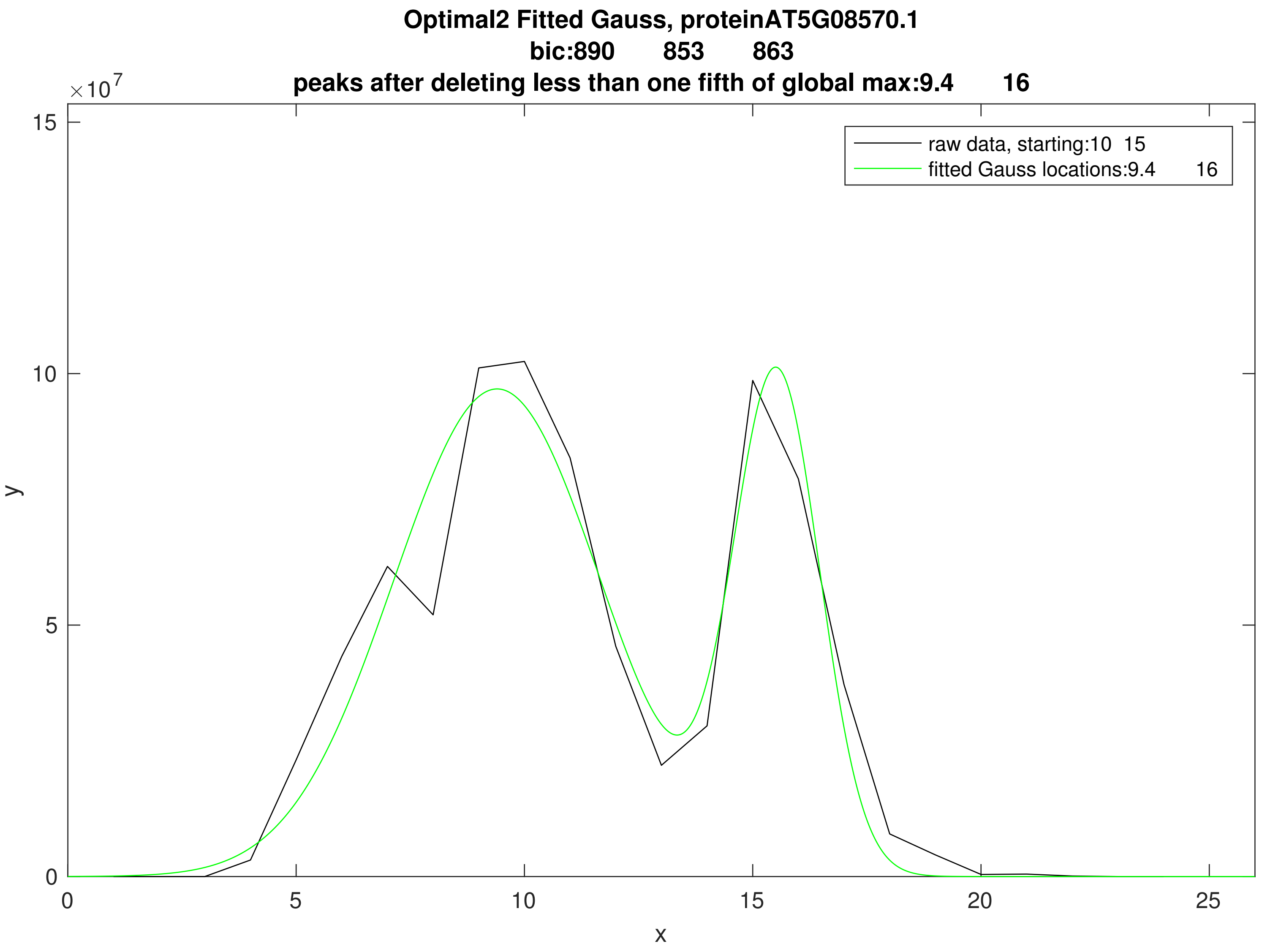

Optimal2 Fitted Gauss, proteinAT5G08570.1
bic:890 853 863
peaks after deleting less than one fifth of global max:9.4 16
×10^7
raw data, starting:10 15
fitted Gauss locations:9.4 16
y
x
0
5
10
15
20
25

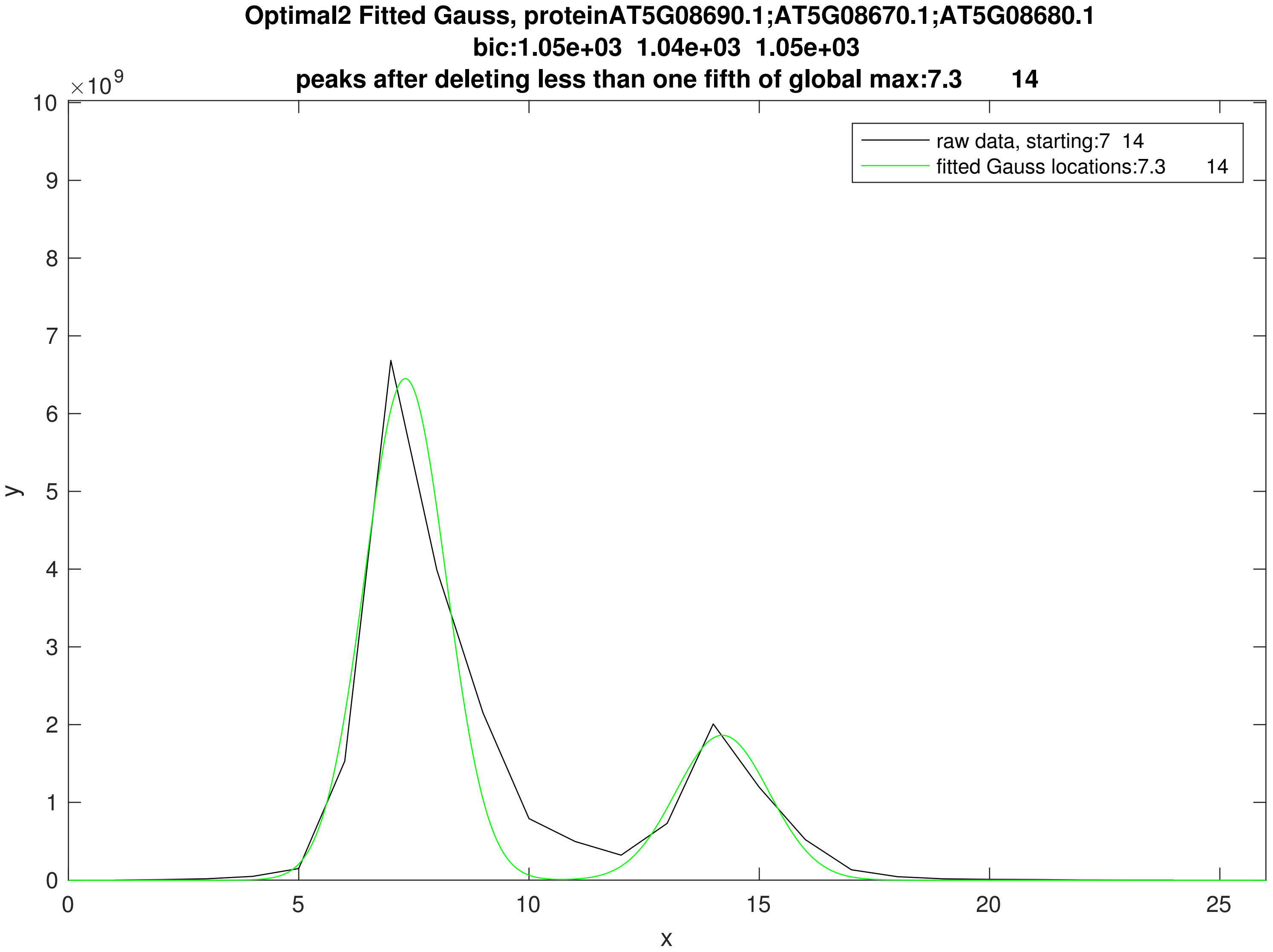
Optimal2 Fitted Gauss, proteinAT5G08690.1;AT5G08670.1;AT5G08680.1
bic:1.05e+03  1.04e+03  1.05e+03
peaks after deleting less than one fifth of global max:7.3 14
raw data, starting:7 14
fitted Gauss locations:7.3 14
×10^9
y
x

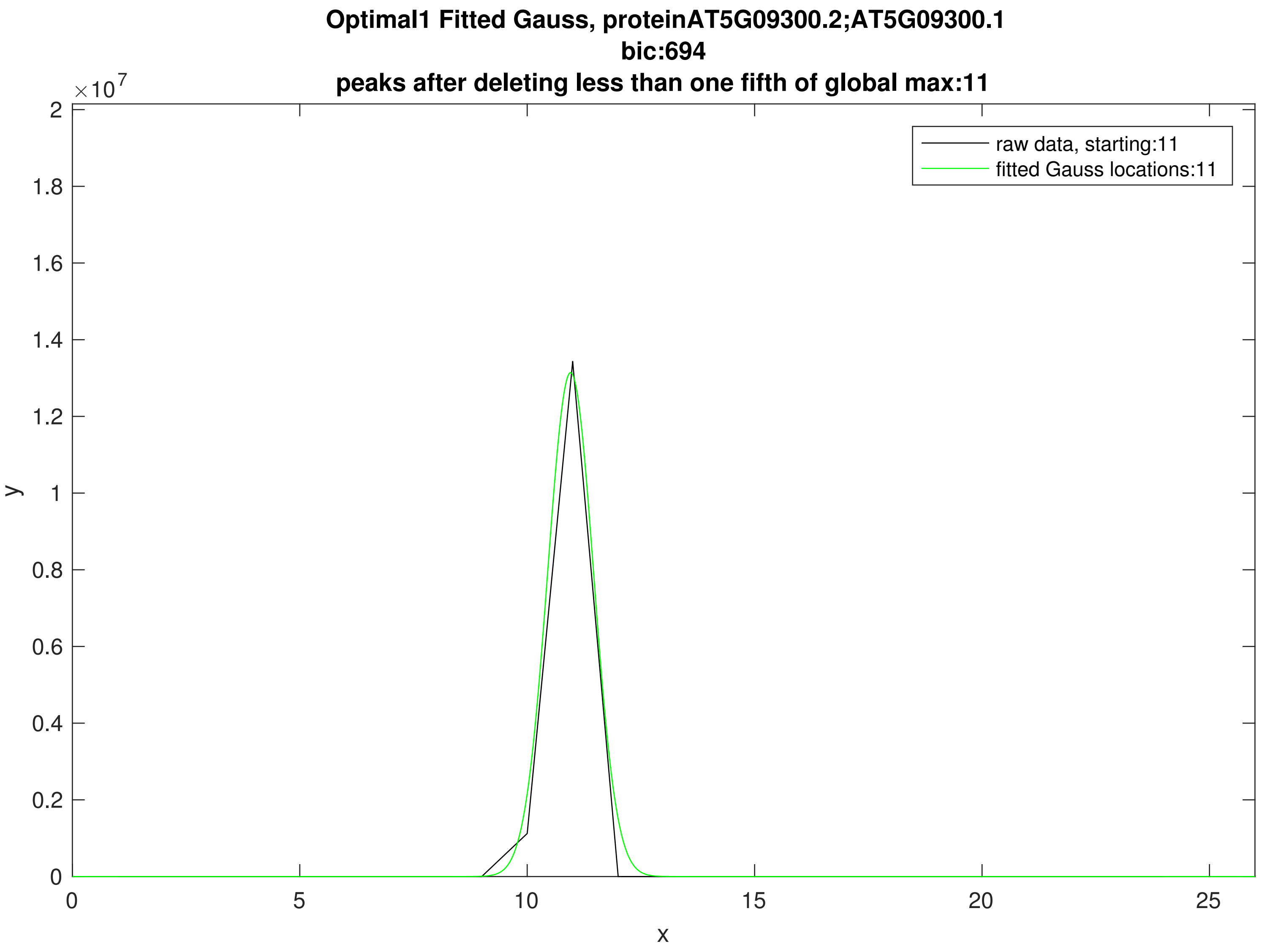

Optimal1 Fitted Gauss, proteinAT5G09300.2;AT5G09300.1
bic:694
peaks after deleting less than one fifth of global max:11
×10^7
raw data, starting:11
fitted Gauss locations:11
y
x
0
0.2
0.4
0.6
0.8
1
1.2
1.4
1.6
1.8
2
0
5
10
15
20
25

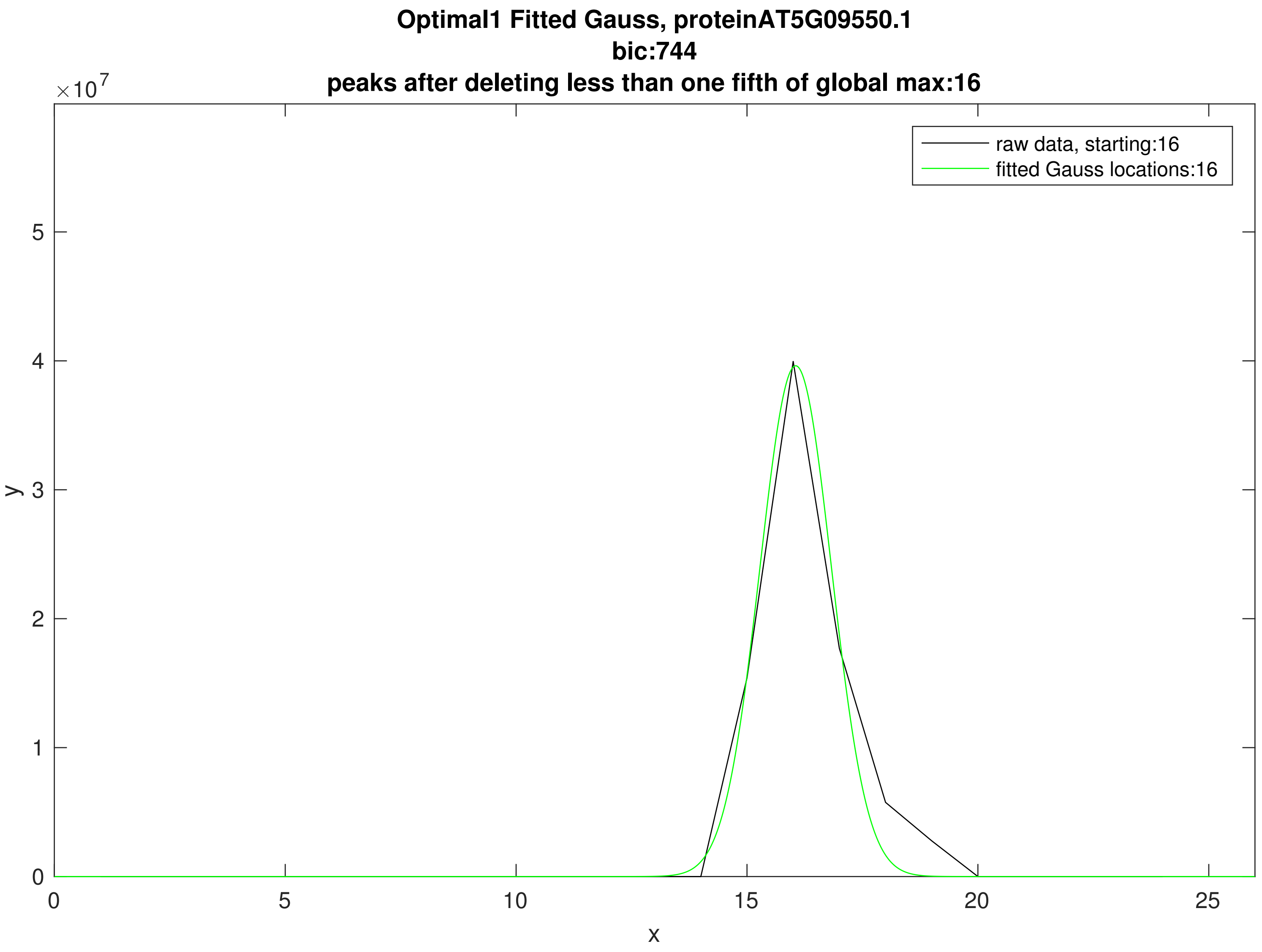

Optimal1 Fitted Gauss, proteinAT5G09550.1
bic:744
peaks after deleting less than one fifth of global max:16
×10^7
raw data, starting:16
fitted Gauss locations:16
y
x
0
1
2
3
4
5
0
5
10
15
20
25

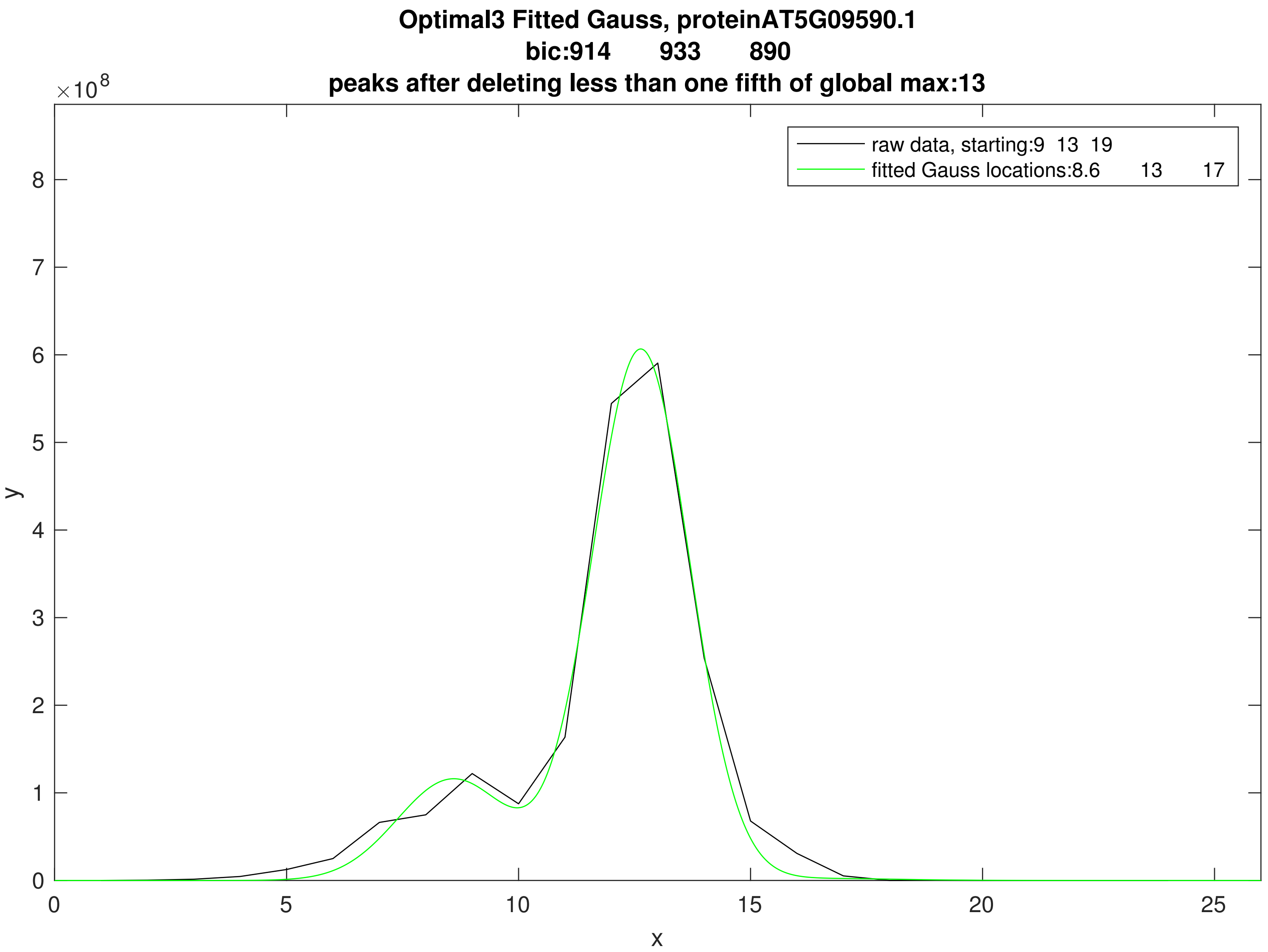

Optimal3 Fitted Gauss, proteinAT5G09590.1
bic:914 933 890
peaks after deleting less than one fifth of global max:13
×10^8
raw data, starting:9 13 19
fitted Gauss locations:8.6 13 17
y
x
0
1
2
3
4
5
6
7
8
0
5
10
15
20
25

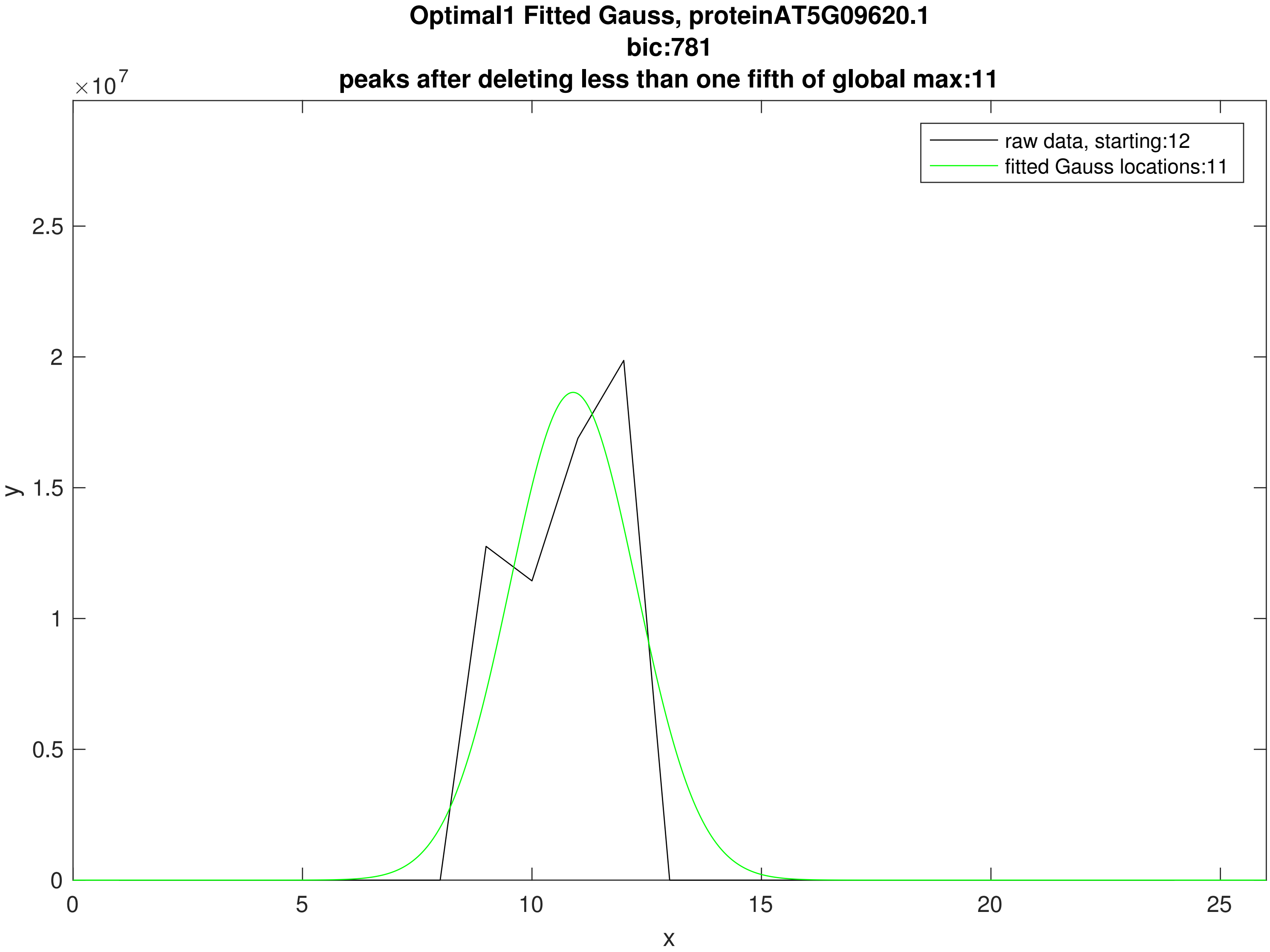

Optimal1 Fitted Gauss, proteinAT5G09620.1
bic:781
peaks after deleting less than one fifth of global max:11
×10^7
raw data, starting:12
fitted Gauss locations:11
2.5
2
1.5
1
0.5
0
y
0
5
10
15
20
25
x

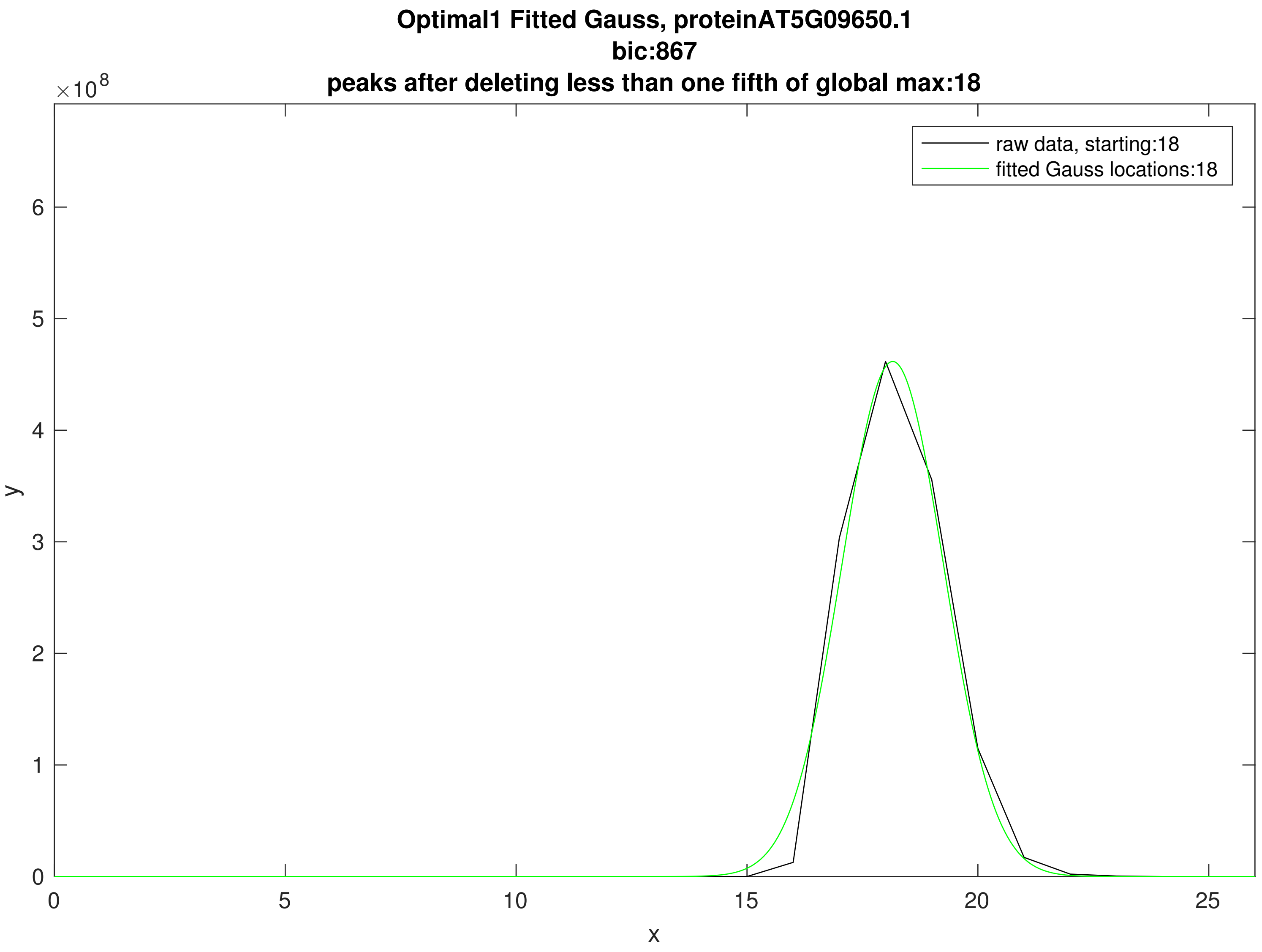

Optimal1 Fitted Gauss, proteinAT5G09650.1
bic:867
peaks after deleting less than one fifth of global max:18
×10^8
raw data, starting:18
fitted Gauss locations:18
y
x
0
1
2
3
4
5
6
0
5
10
15
20
25

Optimal1 Fitted Gauss, proteinAT5G09760.1
bic:785
peaks after deleting less than one fifth of global max:16

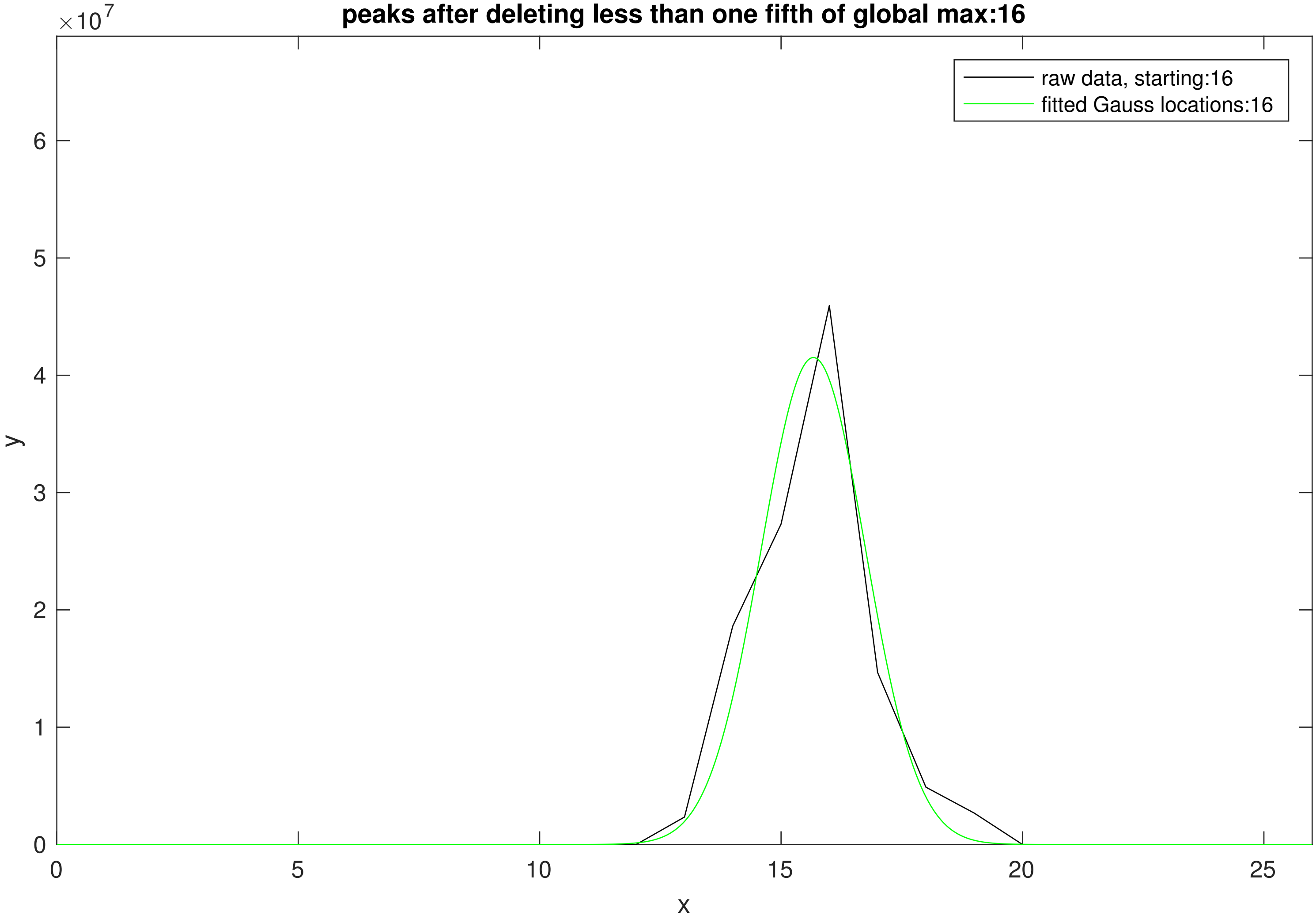

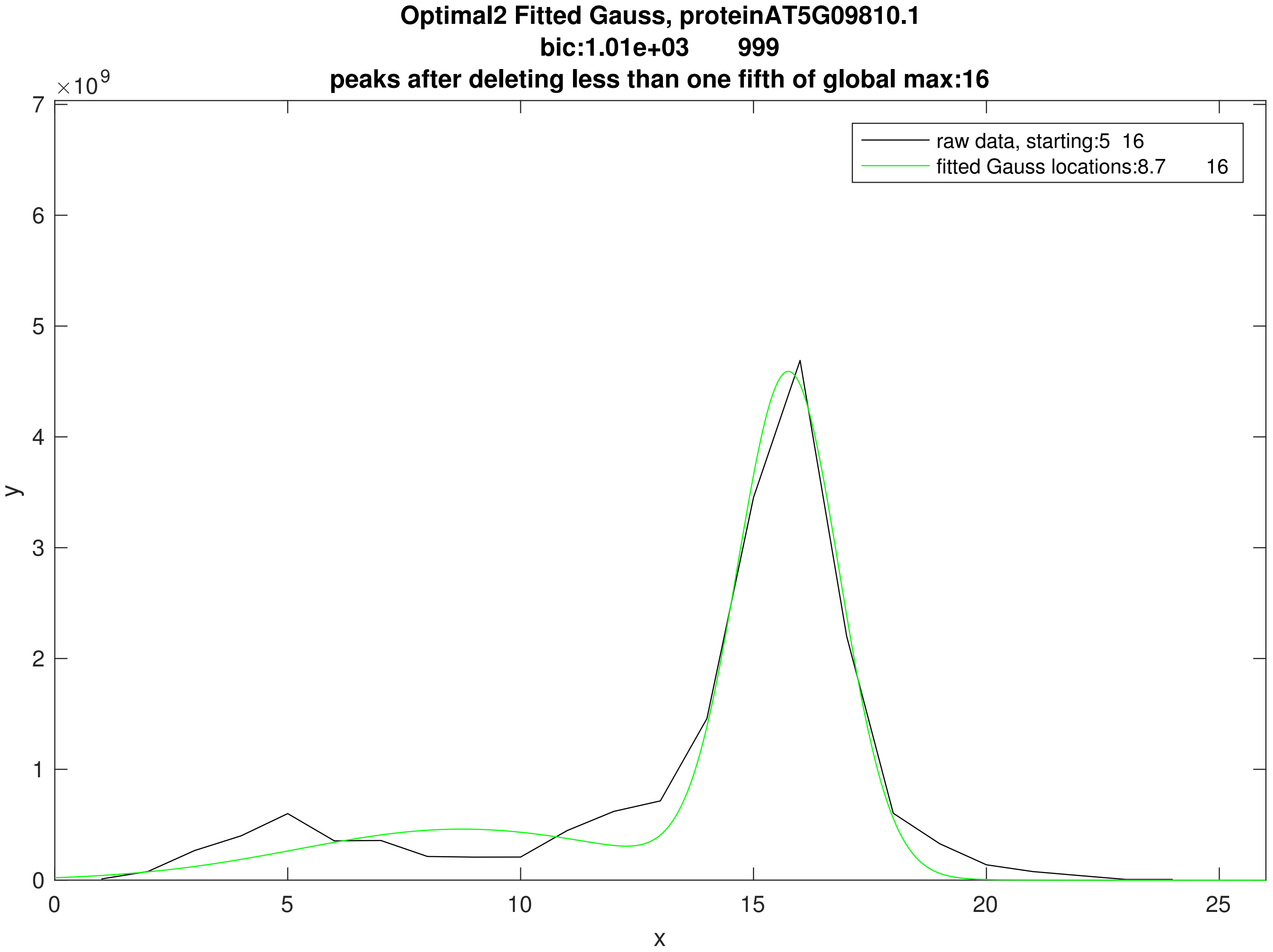
Optimal2 Fitted Gauss, proteinAT5G09810.1
bic:1.01e+03 999
peaks after deleting less than one fifth of global max:16
×10^9
raw data, starting:5 16
fitted Gauss locations:8.7 16
y
x
0
1
2
3
4
5
6
7
0
5
10
15
20
25

#### Optimal1 Fitted Gauss, proteinAT5G09830.1
#### bic:722
#### peaks after deleting less than one fifth of global max:20

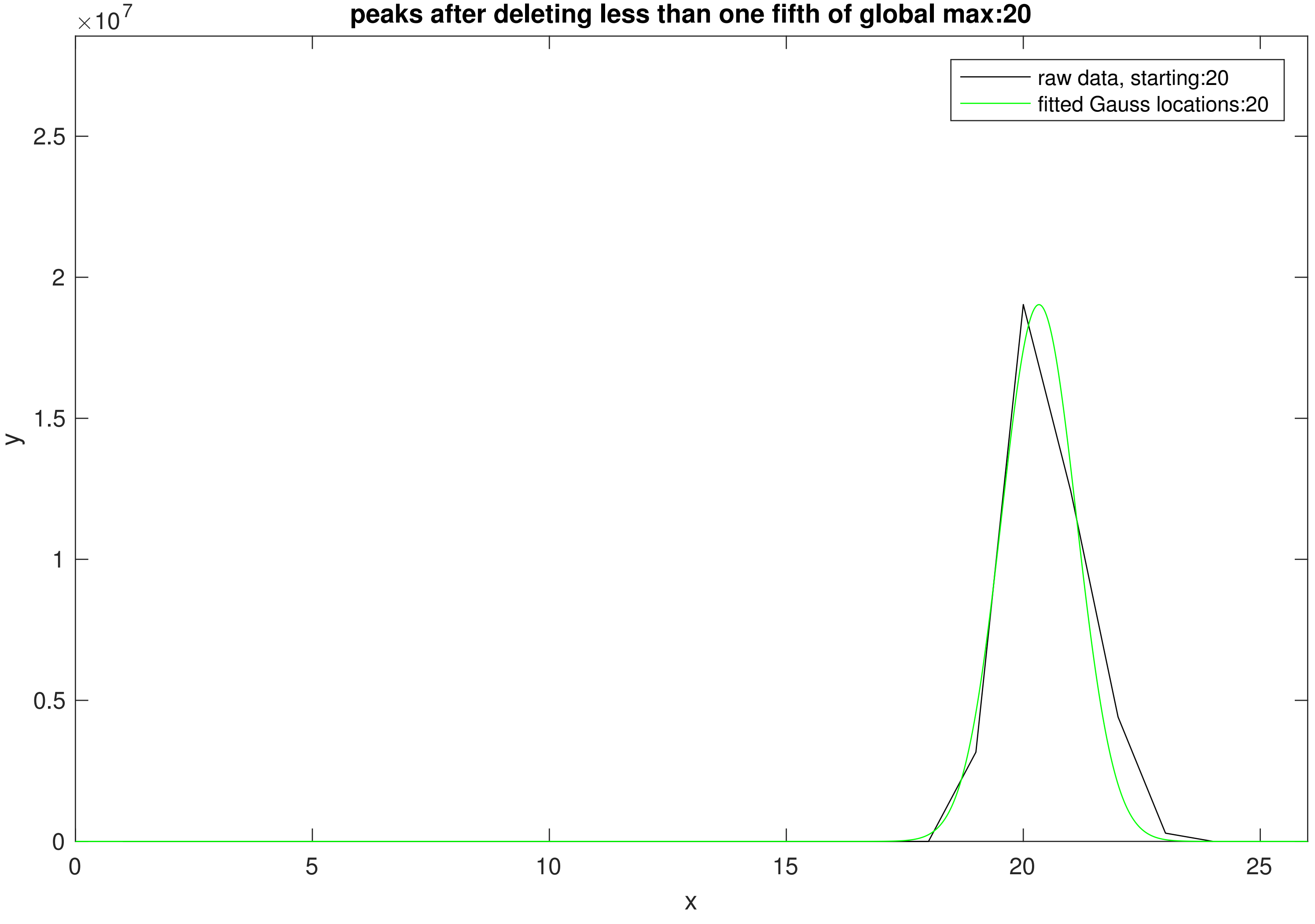

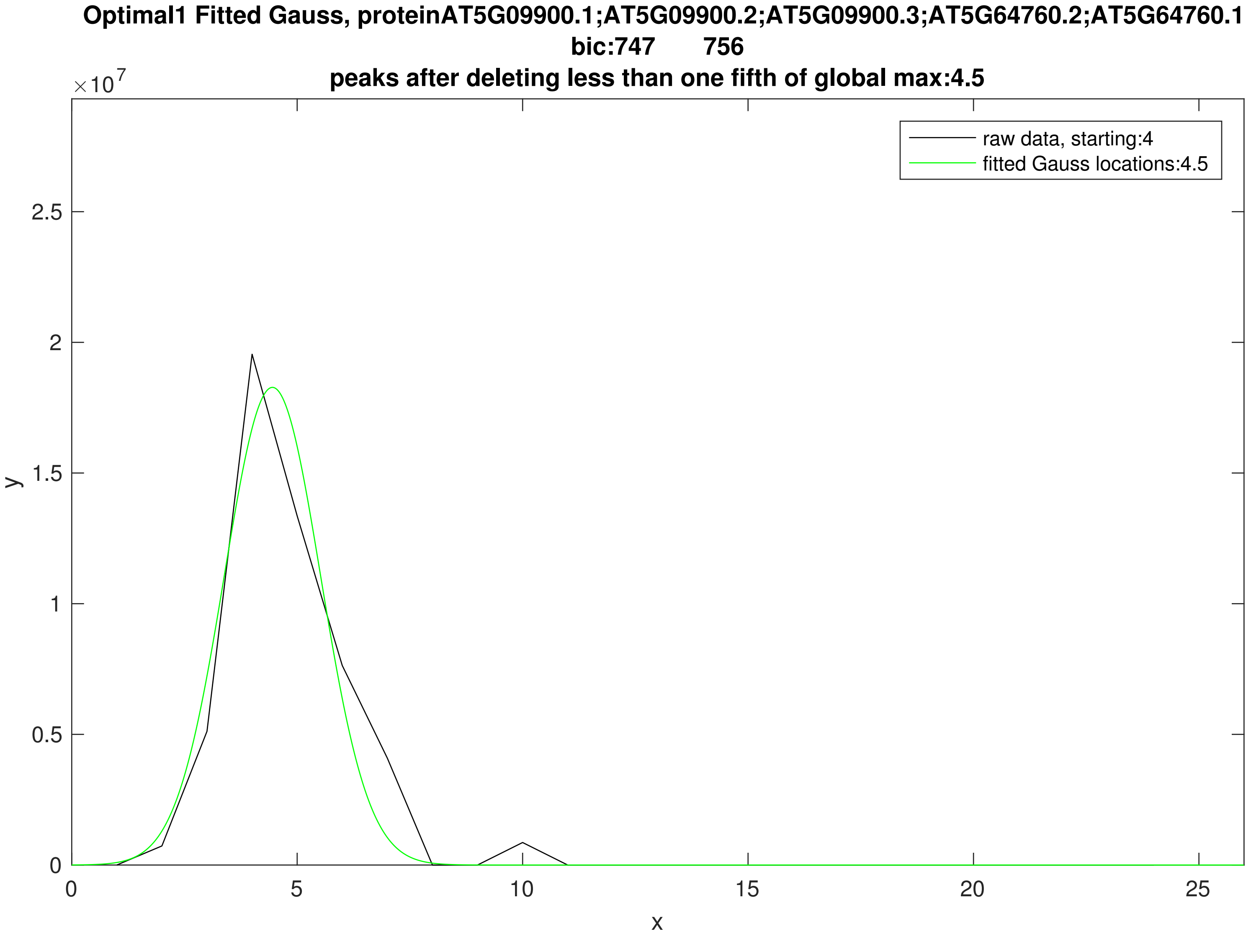
Optimal1 Fitted Gauss, proteinAT5G09900.1;AT5G09900.2;AT5G09900.3;AT5G64760.2;AT5G64760.1
bic:747 756
peaks after deleting less than one fifth of global max:4.5
raw data, starting:4
fitted Gauss locations:4.5
×10^7
y
x
0
0.5
1
1.5
2
2.5
5
10
15
20
25

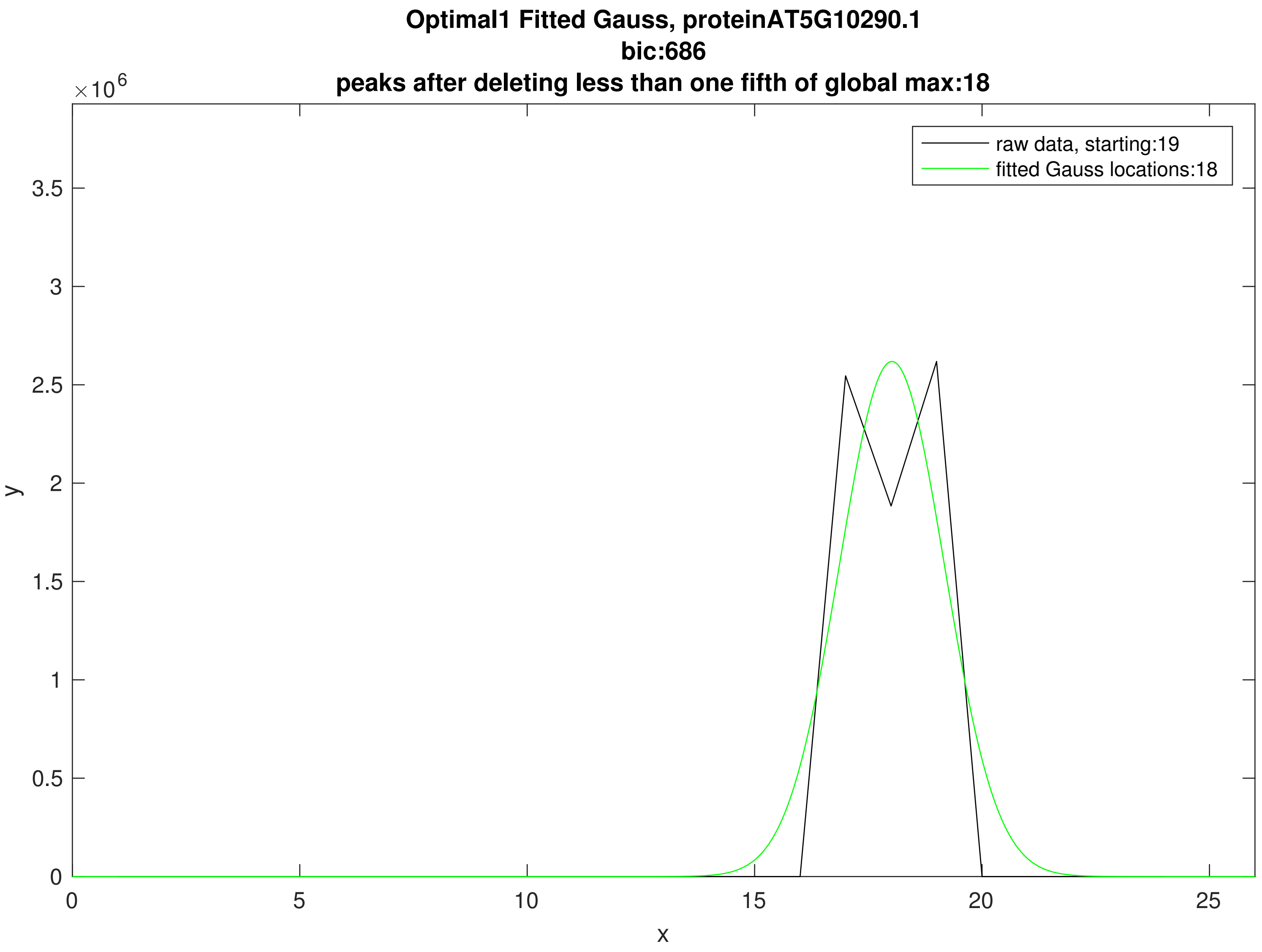
Optimal1 Fitted Gauss, proteinAT5G10290.1
bic:686
peaks after deleting less than one fifth of global max:18
×10^6
raw data, starting:19
fitted Gauss locations:18
y
x
0
0.5
1
1.5
2
2.5
3
3.5
0
5
10
15
20
25

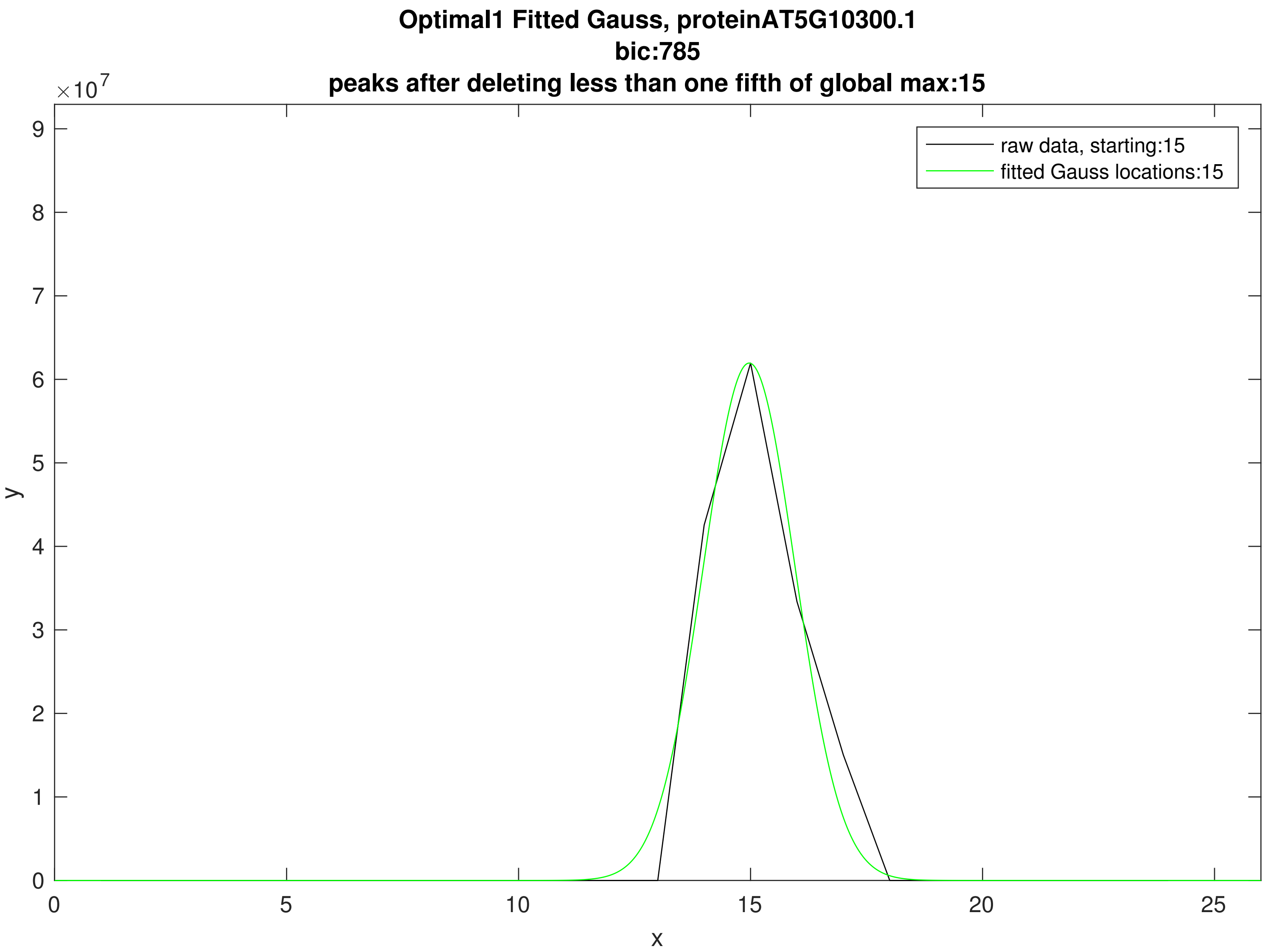

Optimal1 Fitted Gauss, proteinAT5G10300.1
bic:785
peaks after deleting less than one fifth of global max:15
×10^7
raw data, starting:15
fitted Gauss locations:15
y
x
0
1
2
3
4
5
6
7
8
9
0
5
10
15
20
25

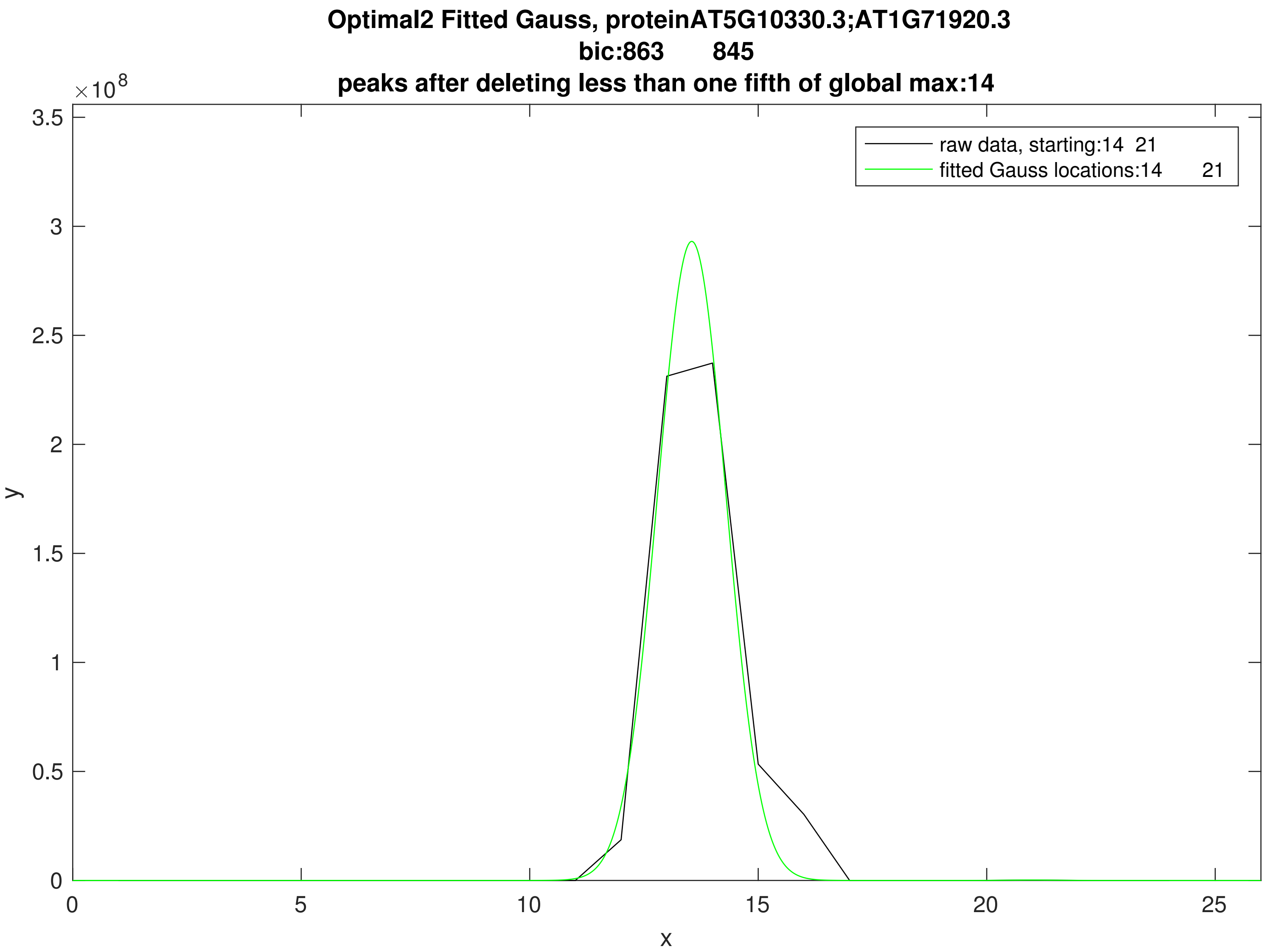

Optimal2 Fitted Gauss, proteinAT5G10330.3;AT1G71920.3
bic:863 845
peaks after deleting less than one fifth of global max:14
raw data, starting:14 21
fitted Gauss locations:14 21
y
x

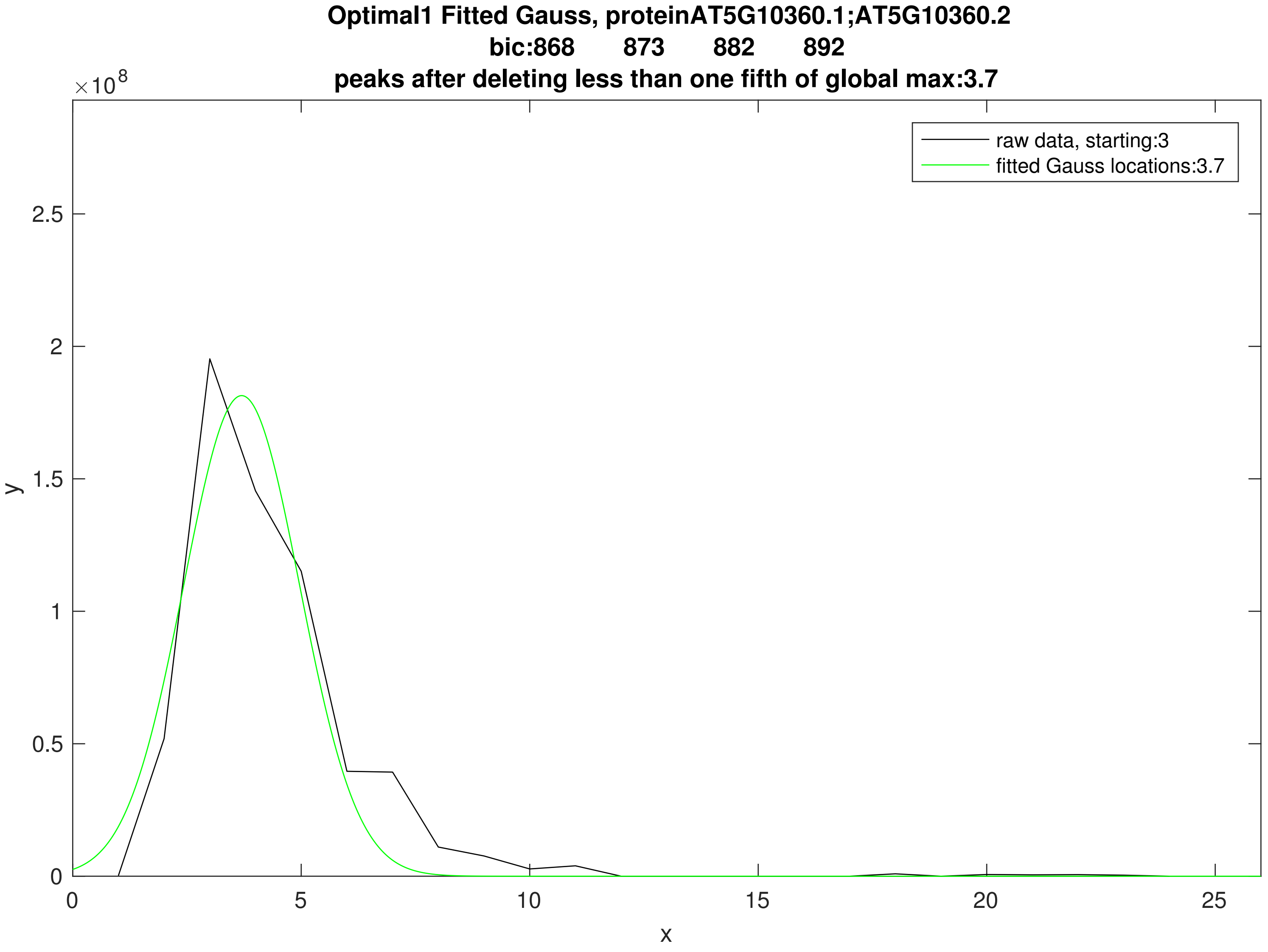
Optimal1 Fitted Gauss, proteinAT5G10360.1;AT5G10360.2
bic:868 873 882 892
peaks after deleting less than one fifth of global max:3.7
×10^8
raw data, starting:3
fitted Gauss locations:3.7
y
x
0
0.5
1
1.5
2
2.5
0
5
10
15
20
25

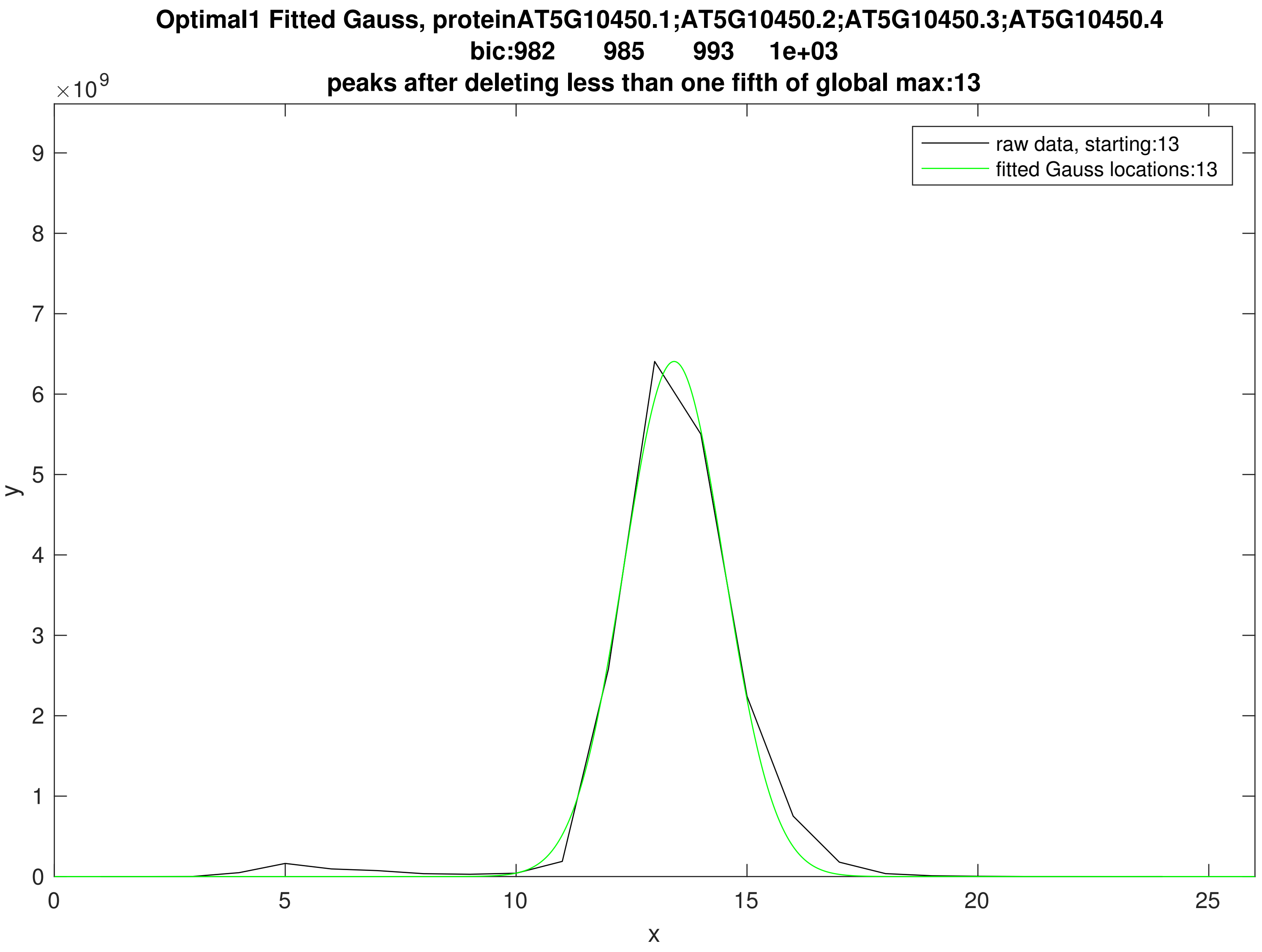
Optimal1 Fitted Gauss, proteinAT5G10450.1;AT5G10450.2;AT5G10450.3;AT5G10450.4
bic:982 985 993 1e+03
peaks after deleting less than one fifth of global max:13
×10^9
raw data, starting:13
fitted Gauss locations:13
y
x
0
1
2
3
4
5
6
7
8
9
0
5
10
15
20
25

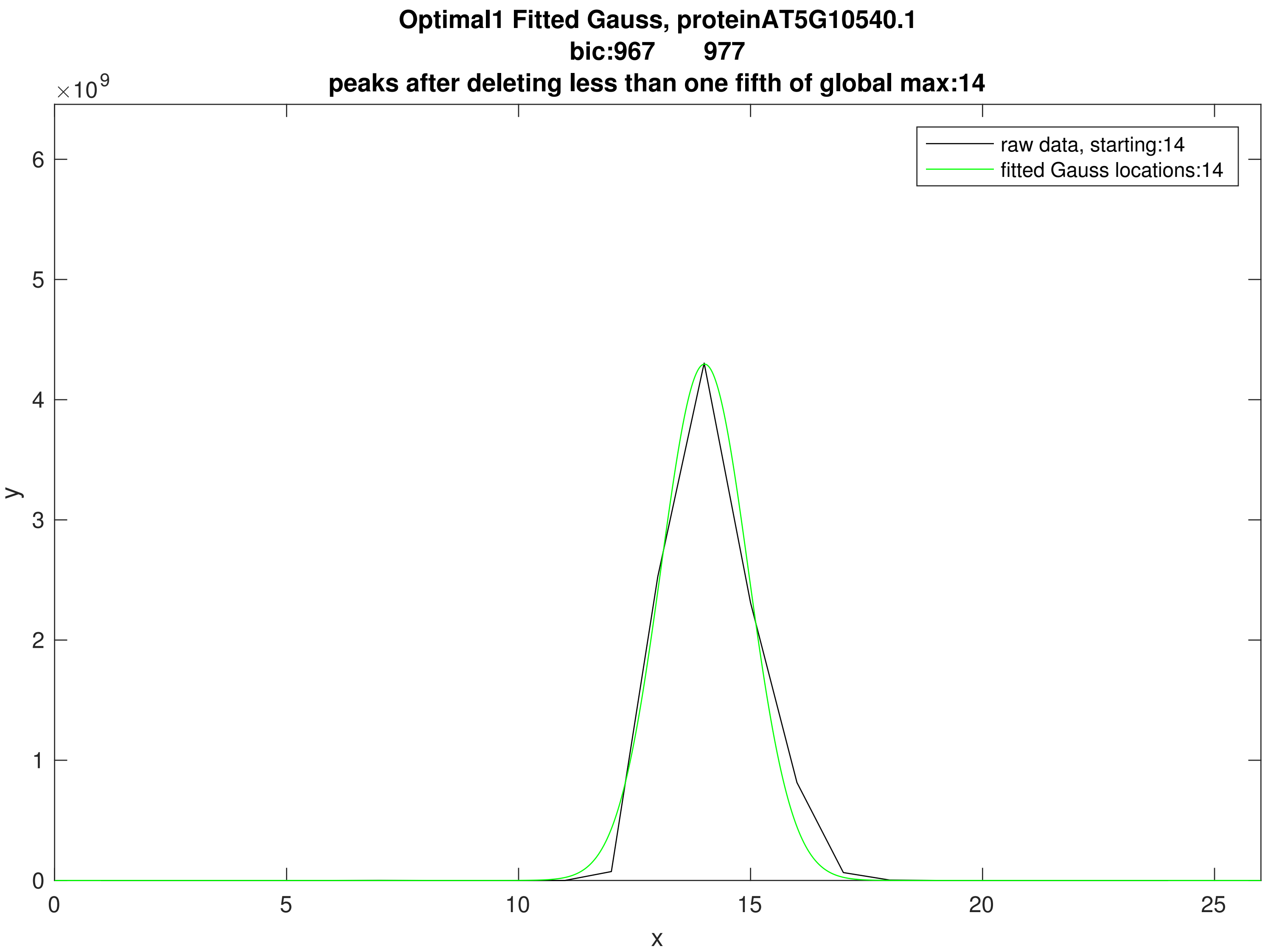

Optimal1 Fitted Gauss, proteinAT5G10540.1
bic:967 977
peaks after deleting less than one fifth of global max:14
×10^9
raw data, starting:14
fitted Gauss locations:14
y
x
0
1
2
3
4
5
6
0
5
10
15
20
25

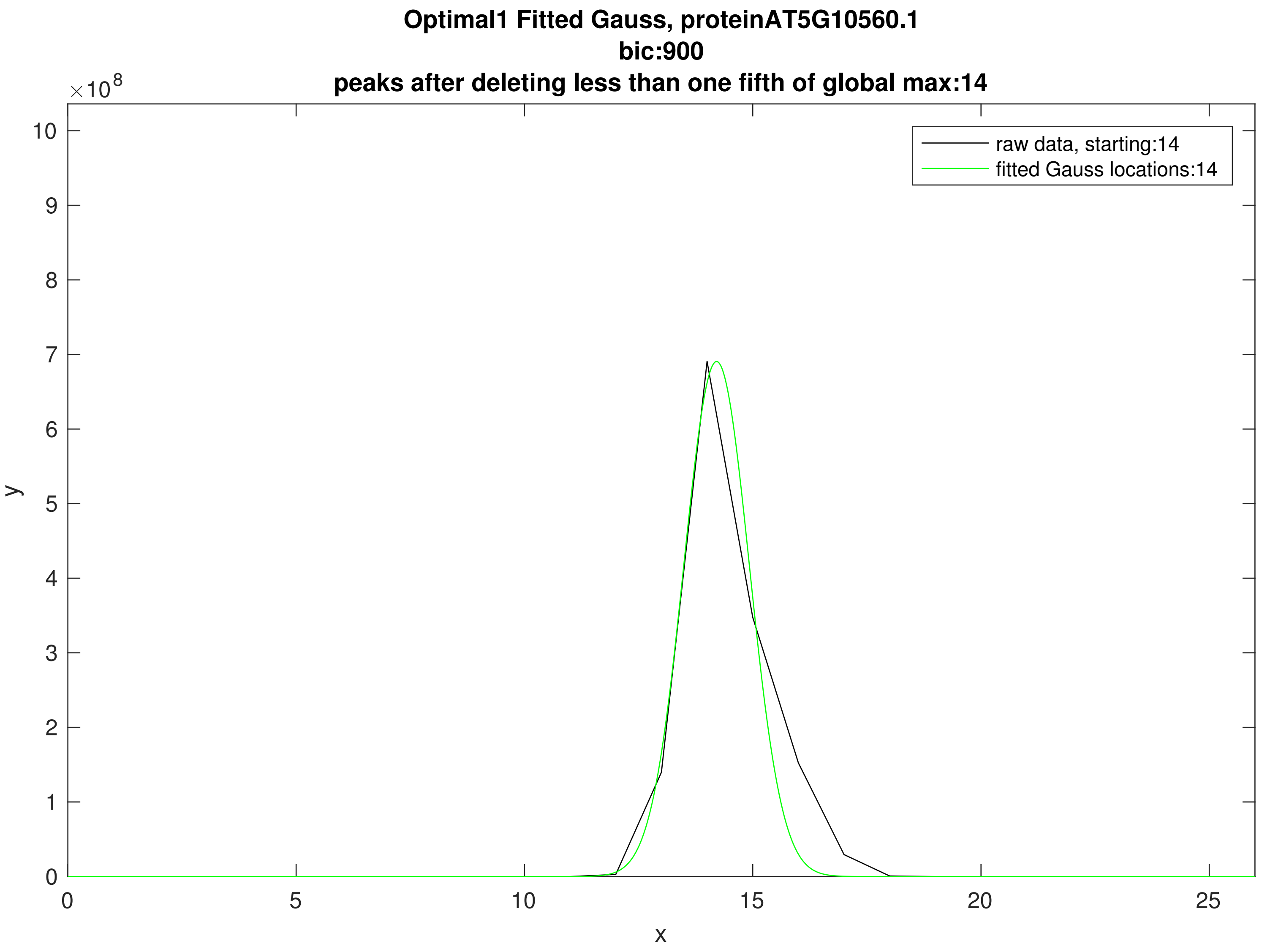

Optimal1 Fitted Gauss, proteinAT5G10560.1
bic:900
peaks after deleting less than one fifth of global max:14
×10^8
raw data, starting:14
fitted Gauss locations:14
y
x
0
1
2
3
4
5
6
7
8
9
10
0
5
10
15
20
25

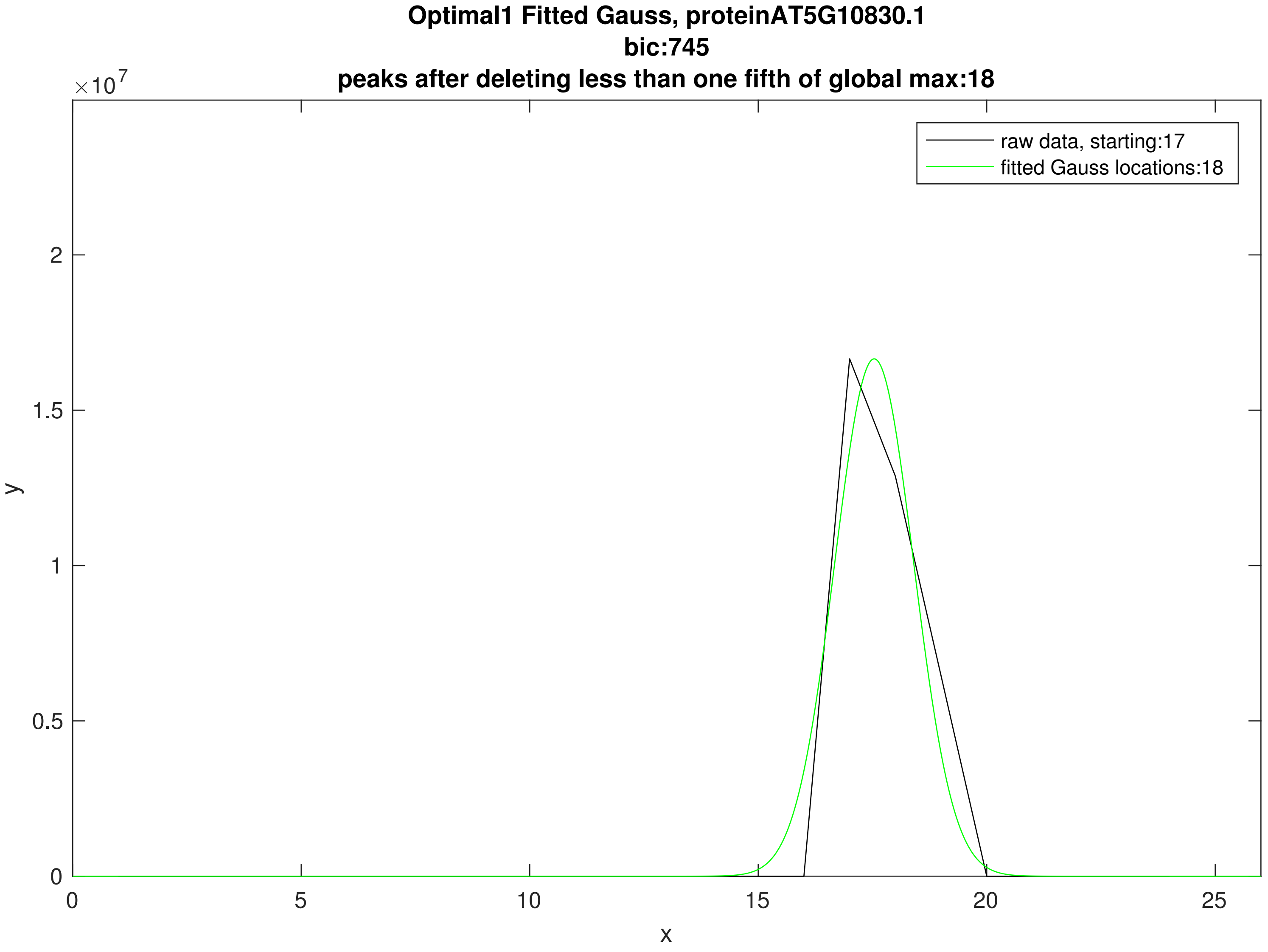
Optimal1 Fitted Gauss, proteinAT5G10830.1
bic:745
peaks after deleting less than one fifth of global max:18
×10^7
raw data, starting:17
fitted Gauss locations:18
2
1.5
1
0.5
0
y
0
5
10
15
20
25
x

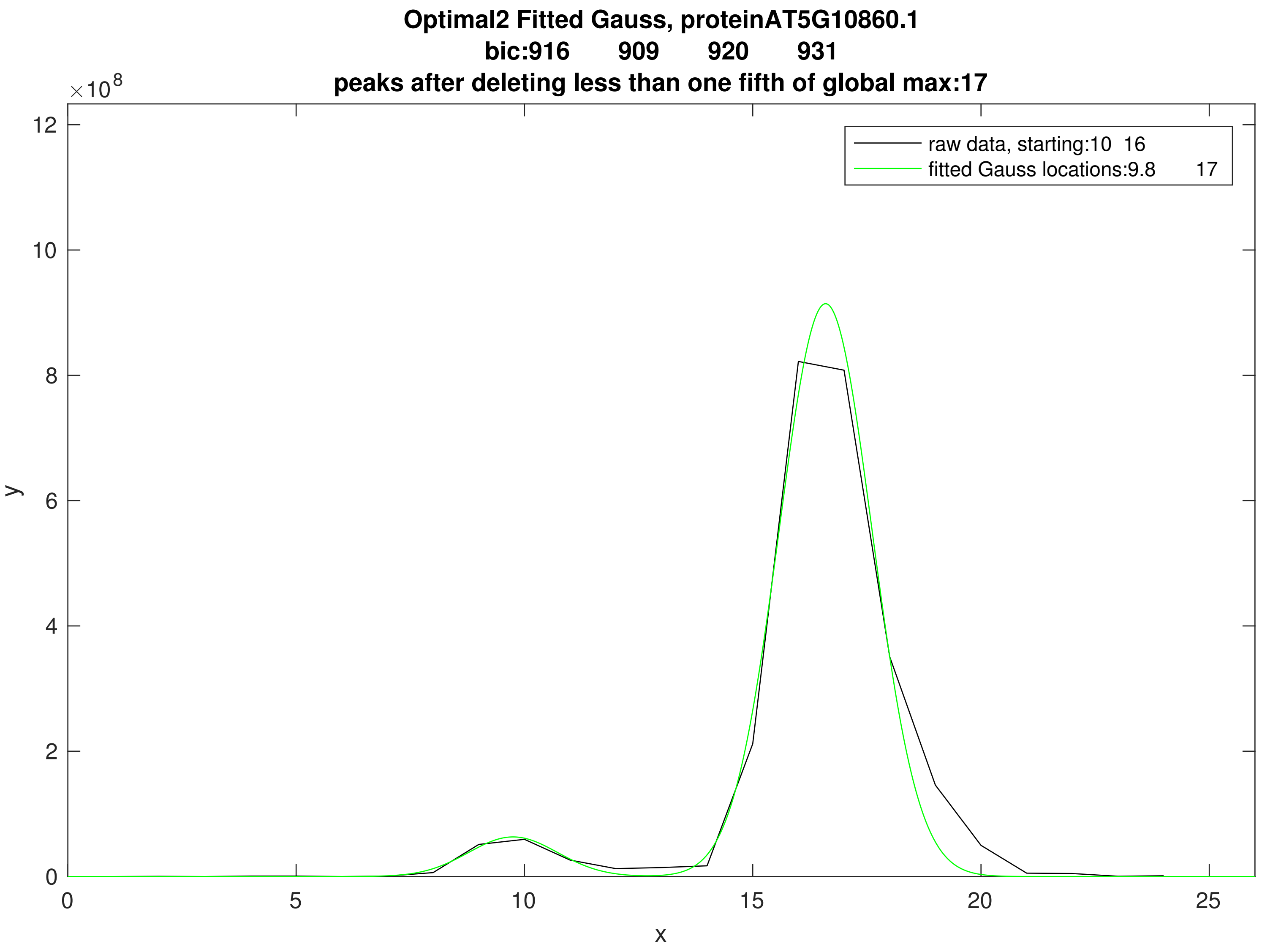
Optimal2 Fitted Gauss, proteinAT5G10860.1
bic:916 909 920 931
peaks after deleting less than one fifth of global max:17
×10^8
raw data, starting:10 16
fitted Gauss locations:9.8 17
y
x
0
2
4
6
8
10
12
0
5
10
15
20
25

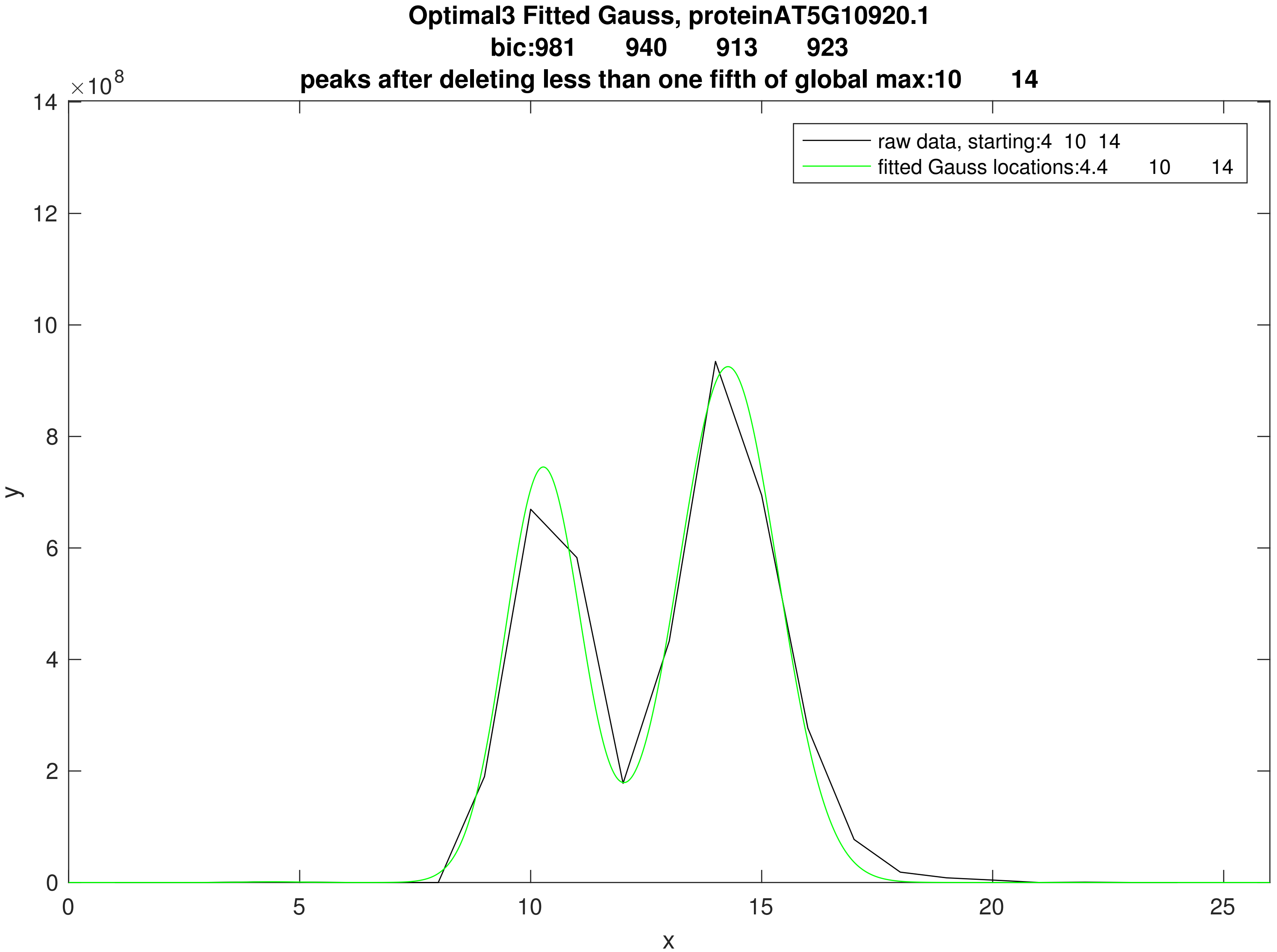
Optimal3 Fitted Gauss, proteinAT5G10920.1
bic:981 940 913 923
peaks after deleting less than one fifth of global max:10 14
×10^8
raw data, starting:4 10 14
fitted Gauss locations:4.4 10 14
y
x
0
2
4
6
8
10
12
14
0
5
10
15
20
25

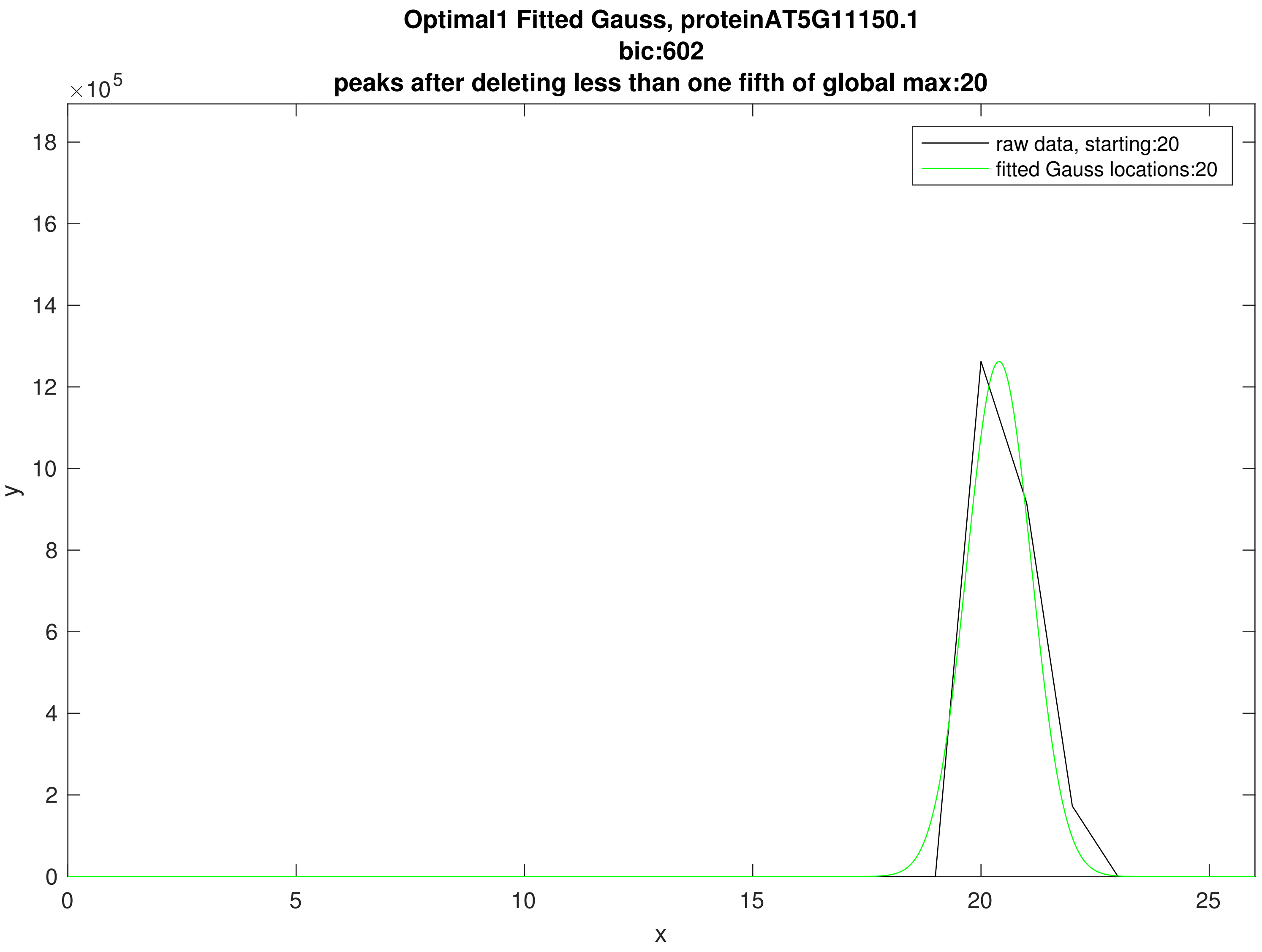
Optimal1 Fitted Gauss, proteinAT5G11150.1
bic:602
peaks after deleting less than one fifth of global max:20
×10^5
raw data, starting:20
fitted Gauss locations:20
y
x
0
2
4
6
8
10
12
14
16
18
0
5
10
15
20
25

**Optimal1 Fitted Gauss, proteinAT5G11200.1;AT5G11170.1;AT5G11200.3;AT5G11200.2;AT5G11170.2**
**bic:921**
**peaks after deleting less than one fifth of global max:14**

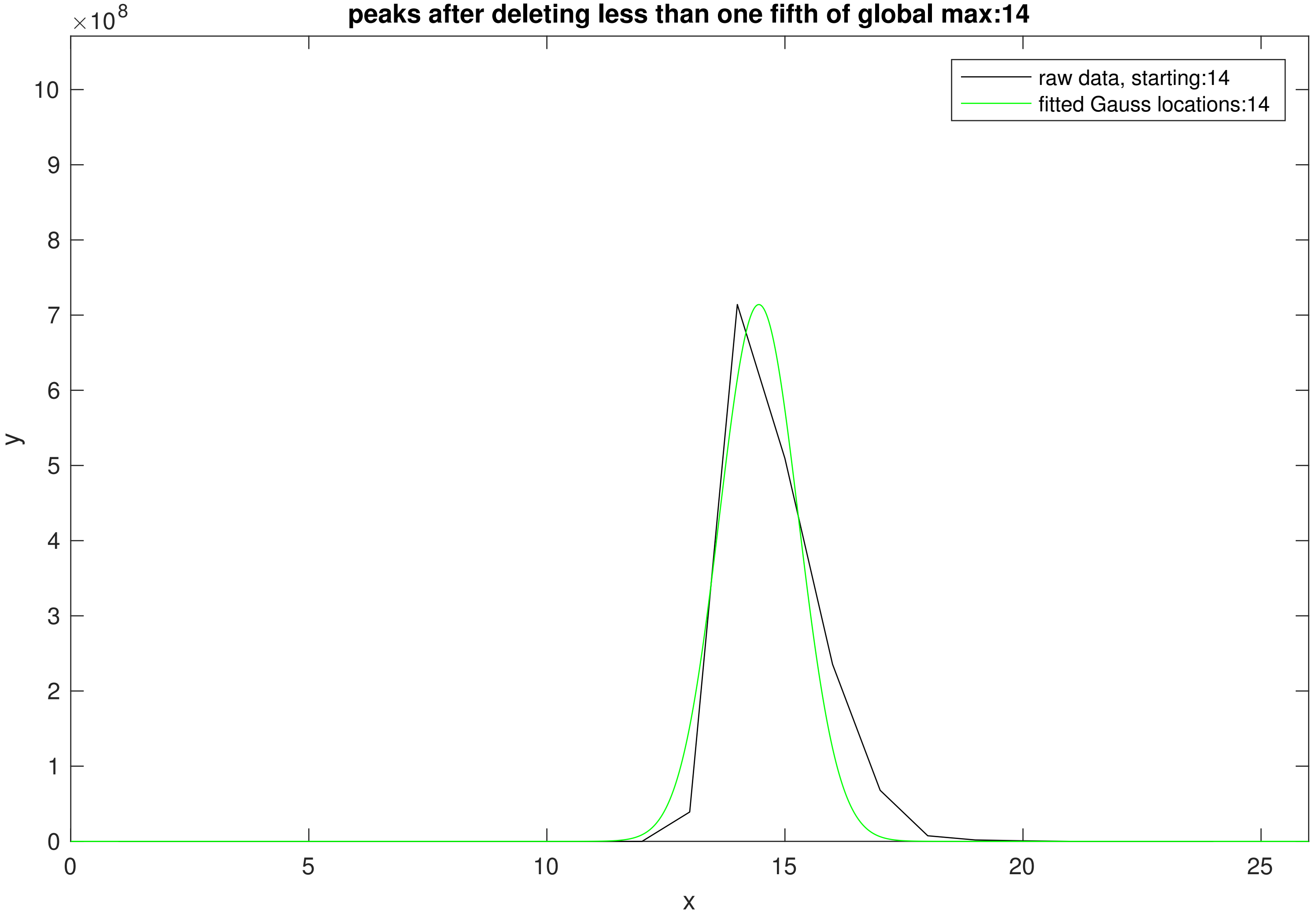

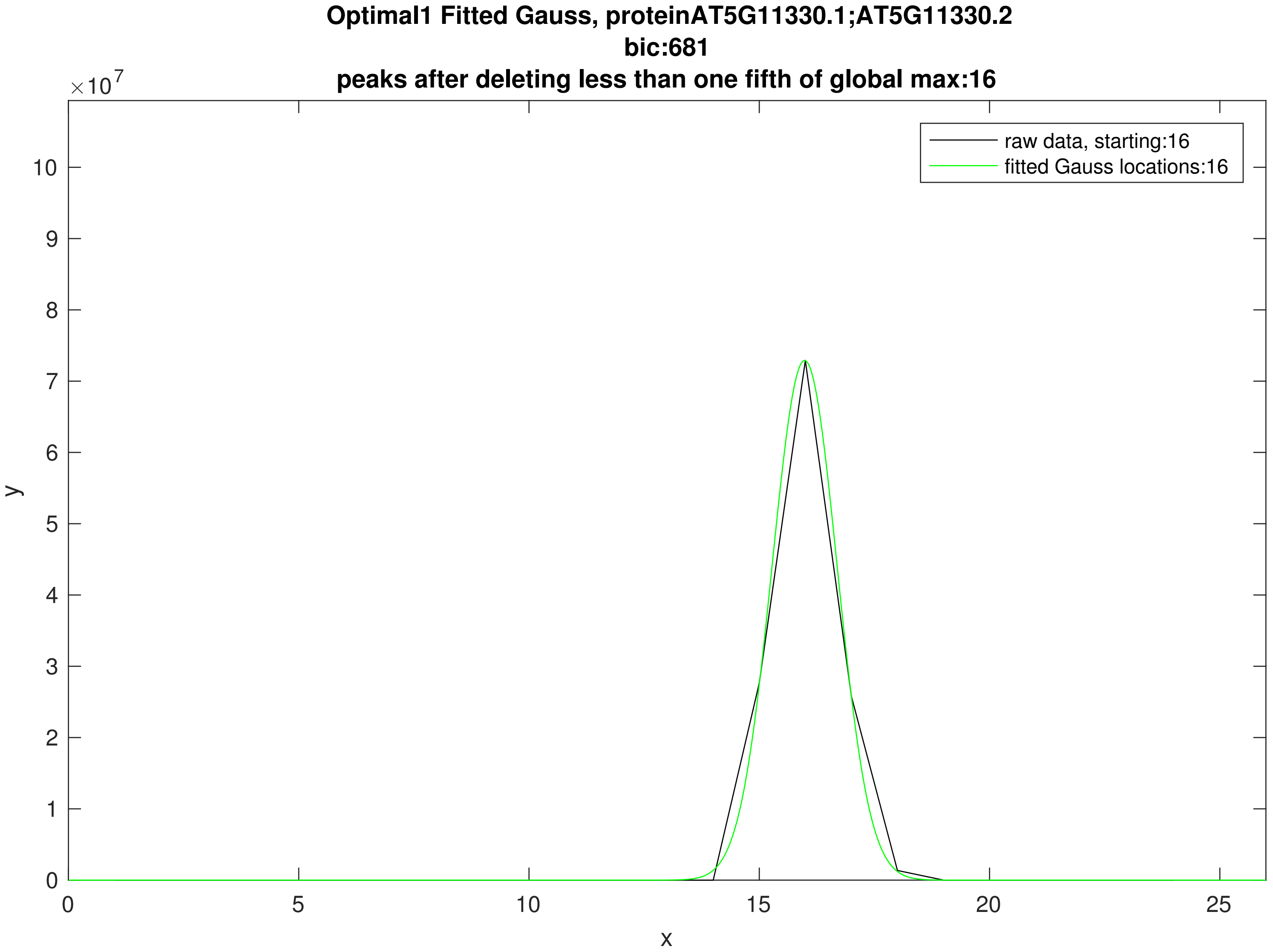
Optimal1 Fitted Gauss, proteinAT5G11330.1;AT5G11330.2
bic:681
peaks after deleting less than one fifth of global max:16
×10^7
raw data, starting:16
fitted Gauss locations:16
y
x
0
1
2
3
4
5
6
7
8
9
10
0
5
10
15
20
25

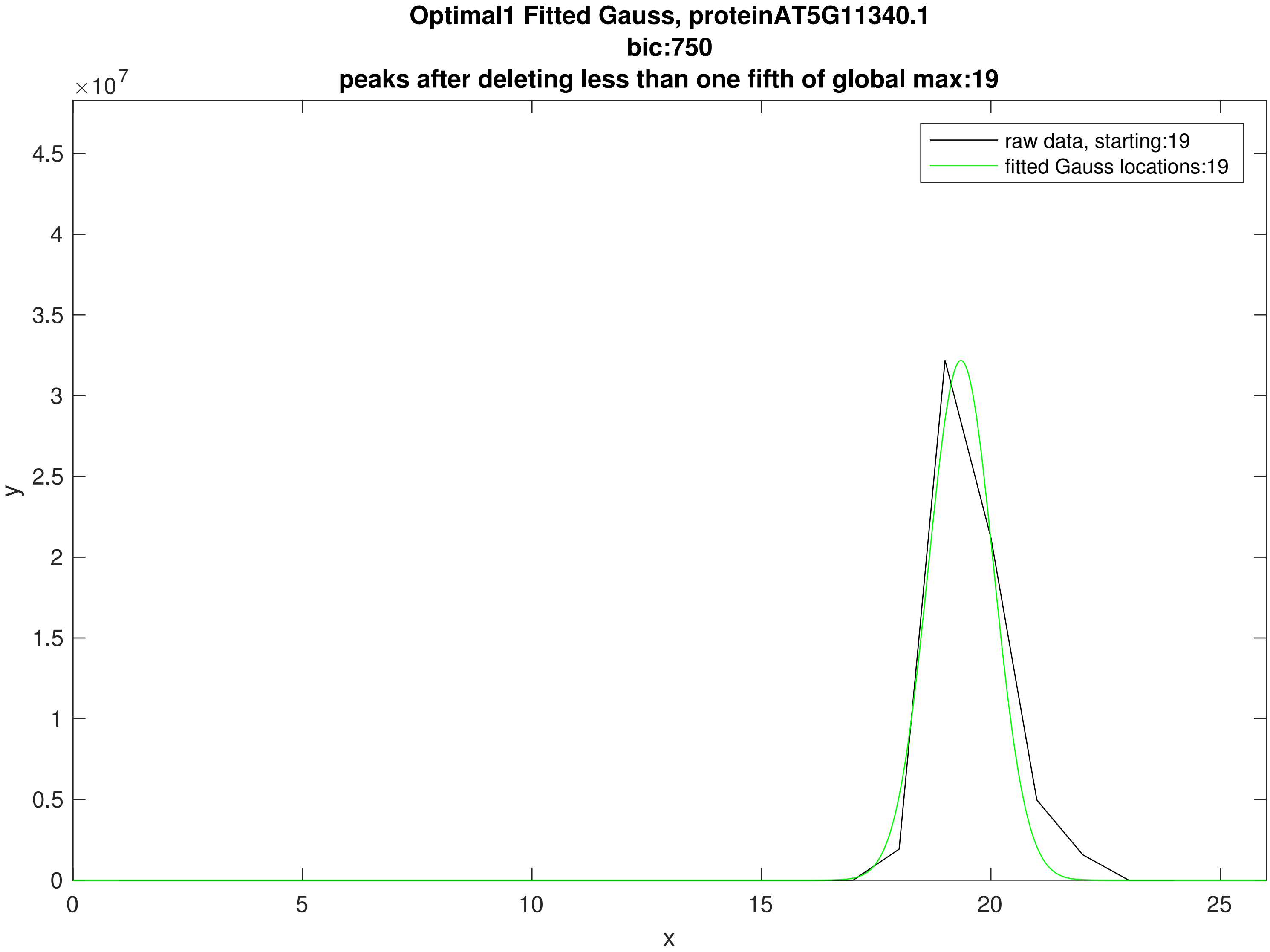
Optimal1 Fitted Gauss, proteinAT5G11340.1
bic:750
peaks after deleting less than one fifth of global max:19
×10^7
4.5
4
3.5
3
2.5
2
1.5
1
0.5
0
y
0
5
10
15
20
25
x
raw data, starting:19
fitted Gauss locations:19

**Optimal1 Fitted Gauss, proteinAT5G11450.1**
**bic:679**
**peaks after deleting less than one fifth of global max:21**

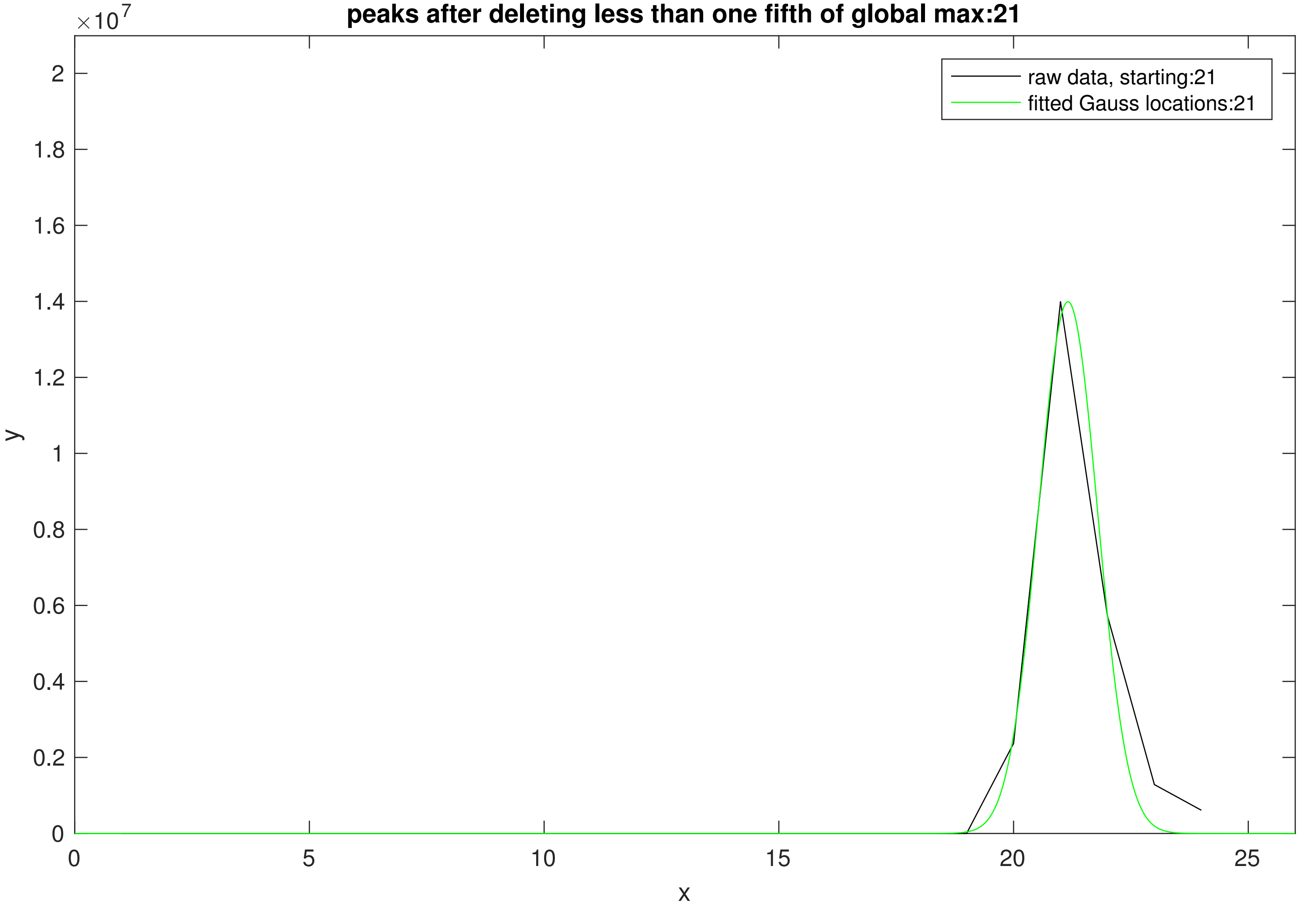

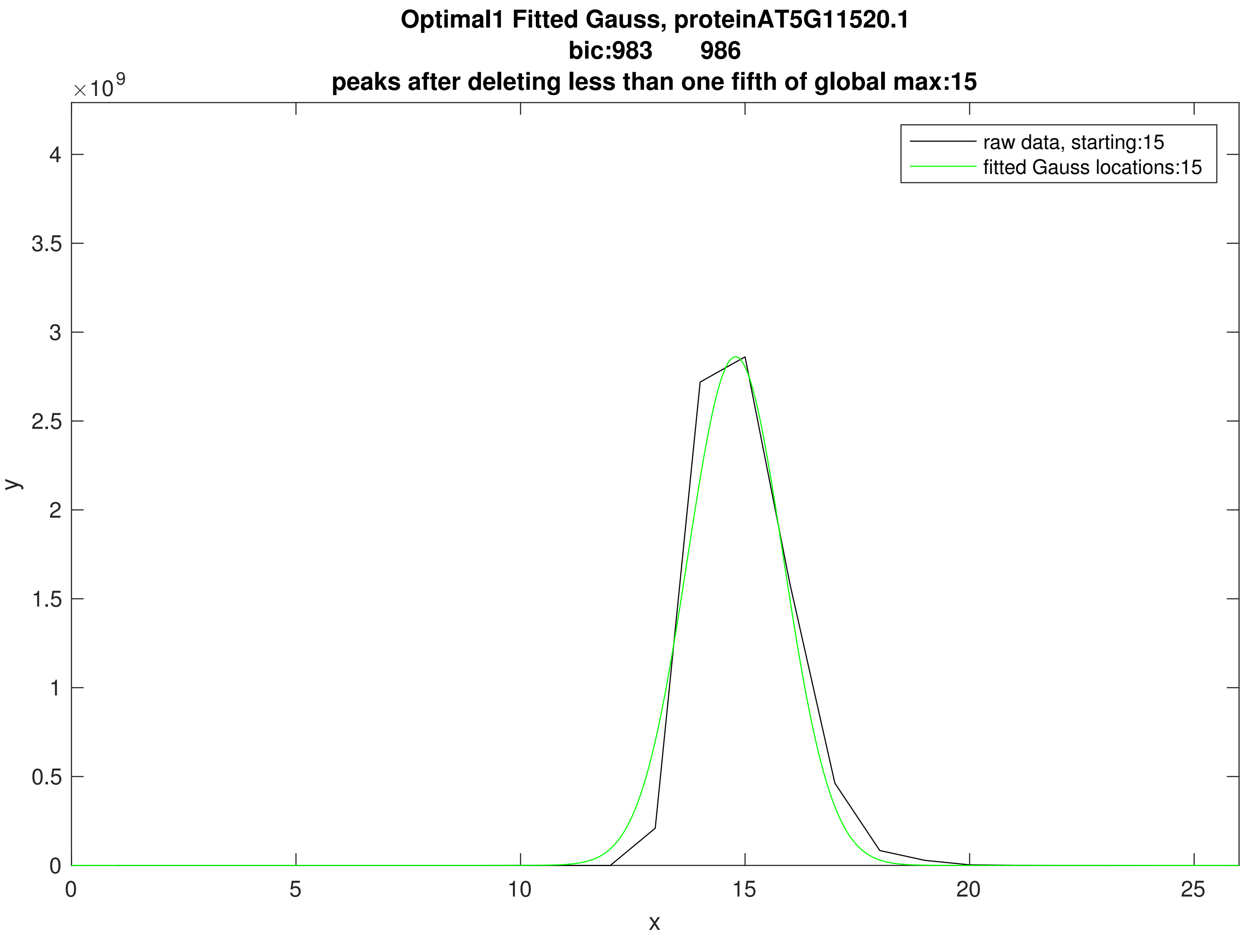

Optimal1 Fitted Gauss, proteinAT5G11520.1
bic:983 986
peaks after deleting less than one fifth of global max:15
×10^9
4
3.5
3
2.5
2
1.5
1
0.5
0
y
0
5
10
15
20
25
x
raw data, starting:15
fitted Gauss locations:15

Optimal1 Fitted Gauss, proteinAT5G11580.1
bic:603
peaks after deleting less than one fifth of global max:5.1

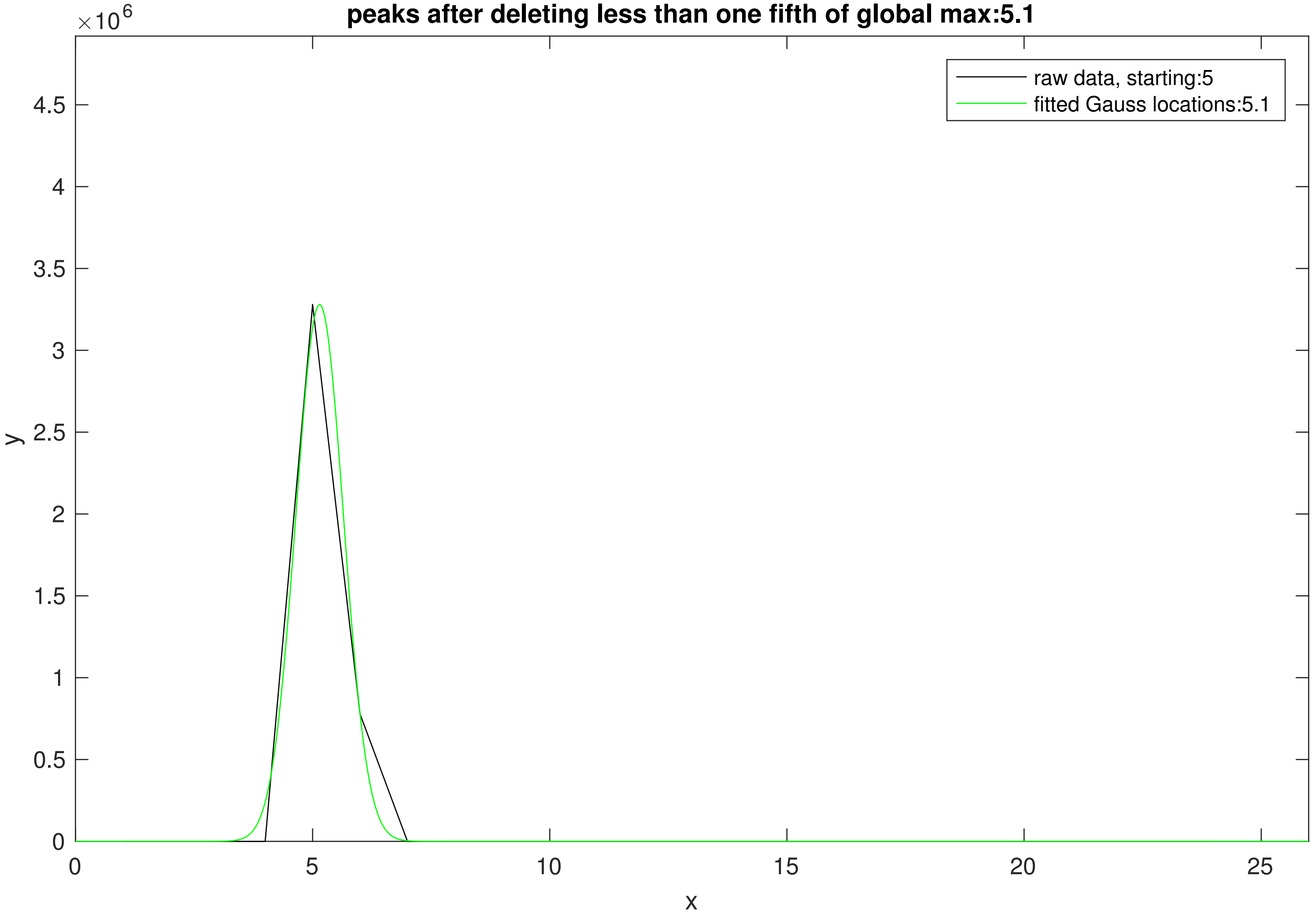

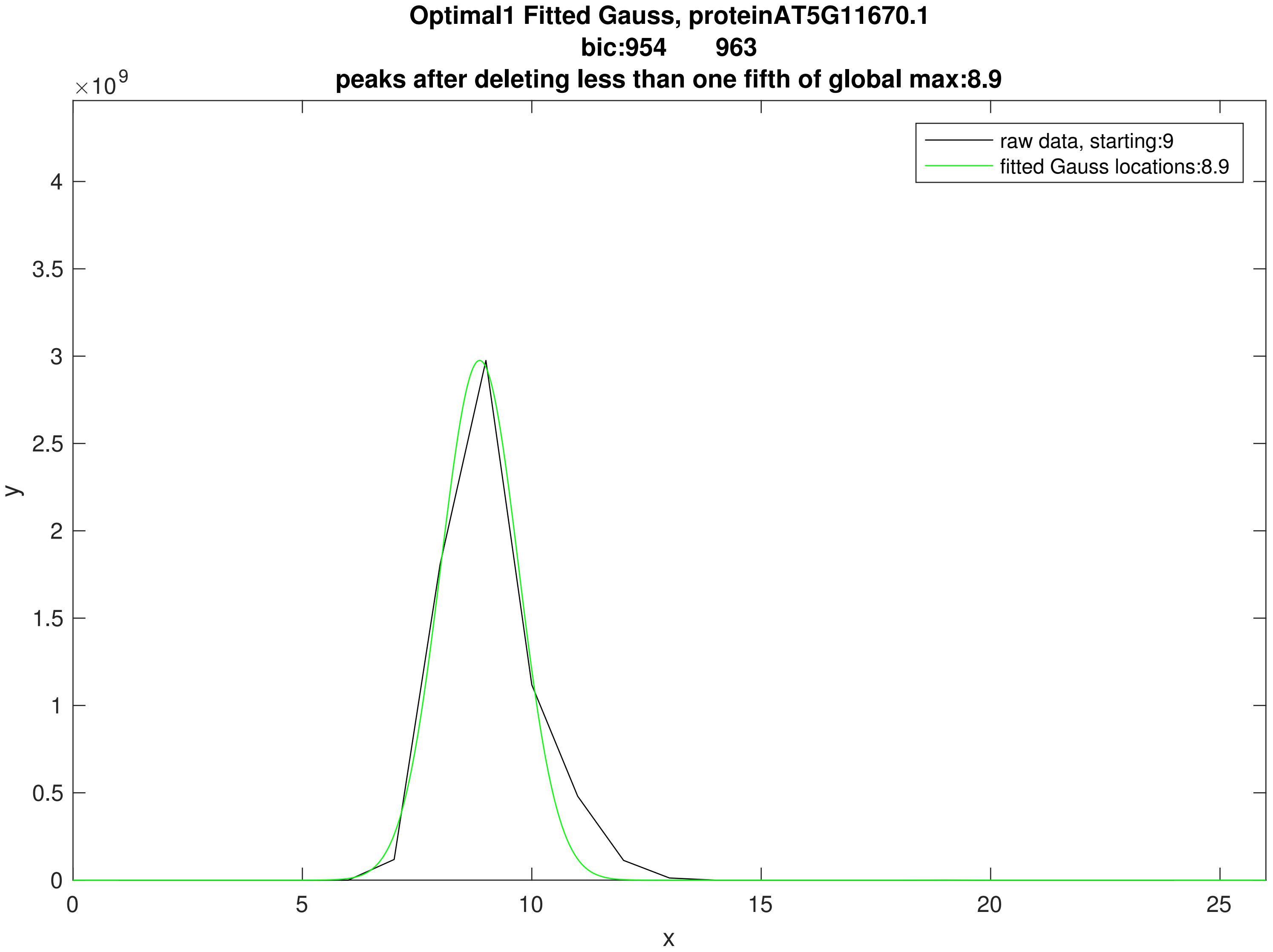

Optimal1 Fitted Gauss, proteinAT5G11670.1
bic:954 963
peaks after deleting less than one fifth of global max:8.9
×10^9
raw data, starting:9
fitted Gauss locations:8.9
y
x
0
0.5
1
1.5
2
2.5
3
3.5
4
5
10
15
20
25

Optimal1 Fitted Gauss, proteinAT5G11680.1
bic:659
peaks after deleting less than one fifth of global max:20

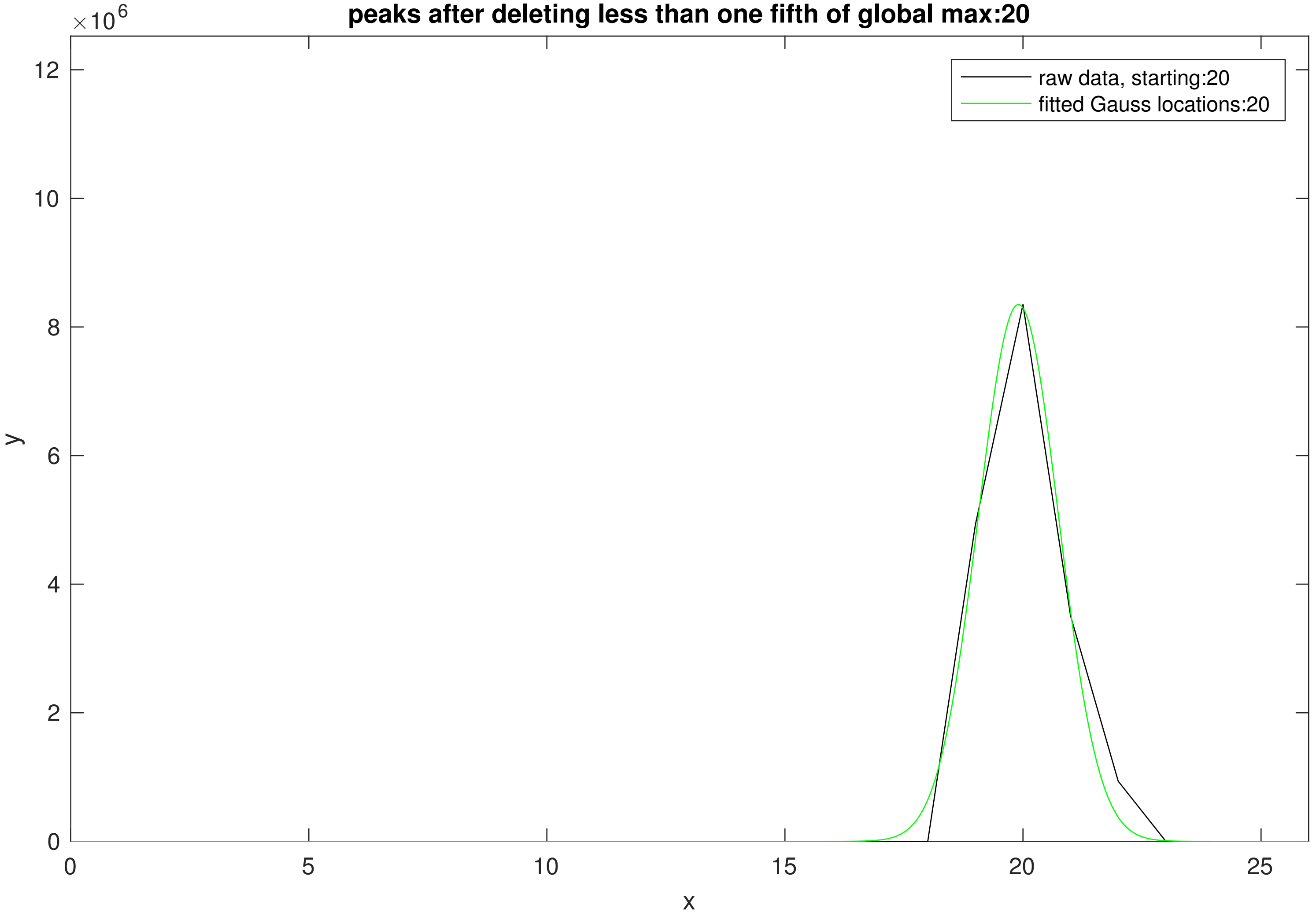

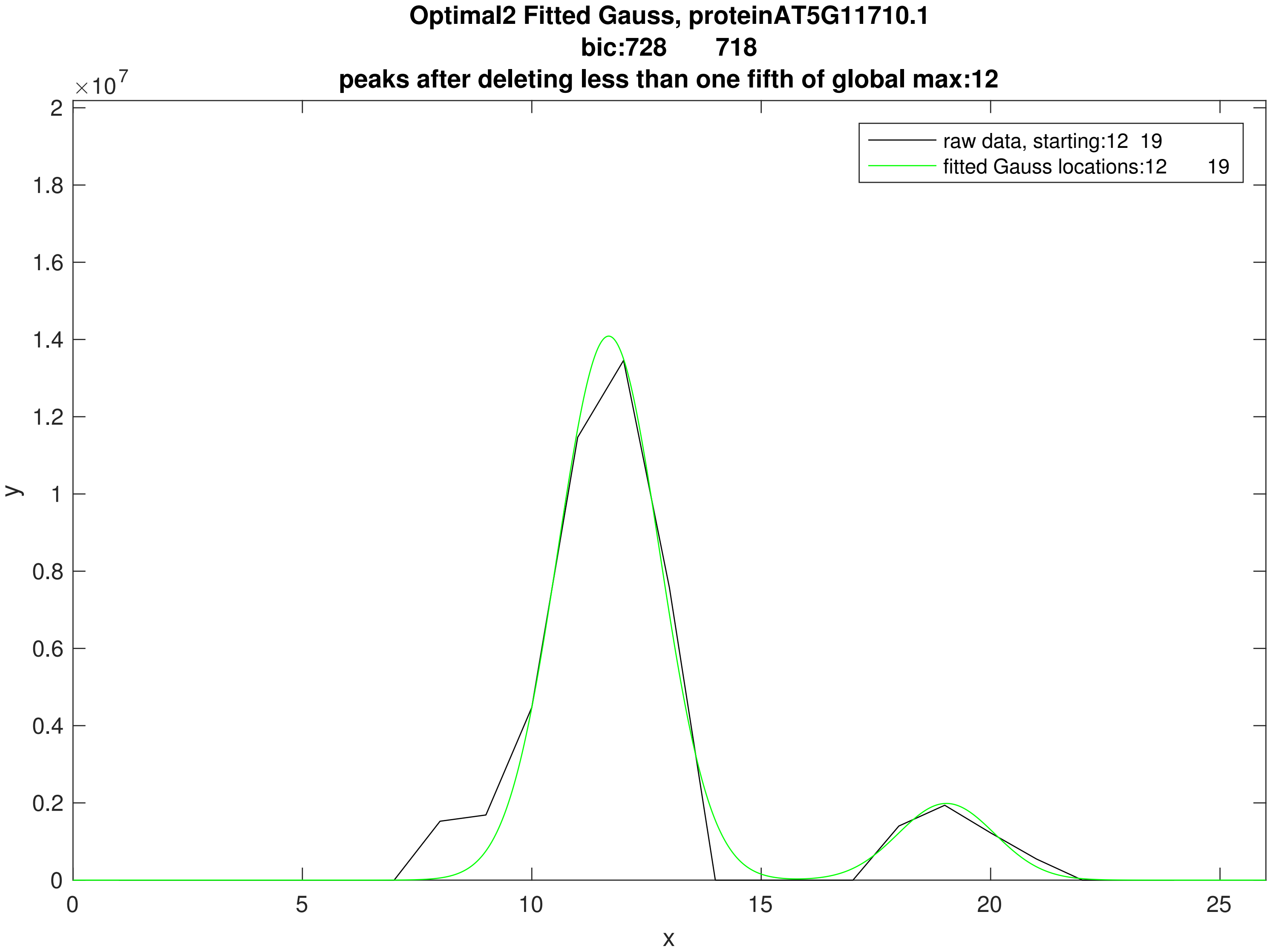
Optimal2 Fitted Gauss, proteinAT5G11710.1
bic:728 718
peaks after deleting less than one fifth of global max:12
×10^7
raw data, starting:12 19
fitted Gauss locations:12 19
y
x
0
0.2
0.4
0.6
0.8
1
1.2
1.4
1.6
1.8
2
5
10
15
20
25

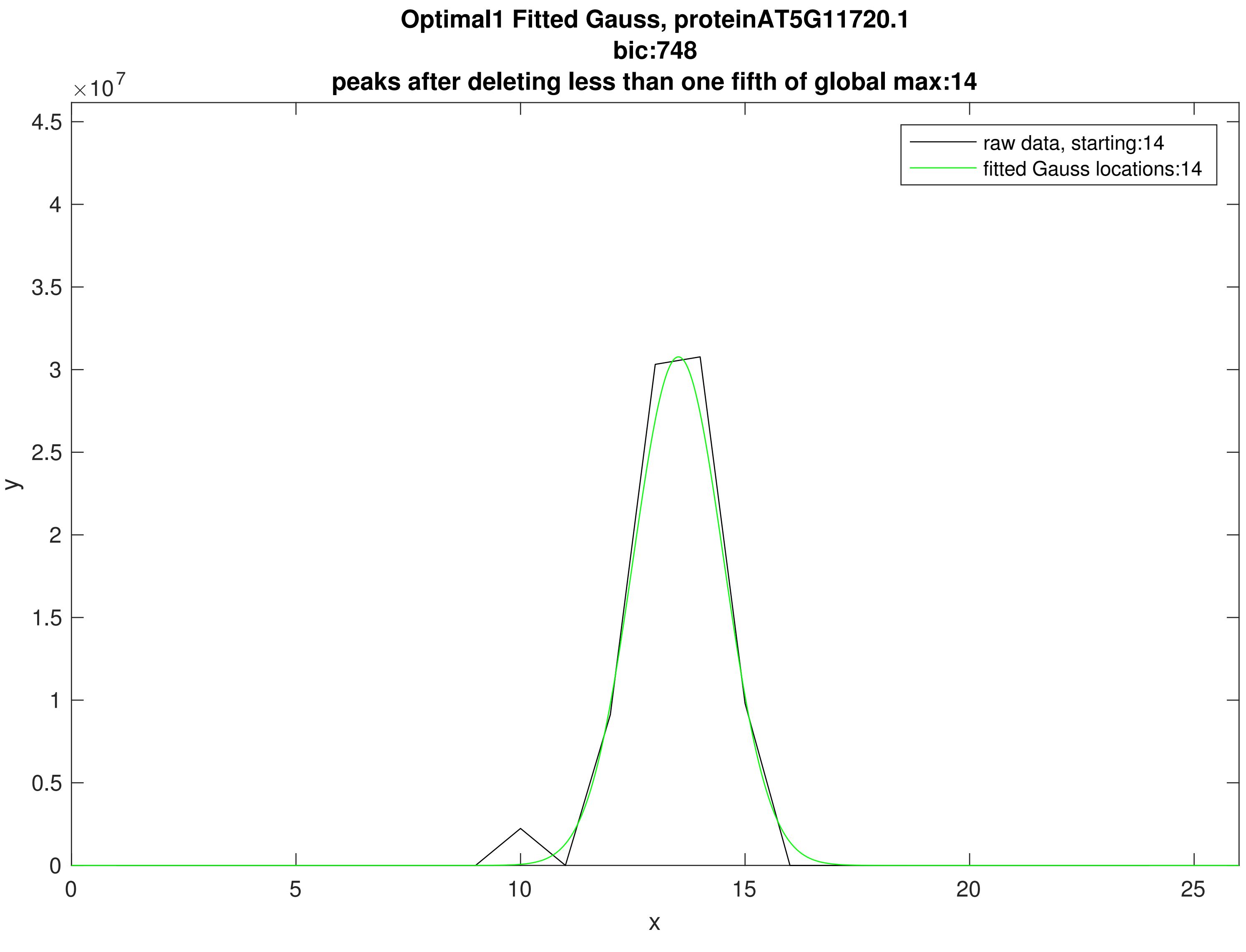

Optimal1 Fitted Gauss, proteinAT5G11720.1
bic:748
peaks after deleting less than one fifth of global max:14
×10^7
4.5
4
3.5
3
2.5
2
1.5
1
0.5
0
y
0
5
10
15
20
25
x
raw data, starting:14
fitted Gauss locations:14

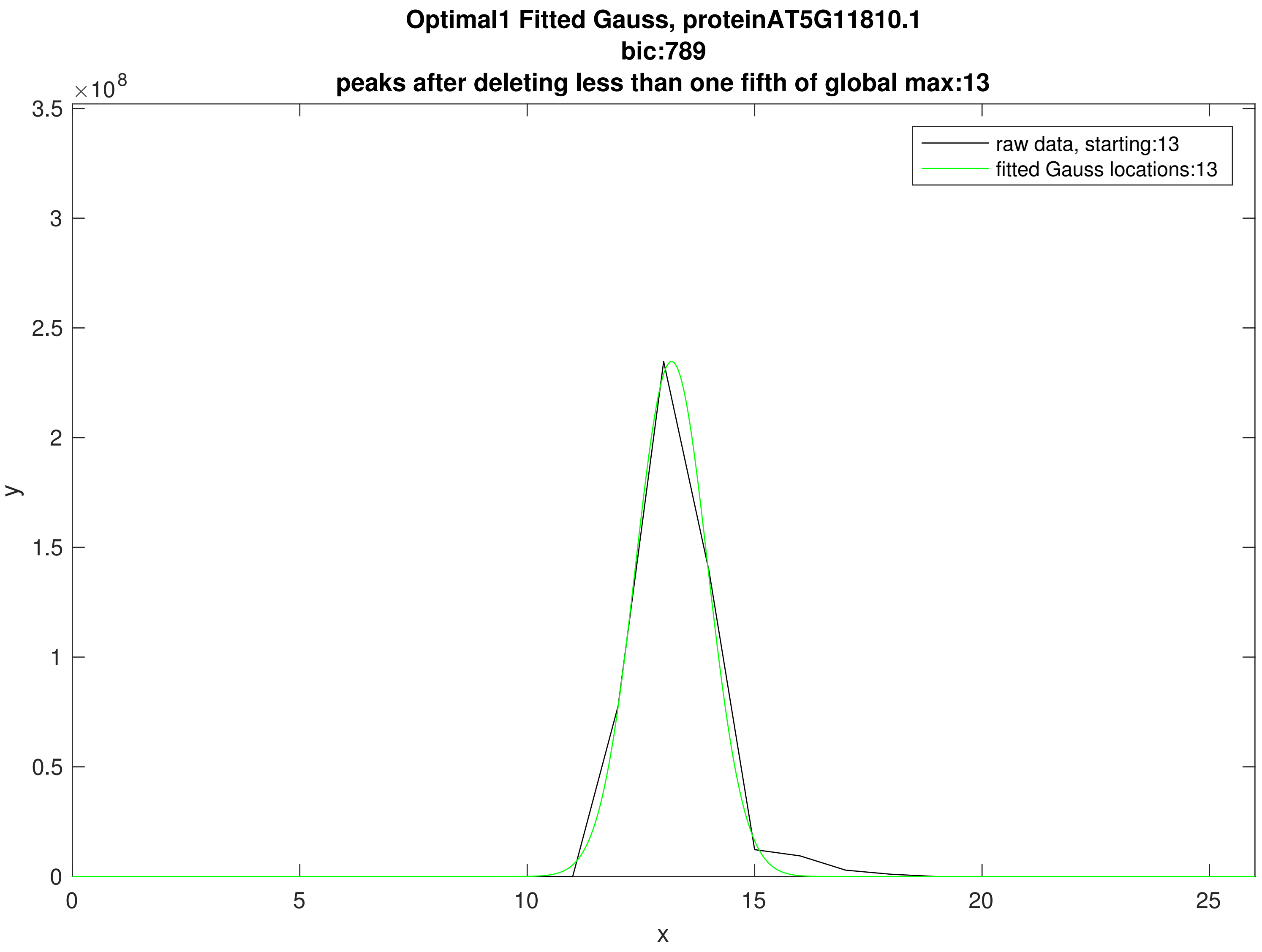

Optimal1 Fitted Gauss, proteinAT5G11810.1
bic:789
peaks after deleting less than one fifth of global max:13
×10^8
3.5
3
2.5
2
1.5
1
0.5
0
y
0
5
10
15
20
25
x
raw data, starting:13
fitted Gauss locations:13

**Optimal1 Fitted Gauss, proteinAT5G11880.1**
**bic:921**
**peaks after deleting less than one fifth of global max:13**

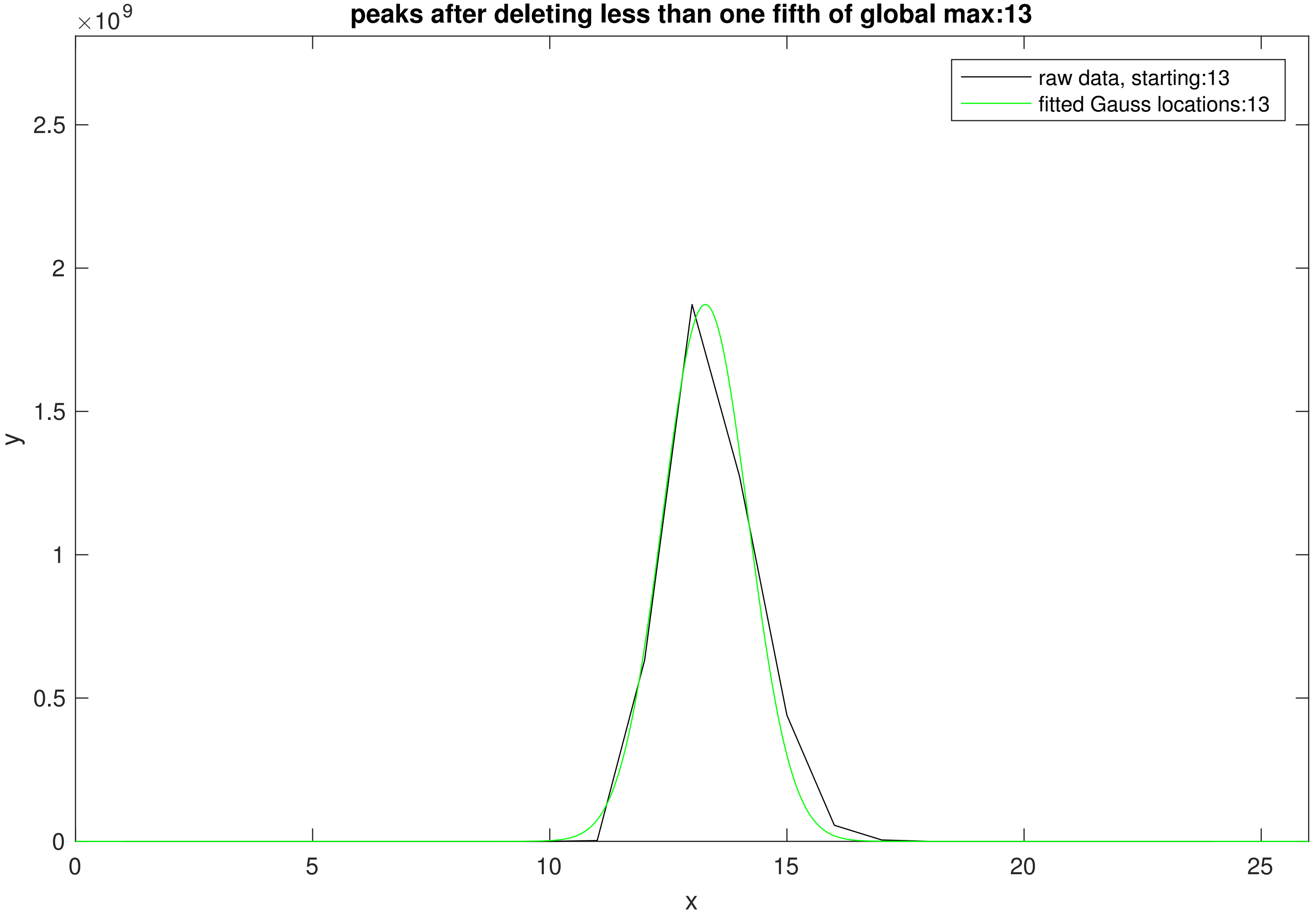

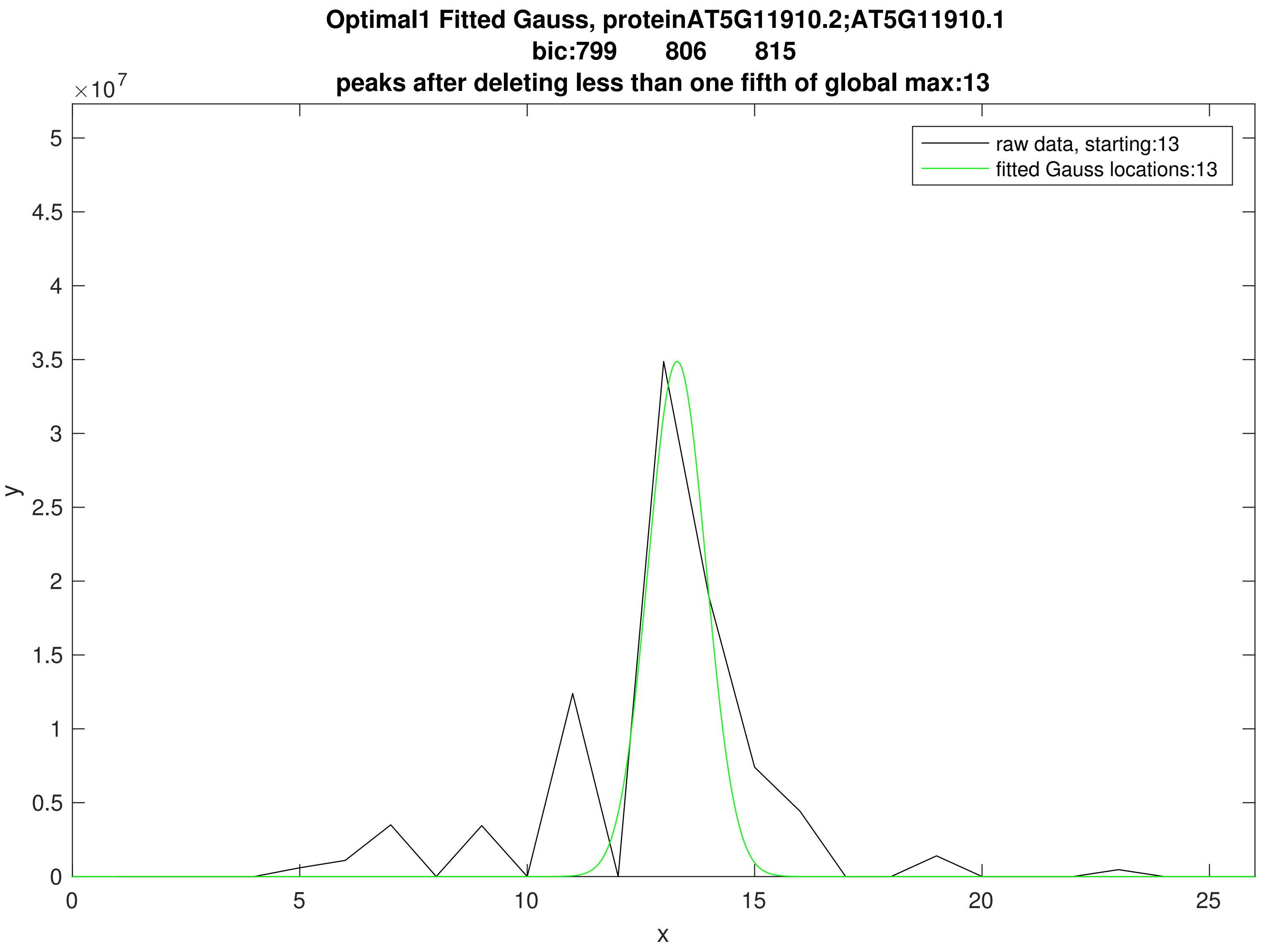

Optimal1 Fitted Gauss, proteinAT5G11910.2;AT5G11910.1
bic:799 806 815
peaks after deleting less than one fifth of global max:13
×10^7
raw data, starting:13
fitted Gauss locations:13
y
x
0
0.5
1
1.5
2
2.5
3
3.5
4
4.5
5
0
5
10
15
20
25

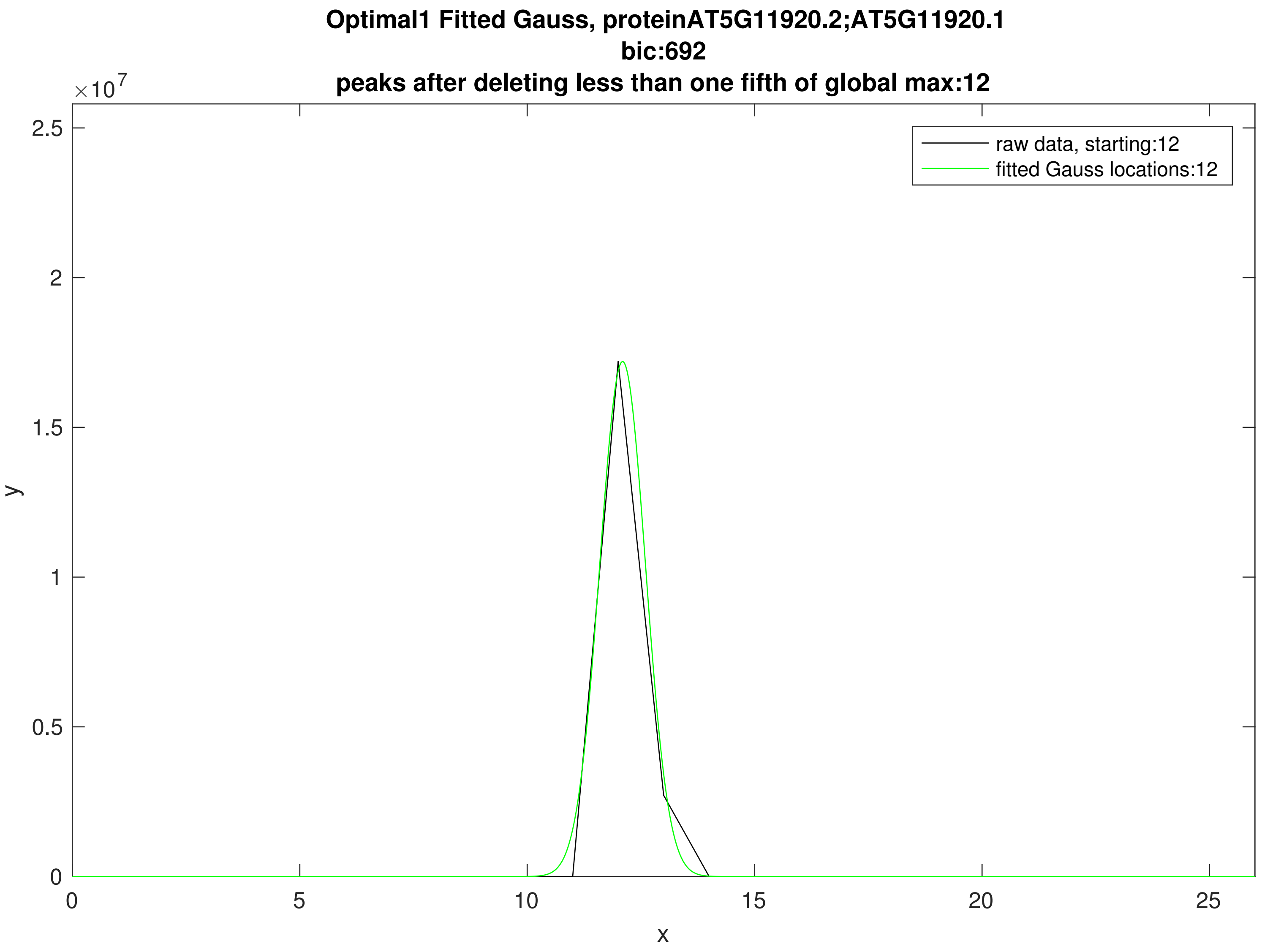

Optimal1 Fitted Gauss, proteinAT5G11920.2;AT5G11920.1
bic:692
peaks after deleting less than one fifth of global max:12
×10^7
2.5
2
1.5
1
0.5
0
y
0
5
10
15
20
25
x
raw data, starting:12
fitted Gauss locations:12

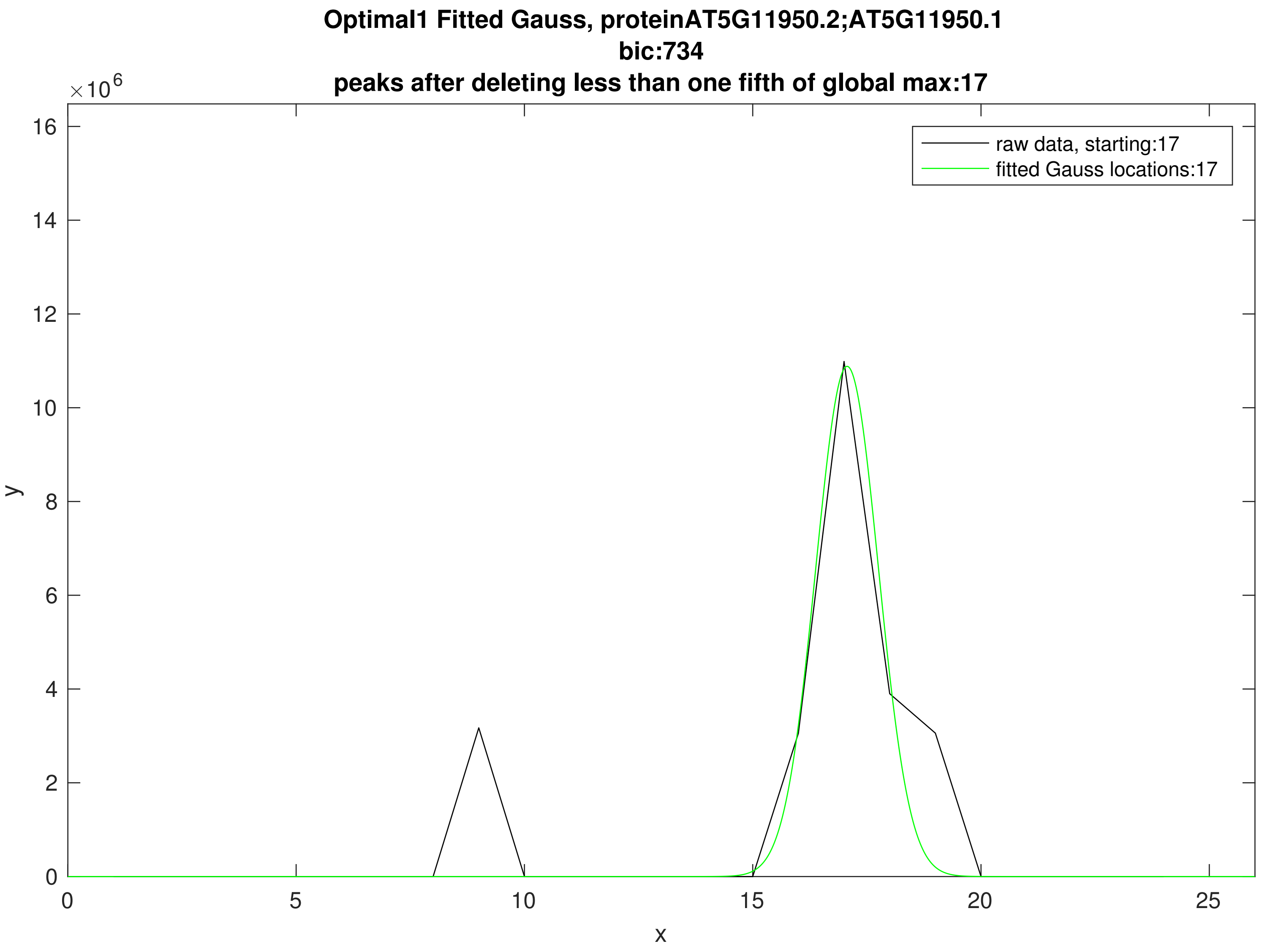
Optimal1 Fitted Gauss, proteinAT5G11950.2;AT5G11950.1
bic:734
peaks after deleting less than one fifth of global max:17
×10^6
raw data, starting:17
fitted Gauss locations:17
y
x
0
2
4
6
8
10
12
14
16
0
5
10
15
20
25

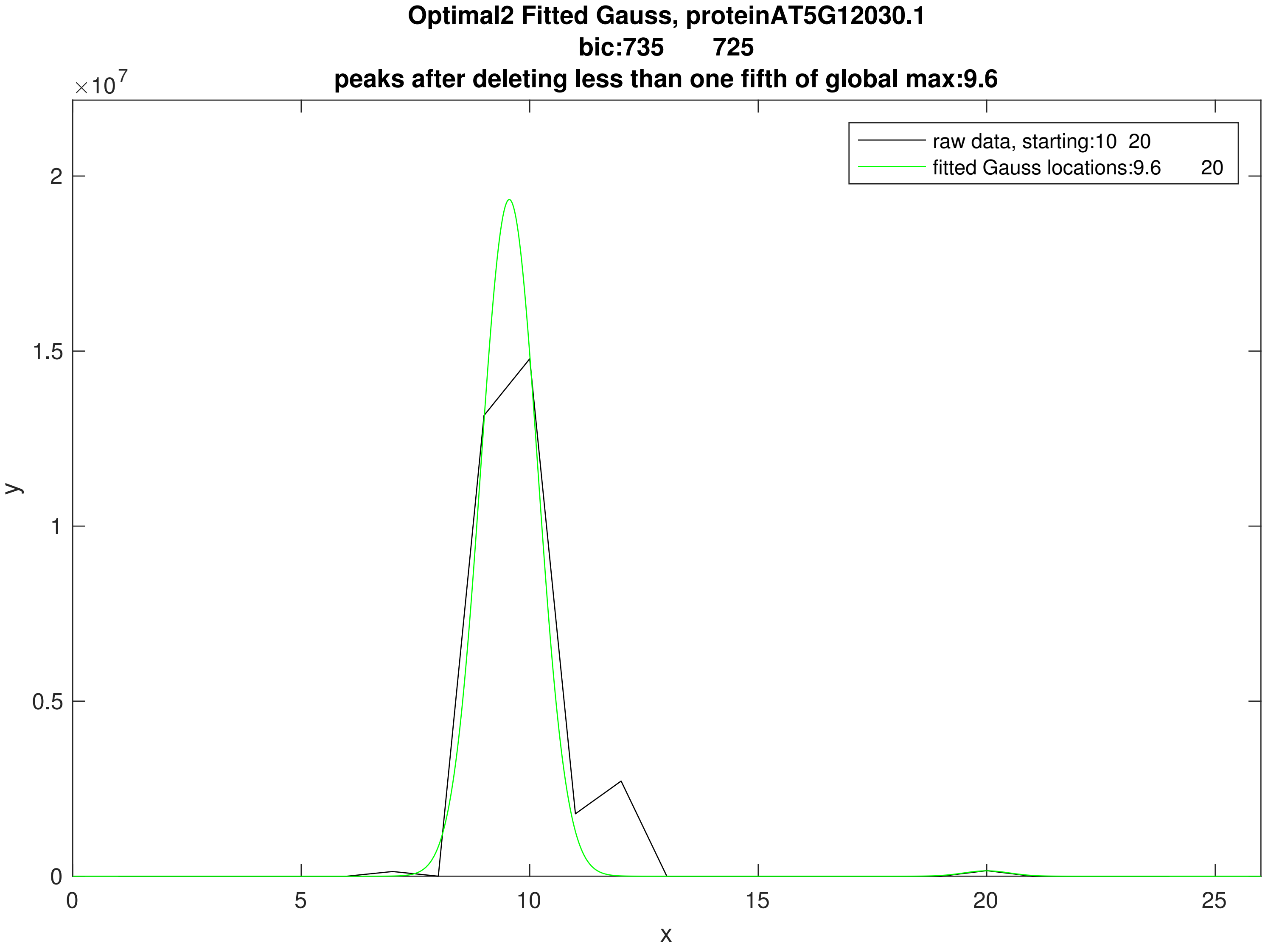

Optimal2 Fitted Gauss, proteinAT5G12030.1
bic:735 725
peaks after deleting less than one fifth of global max:9.6
×10^7
raw data, starting:10 20
fitted Gauss locations:9.6 20
y
x
0
0.5
1
1.5
2
0
5
10
15
20
25

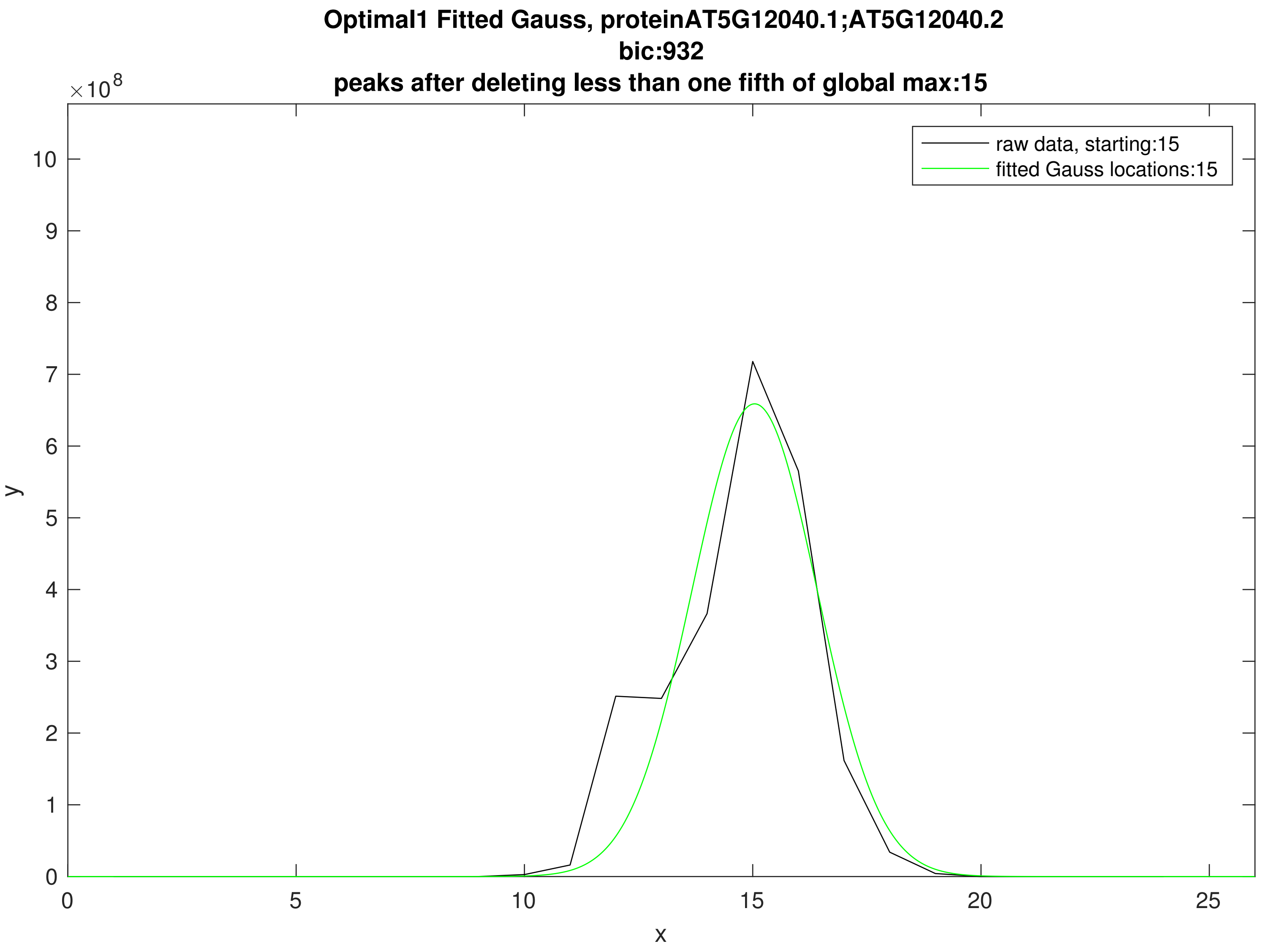

Optimal1 Fitted Gauss, proteinAT5G12040.1;AT5G12040.2
bic:932
peaks after deleting less than one fifth of global max:15
×10^8
raw data, starting:15
fitted Gauss locations:15
y
x
0
1
2
3
4
5
6
7
8
9
10
0
5
10
15
20
25

**Optimal1 Fitted Gauss, proteinAT5G12110.1**
**bic:754 761**
**peaks after deleting less than one fifth of global max:8.6**

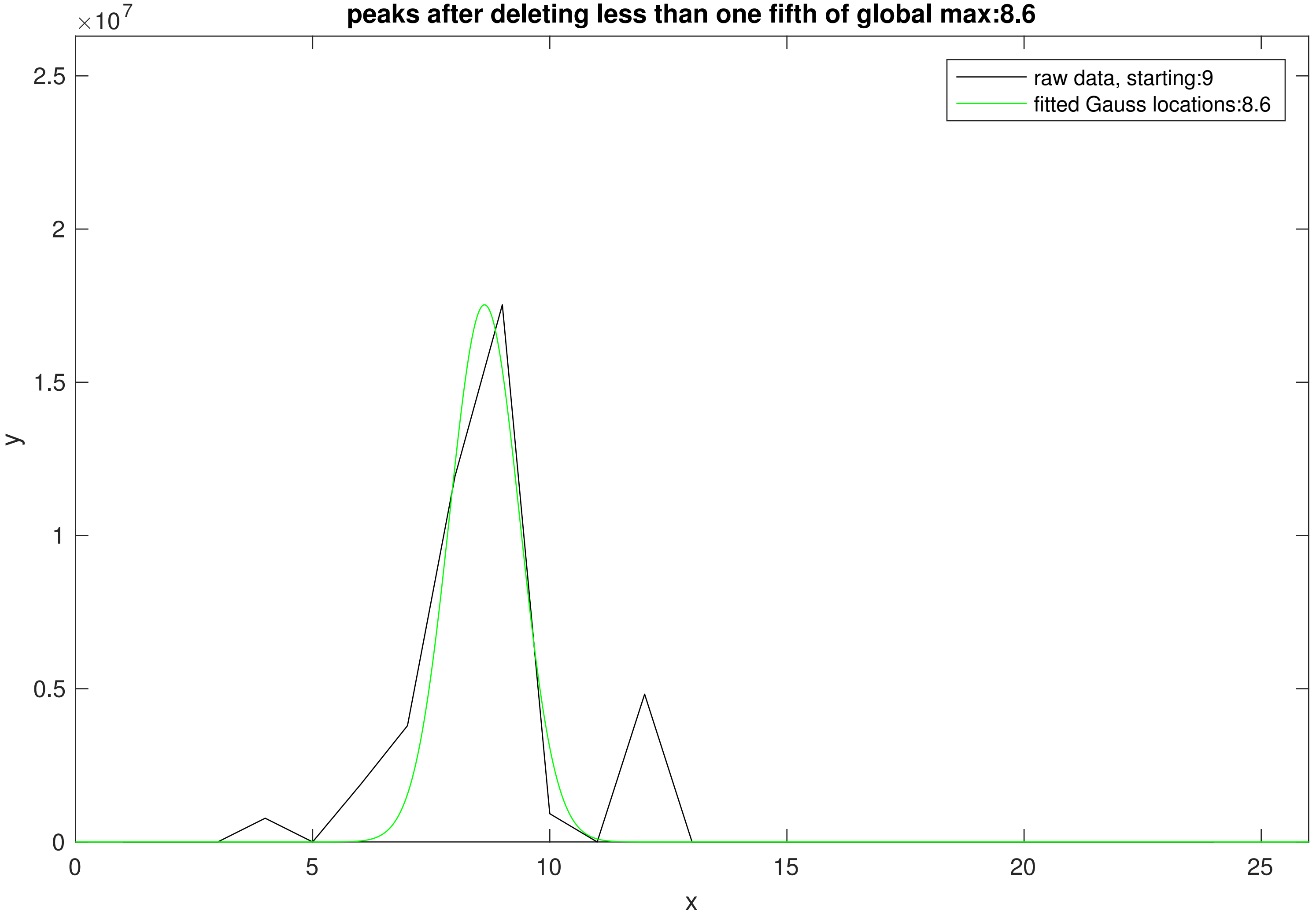

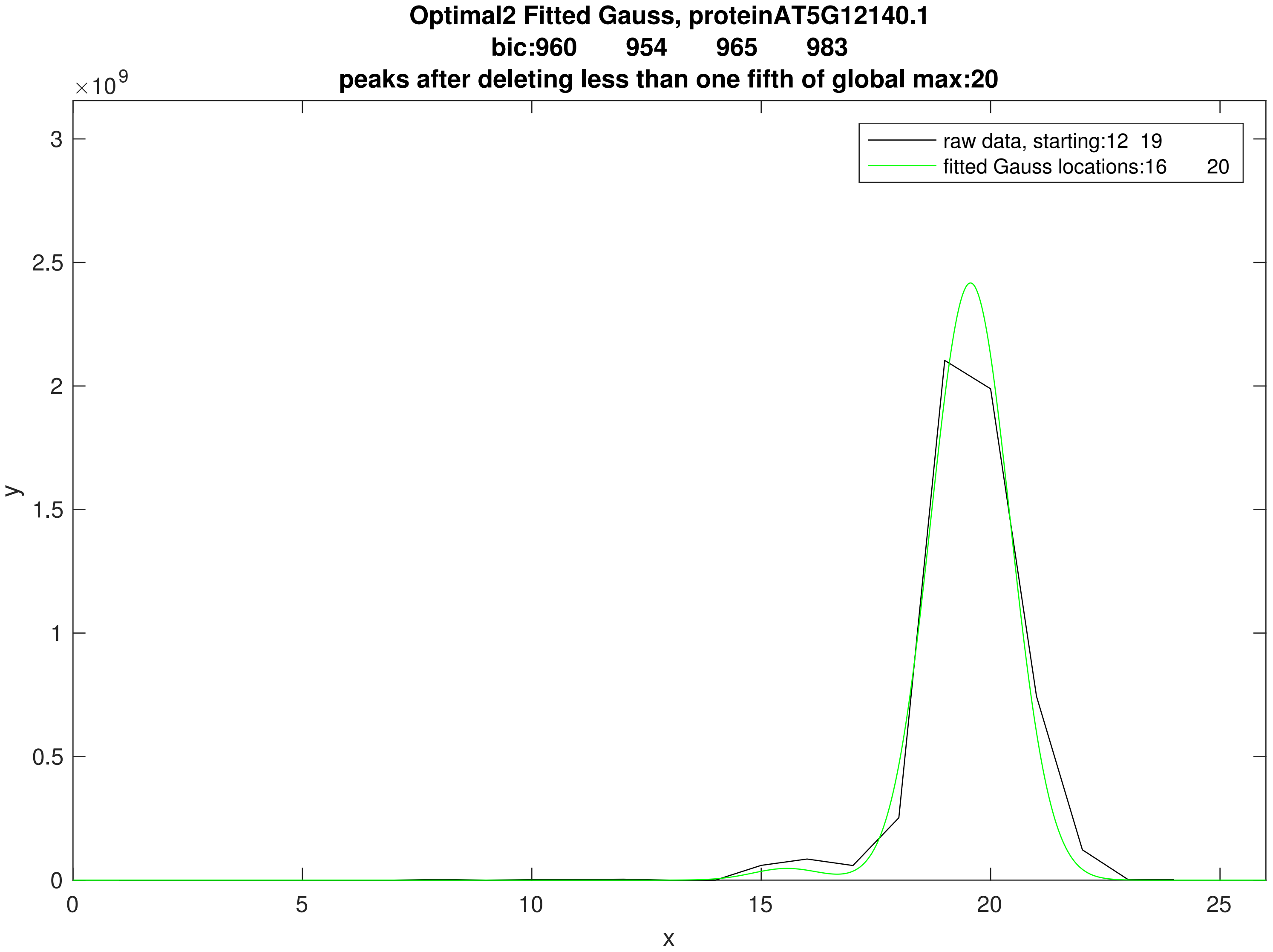

Optimal2 Fitted Gauss, proteinAT5G12140.1
bic:960 954 965 983
peaks after deleting less than one fifth of global max:20
×10^9
raw data, starting:12 19
fitted Gauss locations:16 20
3
2.5
2
1.5
1
0.5
0
y
0
5
10
15
20
25
x

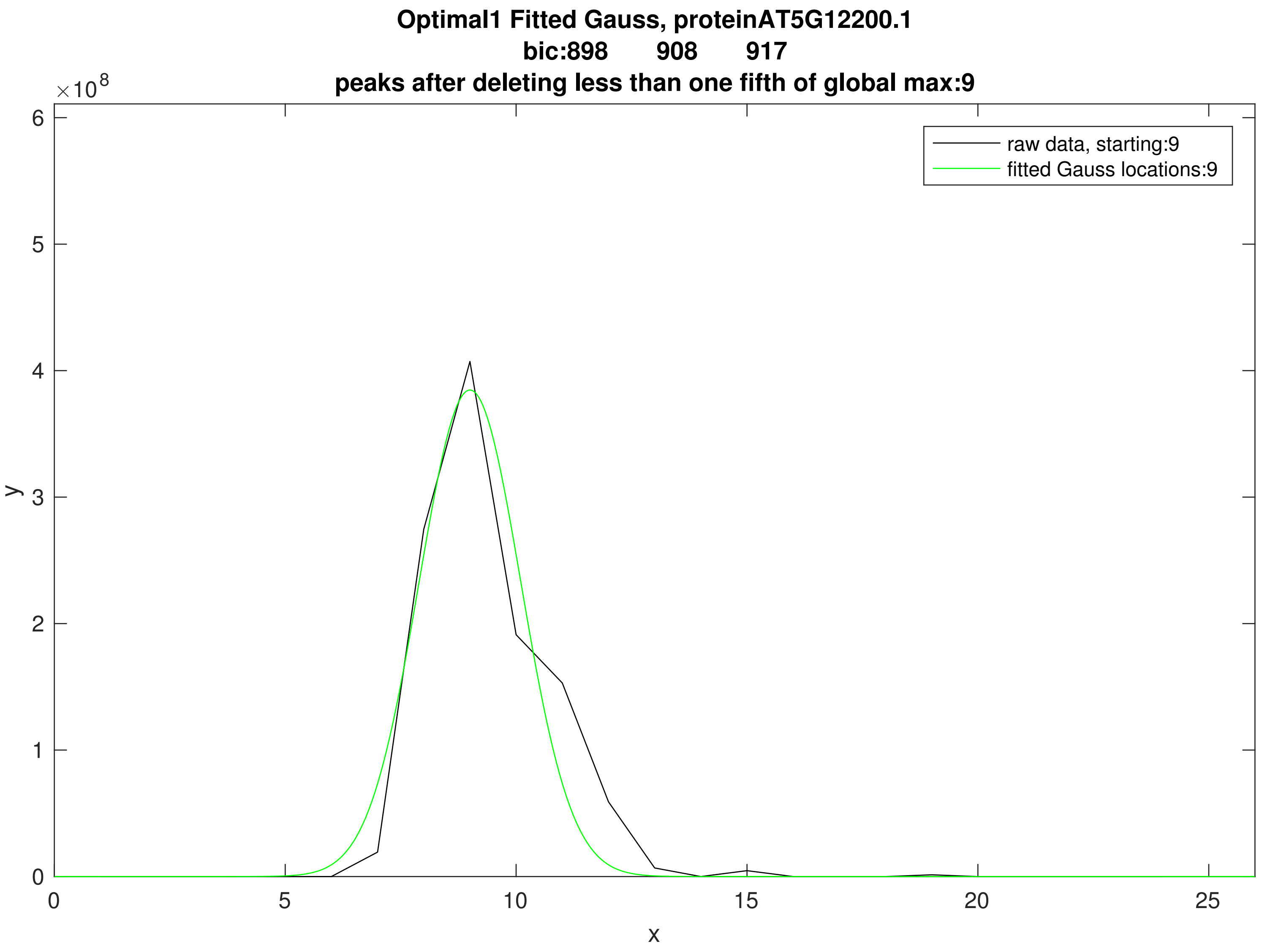

Optimal1 Fitted Gauss, proteinAT5G12200.1
bic:898 908 917
peaks after deleting less than one fifth of global max:9
×10^8
6
5
4
3
2
1
0
y
0
5
10
15
20
25
x
raw data, starting:9
fitted Gauss locations:9

**Optimal1 Fitted Gauss, proteinAT5G12210.2;AT5G12210.1**
**bic:793**
**peaks after deleting less than one fifth of global max:12**

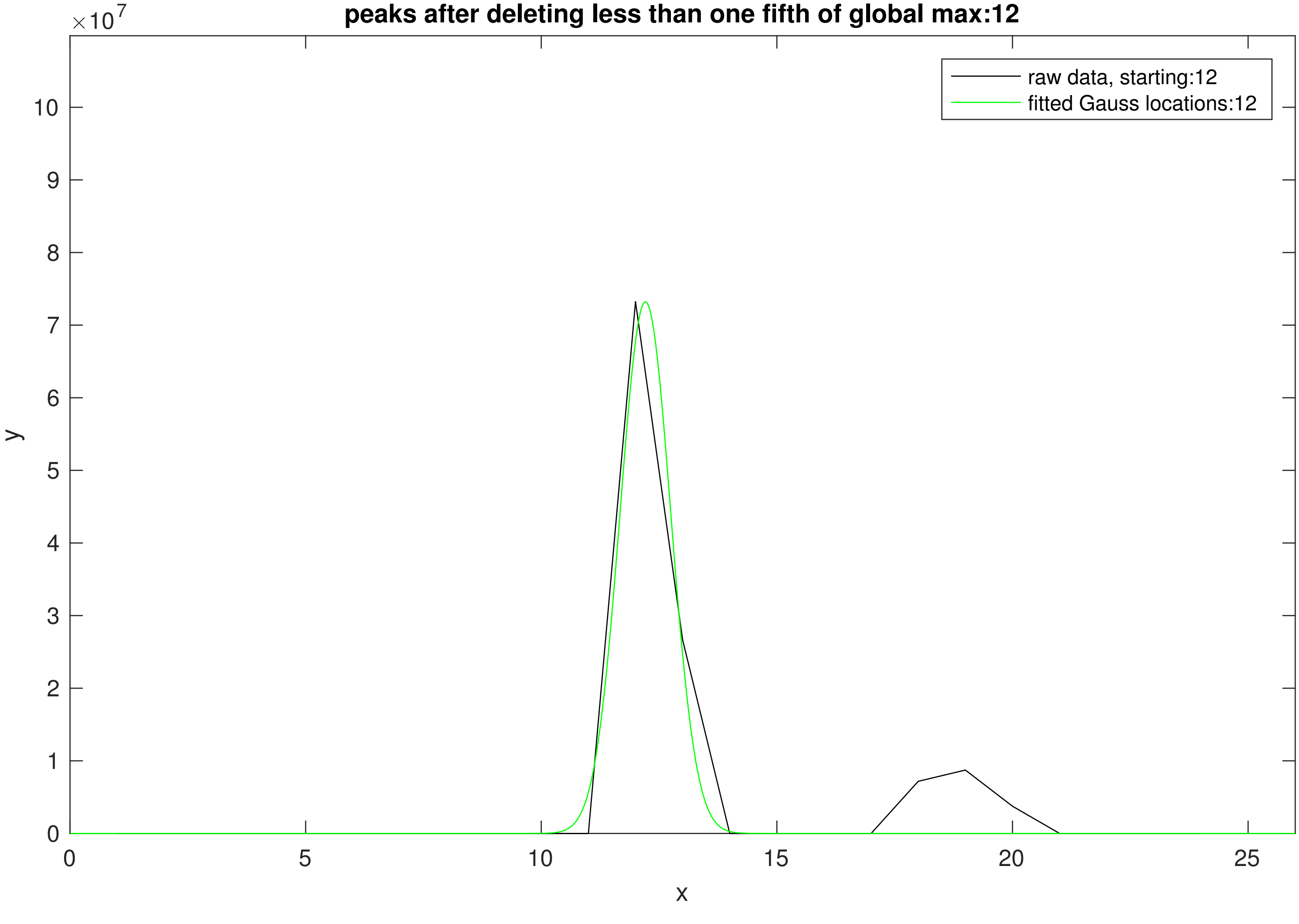

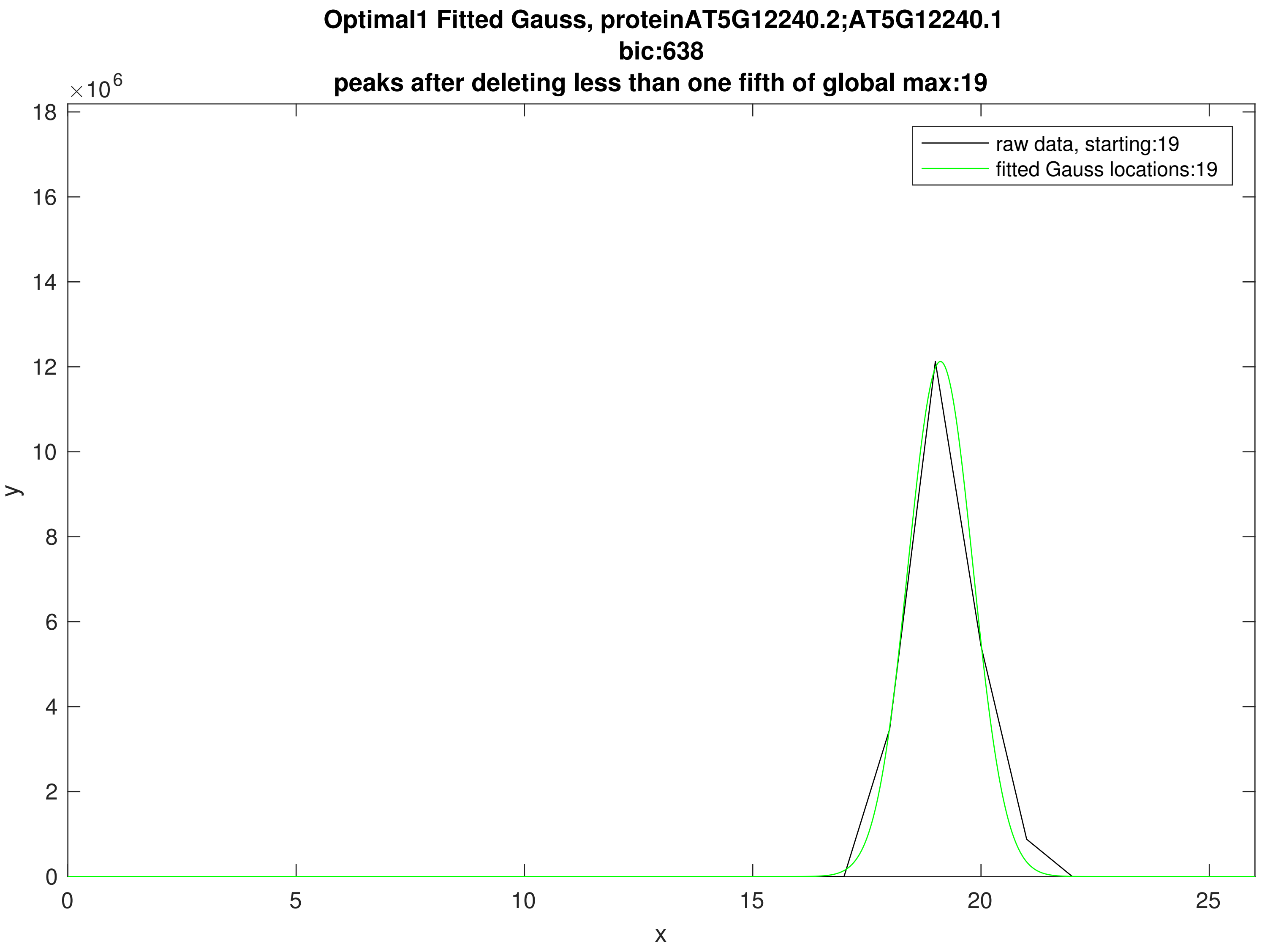

Optimal1 Fitted Gauss, proteinAT5G12240.2;AT5G12240.1
bic:638
peaks after deleting less than one fifth of global max:19
×10^6
18
16
14
12
10
8
6
4
2
0
y
0
5
10
15
20
25
x
raw data, starting:19
fitted Gauss locations:19

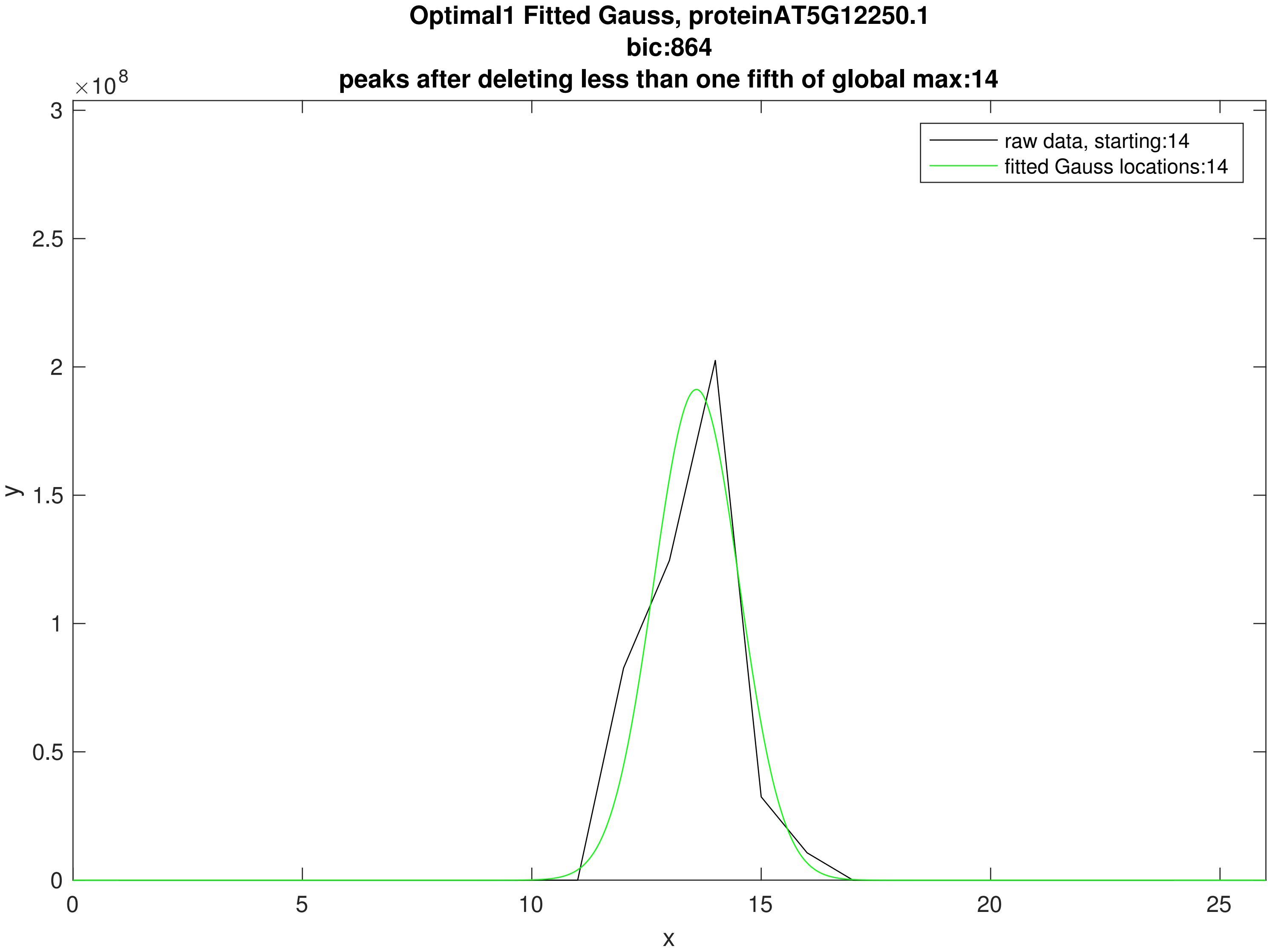
Optimal1 Fitted Gauss, proteinAT5G12250.1
bic:864
peaks after deleting less than one fifth of global max:14
×10^8
3
2.5
2
1.5
1
0.5
0
y
0
5
10
15
20
25
x
raw data, starting:14
fitted Gauss locations:14

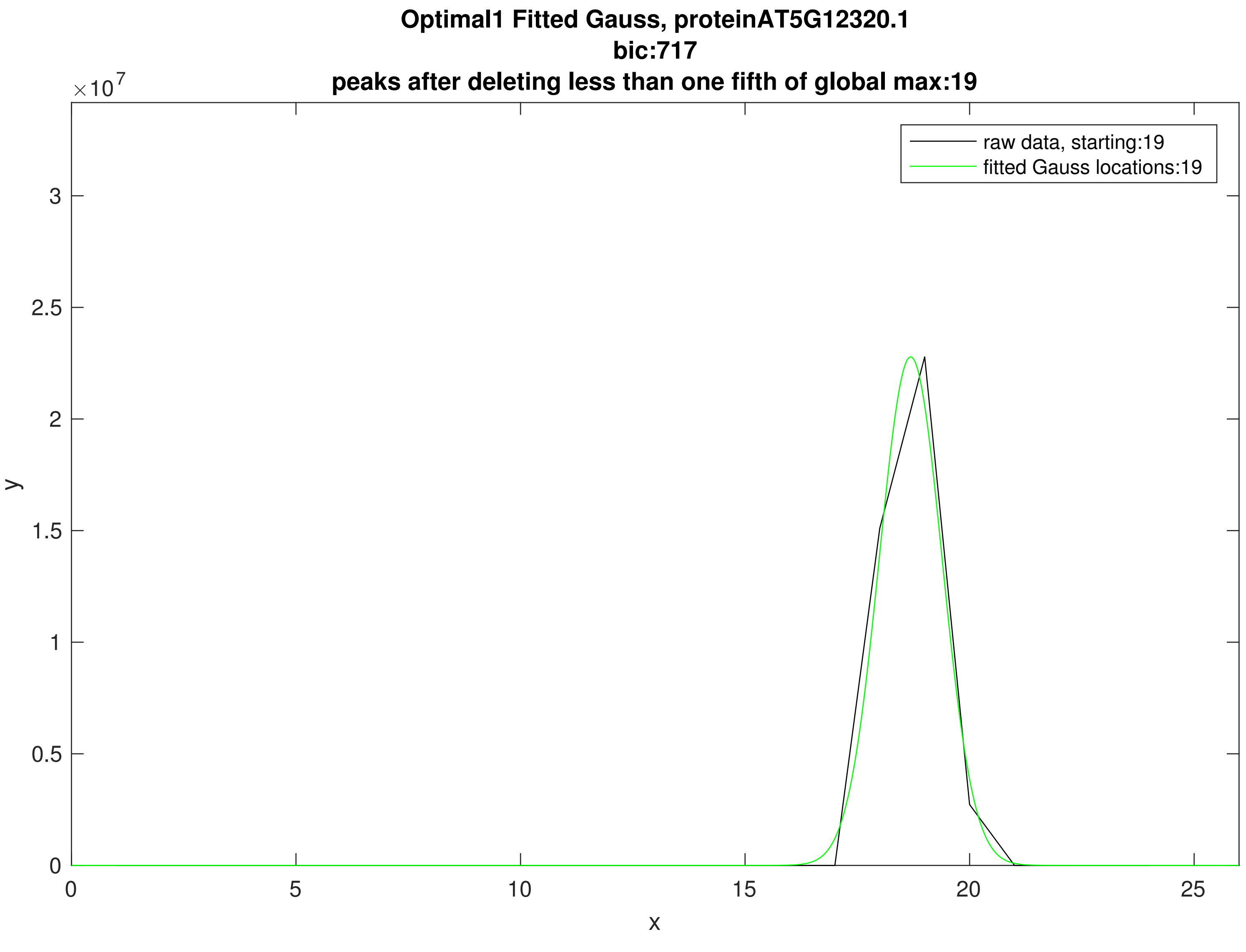

Optimal1 Fitted Gauss, proteinAT5G12320.1
bic:717
peaks after deleting less than one fifth of global max:19
×10^7
raw data, starting:19
fitted Gauss locations:19
3
2.5
2
1.5
1
0.5
0
y
0
5
10
15
20
25
x

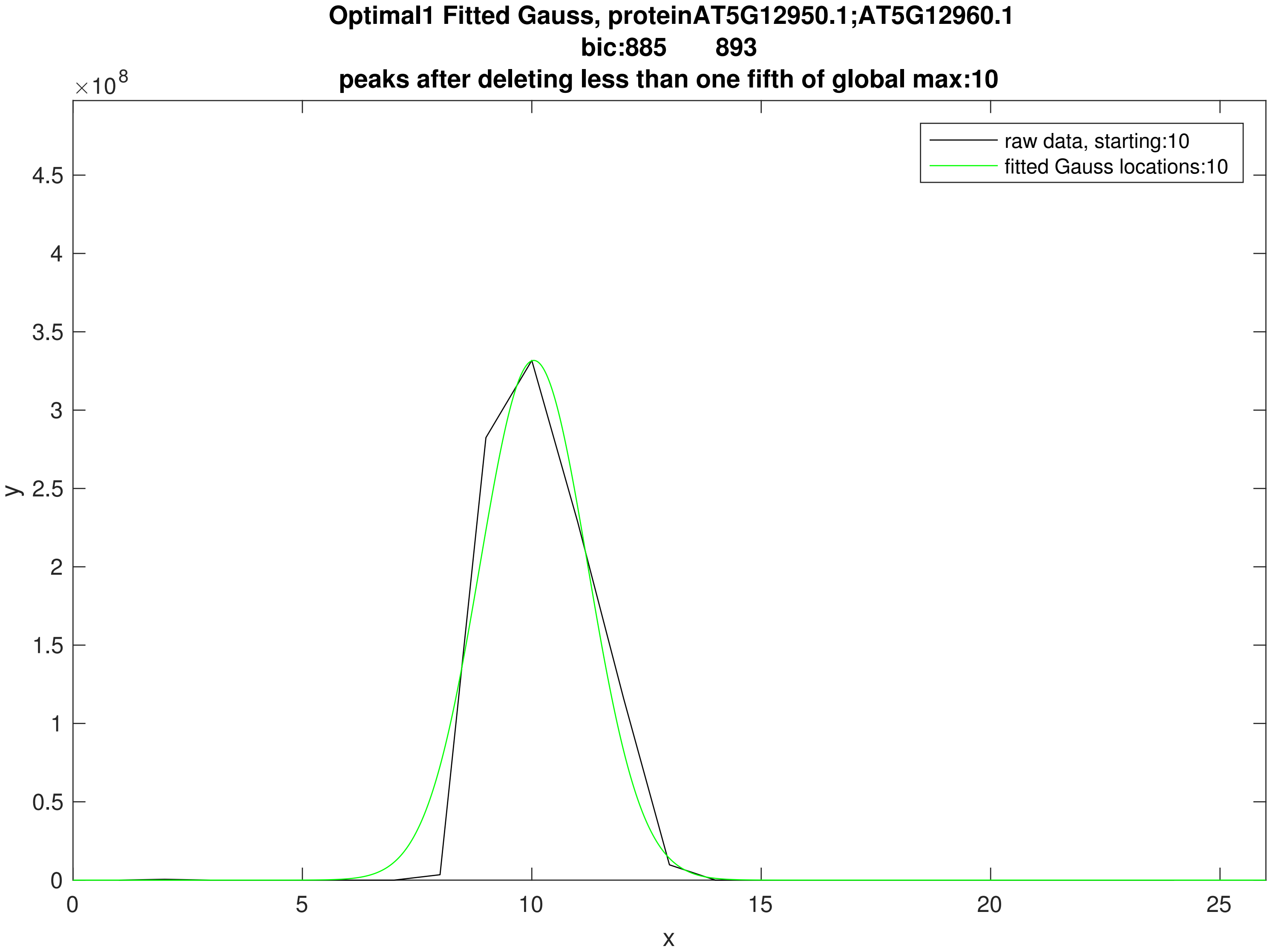
Optimal1 Fitted Gauss, proteinAT5G12950.1;AT5G12960.1
bic:885 893
peaks after deleting less than one fifth of global max:10
×10^8
raw data, starting:10
fitted Gauss locations:10
y
x
0
0.5
1
1.5
2
2.5
3
3.5
4
4.5
5
10
15
20
25

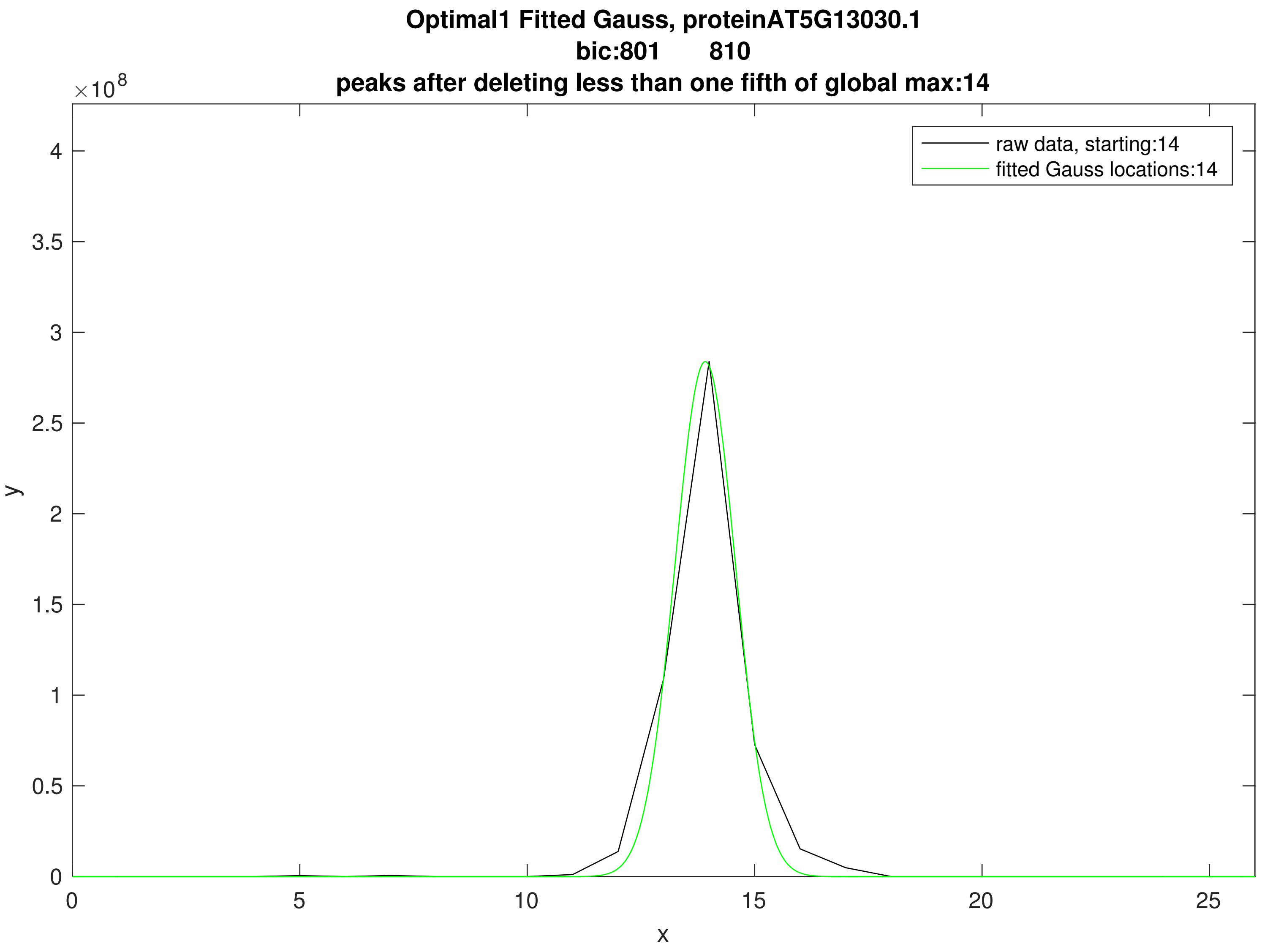

Optimal1 Fitted Gauss, proteinAT5G13030.1
bic:801 810
peaks after deleting less than one fifth of global max:14
×10^8
raw data, starting:14
fitted Gauss locations:14
y
x
0
0.5
1
1.5
2
2.5
3
3.5
4
0
5
10
15
20
25

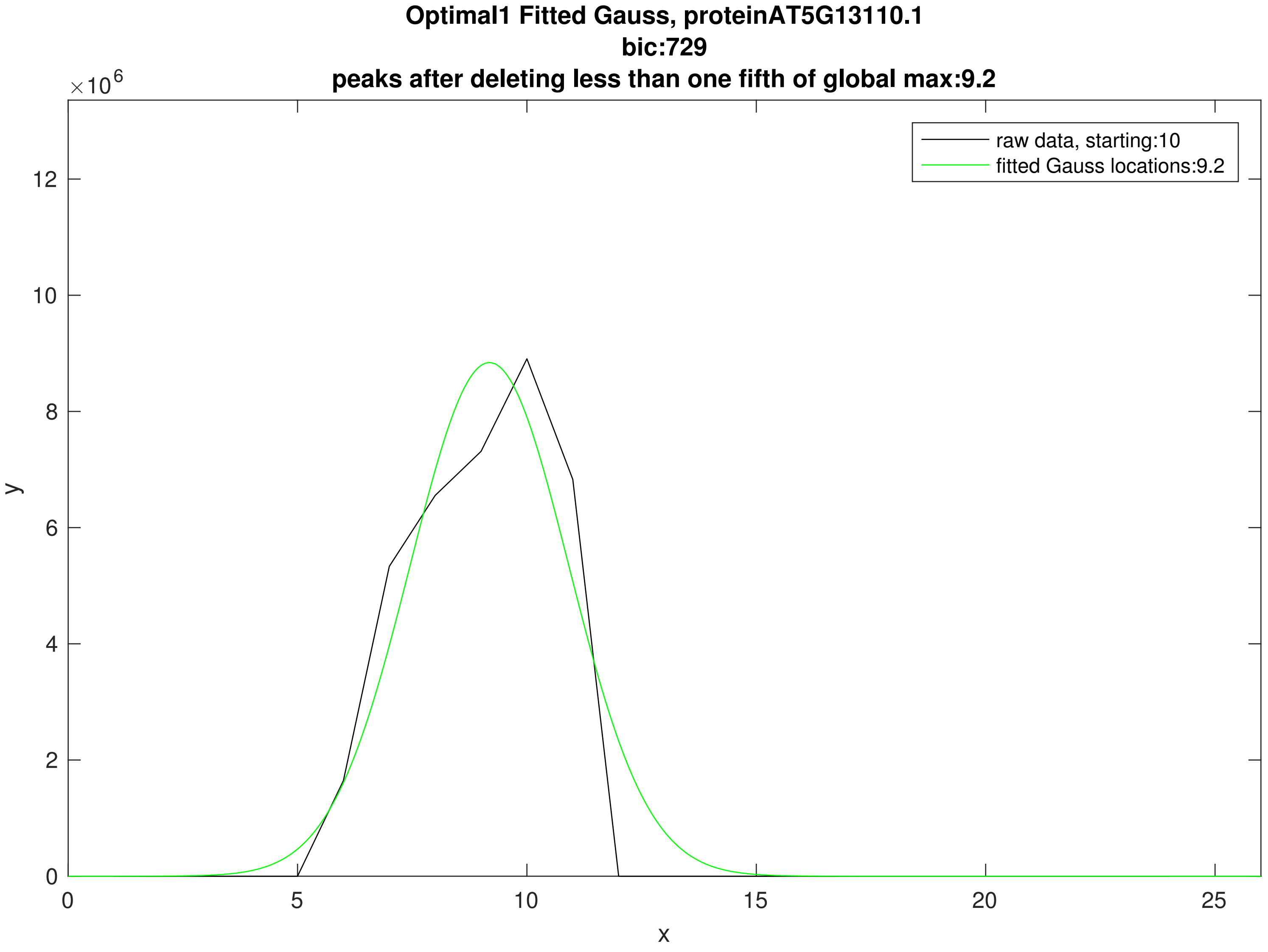

Optimal1 Fitted Gauss, proteinAT5G13110.1
bic:729
peaks after deleting less than one fifth of global max:9.2
×10^6
raw data, starting:10
fitted Gauss locations:9.2
y
x
0
2
4
6
8
10
12
5
15
20
25

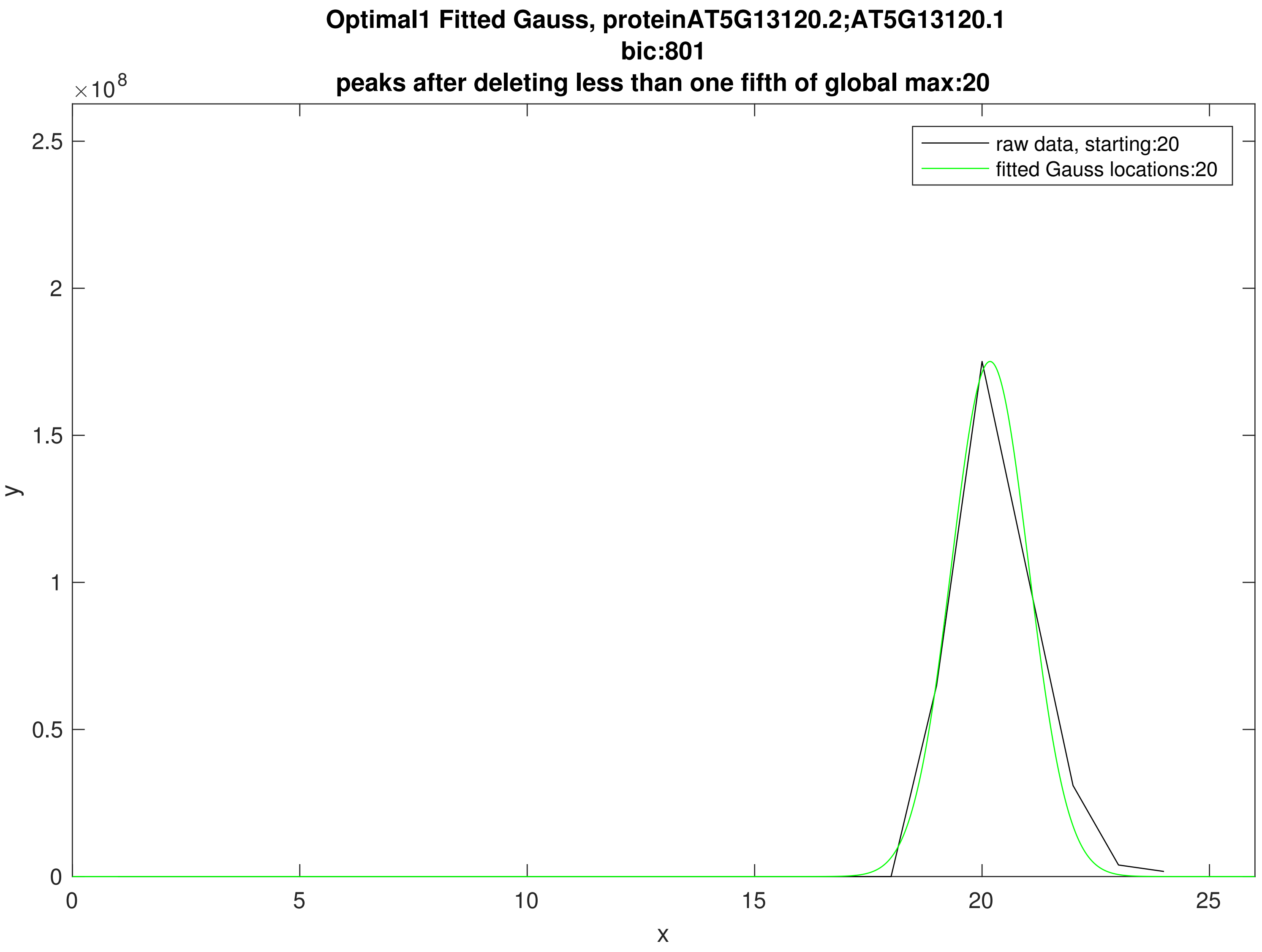

Optimal1 Fitted Gauss, proteinAT5G13120.2;AT5G13120.1
bic:801
peaks after deleting less than one fifth of global max:20
×10^8
2.5
2
1.5
1
0.5
0
y
0
5
10
15
20
25
x
raw data, starting:20
fitted Gauss locations:20

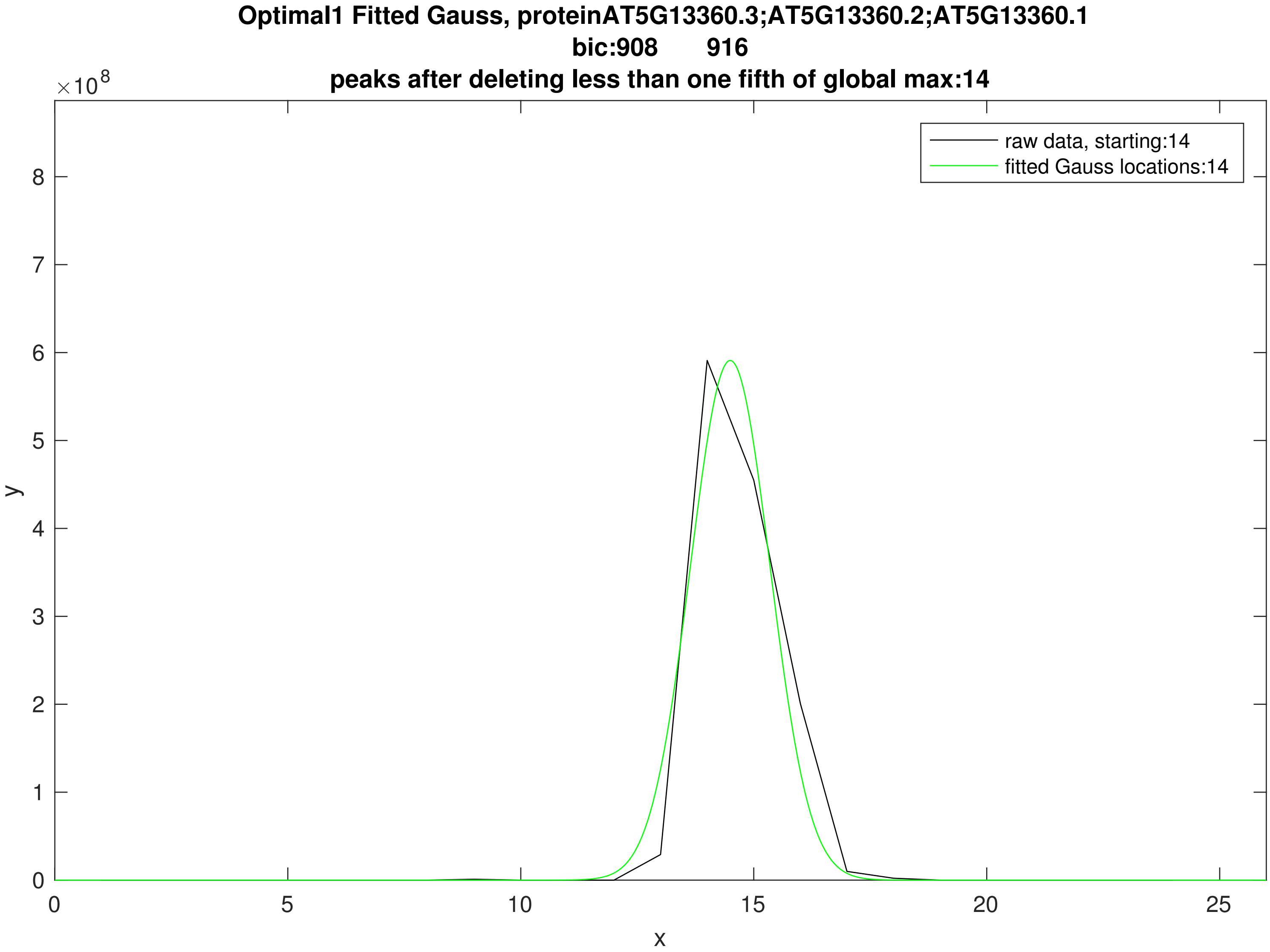

Optimal1 Fitted Gauss, proteinAT5G13360.3;AT5G13360.2;AT5G13360.1
bic:908 916
peaks after deleting less than one fifth of global max:14
×10^8
raw data, starting:14
fitted Gauss locations:14
y
x
0
1
2
3
4
5
6
7
8
0
5
10
15
20
25

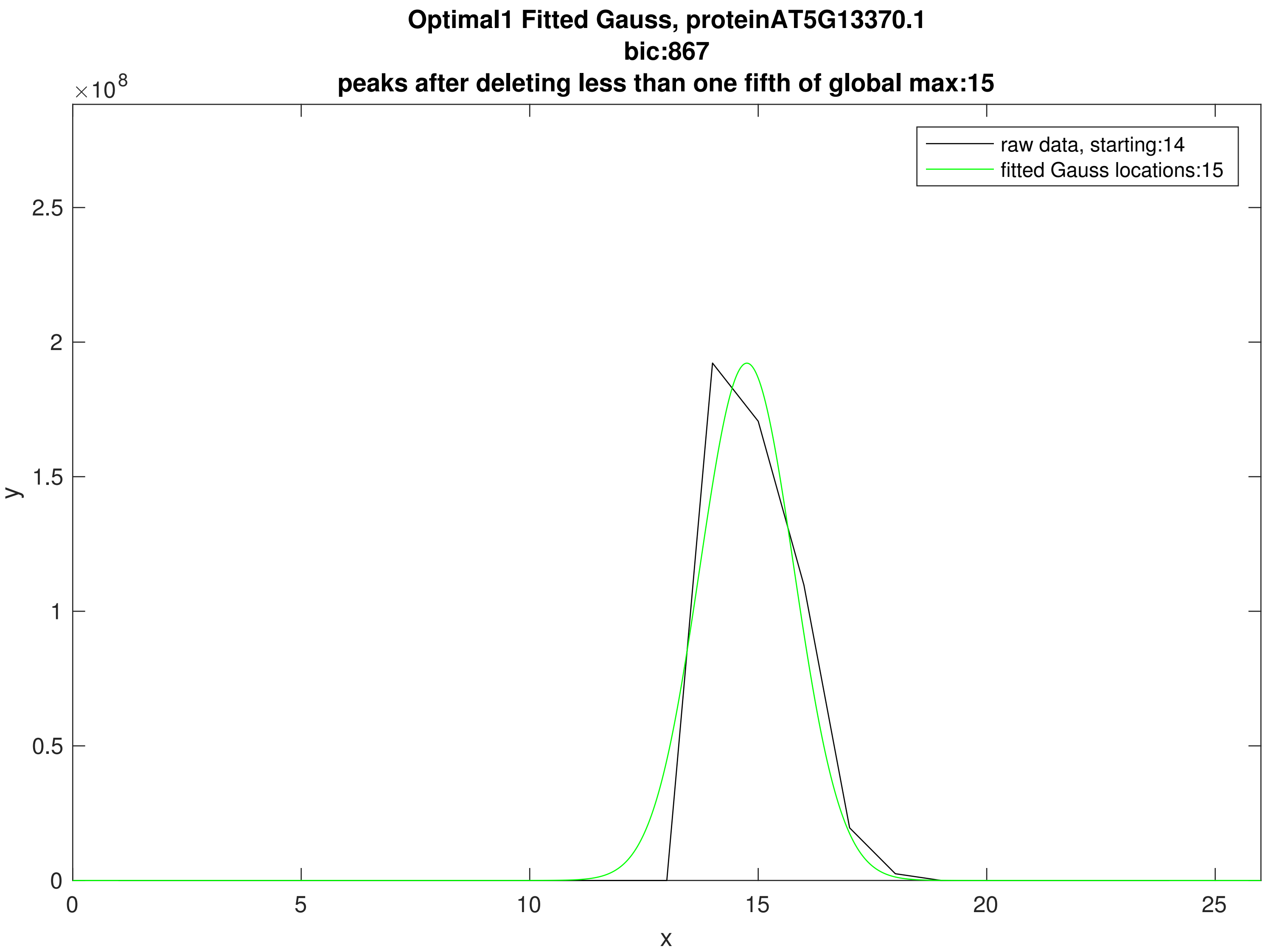
Optimal1 Fitted Gauss, proteinAT5G13370.1
bic:867
peaks after deleting less than one fifth of global max:15
×10^8
2.5
2
1.5
1
0.5
0
y
0
5
10
15
20
25
x
raw data, starting:14
fitted Gauss locations:15

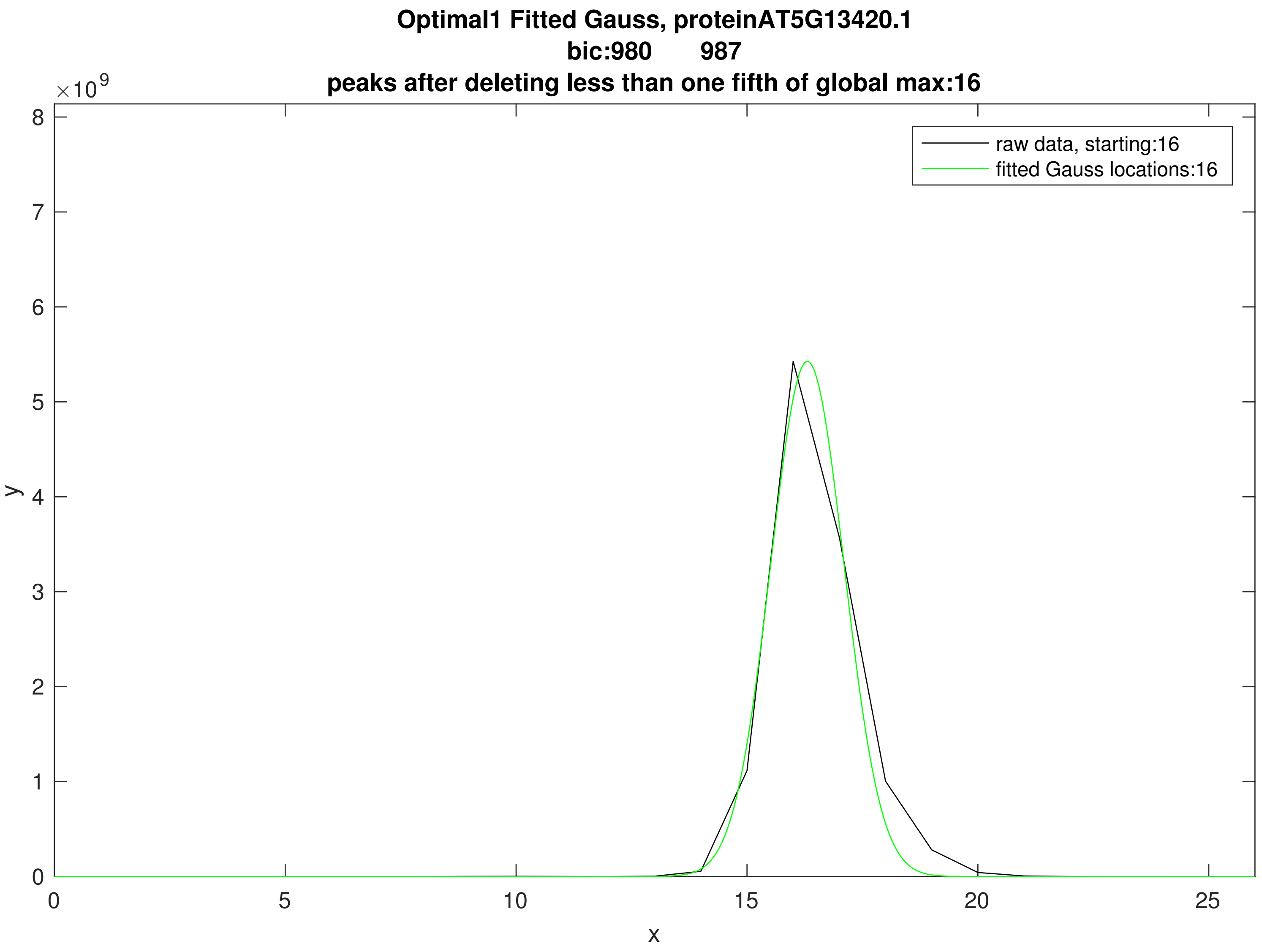
Optimal1 Fitted Gauss, proteinAT5G13420.1
bic:980 987
peaks after deleting less than one fifth of global max:16
×10^9
raw data, starting:16
fitted Gauss locations:16
y
x
0
1
2
3
4
5
6
7
8
0
5
10
15
20
25

**Optimal1 Fitted Gauss, proteinAT5G13450.1;AT5G13450.2**
**bic:754 800**
**peaks after deleting less than one fifth of global max:7.1**

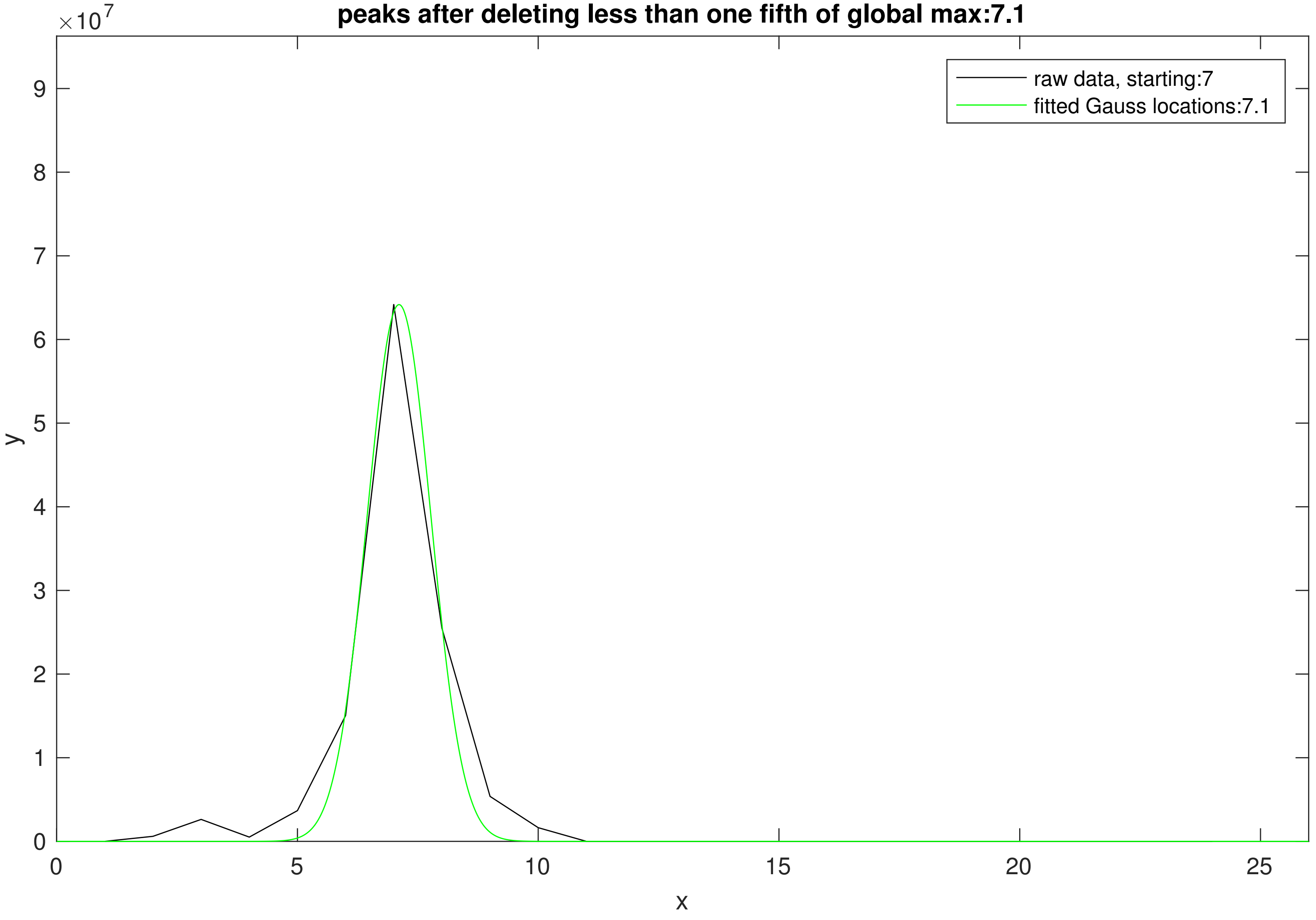

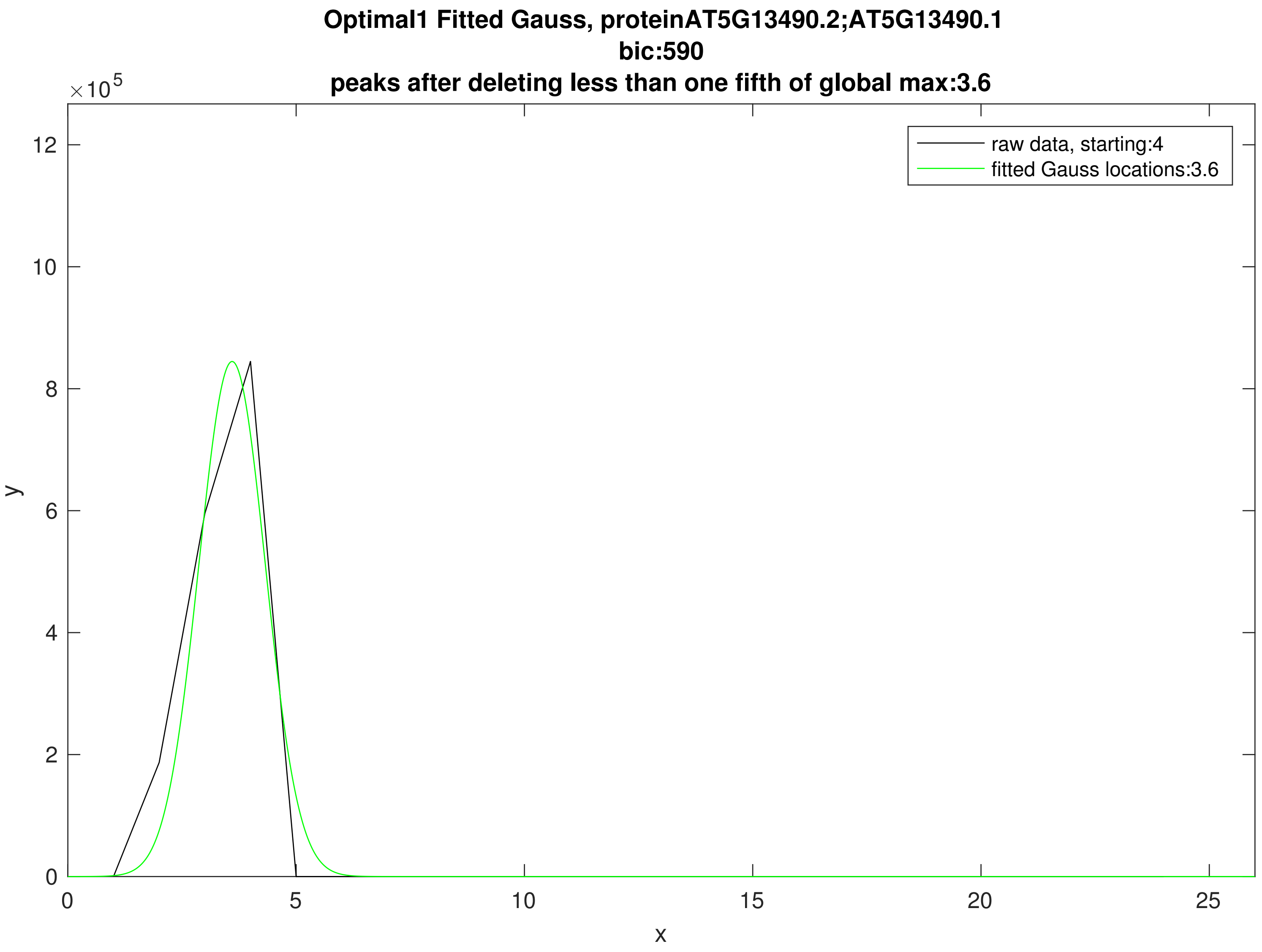

Optimal1 Fitted Gauss, proteinAT5G13490.2;AT5G13490.1
bic:590
peaks after deleting less than one fifth of global max:3.6
×10^5
raw data, starting:4
fitted Gauss locations:3.6
y
x
0
2
4
6
8
10
12
0
5
10
15
20
25

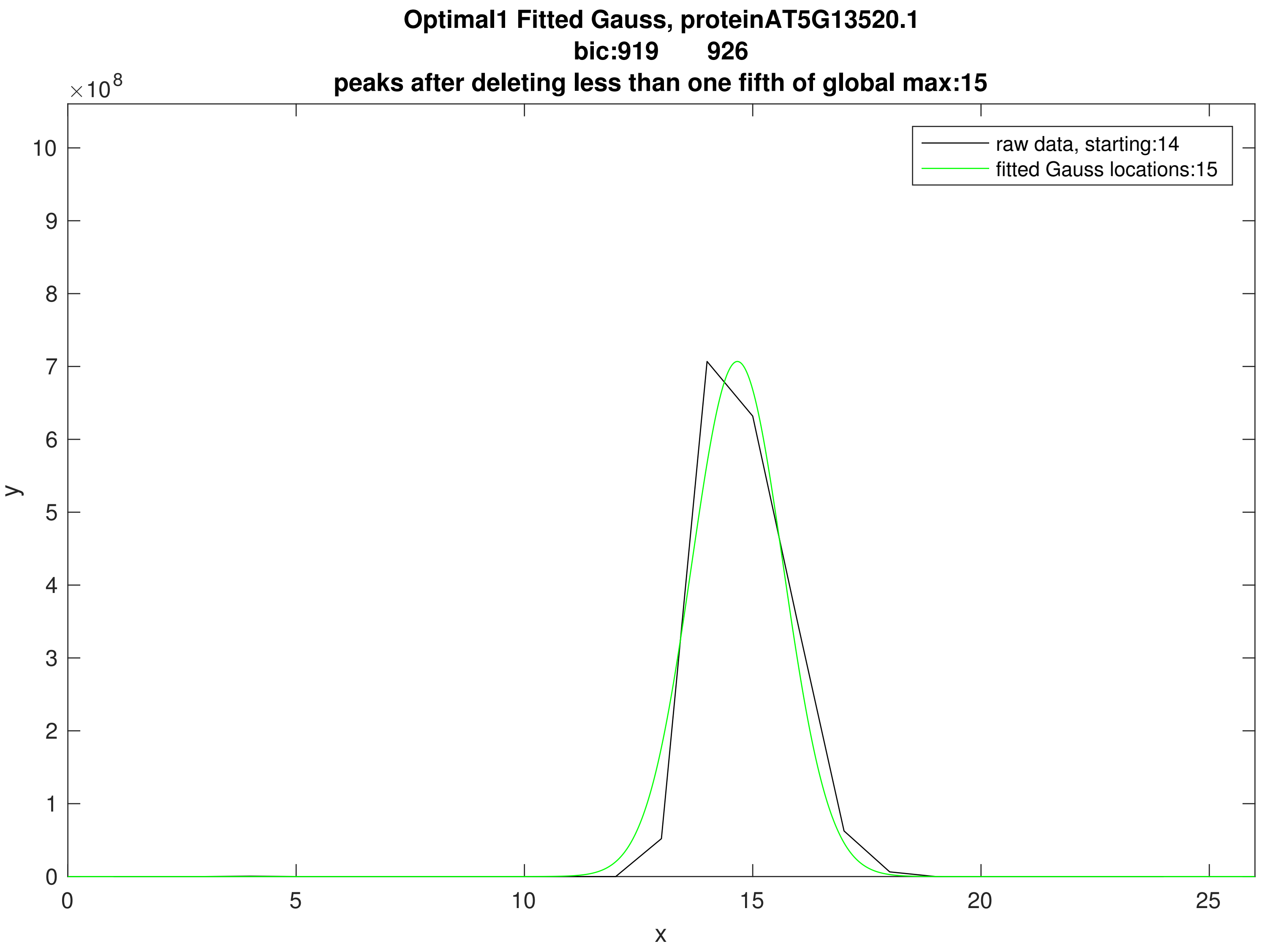
Optimal1 Fitted Gauss, proteinAT5G13520.1
bic:919 926
peaks after deleting less than one fifth of global max:15
×10^8
10
9
8
7
6
5
4
3
2
1
0
y
0
5
10
15
20
25
x
raw data, starting:14
fitted Gauss locations:15

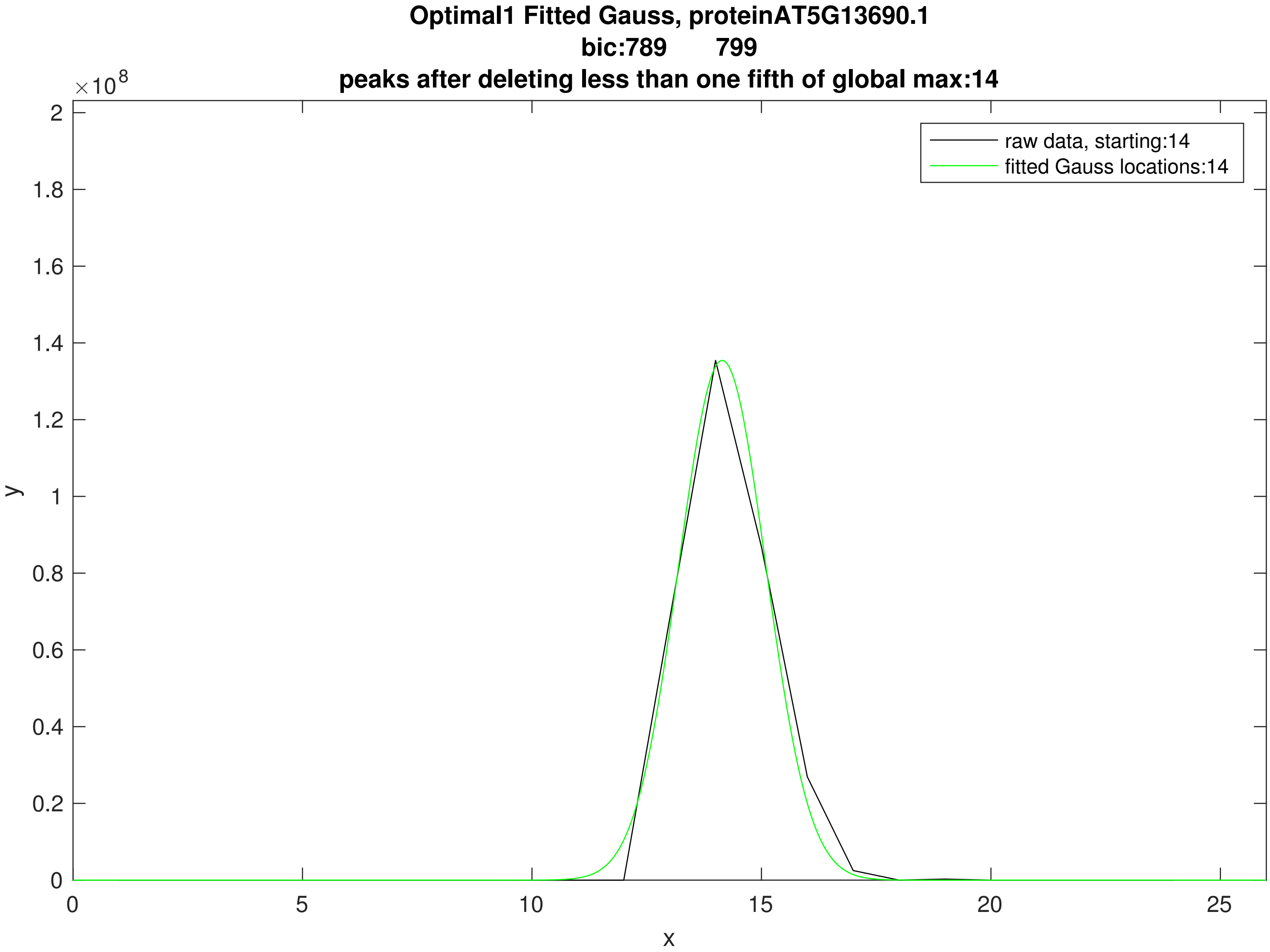
Optimal1 Fitted Gauss, proteinAT5G13690.1
bic:789 799
peaks after deleting less than one fifth of global max:14
×10^8
raw data, starting:14
fitted Gauss locations:14
y
x
0
0.2
0.4
0.6
0.8
1
1.2
1.4
1.6
1.8
2
0
5
10
15
20
25

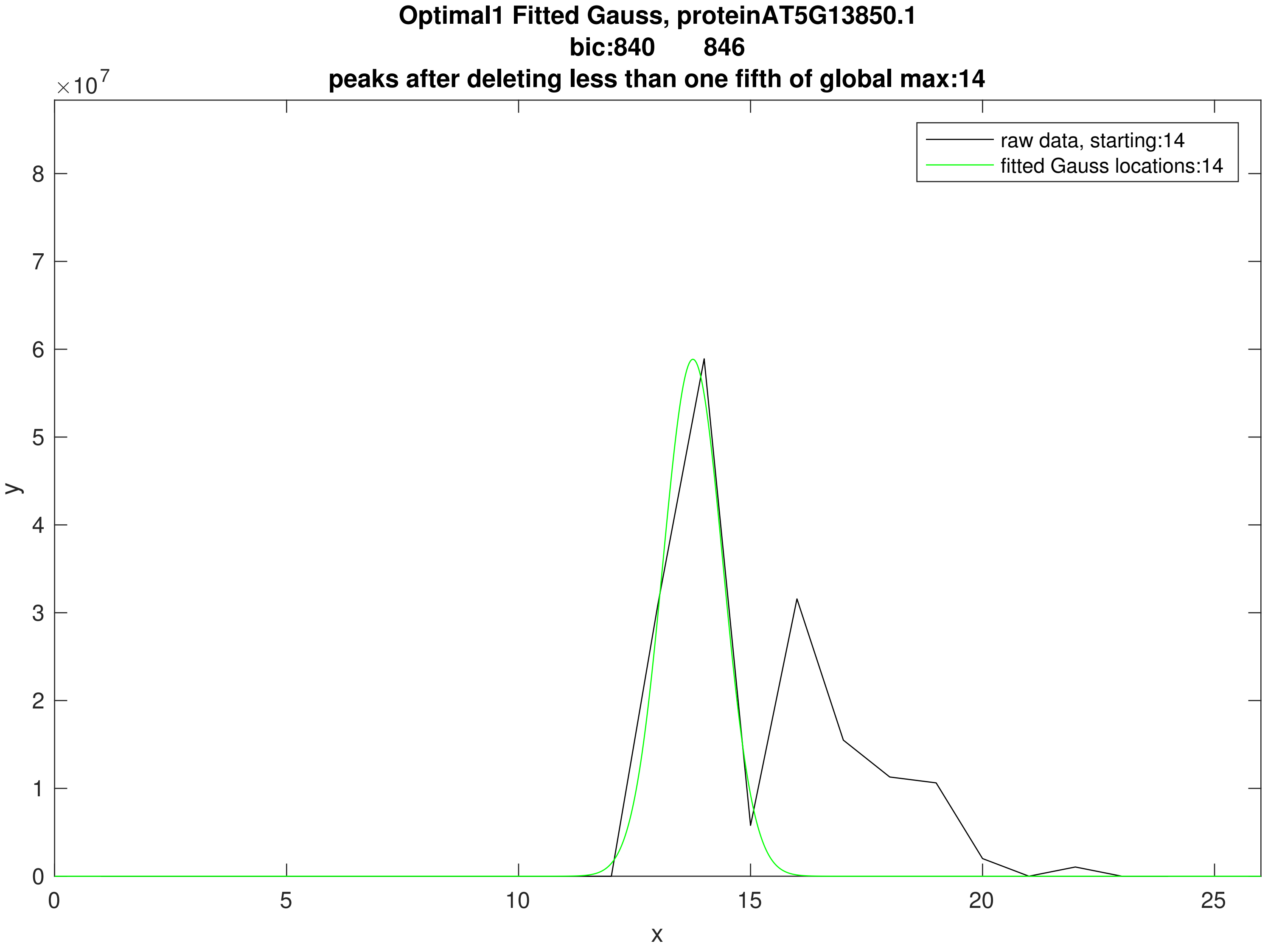

Optimal1 Fitted Gauss, proteinAT5G13850.1
bic:840 846
peaks after deleting less than one fifth of global max:14
×10^7
raw data, starting:14
fitted Gauss locations:14
y
x
0
1
2
3
4
5
6
7
8
0
5
10
15
20
25

Optimal1 Fitted Gauss, proteinAT5G13980.1;AT5G13980.2;AT5G13980.3
bic:893
peaks after deleting less than one fifth of global max:9.1

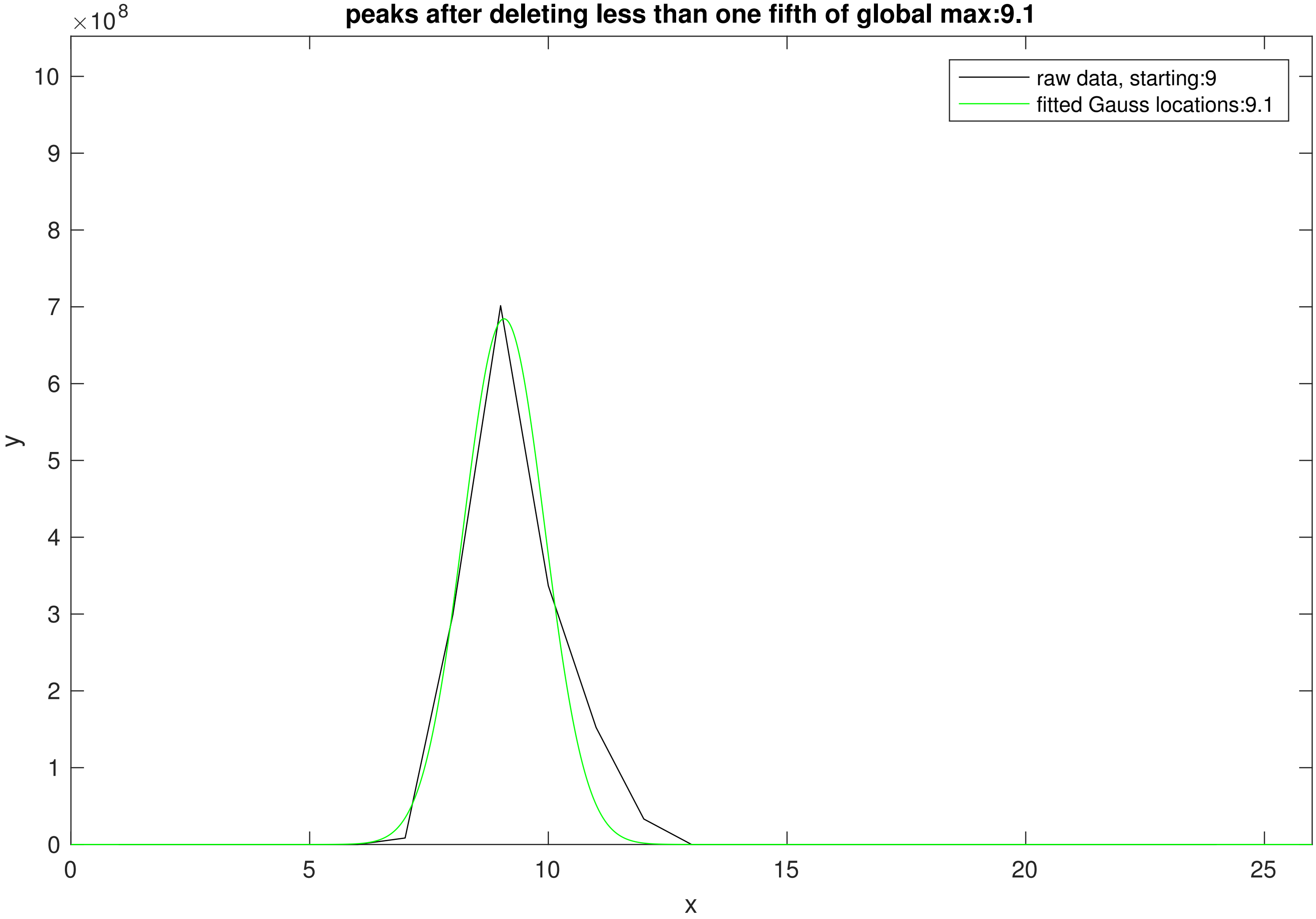

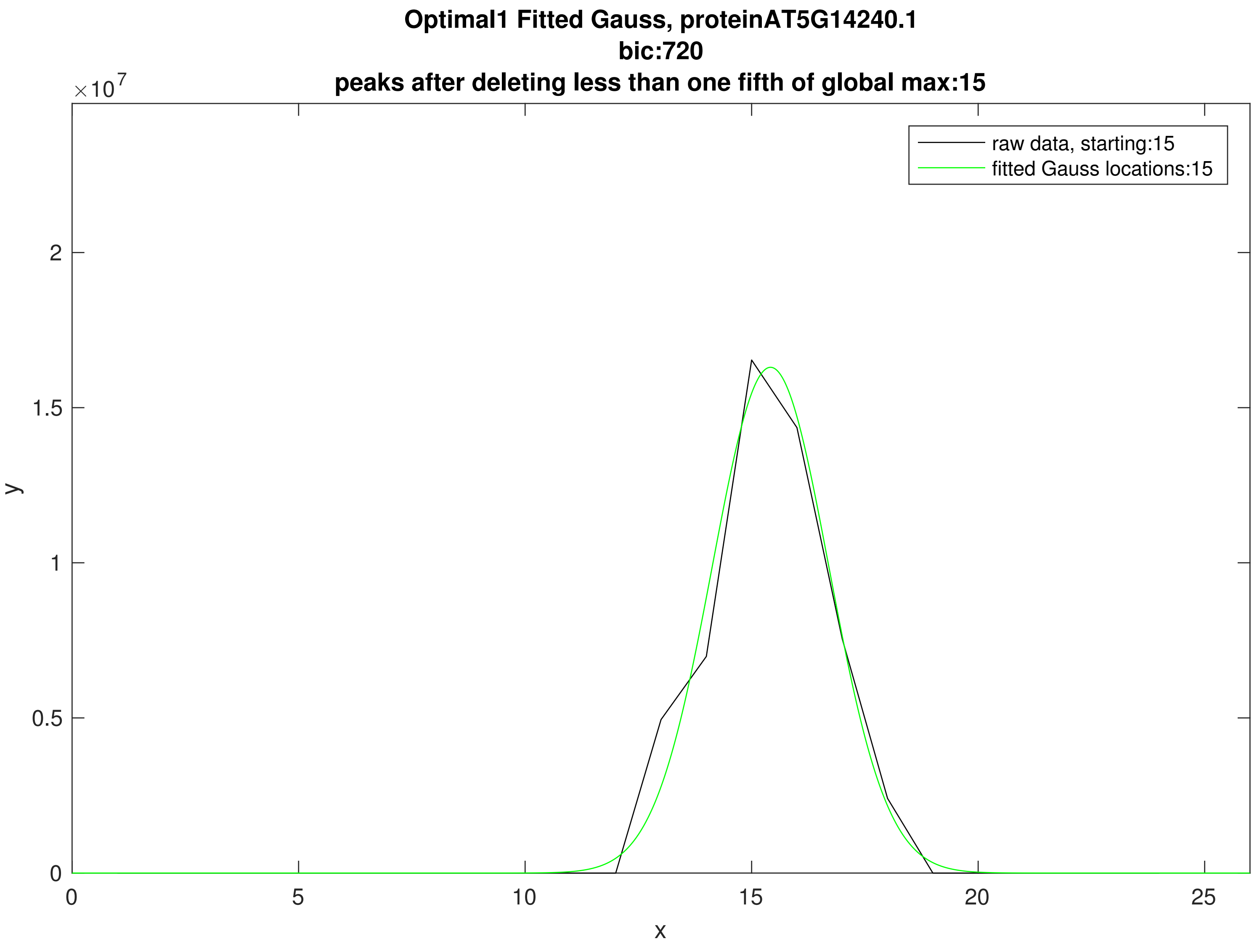

Optimal1 Fitted Gauss, proteinAT5G14240.1
bic:720
peaks after deleting less than one fifth of global max:15
×10^7
raw data, starting:15
fitted Gauss locations:15
y
x
0
0.5
1
1.5
2
0
5
10
15
20
25

Optimal1 Fitted Gauss, proteinAT5G14320.2;AT5G14320.1
bic:676
peaks after deleting less than one fifth of global max:5.2

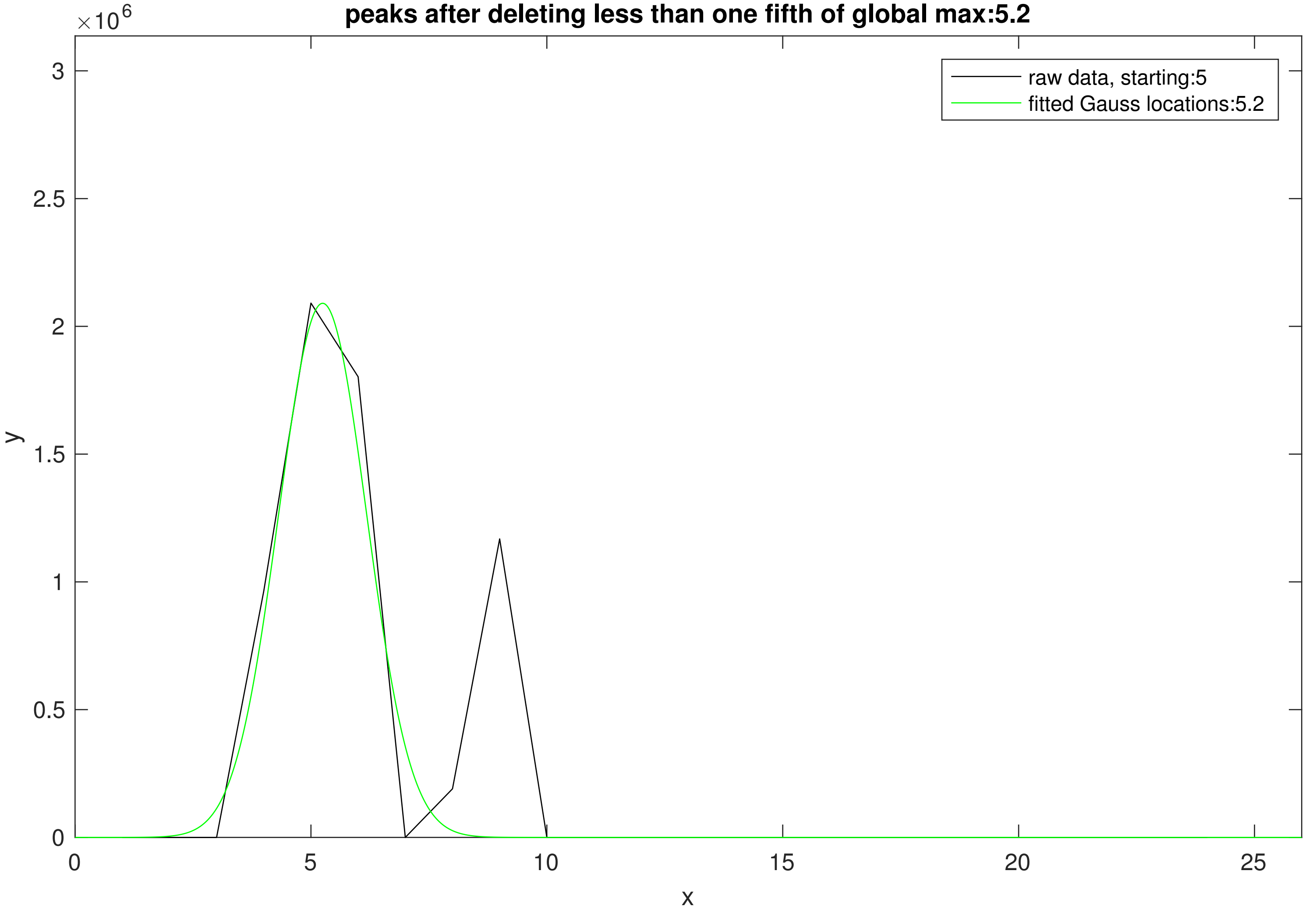

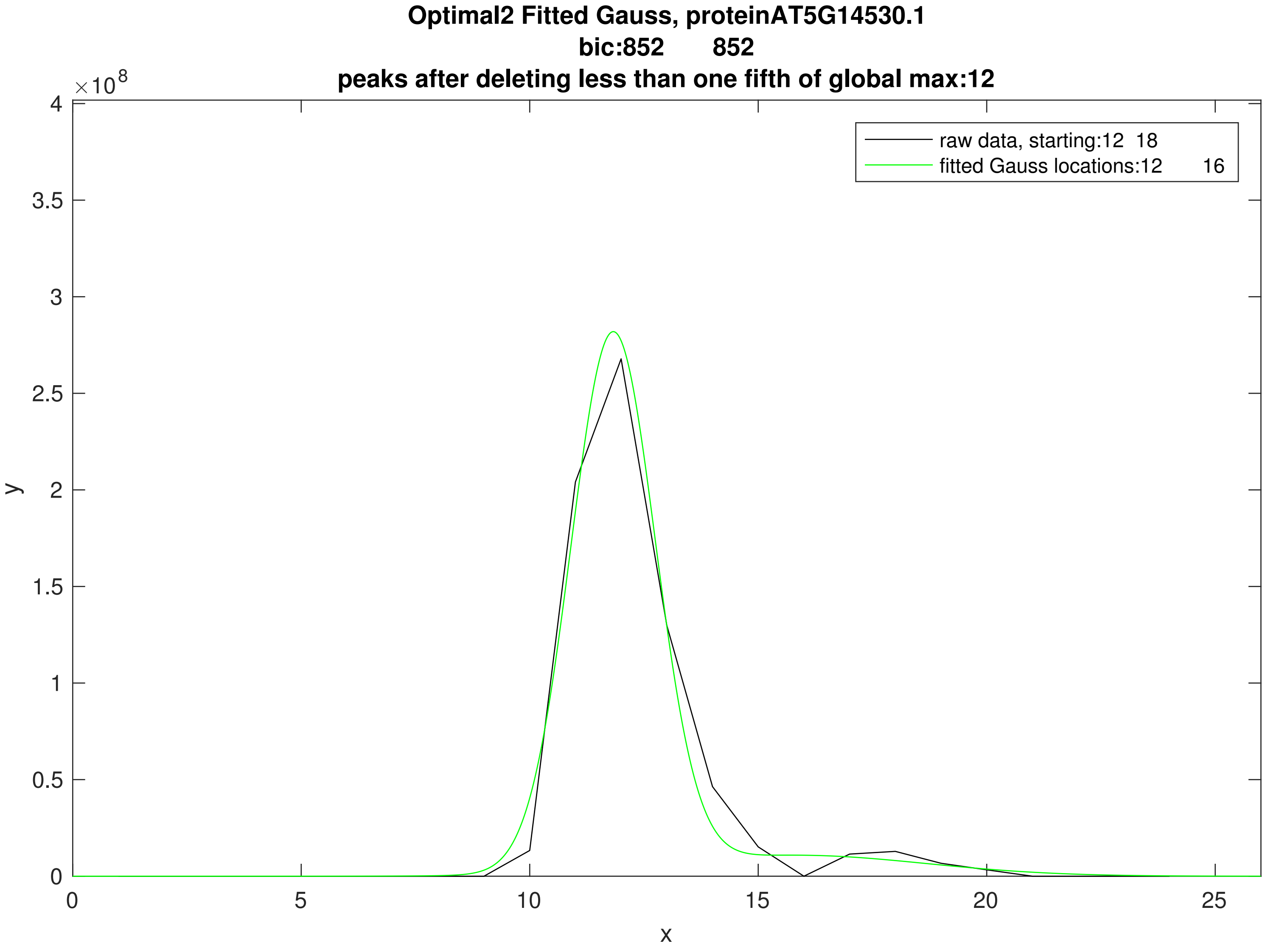

Optimal2 Fitted Gauss, proteinAT5G14530.1
bic:852 852
peaks after deleting less than one fifth of global max:12
×10^8
4
3.5
3
2.5
2
1.5
1
0.5
0
y
0
5
10
15
20
25
x
raw data, starting:12 18
fitted Gauss locations:12 16

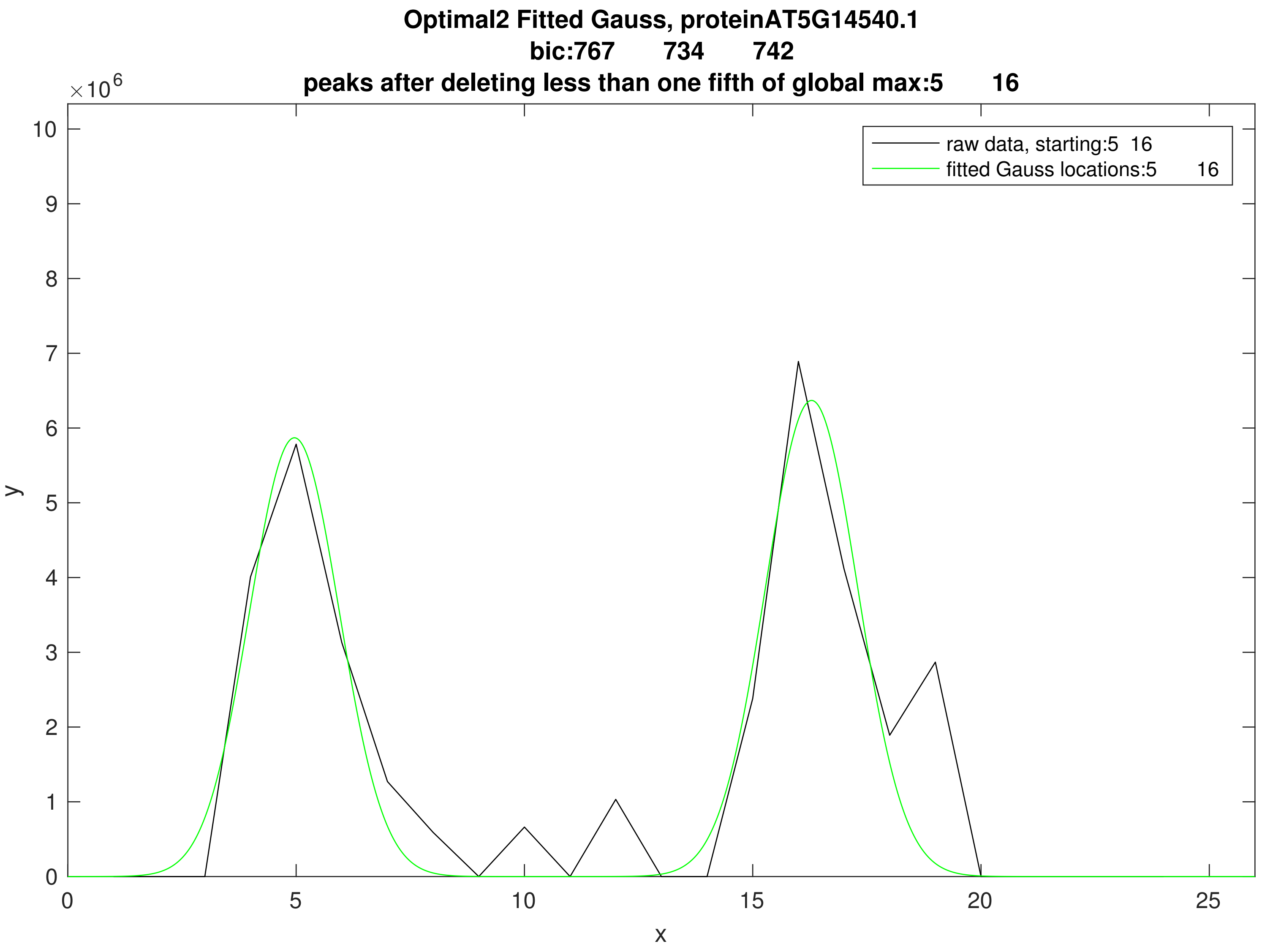
Optimal2 Fitted Gauss, proteinAT5G14540.1
bic:767 734 742
peaks after deleting less than one fifth of global max:5 16
×10^6
raw data, starting:5 16
fitted Gauss locations:5 16
y
x
0
1
2
3
4
5
6
7
8
9
10
0
5
10
15
20
25

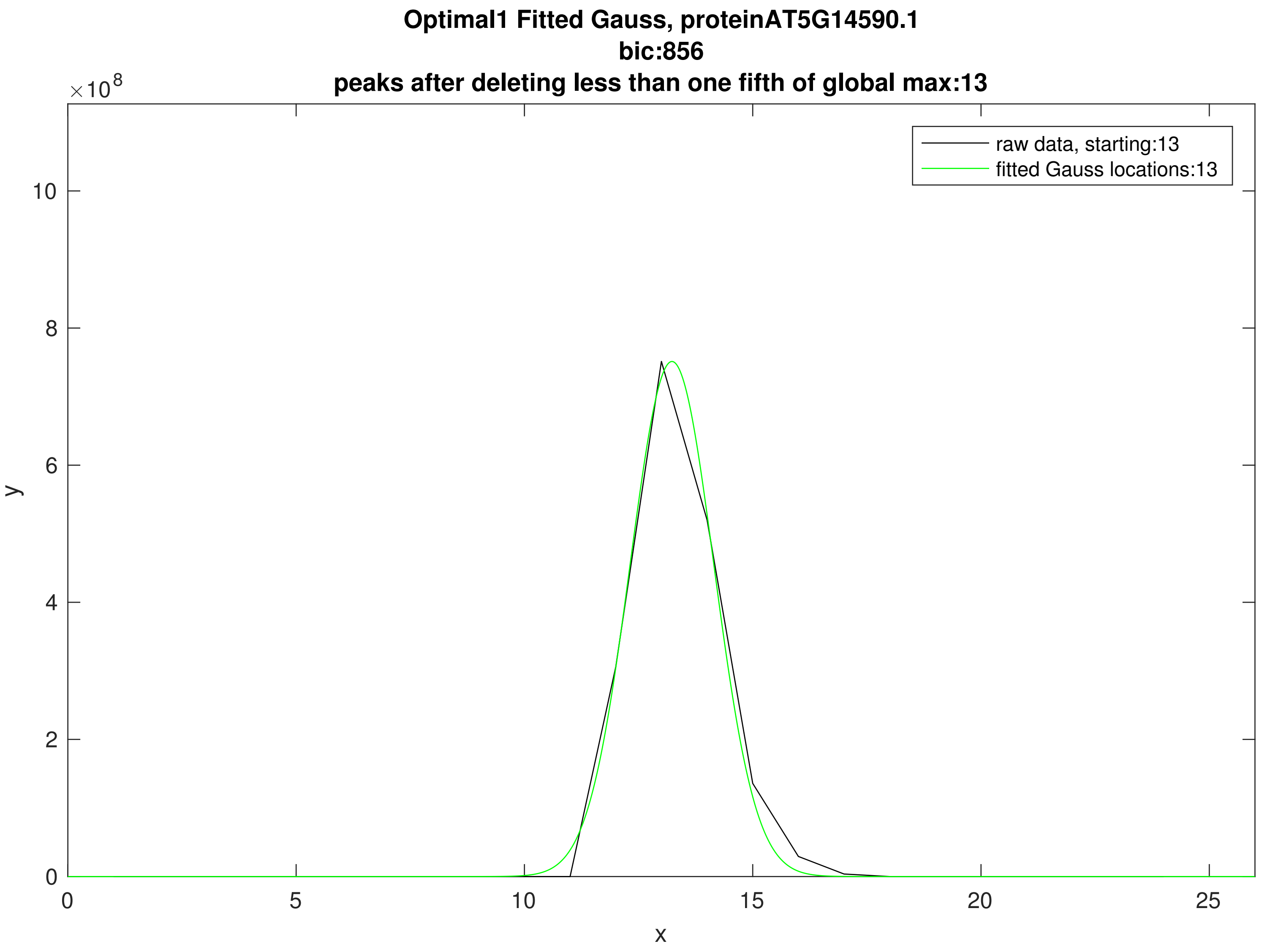
Optimal1 Fitted Gauss, proteinAT5G14590.1
bic:856
peaks after deleting less than one fifth of global max:13
×10^8
raw data, starting:13
fitted Gauss locations:13
y
x
0
2
4
6
8
10
0
5
10
15
20
25

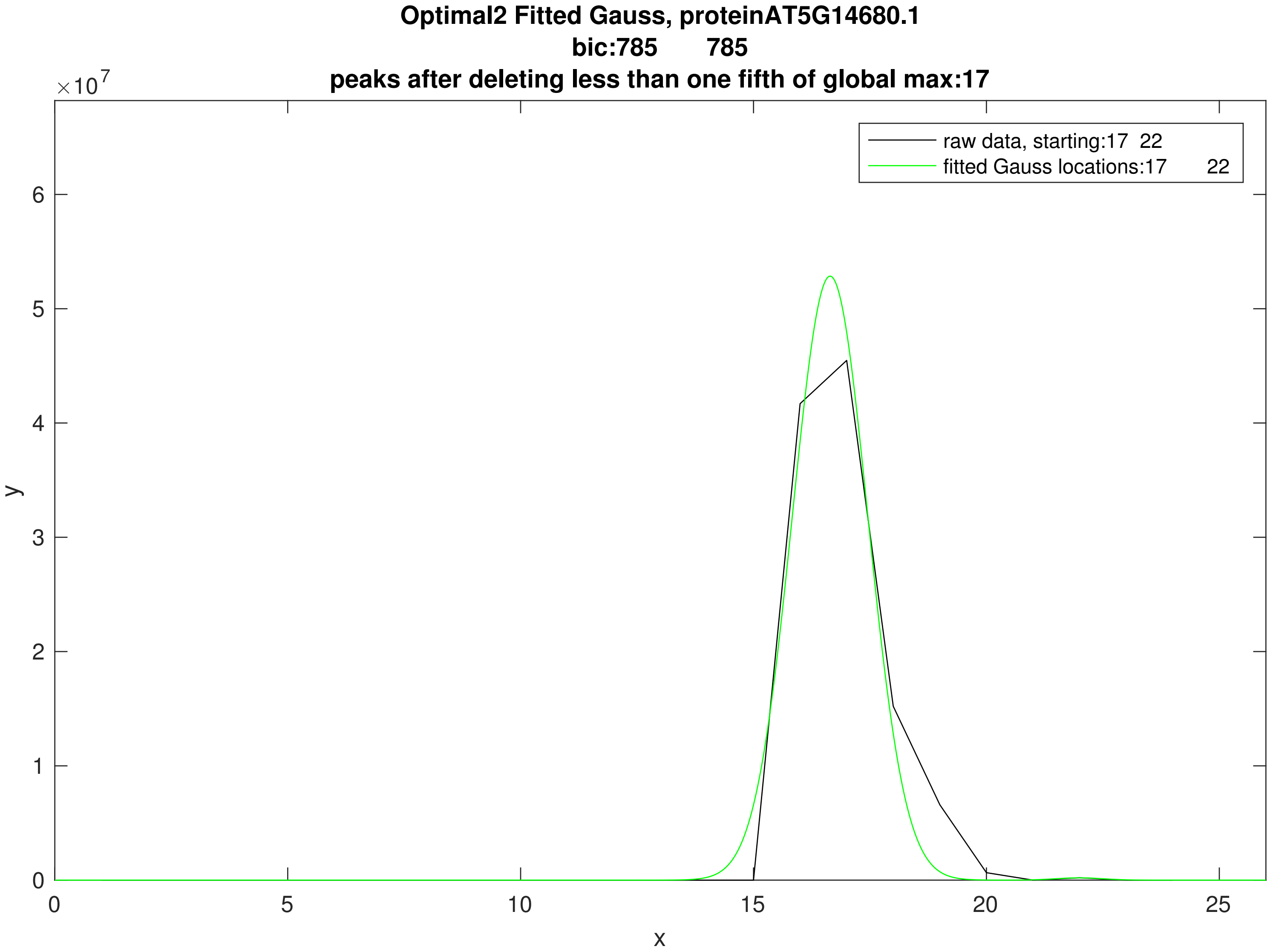

Optimal2 Fitted Gauss, proteinAT5G14680.1
bic:785 785
peaks after deleting less than one fifth of global max:17
×10^7
raw data, starting:17 22
fitted Gauss locations:17 22
y
x
0
1
2
3
4
5
6
0
5
10
15
20
25

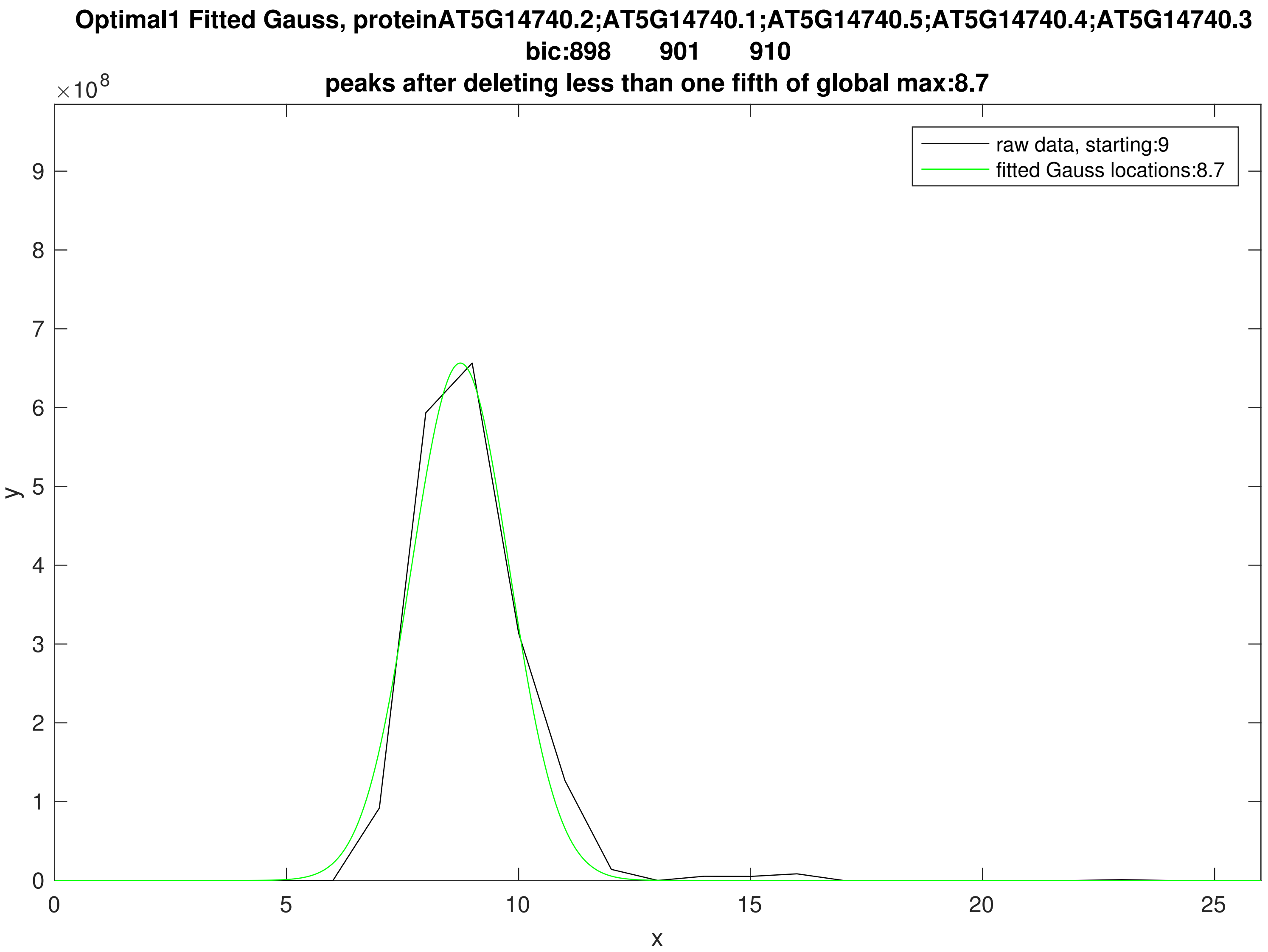

Optimal1 Fitted Gauss, proteinAT5G14740.2;AT5G14740.1;AT5G14740.5;AT5G14740.4;AT5G14740.3
bic:898 901 910
peaks after deleting less than one fifth of global max:8.7
×10^8
raw data, starting:9
fitted Gauss locations:8.7
y
x
0
1
2
3
4
5
6
7
8
9
0
5
10
15
20
25

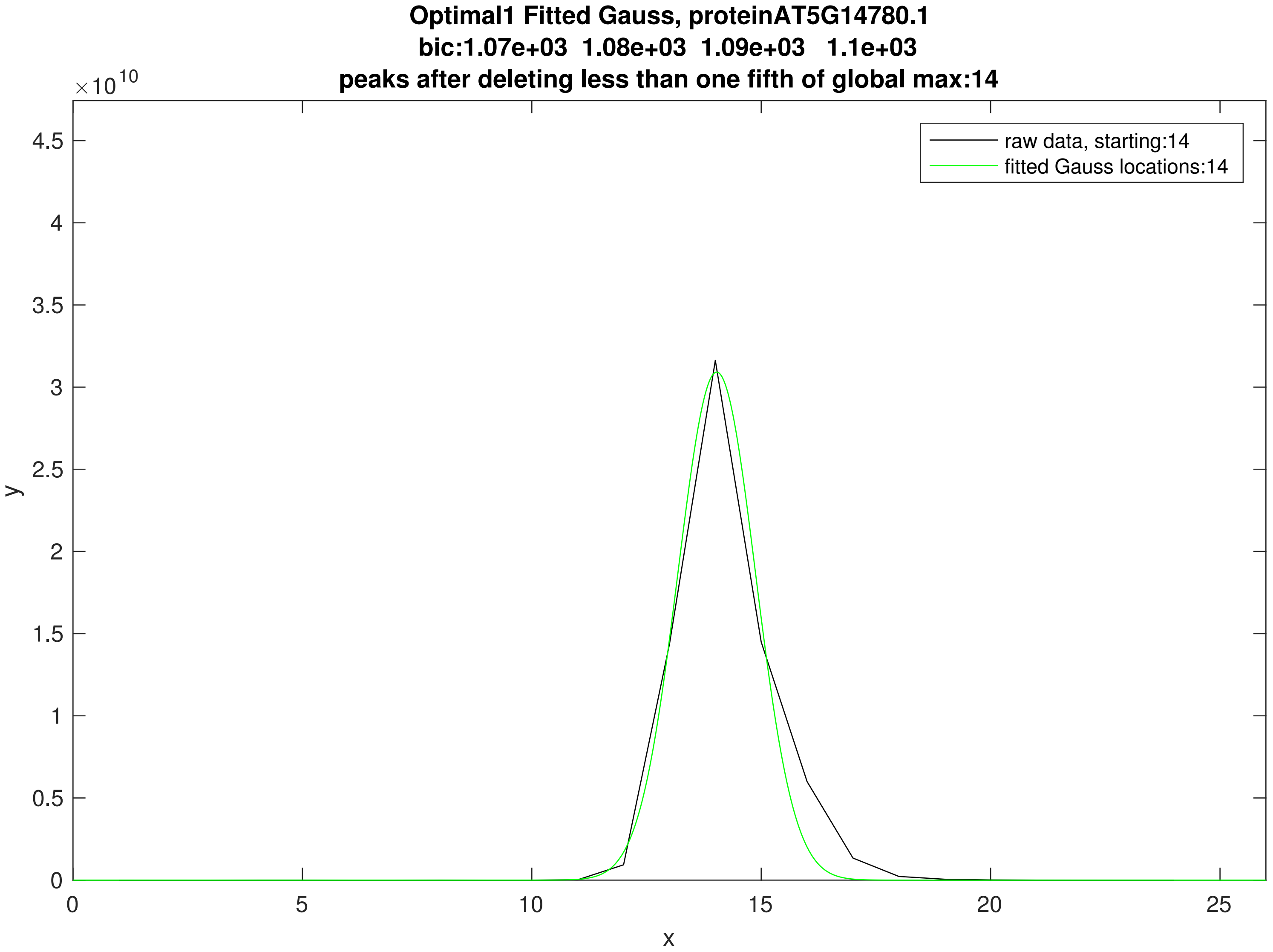

Optimal1 Fitted Gauss, proteinAT5G14780.1
bic:1.07e+03  1.08e+03  1.09e+03   1.1e+03
peaks after deleting less than one fifth of global max:14
×10^10
4.5
4
3.5
3
2.5
2
1.5
1
0.5
0
y
0
5
10
15
20
25
x
raw data, starting:14
fitted Gauss locations:14

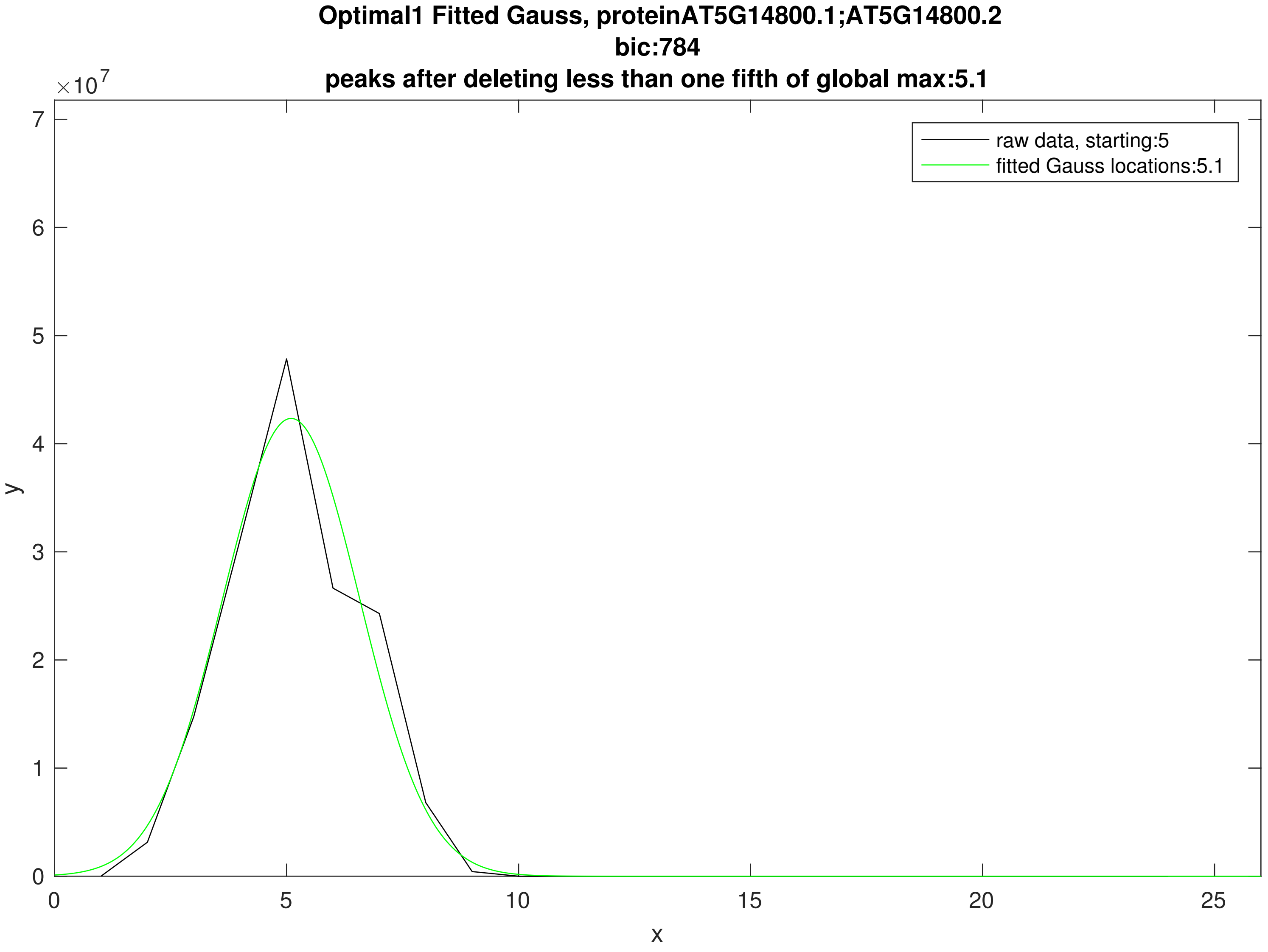

Optimal1 Fitted Gauss, proteinAT5G14800.1;AT5G14800.2
bic:784
peaks after deleting less than one fifth of global max:5.1
×10^7
raw data, starting:5
fitted Gauss locations:5.1
y
x
0
1
2
3
4
5
6
7
0
5
10
15
20
25

**Optimal2 Fitted Gauss, proteinAT5G14910.1**
**bic:740       737**
**peaks after deleting less than one fifth of global max:16**

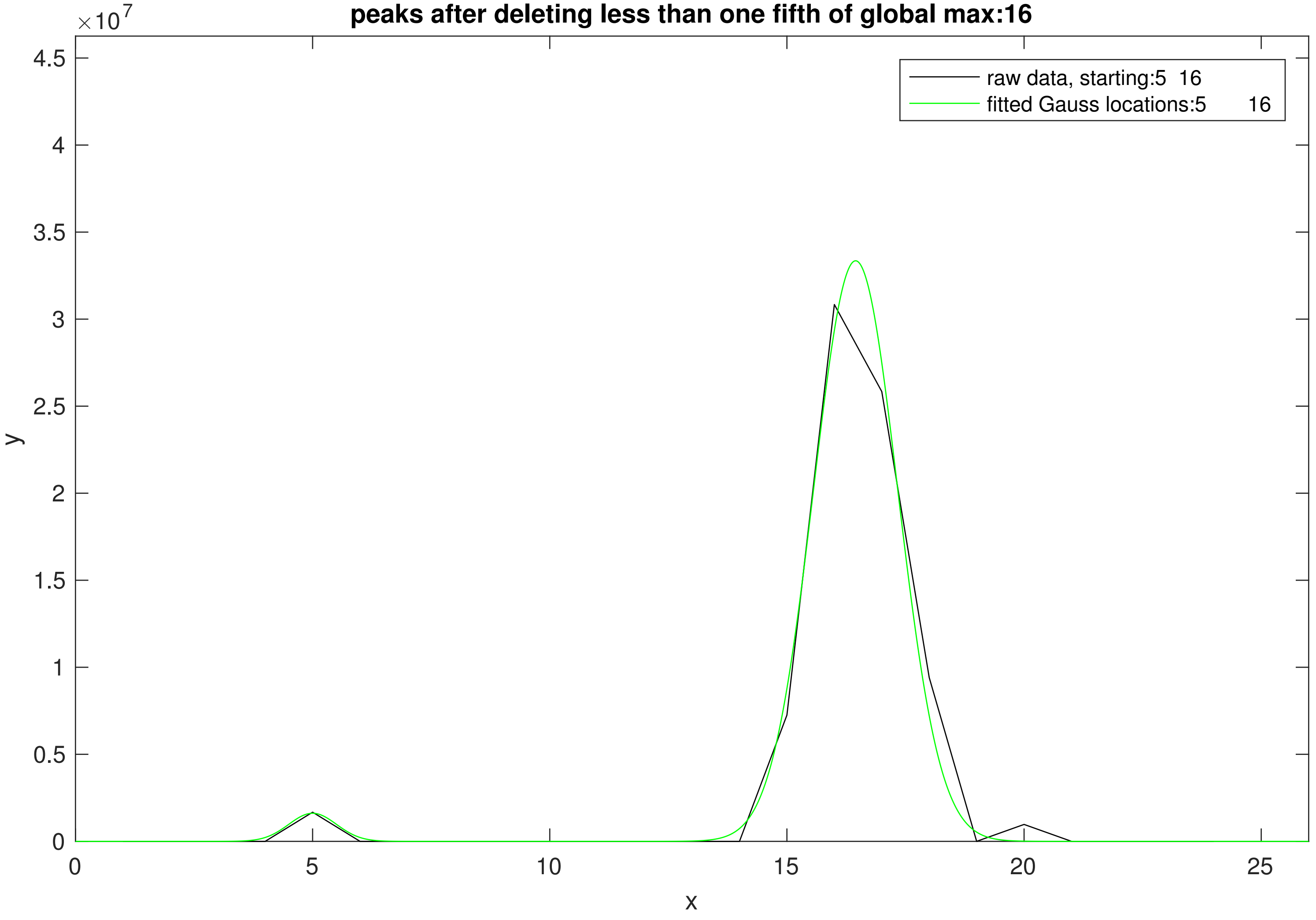

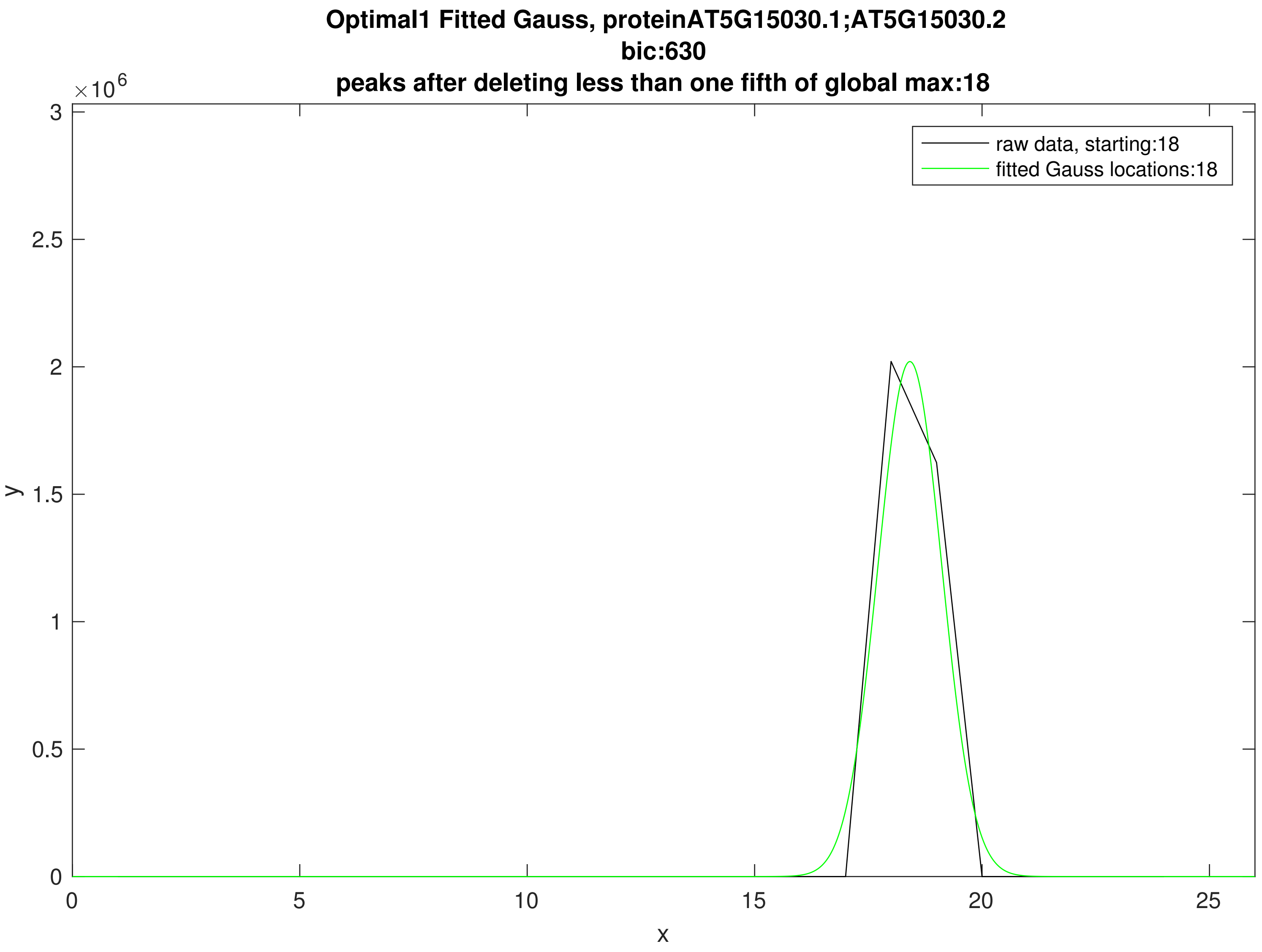

Optimal1 Fitted Gauss, proteinAT5G15030.1;AT5G15030.2
bic:630
peaks after deleting less than one fifth of global max:18
×10^6
3
2.5
2
1.5
1
0.5
0
y
0
5
10
15
20
25
x
raw data, starting:18
fitted Gauss locations:18

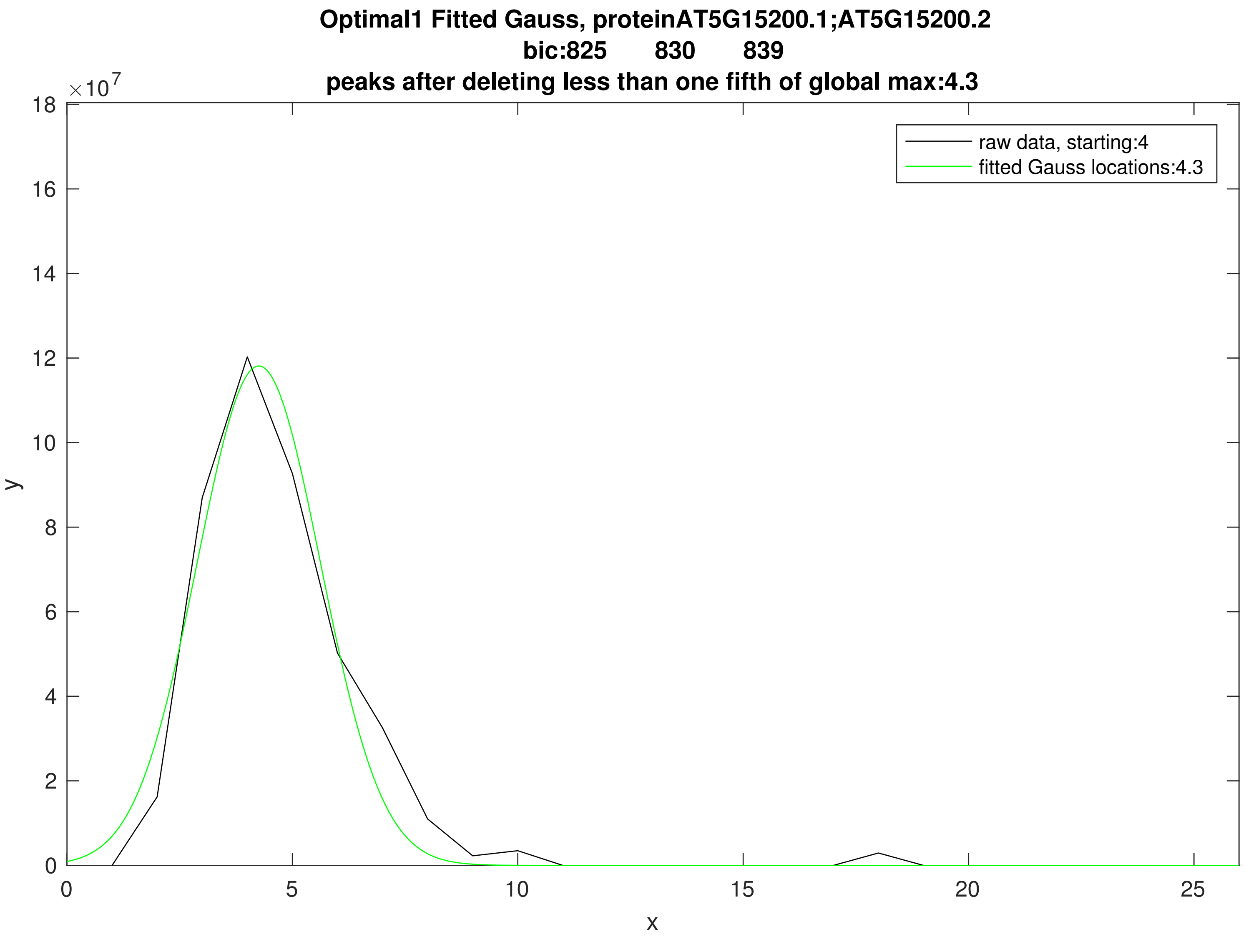

Optimal1 Fitted Gauss, proteinAT5G15200.1;AT5G15200.2
bic:825 830 839
peaks after deleting less than one fifth of global max:4.3
×10^7
raw data, starting:4
fitted Gauss locations:4.3
y
x
0
2
4
6
8
10
12
14
16
18
0
5
10
15
20
25

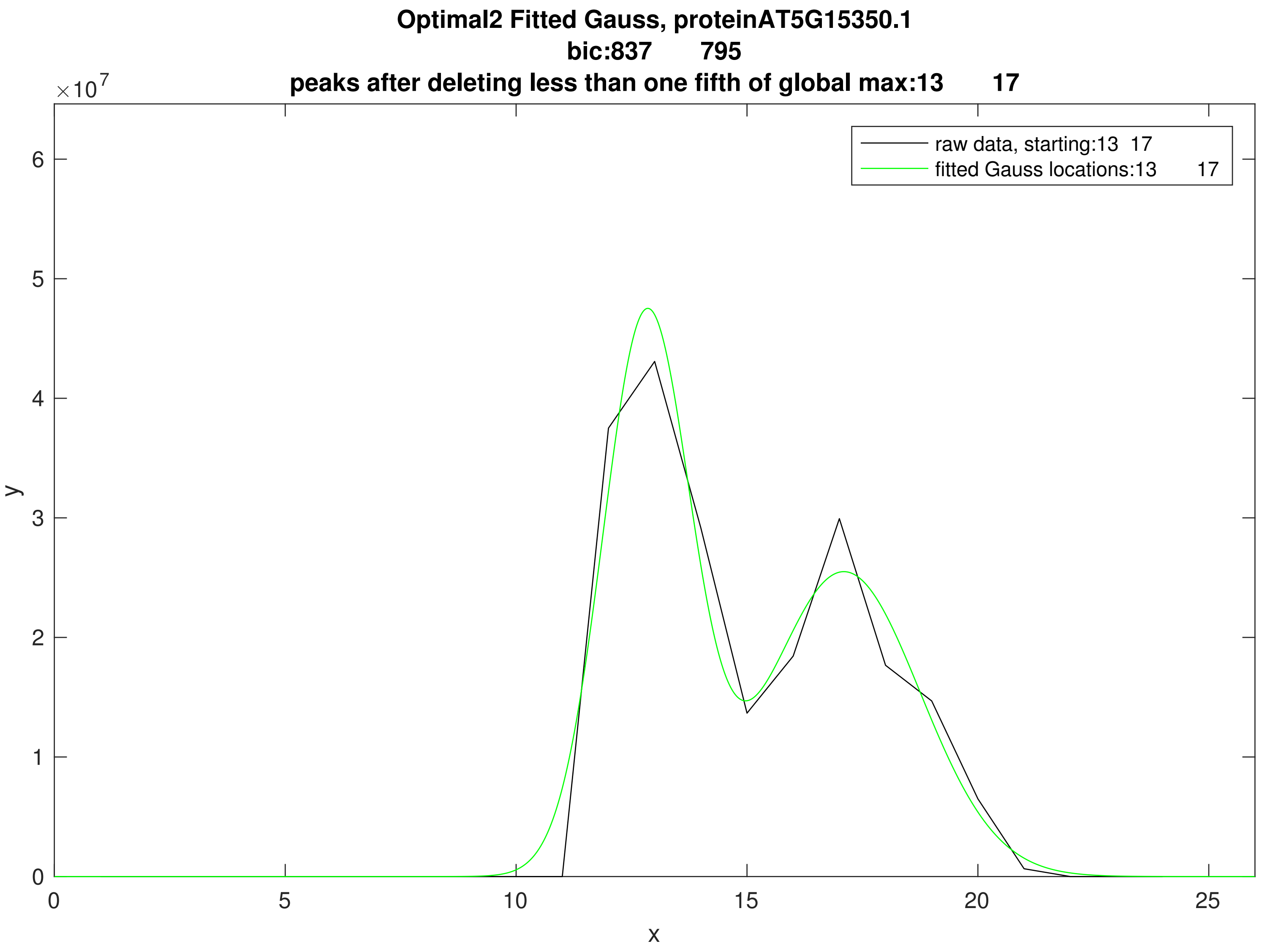
Optimal2 Fitted Gauss, proteinAT5G15350.1
bic:837 795
peaks after deleting less than one fifth of global max:13 17
×10^7
raw data, starting:13 17
fitted Gauss locations:13 17
y
x
0
1
2
3
4
5
6
0
5
10
15
20
25

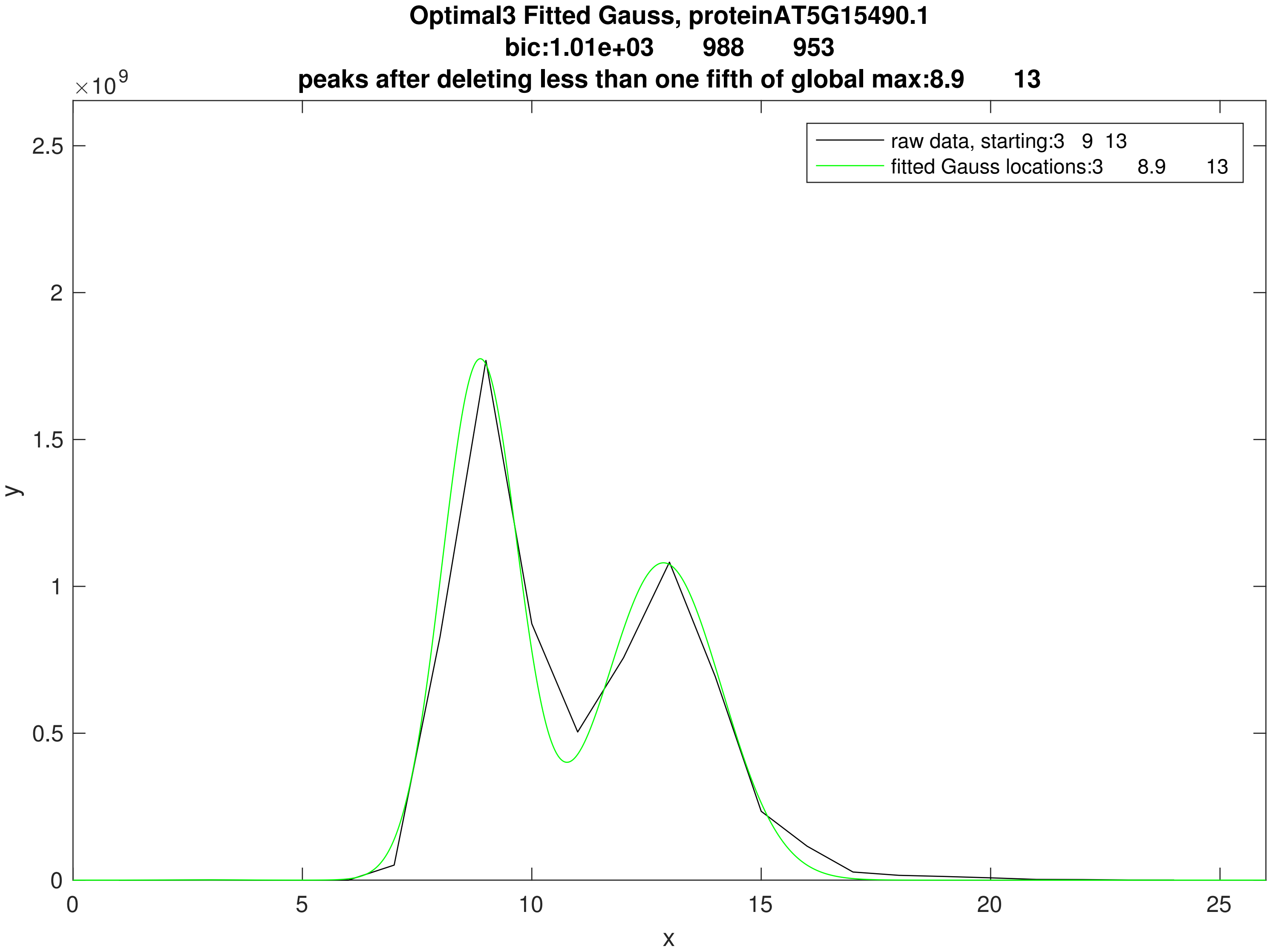
Optimal3 Fitted Gauss, proteinAT5G15490.1
bic:1.01e+03 988 953
peaks after deleting less than one fifth of global max:8.9 13
×10^9
raw data, starting:3 9 13
fitted Gauss locations:3 8.9 13
y
x
0
0.5
1
1.5
2
2.5
5
10
15
20
25

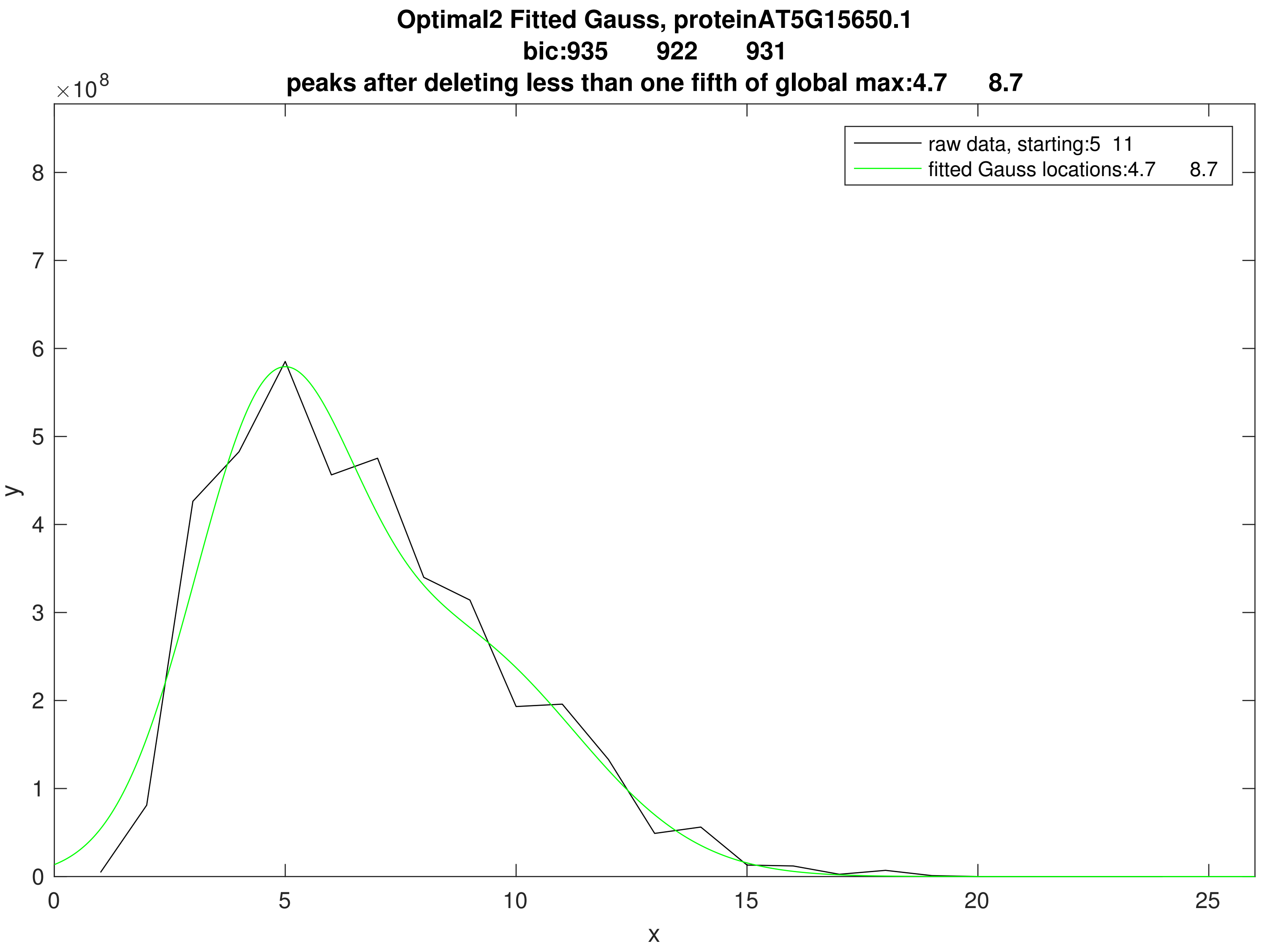

Optimal2 Fitted Gauss, proteinAT5G15650.1
bic:935 922 931
peaks after deleting less than one fifth of global max:4.7 8.7
×10^8
raw data, starting:5 11
fitted Gauss locations:4.7 8.7
y
x

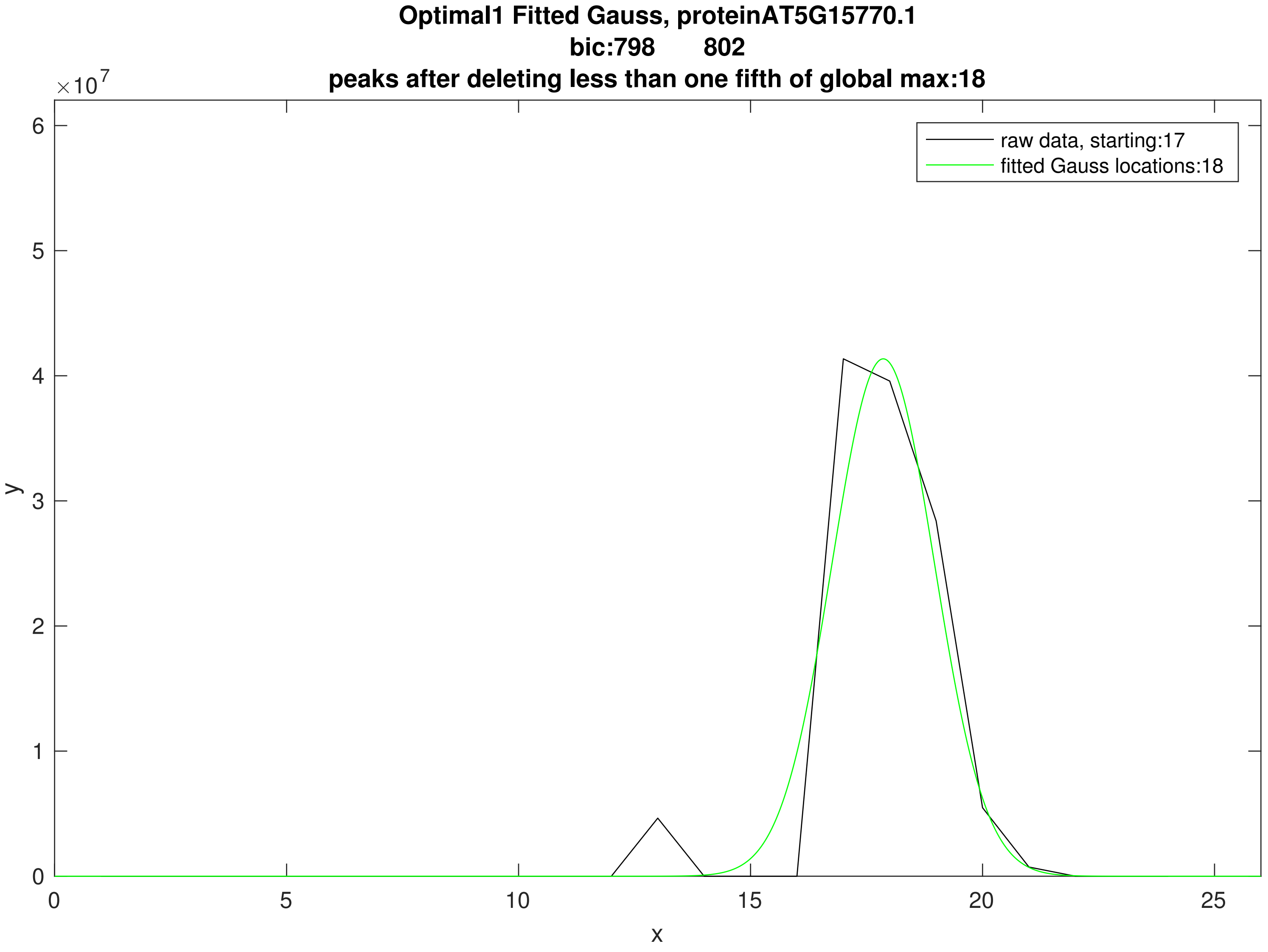
Optimal1 Fitted Gauss, proteinAT5G15770.1
bic:798 802
peaks after deleting less than one fifth of global max:18
×10^7
raw data, starting:17
fitted Gauss locations:18
y
x
0
1
2
3
4
5
6
0
5
10
15
20
25

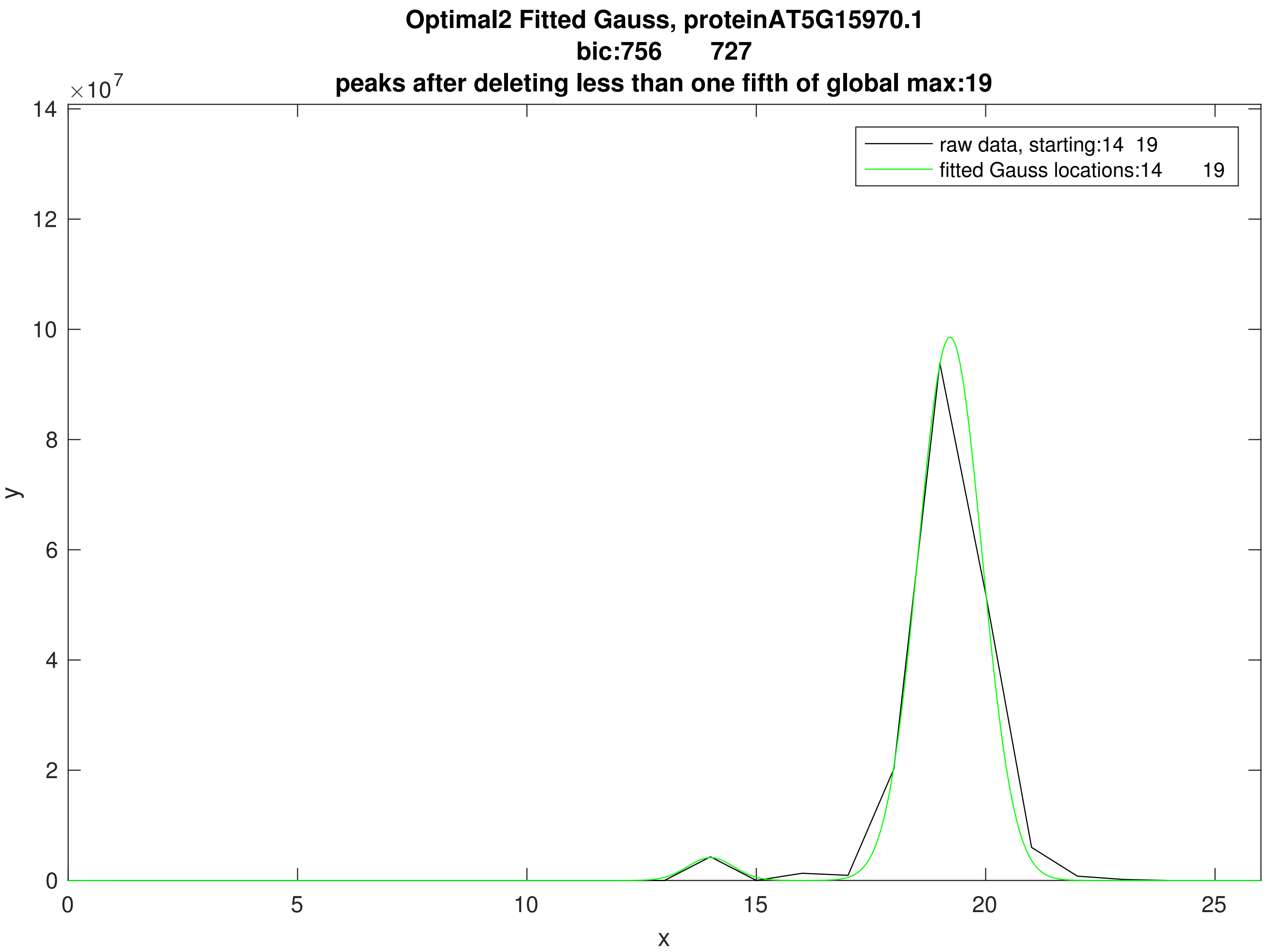
Optimal2 Fitted Gauss, proteinAT5G15970.1
bic:756 727
peaks after deleting less than one fifth of global max:19
×10^7
raw data, starting:14 19
fitted Gauss locations:14 19
y
x
0
2
4
6
8
10
12
14
5
15
20
25

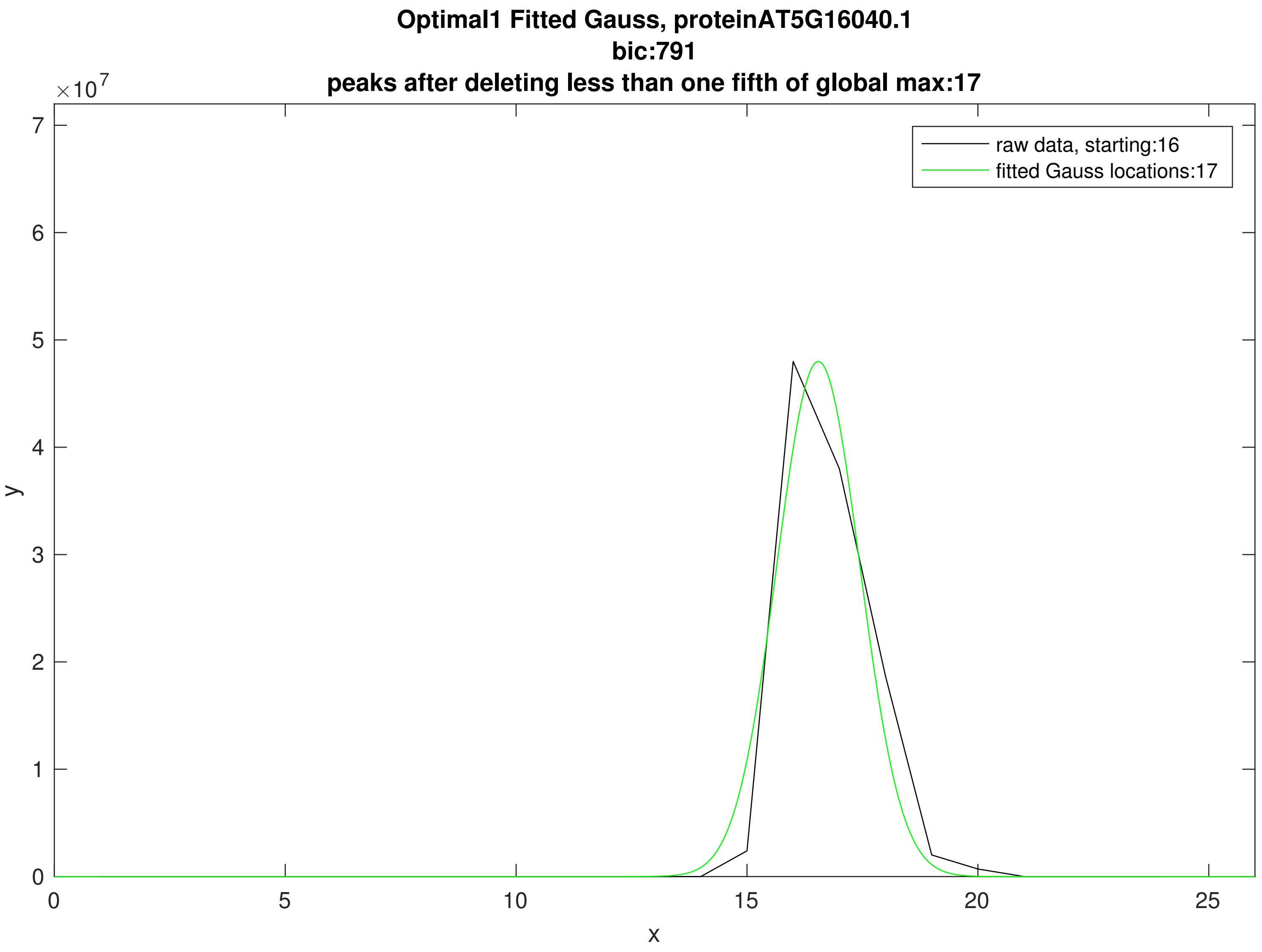
Optimal1 Fitted Gauss, proteinAT5G16040.1
bic:791
peaks after deleting less than one fifth of global max:17
×10^7
7
6
5
4
3
2
1
0
y
0
5
10
15
20
25
x
raw data, starting:16
fitted Gauss locations:17

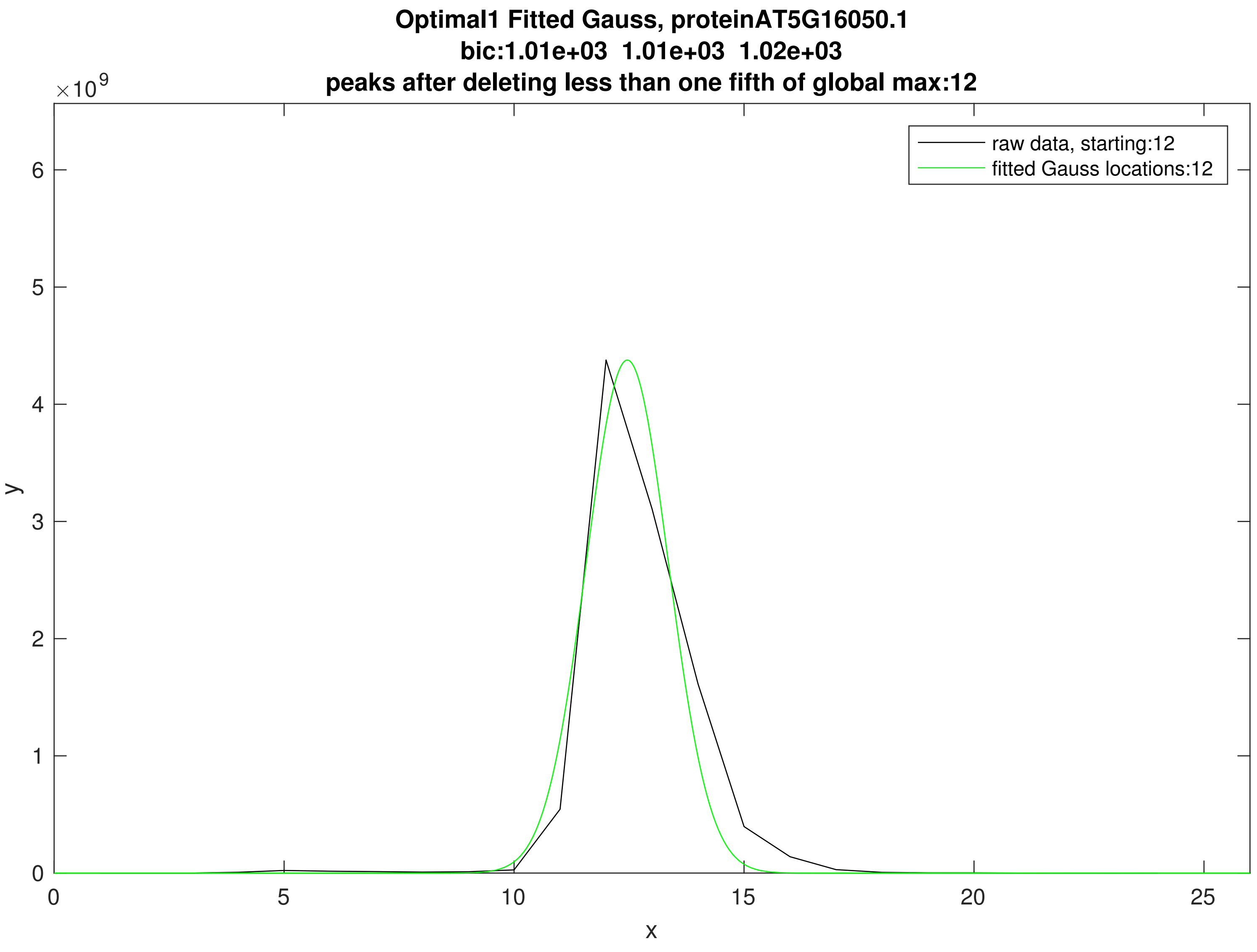

Optimal1 Fitted Gauss, proteinAT5G16050.1
bic:1.01e+03  1.01e+03  1.02e+03
peaks after deleting less than one fifth of global max:12
×10^9
raw data, starting:12
fitted Gauss locations:12
y
x
0
1
2
3
4
5
6
0
5
10
15
20
25

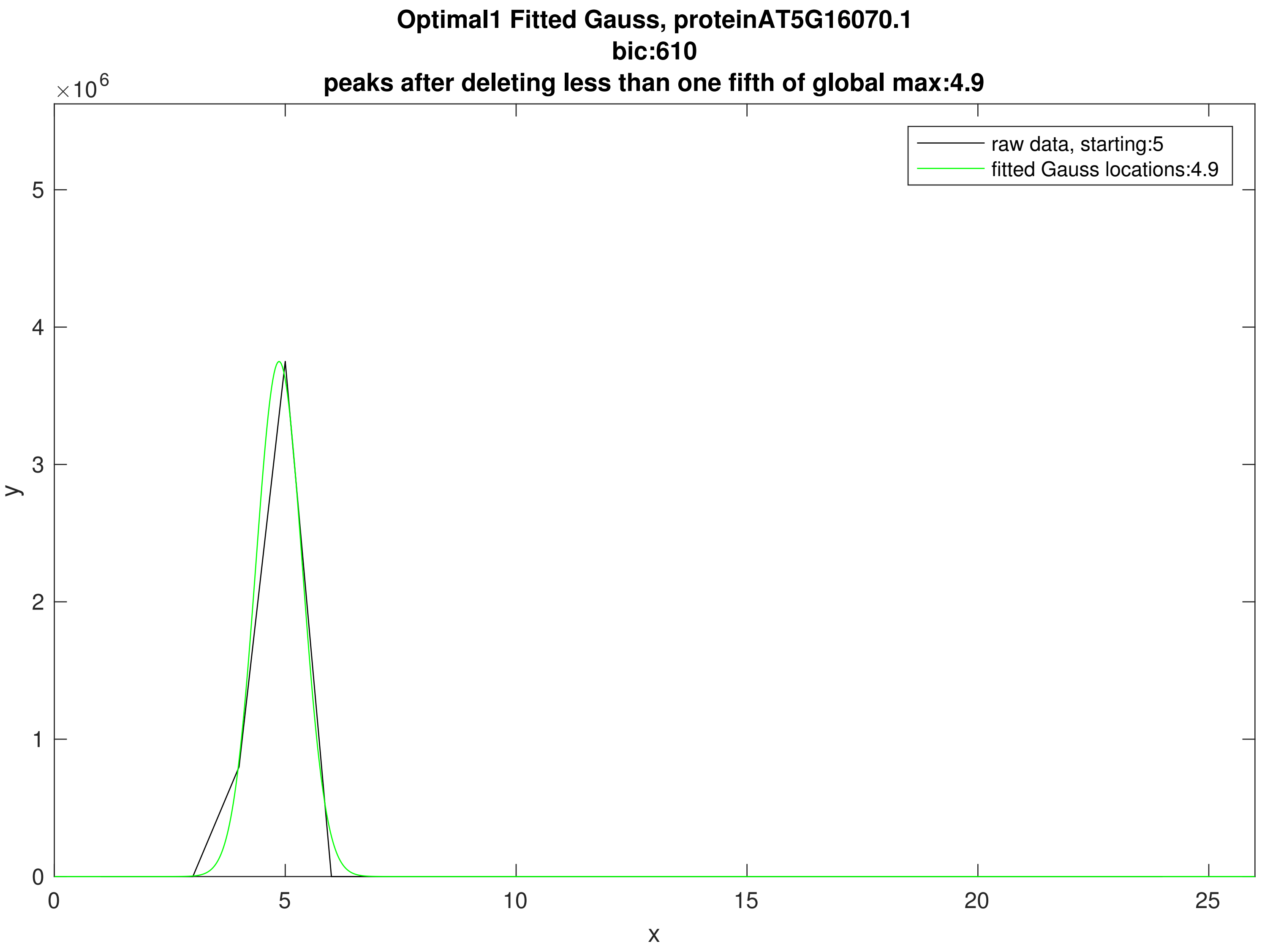
Optimal1 Fitted Gauss, proteinAT5G16070.1
bic:610
peaks after deleting less than one fifth of global max:4.9
×10^6
raw data, starting:5
fitted Gauss locations:4.9
y
x
0
1
2
3
4
5
0
5
10
15
20
25

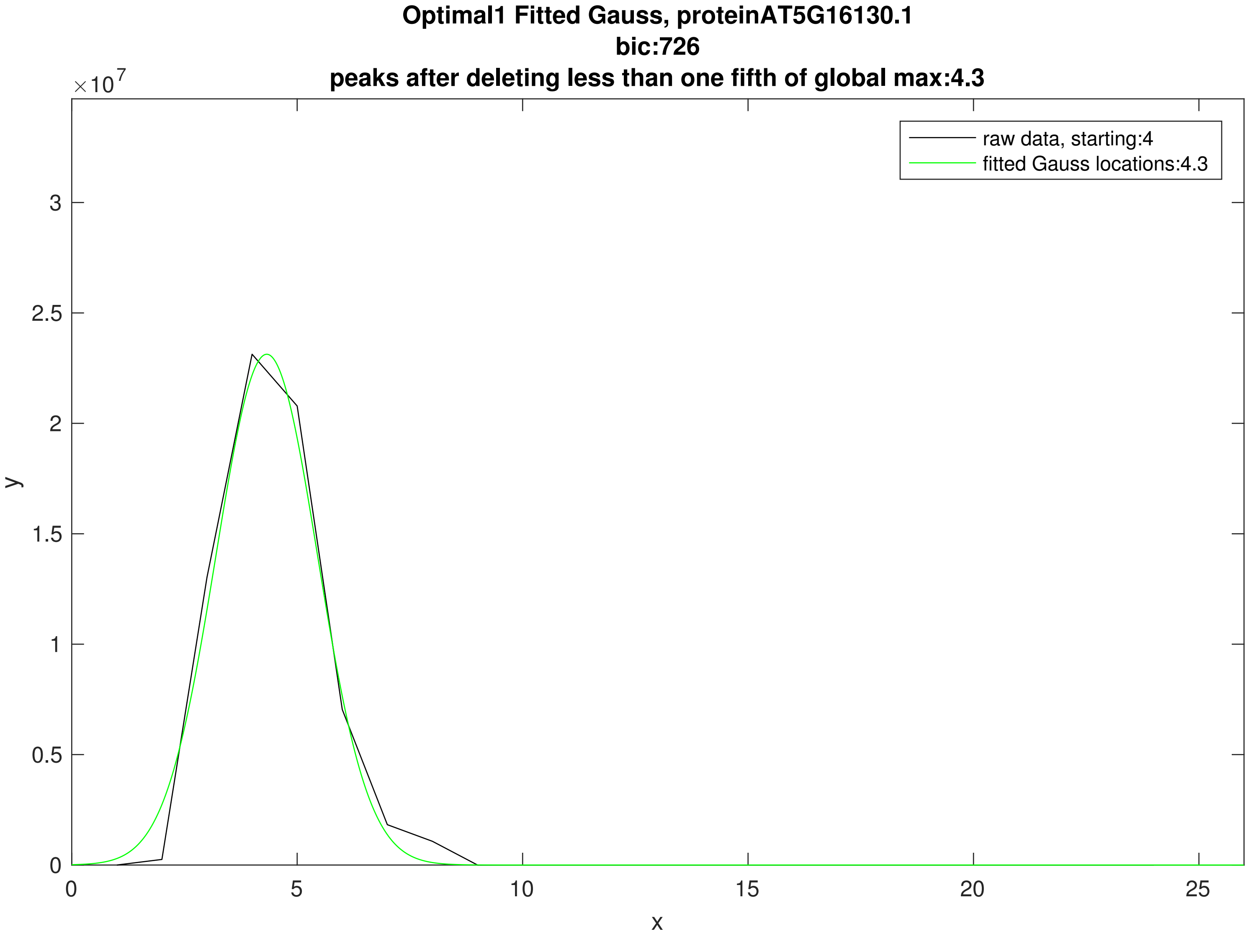
Optimal1 Fitted Gauss, proteinAT5G16130.1
bic:726
peaks after deleting less than one fifth of global max:4.3
×10^7
raw data, starting:4
fitted Gauss locations:4.3
y
x
0
0.5
1
1.5
2
2.5
3
5
10
15
20
25

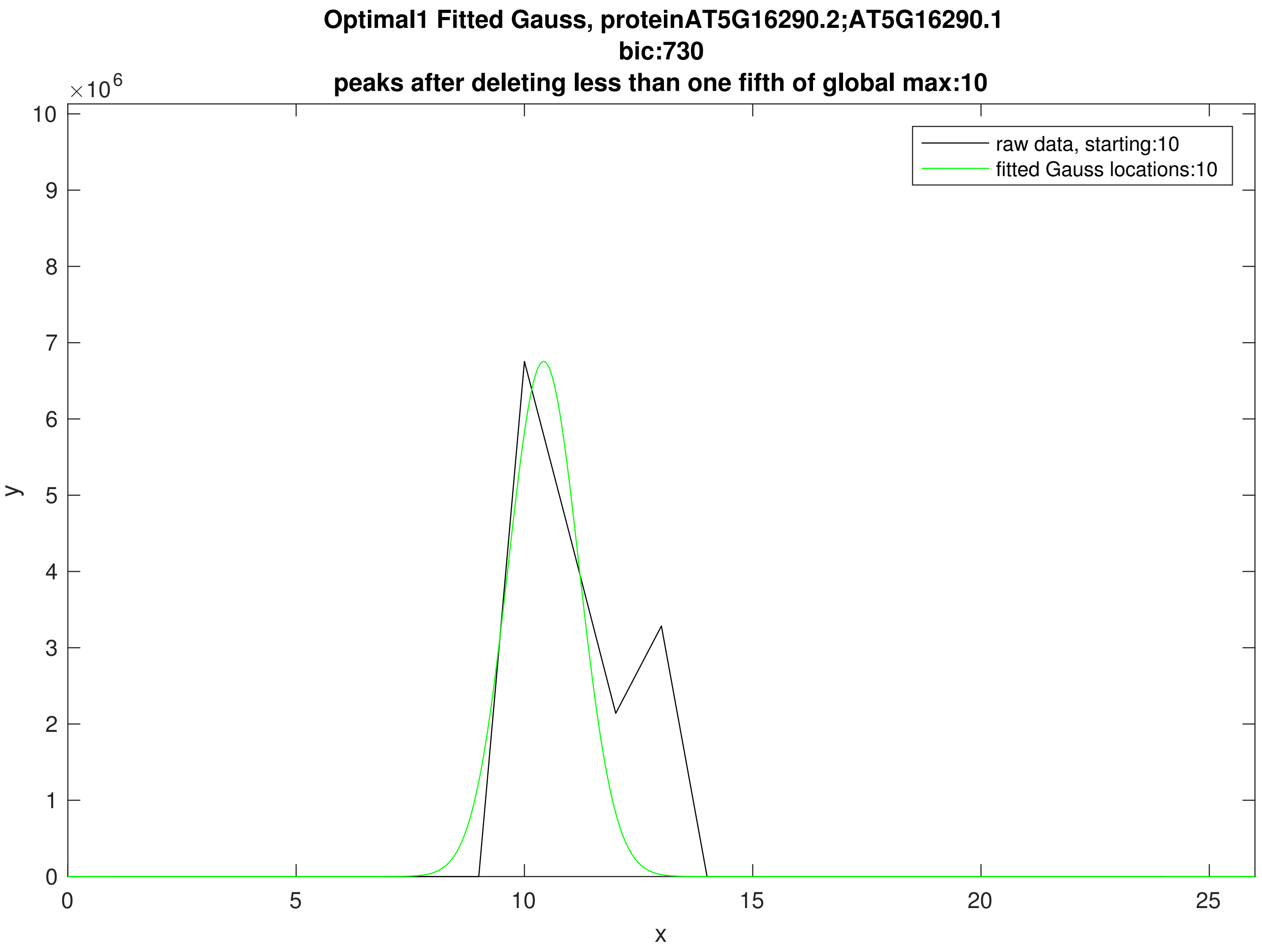
Optimal1 Fitted Gauss, proteinAT5G16290.2;AT5G16290.1
bic:730
peaks after deleting less than one fifth of global max:10
$\times 10^6$
10
9
8
7
6
5
4
3
2
1
0
y
0
5
10
15
20
25
x
raw data, starting:10
fitted Gauss locations:10

Optimal1 Fitted Gauss, proteinAT5G16340.1;AT5G16370.1
bic:677
peaks after deleting less than one fifth of global max:11

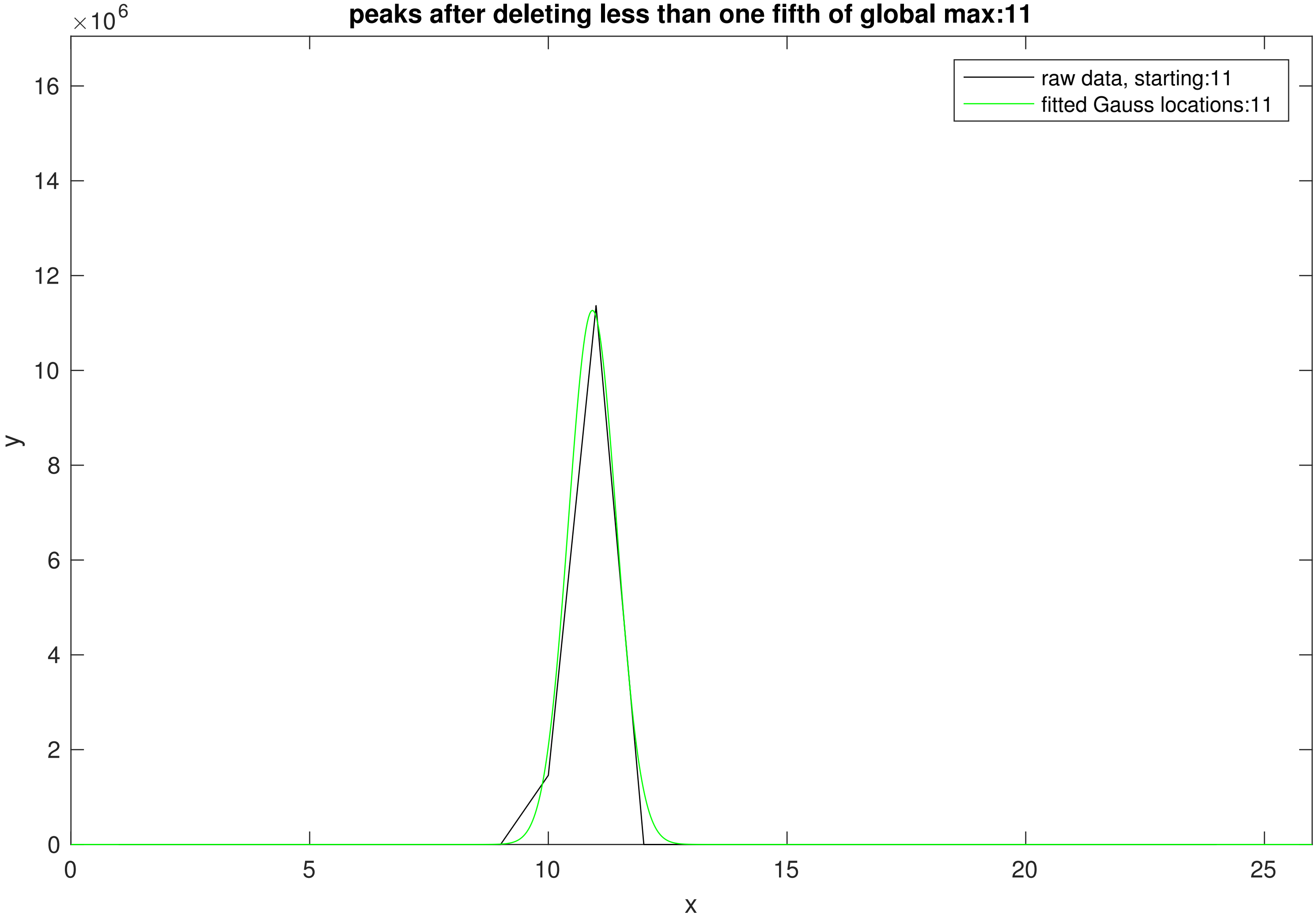

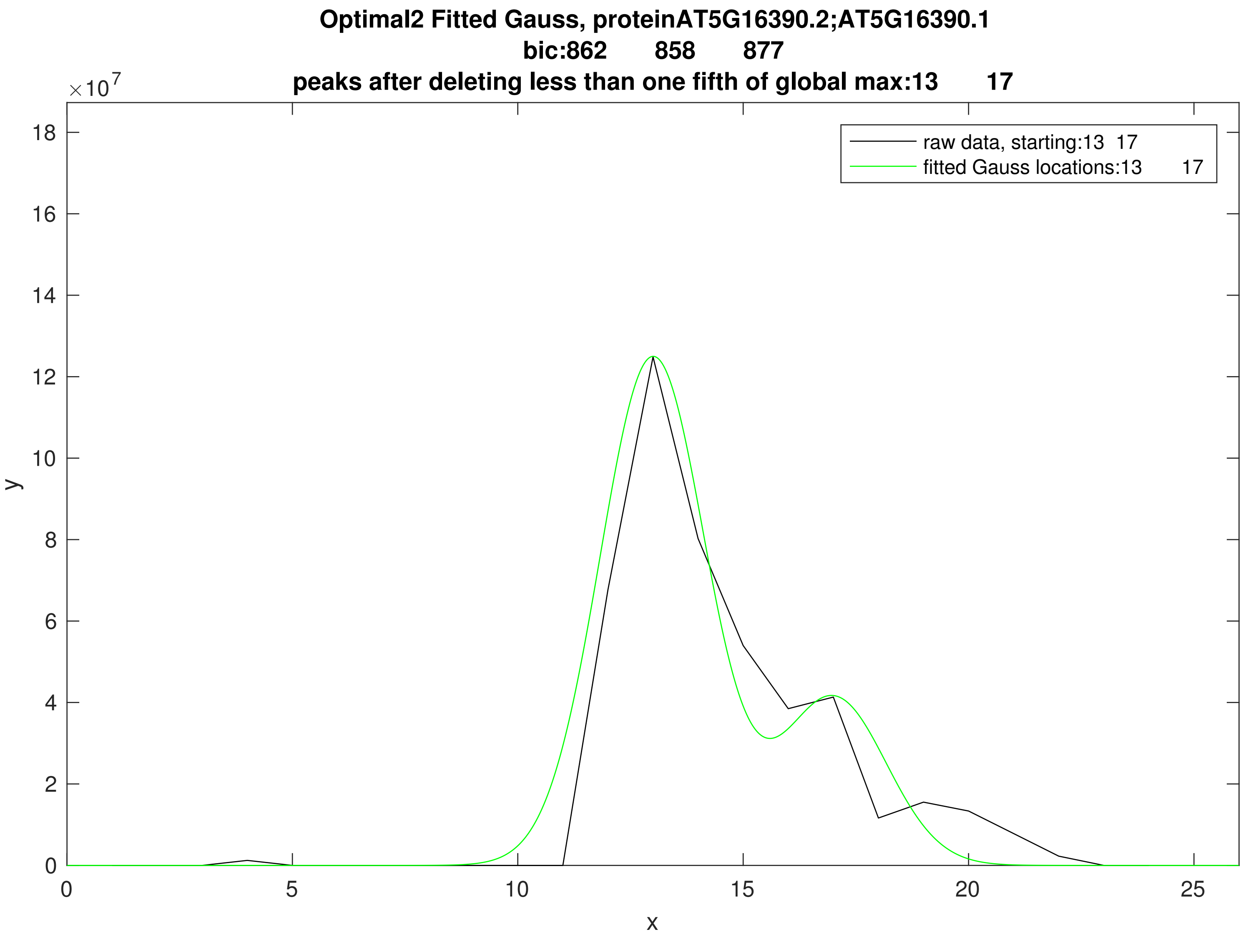

Optimal2 Fitted Gauss, proteinAT5G16390.2;AT5G16390.1
bic:862 858 877
peaks after deleting less than one fifth of global max:13 17
×10^7
raw data, starting:13 17
fitted Gauss locations:13 17
y
x
0
2
4
6
8
10
12
14
16
18
0
5
10
15
20
25

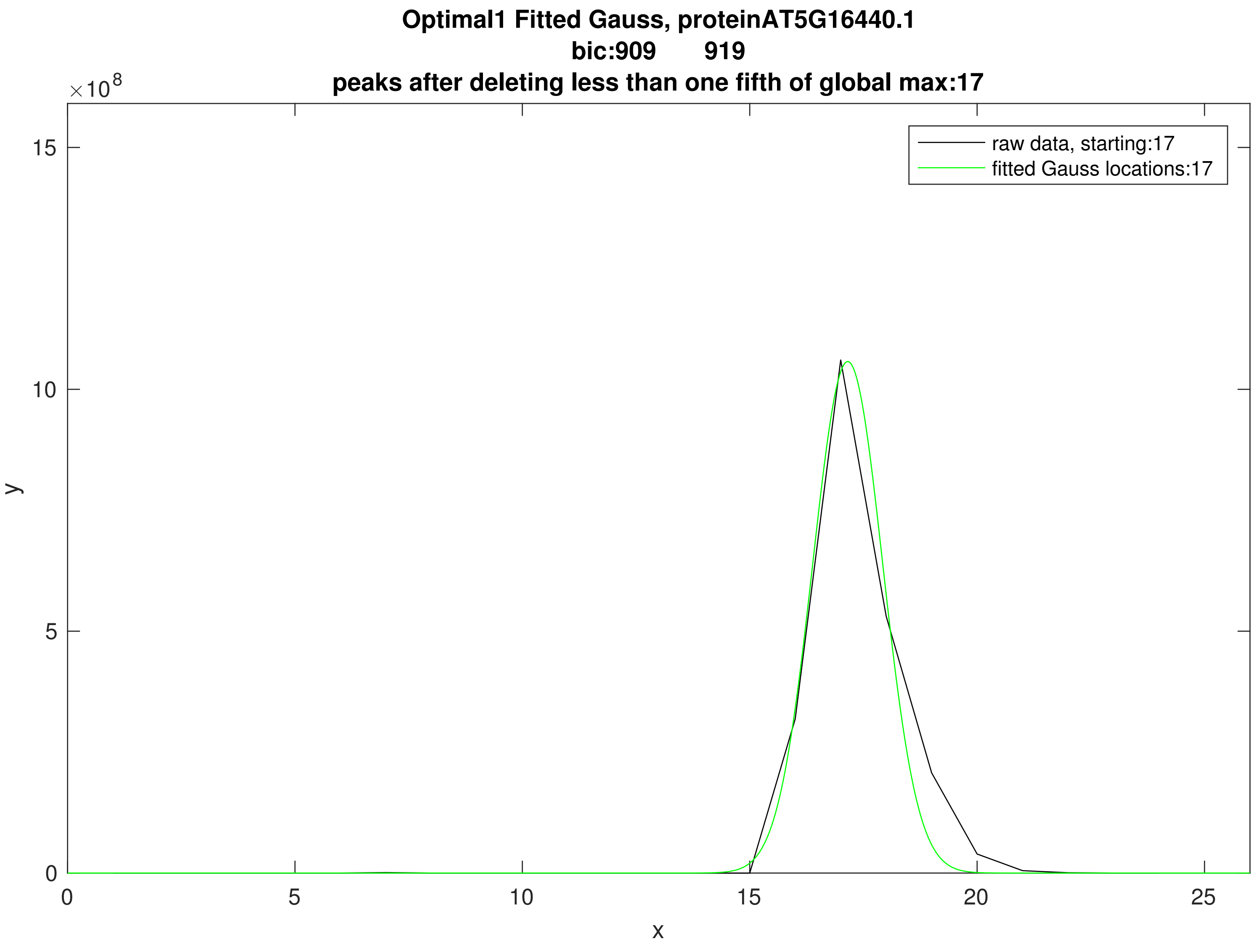
Optimal1 Fitted Gauss, proteinAT5G16440.1
bic:909 919
peaks after deleting less than one fifth of global max:17
×10^8
15
10
5
0
y
0
5
10
15
20
25
x
raw data, starting:17
fitted Gauss locations:17

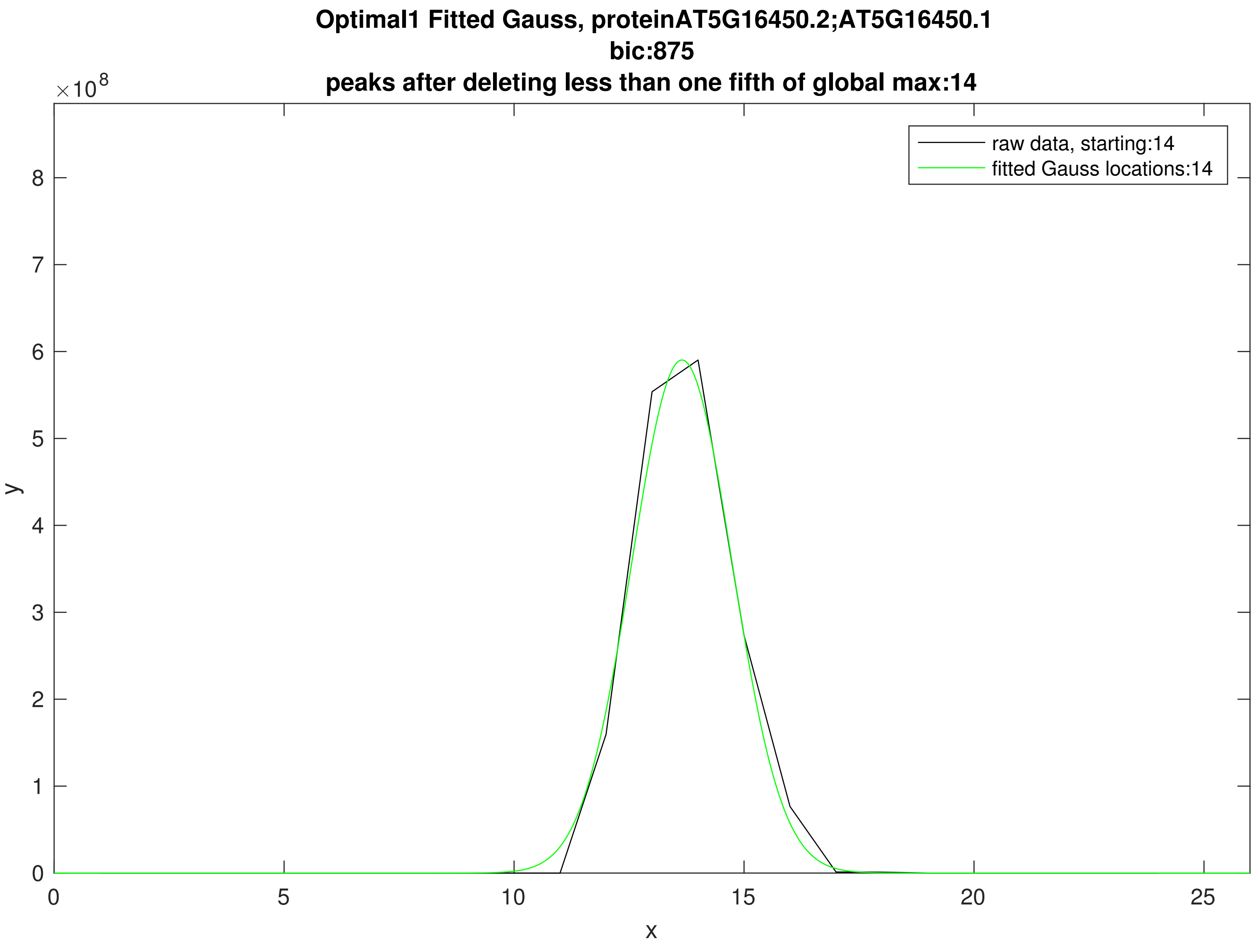
Optimal1 Fitted Gauss, proteinAT5G16450.2;AT5G16450.1
bic:875
peaks after deleting less than one fifth of global max:14
×10^8
raw data, starting:14
fitted Gauss locations:14
y
x
0
1
2
3
4
5
6
7
8
0
5
10
15
20
25

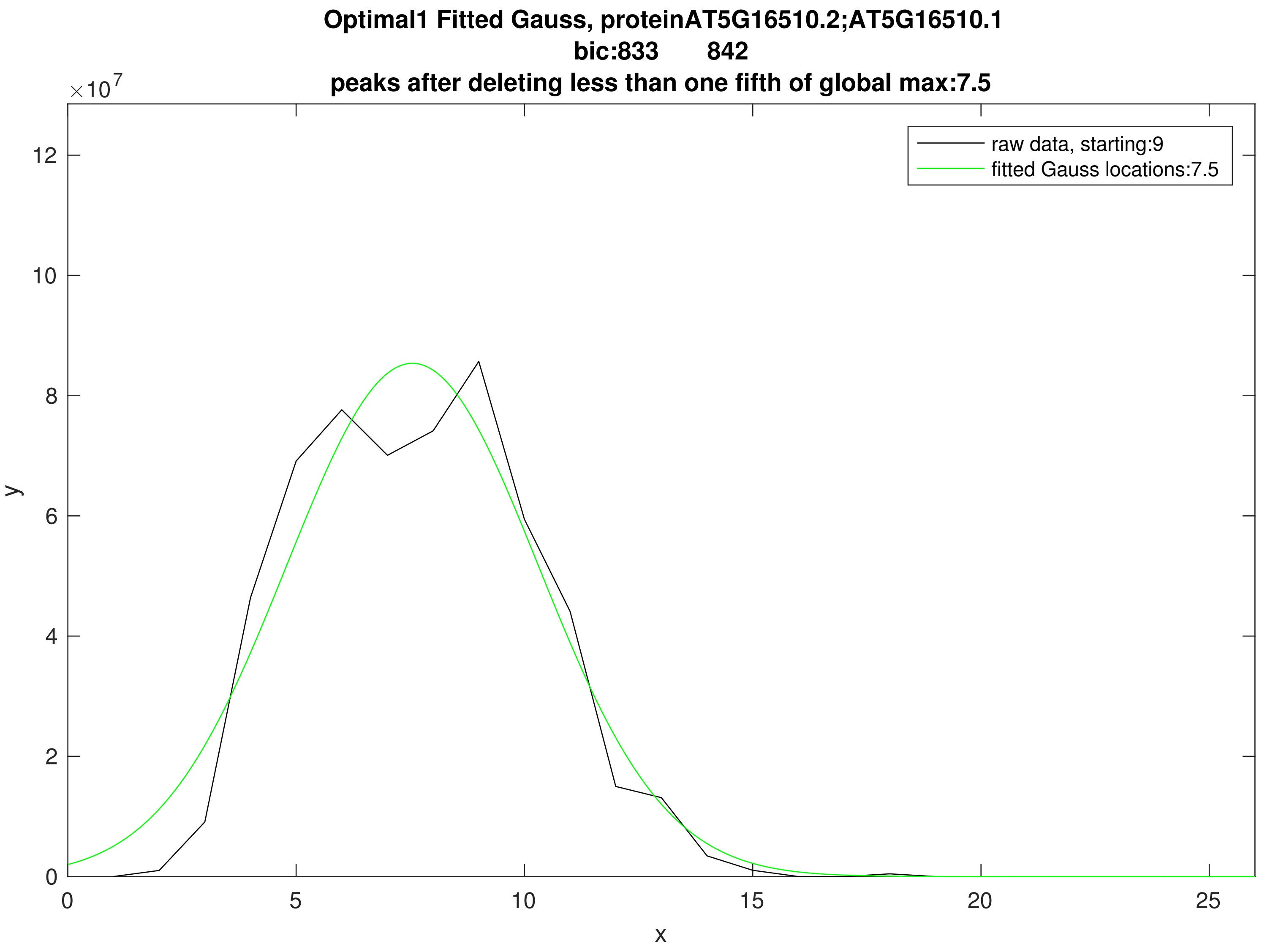

Optimal1 Fitted Gauss, proteinAT5G16510.2;AT5G16510.1
bic:833 842
peaks after deleting less than one fifth of global max:7.5
×10^7
raw data, starting:9
fitted Gauss locations:7.5
y
x
0
2
4
6
8
10
12
0
5
10
15
20
25

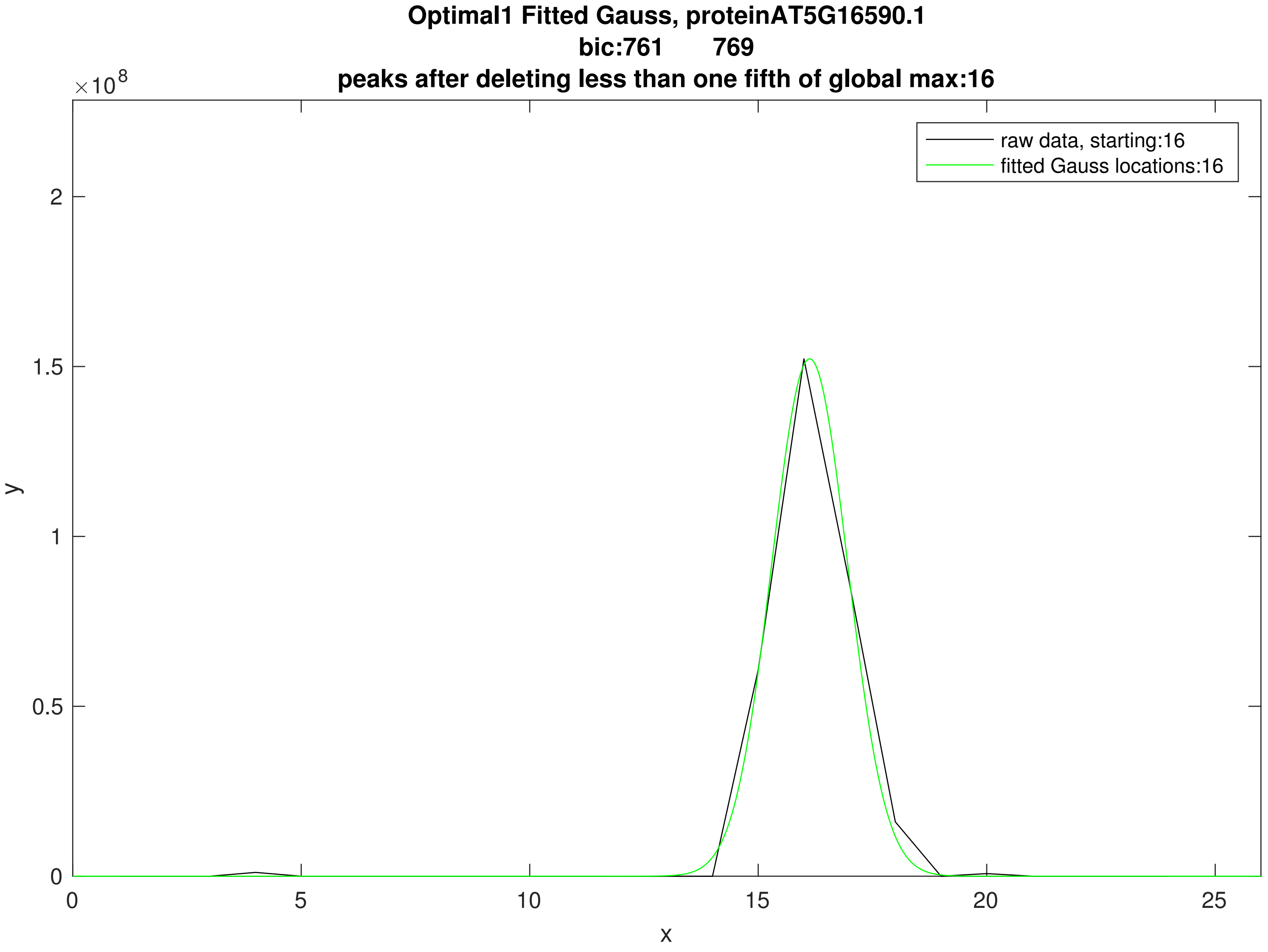

Optimal1 Fitted Gauss, proteinAT5G16590.1
bic:761 769
peaks after deleting less than one fifth of global max:16
×10^8
raw data, starting:16
fitted Gauss locations:16
y
x
0
0.5
1
1.5
2
0
5
10
15
20
25

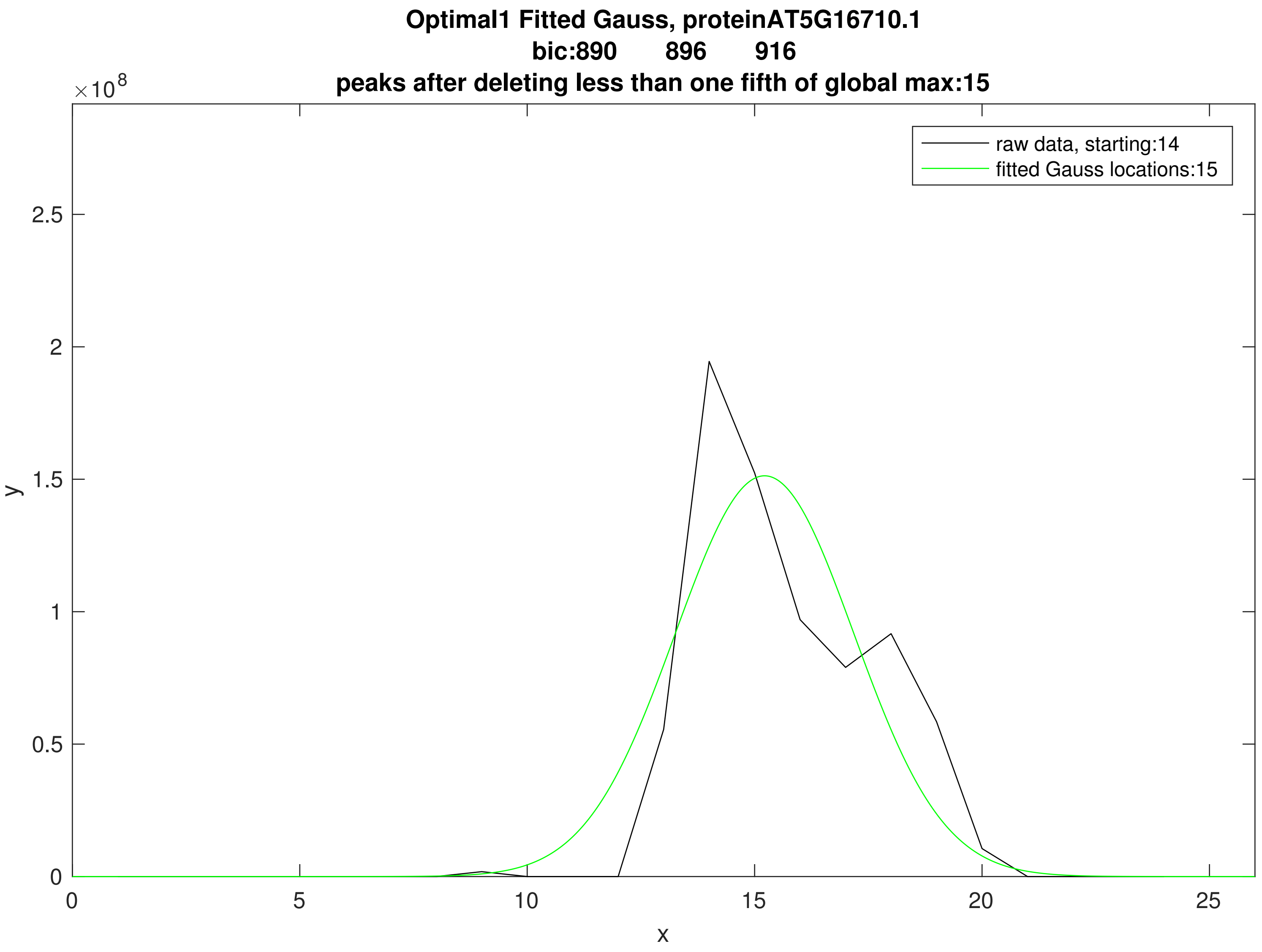
Optimal1 Fitted Gauss, proteinAT5G16710.1
bic:890 896 916
peaks after deleting less than one fifth of global max:15
×10^8
raw data, starting:14
fitted Gauss locations:15
y
x
0
0.5
1
1.5
2
2.5
0
5
10
15
20
25

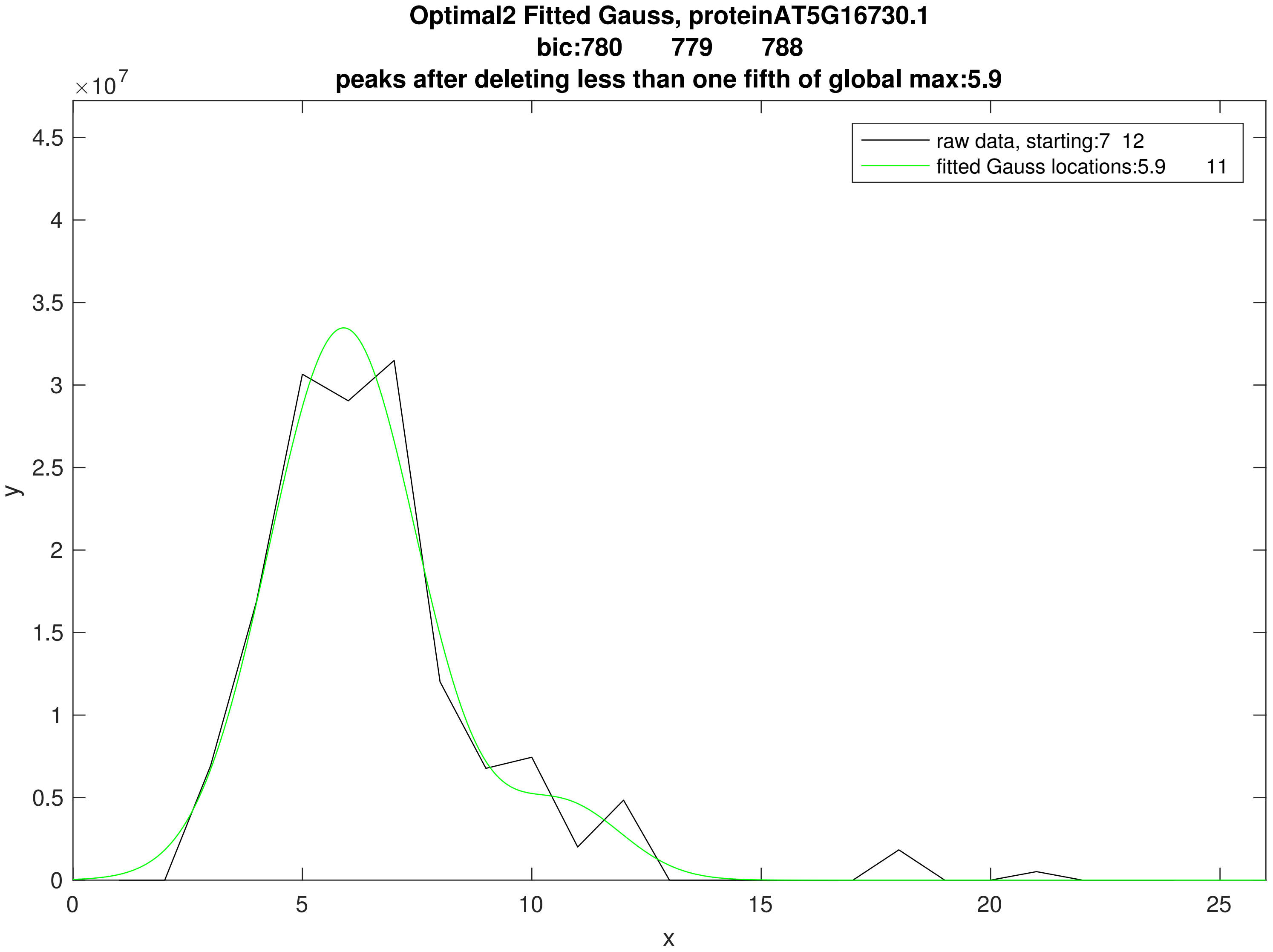
Optimal2 Fitted Gauss, proteinAT5G16730.1
bic:780 779 788
peaks after deleting less than one fifth of global max:5.9
×10^7
raw data, starting:7 12
fitted Gauss locations:5.9 11
y
x
0
0.5
1
1.5
2
2.5
3
3.5
4
4.5
5
10
15
20
25

Optimal2 Fitted Gauss, proteinAT5G16840.3;AT5G16840.1;AT5G16840.2
bic:872 867 877
peaks after deleting less than one fifth of global max:16

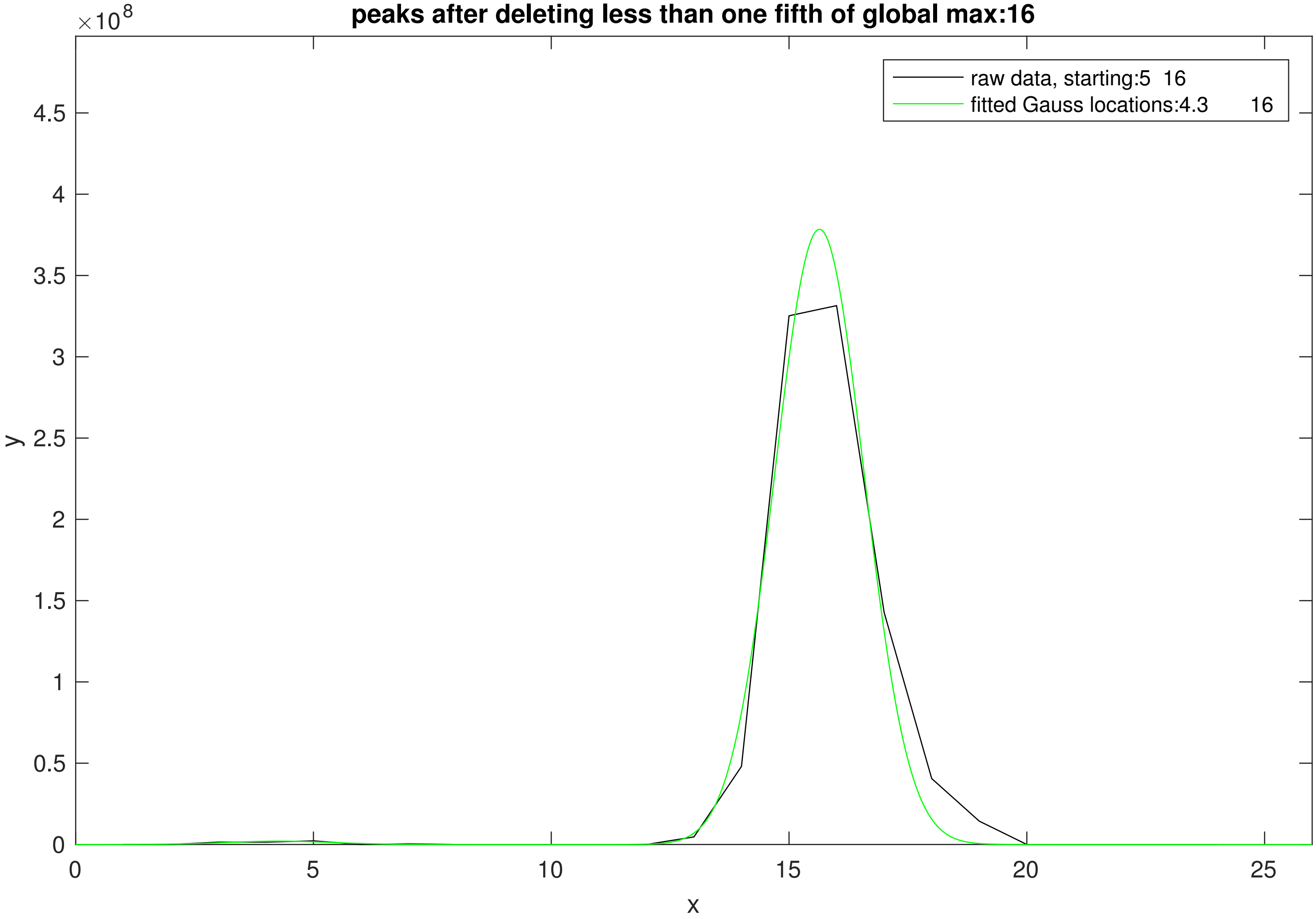

**Optimal1 Fitted Gauss, proteinAT5G16880.1;AT5G16880.2;AT5G16880.3**
**bic:834 841 851**
**peaks after deleting less than one fifth of global max:12**

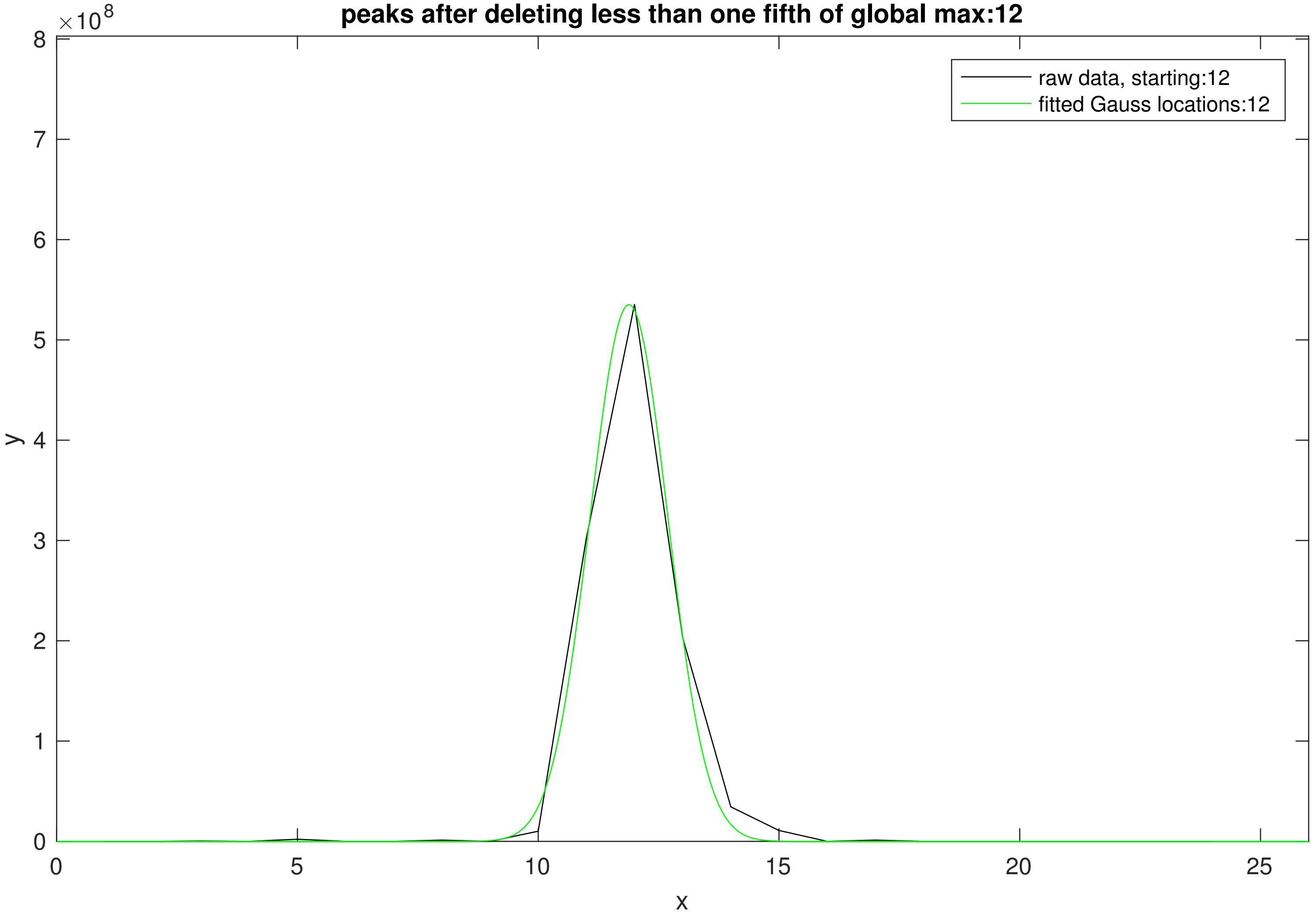

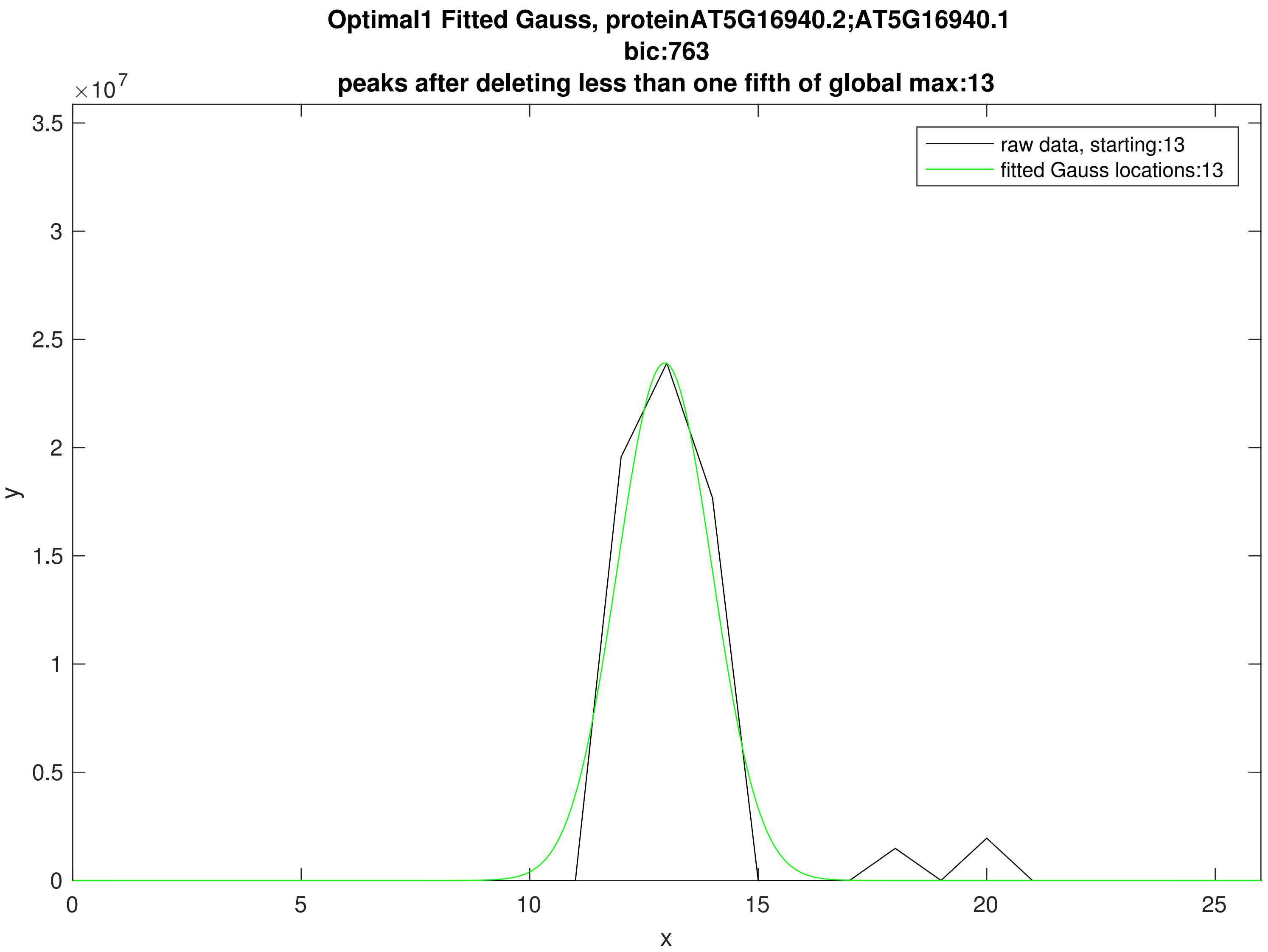

Optimal1 Fitted Gauss, proteinAT5G16940.2;AT5G16940.1
bic:763
peaks after deleting less than one fifth of global max:13
×10^7
3.5
3
2.5
2
1.5
1
0.5
0
y
0
5
10
15
20
25
x
raw data, starting:13
fitted Gauss locations:13

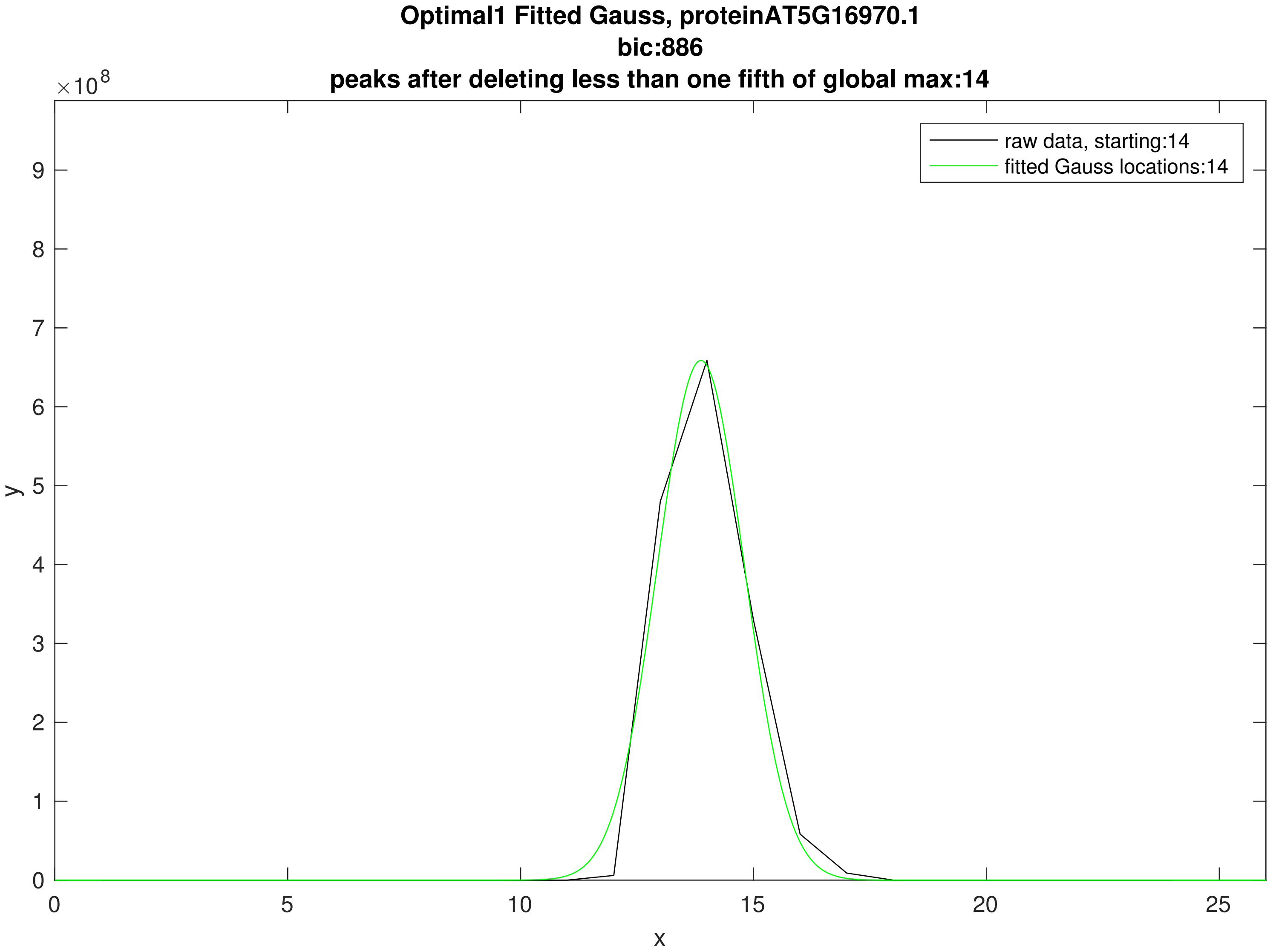
Optimal1 Fitted Gauss, proteinAT5G16970.1
bic:886
peaks after deleting less than one fifth of global max:14
×10^8
raw data, starting:14
fitted Gauss locations:14
y
x
0
1
2
3
4
5
6
7
8
9
0
5
10
15
20
25

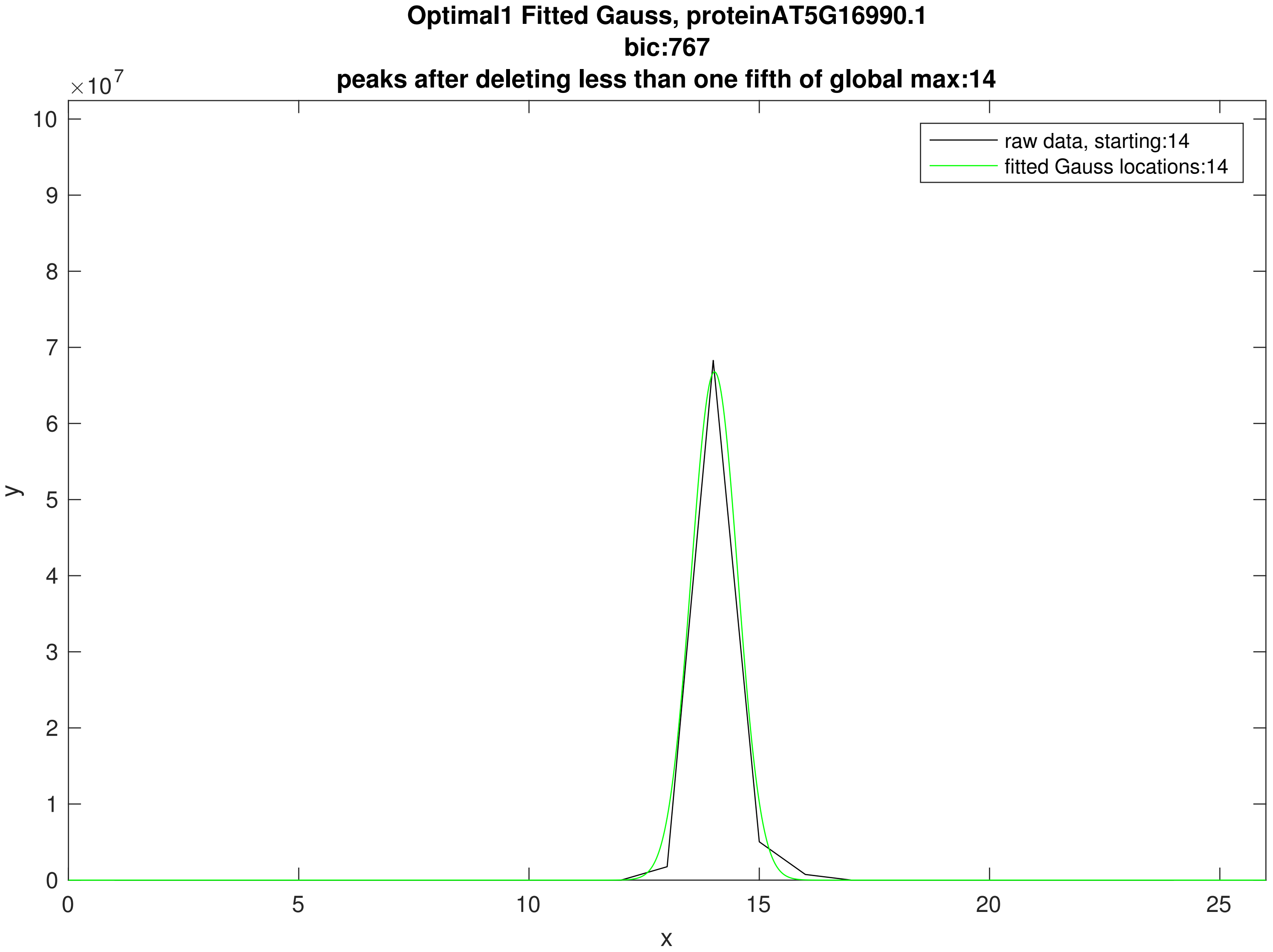
Optimal1 Fitted Gauss, proteinAT5G16990.1
bic:767
peaks after deleting less than one fifth of global max:14
×10^7
y
x
raw data, starting:14
fitted Gauss locations:14
0
1
2
3
4
5
6
7
8
9
10
0
5
10
15
20
25

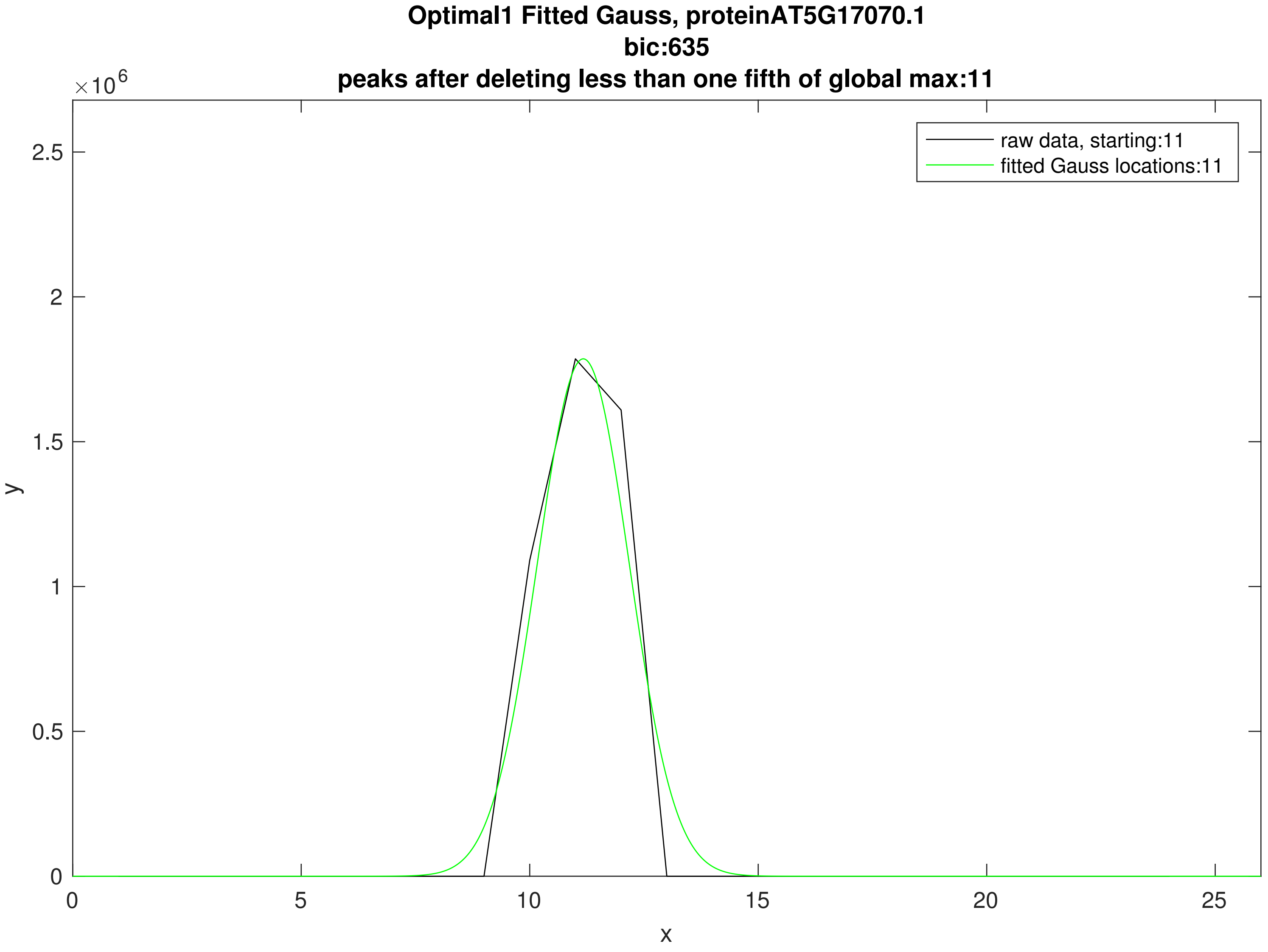
Optimal1 Fitted Gauss, proteinAT5G17070.1
bic:635
peaks after deleting less than one fifth of global max:11
×10^6
2.5
2
1.5
1
0.5
0
y
0
5
10
15
20
25
x
raw data, starting:11
fitted Gauss locations:11

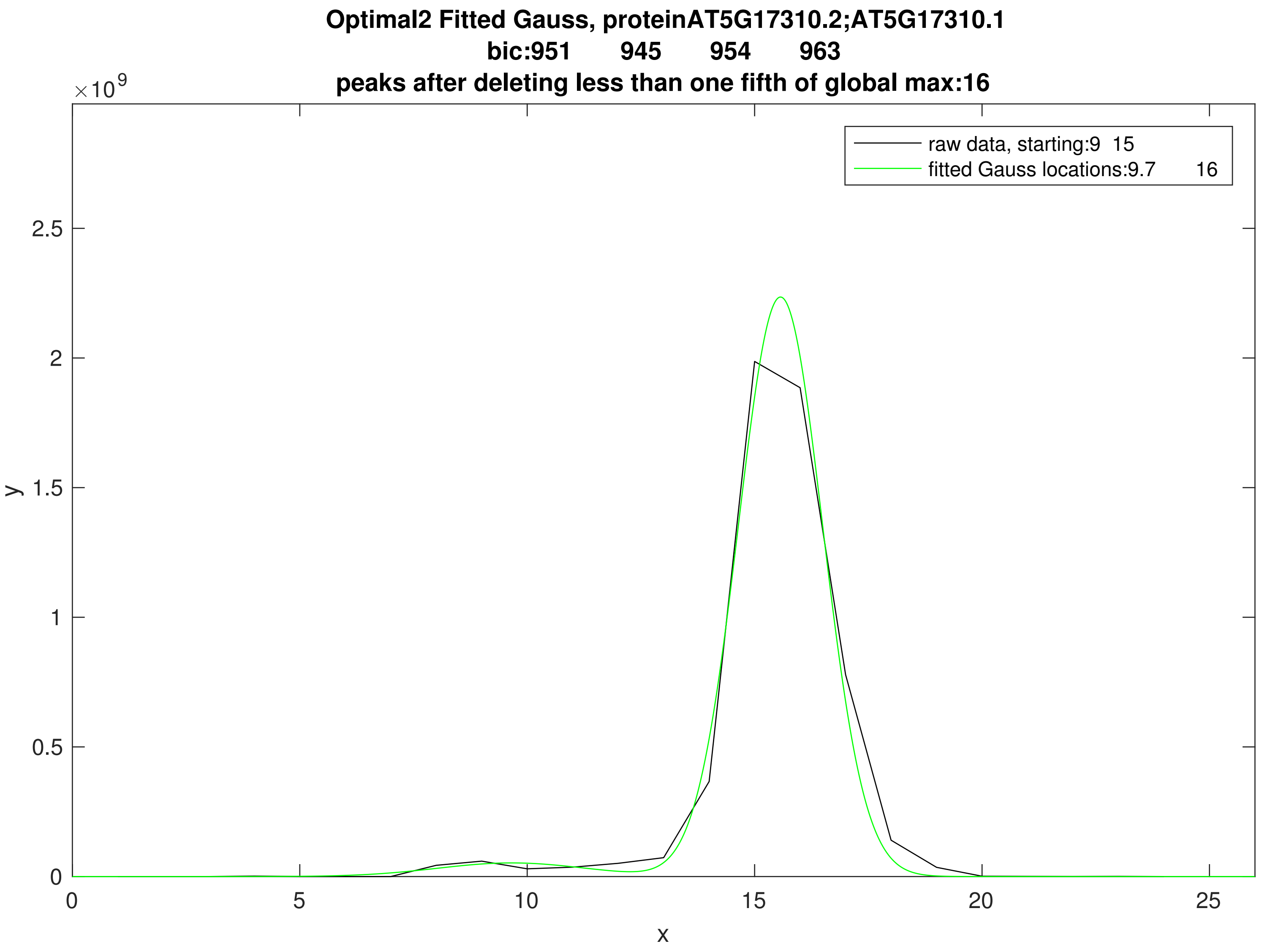
Optimal2 Fitted Gauss, proteinAT5G17310.2;AT5G17310.1
bic:951 945 954 963
peaks after deleting less than one fifth of global max:16
×10^9
raw data, starting:9 15
fitted Gauss locations:9.7 16
2.5
2
1.5
1
0.5
0
y
0
5
10
15
20
25
x

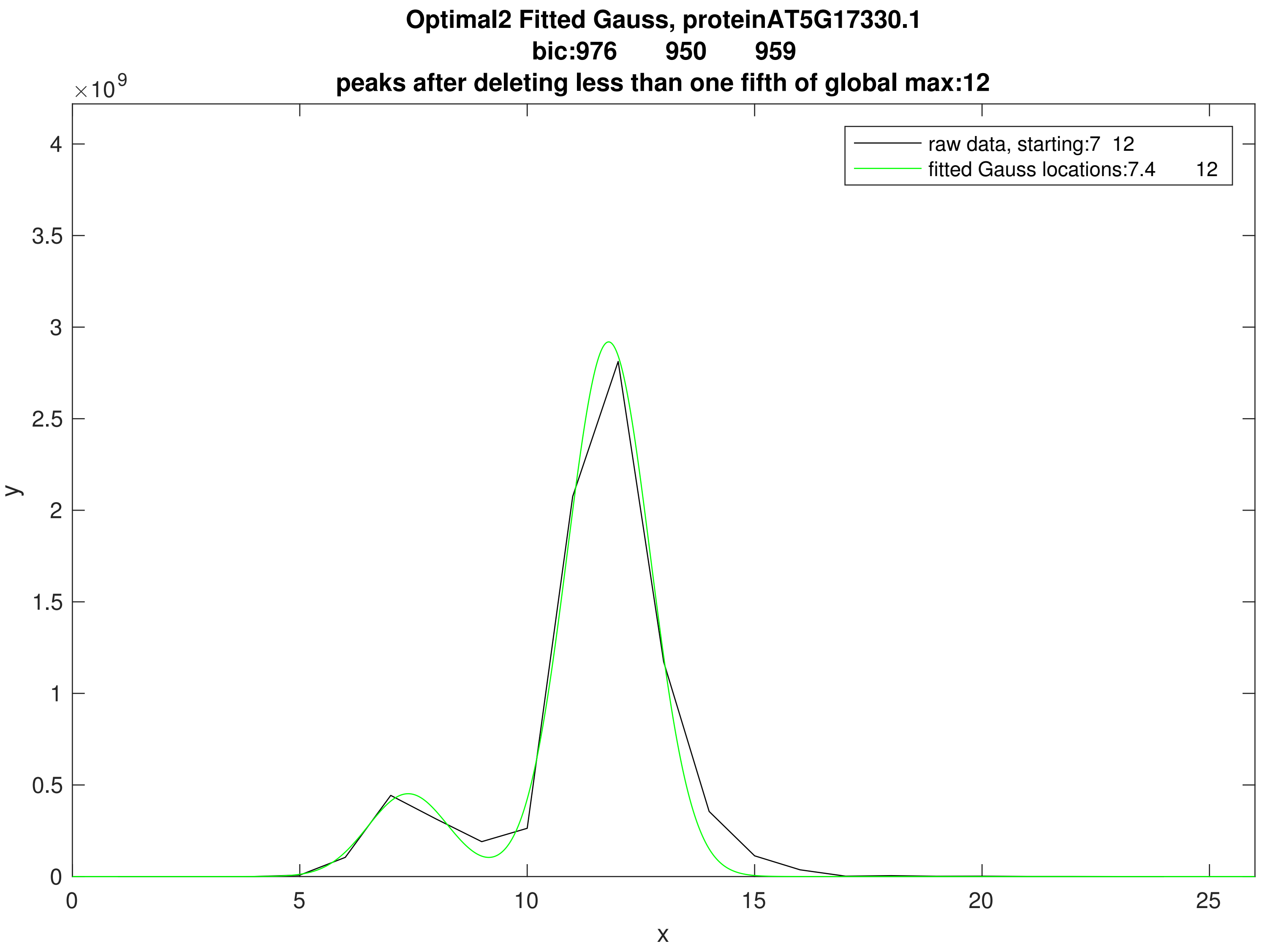
Optimal2 Fitted Gauss, proteinAT5G17330.1
bic:976 950 959
peaks after deleting less than one fifth of global max:12
×10^9
4
3.5
3
2.5
2
1.5
1
0.5
0
y
0
5
10
15
20
25
x
raw data, starting:7 12
fitted Gauss locations:7.4 12

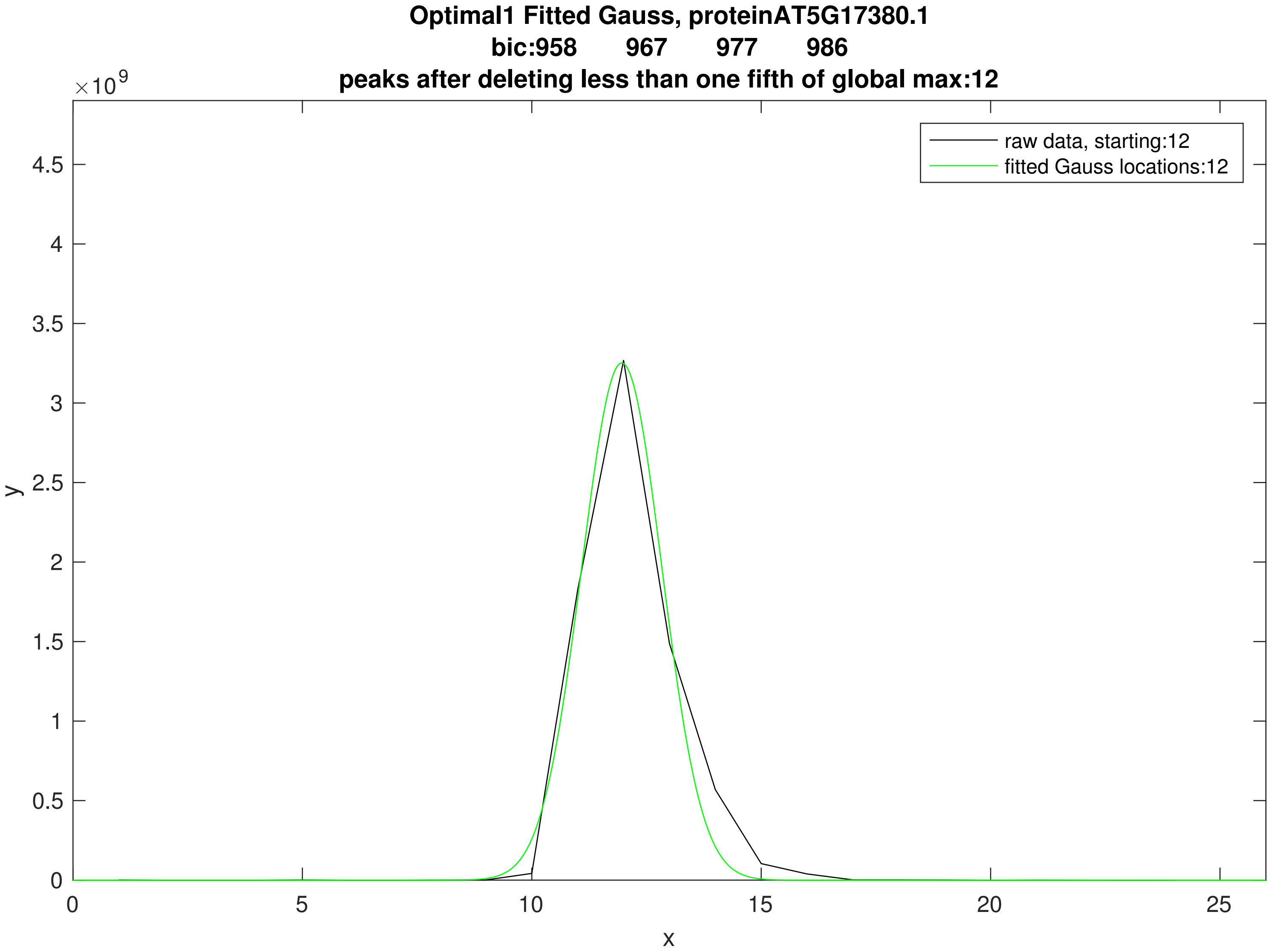
Optimal1 Fitted Gauss, proteinAT5G17380.1
bic:958 967 977 986
peaks after deleting less than one fifth of global max:12
×10^9
raw data, starting:12
fitted Gauss locations:12
4.5
4
3.5
3
2.5
2
1.5
1
0.5
0
y
0
5
10
15
20
25
x

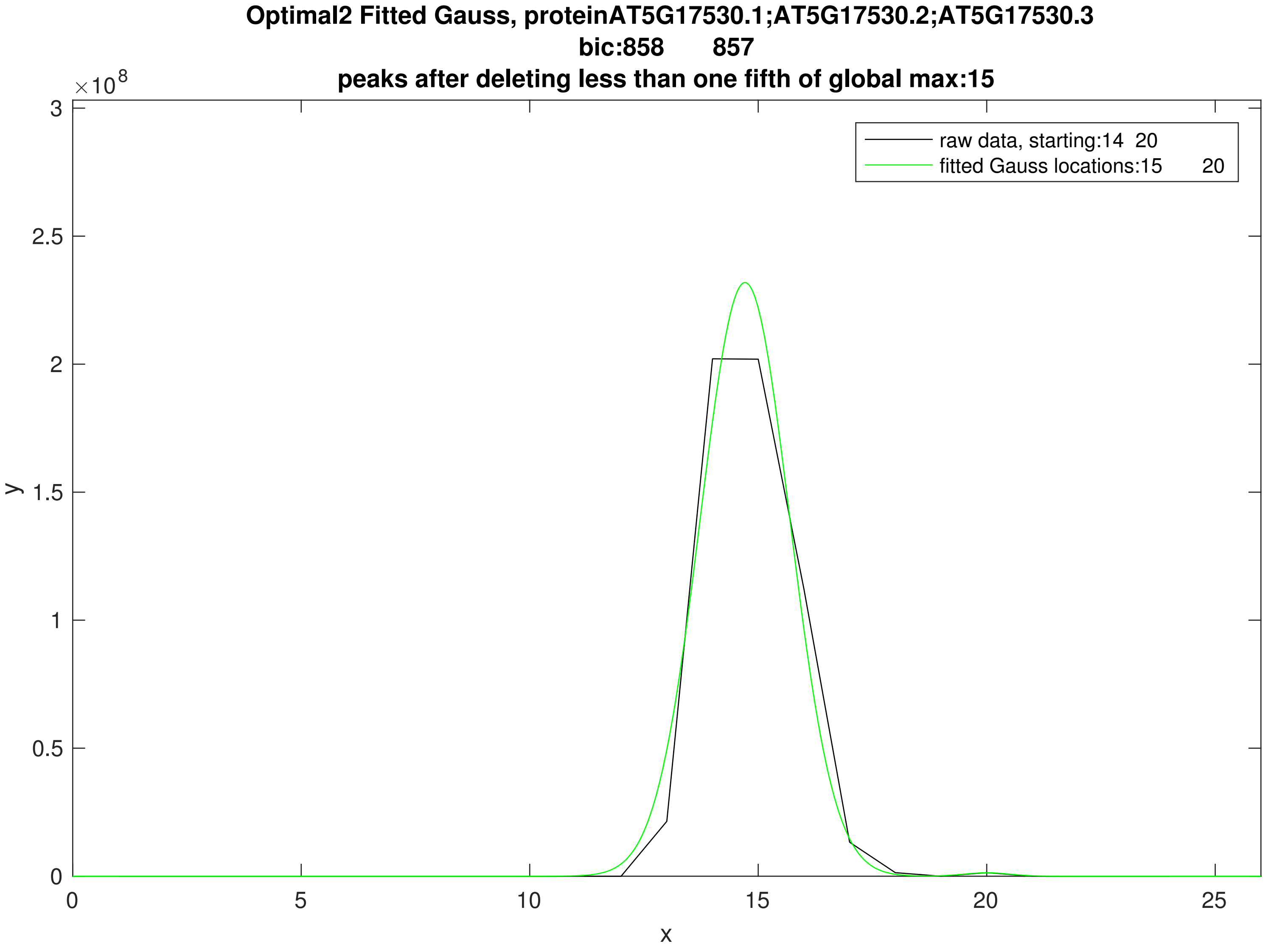
Optimal2 Fitted Gauss, proteinAT5G17530.1;AT5G17530.2;AT5G17530.3
bic:858 857
peaks after deleting less than one fifth of global max:15
×10^8
raw data, starting:14 20
fitted Gauss locations:15 20
3
2.5
2
1.5
1
0.5
0
y
0
5
10
15
20
25
x

Optimal1 Fitted Gauss, proteinAT5G17560.1
bic:687
peaks after deleting less than one fifth of global max:20

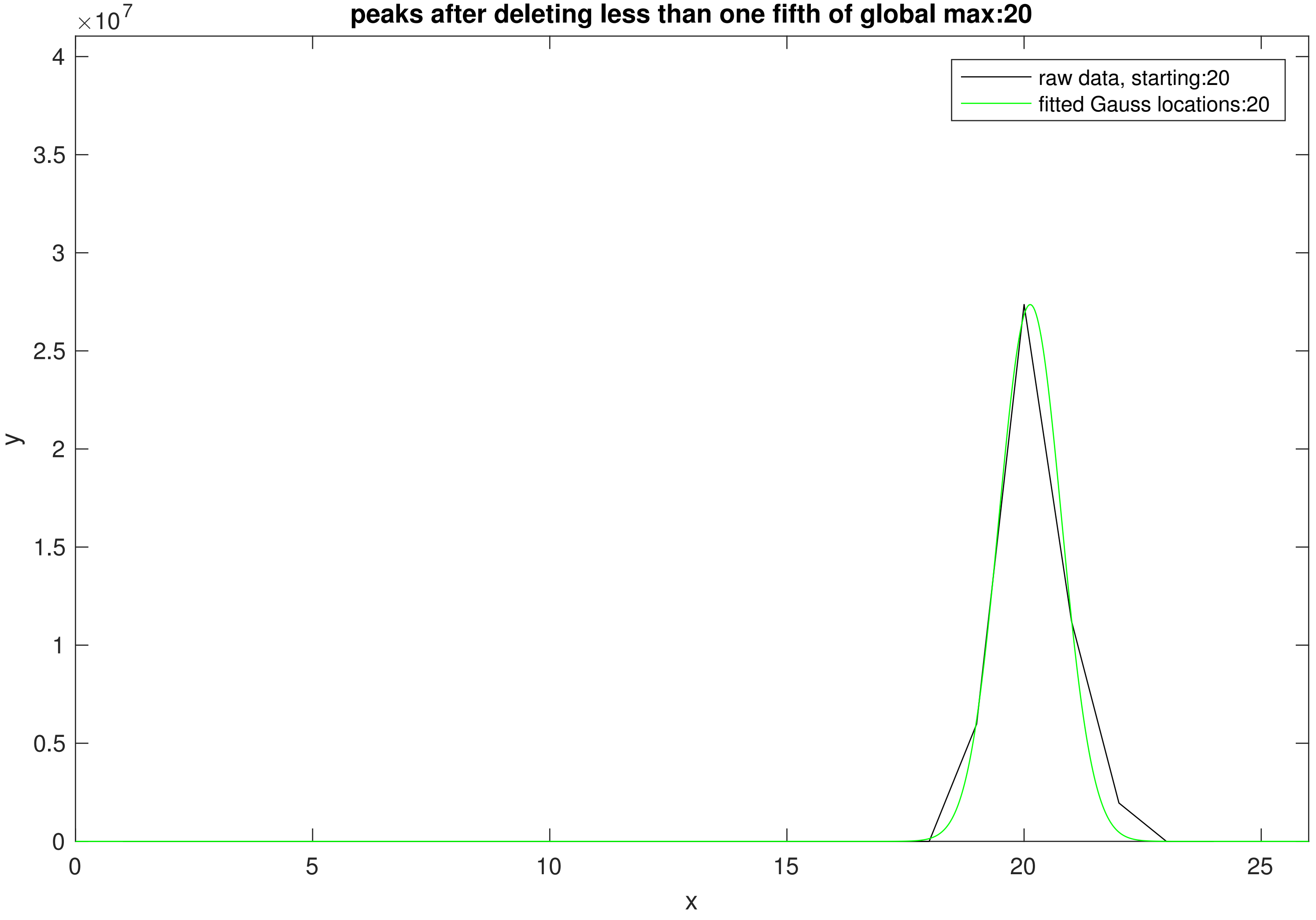

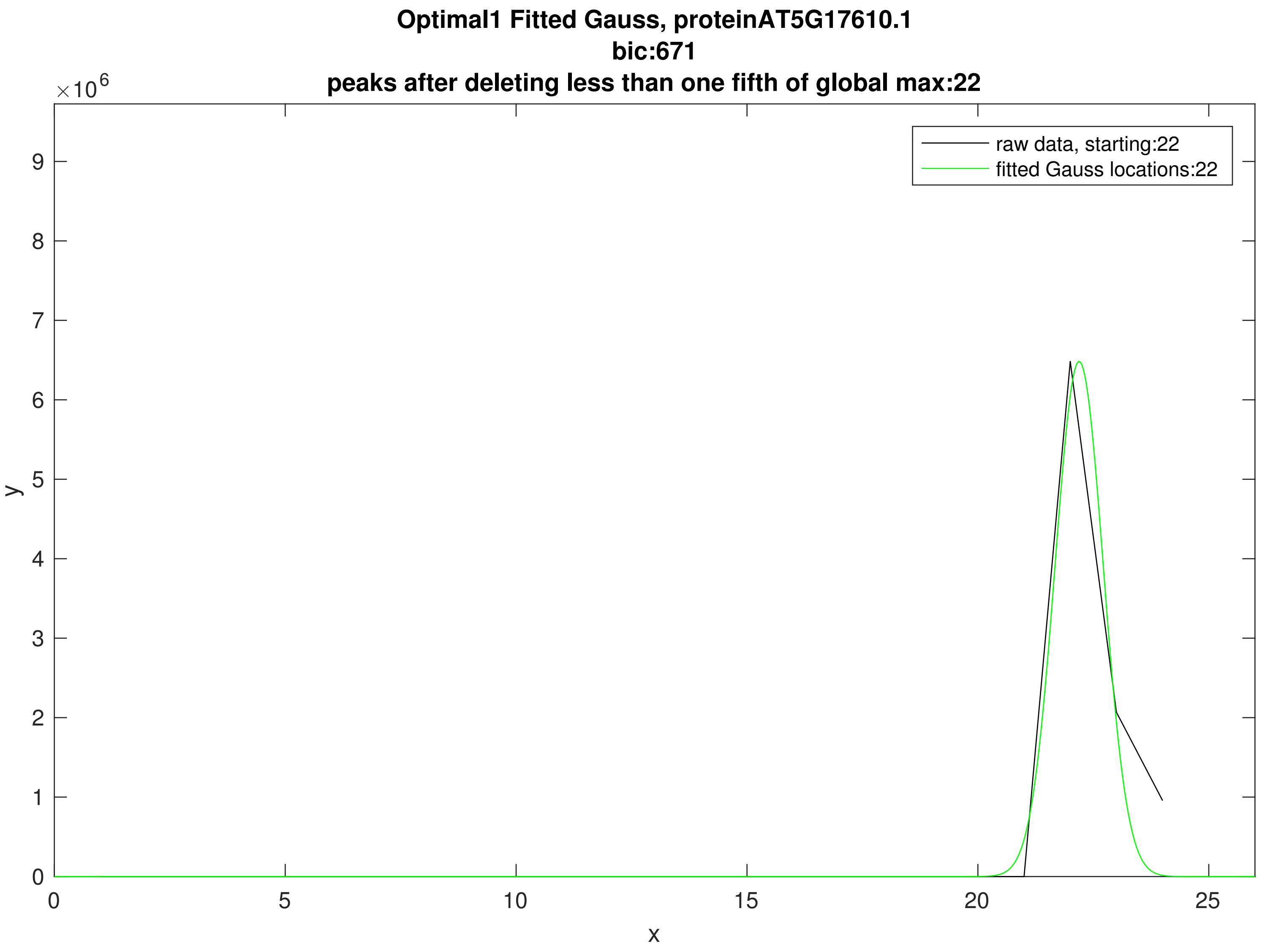
Optimal1 Fitted Gauss, proteinAT5G17610.1
bic:671
peaks after deleting less than one fifth of global max:22
×10^6
raw data, starting:22
fitted Gauss locations:22
y
x
0
1
2
3
4
5
6
7
8
9
0
5
10
15
20
25

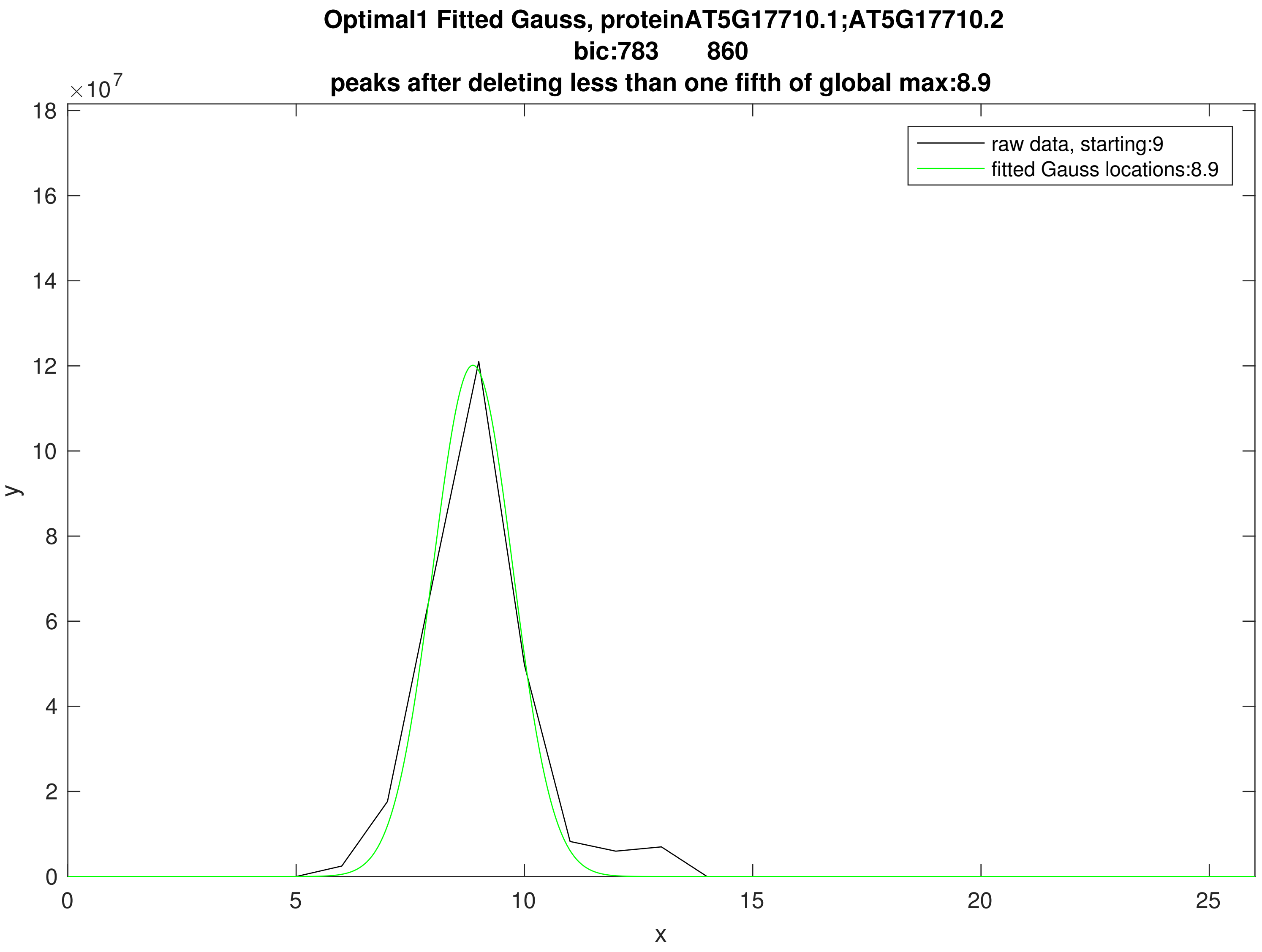

Optimal1 Fitted Gauss, proteinAT5G17710.1;AT5G17710.2
bic:783 860
peaks after deleting less than one fifth of global max:8.9
×10^7
raw data, starting:9
fitted Gauss locations:8.9
y
x
0
2
4
6
8
10
12
14
16
18
0
5
10
15
20
25

**Optimal1 Fitted Gauss, proteinAT5G17910.2;AT5G17910.1**
**bic:830 834 848**
**peaks after deleting less than one fifth of global max:5.3**

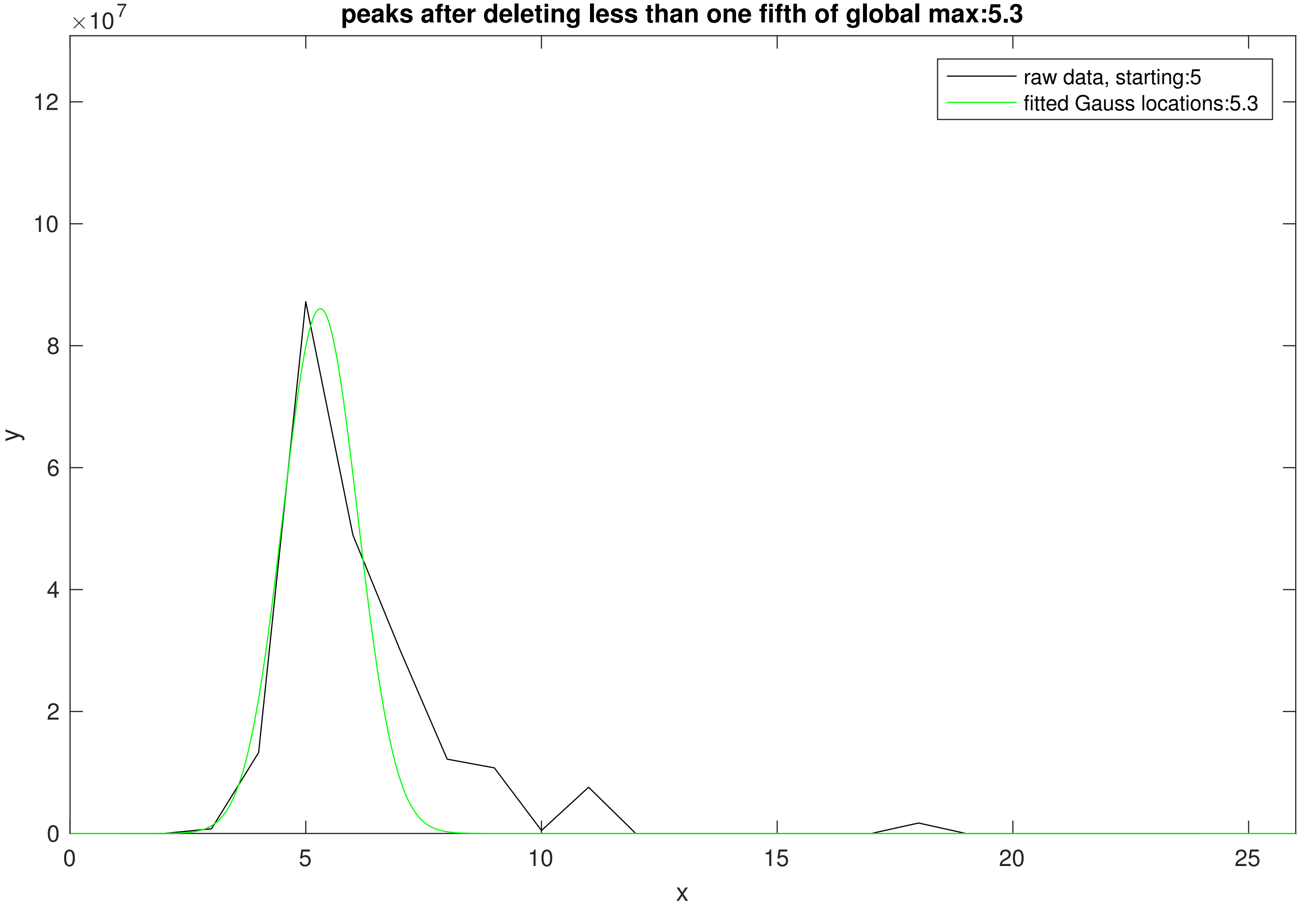

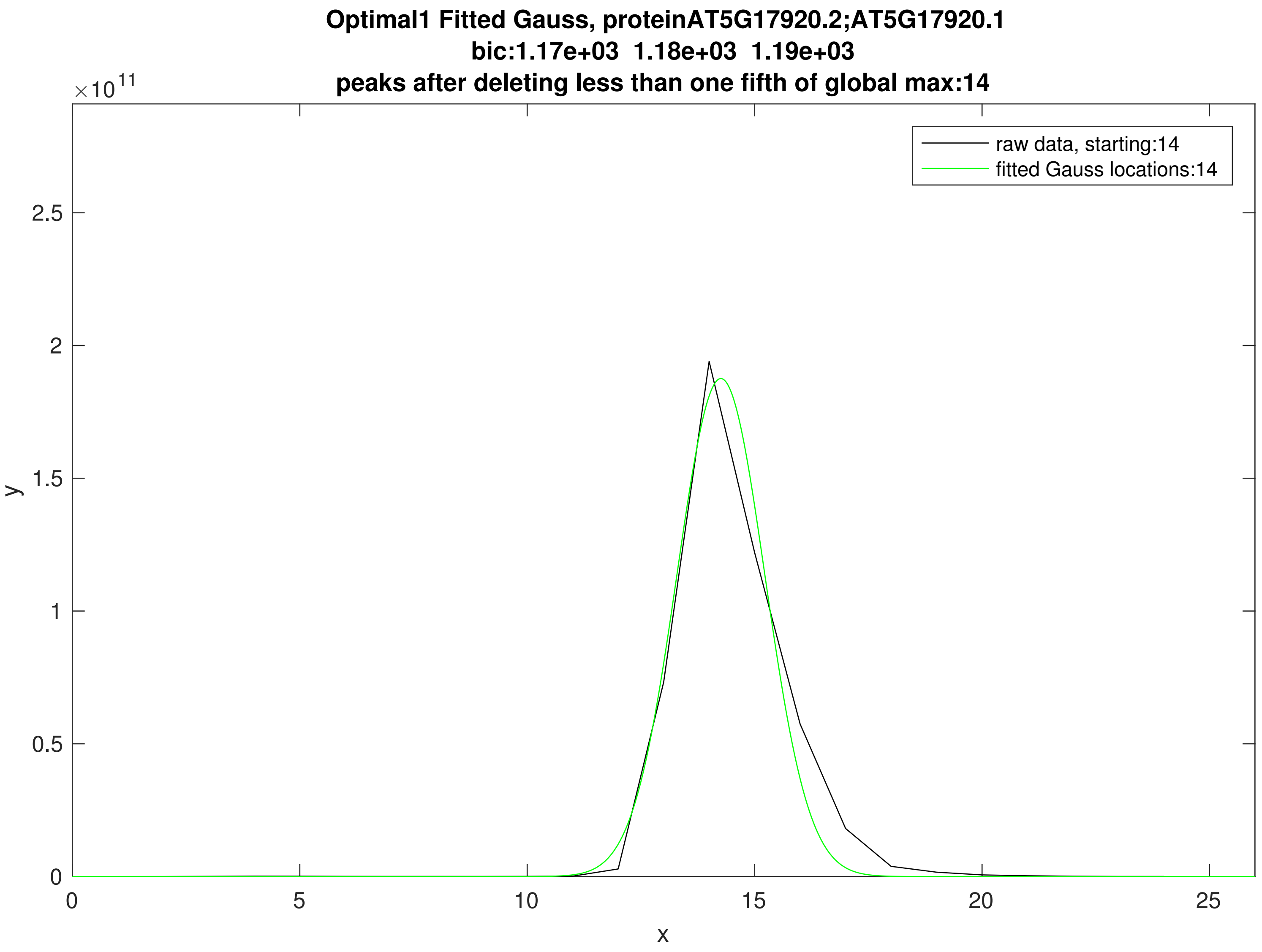

Optimal1 Fitted Gauss, proteinAT5G17920.2;AT5G17920.1
bic:1.17e+03 1.18e+03 1.19e+03
peaks after deleting less than one fifth of global max:14
×10^11
raw data, starting:14
fitted Gauss locations:14
y
x
0
0.5
1
1.5
2
2.5
5
10
15
20
25

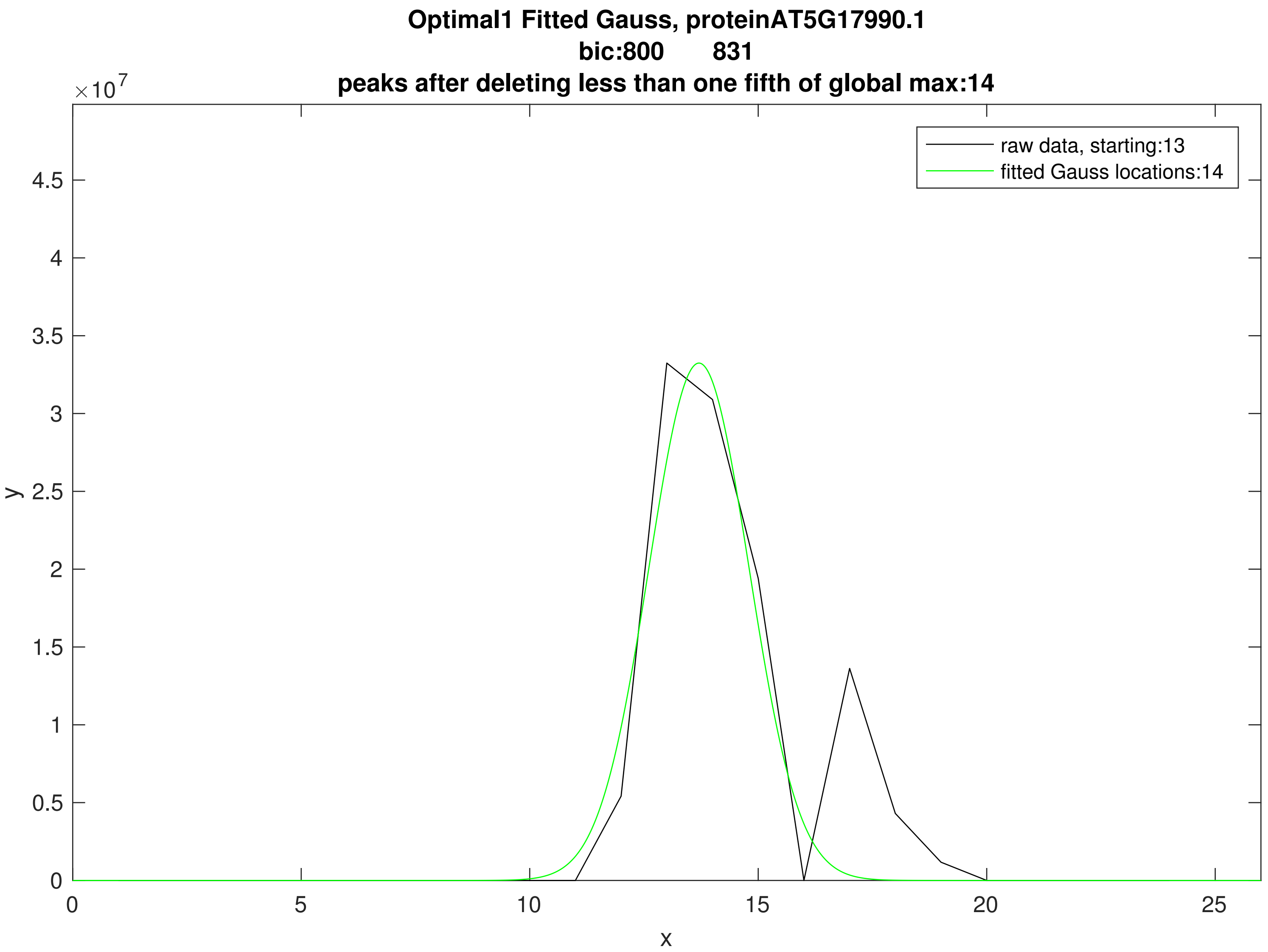
Optimal1 Fitted Gauss, proteinAT5G17990.1
bic:800 831
peaks after deleting less than one fifth of global max:14
×10^7
raw data, starting:13
fitted Gauss locations:14
y
x
4.5
4
3.5
3
2.5
2
1.5
1
0.5
0
5
10
15
20
25

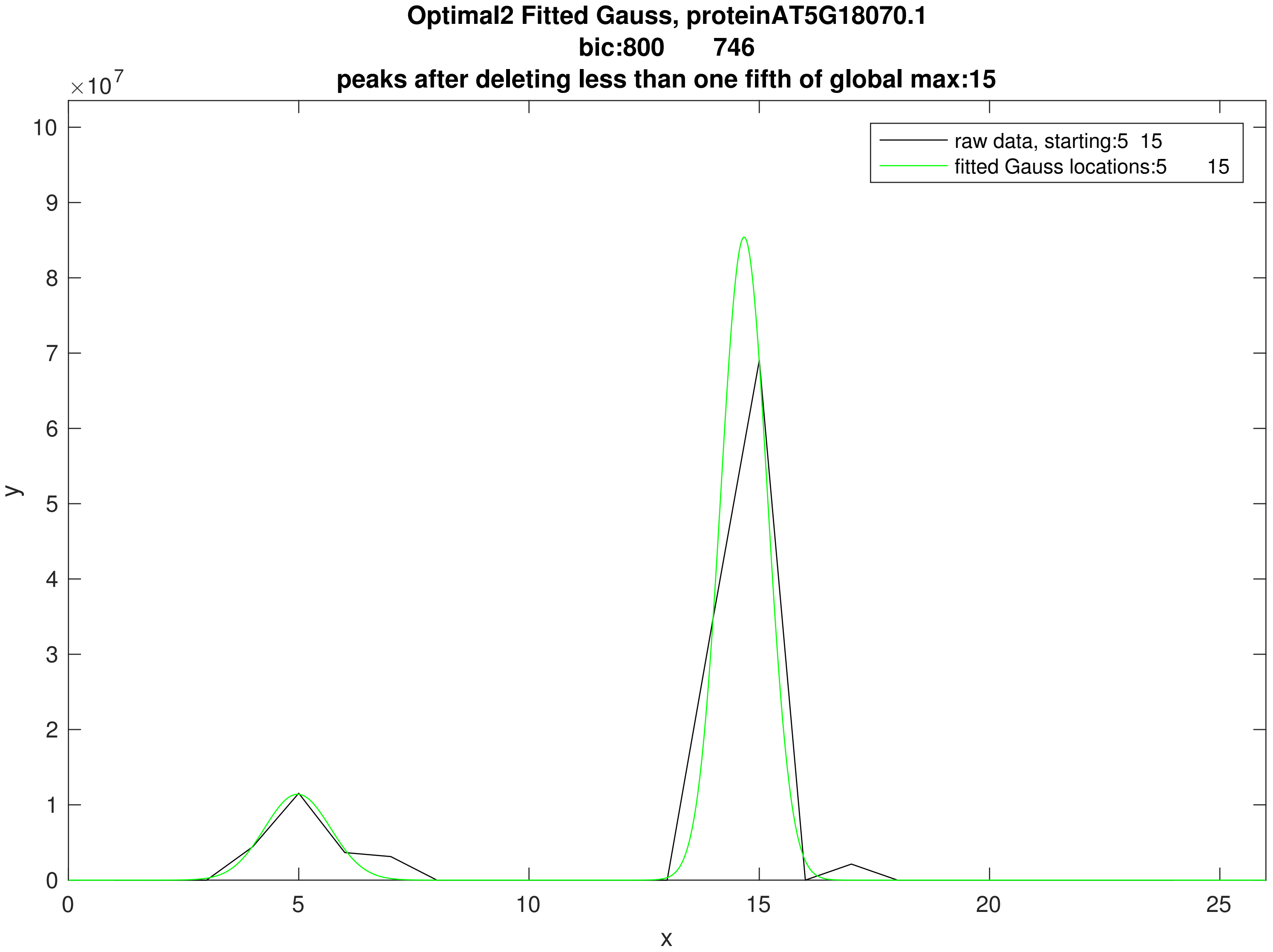

Optimal2 Fitted Gauss, proteinAT5G18070.1
bic:800 746
peaks after deleting less than one fifth of global max:15
×10^7
raw data, starting:5 15
fitted Gauss locations:5 15
y
x
0
1
2
3
4
5
6
7
8
9
10
0
5
10
15
20
25

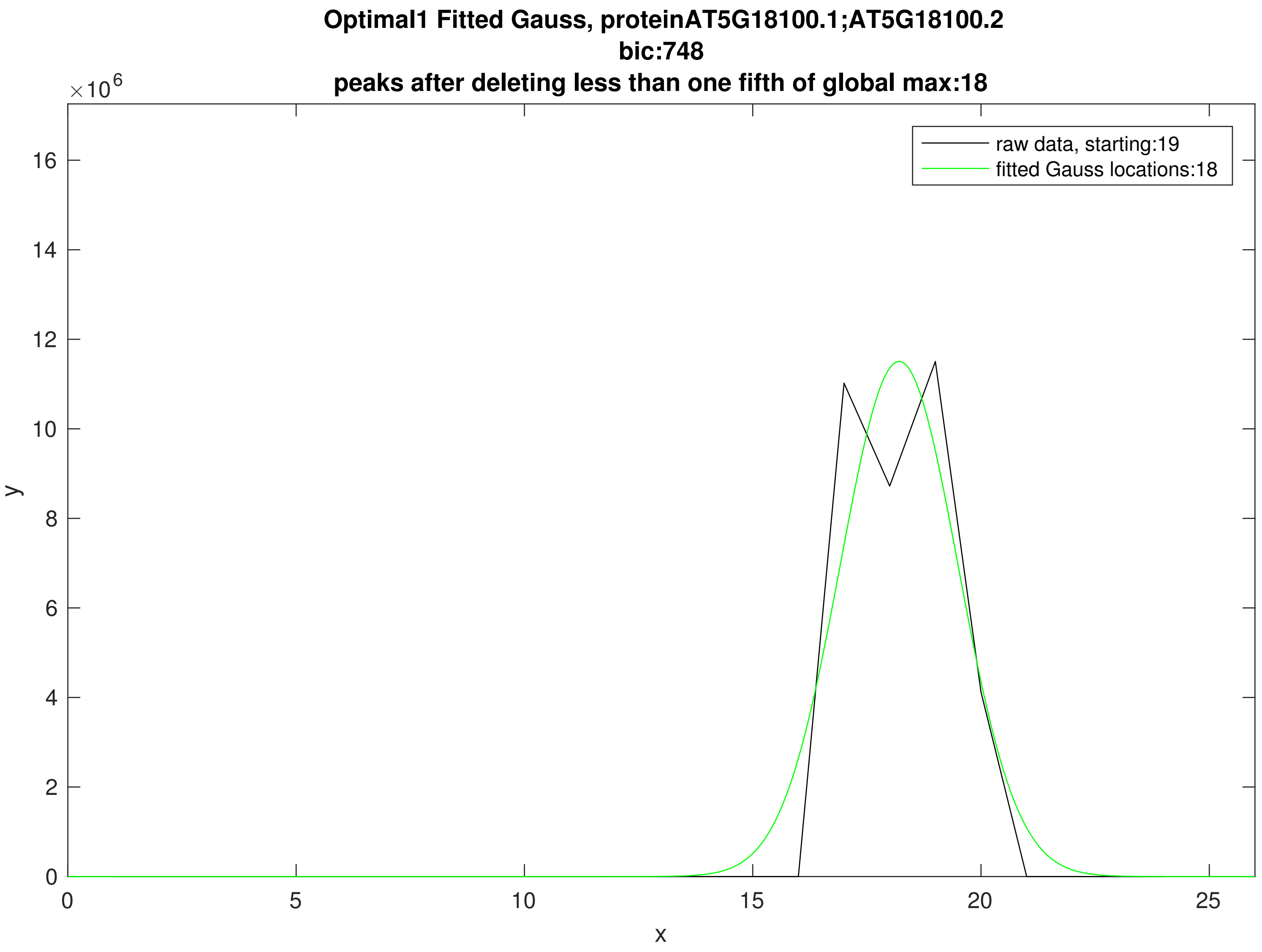
Optimal1 Fitted Gauss, proteinAT5G18100.1;AT5G18100.2
bic:748
peaks after deleting less than one fifth of global max:18
×10^6
raw data, starting:19
fitted Gauss locations:18
y
x
0
2
4
6
8
10
12
14
16
0
5
10
15
20
25

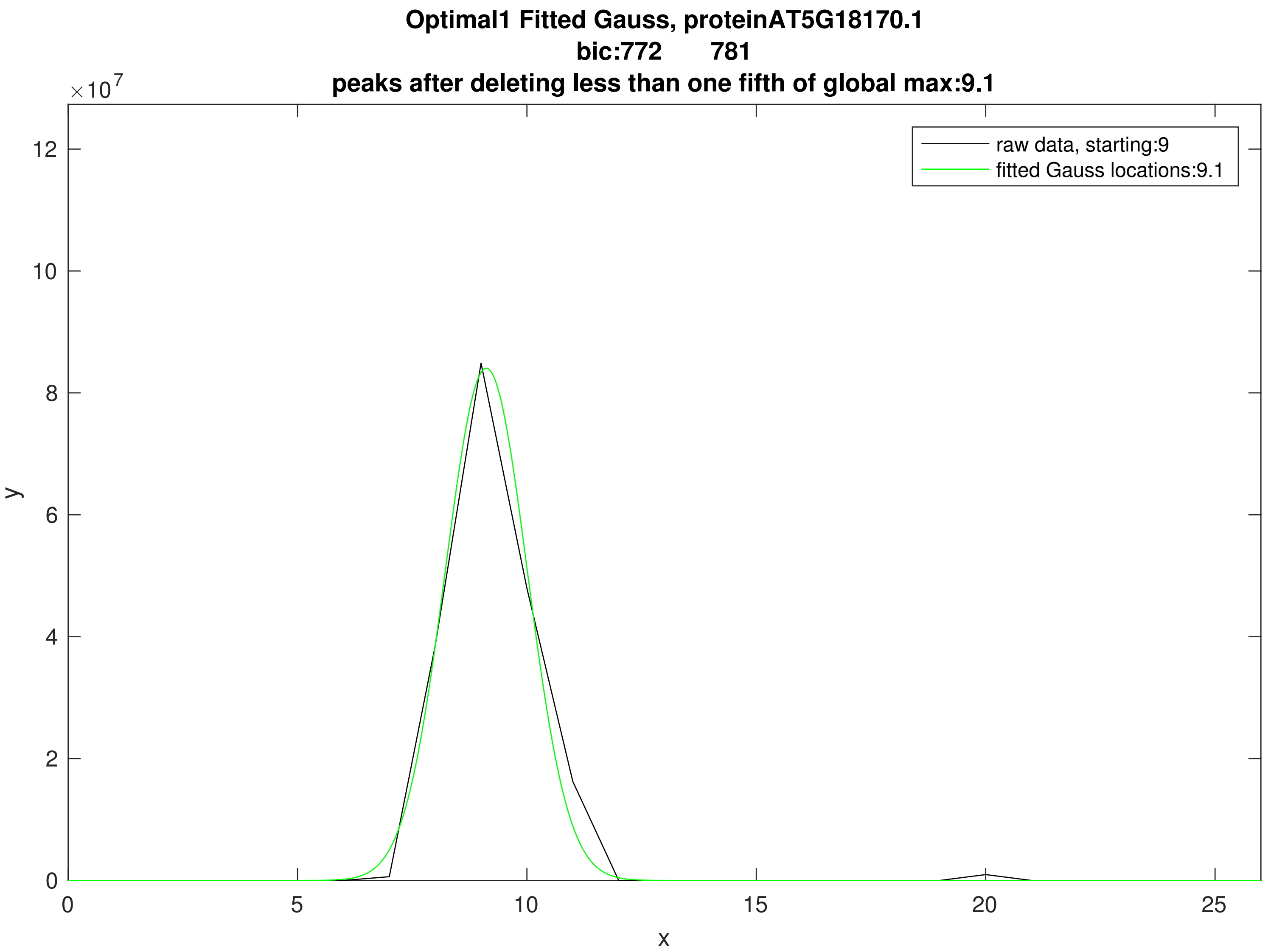

Optimal1 Fitted Gauss, proteinAT5G18170.1
bic:772 781
peaks after deleting less than one fifth of global max:9.1
×10^7
raw data, starting:9
fitted Gauss locations:9.1
y
x
0
2
4
6
8
10
12
0
5
10
15
20
25

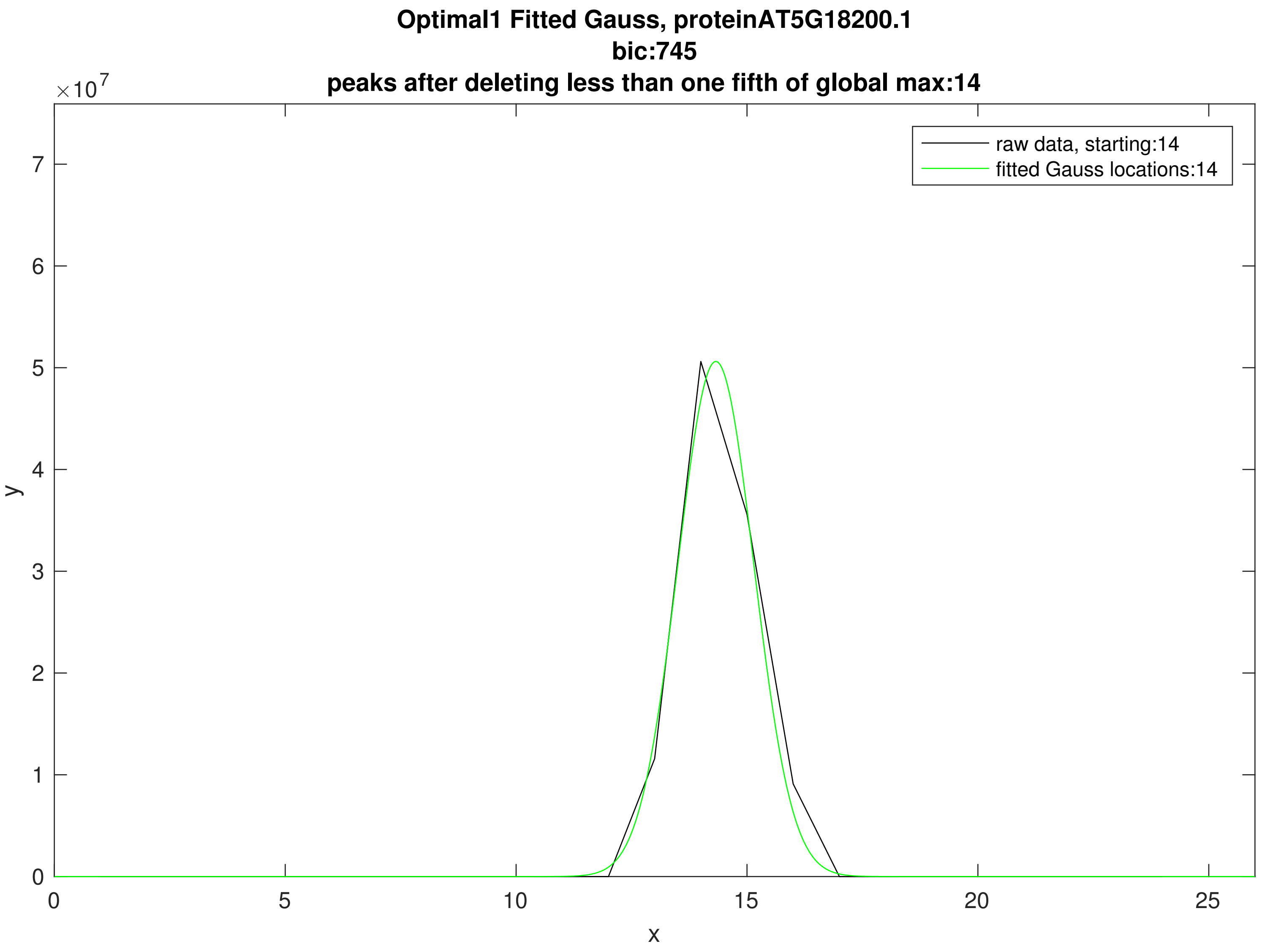

Optimal1 Fitted Gauss, proteinAT5G18200.1
bic:745
peaks after deleting less than one fifth of global max:14
×10^7
raw data, starting:14
fitted Gauss locations:14
y
x
0
1
2
3
4
5
6
7
0
5
10
15
20
25

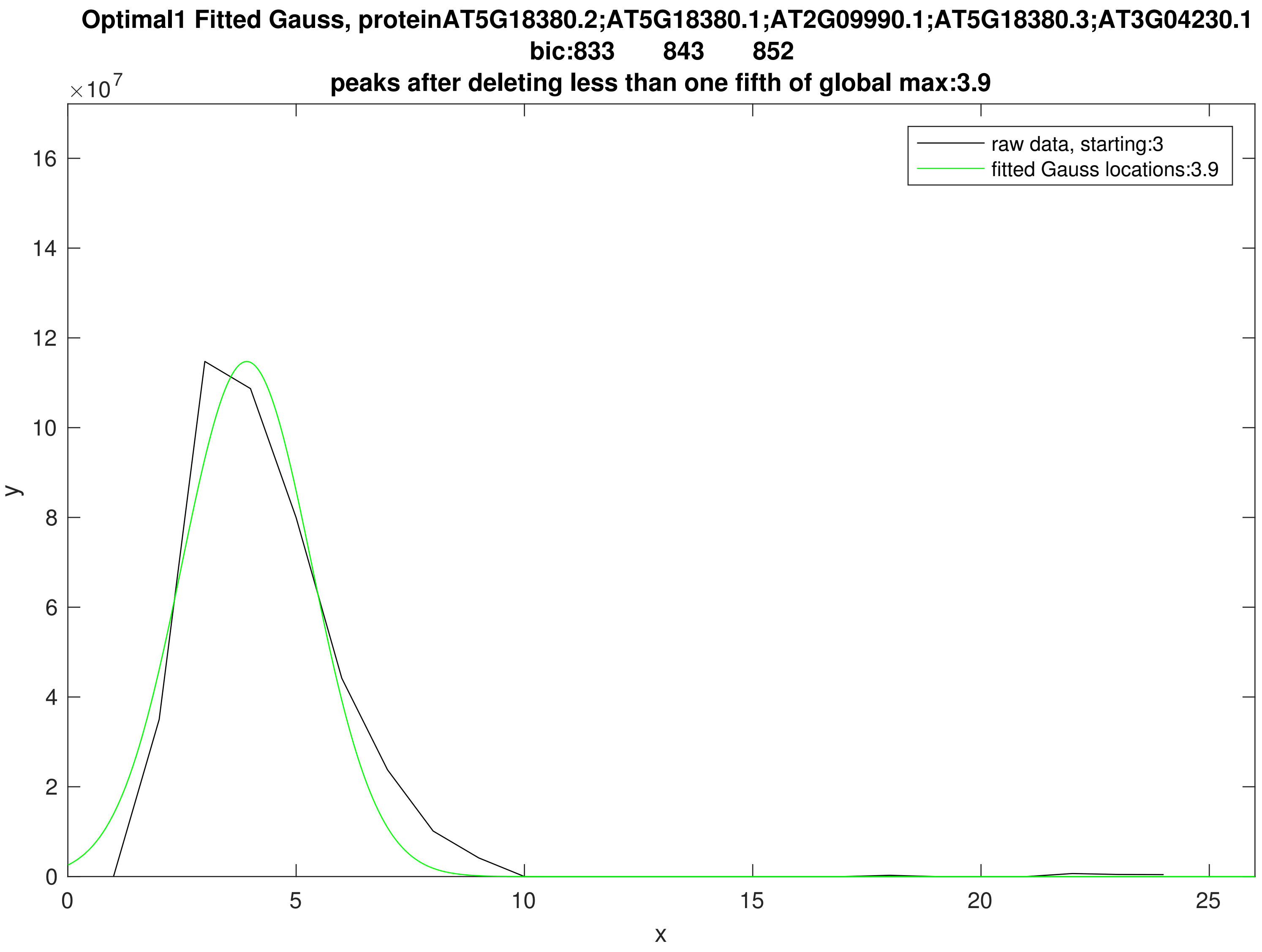
Optimal1 Fitted Gauss, proteinAT5G18380.2;AT5G18380.1;AT2G09990.1;AT5G18380.3;AT3G04230.1
bic:833 843 852
peaks after deleting less than one fifth of global max:3.9
×10^7
raw data, starting:3
fitted Gauss locations:3.9
y
x
0
2
4
6
8
10
12
14
16
5
15
20
25

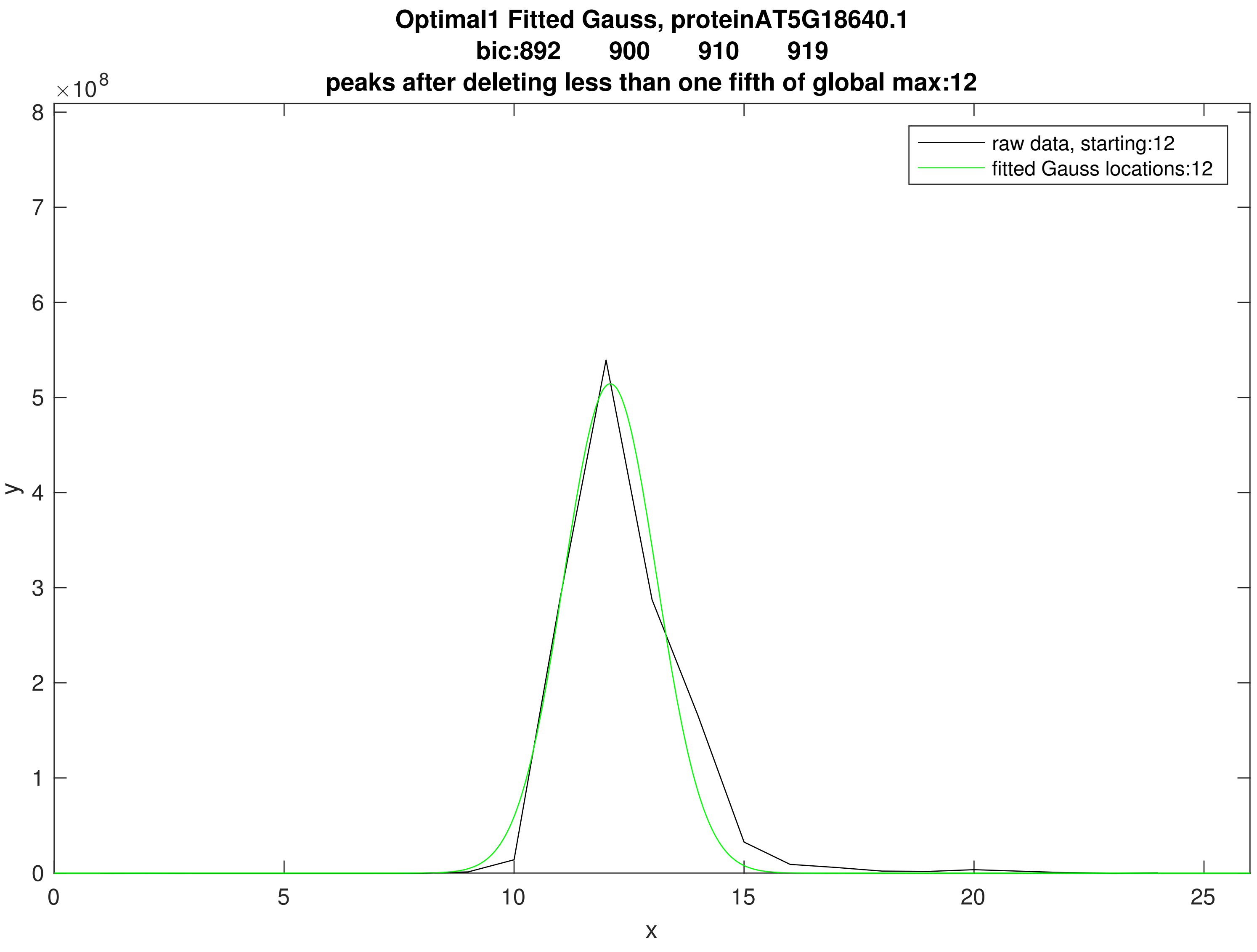
Optimal1 Fitted Gauss, proteinAT5G18640.1
bic:892 900 910 919
peaks after deleting less than one fifth of global max:12
×10^8
raw data, starting:12
fitted Gauss locations:12
y
x
0
1
2
3
4
5
6
7
8
0
5
10
15
20
25

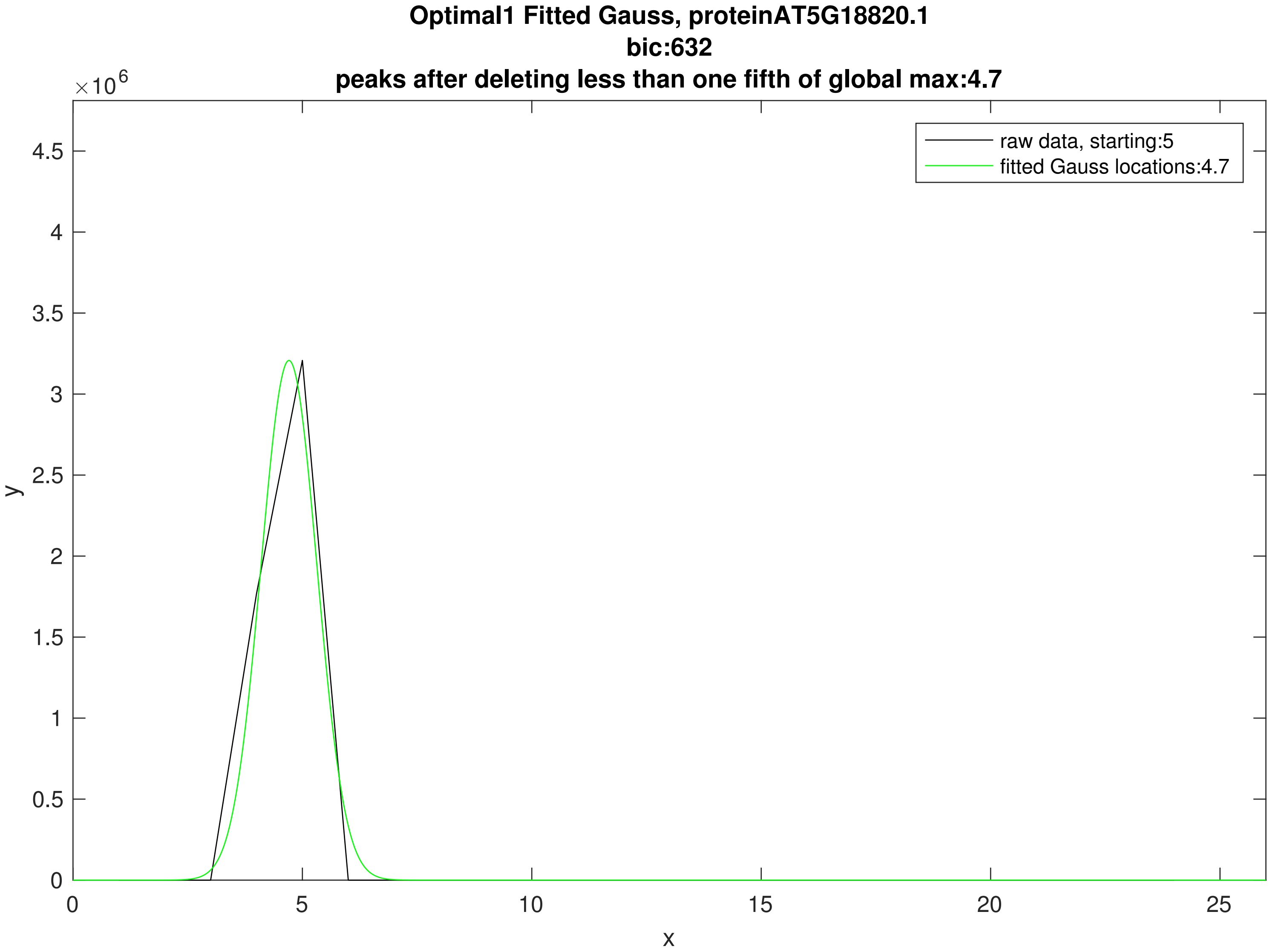
Optimal1 Fitted Gauss, proteinAT5G18820.1
bic:632
peaks after deleting less than one fifth of global max:4.7
×10^6
raw data, starting:5
fitted Gauss locations:4.7
y
x
0
0.5
1
1.5
2
2.5
3
3.5
4
4.5
5
10
15
20
25

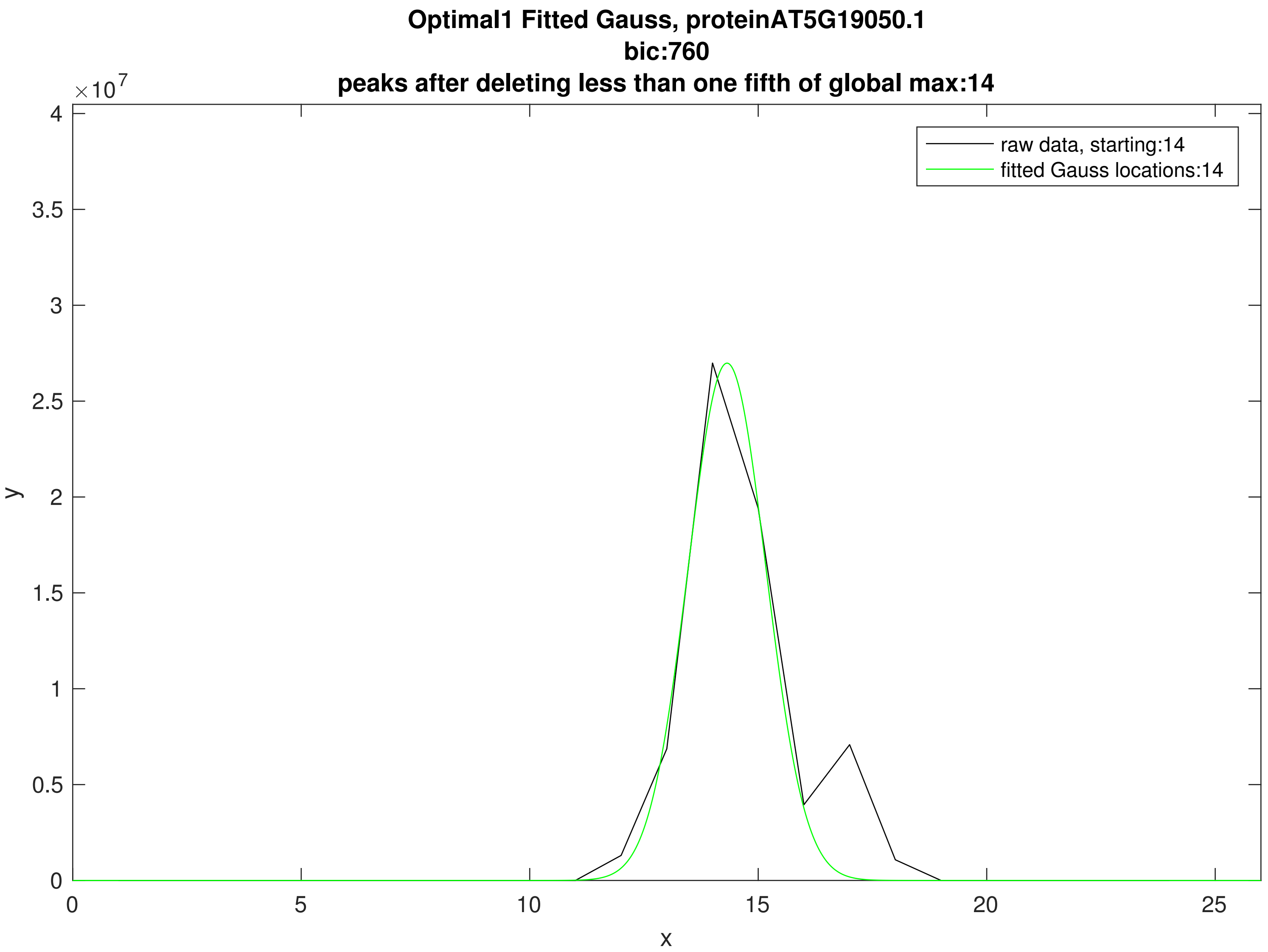
Optimal1 Fitted Gauss, proteinAT5G19050.1
bic:760
peaks after deleting less than one fifth of global max:14
×10^7
4
3.5
3
2.5
2
1.5
1
0.5
0
y
0
5
10
15
20
25
x
raw data, starting:14
fitted Gauss locations:14

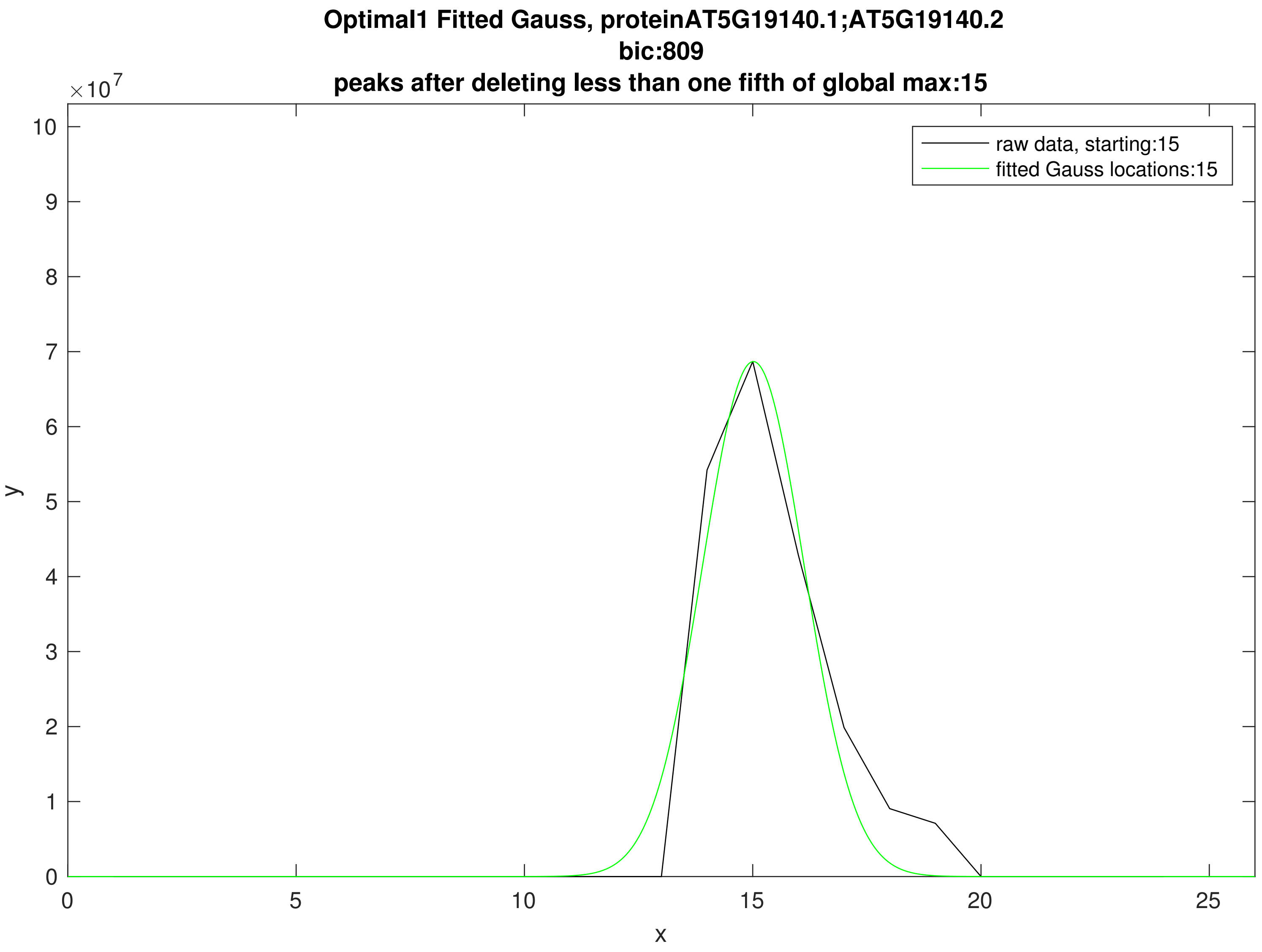

Optimal1 Fitted Gauss, proteinAT5G19140.1;AT5G19140.2
bic:809
peaks after deleting less than one fifth of global max:15
×10^7
raw data, starting:15
fitted Gauss locations:15
y
x
0
1
2
3
4
5
6
7
8
9
10
0
5
10
15
20
25

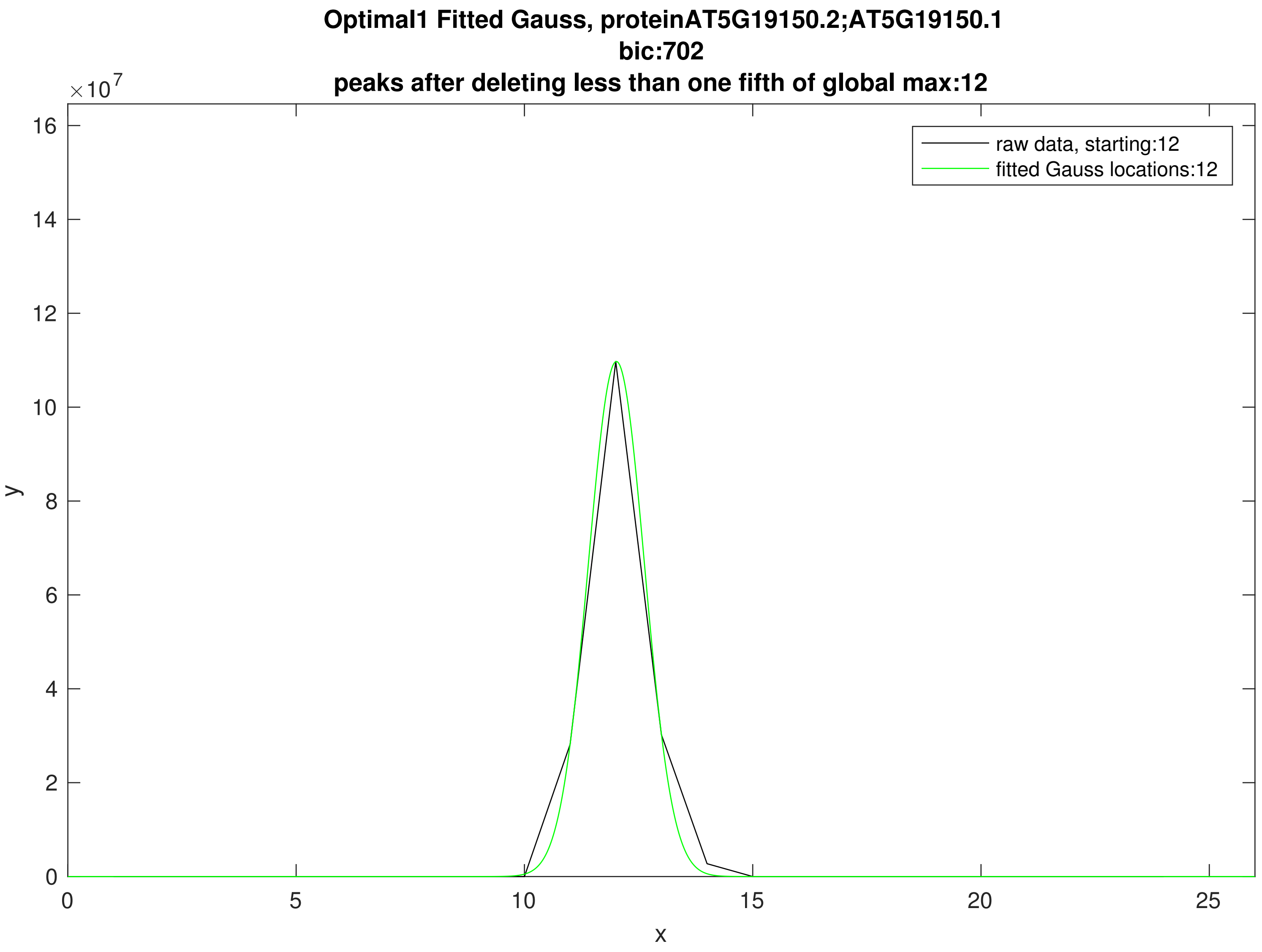
Optimal1 Fitted Gauss, proteinAT5G19150.2;AT5G19150.1
bic:702
peaks after deleting less than one fifth of global max:12
×10^7
raw data, starting:12
fitted Gauss locations:12
y
x
0
2
4
6
8
10
12
14
16
0
5
10
15
20
25

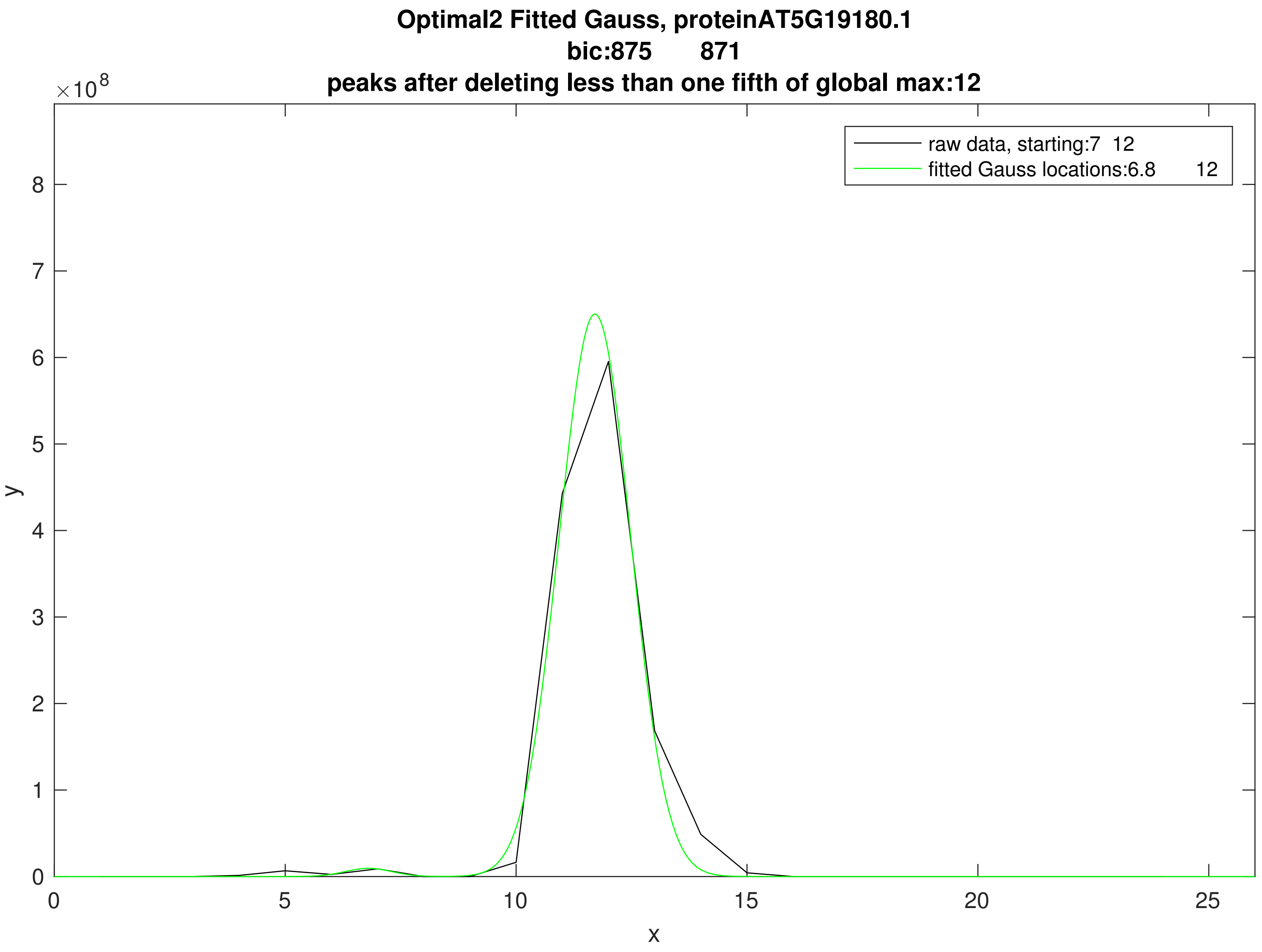
Optimal2 Fitted Gauss, proteinAT5G19180.1
bic:875 871
peaks after deleting less than one fifth of global max:12
×10^8
raw data, starting:7 12
fitted Gauss locations:6.8 12
y
x
0
1
2
3
4
5
6
7
8
0
5
10
15
20
25

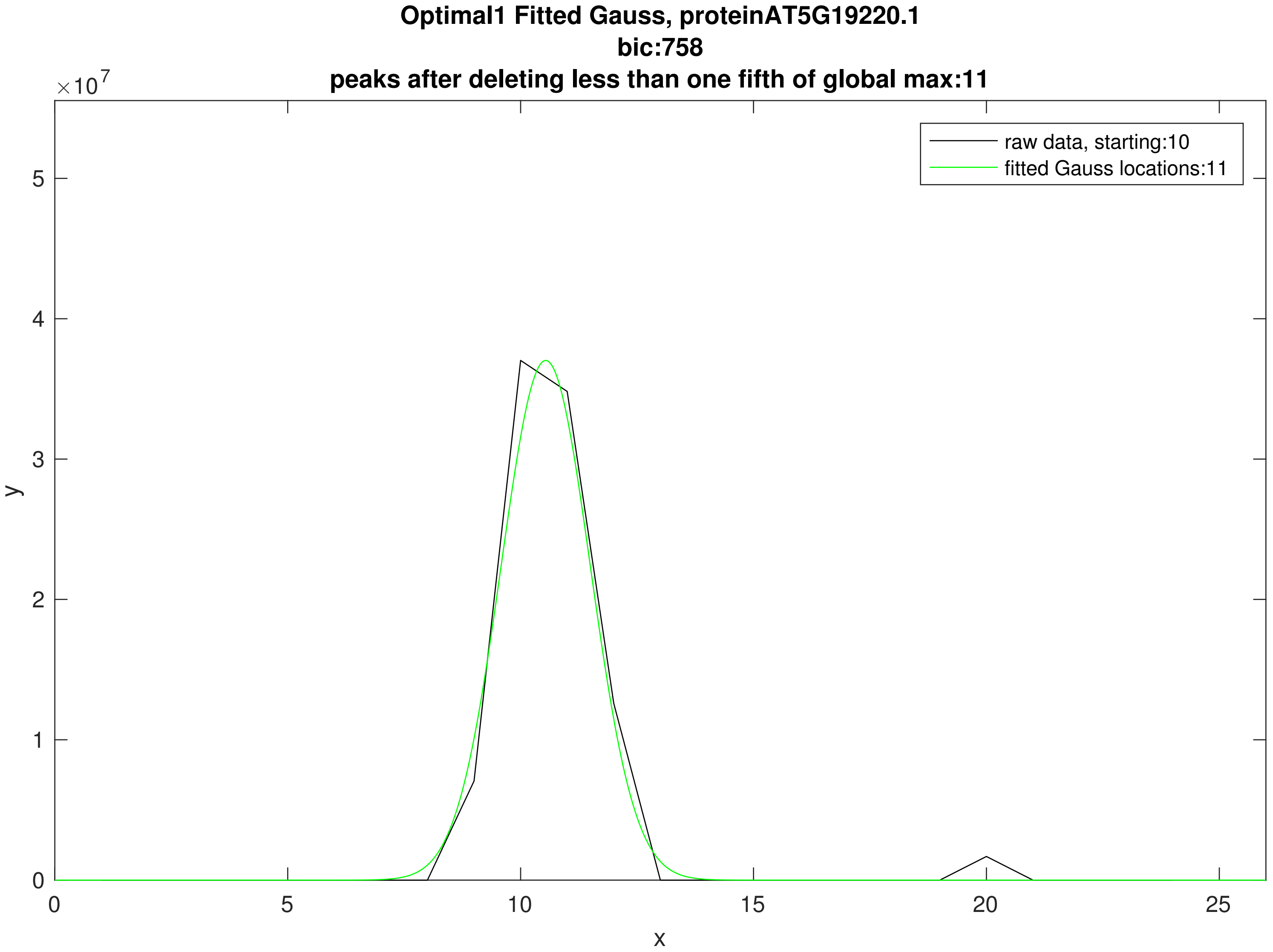
Optimal1 Fitted Gauss, proteinAT5G19220.1
bic:758
peaks after deleting less than one fifth of global max:11
×10^7
raw data, starting:10
fitted Gauss locations:11
y
x
0
1
2
3
4
5
0
5
10
15
20
25

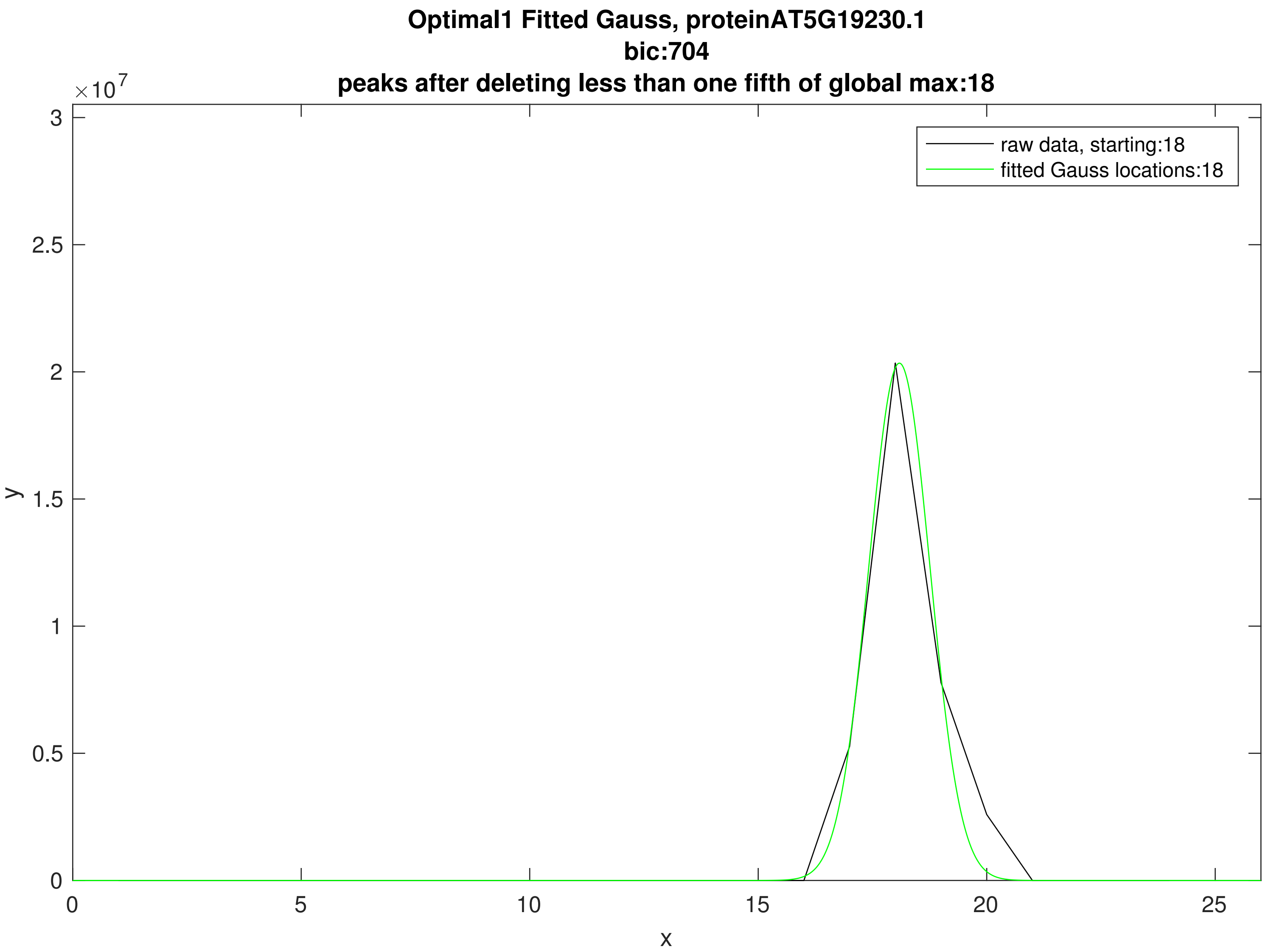

Optimal1 Fitted Gauss, proteinAT5G19230.1
bic:704
peaks after deleting less than one fifth of global max:18
×10^7
raw data, starting:18
fitted Gauss locations:18
3
2.5
2
1.5
1
0.5
0
y
0
5
10
15
20
25
x

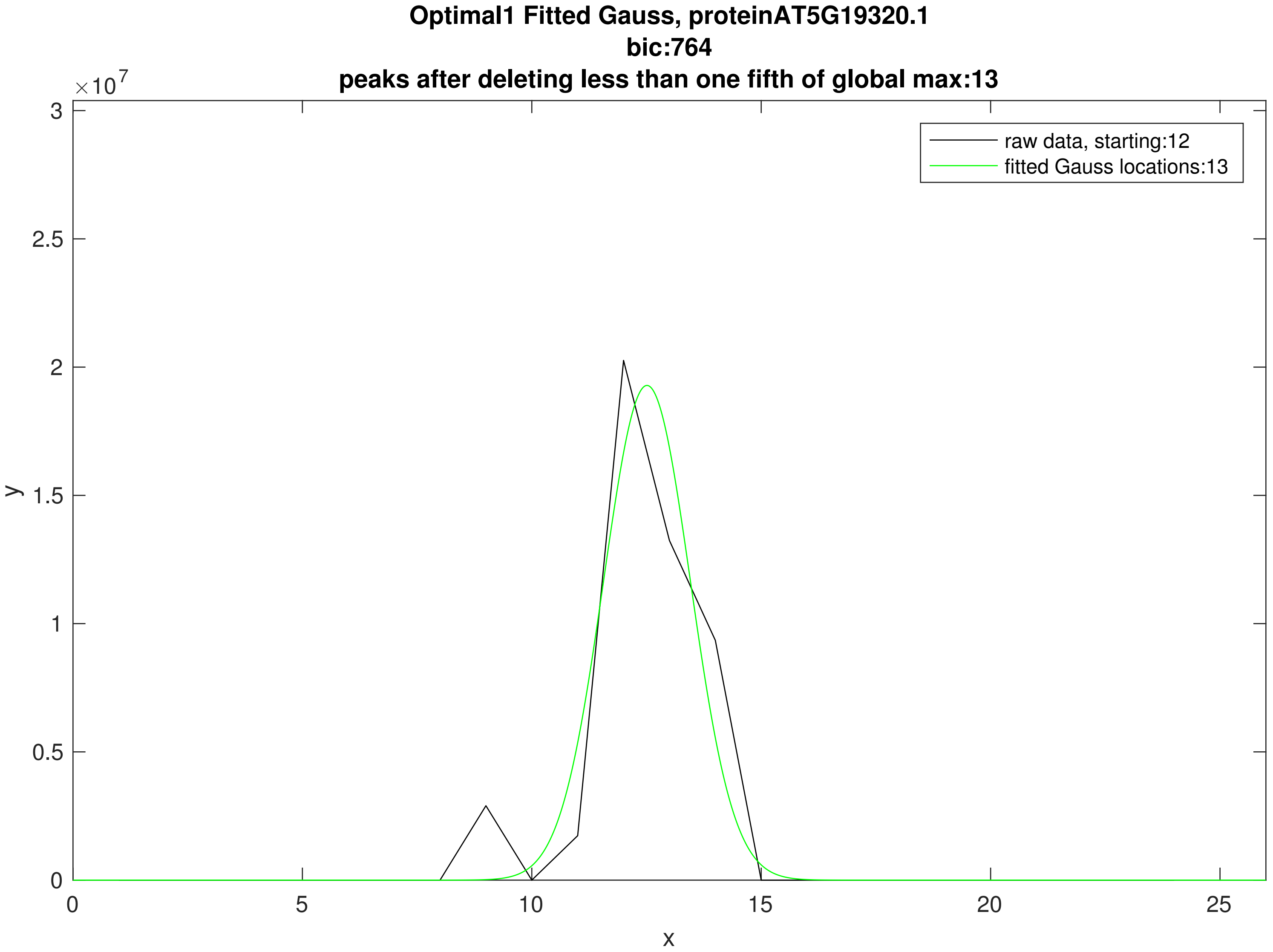

Optimal1 Fitted Gauss, proteinAT5G19320.1
bic:764
peaks after deleting less than one fifth of global max:13
×10^7
3
2.5
2
1.5
1
0.5
0
y
0
5
10
15
20
25
x
raw data, starting:12
fitted Gauss locations:13

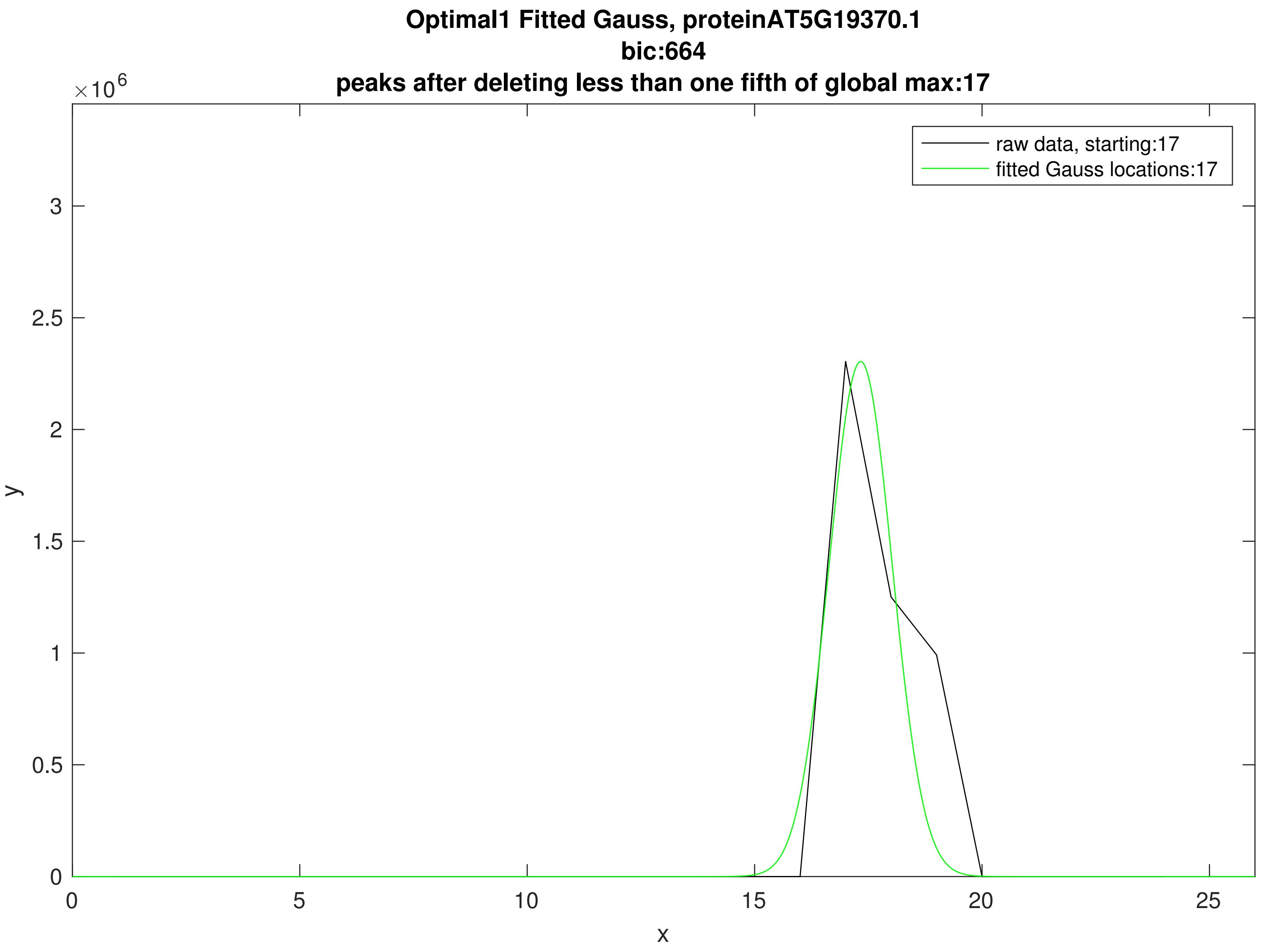
Optimal1 Fitted Gauss, proteinAT5G19370.1
bic:664
peaks after deleting less than one fifth of global max:17
×10^6
raw data, starting:17
fitted Gauss locations:17
y
x
0
0.5
1
1.5
2
2.5
3
0
5
10
15
20
25

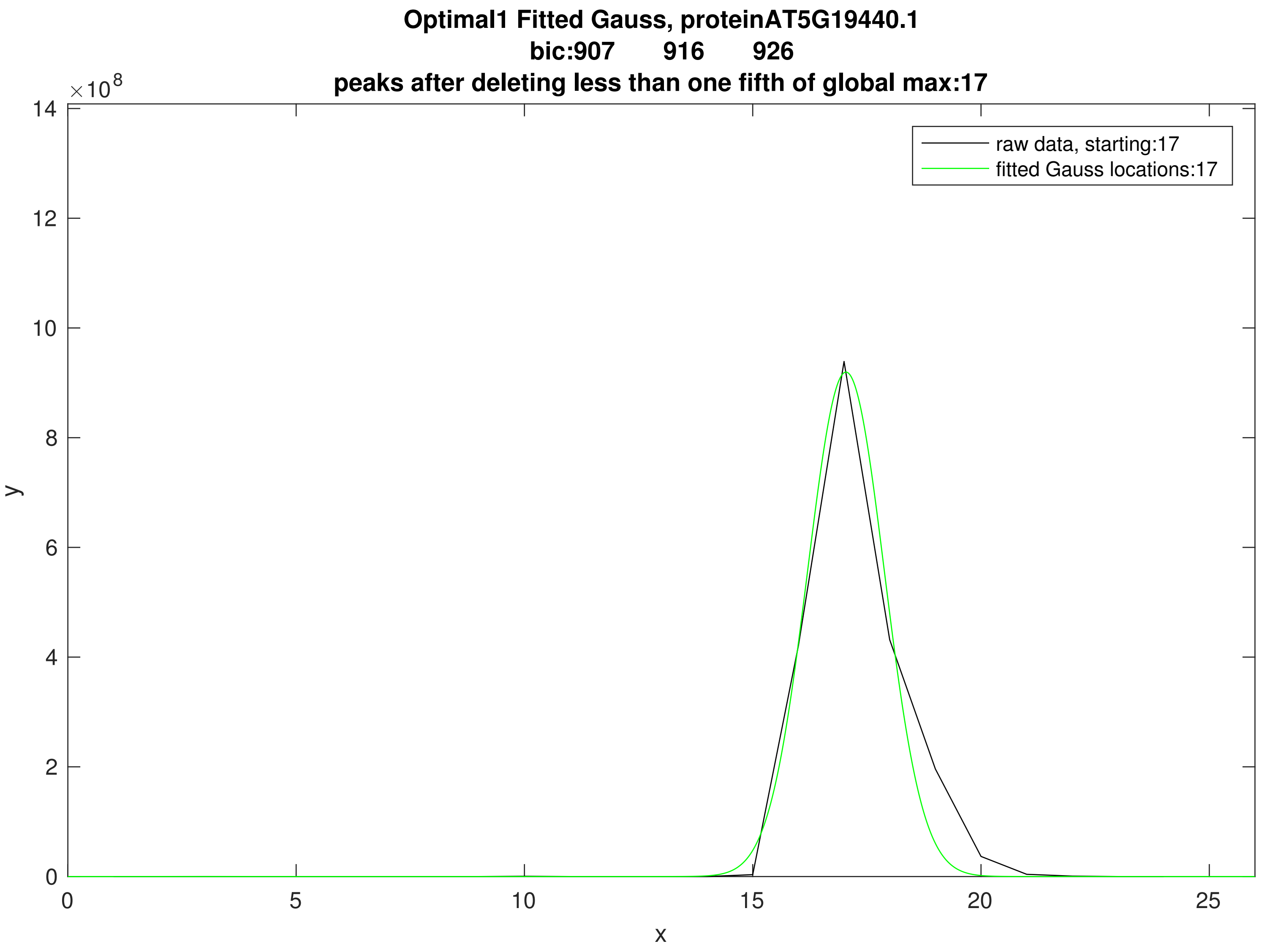

Optimal1 Fitted Gauss, proteinAT5G19440.1
bic:907 916 926
peaks after deleting less than one fifth of global max:17
×10^8
14
12
10
8
6
4
2
0
y
0
5
10
15
20
25
x
raw data, starting:17
fitted Gauss locations:17

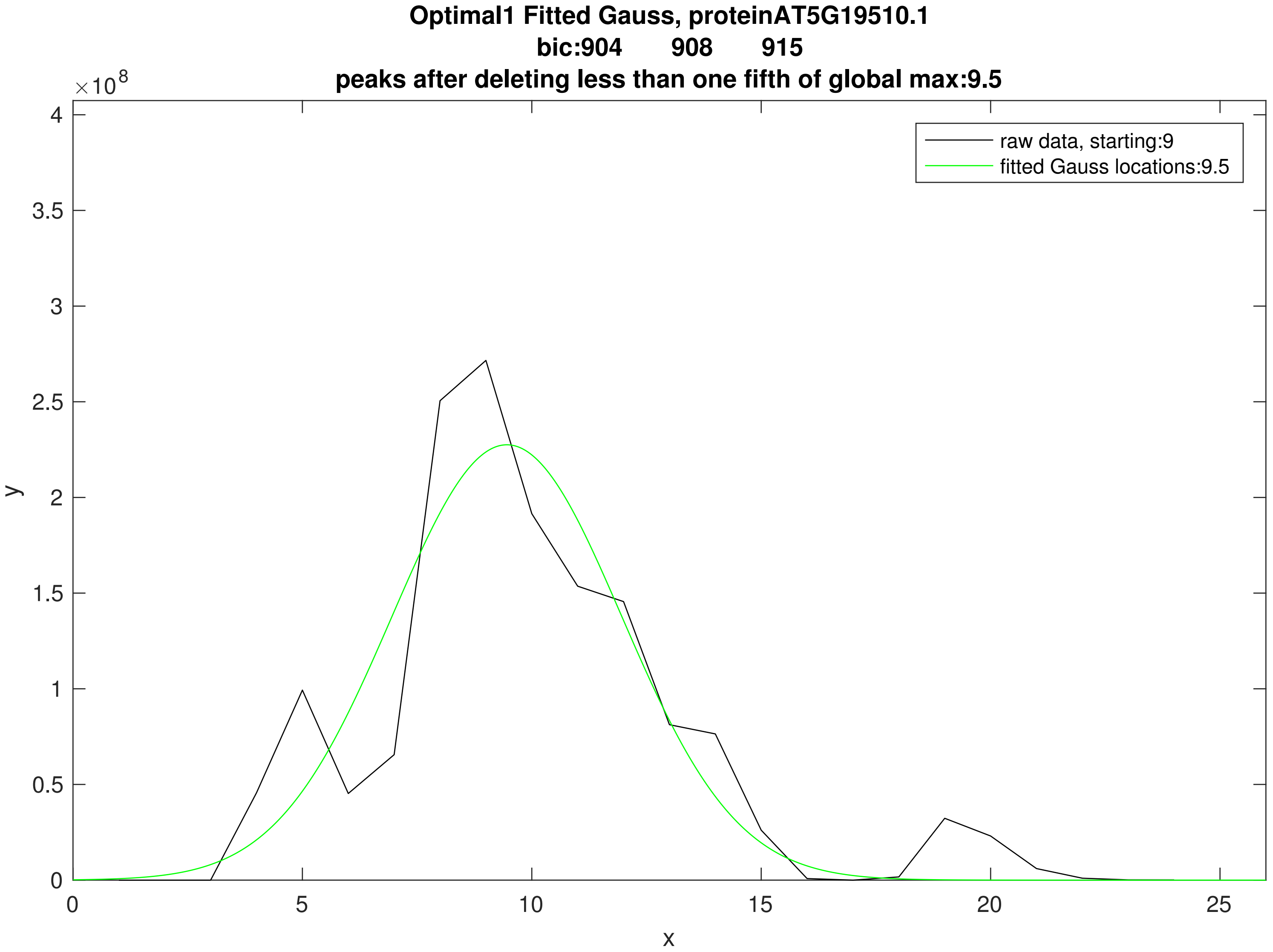
Optimal1 Fitted Gauss, proteinAT5G19510.1
bic:904 908 915
peaks after deleting less than one fifth of global max:9.5
×10^8
4
3.5
3
2.5
2
1.5
1
0.5
0
y
0
5
10
15
20
25
x
raw data, starting:9
fitted Gauss locations:9.5

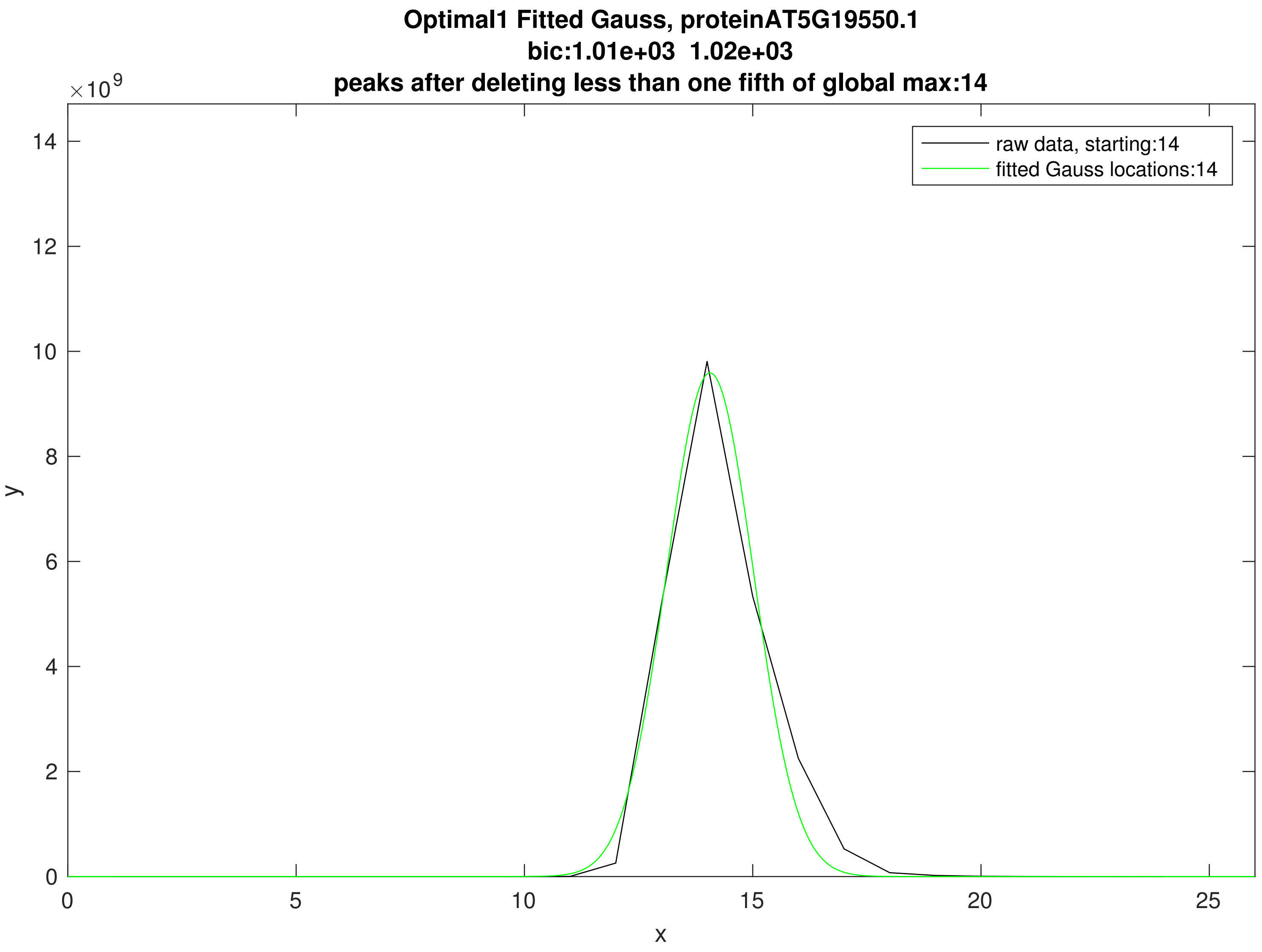

Optimal1 Fitted Gauss, proteinAT5G19550.1
bic:1.01e+03  1.02e+03
peaks after deleting less than one fifth of global max:14
×10^9
14
12
10
8
6
4
2
0
y
0
5
10
15
20
25
x
raw data, starting:14
fitted Gauss locations:14

**Optimal1 Fitted Gauss, proteinAT5G19680.1**
**bic:773**
**peaks after deleting less than one fifth of global max:15**

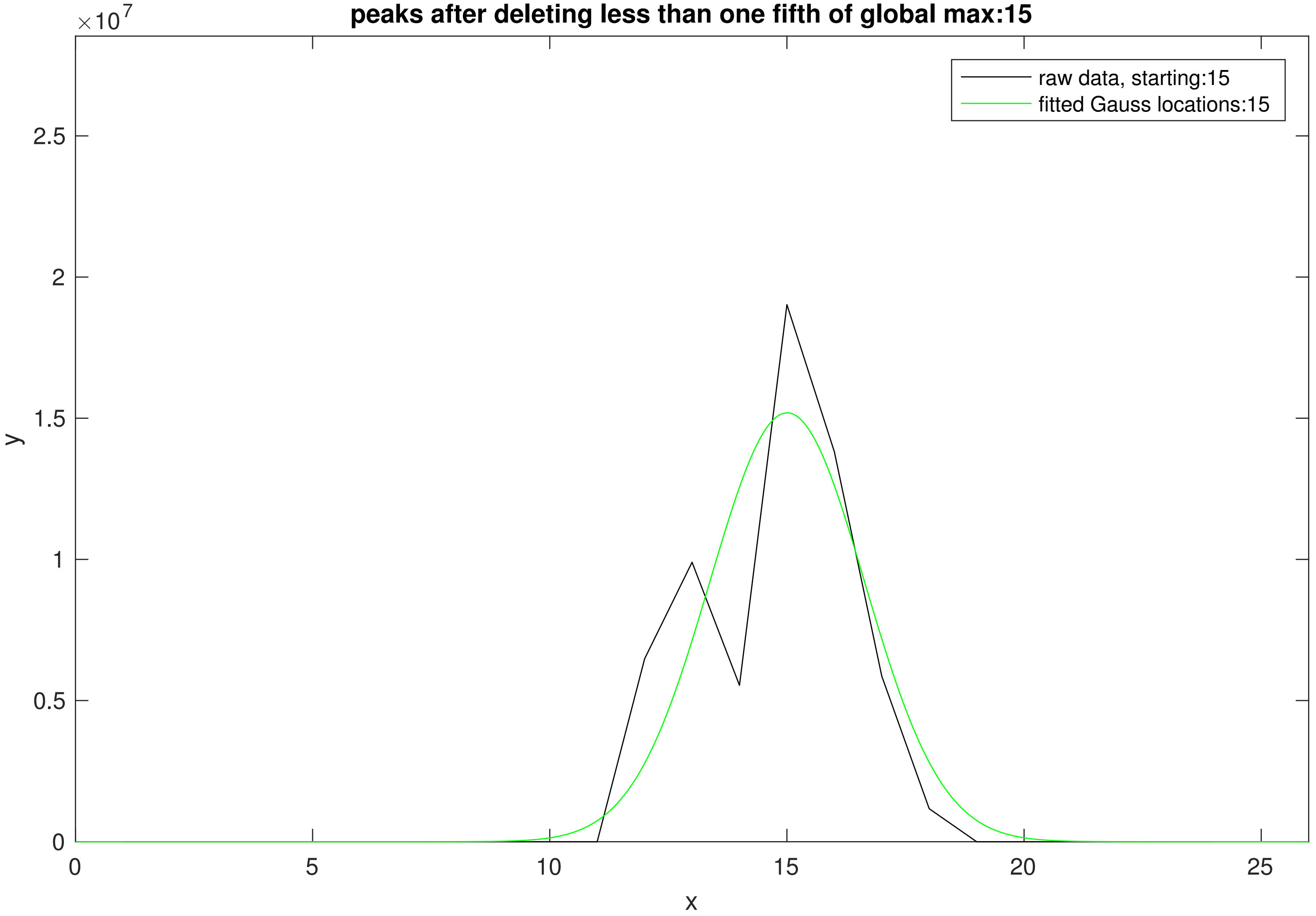

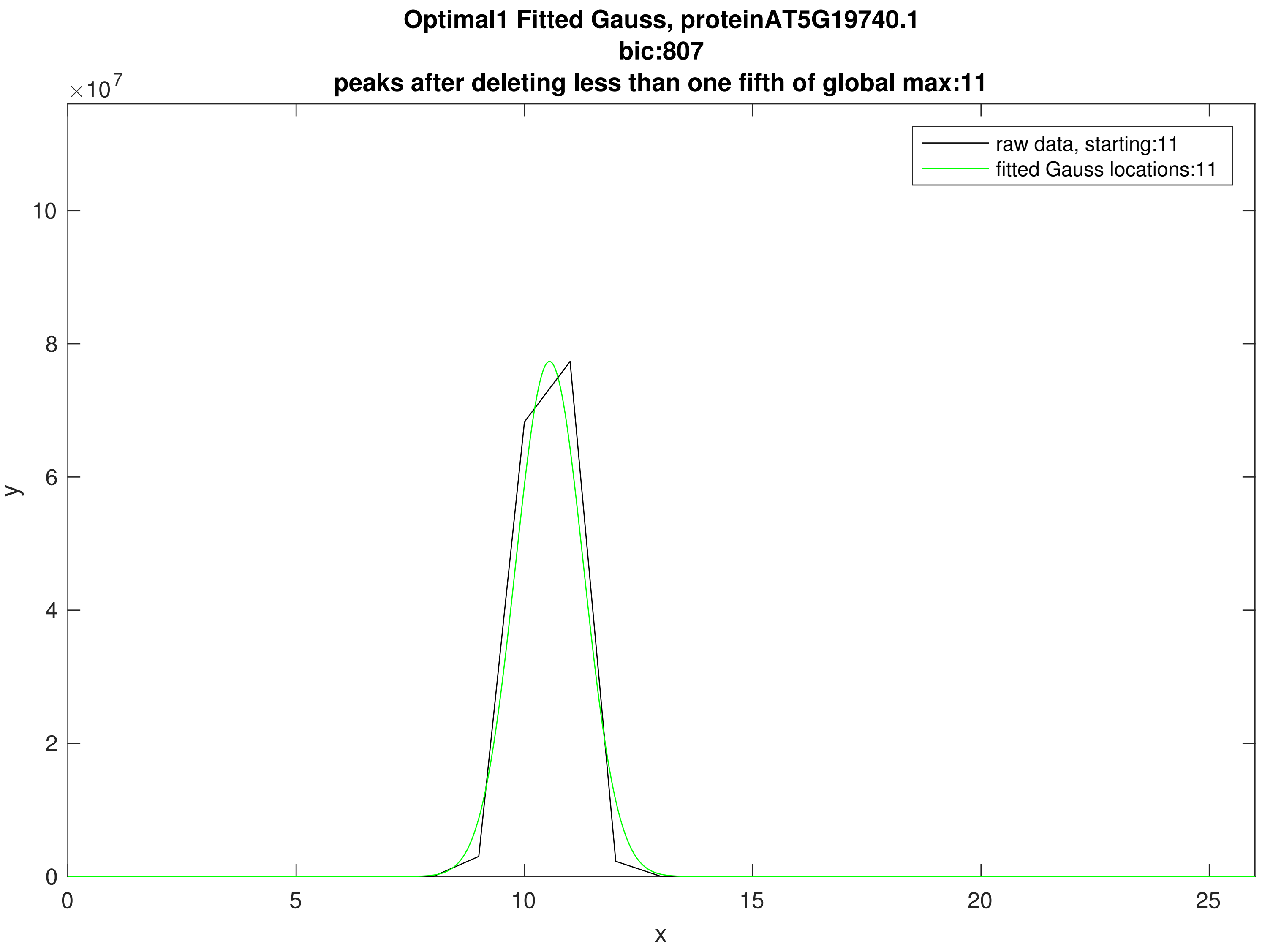

Optimal1 Fitted Gauss, proteinAT5G19740.1
bic:807
peaks after deleting less than one fifth of global max:11
×10^7
raw data, starting:11
fitted Gauss locations:11
y
x
0
2
4
6
8
10
0
5
10
15
20
25

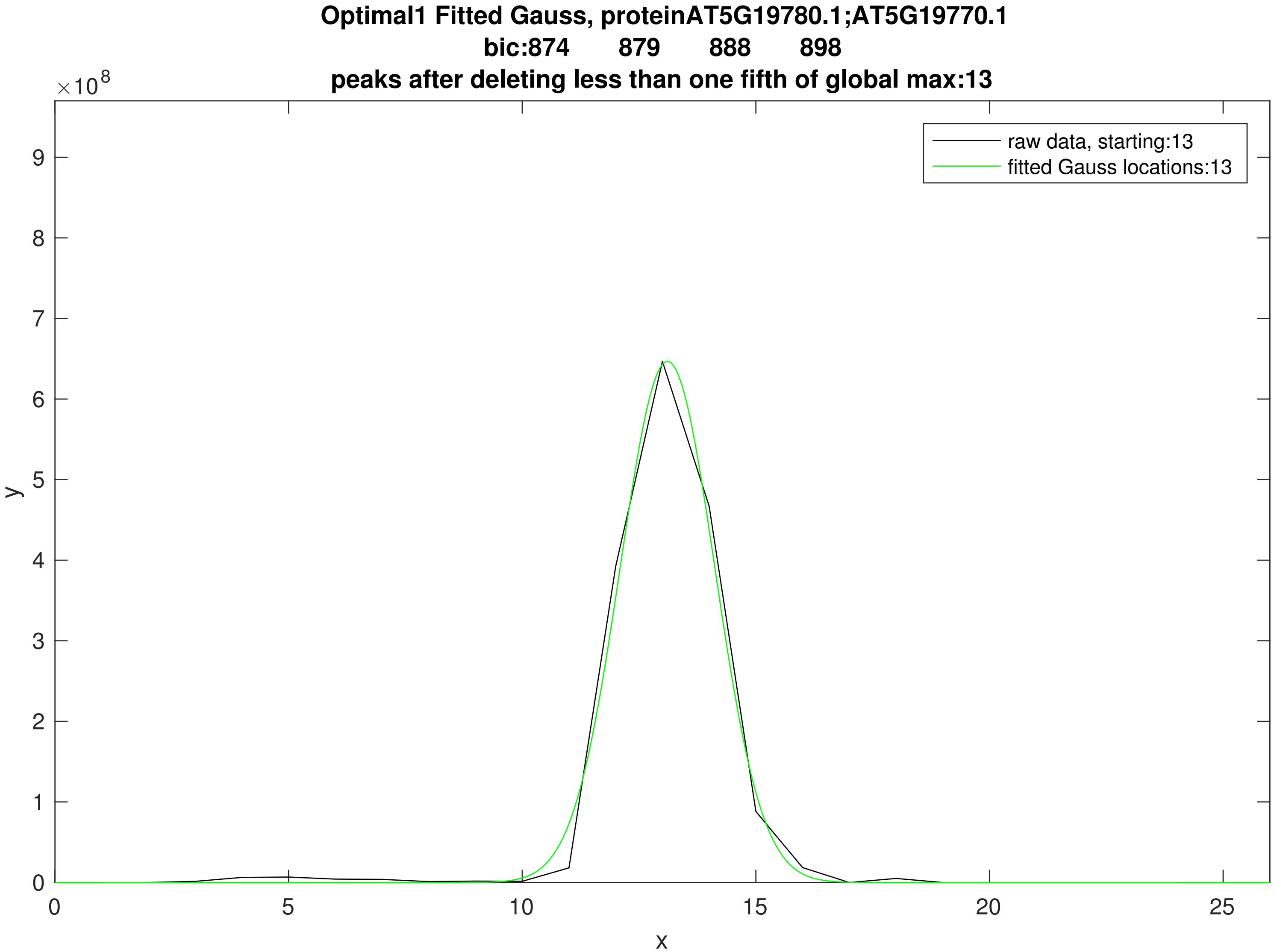
Optimal1 Fitted Gauss, proteinAT5G19780.1;AT5G19770.1
bic:874 879 888 898
peaks after deleting less than one fifth of global max:13
×10^8
raw data, starting:13
fitted Gauss locations:13
y
x
0
1
2
3
4
5
6
7
8
9
0
5
10
15
20
25

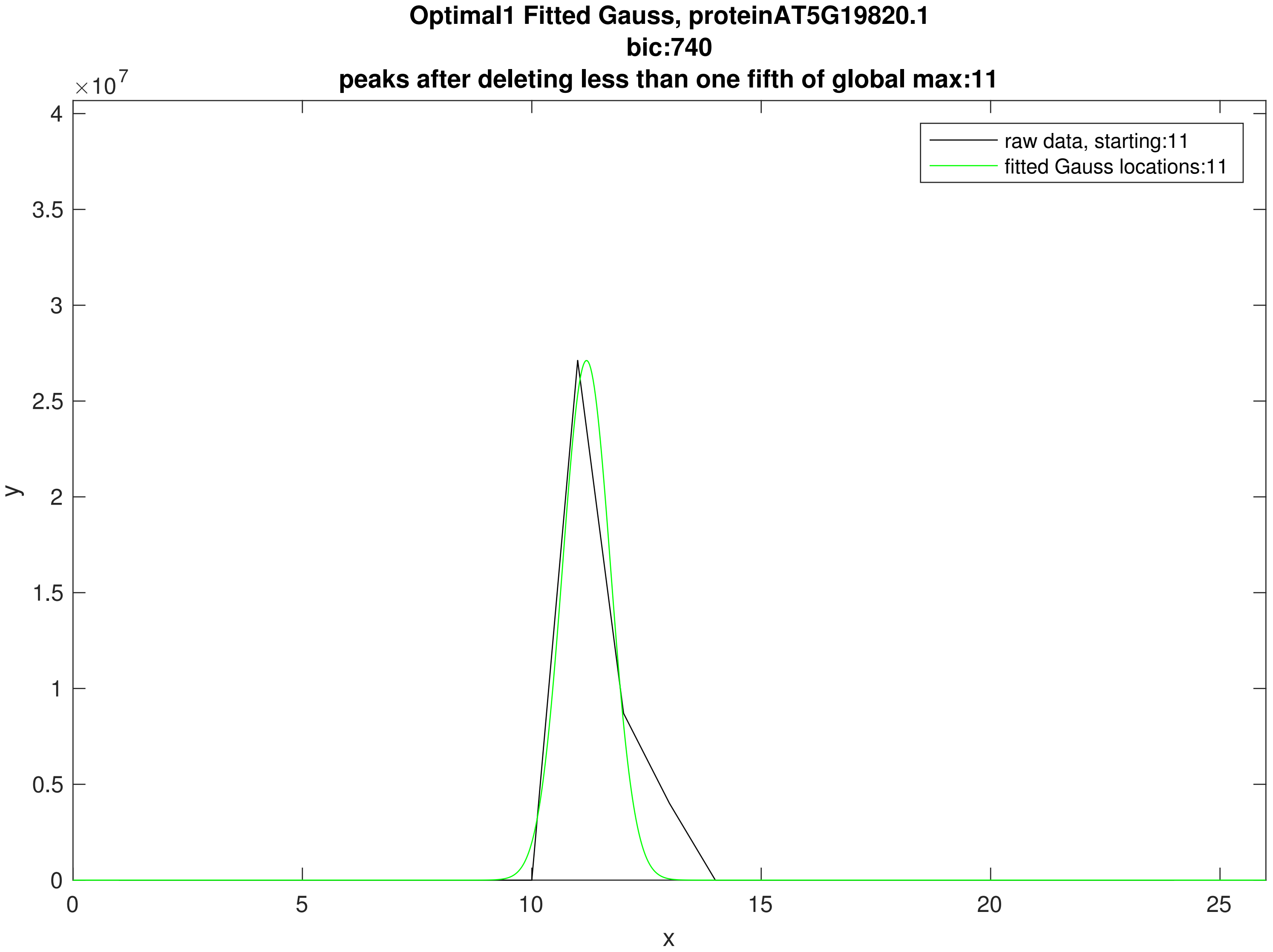
Optimal1 Fitted Gauss, proteinAT5G19820.1
bic:740
peaks after deleting less than one fifth of global max:11
×10^7
raw data, starting:11
fitted Gauss locations:11
y
x
0
0.5
1
1.5
2
2.5
3
3.5
4
5
10
15
20
25

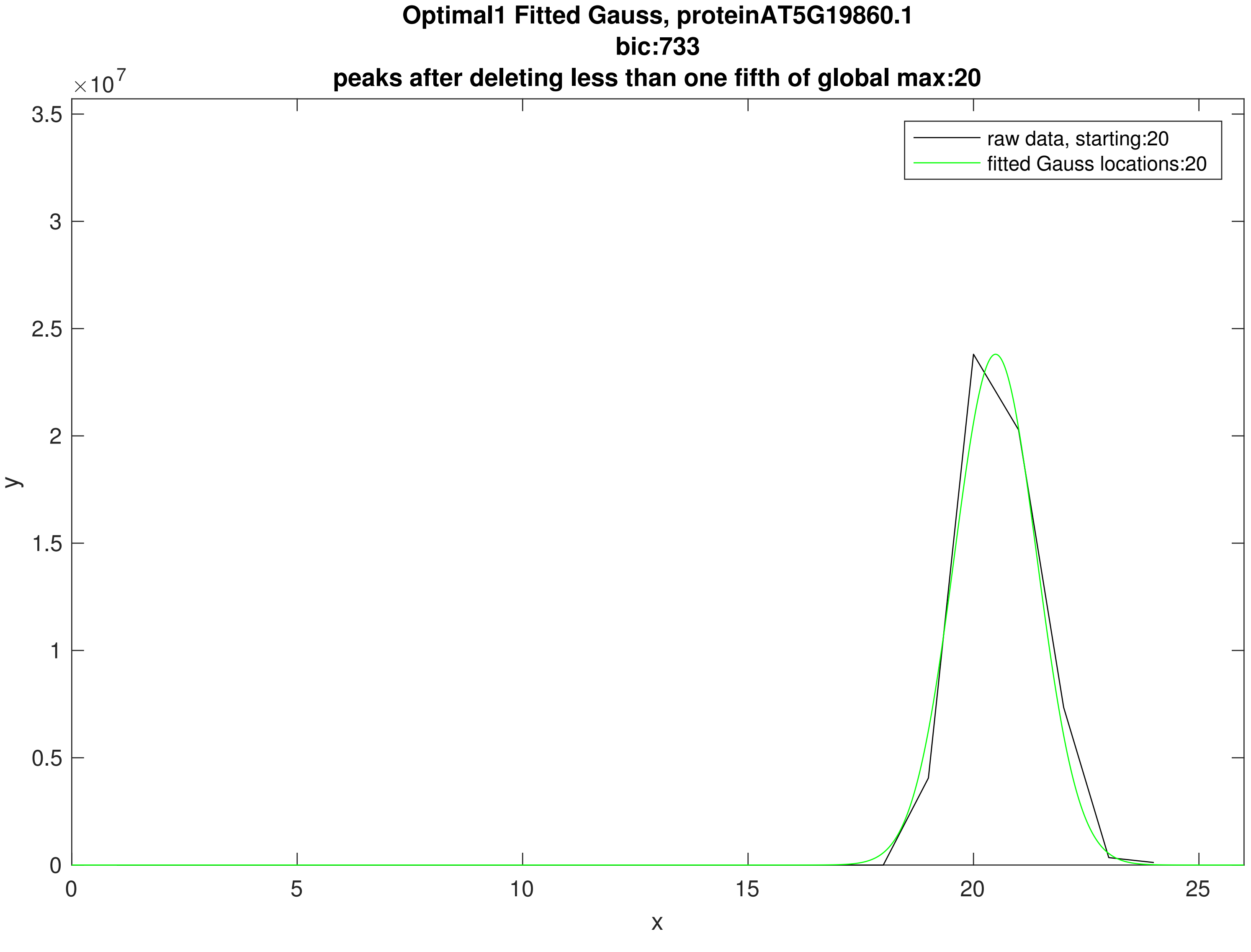
Optimal1 Fitted Gauss, proteinAT5G19860.1
bic:733
peaks after deleting less than one fifth of global max:20
×10^7
raw data, starting:20
fitted Gauss locations:20
y
x
0
0.5
1
1.5
2
2.5
3
3.5
0
5
10
15
20
25

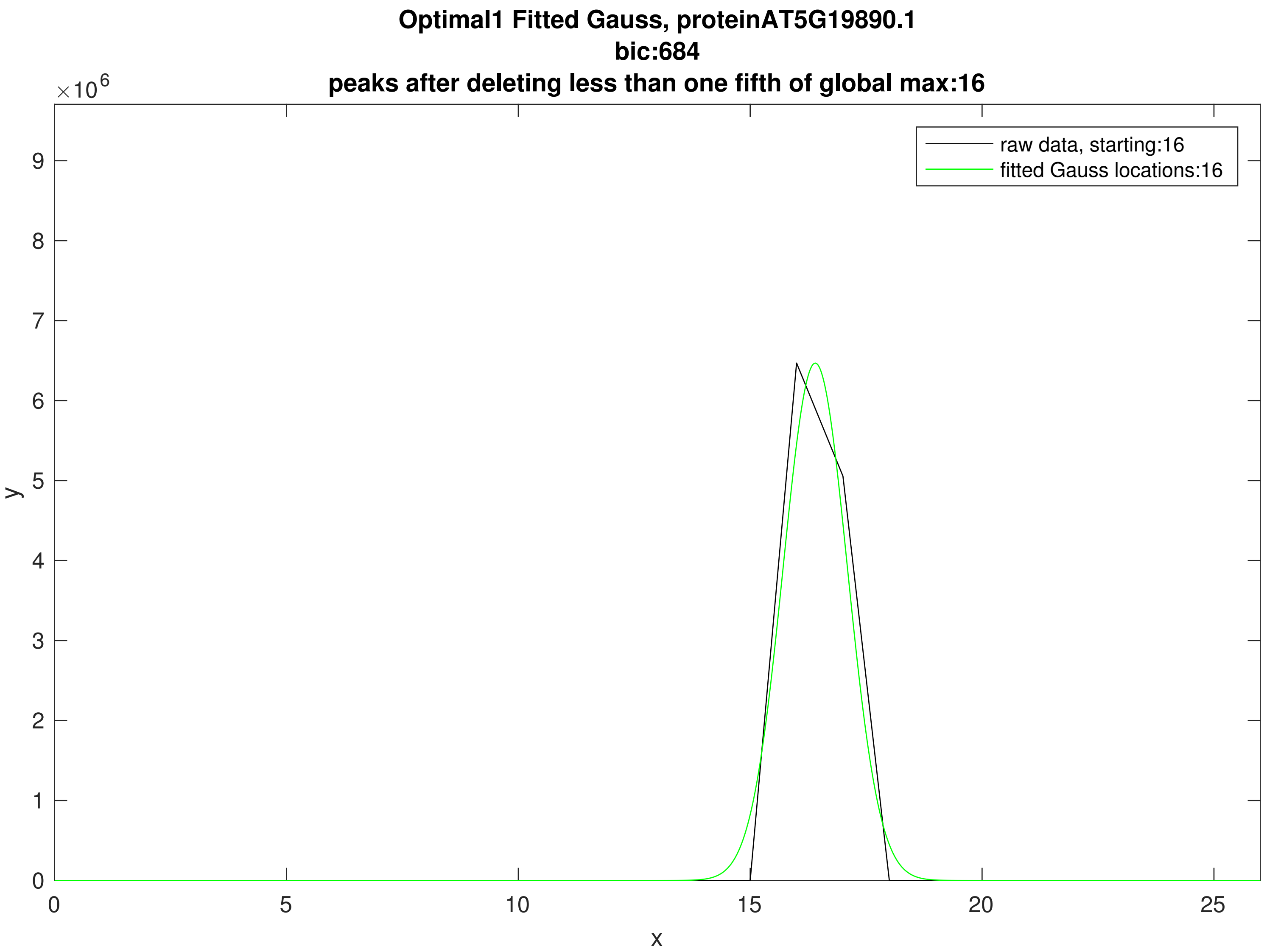
Optimal1 Fitted Gauss, proteinAT5G19890.1
bic:684
peaks after deleting less than one fifth of global max:16
×10^6
raw data, starting:16
fitted Gauss locations:16
y
x
0
1
2
3
4
5
6
7
8
9
0
5
10
15
20
25

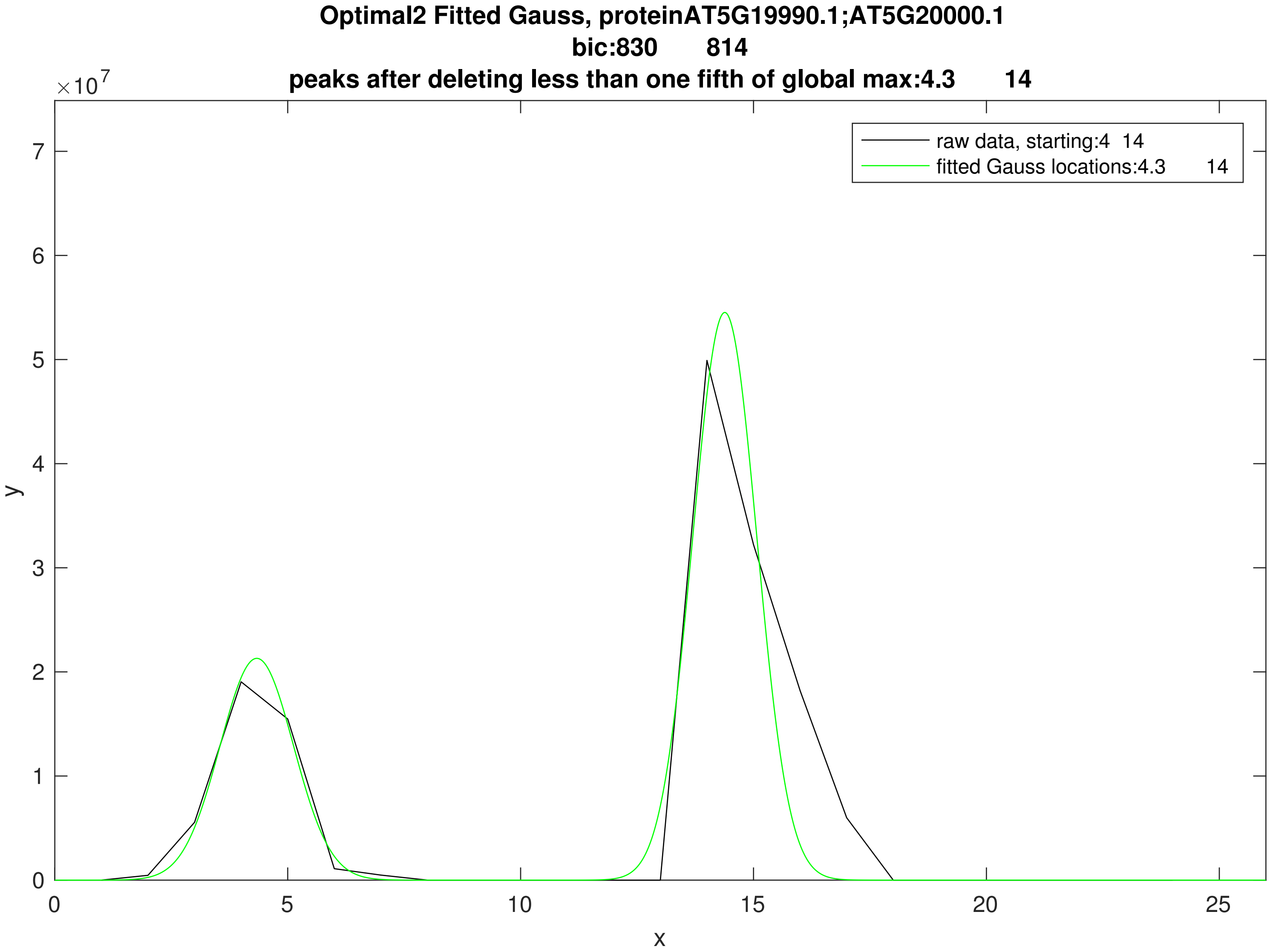

Optimal2 Fitted Gauss, proteinAT5G19990.1;AT5G20000.1
bic:830 814
peaks after deleting less than one fifth of global max:4.3 14
×10^7
raw data, starting:4 14
fitted Gauss locations:4.3 14
y
x
0
1
2
3
4
5
6
7
10
15
20
25

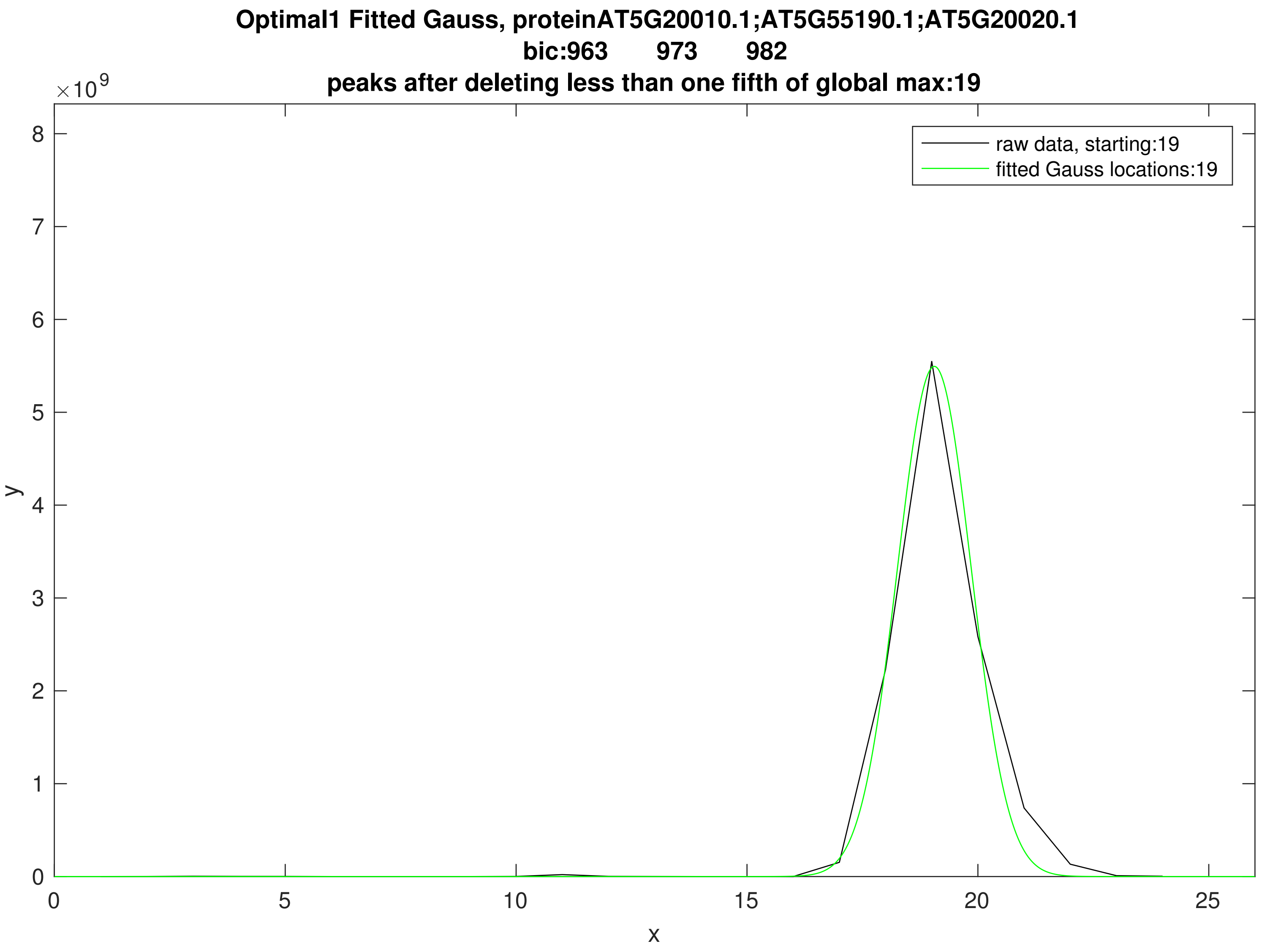
Optimal1 Fitted Gauss, proteinAT5G20010.1;AT5G55190.1;AT5G20020.1
bic:963 973 982
peaks after deleting less than one fifth of global max:19
×10^9
raw data, starting:19
fitted Gauss locations:19
y
x
0
1
2
3
4
5
6
7
8
0
5
10
15
20
25

Optimal1 Fitted Gauss, proteinAT5G20060.3;AT5G20060.2;AT5G20060.1
bic:805
peaks after deleting less than one fifth of global max:19

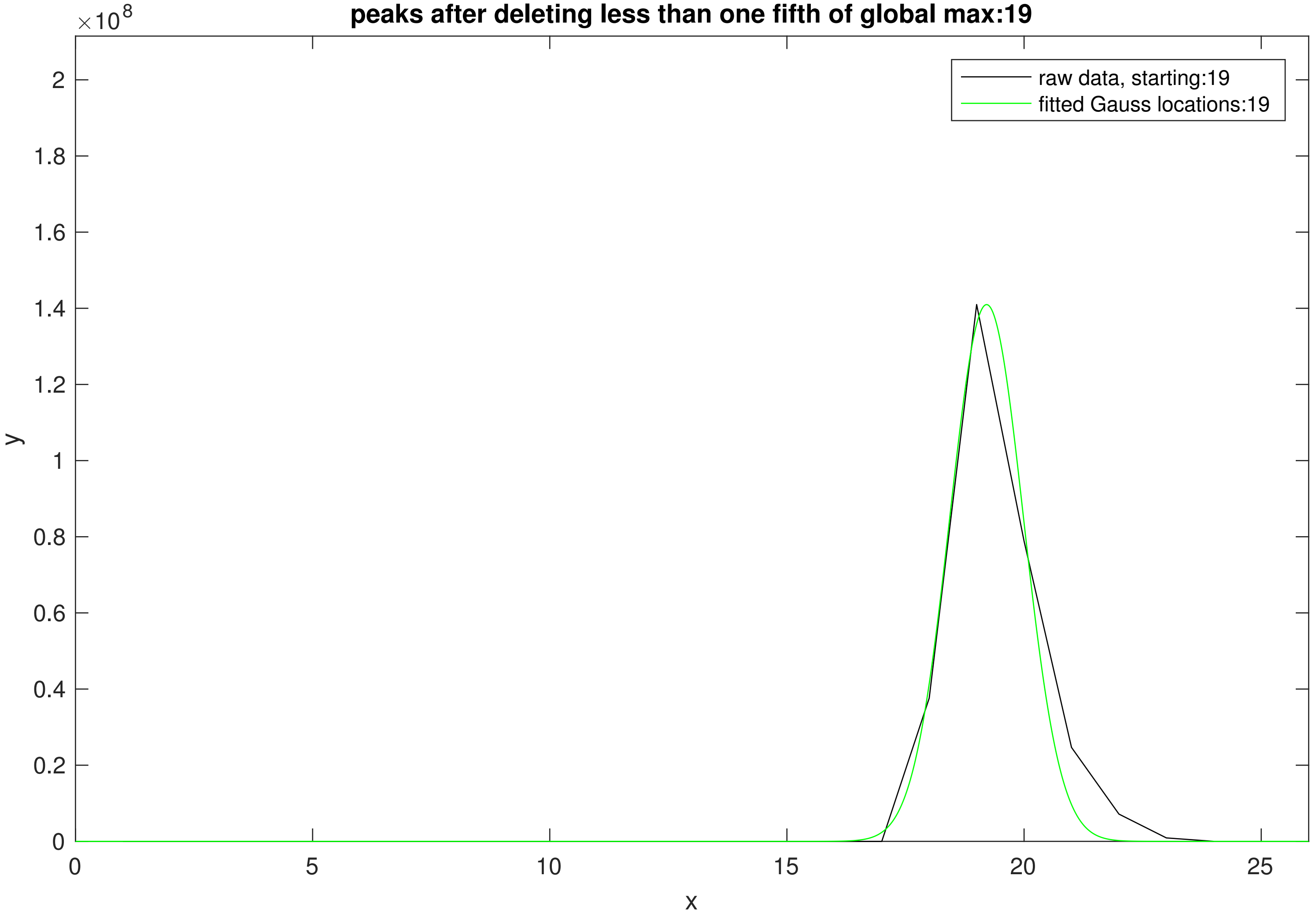

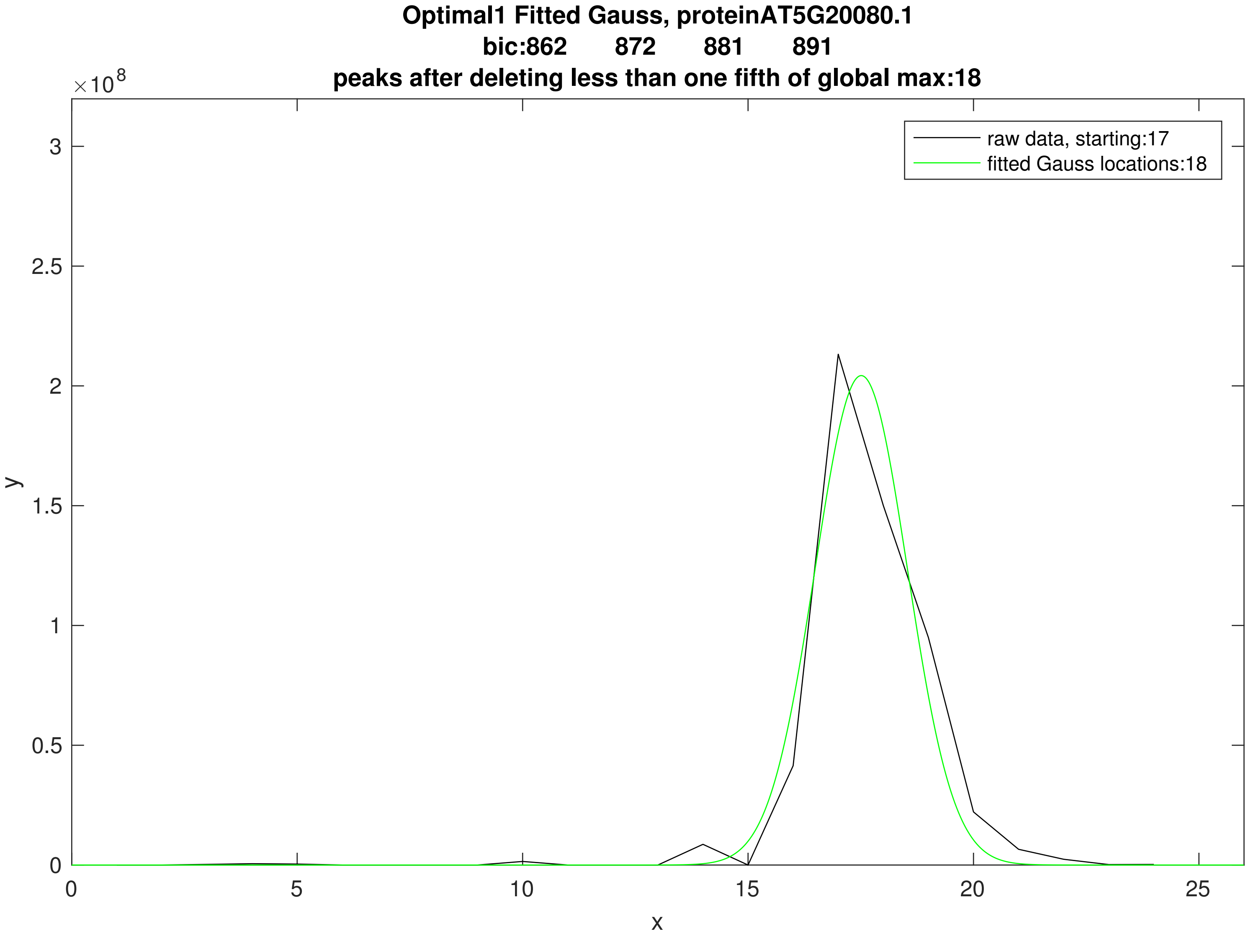

Optimal1 Fitted Gauss, proteinAT5G20080.1
bic:862 872 881 891
peaks after deleting less than one fifth of global max:18
×10^8
raw data, starting:17
fitted Gauss locations:18
y
x
0
0.5
1
1.5
2
2.5
3
0
5
10
15
20
25

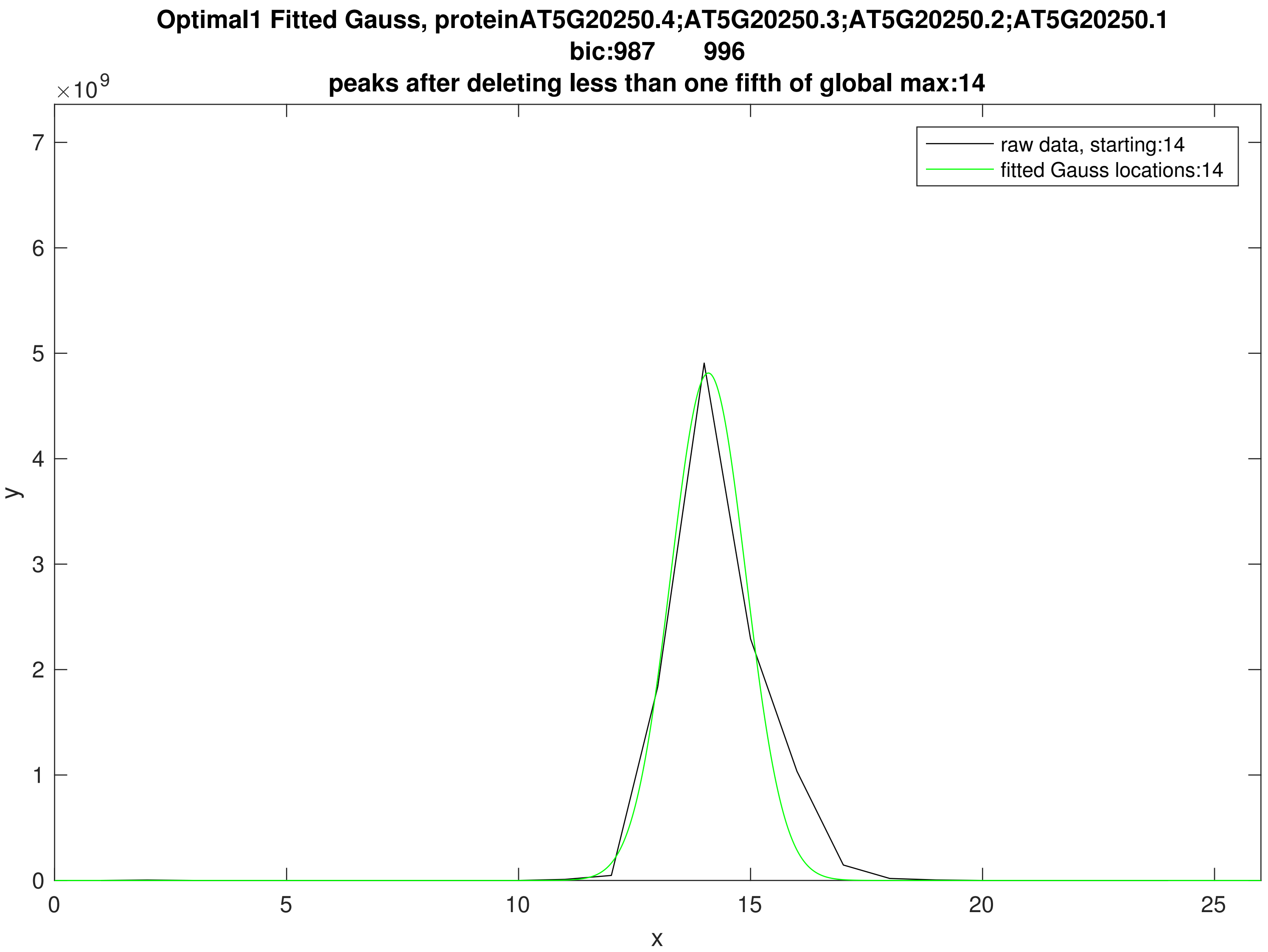
Optimal1 Fitted Gauss, proteinAT5G20250.4;AT5G20250.3;AT5G20250.2;AT5G20250.1
bic:987 996
peaks after deleting less than one fifth of global max:14
×10^9
raw data, starting:14
fitted Gauss locations:14
y
x

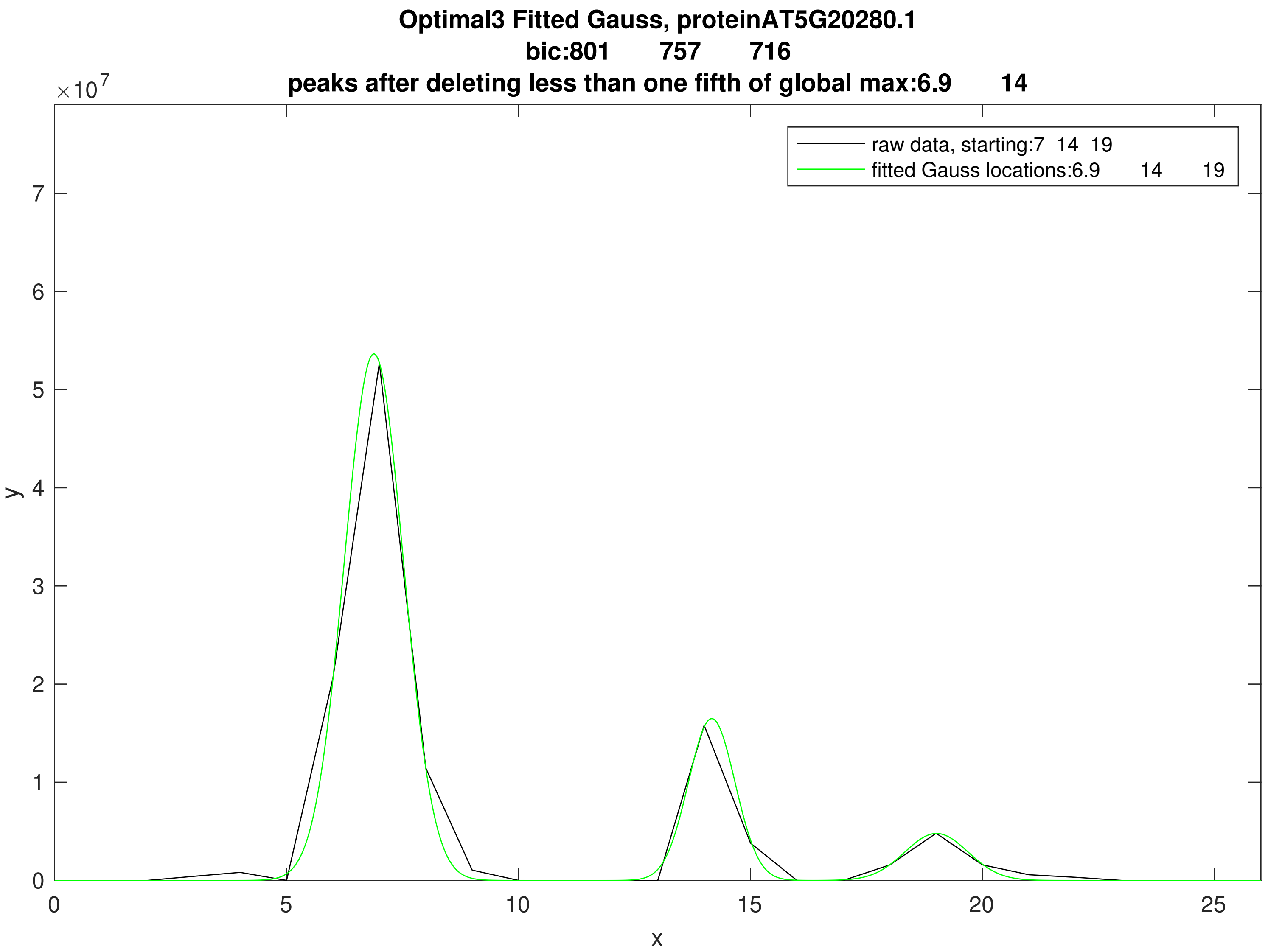
Optimal3 Fitted Gauss, proteinAT5G20280.1
bic:801 757 716
peaks after deleting less than one fifth of global max:6.9 14
×10^7
raw data, starting:7 14 19
fitted Gauss locations:6.9 14 19
0
1
2
3
4
5
6
7
y
0
5
10
15
20
25
x

**Optimal1 Fitted Gauss, proteinAT5G20290.1**
**bic:874 874 884**
**peaks after deleting less than one fifth of global max:4.2**

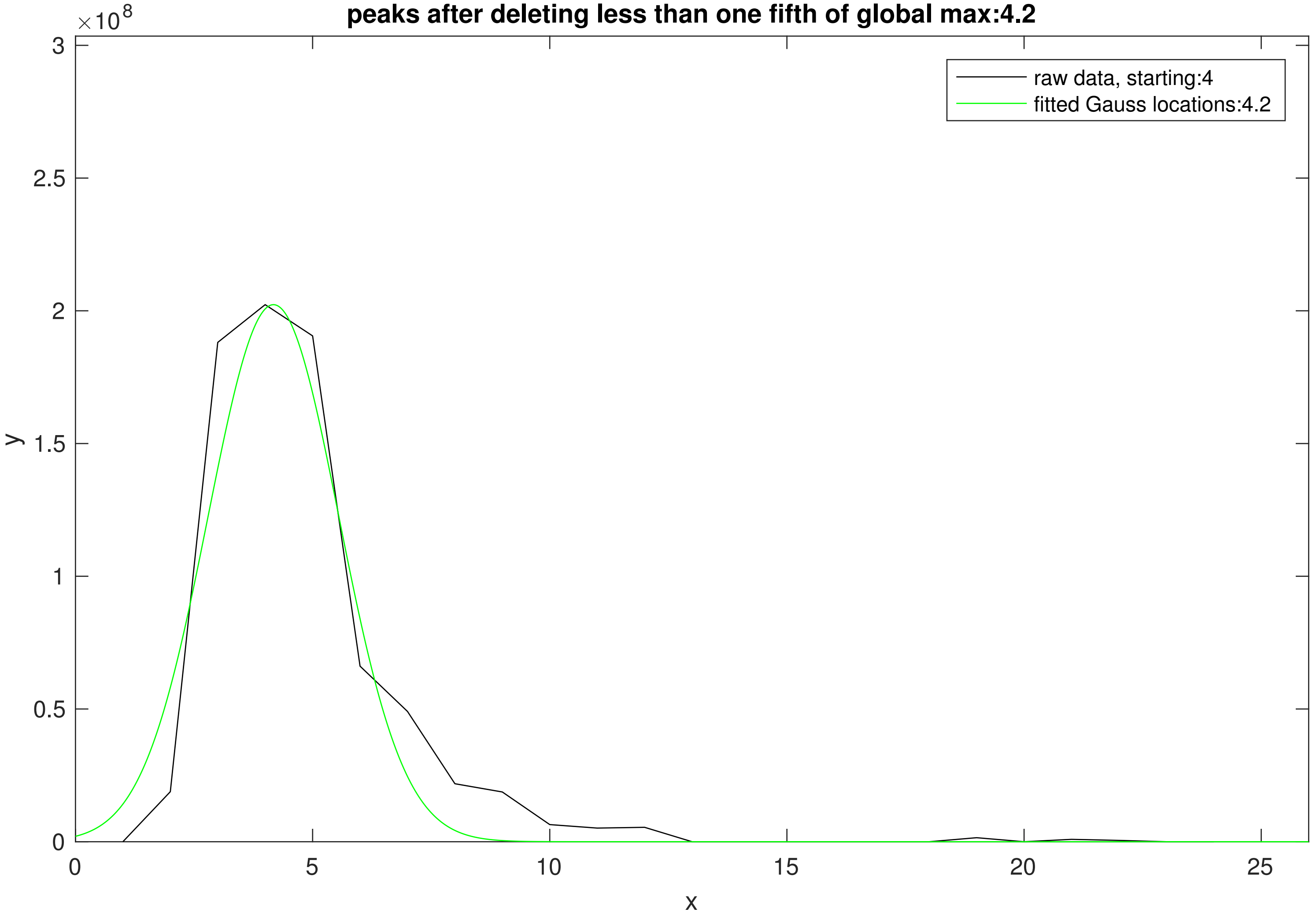

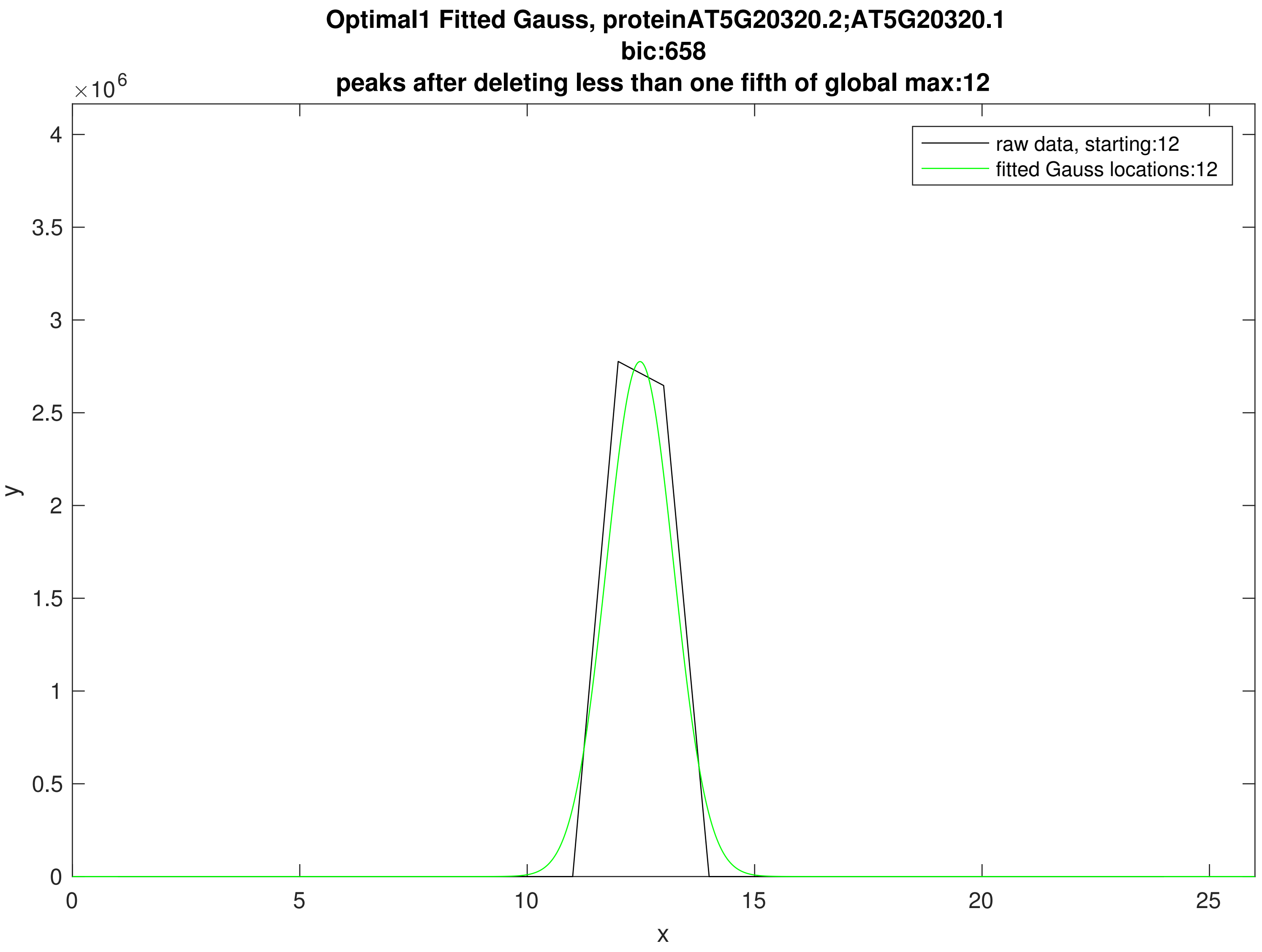
Optimal1 Fitted Gauss, proteinAT5G20320.2;AT5G20320.1
bic:658
peaks after deleting less than one fifth of global max:12
×10^6
4
3.5
3
2.5
2
1.5
1
0.5
0
y
0
5
10
15
20
25
x
raw data, starting:12
fitted Gauss locations:12

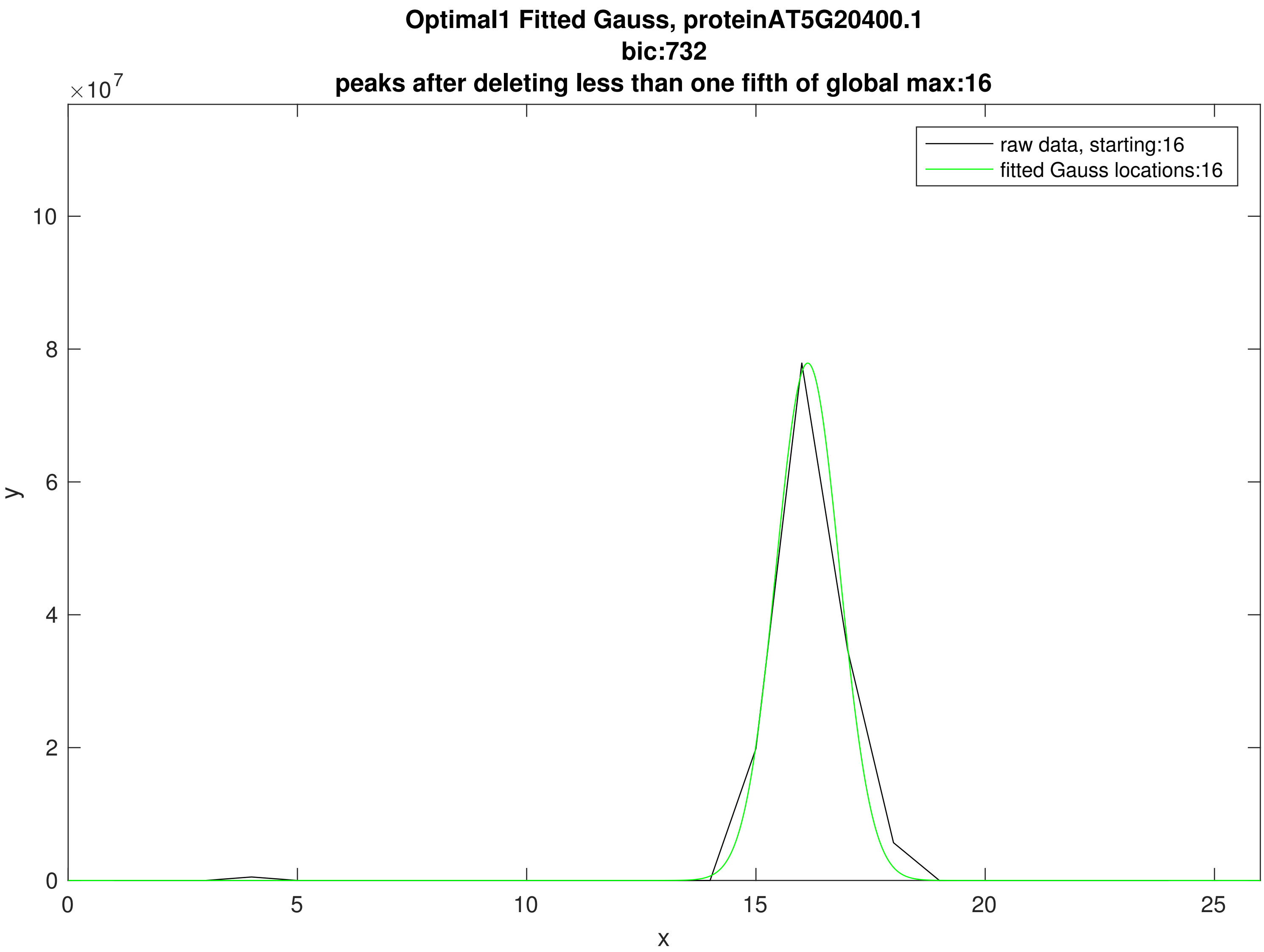
Optimal1 Fitted Gauss, proteinAT5G20400.1
bic:732
peaks after deleting less than one fifth of global max:16
×10^7
raw data, starting:16
fitted Gauss locations:16
y
x
0
2
4
6
8
10
0
5
10
15
20
25

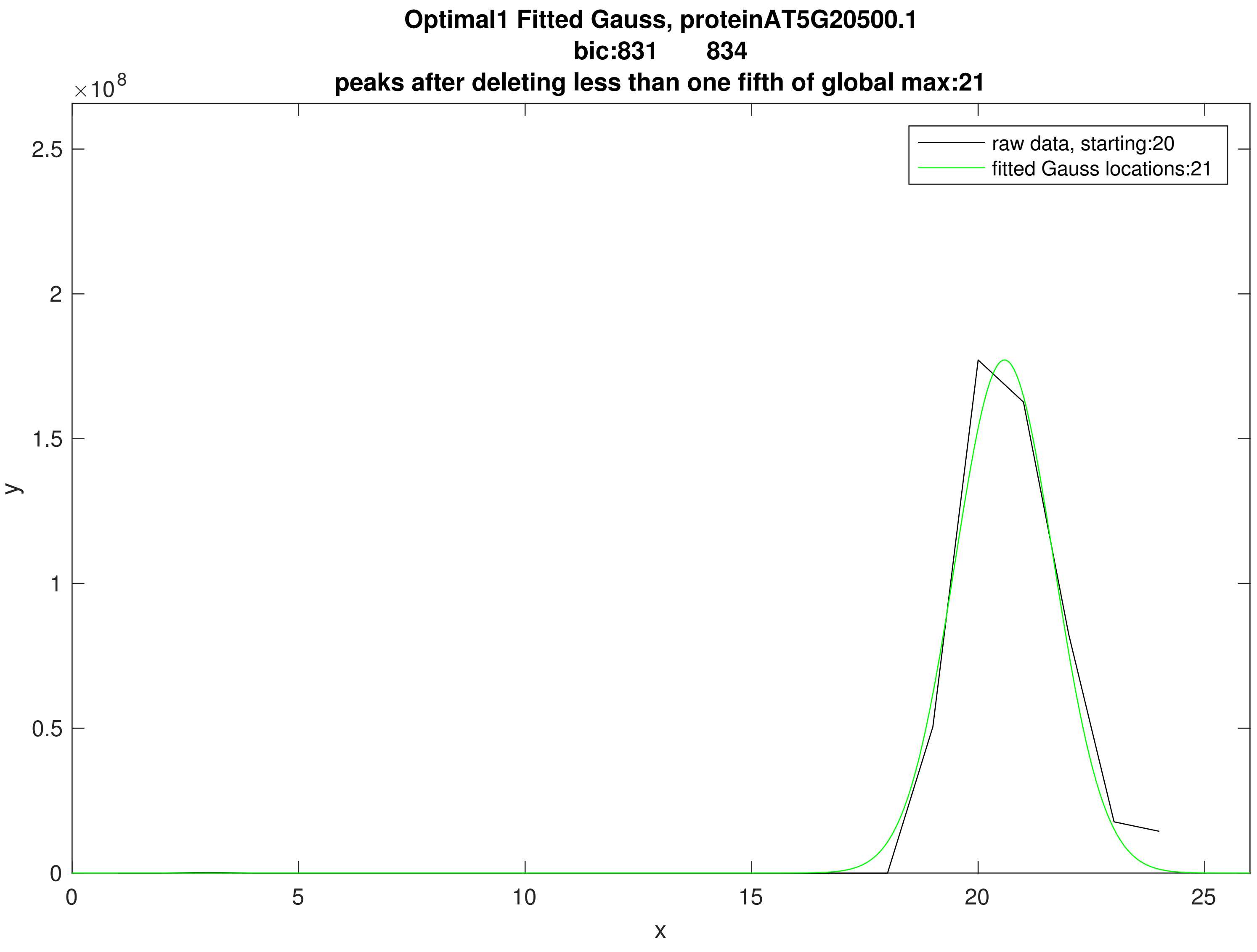

Optimal1 Fitted Gauss, proteinAT5G20500.1
bic:831 834
peaks after deleting less than one fifth of global max:21
×10^8
2.5
2
1.5
1
0.5
0
y
0
5
10
15
20
25
x
raw data, starting:20
fitted Gauss locations:21

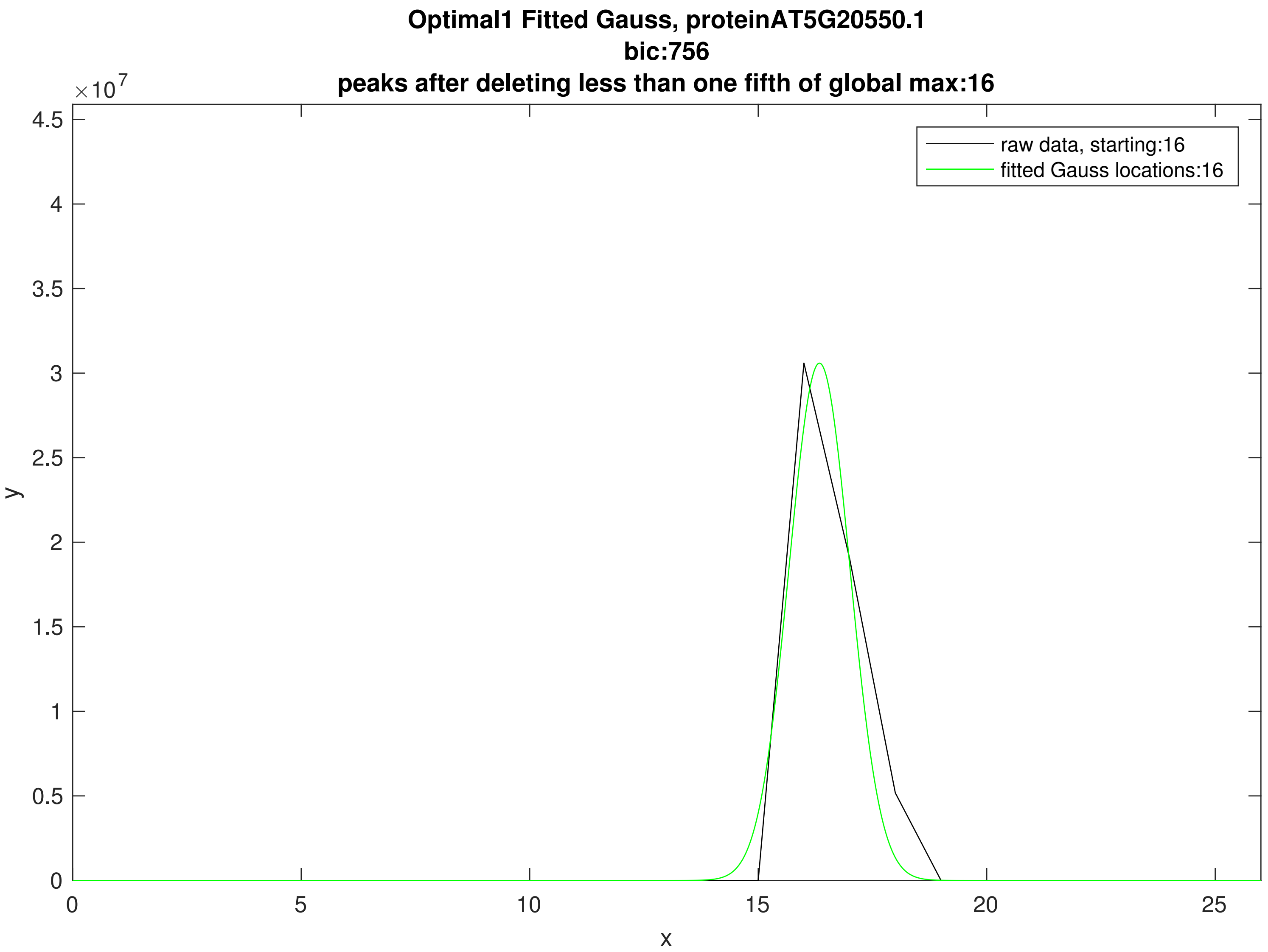
Optimal1 Fitted Gauss, proteinAT5G20550.1
bic:756
peaks after deleting less than one fifth of global max:16
×10^7
4.5
4
3.5
3
2.5
2
1.5
1
0.5
0
y
0
5
10
15
20
25
x
raw data, starting:16
fitted Gauss locations:16

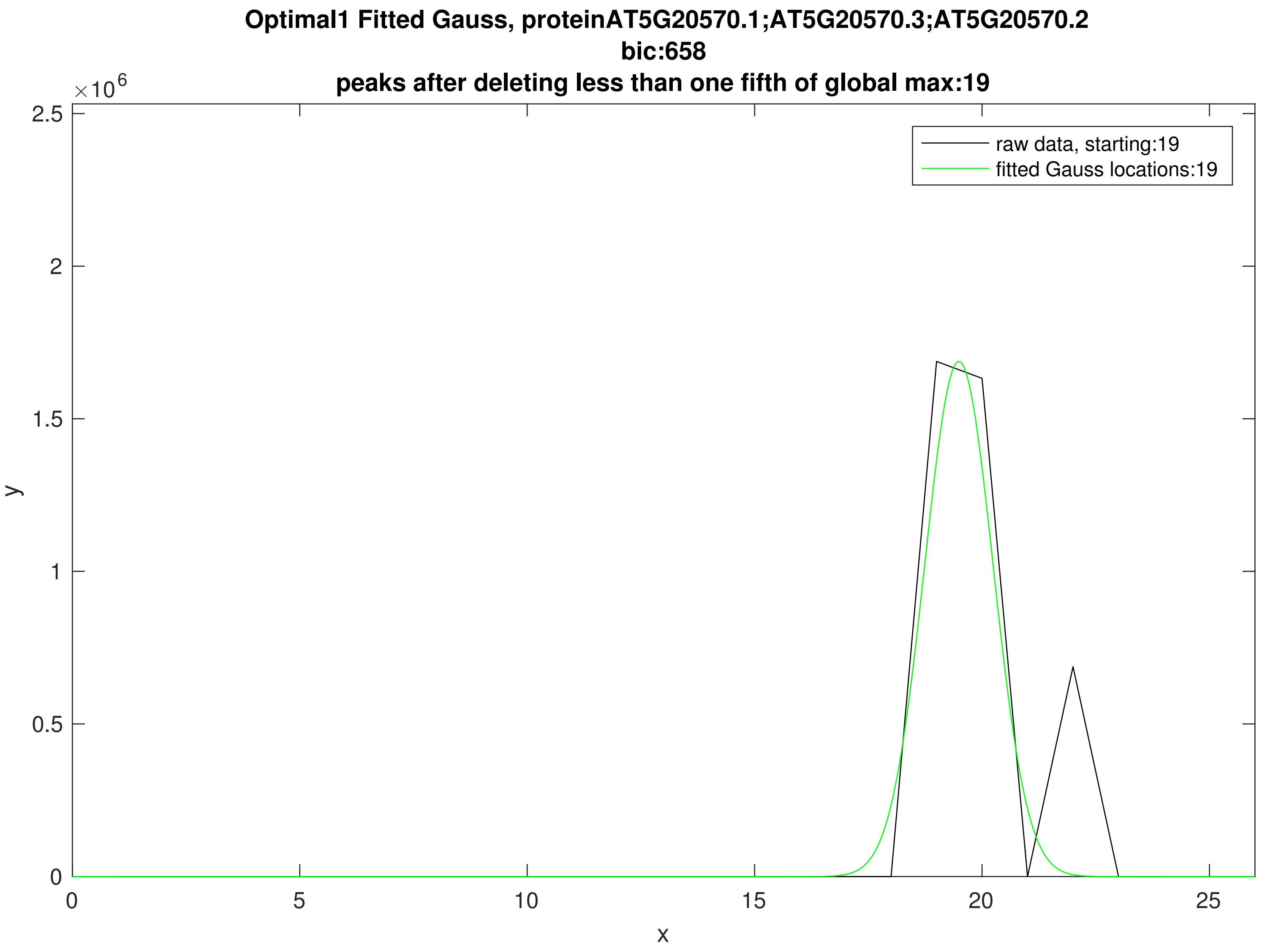

Optimal1 Fitted Gauss, proteinAT5G20570.1;AT5G20570.3;AT5G20570.2
bic:658
peaks after deleting less than one fifth of global max:19
×10^6
2.5
2
1.5
1
0.5
0
y
0
5
10
15
20
25
x
raw data, starting:19
fitted Gauss locations:19

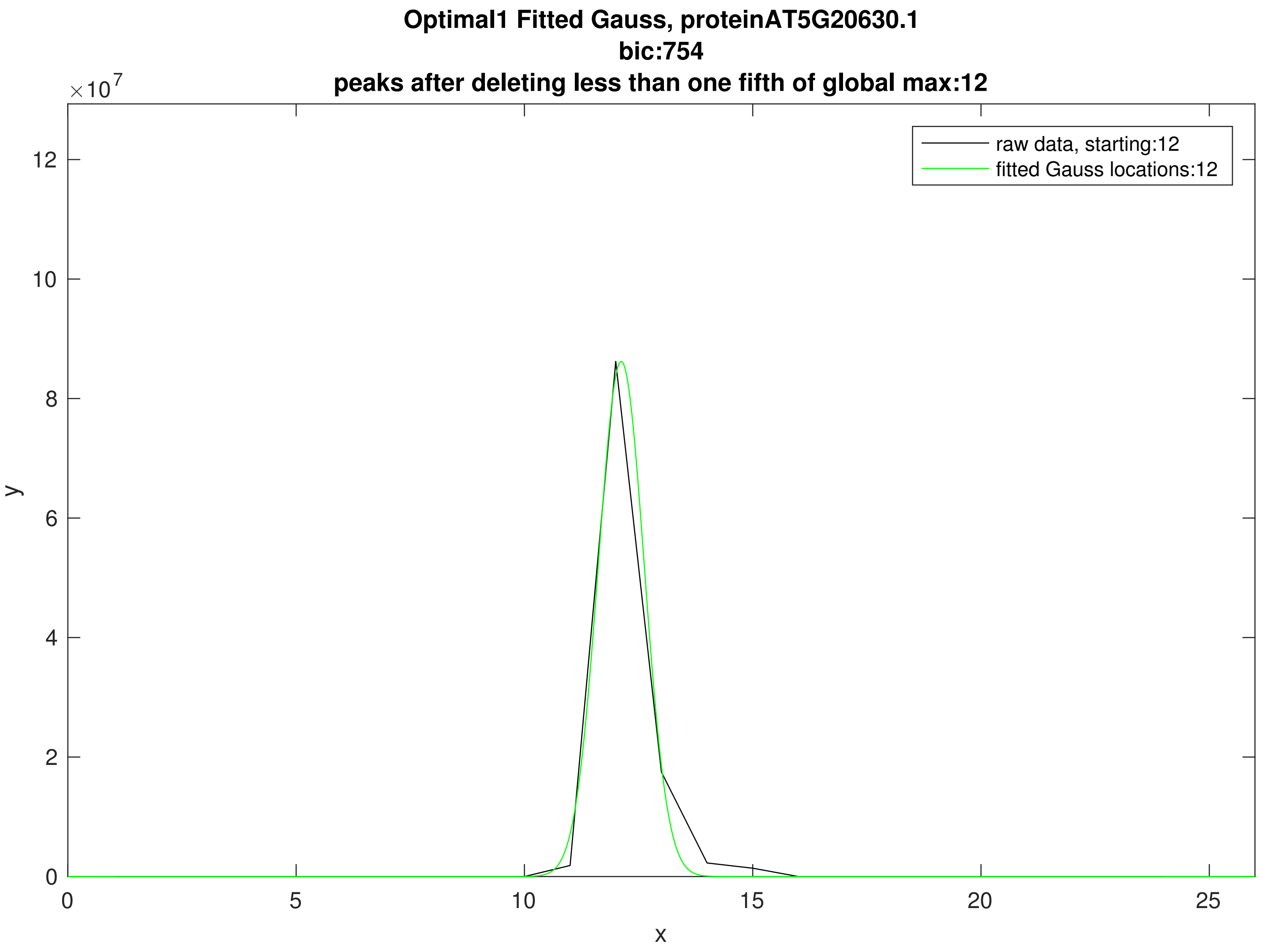

Optimal1 Fitted Gauss, proteinAT5G20630.1
bic:754
peaks after deleting less than one fifth of global max:12
×10^7
raw data, starting:12
fitted Gauss locations:12
y
x
0
2
4
6
8
10
12
0
5
10
15
20
25

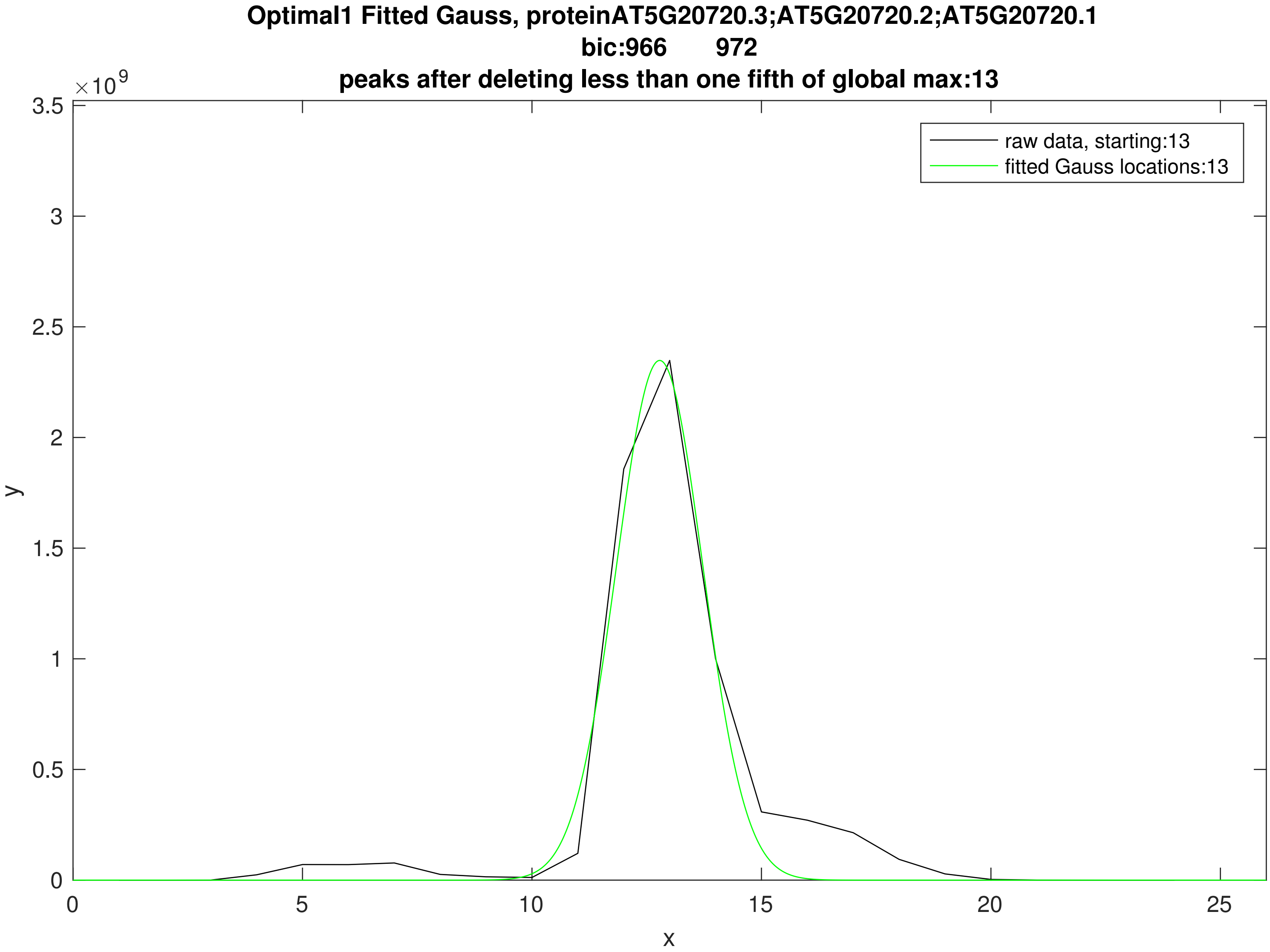

Optimal1 Fitted Gauss, proteinAT5G20720.3;AT5G20720.2;AT5G20720.1
bic:966 972
peaks after deleting less than one fifth of global max:13
×10^9
3.5
3
2.5
2
1.5
1
0.5
0
y
0
5
10
15
20
25
x
raw data, starting:13
fitted Gauss locations:13

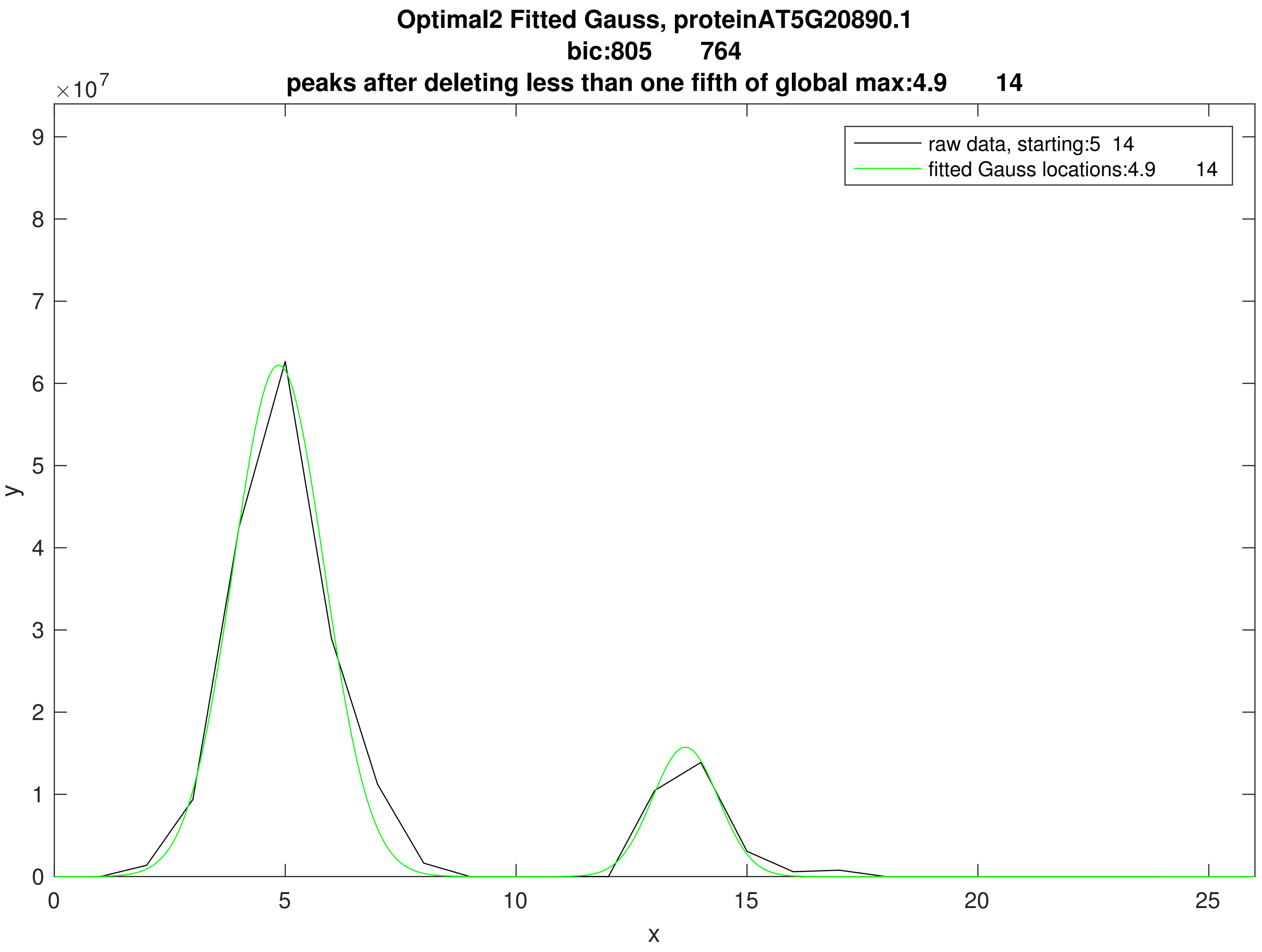
Optimal2 Fitted Gauss, proteinAT5G20890.1
bic:805 764
peaks after deleting less than one fifth of global max:4.9 14
×10^7
raw data, starting:5 14
fitted Gauss locations:4.9 14
y
x

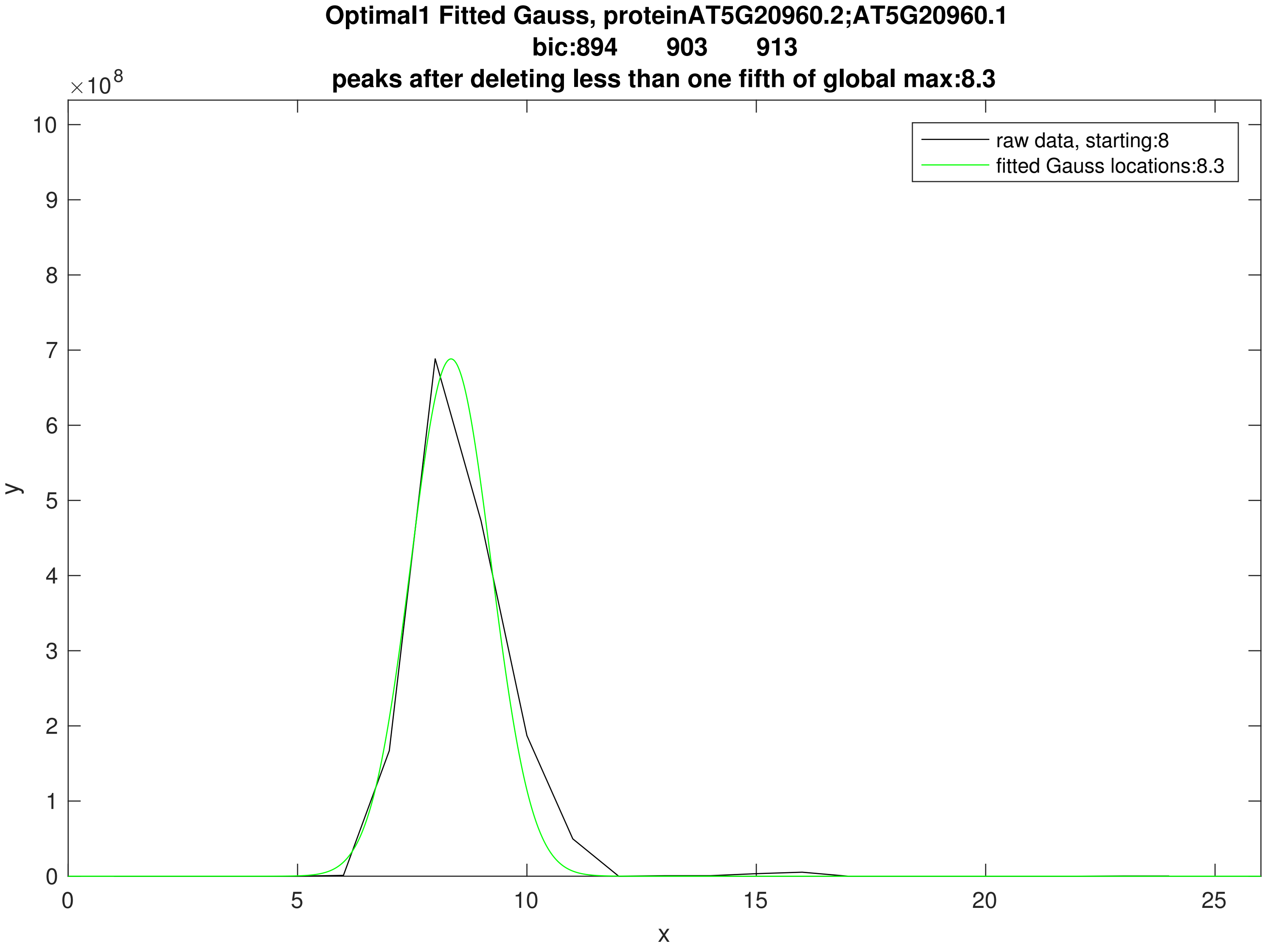
Optimal1 Fitted Gauss, proteinAT5G20960.2;AT5G20960.1
bic:894 903 913
peaks after deleting less than one fifth of global max:8.3
×10^8
raw data, starting:8
fitted Gauss locations:8.3
y
x
0
1
2
3
4
5
6
7
8
9
10
0
5
10
15
20
25

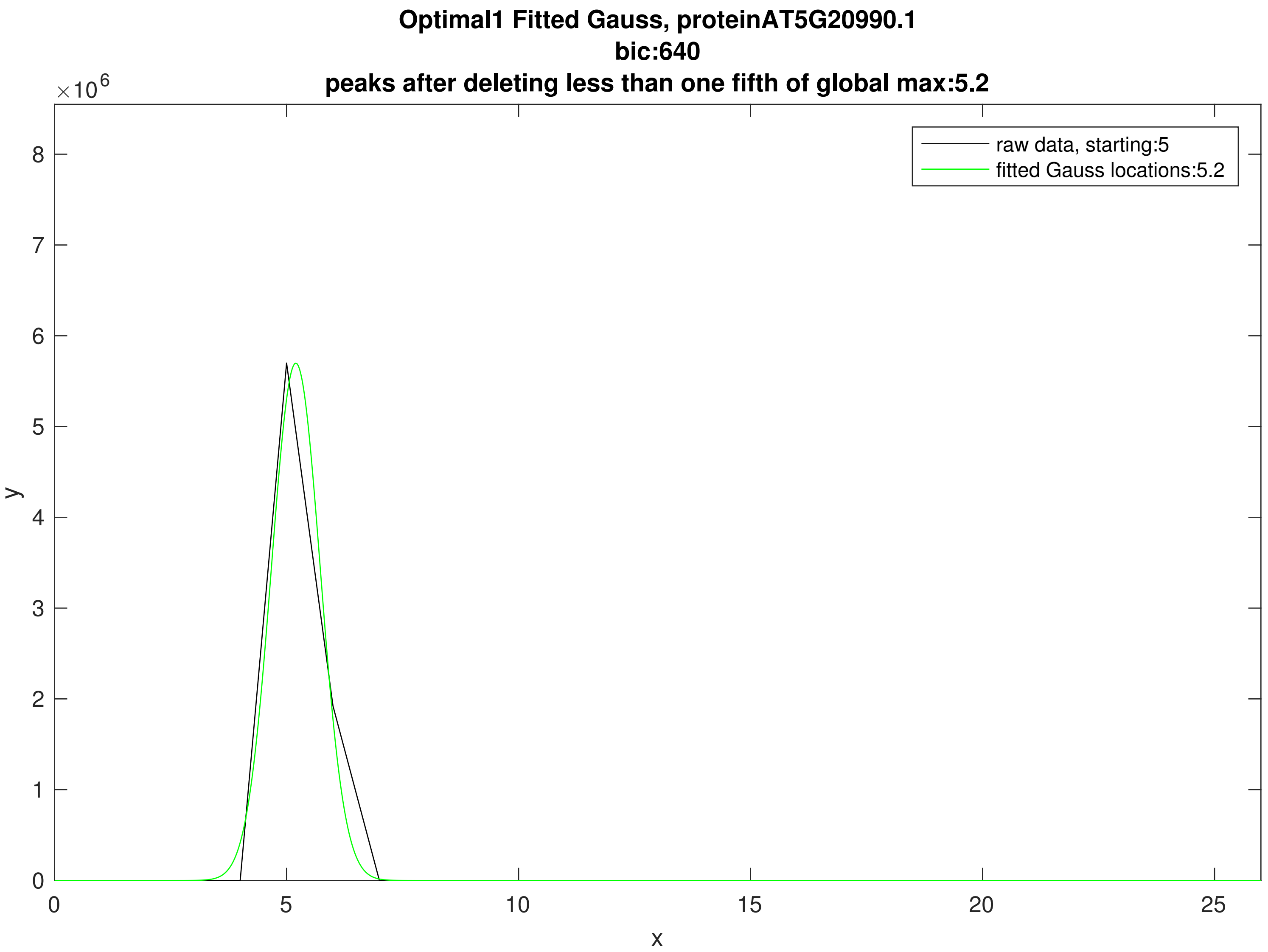
Optimal1 Fitted Gauss, proteinAT5G20990.1
bic:640
peaks after deleting less than one fifth of global max:5.2
×10^6
raw data, starting:5
fitted Gauss locations:5.2
y
x
0
1
2
3
4
5
6
7
8
0
5
10
15
20
25

**Optimal1 Fitted Gauss, proteinAT5G21060.1;AT5G21060.2;AT5G21060.3**
**bic:765**
**peaks after deleting less than one fifth of global max:14**

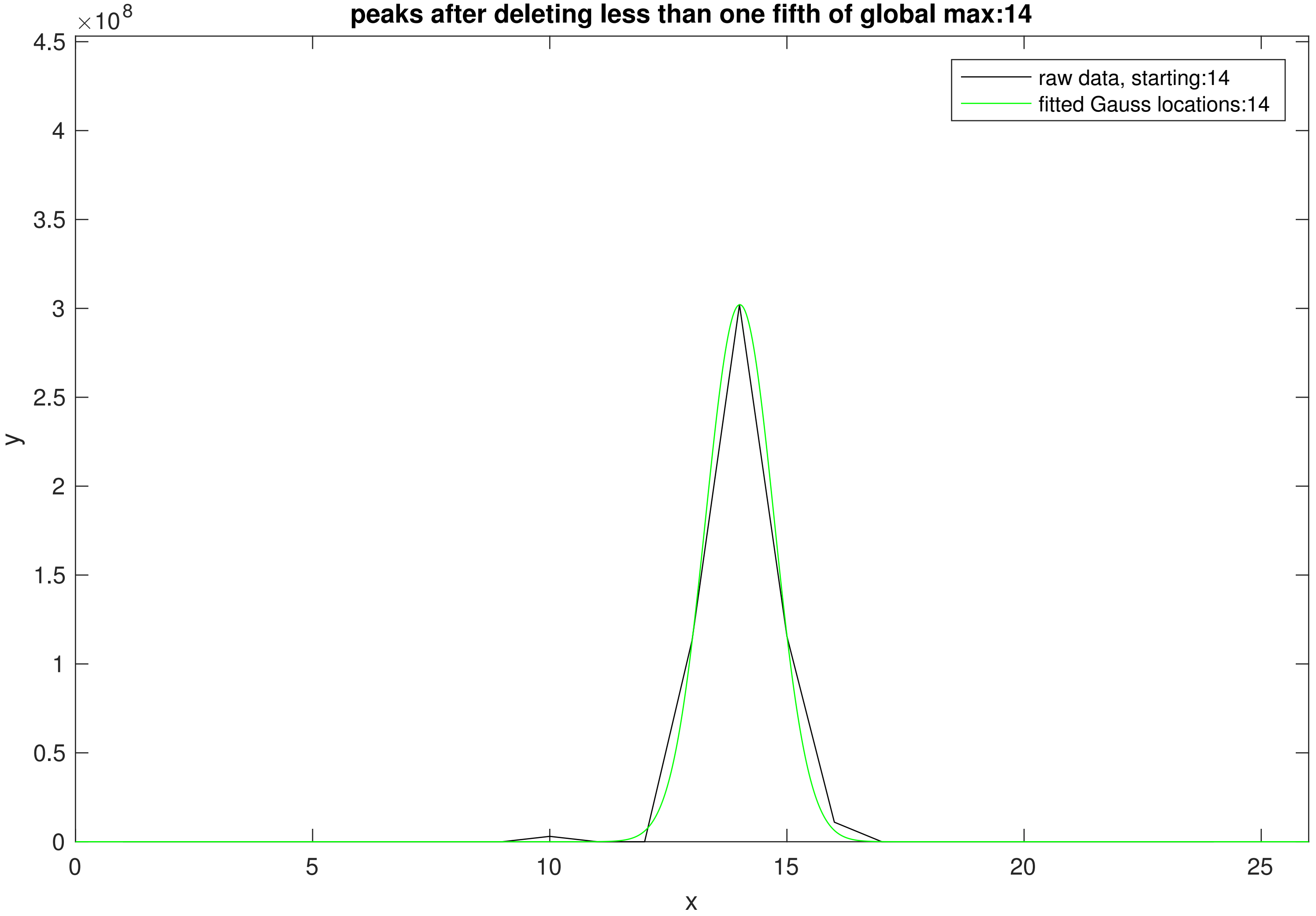

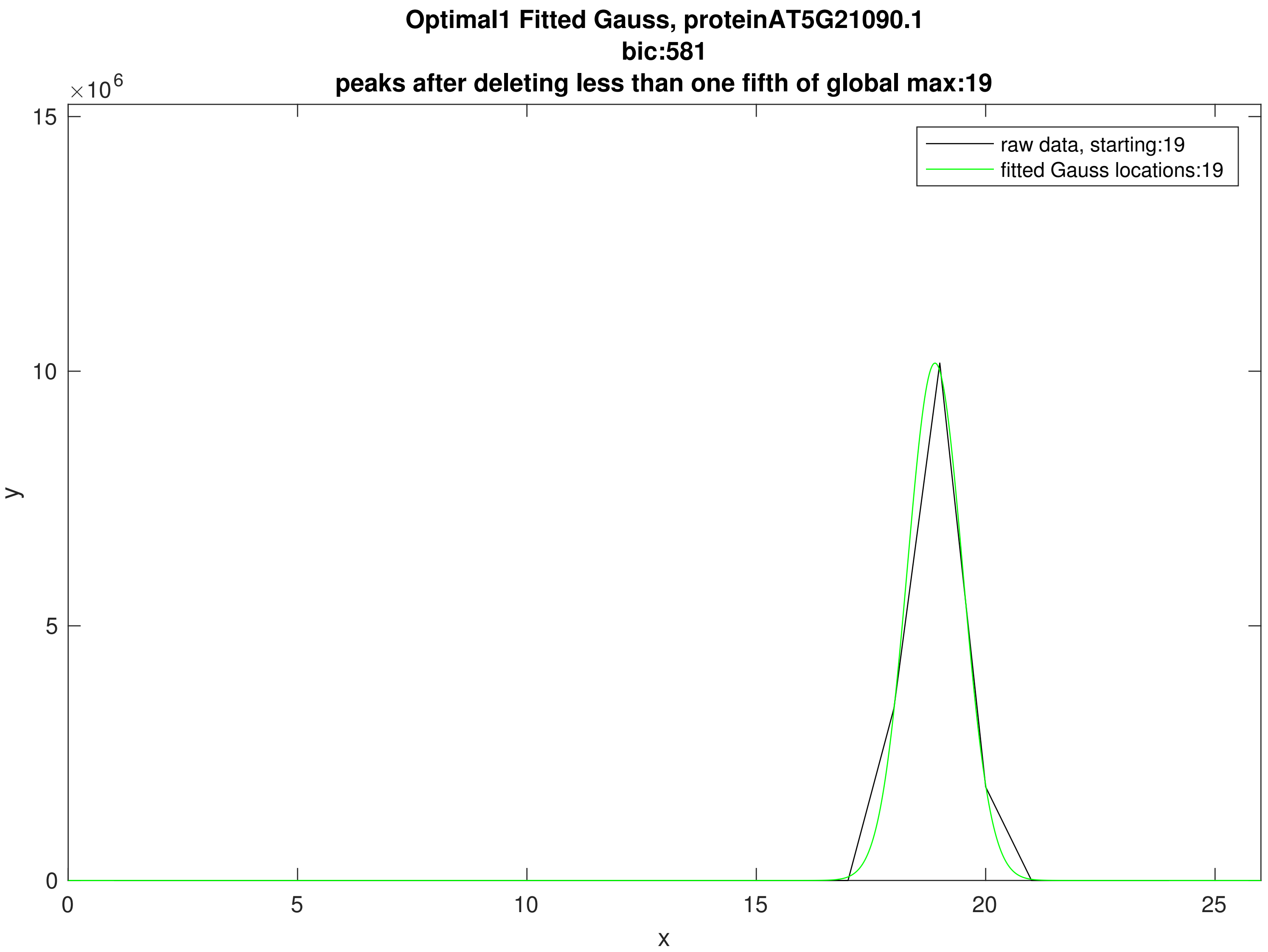
Optimal1 Fitted Gauss, proteinAT5G21090.1
bic:581
peaks after deleting less than one fifth of global max:19
×10^6
15
10
5
0
y
0
5
10
15
20
25
x
raw data, starting:19
fitted Gauss locations:19

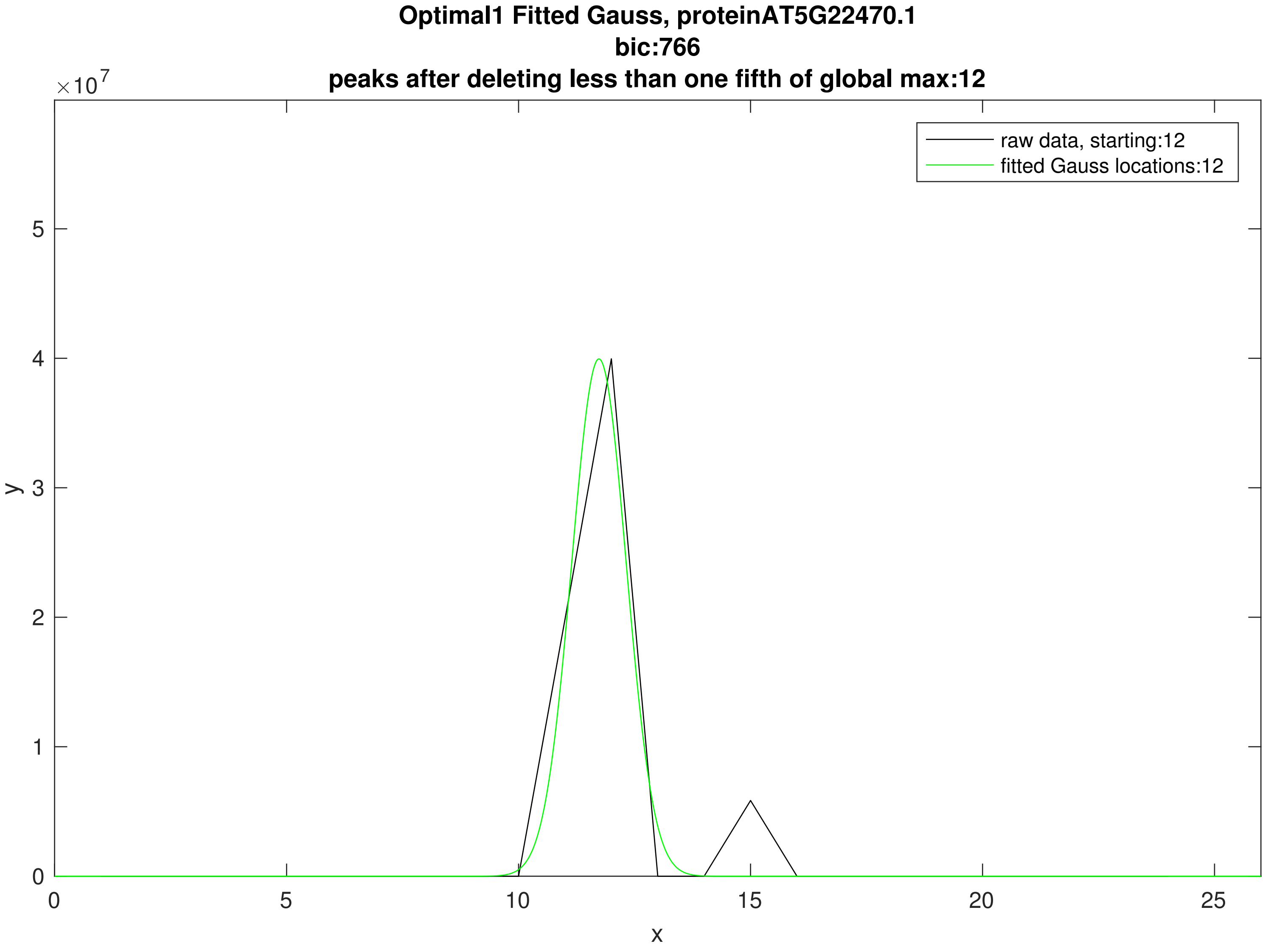
Optimal1 Fitted Gauss, proteinAT5G22470.1
bic:766
peaks after deleting less than one fifth of global max:12
$\times 10^7$
raw data, starting:12
fitted Gauss locations:12
y
x
0
1
2
3
4
5
0
5
10
15
20
25

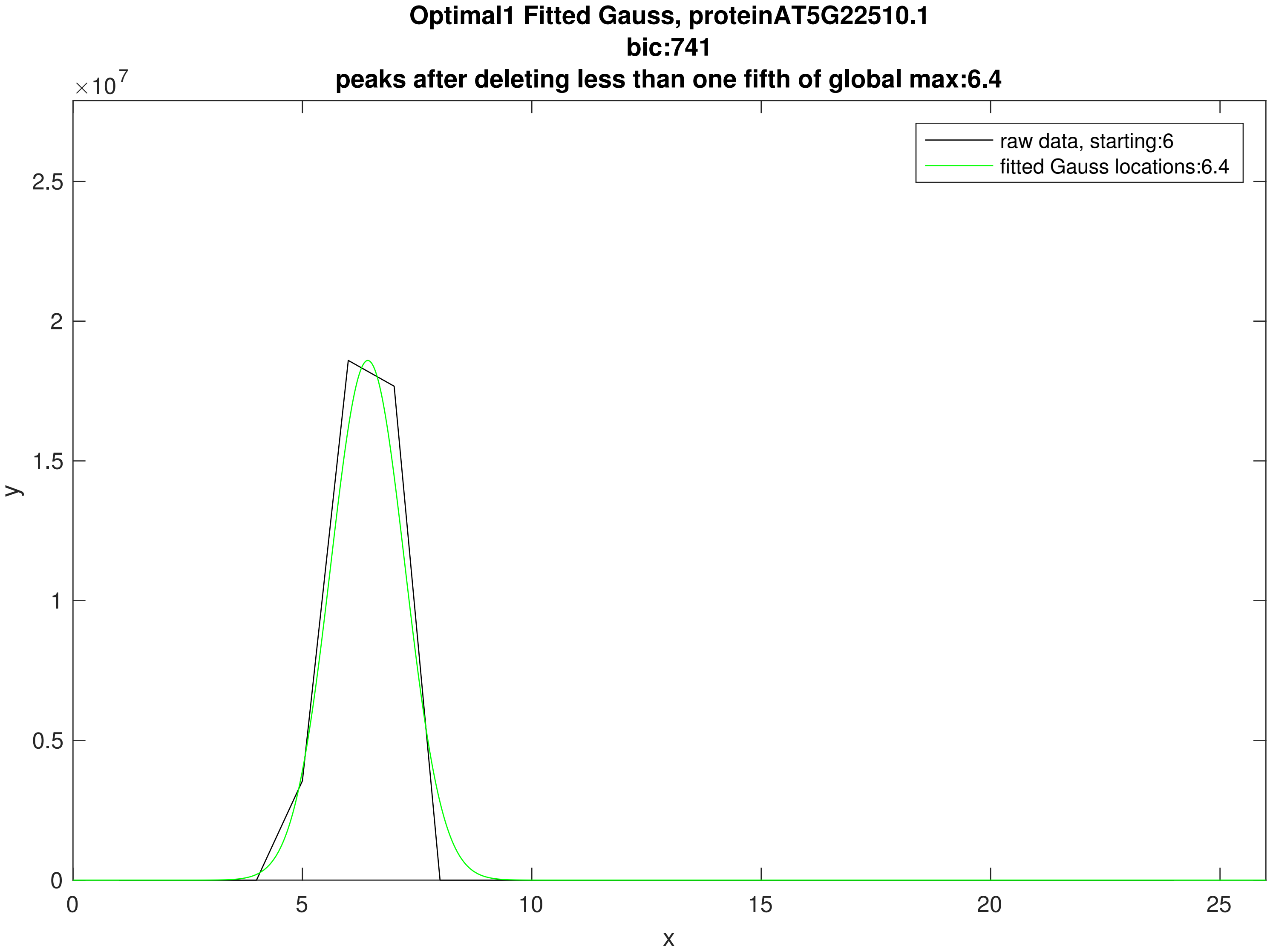

Optimal1 Fitted Gauss, proteinAT5G22510.1
bic:741
peaks after deleting less than one fifth of global max:6.4
×10^7
raw data, starting:6
fitted Gauss locations:6.4
y
x
0
0.5
1
1.5
2
2.5
5
10
15
20
25

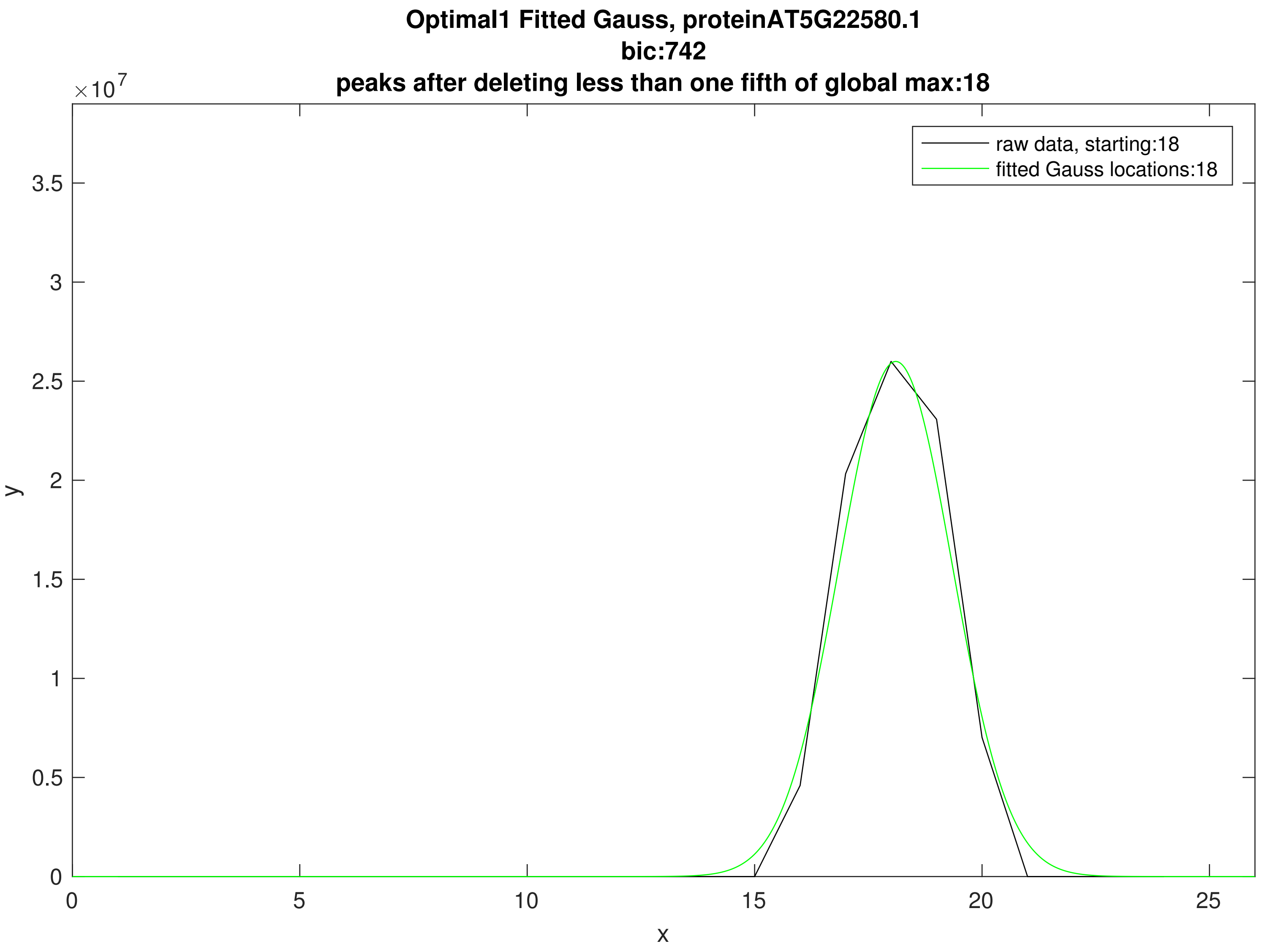
Optimal1 Fitted Gauss, proteinAT5G22580.1
bic:742
peaks after deleting less than one fifth of global max:18
×10^7
3.5
3
2.5
2
1.5
1
0.5
0
y
0
5
10
15
20
25
x
raw data, starting:18
fitted Gauss locations:18

**Optimal1 Fitted Gauss, proteinAT5G22650.2;AT5G22650.1**
**bic:665**
**peaks after deleting less than one fifth of global max:17**

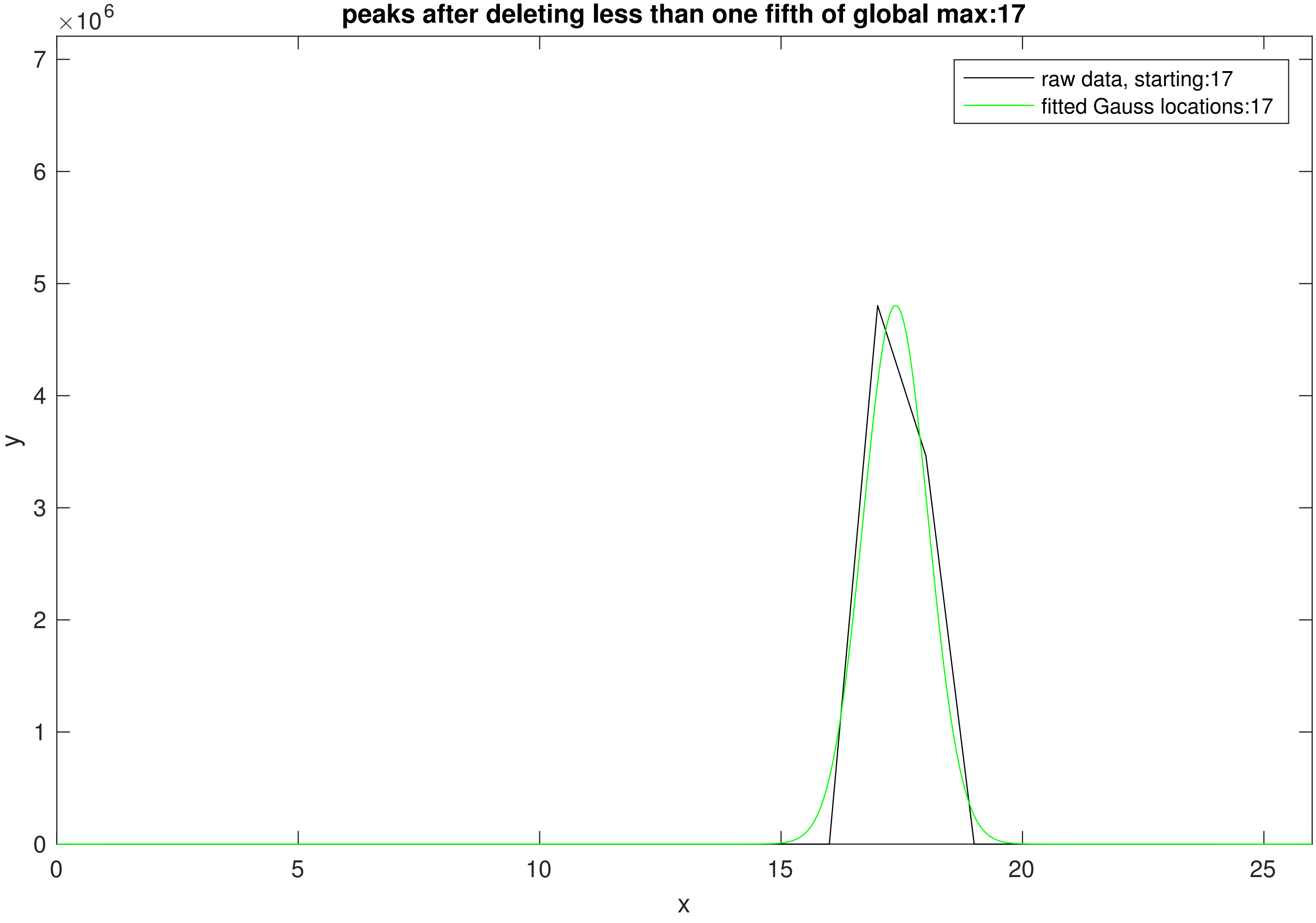

**Optimal1 Fitted Gauss, proteinAT5G22800.1**
**bic:817**
**peaks after deleting less than one fifth of global max:13**

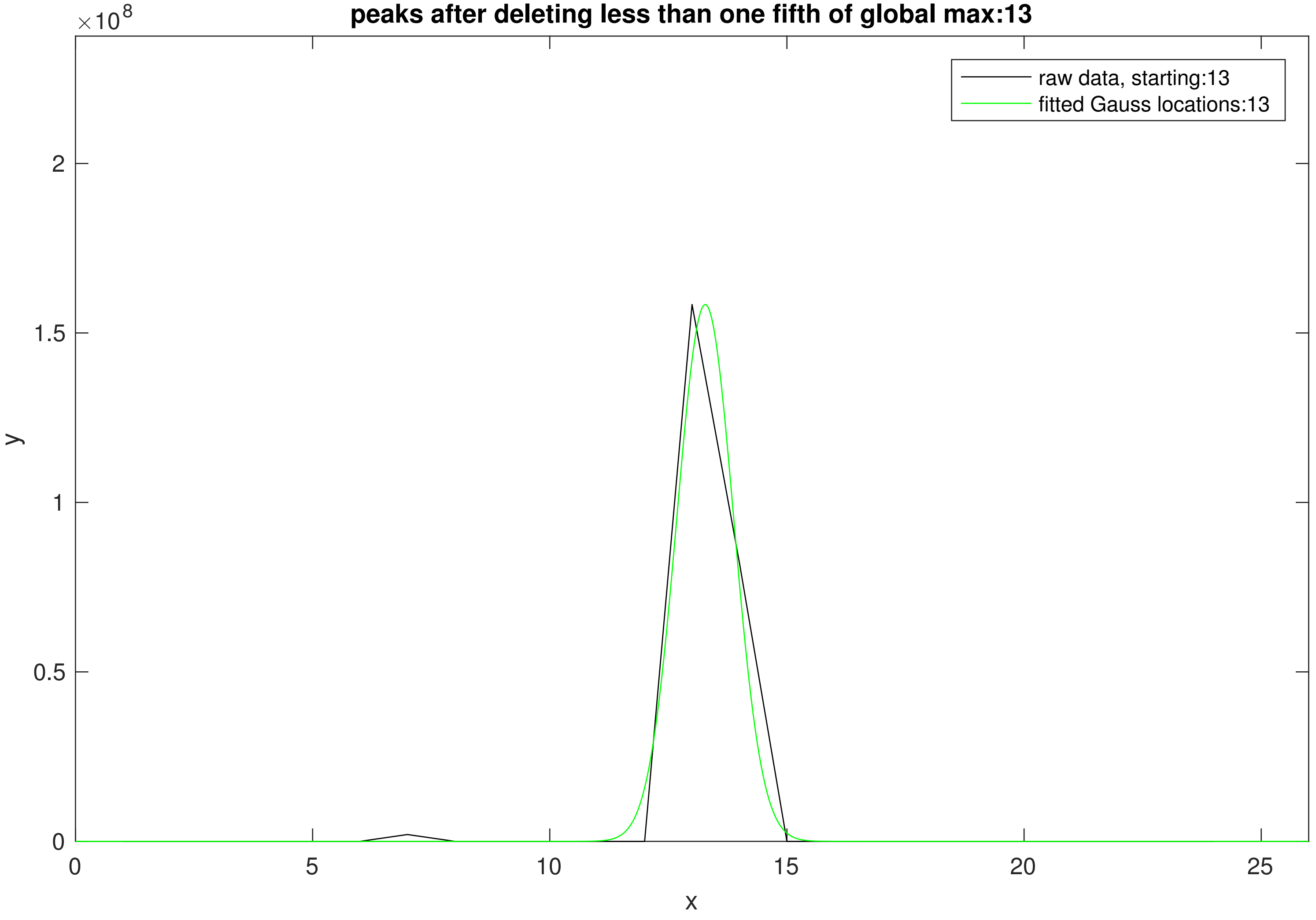

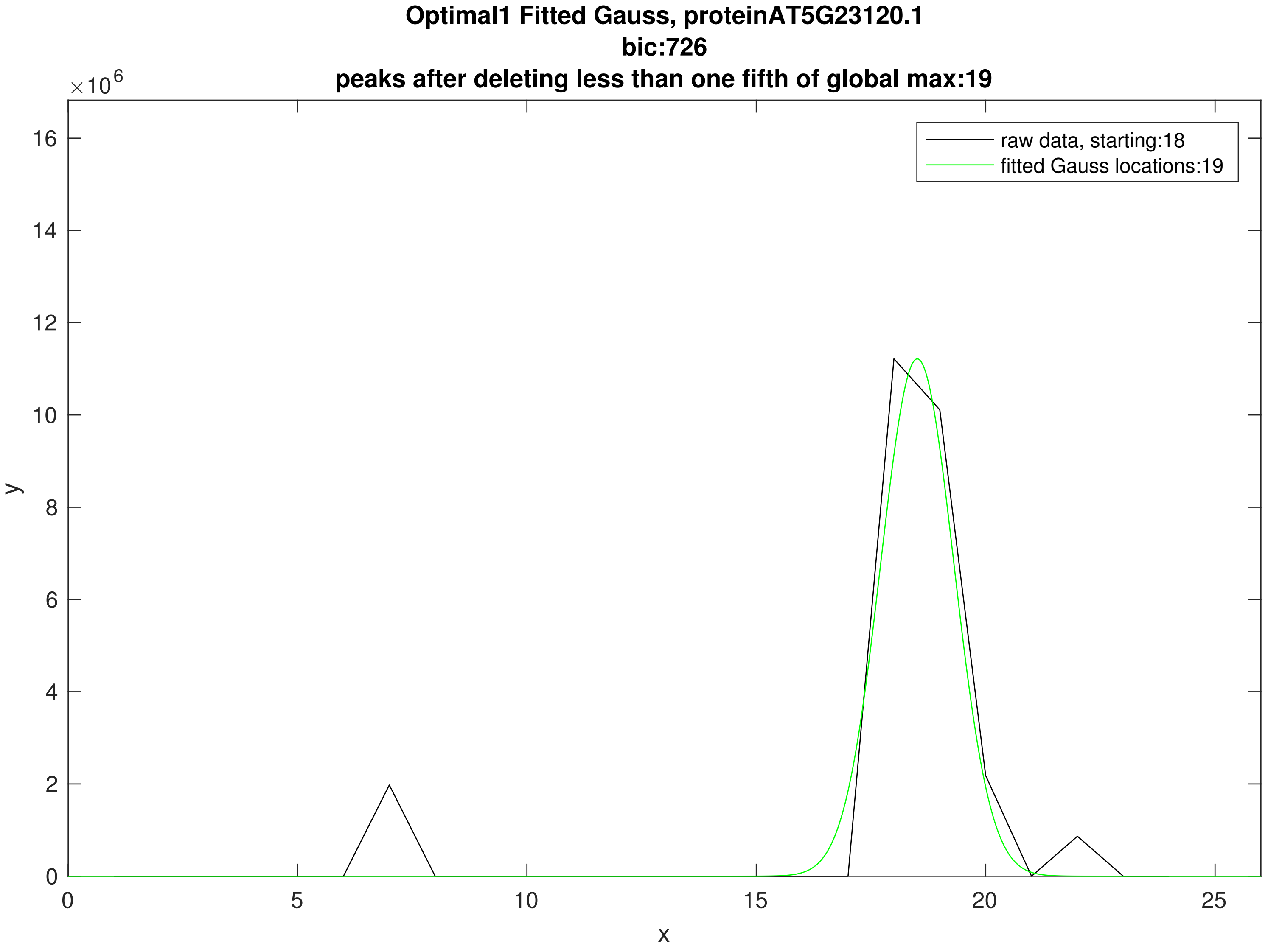
Optimal1 Fitted Gauss, proteinAT5G23120.1
bic:726
peaks after deleting less than one fifth of global max:19
×10^6
raw data, starting:18
fitted Gauss locations:19
y
x
0
2
4
6
8
10
12
14
16
0
5
10
15
20
25

**Optimal1 Fitted Gauss, proteinAT5G23140.1**
**bic:853 863**
**peaks after deleting less than one fifth of global max:7.2**

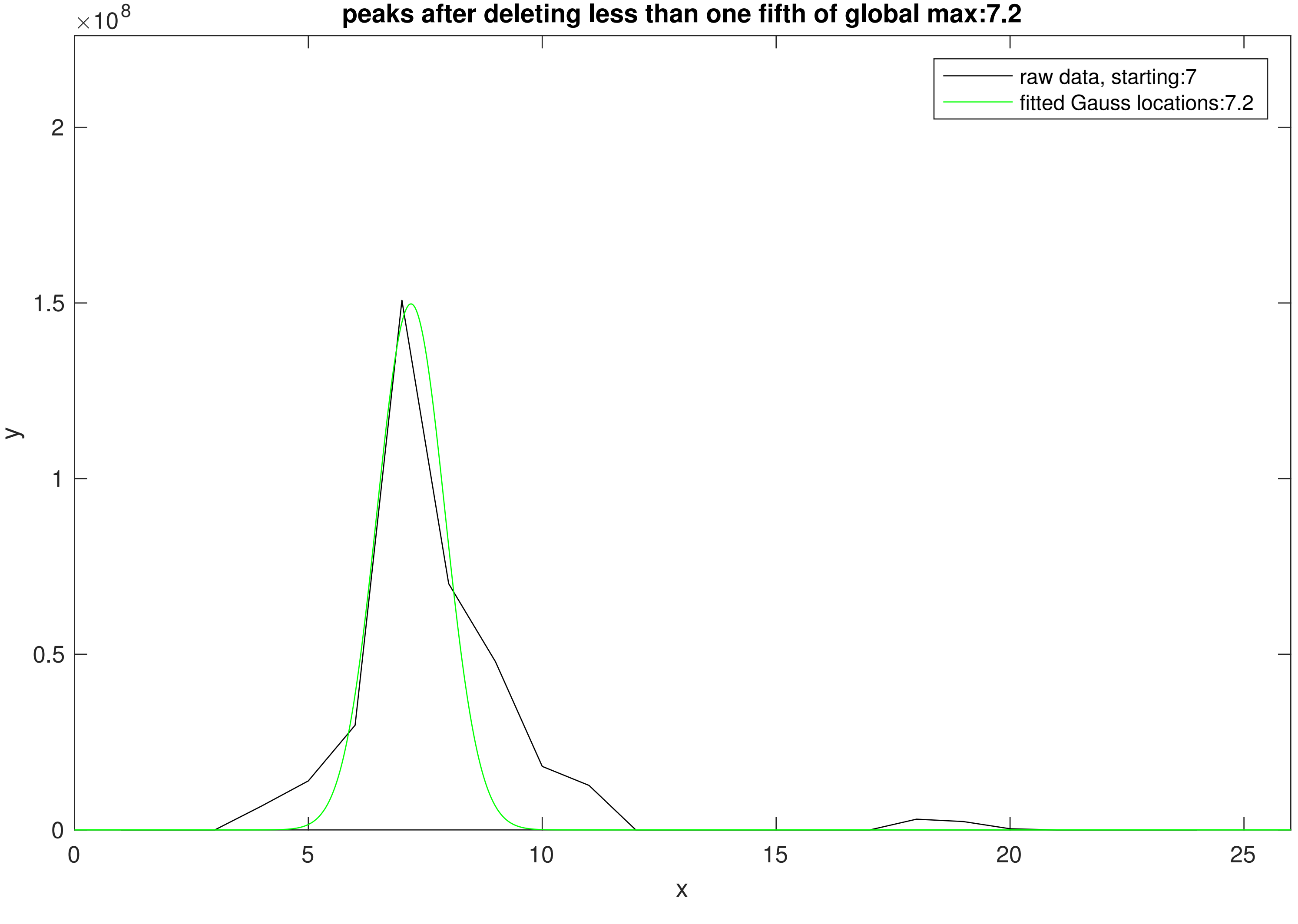

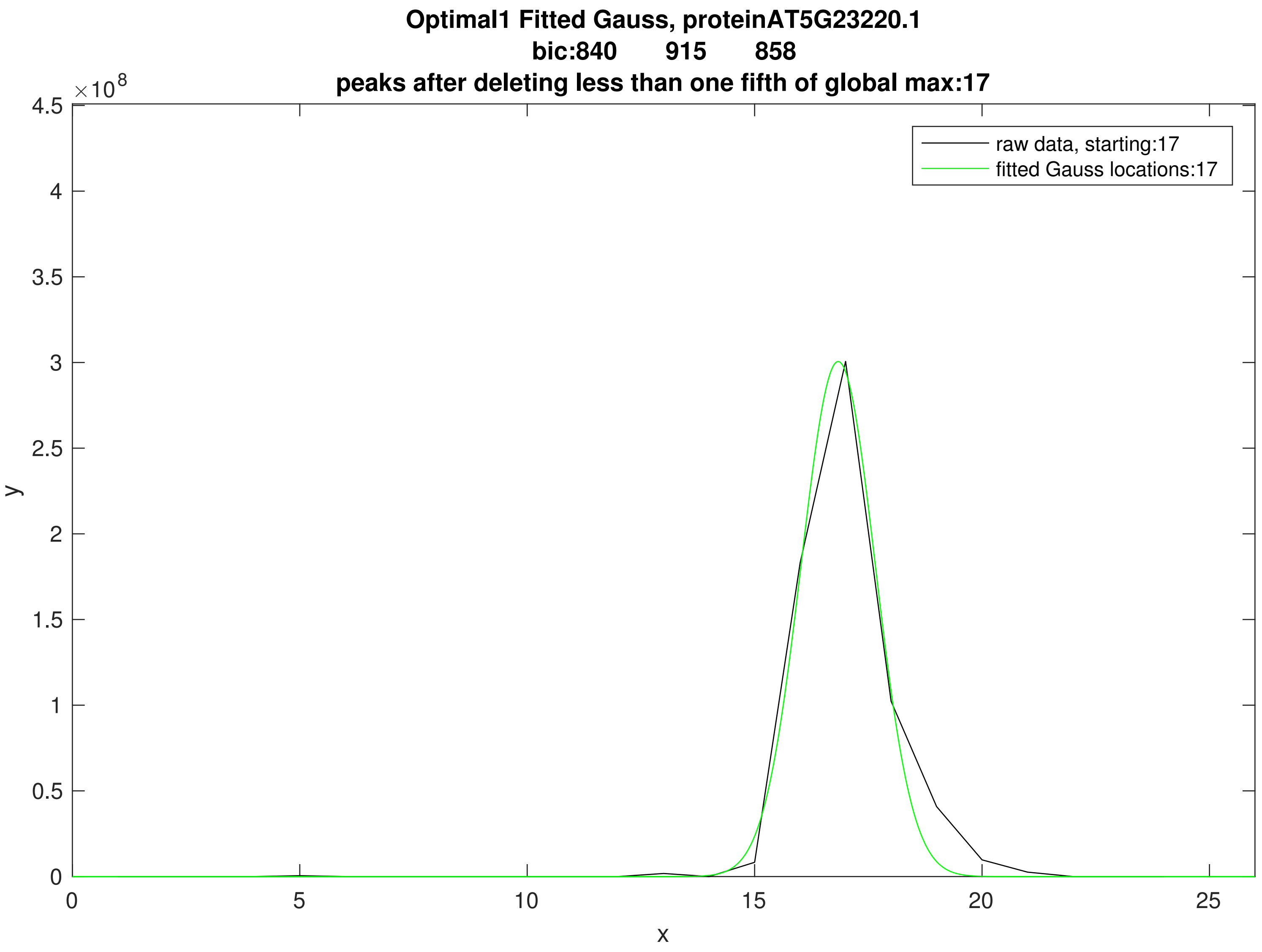
Optimal1 Fitted Gauss, proteinAT5G23220.1
bic:840 915 858
peaks after deleting less than one fifth of global max:17
raw data, starting:17
fitted Gauss locations:17
×10^8
y
x

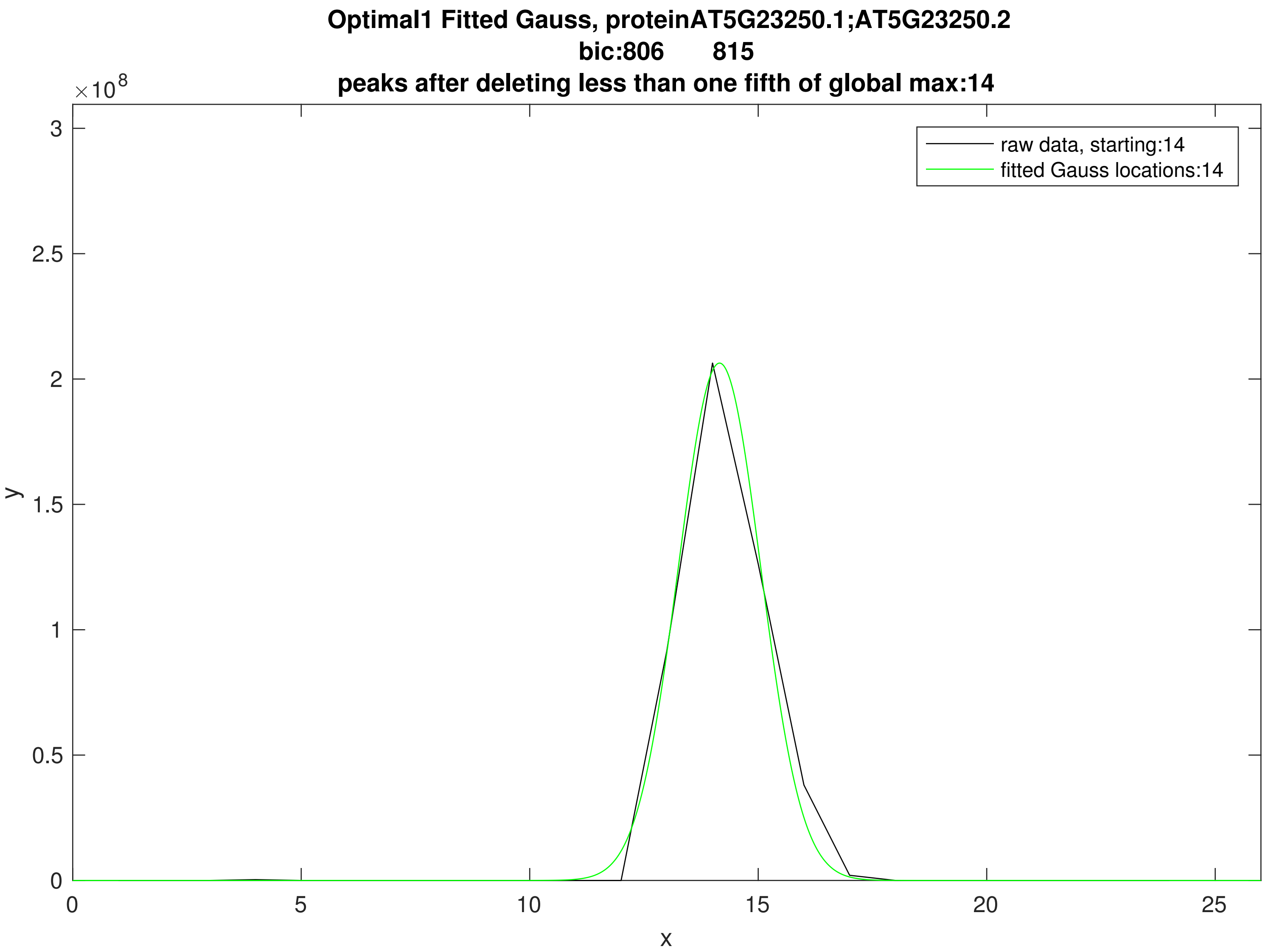
Optimal1 Fitted Gauss, proteinAT5G23250.1;AT5G23250.2
bic:806 815
peaks after deleting less than one fifth of global max:14
×10^8
raw data, starting:14
fitted Gauss locations:14
y
x
0
0.5
1
1.5
2
2.5
3
5
10
15
20
25

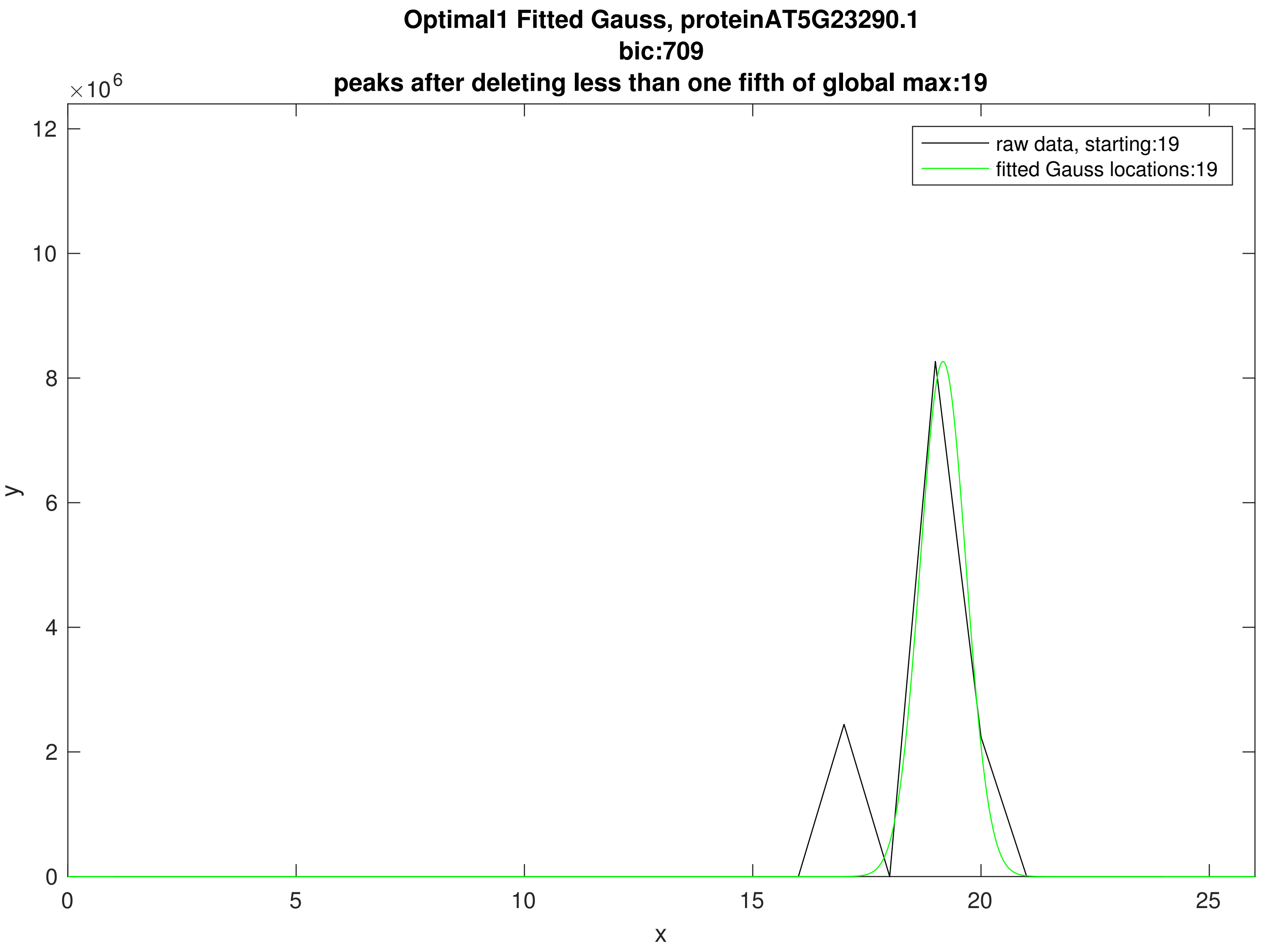
Optimal1 Fitted Gauss, proteinAT5G23290.1
bic:709
peaks after deleting less than one fifth of global max:19
×10^6
12
10
8
6
4
2
0
y
0
5
10
15
20
25
x
raw data, starting:19
fitted Gauss locations:19

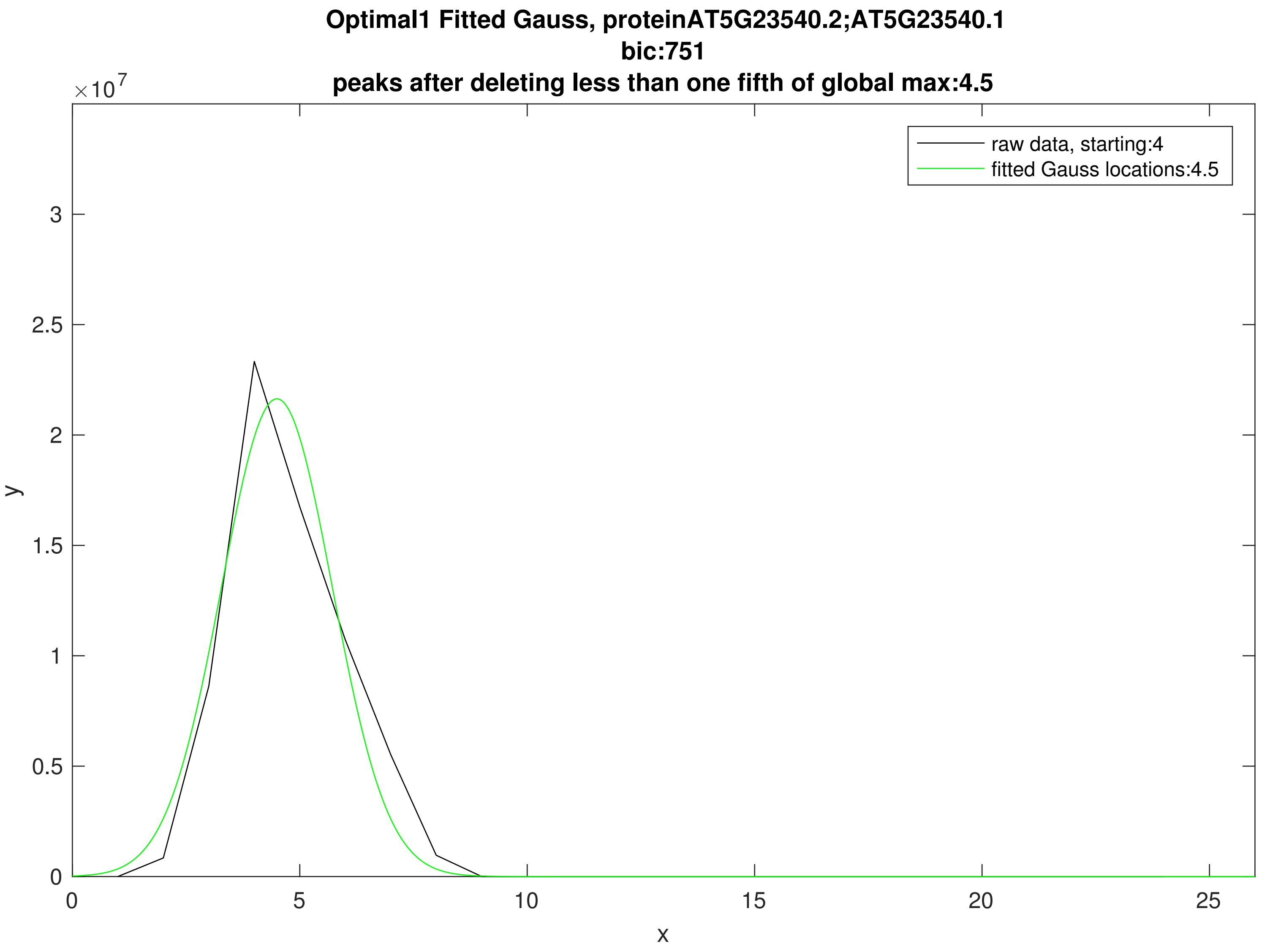
Optimal1 Fitted Gauss, proteinAT5G23540.2;AT5G23540.1
bic:751
peaks after deleting less than one fifth of global max:4.5
×10^7
raw data, starting:4
fitted Gauss locations:4.5
y
x
0
0.5
1
1.5
2
2.5
3
5
10
15
20
25

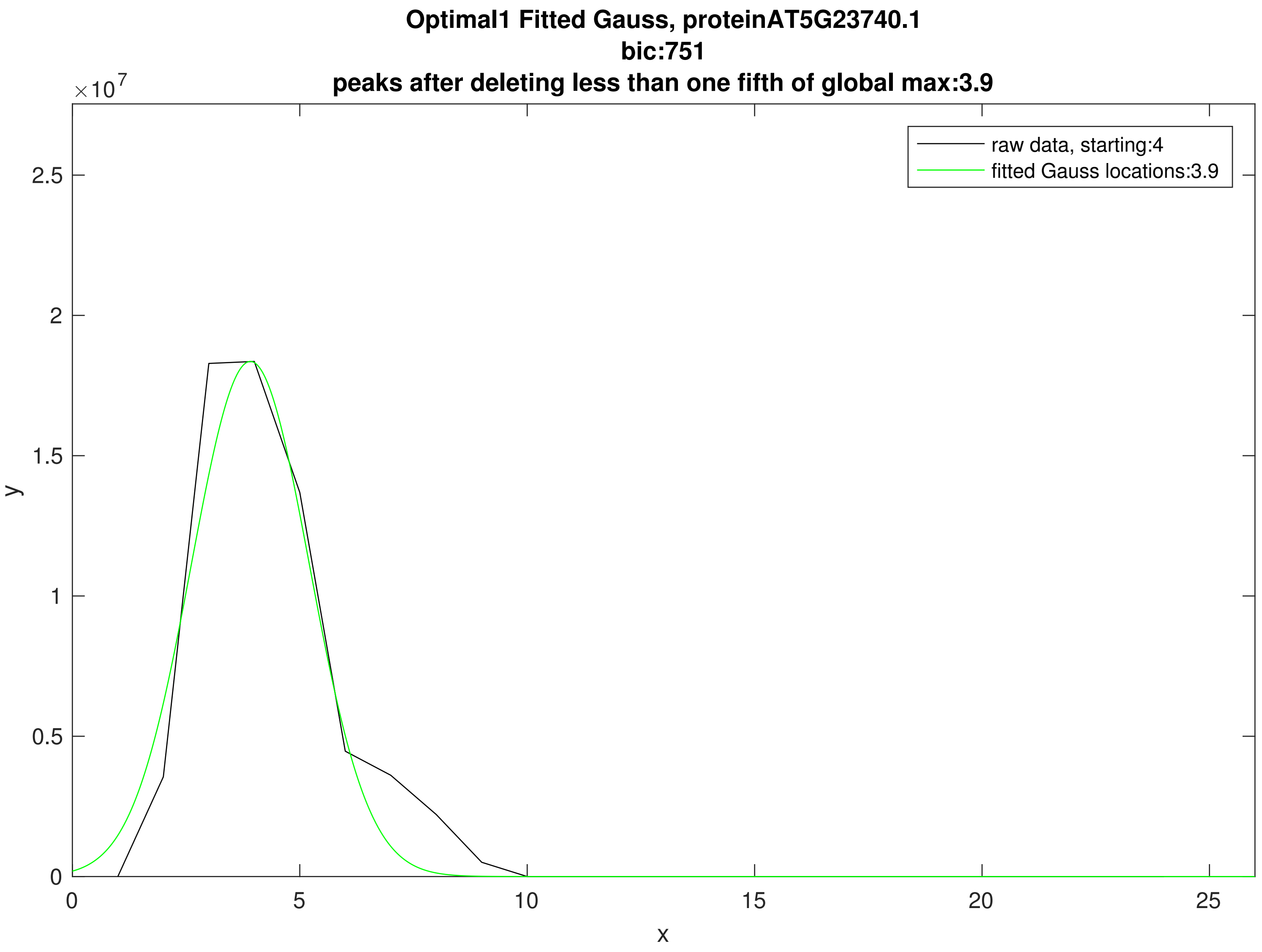

Optimal1 Fitted Gauss, proteinAT5G23740.1
bic:751
peaks after deleting less than one fifth of global max:3.9
×10^7
2.5
2
1.5
1
0.5
0
y
0
5
10
15
20
25
x
raw data, starting:4
fitted Gauss locations:3.9

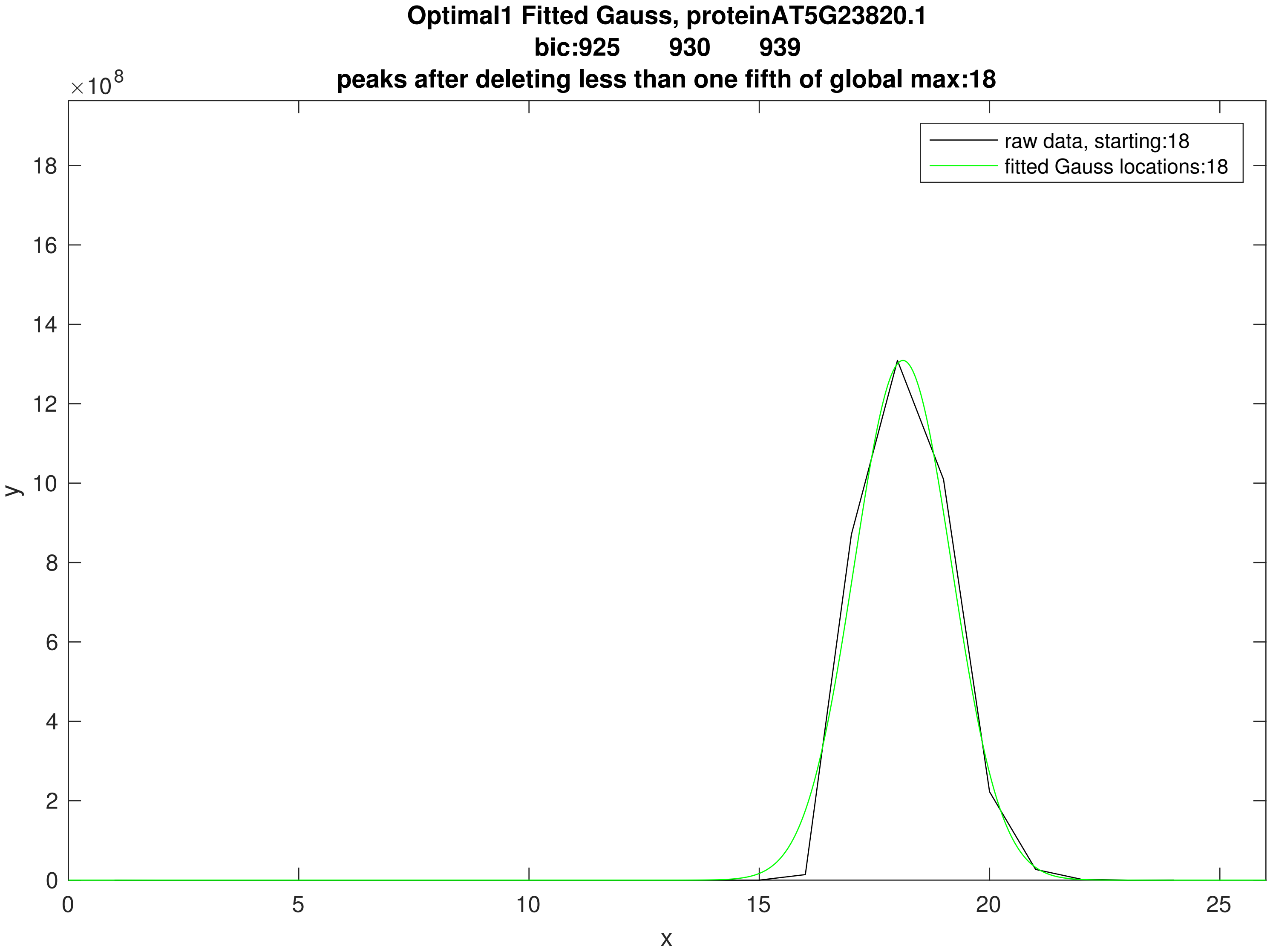
Optimal1 Fitted Gauss, proteinAT5G23820.1
bic:925 930 939
peaks after deleting less than one fifth of global max:18
×10^8
raw data, starting:18
fitted Gauss locations:18
18
16
14
12
10
8
6
4
2
0
y
0
5
10
15
20
25
x

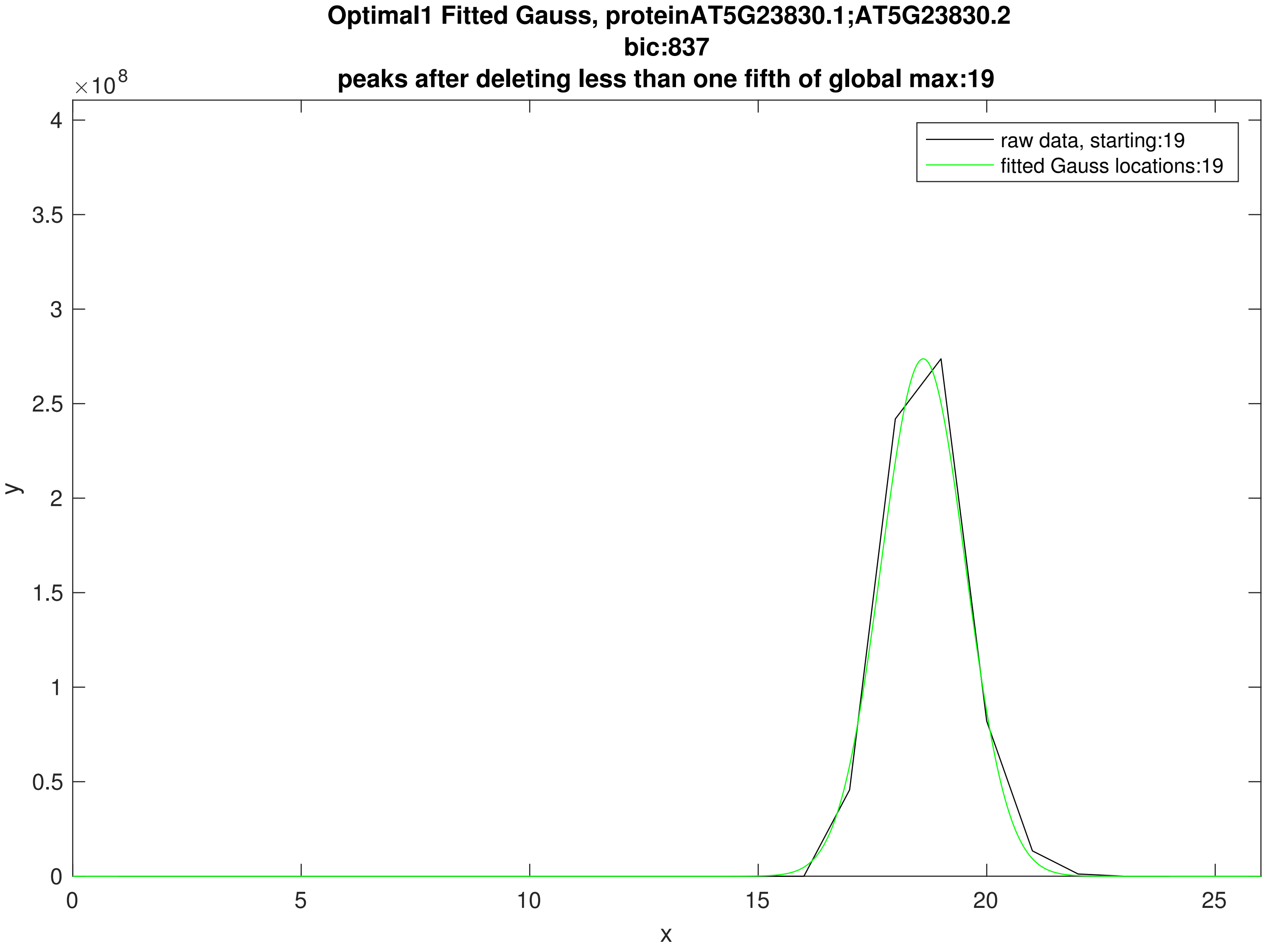
Optimal1 Fitted Gauss, proteinAT5G23830.1;AT5G23830.2
bic:837
peaks after deleting less than one fifth of global max:19
raw data, starting:19
fitted Gauss locations:19
×10^8
y
x

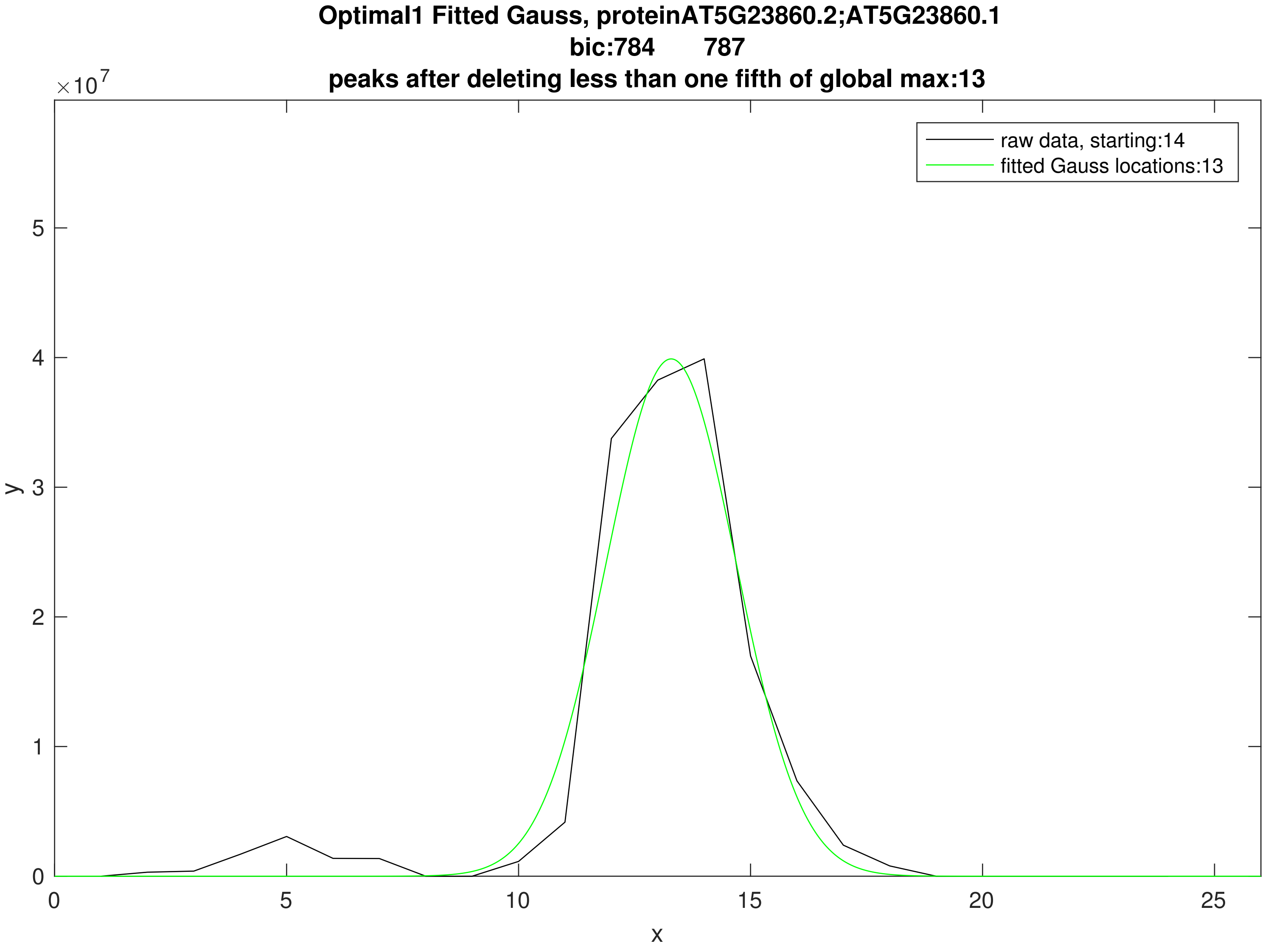

Optimal1 Fitted Gauss, proteinAT5G23860.2;AT5G23860.1
bic:784 787
peaks after deleting less than one fifth of global max:13
×10^7
raw data, starting:14
fitted Gauss locations:13
y
x
0
1
2
3
4
5
0
5
10
15
20
25

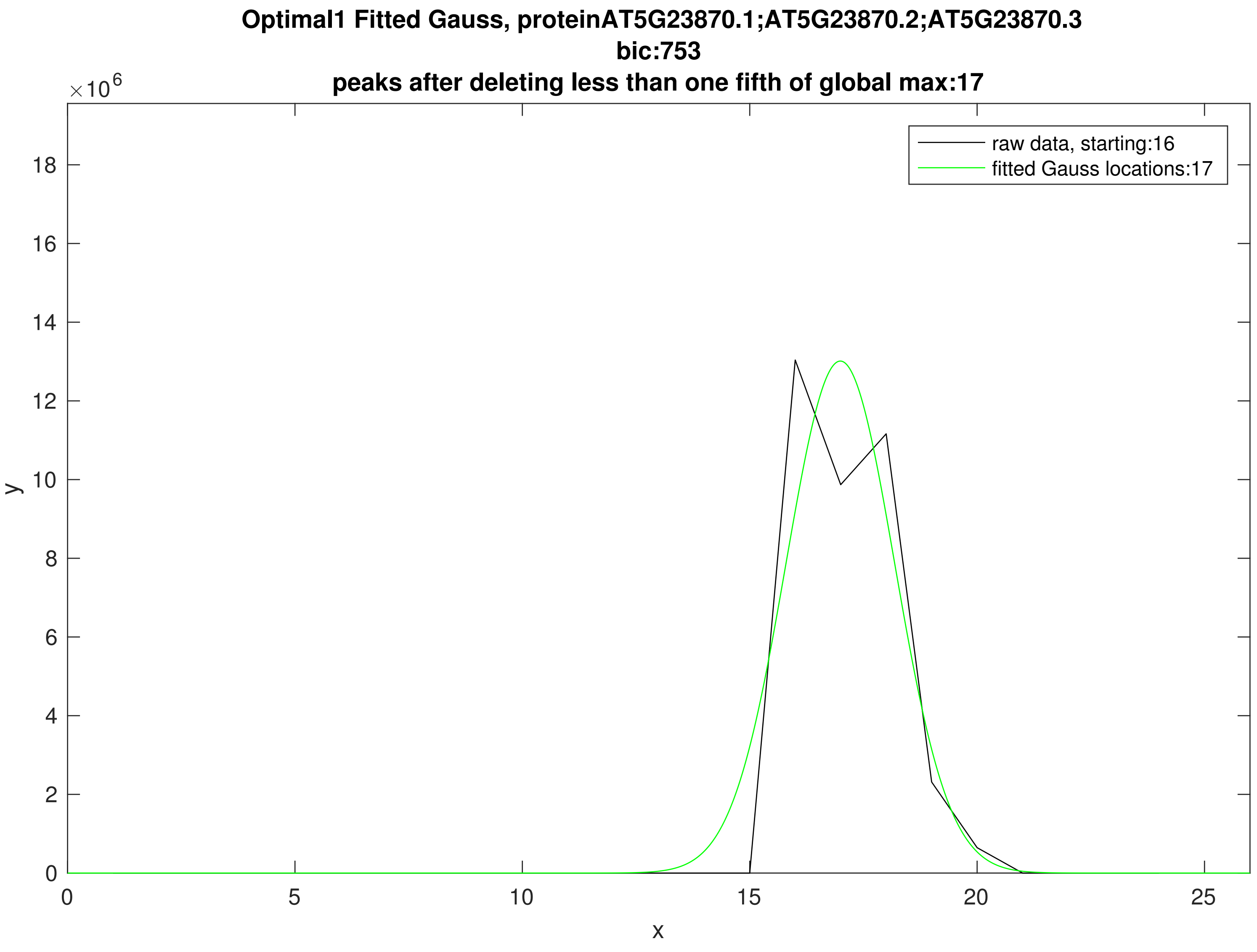
Optimal1 Fitted Gauss, proteinAT5G23870.1;AT5G23870.2;AT5G23870.3
bic:753
peaks after deleting less than one fifth of global max:17
×10^6
raw data, starting:16
fitted Gauss locations:17
y
x
0
2
4
6
8
10
12
14
16
18
0
5
10
15
20
25

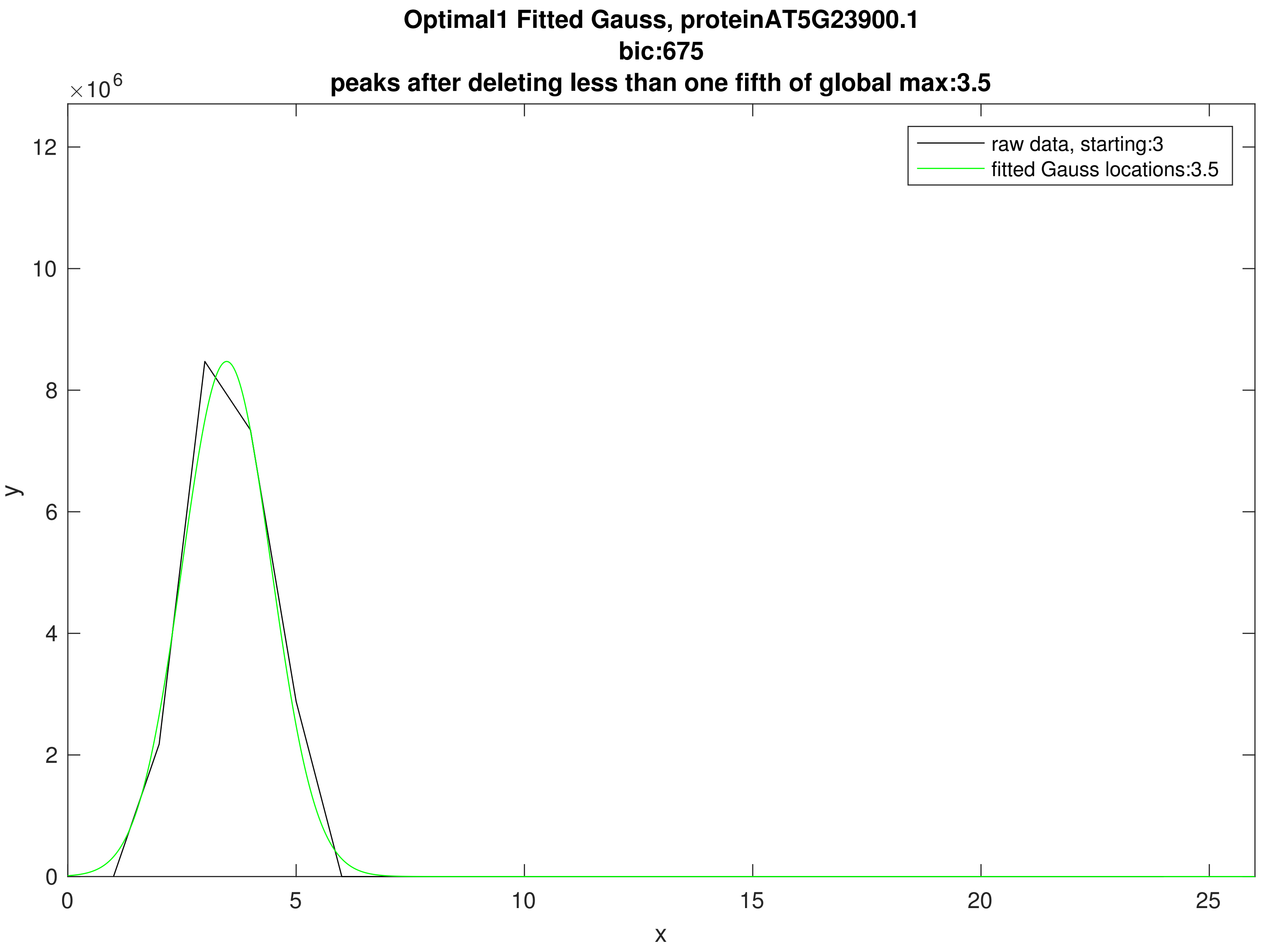
Optimal1 Fitted Gauss, proteinAT5G23900.1
bic:675
peaks after deleting less than one fifth of global max:3.5
×10^6
12
10
8
6
4
2
0
y
0
5
10
15
20
25
x
raw data, starting:3
fitted Gauss locations:3.5

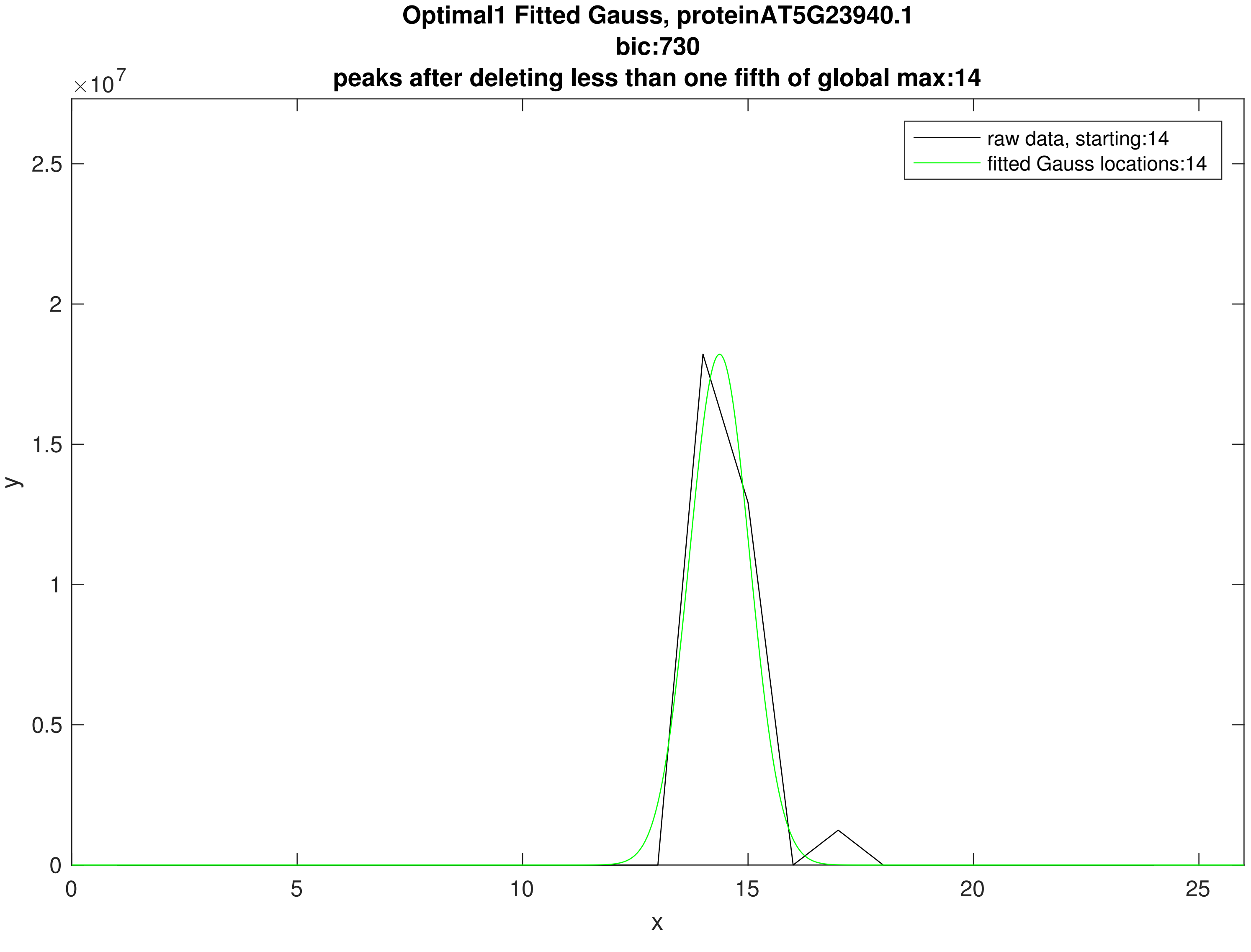
Optimal1 Fitted Gauss, proteinAT5G23940.1
bic:730
peaks after deleting less than one fifth of global max:14
×10^7
raw data, starting:14
fitted Gauss locations:14
2.5
2
1.5
1
0.5
0
y
0
5
10
15
20
25
x

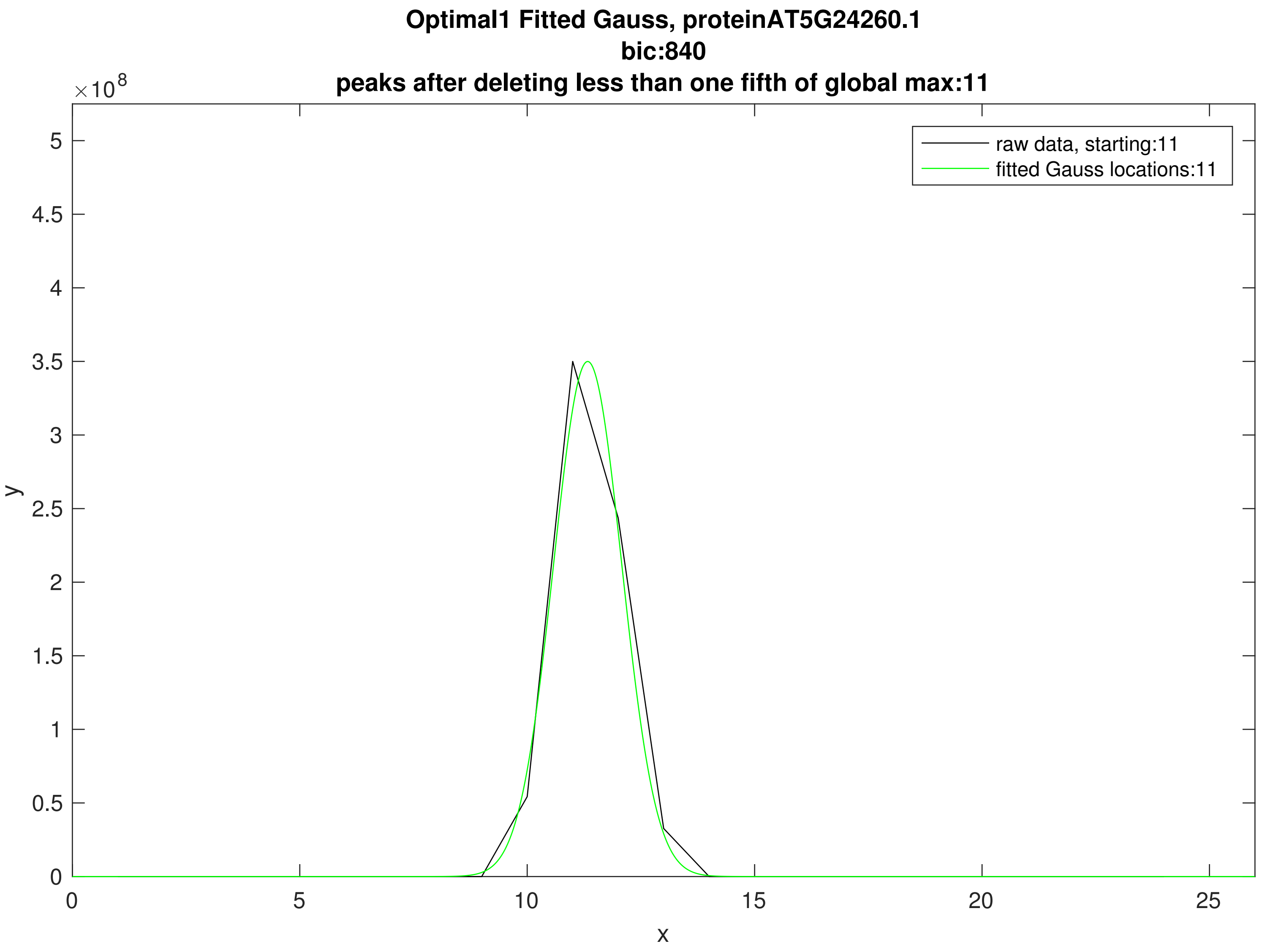

Optimal1 Fitted Gauss, proteinAT5G24260.1
bic:840
peaks after deleting less than one fifth of global max:11
×10^8
5
4.5
4
3.5
3
2.5
2
1.5
1
0.5
0
y
0
5
10
15
20
25
x
raw data, starting:11
fitted Gauss locations:11

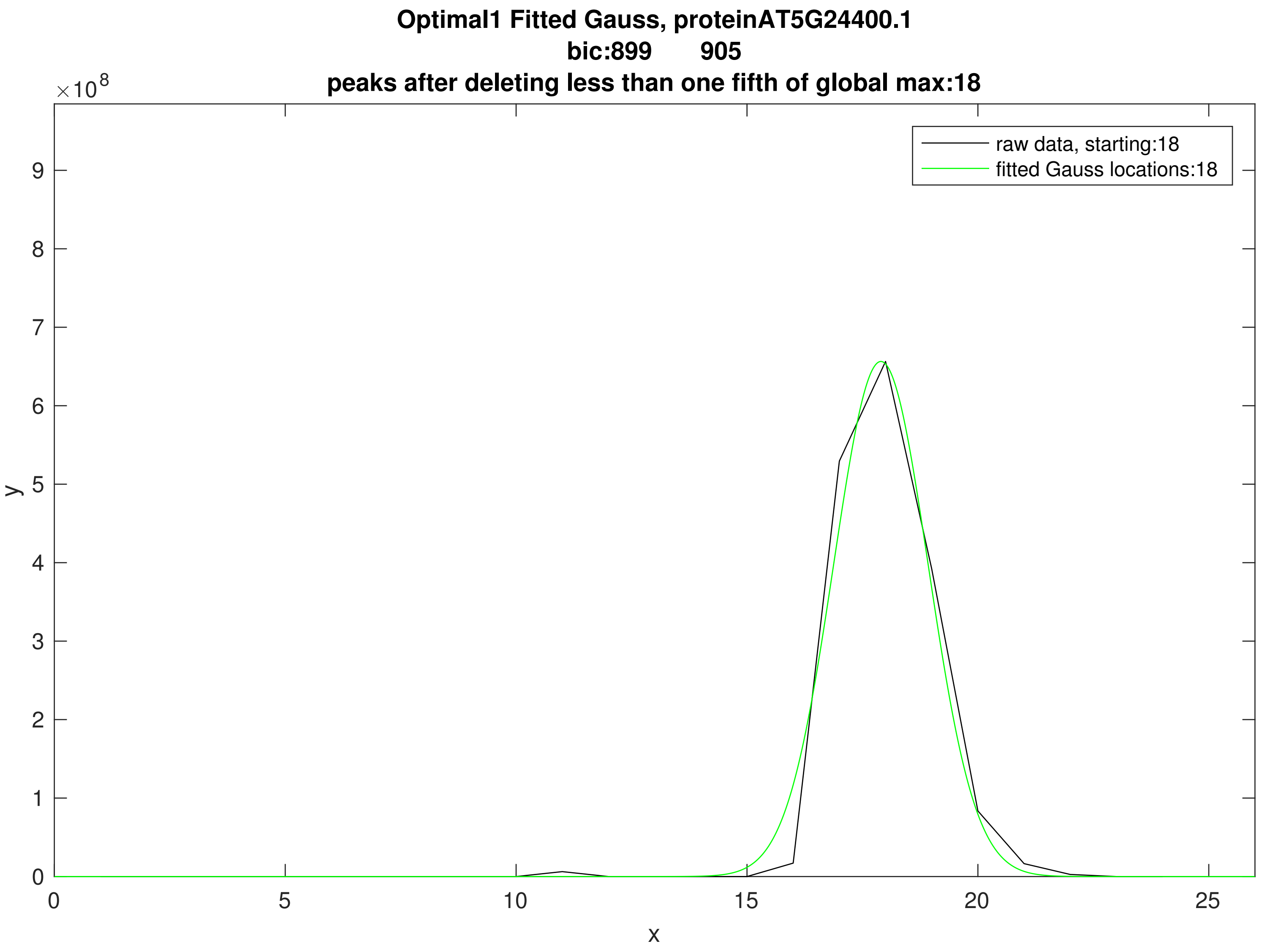
Optimal1 Fitted Gauss, proteinAT5G24400.1
bic:899 905
peaks after deleting less than one fifth of global max:18
×10^8
raw data, starting:18
fitted Gauss locations:18
y
x
0
1
2
3
4
5
6
7
8
9
0
5
10
15
20
25

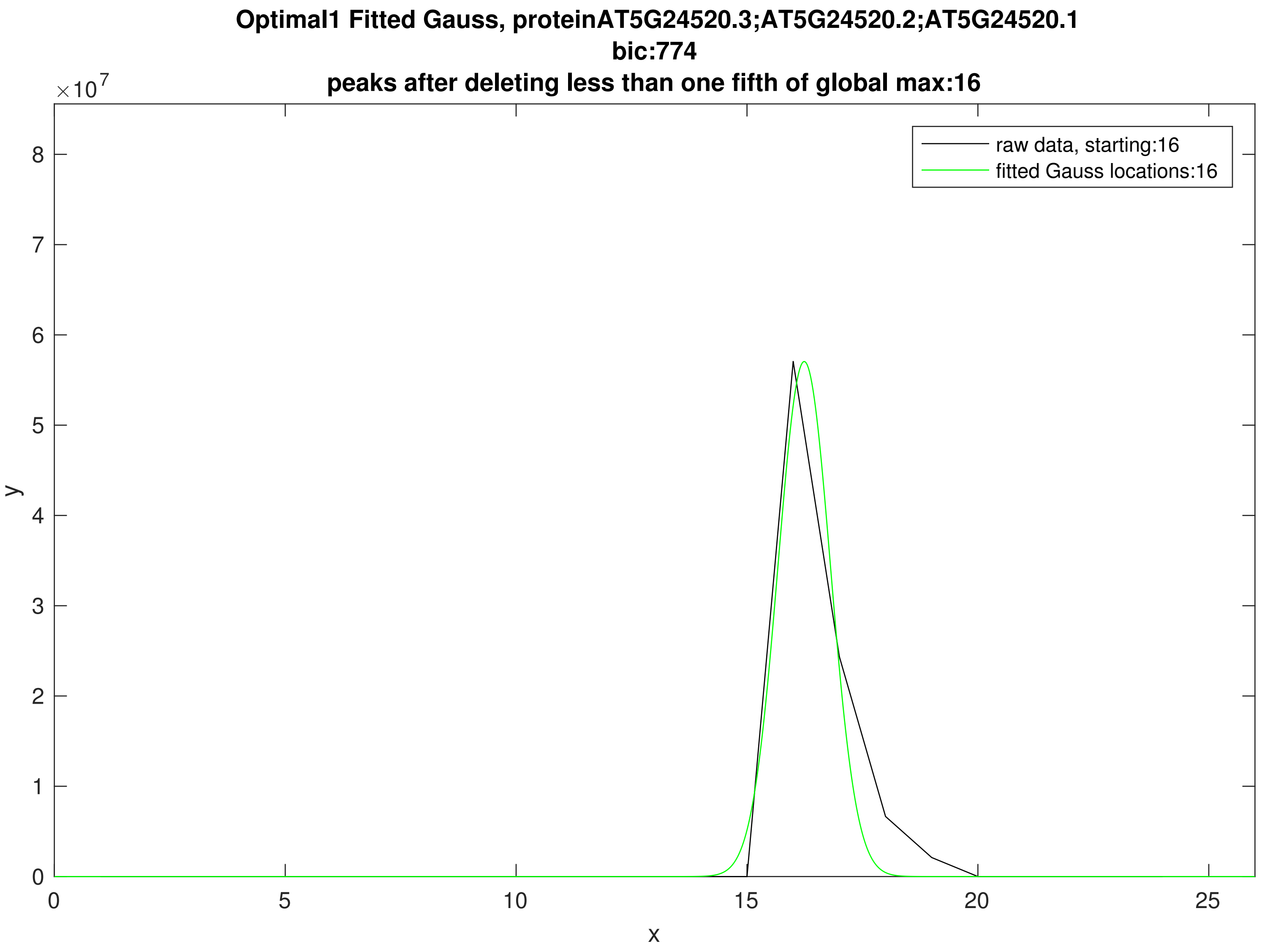
Optimal1 Fitted Gauss, proteinAT5G24520.3;AT5G24520.2;AT5G24520.1
bic:774
peaks after deleting less than one fifth of global max:16
×10^7
raw data, starting:16
fitted Gauss locations:16
y
x

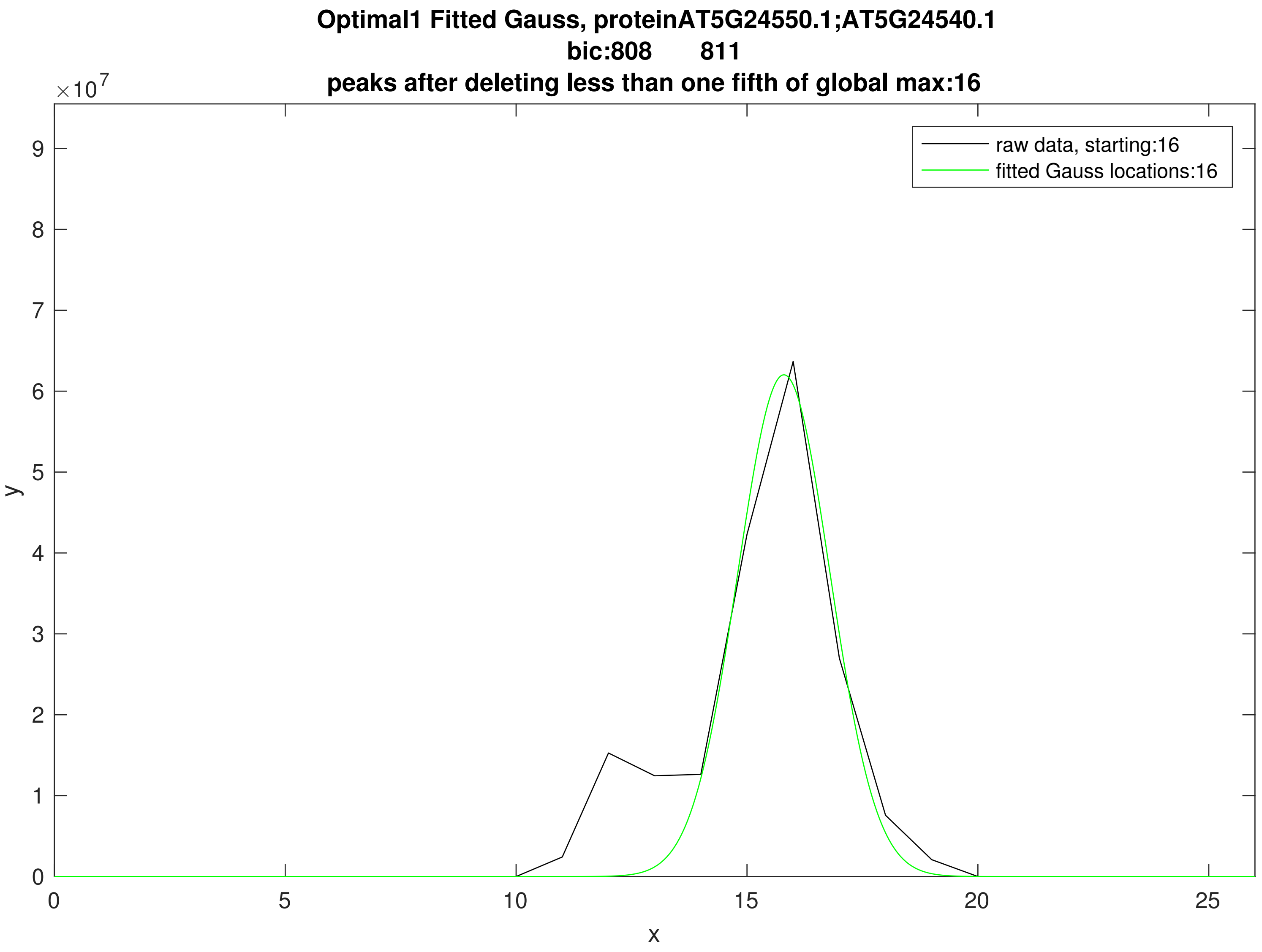
Optimal1 Fitted Gauss, proteinAT5G24550.1;AT5G24540.1
bic:808 811
peaks after deleting less than one fifth of global max:16
×10^7
raw data, starting:16
fitted Gauss locations:16
y
x
0
1
2
3
4
5
6
7
8
9
0
5
10
15
20
25

Optimal1 Fitted Gauss, proteinAT5G24760.3;AT5G24760.2;AT5G24760.1
bic:765
peaks after deleting less than one fifth of global max:14

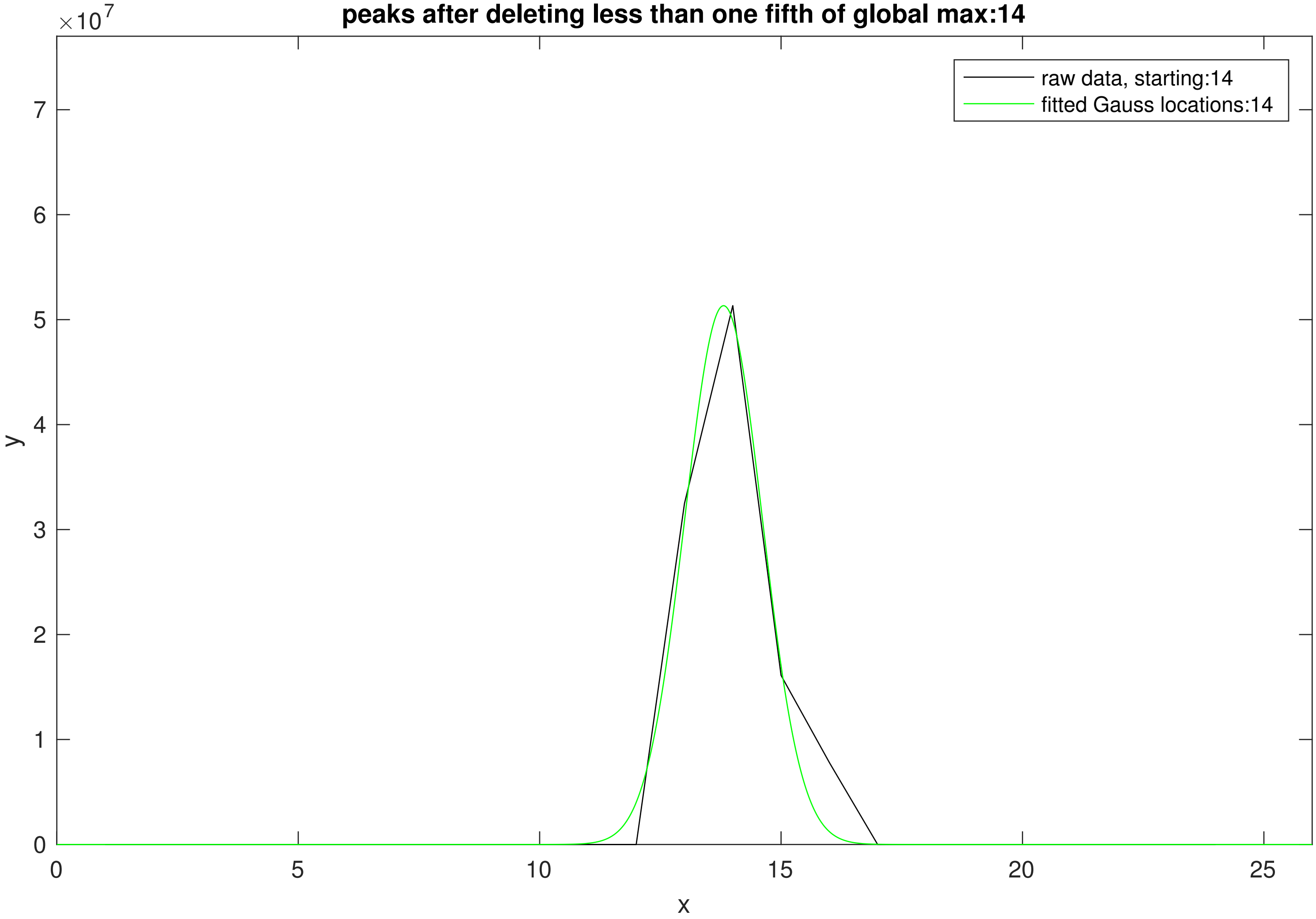

**Optimal1 Fitted Gauss, proteinAT5G25770.1;AT5G25770.2;AT5G25770.3**
**bic:874**
**peaks after deleting less than one fifth of global max:17**

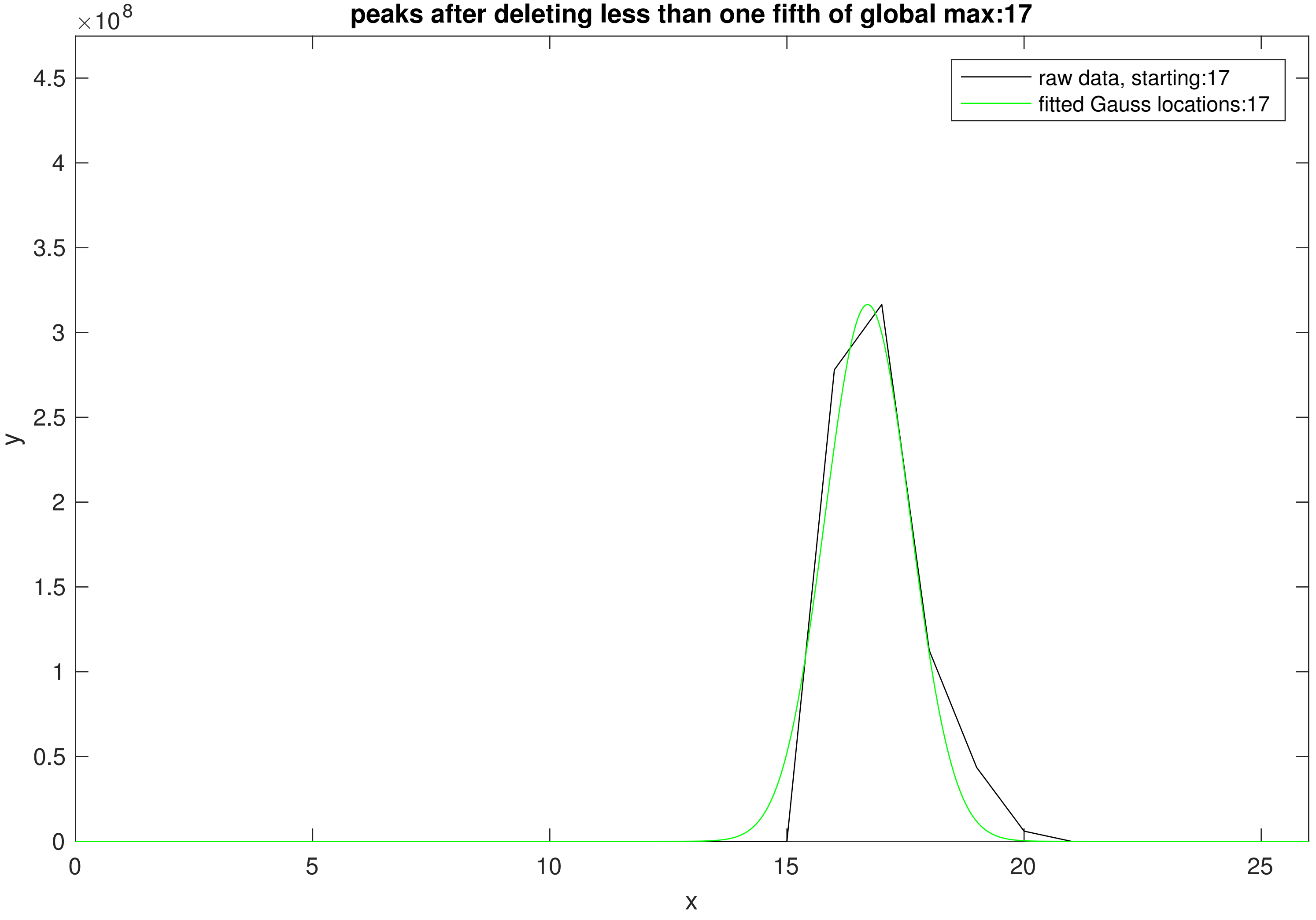

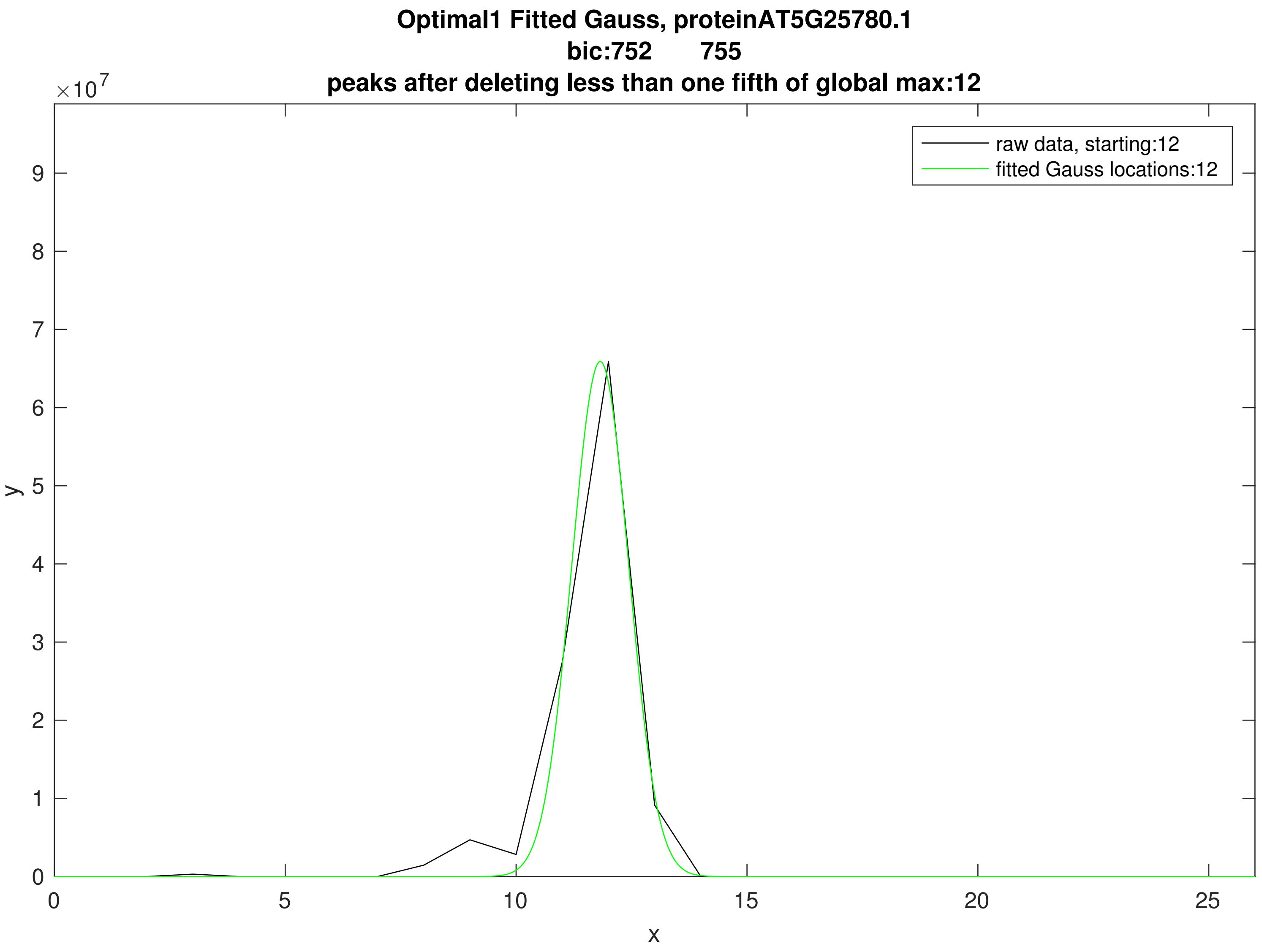
Optimal1 Fitted Gauss, proteinAT5G25780.1
bic:752 755
peaks after deleting less than one fifth of global max:12
×10^7
raw data, starting:12
fitted Gauss locations:12
y
x
0
1
2
3
4
5
6
7
8
9
10
15
20
25

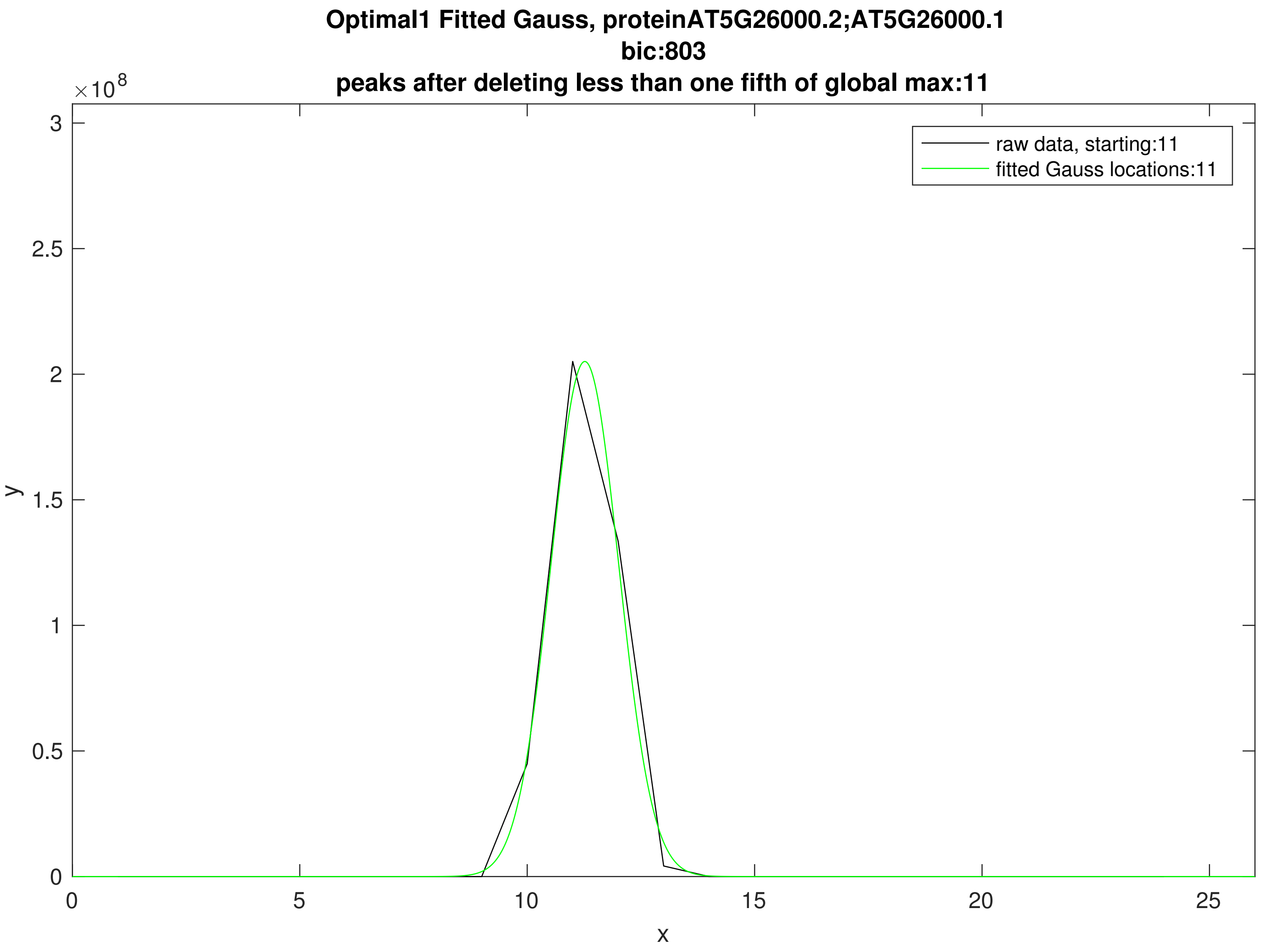
Optimal1 Fitted Gauss, proteinAT5G26000.2;AT5G26000.1
bic:803
peaks after deleting less than one fifth of global max:11
×10^8
raw data, starting:11
fitted Gauss locations:11
y
x
0
0.5
1
1.5
2
2.5
3
0
5
10
15
20
25

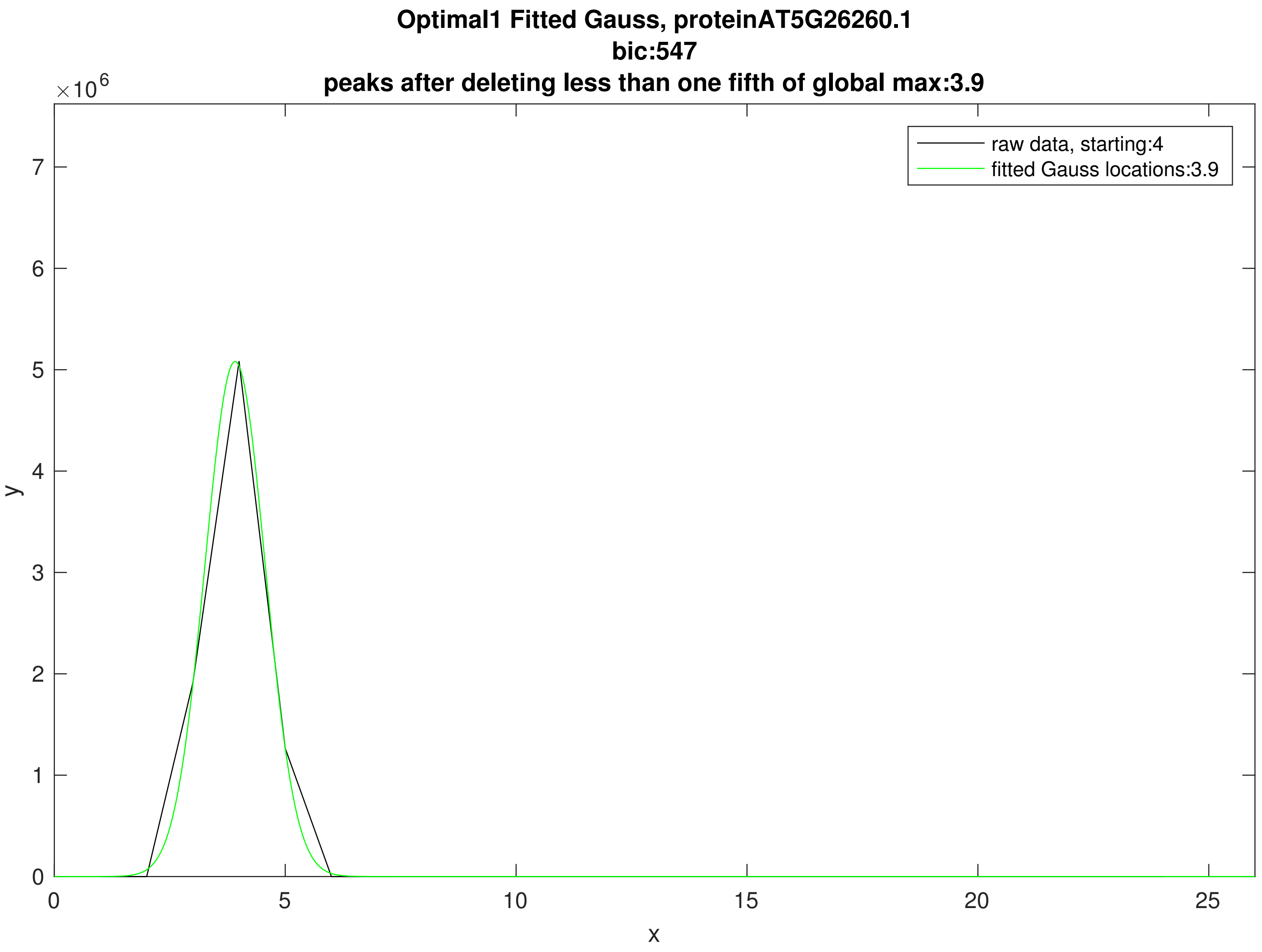
Optimal1 Fitted Gauss, proteinAT5G26260.1
bic:547
peaks after deleting less than one fifth of global max:3.9
×10^6
raw data, starting:4
fitted Gauss locations:3.9
7
6
5
4
3
2
1
0
y
0
5
10
15
20
25
x

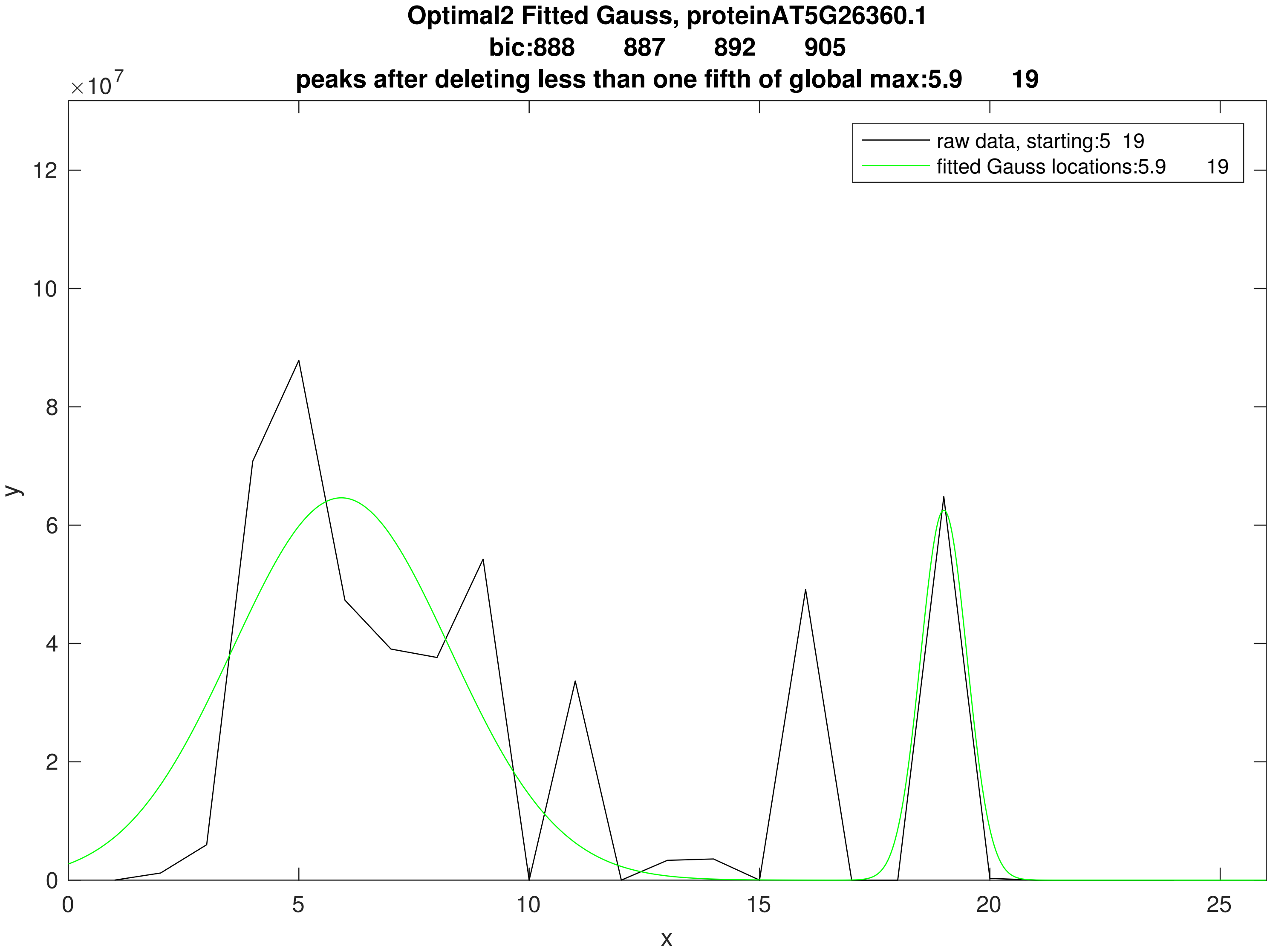

Optimal2 Fitted Gauss, proteinAT5G26360.1
bic:888 887 892 905
peaks after deleting less than one fifth of global max:5.9 19
×10^7
raw data, starting:5 19
fitted Gauss locations:5.9 19
12
10
8
6
4
2
0
y
0
5
10
15
20
25
x

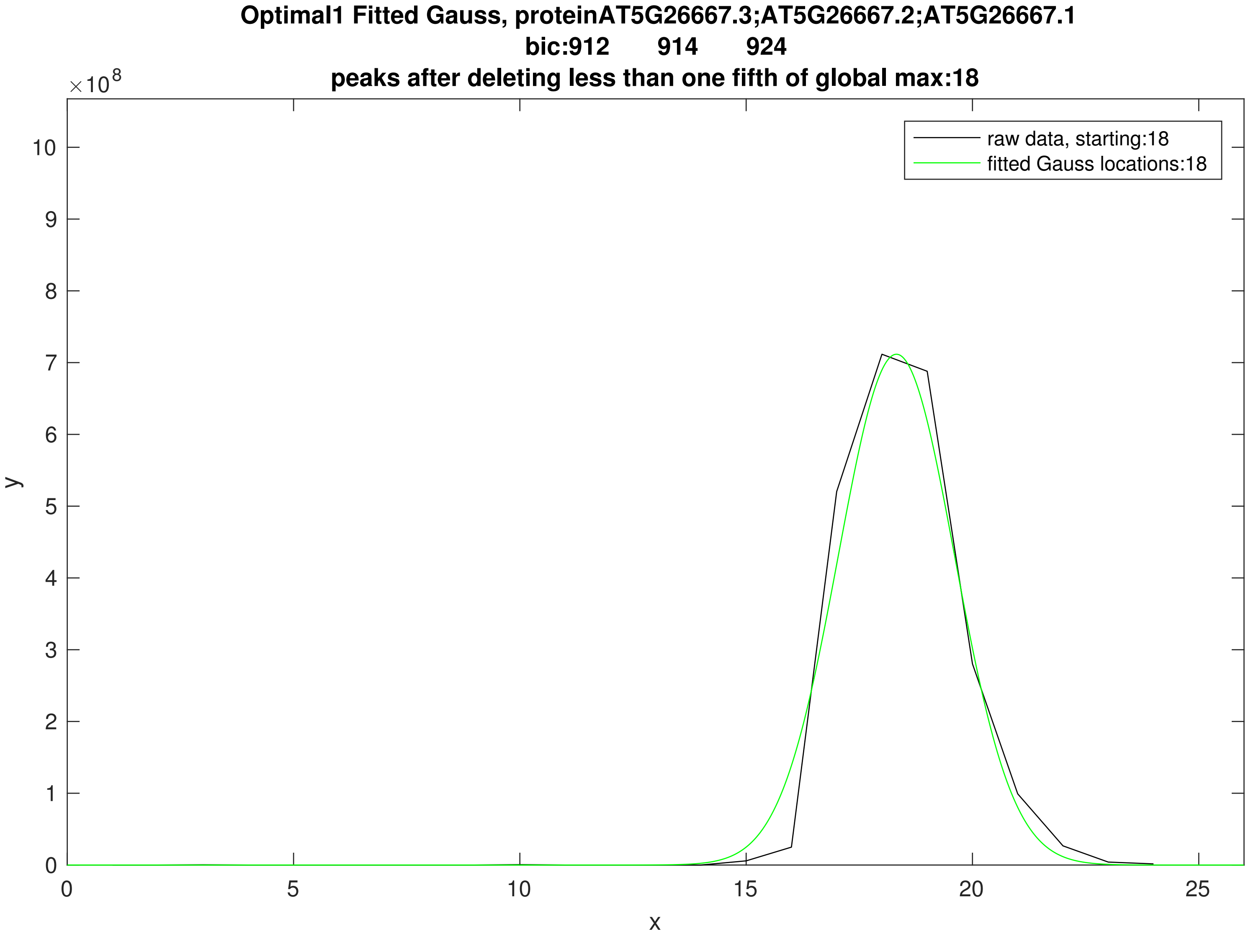
Optimal1 Fitted Gauss, proteinAT5G26667.3;AT5G26667.2;AT5G26667.1
bic:912 914 924
peaks after deleting less than one fifth of global max:18
×10^8
raw data, starting:18
fitted Gauss locations:18
y
x
0
1
2
3
4
5
6
7
8
9
10
0
5
10
15
20
25

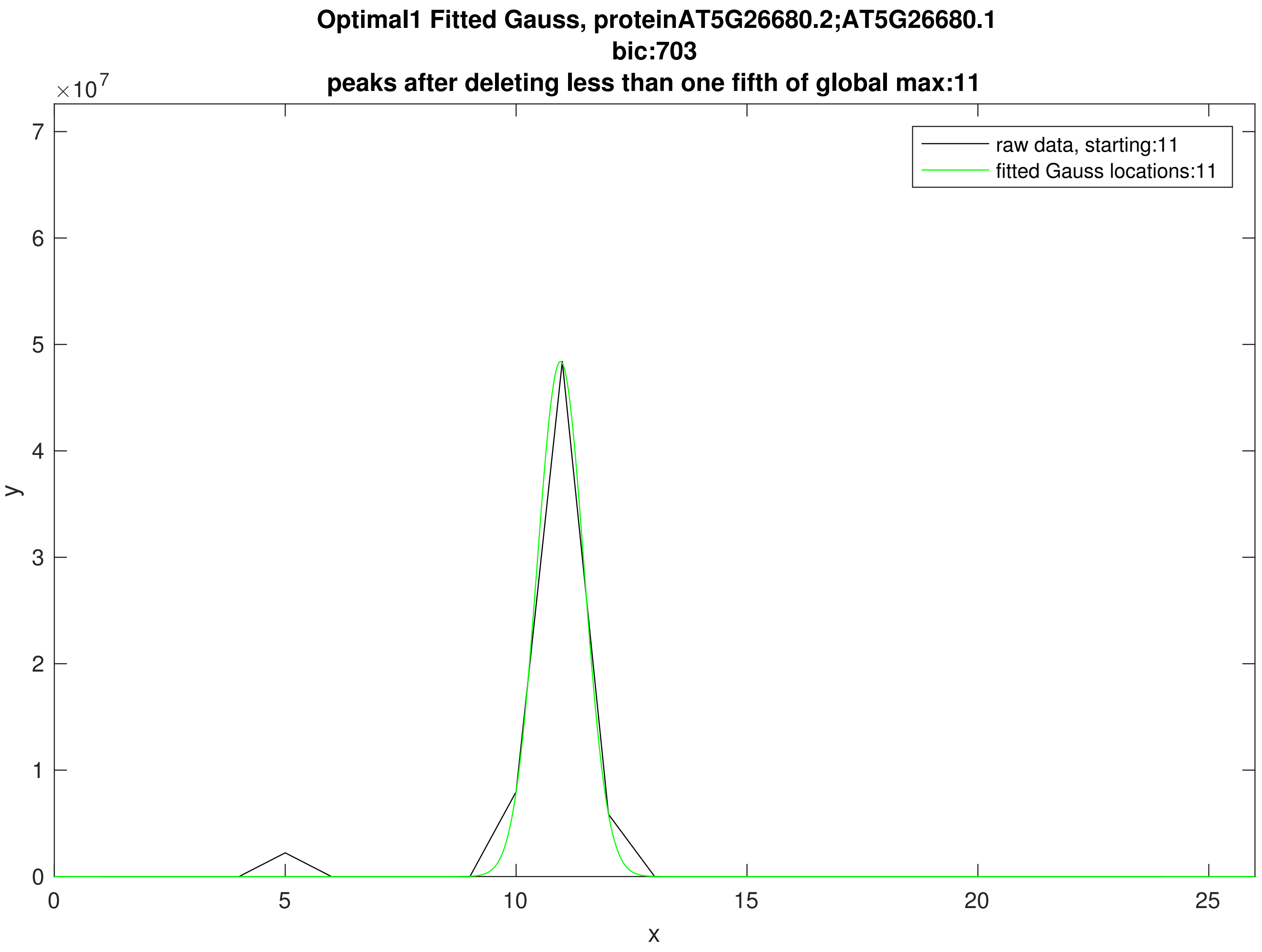

Optimal1 Fitted Gauss, proteinAT5G26680.2;AT5G26680.1
bic:703
peaks after deleting less than one fifth of global max:11
×10^7
raw data, starting:11
fitted Gauss locations:11
y
x
0
1
2
3
4
5
6
7
0
5
10
15
20
25

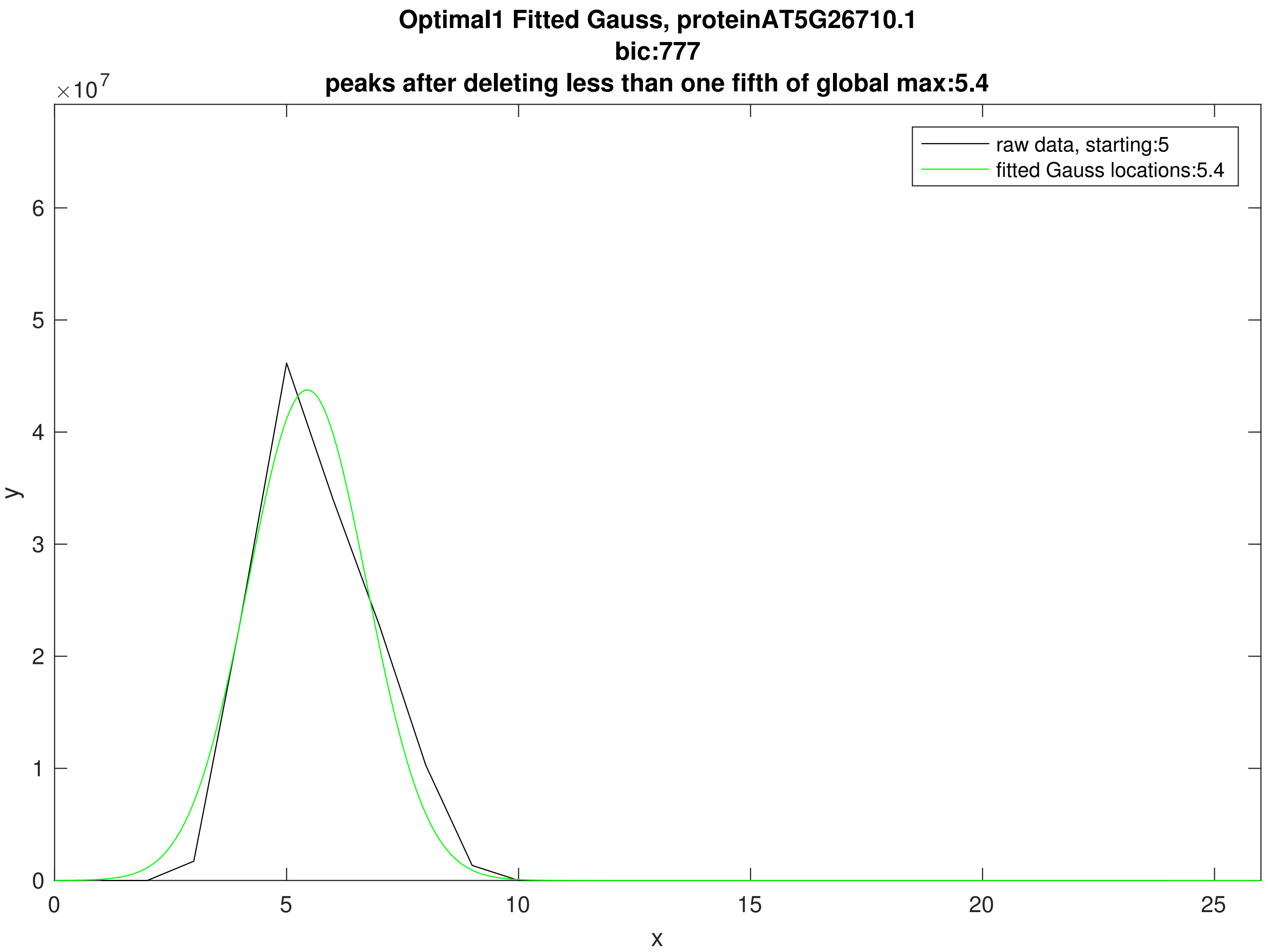
Optimal1 Fitted Gauss, proteinAT5G26710.1
bic:777
peaks after deleting less than one fifth of global max:5.4
×10^7
raw data, starting:5
fitted Gauss locations:5.4
y
x
0
1
2
3
4
5
6
0
5
10
15
20
25

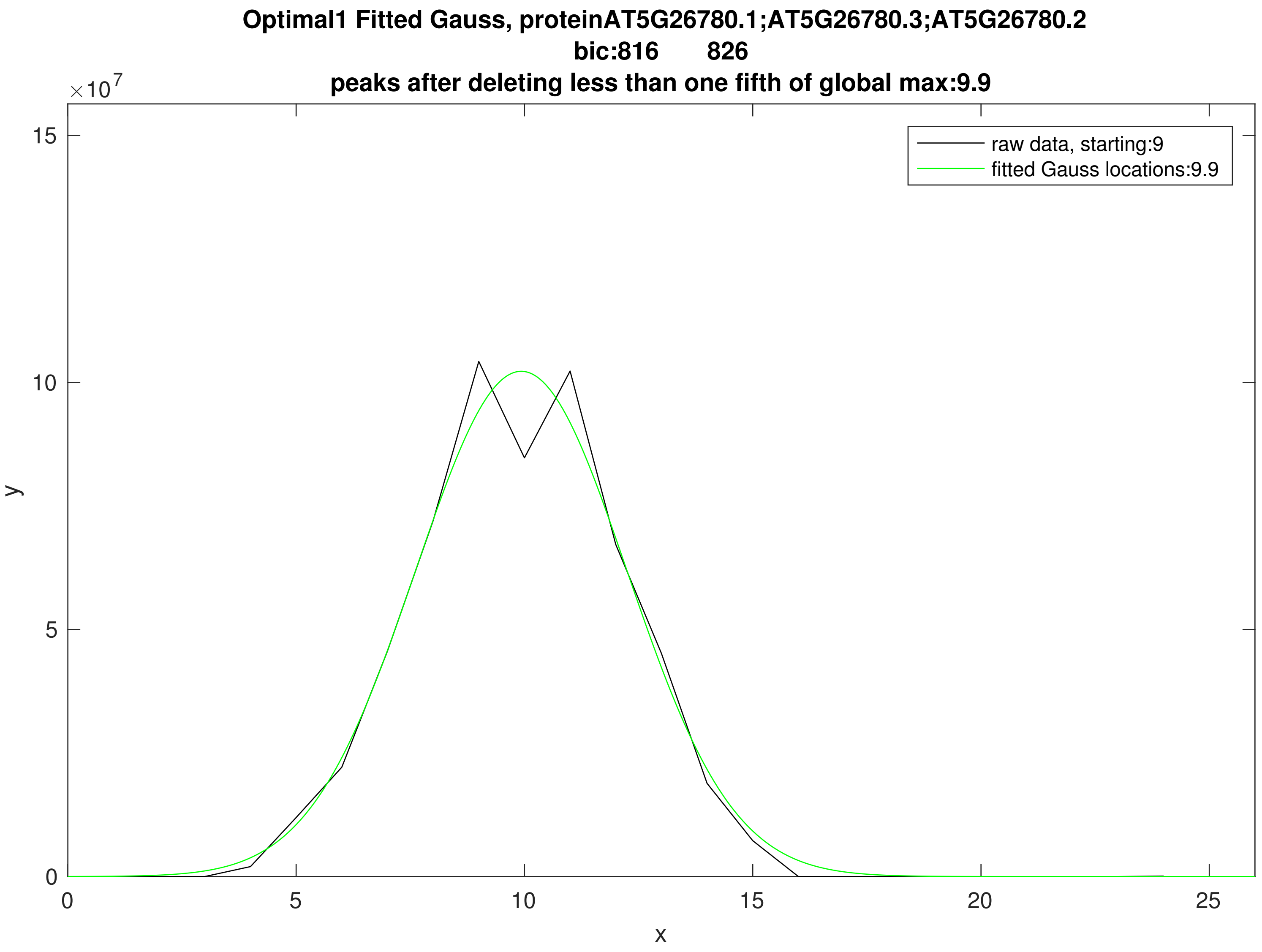

Optimal1 Fitted Gauss, proteinAT5G26780.1;AT5G26780.3;AT5G26780.2
bic:816 826
peaks after deleting less than one fifth of global max:9.9
×10^7
15
10
5
0
y
0
5
10
15
20
25
x
raw data, starting:9
fitted Gauss locations:9.9

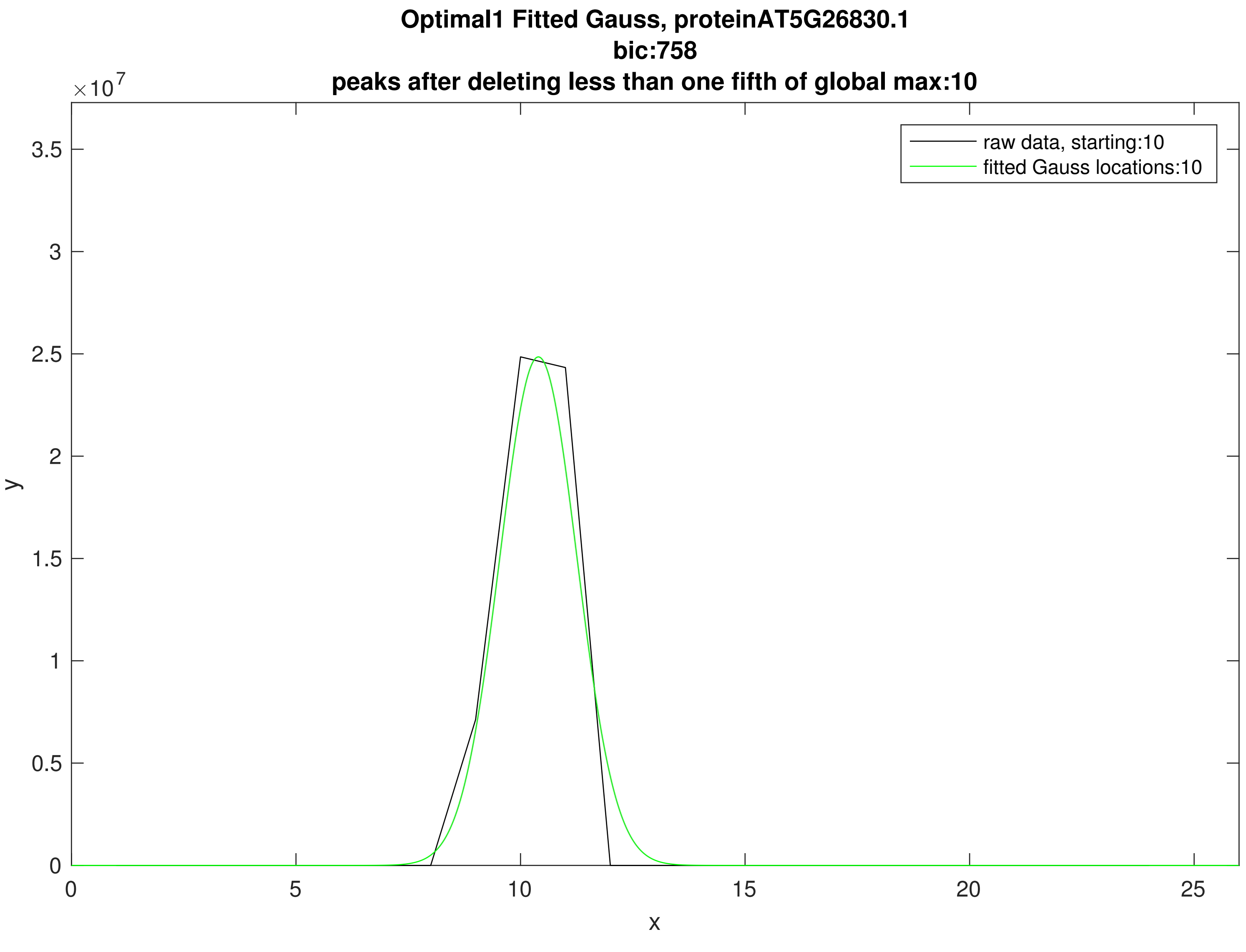
Optimal1 Fitted Gauss, proteinAT5G26830.1
bic:758
peaks after deleting less than one fifth of global max:10
×10^7
raw data, starting:10
fitted Gauss locations:10
3.5
3
2.5
2
1.5
1
0.5
0
y
0
5
10
15
20
25
x

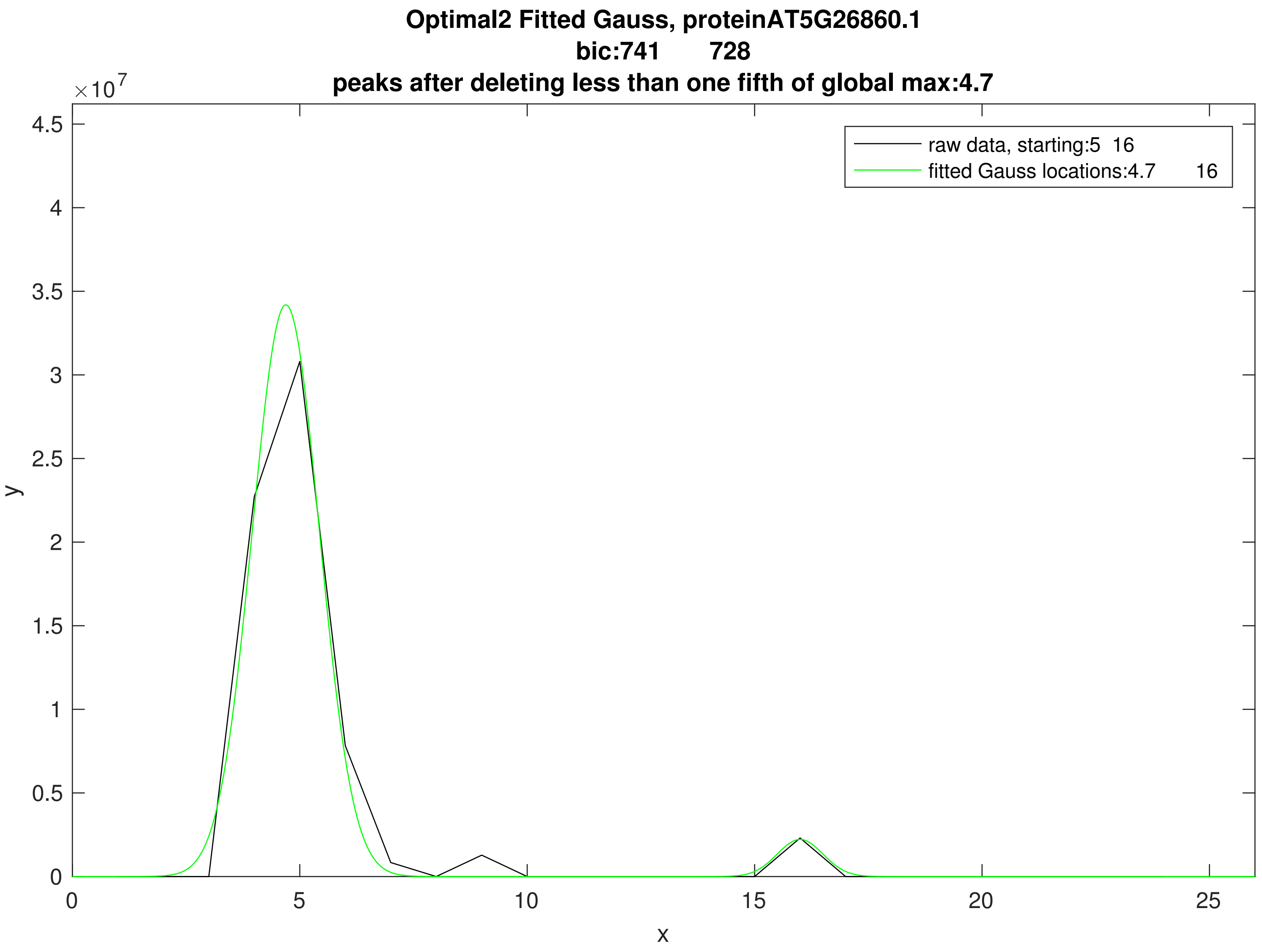

Optimal2 Fitted Gauss, proteinAT5G26860.1
bic:741 728
peaks after deleting less than one fifth of global max:4.7
×10^7
raw data, starting:5 16
fitted Gauss locations:4.7 16
y
x
0
0.5
1
1.5
2
2.5
3
3.5
4
4.5
0
5
10
15
20
25

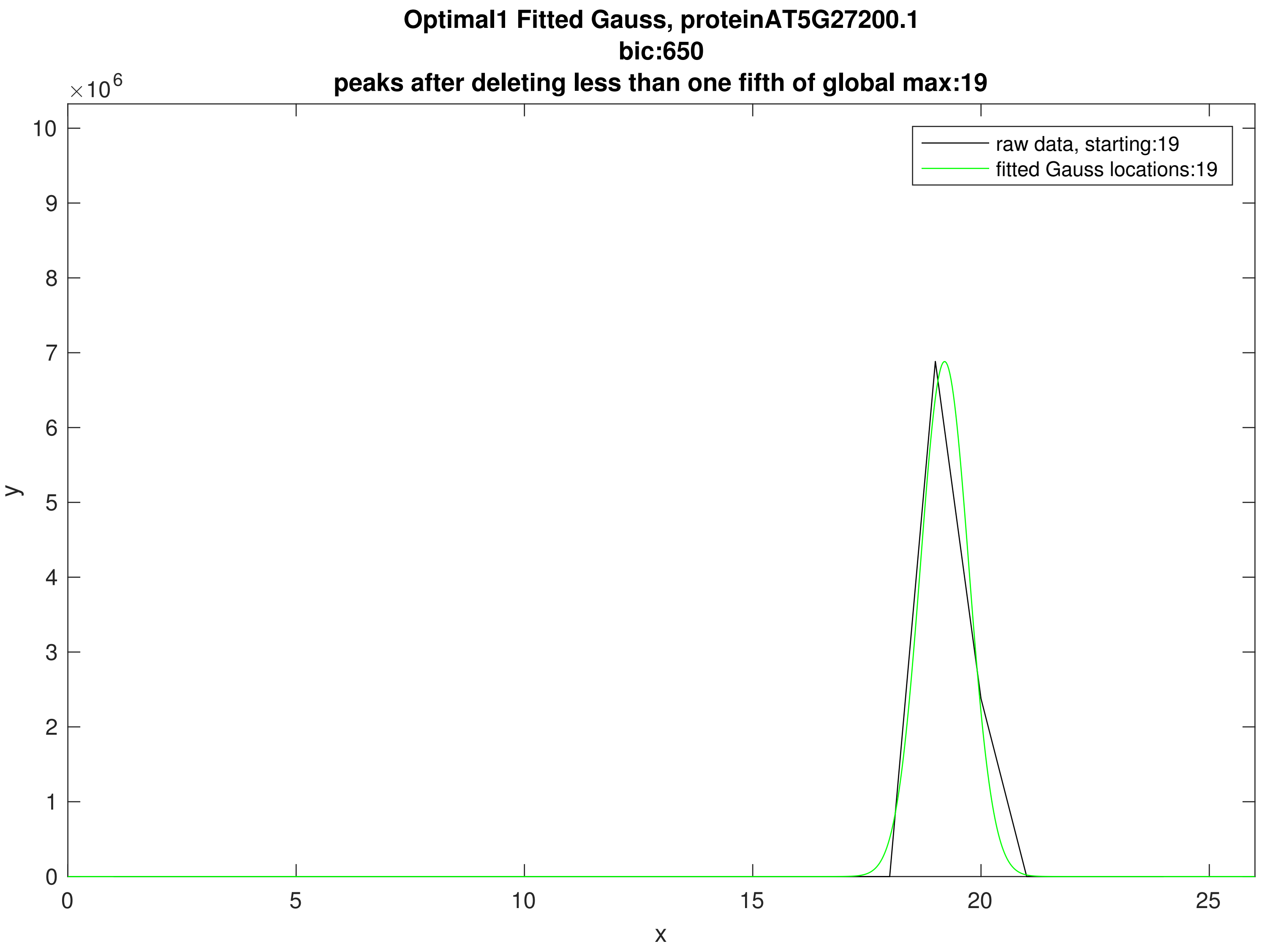
Optimal1 Fitted Gauss, proteinAT5G27200.1
bic:650
peaks after deleting less than one fifth of global max:19
×10^6
raw data, starting:19
fitted Gauss locations:19
y
x
0
1
2
3
4
5
6
7
8
9
10
0
5
10
15
20
25

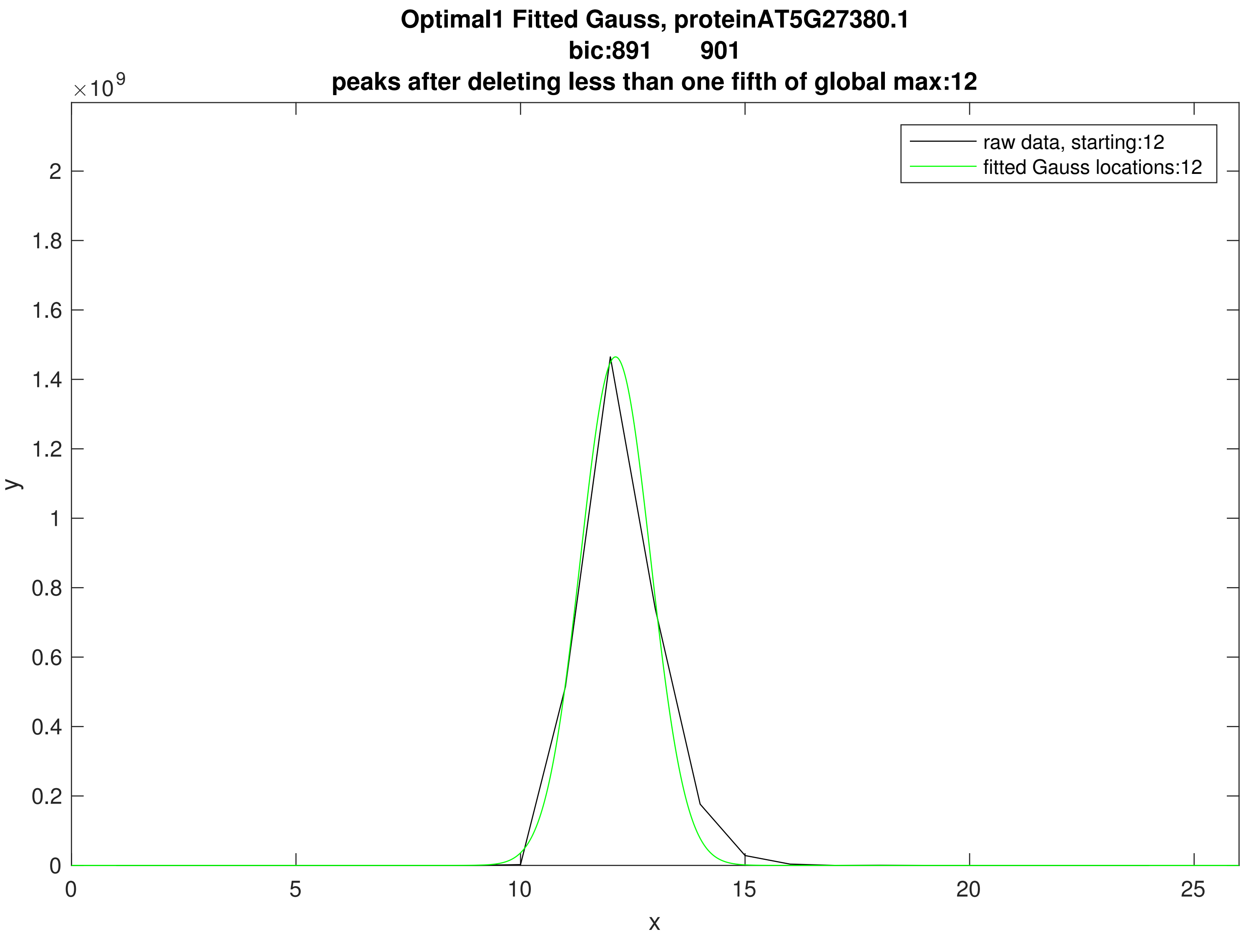

Optimal1 Fitted Gauss, proteinAT5G27380.1
bic:891 901
peaks after deleting less than one fifth of global max:12
×10^9
2
1.8
1.6
1.4
1.2
1
0.8
0.6
0.4
0.2
0
y
0
5
10
15
20
25
x
raw data, starting:12
fitted Gauss locations:12

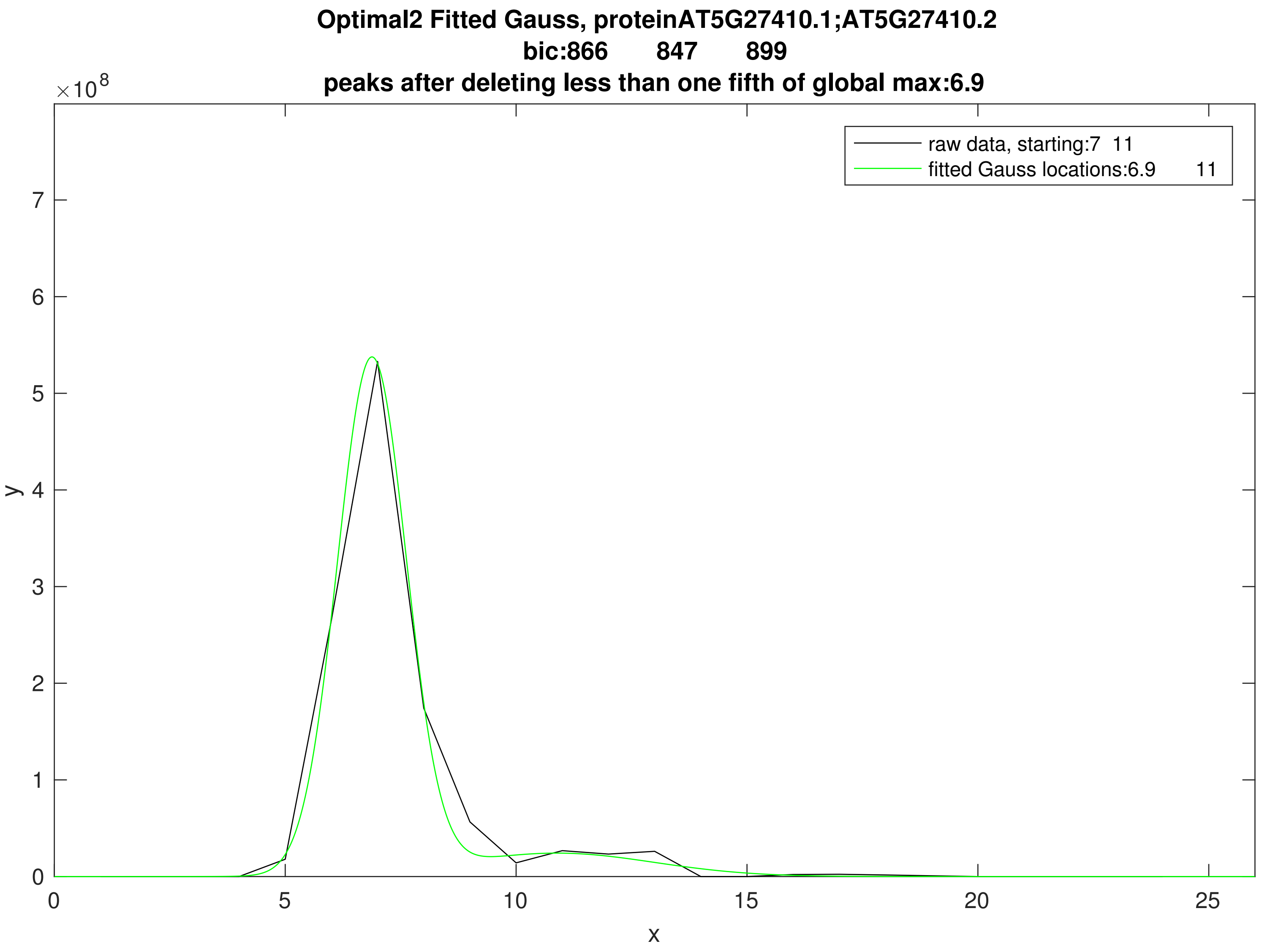

Optimal2 Fitted Gauss, proteinAT5G27410.1;AT5G27410.2
bic:866 847 899
peaks after deleting less than one fifth of global max:6.9
×10^8
raw data, starting:7 11
fitted Gauss locations:6.9 11
y
x
0
1
2
3
4
5
6
7
0
5
10
15
20
25

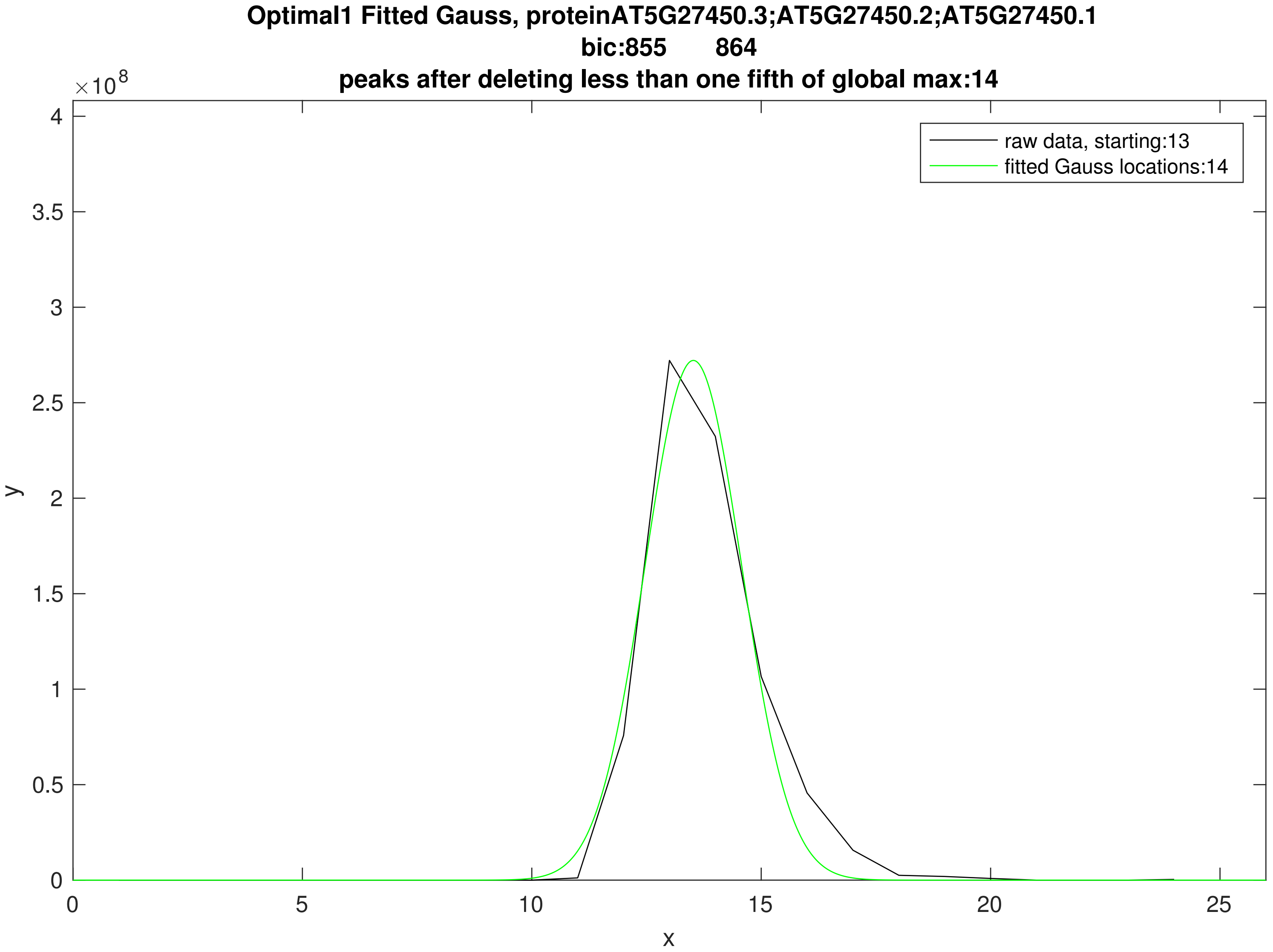
Optimal1 Fitted Gauss, proteinAT5G27450.3;AT5G27450.2;AT5G27450.1
bic:855 864
peaks after deleting less than one fifth of global max:14
×10^8
raw data, starting:13
fitted Gauss locations:14
y
x
0
0.5
1
1.5
2
2.5
3
3.5
4
0
5
10
15
20
25

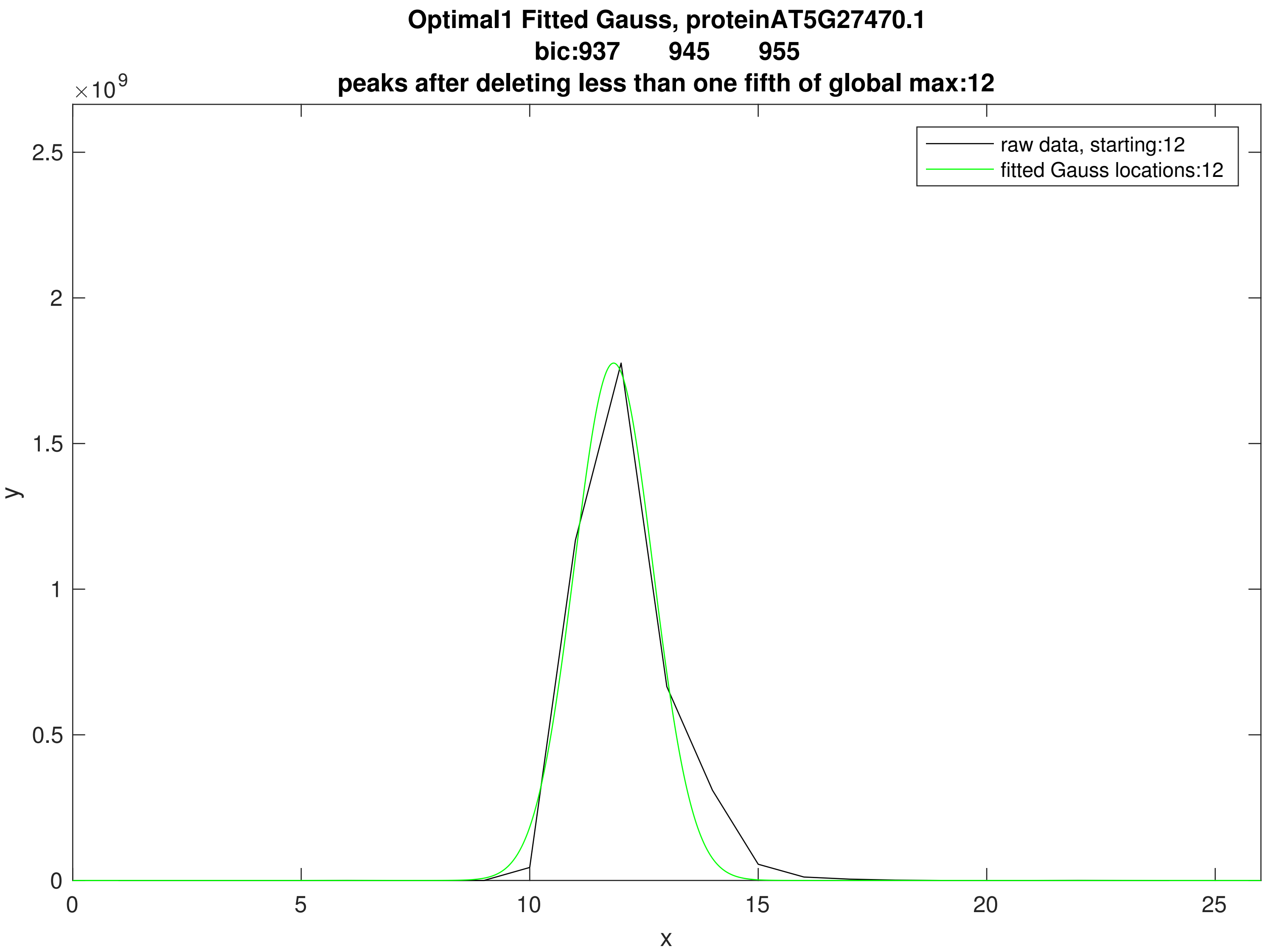
Optimal1 Fitted Gauss, proteinAT5G27470.1
bic:937 945 955
peaks after deleting less than one fifth of global max:12
×10^9
2.5
2
1.5
1
0.5
0
y
0
5
10
15
20
25
x
raw data, starting:12
fitted Gauss locations:12

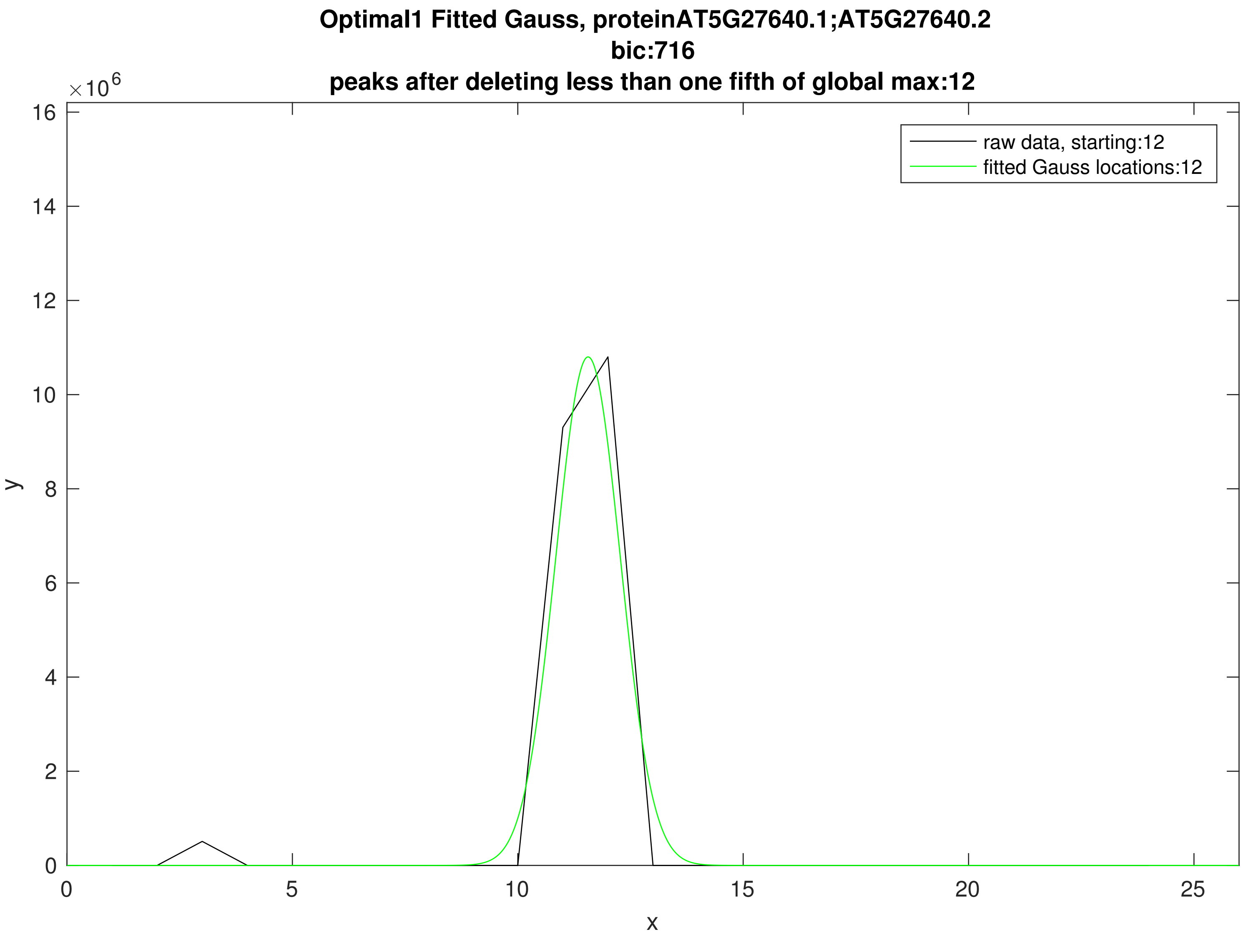

Optimal1 Fitted Gauss, proteinAT5G27640.1;AT5G27640.2
bic:716
peaks after deleting less than one fifth of global max:12
×10^6
16
14
12
10
8
6
4
2
0
y
0
5
10
15
20
25
x
raw data, starting:12
fitted Gauss locations:12

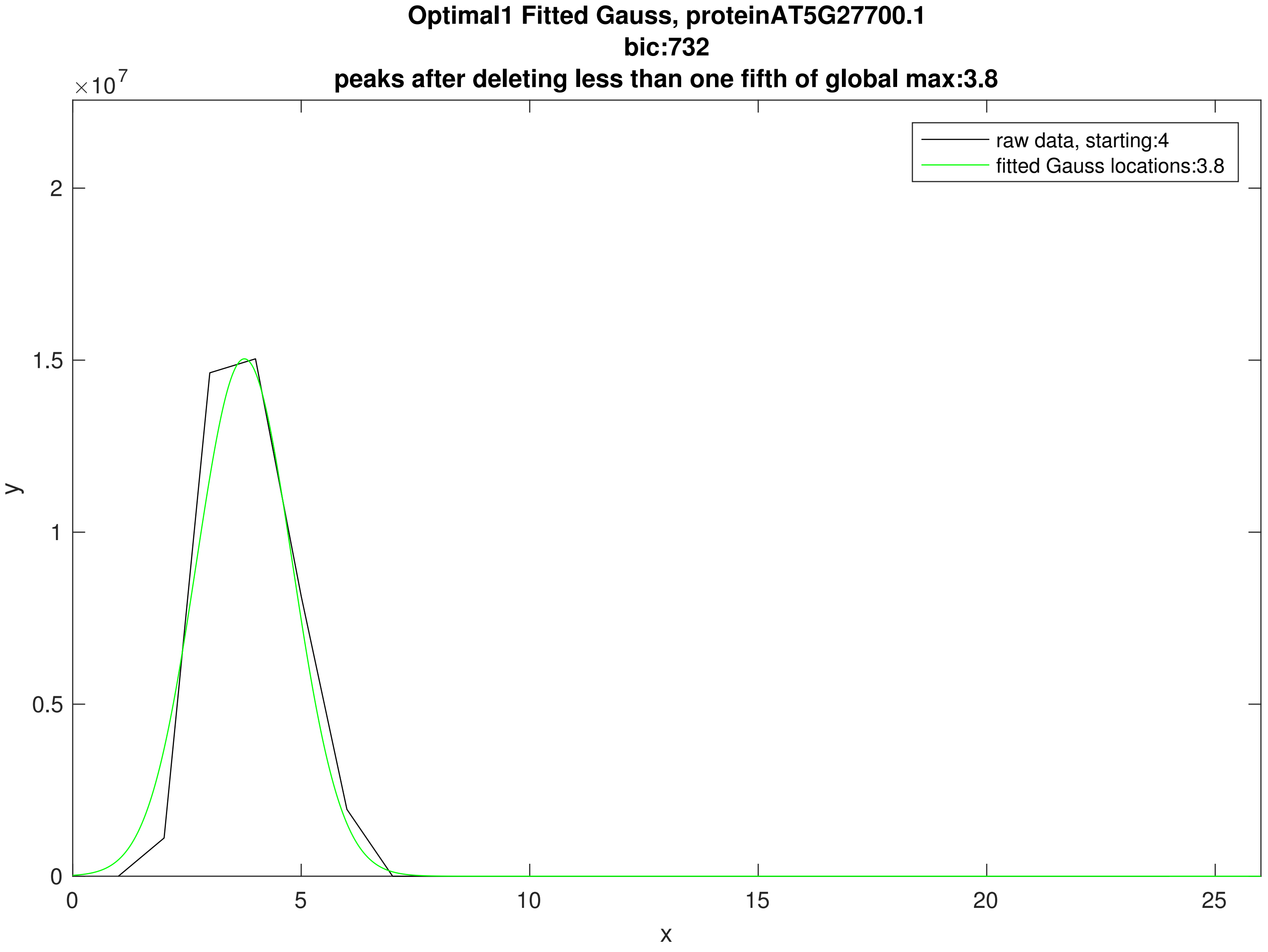
Optimal1 Fitted Gauss, proteinAT5G27700.1
bic:732
peaks after deleting less than one fifth of global max:3.8
×10^7
raw data, starting:4
fitted Gauss locations:3.8
y
x
0
0.5
1
1.5
2
5
10
15
20
25

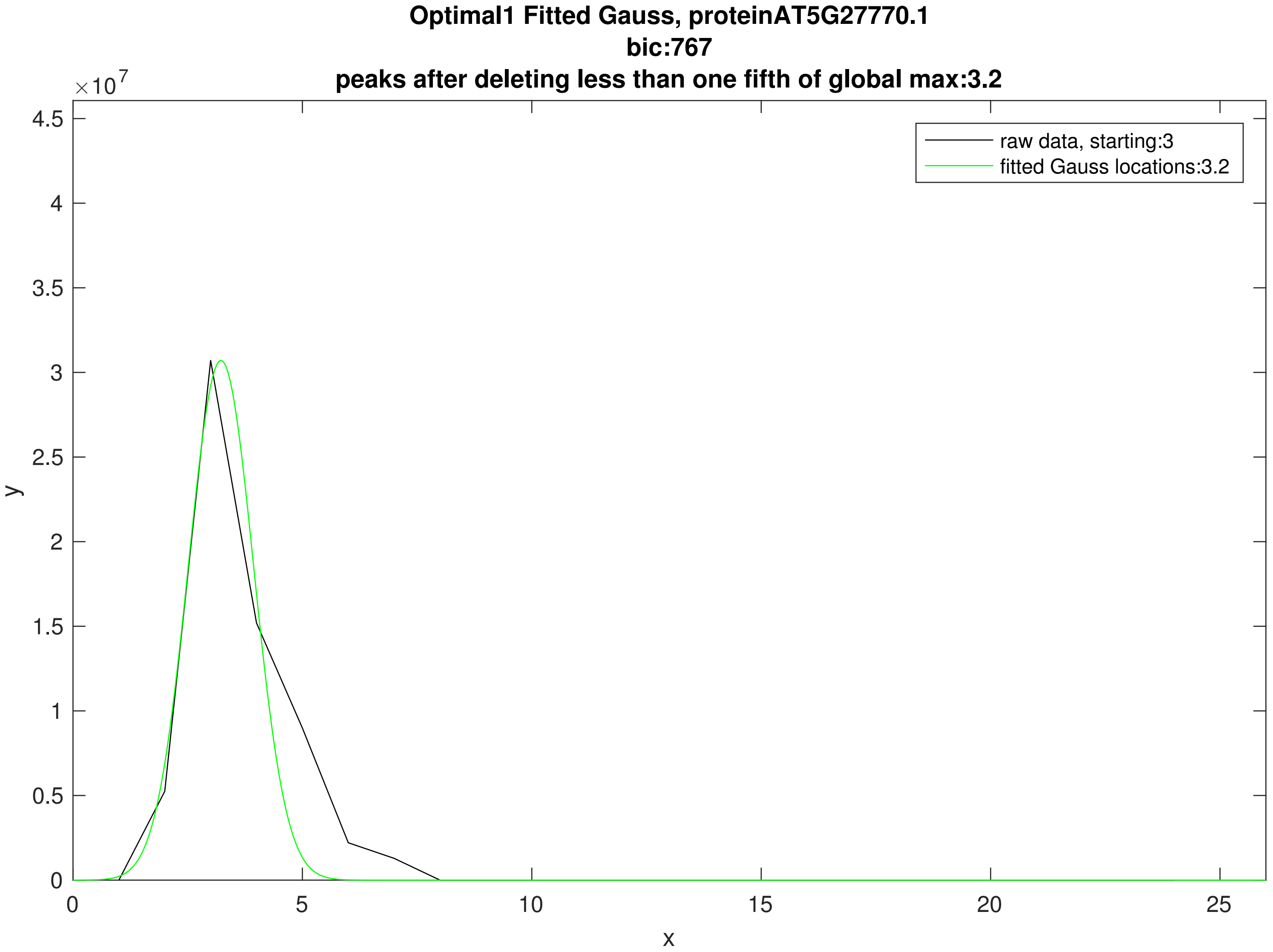
Optimal1 Fitted Gauss, proteinAT5G27770.1
bic:767
peaks after deleting less than one fifth of global max:3.2
×10^7
4.5
4
3.5
3
2.5
2
1.5
1
0.5
0
y
0
5
10
15
20
25
x
raw data, starting:3
fitted Gauss locations:3.2

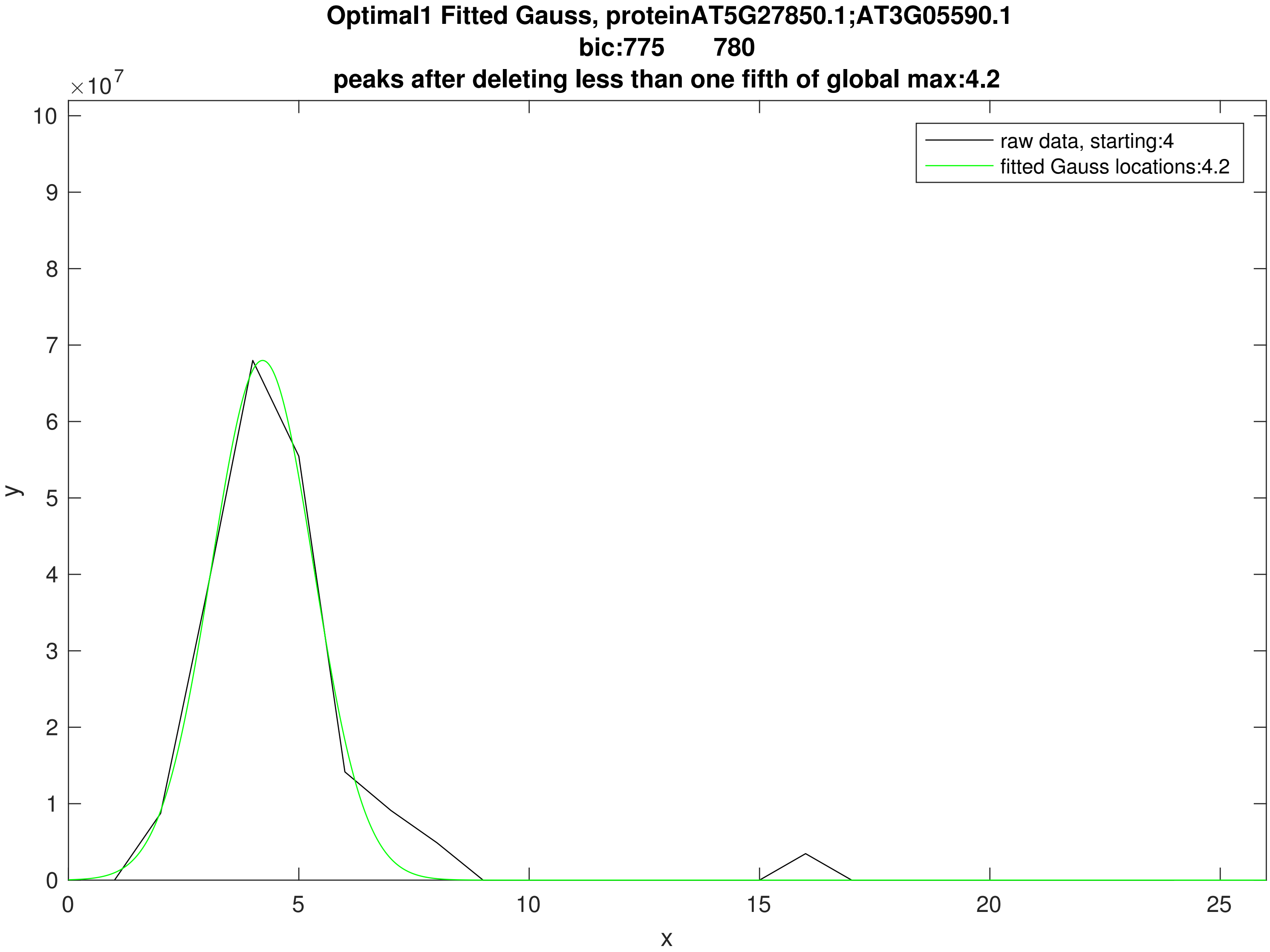

Optimal1 Fitted Gauss, proteinAT5G27850.1;AT3G05590.1
bic:775 780
peaks after deleting less than one fifth of global max:4.2
×10^7
raw data, starting:4
fitted Gauss locations:4.2
y
x

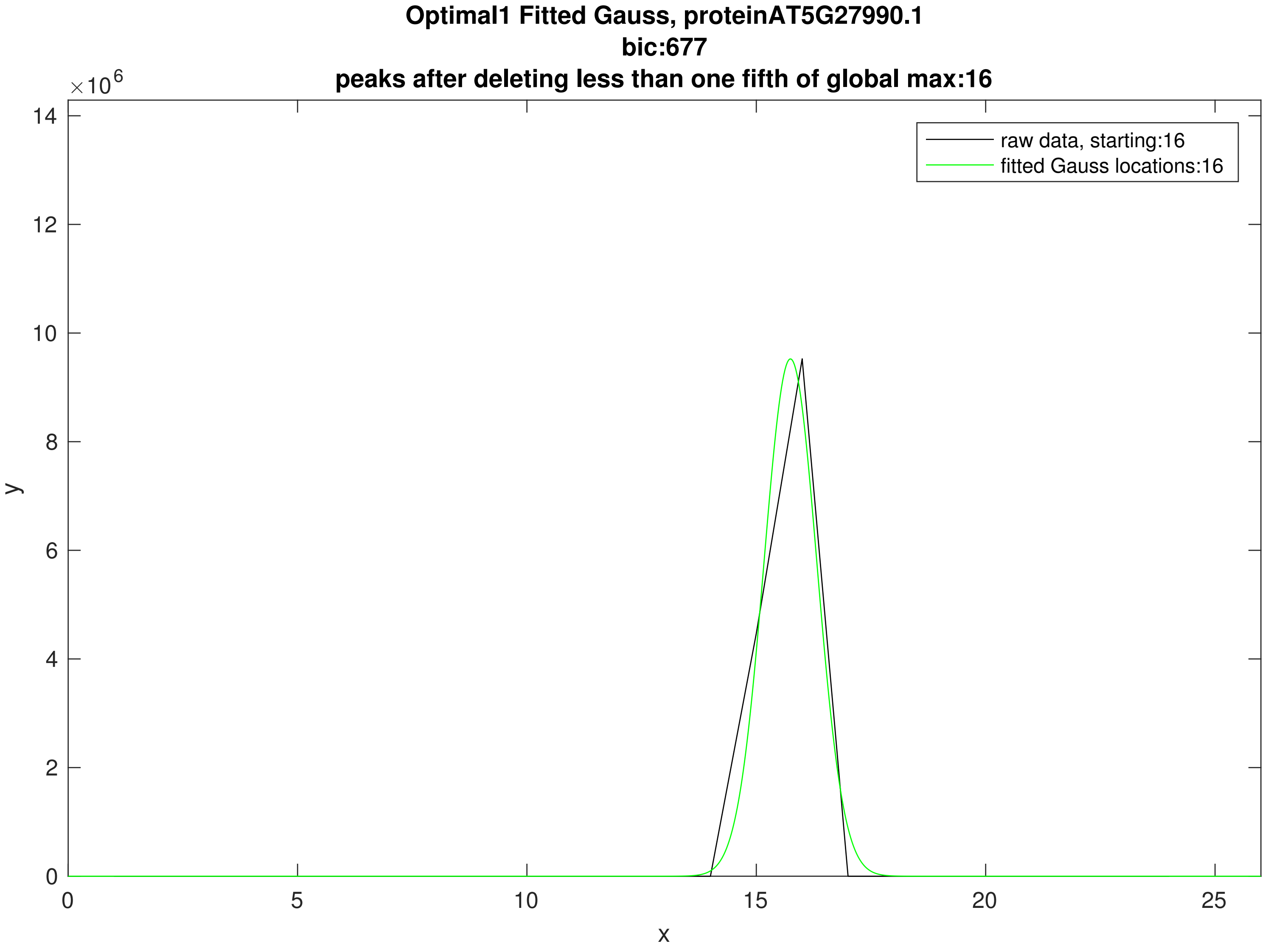

Optimal1 Fitted Gauss, proteinAT5G27990.1
bic:677
peaks after deleting less than one fifth of global max:16
×10^6
14
12
10
8
6
4
2
0
y
0
5
10
15
20
25
x
raw data, starting:16
fitted Gauss locations:16

**Optimal1 Fitted Gauss, proteinAT5G28020.6;AT5G28020.4;AT5G28020.3;AT5G28020.2;AT5G28020.1;AT5G28020.**
**bic:858**
**peaks after deleting less than one fifth of global max:15**

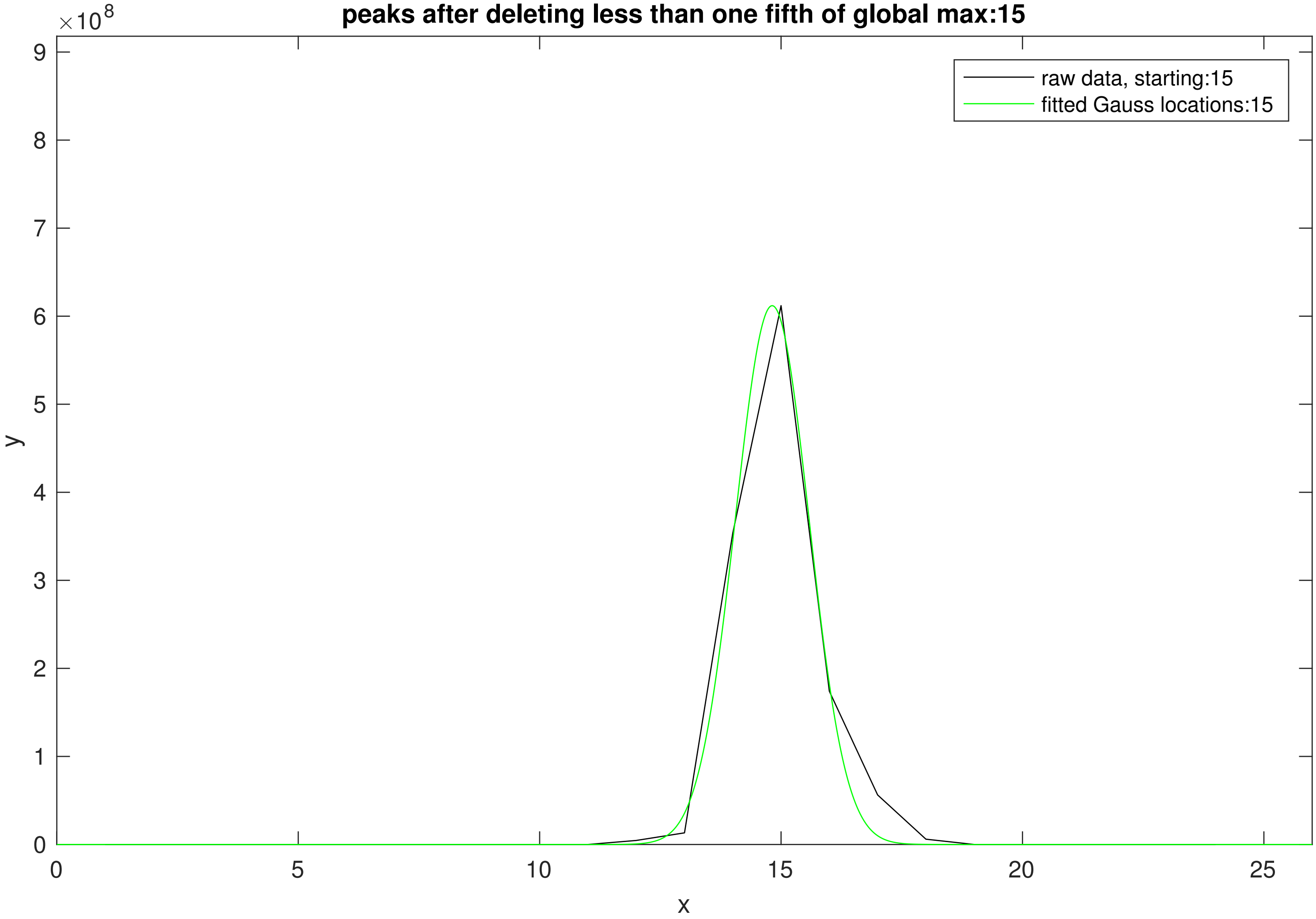

**Optimal1 Fitted Gauss, proteinAT5G28050.1;AT5G28050.2**
**bic:866 875 884**
**peaks after deleting less than one fifth of global max:15**

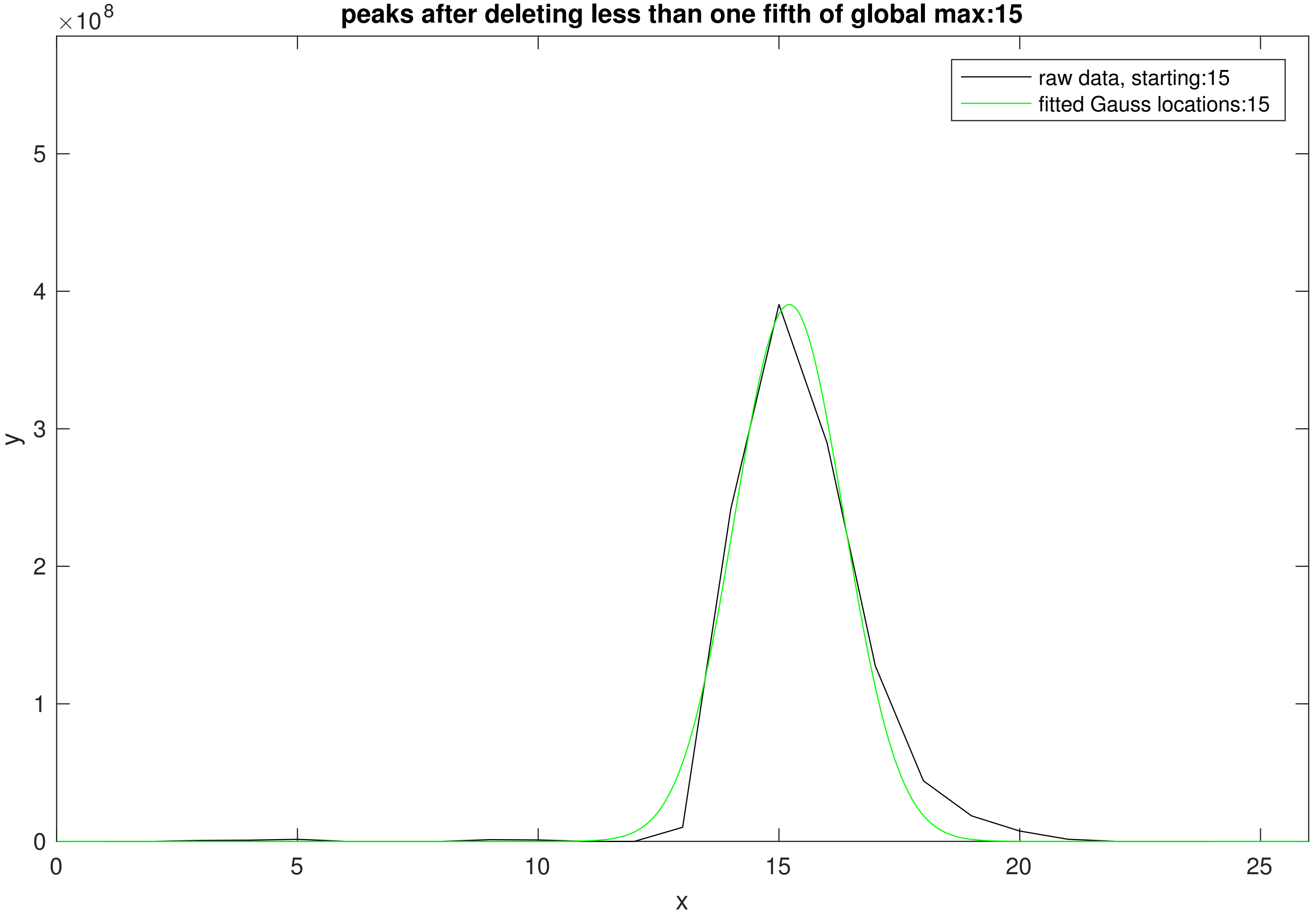

#### Optimal1 Fitted Gauss, proteinAT5G28060.1
#### bic:795
#### peaks after deleting less than one fifth of global max:3.9

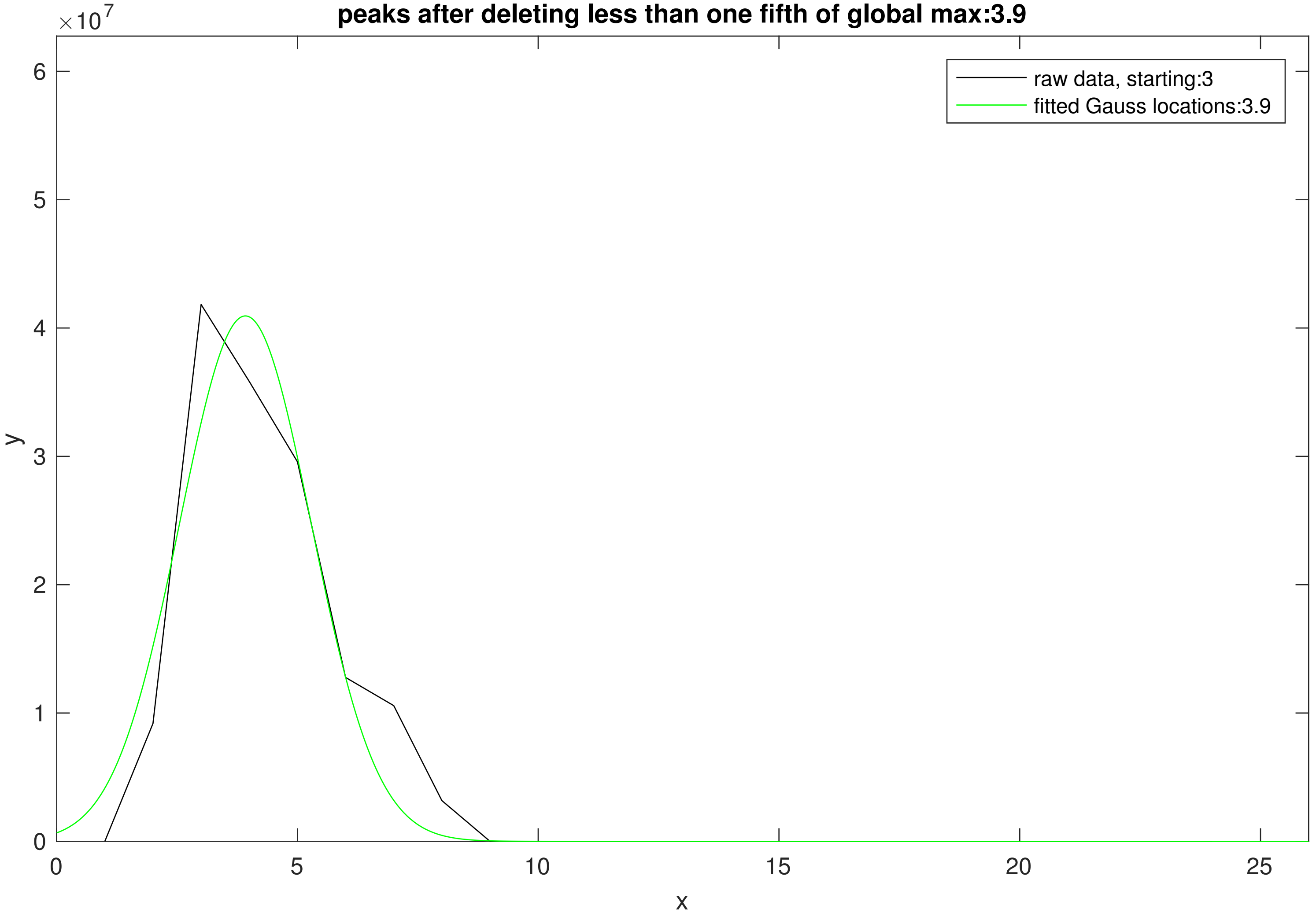

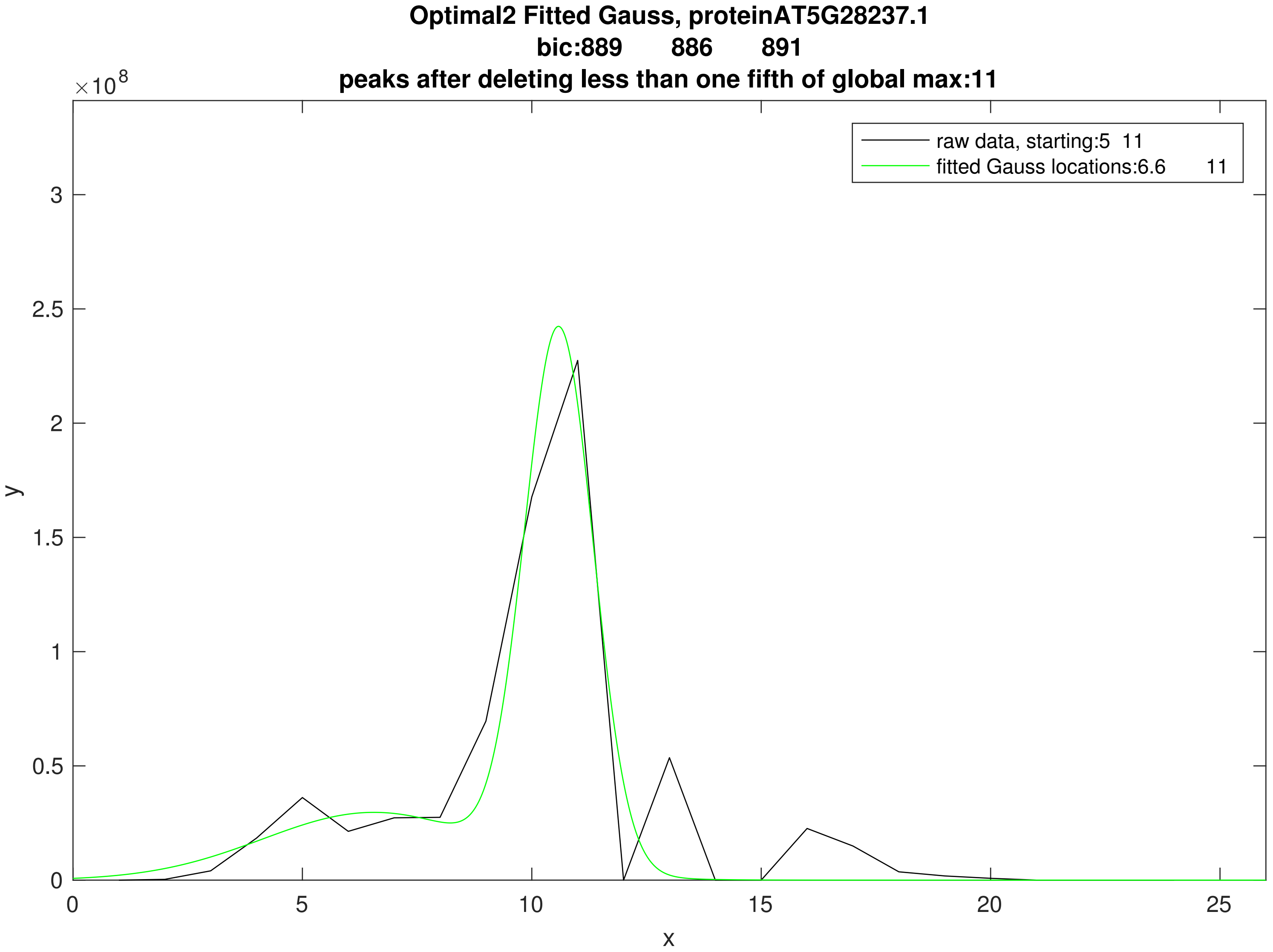

Optimal2 Fitted Gauss, proteinAT5G28237.1
bic:889 886 891
peaks after deleting less than one fifth of global max:11
×10^8
raw data, starting:5 11
fitted Gauss locations:6.6 11
y
x
0
0.5
1
1.5
2
2.5
3
0
5
10
15
20
25

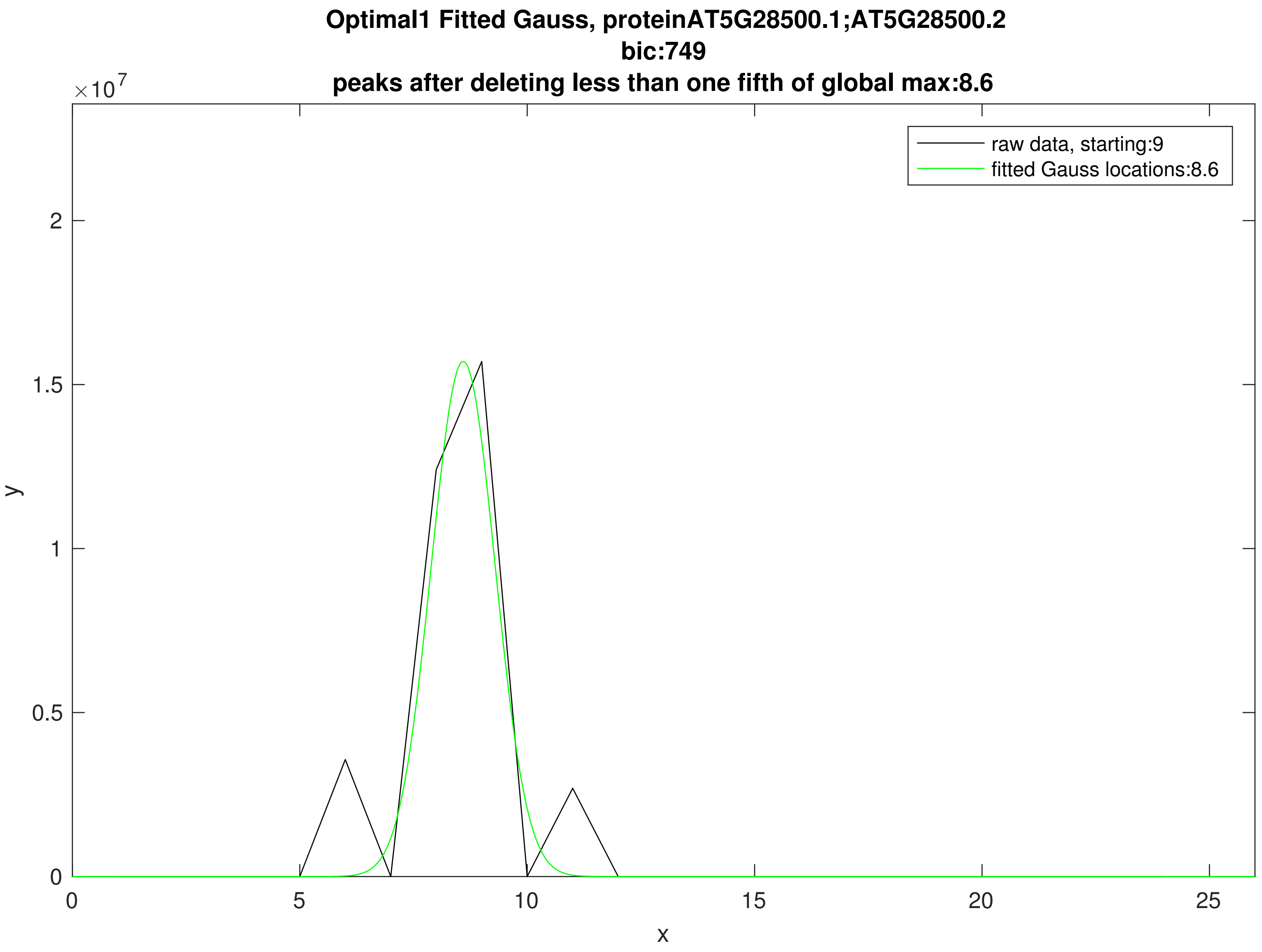

Optimal1 Fitted Gauss, proteinAT5G28500.1;AT5G28500.2
bic:749
peaks after deleting less than one fifth of global max:8.6
×10^7
raw data, starting:9
fitted Gauss locations:8.6
y
x
0
0.5
1
1.5
2
0
5
10
15
20
25

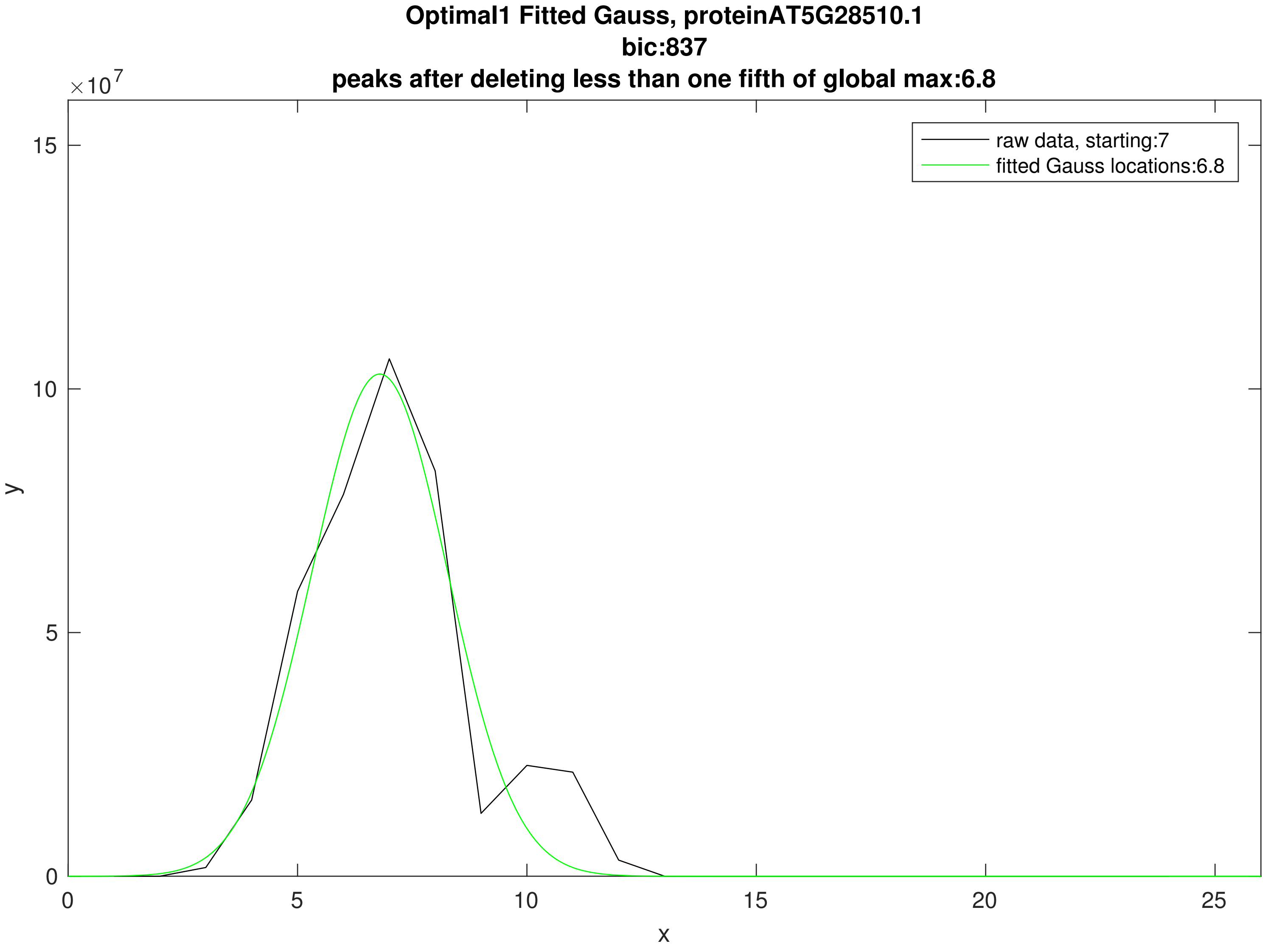
Optimal1 Fitted Gauss, proteinAT5G28510.1
bic:837
peaks after deleting less than one fifth of global max:6.8
×10^7
raw data, starting:7
fitted Gauss locations:6.8
15
10
5
0
y
0
5
10
15
20
25
x

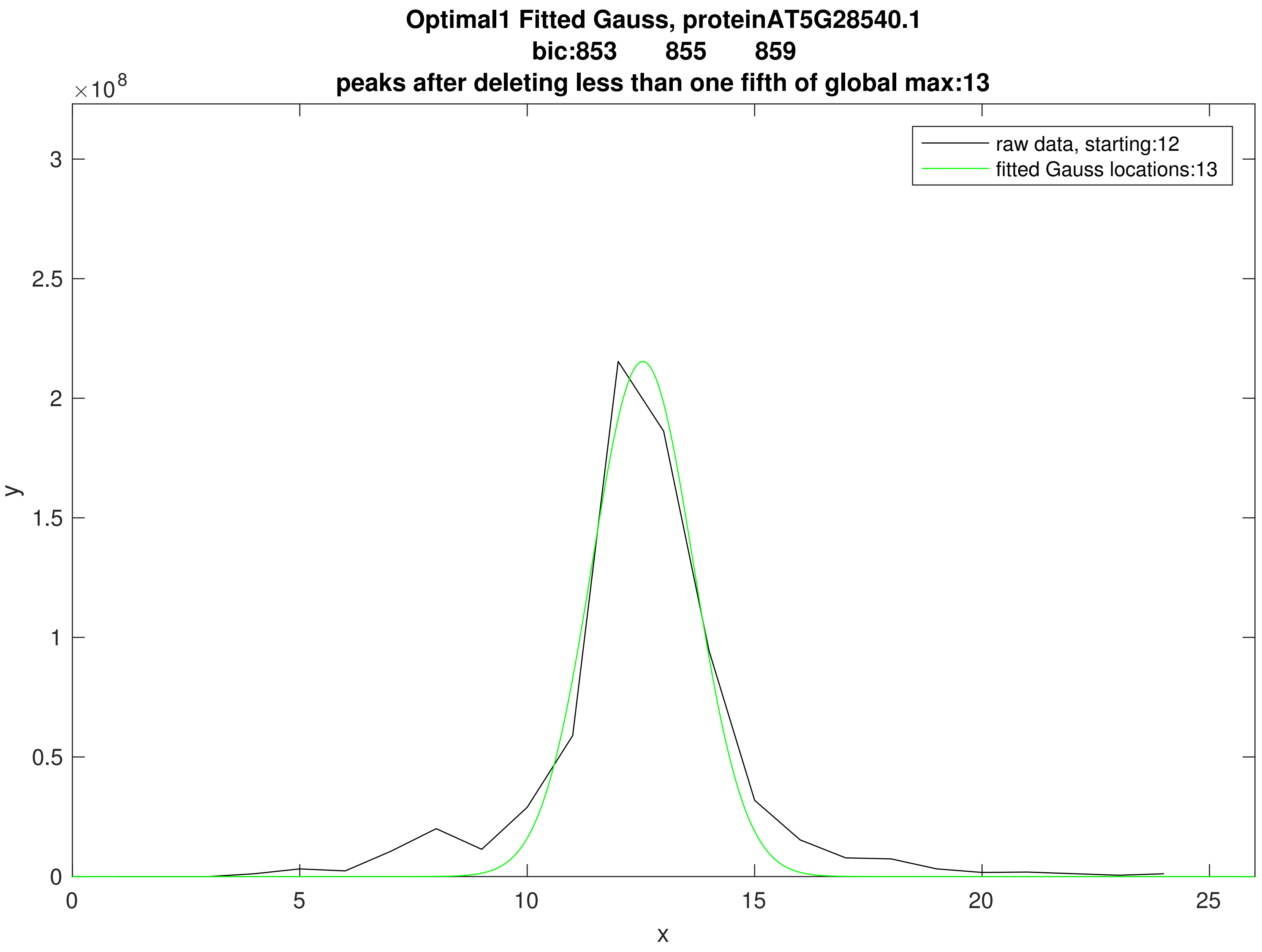

Optimal1 Fitted Gauss, proteinAT5G28540.1
bic:853 855 859
peaks after deleting less than one fifth of global max:13
×10^8
raw data, starting:12
fitted Gauss locations:13
y
x
0
0.5
1
1.5
2
2.5
3
0
5
10
15
20
25

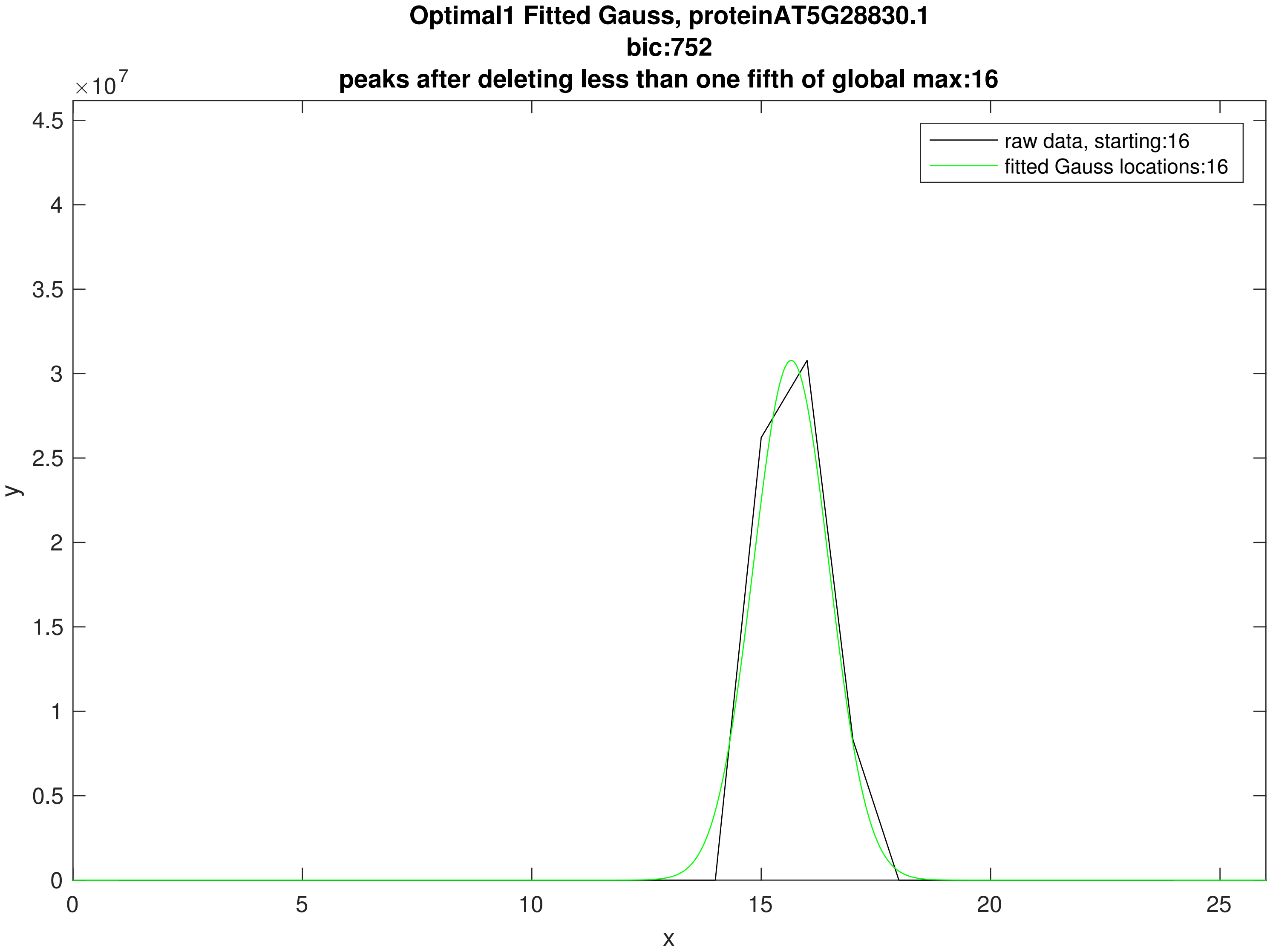
Optimal1 Fitted Gauss, proteinAT5G28830.1
bic:752
peaks after deleting less than one fifth of global max:16
×10^7
4.5
4
3.5
3
2.5
2
1.5
1
0.5
0
y
0
5
10
15
20
25
x
raw data, starting:16
fitted Gauss locations:16

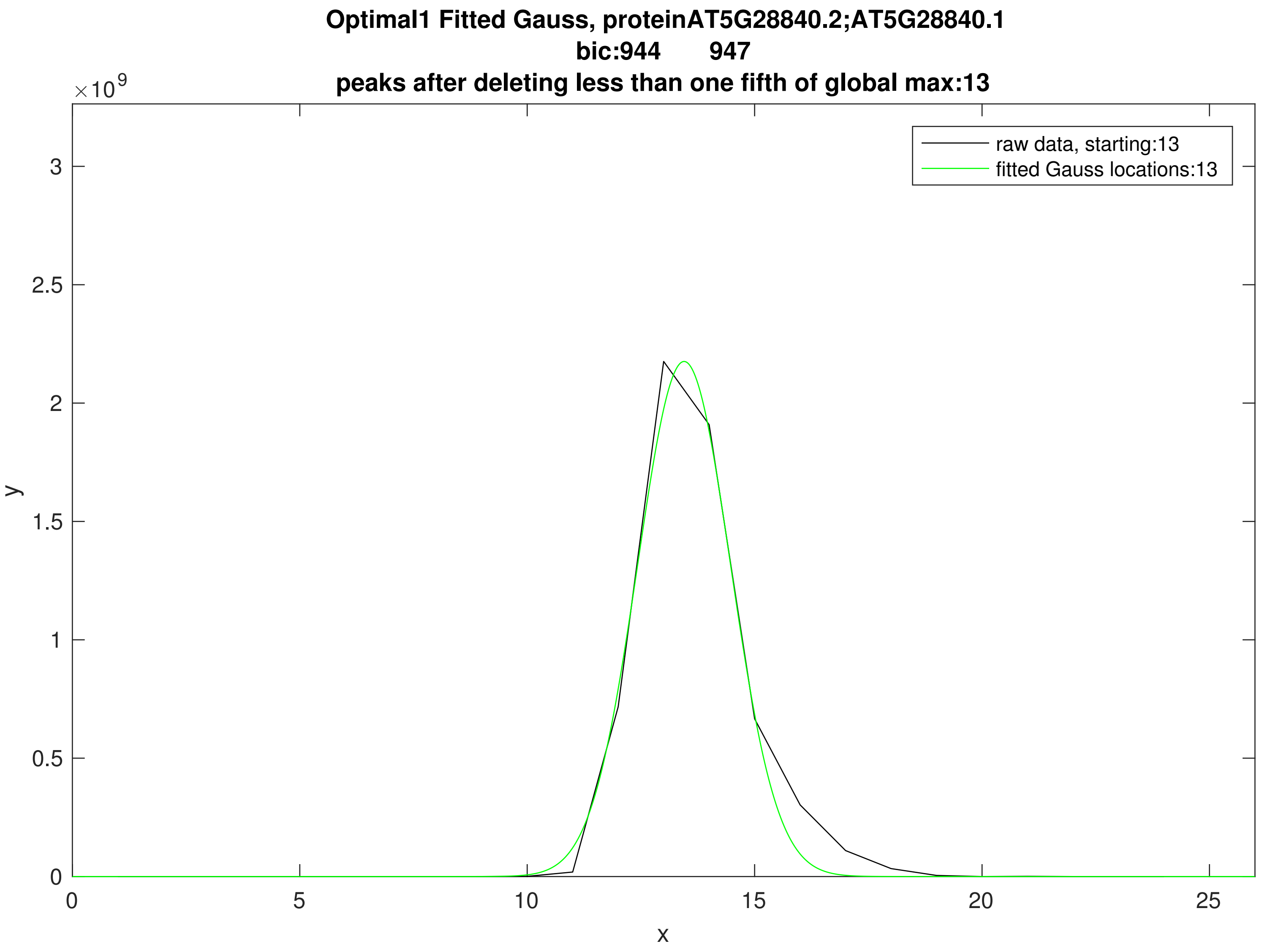
Optimal1 Fitted Gauss, proteinAT5G28840.2;AT5G28840.1
bic:944 947
peaks after deleting less than one fifth of global max:13
×10^9
raw data, starting:13
fitted Gauss locations:13
y
x
0
0.5
1
1.5
2
2.5
3
0
5
10
15
20
25

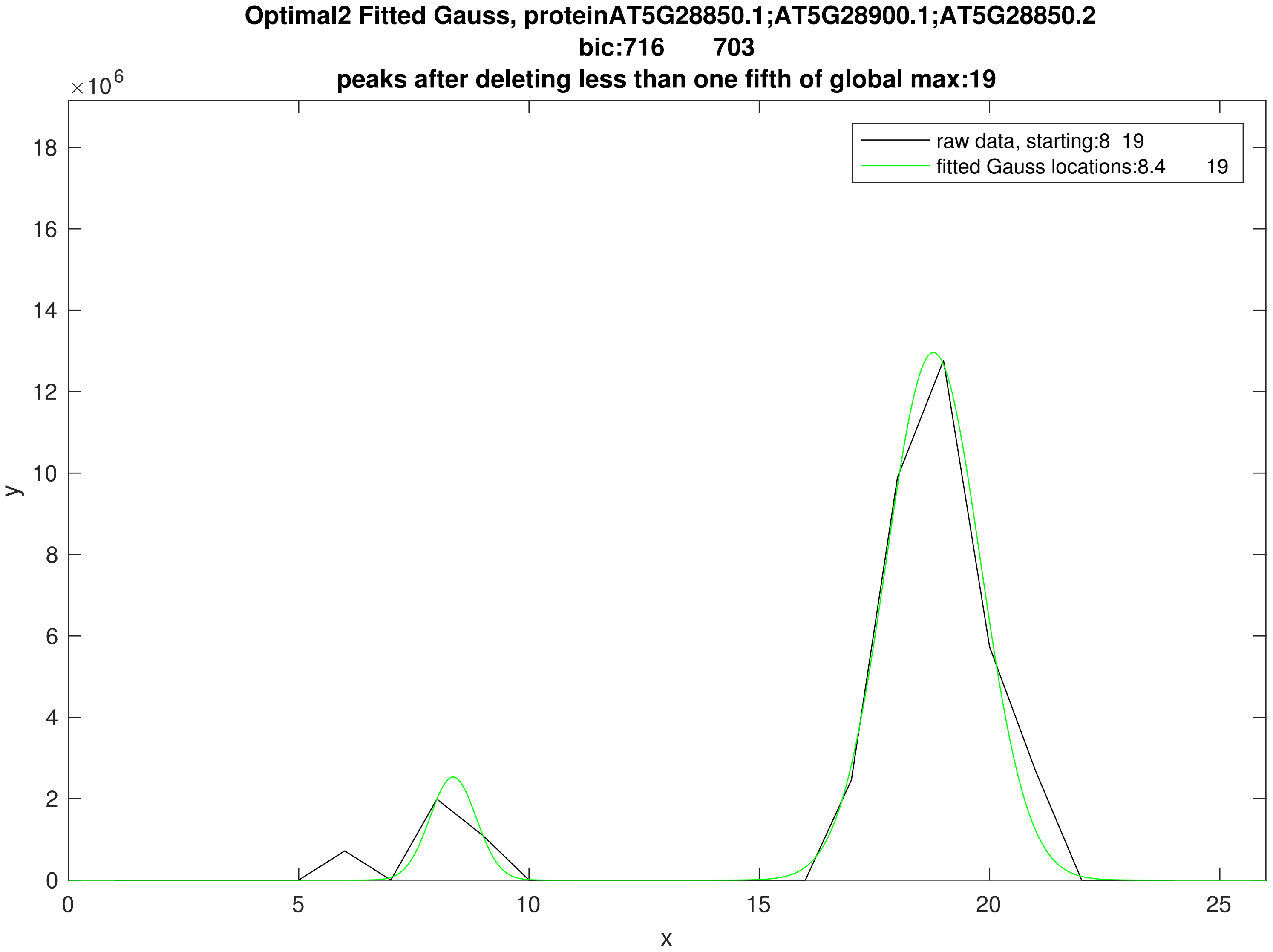

Optimal2 Fitted Gauss, proteinAT5G28850.1;AT5G28900.1;AT5G28850.2
bic:716 703
peaks after deleting less than one fifth of global max:19
×10^6
raw data, starting:8 19
fitted Gauss locations:8.4 19
y
x

**Optimal1 Fitted Gauss, proteinAT5G34850.1**
**bic:987 992 1e+03**
**peaks after deleting less than one fifth of global max:13**

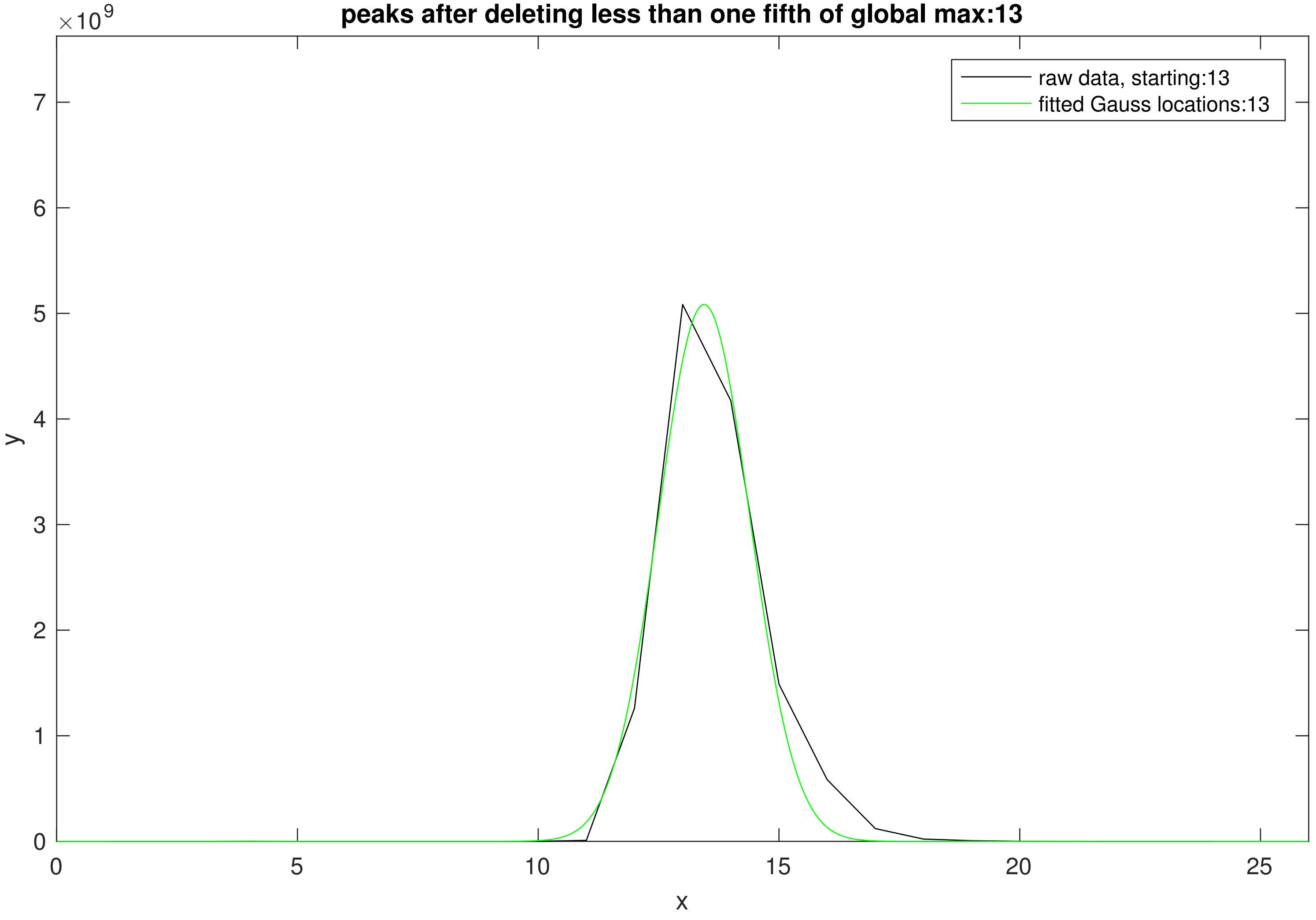

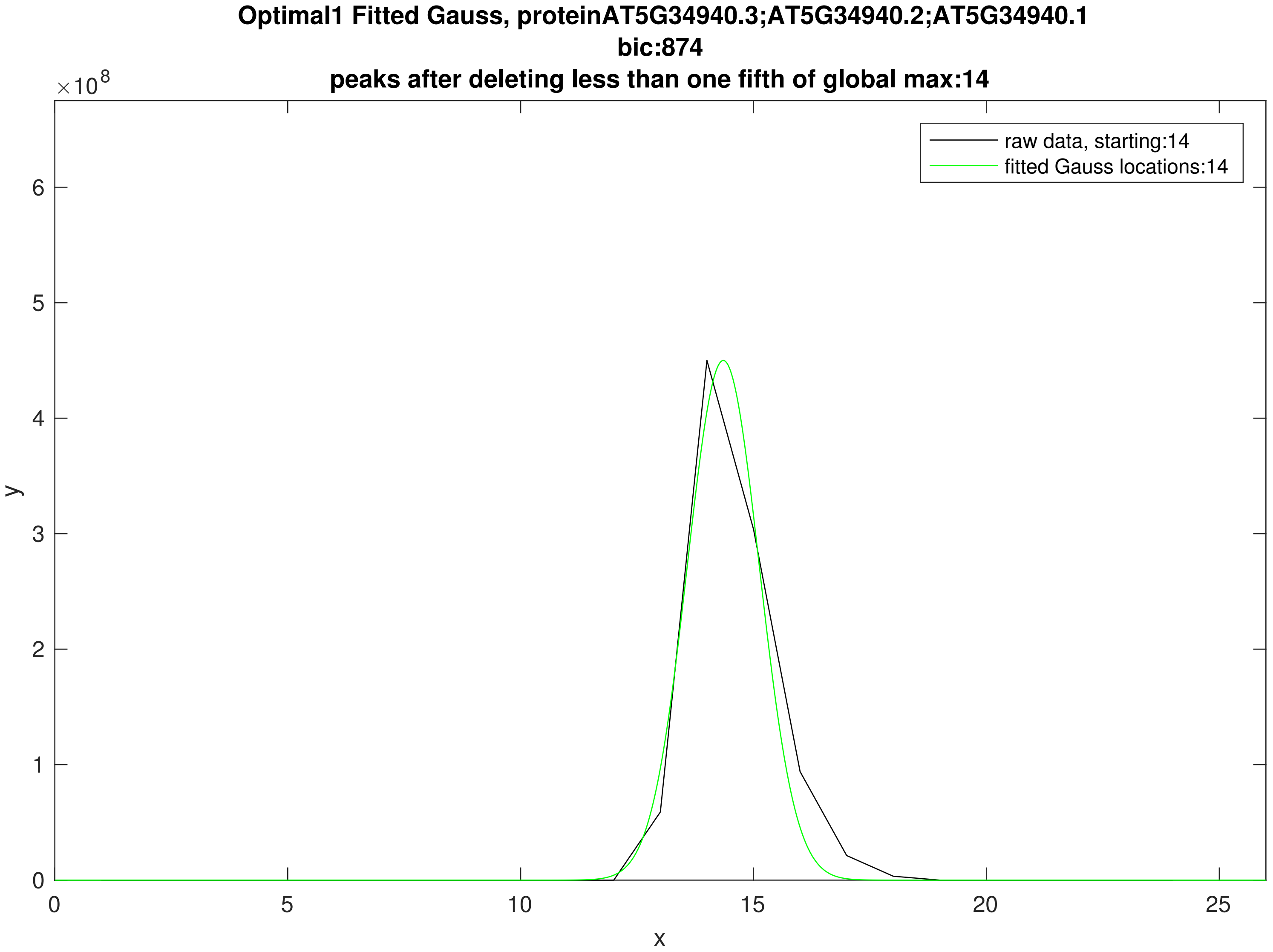
Optimal1 Fitted Gauss, proteinAT5G34940.3;AT5G34940.2;AT5G34940.1
bic:874
peaks after deleting less than one fifth of global max:14
×10^8
raw data, starting:14
fitted Gauss locations:14
y
x
0
1
2
3
4
5
6
0
5
10
15
20
25

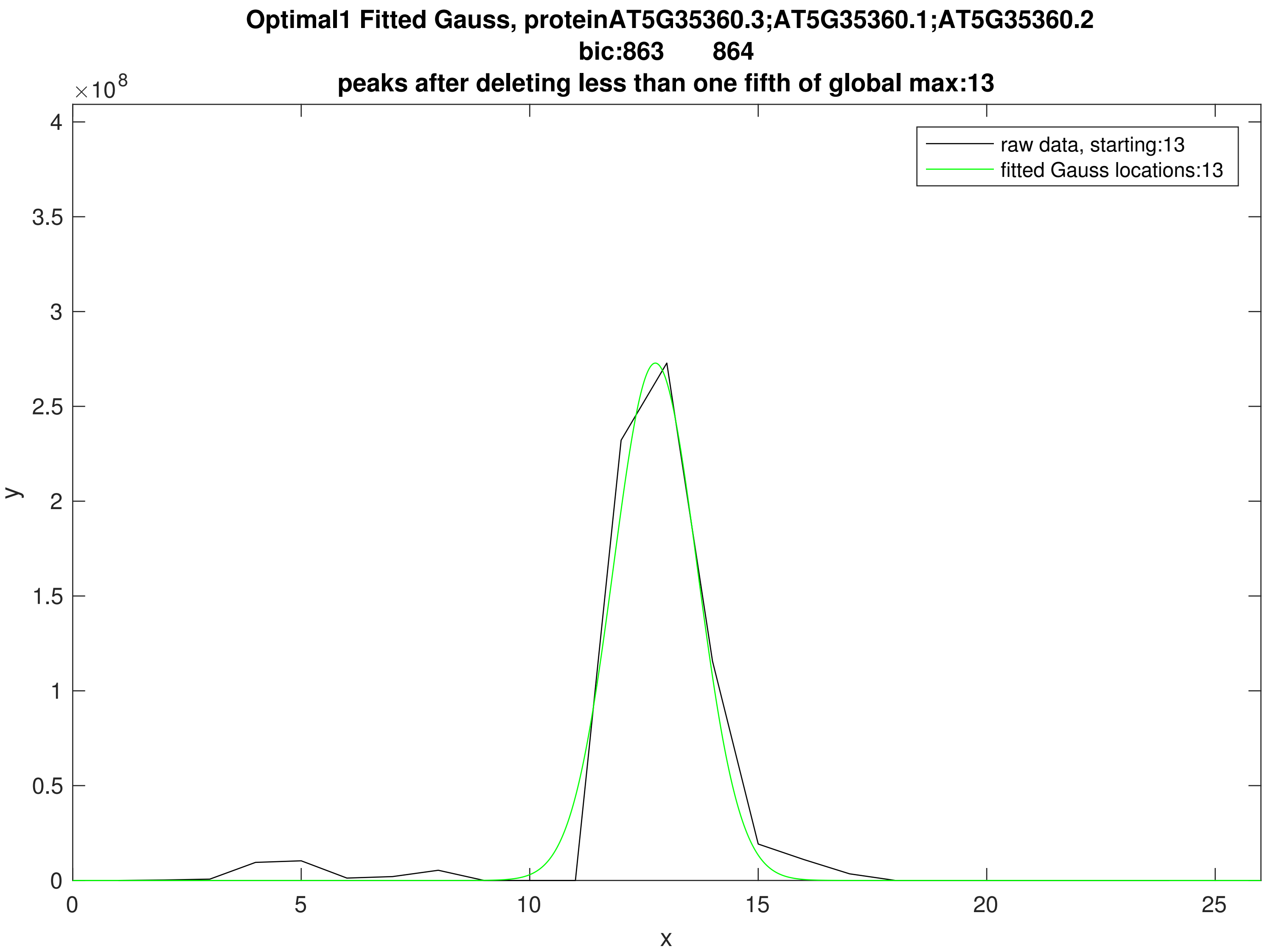
Optimal1 Fitted Gauss, proteinAT5G35360.3;AT5G35360.1;AT5G35360.2
bic:863 864
peaks after deleting less than one fifth of global max:13
×10^8
4
3.5
3
2.5
2
1.5
1
0.5
0
y
0
5
10
15
20
25
x
raw data, starting:13
fitted Gauss locations:13

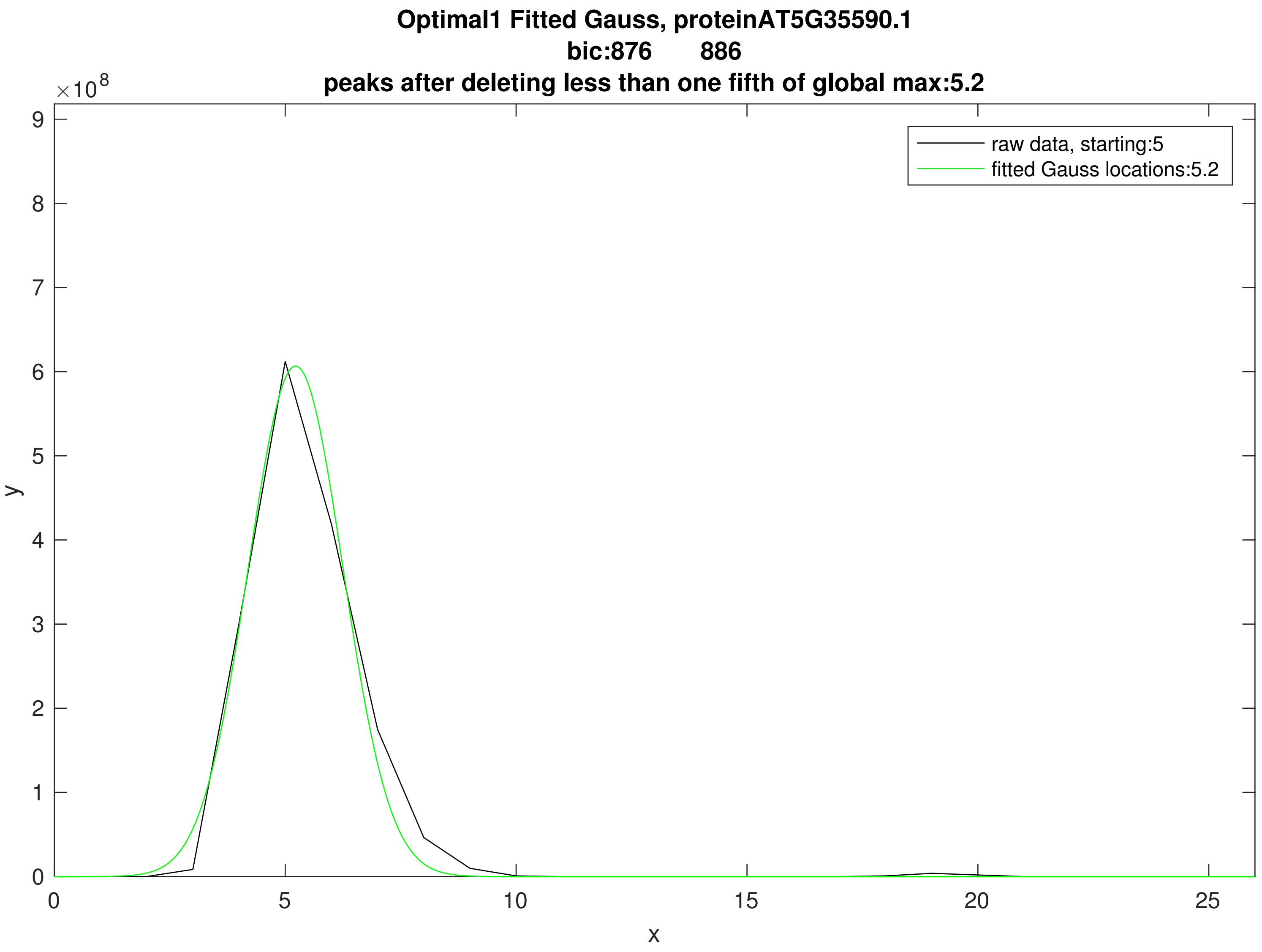
Optimal1 Fitted Gauss, proteinAT5G35590.1
bic:876 886
peaks after deleting less than one fifth of global max:5.2
×10^8
9
8
7
6
5
4
3
2
1
0
y
0
5
10
15
20
25
x
raw data, starting:5
fitted Gauss locations:5.2

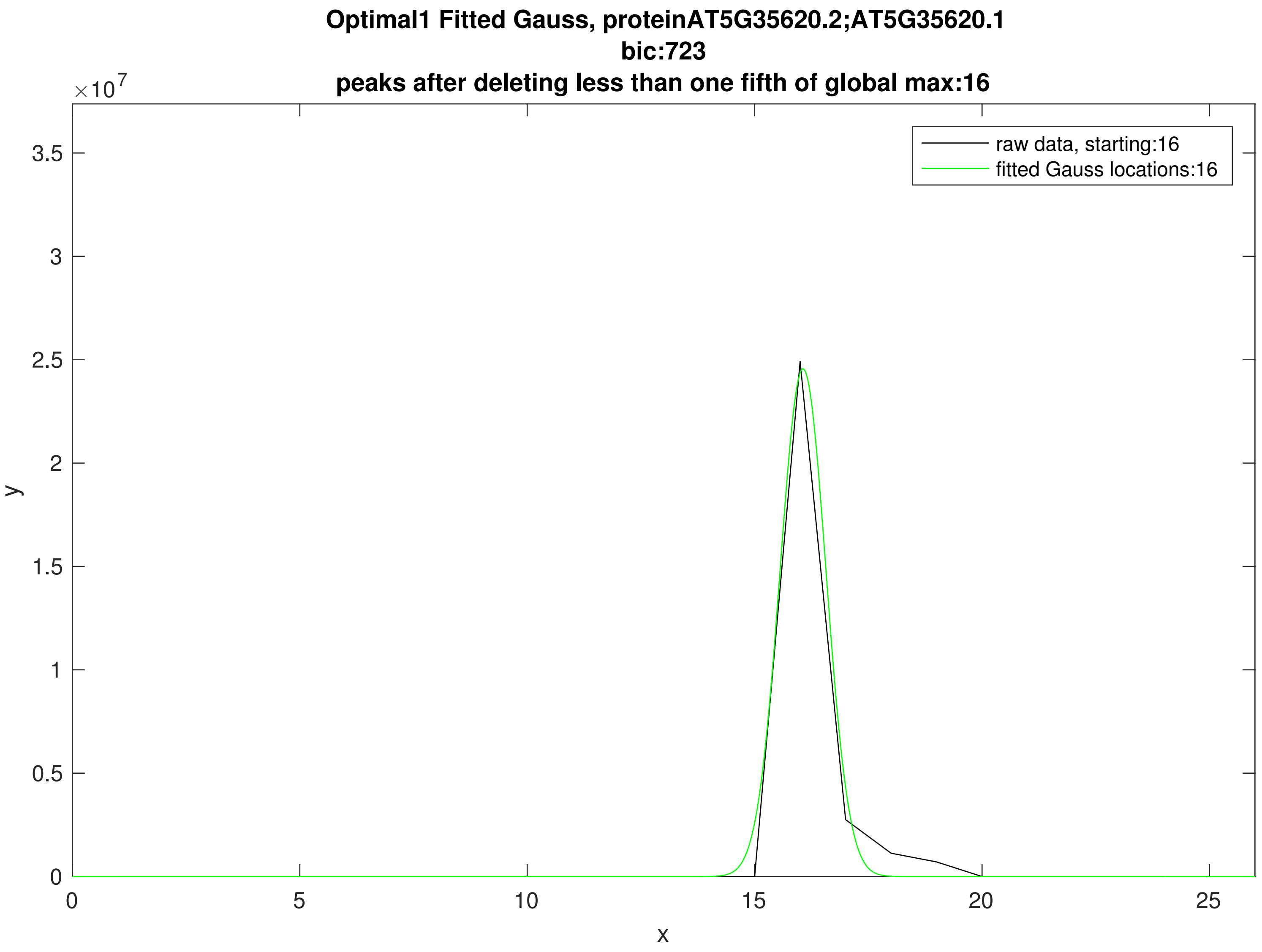

Optimal1 Fitted Gauss, proteinAT5G35620.2;AT5G35620.1
bic:723
peaks after deleting less than one fifth of global max:16
×10^7
3.5
3
2.5
2
1.5
1
0.5
0
y
0
5
10
15
20
25
x
raw data, starting:16
fitted Gauss locations:16

#### Optimal1 Fitted Gauss, proteinAT5G35630.3;AT5G35630.2;AT5G35630.1
#### bic:953 963
#### peaks after deleting less than one fifth of global max:7.1

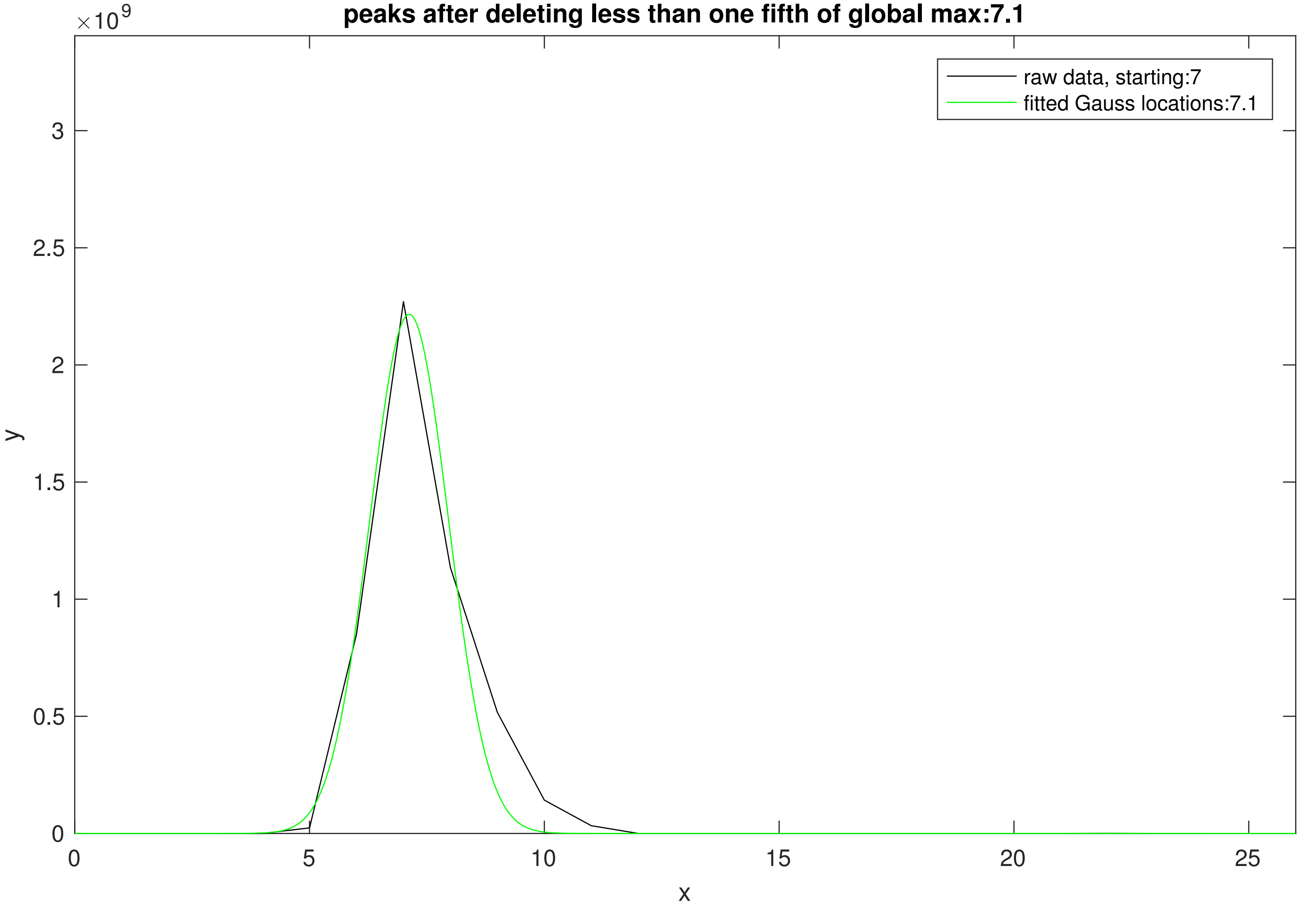

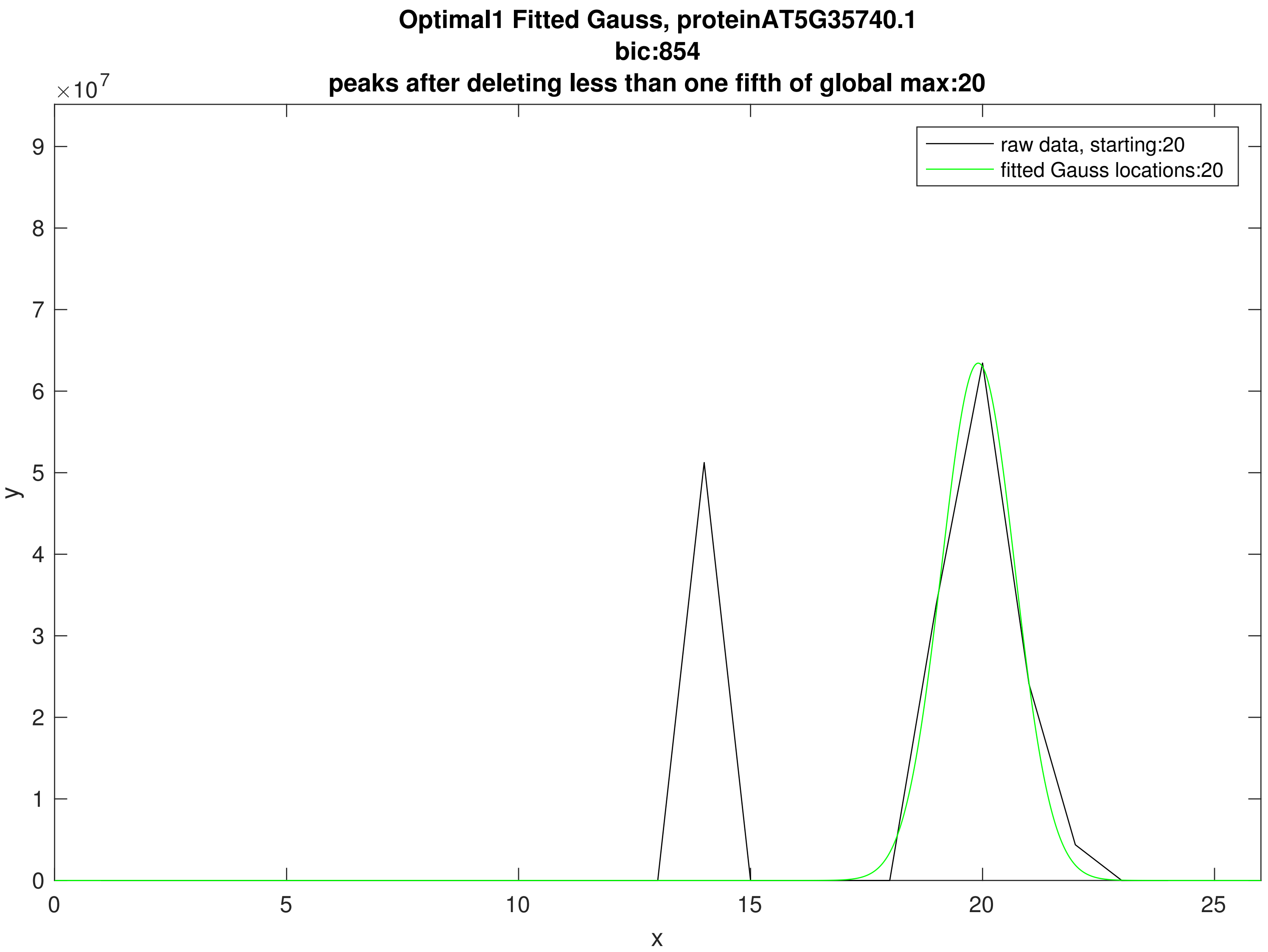
Optimal1 Fitted Gauss, proteinAT5G35740.1
bic:854
peaks after deleting less than one fifth of global max:20
×10^7
raw data, starting:20
fitted Gauss locations:20
y
x
0
1
2
3
4
5
6
7
8
9
0
5
10
15
20
25

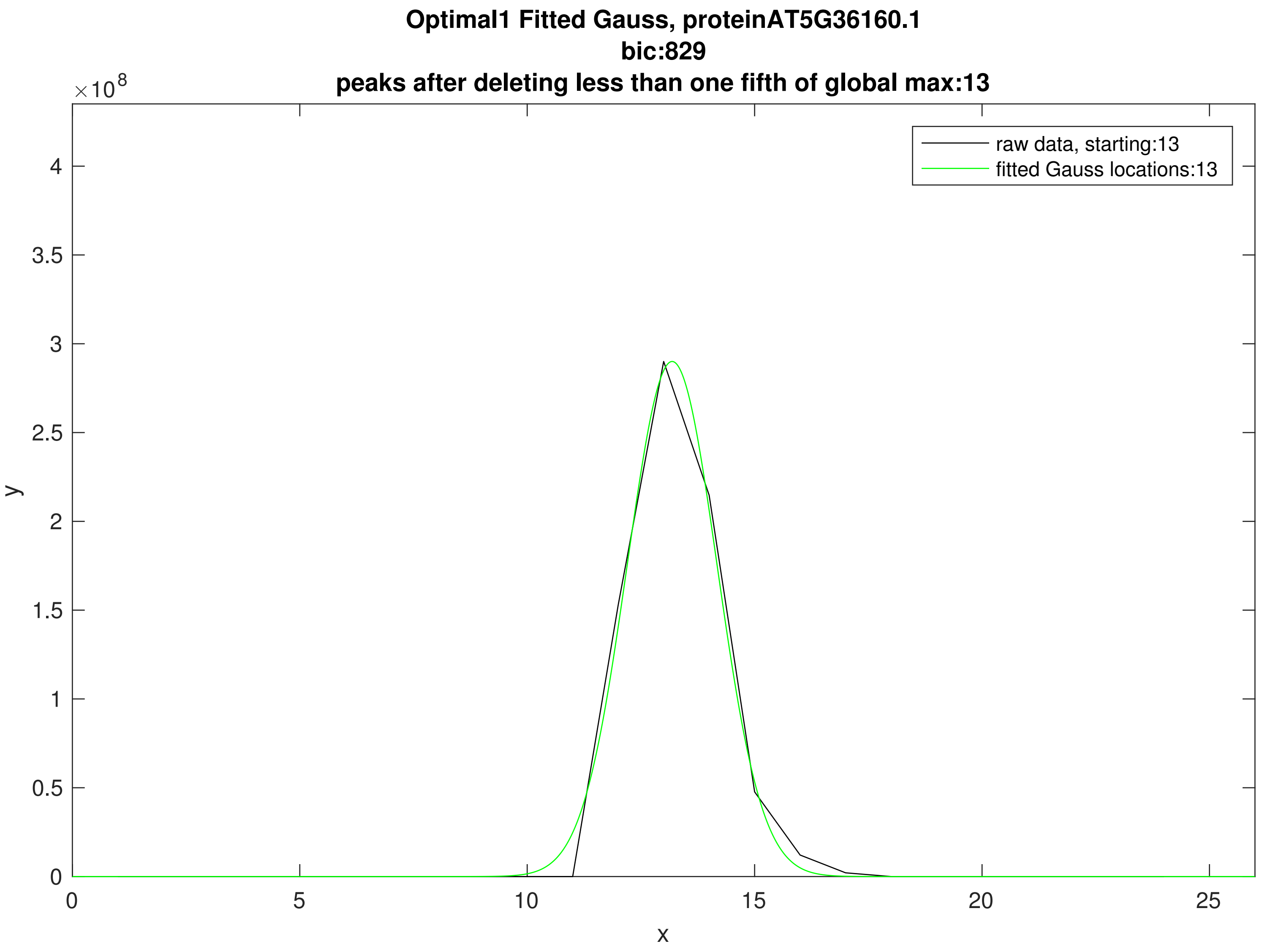

Optimal1 Fitted Gauss, proteinAT5G36160.1
bic:829
peaks after deleting less than one fifth of global max:13
×10^8
4
3.5
3
2.5
2
1.5
1
0.5
0
y
0
5
10
15
20
25
x
raw data, starting:13
fitted Gauss locations:13

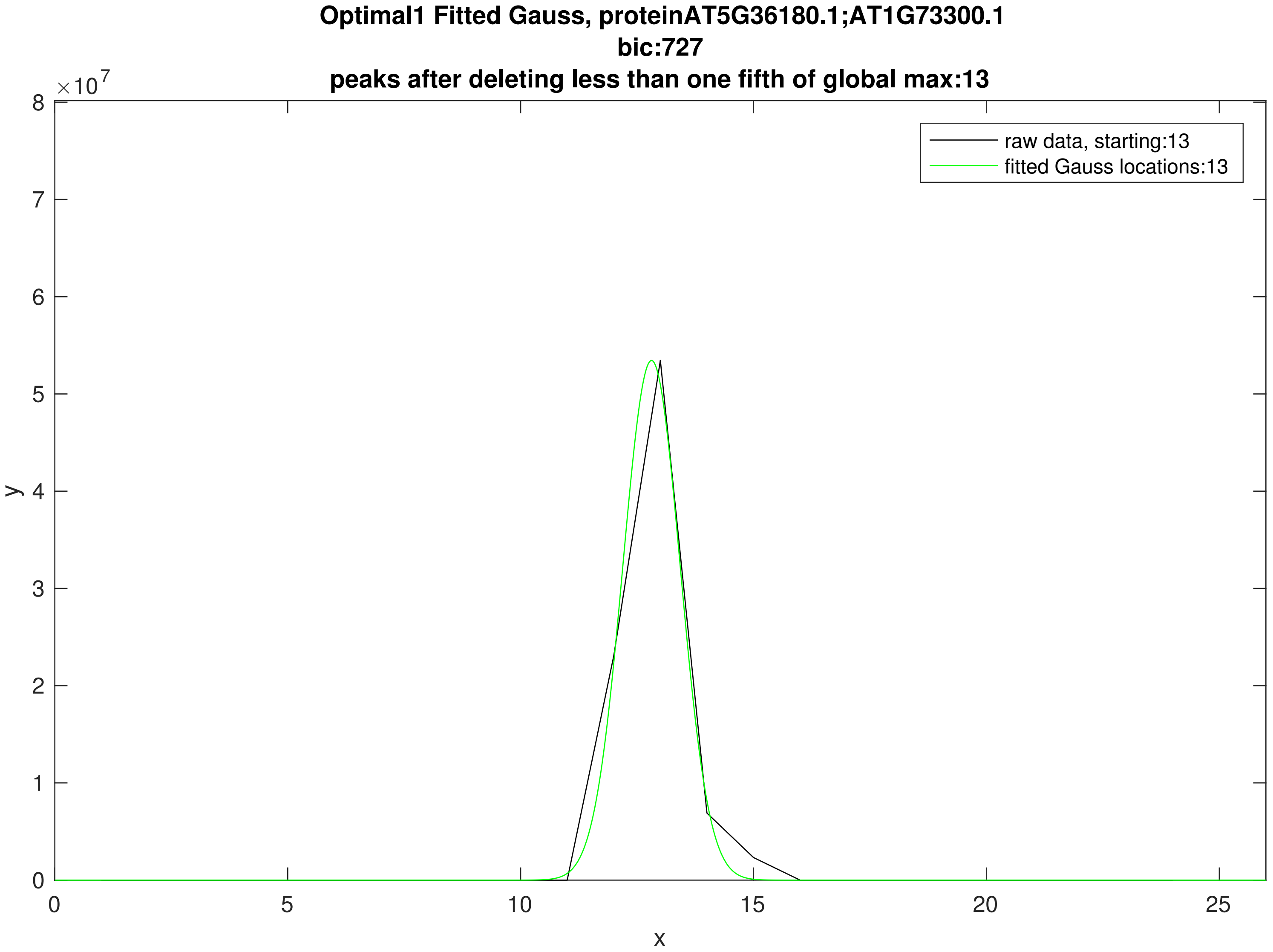
Optimal1 Fitted Gauss, proteinAT5G36180.1;AT1G73300.1
bic:727
peaks after deleting less than one fifth of global max:13
×10^7
8
7
6
5
4
3
2
1
0
y
0
5
10
15
20
25
x
raw data, starting:13
fitted Gauss locations:13

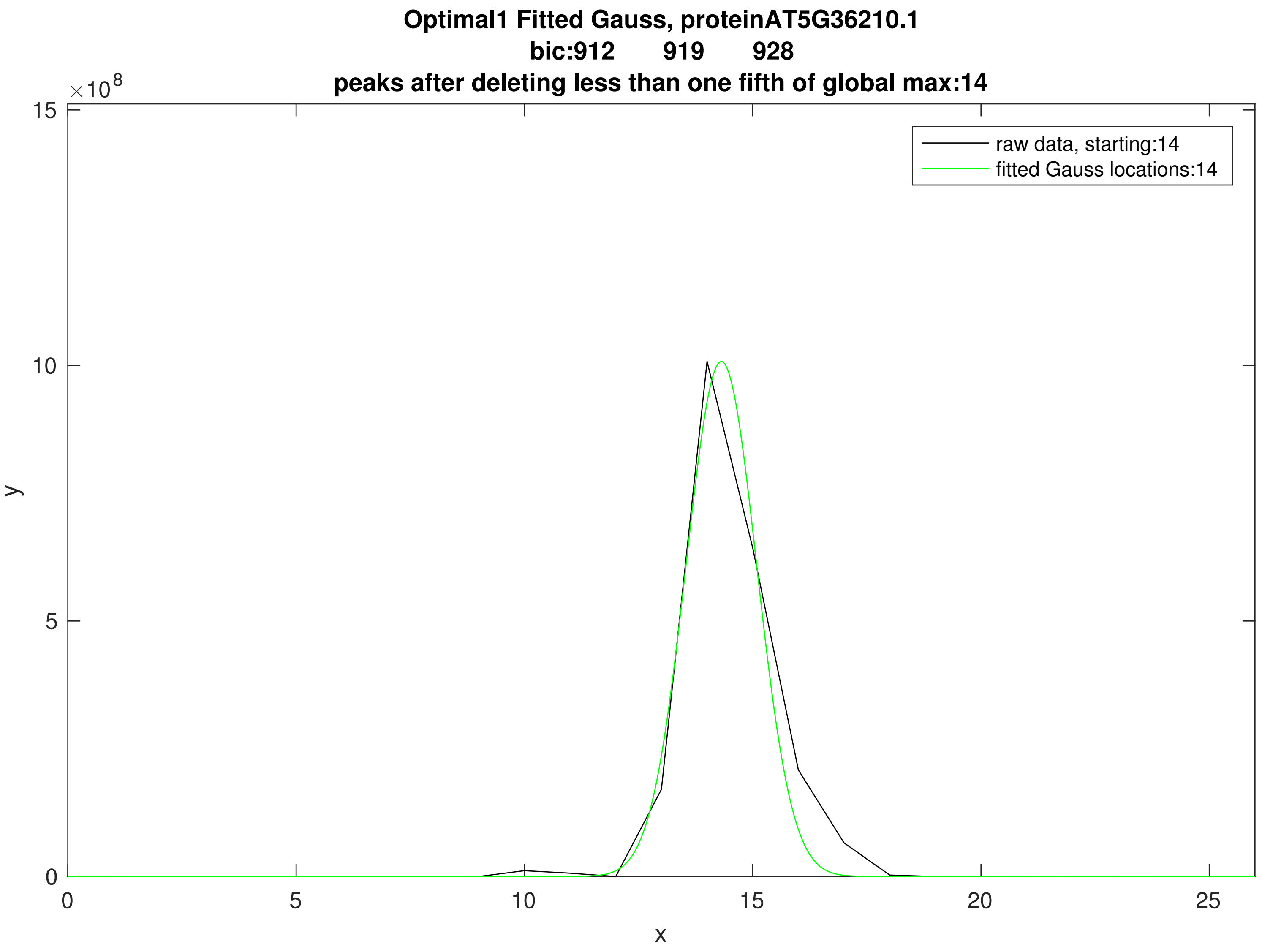

Optimal1 Fitted Gauss, proteinAT5G36210.1
bic:912 919 928
peaks after deleting less than one fifth of global max:14
×10^8
raw data, starting:14
fitted Gauss locations:14
y
x
0
5
10
15
20
25

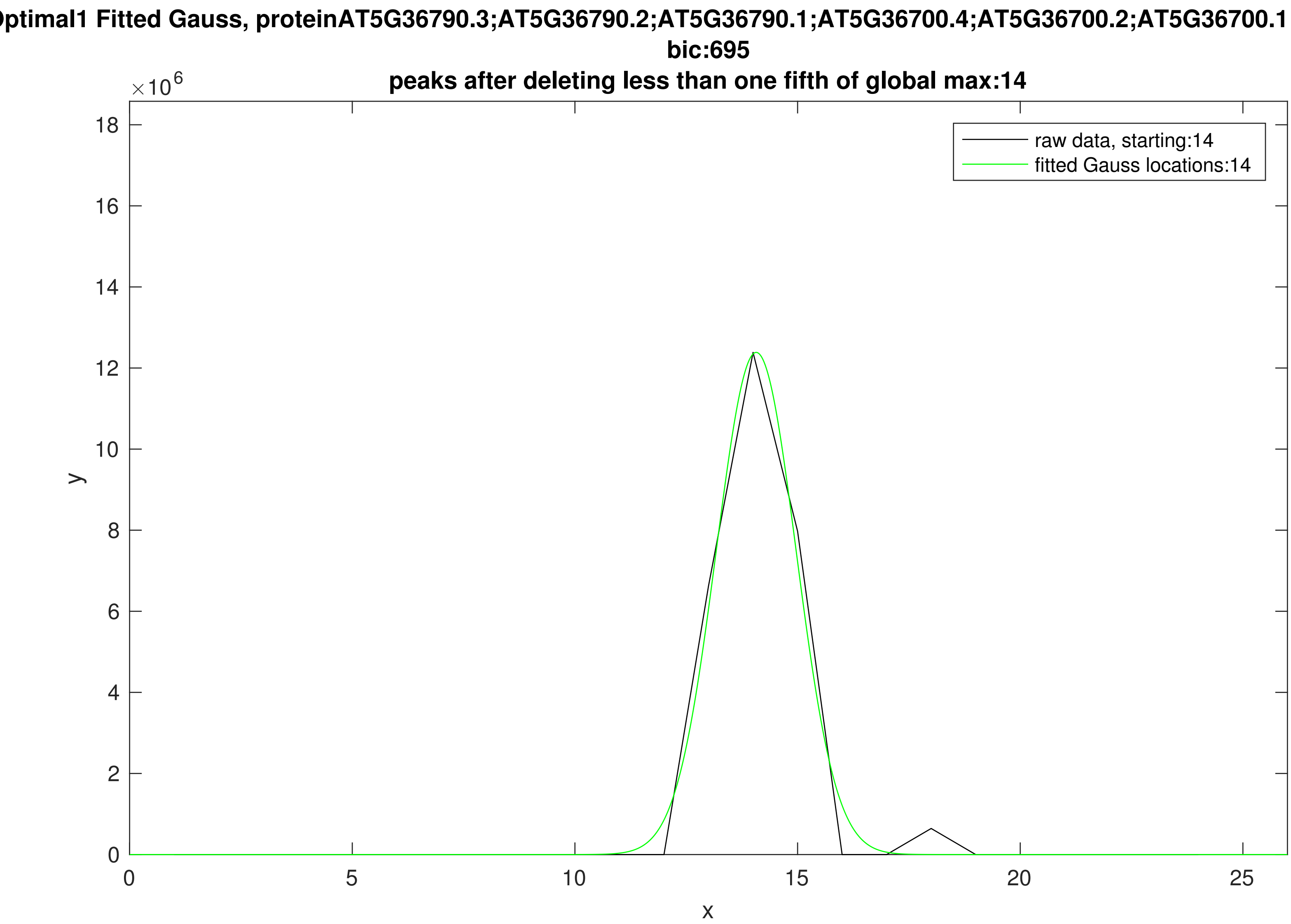

Optimal1 Fitted Gauss, proteinAT5G36790.3;AT5G36790.2;AT5G36790.1;AT5G36700.4;AT5G36700.2;AT5G36700.1;AT5
bic:695
peaks after deleting less than one fifth of global max:14
×10^6
raw data, starting:14
fitted Gauss locations:14
y
x
0
2
4
6
8
10
12
14
16
18
0
5
10
15
20
25

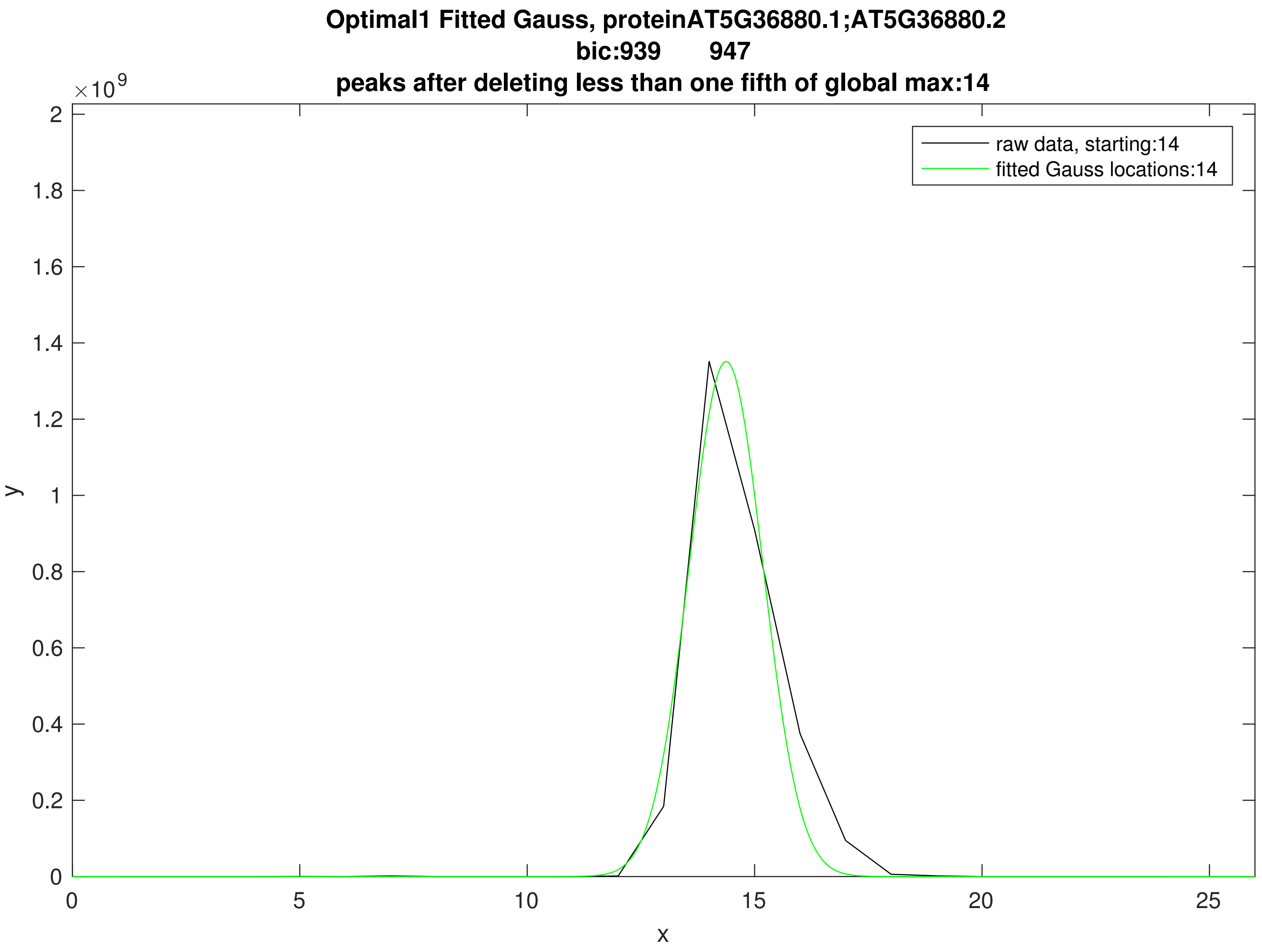
Optimal1 Fitted Gauss, proteinAT5G36880.1;AT5G36880.2
bic:939 947
peaks after deleting less than one fifth of global max:14
×10^9
2
1.8
1.6
1.4
1.2
1
0.8
0.6
0.4
0.2
0
y
0
5
10
15
20
25
x
raw data, starting:14
fitted Gauss locations:14

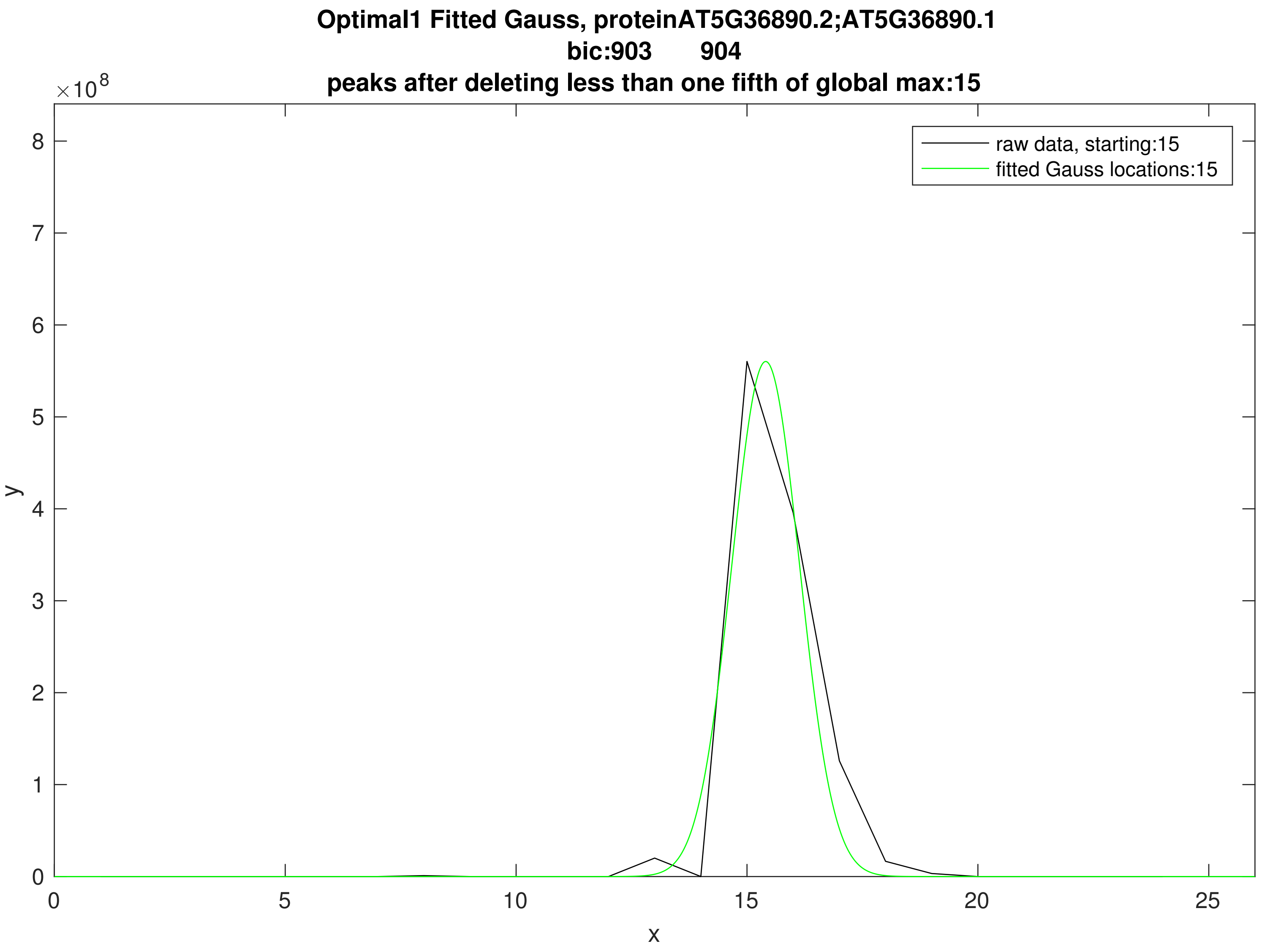
Optimal1 Fitted Gauss, proteinAT5G36890.2;AT5G36890.1
bic:903 904
peaks after deleting less than one fifth of global max:15
×10^8
raw data, starting:15
fitted Gauss locations:15
y
x
0
1
2
3
4
5
6
7
8
0
5
10
15
20
25

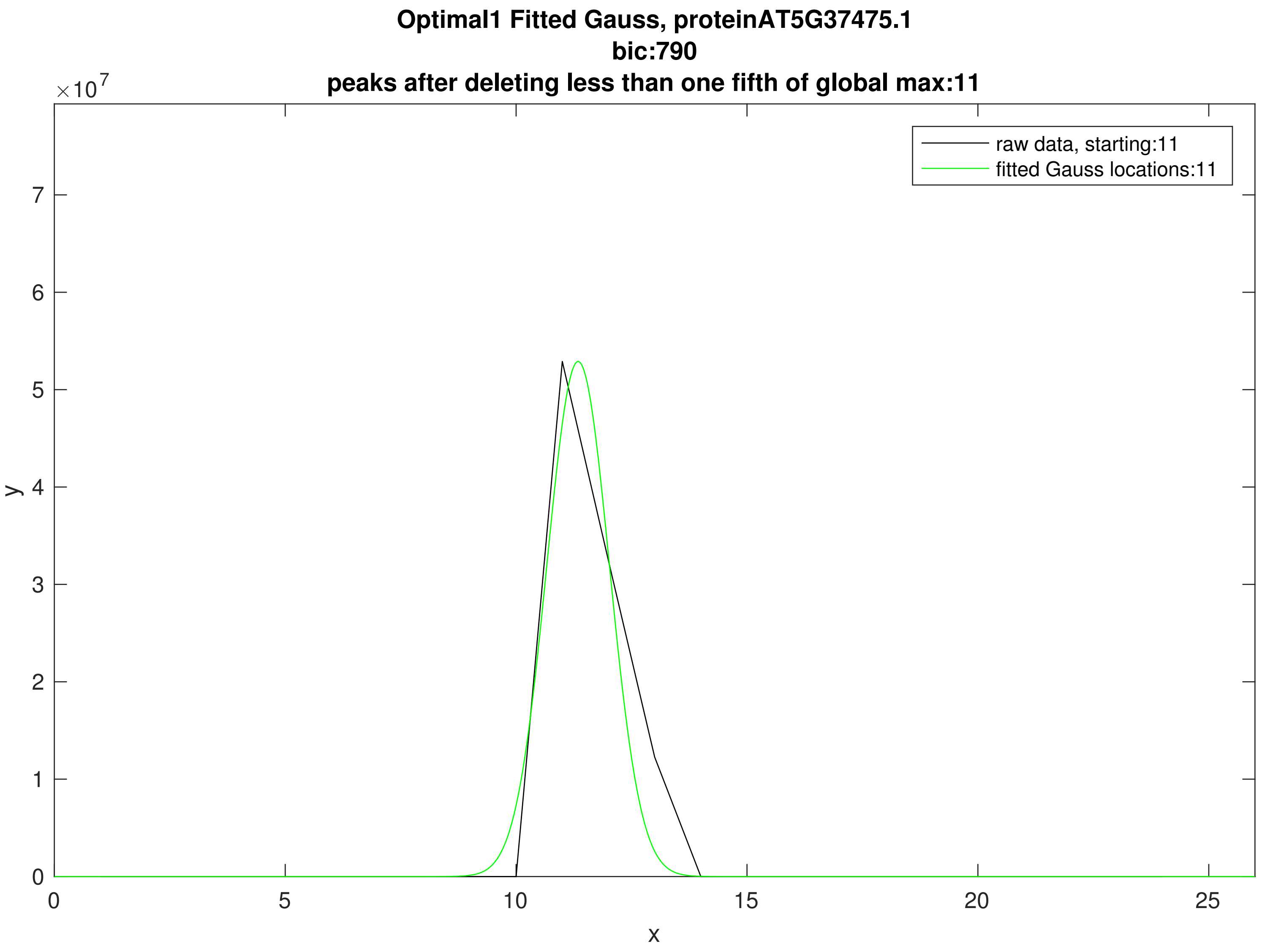
Optimal1 Fitted Gauss, proteinAT5G37475.1
bic:790
peaks after deleting less than one fifth of global max:11
×10^7
raw data, starting:11
fitted Gauss locations:11
y
x
0
1
2
3
4
5
6
7
0
5
10
15
20
25

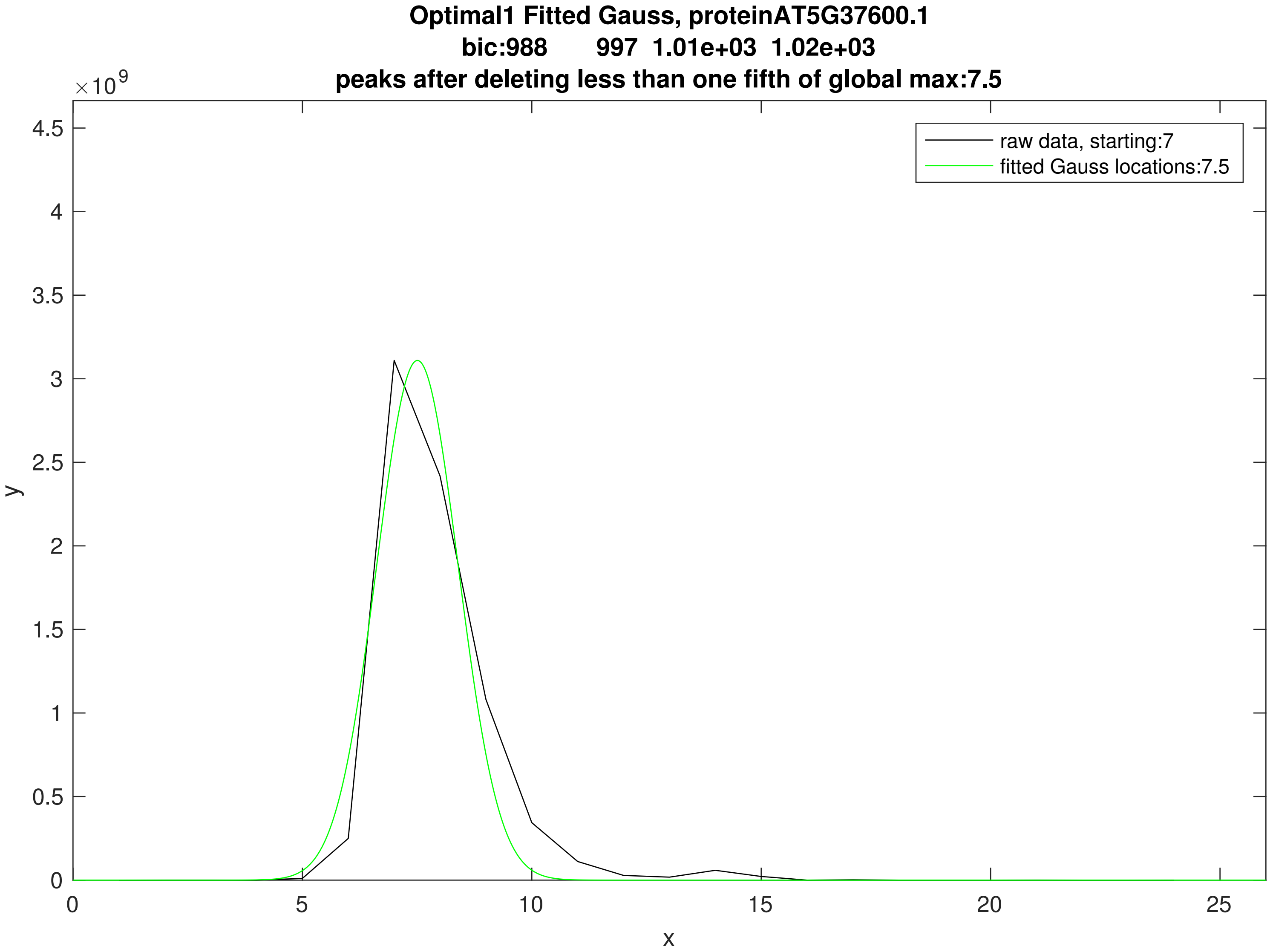

Optimal1 Fitted Gauss, proteinAT5G37600.1
bic:988 997 1.01e+03 1.02e+03
peaks after deleting less than one fifth of global max:7.5
×10^9
raw data, starting:7
fitted Gauss locations:7.5
y
x
0
0.5
1
1.5
2
2.5
3
3.5
4
4.5
5
10
15
20
25

**Optimal1 Fitted Gauss, proteinAT5G37660.1;AT5G37660.2**
**bic:633**
**peaks after deleting less than one fifth of global max:18**

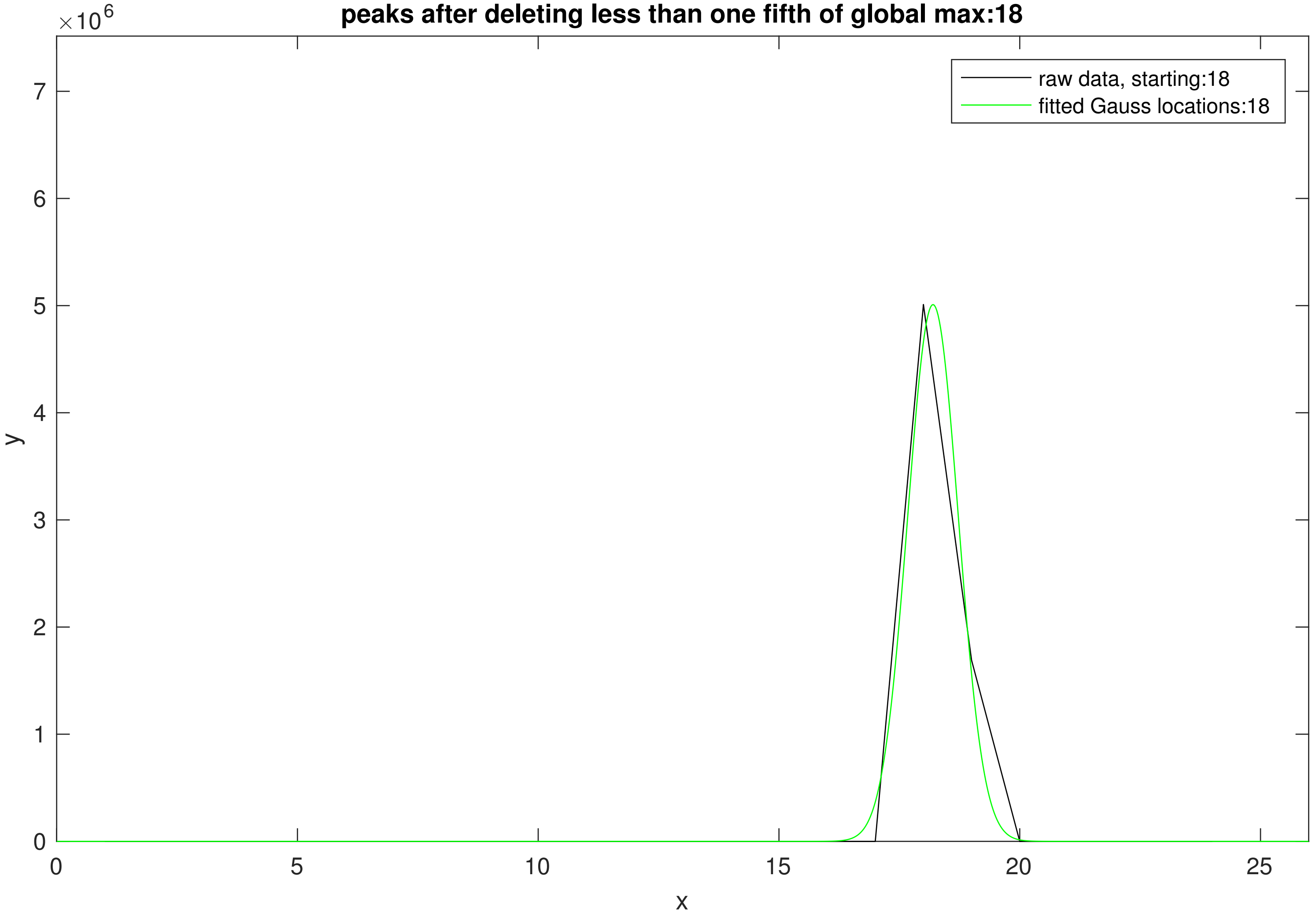

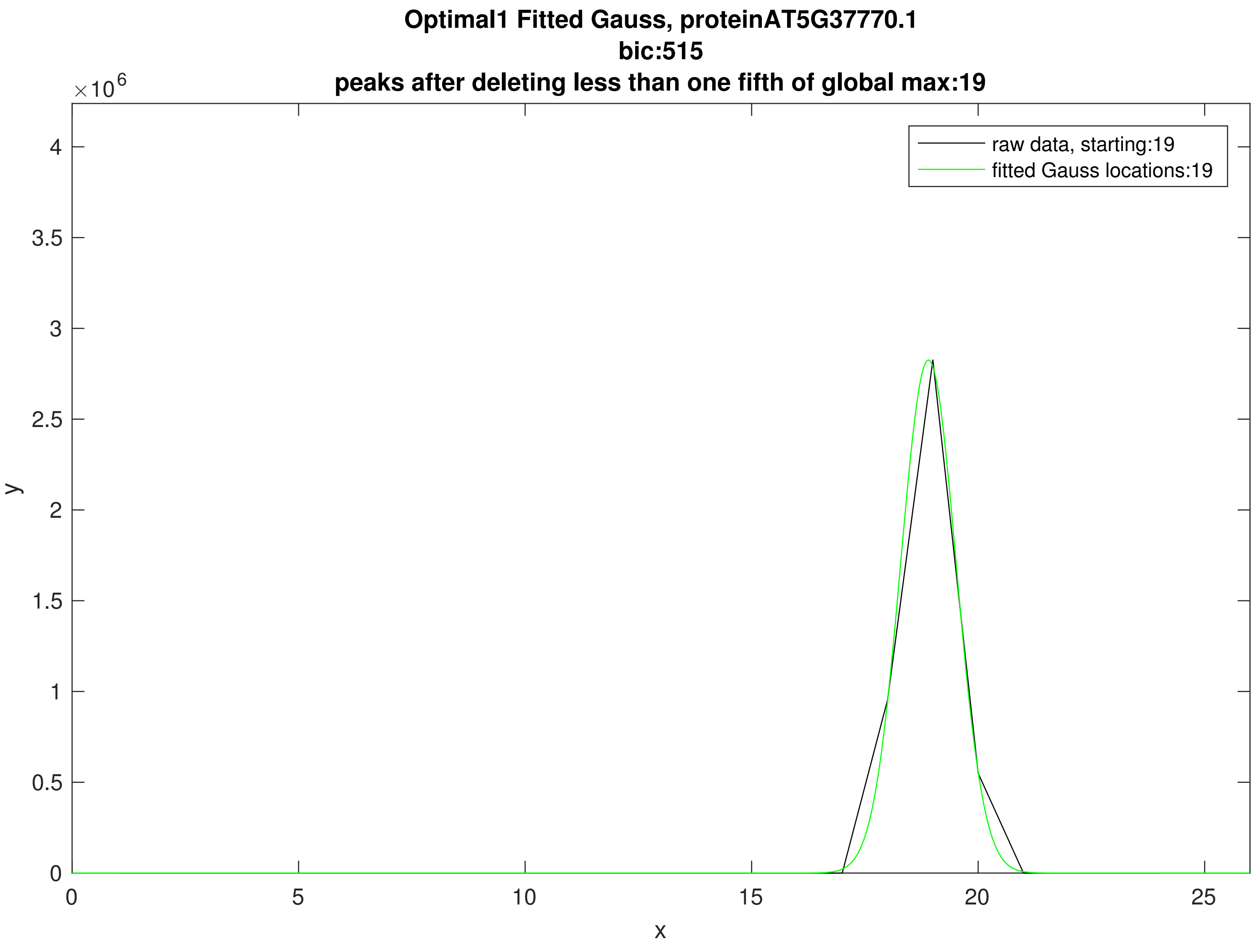

Optimal1 Fitted Gauss, proteinAT5G37770.1
bic:515
peaks after deleting less than one fifth of global max:19
×10^6
raw data, starting:19
fitted Gauss locations:19
y
x
0
0.5
1
1.5
2
2.5
3
3.5
4
0
5
10
15
20
25

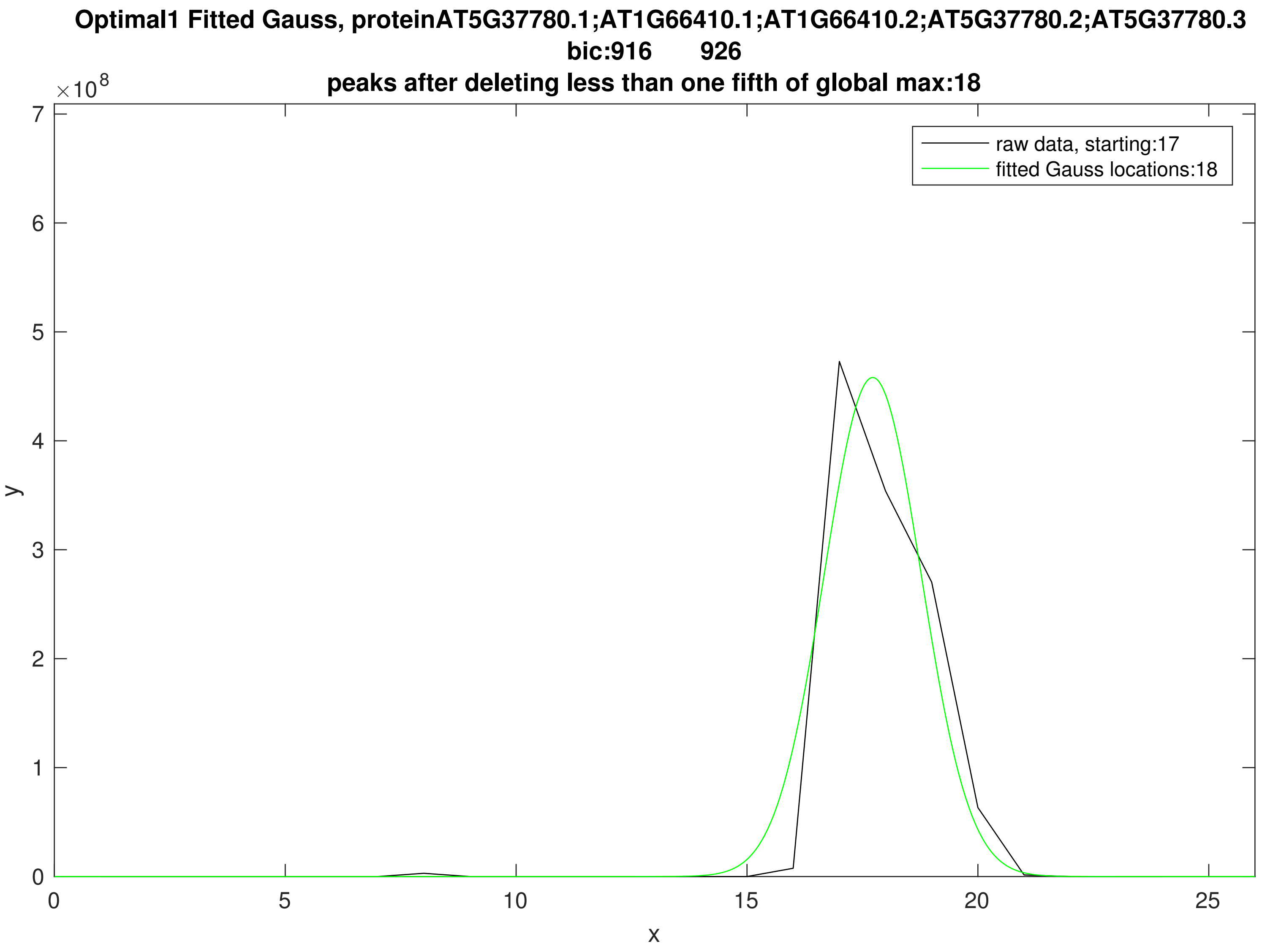
Optimal1 Fitted Gauss, proteinAT5G37780.1;AT1G66410.1;AT1G66410.2;AT5G37780.2;AT5G37780.3
bic:916 926
peaks after deleting less than one fifth of global max:18
×10^8
raw data, starting:17
fitted Gauss locations:18
y
x
0
1
2
3
4
5
6
7
10
15
20
25

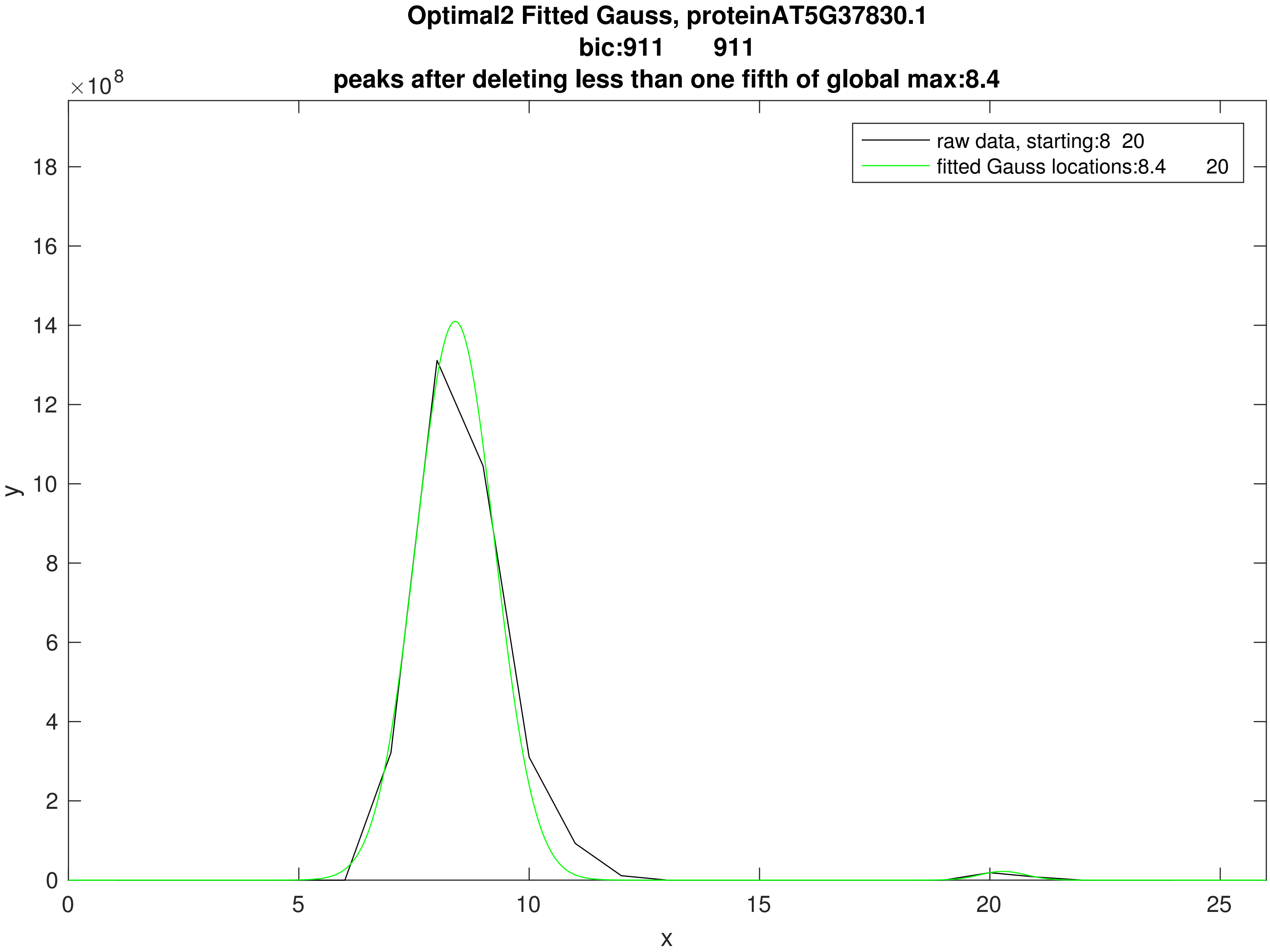
Optimal2 Fitted Gauss, proteinAT5G37830.1
bic:911 911
peaks after deleting less than one fifth of global max:8.4
×10^8
raw data, starting:8 20
fitted Gauss locations:8.4 20
y
x
0
2
4
6
8
10
12
14
16
18
0
5
10
15
20
25

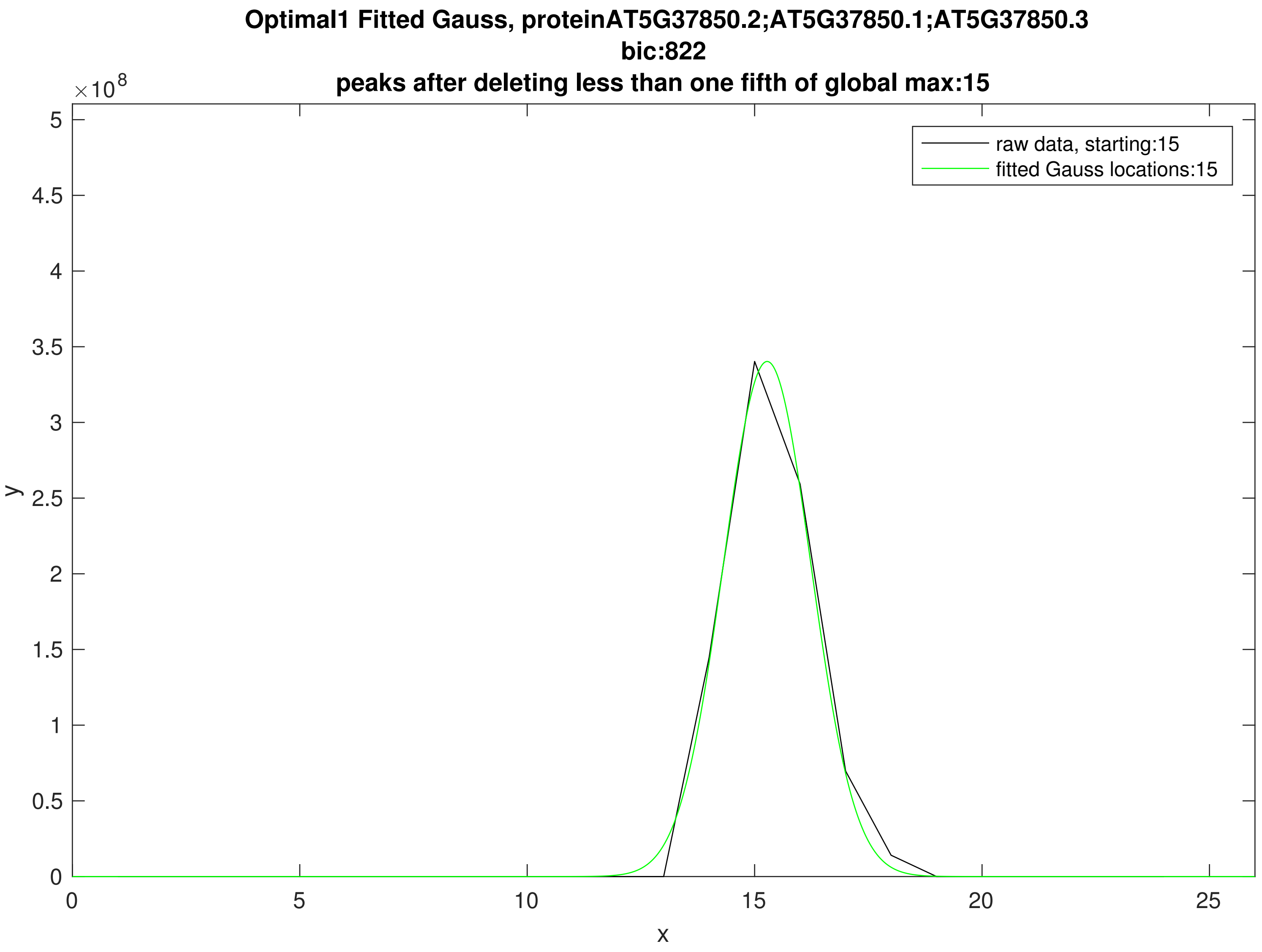
Optimal1 Fitted Gauss, proteinAT5G37850.2;AT5G37850.1;AT5G37850.3
bic:822
peaks after deleting less than one fifth of global max:15
×10^8
raw data, starting:15
fitted Gauss locations:15
y
x
0
0.5
1
1.5
2
2.5
3
3.5
4
4.5
5
0
5
10
15
20
25

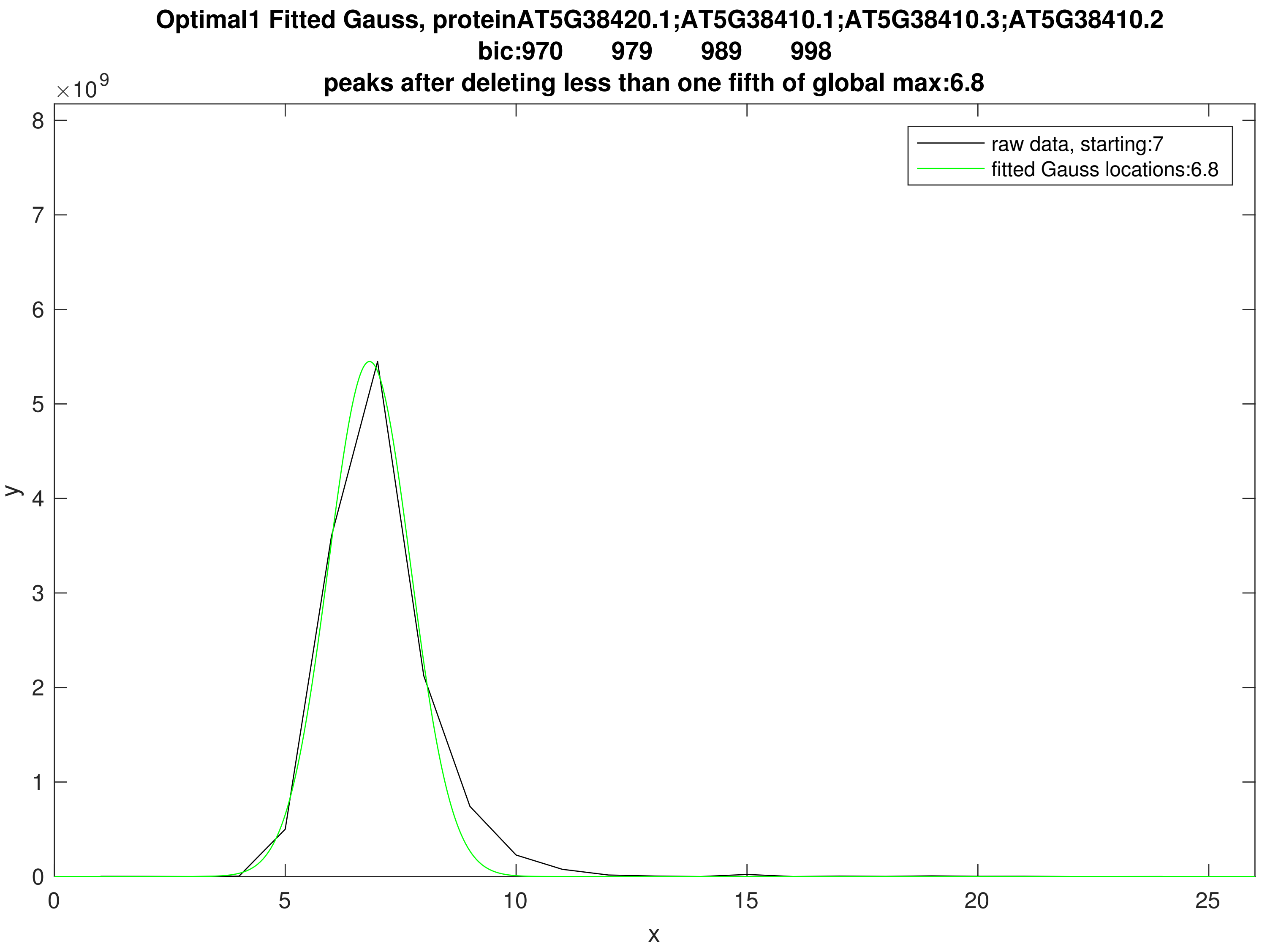

Optimal1 Fitted Gauss, proteinAT5G38420.1;AT5G38410.1;AT5G38410.3;AT5G38410.2
bic:970 979 989 998
peaks after deleting less than one fifth of global max:6.8
×10^9
raw data, starting:7
fitted Gauss locations:6.8
y
x
0
1
2
3
4
5
6
7
8
0
5
10
15
20
25

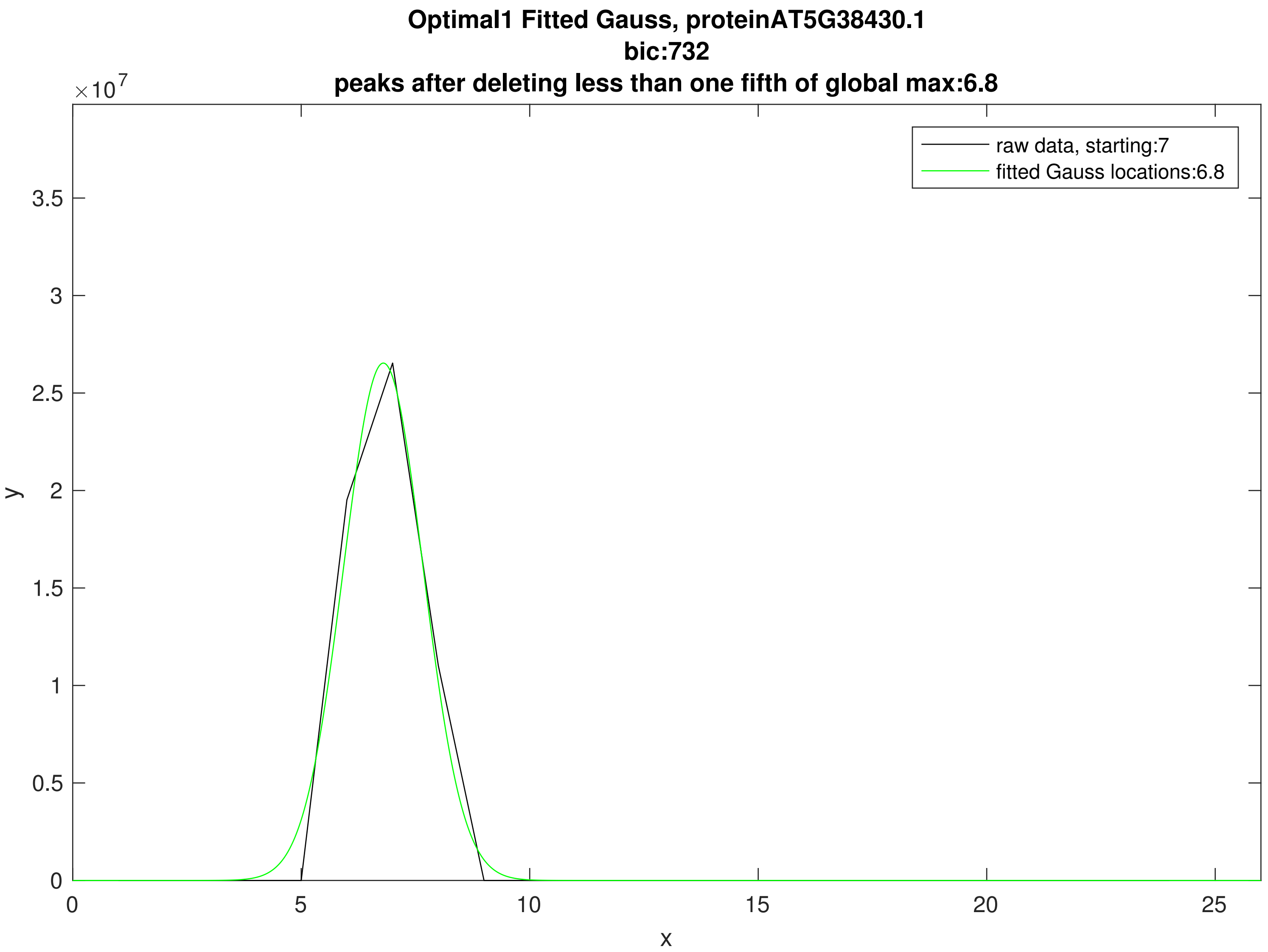
Optimal1 Fitted Gauss, proteinAT5G38430.1
bic:732
peaks after deleting less than one fifth of global max:6.8
×10^7
raw data, starting:7
fitted Gauss locations:6.8
3.5
3
2.5
2
1.5
1
0.5
0
y
0
5
10
15
20
25
x

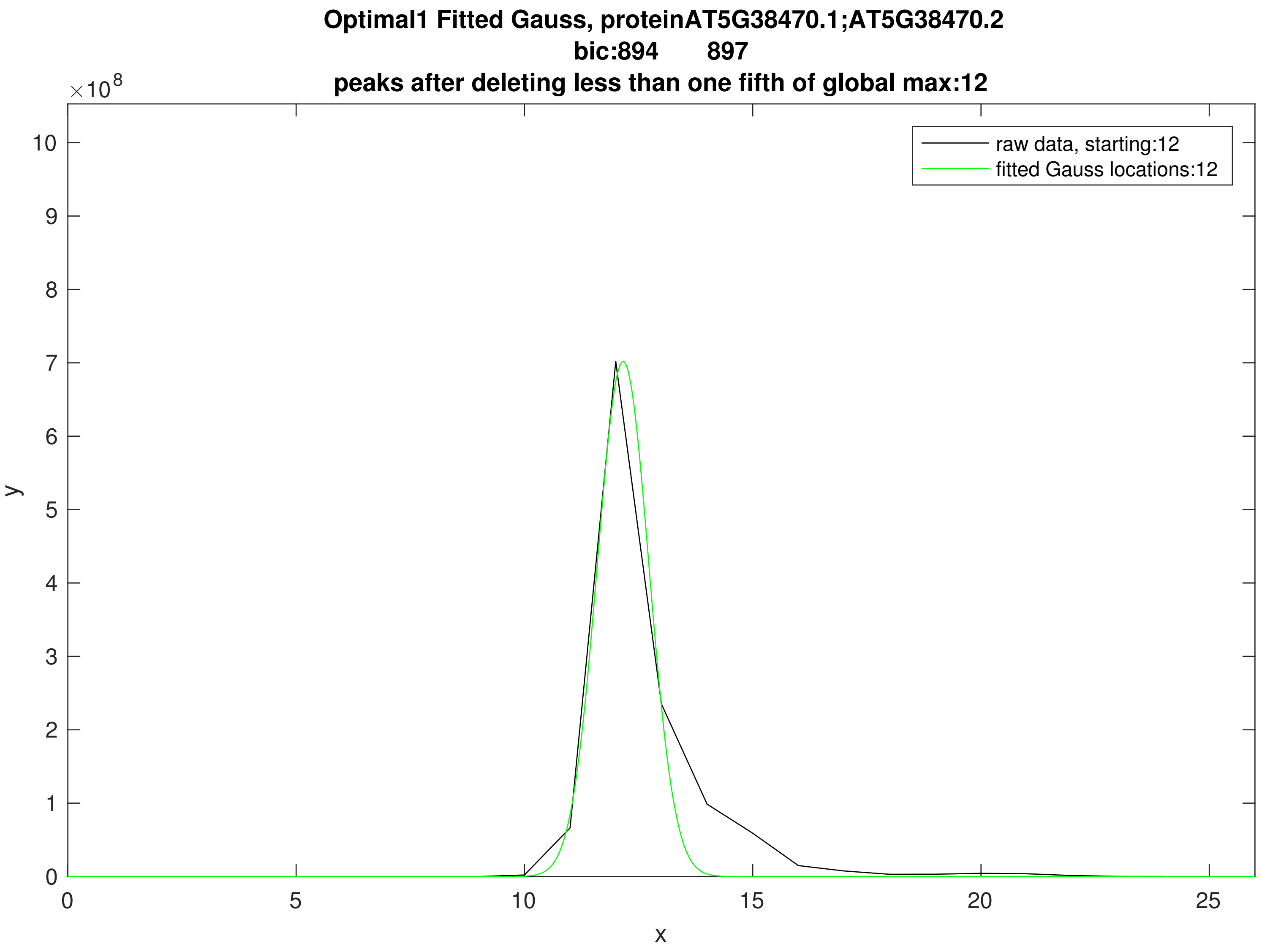

Optimal1 Fitted Gauss, proteinAT5G38470.1;AT5G38470.2
bic:894 897
peaks after deleting less than one fifth of global max:12
×10^8
raw data, starting:12
fitted Gauss locations:12
y
x

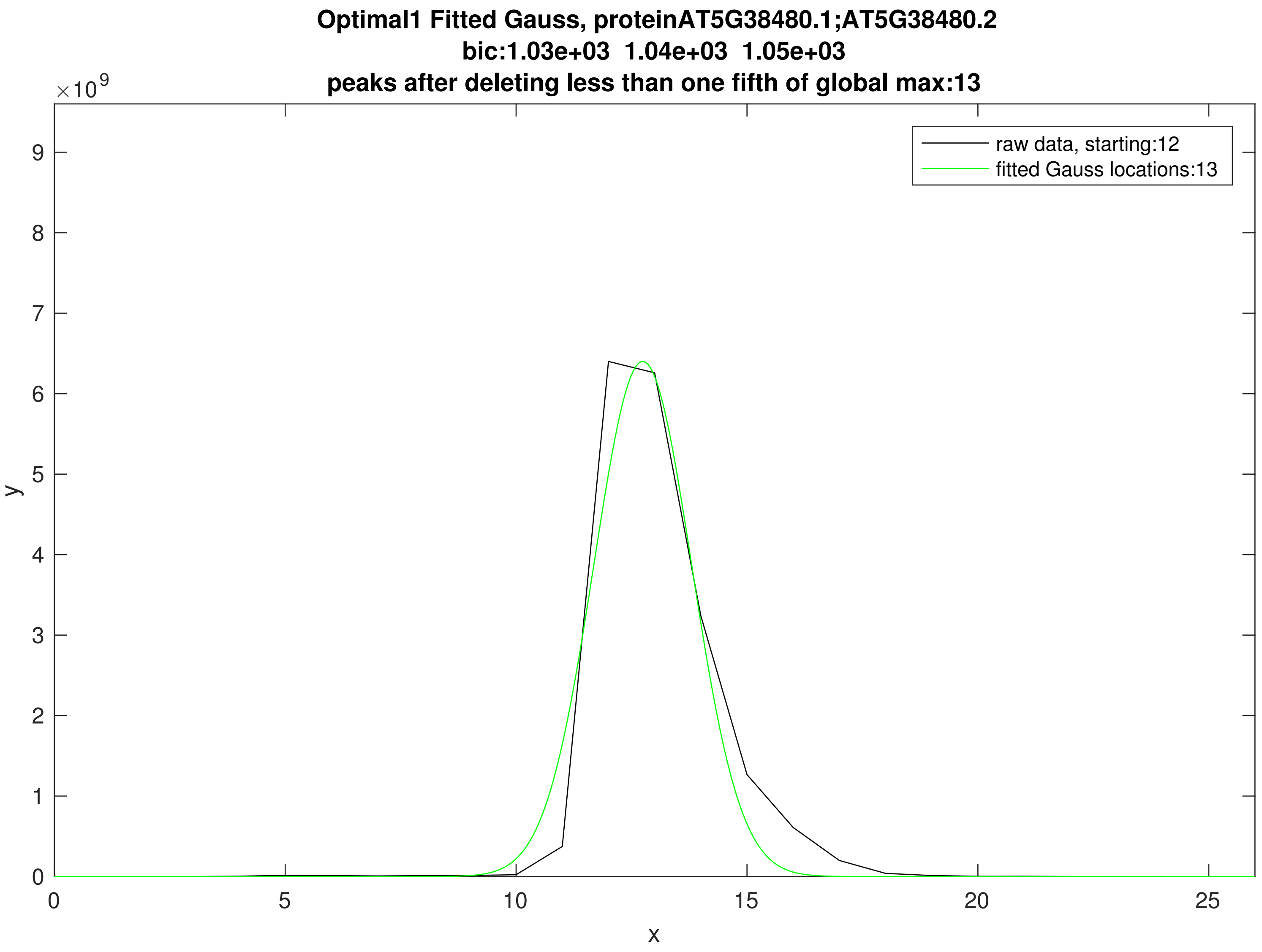

Optimal1 Fitted Gauss, proteinAT5G38480.1;AT5G38480.2
bic:1.03e+03 1.04e+03 1.05e+03
peaks after deleting less than one fifth of global max:13
×10^9
raw data, starting:12
fitted Gauss locations:13
y
x
0
1
2
3
4
5
6
7
8
9
0
5
10
15
20
25

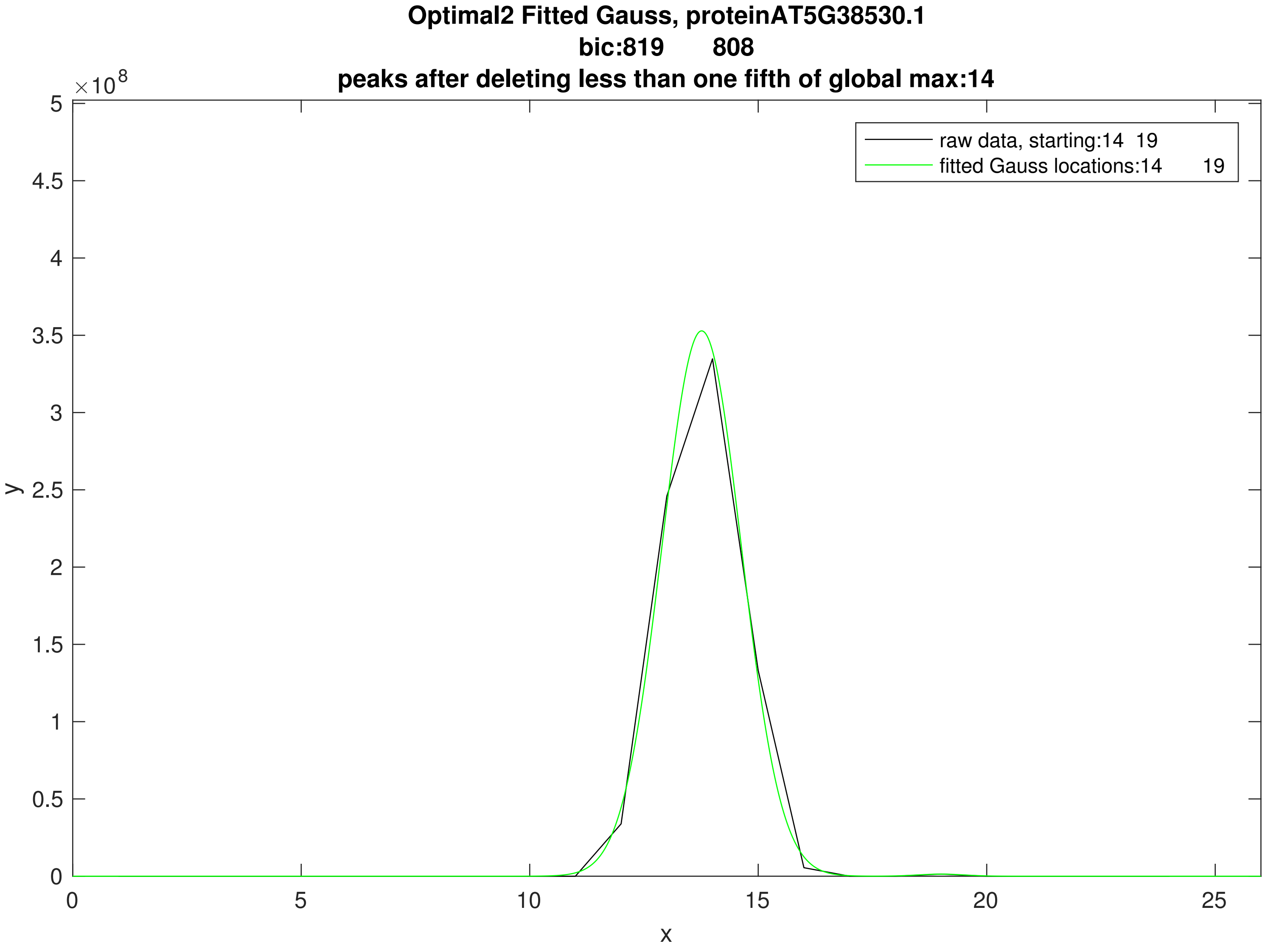

Optimal2 Fitted Gauss, proteinAT5G38530.1
bic:819 808
peaks after deleting less than one fifth of global max:14
×10^8
raw data, starting:14 19
fitted Gauss locations:14 19
y
x
0
0.5
1
1.5
2
2.5
3
3.5
4
4.5
5
0
5
10
15
20
25

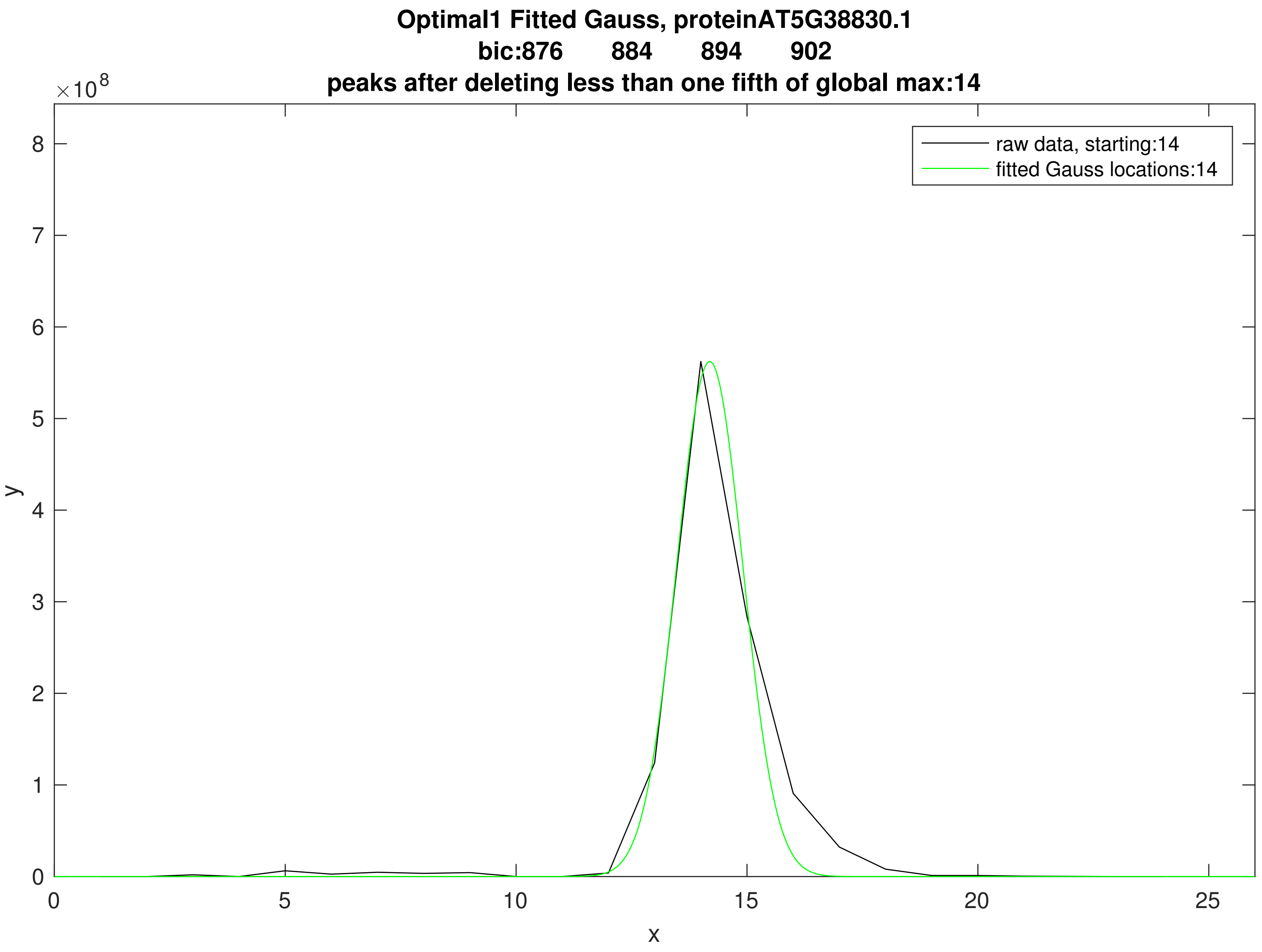

Optimal1 Fitted Gauss, proteinAT5G38830.1
bic:876 884 894 902
peaks after deleting less than one fifth of global max:14
×10^8
raw data, starting:14
fitted Gauss locations:14
y
x
0
1
2
3
4
5
6
7
8
0
5
10
15
20
25

**Optimal1 Fitted Gauss, proteinAT5G38860.1**
**bic:808**
**peaks after deleting less than one fifth of global max:12**

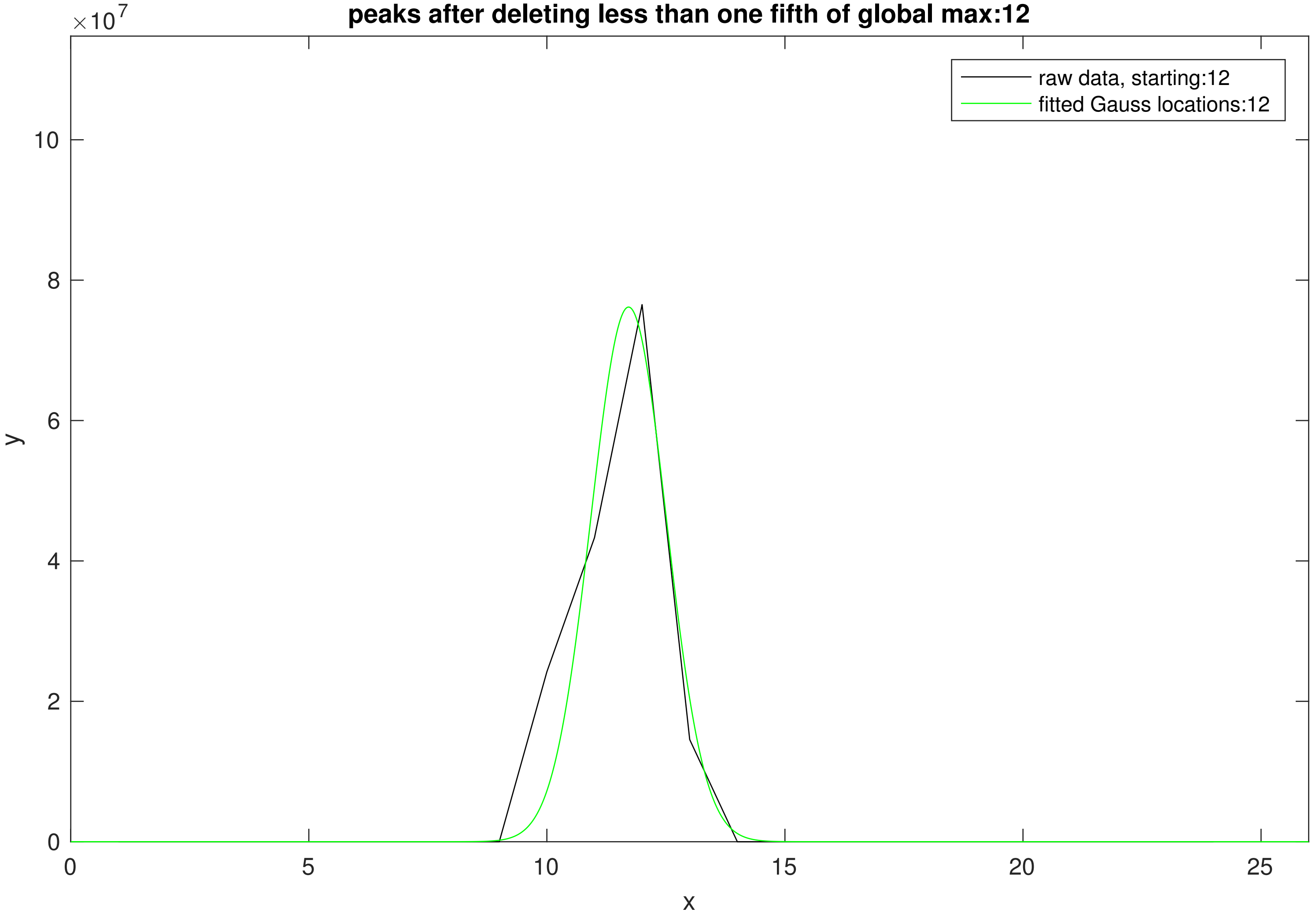

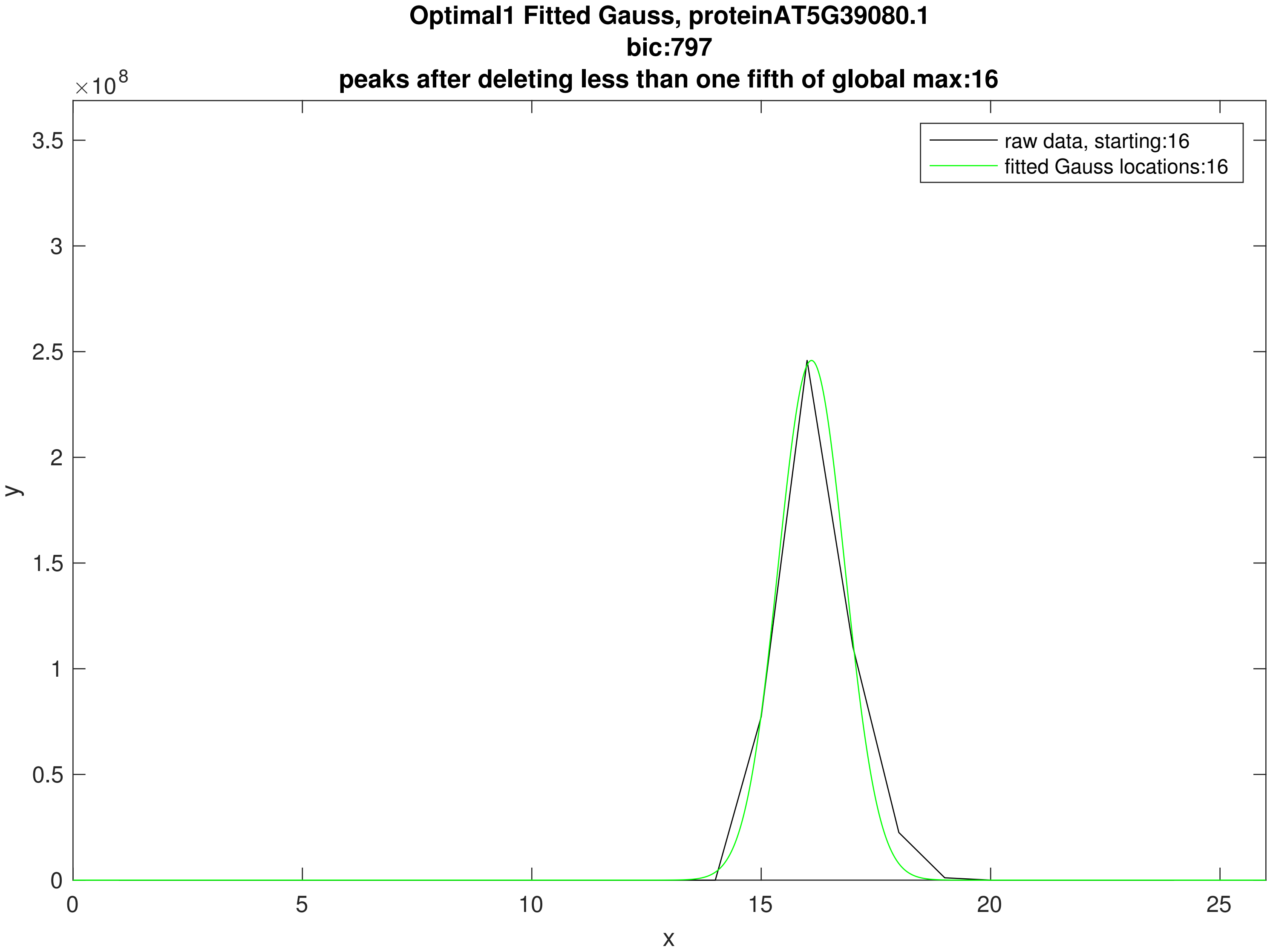

Optimal1 Fitted Gauss, proteinAT5G39080.1
bic:797
peaks after deleting less than one fifth of global max:16
×10^8
3.5
3
2.5
2
1.5
1
0.5
0
y
0
5
10
15
20
25
x
raw data, starting:16
fitted Gauss locations:16

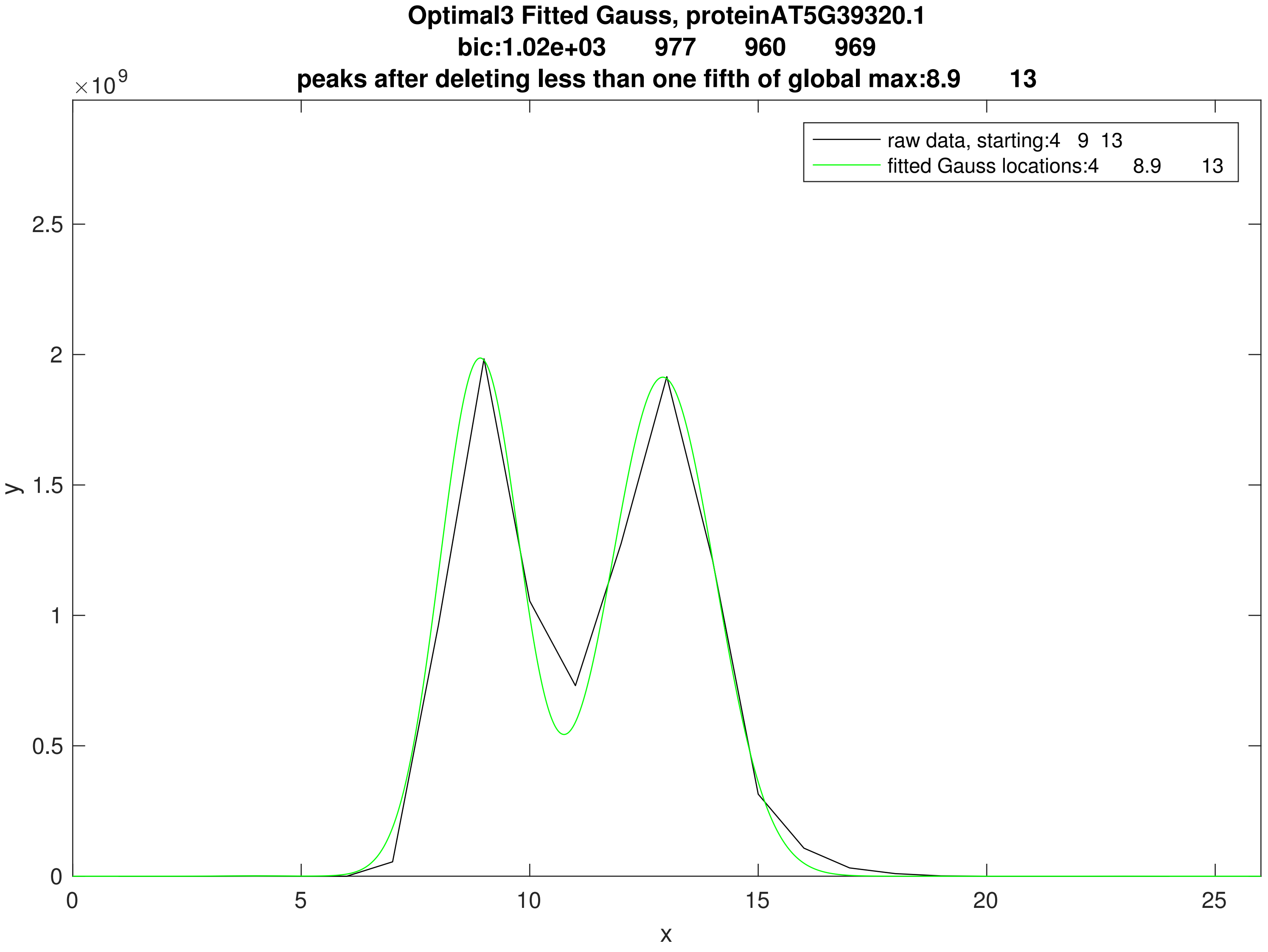
Optimal3 Fitted Gauss, proteinAT5G39320.1
bic:1.02e+03 977 960 969
peaks after deleting less than one fifth of global max:8.9 13
×10^9
raw data, starting:4 9 13
fitted Gauss locations:4 8.9 13
y
x
0
0.5
1
1.5
2
2.5
0
5
10
15
20
25

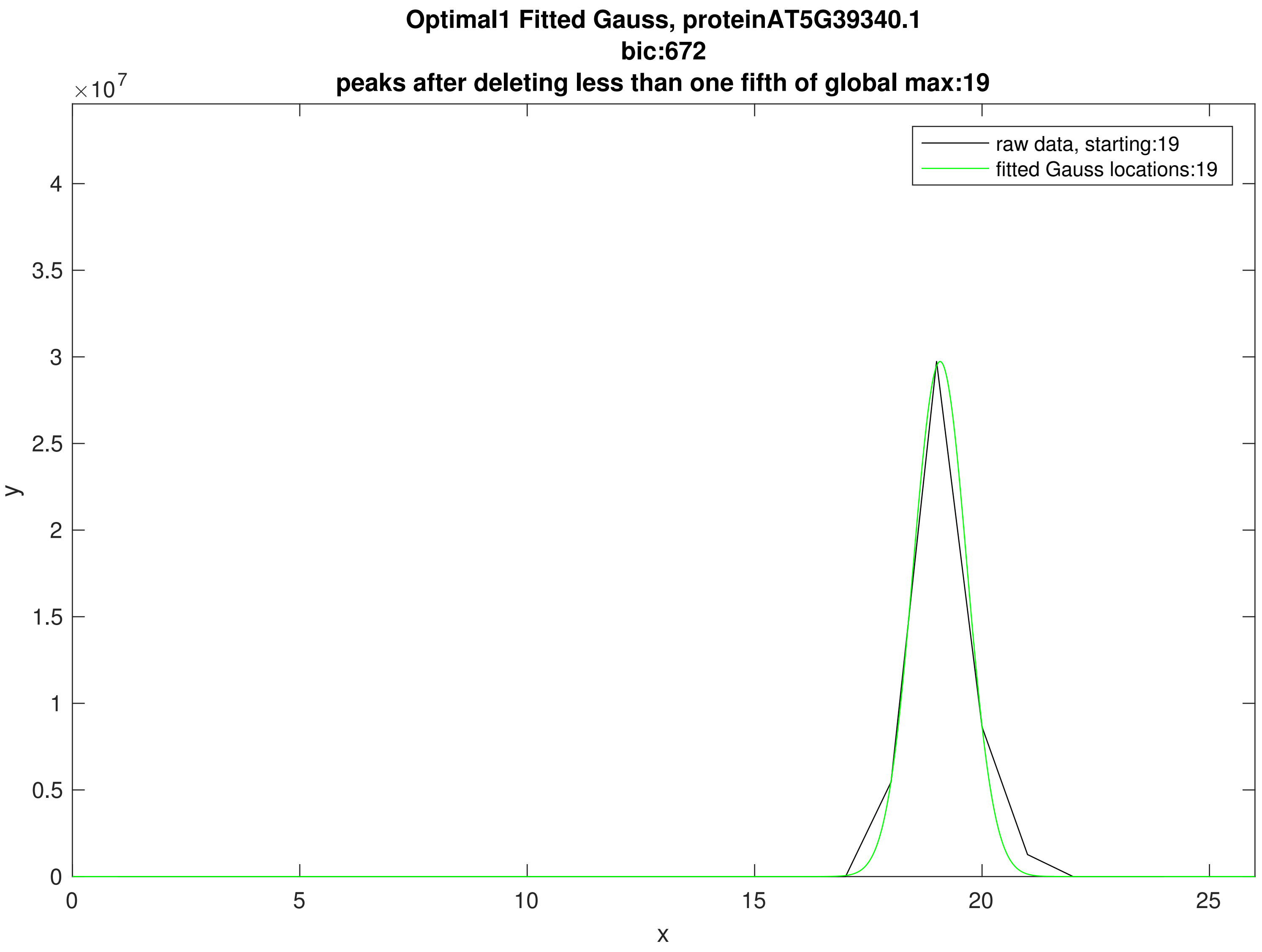
Optimal1 Fitted Gauss, proteinAT5G39340.1
bic:672
peaks after deleting less than one fifth of global max:19
×10^7
raw data, starting:19
fitted Gauss locations:19
4
3.5
3
2.5
2
1.5
1
0.5
0
y
0
5
10
15
20
25
x

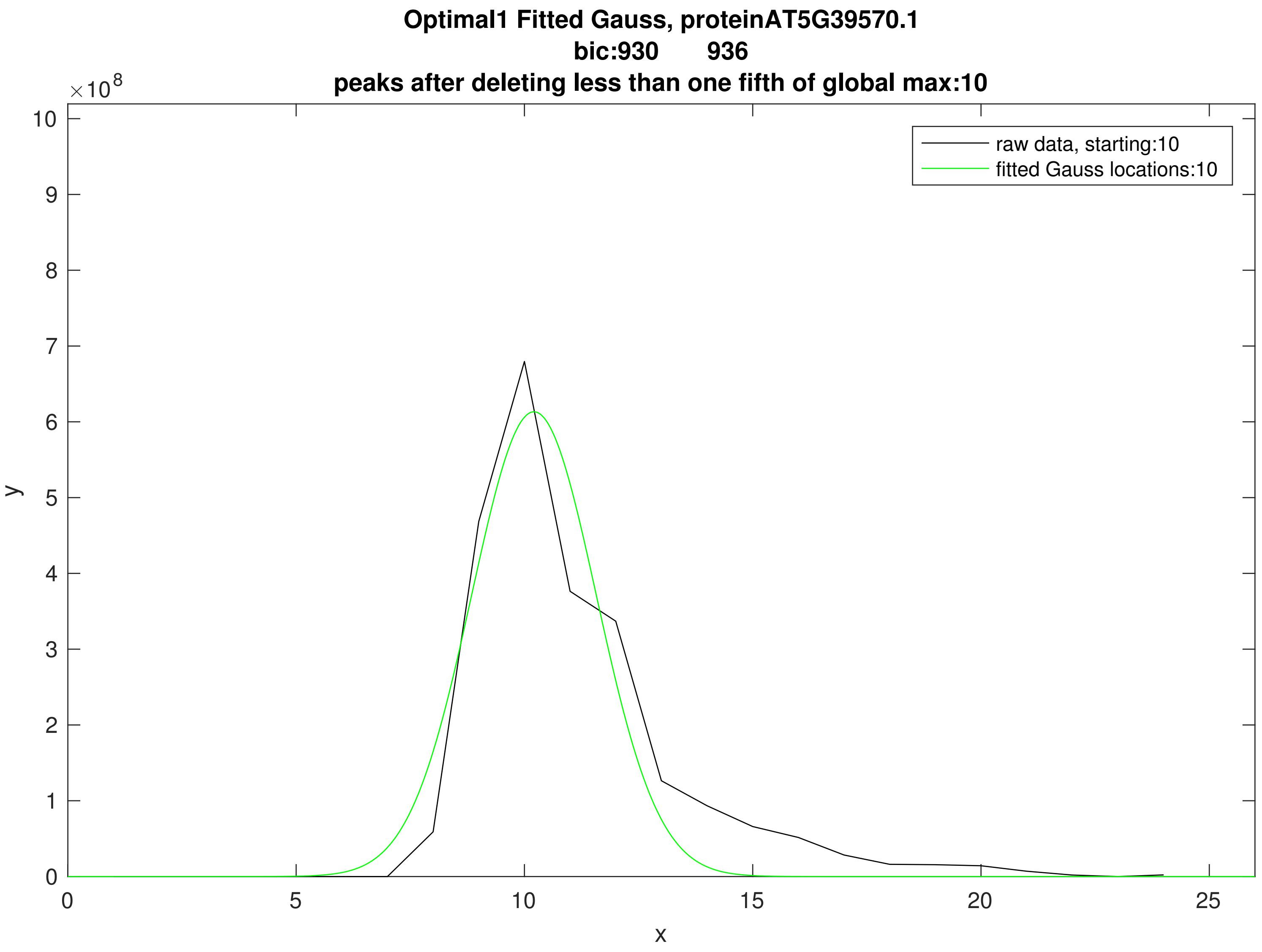

Optimal1 Fitted Gauss, proteinAT5G39570.1
bic:930 936
peaks after deleting less than one fifth of global max:10
×10^8
raw data, starting:10
fitted Gauss locations:10
y
x
0
1
2
3
4
5
6
7
8
9
10
0
5
10
15
20
25

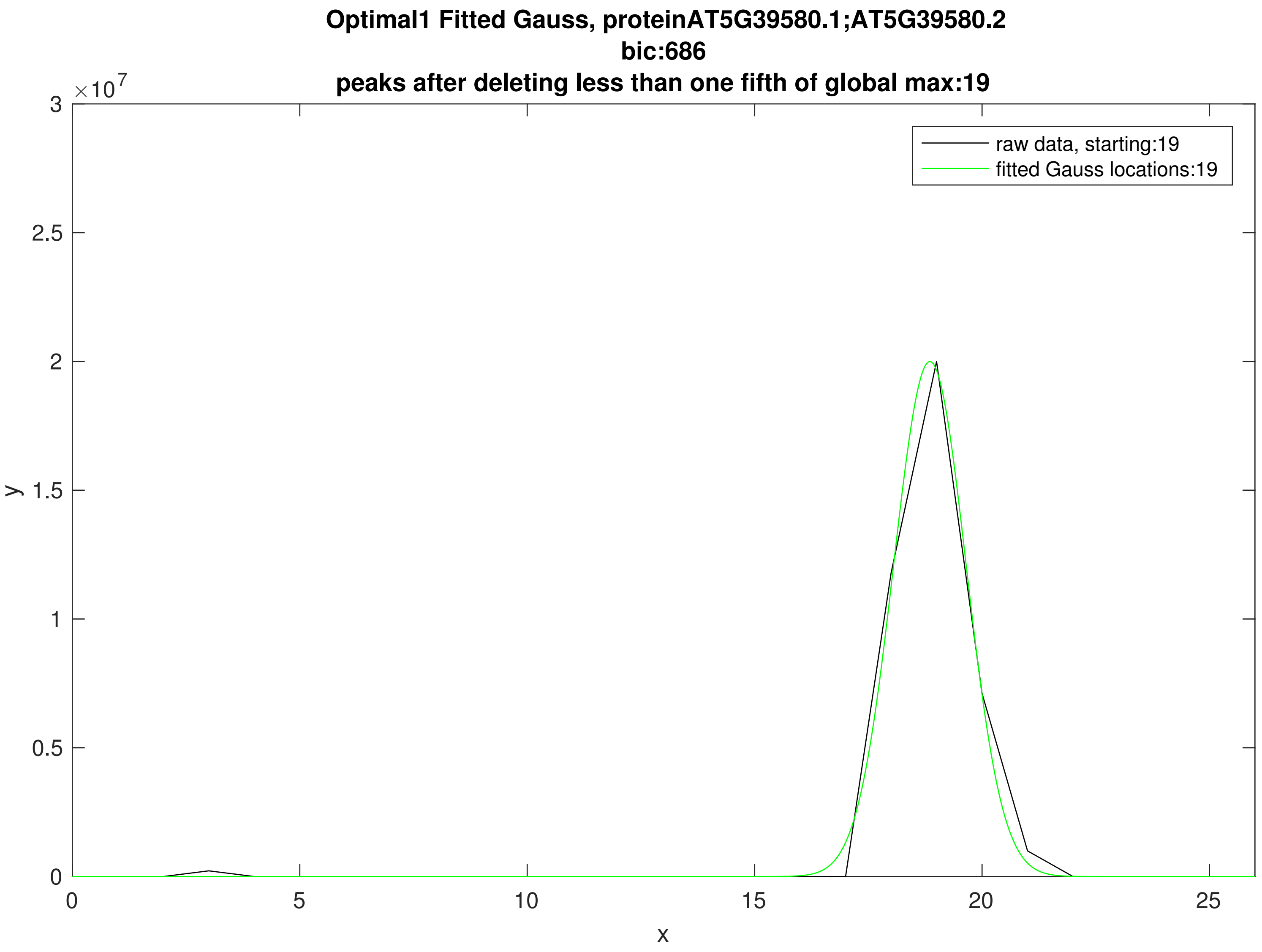
Optimal1 Fitted Gauss, proteinAT5G39580.1;AT5G39580.2
bic:686
peaks after deleting less than one fifth of global max:19
raw data, starting:19
fitted Gauss locations:19
$\times 10^7$
3
2.5
2
1.5
1
0.5
0
y
0
5
10
15
20
25
x

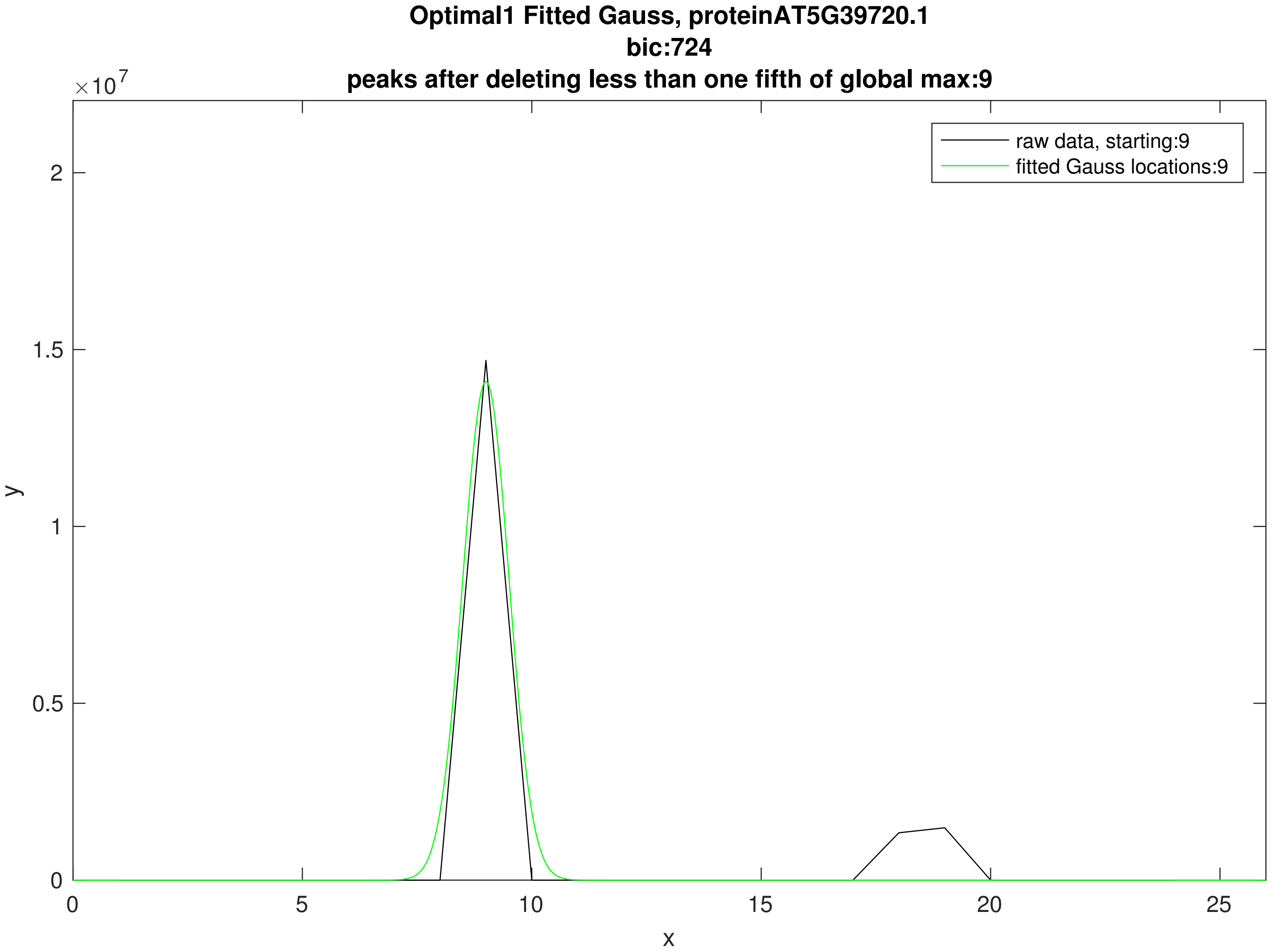
Optimal1 Fitted Gauss, proteinAT5G39720.1
bic:724
peaks after deleting less than one fifth of global max:9
×10^7
raw data, starting:9
fitted Gauss locations:9
y
x
0
0.5
1
1.5
2
0
5
10
15
20
25

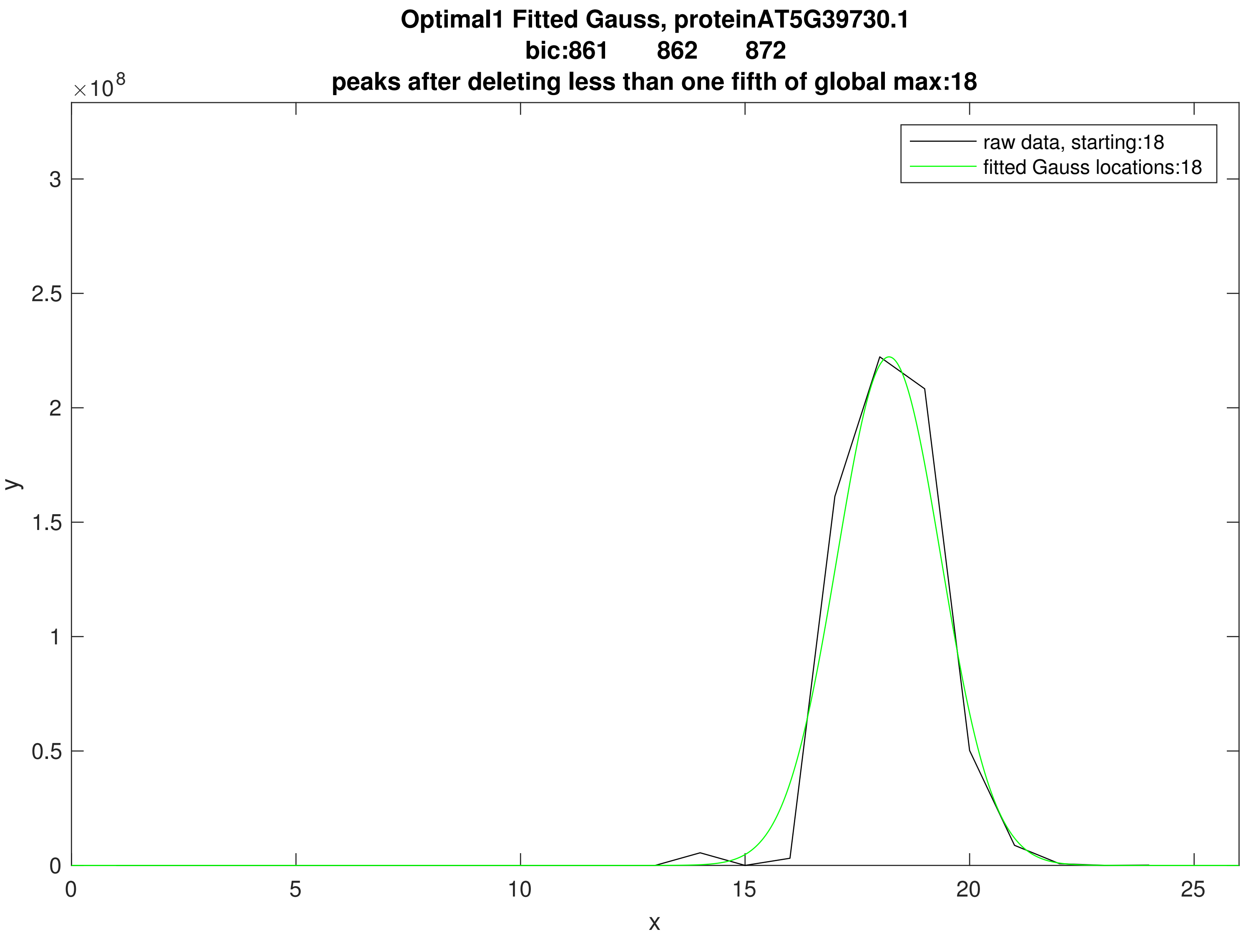
Optimal1 Fitted Gauss, proteinAT5G39730.1
bic:861 862 872
peaks after deleting less than one fifth of global max:18
×10^8
3
2.5
2
1.5
1
0.5
0
y
0
5
10
15
20
25
x
raw data, starting:18
fitted Gauss locations:18

**Optimal1 Fitted Gauss, proteinAT5G39740.2;AT5G39740.1**
**bic:609**
**peaks after deleting less than one fifth of global max:3.8**

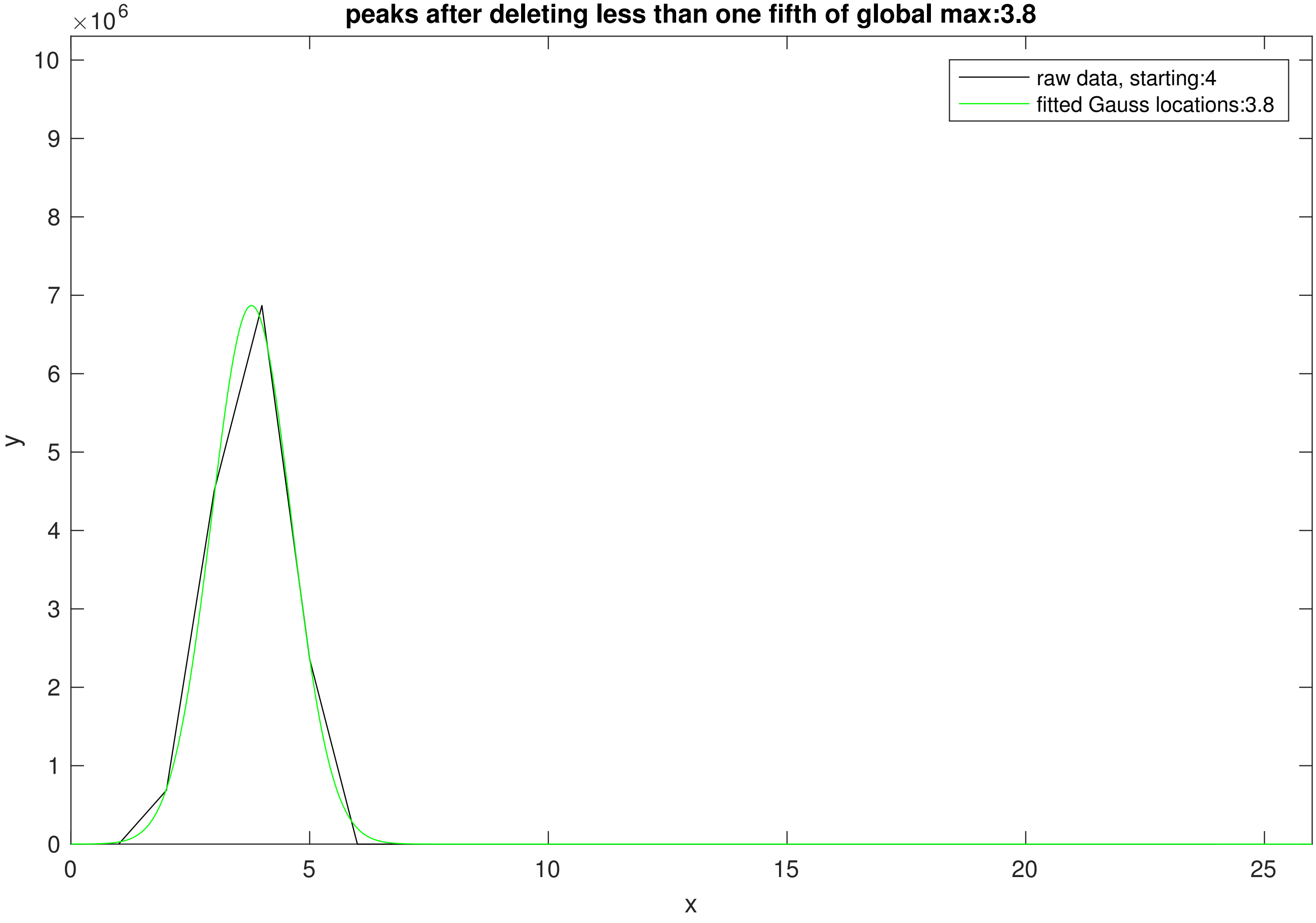

**Optimal1 Fitted Gauss, proteinAT5G40060.1**
**bic:615**
**peaks after deleting less than one fifth of global max:19**

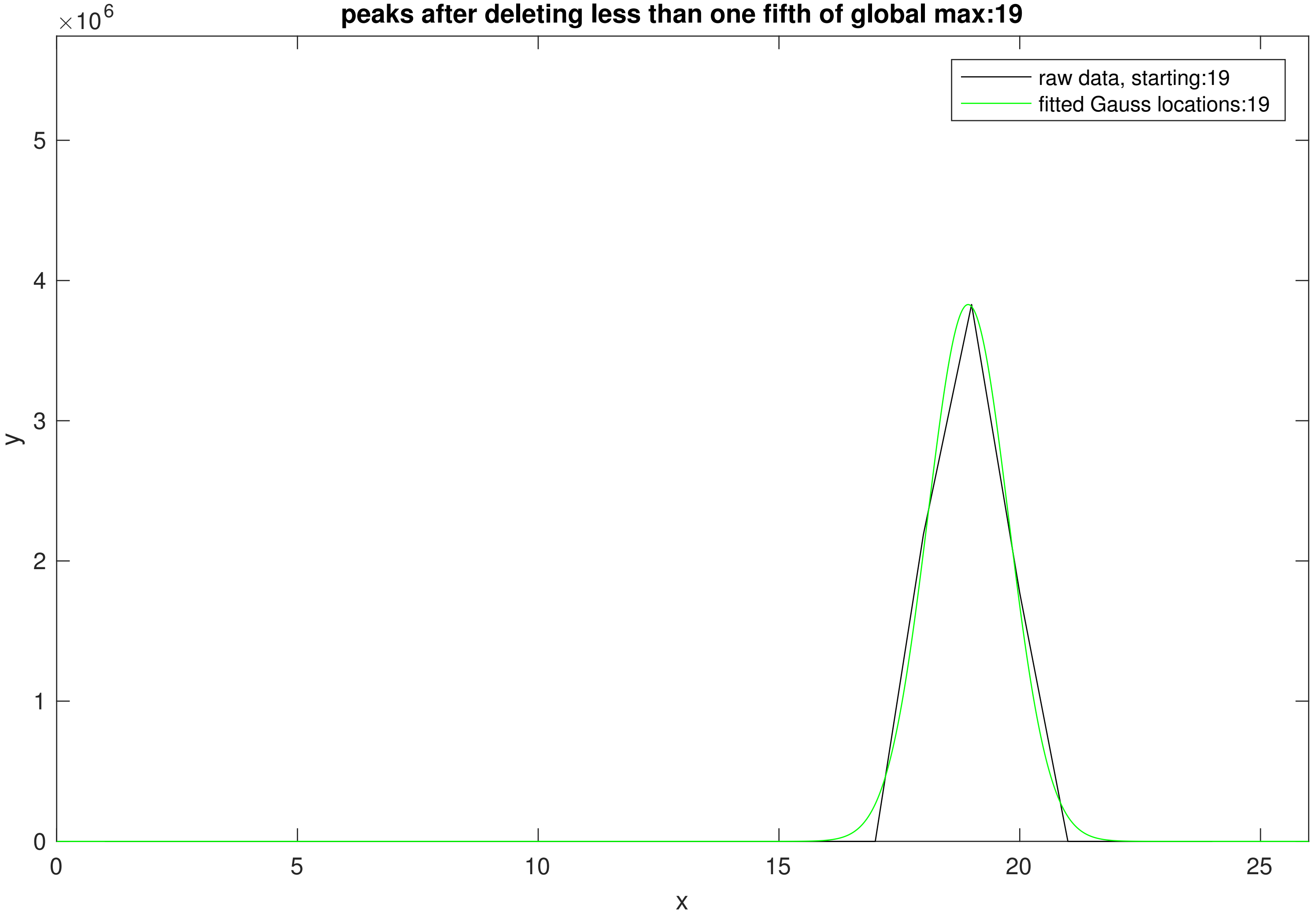

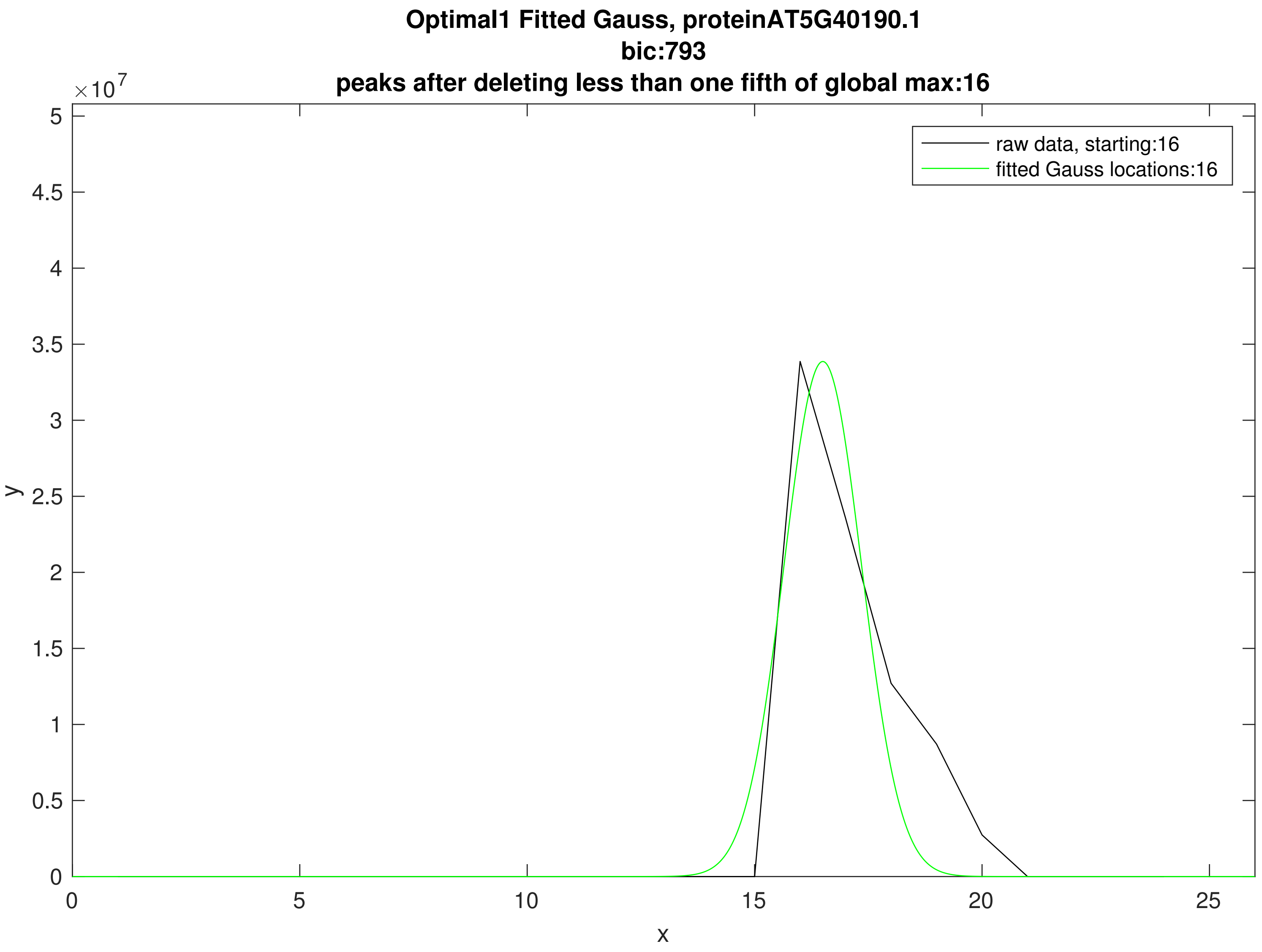

Optimal1 Fitted Gauss, proteinAT5G40190.1
bic:793
peaks after deleting less than one fifth of global max:16
×10^7
raw data, starting:16
fitted Gauss locations:16
y
x
0
0.5
1
1.5
2
2.5
3
3.5
4
4.5
5
0
5
10
15
20
25

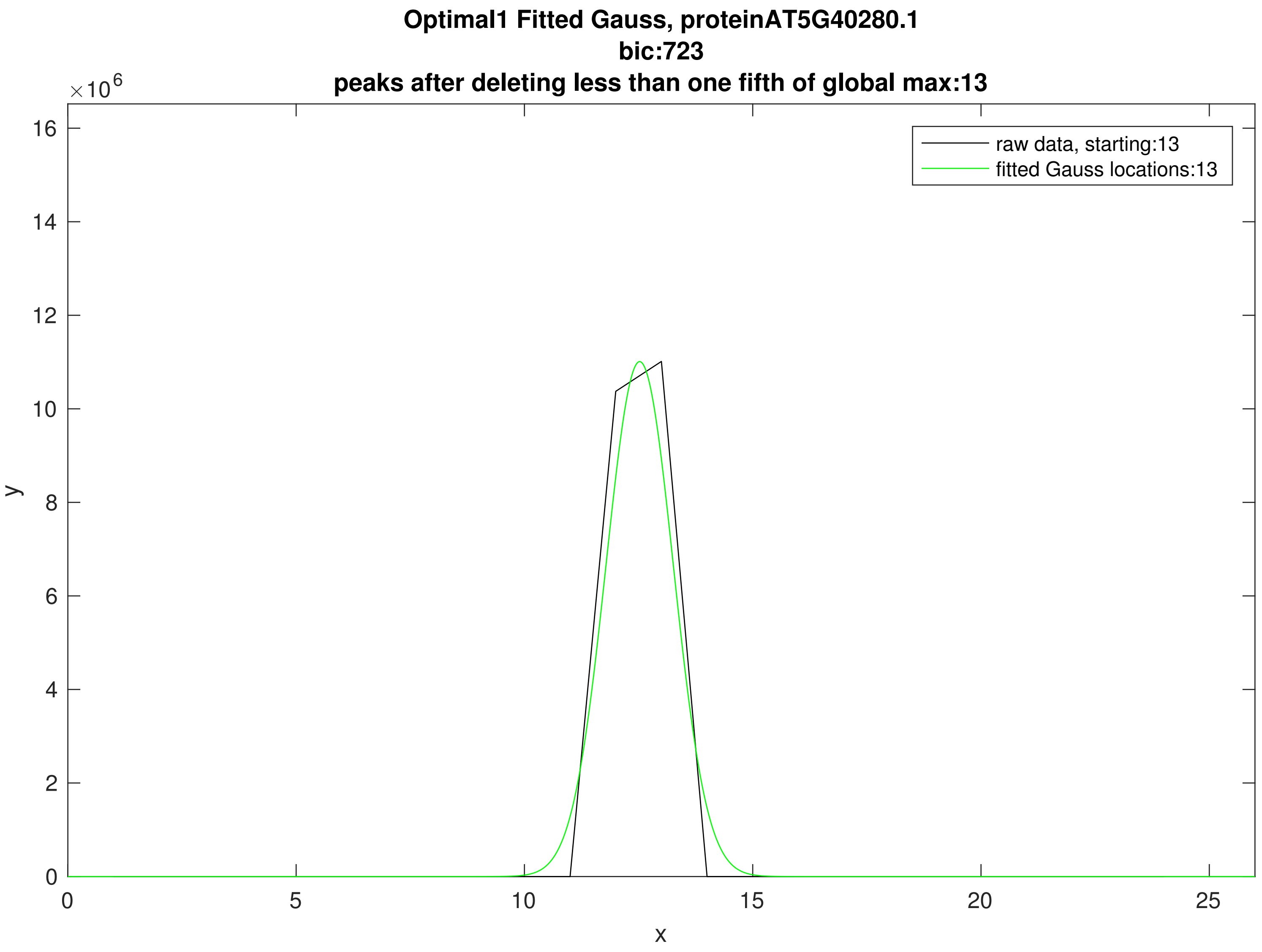

Optimal1 Fitted Gauss, proteinAT5G40280.1
bic:723
peaks after deleting less than one fifth of global max:13
×10^6
y
x
raw data, starting:13
fitted Gauss locations:13
0
2
4
6
8
10
12
14
16
0
5
10
15
20
25

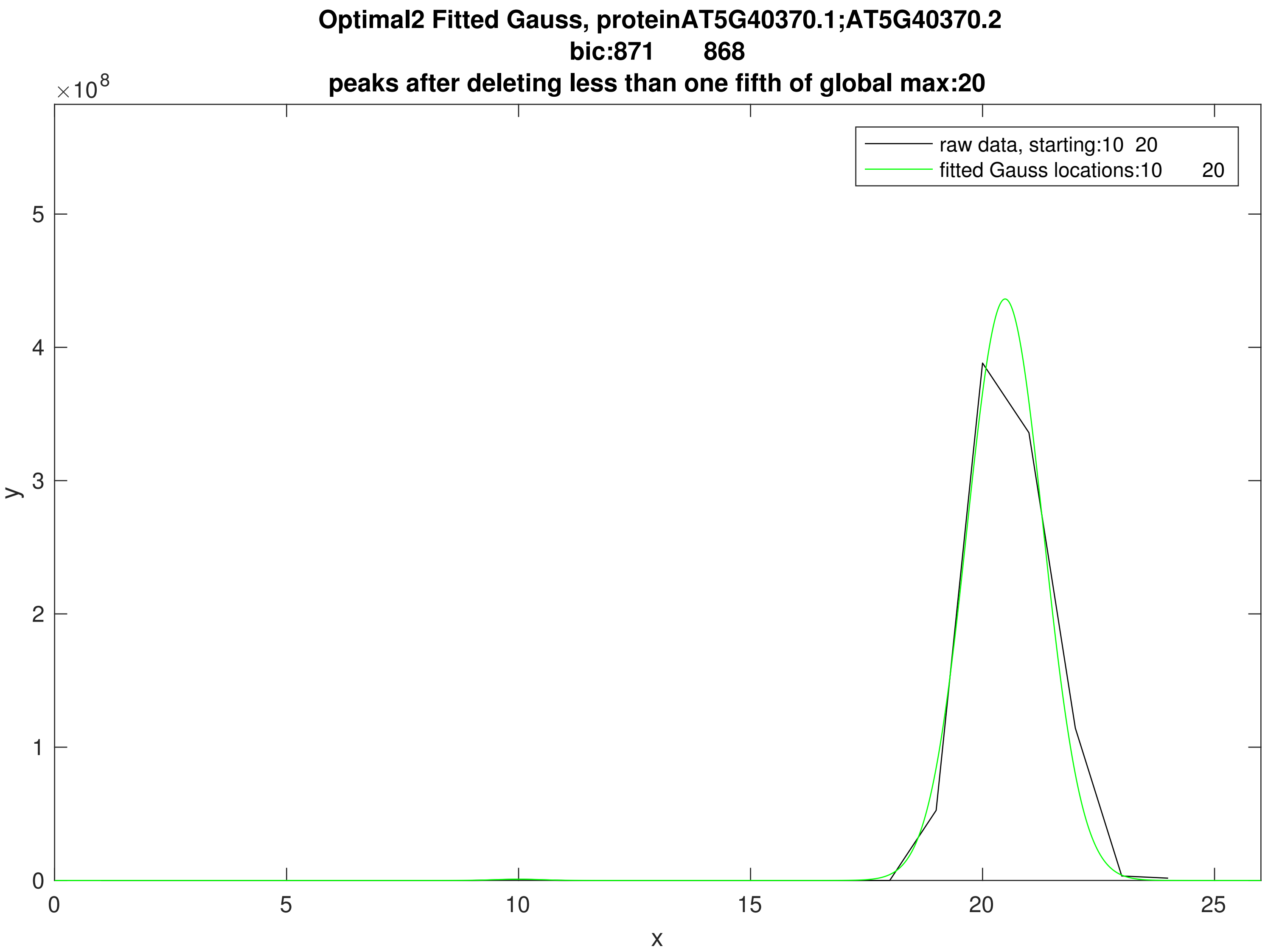
Optimal2 Fitted Gauss, proteinAT5G40370.1;AT5G40370.2
bic:871 868
peaks after deleting less than one fifth of global max:20
×10^8
raw data, starting:10 20
fitted Gauss locations:10 20
y
x
0
1
2
3
4
5
0
5
10
15
20
25

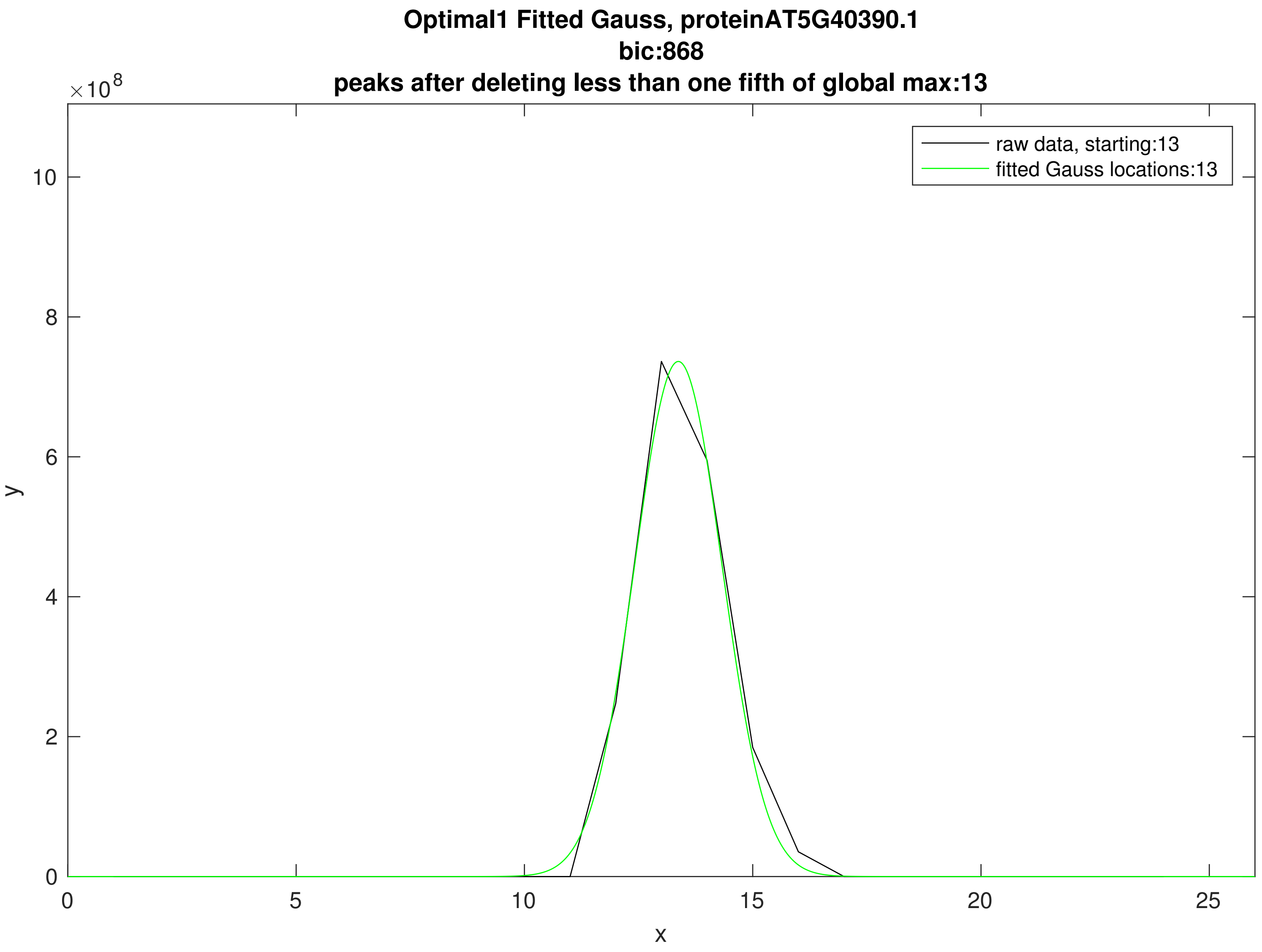

Optimal1 Fitted Gauss, proteinAT5G40390.1
bic:868
peaks after deleting less than one fifth of global max:13
×10^8
10
8
6
4
2
0
y
0
5
10
15
20
25
x
raw data, starting:13
fitted Gauss locations:13

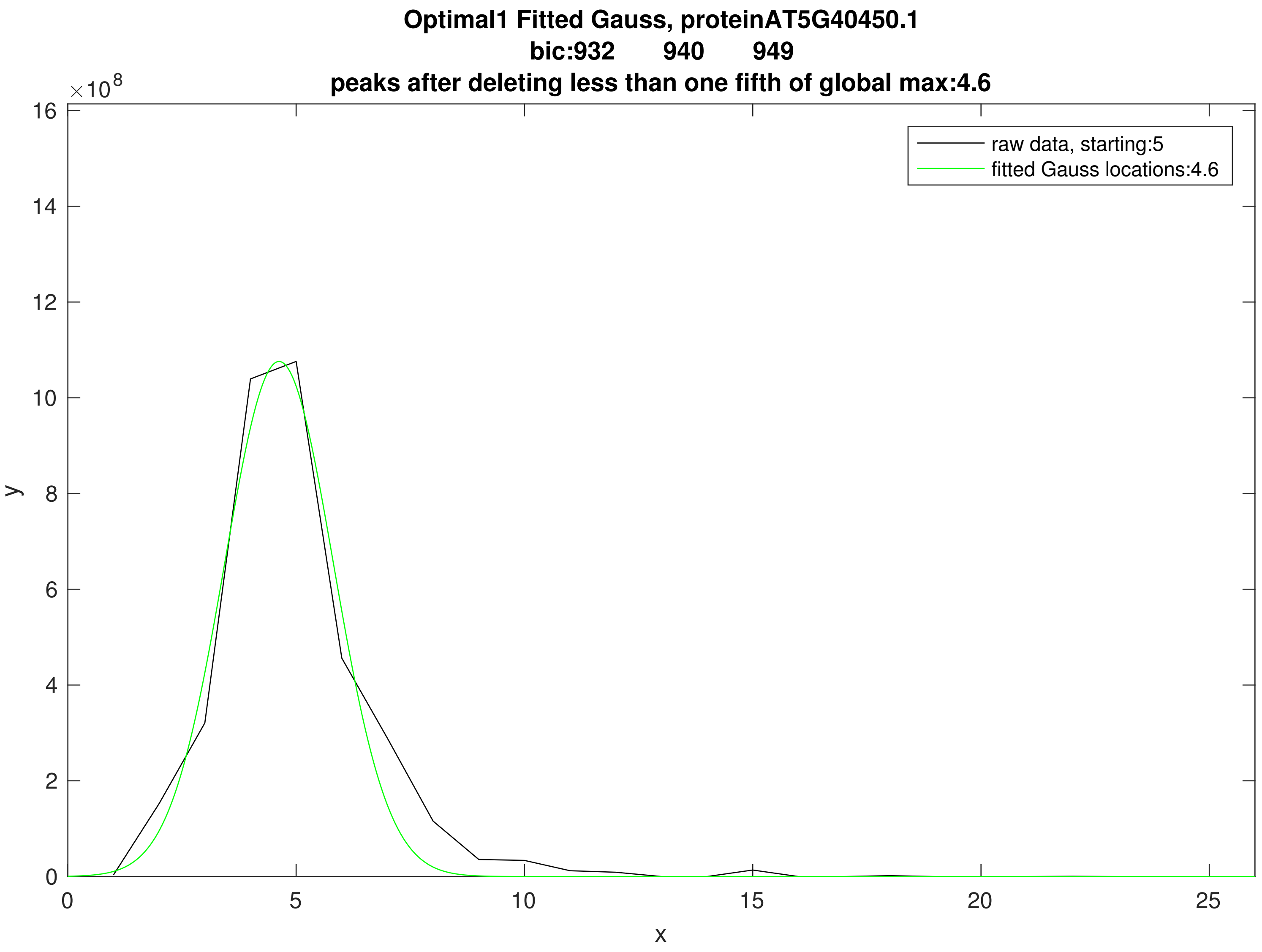
Optimal1 Fitted Gauss, proteinAT5G40450.1
bic:932 940 949
peaks after deleting less than one fifth of global max:4.6
×10^8
raw data, starting:5
fitted Gauss locations:4.6
y
x
0
2
4
6
8
10
12
14
16
0
5
10
15
20
25

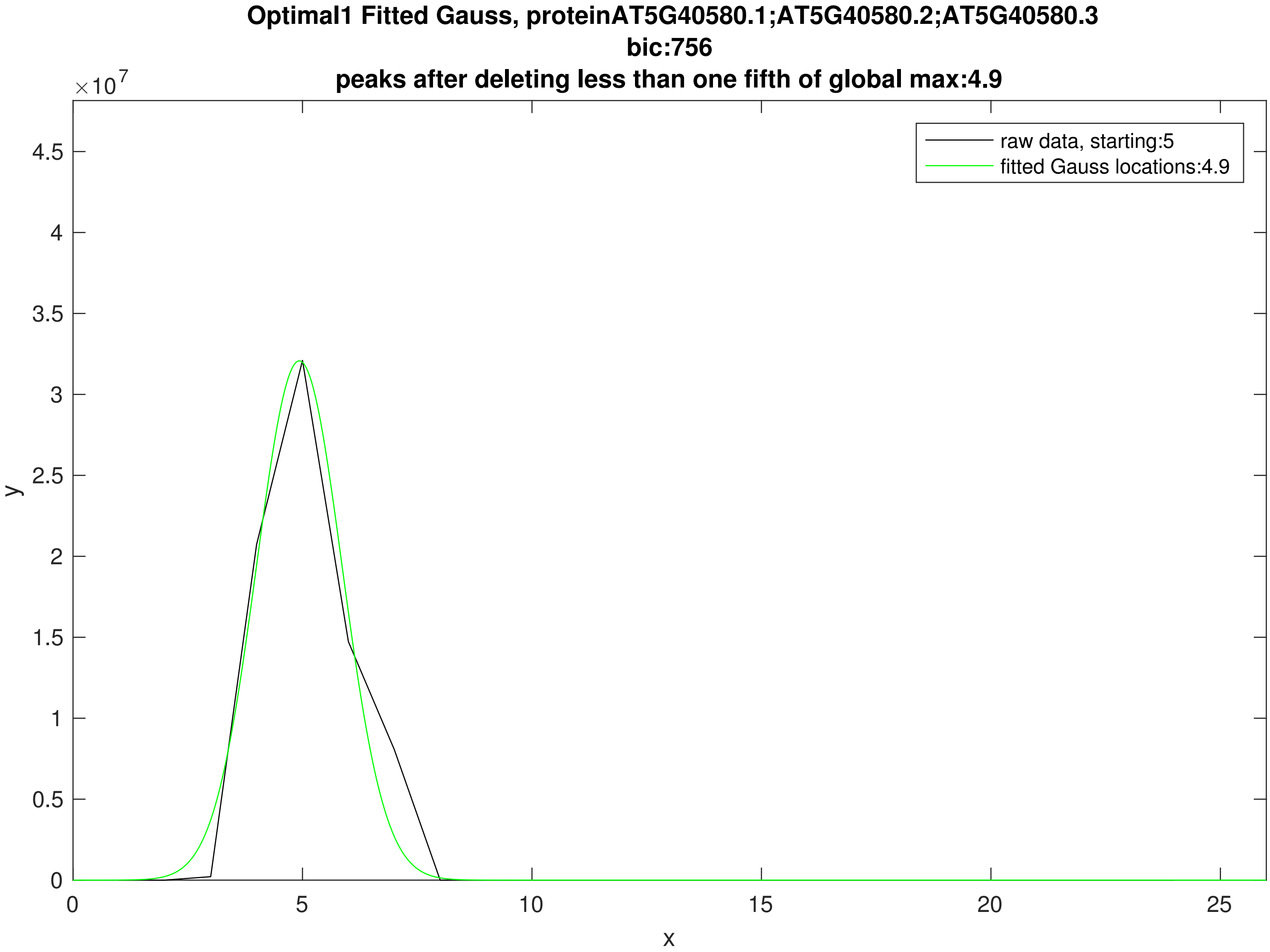

Optimal1 Fitted Gauss, proteinAT5G40580.1;AT5G40580.2;AT5G40580.3
bic:756
peaks after deleting less than one fifth of global max:4.9
×10^7
4.5
4
3.5
3
2.5
2
1.5
1
0.5
0
y
0
5
10
15
20
25
x
raw data, starting:5
fitted Gauss locations:4.9

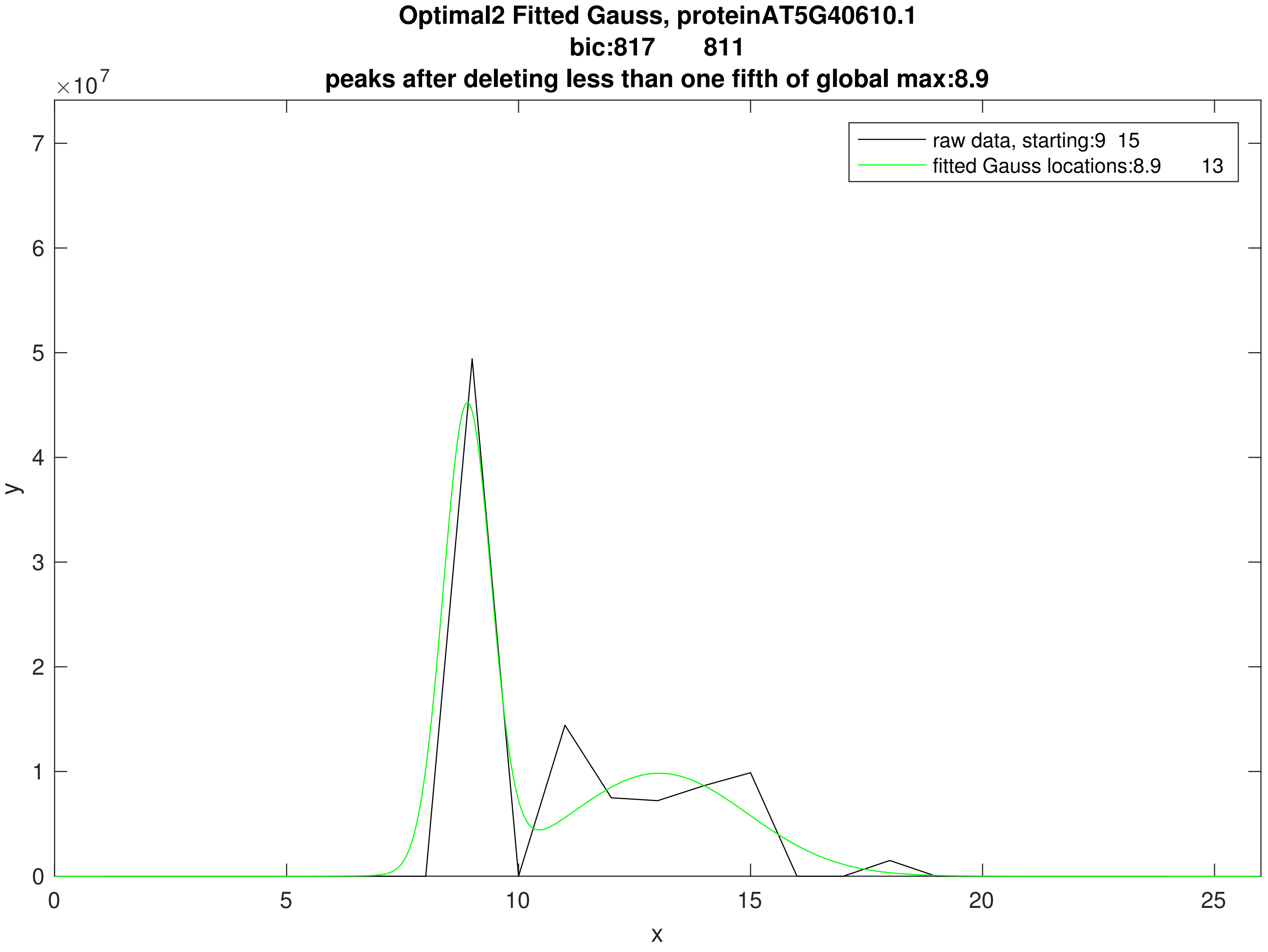
Optimal2 Fitted Gauss, proteinAT5G40610.1
bic:817 811
peaks after deleting less than one fifth of global max:8.9
×10^7
raw data, starting:9 15
fitted Gauss locations:8.9 13
y
x
0
1
2
3
4
5
6
7
0
5
10
15
20
25

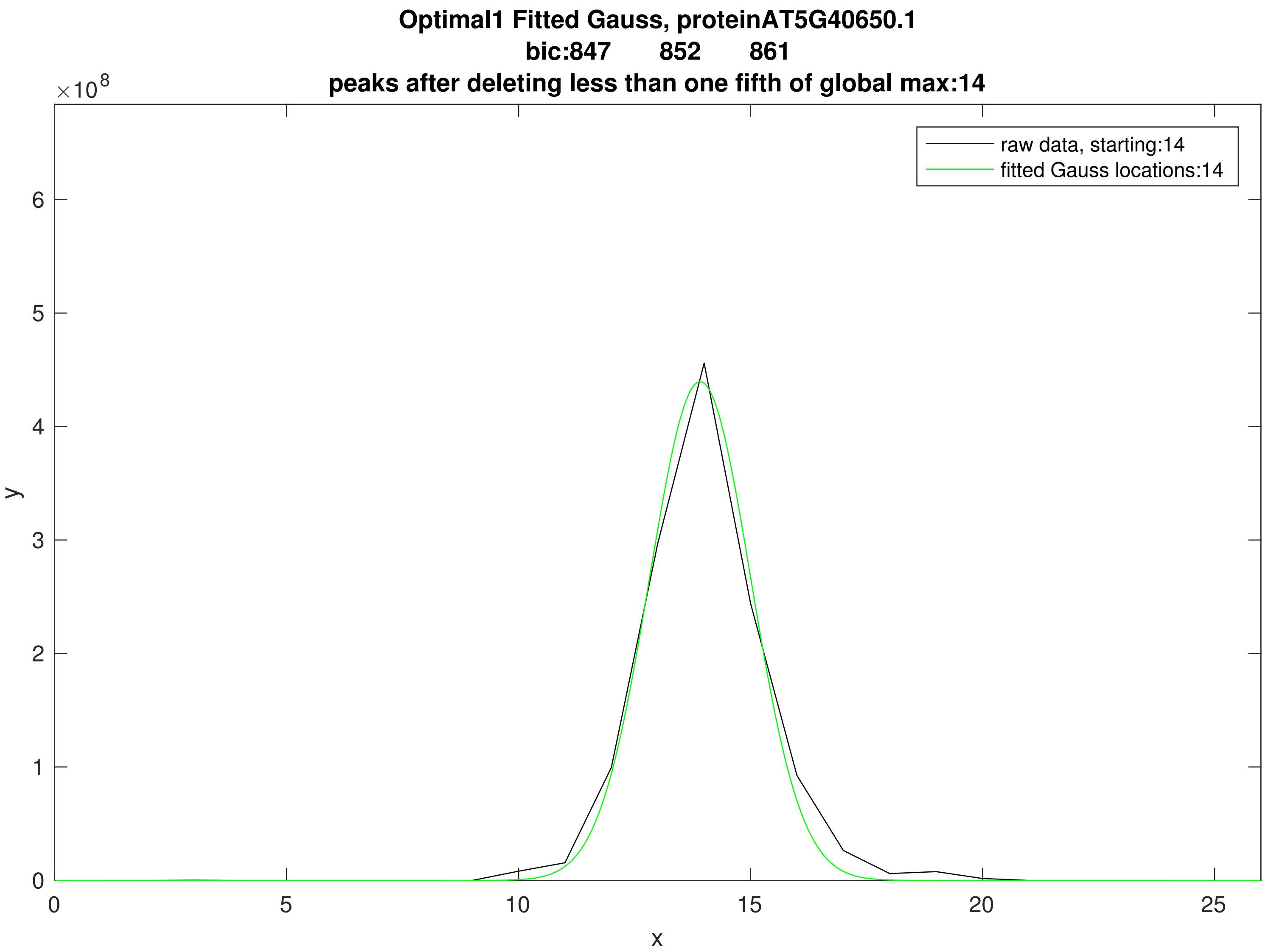

Optimal1 Fitted Gauss, proteinAT5G40650.1
bic:847 852 861
peaks after deleting less than one fifth of global max:14
×10^8
raw data, starting:14
fitted Gauss locations:14
y
x
0
1
2
3
4
5
6
0
5
10
15
20
25

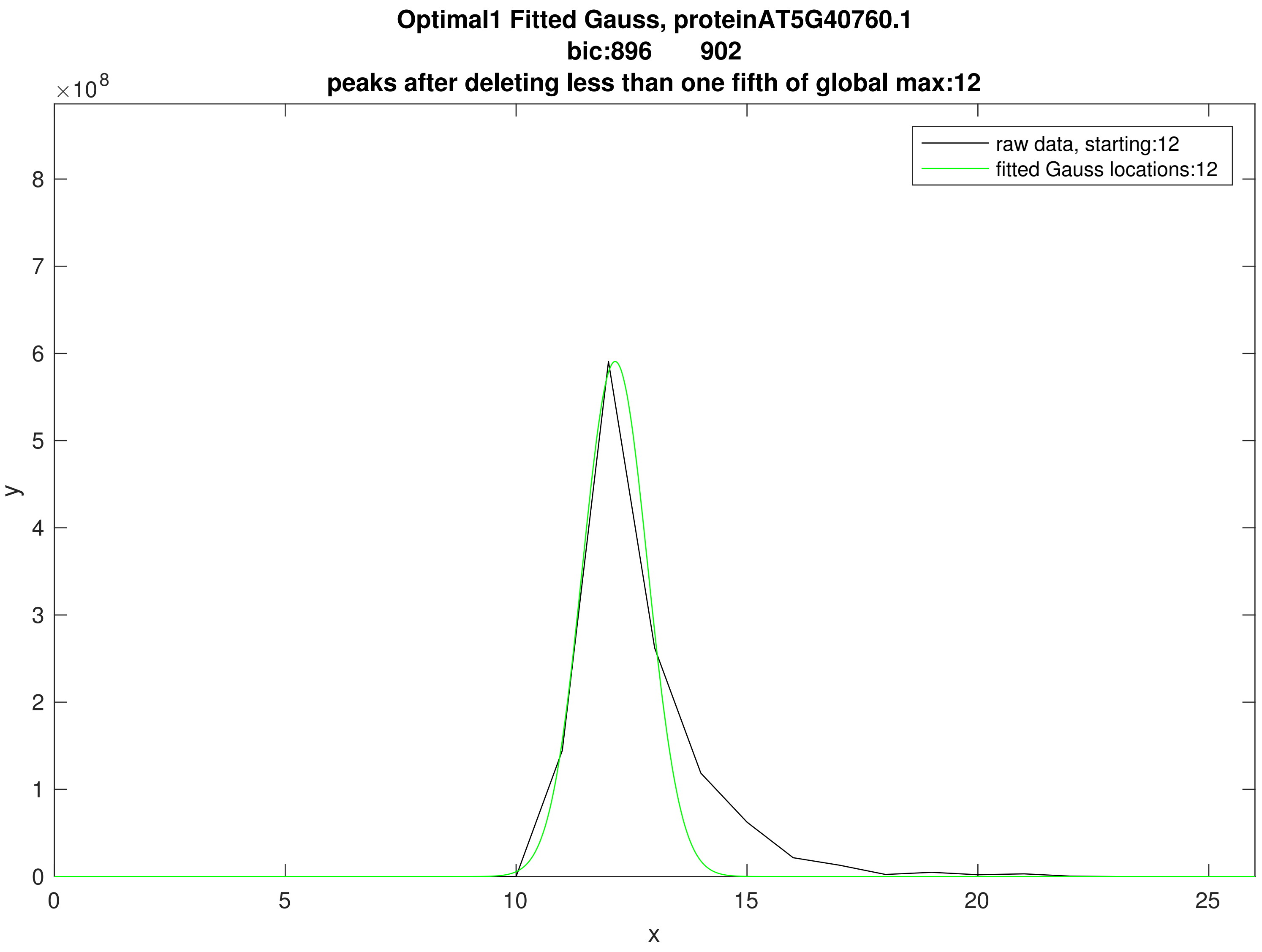

Optimal1 Fitted Gauss, proteinAT5G40760.1
bic:896 902
peaks after deleting less than one fifth of global max:12
×10^8
raw data, starting:12
fitted Gauss locations:12
y
x
0
1
2
3
4
5
6
7
8
0
5
10
15
20
25

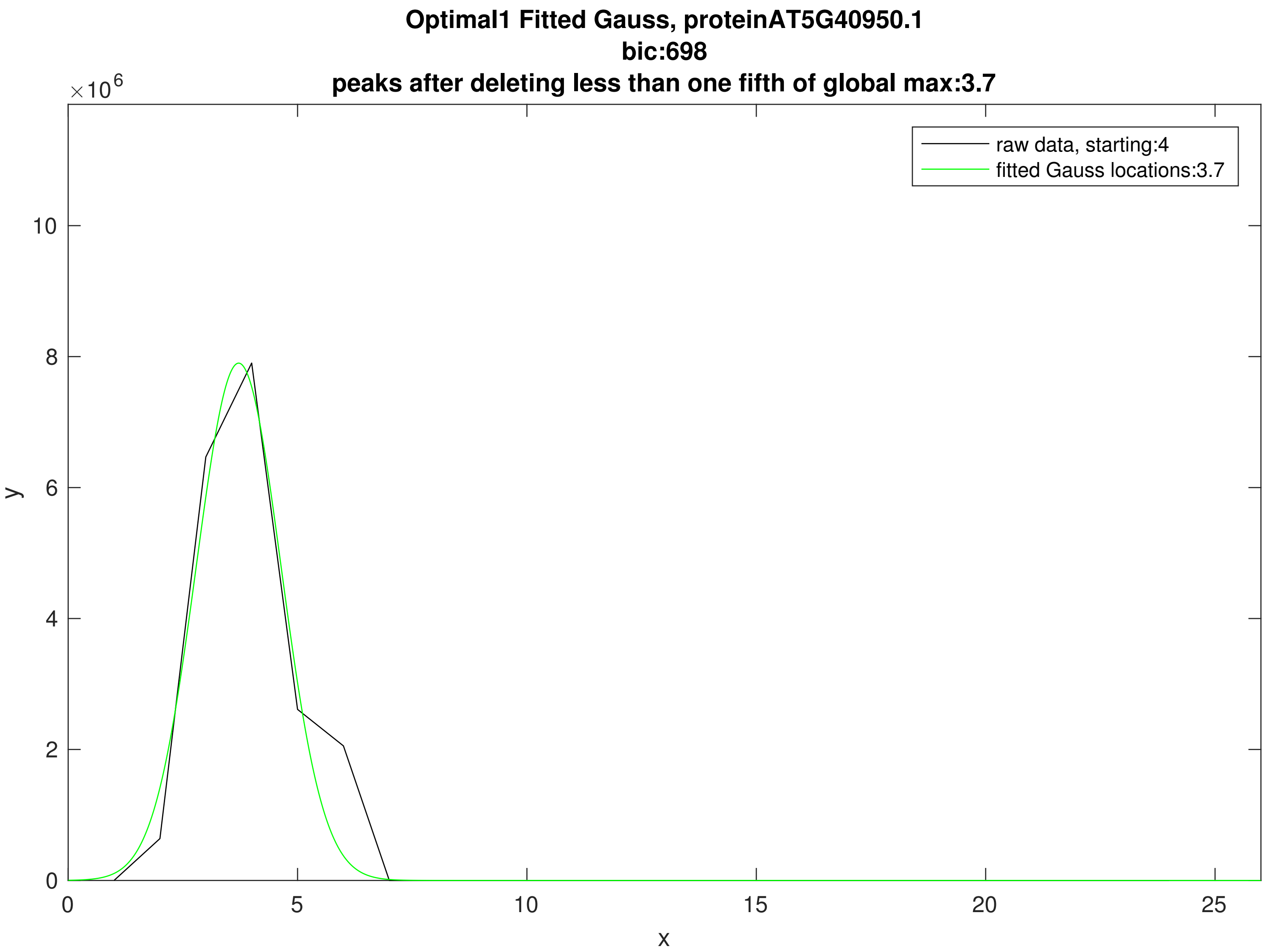
Optimal1 Fitted Gauss, proteinAT5G40950.1
bic:698
peaks after deleting less than one fifth of global max:3.7
×10^6
raw data, starting:4
fitted Gauss locations:3.7
y
0
2
4
6
8
10
0
5
10
15
20
25
x

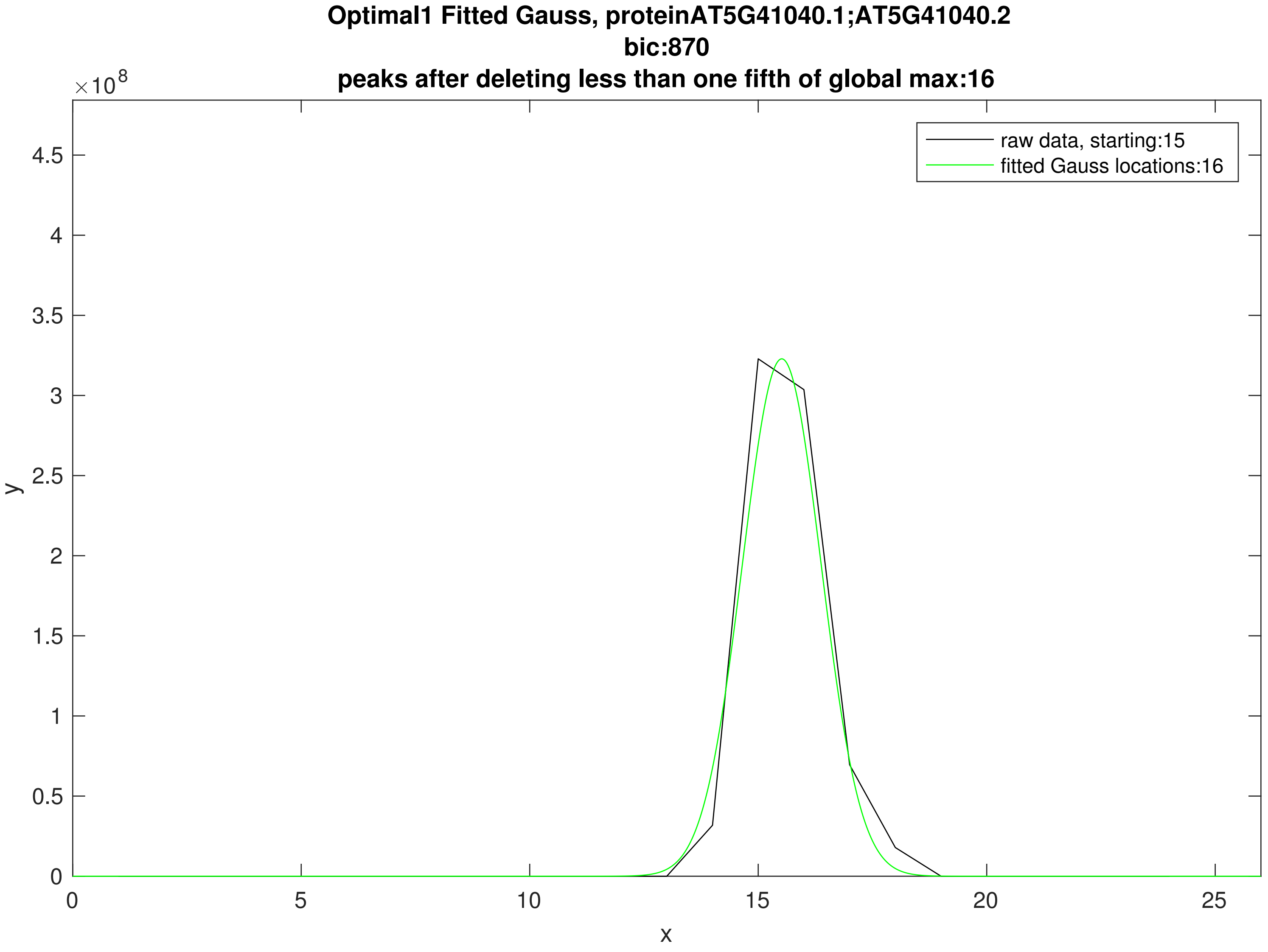

Optimal1 Fitted Gauss, proteinAT5G41040.1;AT5G41040.2
bic:870
peaks after deleting less than one fifth of global max:16
×10^8
raw data, starting:15
fitted Gauss locations:16
4.5
4
3.5
3
2.5
2
1.5
1
0.5
0
y
0
5
10
15
20
25
x

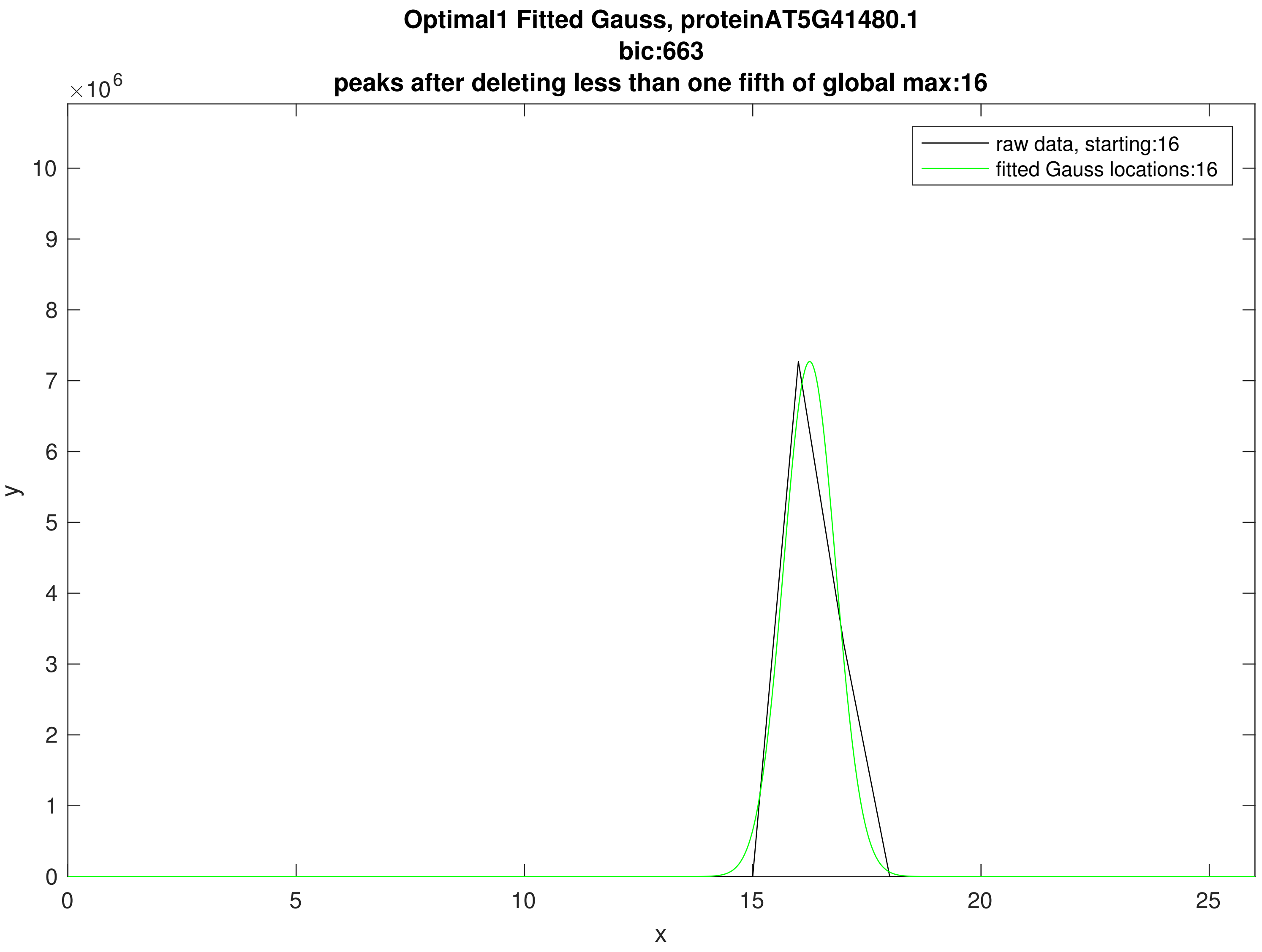

Optimal1 Fitted Gauss, proteinAT5G41480.1
bic:663
peaks after deleting less than one fifth of global max:16
×10^6
raw data, starting:16
fitted Gauss locations:16
y
x
0
1
2
3
4
5
6
7
8
9
10
0
5
10
15
20
25

Optimal1 Fitted Gauss, proteinAT5G41520.1;AT5G41520.2
bic:530
peaks after deleting less than one fifth of global max:3.1

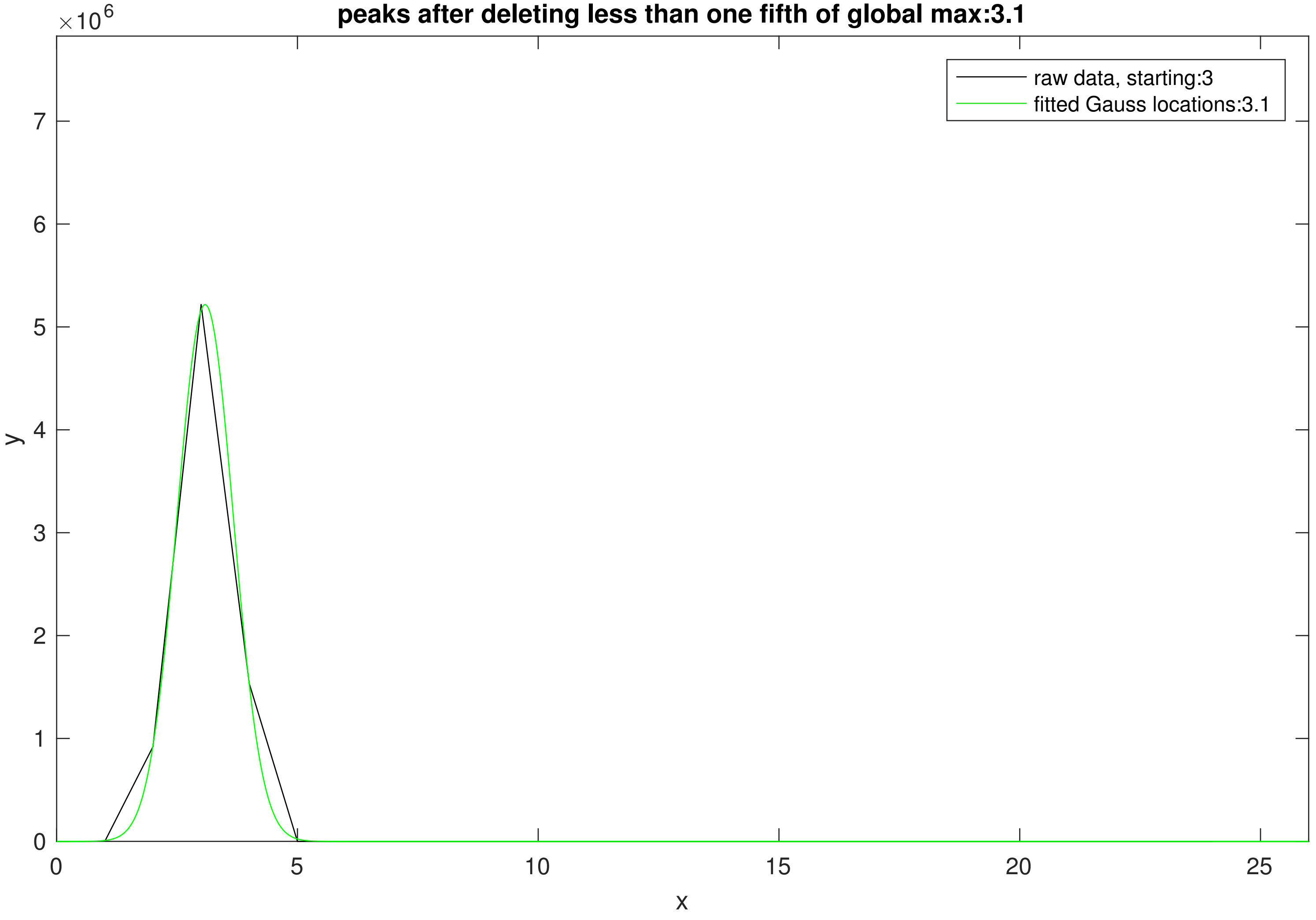

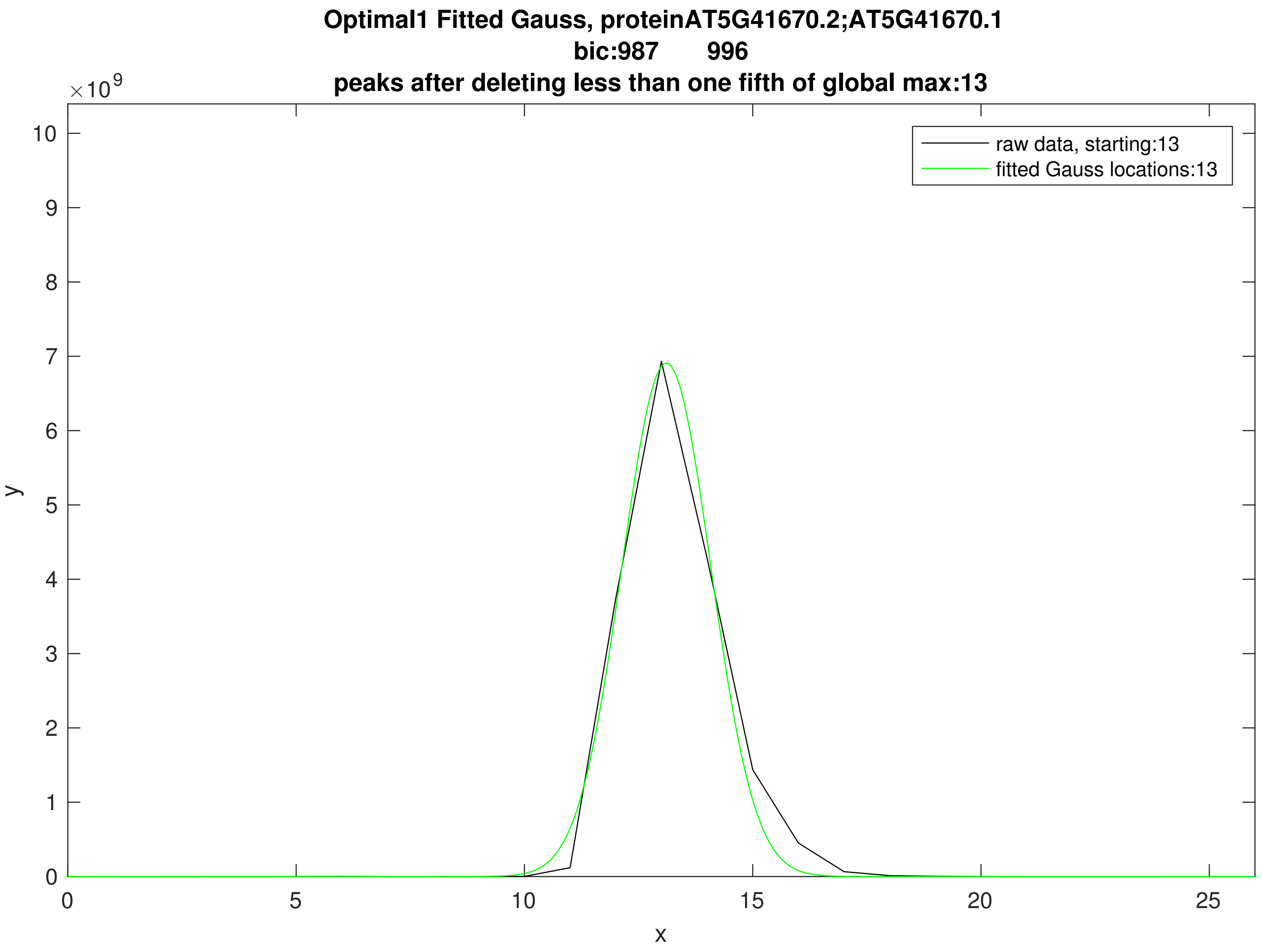
Optimal1 Fitted Gauss, proteinAT5G41670.2;AT5G41670.1
bic:987 996
peaks after deleting less than one fifth of global max:13
×10^9
raw data, starting:13
fitted Gauss locations:13
y
x
0
1
2
3
4
5
6
7
8
9
10
0
5
10
15
20
25

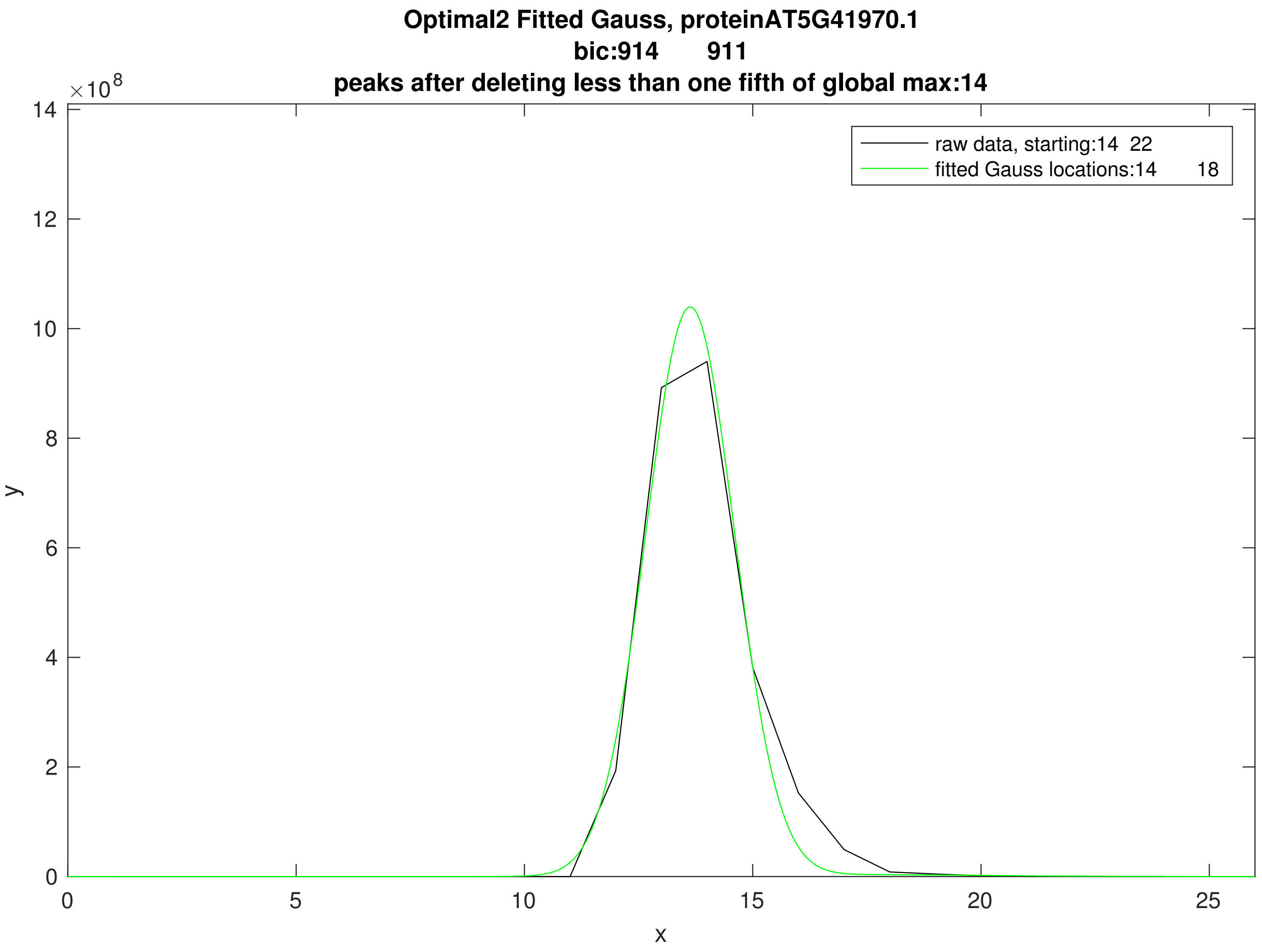
Optimal2 Fitted Gauss, proteinAT5G41970.1
bic:914 911
peaks after deleting less than one fifth of global max:14
raw data, starting:14 22
fitted Gauss locations:14 18
×10^8
y
x
0
2
4
6
8
10
12
14
0
5
10
15
20
25

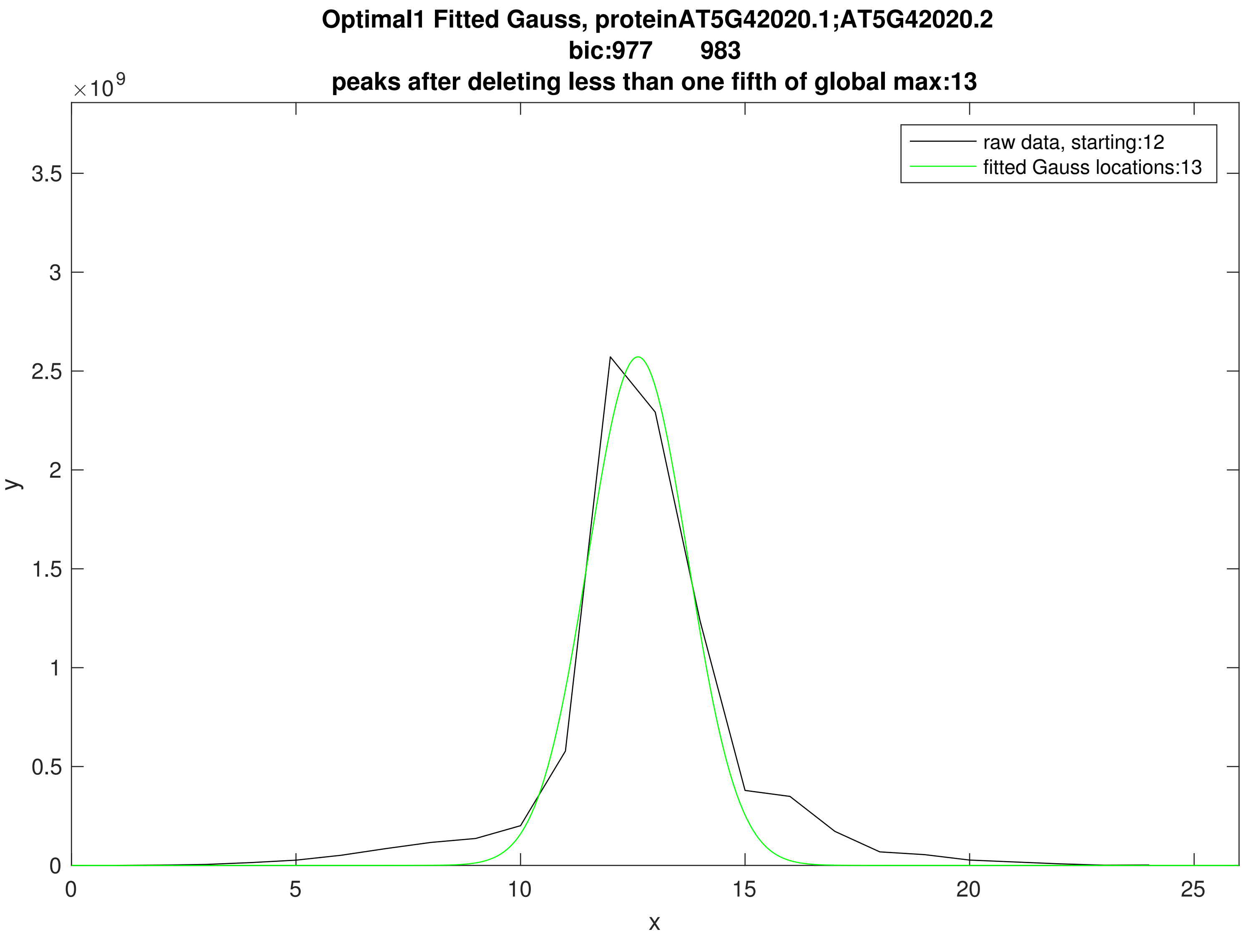
Optimal1 Fitted Gauss, proteinAT5G42020.1;AT5G42020.2
bic:977 983
peaks after deleting less than one fifth of global max:13
×10^9
raw data, starting:12
fitted Gauss locations:13
y
x
0
0.5
1
1.5
2
2.5
3
3.5
0
5
10
15
20
25

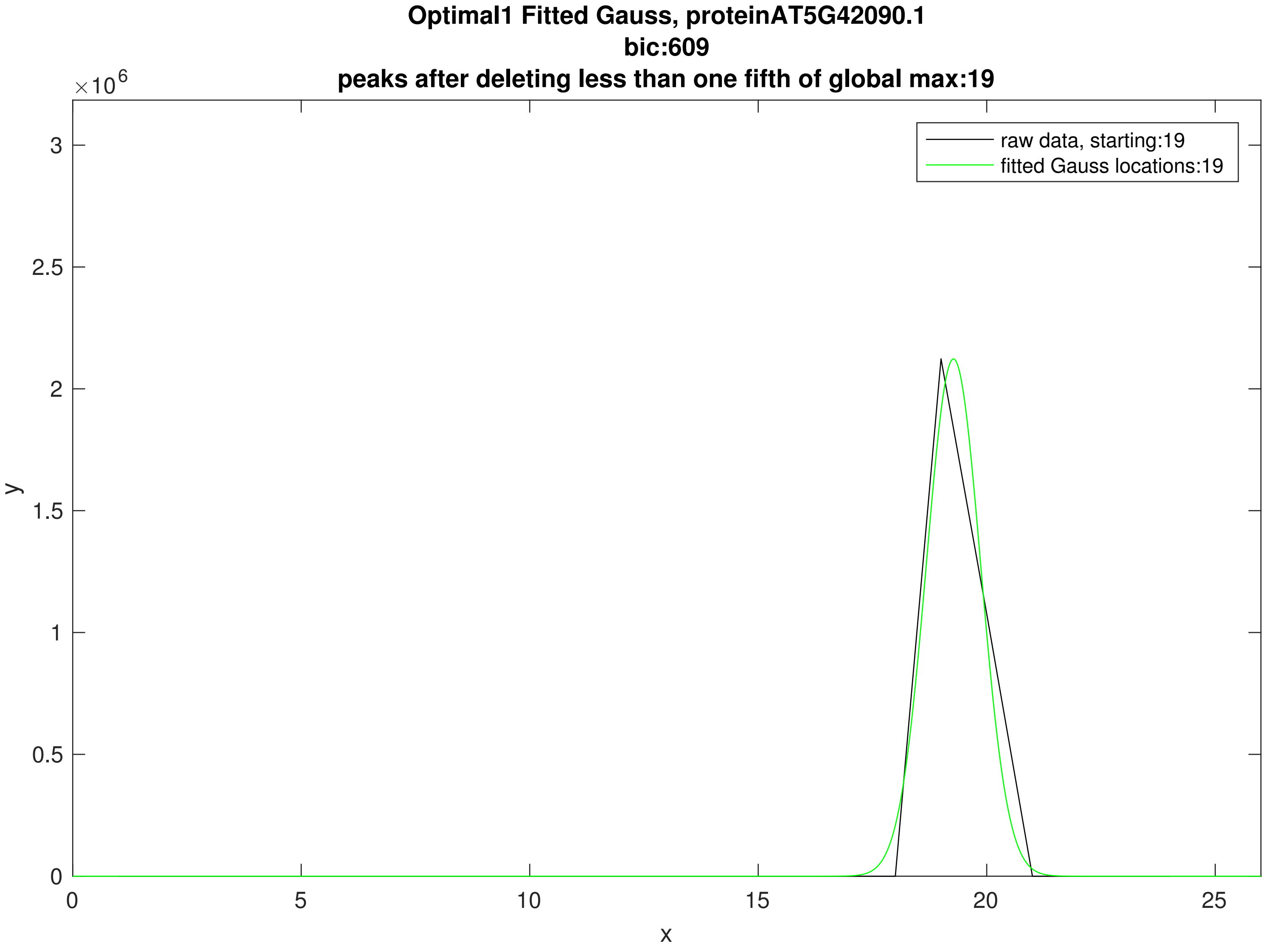
Optimal1 Fitted Gauss, proteinAT5G42090.1
bic:609
peaks after deleting less than one fifth of global max:19
×10^6
raw data, starting:19
fitted Gauss locations:19
y
3
2.5
2
1.5
1
0.5
0
0
5
10
15
20
25
x

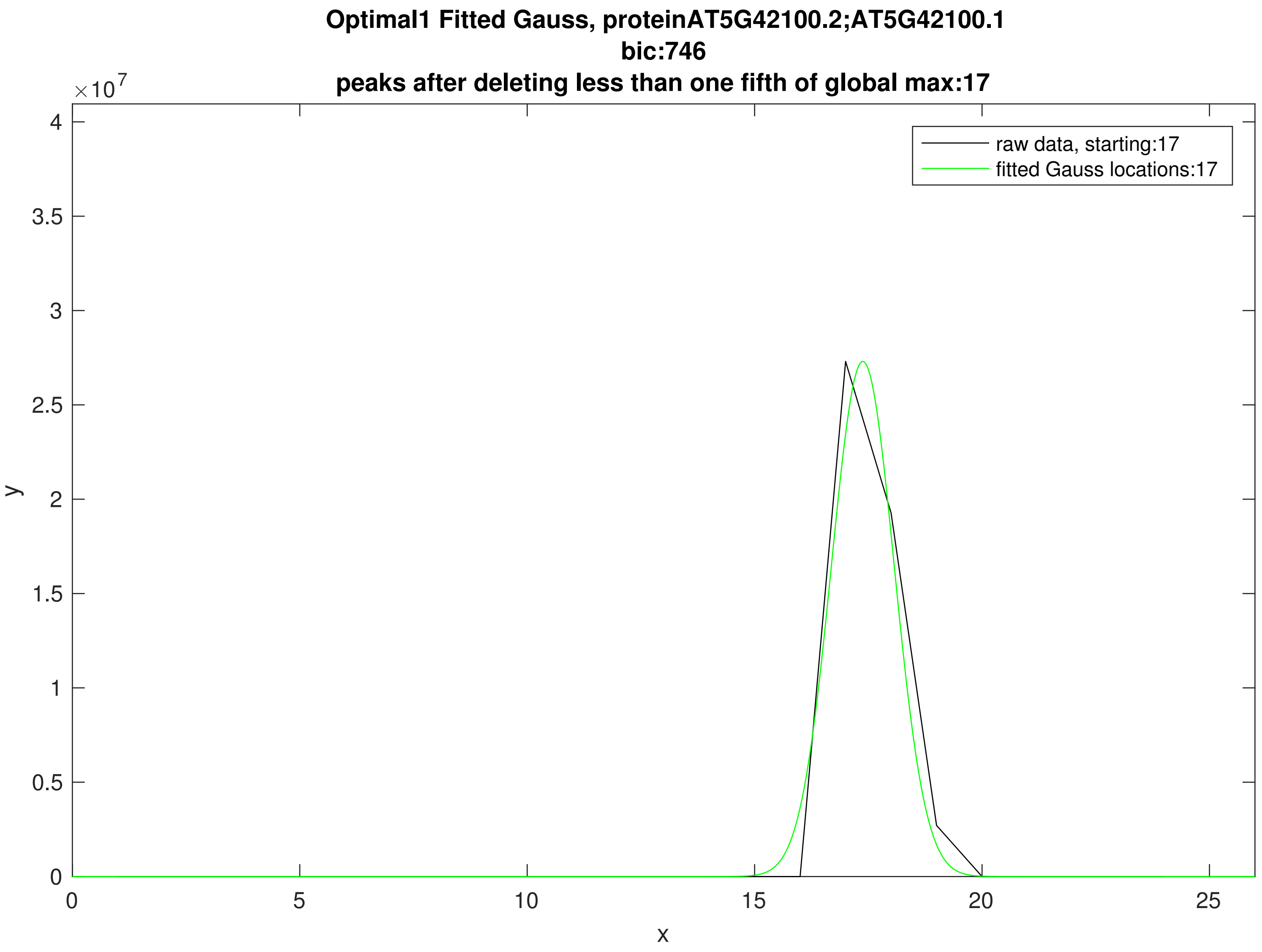

Optimal1 Fitted Gauss, proteinAT5G42100.2;AT5G42100.1
bic:746
peaks after deleting less than one fifth of global max:17
×10^7
raw data, starting:17
fitted Gauss locations:17
y
x
0
0.5
1
1.5
2
2.5
3
3.5
4
5
10
15
20
25

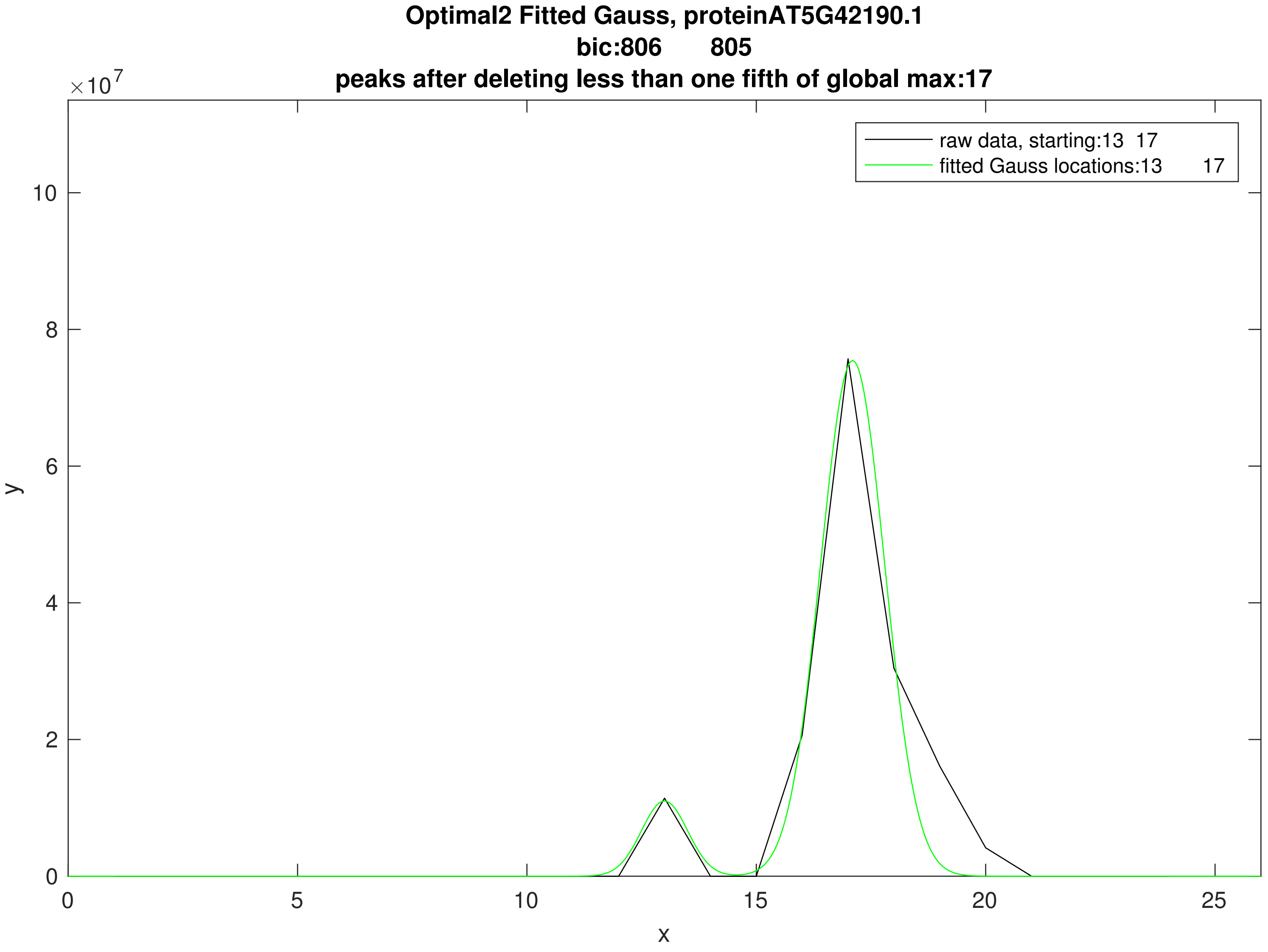

Optimal2 Fitted Gauss, proteinAT5G42190.1
bic:806 805
peaks after deleting less than one fifth of global max:17
×10^7
raw data, starting:13 17
fitted Gauss locations:13 17
y
x
0
2
4
6
8
10
0
5
10
15
20
25

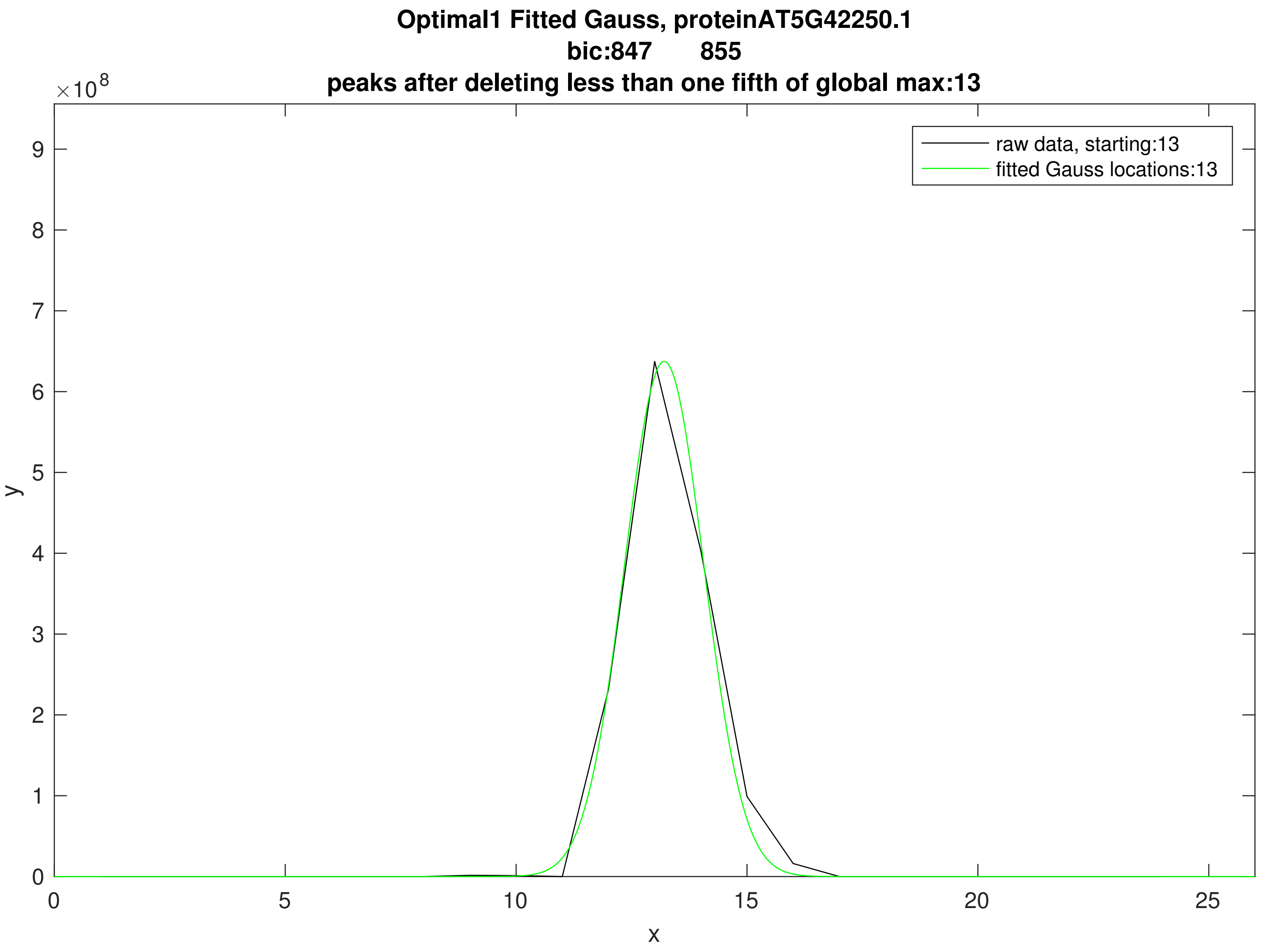
Optimal1 Fitted Gauss, proteinAT5G42250.1
bic:847 855
peaks after deleting less than one fifth of global max:13
×10^8
9
8
7
6
5
4
3
2
1
0
y
0
5
10
15
20
25
x
raw data, starting:13
fitted Gauss locations:13

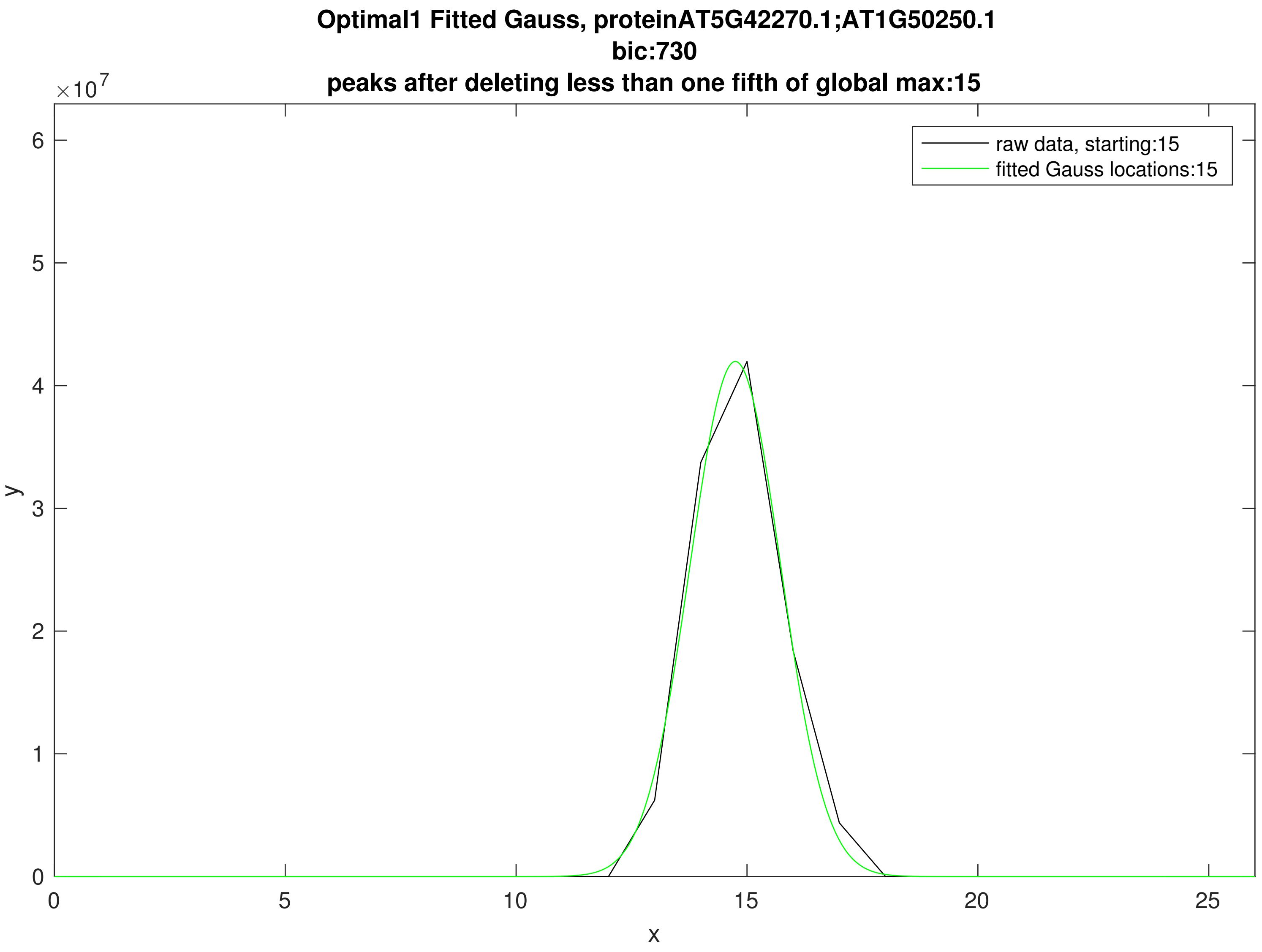
Optimal1 Fitted Gauss, proteinAT5G42270.1;AT1G50250.1
bic:730
peaks after deleting less than one fifth of global max:15
×10^7
raw data, starting:15
fitted Gauss locations:15
y
x
0
1
2
3
4
5
6
0
5
10
15
20
25

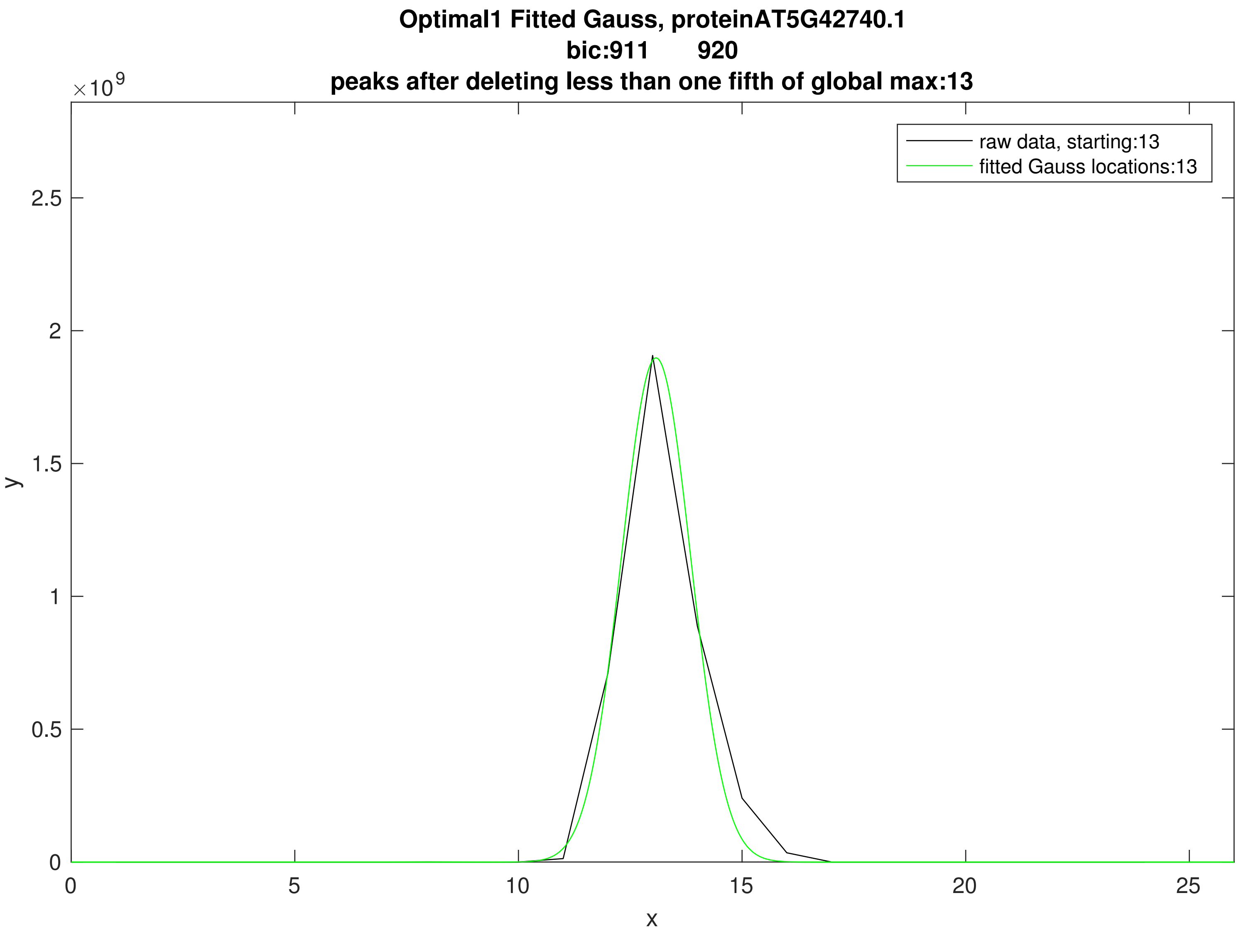
Optimal1 Fitted Gauss, proteinAT5G42740.1
bic:911 920
peaks after deleting less than one fifth of global max:13
×10^9
raw data, starting:13
fitted Gauss locations:13
2.5
2
1.5
1
0.5
0
y
0
5
10
15
20
25
x

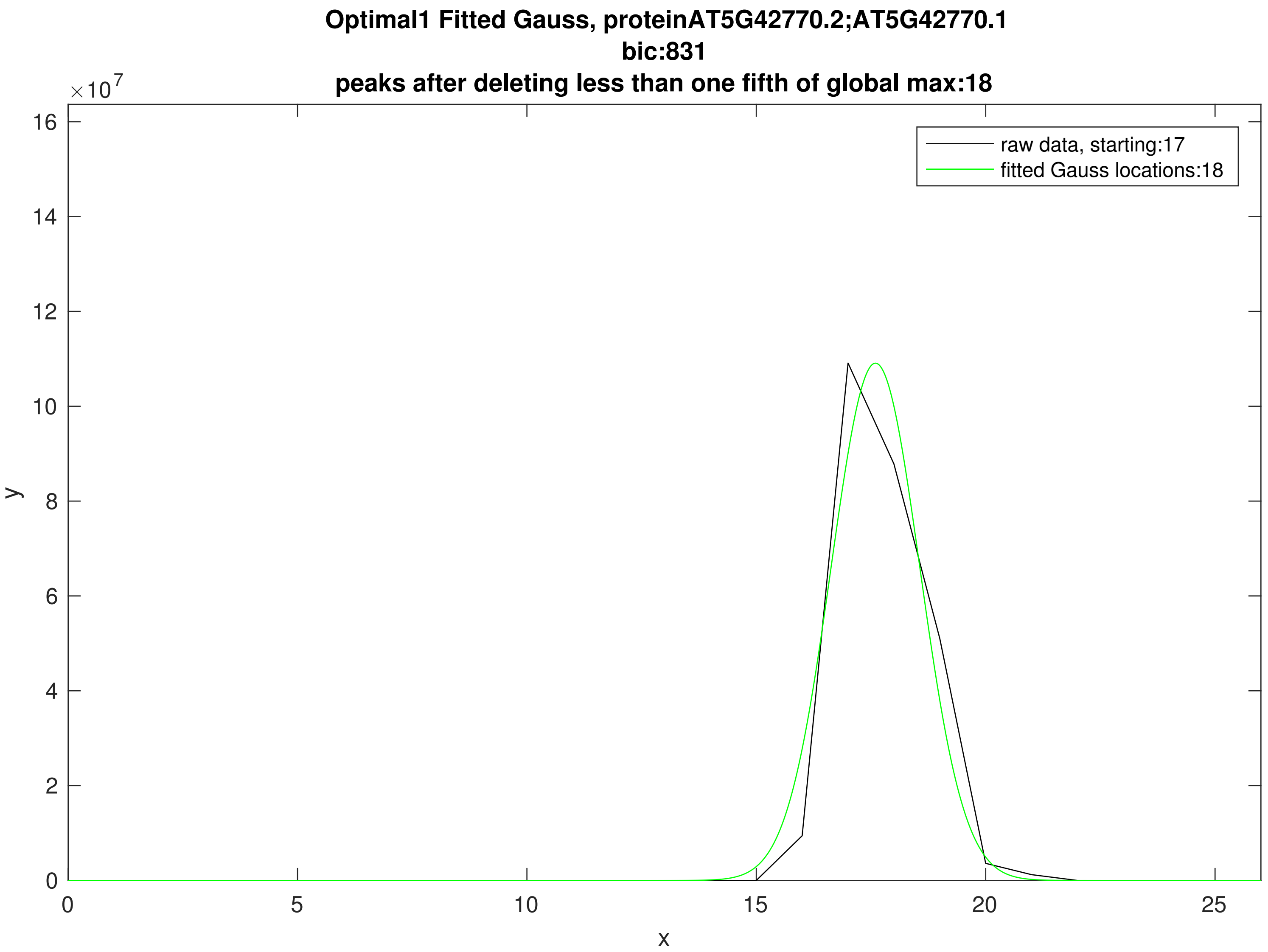
Optimal1 Fitted Gauss, proteinAT5G42770.2;AT5G42770.1
bic:831
peaks after deleting less than one fifth of global max:18
×10^7
raw data, starting:17
fitted Gauss locations:18
y
x
0
2
4
6
8
10
12
14
16
0
5
10
15
20
25

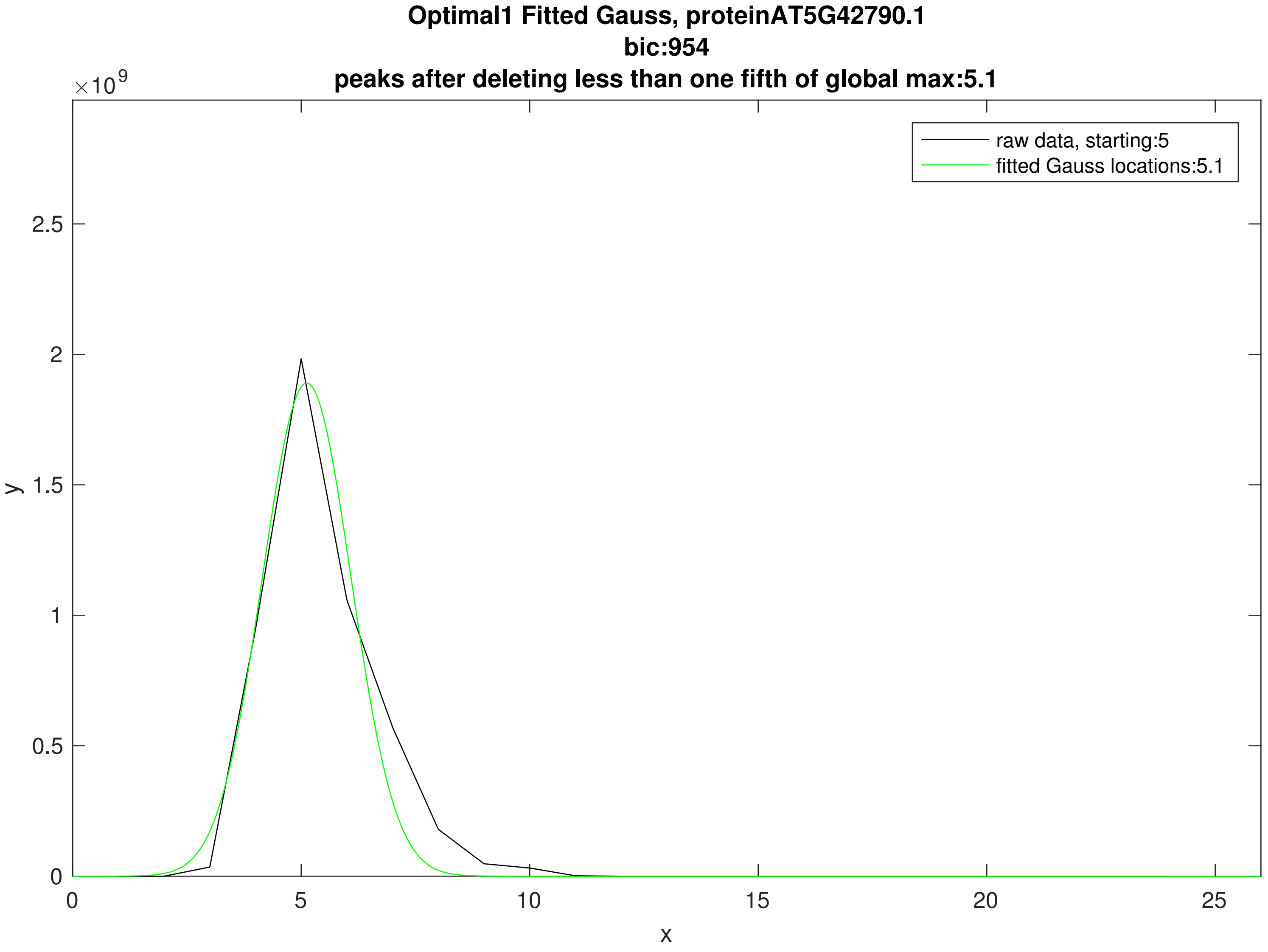

Optimal1 Fitted Gauss, proteinAT5G42790.1
bic:954
peaks after deleting less than one fifth of global max:5.1
×10^9
raw data, starting:5
fitted Gauss locations:5.1
y
x
0
0.5
1
1.5
2
2.5
0
5
10
15
20
25

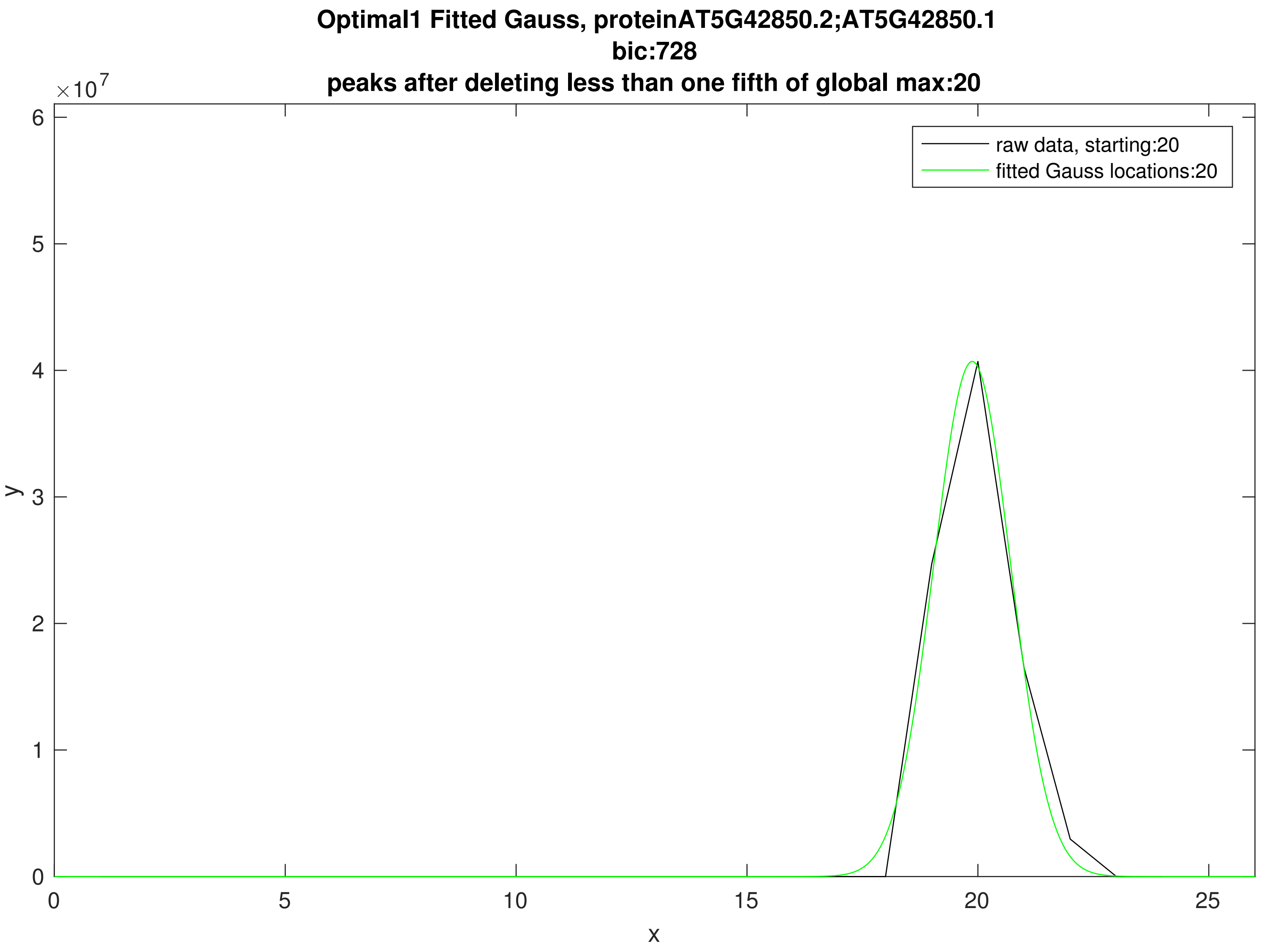
Optimal1 Fitted Gauss, proteinAT5G42850.2;AT5G42850.1
bic:728
peaks after deleting less than one fifth of global max:20
×10^7
6
5
4
3
2
1
0
y
0
5
10
15
20
25
x
raw data, starting:20
fitted Gauss locations:20

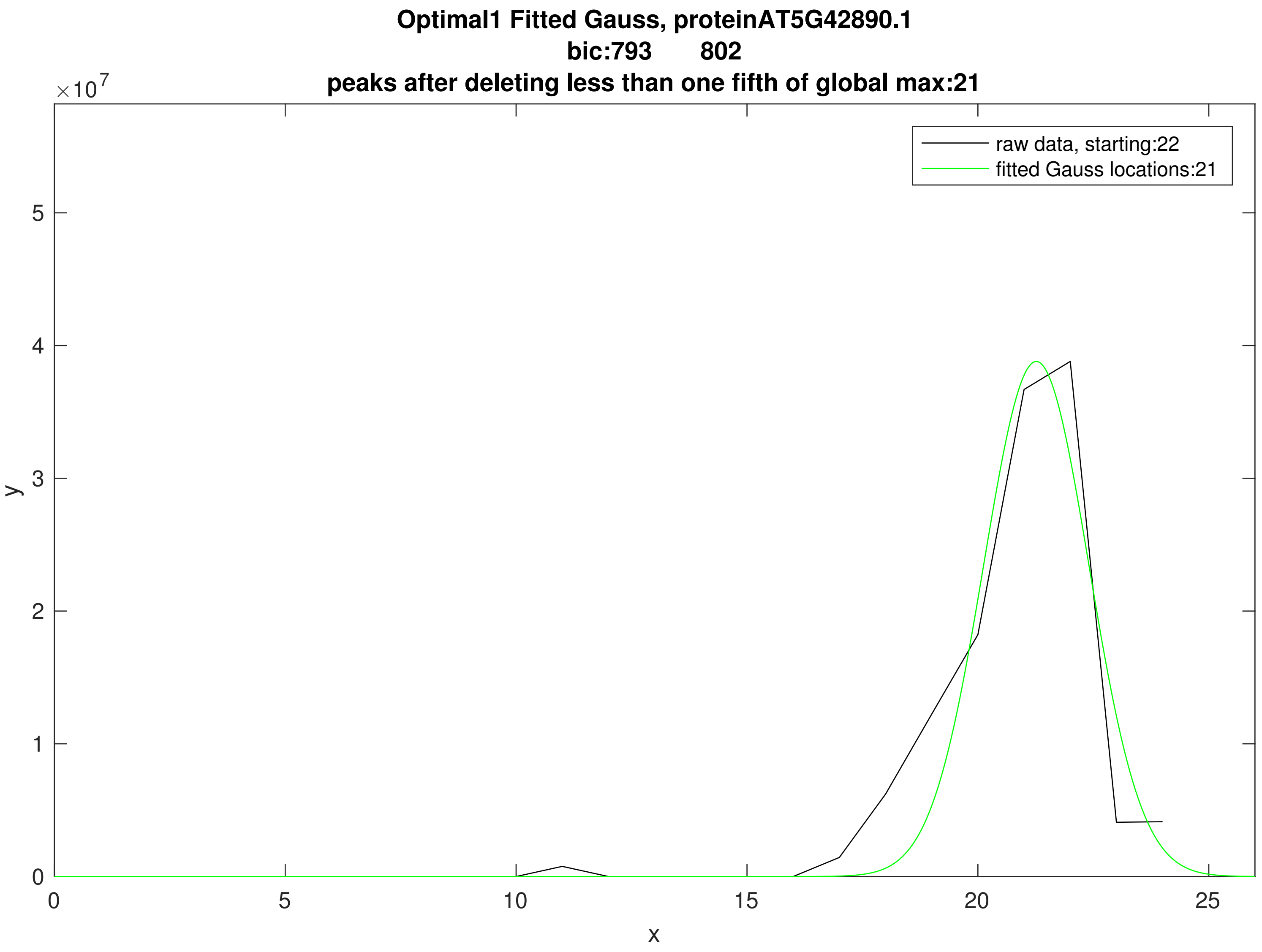
Optimal1 Fitted Gauss, proteinAT5G42890.1
bic:793 802
peaks after deleting less than one fifth of global max:21
×10^7
raw data, starting:22
fitted Gauss locations:21
y
x
0
1
2
3
4
5
0
5
10
15
20
25

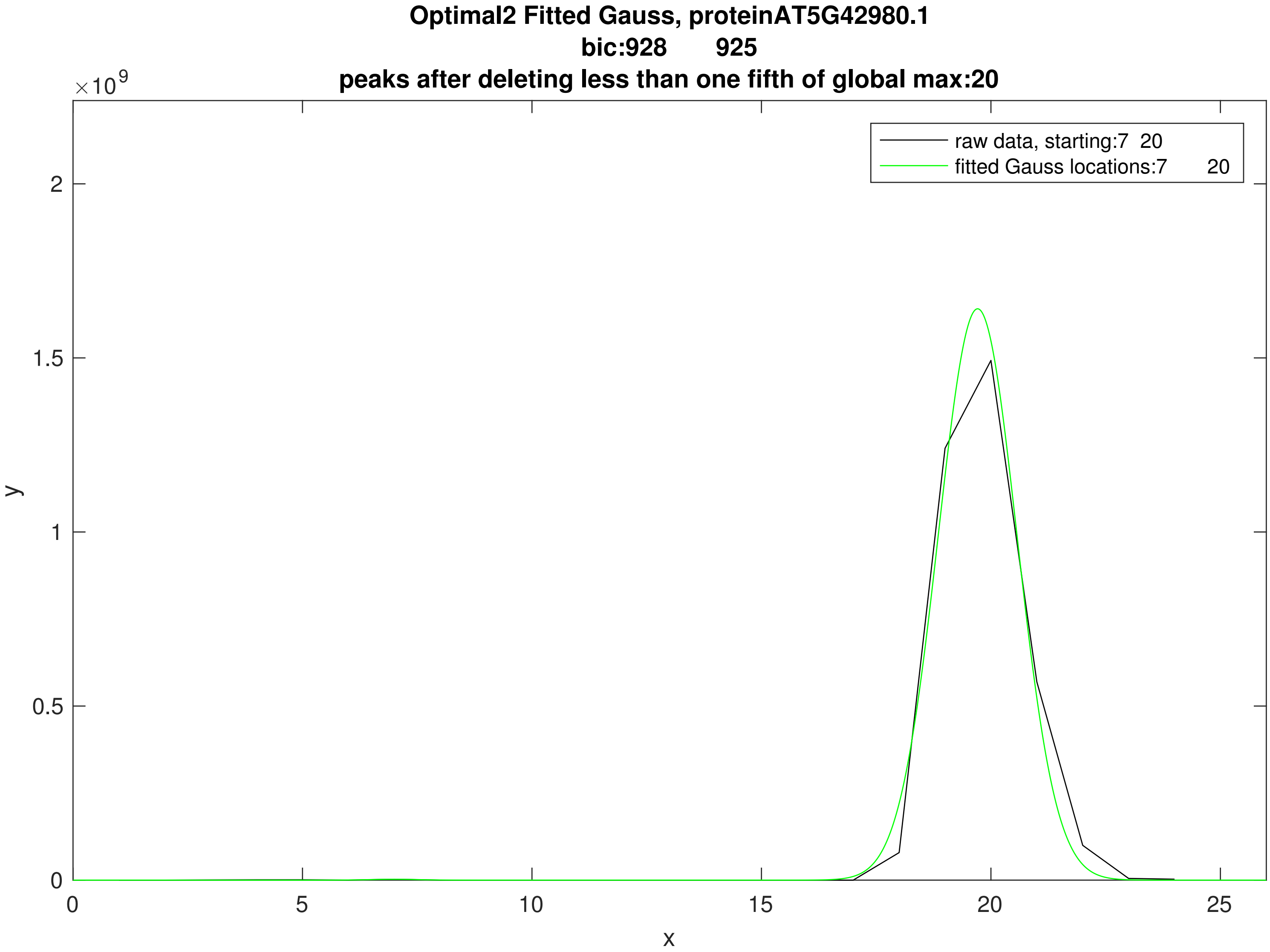

Optimal2 Fitted Gauss, proteinAT5G42980.1
bic:928 925
peaks after deleting less than one fifth of global max:20
×10^9
raw data, starting:7 20
fitted Gauss locations:7 20
y
x
0
0.5
1
1.5
2
0
5
10
15
20
25

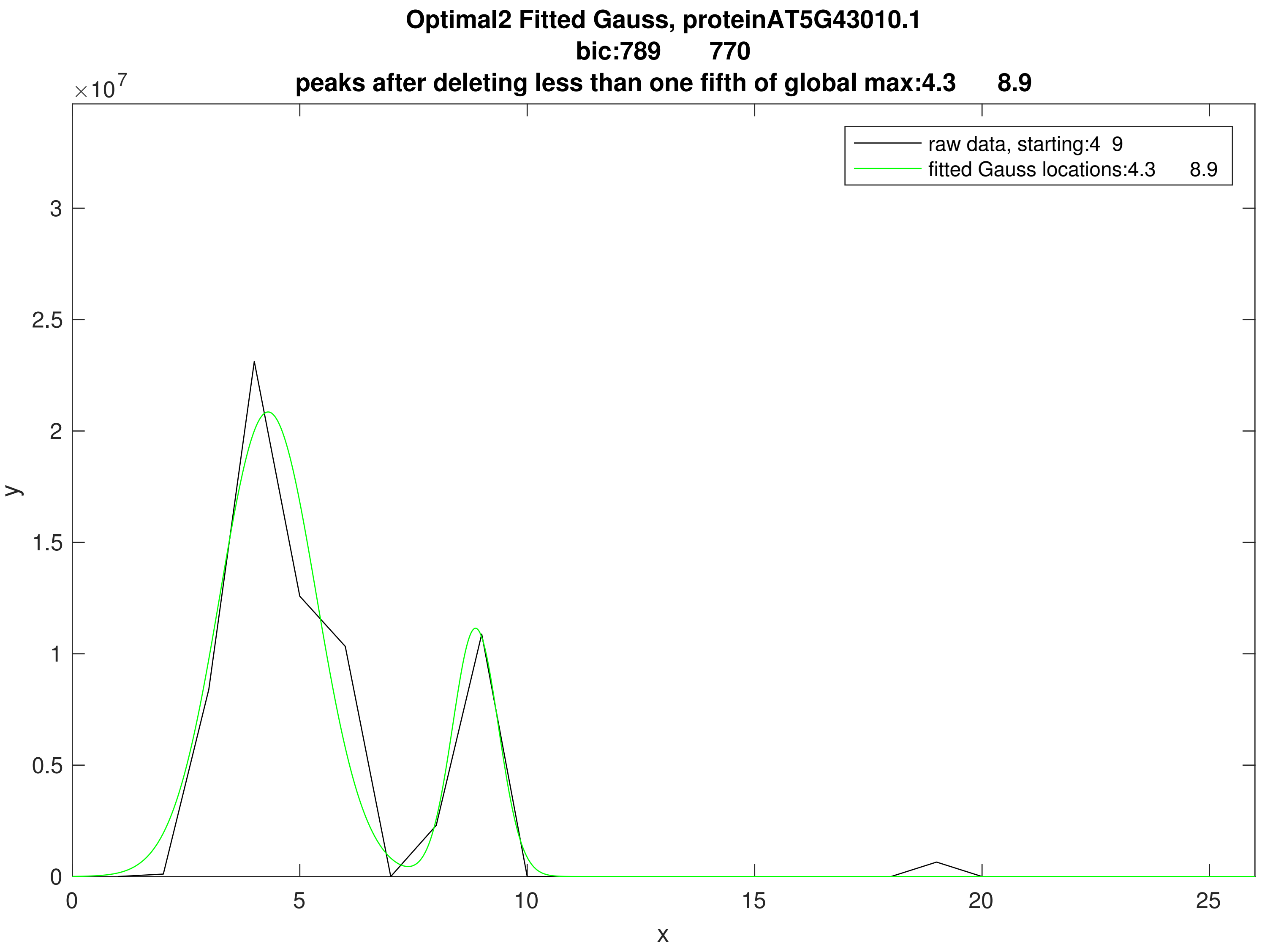
Optimal2 Fitted Gauss, proteinAT5G43010.1
bic:789 770
peaks after deleting less than one fifth of global max:4.3 8.9
×10^7
raw data, starting:4 9
fitted Gauss locations:4.3 8.9
3
2.5
2
1.5
1
0.5
0
y
0
5
10
15
20
25
x

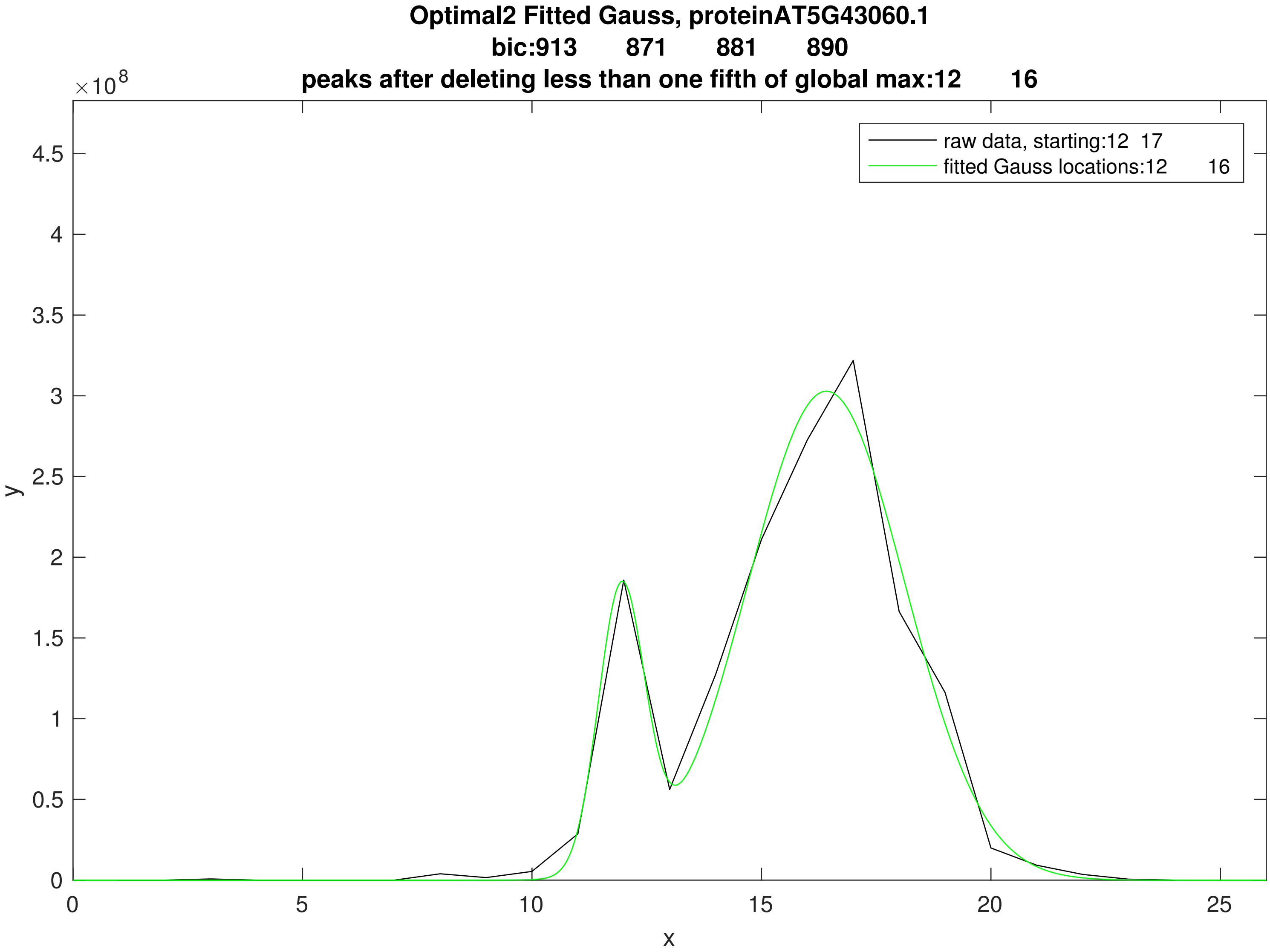
Optimal2 Fitted Gauss, proteinAT5G43060.1
bic:913 871 881 890
peaks after deleting less than one fifth of global max:12 16
×10^8
raw data, starting:12 17
fitted Gauss locations:12 16
0
0.5
1
1.5
2
2.5
3
3.5
4
4.5
y
0
5
10
15
20
25
x

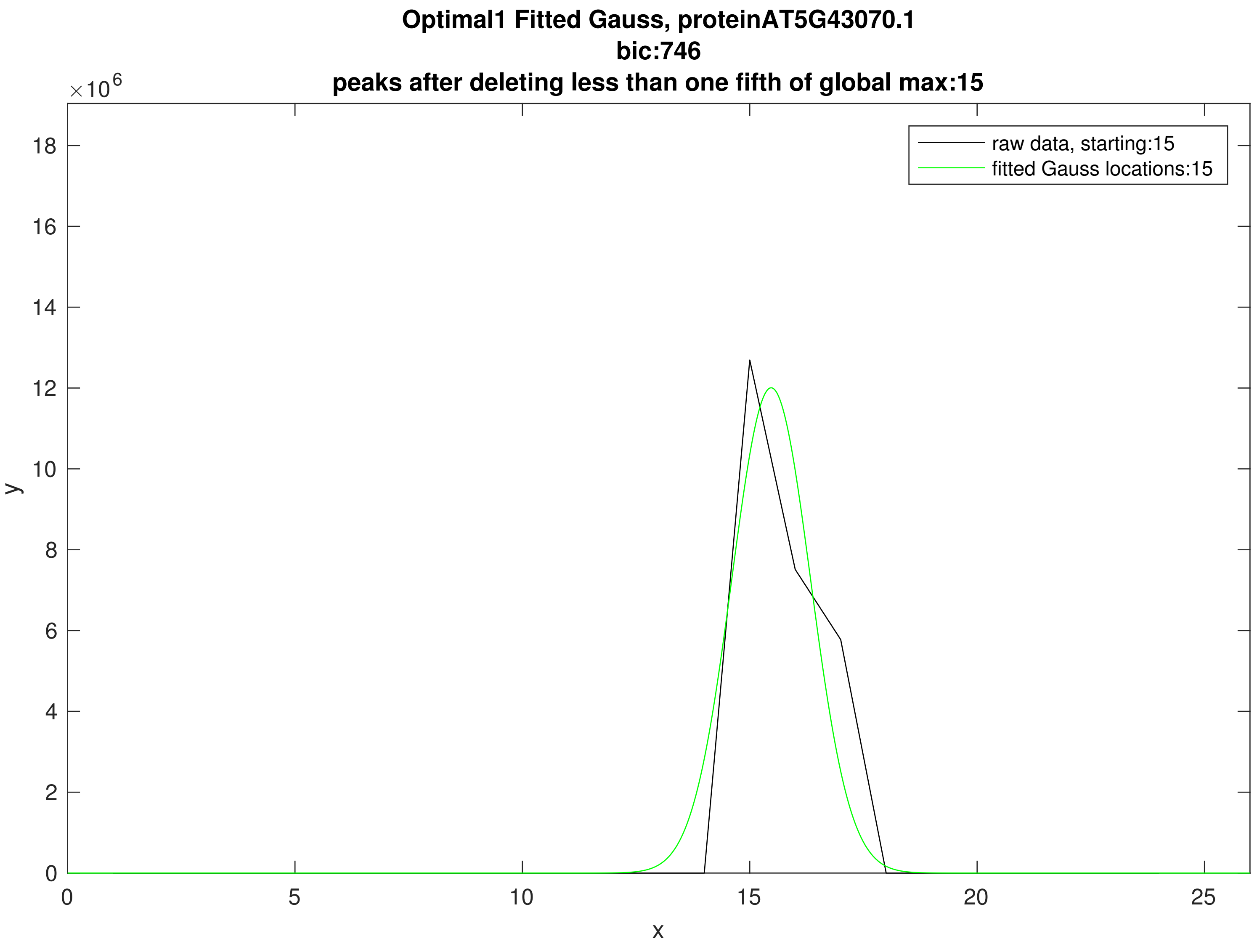

Optimal1 Fitted Gauss, proteinAT5G43070.1
bic:746
peaks after deleting less than one fifth of global max:15
×10^6
raw data, starting:15
fitted Gauss locations:15
y
x
0
2
4
6
8
10
12
14
16
18
0
5
10
15
20
25

**Optimal1 Fitted Gauss, proteinAT5G43100.1**
**bic:659**
**peaks after deleting less than one fifth of global max:16**

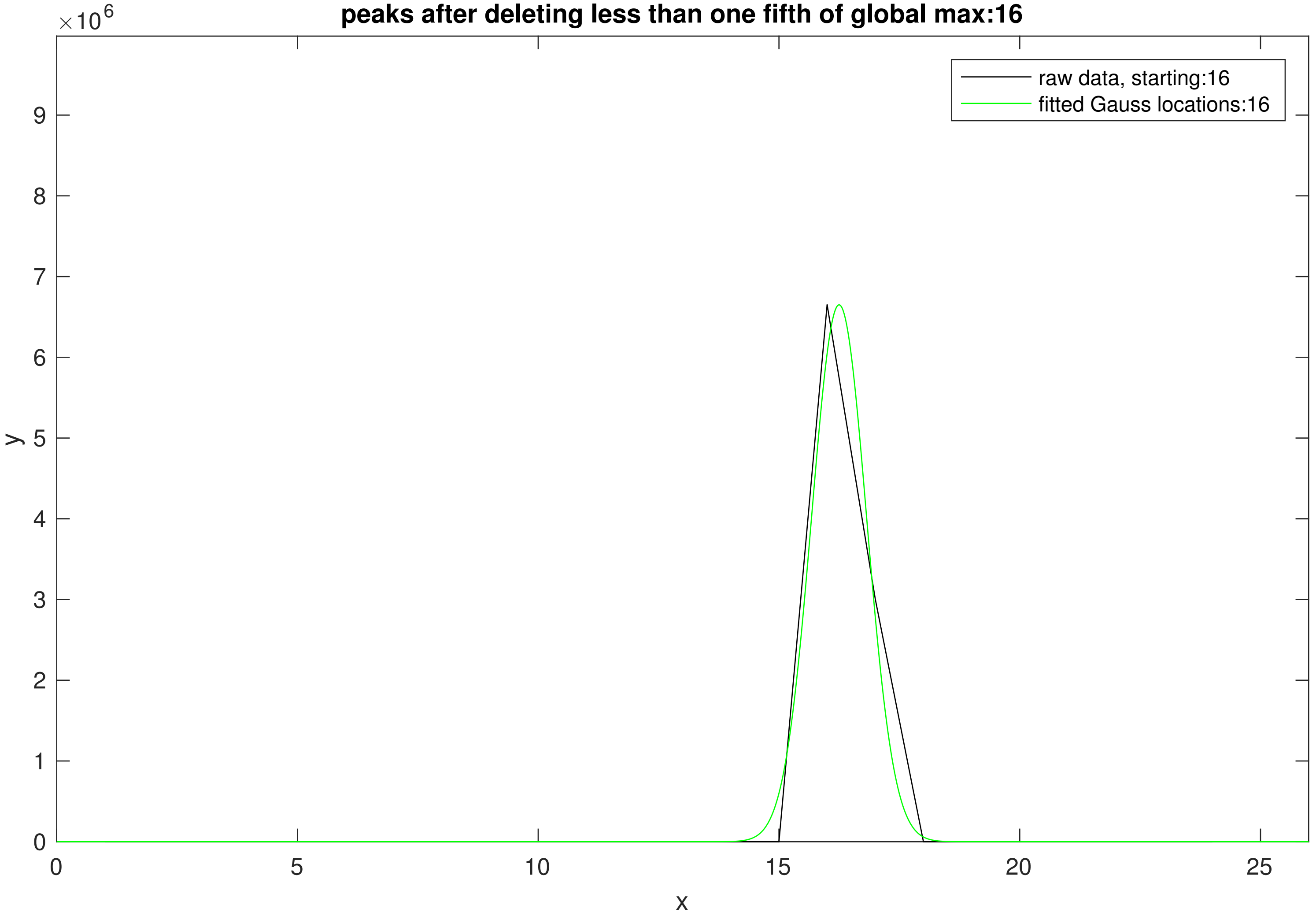

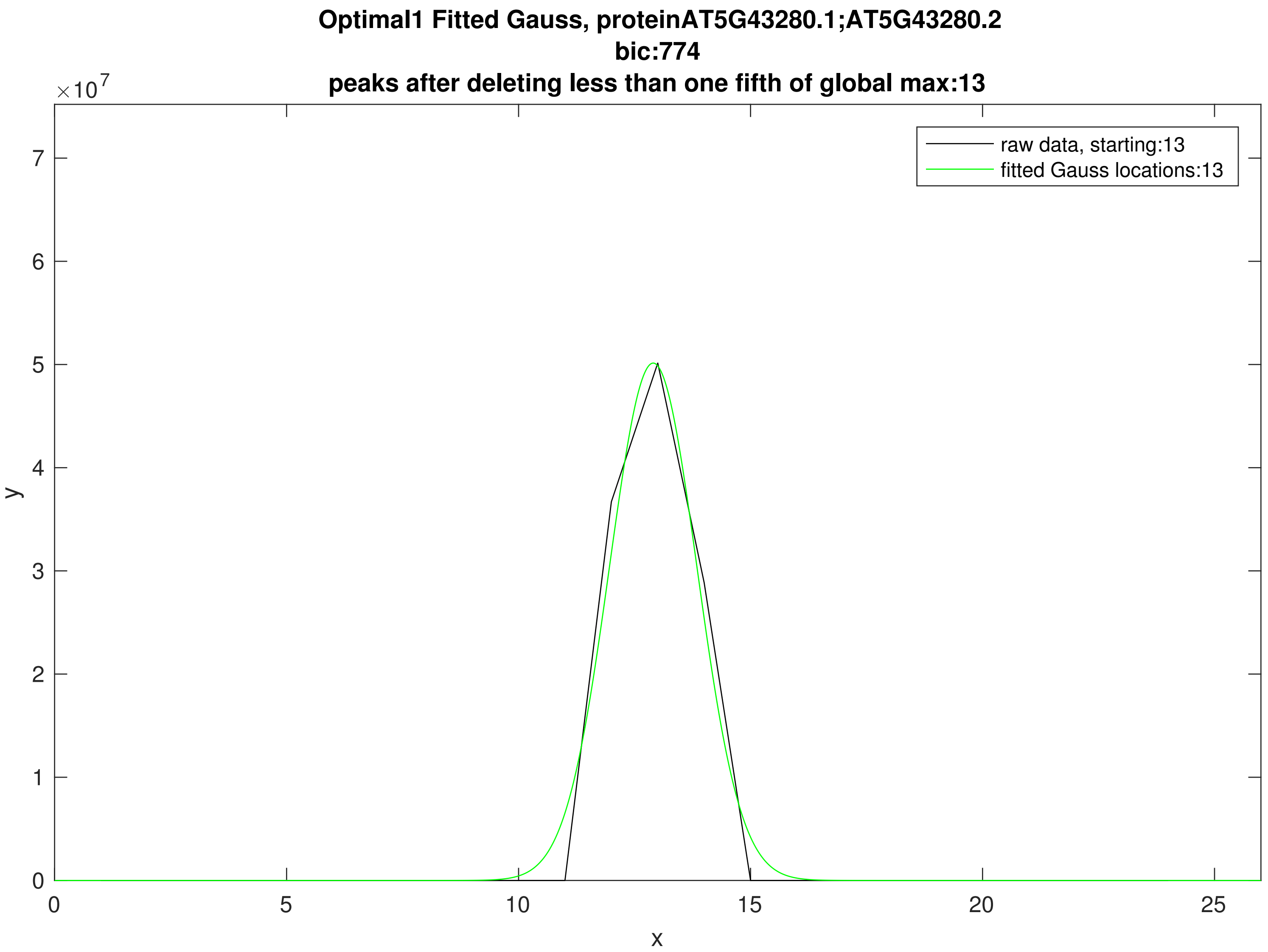
Optimal1 Fitted Gauss, proteinAT5G43280.1;AT5G43280.2
bic:774
peaks after deleting less than one fifth of global max:13
×10^7
raw data, starting:13
fitted Gauss locations:13
y
x
0
1
2
3
4
5
6
7
0
5
10
15
20
25

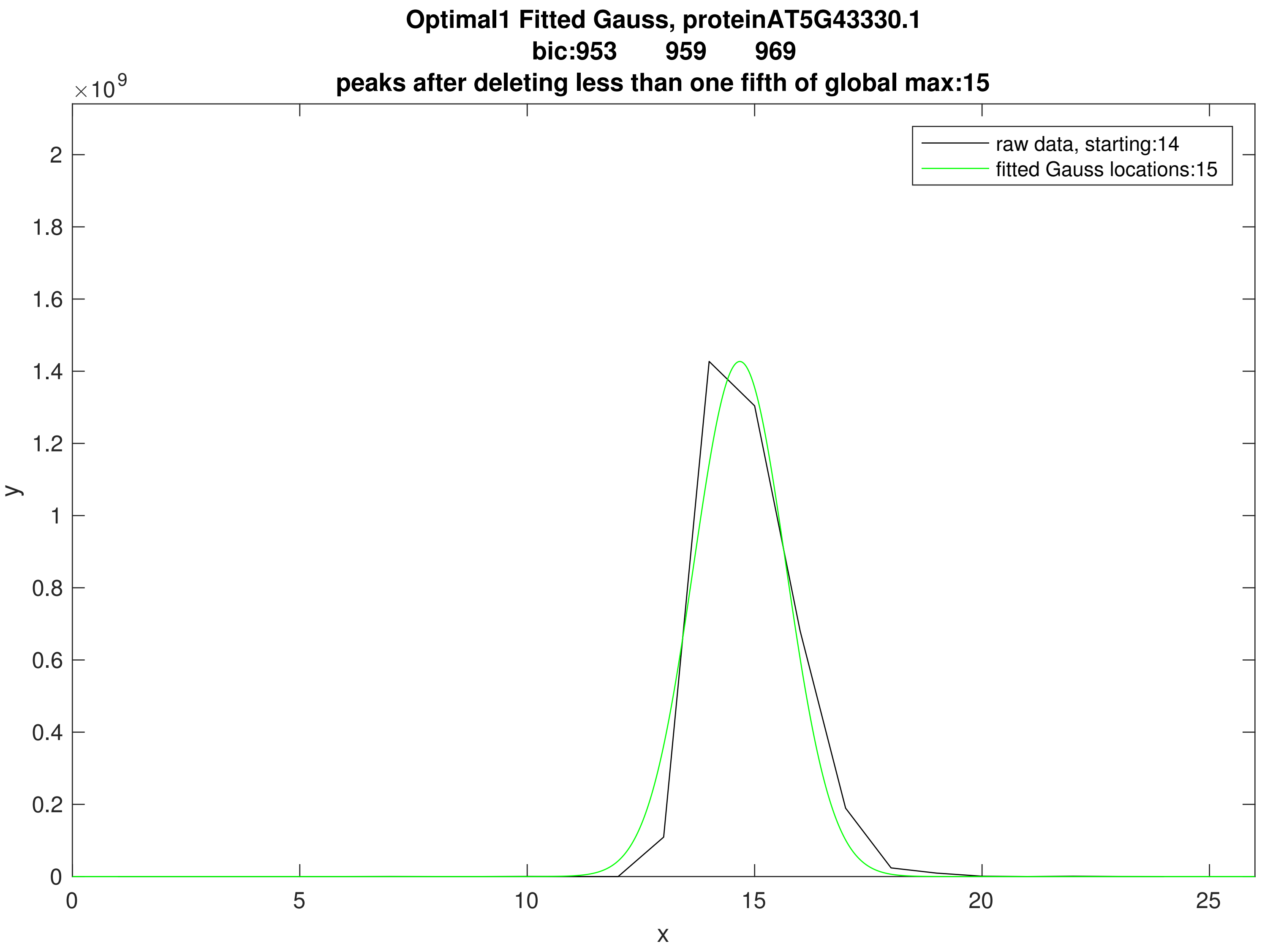

Optimal1 Fitted Gauss, proteinAT5G43330.1
bic:953 959 969
peaks after deleting less than one fifth of global max:15
×10^9
raw data, starting:14
fitted Gauss locations:15
2
1.8
1.6
1.4
1.2
1
0.8
0.6
0.4
0.2
0
y
0
5
10
15
20
25
x

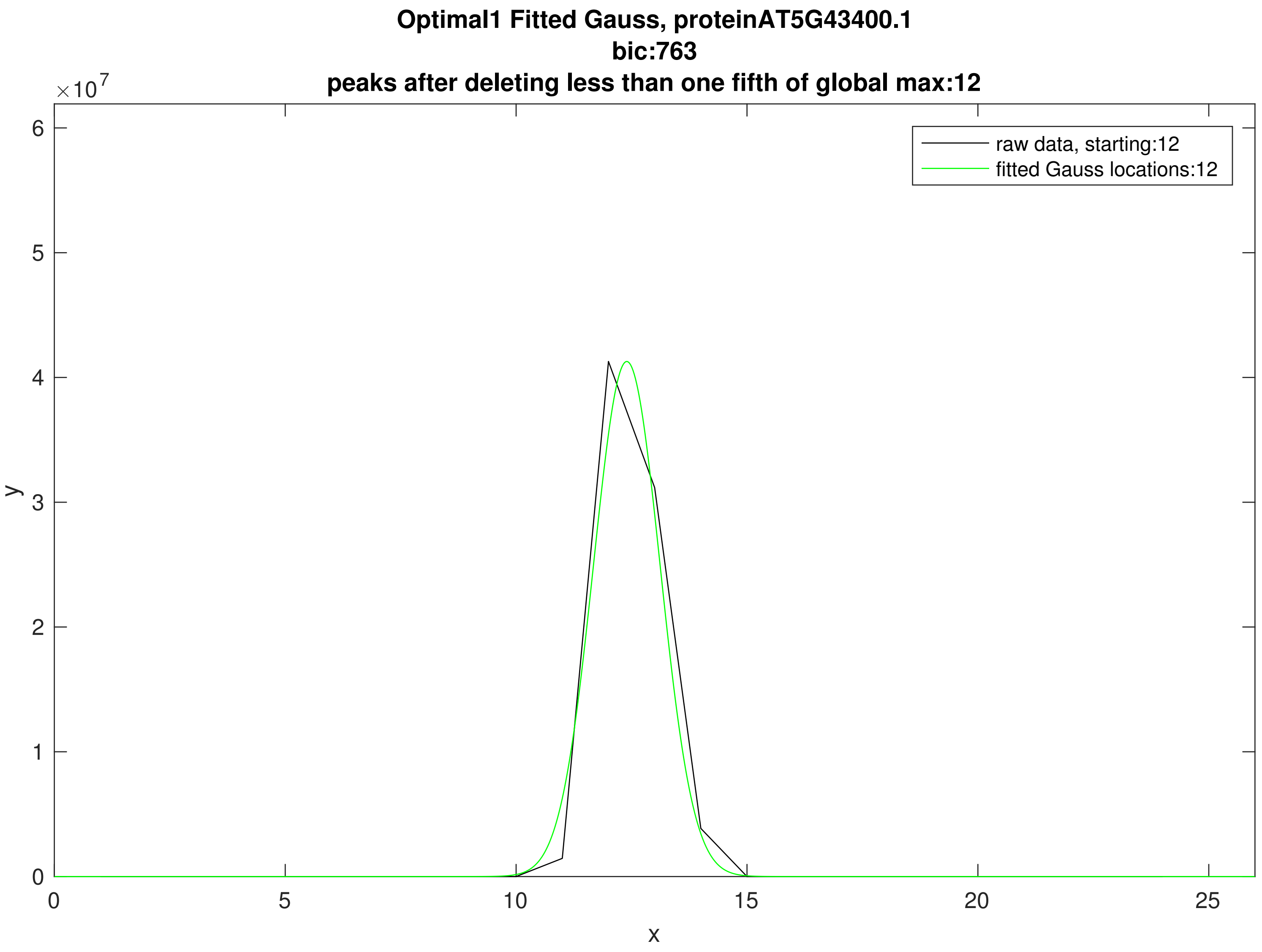
Optimal1 Fitted Gauss, proteinAT5G43400.1
bic:763
peaks after deleting less than one fifth of global max:12
×10^7
raw data, starting:12
fitted Gauss locations:12
y
x
0
1
2
3
4
5
6
0
5
10
15
20
25

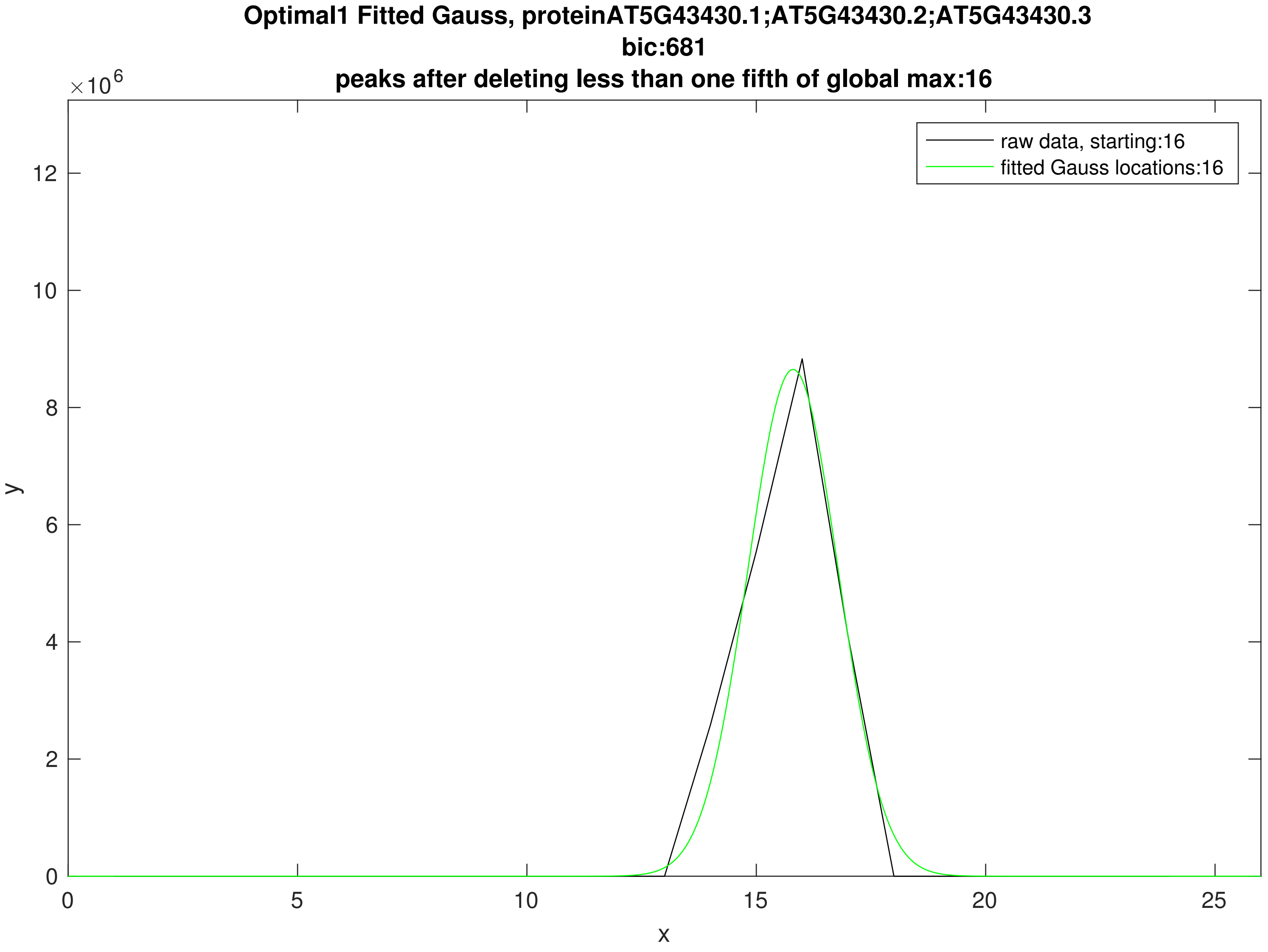

Optimal1 Fitted Gauss, proteinAT5G43430.1;AT5G43430.2;AT5G43430.3
bic:681
peaks after deleting less than one fifth of global max:16
×10^6
raw data, starting:16
fitted Gauss locations:16
12
10
8
6
4
2
0
y
0
5
10
15
20
25
x

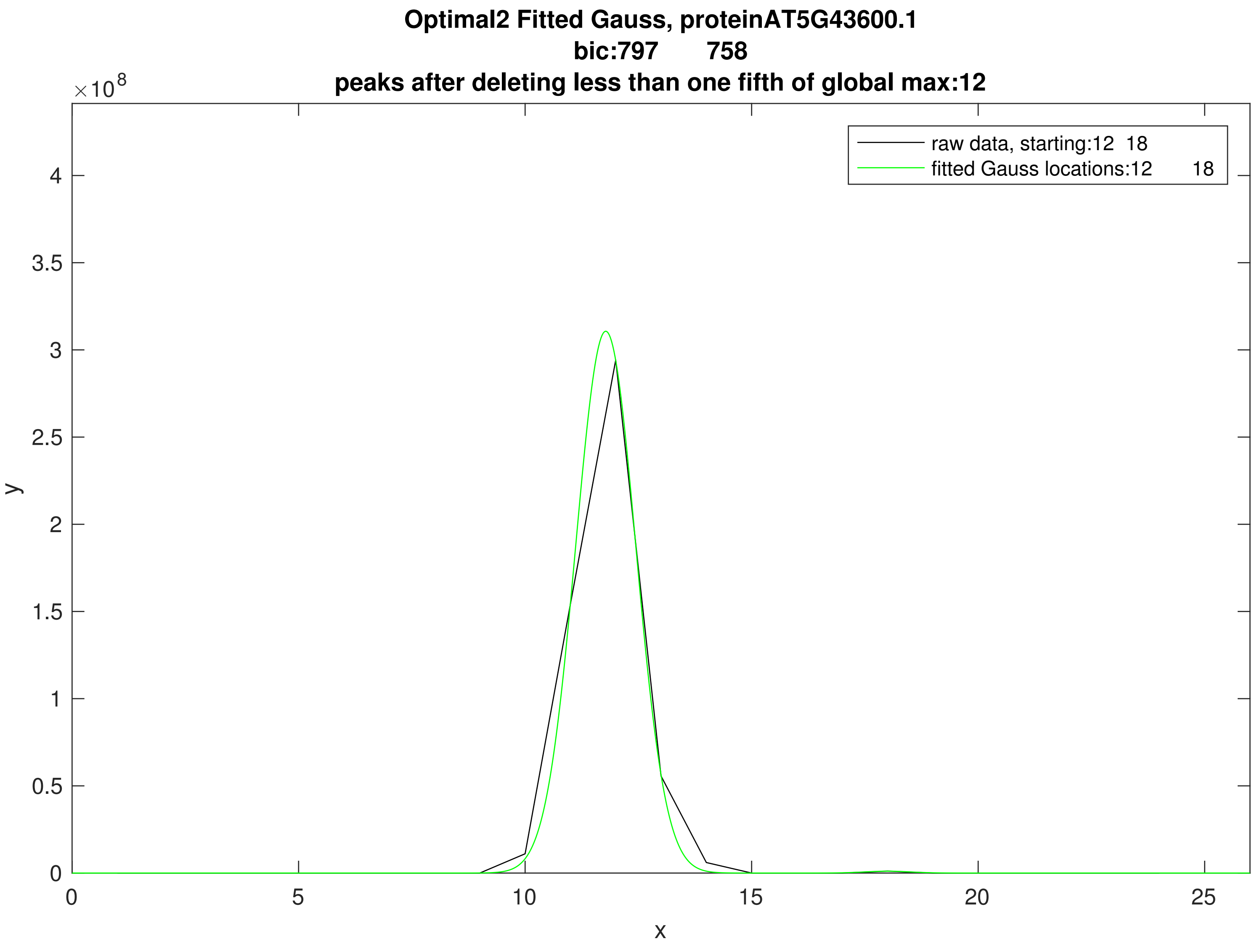
Optimal2 Fitted Gauss, proteinAT5G43600.1
bic:797 758
peaks after deleting less than one fifth of global max:12
×10^8
4
3.5
3
2.5
2
1.5
1
0.5
0
y
0
5
10
15
20
25
x
raw data, starting:12 18
fitted Gauss locations:12 18

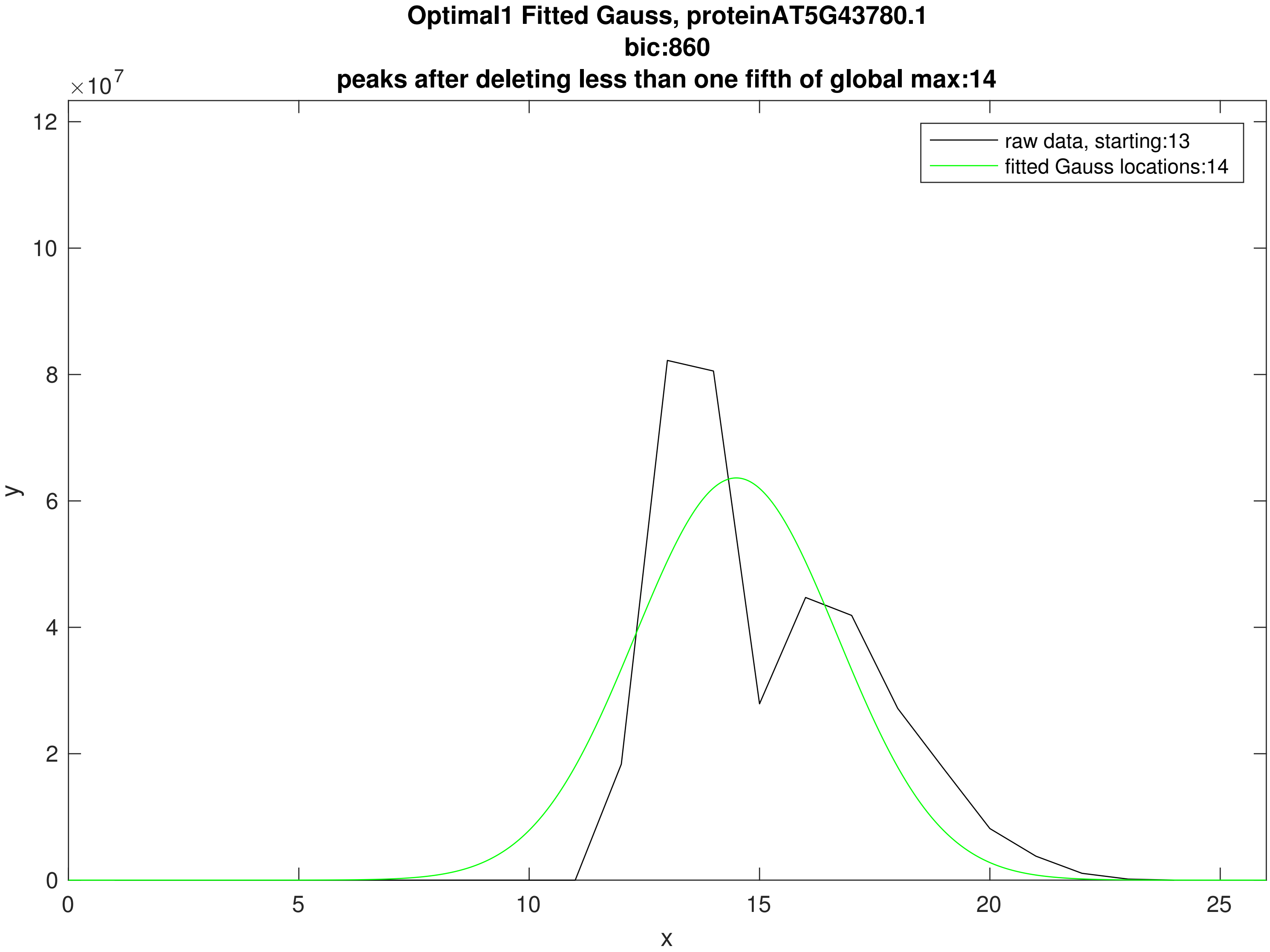

Optimal1 Fitted Gauss, proteinAT5G43780.1
bic:860
peaks after deleting less than one fifth of global max:14
×10^7
12
10
8
6
4
2
0
y
0
5
10
15
20
25
x
raw data, starting:13
fitted Gauss locations:14

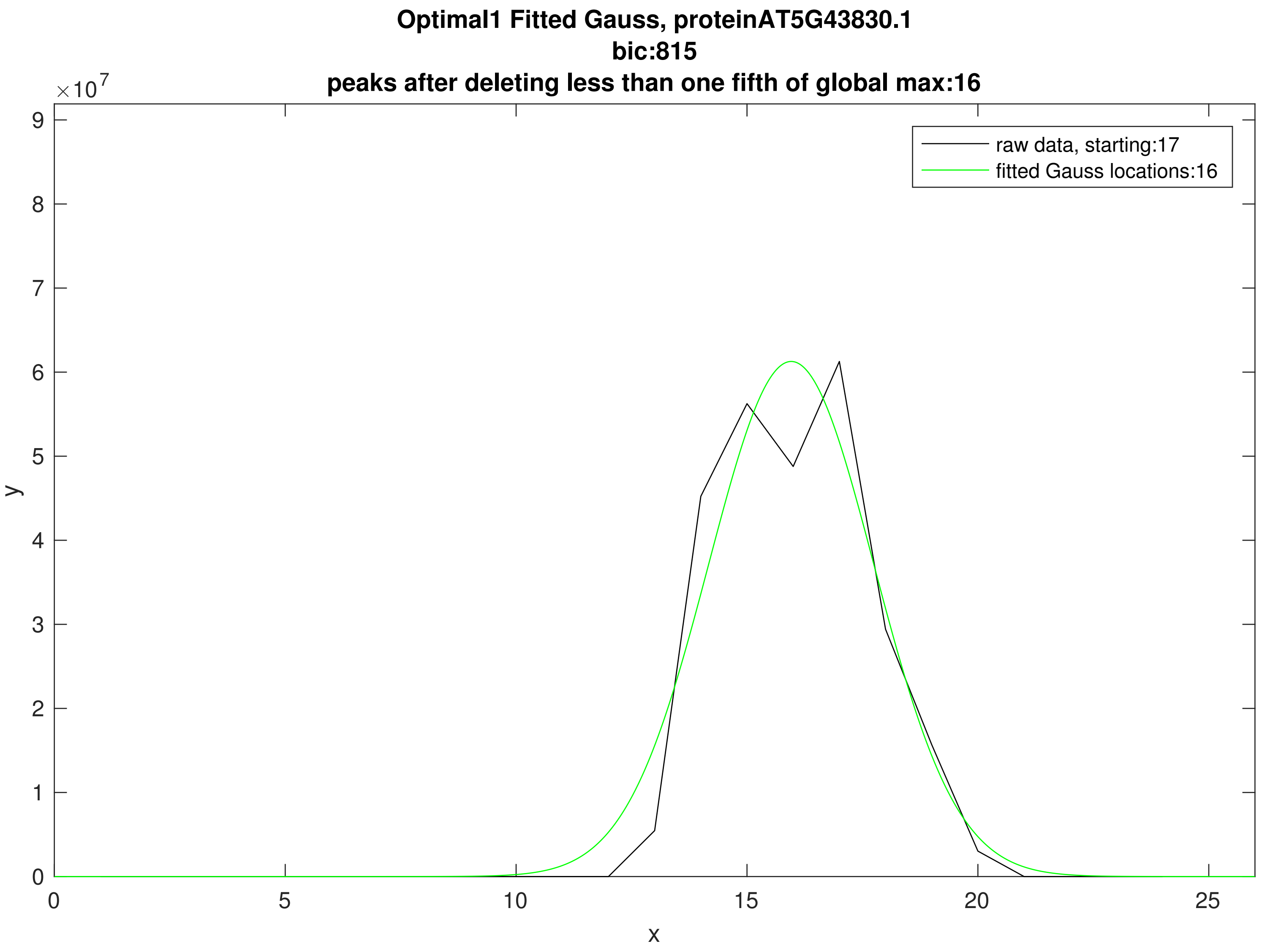
Optimal1 Fitted Gauss, proteinAT5G43830.1
bic:815
peaks after deleting less than one fifth of global max:16
×10^7
raw data, starting:17
fitted Gauss locations:16
y
x
0
1
2
3
4
5
6
7
8
9
0
5
10
15
20
25

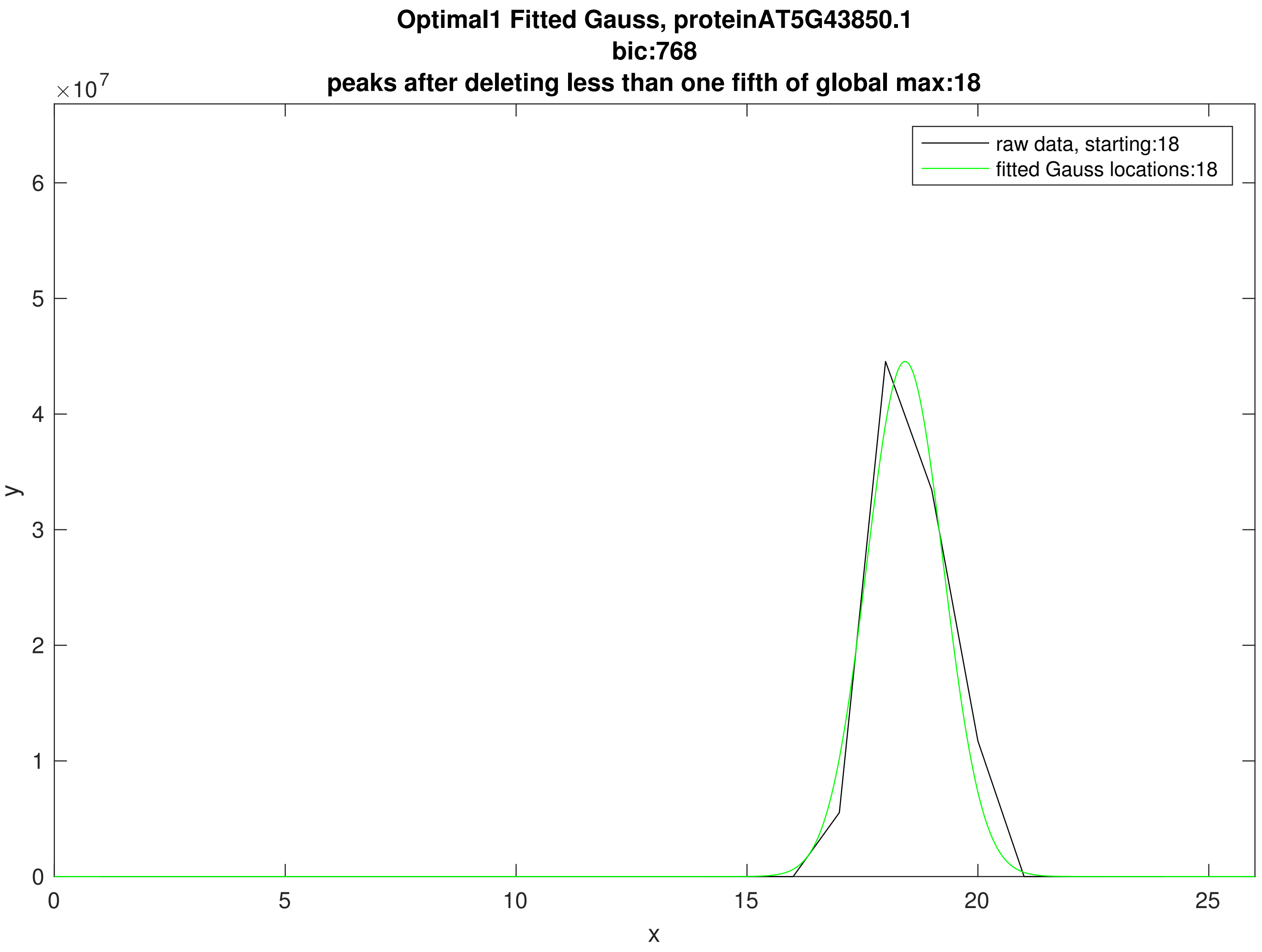
Optimal1 Fitted Gauss, proteinAT5G43850.1
bic:768
peaks after deleting less than one fifth of global max:18
×10^7
raw data, starting:18
fitted Gauss locations:18
y
x
0
1
2
3
4
5
6
0
5
10
15
20
25

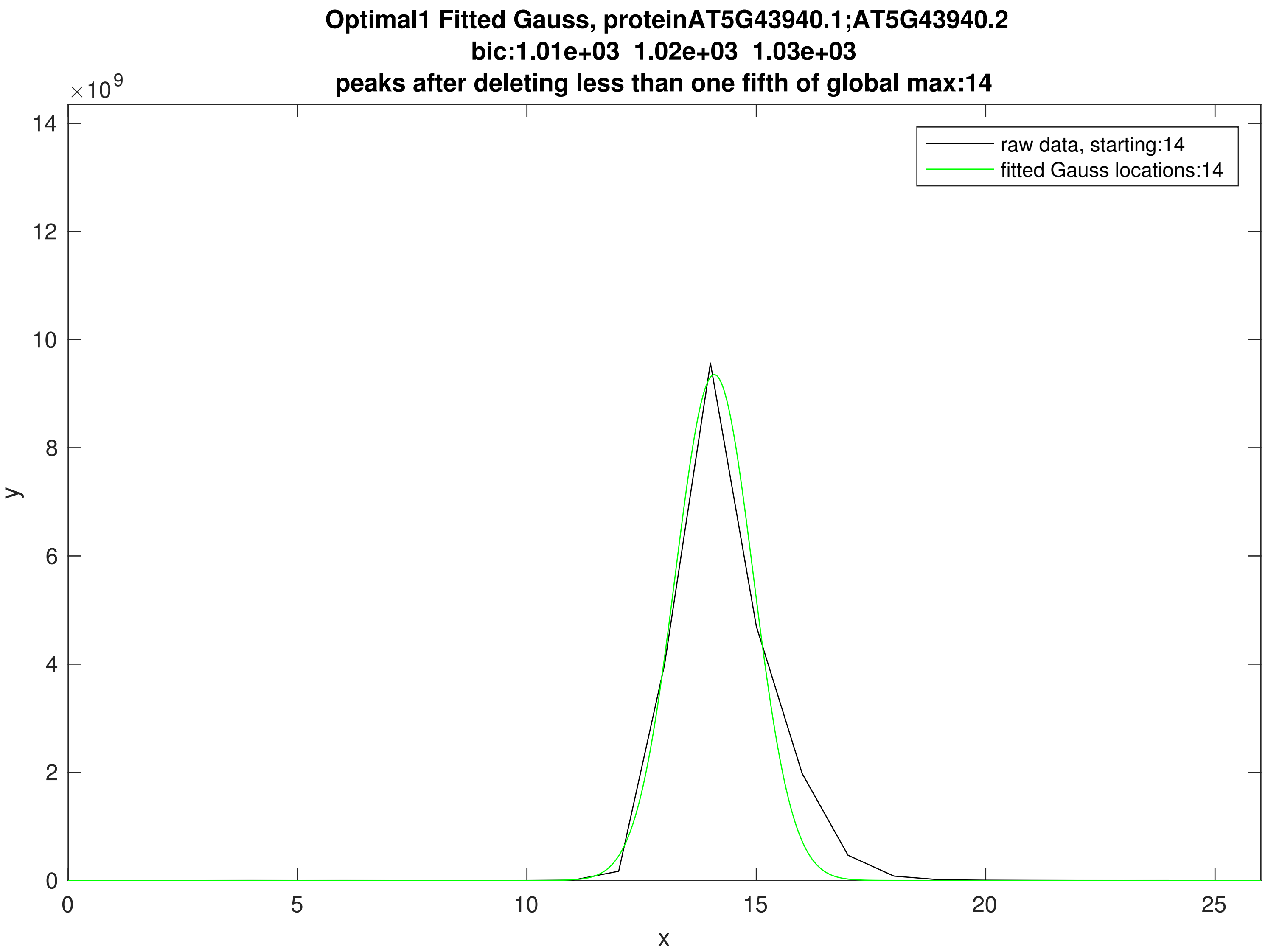
Optimal1 Fitted Gauss, proteinAT5G43940.1;AT5G43940.2
bic:1.01e+03 1.02e+03 1.03e+03
peaks after deleting less than one fifth of global max:14
×10^9
y
x
raw data, starting:14
fitted Gauss locations:14
0
2
4
6
8
10
12
14
0
5
10
15
20
25

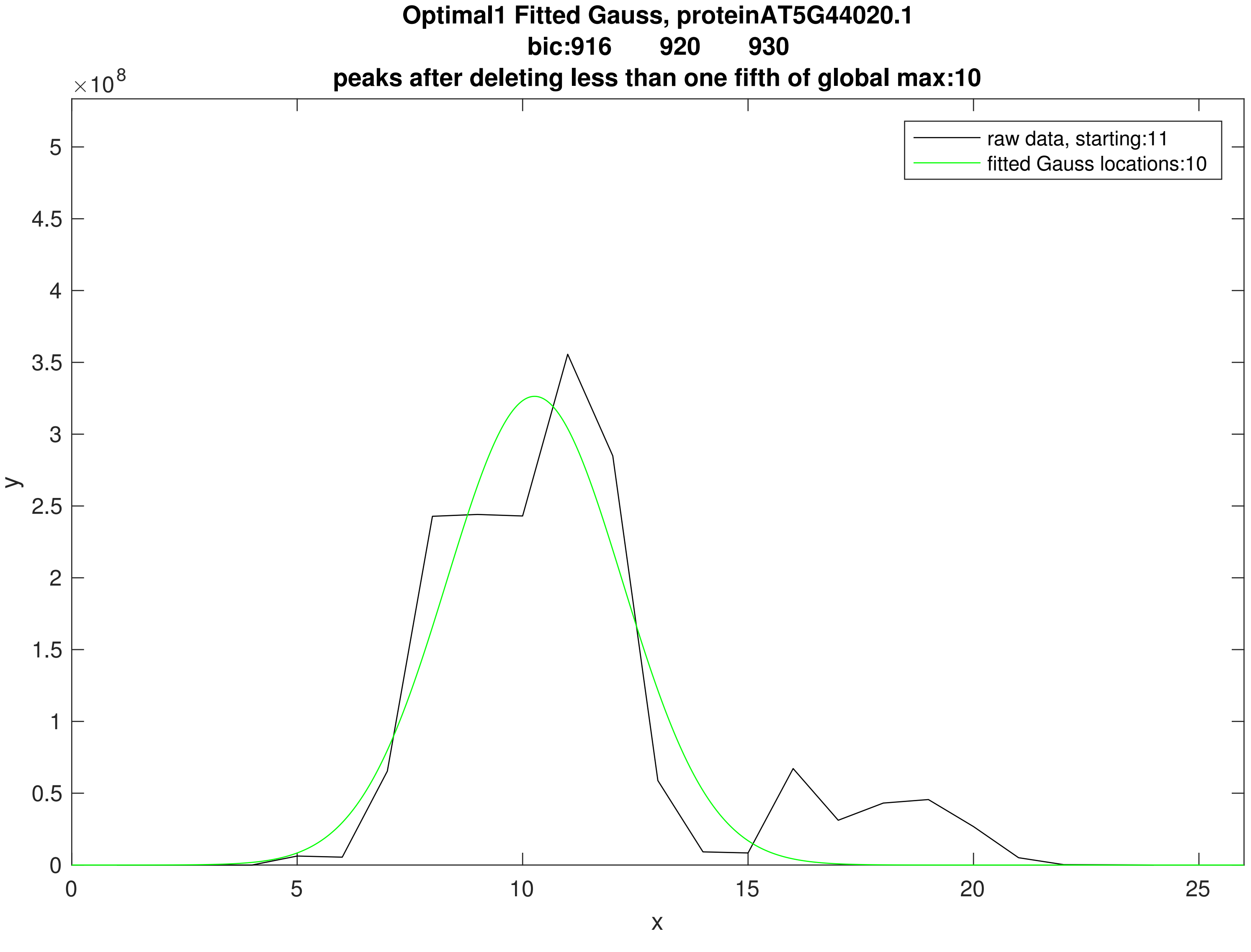

Optimal1 Fitted Gauss, proteinAT5G44020.1
bic:916 920 930
peaks after deleting less than one fifth of global max:10
×10^8
raw data, starting:11
fitted Gauss locations:10
y
x
0
0.5
1
1.5
2
2.5
3
3.5
4
4.5
5
0
5
10
15
20
25

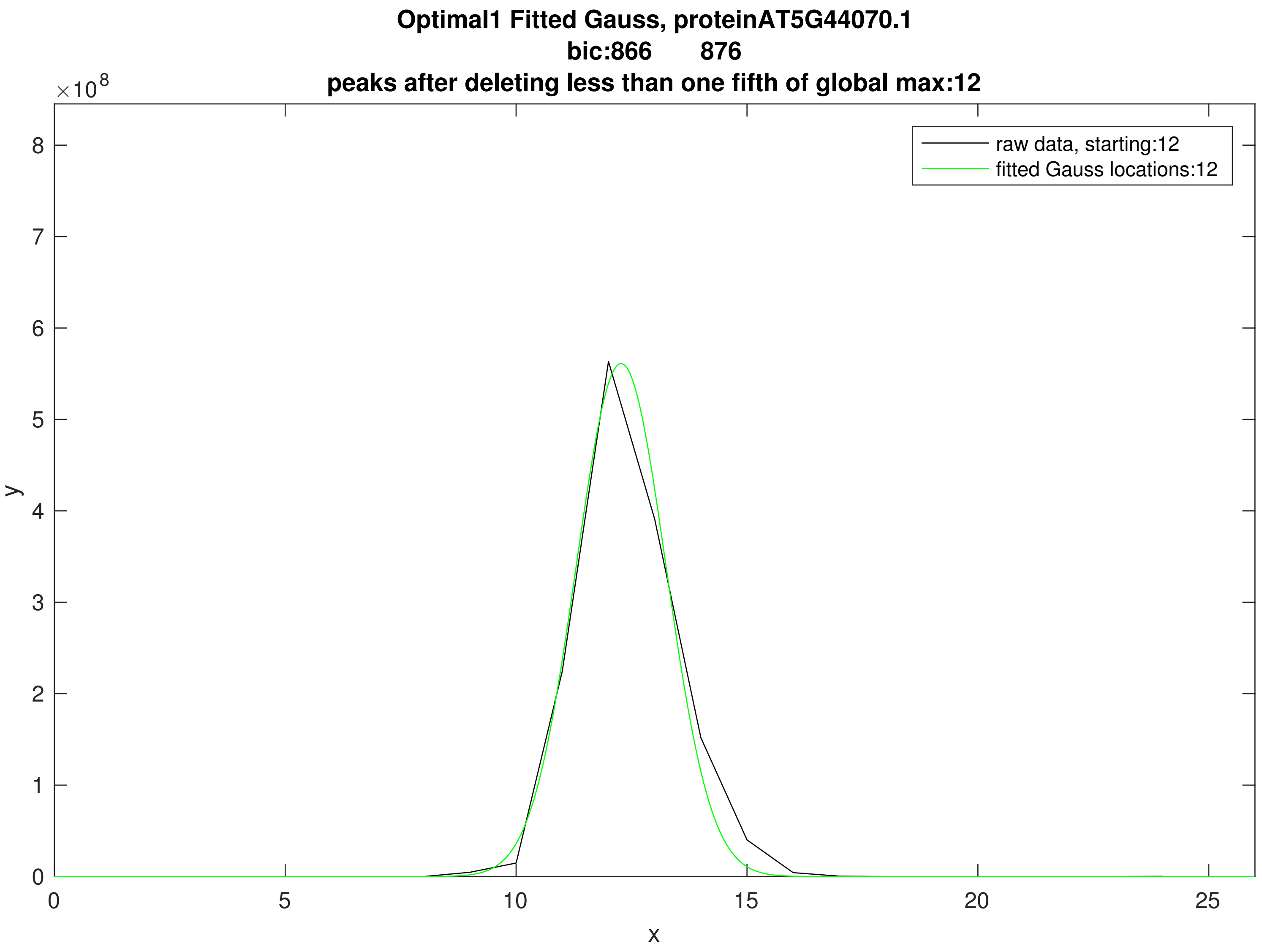
Optimal1 Fitted Gauss, proteinAT5G44070.1
bic:866 876
peaks after deleting less than one fifth of global max:12
×10^8
raw data, starting:12
fitted Gauss locations:12
y
x

**Optimal1 Fitted Gauss, proteinAT5G44110.1;AT5G44110.2;AT5G44110.3**
**bic:777**
**peaks after deleting less than one fifth of global max:8.6**

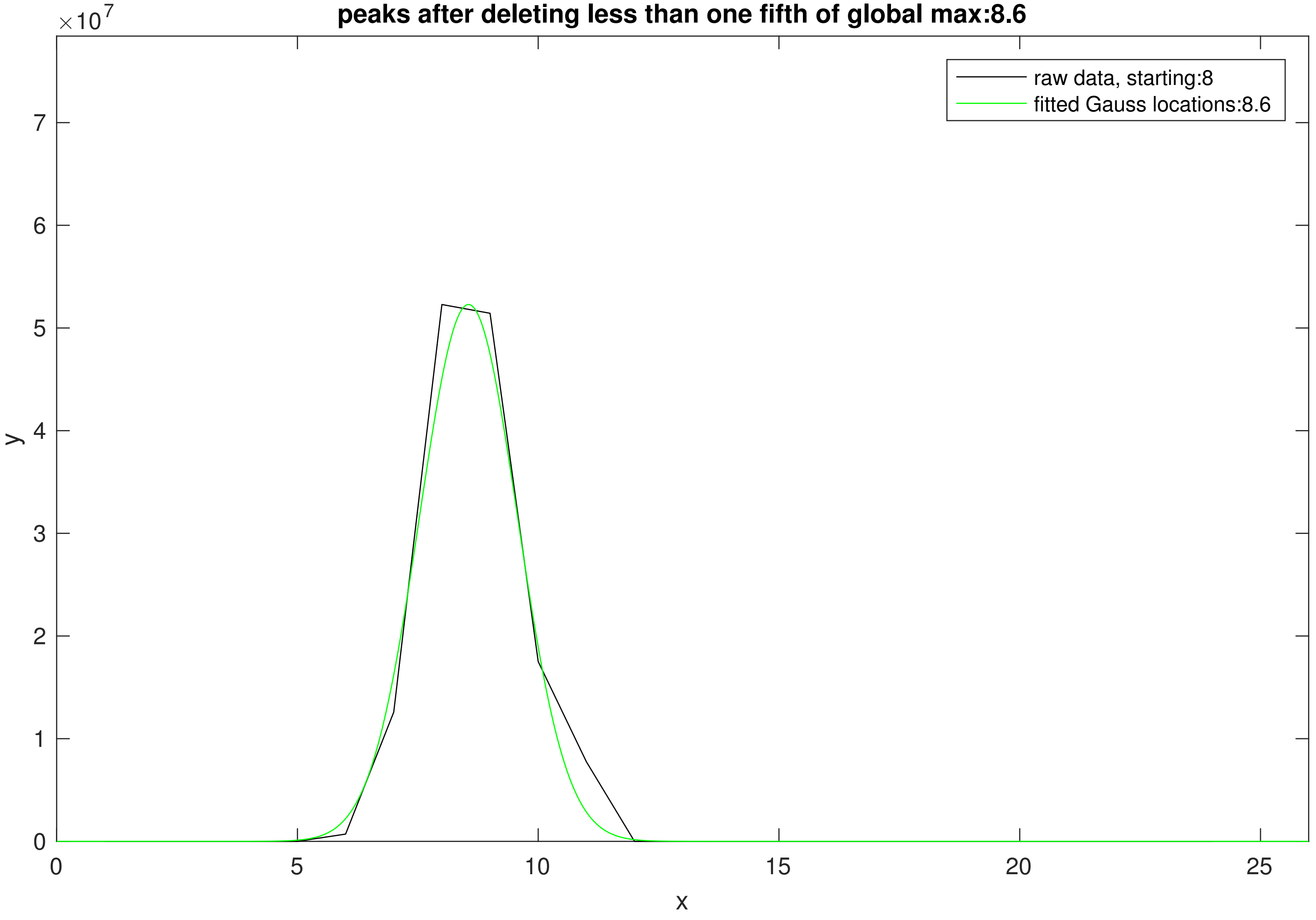

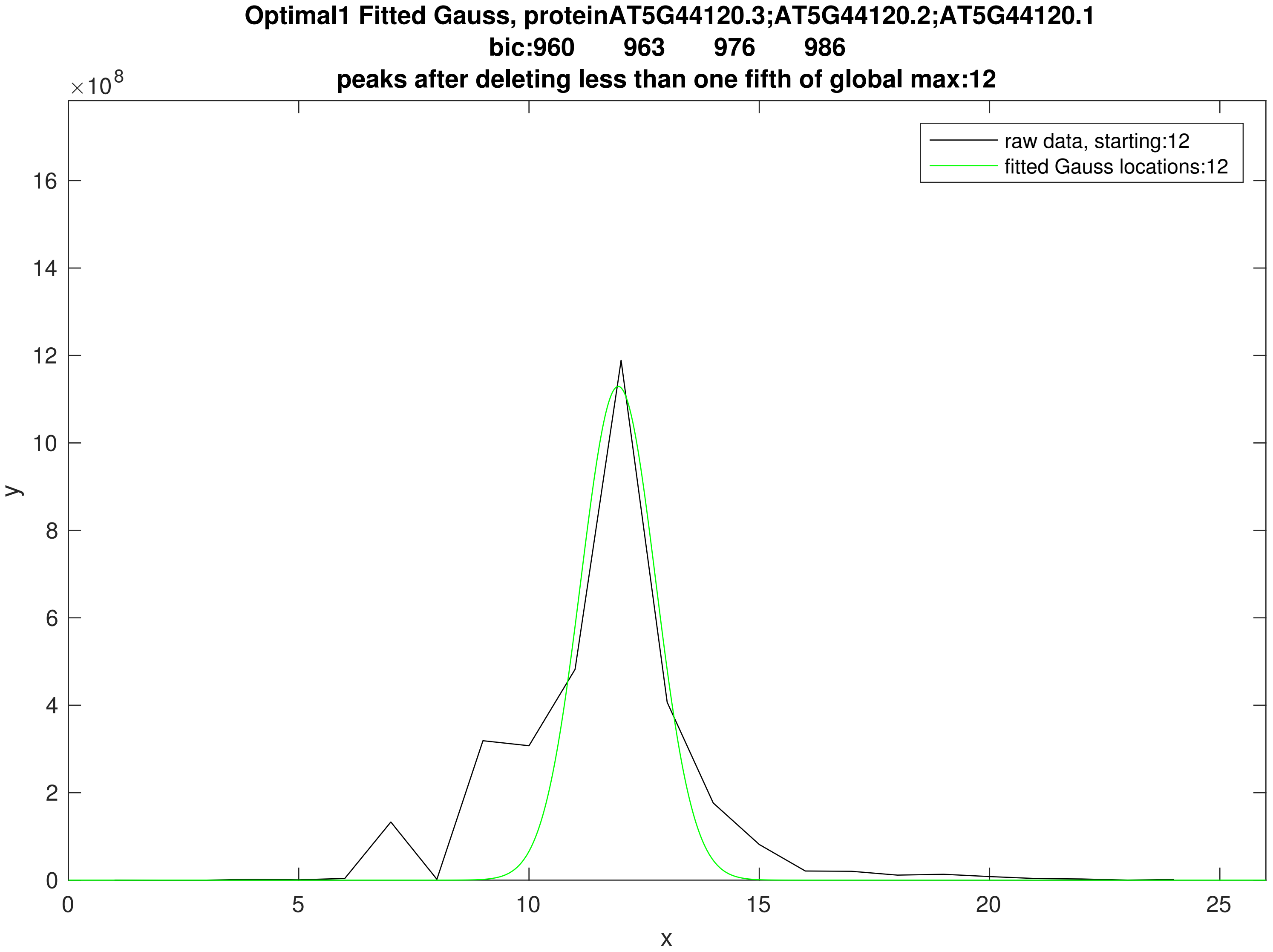
Optimal1 Fitted Gauss, proteinAT5G44120.3;AT5G44120.2;AT5G44120.1
bic:960 963 976 986
peaks after deleting less than one fifth of global max:12
×10^8
raw data, starting:12
fitted Gauss locations:12
y
x
0
2
4
6
8
10
12
14
16
0
5
10
15
20
25

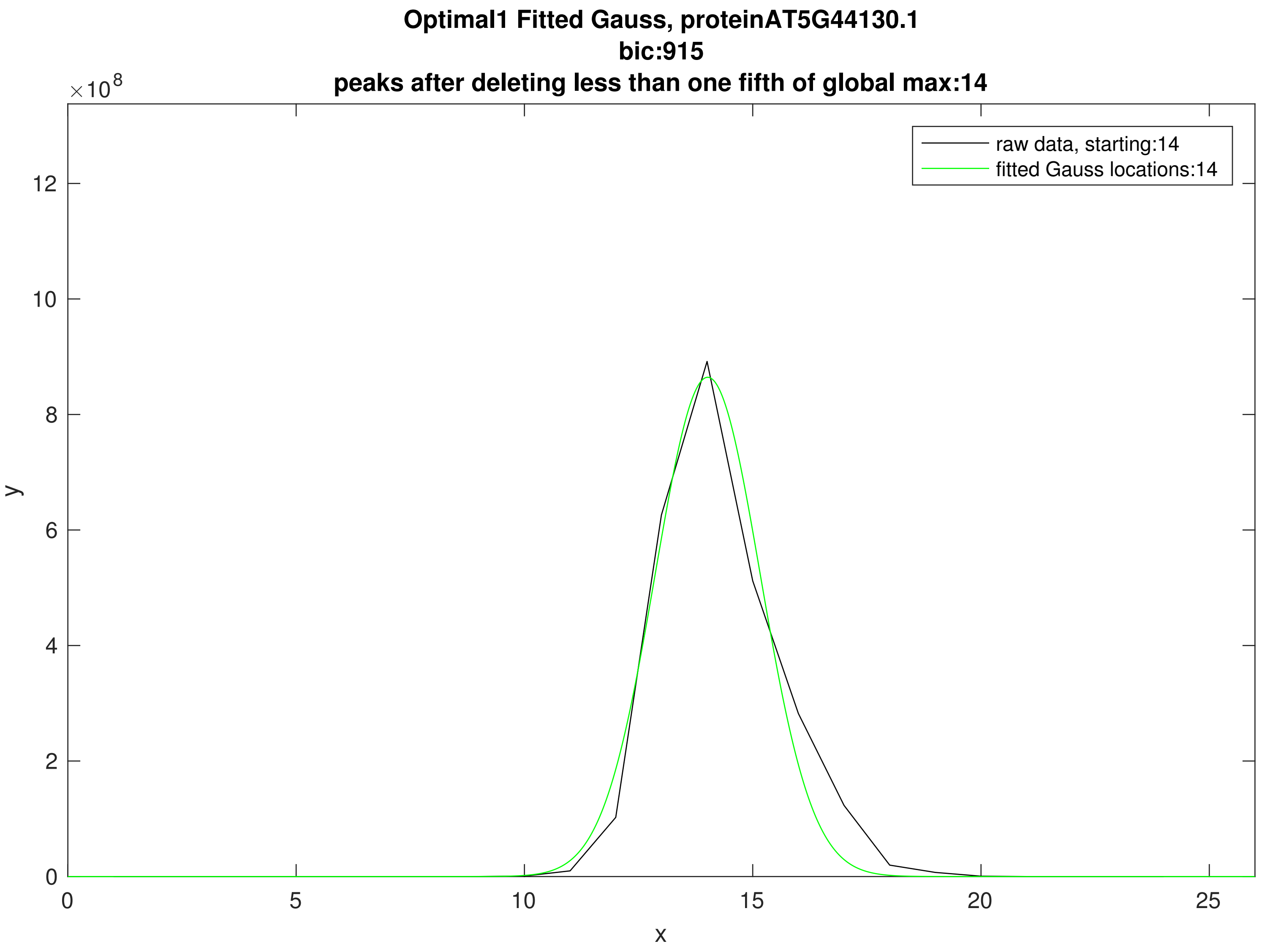
Optimal1 Fitted Gauss, proteinAT5G44130.1
bic:915
peaks after deleting less than one fifth of global max:14
×10^8
raw data, starting:14
fitted Gauss locations:14
y
x
0
2
4
6
8
10
12
0
5
10
15
20
25

**Optimal1 Fitted Gauss, proteinAT5G44310.2;AT5G44310.1**
**bic:659**
**peaks after deleting less than one fifth of global max:4.4**

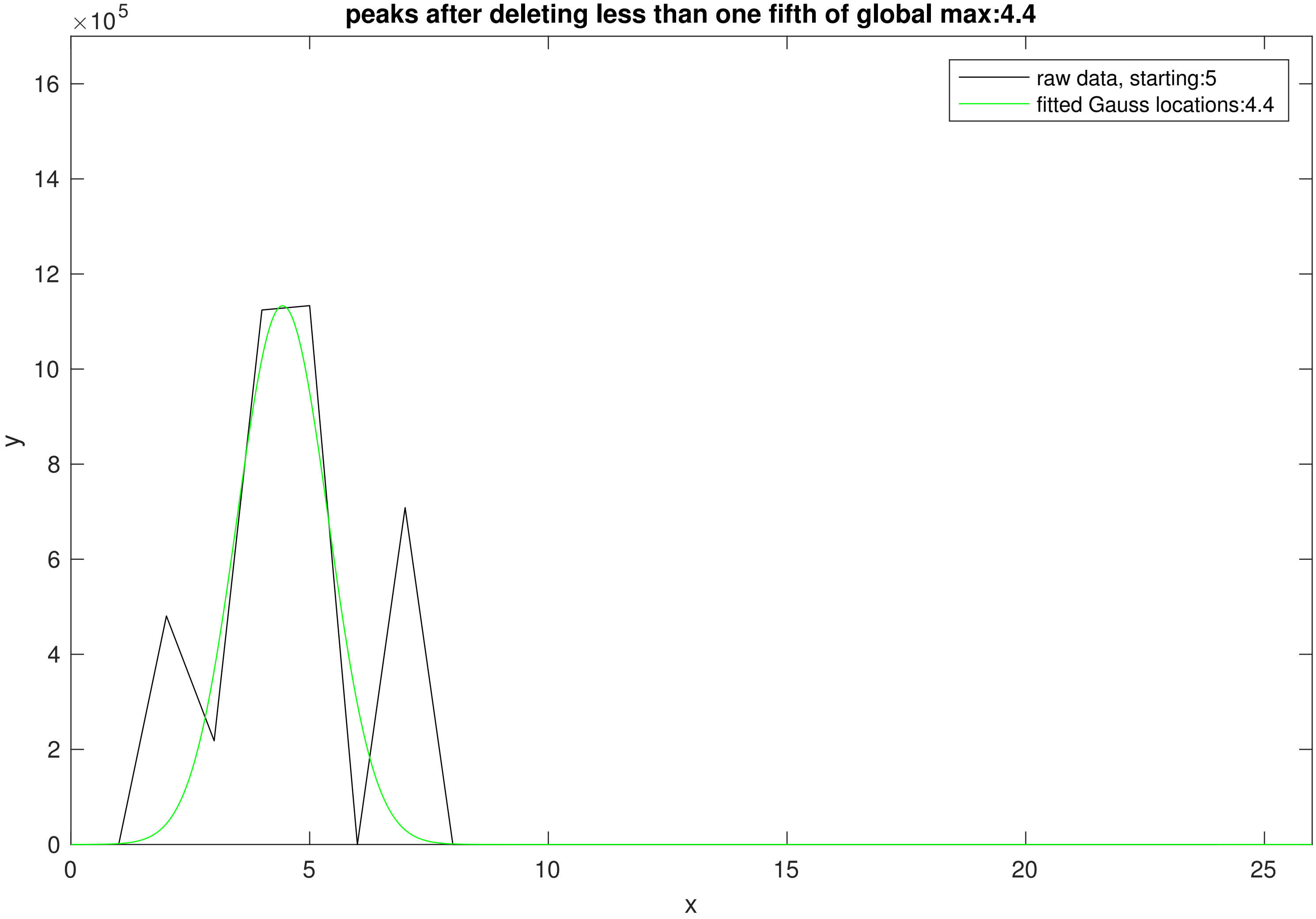

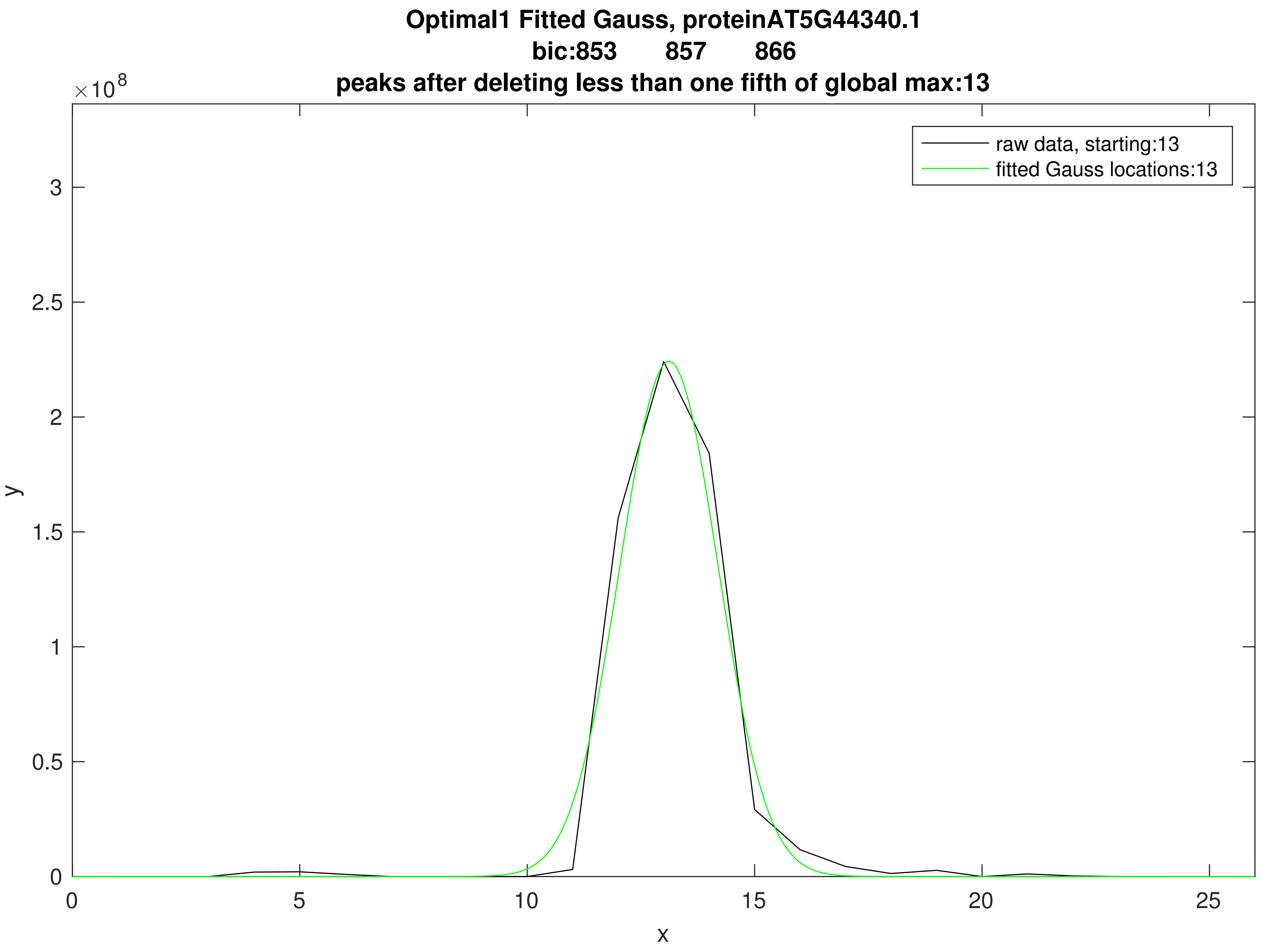
Optimal1 Fitted Gauss, proteinAT5G44340.1
bic:853 857 866
peaks after deleting less than one fifth of global max:13
×10^8
raw data, starting:13
fitted Gauss locations:13
y
x
0
0.5
1
1.5
2
2.5
3
0
5
10
15
20
25

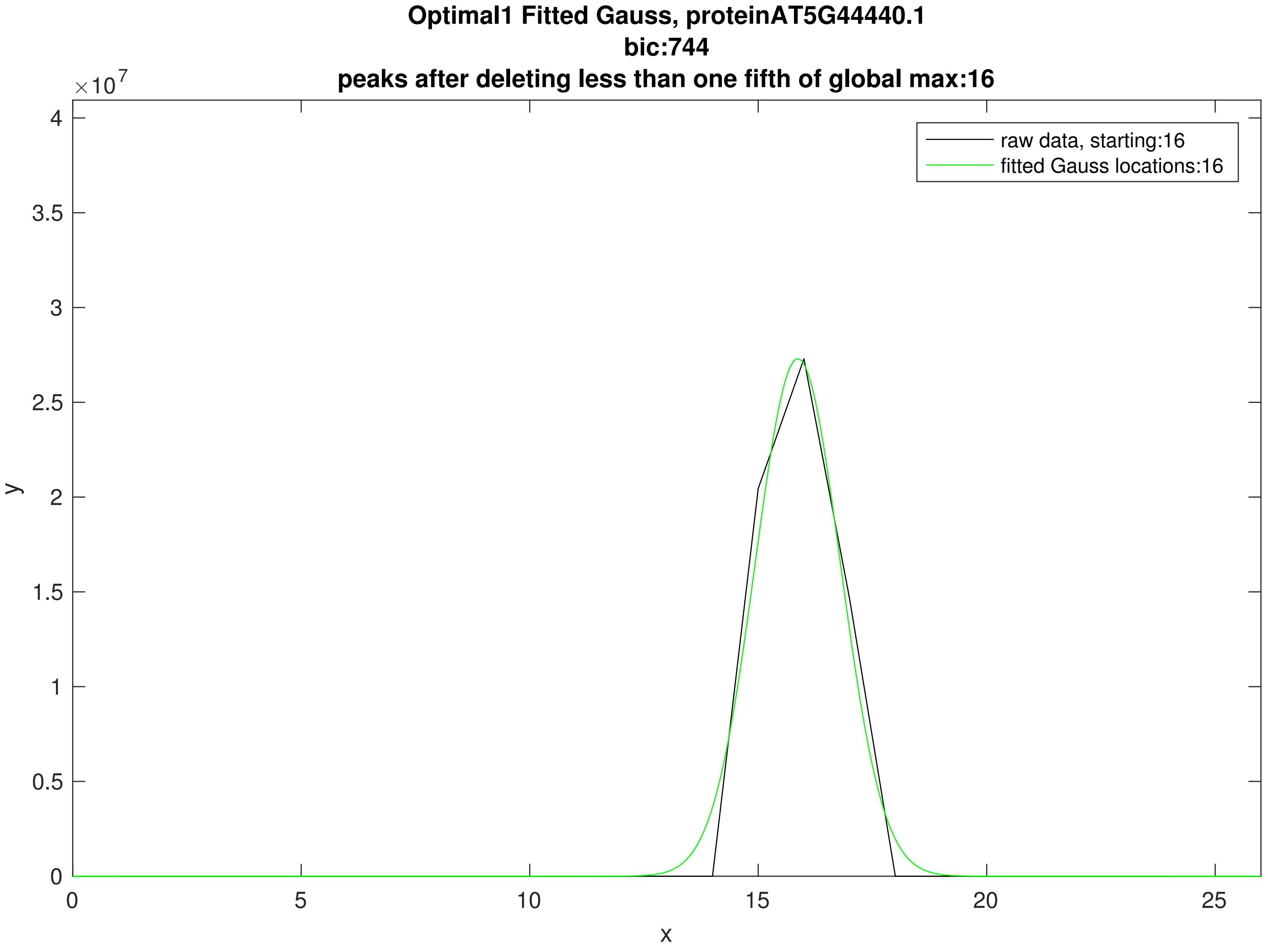
Optimal1 Fitted Gauss, proteinAT5G44440.1
bic:744
peaks after deleting less than one fifth of global max:16
×10^7
4
3.5
3
2.5
2
1.5
1
0.5
0
y
0
5
10
15
20
25
x
raw data, starting:16
fitted Gauss locations:16

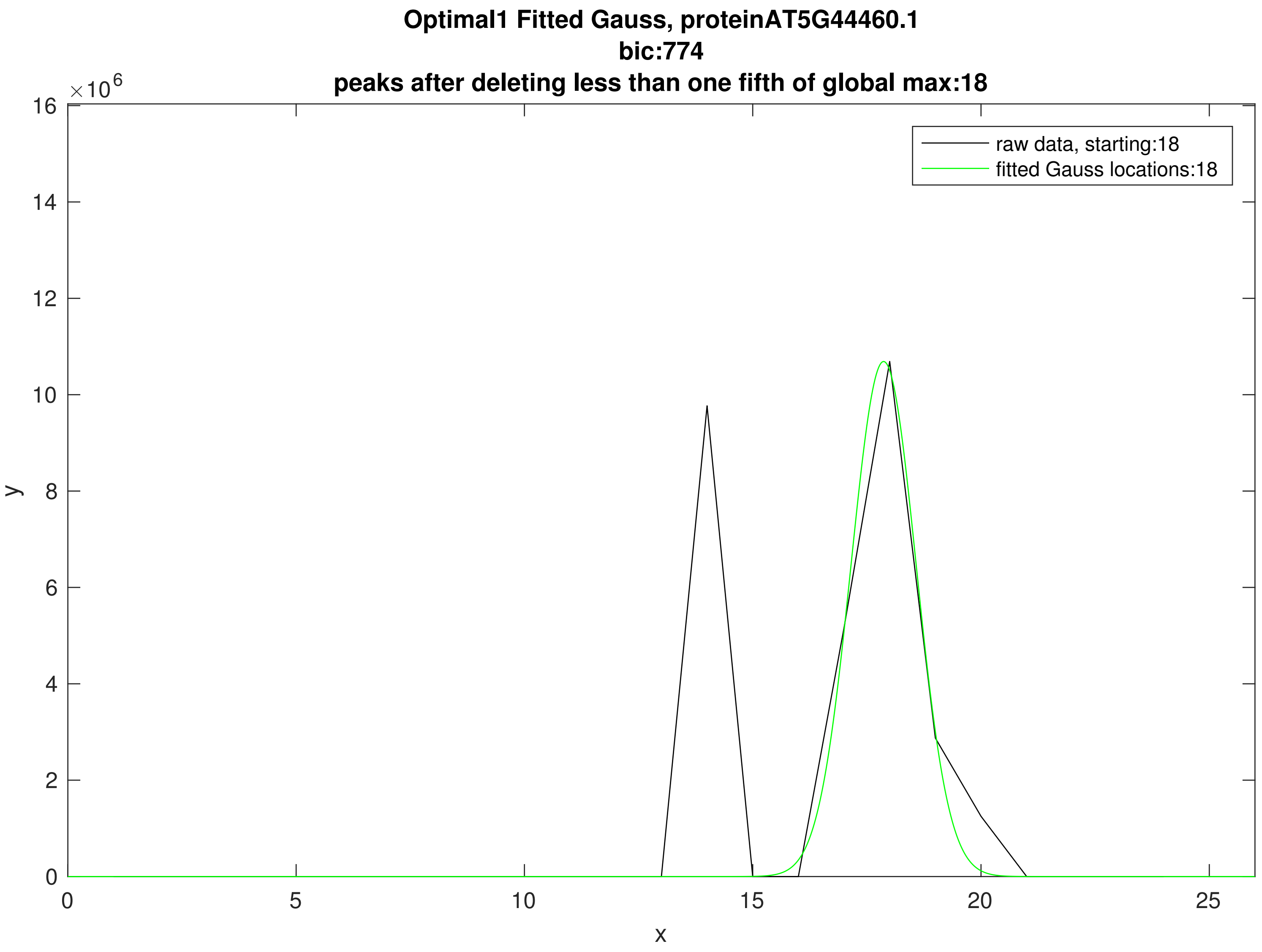
Optimal1 Fitted Gauss, proteinAT5G44460.1
bic:774
peaks after deleting less than one fifth of global max:18
×10^6
16
14
12
10
8
6
4
2
0
y
0
5
10
15
20
25
x
raw data, starting:18
fitted Gauss locations:18

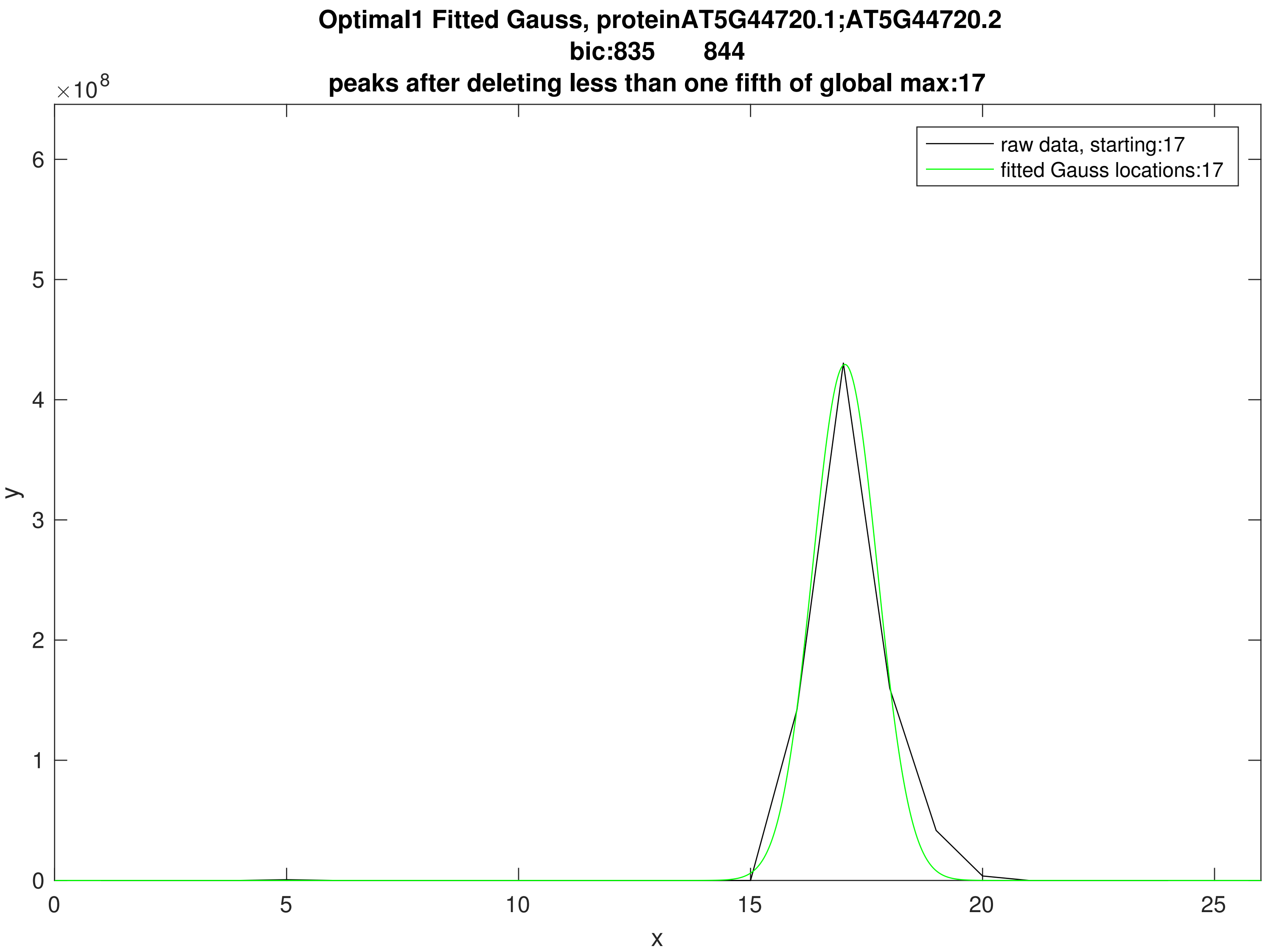
Optimal1 Fitted Gauss, proteinAT5G44720.1;AT5G44720.2
bic:835 844
peaks after deleting less than one fifth of global max:17
×10^8
raw data, starting:17
fitted Gauss locations:17
y
x
0
1
2
3
4
5
6
0
5
10
15
20
25

**Optimal1 Fitted Gauss, proteinAT5G45020.2;AT5G45020.1**
**bic:747**
**peaks after deleting less than one fifth of global max:14**

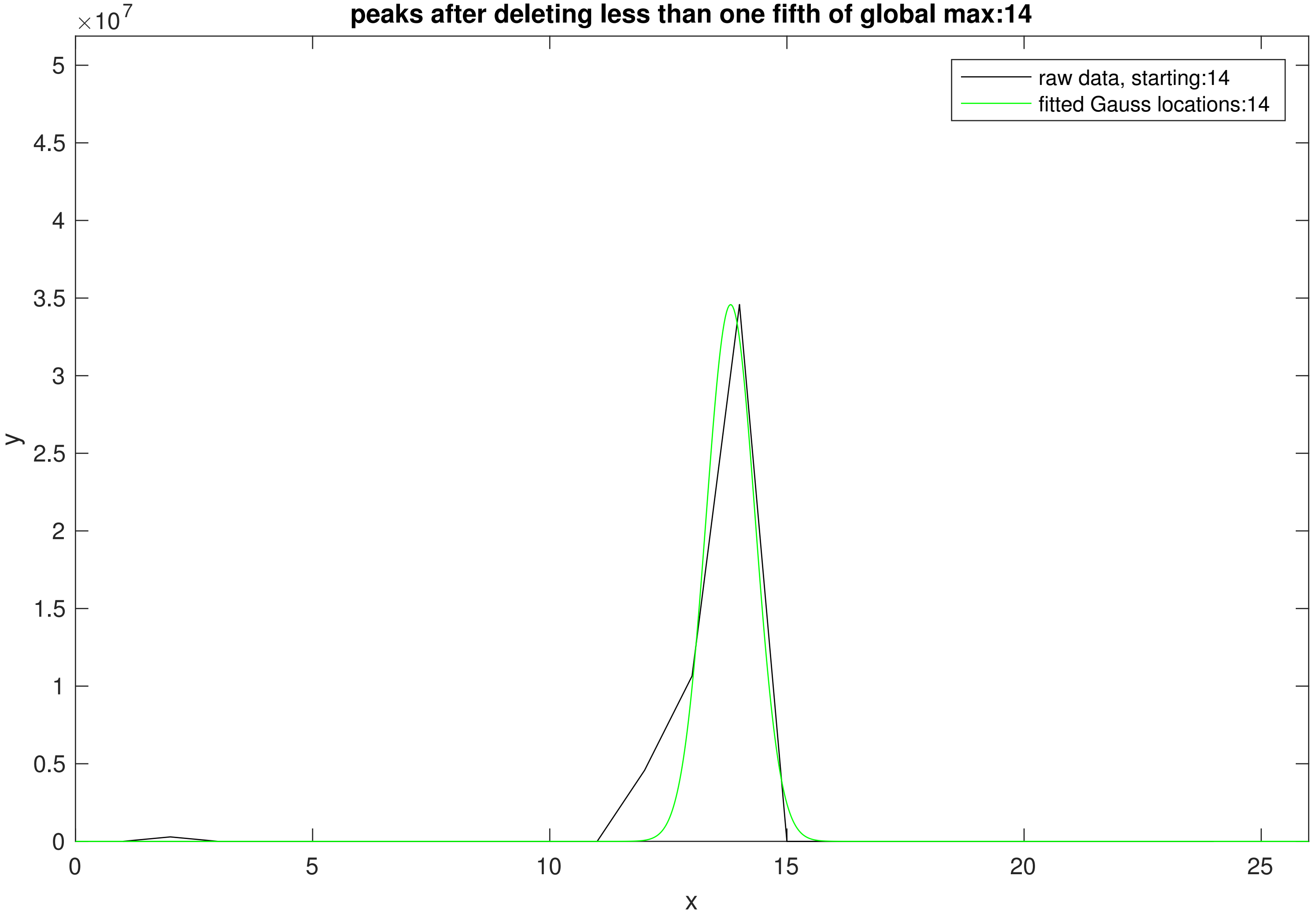

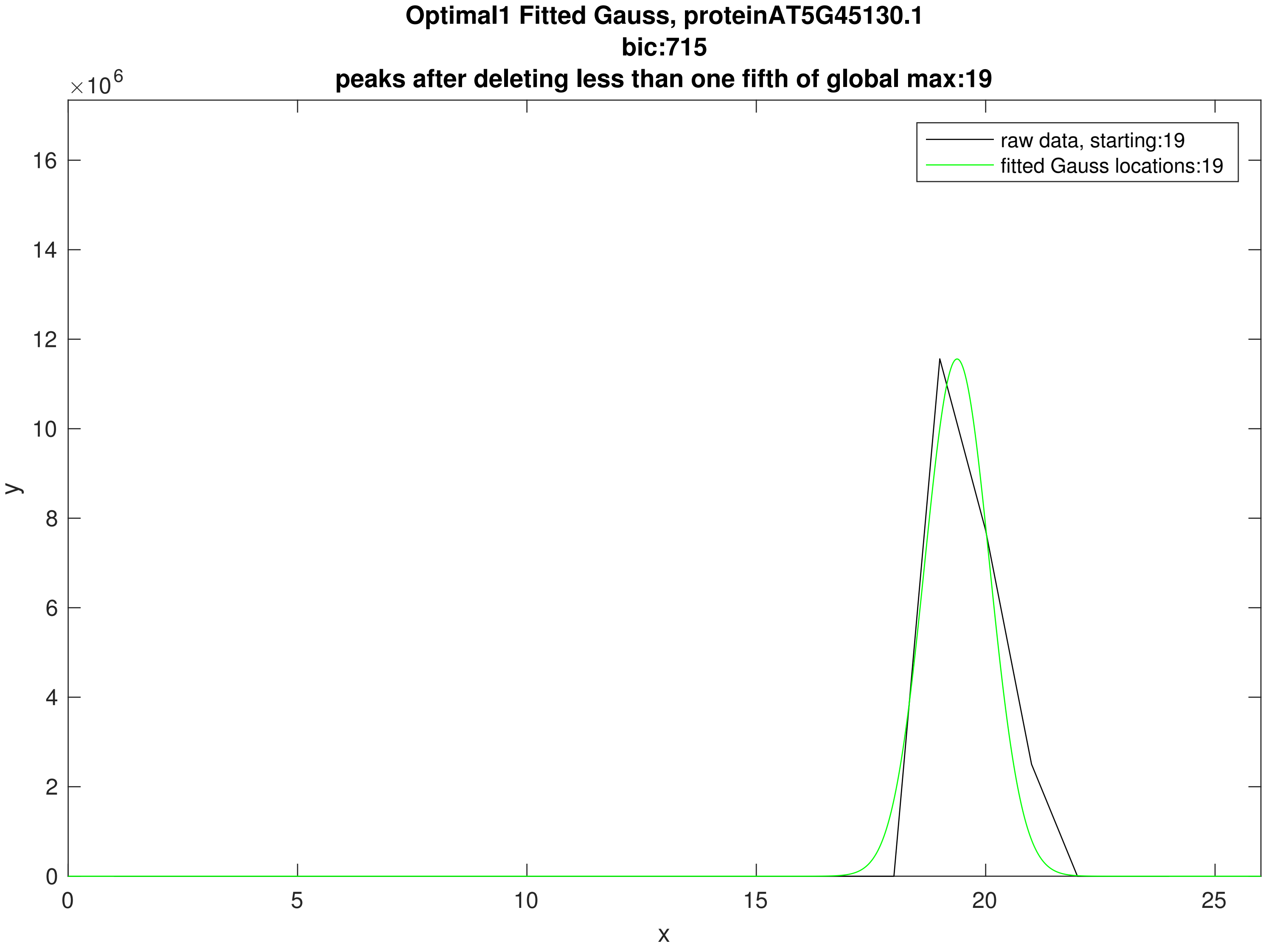

Optimal1 Fitted Gauss, proteinAT5G45130.1
bic:715
peaks after deleting less than one fifth of global max:19
×10^6
raw data, starting:19
fitted Gauss locations:19
y
x
0
2
4
6
8
10
12
14
16
0
5
10
15
20
25

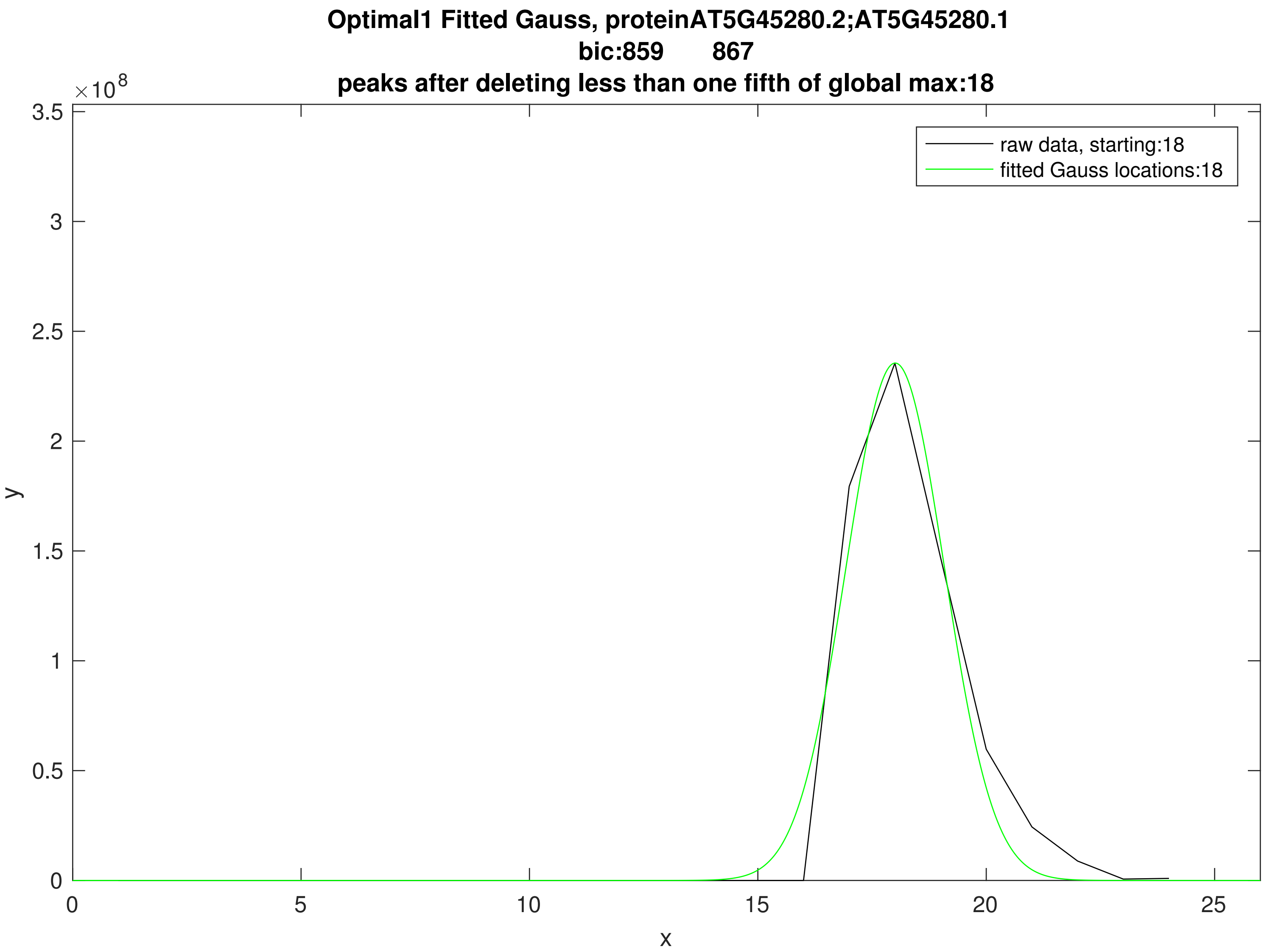
Optimal1 Fitted Gauss, proteinAT5G45280.2;AT5G45280.1
bic:859 867
peaks after deleting less than one fifth of global max:18
×10^8
3.5
3
2.5
2
1.5
1
0.5
0
y
0
5
10
15
20
25
x
raw data, starting:18
fitted Gauss locations:18

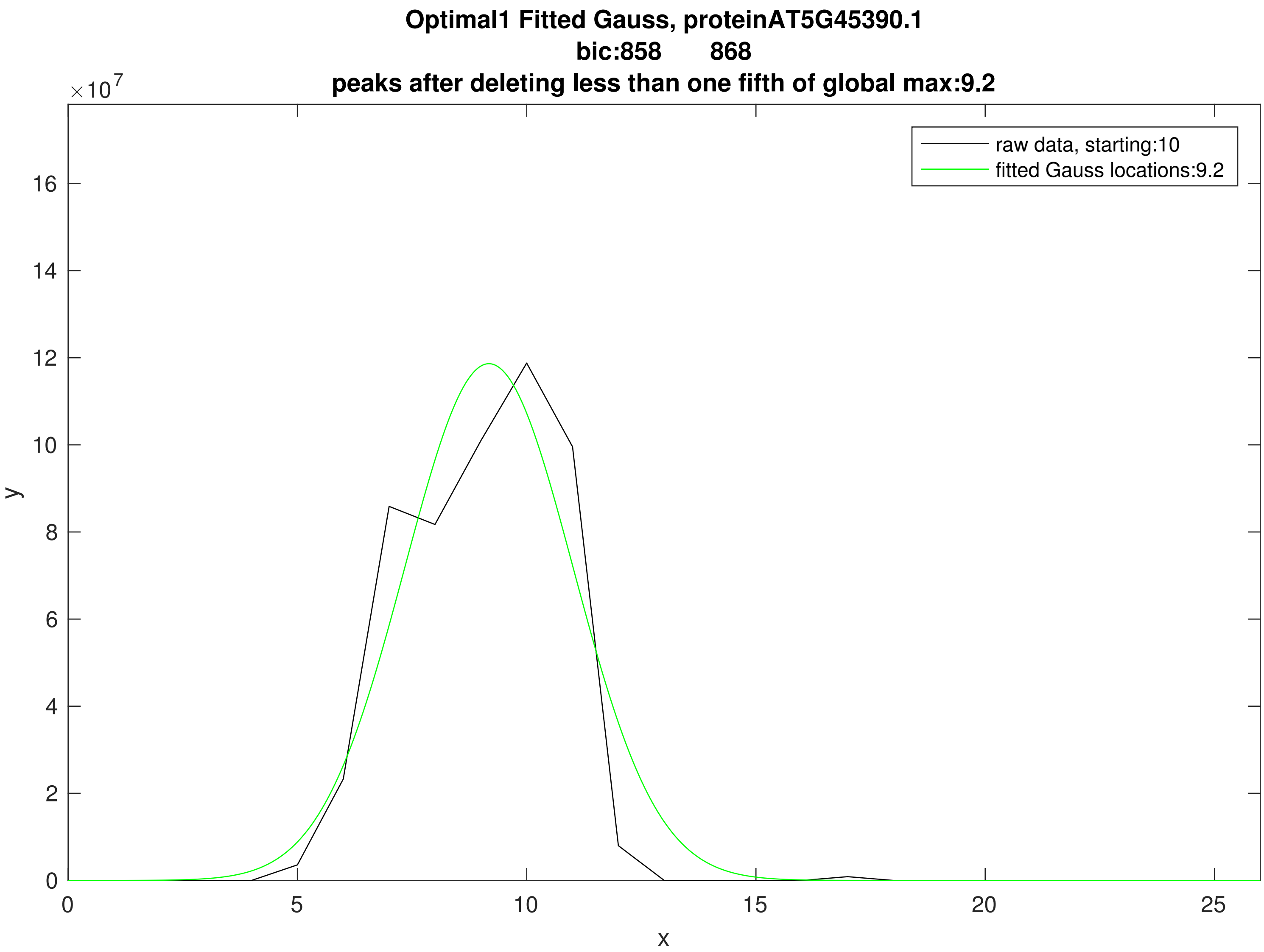
Optimal1 Fitted Gauss, proteinAT5G45390.1
bic:858 868
peaks after deleting less than one fifth of global max:9.2
×10^7
raw data, starting:10
fitted Gauss locations:9.2
y
x
0
2
4
6
8
10
12
14
16
0
5
10
15
20
25

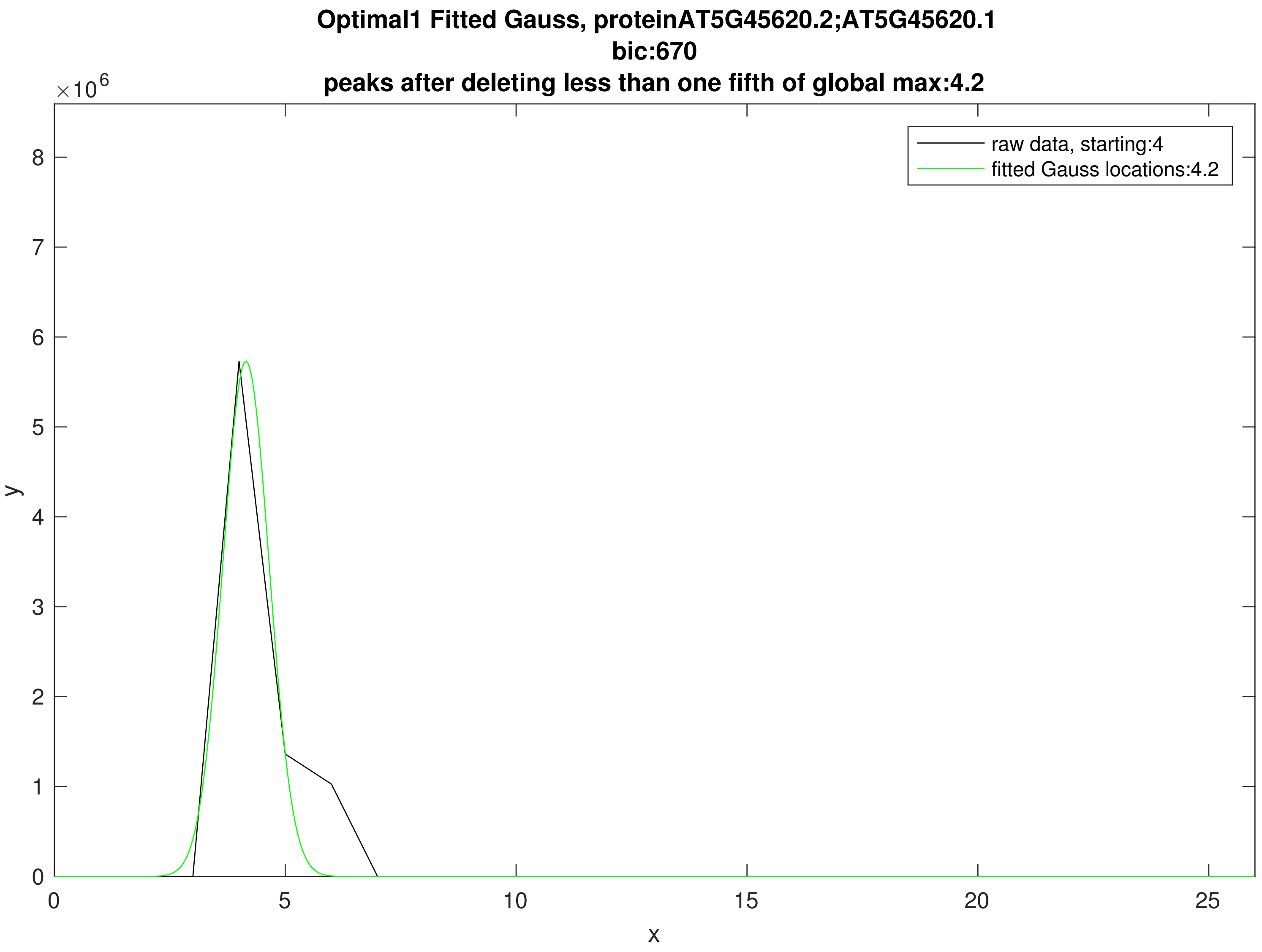
Optimal1 Fitted Gauss, proteinAT5G45620.2;AT5G45620.1
bic:670
peaks after deleting less than one fifth of global max:4.2
×10^6
raw data, starting:4
fitted Gauss locations:4.2
y
x
0
1
2
3
4
5
6
7
8
0
5
10
15
20
25

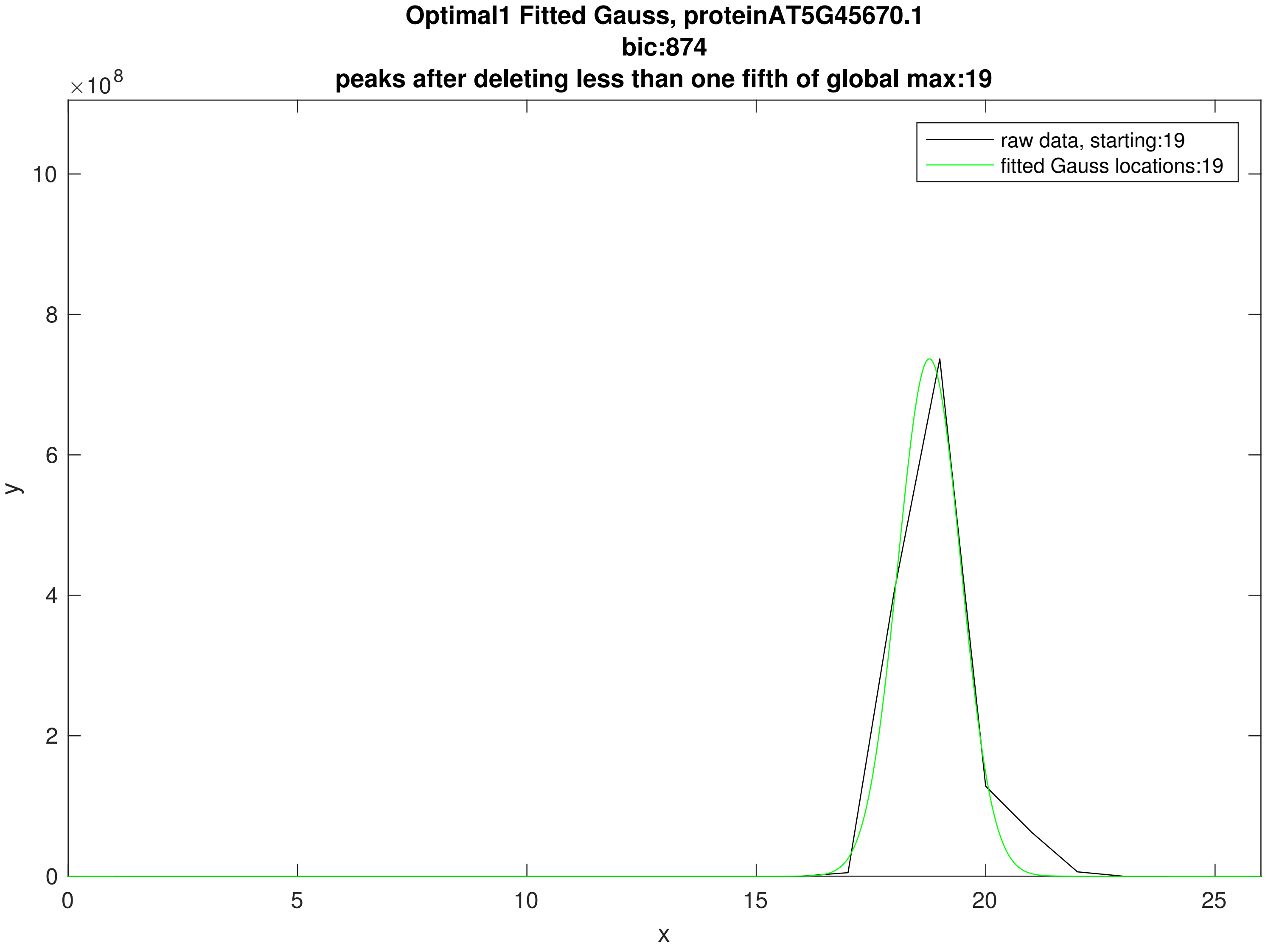
Optimal1 Fitted Gauss, proteinAT5G45670.1
bic:874
peaks after deleting less than one fifth of global max:19
×10^8
raw data, starting:19
fitted Gauss locations:19
y
x
0
2
4
6
8
10
0
5
10
15
20
25

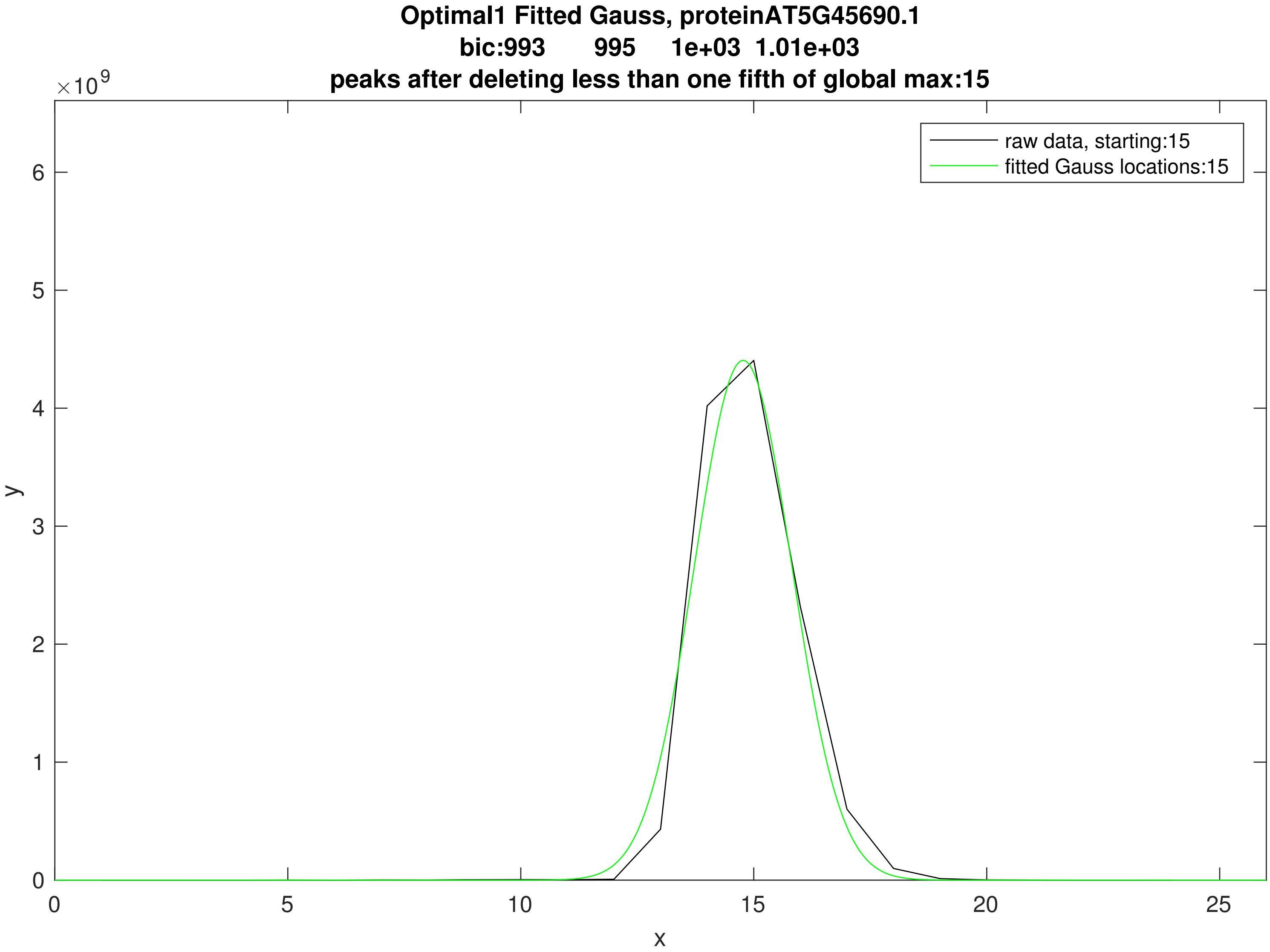

Optimal1 Fitted Gauss, proteinAT5G45690.1
bic:993 995 1e+03 1.01e+03
peaks after deleting less than one fifth of global max:15
×10^9
raw data, starting:15
fitted Gauss locations:15
y
x
0
1
2
3
4
5
6
0
5
10
15
20
25

**Optimal1 Fitted Gauss, proteinAT5G45775.1;AT5G45775.2;AT4G18730.1;AT3G58700.1;AT2G42740.1**
**bic:793**
**peaks after deleting less than one fifth of global max:3.7**

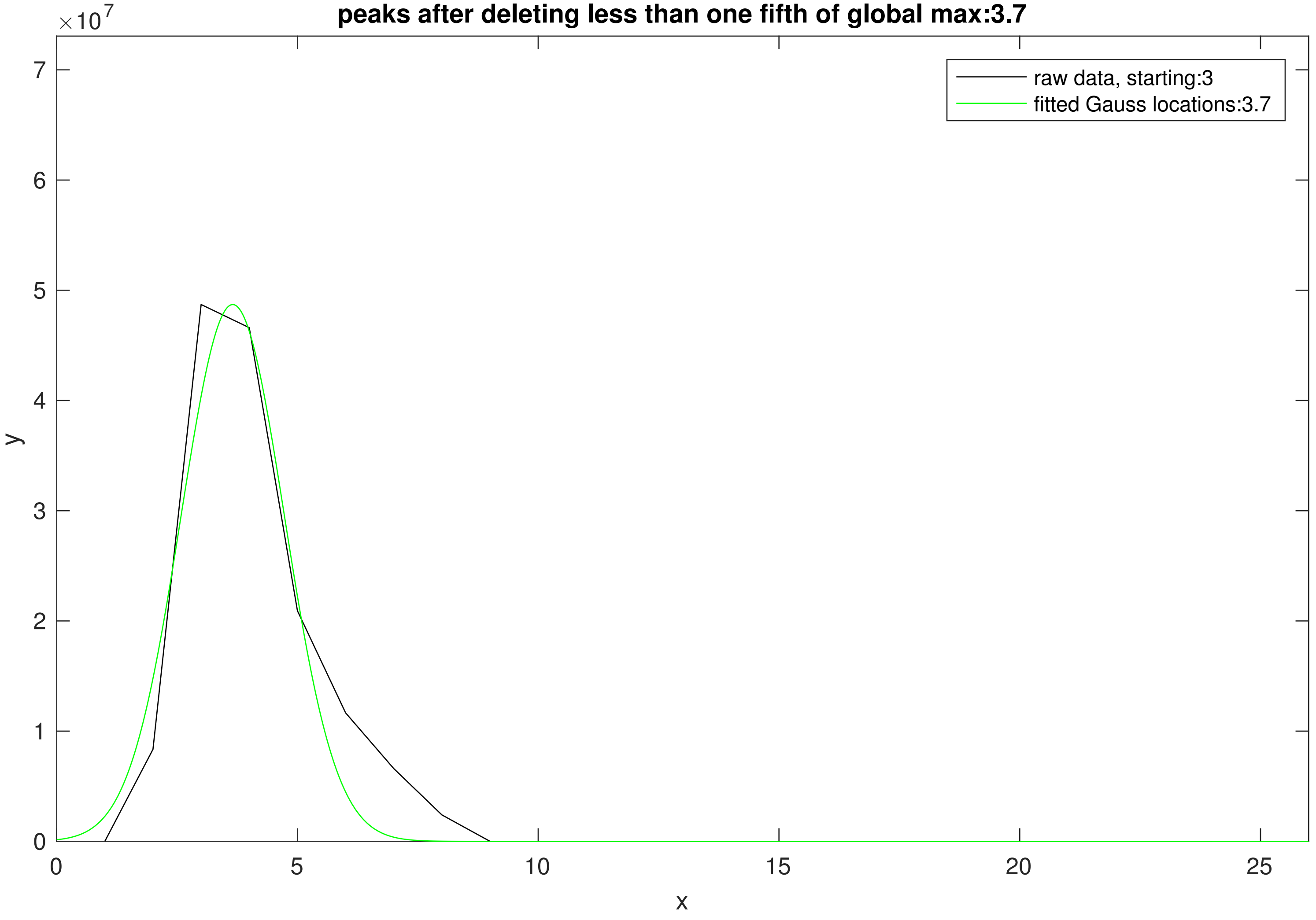

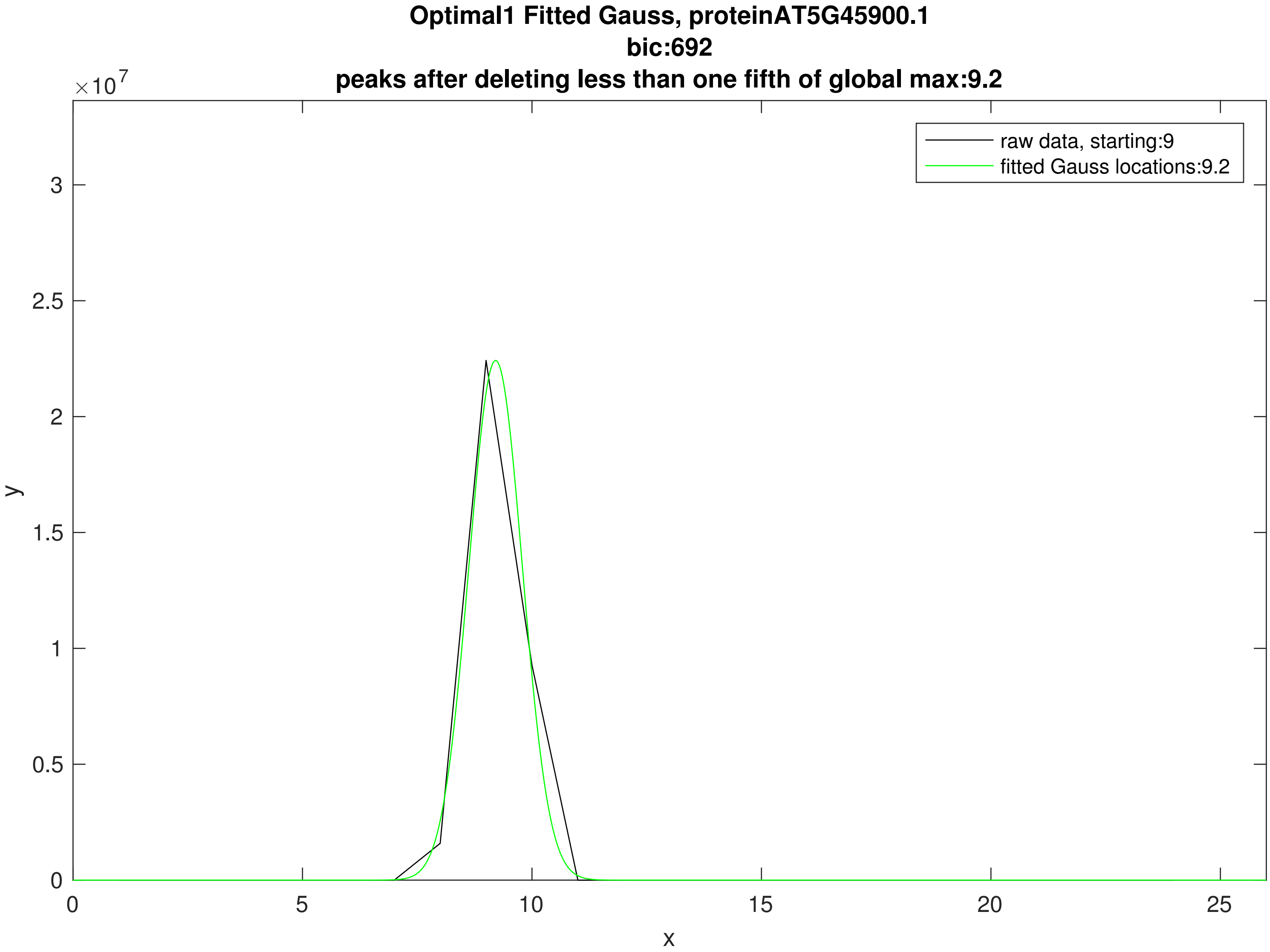
Optimal1 Fitted Gauss, proteinAT5G45900.1
bic:692
peaks after deleting less than one fifth of global max:9.2
×10^7
raw data, starting:9
fitted Gauss locations:9.2
y
x
0
0.5
1
1.5
2
2.5
3
0
5
10
15
20
25

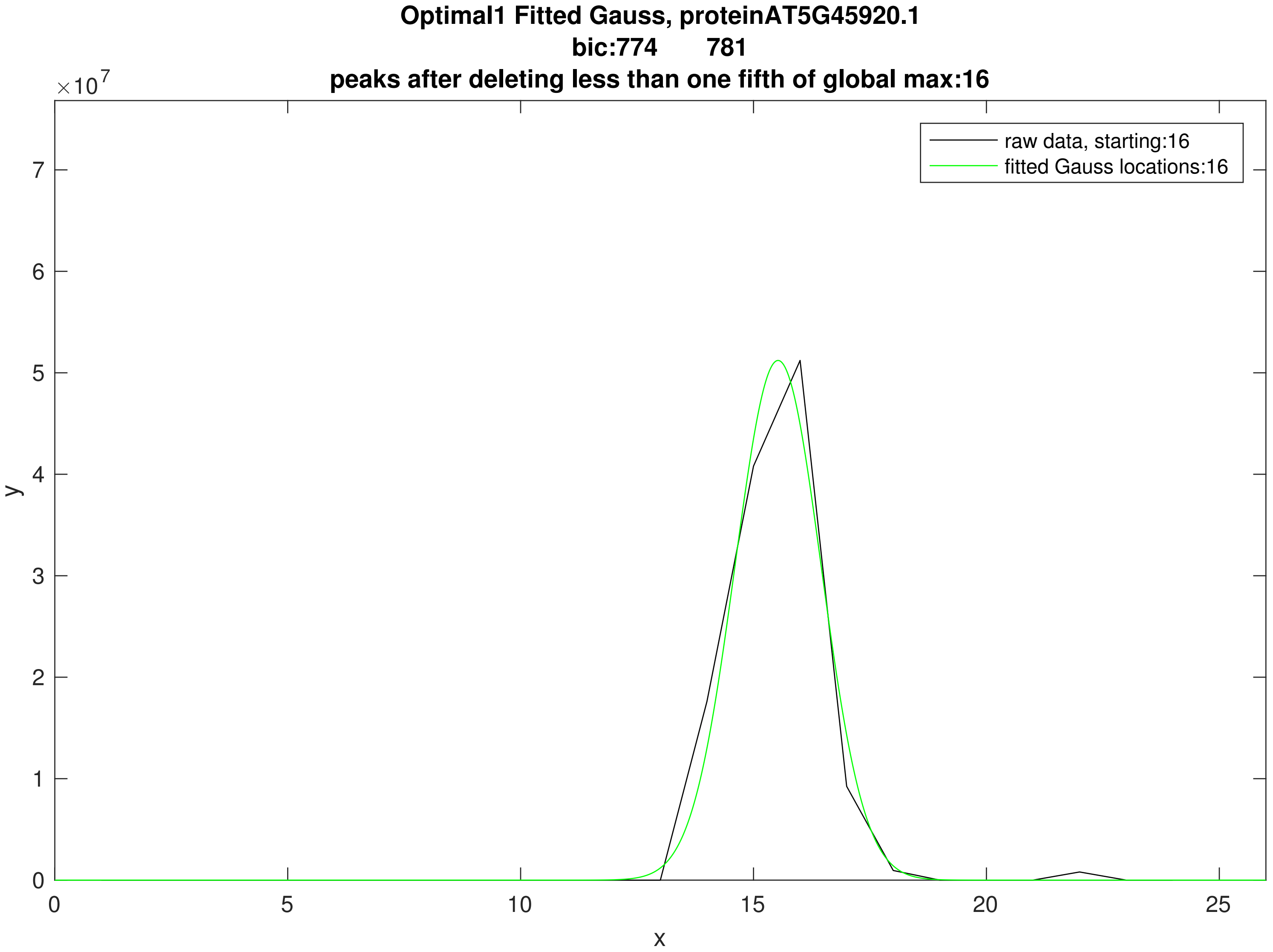
Optimal1 Fitted Gauss, proteinAT5G45920.1
bic:774 781
peaks after deleting less than one fifth of global max:16
×10^7
raw data, starting:16
fitted Gauss locations:16
y
x
0
1
2
3
4
5
6
7
0
5
10
15
20
25

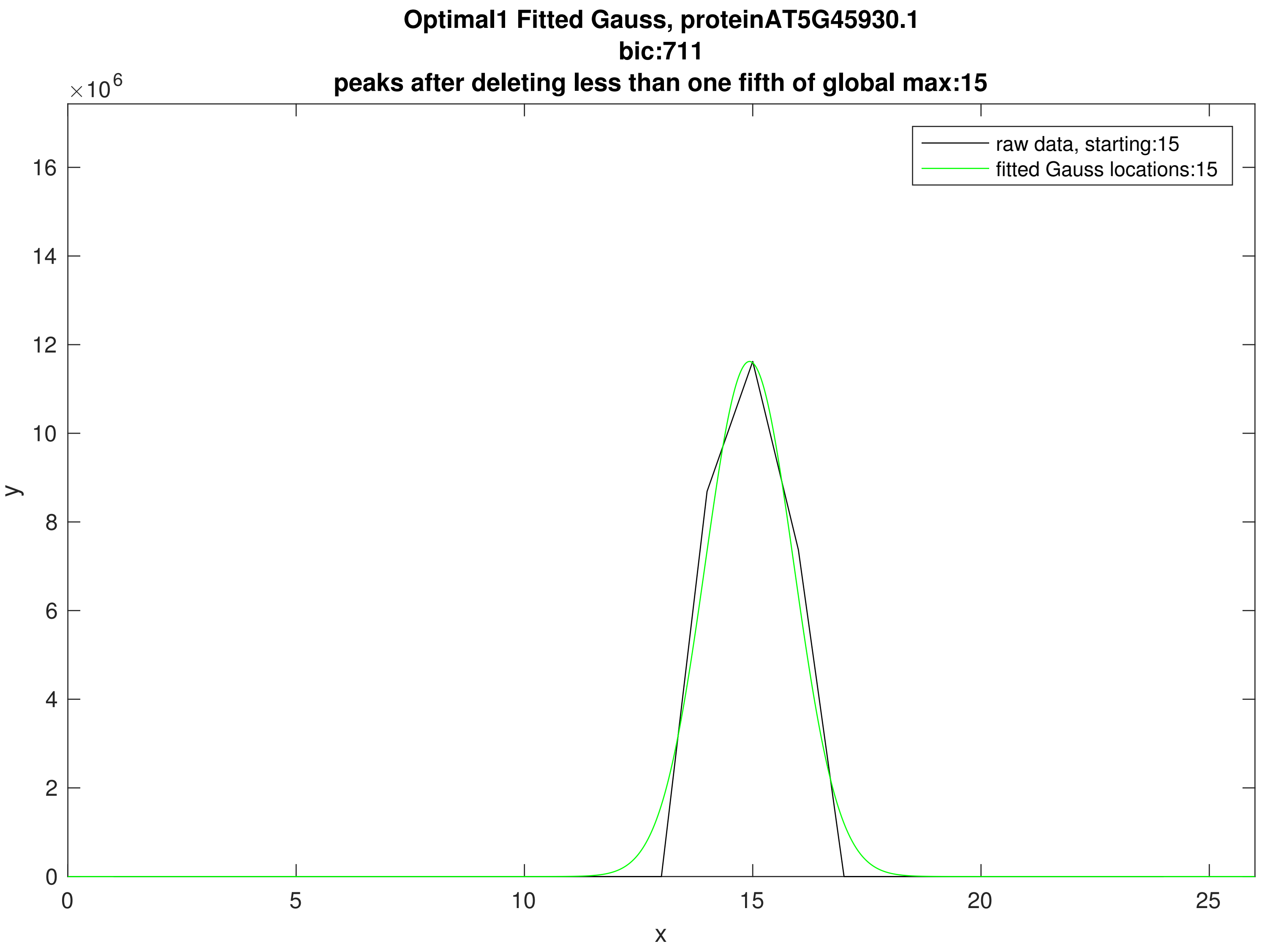

Optimal1 Fitted Gauss, proteinAT5G45930.1
bic:711
peaks after deleting less than one fifth of global max:15
×10^6
raw data, starting:15
fitted Gauss locations:15
16
14
12
10
8
6
4
2
0
y
0
5
10
15
20
25
x

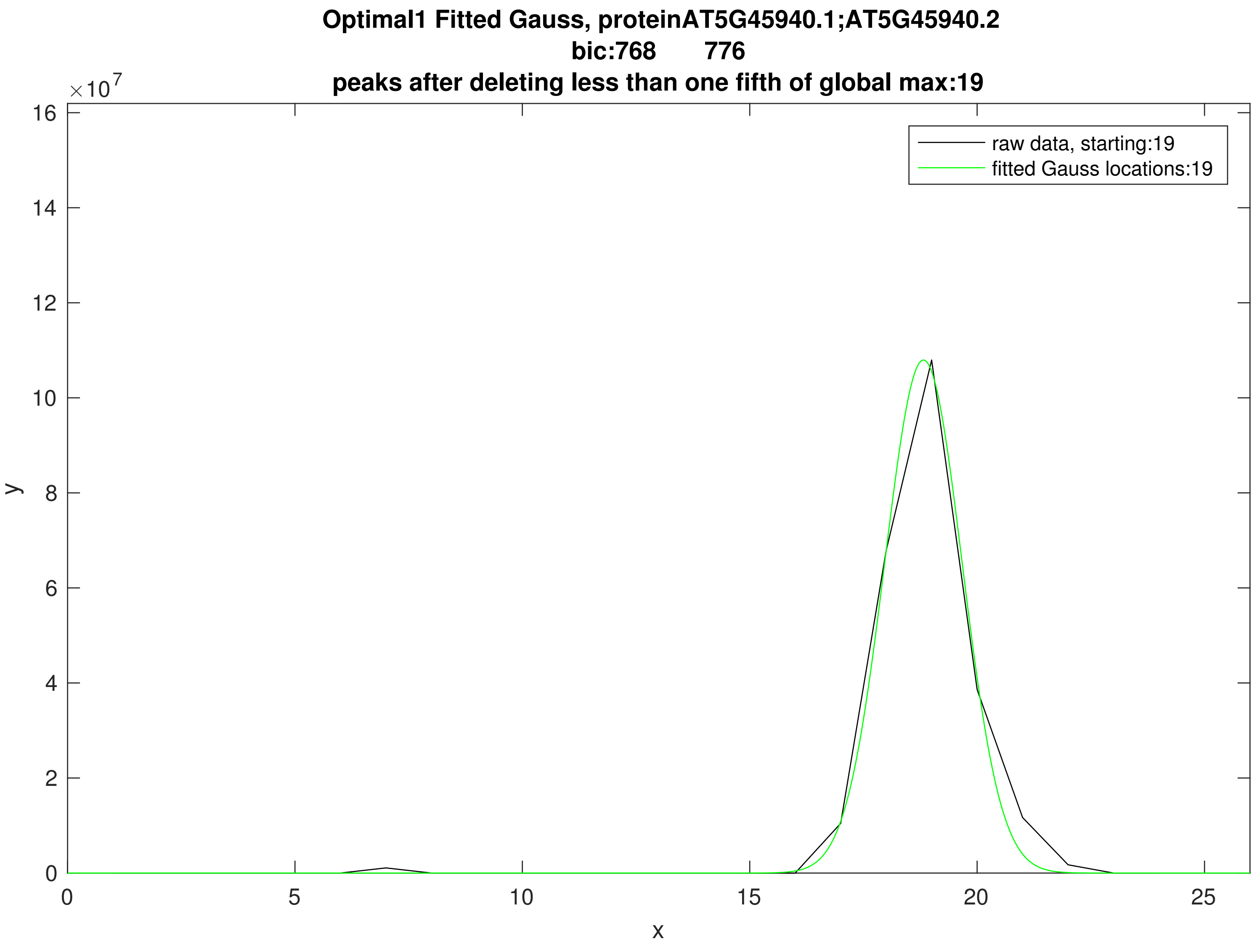
Optimal1 Fitted Gauss, proteinAT5G45940.1;AT5G45940.2
bic:768 776
peaks after deleting less than one fifth of global max:19
×10^7
raw data, starting:19
fitted Gauss locations:19
y
x
0
2
4
6
8
10
12
14
16
0
5
10
15
20
25

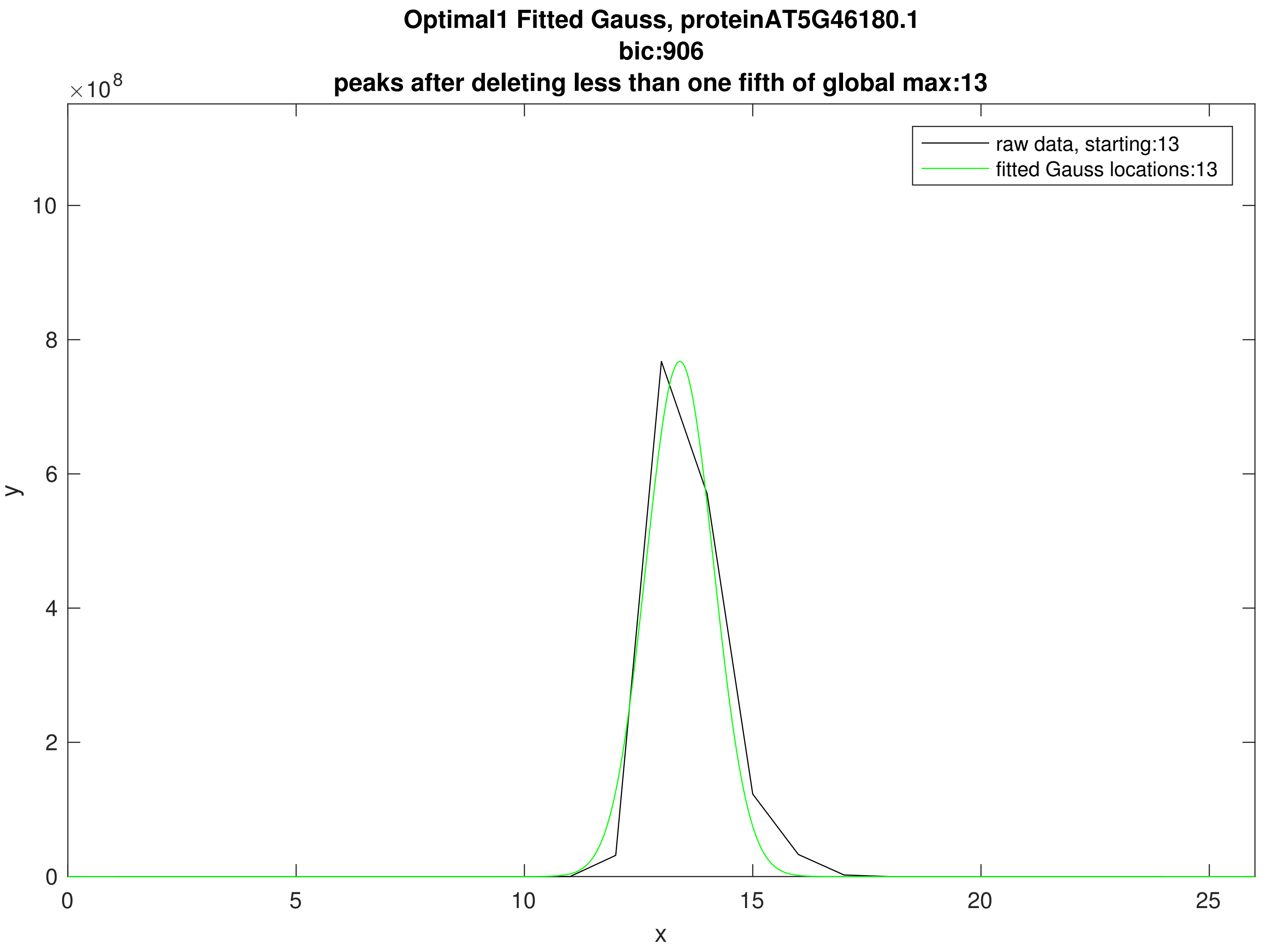

Optimal1 Fitted Gauss, proteinAT5G46180.1
bic:906
peaks after deleting less than one fifth of global max:13
×10^8
raw data, starting:13
fitted Gauss locations:13
y
x
0
2
4
6
8
10
0
5
10
15
20
25

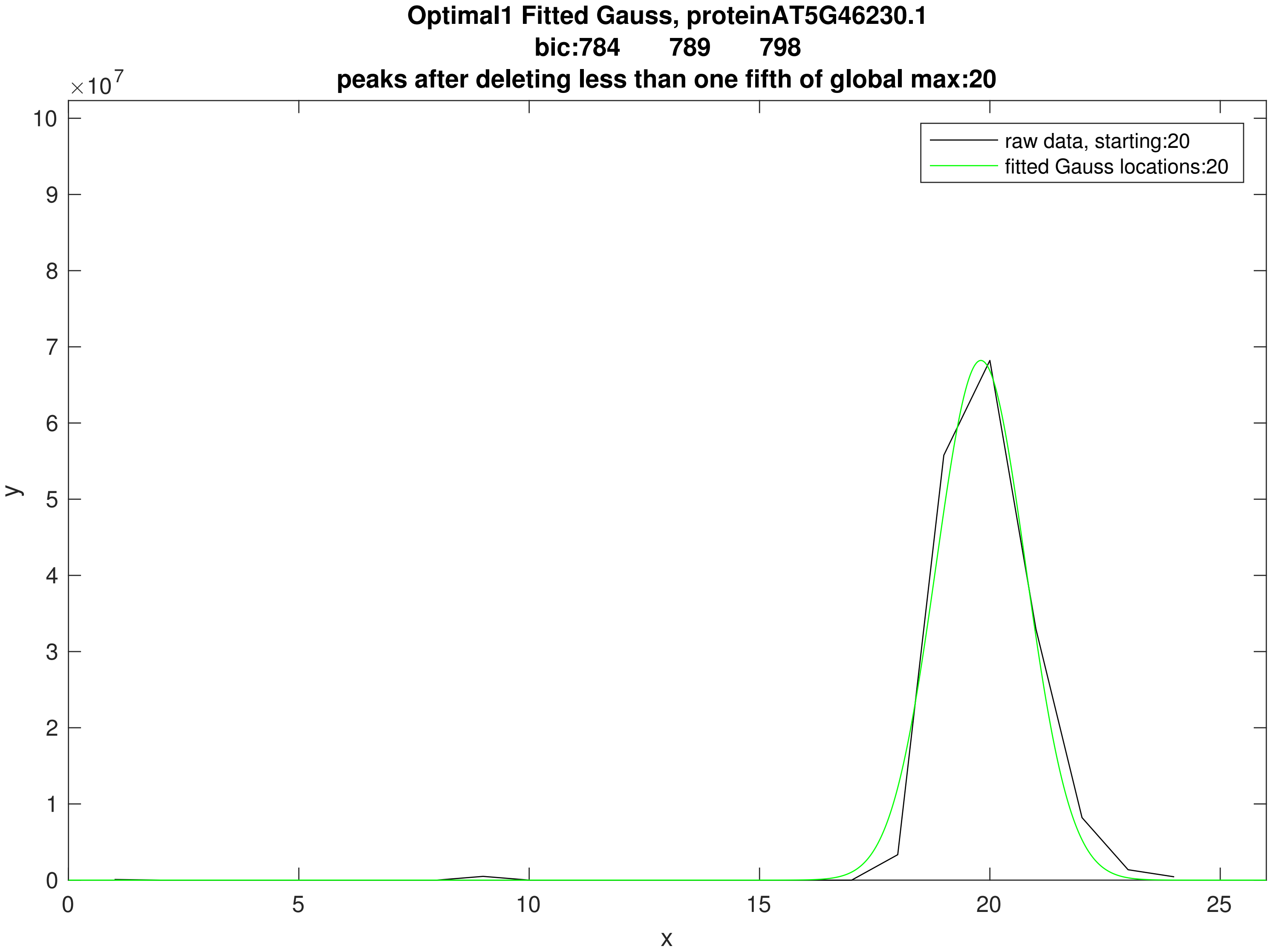

Optimal1 Fitted Gauss, proteinAT5G46230.1
bic:784 789 798
peaks after deleting less than one fifth of global max:20
×10^7
raw data, starting:20
fitted Gauss locations:20
y
x
0
1
2
3
4
5
6
7
8
9
10
0
5
10
15
20
25

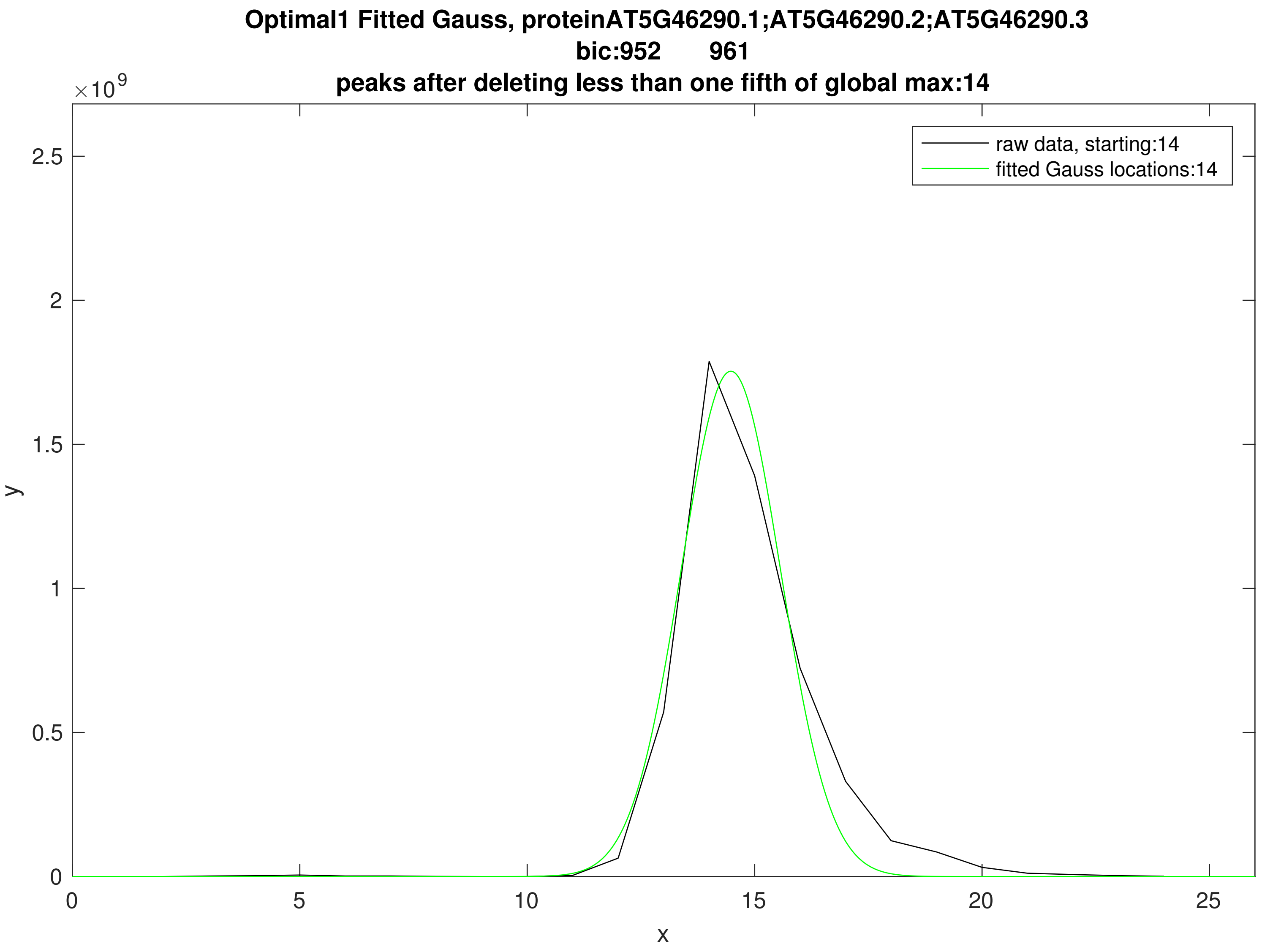
Optimal1 Fitted Gauss, proteinAT5G46290.1;AT5G46290.2;AT5G46290.3
bic:952 961
peaks after deleting less than one fifth of global max:14
×10^9
2.5
2
1.5
1
0.5
0
y
0
5
10
15
20
25
x
raw data, starting:14
fitted Gauss locations:14

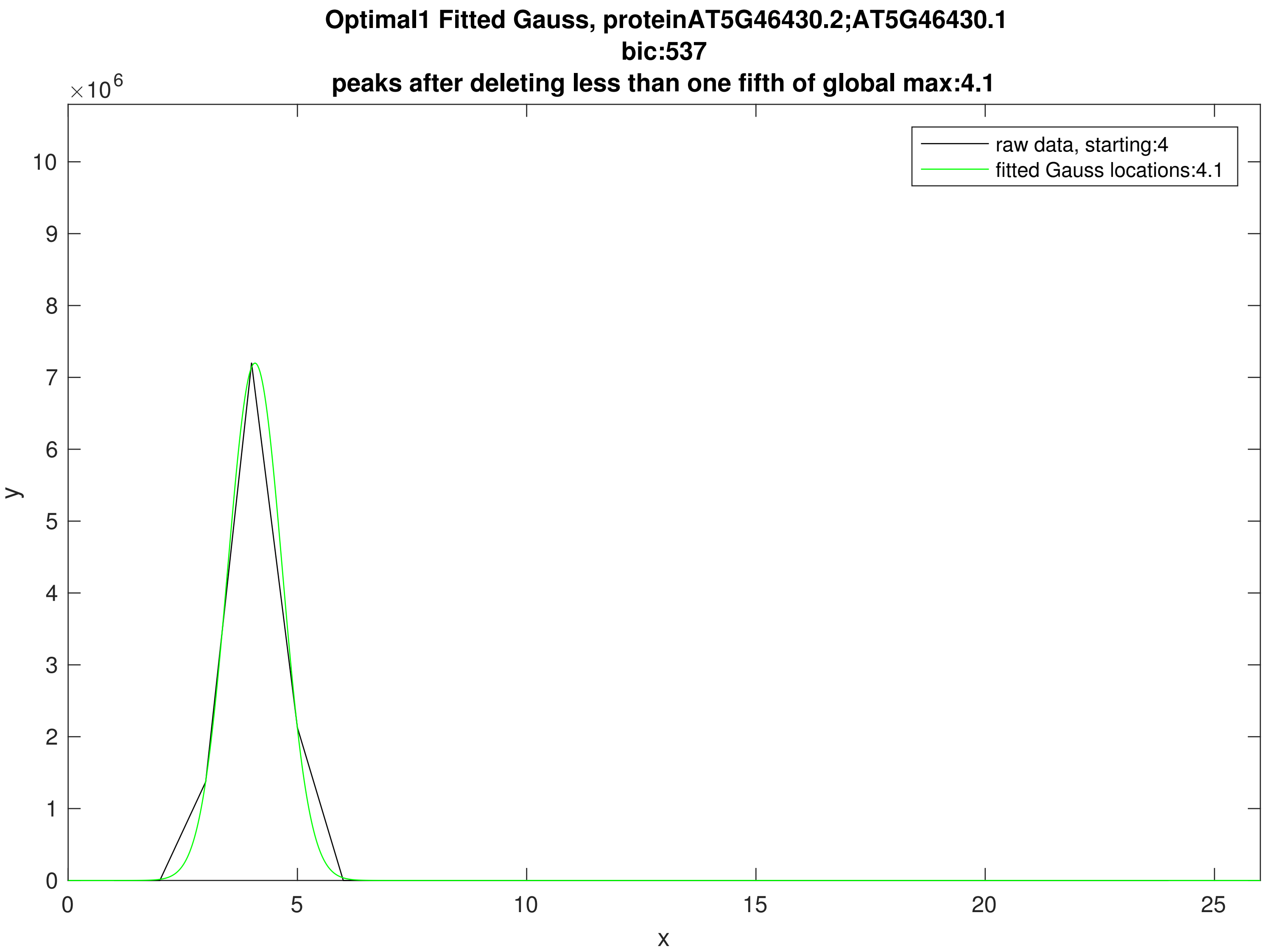

Optimal1 Fitted Gauss, proteinAT5G46430.2;AT5G46430.1
bic:537
peaks after deleting less than one fifth of global max:4.1
×10^6
10
9
8
7
6
5
4
3
2
1
0
y
0
5
10
15
20
25
x
raw data, starting:4
fitted Gauss locations:4.1

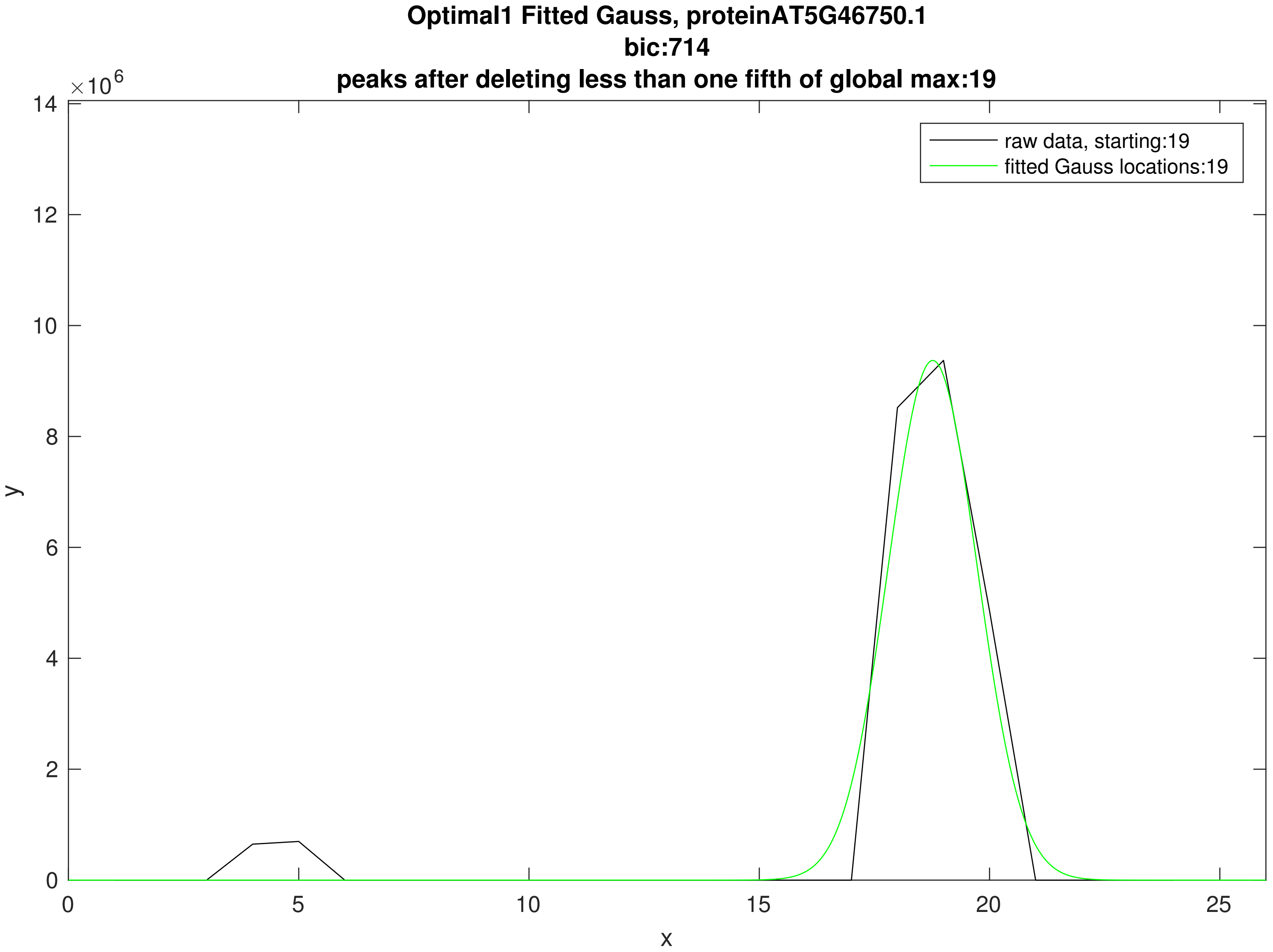

Optimal1 Fitted Gauss, proteinAT5G46750.1
bic:714
peaks after deleting less than one fifth of global max:19
×10^6
14
12
10
8
6
4
2
0
y
0
5
10
15
20
25
x
raw data, starting:19
fitted Gauss locations:19

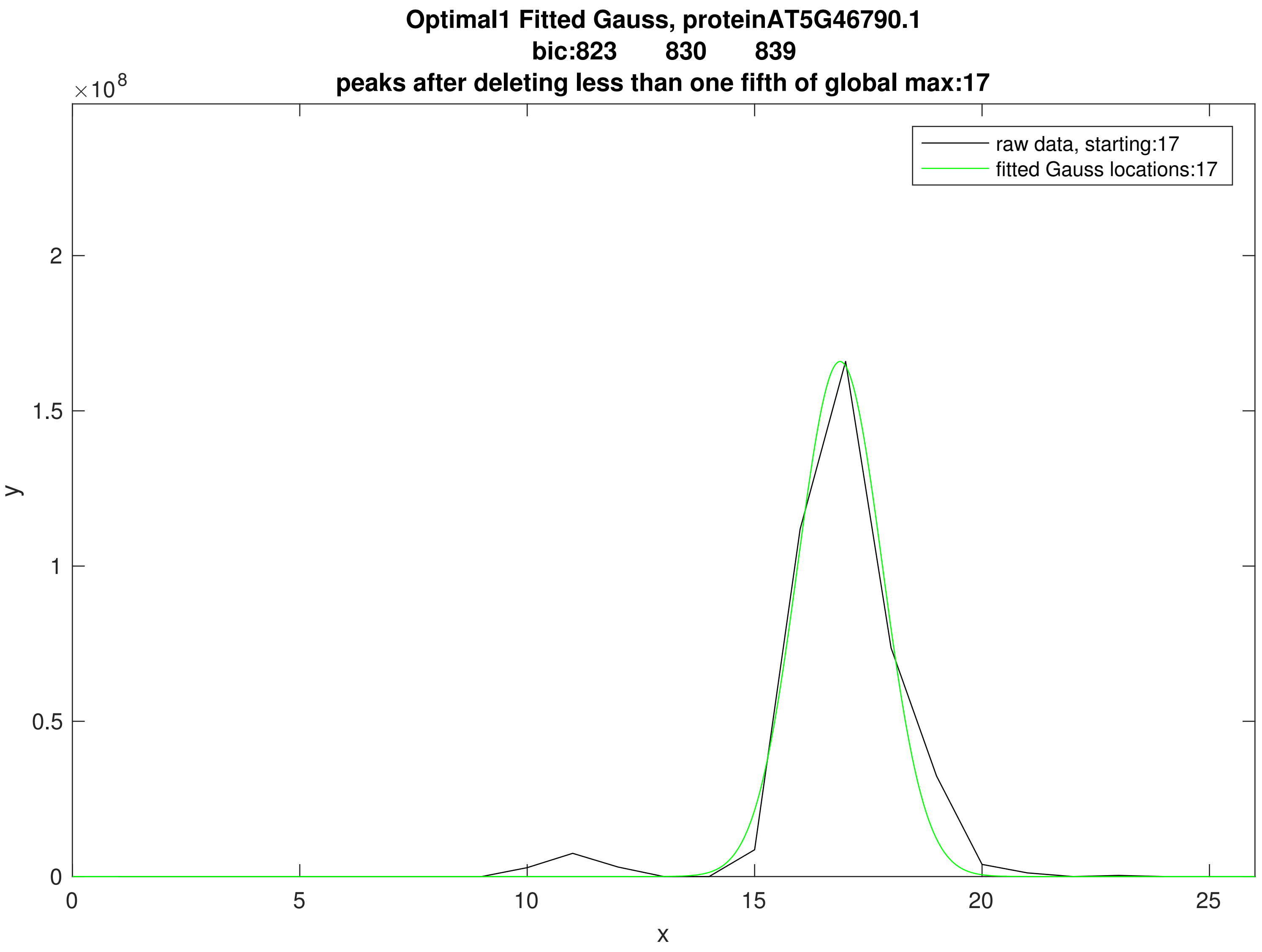
Optimal1 Fitted Gauss, proteinAT5G46790.1
bic:823 830 839
peaks after deleting less than one fifth of global max:17
×10^8
raw data, starting:17
fitted Gauss locations:17
2
1.5
1
0.5
0
y
0
5
10
15
20
25
x

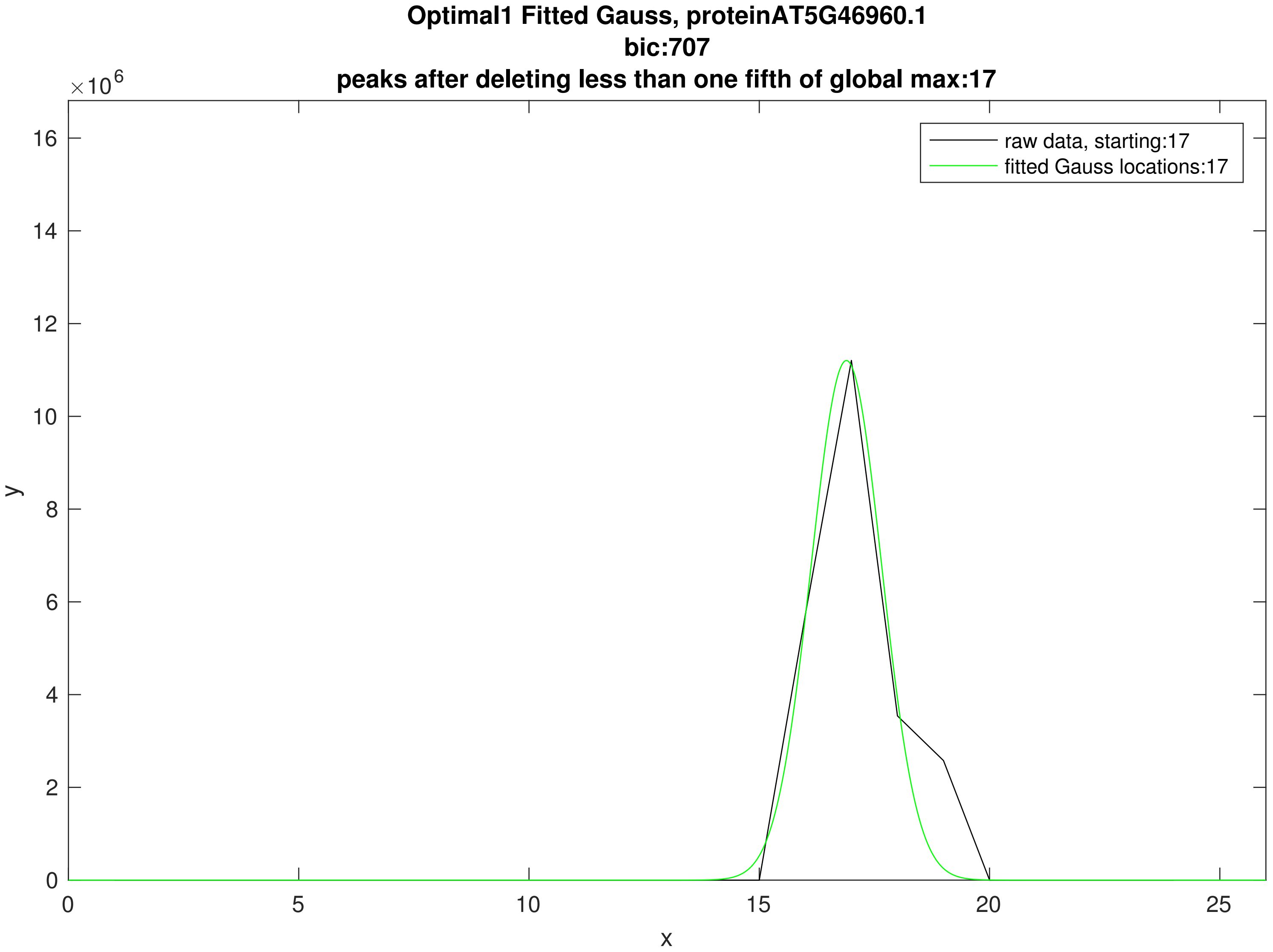
Optimal1 Fitted Gauss, proteinAT5G46960.1
bic:707
peaks after deleting less than one fifth of global max:17
×10^6
raw data, starting:17
fitted Gauss locations:17
y
x
0
2
4
6
8
10
12
14
16
0
5
10
15
20
25

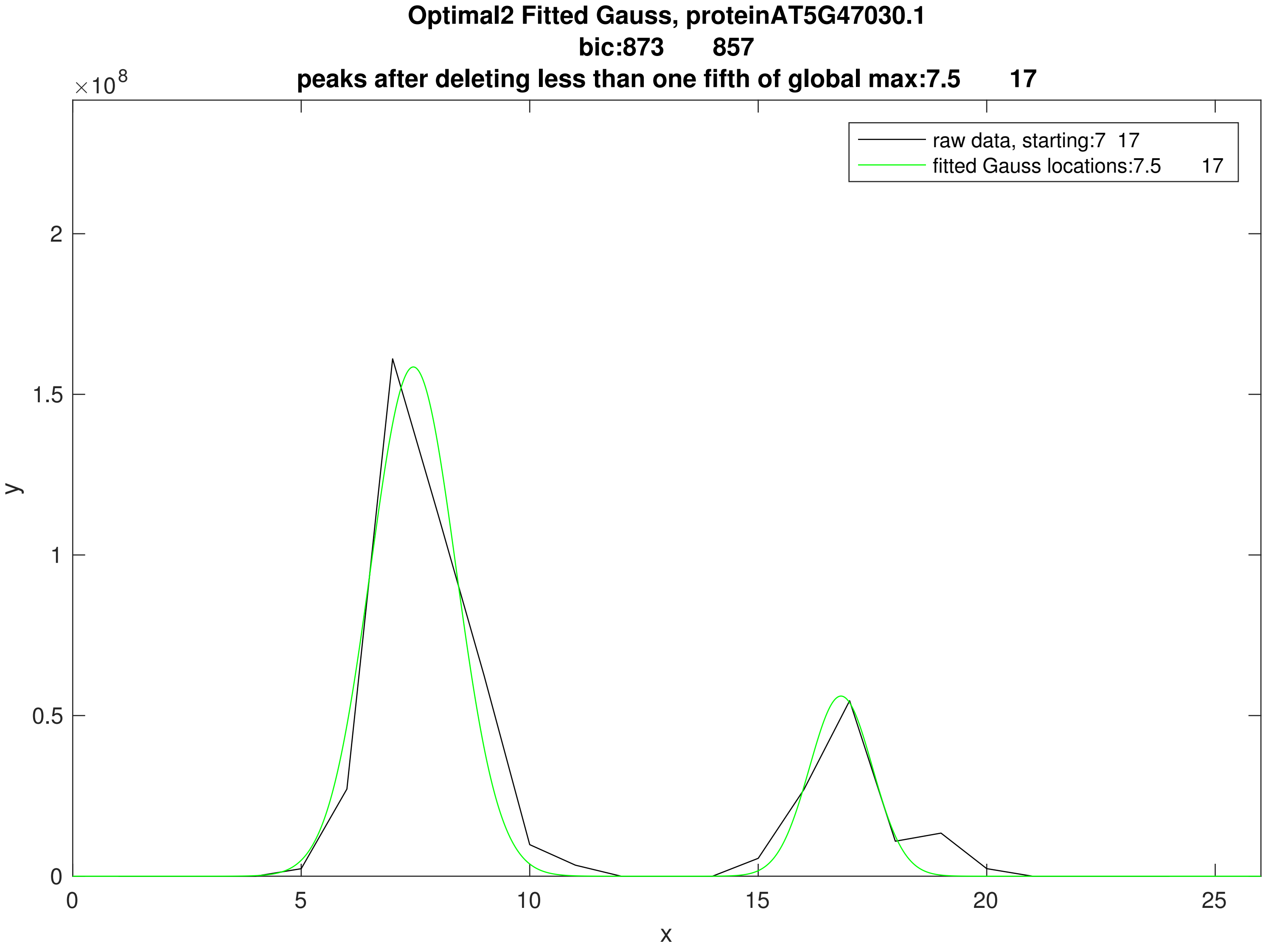
Optimal2 Fitted Gauss, proteinAT5G47030.1
bic:873 857
peaks after deleting less than one fifth of global max:7.5 17
×10^8
raw data, starting:7 17
fitted Gauss locations:7.5 17
y
x
0
0.5
1
1.5
2
0
5
10
15
20
25

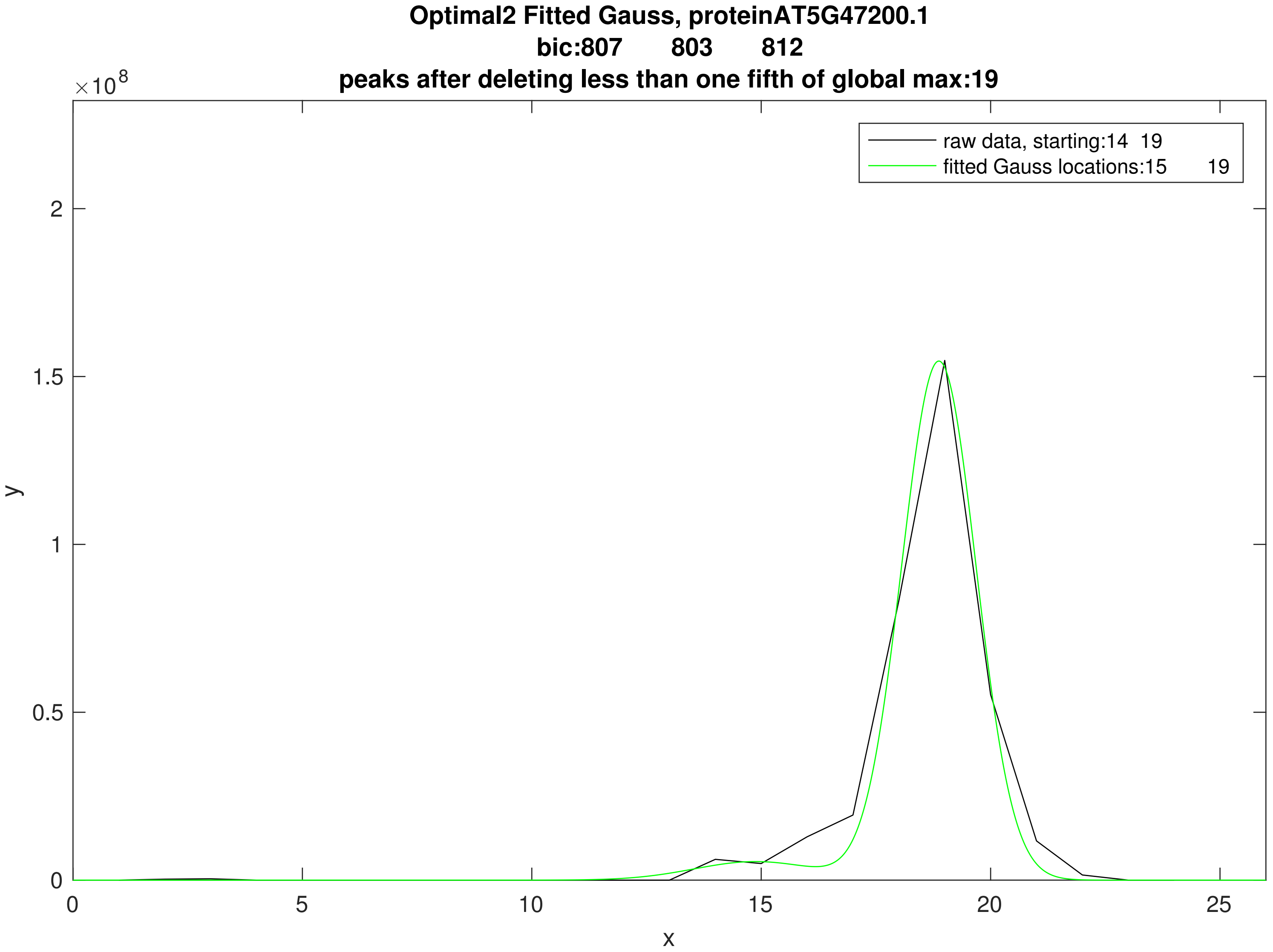
Optimal2 Fitted Gauss, proteinAT5G47200.1
bic:807 803 812
peaks after deleting less than one fifth of global max:19
×10^8
raw data, starting:14 19
fitted Gauss locations:15 19
y
x
0
0.5
1
1.5
2
0
5
10
15
20
25

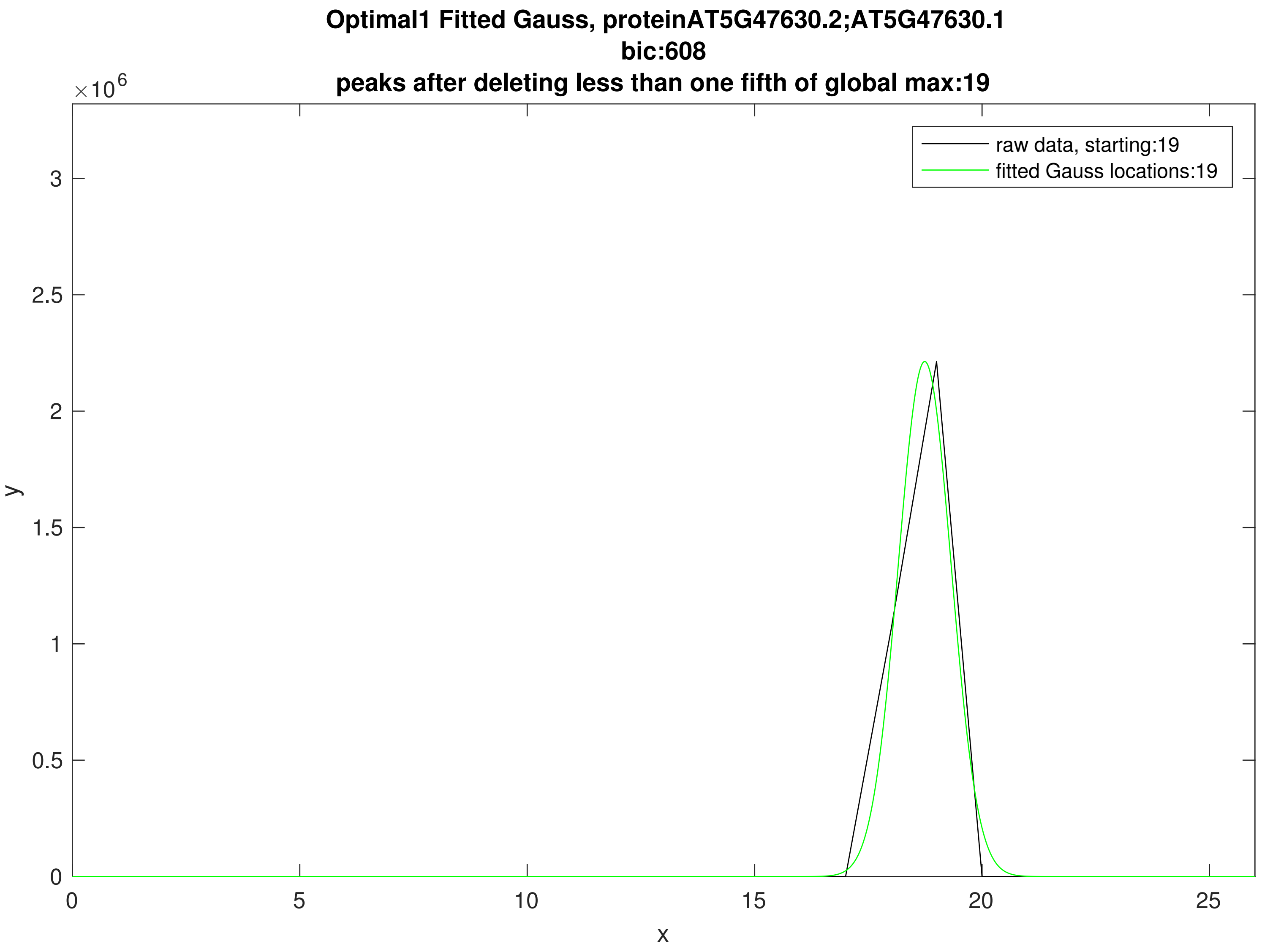
Optimal1 Fitted Gauss, proteinAT5G47630.2;AT5G47630.1
bic:608
peaks after deleting less than one fifth of global max:19
×10^6
raw data, starting:19
fitted Gauss locations:19
y
x
0
0.5
1
1.5
2
2.5
3
0
5
10
15
20
25

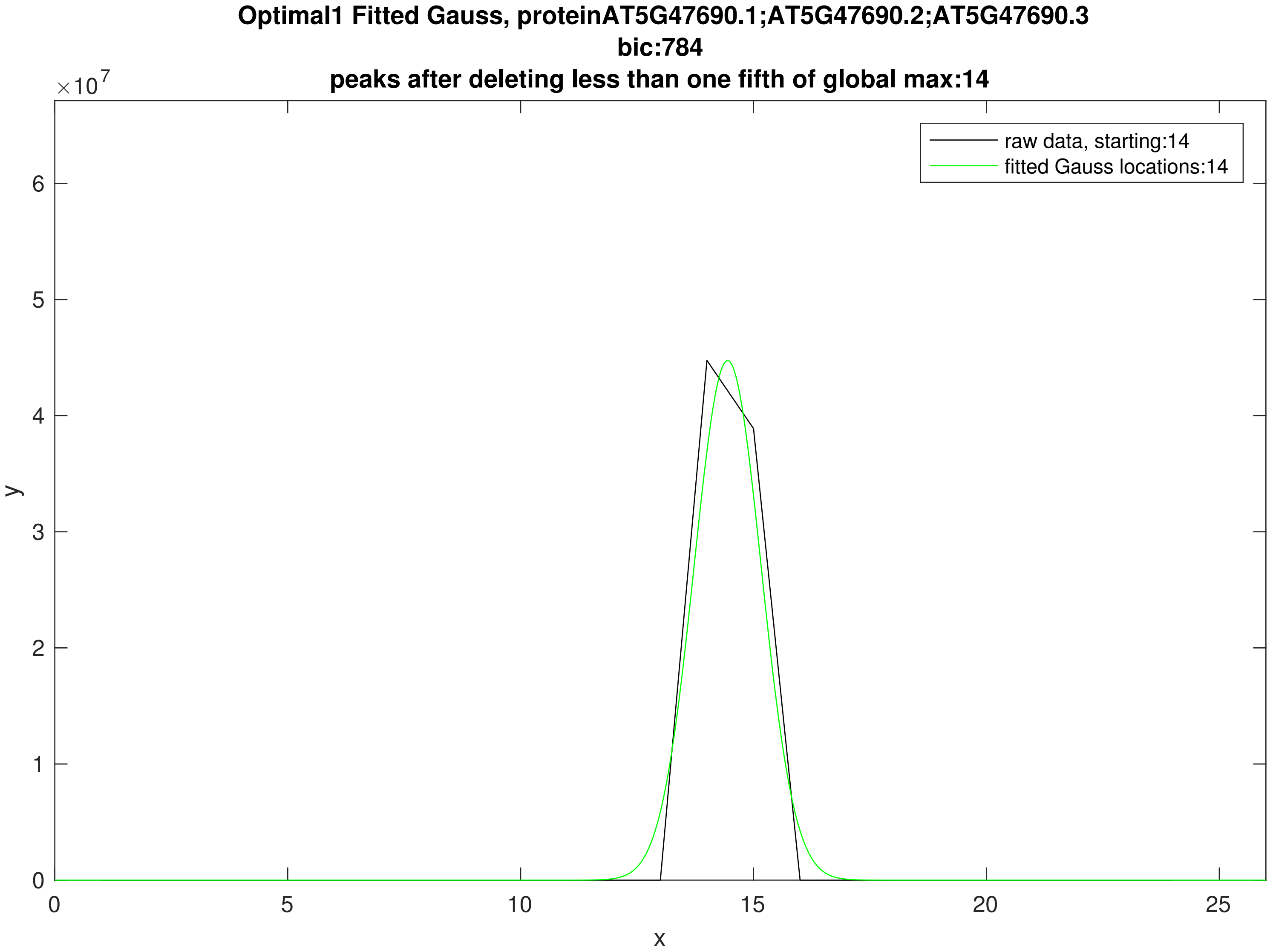
Optimal1 Fitted Gauss, proteinAT5G47690.1;AT5G47690.2;AT5G47690.3
bic:784
peaks after deleting less than one fifth of global max:14
×10^7
raw data, starting:14
fitted Gauss locations:14
y
x
0
1
2
3
4
5
6
0
5
10
15
20
25

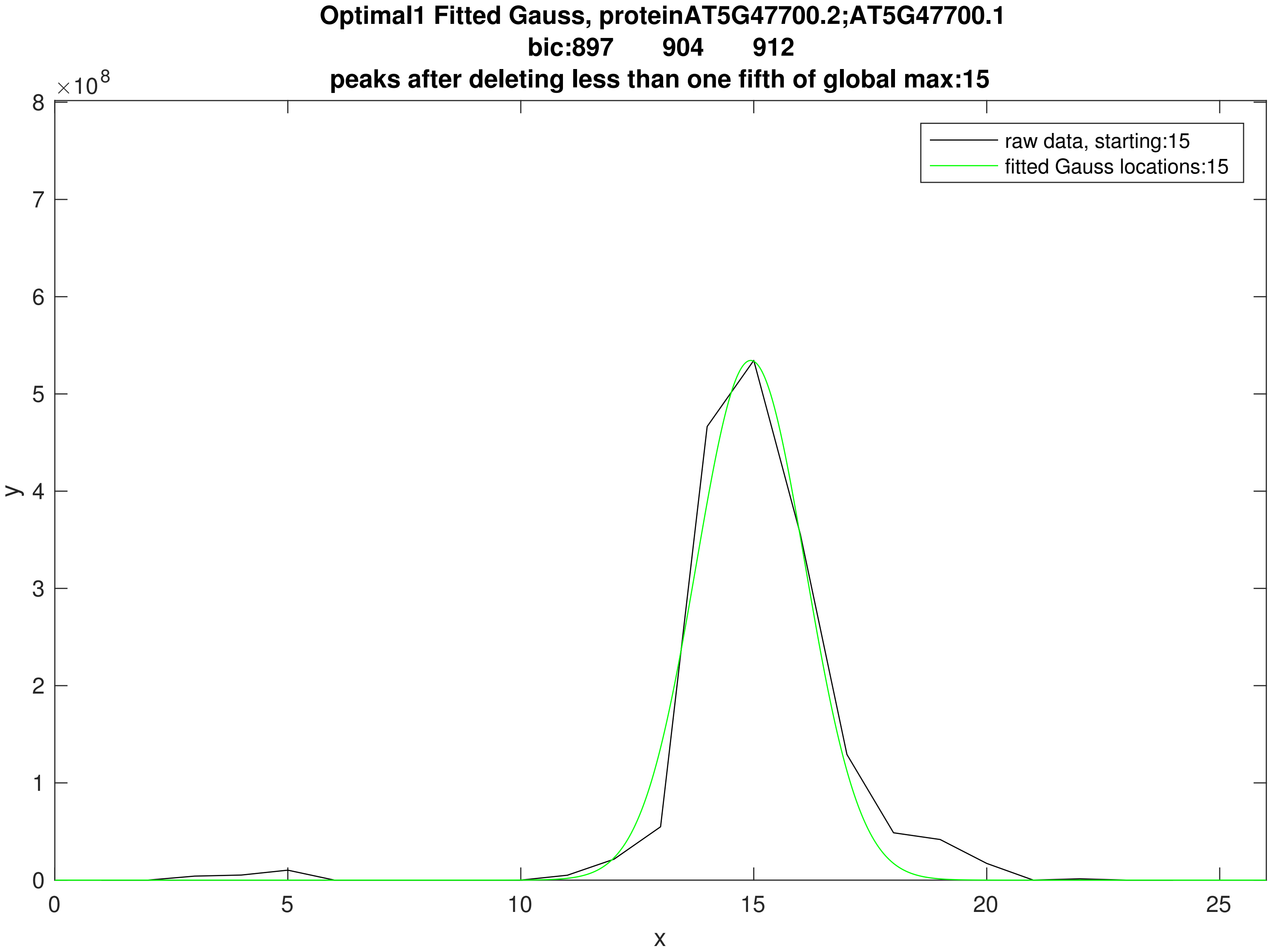

Optimal1 Fitted Gauss, proteinAT5G47700.2;AT5G47700.1
bic:897 904 912
peaks after deleting less than one fifth of global max:15
×10^8
raw data, starting:15
fitted Gauss locations:15
y
x
0
1
2
3
4
5
6
7
8
0
5
10
15
20
25

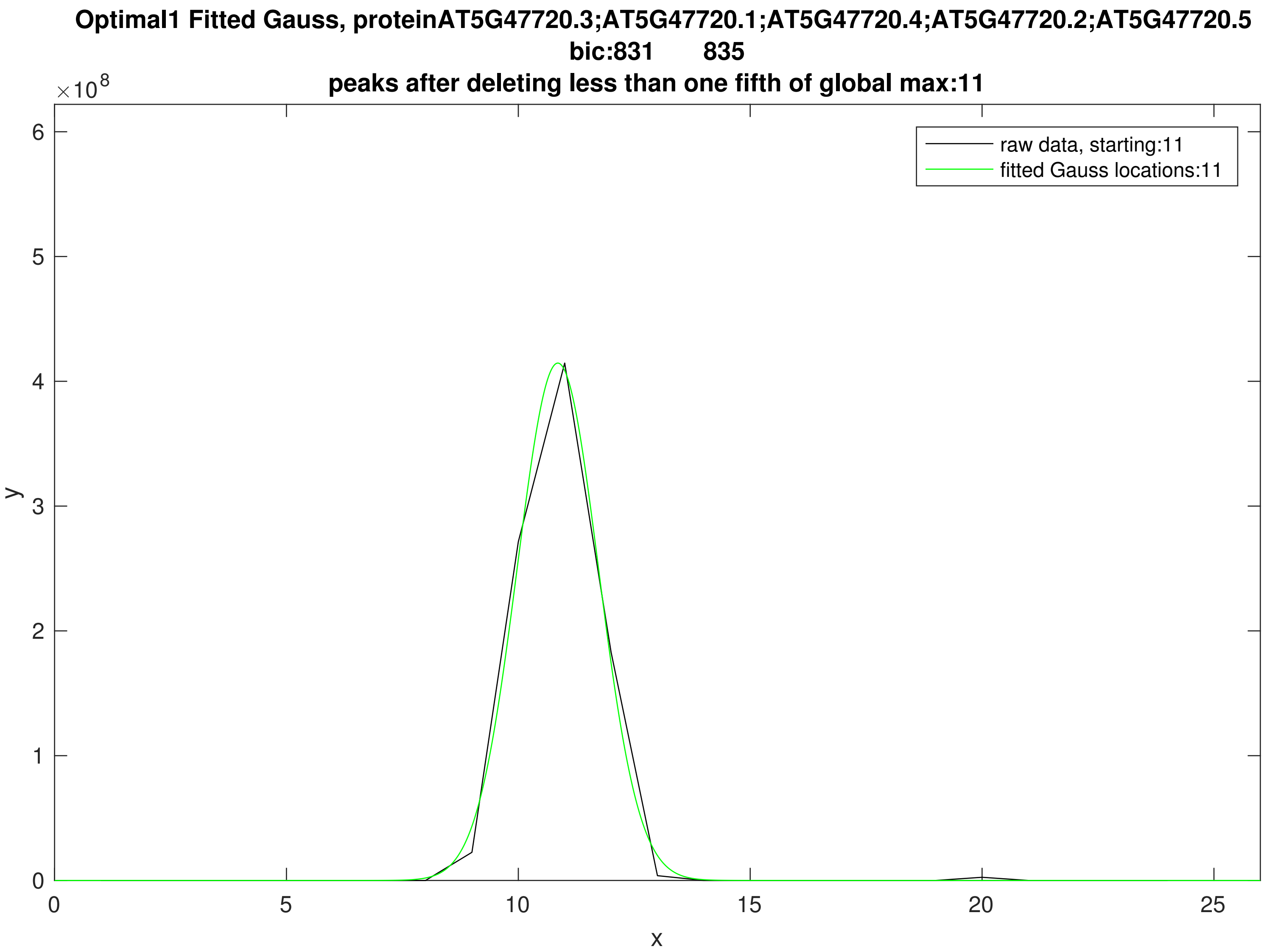
Optimal1 Fitted Gauss, proteinAT5G47720.3;AT5G47720.1;AT5G47720.4;AT5G47720.2;AT5G47720.5
bic:831 835
peaks after deleting less than one fifth of global max:11
×10^8
raw data, starting:11
fitted Gauss locations:11
y
x
0
1
2
3
4
5
6
0
5
10
15
20
25

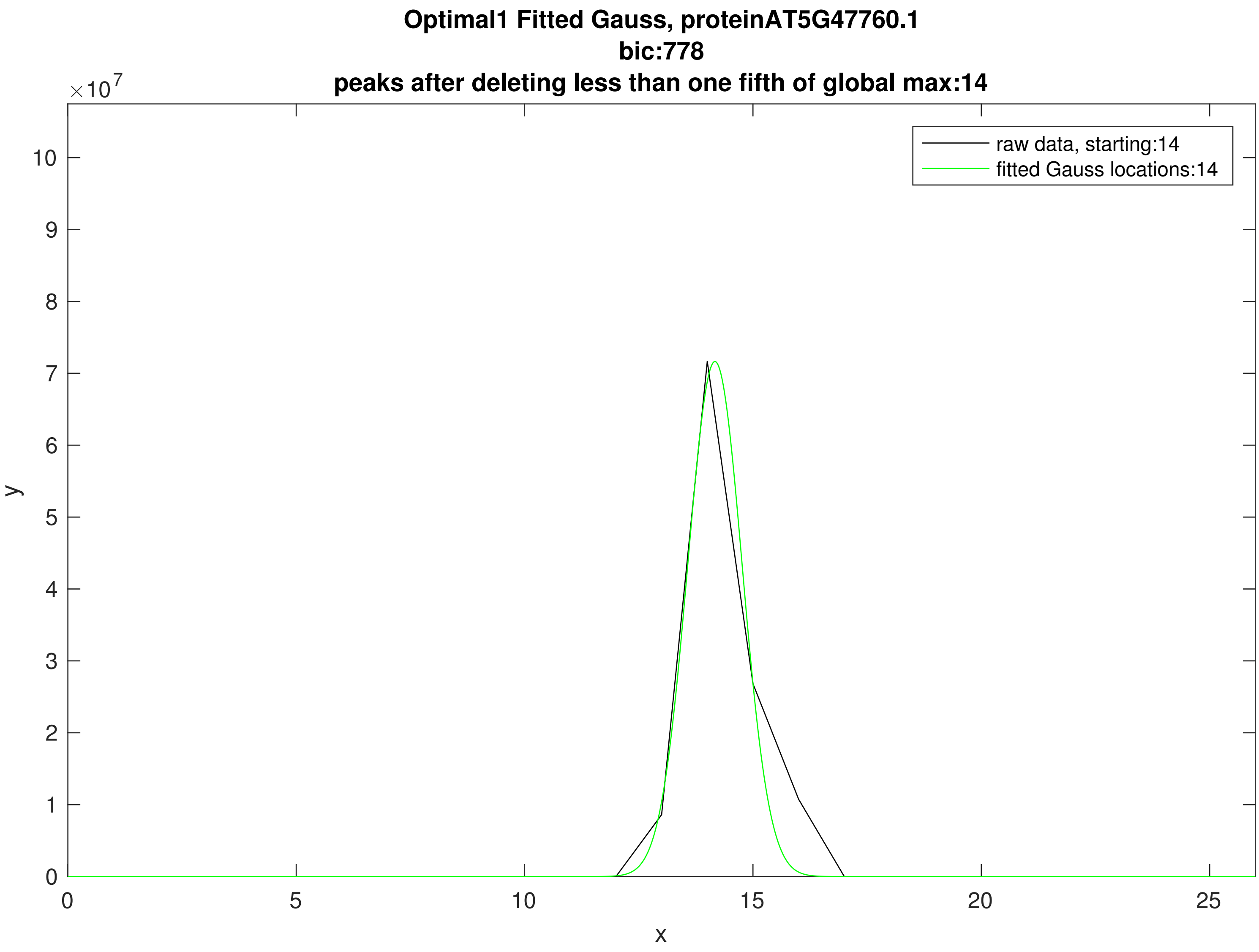

Optimal1 Fitted Gauss, proteinAT5G47760.1
bic:778
peaks after deleting less than one fifth of global max:14
×10^7
raw data, starting:14
fitted Gauss locations:14
y
x
0
1
2
3
4
5
6
7
8
9
10
0
5
10
15
20
25

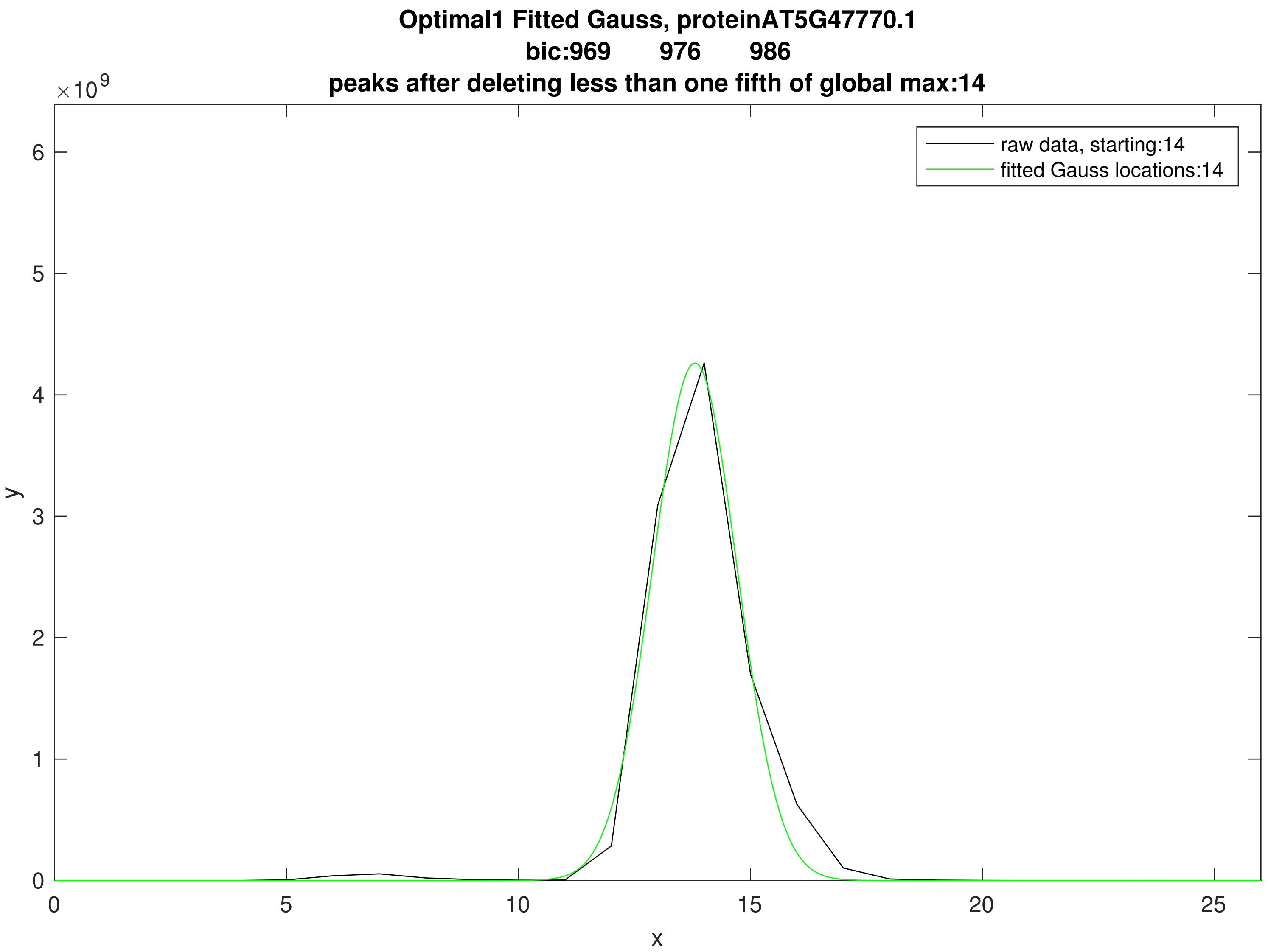
Optimal1 Fitted Gauss, proteinAT5G47770.1
bic:969 976 986
peaks after deleting less than one fifth of global max:14
×10^9
raw data, starting:14
fitted Gauss locations:14
y
x
0
1
2
3
4
5
6
0
5
10
15
20
25

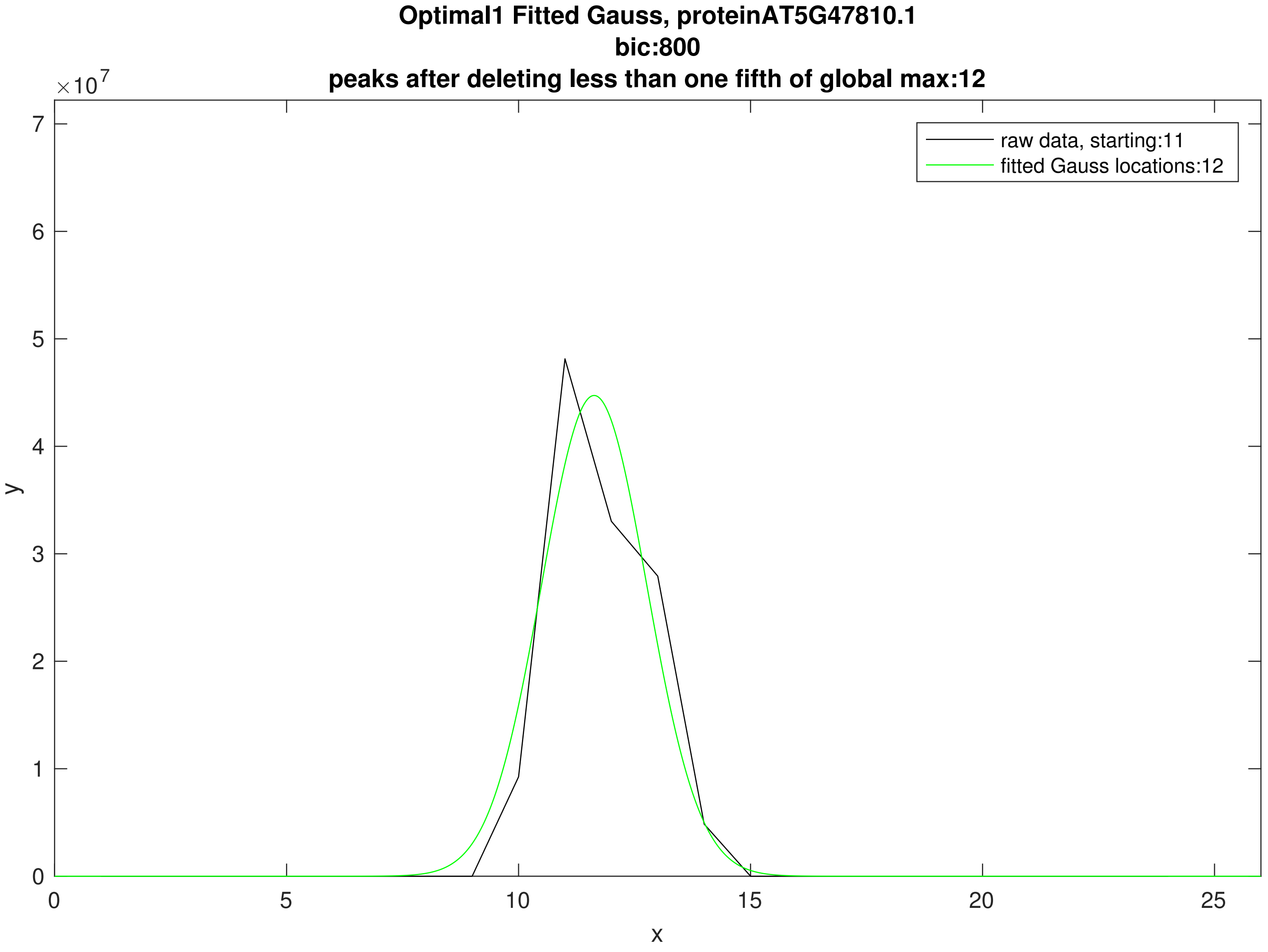

Optimal1 Fitted Gauss, proteinAT5G47810.1
bic:800
peaks after deleting less than one fifth of global max:12
×10^7
raw data, starting:11
fitted Gauss locations:12
y
x
0
1
2
3
4
5
6
7
0
5
10
15
20
25

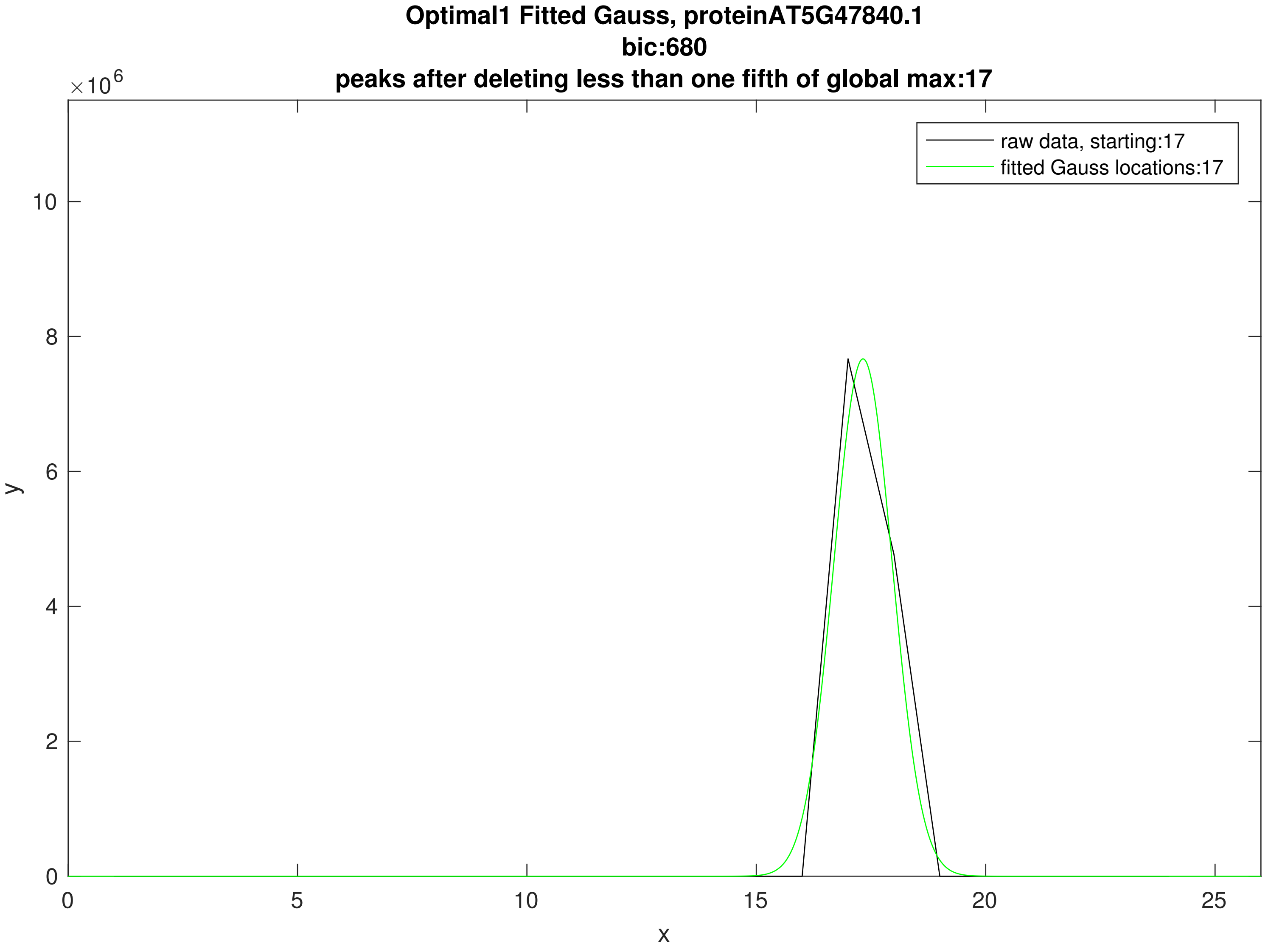
Optimal1 Fitted Gauss, proteinAT5G47840.1
bic:680
peaks after deleting less than one fifth of global max:17
×10^6
raw data, starting:17
fitted Gauss locations:17
10
8
6
4
2
0
y
0
5
10
15
20
25
x

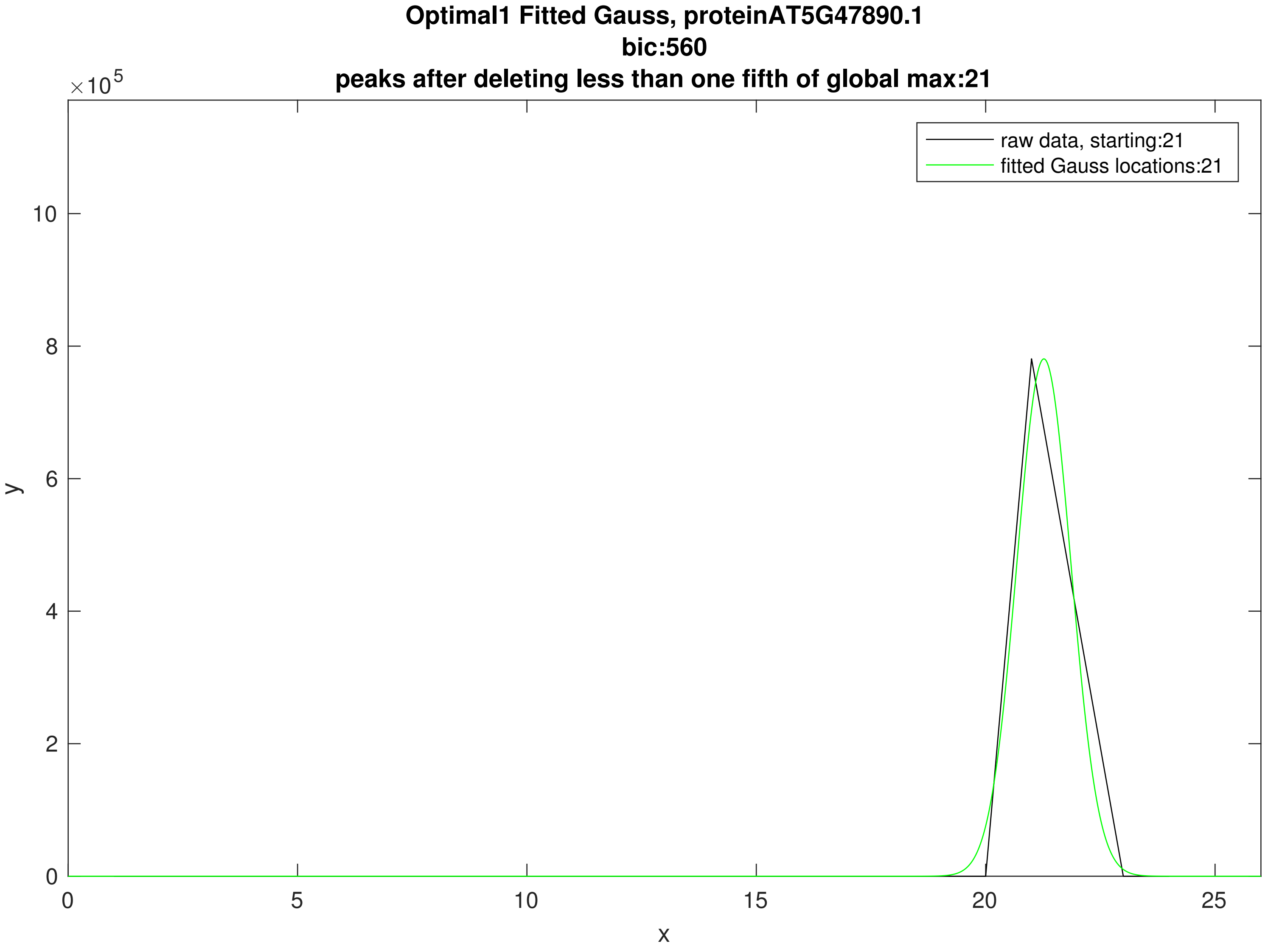

Optimal1 Fitted Gauss, proteinAT5G47890.1
bic:560
peaks after deleting less than one fifth of global max:21
×10^5
raw data, starting:21
fitted Gauss locations:21
10
8
6
4
2
0
y
0
5
10
15
20
25
x

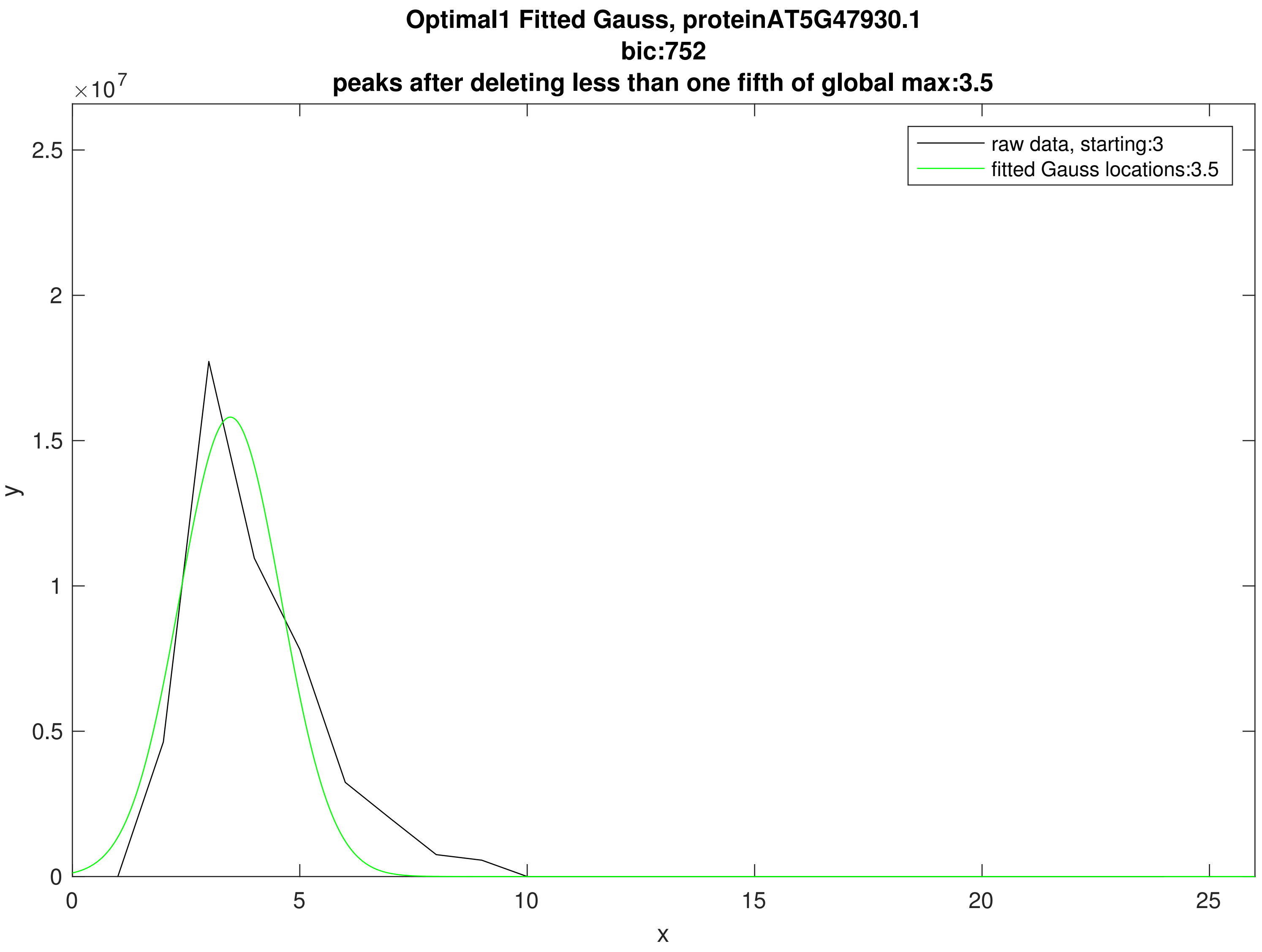
Optimal1 Fitted Gauss, proteinAT5G47930.1
bic:752
peaks after deleting less than one fifth of global max:3.5
×10^7
2.5
2
1.5
1
0.5
0
y
0
5
10
15
20
25
x
raw data, starting:3
fitted Gauss locations:3.5

Optimal1 Fitted Gauss, proteinAT5G48020.1
bic:869
peaks after deleting less than one fifth of global max:16

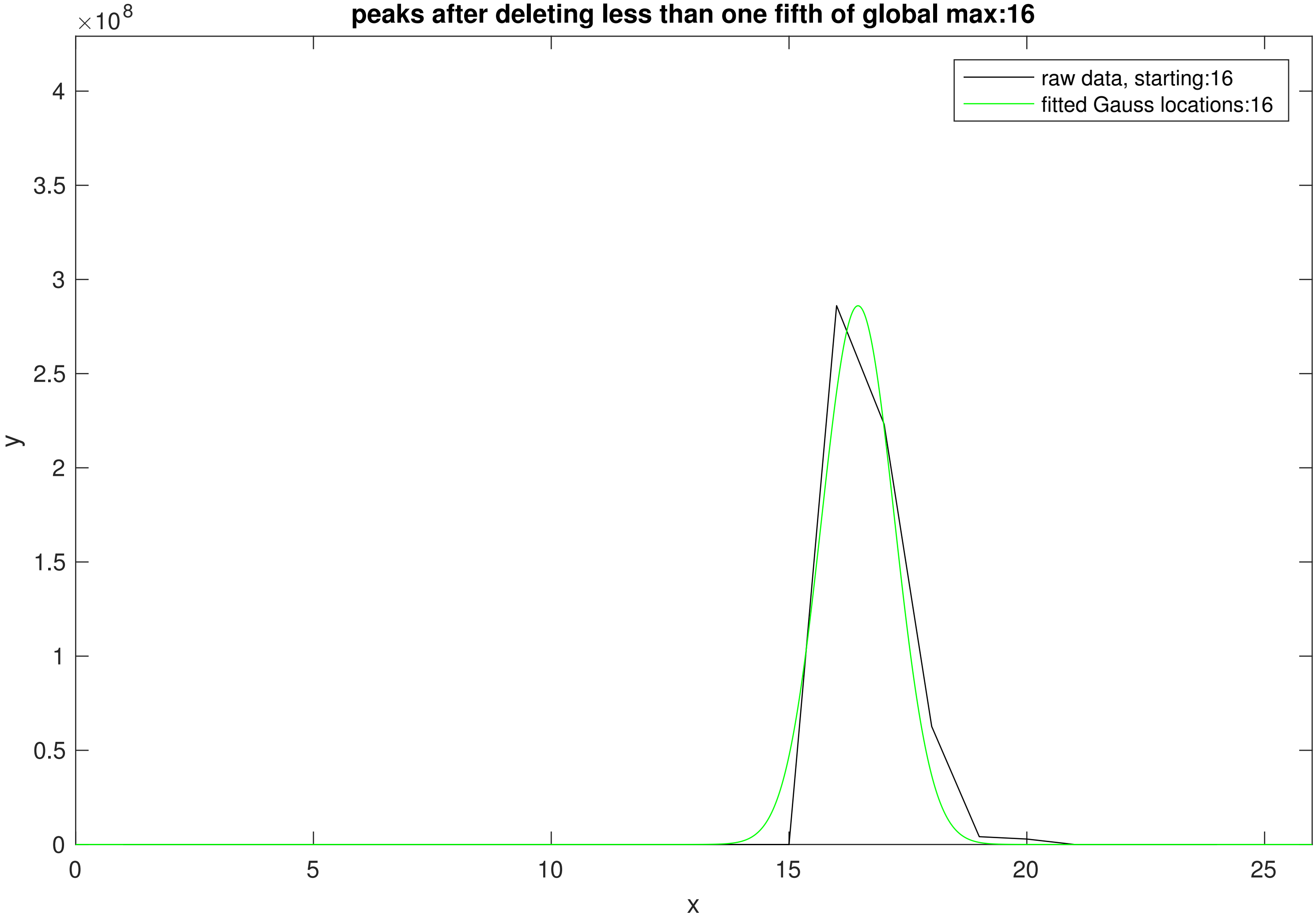

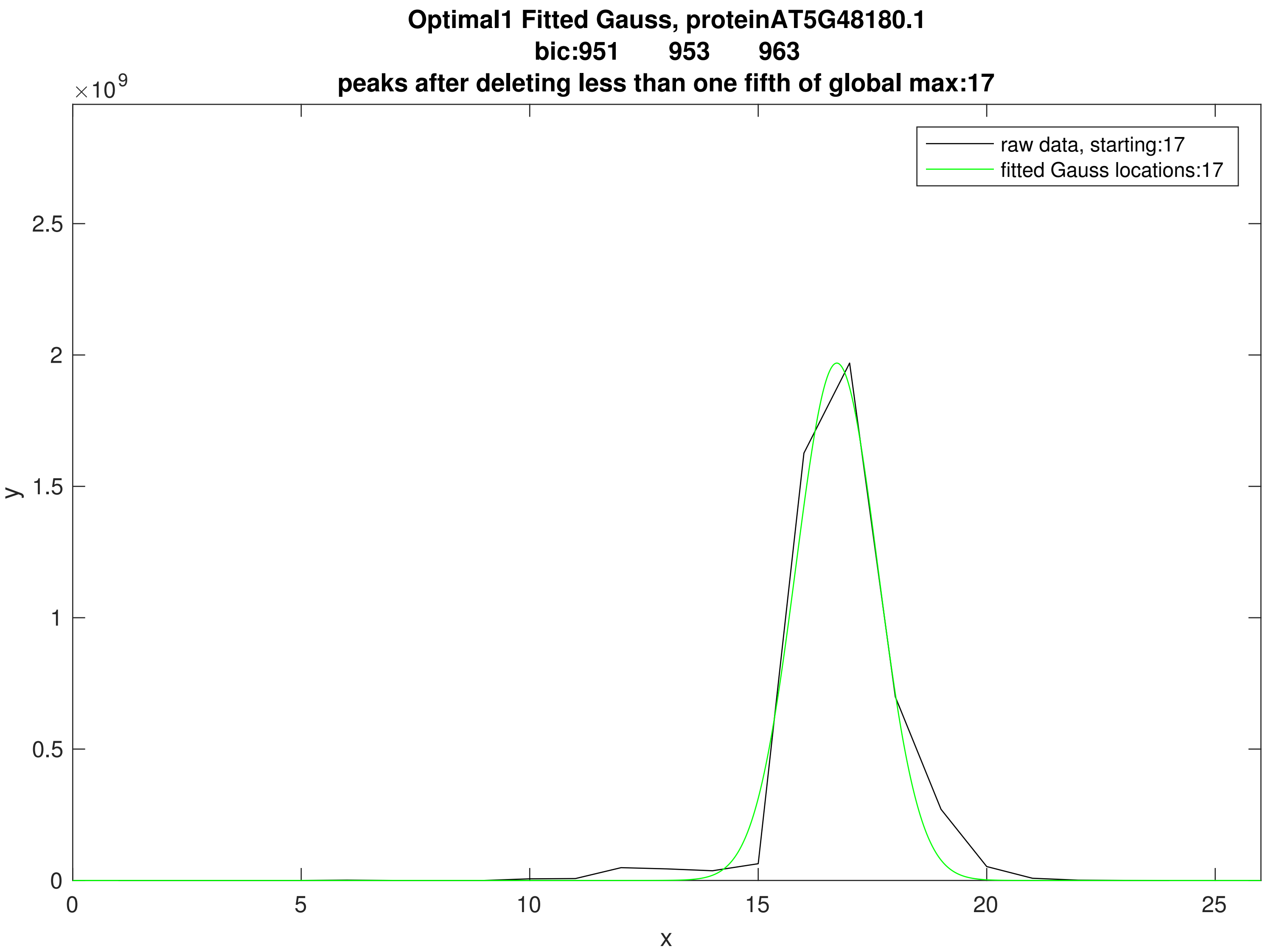
Optimal1 Fitted Gauss, proteinAT5G48180.1
bic:951 953 963
peaks after deleting less than one fifth of global max:17
×10^9
raw data, starting:17
fitted Gauss locations:17
y
x
0
0.5
1
1.5
2
2.5
0
5
10
15
20
25

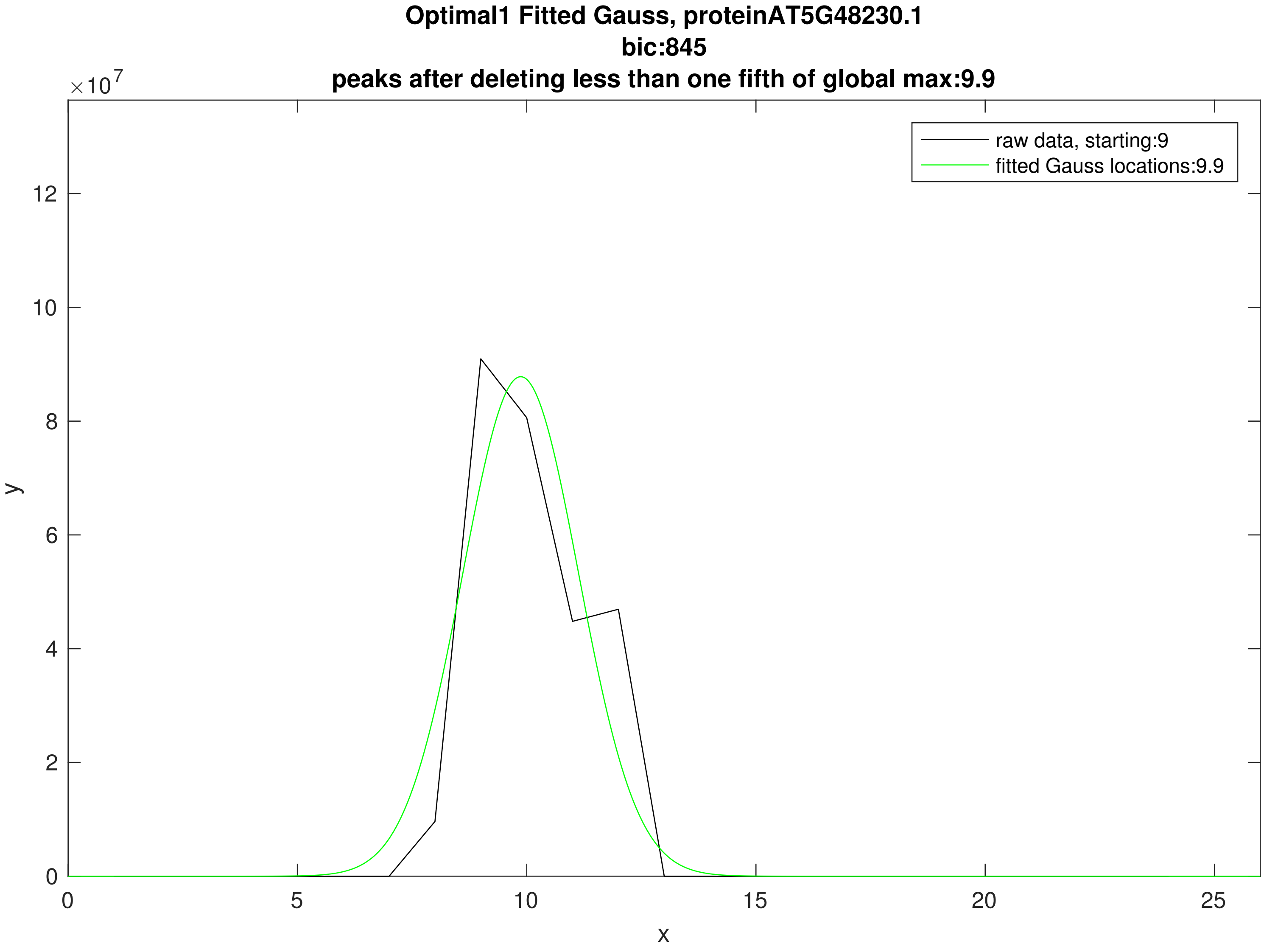

Optimal1 Fitted Gauss, proteinAT5G48230.1
bic:845
peaks after deleting less than one fifth of global max:9.9
×10^7
raw data, starting:9
fitted Gauss locations:9.9
y
x
0
2
4
6
8
10
12
0
5
10
15
20
25

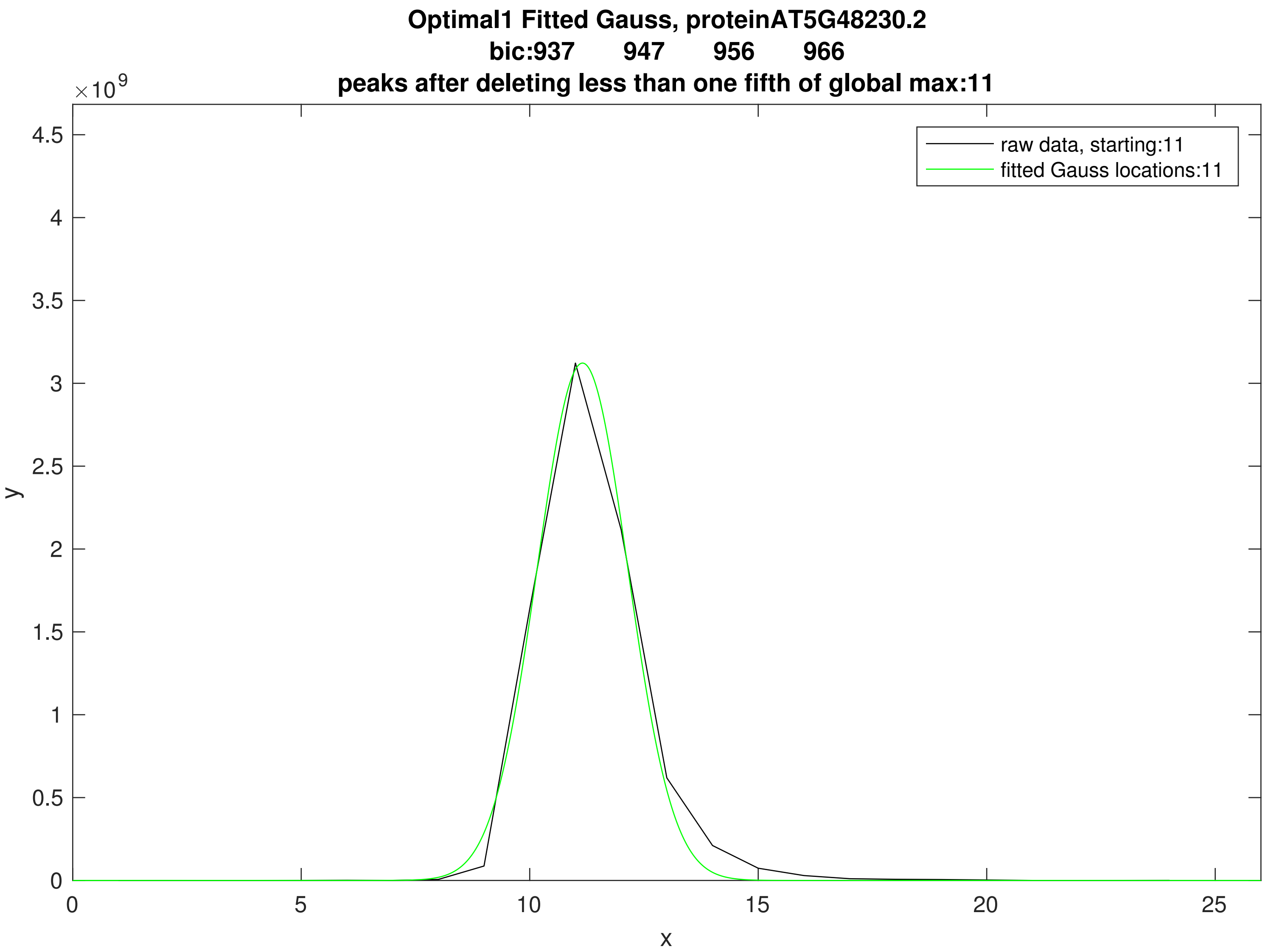
Optimal1 Fitted Gauss, proteinAT5G48230.2
bic:937 947 956 966
peaks after deleting less than one fifth of global max:11
×10^9
4.5
4
3.5
3
2.5
2
1.5
1
0.5
0
y
0
5
10
15
20
25
x
raw data, starting:11
fitted Gauss locations:11

**Optimal1 Fitted Gauss, proteinAT5G48300.1**
**bic:882**
**peaks after deleting less than one fifth of global max:11**

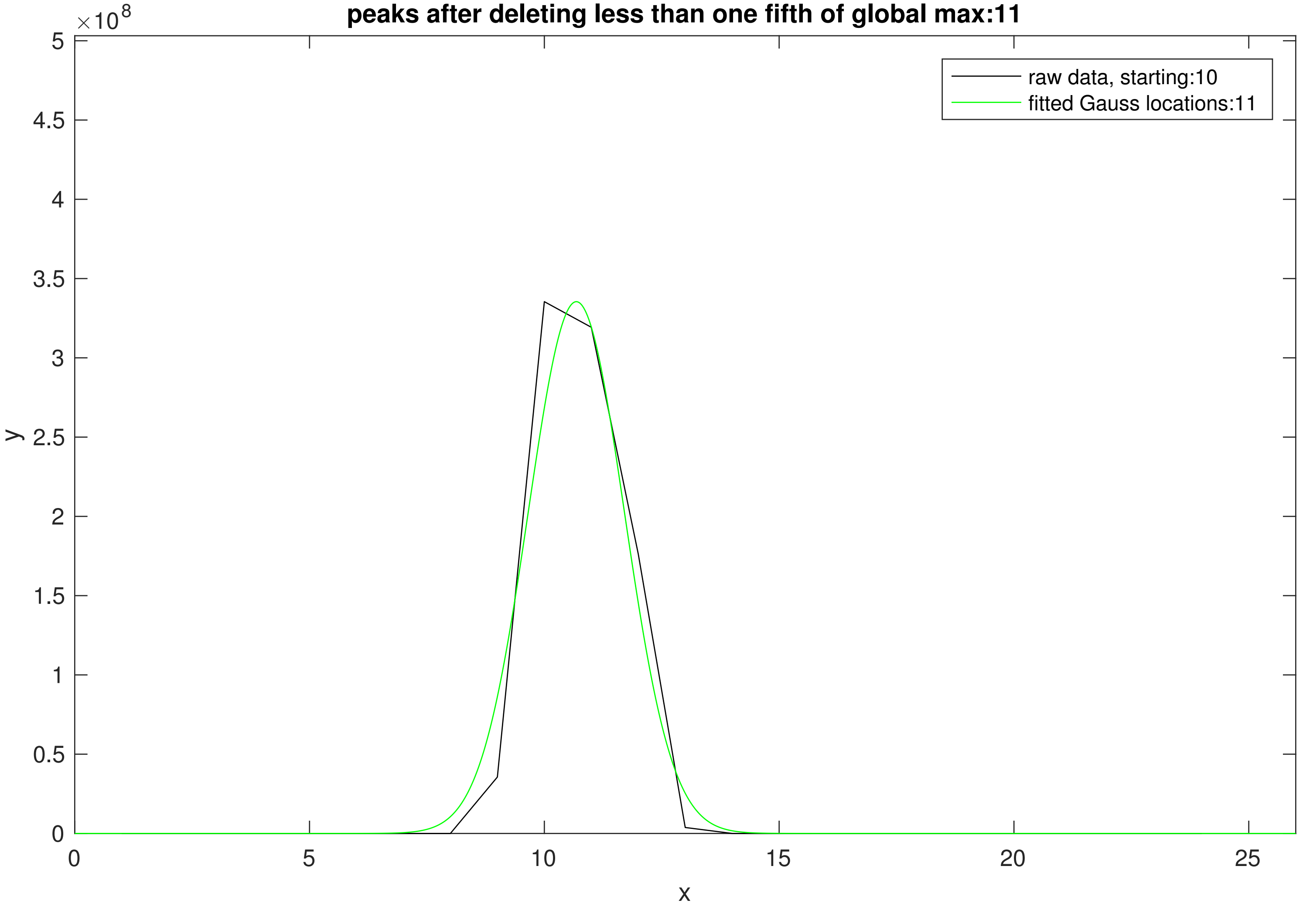

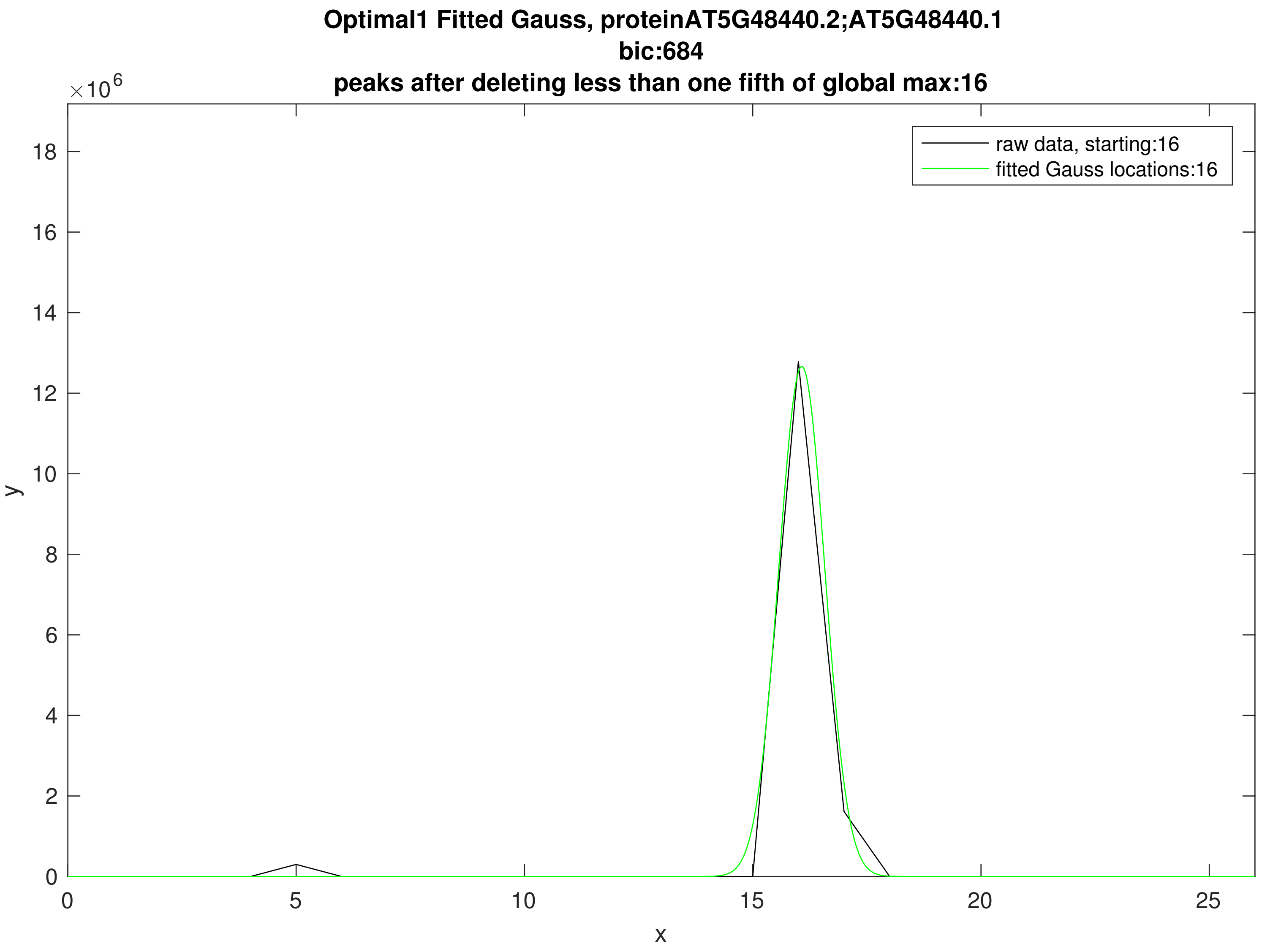

Optimal1 Fitted Gauss, proteinAT5G48440.2;AT5G48440.1
bic:684
peaks after deleting less than one fifth of global max:16
×10^6
raw data, starting:16
fitted Gauss locations:16
y
x
0
2
4
6
8
10
12
14
16
18
0
5
10
15
20
25

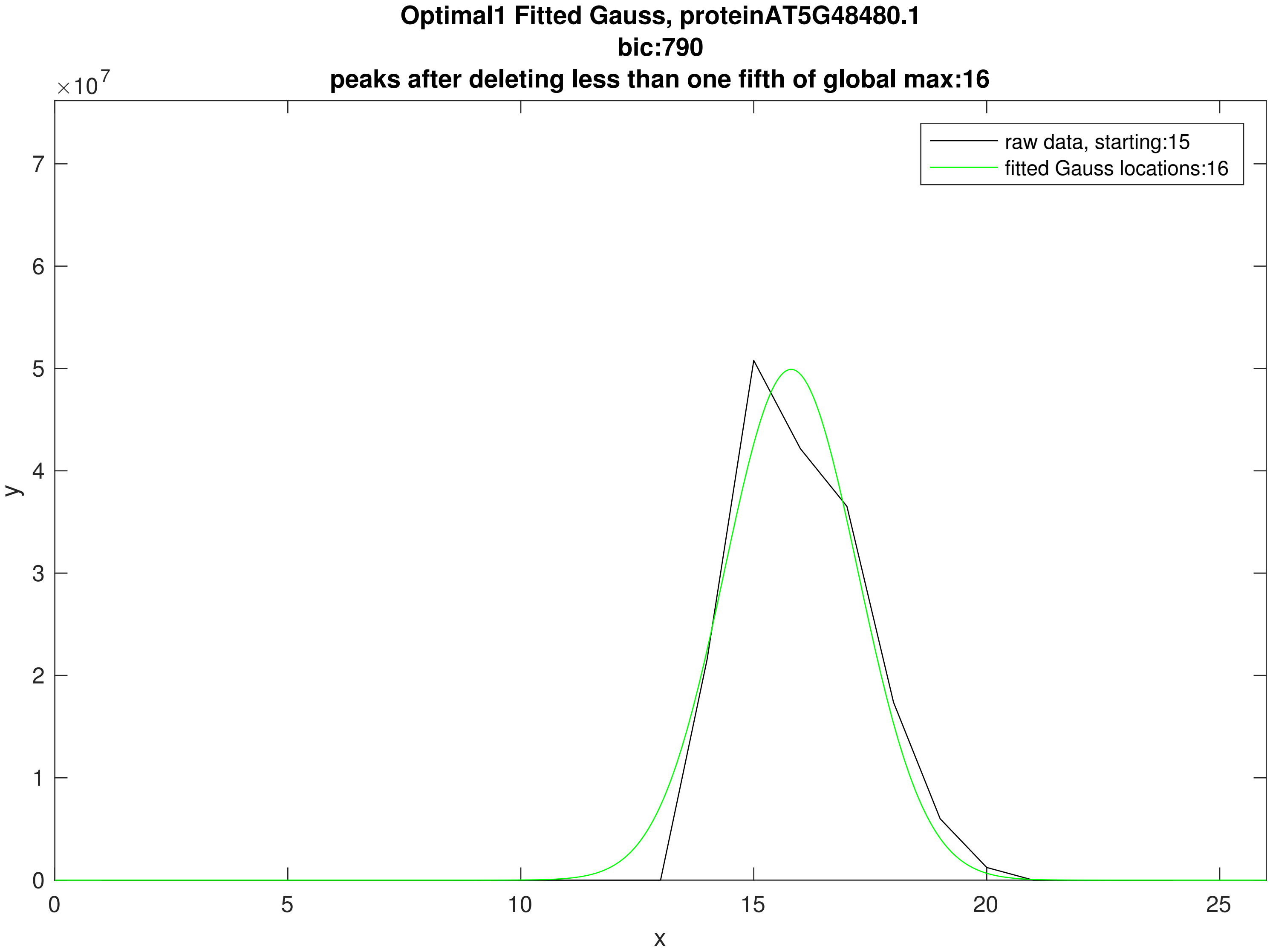

Optimal1 Fitted Gauss, proteinAT5G48480.1
bic:790
peaks after deleting less than one fifth of global max:16
×10^7
7
6
5
4
3
2
1
0
y
0
5
10
15
20
25
x
raw data, starting:15
fitted Gauss locations:16

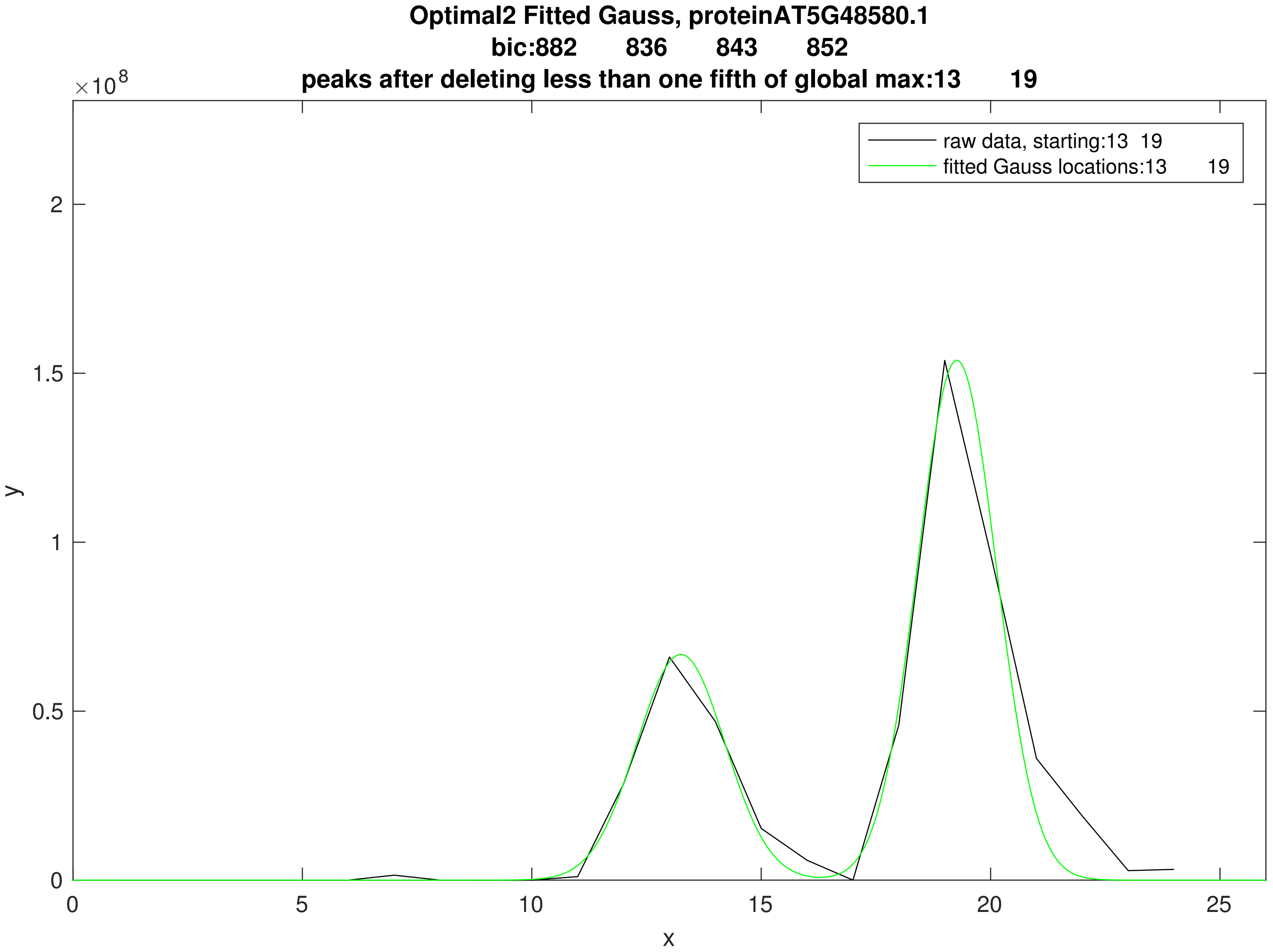
Optimal2 Fitted Gauss, proteinAT5G48580.1
bic:882 836 843 852
peaks after deleting less than one fifth of global max:13 19
×10^8
raw data, starting:13 19
fitted Gauss locations:13 19
y
x
0
0.5
1
1.5
2
5
10
15
20
25

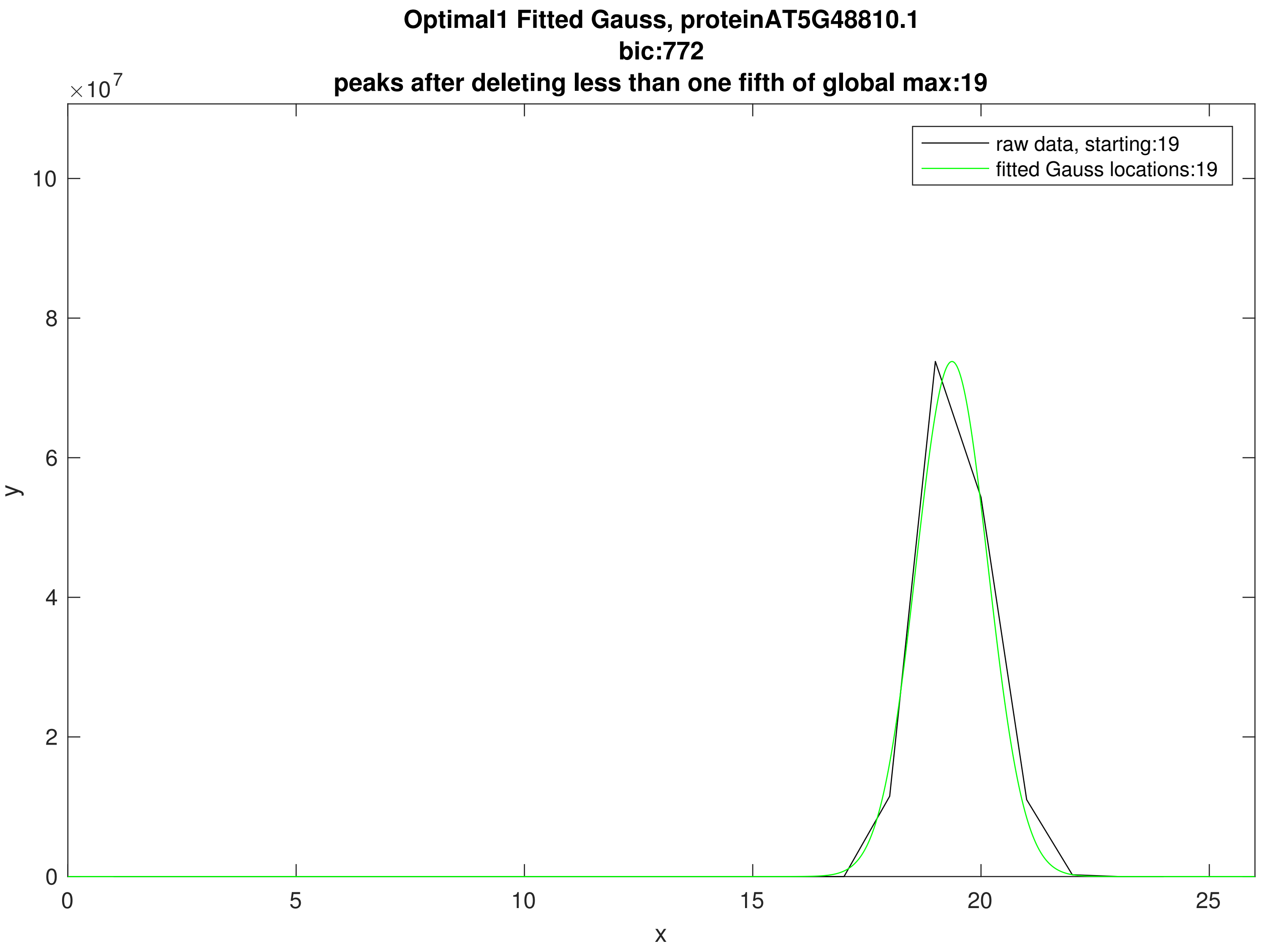

Optimal1 Fitted Gauss, proteinAT5G48810.1
bic:772
peaks after deleting less than one fifth of global max:19
×10^7
raw data, starting:19
fitted Gauss locations:19
10
8
6
4
2
0
y
0
5
10
15
20
25
x

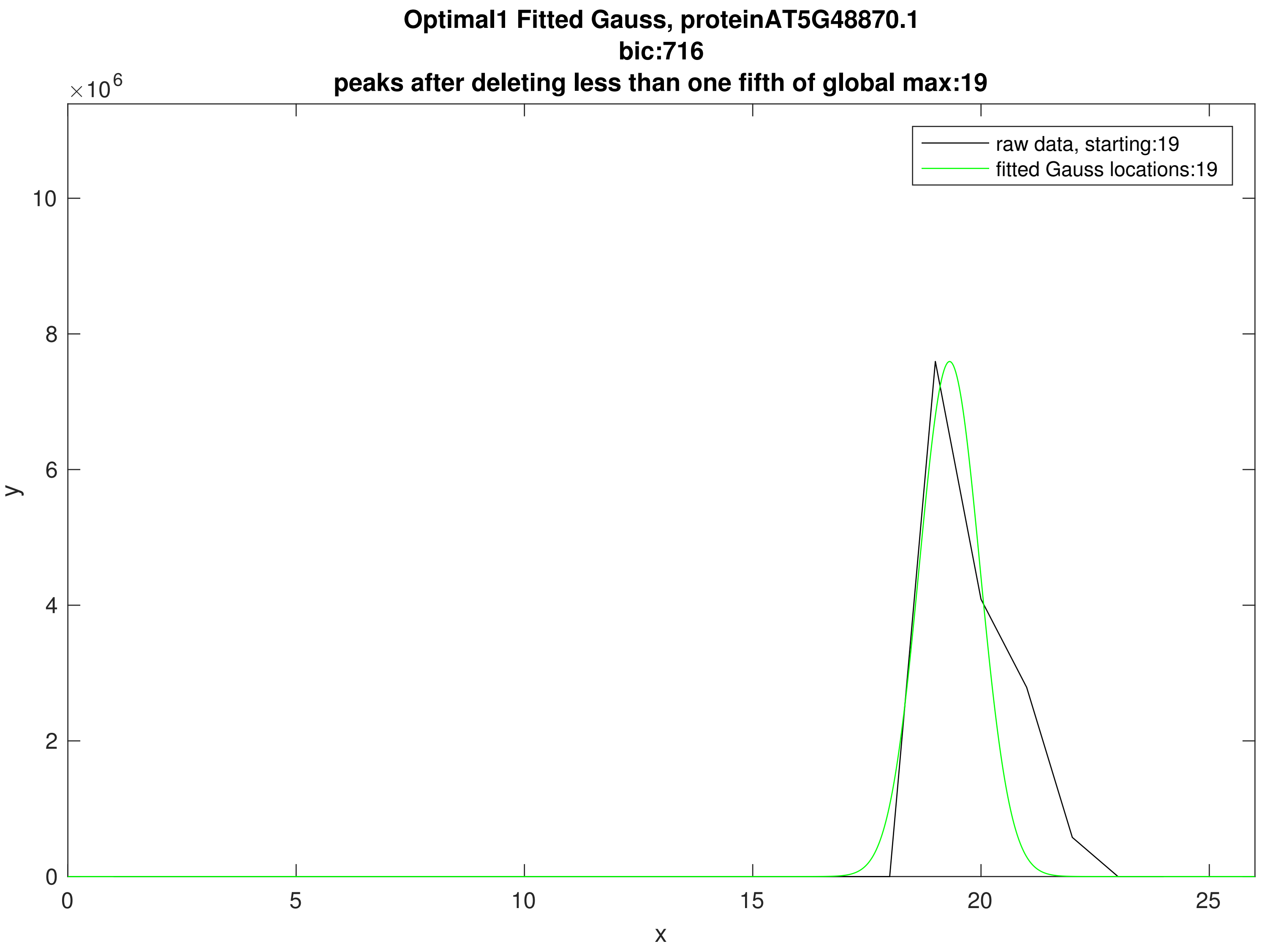

Optimal1 Fitted Gauss, proteinAT5G48870.1
bic:716
peaks after deleting less than one fifth of global max:19
×10^6
raw data, starting:19
fitted Gauss locations:19
10
8
6
4
2
0
y
0
5
10
15
20
25
x

Optimal1 Fitted Gauss, proteinAT5G48880.1;AT5G48880.3;AT5G48880.2
bic:707
peaks after deleting less than one fifth of global max:14

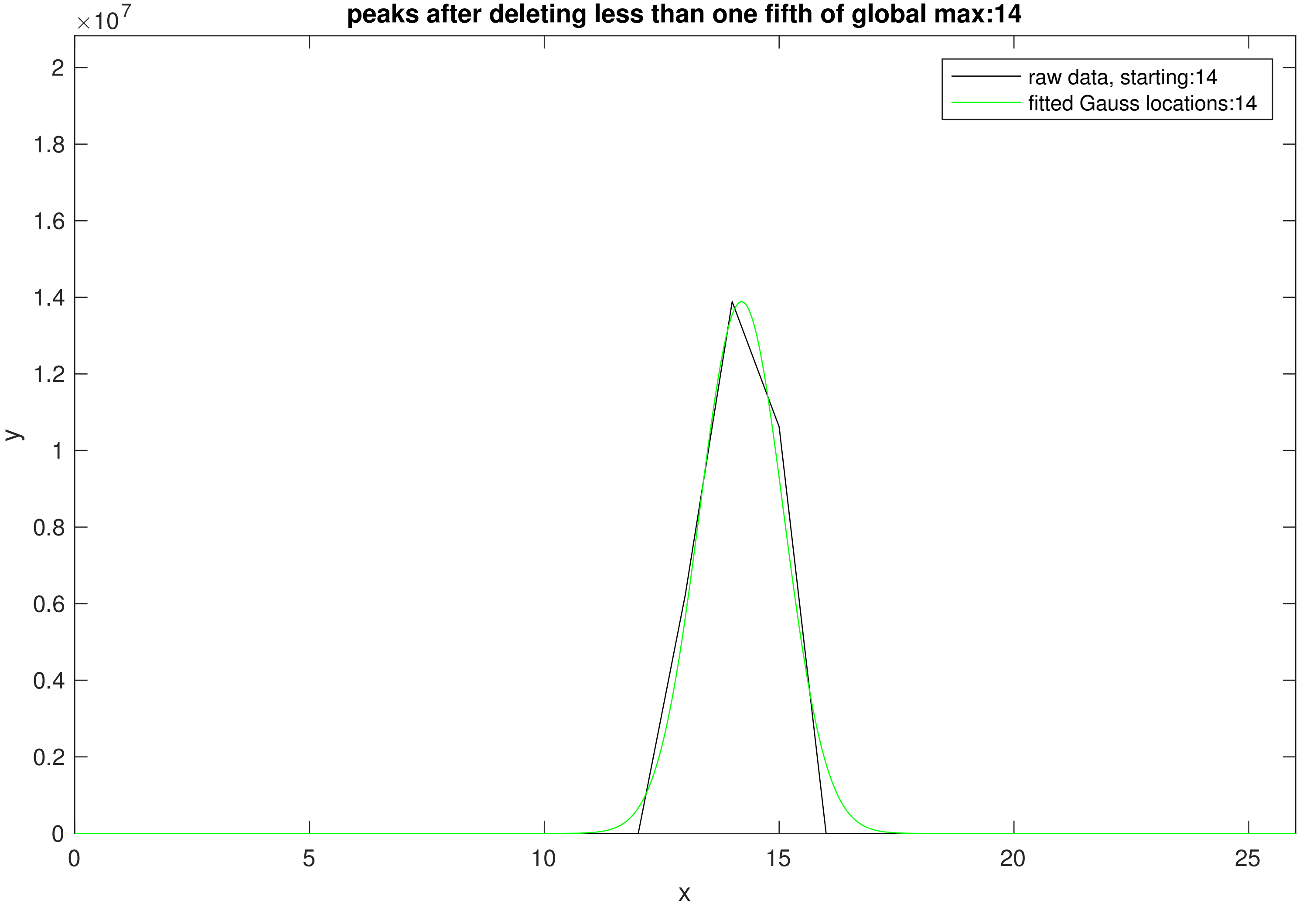

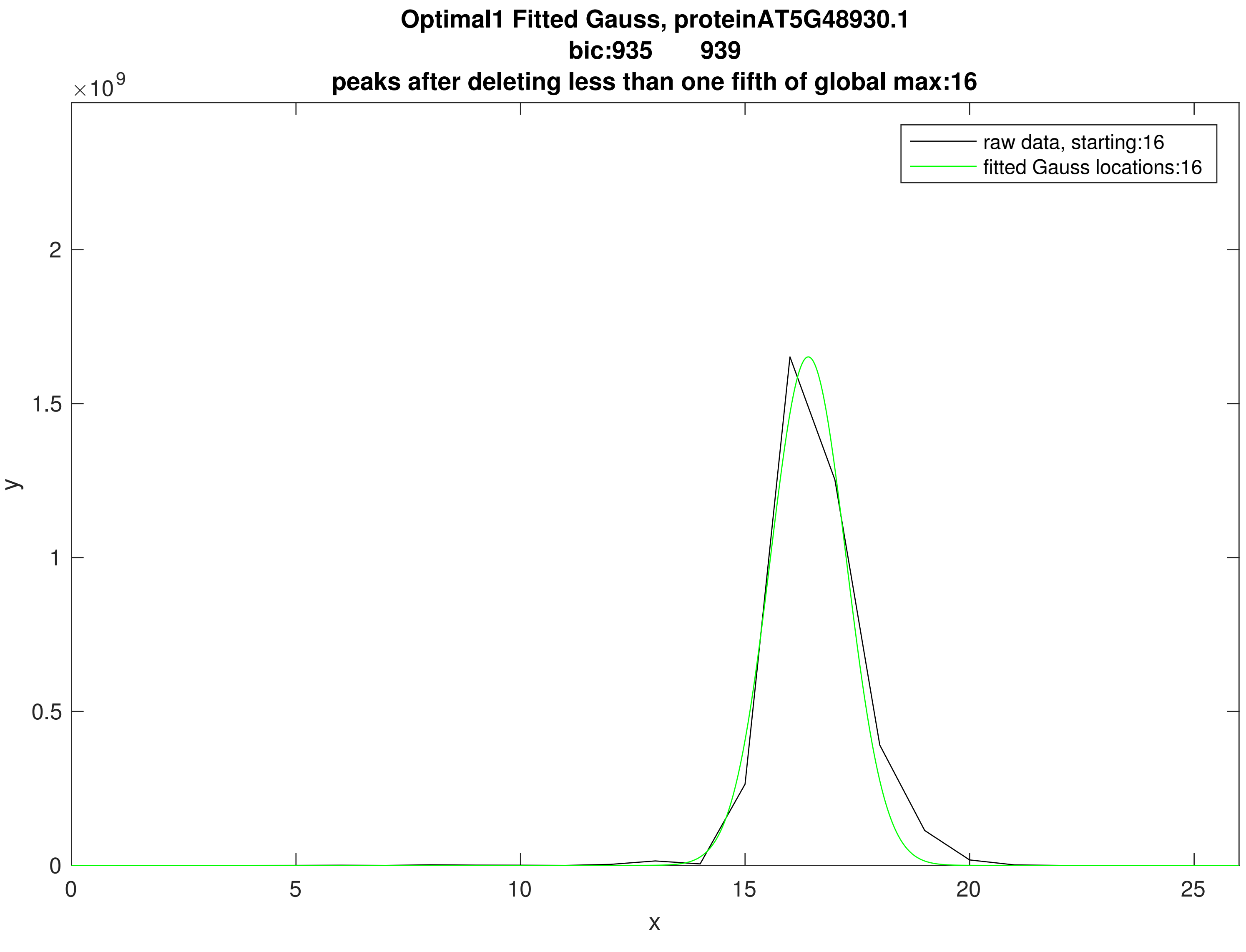
Optimal1 Fitted Gauss, proteinAT5G48930.1
bic:935 939
peaks after deleting less than one fifth of global max:16
×10^9
raw data, starting:16
fitted Gauss locations:16
y
x
0
0.5
1
1.5
2
0
5
10
15
20
25

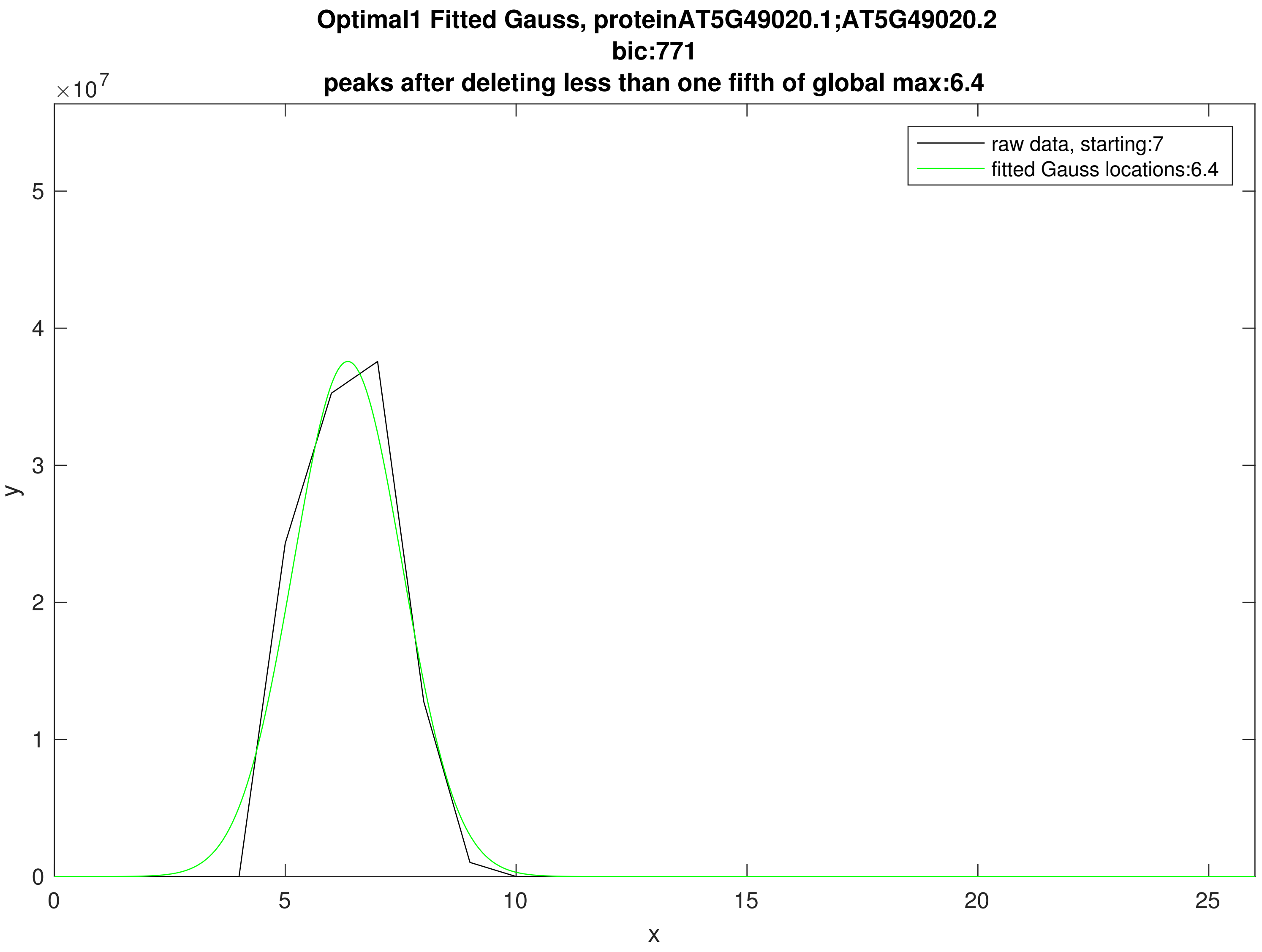
Optimal1 Fitted Gauss, proteinAT5G49020.1;AT5G49020.2
bic:771
peaks after deleting less than one fifth of global max:6.4
×10^7
raw data, starting:7
fitted Gauss locations:6.4
y
x
0
1
2
3
4
5
0
5
10
15
20
25

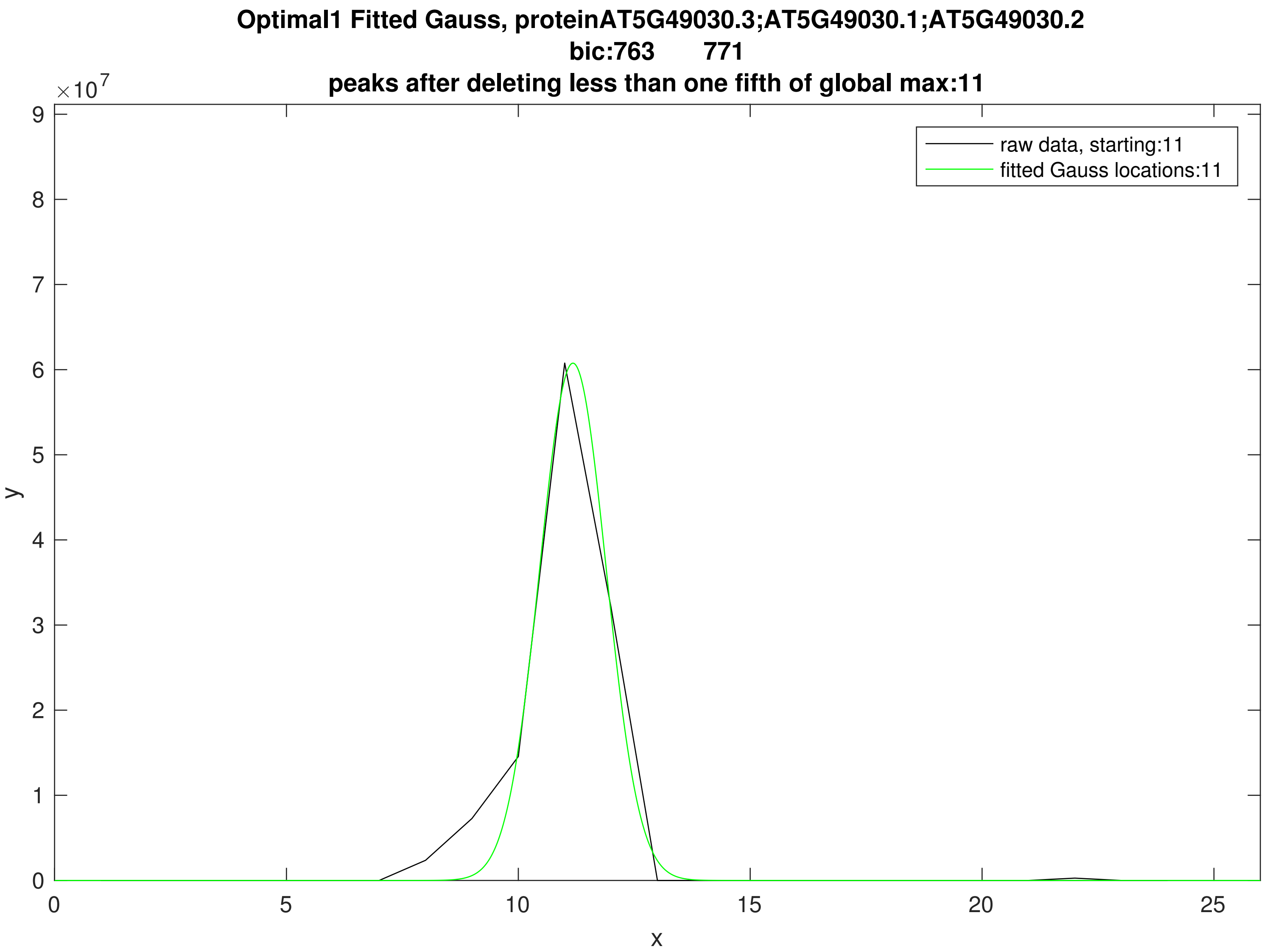
Optimal1 Fitted Gauss, proteinAT5G49030.3;AT5G49030.1;AT5G49030.2
bic:763 771
peaks after deleting less than one fifth of global max:11
×10^7
raw data, starting:11
fitted Gauss locations:11
y
x
0
1
2
3
4
5
6
7
8
9
0
5
10
15
20
25

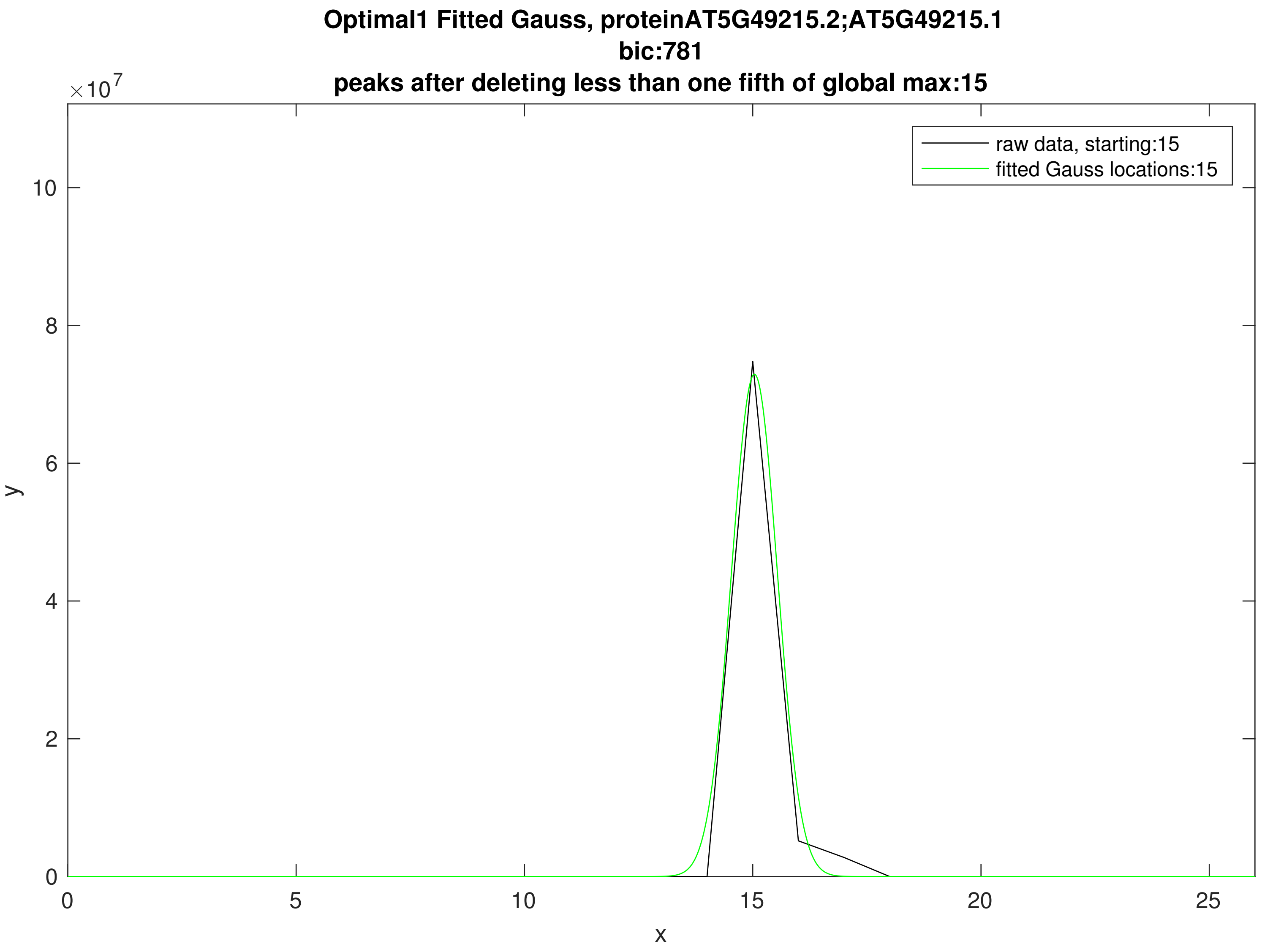
Optimal1 Fitted Gauss, proteinAT5G49215.2;AT5G49215.1
bic:781
peaks after deleting less than one fifth of global max:15
×10^7
raw data, starting:15
fitted Gauss locations:15
10
8
6
4
2
0
y
0
5
10
15
20
25
x

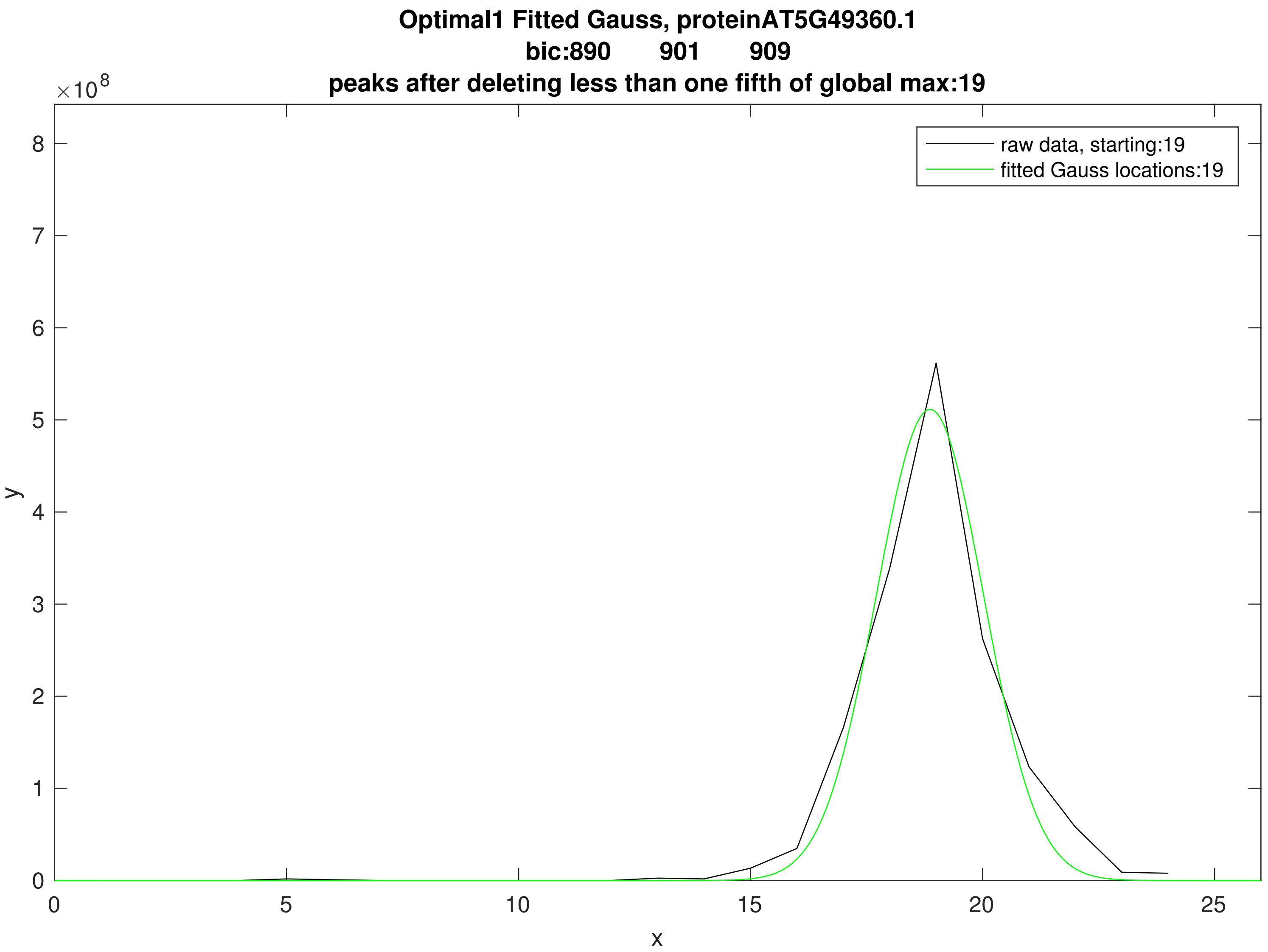
Optimal1 Fitted Gauss, proteinAT5G49360.1
bic:890 901 909
peaks after deleting less than one fifth of global max:19
×10^8
raw data, starting:19
fitted Gauss locations:19
y
x
0
1
2
3
4
5
6
7
8
10
15
20
25

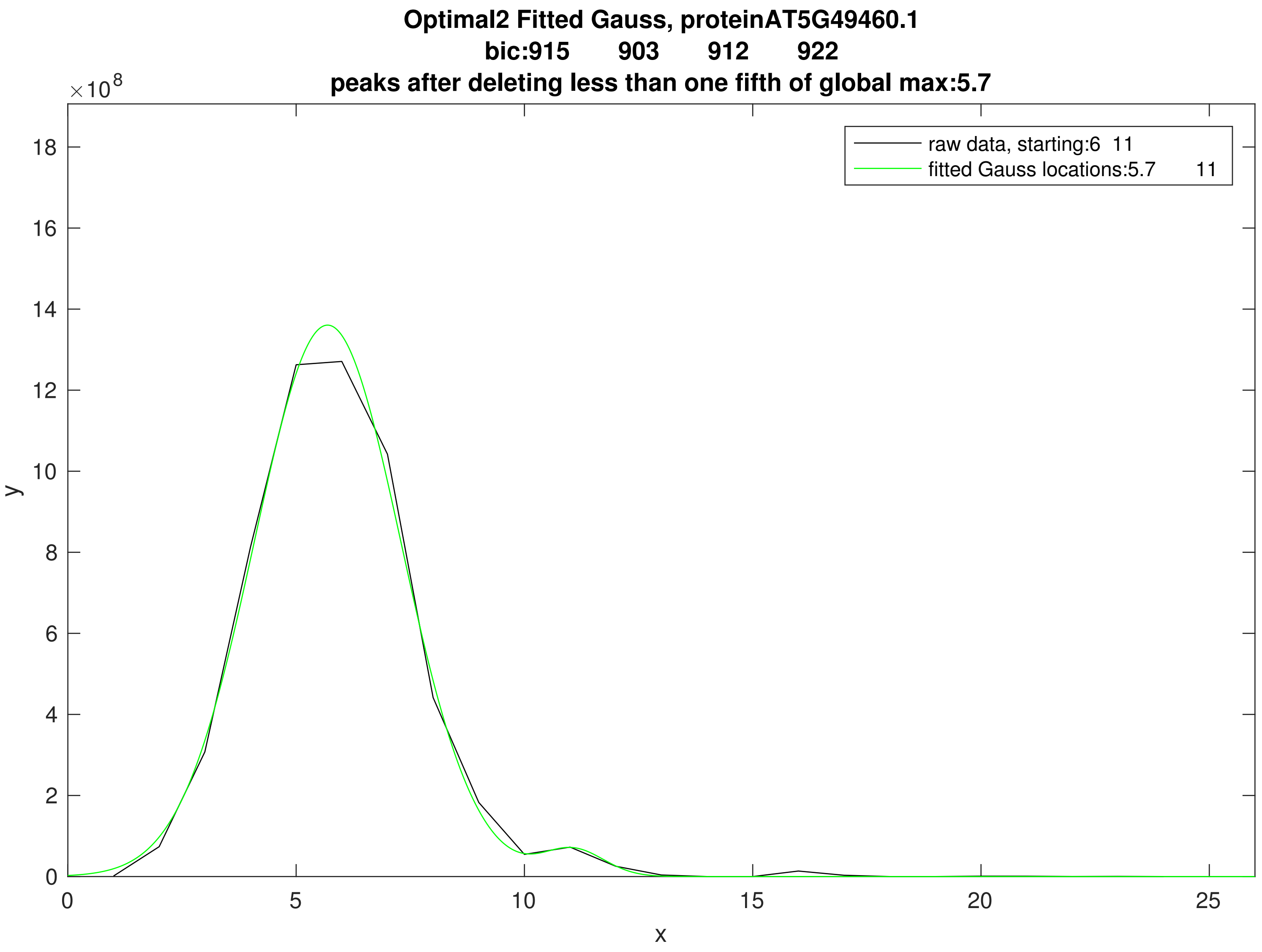
Optimal2 Fitted Gauss, proteinAT5G49460.1
bic:915 903 912 922
peaks after deleting less than one fifth of global max:5.7
×10^8
raw data, starting:6 11
fitted Gauss locations:5.7 11
y
x
0
2
4
6
8
10
12
14
16
18
0
5
10
15
20
25

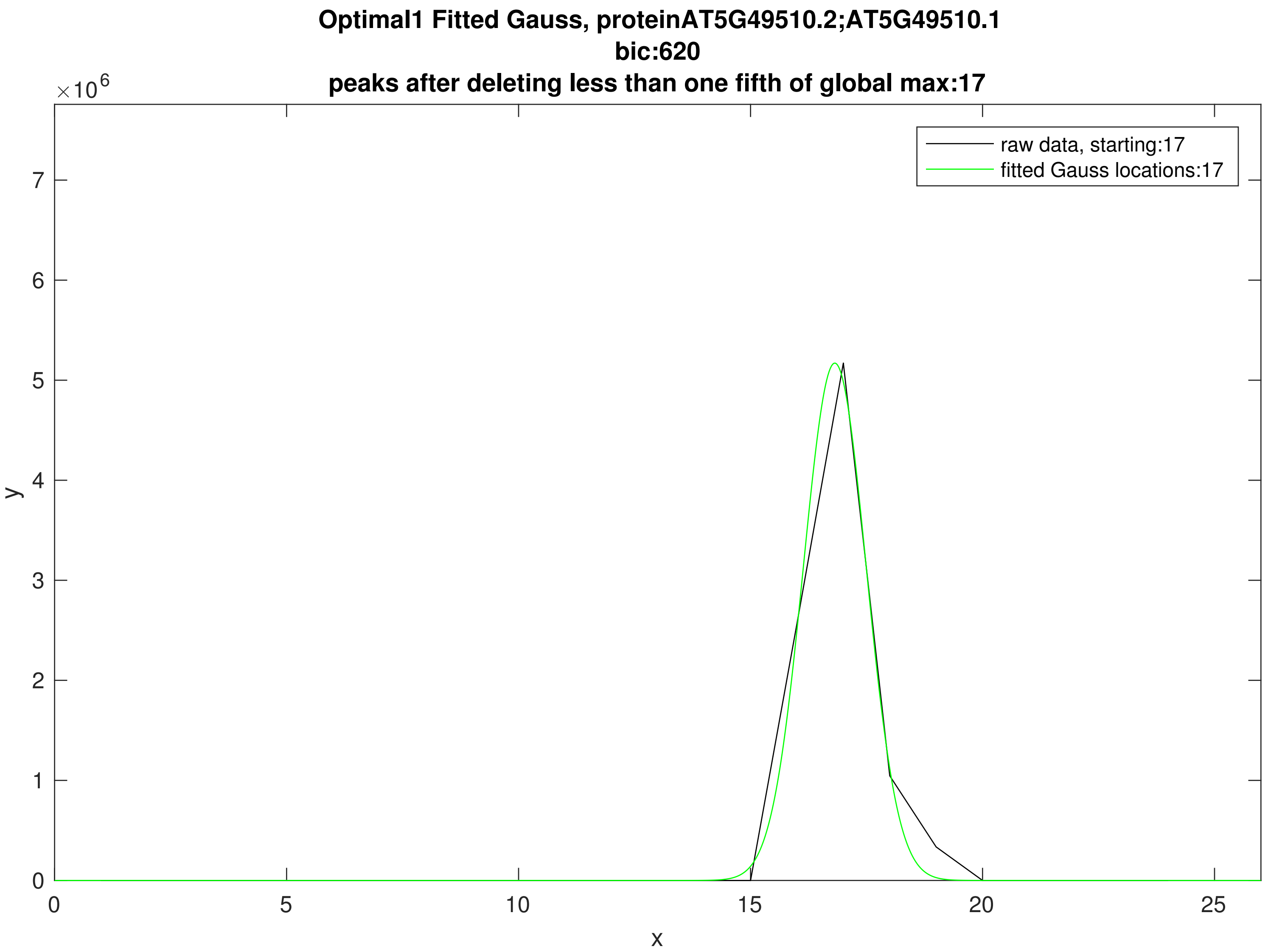
Optimal1 Fitted Gauss, proteinAT5G49510.2;AT5G49510.1
bic:620
peaks after deleting less than one fifth of global max:17
×10^6
raw data, starting:17
fitted Gauss locations:17
y
x
0
1
2
3
4
5
6
7
0
5
10
15
20
25

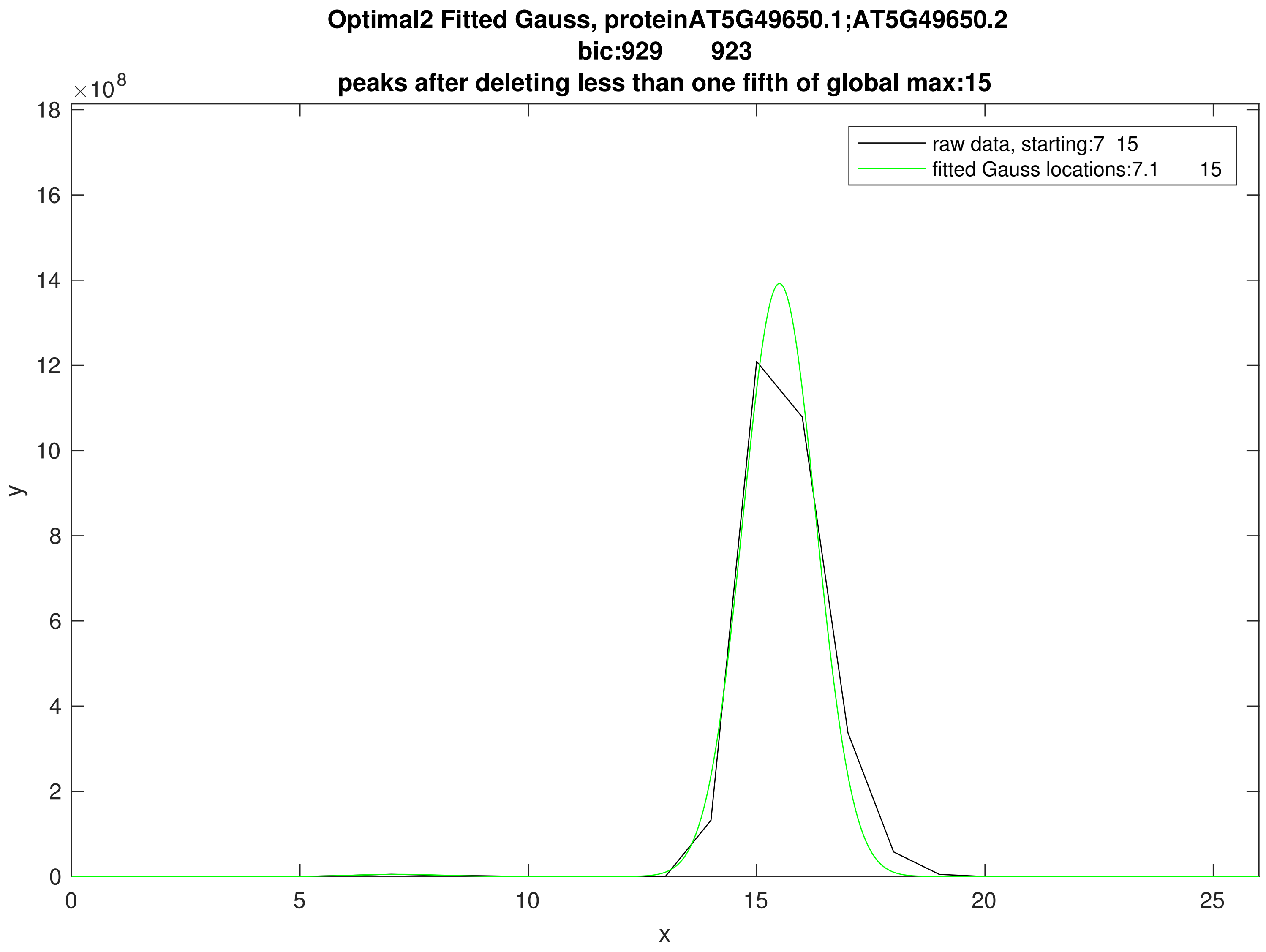
Optimal2 Fitted Gauss, proteinAT5G49650.1;AT5G49650.2
bic:929 923
peaks after deleting less than one fifth of global max:15
×10^8
raw data, starting:7 15
fitted Gauss locations:7.1 15
y
x
0
2
4
6
8
10
12
14
16
18
0
5
10
15
20
25

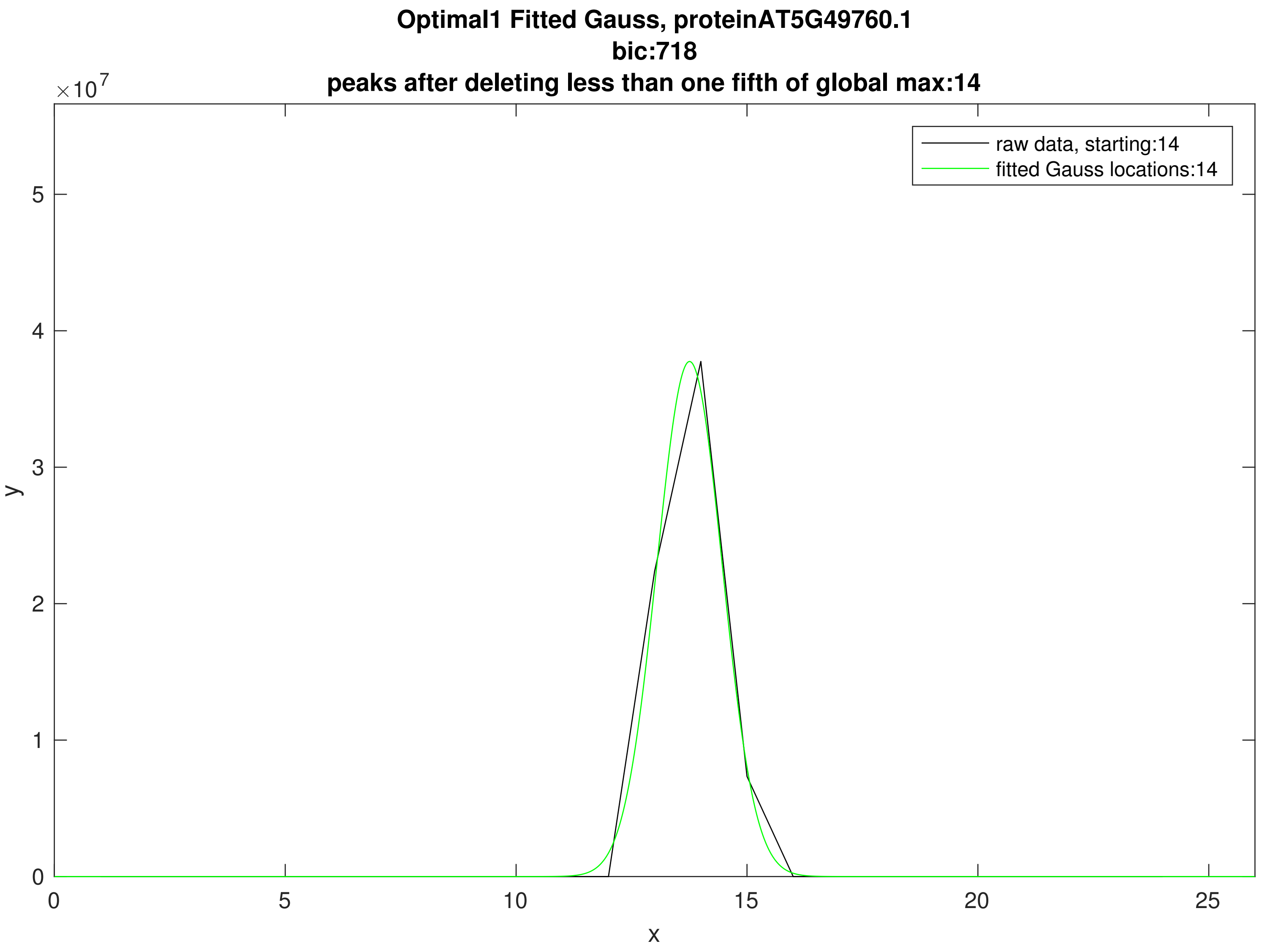

Optimal1 Fitted Gauss, proteinAT5G49760.1
bic:718
peaks after deleting less than one fifth of global max:14
×10^7
raw data, starting:14
fitted Gauss locations:14
y
x
0
1
2
3
4
5
0
5
10
15
20
25

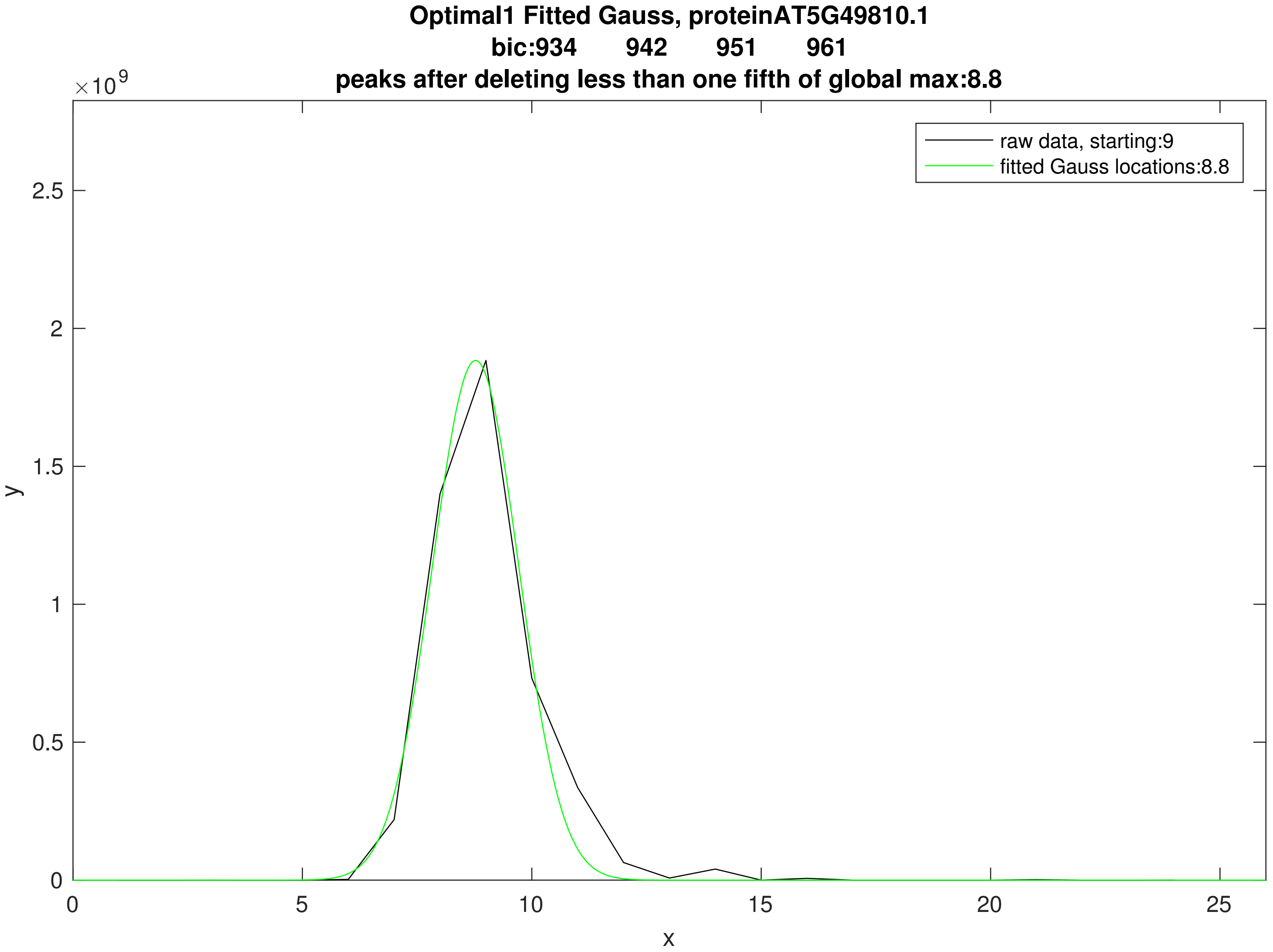
Optimal1 Fitted Gauss, proteinAT5G49810.1
bic:934 942 951 961
peaks after deleting less than one fifth of global max:8.8
×10^9
raw data, starting:9
fitted Gauss locations:8.8
2.5
2
1.5
1
0.5
0
y
0
5
10
15
20
25
x

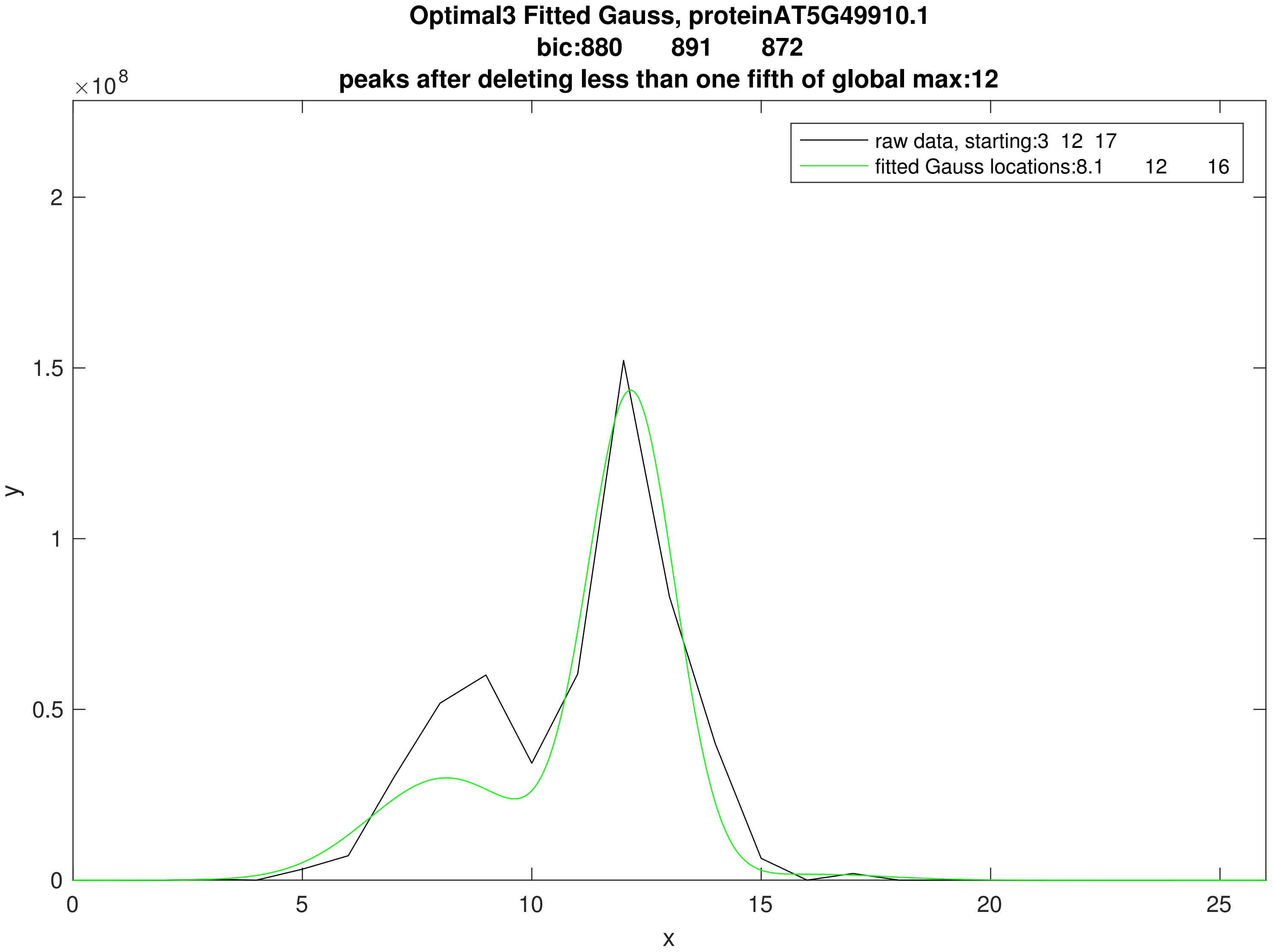

Optimal3 Fitted Gauss, proteinAT5G49910.1
bic:880 891 872
peaks after deleting less than one fifth of global max:12
×10^8
raw data, starting:3 12 17
fitted Gauss locations:8.1 12 16
y
x
0
0.5
1
1.5
2
5
10
15
20
25

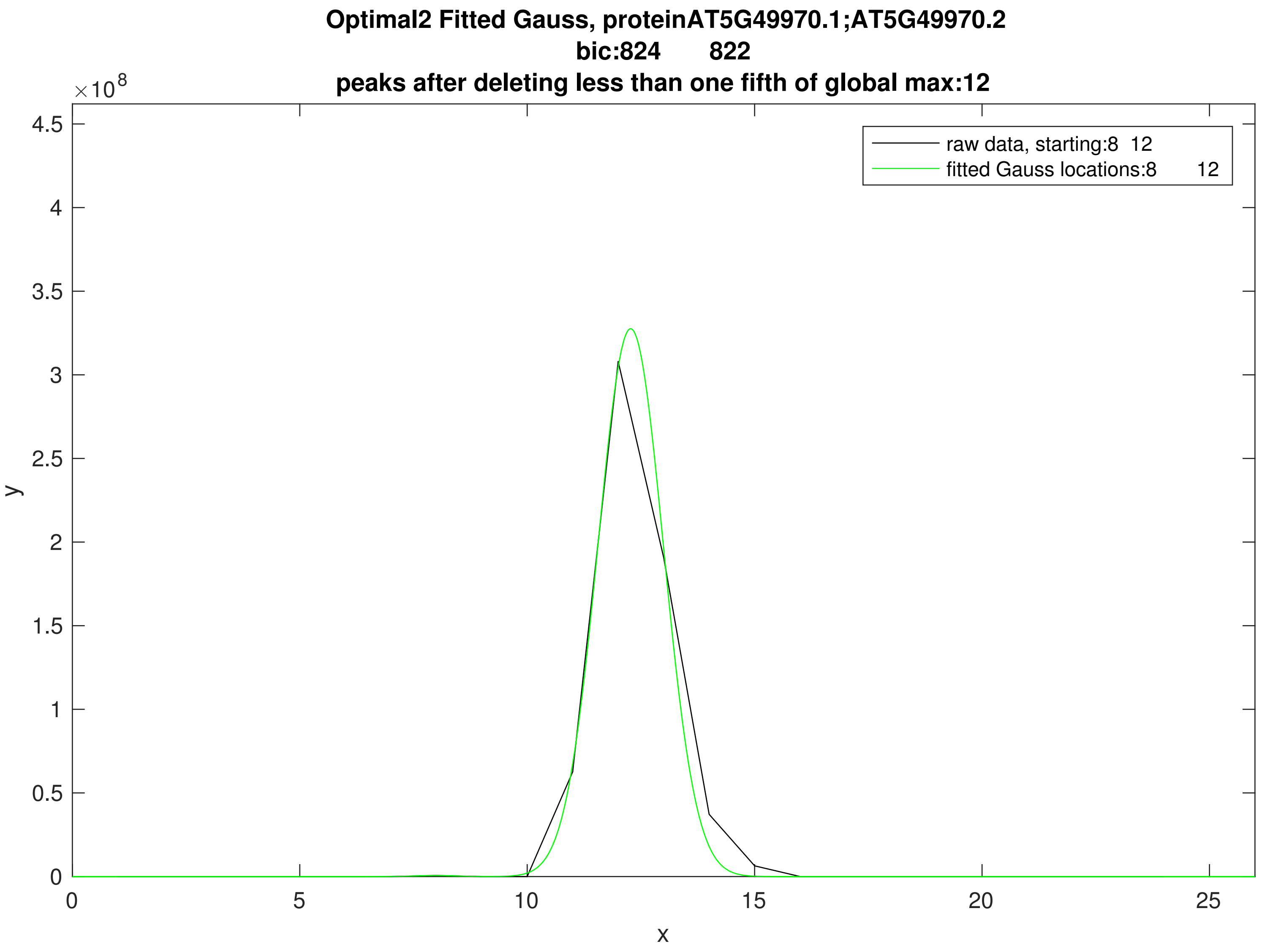

Optimal2 Fitted Gauss, proteinAT5G49970.1;AT5G49970.2
bic:824 822
peaks after deleting less than one fifth of global max:12
×10^8
raw data, starting:8 12
fitted Gauss locations:8 12
y
x
0
0.5
1
1.5
2
2.5
3
3.5
4
4.5
5
10
15
20
25

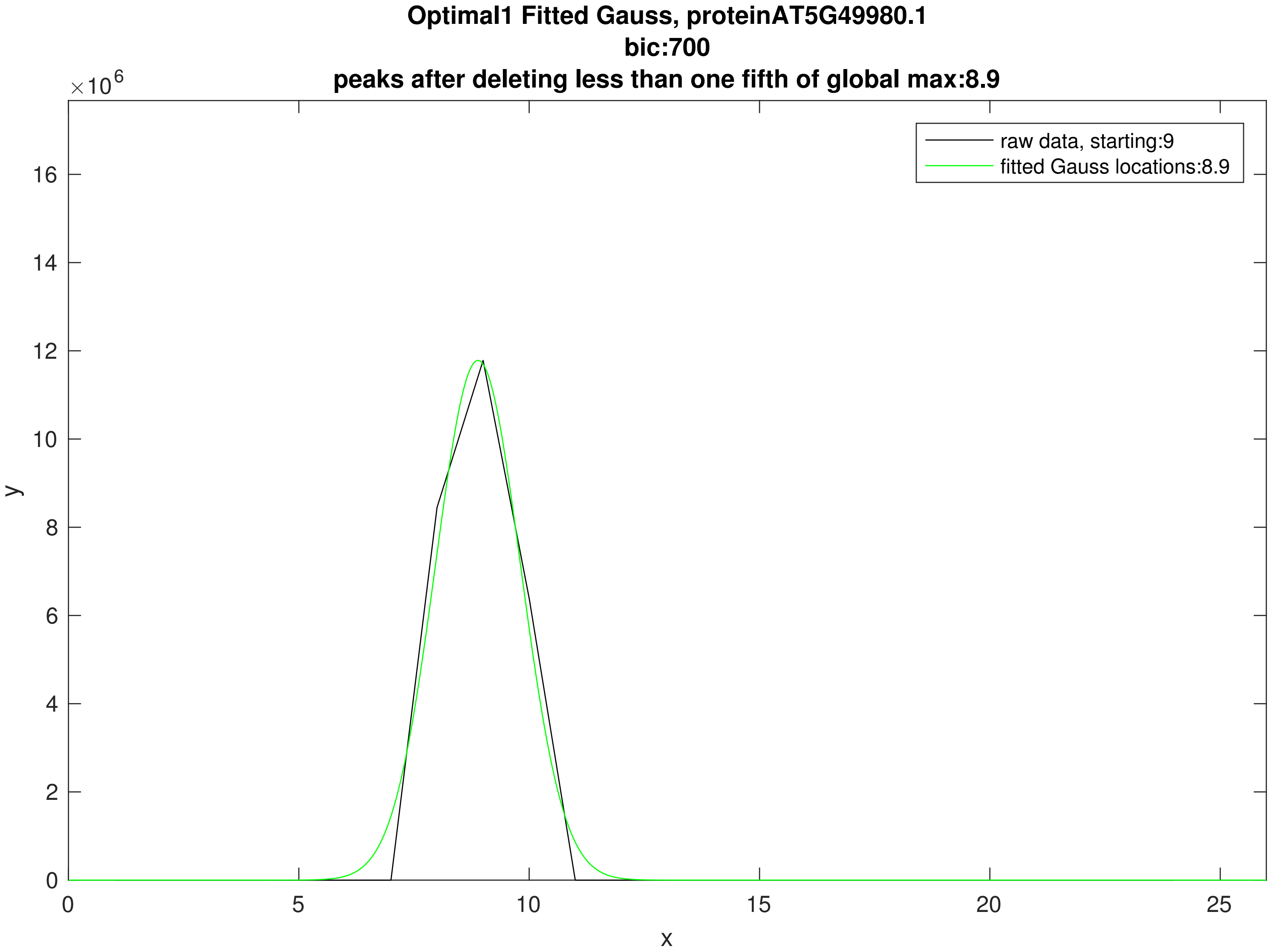

Optimal1 Fitted Gauss, proteinAT5G49980.1
bic:700
peaks after deleting less than one fifth of global max:8.9
×10^6
raw data, starting:9
fitted Gauss locations:8.9
y
x
0
2
4
6
8
10
12
14
16
0
5
10
15
20
25

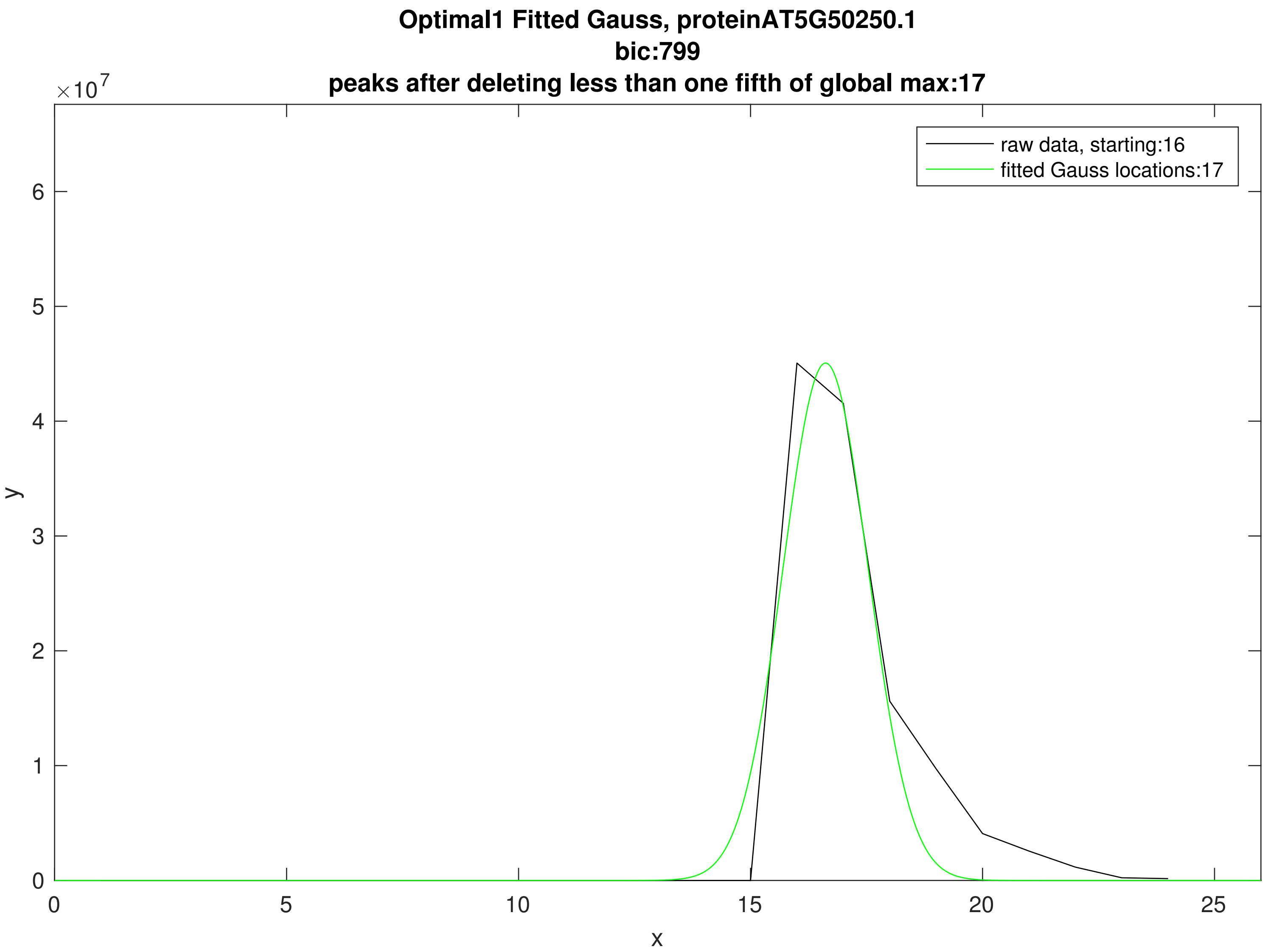
Optimal1 Fitted Gauss, proteinAT5G50250.1
bic:799
peaks after deleting less than one fifth of global max:17
×10^7
raw data, starting:16
fitted Gauss locations:17
y
x
0
1
2
3
4
5
6
0
5
10
15
20
25

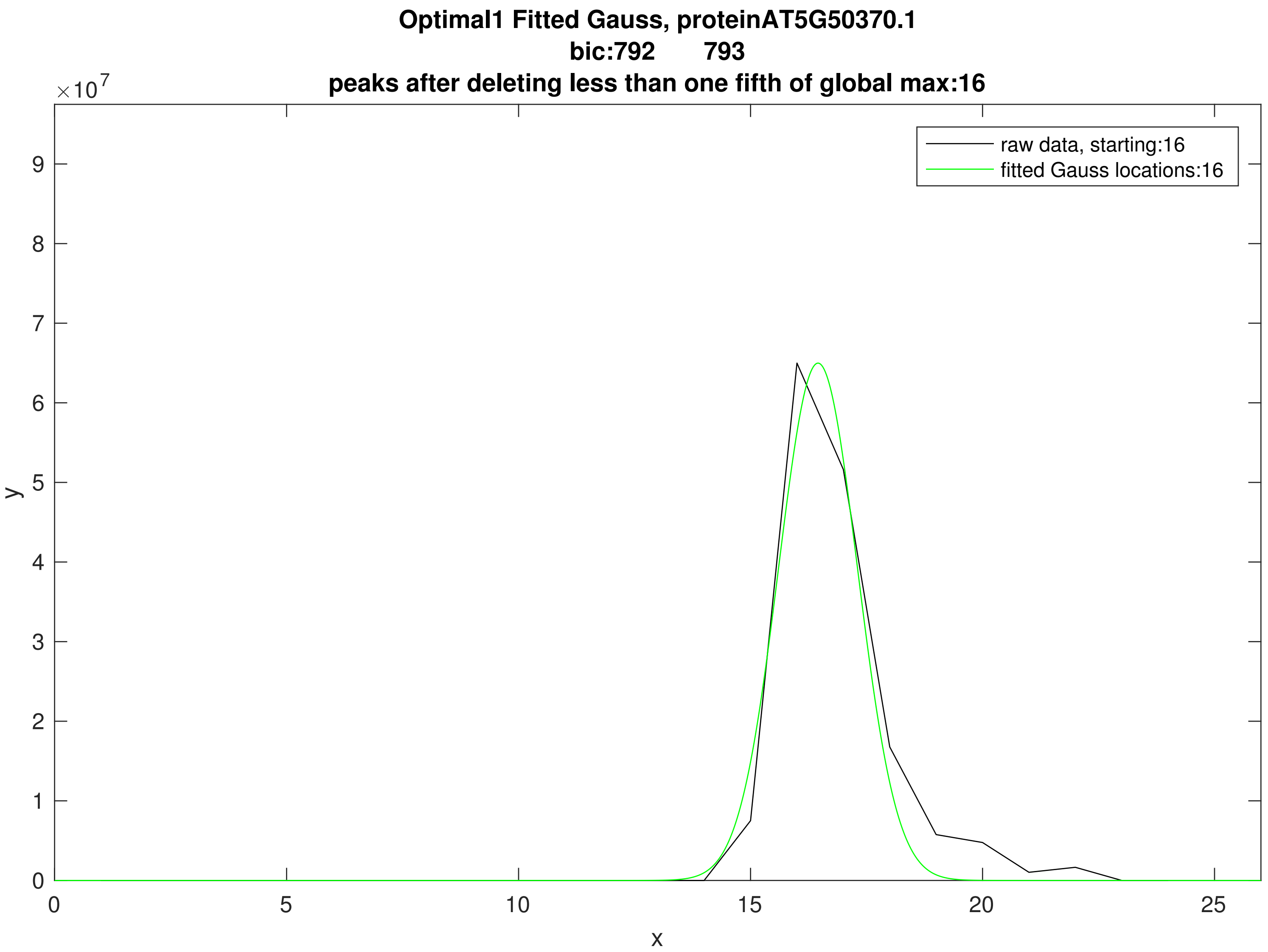
Optimal1 Fitted Gauss, proteinAT5G50370.1
bic:792 793
peaks after deleting less than one fifth of global max:16
×10^7
raw data, starting:16
fitted Gauss locations:16
y
x
0
1
2
3
4
5
6
7
8
9
0
5
10
15
20
25

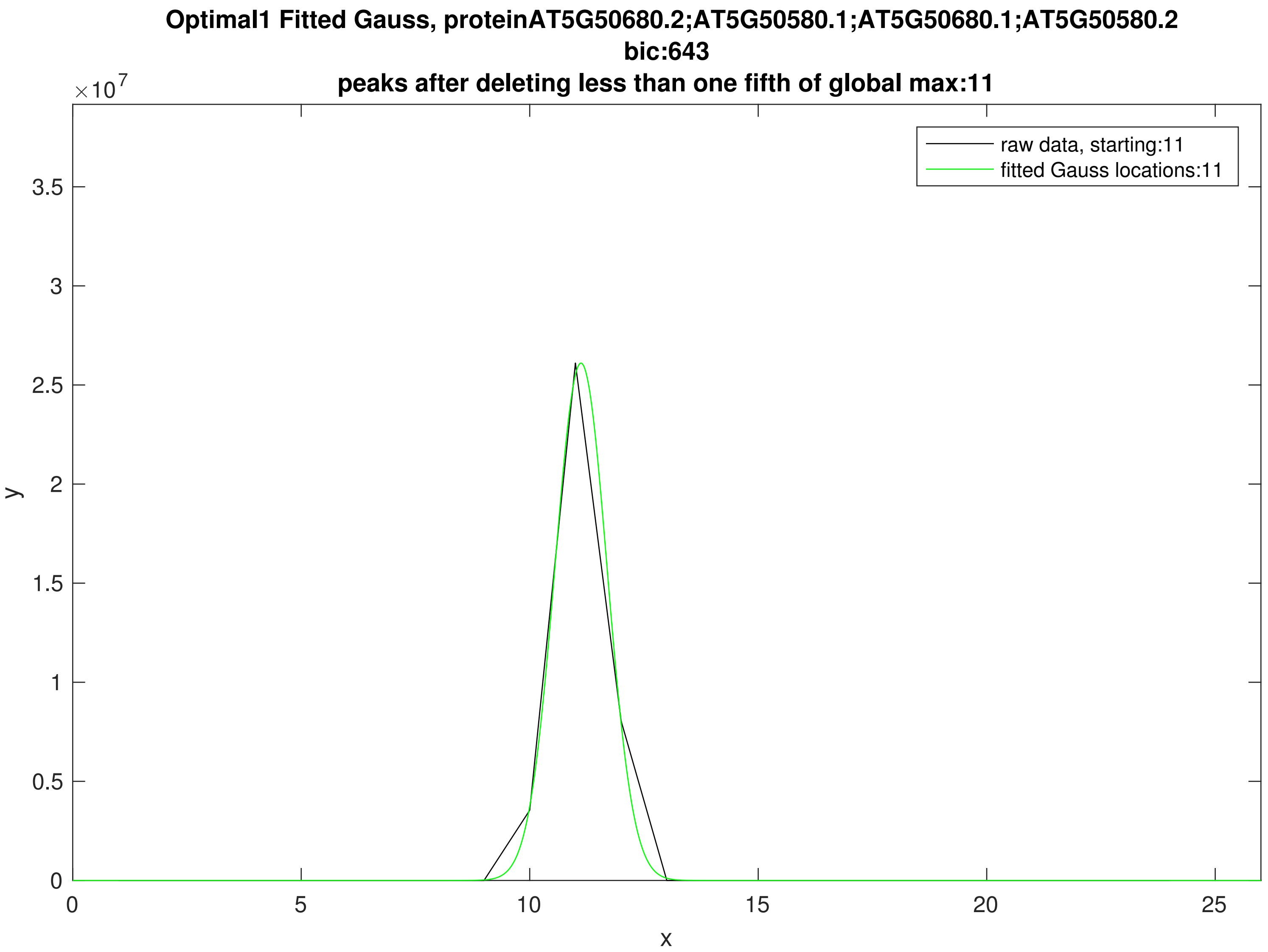
Optimal1 Fitted Gauss, proteinAT5G50680.2;AT5G50580.1;AT5G50680.1;AT5G50580.2
bic:643
peaks after deleting less than one fifth of global max:11
×10^7
raw data, starting:11
fitted Gauss locations:11
3.5
3
2.5
2
1.5
1
0.5
0
y
0
5
10
15
20
25
x

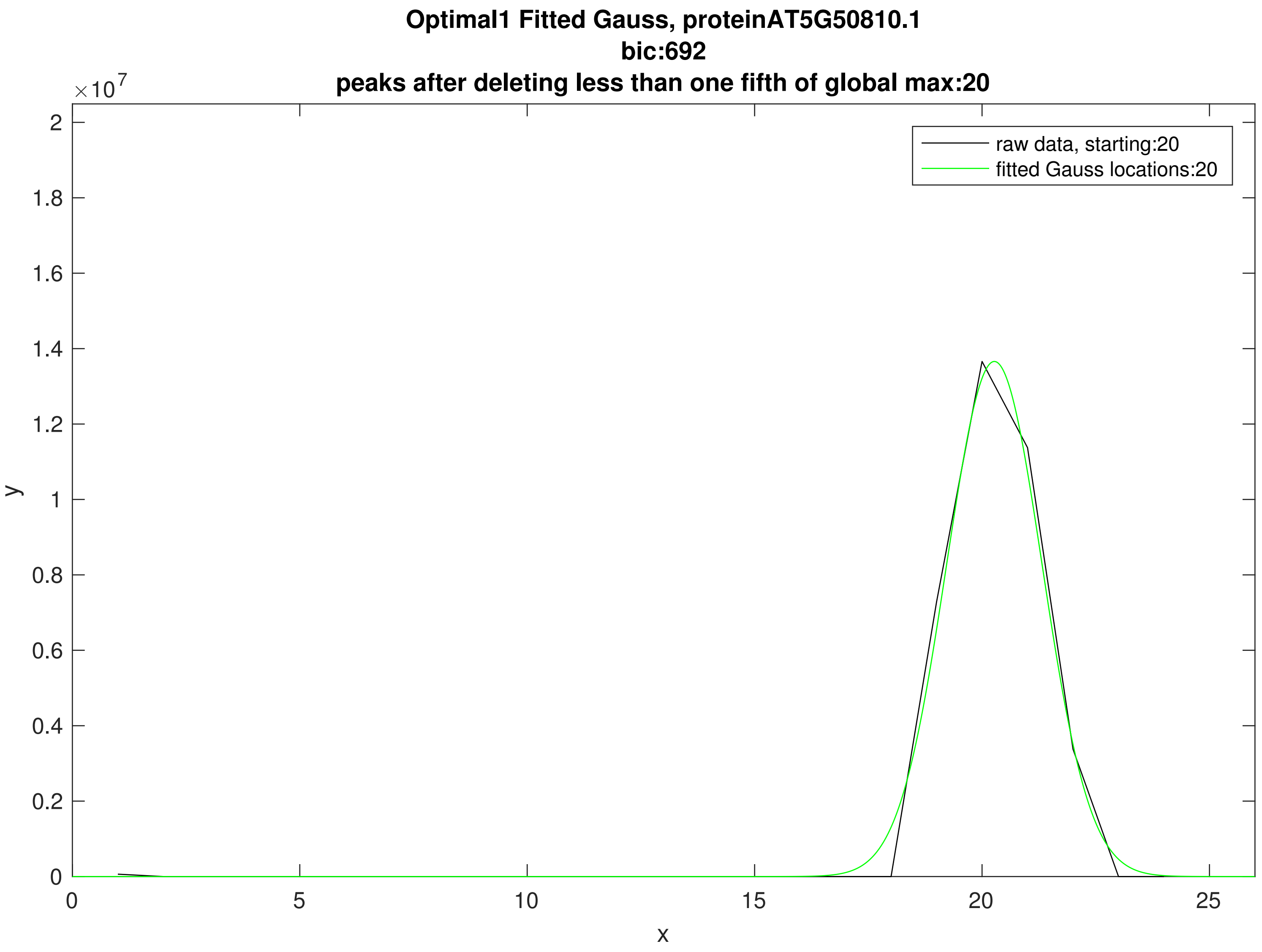
Optimal1 Fitted Gauss, proteinAT5G50810.1
bic:692
peaks after deleting less than one fifth of global max:20
×10^7
2
1.8
1.6
1.4
1.2
1
0.8
0.6
0.4
0.2
0
y
0
5
10
15
20
25
x
raw data, starting:20
fitted Gauss locations:20

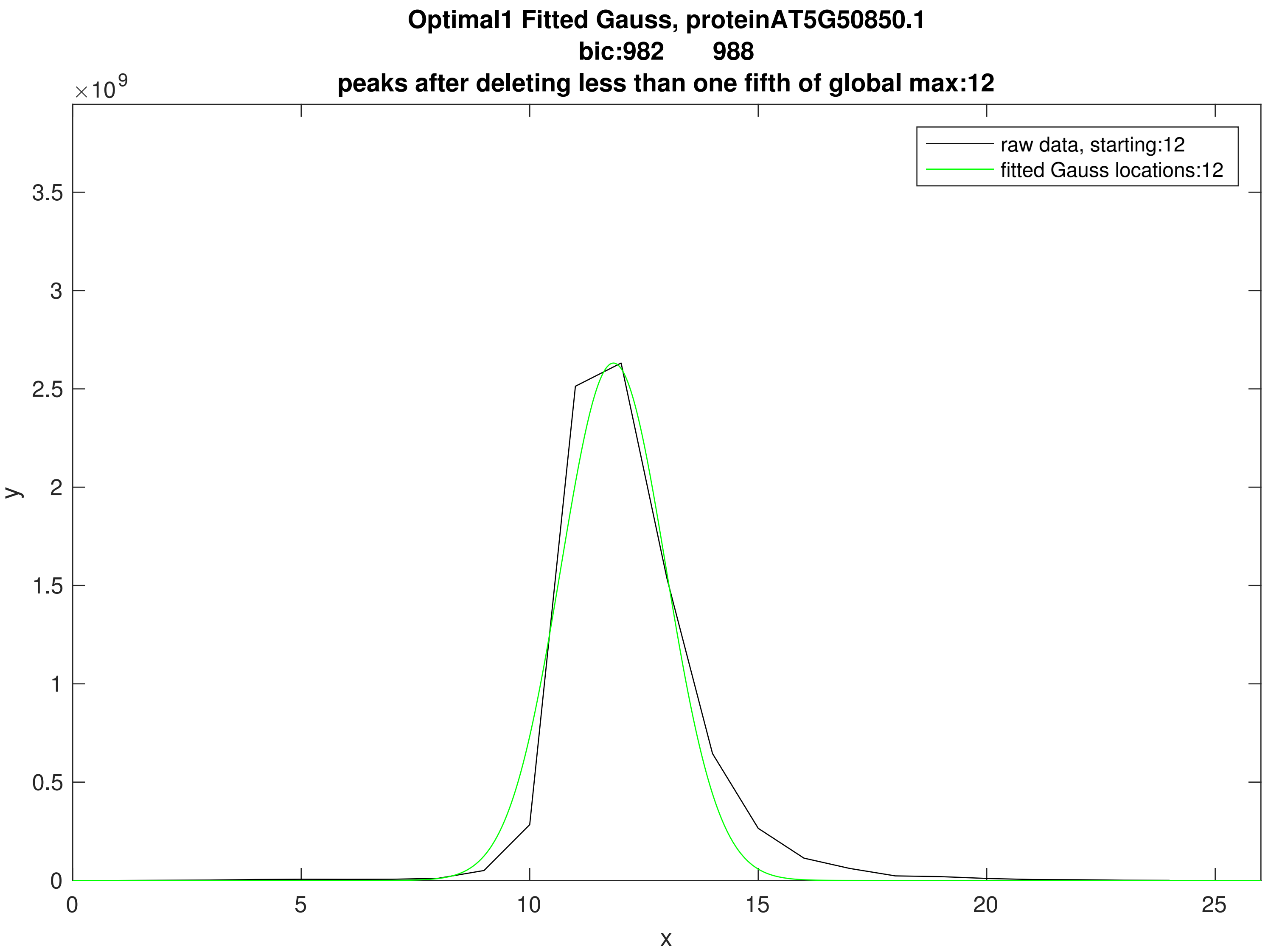
Optimal1 Fitted Gauss, proteinAT5G50850.1
bic:982 988
peaks after deleting less than one fifth of global max:12
×10^9
raw data, starting:12
fitted Gauss locations:12
3.5
3
2.5
2
1.5
1
0.5
0
y
0
5
10
15
20
25
x

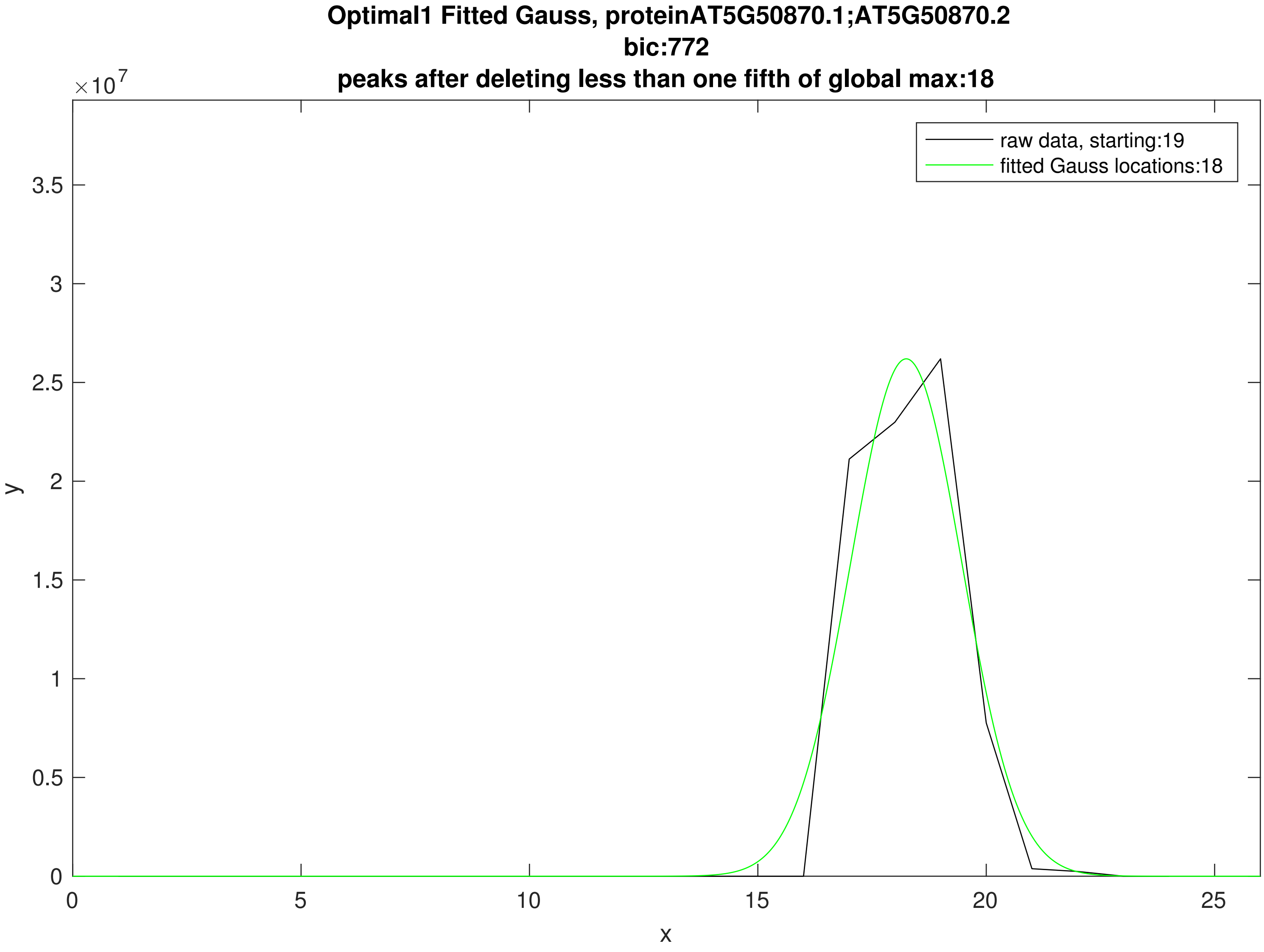

Optimal1 Fitted Gauss, proteinAT5G50870.1;AT5G50870.2
bic:772
peaks after deleting less than one fifth of global max:18
×10^7
raw data, starting:19
fitted Gauss locations:18
3.5
3
2.5
2
1.5
1
0.5
0
y
0
5
10
15
20
25
x

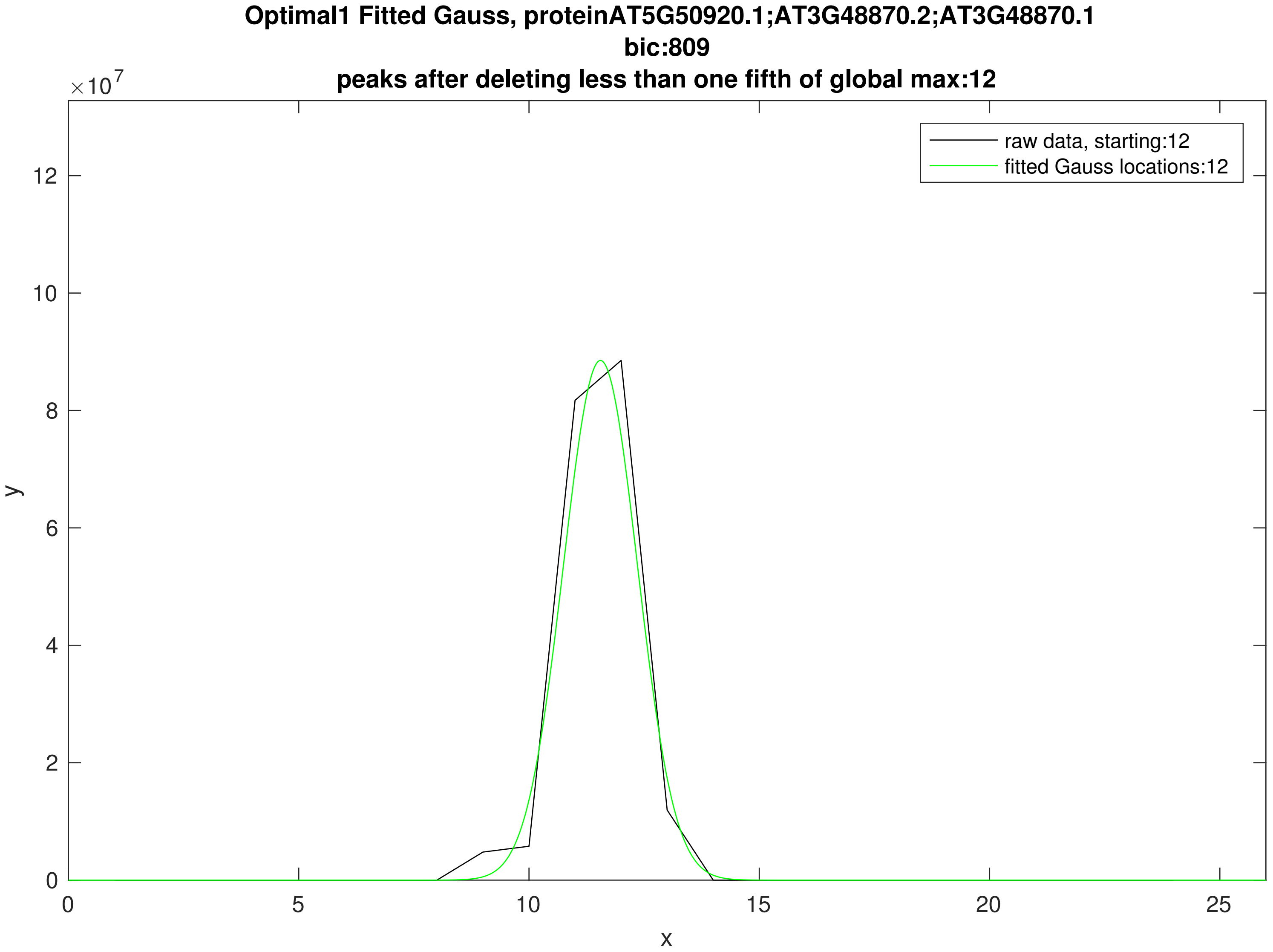
Optimal1 Fitted Gauss, proteinAT5G50920.1;AT3G48870.2;AT3G48870.1
bic:809
peaks after deleting less than one fifth of global max:12
×10^7
raw data, starting:12
fitted Gauss locations:12
y
x
0
2
4
6
8
10
12
0
5
10
15
20
25

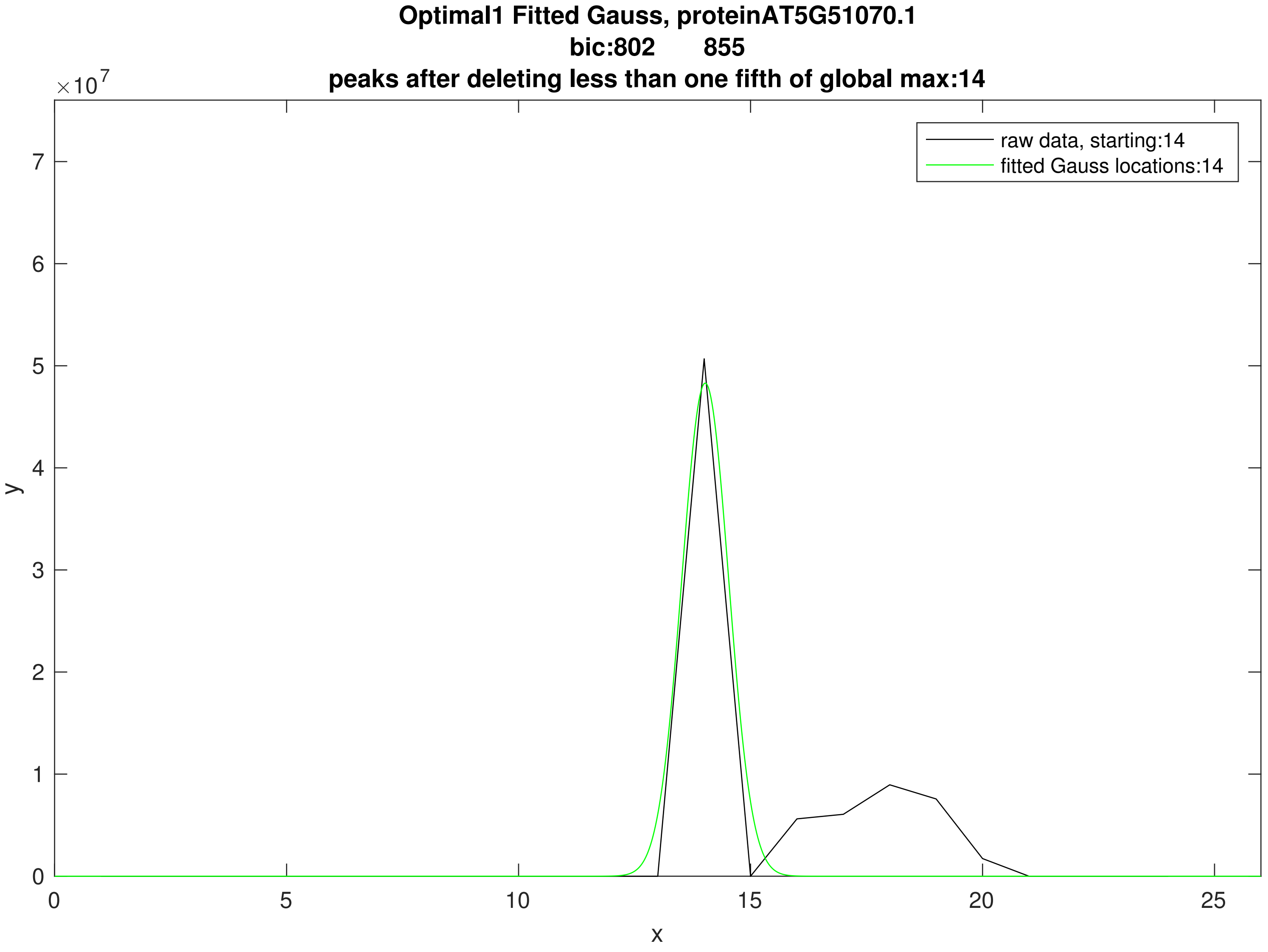
Optimal1 Fitted Gauss, proteinAT5G51070.1
bic:802 855
peaks after deleting less than one fifth of global max:14
raw data, starting:14
fitted Gauss locations:14
×10^7
y
x
0
1
2
3
4
5
6
7
0
5
10
15
20
25

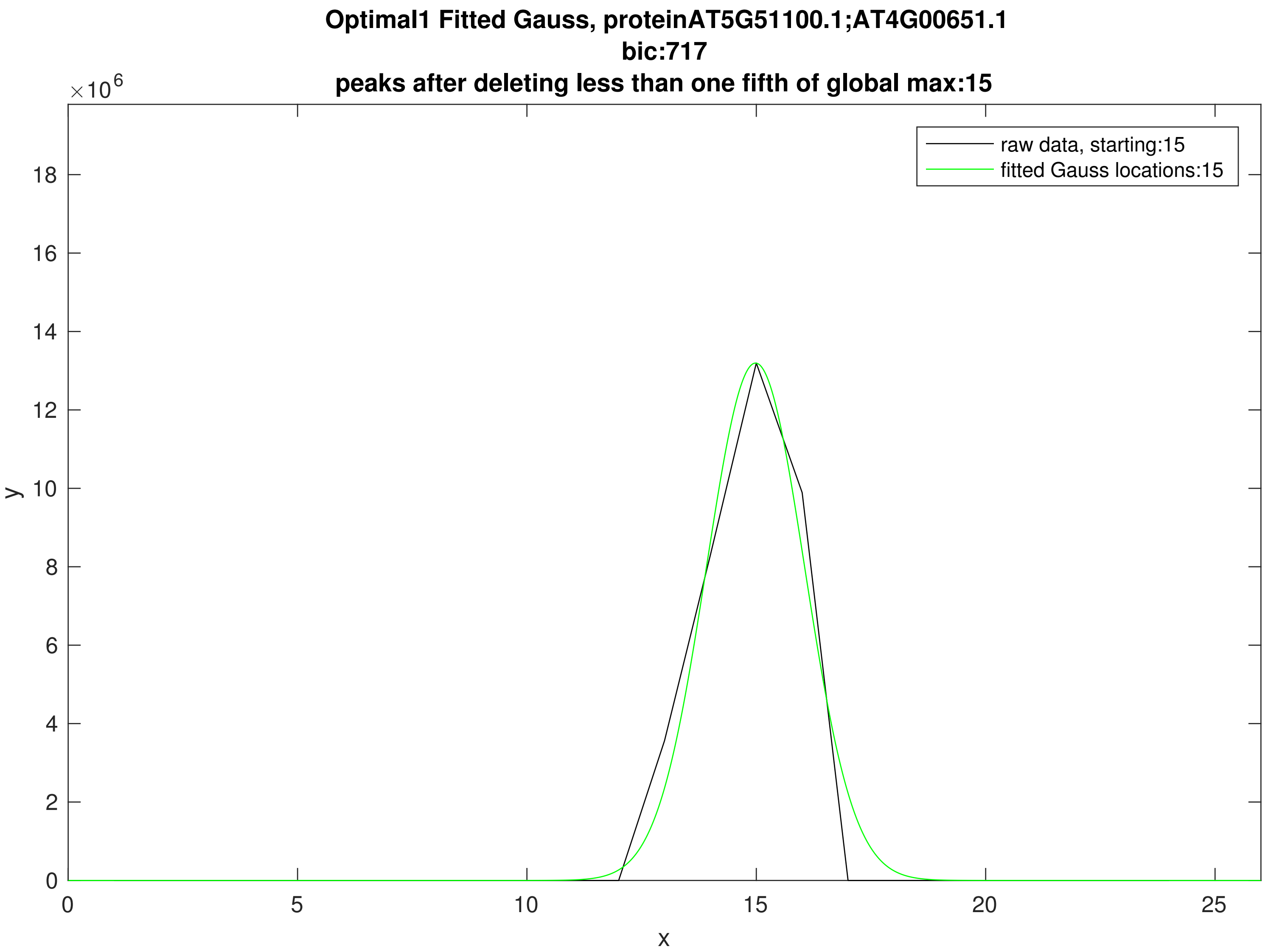

Optimal1 Fitted Gauss, proteinAT5G51100.1;AT4G00651.1
bic:717
peaks after deleting less than one fifth of global max:15
×10^6
raw data, starting:15
fitted Gauss locations:15
y
x
0
2
4
6
8
10
12
14
16
18
0
5
10
15
20
25

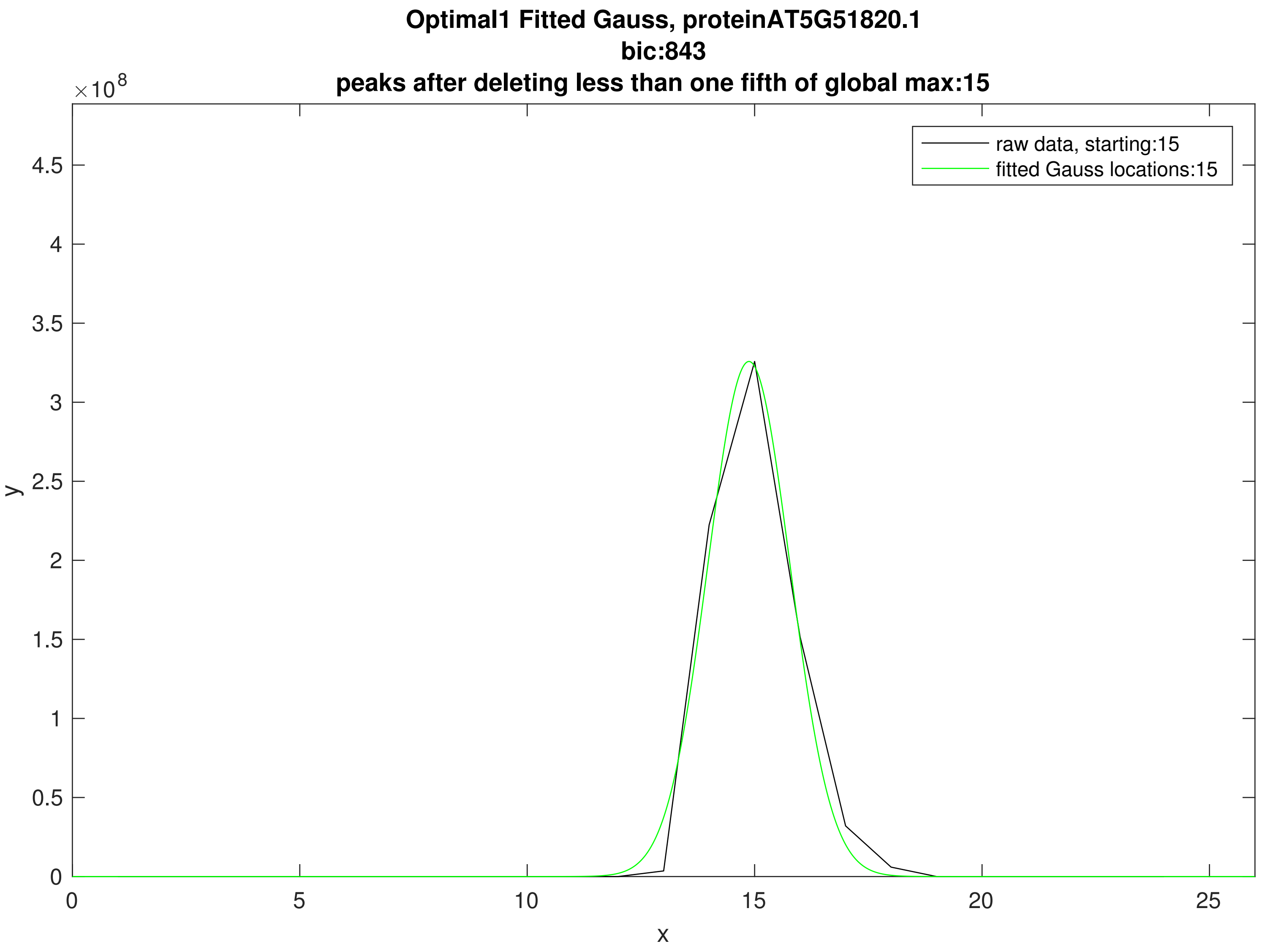
Optimal1 Fitted Gauss, proteinAT5G51820.1
bic:843
peaks after deleting less than one fifth of global max:15
×10^8
4.5
4
3.5
3
2.5
2
1.5
1
0.5
0
y
0
5
10
15
20
25
x
raw data, starting:15
fitted Gauss locations:15

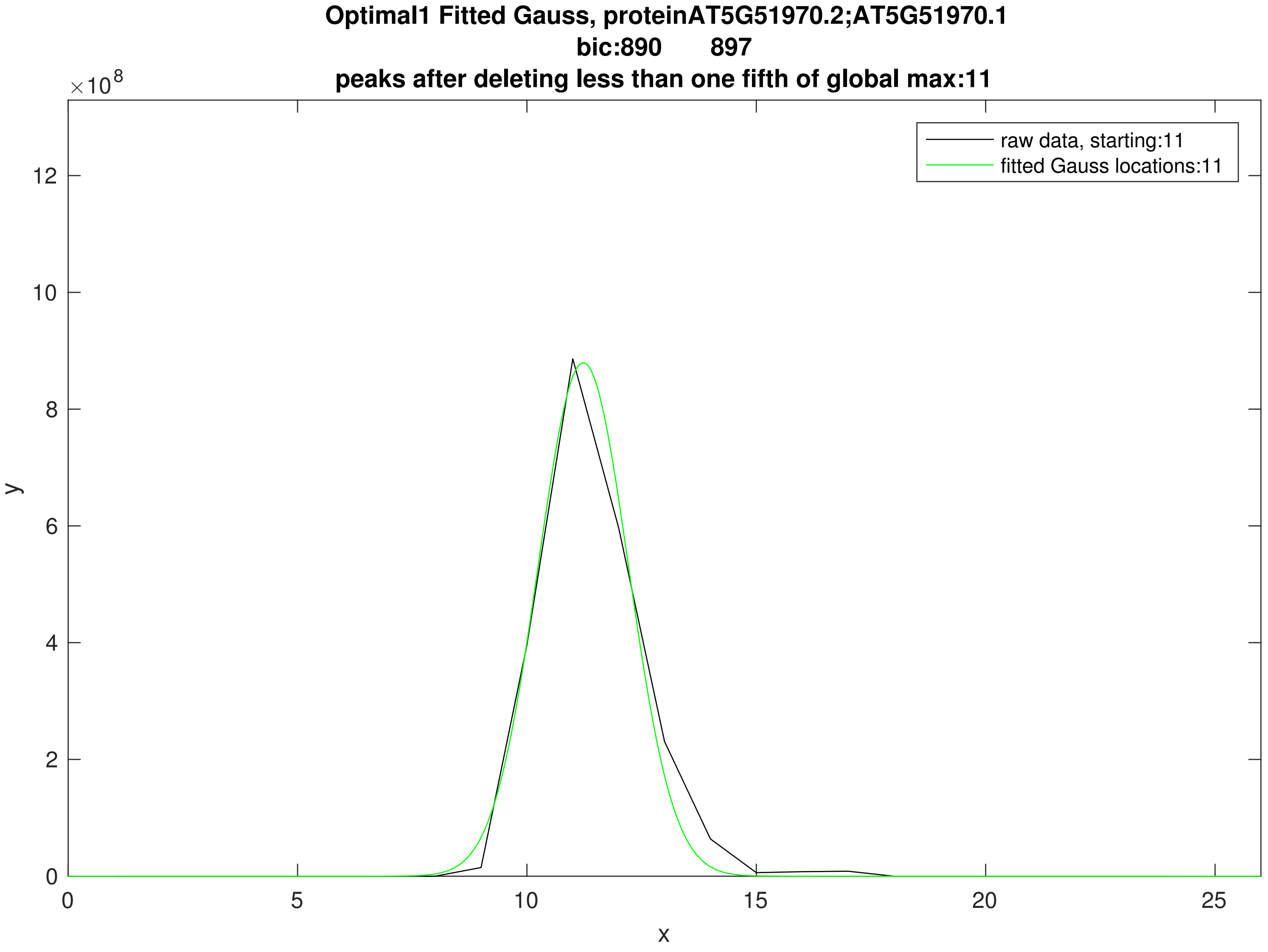
Optimal1 Fitted Gauss, proteinAT5G51970.2;AT5G51970.1
bic:890 897
peaks after deleting less than one fifth of global max:11
×10^8
raw data, starting:11
fitted Gauss locations:11
y
x
0
2
4
6
8
10
12
0
5
10
15
20
25

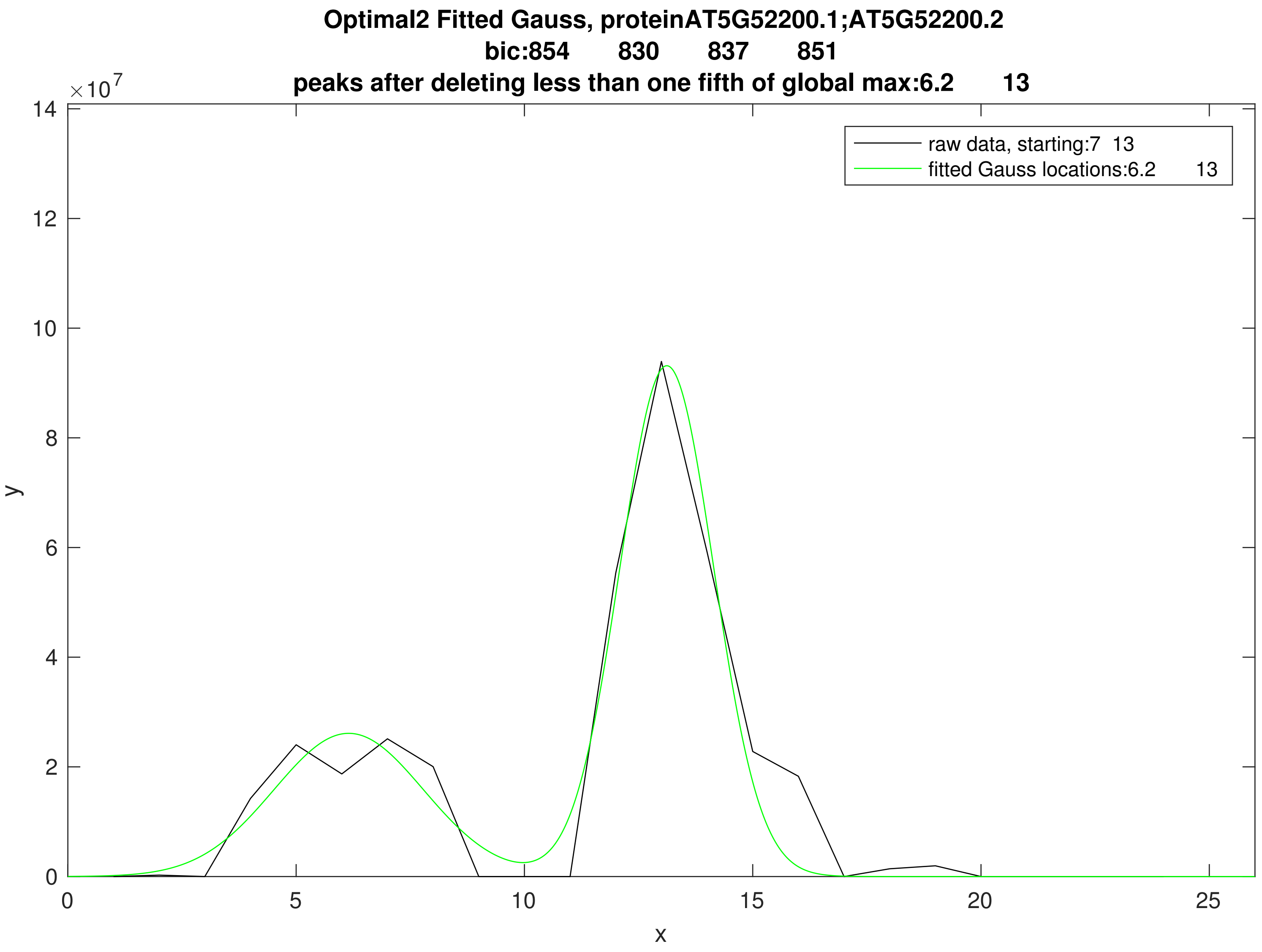
Optimal2 Fitted Gauss, proteinAT5G52200.1;AT5G52200.2
bic:854 830 837 851
peaks after deleting less than one fifth of global max:6.2 13
×10^7
raw data, starting:7 13
fitted Gauss locations:6.2 13
y
x
0
2
4
6
8
10
12
14
0
5
10
15
20
25

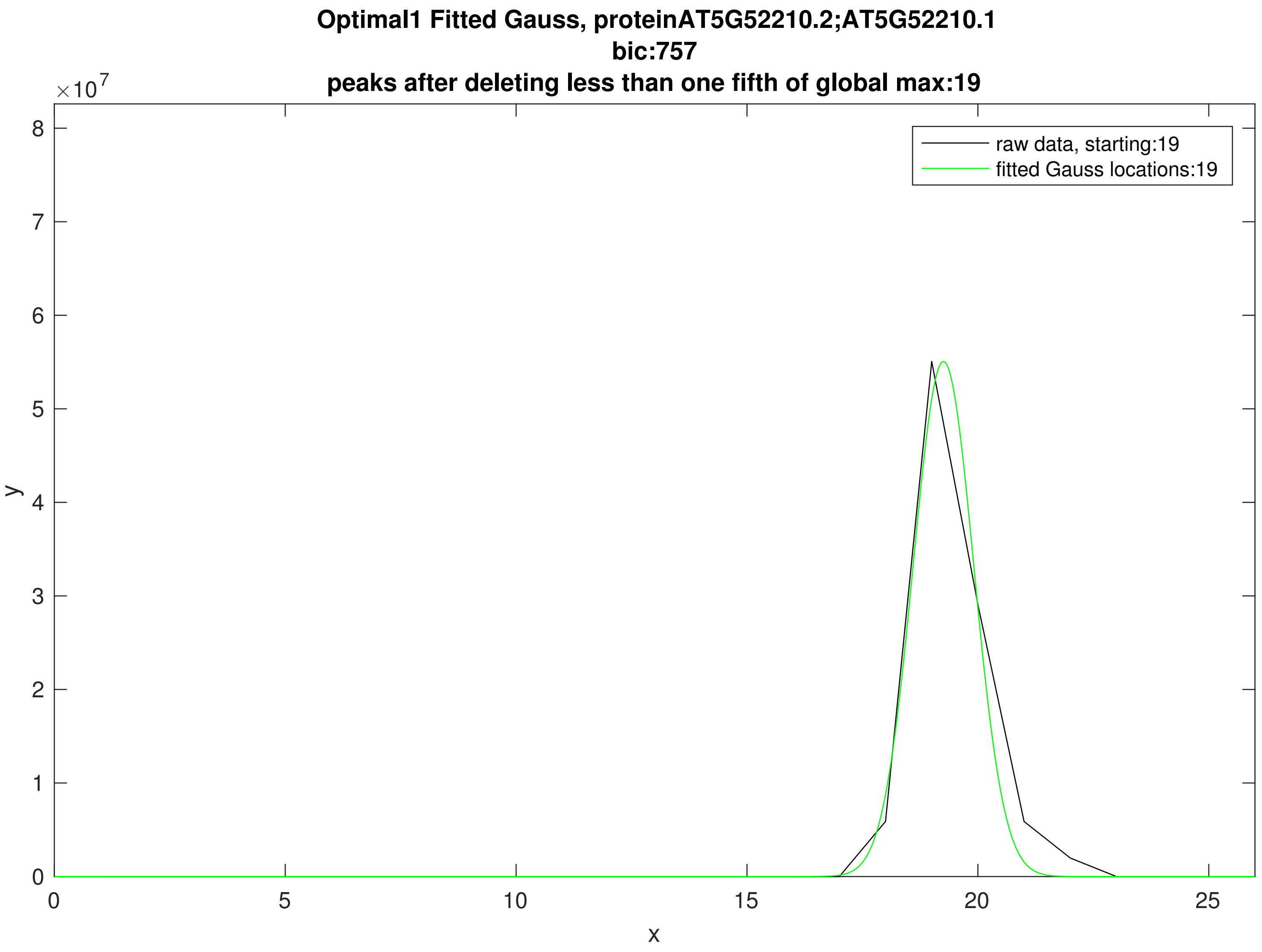

Optimal1 Fitted Gauss, proteinAT5G52210.2;AT5G52210.1
bic:757
peaks after deleting less than one fifth of global max:19
×10^7
raw data, starting:19
fitted Gauss locations:19
y
x
0
1
2
3
4
5
6
7
8
0
5
10
15
20
25

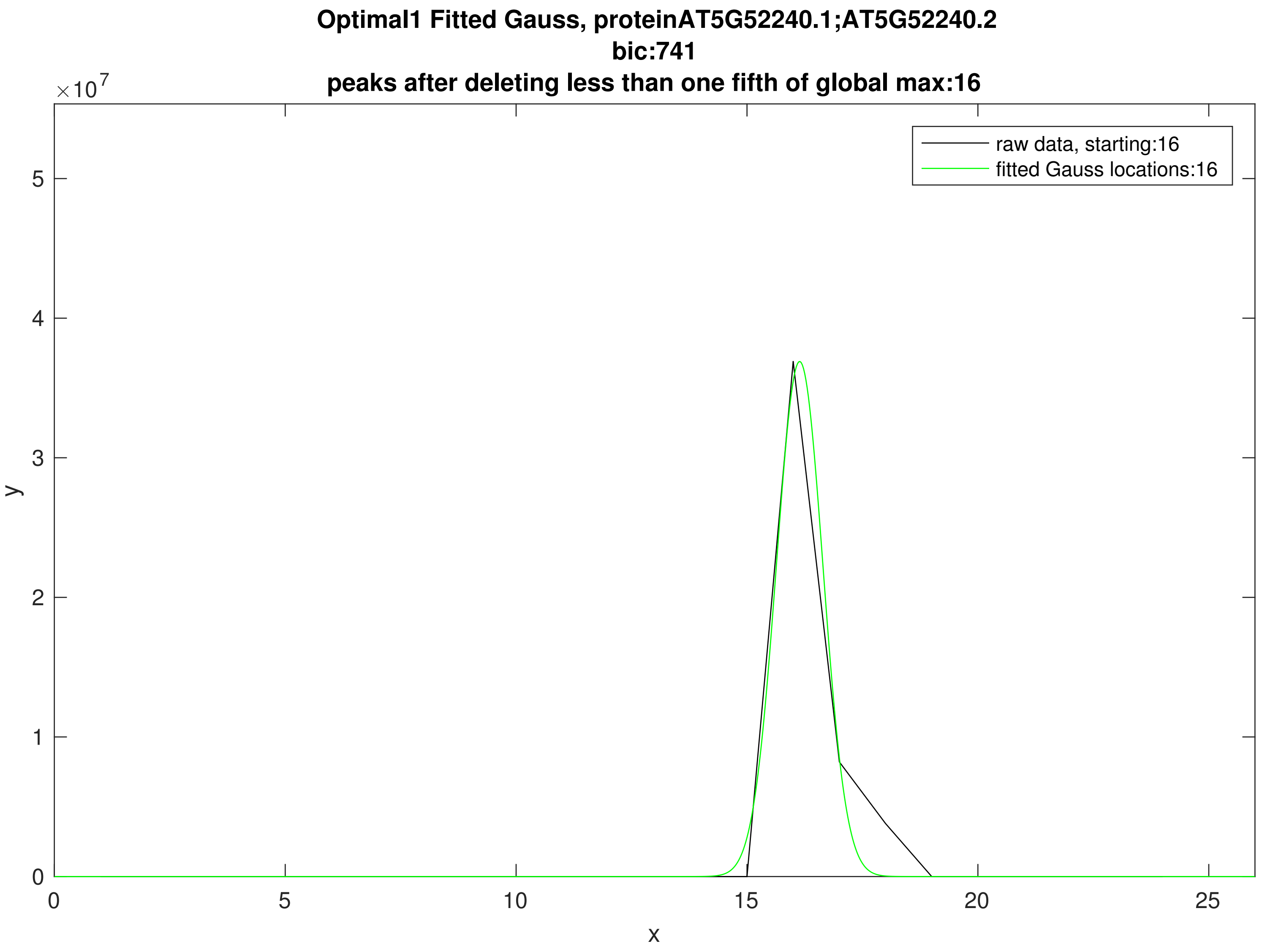
Optimal1 Fitted Gauss, proteinAT5G52240.1;AT5G52240.2
bic:741
peaks after deleting less than one fifth of global max:16
×10^7
raw data, starting:16
fitted Gauss locations:16
y
x
0
1
2
3
4
5
0
5
10
15
20
25

**Optimal1 Fitted Gauss, proteinAT5G52300.2;AT5G52300.1**
**bic:729**
**peaks after deleting less than one fifth of global max:7.3**

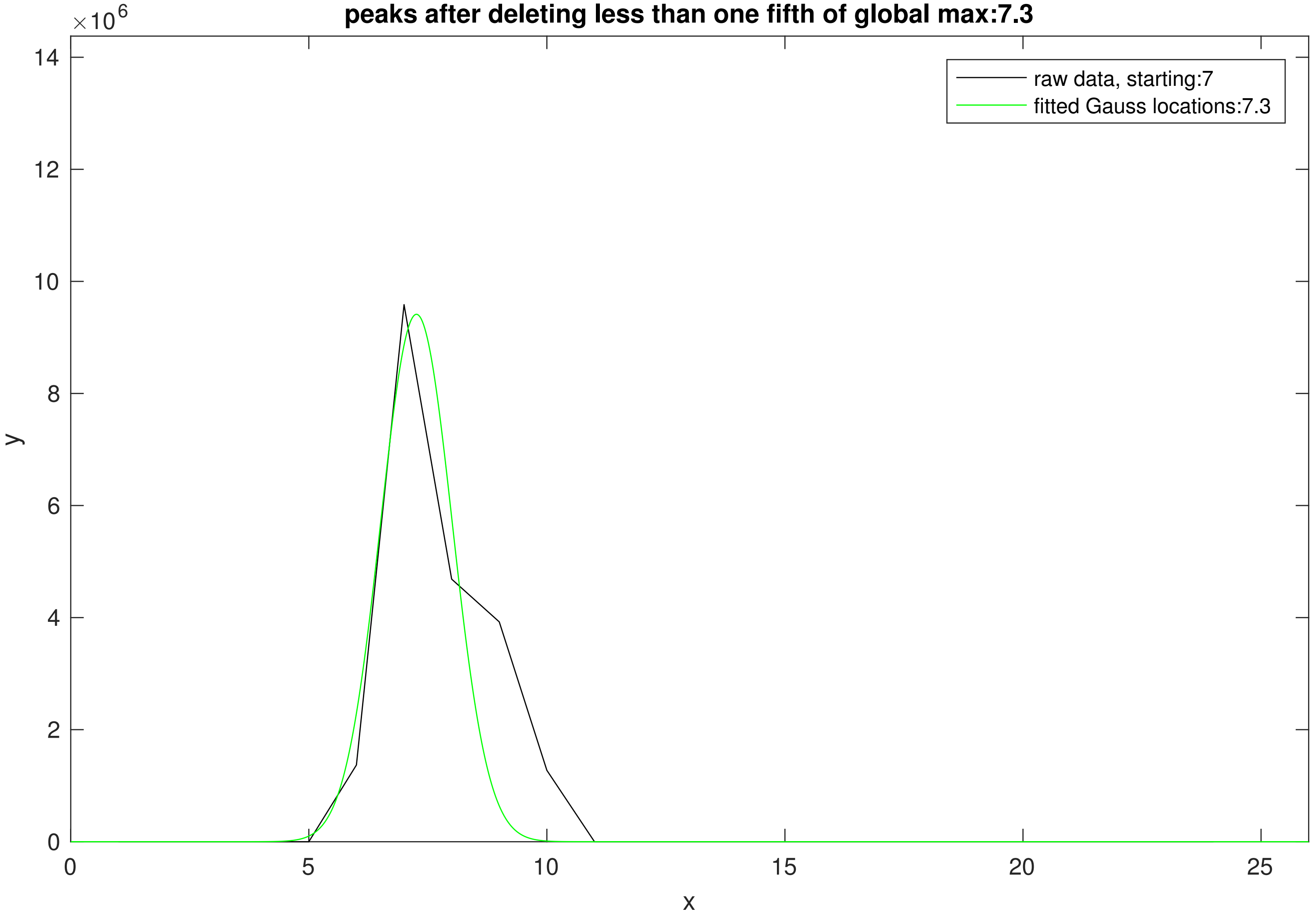

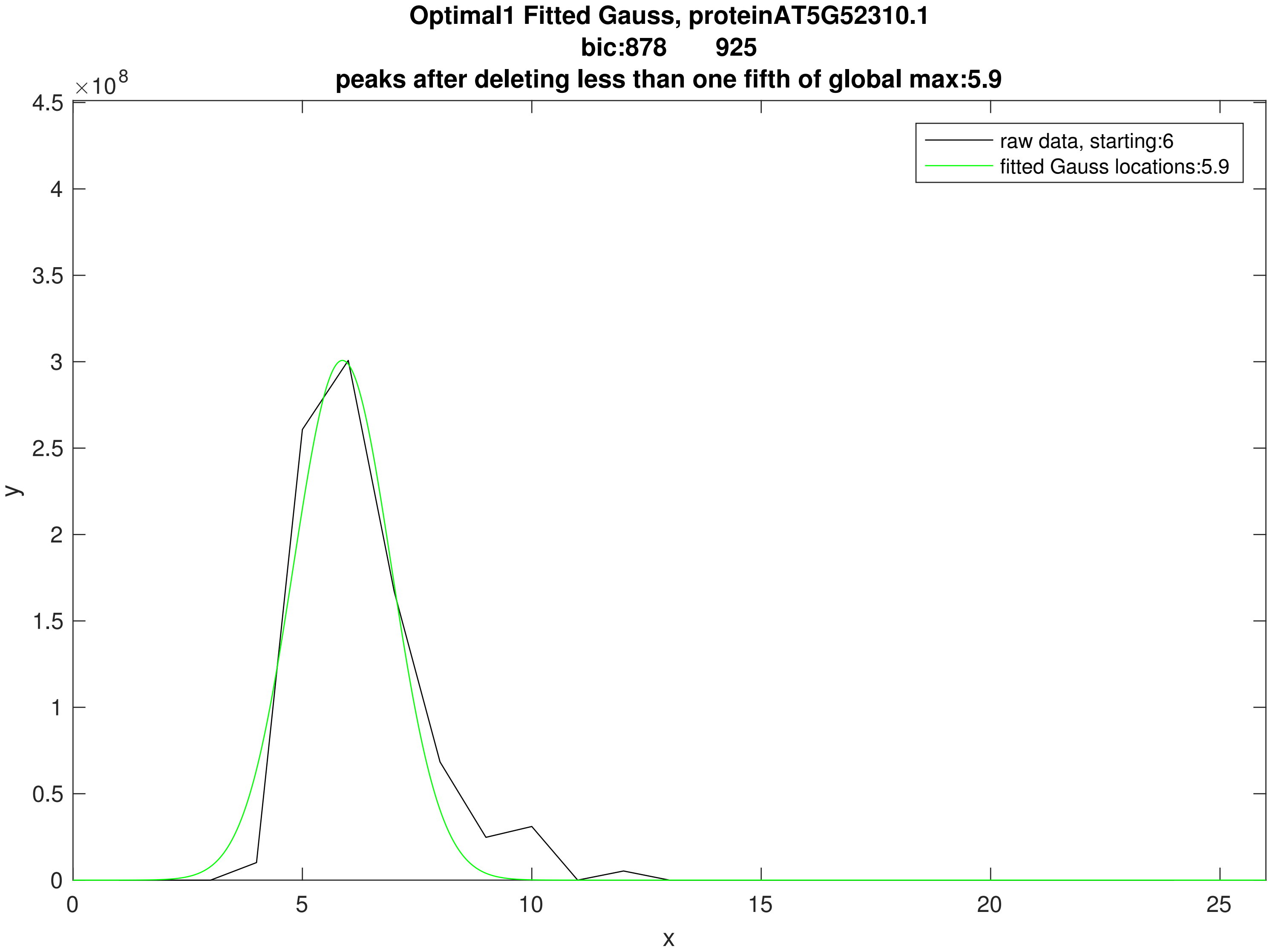

Optimal1 Fitted Gauss, proteinAT5G52310.1
bic:878 925
peaks after deleting less than one fifth of global max:5.9
raw data, starting:6
fitted Gauss locations:5.9
y
x

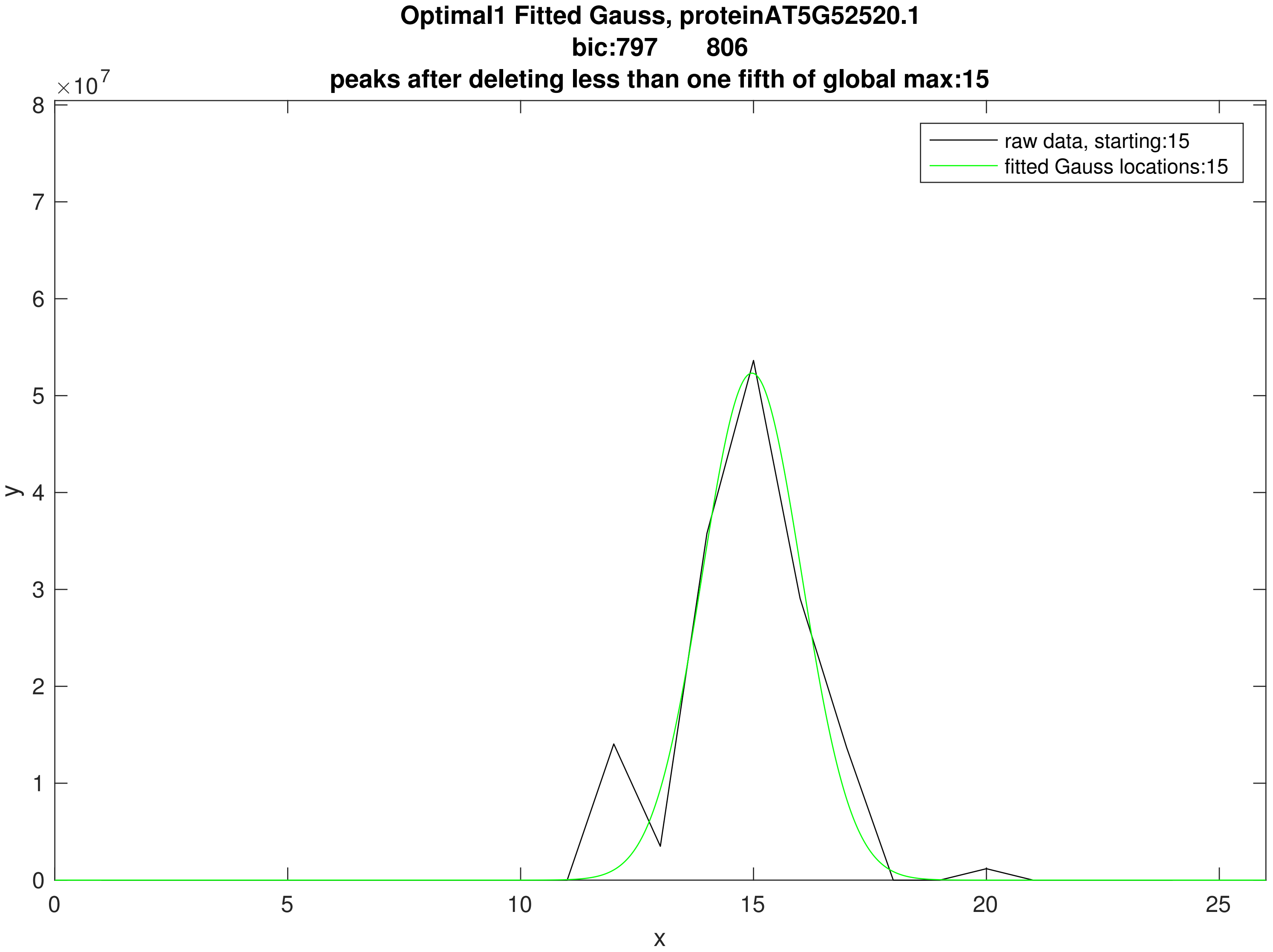
Optimal1 Fitted Gauss, proteinAT5G52520.1
bic:797 806
peaks after deleting less than one fifth of global max:15
×10^7
raw data, starting:15
fitted Gauss locations:15
y
x
0
1
2
3
4
5
6
7
8
0
5
10
15
20
25

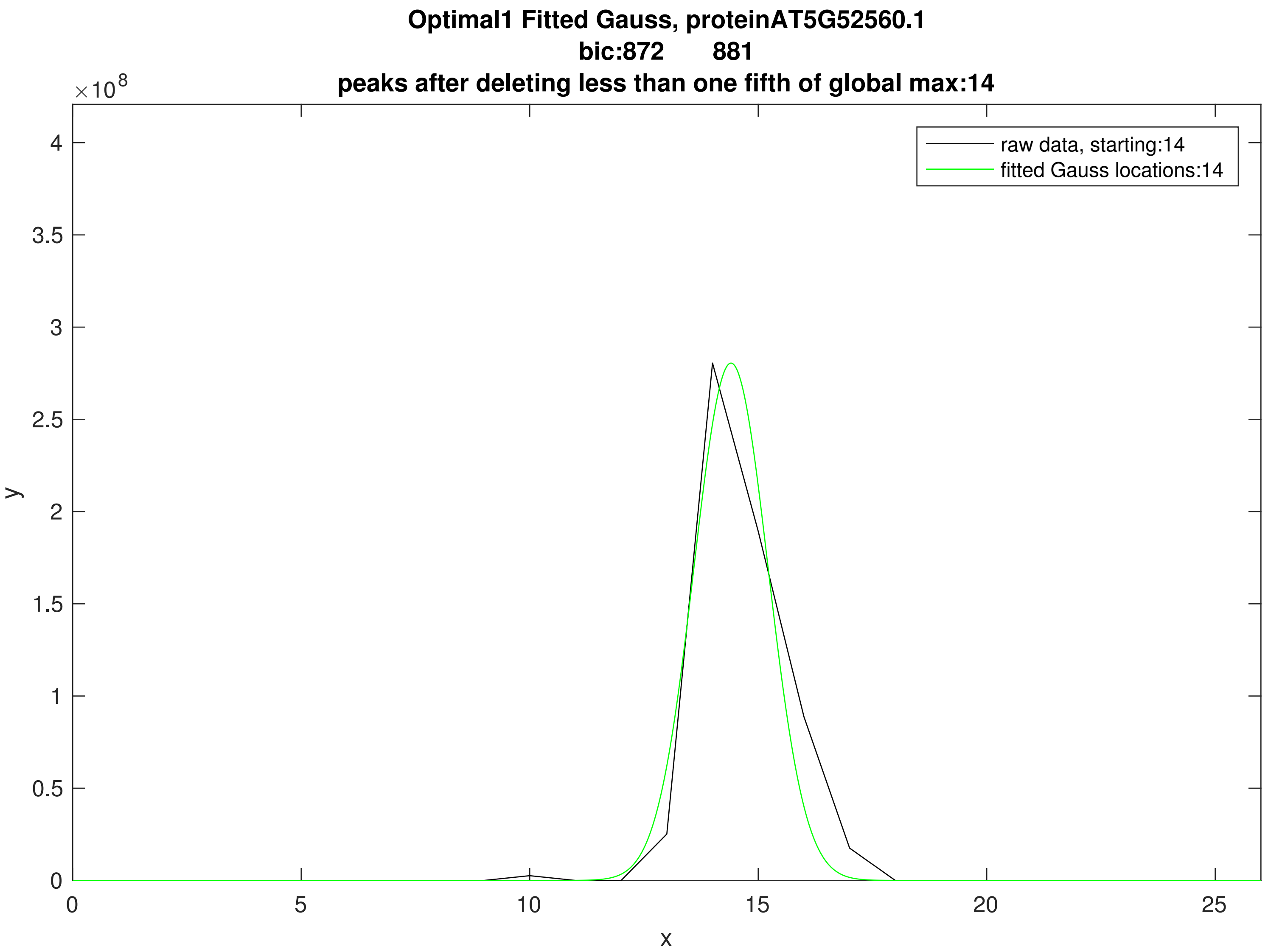
Optimal1 Fitted Gauss, proteinAT5G52560.1
bic:872 881
peaks after deleting less than one fifth of global max:14
×10^8
raw data, starting:14
fitted Gauss locations:14
4
3.5
3
2.5
2
1.5
1
0.5
0
y
0
5
10
15
20
25
x

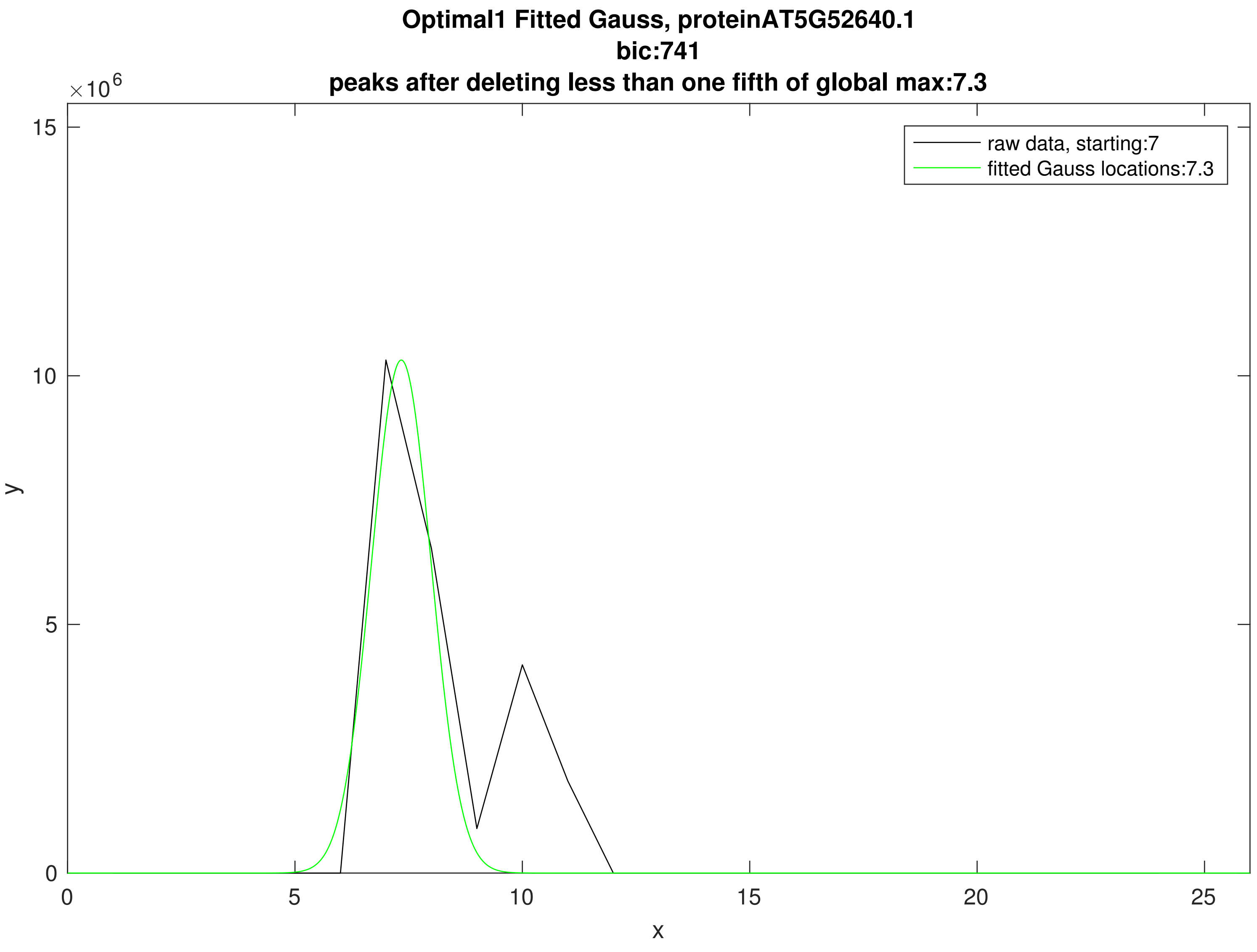
Optimal1 Fitted Gauss, proteinAT5G52640.1
bic:741
peaks after deleting less than one fifth of global max:7.3
×10^6
raw data, starting:7
fitted Gauss locations:7.3
y
x
0
5
10
15
20
25

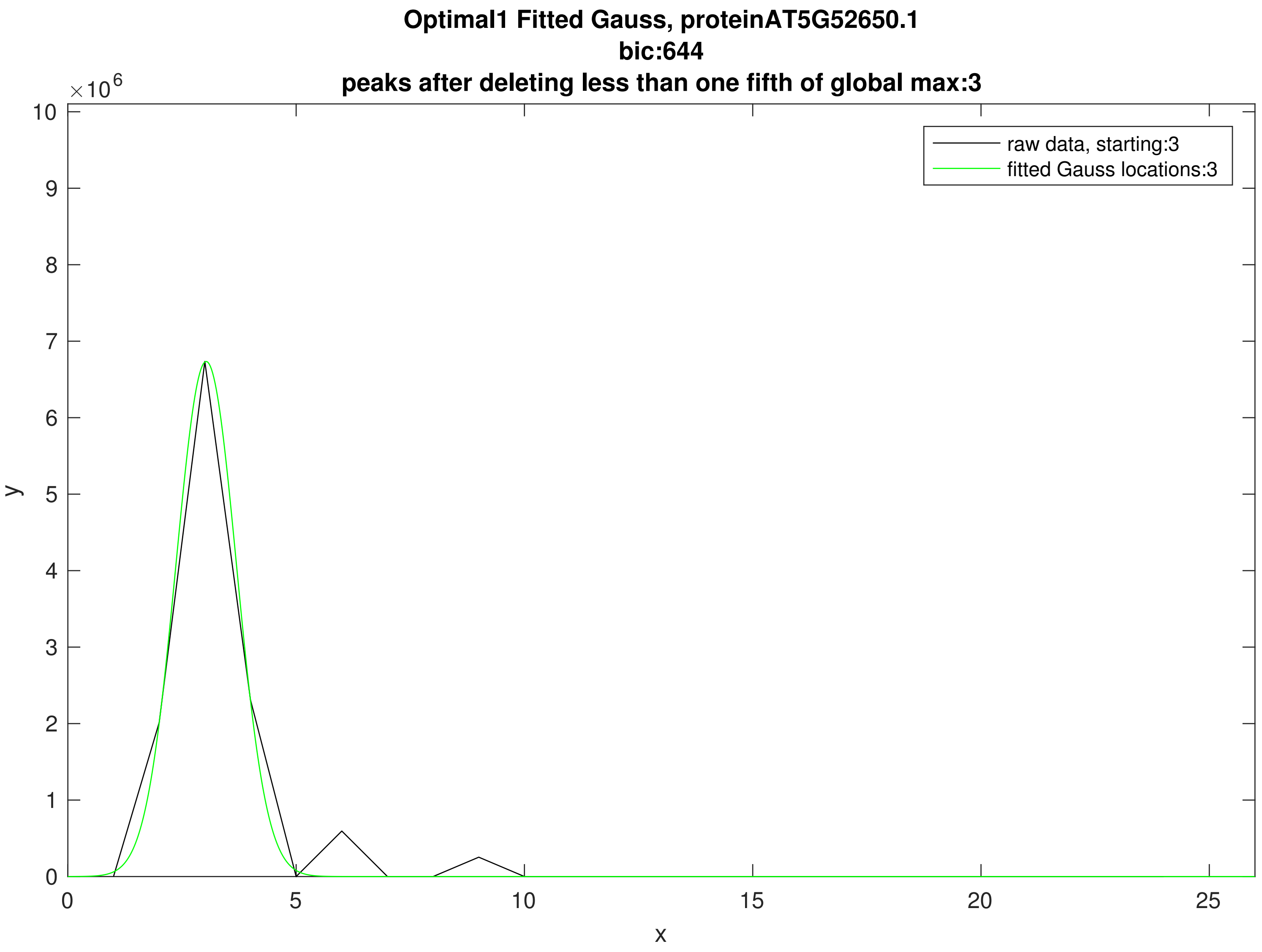
Optimal1 Fitted Gauss, proteinAT5G52650.1
bic:644
peaks after deleting less than one fifth of global max:3
×10^6
10
9
8
7
6
5
4
3
2
1
0
y
0
5
10
15
20
25
x
raw data, starting:3
fitted Gauss locations:3

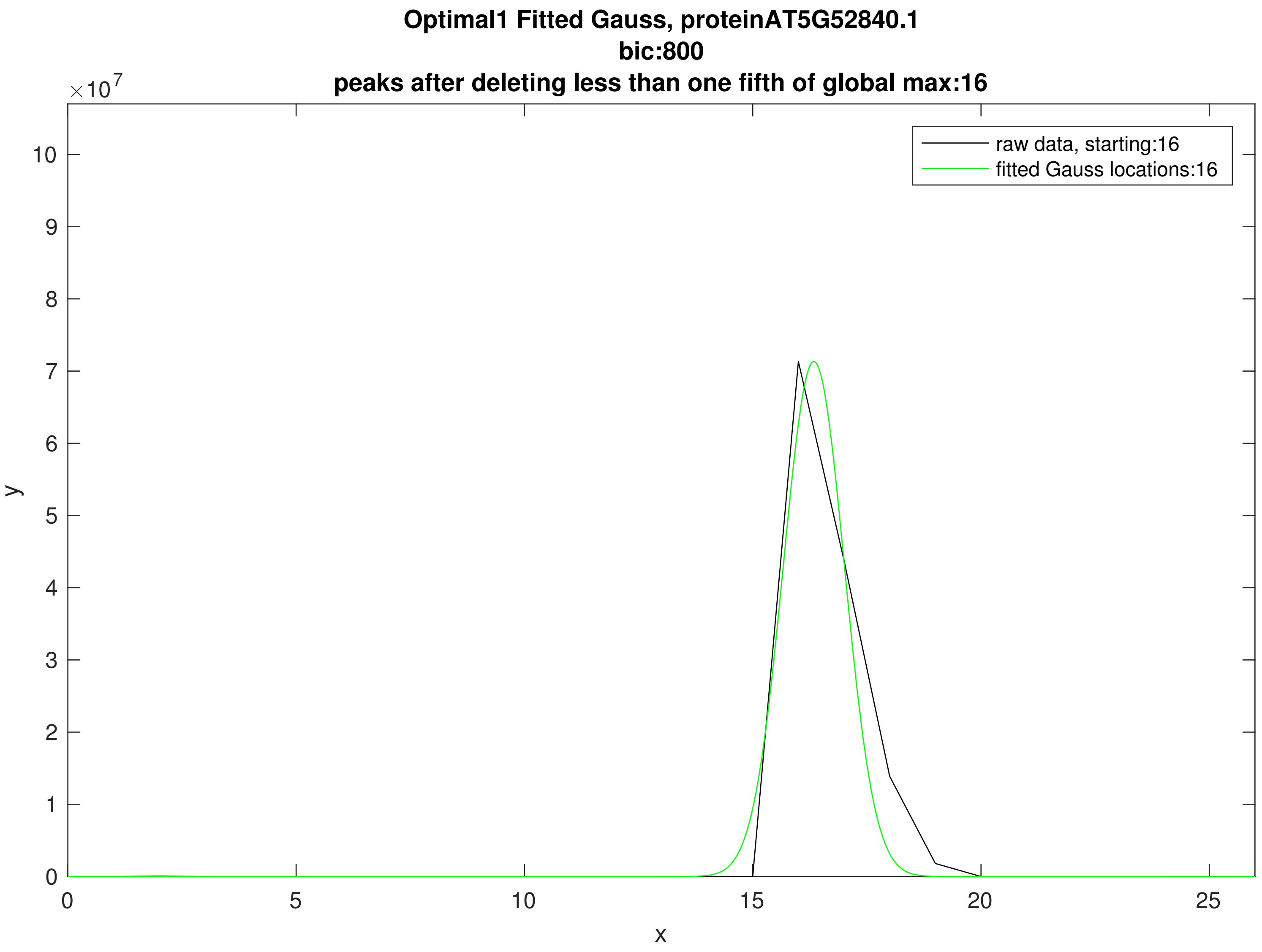
Optimal1 Fitted Gauss, proteinAT5G52840.1
bic:800
peaks after deleting less than one fifth of global max:16
×10^7
raw data, starting:16
fitted Gauss locations:16
y
x
0
1
2
3
4
5
6
7
8
9
10
0
5
10
15
20
25

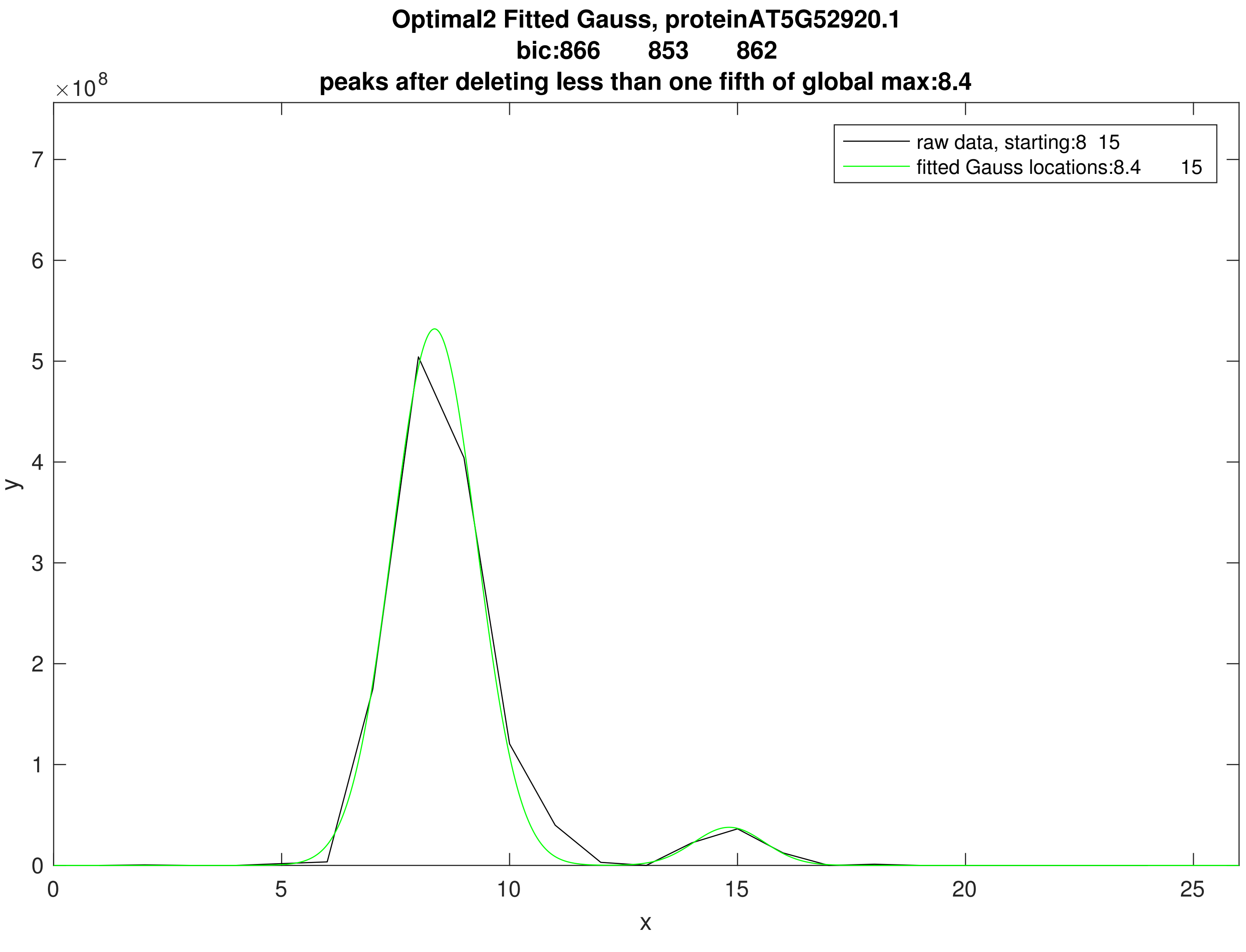
Optimal2 Fitted Gauss, proteinAT5G52920.1
bic:866 853 862
peaks after deleting less than one fifth of global max:8.4
×10^8
raw data, starting:8 15
fitted Gauss locations:8.4 15
y
x
0
1
2
3
4
5
6
7
0
5
10
15
20
25

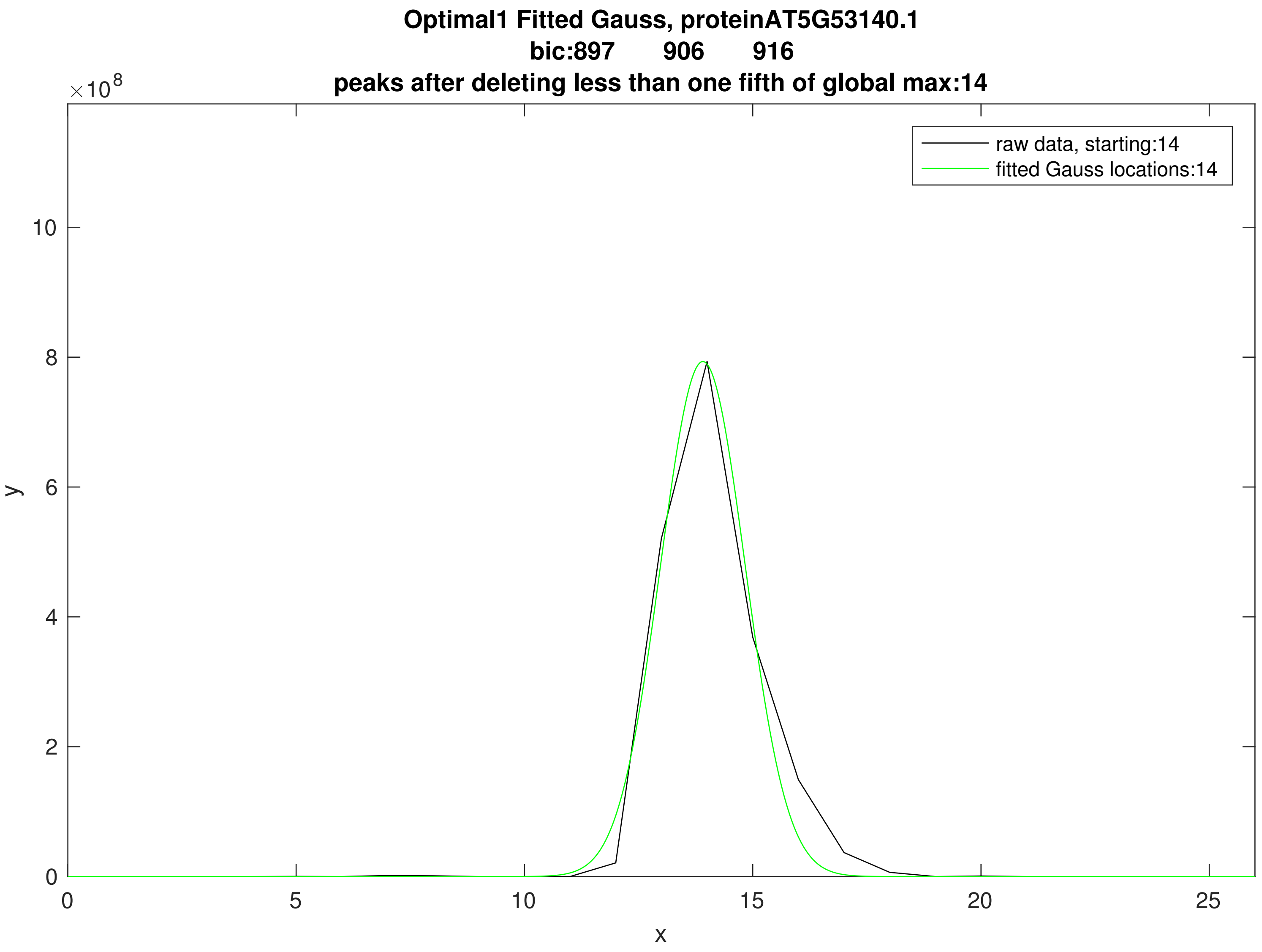
Optimal1 Fitted Gauss, proteinAT5G53140.1
bic:897 906 916
peaks after deleting less than one fifth of global max:14
×10^8
raw data, starting:14
fitted Gauss locations:14
y
x
0
2
4
6
8
10
0
5
10
15
20
25

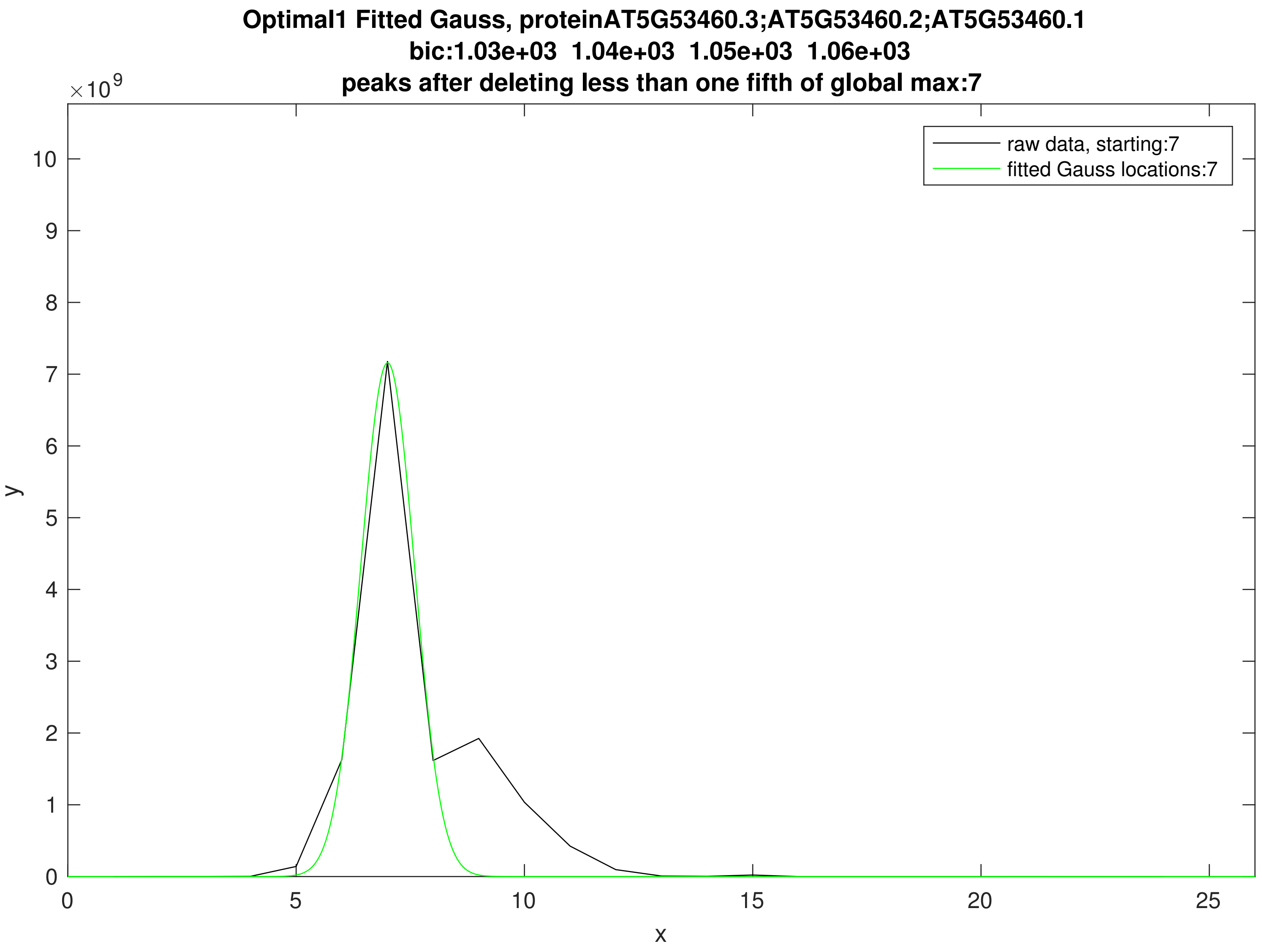
Optimal1 Fitted Gauss, proteinAT5G53460.3;AT5G53460.2;AT5G53460.1
bic:1.03e+03 1.04e+03 1.05e+03 1.06e+03
peaks after deleting less than one fifth of global max:7
×10^9
raw data, starting:7
fitted Gauss locations:7
y
x
0
1
2
3
4
5
6
7
8
9
10
0
5
10
15
20
25

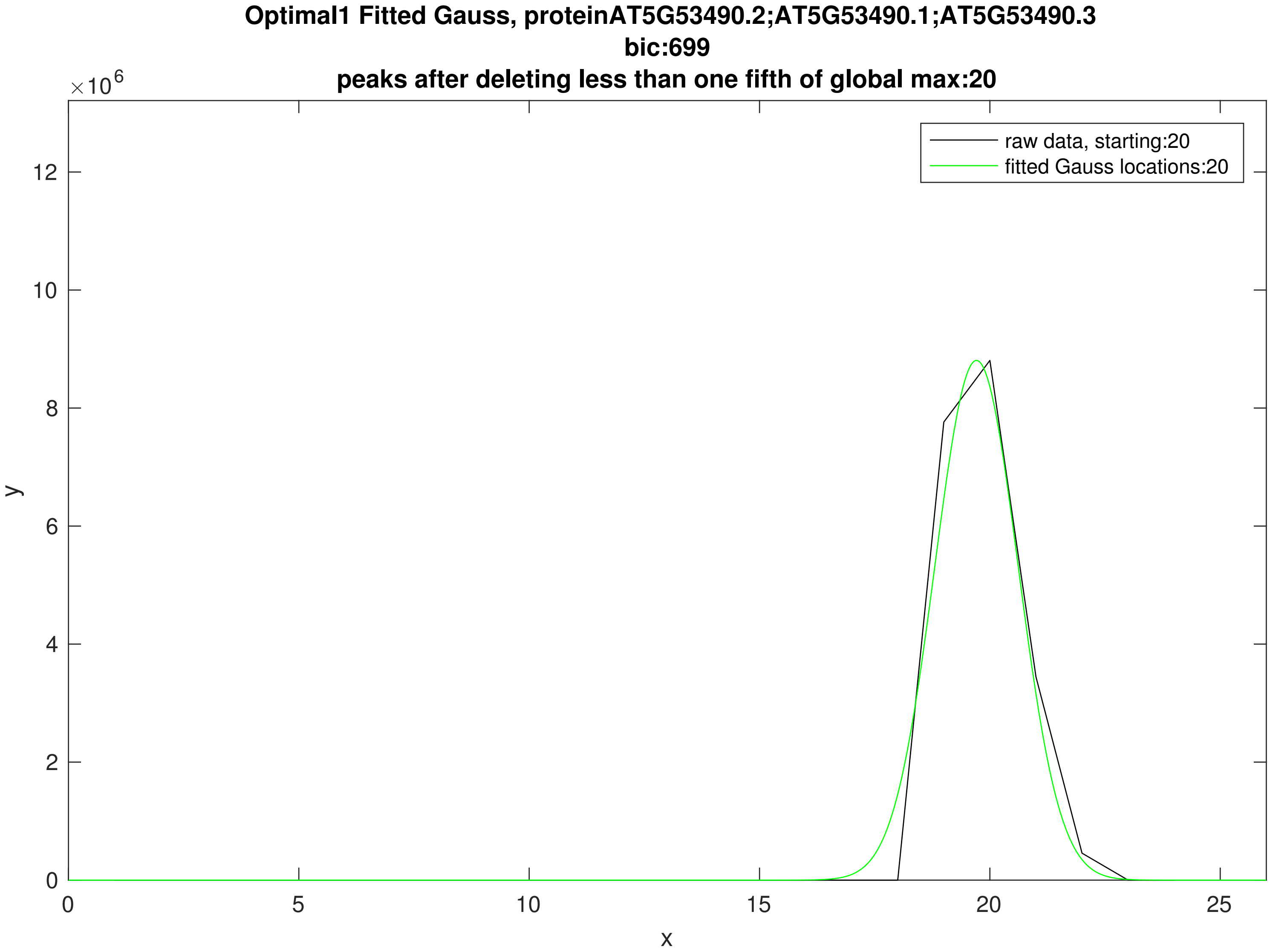

Optimal1 Fitted Gauss, proteinAT5G53490.2;AT5G53490.1;AT5G53490.3
bic:699
peaks after deleting less than one fifth of global max:20
×10^6
raw data, starting:20
fitted Gauss locations:20
y
x
0
2
4
6
8
10
12
0
5
10
15
20
25

**Optimal1 Fitted Gauss, proteinAT5G53530.1**
**bic:837**
**peaks after deleting less than one fifth of global max:17**

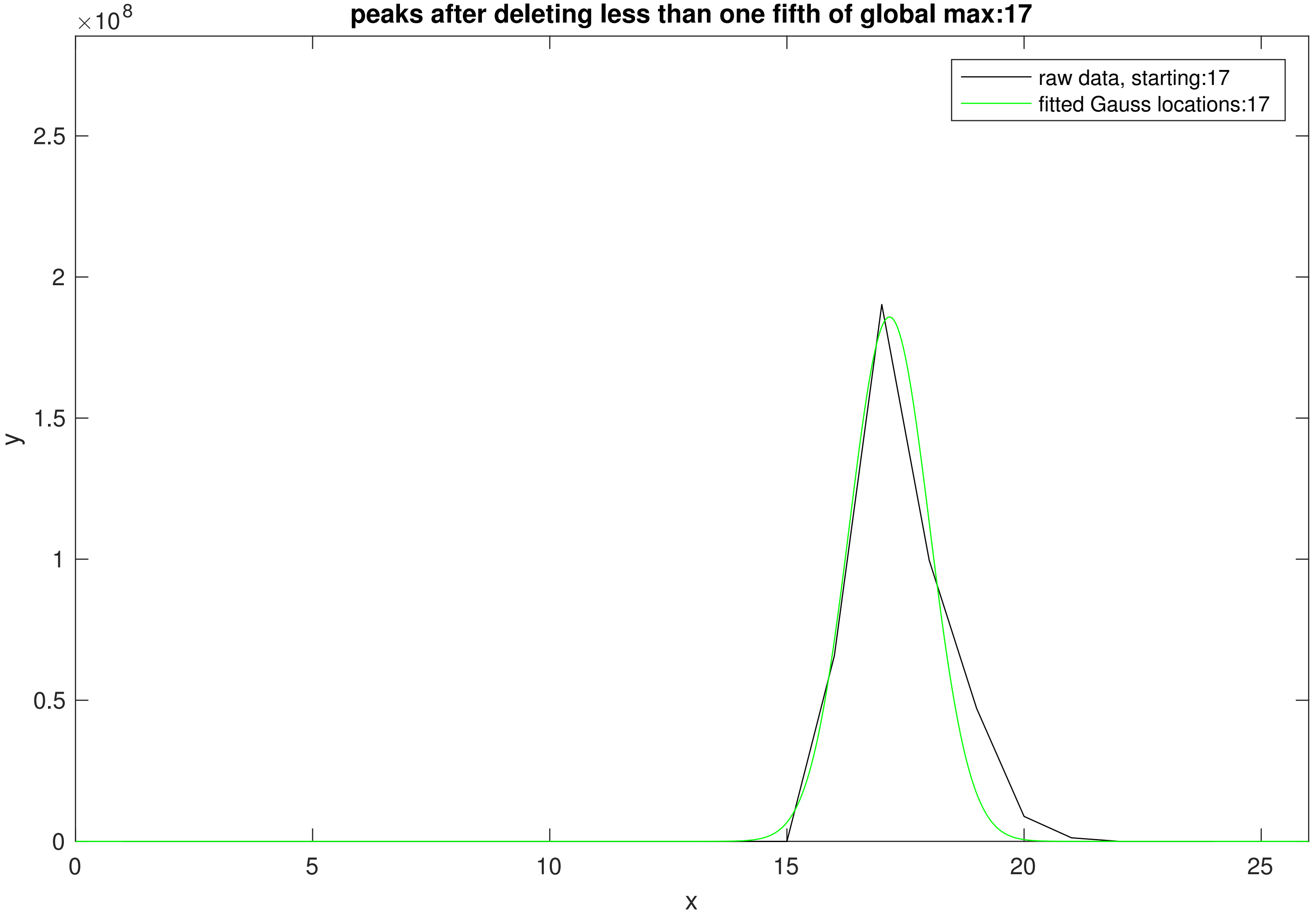

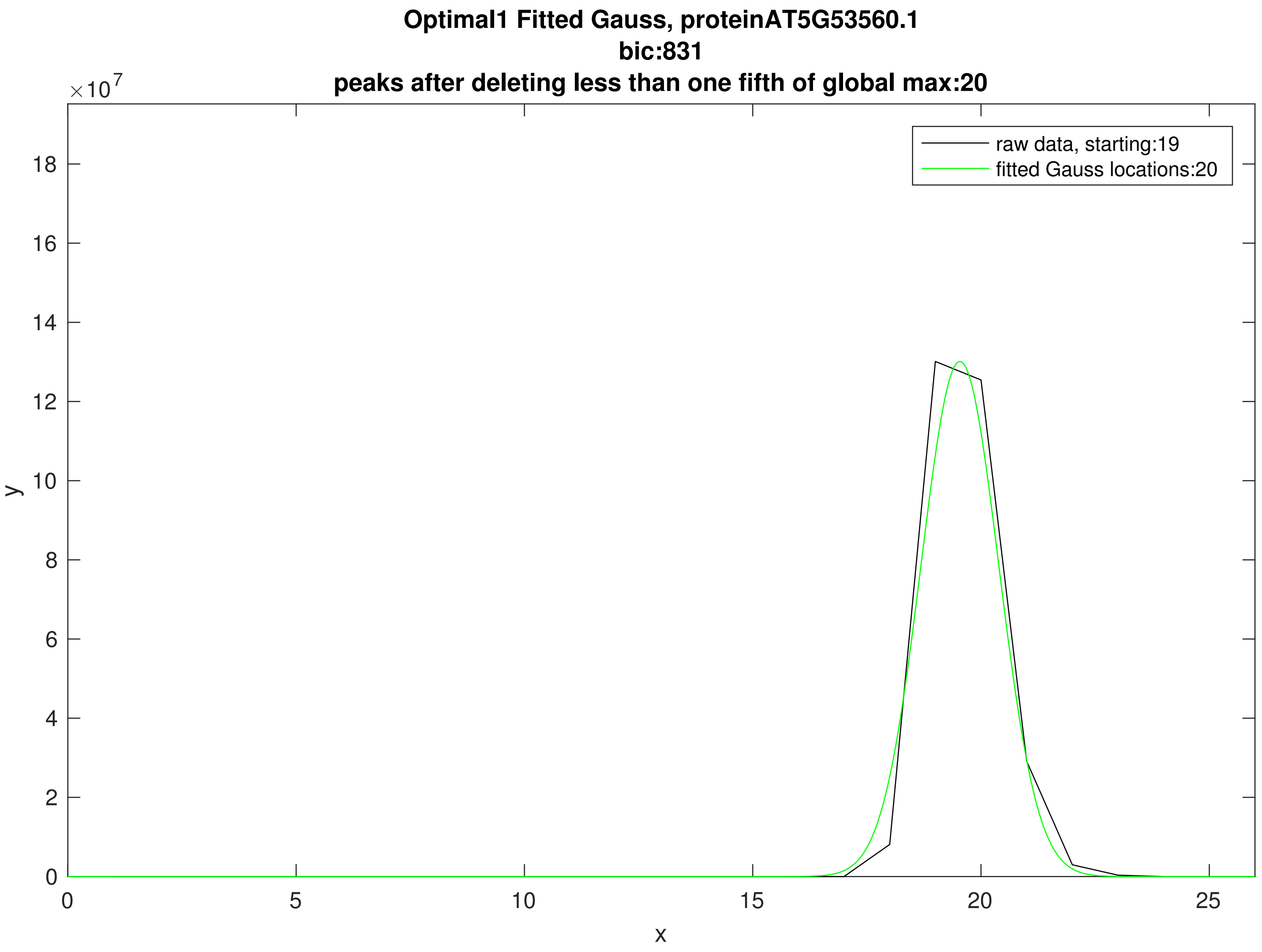

Optimal1 Fitted Gauss, proteinAT5G53560.1
bic:831
peaks after deleting less than one fifth of global max:20
×10^7
raw data, starting:19
fitted Gauss locations:20
18
16
14
12
10
8
6
4
2
0
y
0
5
10
15
20
25
x

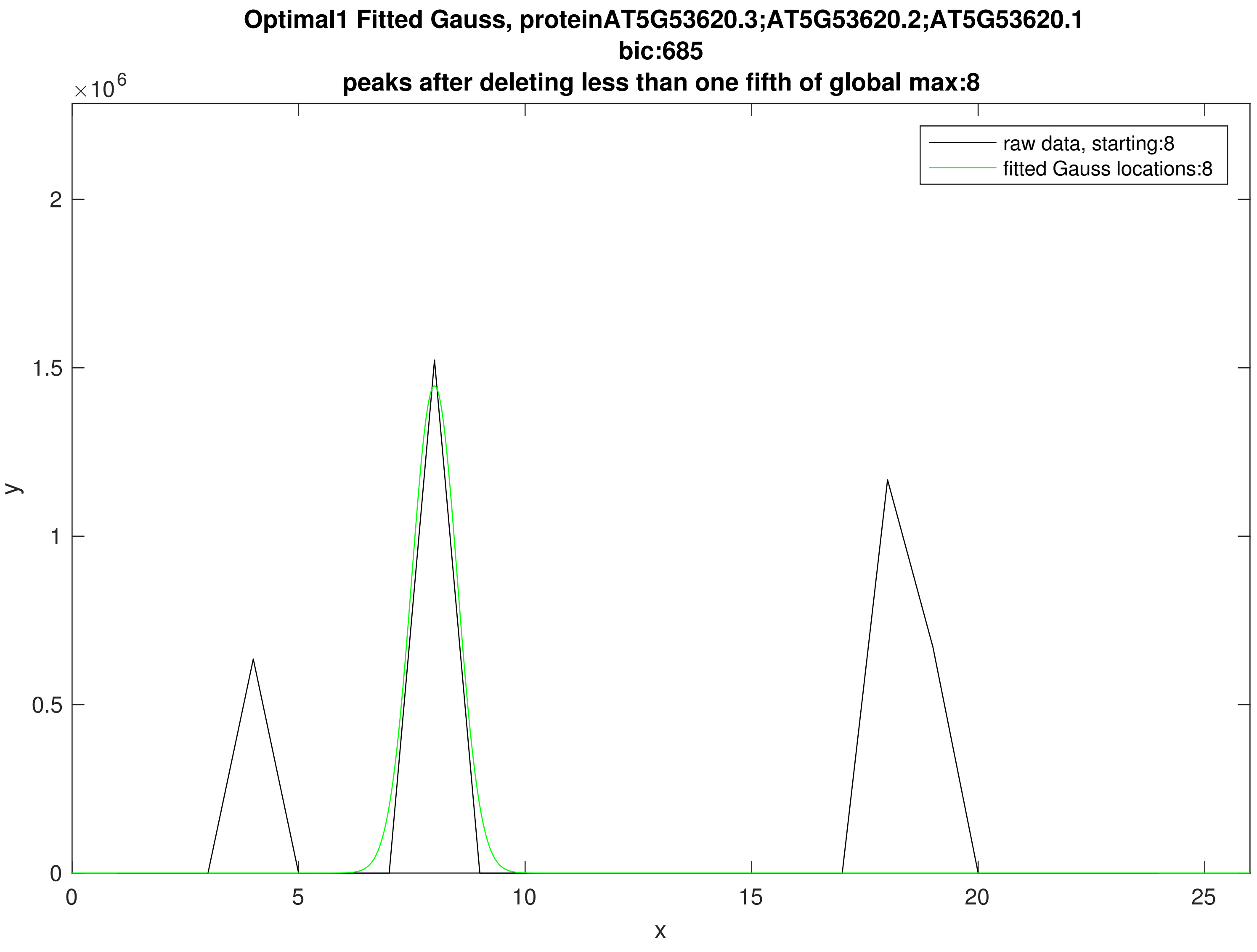

Optimal1 Fitted Gauss, proteinAT5G53620.3;AT5G53620.2;AT5G53620.1
bic:685
peaks after deleting less than one fifth of global max:8
×10^6
raw data, starting:8
fitted Gauss locations:8
y
x
0
0.5
1
1.5
2
0
5
10
15
20
25

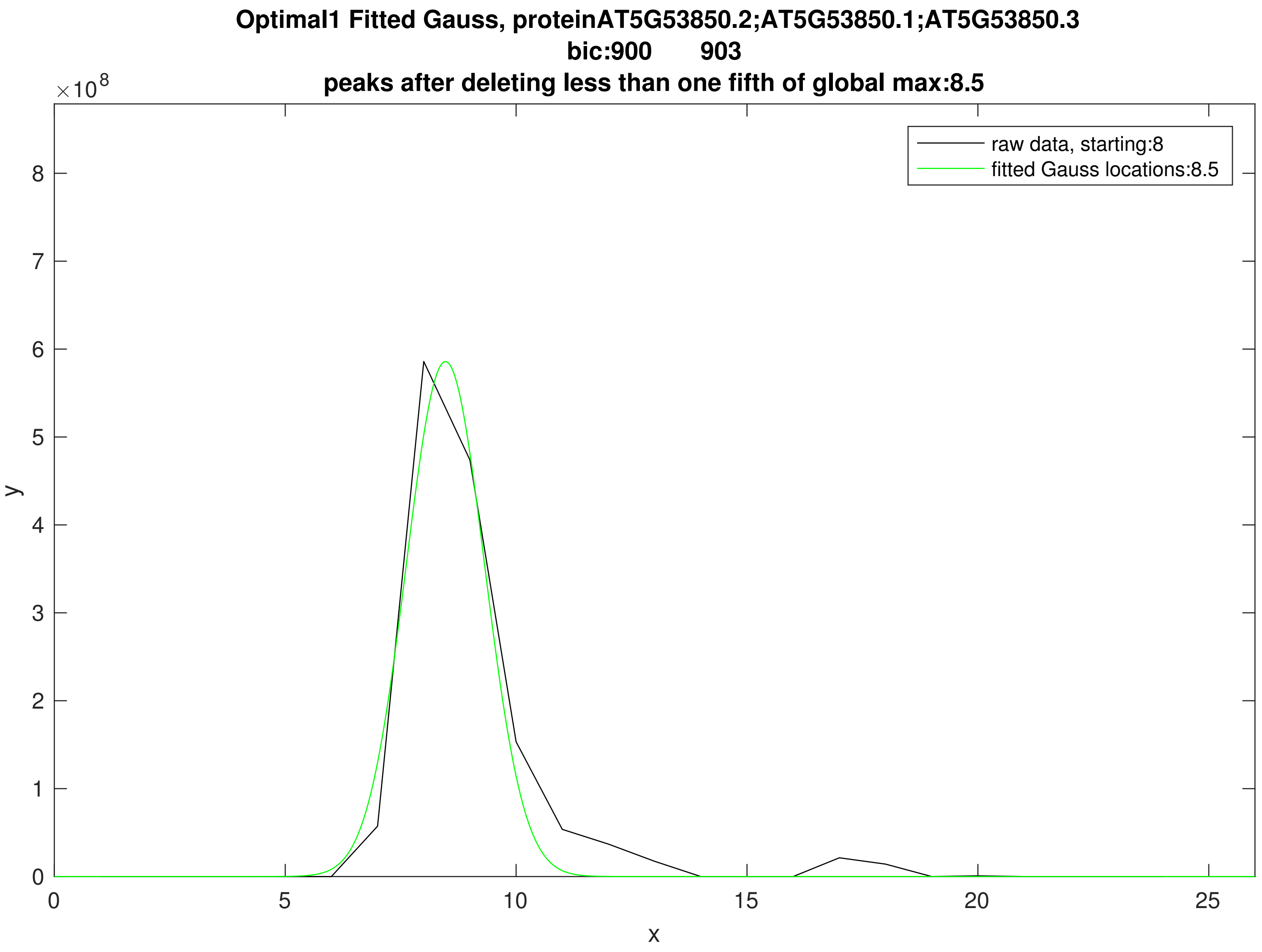
Optimal1 Fitted Gauss, proteinAT5G53850.2;AT5G53850.1;AT5G53850.3
bic:900 903
peaks after deleting less than one fifth of global max:8.5
×10^8
raw data, starting:8
fitted Gauss locations:8.5
y
x
0
1
2
3
4
5
6
7
8
0
5
10
15
20
25

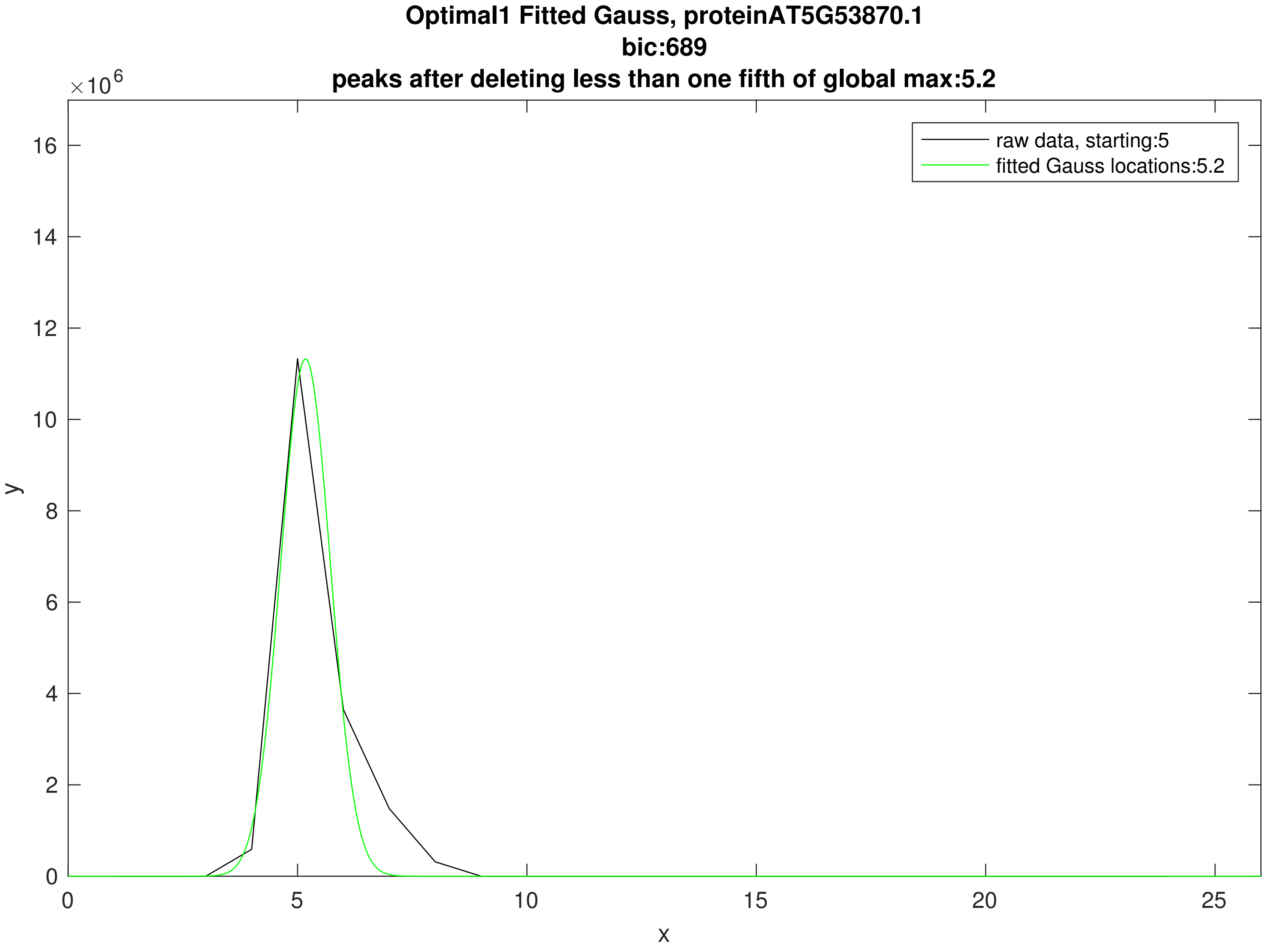
Optimal1 Fitted Gauss, proteinAT5G53870.1
bic:689
peaks after deleting less than one fifth of global max:5.2
×10^6
raw data, starting:5
fitted Gauss locations:5.2
y
x
0
2
4
6
8
10
12
14
16
0
5
10
15
20
25

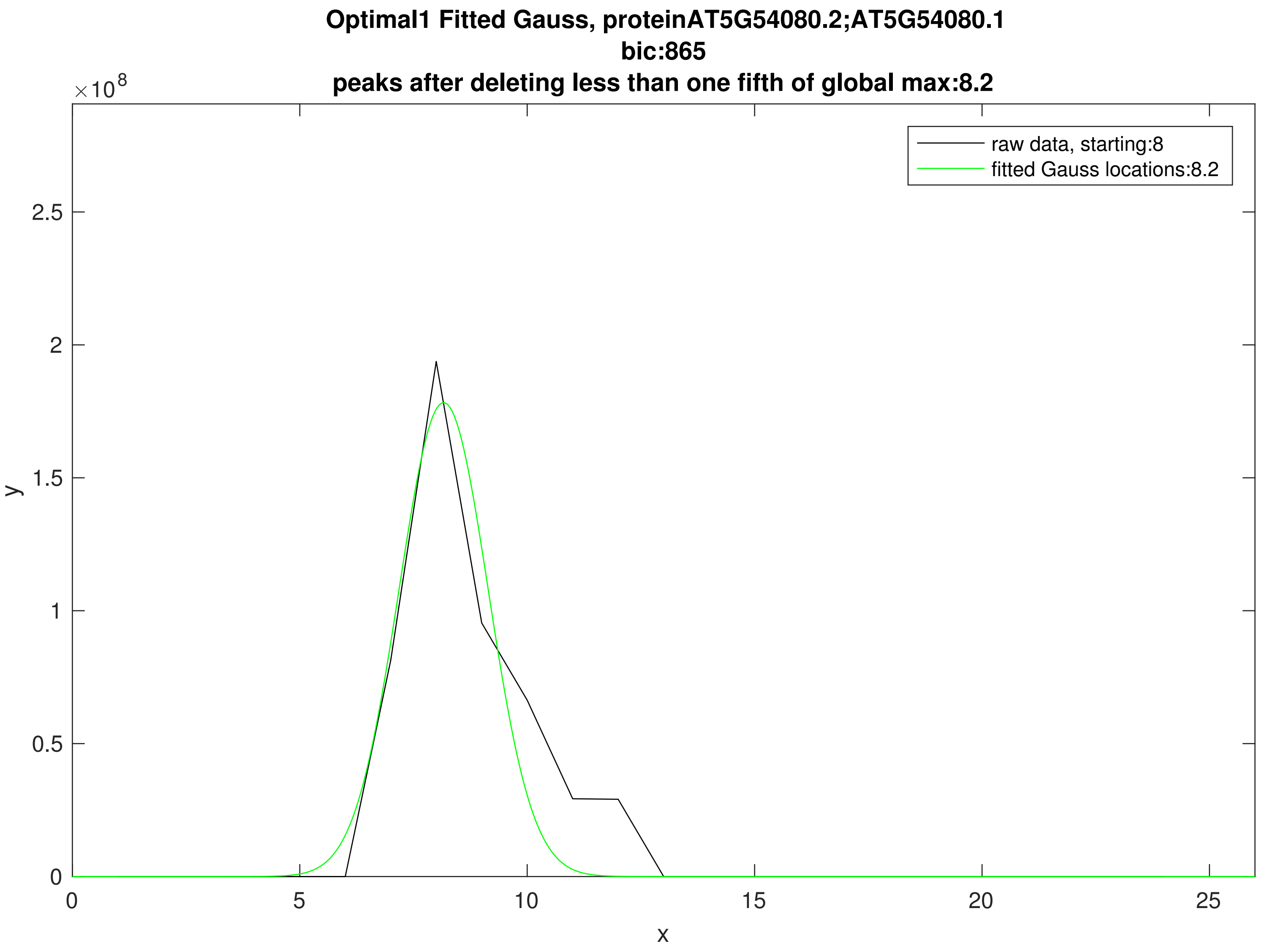

Optimal1 Fitted Gauss, proteinAT5G54080.2;AT5G54080.1
bic:865
peaks after deleting less than one fifth of global max:8.2
×10^8
raw data, starting:8
fitted Gauss locations:8.2
y
x
0
0.5
1
1.5
2
2.5
5
10
15
20
25

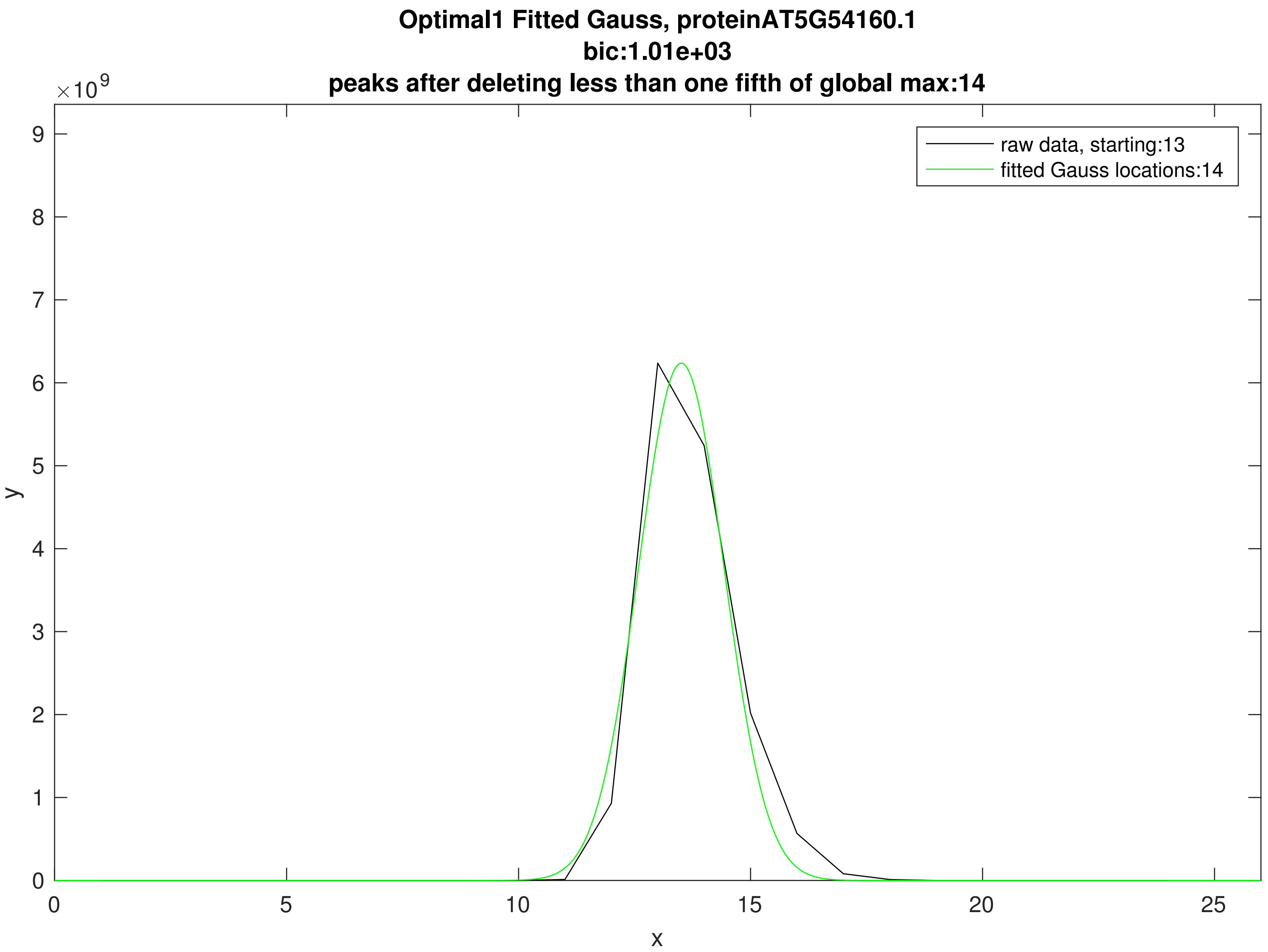
Optimal1 Fitted Gauss, proteinAT5G54160.1
bic:1.01e+03
peaks after deleting less than one fifth of global max:14
×10^9
raw data, starting:13
fitted Gauss locations:14
y
x
0
5
10
15
20
25
1
2
3
4
6
7
8
9

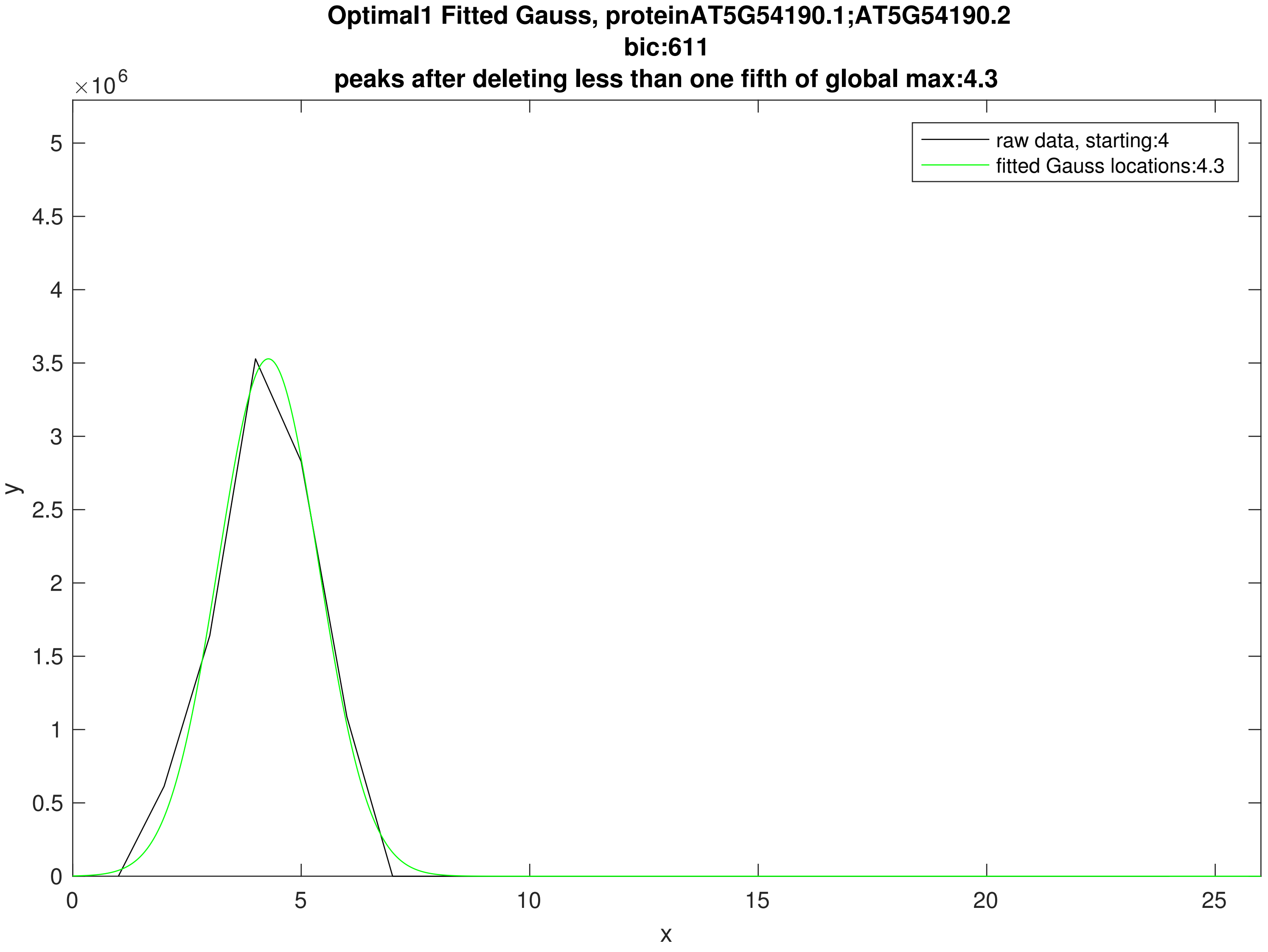

Optimal1 Fitted Gauss, proteinAT5G54190.1;AT5G54190.2
bic:611
peaks after deleting less than one fifth of global max:4.3
×10^6
raw data, starting:4
fitted Gauss locations:4.3
y
x
0
0.5
1
1.5
2
2.5
3
3.5
4
4.5
5
0
5
10
15
20
25

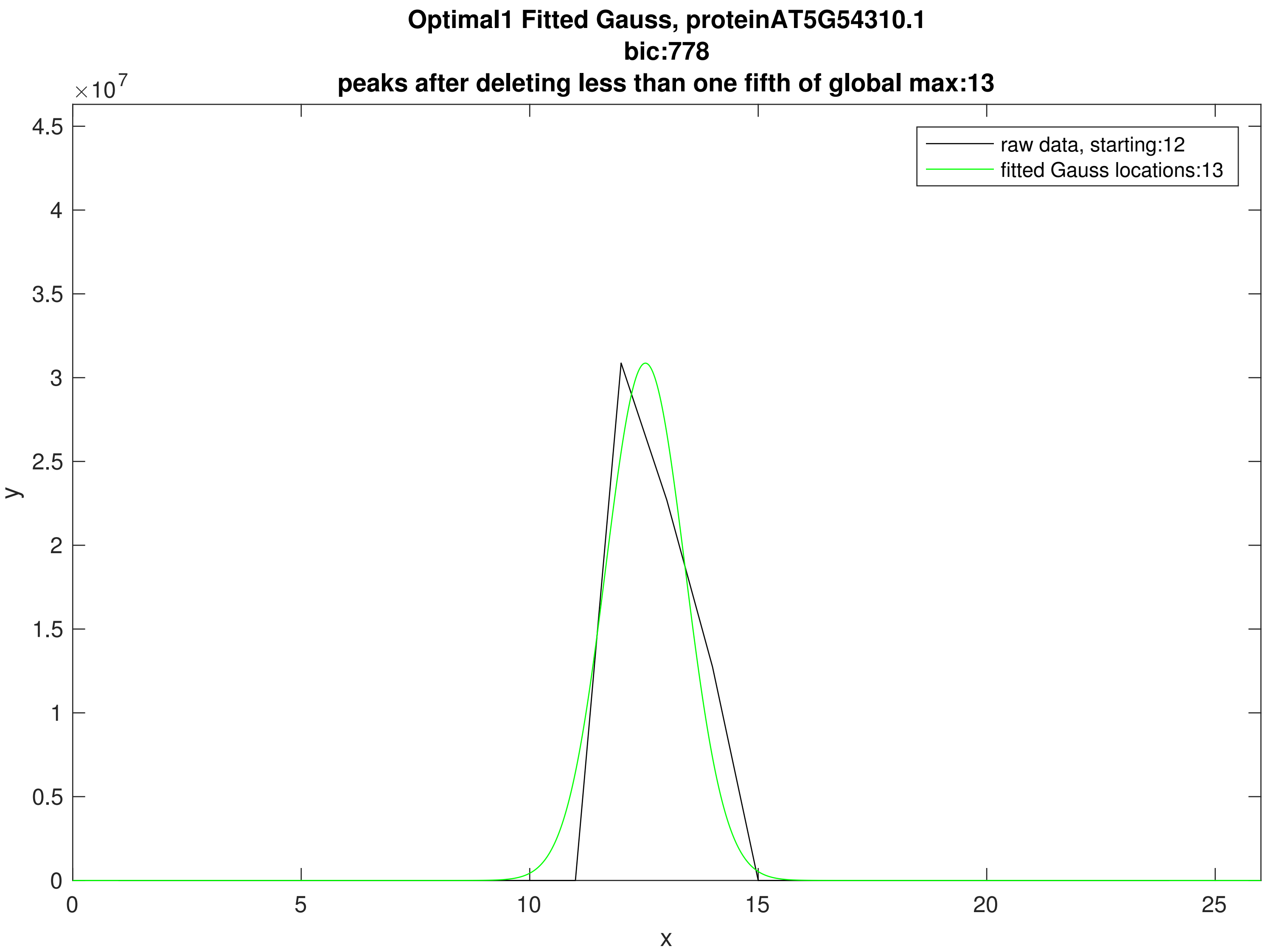
Optimal1 Fitted Gauss, proteinAT5G54310.1
bic:778
peaks after deleting less than one fifth of global max:13
×10^7
4.5
4
3.5
3
2.5
2
1.5
1
0.5
0
y
0
5
10
15
20
25
x
raw data, starting:12
fitted Gauss locations:13

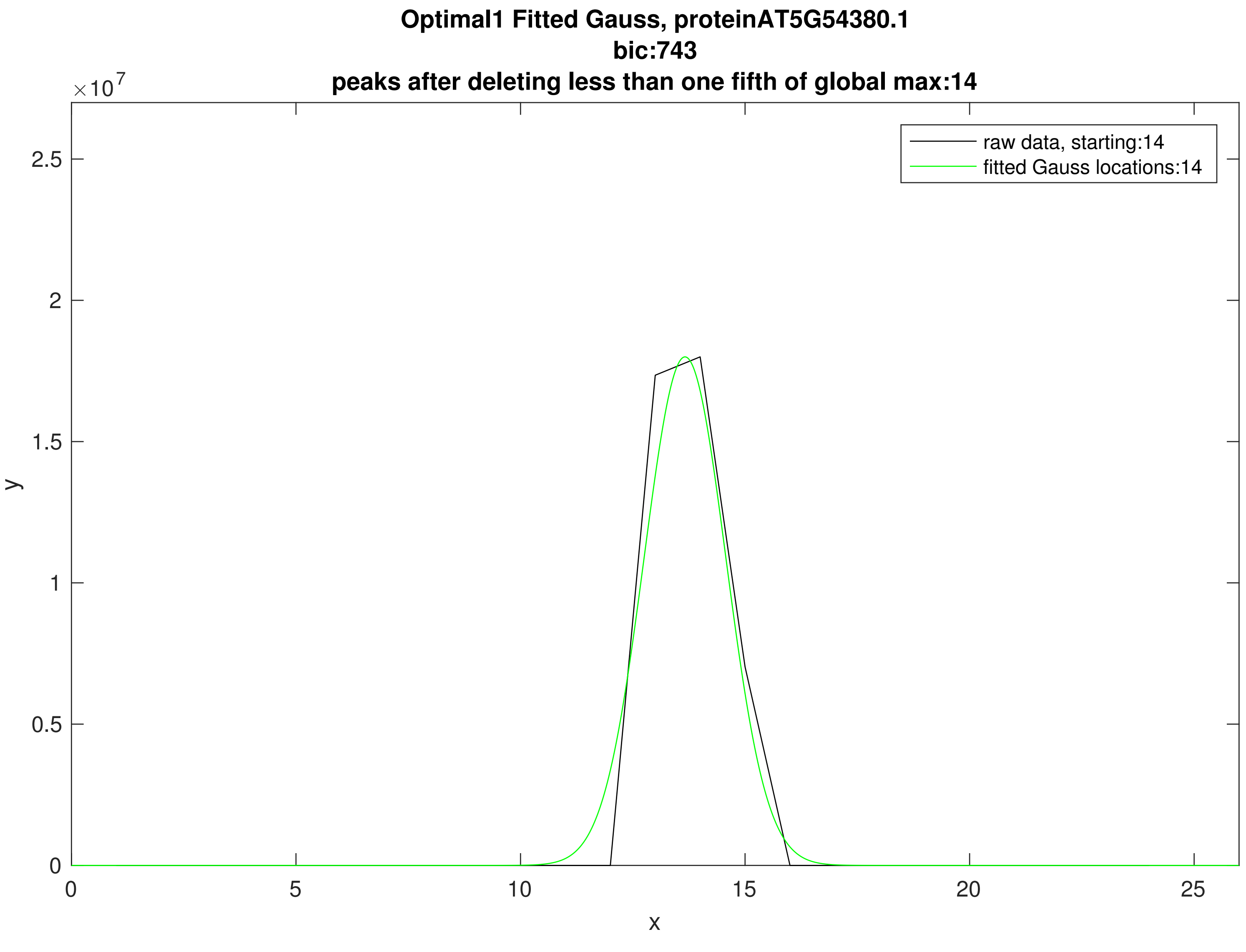
Optimal1 Fitted Gauss, proteinAT5G54380.1
bic:743
peaks after deleting less than one fifth of global max:14
$\times 10^7$
raw data, starting:14
fitted Gauss locations:14
y
x
0
0.5
1
1.5
2
2.5
0
5
10
15
20
25

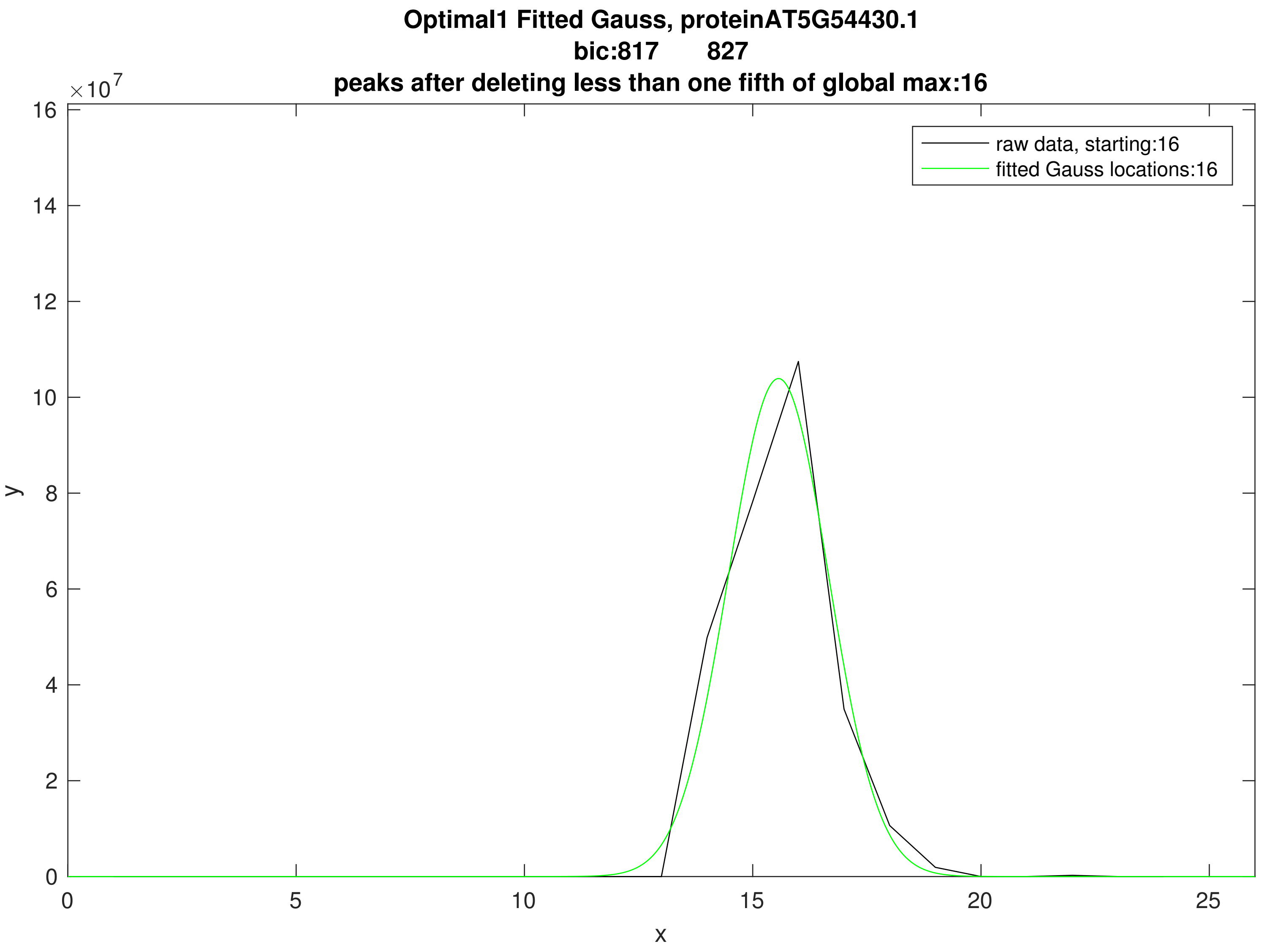
Optimal1 Fitted Gauss, proteinAT5G54430.1
bic:817 827
peaks after deleting less than one fifth of global max:16
×10^7
raw data, starting:16
fitted Gauss locations:16
y
x
0
2
4
6
8
10
12
14
16
0
5
10
15
20
25

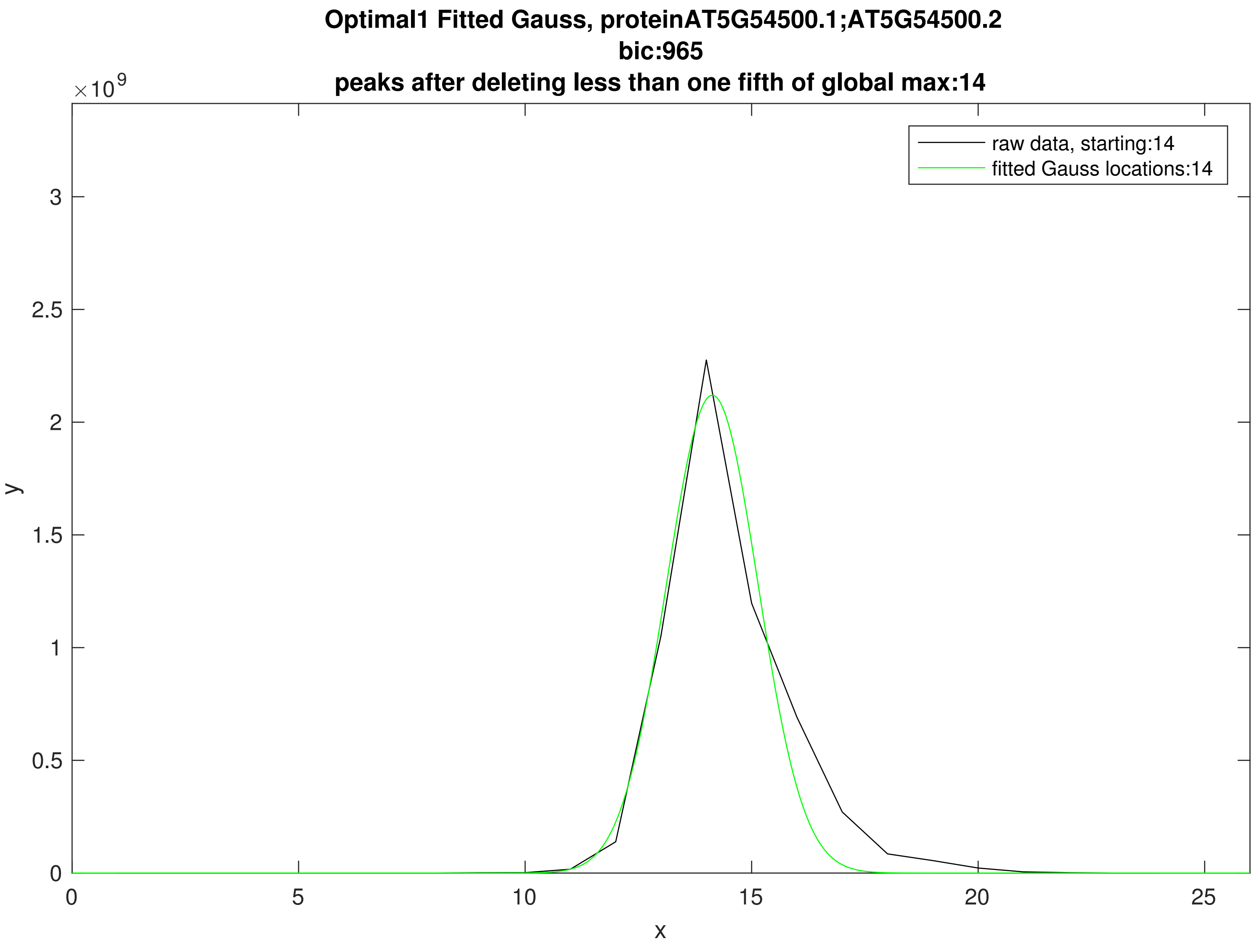
Optimal1 Fitted Gauss, proteinAT5G54500.1;AT5G54500.2
bic:965
peaks after deleting less than one fifth of global max:14
×10^9
raw data, starting:14
fitted Gauss locations:14
3
2.5
2
1.5
1
0.5
0
y
0
5
10
15
20
25
x

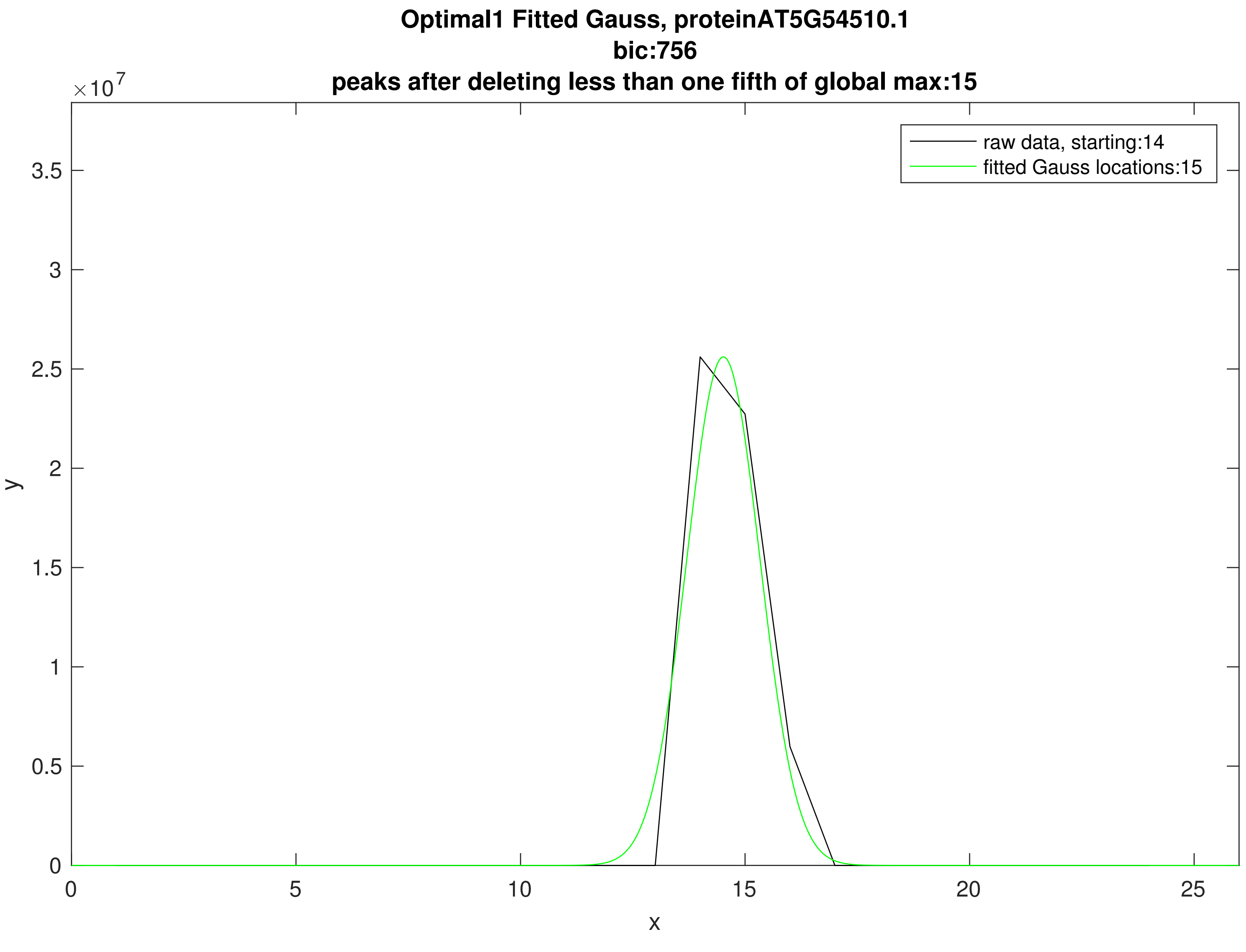
Optimal1 Fitted Gauss, proteinAT5G54510.1
bic:756
peaks after deleting less than one fifth of global max:15
×10^7
raw data, starting:14
fitted Gauss locations:15
3.5
3
2.5
2
1.5
1
0.5
0
y
0
5
10
15
20
25
x

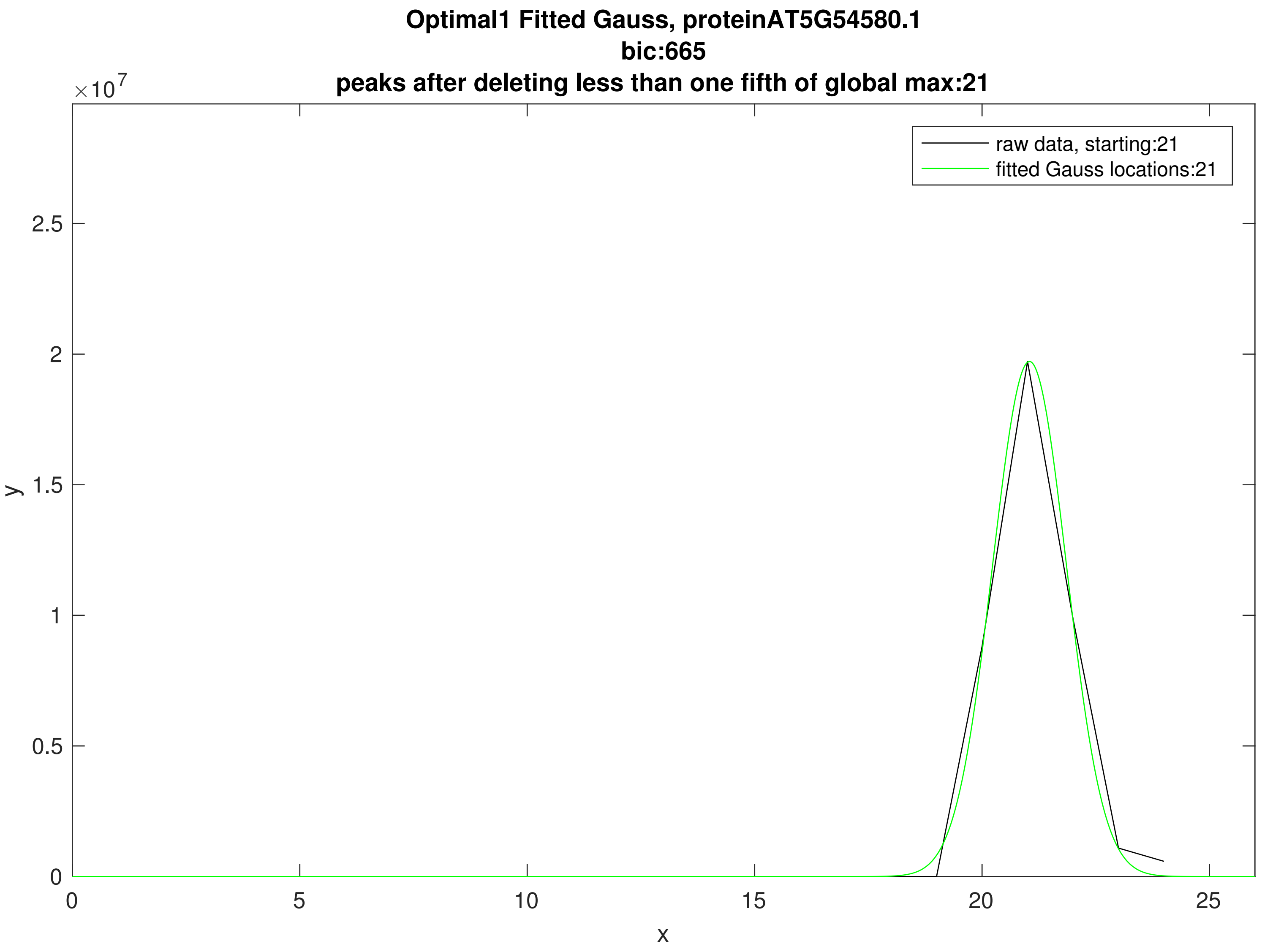
Optimal1 Fitted Gauss, proteinAT5G54580.1
bic:665
peaks after deleting less than one fifth of global max:21
×10^7
raw data, starting:21
fitted Gauss locations:21
2.5
2
1.5
1
0.5
0
y
0
5
10
15
20
25
x

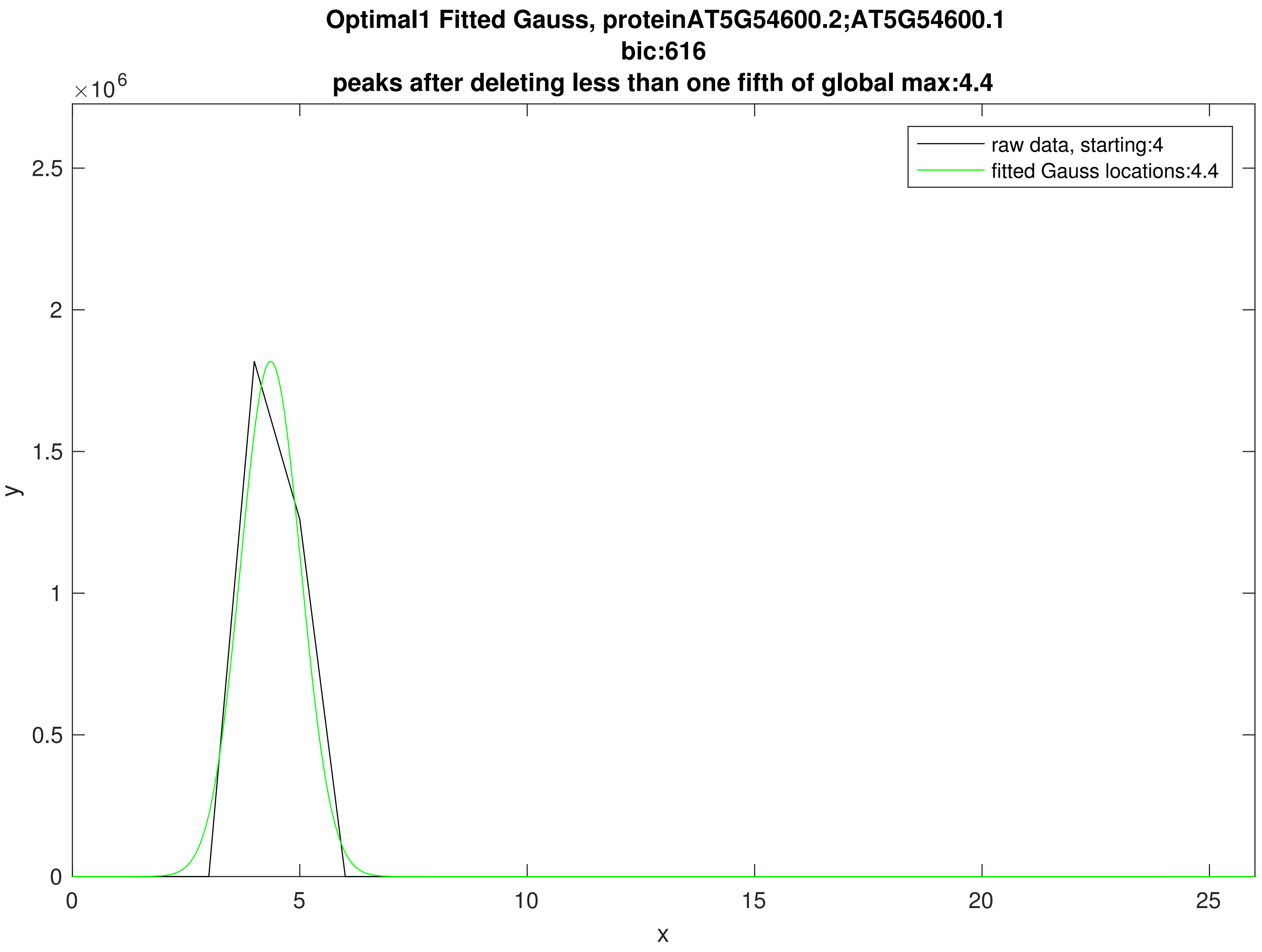

Optimal1 Fitted Gauss, proteinAT5G54600.2;AT5G54600.1
bic:616
peaks after deleting less than one fifth of global max:4.4
×10^6
raw data, starting:4
fitted Gauss locations:4.4
2.5
2
1.5
1
0.5
0
y
0
5
10
15
20
25
x

**Optimal1 Fitted Gauss, proteinAT5G54750.1;AT5G54750.2**
**bic:828 837 858**
**peaks after deleting less than one fifth of global max:13**

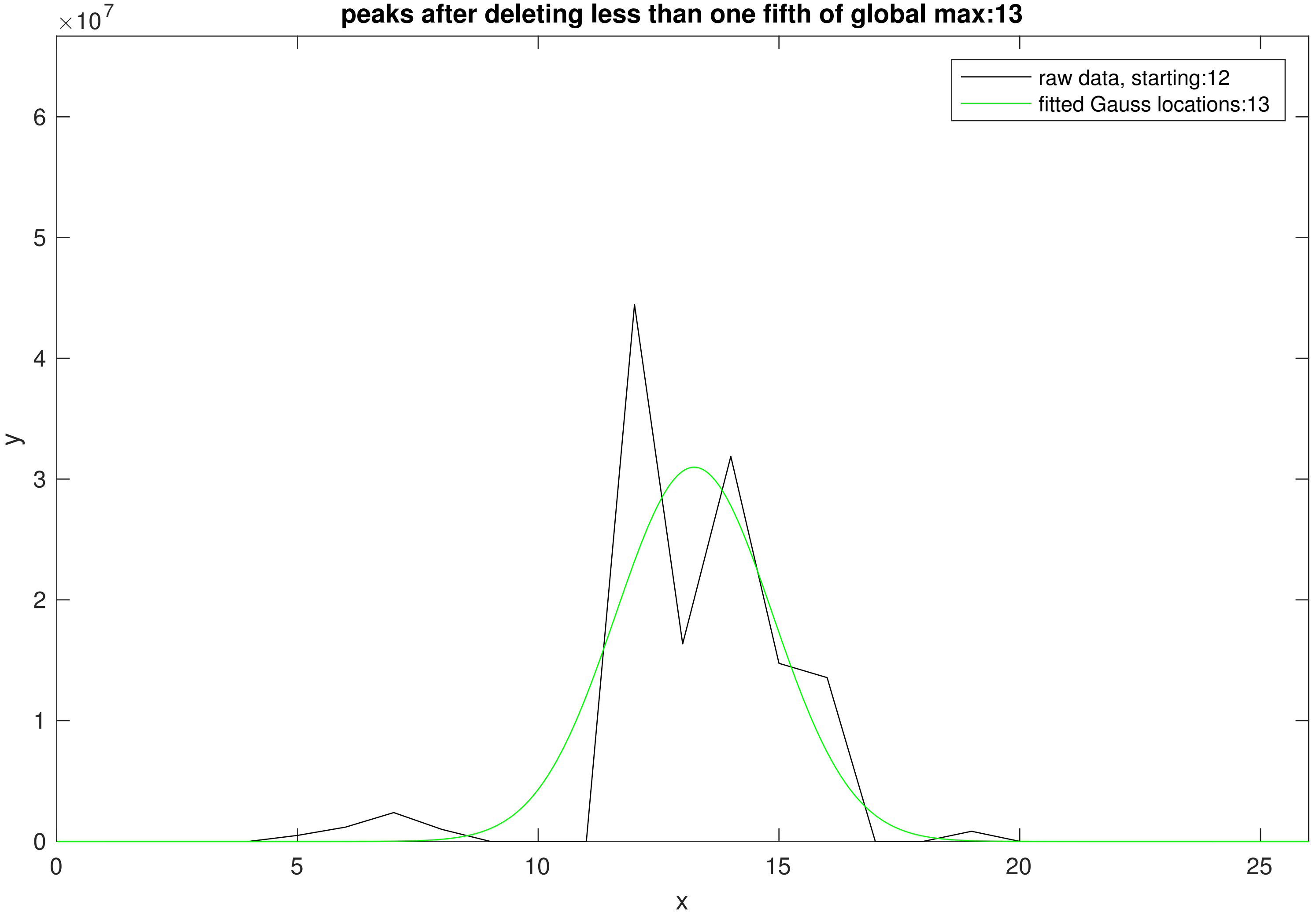

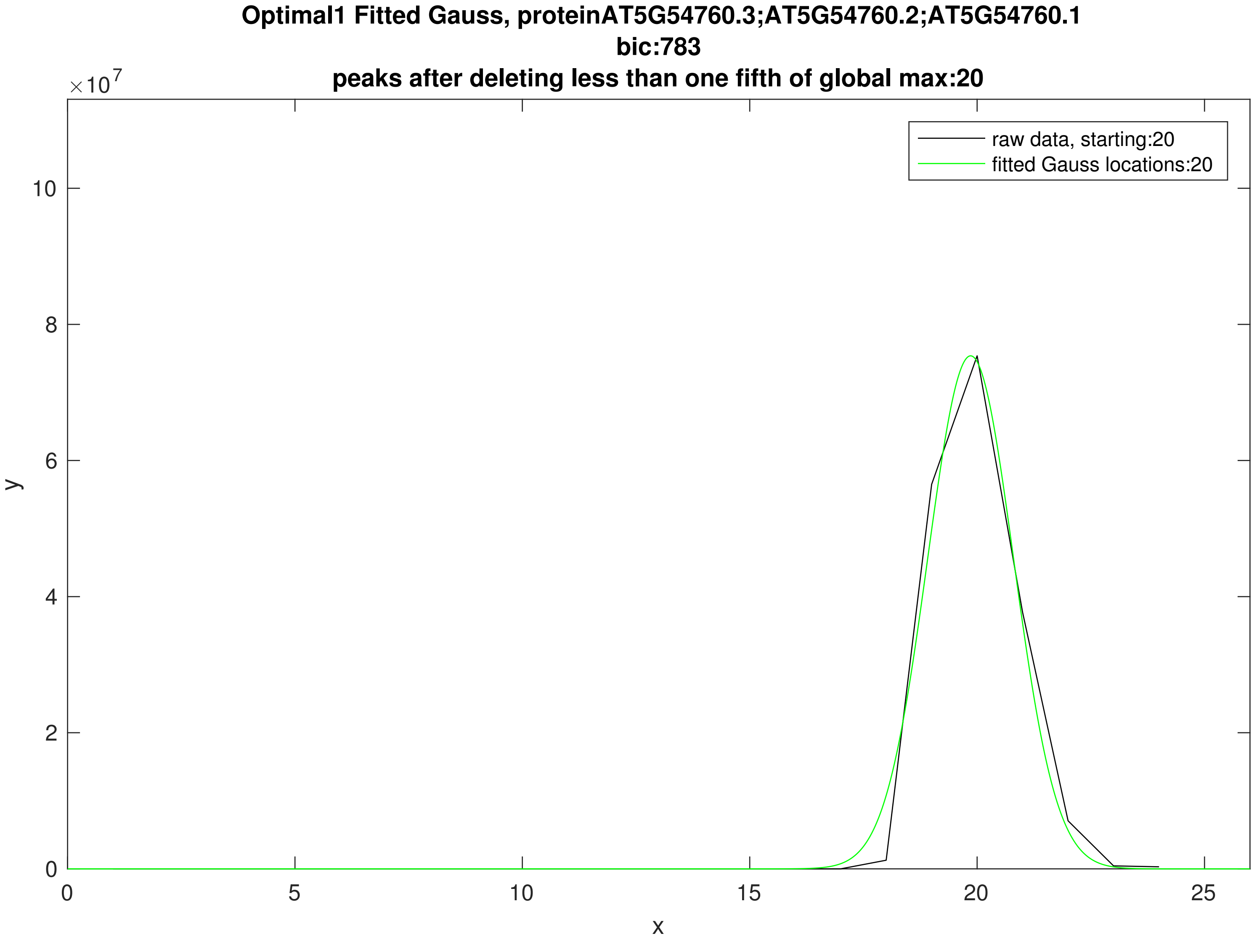

Optimal1 Fitted Gauss, proteinAT5G54760.3;AT5G54760.2;AT5G54760.1
bic:783
peaks after deleting less than one fifth of global max:20
×10^7
raw data, starting:20
fitted Gauss locations:20
y
x
0
2
4
6
8
10
5
15
20
25

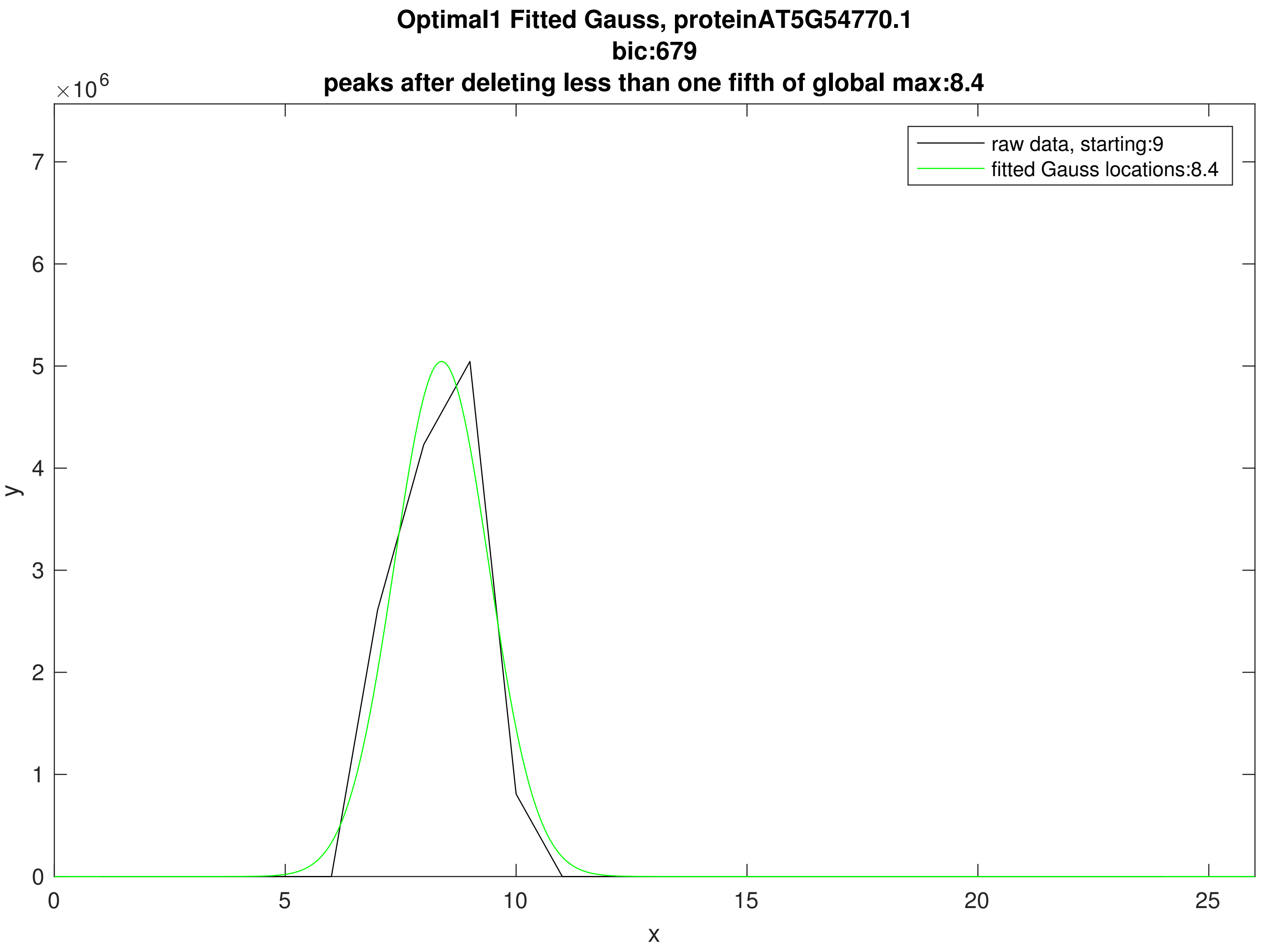
Optimal1 Fitted Gauss, proteinAT5G54770.1
bic:679
peaks after deleting less than one fifth of global max:8.4
×10^6
raw data, starting:9
fitted Gauss locations:8.4
y
x
0
1
2
3
4
5
6
7
0
5
10
15
20
25

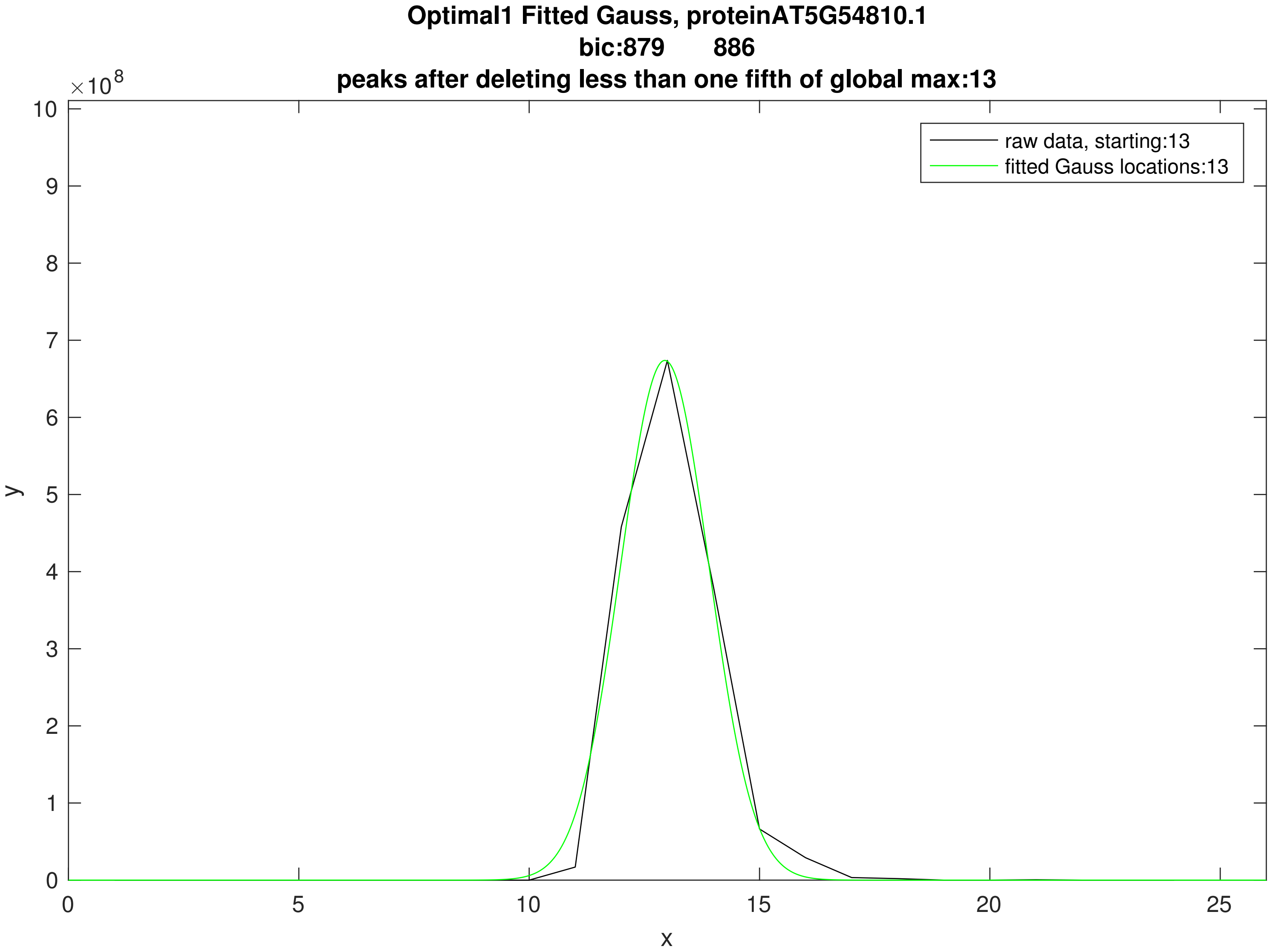

Optimal1 Fitted Gauss, proteinAT5G54810.1
bic:879 886
peaks after deleting less than one fifth of global max:13
×10^8
10
9
8
7
6
5
4
3
2
1
0
y
0
5
10
15
20
25
x
raw data, starting:13
fitted Gauss locations:13

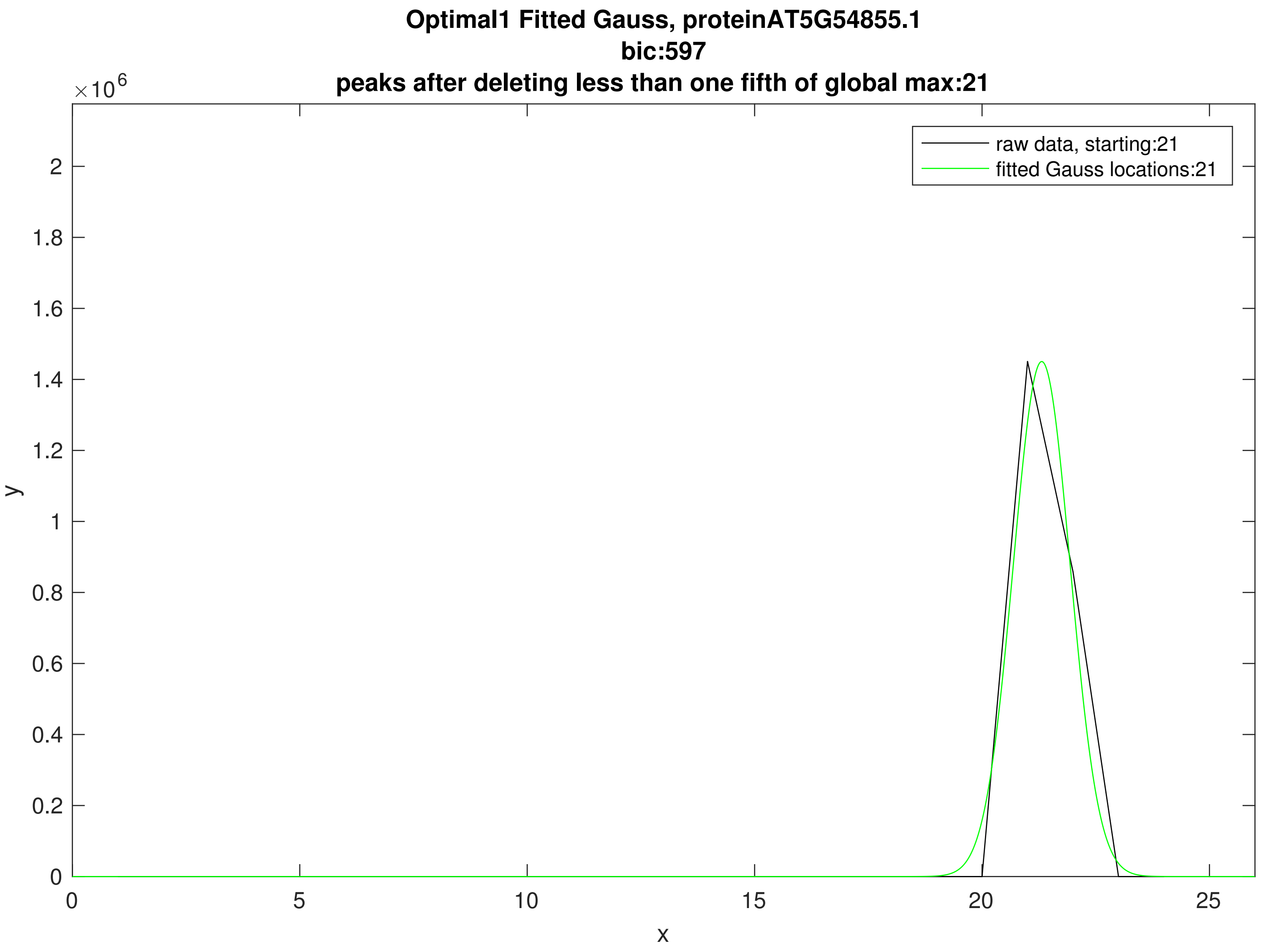

Optimal1 Fitted Gauss, proteinAT5G54855.1
bic:597
peaks after deleting less than one fifth of global max:21
×10^6
raw data, starting:21
fitted Gauss locations:21
y
x
0
0.2
0.4
0.6
0.8
1
1.2
1.4
1.6
1.8
2
5
10
15
20
25

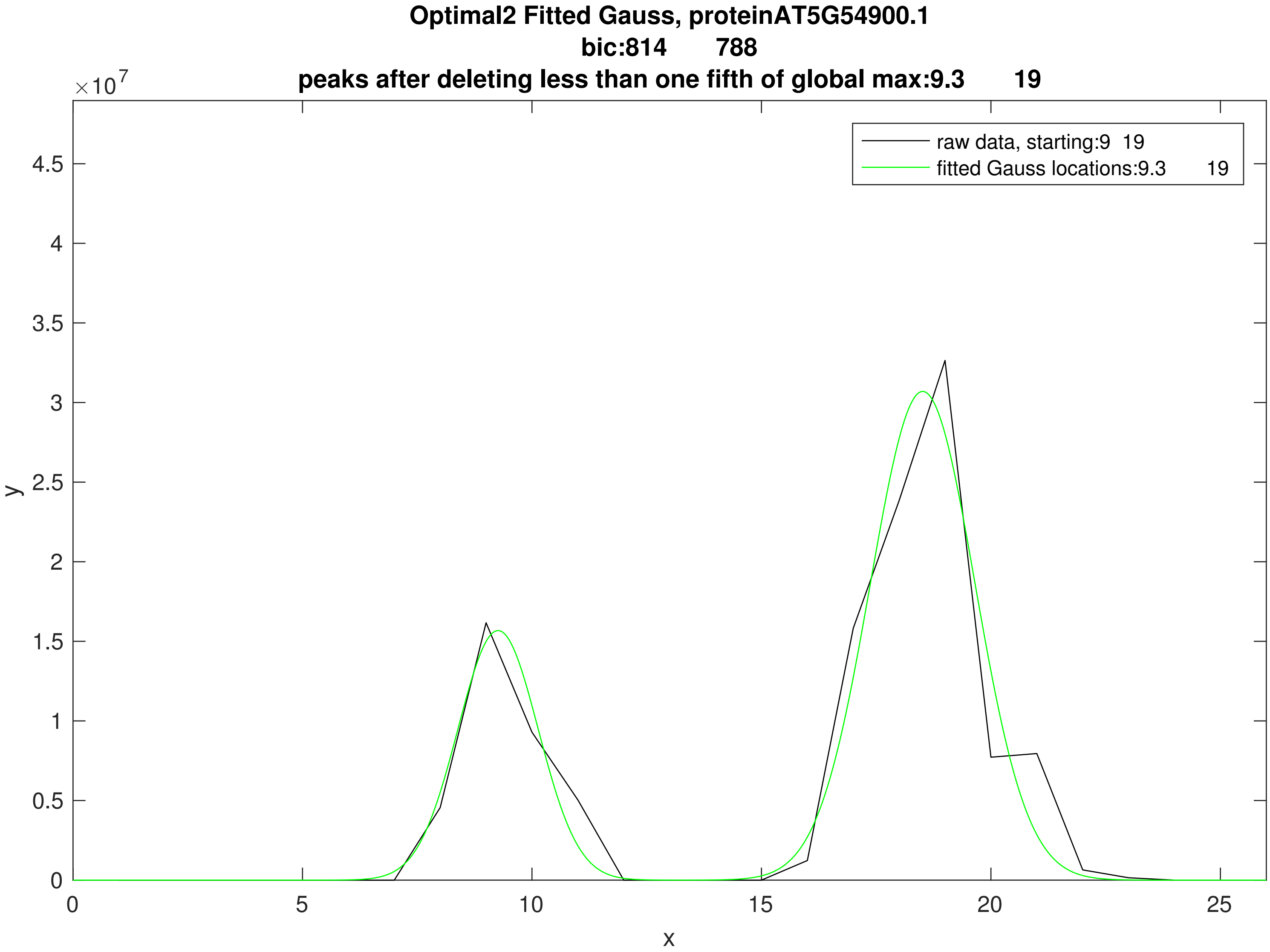
Optimal2 Fitted Gauss, proteinAT5G54900.1
bic:814 788
peaks after deleting less than one fifth of global max:9.3 19
×10^7
4.5
4
3.5
3
2.5
2
1.5
1
0.5
0
y
0
5
10
15
20
25
x
raw data, starting:9 19
fitted Gauss locations:9.3 19

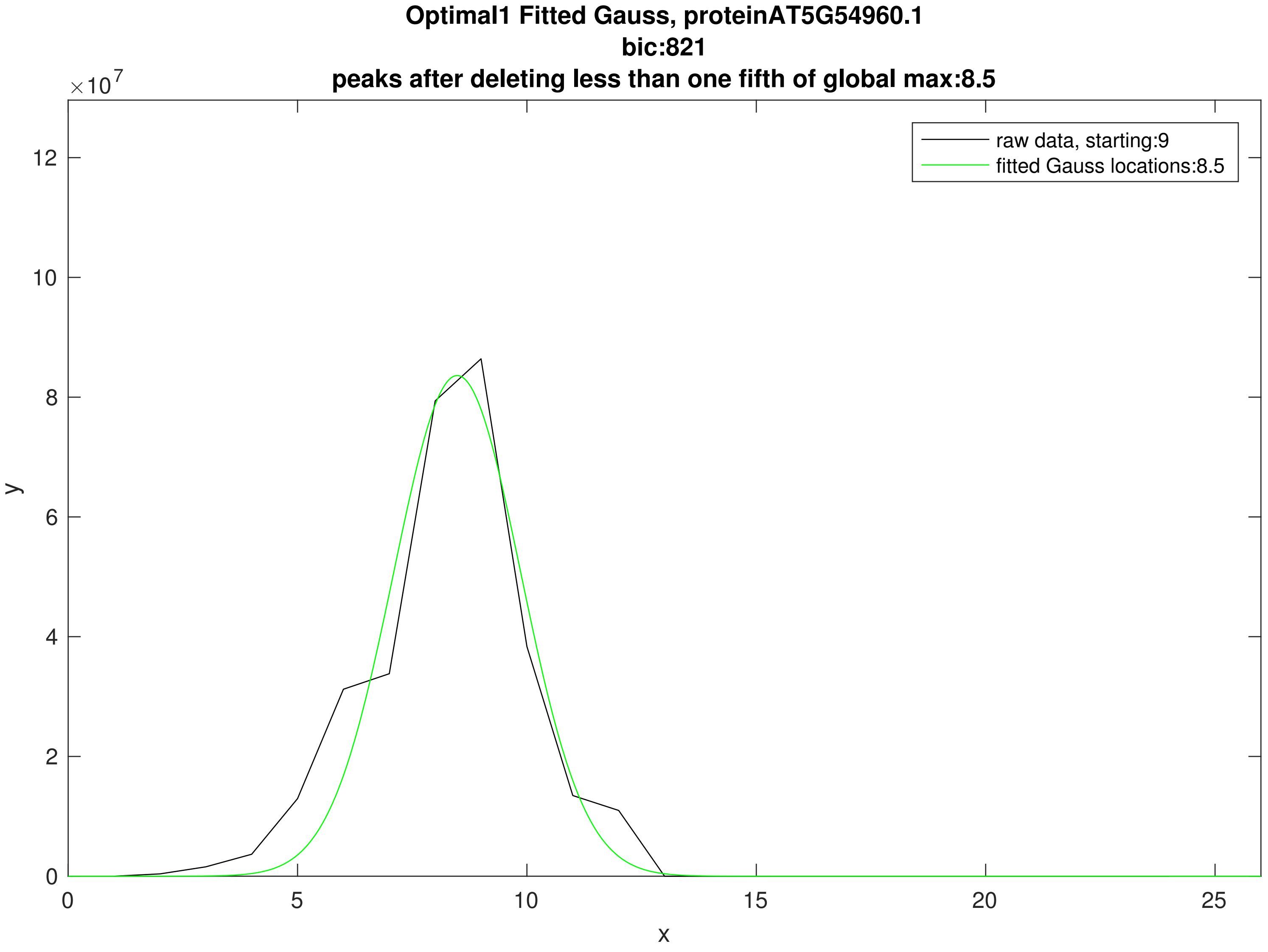

Optimal1 Fitted Gauss, proteinAT5G54960.1
bic:821
peaks after deleting less than one fifth of global max:8.5
×10^7
raw data, starting:9
fitted Gauss locations:8.5
12
10
8
6
4
2
0
y
0
5
10
15
20
25
x

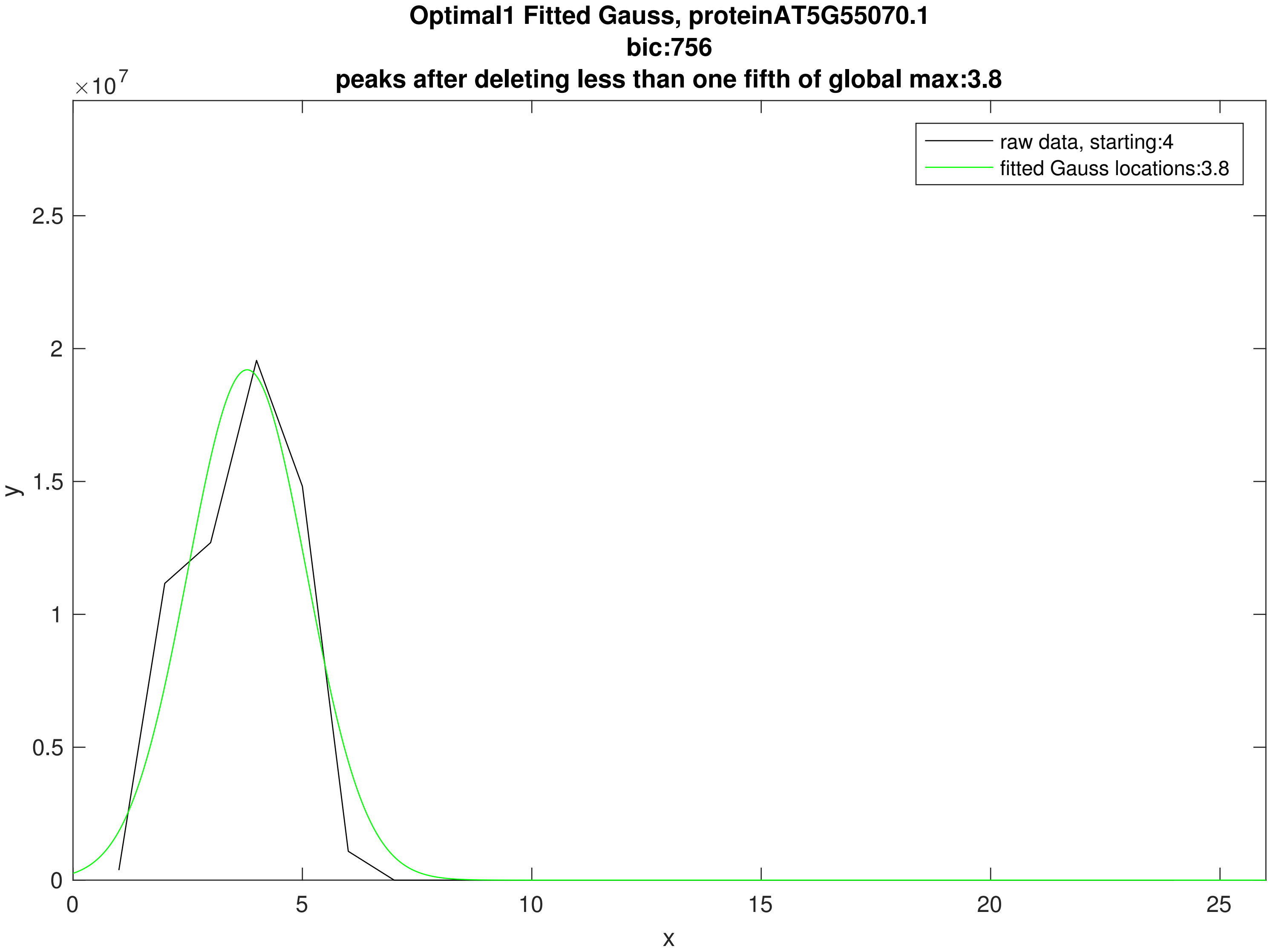
Optimal1 Fitted Gauss, proteinAT5G55070.1
bic:756
peaks after deleting less than one fifth of global max:3.8
raw data, starting:4
fitted Gauss locations:3.8
$\times 10^7$
2.5
2
1.5
1
0.5
0
y
0
5
10
15
20
25
x

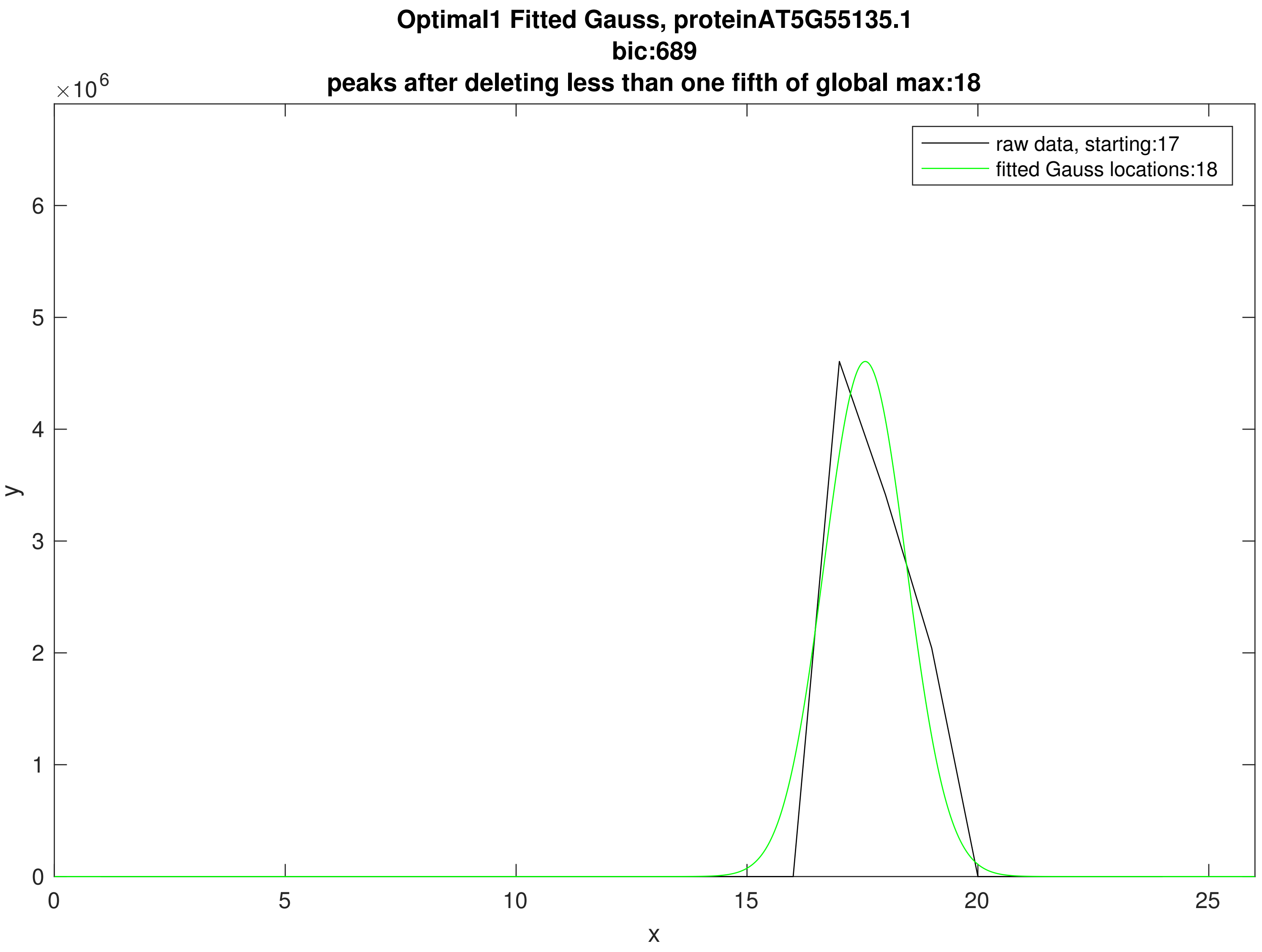
Optimal1 Fitted Gauss, proteinAT5G55135.1
bic:689
peaks after deleting less than one fifth of global max:18
×10^6
raw data, starting:17
fitted Gauss locations:18
y
x
0
1
2
3
4
5
6
0
5
10
15
20
25

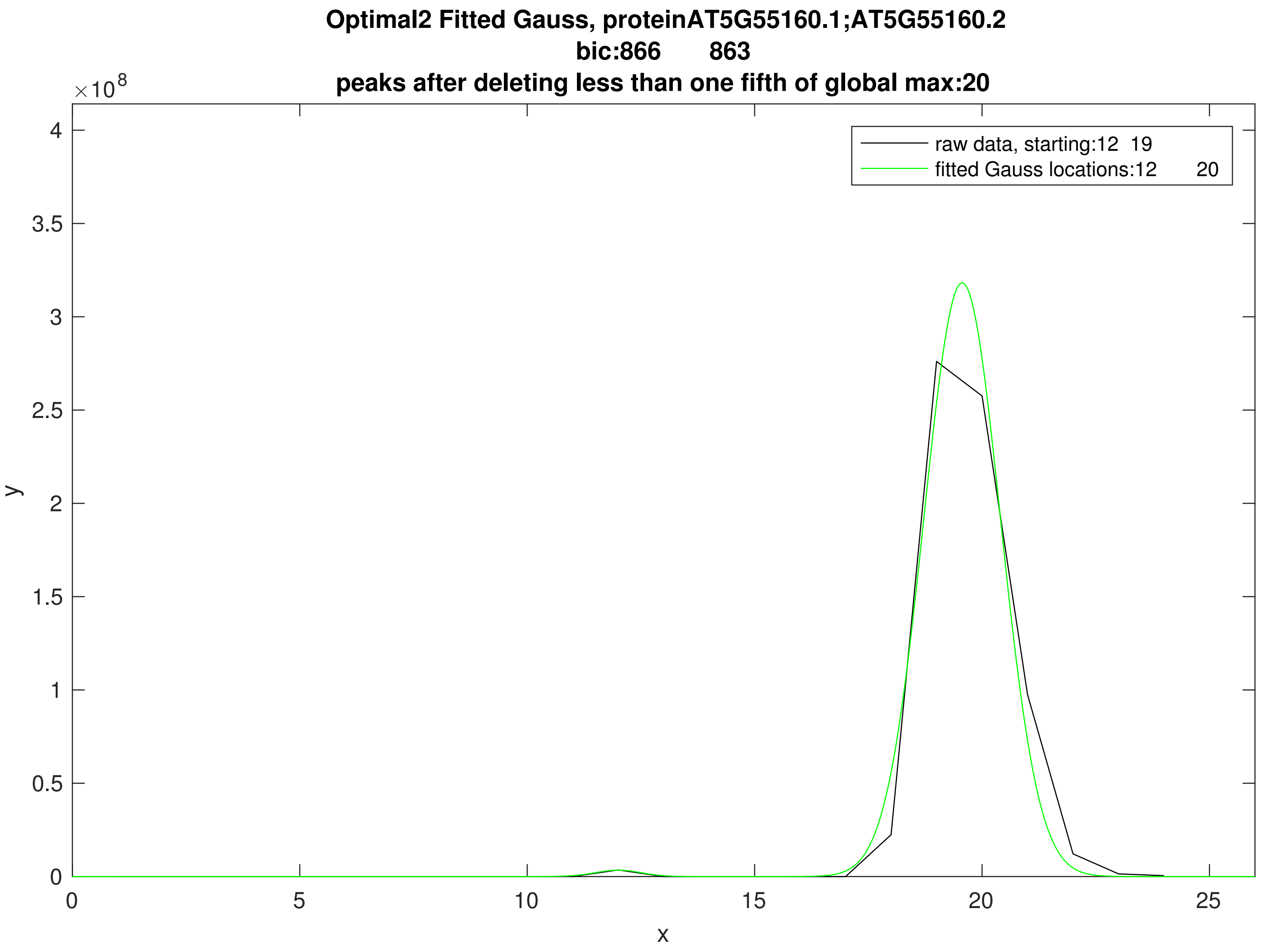
Optimal2 Fitted Gauss, proteinAT5G55160.1;AT5G55160.2
bic:866 863
peaks after deleting less than one fifth of global max:20
×10^8
raw data, starting:12 19
fitted Gauss locations:12 20
y
x
0
0.5
1
1.5
2
2.5
3
3.5
4
0
5
10
15
20
25

**Optimal1 Fitted Gauss, proteinAT5G55200.1**
**bic:640**
**peaks after deleting less than one fifth of global max:9.2**

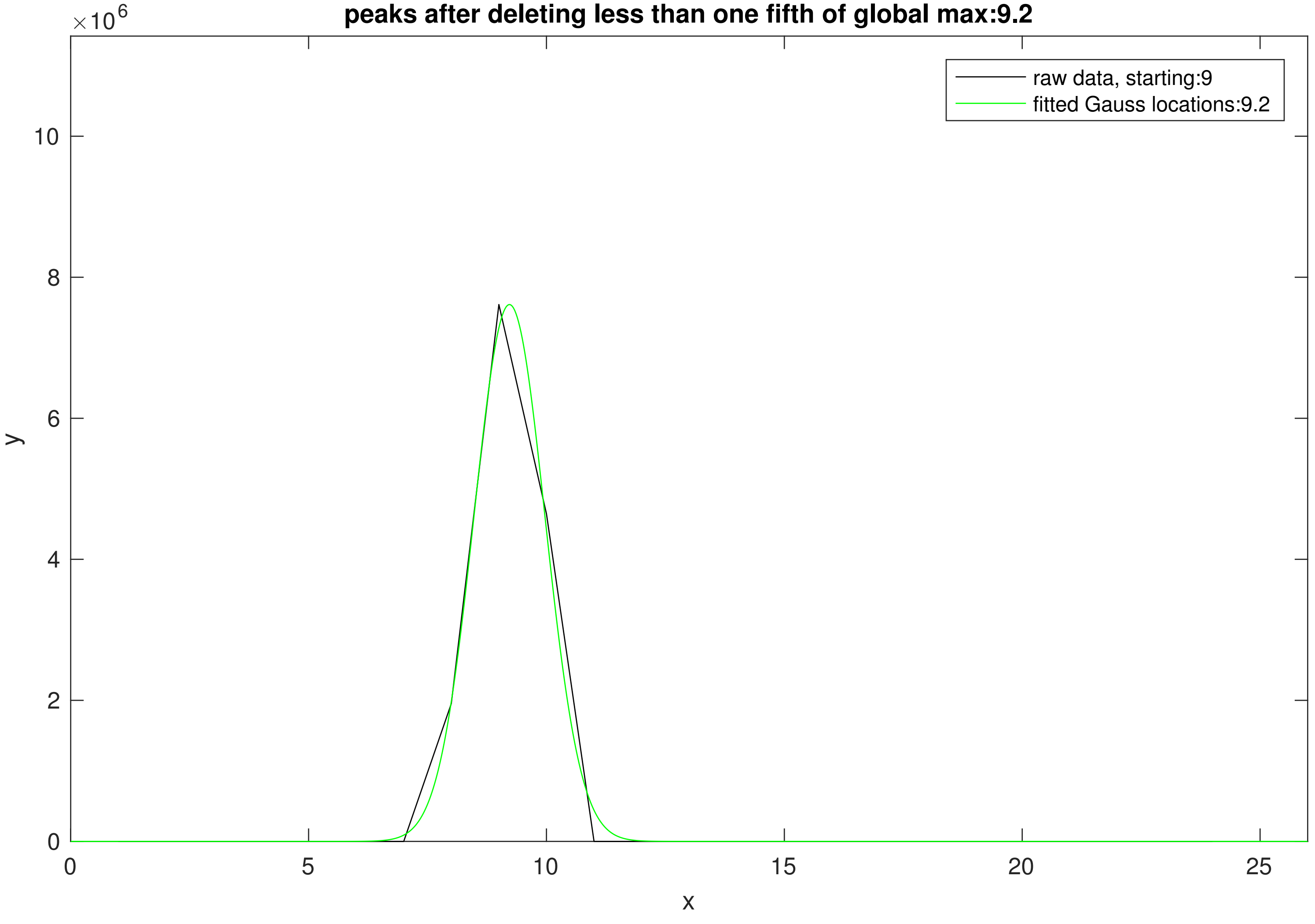

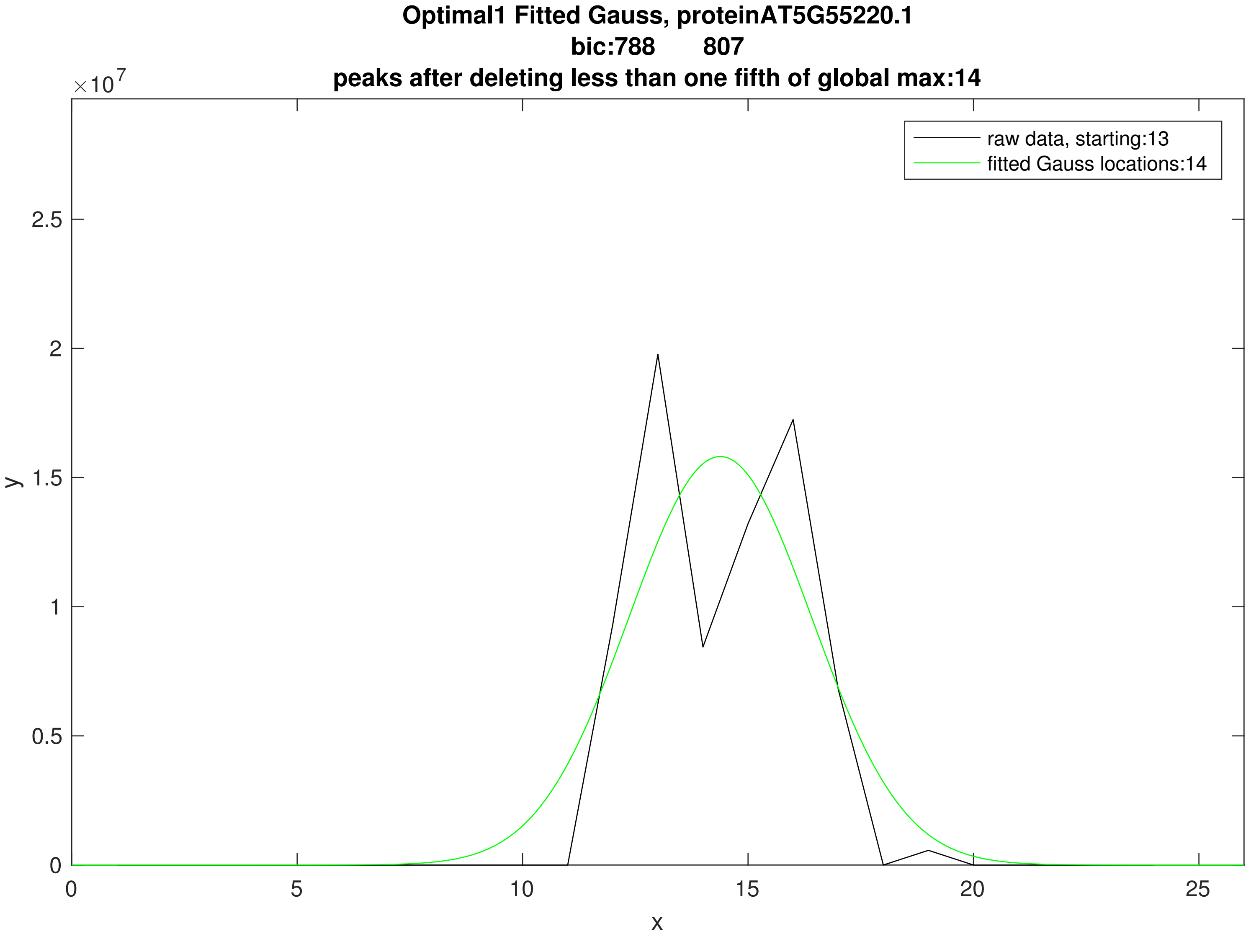

Optimal1 Fitted Gauss, proteinAT5G55220.1
bic:788 807
peaks after deleting less than one fifth of global max:14
×10^7
raw data, starting:13
fitted Gauss locations:14
2.5
2
1.5
1
0.5
0
y
0
5
10
15
20
25
x

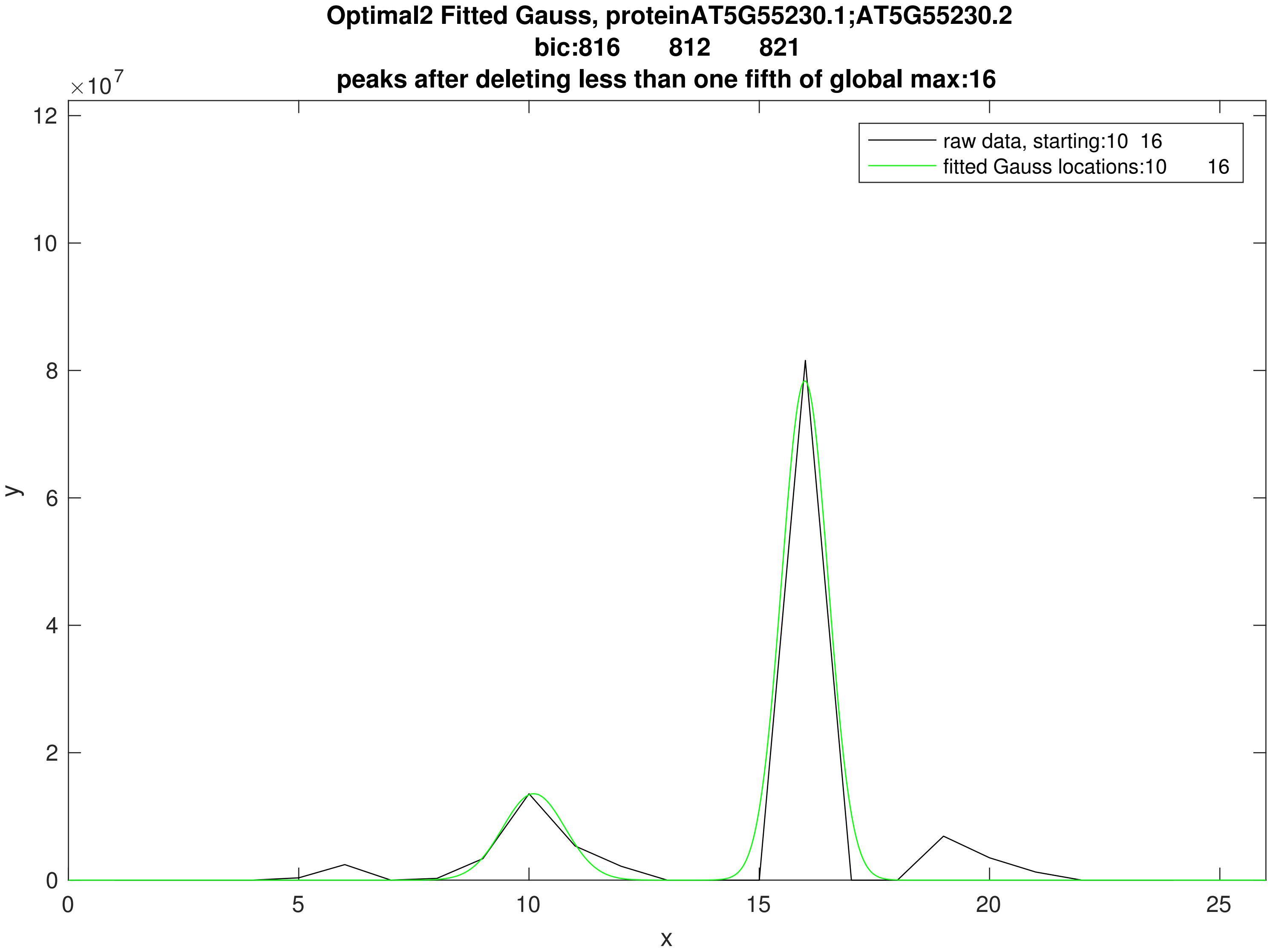
Optimal2 Fitted Gauss, proteinAT5G55230.1;AT5G55230.2
bic:816 812 821
peaks after deleting less than one fifth of global max:16
×10^7
12
10
8
6
4
2
0
y
0
5
10
15
20
25
x
raw data, starting:10 16
fitted Gauss locations:10 16

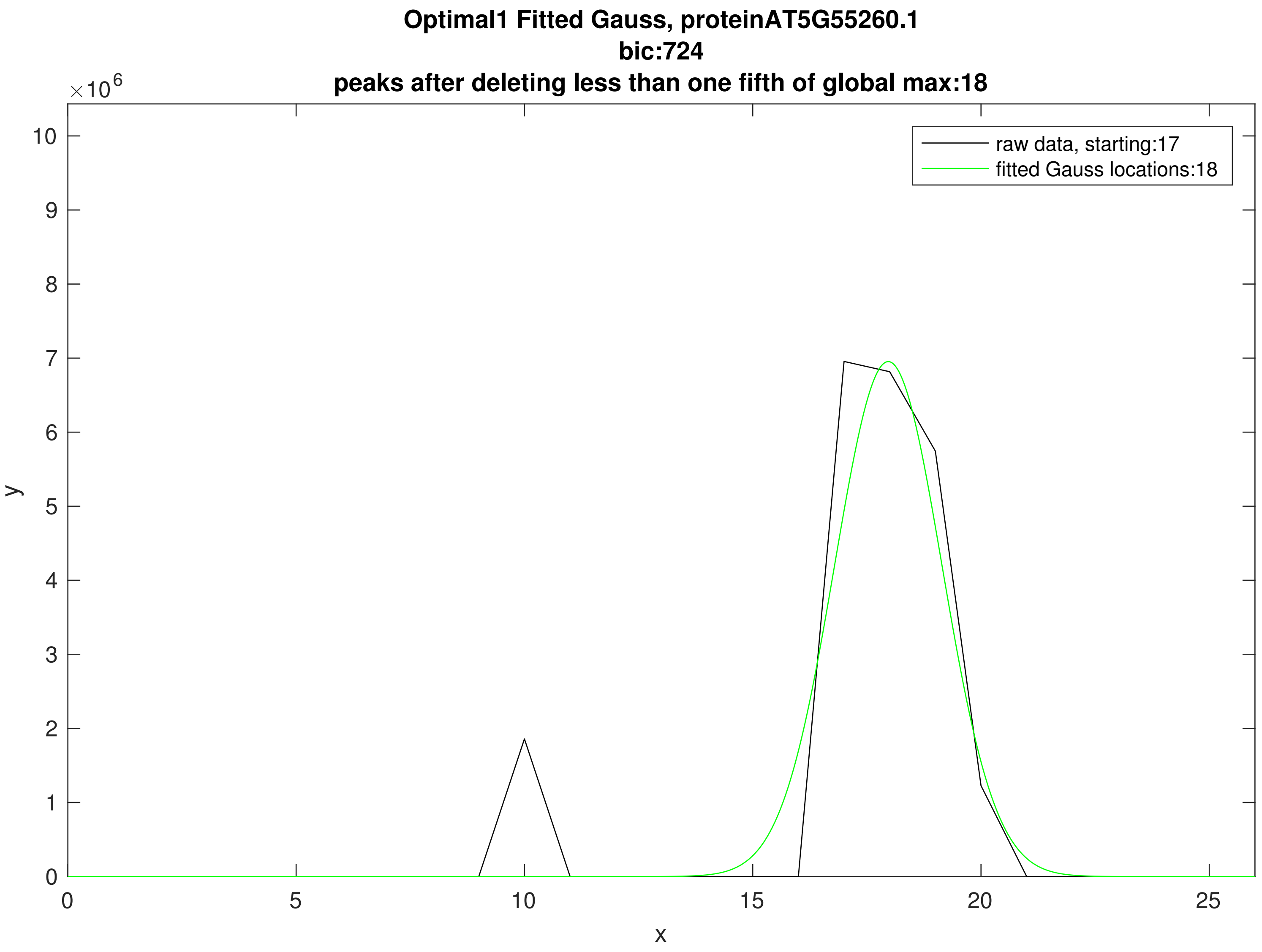
Optimal1 Fitted Gauss, proteinAT5G55260.1
bic:724
peaks after deleting less than one fifth of global max:18
×10^6
raw data, starting:17
fitted Gauss locations:18
y
x
0
1
2
3
4
5
6
7
8
9
10
0
5
10
15
20
25

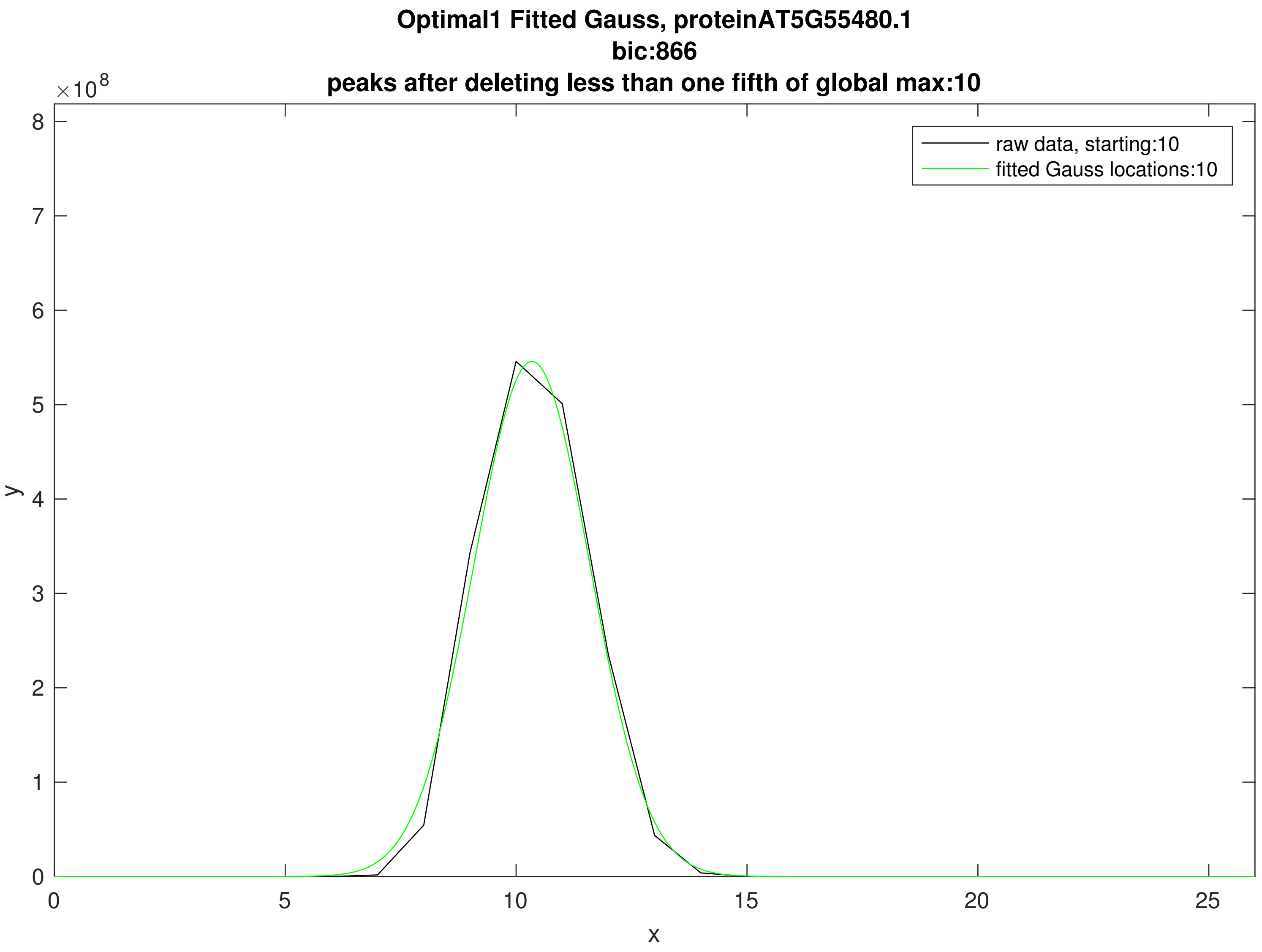
Optimal1 Fitted Gauss, proteinAT5G55480.1
bic:866
peaks after deleting less than one fifth of global max:10
×10^8
raw data, starting:10
fitted Gauss locations:10
y
x
0
1
2
3
4
5
6
7
8
0
5
10
15
20
25

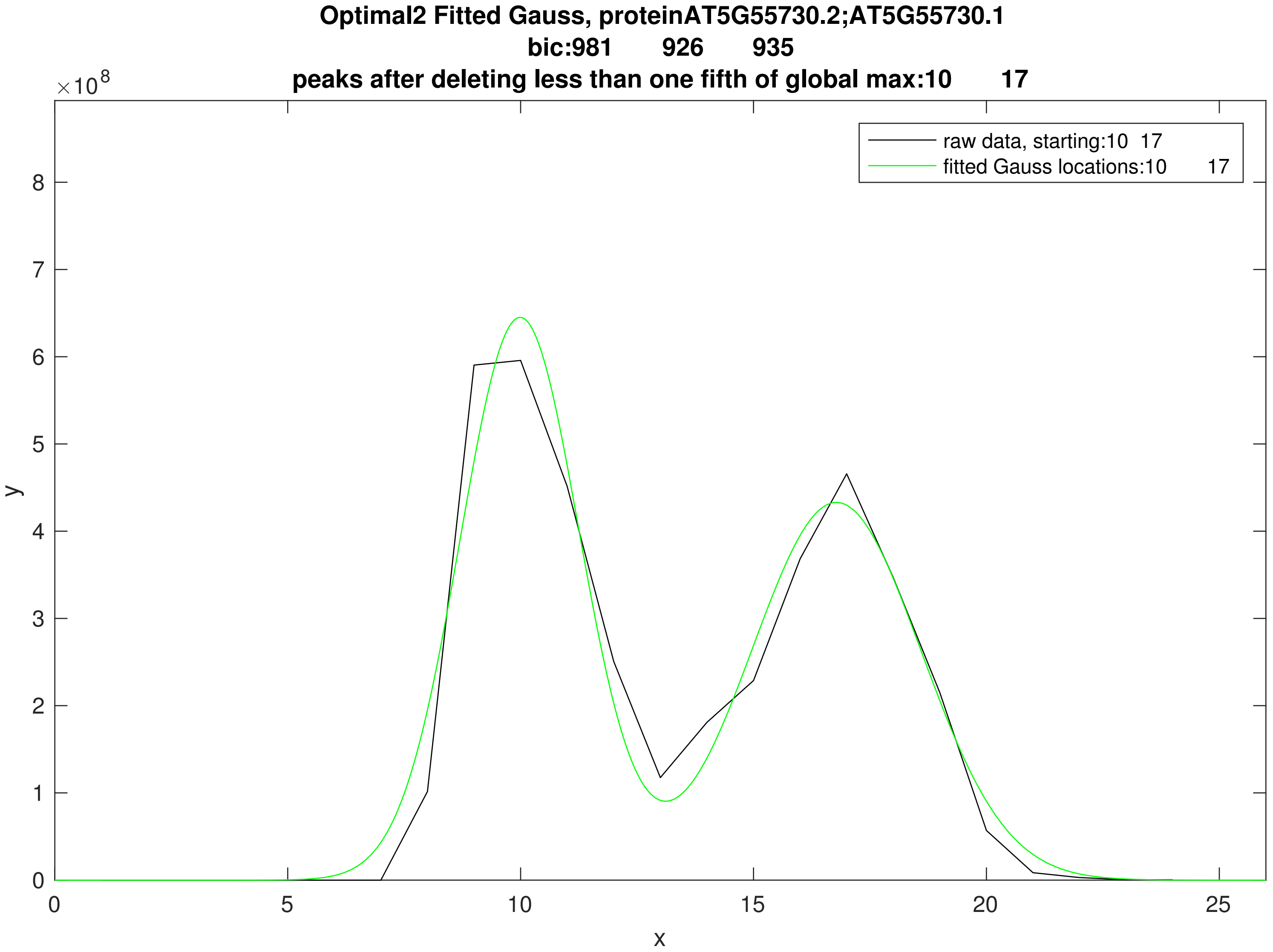
Optimal2 Fitted Gauss, proteinAT5G55730.2;AT5G55730.1
bic:981 926 935
peaks after deleting less than one fifth of global max:10 17
×10^8
raw data, starting:10 17
fitted Gauss locations:10 17
y
x
0
1
2
3
4
5
6
7
8
0
5
10
15
20
25

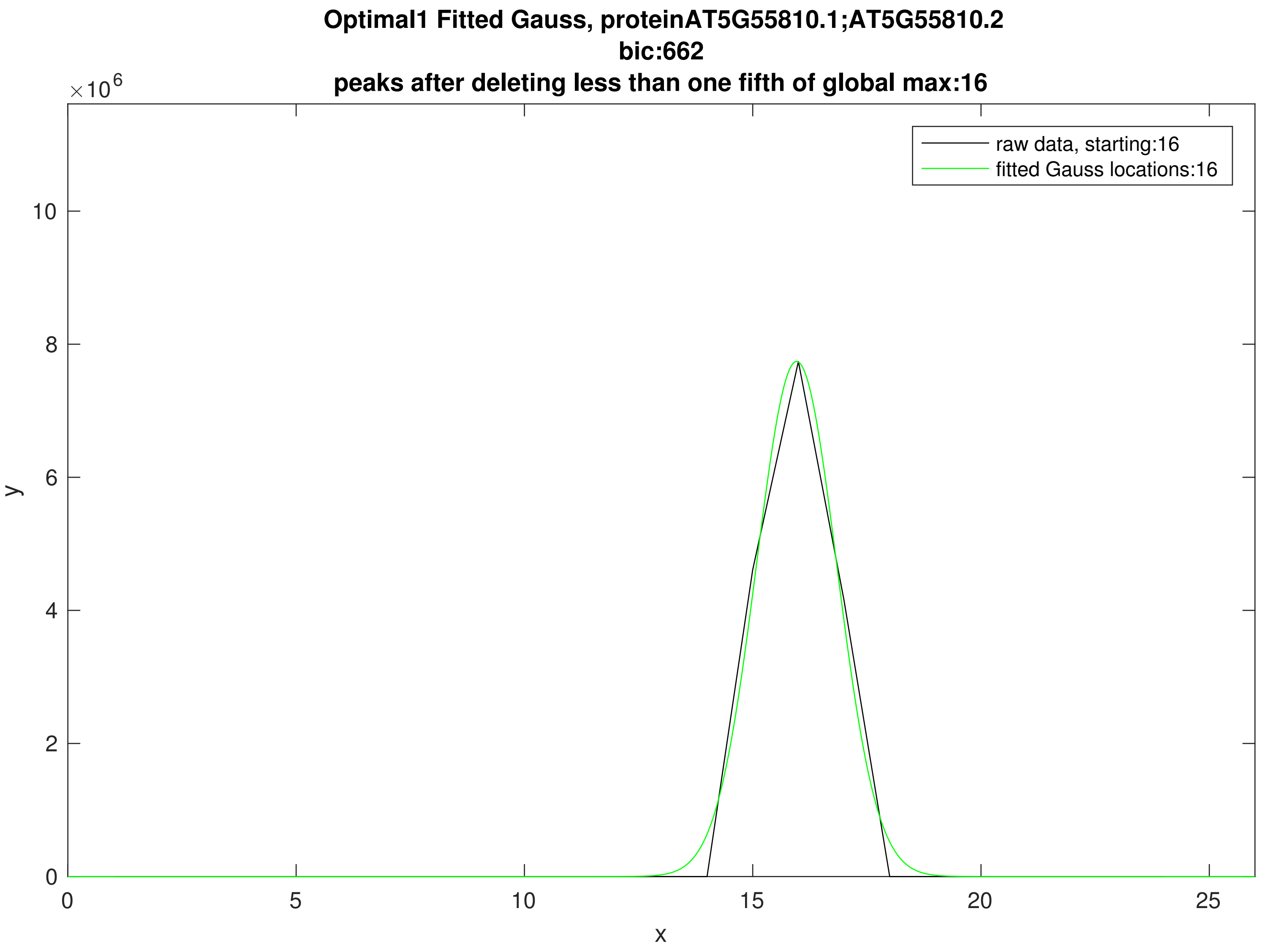

Optimal1 Fitted Gauss, proteinAT5G55810.1;AT5G55810.2
bic:662
peaks after deleting less than one fifth of global max:16
×10^6
raw data, starting:16
fitted Gauss locations:16
10
8
6
4
2
0
y
0
5
10
15
20
25
x

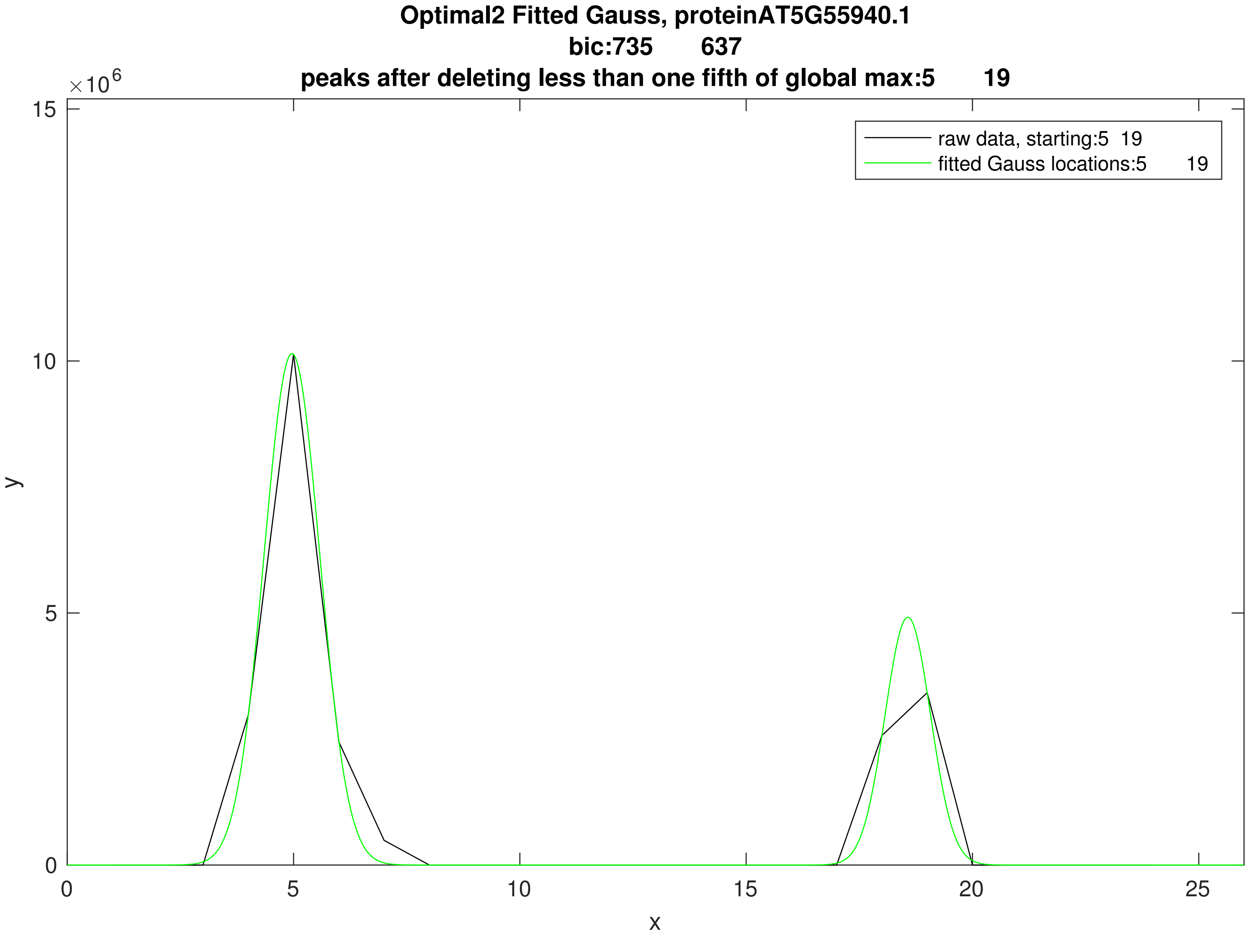
Optimal2 Fitted Gauss, proteinAT5G55940.1
bic:735 637
peaks after deleting less than one fifth of global max:5 19
×10^6
15
10
5
0
y
0
5
10
15
20
25
x
raw data, starting:5 19
fitted Gauss locations:5 19

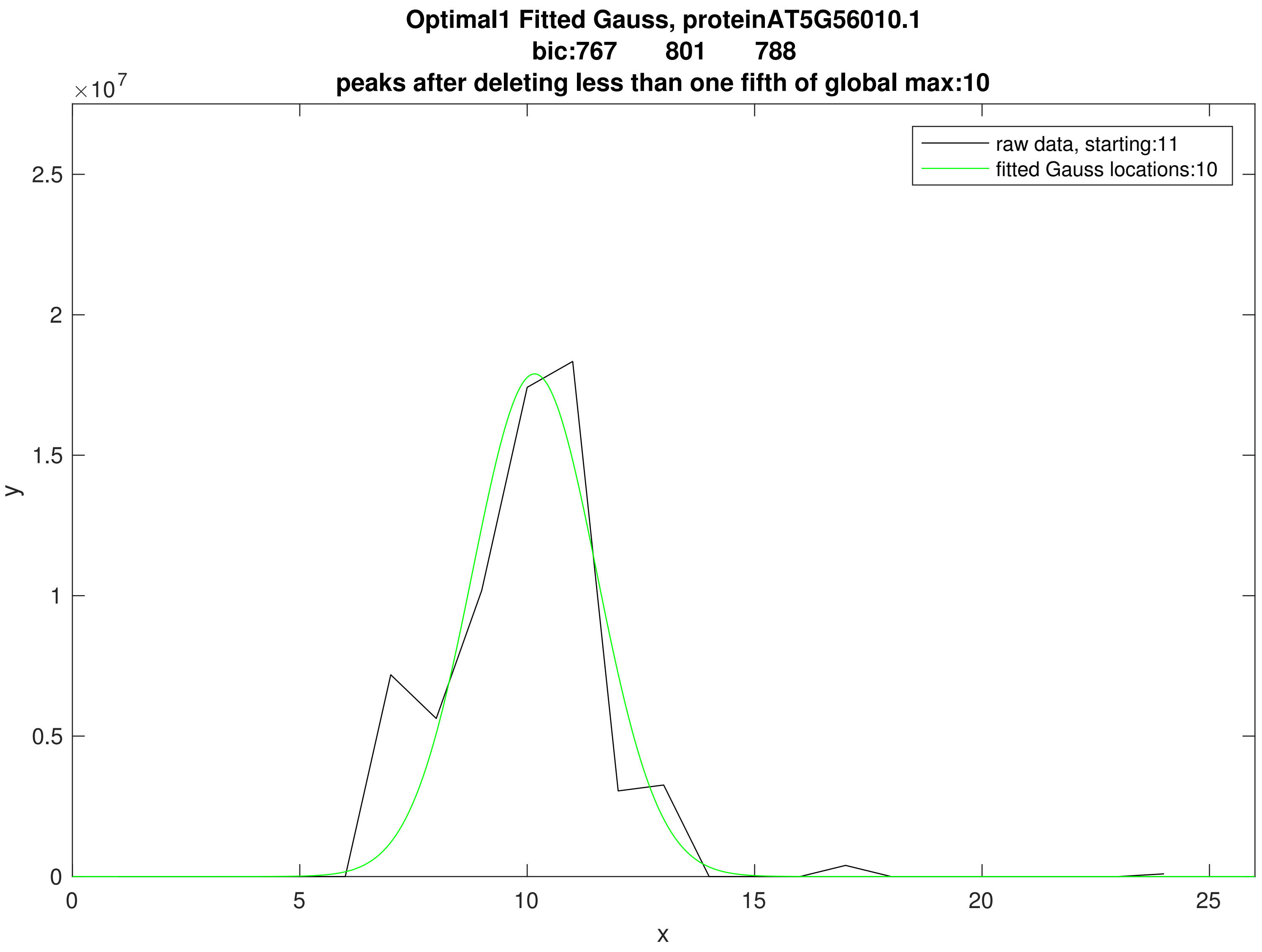

Optimal1 Fitted Gauss, proteinAT5G56010.1
bic:767 801 788
peaks after deleting less than one fifth of global max:10
×10^7
raw data, starting:11
fitted Gauss locations:10
2.5
2
1.5
1
0.5
0
y
0
5
10
15
20
25
x

#### Optimal3 Fitted Gauss, proteinAT5G56030.1;AT5G56030.2
#### bic:969 960 957
#### peaks after deleting less than one fifth of global max:6.9 11

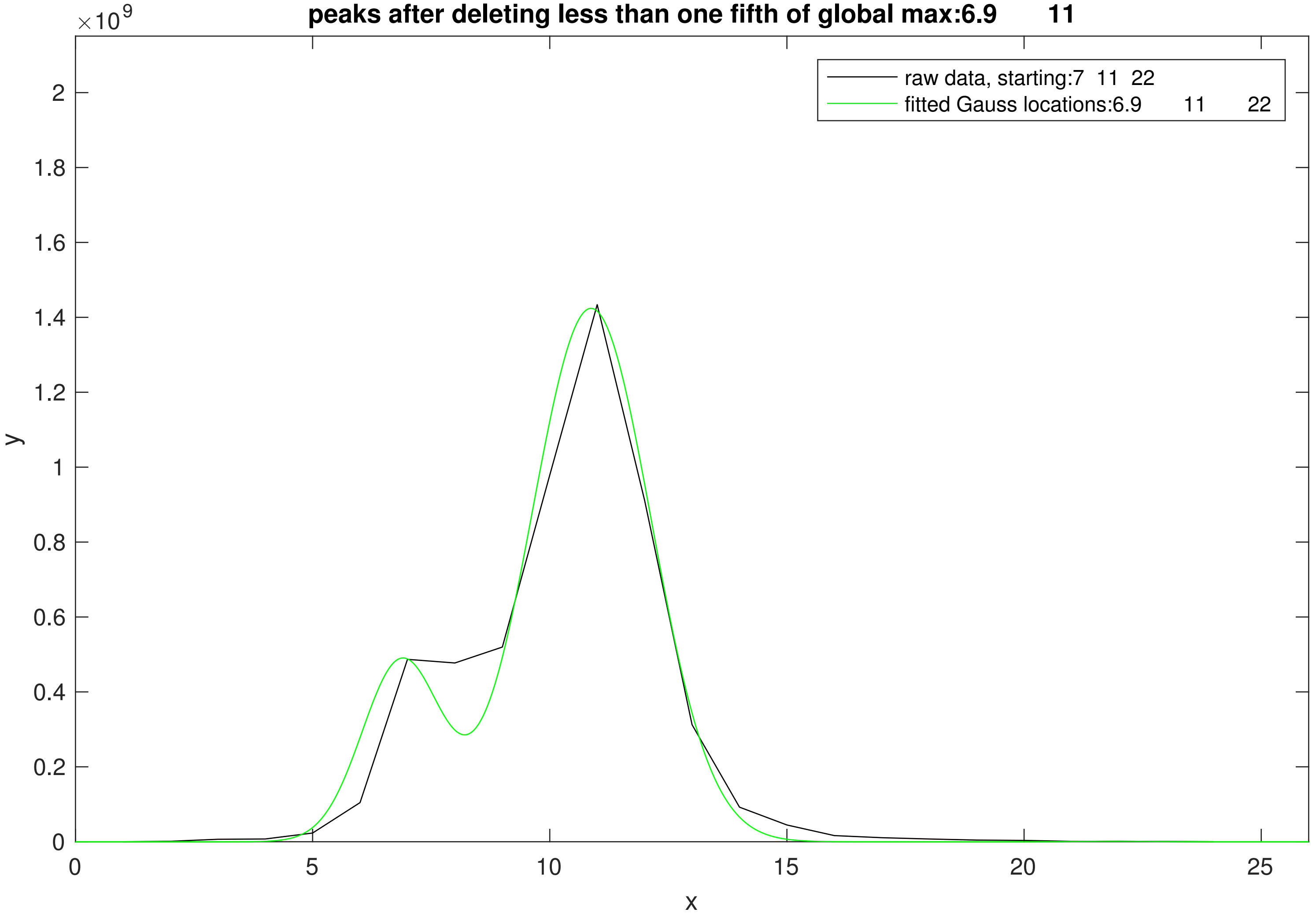

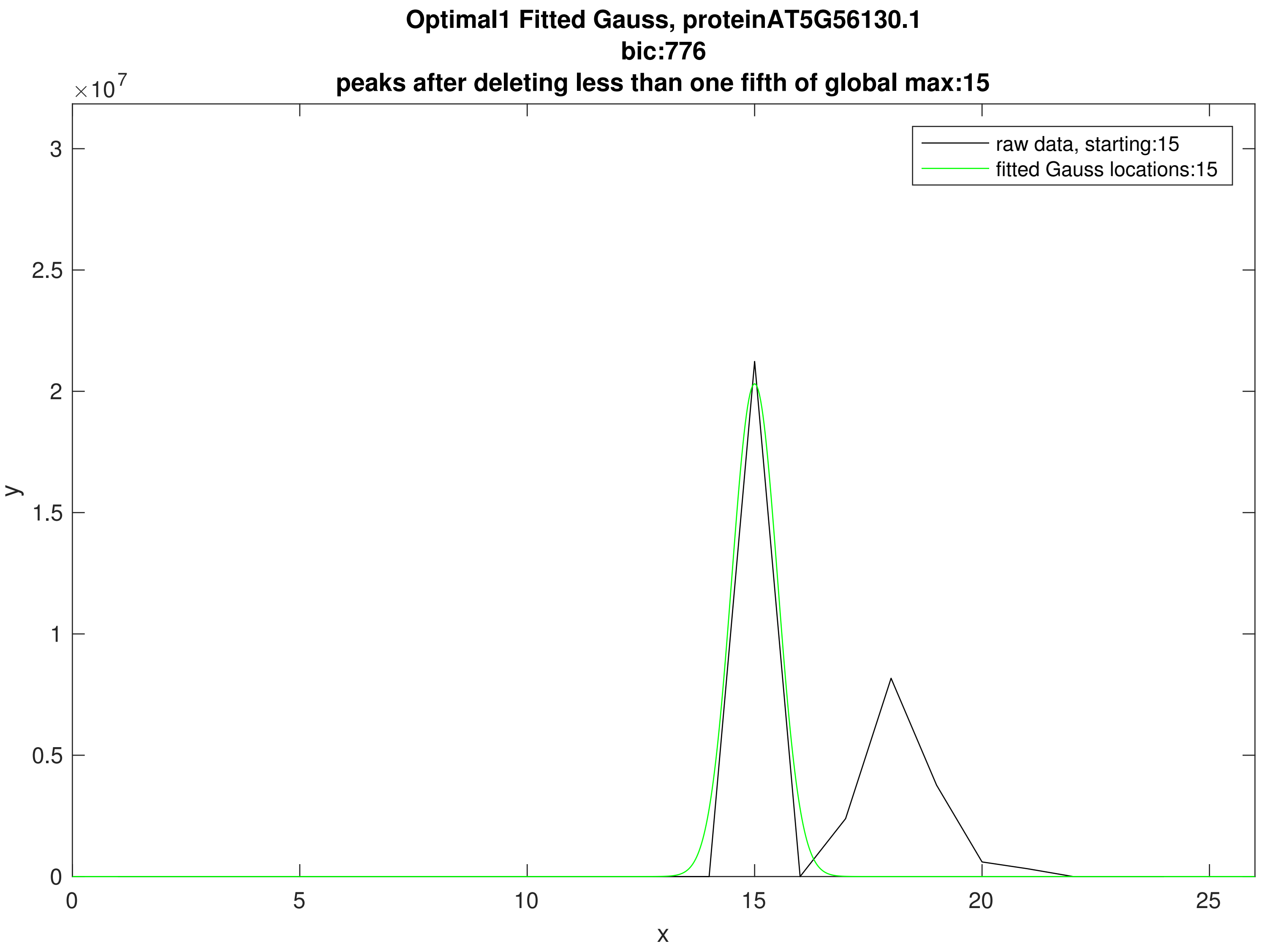
Optimal1 Fitted Gauss, proteinAT5G56130.1
bic:776
peaks after deleting less than one fifth of global max:15
×10^7
3
2.5
2
1.5
1
0.5
0
y
0
5
10
15
20
25
x
raw data, starting:15
fitted Gauss locations:15

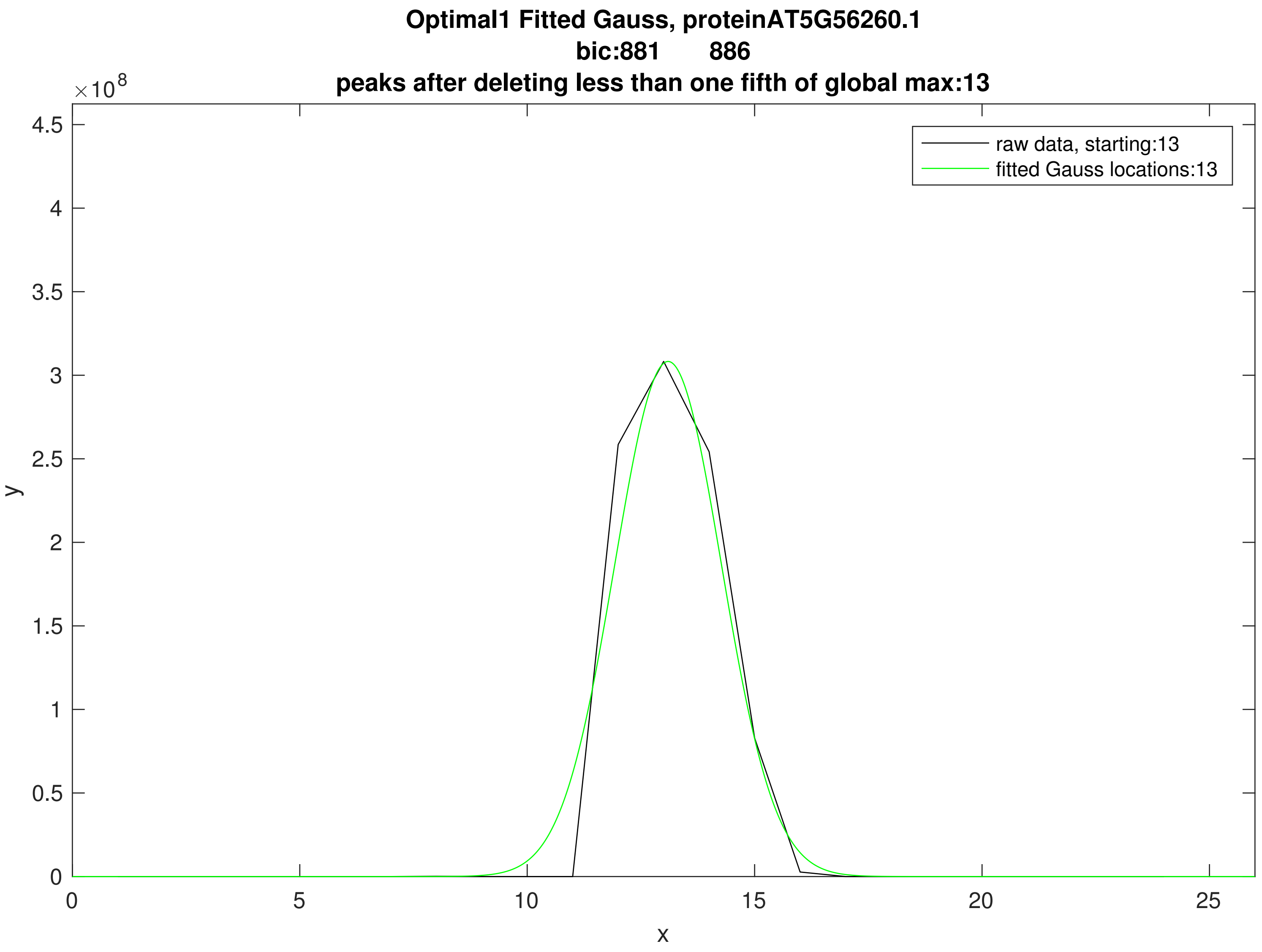

Optimal1 Fitted Gauss, proteinAT5G56260.1
bic:881 886
peaks after deleting less than one fifth of global max:13
×10^8
4.5
4
3.5
3
2.5
2
1.5
1
0.5
0
y
0
5
10
15
20
25
x
raw data, starting:13
fitted Gauss locations:13

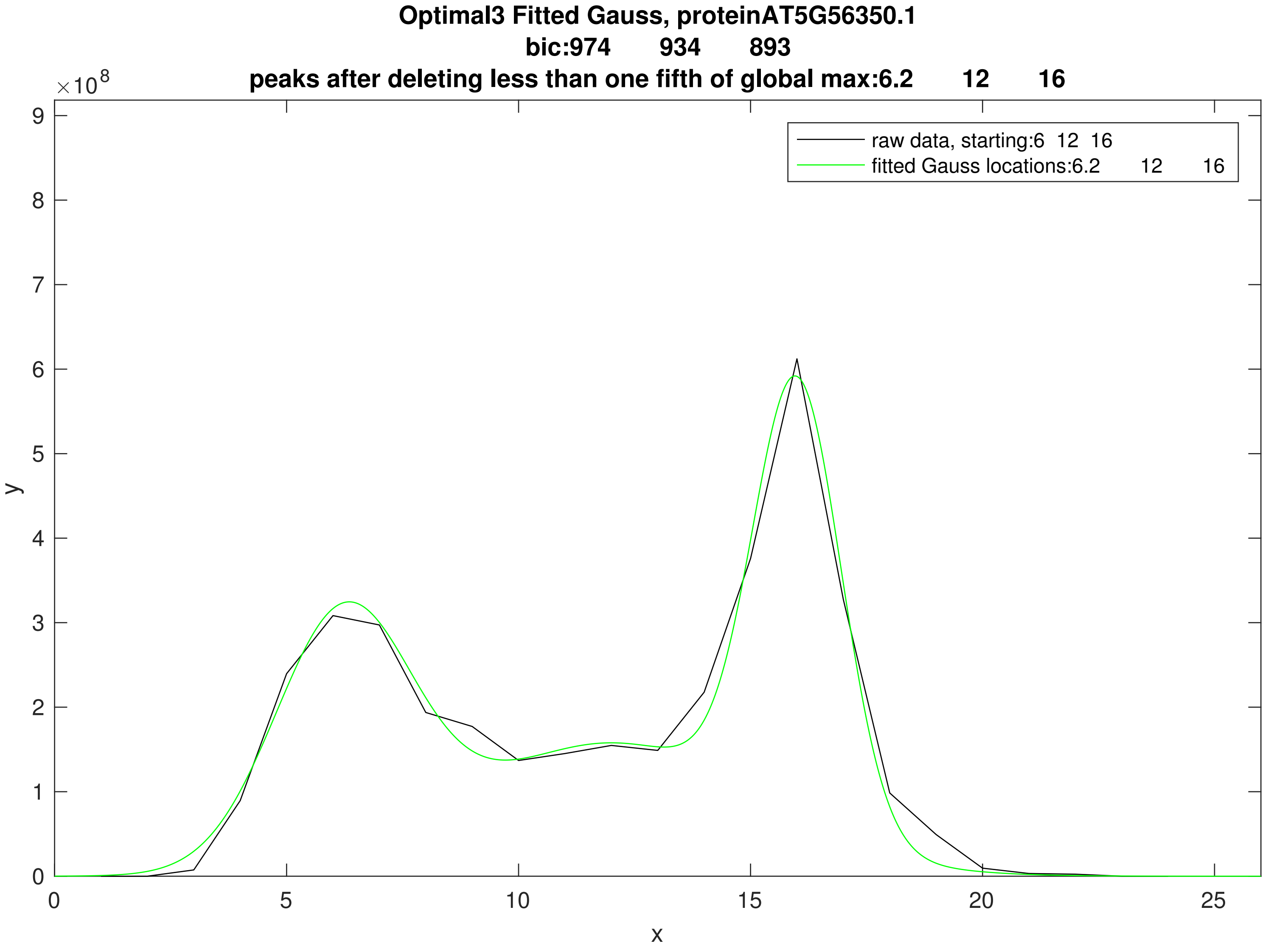

Optimal3 Fitted Gauss, proteinAT5G56350.1
bic:974 934 893
peaks after deleting less than one fifth of global max:6.2 12 16
×10^8
raw data, starting:6 12 16
fitted Gauss locations:6.2 12 16
y
x
0
1
2
3
4
5
6
7
8
9
0
5
10
15
20
25

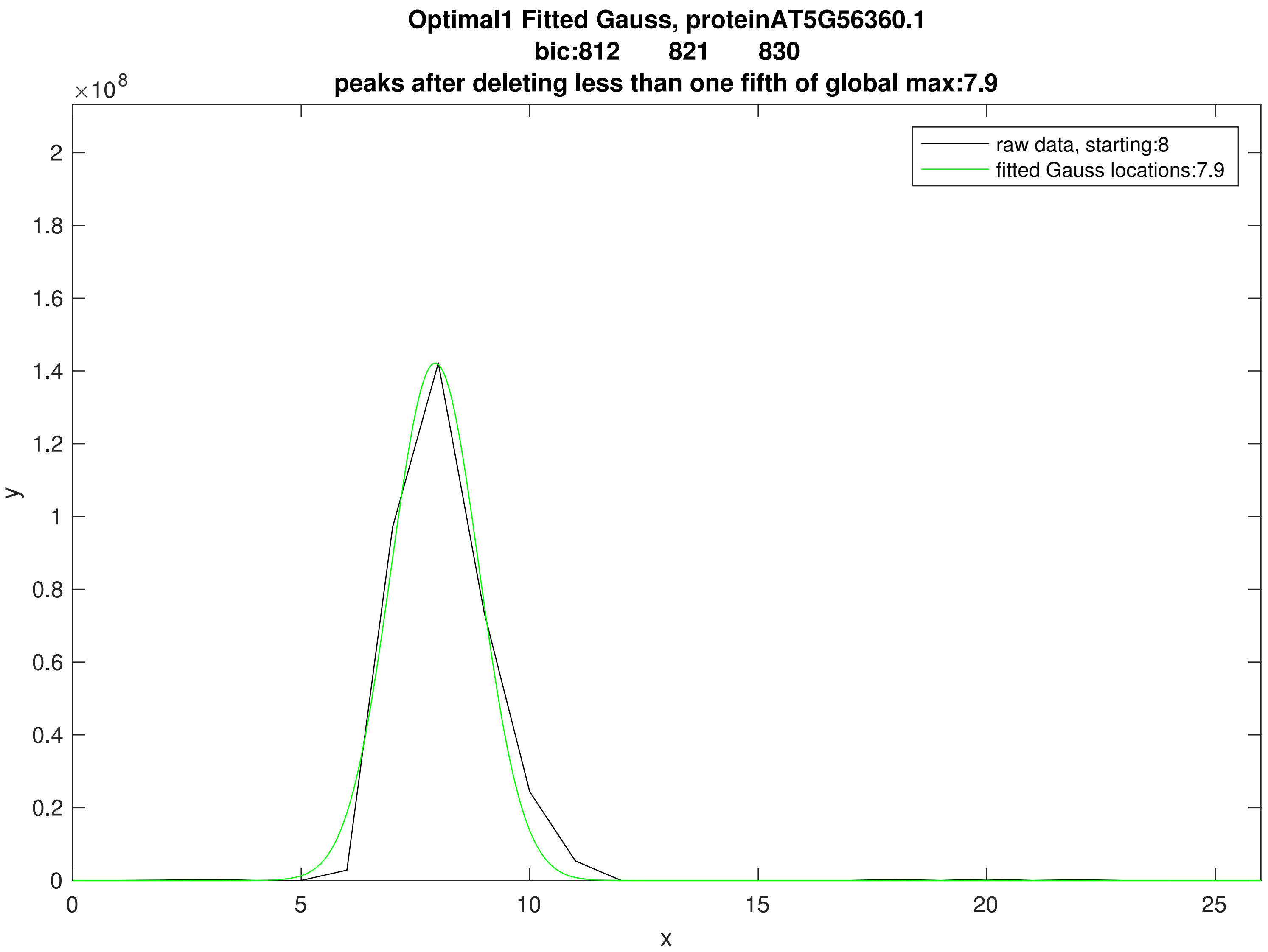

Optimal1 Fitted Gauss, proteinAT5G56360.1
bic:812 821 830
peaks after deleting less than one fifth of global max:7.9
×10^8
raw data, starting:8
fitted Gauss locations:7.9
y
x
0
0.2
0.4
0.6
0.8
1
1.2
1.4
1.6
1.8
2
5
10
15
20
25

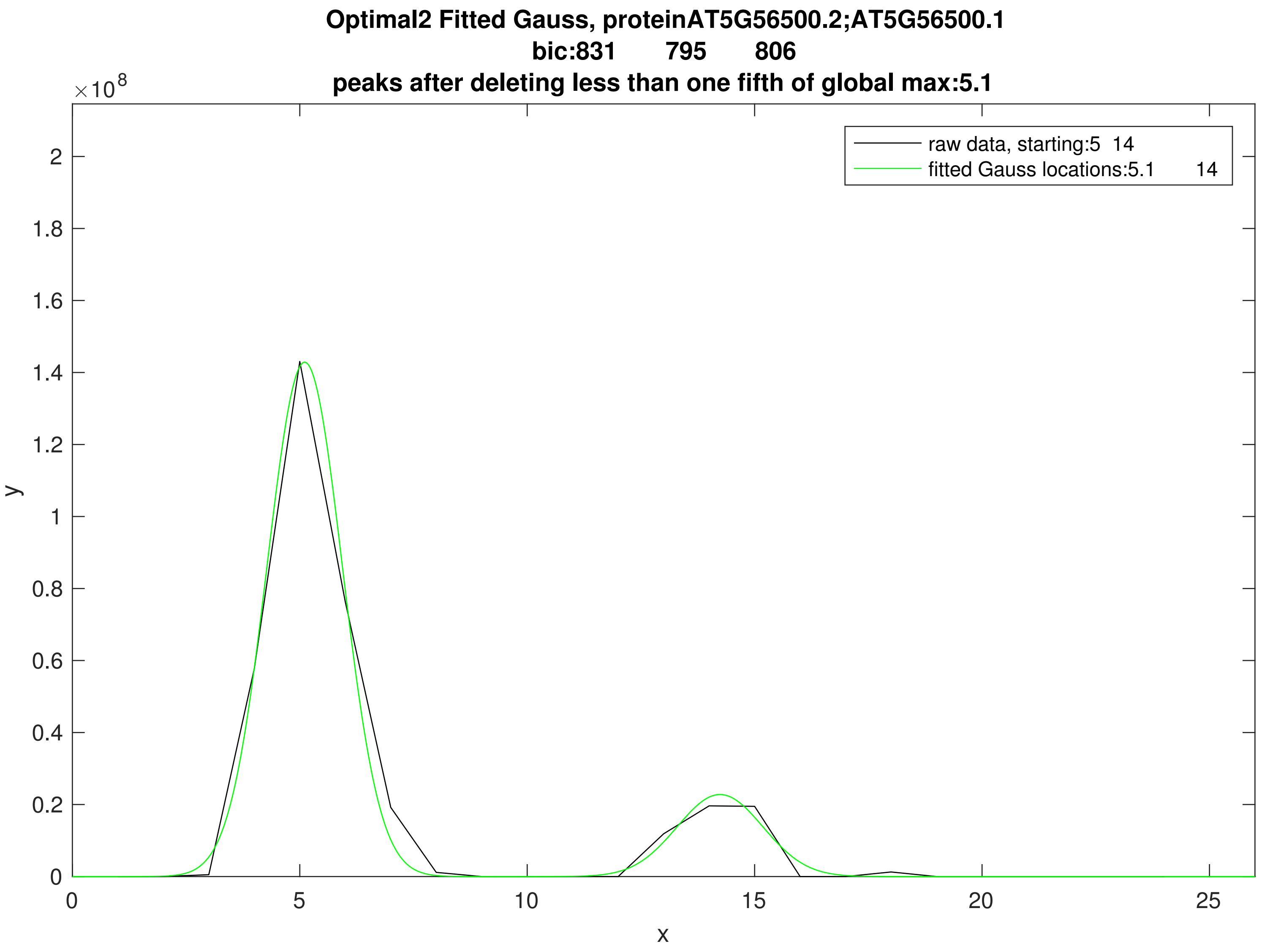
Optimal2 Fitted Gauss, proteinAT5G56500.2;AT5G56500.1
bic:831 795 806
peaks after deleting less than one fifth of global max:5.1
×10^8
raw data, starting:5 14
fitted Gauss locations:5.1 14
y
x
0
0.2
0.4
0.6
0.8
1
1.2
1.4
1.6
1.8
2
5
10
15
20
25

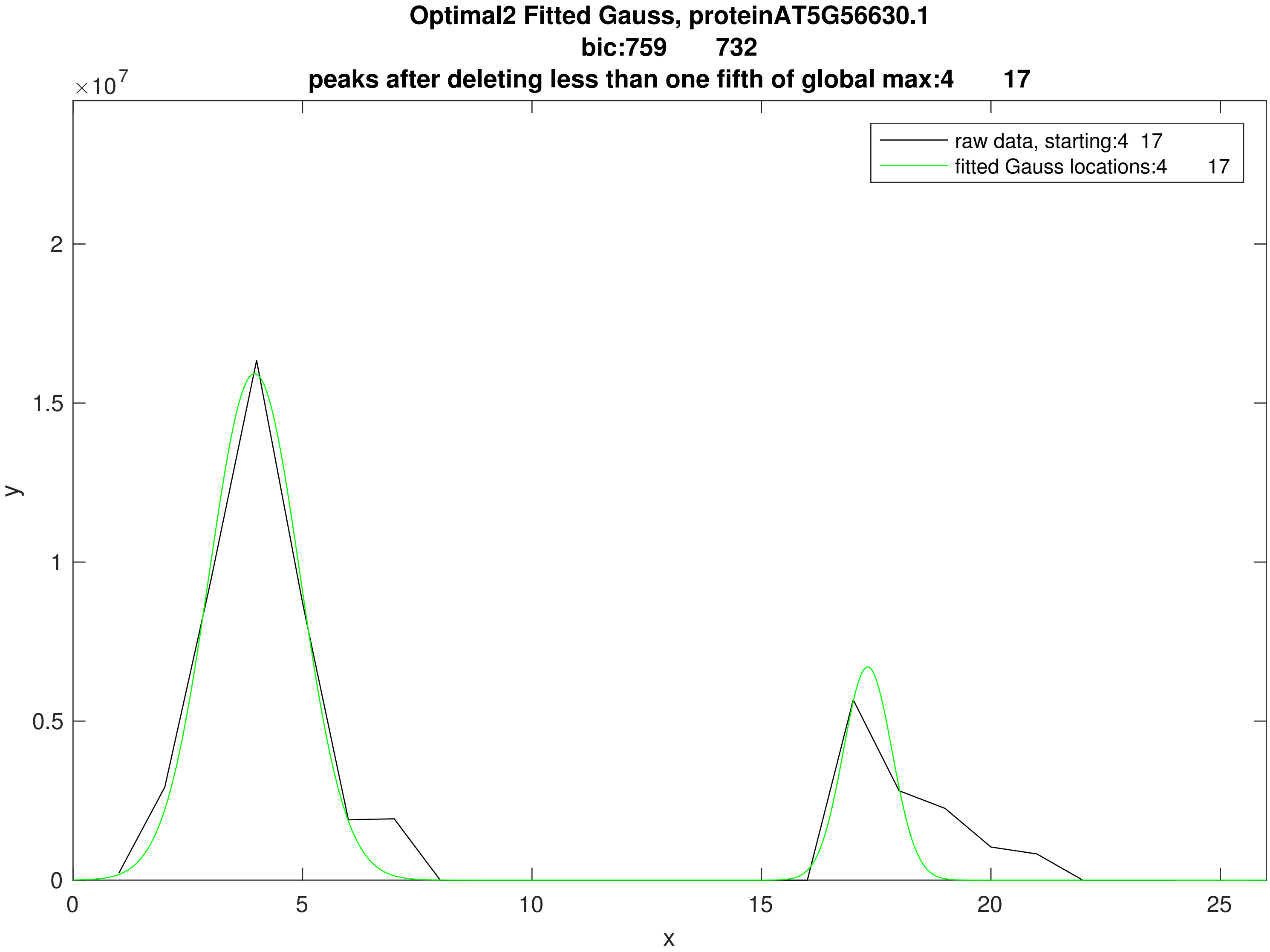
Optimal2 Fitted Gauss, proteinAT5G56630.1
bic:759 732
peaks after deleting less than one fifth of global max:4 17
×10^7
raw data, starting:4 17
fitted Gauss locations:4 17
y
x
0
0.5
1
1.5
2
5
10
15
20
25

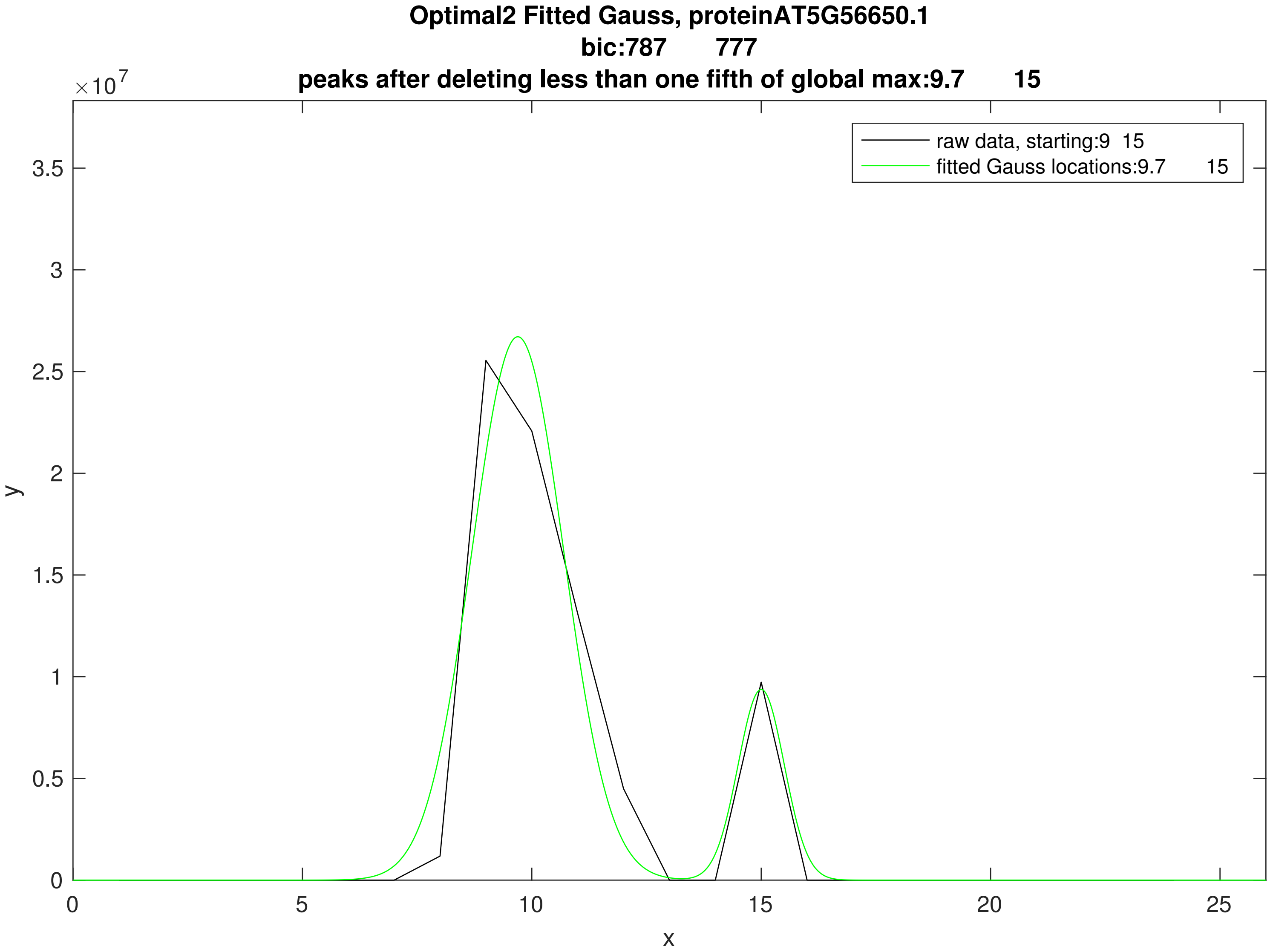

Optimal2 Fitted Gauss, proteinAT5G56650.1
bic:787 777
peaks after deleting less than one fifth of global max:9.7 15
×10^7
raw data, starting:9 15
fitted Gauss locations:9.7 15
3.5
3
2.5
2
1.5
1
0.5
0
y
0
5
10
15
20
25
x

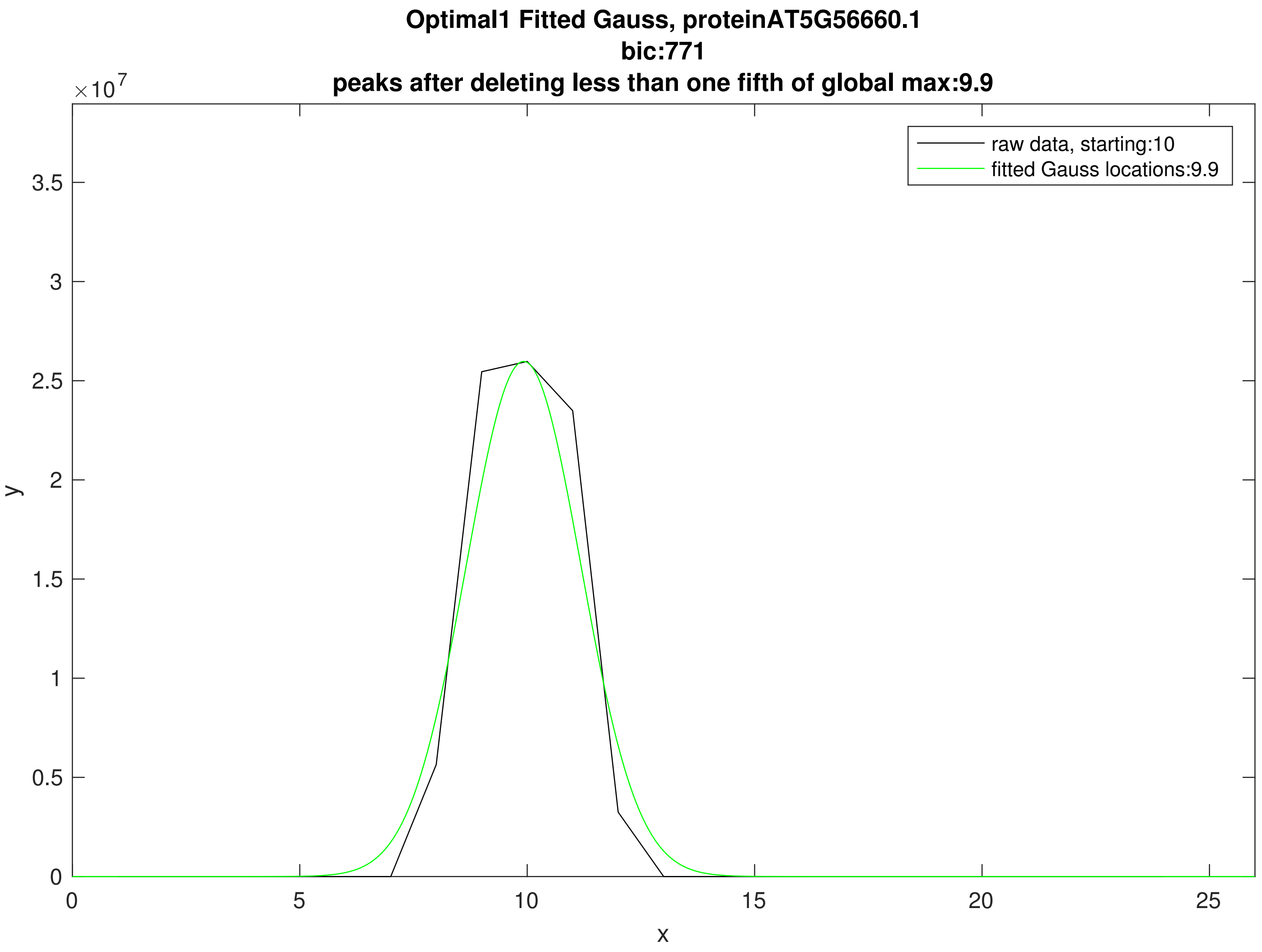

Optimal1 Fitted Gauss, proteinAT5G56660.1
bic:771
peaks after deleting less than one fifth of global max:9.9
×10^7
raw data, starting:10
fitted Gauss locations:9.9
3.5
3
2.5
2
1.5
1
0.5
0
y
0
5
10
15
20
25
x

**Optimal1 Fitted Gauss, proteinAT5G56670.1;AT4G29390.1;AT2G19750.1**
**bic:720**
**peaks after deleting less than one fifth of global max:3.9**

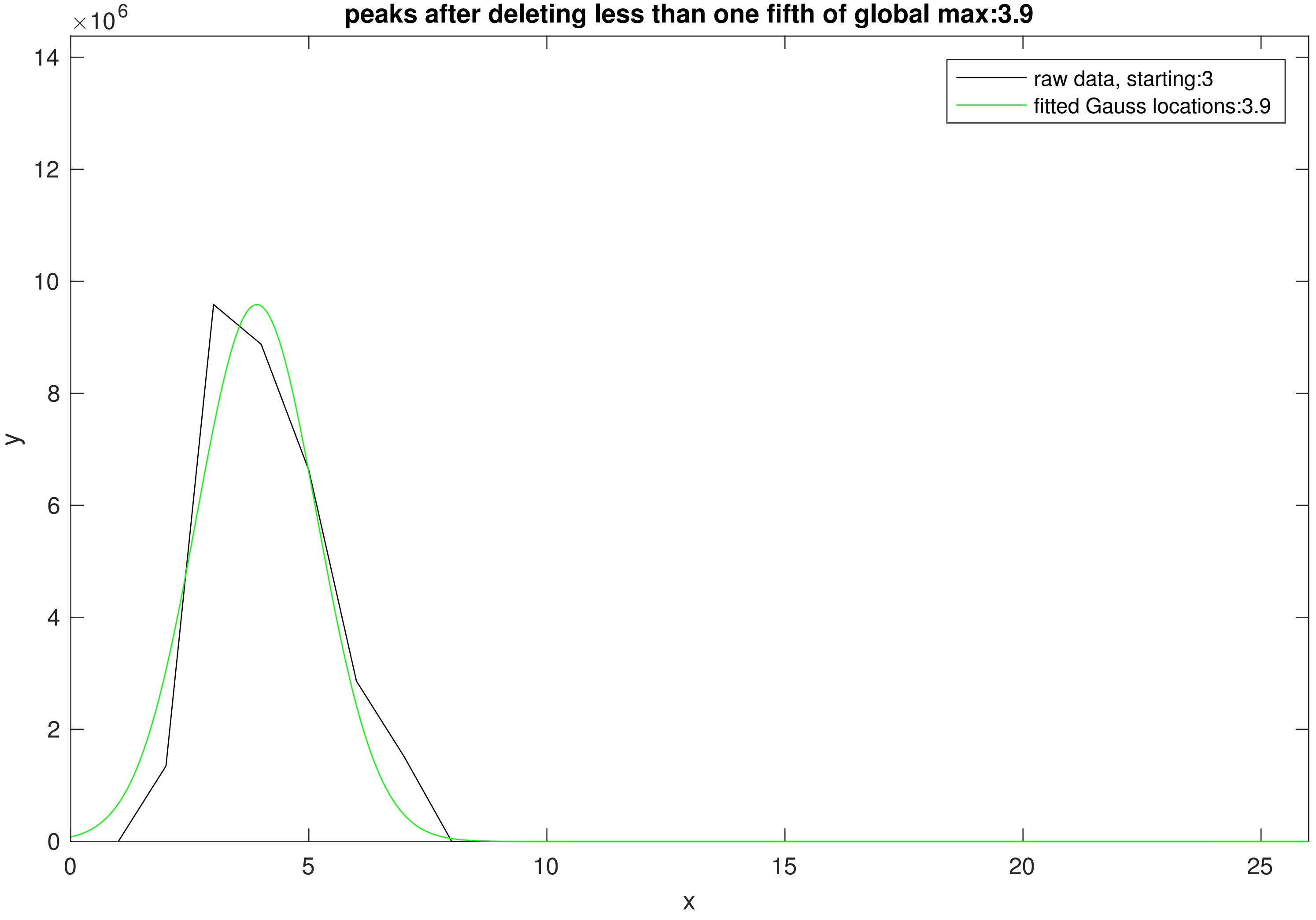

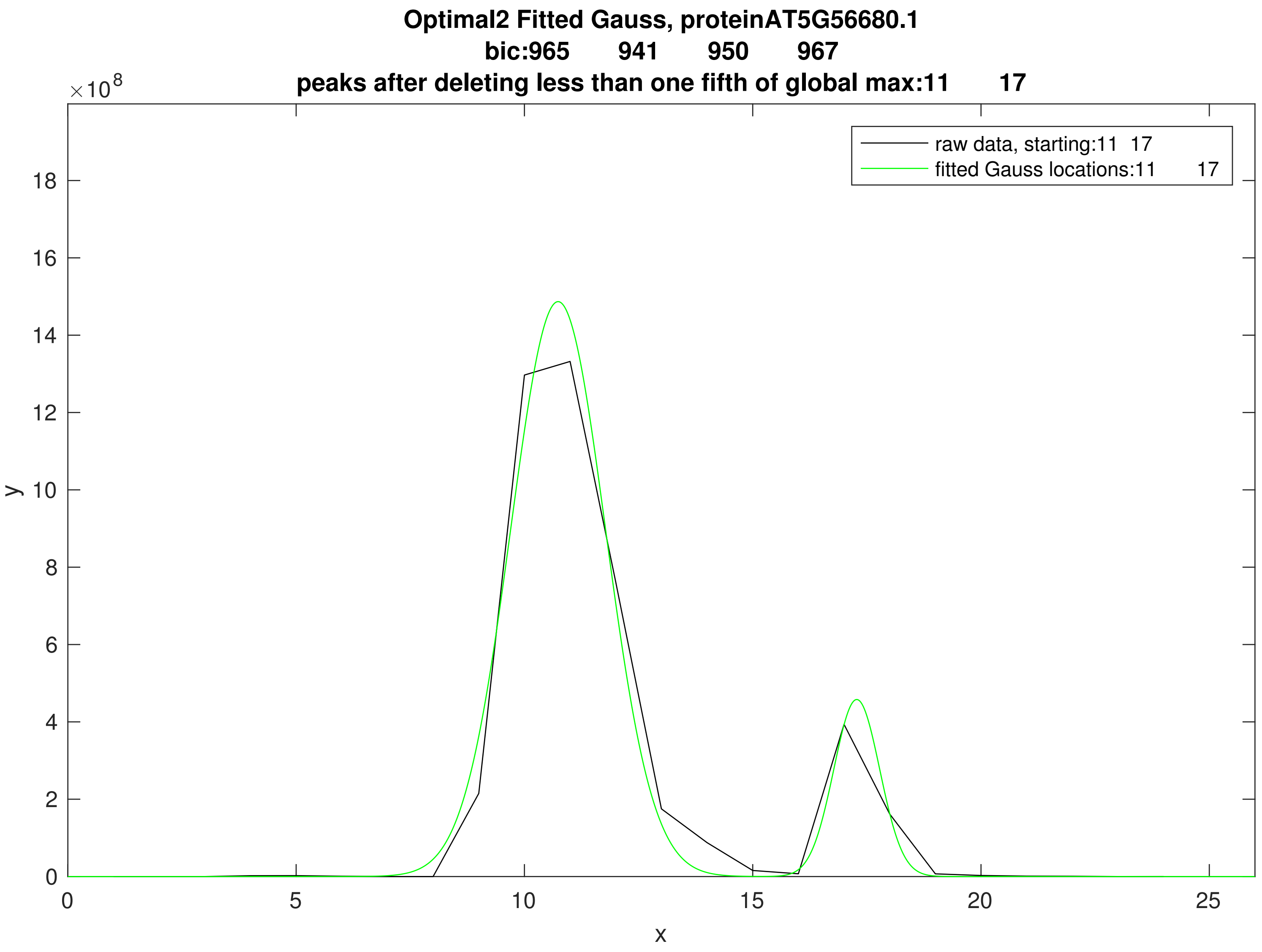

Optimal2 Fitted Gauss, proteinAT5G56680.1
bic:965 941 950 967
peaks after deleting less than one fifth of global max:11 17
×10^8
raw data, starting:11 17
fitted Gauss locations:11 17
y
x
0
2
4
6
8
10
12
14
16
18
0
5
10
15
20
25

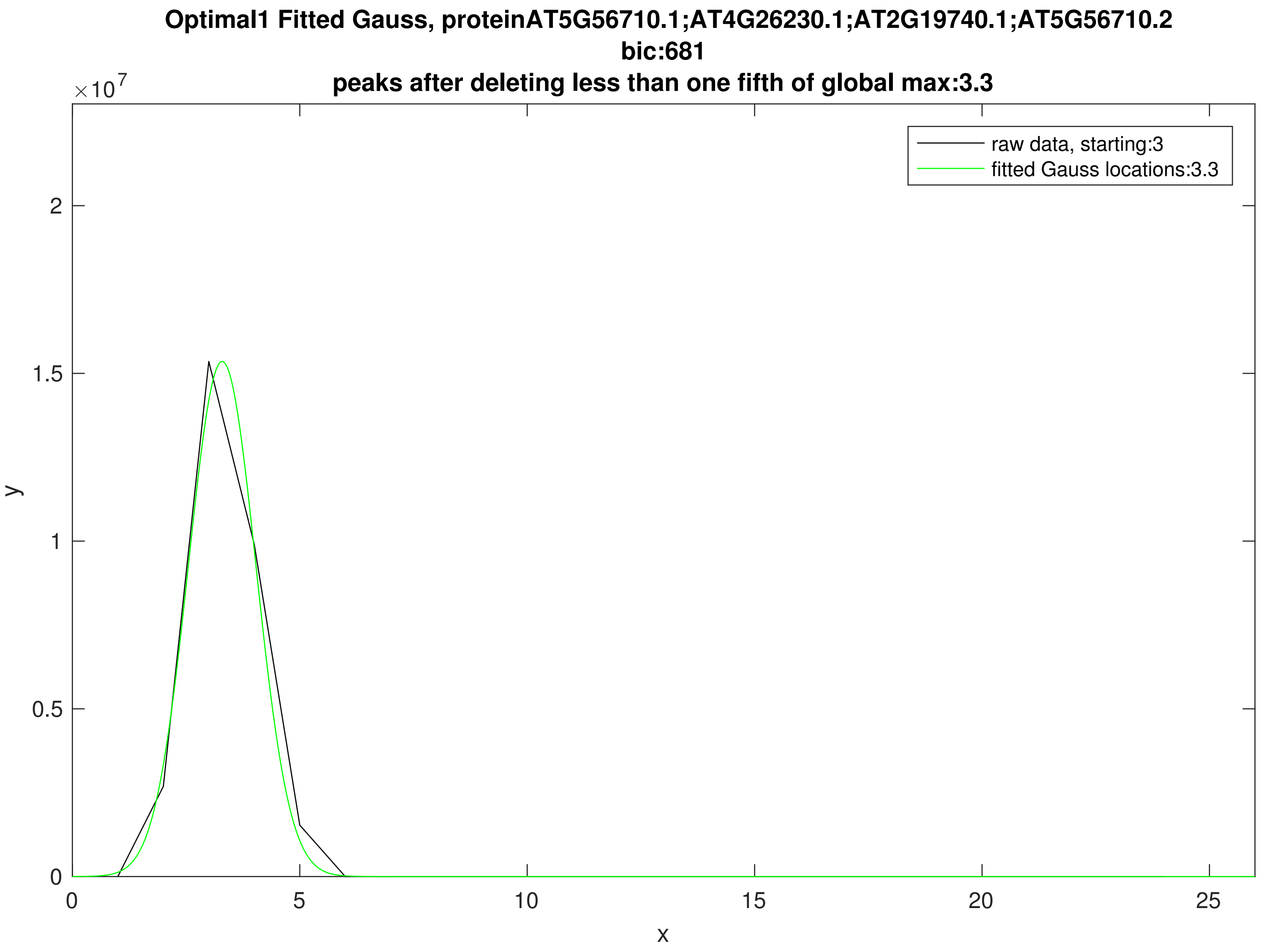
Optimal1 Fitted Gauss, proteinAT5G56710.1;AT4G26230.1;AT2G19740.1;AT5G56710.2
bic:681
peaks after deleting less than one fifth of global max:3.3
×10^7
raw data, starting:3
fitted Gauss locations:3.3
y
x
0
0.5
1
1.5
2
5
10
15
20
25

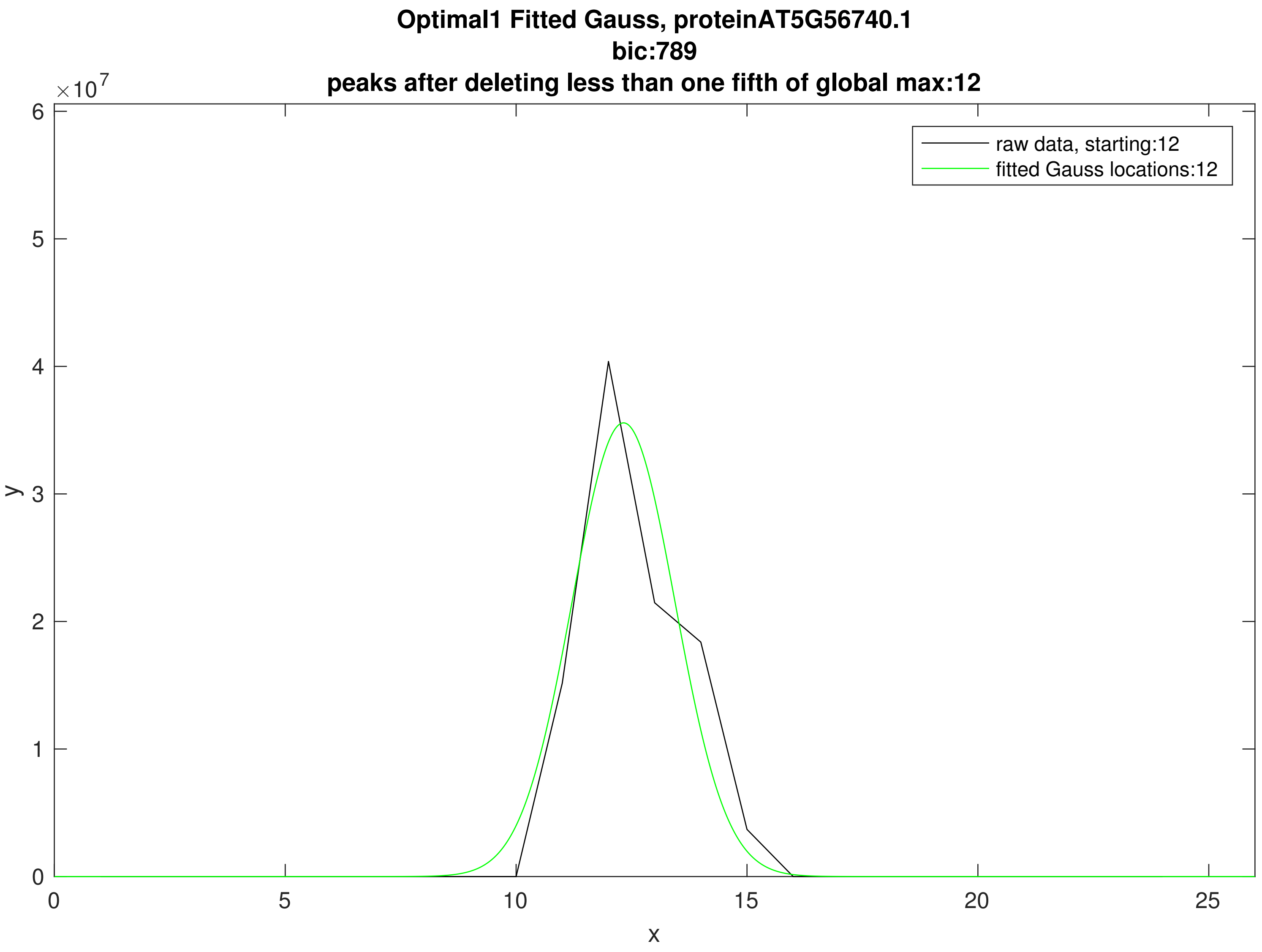

Optimal1 Fitted Gauss, proteinAT5G56740.1
bic:789
peaks after deleting less than one fifth of global max:12
×10^7
6
5
4
3
2
1
0
y
0
5
10
15
20
25
x
raw data, starting:12
fitted Gauss locations:12

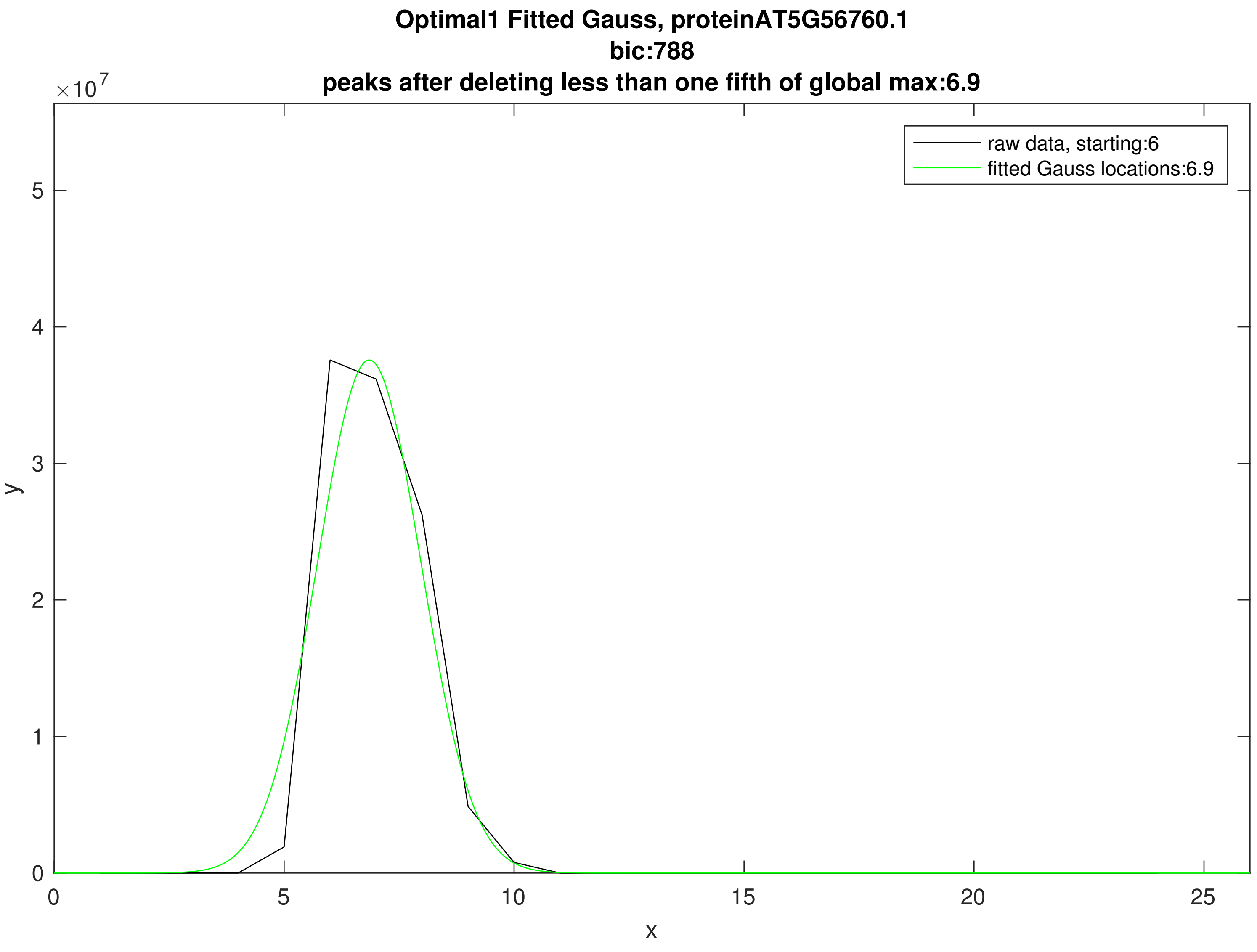

Optimal1 Fitted Gauss, proteinAT5G56760.1
bic:788
peaks after deleting less than one fifth of global max:6.9
×10^7
raw data, starting:6
fitted Gauss locations:6.9
y
x
0
1
2
3
4
5
0
5
10
15
20
25

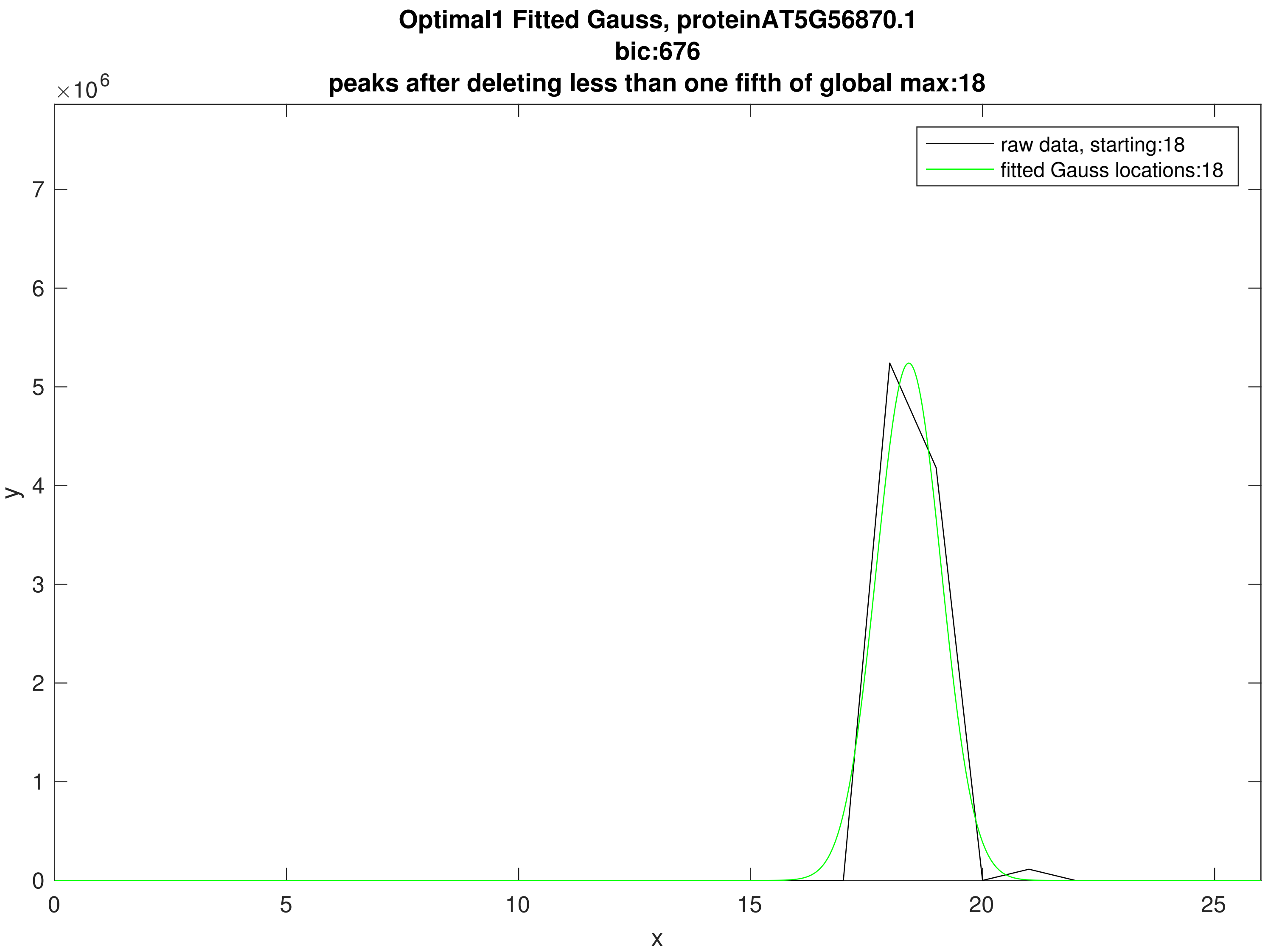
Optimal1 Fitted Gauss, proteinAT5G56870.1
bic:676
peaks after deleting less than one fifth of global max:18
×10^6
raw data, starting:18
fitted Gauss locations:18
y
x
0
1
2
3
4
5
6
7
0
5
10
15
20
25

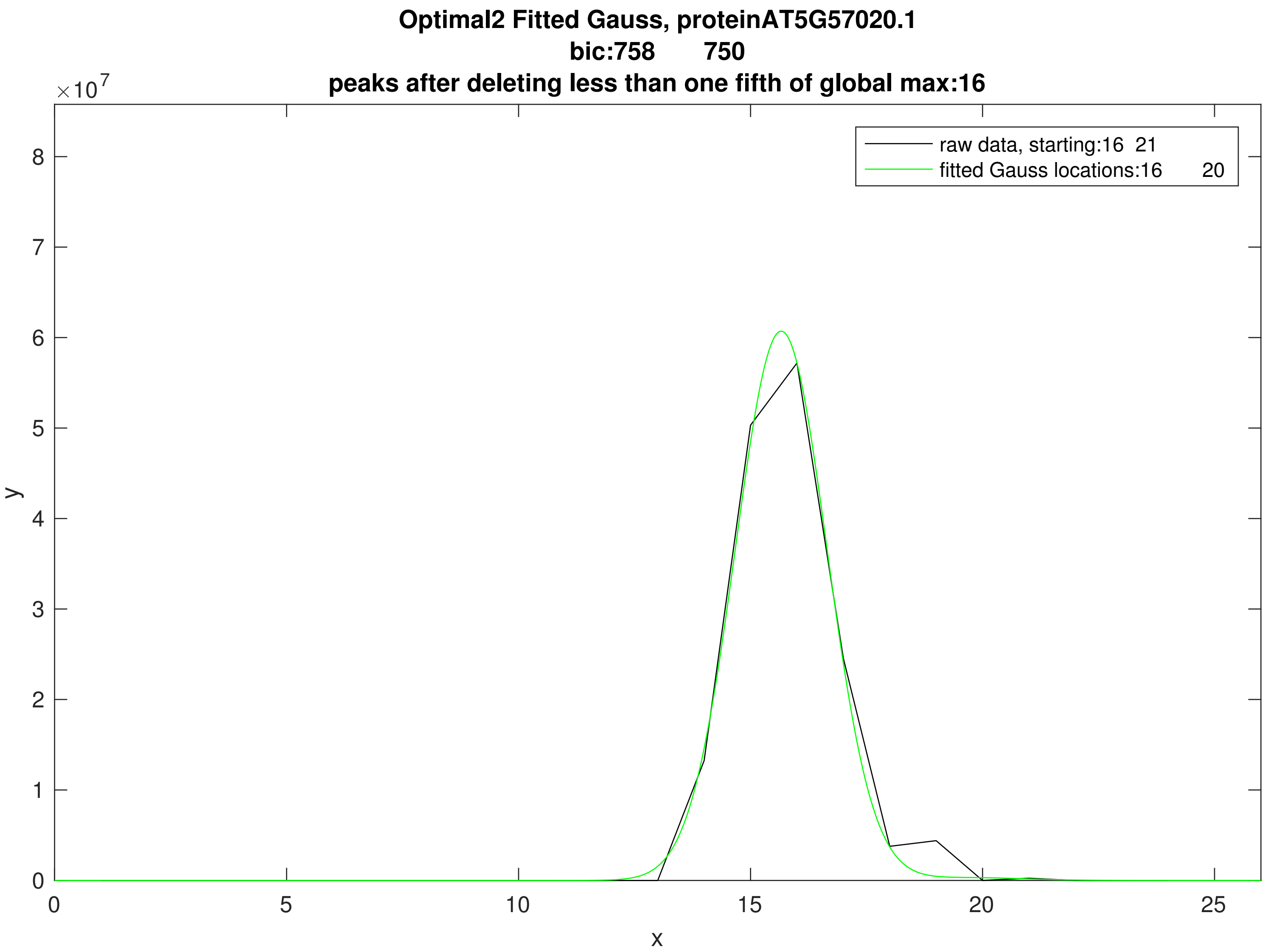
Optimal2 Fitted Gauss, proteinAT5G57020.1
bic:758 750
peaks after deleting less than one fifth of global max:16
×10^7
raw data, starting:16 21
fitted Gauss locations:16 20
y
x
0
1
2
3
4
5
6
7
8
0
5
10
15
20
25

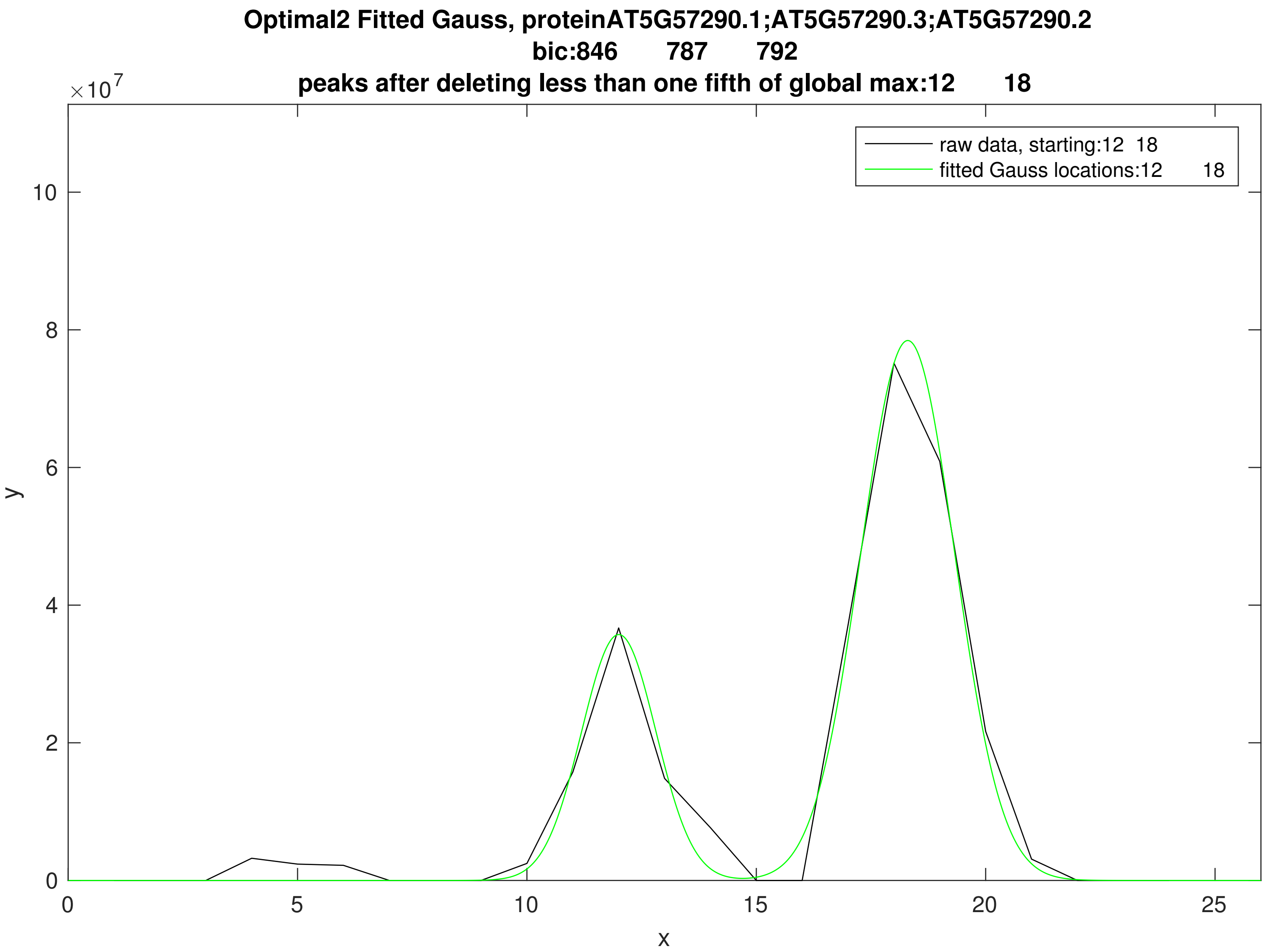
Optimal2 Fitted Gauss, proteinAT5G57290.1;AT5G57290.3;AT5G57290.2
bic:846 787 792
peaks after deleting less than one fifth of global max:12 18
×10^7
raw data, starting:12 18
fitted Gauss locations:12 18
y
x
0
2
4
6
8
10
0
5
10
15
20
25

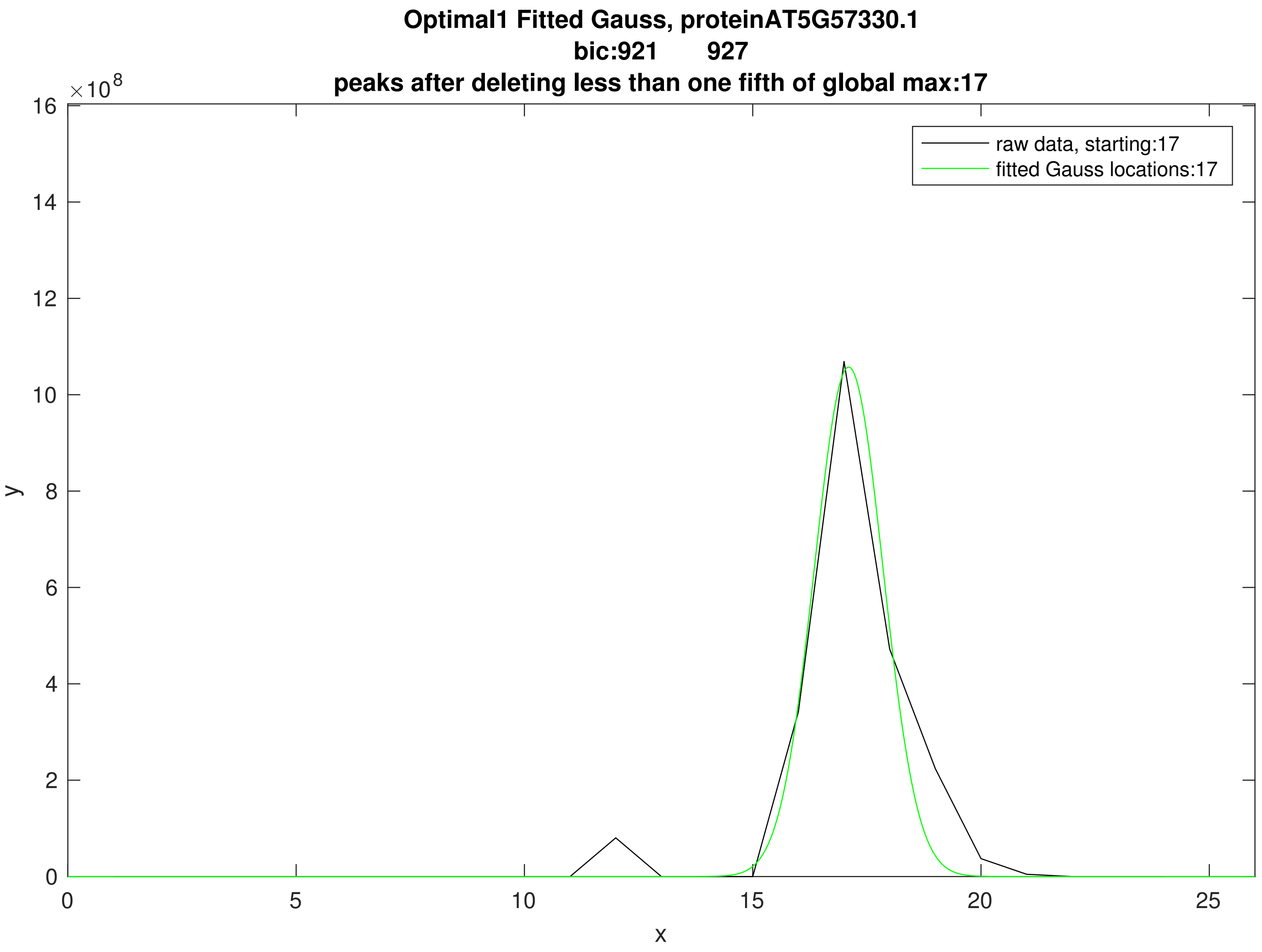
Optimal1 Fitted Gauss, proteinAT5G57330.1
bic:921 927
peaks after deleting less than one fifth of global max:17
×10^8
16
14
12
10
8
6
4
2
0
y
0
5
10
15
20
25
x
raw data, starting:17
fitted Gauss locations:17

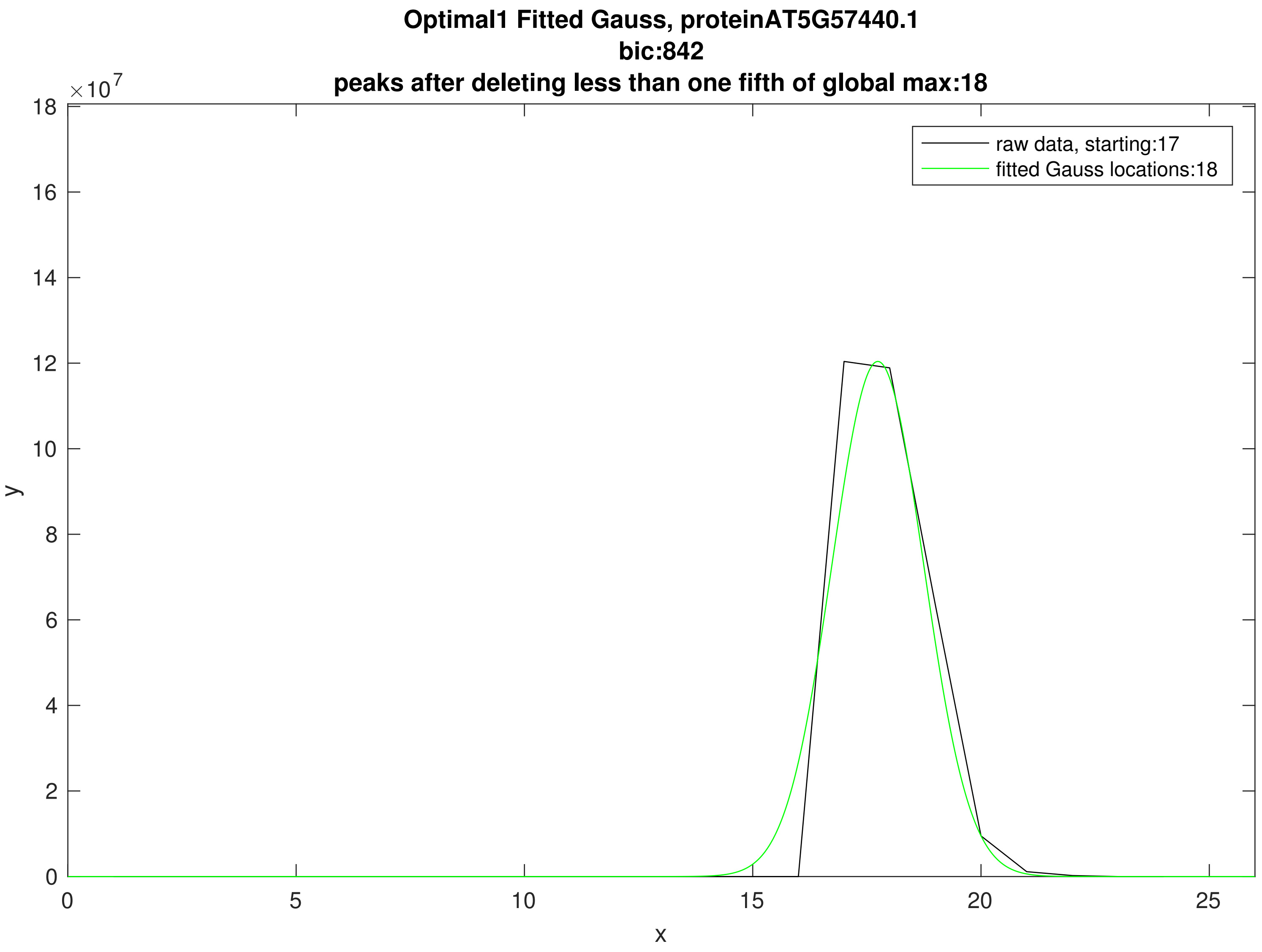

Optimal1 Fitted Gauss, proteinAT5G57440.1
bic:842
peaks after deleting less than one fifth of global max:18
×10^7
18
16
14
12
10
8
6
4
2
0
y
0
5
10
15
20
25
x
raw data, starting:17
fitted Gauss locations:18

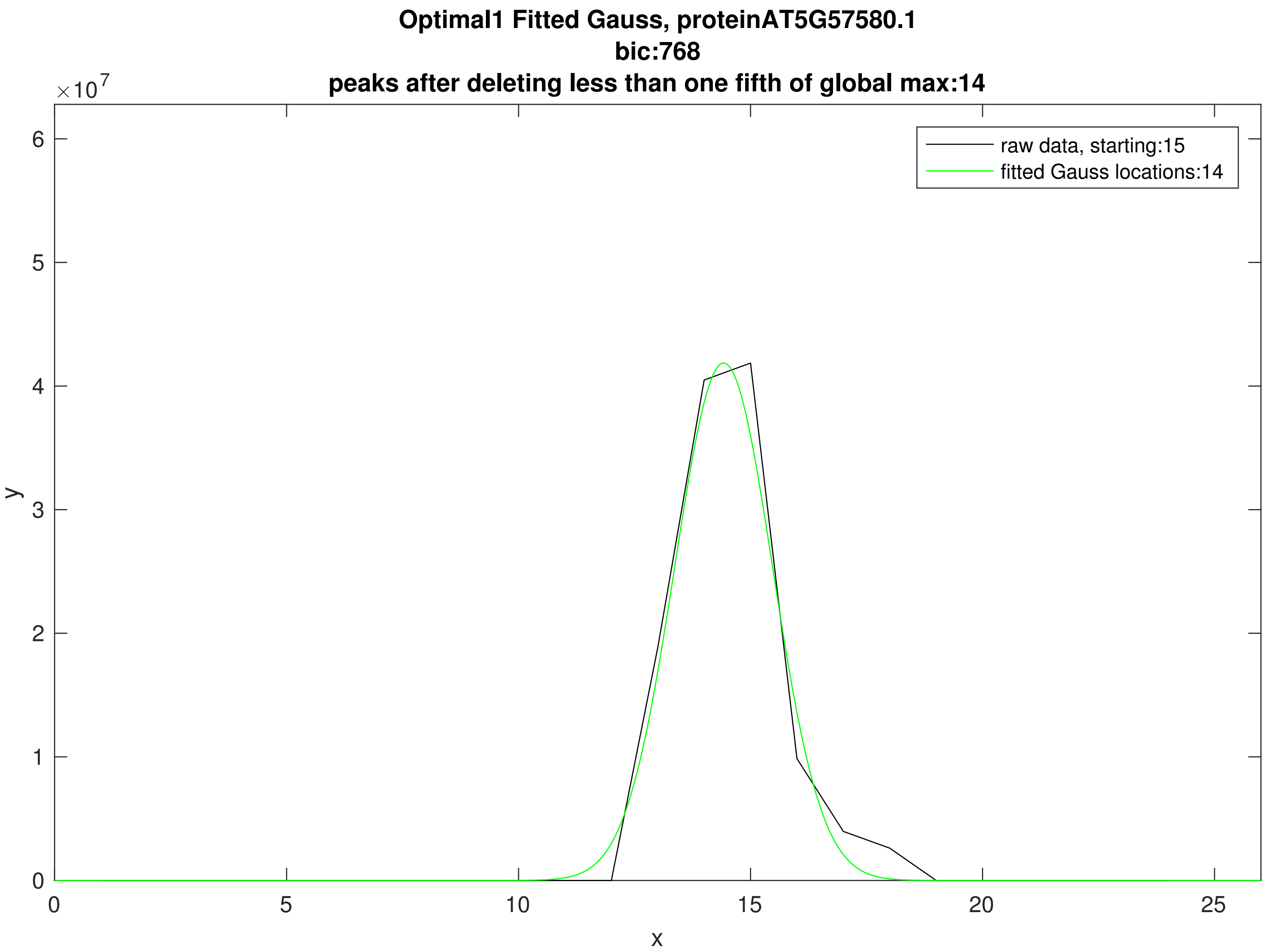

Optimal1 Fitted Gauss, proteinAT5G57580.1
bic:768
peaks after deleting less than one fifth of global max:14
×10^7
raw data, starting:15
fitted Gauss locations:14
y
x
0
1
2
3
4
5
6
0
5
10
15
20
25

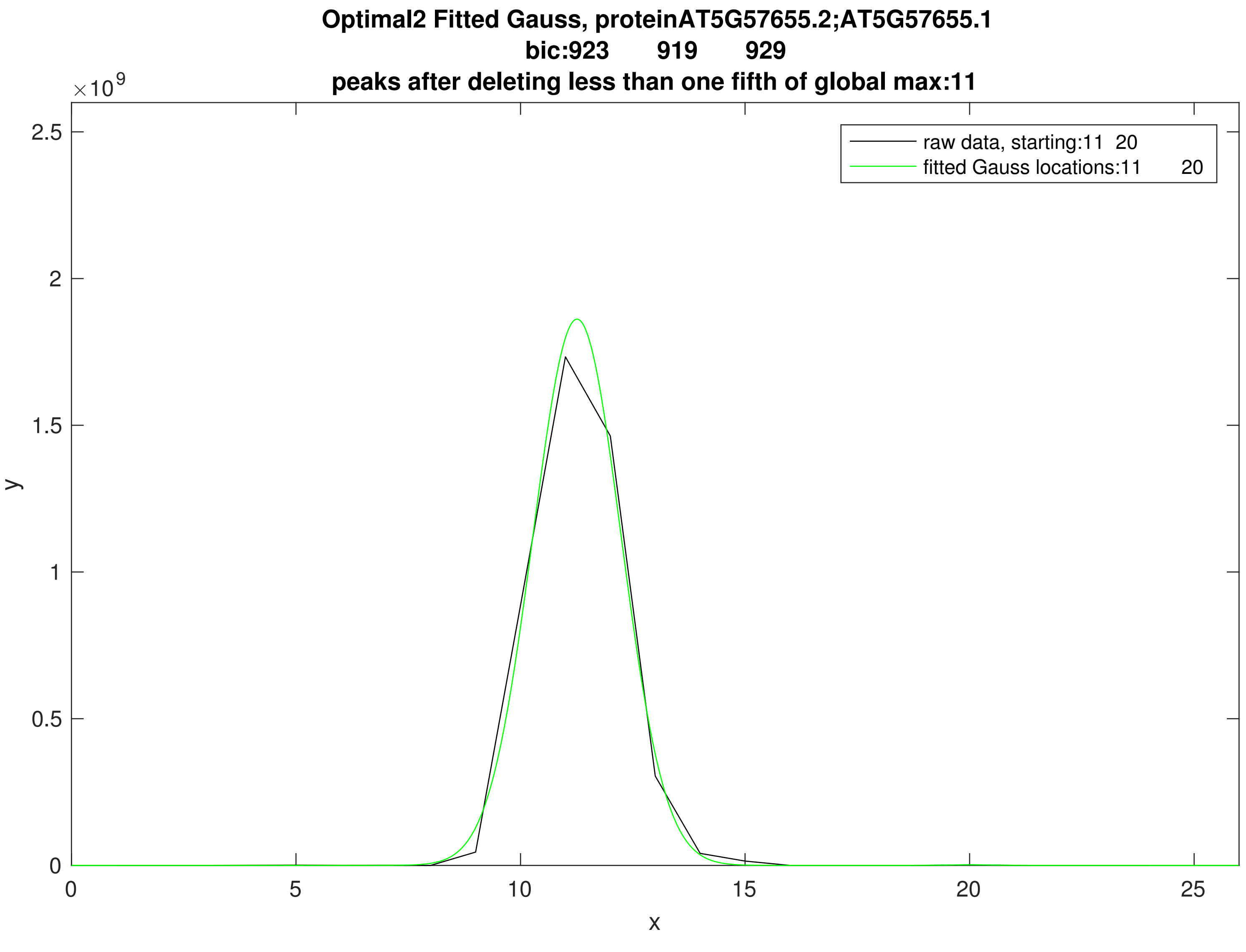
Optimal2 Fitted Gauss, proteinAT5G57655.2;AT5G57655.1
bic:923 919 929
peaks after deleting less than one fifth of global max:11
×10^9
raw data, starting:11 20
fitted Gauss locations:11 20
y
x
0
0.5
1
1.5
2
2.5
5
10
15
20
25

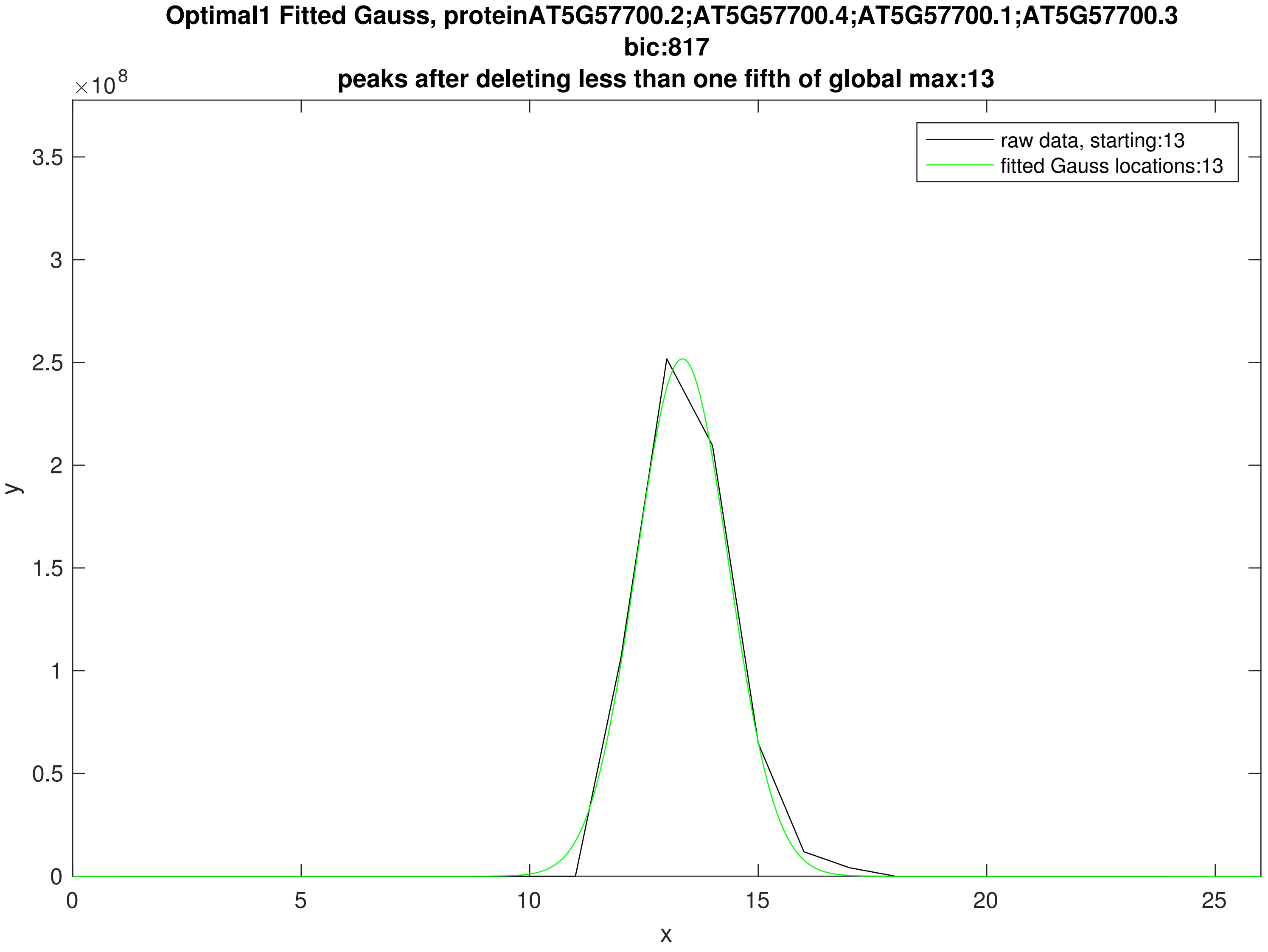
Optimal1 Fitted Gauss, proteinAT5G57700.2;AT5G57700.4;AT5G57700.1;AT5G57700.3
bic:817
peaks after deleting less than one fifth of global max:13
×10^8
raw data, starting:13
fitted Gauss locations:13
y
x
0
0.5
1
1.5
2
2.5
3
3.5
5
10
15
20
25

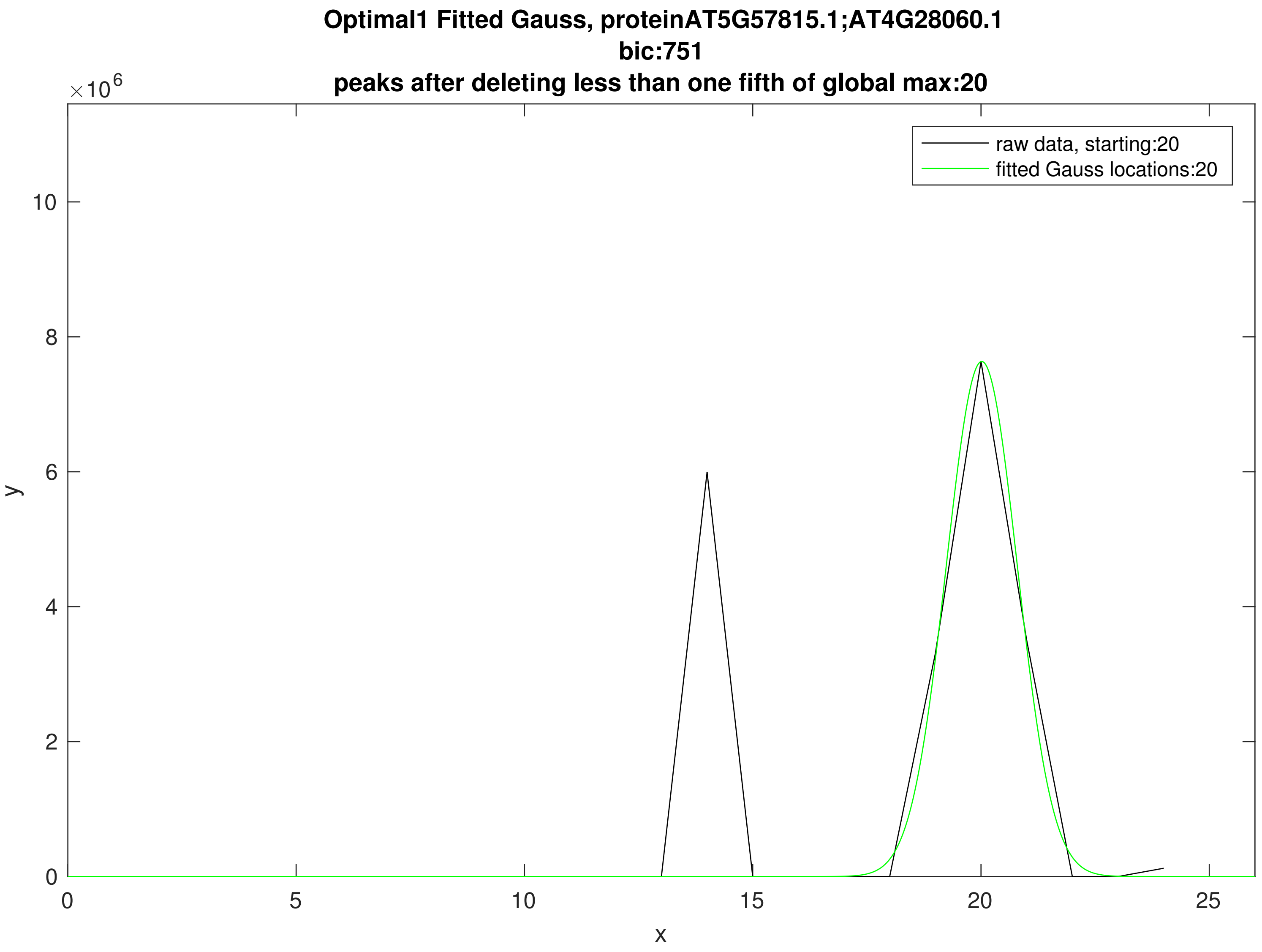
Optimal1 Fitted Gauss, proteinAT5G57815.1;AT4G28060.1
bic:751
peaks after deleting less than one fifth of global max:20
×10^6
raw data, starting:20
fitted Gauss locations:20
10
8
6
4
2
0
y
0
5
10
15
20
25
x

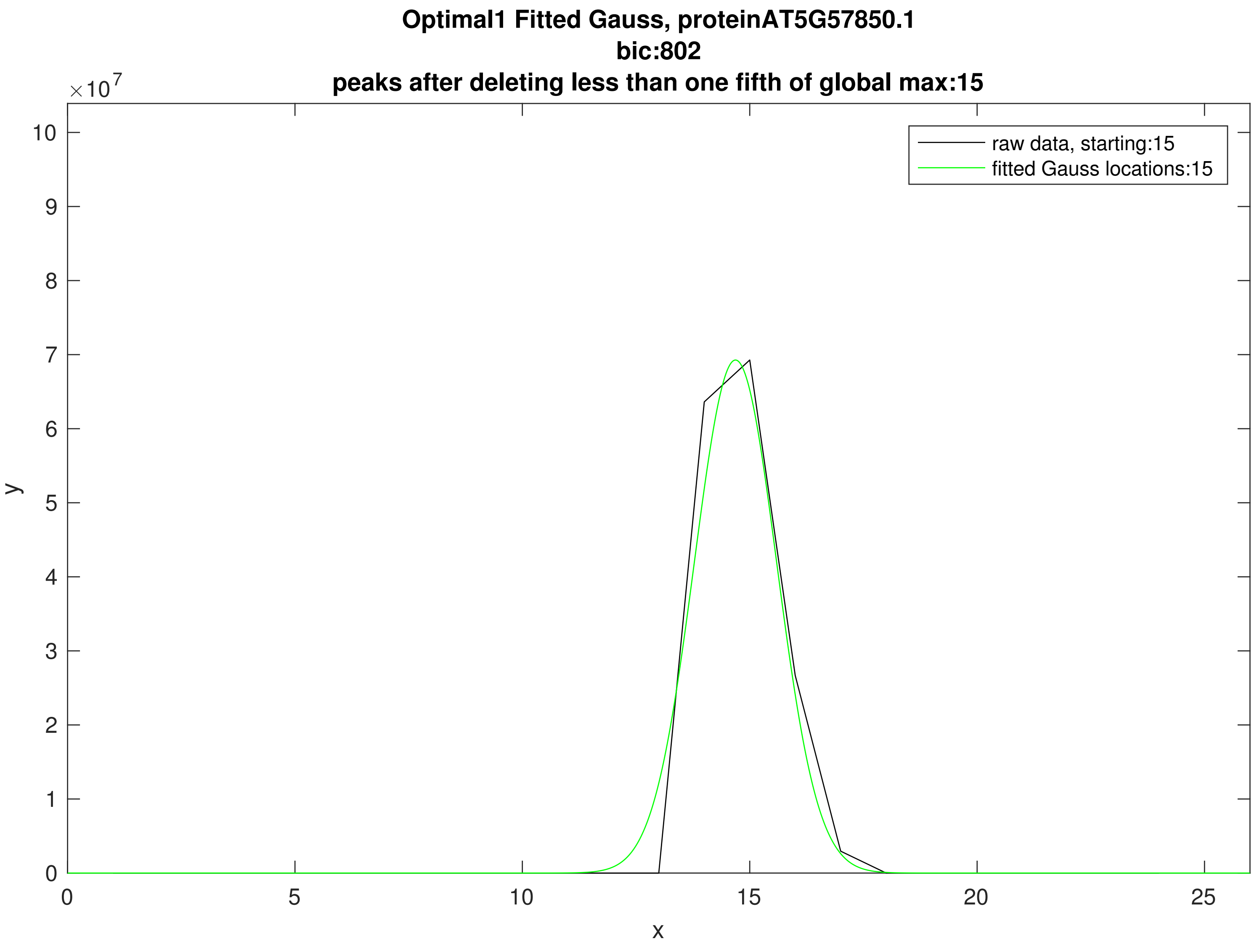

Optimal1 Fitted Gauss, proteinAT5G57850.1
bic:802
peaks after deleting less than one fifth of global max:15
×10^7
raw data, starting:15
fitted Gauss locations:15
y
x
0
1
2
3
4
5
6
7
8
9
10
0
5
10
15
20
25

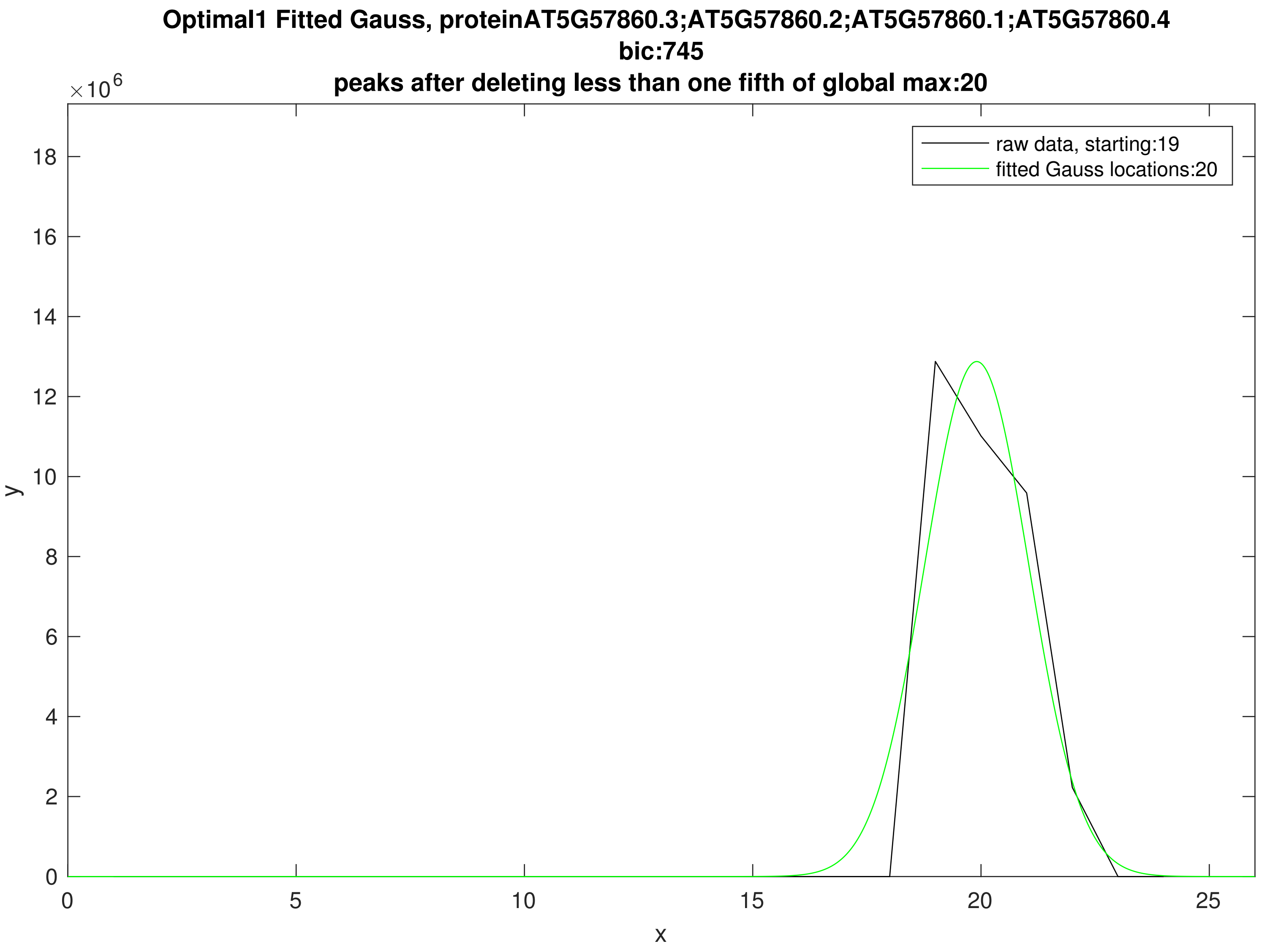
Optimal1 Fitted Gauss, proteinAT5G57860.3;AT5G57860.2;AT5G57860.1;AT5G57860.4
bic:745
peaks after deleting less than one fifth of global max:20
×10^6
raw data, starting:19
fitted Gauss locations:20
y
x
0
2
4
6
8
10
12
14
16
18
0
5
10
15
20
25

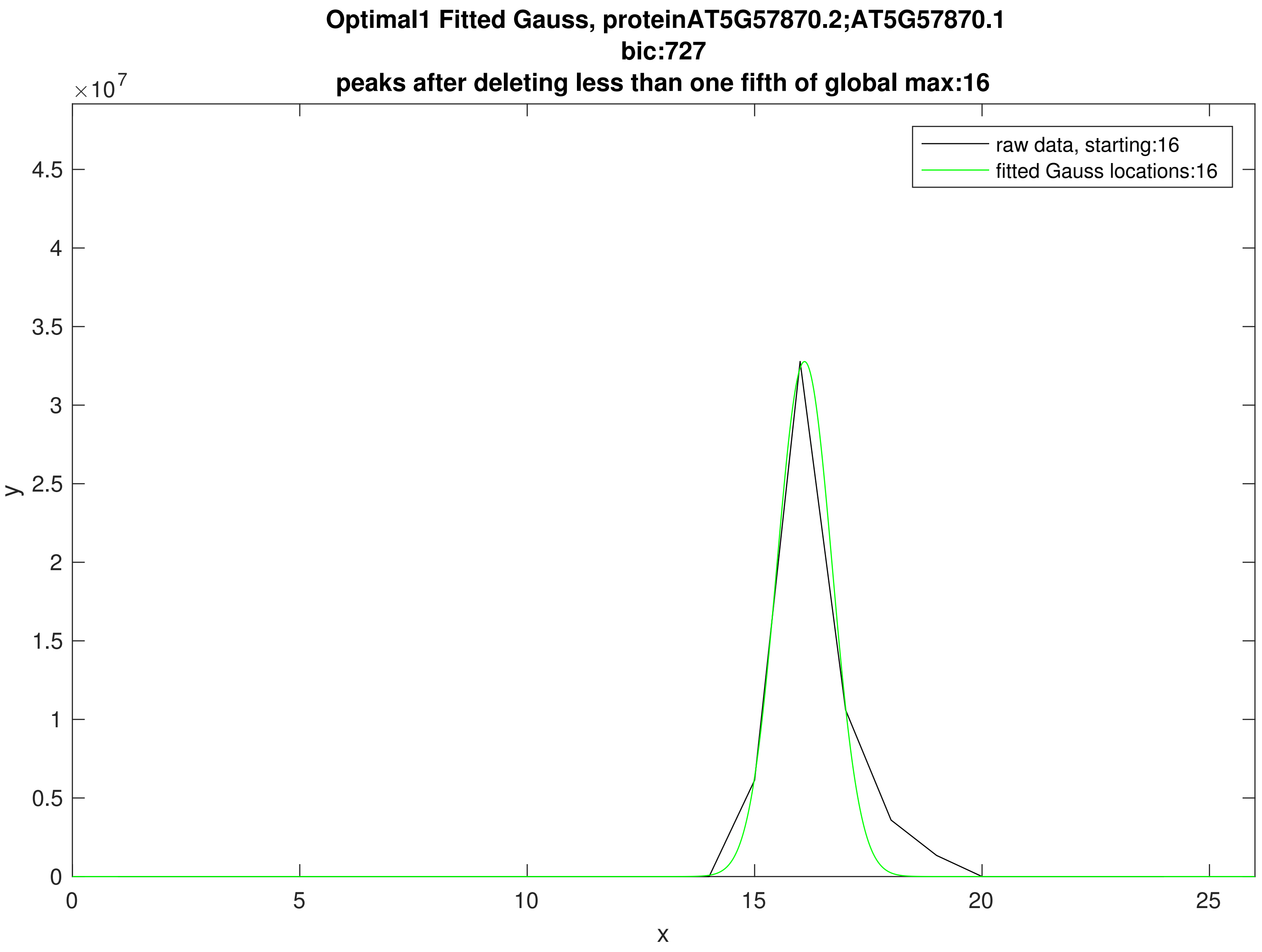
Optimal1 Fitted Gauss, proteinAT5G57870.2;AT5G57870.1
bic:727
peaks after deleting less than one fifth of global max:16
×10^7
raw data, starting:16
fitted Gauss locations:16
y
x
0
0.5
1
1.5
2
2.5
3
3.5
4
4.5
5
10
15
20
25

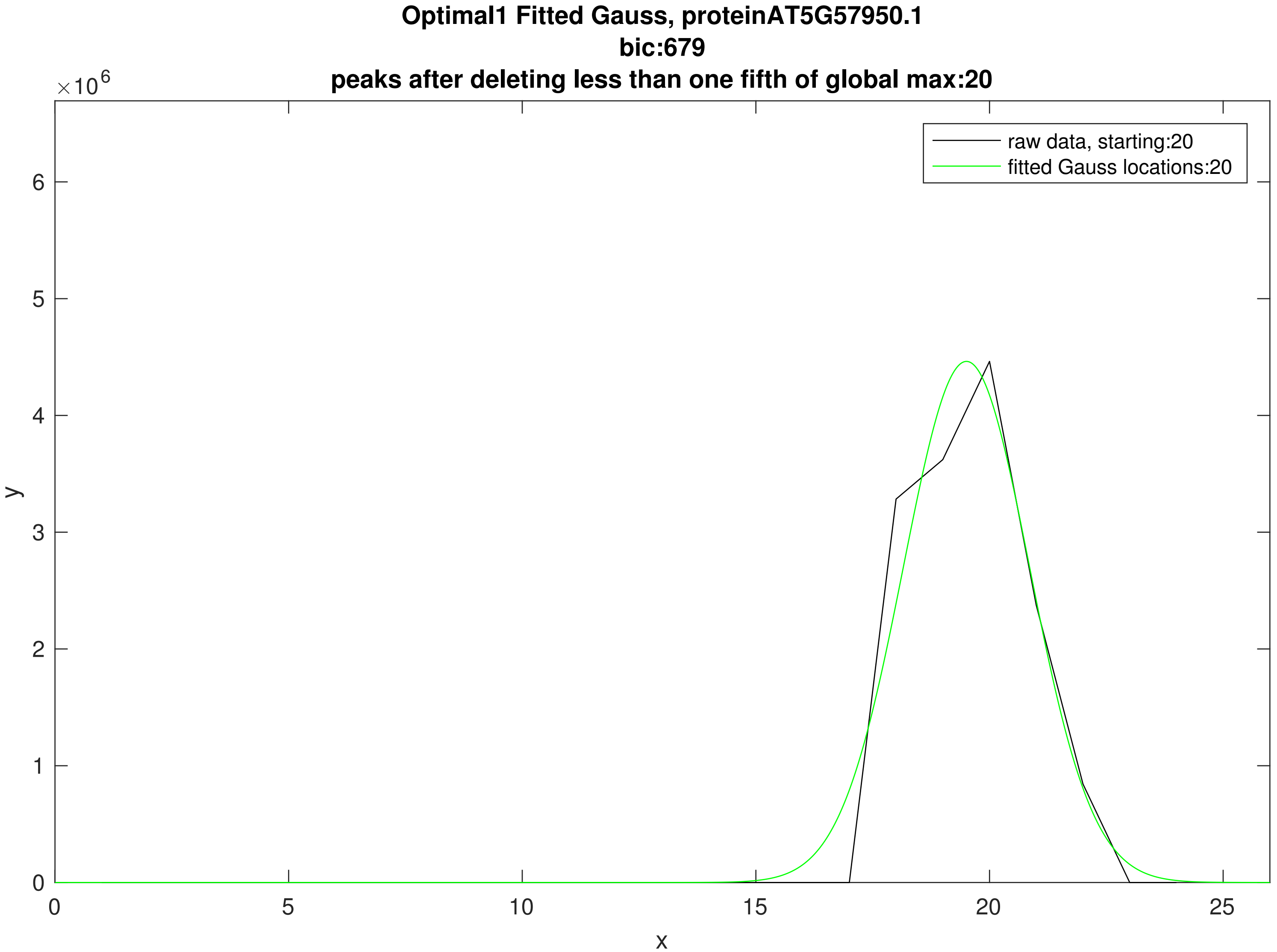

Optimal1 Fitted Gauss, proteinAT5G57950.1
bic:679
peaks after deleting less than one fifth of global max:20
×10^6
raw data, starting:20
fitted Gauss locations:20
y
x
0
1
2
3
4
5
6
0
5
10
15
20
25

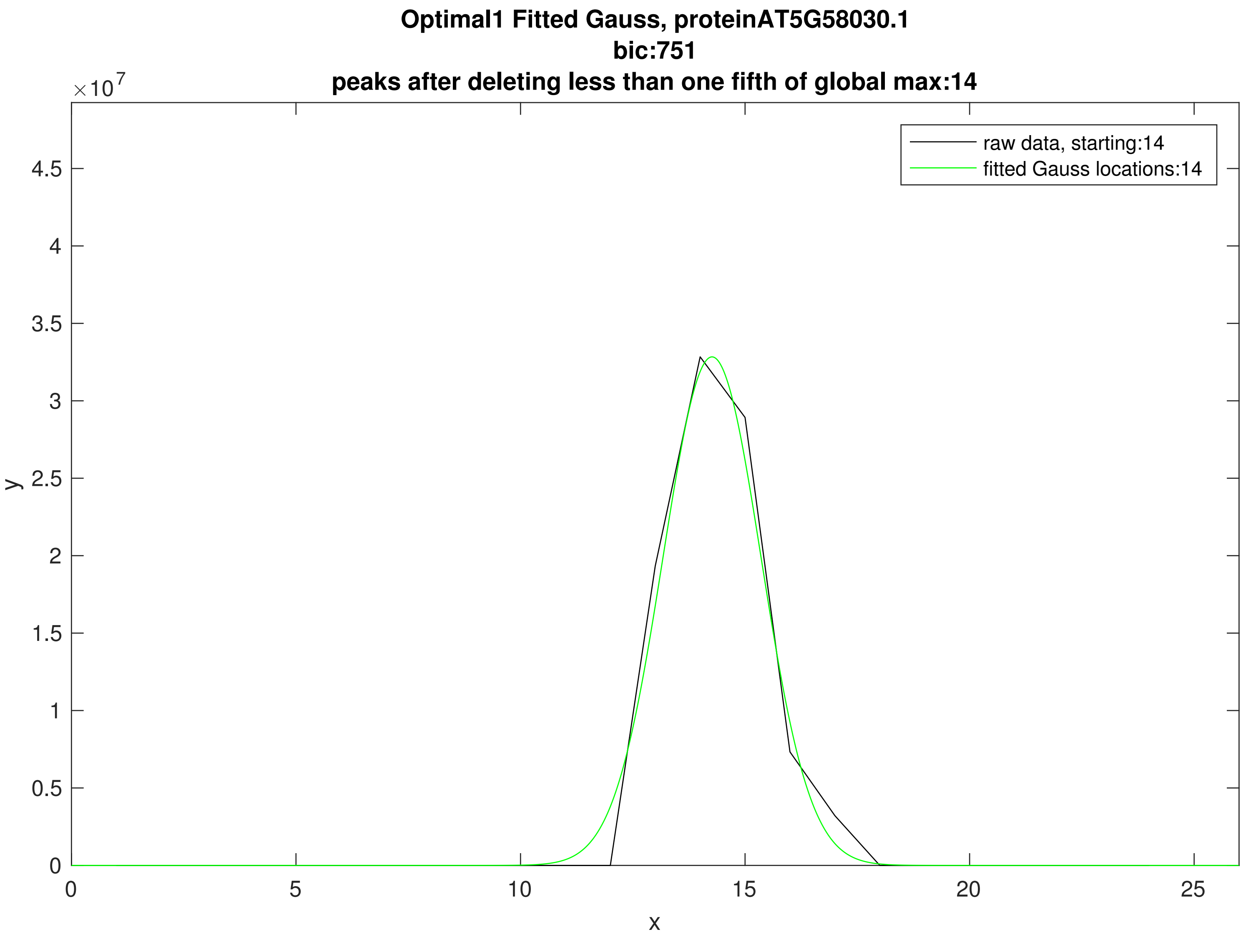
Optimal1 Fitted Gauss, proteinAT5G58030.1
bic:751
peaks after deleting less than one fifth of global max:14
×10^7
raw data, starting:14
fitted Gauss locations:14
4.5
4
3.5
3
2.5
2
1.5
1
0.5
0
y
0
5
10
15
20
25
x

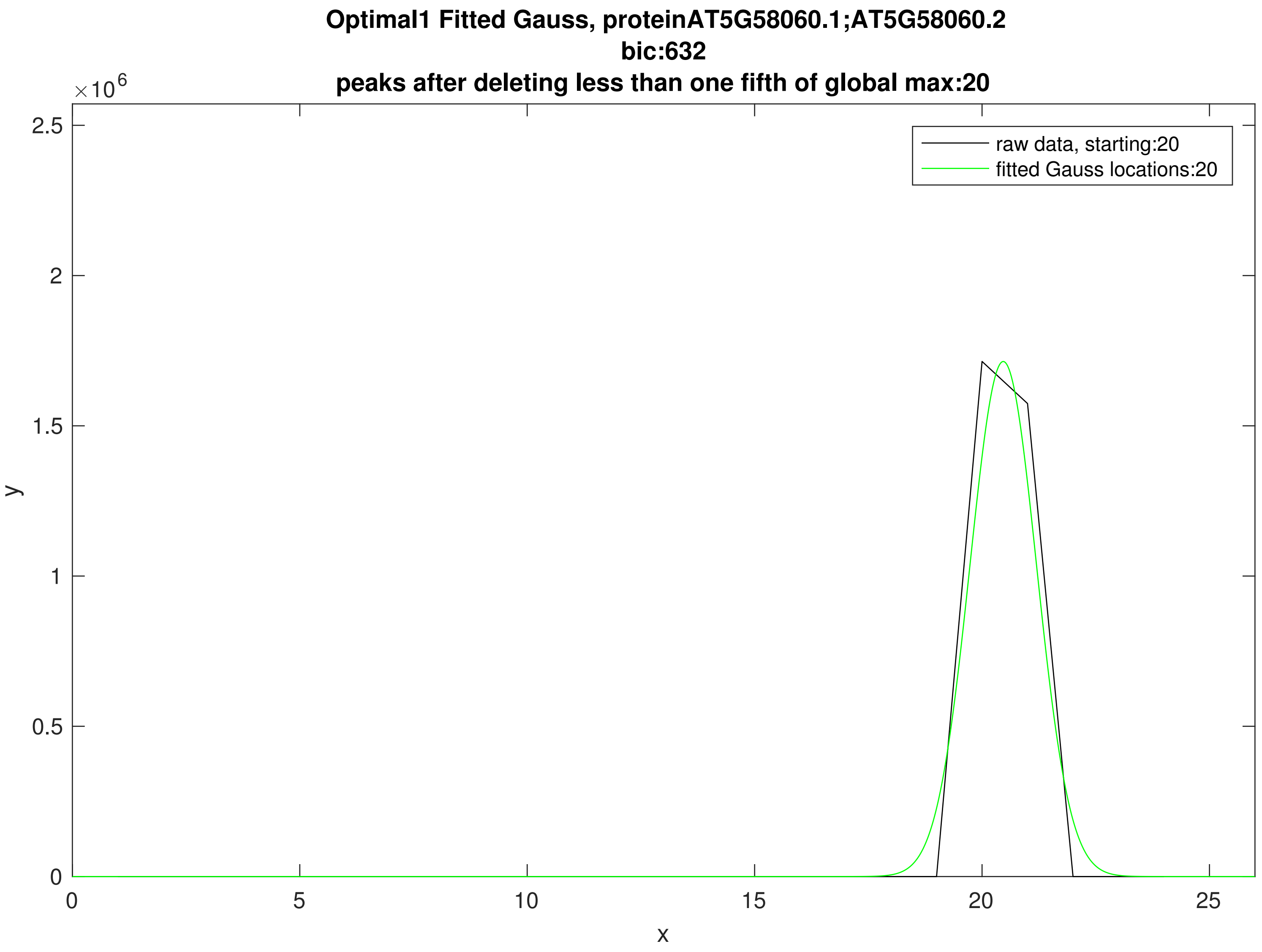
Optimal1 Fitted Gauss, proteinAT5G58060.1;AT5G58060.2
bic:632
peaks after deleting less than one fifth of global max:20
×10^6
raw data, starting:20
fitted Gauss locations:20
y
x
0
0.5
1
1.5
2
2.5
0
5
10
15
20
25

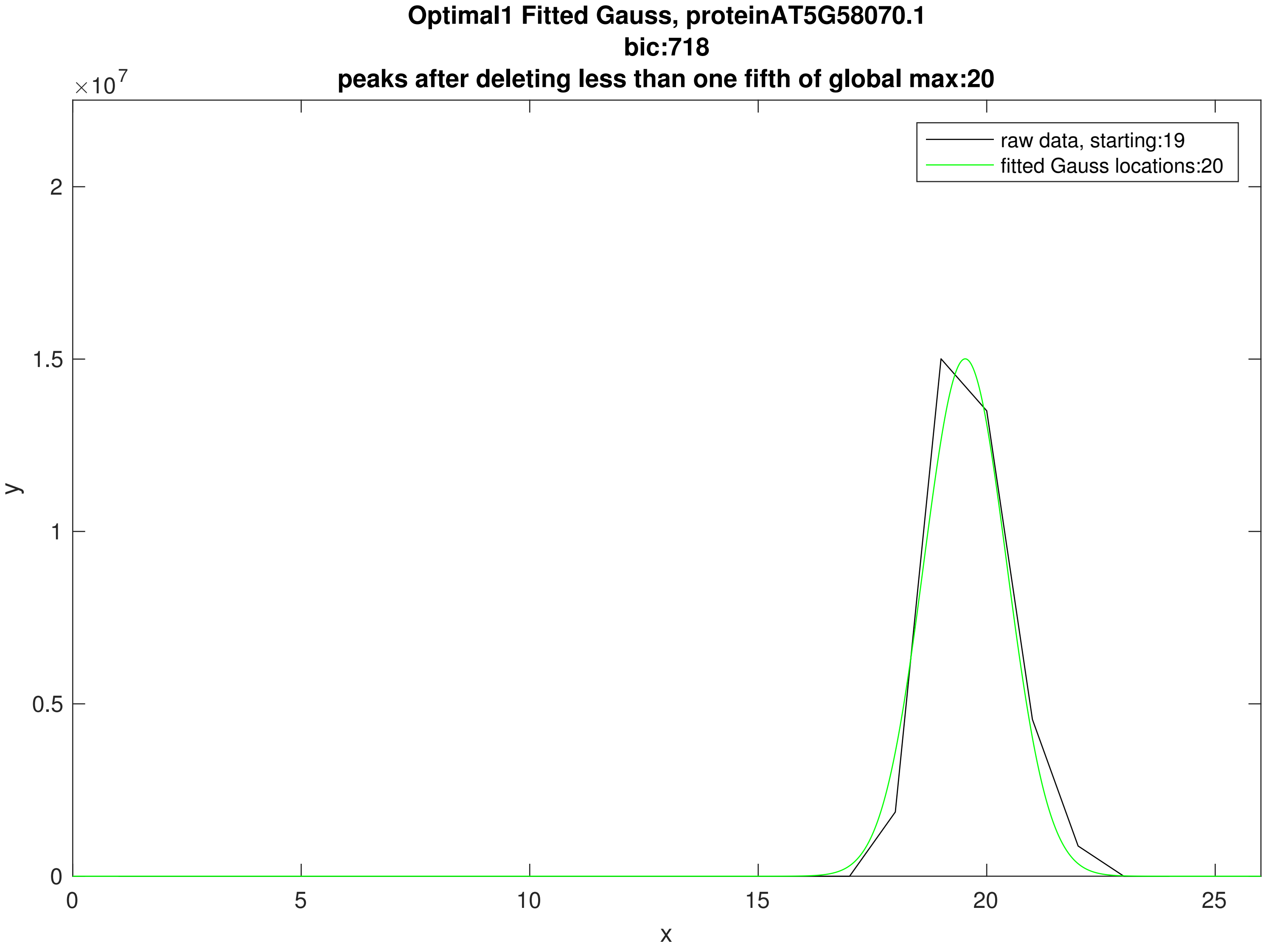
Optimal1 Fitted Gauss, proteinAT5G58070.1
bic:718
peaks after deleting less than one fifth of global max:20
×10^7
raw data, starting:19
fitted Gauss locations:20
y
x
0
0.5
1
1.5
2
0
5
10
15
20
25

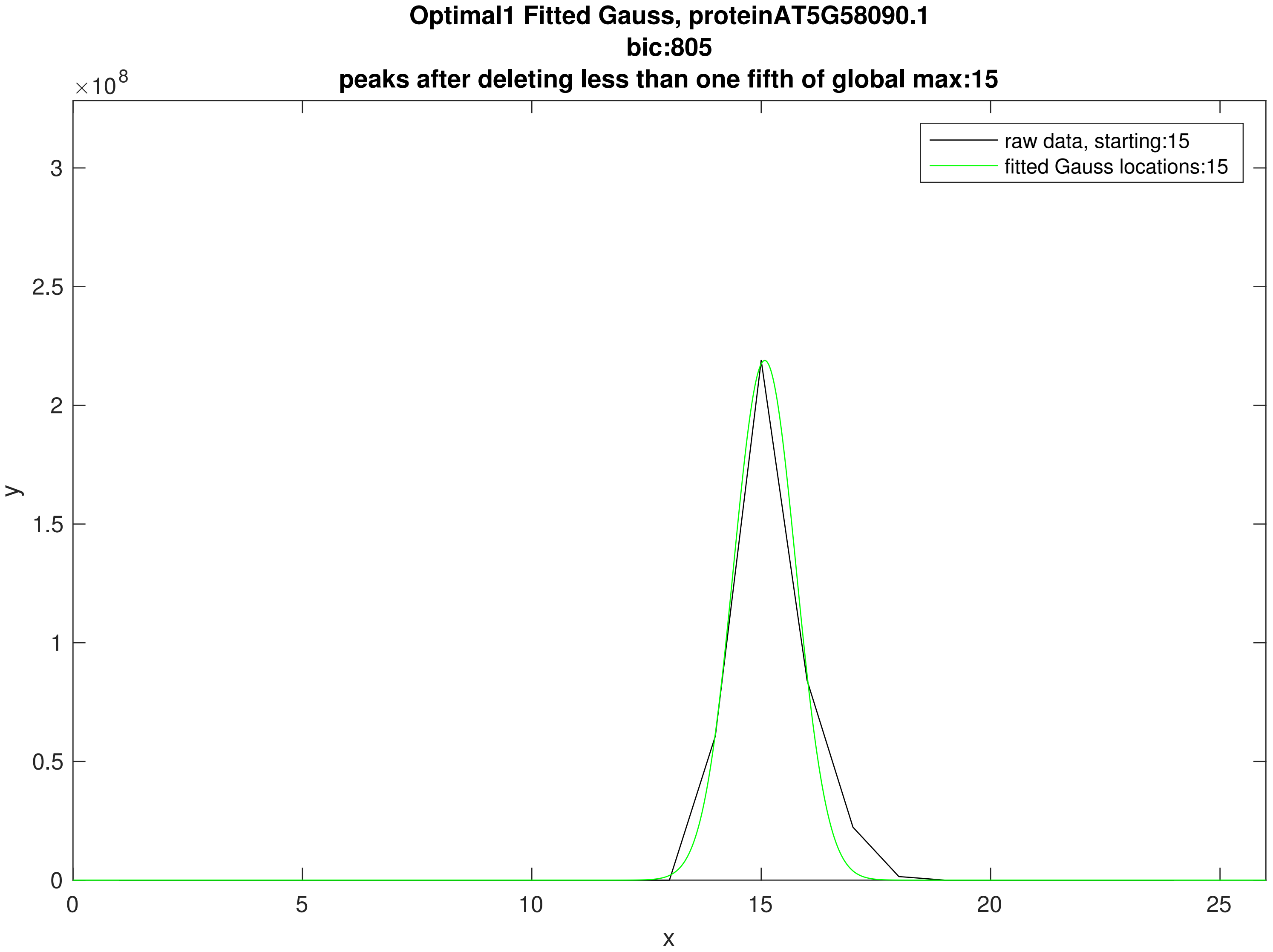
Optimal1 Fitted Gauss, proteinAT5G58090.1
bic:805
peaks after deleting less than one fifth of global max:15
×10^8
raw data, starting:15
fitted Gauss locations:15
y
x
0
0.5
1
1.5
2
2.5
3
0
5
10
15
20
25

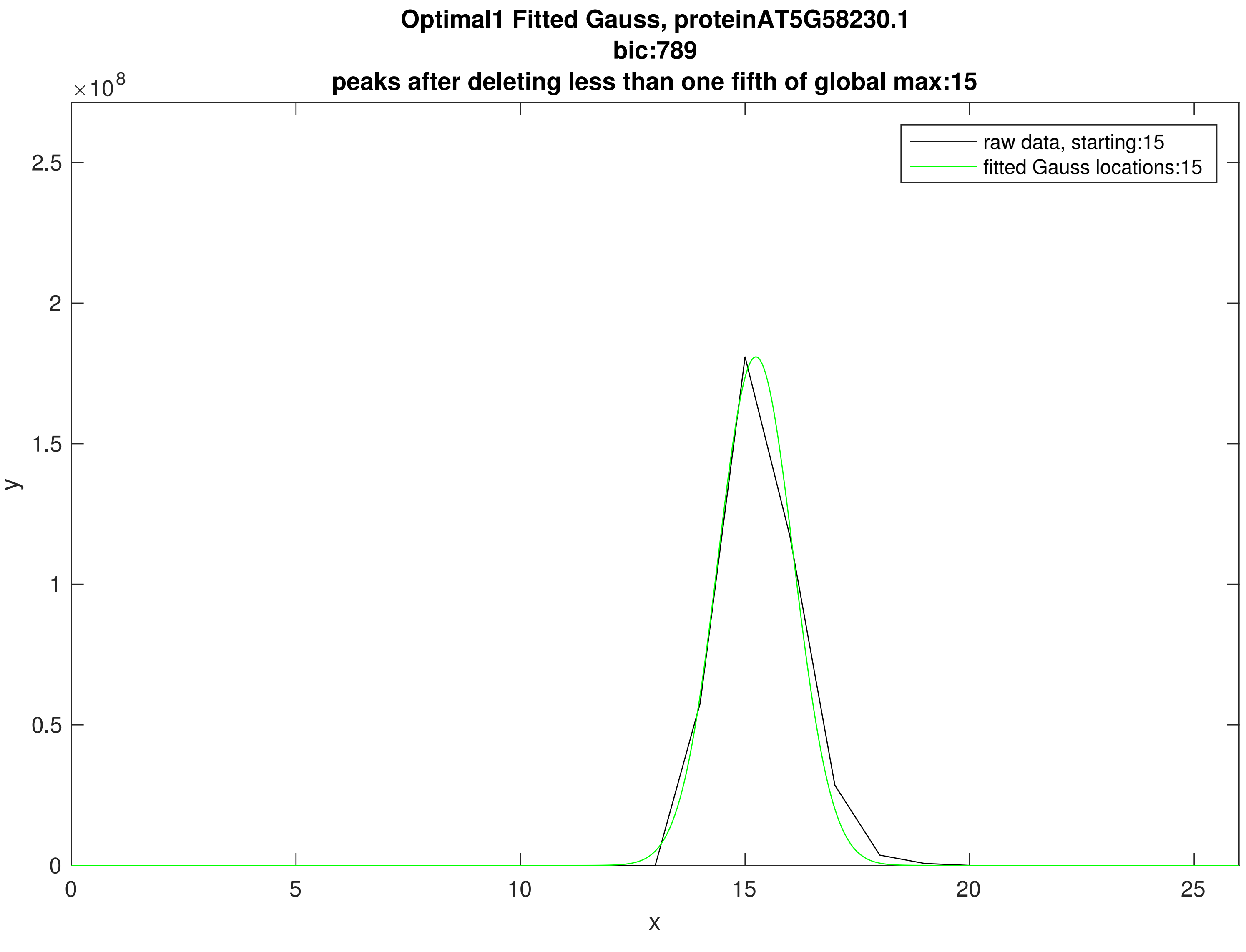

Optimal1 Fitted Gauss, proteinAT5G58230.1
bic:789
peaks after deleting less than one fifth of global max:15
×10^8
raw data, starting:15
fitted Gauss locations:15
2.5
2
1.5
1
0.5
0
y
0
5
10
15
20
25
x

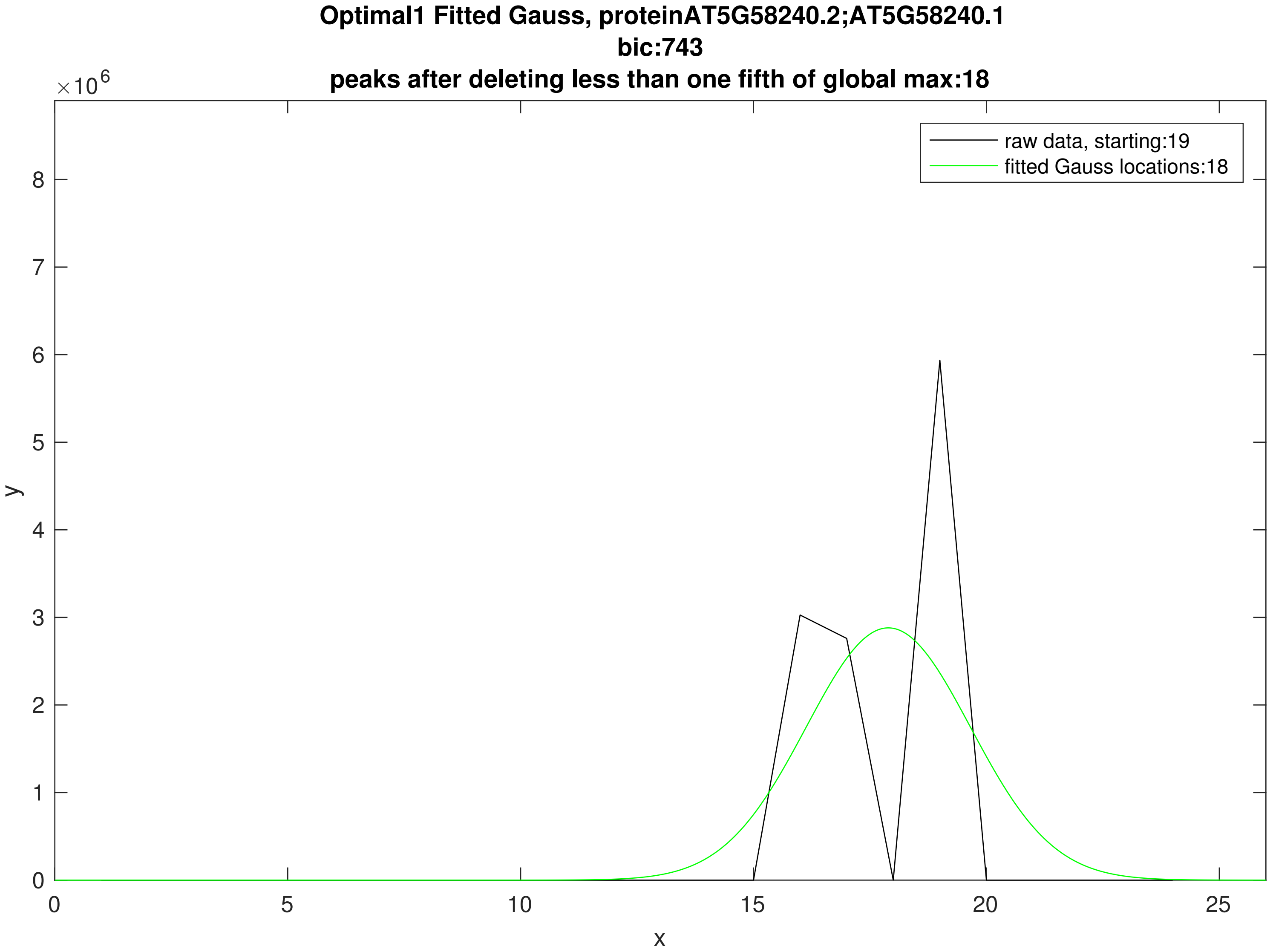
Optimal1 Fitted Gauss, proteinAT5G58240.2;AT5G58240.1
bic:743
peaks after deleting less than one fifth of global max:18
×10^6
raw data, starting:19
fitted Gauss locations:18
y
x
0
1
2
3
4
5
6
7
8
0
5
10
15
20
25

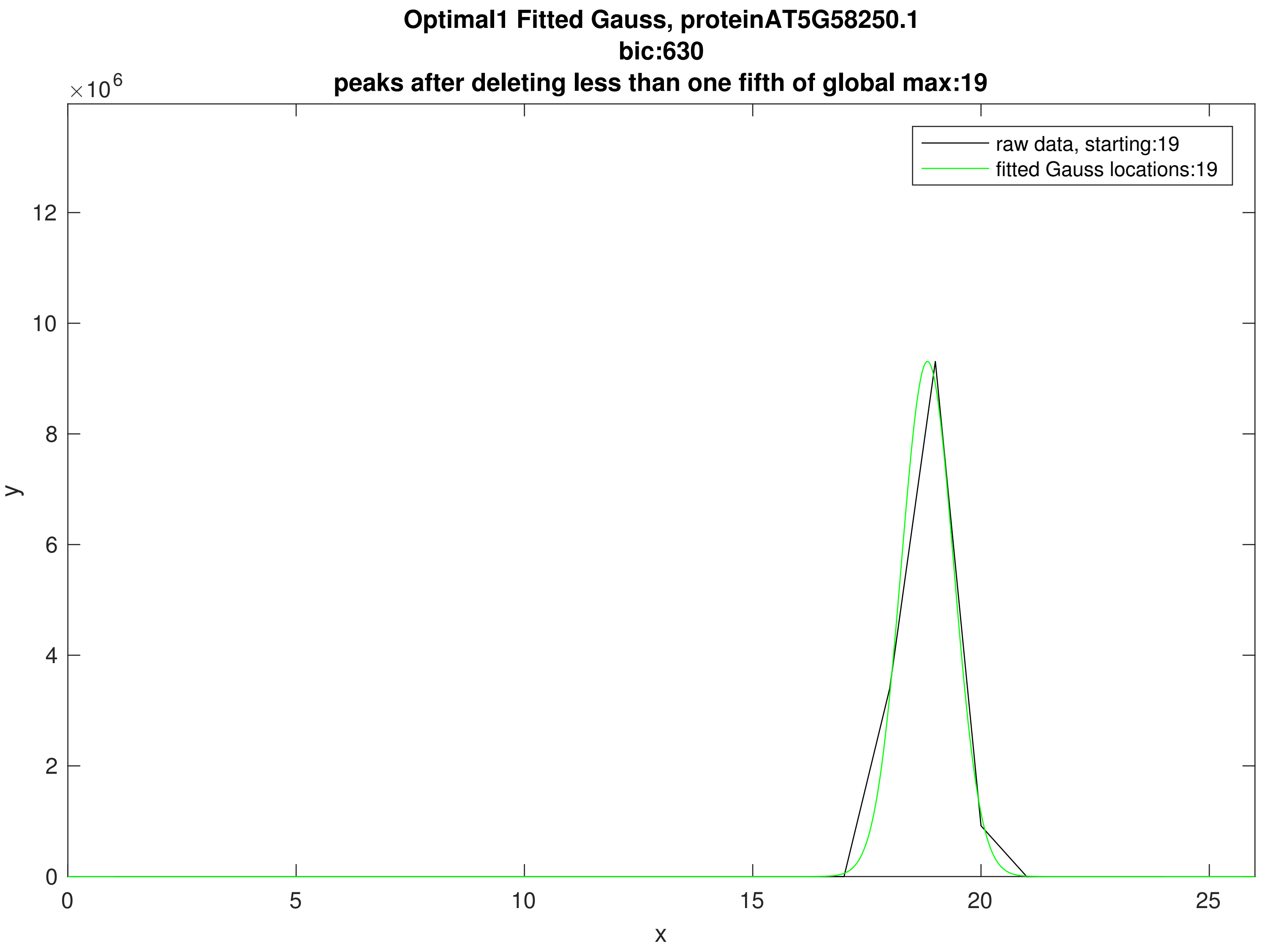
Optimal1 Fitted Gauss, proteinAT5G58250.1
bic:630
peaks after deleting less than one fifth of global max:19
×10^6
raw data, starting:19
fitted Gauss locations:19
y
x
0
2
4
6
8
10
12
5
15
20
25

**Optimal1 Fitted Gauss, proteinAT5G58290.1**
**bic:736**
**peaks after deleting less than one fifth of global max:4**

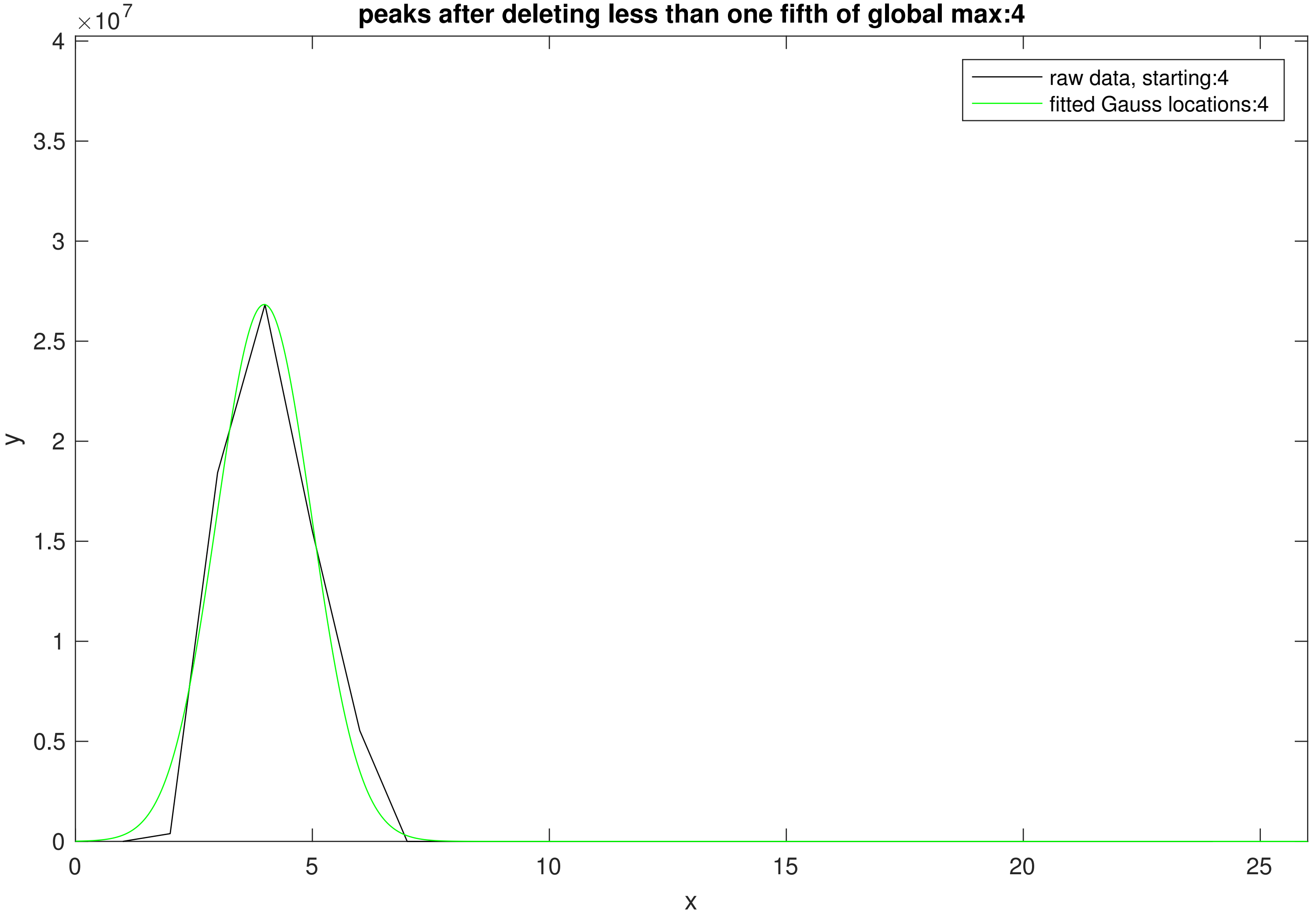

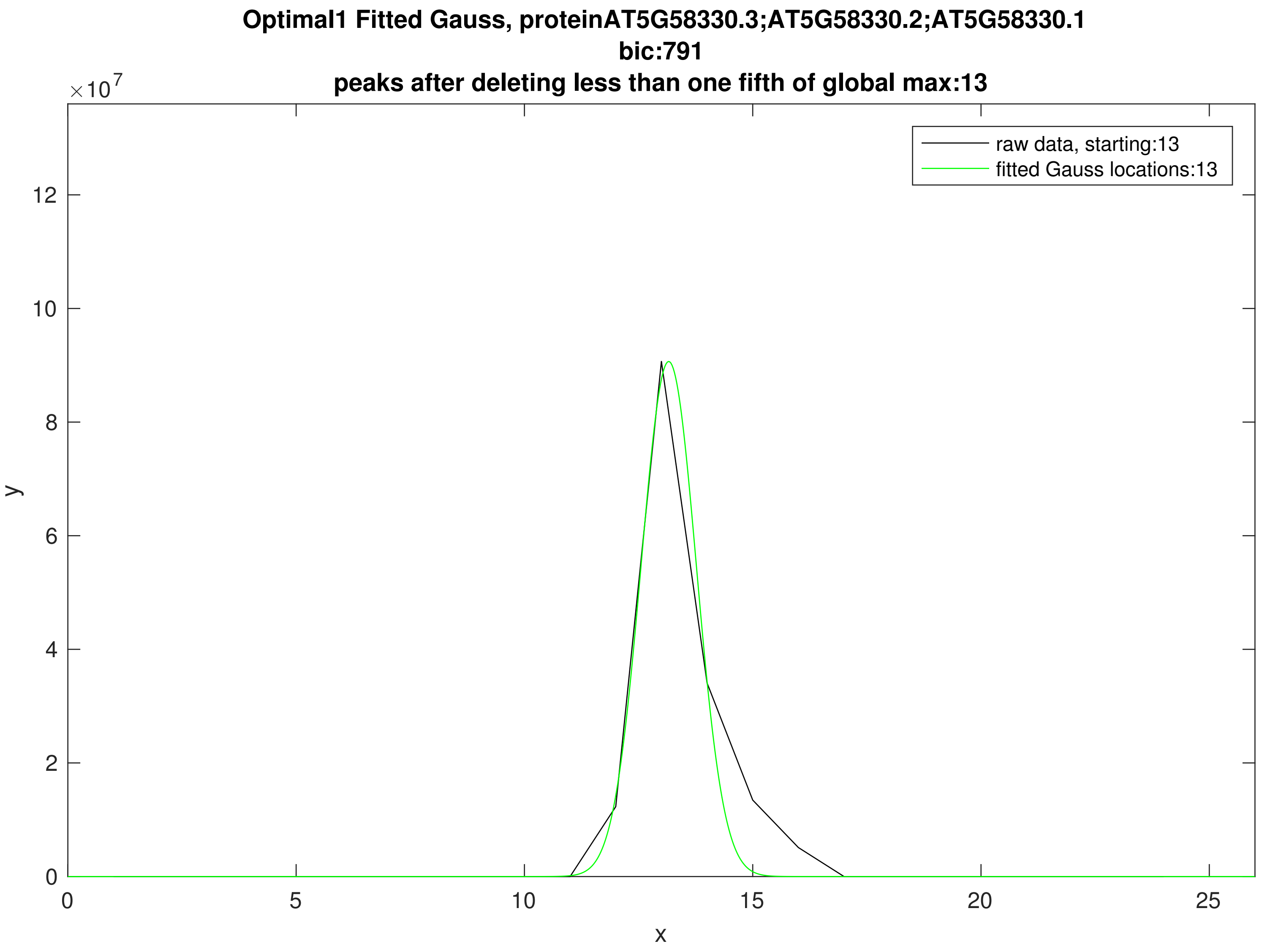

Optimal1 Fitted Gauss, proteinAT5G58330.3;AT5G58330.2;AT5G58330.1
bic:791
peaks after deleting less than one fifth of global max:13
×10^7
raw data, starting:13
fitted Gauss locations:13
12
10
8
6
4
2
0
y
0
5
10
15
20
25
x

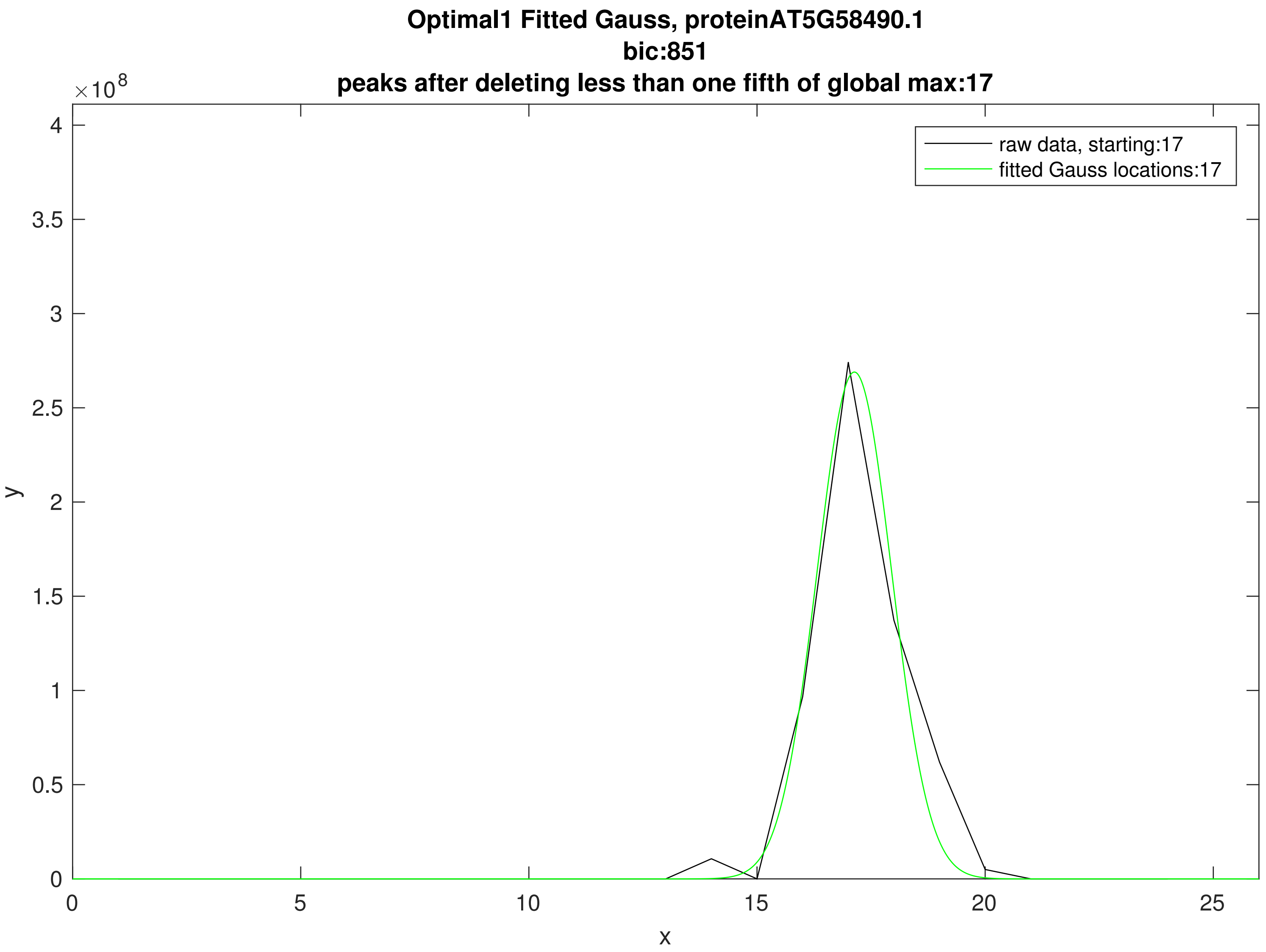

Optimal1 Fitted Gauss, proteinAT5G58490.1
bic:851
peaks after deleting less than one fifth of global max:17
×10^8
raw data, starting:17
fitted Gauss locations:17
y
x
0
0.5
1
1.5
2
2.5
3
3.5
4
0
5
10
15
20
25

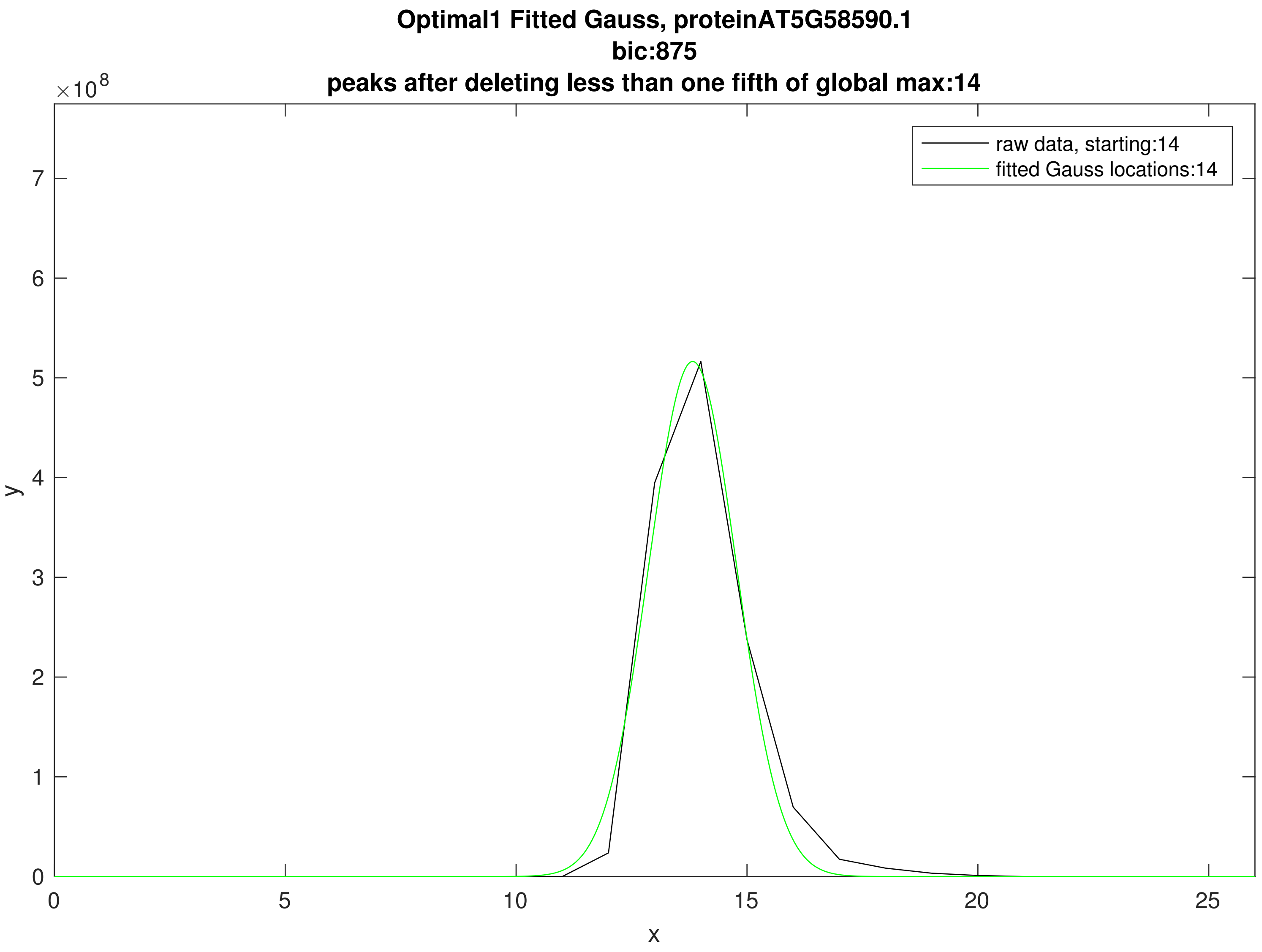
Optimal1 Fitted Gauss, proteinAT5G58590.1
bic:875
peaks after deleting less than one fifth of global max:14
×10^8
raw data, starting:14
fitted Gauss locations:14
y
x
0
1
2
3
4
5
6
7
0
5
10
15
20
25

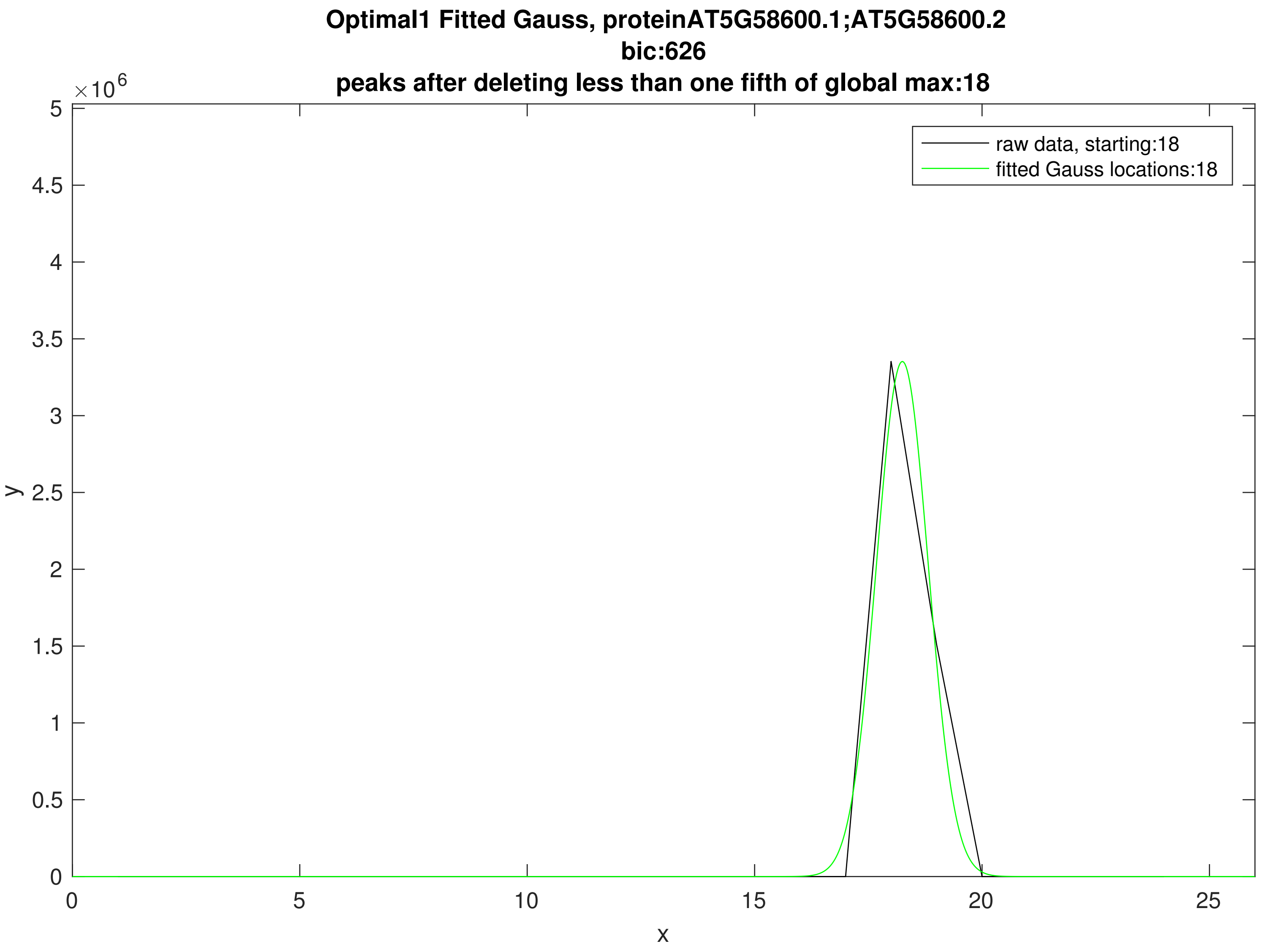
Optimal1 Fitted Gauss, proteinAT5G58600.1;AT5G58600.2
bic:626
peaks after deleting less than one fifth of global max:18
×10^6
5
4.5
4
3.5
3
2.5
2
1.5
1
0.5
0
y
0
5
10
15
20
25
x
raw data, starting:18
fitted Gauss locations:18

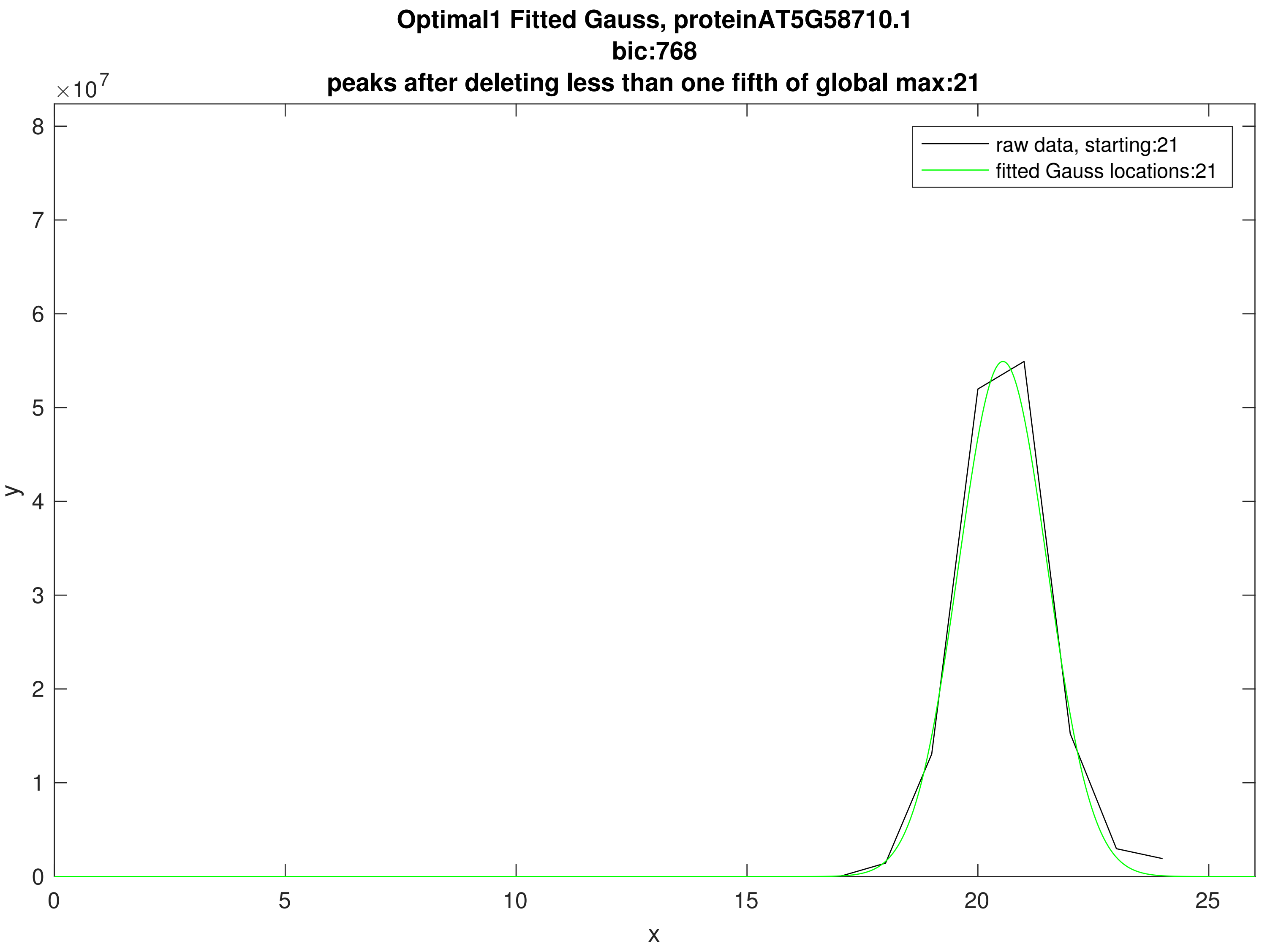
Optimal1 Fitted Gauss, proteinAT5G58710.1
bic:768
peaks after deleting less than one fifth of global max:21
×10^7
raw data, starting:21
fitted Gauss locations:21
y
x
0
1
2
3
4
5
6
7
8
0
5
10
15
20
25

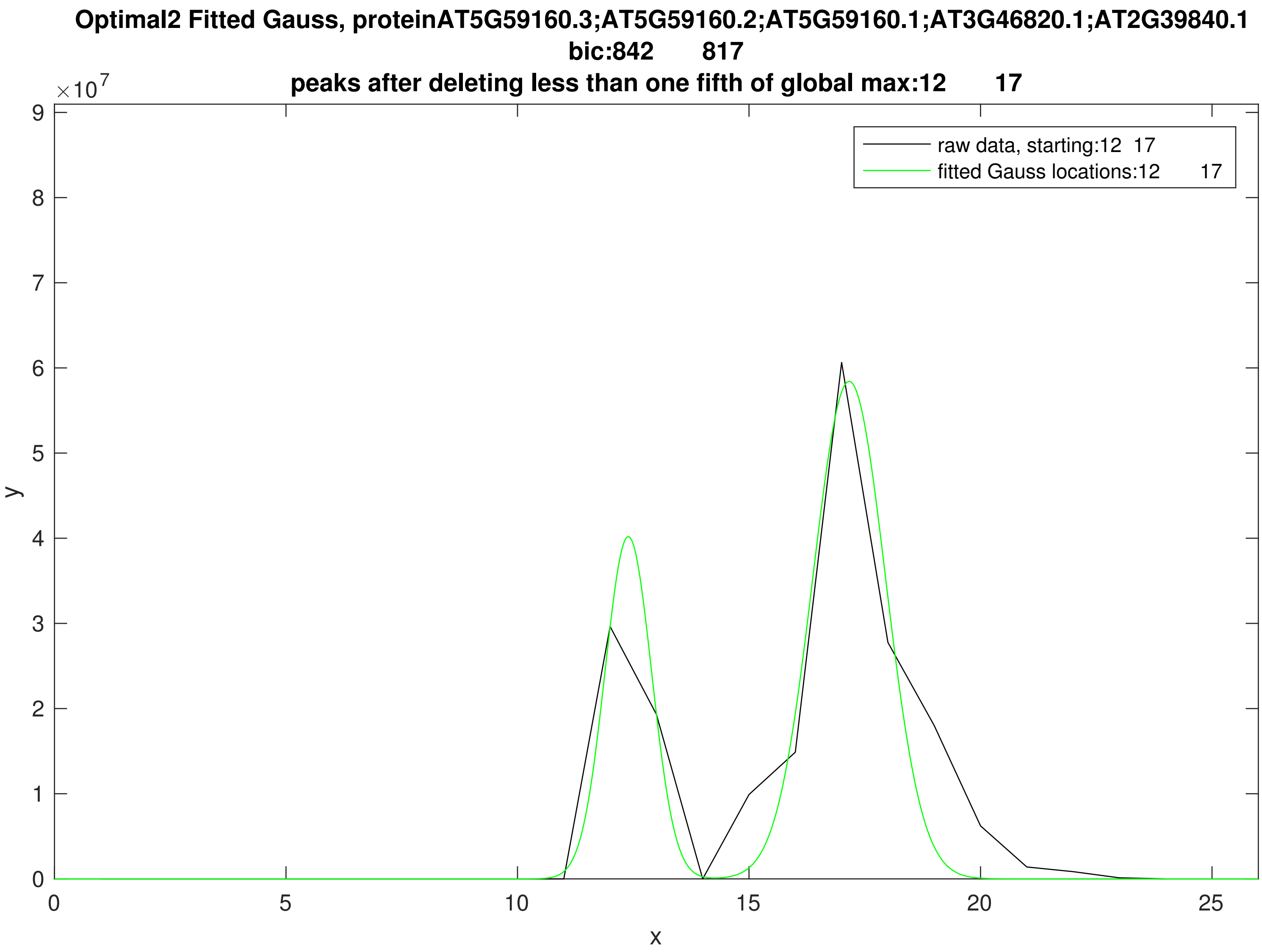

Optimal2 Fitted Gauss, proteinAT5G59160.3;AT5G59160.2;AT5G59160.1;AT3G46820.1;AT2G39840.1
bic:842 817
peaks after deleting less than one fifth of global max:12 17
×10^7
raw data, starting:12 17
fitted Gauss locations:12 17
y
x
0
5
10
15
20
25
1
2
3
4
5
6
7
8
9

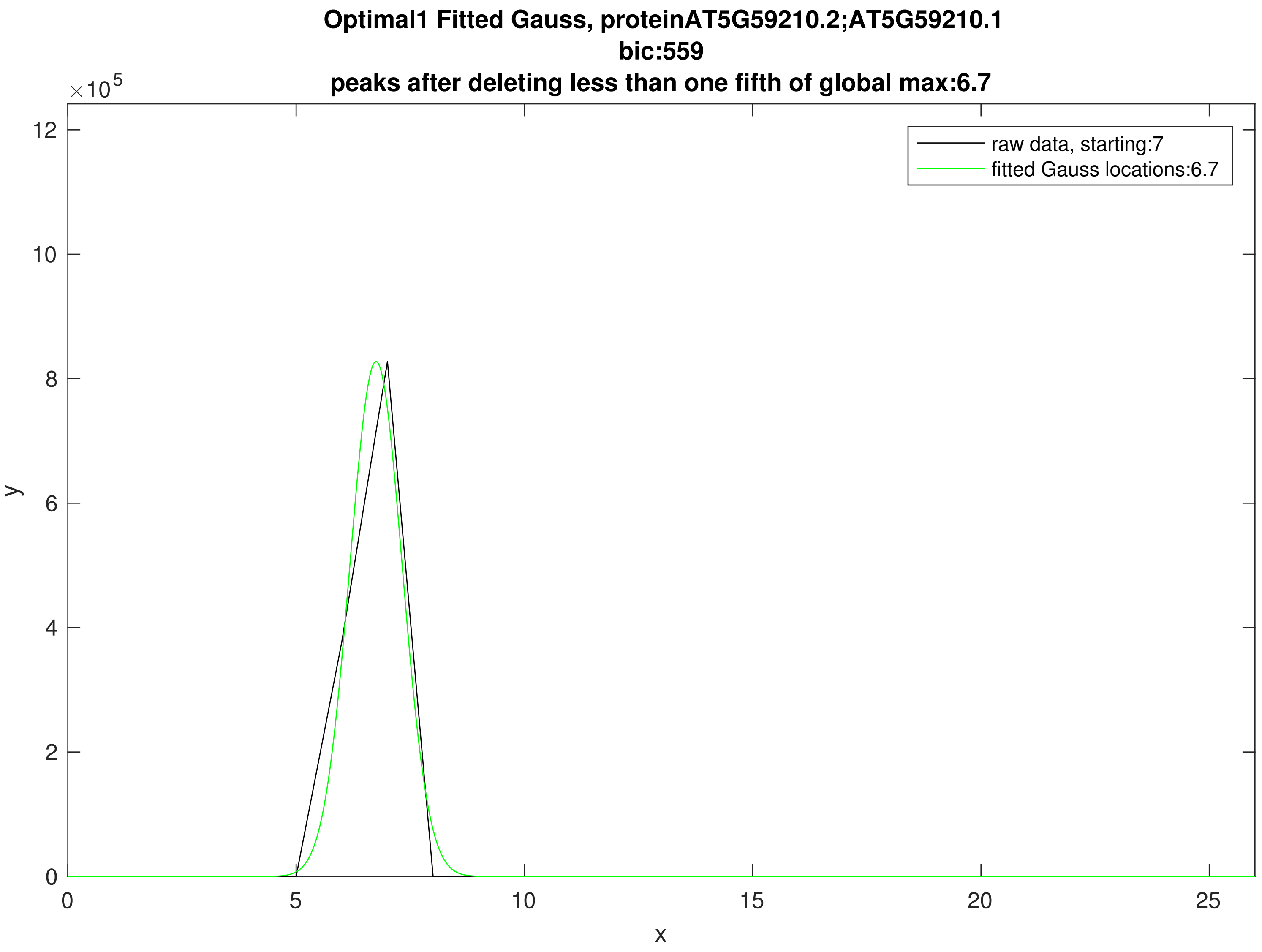

Optimal1 Fitted Gauss, proteinAT5G59210.2;AT5G59210.1
bic:559
peaks after deleting less than one fifth of global max:6.7
×10^5
raw data, starting:7
fitted Gauss locations:6.7
y
x
0
2
4
6
8
10
12
0
5
10
15
20
25

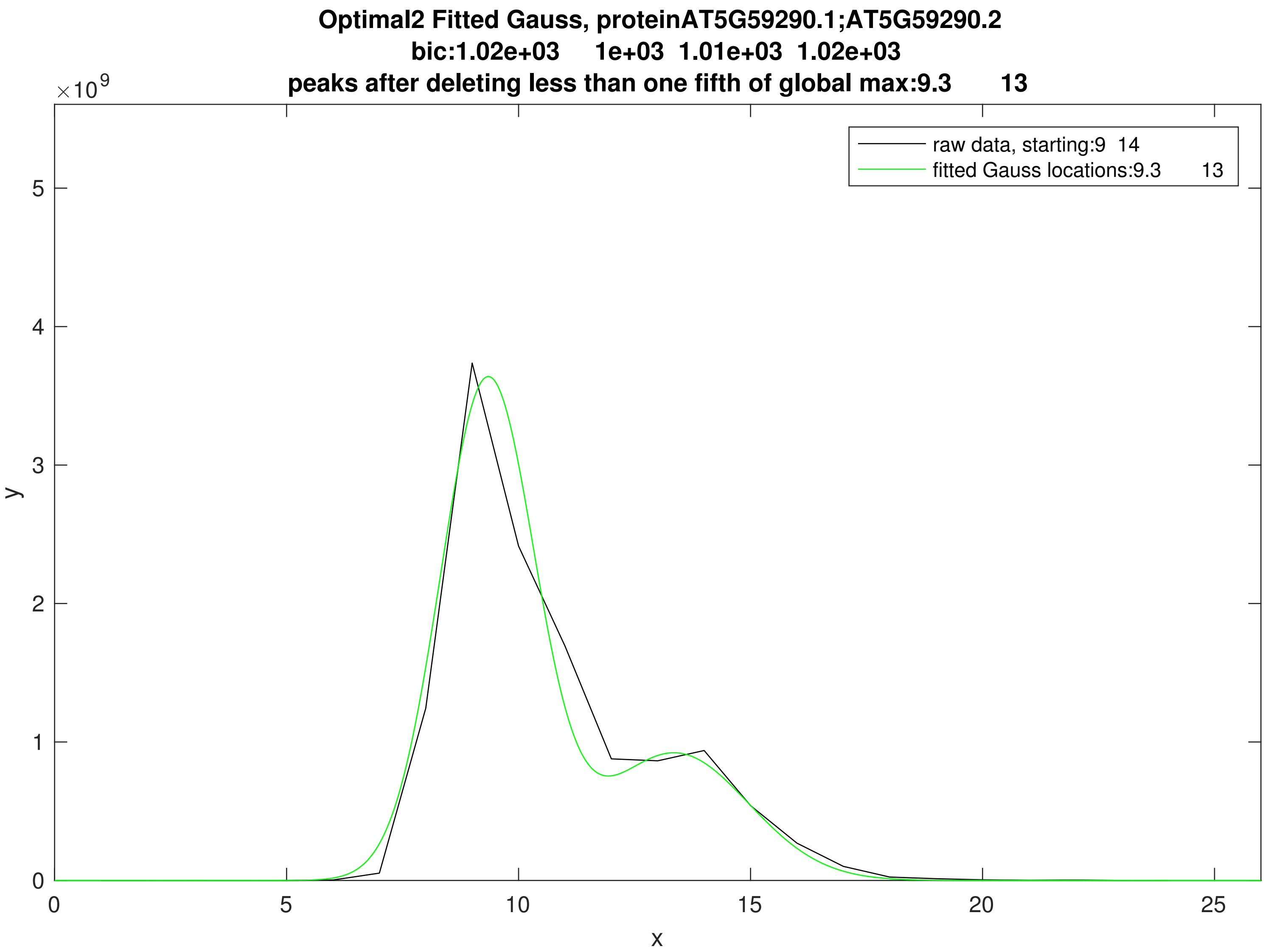

Optimal2 Fitted Gauss, proteinAT5G59290.1;AT5G59290.2
bic:1.02e+03 1e+03 1.01e+03 1.02e+03
peaks after deleting less than one fifth of global max:9.3 13
×10^9
raw data, starting:9 14
fitted Gauss locations:9.3 13
y
x
0
1
2
3
4
5
0
5
10
15
20
25

**Optimal1 Fitted Gauss, proteinAT5G59300.1**
**bic:730**
**peaks after deleting less than one fifth of global max:18**

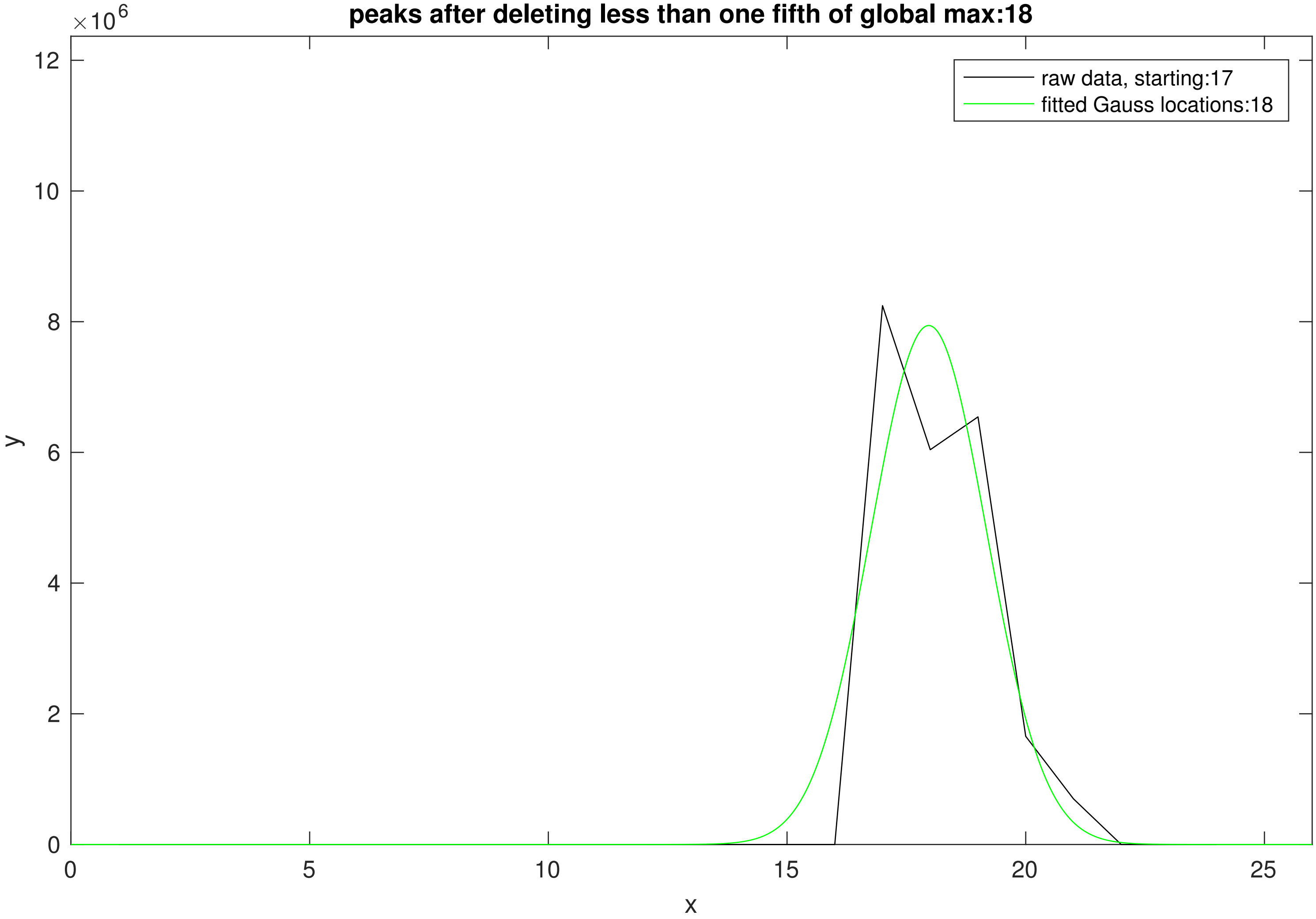

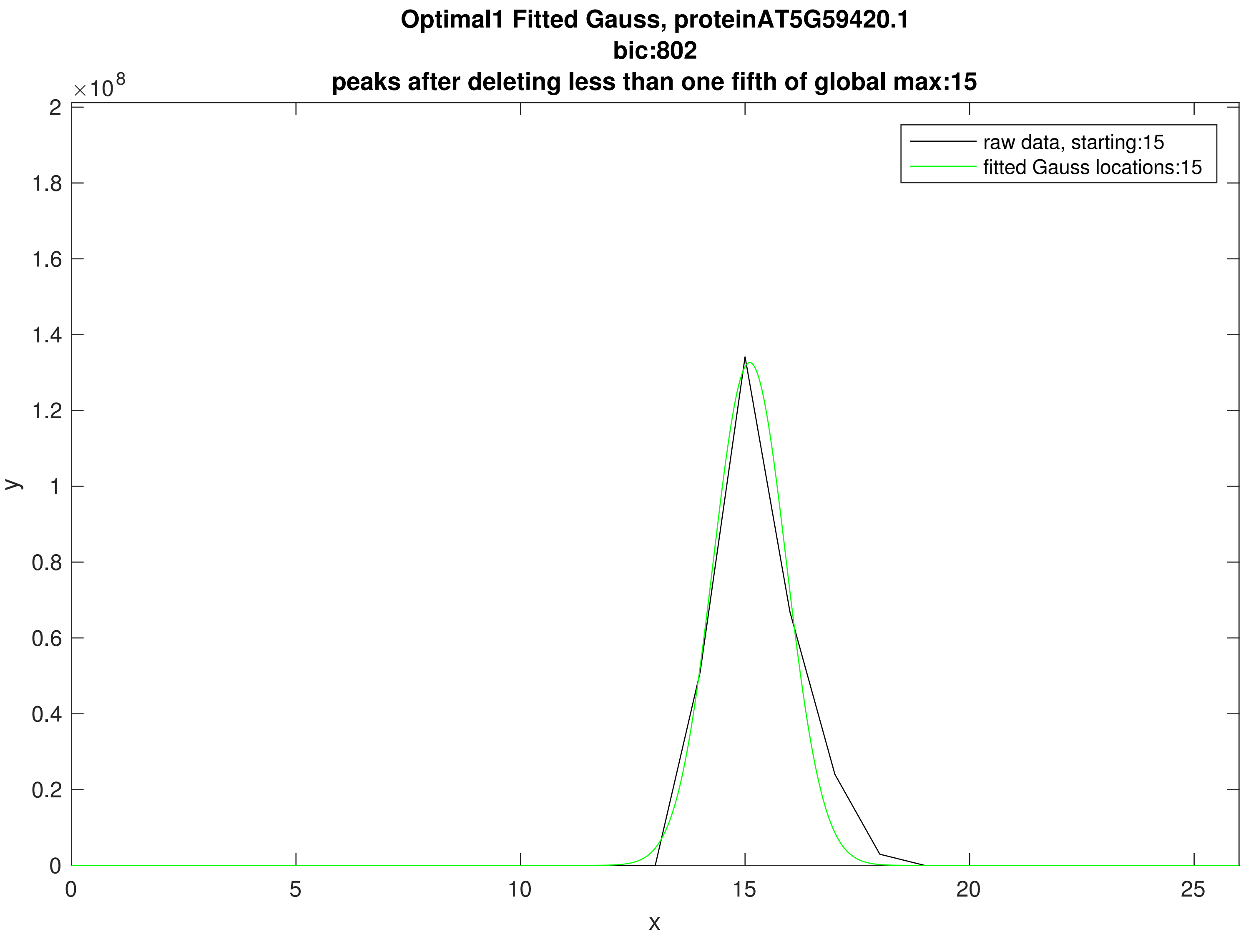
Optimal1 Fitted Gauss, proteinAT5G59420.1
bic:802
peaks after deleting less than one fifth of global max:15
×10^8
2
1.8
1.6
1.4
1.2
1
0.8
0.6
0.4
0.2
0
y
0
5
10
15
20
25
x
raw data, starting:15
fitted Gauss locations:15

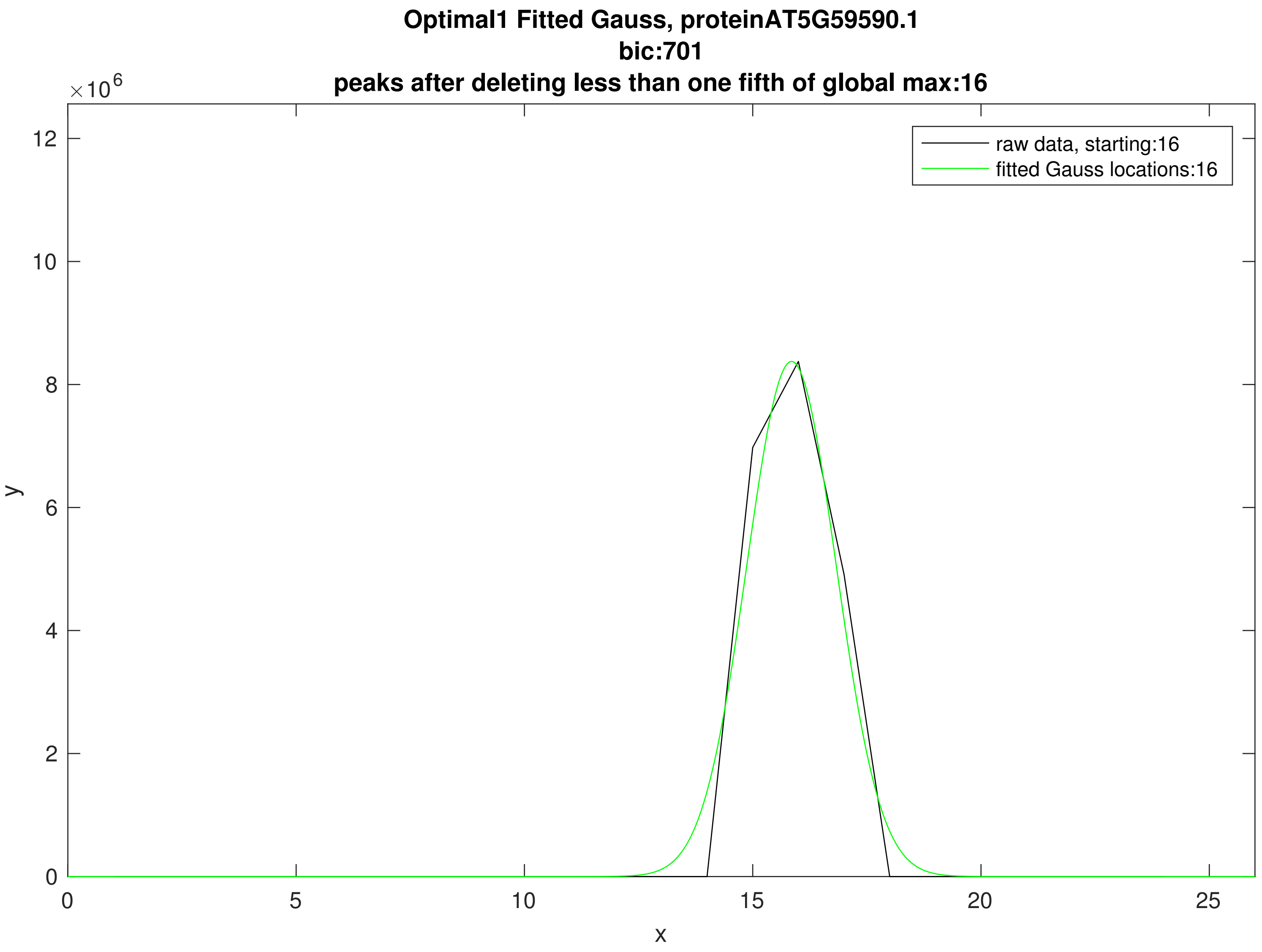
Optimal1 Fitted Gauss, proteinAT5G59590.1
bic:701
peaks after deleting less than one fifth of global max:16
×10^6
12
10
8
6
4
2
0
y
0
5
10
15
20
25
x
raw data, starting:16
fitted Gauss locations:16

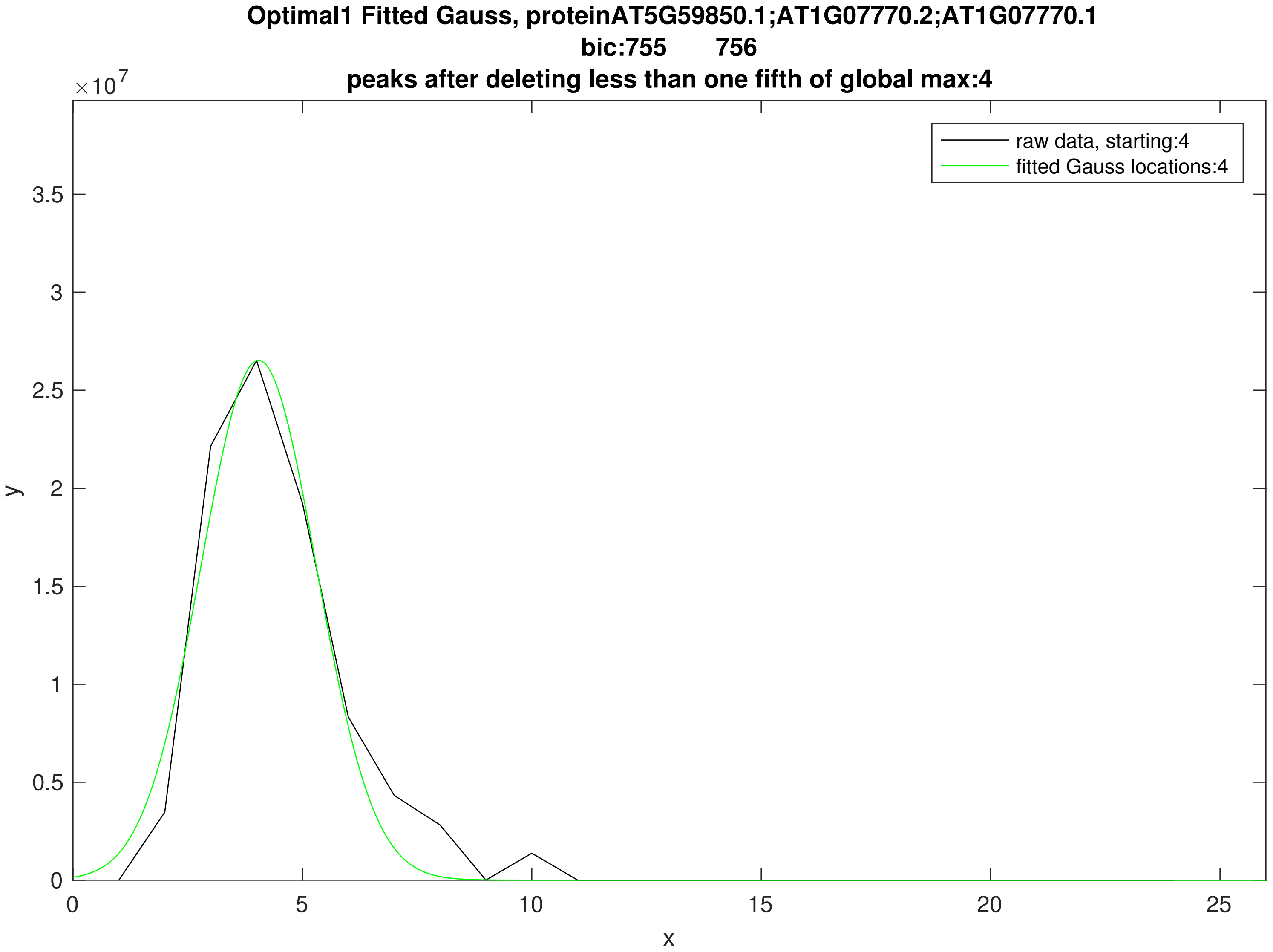

Optimal1 Fitted Gauss, proteinAT5G59850.1;AT1G07770.2;AT1G07770.1
bic:755 756
peaks after deleting less than one fifth of global max:4
×10^7
raw data, starting:4
fitted Gauss locations:4
3.5
3
2.5
2
1.5
1
0.5
0
y
0
5
10
15
20
25
x

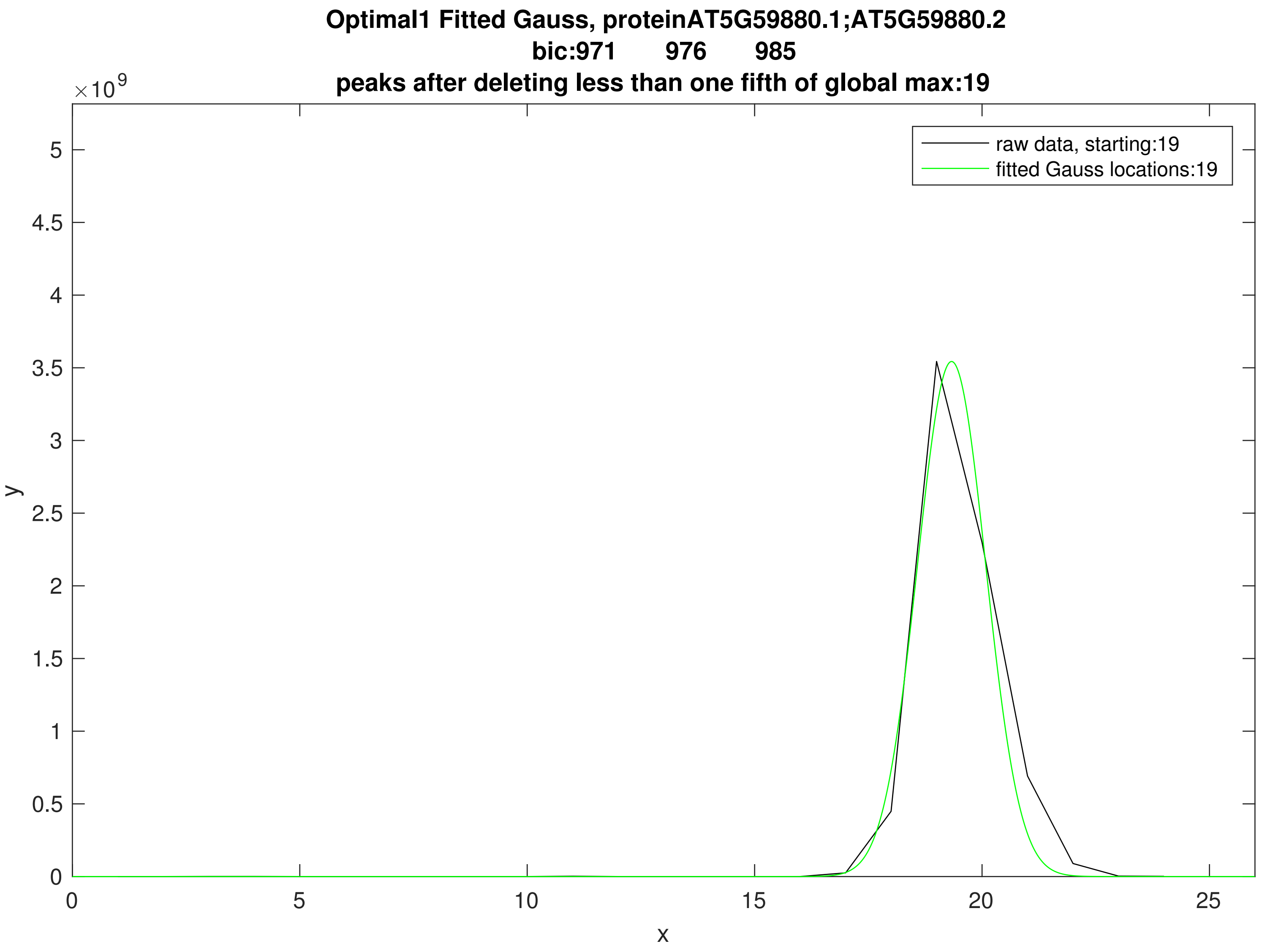

Optimal1 Fitted Gauss, proteinAT5G59880.1;AT5G59880.2
bic:971 976 985
peaks after deleting less than one fifth of global max:19
×10^9
raw data, starting:19
fitted Gauss locations:19
y
x
0
0.5
1
1.5
2
2.5
3
3.5
4
4.5
5
0
5
10
15
20
25

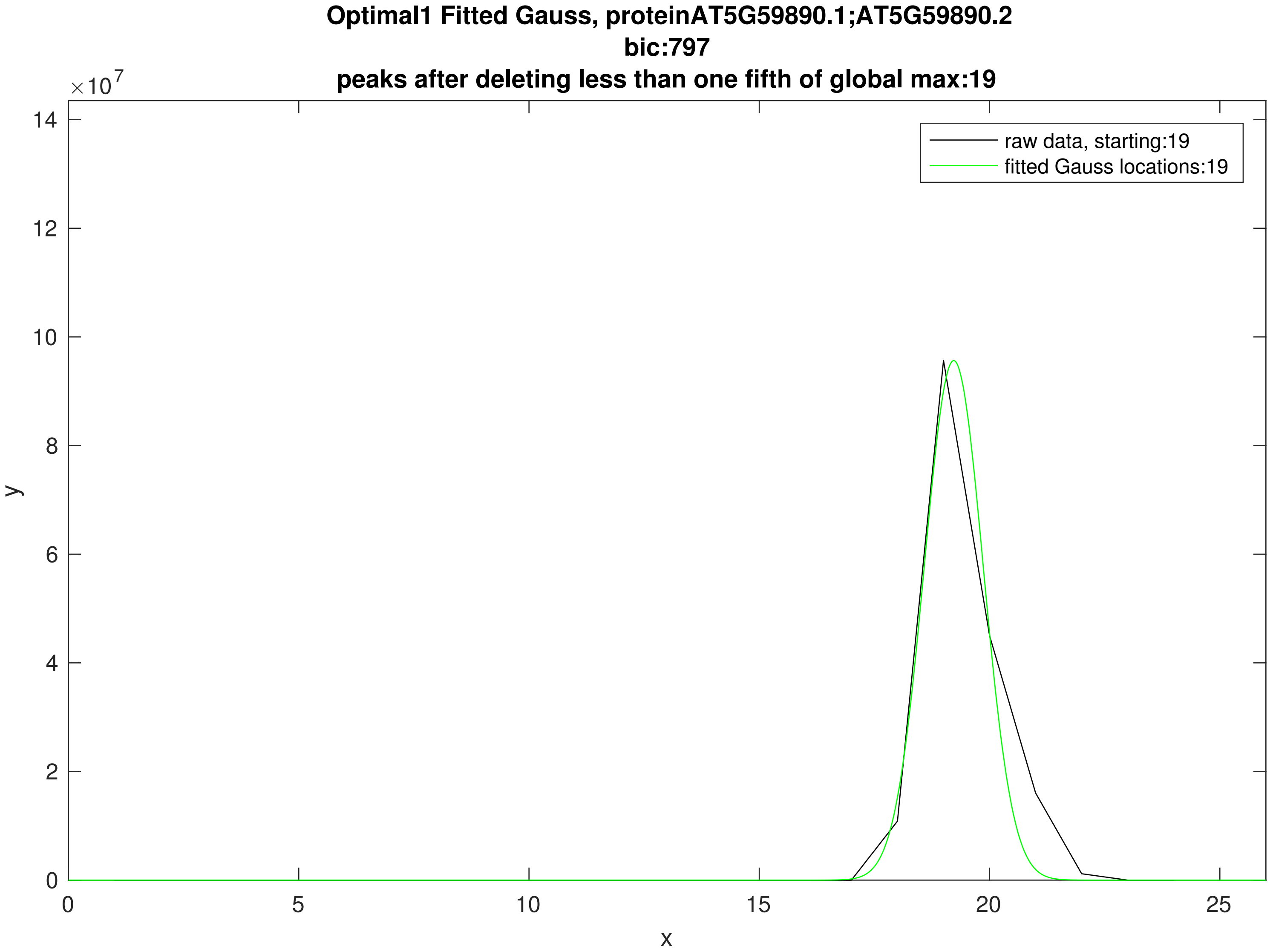
Optimal1 Fitted Gauss, proteinAT5G59890.1;AT5G59890.2
bic:797
peaks after deleting less than one fifth of global max:19
×10^7
raw data, starting:19
fitted Gauss locations:19
y
x
0
2
4
6
8
10
12
14
0
5
10
15
20
25

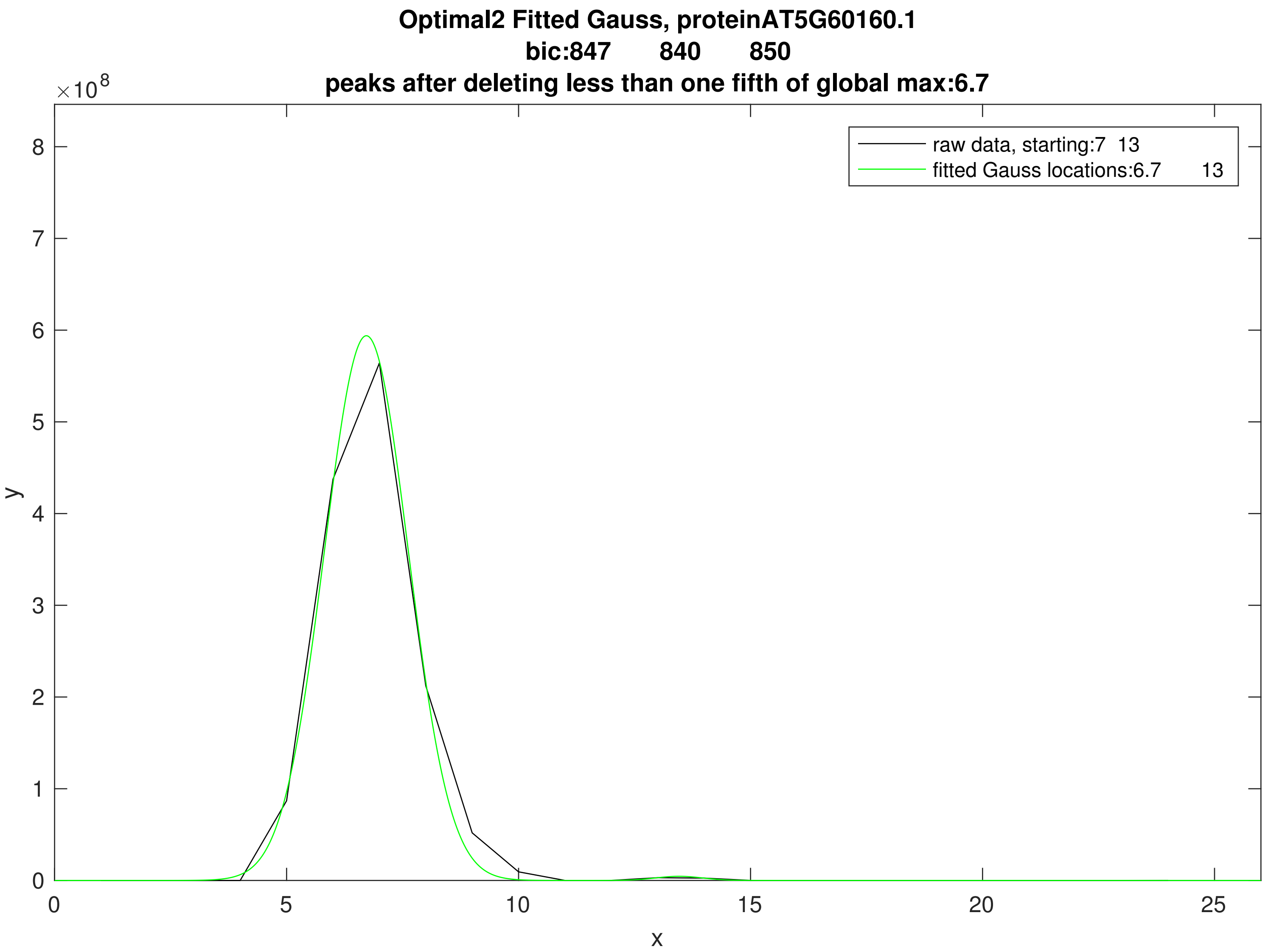

Optimal2 Fitted Gauss, proteinAT5G60160.1
bic:847 840 850
peaks after deleting less than one fifth of global max:6.7
×10^8
raw data, starting:7 13
fitted Gauss locations:6.7 13
y
x
0
1
2
3
4
5
6
7
8
0
5
10
15
20
25

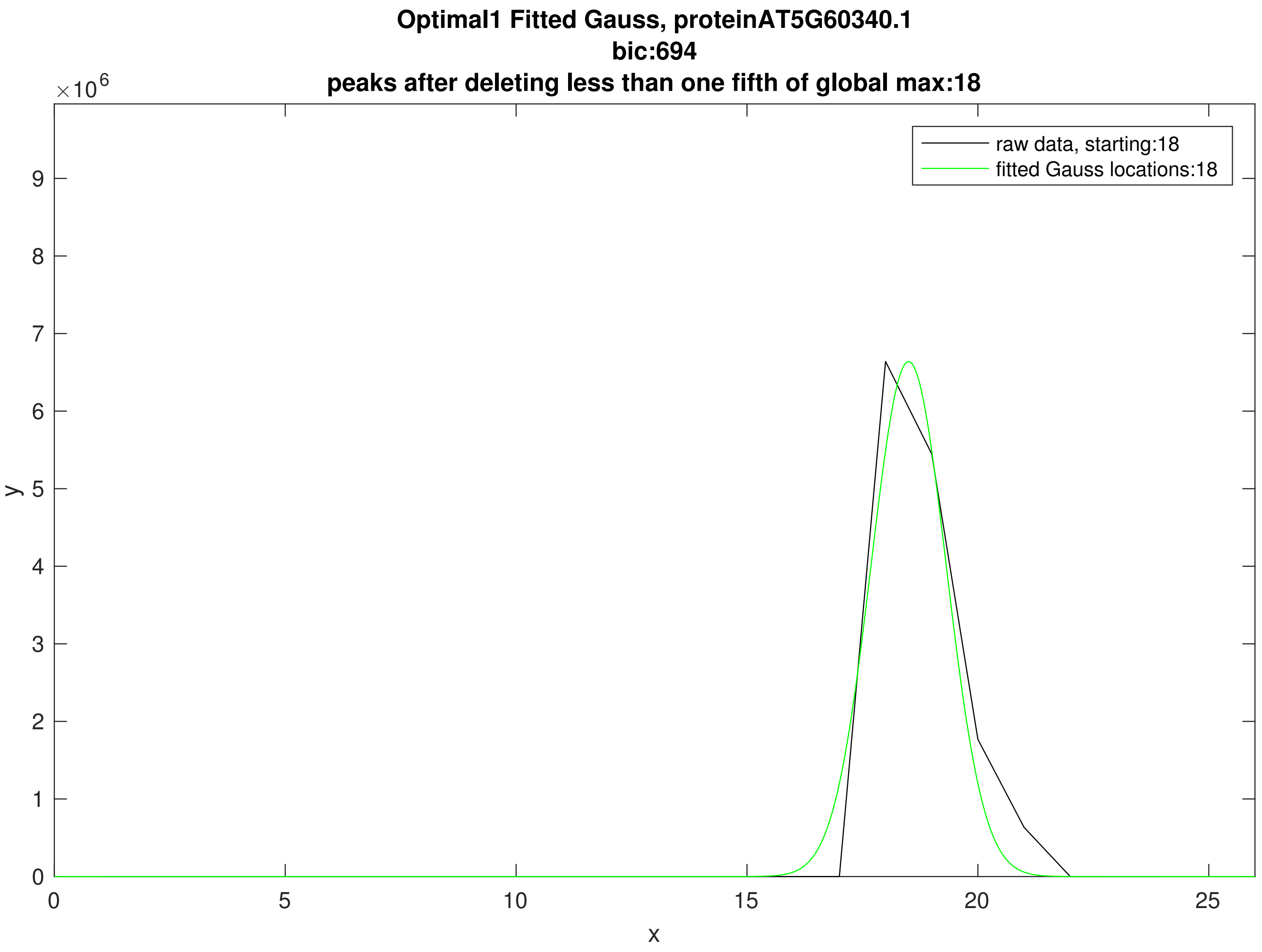
Optimal1 Fitted Gauss, proteinAT5G60340.1
bic:694
peaks after deleting less than one fifth of global max:18
×10^6
raw data, starting:18
fitted Gauss locations:18
y
x
0
1
2
3
4
5
6
7
8
9
0
5
10
15
20
25

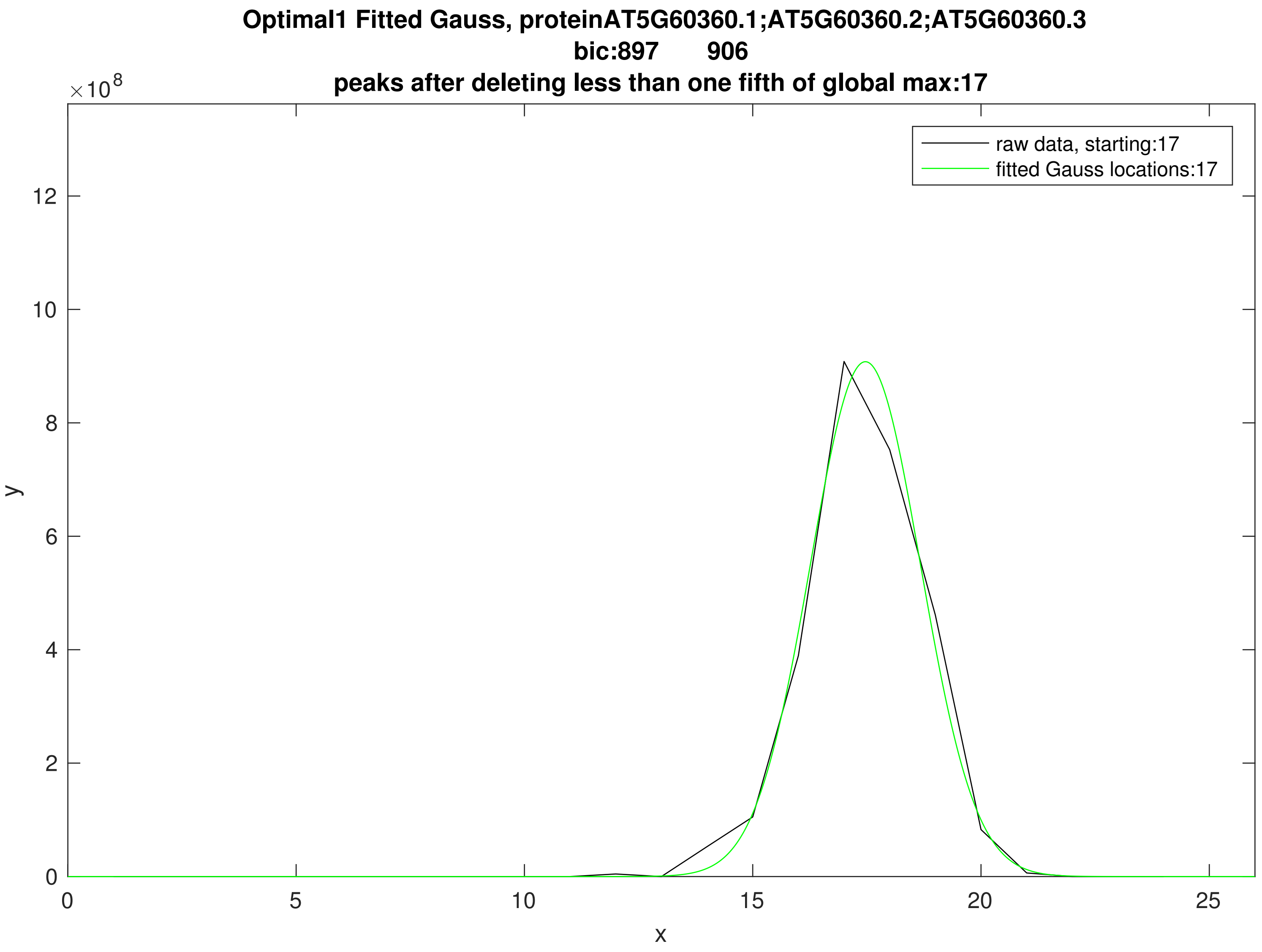

Optimal1 Fitted Gauss, proteinAT5G60360.1;AT5G60360.2;AT5G60360.3
bic:897 906
peaks after deleting less than one fifth of global max:17
×10^8
raw data, starting:17
fitted Gauss locations:17
y
x
0
2
4
6
8
10
12
0
5
10
15
20
25

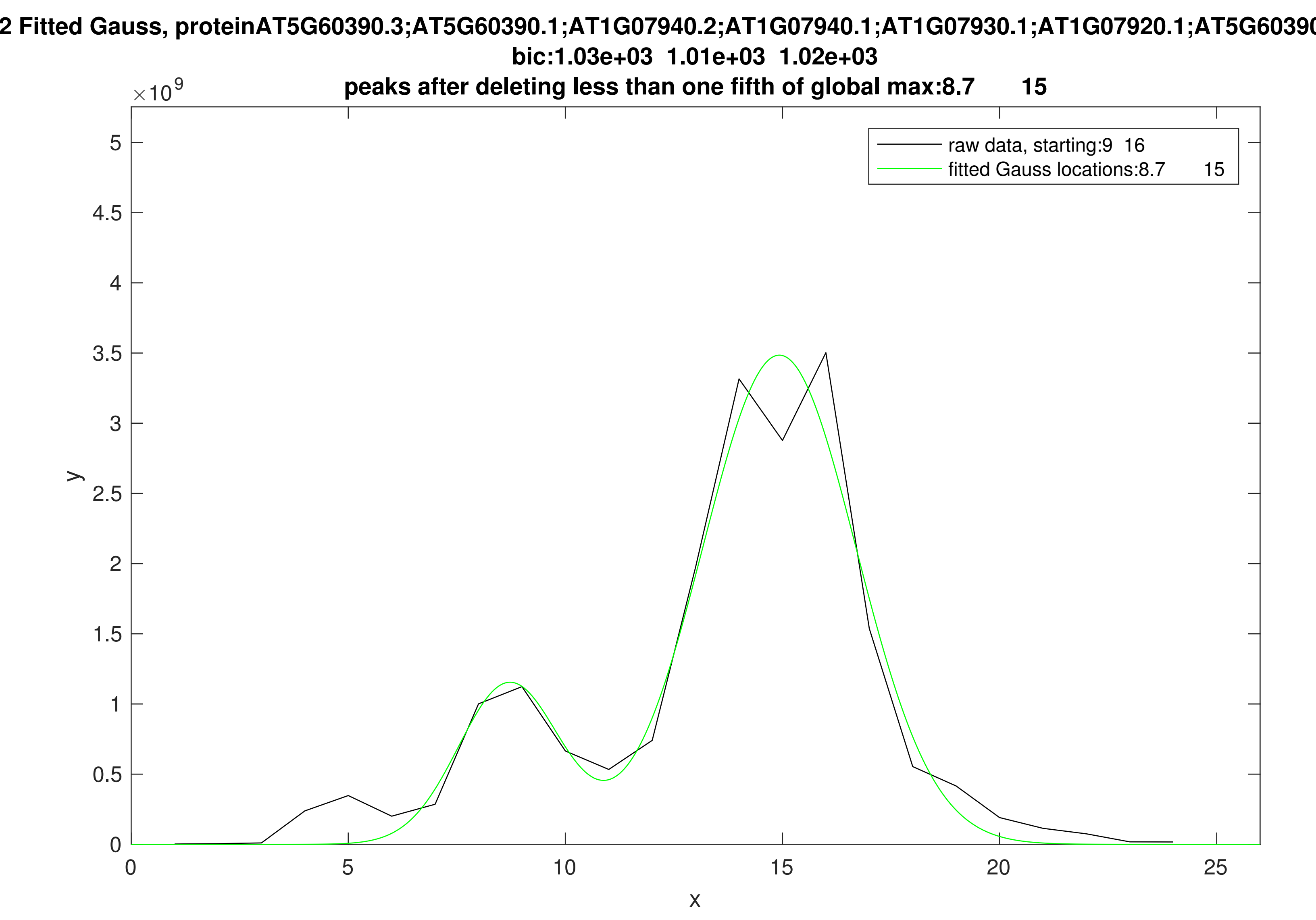
2 Fitted Gauss, proteinAT5G60390.3;AT5G60390.1;AT1G07940.2;AT1G07940.1;AT1G07930.1;AT1G07920.1;AT5G6039
bic:1.03e+03 1.01e+03 1.02e+03
peaks after deleting less than one fifth of global max:8.7 15
×10^9
raw data, starting:9 16
fitted Gauss locations:8.7 15
y
x
0
0.5
1
1.5
2
2.5
3
3.5
4
4.5
5
0
5
10
15
20
25

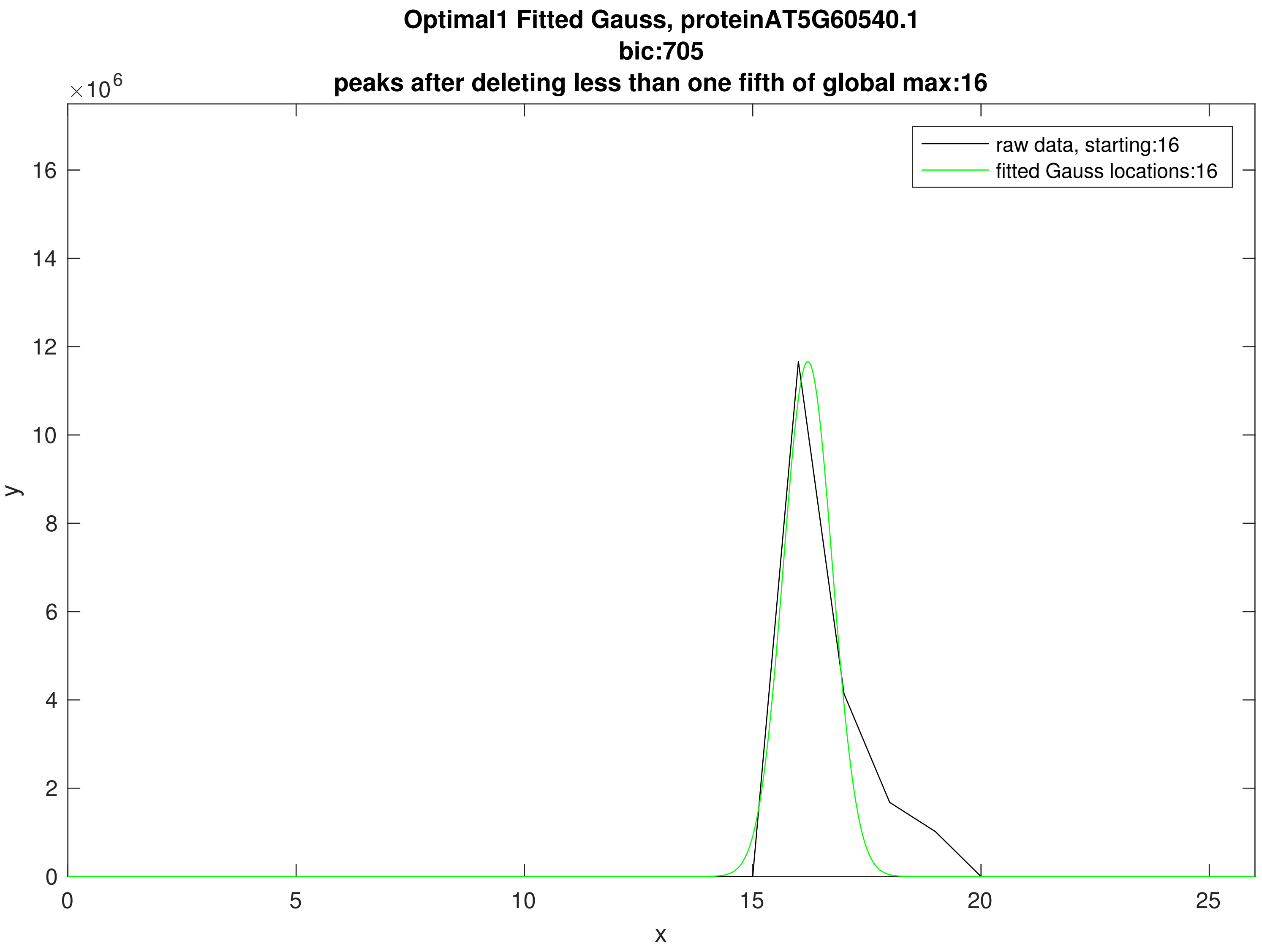
Optimal1 Fitted Gauss, proteinAT5G60540.1
bic:705
peaks after deleting less than one fifth of global max:16
×10^6
raw data, starting:16
fitted Gauss locations:16
16
14
12
10
8
6
4
2
0
y
0
5
10
15
20
25
x

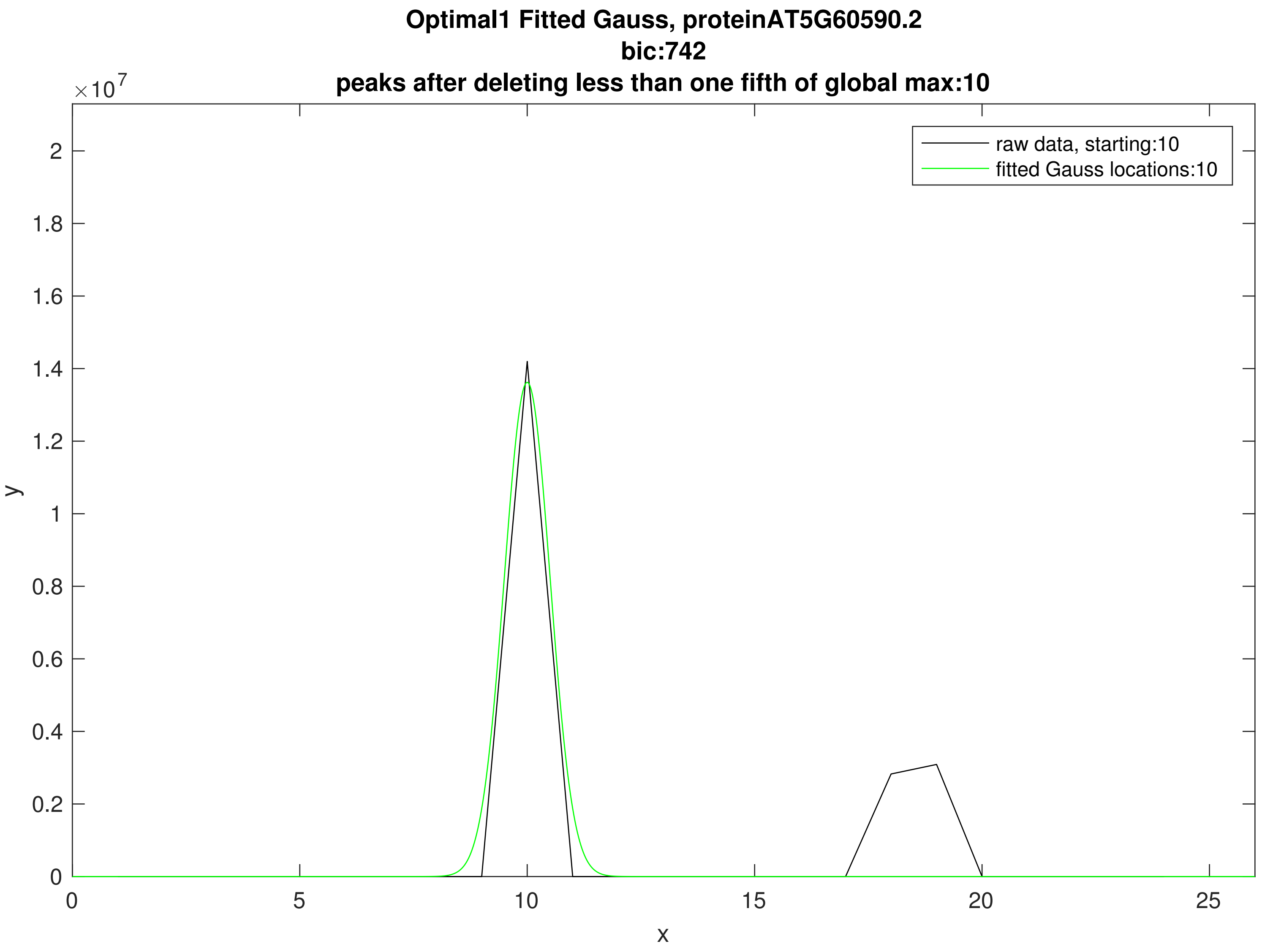
Optimal1 Fitted Gauss, proteinAT5G60590.2
bic:742
peaks after deleting less than one fifth of global max:10
×10^7
raw data, starting:10
fitted Gauss locations:10
2
1.8
1.6
1.4
1.2
1
0.8
0.6
0.4
0.2
0
y
0
5
10
15
20
25
x

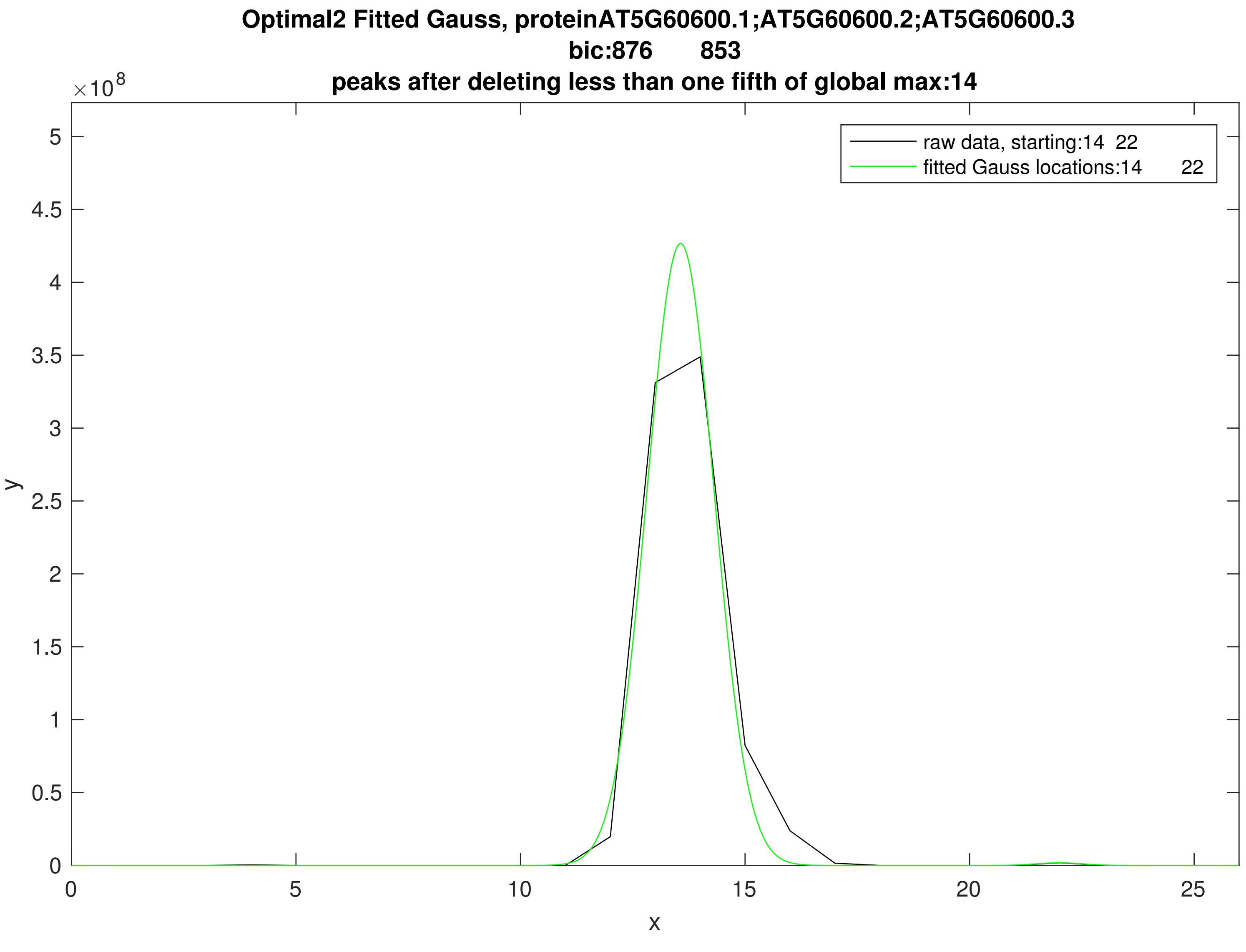

Optimal2 Fitted Gauss, proteinAT5G60600.1;AT5G60600.2;AT5G60600.3
bic:876 853
peaks after deleting less than one fifth of global max:14
×10^8
raw data, starting:14 22
fitted Gauss locations:14 22
y
x
0
0.5
1
1.5
2
2.5
3
3.5
4
4.5
5
0
5
10
15
20
25

Optimal2 Fitted Gauss, proteinAT5G60640.1;AT5G60640.2;AT5G60640.3
bic:1.01e+03 997 1.01e+03 1.01e+03
peaks after deleting less than one fifth of global max:11 16

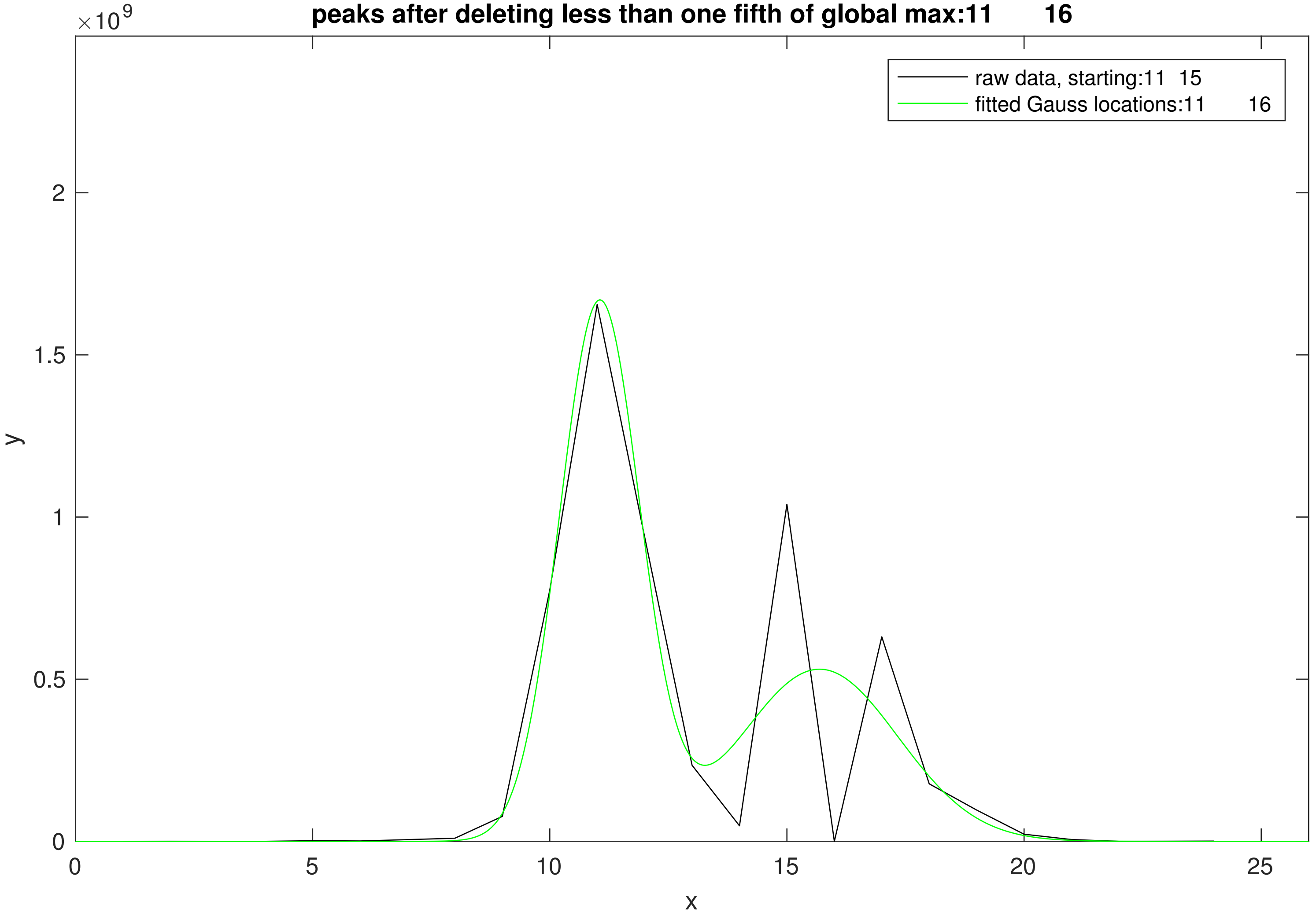

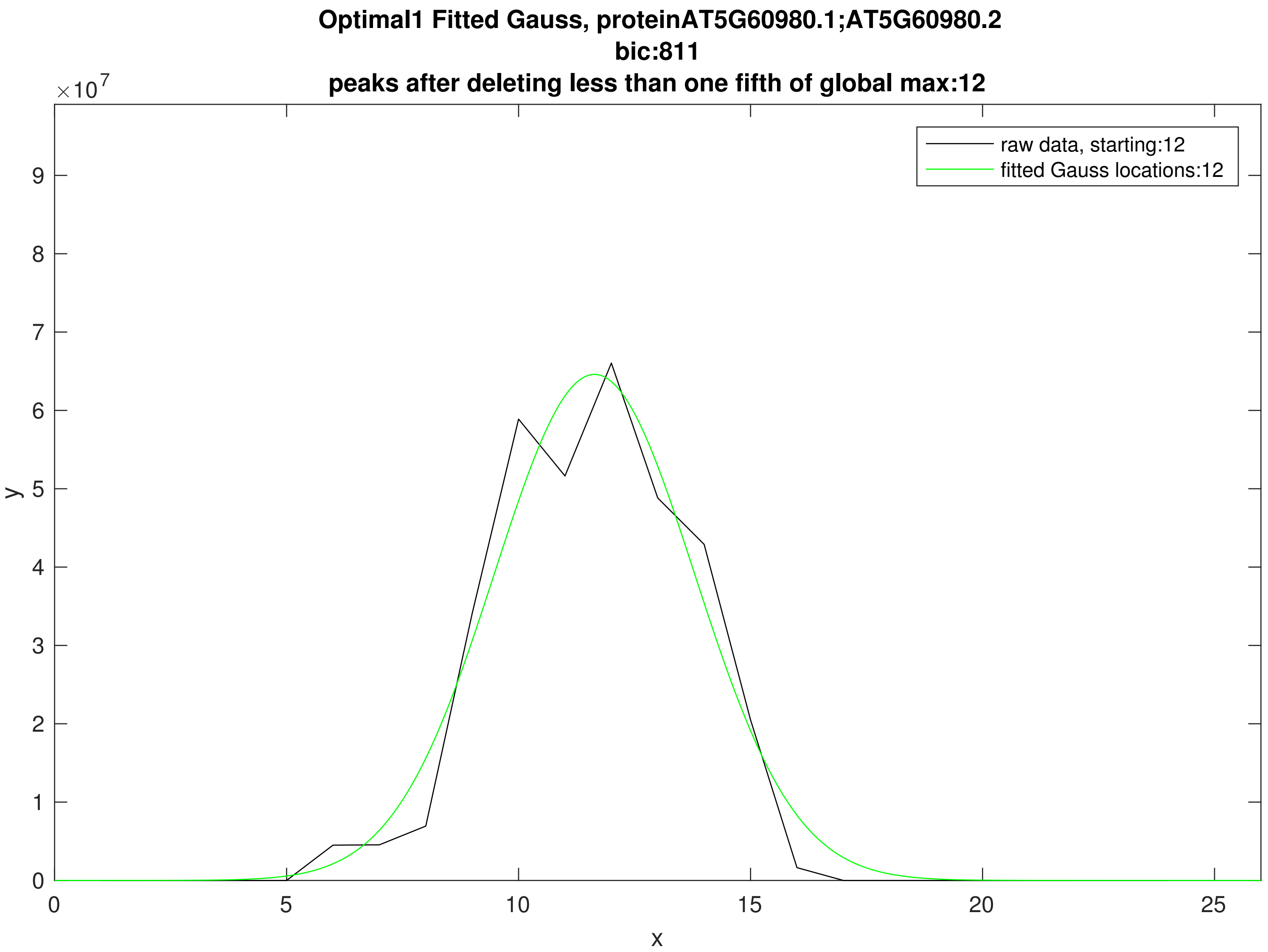
Optimal1 Fitted Gauss, proteinAT5G60980.1;AT5G60980.2
bic:811
peaks after deleting less than one fifth of global max:12
×10^7
raw data, starting:12
fitted Gauss locations:12
y
x
0
1
2
3
4
5
6
7
8
9
0
5
10
15
20
25

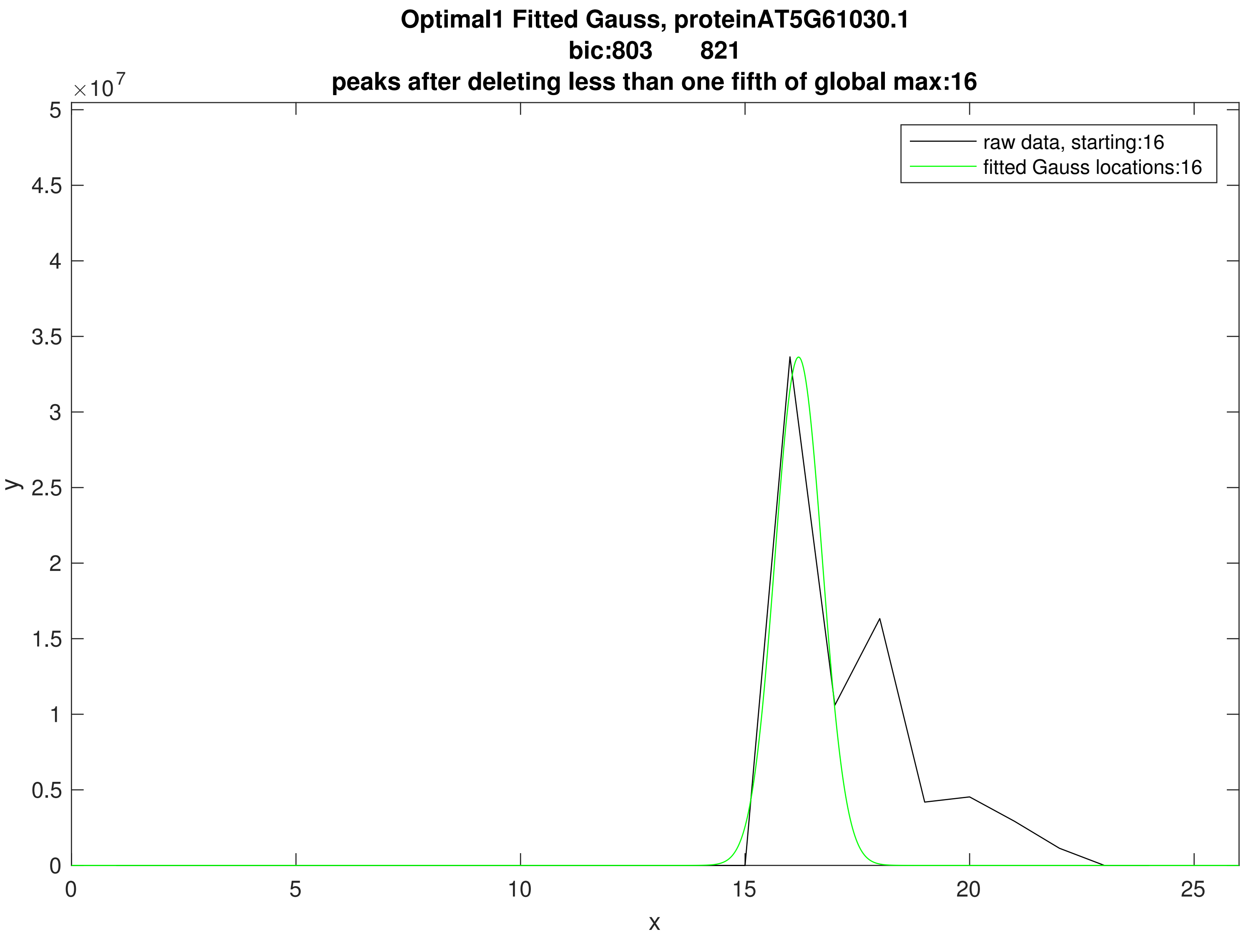

Optimal1 Fitted Gauss, proteinAT5G61030.1
bic:803 821
peaks after deleting less than one fifth of global max:16
×10^7
raw data, starting:16
fitted Gauss locations:16
y
x

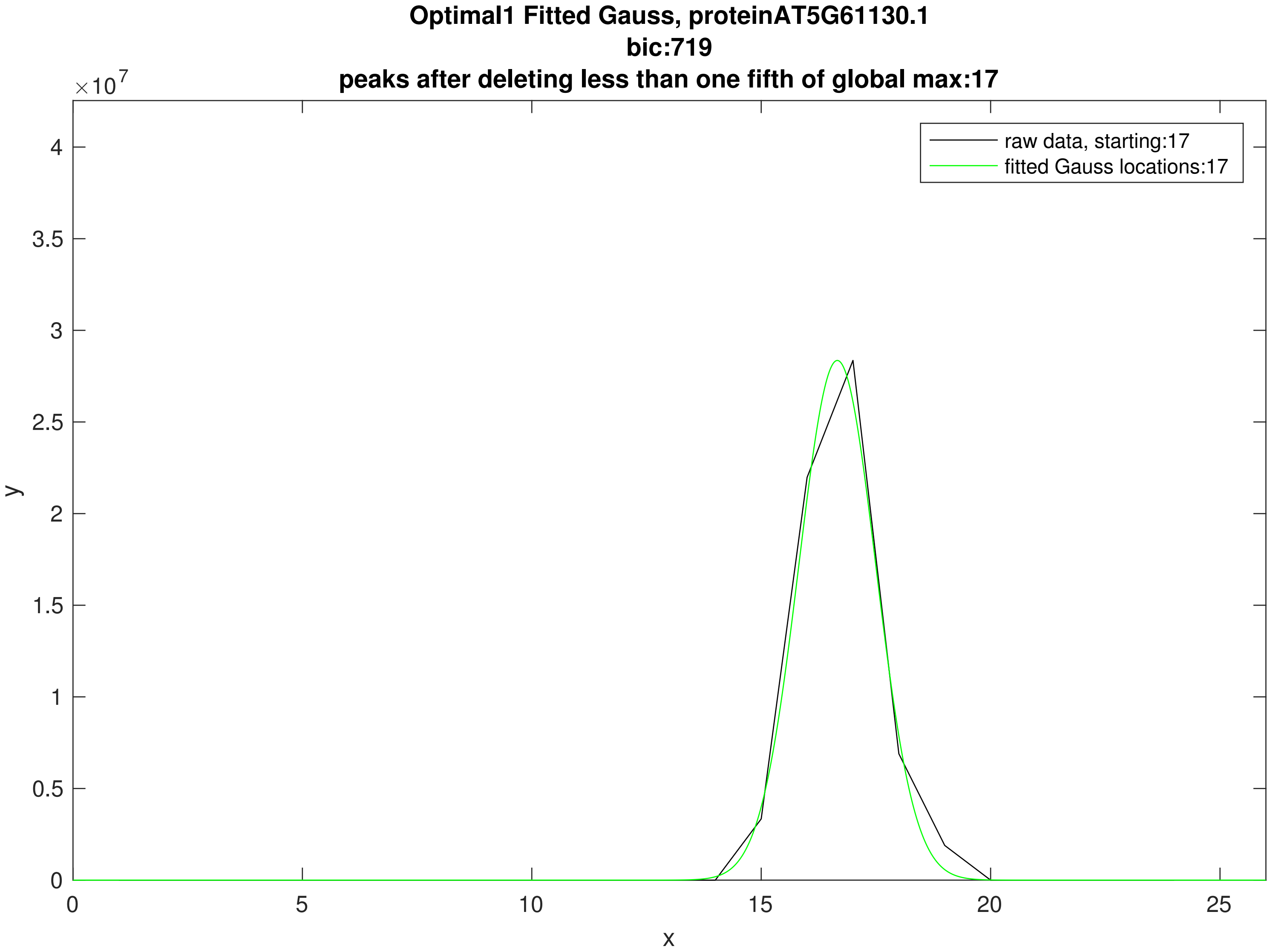

Optimal1 Fitted Gauss, proteinAT5G61130.1
bic:719
peaks after deleting less than one fifth of global max:17
×10^7
raw data, starting:17
fitted Gauss locations:17
y
x
0
0.5
1
1.5
2
2.5
3
3.5
4
5
10
15
20
25

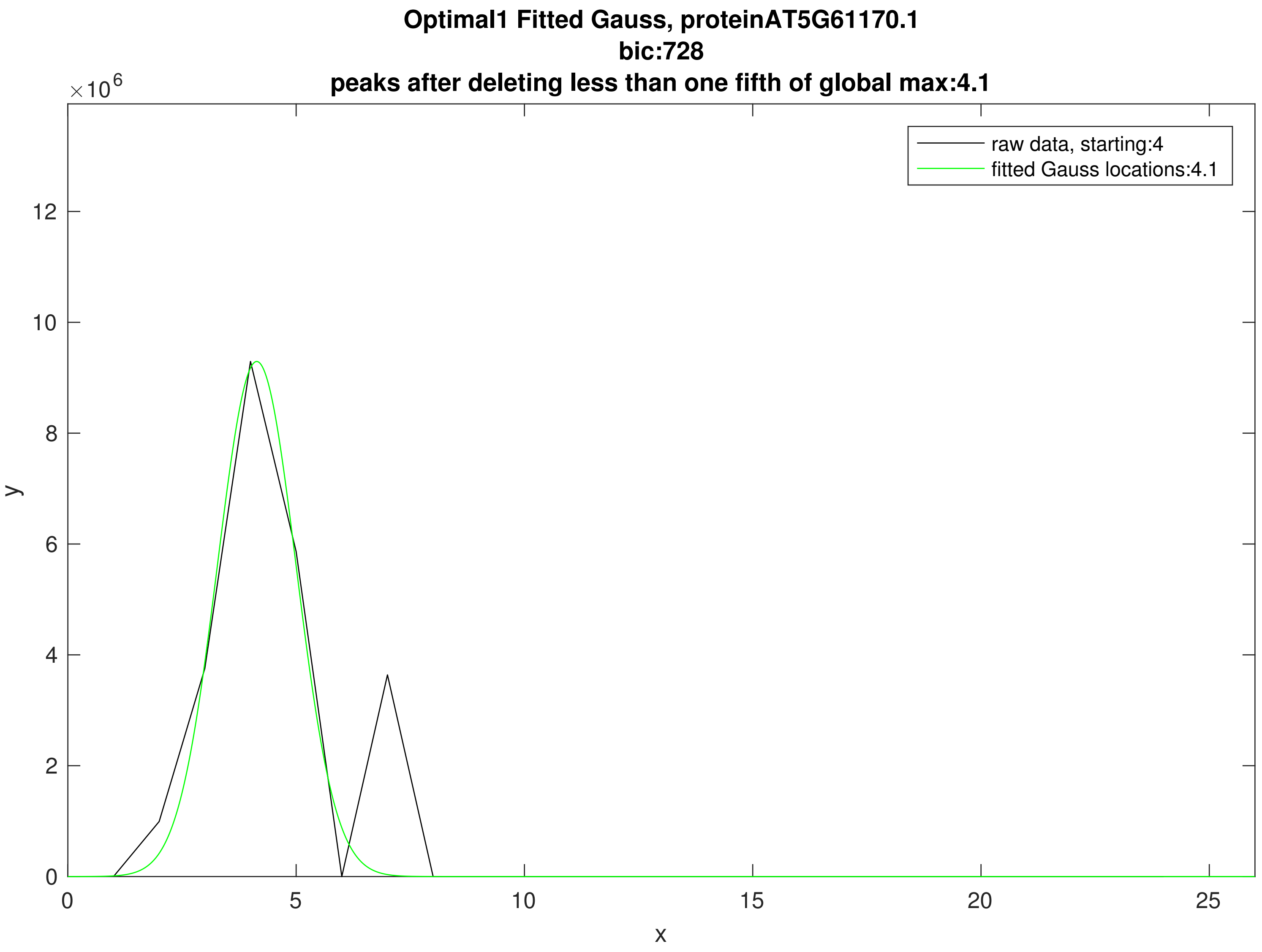
Optimal1 Fitted Gauss, proteinAT5G61170.1
bic:728
peaks after deleting less than one fifth of global max:4.1
×10^6
raw data, starting:4
fitted Gauss locations:4.1
y
x
0
2
4
6
8
10
12
0
5
10
15
20
25

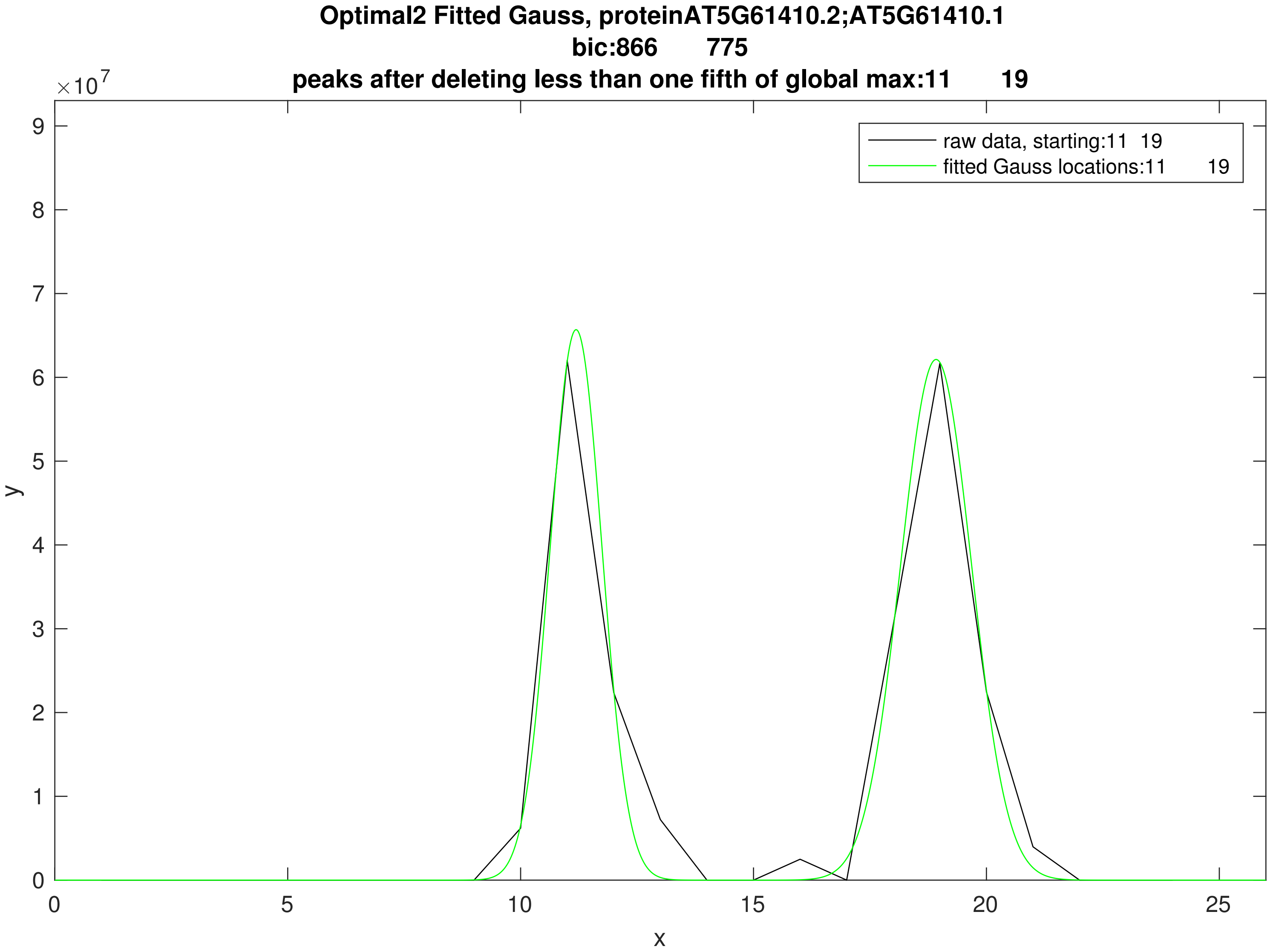

Optimal2 Fitted Gauss, proteinAT5G61410.2;AT5G61410.1
bic:866 775
peaks after deleting less than one fifth of global max:11 19
×10^7
raw data, starting:11 19
fitted Gauss locations:11 19
y
x
0
1
2
3
4
5
6
7
8
9
0
5
10
15
20
25

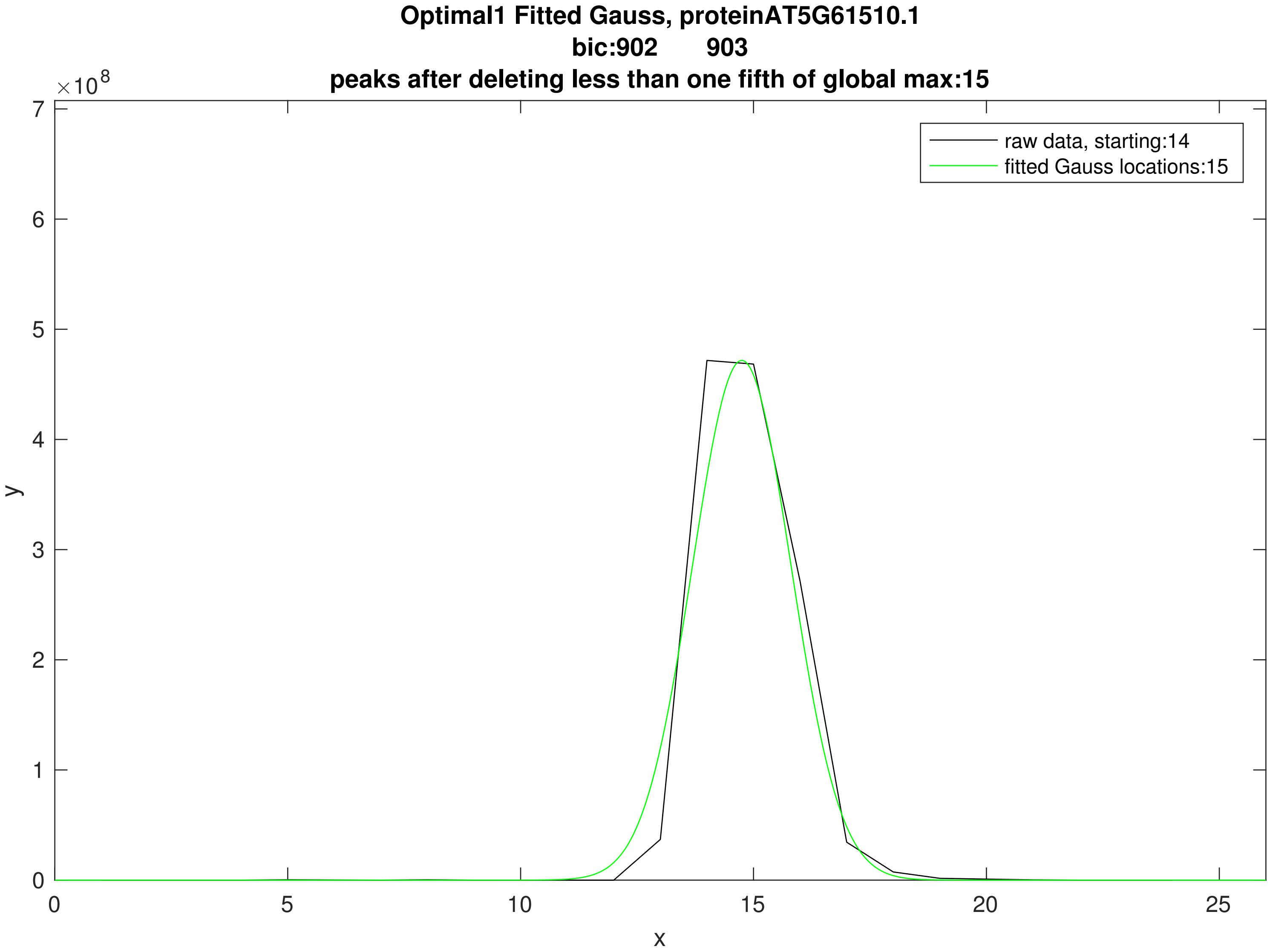
Optimal1 Fitted Gauss, proteinAT5G61510.1
bic:902 903
peaks after deleting less than one fifth of global max:15
×10^8
raw data, starting:14
fitted Gauss locations:15
y
x
0
1
2
3
4
5
6
7
0
5
10
15
20
25

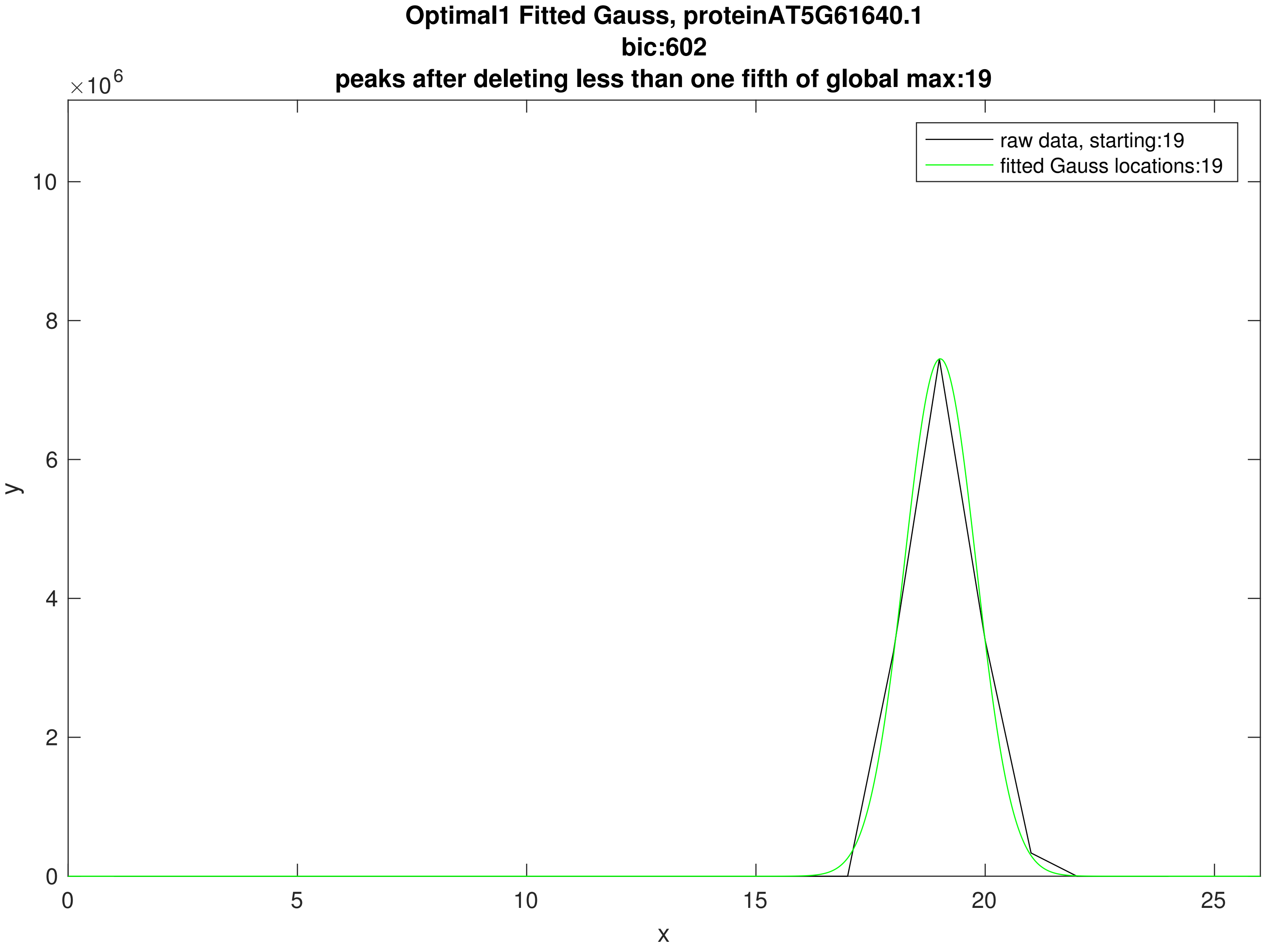
Optimal1 Fitted Gauss, proteinAT5G61640.1
bic:602
peaks after deleting less than one fifth of global max:19
×10^6
raw data, starting:19
fitted Gauss locations:19
y
x
0
2
4
6
8
10
0
5
10
15
20
25

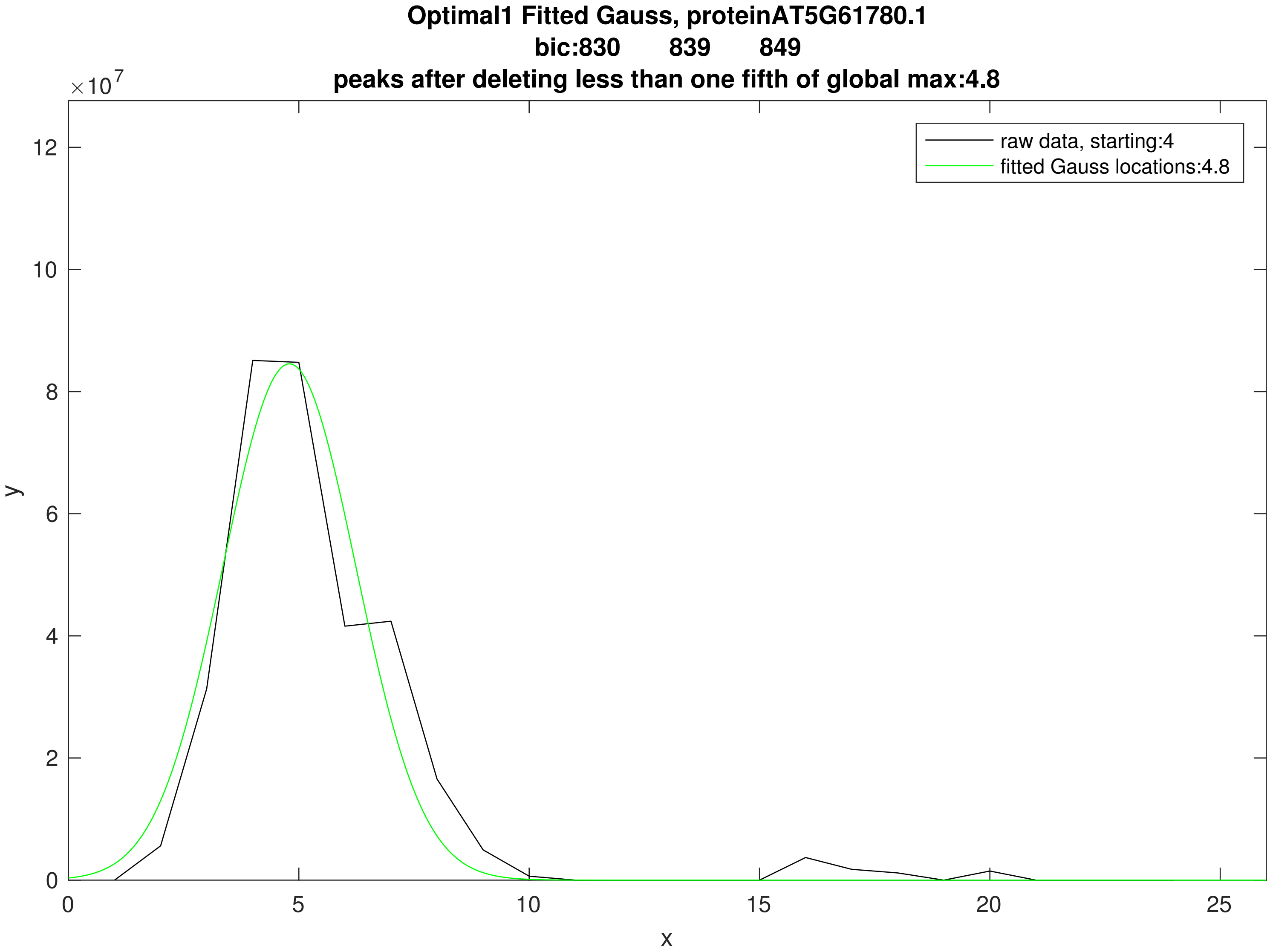
Optimal1 Fitted Gauss, proteinAT5G61780.1
bic:830 839 849
peaks after deleting less than one fifth of global max:4.8
×10^7
raw data, starting:4
fitted Gauss locations:4.8
y
x
0
2
4
6
8
10
12
0
5
10
15
20
25

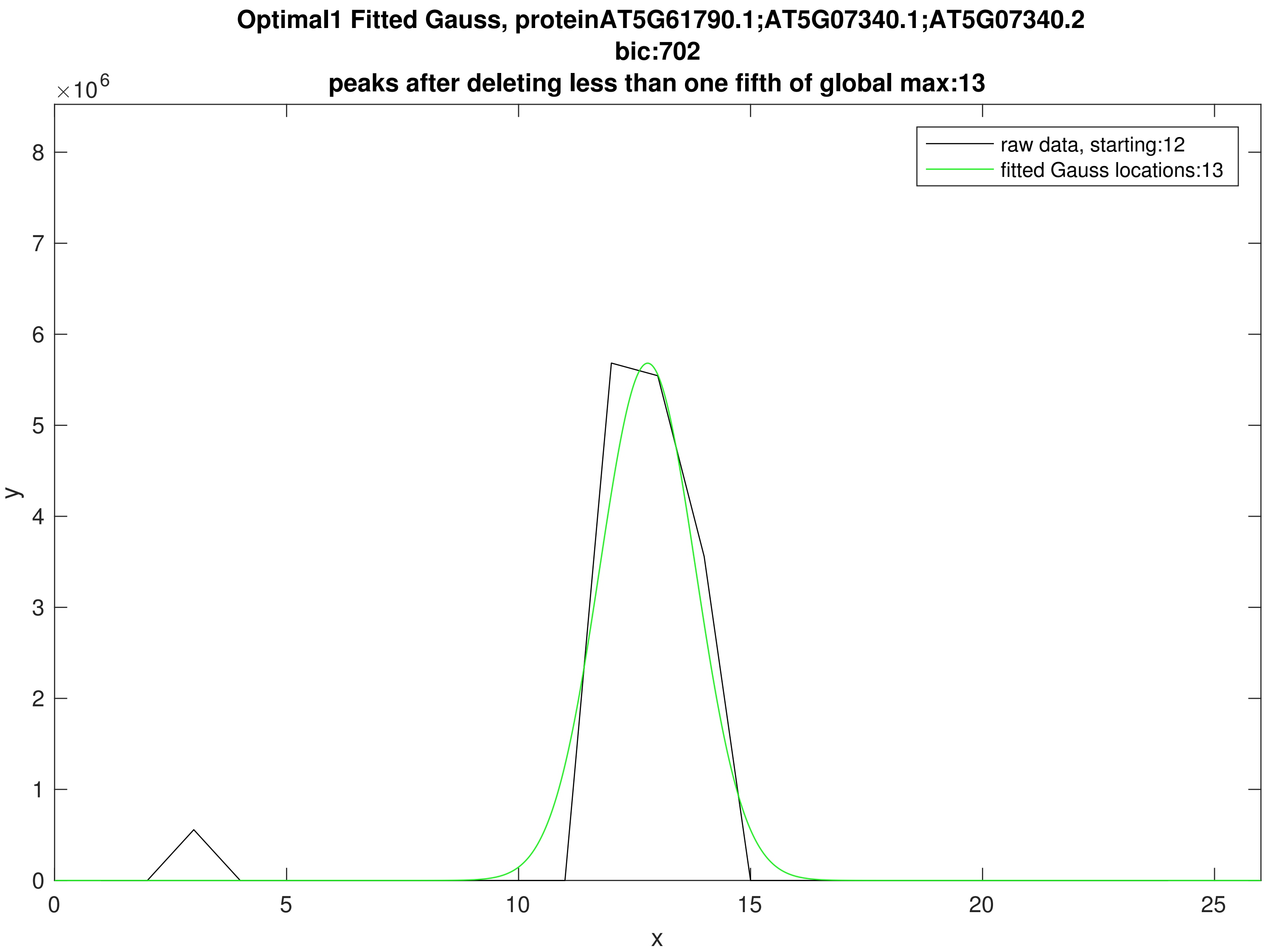
Optimal1 Fitted Gauss, proteinAT5G61790.1;AT5G07340.1;AT5G07340.2
bic:702
peaks after deleting less than one fifth of global max:13
×10^6
8
7
6
5
4
3
2
1
0
y
0
5
10
15
20
25
x
raw data, starting:12
fitted Gauss locations:13

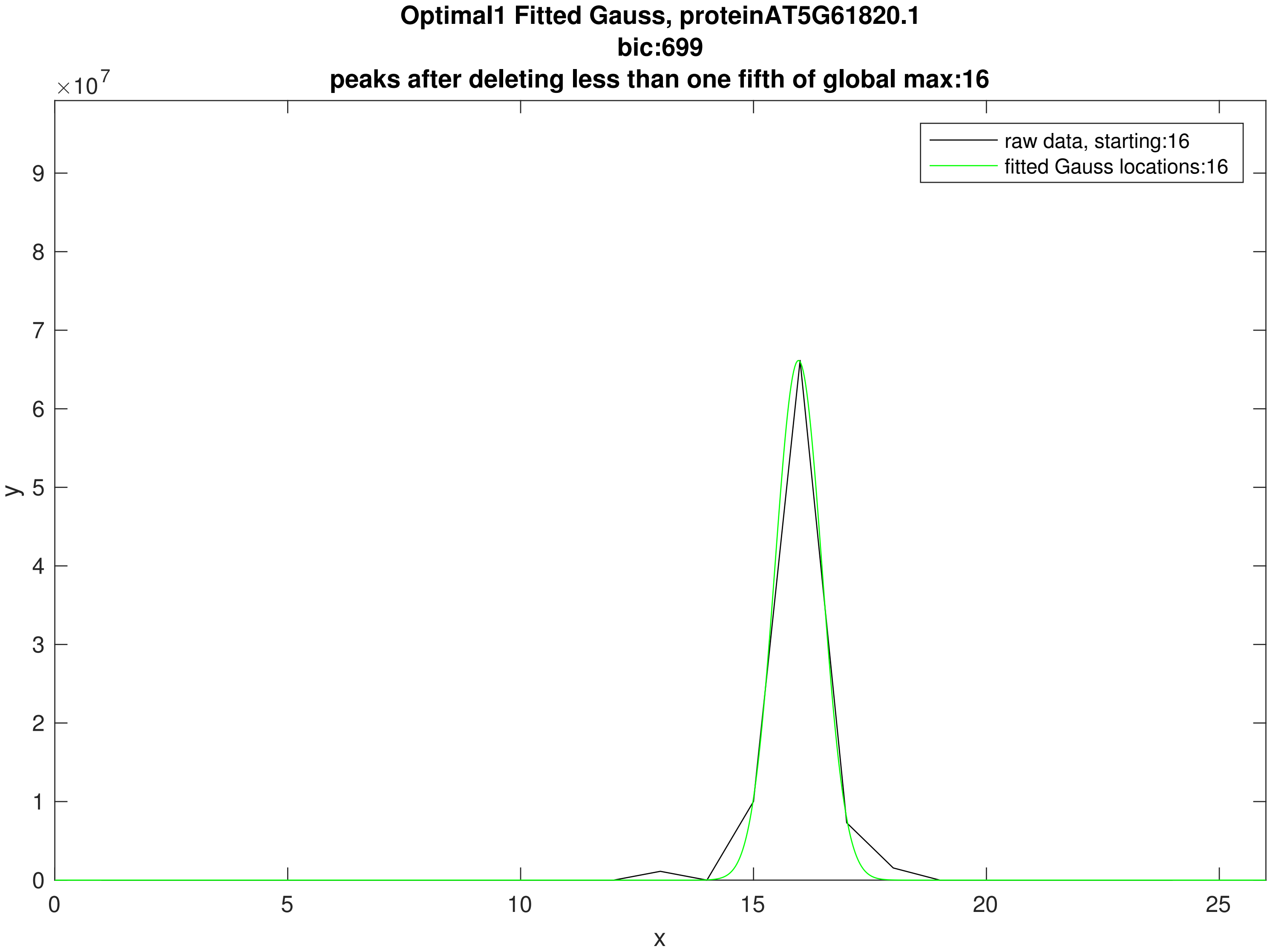
Optimal1 Fitted Gauss, proteinAT5G61820.1
bic:699
peaks after deleting less than one fifth of global max:16
×10^7
raw data, starting:16
fitted Gauss locations:16
y
x
0
1
2
3
4
5
6
7
8
9
0
5
10
15
20
25

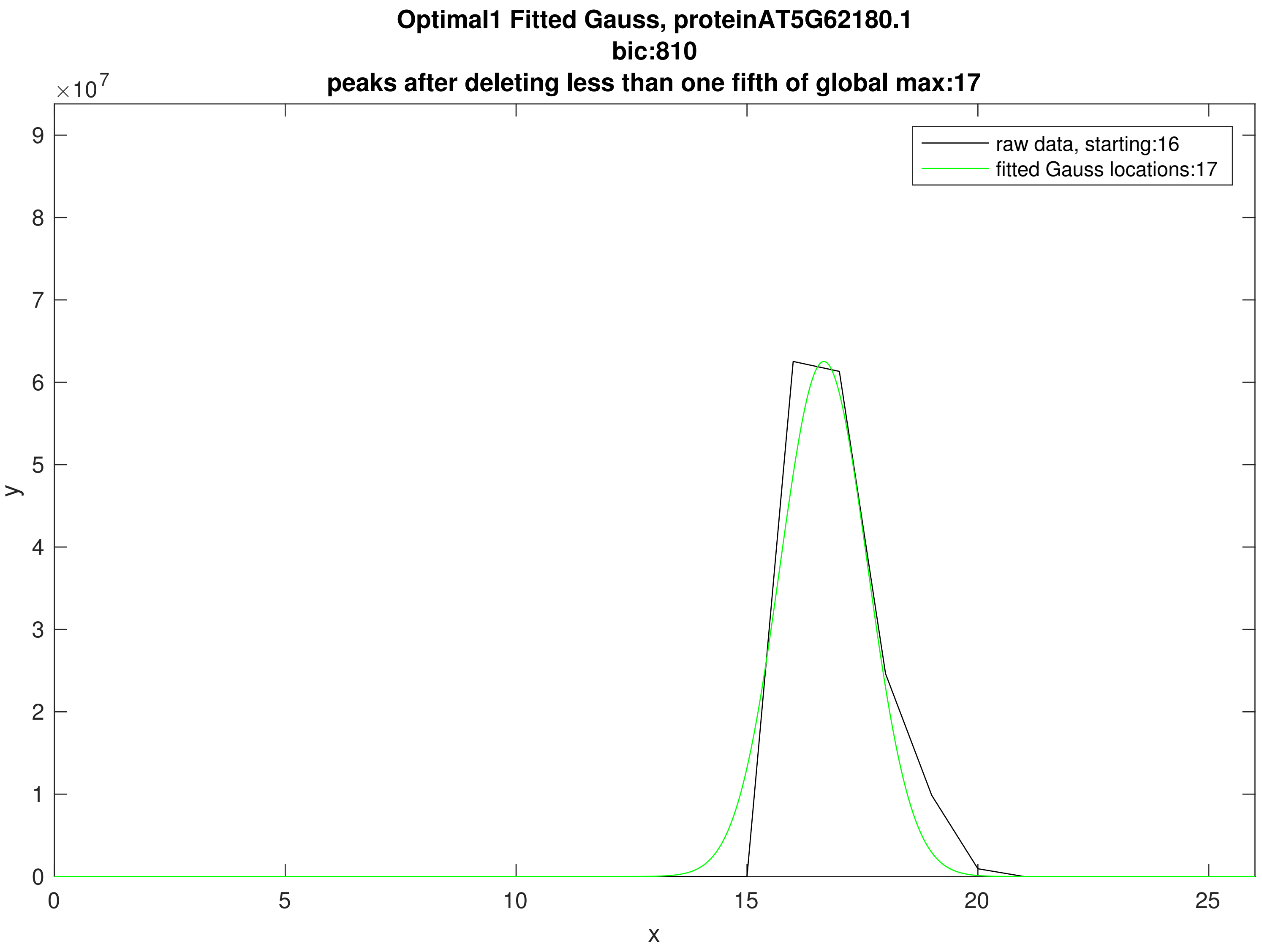
Optimal1 Fitted Gauss, proteinAT5G62180.1
bic:810
peaks after deleting less than one fifth of global max:17
×10^7
raw data, starting:16
fitted Gauss locations:17
y
x
0
1
2
3
4
5
6
7
8
9
0
5
10
15
20
25

Optimal1 Fitted Gauss, proteinAT5G62300.2;AT5G62300.1;AT3G45030.1
bic:806
peaks after deleting less than one fifth of global max:4.2

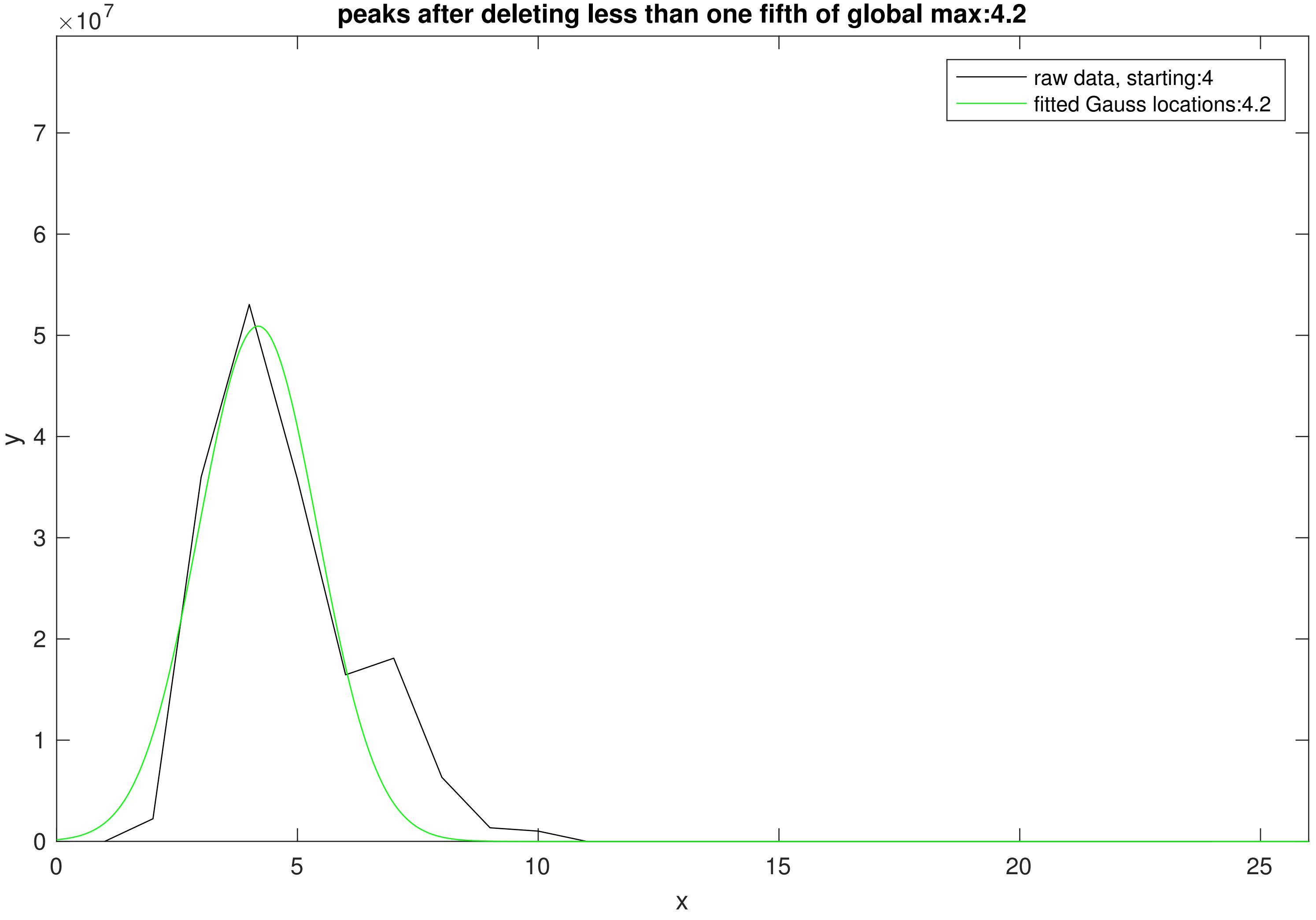

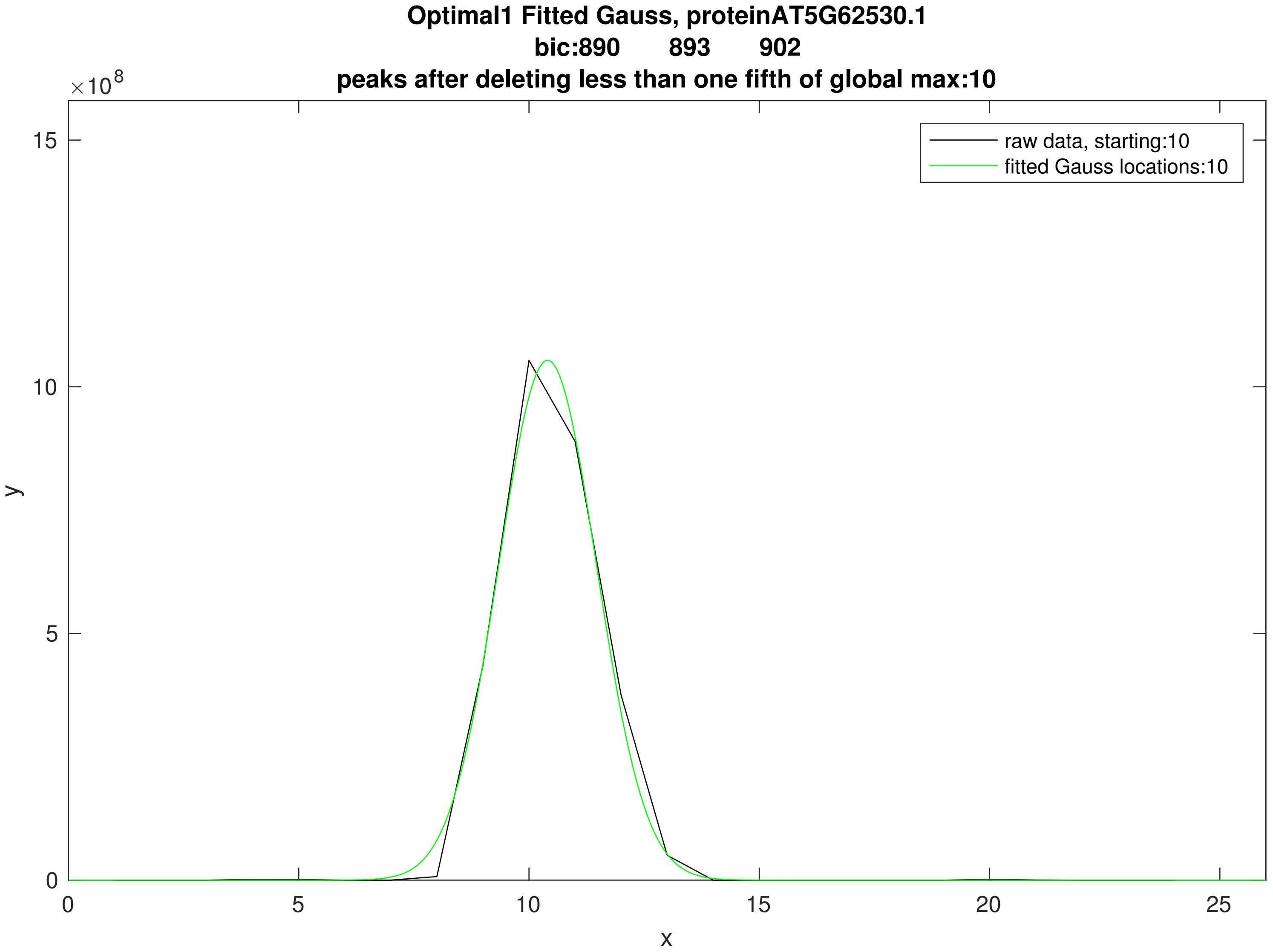

Optimal1 Fitted Gauss, proteinAT5G62530.1
bic:890 893 902
peaks after deleting less than one fifth of global max:10
×10^8
15
10
5
0
y
0
5
10
15
20
25
x
raw data, starting:10
fitted Gauss locations:10

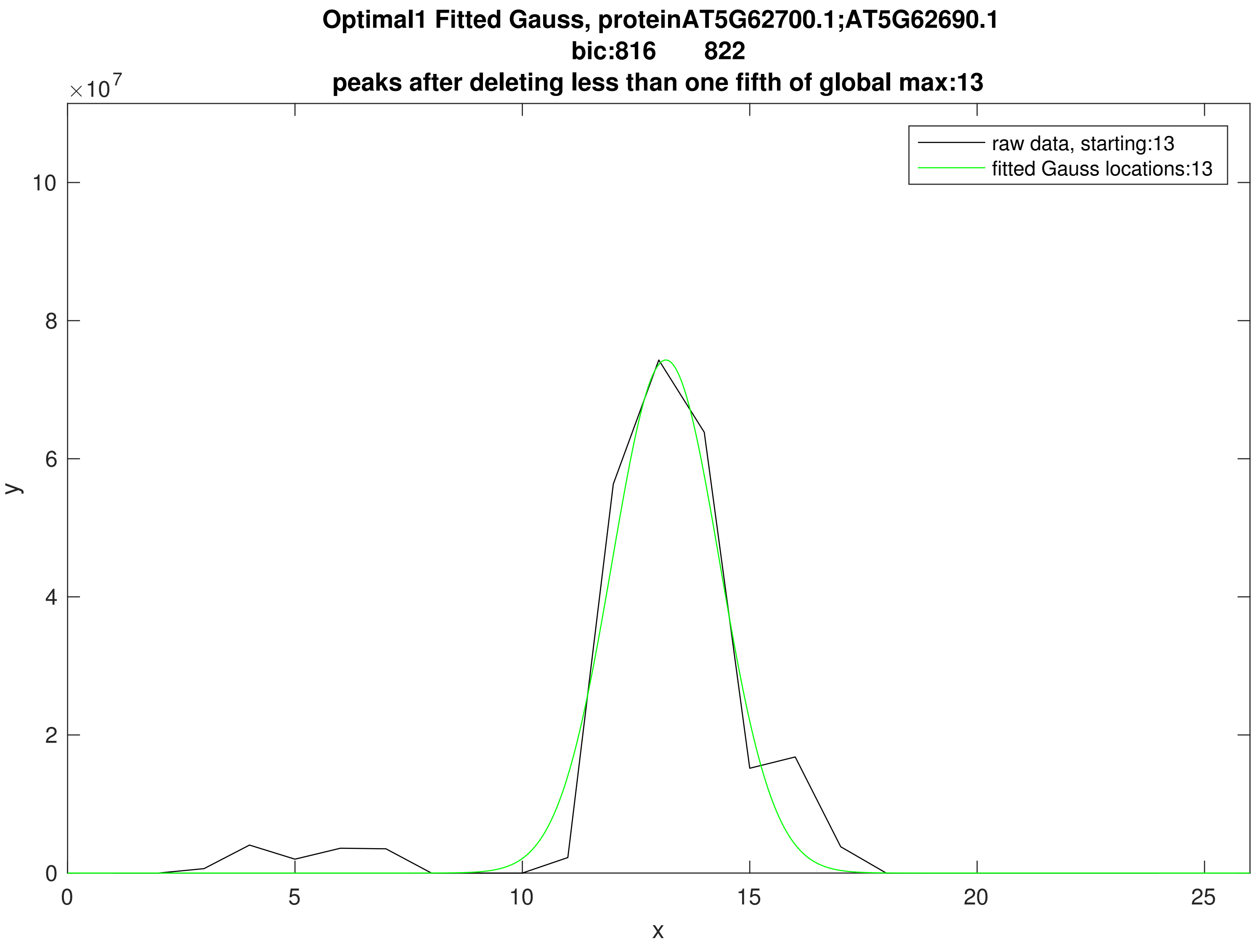
Optimal1 Fitted Gauss, proteinAT5G62700.1;AT5G62690.1
bic:816 822
peaks after deleting less than one fifth of global max:13
×10^7
raw data, starting:13
fitted Gauss locations:13
y
x
0
2
4
6
8
10
0
5
10
15
20
25

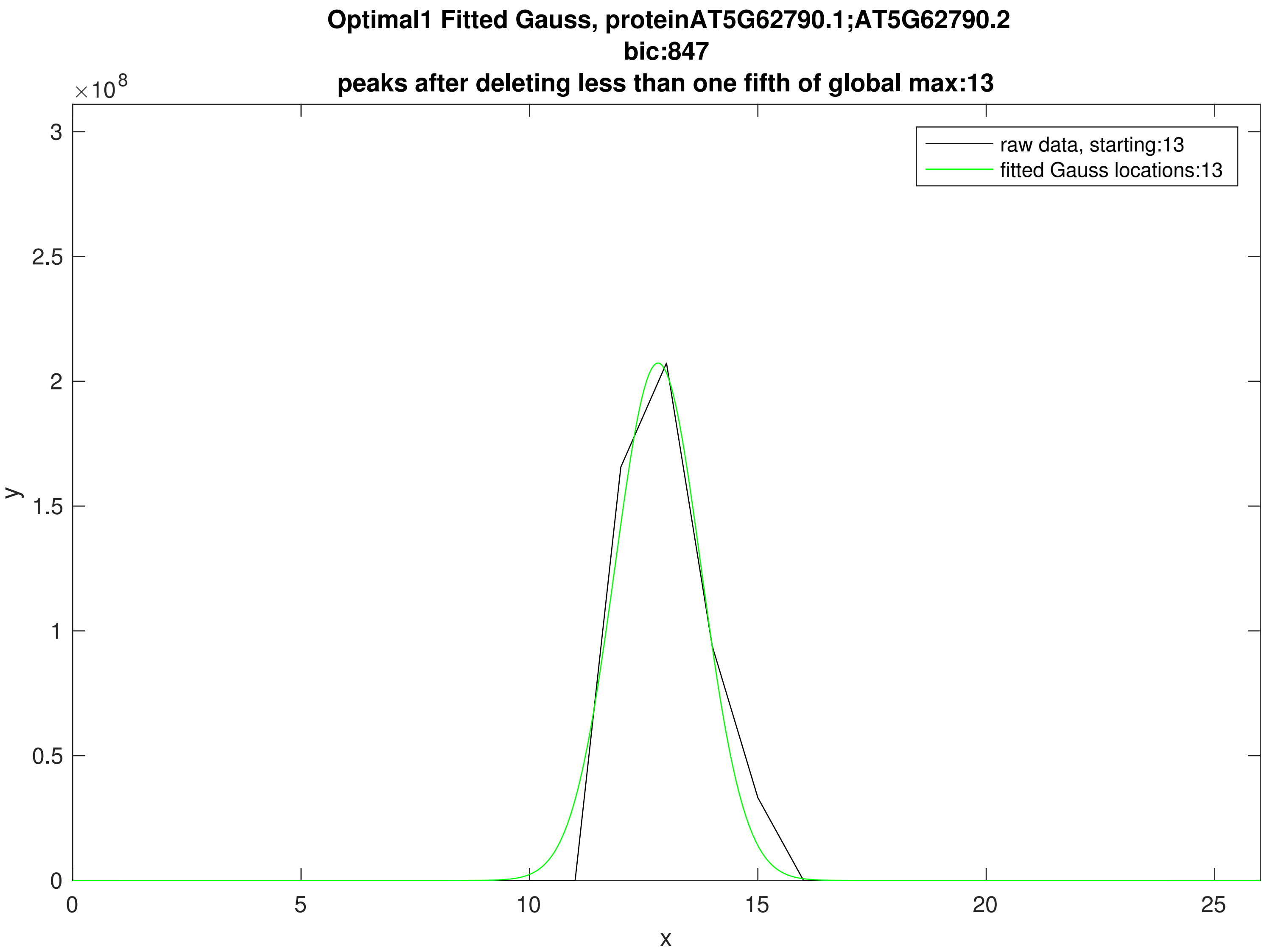
Optimal1 Fitted Gauss, proteinAT5G62790.1;AT5G62790.2
bic:847
peaks after deleting less than one fifth of global max:13
×10^8
3
2.5
2
1.5
1
0.5
0
y
0
5
10
15
20
25
x
raw data, starting:13
fitted Gauss locations:13

Optimal1 Fitted Gauss, proteinAT5G62810.1
bic:699
peaks after deleting less than one fifth of global max:12

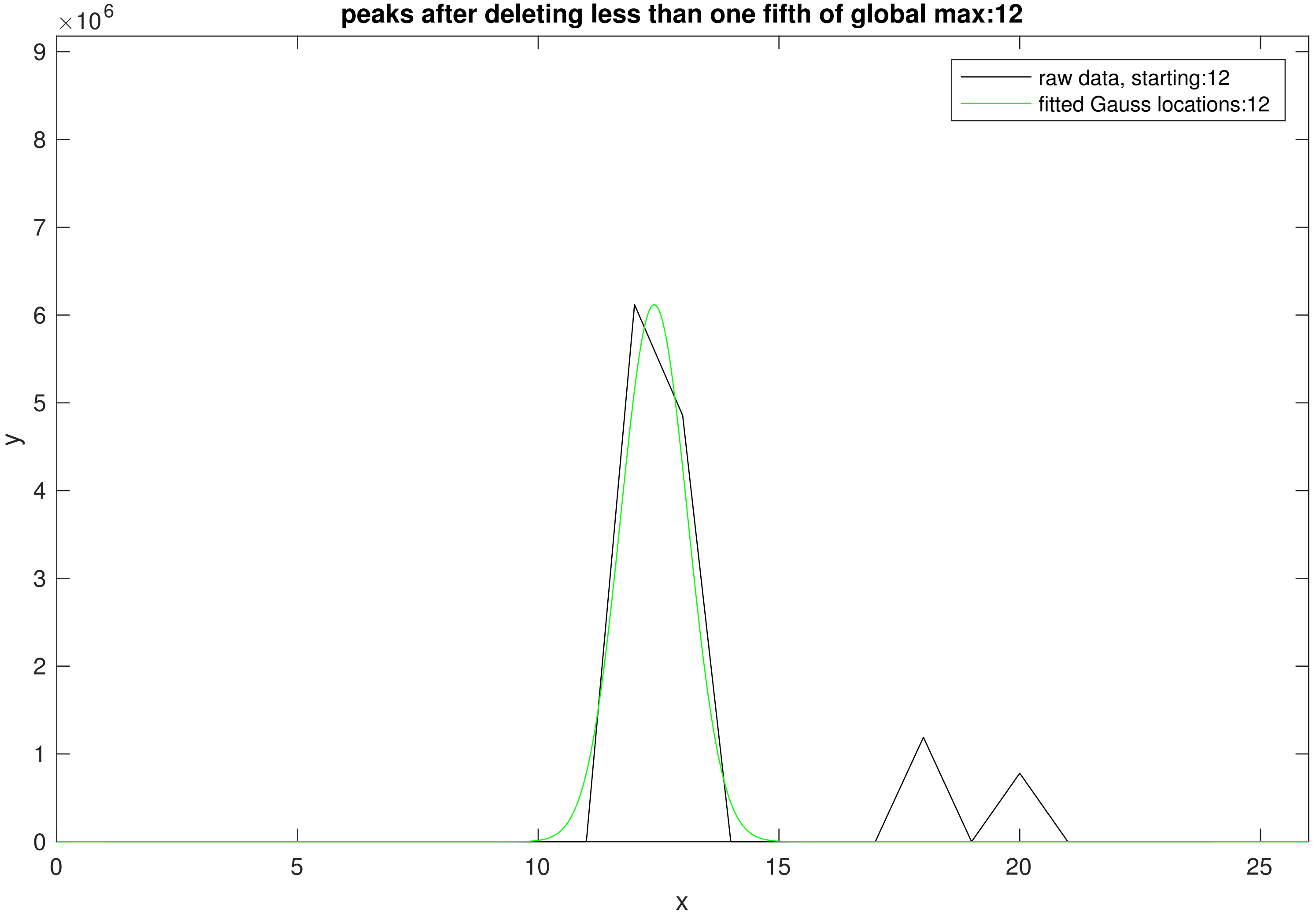

**Optimal1 Fitted Gauss, proteinAT5G62930.1**
**bic:879 884 893**
**peaks after deleting less than one fifth of global max:15**

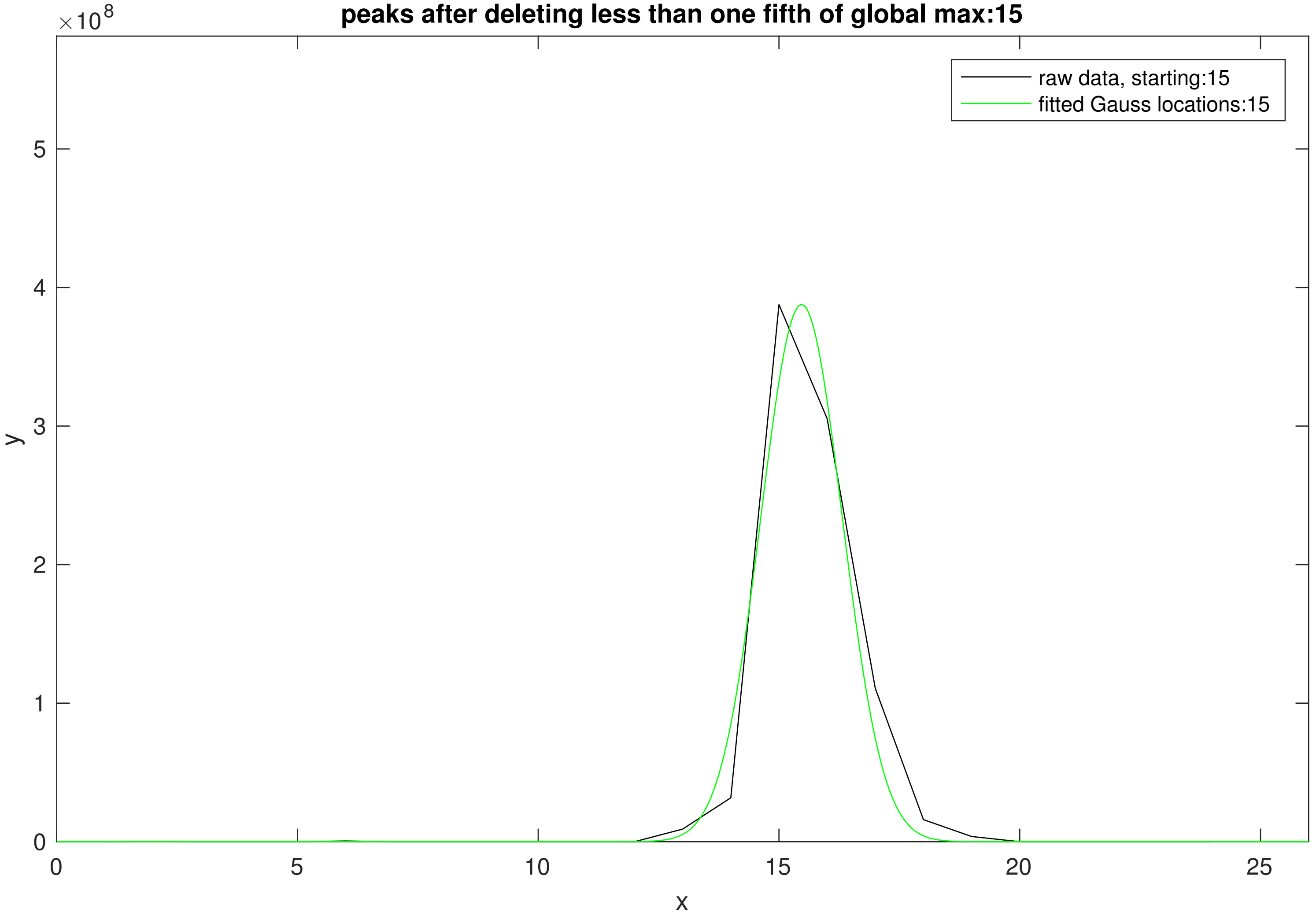

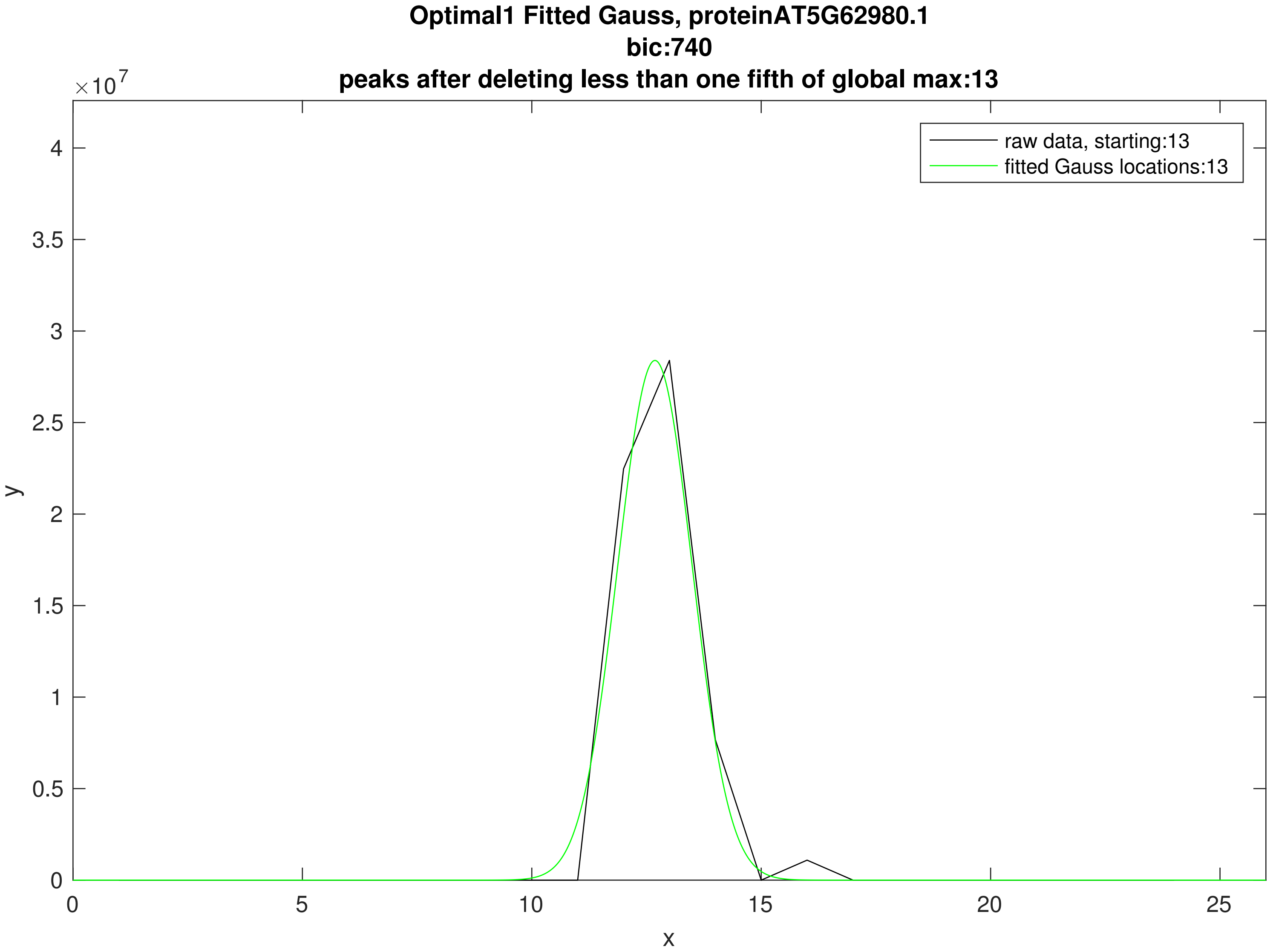
Optimal1 Fitted Gauss, proteinAT5G62980.1
bic:740
peaks after deleting less than one fifth of global max:13
×10^7
4
3.5
3
2.5
2
1.5
1
0.5
0
y
0
5
10
15
20
25
x
raw data, starting:13
fitted Gauss locations:13

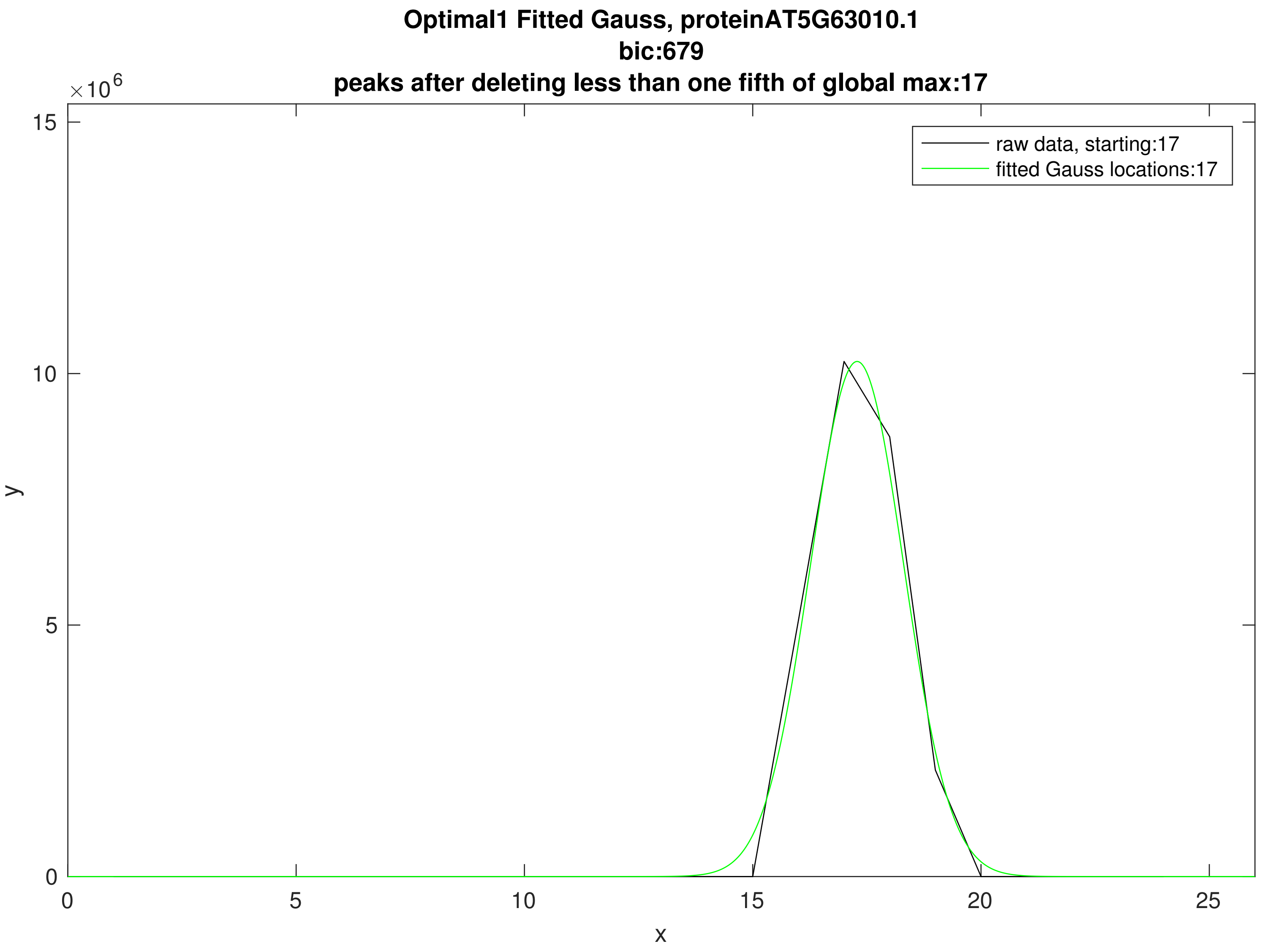
Optimal1 Fitted Gauss, proteinAT5G63010.1
bic:679
peaks after deleting less than one fifth of global max:17
×10^6
15
10
5
0
y
0
5
10
15
20
25
x
raw data, starting:17
fitted Gauss locations:17

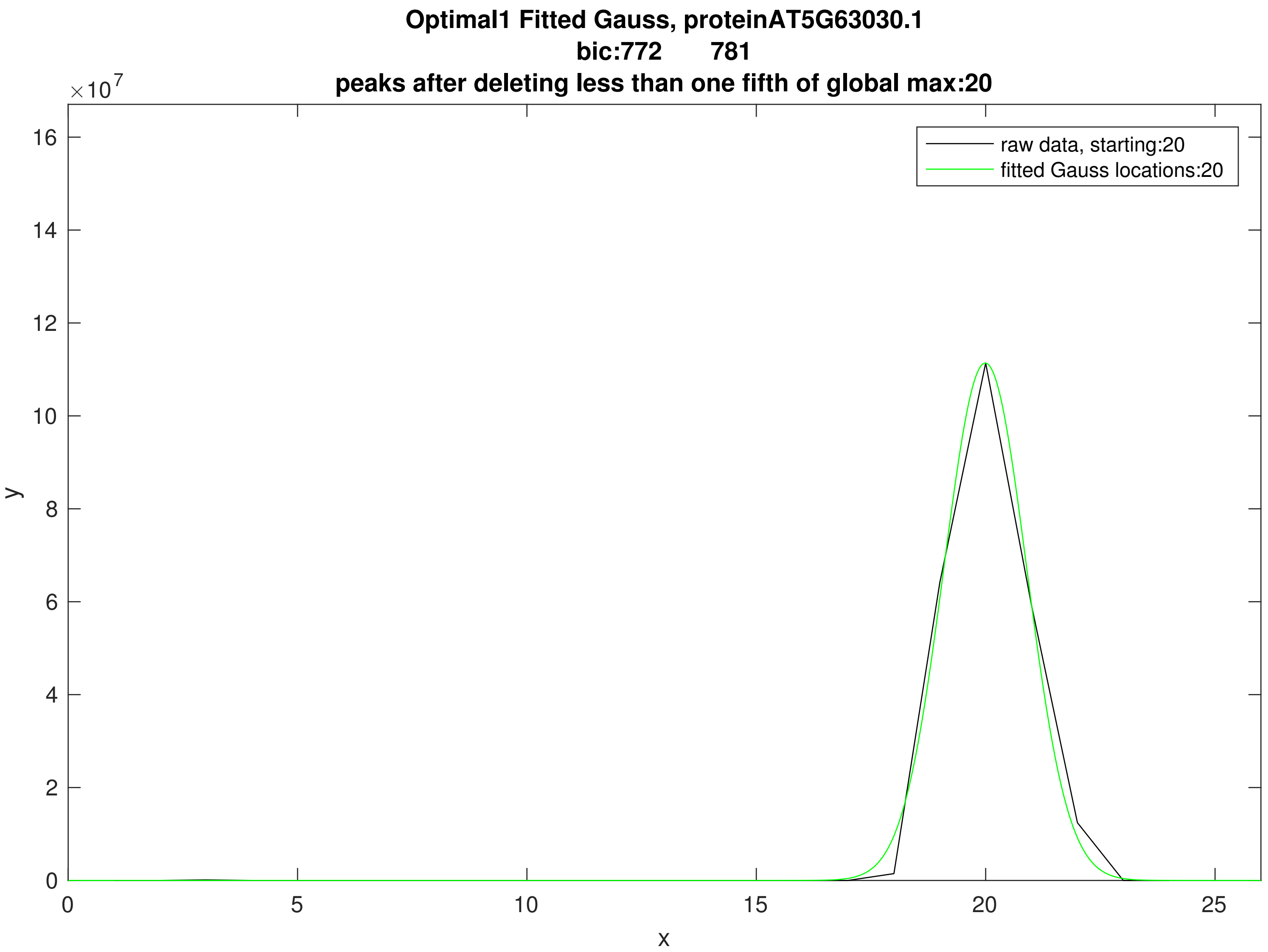

Optimal1 Fitted Gauss, proteinAT5G63030.1
bic:772 781
peaks after deleting less than one fifth of global max:20
×10^7
16
14
12
10
8
6
4
2
0
y
0
5
10
15
20
25
x
raw data, starting:20
fitted Gauss locations:20

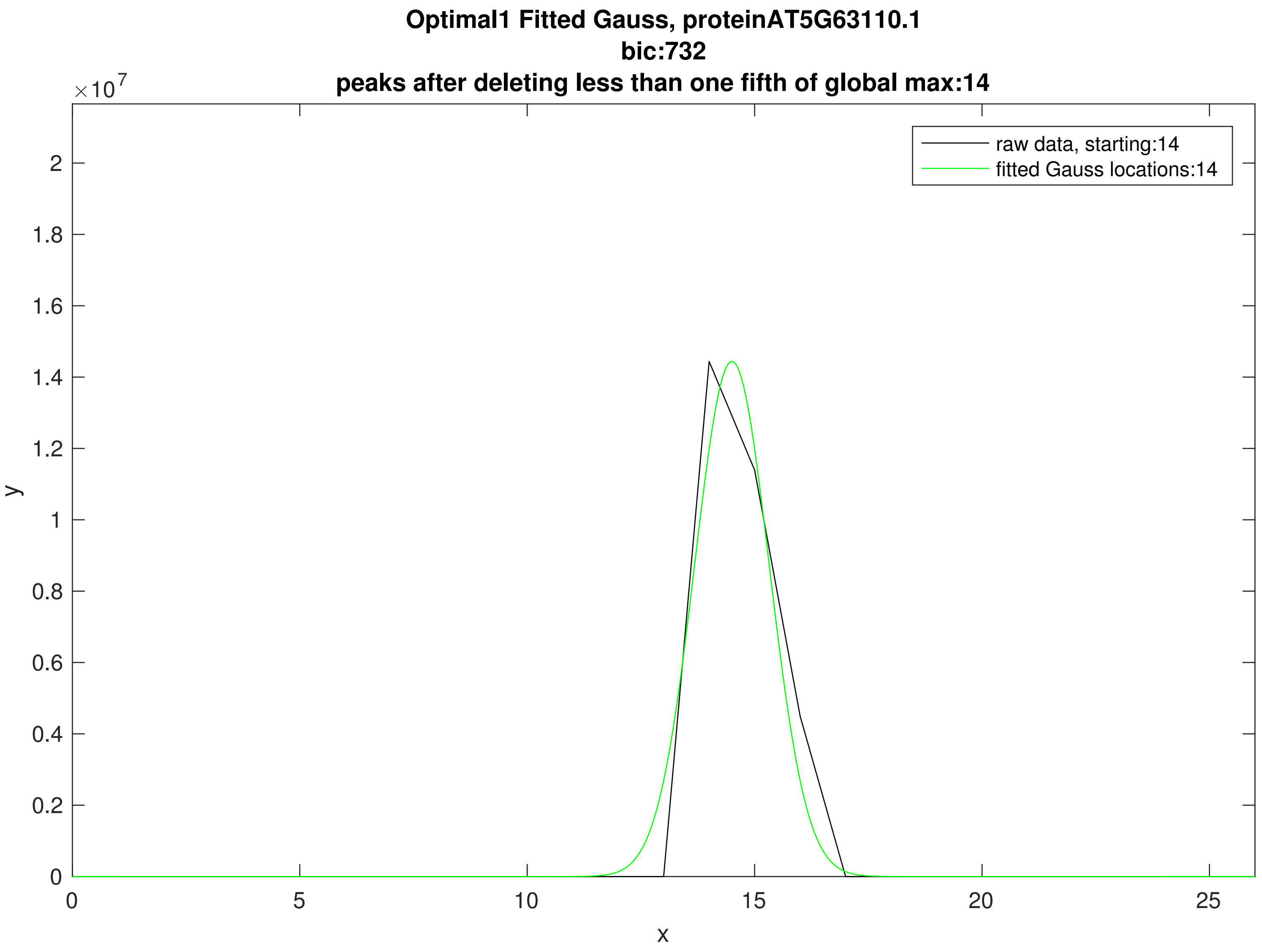
Optimal1 Fitted Gauss, proteinAT5G63110.1
bic:732
peaks after deleting less than one fifth of global max:14
×10^7
raw data, starting:14
fitted Gauss locations:14
2
1.8
1.6
1.4
1.2
1
0.8
0.6
0.4
0.2
0
y
0
5
10
15
20
25
x

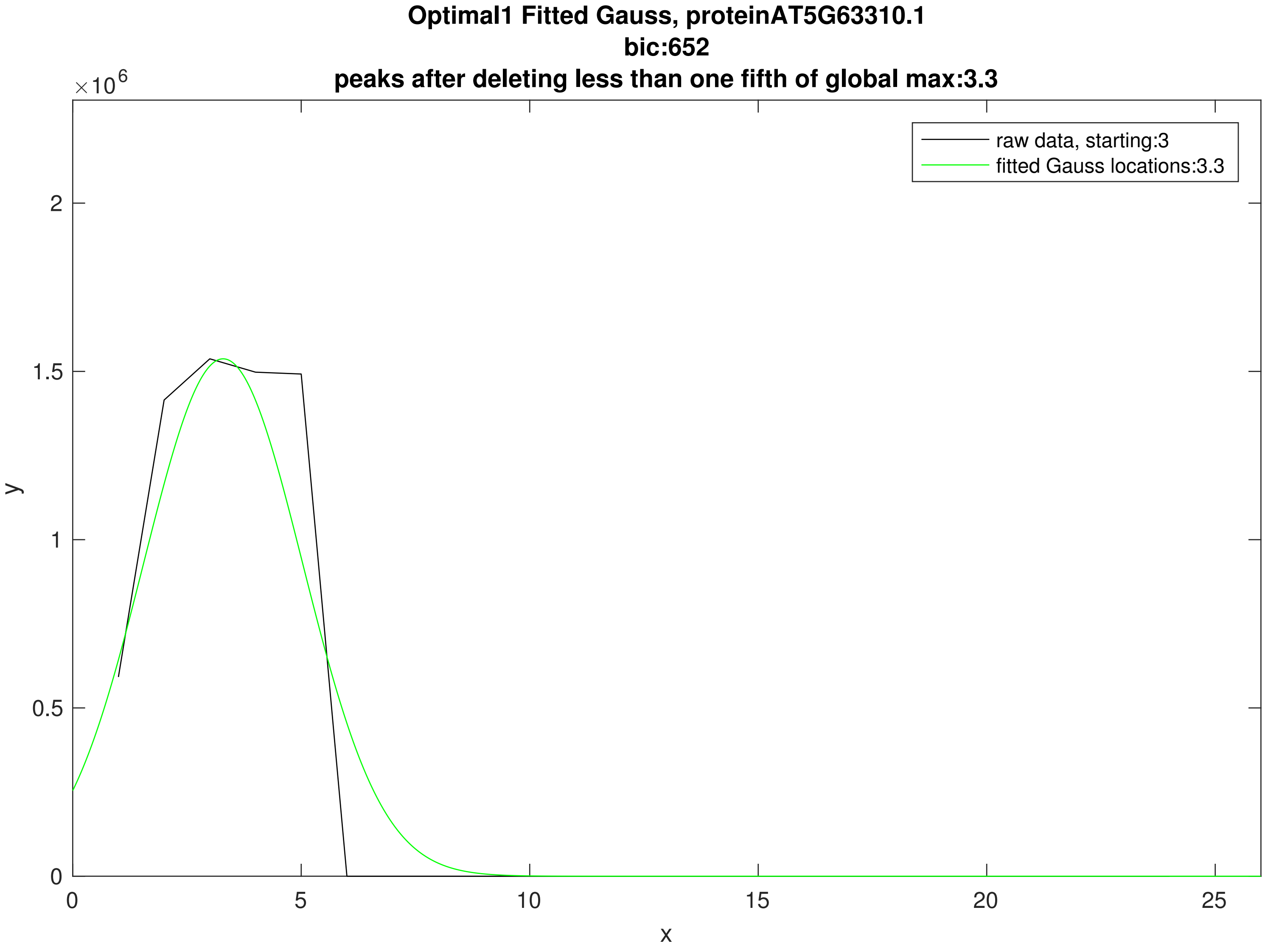
Optimal1 Fitted Gauss, proteinAT5G63310.1
bic:652
peaks after deleting less than one fifth of global max:3.3
×10^6
raw data, starting:3
fitted Gauss locations:3.3
2
1.5
1
0.5
0
y
0
5
10
15
20
25
x

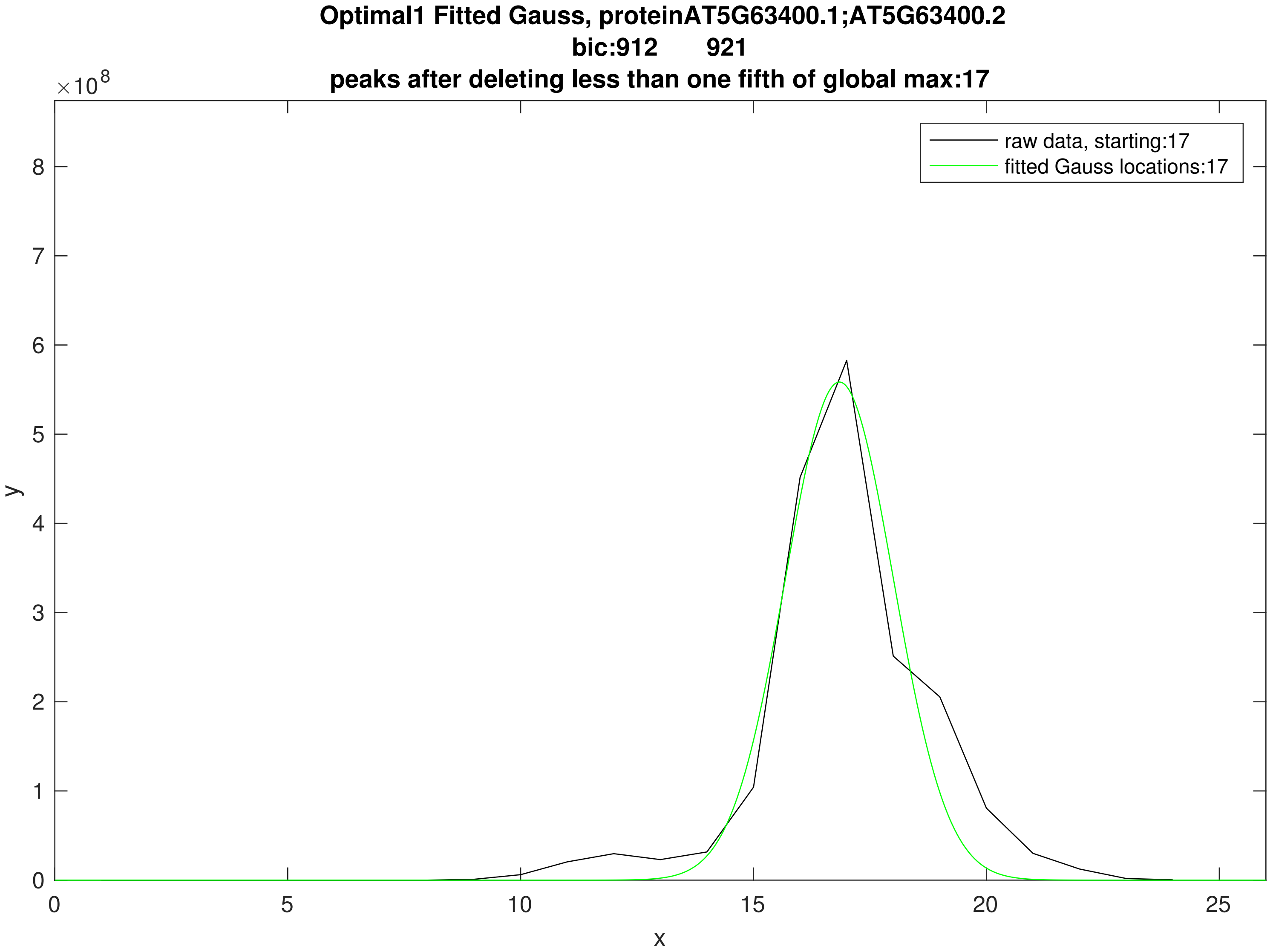
Optimal1 Fitted Gauss, proteinAT5G63400.1;AT5G63400.2
bic:912 921
peaks after deleting less than one fifth of global max:17
×10^8
raw data, starting:17
fitted Gauss locations:17
y
x
0
1
2
3
4
5
6
7
8
0
5
10
15
20
25

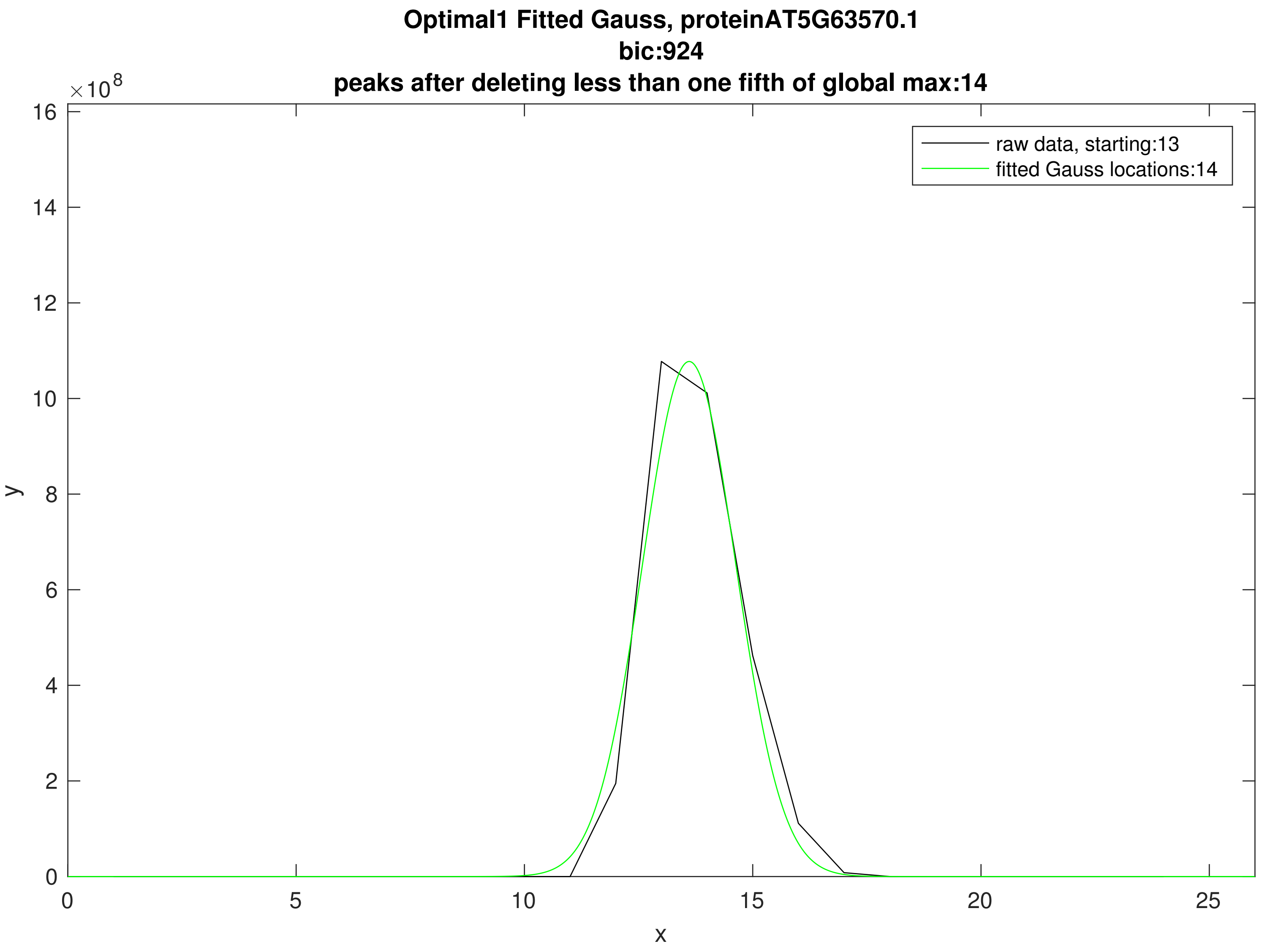
Optimal1 Fitted Gauss, proteinAT5G63570.1
bic:924
peaks after deleting less than one fifth of global max:14
×10^8
raw data, starting:13
fitted Gauss locations:14
y
x
0
2
4
6
8
10
12
14
16
0
5
10
15
20
25

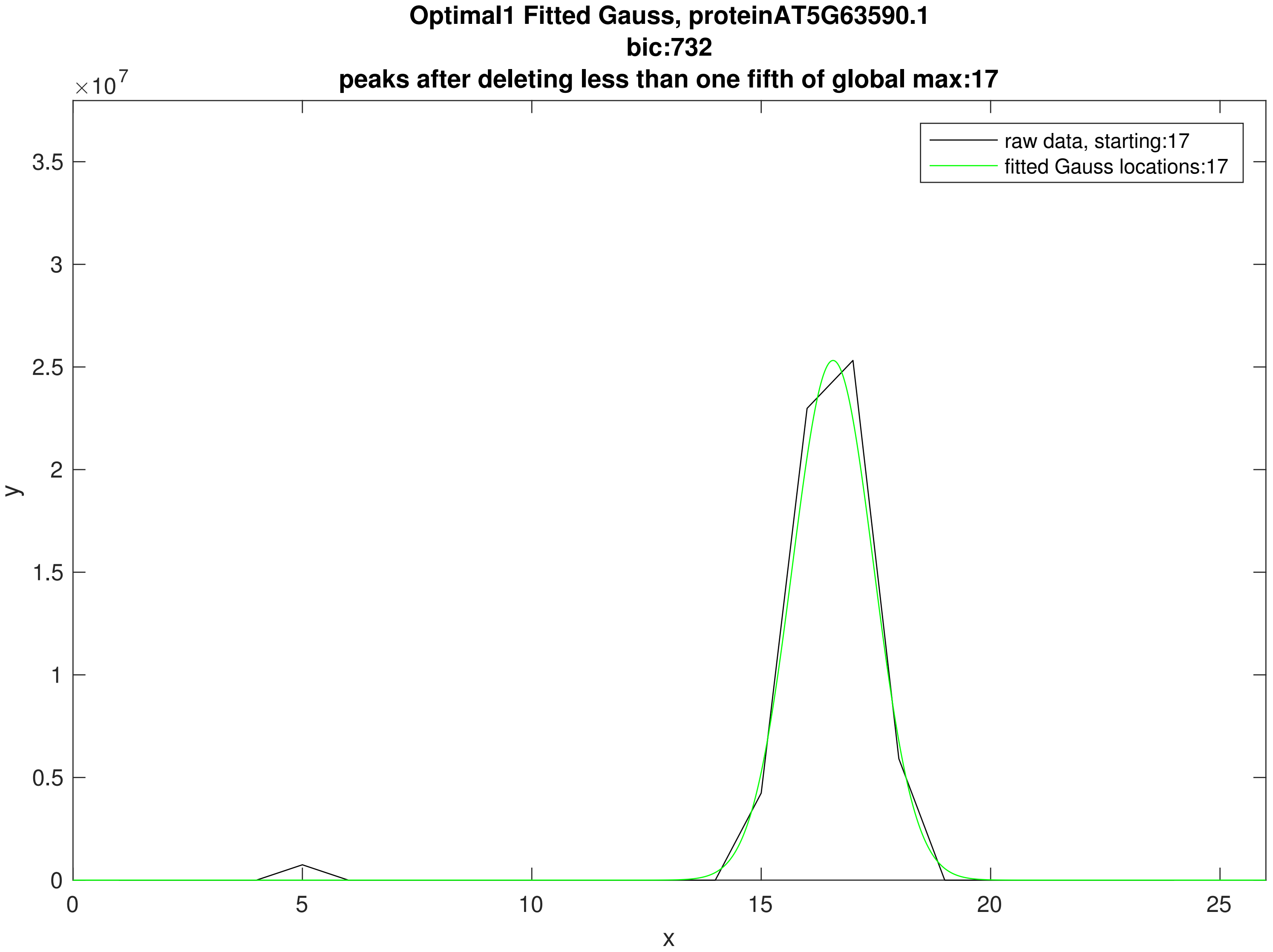
Optimal1 Fitted Gauss, proteinAT5G63590.1
bic:732
peaks after deleting less than one fifth of global max:17
×10^7
raw data, starting:17
fitted Gauss locations:17
y
x
3.5
3
2.5
2
1.5
1
0.5
0
5
10
15
20
25

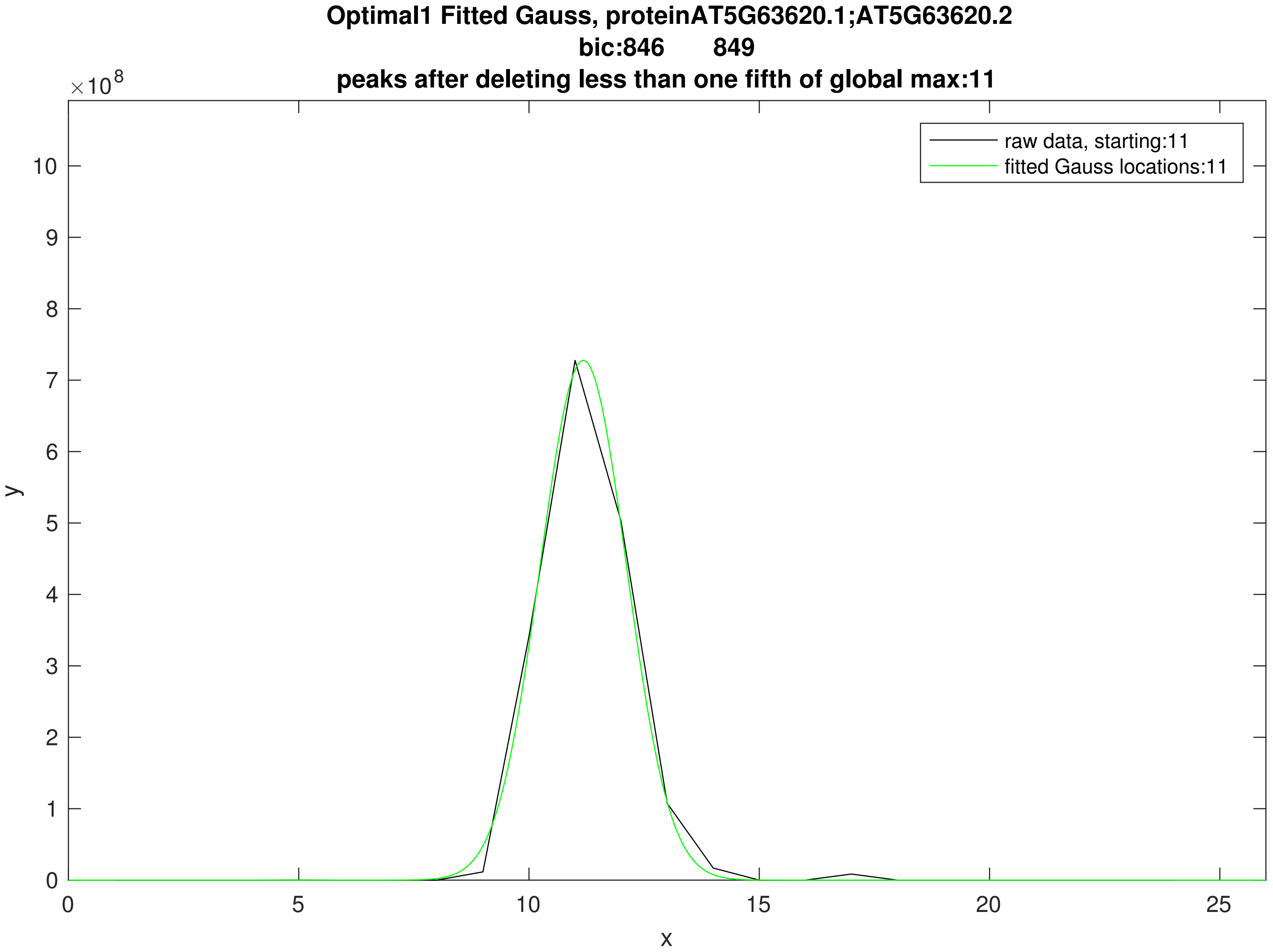

Optimal1 Fitted Gauss, proteinAT5G63620.1;AT5G63620.2
bic:846 849
peaks after deleting less than one fifth of global max:11
×10^8
raw data, starting:11
fitted Gauss locations:11
y
x
0
1
2
3
4
5
6
7
8
9
10
0
5
10
15
20
25

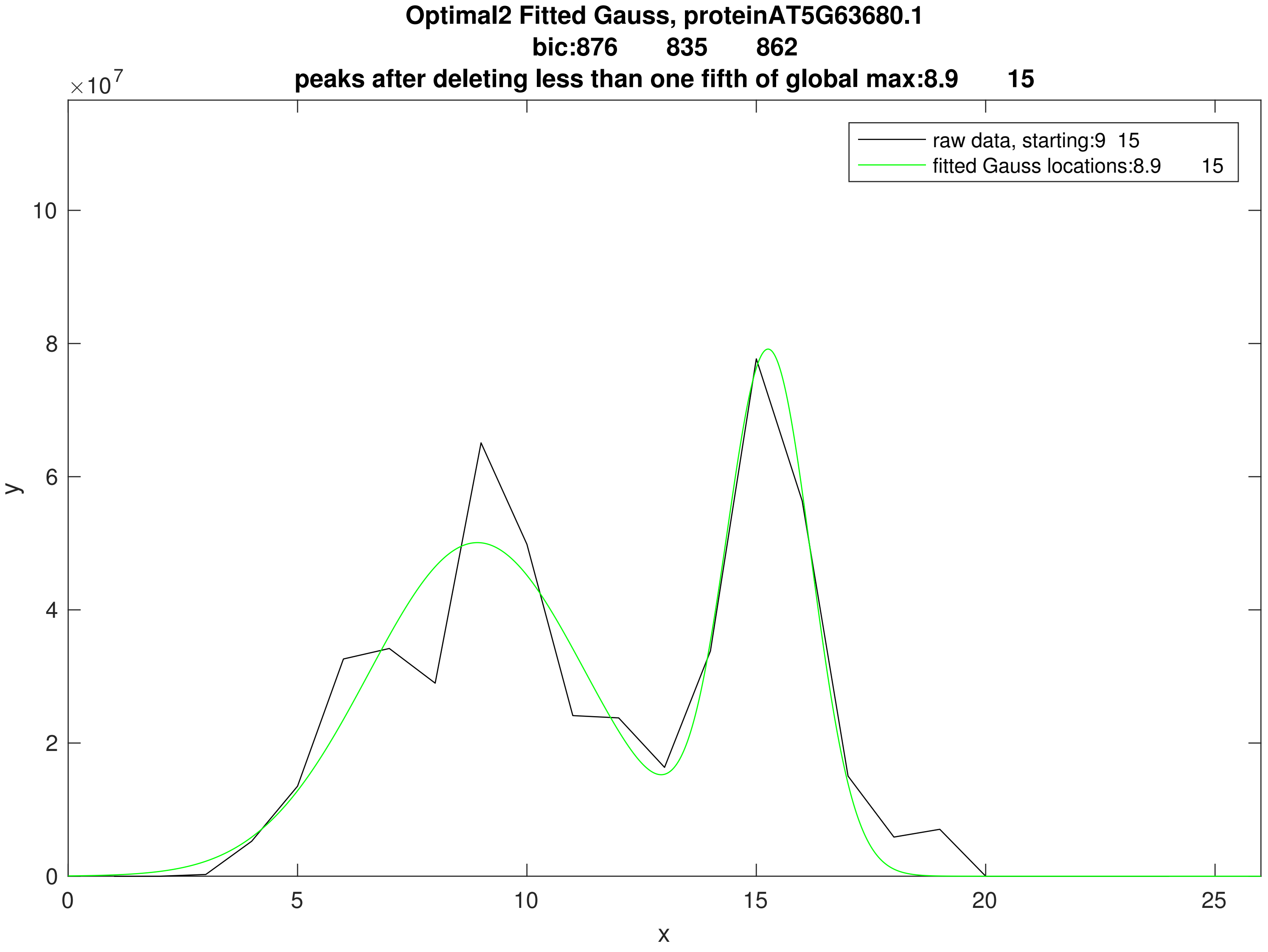

Optimal2 Fitted Gauss, proteinAT5G63680.1
bic:876 835 862
peaks after deleting less than one fifth of global max:8.9 15
×10^7
raw data, starting:9 15
fitted Gauss locations:8.9 15
y
x
0
2
4
6
8
10
0
5
10
15
20
25

#### Optimal1 Fitted Gauss, proteinAT5G63750.1
#### bic:811 819
#### peaks after deleting less than one fifth of global max:12

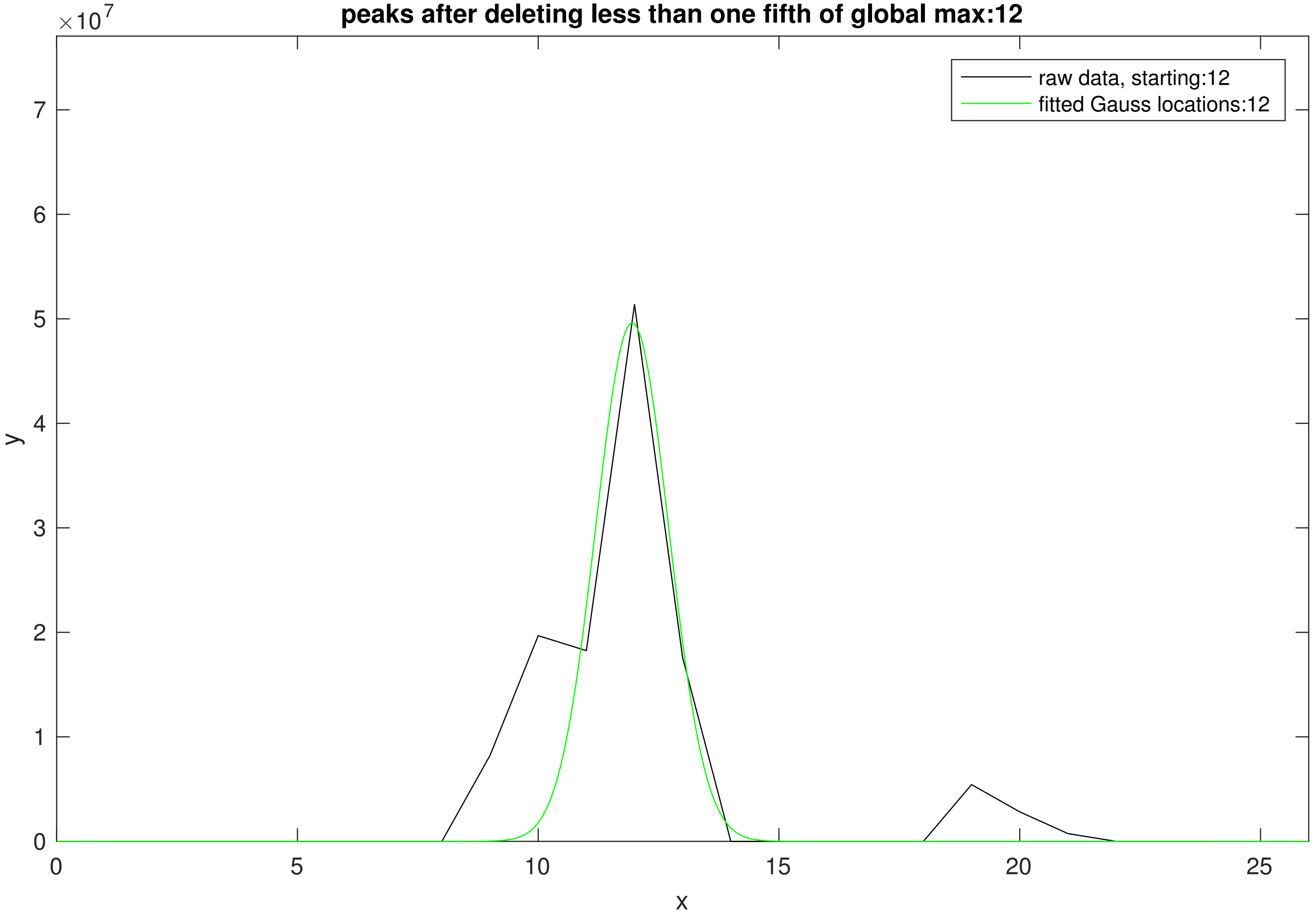

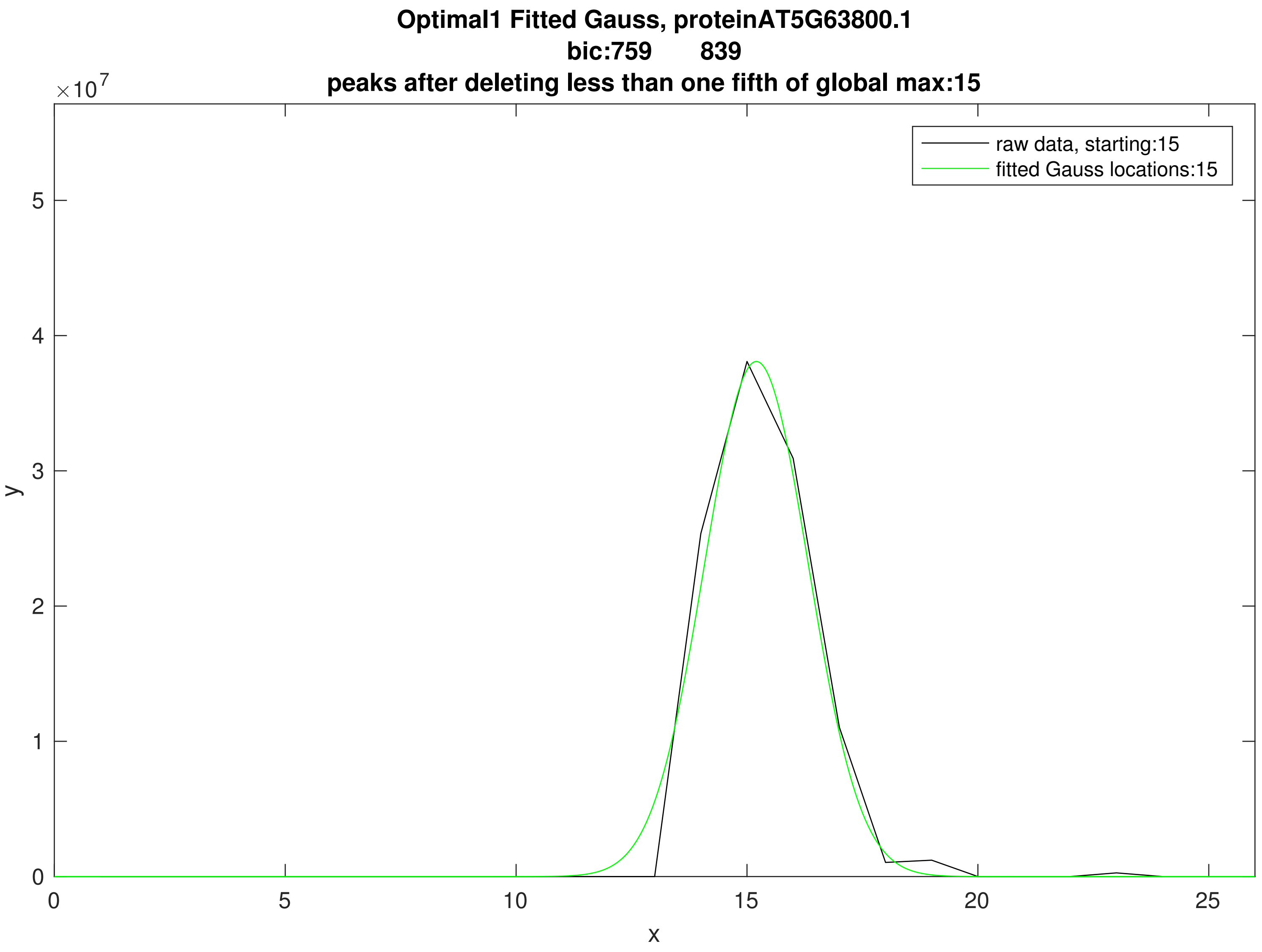
Optimal1 Fitted Gauss, proteinAT5G63800.1
bic:759 839
peaks after deleting less than one fifth of global max:15
×10^7
raw data, starting:15
fitted Gauss locations:15
y
x
0
1
2
3
4
5
0
5
10
15
20
25

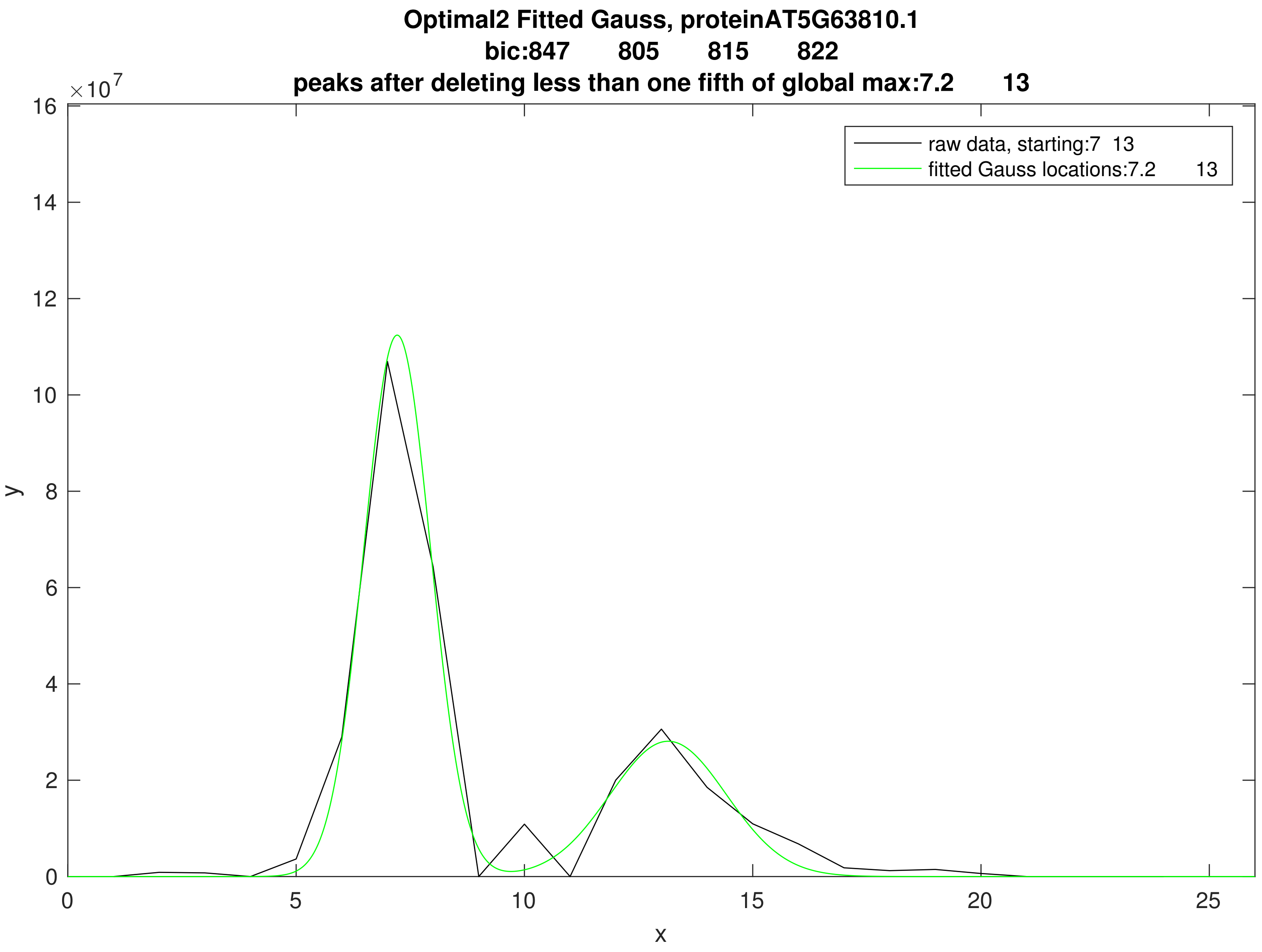

Optimal2 Fitted Gauss, proteinAT5G63810.1
bic:847 805 815 822
peaks after deleting less than one fifth of global max:7.2 13
×10^7
16
14
12
10
8
6
4
2
0
y
0
5
10
15
20
25
x
raw data, starting:7 13
fitted Gauss locations:7.2 13

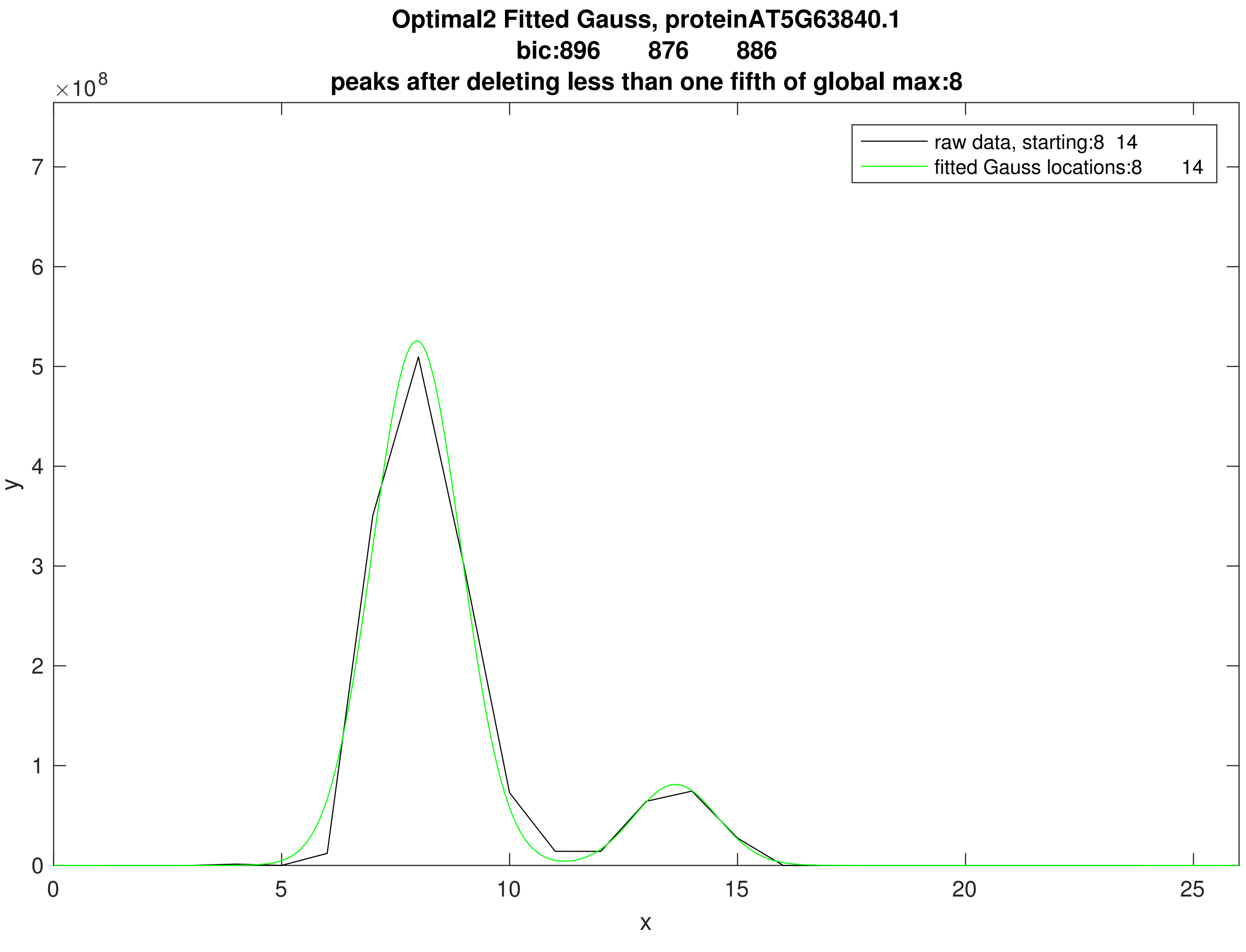
Optimal2 Fitted Gauss, proteinAT5G63840.1
bic:896 876 886
peaks after deleting less than one fifth of global max:8
×10^8
raw data, starting:8 14
fitted Gauss locations:8 14
y
x
0
1
2
3
4
5
6
7
5
10
15
20
25

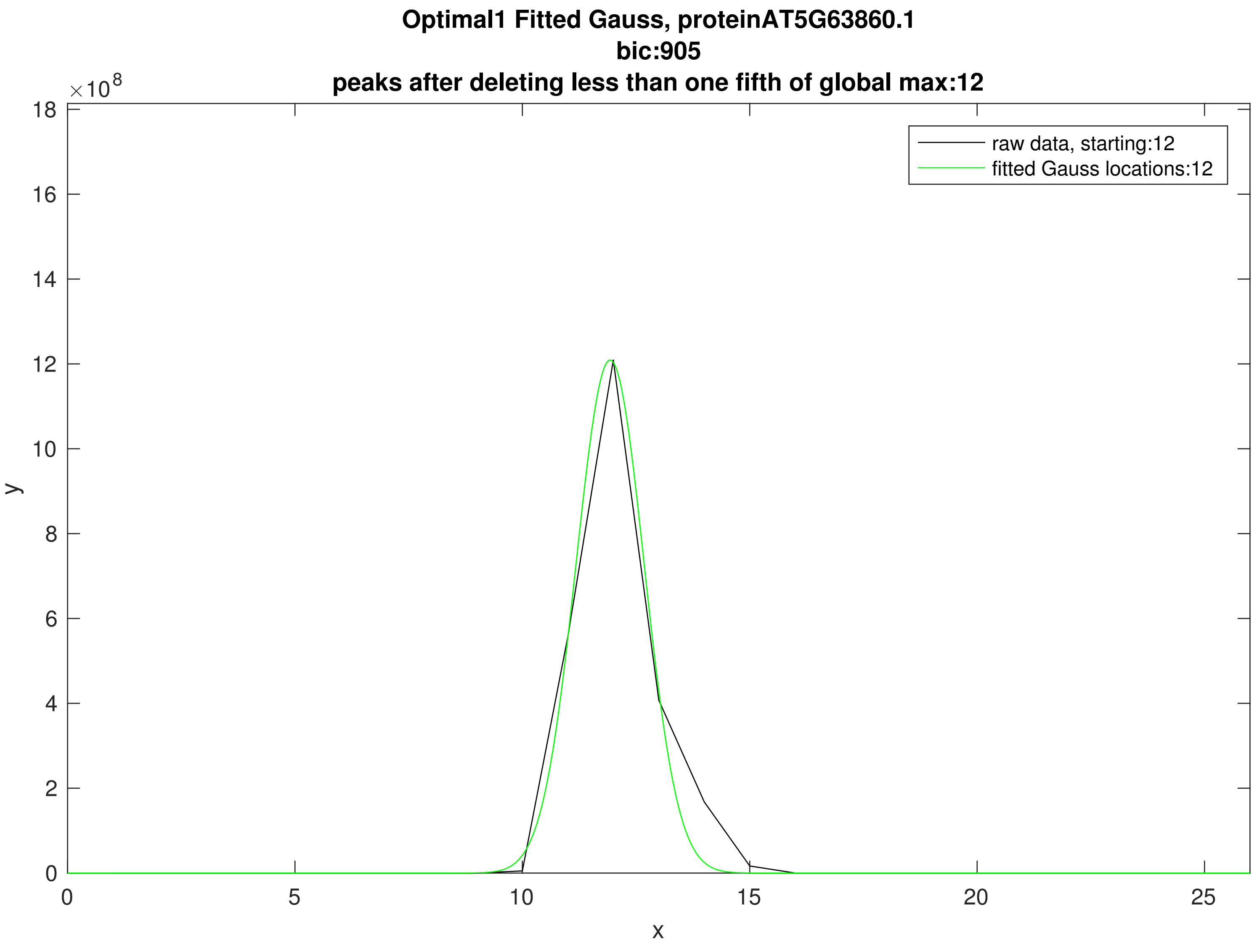

Optimal1 Fitted Gauss, proteinAT5G63860.1
bic:905
peaks after deleting less than one fifth of global max:12
×10^8
18
16
14
12
10
8
6
4
2
0
y
0
5
10
15
20
25
x
raw data, starting:12
fitted Gauss locations:12

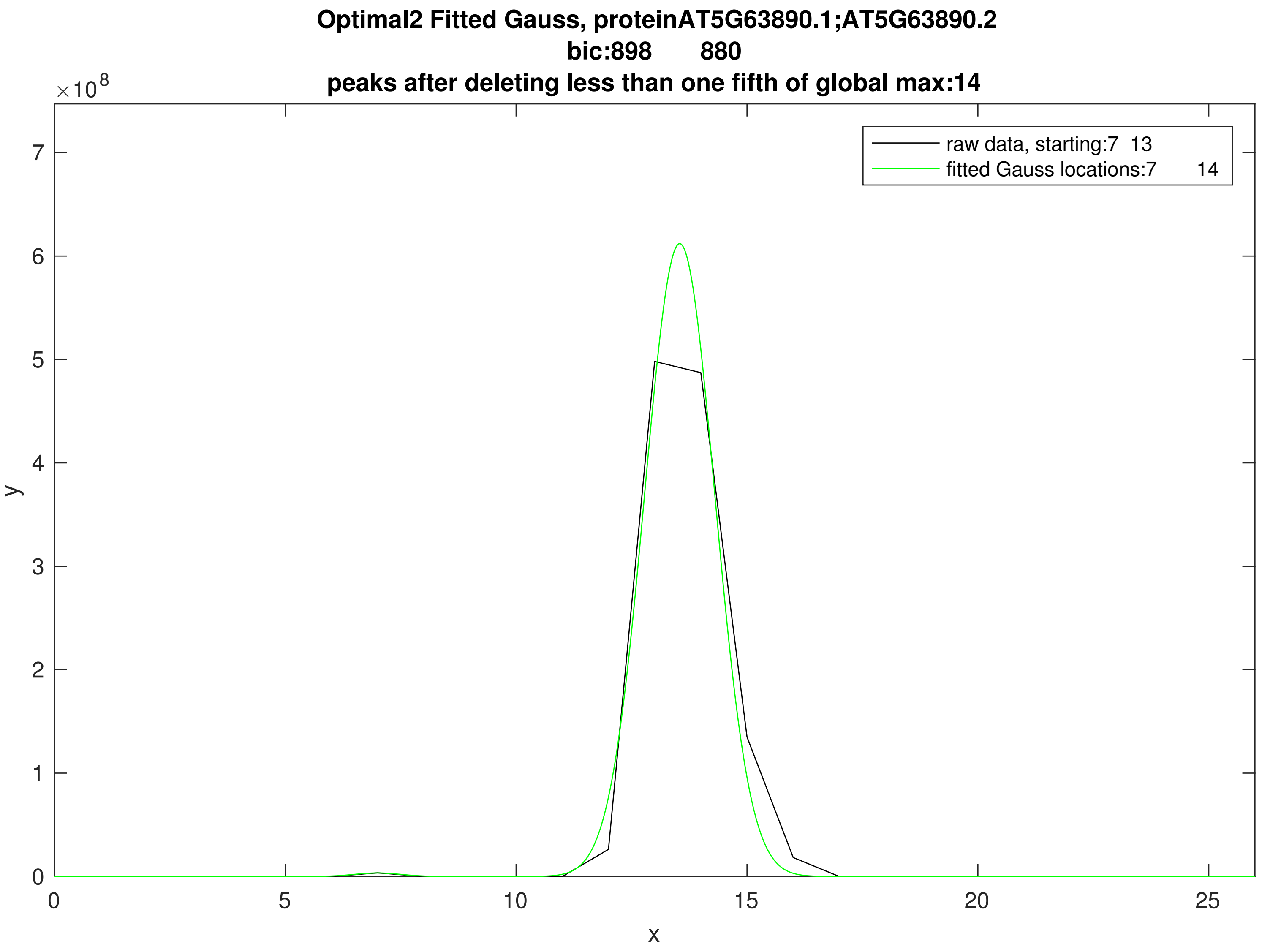

Optimal2 Fitted Gauss, proteinAT5G63890.1;AT5G63890.2
bic:898 880
peaks after deleting less than one fifth of global max:14
×10^8
raw data, starting:7 13
fitted Gauss locations:7 14
y
x
0
1
2
3
4
5
6
7
0
5
10
15
20
25

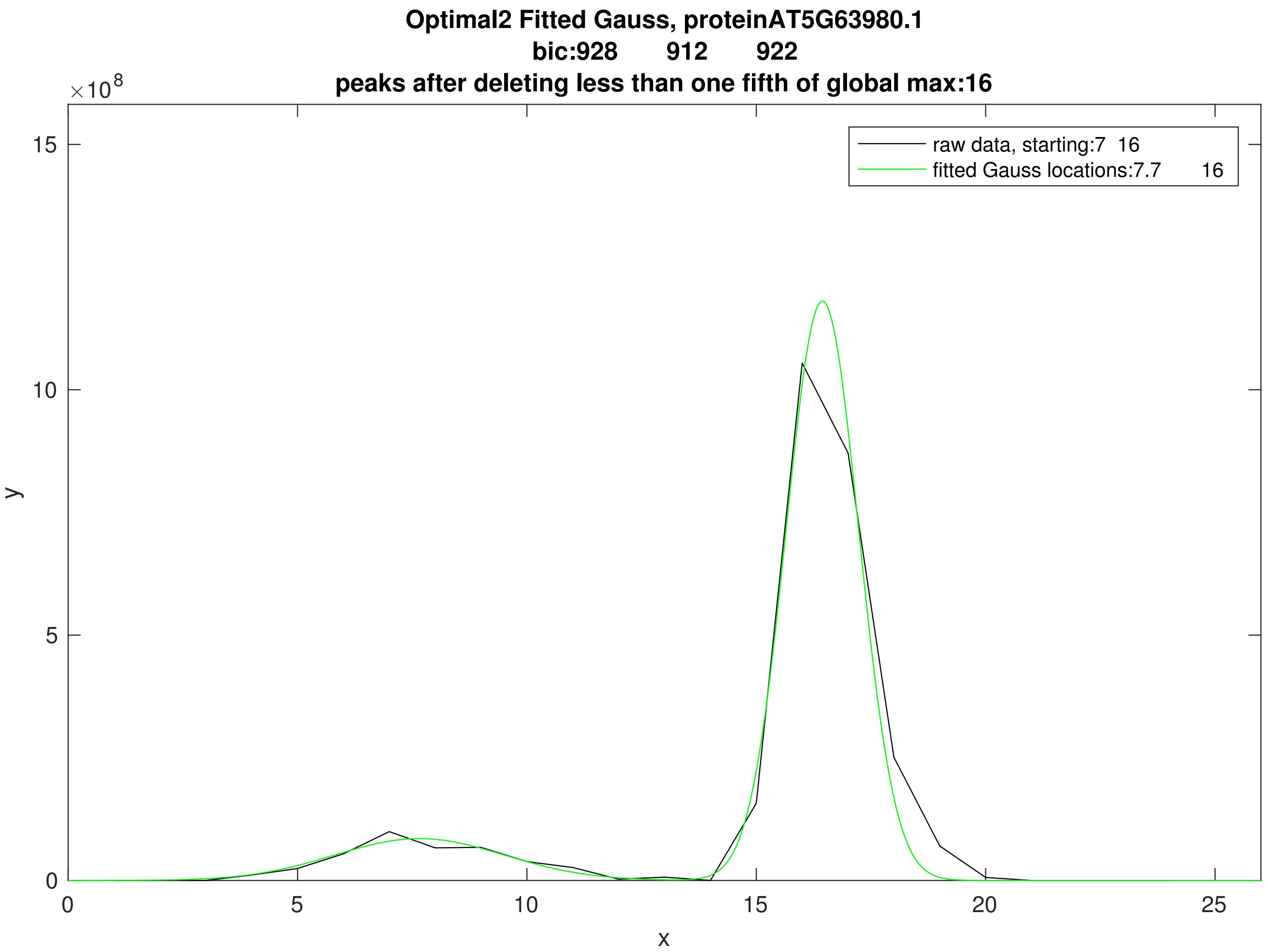
Optimal2 Fitted Gauss, proteinAT5G63980.1
bic:928 912 922
peaks after deleting less than one fifth of global max:16
×10^8
15
10
5
0
y
0
5
10
15
20
25
x
raw data, starting:7 16
fitted Gauss locations:7.7 16

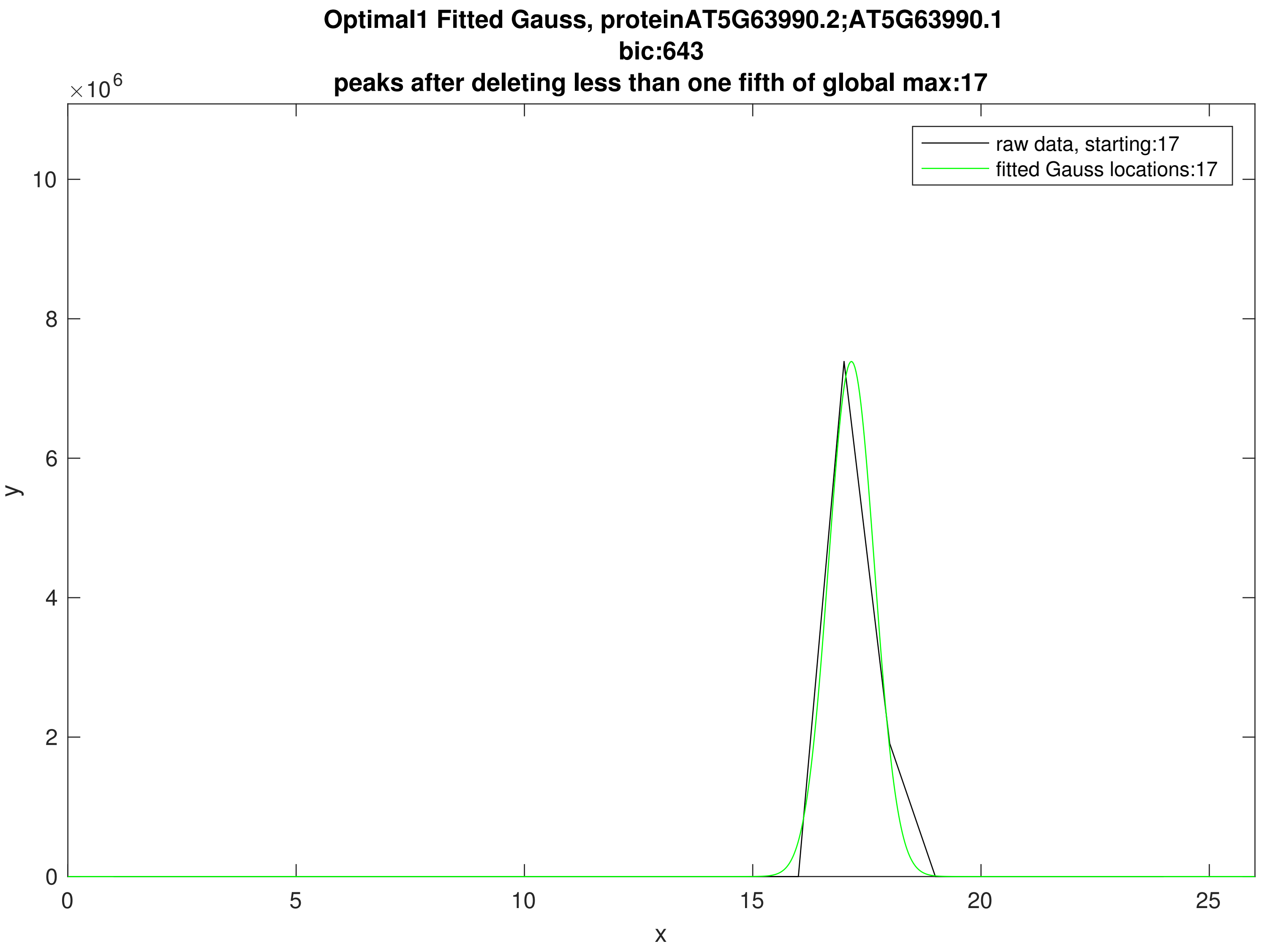

Optimal1 Fitted Gauss, proteinAT5G63990.2;AT5G63990.1
bic:643
peaks after deleting less than one fifth of global max:17
×10^6
raw data, starting:17
fitted Gauss locations:17
y
x
0
2
4
6
8
10
0
5
10
15
20
25

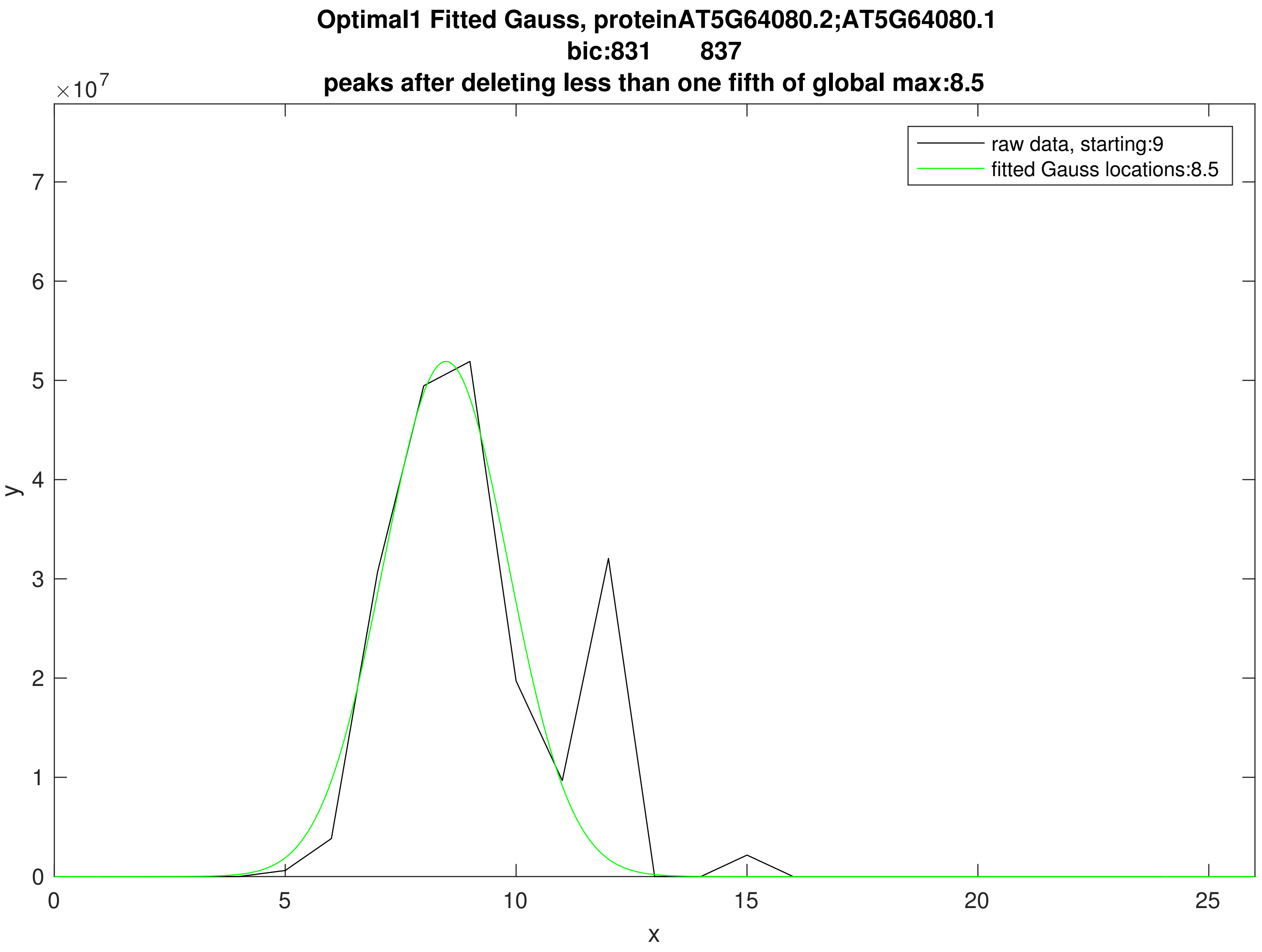
Optimal1 Fitted Gauss, proteinAT5G64080.2;AT5G64080.1
bic:831 837
peaks after deleting less than one fifth of global max:8.5
×10^7
raw data, starting:9
fitted Gauss locations:8.5
y
x
0
1
2
3
4
5
6
7
0
5
10
15
20
25

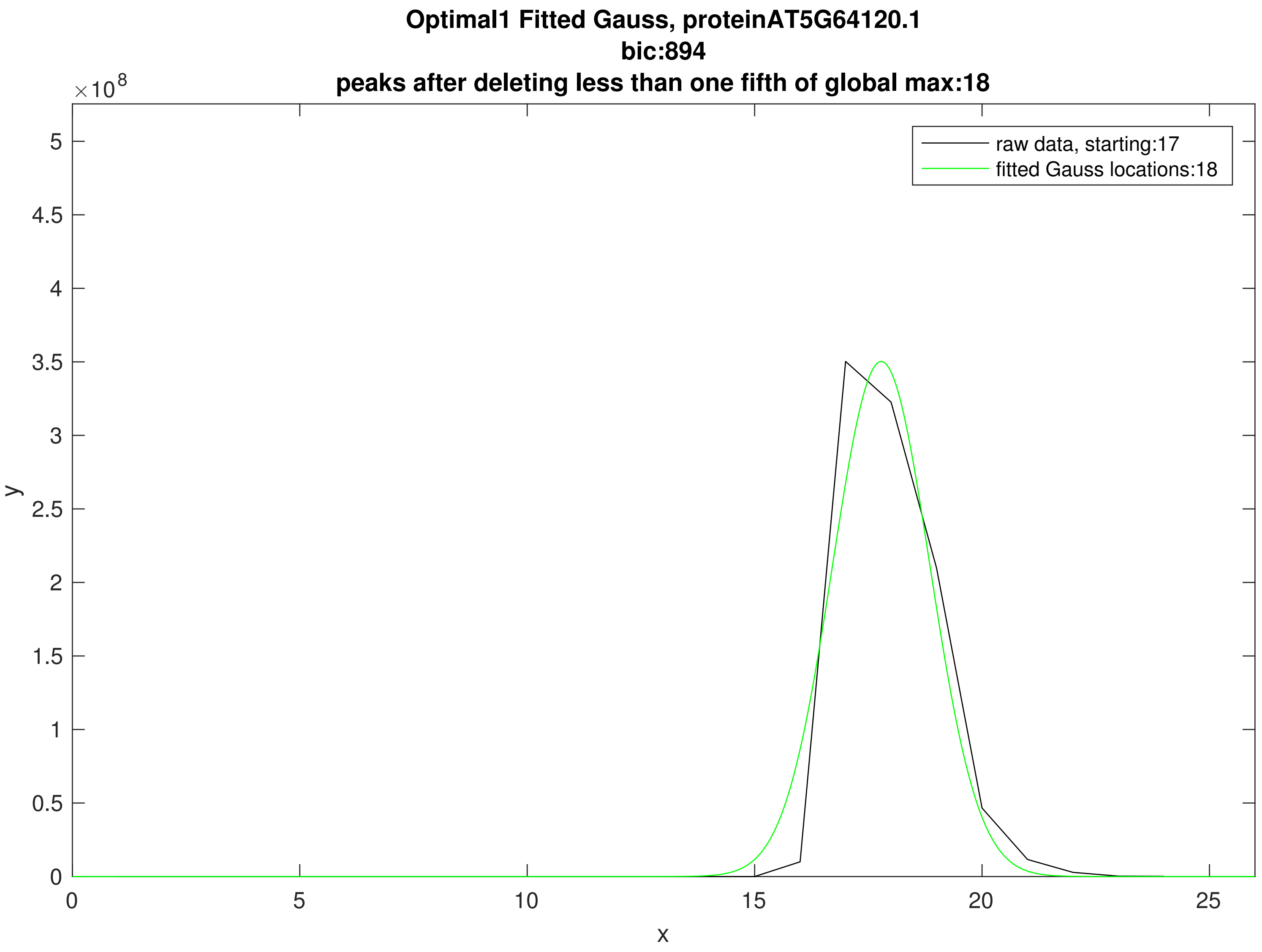
Optimal1 Fitted Gauss, proteinAT5G64120.1
bic:894
peaks after deleting less than one fifth of global max:18
×10^8
5
4.5
4
3.5
3
2.5
2
1.5
1
0.5
0
y
0
5
10
15
20
25
x
raw data, starting:17
fitted Gauss locations:18

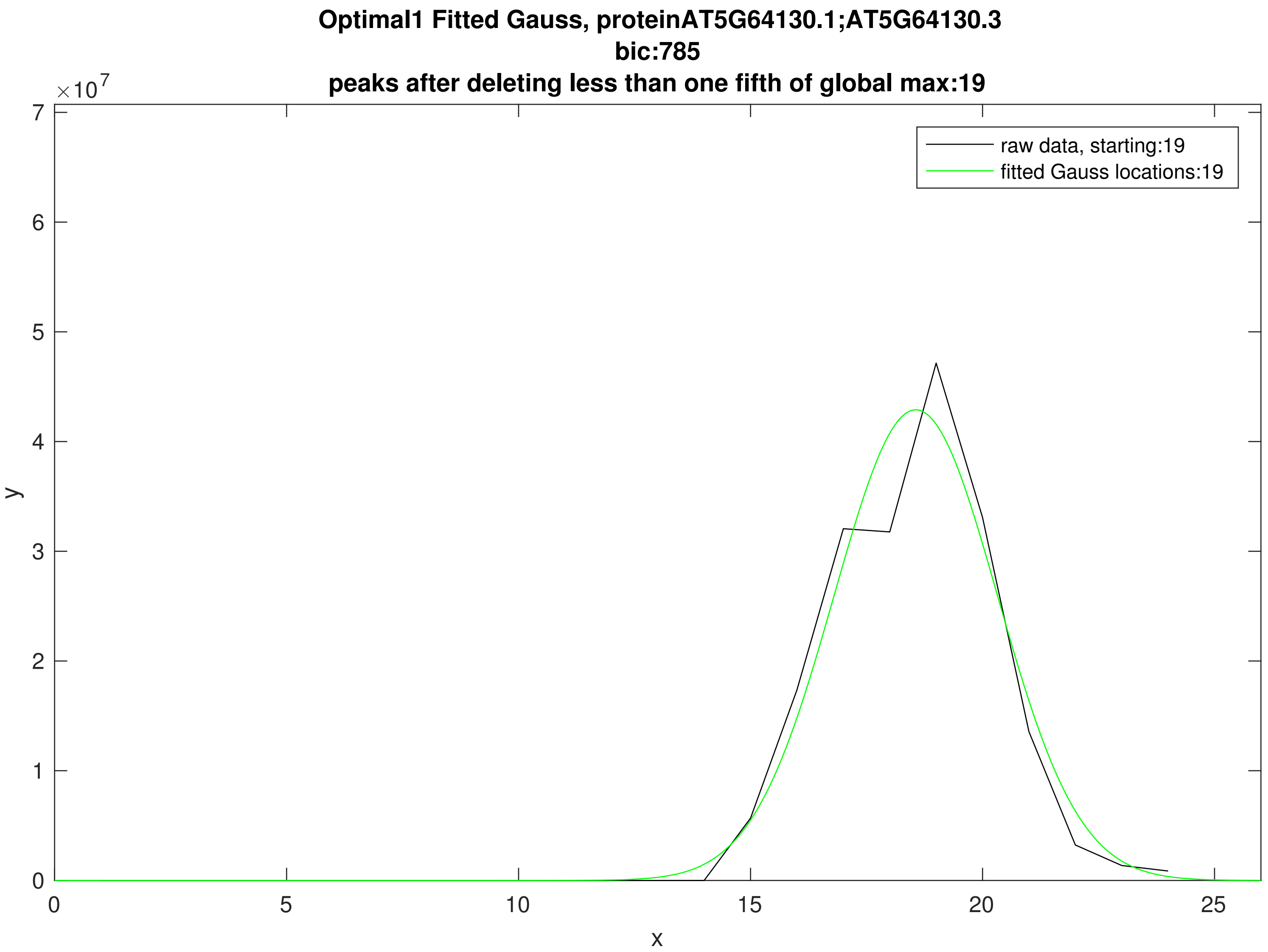
Optimal1 Fitted Gauss, proteinAT5G64130.1;AT5G64130.3
bic:785
peaks after deleting less than one fifth of global max:19
×10^7
raw data, starting:19
fitted Gauss locations:19
y
x
0
1
2
3
4
5
6
7
0
5
10
15
20
25

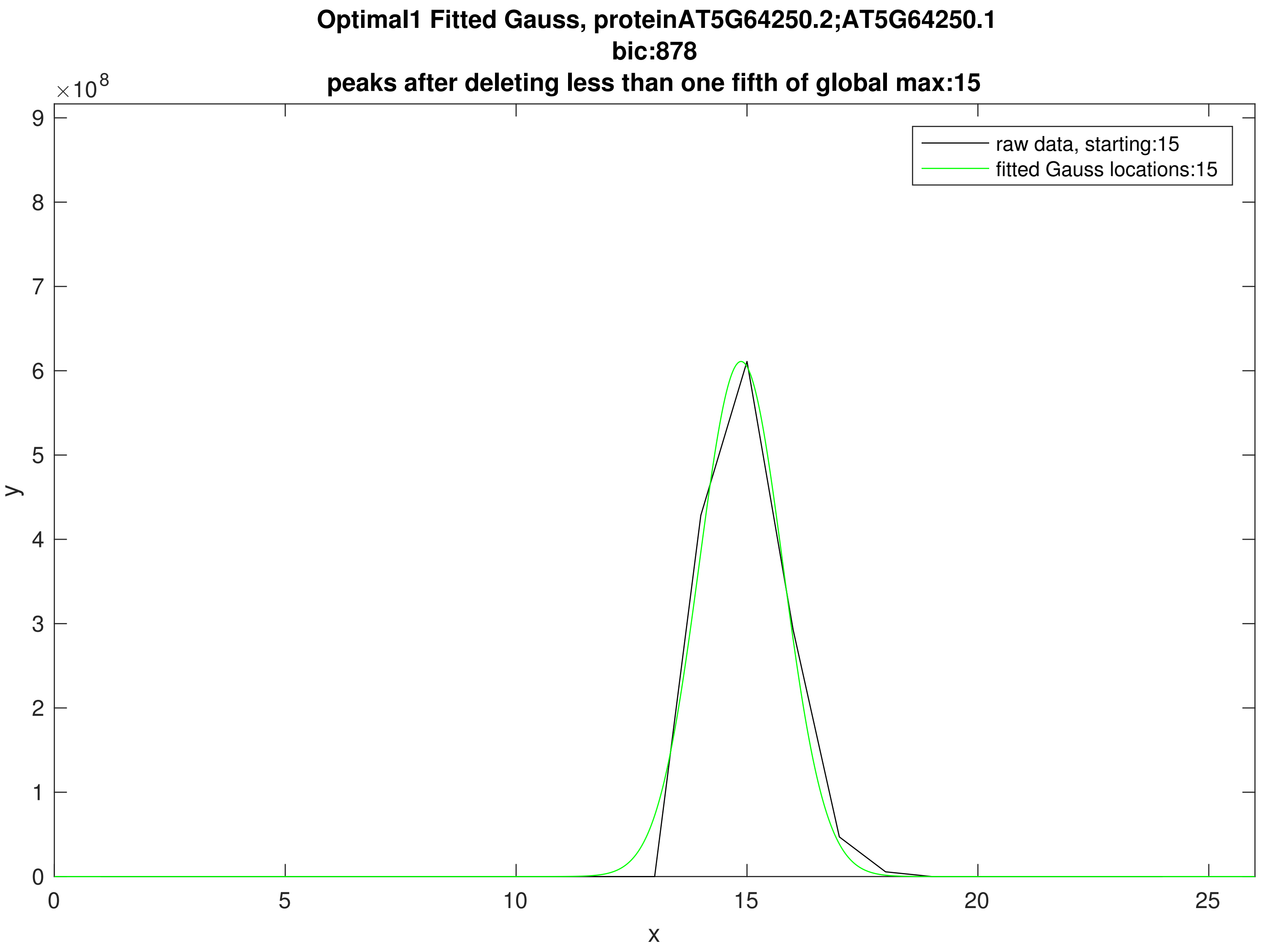

Optimal1 Fitted Gauss, proteinAT5G64250.2;AT5G64250.1
bic:878
peaks after deleting less than one fifth of global max:15
×10^8
raw data, starting:15
fitted Gauss locations:15
y
x
0
1
2
3
4
5
6
7
8
9
0
5
10
15
20
25

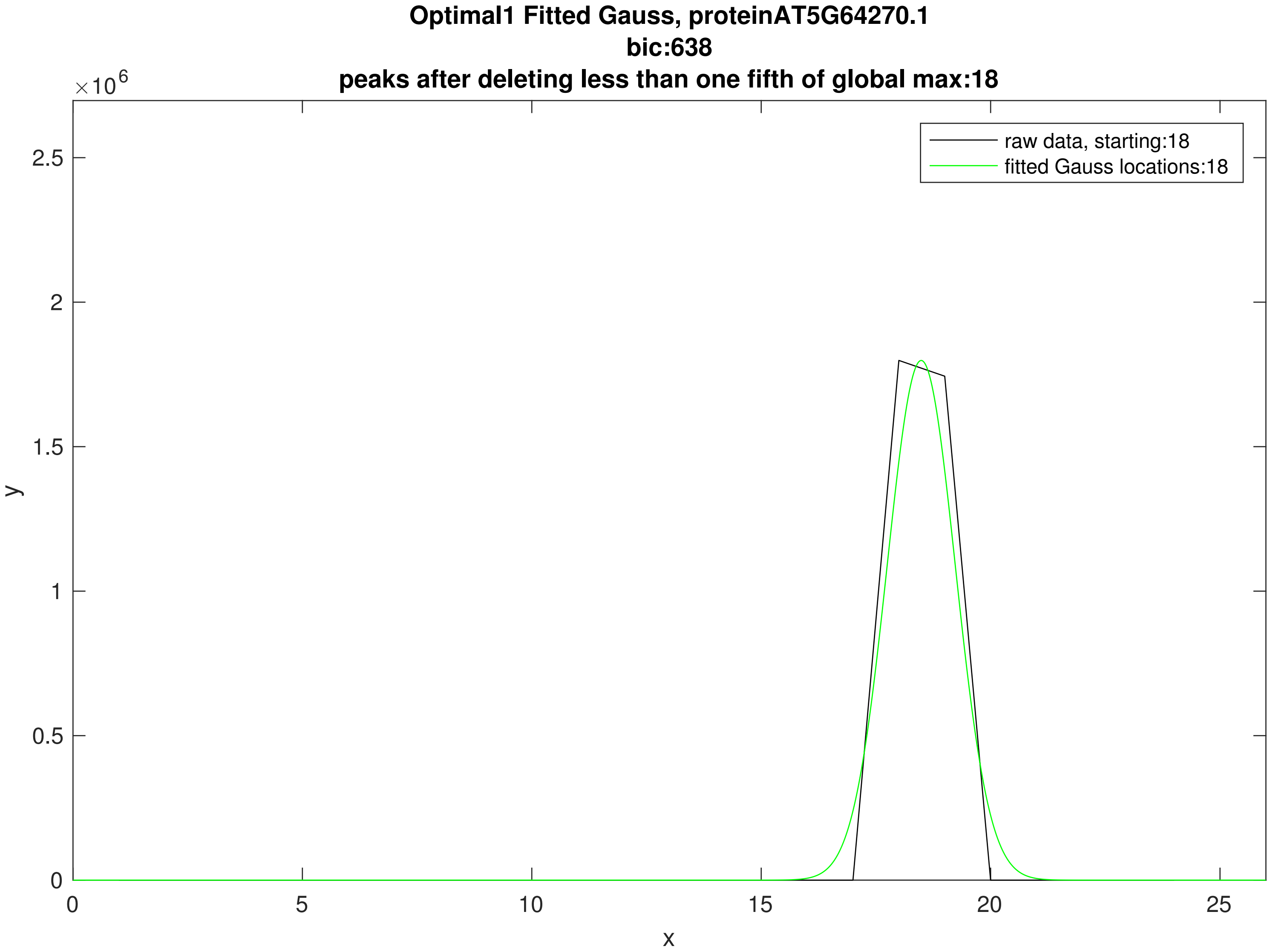
Optimal1 Fitted Gauss, proteinAT5G64270.1
bic:638
peaks after deleting less than one fifth of global max:18
×10^6
raw data, starting:18
fitted Gauss locations:18
y
x
0
0.5
1
1.5
2
2.5
0
5
10
15
20
25

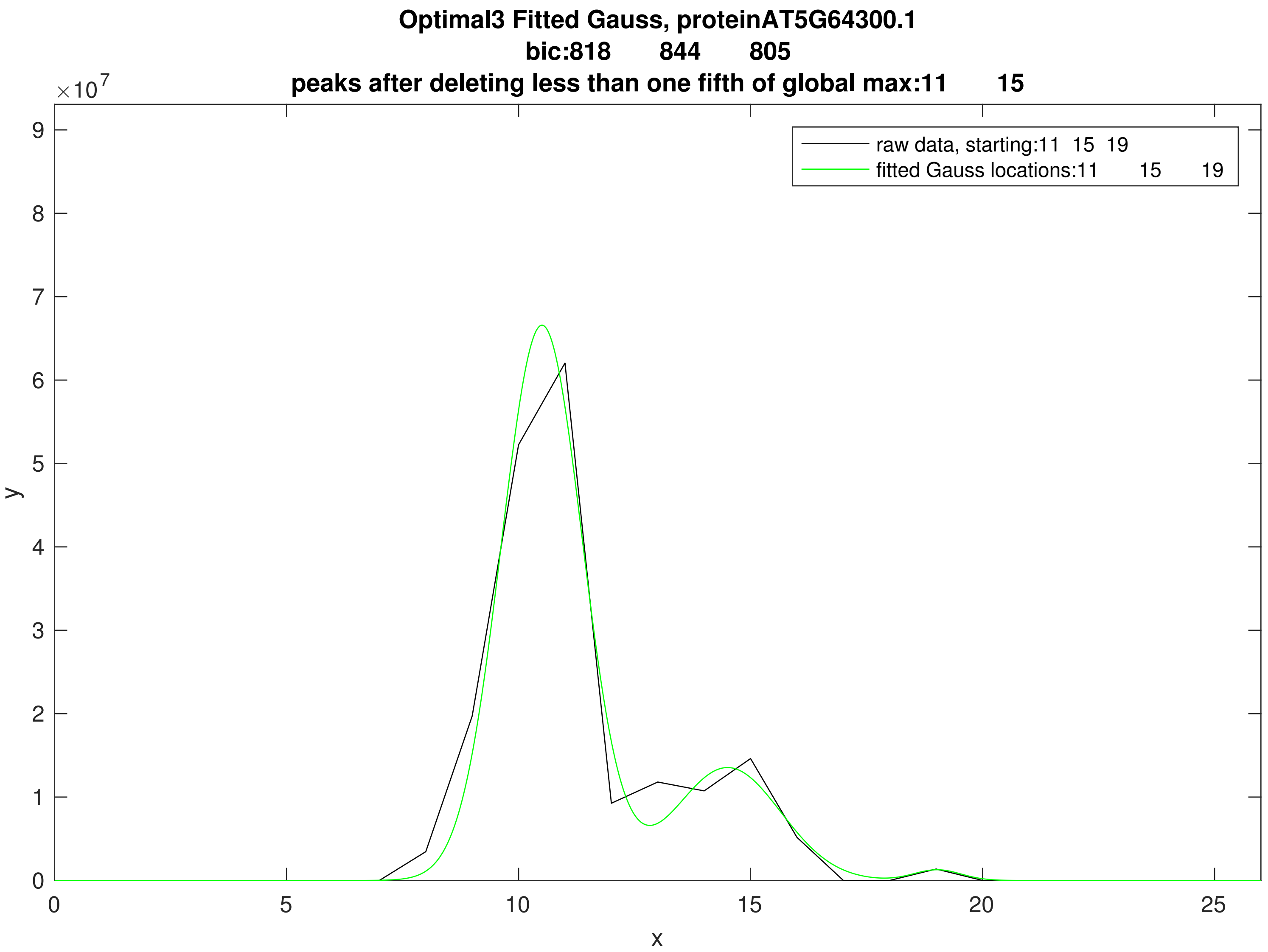
Optimal3 Fitted Gauss, proteinAT5G64300.1
bic:818 844 805
peaks after deleting less than one fifth of global max:11 15
raw data, starting:11 15 19
fitted Gauss locations:11 15 19
×10^7
y
x

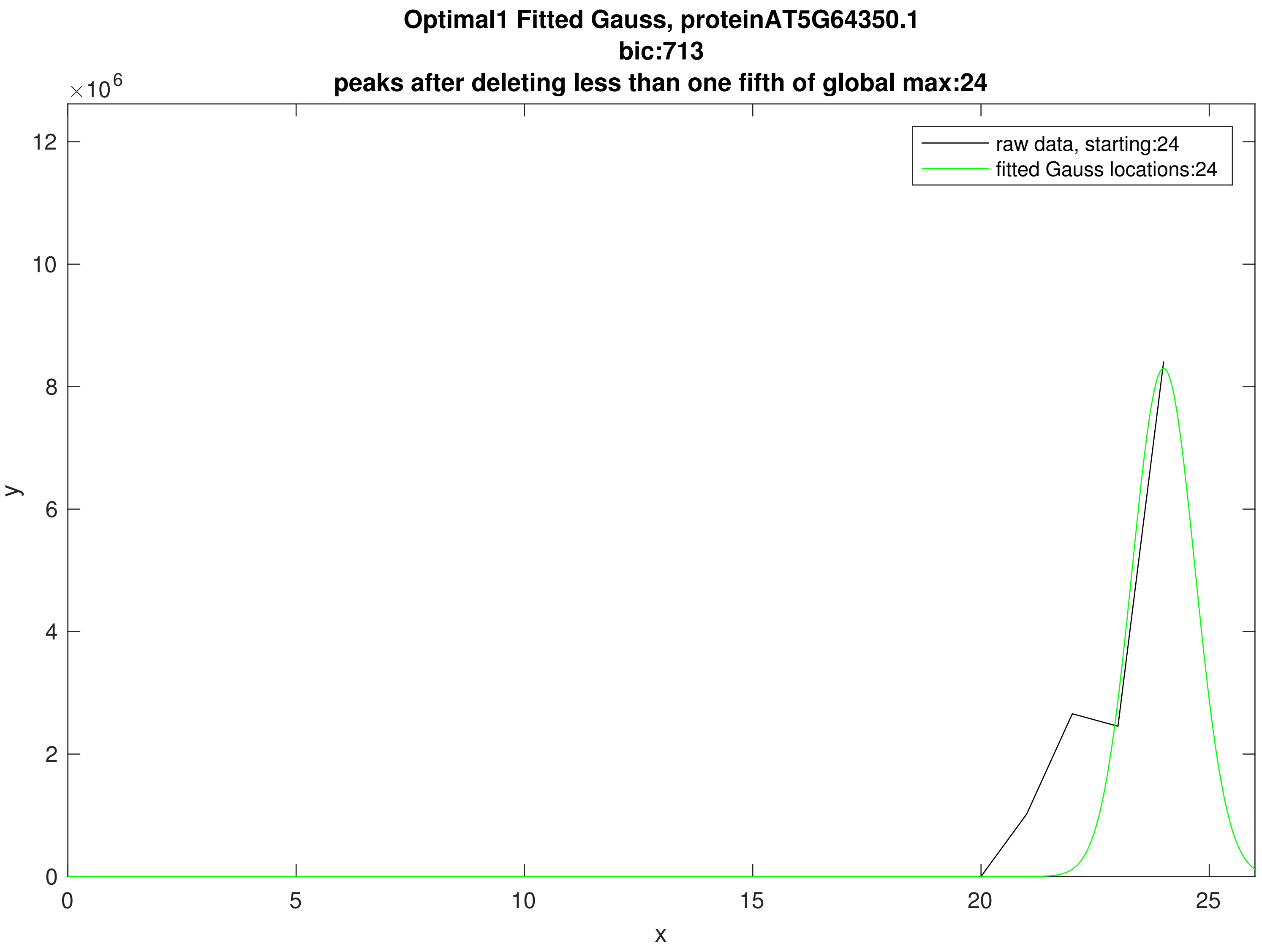
Optimal1 Fitted Gauss, proteinAT5G64350.1
bic:713
peaks after deleting less than one fifth of global max:24
×10^6
raw data, starting:24
fitted Gauss locations:24
12
10
8
6
4
2
0
0
5
10
15
20
25
x
y

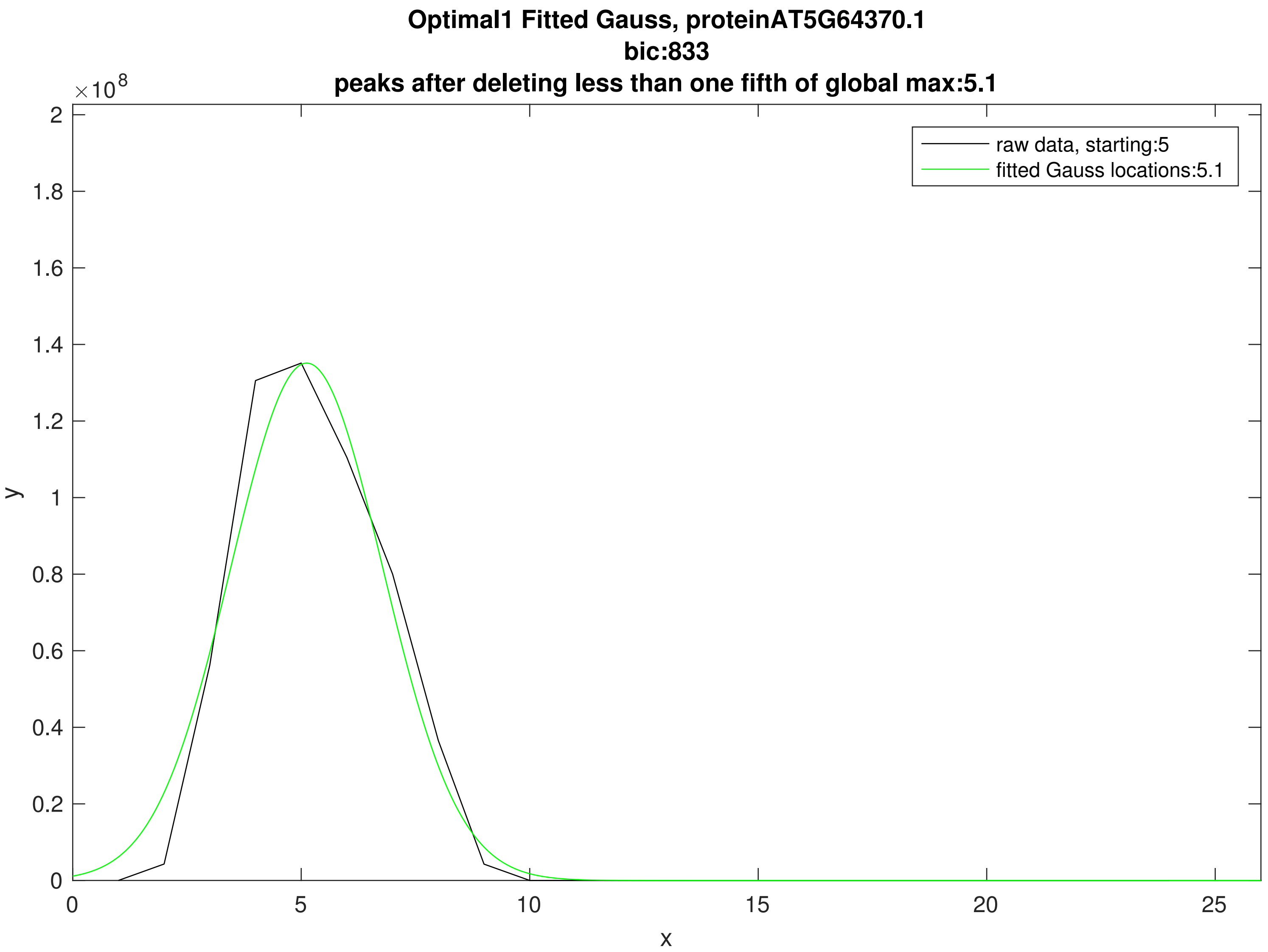

Optimal1 Fitted Gauss, proteinAT5G64370.1
bic:833
peaks after deleting less than one fifth of global max:5.1
×10^8
2
1.8
1.6
1.4
1.2
1
0.8
0.6
0.4
0.2
0
y
0
5
10
15
20
25
x
raw data, starting:5
fitted Gauss locations:5.1

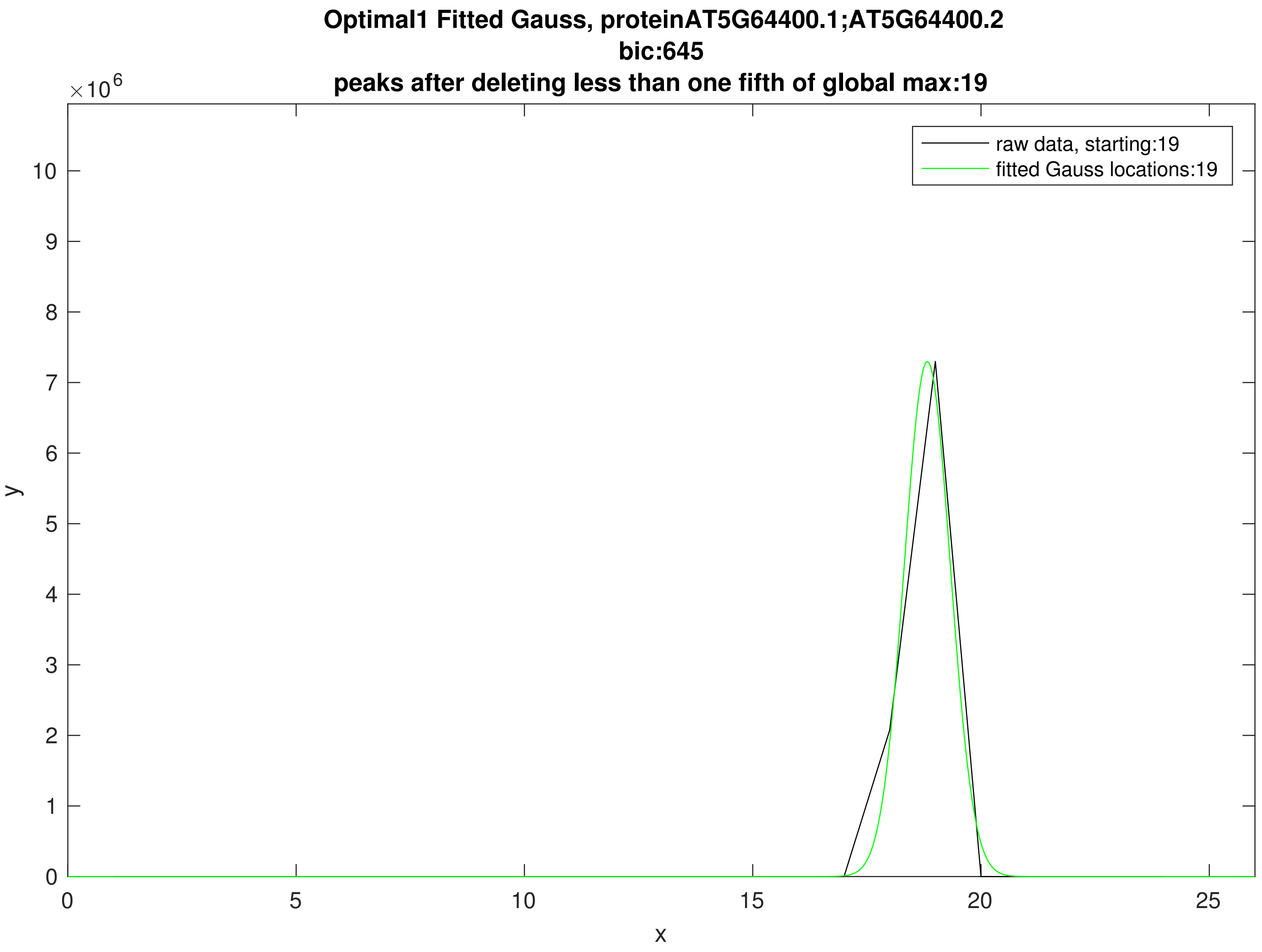
Optimal1 Fitted Gauss, proteinAT5G64400.1;AT5G64400.2
bic:645
peaks after deleting less than one fifth of global max:19
×10^6
raw data, starting:19
fitted Gauss locations:19
y
x
0
1
2
3
4
5
6
7
8
9
10
0
5
10
15
20
25

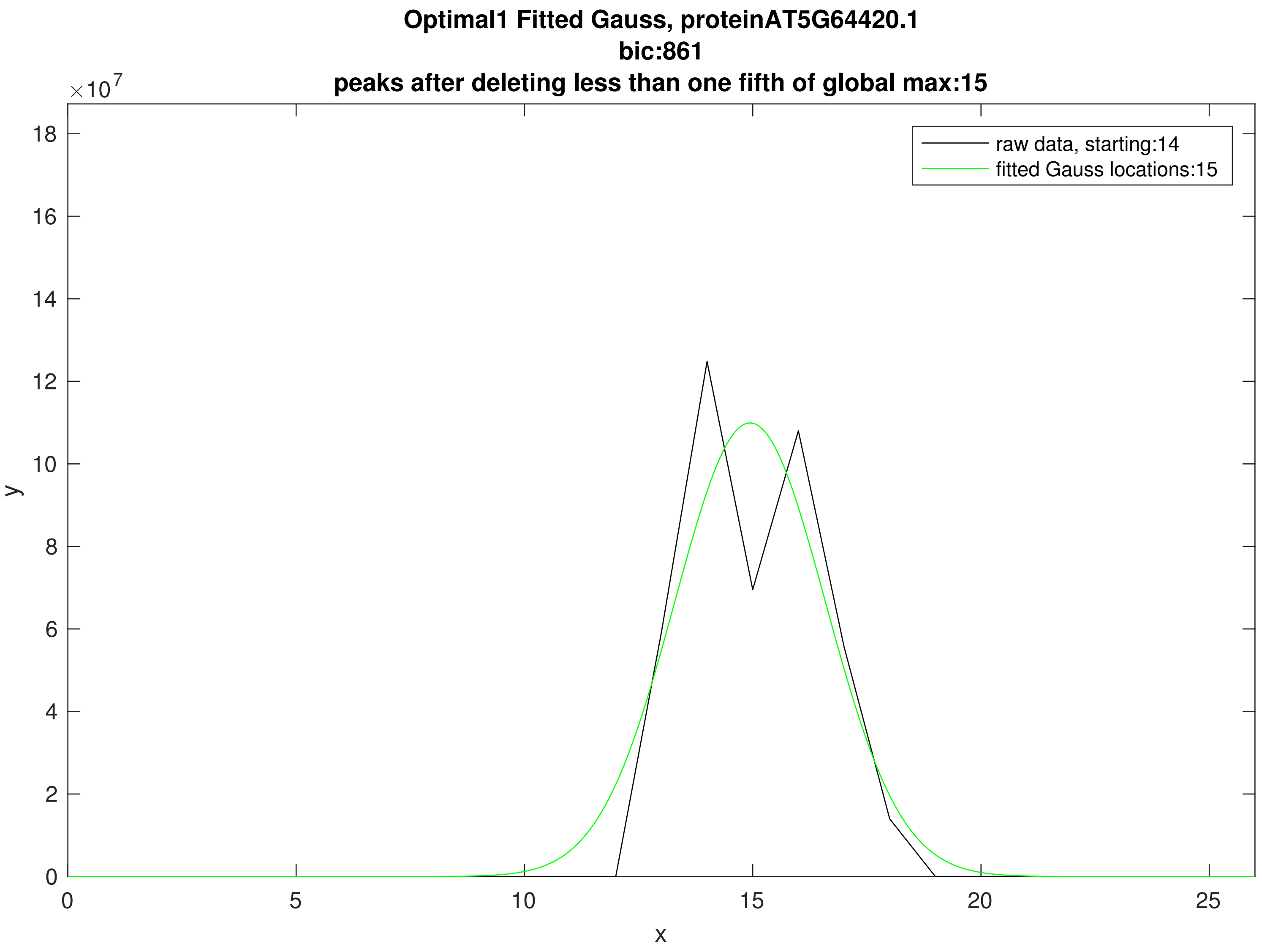
Optimal1 Fitted Gauss, proteinAT5G64420.1
bic:861
peaks after deleting less than one fifth of global max:15
×10^7
raw data, starting:14
fitted Gauss locations:15
y
x

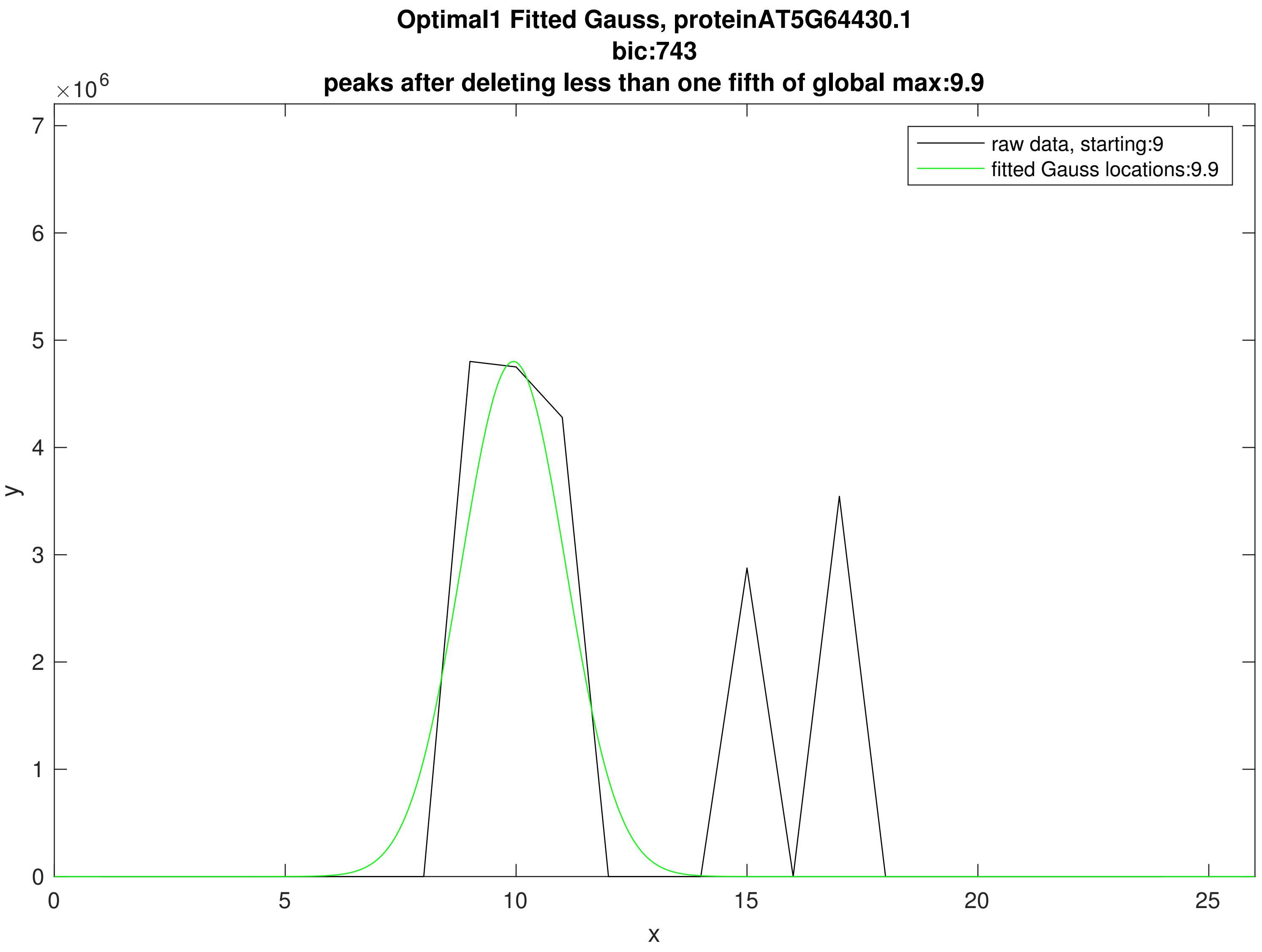

Optimal1 Fitted Gauss, proteinAT5G64430.1
bic:743
peaks after deleting less than one fifth of global max:9.9
×10^6
raw data, starting:9
fitted Gauss locations:9.9
y
x
0
1
2
3
4
5
6
7
0
5
10
15
20
25

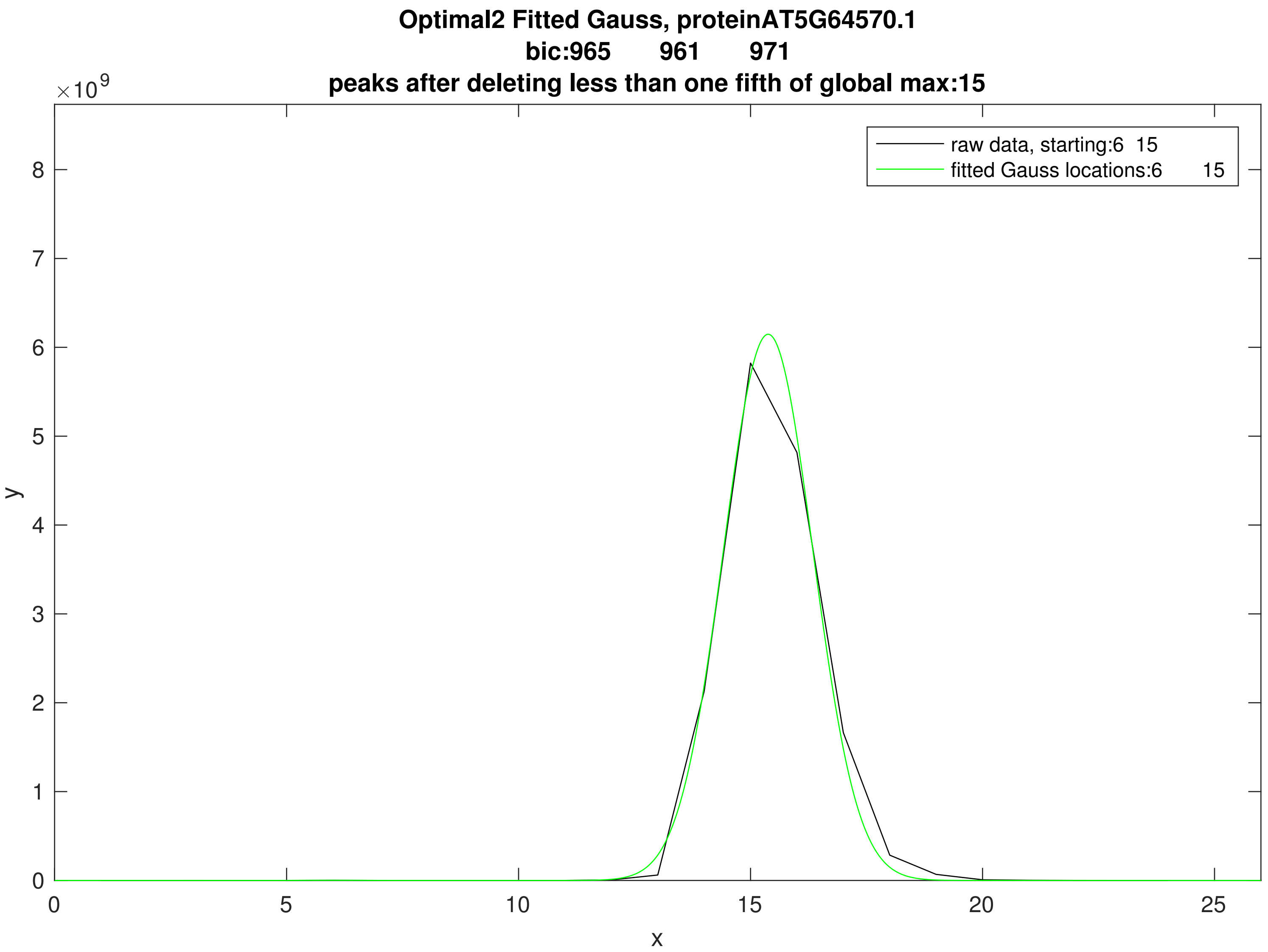
Optimal2 Fitted Gauss, proteinAT5G64570.1
bic:965 961 971
peaks after deleting less than one fifth of global max:15
×10^9
raw data, starting:6 15
fitted Gauss locations:6 15
y
x
0
1
2
3
4
5
6
7
8
0
5
10
15
20
25

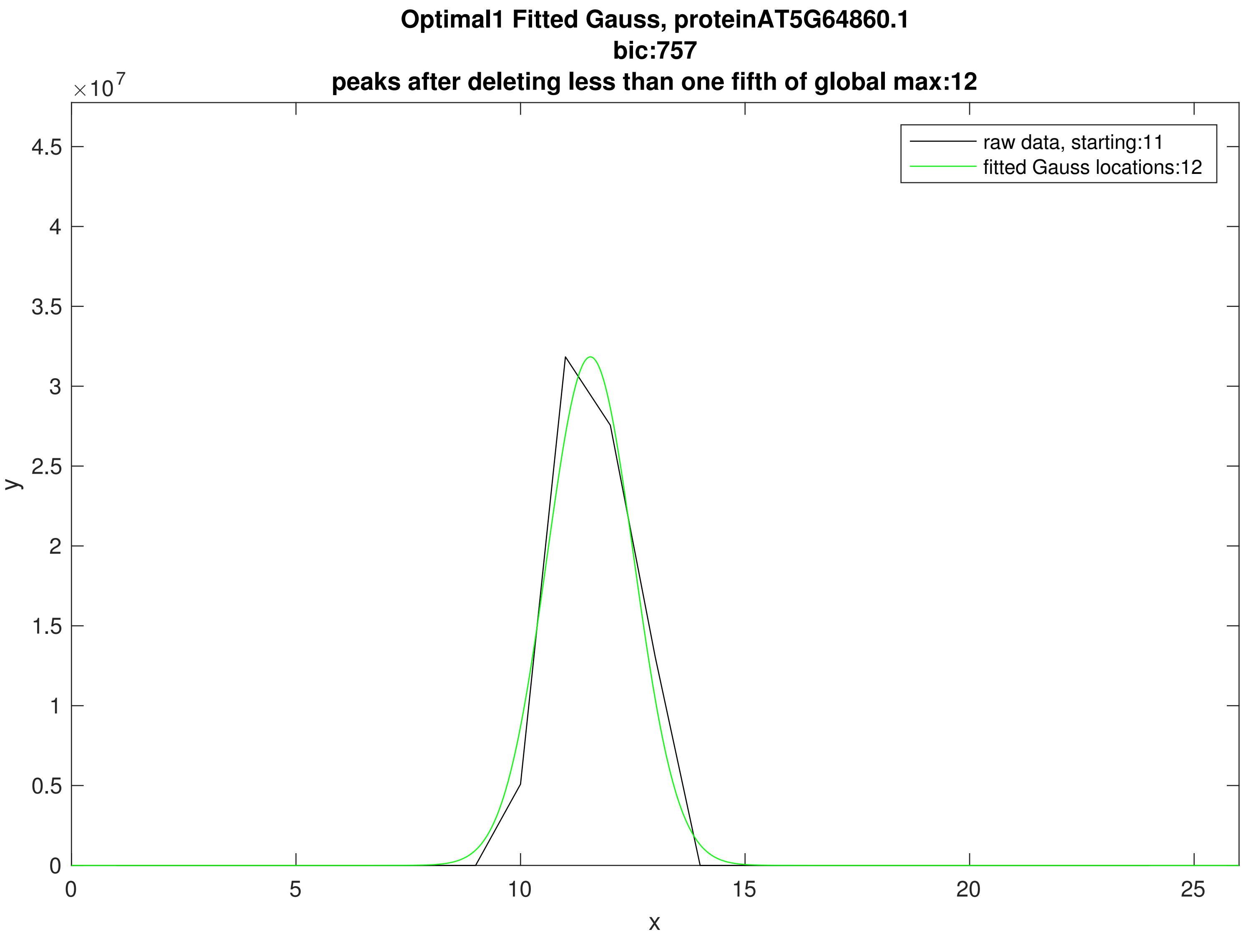

Optimal1 Fitted Gauss, proteinAT5G64860.1
bic:757
peaks after deleting less than one fifth of global max:12
×10^7
raw data, starting:11
fitted Gauss locations:12
4.5
4
3.5
3
2.5
2
1.5
1
0.5
0
y
0
5
10
15
20
25
x

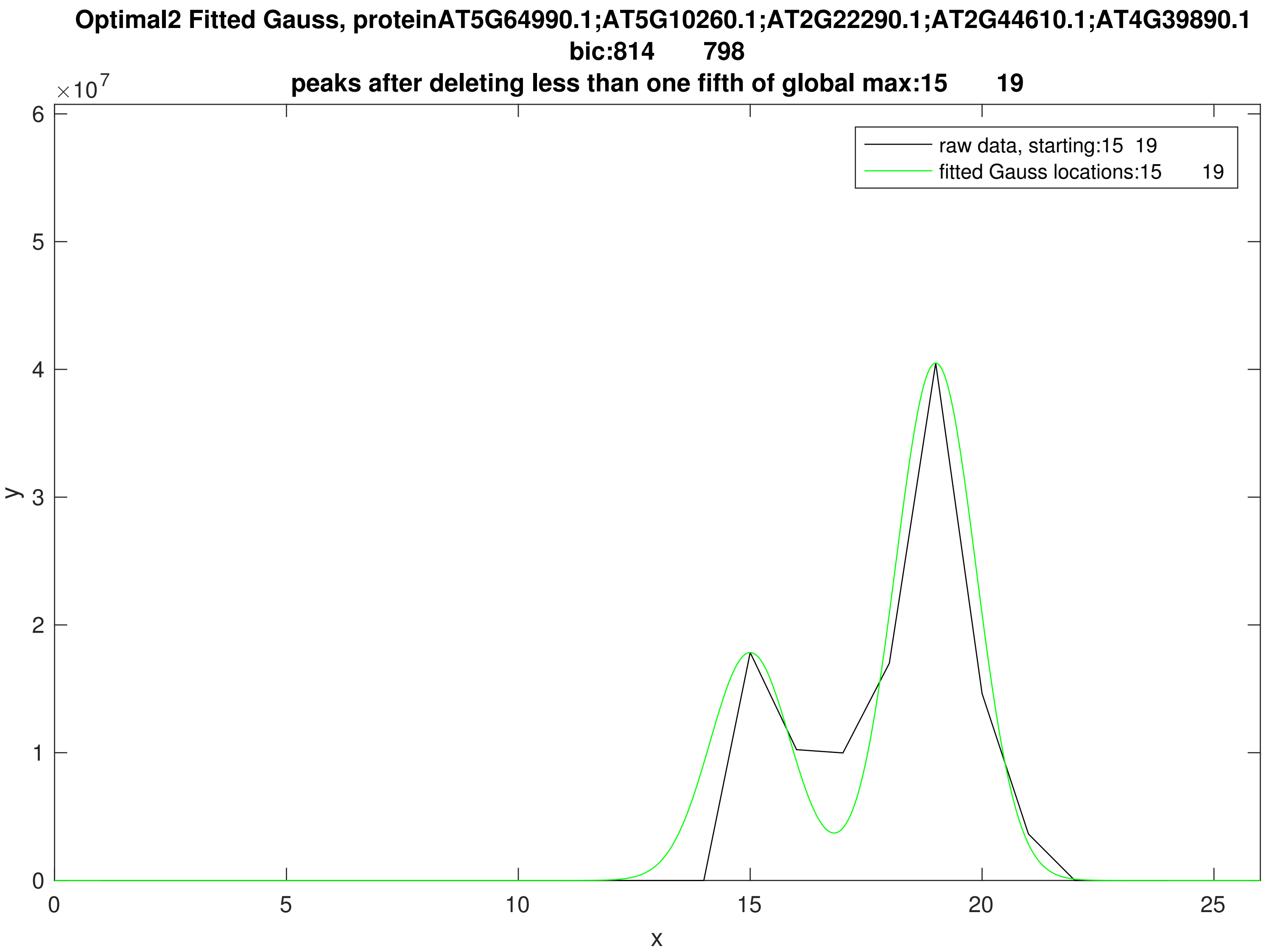

Optimal2 Fitted Gauss, proteinAT5G64990.1;AT5G10260.1;AT2G22290.1;AT2G44610.1;AT4G39890.1
bic:814 798
peaks after deleting less than one fifth of global max:15 19
×10^7
raw data, starting:15 19
fitted Gauss locations:15 19
y
x
0
1
2
3
4
5
6
0
5
10
15
20
25

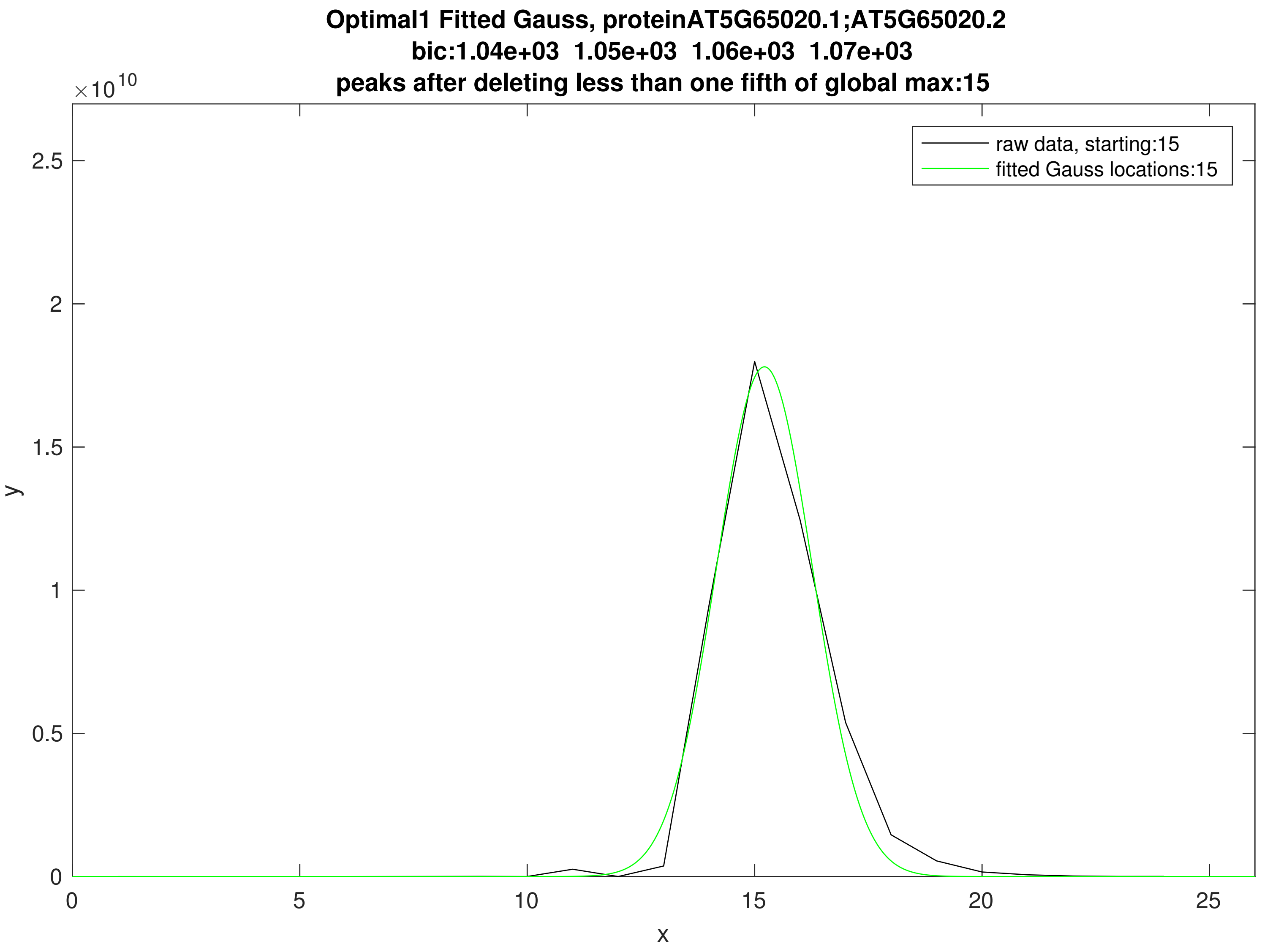

Optimal1 Fitted Gauss, proteinAT5G65020.1;AT5G65020.2
bic:1.04e+03 1.05e+03 1.06e+03 1.07e+03
peaks after deleting less than one fifth of global max:15
×10^10
raw data, starting:15
fitted Gauss locations:15
y
x
0
0.5
1
1.5
2
2.5
5
10
15
20
25

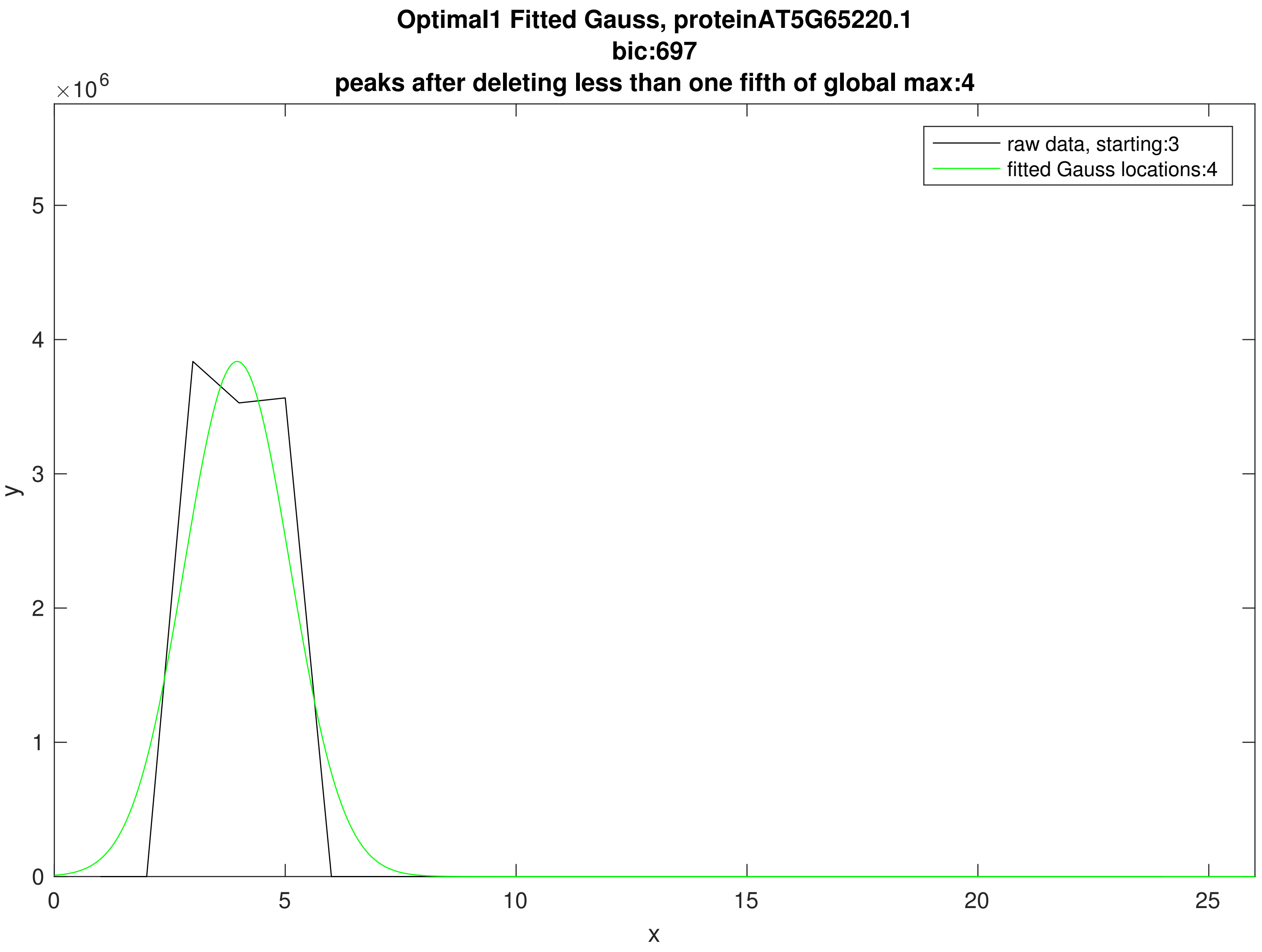

Optimal1 Fitted Gauss, proteinAT5G65220.1
bic:697
peaks after deleting less than one fifth of global max:4
×10^6
raw data, starting:3
fitted Gauss locations:4
y
x
0
1
2
3
4
5
0
5
10
15
20
25

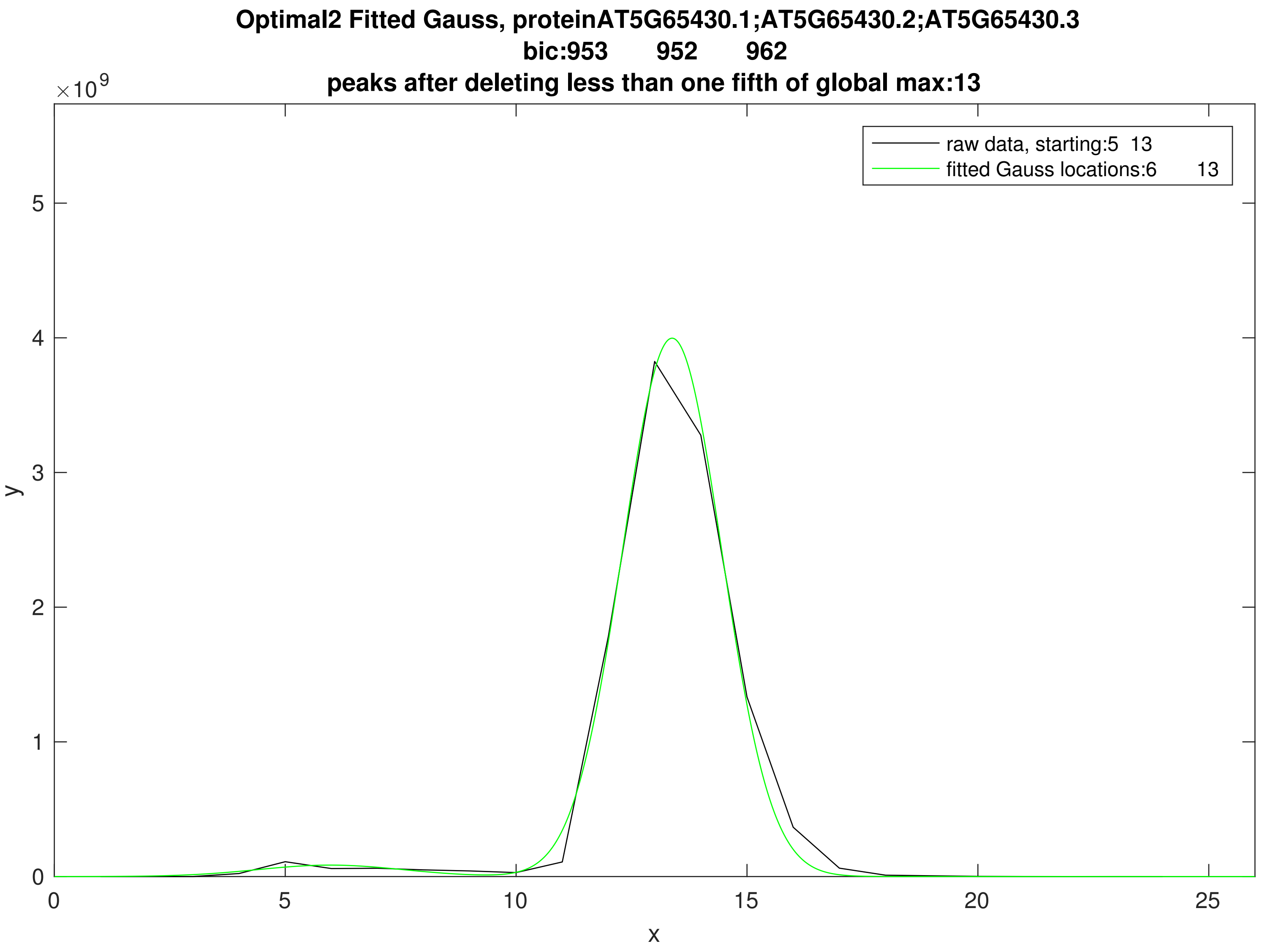

Optimal2 Fitted Gauss, proteinAT5G65430.1;AT5G65430.2;AT5G65430.3
bic:953 952 962
peaks after deleting less than one fifth of global max:13
×10^9
raw data, starting:5 13
fitted Gauss locations:6 13
y
x
0
1
2
3
4
5
10
15
20
25

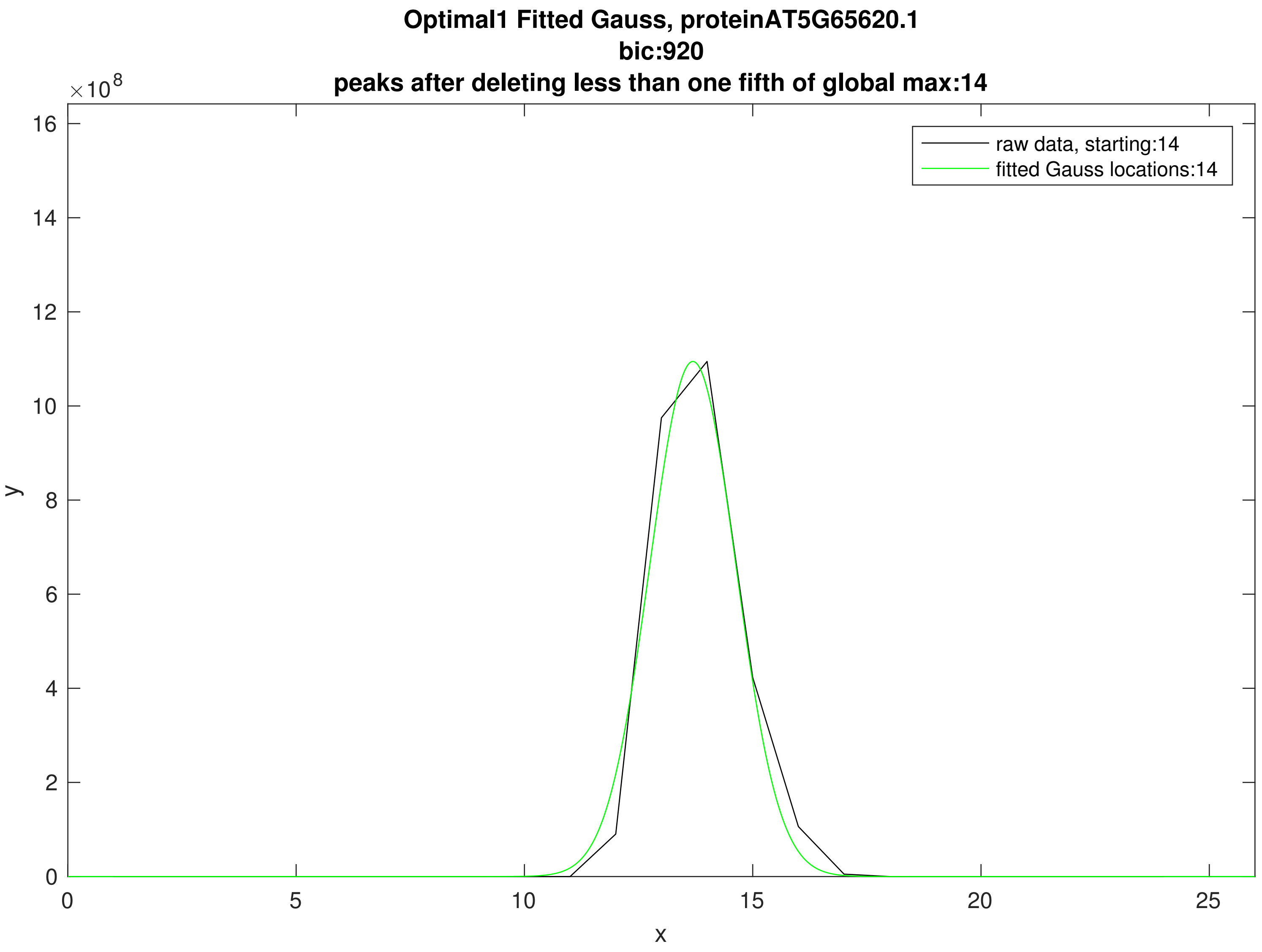
Optimal1 Fitted Gauss, proteinAT5G65620.1
bic:920
peaks after deleting less than one fifth of global max:14
×10^8
16
14
12
10
8
6
4
2
0
y
0
5
10
15
20
25
x
raw data, starting:14
fitted Gauss locations:14

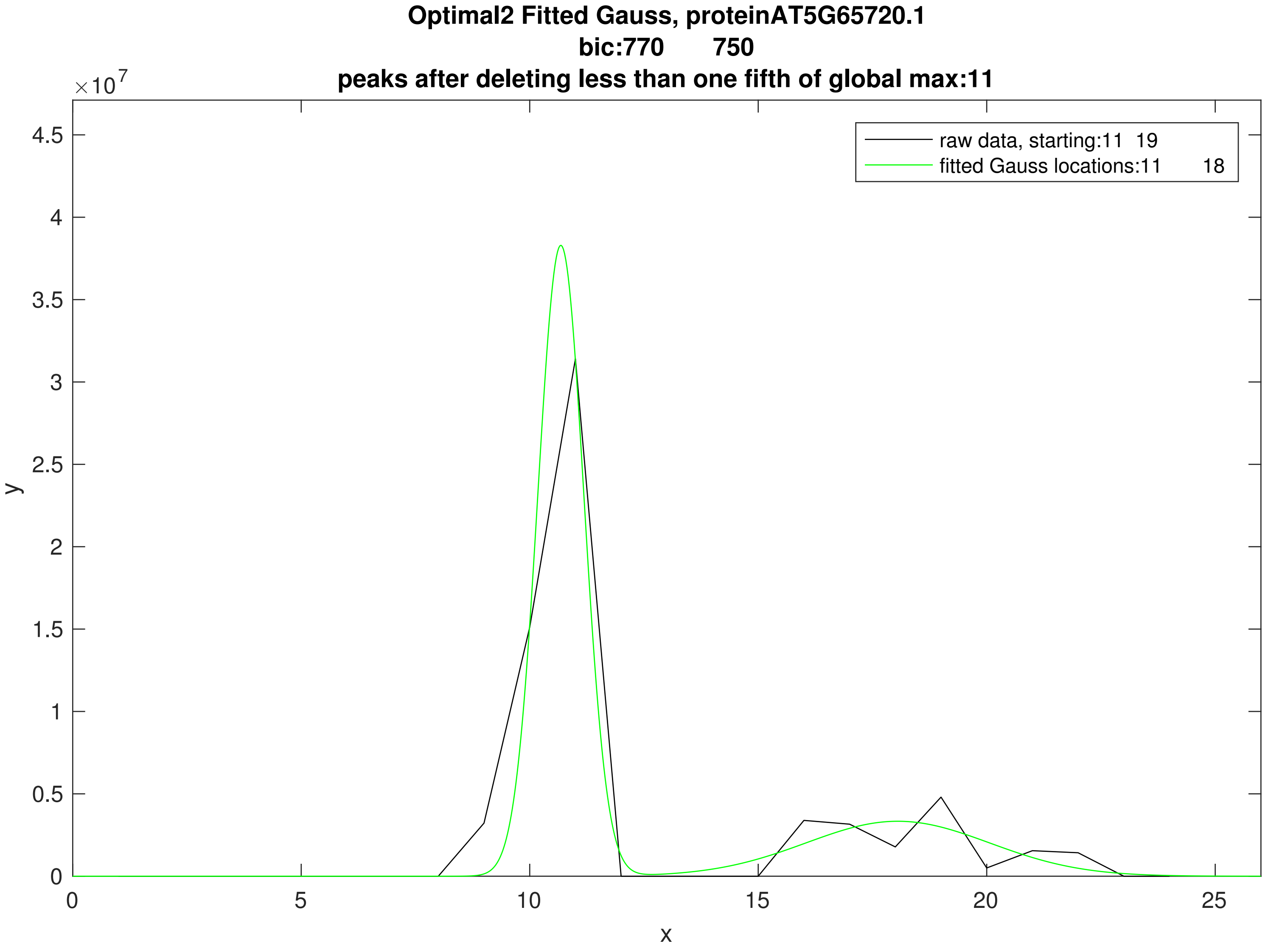
Optimal2 Fitted Gauss, proteinAT5G65720.1
bic:770 750
peaks after deleting less than one fifth of global max:11
×10^7
raw data, starting:11 19
fitted Gauss locations:11 18
4.5
4
3.5
3
2.5
2
1.5
1
0.5
0
y
0
5
10
15
20
25
x

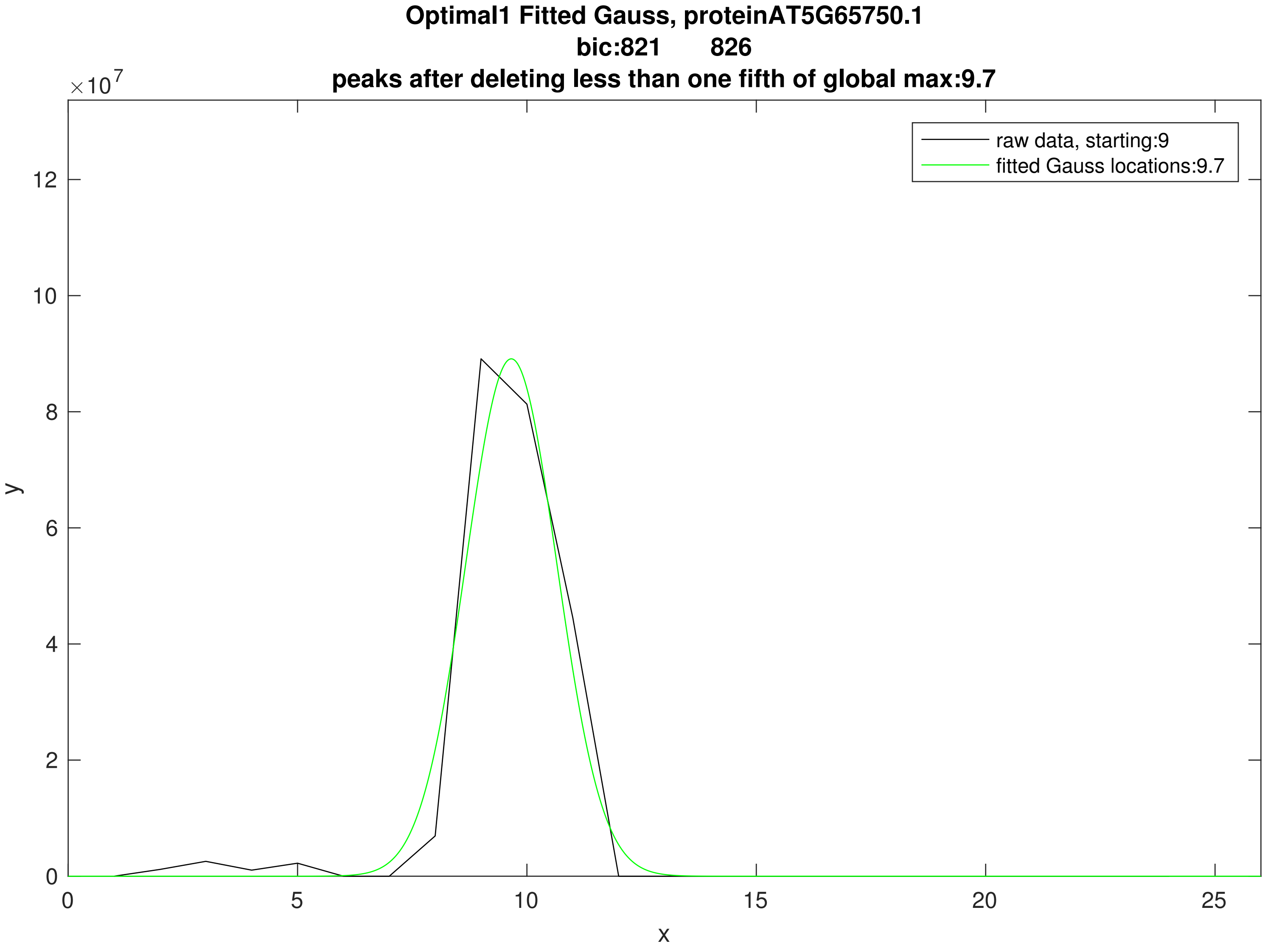
Optimal1 Fitted Gauss, proteinAT5G65750.1
bic:821 826
peaks after deleting less than one fifth of global max:9.7
×10^7
raw data, starting:9
fitted Gauss locations:9.7
y
x
0
2
4
6
8
10
12
0
5
10
15
20
25

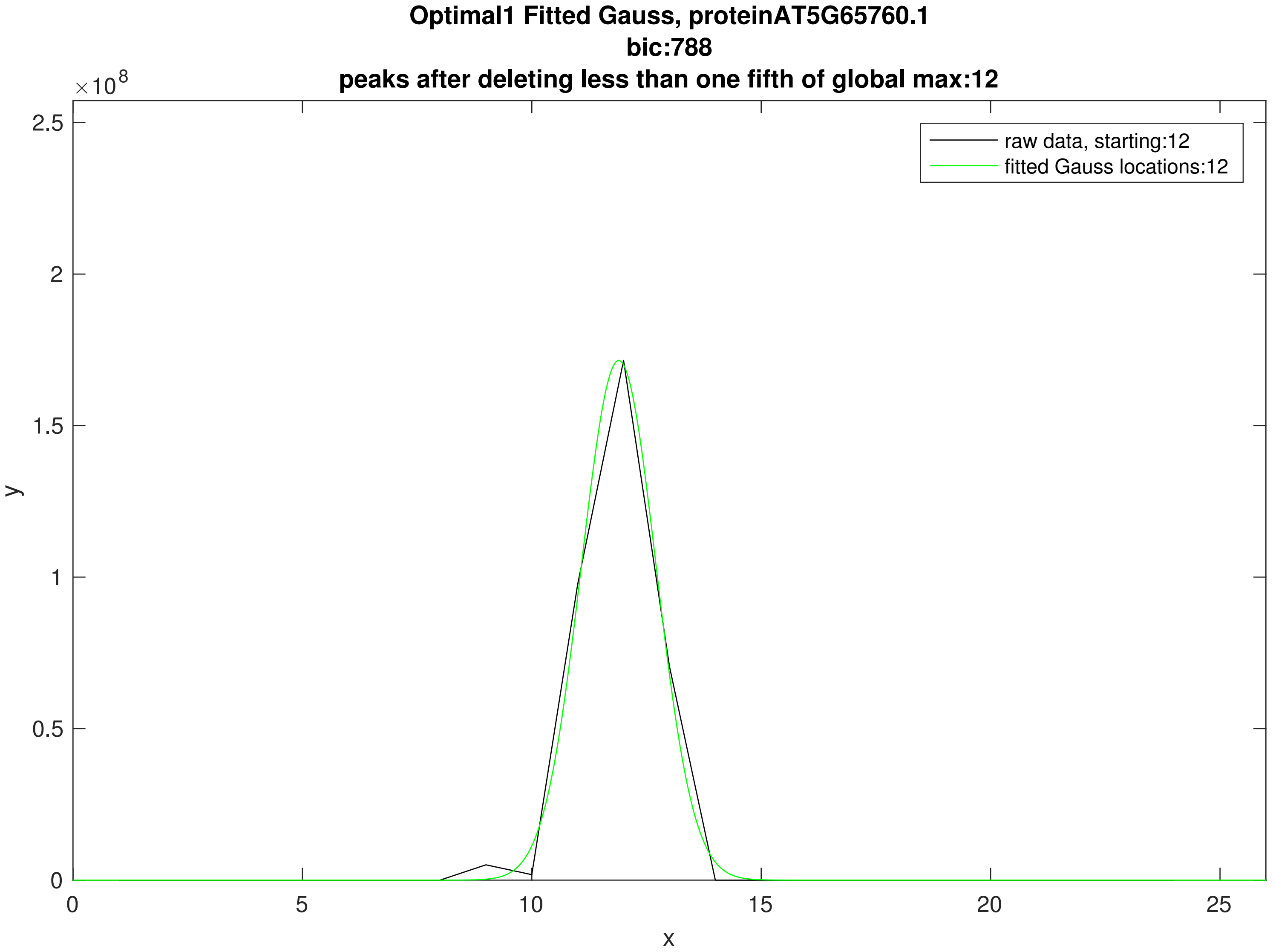

Optimal1 Fitted Gauss, proteinAT5G65760.1
bic:788
peaks after deleting less than one fifth of global max:12
×10^8
2.5
2
1.5
1
0.5
0
y
0
5
10
15
20
25
x
raw data, starting:12
fitted Gauss locations:12

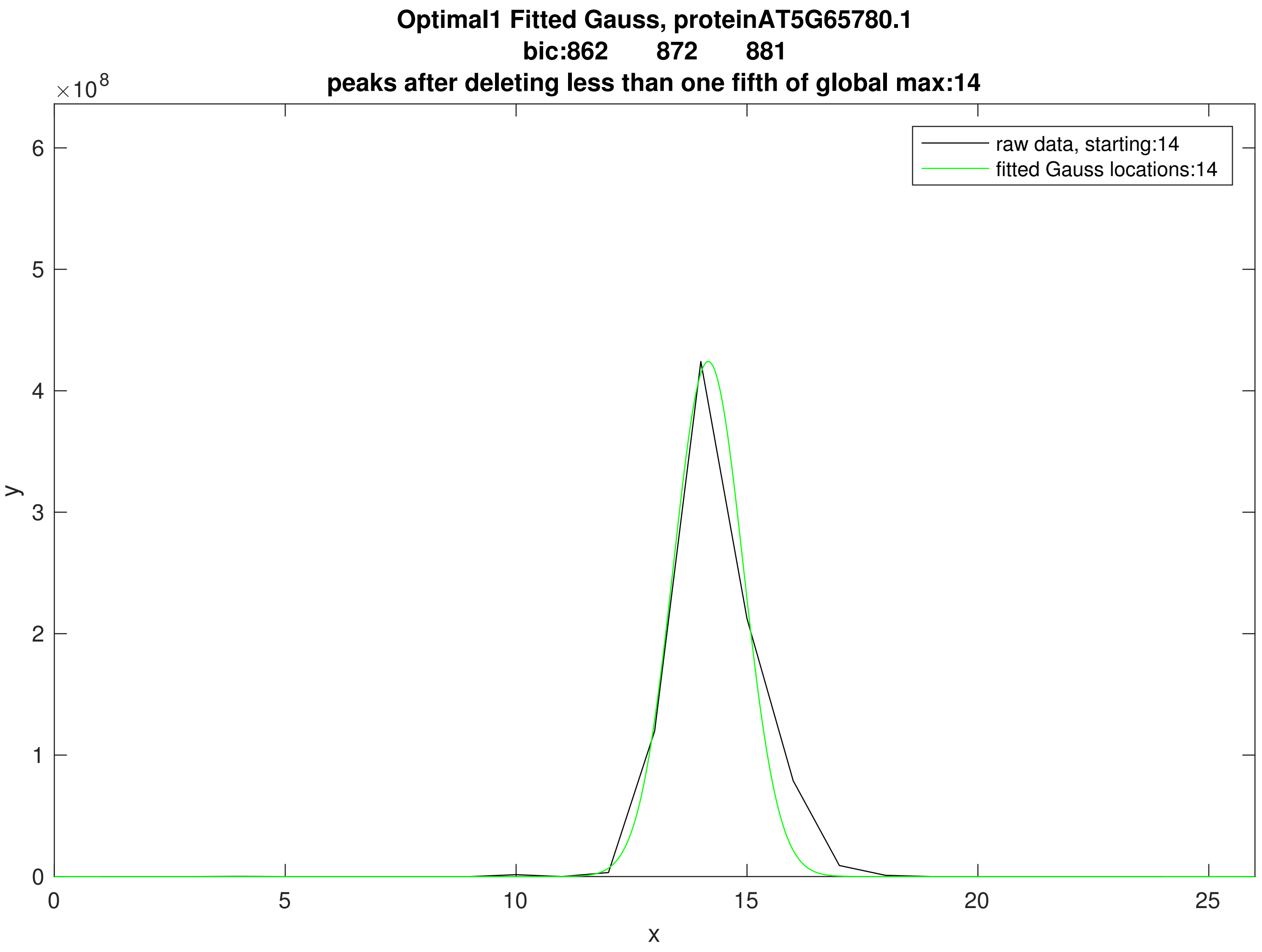

Optimal1 Fitted Gauss, proteinAT5G65780.1
bic:862 872 881
peaks after deleting less than one fifth of global max:14
×10^8
raw data, starting:14
fitted Gauss locations:14
6
5
4
3
2
1
0
y
0
5
10
15
20
25
x

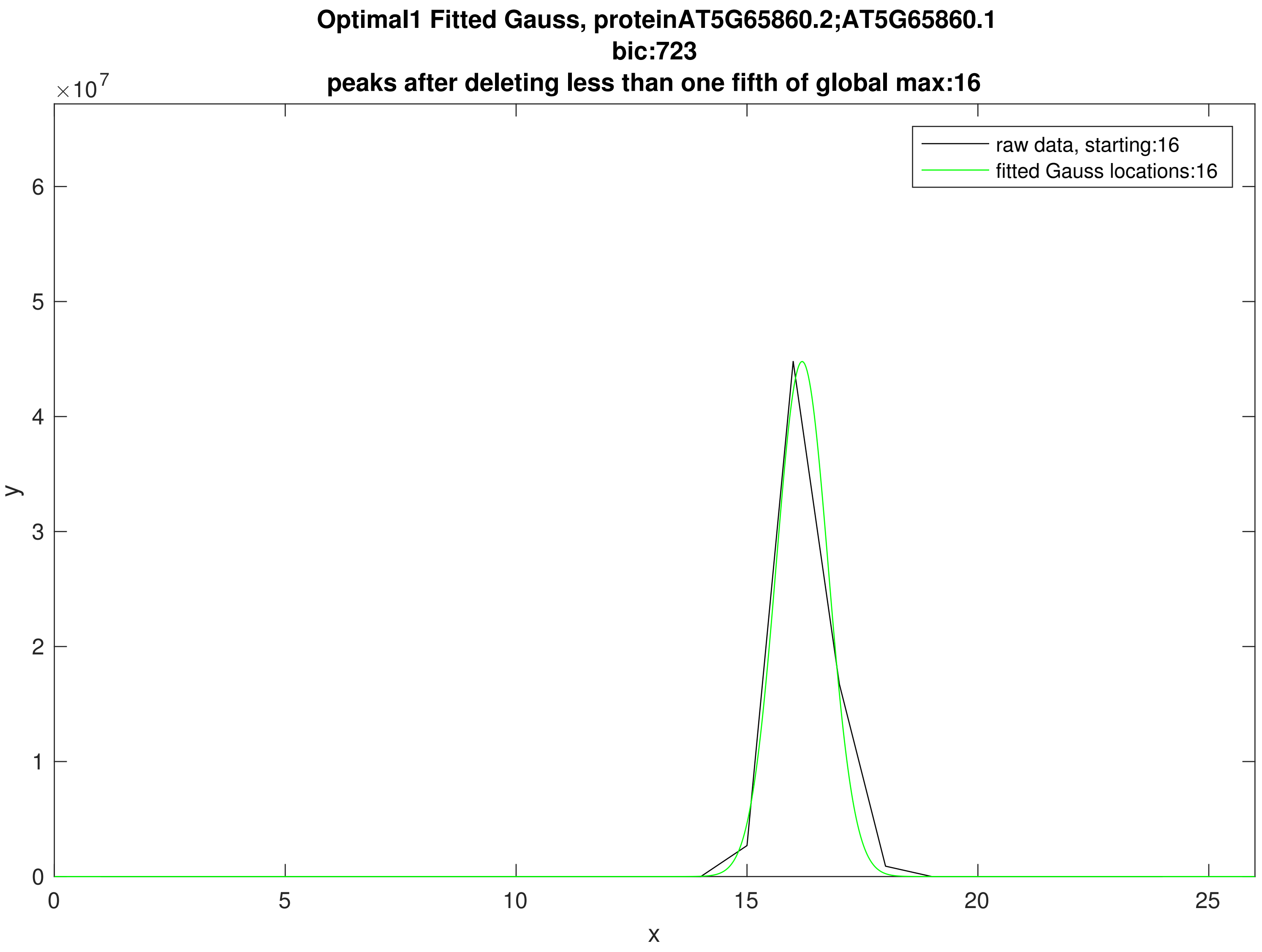
Optimal1 Fitted Gauss, proteinAT5G65860.2;AT5G65860.1
bic:723
peaks after deleting less than one fifth of global max:16
×10^7
raw data, starting:16
fitted Gauss locations:16
y
x
0
1
2
3
4
5
6
0
5
10
15
20
25

**Optimal1 Fitted Gauss, proteinAT5G65940.3;AT5G65940.1;AT5G65940.2**
**bic:663**
**peaks after deleting less than one fifth of global max:19**

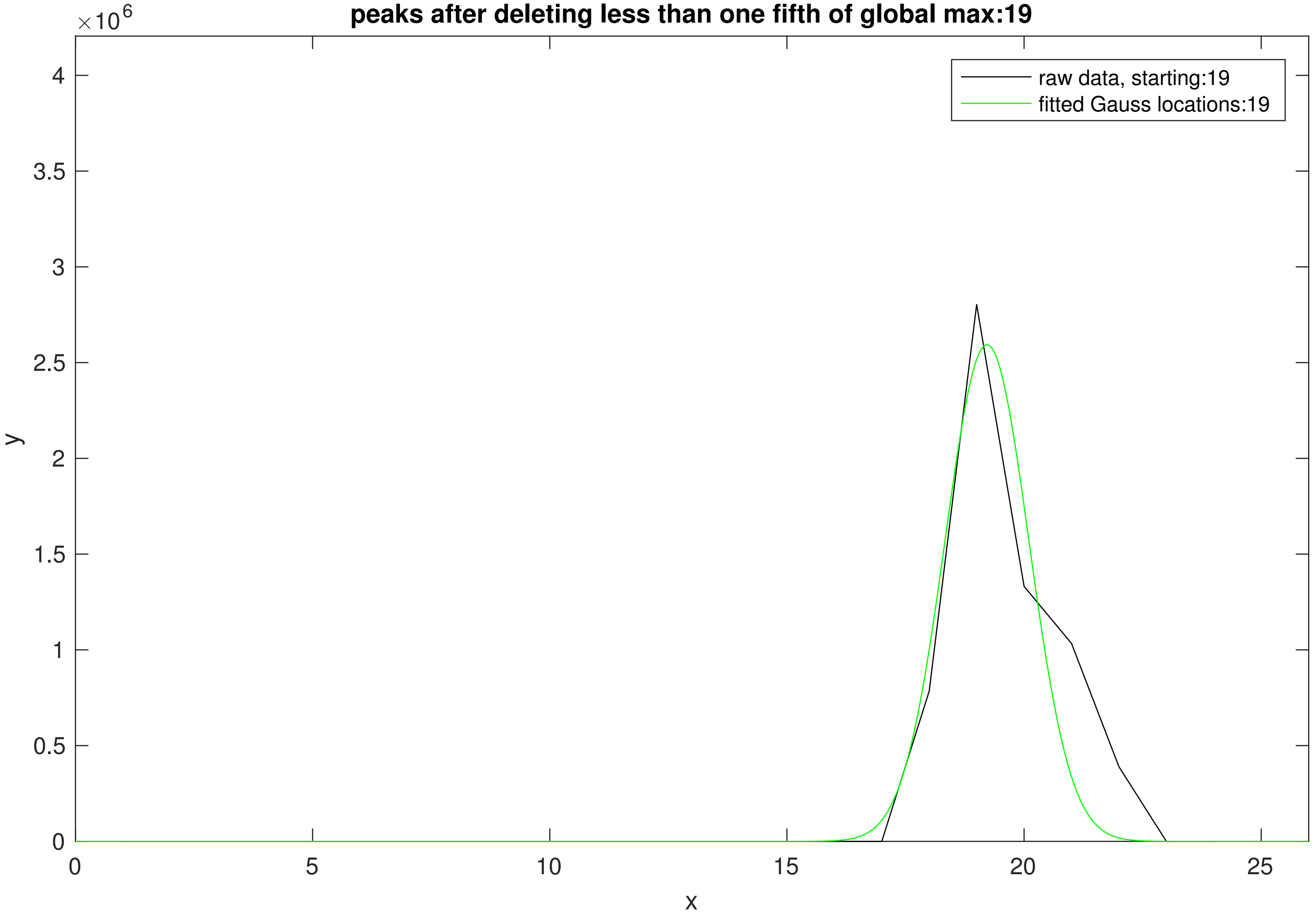

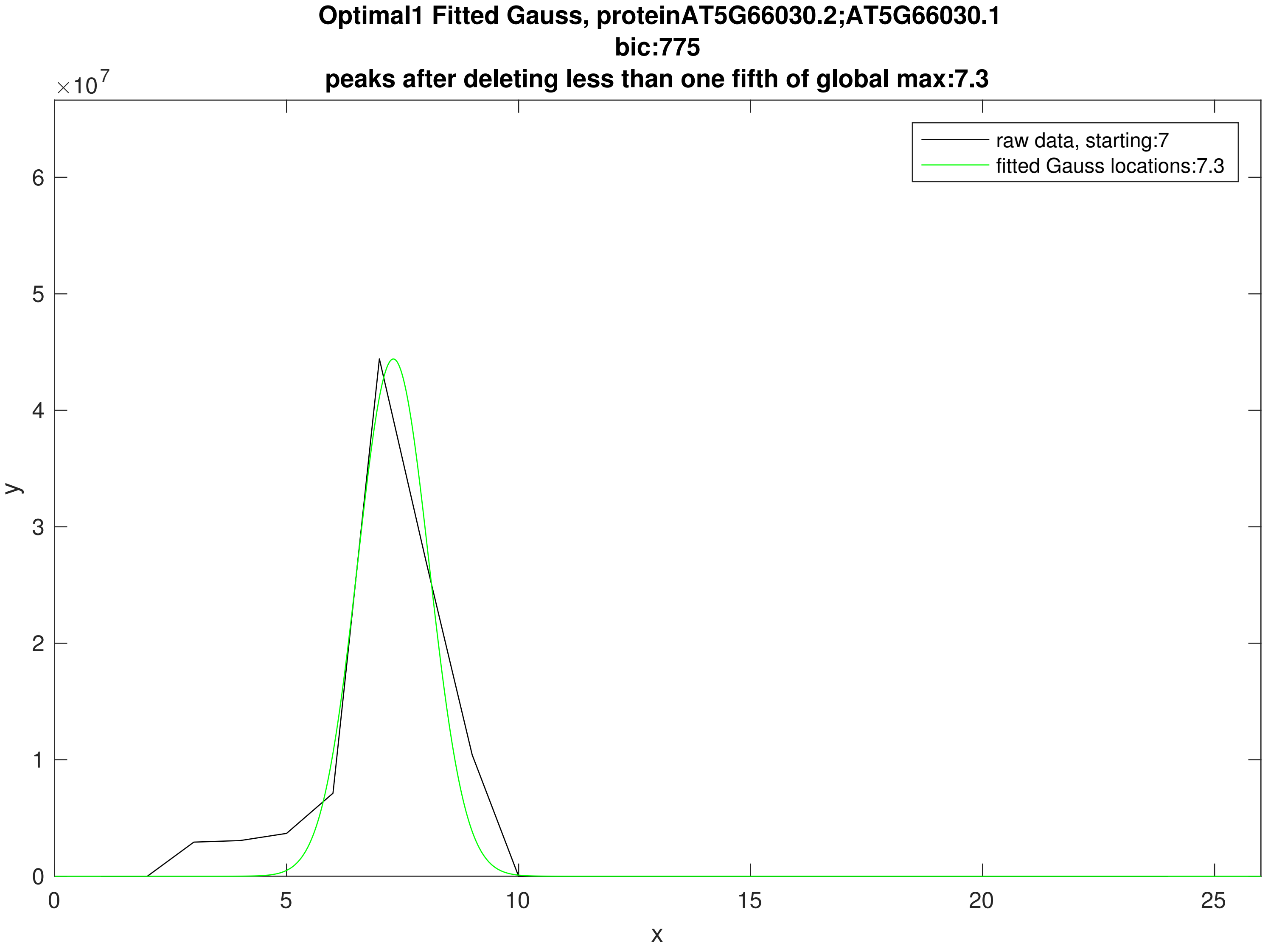

Optimal1 Fitted Gauss, proteinAT5G66030.2;AT5G66030.1
bic:775
peaks after deleting less than one fifth of global max:7.3
×10^7
6
5
4
3
2
1
0
y
0
5
10
15
20
25
x
raw data, starting:7
fitted Gauss locations:7.3

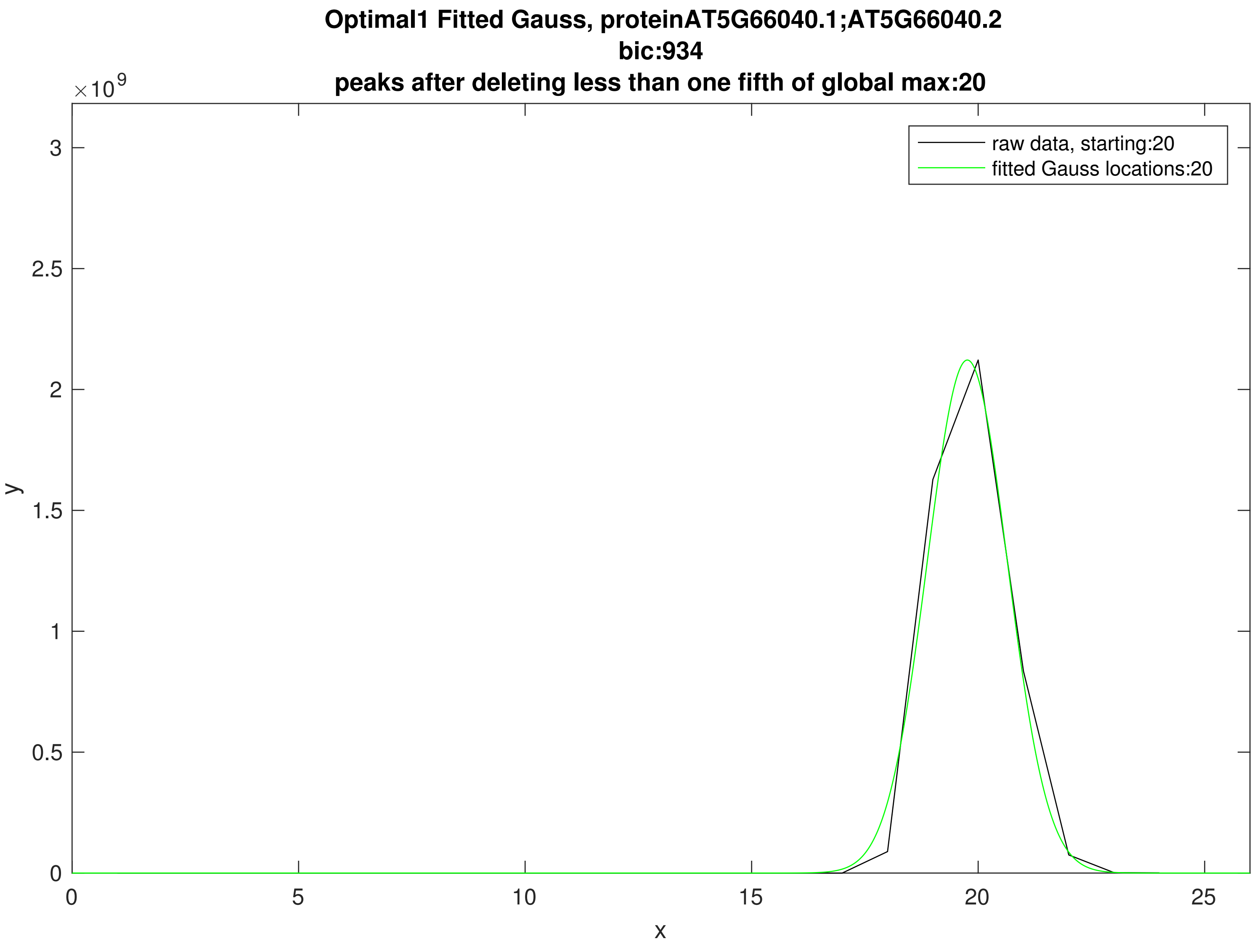

Optimal1 Fitted Gauss, proteinAT5G66040.1;AT5G66040.2
bic:934
peaks after deleting less than one fifth of global max:20
×10^9
raw data, starting:20
fitted Gauss locations:20
y
x
0
0.5
1
1.5
2
2.5
3
0
5
10
15
20
25

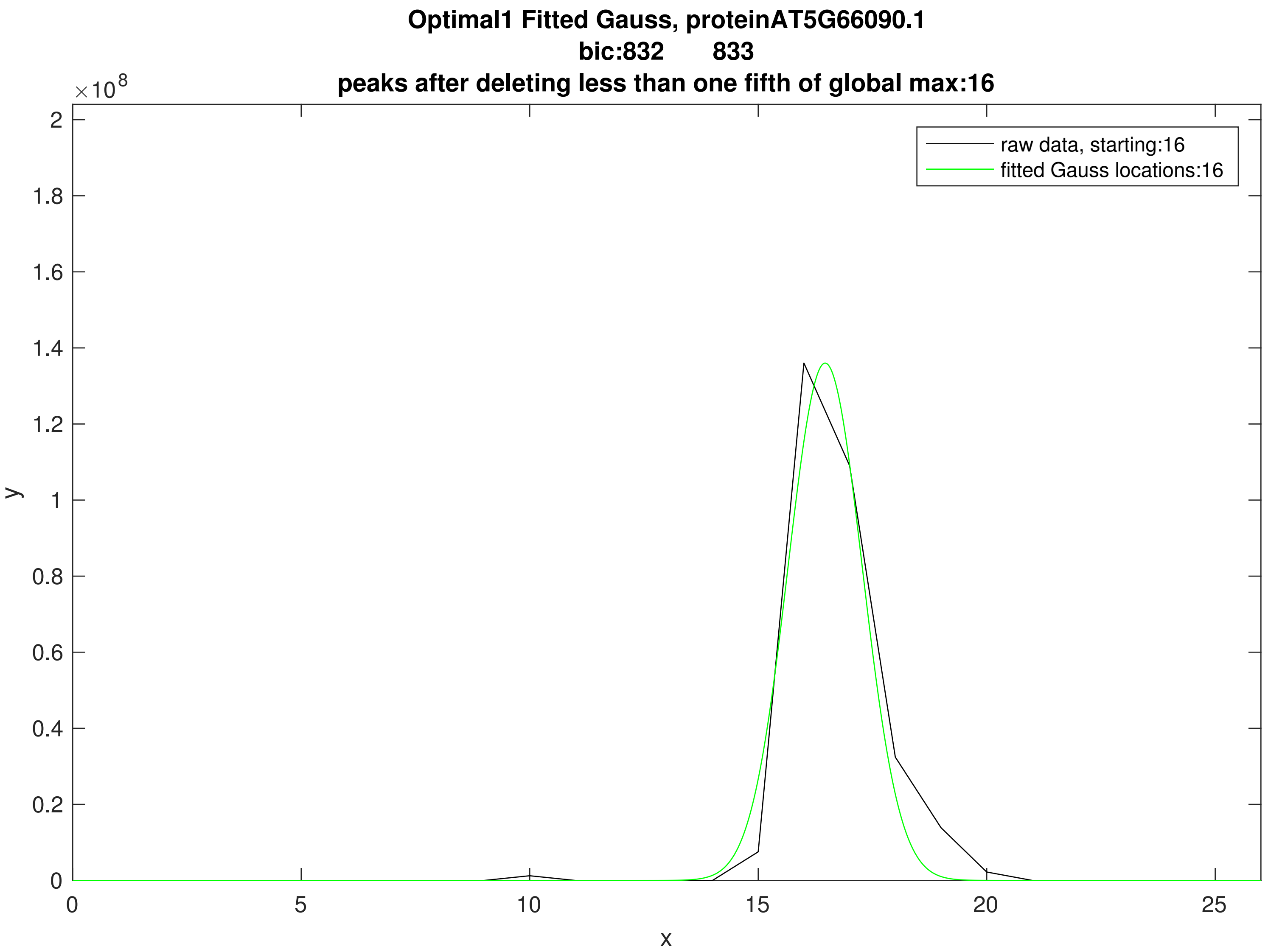
Optimal1 Fitted Gauss, proteinAT5G66090.1
bic:832 833
peaks after deleting less than one fifth of global max:16
×10^8
2
1.8
1.6
1.4
1.2
1
0.8
0.6
0.4
0.2
0
y
0
5
10
15
20
25
x
raw data, starting:16
fitted Gauss locations:16

Optimal1 Fitted Gauss, proteinAT5G66120.2;AT5G66120.1
bic:915
peaks after deleting less than one fifth of global max:14

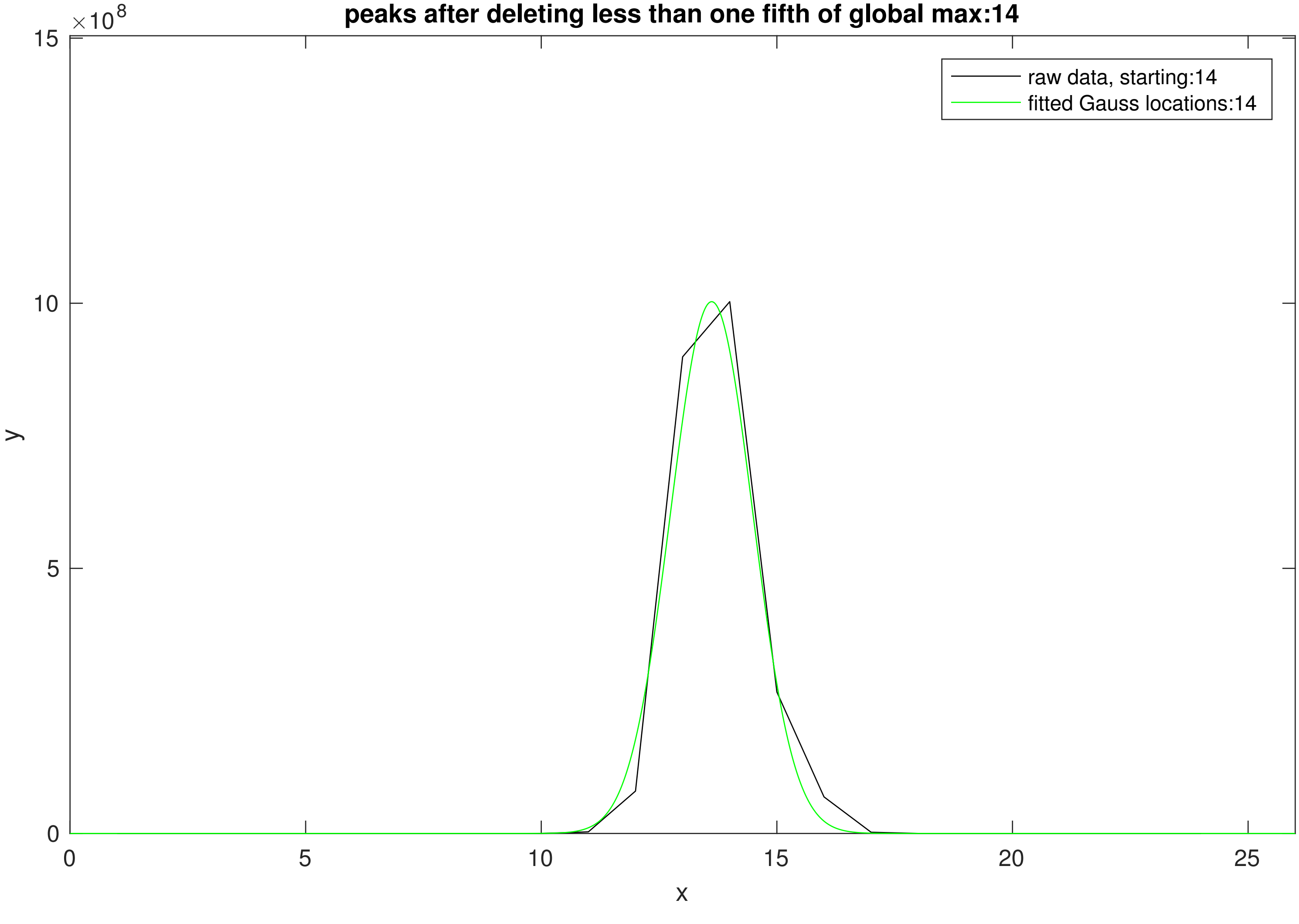

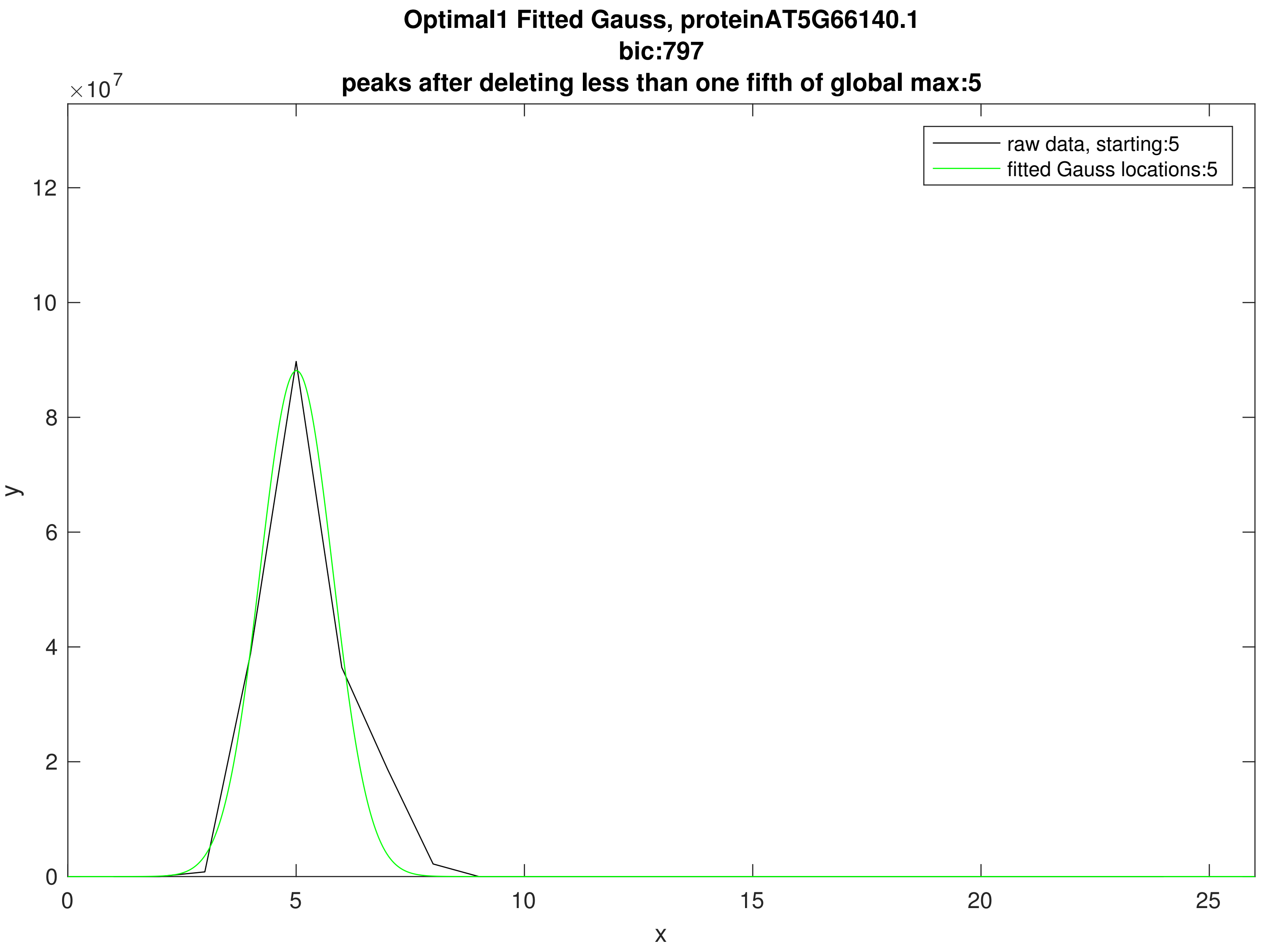

Optimal1 Fitted Gauss, proteinAT5G66140.1
bic:797
peaks after deleting less than one fifth of global max:5
×10^7
12
10
8
6
4
2
0
y
0
5
10
15
20
25
x
raw data, starting:5
fitted Gauss locations:5

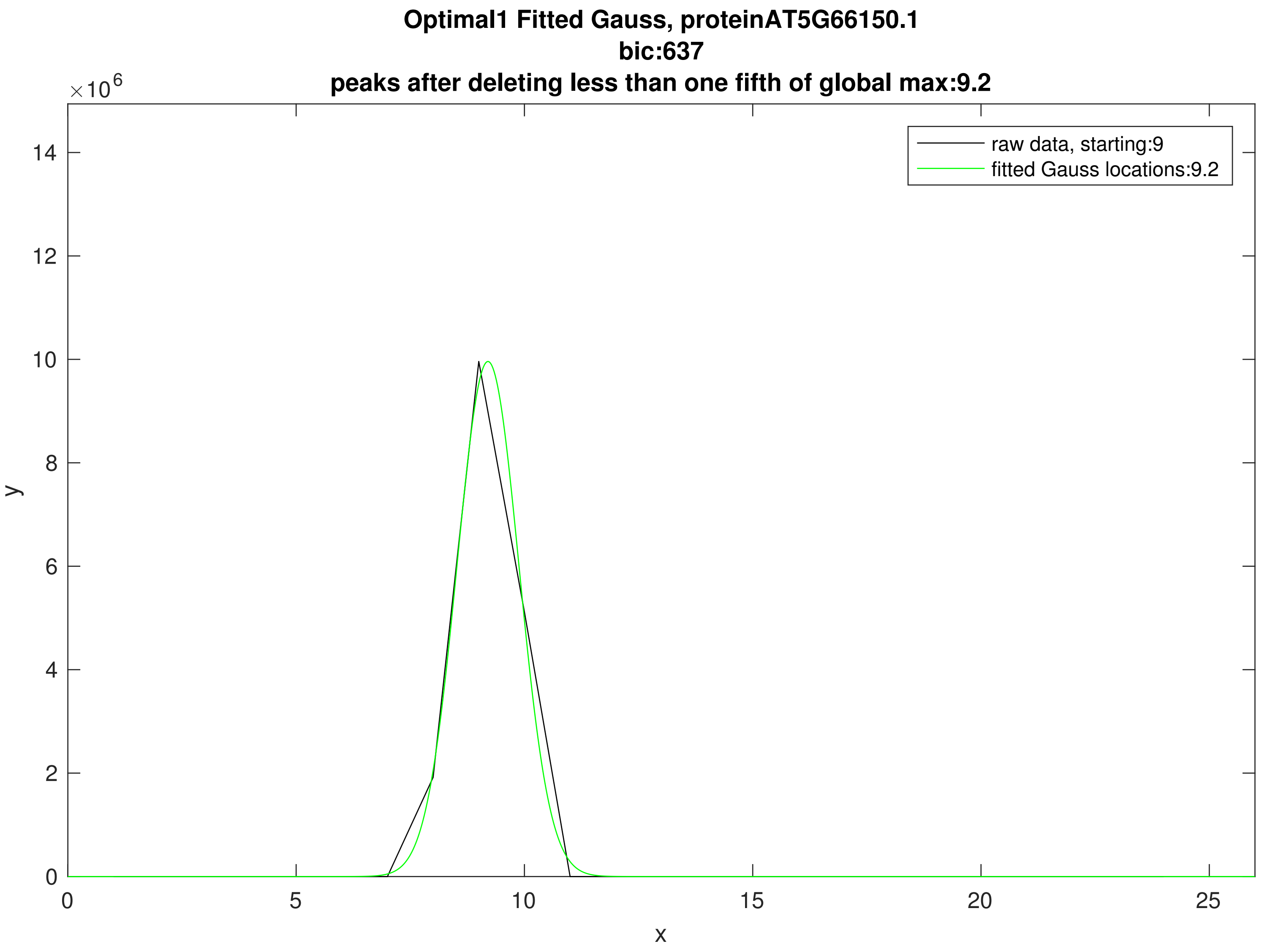
Optimal1 Fitted Gauss, proteinAT5G66150.1
bic:637
peaks after deleting less than one fifth of global max:9.2
×10^6
raw data, starting:9
fitted Gauss locations:9.2
y
x
0
2
4
6
8
10
12
14
0
5
10
15
20
25

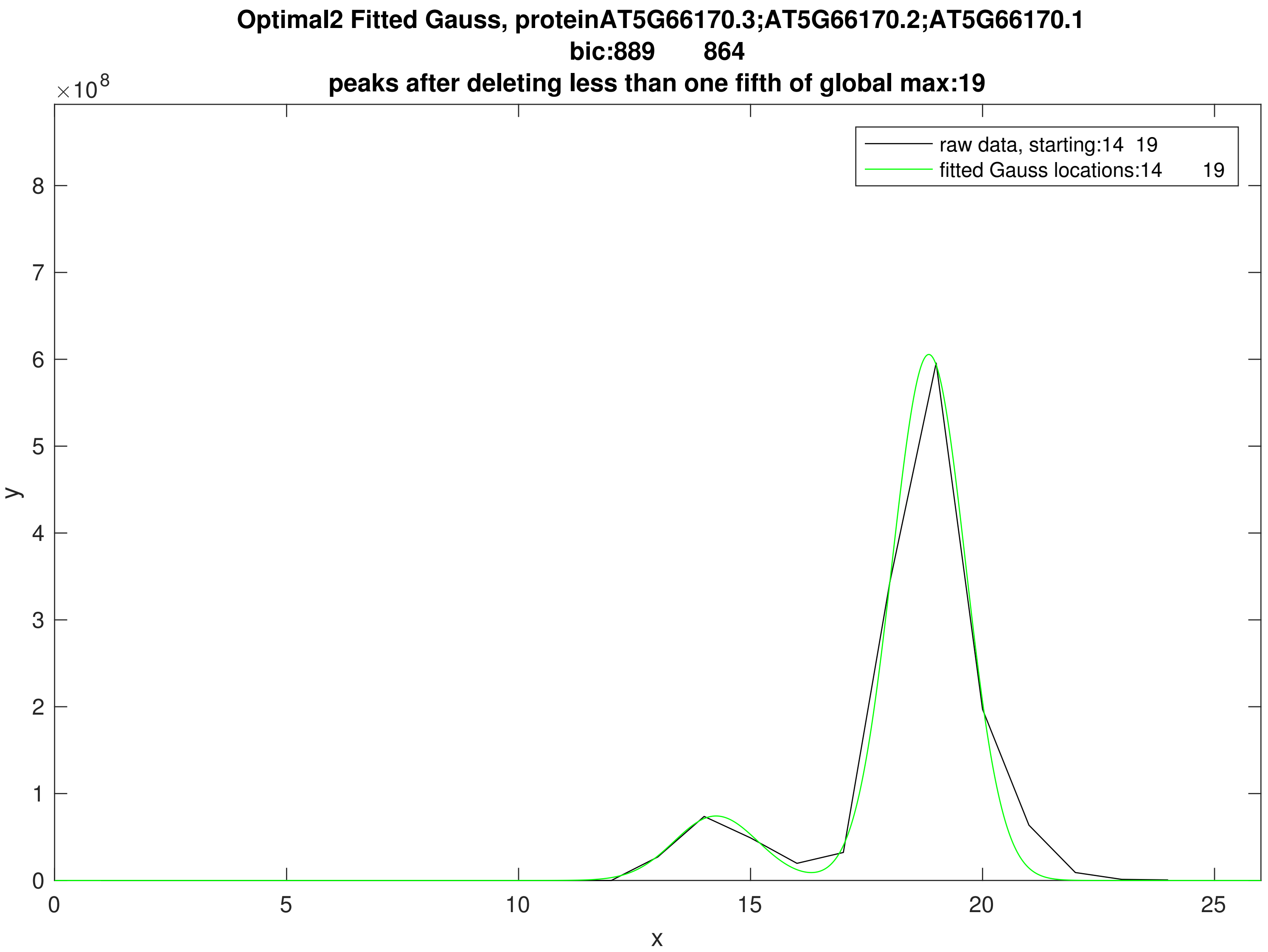
Optimal2 Fitted Gauss, proteinAT5G66170.3;AT5G66170.2;AT5G66170.1
bic:889 864
peaks after deleting less than one fifth of global max:19
×10^8
raw data, starting:14 19
fitted Gauss locations:14 19
y
x
0
1
2
3
4
5
6
7
8
0
5
10
15
20
25

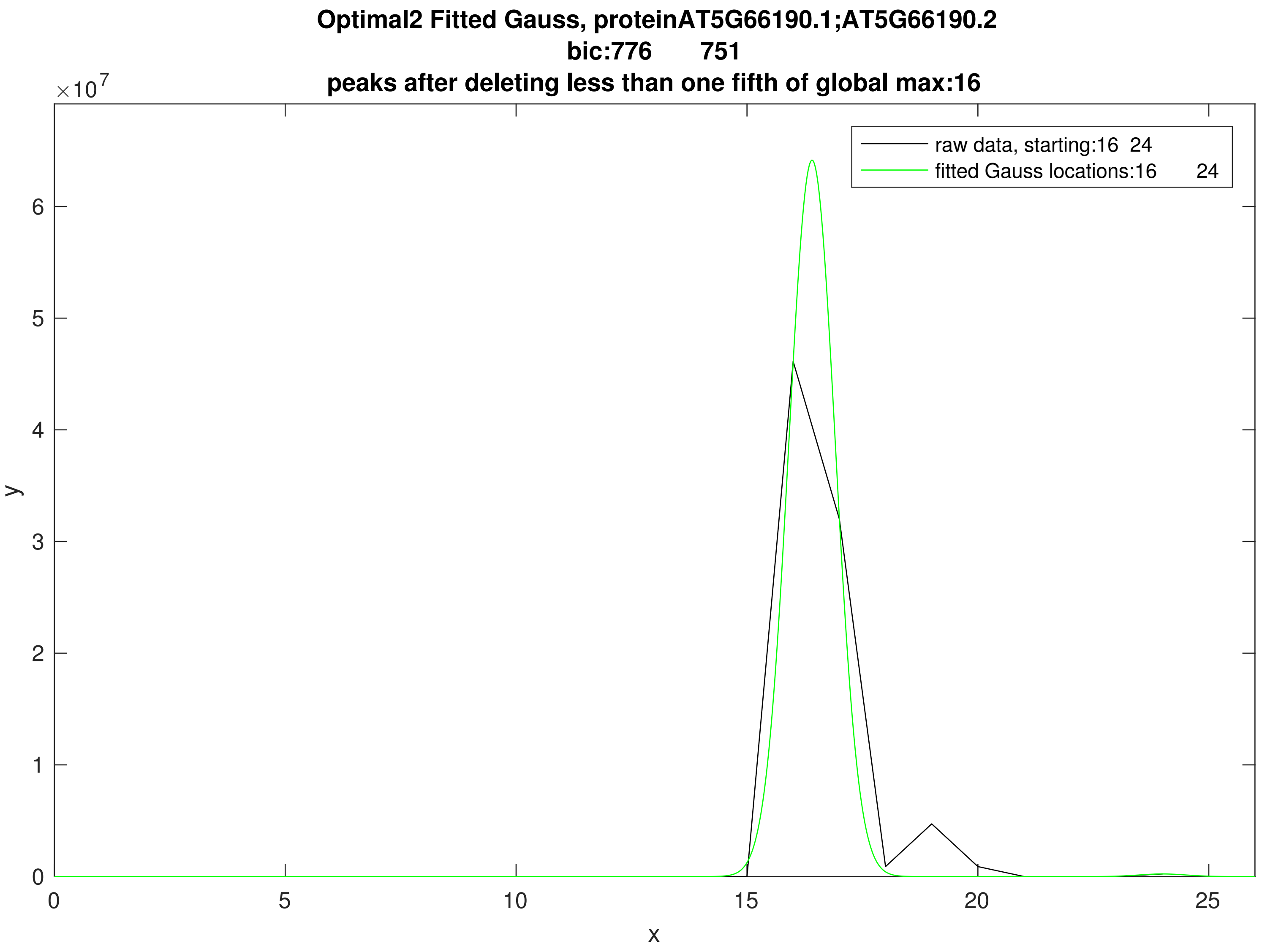
Optimal2 Fitted Gauss, proteinAT5G66190.1;AT5G66190.2
bic:776 751
peaks after deleting less than one fifth of global max:16
×10^7
raw data, starting:16 24
fitted Gauss locations:16 24
y
x
0
1
2
3
4
5
6
0
5
10
15
20
25

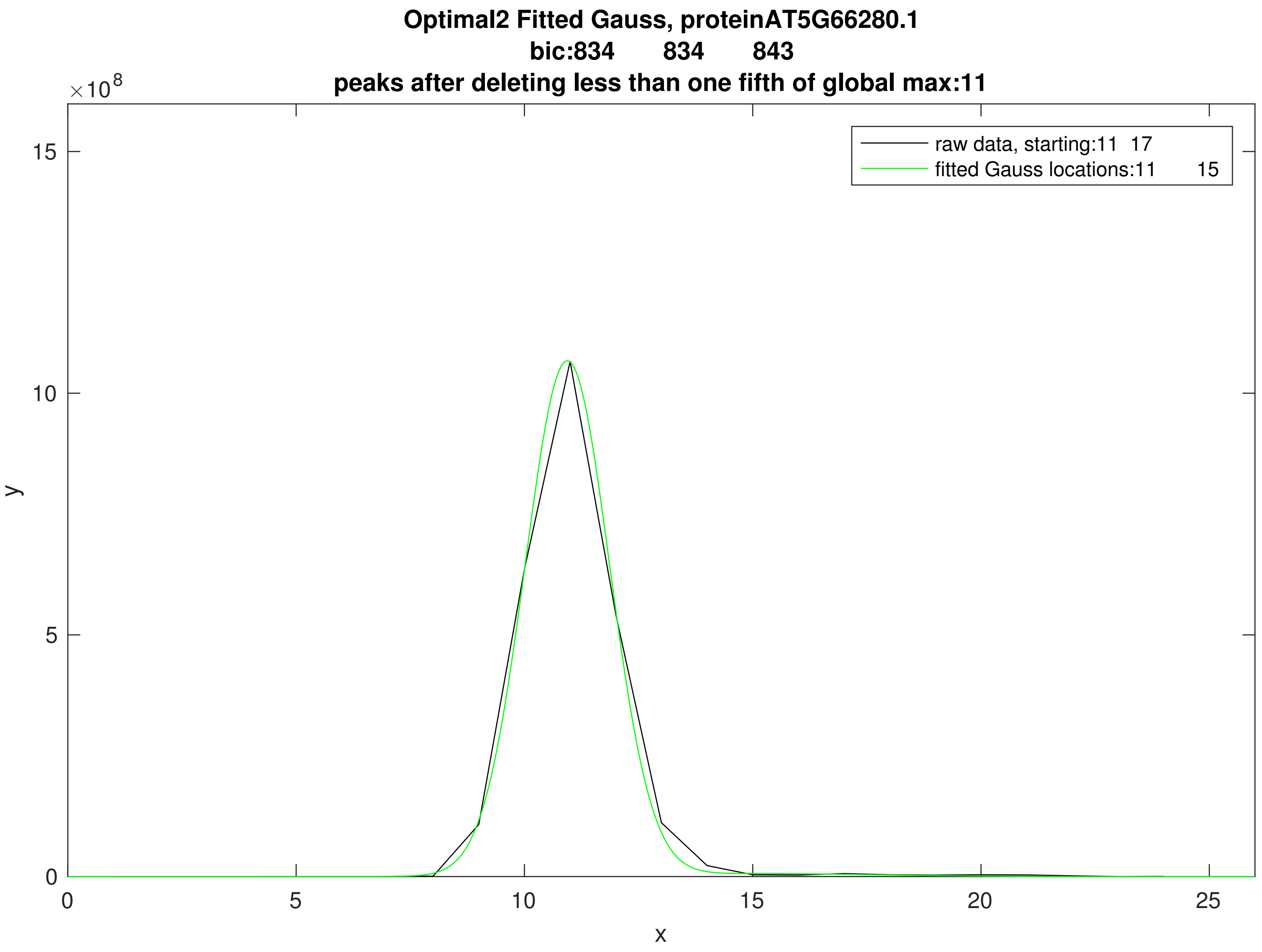
Optimal2 Fitted Gauss, proteinAT5G66280.1
bic:834 834 843
peaks after deleting less than one fifth of global max:11
×10^8
raw data, starting:11 17
fitted Gauss locations:11 15
y
x
0
5
10
15
20
25

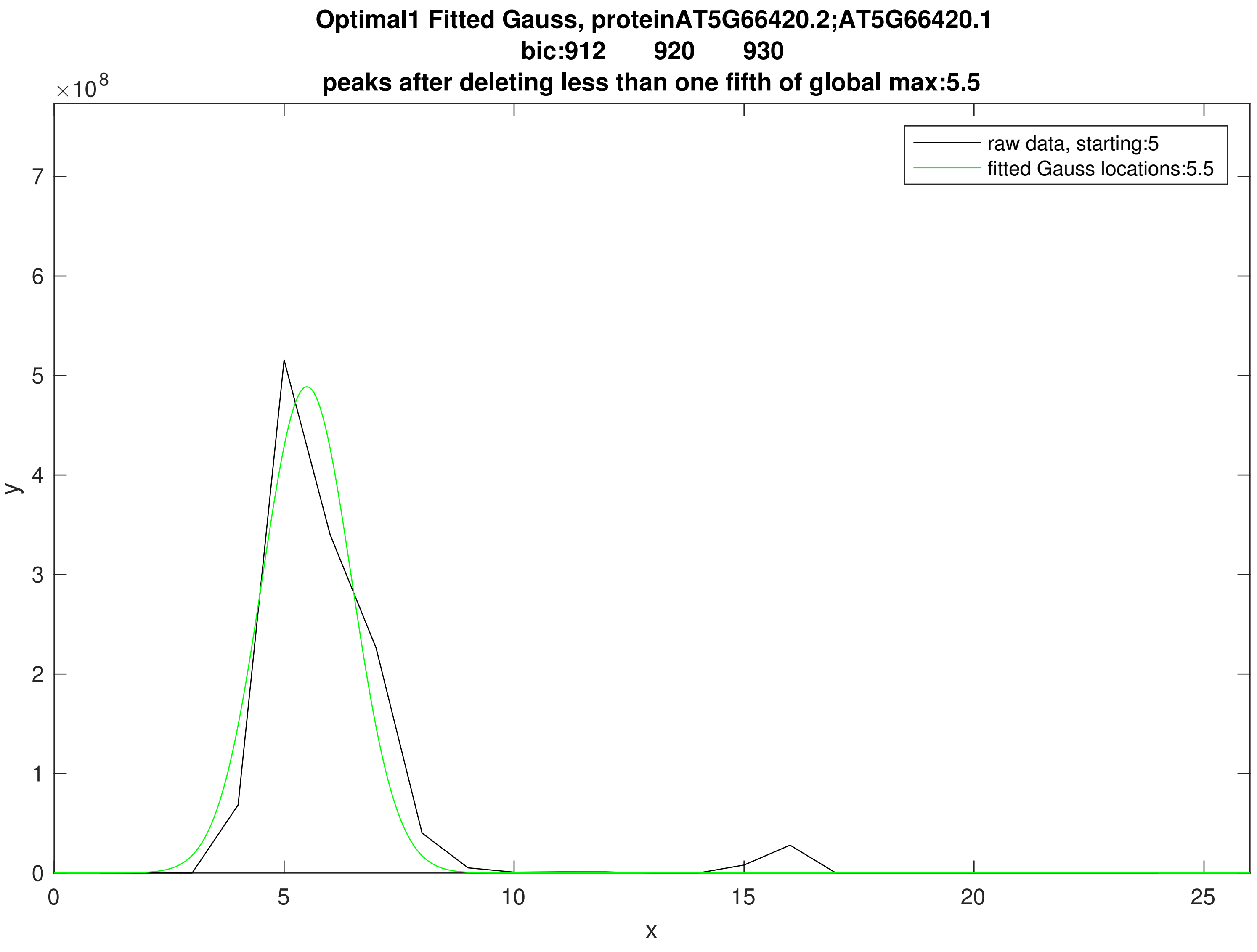
Optimal1 Fitted Gauss, proteinAT5G66420.2;AT5G66420.1
bic:912 920 930
peaks after deleting less than one fifth of global max:5.5
×10^8
raw data, starting:5
fitted Gauss locations:5.5
y
x
0
1
2
3
4
5
6
7
0
5
10
15
20
25

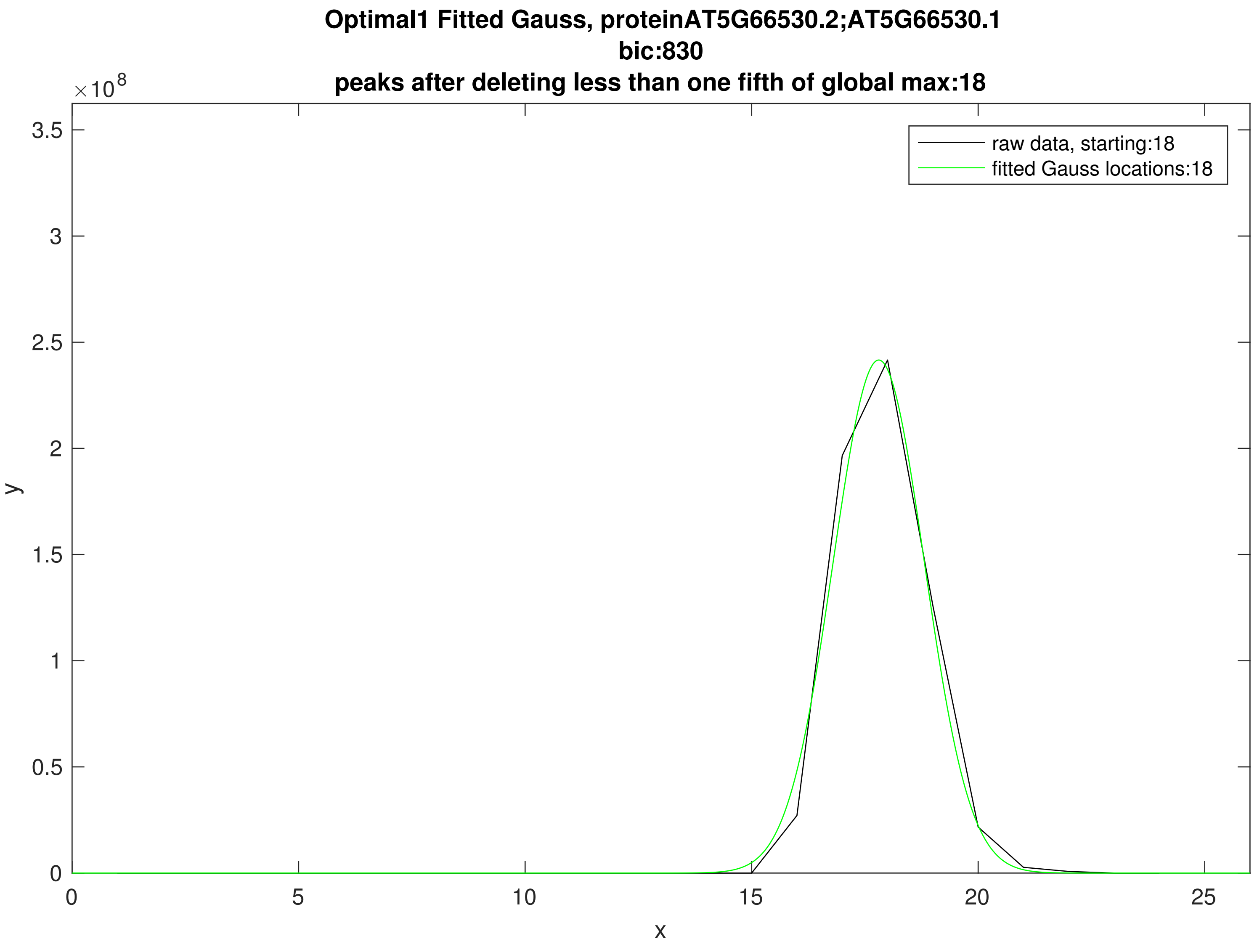
Optimal1 Fitted Gauss, proteinAT5G66530.2;AT5G66530.1
bic:830
peaks after deleting less than one fifth of global max:18
×10^8
3.5
3
2.5
2
1.5
1
0.5
0
y
0
5
10
15
20
25
x
raw data, starting:18
fitted Gauss locations:18

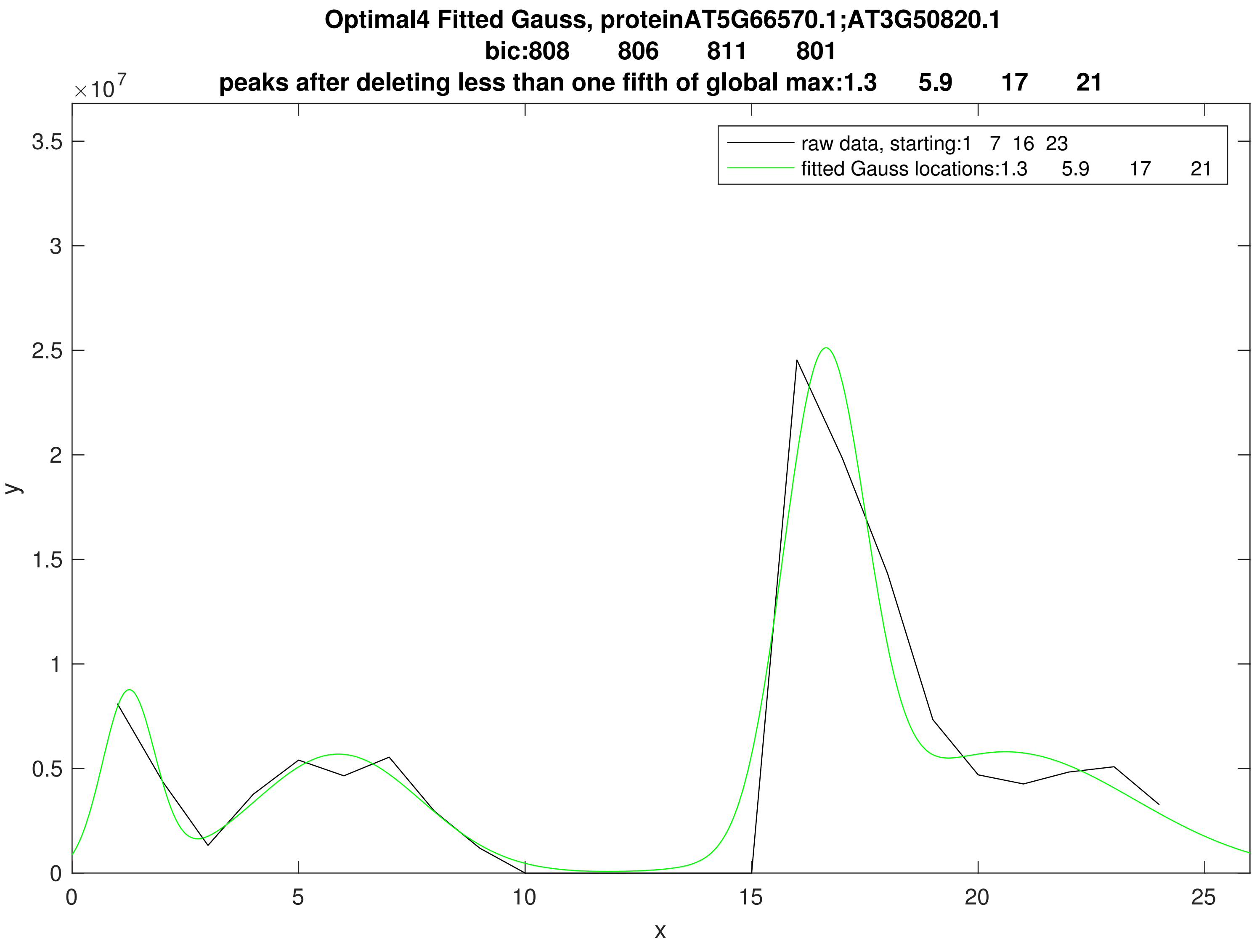

Optimal4 Fitted Gauss, proteinAT5G66570.1;AT3G50820.1
bic:808 806 811 801
peaks after deleting less than one fifth of global max:1.3 5.9 17 21
×10^7
raw data, starting:1 7 16 23
fitted Gauss locations:1.3 5.9 17 21
3.5
3
2.5
2
1.5
1
0.5
0
y
0
5
10
15
20
25
x

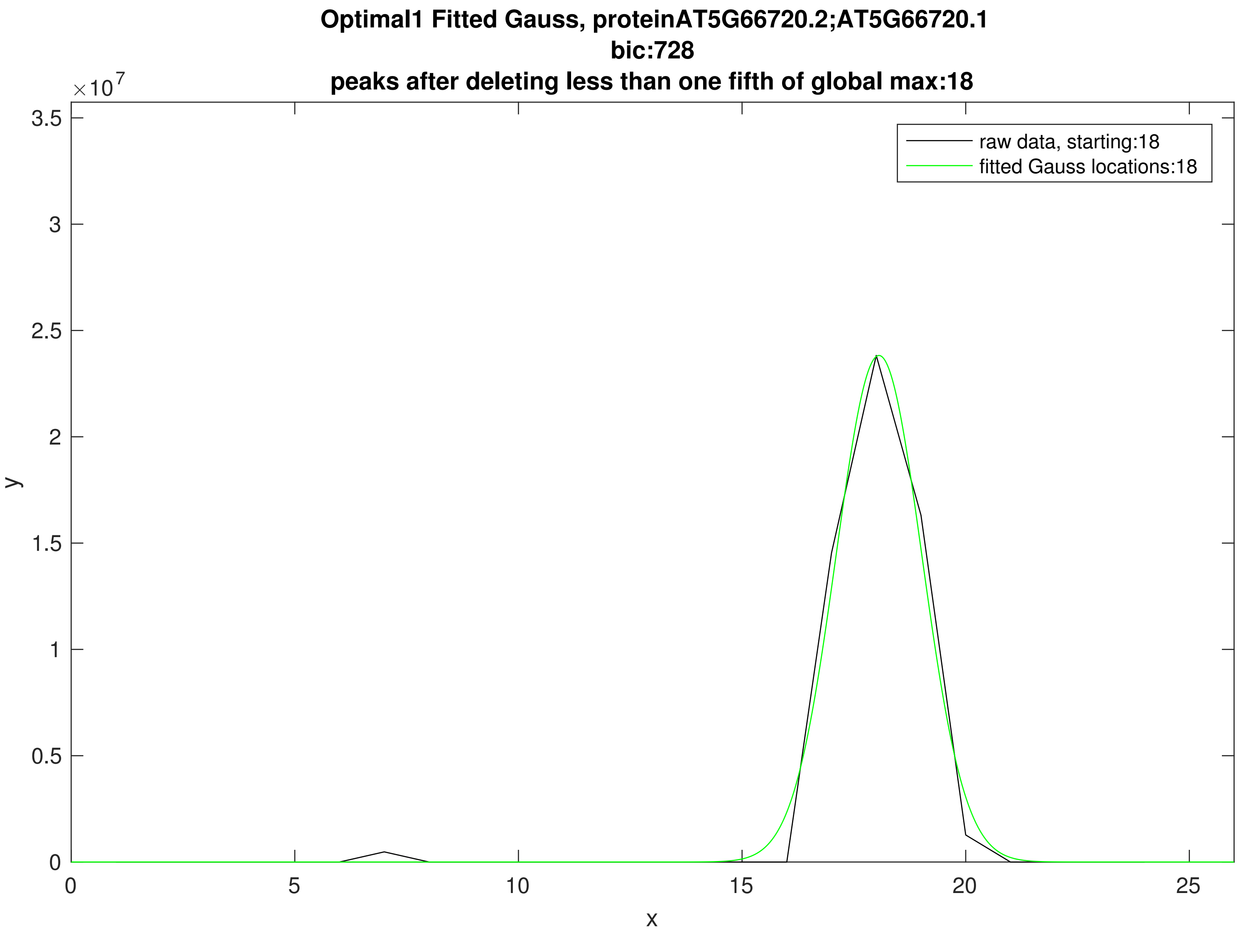
Optimal1 Fitted Gauss, proteinAT5G66720.2;AT5G66720.1
bic:728
peaks after deleting less than one fifth of global max:18
×10^7
raw data, starting:18
fitted Gauss locations:18
y
x
0
0.5
1
1.5
2
2.5
3
3.5
0
5
10
15
20
25

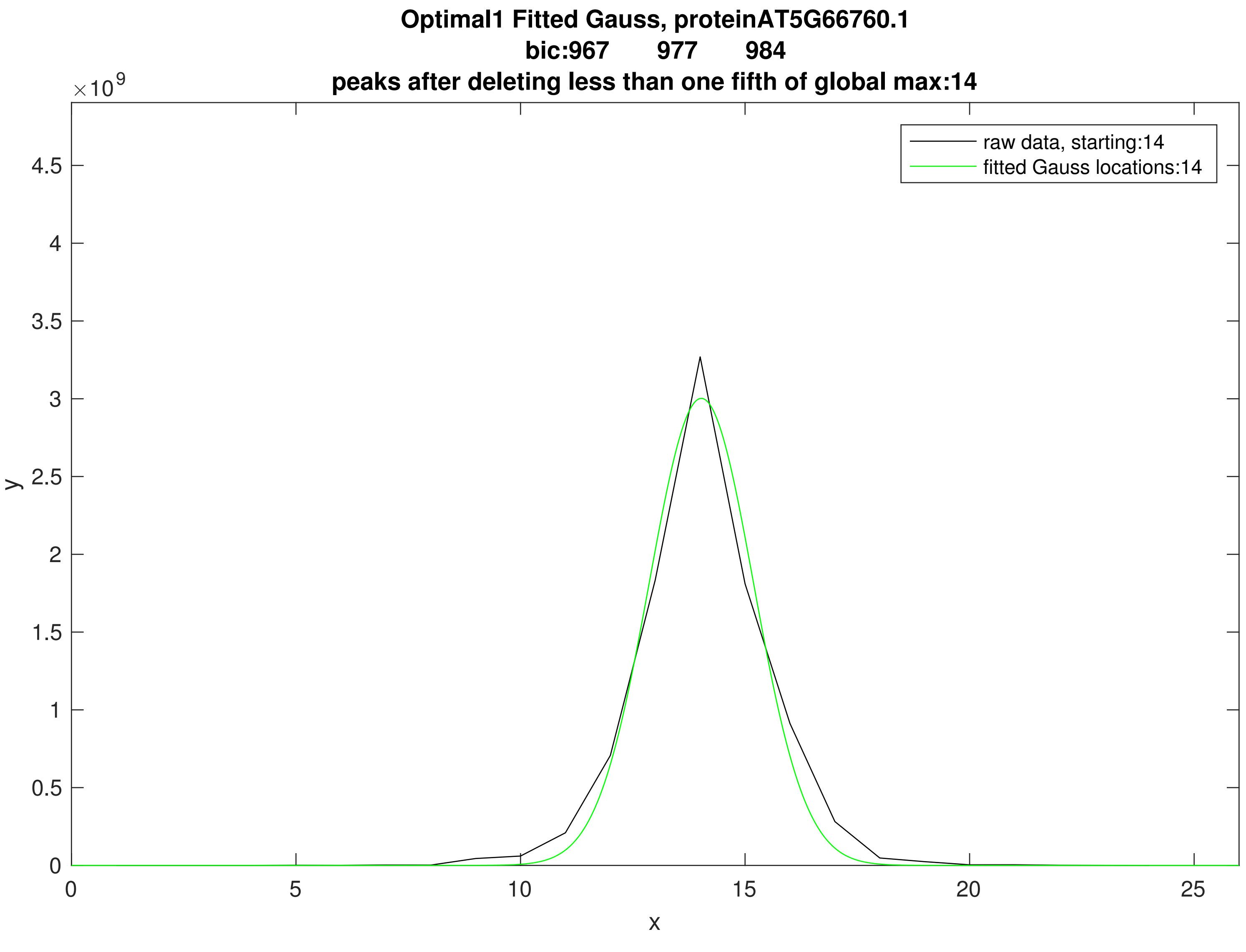
Optimal1 Fitted Gauss, proteinAT5G66760.1
bic:967 977 984
peaks after deleting less than one fifth of global max:14
×10^9
raw data, starting:14
fitted Gauss locations:14
y
x
0
0.5
1
1.5
2
2.5
3
3.5
4
4.5
0
5
10
15
20
25

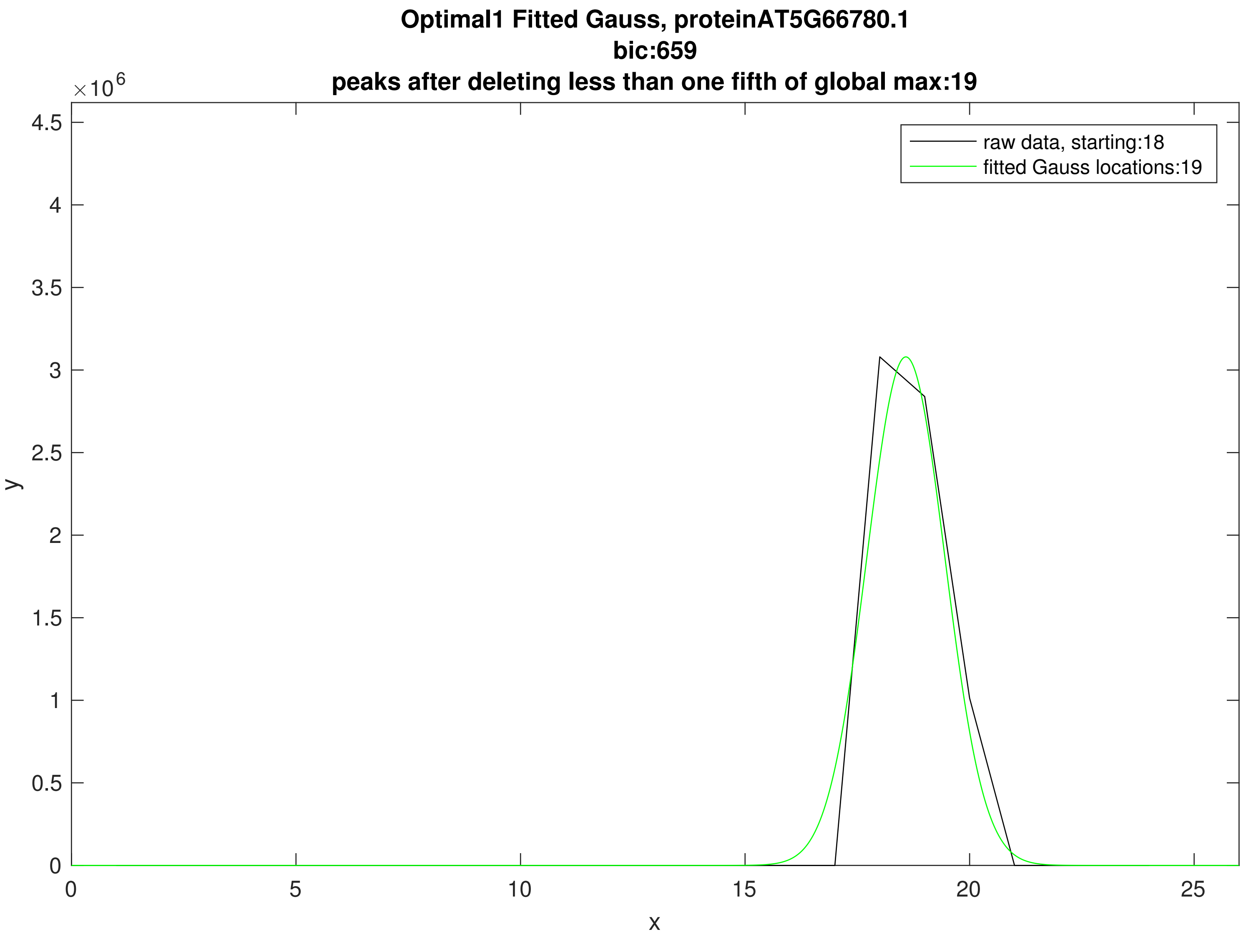
Optimal1 Fitted Gauss, proteinAT5G66780.1
bic:659
peaks after deleting less than one fifth of global max:19
×10^6
4.5
4
3.5
3
2.5
2
1.5
1
0.5
0
y
0
5
10
15
20
25
x
raw data, starting:18
fitted Gauss locations:19

**Optimal1 Fitted Gauss, proteinAT5G66920.1**
**bic:828**
**peaks after deleting less than one fifth of global max:17**

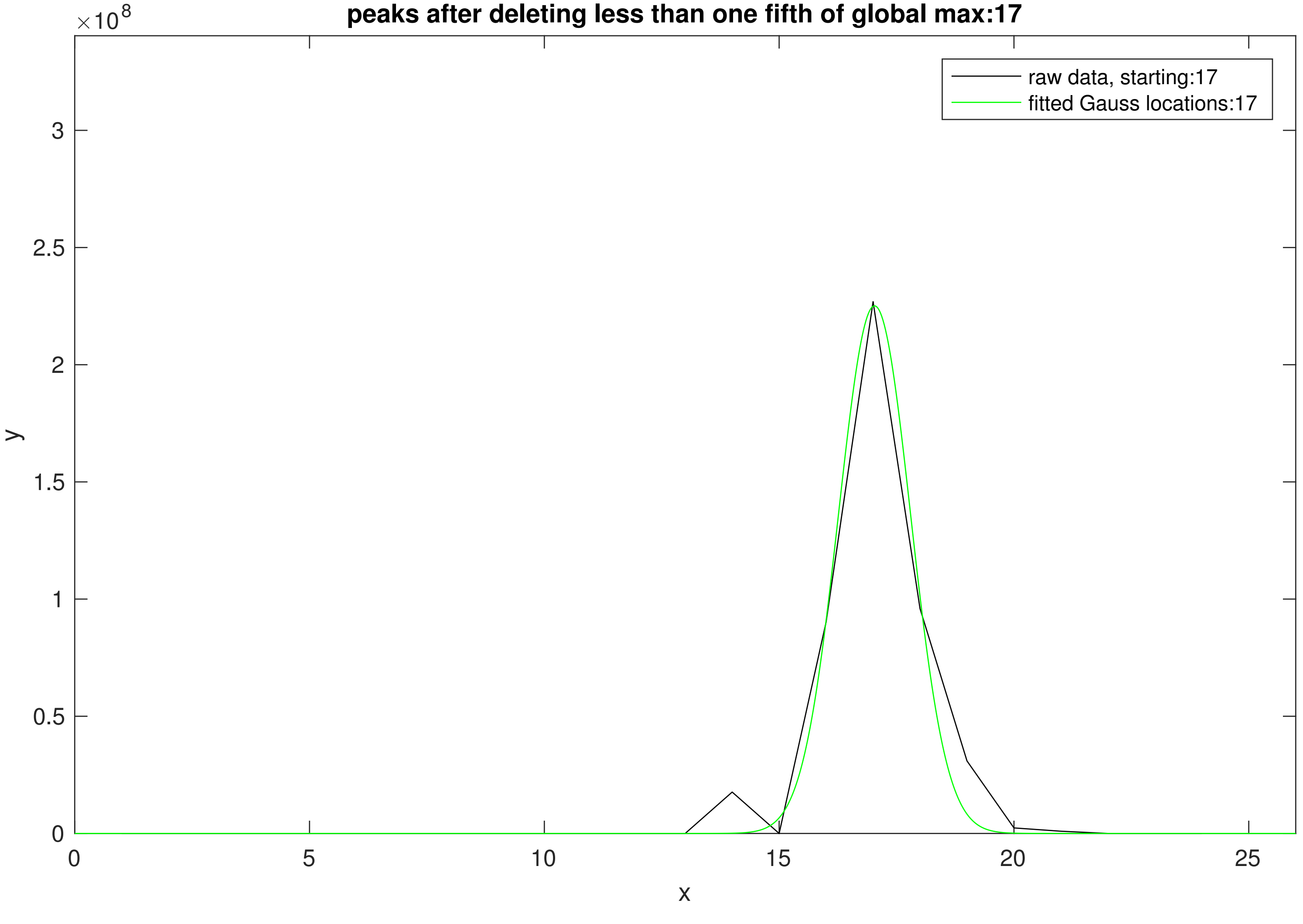

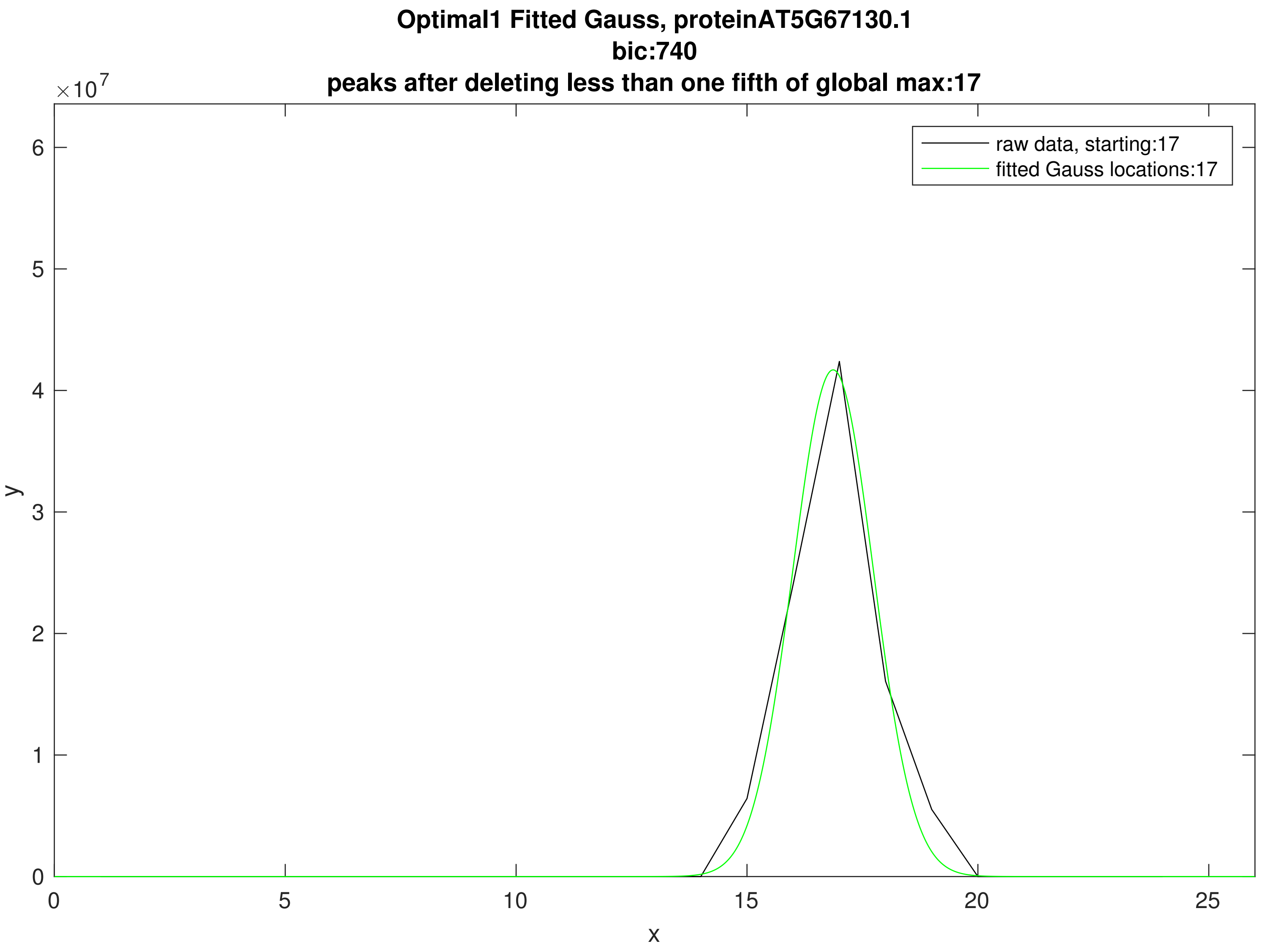
Optimal1 Fitted Gauss, proteinAT5G67130.1
bic:740
peaks after deleting less than one fifth of global max:17
×10^7
raw data, starting:17
fitted Gauss locations:17
y
x
0
1
2
3
4
5
6
0
5
10
15
20
25

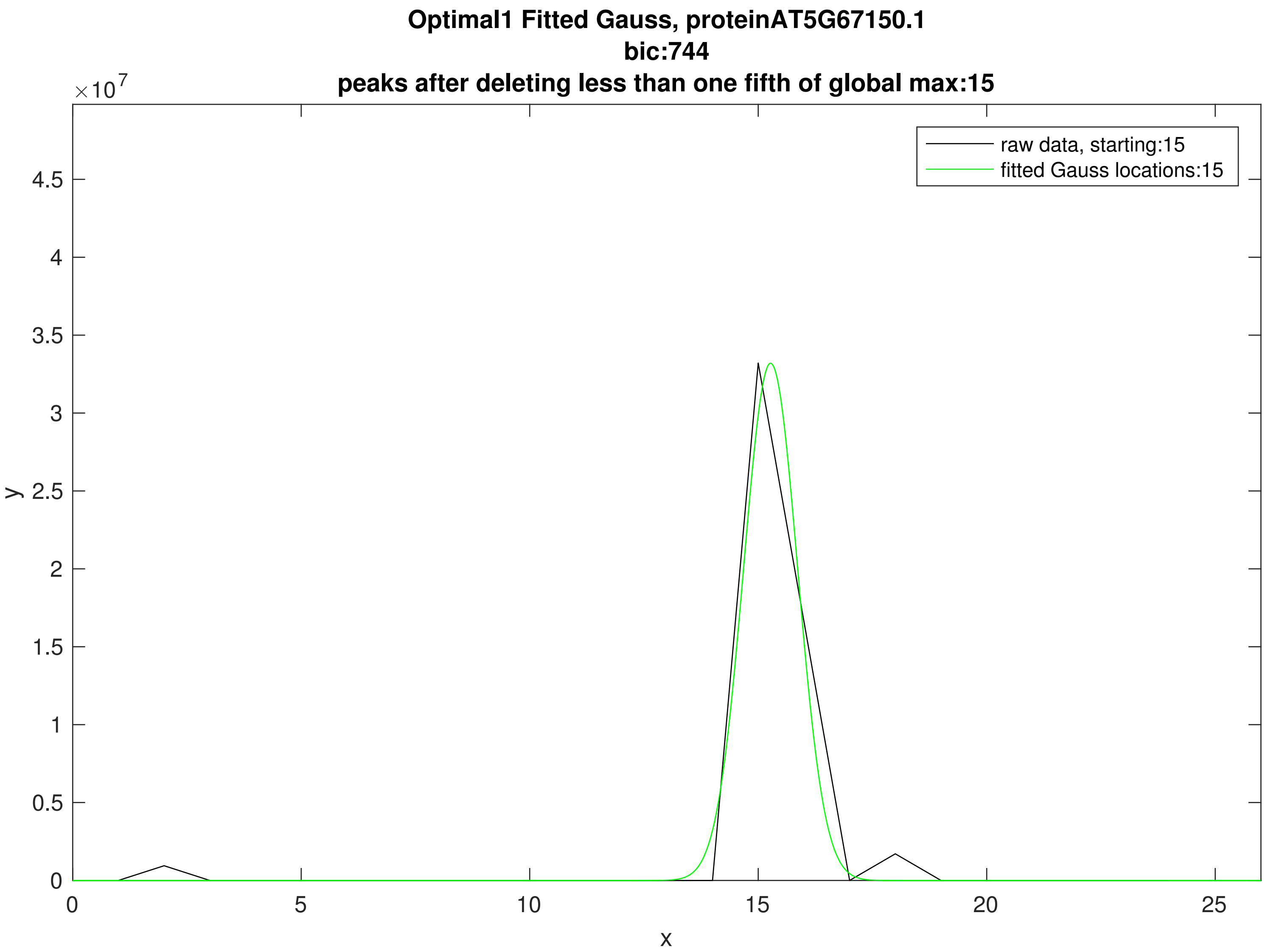

Optimal1 Fitted Gauss, proteinAT5G67150.1
bic:744
peaks after deleting less than one fifth of global max:15
×10^7
raw data, starting:15
fitted Gauss locations:15
4.5
4
3.5
3
2.5
2
1.5
1
0.5
0
y
0
5
10
15
20
25
x

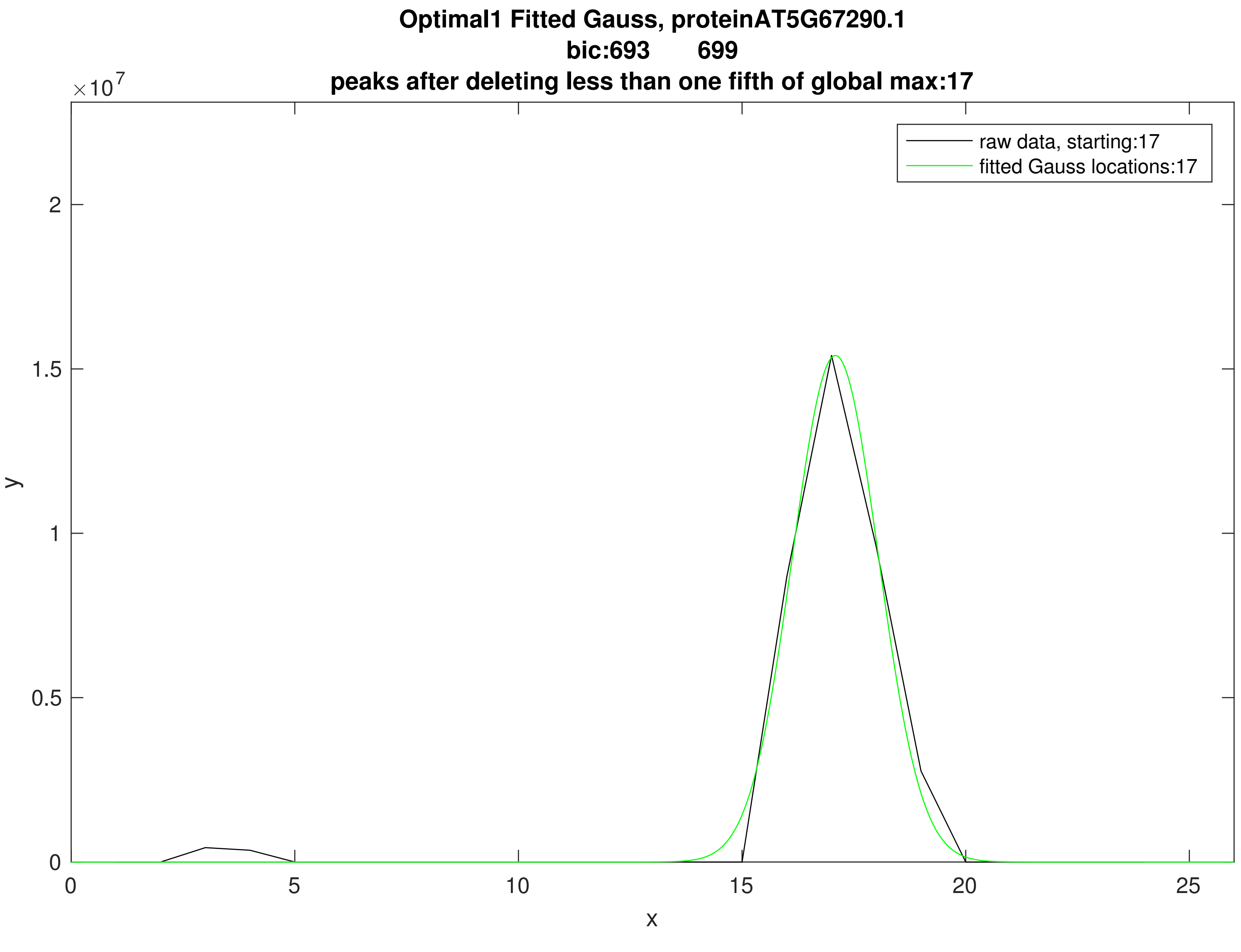
Optimal1 Fitted Gauss, proteinAT5G67290.1
bic:693 699
peaks after deleting less than one fifth of global max:17
×10^7
raw data, starting:17
fitted Gauss locations:17
2
1.5
1
0.5
0
y
0
5
10
15
20
25
x

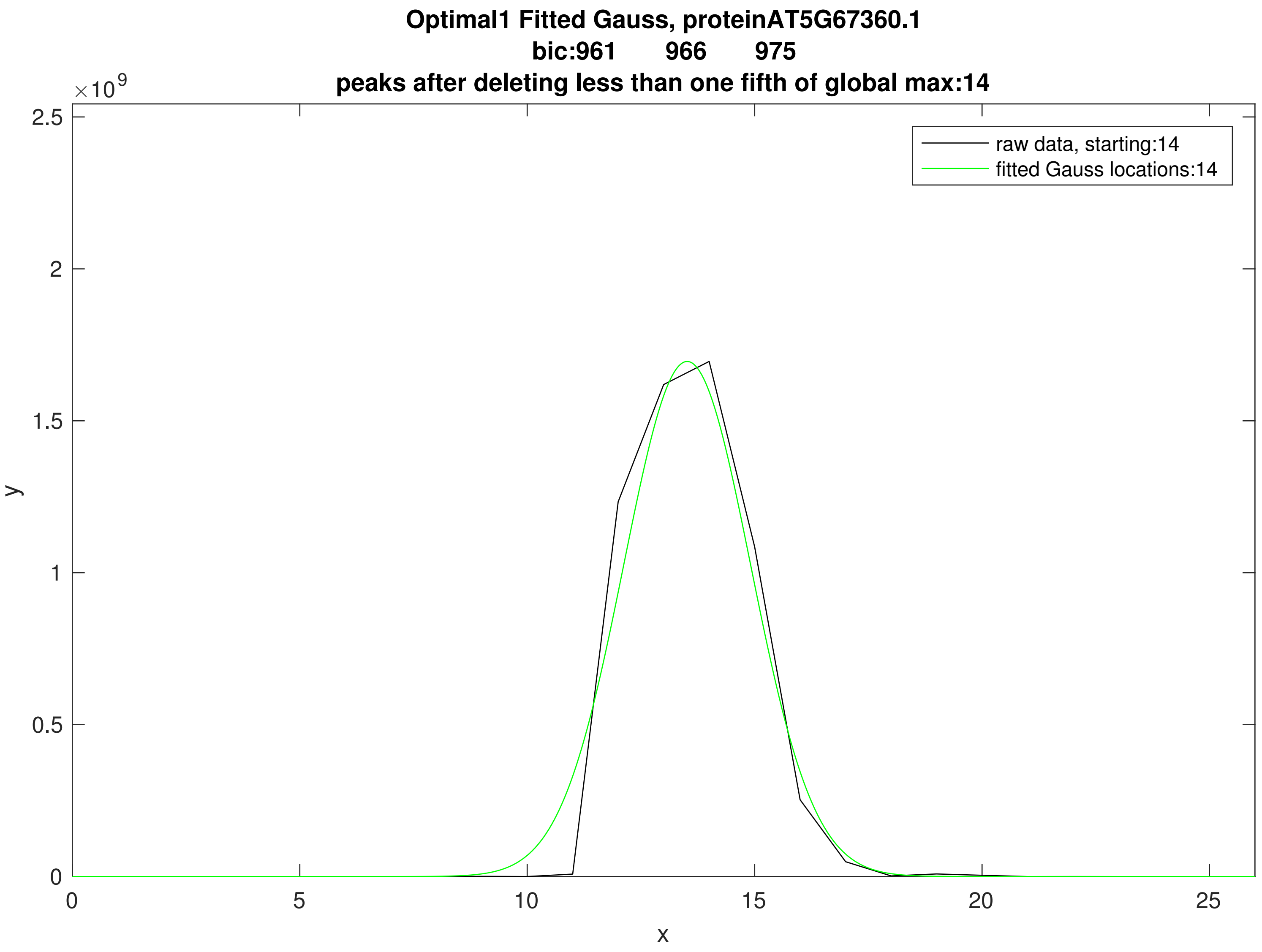

Optimal1 Fitted Gauss, proteinAT5G67360.1
bic:961 966 975
peaks after deleting less than one fifth of global max:14
×10^9
2.5
2
1.5
1
0.5
0
y
0
5
10
15
20
25
x
raw data, starting:14
fitted Gauss locations:14

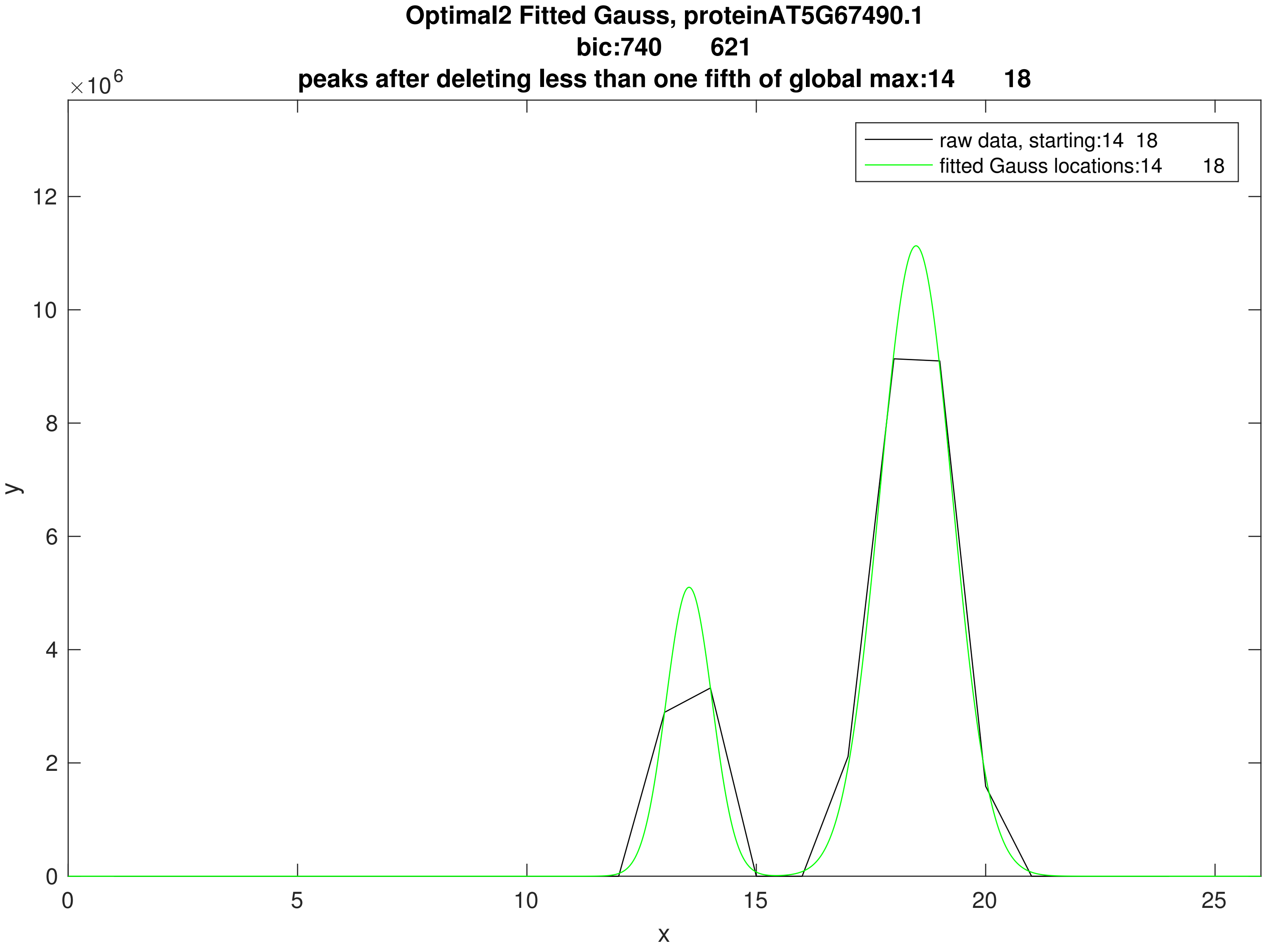

Optimal2 Fitted Gauss, proteinAT5G67490.1
bic:740 621
peaks after deleting less than one fifth of global max:14 18
×10^6
raw data, starting:14 18
fitted Gauss locations:14 18
y
x
0
2
4
6
8
10
12
0
5
10
15
20
25

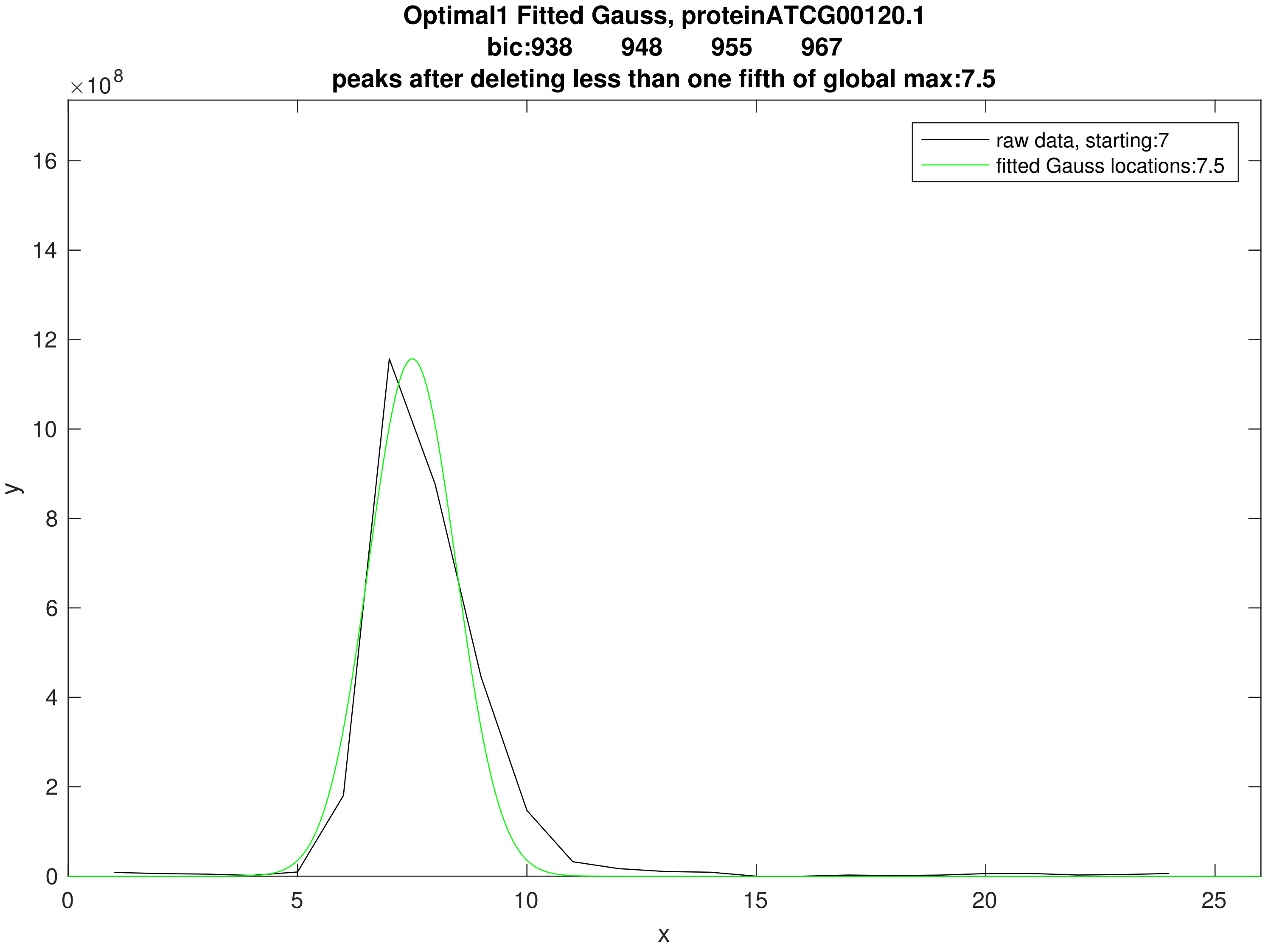

Optimal1 Fitted Gauss, proteinATCG00120.1
bic:938 948 955 967
peaks after deleting less than one fifth of global max:7.5
×10^8
16
14
12
10
8
6
4
2
0
y
0
5
10
15
20
25
x
raw data, starting:7
fitted Gauss locations:7.5

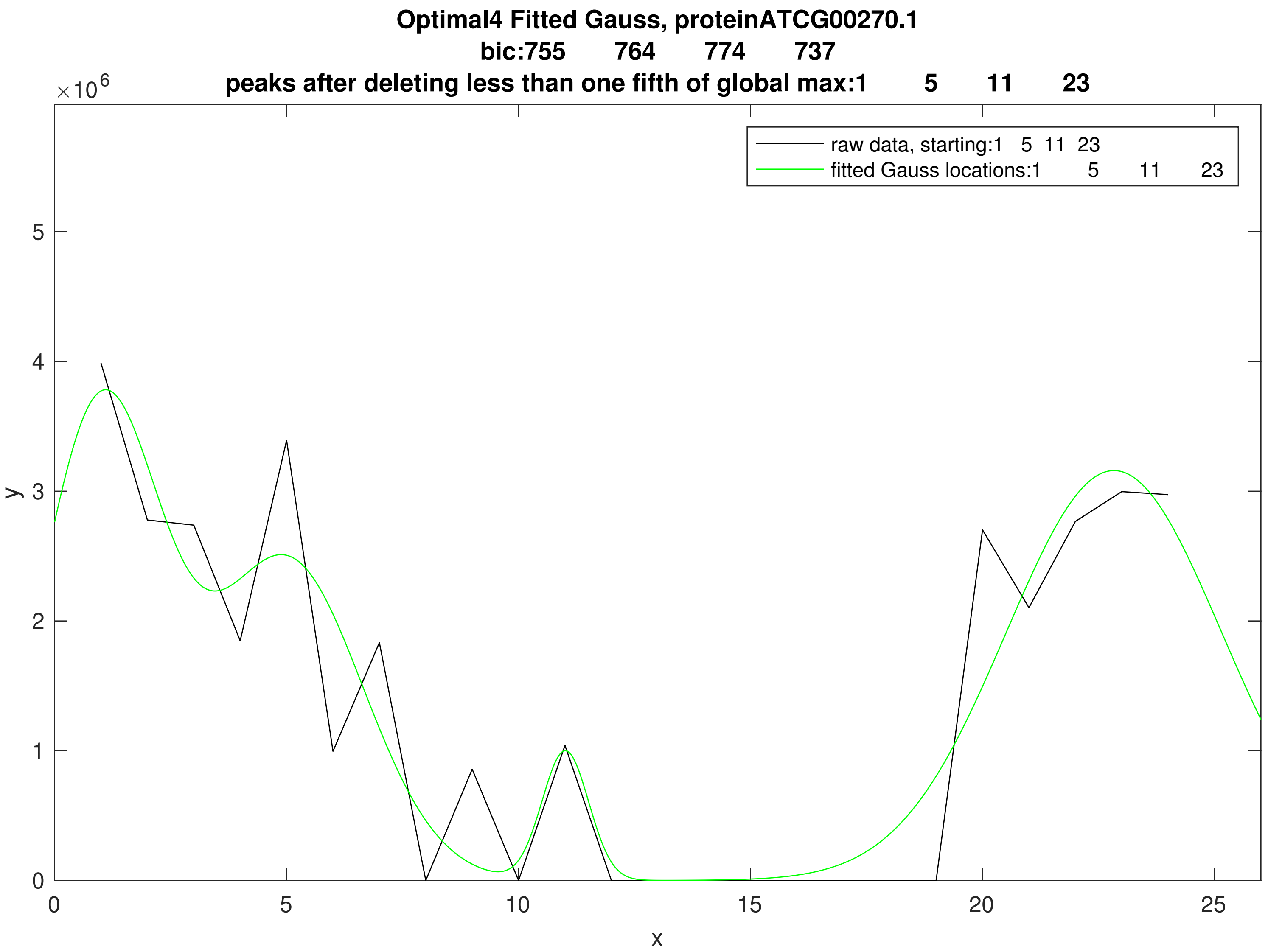
Optimal4 Fitted Gauss, proteinATCG00270.1
bic:755 764 774 737
peaks after deleting less than one fifth of global max:1 5 11 23
×10^6
raw data, starting:1 5 11 23
fitted Gauss locations:1 5 11 23
y
x
0
1
2
3
4
5
0
5
10
15
20
25

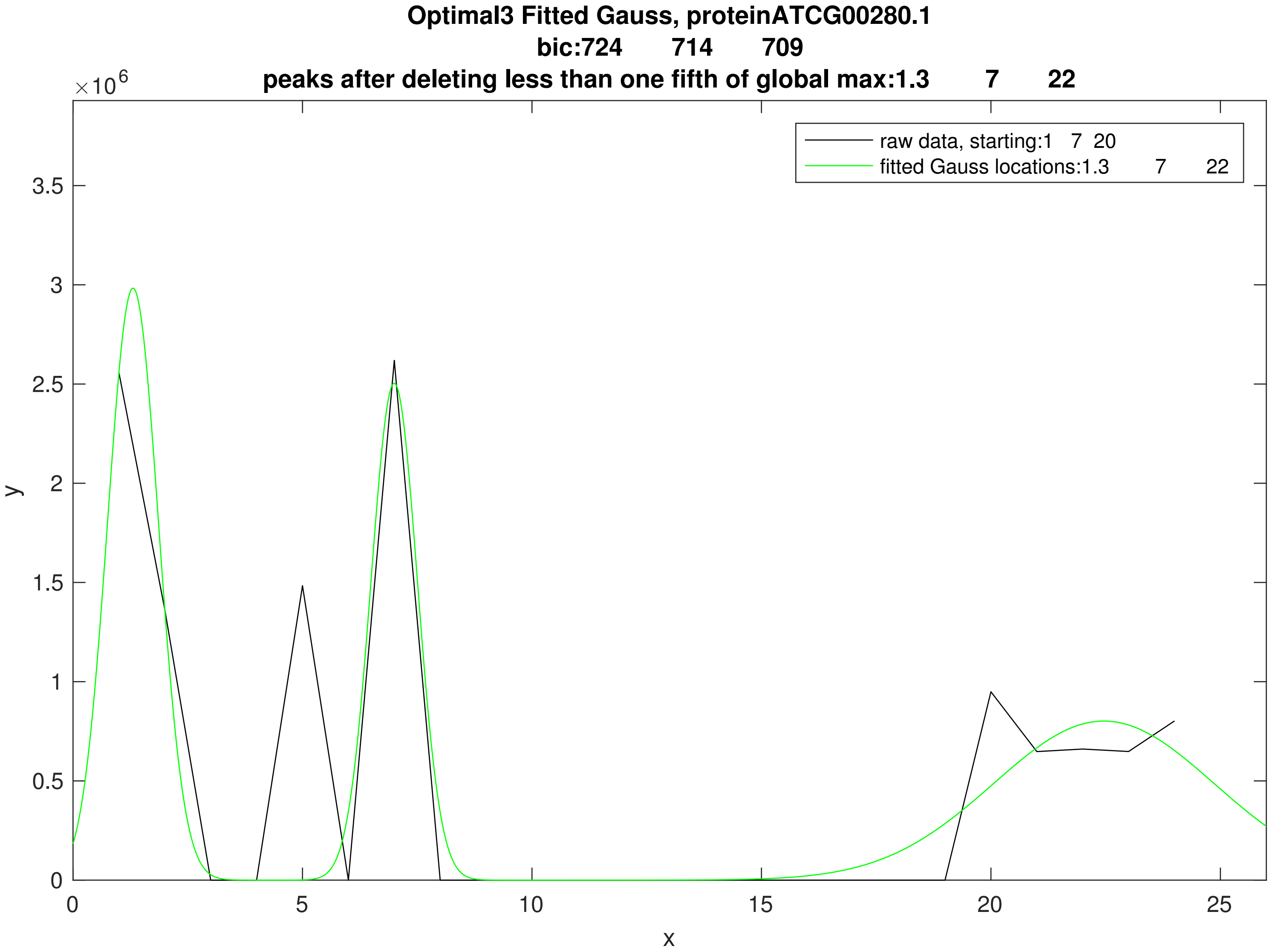
Optimal3 Fitted Gauss, proteinATCG00280.1
bic:724 714 709
peaks after deleting less than one fifth of global max:1.3 7 22
×10^6
raw data, starting:1 7 20
fitted Gauss locations:1.3 7 22
0
0.5
1
1.5
2
2.5
3
3.5
y
0
5
10
15
20
25
x

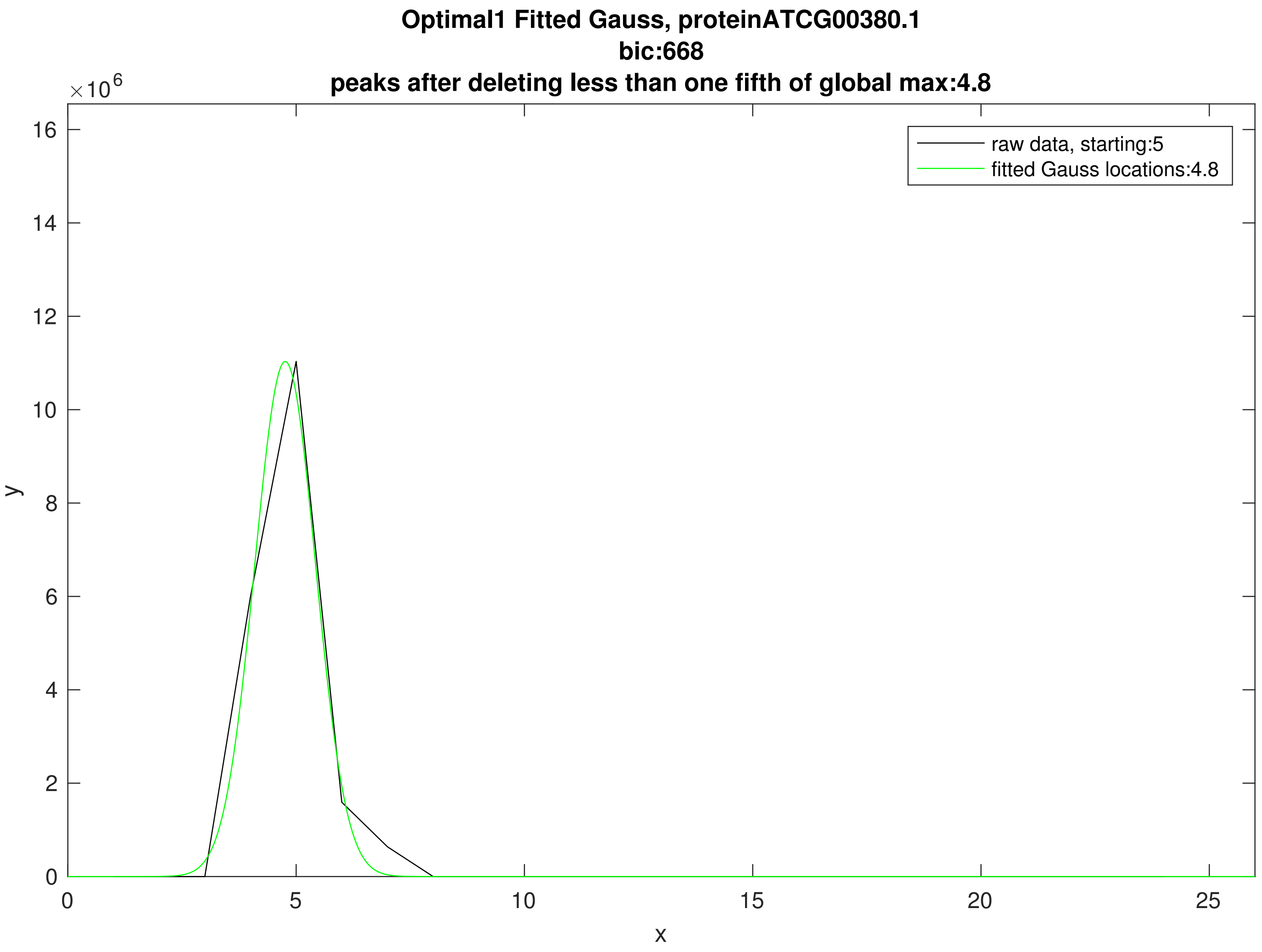
Optimal1 Fitted Gauss, proteinATCG00380.1
bic:668
peaks after deleting less than one fifth of global max:4.8
×10^6
raw data, starting:5
fitted Gauss locations:4.8
y
x
0
2
4
6
8
10
12
14
16
0
5
10
15
20
25

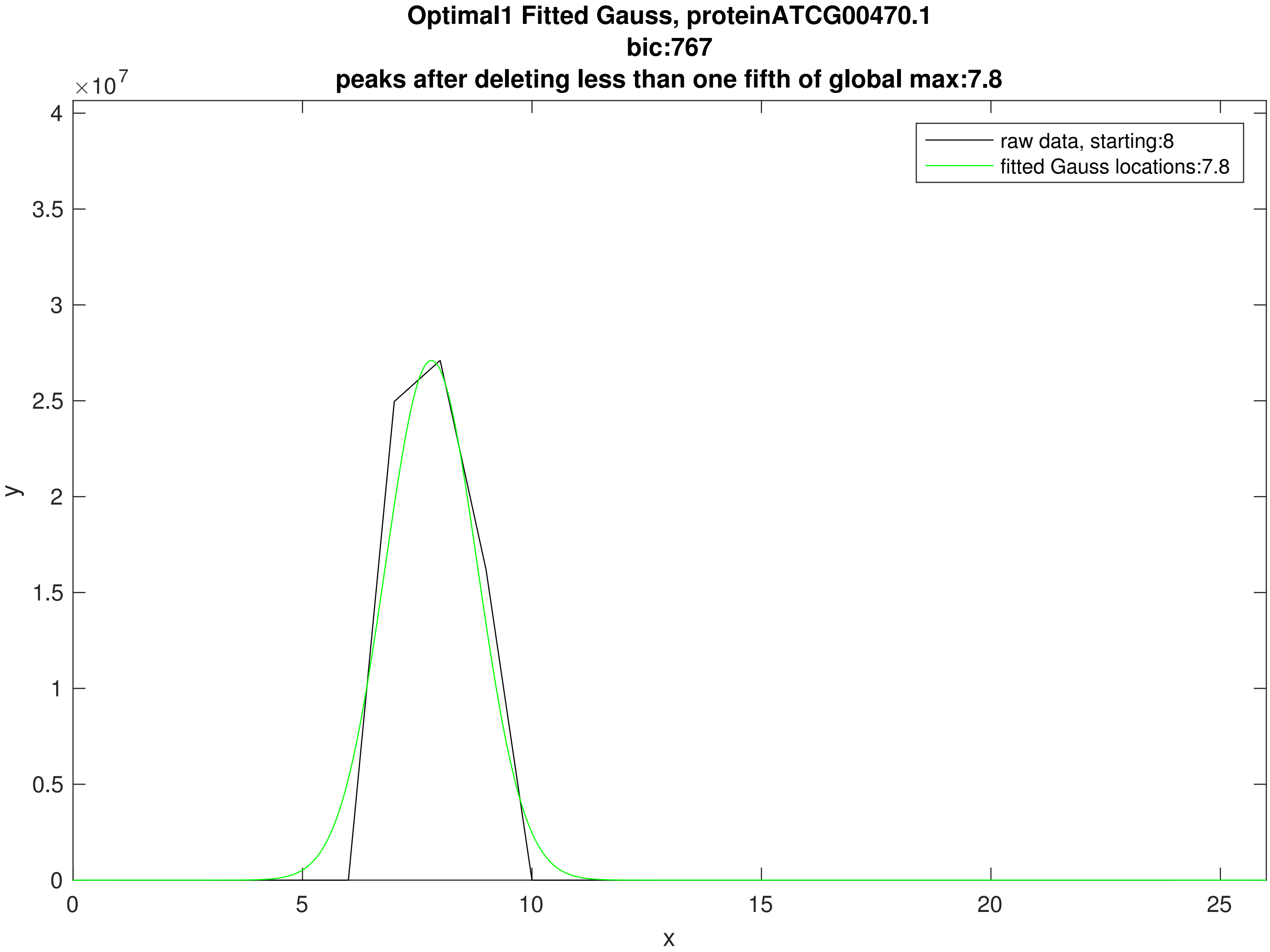
Optimal1 Fitted Gauss, proteinATCG00470.1
bic:767
peaks after deleting less than one fifth of global max:7.8
×10^7
4
3.5
3
2.5
2
1.5
1
0.5
0
y
0
5
10
15
20
25
x
raw data, starting:8
fitted Gauss locations:7.8

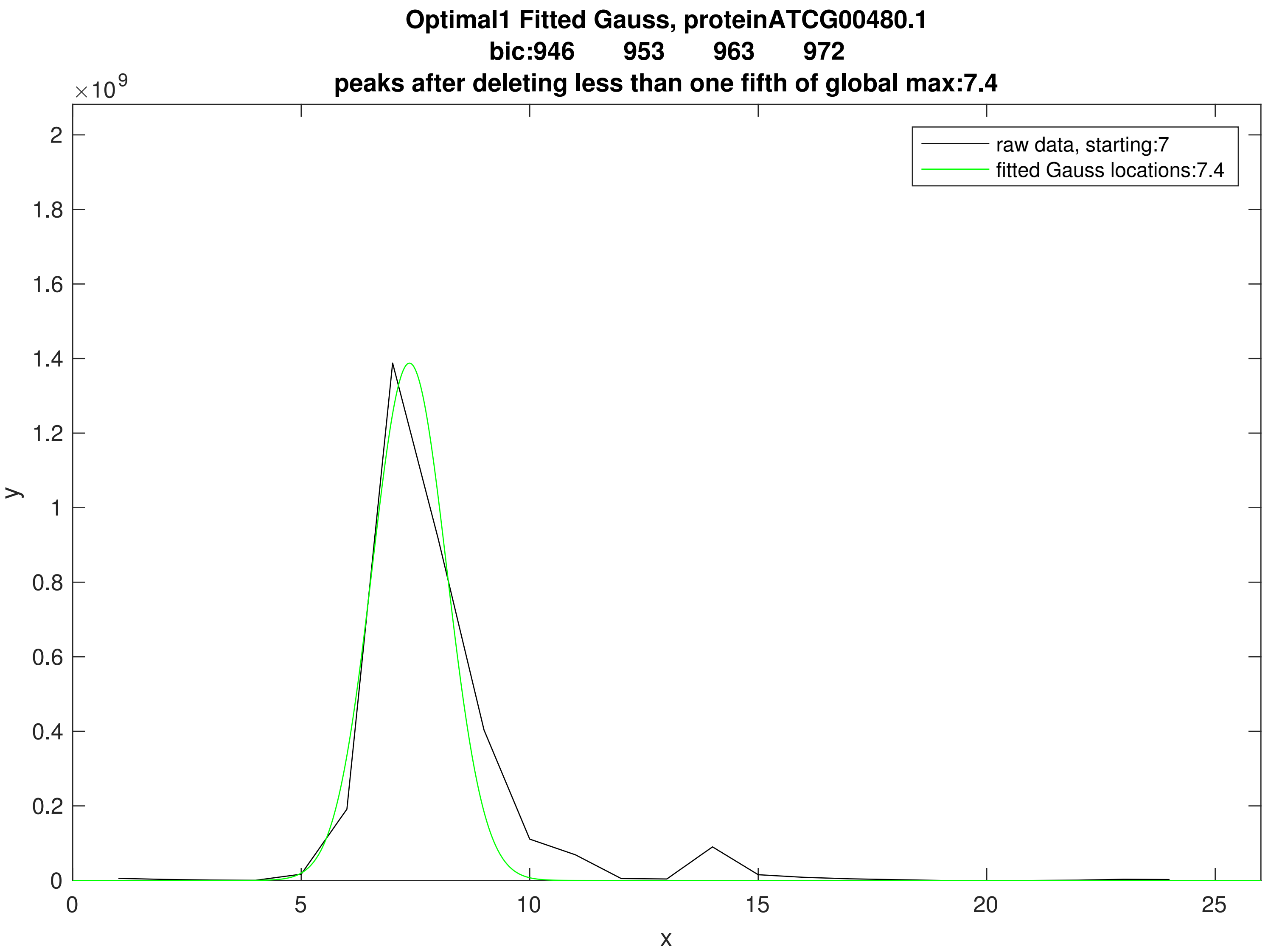
Optimal1 Fitted Gauss, proteinATCG00480.1
bic:946 953 963 972
peaks after deleting less than one fifth of global max:7.4
×10^9
raw data, starting:7
fitted Gauss locations:7.4
y
x

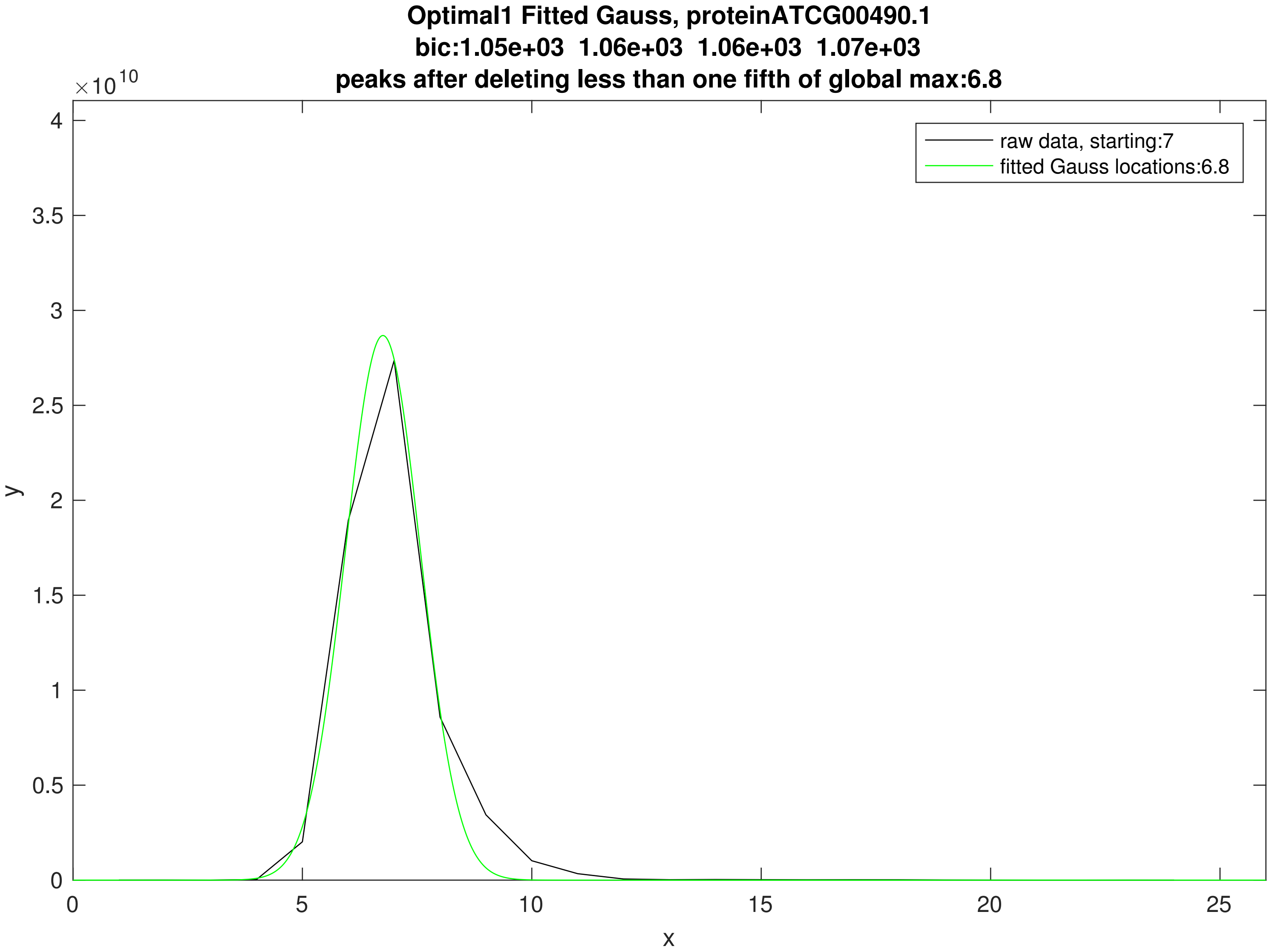

Optimal1 Fitted Gauss, proteinATCG00490.1
bic:1.05e+03  1.06e+03  1.06e+03  1.07e+03
peaks after deleting less than one fifth of global max:6.8
×10^10
4
3.5
3
2.5
2
1.5
1
0.5
0
y
0
5
10
15
20
25
x
raw data, starting:7
fitted Gauss locations:6.8

#### Optimal1 Fitted Gauss, proteinATCG00500.1
#### bic:575
#### peaks after deleting less than one fifth of global max:5.1

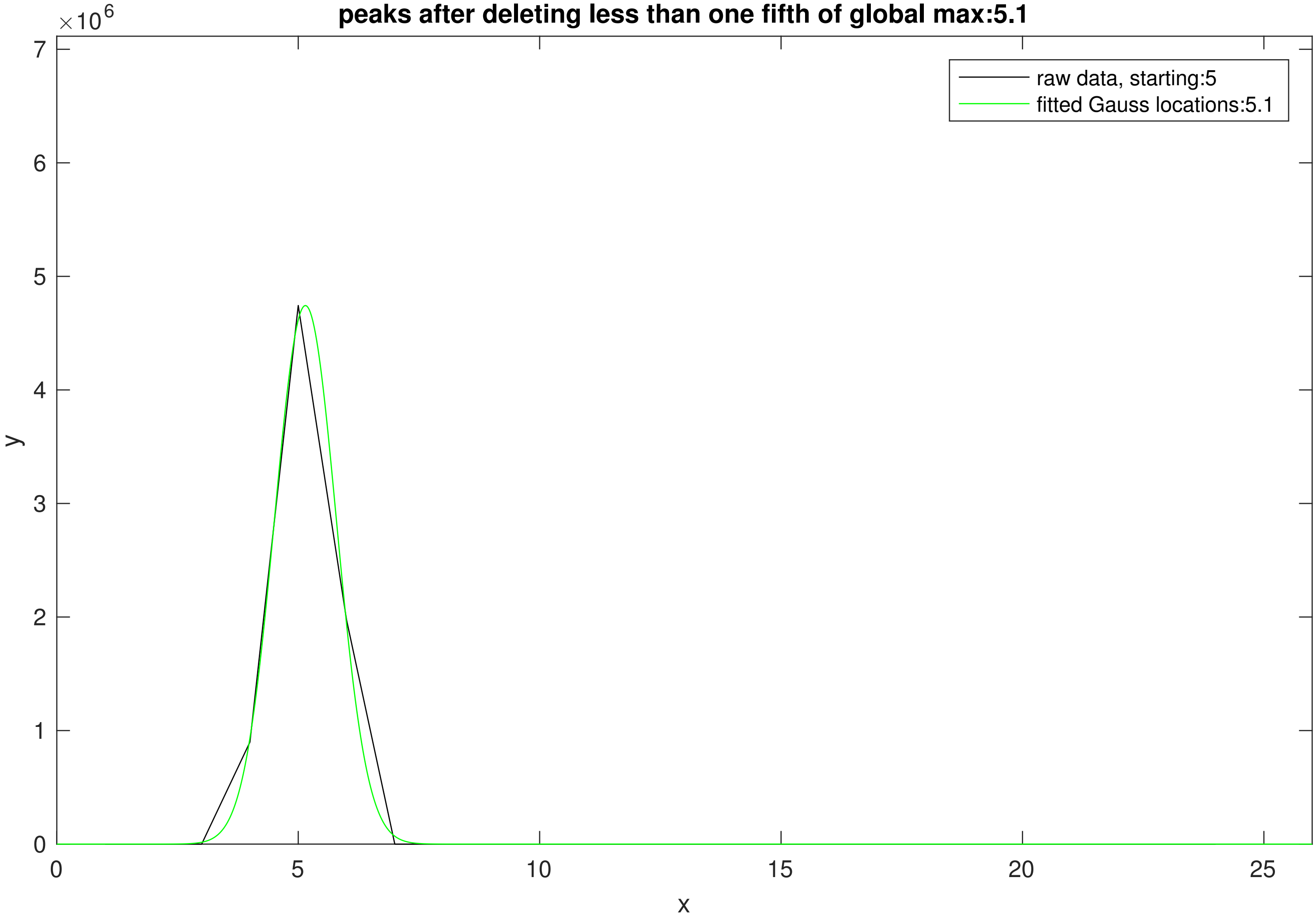

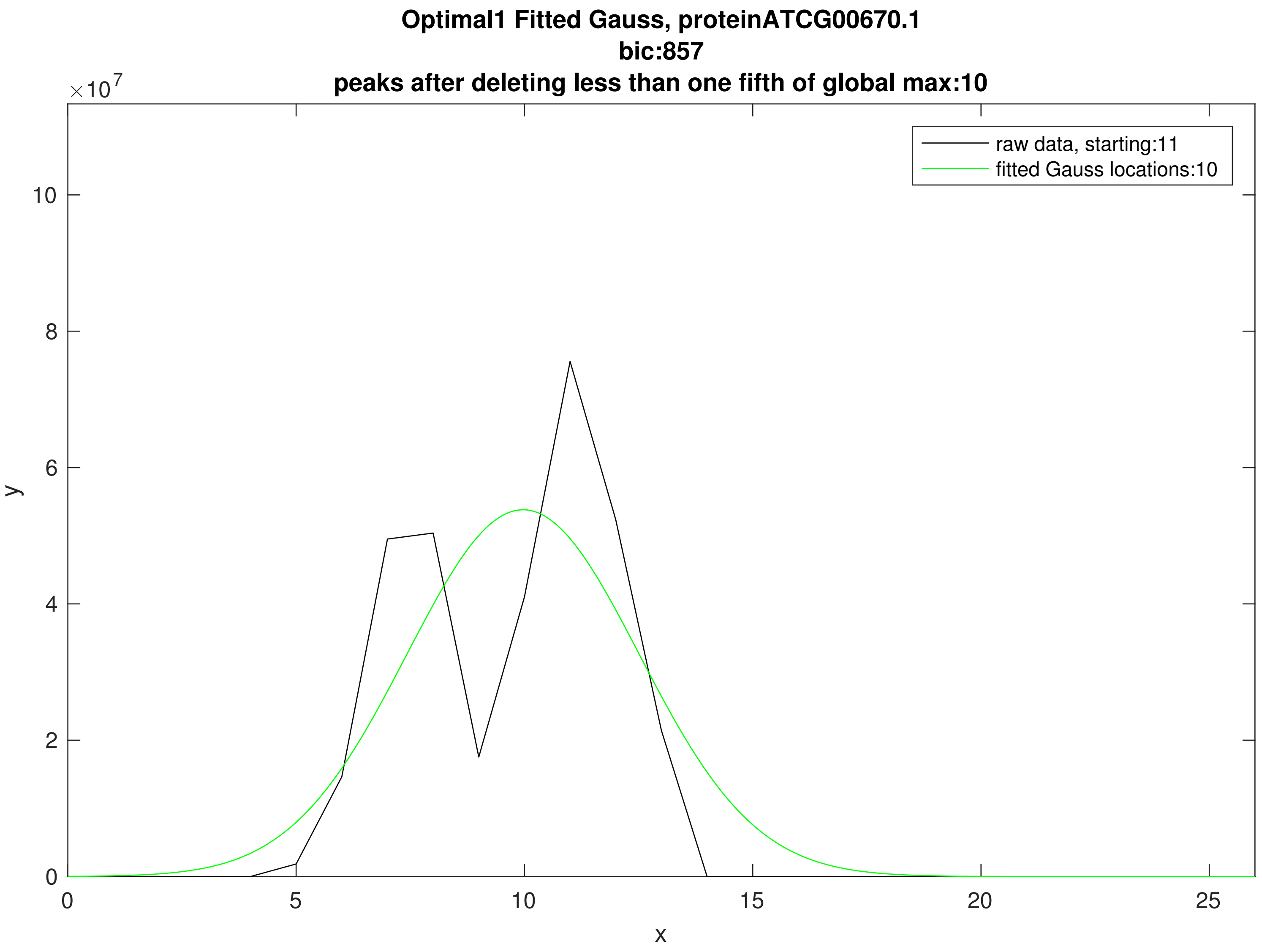
Optimal1 Fitted Gauss, proteinATCG00670.1
bic:857
peaks after deleting less than one fifth of global max:10
×10^7
raw data, starting:11
fitted Gauss locations:10
y
x
0
2
4
6
8
10
0
5
10
15
20
25

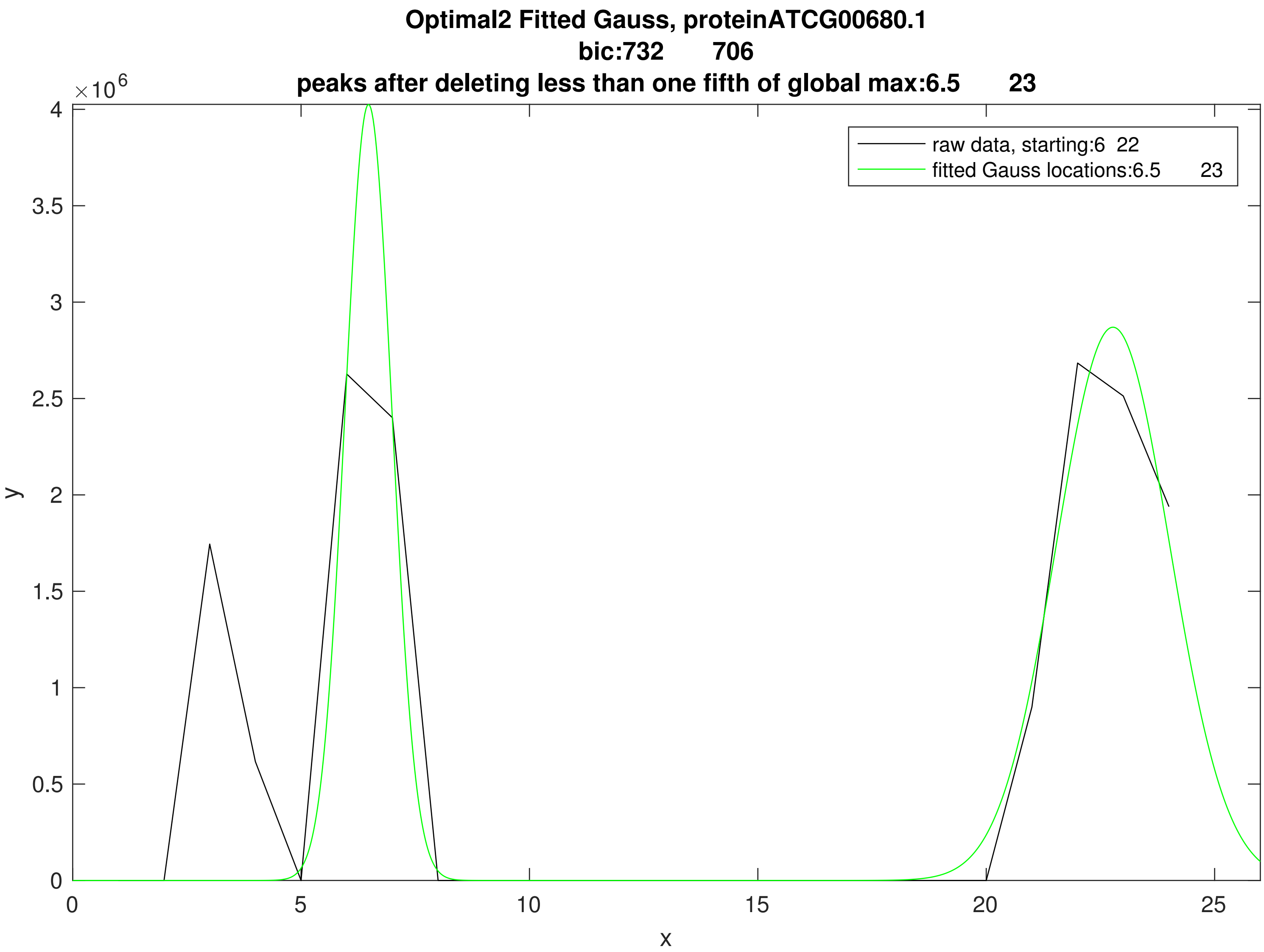

Optimal2 Fitted Gauss, proteinATCG00680.1
bic:732 706
peaks after deleting less than one fifth of global max:6.5 23
×10^6
raw data, starting:6 22
fitted Gauss locations:6.5 23
y
x
0
0.5
1
1.5
2
2.5
3
3.5
4
0
5
10
15
20
25

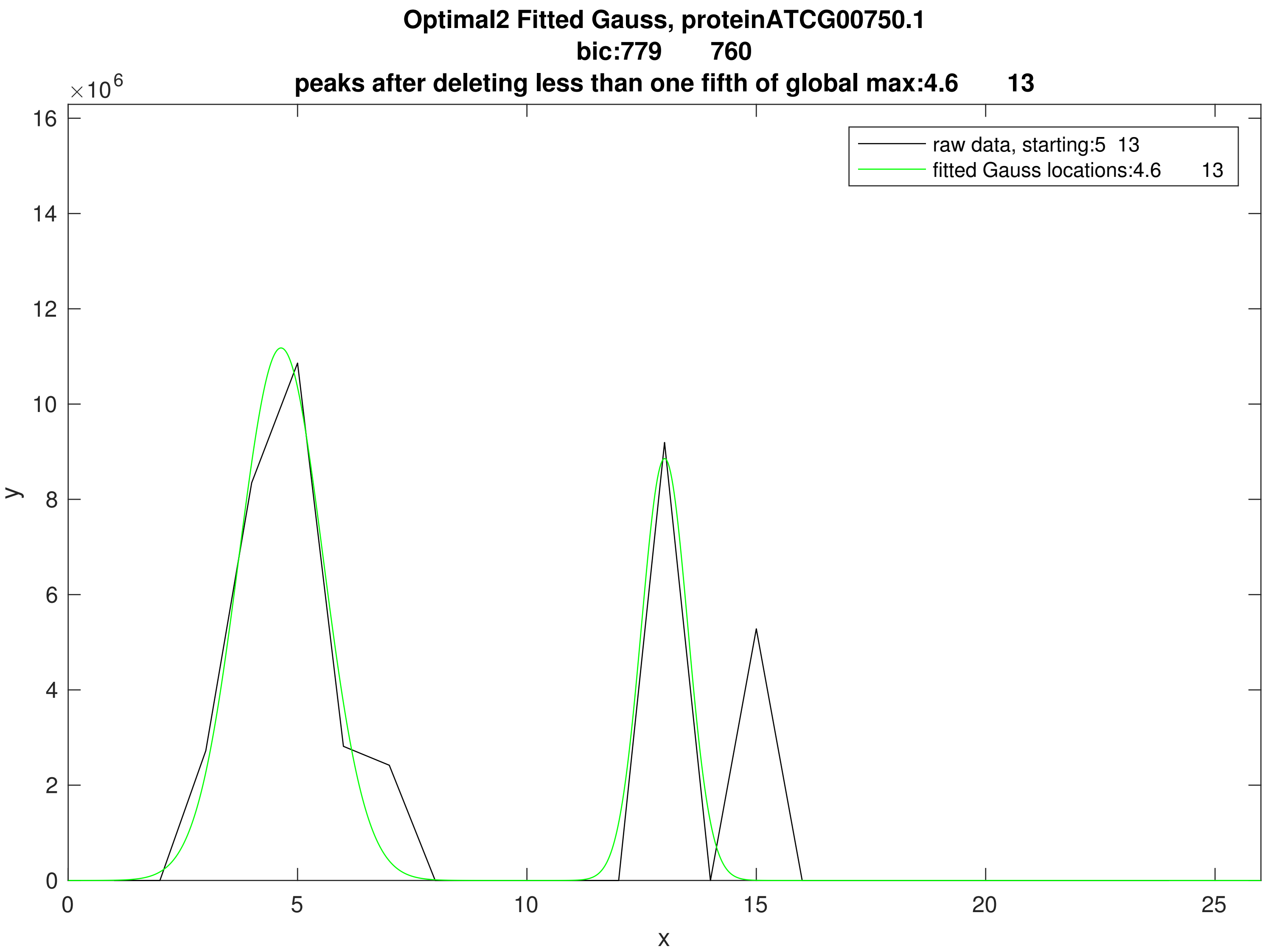

Optimal2 Fitted Gauss, proteinATCG00750.1
bic:779 760
peaks after deleting less than one fifth of global max:4.6 13
×10^6
raw data, starting:5 13
fitted Gauss locations:4.6 13
y
x
0
2
4
6
8
10
12
14
16
0
5
10
15
20
25

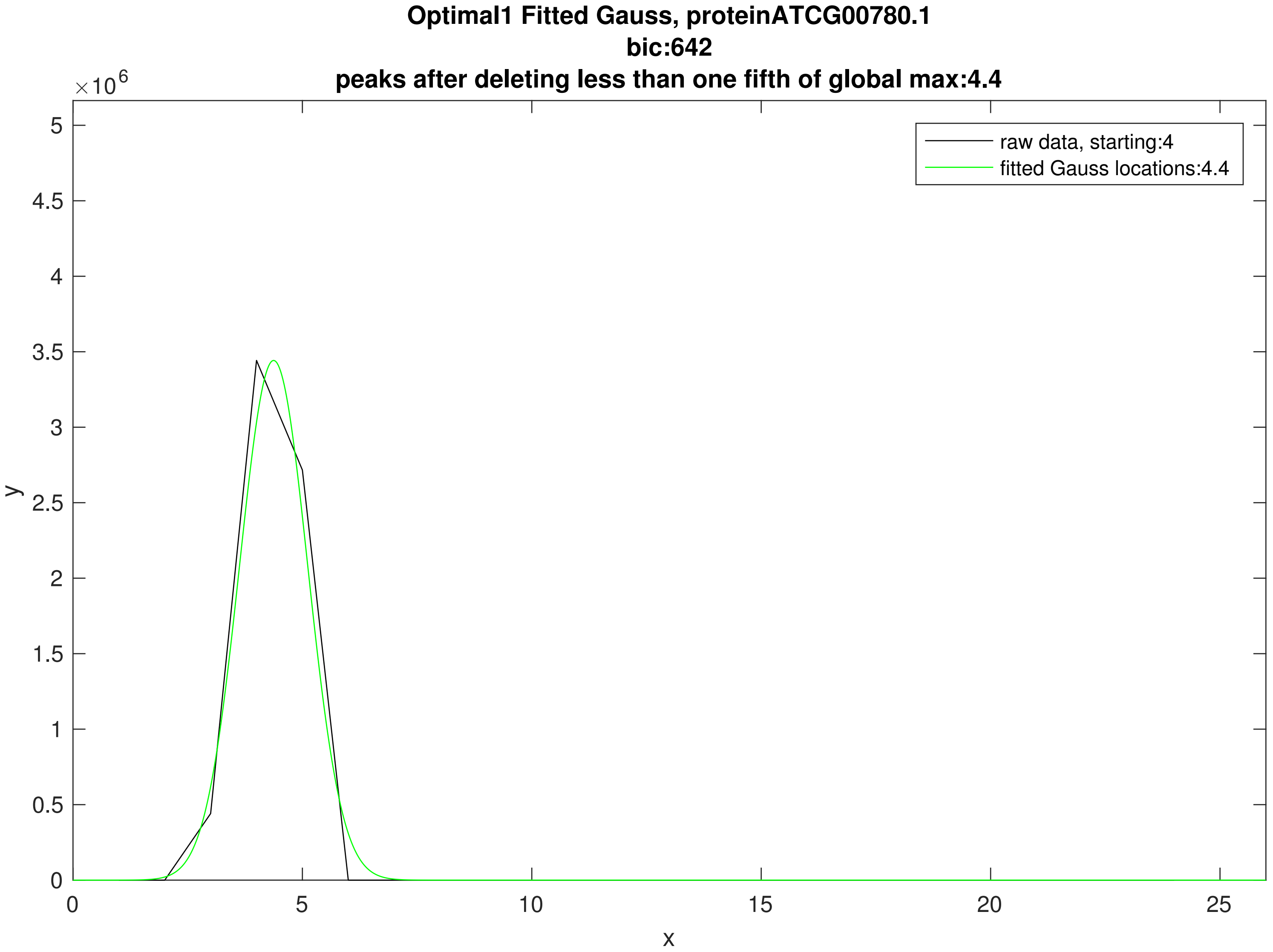
Optimal1 Fitted Gauss, proteinATCG00780.1
bic:642
peaks after deleting less than one fifth of global max:4.4
×10^6
5
4.5
4
3.5
3
2.5
2
1.5
1
0.5
0
y
0
5
10
15
20
25
x
raw data, starting:4
fitted Gauss locations:4.4

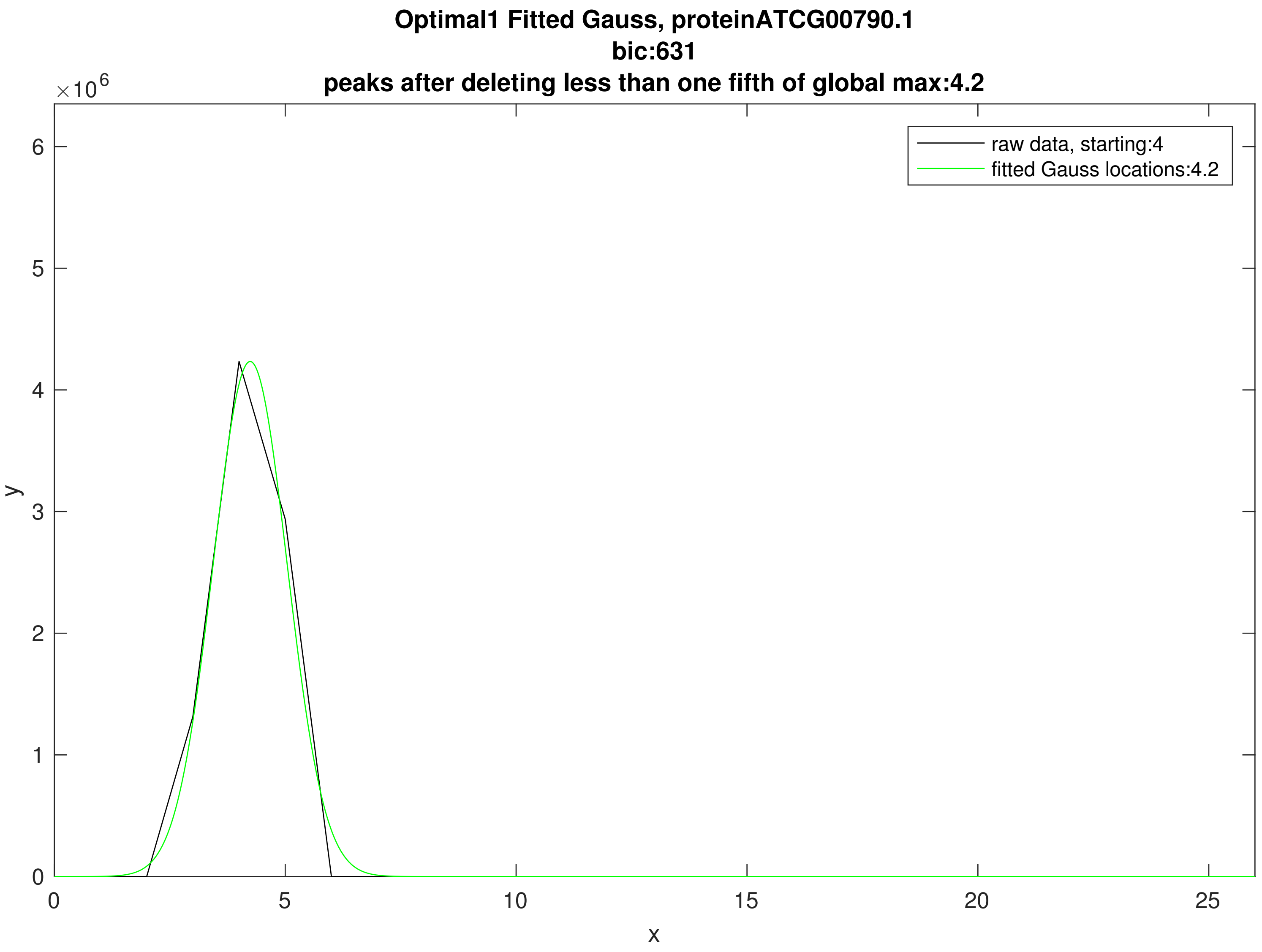

Optimal1 Fitted Gauss, proteinATCG00790.1
bic:631
peaks after deleting less than one fifth of global max:4.2
×10^6
6
5
4
3
2
1
0
y
0
5
10
15
20
25
x
raw data, starting:4
fitted Gauss locations:4.2

Optimal1 Fitted Gauss, proteinATCG00800.1
bic:619
peaks after deleting less than one fifth of global max:4.8

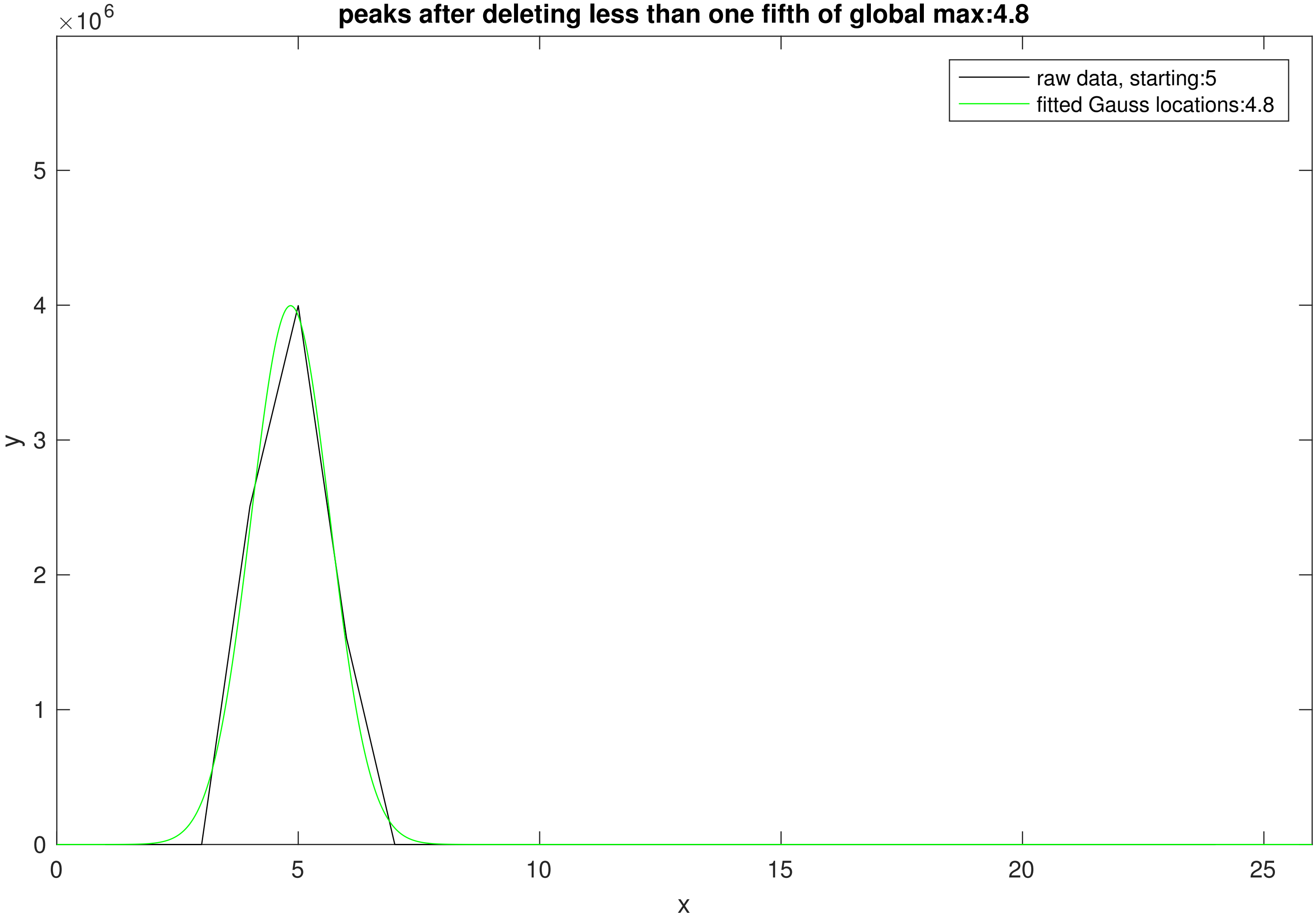

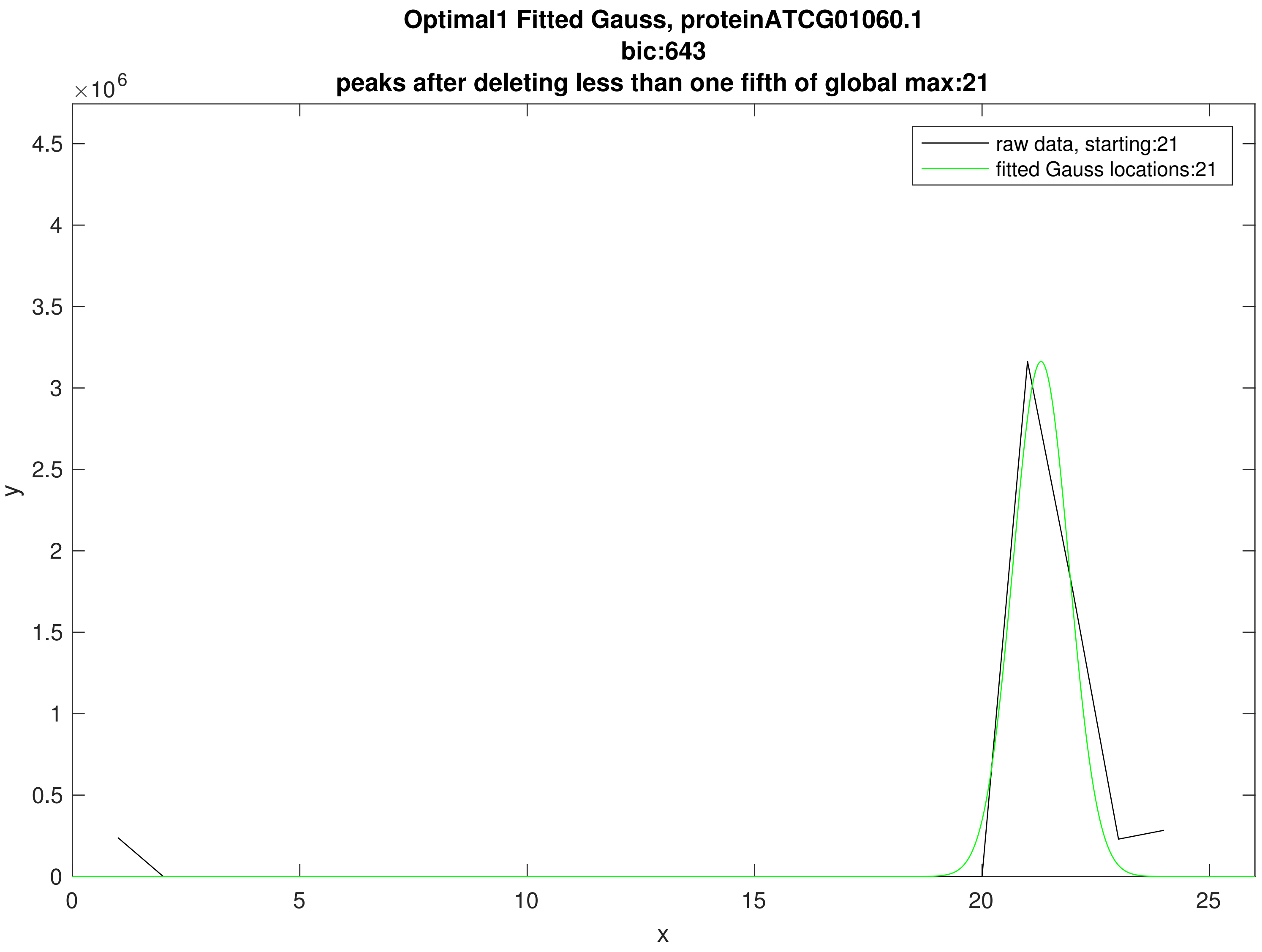

Optimal1 Fitted Gauss, proteinATCG01060.1
bic:643
peaks after deleting less than one fifth of global max:21
×10^6
raw data, starting:21
fitted Gauss locations:21
4.5
4
3.5
3
2.5
2
1.5
1
0.5
0
y
0
5
10
15
20
25
x

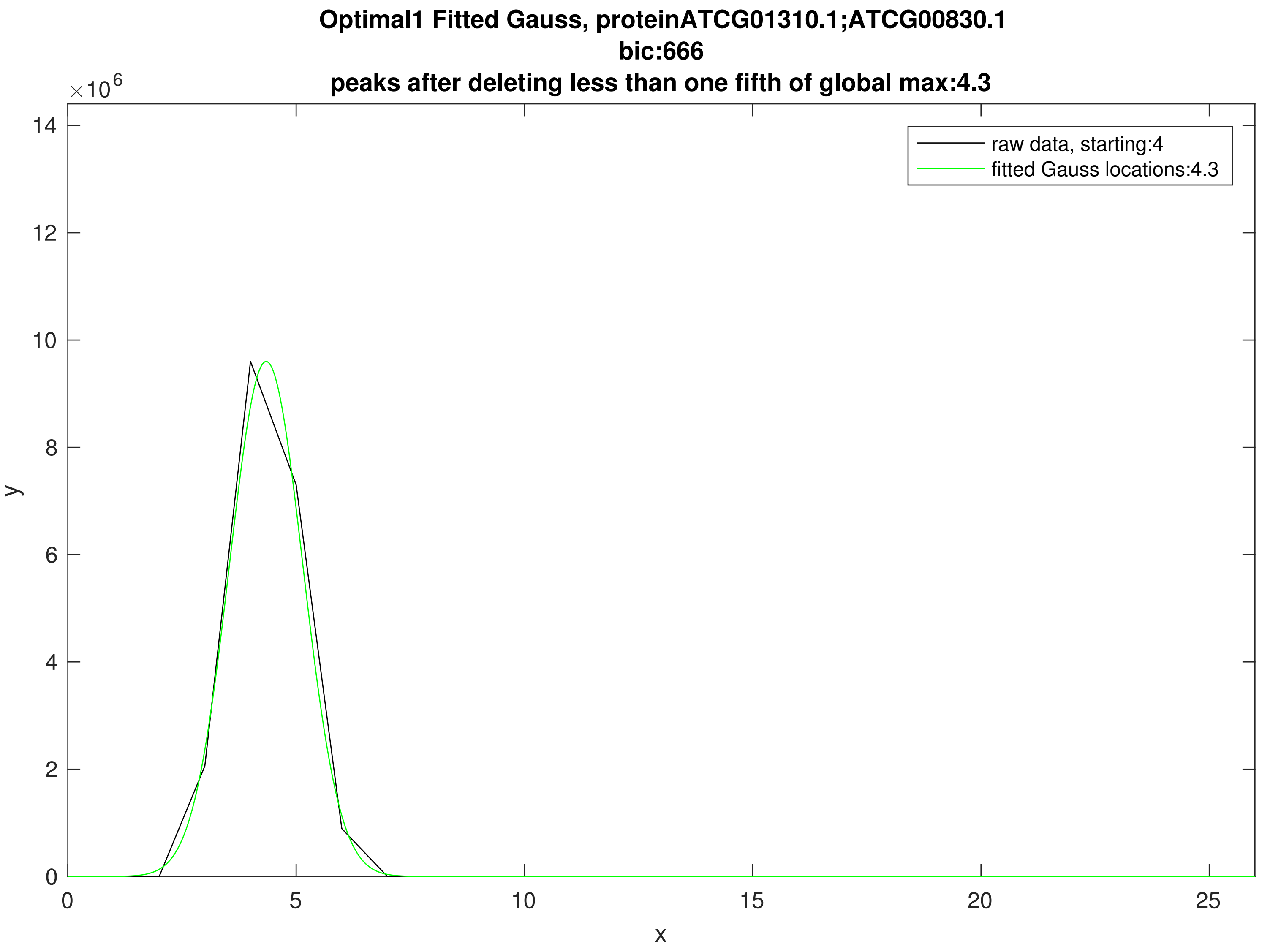
Optimal1 Fitted Gauss, proteinATCG01310.1;ATCG00830.1
bic:666
peaks after deleting less than one fifth of global max:4.3
×10^6
14
12
10
8
6
4
2
0
y
0
5
10
15
20
25
x
raw data, starting:4
fitted Gauss locations:4.3

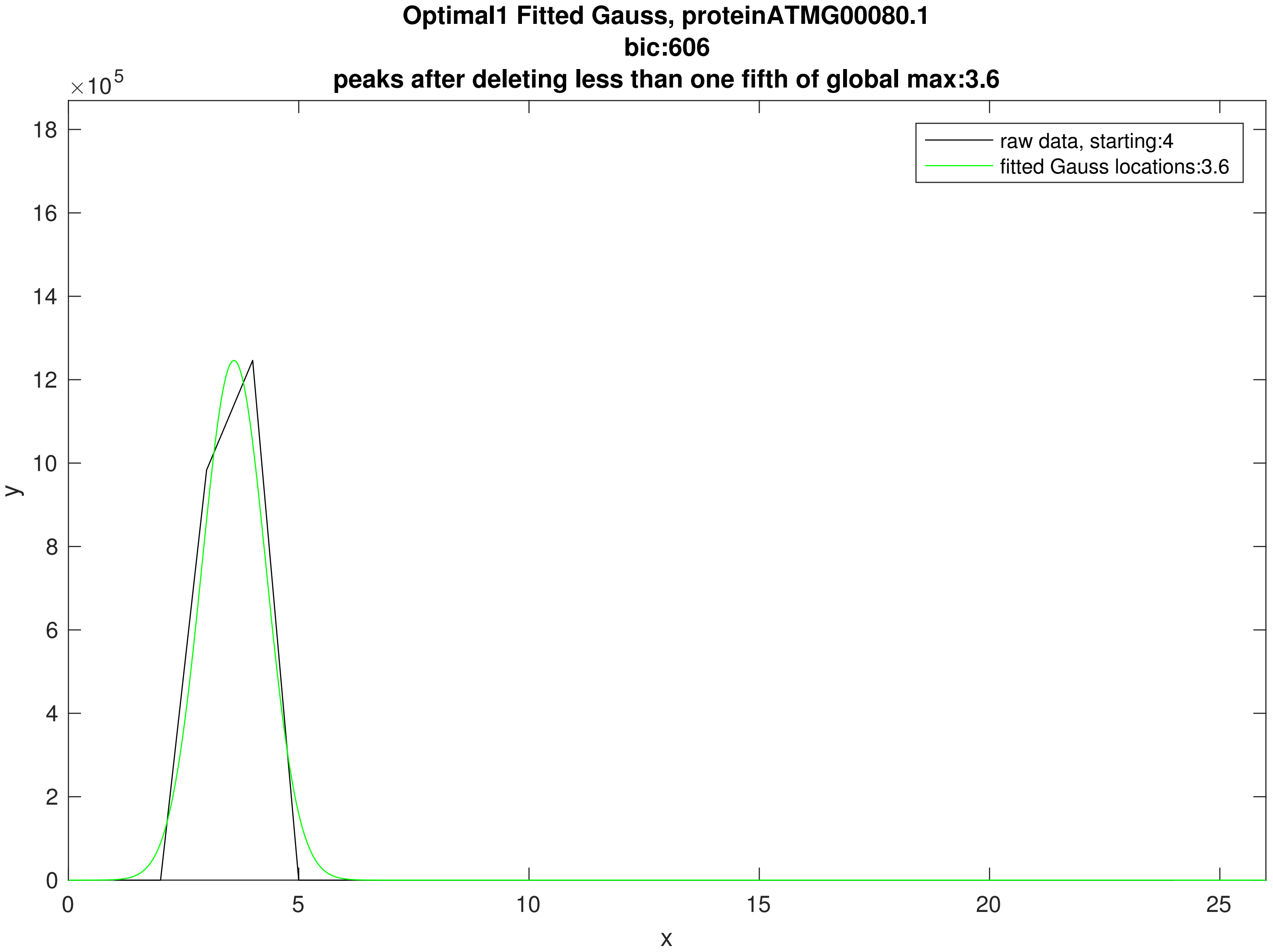

Optimal1 Fitted Gauss, proteinATMG00080.1
bic:606
peaks after deleting less than one fifth of global max:3.6
×10^5
raw data, starting:4
fitted Gauss locations:3.6
18
16
14
12
10
8
6
4
2
0
y
0
5
10
15
20
25
x

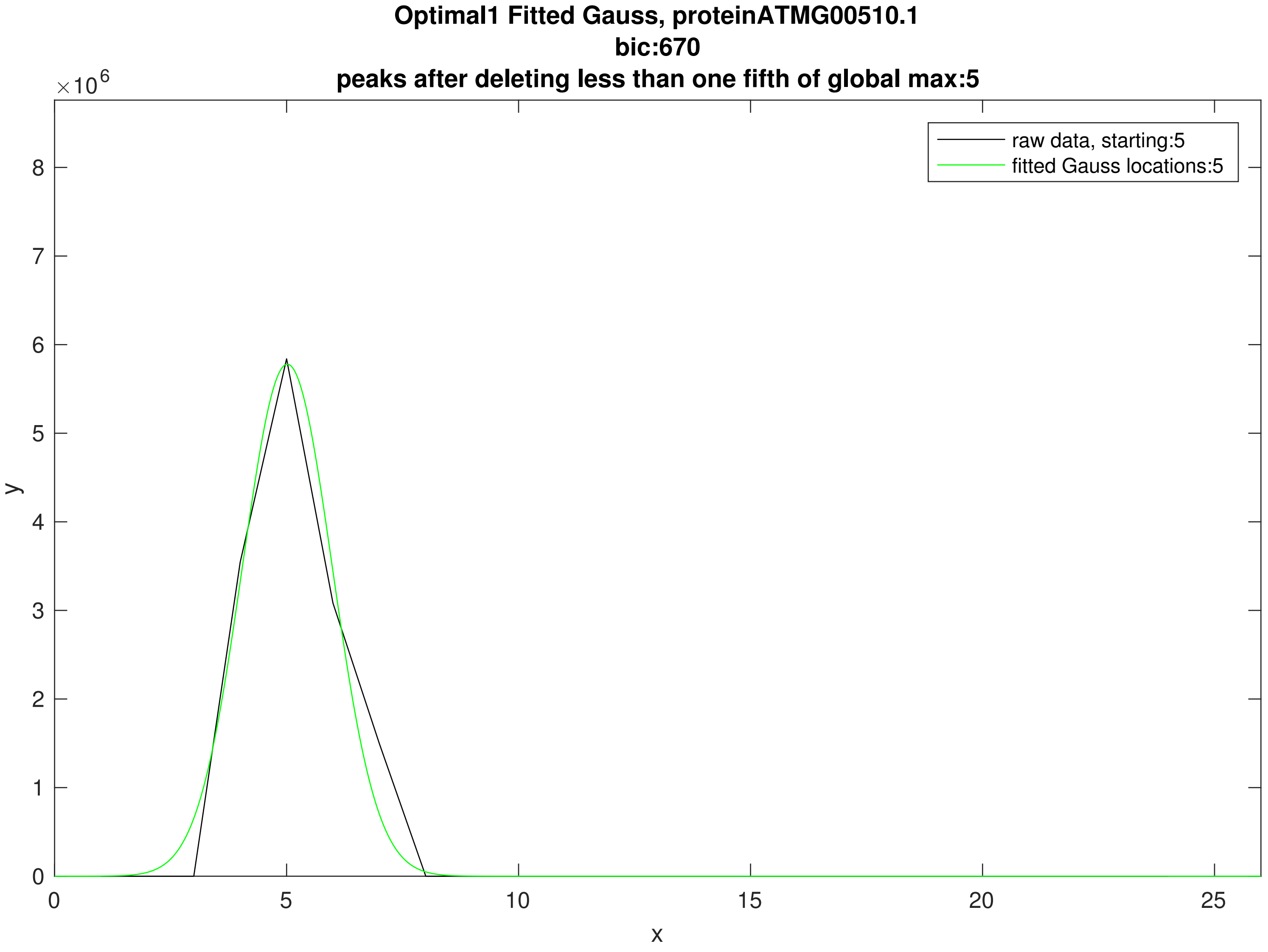

Optimal1 Fitted Gauss, proteinATMG00510.1
bic:670
peaks after deleting less than one fifth of global max:5
×10^6
raw data, starting:5
fitted Gauss locations:5
y
x
0
1
2
3
4
5
6
7
8
0
5
10
15
20
25

$SEC_{Ethylene}$ replicate 2 elution profiles

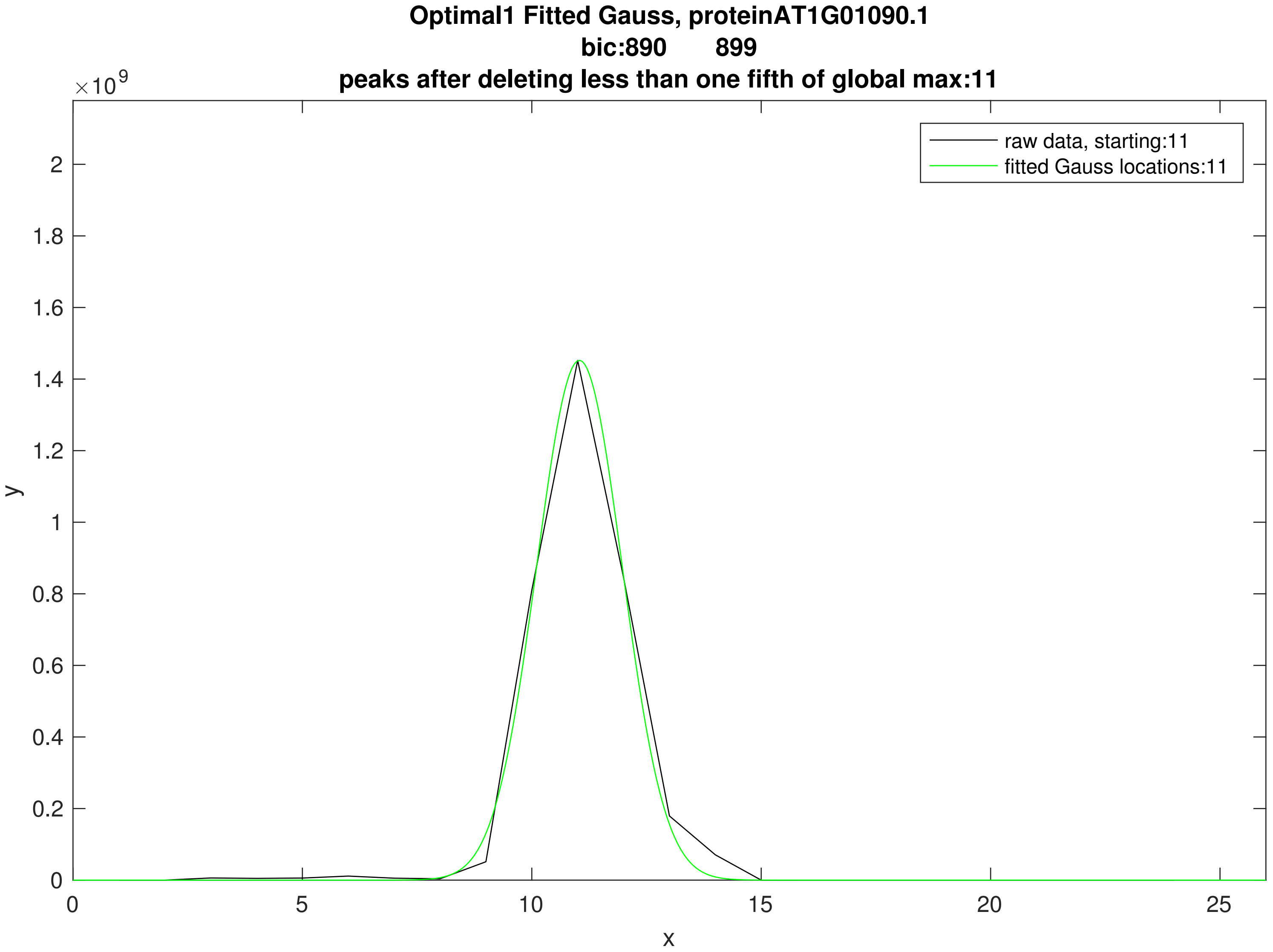
Optimal1 Fitted Gauss, proteinAT1G01090.1
bic:890 899
peaks after deleting less than one fifth of global max:11
×10^9
raw data, starting:11
fitted Gauss locations:11
y
x
0
0.2
0.4
0.6
0.8
1
1.2
1.4
1.6
1.8
2
0
5
10
15
20
25

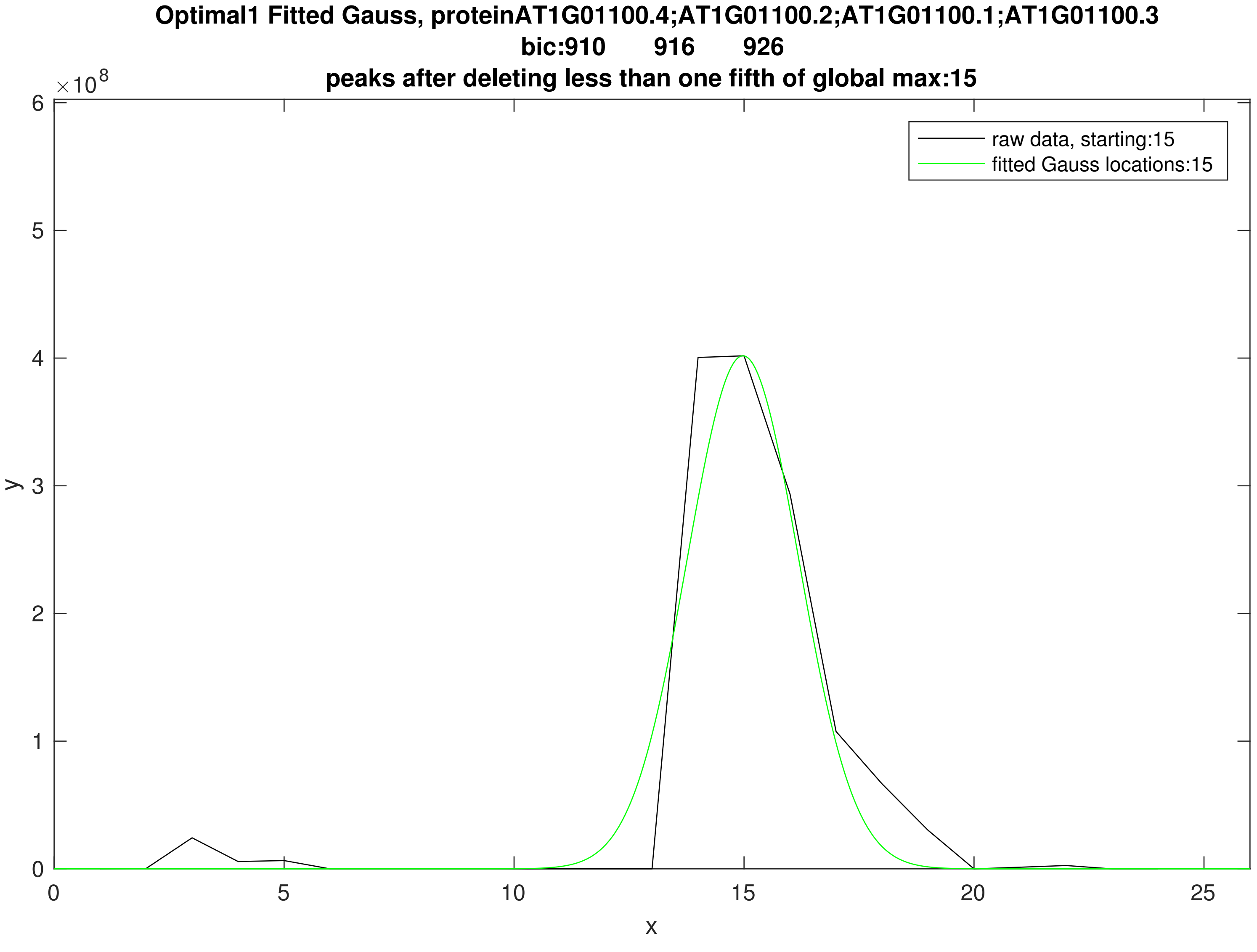

Optimal1 Fitted Gauss, proteinAT1G01100.4;AT1G01100.2;AT1G01100.1;AT1G01100.3
bic:910 916 926
peaks after deleting less than one fifth of global max:15
×10^8
6
5
4
3
2
1
0
y
0
5
10
15
20
25
x
raw data, starting:15
fitted Gauss locations:15

ptimal1 Fitted Gauss, proteinAT1G01200.1;AT2G43130.1;AT2G31680.1;AT3G12160.1;AT5G47960.1;AT5G65270.1;AT1G
bic:673
peaks after deleting less than one fifth of global max:18

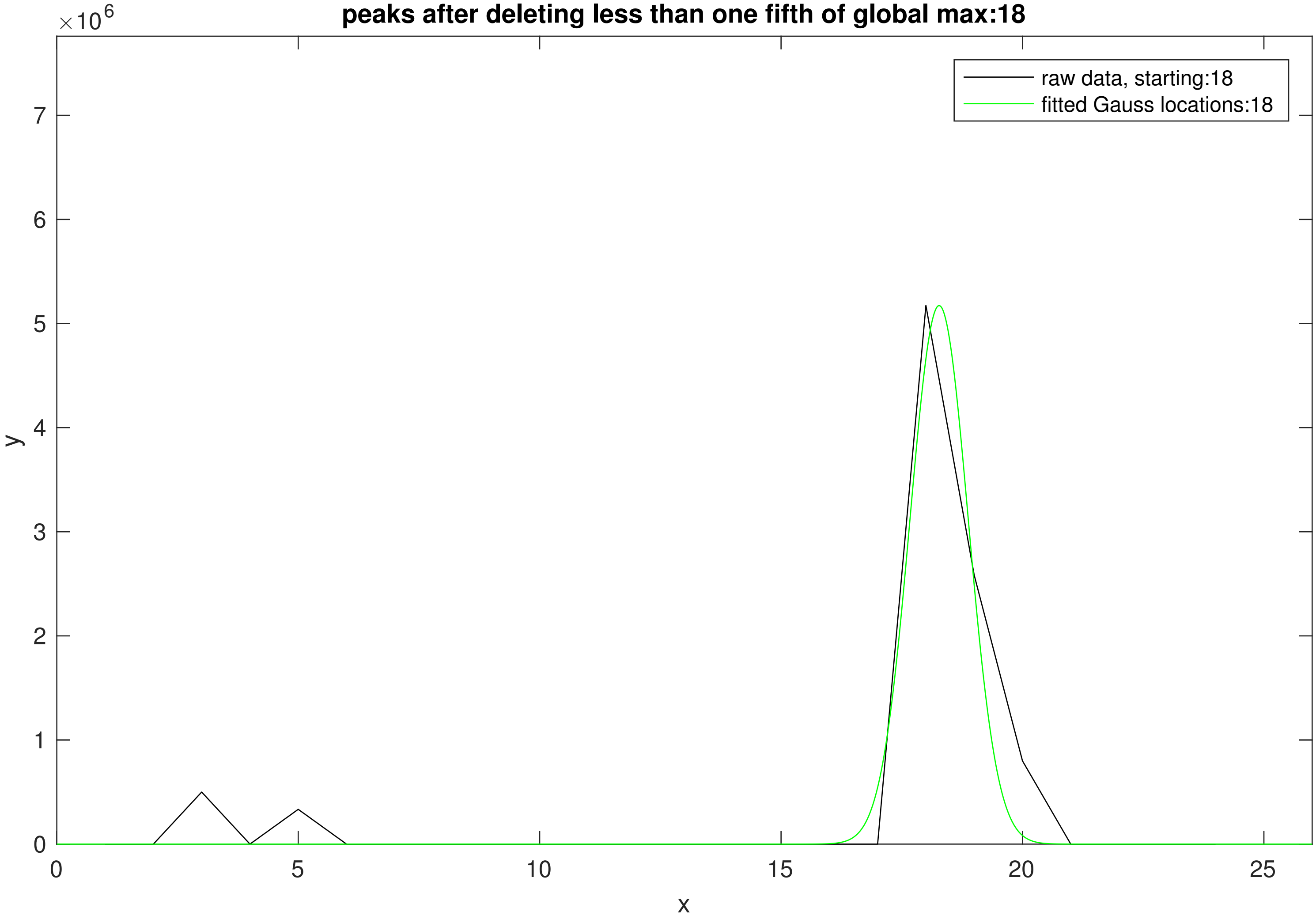

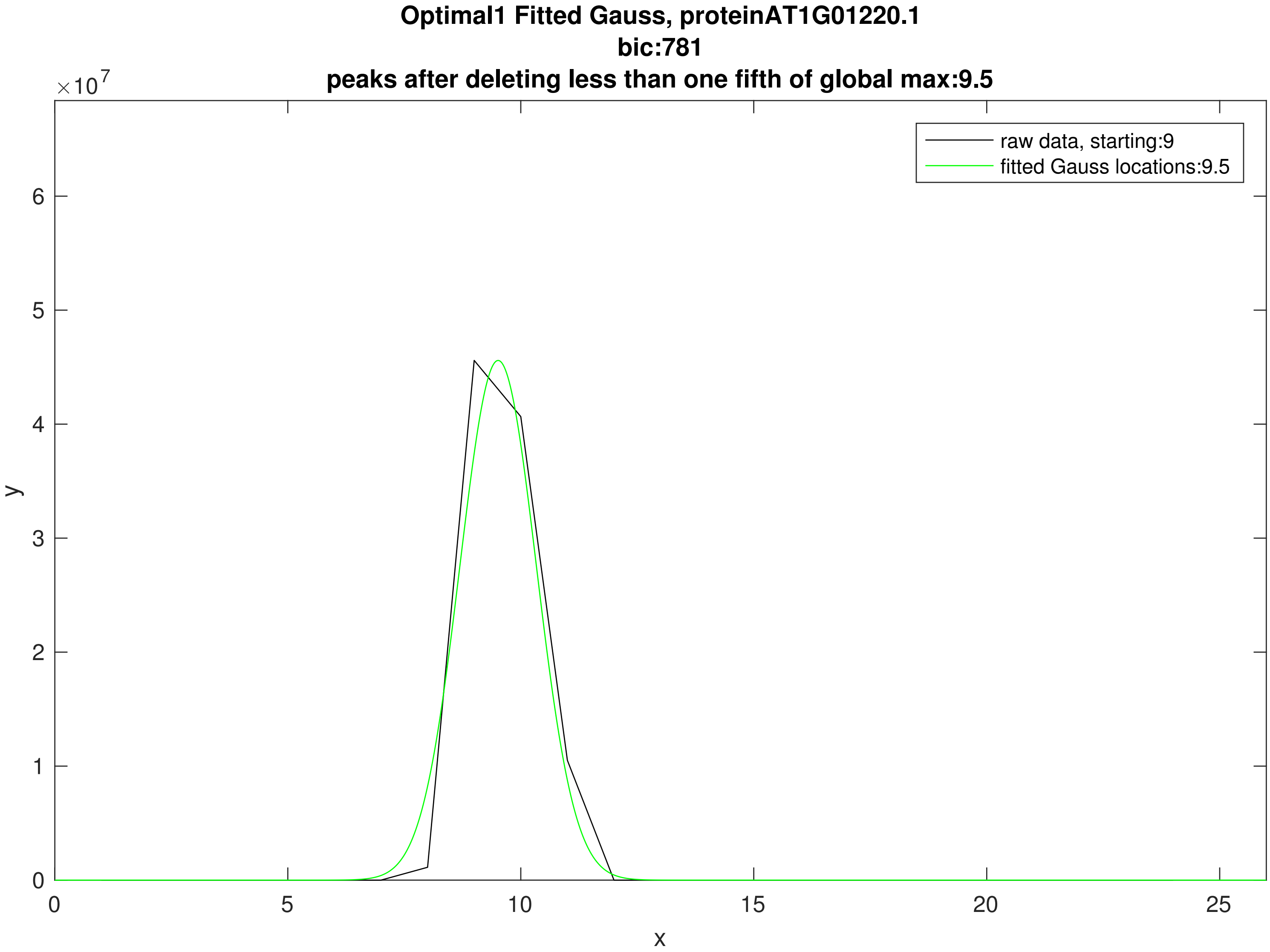

Optimal1 Fitted Gauss, proteinAT1G01220.1
bic:781
peaks after deleting less than one fifth of global max:9.5
×10^7
raw data, starting:9
fitted Gauss locations:9.5
6
5
4
3
2
1
0
y
0
5
10
15
20
25
x

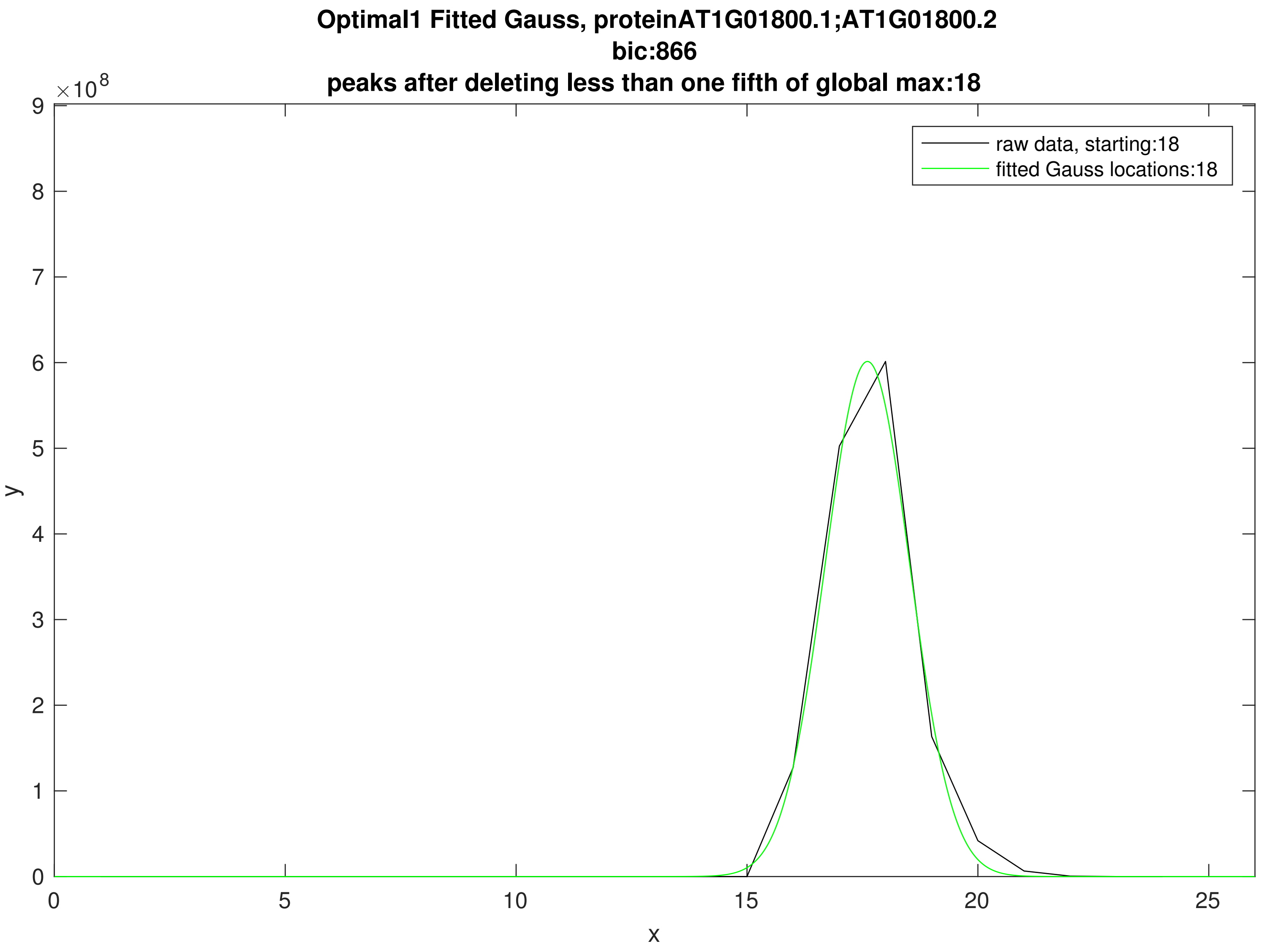

Optimal1 Fitted Gauss, proteinAT1G01800.1;AT1G01800.2
bic:866
peaks after deleting less than one fifth of global max:18
×10^8
raw data, starting:18
fitted Gauss locations:18
y
x
0
1
2
3
4
5
6
7
8
9
0
5
10
15
20
25

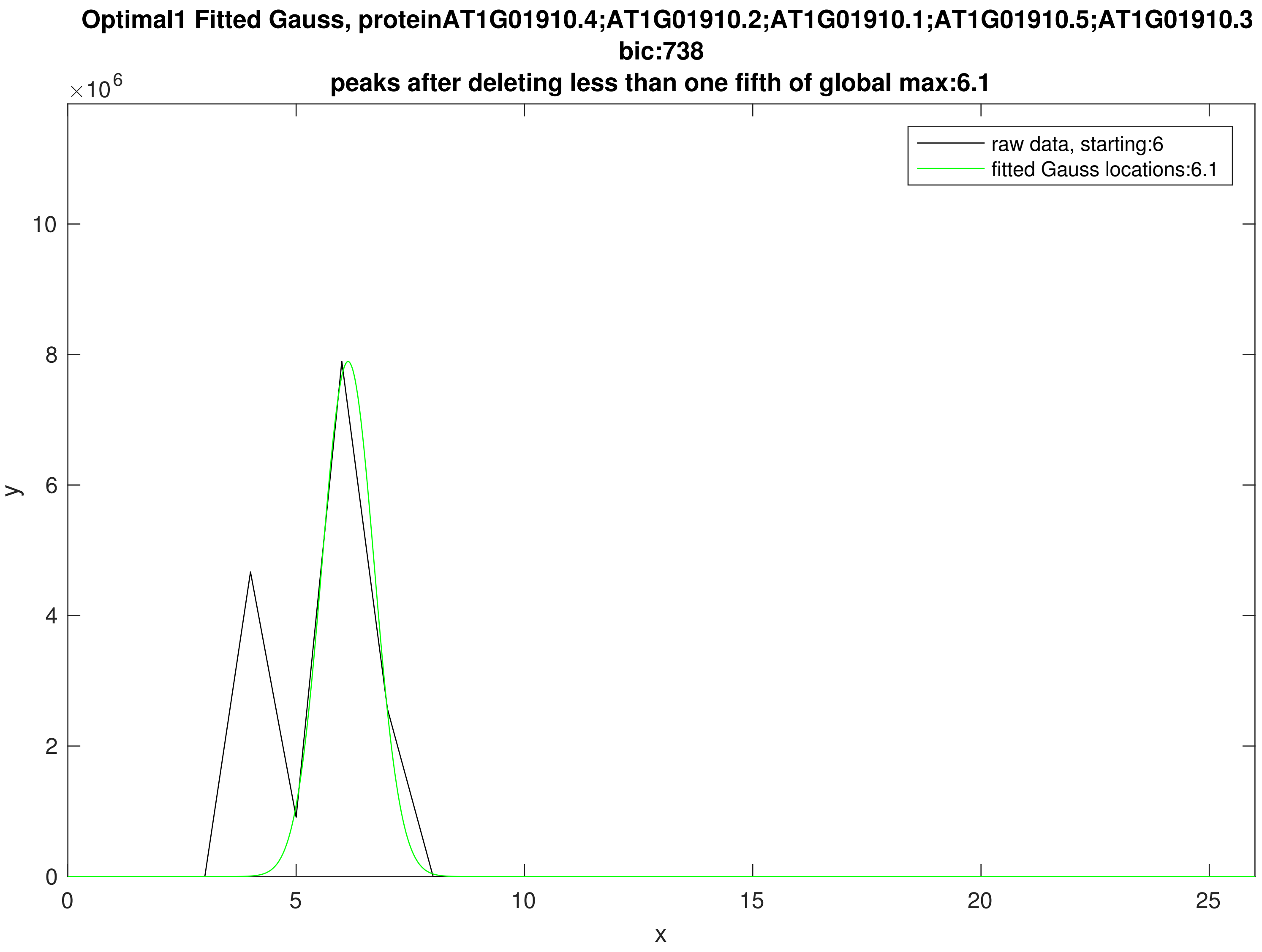
Optimal1 Fitted Gauss, proteinAT1G01910.4;AT1G01910.2;AT1G01910.1;AT1G01910.5;AT1G01910.3
bic:738
peaks after deleting less than one fifth of global max:6.1
×10^6
raw data, starting:6
fitted Gauss locations:6.1
y
x
0
2
4
6
8
10
0
5
10
15
20
25

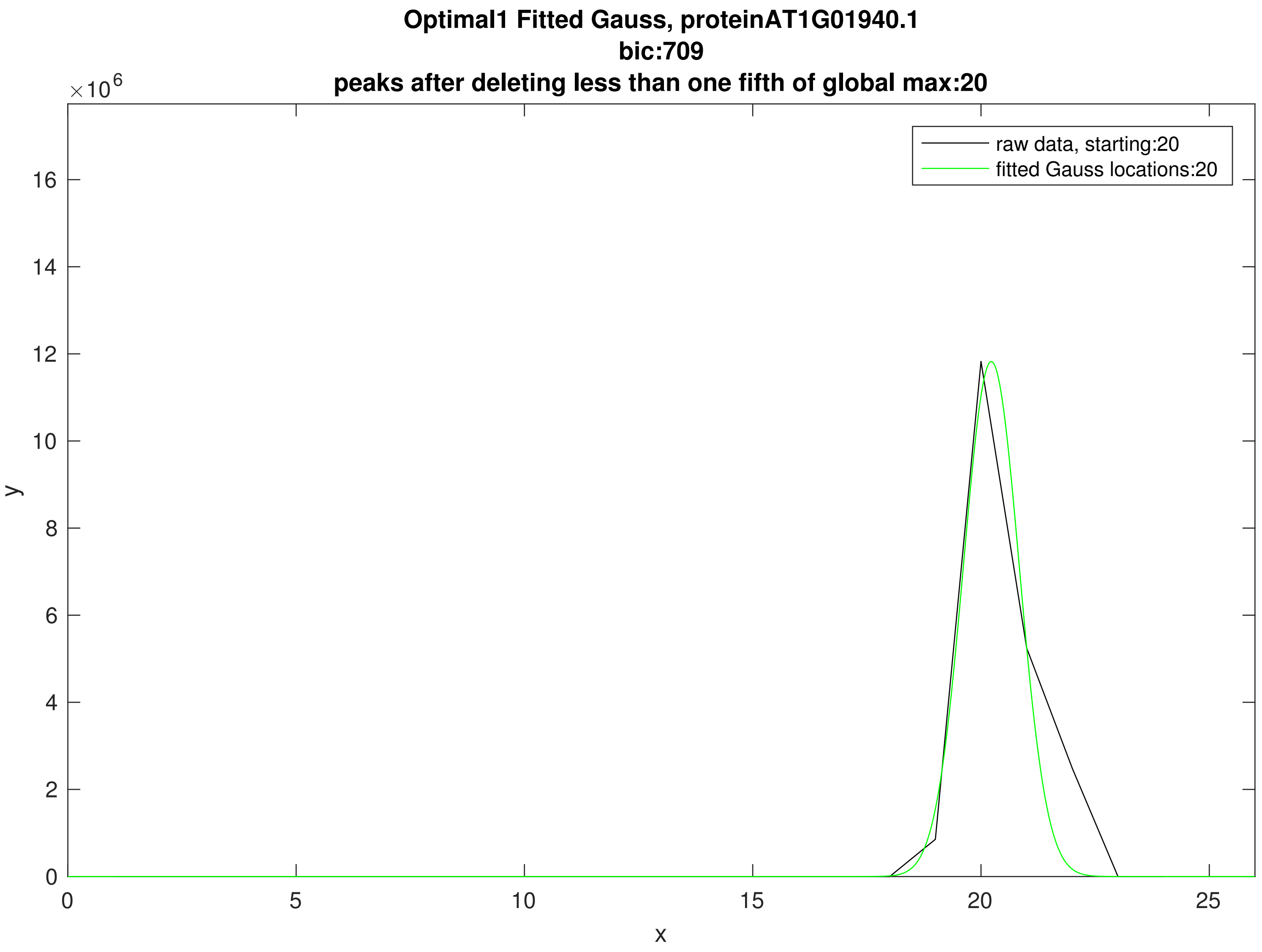
Optimal1 Fitted Gauss, proteinAT1G01940.1
bic:709
peaks after deleting less than one fifth of global max:20
×10^6
16
14
12
10
8
6
4
2
0
y
0
5
10
15
20
25
x
raw data, starting:20
fitted Gauss locations:20

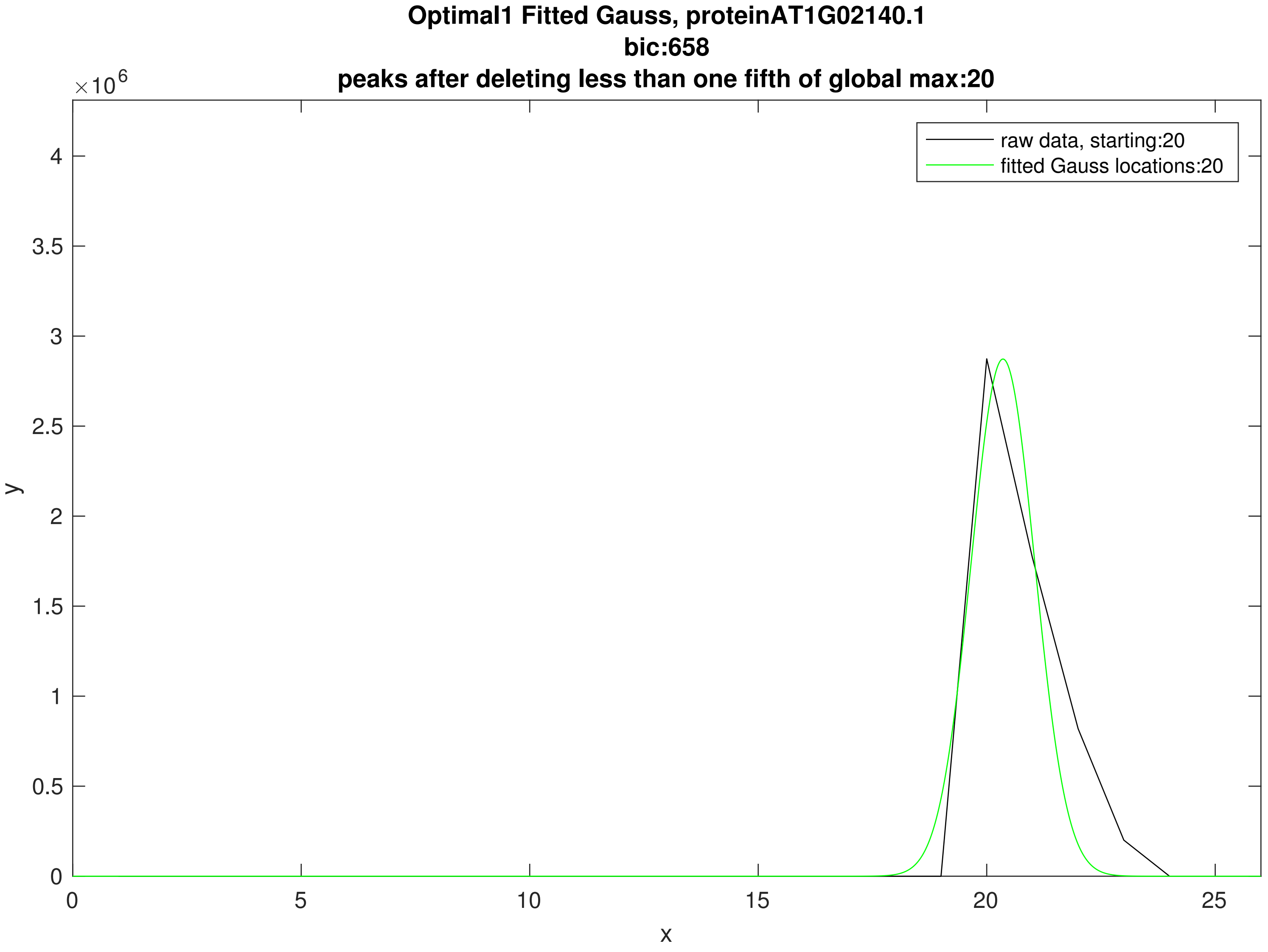
Optimal1 Fitted Gauss, proteinAT1G02140.1
bic:658
peaks after deleting less than one fifth of global max:20
×10^6
raw data, starting:20
fitted Gauss locations:20
y
x
0
0.5
1
1.5
2
2.5
3
3.5
4
5
10
15
20
25

Optimal1 Fitted Gauss, proteinAT1G02360.1
bic:666
peaks after deleting less than one fifth of global max:18

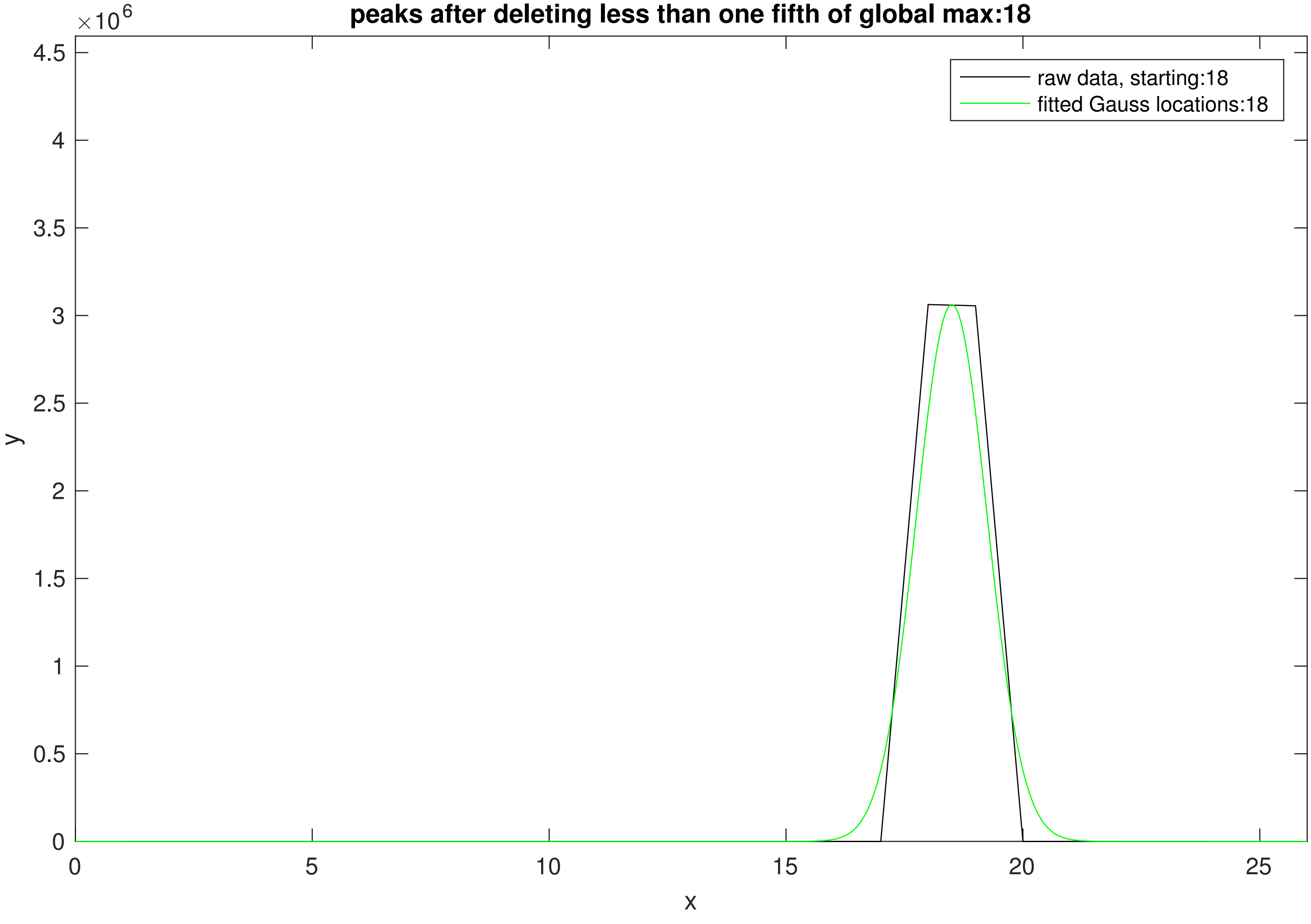

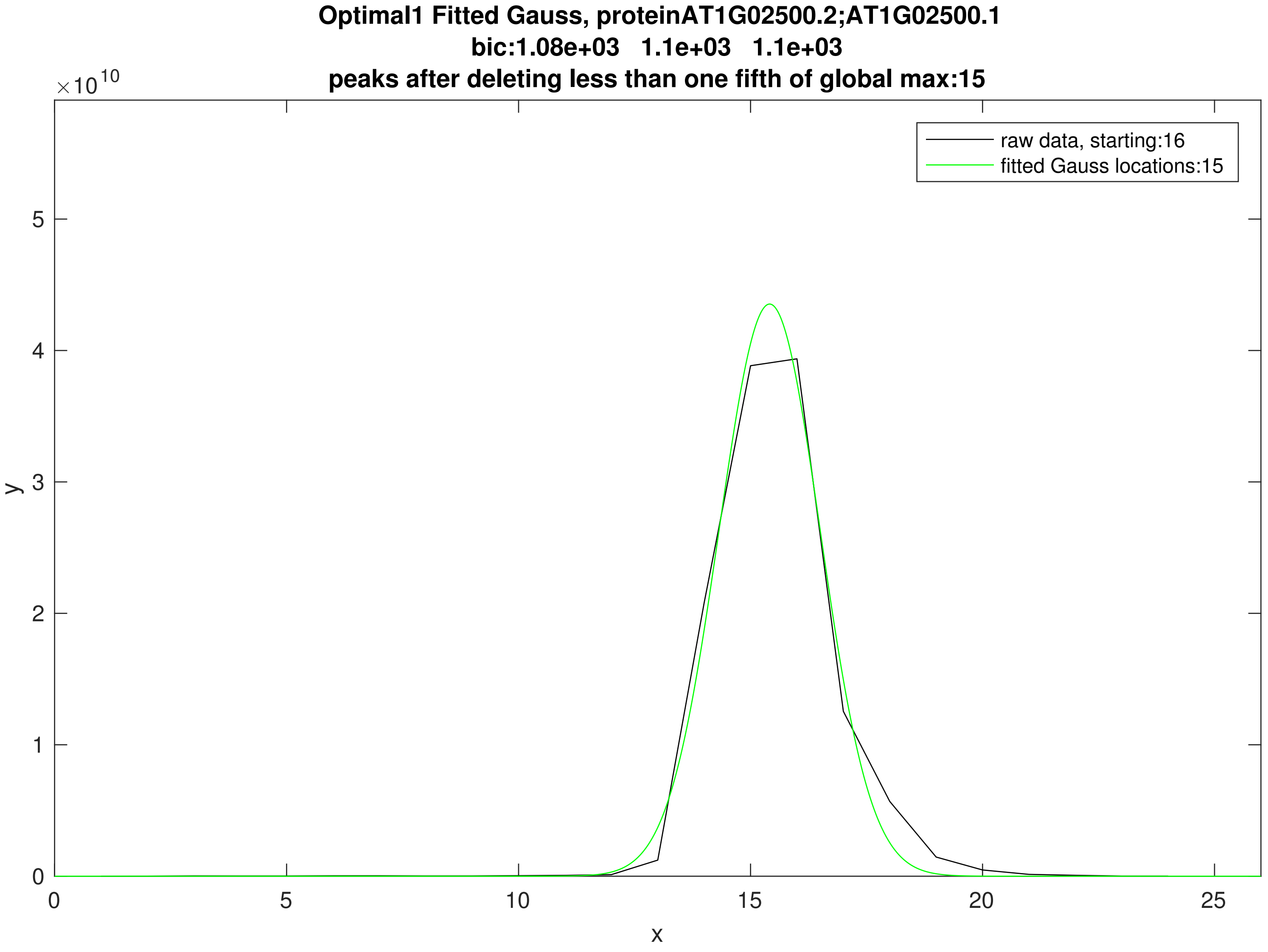
Optimal1 Fitted Gauss, proteinAT1G02500.2;AT1G02500.1
bic:1.08e+03 1.1e+03 1.1e+03
peaks after deleting less than one fifth of global max:15
×10^10
raw data, starting:16
fitted Gauss locations:15
y
x
0
1
2
3
4
5
0
5
10
15
20
25

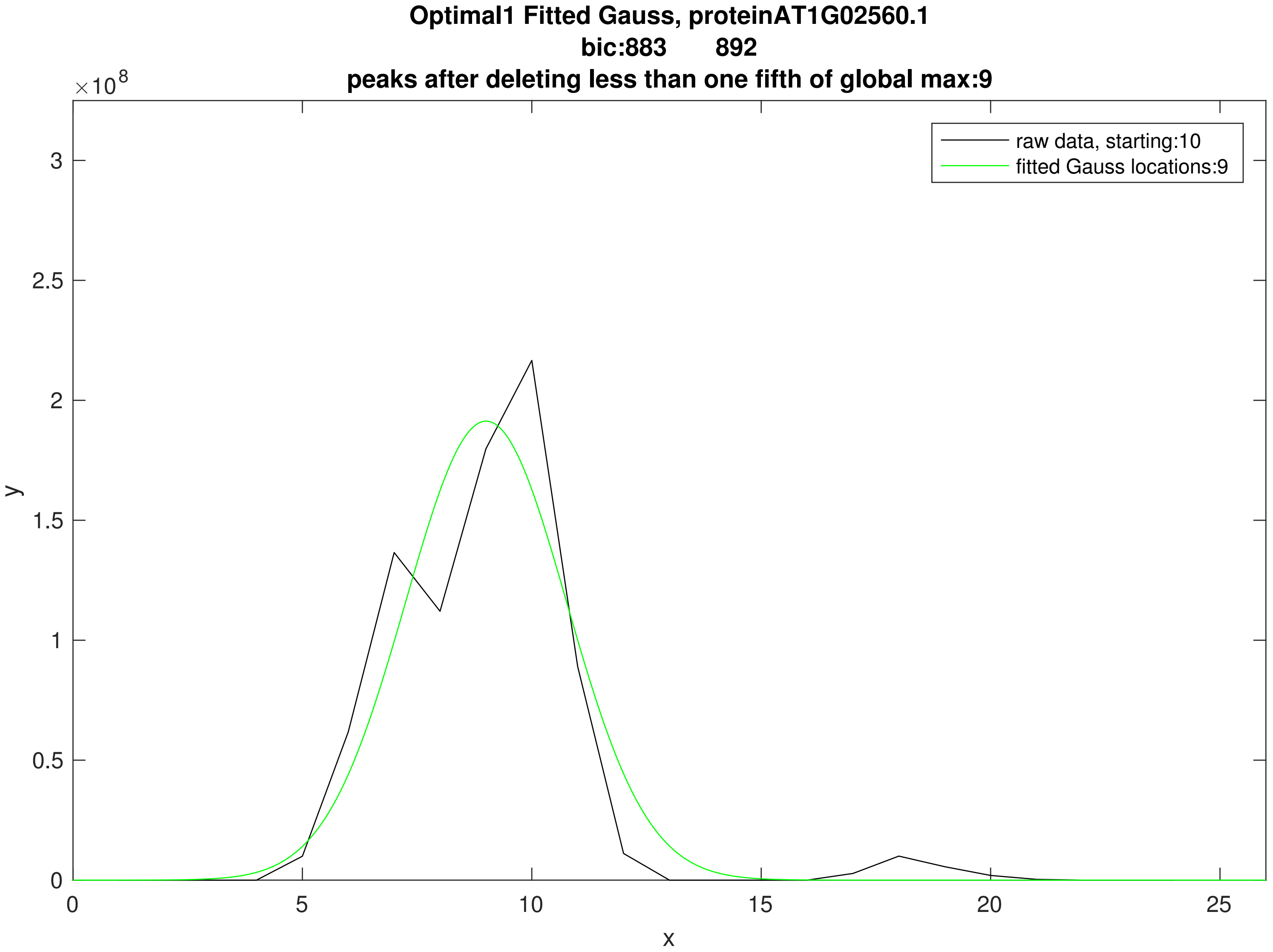
Optimal1 Fitted Gauss, proteinAT1G02560.1
bic:883 892
peaks after deleting less than one fifth of global max:9
×10^8
raw data, starting:10
fitted Gauss locations:9
y
x
0
0.5
1
1.5
2
2.5
3
0
5
10
15
20
25

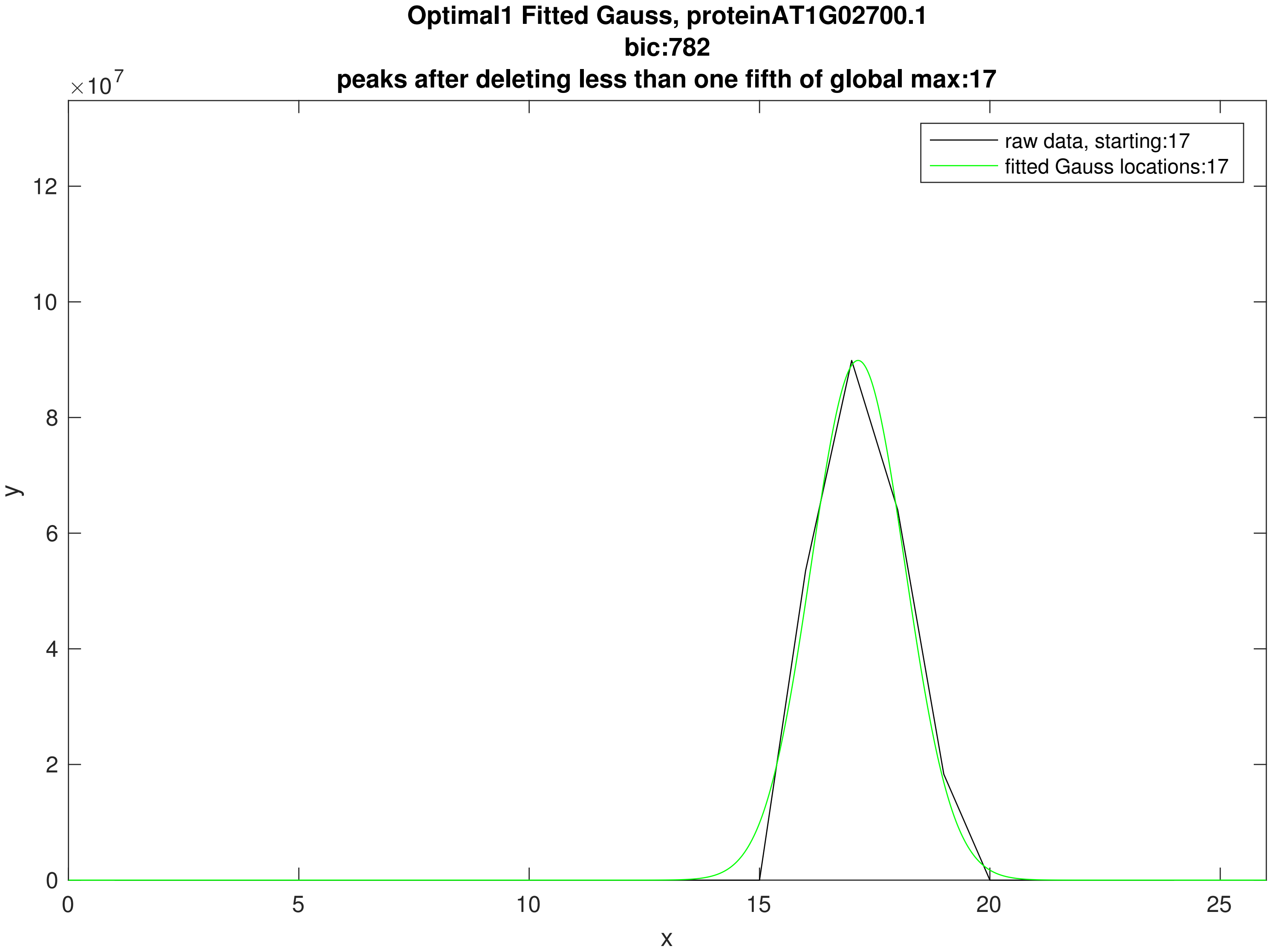

Optimal1 Fitted Gauss, proteinAT1G02700.1
bic:782
peaks after deleting less than one fifth of global max:17
×10^7
raw data, starting:17
fitted Gauss locations:17
12
10
8
6
4
2
0
y
0
5
10
15
20
25
x

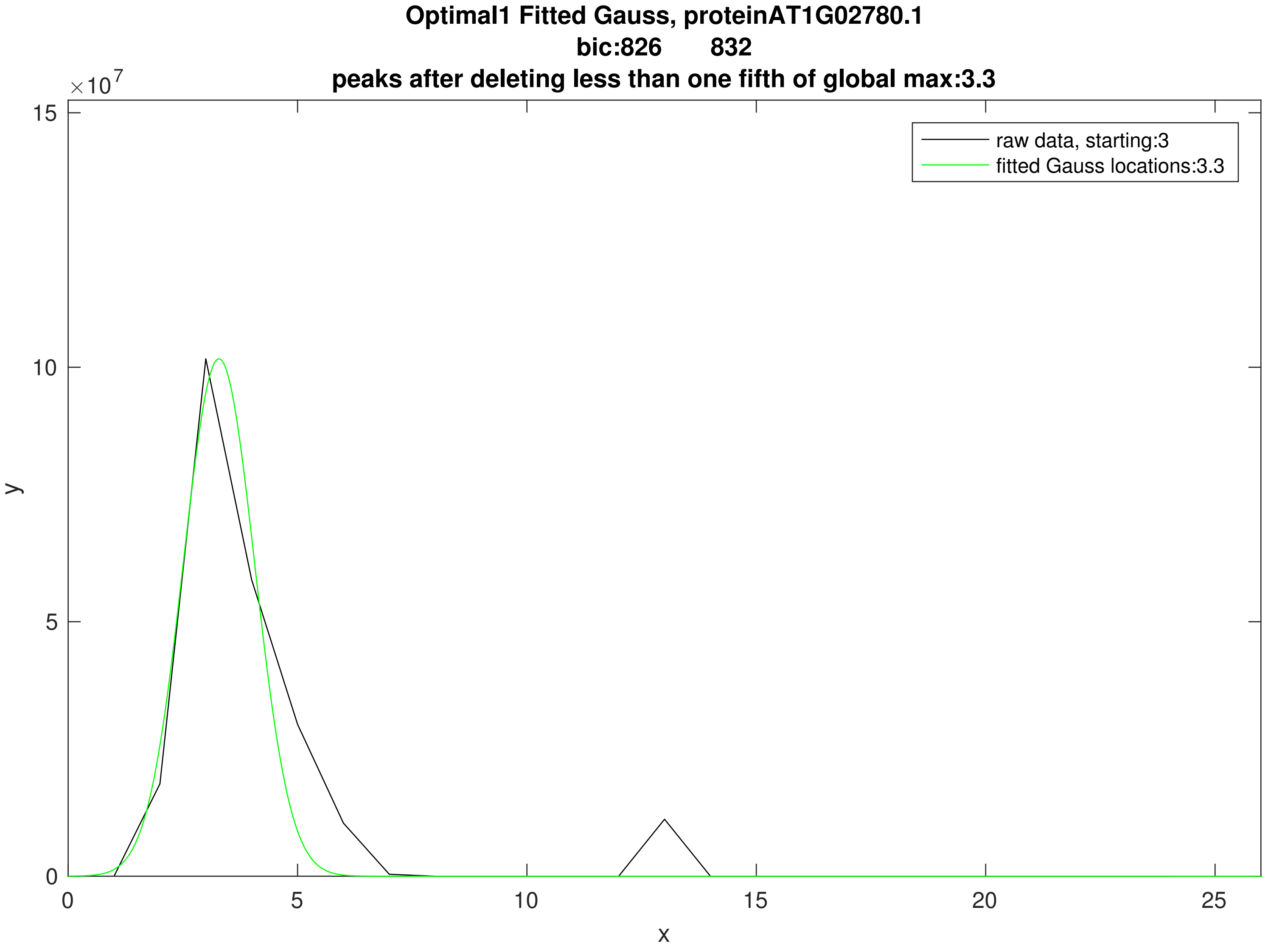

Optimal1 Fitted Gauss, proteinAT1G02780.1
bic:826 832
peaks after deleting less than one fifth of global max:3.3
×10^7
15
10
5
0
y
0
5
10
15
20
25
x
raw data, starting:3
fitted Gauss locations:3.3

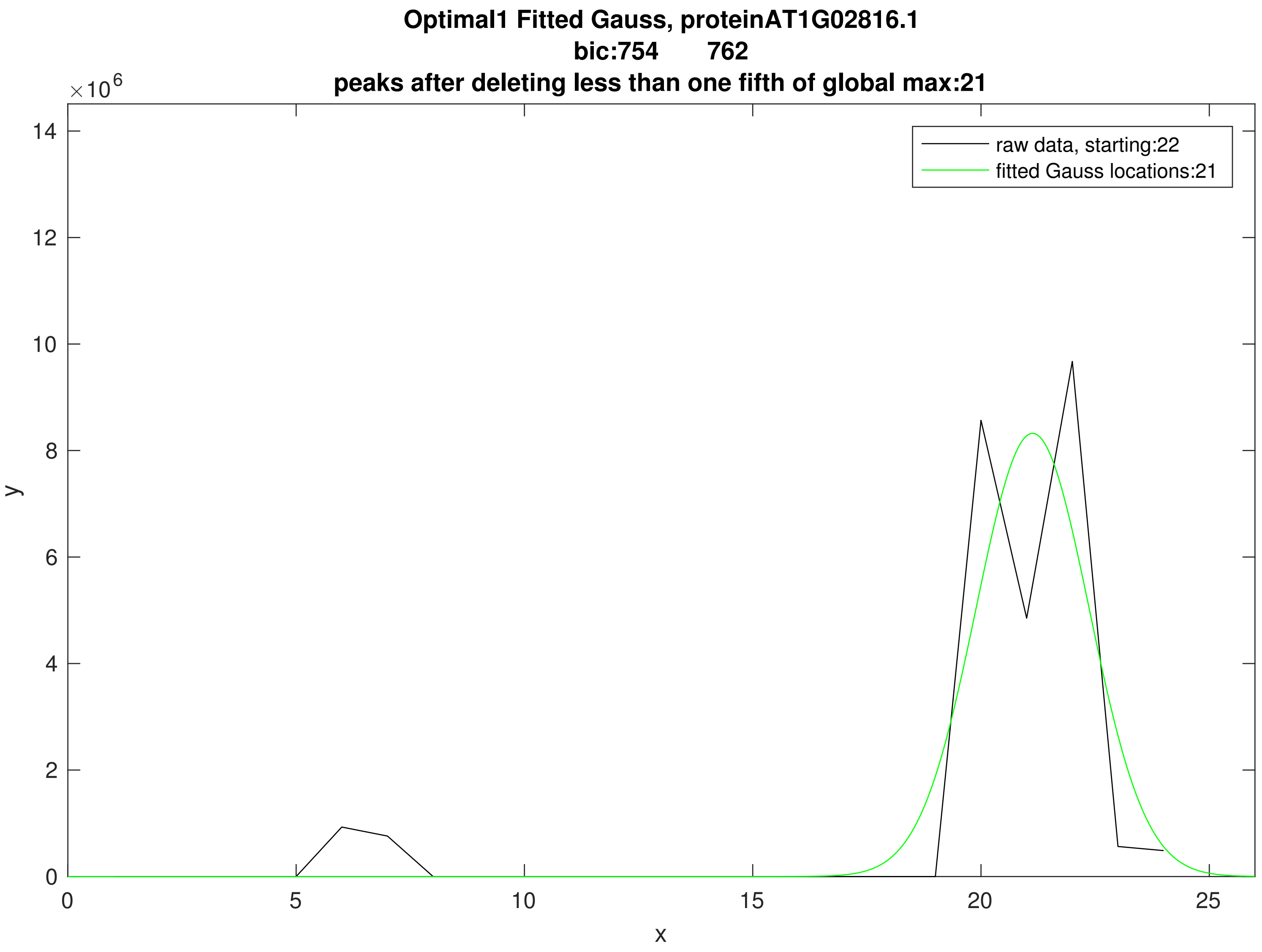
Optimal1 Fitted Gauss, proteinAT1G02816.1
bic:754 762
peaks after deleting less than one fifth of global max:21
×10^6
14
12
10
8
6
4
2
0
y
0
5
10
15
20
25
x
raw data, starting:22
fitted Gauss locations:21

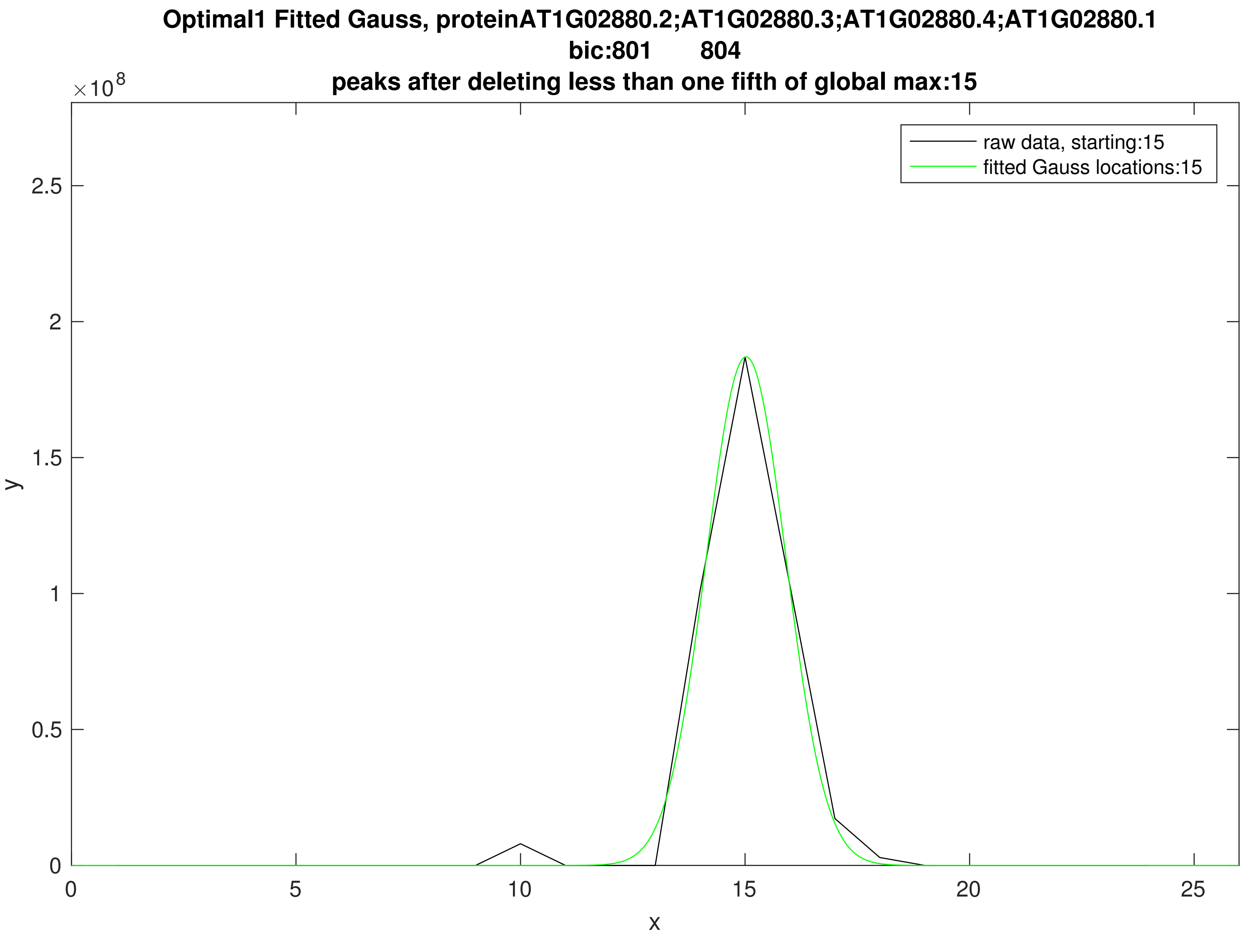
Optimal1 Fitted Gauss, proteinAT1G02880.2;AT1G02880.3;AT1G02880.4;AT1G02880.1
bic:801 804
peaks after deleting less than one fifth of global max:15
×10^8
2.5
2
1.5
1
0.5
0
y
0
5
10
15
20
25
x
raw data, starting:15
fitted Gauss locations:15

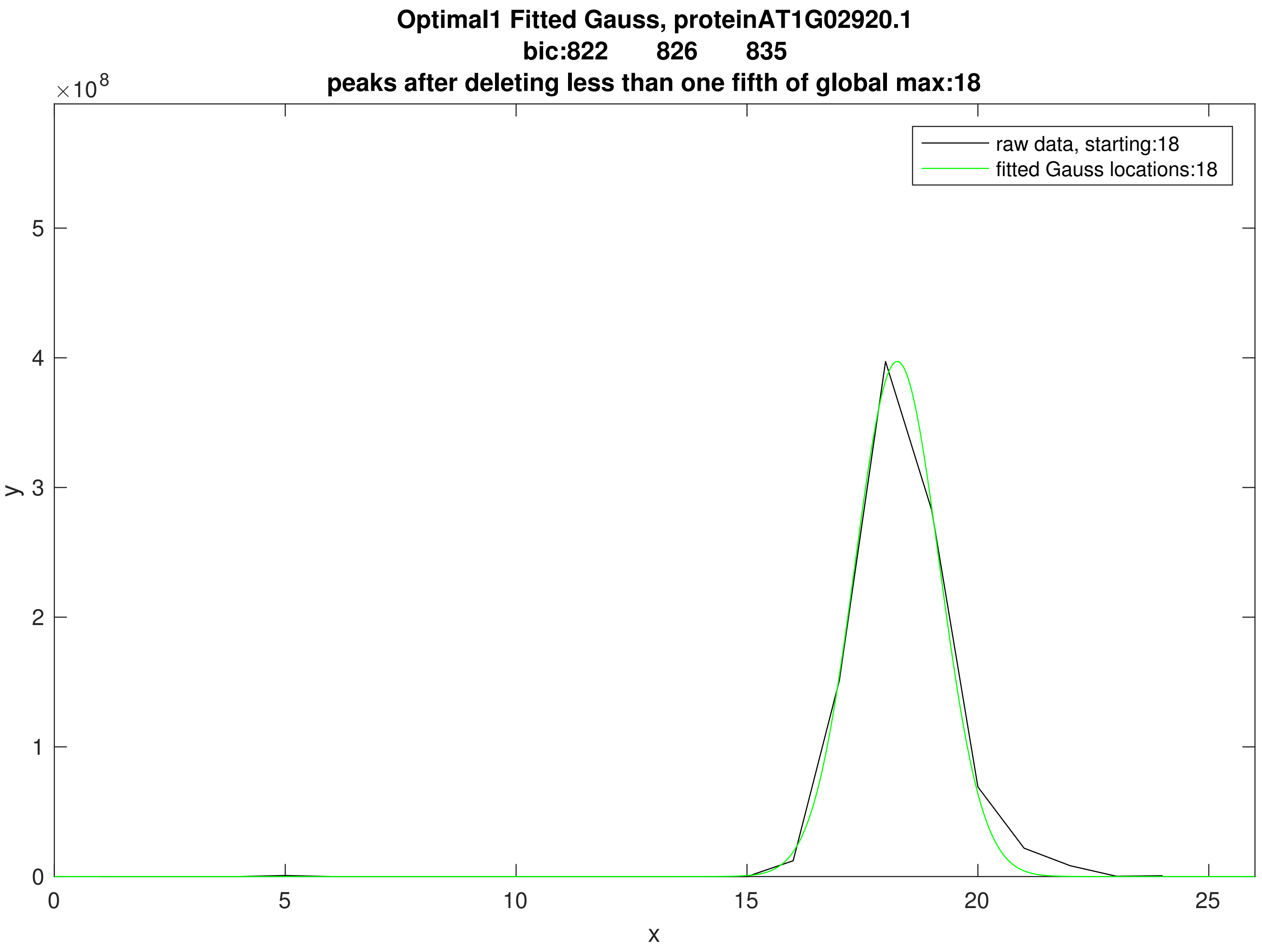
Optimal1 Fitted Gauss, proteinAT1G02920.1
bic:822 826 835
peaks after deleting less than one fifth of global max:18
×10^8
y
x
raw data, starting:18
fitted Gauss locations:18
0
1
2
3
4
5
5
10
15
20
25

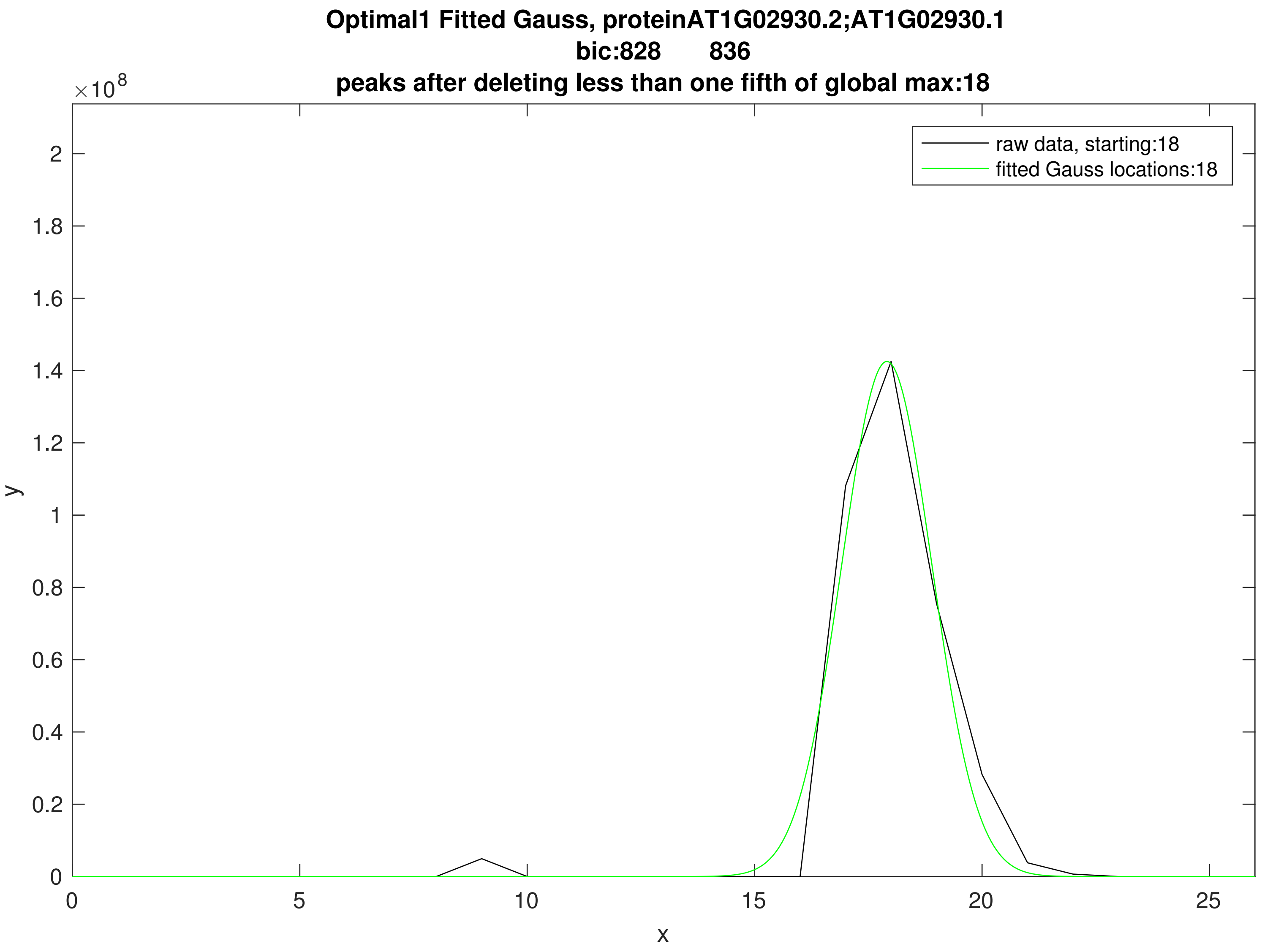

Optimal1 Fitted Gauss, proteinAT1G02930.2;AT1G02930.1
bic:828 836
peaks after deleting less than one fifth of global max:18
×10^8
2
1.8
1.6
1.4
1.2
1
0.8
0.6
0.4
0.2
0
y
0
5
10
15
20
25
x
raw data, starting:18
fitted Gauss locations:18

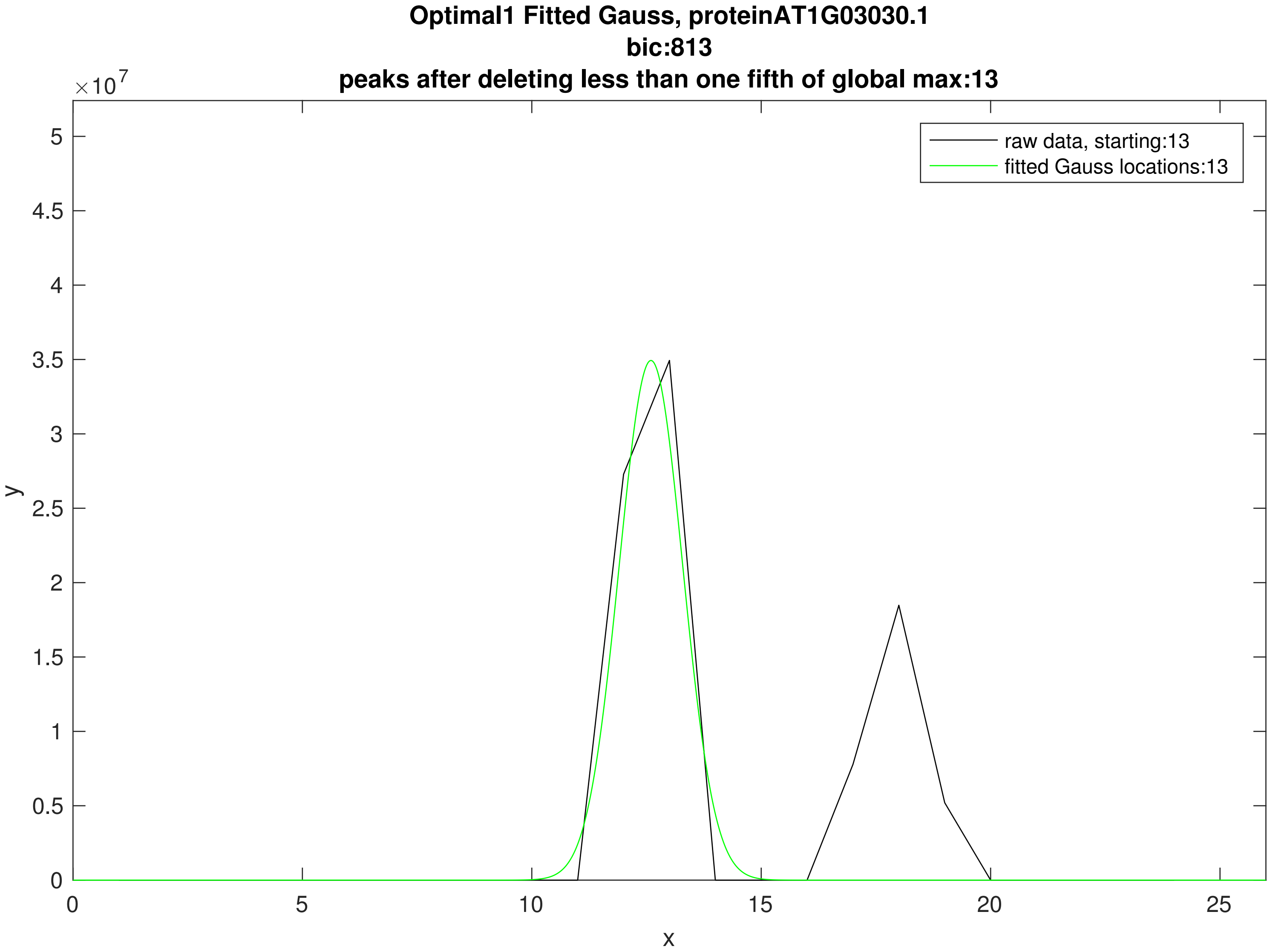

Optimal1 Fitted Gauss, proteinAT1G03030.1
bic:813
peaks after deleting less than one fifth of global max:13
×10^7
raw data, starting:13
fitted Gauss locations:13
y
x
0
0.5
1
1.5
2
2.5
3
3.5
4
4.5
5
0
5
10
15
20
25

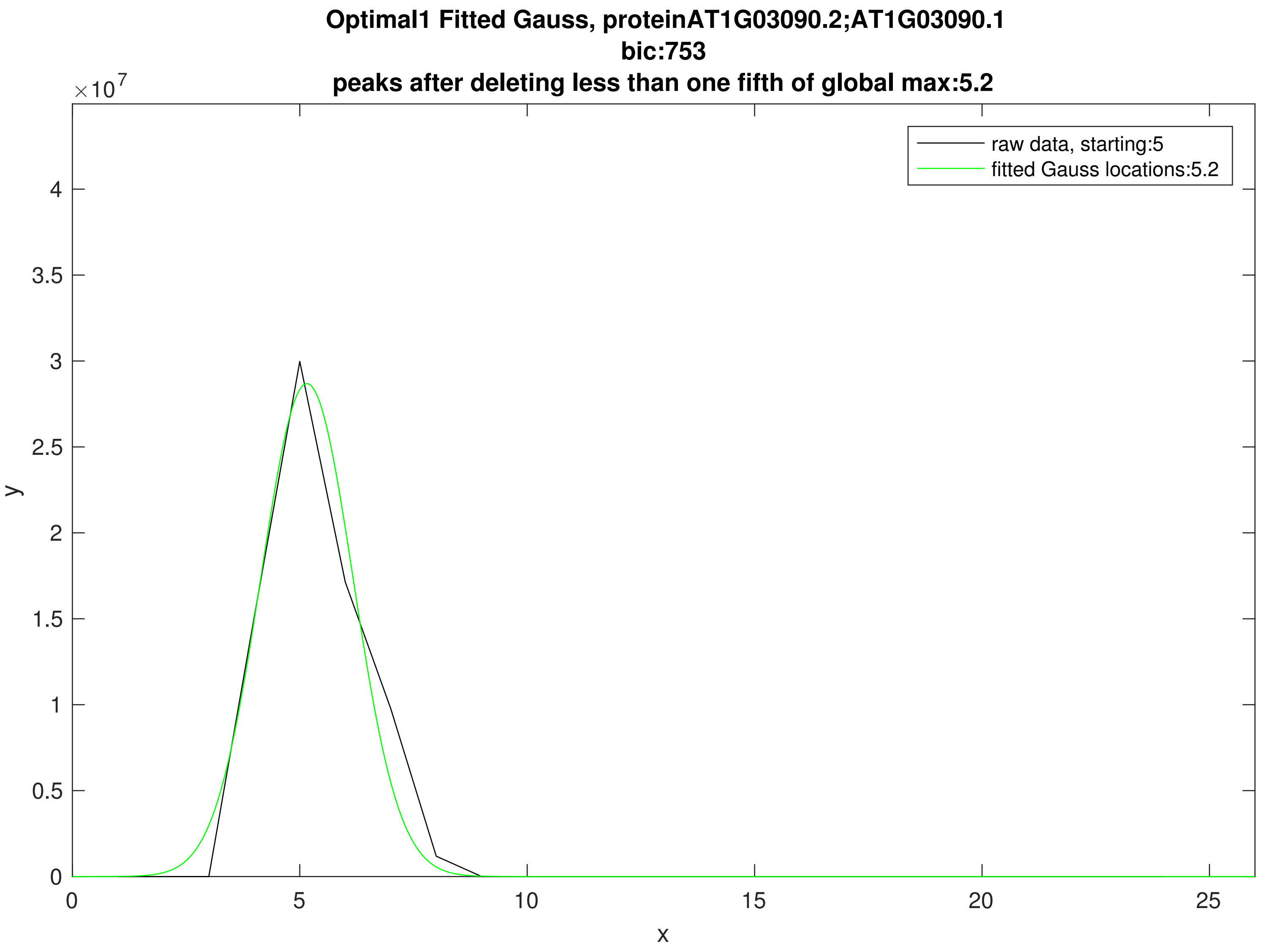

Optimal1 Fitted Gauss, proteinAT1G03090.2;AT1G03090.1
bic:753
peaks after deleting less than one fifth of global max:5.2
×10^7
raw data, starting:5
fitted Gauss locations:5.2
y
x
0
0.5
1
1.5
2
2.5
3
3.5
4
0
5
10
15
20
25

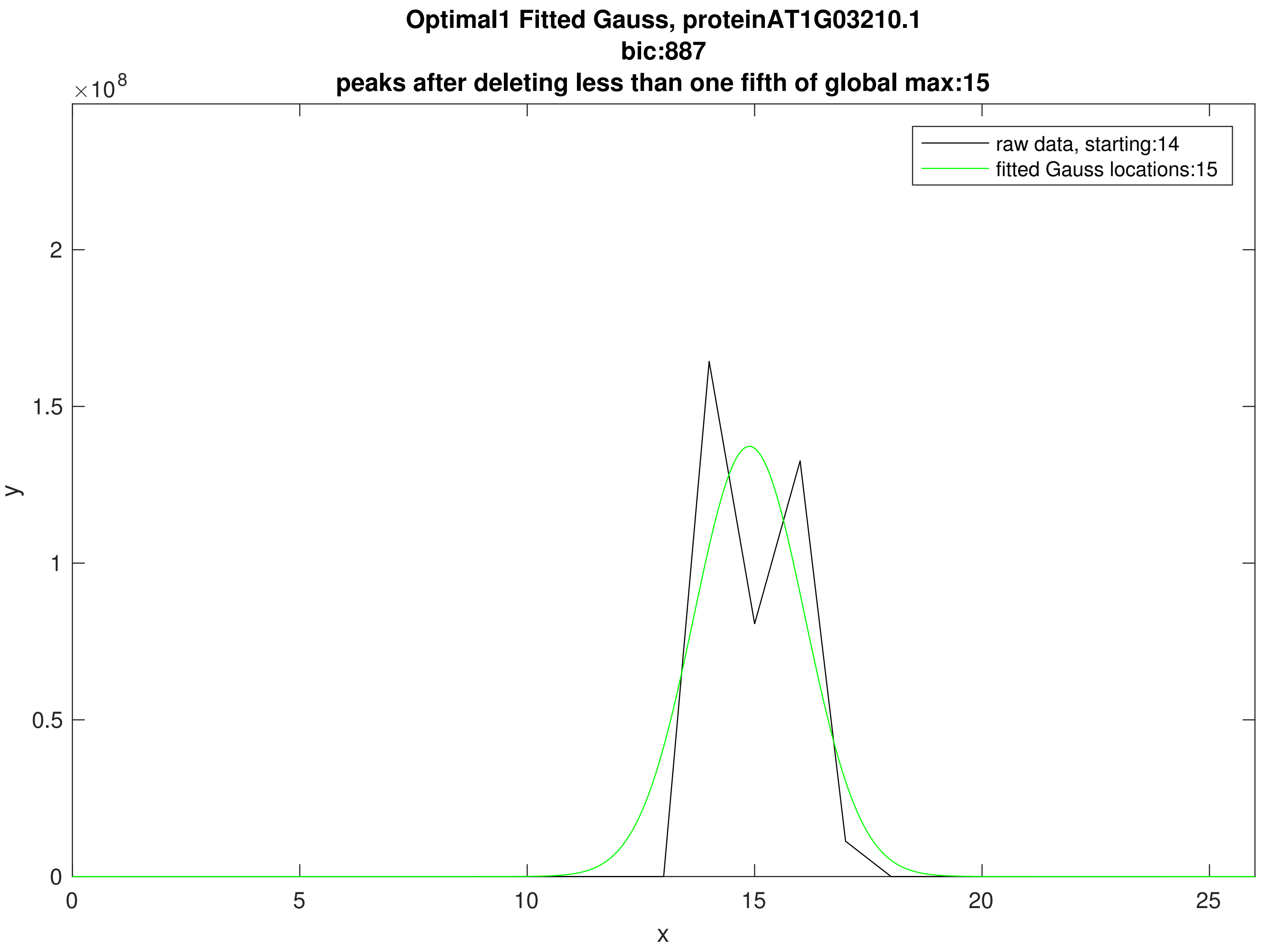

Optimal1 Fitted Gauss, proteinAT1G03210.1
bic:887
peaks after deleting less than one fifth of global max:15
×10^8
raw data, starting:14
fitted Gauss locations:15
y
x
0
0.5
1
1.5
2
0
5
10
15
20
25

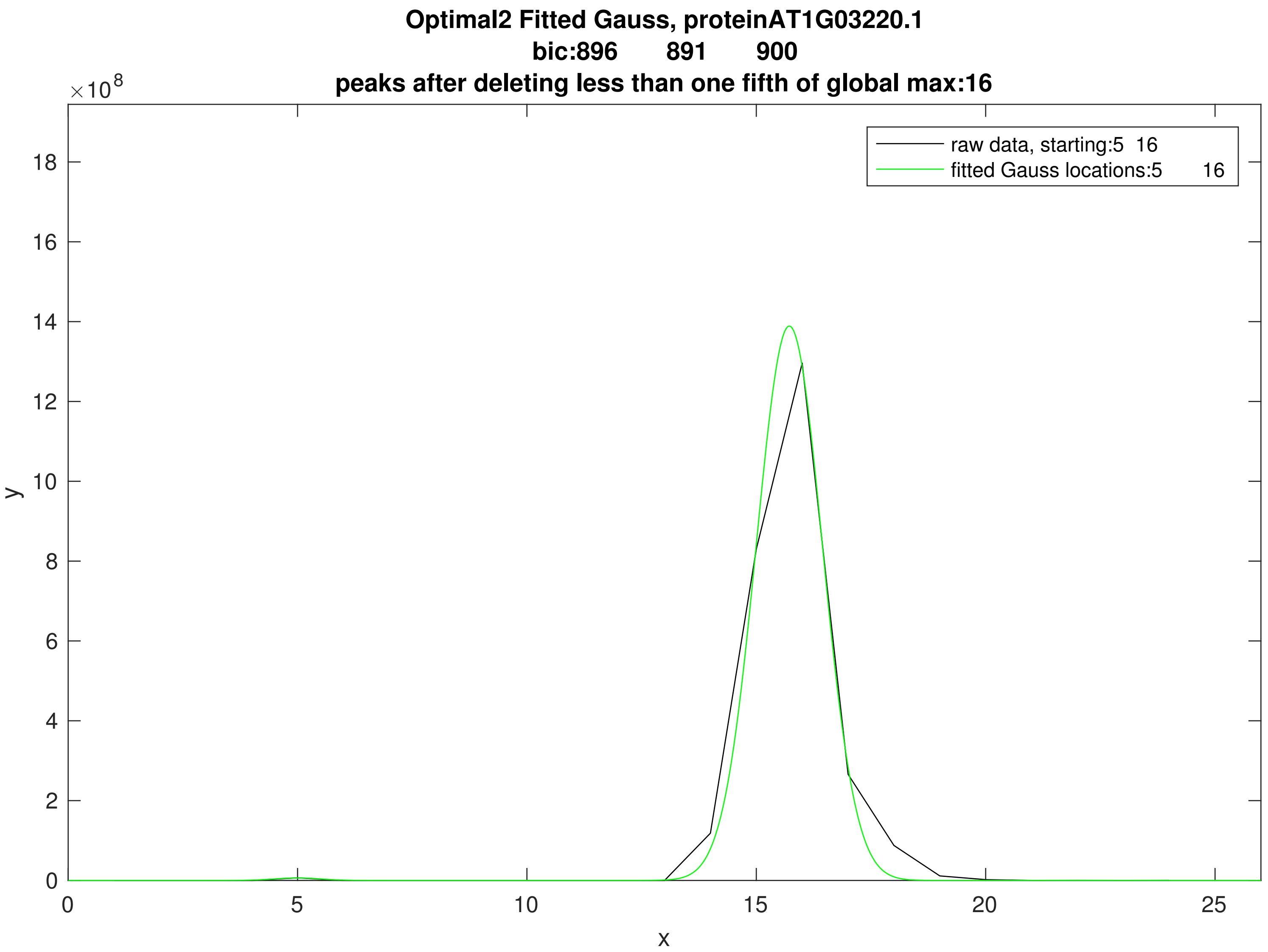

Optimal2 Fitted Gauss, proteinAT1G03220.1
bic:896 891 900
peaks after deleting less than one fifth of global max:16
×10^8
raw data, starting:5 16
fitted Gauss locations:5 16
y
x
0
2
4
6
8
10
12
14
16
18
0
5
10
15
20
25

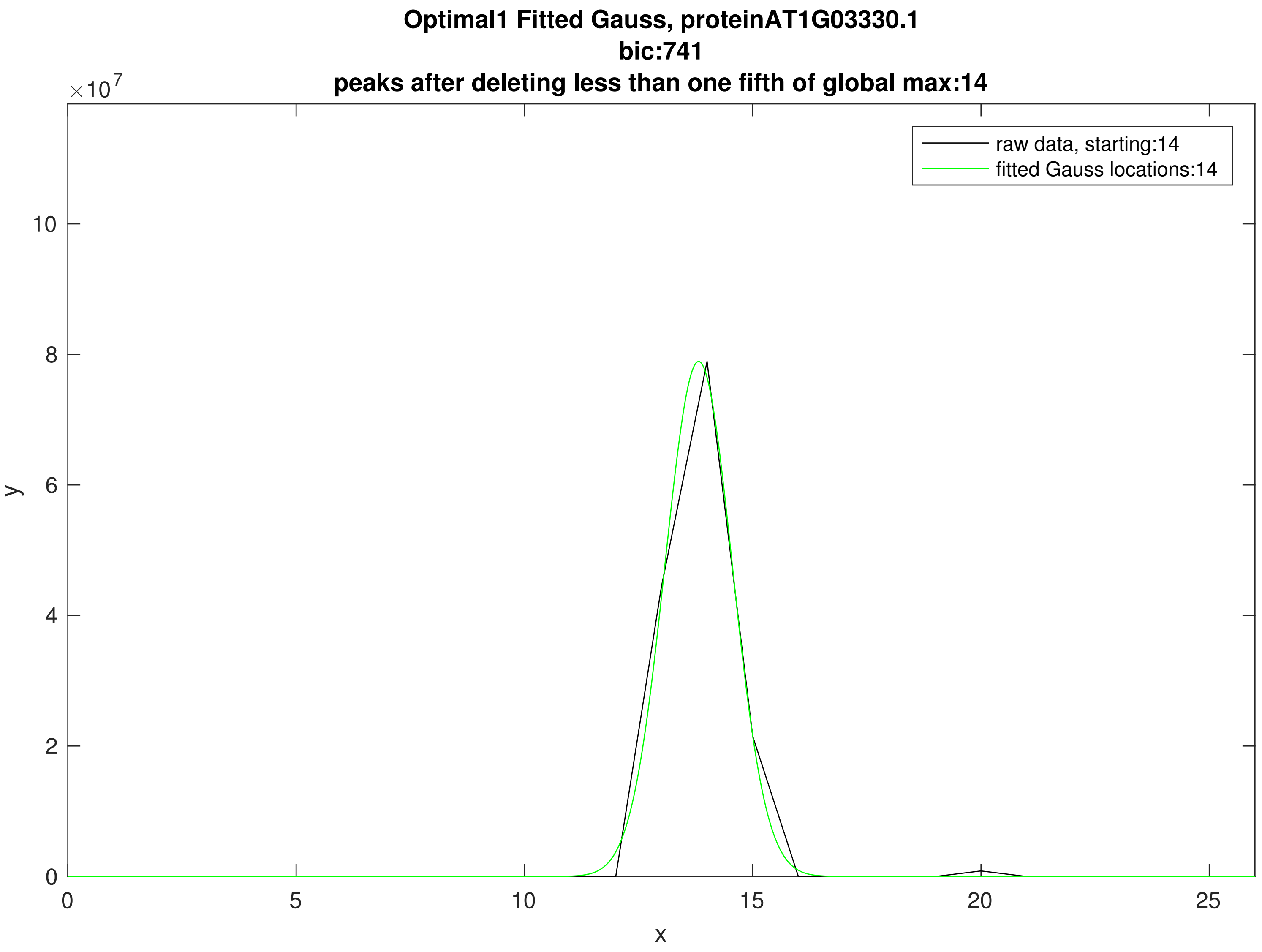
Optimal1 Fitted Gauss, proteinAT1G03330.1
bic:741
peaks after deleting less than one fifth of global max:14
×10^7
raw data, starting:14
fitted Gauss locations:14
y
x
0
2
4
6
8
10
0
5
10
15
20
25

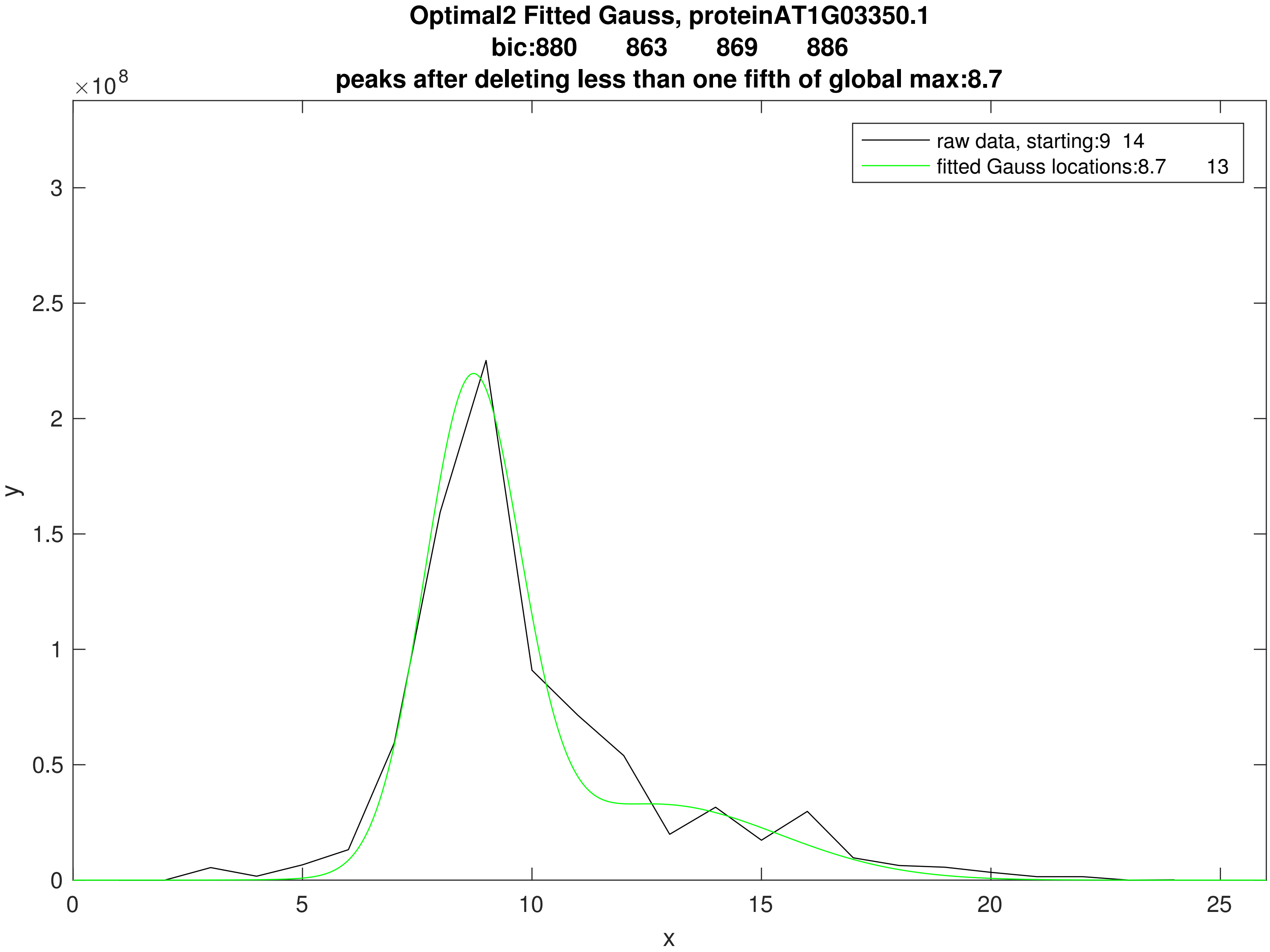

Optimal2 Fitted Gauss, proteinAT1G03350.1
bic:880 863 869 886
peaks after deleting less than one fifth of global max:8.7
×10^8
raw data, starting:9 14
fitted Gauss locations:8.7 13
3
2.5
2
1.5
1
0.5
0
y
0
5
10
15
20
25
x

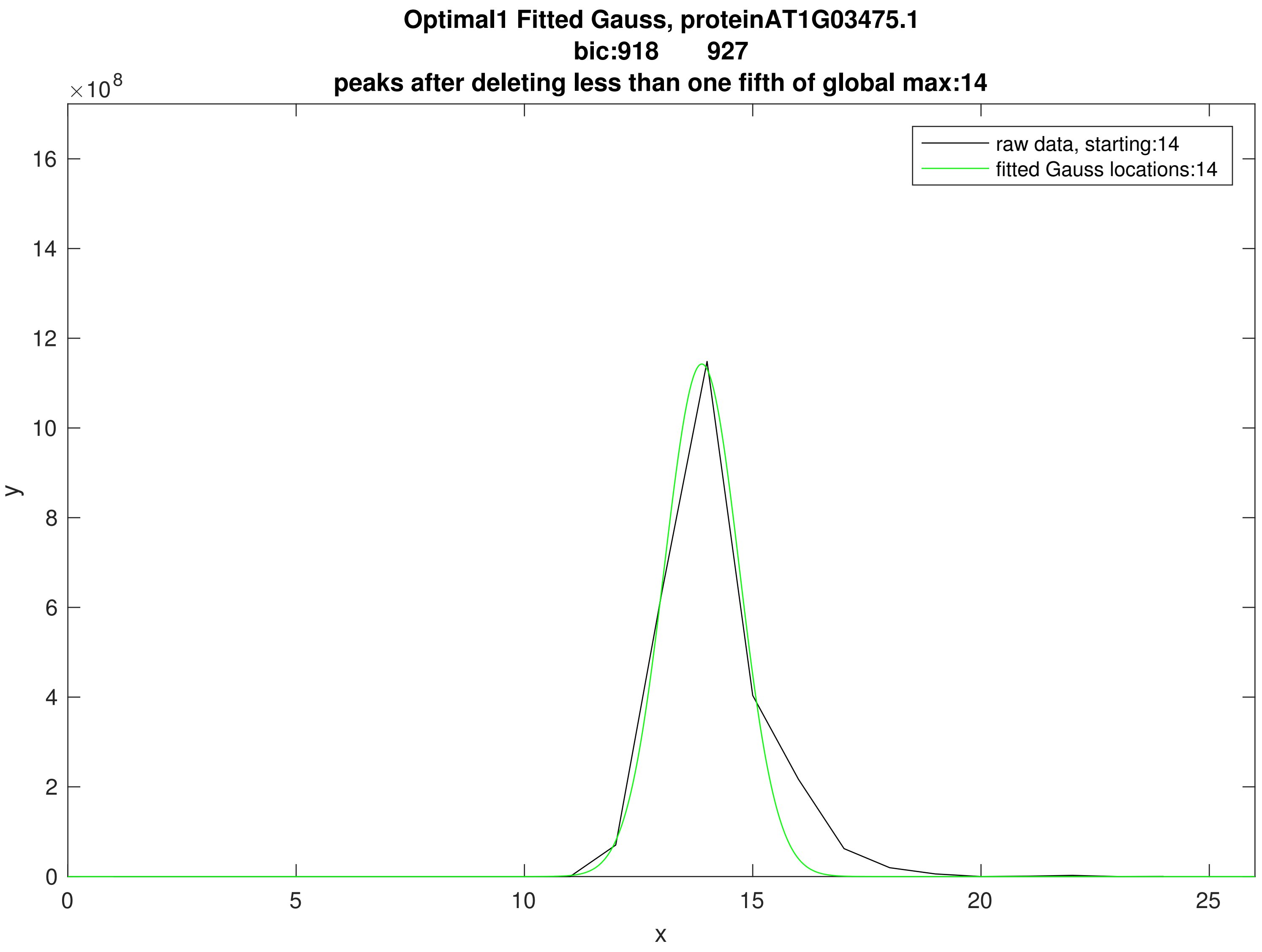

Optimal1 Fitted Gauss, proteinAT1G03475.1
bic:918 927
peaks after deleting less than one fifth of global max:14
×10^8
16
14
12
10
8
6
4
2
0
y
0
5
10
15
20
25
x
raw data, starting:14
fitted Gauss locations:14

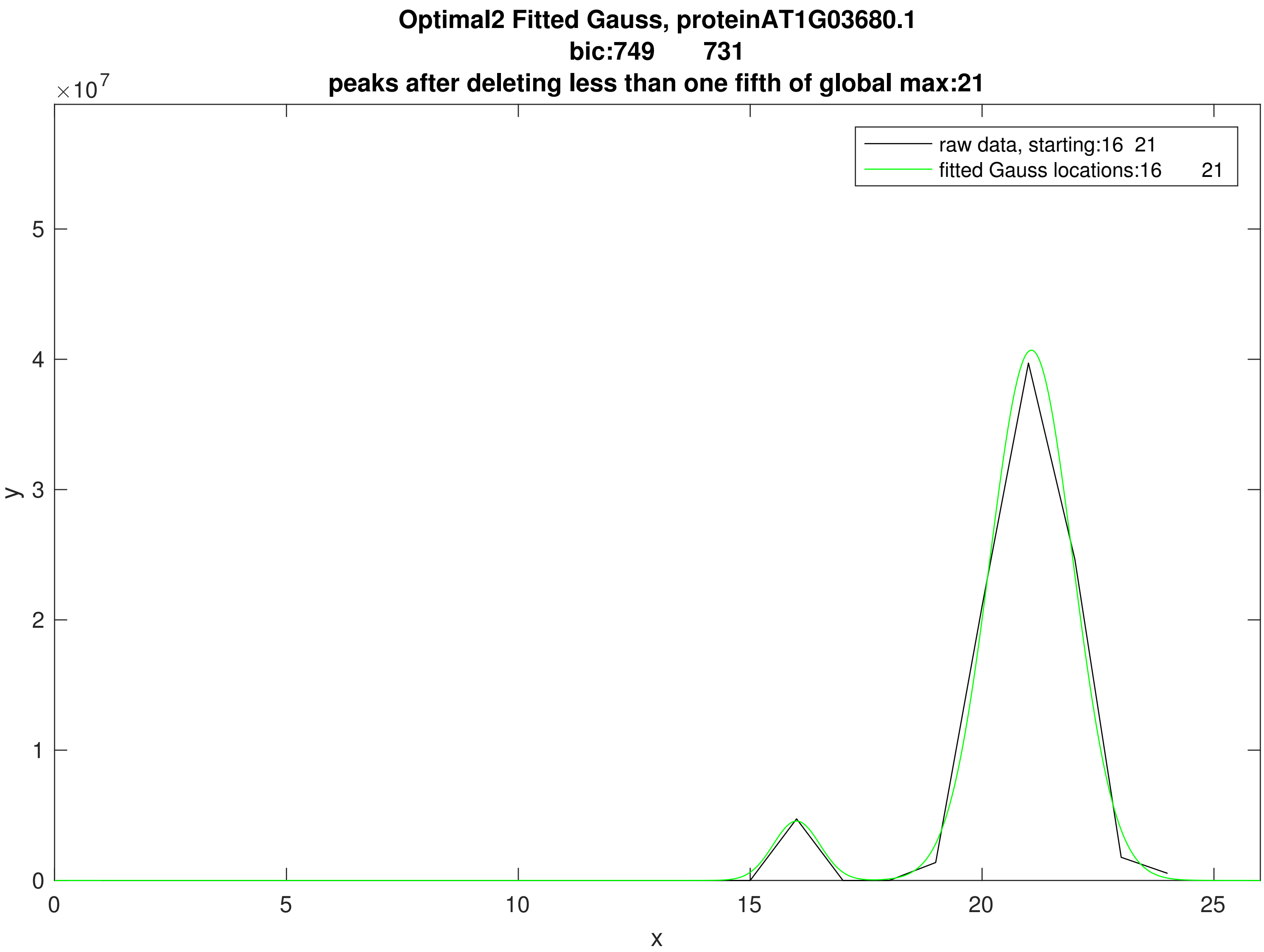
Optimal2 Fitted Gauss, proteinAT1G03680.1
bic:749 731
peaks after deleting less than one fifth of global max:21
×10^7
raw data, starting:16 21
fitted Gauss locations:16 21
y
x
0
1
2
3
4
5
0
5
10
15
20
25

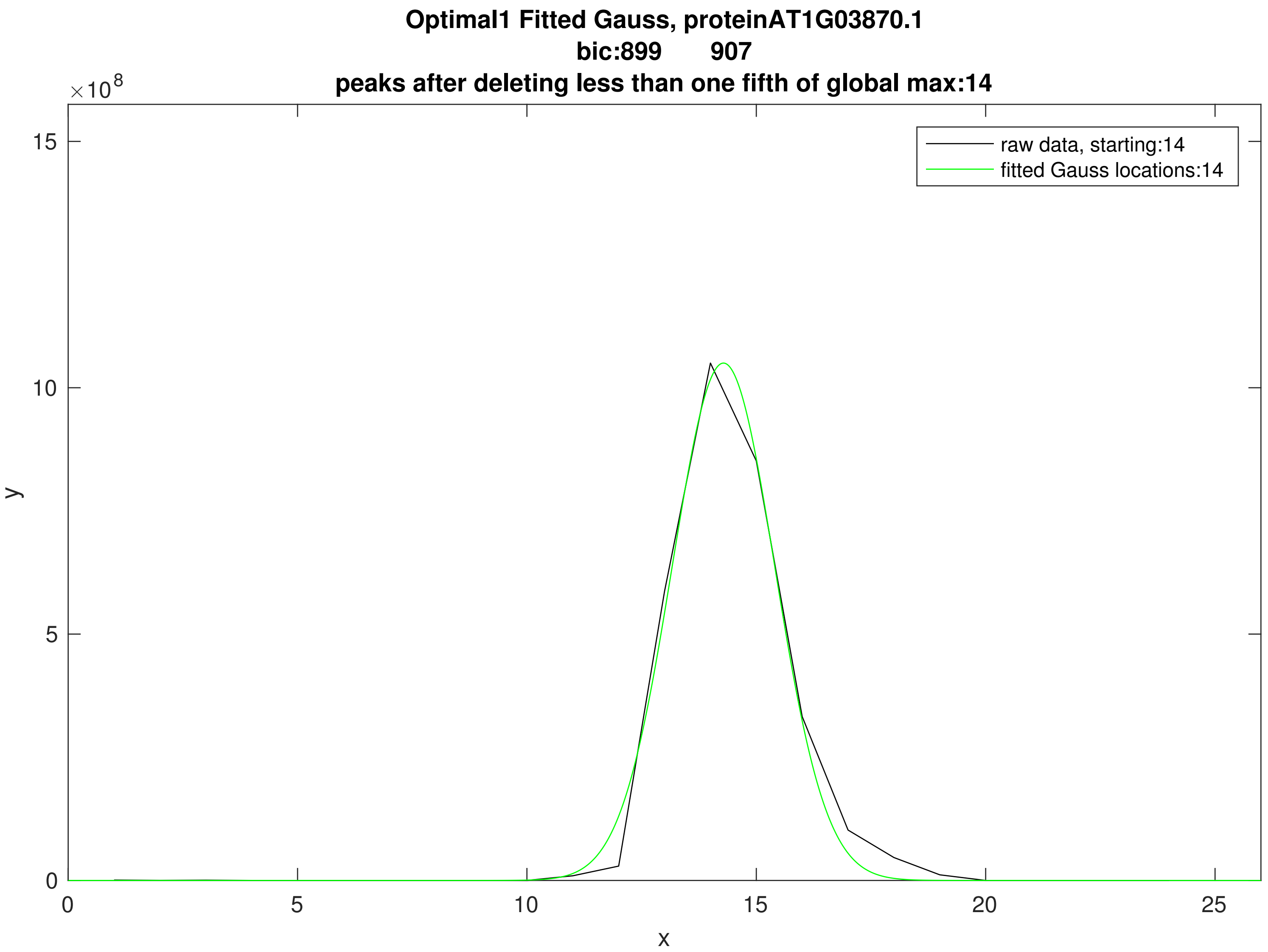
Optimal1 Fitted Gauss, proteinAT1G03870.1
bic:899 907
peaks after deleting less than one fifth of global max:14
×10^8
15
10
5
0
y
0
5
10
15
20
25
x
raw data, starting:14
fitted Gauss locations:14

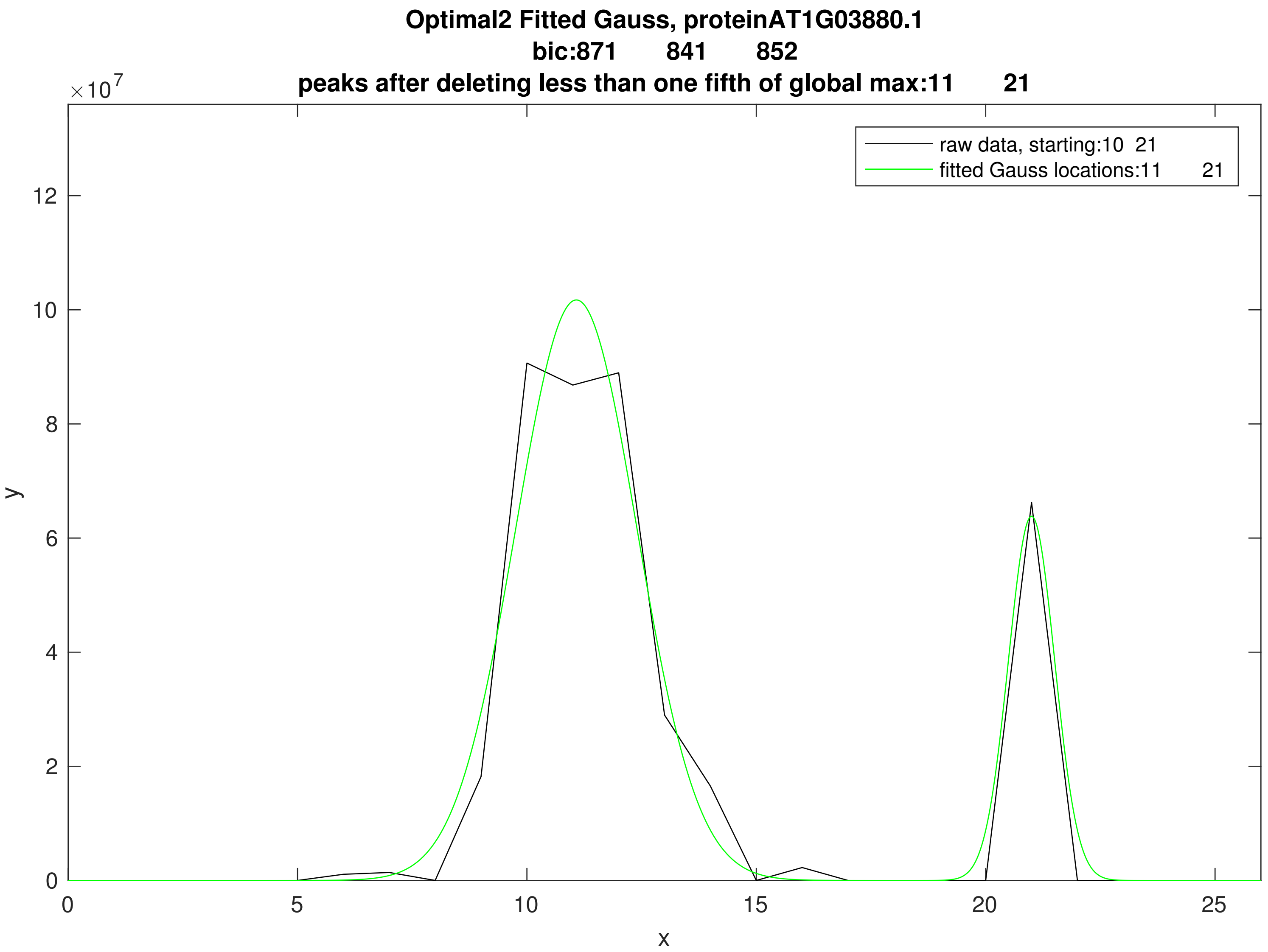

Optimal2 Fitted Gauss, proteinAT1G03880.1
bic:871 841 852
peaks after deleting less than one fifth of global max:11 21
×10^7
raw data, starting:10 21
fitted Gauss locations:11 21
12
10
8
6
4
2
0
y
0
5
10
15
20
25
x

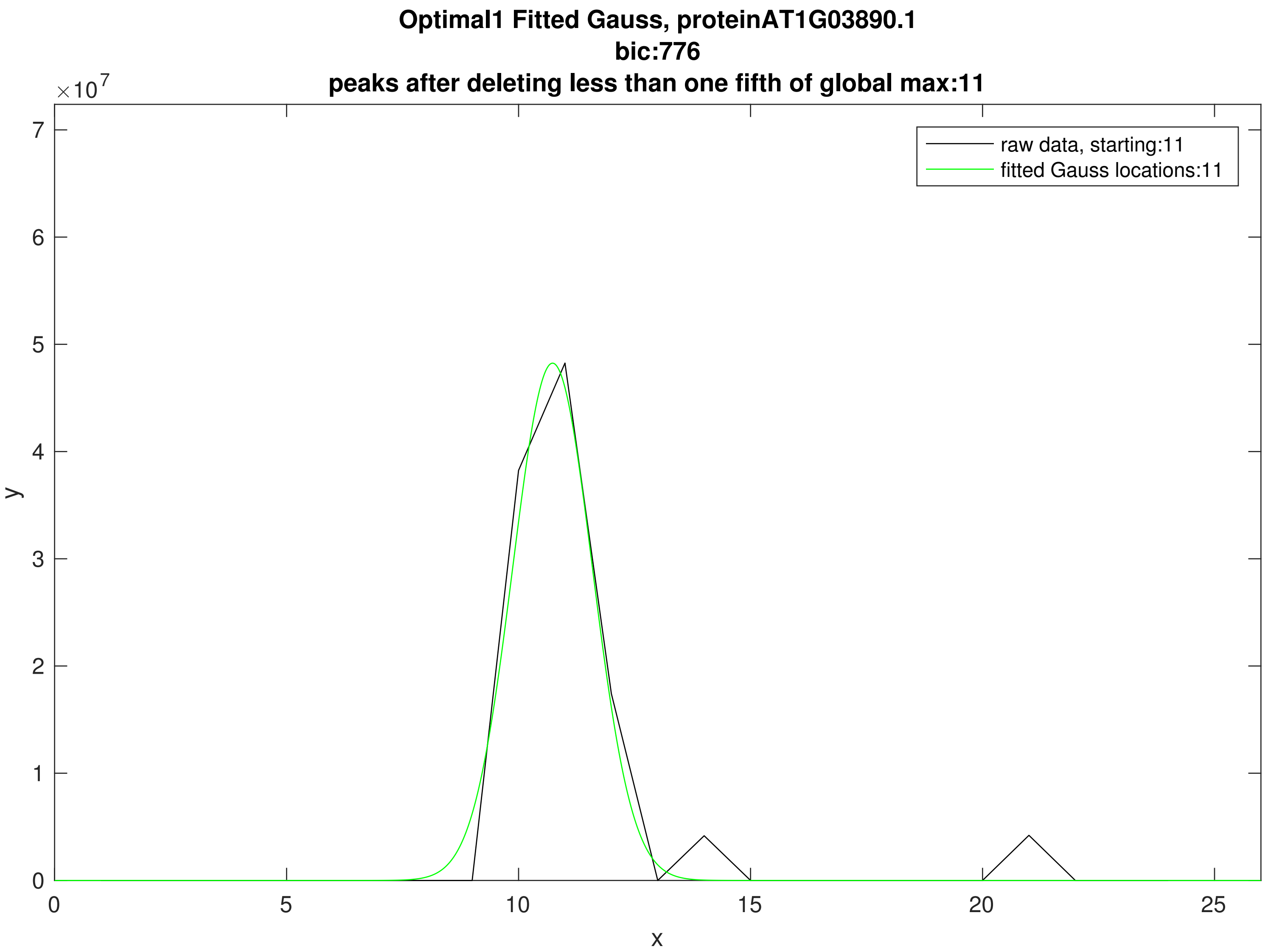
Optimal1 Fitted Gauss, proteinAT1G03890.1
bic:776
peaks after deleting less than one fifth of global max:11
×10^7
7
6
5
4
3
2
1
0
y
0
5
10
15
20
25
x
raw data, starting:11
fitted Gauss locations:11

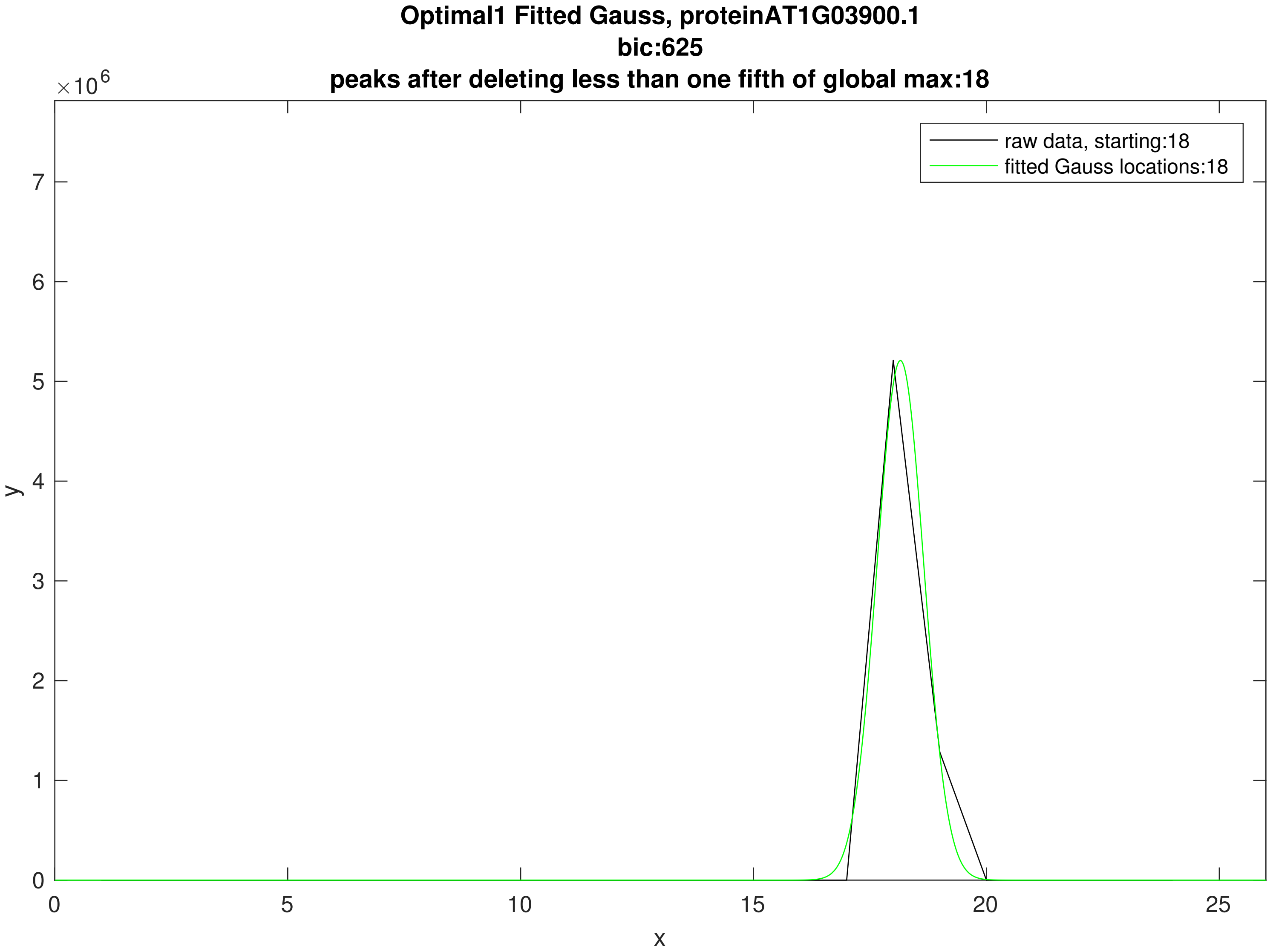

Optimal1 Fitted Gauss, proteinAT1G03900.1
bic:625
peaks after deleting less than one fifth of global max:18
×10^6
raw data, starting:18
fitted Gauss locations:18
y
x
0
1
2
3
4
5
6
7
0
5
10
15
20
25

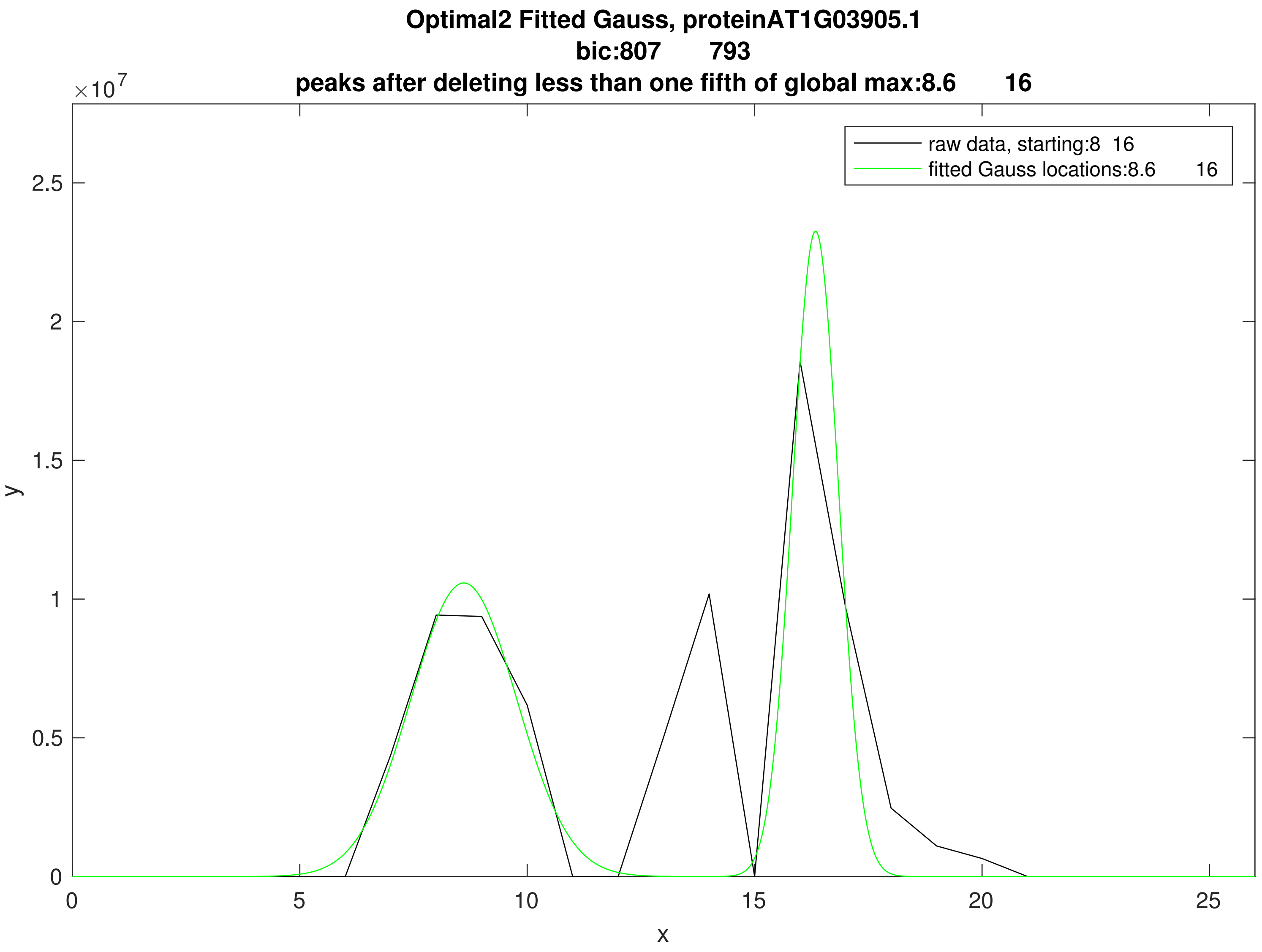
Optimal2 Fitted Gauss, proteinAT1G03905.1
bic:807 793
peaks after deleting less than one fifth of global max:8.6 16
×10^7
raw data, starting:8 16
fitted Gauss locations:8.6 16
y
x
0
0.5
1
1.5
2
2.5
0
5
10
15
20
25

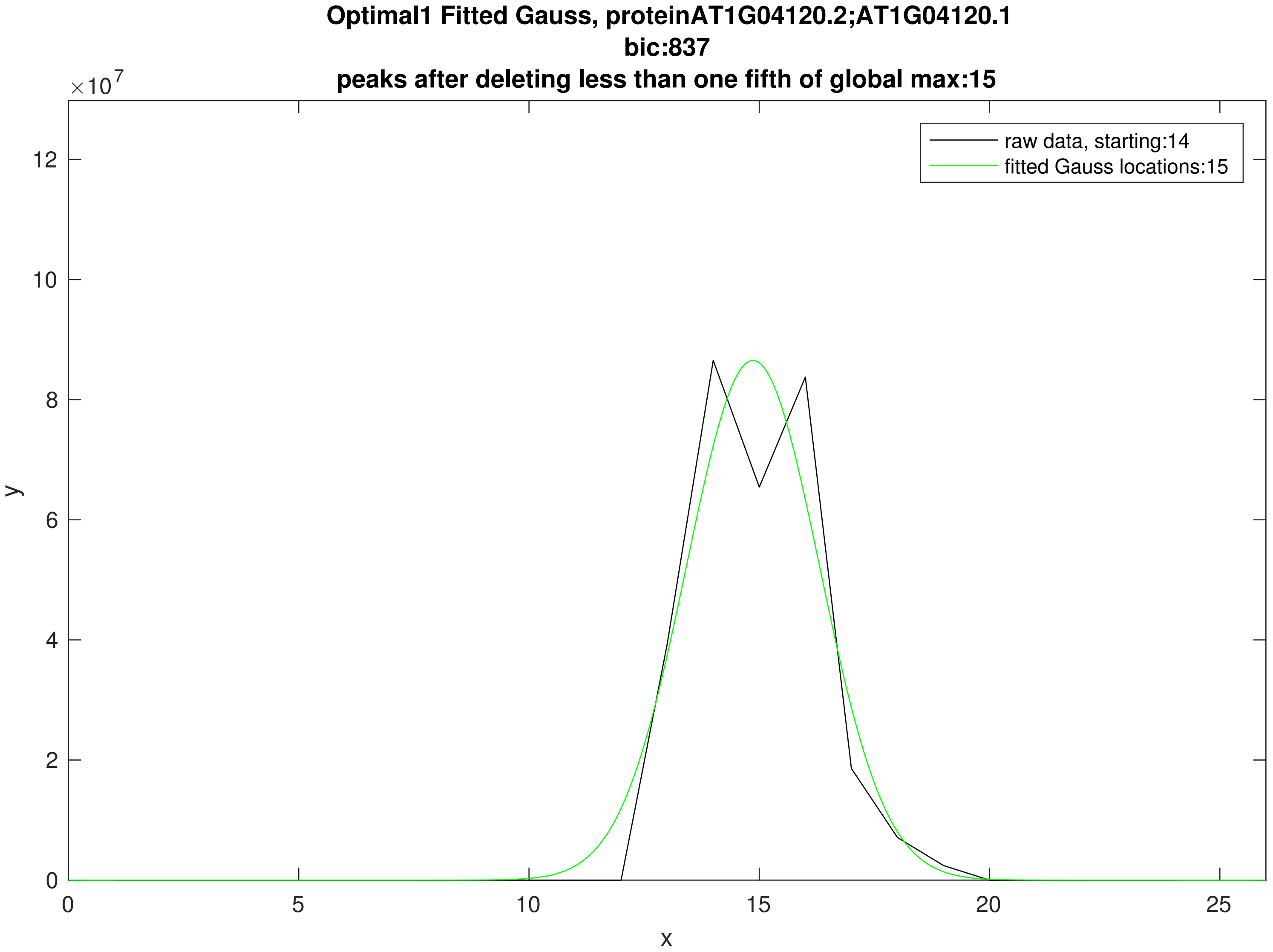
Optimal1 Fitted Gauss, proteinAT1G04120.2;AT1G04120.1
bic:837
peaks after deleting less than one fifth of global max:15
×10^7
raw data, starting:14
fitted Gauss locations:15
y
x
0
2
4
6
8
10
12
0
5
10
15
20
25

Optimal1 Fitted Gauss, proteinAT1G04270.1;AT1G04270.2
bic:851 861
peaks after deleting less than one fifth of global max:3.4

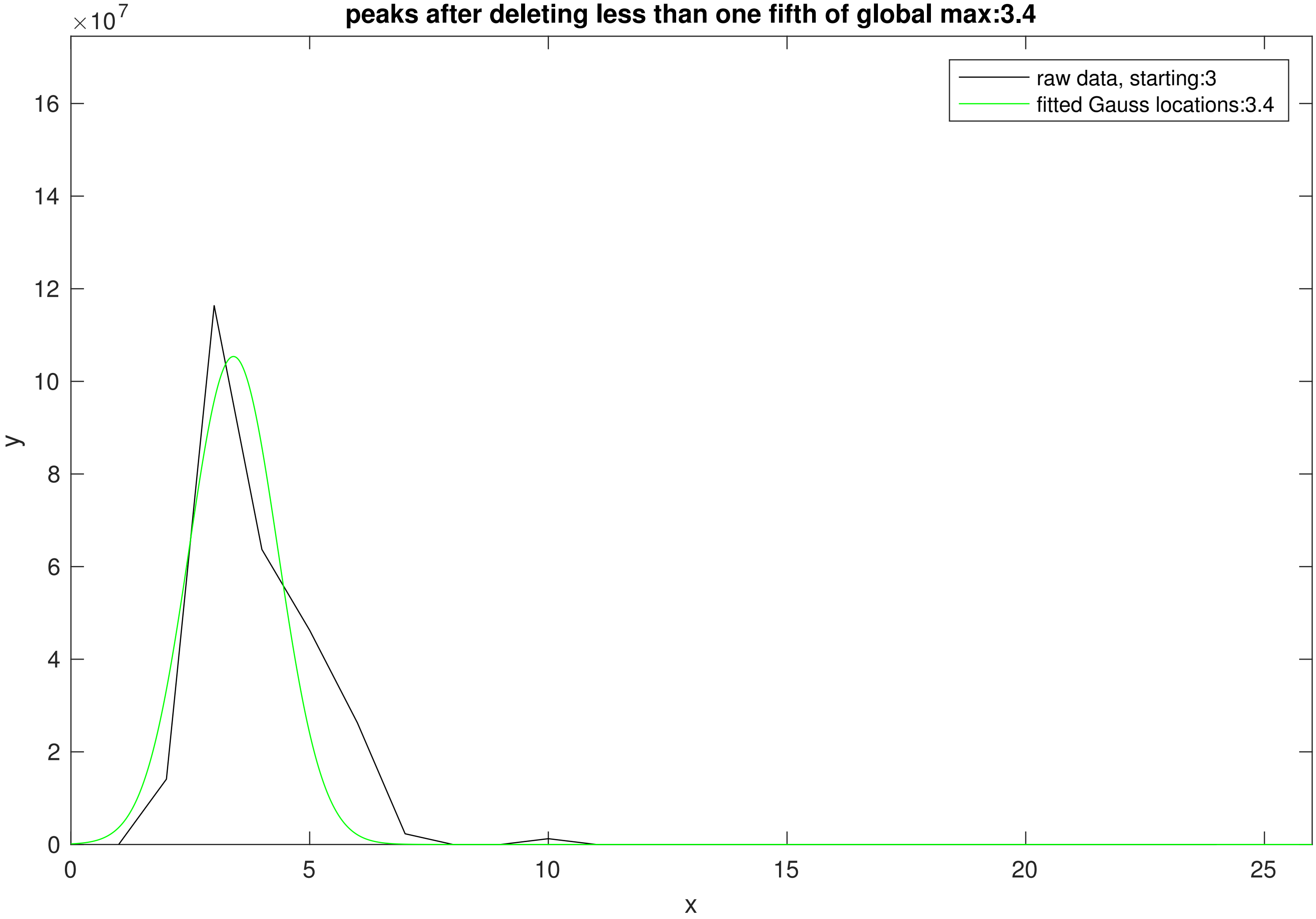

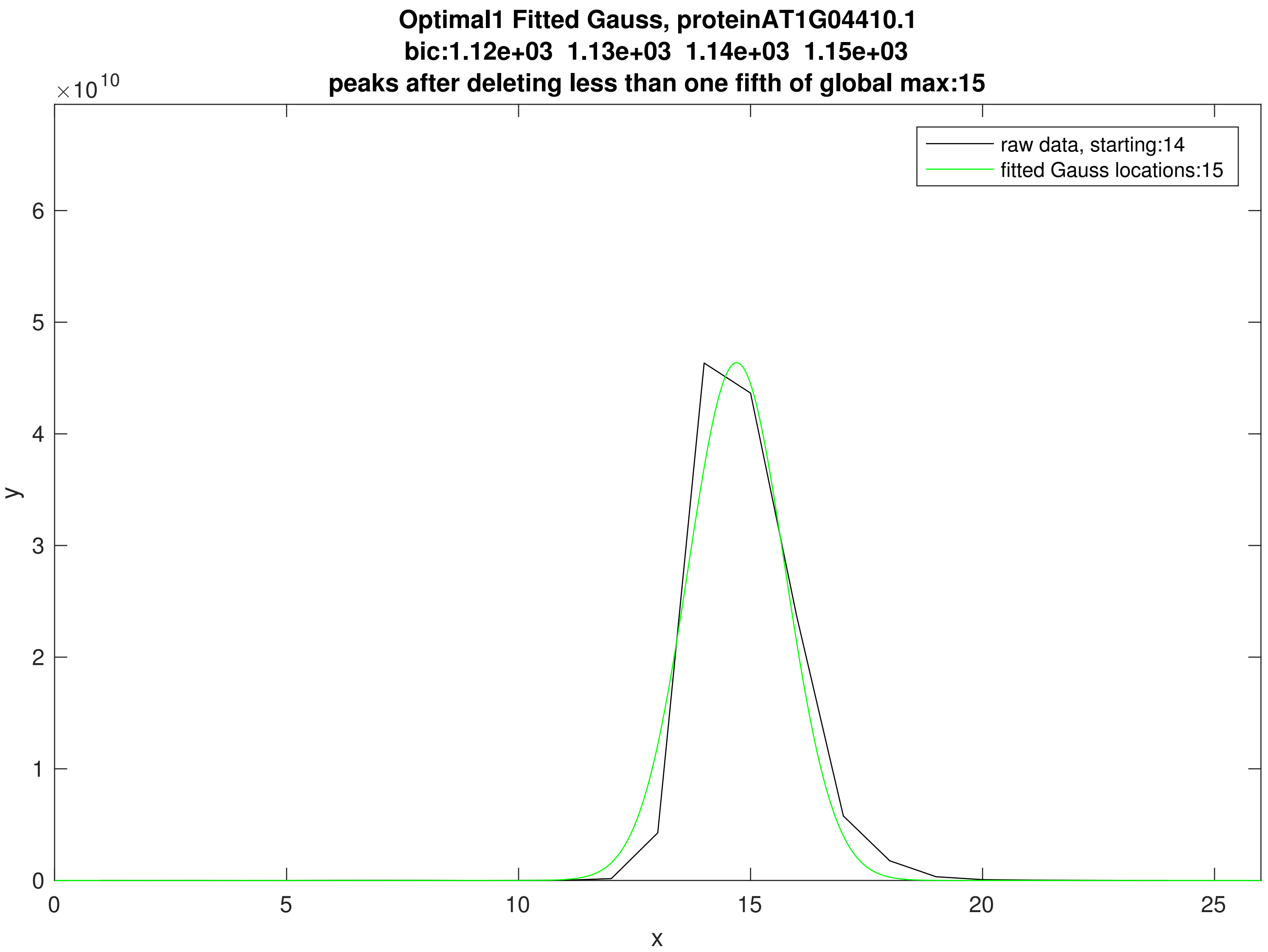
Optimal1 Fitted Gauss, proteinAT1G04410.1
bic:1.12e+03 1.13e+03 1.14e+03 1.15e+03
peaks after deleting less than one fifth of global max:15
×10^10
raw data, starting:14
fitted Gauss locations:15
y
x
0
1
2
3
4
5
6
0
5
10
15
20
25

**Optimal1 Fitted Gauss, proteinAT1G04420.1**
**bic:585**
**peaks after deleting less than one fifth of global max:14**

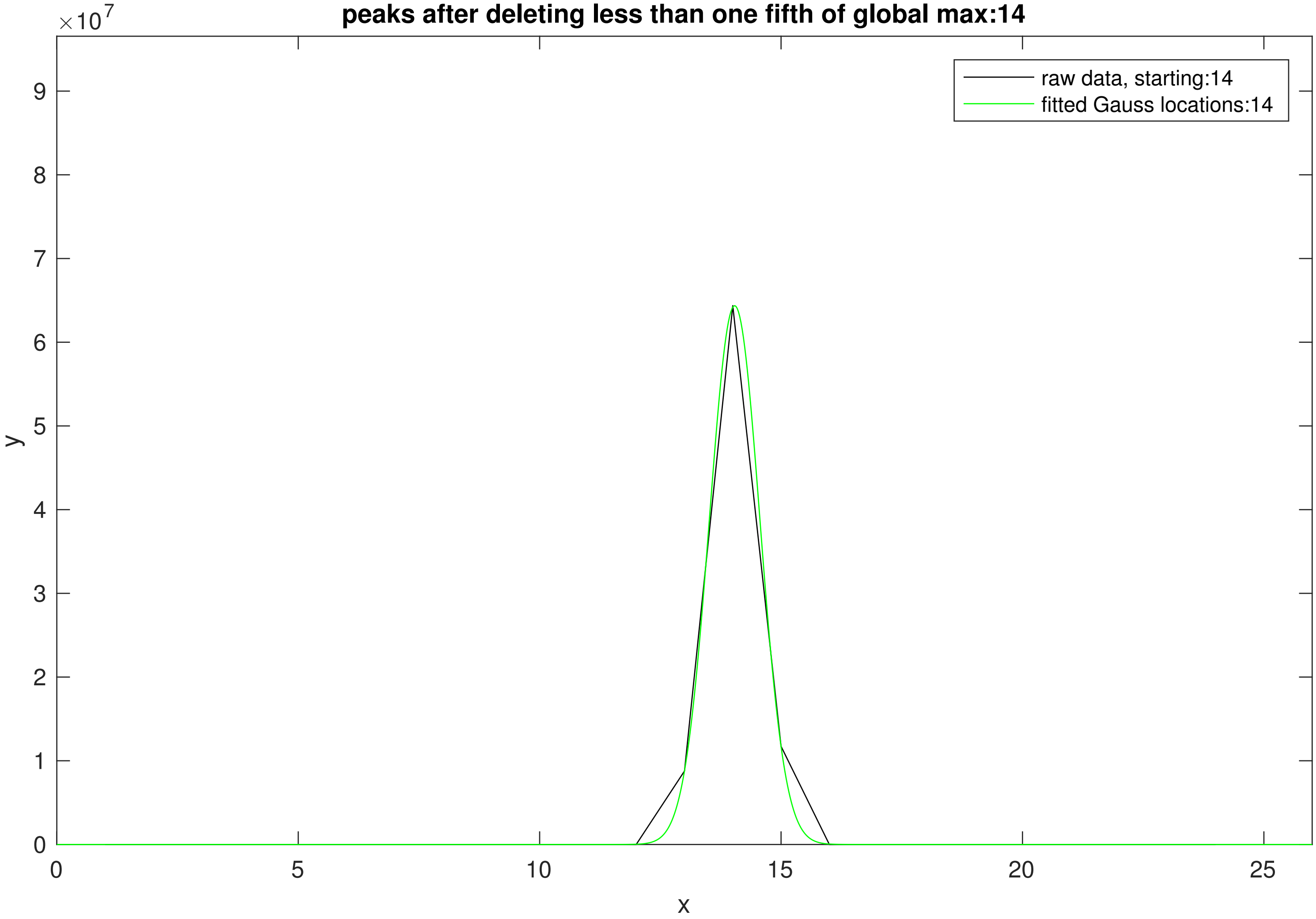

**Optimal1 Fitted Gauss, proteinAT1G04750.2;AT1G04750.1;AT2G33120.1;AT2G33120.2**
**bic:658**
**peaks after deleting less than one fifth of global max:20**

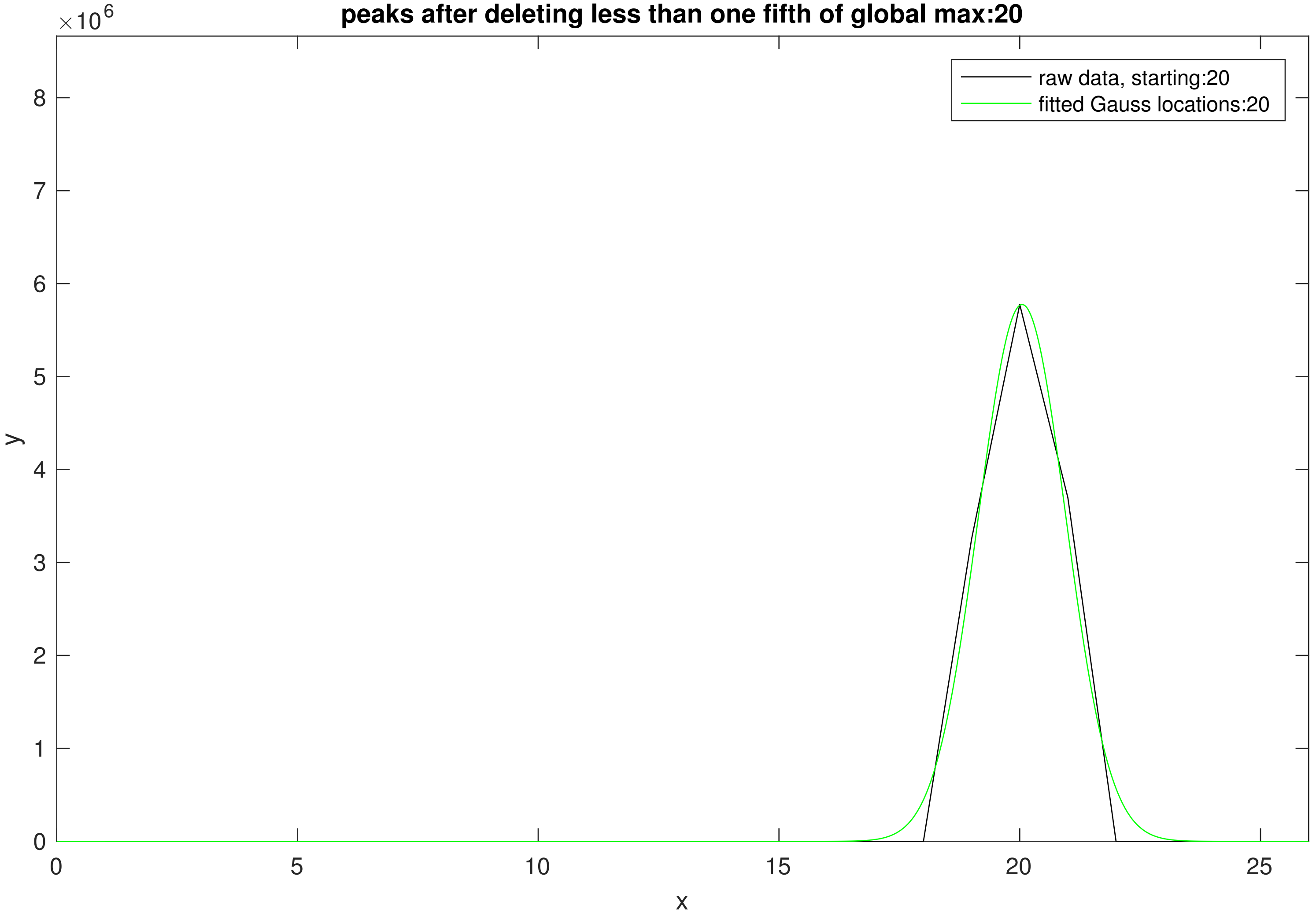

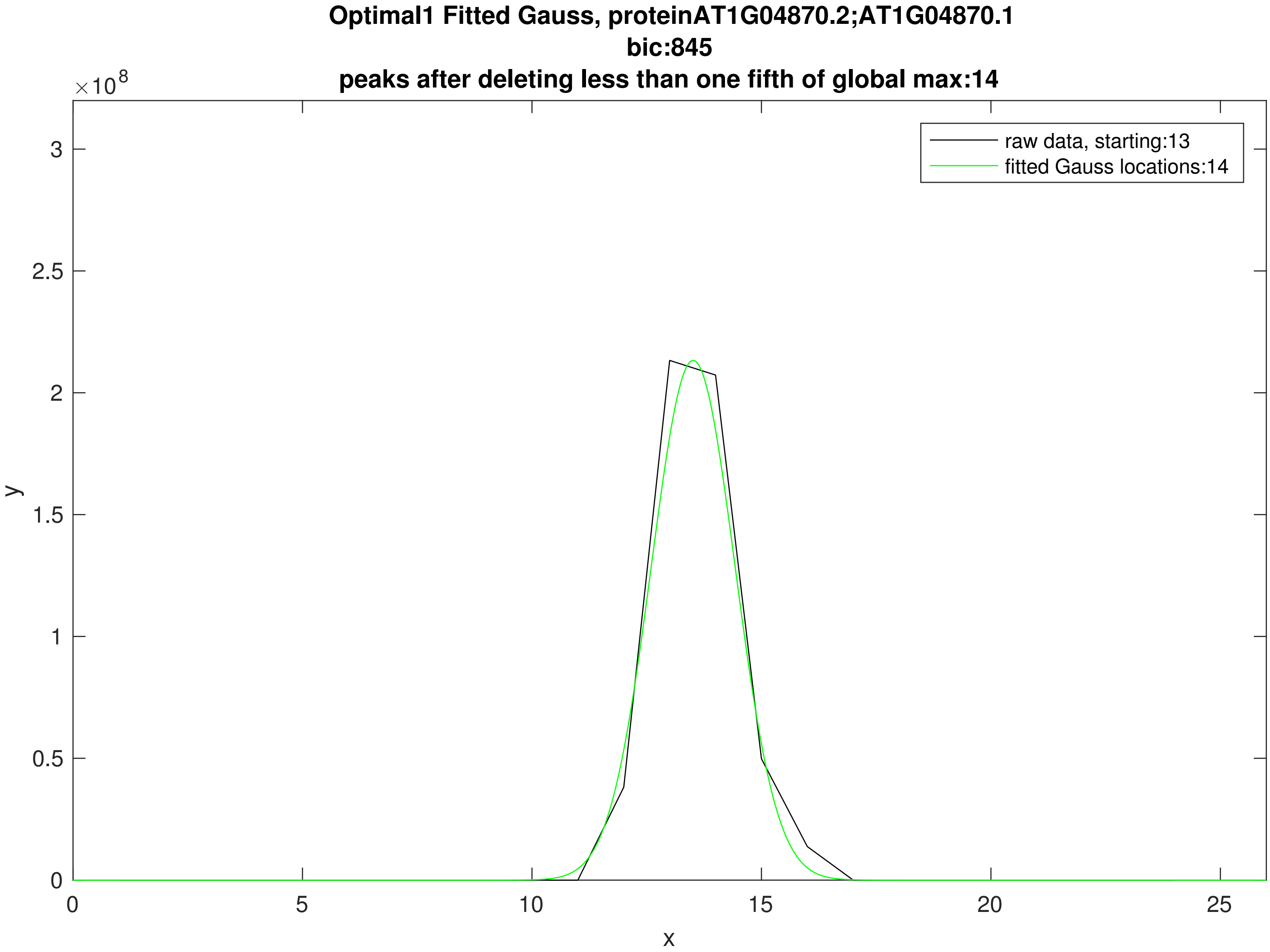

Optimal1 Fitted Gauss, proteinAT1G04870.2;AT1G04870.1
bic:845
peaks after deleting less than one fifth of global max:14
×10^8
raw data, starting:13
fitted Gauss locations:14
y
x
0
0.5
1
1.5
2
2.5
3
5
10
15
20
25

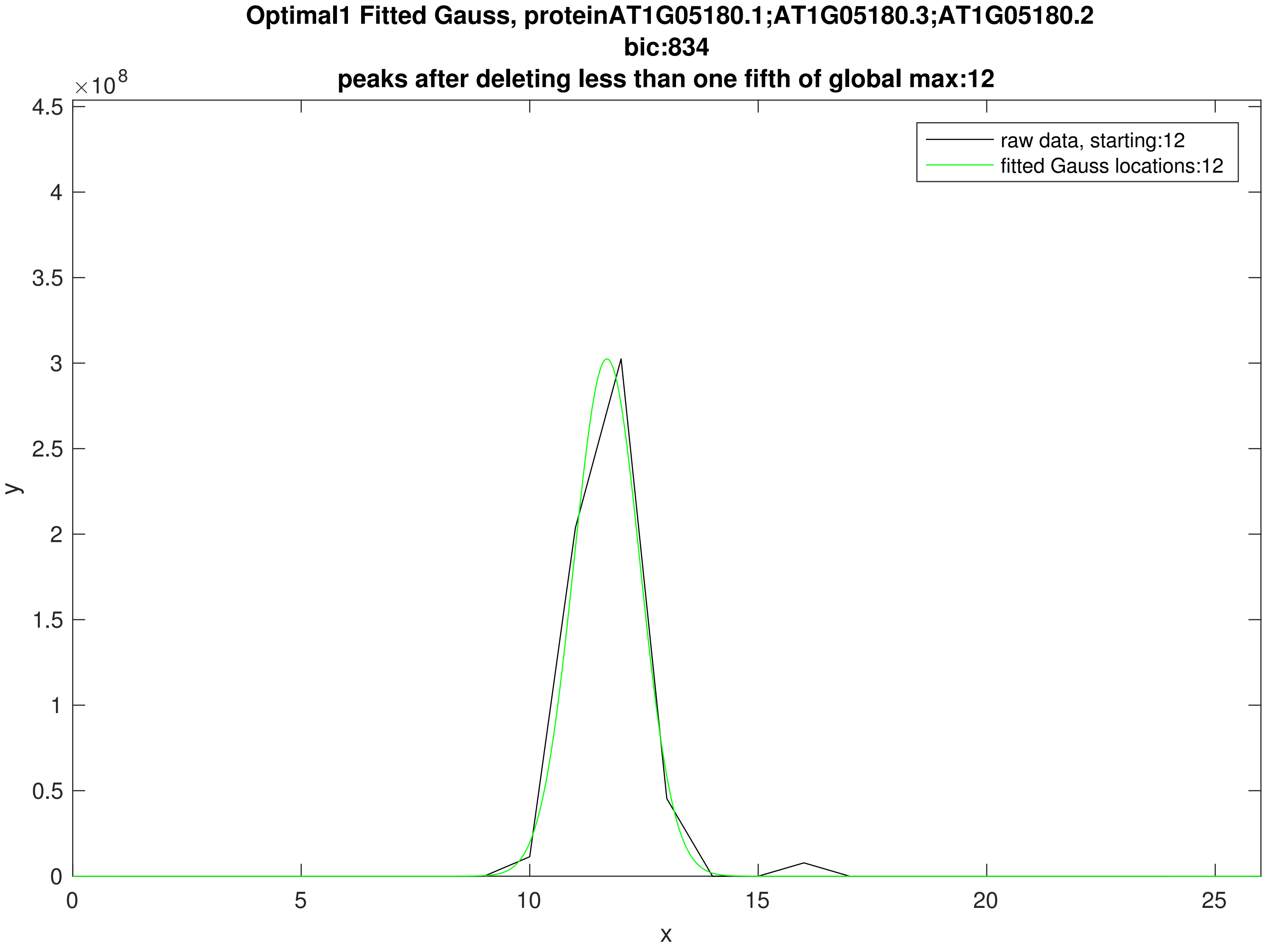
Optimal1 Fitted Gauss, proteinAT1G05180.1;AT1G05180.3;AT1G05180.2
bic:834
peaks after deleting less than one fifth of global max:12
×10^8
4.5
4
3.5
3
2.5
2
1.5
1
0.5
0
y
0
5
10
15
20
25
x
raw data, starting:12
fitted Gauss locations:12

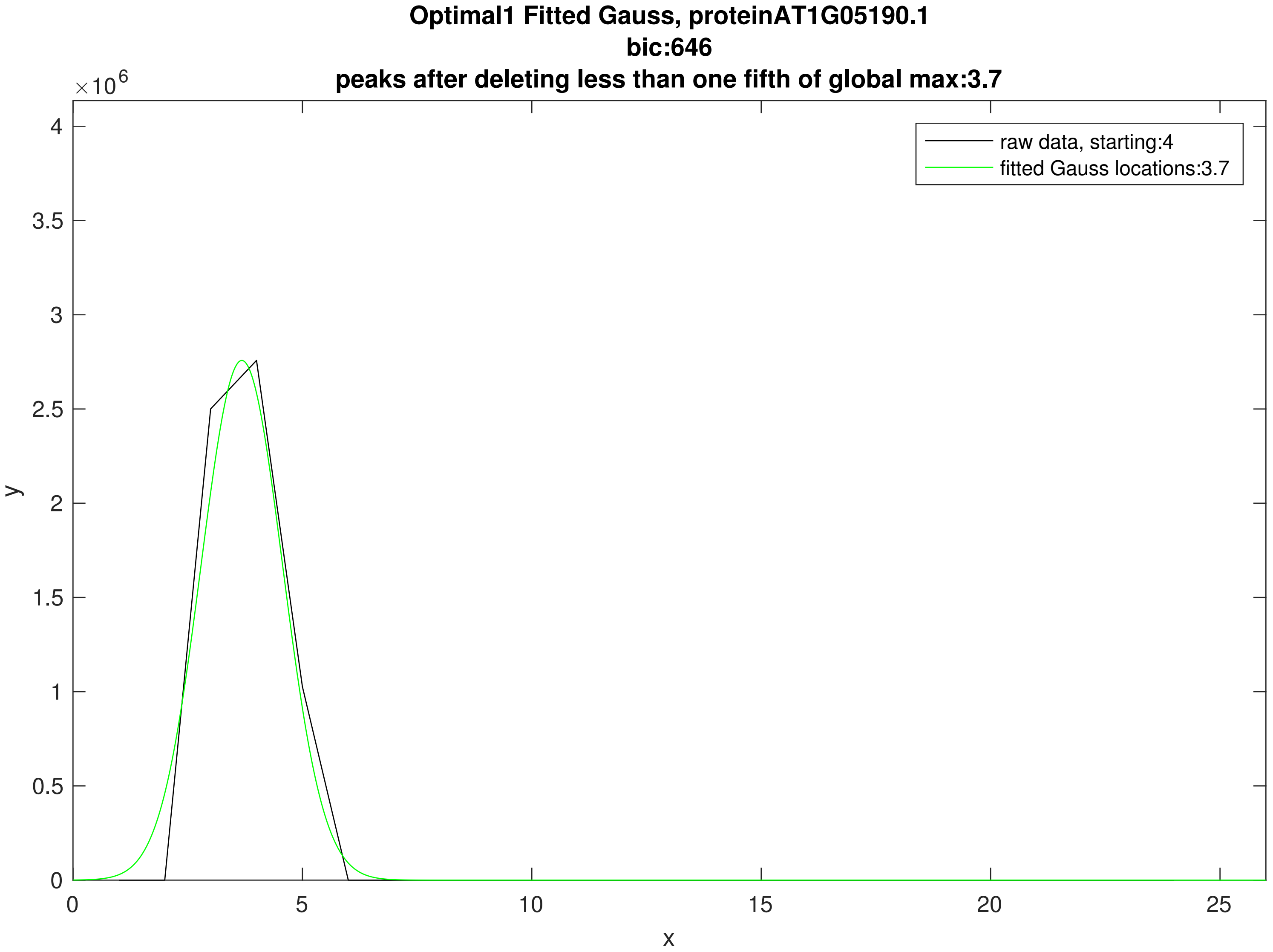

Optimal1 Fitted Gauss, proteinAT1G05190.1
bic:646
peaks after deleting less than one fifth of global max:3.7
×10^6
raw data, starting:4
fitted Gauss locations:3.7
y
x
0
0.5
1
1.5
2
2.5
3
3.5
4
5
10
15
20
25

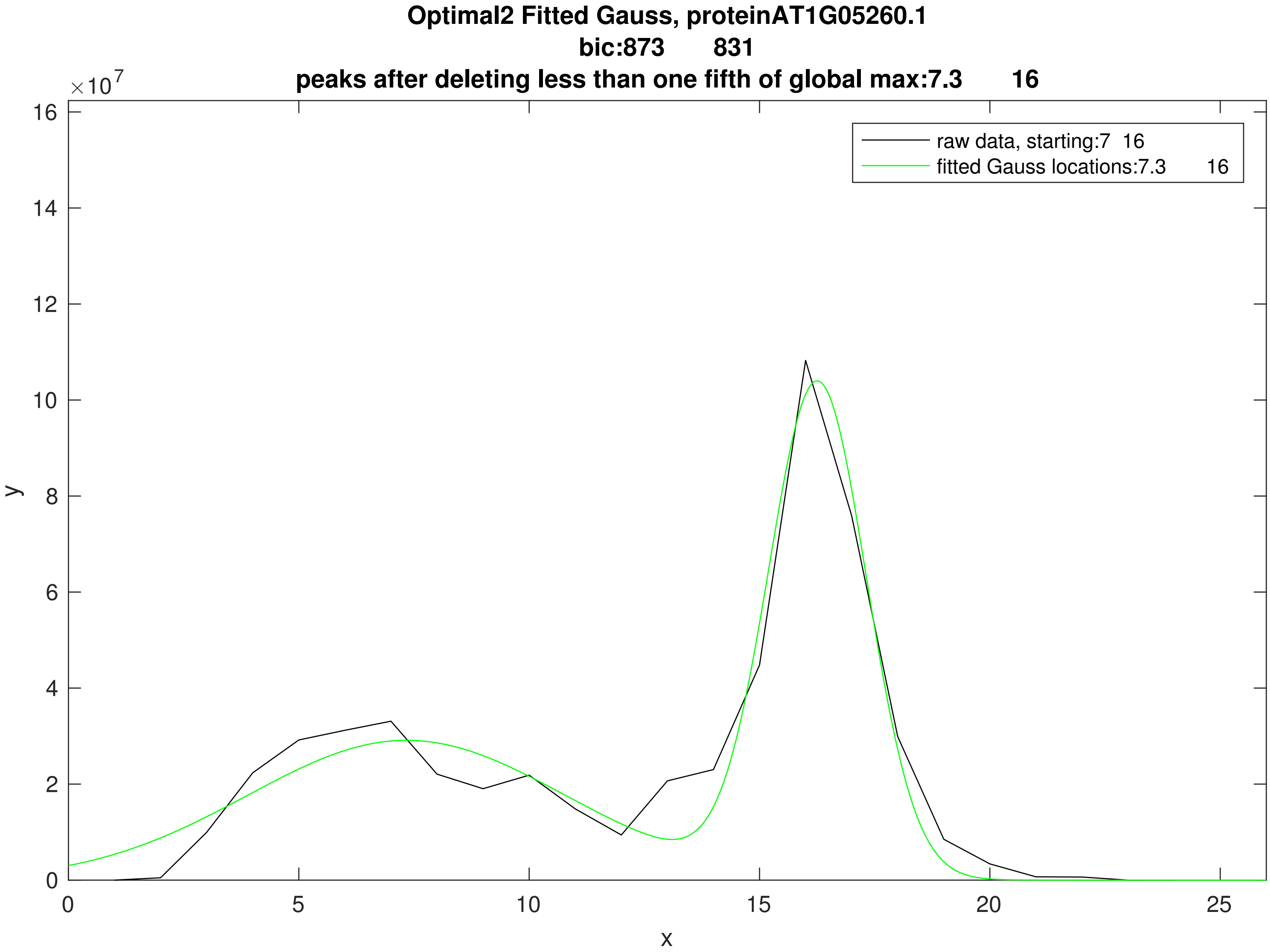

Optimal2 Fitted Gauss, proteinAT1G05260.1
bic:873 831
peaks after deleting less than one fifth of global max:7.3 16
×10^7
raw data, starting:7 16
fitted Gauss locations:7.3 16
y
x
0
2
4
6
8
10
12
14
16
0
5
10
15
20
25

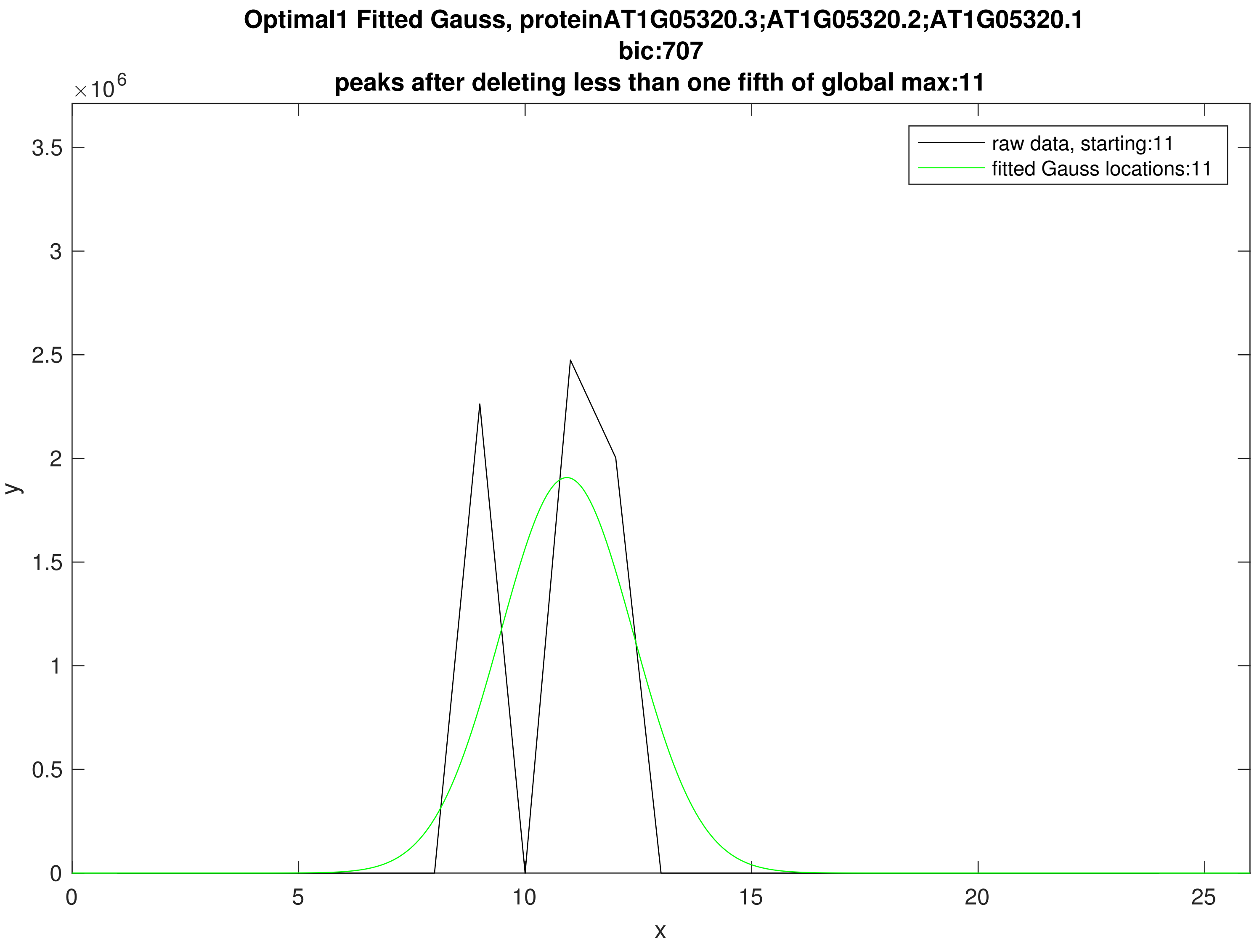

Optimal1 Fitted Gauss, proteinAT1G05320.3;AT1G05320.2;AT1G05320.1
bic:707
peaks after deleting less than one fifth of global max:11
×10^6
3.5
3
2.5
2
1.5
1
0.5
0
y
0
5
10
15
20
25
x
raw data, starting:11
fitted Gauss locations:11

**Optimal1 Fitted Gauss, proteinAT1G05350.1**
**bic:753**
**peaks after deleting less than one fifth of global max:9.3**

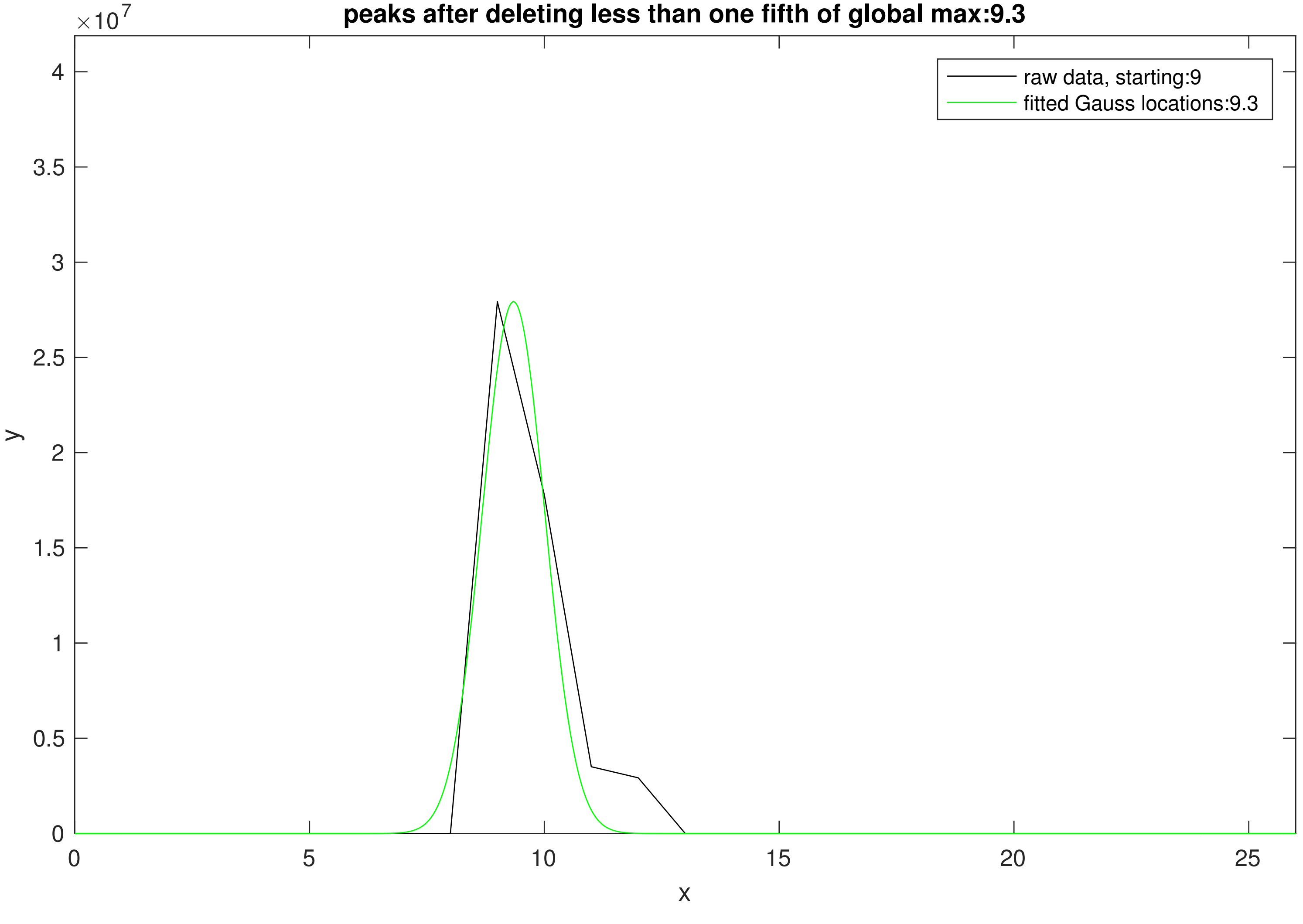

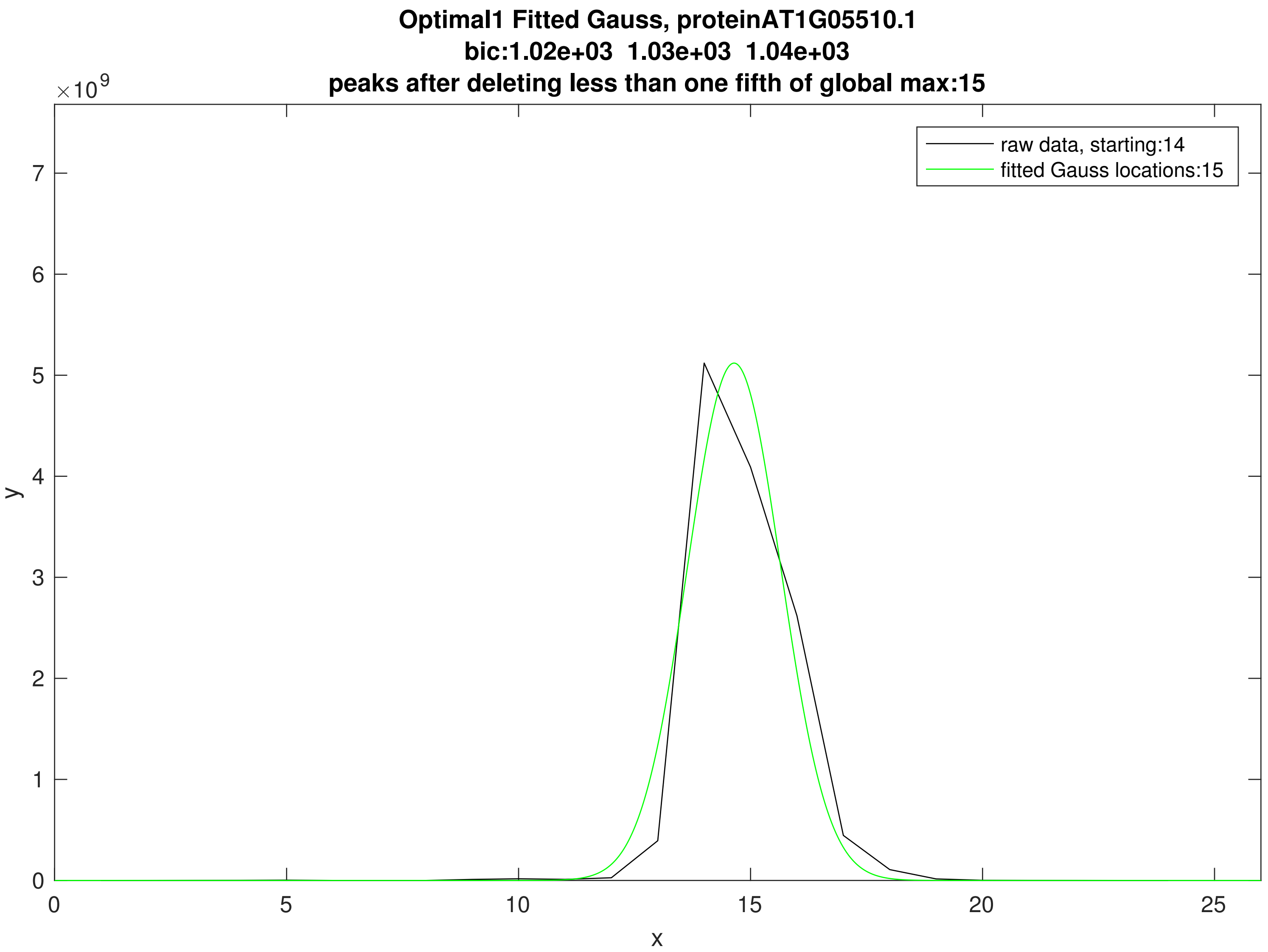

Optimal1 Fitted Gauss, proteinAT1G05510.1
bic:1.02e+03 1.03e+03 1.04e+03
peaks after deleting less than one fifth of global max:15
×10^9
raw data, starting:14
fitted Gauss locations:15
y
x
0
1
2
3
4
5
6
7
0
5
10
15
20
25

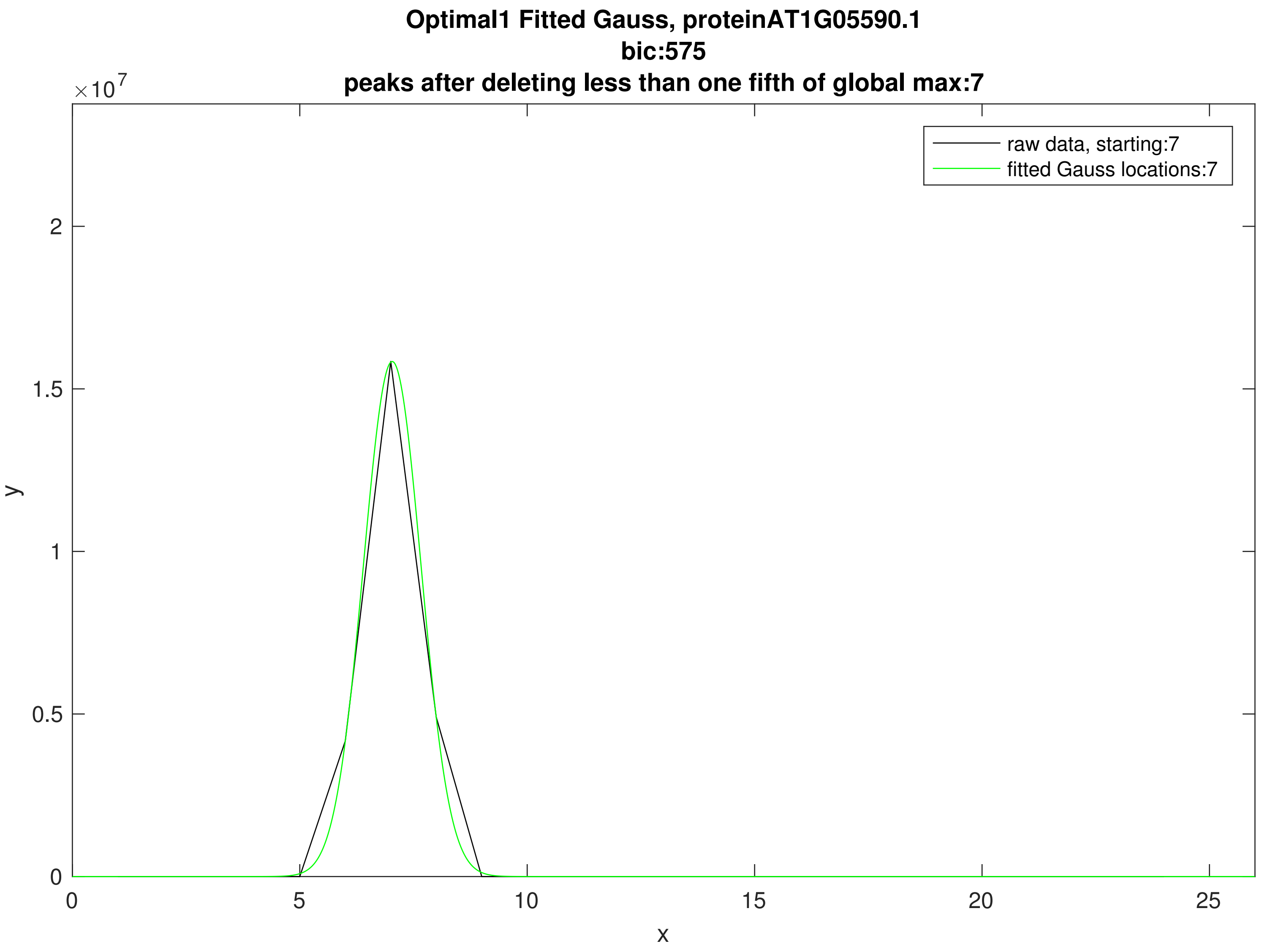
Optimal1 Fitted Gauss, proteinAT1G05590.1
bic:575
peaks after deleting less than one fifth of global max:7
×10^7
raw data, starting:7
fitted Gauss locations:7
2
1.5
1
0.5
0
y
0
5
10
15
20
25
x

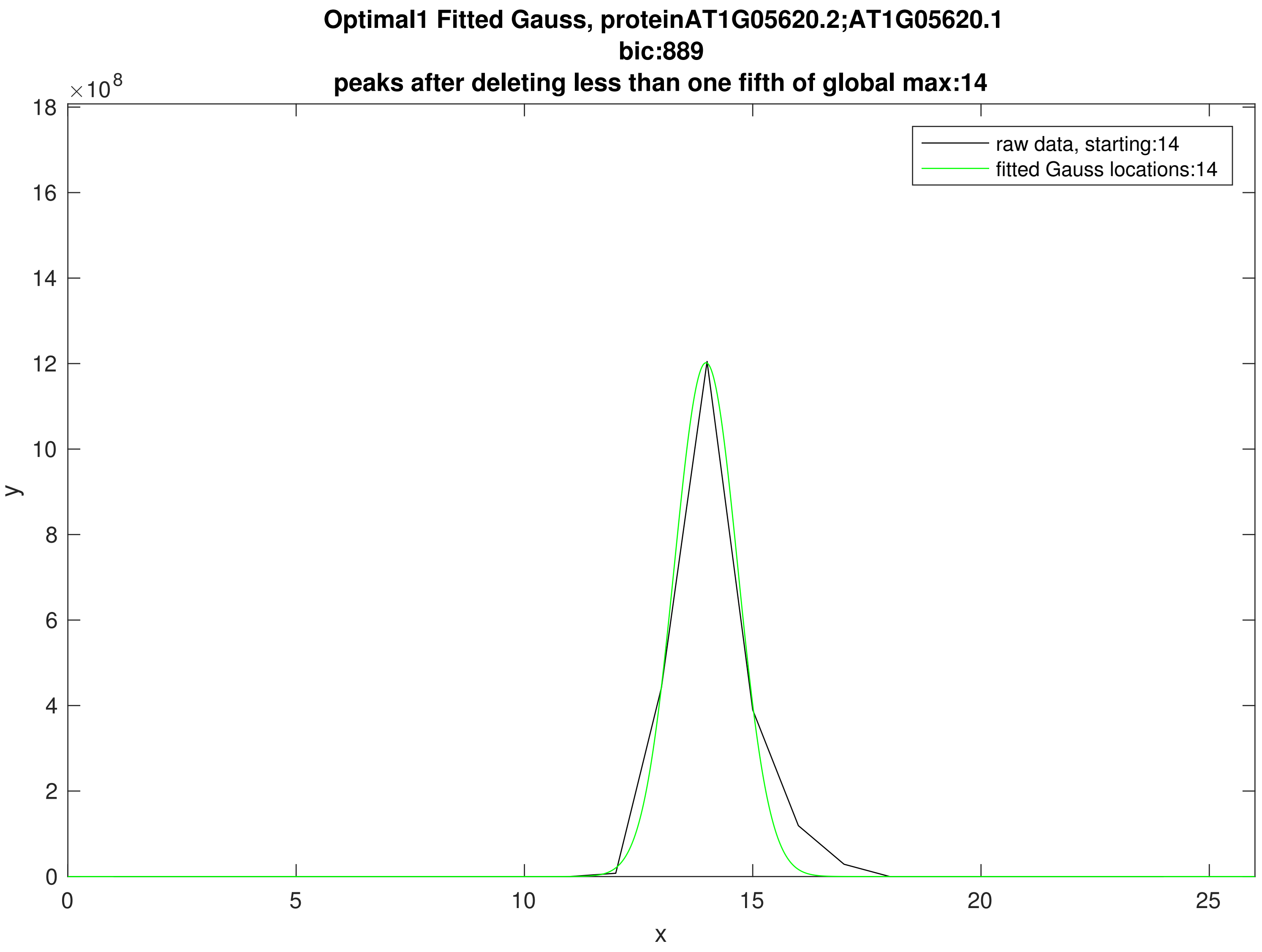
Optimal1 Fitted Gauss, proteinAT1G05620.2;AT1G05620.1
bic:889
peaks after deleting less than one fifth of global max:14
×10^8
raw data, starting:14
fitted Gauss locations:14
y
x
0
2
4
6
8
10
12
14
16
18
5
15
20
25

**Optimal1 Fitted Gauss, proteinAT1G05680.1**
**bic:679**
**peaks after deleting less than one fifth of global max:14**

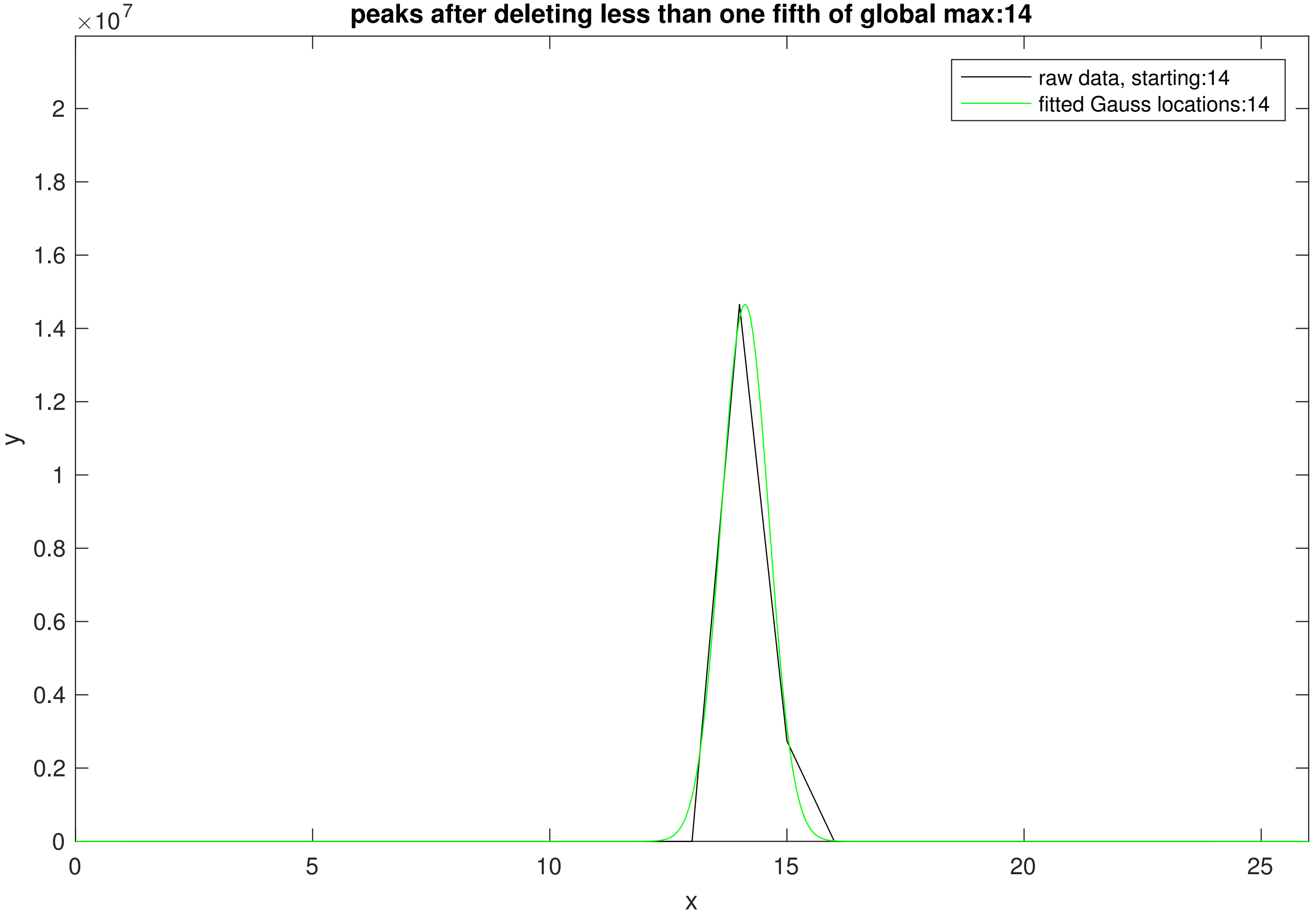

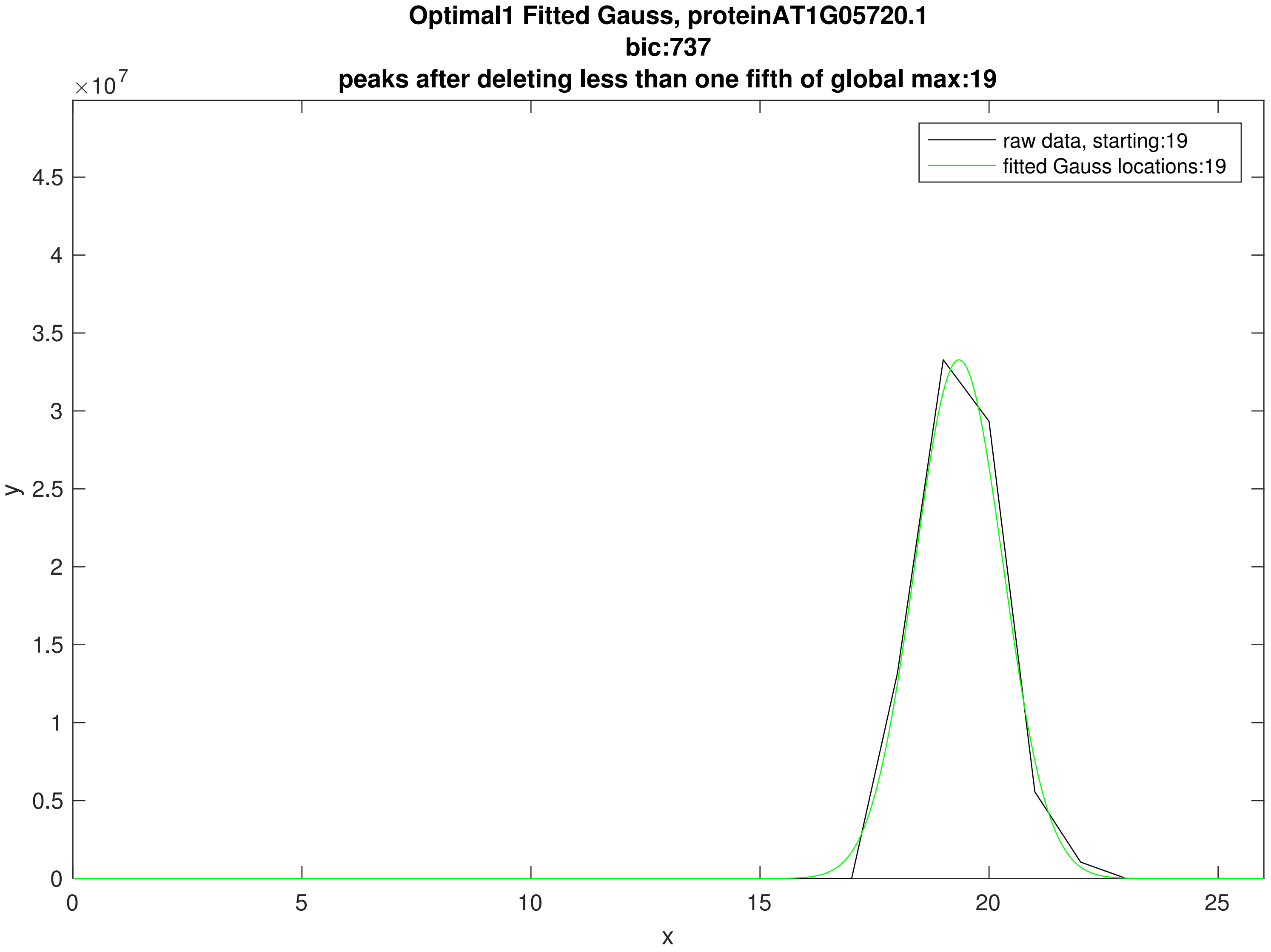
Optimal1 Fitted Gauss, proteinAT1G05720.1
bic:737
peaks after deleting less than one fifth of global max:19
×10^7
raw data, starting:19
fitted Gauss locations:19
y
x
0
0.5
1
1.5
2
2.5
3
3.5
4
4.5
5
10
15
20
25

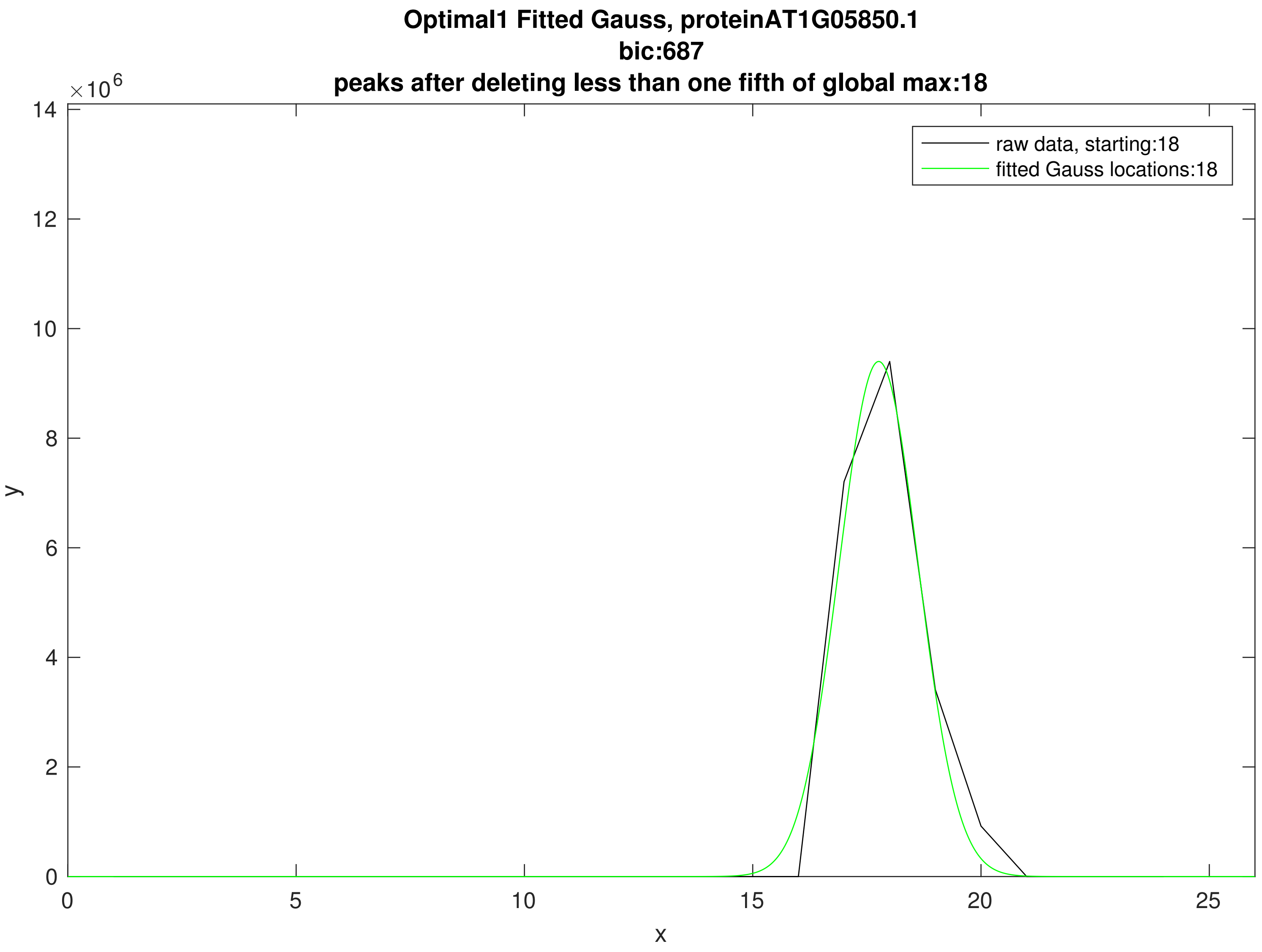
Optimal1 Fitted Gauss, proteinAT1G05850.1
bic:687
peaks after deleting less than one fifth of global max:18
×10^6
14
12
10
8
6
4
2
0
y
0
5
10
15
20
25
x
raw data, starting:18
fitted Gauss locations:18

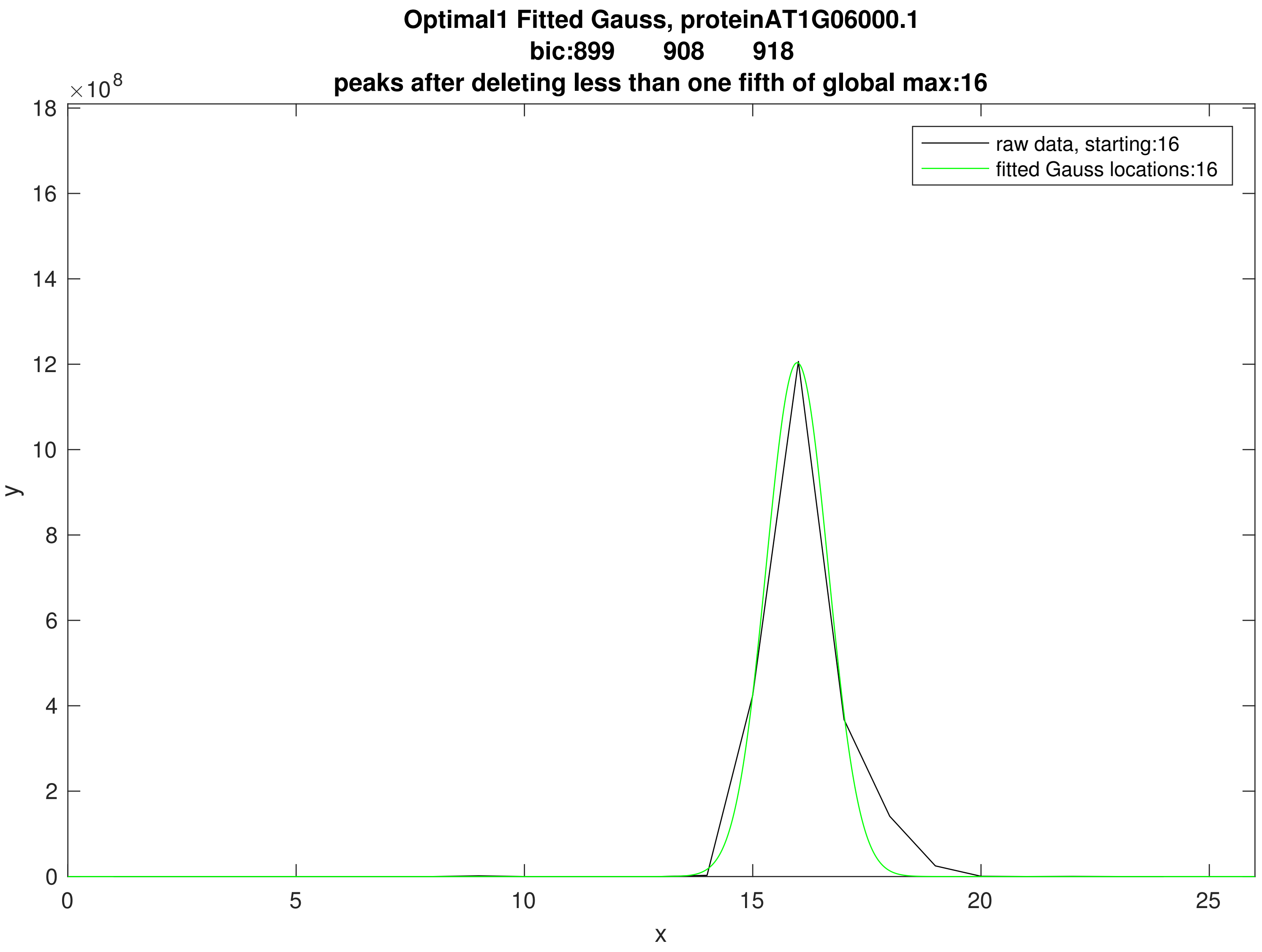
Optimal1 Fitted Gauss, proteinAT1G06000.1
bic:899 908 918
peaks after deleting less than one fifth of global max:16
×10^8
18
16
14
12
10
8
6
4
2
0
y
0
5
10
15
20
25
x
raw data, starting:16
fitted Gauss locations:16

**Optimal1 Fitted Gauss, proteinAT1G06110.1**
**bic:696**
**peaks after deleting less than one fifth of global max:14**

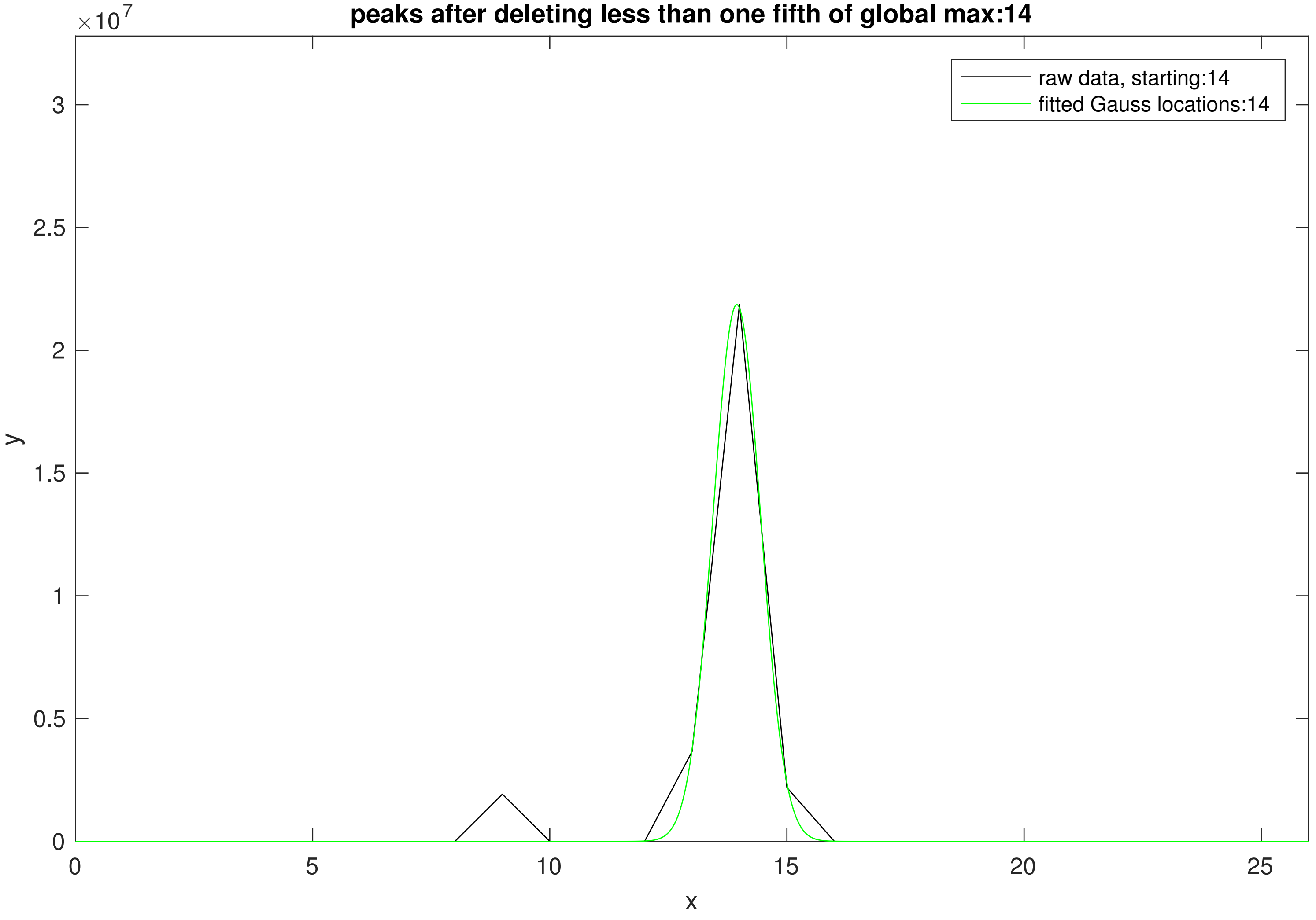

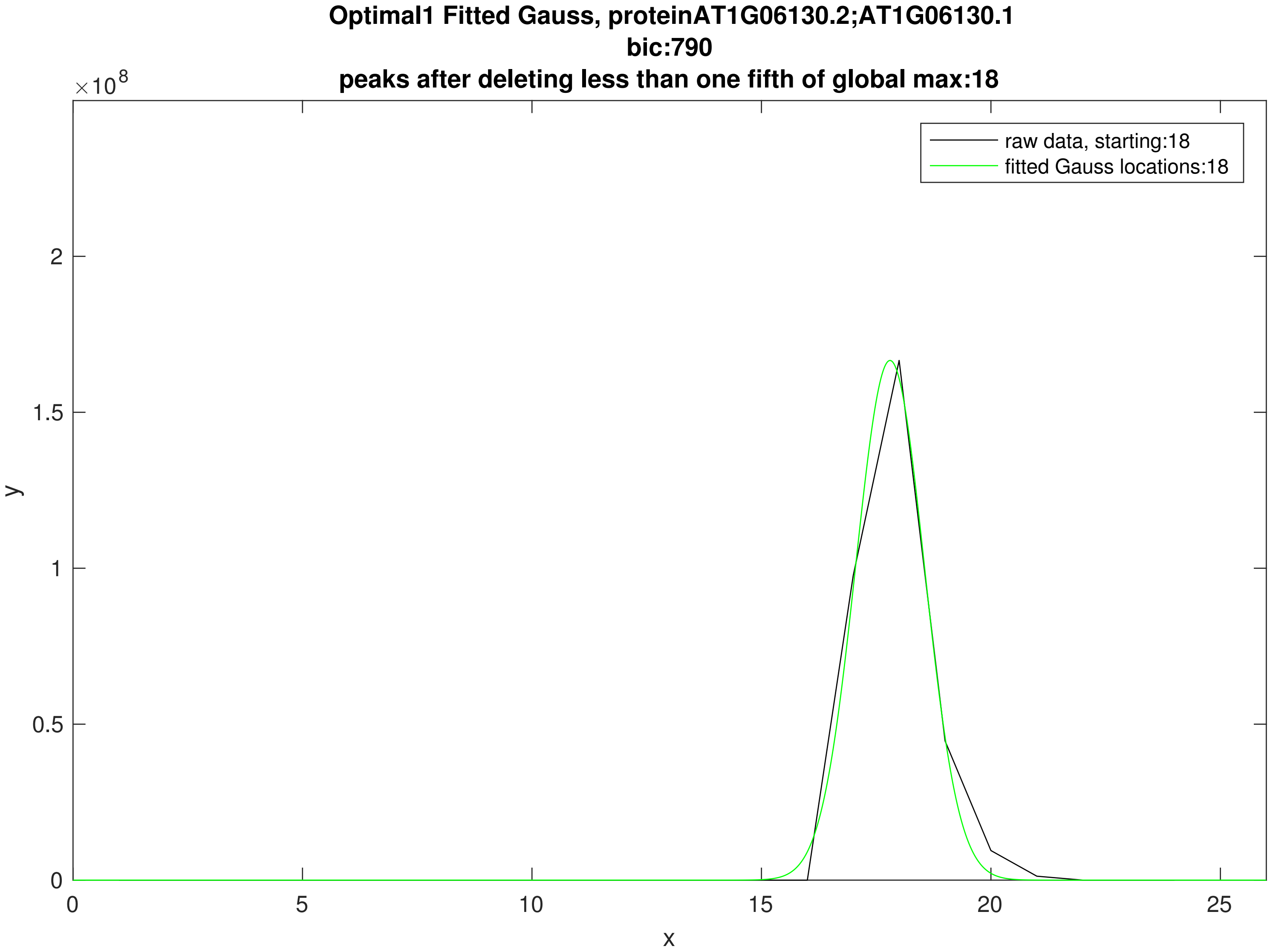
Optimal1 Fitted Gauss, proteinAT1G06130.2;AT1G06130.1
bic:790
peaks after deleting less than one fifth of global max:18
×10^8
raw data, starting:18
fitted Gauss locations:18
2
1.5
1
0.5
0
y
0
5
10
15
20
25
x

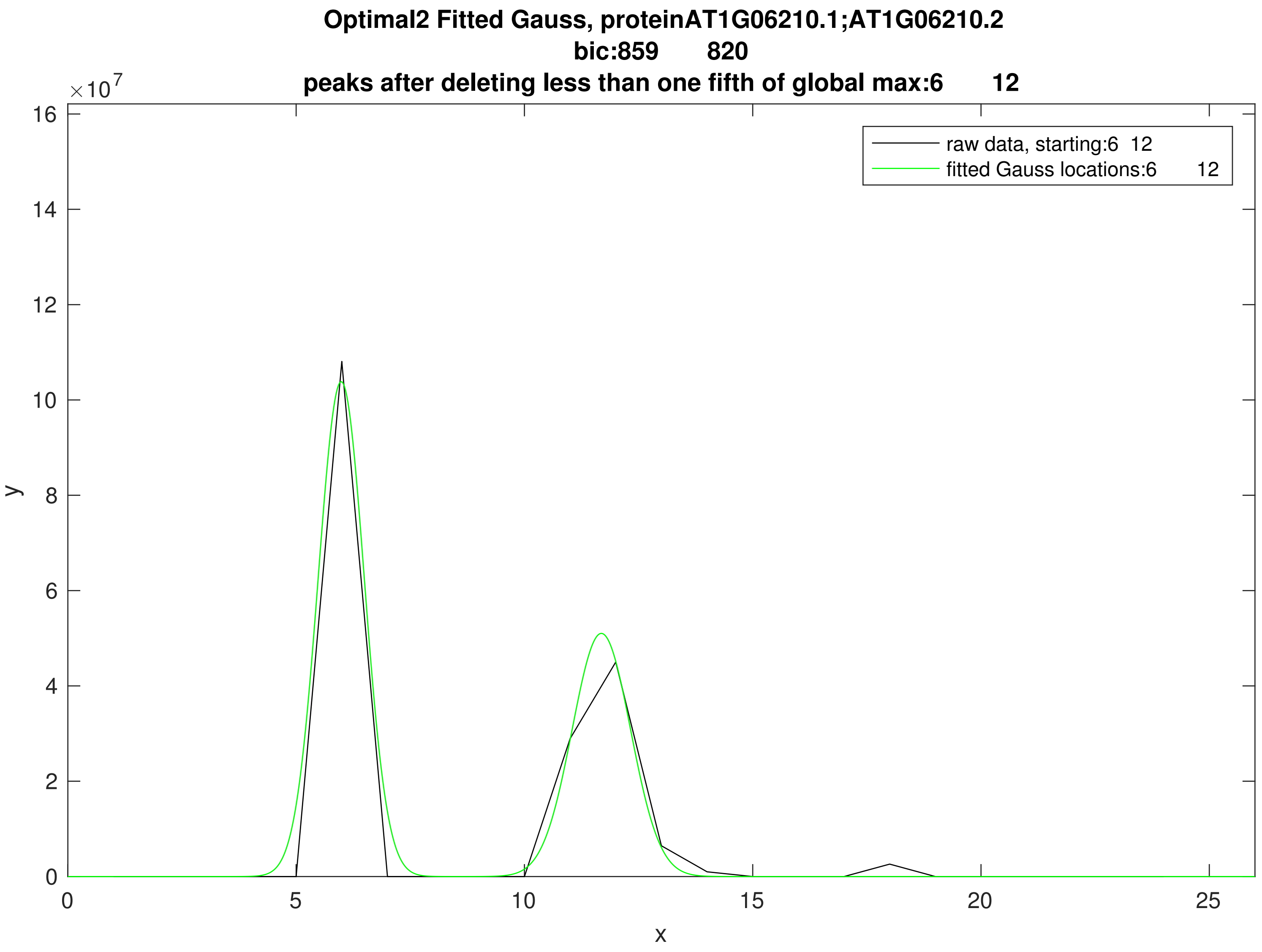

Optimal2 Fitted Gauss, proteinAT1G06210.1;AT1G06210.2
bic:859 820
peaks after deleting less than one fifth of global max:6 12
×10^7
16
14
12
10
8
6
4
2
0
y
0
5
10
15
20
25
x
raw data, starting:6 12
fitted Gauss locations:6 12

#### Optimal1 Fitted Gauss, proteinAT1G06290.1
#### bic:976 978 988
#### peaks after deleting less than one fifth of global max:13

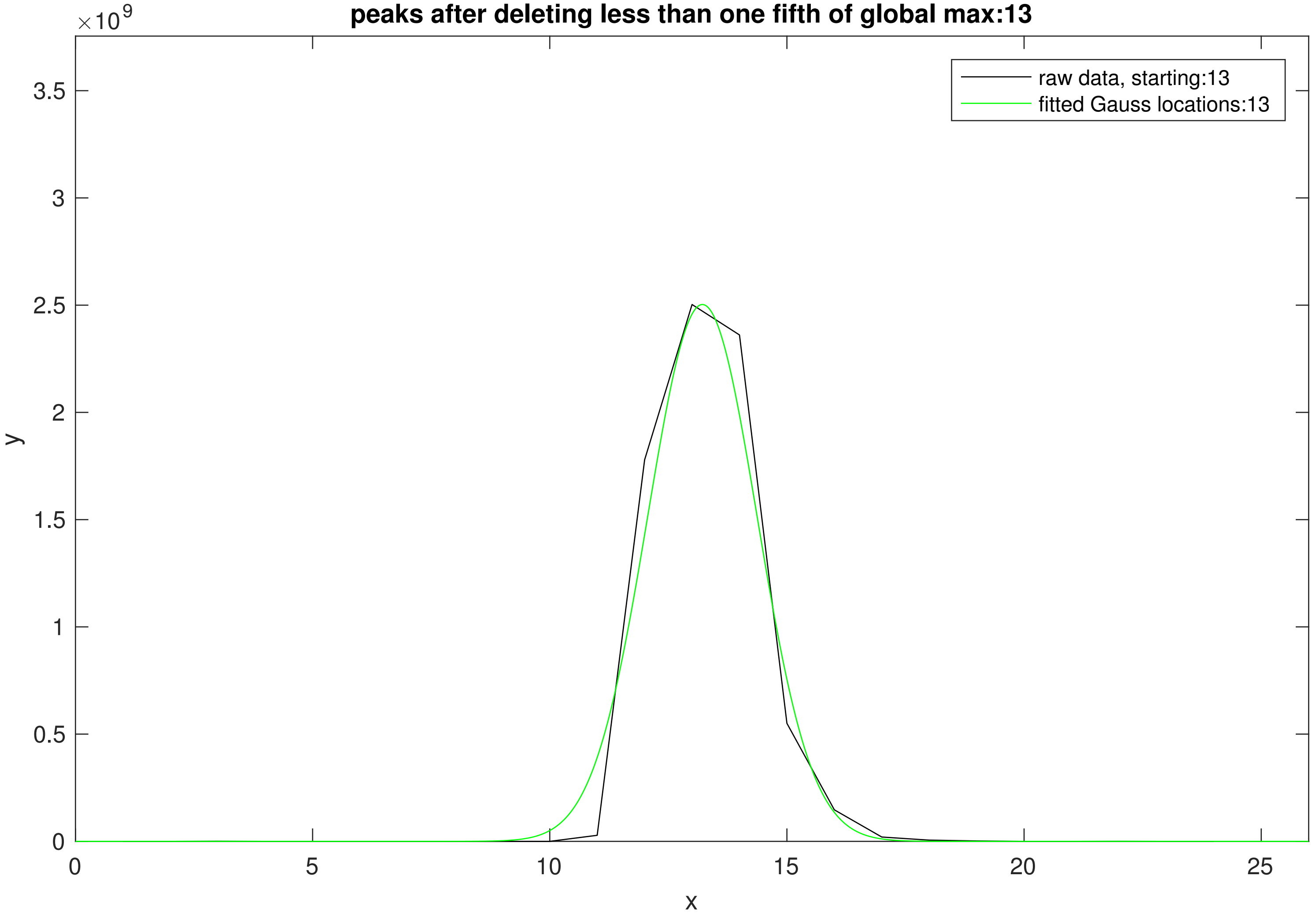

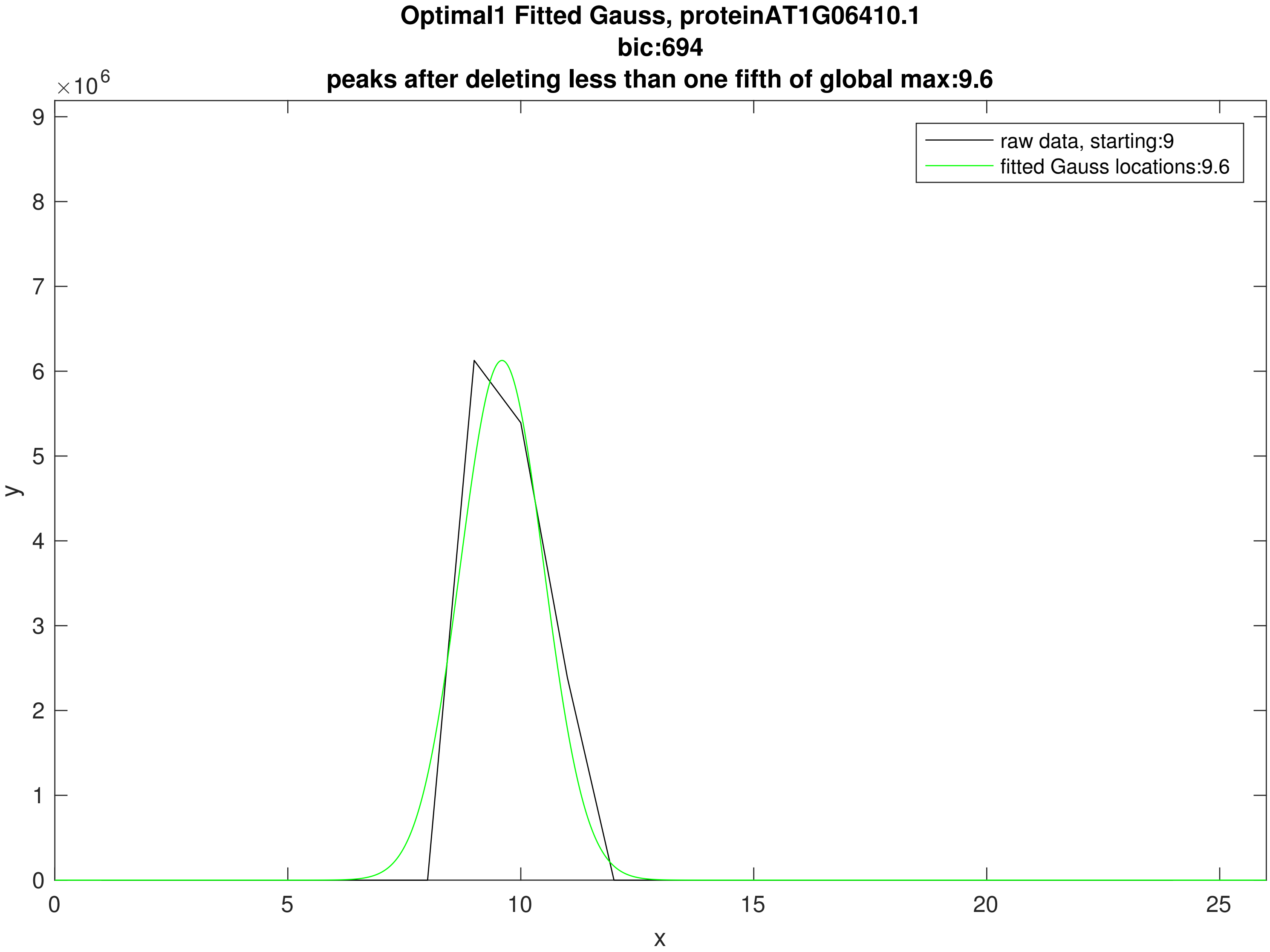
Optimal1 Fitted Gauss, proteinAT1G06410.1
bic:694
peaks after deleting less than one fifth of global max:9.6
×10^6
raw data, starting:9
fitted Gauss locations:9.6
y
x
0
1
2
3
4
5
6
7
8
9
0
5
10
15
20
25

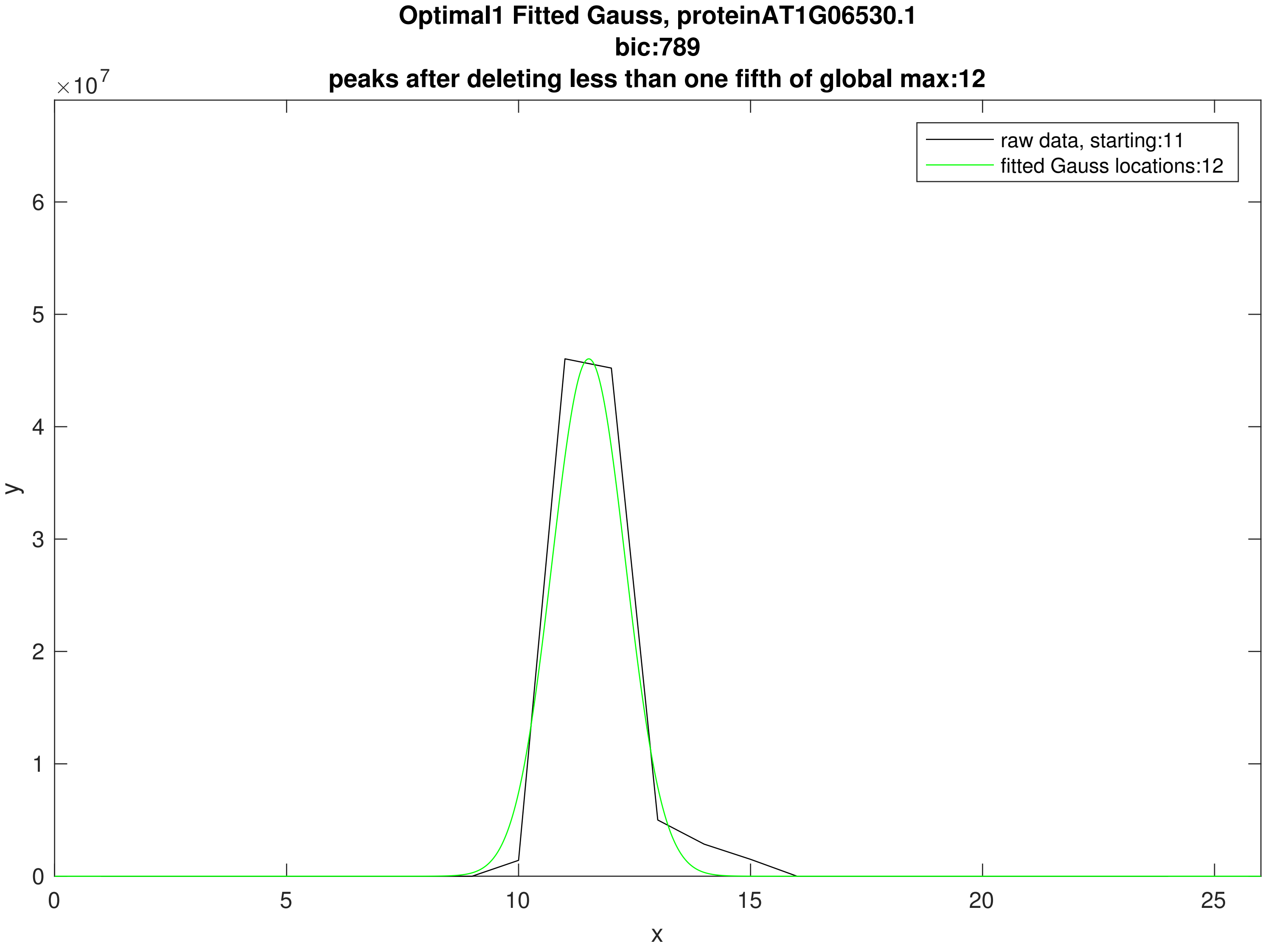
Optimal1 Fitted Gauss, proteinAT1G06530.1
bic:789
peaks after deleting less than one fifth of global max:12
×10^7
raw data, starting:11
fitted Gauss locations:12
y
x
0
1
2
3
4
5
6
0
5
10
15
20
25

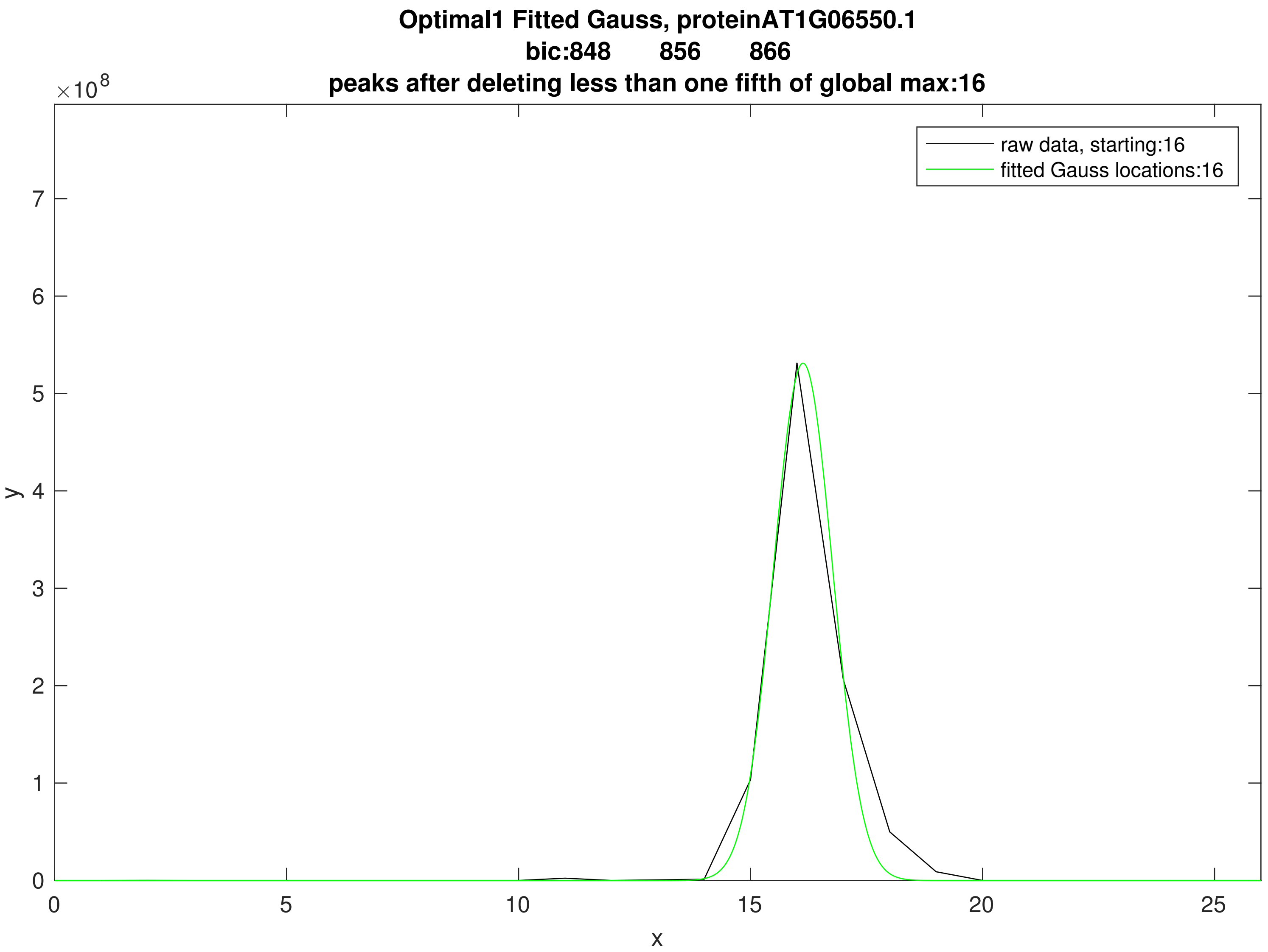

Optimal1 Fitted Gauss, proteinAT1G06550.1
bic:848 856 866
peaks after deleting less than one fifth of global max:16
×10^8
raw data, starting:16
fitted Gauss locations:16
y
x
0
1
2
3
4
5
6
7
0
5
10
15
20
25

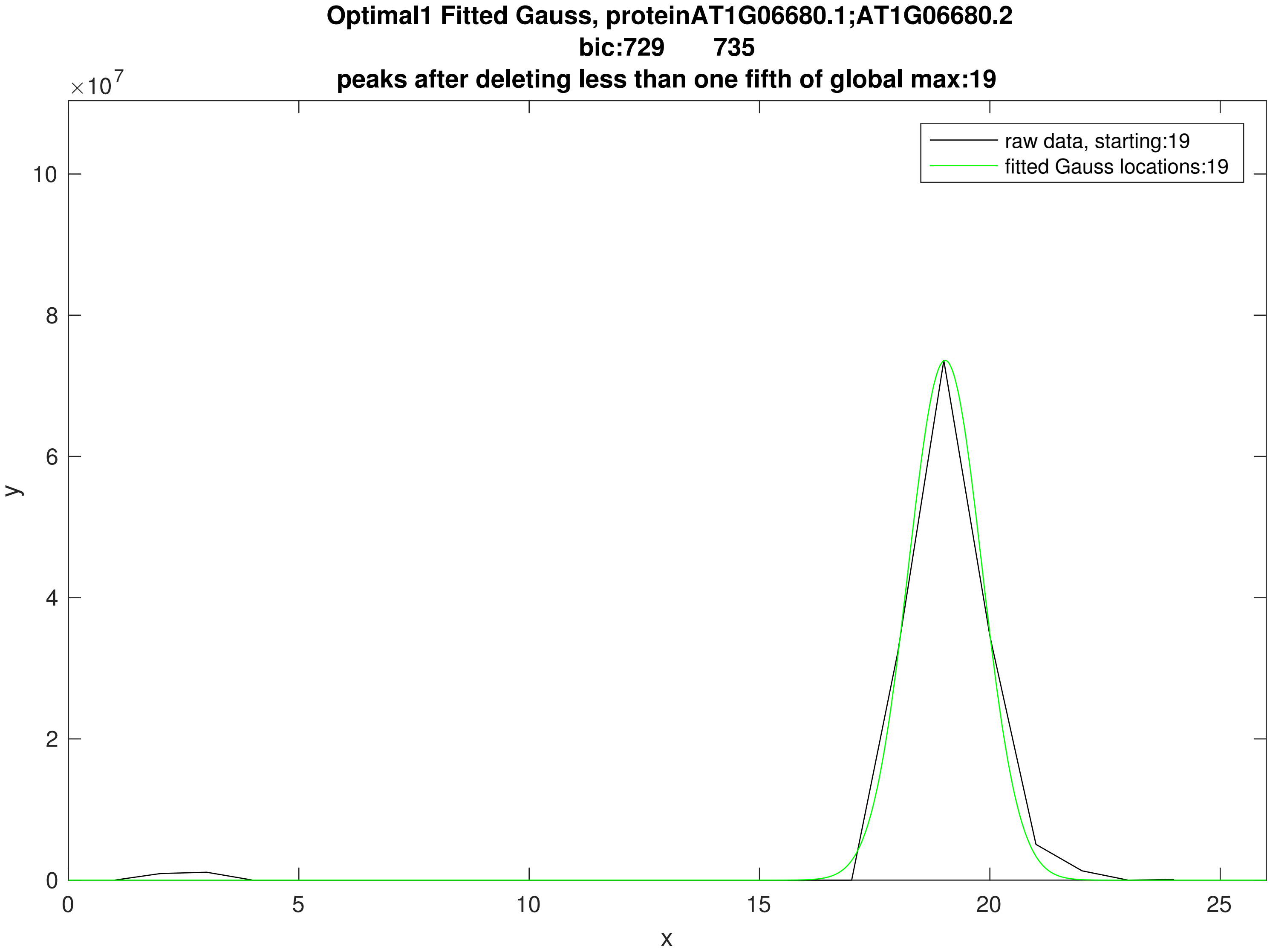

Optimal1 Fitted Gauss, proteinAT1G06680.1;AT1G06680.2
bic:729 735
peaks after deleting less than one fifth of global max:19
×10^7
raw data, starting:19
fitted Gauss locations:19
y
x
0
2
4
6
8
10
5
15
20
25

**Optimal1 Fitted Gauss, proteinAT1G06900.1**
**bic:874 965**
**peaks after deleting less than one fifth of global max:12**

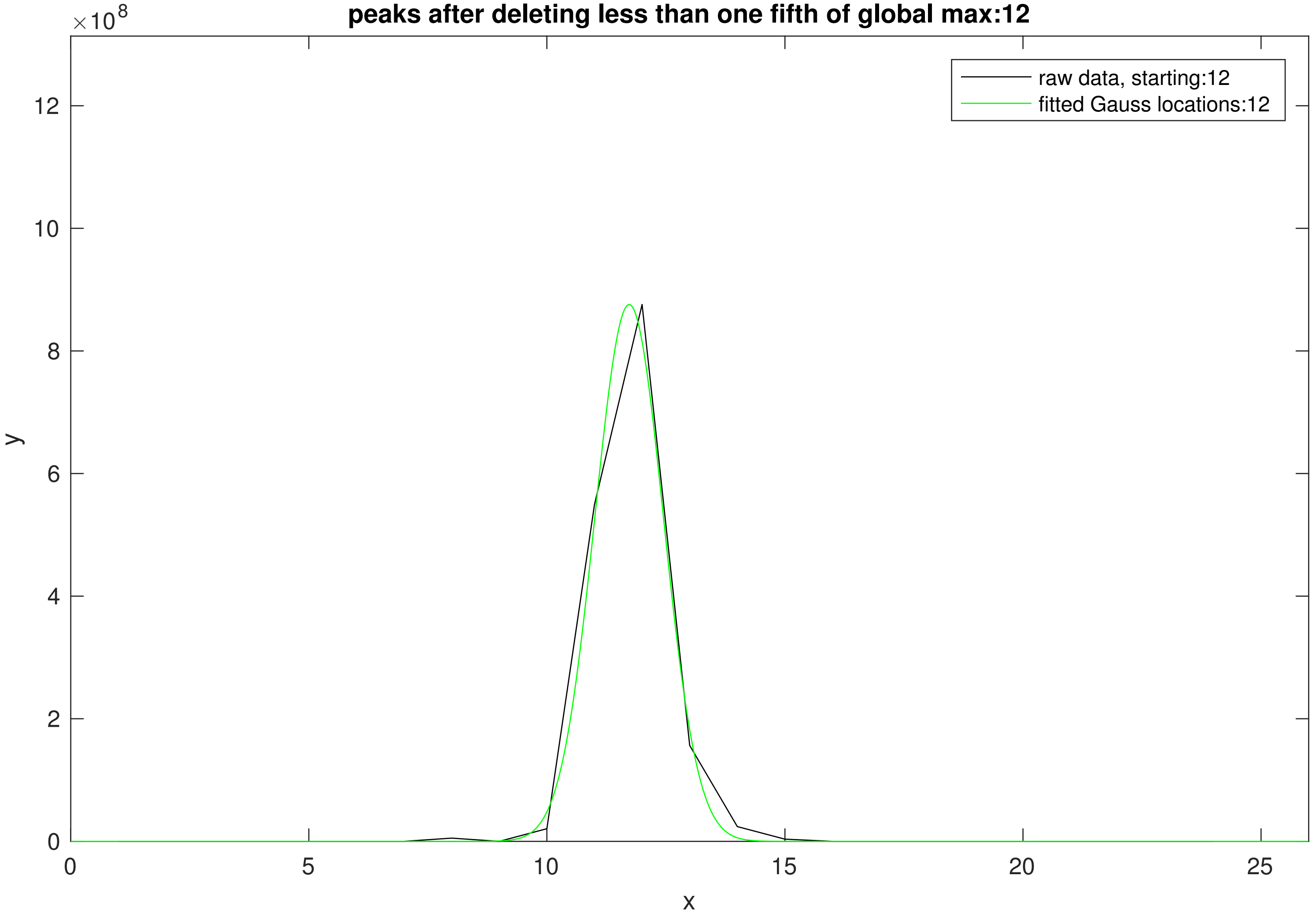

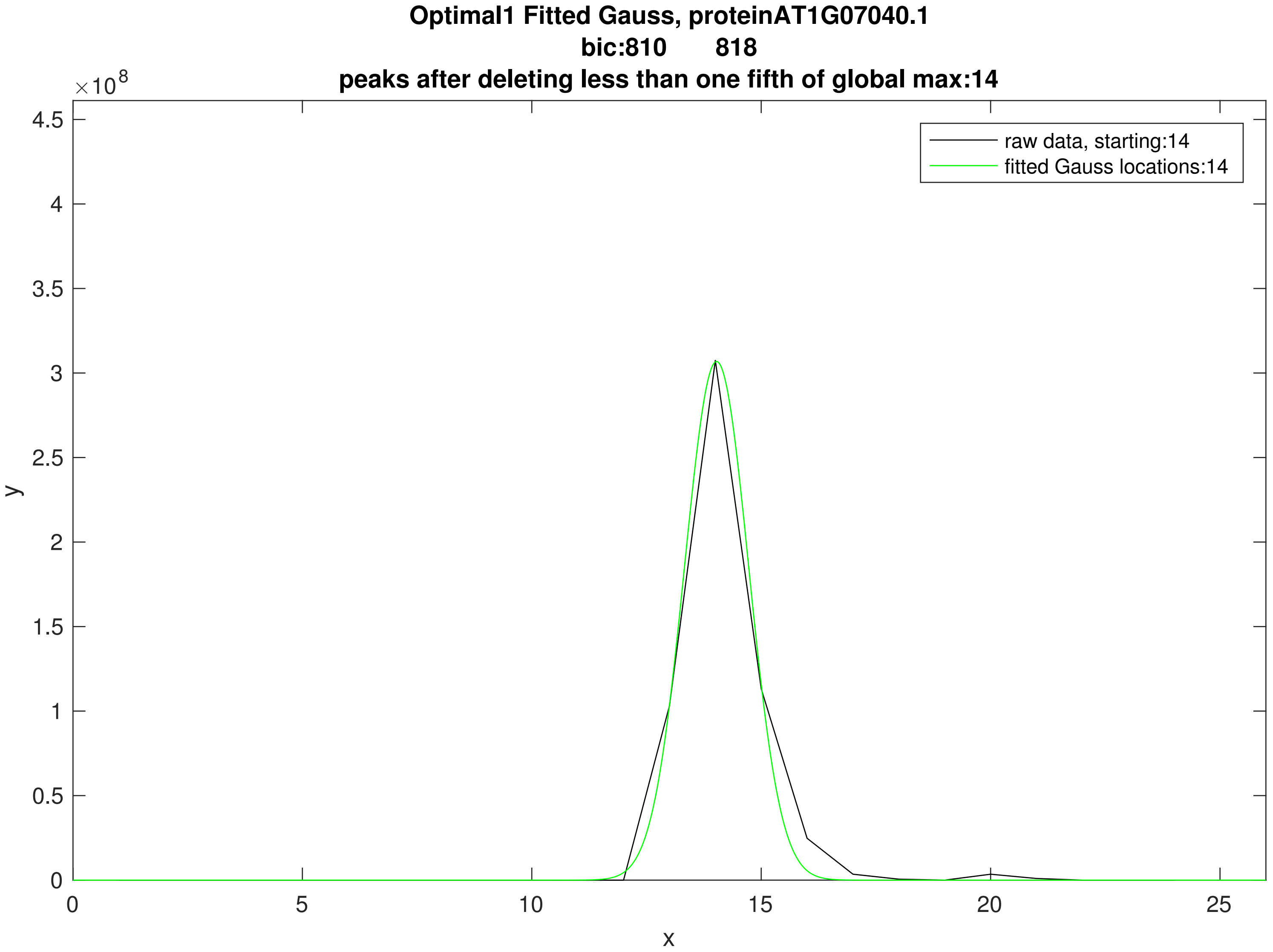

Optimal1 Fitted Gauss, proteinAT1G07040.1
bic:810 818
peaks after deleting less than one fifth of global max:14
×10^8
4.5
4
3.5
3
2.5
2
1.5
1
0.5
0
y
0
5
10
15
20
25
x
raw data, starting:14
fitted Gauss locations:14

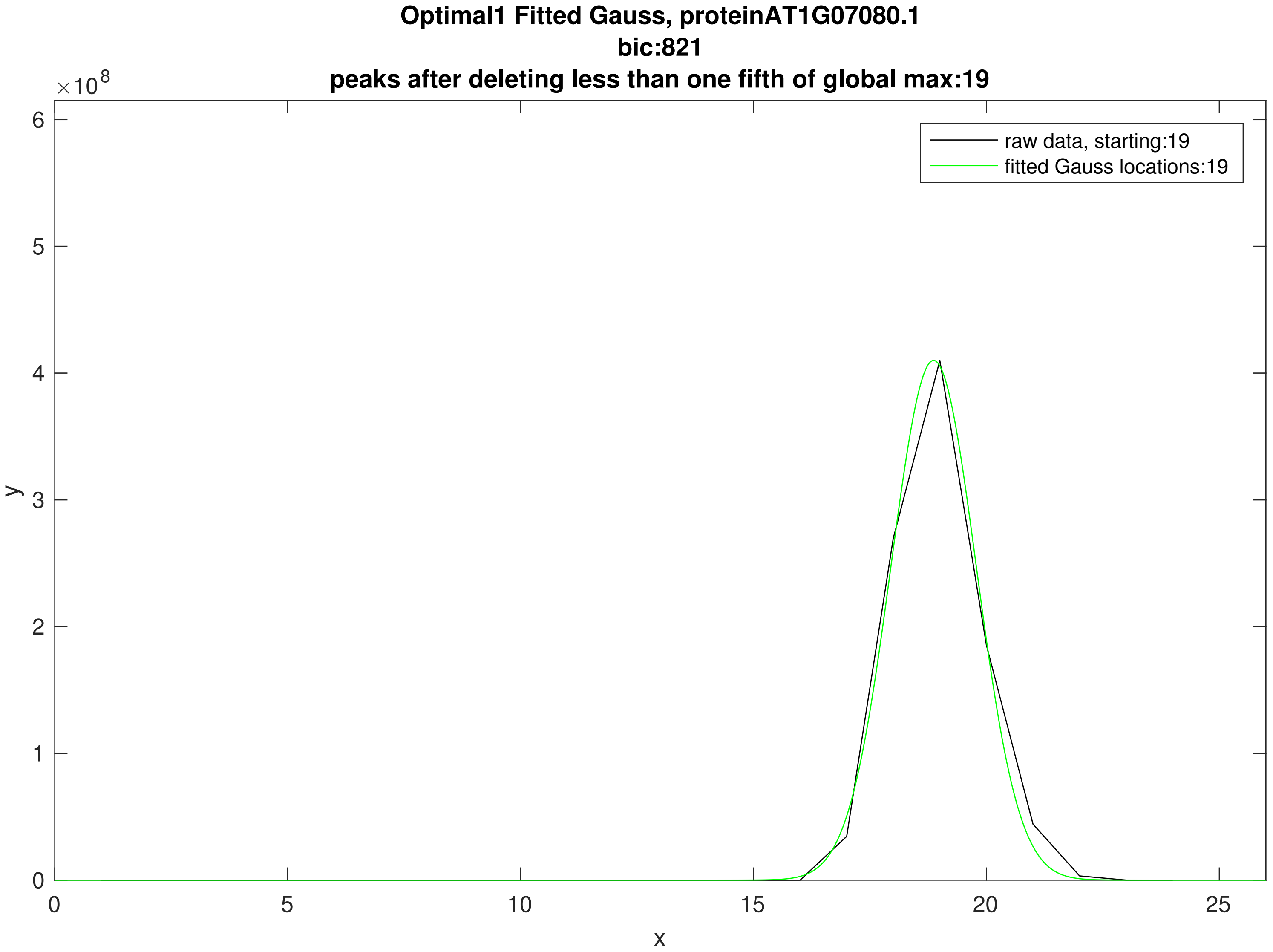
Optimal1 Fitted Gauss, proteinAT1G07080.1
bic:821
peaks after deleting less than one fifth of global max:19
×10^8
6
5
4
3
2
1
0
y
0
5
10
15
20
25
x
raw data, starting:19
fitted Gauss locations:19

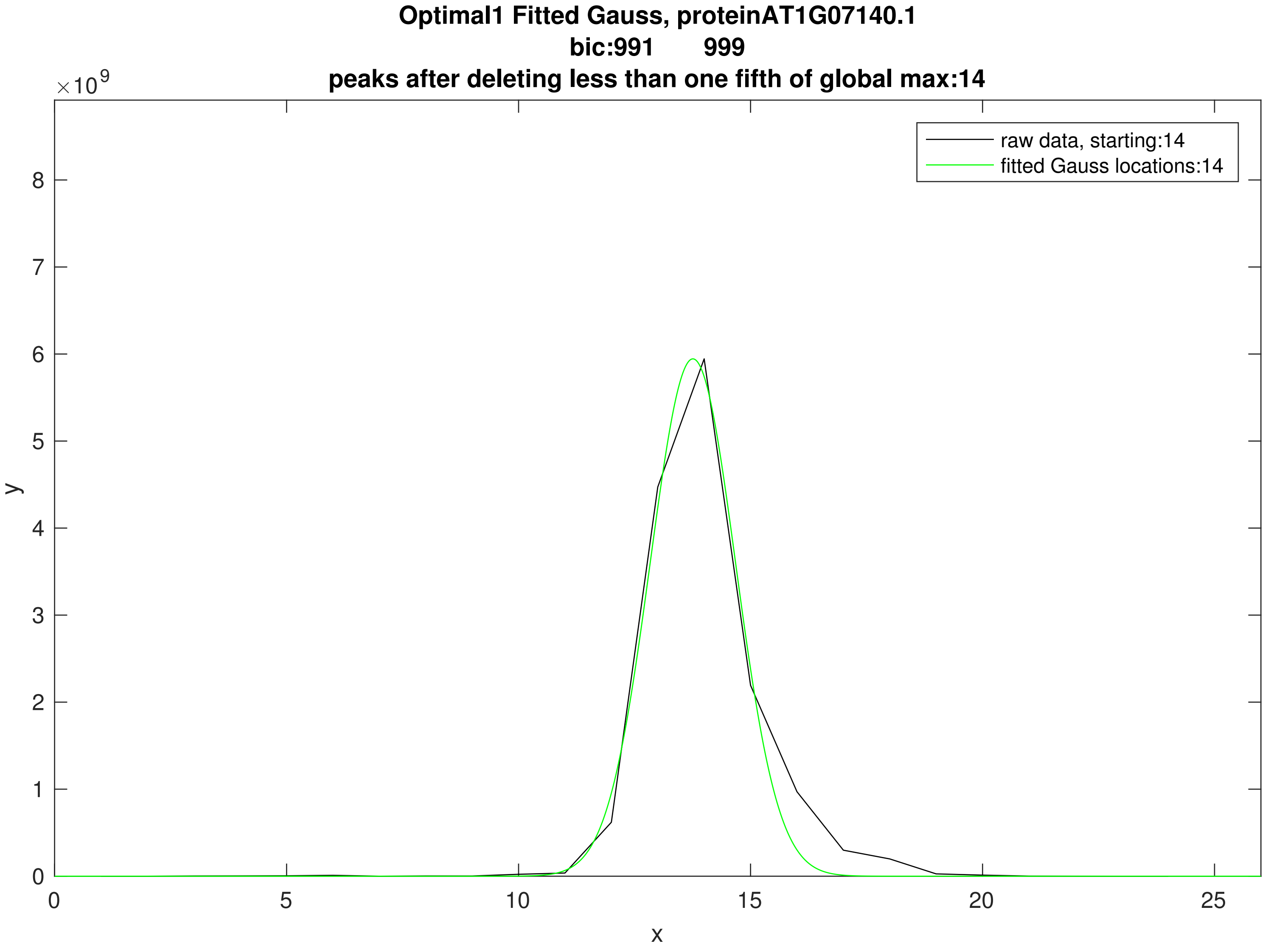
Optimal1 Fitted Gauss, proteinAT1G07140.1
bic:991 999
peaks after deleting less than one fifth of global max:14
×10^9
raw data, starting:14
fitted Gauss locations:14
y
x
0
1
2
3
4
5
6
7
8
0
5
10
15
20
25

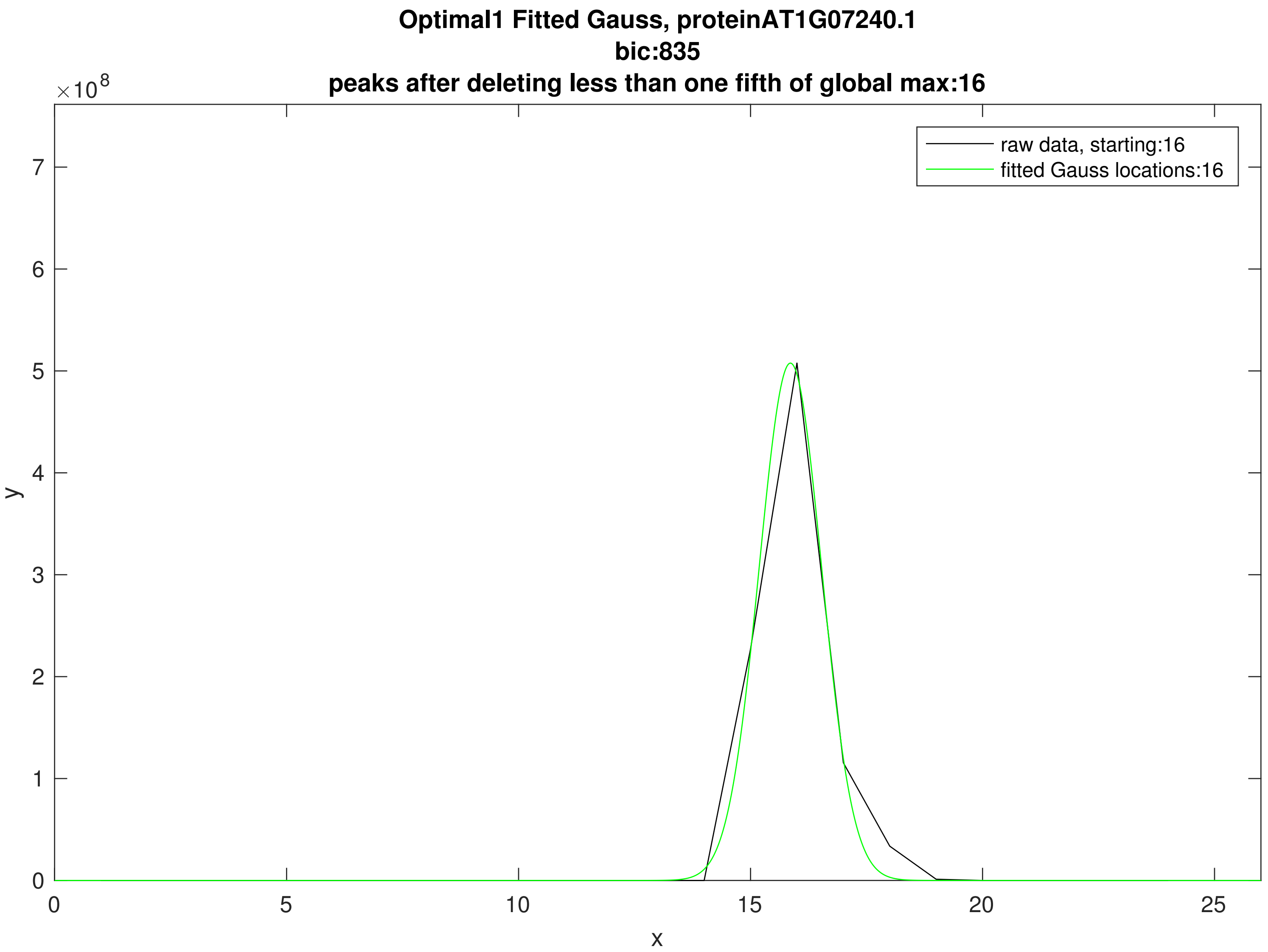
Optimal1 Fitted Gauss, proteinAT1G07240.1
bic:835
peaks after deleting less than one fifth of global max:16
×10^8
raw data, starting:16
fitted Gauss locations:16
y
x
0
1
2
3
4
5
6
7
0
5
10
15
20
25

**Optimal1 Fitted Gauss, proteinAT1G07250.1**
**bic:851       860**
**peaks after deleting less than one fifth of global max:15**

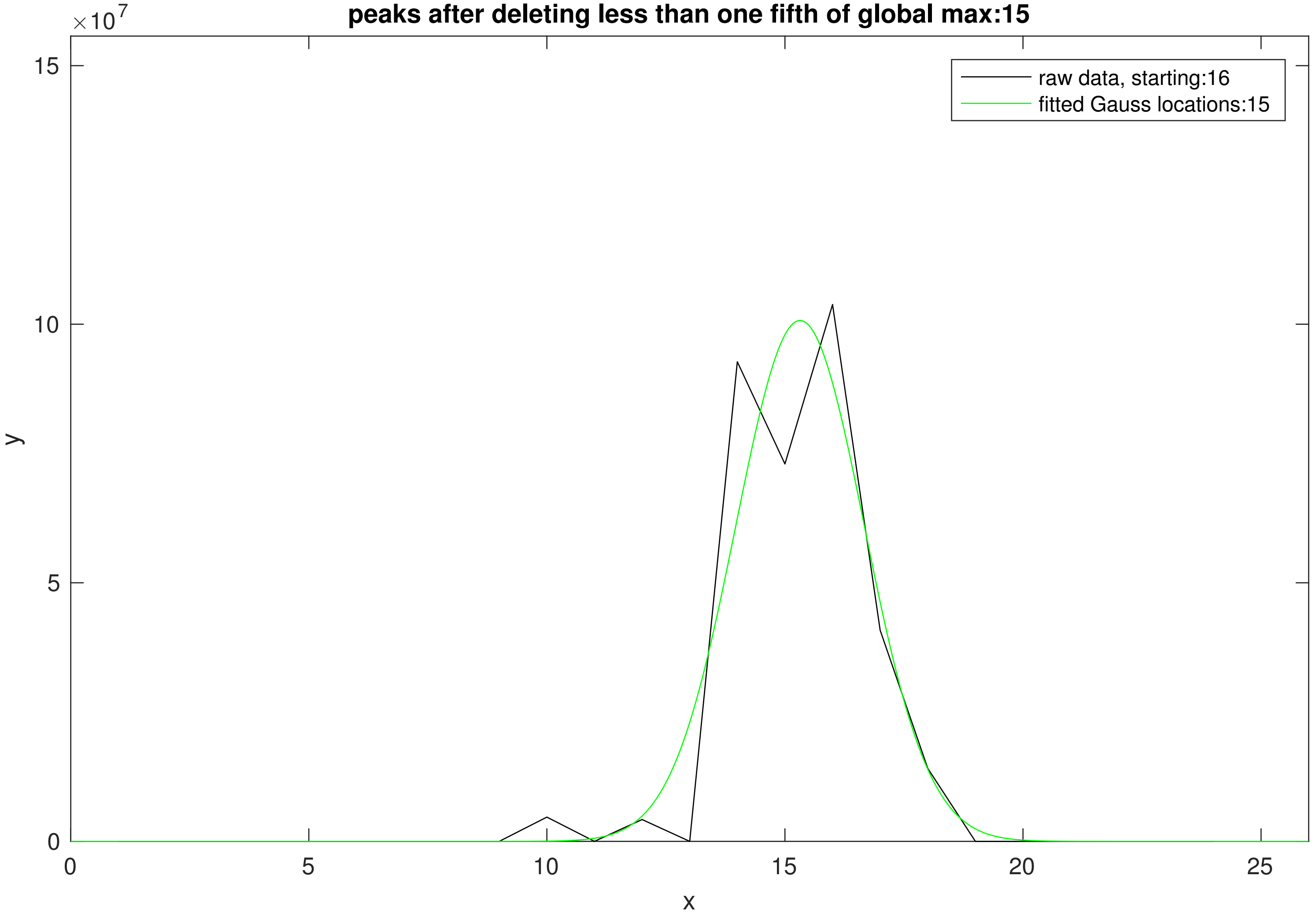

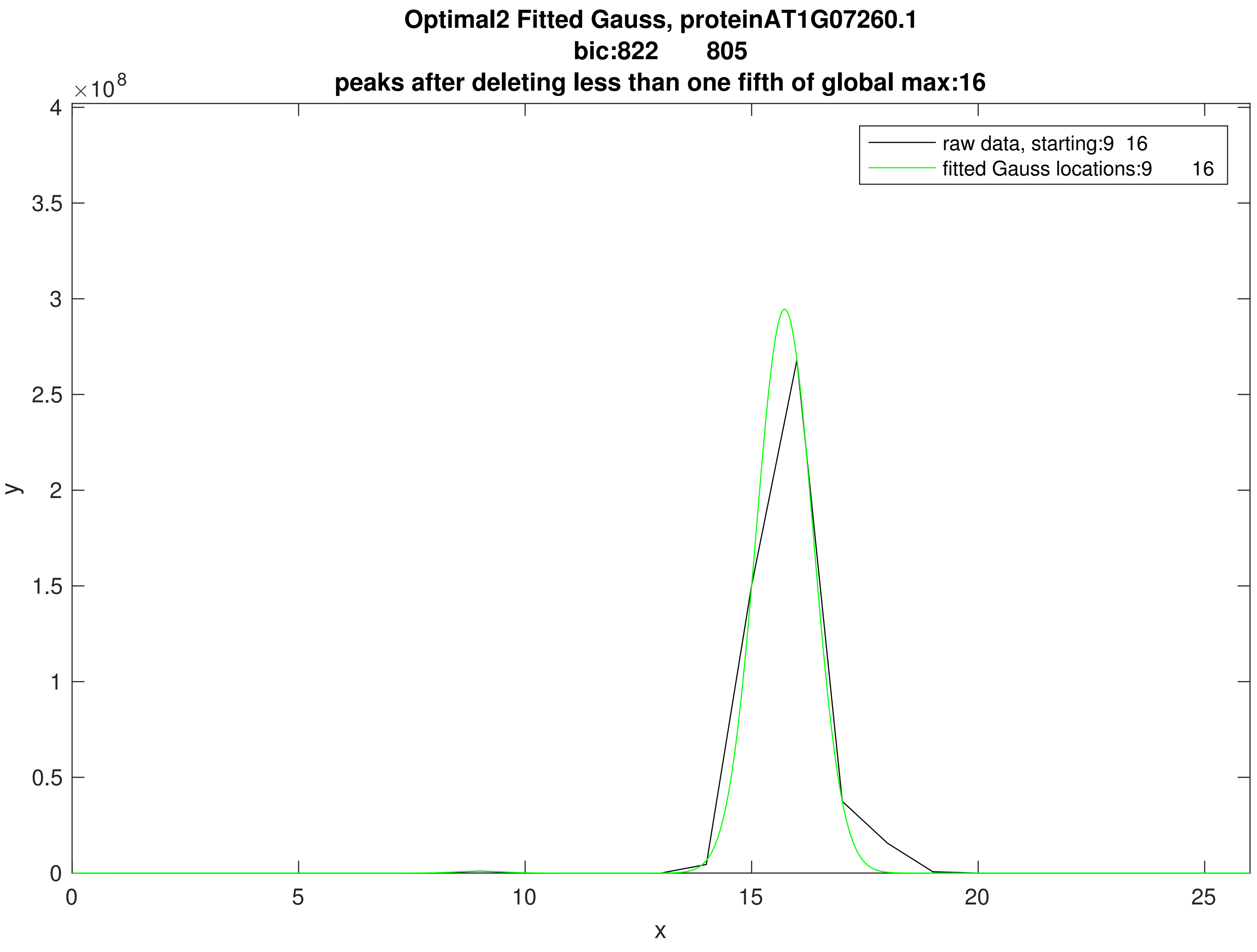
Optimal2 Fitted Gauss, proteinAT1G07260.1
bic:822 805
peaks after deleting less than one fifth of global max:16
raw data, starting:9 16
fitted Gauss locations:9 16
×10^8
4
3.5
3
2.5
2
1.5
1
0.5
0
y
0
5
10
15
20
25
x

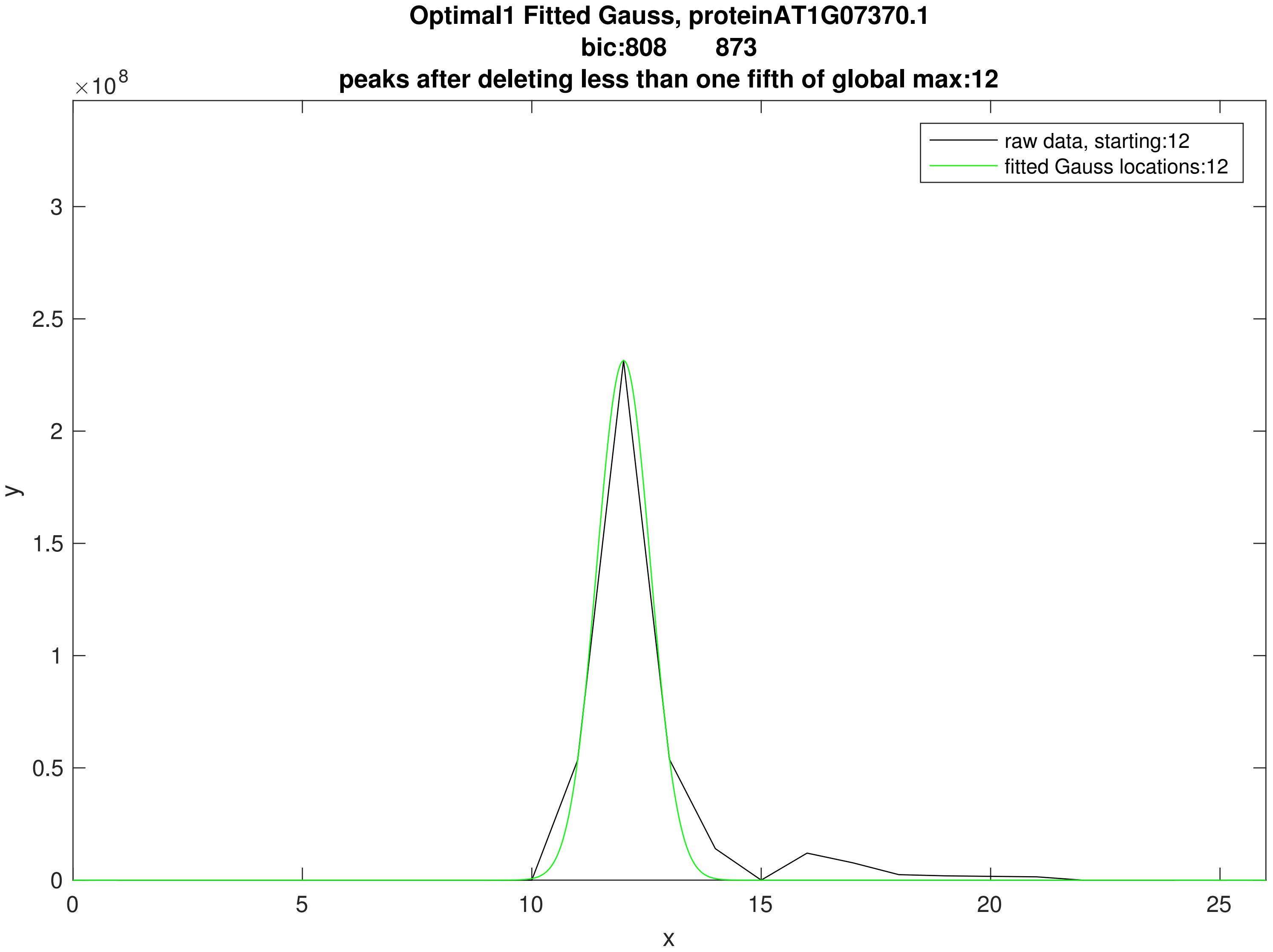
Optimal1 Fitted Gauss, proteinAT1G07370.1
bic:808 873
peaks after deleting less than one fifth of global max:12
×10^8
3
2.5
2
1.5
1
0.5
0
y
0
5
10
15
20
25
x
raw data, starting:12
fitted Gauss locations:12

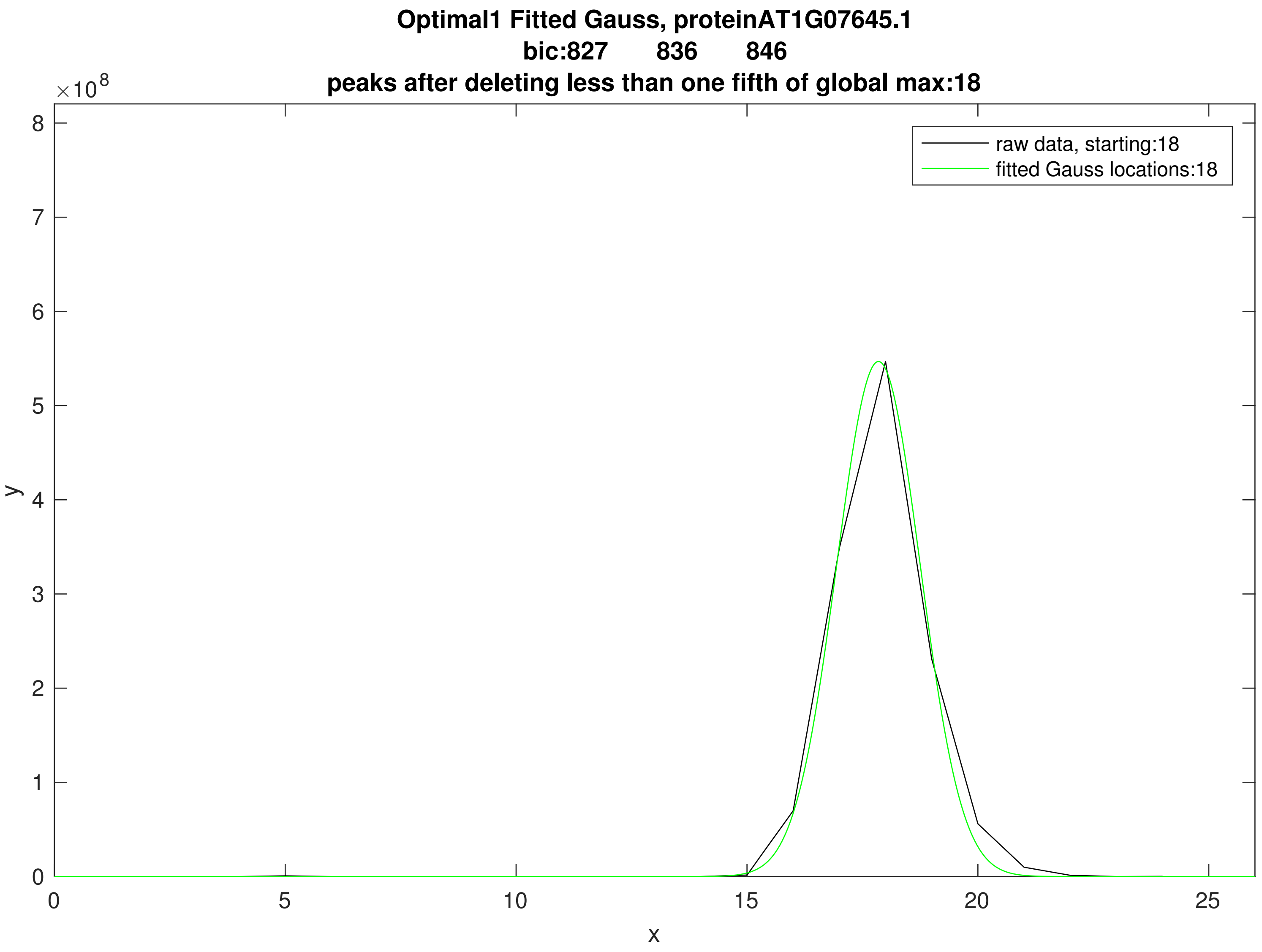
Optimal1 Fitted Gauss, proteinAT1G07645.1
bic:827 836 846
peaks after deleting less than one fifth of global max:18
×10^8
raw data, starting:18
fitted Gauss locations:18
y
x
0
1
2
3
4
5
6
7
8
0
5
10
15
20
25

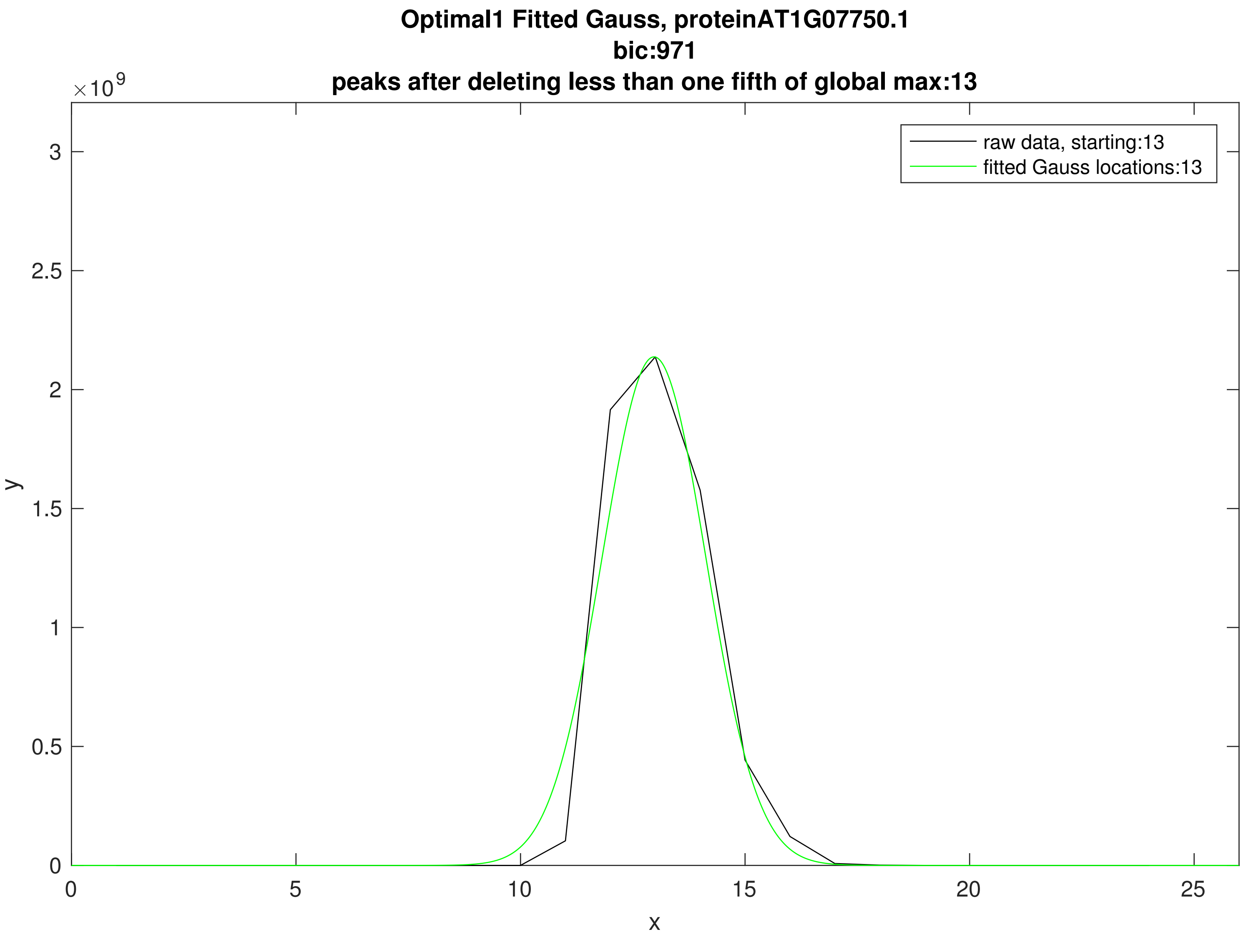

Optimal1 Fitted Gauss, proteinAT1G07750.1
bic:971
peaks after deleting less than one fifth of global max:13
×10^9
3
2.5
2
1.5
1
0.5
0
y
0
5
10
15
20
25
x
raw data, starting:13
fitted Gauss locations:13

Optimal1 Fitted Gauss, proteinAT1G07780.5;AT1G07780.3;AT1G07780.2;AT1G07780.1;AT1G07780.4;AT1G29410.1;AT1

bic:723

peaks after deleting less than one fifth of global max:19

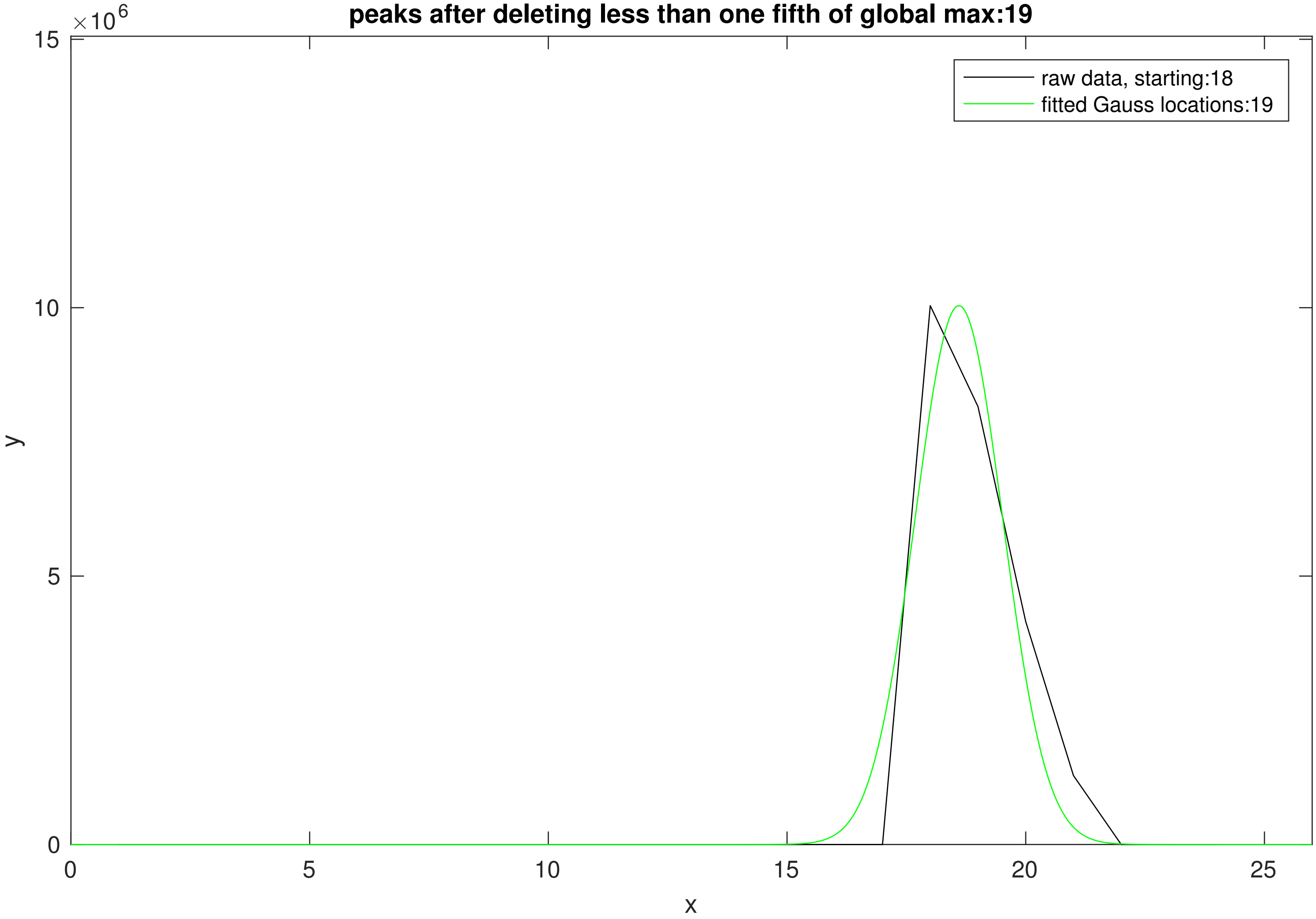

**1 Fitted Gauss, proteinAT1G07890.8;AT1G07890.7;AT1G07890.5;AT1G07890.4;AT1G07890.3;AT1G07890.2;AT1G0789[illegible]**
**bic:1.07e+03  1.08e+03  1.09e+03**
**peaks after deleting less than one fifth of global max:16**

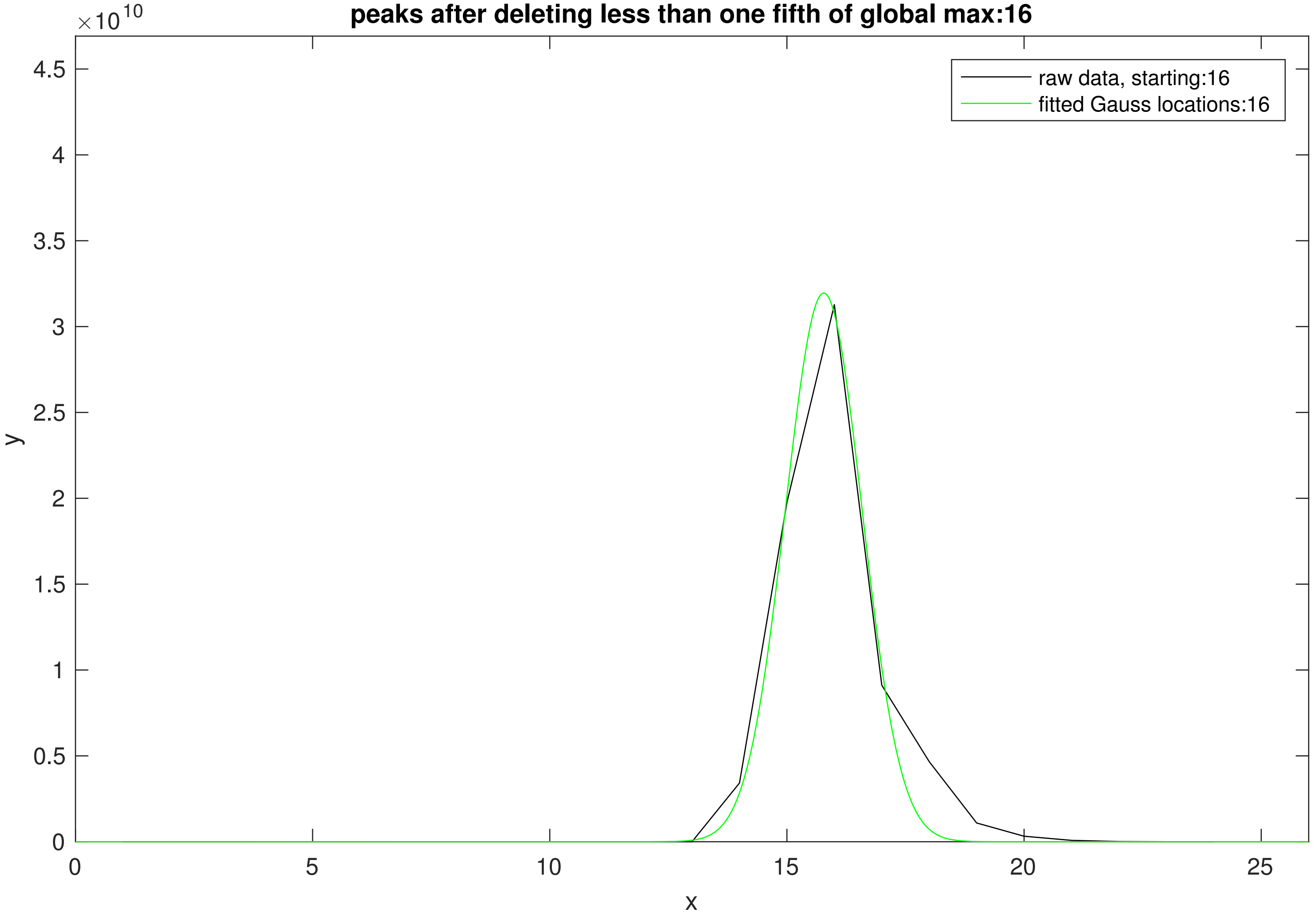

**Optimal1 Fitted Gauss, proteinAT1G07960.3;AT1G07960.2;AT1G07960.1**
**bic:676 700**
**peaks after deleting less than one fifth of global max:20**

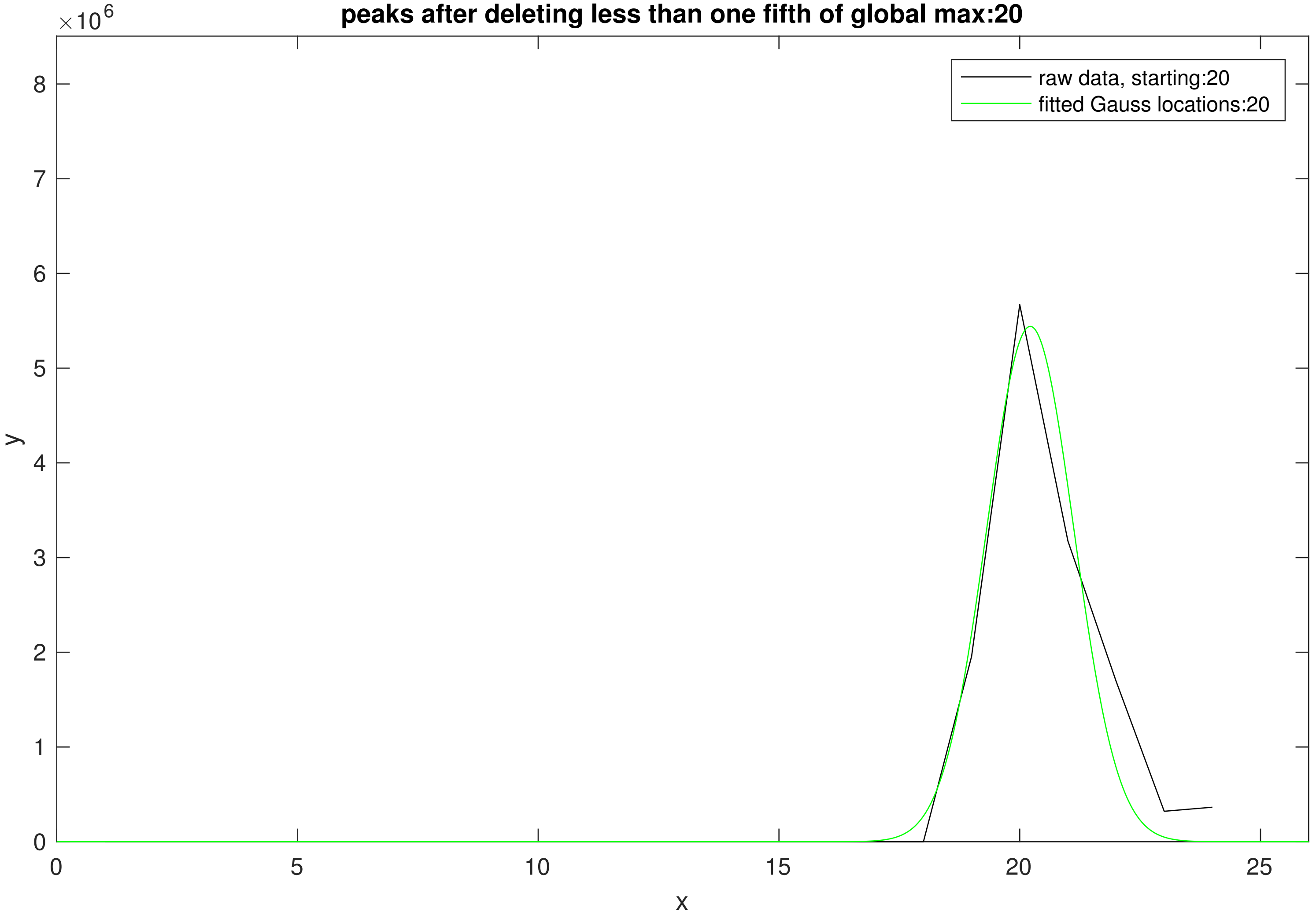

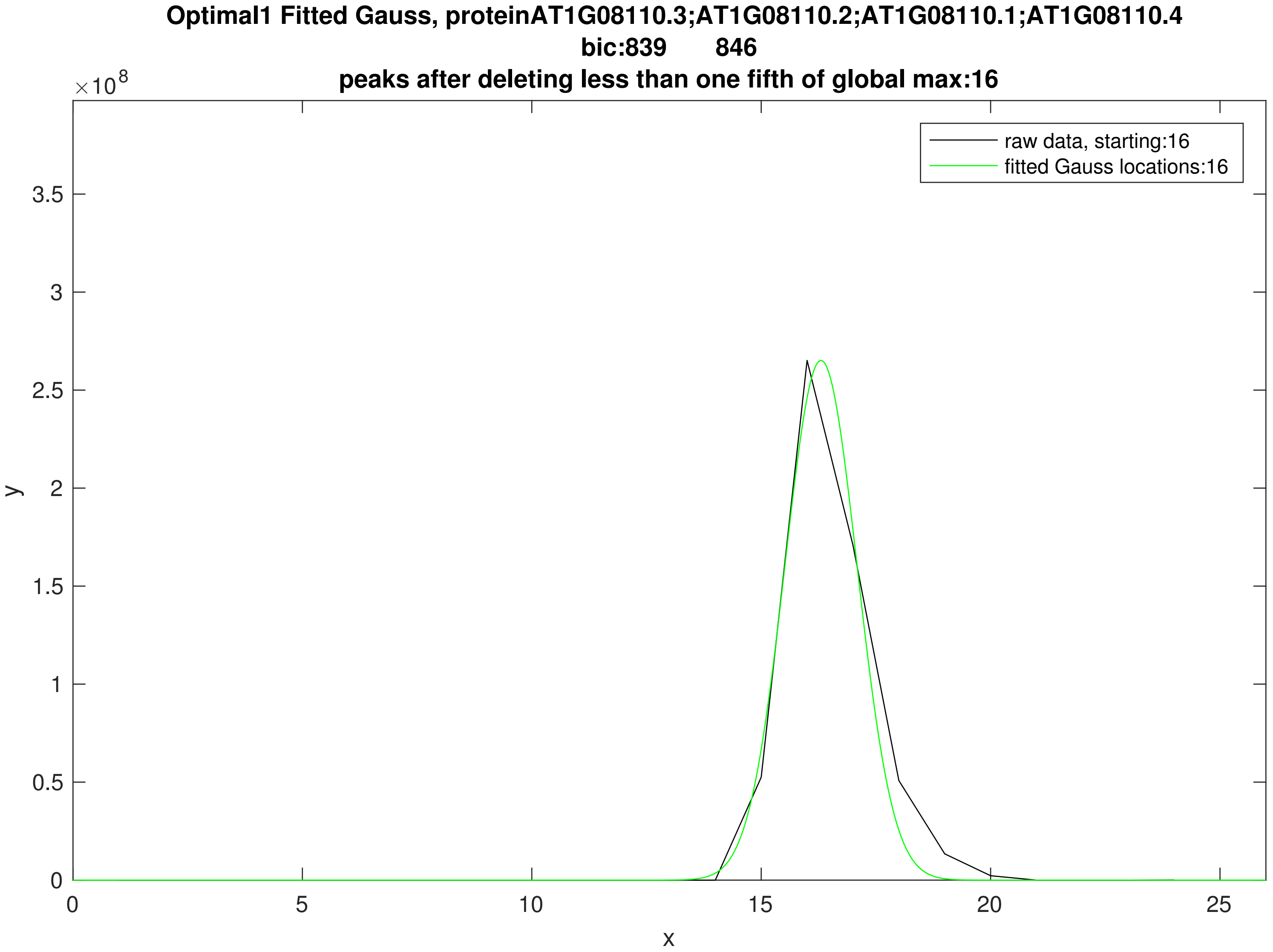
Optimal1 Fitted Gauss, proteinAT1G08110.3;AT1G08110.2;AT1G08110.1;AT1G08110.4
bic:839 846
peaks after deleting less than one fifth of global max:16
×10^8
raw data, starting:16
fitted Gauss locations:16
3.5
3
2.5
2
1.5
1
0.5
0
y
0
5
10
15
20
25
x

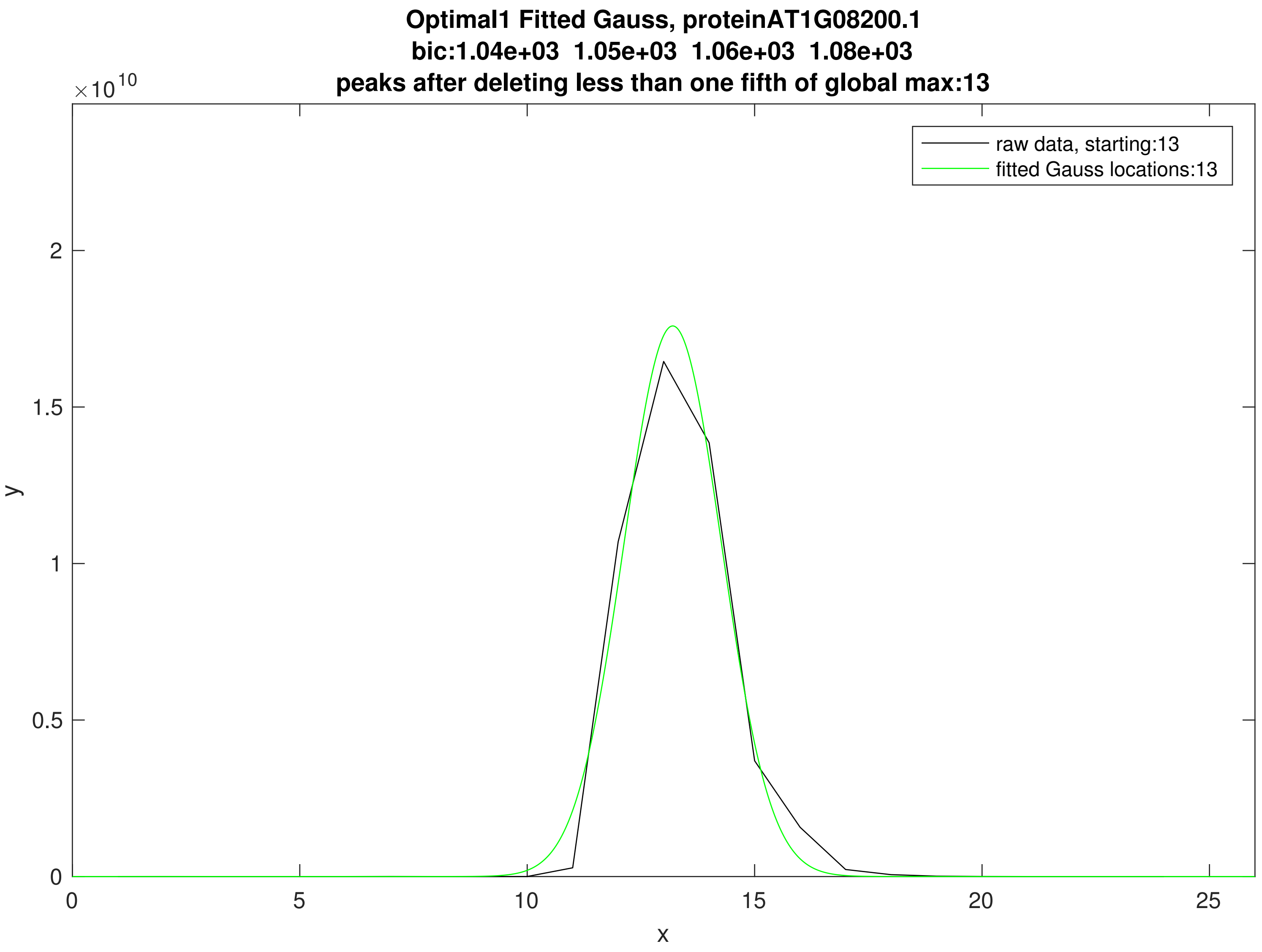

Optimal1 Fitted Gauss, proteinAT1G08200.1
bic:1.04e+03 1.05e+03 1.06e+03 1.08e+03
peaks after deleting less than one fifth of global max:13
×10^10
raw data, starting:13
fitted Gauss locations:13
y
x
0
0.5
1
1.5
2
0
5
10
15
20
25

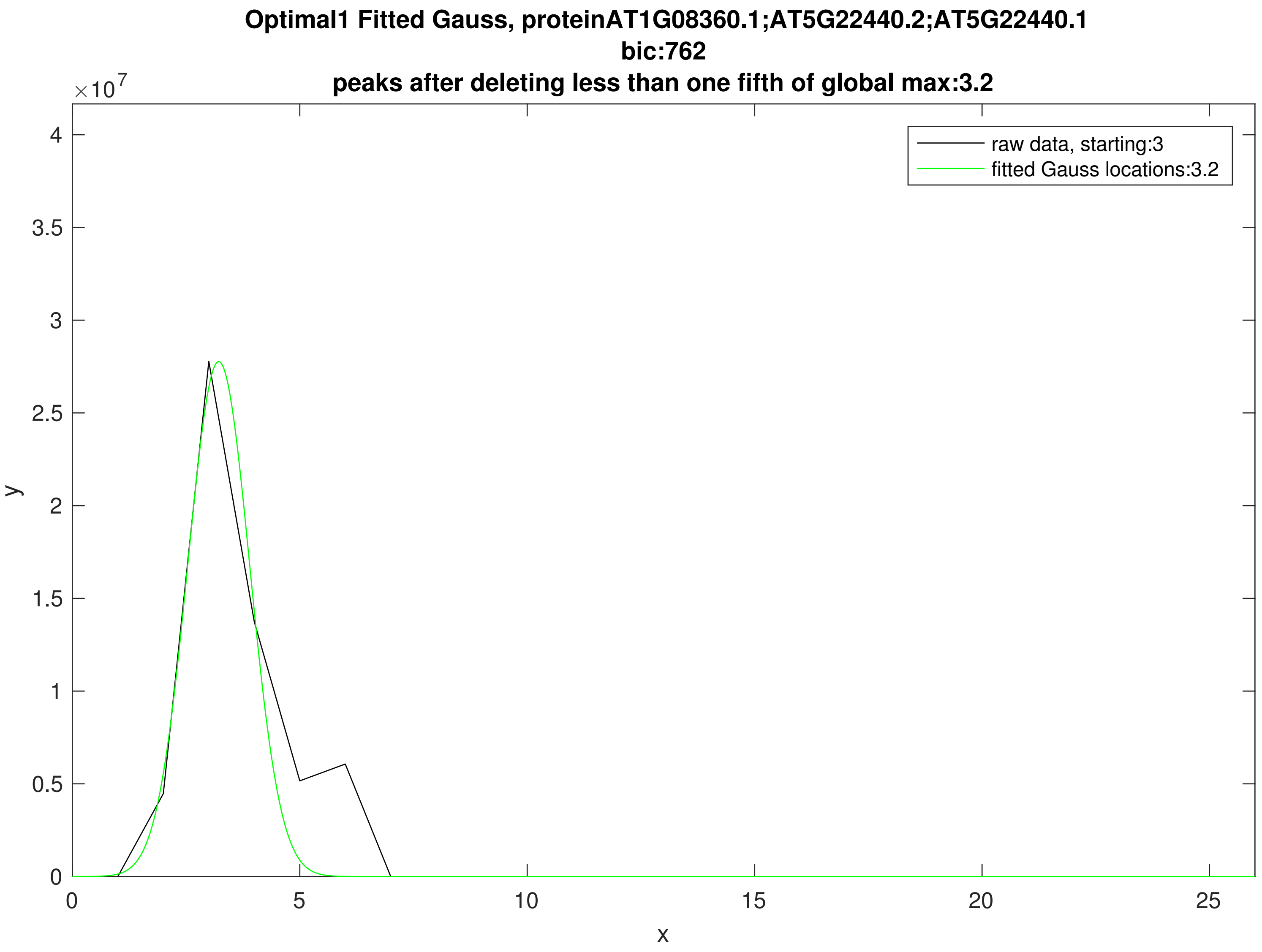

Optimal1 Fitted Gauss, proteinAT1G08360.1;AT5G22440.2;AT5G22440.1
bic:762
peaks after deleting less than one fifth of global max:3.2
×10^7
4
3.5
3
2.5
2
1.5
1
0.5
0
y
0
5
10
15
20
25
x
raw data, starting:3
fitted Gauss locations:3.2

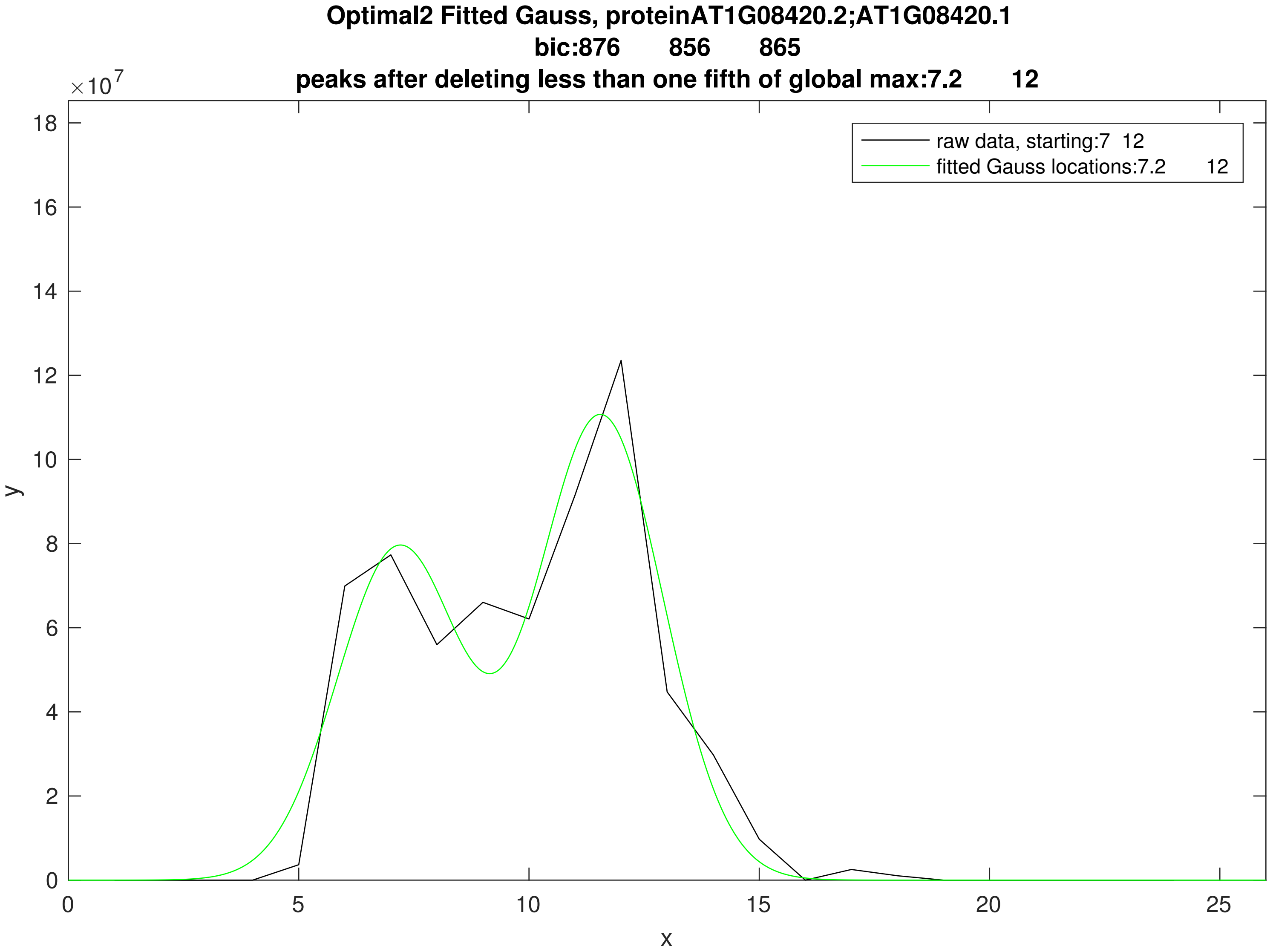
Optimal2 Fitted Gauss, proteinAT1G08420.2;AT1G08420.1
bic:876 856 865
peaks after deleting less than one fifth of global max:7.2 12
×10^7
raw data, starting:7 12
fitted Gauss locations:7.2 12
y
x
0
2
4
6
8
10
12
14
16
18
5
15
20
25

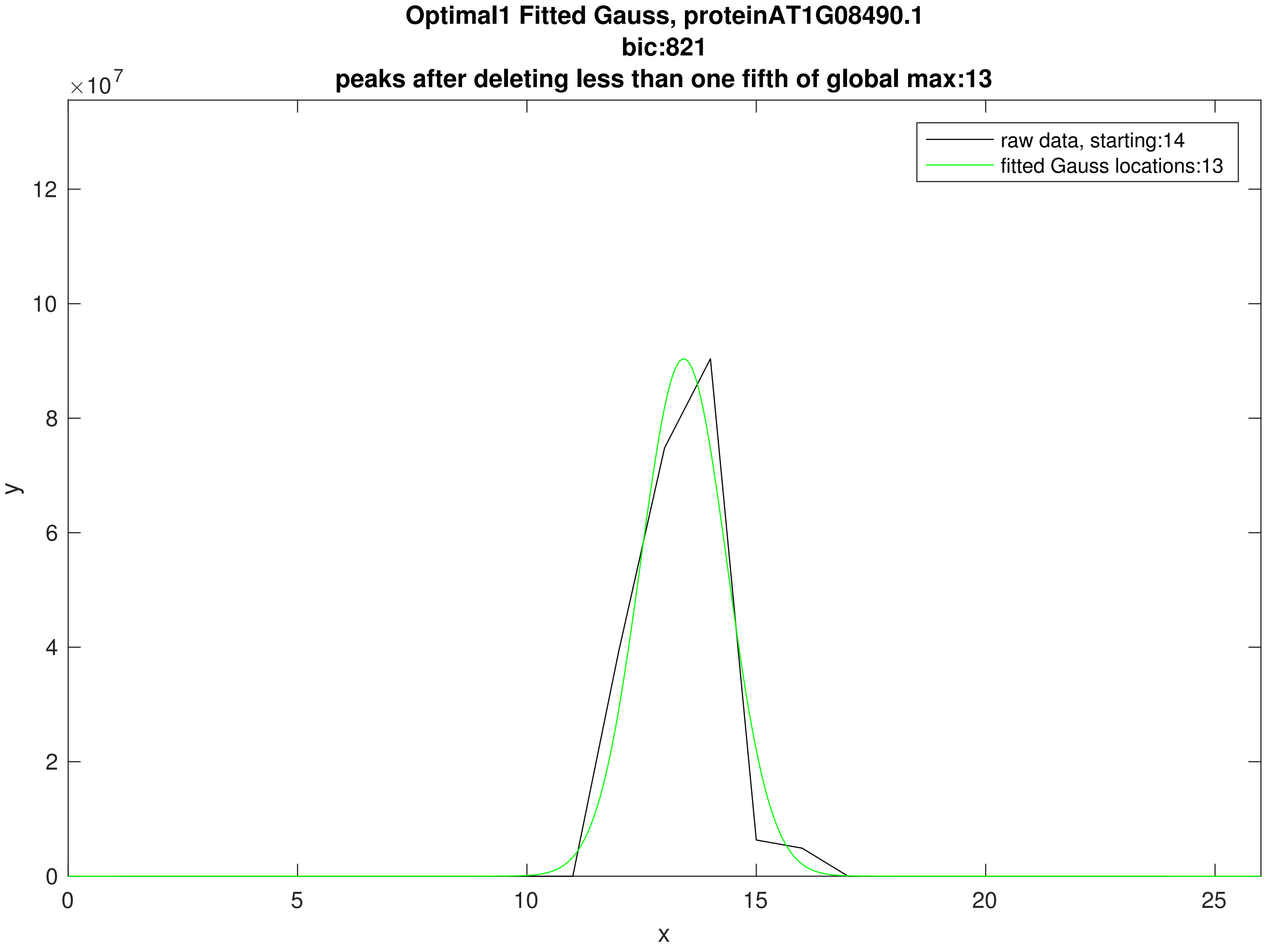

Optimal1 Fitted Gauss, proteinAT1G08490.1
bic:821
peaks after deleting less than one fifth of global max:13
×10^7
raw data, starting:14
fitted Gauss locations:13
12
10
8
6
4
2
0
y
0
5
10
15
20
25
x

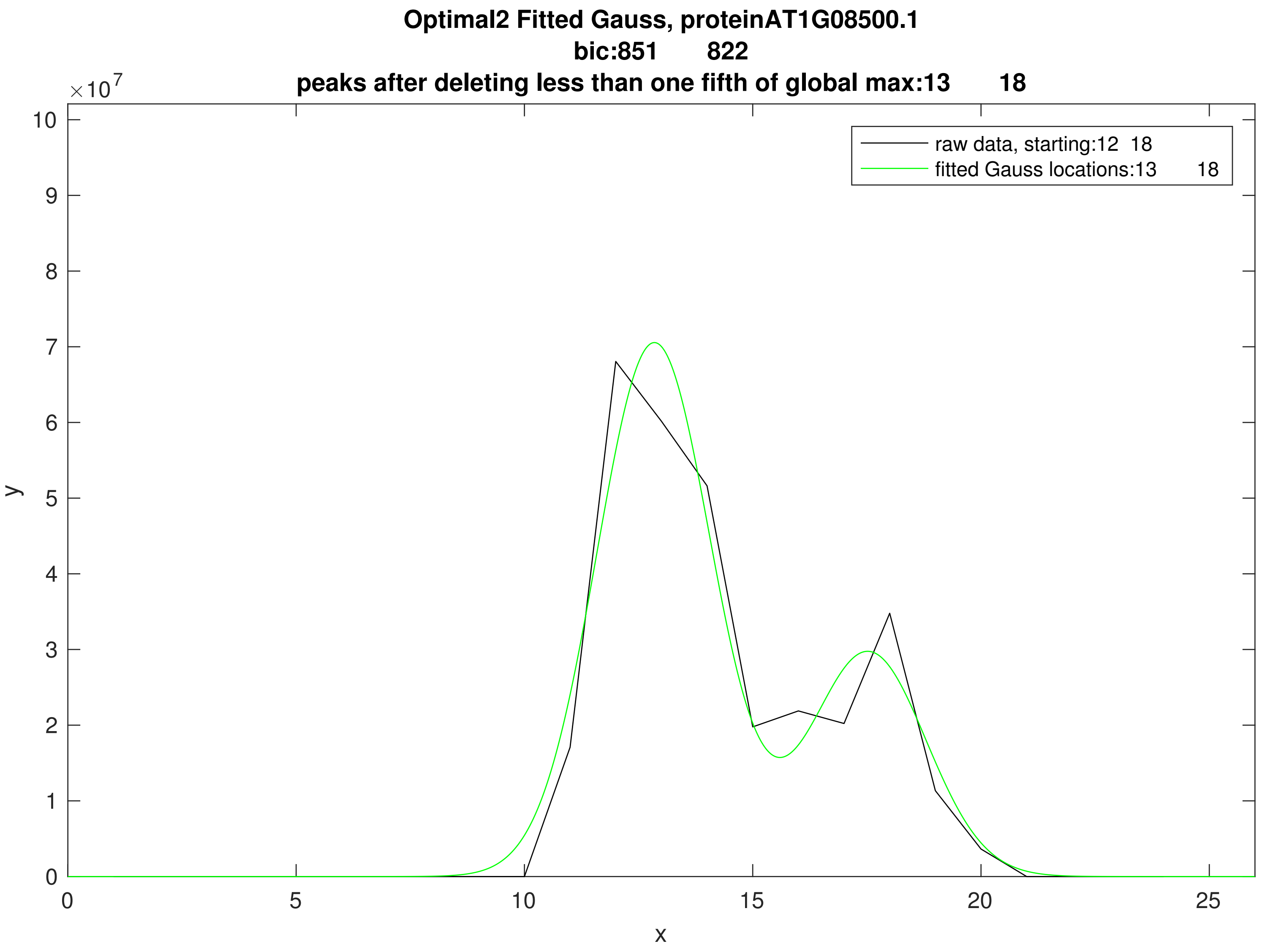
Optimal2 Fitted Gauss, proteinAT1G08500.1
bic:851 822
peaks after deleting less than one fifth of global max:13 18
×10^7
raw data, starting:12 18
fitted Gauss locations:13 18
y
x
0
1
2
3
4
5
6
7
8
9
10
0
5
10
15
20
25

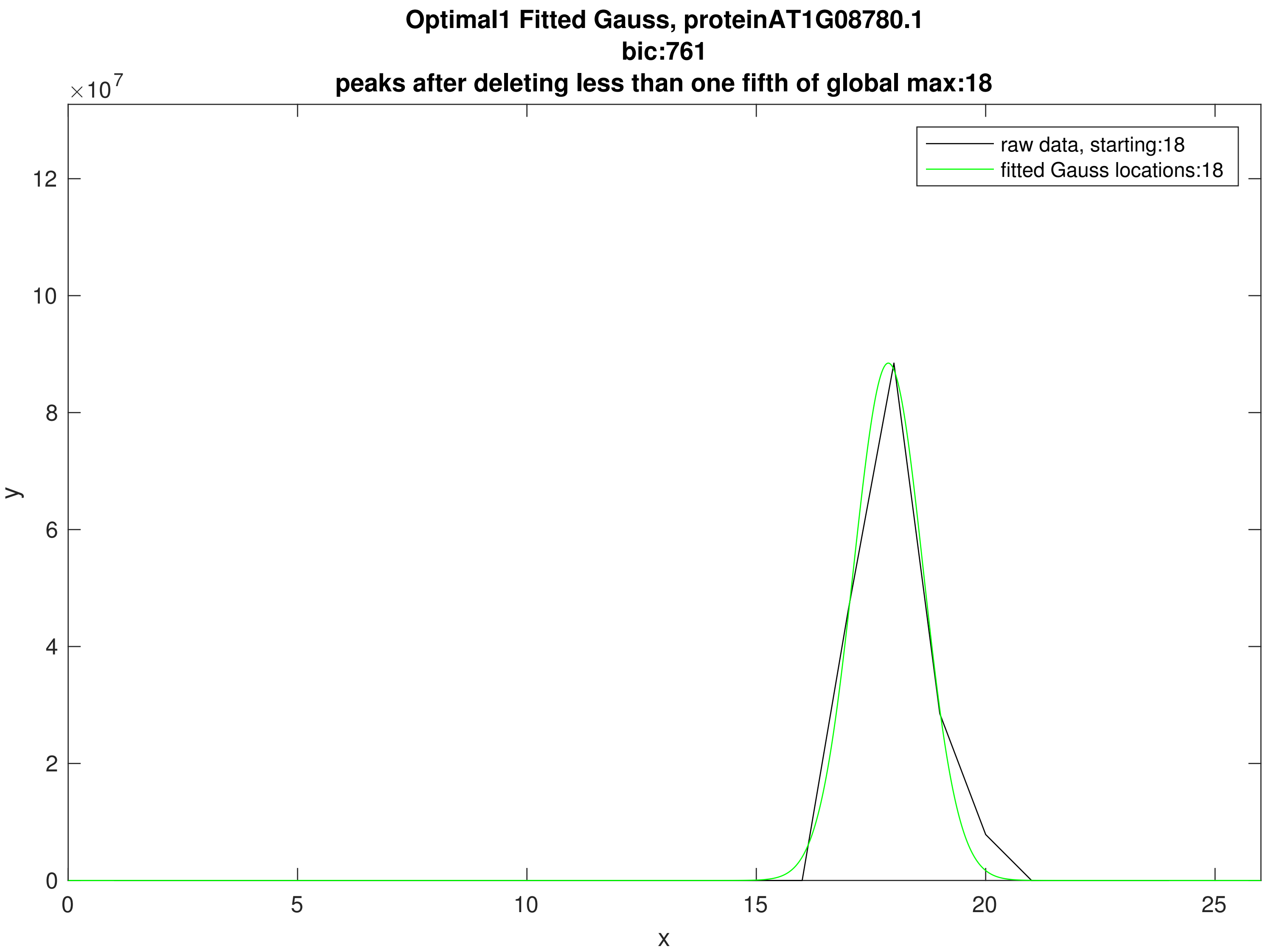
Optimal1 Fitted Gauss, proteinAT1G08780.1
bic:761
peaks after deleting less than one fifth of global max:18
×10^7
raw data, starting:18
fitted Gauss locations:18
y
x
0
2
4
6
8
10
12
0
5
10
15
20
25

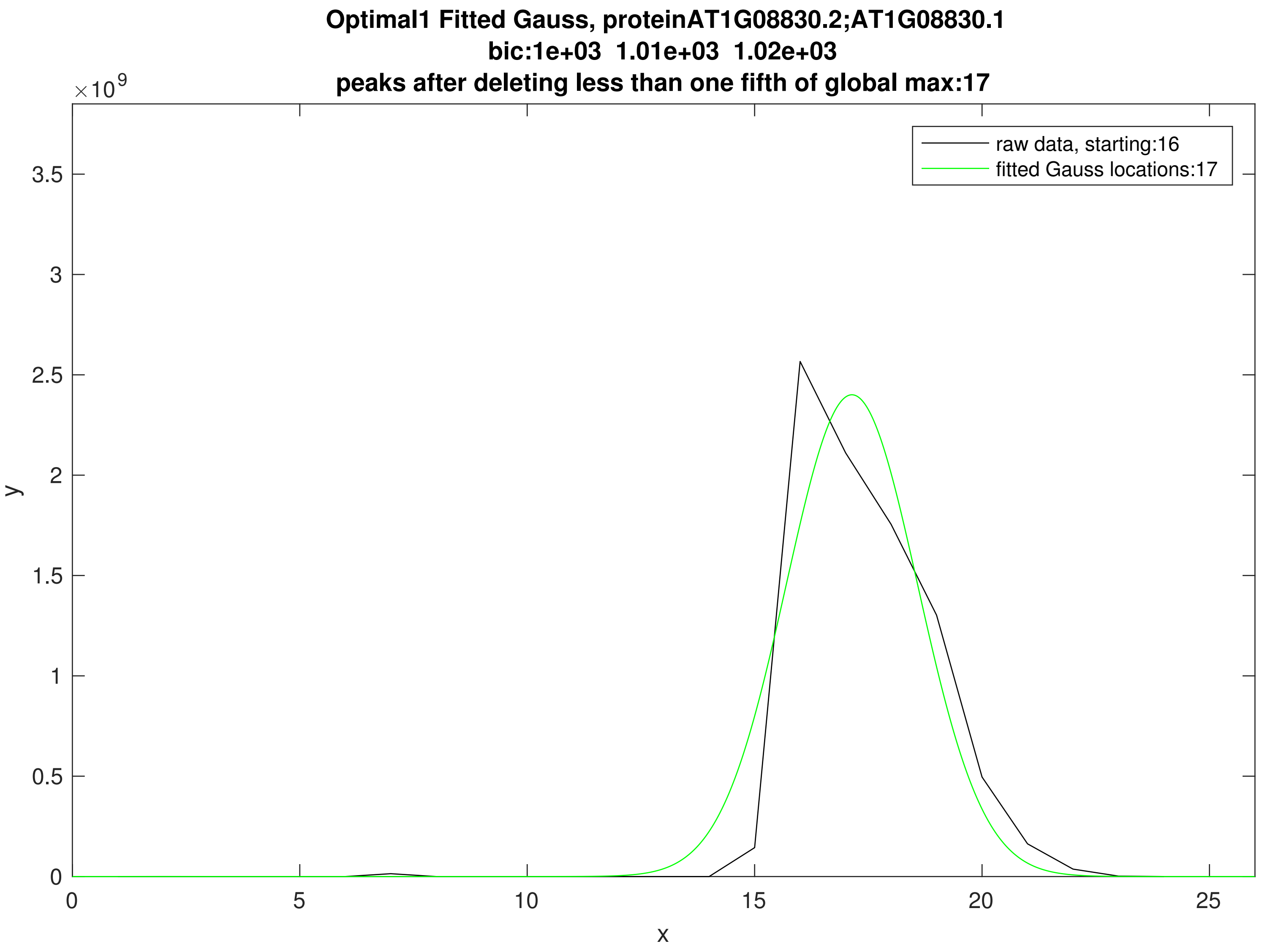

Optimal1 Fitted Gauss, proteinAT1G08830.2;AT1G08830.1
bic:1e+03 1.01e+03 1.02e+03
peaks after deleting less than one fifth of global max:17
×10^9
raw data, starting:16
fitted Gauss locations:17
3.5
3
2.5
2
1.5
1
0.5
0
y
0
5
10
15
20
25
x

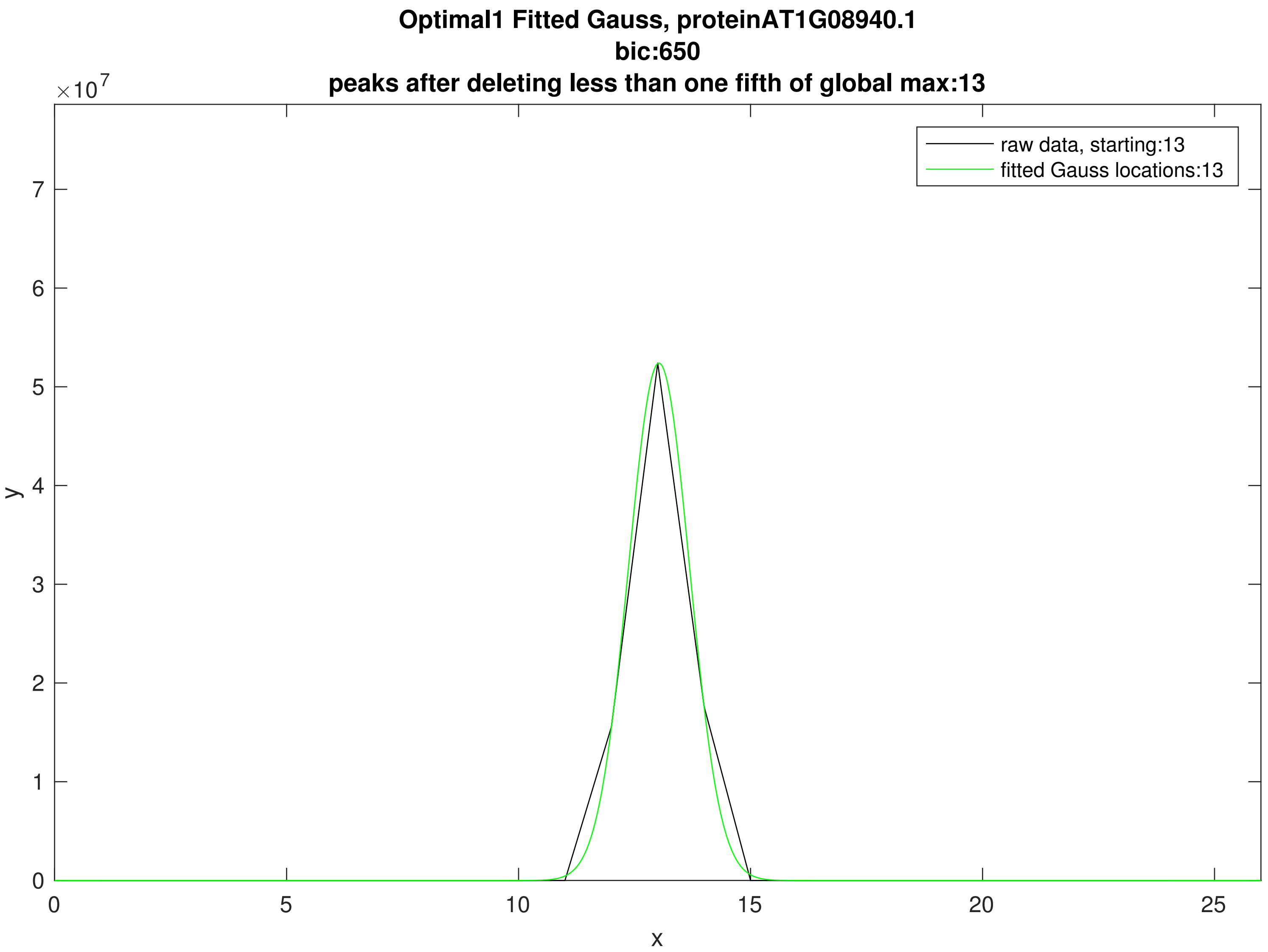
Optimal1 Fitted Gauss, proteinAT1G08940.1
bic:650
peaks after deleting less than one fifth of global max:13
×10^7
raw data, starting:13
fitted Gauss locations:13
y
x
0
1
2
3
4
5
6
7
0
5
10
15
20
25

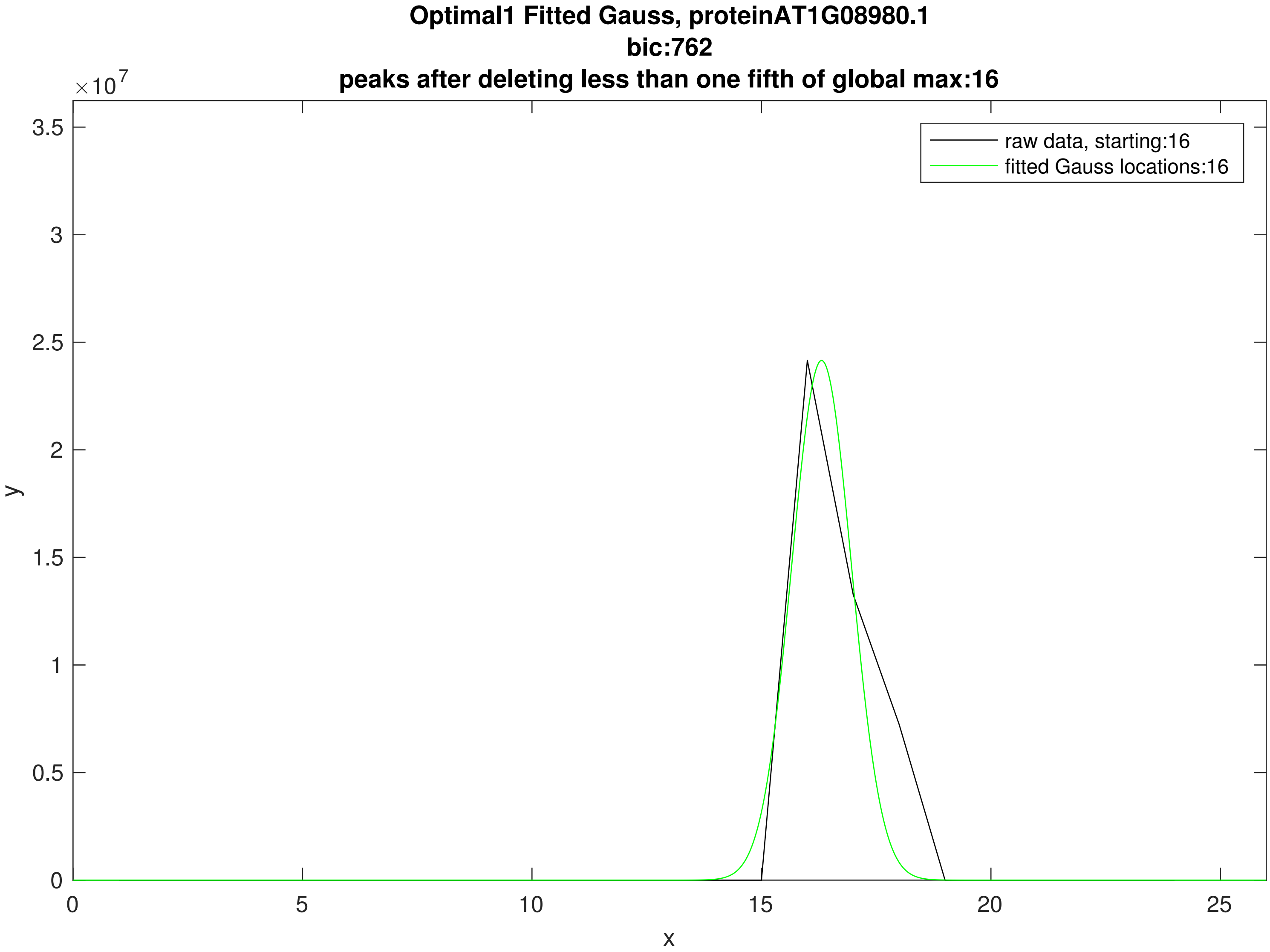

Optimal1 Fitted Gauss, proteinAT1G08980.1
bic:762
peaks after deleting less than one fifth of global max:16
×10^7
3.5
3
2.5
2
1.5
1
0.5
0
y
0
5
10
15
20
25
x
raw data, starting:16
fitted Gauss locations:16

**Optimal1 Fitted Gauss, proteinAT1G09010.1**
**bic:824 834**
**peaks after deleting less than one fifth of global max:14**

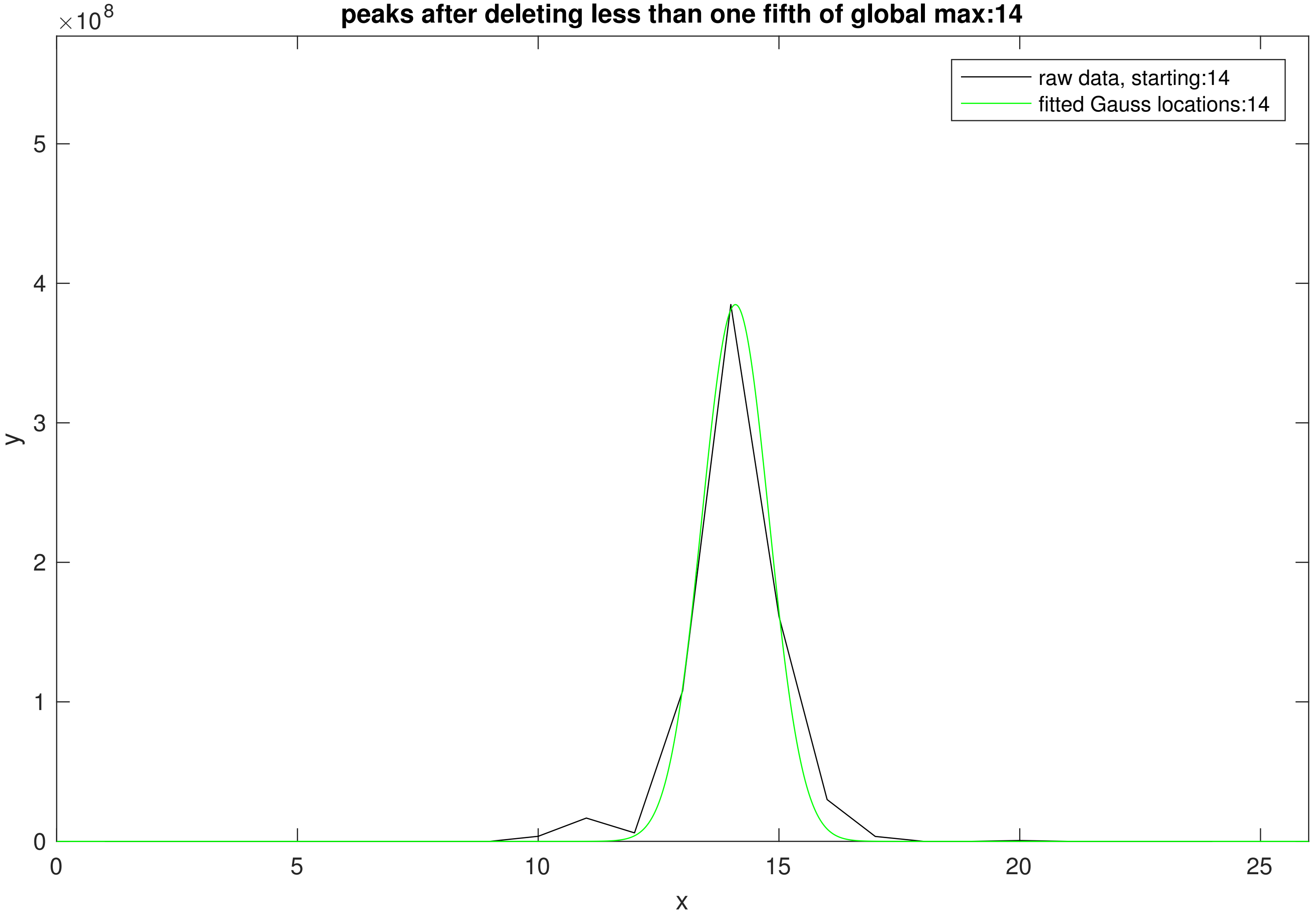

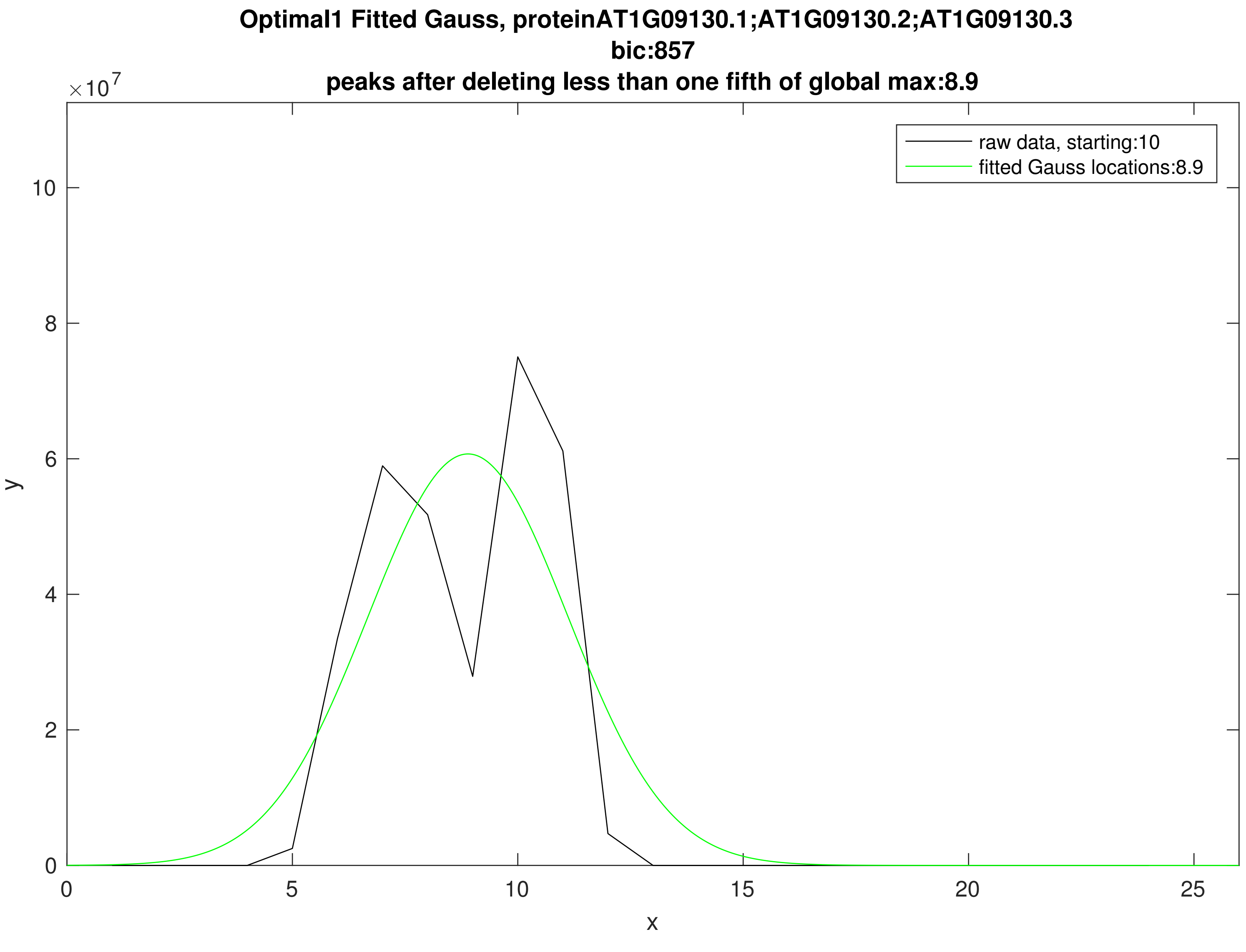
Optimal1 Fitted Gauss, proteinAT1G09130.1;AT1G09130.2;AT1G09130.3
bic:857
peaks after deleting less than one fifth of global max:8.9
×10^7
raw data, starting:10
fitted Gauss locations:8.9
y
x
0
2
4
6
8
10
5
15
20
25

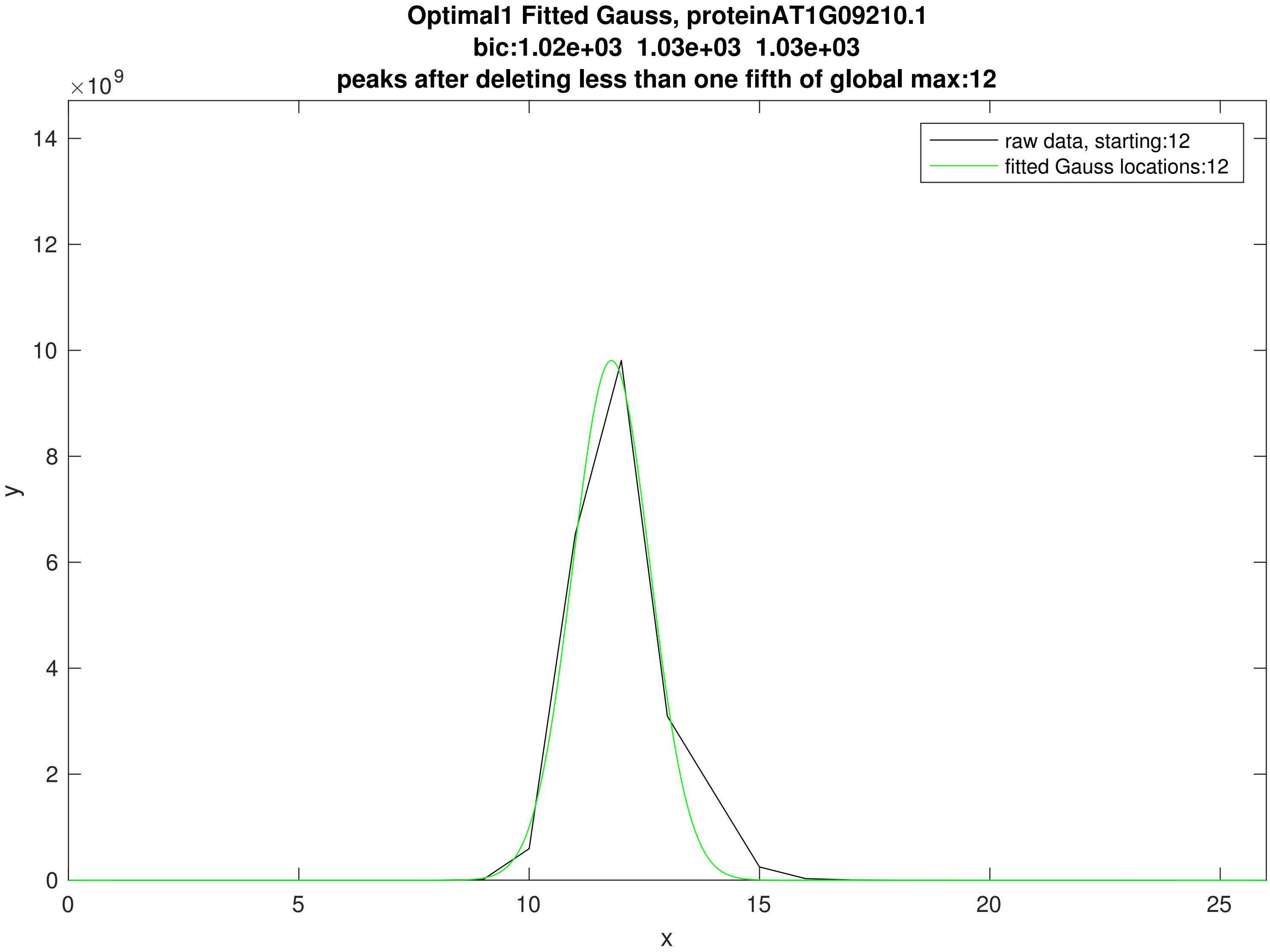
Optimal1 Fitted Gauss, proteinAT1G09210.1
bic:1.02e+03 1.03e+03 1.03e+03
peaks after deleting less than one fifth of global max:12
×10^9
raw data, starting:12
fitted Gauss locations:12
14
12
10
8
6
4
2
0
y
0
5
10
15
20
25
x

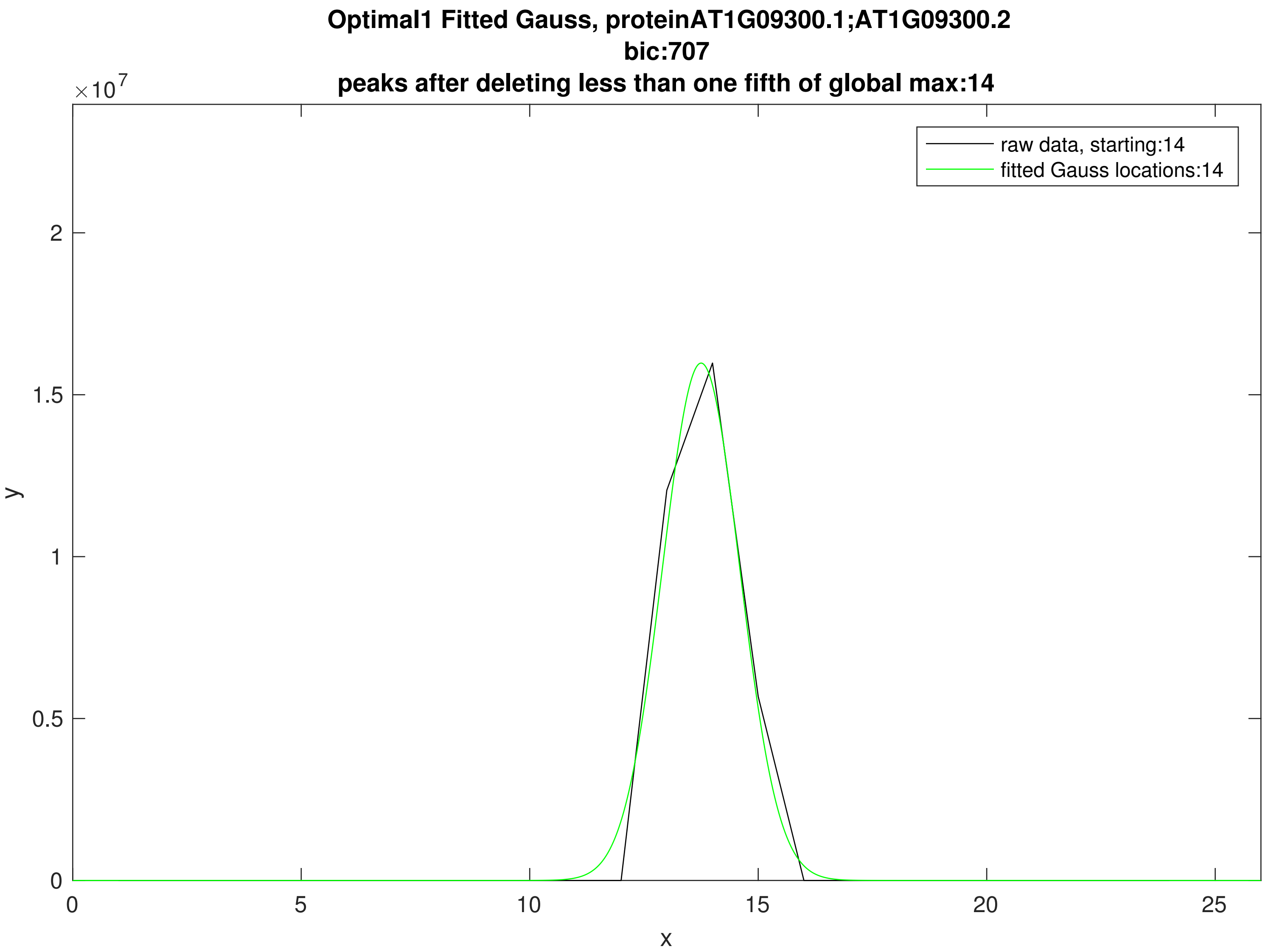
Optimal1 Fitted Gauss, proteinAT1G09300.1;AT1G09300.2
bic:707
peaks after deleting less than one fifth of global max:14
×10^7
raw data, starting:14
fitted Gauss locations:14
y
x
0
0.5
1
1.5
2
0
5
10
15
20
25

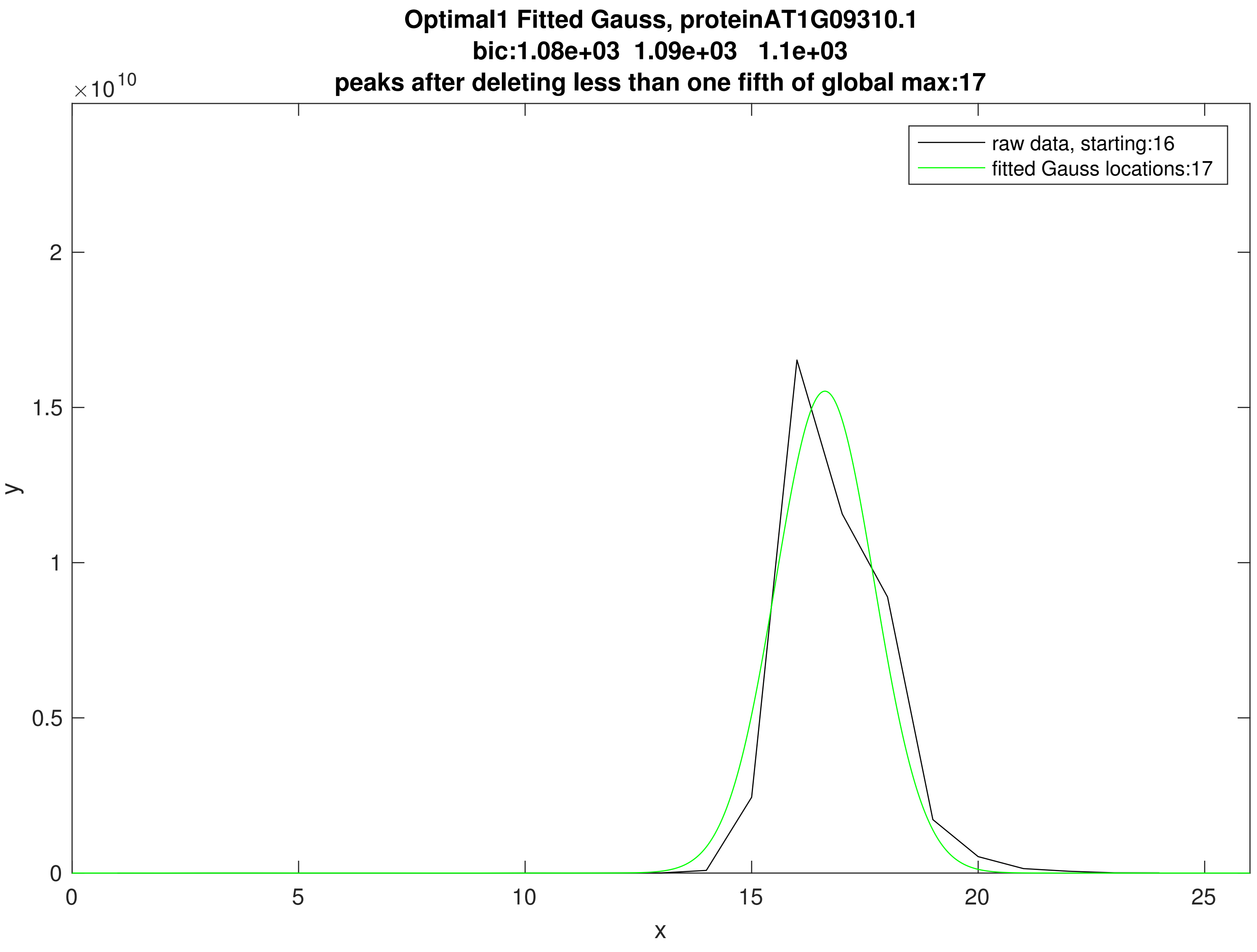

Optimal1 Fitted Gauss, proteinAT1G09310.1
bic:1.08e+03 1.09e+03 1.1e+03
peaks after deleting less than one fifth of global max:17
×10^10
raw data, starting:16
fitted Gauss locations:17
2
1.5
1
0.5
0
y
0
5
10
15
20
25
x

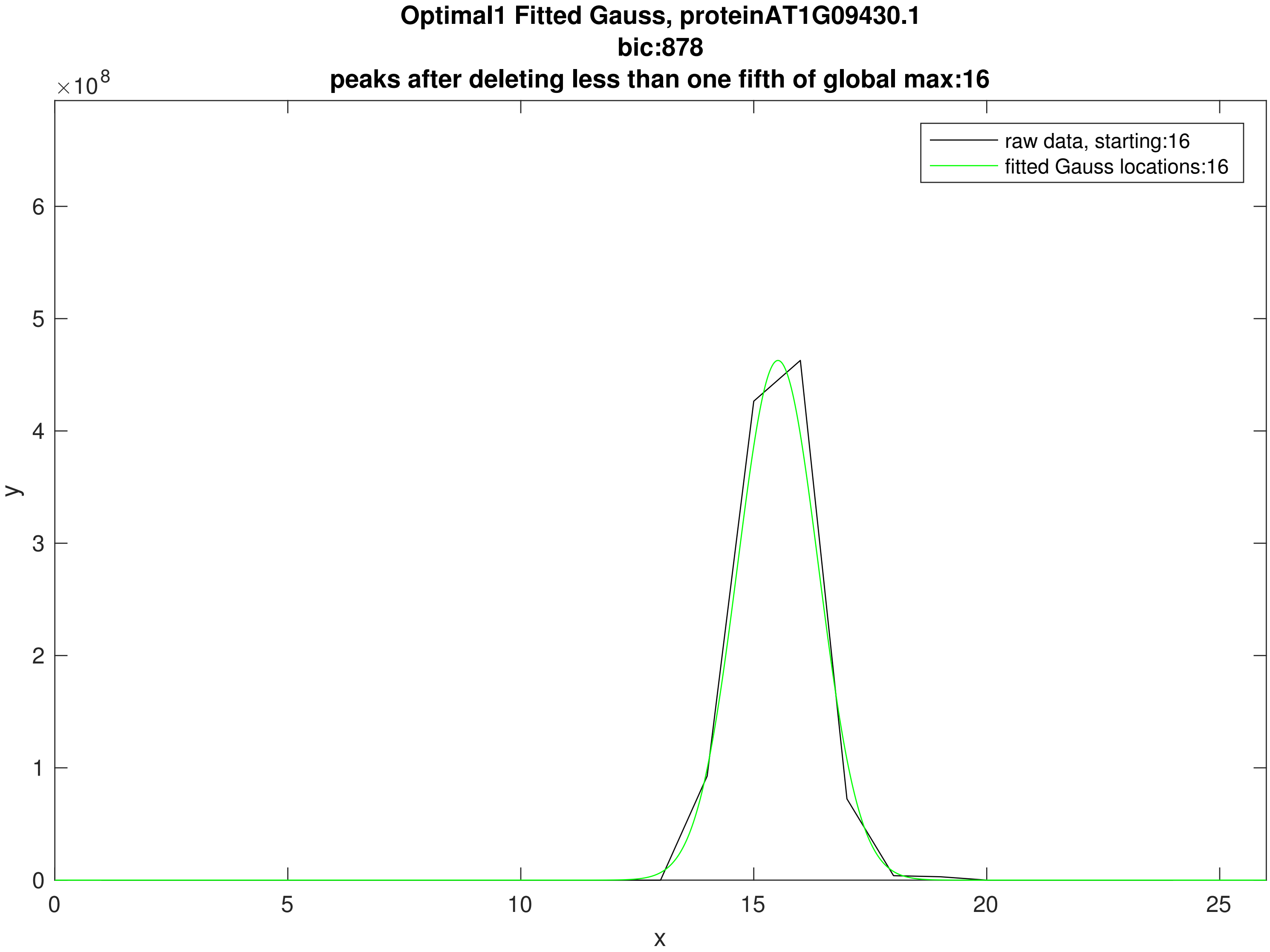
Optimal1 Fitted Gauss, proteinAT1G09430.1
bic:878
peaks after deleting less than one fifth of global max:16
×10^8
raw data, starting:16
fitted Gauss locations:16
y
x
0
1
2
3
4
5
6
0
5
10
15
20
25

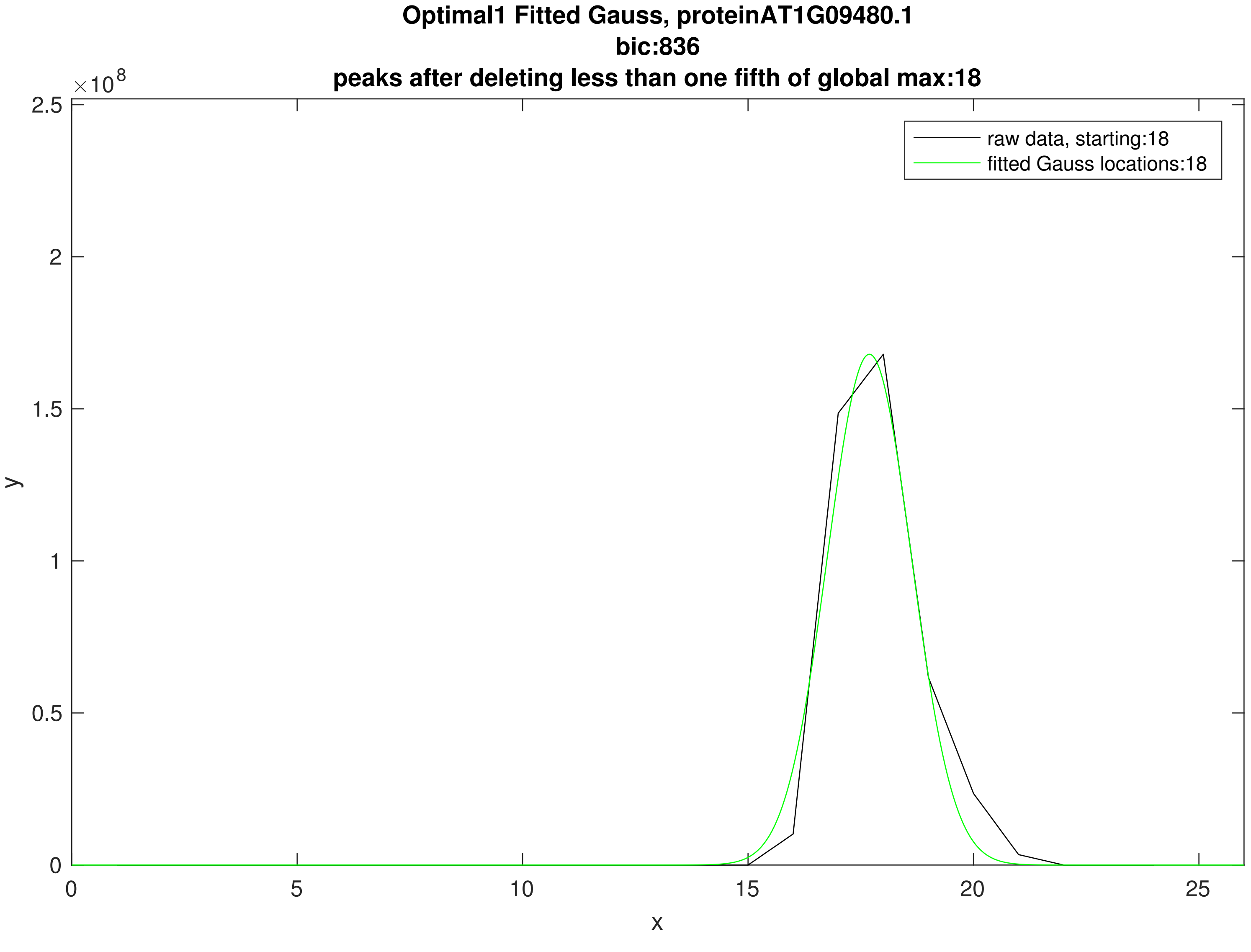
Optimal1 Fitted Gauss, proteinAT1G09480.1
bic:836
peaks after deleting less than one fifth of global max:18
×10^8
2.5
2
1.5
1
0.5
0
y
0
5
10
15
20
25
x
raw data, starting:18
fitted Gauss locations:18

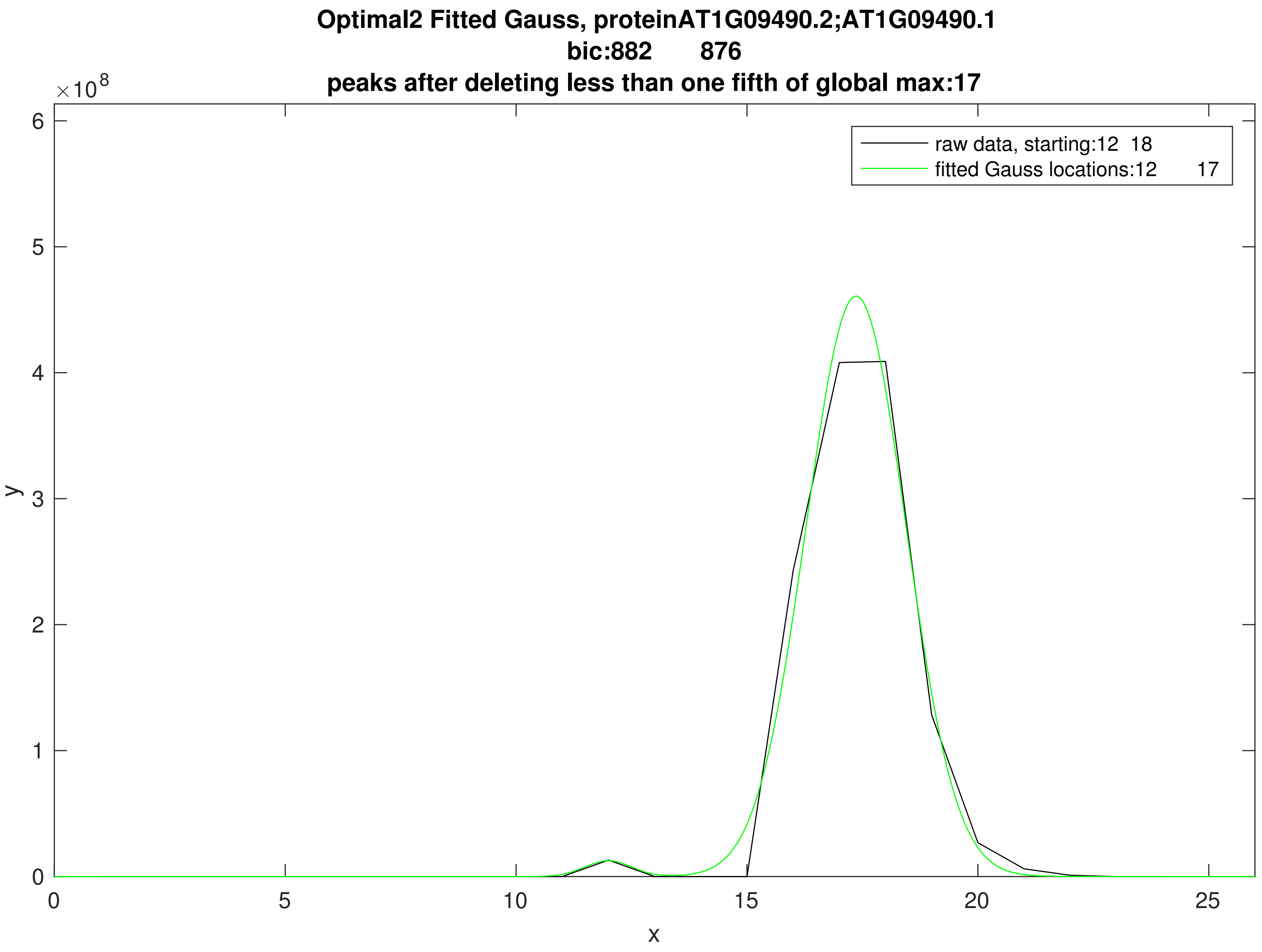

Optimal2 Fitted Gauss, proteinAT1G09490.2;AT1G09490.1
bic:882 876
peaks after deleting less than one fifth of global max:17
×10^8
6
5
4
3
2
1
0
y
0
5
10
15
20
25
x
raw data, starting:12 18
fitted Gauss locations:12 17

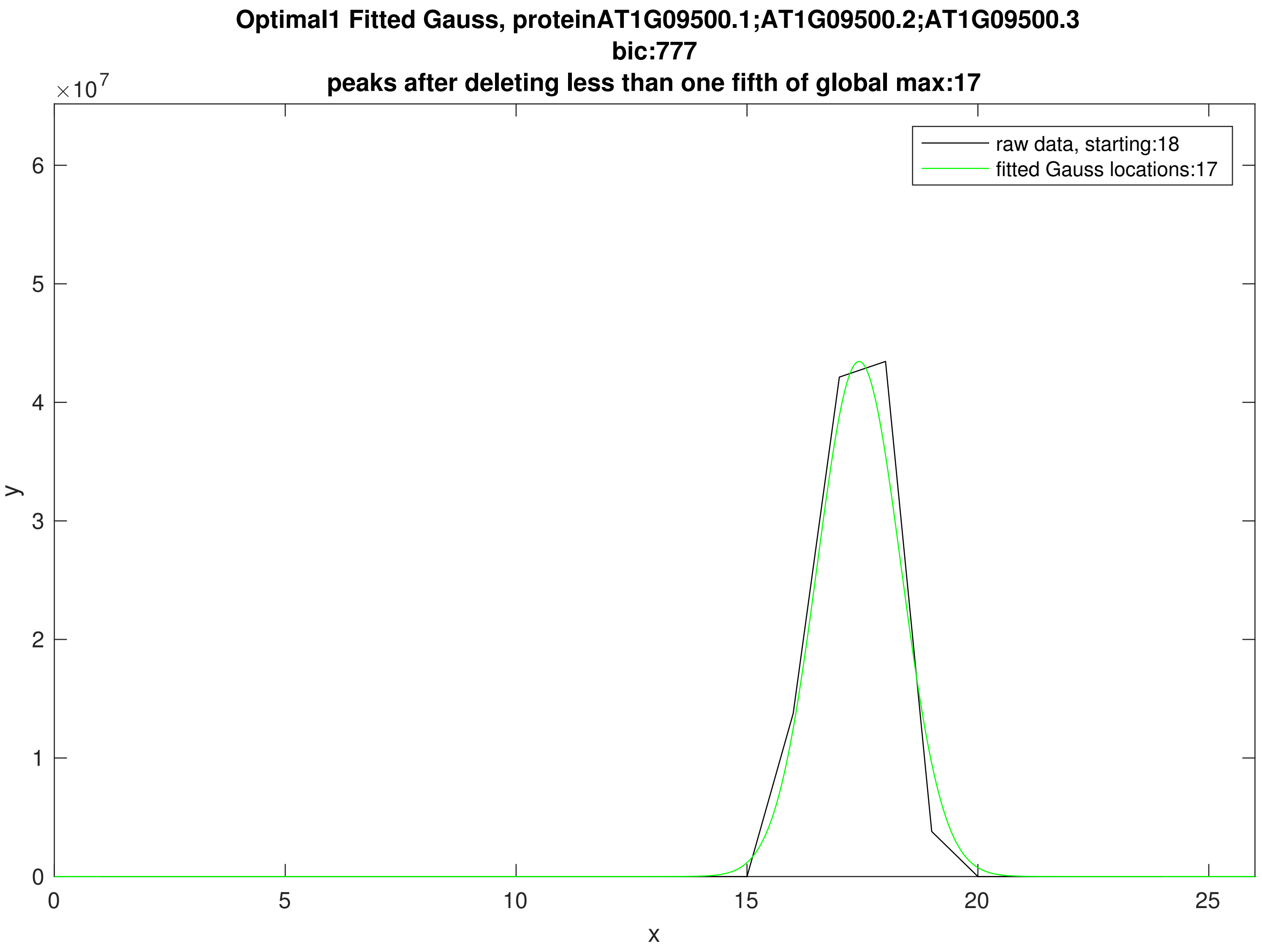

Optimal1 Fitted Gauss, proteinAT1G09500.1;AT1G09500.2;AT1G09500.3
bic:777
peaks after deleting less than one fifth of global max:17
×10^7
raw data, starting:18
fitted Gauss locations:17
y
x
0
1
2
3
4
5
6
0
5
10
15
20
25

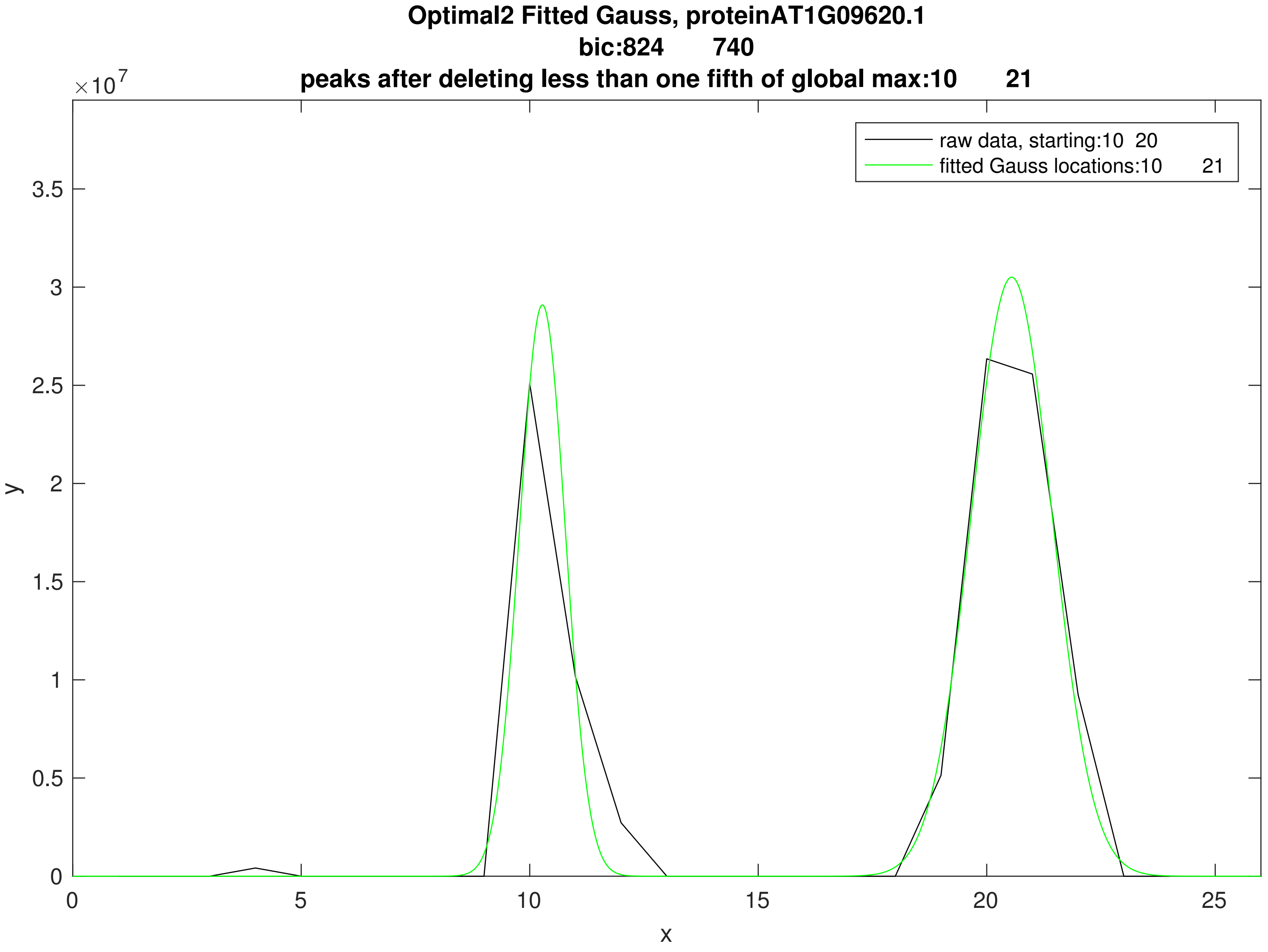
Optimal2 Fitted Gauss, proteinAT1G09620.1
bic:824 740
peaks after deleting less than one fifth of global max:10 21
×10^7
raw data, starting:10 20
fitted Gauss locations:10 21
3.5
3
2.5
2
1.5
1
0.5
0
y
0
5
10
15
20
25
x

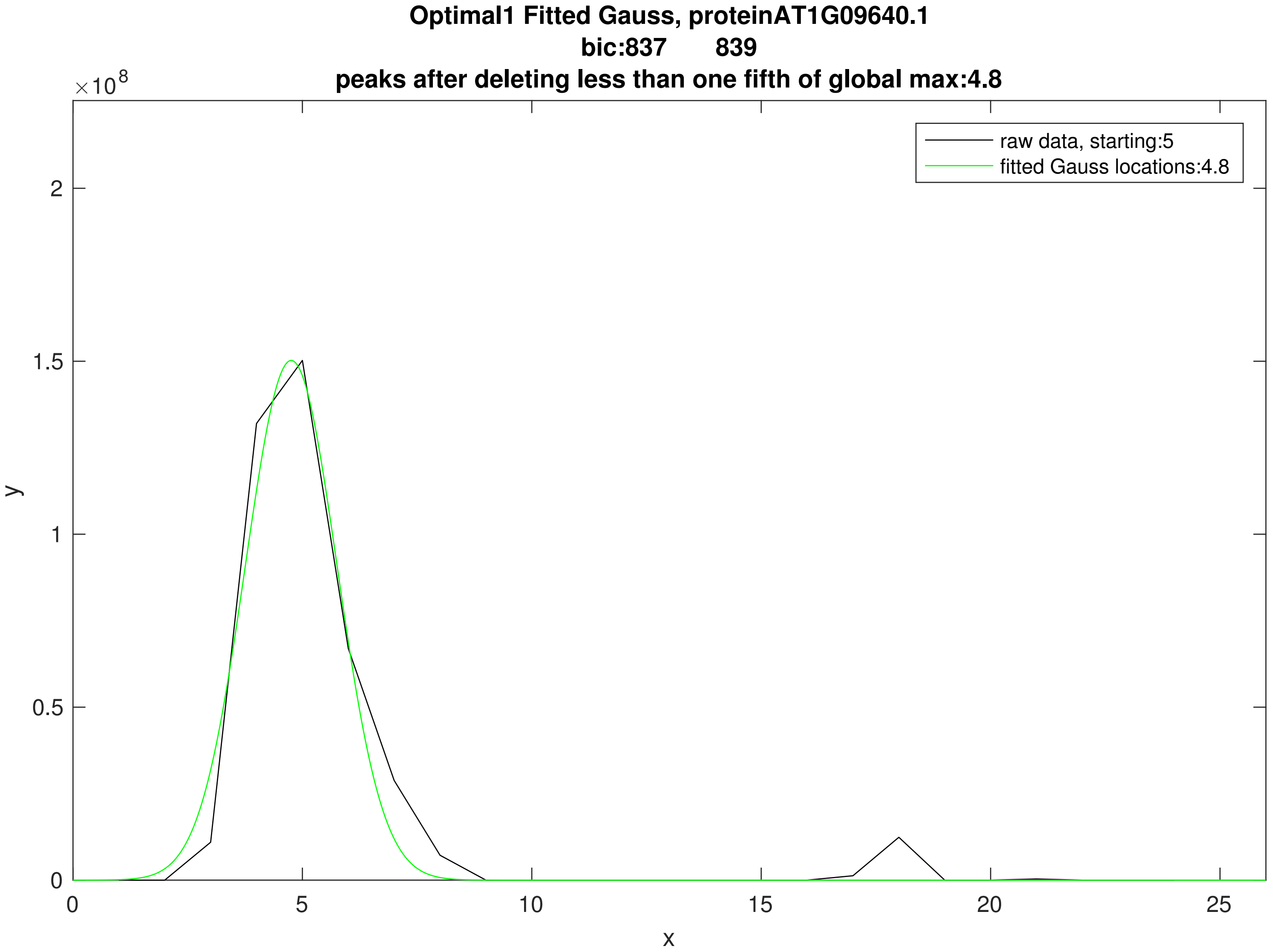
Optimal1 Fitted Gauss, proteinAT1G09640.1
bic:837 839
peaks after deleting less than one fifth of global max:4.8
×10^8
raw data, starting:5
fitted Gauss locations:4.8
y
x
0
0.5
1
1.5
2
0
5
10
15
20
25

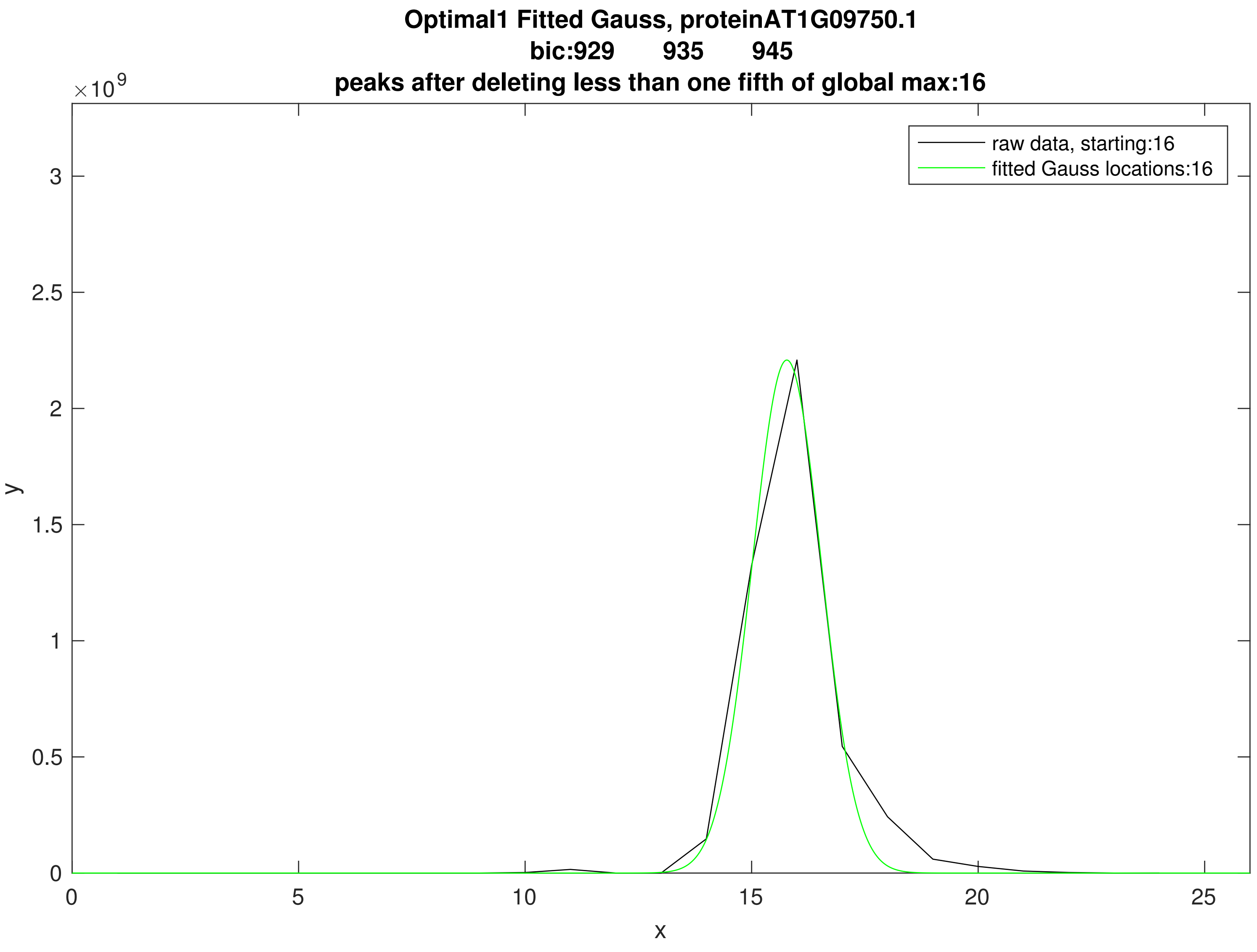

Optimal1 Fitted Gauss, proteinAT1G09750.1
bic:929 935 945
peaks after deleting less than one fifth of global max:16
×10^9
raw data, starting:16
fitted Gauss locations:16
y
x
0
0.5
1
1.5
2
2.5
3
0
5
10
15
20
25

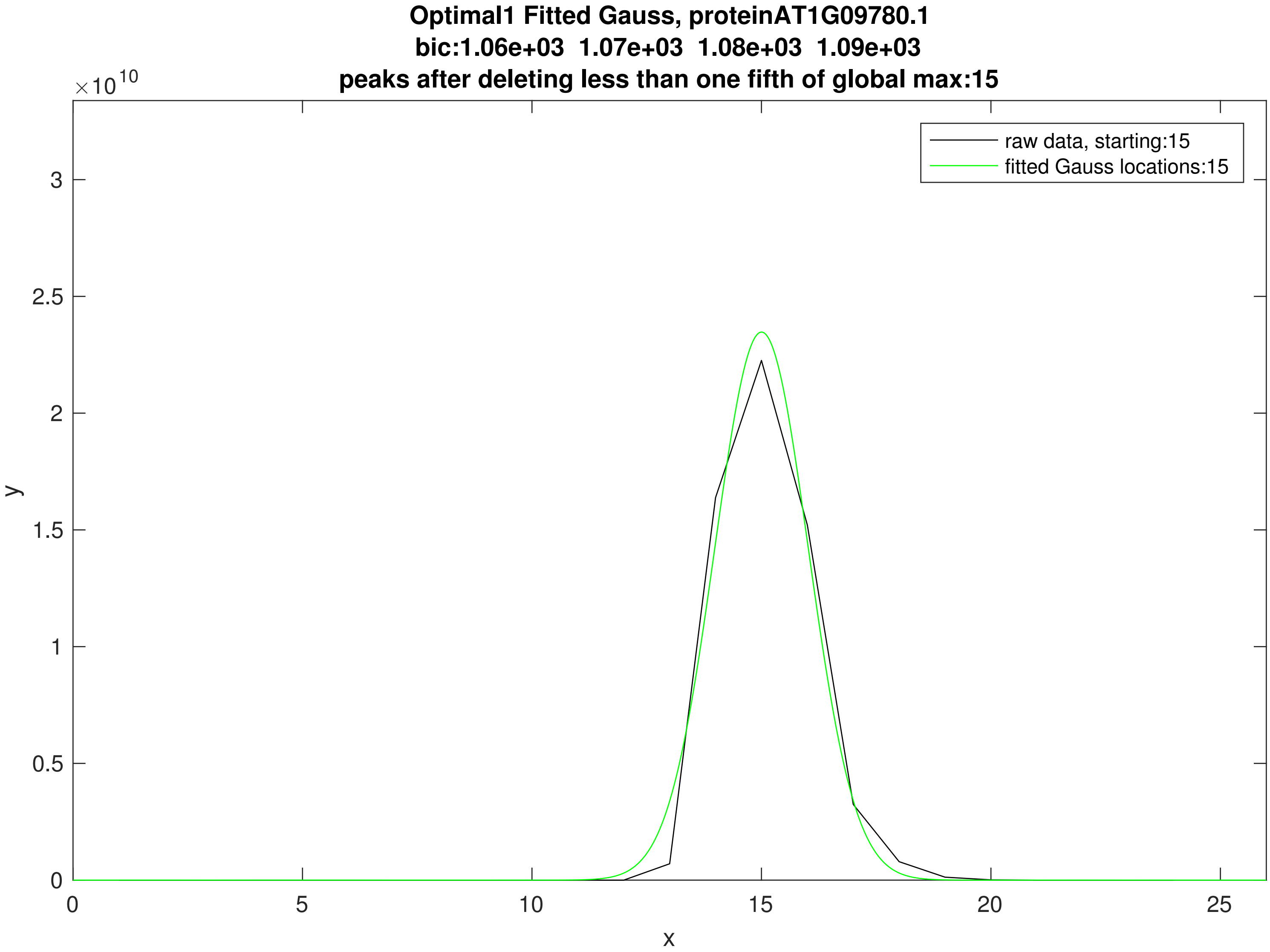

Optimal1 Fitted Gauss, proteinAT1G09780.1
bic:1.06e+03  1.07e+03  1.08e+03  1.09e+03
peaks after deleting less than one fifth of global max:15
×10^10
raw data, starting:15
fitted Gauss locations:15
y
x
0
0.5
1
1.5
2
2.5
3
0
5
10
15
20
25

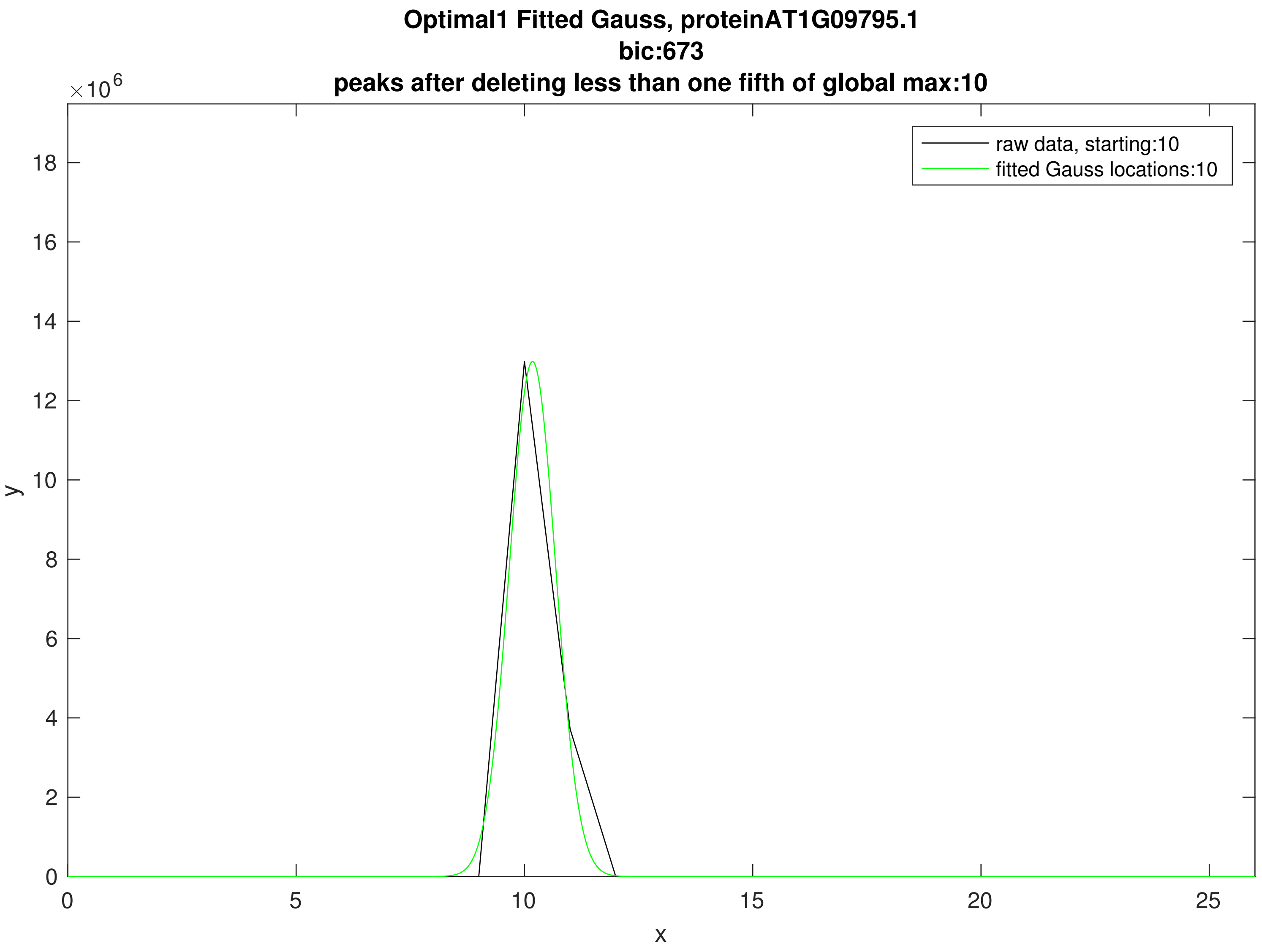
Optimal1 Fitted Gauss, proteinAT1G09795.1
bic:673
peaks after deleting less than one fifth of global max:10
×10^6
raw data, starting:10
fitted Gauss locations:10
18
16
14
12
10
8
6
4
2
0
y
0
5
10
15
20
25
x

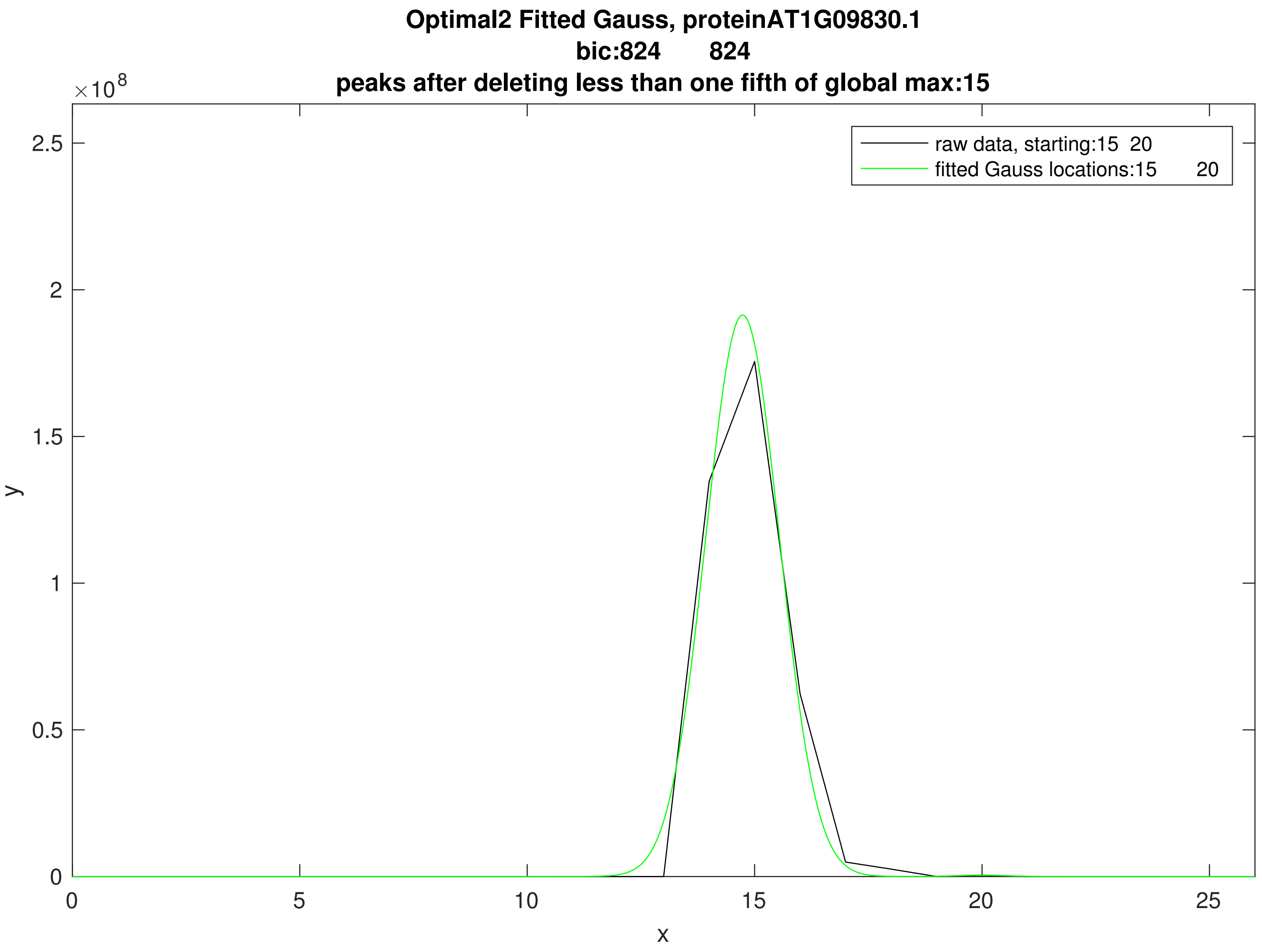
Optimal2 Fitted Gauss, proteinAT1G09830.1
bic:824 824
peaks after deleting less than one fifth of global max:15
×10^8
2.5
2
1.5
1
0.5
0
y
0
5
10
15
20
25
x
raw data, starting:15 20
fitted Gauss locations:15 20

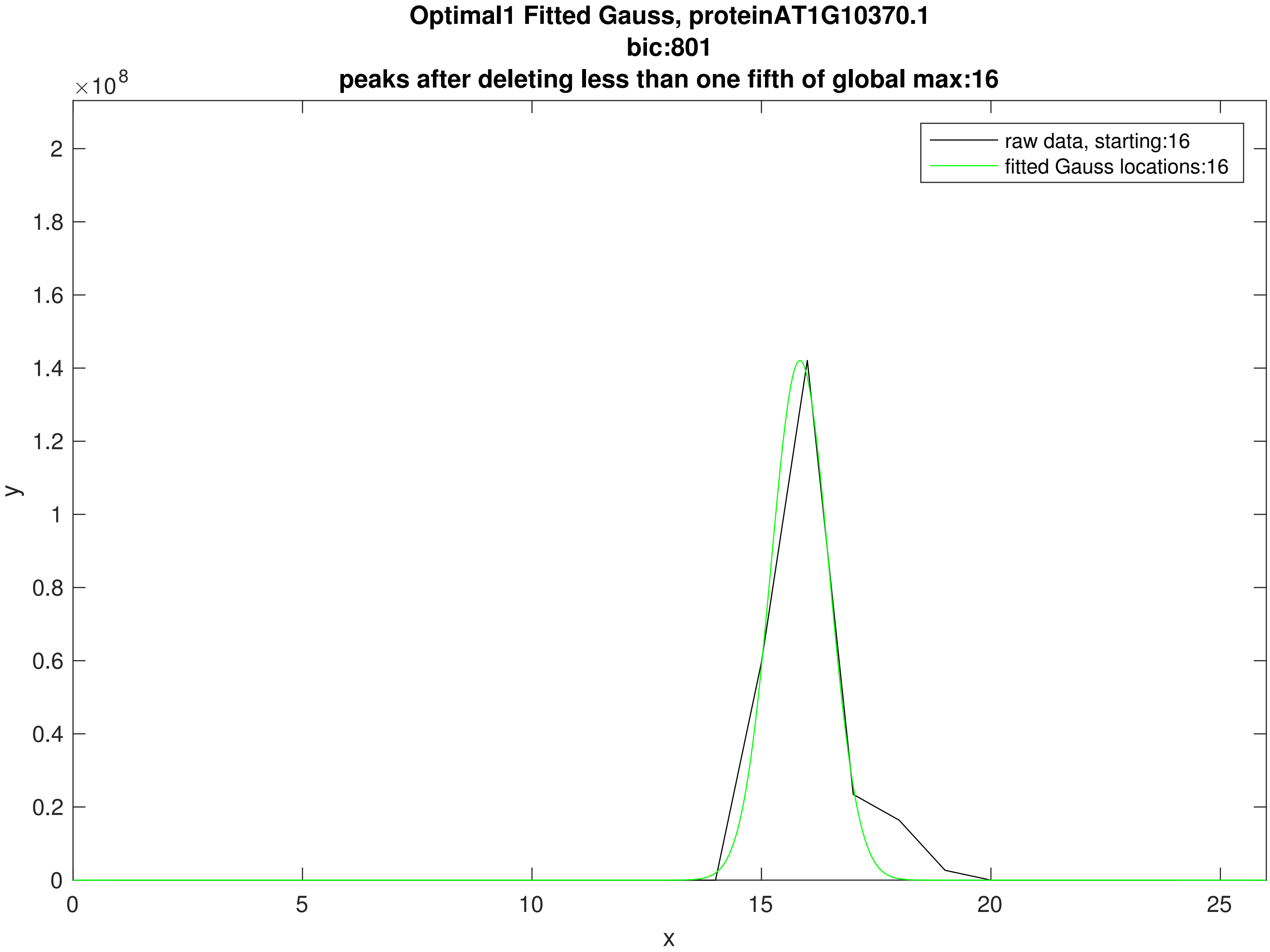
Optimal1 Fitted Gauss, proteinAT1G10370.1
bic:801
peaks after deleting less than one fifth of global max:16
×10^8
raw data, starting:16
fitted Gauss locations:16
2
1.8
1.6
1.4
1.2
1
0.8
0.6
0.4
0.2
0
y
0
5
10
15
20
25
x

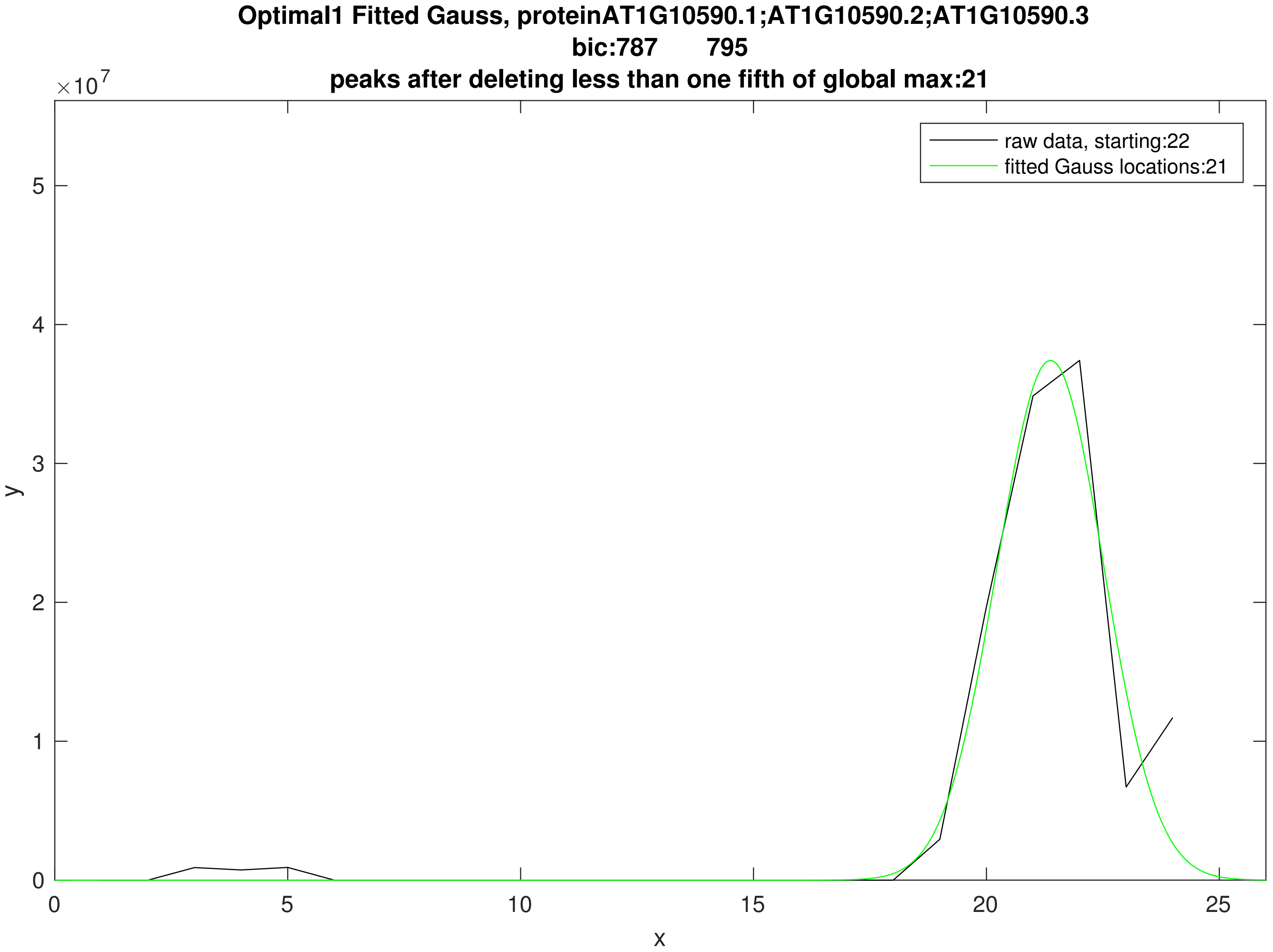
Optimal1 Fitted Gauss, proteinAT1G10590.1;AT1G10590.2;AT1G10590.3
bic:787 795
peaks after deleting less than one fifth of global max:21
×10^7
raw data, starting:22
fitted Gauss locations:21
y
x
0
1
2
3
4
5
0
5
10
15
20
25

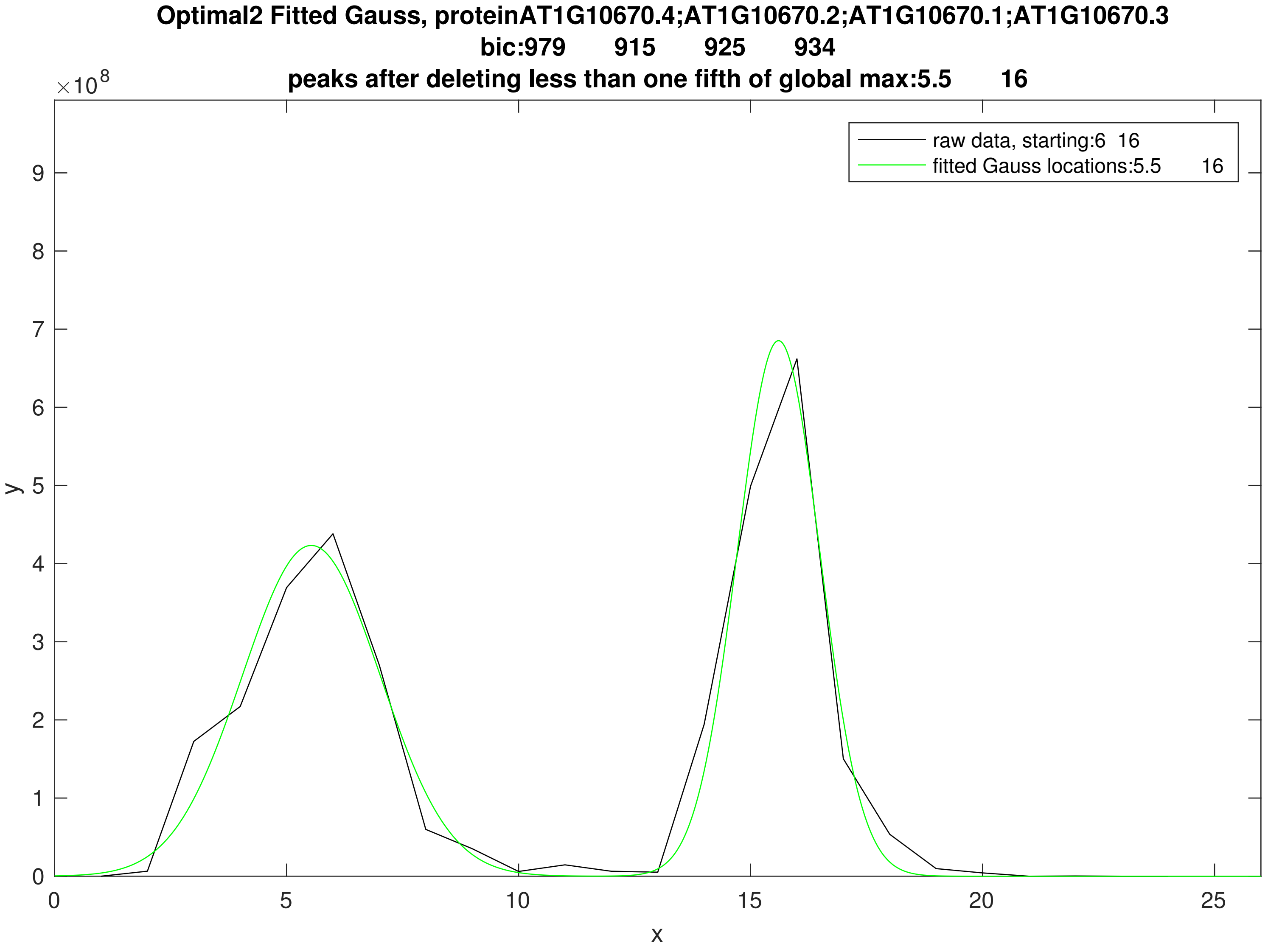

Optimal2 Fitted Gauss, proteinAT1G10670.4;AT1G10670.2;AT1G10670.1;AT1G10670.3
bic:979 915 925 934
peaks after deleting less than one fifth of global max:5.5 16
×10^8
raw data, starting:6 16
fitted Gauss locations:5.5 16
y
x
0
1
2
3
4
5
6
7
8
9
0
5
10
15
20
25

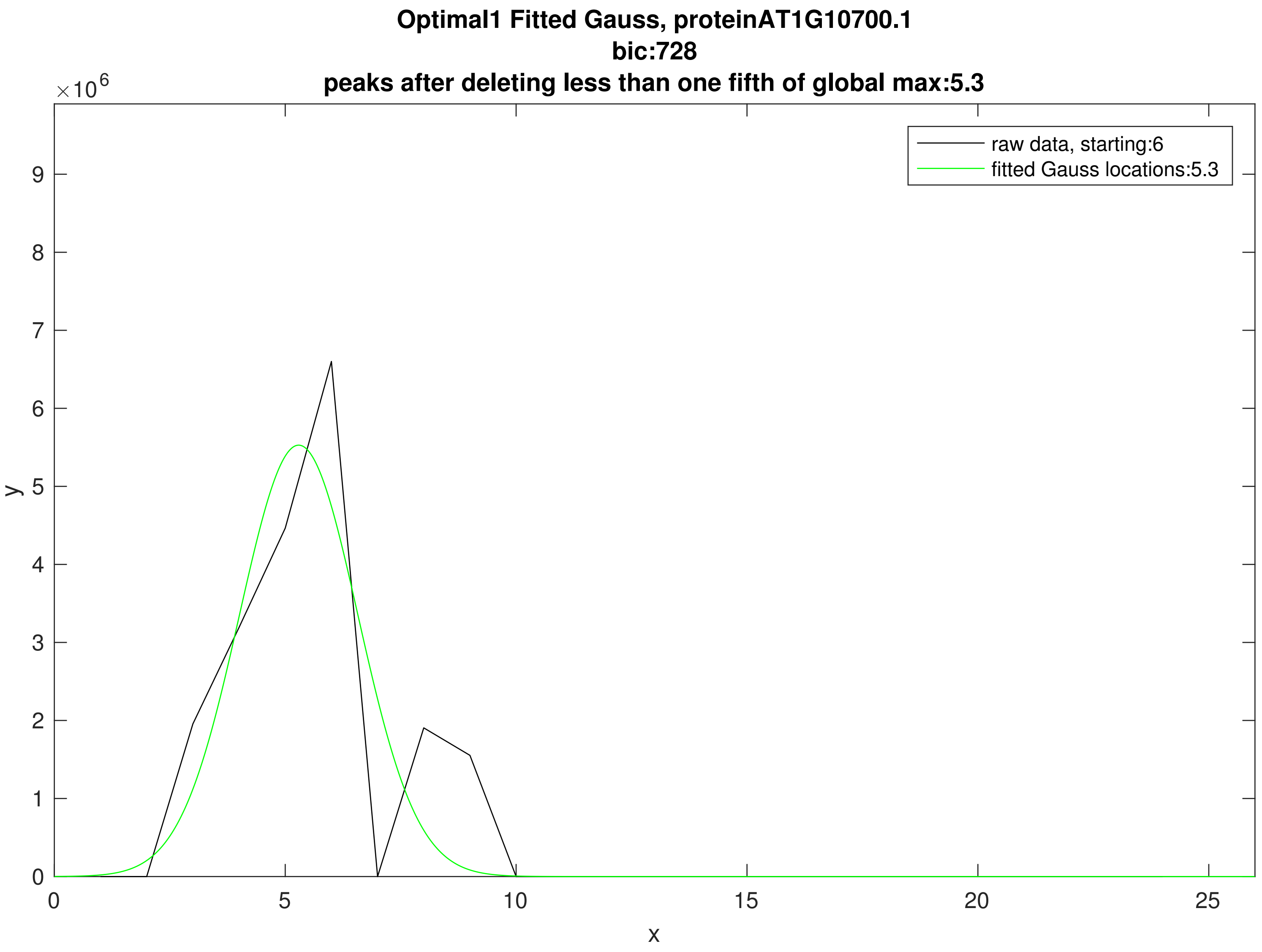

Optimal1 Fitted Gauss, proteinAT1G10700.1
bic:728
peaks after deleting less than one fifth of global max:5.3
×10^6
raw data, starting:6
fitted Gauss locations:5.3
y
x
0
1
2
3
4
5
6
7
8
9
10
15
20
25

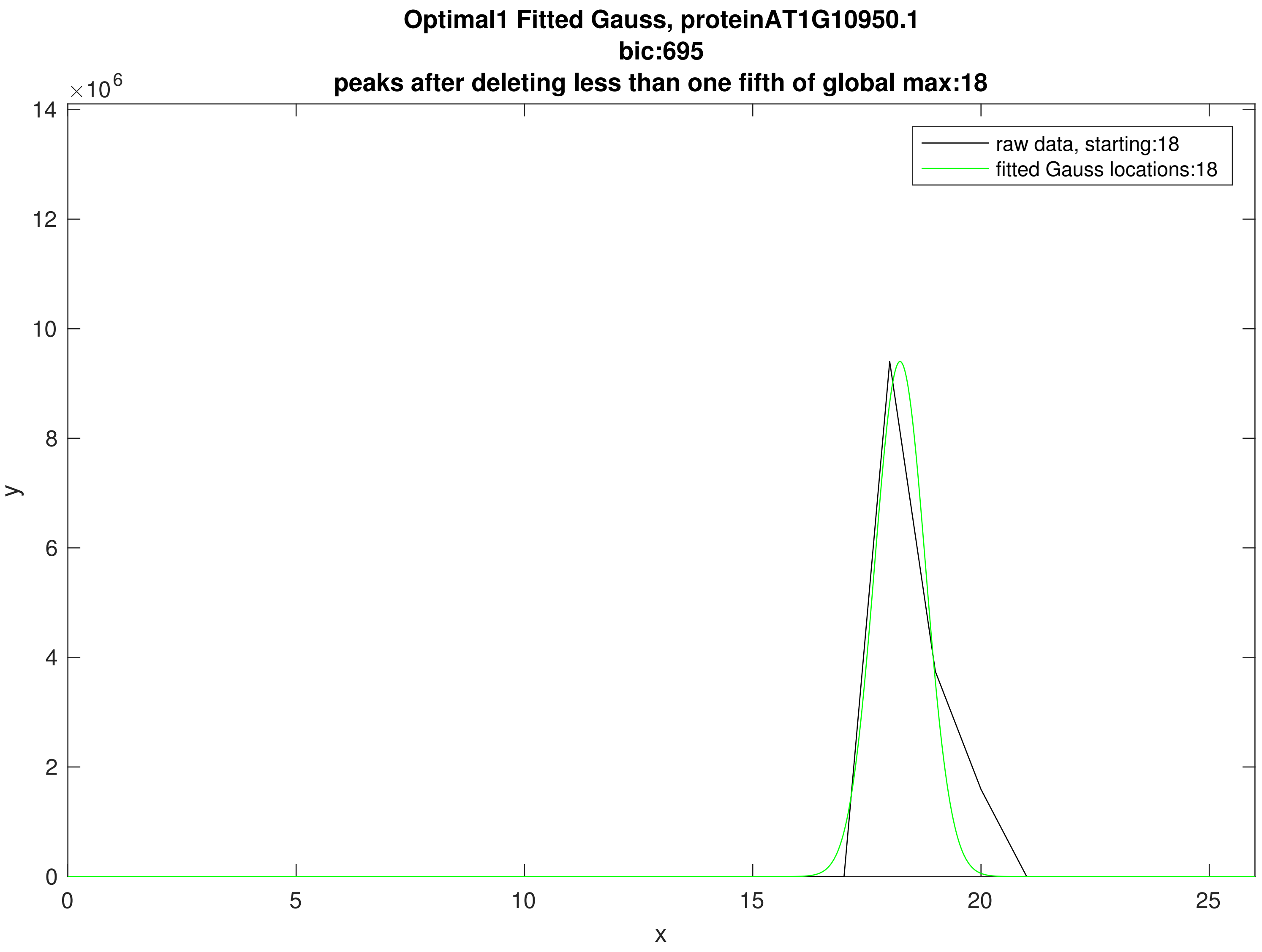
Optimal1 Fitted Gauss, proteinAT1G10950.1
bic:695
peaks after deleting less than one fifth of global max:18
×10^6
14
12
10
8
6
4
2
0
y
0
5
10
15
20
25
x
raw data, starting:18
fitted Gauss locations:18

**Optimal1 Fitted Gauss, proteinAT1G10960.1**
**bic:758**
**peaks after deleting less than one fifth of global max:19**

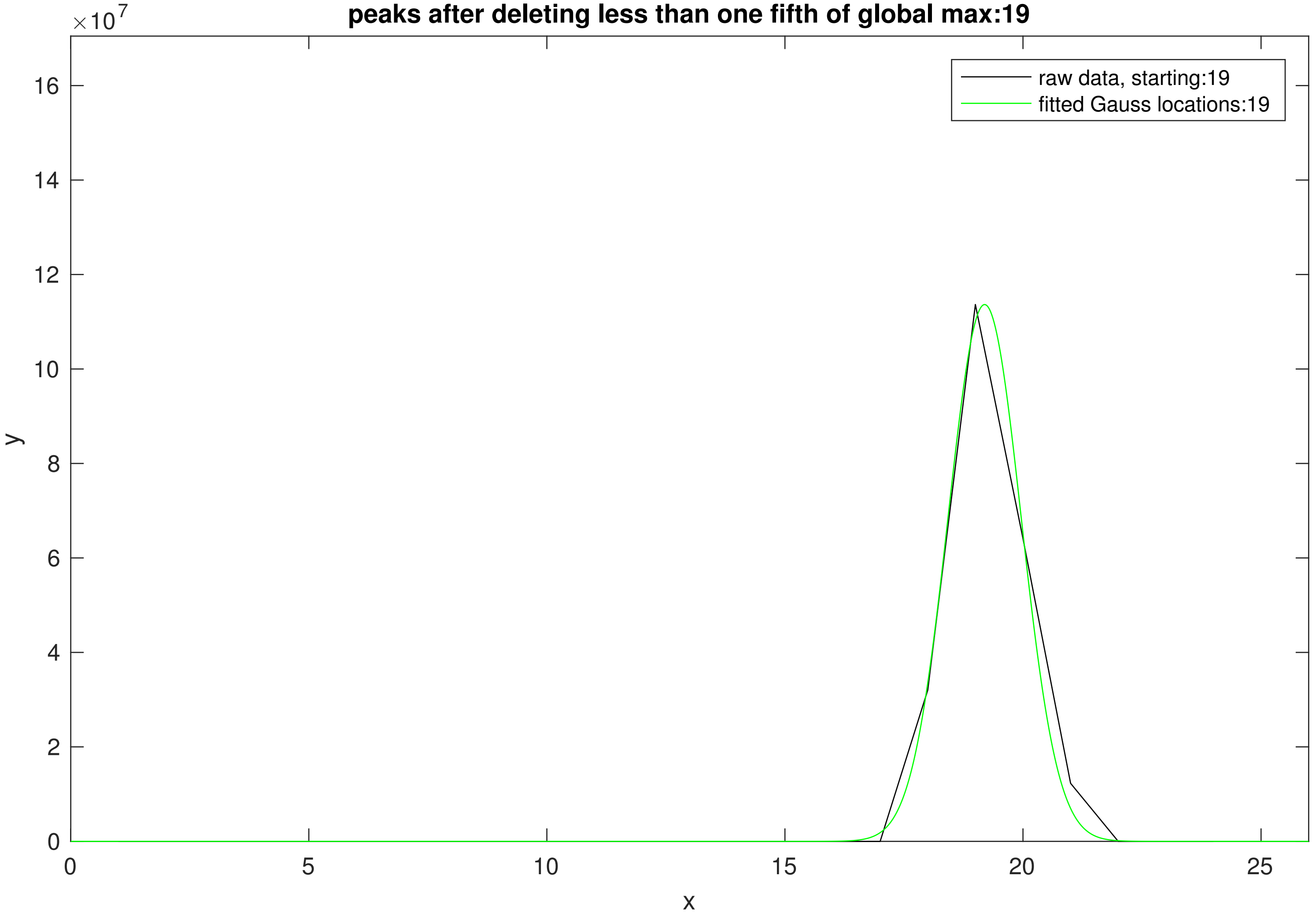

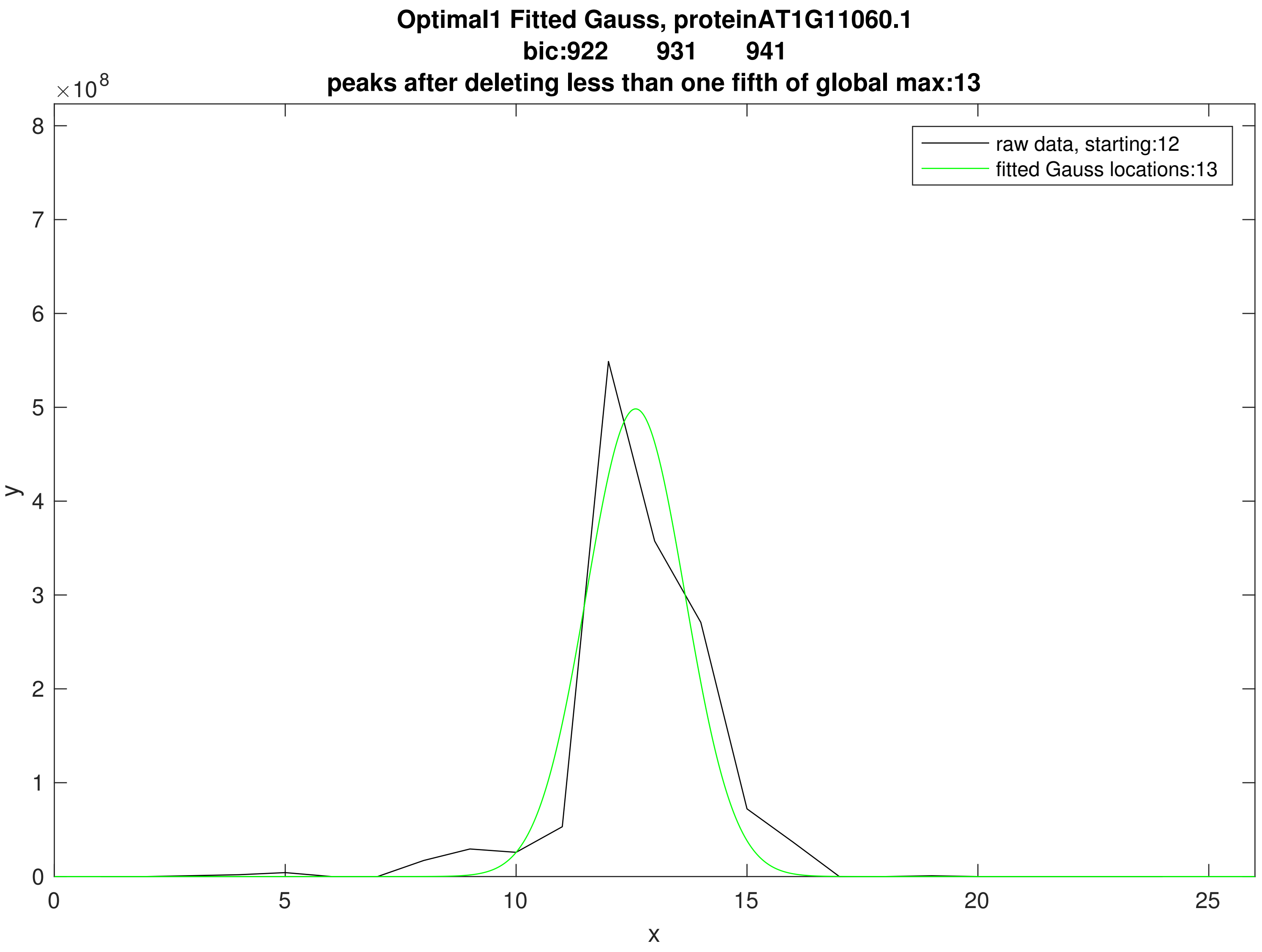

Optimal1 Fitted Gauss, proteinAT1G11060.1
bic:922 931 941
peaks after deleting less than one fifth of global max:13
×10^8
raw data, starting:12
fitted Gauss locations:13
y
x
0
1
2
3
4
5
6
7
8
0
5
10
15
20
25

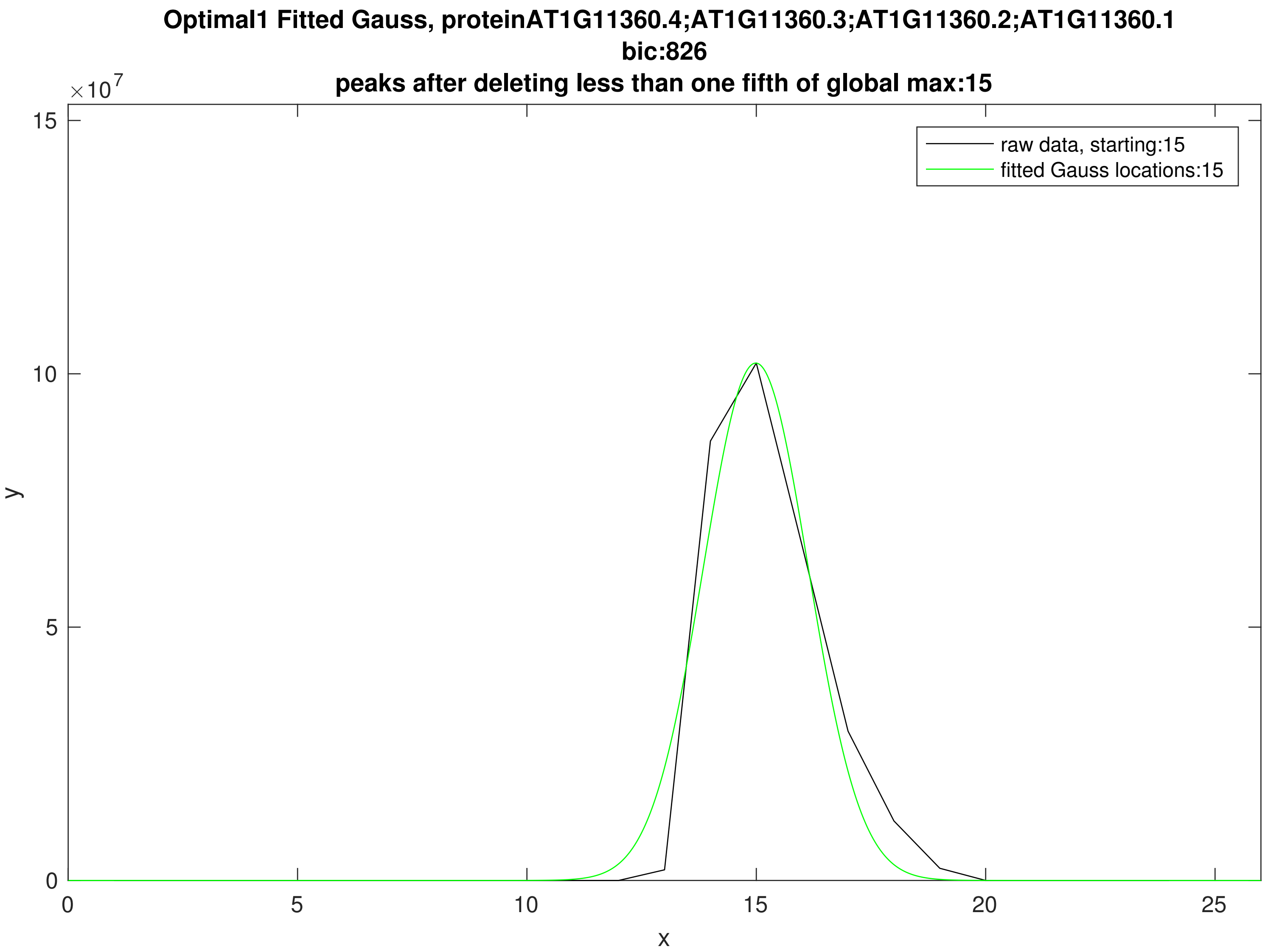
Optimal1 Fitted Gauss, proteinAT1G11360.4;AT1G11360.3;AT1G11360.2;AT1G11360.1
bic:826
peaks after deleting less than one fifth of global max:15
×10^7
15
10
5
0
y
0
5
10
15
20
25
x
raw data, starting:15
fitted Gauss locations:15

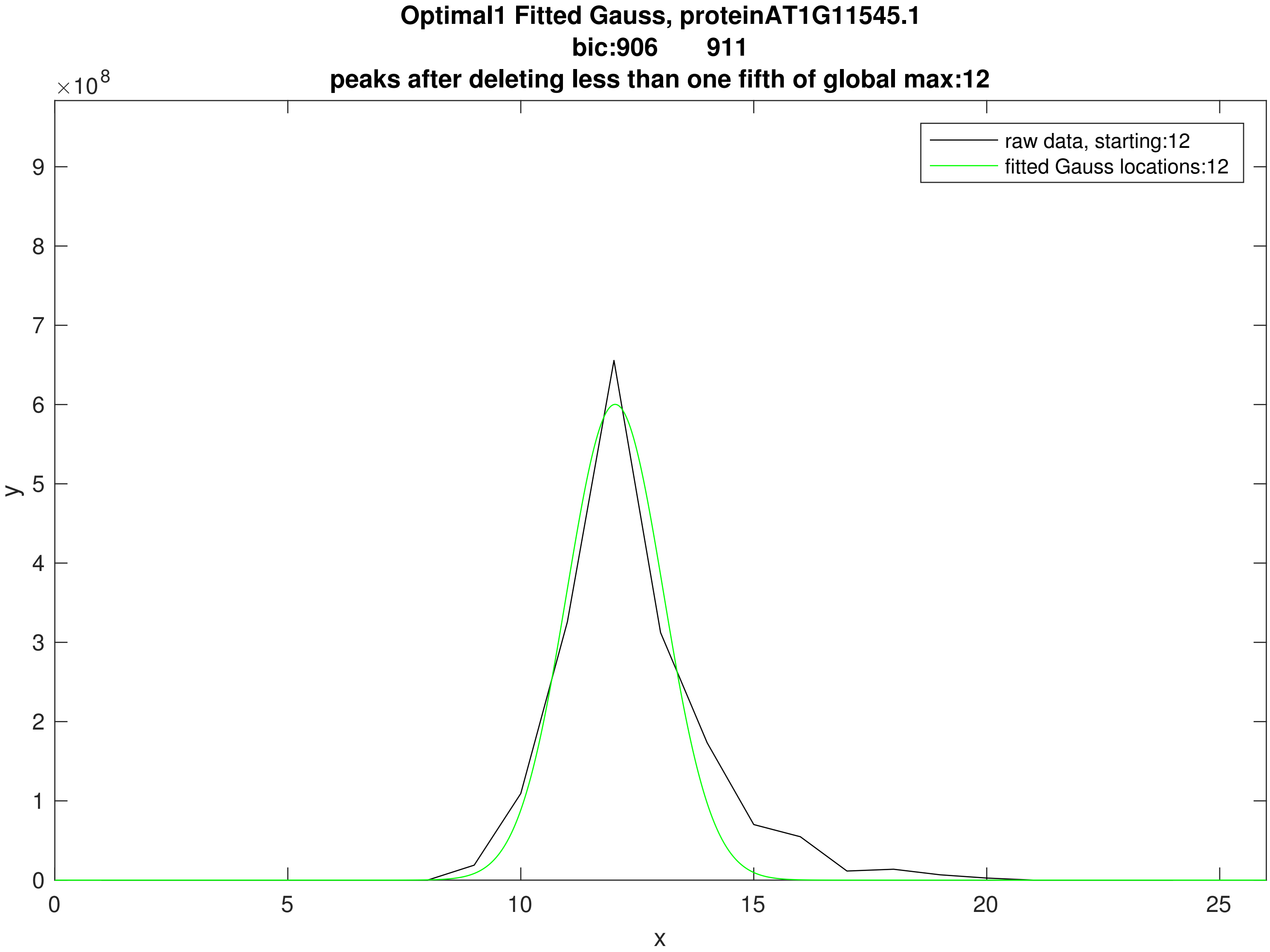
Optimal1 Fitted Gauss, proteinAT1G11545.1
bic:906 911
peaks after deleting less than one fifth of global max:12
×10^8
raw data, starting:12
fitted Gauss locations:12
y
x
0
1
2
3
4
5
6
7
8
9
0
5
10
15
20
25

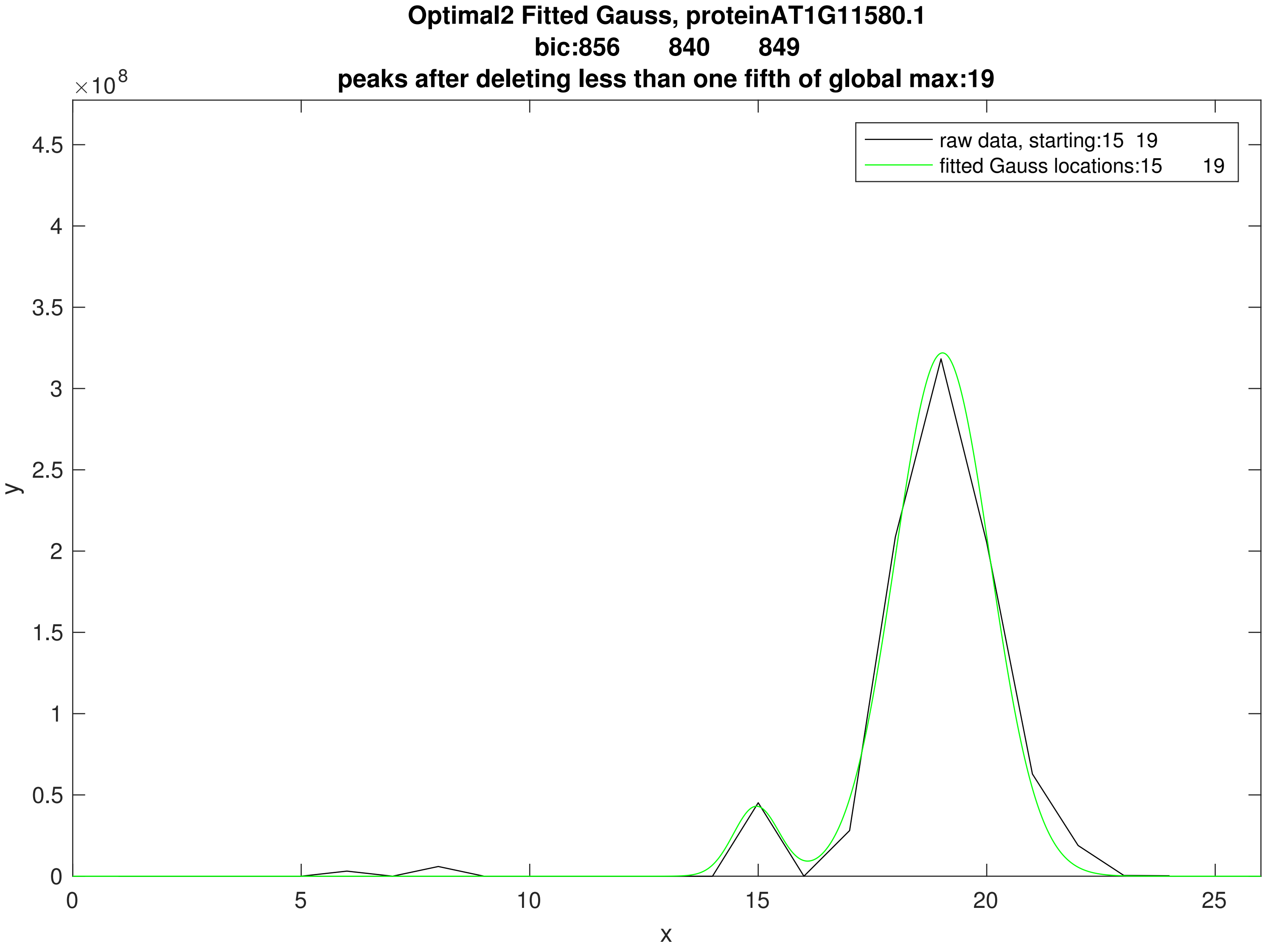
Optimal2 Fitted Gauss, proteinAT1G11580.1
bic:856 840 849
peaks after deleting less than one fifth of global max:19
×10^8
raw data, starting:15 19
fitted Gauss locations:15 19
y
x
0
0.5
1
1.5
2
2.5
3
3.5
4
4.5
5
10
15
20
25

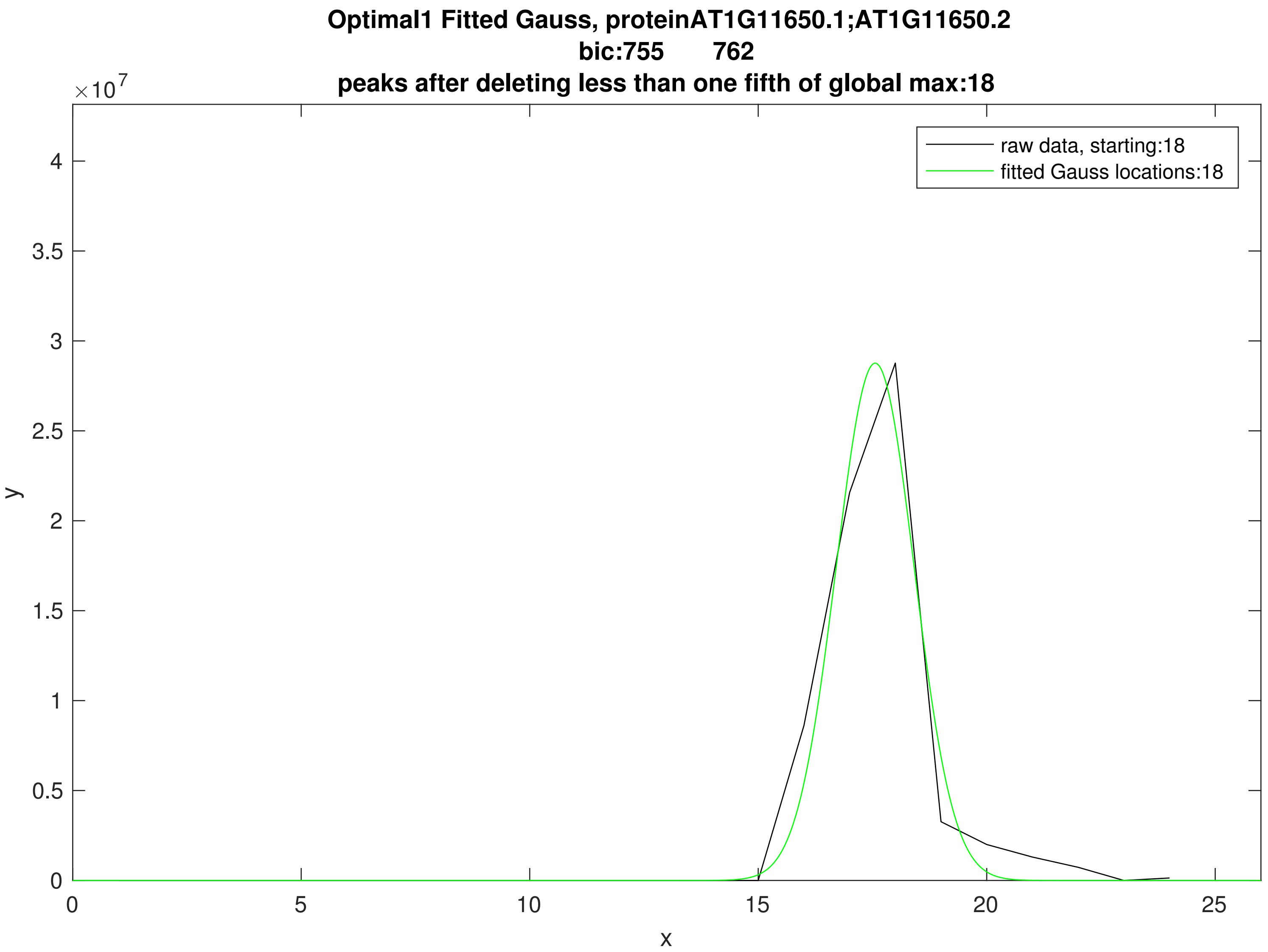

Optimal1 Fitted Gauss, proteinAT1G11650.1;AT1G11650.2
bic:755 762
peaks after deleting less than one fifth of global max:18
×10^7
raw data, starting:18
fitted Gauss locations:18
y
x
0
0.5
1
1.5
2
2.5
3
3.5
4
0
5
10
15
20
25

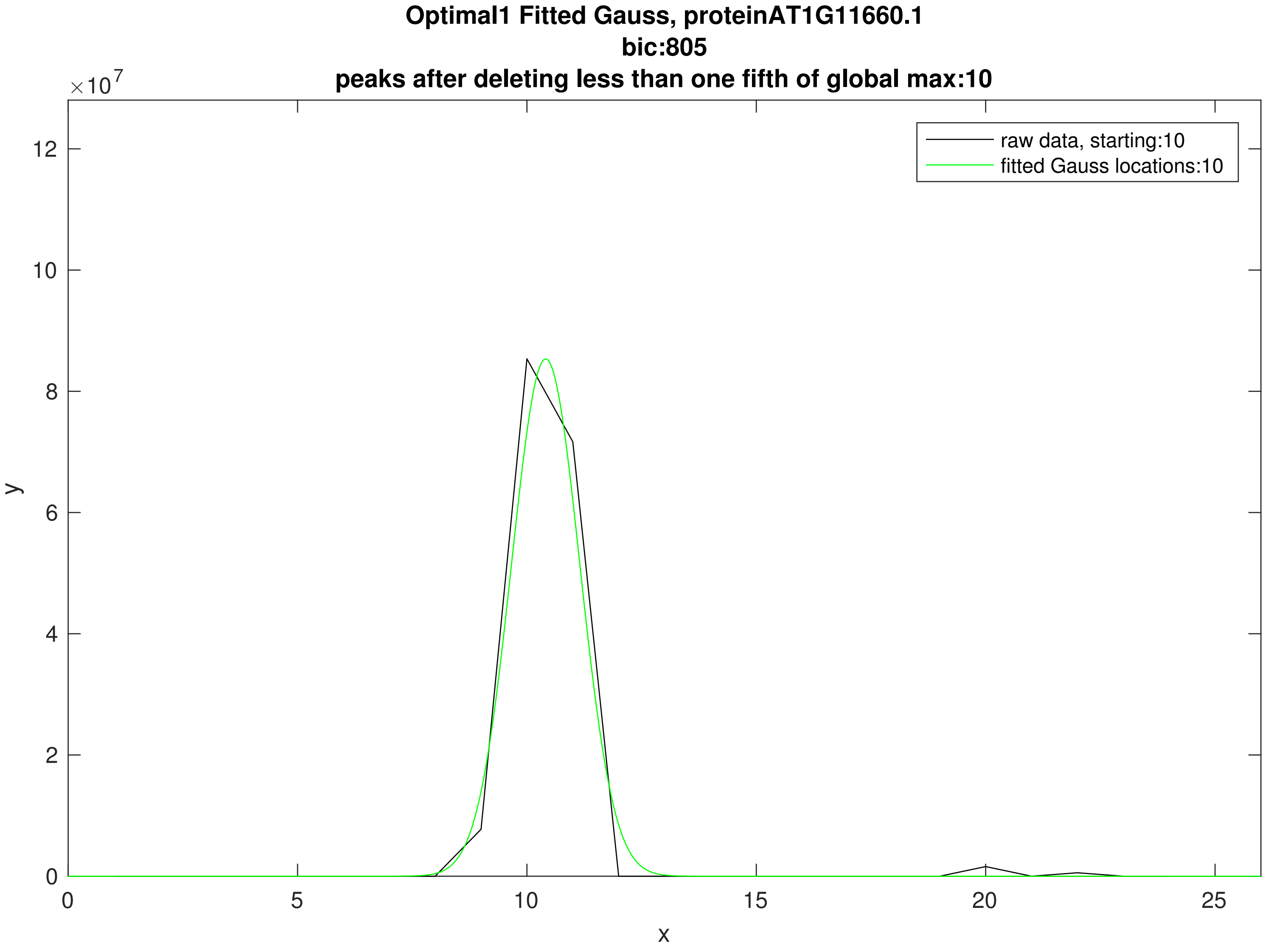

Optimal1 Fitted Gauss, proteinAT1G11660.1
bic:805
peaks after deleting less than one fifth of global max:10
×10^7
12
10
8
6
4
2
0
y
0
5
10
15
20
25
x
raw data, starting:10
fitted Gauss locations:10

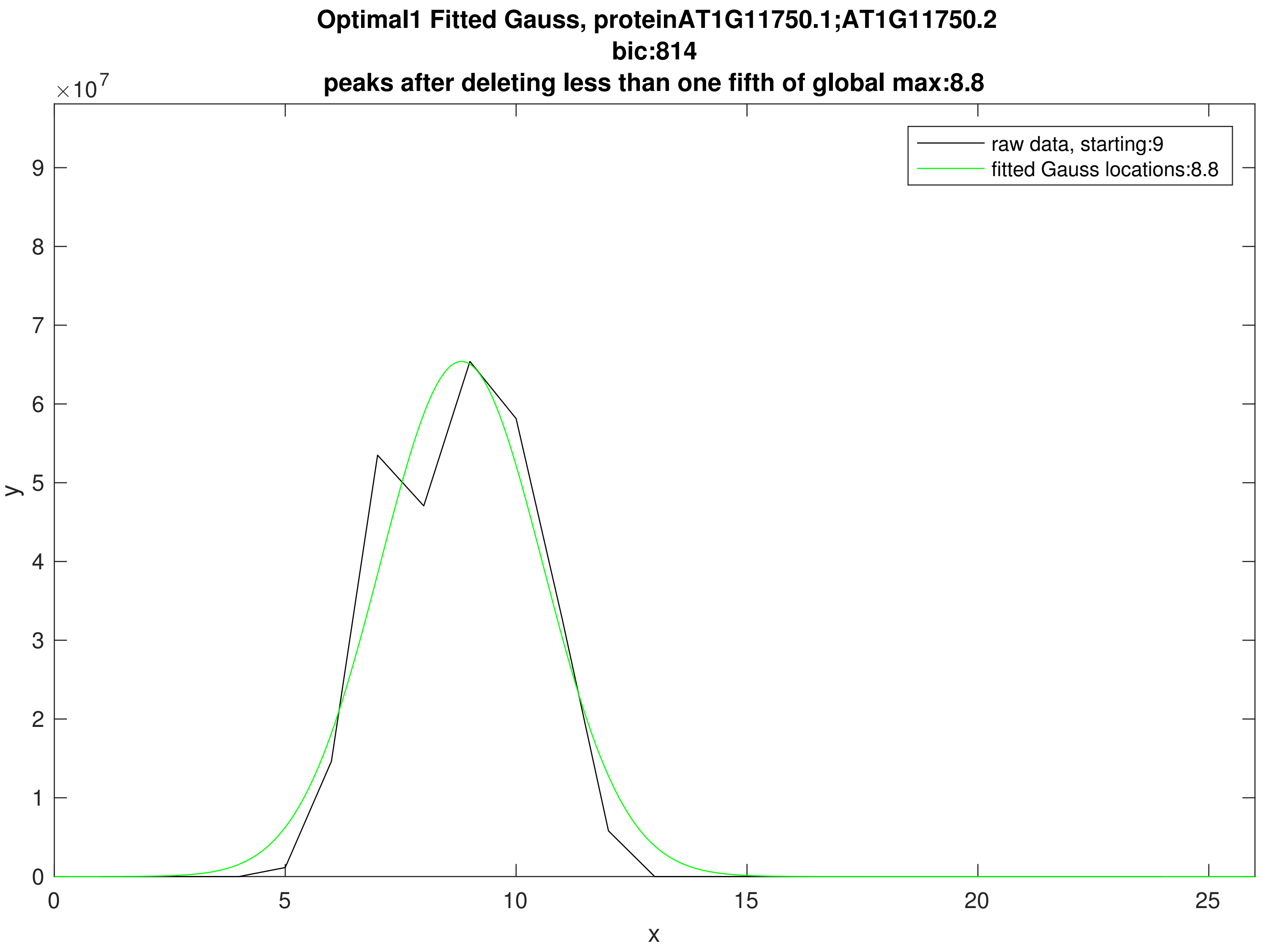

Optimal1 Fitted Gauss, proteinAT1G11750.1;AT1G11750.2
bic:814
peaks after deleting less than one fifth of global max:8.8
×10^7
raw data, starting:9
fitted Gauss locations:8.8
y
x
0
1
2
3
4
5
6
7
8
9
0
5
10
15
20
25

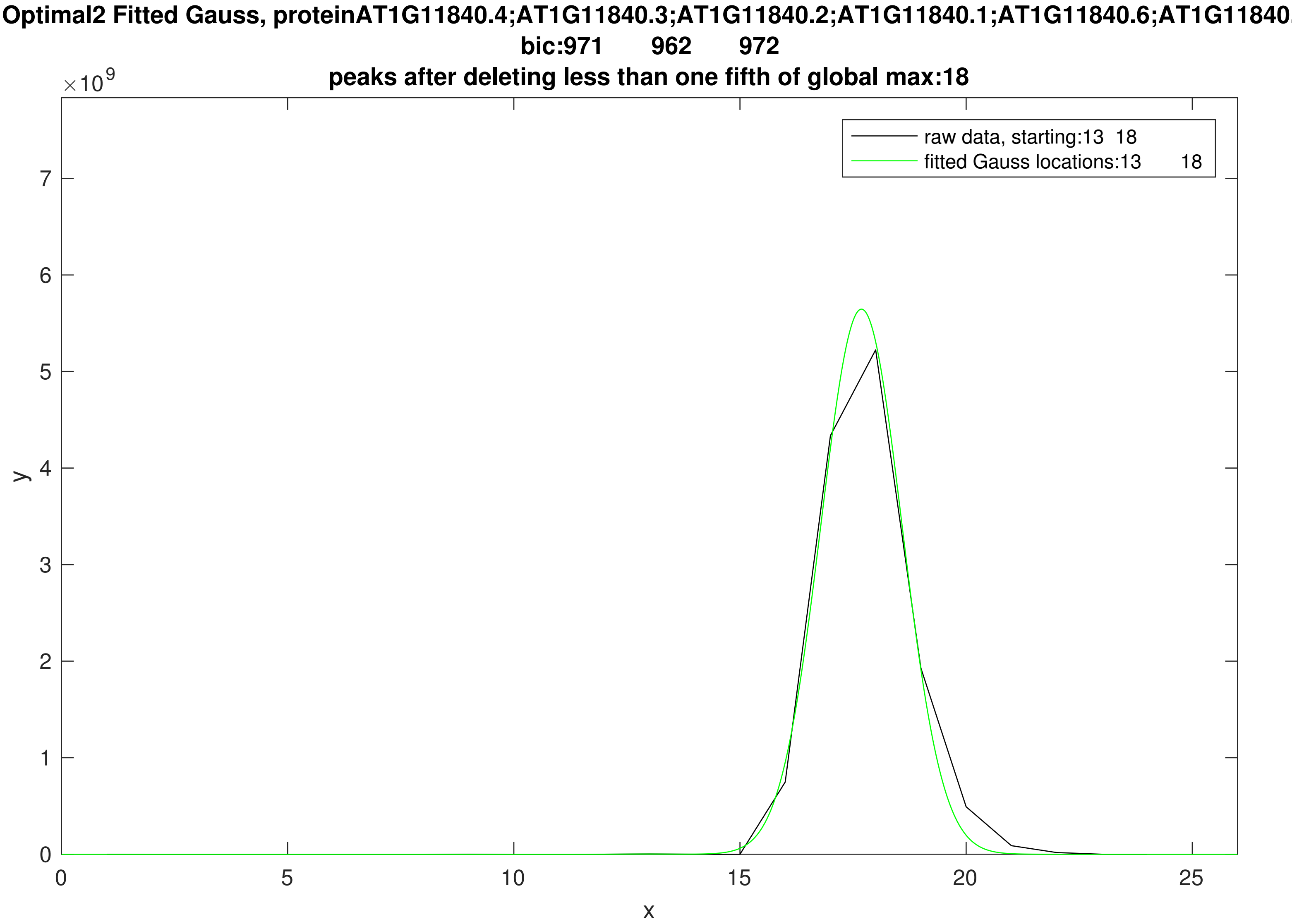
Optimal2 Fitted Gauss, proteinAT1G11840.4;AT1G11840.3;AT1G11840.2;AT1G11840.1;AT1G11840.6;AT1G11840.
bic:971 962 972
peaks after deleting less than one fifth of global max:18
×10^9
raw data, starting:13 18
fitted Gauss locations:13 18
y
x
0
1
2
3
4
5
6
7
0
5
10
15
20
25

Optimal1 Fitted Gauss, proteinAT1G11860.3;AT1G11860.2;AT1G11860.1
bic:930 939 948
peaks after deleting less than one fifth of global max:17

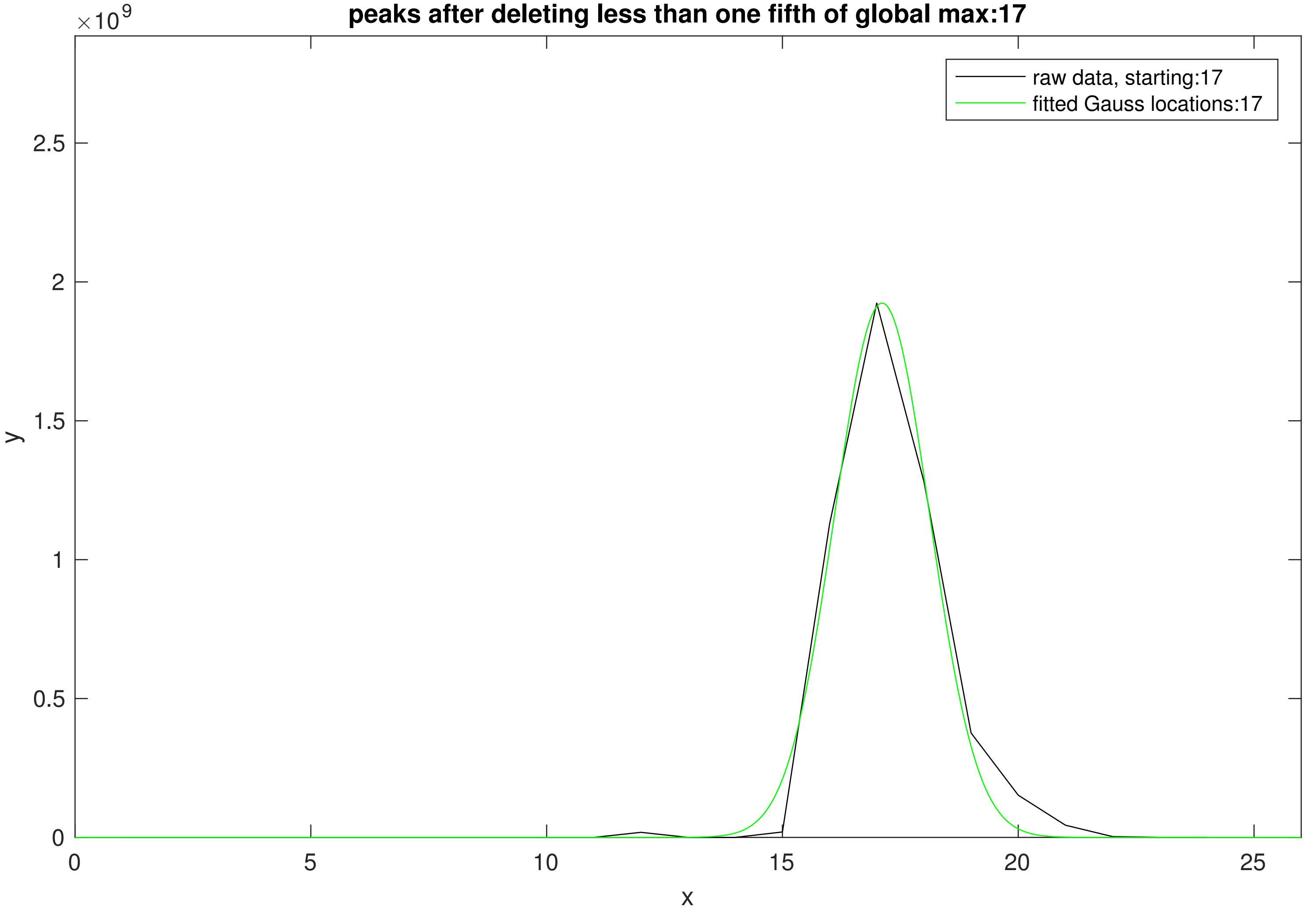

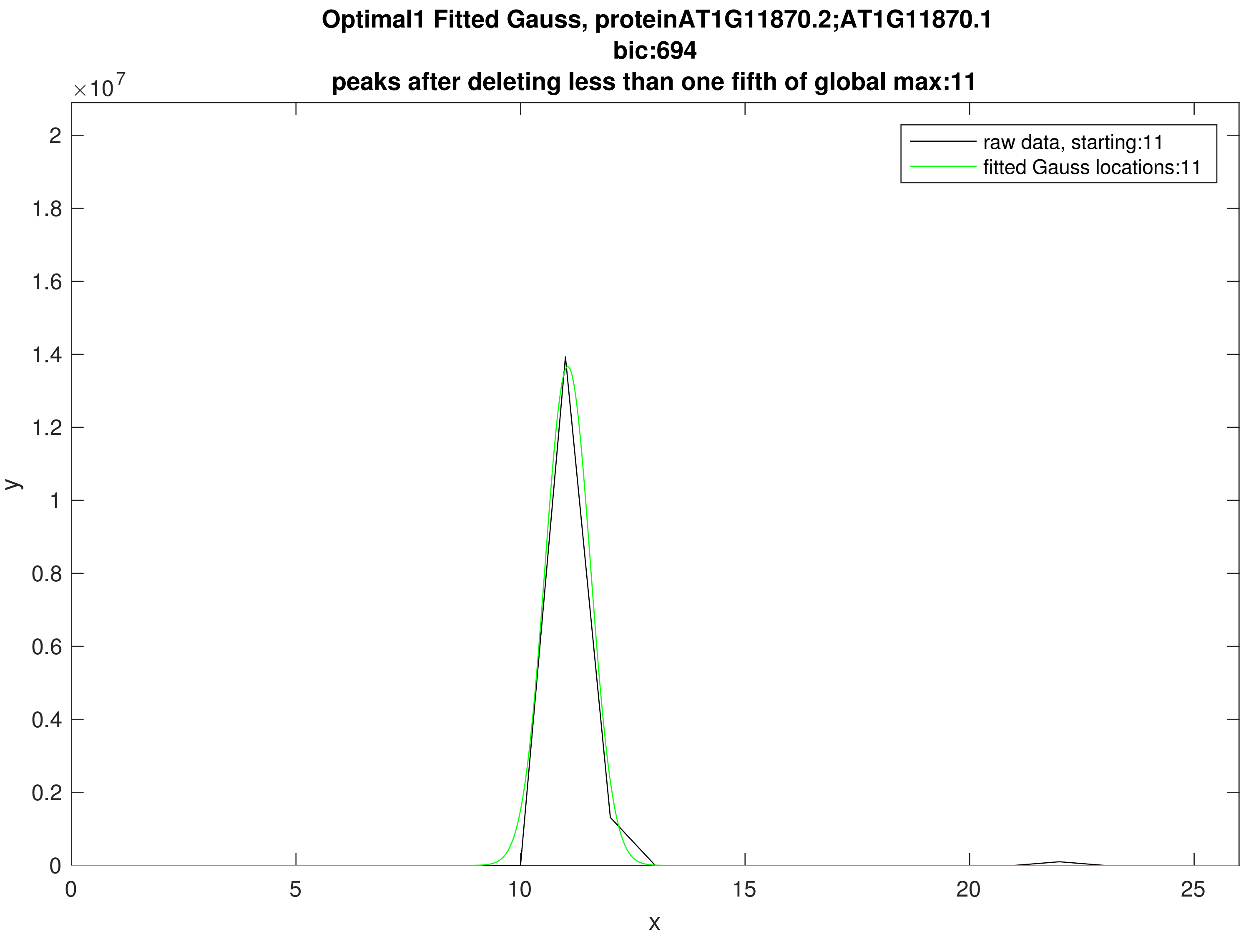
Optimal1 Fitted Gauss, proteinAT1G11870.2;AT1G11870.1
bic:694
peaks after deleting less than one fifth of global max:11
×10^7
2
1.8
1.6
1.4
1.2
1
0.8
0.6
0.4
0.2
0
y
0
5
10
15
20
25
x
raw data, starting:11
fitted Gauss locations:11

Optimal1 Fitted Gauss, proteinAT1G11910.1
bic:796 801
peaks after deleting less than one fifth of global max:5.1

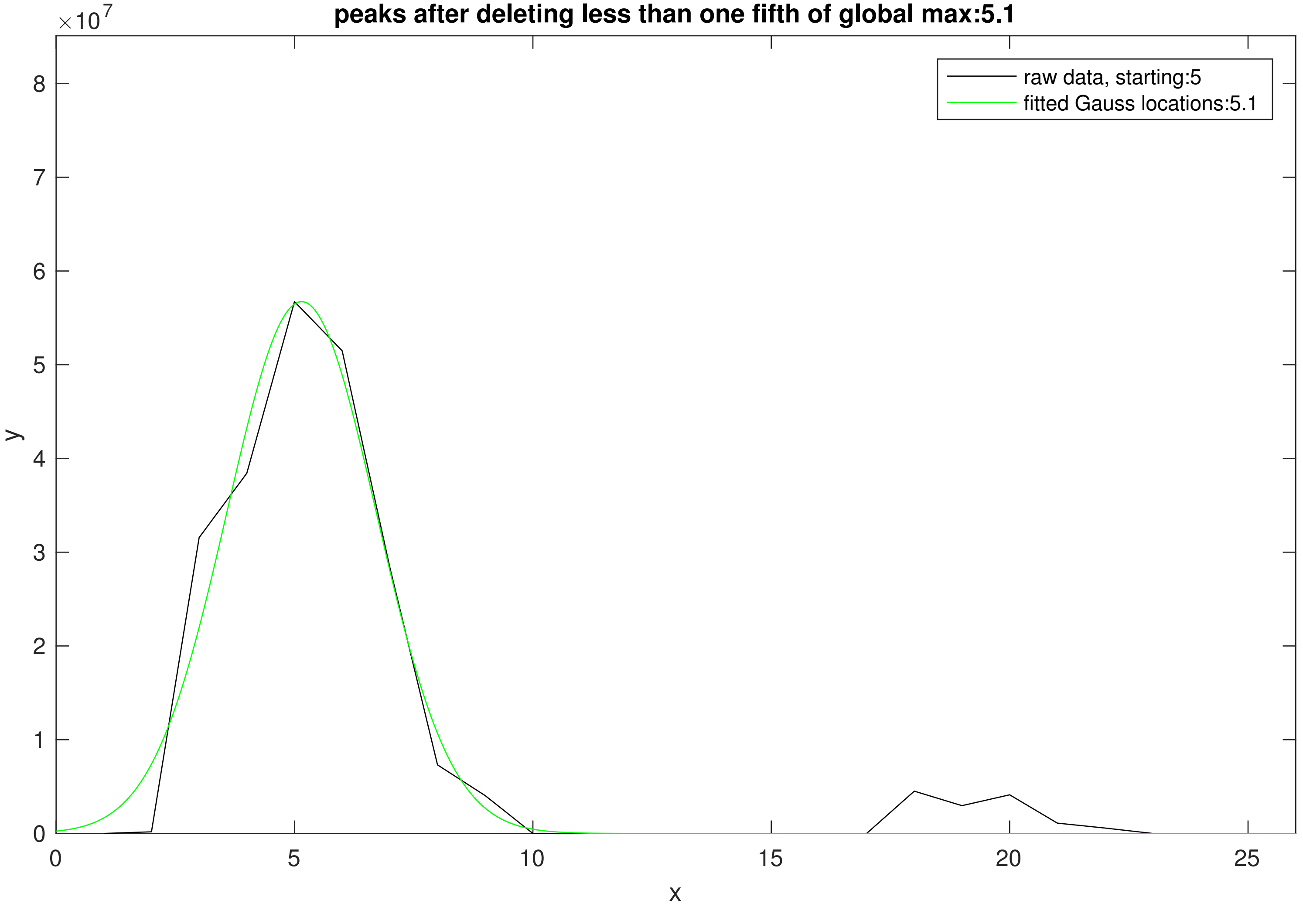

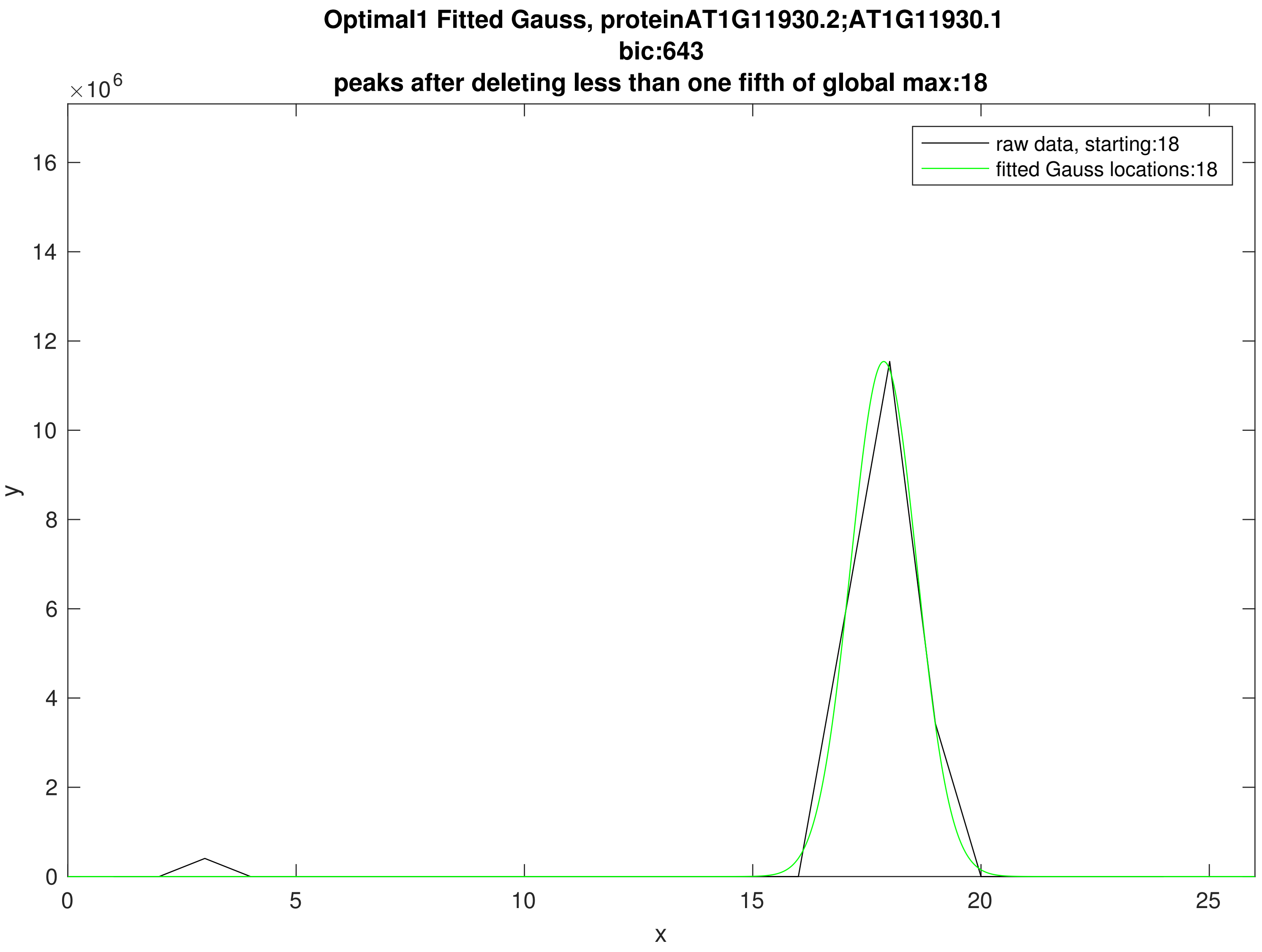
Optimal1 Fitted Gauss, proteinAT1G11930.2;AT1G11930.1
bic:643
peaks after deleting less than one fifth of global max:18
×10^6
raw data, starting:18
fitted Gauss locations:18
y
x
0
2
4
6
8
10
12
14
16
0
5
10
15
20
25

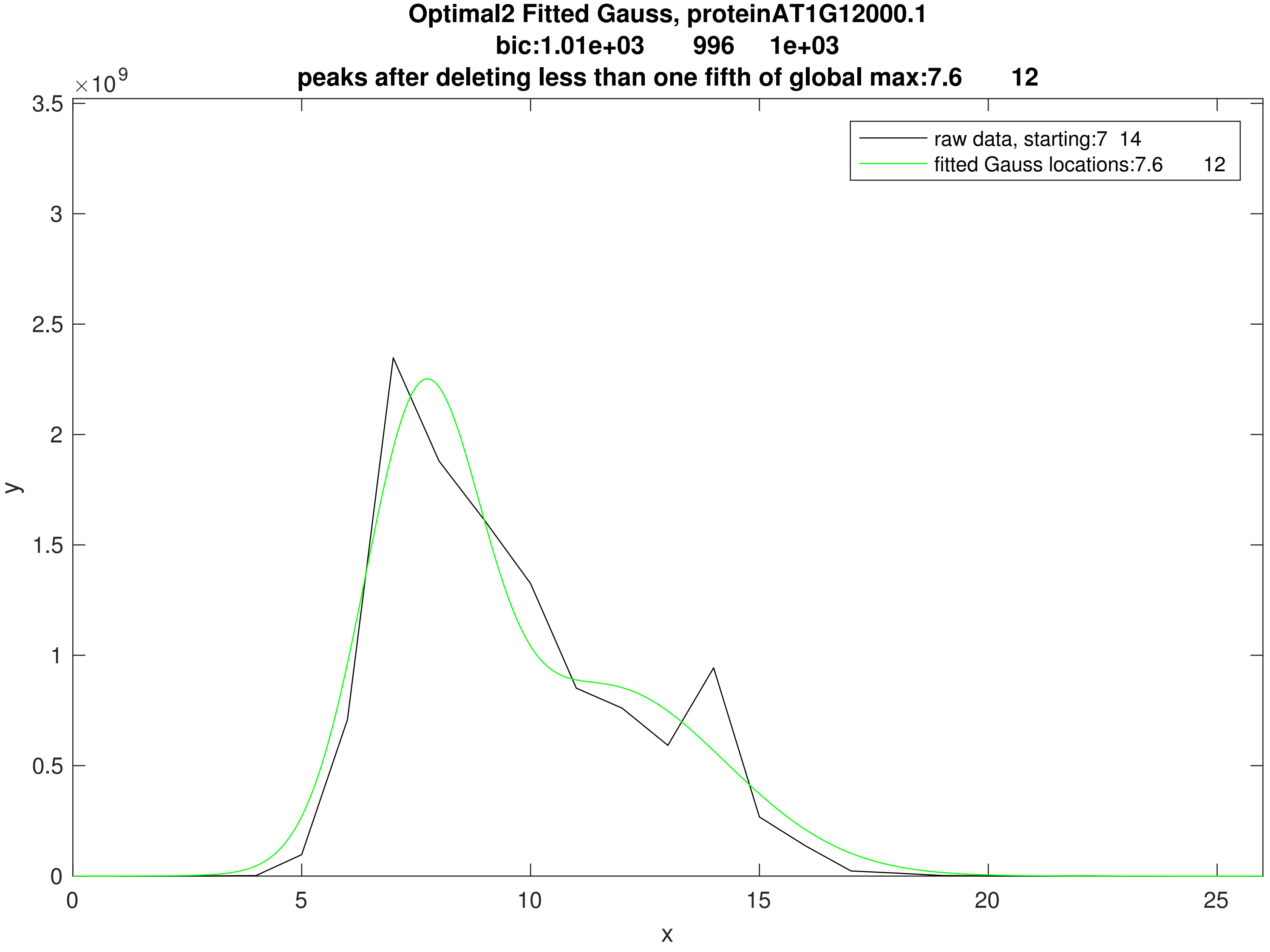
Optimal2 Fitted Gauss, proteinAT1G12000.1
bic:1.01e+03 996 1e+03
peaks after deleting less than one fifth of global max:7.6 12
×10^9
3.5
3
2.5
2
1.5
1
0.5
0
y
0
5
10
15
20
25
x
raw data, starting:7 14
fitted Gauss locations:7.6 12

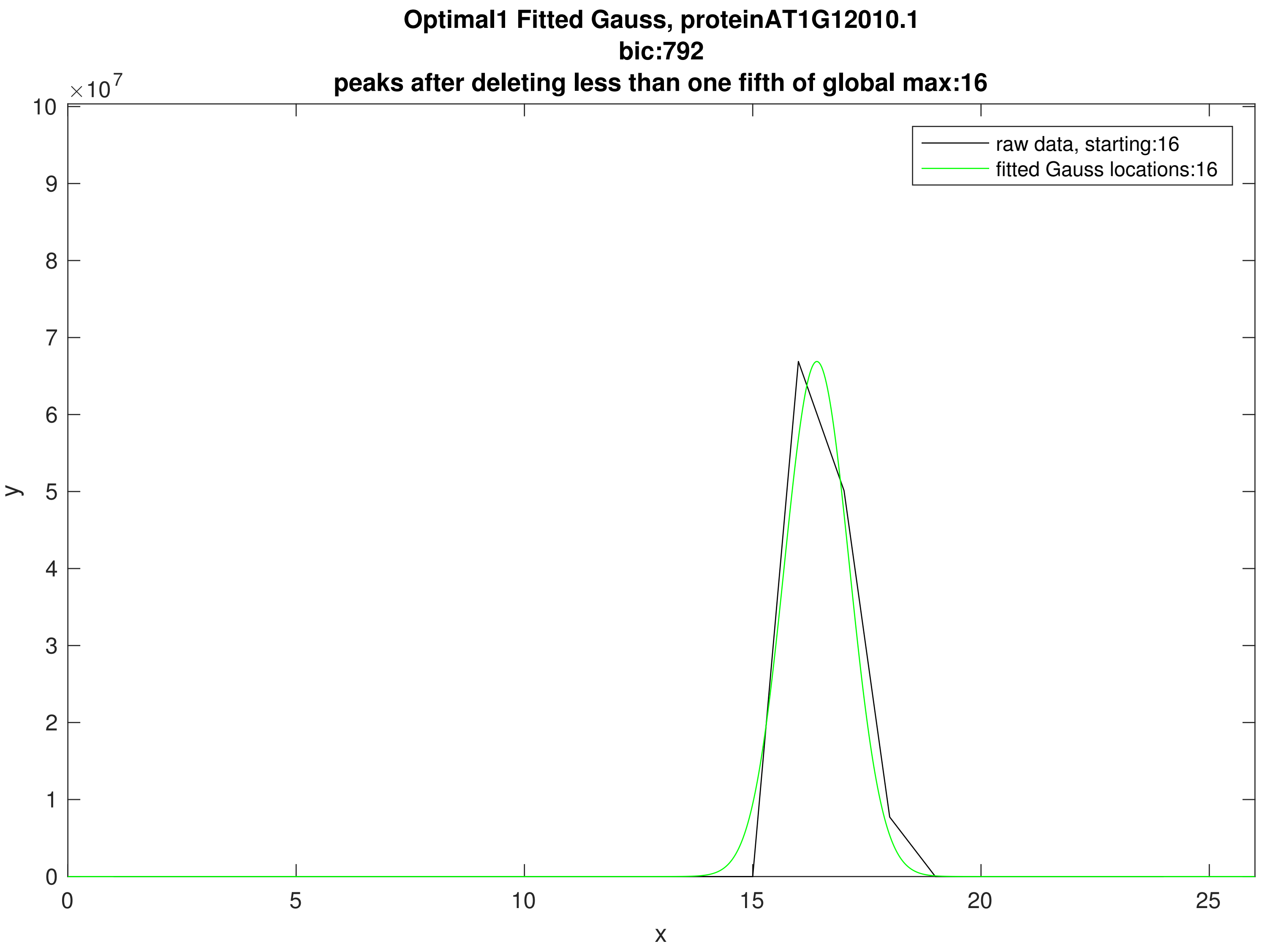
Optimal1 Fitted Gauss, proteinAT1G12010.1
bic:792
peaks after deleting less than one fifth of global max:16
×10^7
10
9
8
7
6
5
4
3
2
1
0
y
0
5
10
15
20
25
x
raw data, starting:16
fitted Gauss locations:16

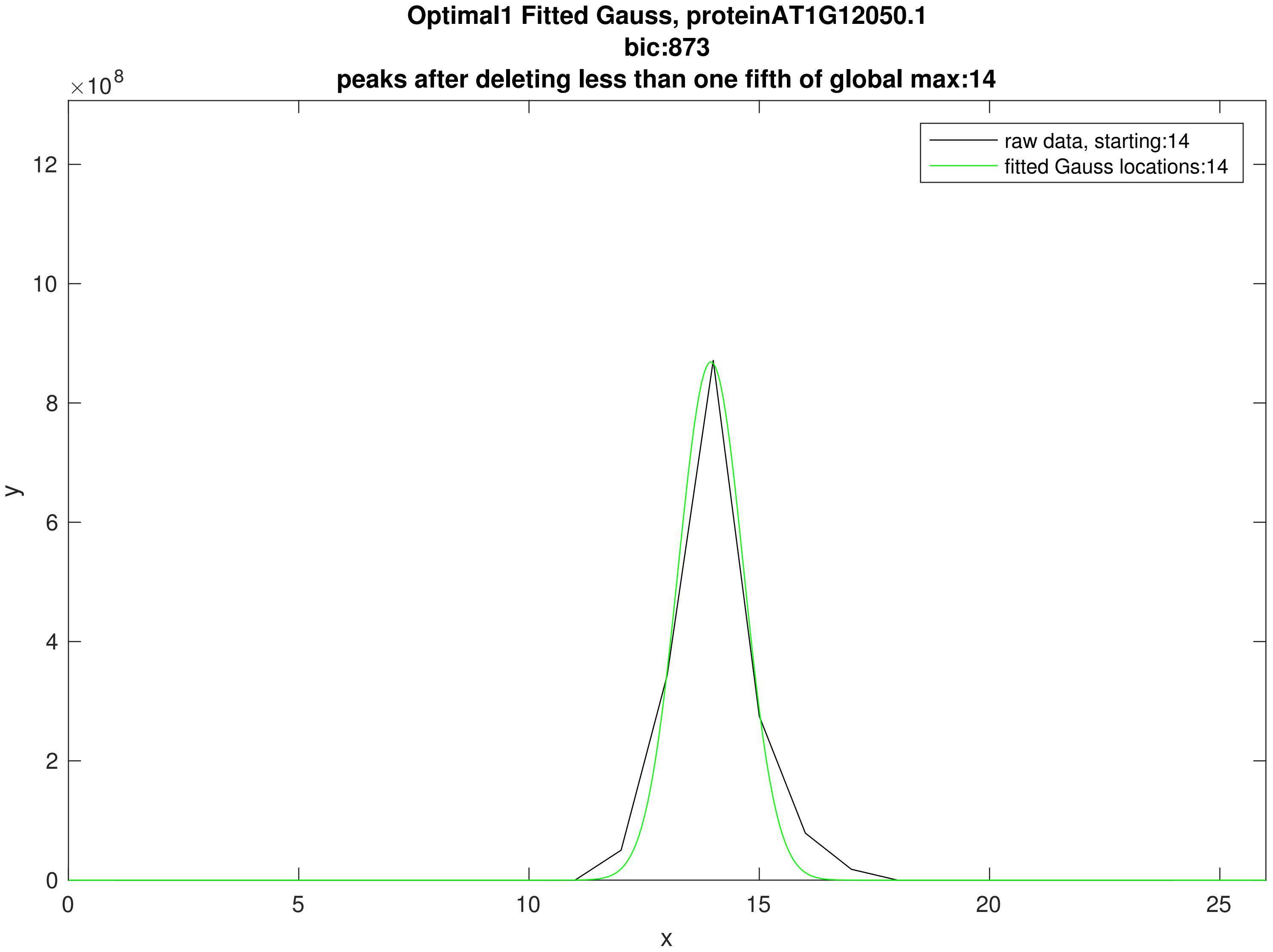

Optimal1 Fitted Gauss, proteinAT1G12050.1
bic:873
peaks after deleting less than one fifth of global max:14
×10^8
raw data, starting:14
fitted Gauss locations:14
y
x
0
2
4
6
8
10
12
0
5
10
15
20
25

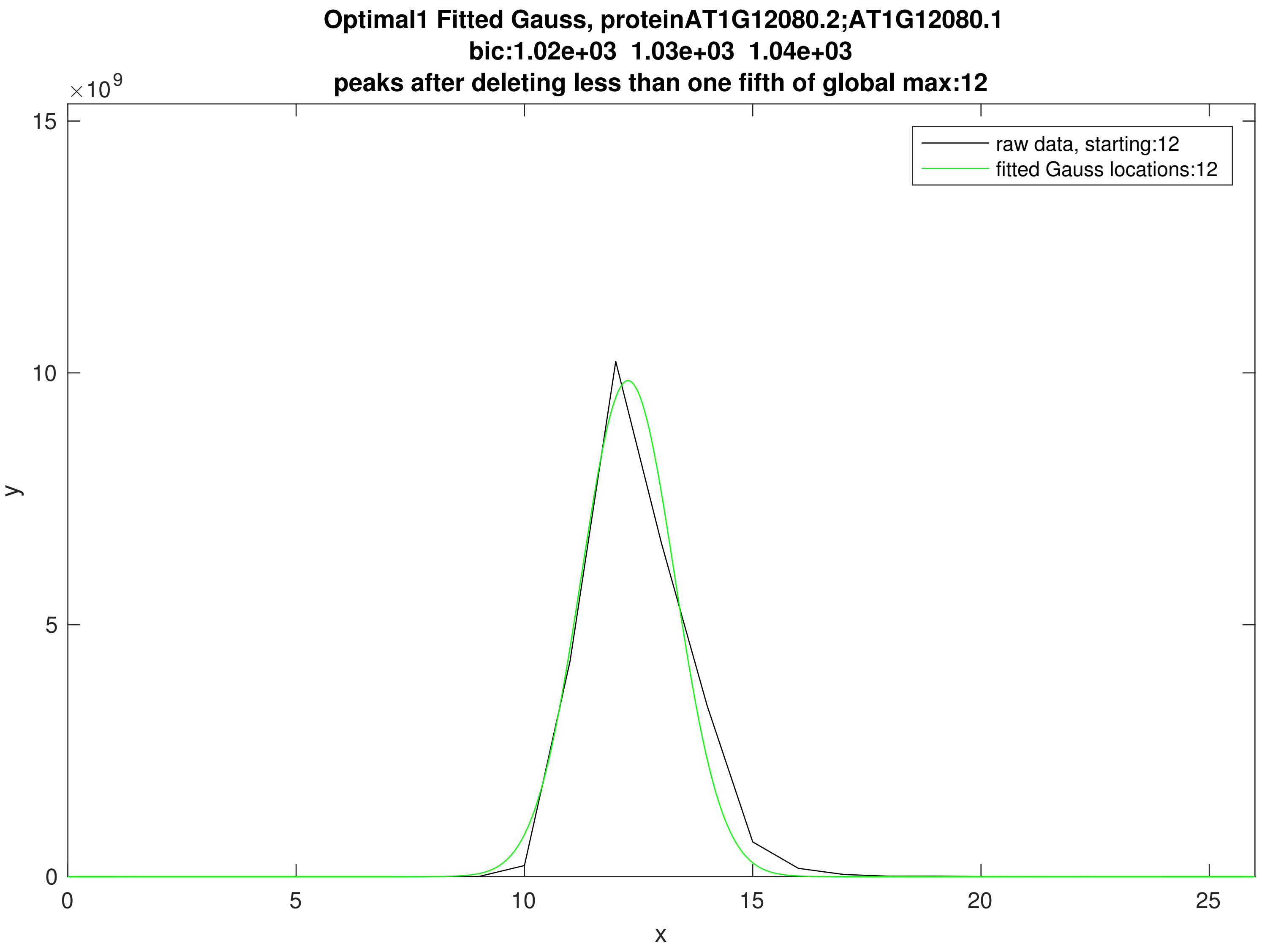
Optimal1 Fitted Gauss, proteinAT1G12080.2;AT1G12080.1
bic:1.02e+03 1.03e+03 1.04e+03
peaks after deleting less than one fifth of global max:12
×10^9
15
10
5
0
y
0
5
10
15
20
25
x
raw data, starting:12
fitted Gauss locations:12

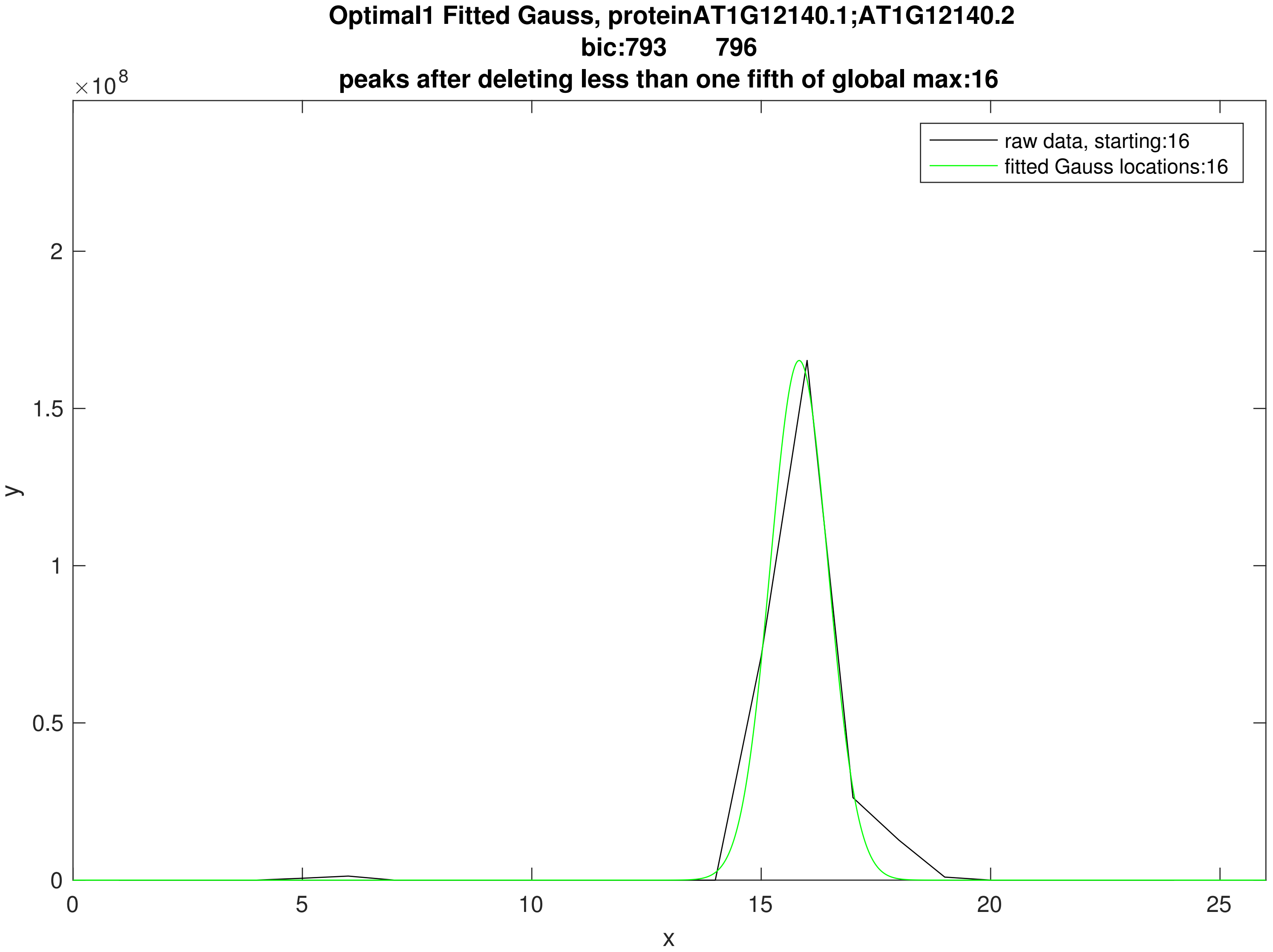
Optimal1 Fitted Gauss, proteinAT1G12140.1;AT1G12140.2
bic:793 796
peaks after deleting less than one fifth of global max:16
×10^8
raw data, starting:16
fitted Gauss locations:16
y
x
0
0.5
1
1.5
2
0
5
10
15
20
25

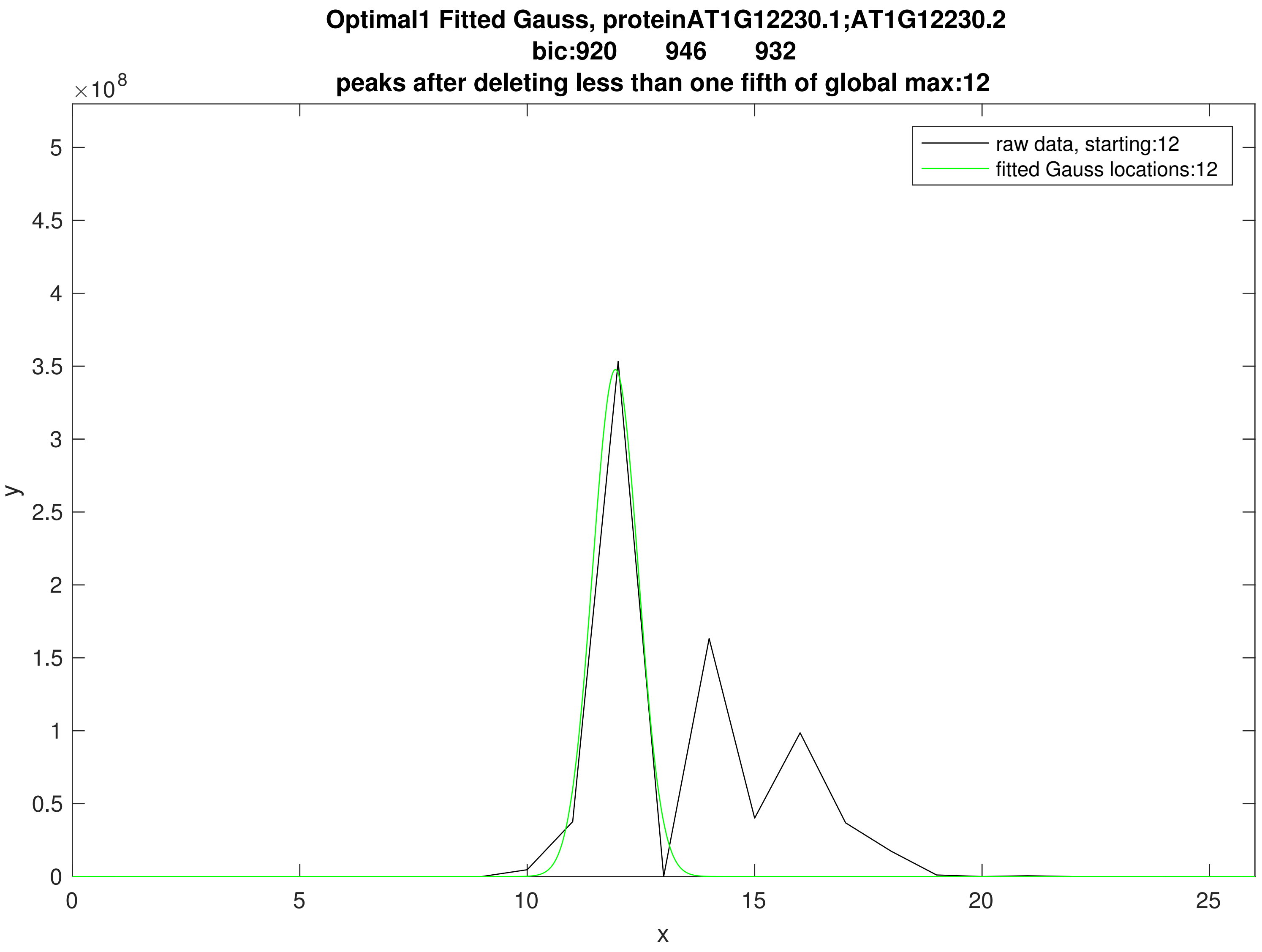
Optimal1 Fitted Gauss, proteinAT1G12230.1;AT1G12230.2
bic:920 946 932
peaks after deleting less than one fifth of global max:12
×10^8
raw data, starting:12
fitted Gauss locations:12
y
x
0
0.5
1
1.5
2
2.5
3
3.5
4
4.5
5
0
5
10
15
20
25

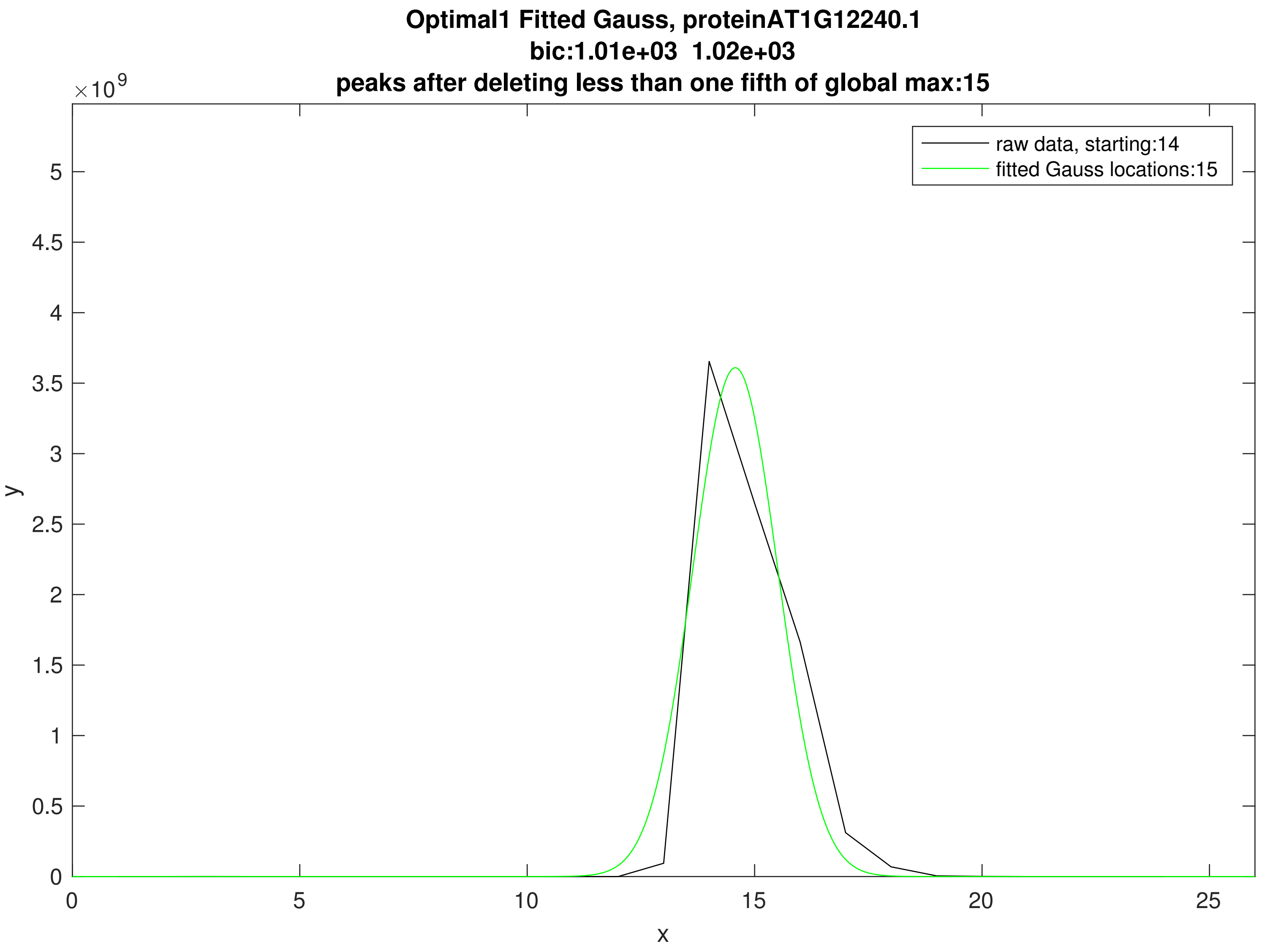

Optimal1 Fitted Gauss, proteinAT1G12240.1
bic:1.01e+03  1.02e+03
peaks after deleting less than one fifth of global max:15
×10^9
raw data, starting:14
fitted Gauss locations:15
y
x
0
0.5
1
1.5
2
2.5
3
3.5
4
4.5
5
0
5
10
15
20
25

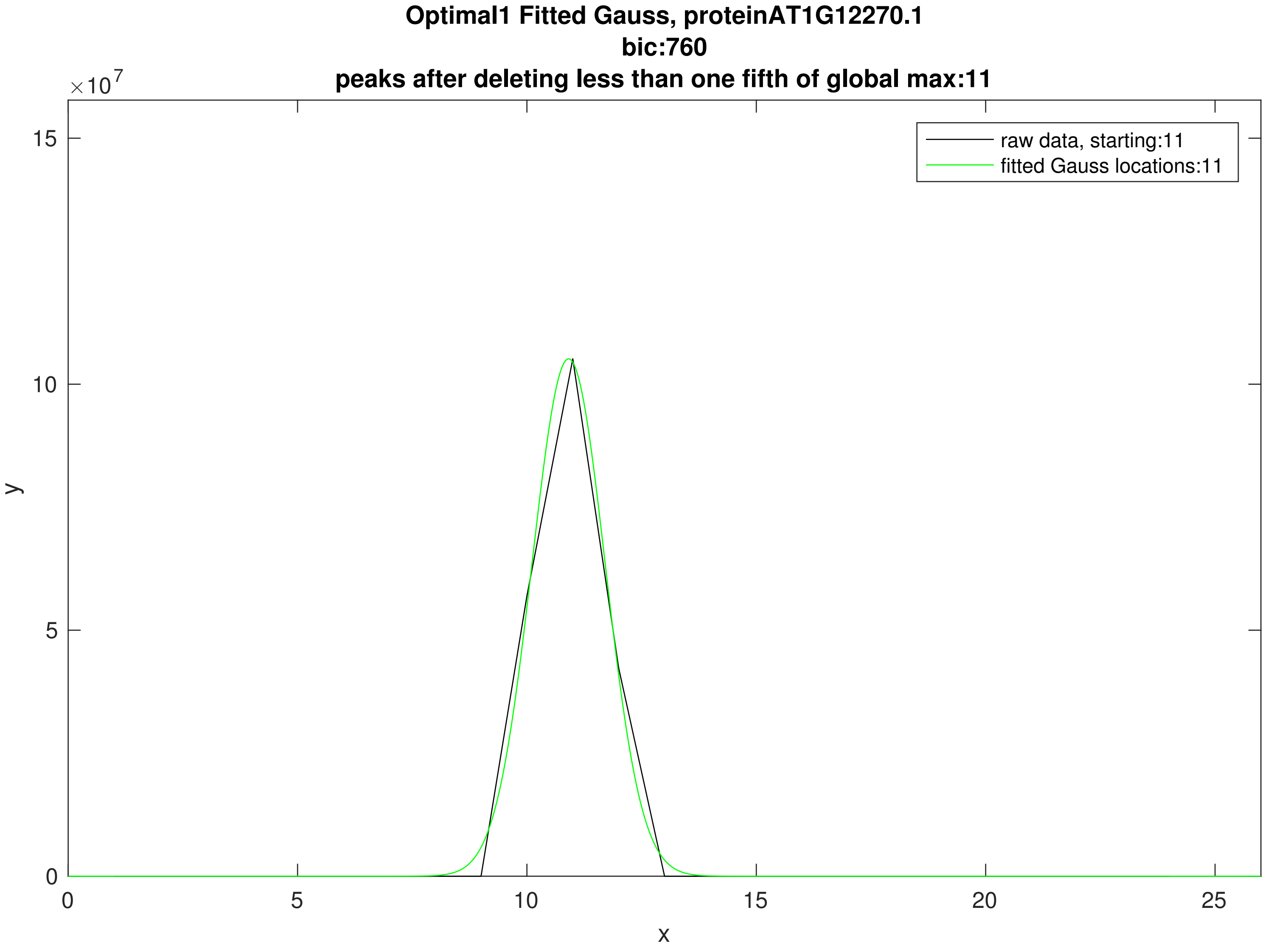
Optimal1 Fitted Gauss, proteinAT1G12270.1
bic:760
peaks after deleting less than one fifth of global max:11
×10^7
15
10
5
0
y
0
5
10
15
20
25
x
raw data, starting:11
fitted Gauss locations:11

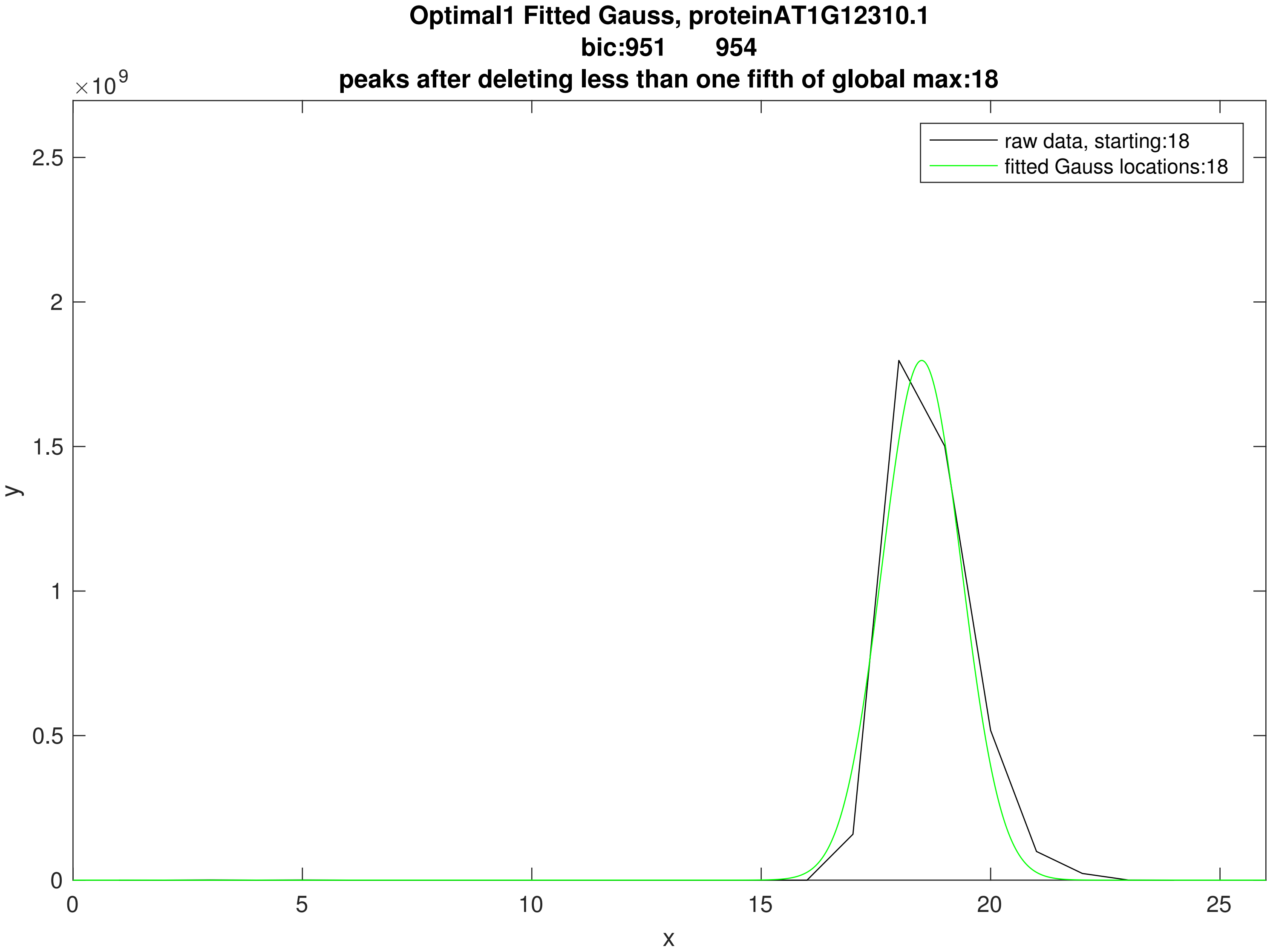

Optimal1 Fitted Gauss, proteinAT1G12310.1
bic:951 954
peaks after deleting less than one fifth of global max:18
×10^9
2.5
2
1.5
1
0.5
0
y
0
5
10
15
20
25
x
raw data, starting:18
fitted Gauss locations:18

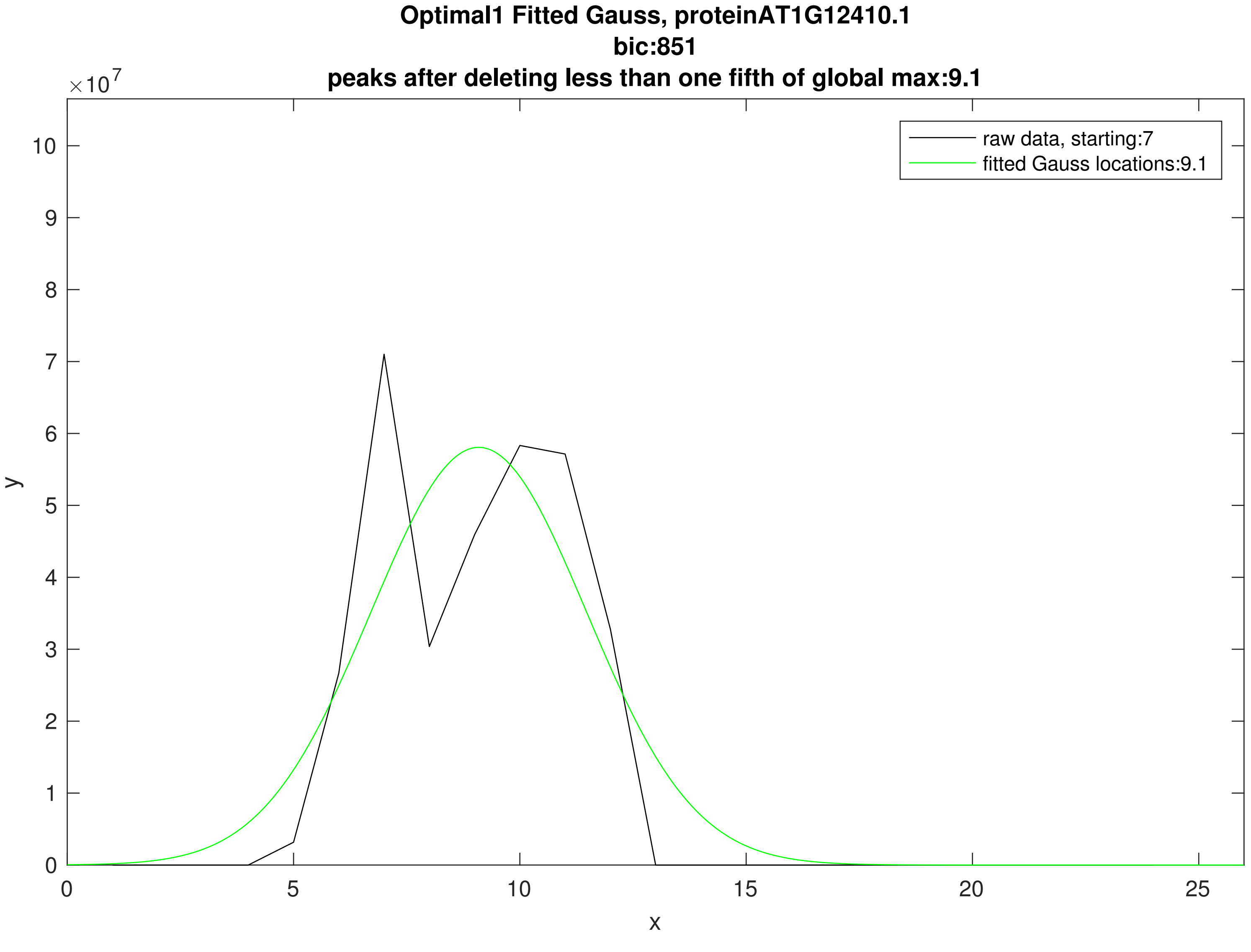

Optimal1 Fitted Gauss, proteinAT1G12410.1
bic:851
peaks after deleting less than one fifth of global max:9.1
×10^7
raw data, starting:7
fitted Gauss locations:9.1
y
x
0
1
2
3
4
5
6
7
8
9
10
0
5
10
15
20
25

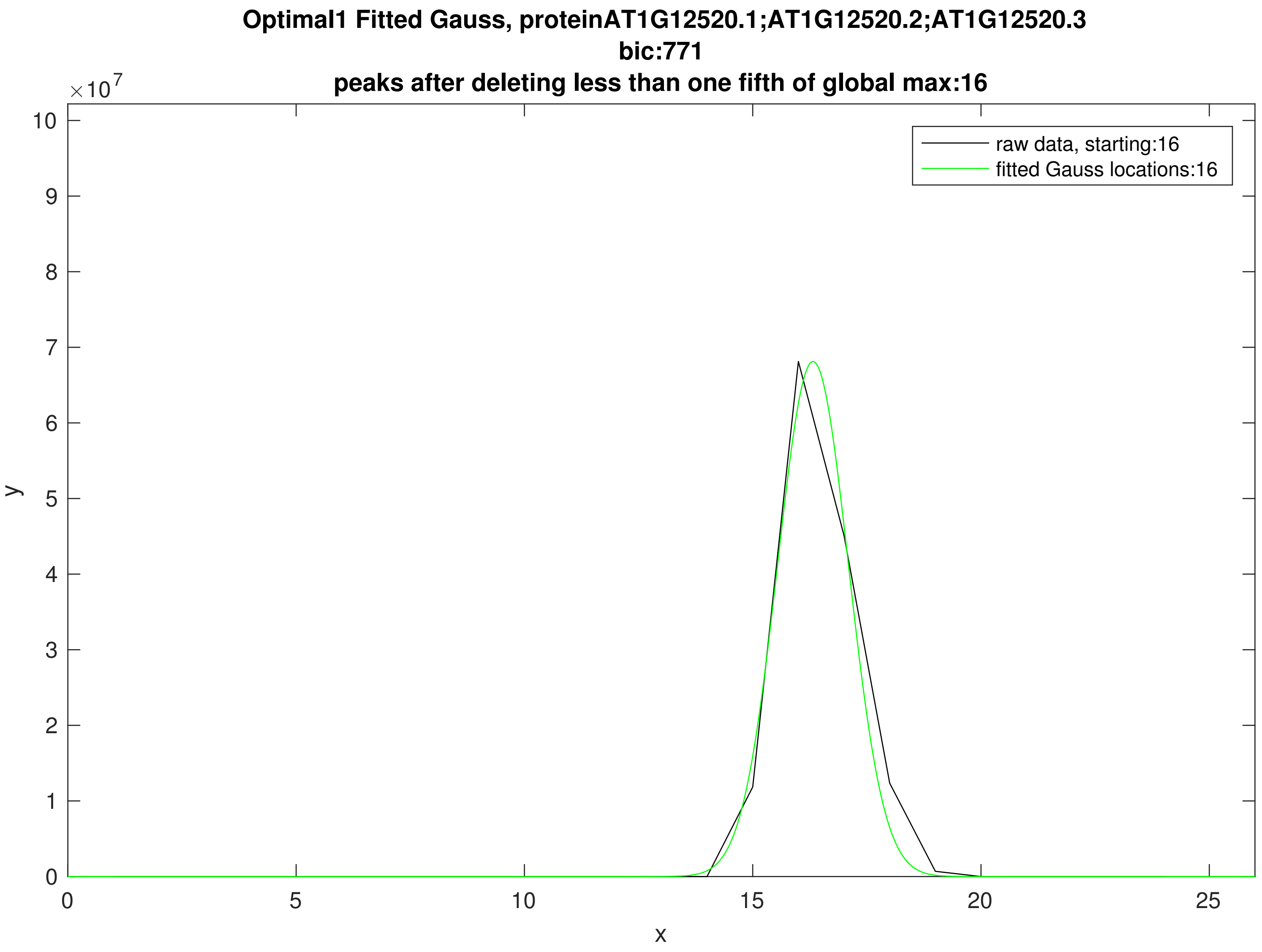
Optimal1 Fitted Gauss, proteinAT1G12520.1;AT1G12520.2;AT1G12520.3
bic:771
peaks after deleting less than one fifth of global max:16
×10^7
raw data, starting:16
fitted Gauss locations:16
y
x
0
1
2
3
4
5
6
7
8
9
10
0
5
10
15
20
25

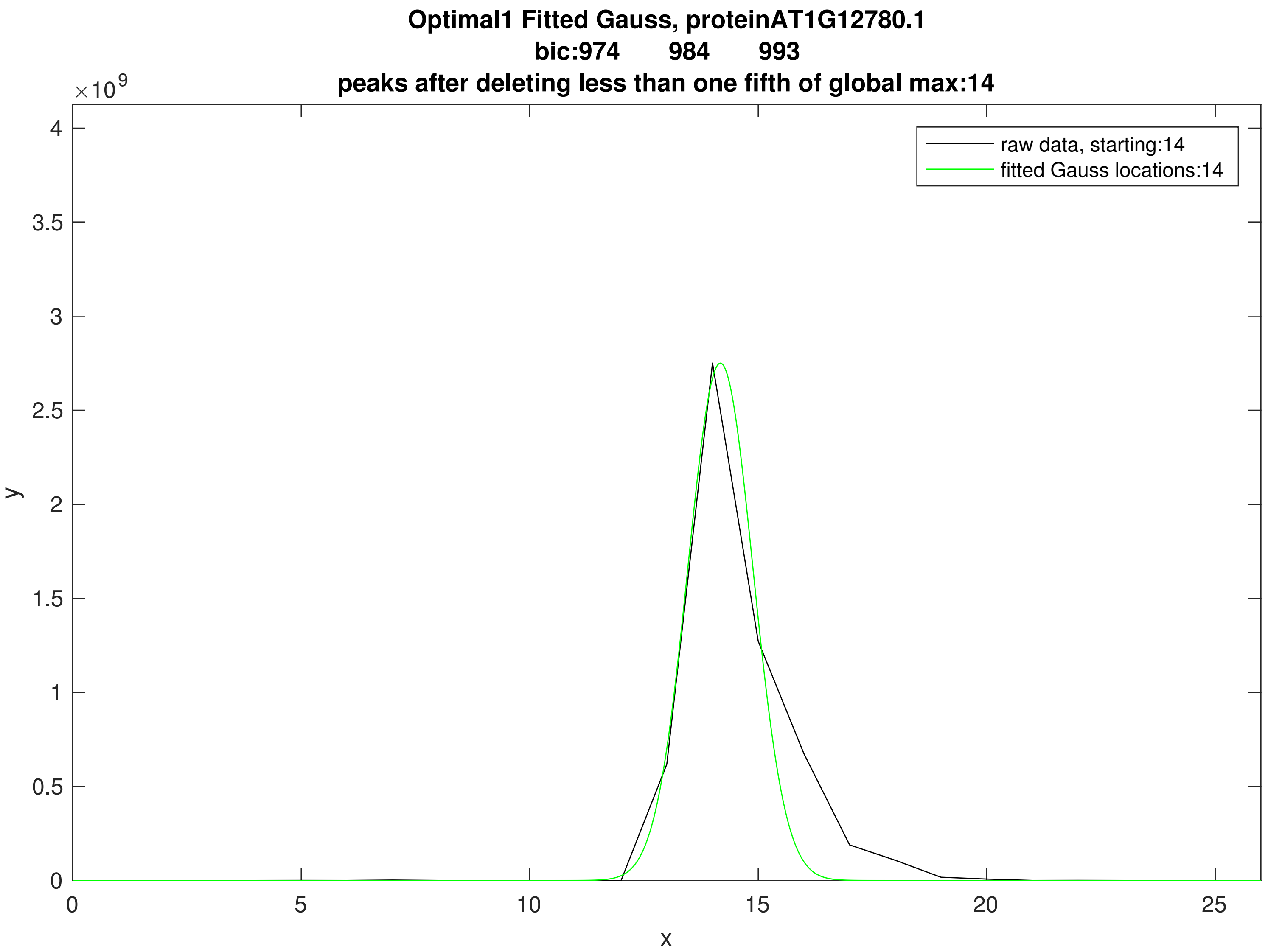

Optimal1 Fitted Gauss, proteinAT1G12780.1
bic:974 984 993
peaks after deleting less than one fifth of global max:14
×10^9
raw data, starting:14
fitted Gauss locations:14
y
x
0
0.5
1
1.5
2
2.5
3
3.5
4
5
10
15
20
25

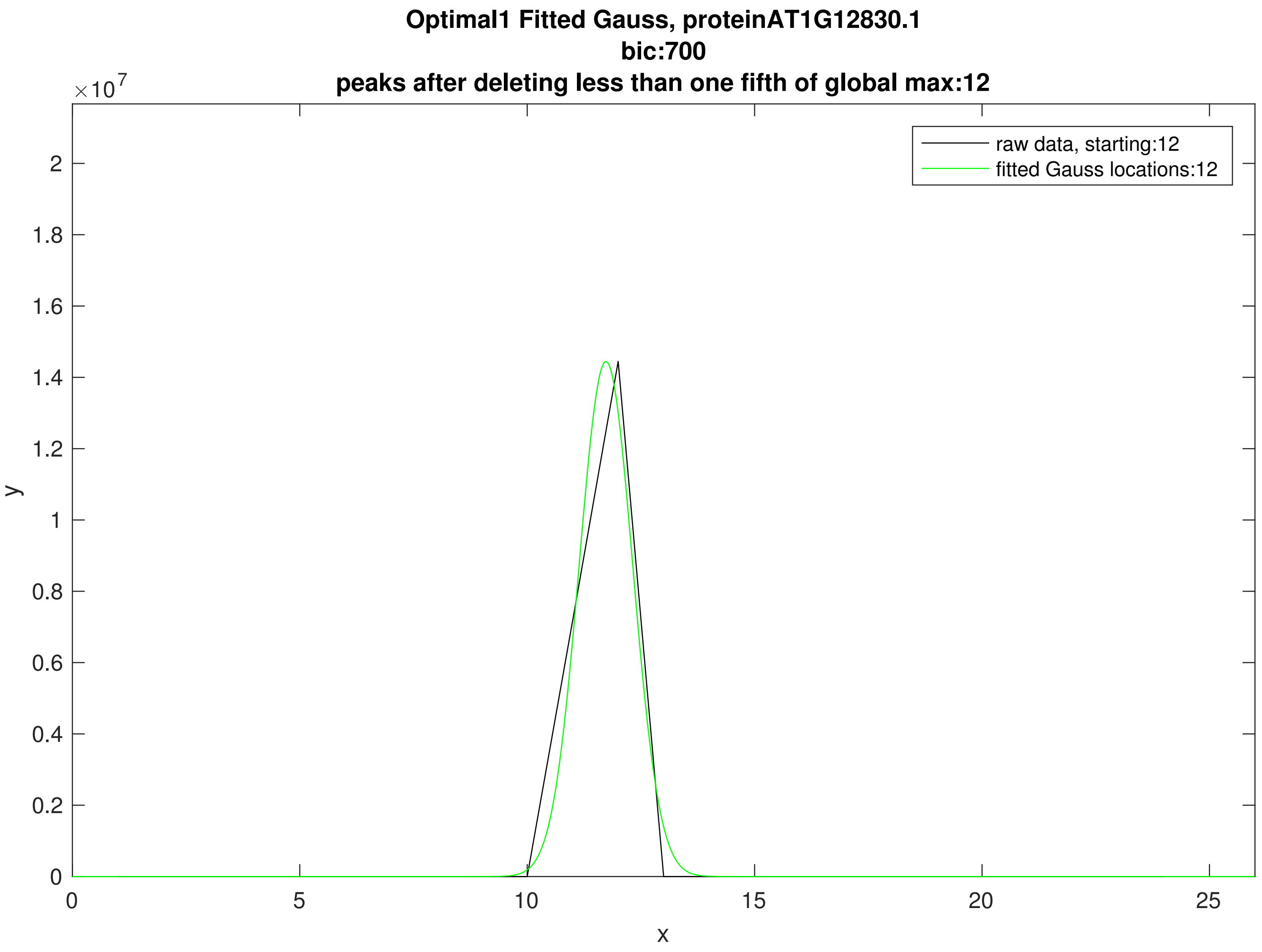
Optimal1 Fitted Gauss, proteinAT1G12830.1
bic:700
peaks after deleting less than one fifth of global max:12
×10^7
raw data, starting:12
fitted Gauss locations:12
y
x
0
0.2
0.4
0.6
0.8
1
1.2
1.4
1.6
1.8
2
0
5
10
15
20
25

**Optimal1 Fitted Gauss, proteinAT1G12840.1**
**bic:781**
**peaks after deleting less than one fifth of global max:16**

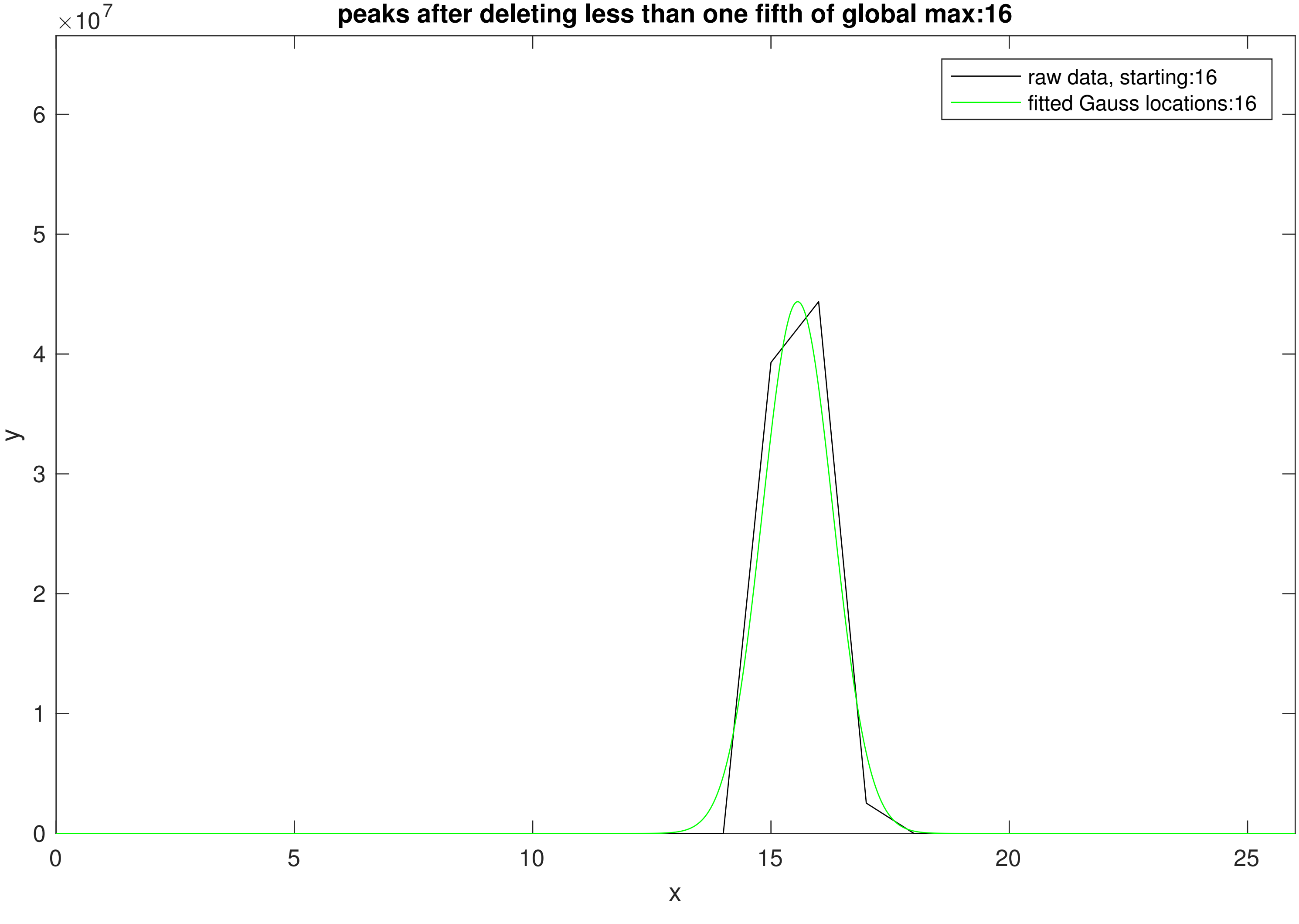

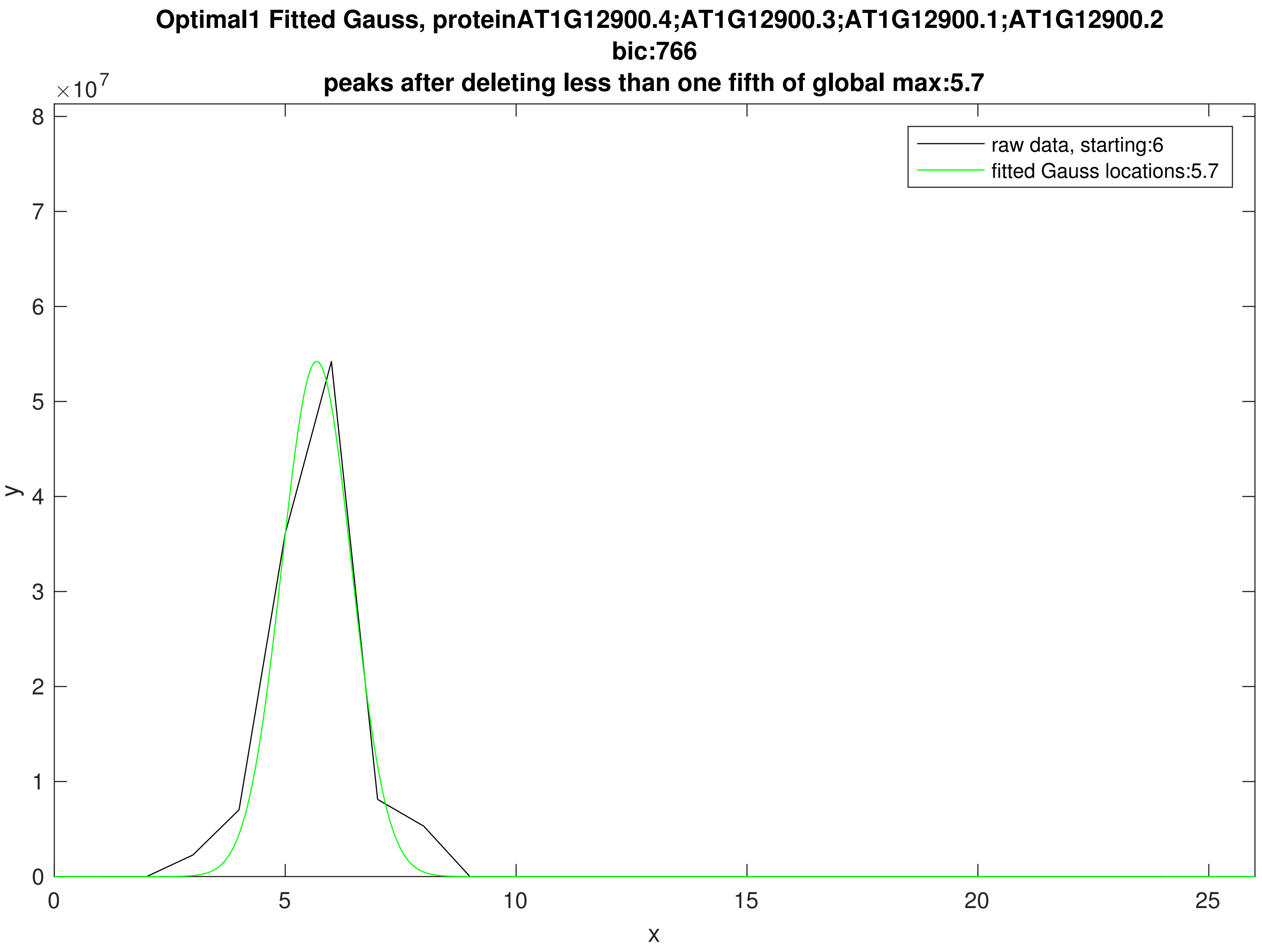

Optimal1 Fitted Gauss, proteinAT1G12900.4;AT1G12900.3;AT1G12900.1;AT1G12900.2
bic:766
peaks after deleting less than one fifth of global max:5.7
×10^7
raw data, starting:6
fitted Gauss locations:5.7
y
x
0
1
2
3
4
5
6
7
8
0
5
10
15
20
25

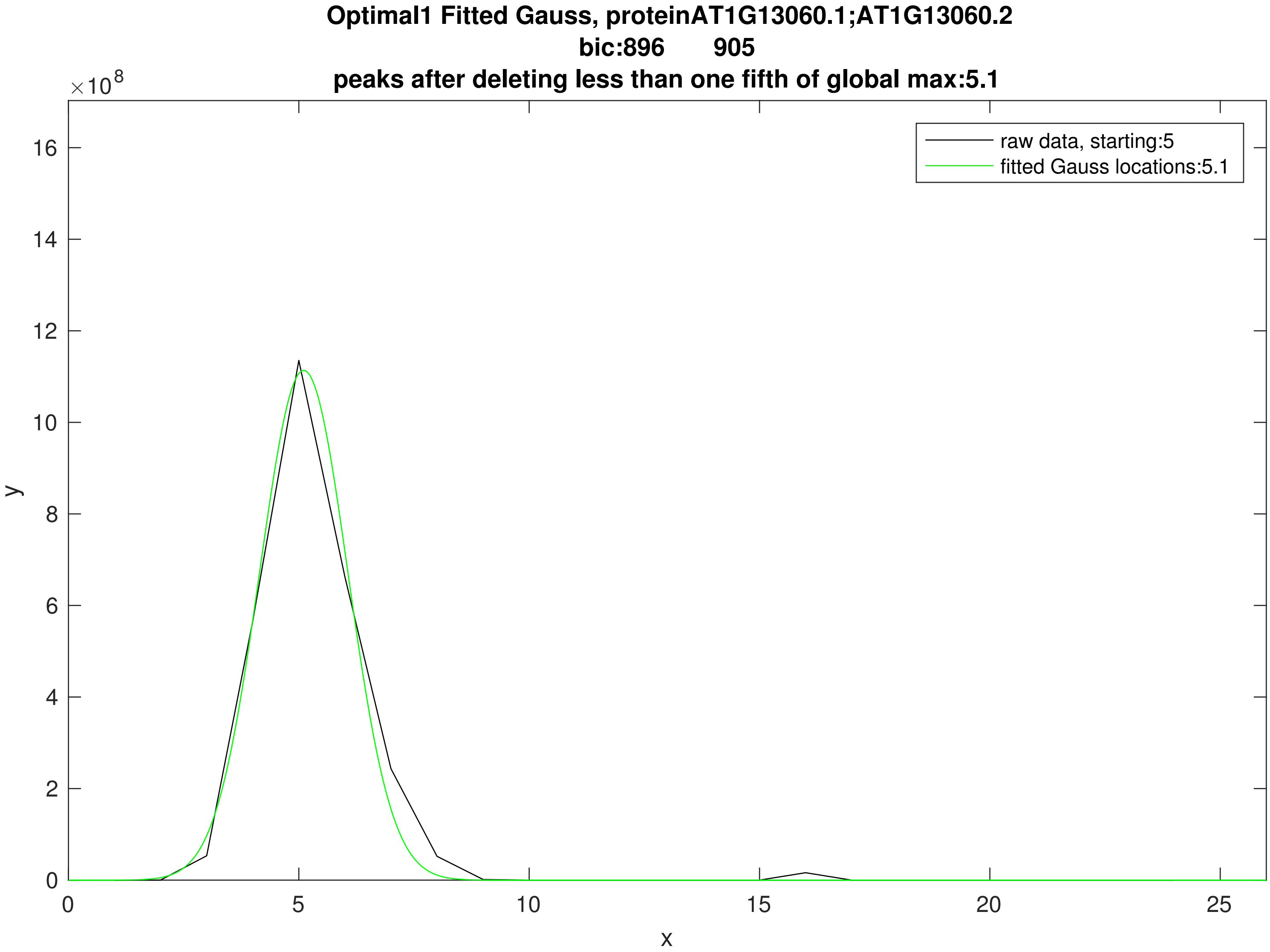

Optimal1 Fitted Gauss, proteinAT1G13060.1;AT1G13060.2
bic:896 905
peaks after deleting less than one fifth of global max:5.1
×10^8
raw data, starting:5
fitted Gauss locations:5.1
y
x
0
2
4
6
8
10
12
14
16
0
5
10
15
20
25

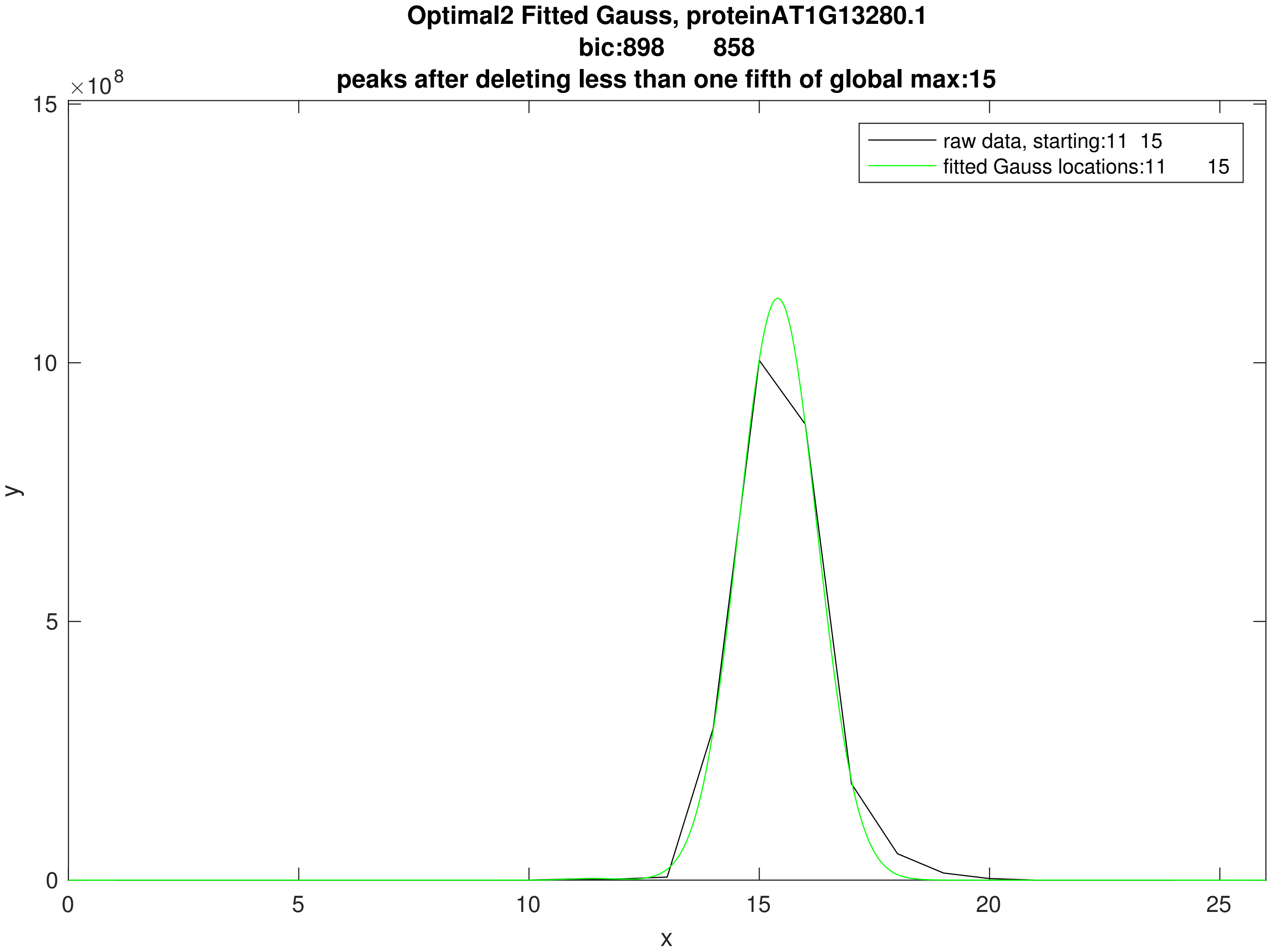

Optimal2 Fitted Gauss, proteinAT1G13280.1
bic:898 858
peaks after deleting less than one fifth of global max:15
×10^8
15
10
5
0
y
0
5
10
15
20
25
x
raw data, starting:11 15
fitted Gauss locations:11 15

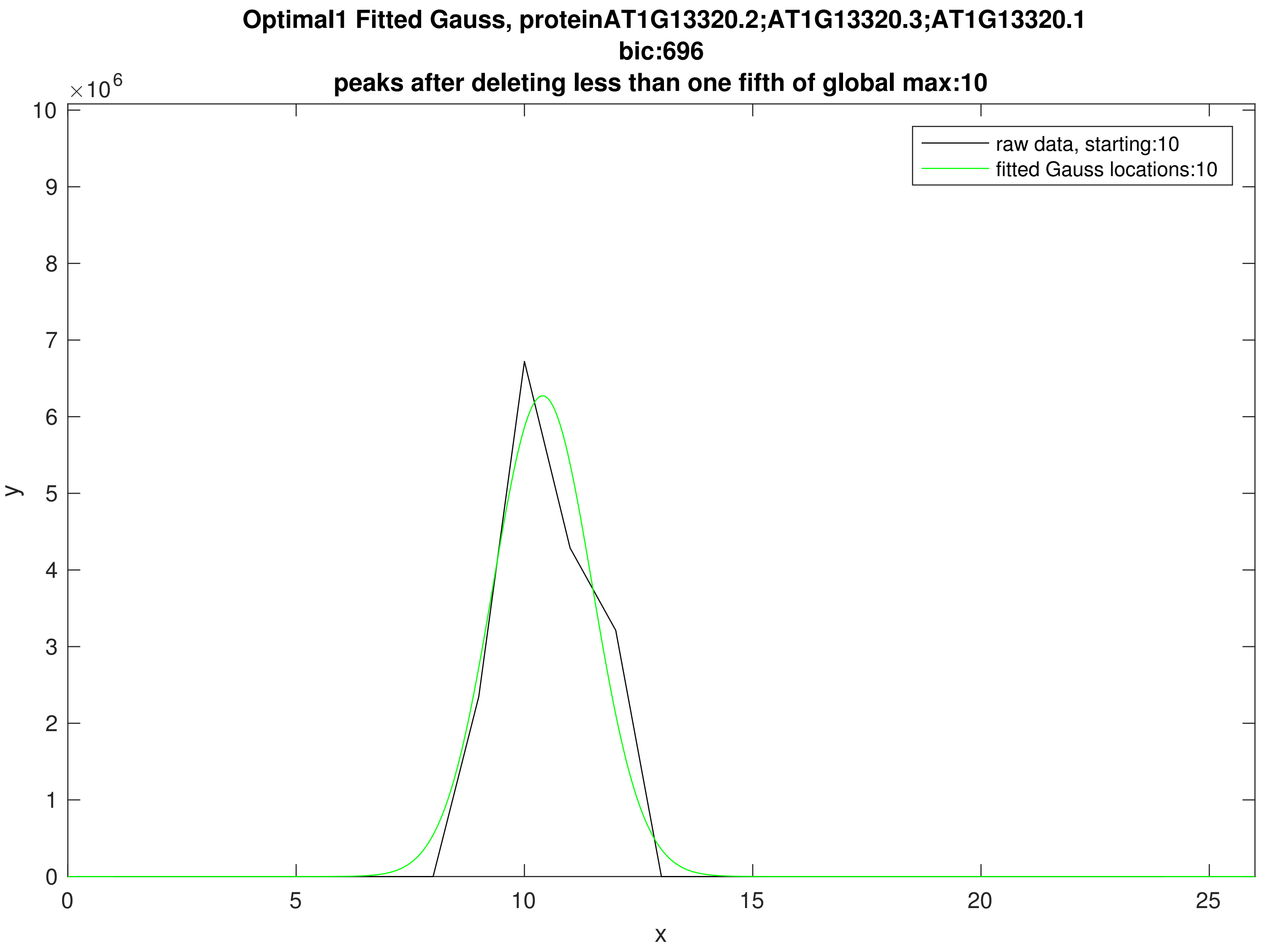
Optimal1 Fitted Gauss, proteinAT1G13320.2;AT1G13320.3;AT1G13320.1
bic:696
peaks after deleting less than one fifth of global max:10
×10^6
raw data, starting:10
fitted Gauss locations:10
y
x
0
1
2
3
4
5
6
7
8
9
10
0
5
10
15
20
25

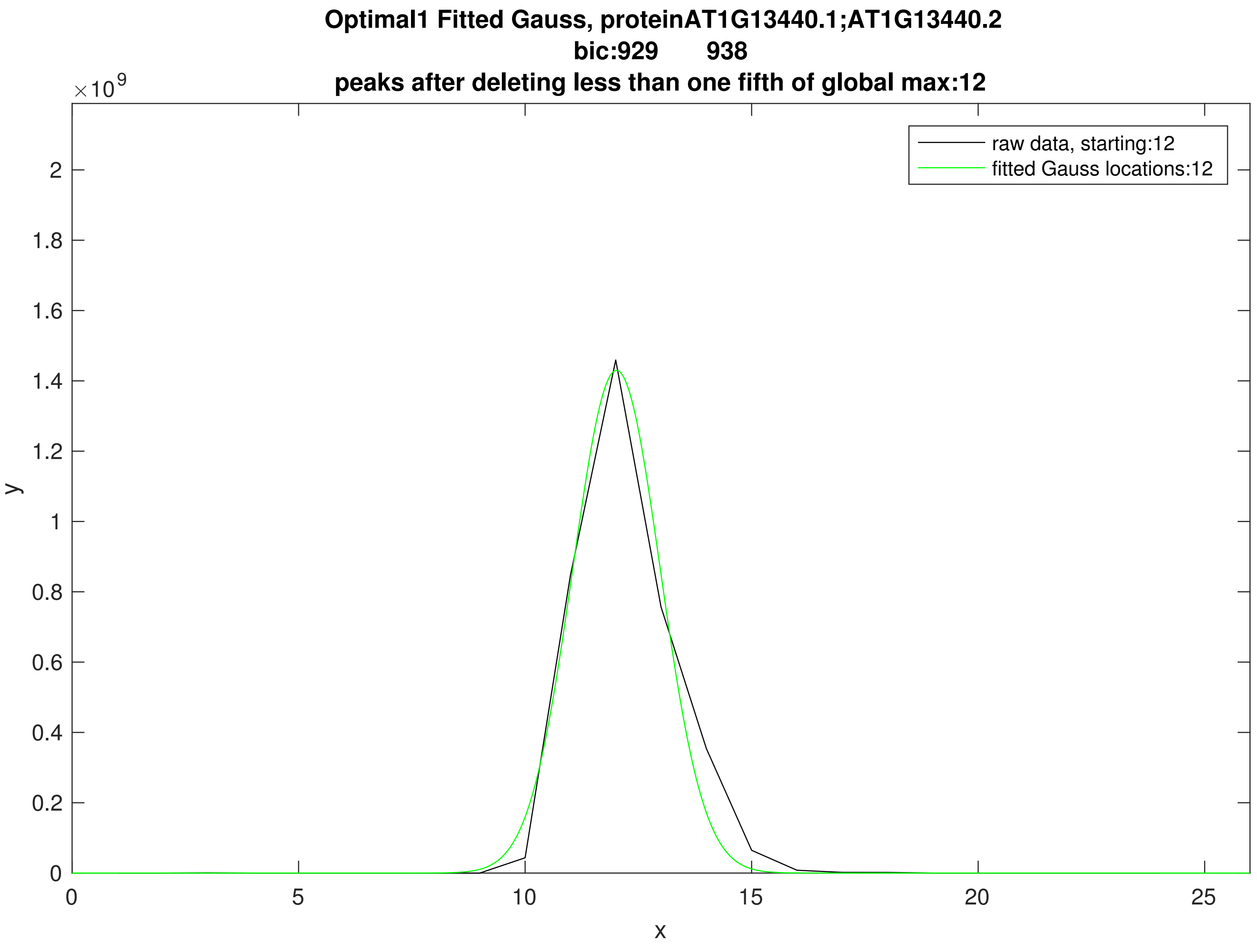
Optimal1 Fitted Gauss, proteinAT1G13440.1;AT1G13440.2
bic:929 938
peaks after deleting less than one fifth of global max:12
×10^9
raw data, starting:12
fitted Gauss locations:12
y
x
0
0.2
0.4
0.6
0.8
1
1.2
1.4
1.6
1.8
2
5
10
15
20
25

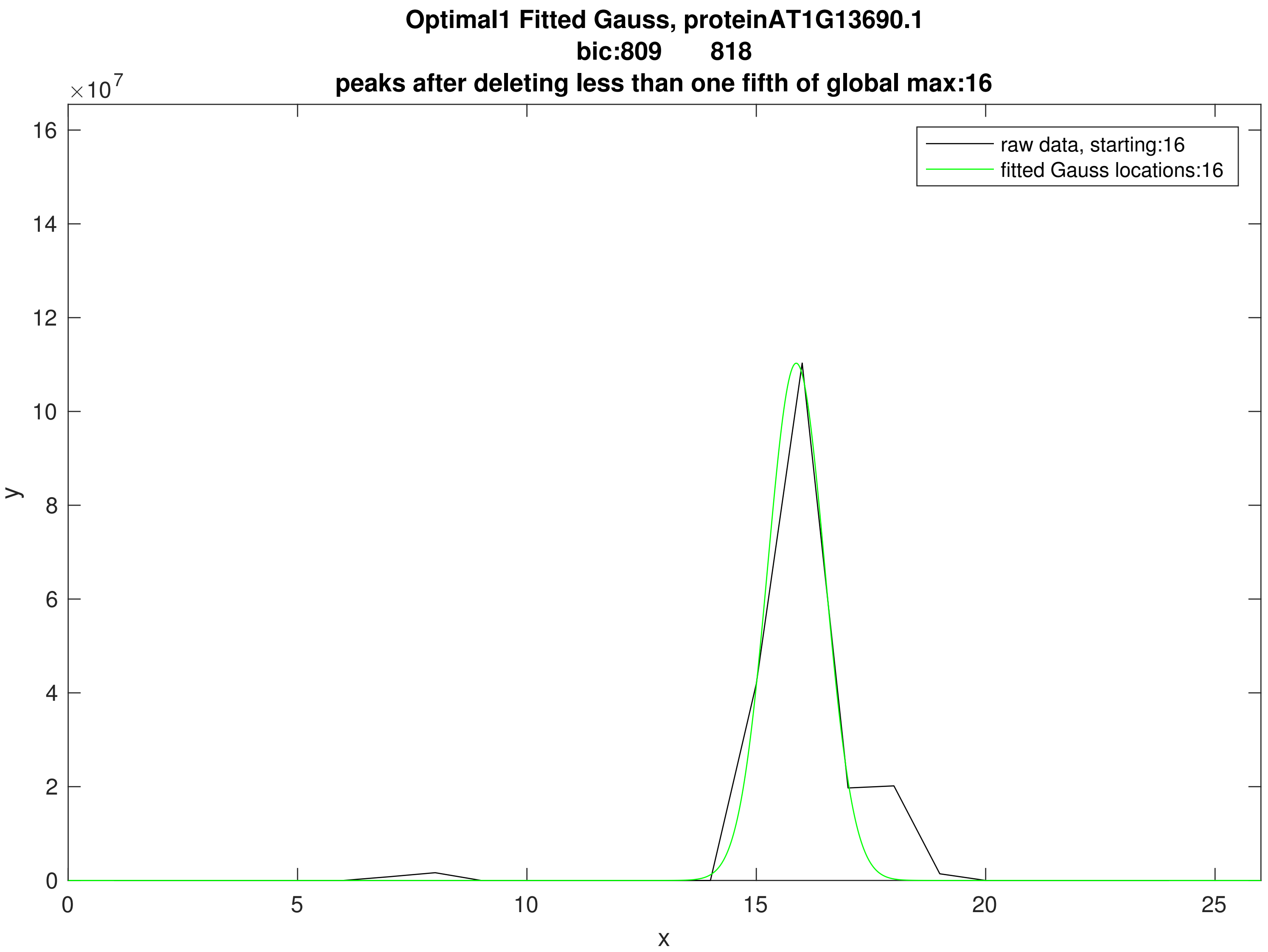
Optimal1 Fitted Gauss, proteinAT1G13690.1
bic:809 818
peaks after deleting less than one fifth of global max:16
×10^7
raw data, starting:16
fitted Gauss locations:16
y
x
0
2
4
6
8
10
12
14
16
0
5
10
15
20
25

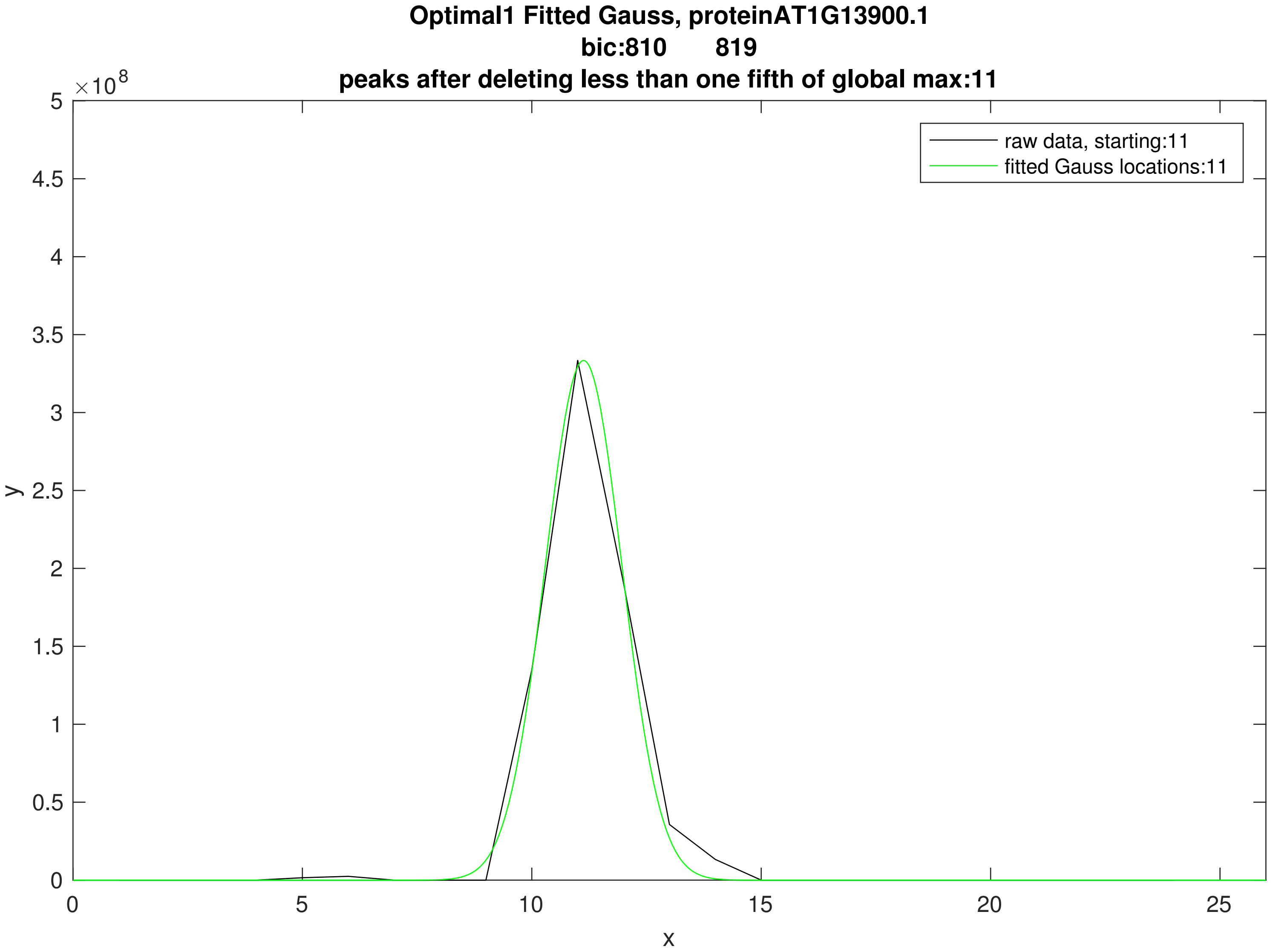

Optimal1 Fitted Gauss, proteinAT1G13900.1
bic:810 819
peaks after deleting less than one fifth of global max:11
×10^8
5
4.5
4
3.5
3
2.5
2
1.5
1
0.5
0
y
0
5
10
15
20
25
x
raw data, starting:11
fitted Gauss locations:11

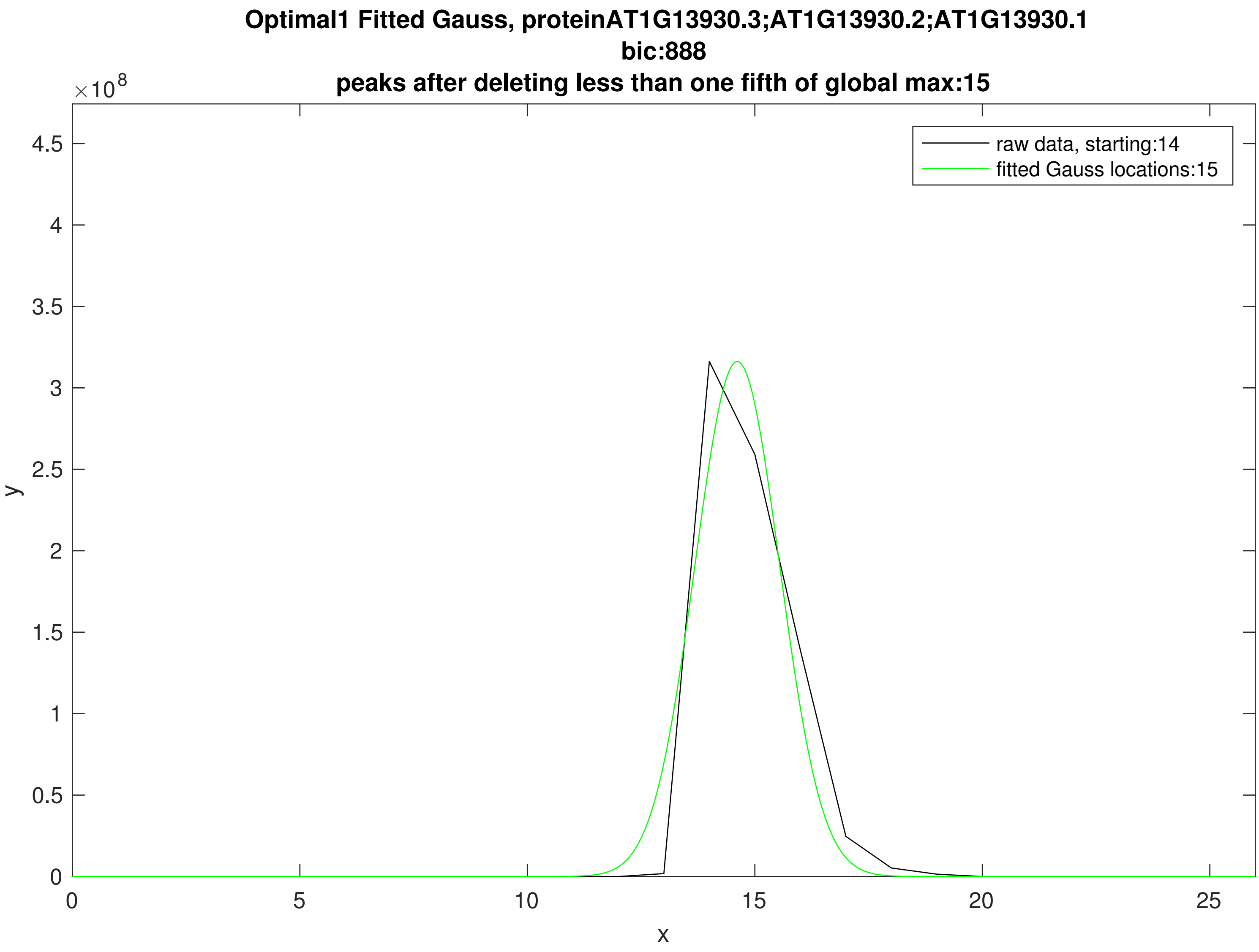

Optimal1 Fitted Gauss, proteinAT1G13930.3;AT1G13930.2;AT1G13930.1
bic:888
peaks after deleting less than one fifth of global max:15
×10^8
4.5
4
3.5
3
2.5
2
1.5
1
0.5
0
y
0
5
10
15
20
25
x
raw data, starting:14
fitted Gauss locations:15

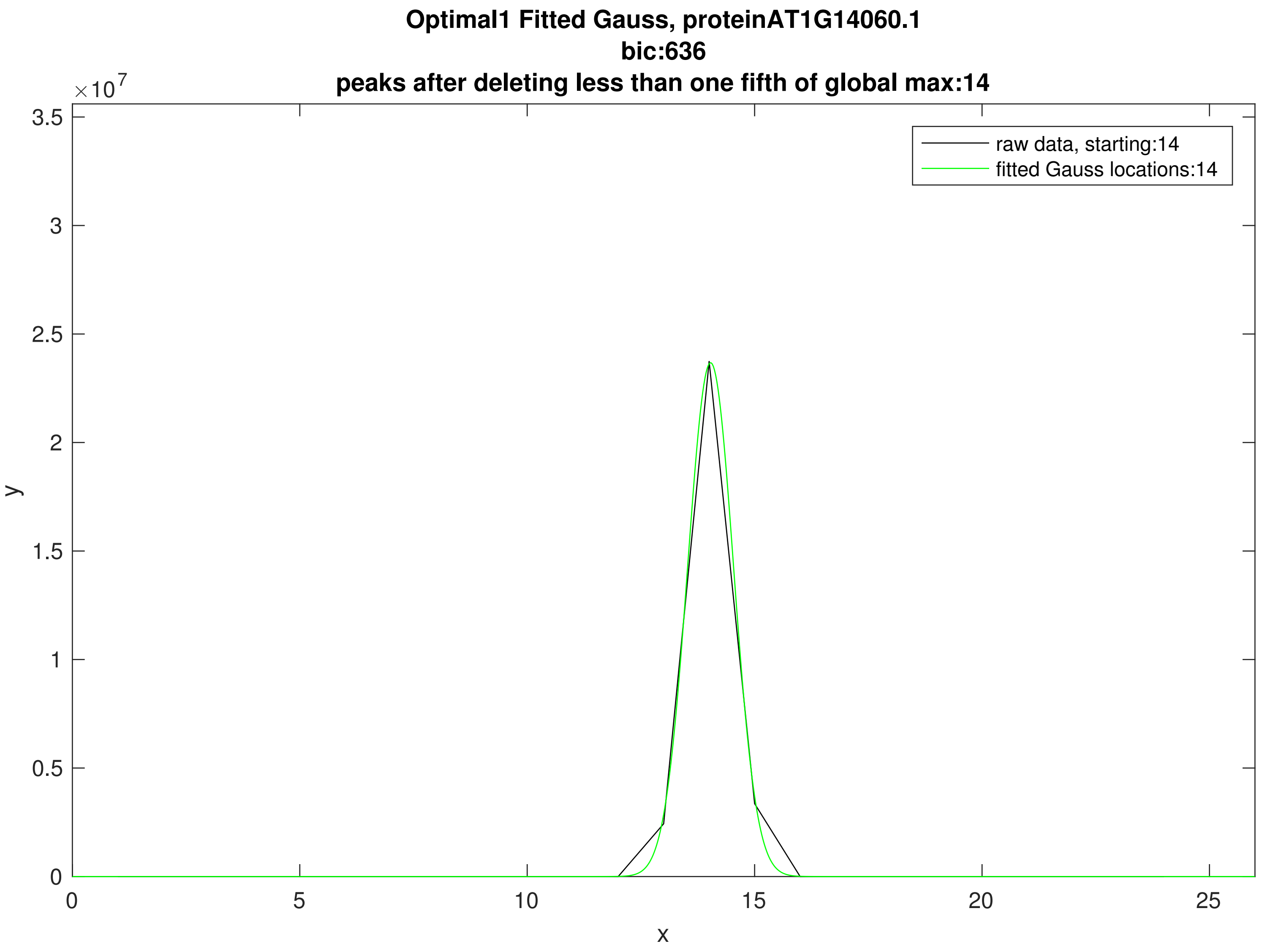
Optimal1 Fitted Gauss, proteinAT1G14060.1
bic:636
peaks after deleting less than one fifth of global max:14
×10^7
3.5
3
2.5
2
1.5
1
0.5
0
y
0
5
10
15
20
25
x
raw data, starting:14
fitted Gauss locations:14

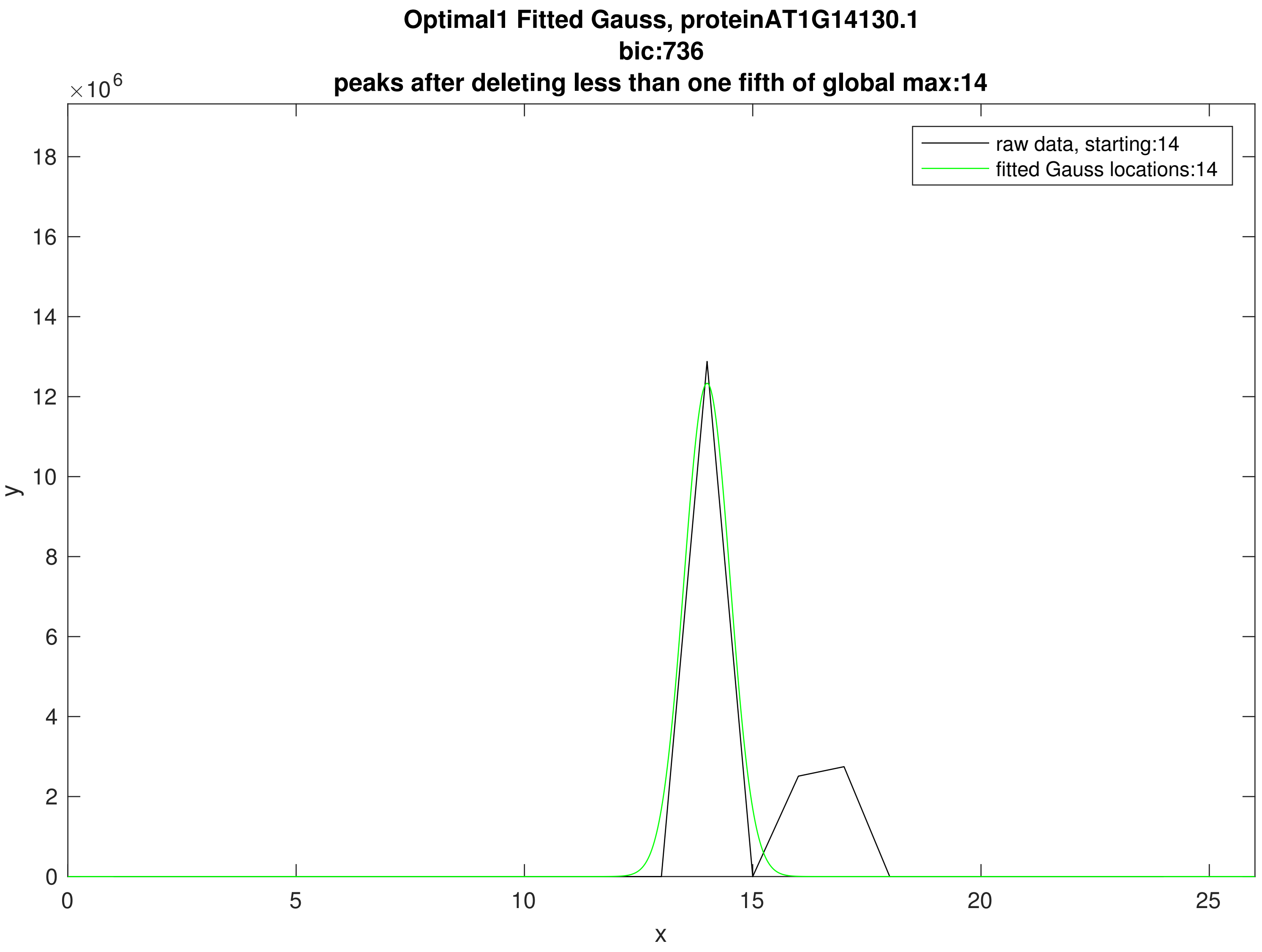
Optimal1 Fitted Gauss, proteinAT1G14130.1
bic:736
peaks after deleting less than one fifth of global max:14
×10^6
raw data, starting:14
fitted Gauss locations:14
y
x
0
2
4
6
8
10
12
14
16
18
0
5
10
15
20
25

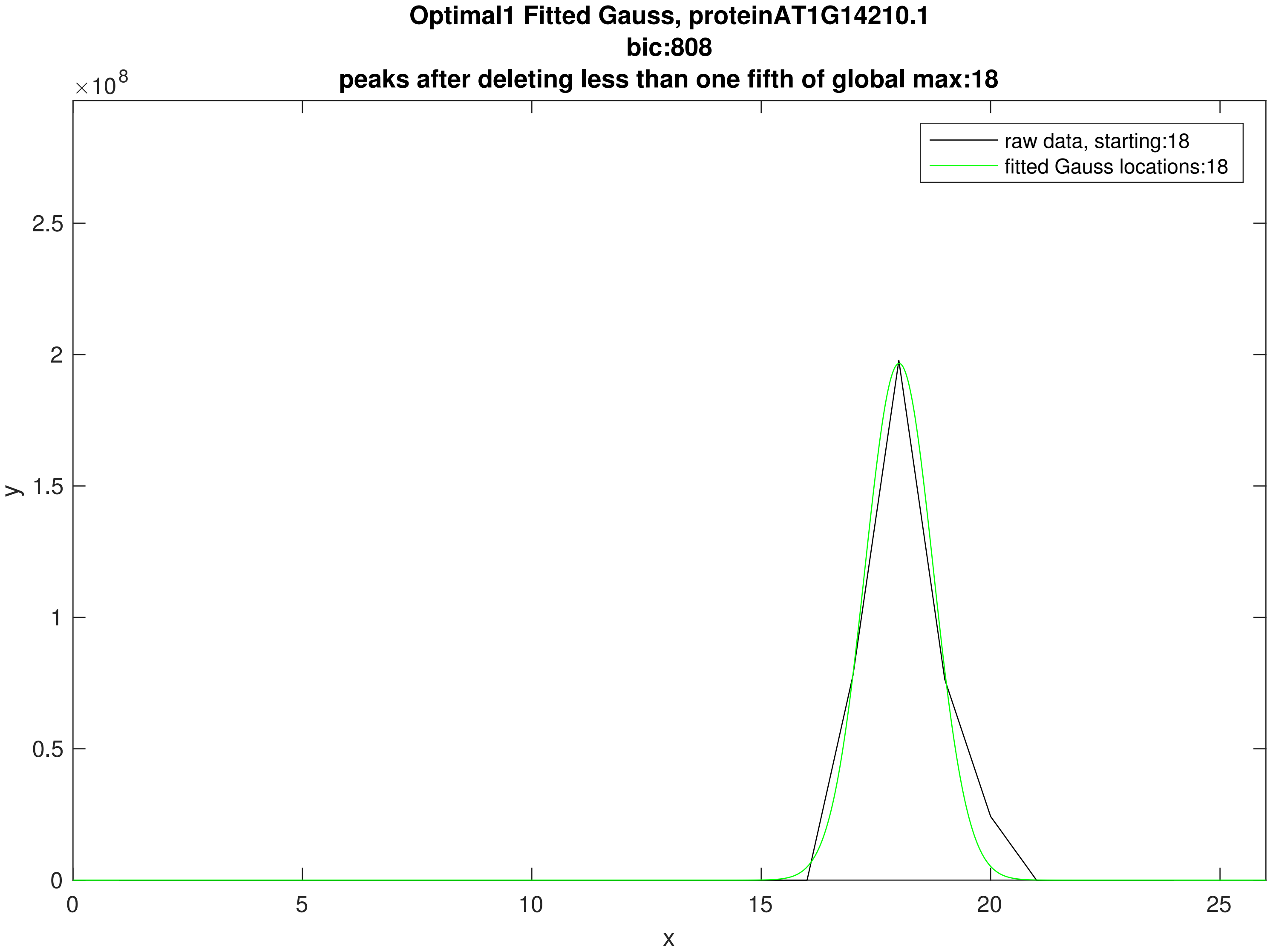
Optimal1 Fitted Gauss, proteinAT1G14210.1
bic:808
peaks after deleting less than one fifth of global max:18
×10^8
raw data, starting:18
fitted Gauss locations:18
y
x
0
0.5
1
1.5
2
2.5
0
5
10
15
20
25

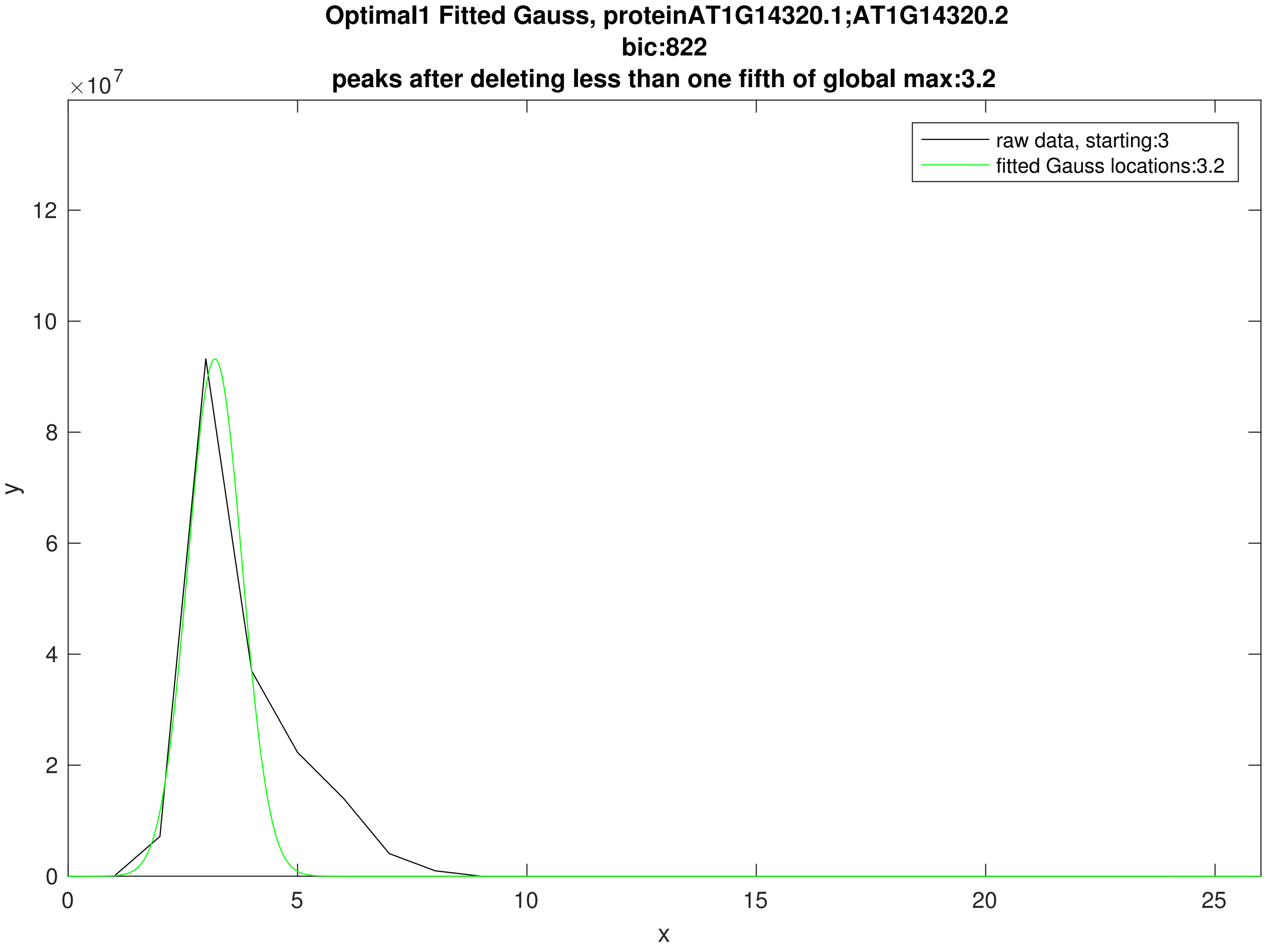

Optimal1 Fitted Gauss, proteinAT1G14320.1;AT1G14320.2
bic:822
peaks after deleting less than one fifth of global max:3.2
×10^7
raw data, starting:3
fitted Gauss locations:3.2
y
x
0
2
4
6
8
10
12
5
10
15
20
25

Optimal1 Fitted Gauss, proteinAT1G14400.2;AT1G14400.1
bic:737 800
peaks after deleting less than one fifth of global max:20

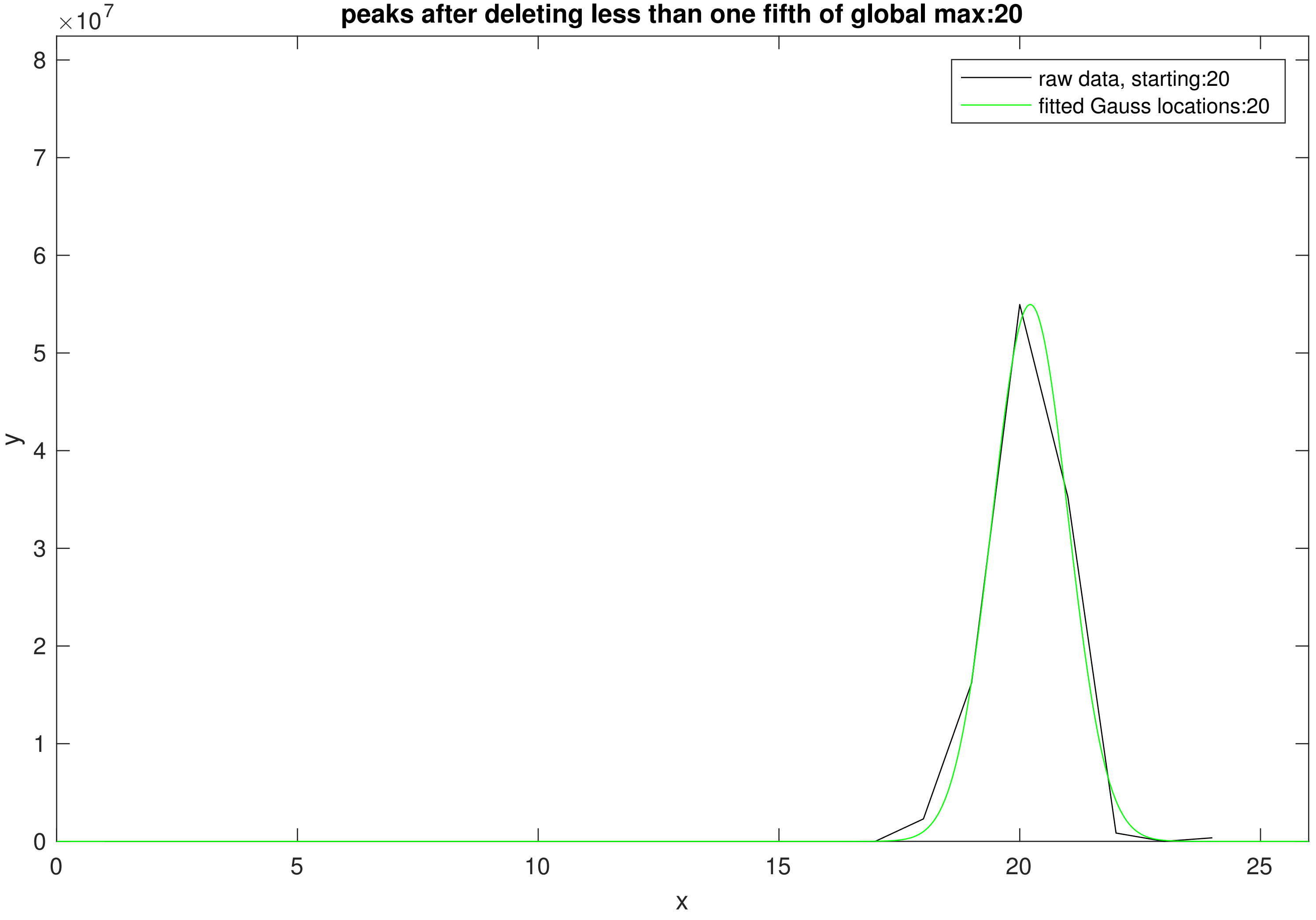

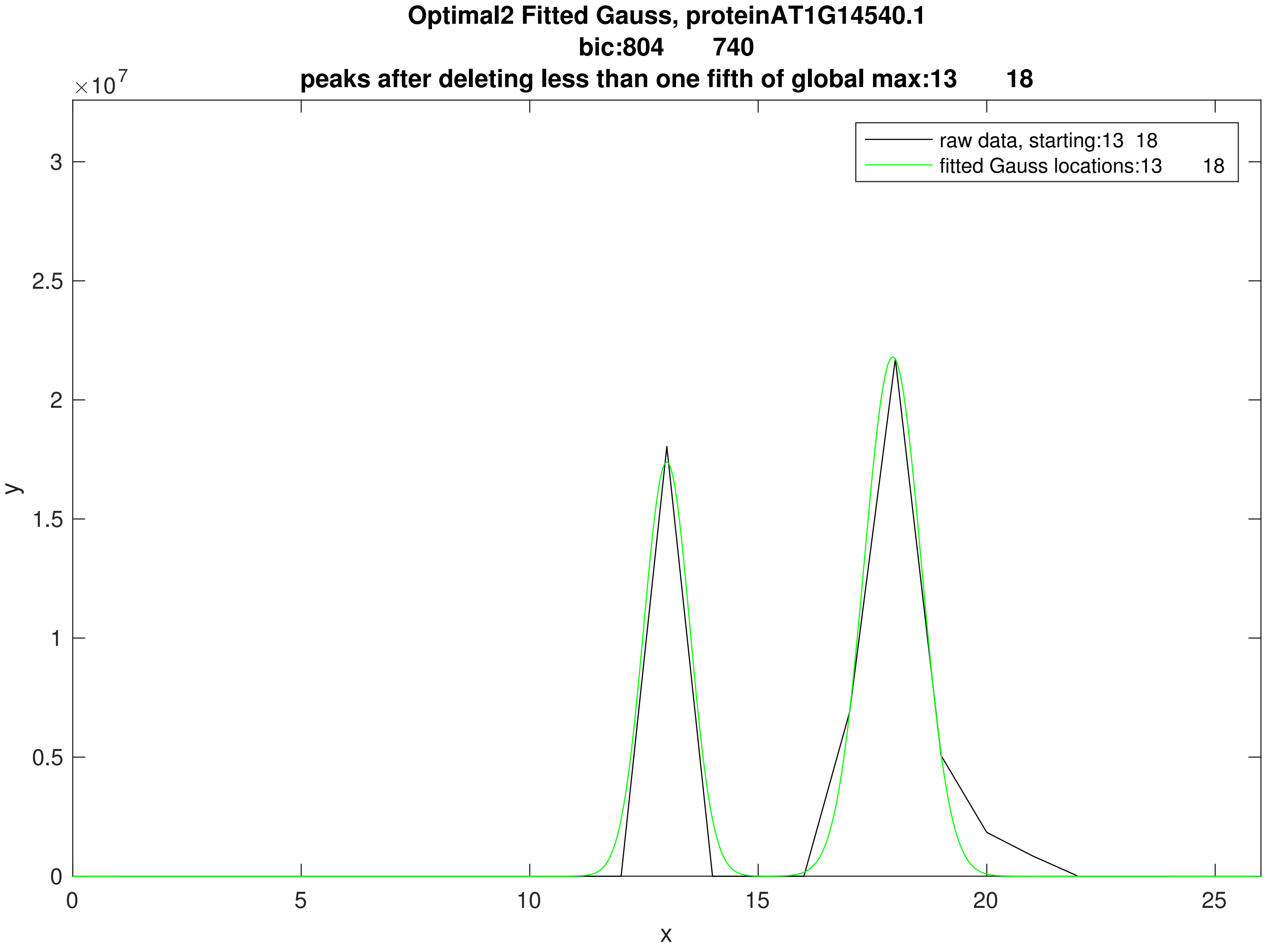
Optimal2 Fitted Gauss, proteinAT1G14540.1
bic:804 740
peaks after deleting less than one fifth of global max:13 18
raw data, starting:13 18
fitted Gauss locations:13 18
×10^7
y
x
0
0.5
1
1.5
2
2.5
3
5
10
15
20
25

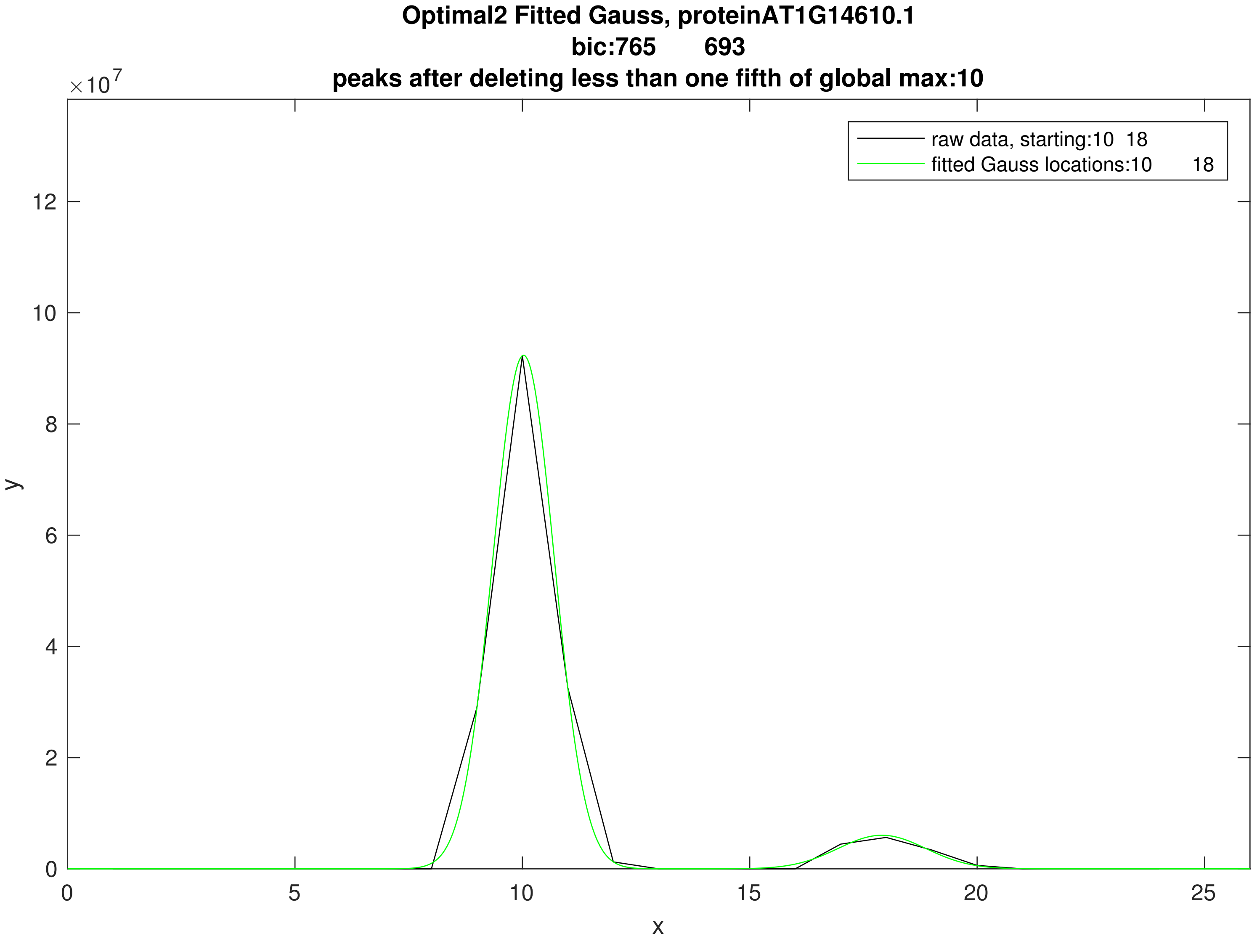

Optimal2 Fitted Gauss, proteinAT1G14610.1
bic:765 693
peaks after deleting less than one fifth of global max:10
×10^7
raw data, starting:10 18
fitted Gauss locations:10 18
12
10
8
6
4
2
0
y
0
5
10
15
20
25
x

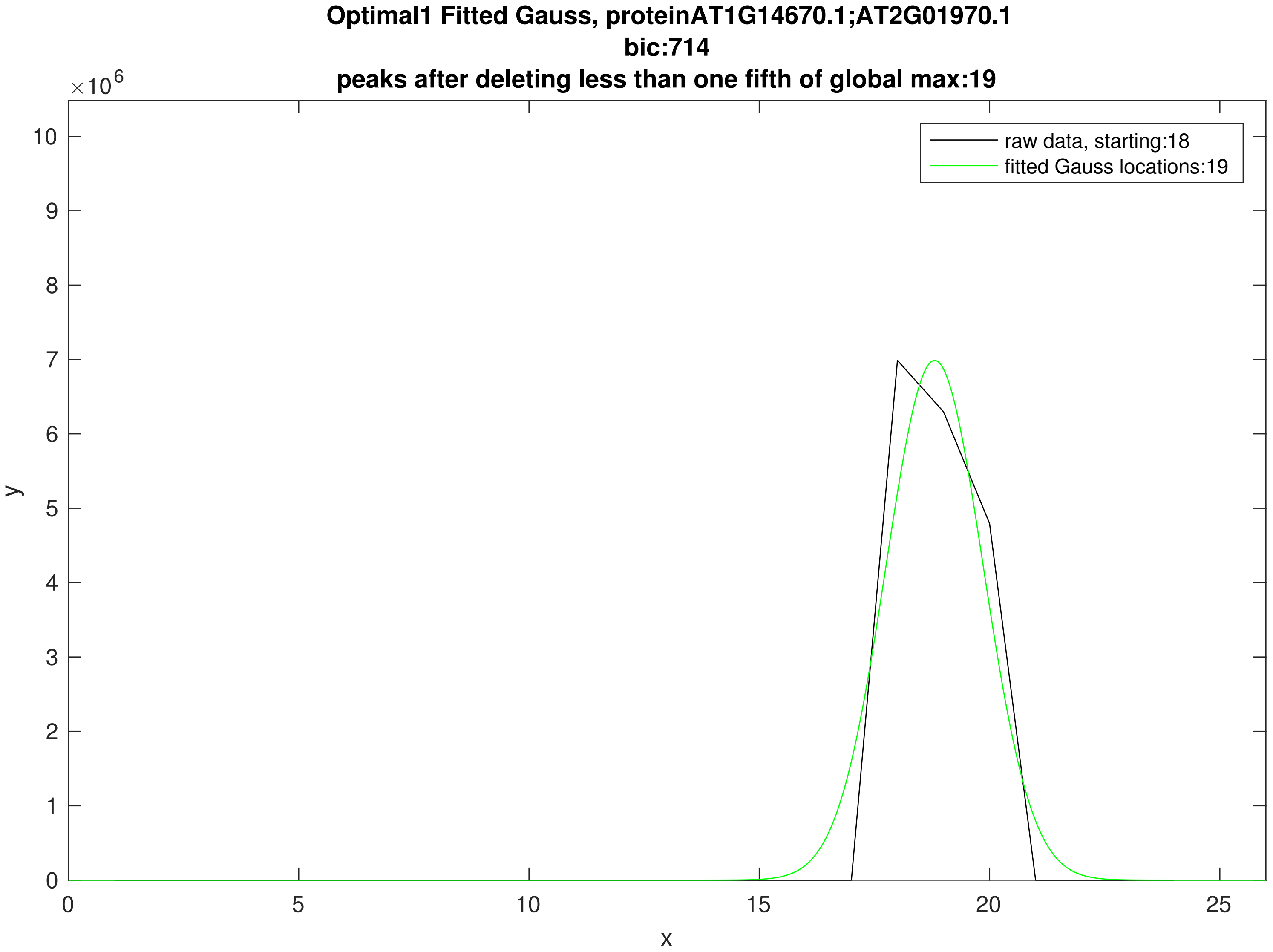
Optimal1 Fitted Gauss, proteinAT1G14670.1;AT2G01970.1
bic:714
peaks after deleting less than one fifth of global max:19
×10^6
raw data, starting:18
fitted Gauss locations:19
y
x
0
1
2
3
4
5
6
7
8
9
10
15
20
25

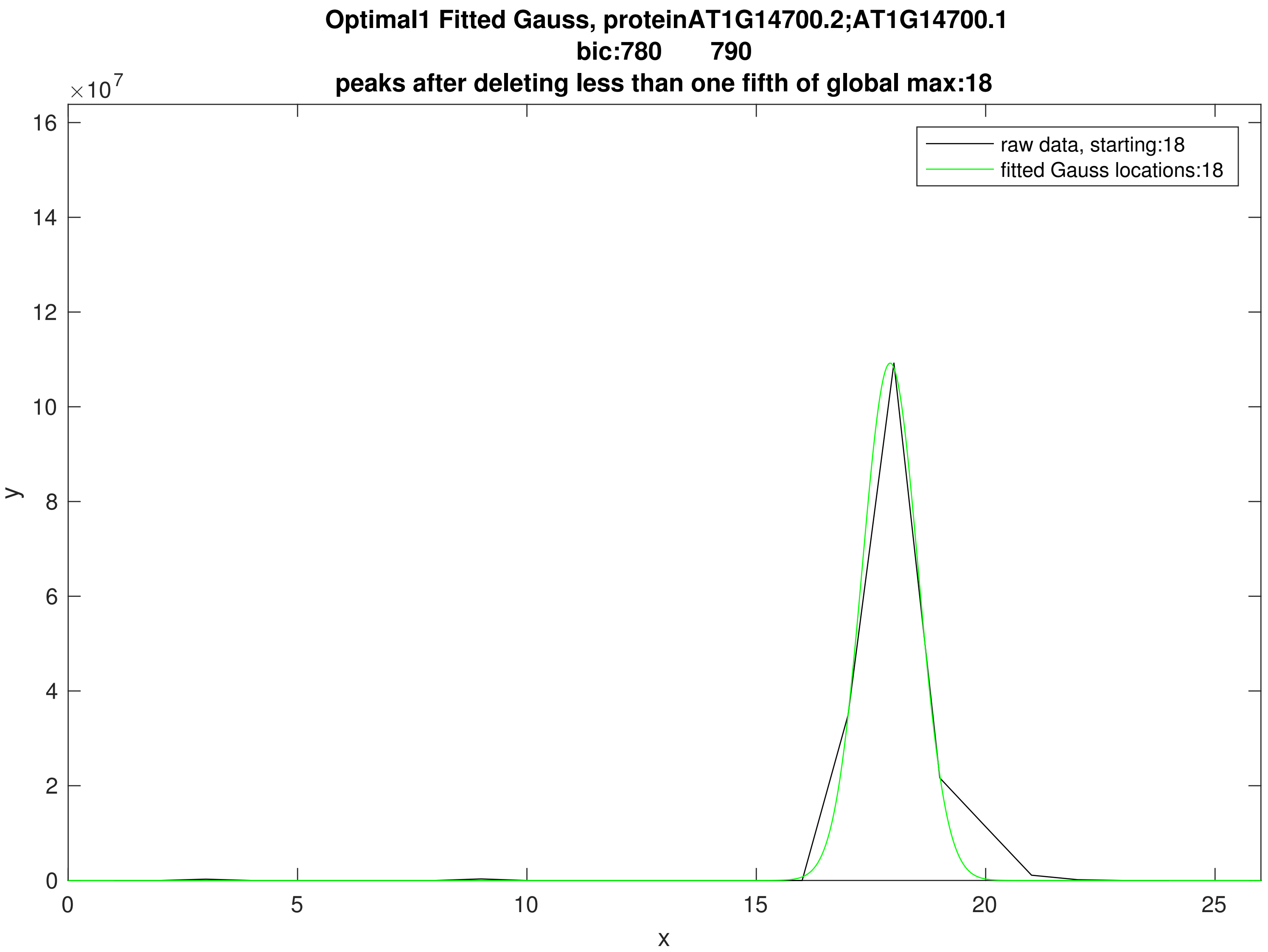
Optimal1 Fitted Gauss, proteinAT1G14700.2;AT1G14700.1
bic:780 790
peaks after deleting less than one fifth of global max:18
×10^7
raw data, starting:18
fitted Gauss locations:18
y
x
0
2
4
6
8
10
12
14
16
0
5
10
15
20
25

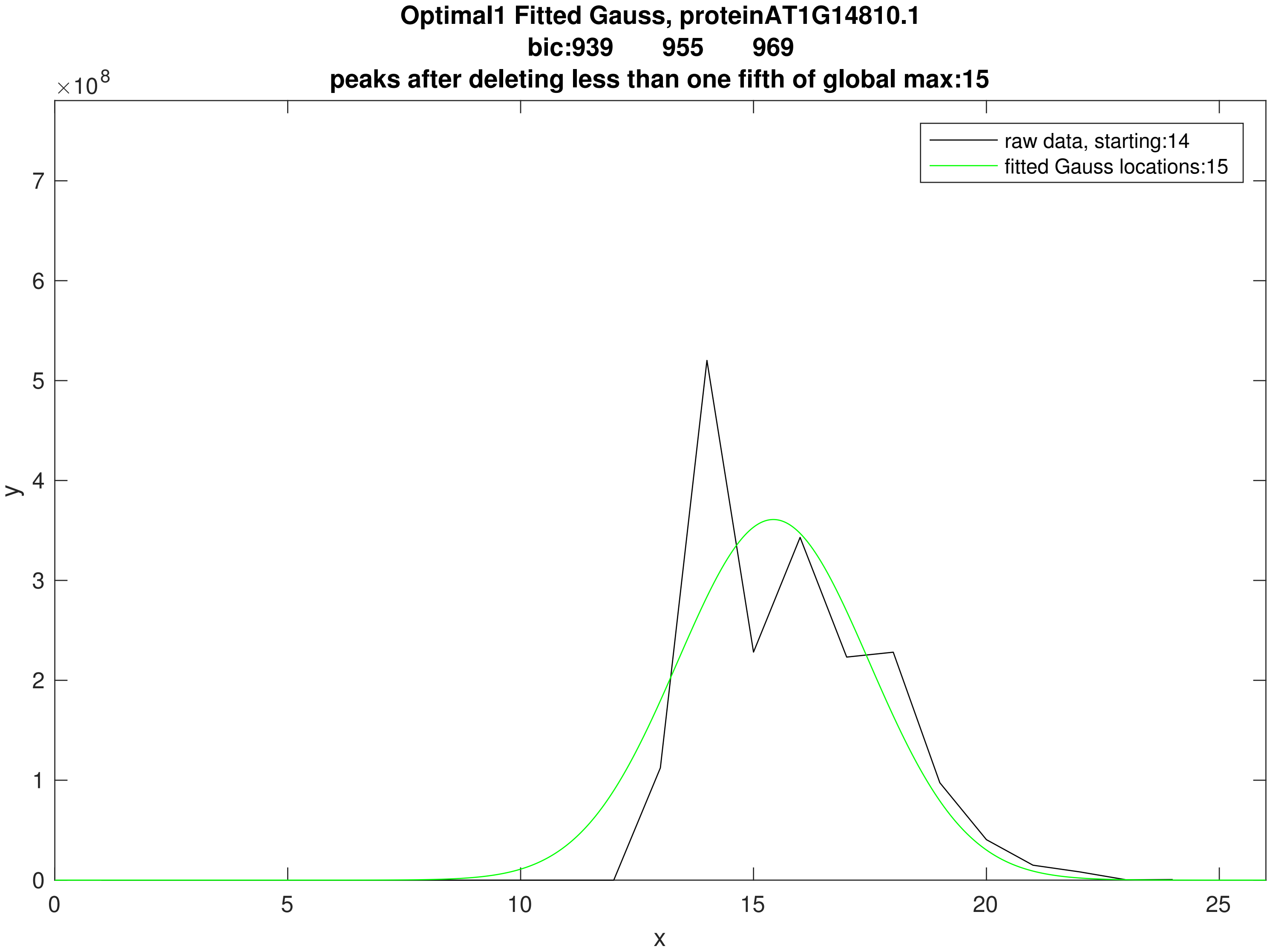

Optimal1 Fitted Gauss, proteinAT1G14810.1
bic:939 955 969
peaks after deleting less than one fifth of global max:15
×10^8
raw data, starting:14
fitted Gauss locations:15
y
x
0
1
2
3
4
5
6
7
0
5
10
15
20
25

Optimal1 Fitted Gauss, proteinAT1G14930.1
bic:789
peaks after deleting less than one fifth of global max:19

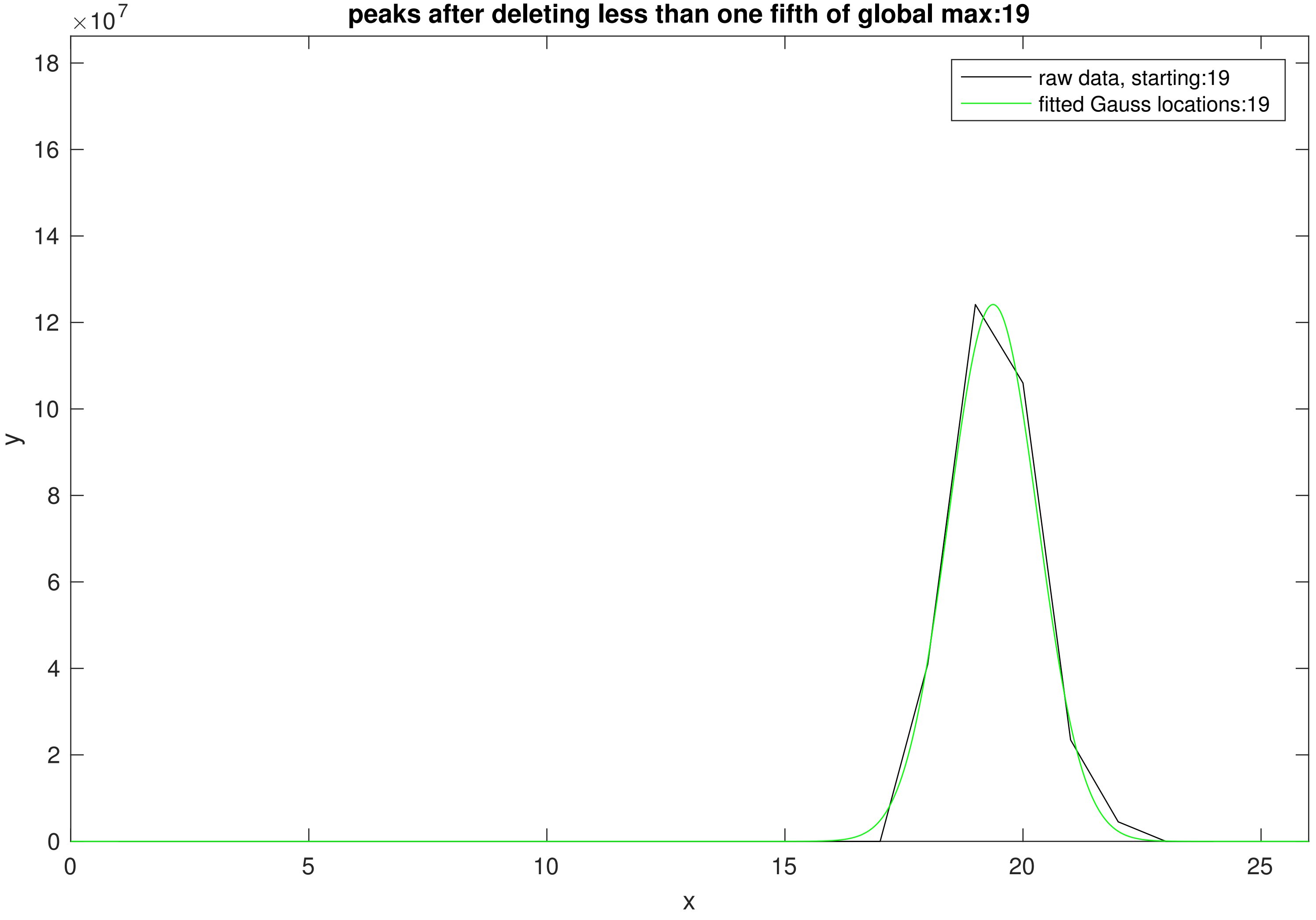

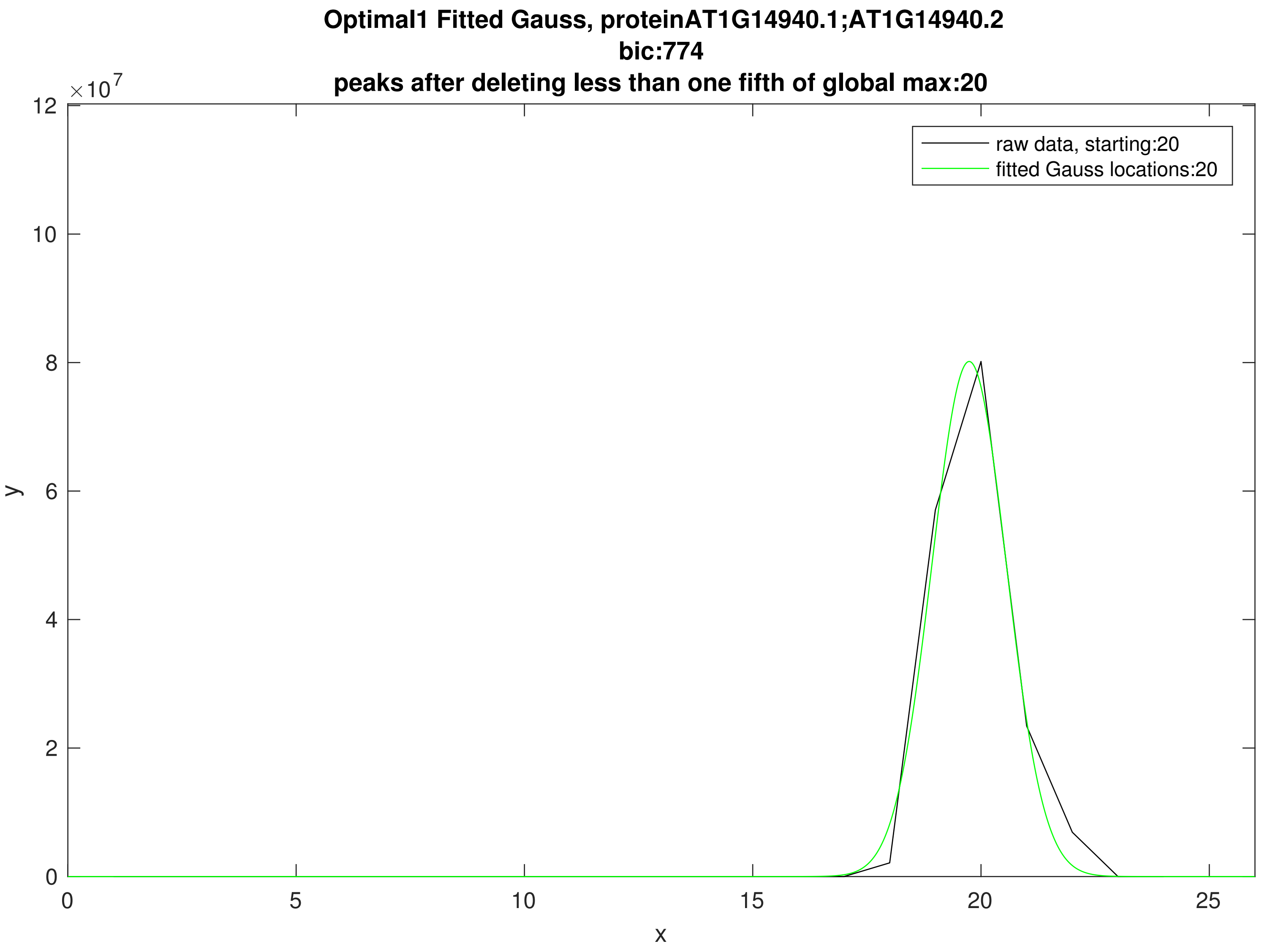
Optimal1 Fitted Gauss, proteinAT1G14940.1;AT1G14940.2
bic:774
peaks after deleting less than one fifth of global max:20
×10^7
12
10
8
6
4
2
0
y
0
5
10
15
20
25
x
raw data, starting:20
fitted Gauss locations:20

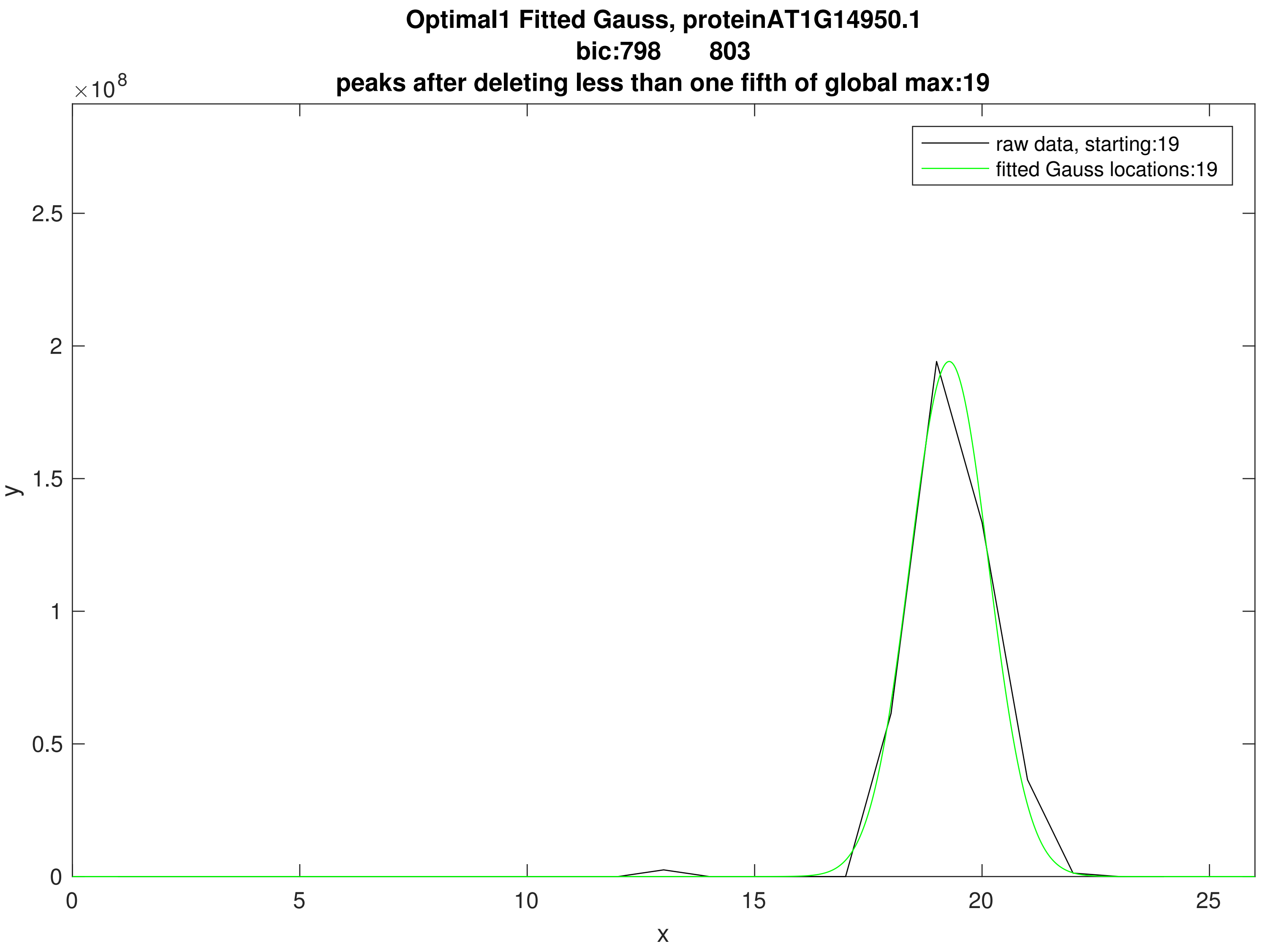

Optimal1 Fitted Gauss, proteinAT1G14950.1
bic:798 803
peaks after deleting less than one fifth of global max:19
×10^8
raw data, starting:19
fitted Gauss locations:19
y
x
0
0.5
1
1.5
2
2.5
5
10
15
20
25

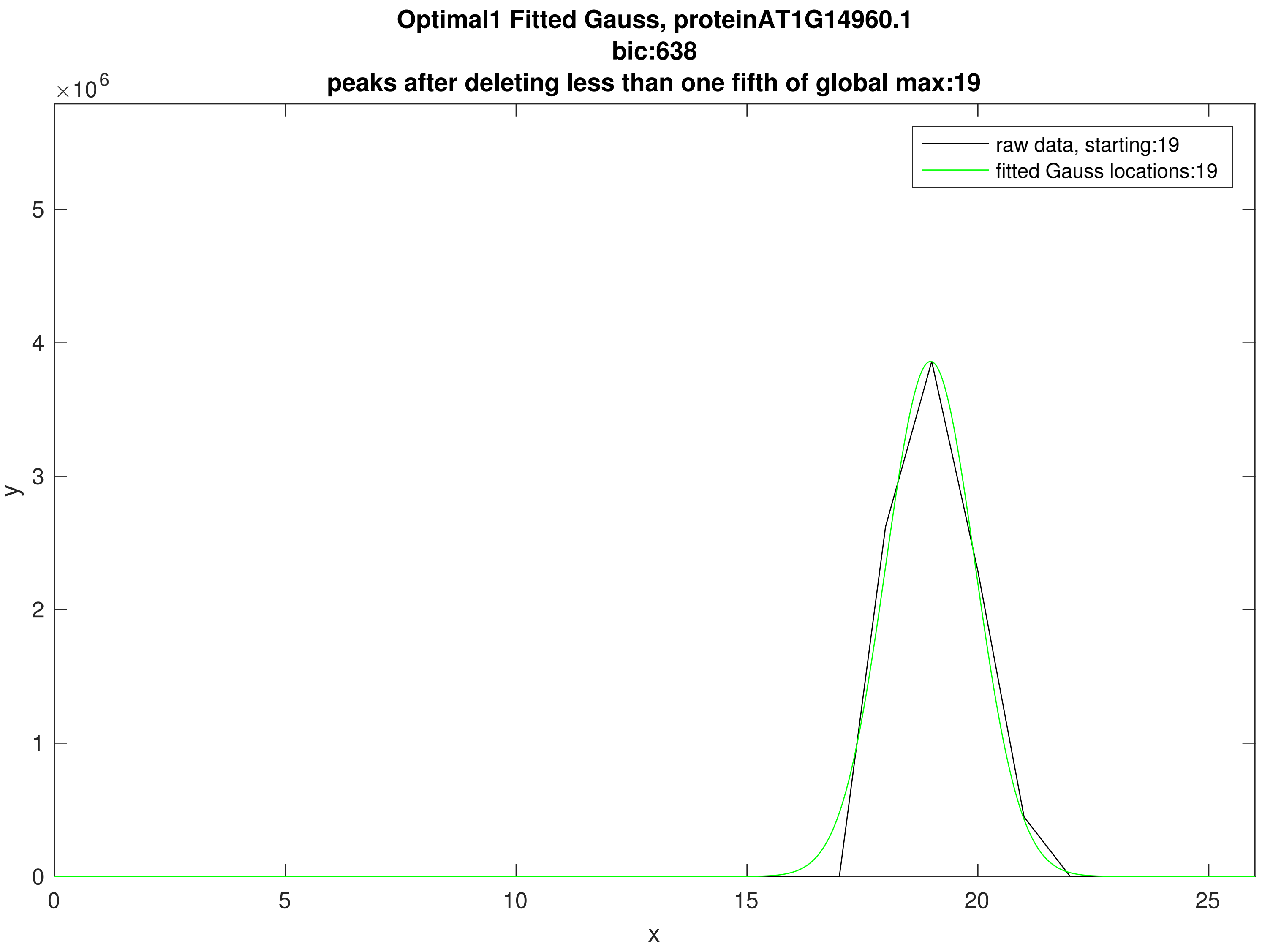

Optimal1 Fitted Gauss, proteinAT1G14960.1
bic:638
peaks after deleting less than one fifth of global max:19
×10^6
raw data, starting:19
fitted Gauss locations:19
y
x
0
1
2
3
4
5
0
5
10
15
20
25

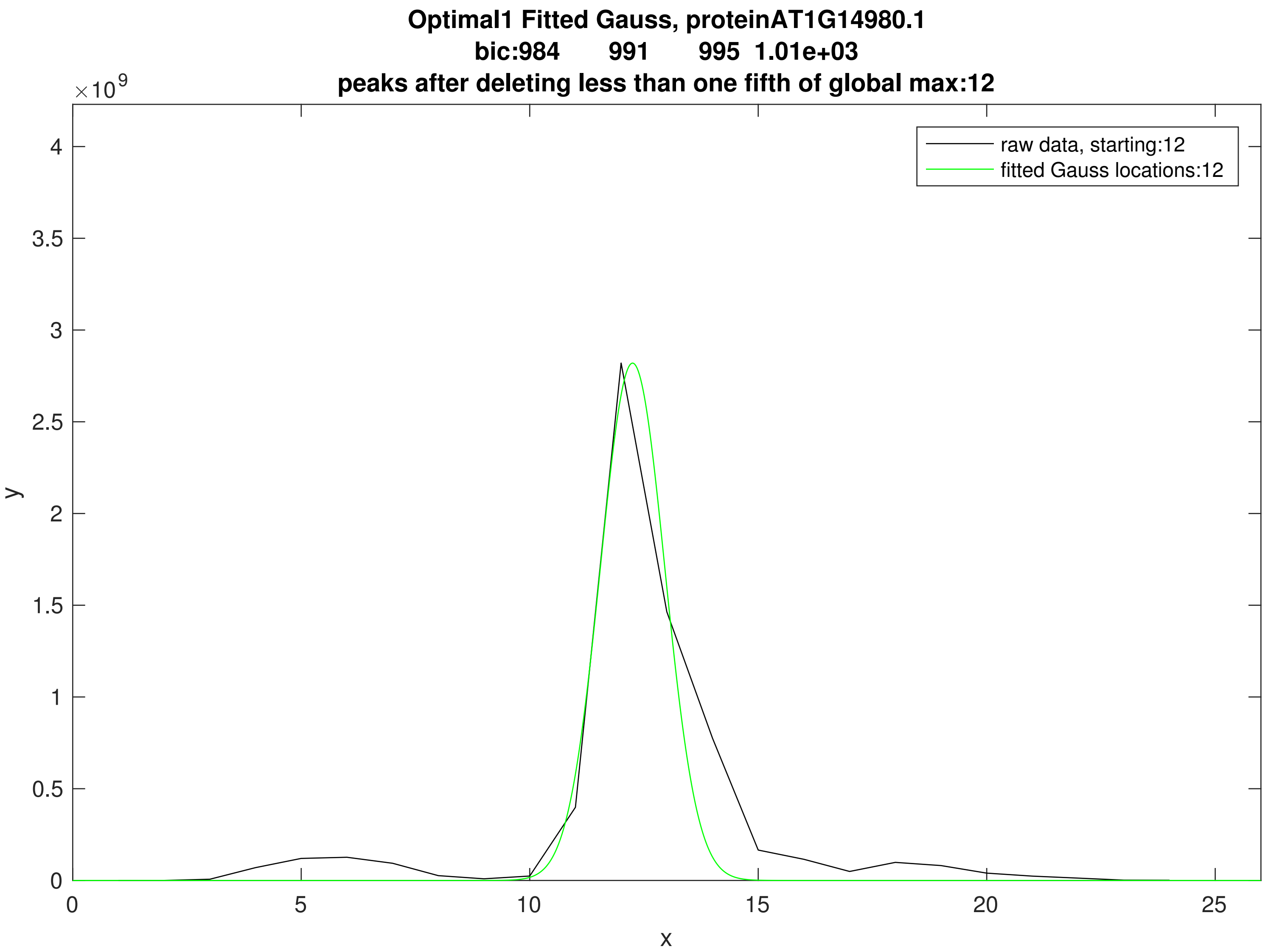
Optimal1 Fitted Gauss, proteinAT1G14980.1
bic:984 991 995 1.01e+03
peaks after deleting less than one fifth of global max:12
×10^9
raw data, starting:12
fitted Gauss locations:12
y
x
0
0.5
1
1.5
2
2.5
3
3.5
4
0
5
10
15
20
25

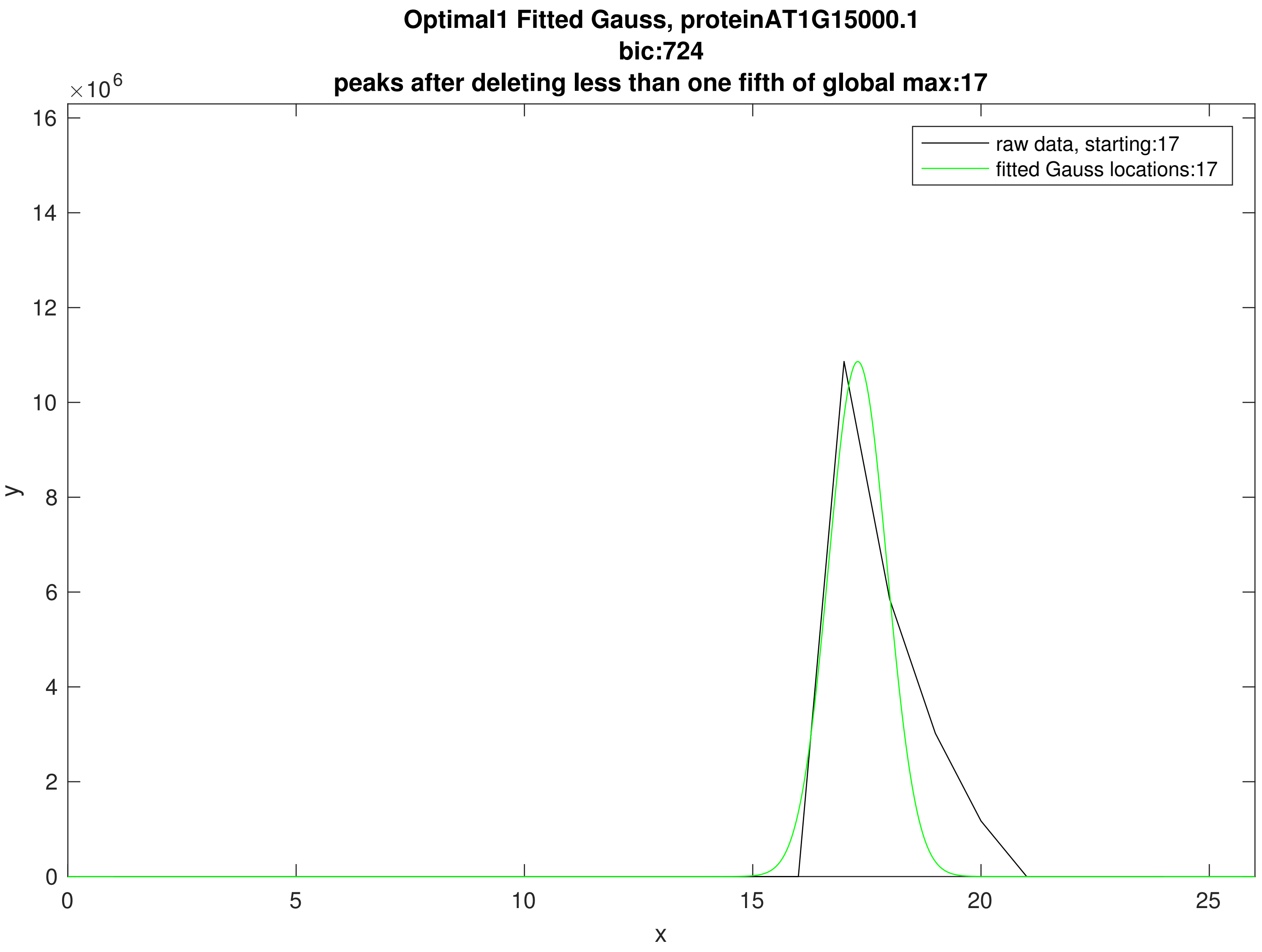
Optimal1 Fitted Gauss, proteinAT1G15000.1
bic:724
peaks after deleting less than one fifth of global max:17
×10^6
16
14
12
10
8
6
4
2
0
y
0
5
10
15
20
25
x
raw data, starting:17
fitted Gauss locations:17

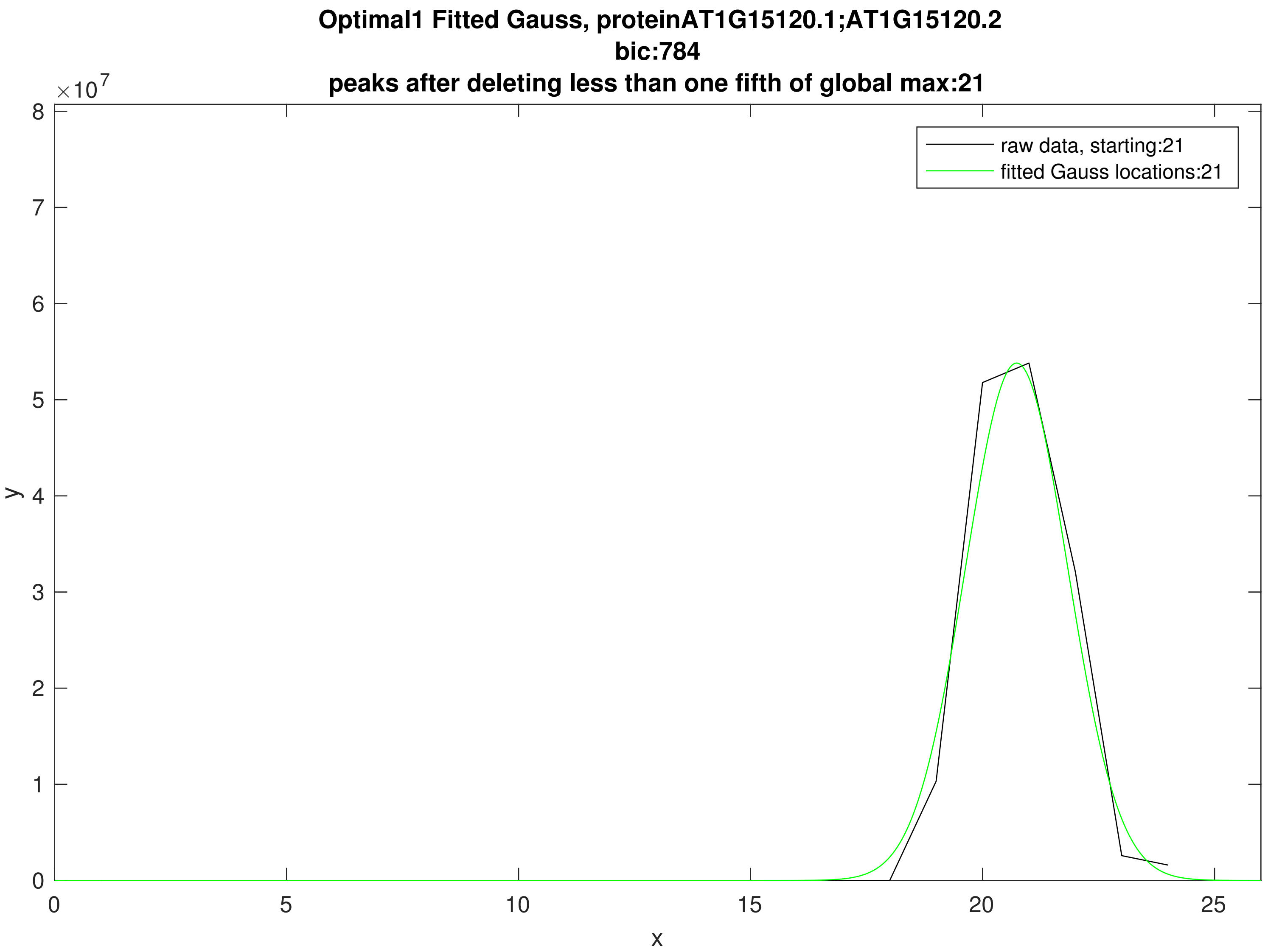
Optimal1 Fitted Gauss, proteinAT1G15120.1;AT1G15120.2
bic:784
peaks after deleting less than one fifth of global max:21
×10^7
raw data, starting:21
fitted Gauss locations:21
y
x
0
1
2
3
4
5
6
7
8
0
5
10
15
20
25

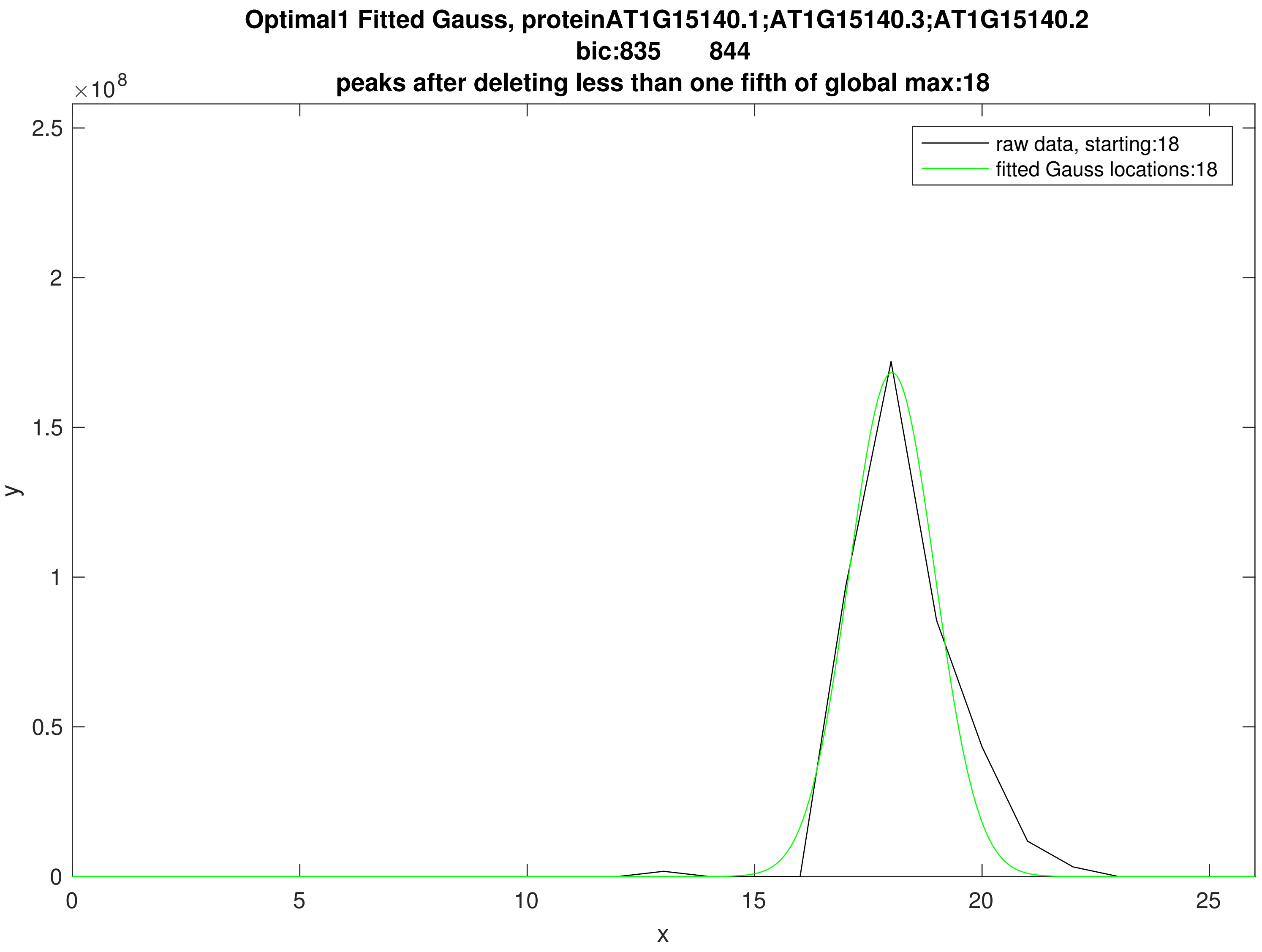

Optimal1 Fitted Gauss, proteinAT1G15140.1;AT1G15140.3;AT1G15140.2
bic:835 844
peaks after deleting less than one fifth of global max:18
×10^8
2.5
2
1.5
1
0.5
0
y
0
5
10
15
20
25
x
raw data, starting:18
fitted Gauss locations:18

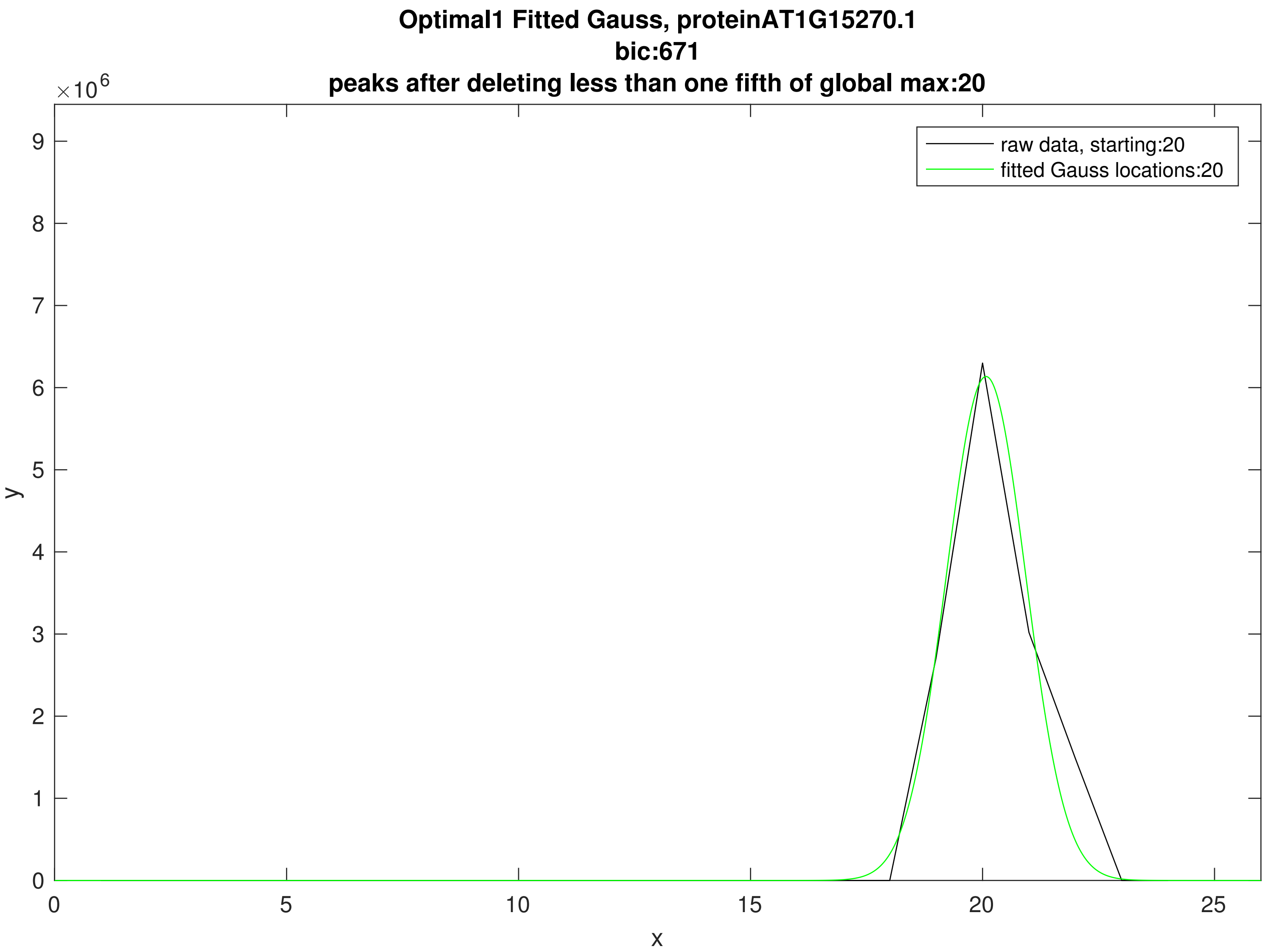
Optimal1 Fitted Gauss, proteinAT1G15270.1
bic:671
peaks after deleting less than one fifth of global max:20
×10^6
raw data, starting:20
fitted Gauss locations:20
y
x
0
1
2
3
4
5
6
7
8
9
0
5
10
15
20
25

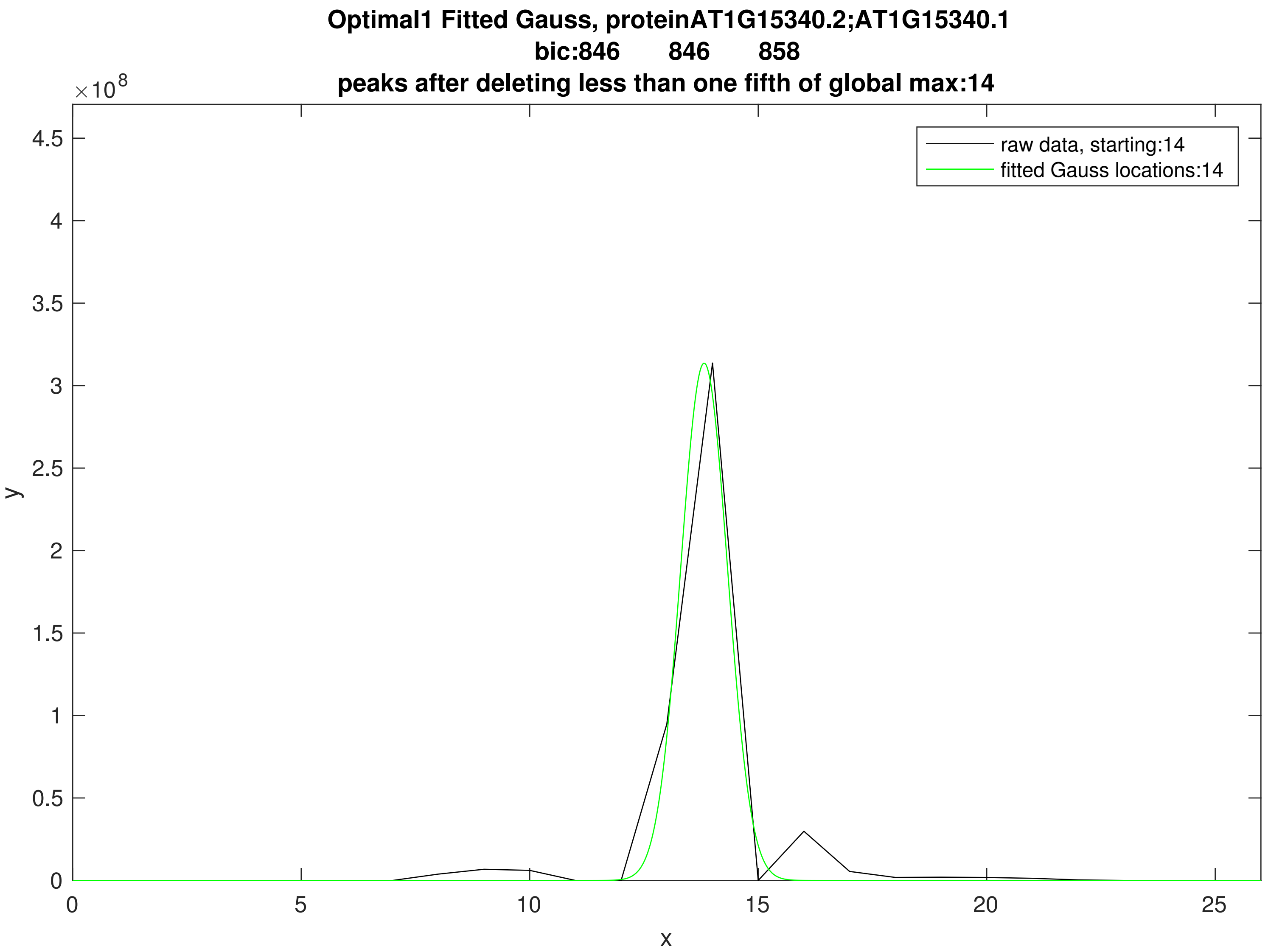
Optimal1 Fitted Gauss, proteinAT1G15340.2;AT1G15340.1
bic:846 846 858
peaks after deleting less than one fifth of global max:14
×10^8
4.5
4
3.5
3
2.5
2
1.5
1
0.5
0
y
0
5
10
15
20
25
x
raw data, starting:14
fitted Gauss locations:14

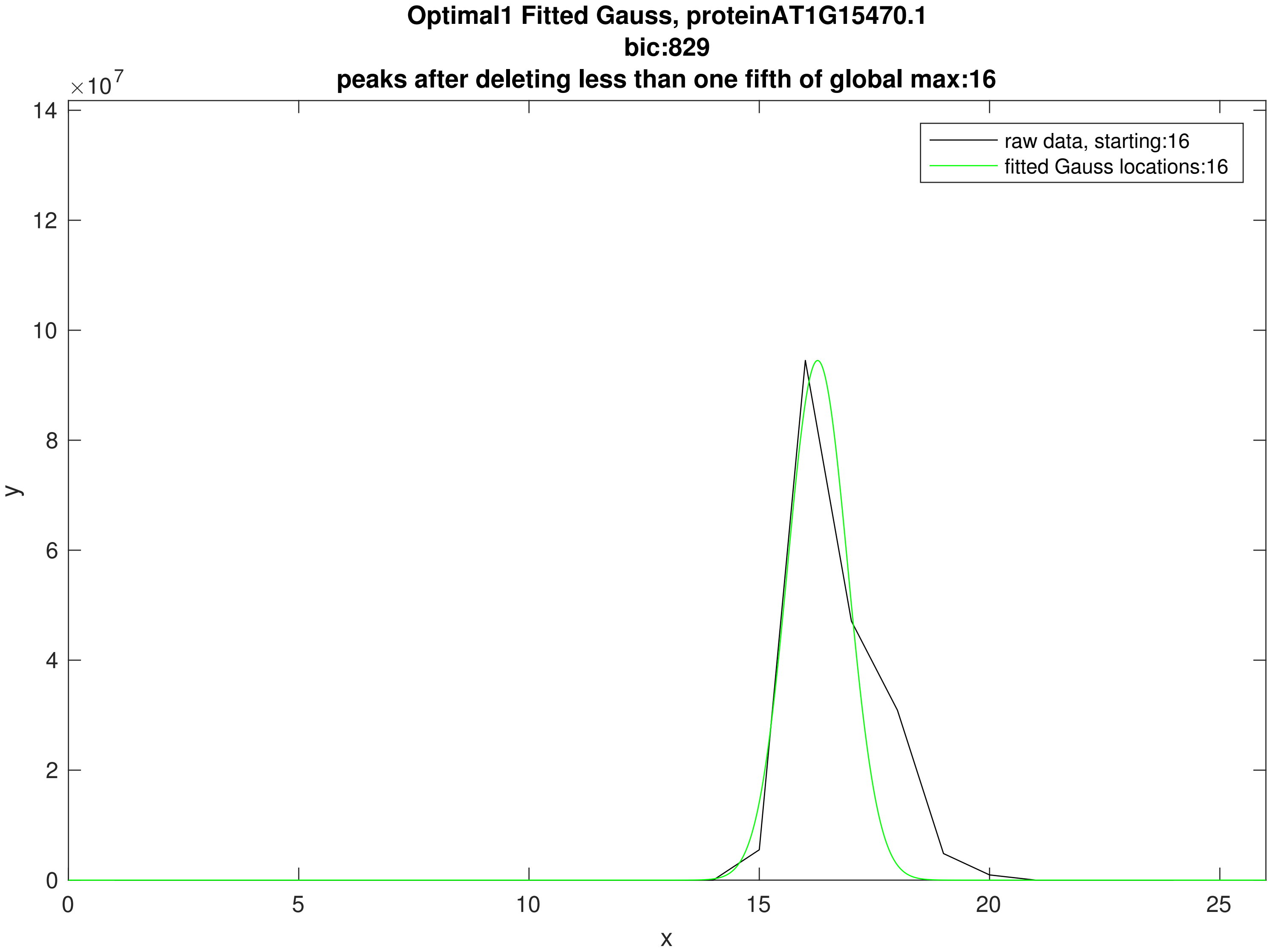

Optimal1 Fitted Gauss, proteinAT1G15470.1
bic:829
peaks after deleting less than one fifth of global max:16
×10^7
14
12
10
8
6
4
2
0
y
0
5
10
15
20
25
x
raw data, starting:16
fitted Gauss locations:16

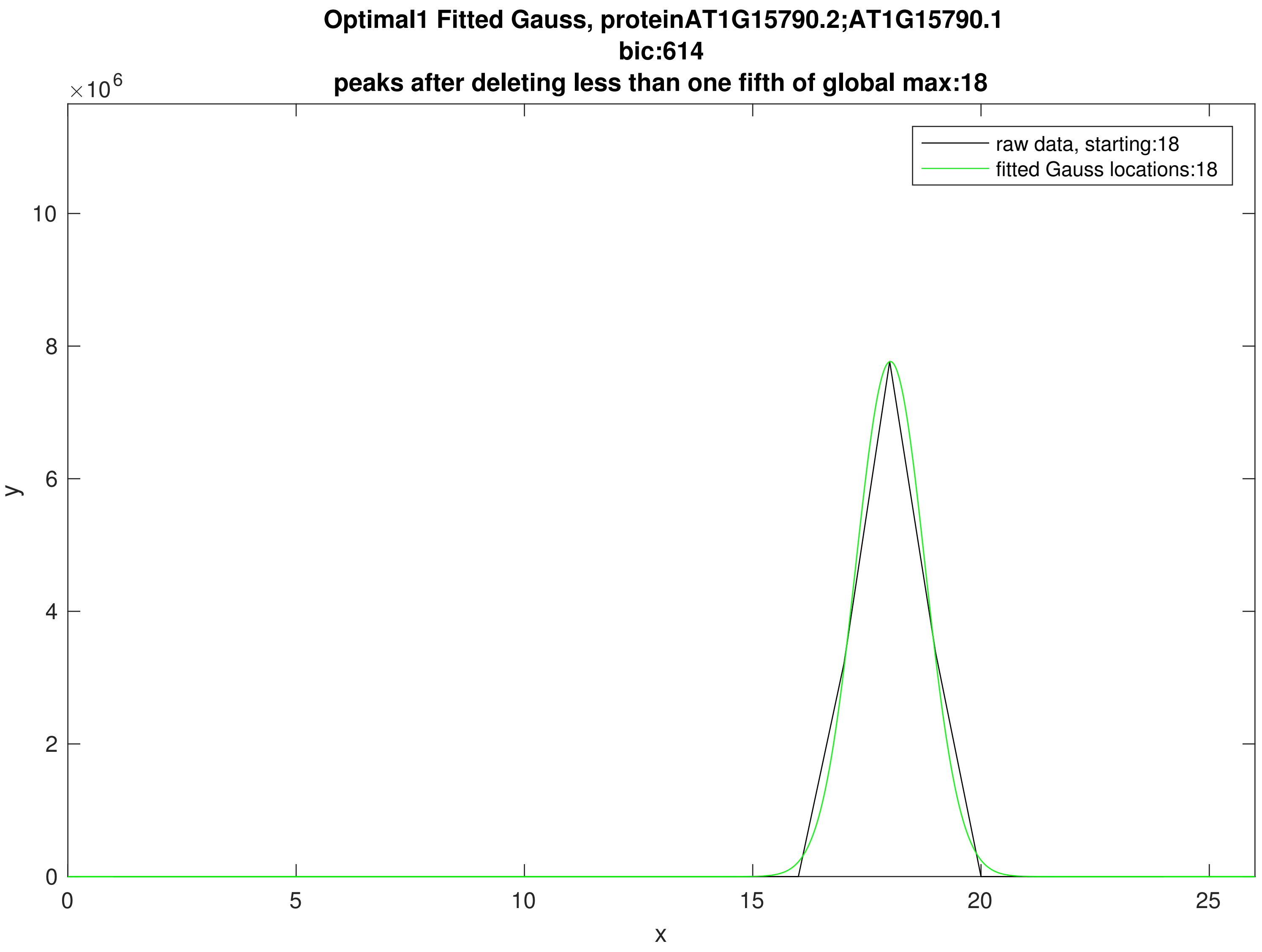
Optimal1 Fitted Gauss, proteinAT1G15790.2;AT1G15790.1
bic:614
peaks after deleting less than one fifth of global max:18
×10^6
raw data, starting:18
fitted Gauss locations:18
10
8
6
4
2
0
y
0
5
10
15
20
25
x

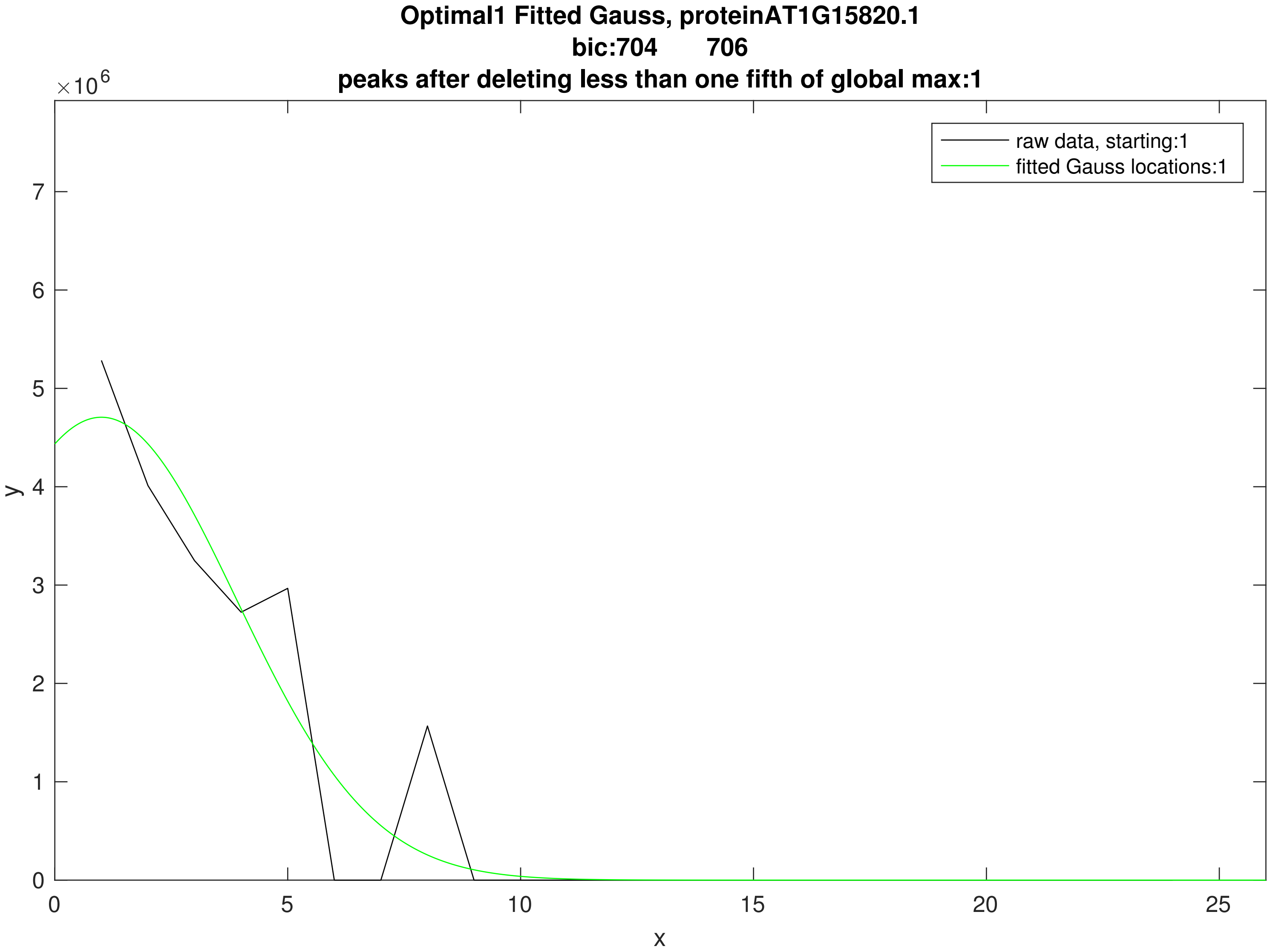
Optimal1 Fitted Gauss, proteinAT1G15820.1
bic:704 706
peaks after deleting less than one fifth of global max:1
×10^6
raw data, starting:1
fitted Gauss locations:1
y
x
0
1
2
3
4
5
6
7
0
5
10
15
20
25

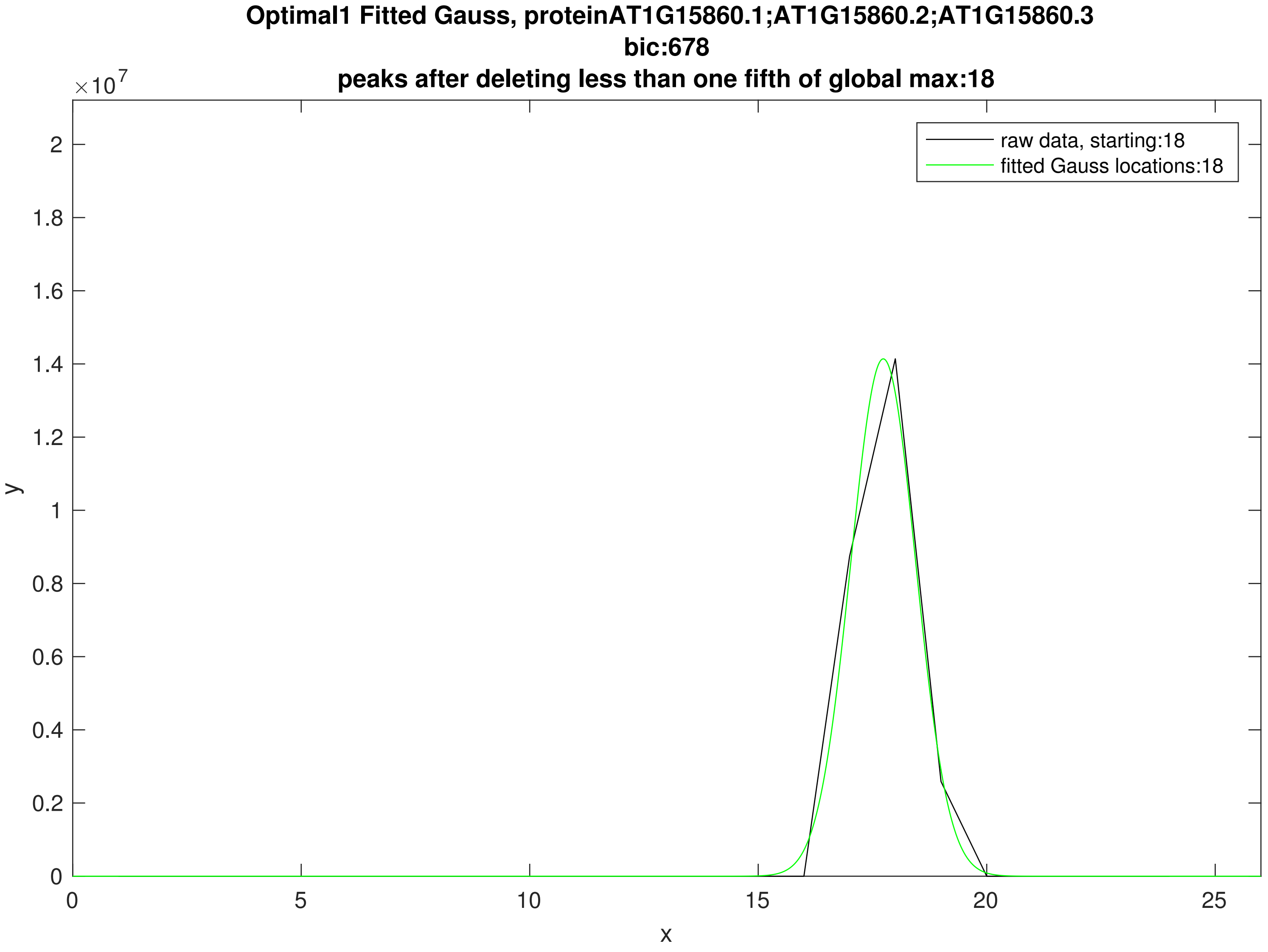

Optimal1 Fitted Gauss, proteinAT1G15860.1;AT1G15860.2;AT1G15860.3
bic:678
peaks after deleting less than one fifth of global max:18
×10^7
raw data, starting:18
fitted Gauss locations:18
y
x
0
0.2
0.4
0.6
0.8
1
1.2
1.4
1.6
1.8
2
0
5
10
15
20
25

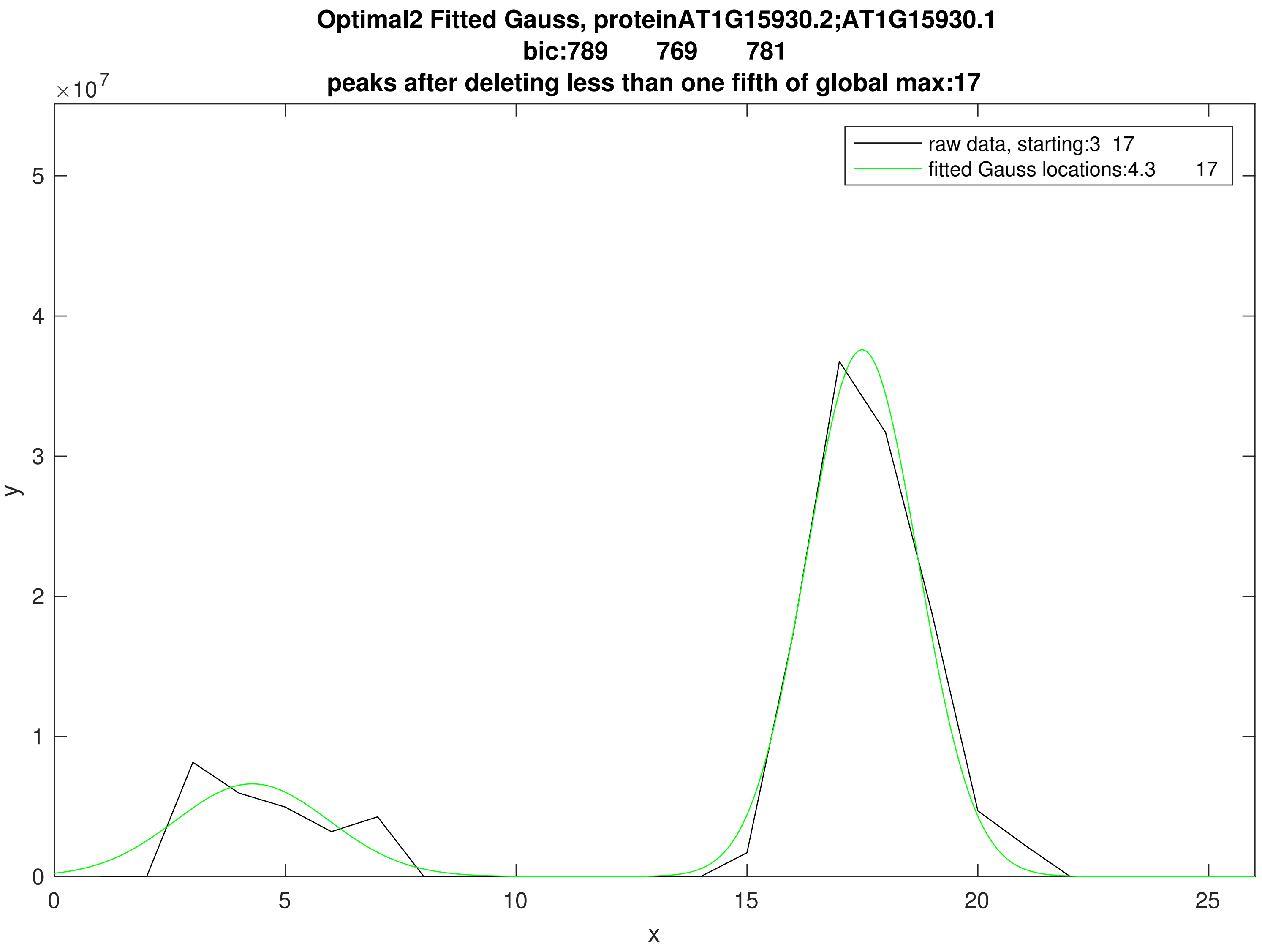
Optimal2 Fitted Gauss, proteinAT1G15930.2;AT1G15930.1
bic:789 769 781
peaks after deleting less than one fifth of global max:17
×10^7
raw data, starting:3 17
fitted Gauss locations:4.3 17
y
x
0
1
2
3
4
5
0
5
10
15
20
25

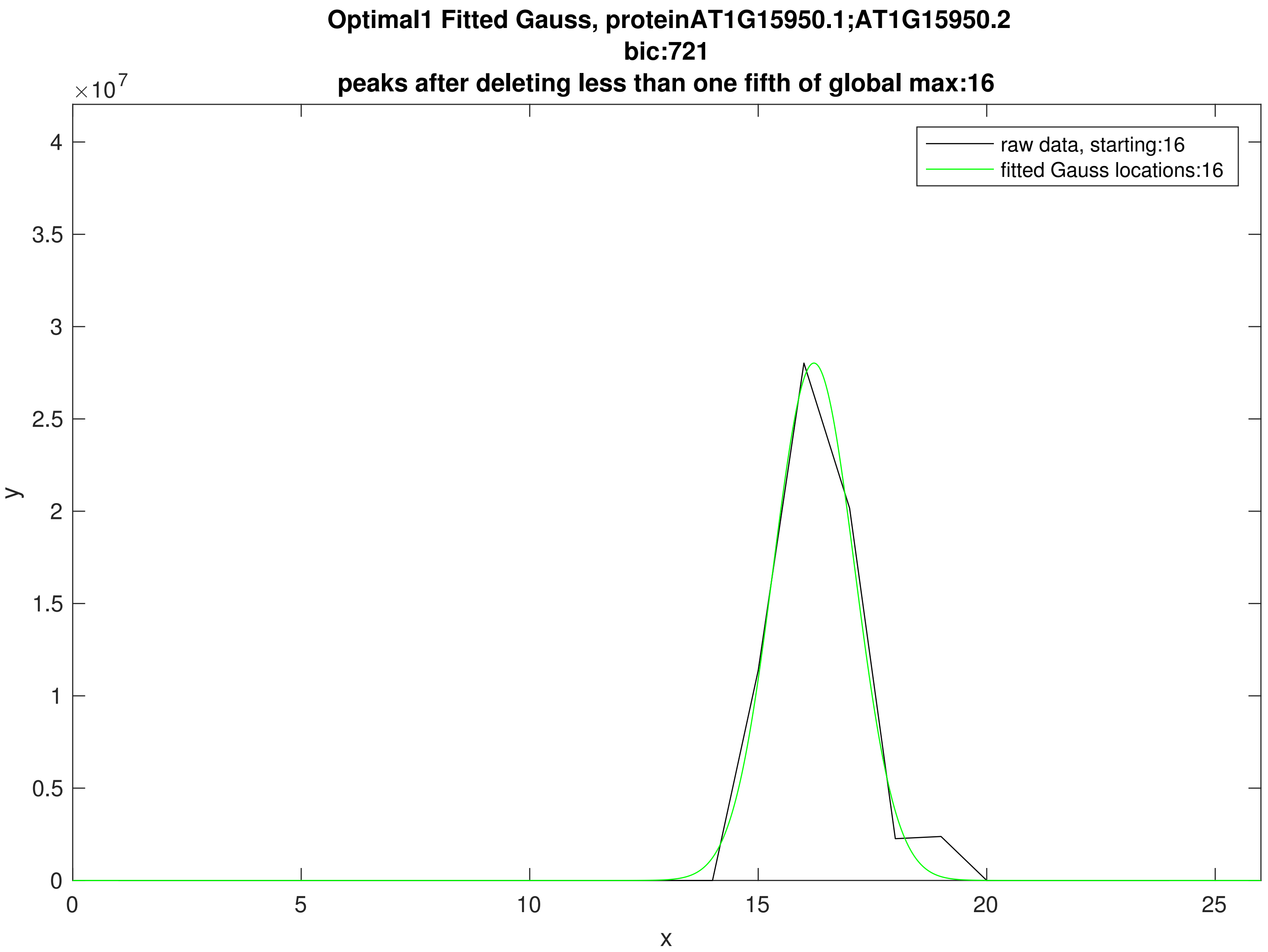

Optimal1 Fitted Gauss, proteinAT1G15950.1;AT1G15950.2
bic:721
peaks after deleting less than one fifth of global max:16
×10^7
raw data, starting:16
fitted Gauss locations:16
y
x
0
0.5
1
1.5
2
2.5
3
3.5
4
0
5
10
15
20
25

Optimal1 Fitted Gauss, proteinAT1G16030.1
bic:703
peaks after deleting less than one fifth of global max:12

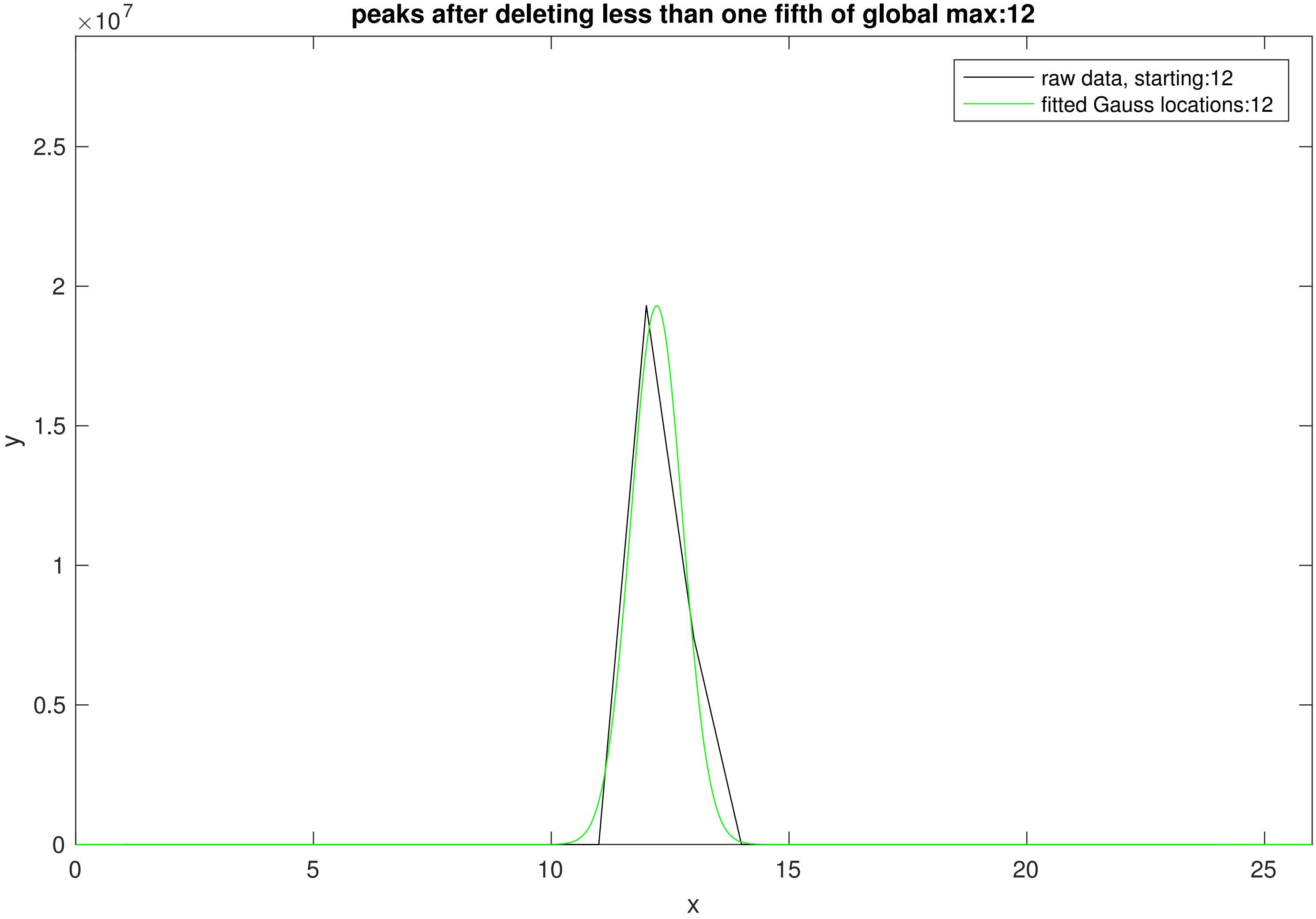

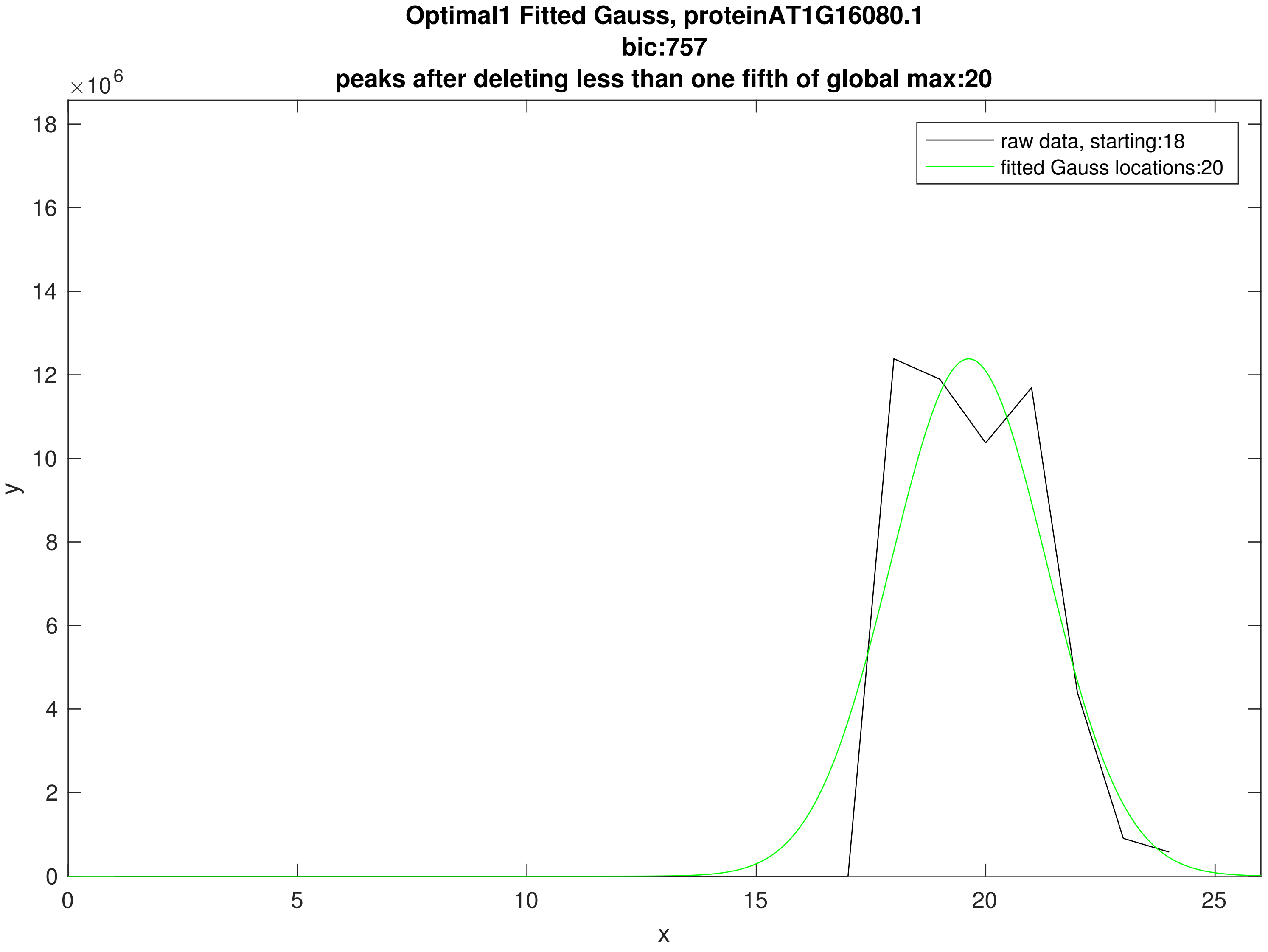
Optimal1 Fitted Gauss, proteinAT1G16080.1
bic:757
peaks after deleting less than one fifth of global max:20
×10^6
raw data, starting:18
fitted Gauss locations:20
y
x
0
2
4
6
8
10
12
14
16
18
0
5
10
15
20
25

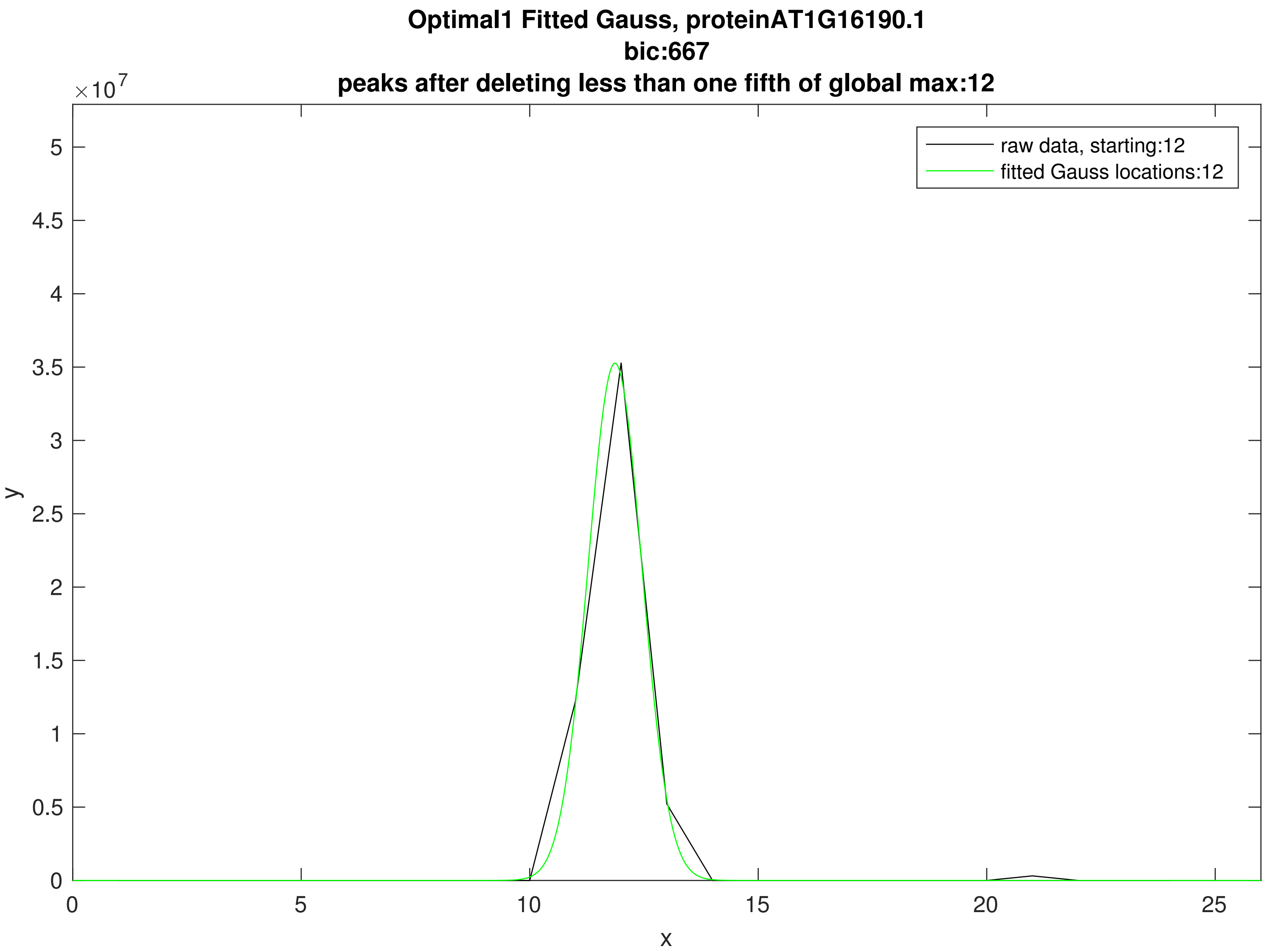
Optimal1 Fitted Gauss, proteinAT1G16190.1
bic:667
peaks after deleting less than one fifth of global max:12
×10^7
raw data, starting:12
fitted Gauss locations:12
y
x
0
0.5
1
1.5
2
2.5
3
3.5
4
4.5
5
0
5
10
15
20
25

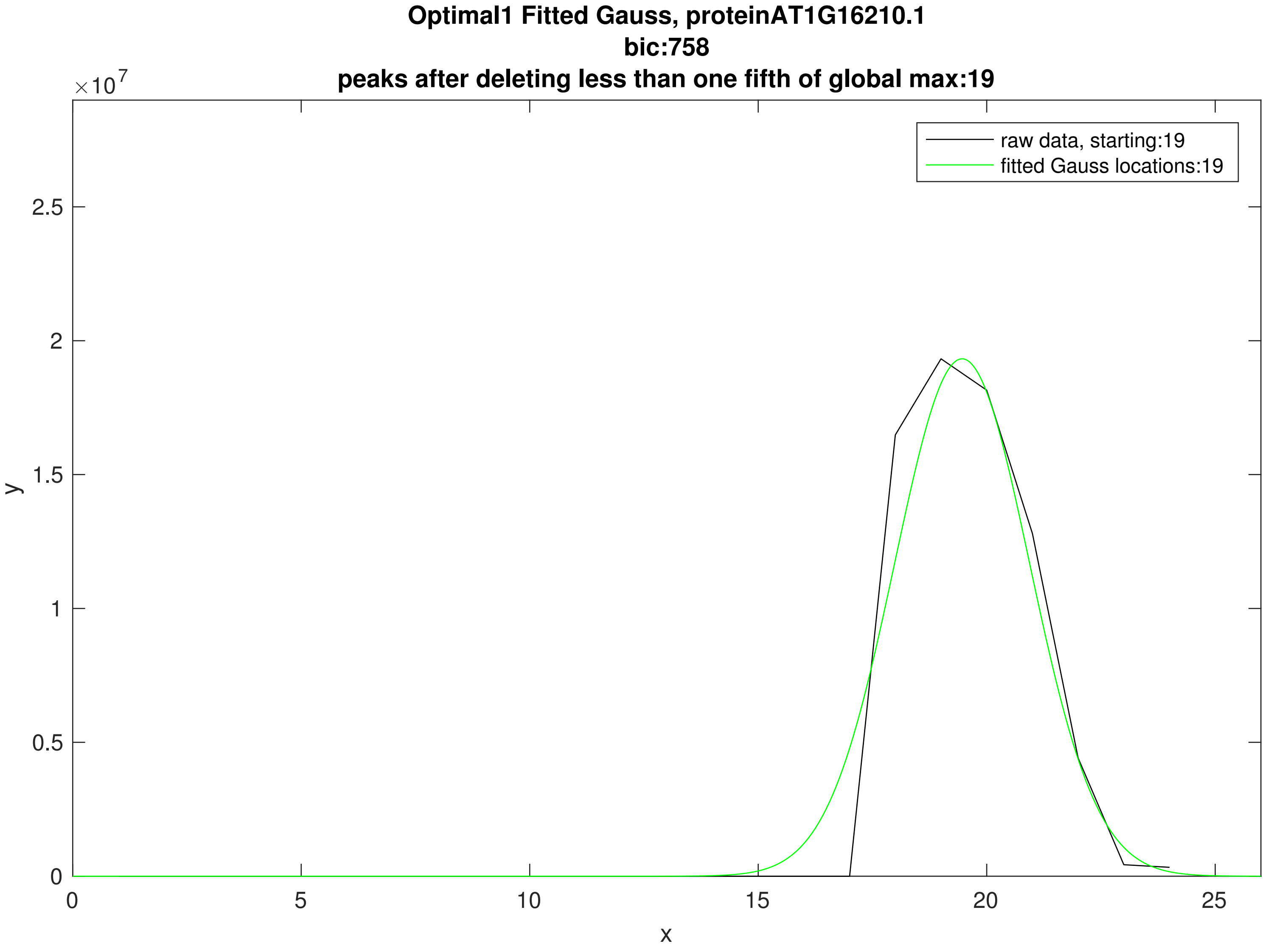
Optimal1 Fitted Gauss, proteinAT1G16210.1
bic:758
peaks after deleting less than one fifth of global max:19
×10^7
raw data, starting:19
fitted Gauss locations:19
y
x
0
0.5
1
1.5
2
2.5
0
5
10
15
20
25

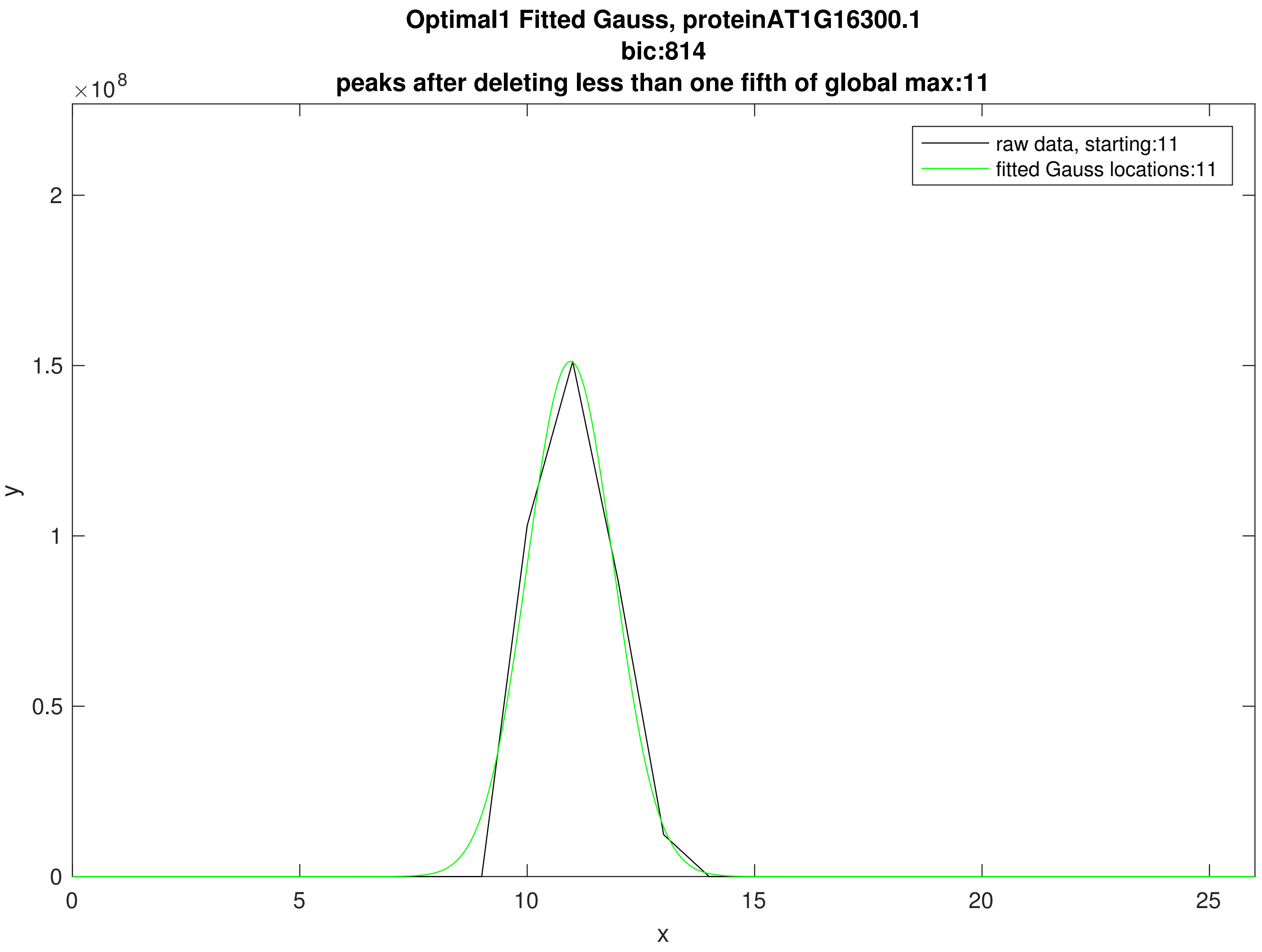
Optimal1 Fitted Gauss, proteinAT1G16300.1
bic:814
peaks after deleting less than one fifth of global max:11
×10^8
raw data, starting:11
fitted Gauss locations:11
y
x
0
0.5
1
1.5
2
0
5
10
15
20
25

**Optimal1 Fitted Gauss, proteinAT1G16340.3;AT1G16340.2;AT1G16340.1;AT1G16340.4**
**bic:755**
**peaks after deleting less than one fifth of global max:12**

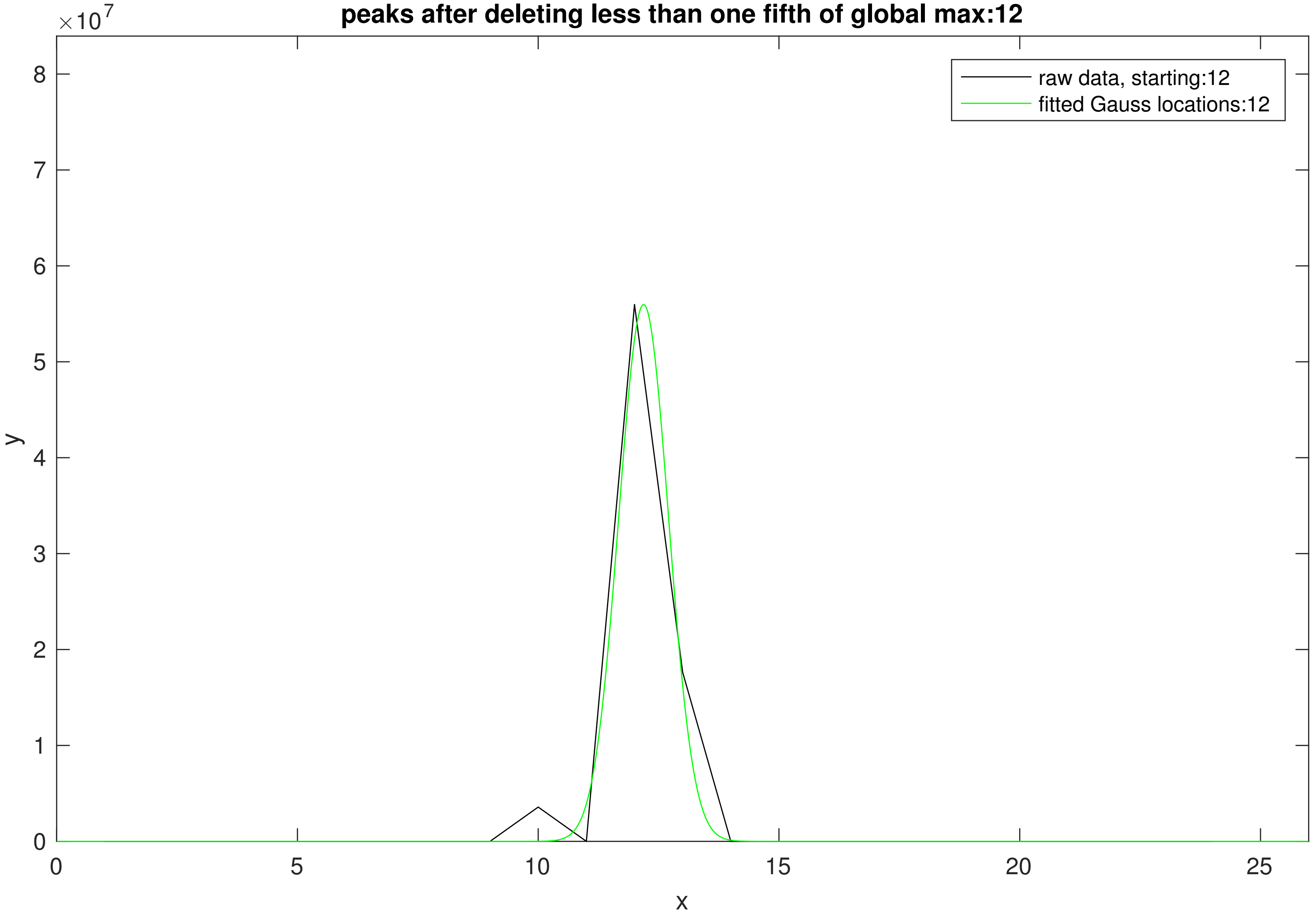

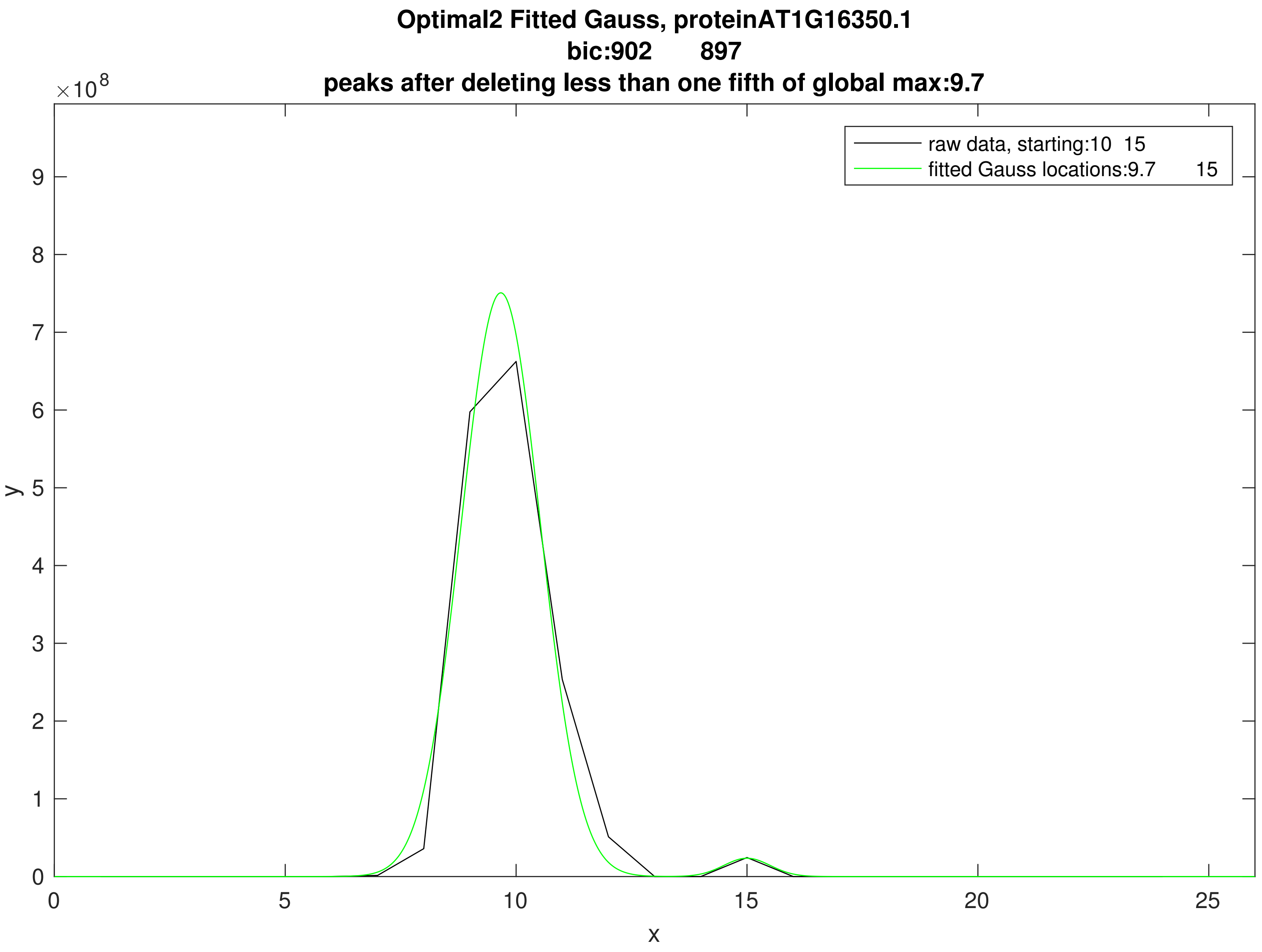
Optimal2 Fitted Gauss, proteinAT1G16350.1
bic:902 897
peaks after deleting less than one fifth of global max:9.7
×10^8
raw data, starting:10 15
fitted Gauss locations:9.7 15
y
x
0
1
2
3
4
5
6
7
8
9
0
5
10
15
20
25

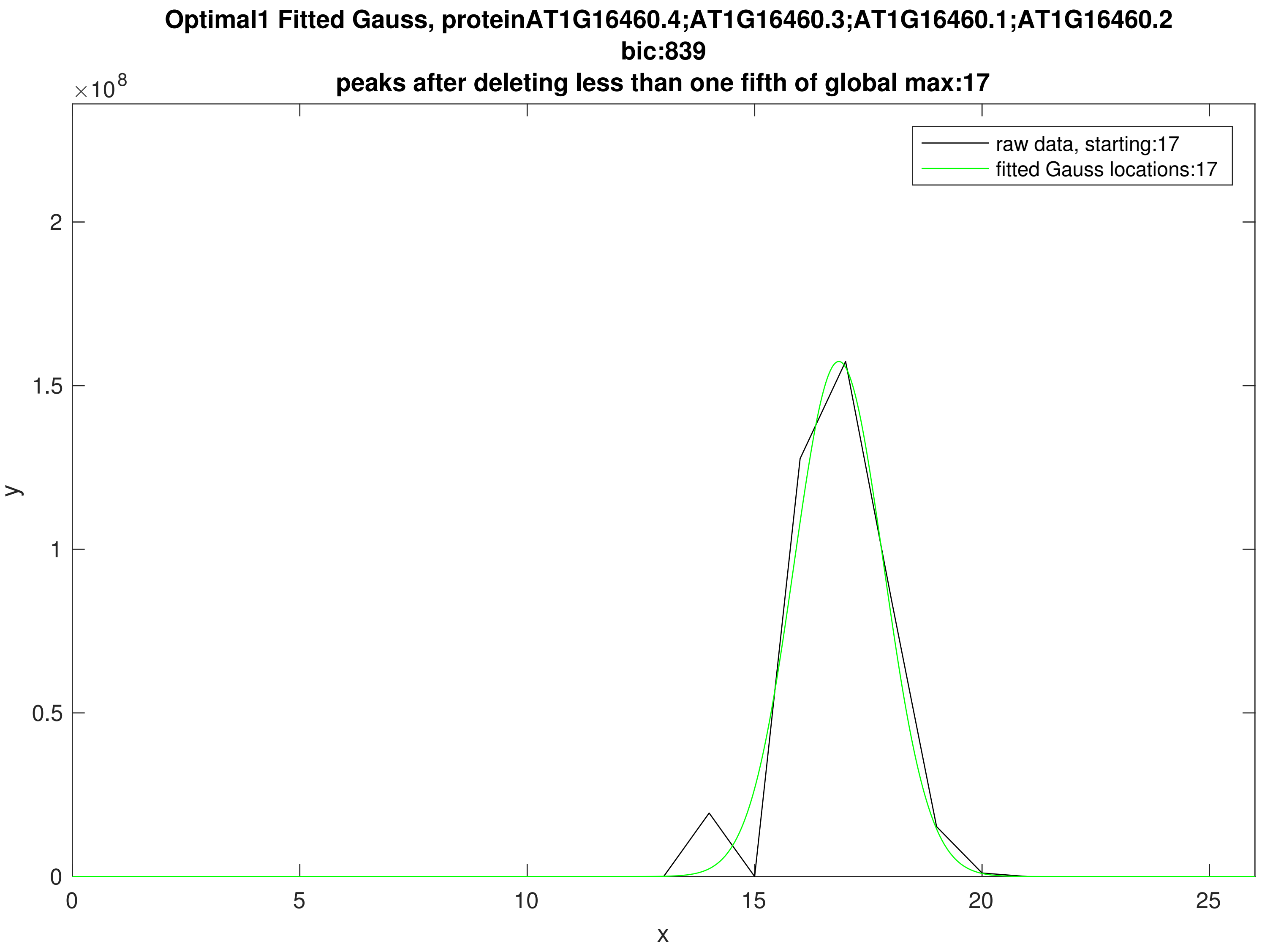

Optimal1 Fitted Gauss, proteinAT1G16460.4;AT1G16460.3;AT1G16460.1;AT1G16460.2
bic:839
peaks after deleting less than one fifth of global max:17
×10^8
raw data, starting:17
fitted Gauss locations:17
y
x
0
0.5
1
1.5
2
0
5
10
15
20
25

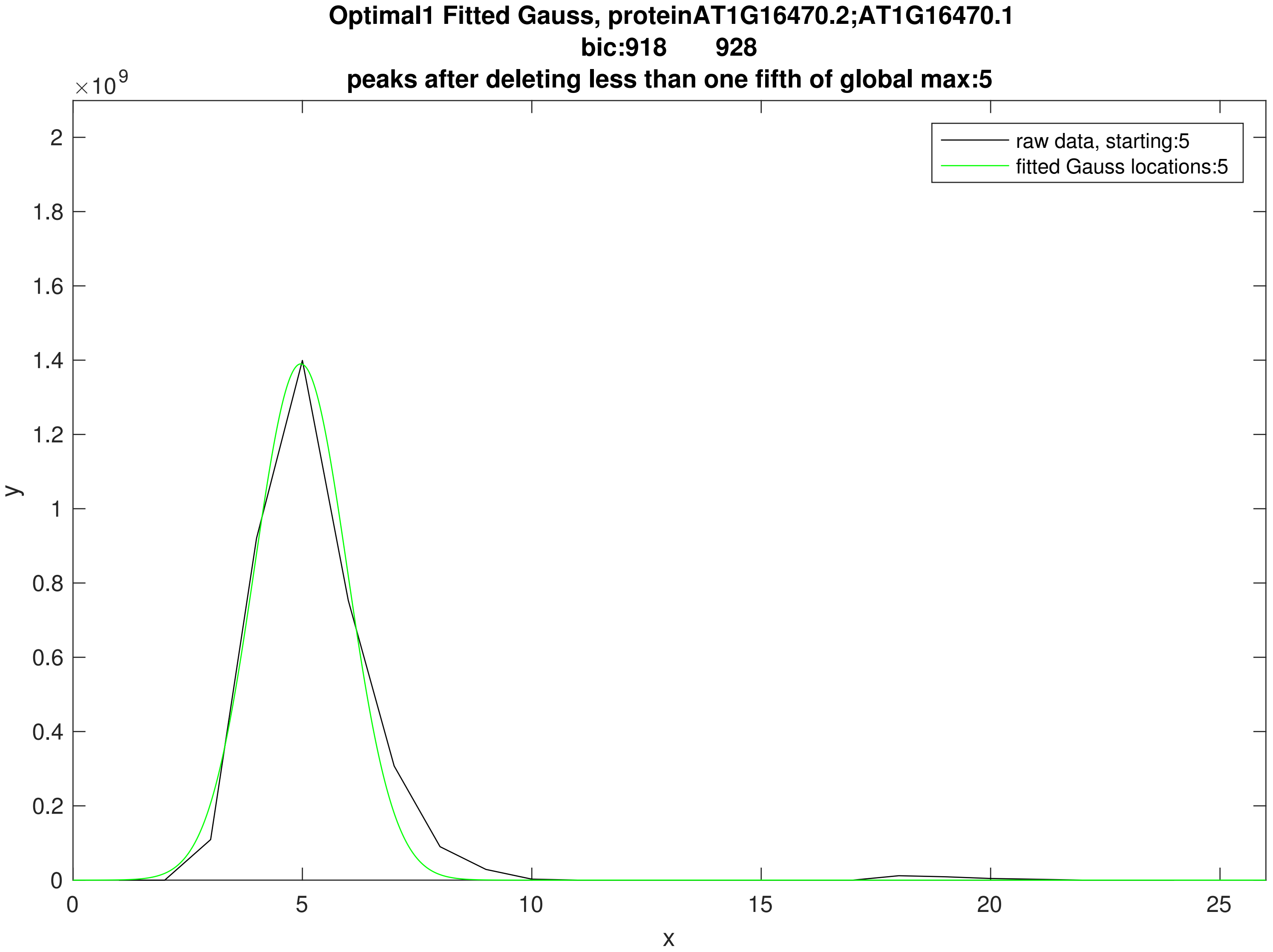
Optimal1 Fitted Gauss, proteinAT1G16470.2;AT1G16470.1
bic:918 928
peaks after deleting less than one fifth of global max:5
×10^9
2
1.8
1.6
1.4
1.2
1
0.8
0.6
0.4
0.2
0
y
0
5
10
15
20
25
x
raw data, starting:5
fitted Gauss locations:5

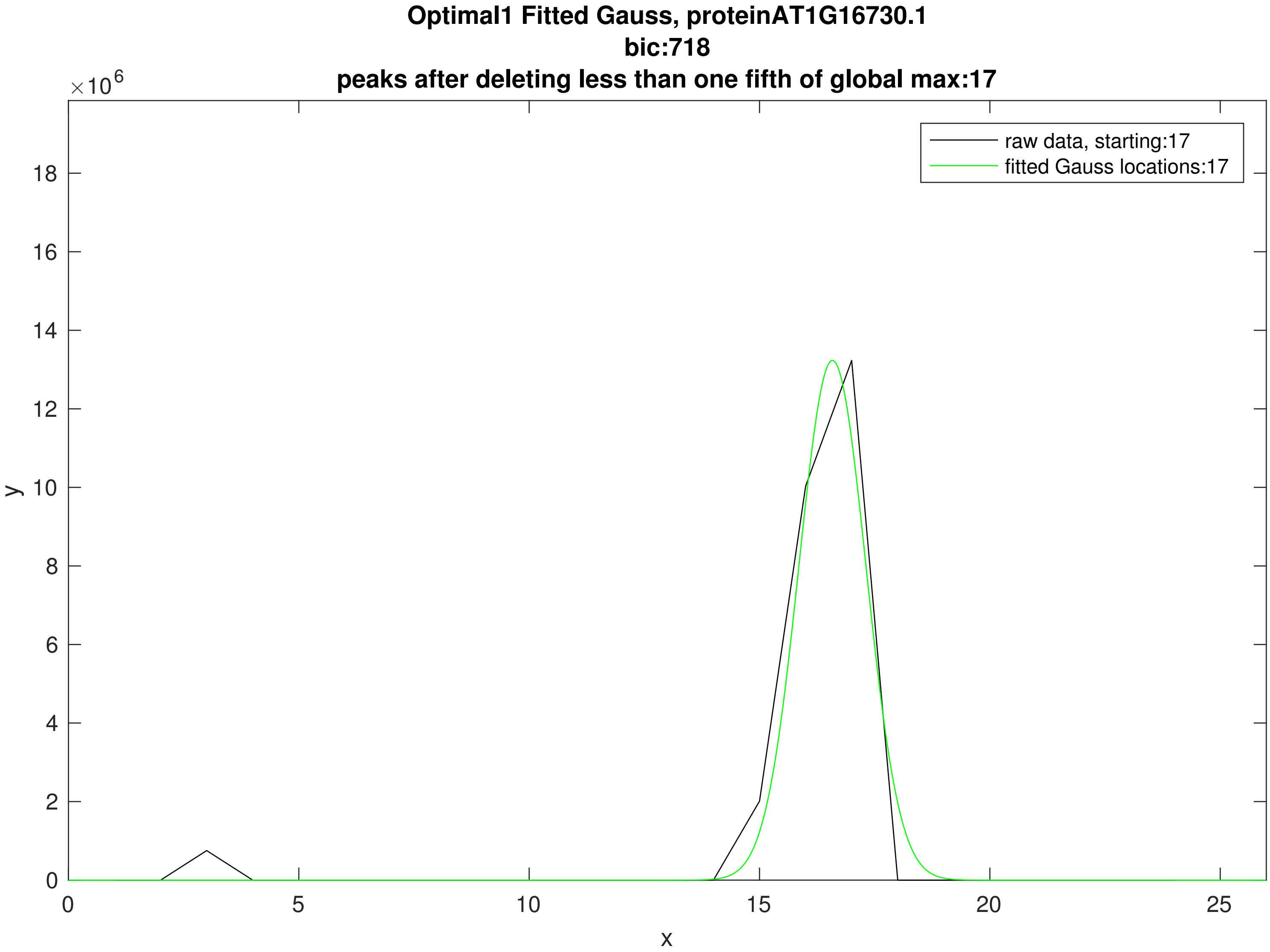

Optimal1 Fitted Gauss, proteinAT1G16730.1
bic:718
peaks after deleting less than one fifth of global max:17
×10^6
raw data, starting:17
fitted Gauss locations:17
y
x
0
2
4
6
8
10
12
14
16
18
0
5
10
15
20
25

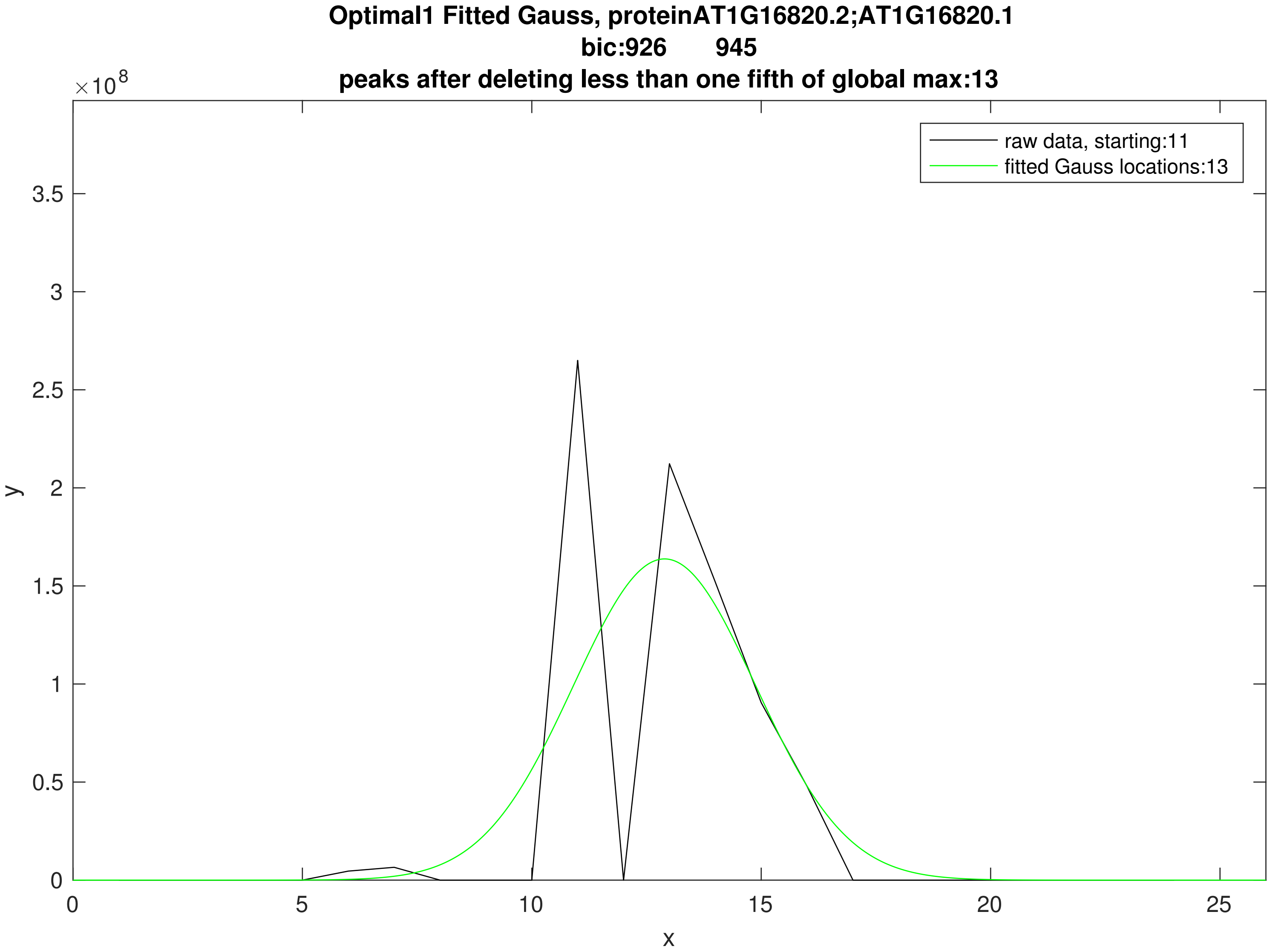

Optimal1 Fitted Gauss, proteinAT1G16820.2;AT1G16820.1
bic:926 945
peaks after deleting less than one fifth of global max:13
×10^8
raw data, starting:11
fitted Gauss locations:13
3.5
3
2.5
2
1.5
1
0.5
0
y
0
5
10
15
20
25
x

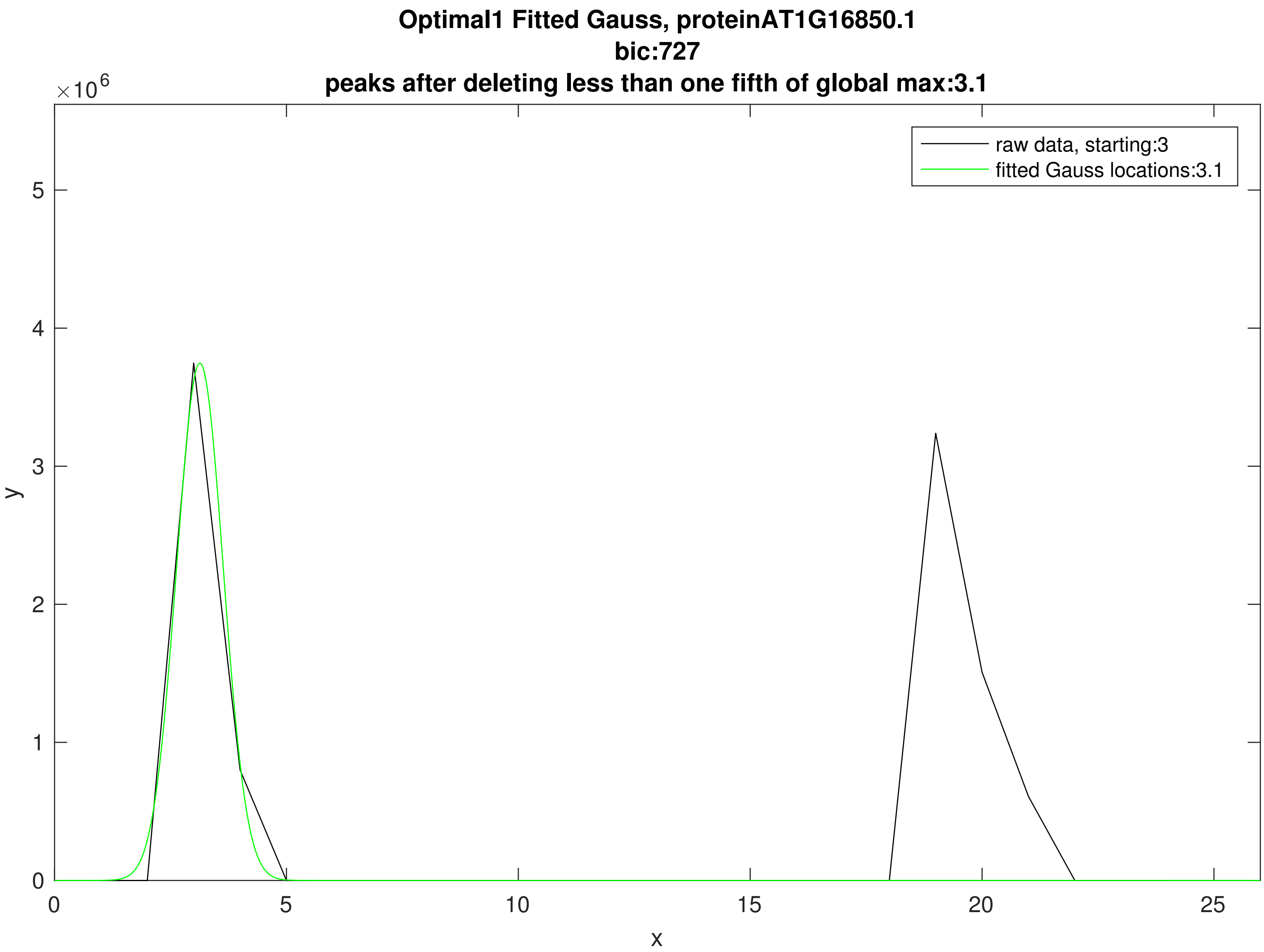
Optimal1 Fitted Gauss, proteinAT1G16850.1
bic:727
peaks after deleting less than one fifth of global max:3.1
×10^6
raw data, starting:3
fitted Gauss locations:3.1
y
x
0
1
2
3
4
5
0
5
10
15
20
25

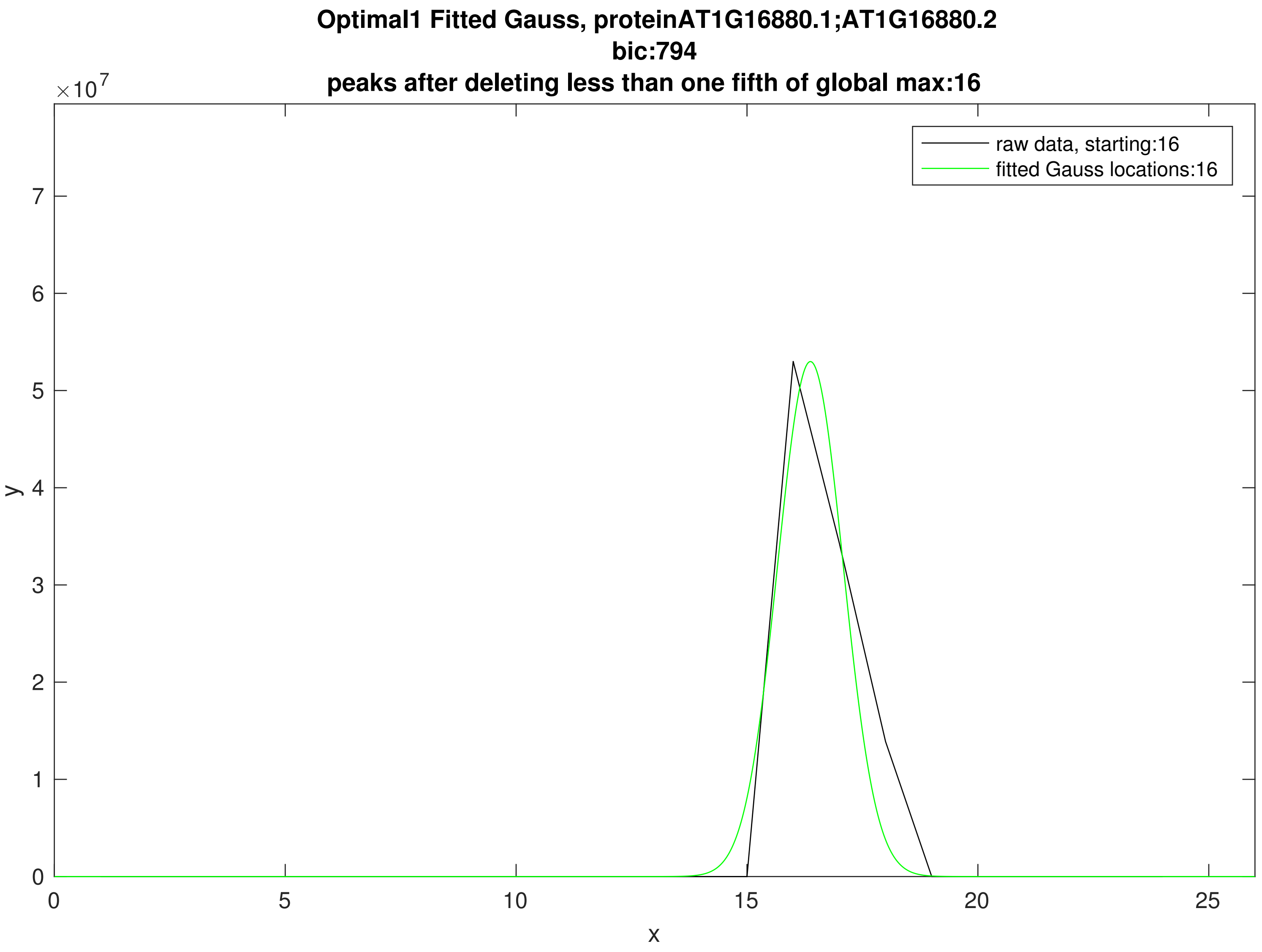
Optimal1 Fitted Gauss, proteinAT1G16880.1;AT1G16880.2
bic:794
peaks after deleting less than one fifth of global max:16
×10^7
raw data, starting:16
fitted Gauss locations:16
y
x
0
1
2
3
4
5
6
7
0
5
10
15
20
25

Optimal1 Fitted Gauss, proteinAT1G16890.2;AT1G16890.3;AT1G16890.1
bic:802 811
peaks after deleting less than one fifth of global max:19

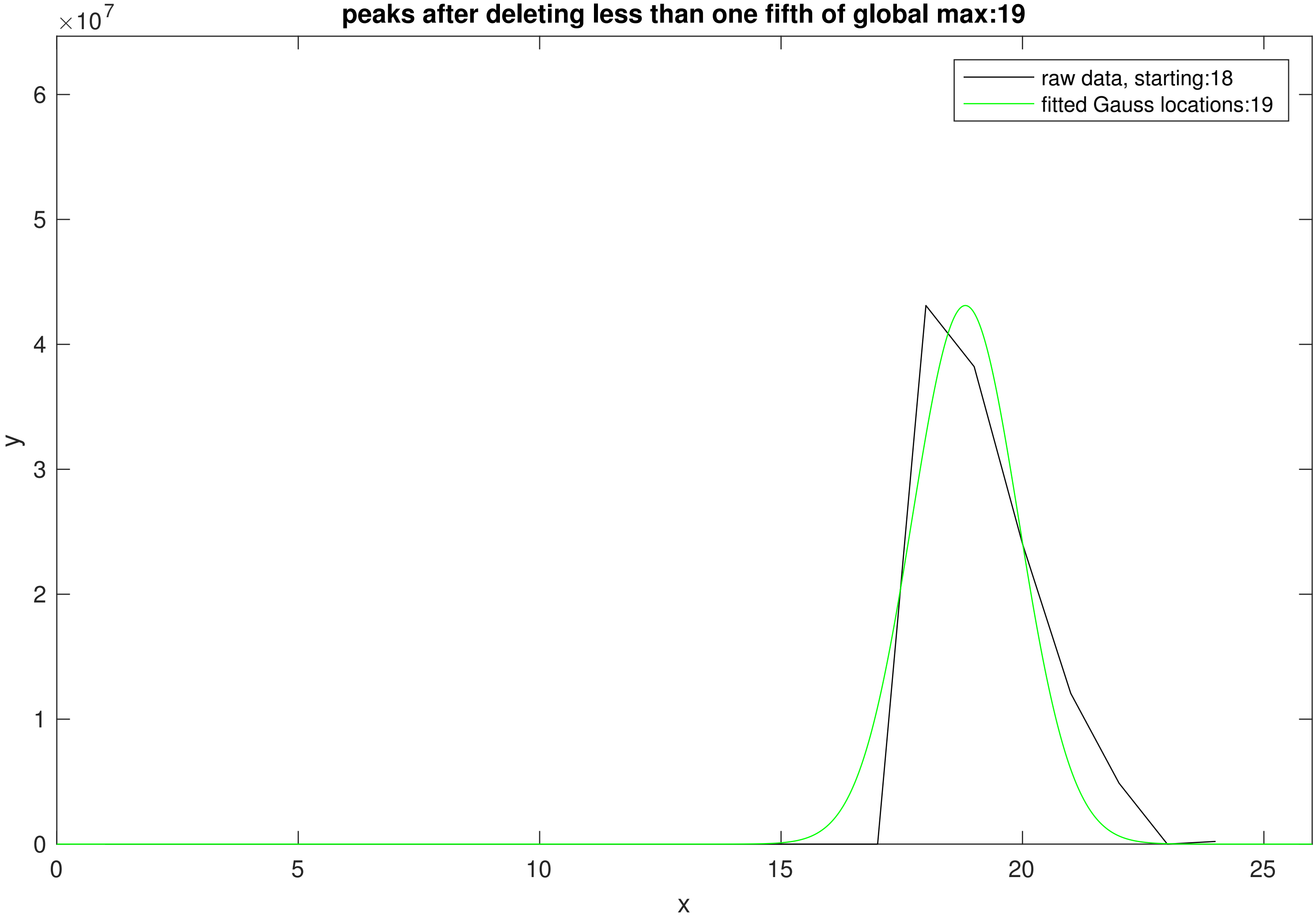

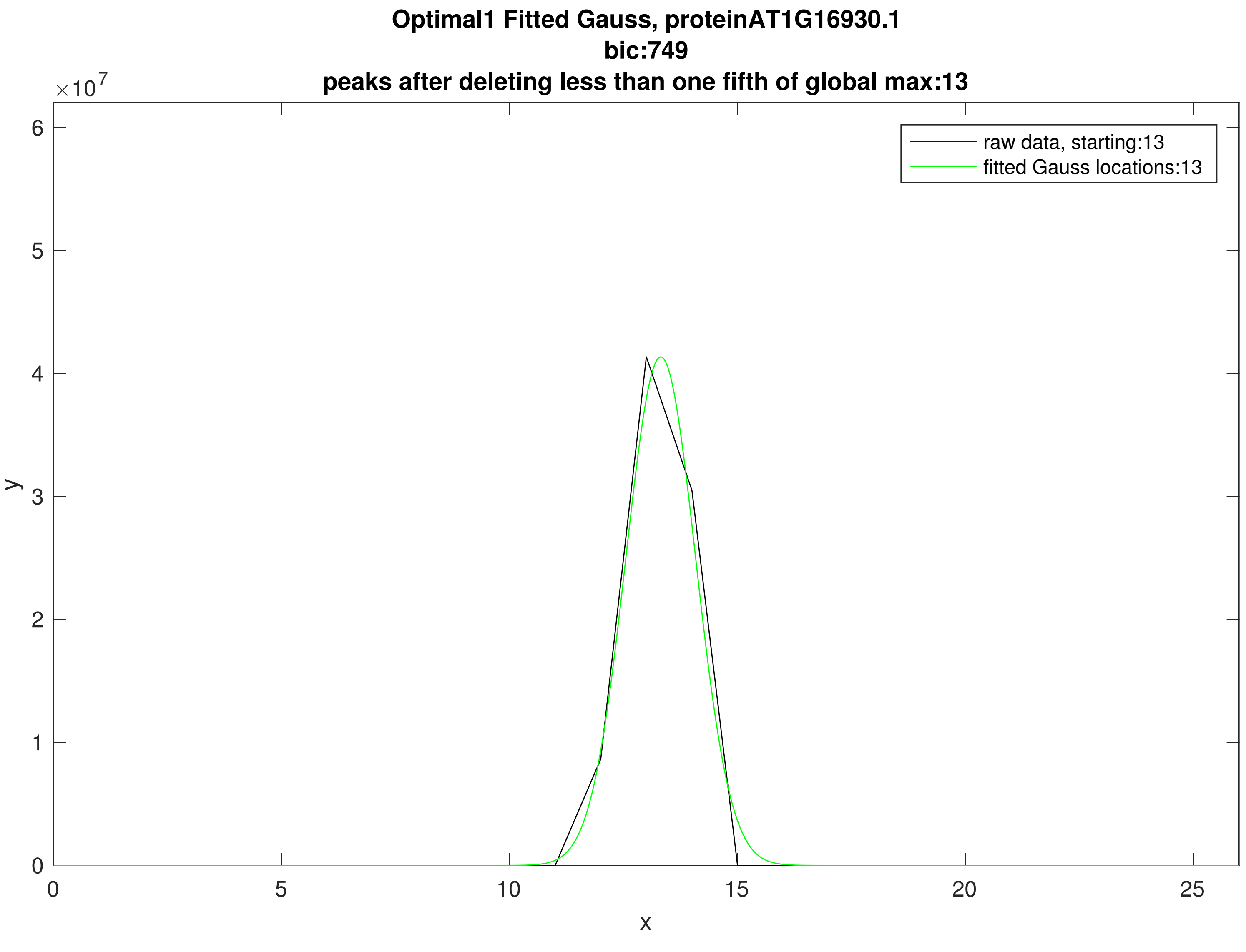
Optimal1 Fitted Gauss, proteinAT1G16930.1
bic:749
peaks after deleting less than one fifth of global max:13
×10^7
y
x
0
1
2
3
4
5
6
0
5
10
15
20
25
raw data, starting:13
fitted Gauss locations:13

#### Optimal1 Fitted Gauss, proteinAT1G17100.1
#### bic:709
#### peaks after deleting less than one fifth of global max:20

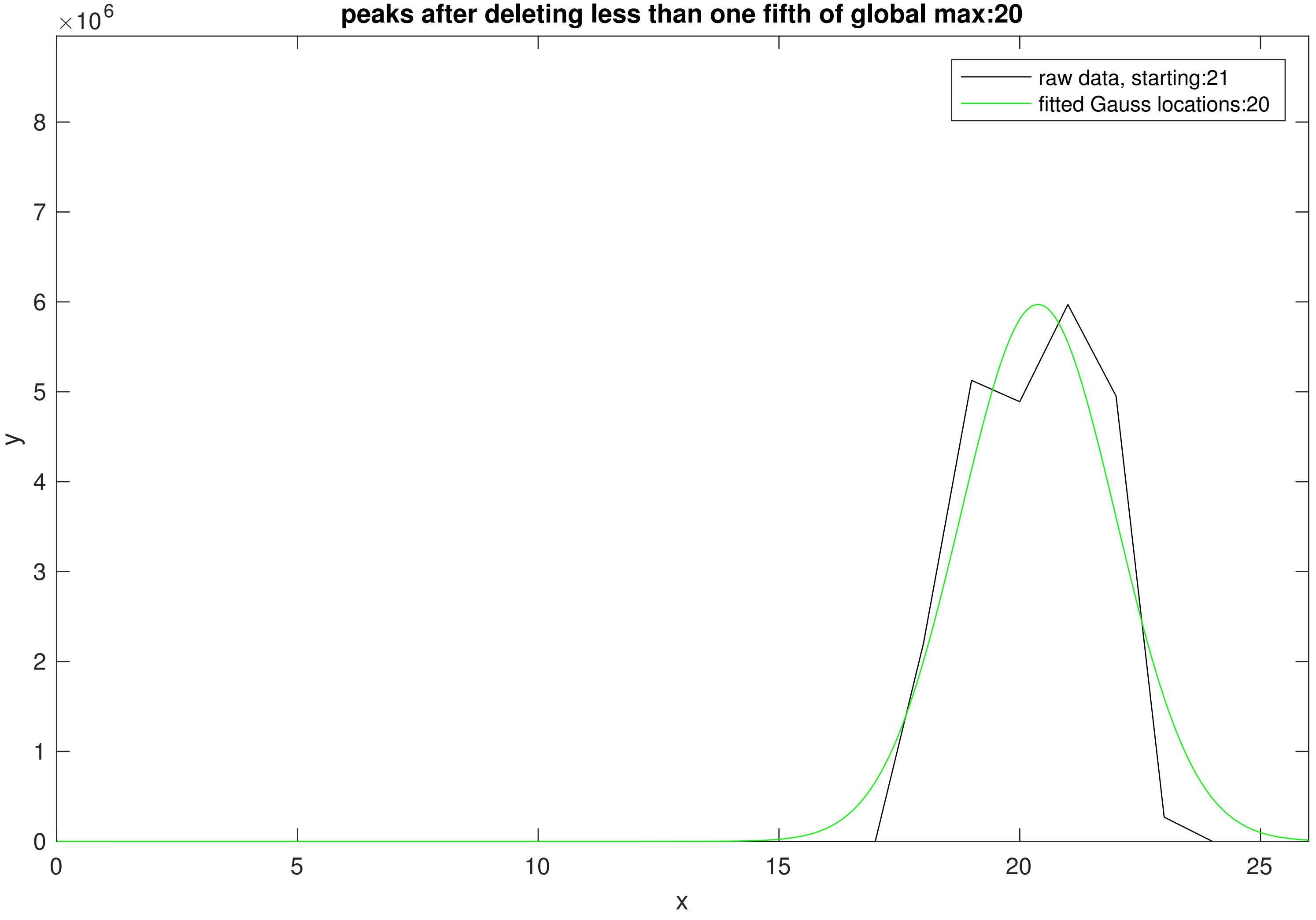

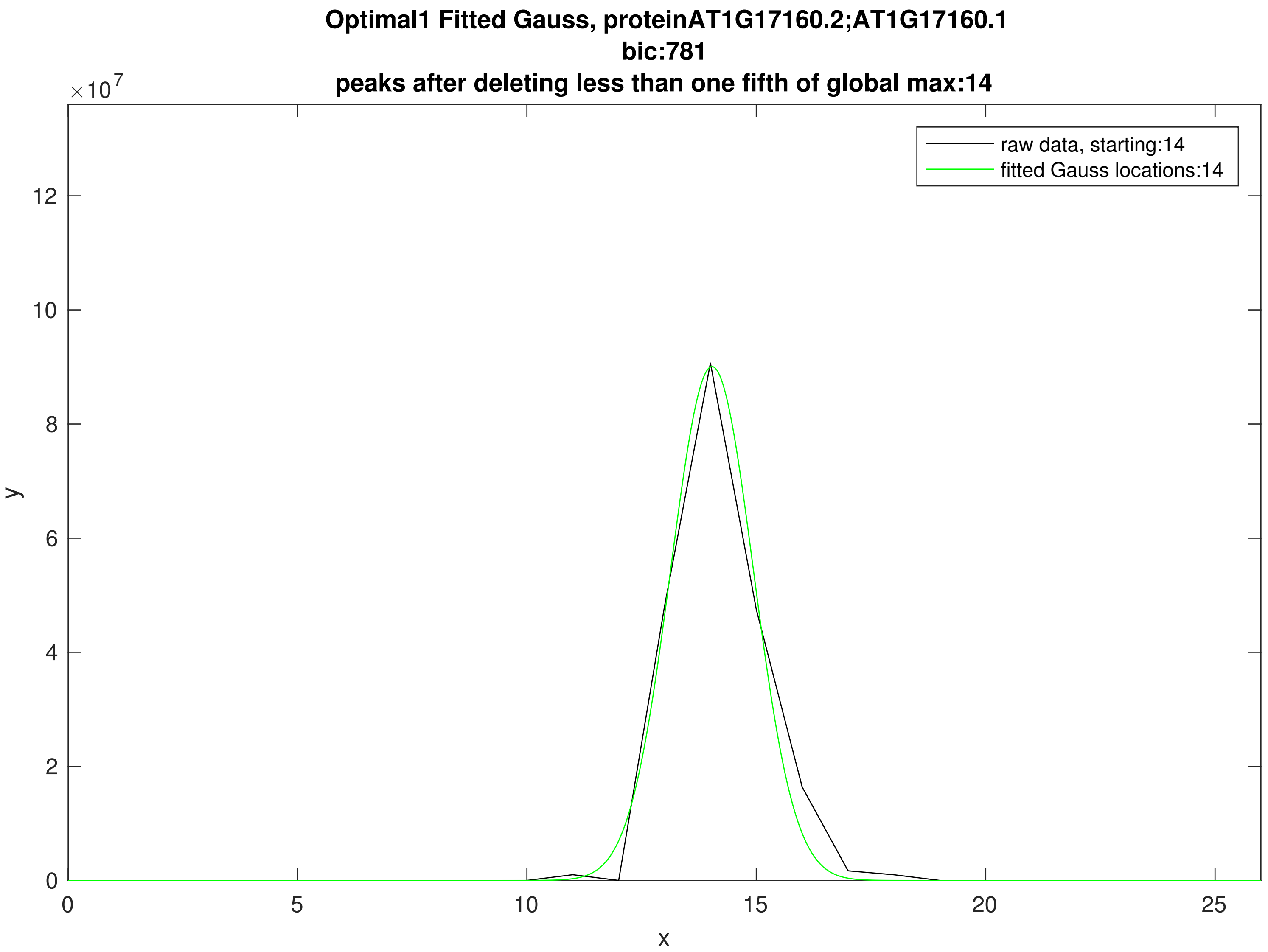
Optimal1 Fitted Gauss, proteinAT1G17160.2;AT1G17160.1
bic:781
peaks after deleting less than one fifth of global max:14
×10^7
raw data, starting:14
fitted Gauss locations:14
y
x
0
2
4
6
8
10
12
0
5
10
15
20
25

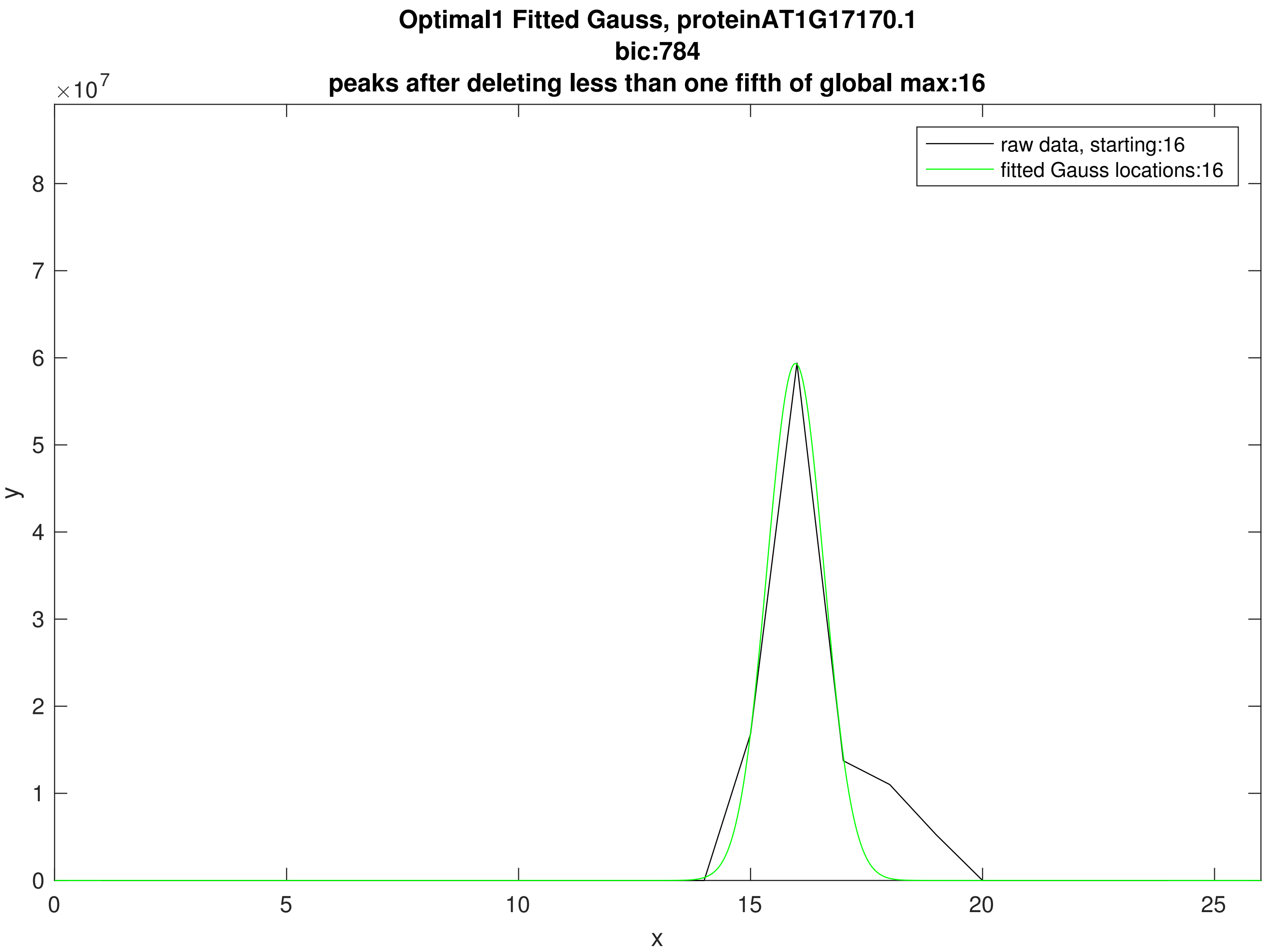
Optimal1 Fitted Gauss, proteinAT1G17170.1
bic:784
peaks after deleting less than one fifth of global max:16
×10^7
raw data, starting:16
fitted Gauss locations:16
y
x
0
1
2
3
4
5
6
7
8
0
5
10
15
20
25

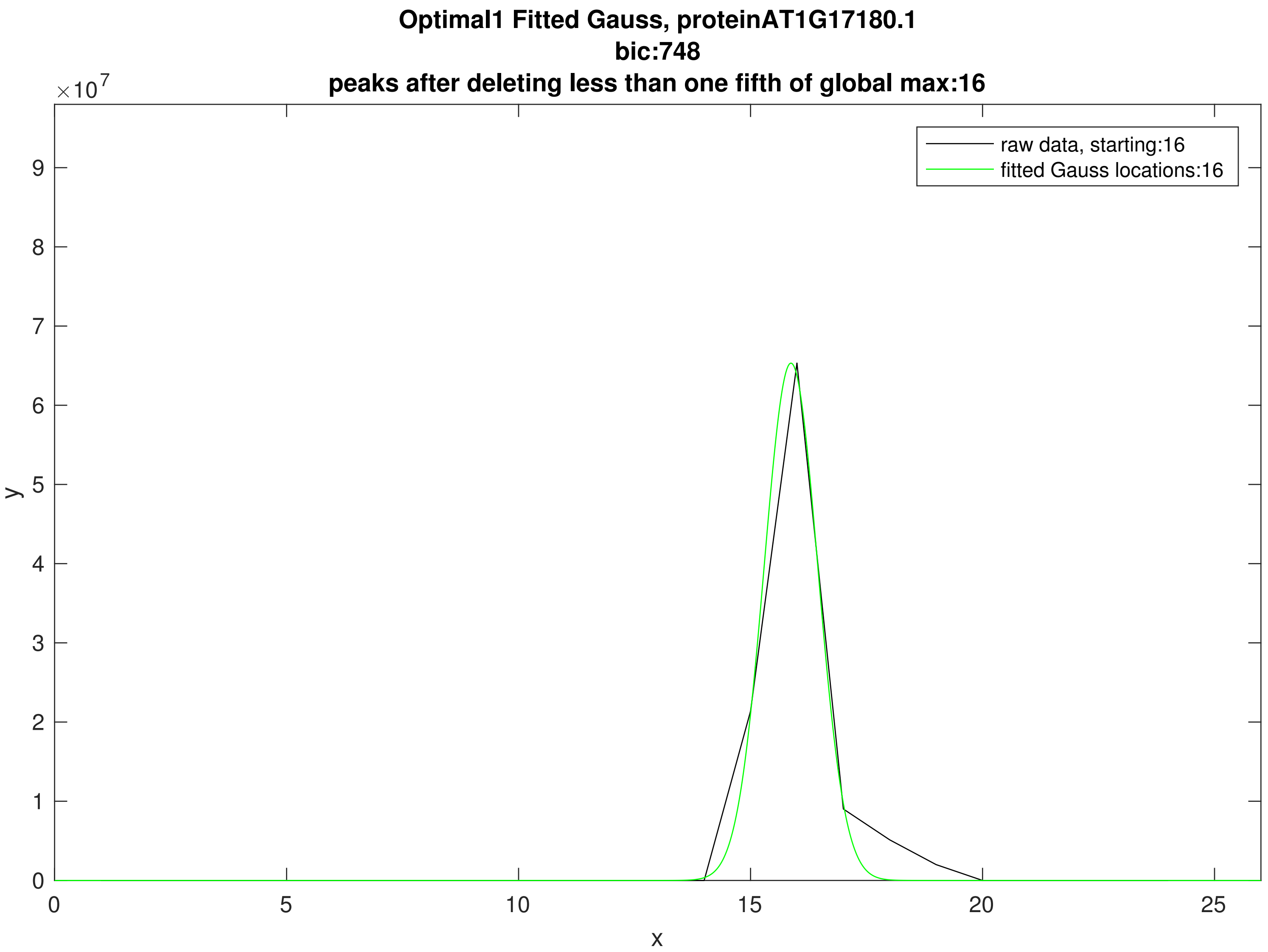
Optimal1 Fitted Gauss, proteinAT1G17180.1
bic:748
peaks after deleting less than one fifth of global max:16
×10^7
raw data, starting:16
fitted Gauss locations:16
y
x
0
1
2
3
4
5
6
7
8
9
0
5
10
15
20
25

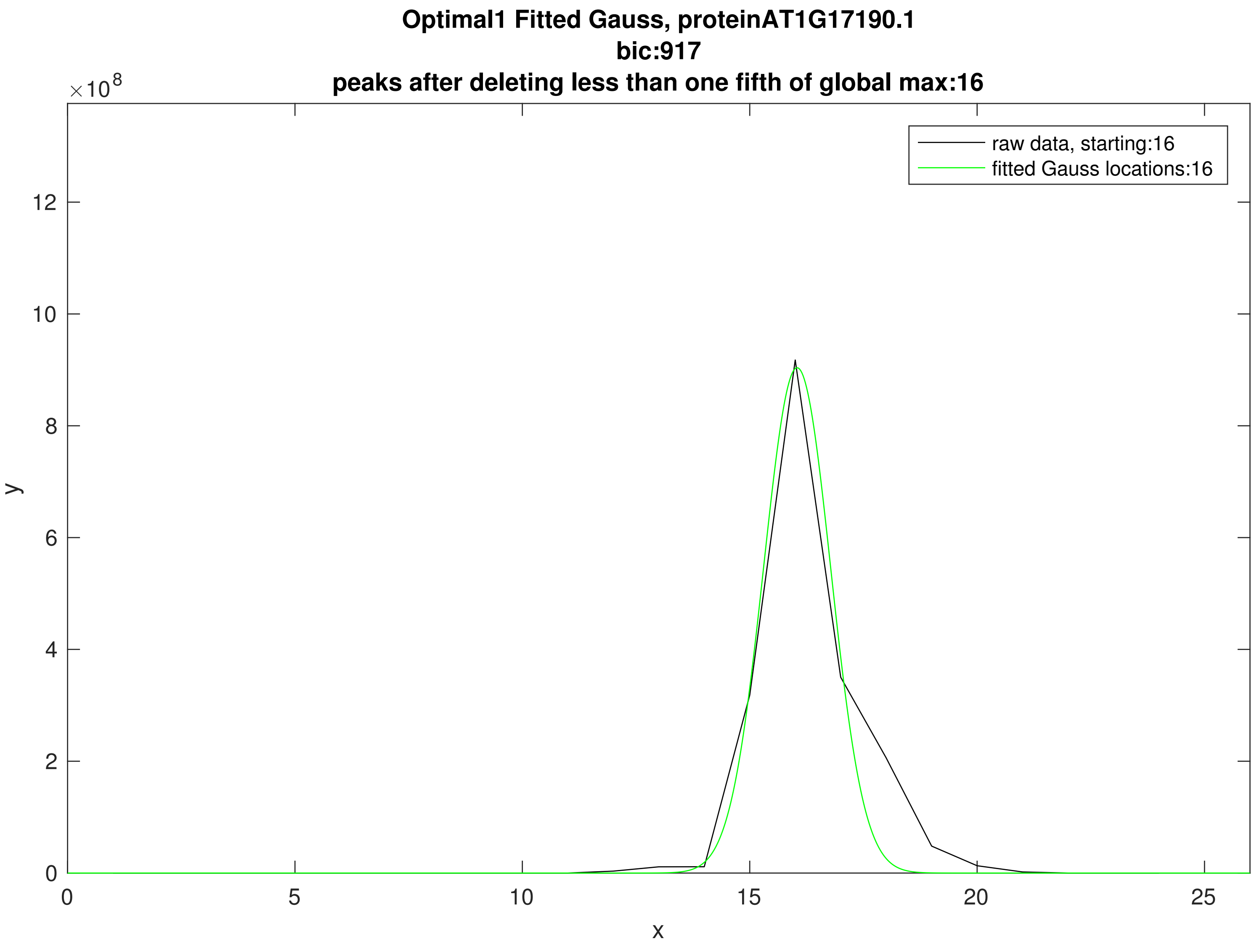

Optimal1 Fitted Gauss, proteinAT1G17190.1
bic:917
peaks after deleting less than one fifth of global max:16
×10^8
raw data, starting:16
fitted Gauss locations:16
0
2
4
6
8
10
12
y
0
5
10
15
20
25
x

**Optimal1 Fitted Gauss, proteinAT1G17290.1**
**bic:1.01e+03**
**peaks after deleting less than one fifth of global max:13**

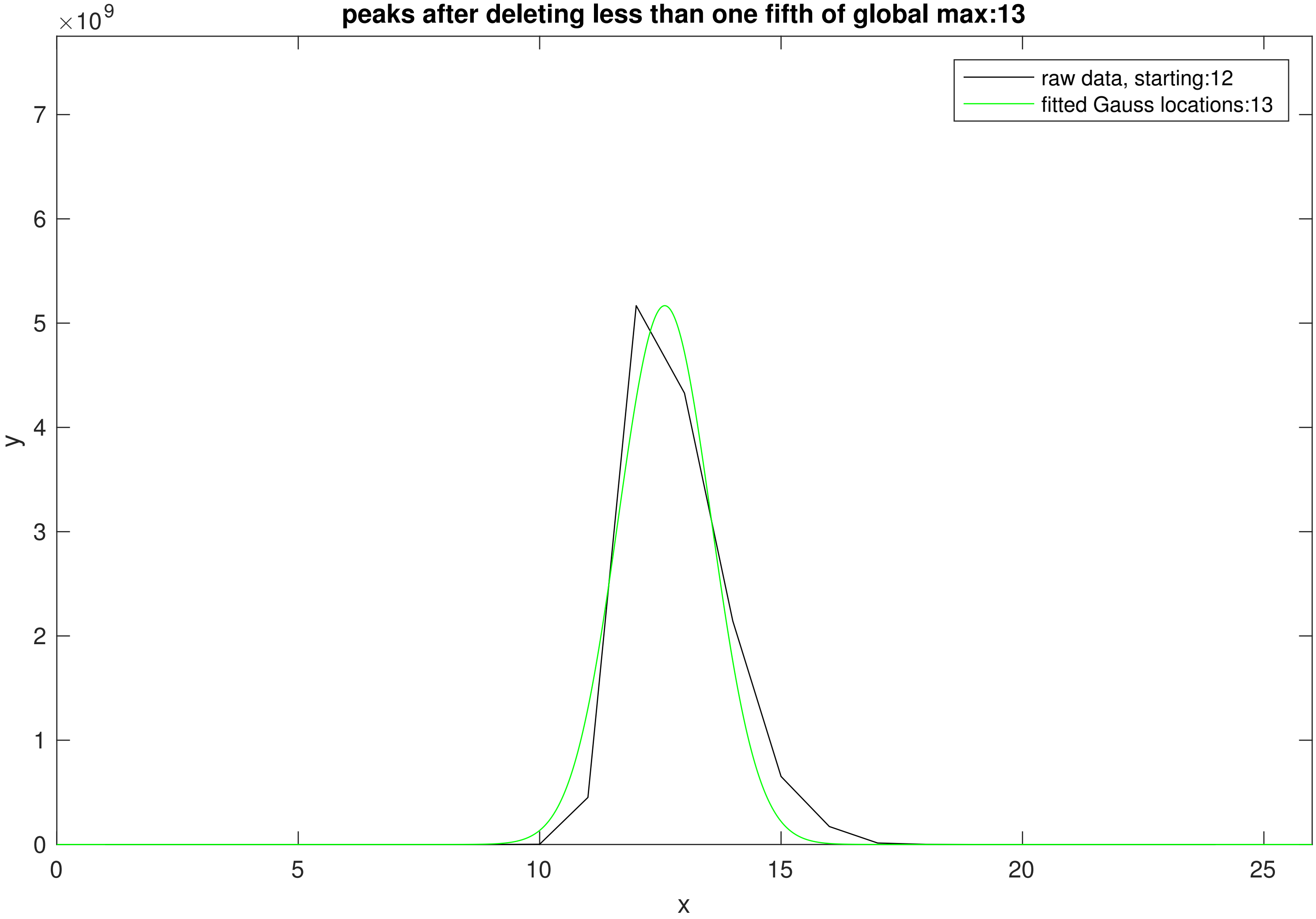

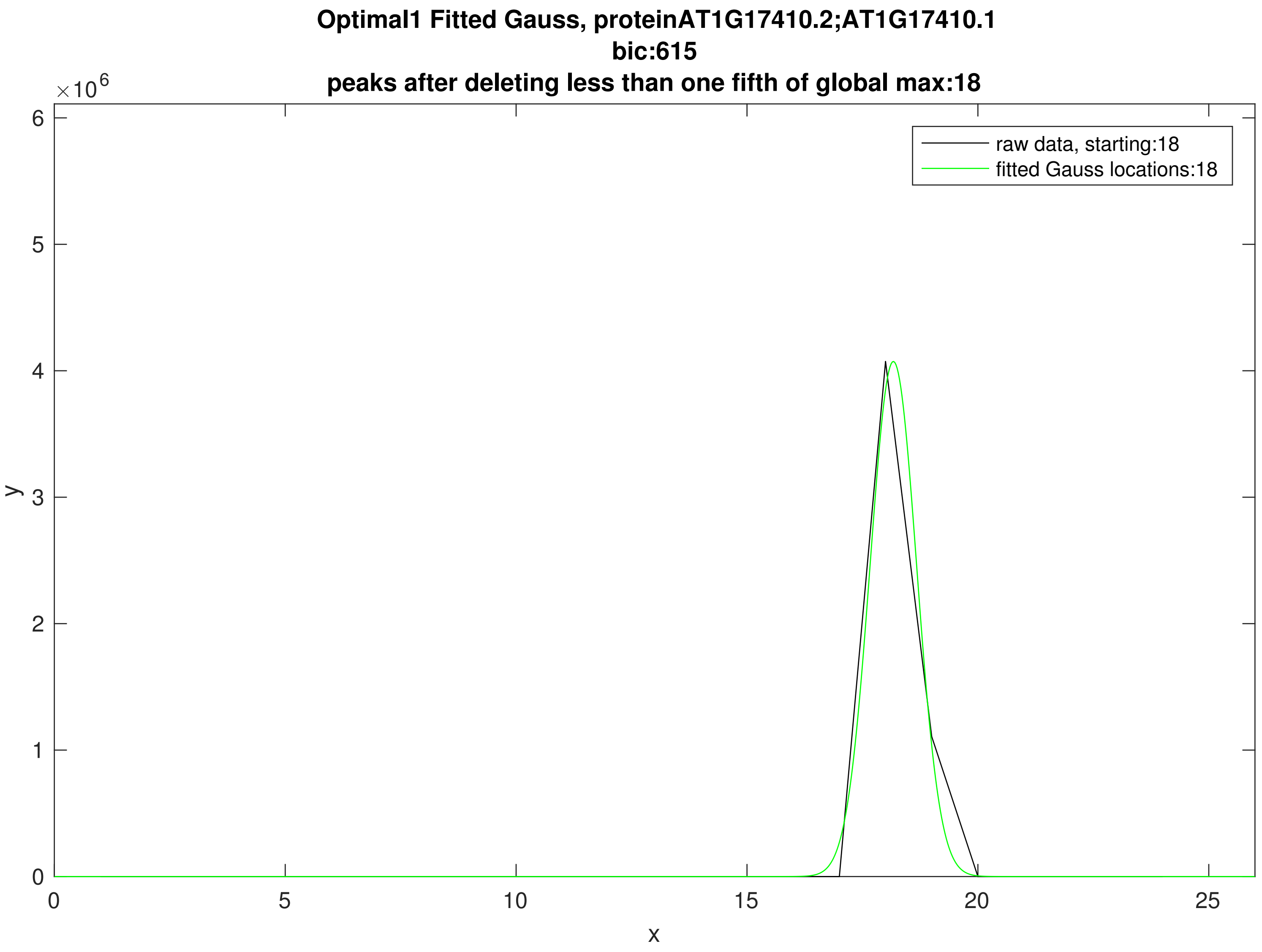

Optimal1 Fitted Gauss, proteinAT1G17410.2;AT1G17410.1
bic:615
peaks after deleting less than one fifth of global max:18
×10^6
raw data, starting:18
fitted Gauss locations:18
y
x
0
1
2
3
4
5
6
0
5
10
15
20
25

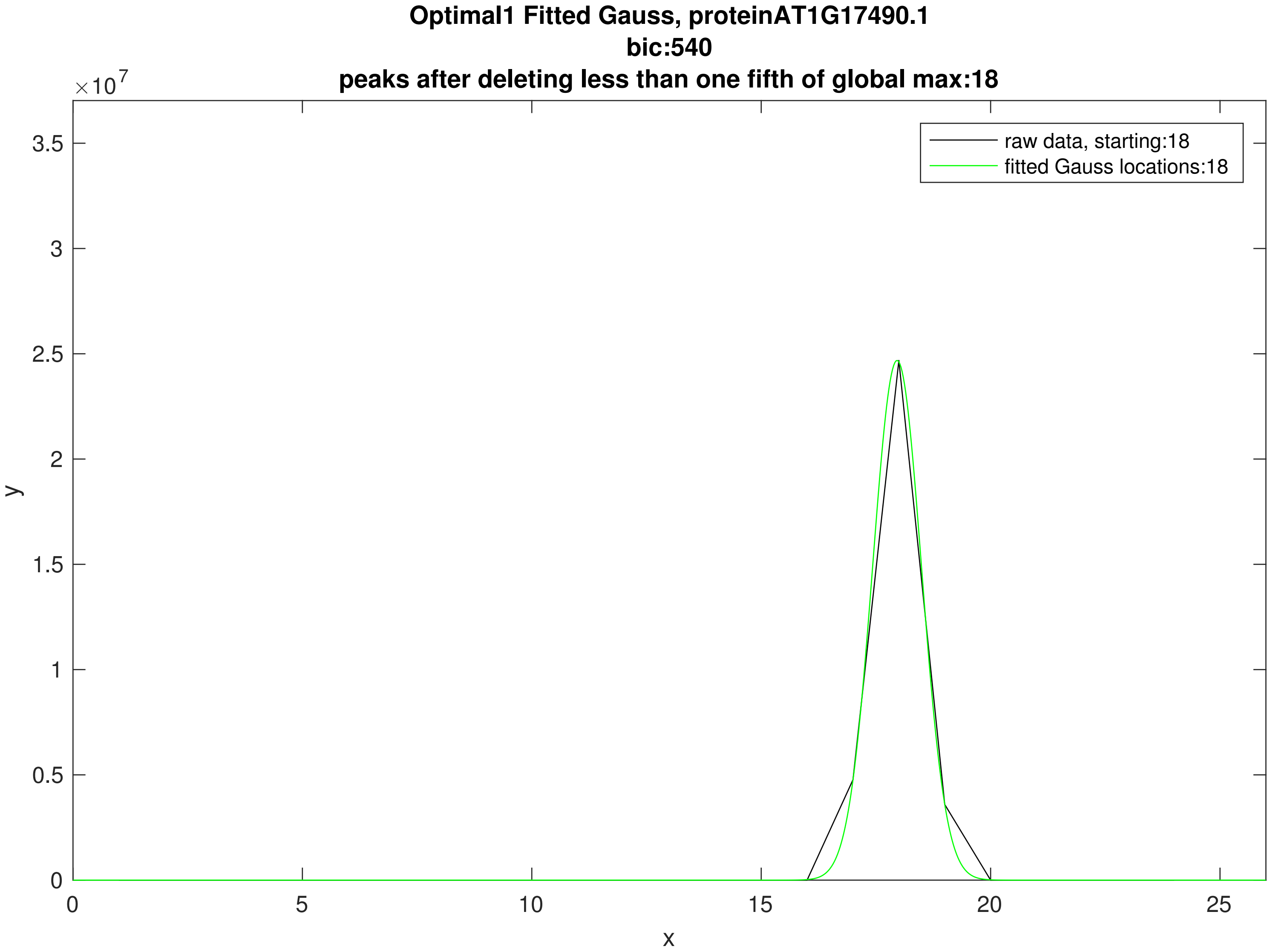

Optimal1 Fitted Gauss, proteinAT1G17490.1
bic:540
peaks after deleting less than one fifth of global max:18
×10^7
3.5
3
2.5
2
1.5
1
0.5
0
y
0
5
10
15
20
25
x
raw data, starting:18
fitted Gauss locations:18

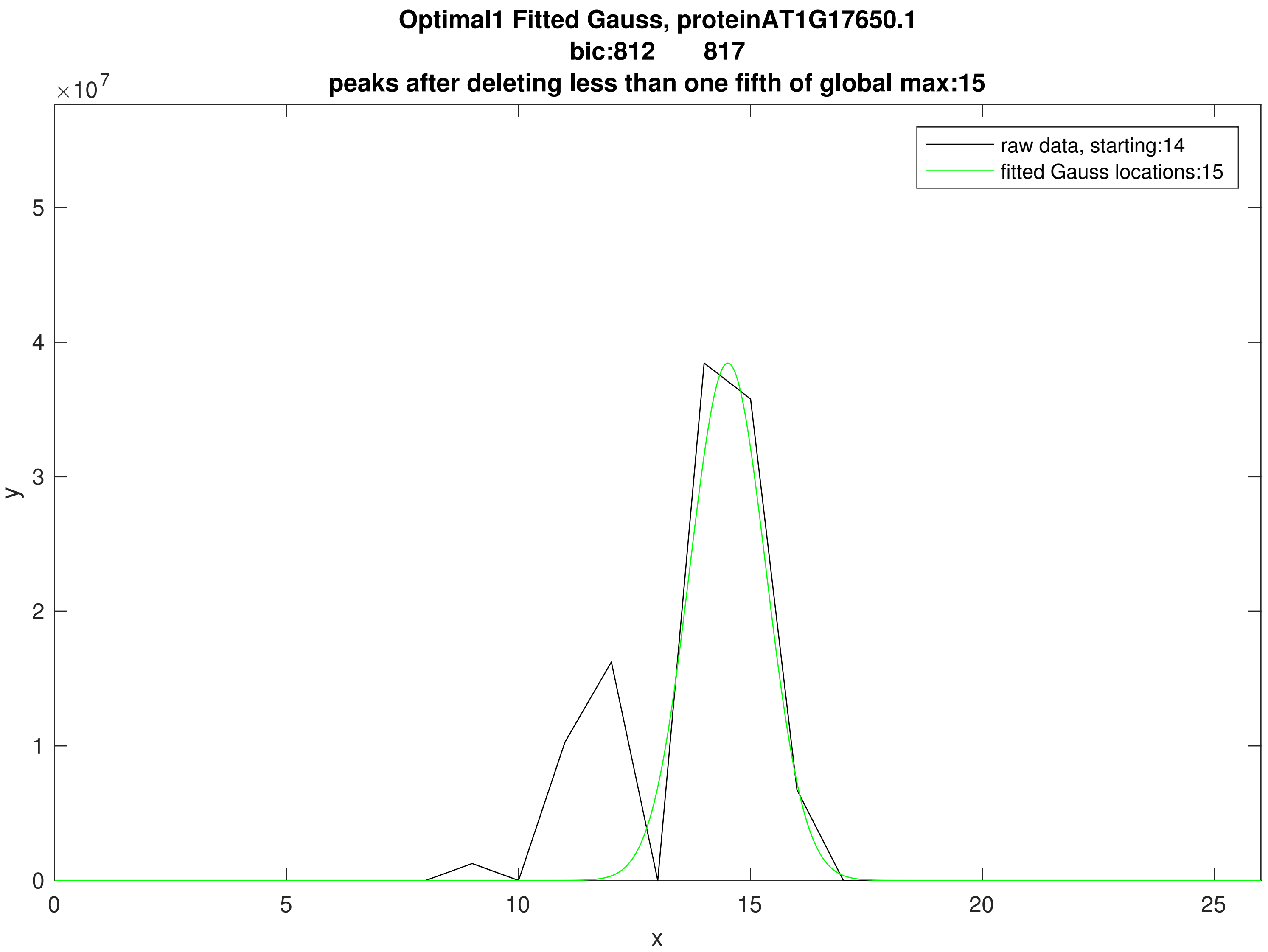

Optimal1 Fitted Gauss, proteinAT1G17650.1
bic:812 817
peaks after deleting less than one fifth of global max:15
×10^7
raw data, starting:14
fitted Gauss locations:15
y
x

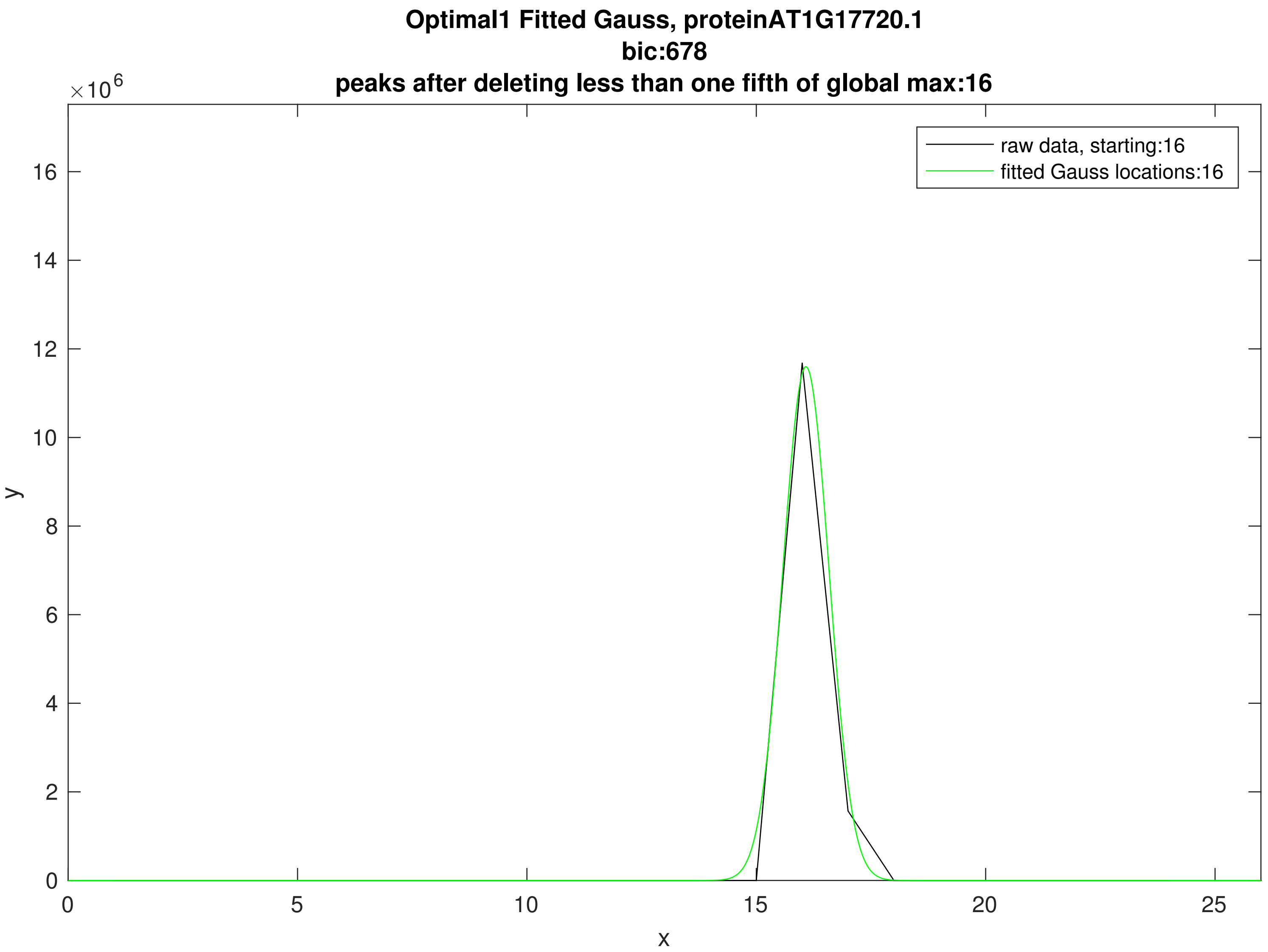

Optimal1 Fitted Gauss, proteinAT1G17720.1
bic:678
peaks after deleting less than one fifth of global max:16
×10^6
raw data, starting:16
fitted Gauss locations:16
y
x
0
2
4
6
8
10
12
14
16
0
5
10
15
20
25

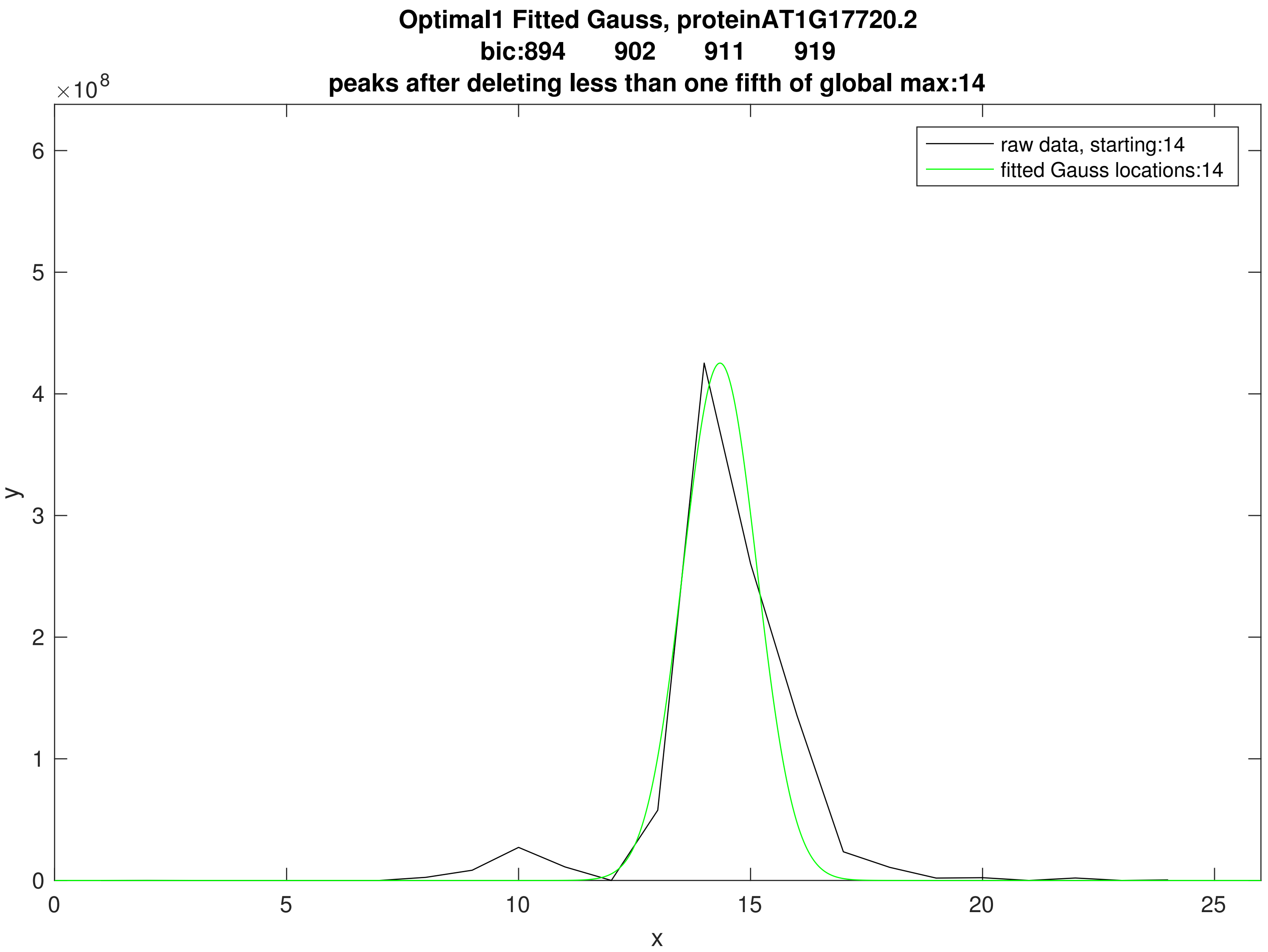

Optimal1 Fitted Gauss, proteinAT1G17720.2
bic:894 902 911 919
peaks after deleting less than one fifth of global max:14
×10^8
6
5
4
3
2
1
0
y
0
5
10
15
20
25
x
raw data, starting:14
fitted Gauss locations:14

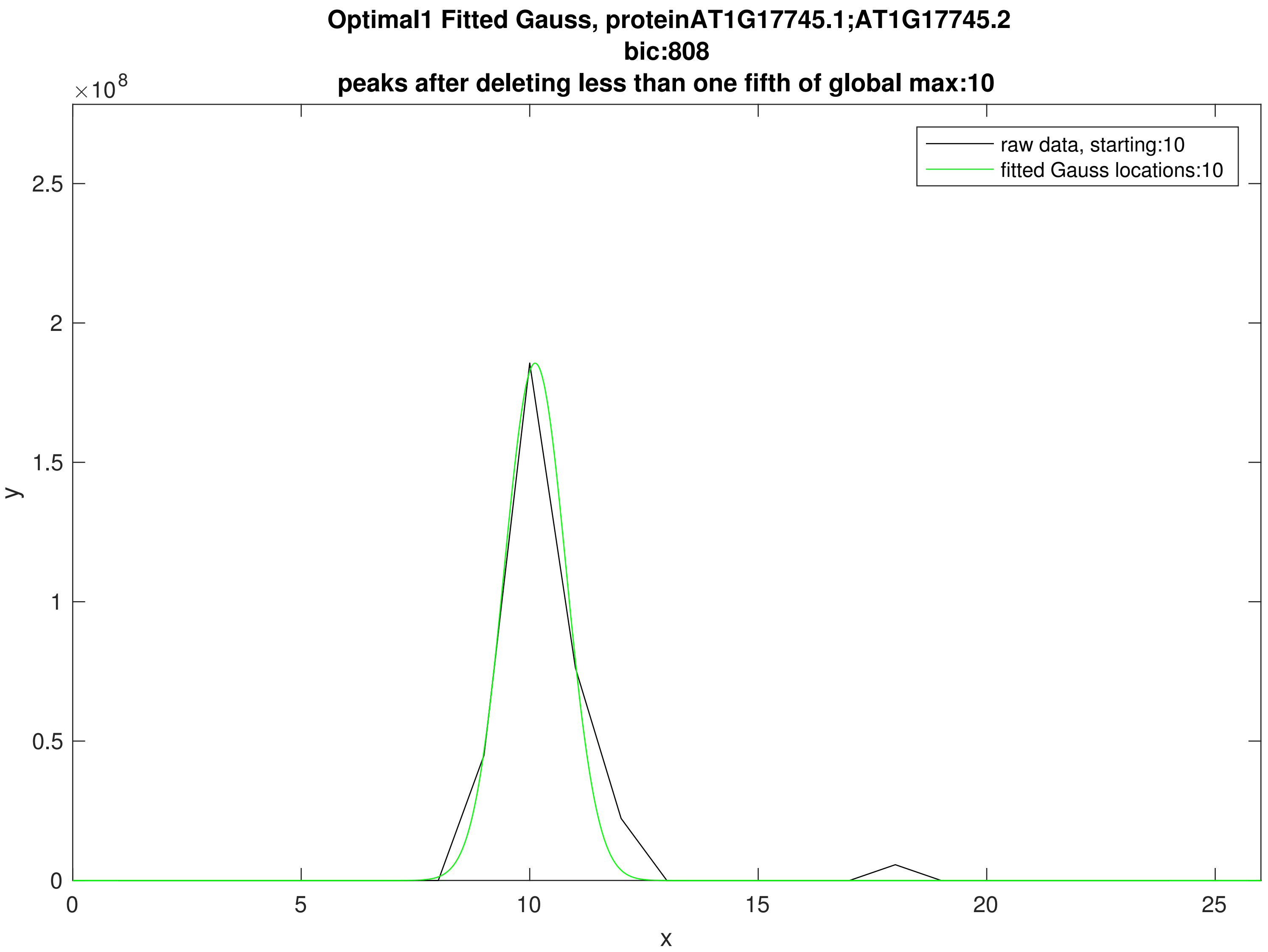
Optimal1 Fitted Gauss, proteinAT1G17745.1;AT1G17745.2
bic:808
peaks after deleting less than one fifth of global max:10
×10^8
raw data, starting:10
fitted Gauss locations:10
y
x
0
0.5
1
1.5
2
2.5
0
5
10
15
20
25

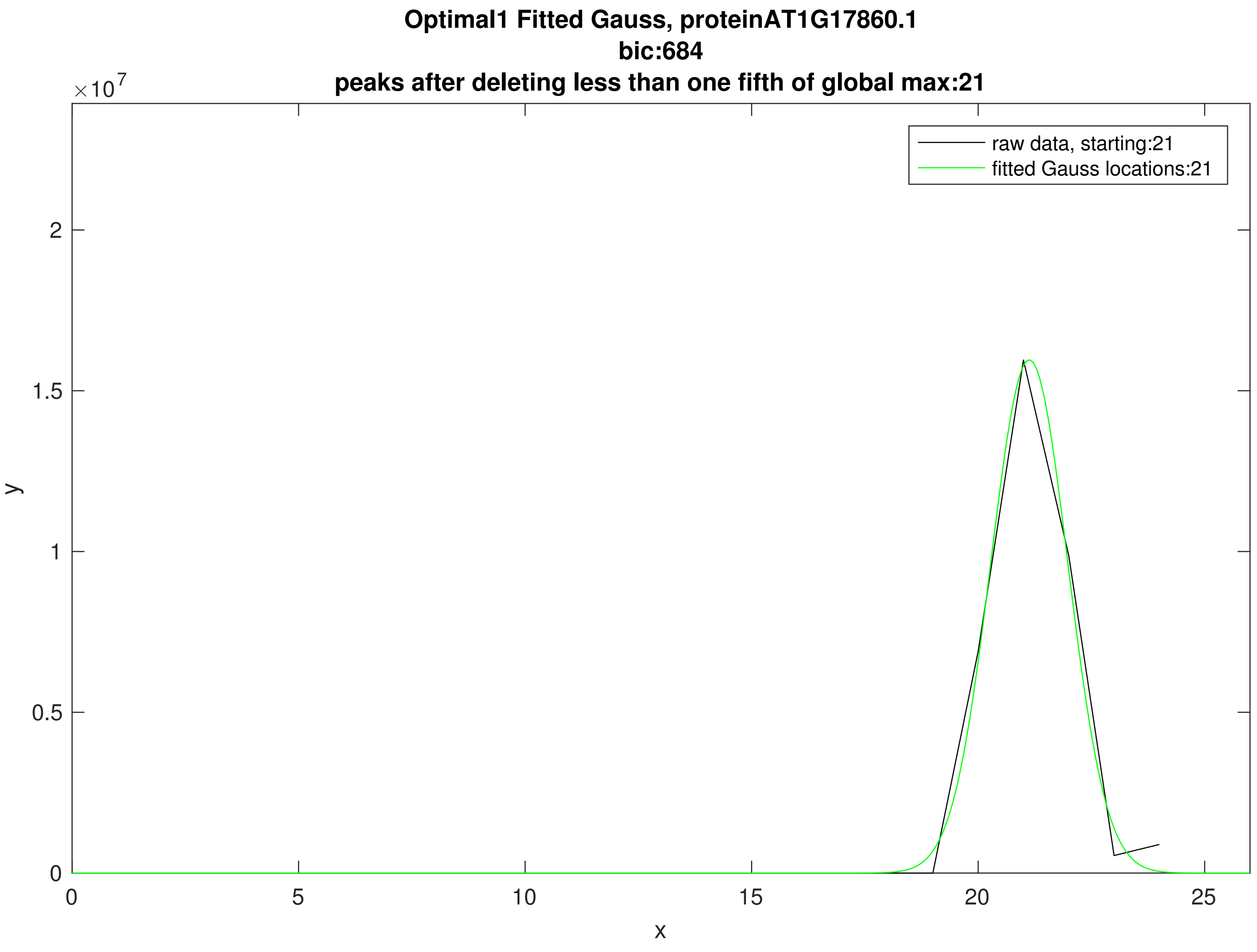
Optimal1 Fitted Gauss, proteinAT1G17860.1
bic:684
peaks after deleting less than one fifth of global max:21
×10^7
raw data, starting:21
fitted Gauss locations:21
y
x
0
0.5
1
1.5
2
0
5
10
15
20
25

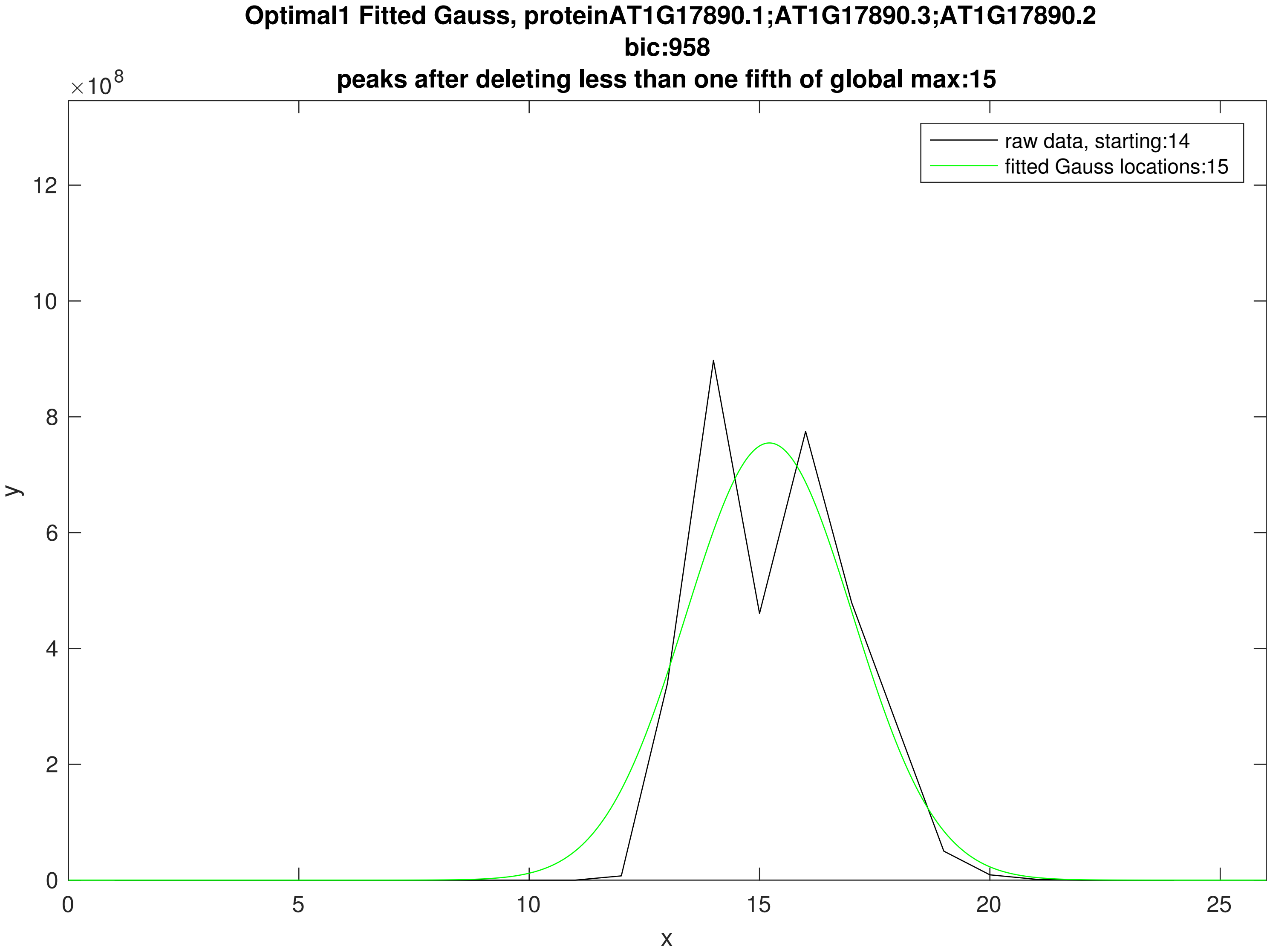

Optimal1 Fitted Gauss, proteinAT1G17890.1;AT1G17890.3;AT1G17890.2
bic:958
peaks after deleting less than one fifth of global max:15
×10^8
raw data, starting:14
fitted Gauss locations:15
y
x
0
2
4
6
8
10
12
0
5
10
15
20
25

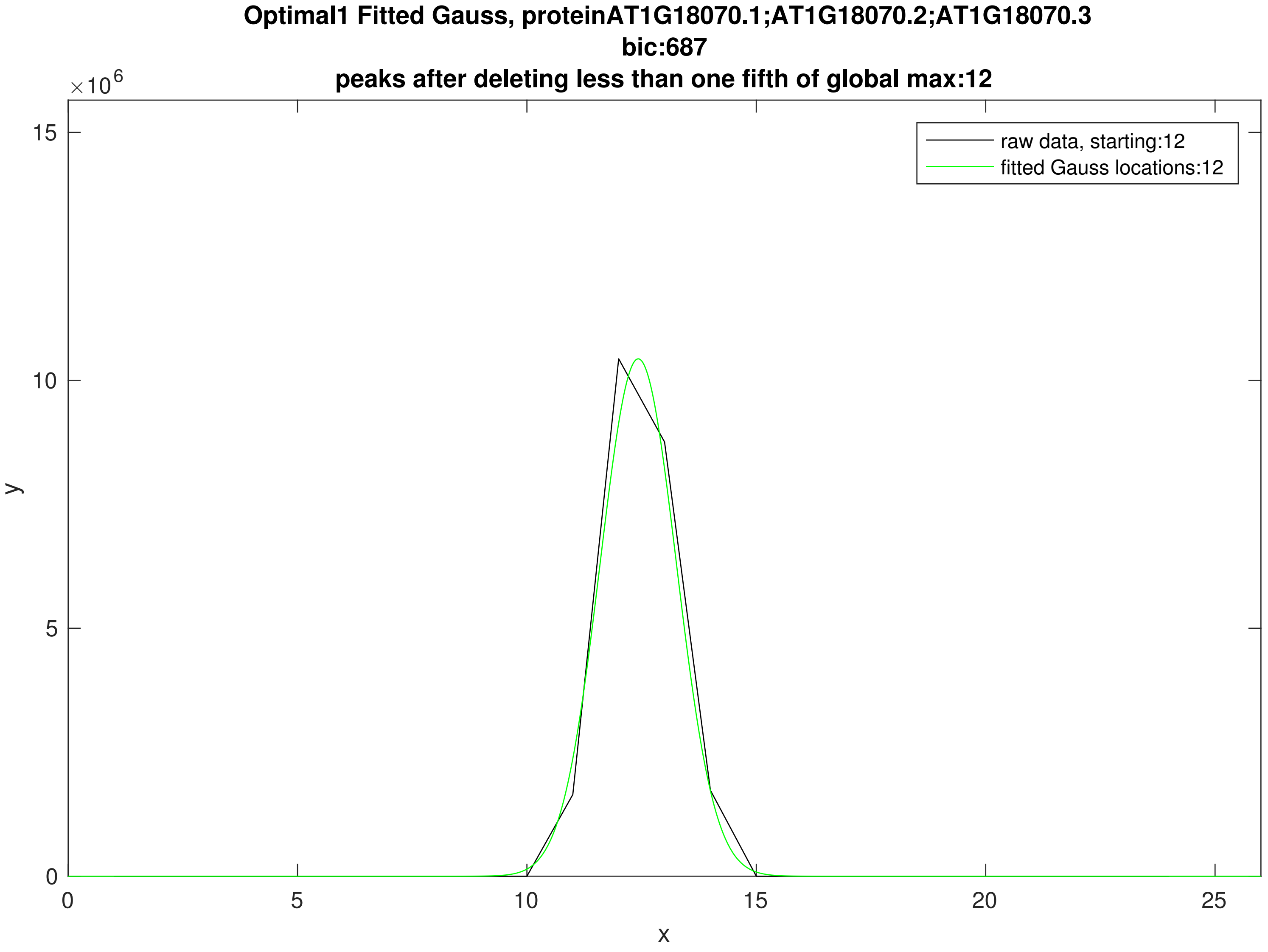
Optimal1 Fitted Gauss, proteinAT1G18070.1;AT1G18070.2;AT1G18070.3
bic:687
peaks after deleting less than one fifth of global max:12
×10^6
15
10
5
0
y
0
5
10
15
20
25
x
raw data, starting:12
fitted Gauss locations:12

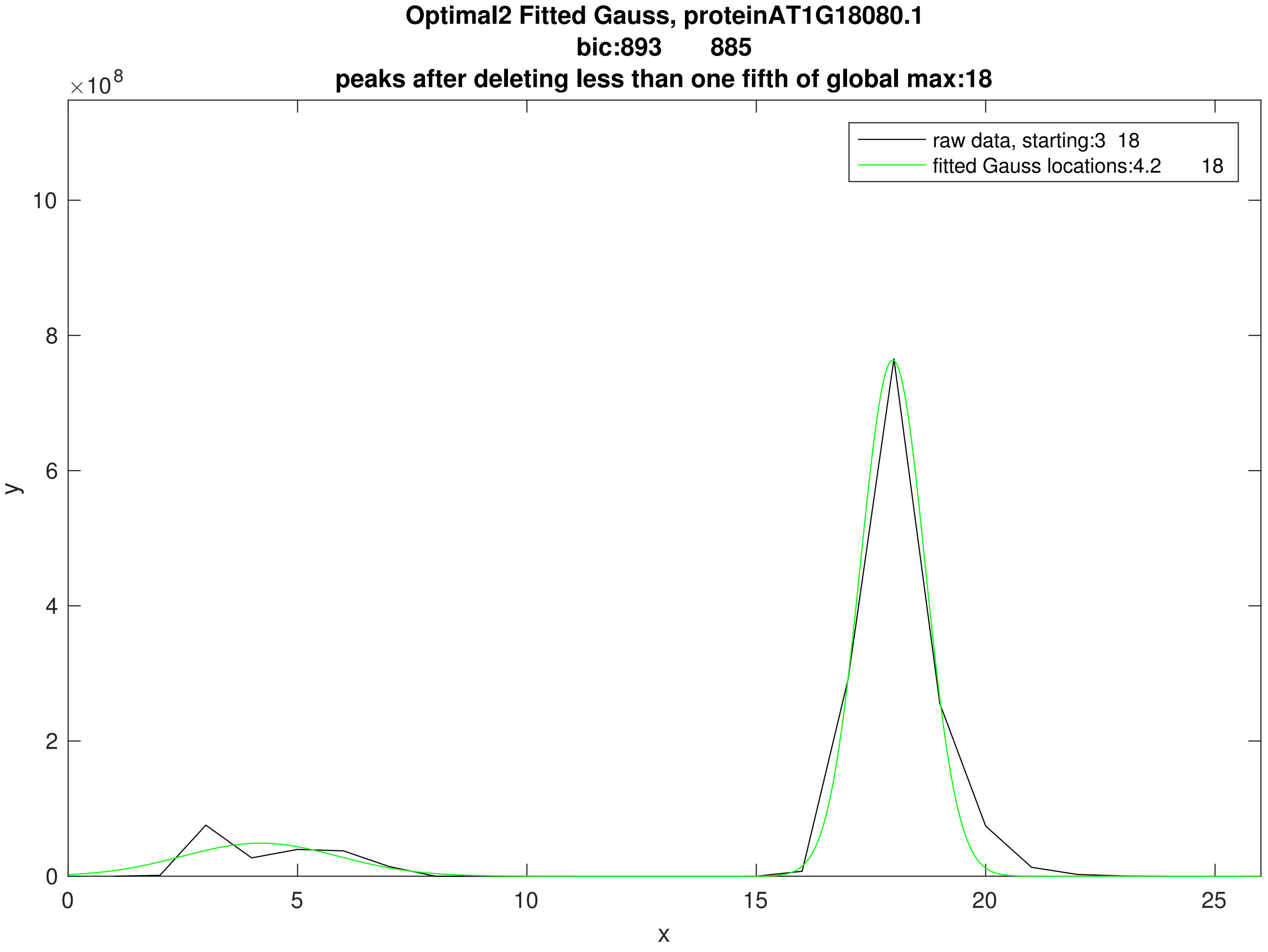

Optimal2 Fitted Gauss, proteinAT1G18080.1
bic:893 885
peaks after deleting less than one fifth of global max:18
×10^8
raw data, starting:3 18
fitted Gauss locations:4.2 18
10
8
6
4
2
0
y
0
5
10
15
20
25
x

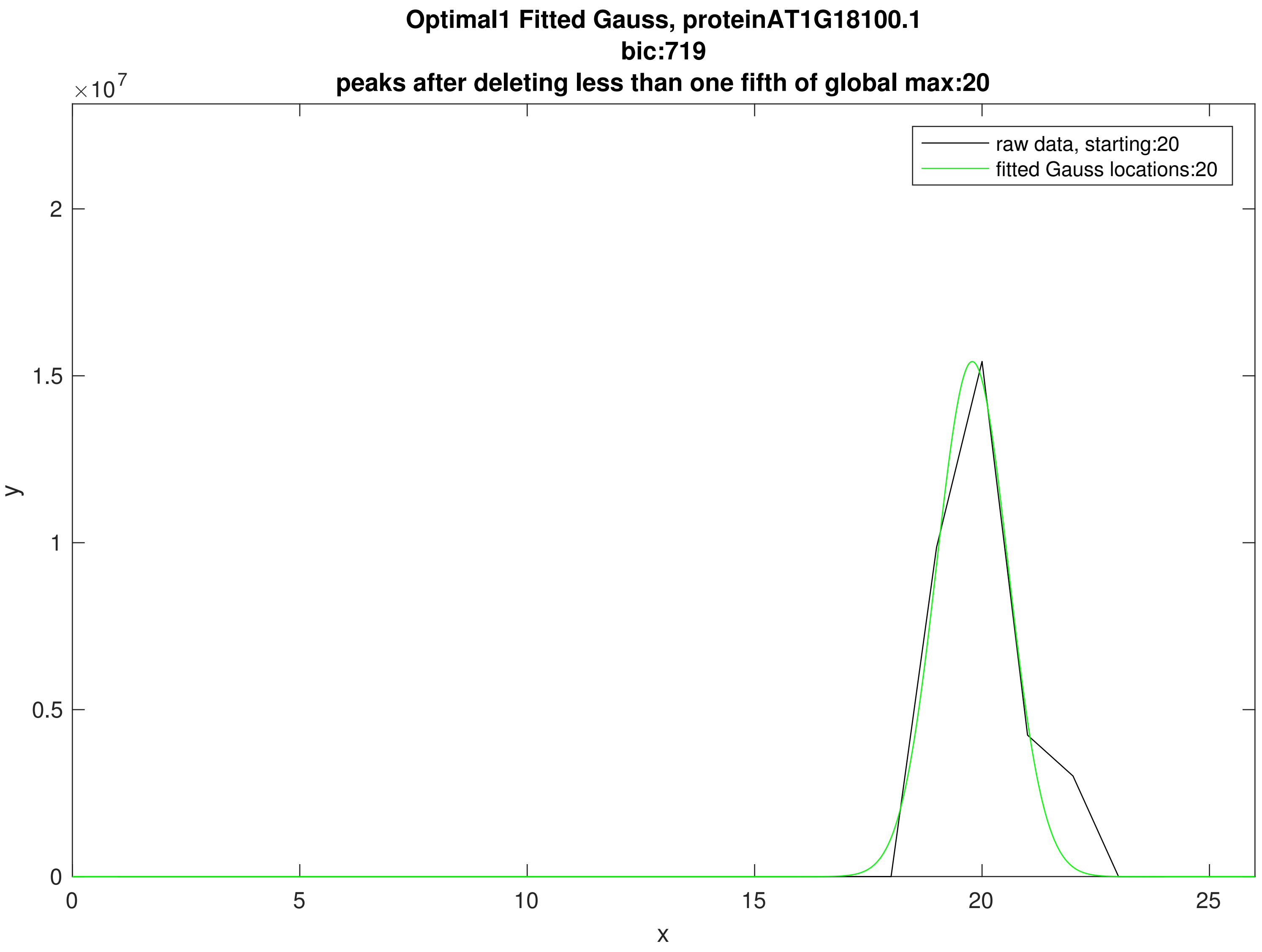
Optimal1 Fitted Gauss, proteinAT1G18100.1
bic:719
peaks after deleting less than one fifth of global max:20
×10^7
2
1.5
1
0.5
0
y
0
5
10
15
20
25
x
raw data, starting:20
fitted Gauss locations:20

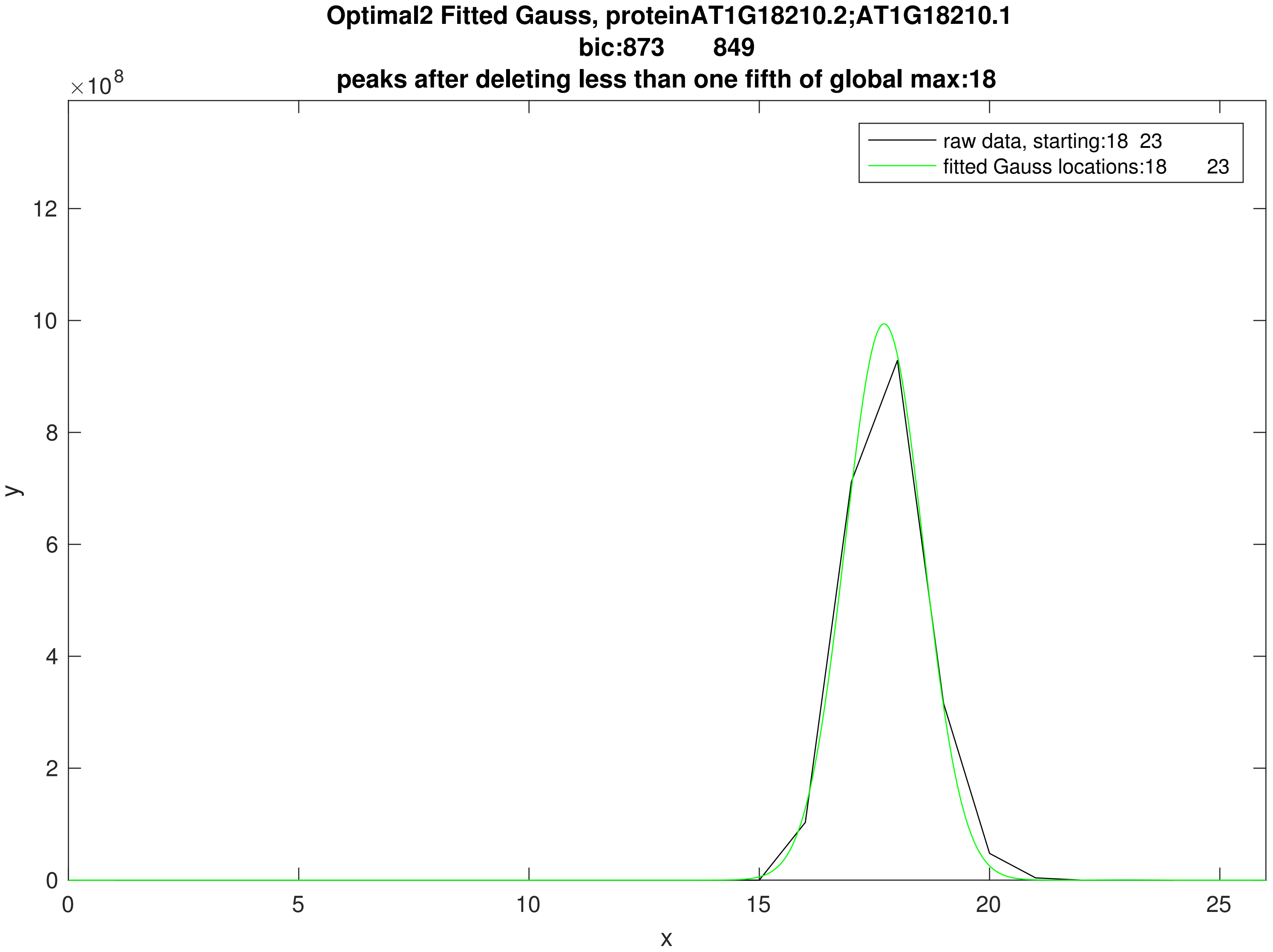
Optimal2 Fitted Gauss, proteinAT1G18210.2;AT1G18210.1
bic:873 849
peaks after deleting less than one fifth of global max:18
×10^8
raw data, starting:18 23
fitted Gauss locations:18 23
y
x
0
2
4
6
8
10
12
0
5
10
15
20
25

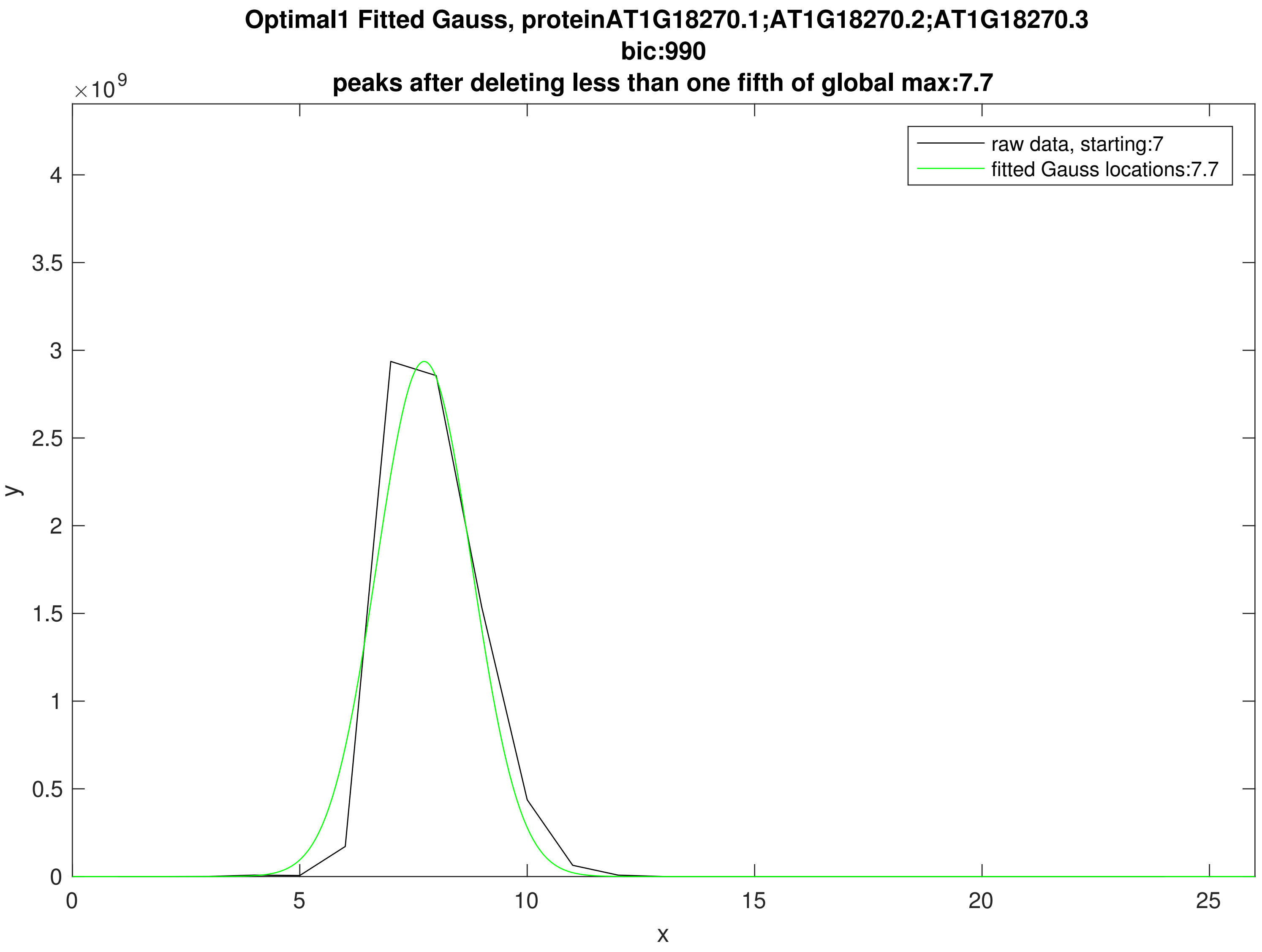

Optimal1 Fitted Gauss, proteinAT1G18270.1;AT1G18270.2;AT1G18270.3
bic:990
peaks after deleting less than one fifth of global max:7.7
×10^9
raw data, starting:7
fitted Gauss locations:7.7
y
x
0
0.5
1
1.5
2
2.5
3
3.5
4
5
10
15
20
25

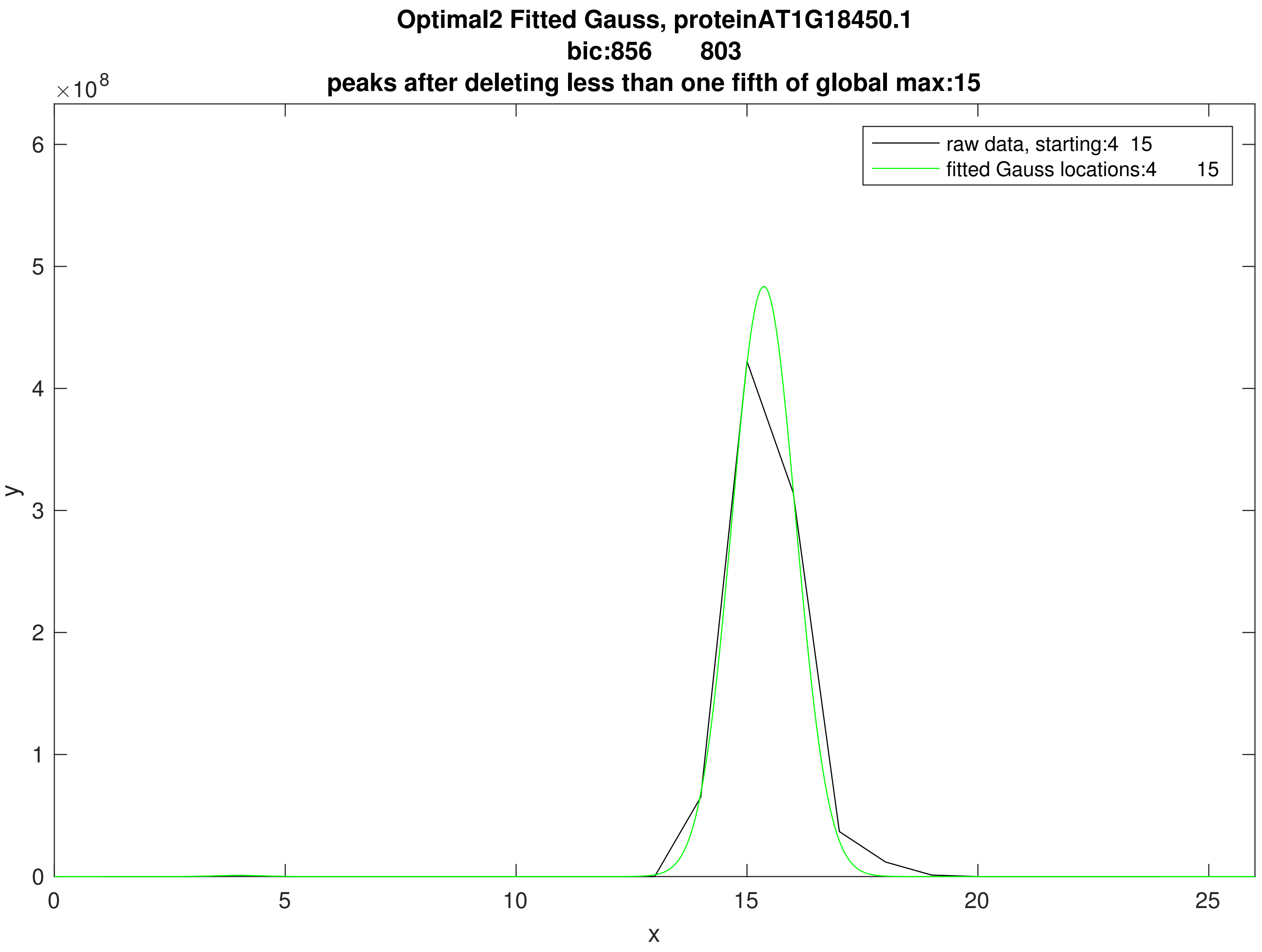
Optimal2 Fitted Gauss, proteinAT1G18450.1
bic:856 803
peaks after deleting less than one fifth of global max:15
×10^8
raw data, starting:4 15
fitted Gauss locations:4 15
y
x
0
1
2
3
4
5
6
0
5
10
15
20
25

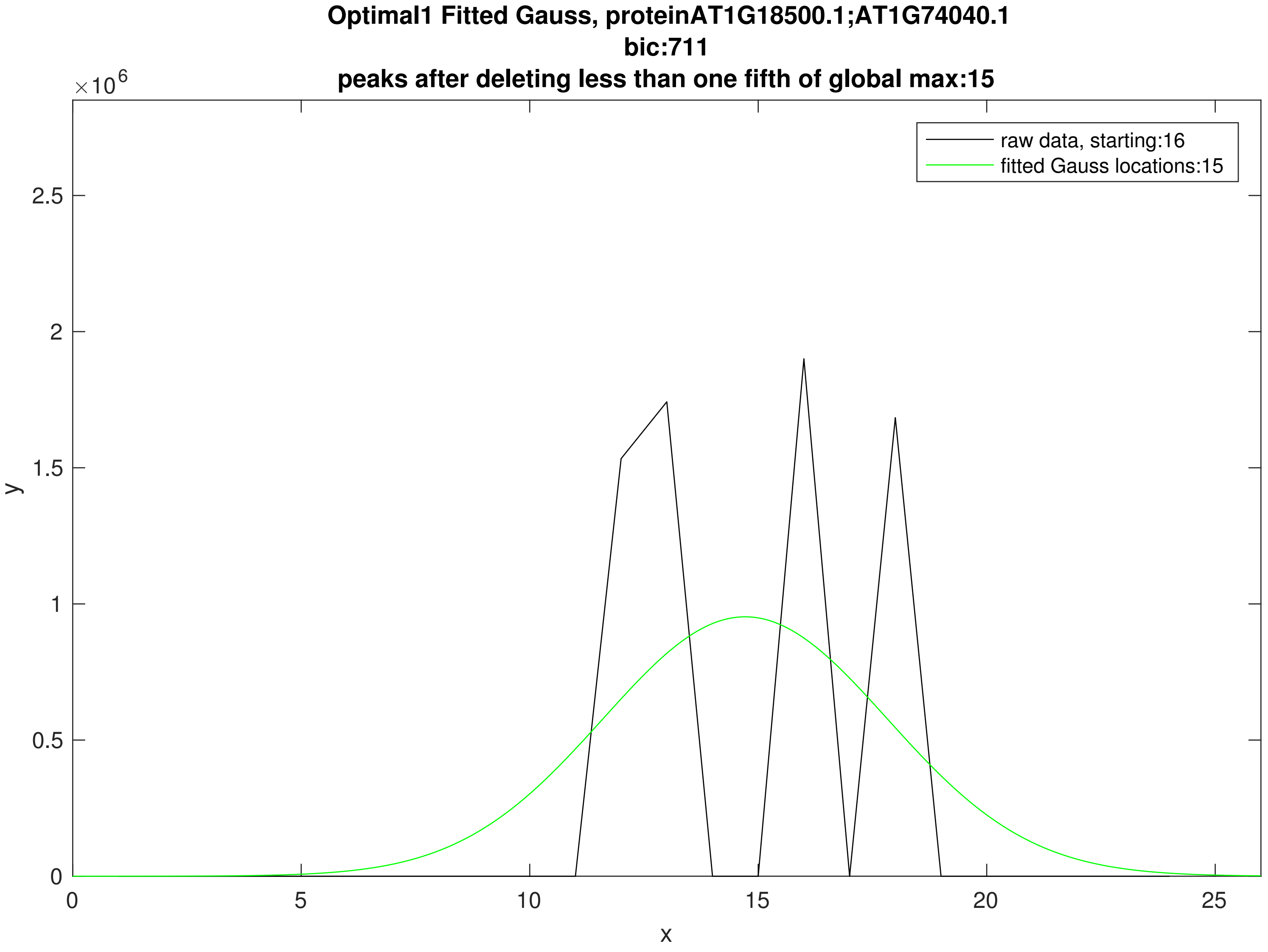
Optimal1 Fitted Gauss, proteinAT1G18500.1;AT1G74040.1
bic:711
peaks after deleting less than one fifth of global max:15
×10^6
raw data, starting:16
fitted Gauss locations:15
y
x
0
0.5
1
1.5
2
2.5
5
10
15
20
25

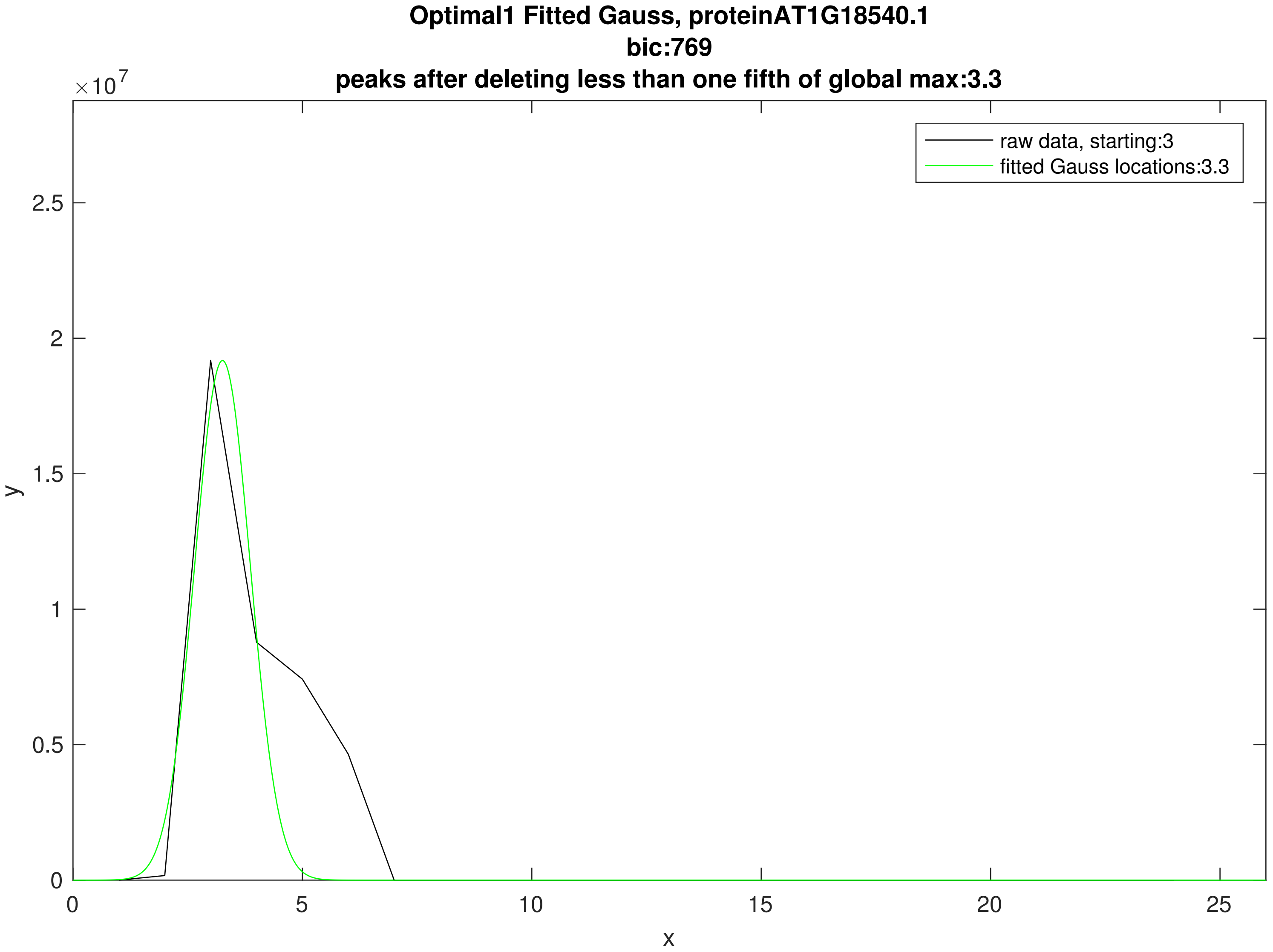
Optimal1 Fitted Gauss, proteinAT1G18540.1
bic:769
peaks after deleting less than one fifth of global max:3.3
×10^7
raw data, starting:3
fitted Gauss locations:3.3
y
0
0.5
1
1.5
2
2.5
x
0
5
10
15
20
25

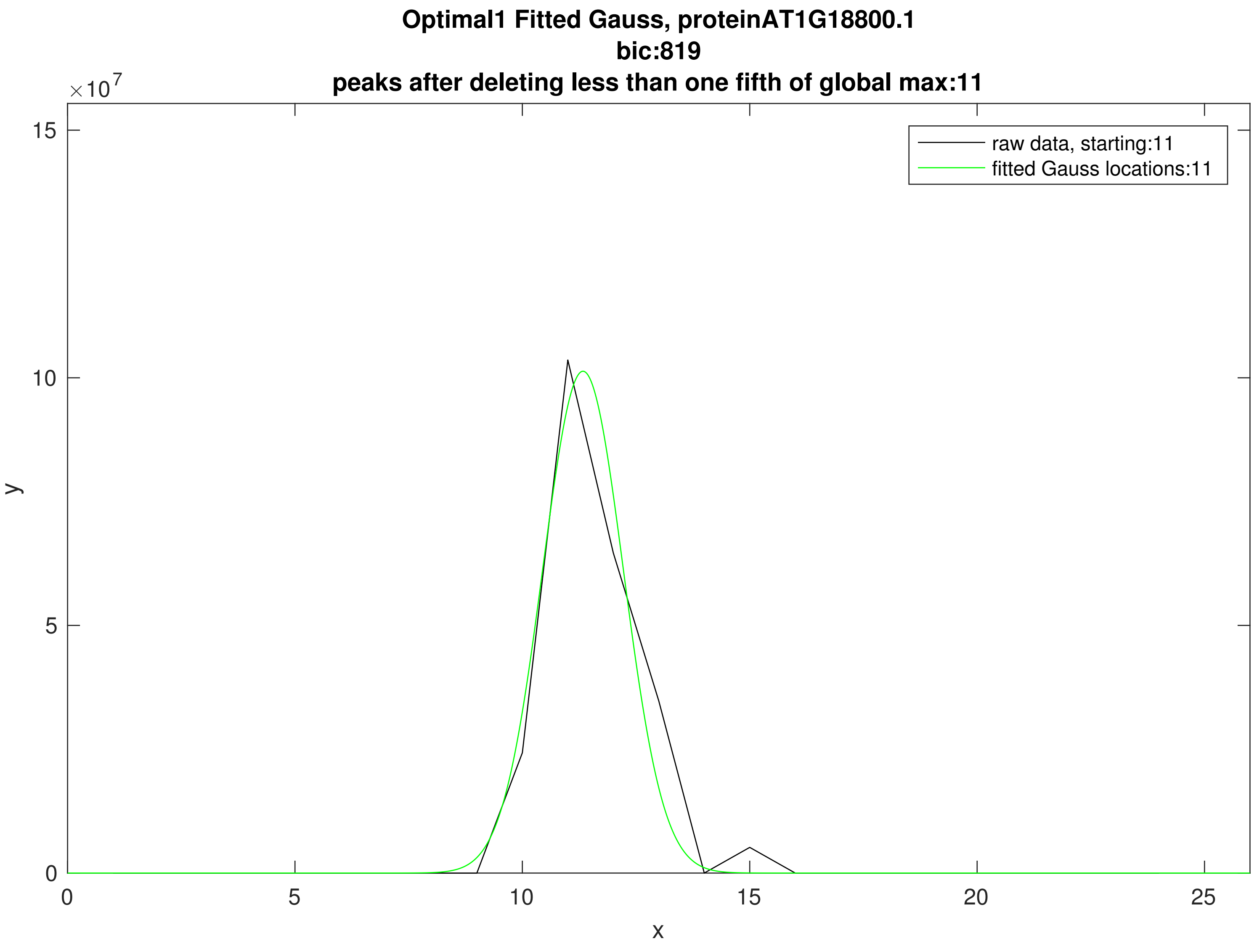
Optimal1 Fitted Gauss, proteinAT1G18800.1
bic:819
peaks after deleting less than one fifth of global max:11
×10^7
raw data, starting:11
fitted Gauss locations:11
y
x
0
5
10
15
20
25

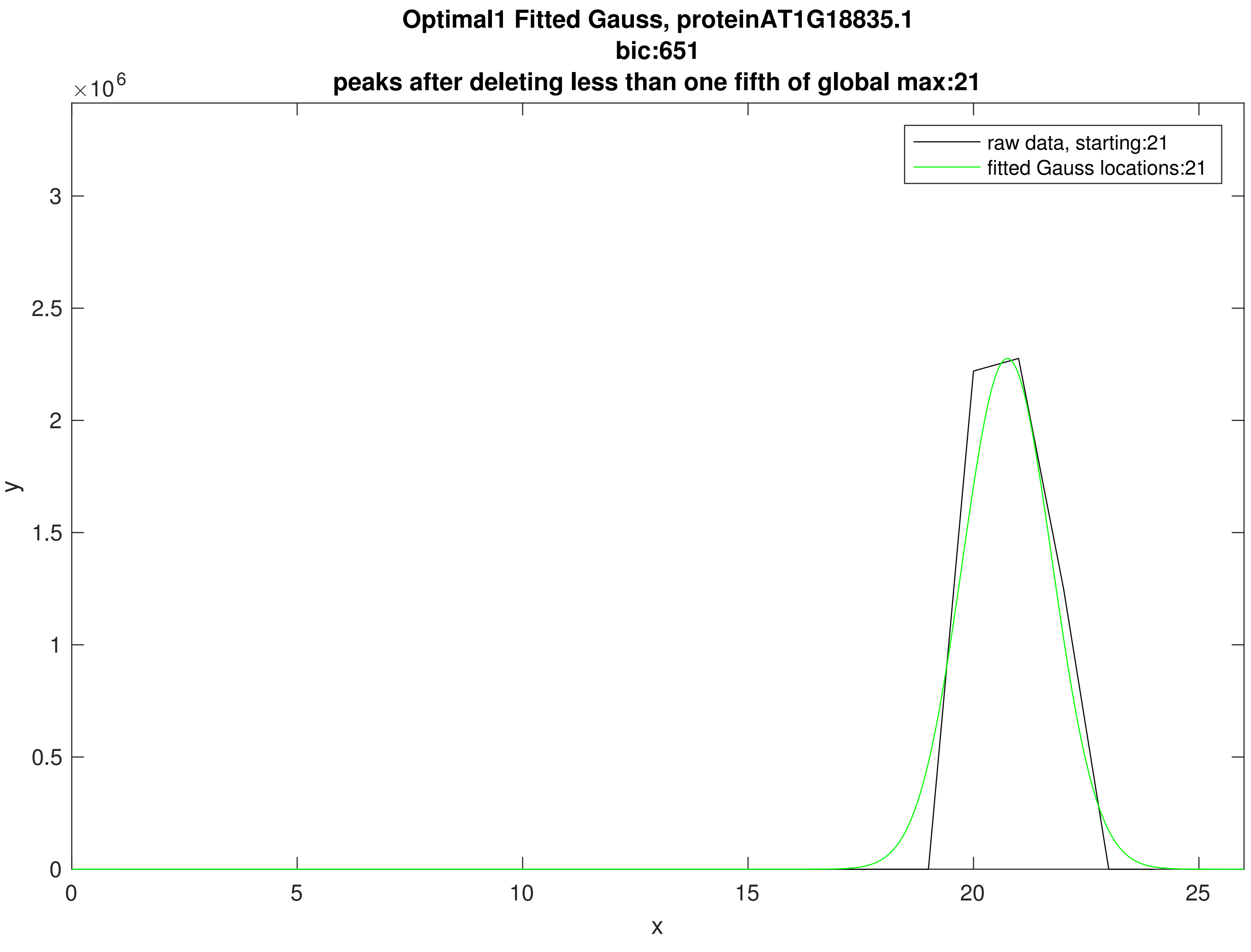
Optimal1 Fitted Gauss, proteinAT1G18835.1
bic:651
peaks after deleting less than one fifth of global max:21
×10^6
raw data, starting:21
fitted Gauss locations:21
y
x
0
0.5
1
1.5
2
2.5
3
0
5
10
15
20
25

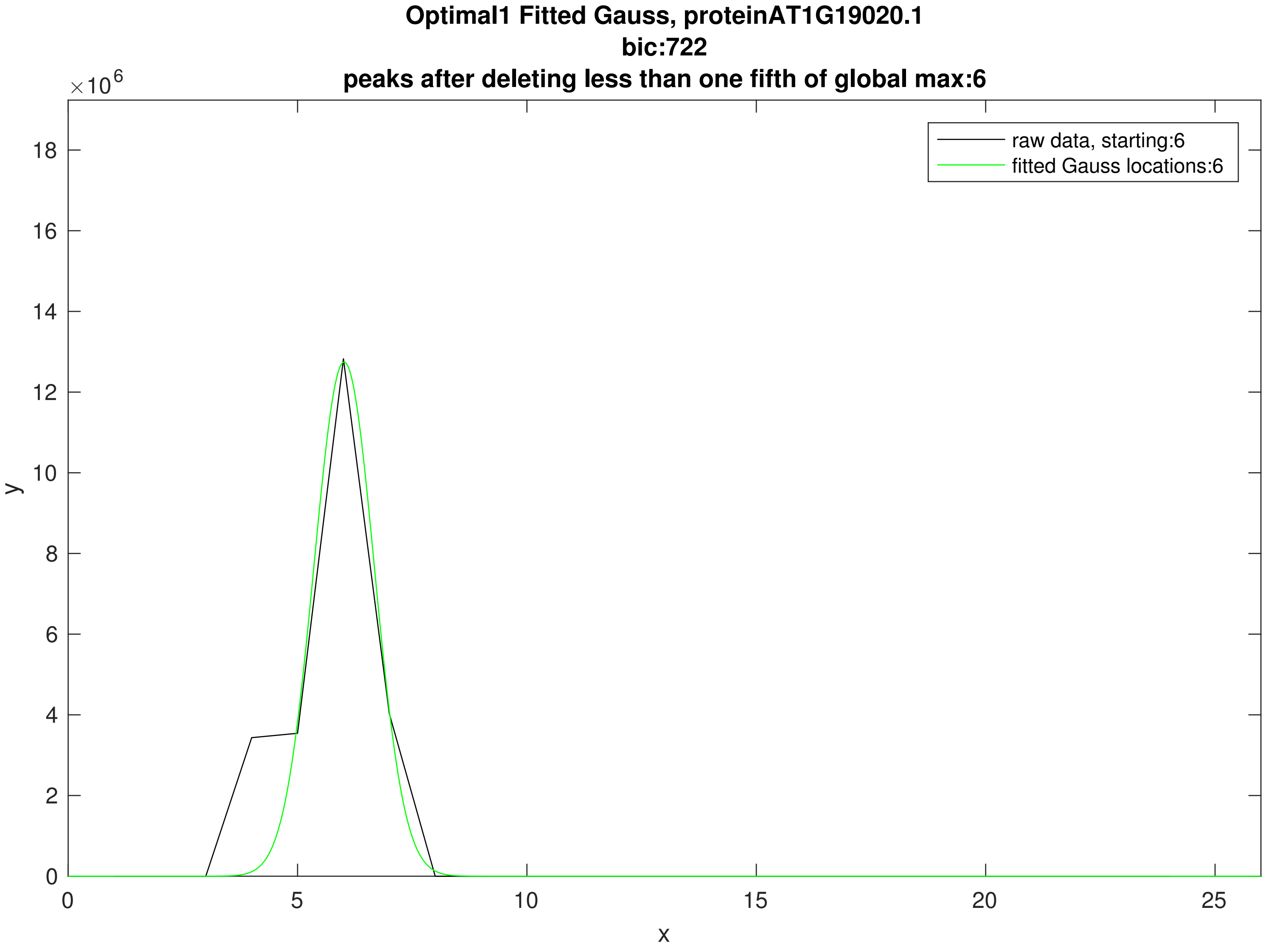
Optimal1 Fitted Gauss, proteinAT1G19020.1
bic:722
peaks after deleting less than one fifth of global max:6
×10^6
raw data, starting:6
fitted Gauss locations:6
y
x
0
2
4
6
8
10
12
14
16
18
0
5
10
15
20
25

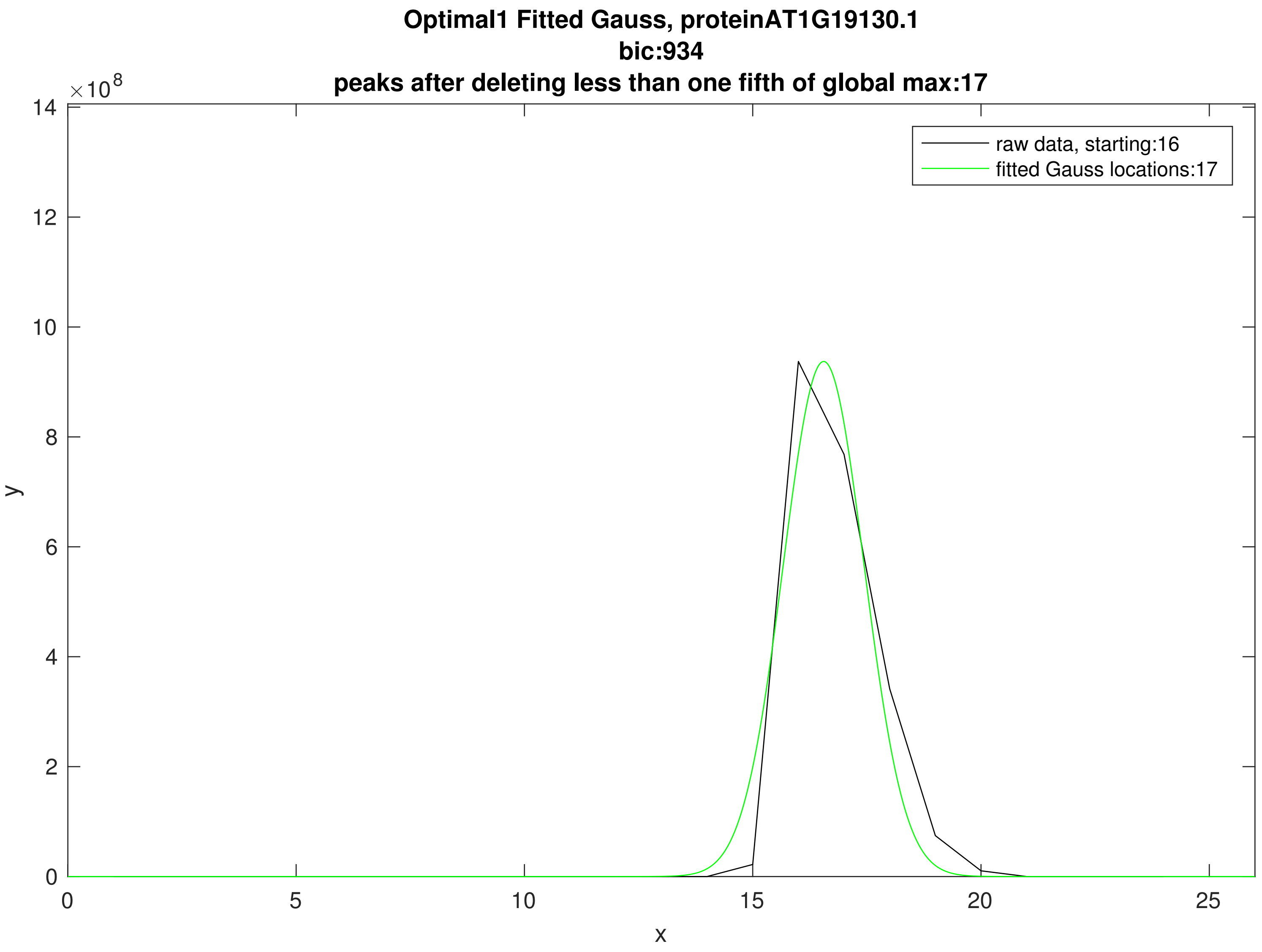

Optimal1 Fitted Gauss, proteinAT1G19130.1
bic:934
peaks after deleting less than one fifth of global max:17
×10^8
14
12
10
8
6
4
2
0
y
0
5
10
15
20
25
x
raw data, starting:16
fitted Gauss locations:17

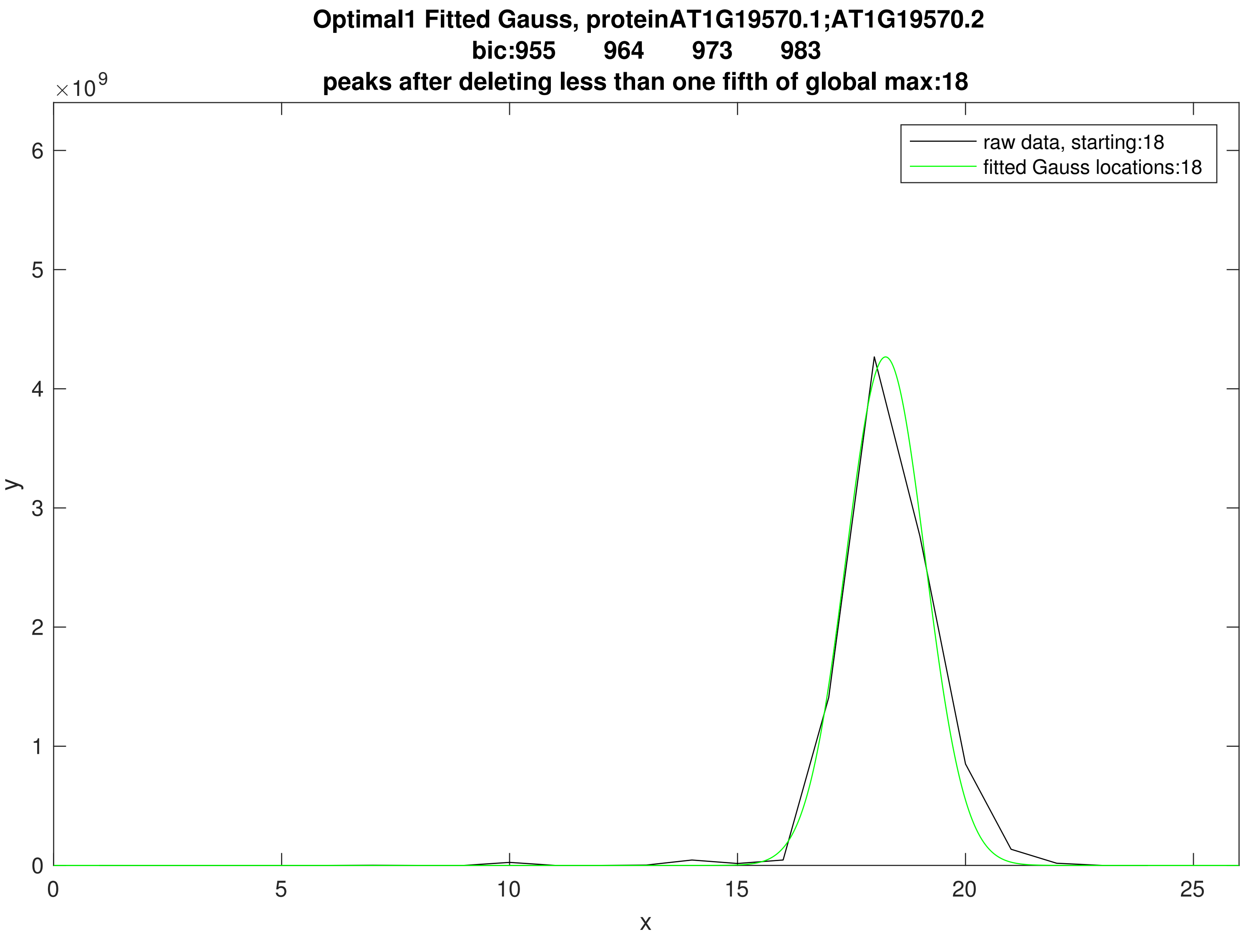

Optimal1 Fitted Gauss, proteinAT1G19570.1;AT1G19570.2
bic:955 964 973 983
peaks after deleting less than one fifth of global max:18
×10^9
raw data, starting:18
fitted Gauss locations:18
y
x
0
1
2
3
4
5
6
0
5
10
15
20
25

**Optimal1 Fitted Gauss, proteinAT1G19730.1**
**bic:709**
**peaks after deleting less than one fifth of global max:20**

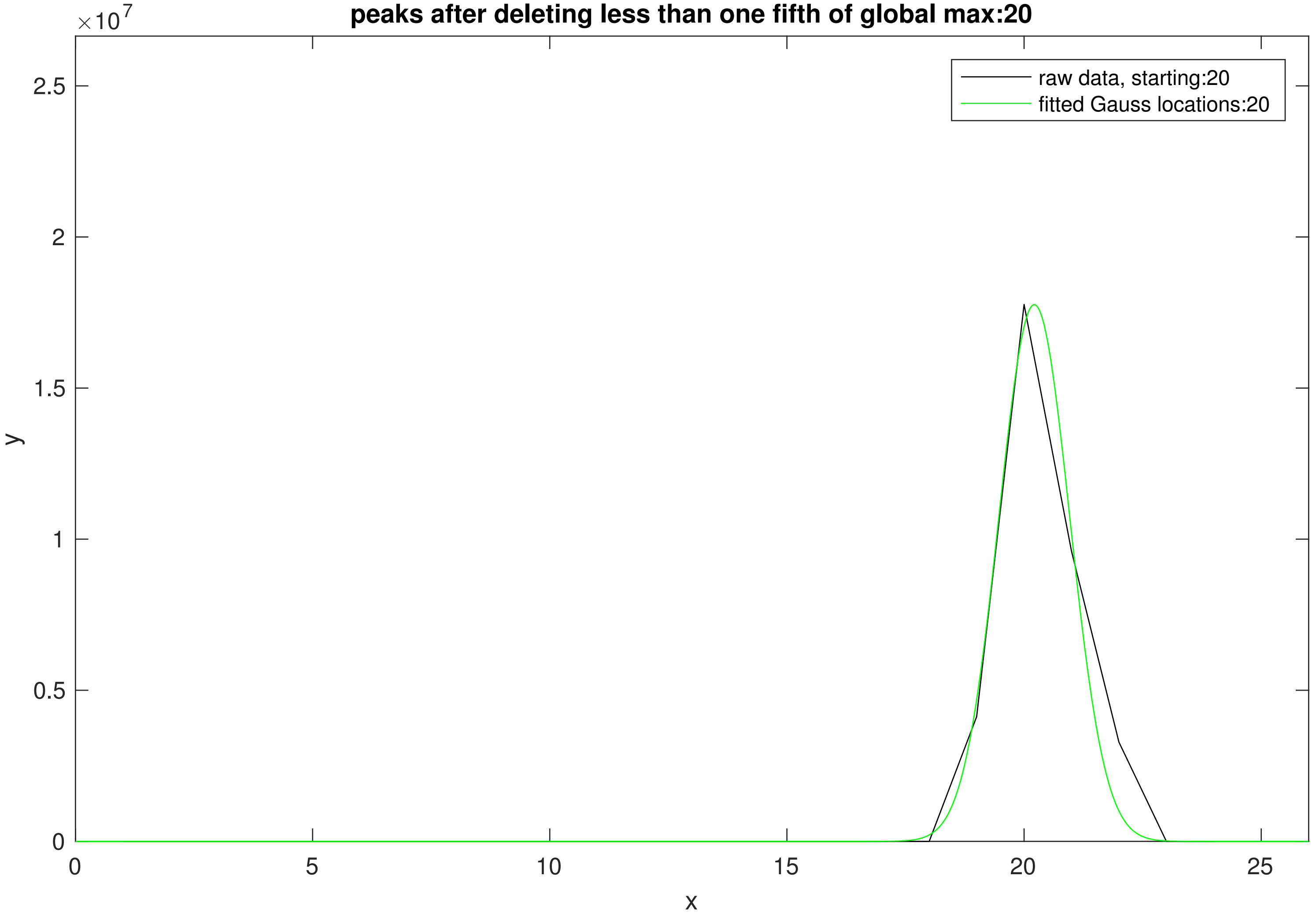

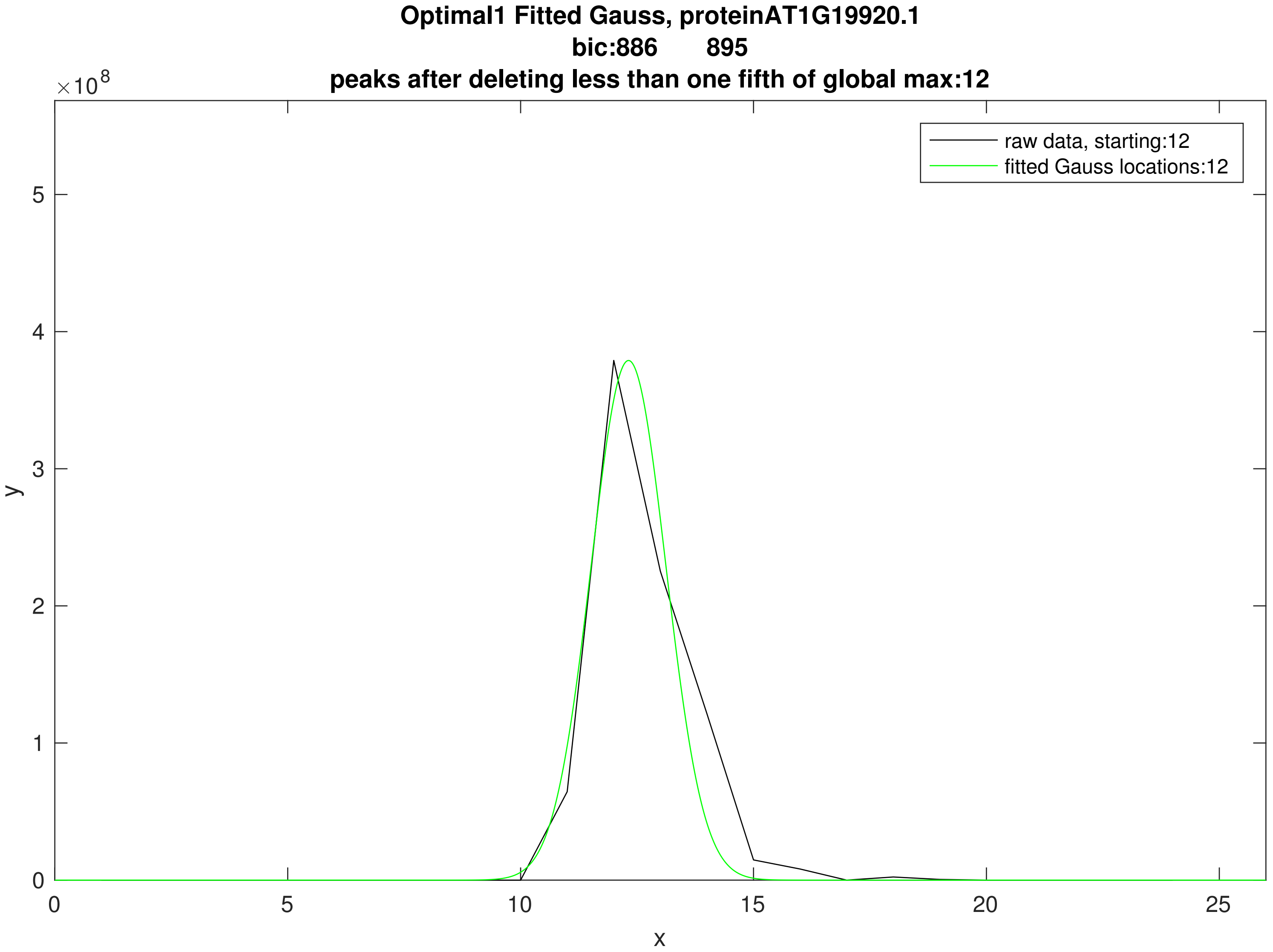

Optimal1 Fitted Gauss, proteinAT1G19920.1
bic:886 895
peaks after deleting less than one fifth of global max:12
×10^8
raw data, starting:12
fitted Gauss locations:12
y
x
0
1
2
3
4
5
0
5
10
15
20
25

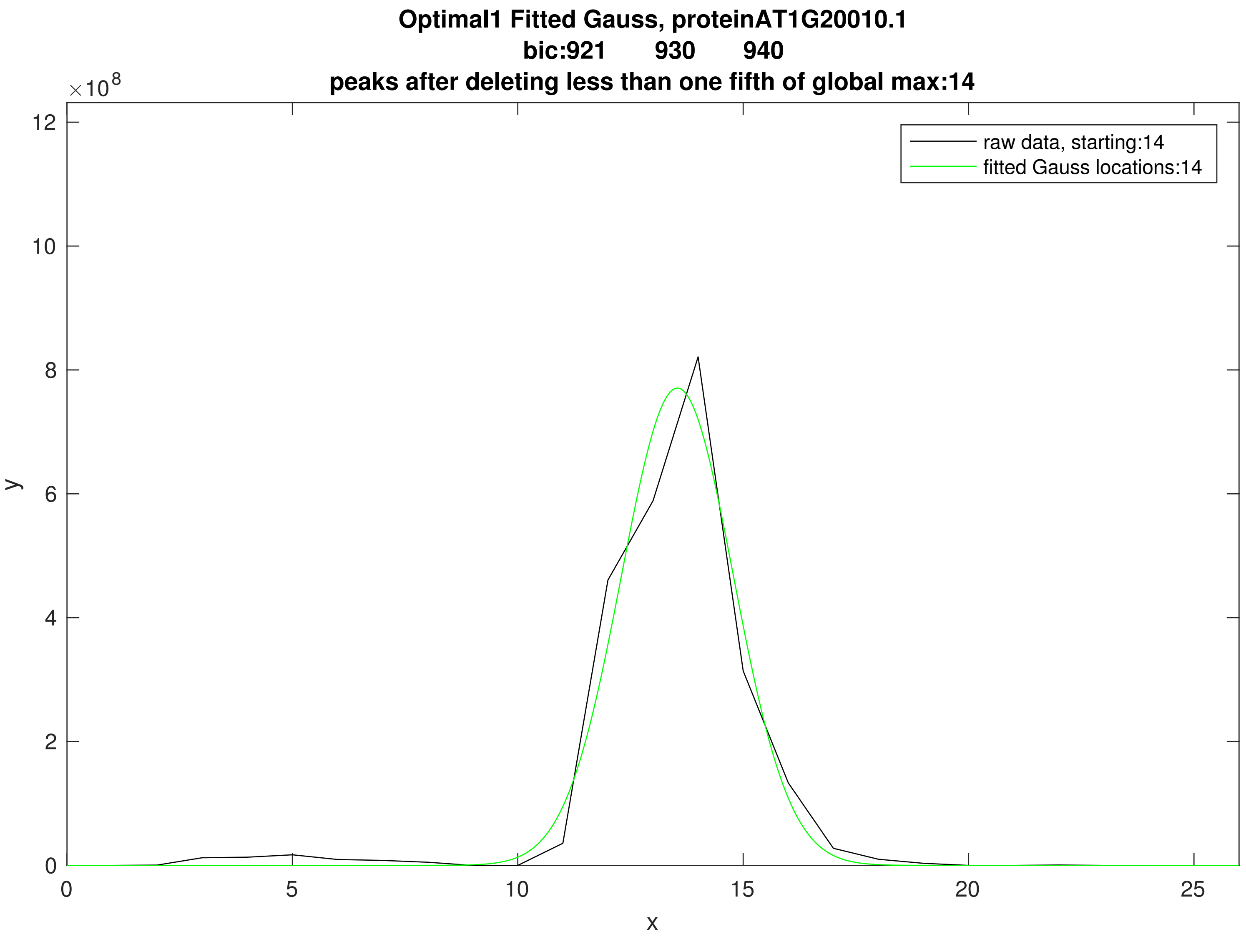
Optimal1 Fitted Gauss, proteinAT1G20010.1
bic:921 930 940
peaks after deleting less than one fifth of global max:14
×10^8
raw data, starting:14
fitted Gauss locations:14
y
x
0
2
4
6
8
10
12
0
5
10
15
20
25

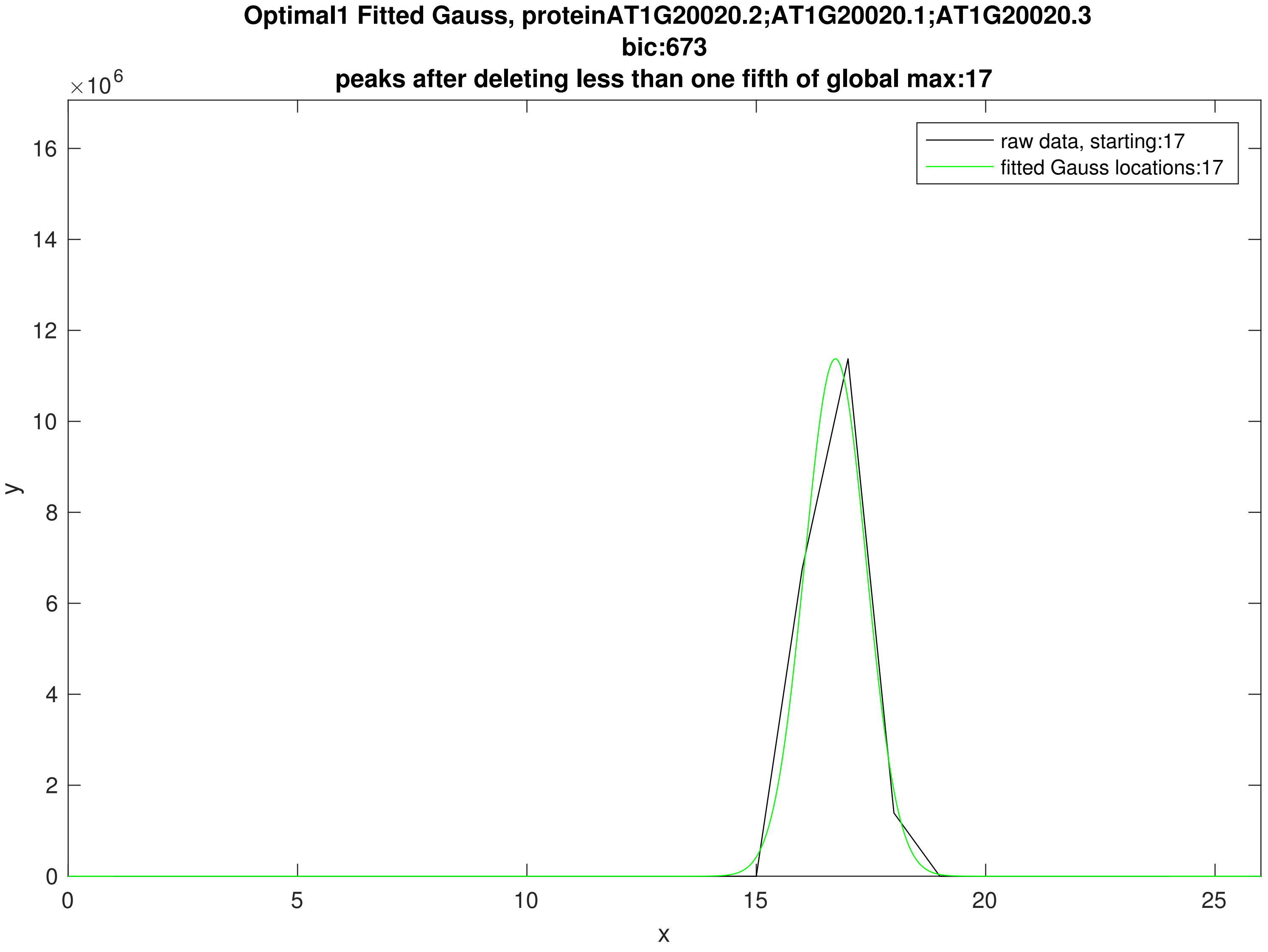

Optimal1 Fitted Gauss, proteinAT1G20020.2;AT1G20020.1;AT1G20020.3
bic:673
peaks after deleting less than one fifth of global max:17
×10^6
raw data, starting:17
fitted Gauss locations:17
y
x
0
2
4
6
8
10
12
14
16
0
5
10
15
20
25

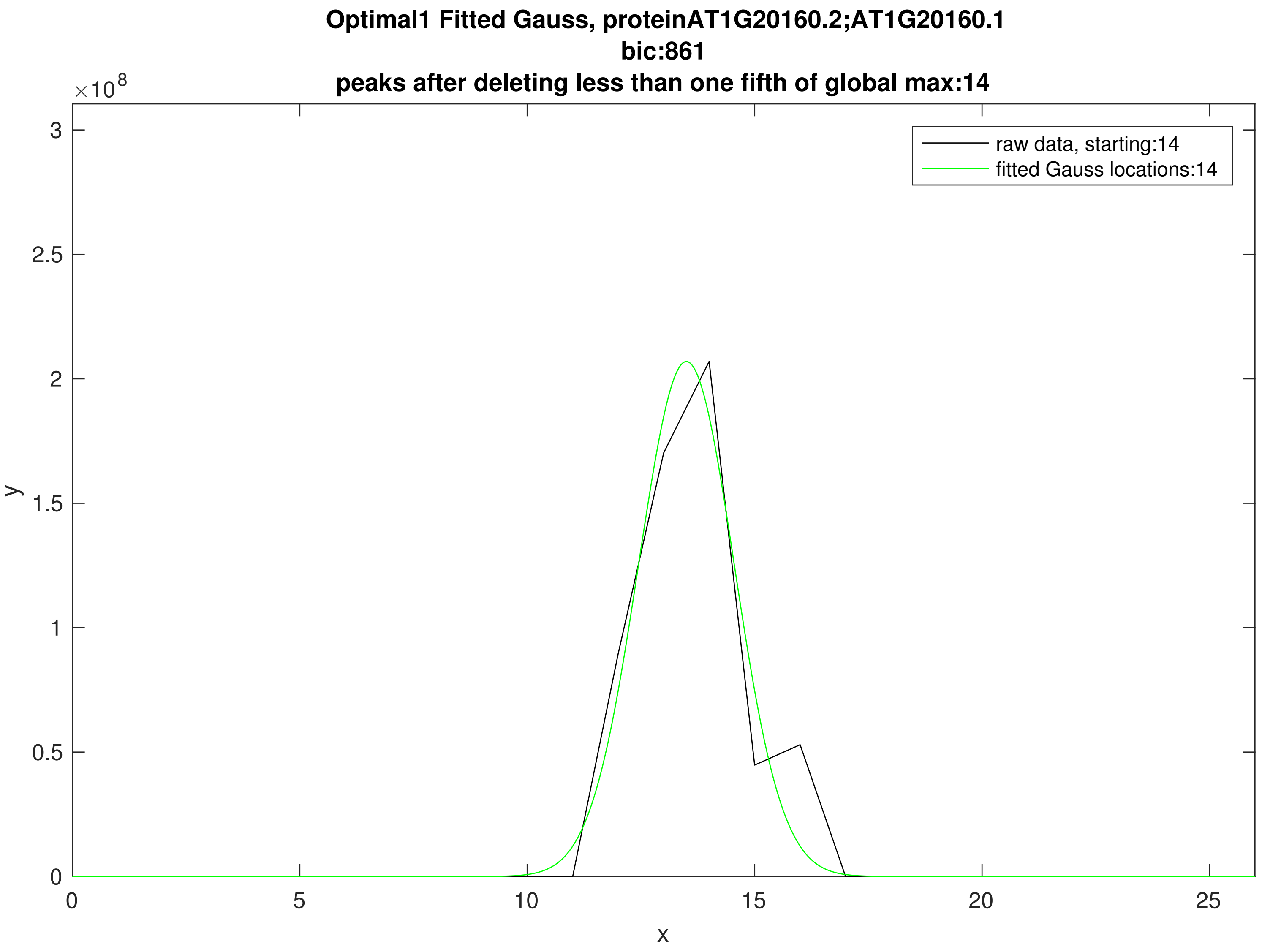

Optimal1 Fitted Gauss, proteinAT1G20160.2;AT1G20160.1
bic:861
peaks after deleting less than one fifth of global max:14
×10^8
3
2.5
2
1.5
1
0.5
0
y
0
5
10
15
20
25
x
raw data, starting:14
fitted Gauss locations:14

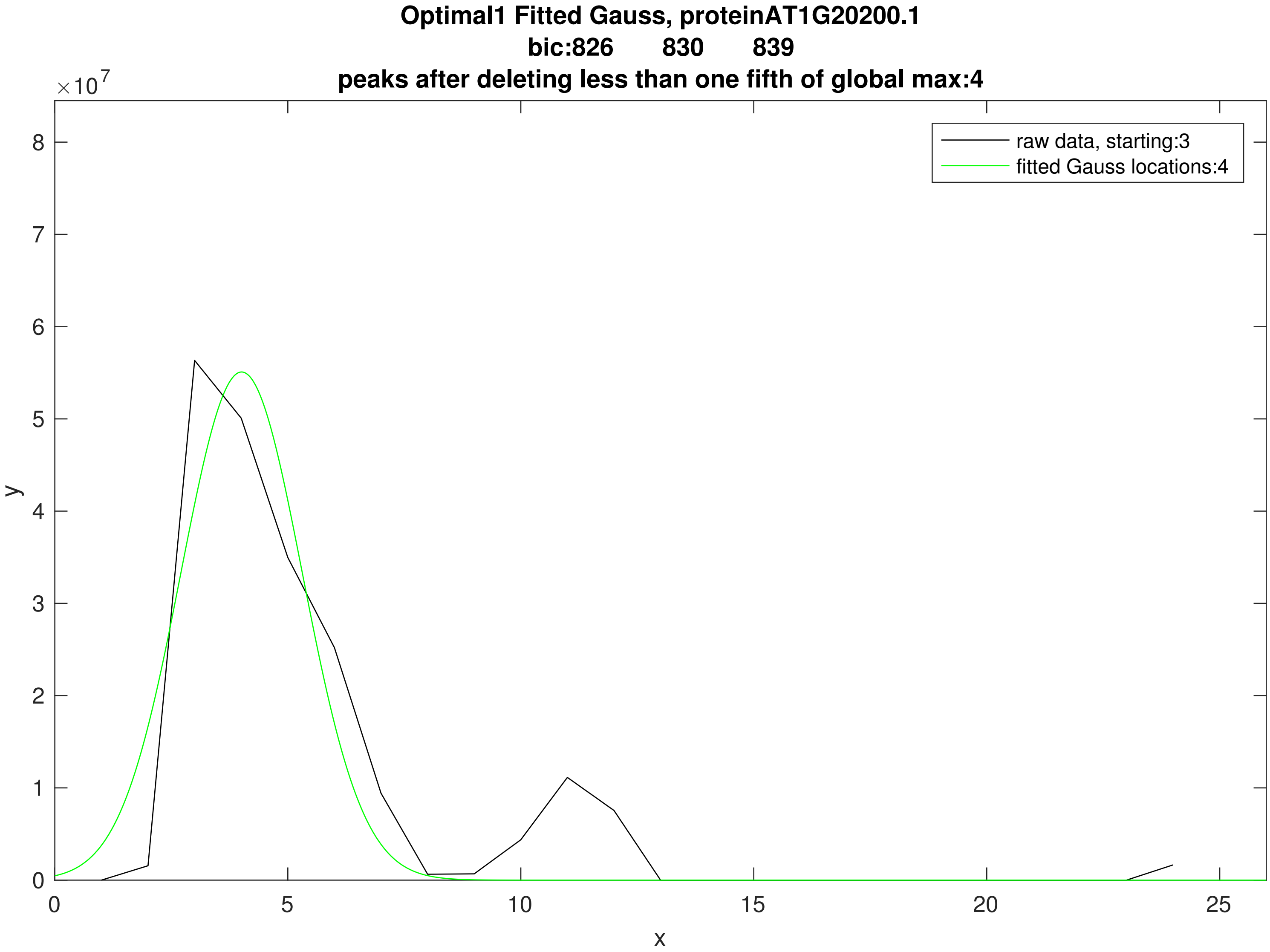

Optimal1 Fitted Gauss, proteinAT1G20200.1
bic:826 830 839
peaks after deleting less than one fifth of global max:4
×10^7
raw data, starting:3
fitted Gauss locations:4
y
x
0
1
2
3
4
5
6
7
8
0
5
10
15
20
25

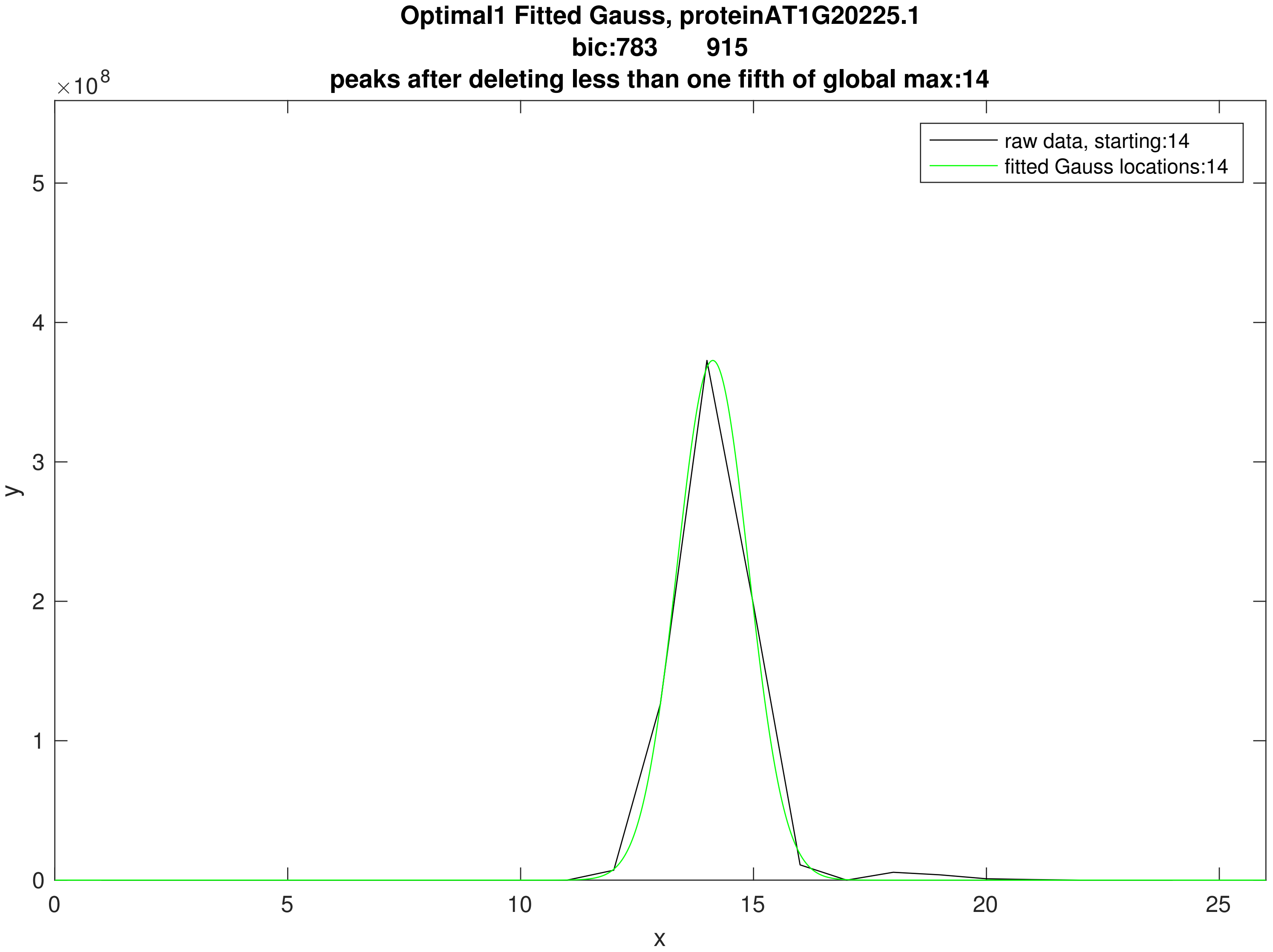

Optimal1 Fitted Gauss, proteinAT1G20225.1
bic:783 915
peaks after deleting less than one fifth of global max:14
×10^8
raw data, starting:14
fitted Gauss locations:14
5
4
3
2
1
0
y
0
5
10
15
20
25
x

**Optimal1 Fitted Gauss, proteinAT1G20260.1**
**bic:1.03e+03  1.04e+03**
**peaks after deleting less than one fifth of global max:13**

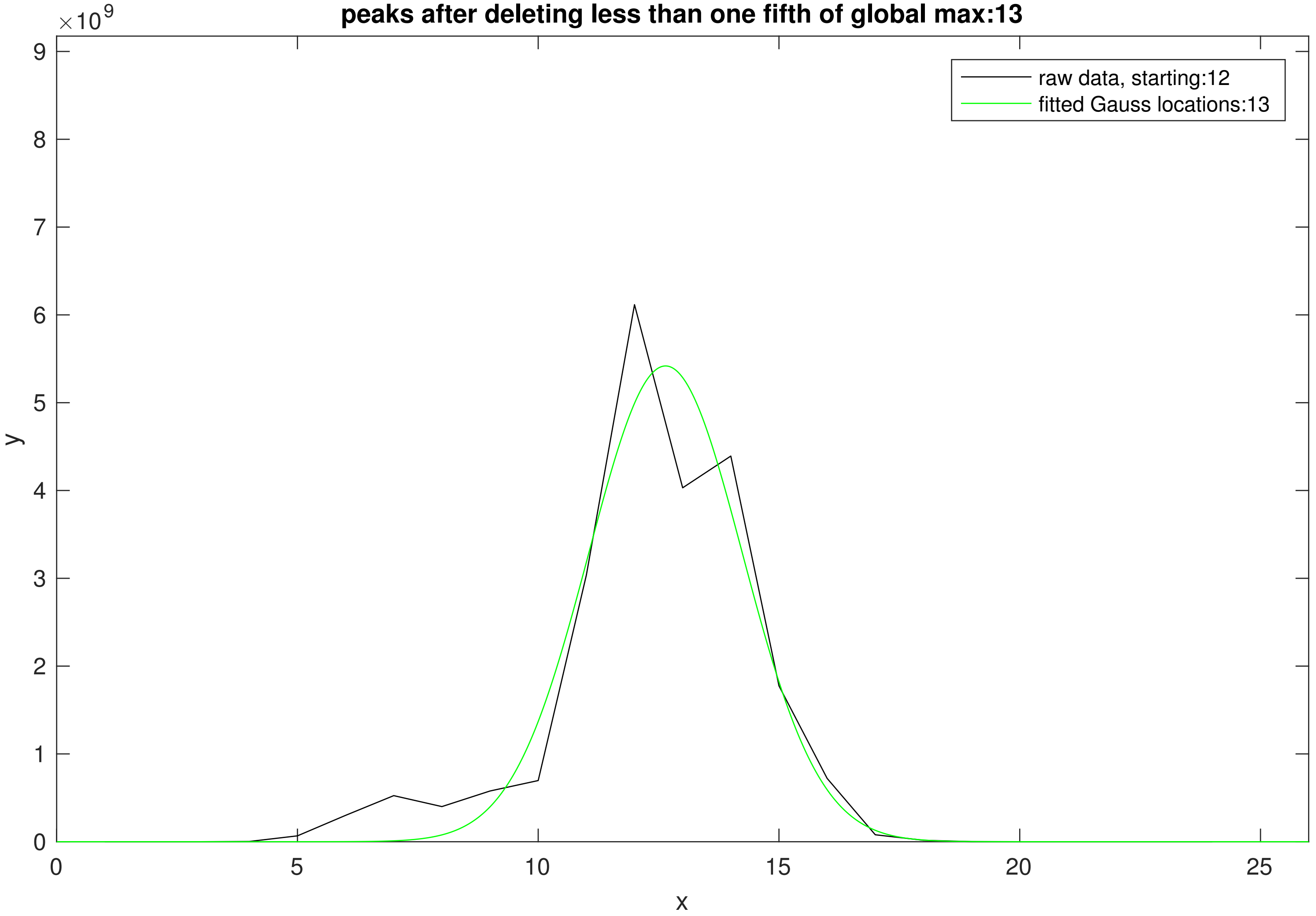

**Optimal1 Fitted Gauss, proteinAT1G20340.1**
**bic:773**
**peaks after deleting less than one fifth of global max:20**

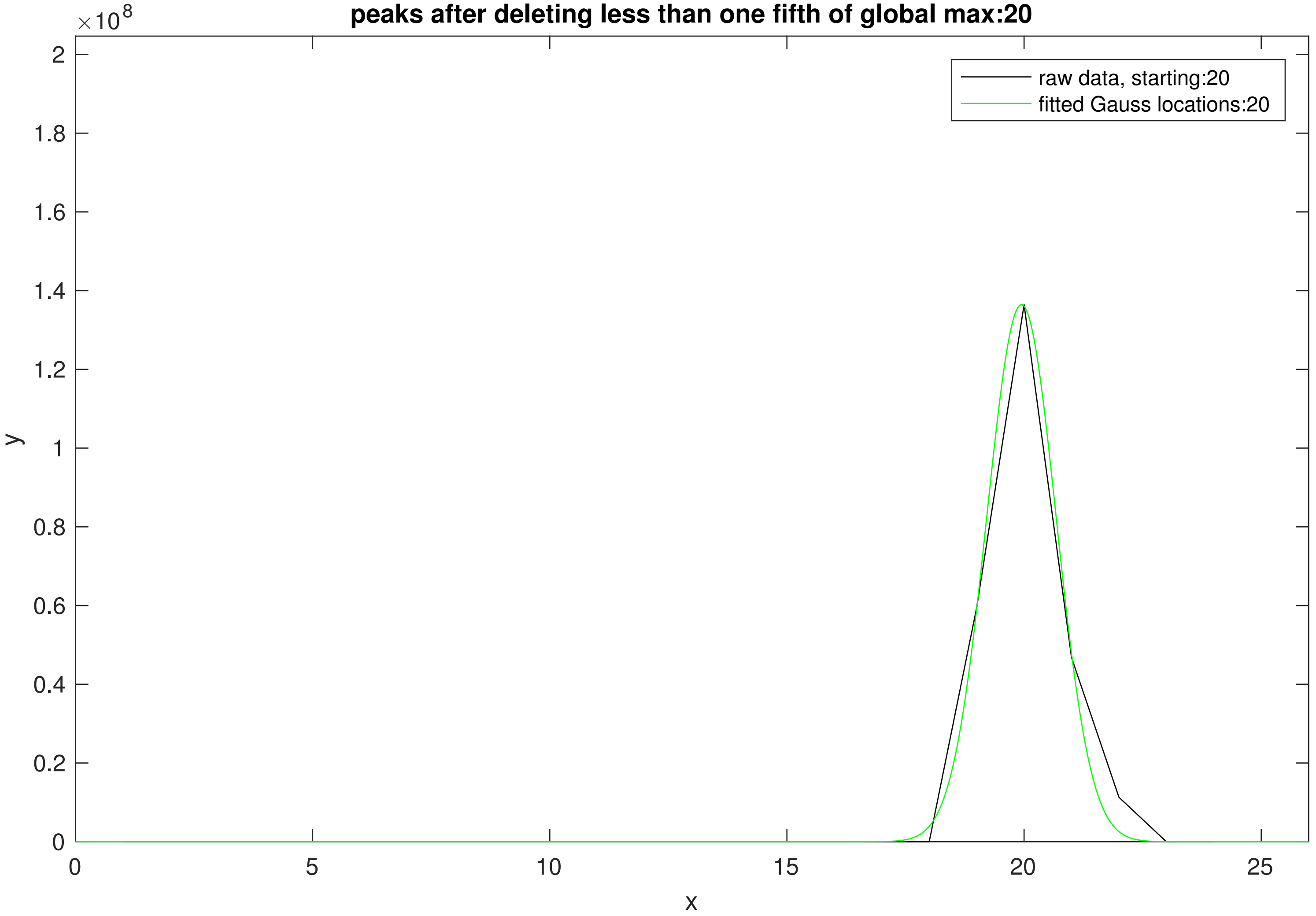

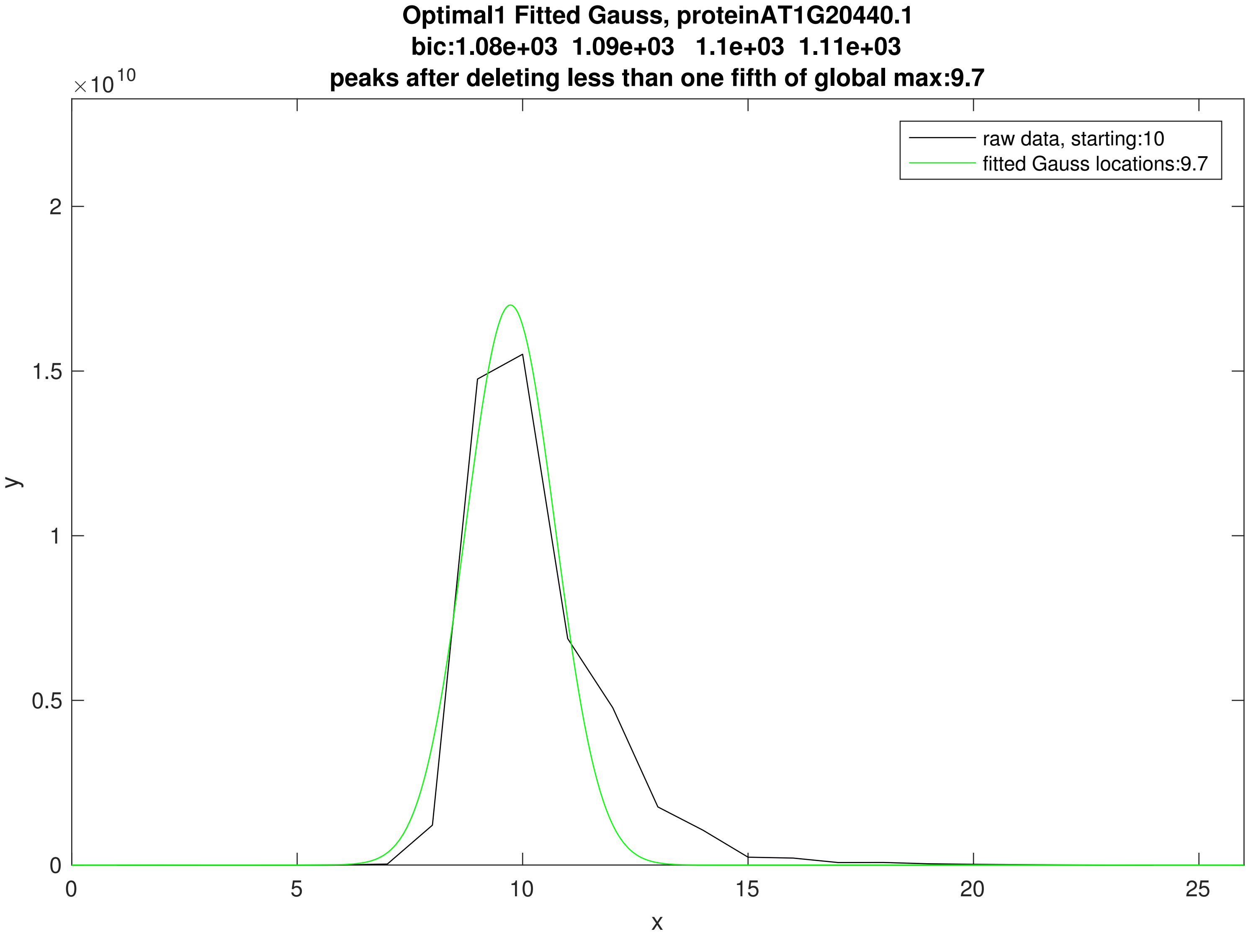
Optimal1 Fitted Gauss, proteinAT1G20440.1
bic:1.08e+03 1.09e+03 1.1e+03 1.11e+03
peaks after deleting less than one fifth of global max:9.7
×10^10
raw data, starting:10
fitted Gauss locations:9.7
y
x
0
0.5
1
1.5
2
0
5
10
15
20
25

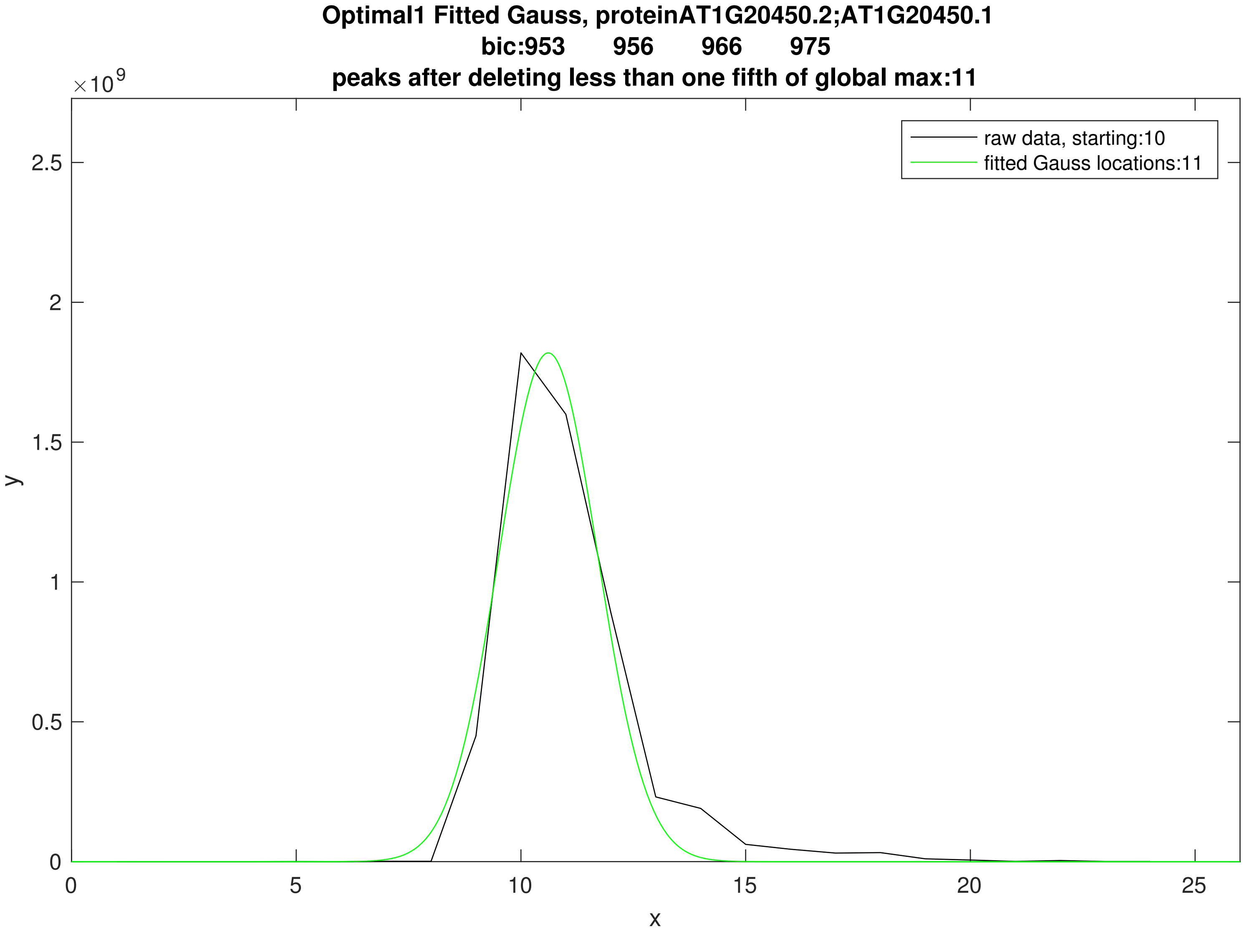

Optimal1 Fitted Gauss, proteinAT1G20450.2;AT1G20450.1
bic:953 956 966 975
peaks after deleting less than one fifth of global max:11
×10^9
raw data, starting:10
fitted Gauss locations:11
y
x
0
0.5
1
1.5
2
2.5
5
10
15
20
25

Optimal1 Fitted Gauss, proteinAT1G20510.2;AT1G20510.1
bic:719
peaks after deleting less than one fifth of global max:12

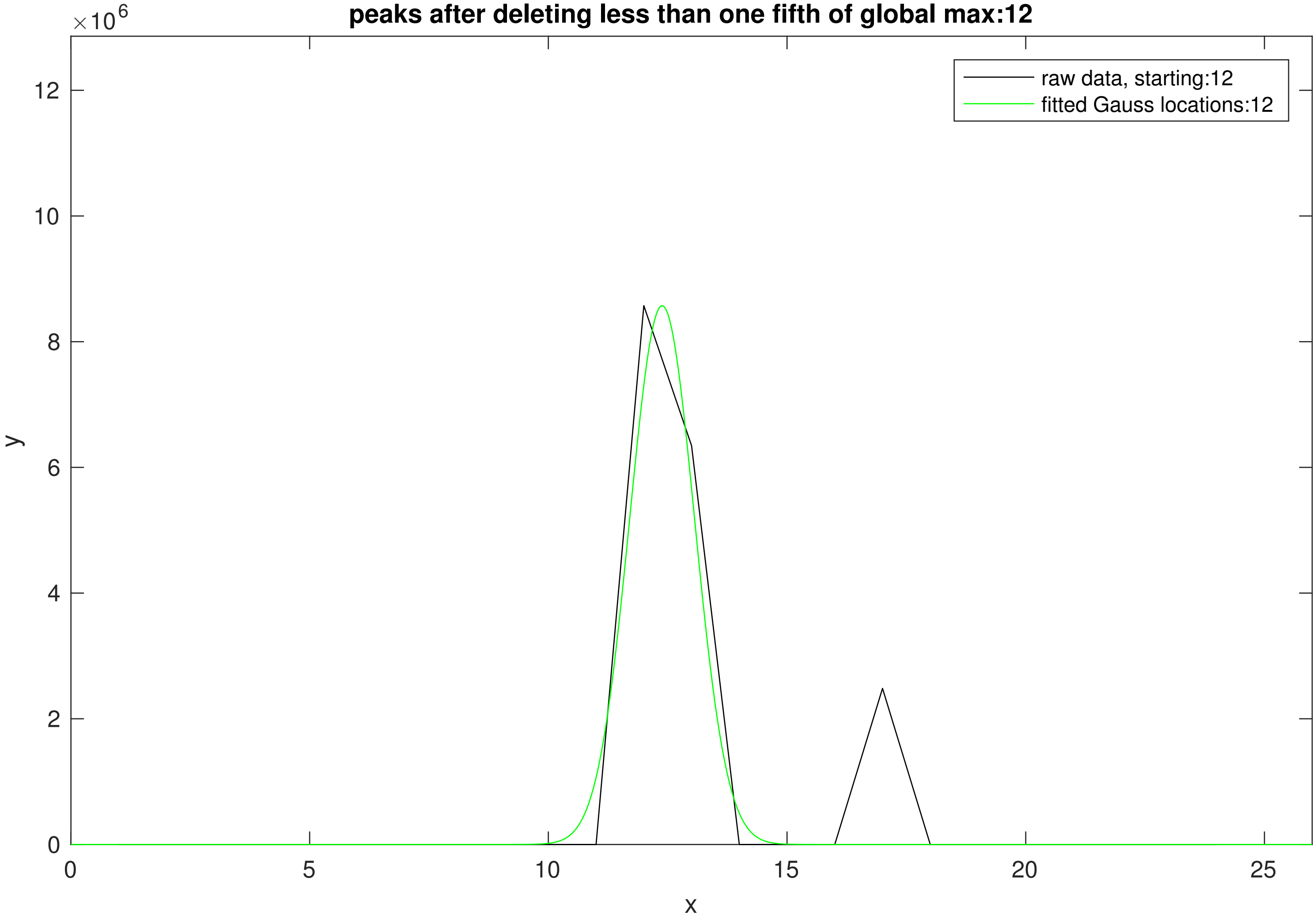

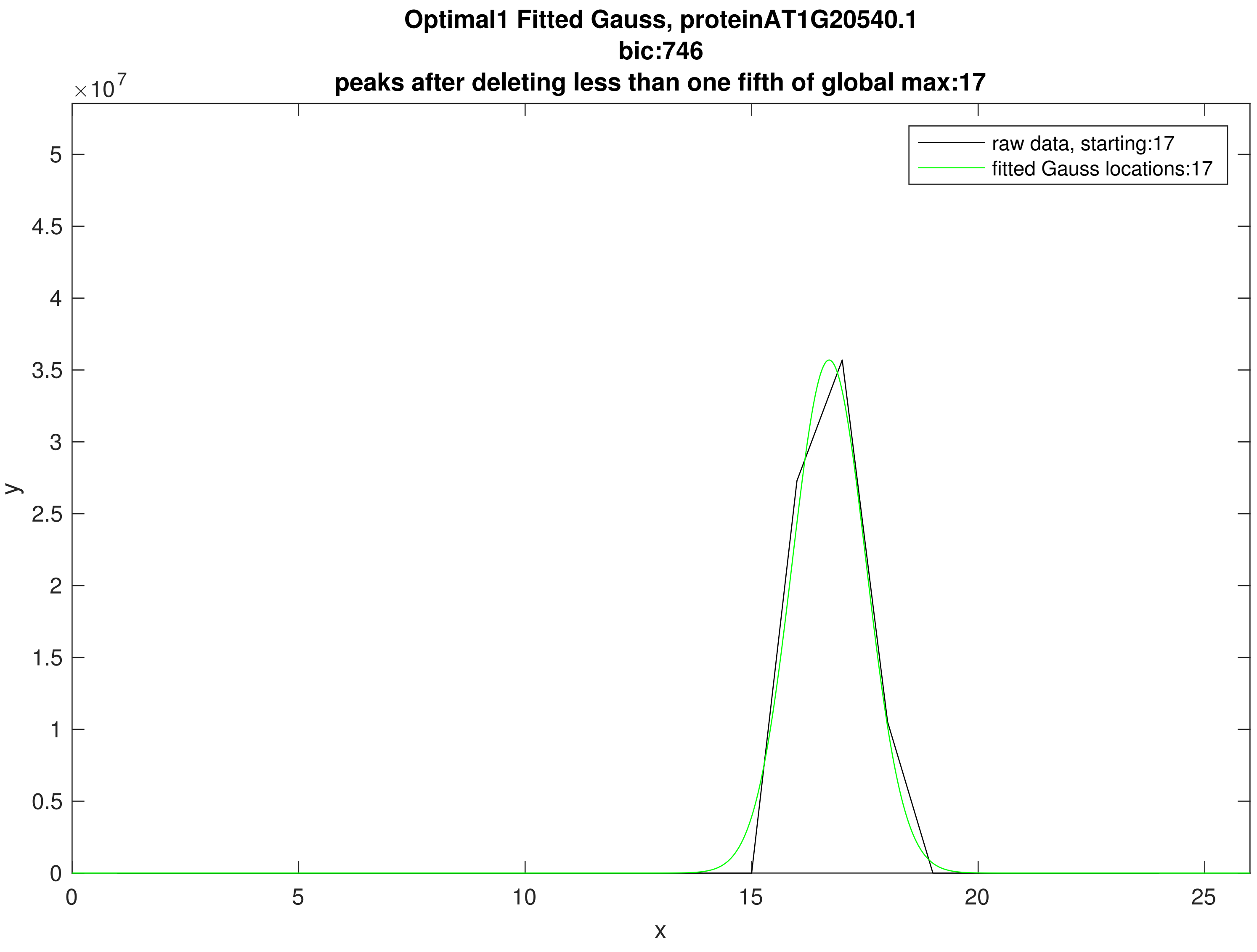

Optimal1 Fitted Gauss, proteinAT1G20540.1
bic:746
peaks after deleting less than one fifth of global max:17
×10^7
raw data, starting:17
fitted Gauss locations:17
y
x
0
0.5
1
1.5
2
2.5
3
3.5
4
4.5
5
0
5
10
15
20
25

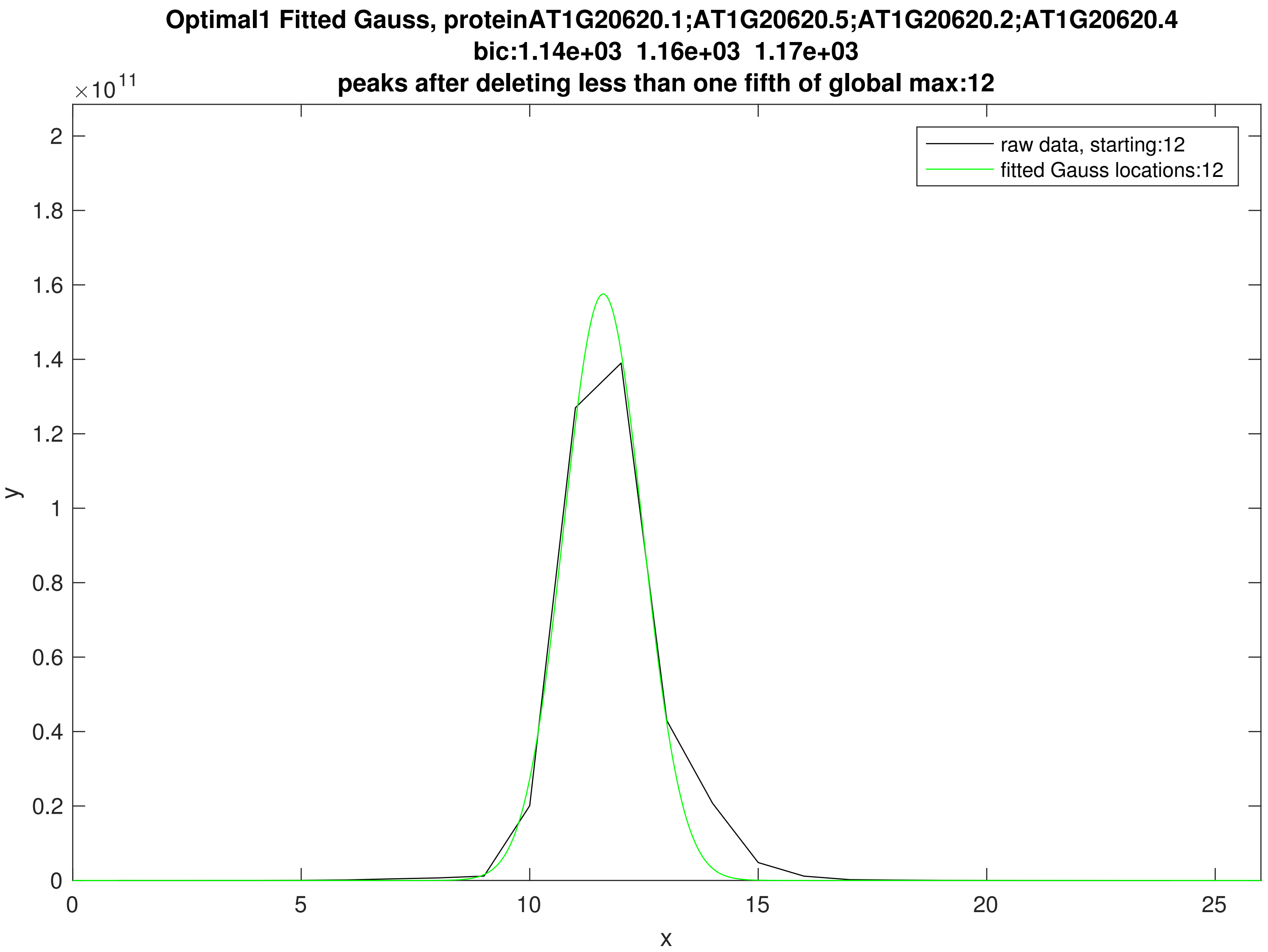
Optimal1 Fitted Gauss, proteinAT1G20620.1;AT1G20620.5;AT1G20620.2;AT1G20620.4
bic:1.14e+03  1.16e+03  1.17e+03
peaks after deleting less than one fifth of global max:12
×10^11
raw data, starting:12
fitted Gauss locations:12
y
x
0
0.2
0.4
0.6
0.8
1
1.2
1.4
1.6
1.8
2
0
5
10
15
20
25

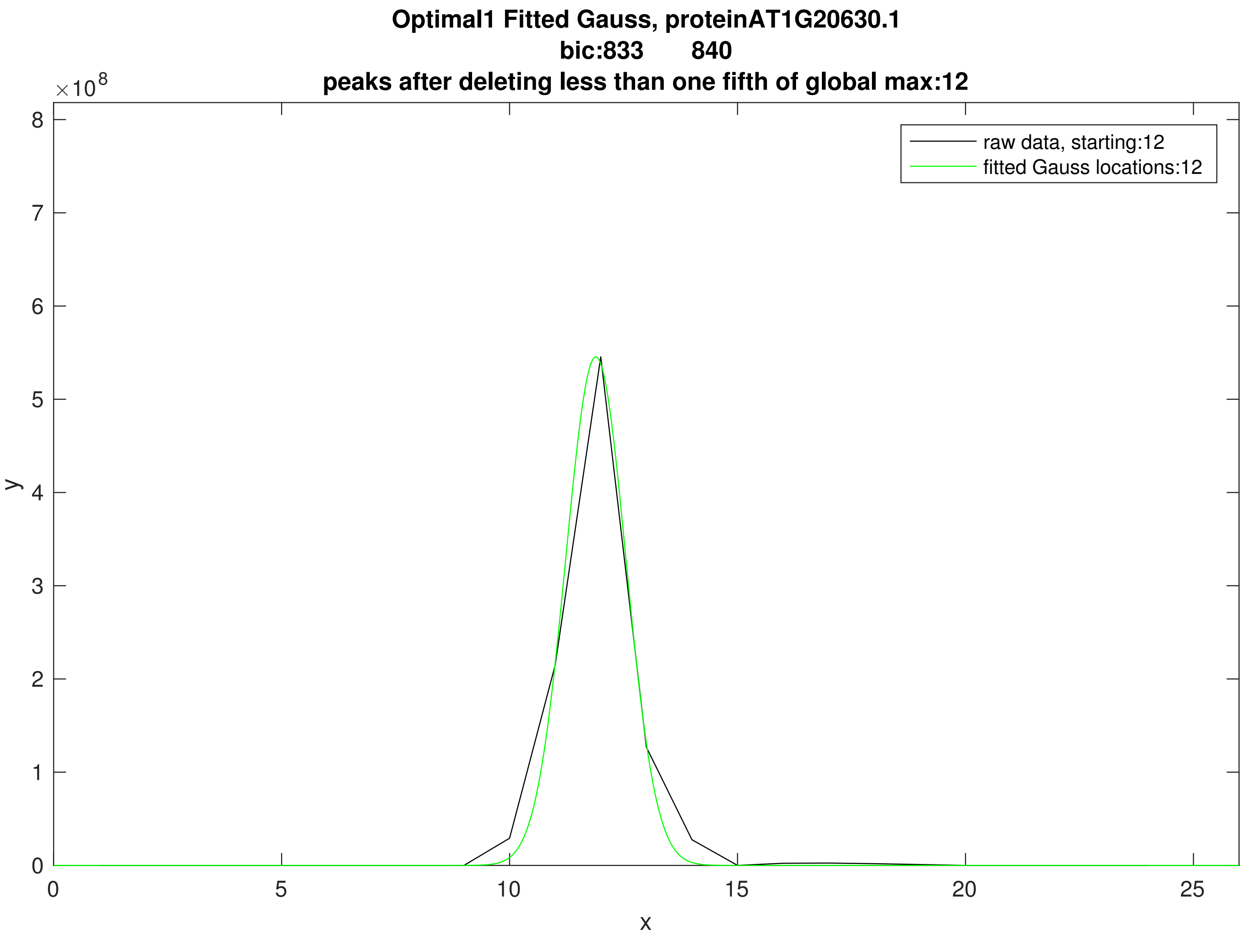
Optimal1 Fitted Gauss, proteinAT1G20630.1
bic:833 840
peaks after deleting less than one fifth of global max:12
×10^8
raw data, starting:12
fitted Gauss locations:12
y
x
0
1
2
3
4
5
6
7
8
0
5
10
15
20
25

**Optimal2 Fitted Gauss, proteinAT1G20760.1**
**bic:754 725**
**peaks after deleting less than one fifth of global max:8.5 19**

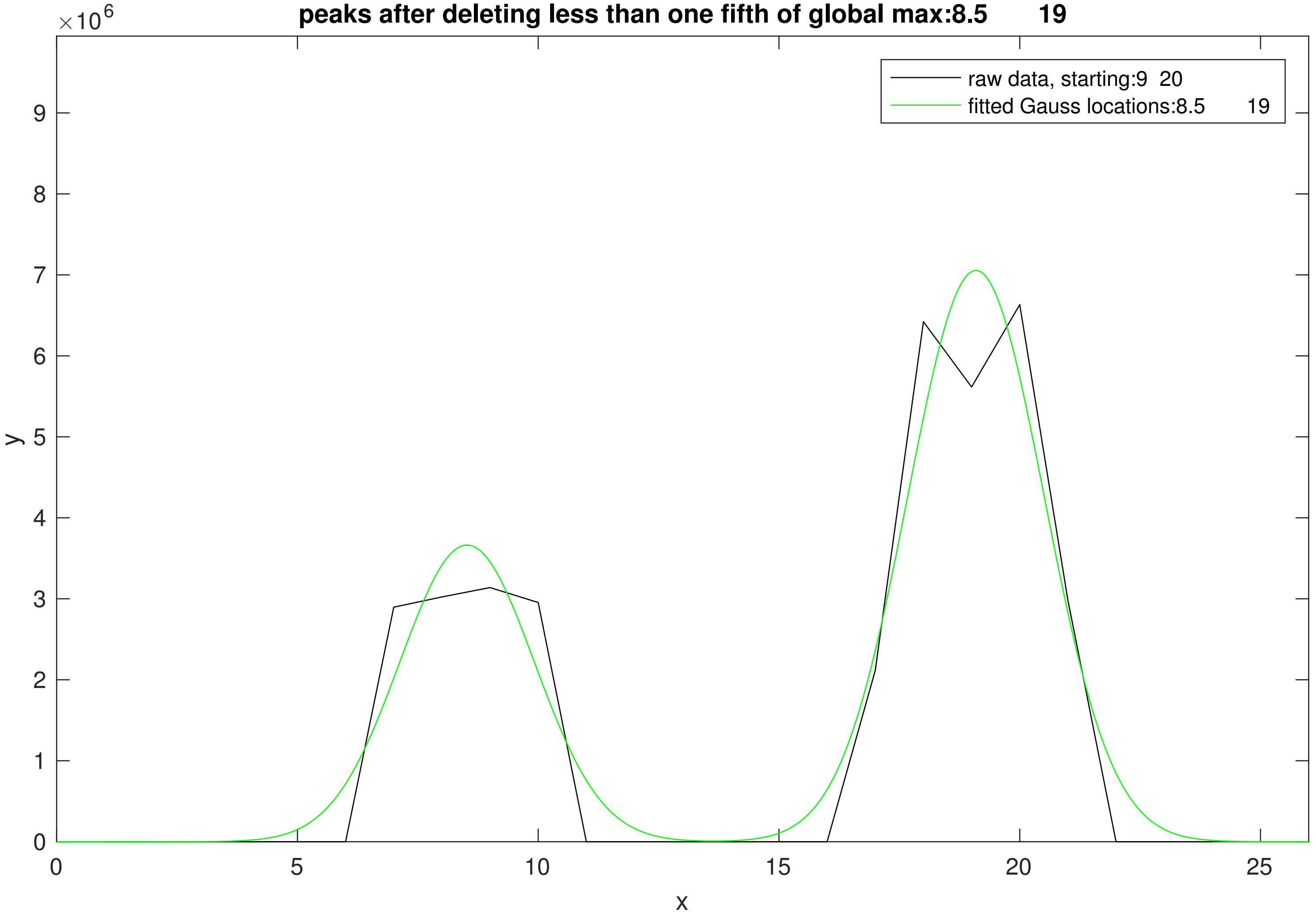

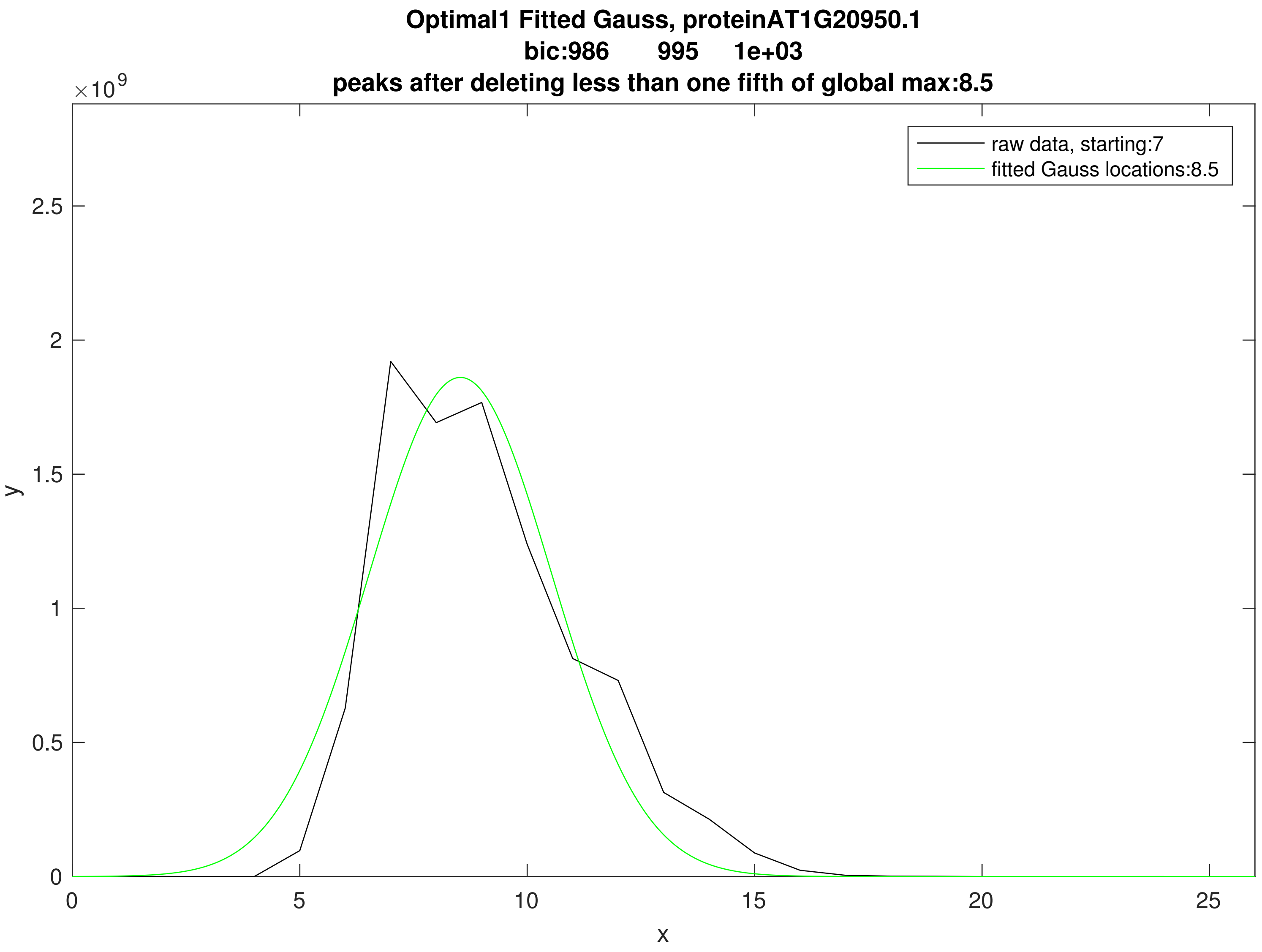

Optimal1 Fitted Gauss, proteinAT1G20950.1
bic:986 995 1e+03
peaks after deleting less than one fifth of global max:8.5
×10^9
raw data, starting:7
fitted Gauss locations:8.5
y
x

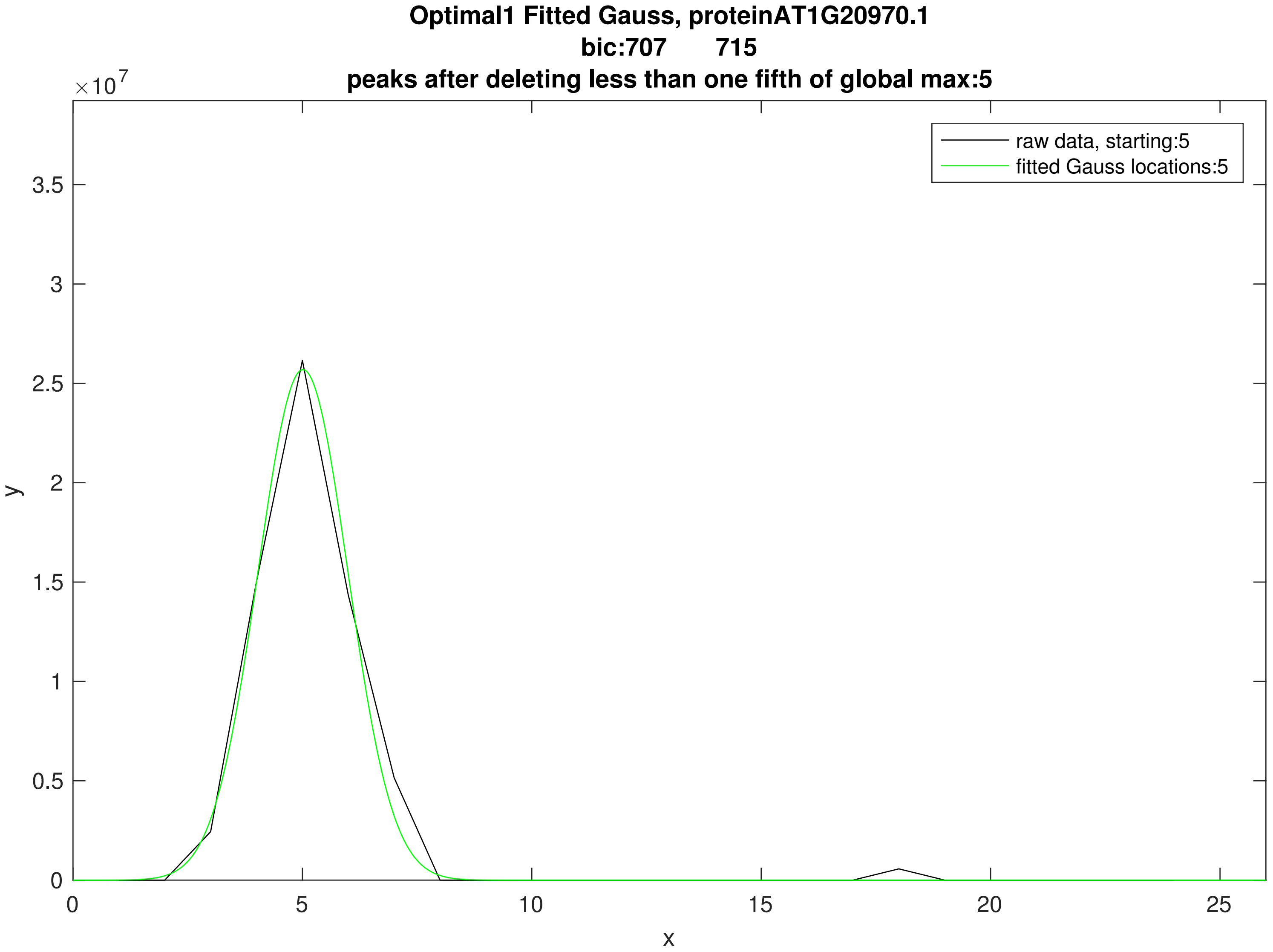
Optimal1 Fitted Gauss, proteinAT1G20970.1
bic:707 715
peaks after deleting less than one fifth of global max:5
×10^7
raw data, starting:5
fitted Gauss locations:5
y
x
0
0.5
1
1.5
2
2.5
3
3.5
5
10
15
20
25

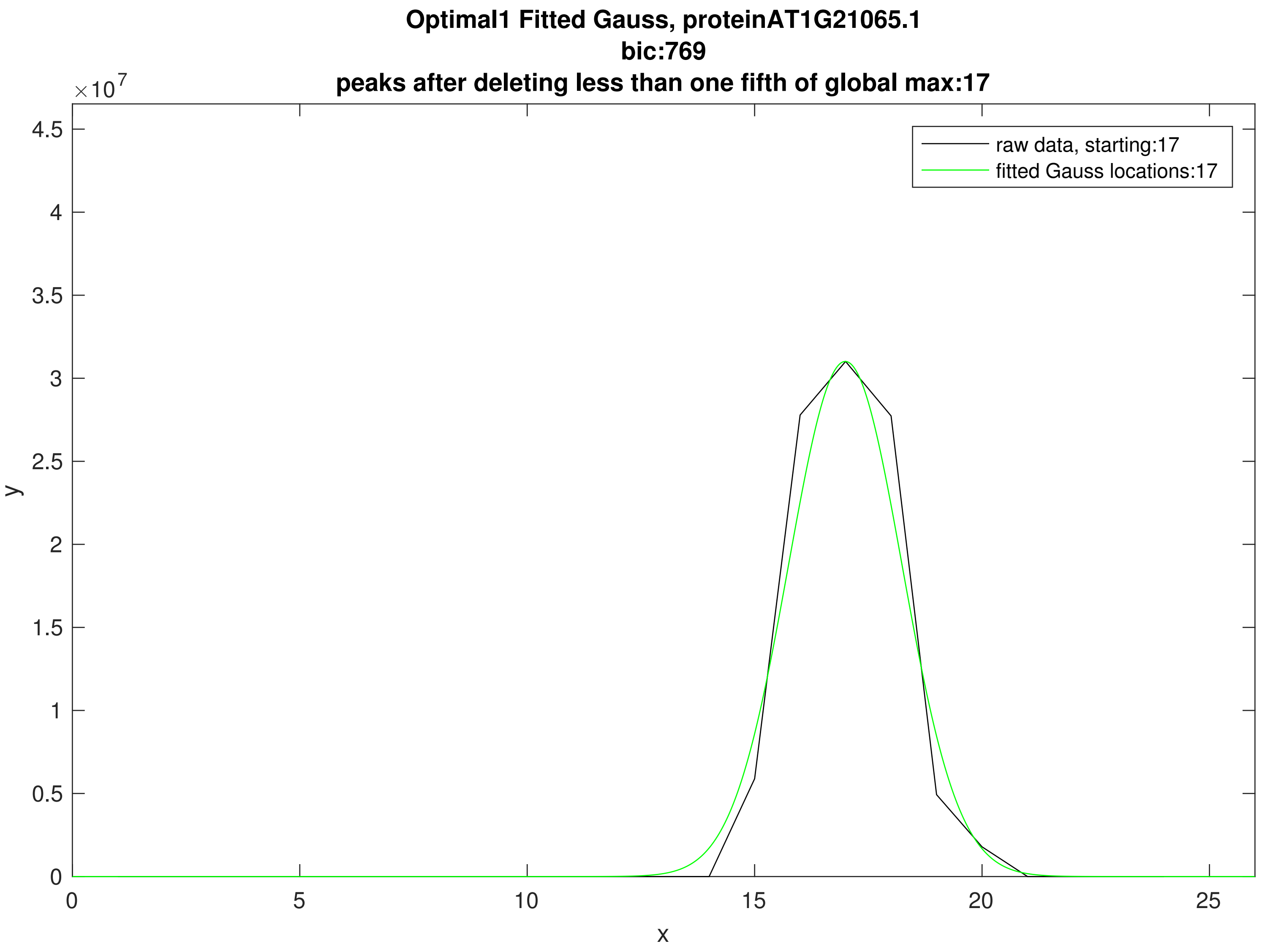

Optimal1 Fitted Gauss, proteinAT1G21065.1
bic:769
peaks after deleting less than one fifth of global max:17
×10^7
4.5
4
3.5
3
2.5
2
1.5
1
0.5
0
y
0
5
10
15
20
25
x
raw data, starting:17
fitted Gauss locations:17

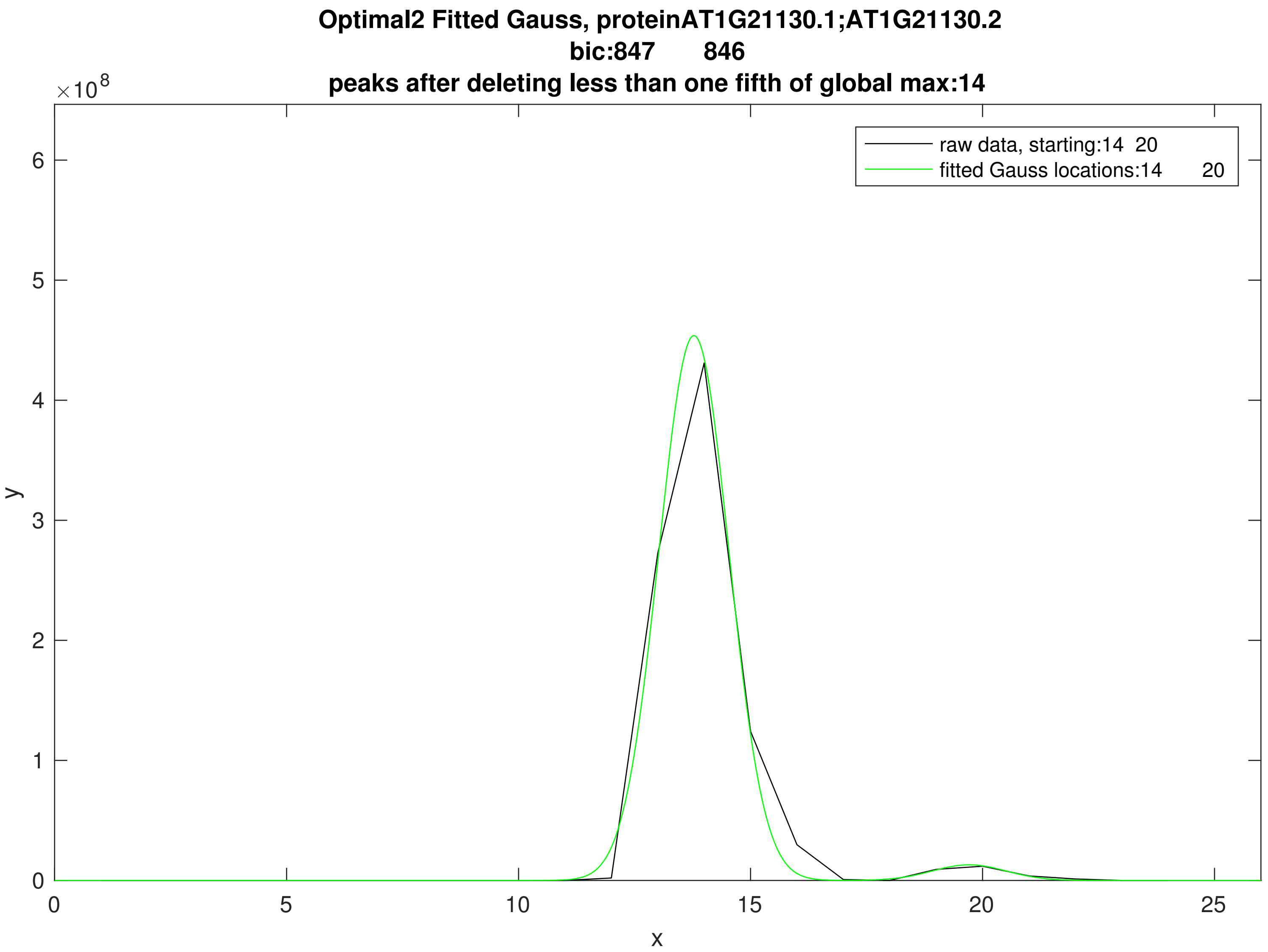

Optimal2 Fitted Gauss, proteinAT1G21130.1;AT1G21130.2
bic:847 846
peaks after deleting less than one fifth of global max:14
×10^8
raw data, starting:14 20
fitted Gauss locations:14 20
y
x
0
1
2
3
4
5
6
0
5
10
15
20
25

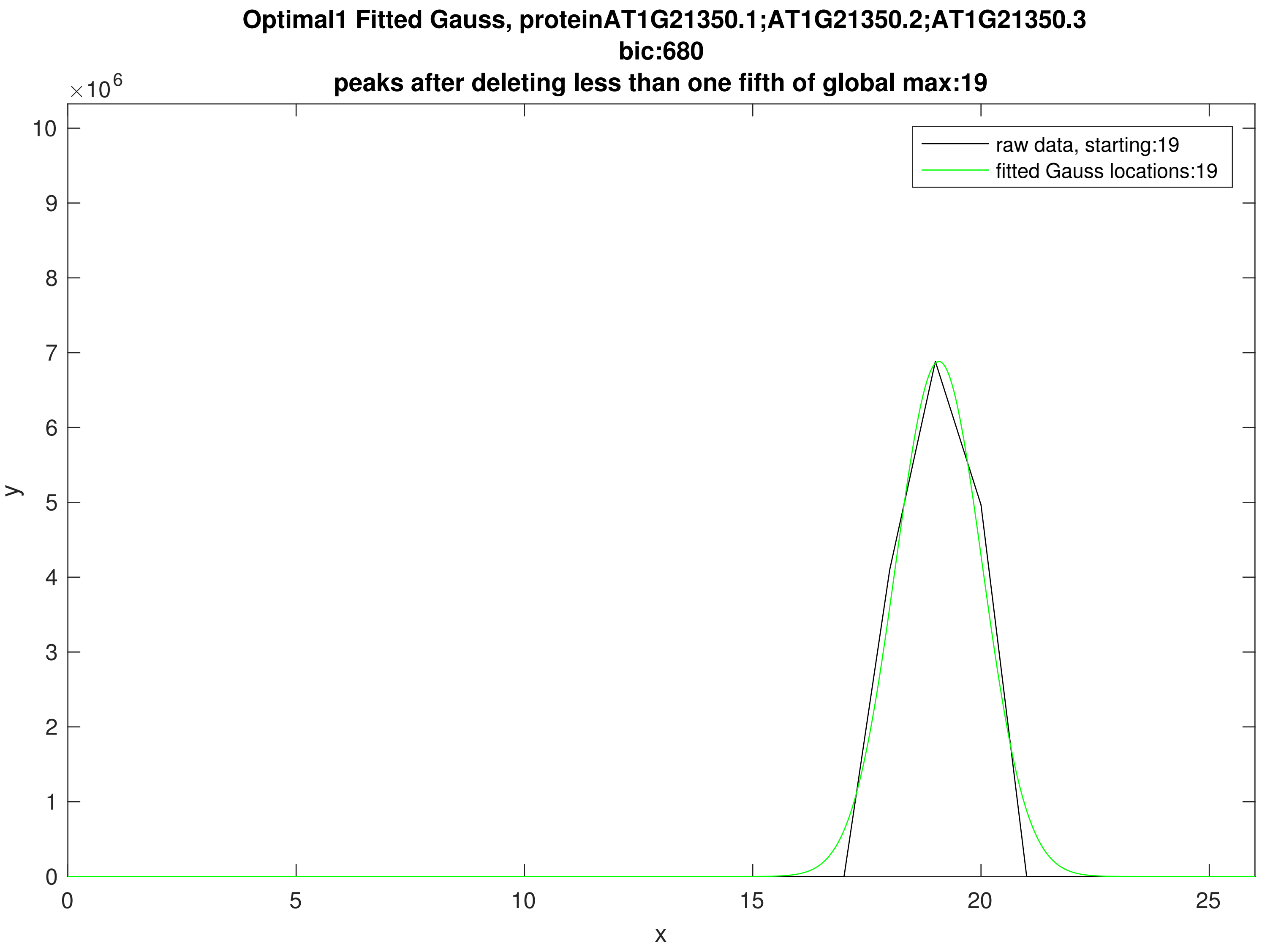

Optimal1 Fitted Gauss, proteinAT1G21350.1;AT1G21350.2;AT1G21350.3
bic:680
peaks after deleting less than one fifth of global max:19
×10^6
y
x
raw data, starting:19
fitted Gauss locations:19
0
1
2
3
4
5
6
7
8
9
10
0
5
10
15
20
25

**Optimal1 Fitted Gauss, proteinAT1G21440.1**
**bic:805**
**peaks after deleting less than one fifth of global max:14**

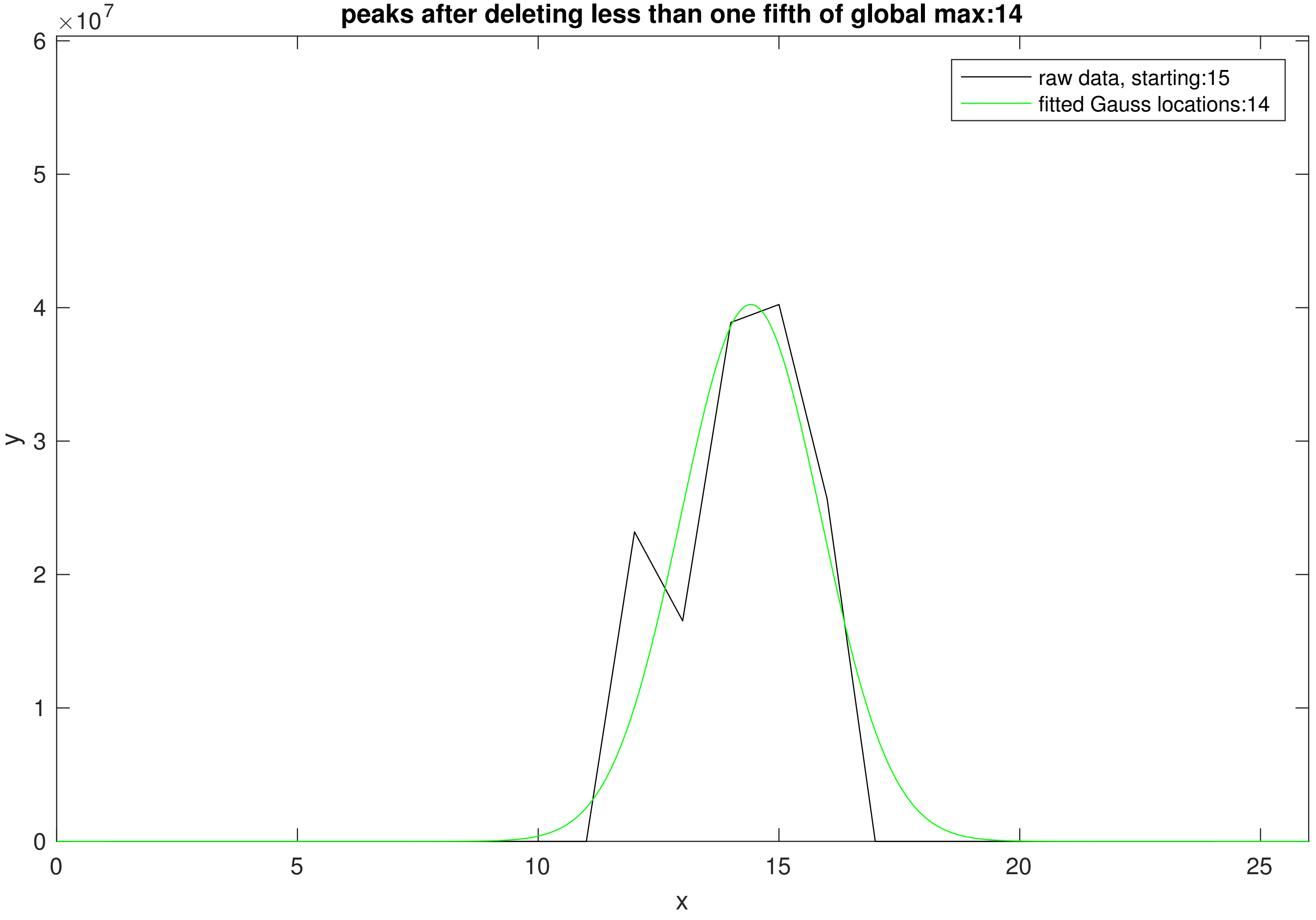

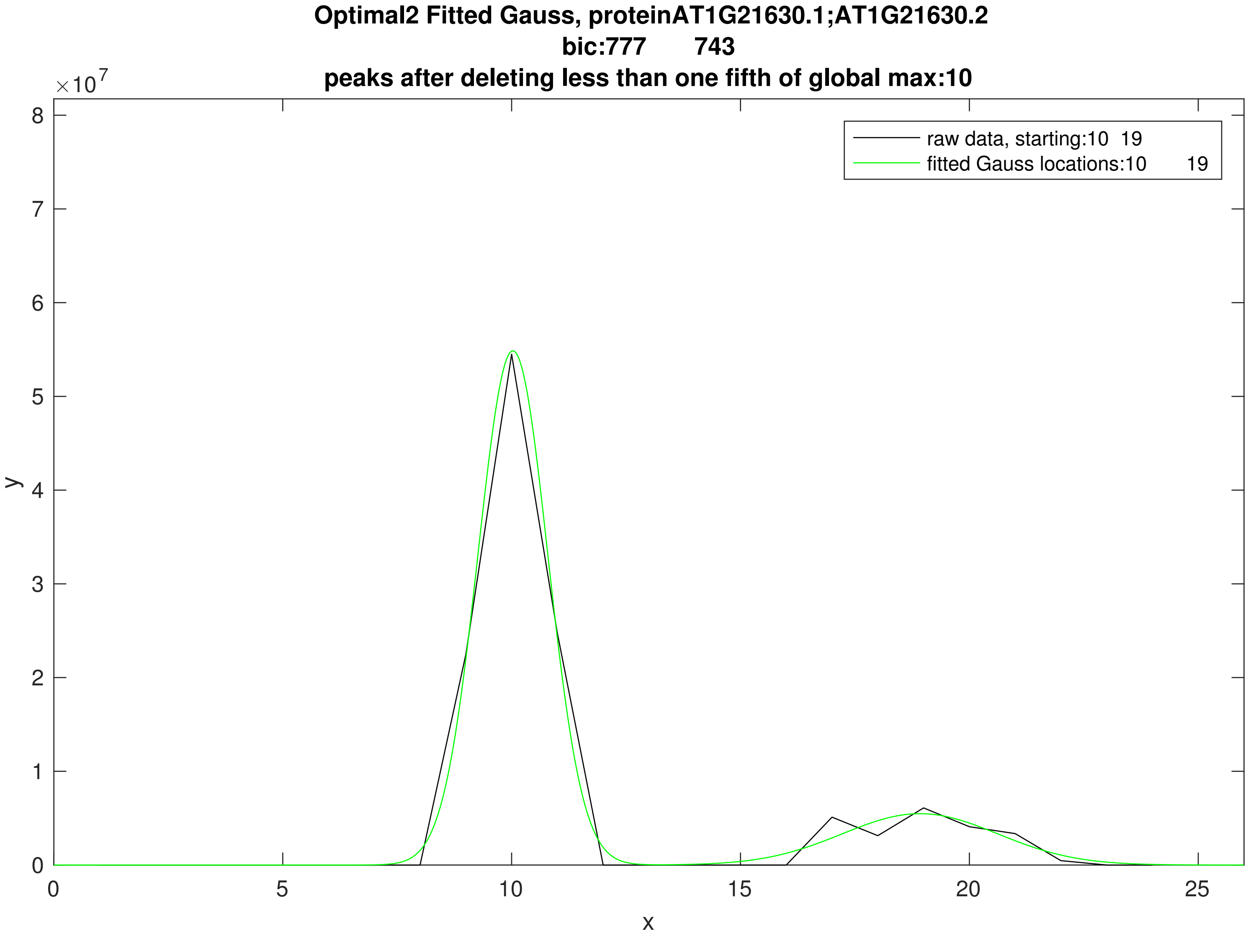

Optimal2 Fitted Gauss, proteinAT1G21630.1;AT1G21630.2
bic:777 743
peaks after deleting less than one fifth of global max:10
×10^7
raw data, starting:10 19
fitted Gauss locations:10 19
y
x
0
1
2
3
4
5
6
7
8
0
5
10
15
20
25

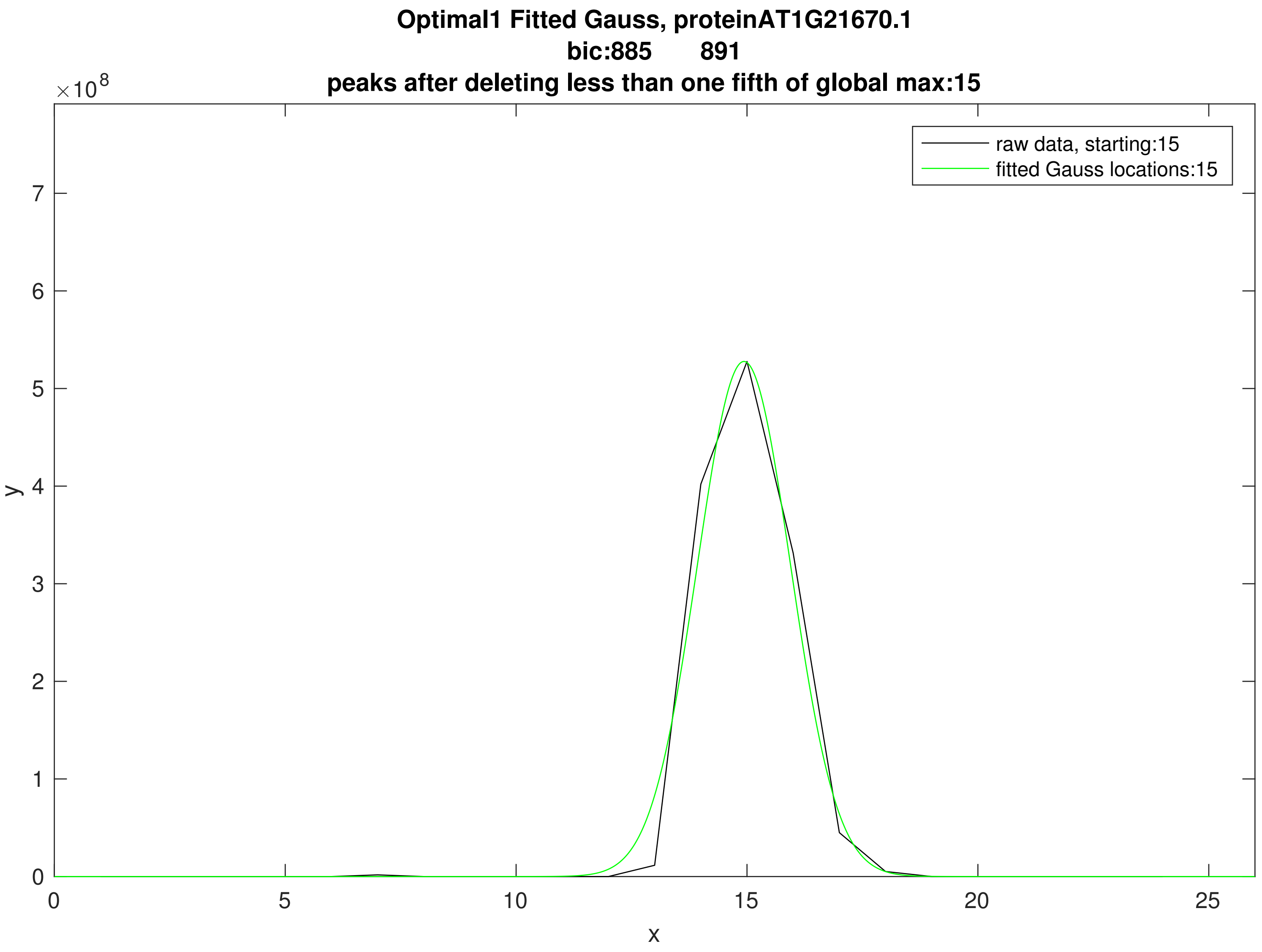

Optimal1 Fitted Gauss, proteinAT1G21670.1
bic:885 891
peaks after deleting less than one fifth of global max:15
×10^8
raw data, starting:15
fitted Gauss locations:15
y
x
0
1
2
3
4
5
6
7
0
5
10
15
20
25

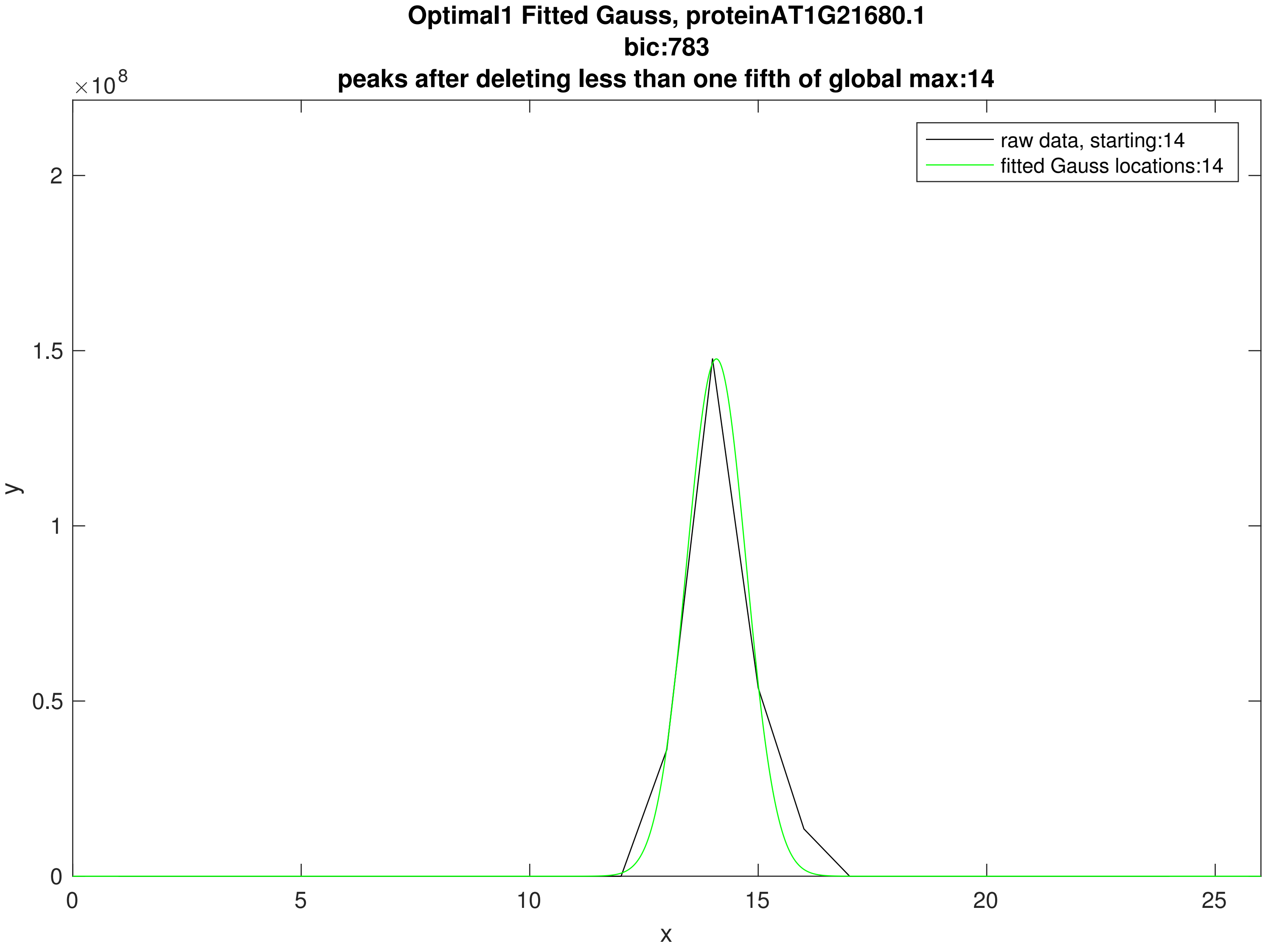
Optimal1 Fitted Gauss, proteinAT1G21680.1
bic:783
peaks after deleting less than one fifth of global max:14
×10^8
raw data, starting:14
fitted Gauss locations:14
y
x
0
0.5
1
1.5
2
0
5
10
15
20
25

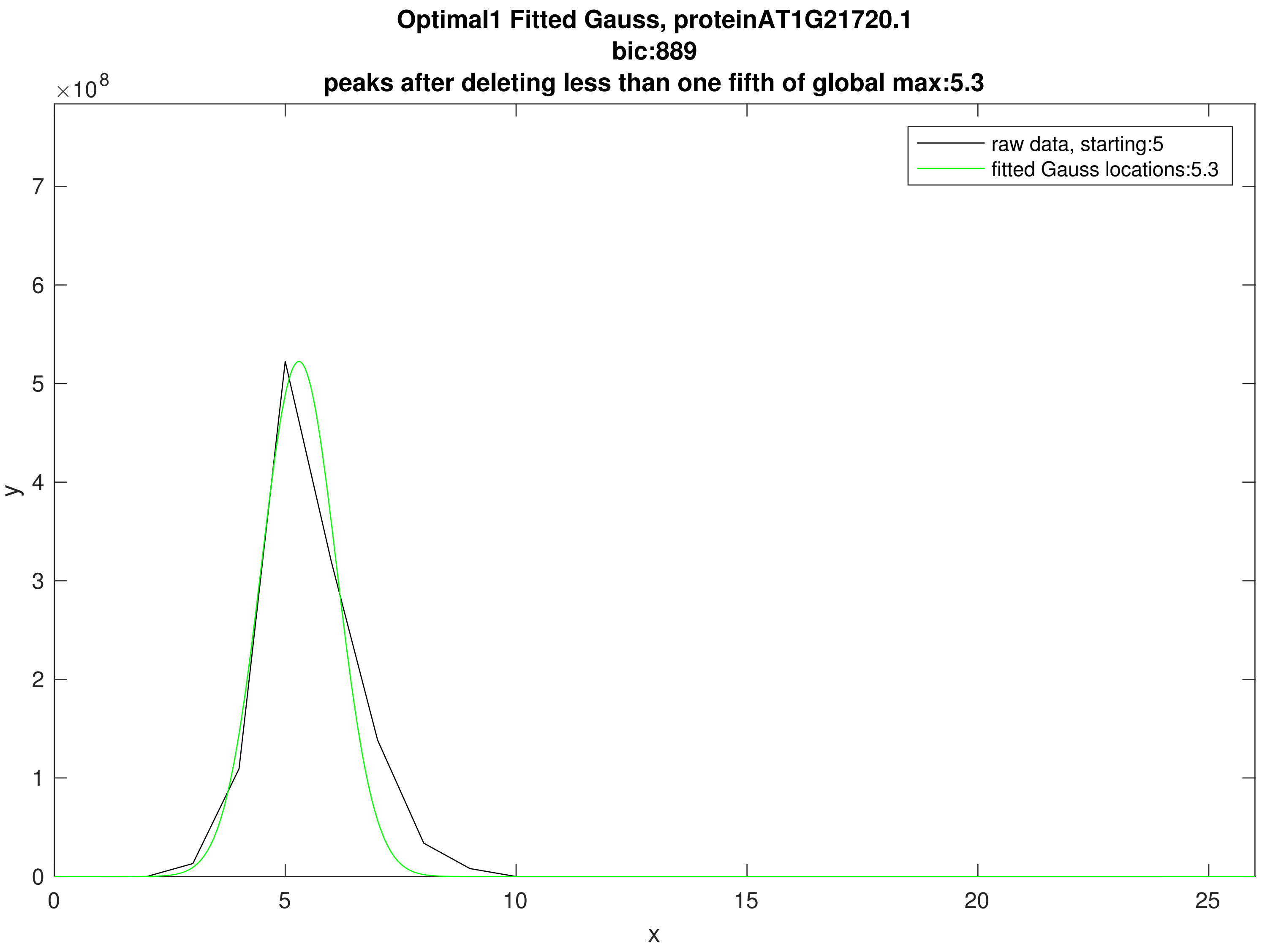
Optimal1 Fitted Gauss, proteinAT1G21720.1
bic:889
peaks after deleting less than one fifth of global max:5.3
×10^8
raw data, starting:5
fitted Gauss locations:5.3
y
x
0
1
2
3
4
5
6
7
10
15
20
25

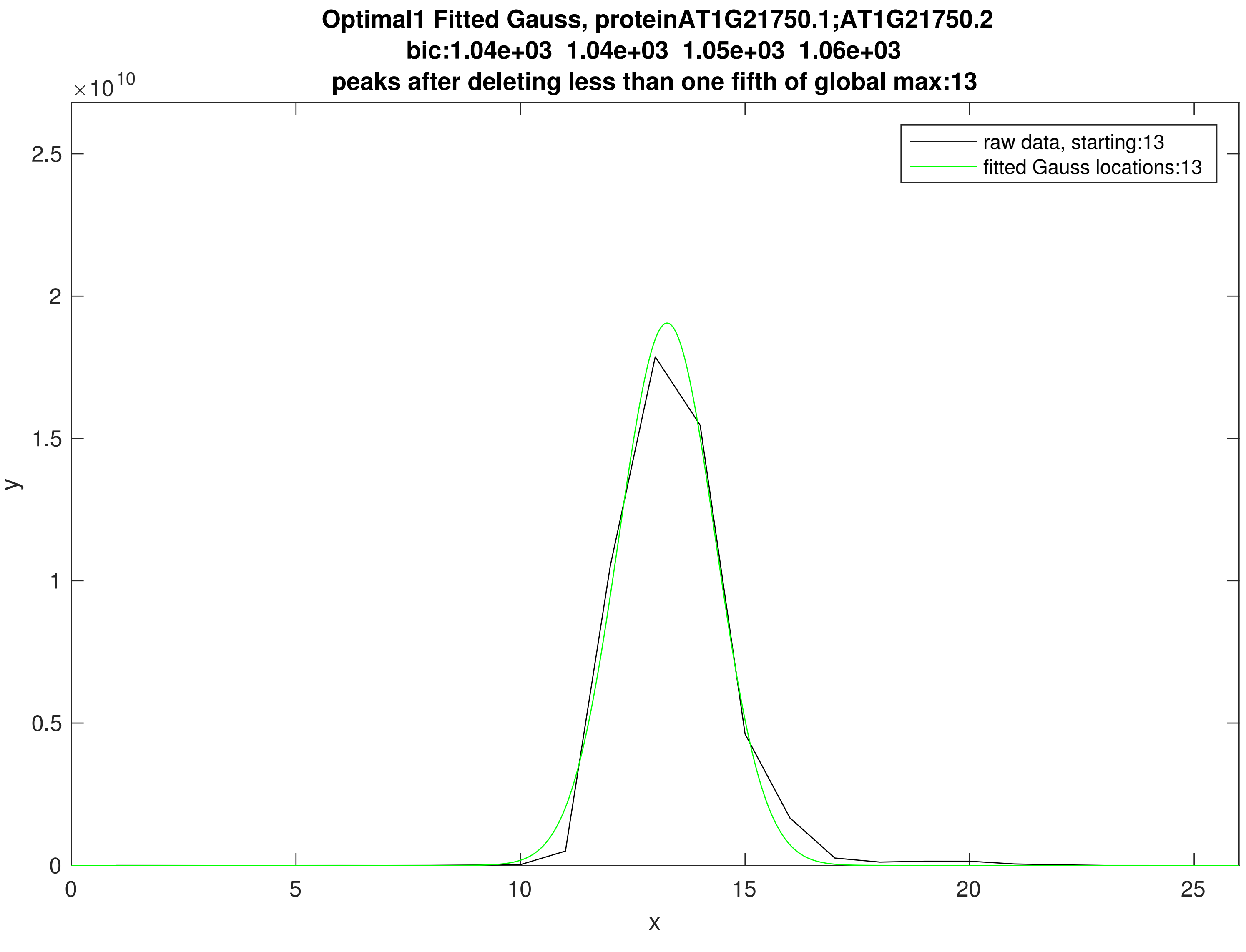
Optimal1 Fitted Gauss, proteinAT1G21750.1;AT1G21750.2
bic:1.04e+03 1.04e+03 1.05e+03 1.06e+03
peaks after deleting less than one fifth of global max:13
×10^10
raw data, starting:13
fitted Gauss locations:13
y
x

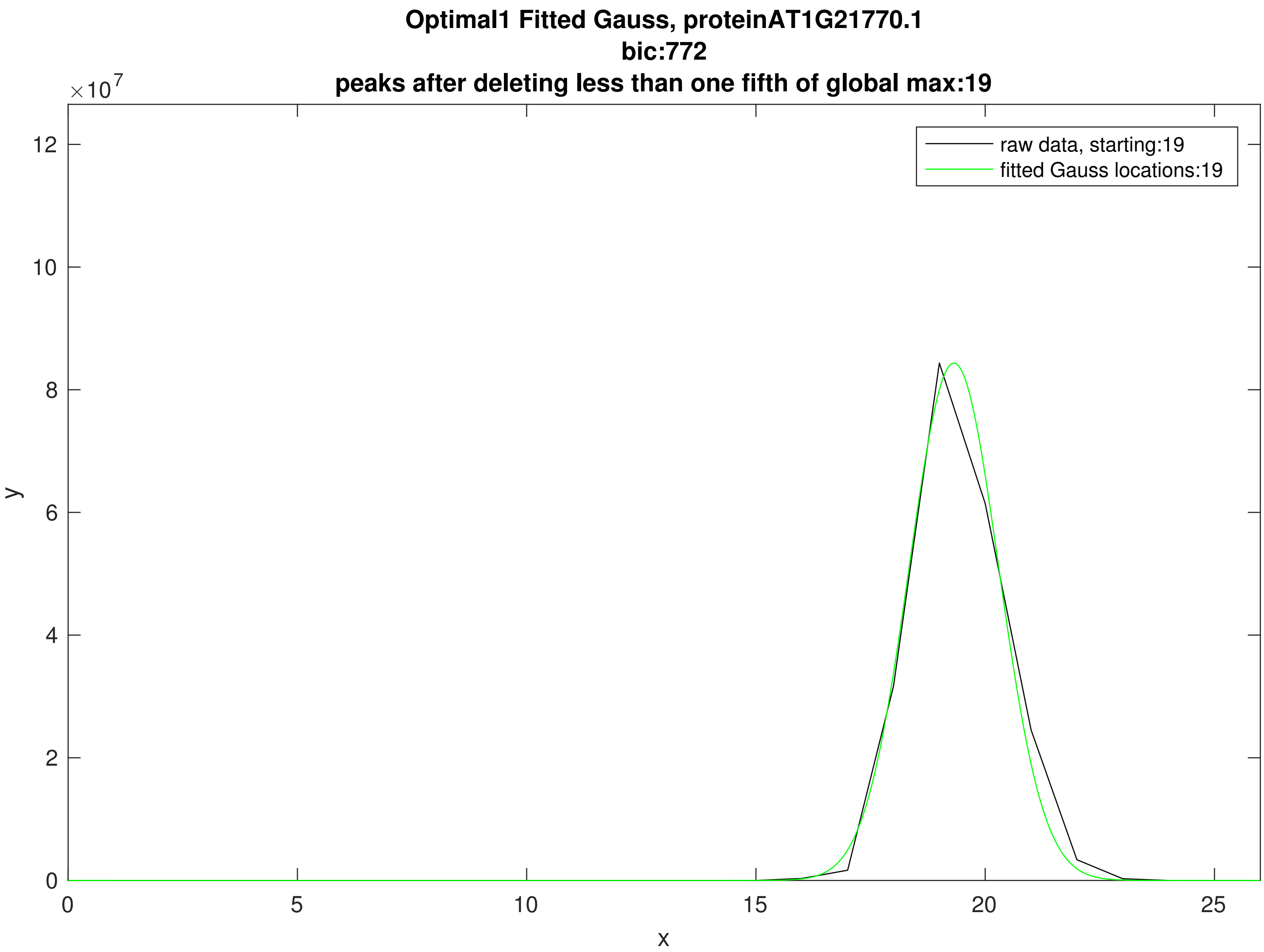
Optimal1 Fitted Gauss, proteinAT1G21770.1
bic:772
peaks after deleting less than one fifth of global max:19
×10^7
12
10
8
6
4
2
0
y
0
5
10
15
20
25
x
raw data, starting:19
fitted Gauss locations:19

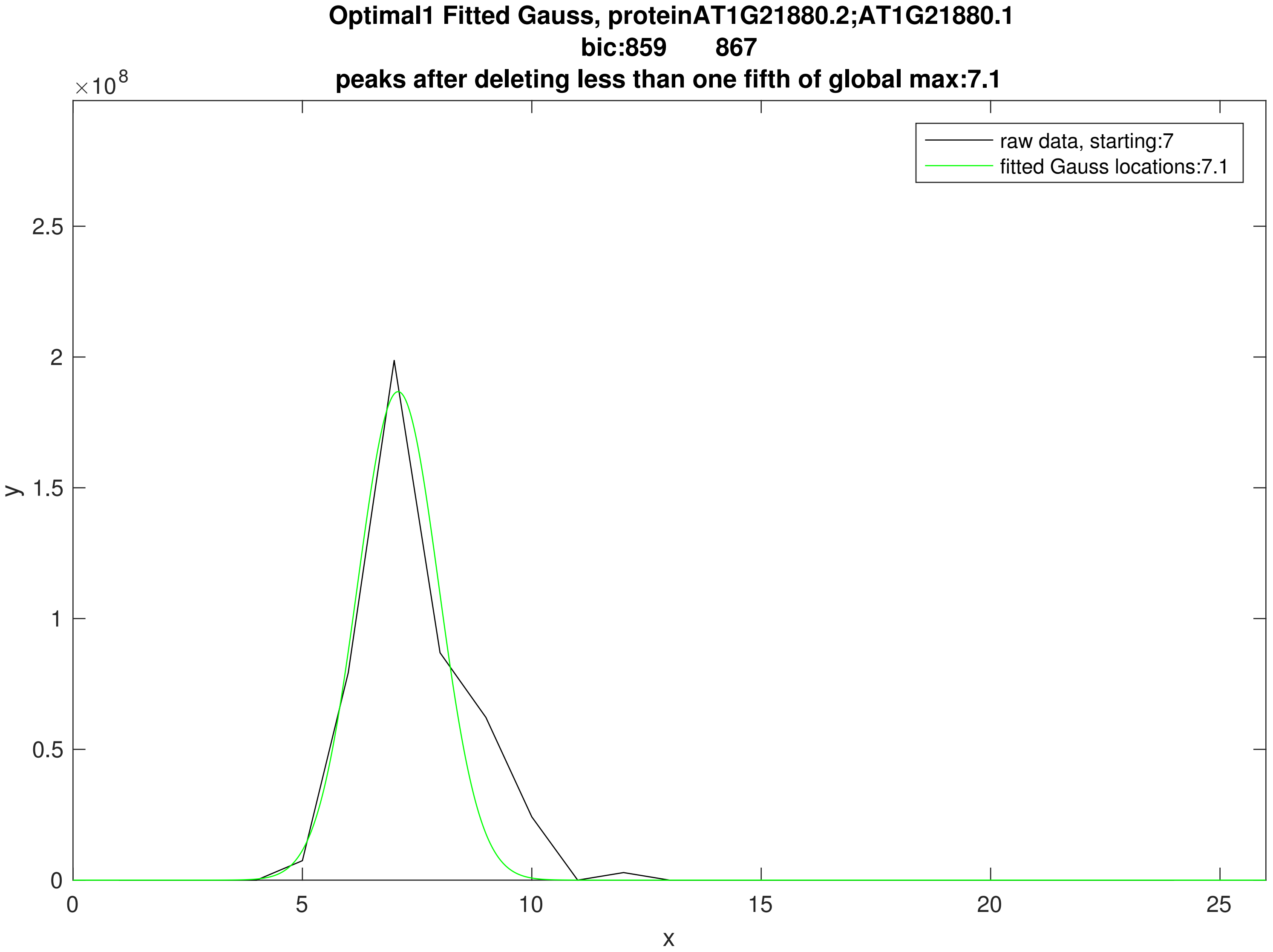
Optimal1 Fitted Gauss, proteinAT1G21880.2;AT1G21880.1
bic:859 867
peaks after deleting less than one fifth of global max:7.1
×10^8
raw data, starting:7
fitted Gauss locations:7.1
y
x
0
0.5
1
1.5
2
2.5
0
5
10
15
20
25

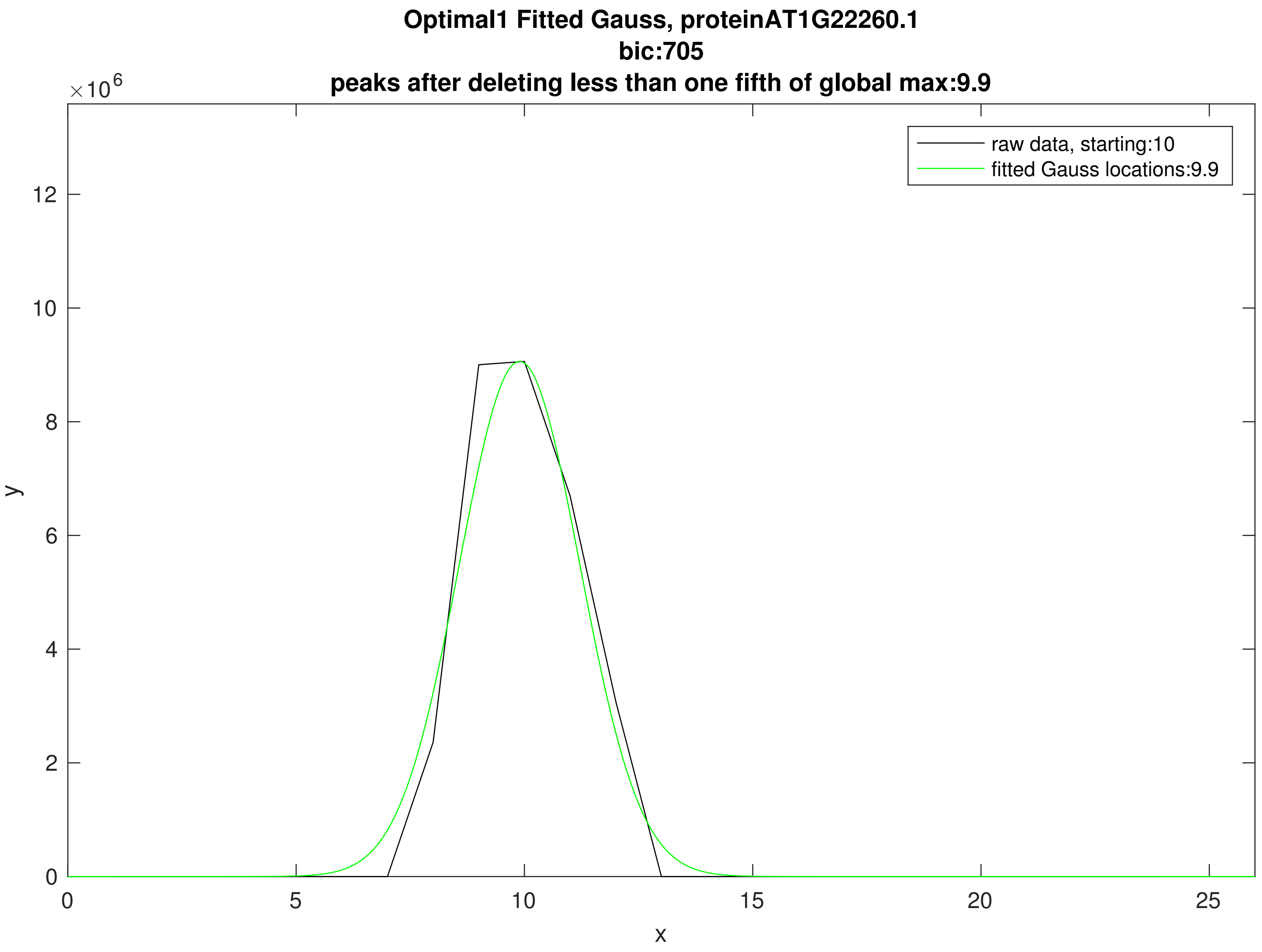

Optimal1 Fitted Gauss, proteinAT1G22260.1
bic:705
peaks after deleting less than one fifth of global max:9.9
×10^6
12
10
8
6
4
2
0
y
0
5
10
15
20
25
x
raw data, starting:10
fitted Gauss locations:9.9

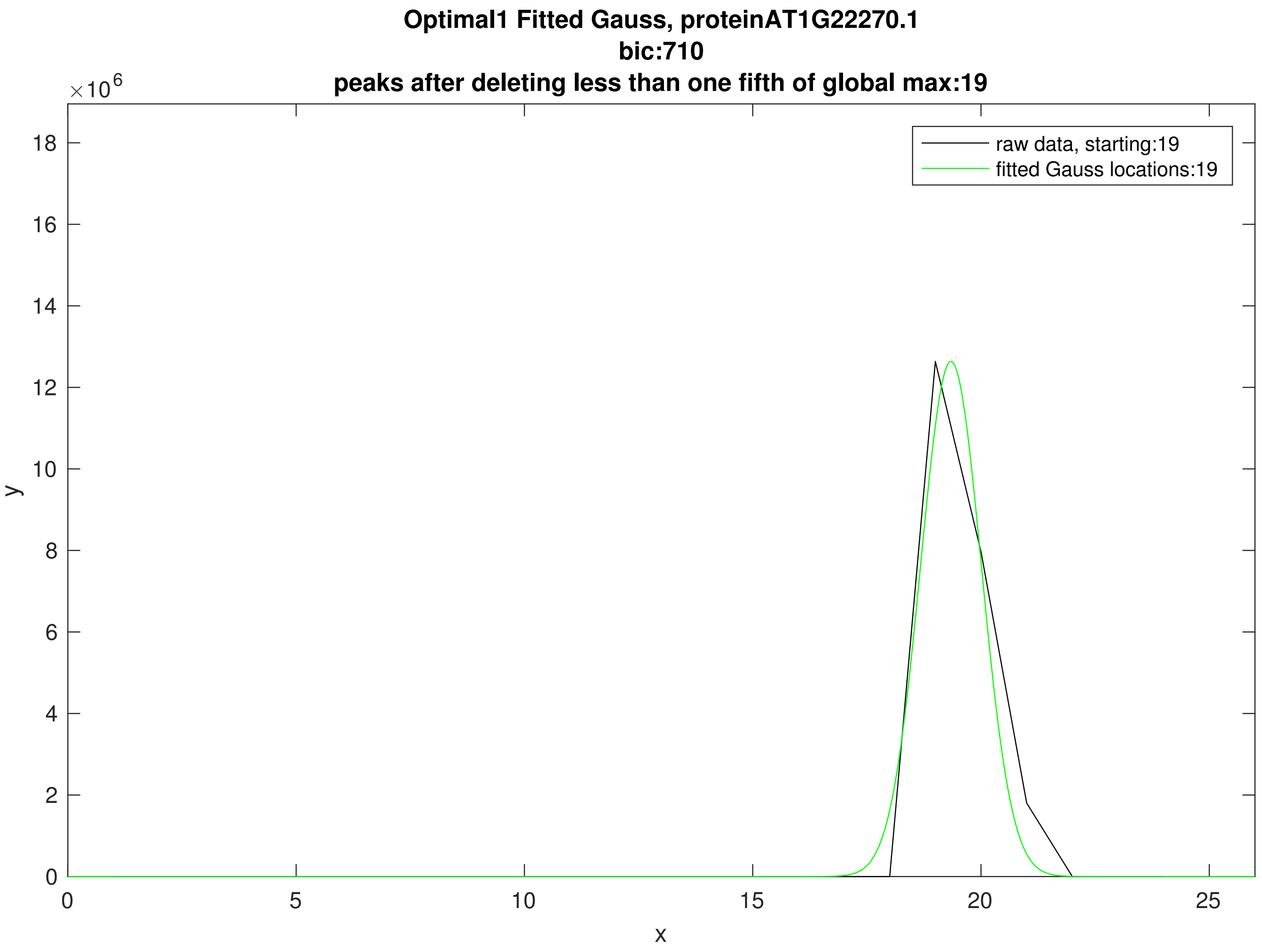
Optimal1 Fitted Gauss, proteinAT1G22270.1
bic:710
peaks after deleting less than one fifth of global max:19
×10^6
raw data, starting:19
fitted Gauss locations:19
y
x

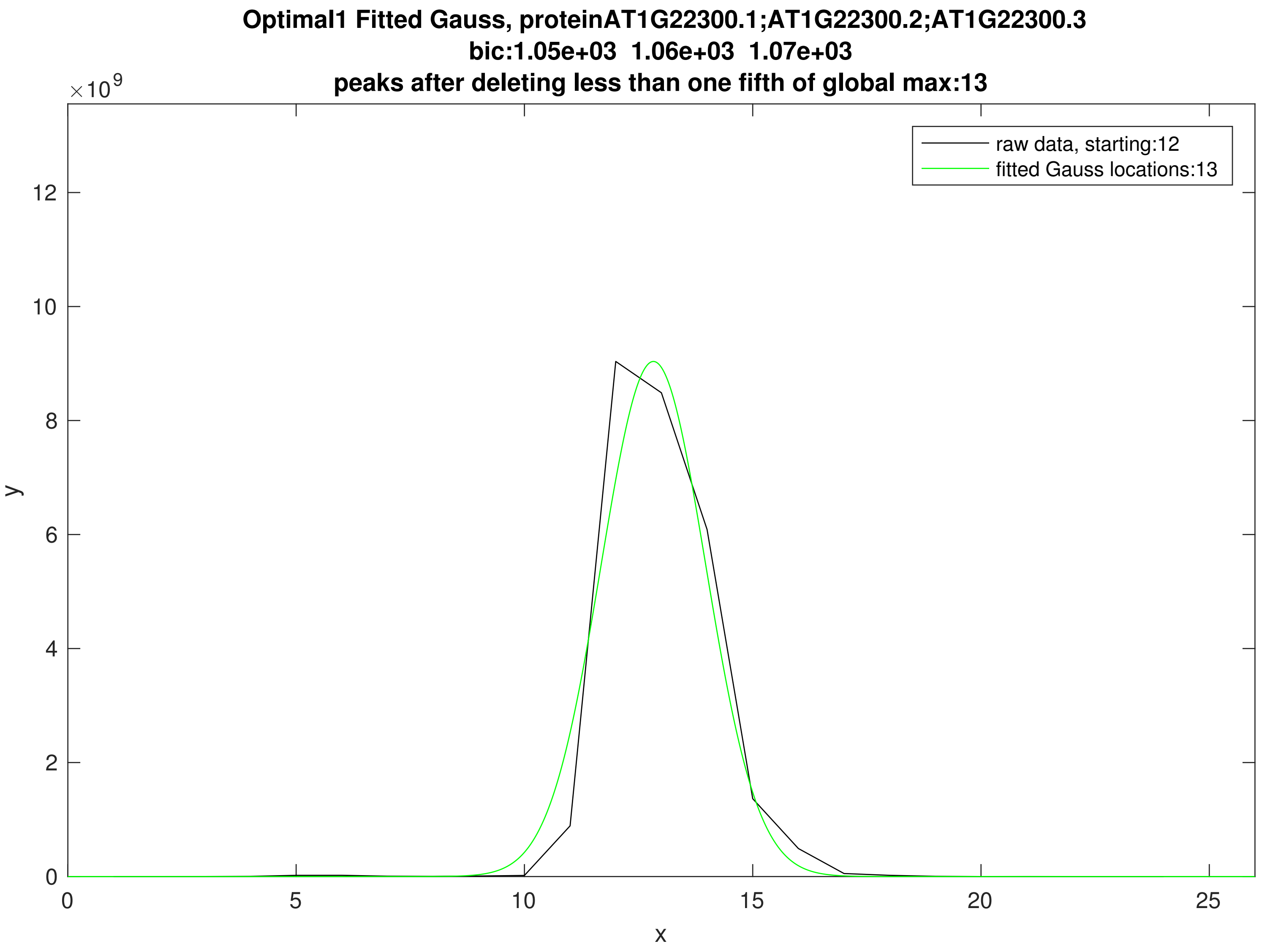
Optimal1 Fitted Gauss, proteinAT1G22300.1;AT1G22300.2;AT1G22300.3
bic:1.05e+03  1.06e+03  1.07e+03
peaks after deleting less than one fifth of global max:13
×10^9
raw data, starting:12
fitted Gauss locations:13
y
x
0
2
4
6
8
10
12
0
5
10
15
20
25

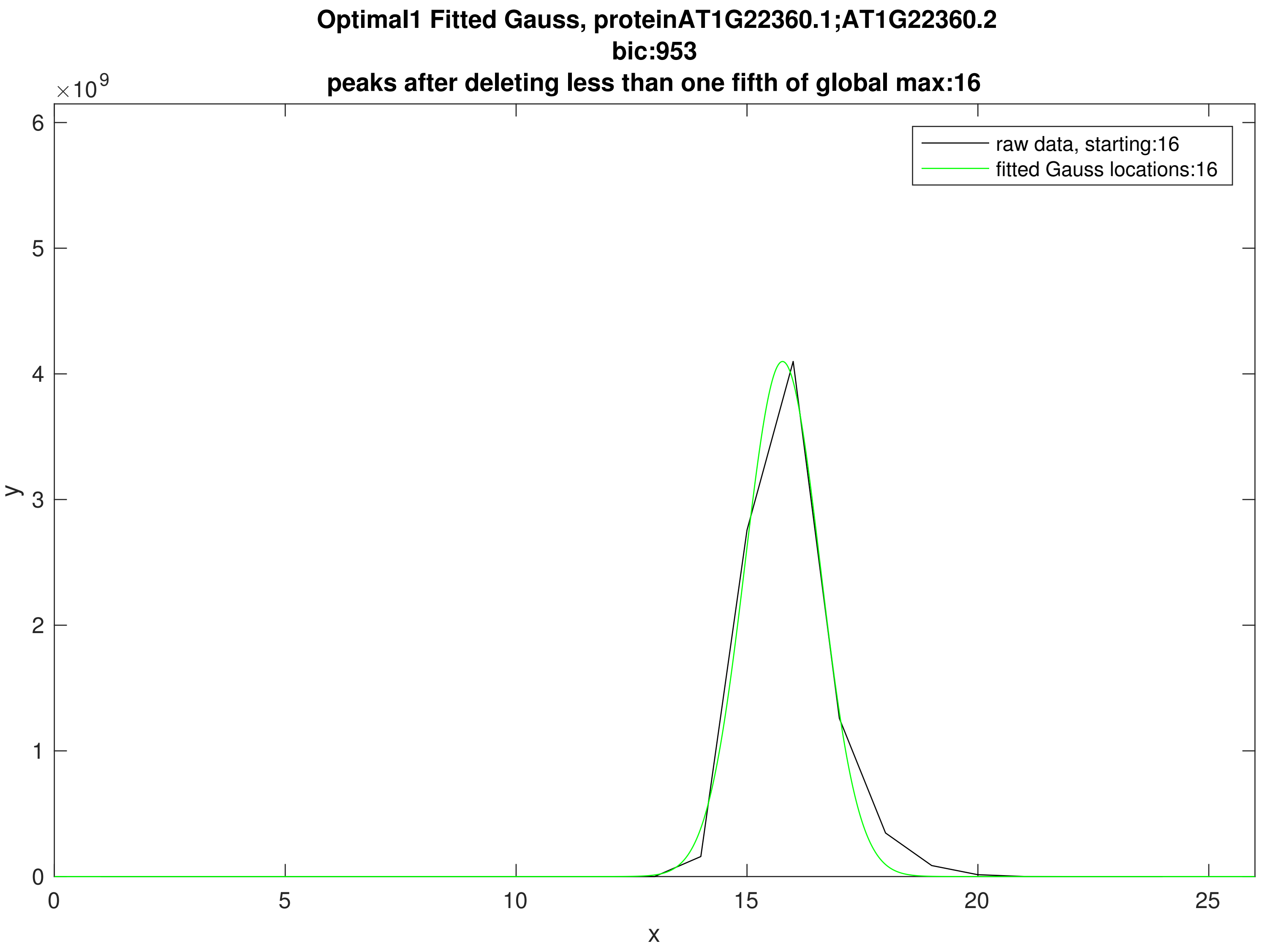
Optimal1 Fitted Gauss, proteinAT1G22360.1;AT1G22360.2
bic:953
peaks after deleting less than one fifth of global max:16
×10^9
raw data, starting:16
fitted Gauss locations:16
y
x
0
1
2
3
4
5
6
0
5
10
15
20
25

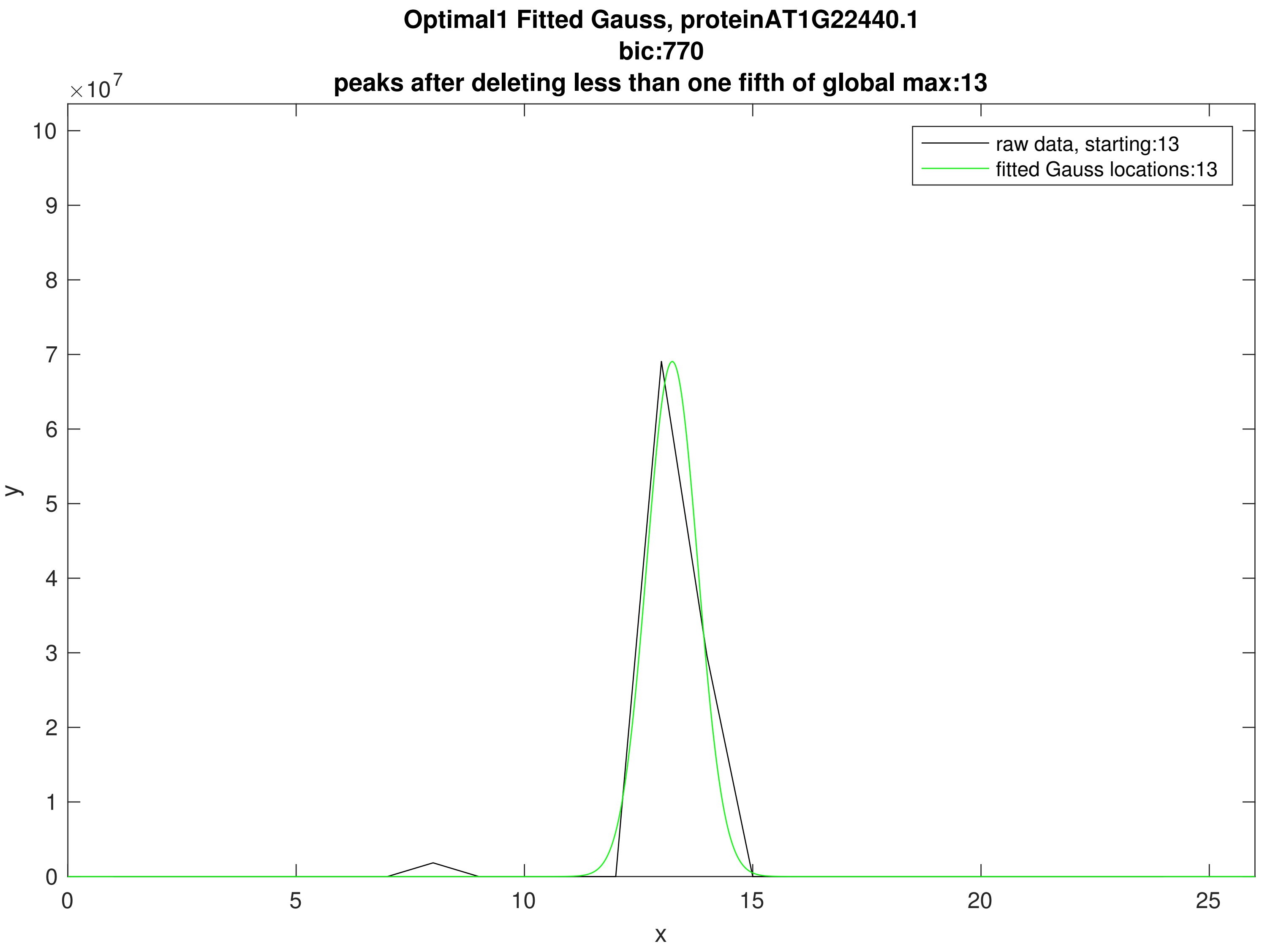
Optimal1 Fitted Gauss, proteinAT1G22440.1
bic:770
peaks after deleting less than one fifth of global max:13
×10^7
raw data, starting:13
fitted Gauss locations:13
y
x
0
1
2
3
4
5
6
7
8
9
10
0
5
10
15
20
25

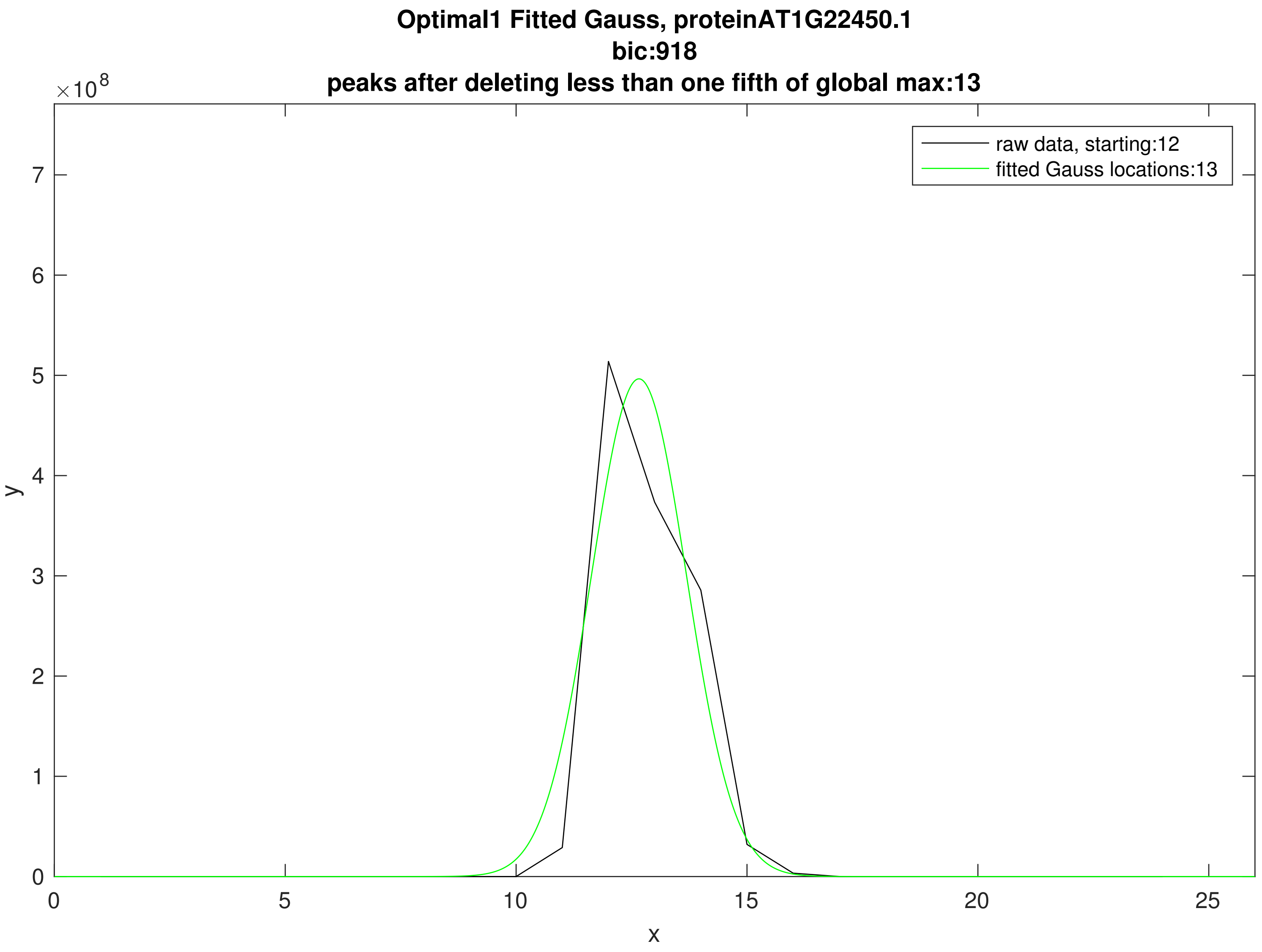

Optimal1 Fitted Gauss, proteinAT1G22450.1
bic:918
peaks after deleting less than one fifth of global max:13
×10^8
raw data, starting:12
fitted Gauss locations:13
y
x
0
1
2
3
4
5
6
7
0
5
10
15
20
25

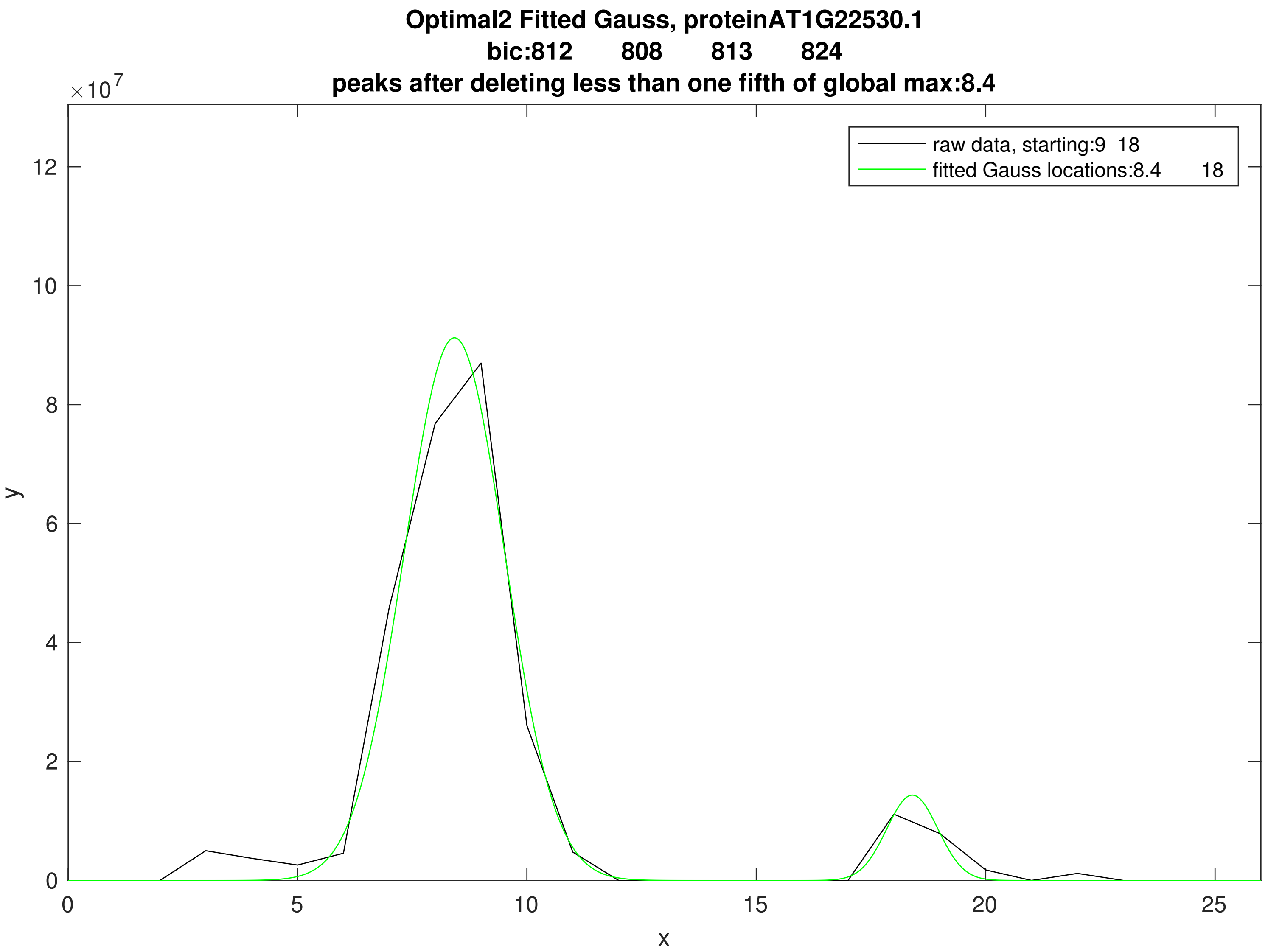

Optimal2 Fitted Gauss, proteinAT1G22530.1
bic:812 808 813 824
peaks after deleting less than one fifth of global max:8.4
×10^7
raw data, starting:9 18
fitted Gauss locations:8.4 18
y
x
0
2
4
6
8
10
12
5
15
20
25

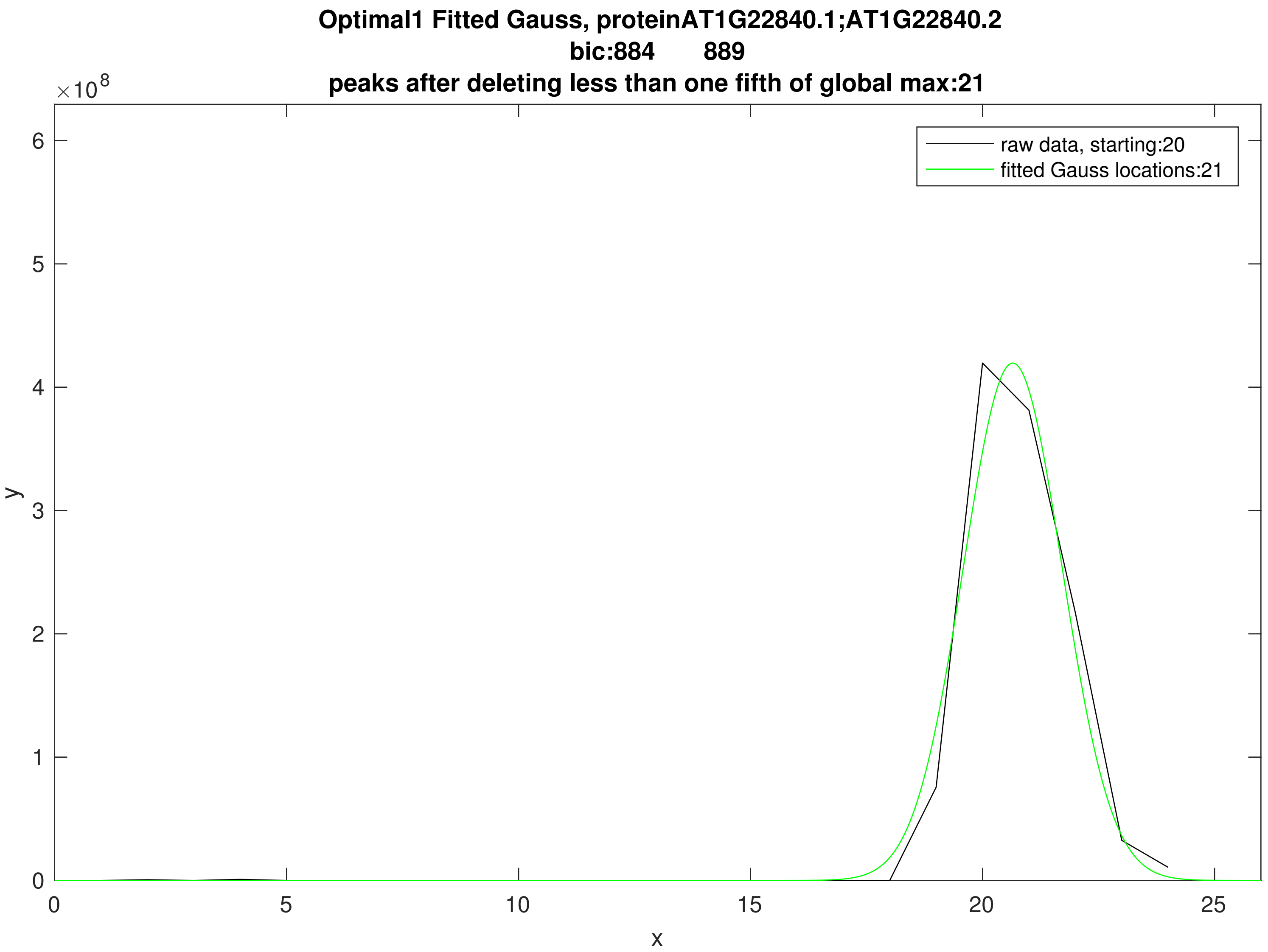

Optimal1 Fitted Gauss, proteinAT1G22840.1;AT1G22840.2
bic:884 889
peaks after deleting less than one fifth of global max:21
×10^8
raw data, starting:20
fitted Gauss locations:21
y
x
0
1
2
3
4
5
6
0
5
10
15
20
25

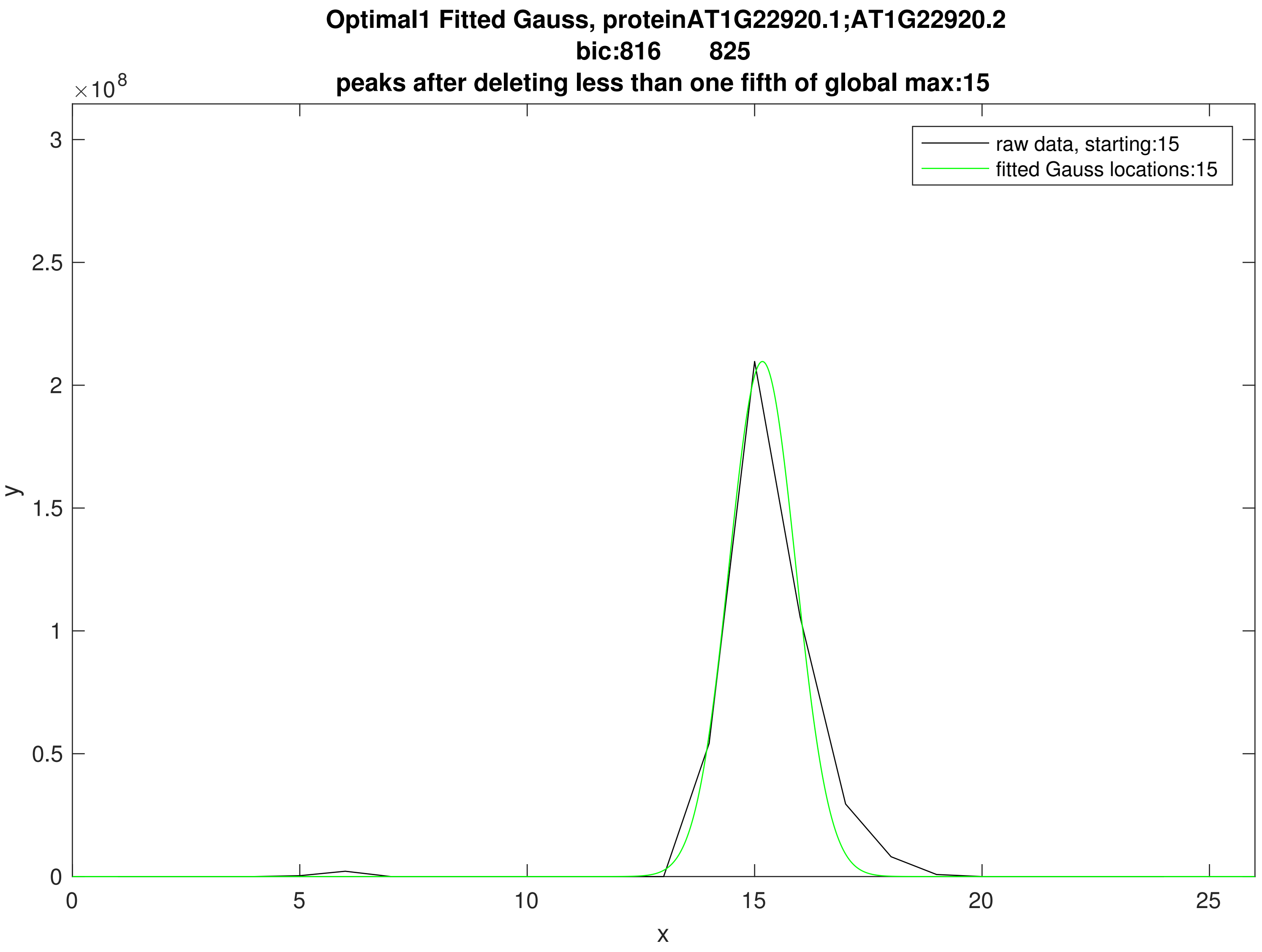
Optimal1 Fitted Gauss, proteinAT1G22920.1;AT1G22920.2
bic:816 825
peaks after deleting less than one fifth of global max:15
×10^8
3
2.5
2
1.5
1
0.5
0
y
0
5
10
15
20
25
x
raw data, starting:15
fitted Gauss locations:15

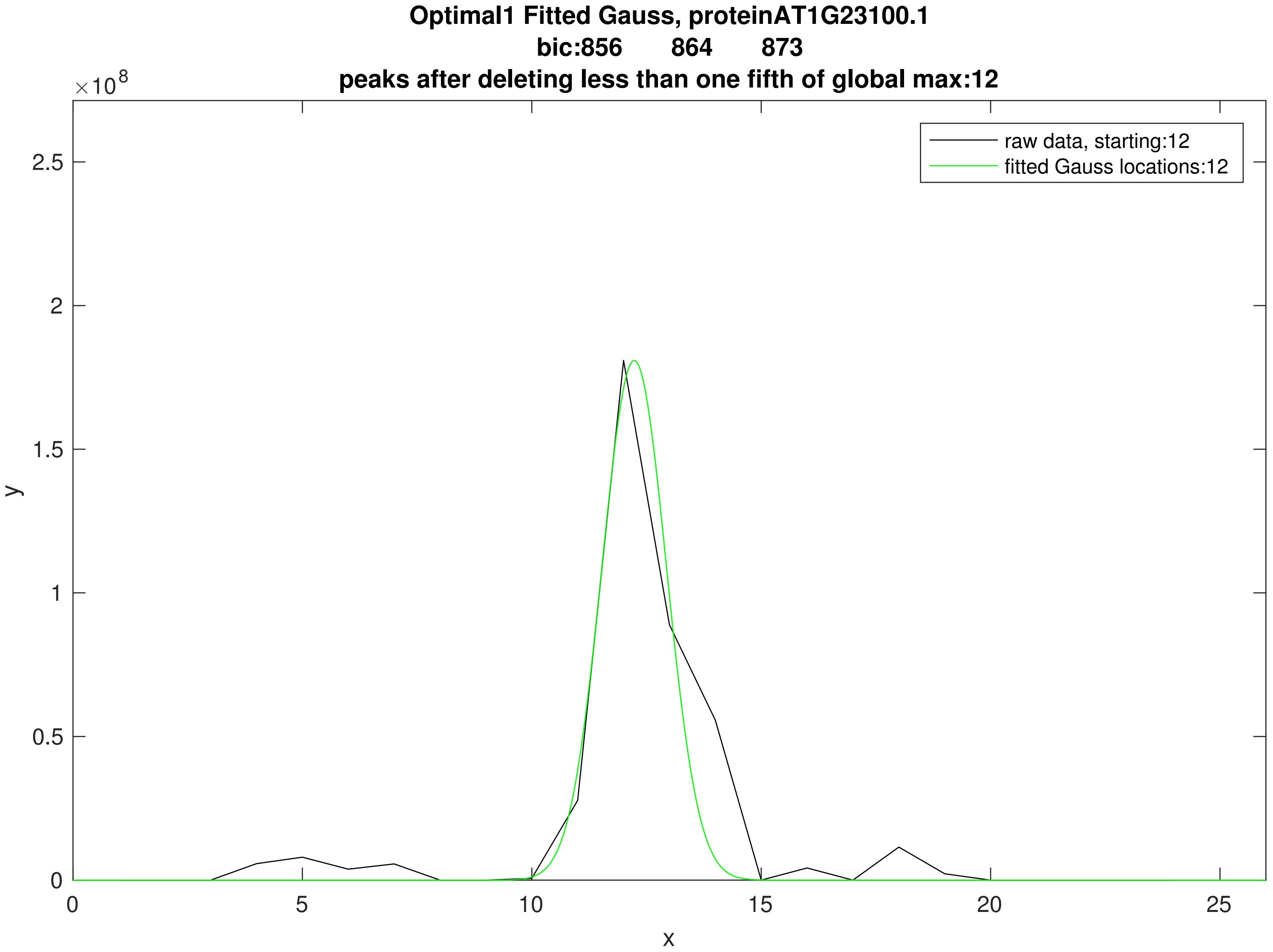

Optimal1 Fitted Gauss, proteinAT1G23100.1
bic:856 864 873
peaks after deleting less than one fifth of global max:12
×10^8
2.5
2
1.5
1
0.5
0
y
0
5
10
15
20
25
x
raw data, starting:12
fitted Gauss locations:12

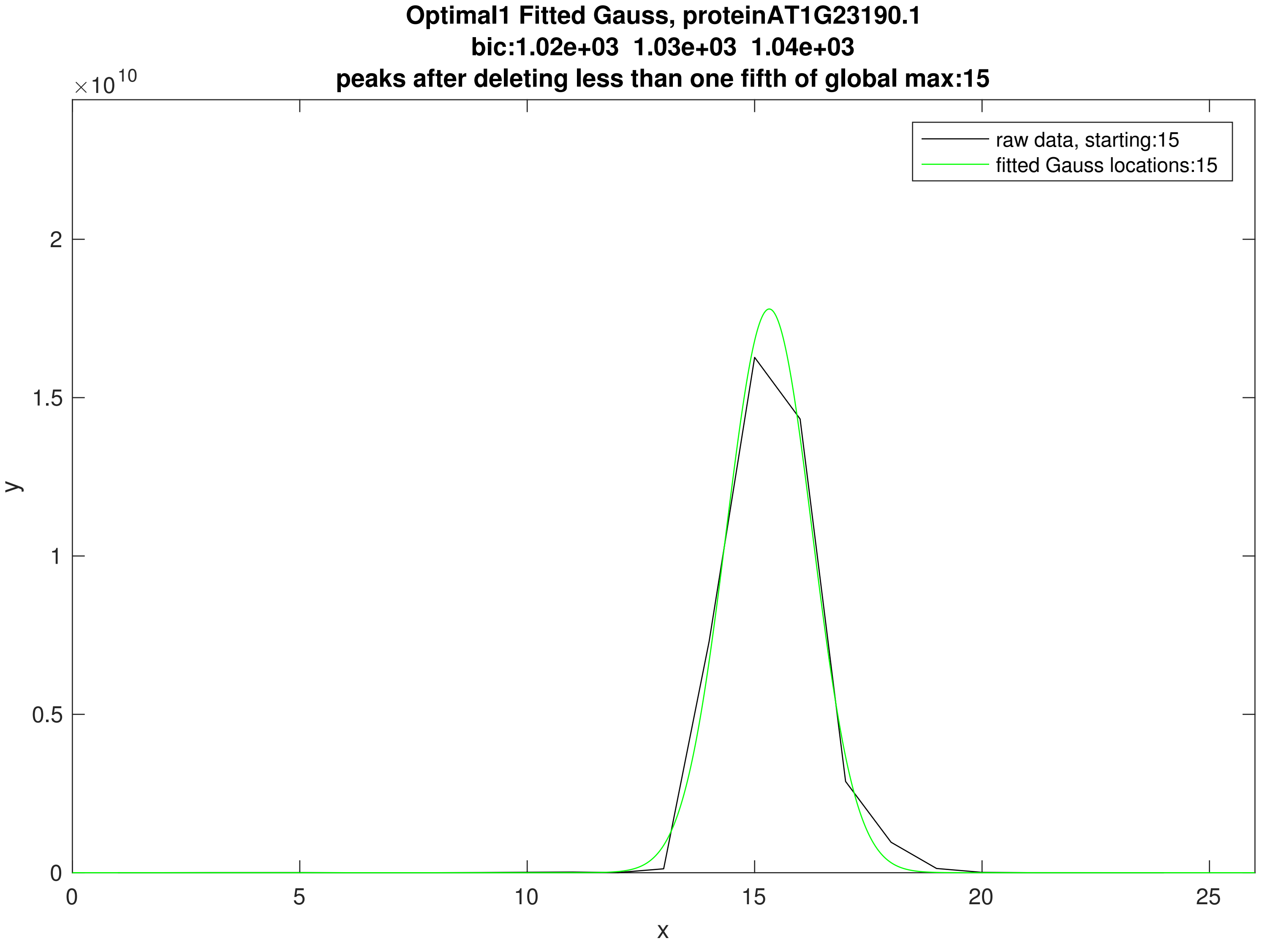

Optimal1 Fitted Gauss, proteinAT1G23190.1
bic:1.02e+03 1.03e+03 1.04e+03
peaks after deleting less than one fifth of global max:15
×10^10
raw data, starting:15
fitted Gauss locations:15
2
1.5
1
0.5
0
y
0
5
10
15
20
25
x

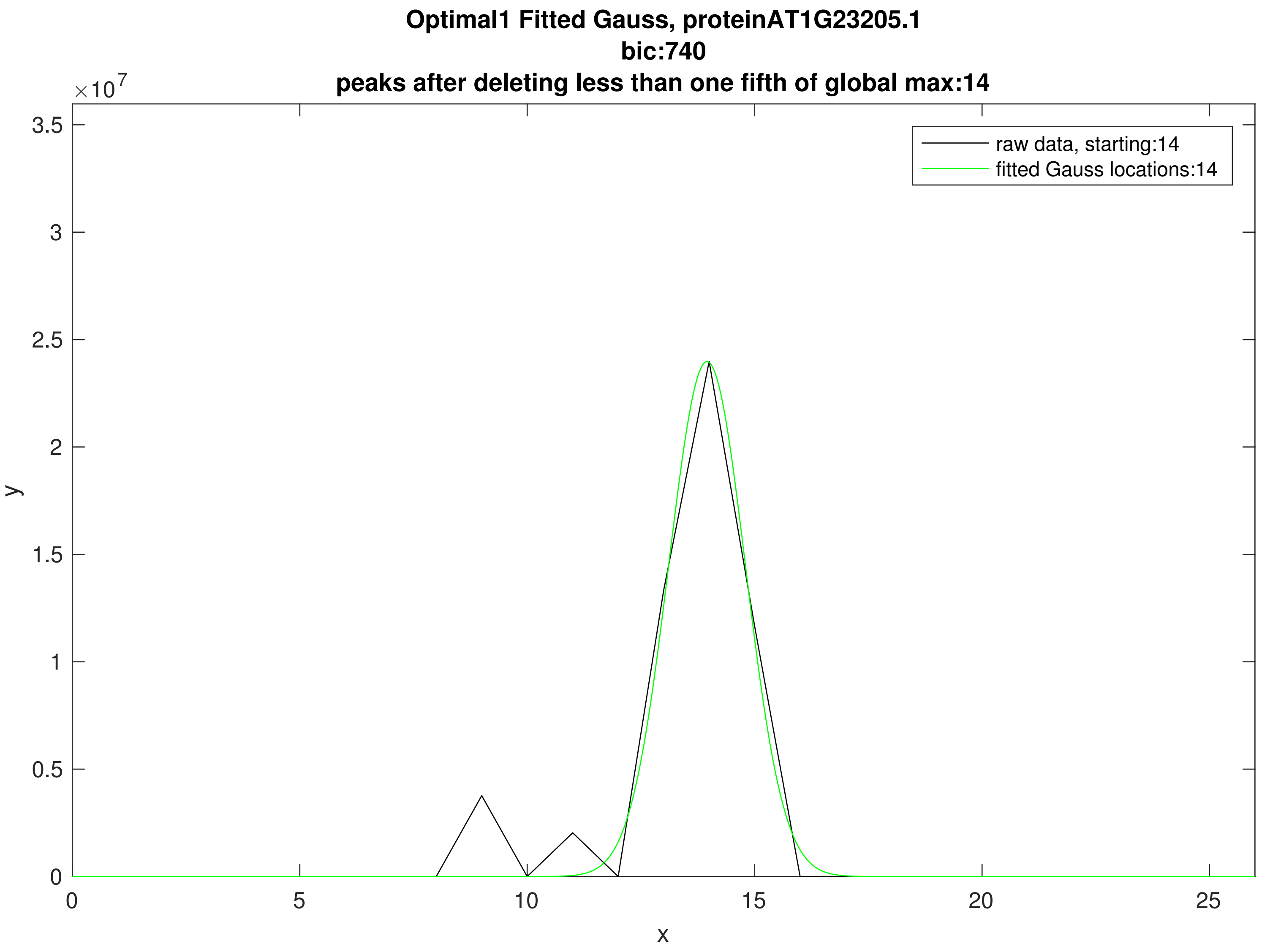
Optimal1 Fitted Gauss, proteinAT1G23205.1
bic:740
peaks after deleting less than one fifth of global max:14
×10^7
3.5
3
2.5
2
1.5
1
0.5
0
y
0
5
10
15
20
25
x
raw data, starting:14
fitted Gauss locations:14

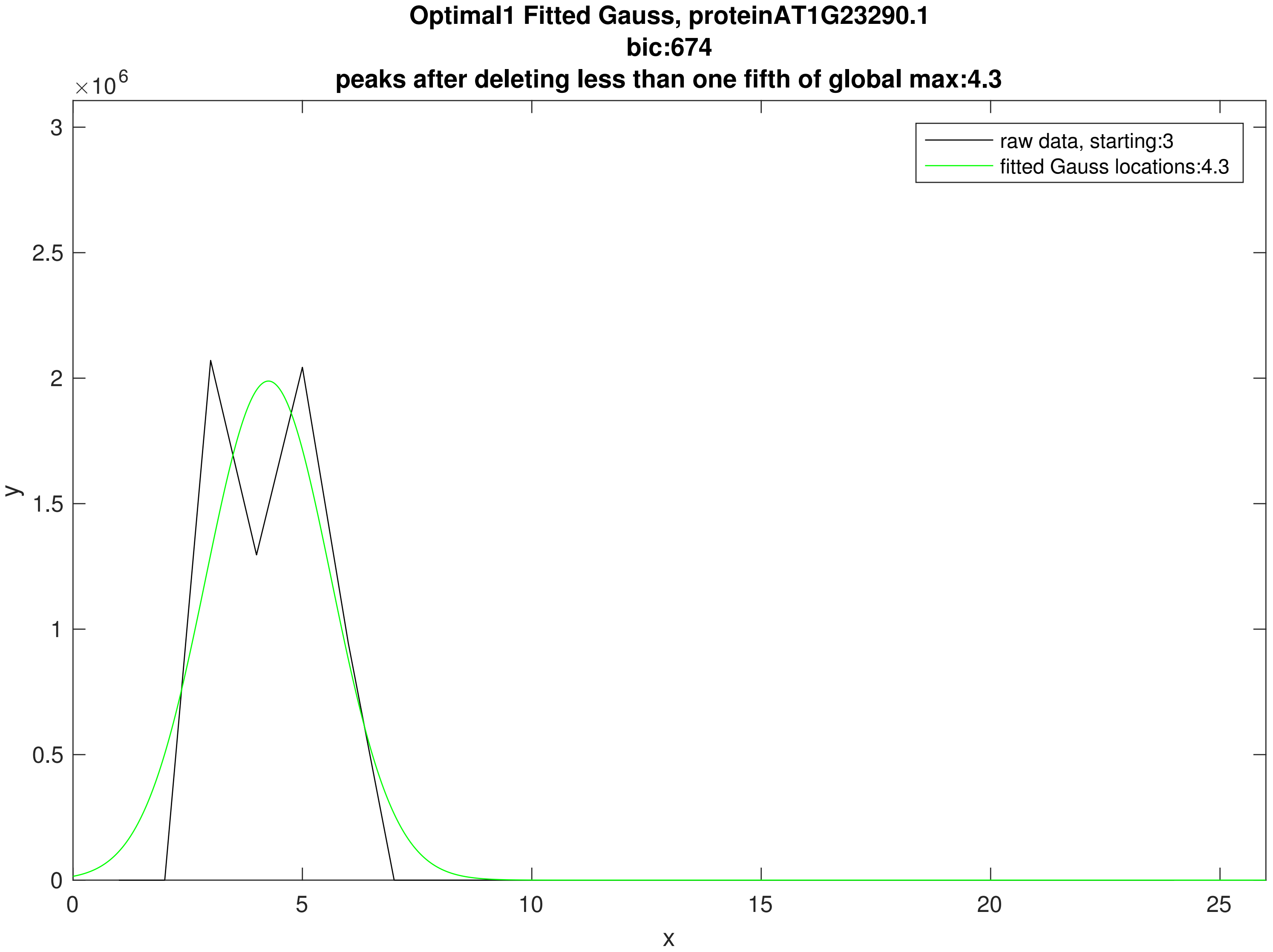
Optimal1 Fitted Gauss, proteinAT1G23290.1
bic:674
peaks after deleting less than one fifth of global max:4.3
×10^6
3
2.5
2
1.5
1
0.5
0
y
0
5
10
15
20
25
x
raw data, starting:3
fitted Gauss locations:4.3

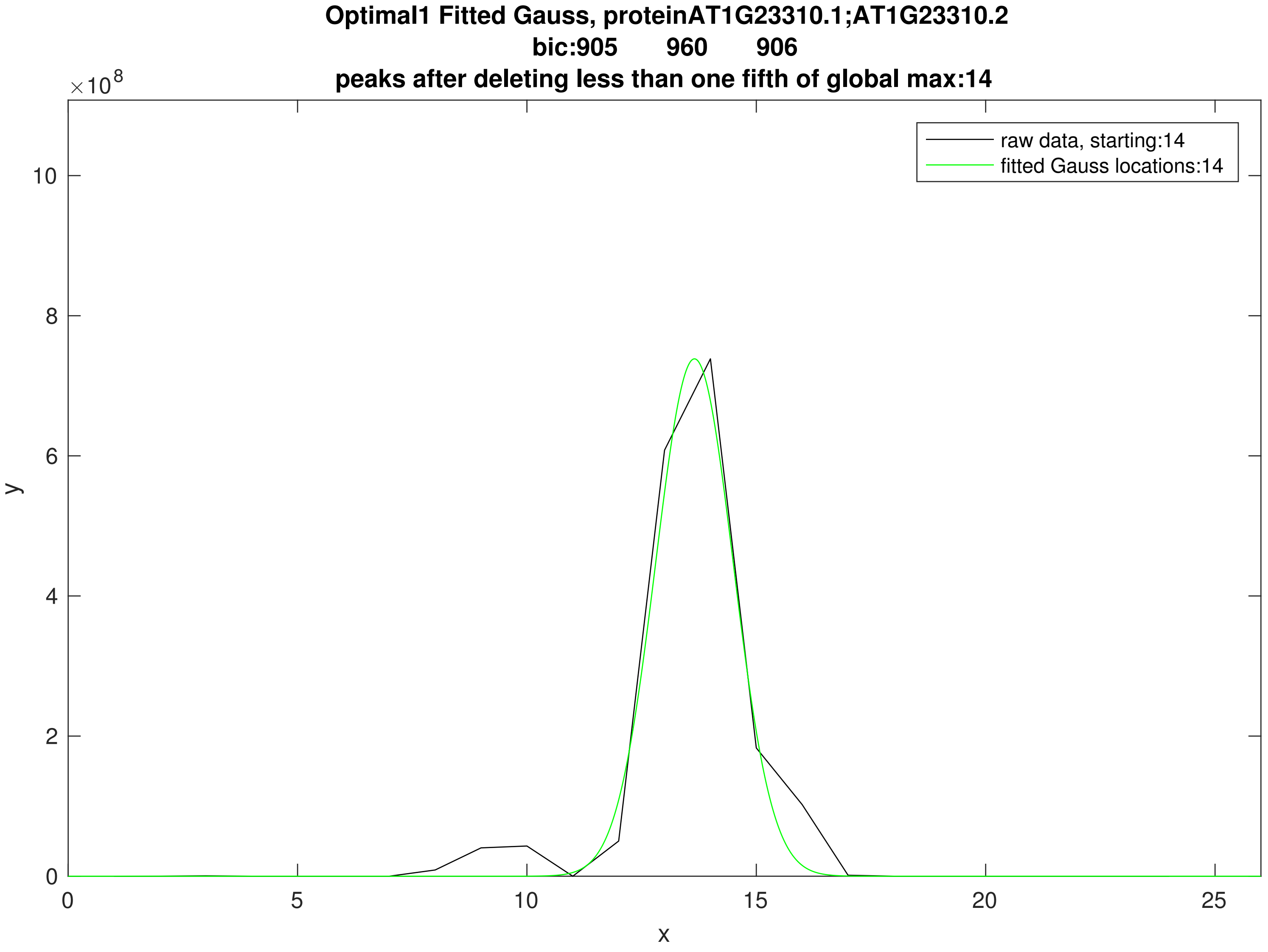
Optimal1 Fitted Gauss, proteinAT1G23310.1;AT1G23310.2
bic:905 960 906
peaks after deleting less than one fifth of global max:14
×10^8
raw data, starting:14
fitted Gauss locations:14
0
2
4
6
8
10
y
0
5
10
15
20
25
x

#### Optimal1 Fitted Gauss, proteinAT1G23440.1
#### bic:884 887
#### peaks after deleting less than one fifth of global max:19

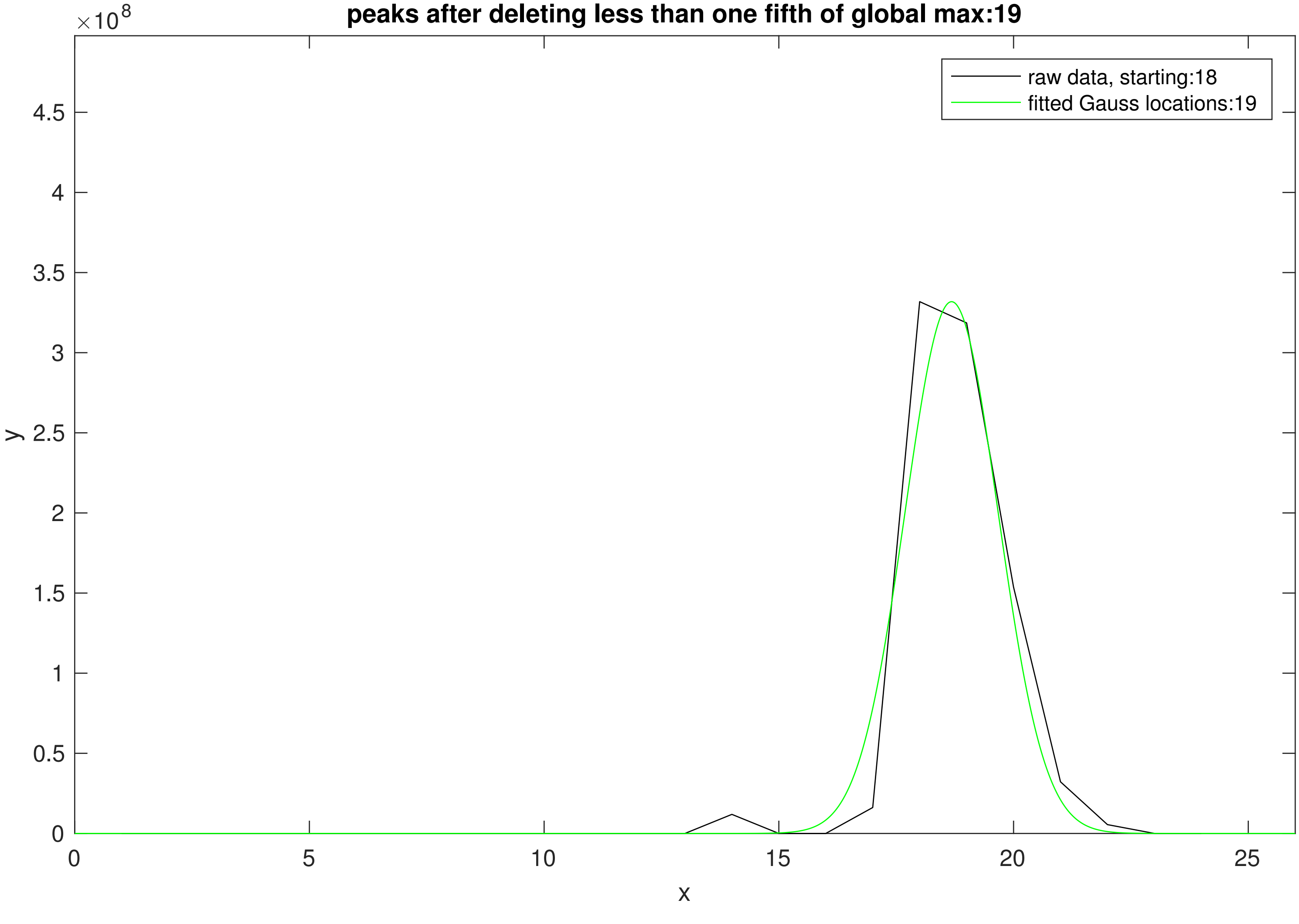

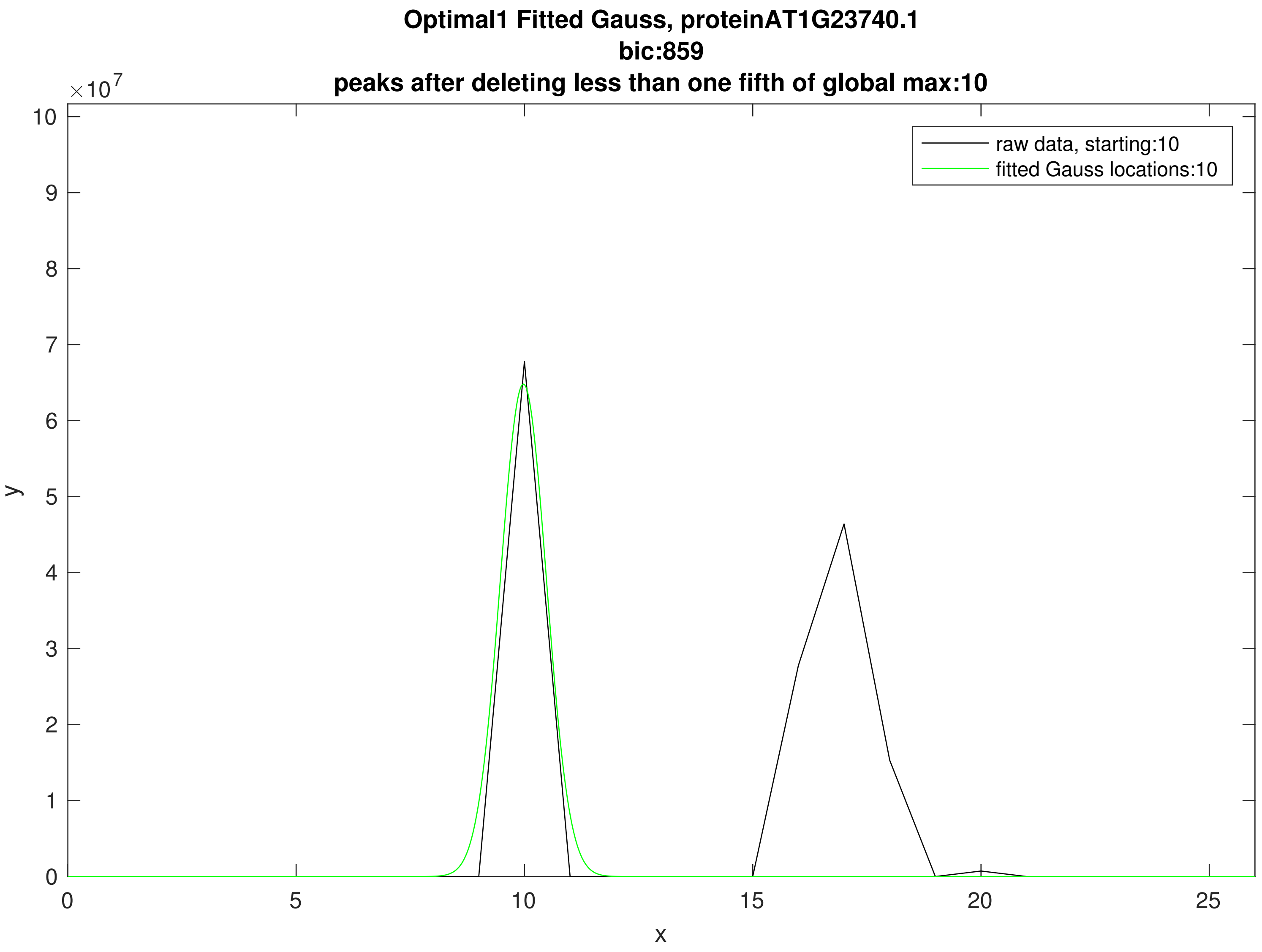

Optimal1 Fitted Gauss, proteinAT1G23740.1
bic:859
peaks after deleting less than one fifth of global max:10
×10^7
10
9
8
7
6
5
4
3
2
1
0
y
0
5
10
15
20
25
x
raw data, starting:10
fitted Gauss locations:10

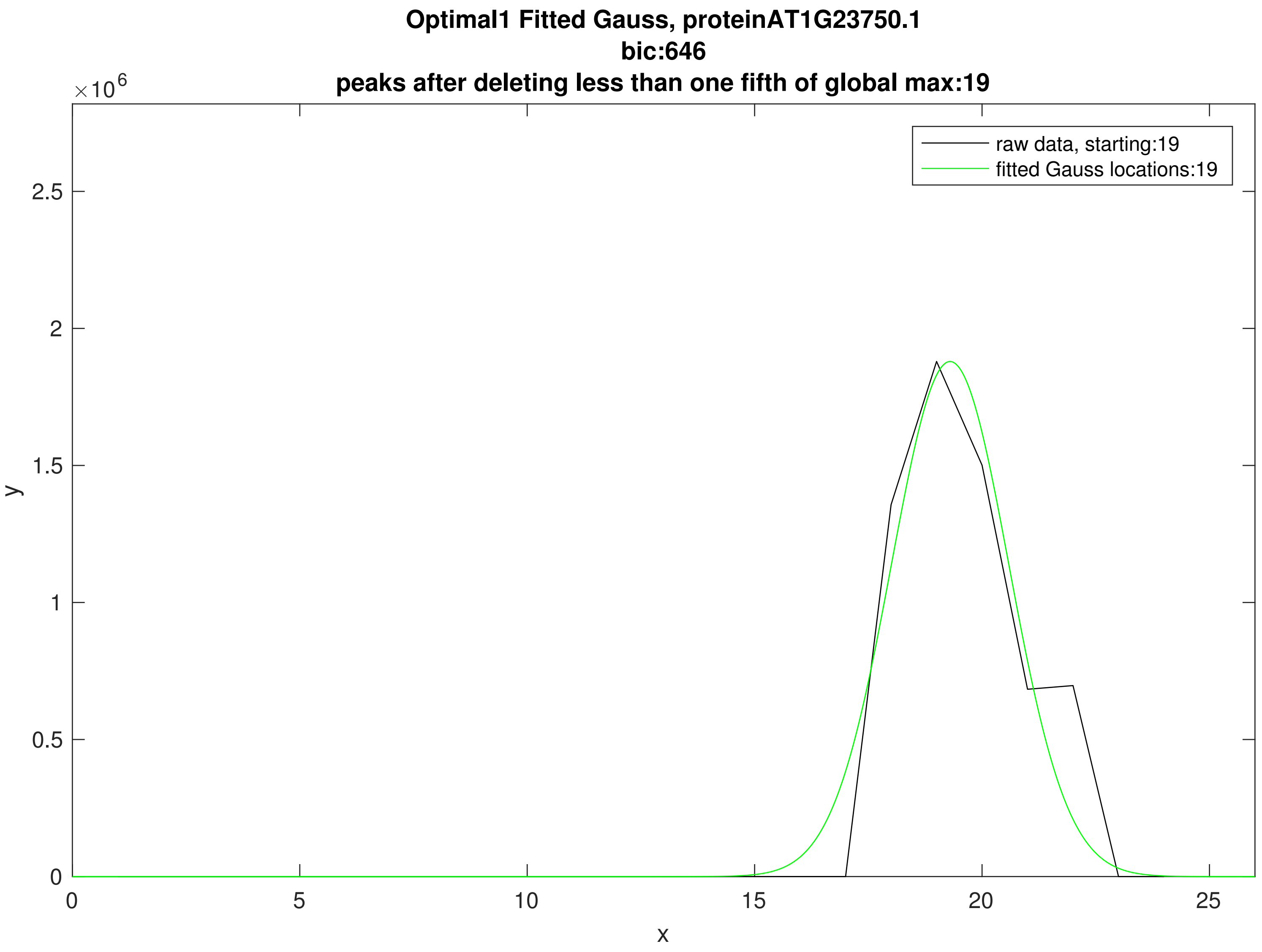
Optimal1 Fitted Gauss, proteinAT1G23750.1
bic:646
peaks after deleting less than one fifth of global max:19
×10^6
raw data, starting:19
fitted Gauss locations:19
2.5
2
1.5
1
0.5
0
y
0
5
10
15
20
25
x

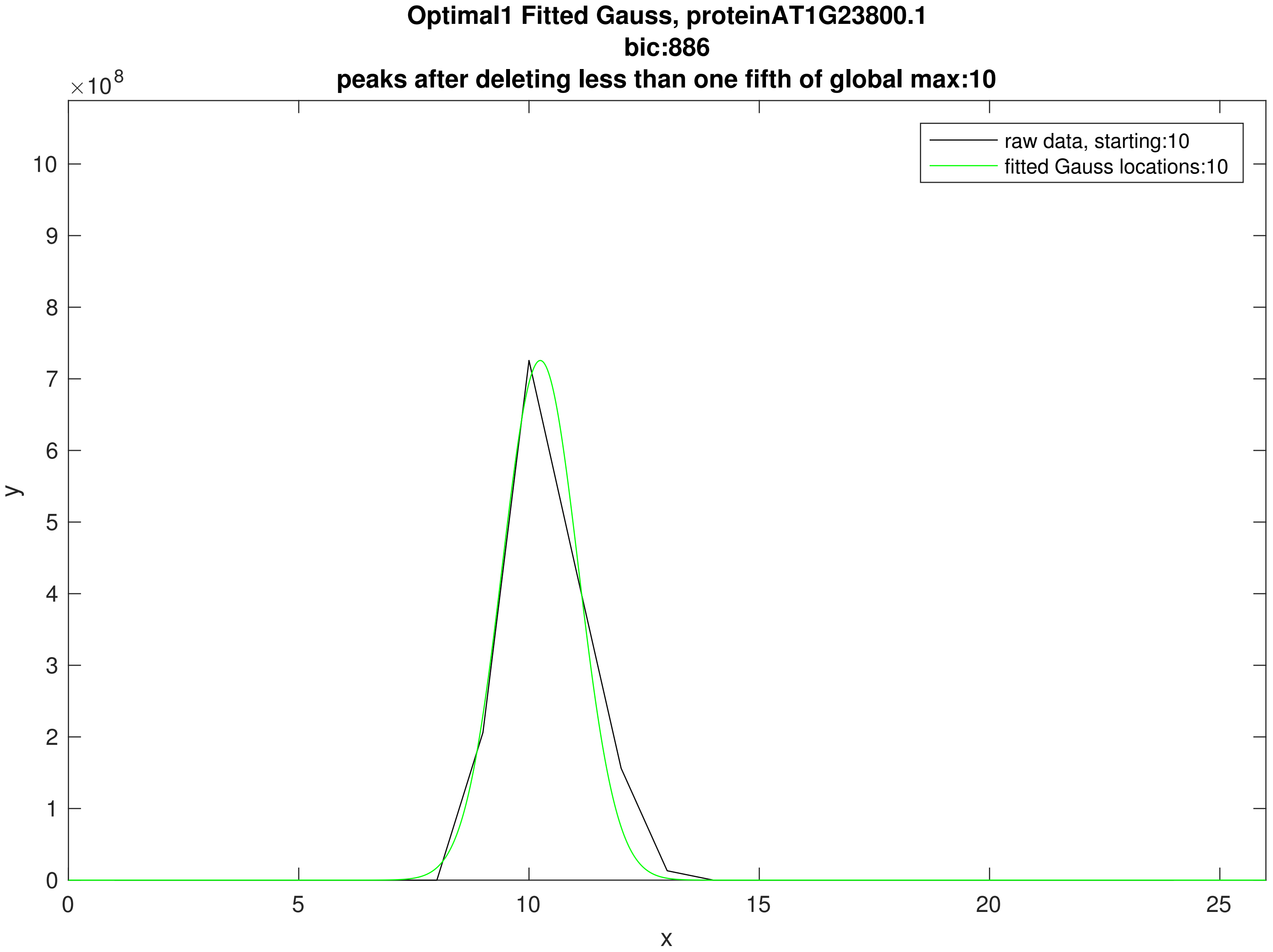

Optimal1 Fitted Gauss, proteinAT1G23800.1
bic:886
peaks after deleting less than one fifth of global max:10
$\times 10^8$
raw data, starting:10
fitted Gauss locations:10
y
x
0
1
2
3
4
5
6
7
8
9
10
0
5
10
15
20
25

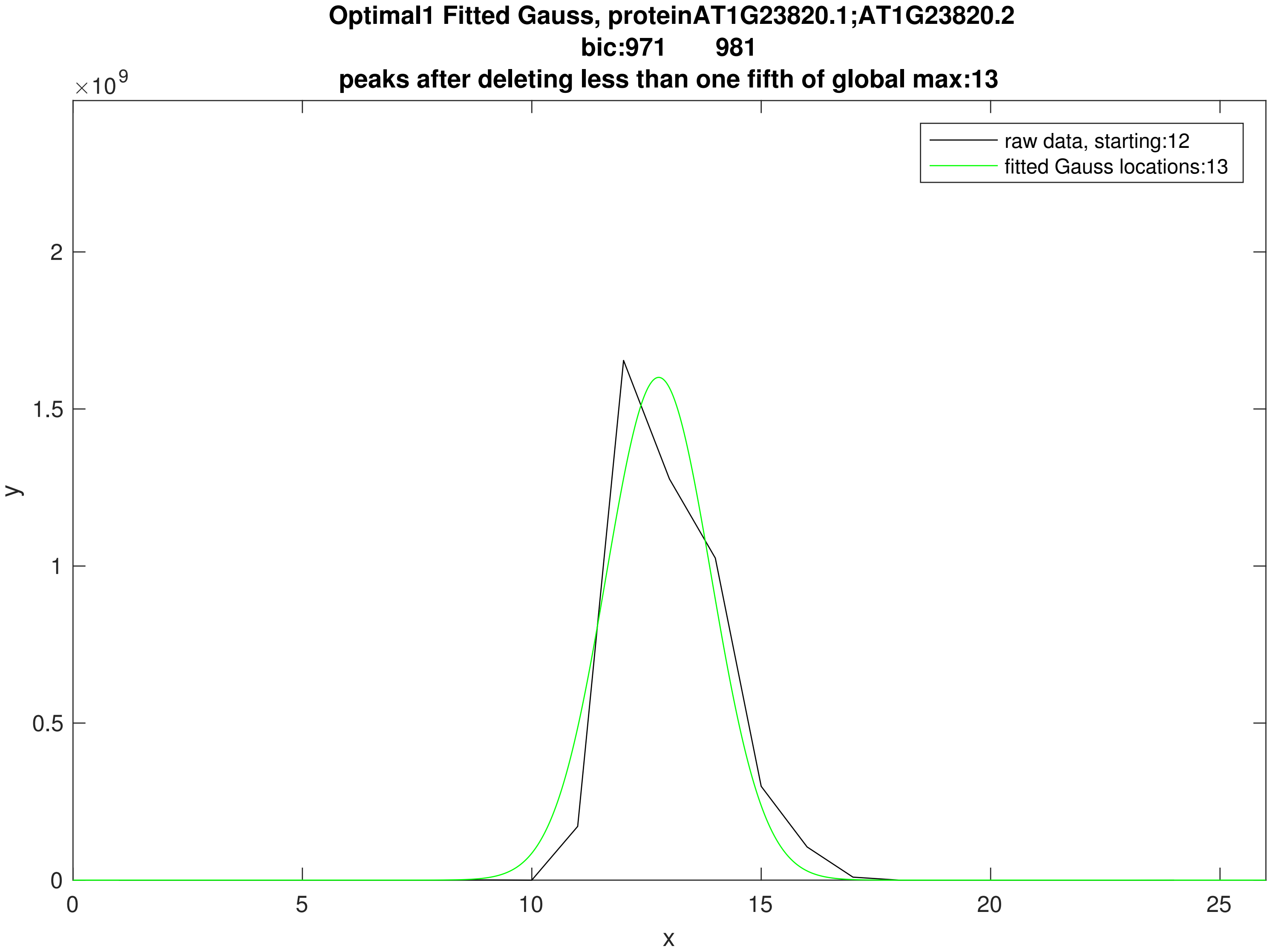
Optimal1 Fitted Gauss, proteinAT1G23820.1;AT1G23820.2
bic:971 981
peaks after deleting less than one fifth of global max:13
$\times 10^9$
raw data, starting:12
fitted Gauss locations:13
y
x
0
0.5
1
1.5
2
5
10
15
20
25

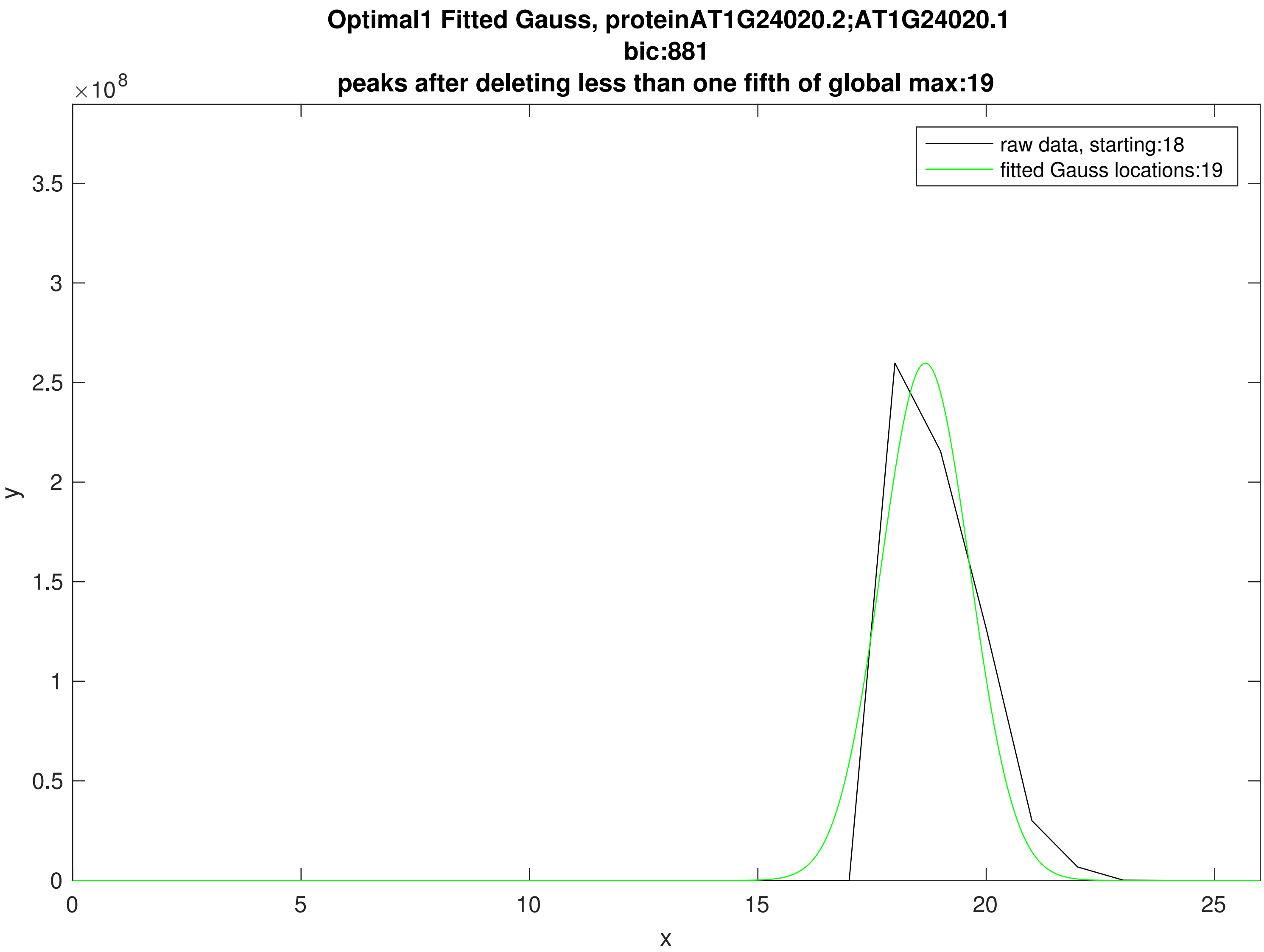

Optimal1 Fitted Gauss, proteinAT1G24020.2;AT1G24020.1
bic:881
peaks after deleting less than one fifth of global max:19
×10^8
raw data, starting:18
fitted Gauss locations:19
y
x
0
0.5
1
1.5
2
2.5
3
3.5
0
5
10
15
20
25

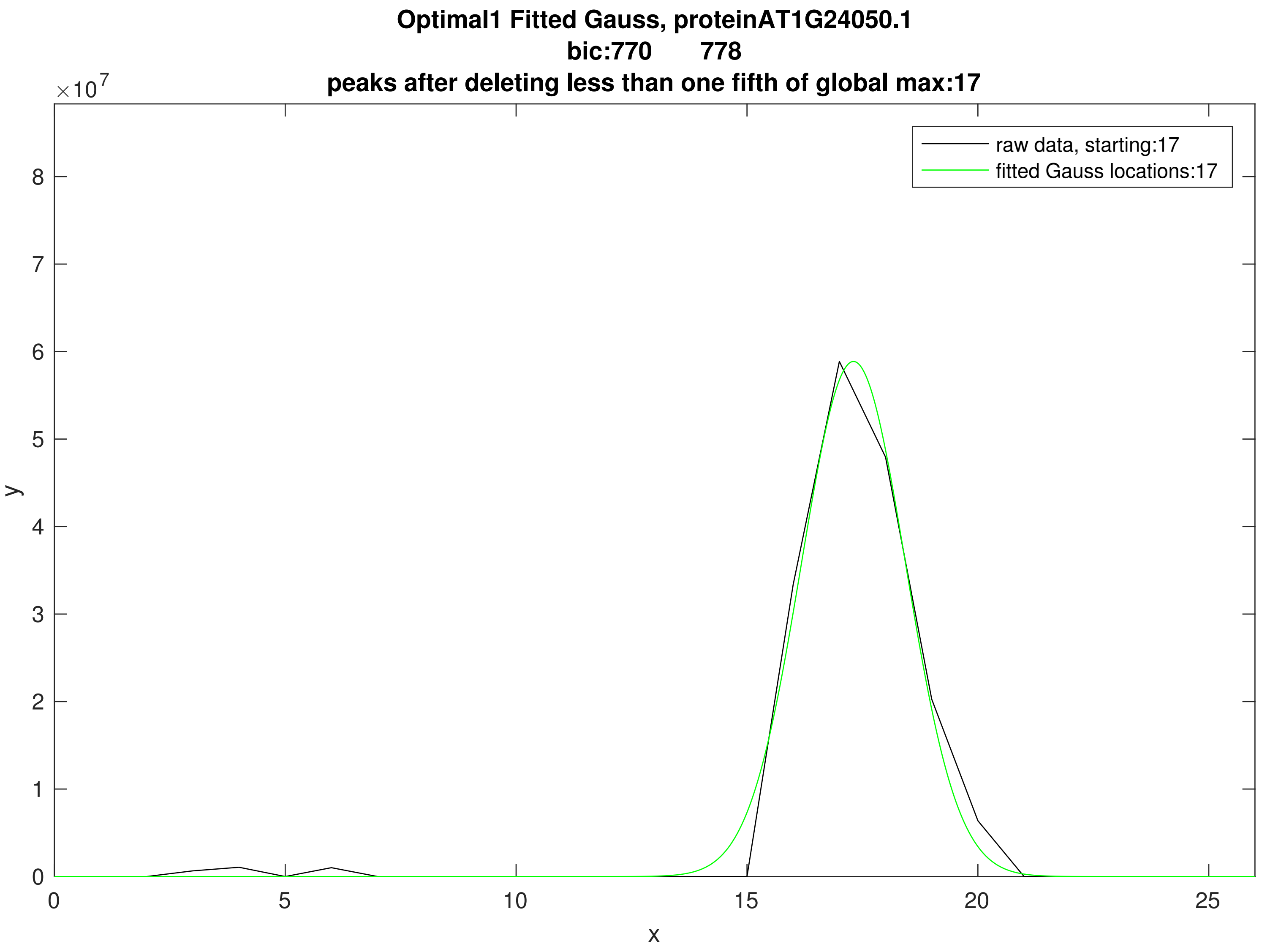

Optimal1 Fitted Gauss, proteinAT1G24050.1
bic:770 778
peaks after deleting less than one fifth of global max:17
×10^7
raw data, starting:17
fitted Gauss locations:17
y
x
0
1
2
3
4
5
6
7
8
0
5
10
15
20
25

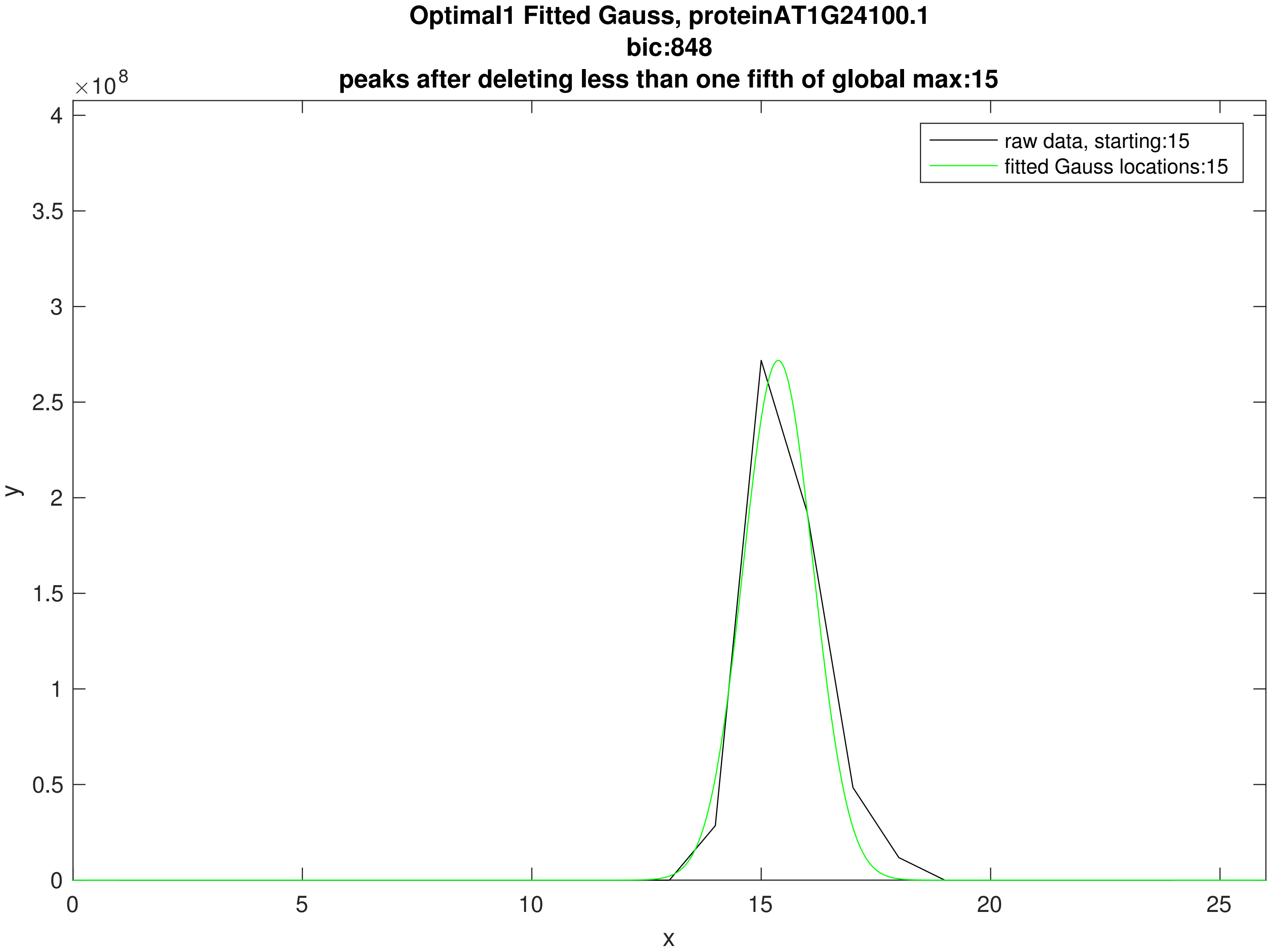

Optimal1 Fitted Gauss, proteinAT1G24100.1
bic:848
peaks after deleting less than one fifth of global max:15
×10^8
4
3.5
3
2.5
2
1.5
1
0.5
0
y
0
5
10
15
20
25
x
raw data, starting:15
fitted Gauss locations:15

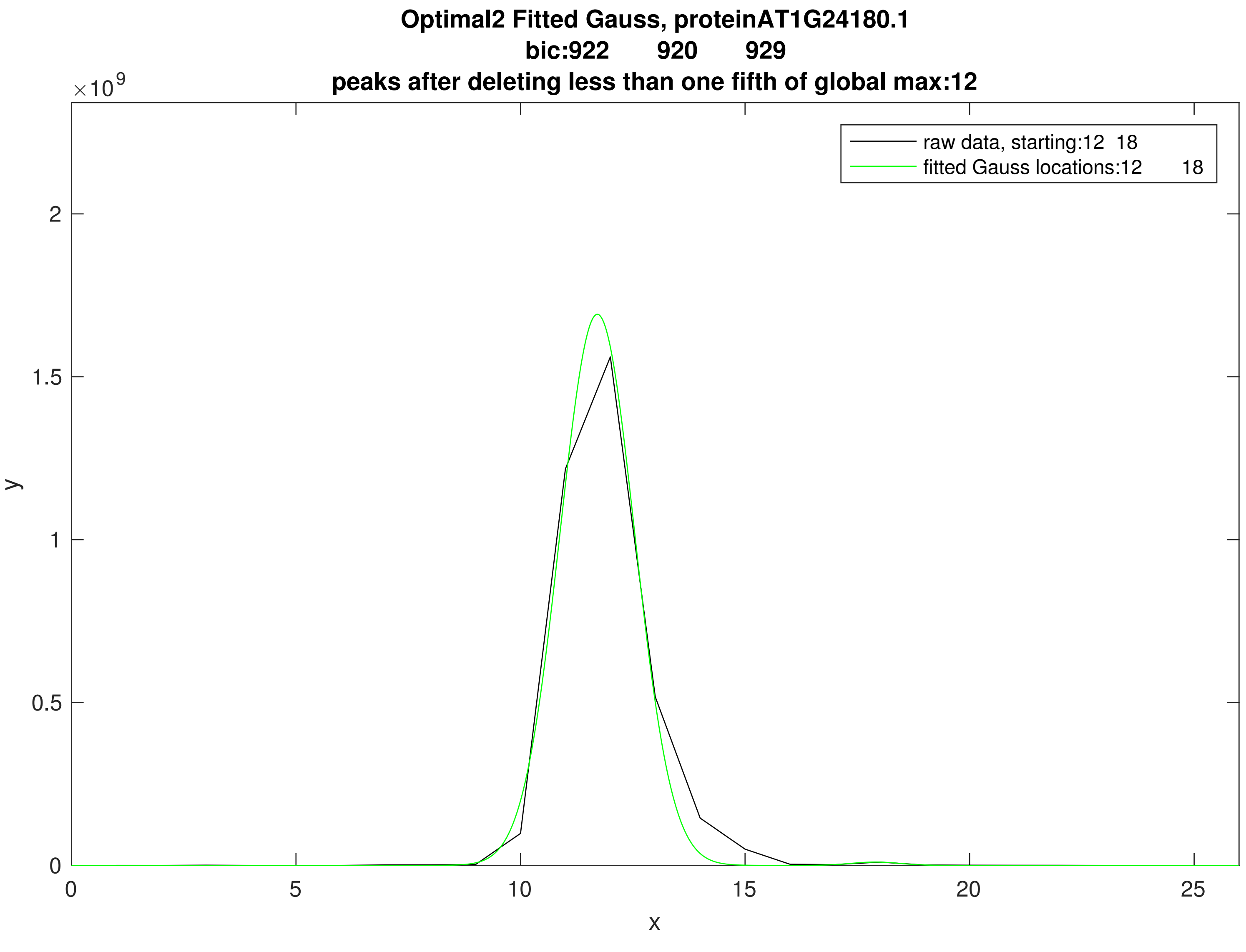
Optimal2 Fitted Gauss, proteinAT1G24180.1
bic:922 920 929
peaks after deleting less than one fifth of global max:12
×10^9
raw data, starting:12 18
fitted Gauss locations:12 18
y
x
0
0.5
1
1.5
2
0
5
10
15
20
25

**Optimal2 Fitted Gauss, proteinAT1G24510.1;AT1G24510.2**
**bic:854 794**
**peaks after deleting less than one fifth of global max:4.7 14**

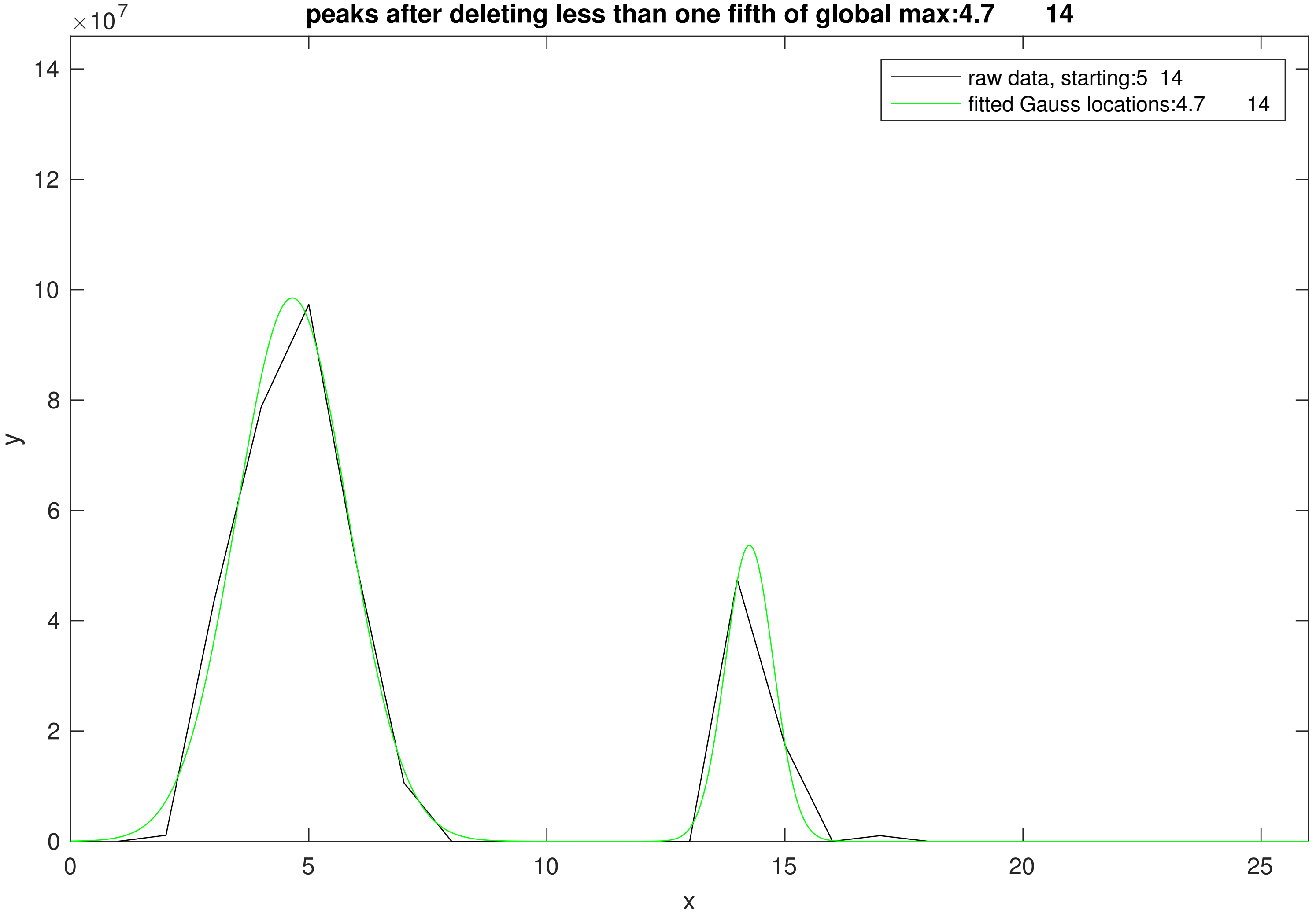

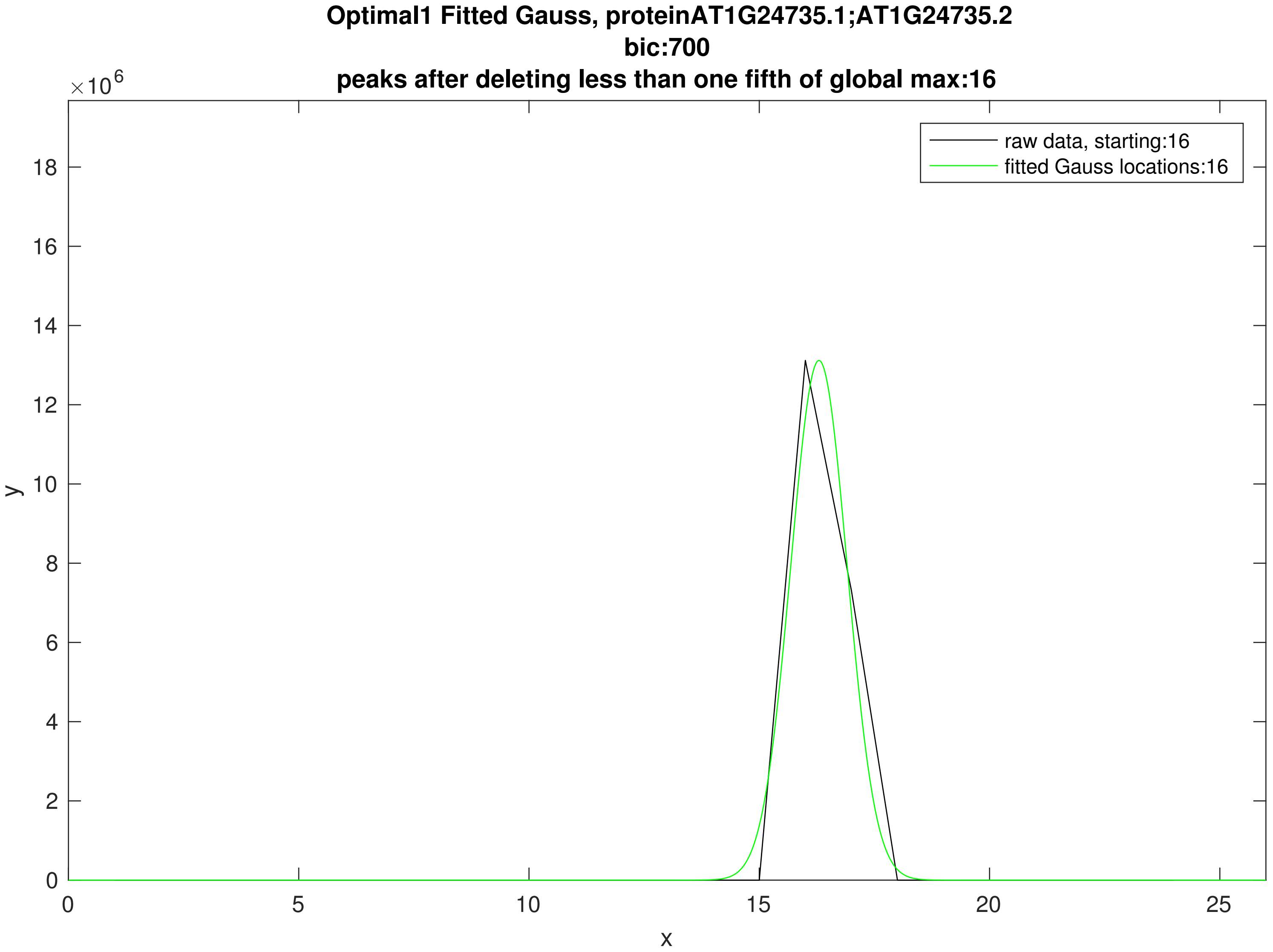
Optimal1 Fitted Gauss, proteinAT1G24735.1;AT1G24735.2
bic:700
peaks after deleting less than one fifth of global max:16
×10^6
raw data, starting:16
fitted Gauss locations:16
y
x
0
2
4
6
8
10
12
14
16
18
0
5
10
15
20
25

Optimal1 Fitted Gauss, proteinAT1G25155.1;AT1G25083.1;AT1G24909.1;AT1G24807.1;AT5G57890.1;AT1G25220.1;AT1

bic:736

peaks after deleting less than one fifth of global max:18

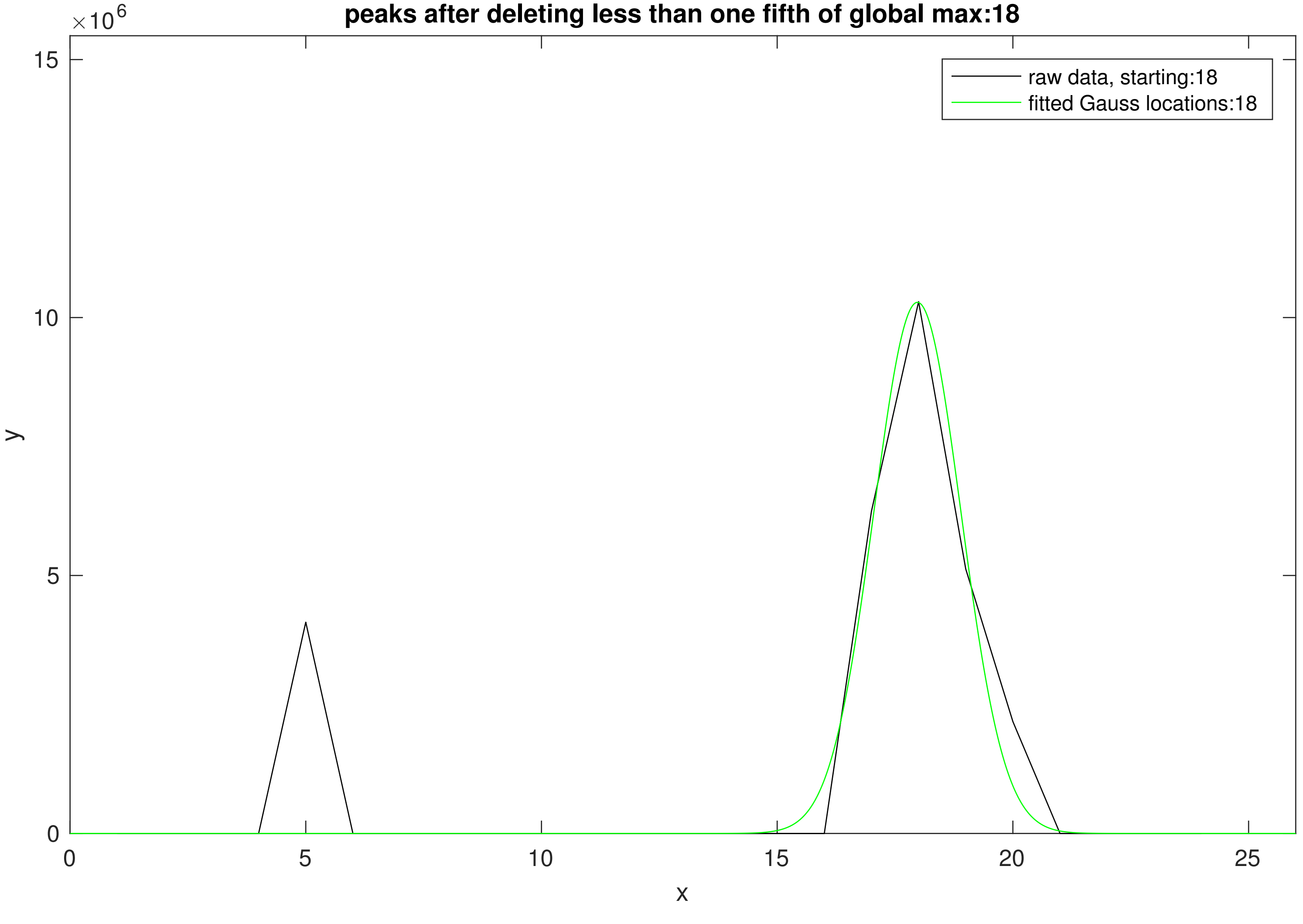

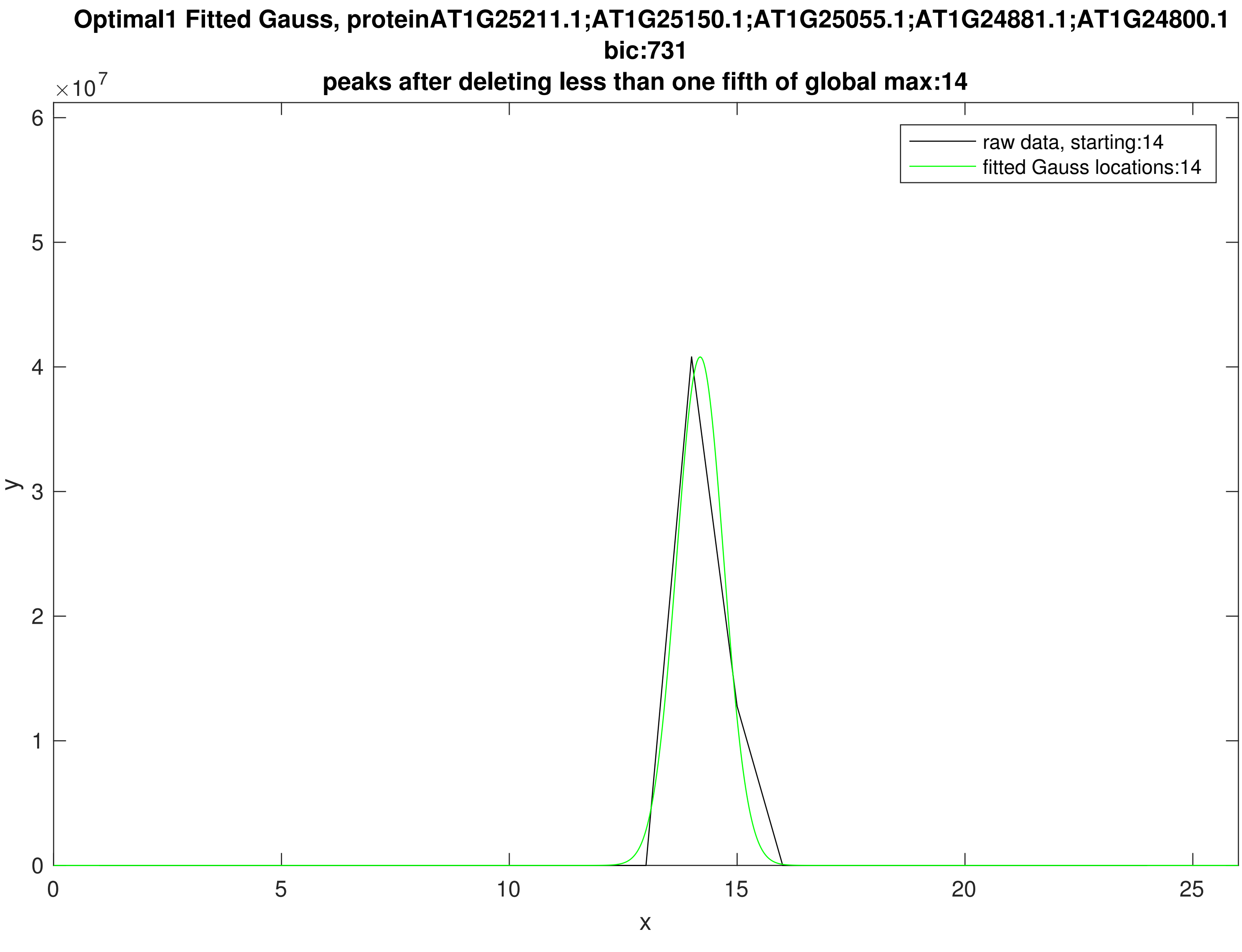
Optimal1 Fitted Gauss, proteinAT1G25211.1;AT1G25150.1;AT1G25055.1;AT1G24881.1;AT1G24800.1
bic:731
peaks after deleting less than one fifth of global max:14
×10^7
6
5
4
3
2
1
0
y
0
5
10
15
20
25
x
raw data, starting:14
fitted Gauss locations:14

Optimal1 Fitted Gauss, proteinAT1G25230.1
bic:874
peaks after deleting less than one fifth of global max:17

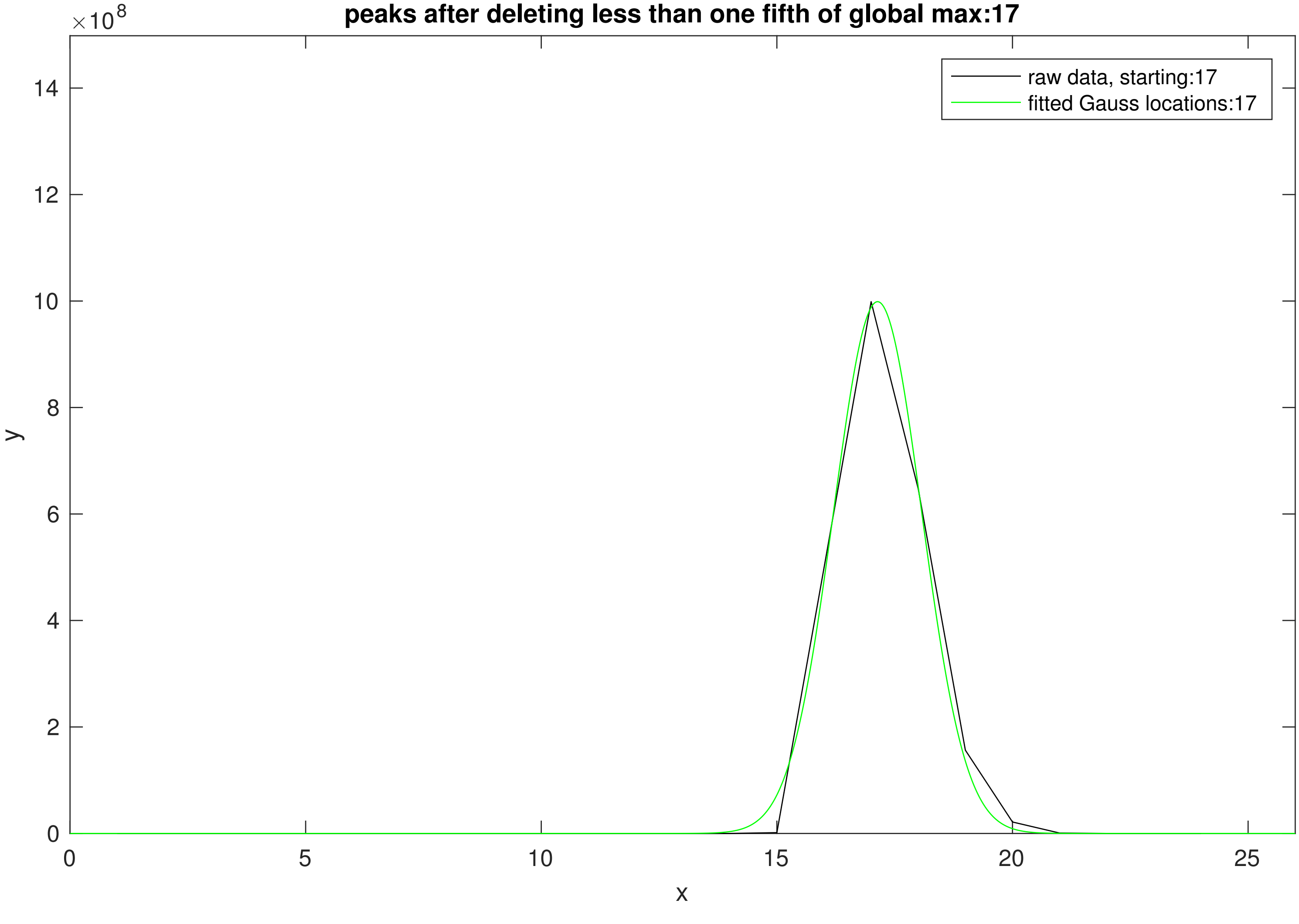

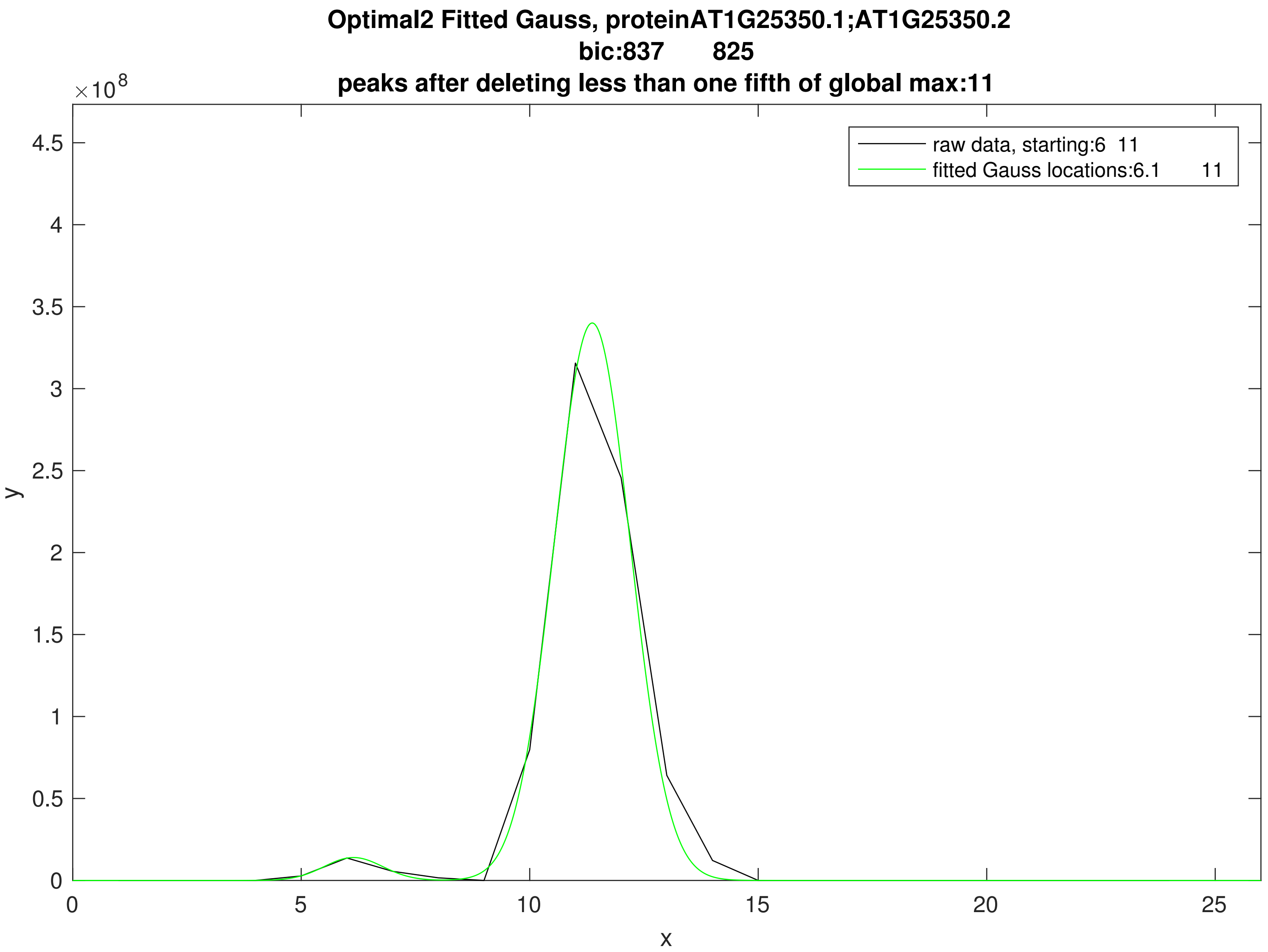
Optimal2 Fitted Gauss, proteinAT1G25350.1;AT1G25350.2
bic:837 825
peaks after deleting less than one fifth of global max:11
×10^8
4.5
4
3.5
3
2.5
2
1.5
1
0.5
0
y
0
5
10
15
20
25
x
raw data, starting:6 11
fitted Gauss locations:6.1 11

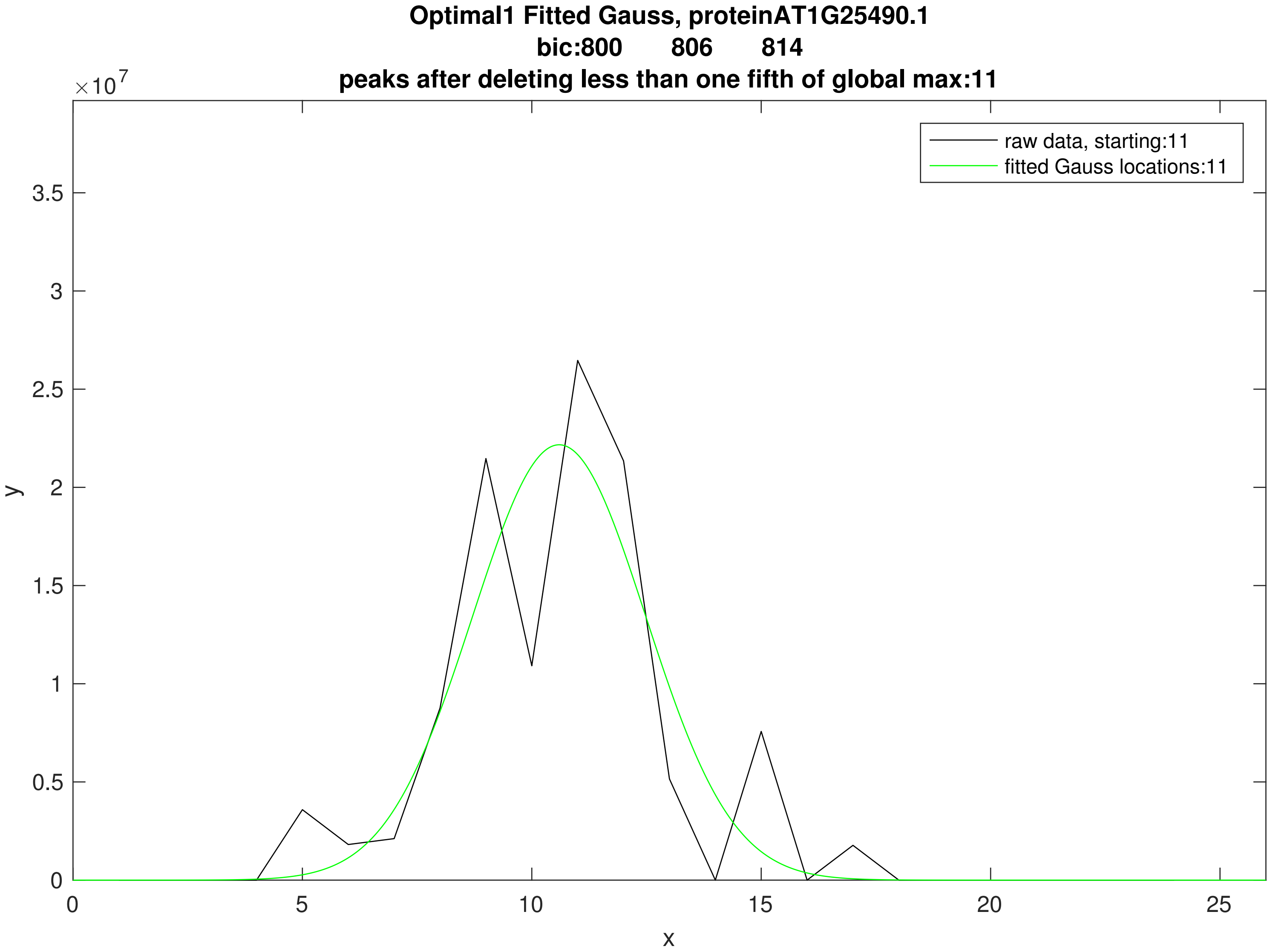
Optimal1 Fitted Gauss, proteinAT1G25490.1
bic:800 806 814
peaks after deleting less than one fifth of global max:11
×10^7
raw data, starting:11
fitted Gauss locations:11
3.5
3
2.5
2
1.5
1
0.5
0
y
0
5
10
15
20
25
x

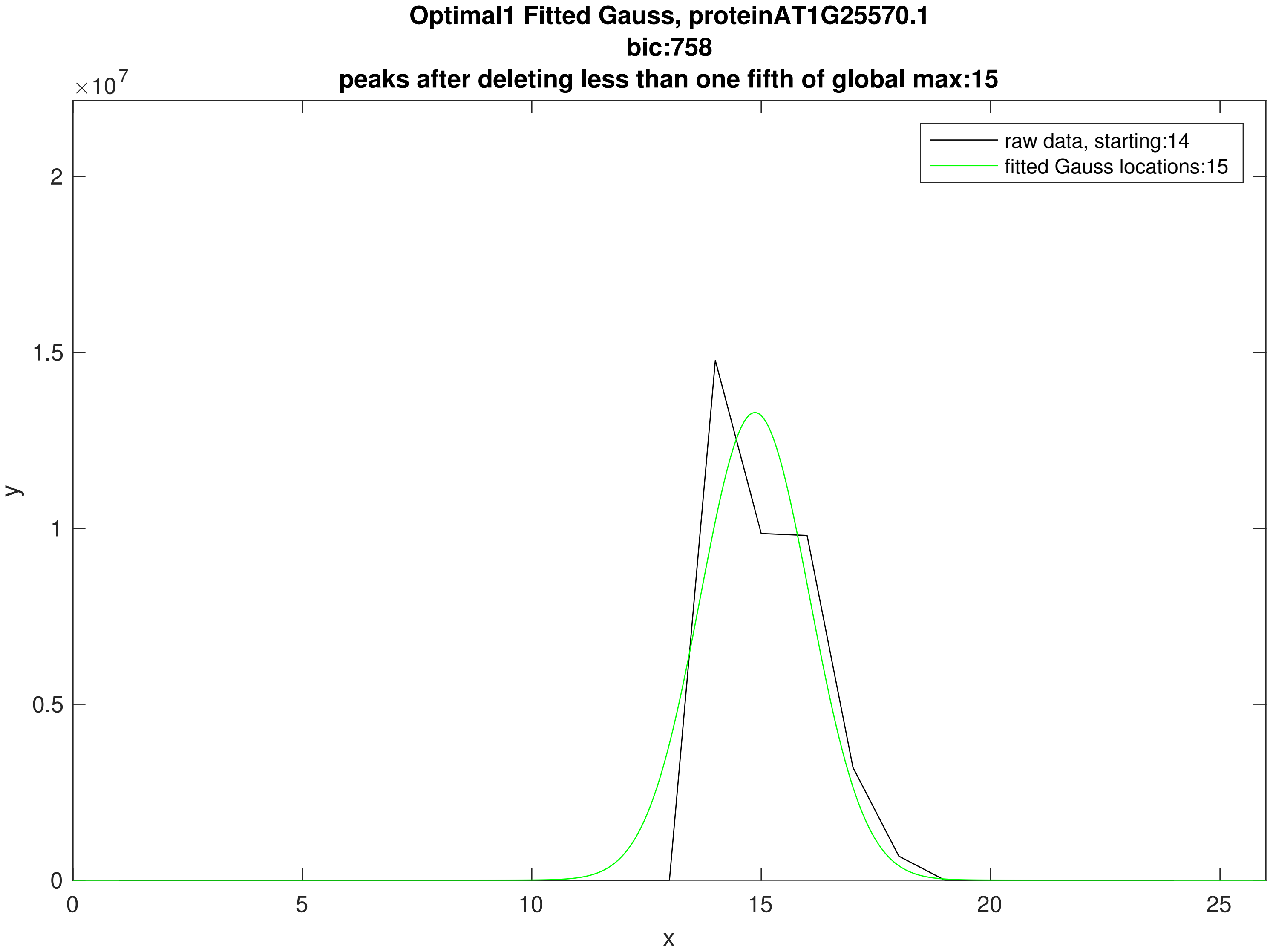

Optimal1 Fitted Gauss, proteinAT1G25570.1
bic:758
peaks after deleting less than one fifth of global max:15
×10^7
raw data, starting:14
fitted Gauss locations:15
2
1.5
1
0.5
0
y
0
5
10
15
20
25
x

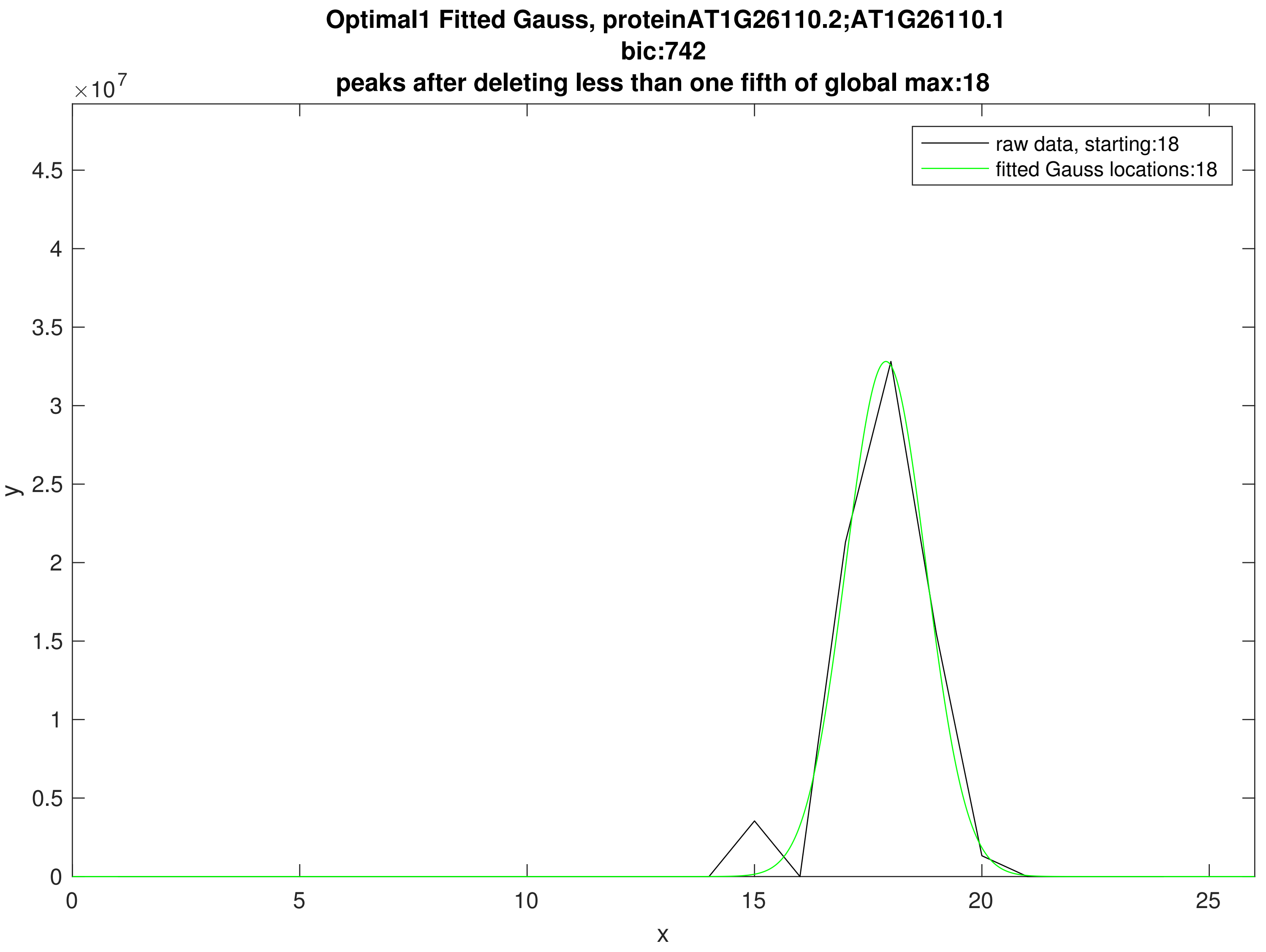

Optimal1 Fitted Gauss, proteinAT1G26110.2;AT1G26110.1
bic:742
peaks after deleting less than one fifth of global max:18
×10^7
raw data, starting:18
fitted Gauss locations:18
y
x
0
0.5
1
1.5
2
2.5
3
3.5
4
4.5
5
10
15
20
25

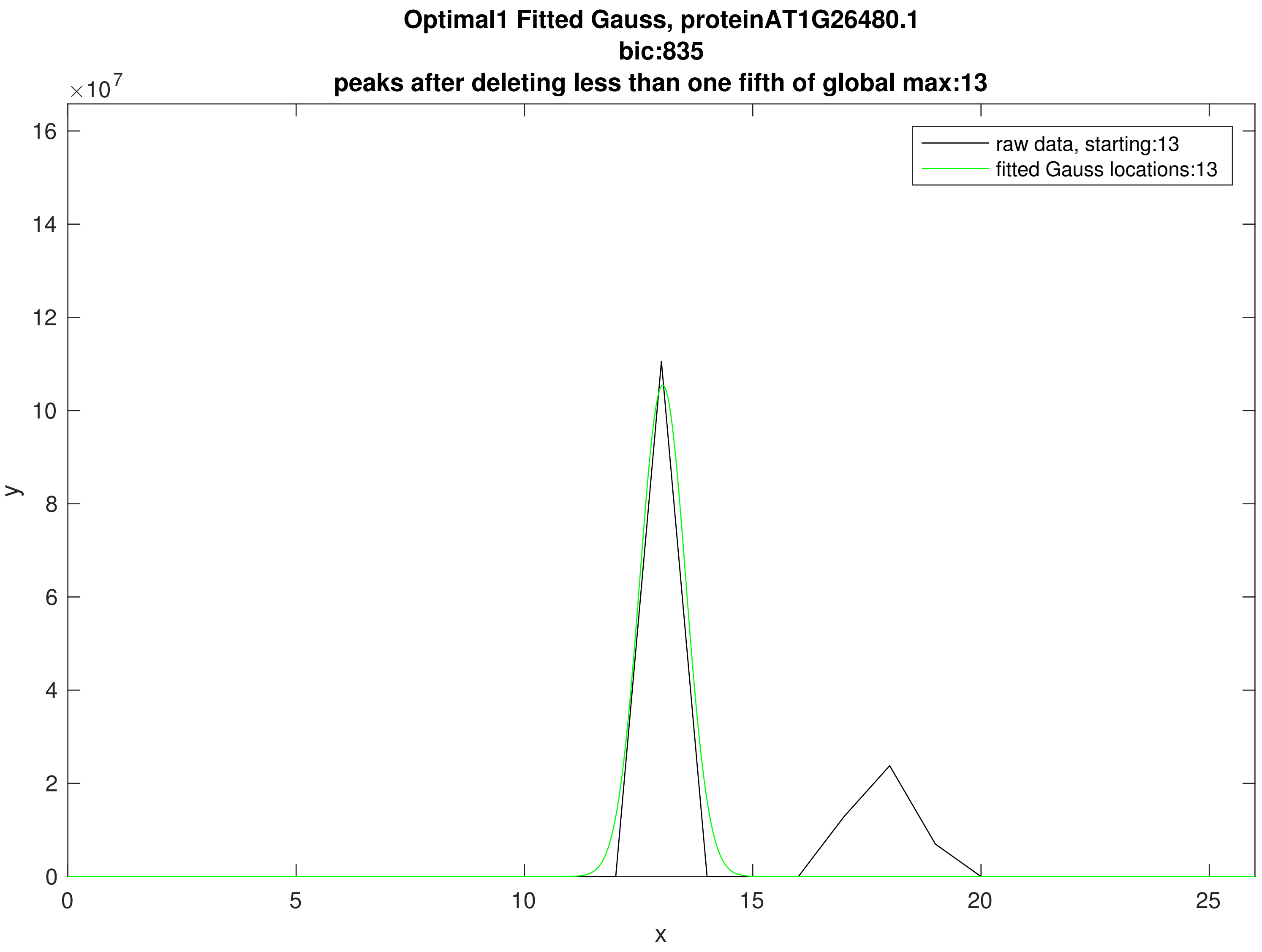
Optimal1 Fitted Gauss, proteinAT1G26480.1
bic:835
peaks after deleting less than one fifth of global max:13
×10^7
16
14
12
10
8
6
4
2
0
y
0
5
10
15
20
25
x
raw data, starting:13
fitted Gauss locations:13

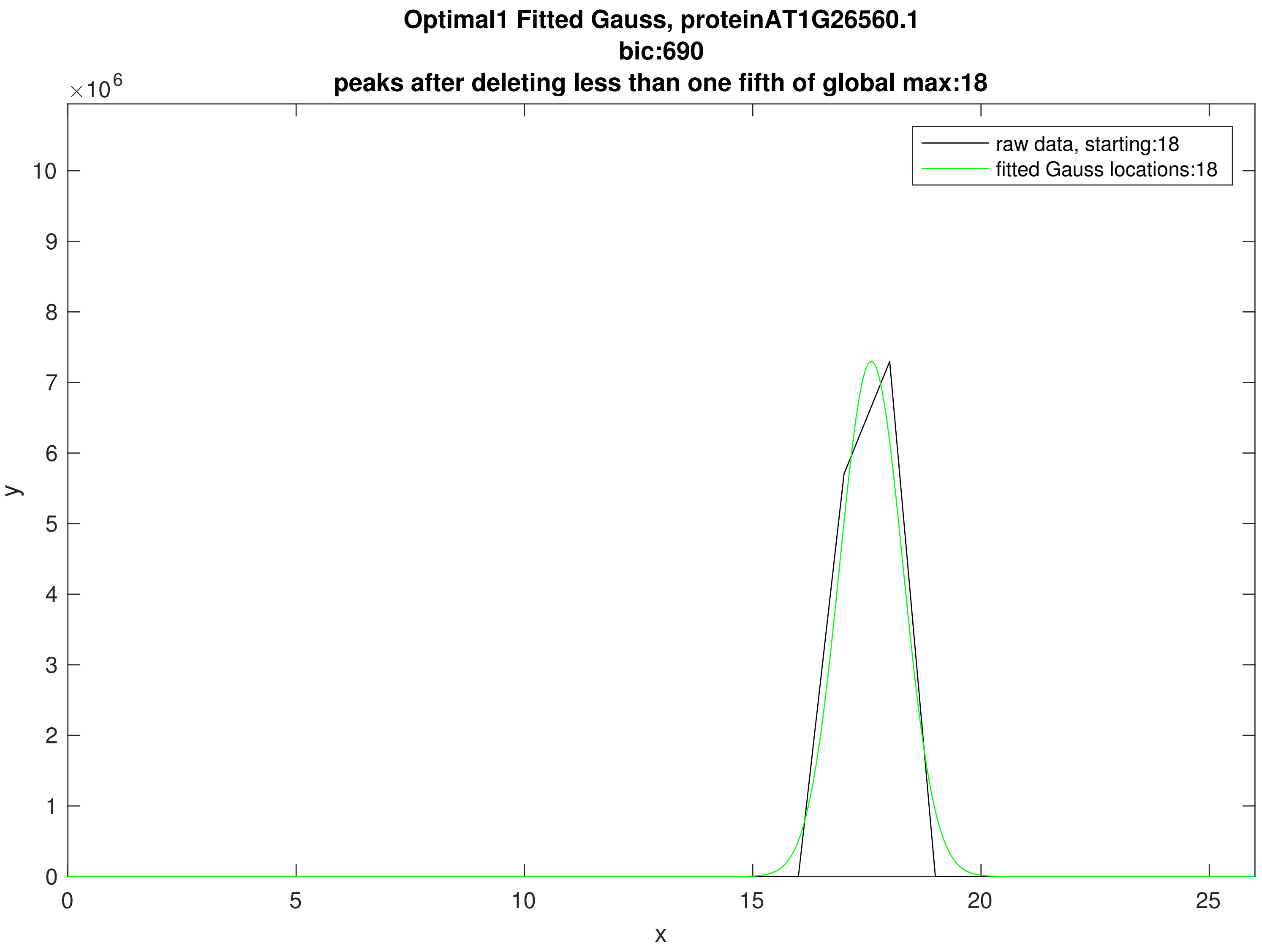

Optimal1 Fitted Gauss, proteinAT1G26560.1
bic:690
peaks after deleting less than one fifth of global max:18
×10^6
raw data, starting:18
fitted Gauss locations:18
y
x

**Optimal2 Fitted Gauss, proteinAT1G26570.1**
**bic:849 711**
**peaks after deleting less than one fifth of global max:9 13**

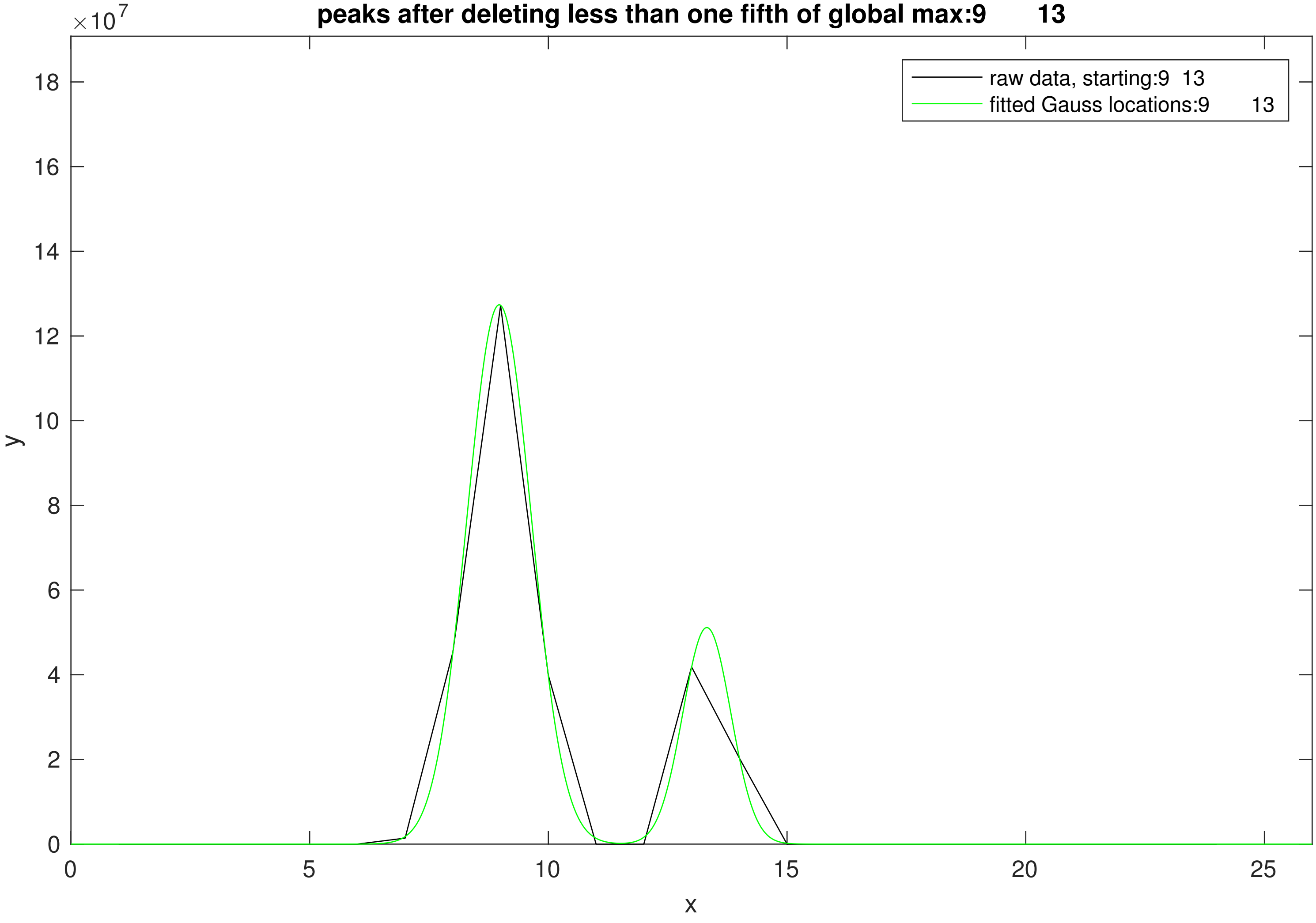

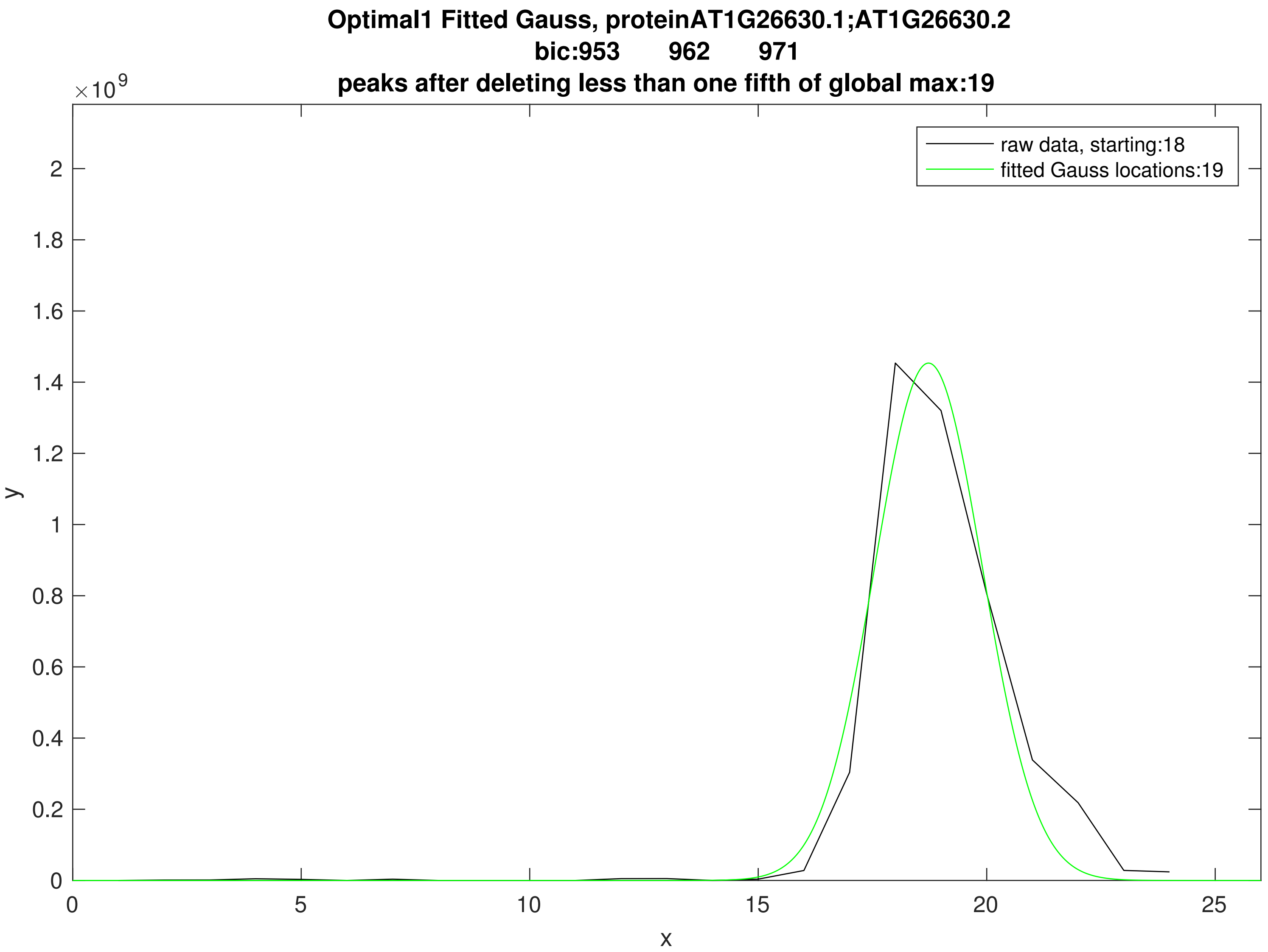
Optimal1 Fitted Gauss, proteinAT1G26630.1;AT1G26630.2
bic:953 962 971
peaks after deleting less than one fifth of global max:19
raw data, starting:18
fitted Gauss locations:19
y
x
$\times 10^9$
0
0.2
0.4
0.6
0.8
1
1.2
1.4
1.6
1.8
2
0
5
10
15
20
25

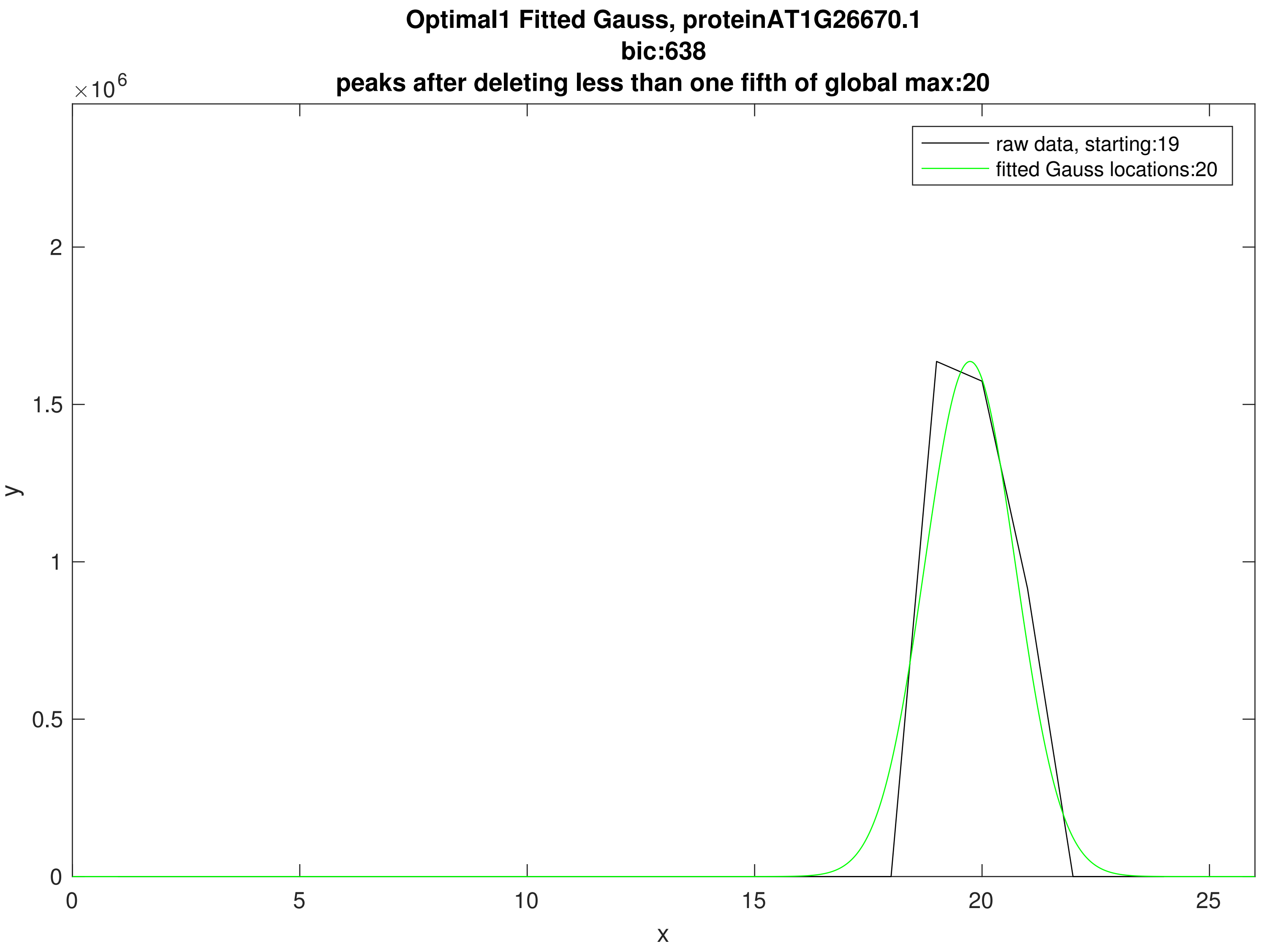
Optimal1 Fitted Gauss, proteinAT1G26670.1
bic:638
peaks after deleting less than one fifth of global max:20
×10^6
raw data, starting:19
fitted Gauss locations:20
2
1.5
1
0.5
0
y
0
5
10
15
20
25
x

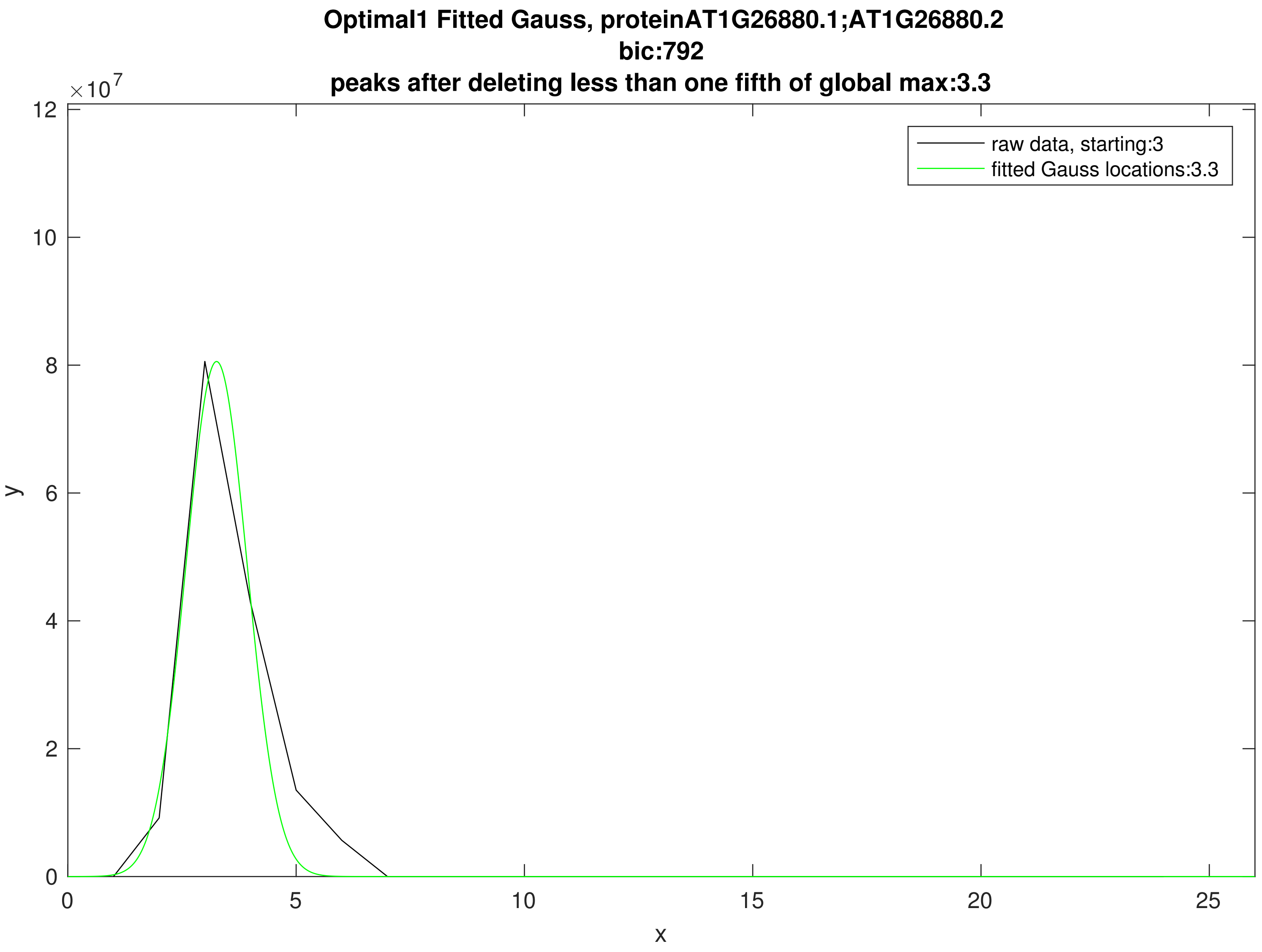

Optimal1 Fitted Gauss, proteinAT1G26880.1;AT1G26880.2
bic:792
peaks after deleting less than one fifth of global max:3.3
×10^7
raw data, starting:3
fitted Gauss locations:3.3
y
x
0
2
4
6
8
10
12
0
5
10
15
20
25

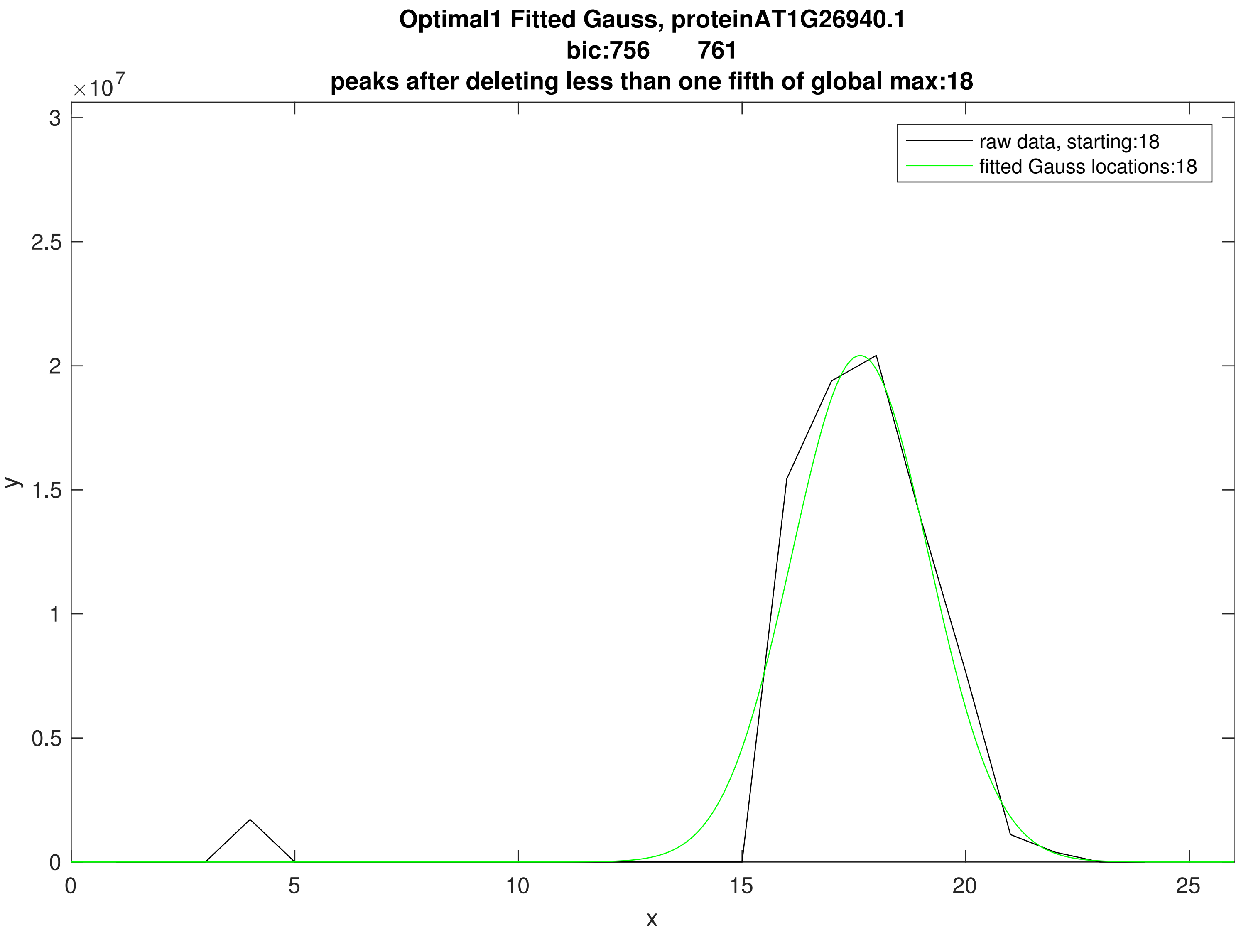

Optimal1 Fitted Gauss, proteinAT1G26940.1
bic:756 761
peaks after deleting less than one fifth of global max:18
×10^7
3
2.5
2
1.5
1
0.5
0
y
0
5
10
15
20
25
x
raw data, starting:18
fitted Gauss locations:18

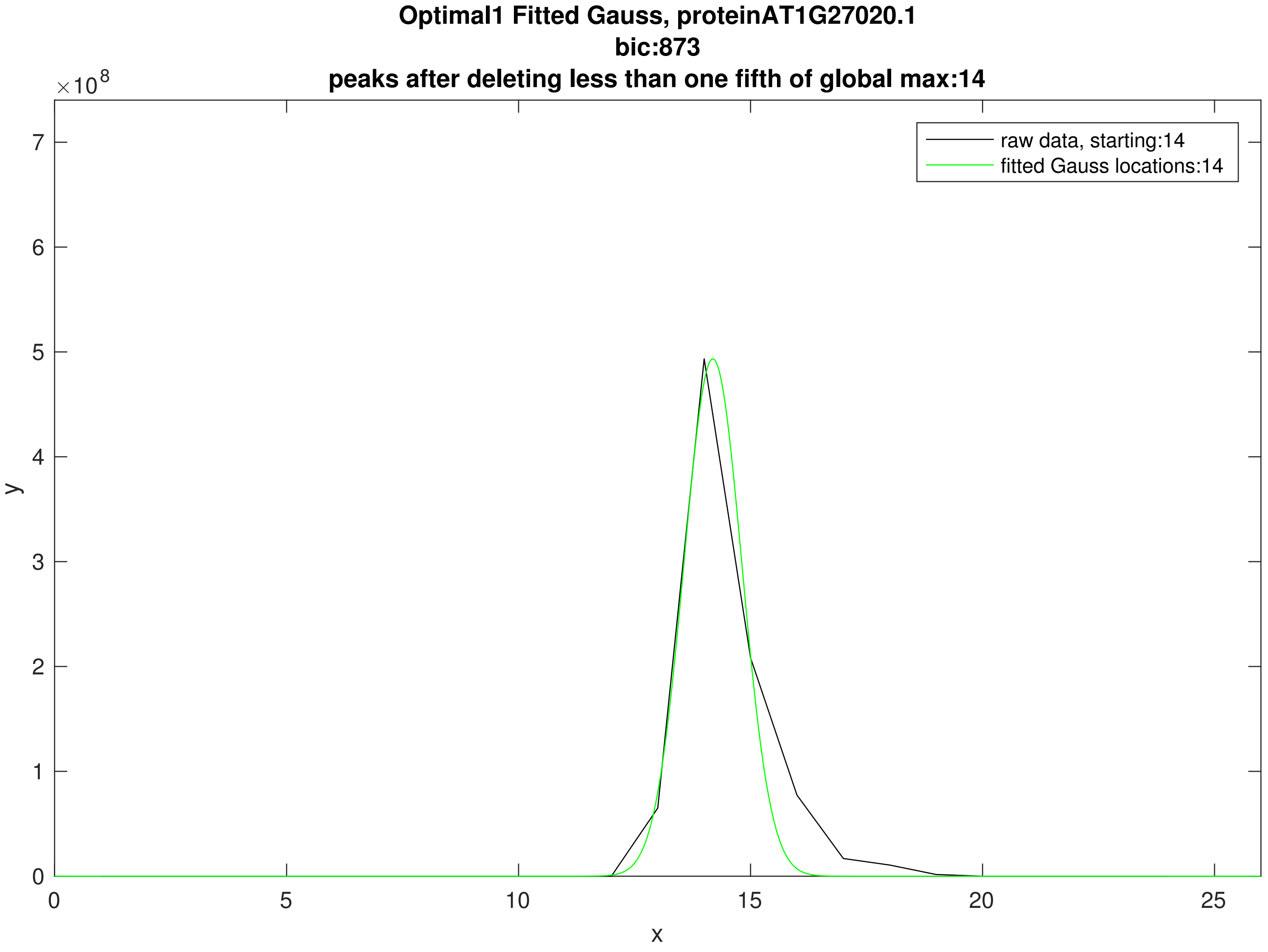
Optimal1 Fitted Gauss, proteinAT1G27020.1
bic:873
peaks after deleting less than one fifth of global max:14
×10^8
raw data, starting:14
fitted Gauss locations:14
y
x
0
1
2
3
4
5
6
7
0
5
10
15
20
25

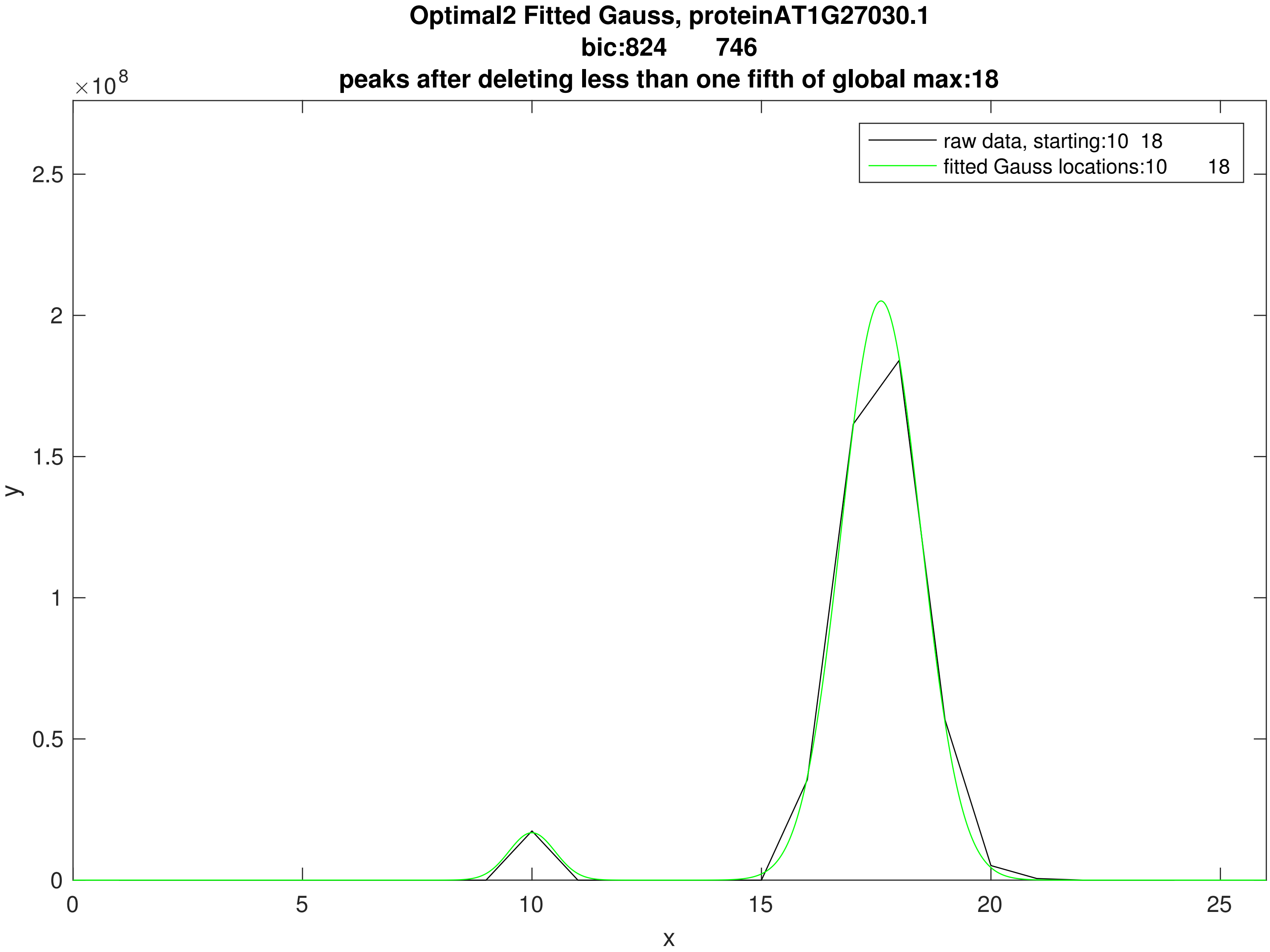

Optimal2 Fitted Gauss, proteinAT1G27030.1
bic:824 746
peaks after deleting less than one fifth of global max:18
×10^8
2.5
2
1.5
1
0.5
0
y
0
5
10
15
20
25
x
raw data, starting:10 18
fitted Gauss locations:10 18

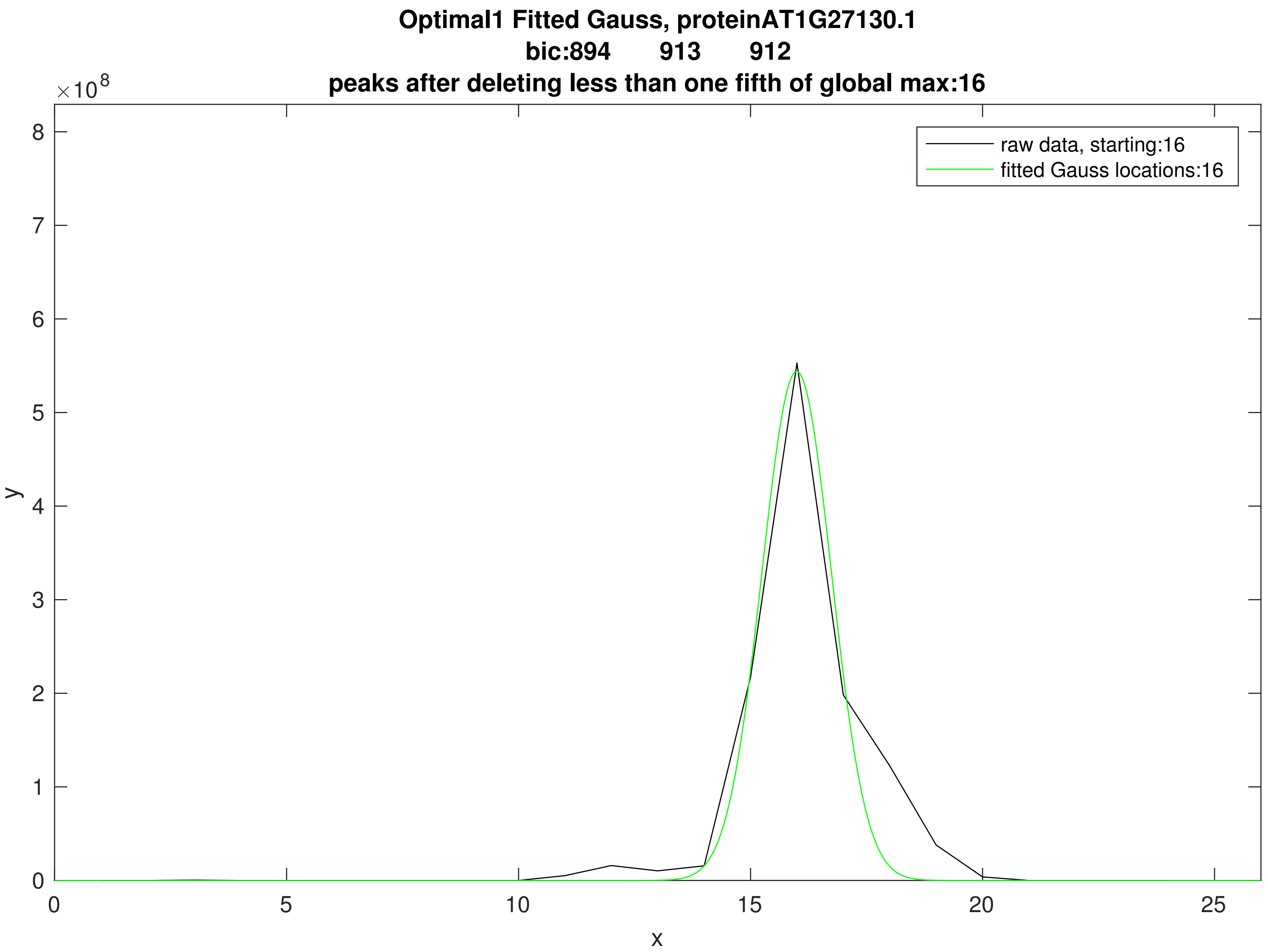
Optimal1 Fitted Gauss, proteinAT1G27130.1
bic:894 913 912
peaks after deleting less than one fifth of global max:16
×10^8
raw data, starting:16
fitted Gauss locations:16
y
x
0
1
2
3
4
5
6
7
8
0
5
10
15
20
25

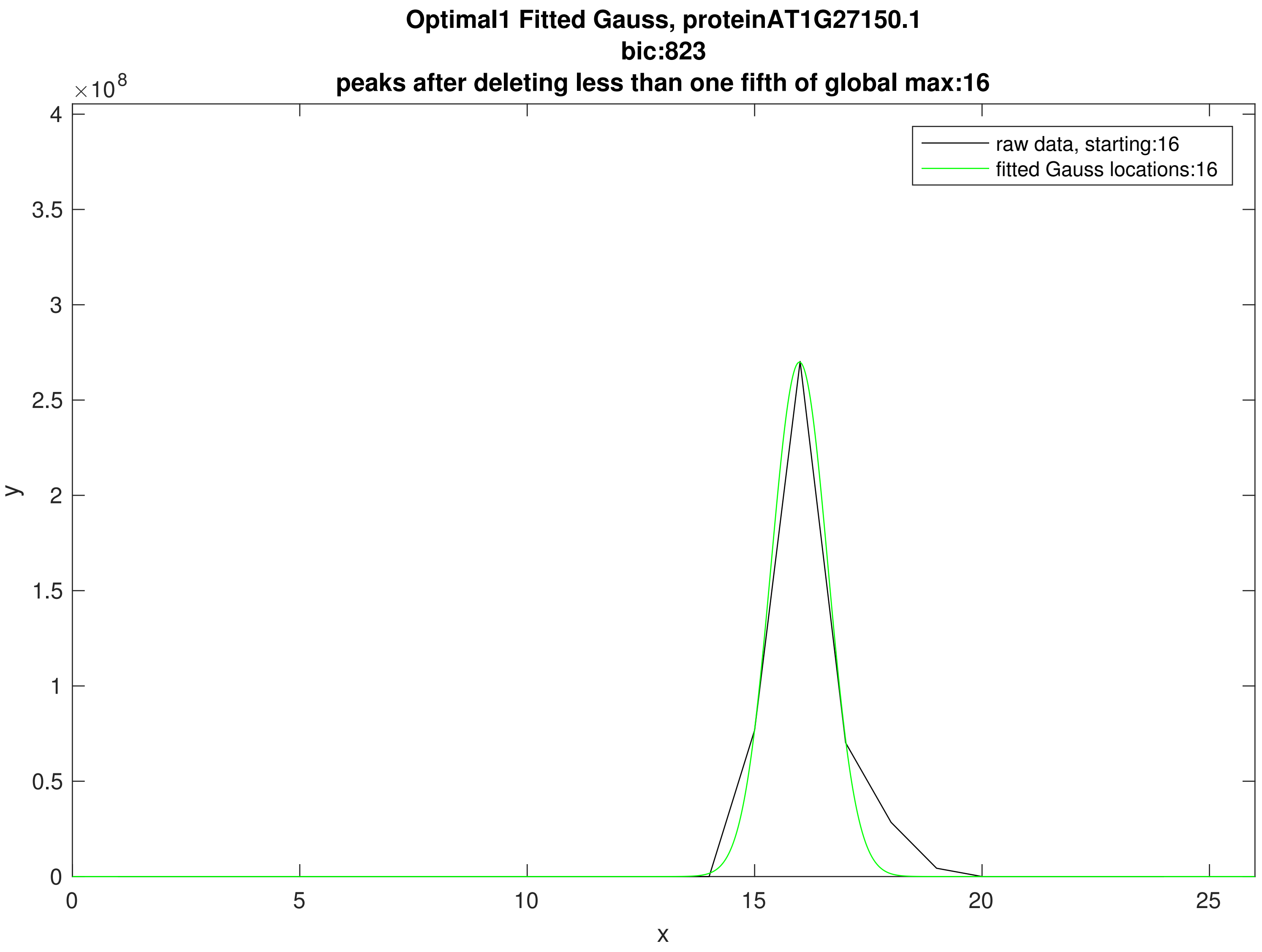
Optimal1 Fitted Gauss, proteinAT1G27150.1
bic:823
peaks after deleting less than one fifth of global max:16
×10^8
raw data, starting:16
fitted Gauss locations:16
y
x
0
0.5
1
1.5
2
2.5
3
3.5
4
0
5
10
15
20
25

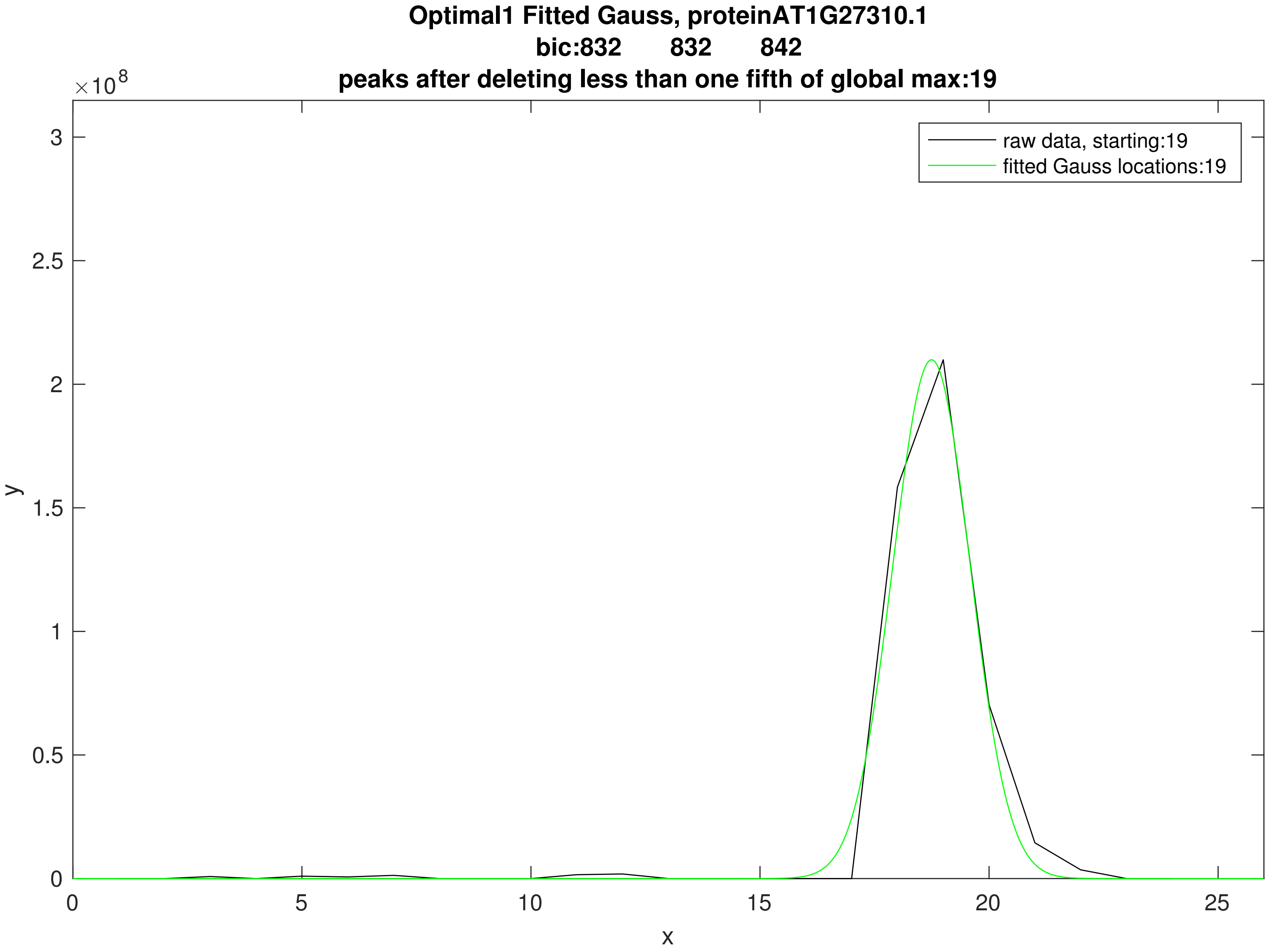

Optimal1 Fitted Gauss, proteinAT1G27310.1
bic:832 832 842
peaks after deleting less than one fifth of global max:19
×10^8
raw data, starting:19
fitted Gauss locations:19
y
x
0
0.5
1
1.5
2
2.5
3
5
10
15
20
25

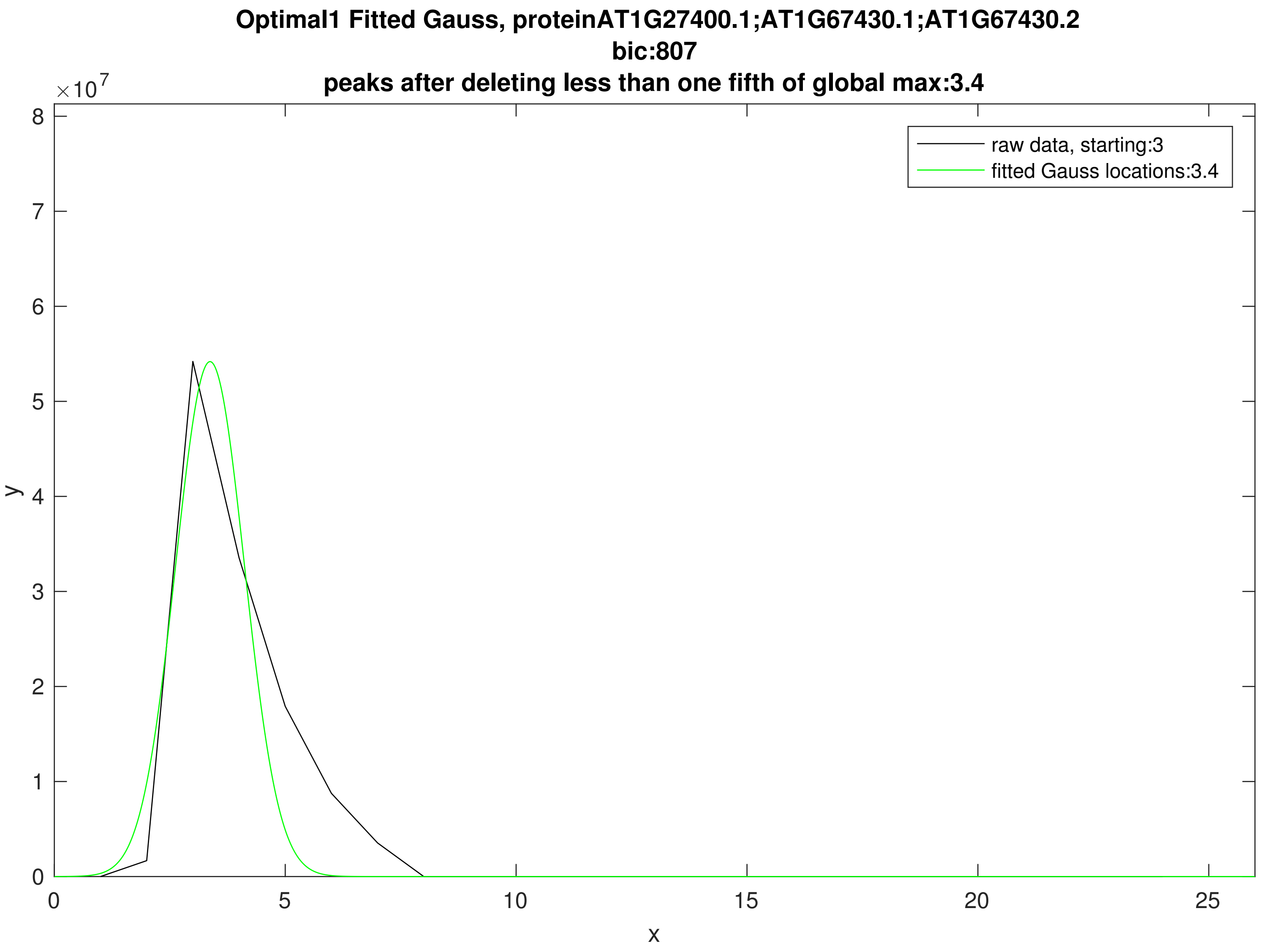
Optimal1 Fitted Gauss, proteinAT1G27400.1;AT1G67430.1;AT1G67430.2
bic:807
peaks after deleting less than one fifth of global max:3.4
×10^7
raw data, starting:3
fitted Gauss locations:3.4
y
x
0
1
2
3
4
5
6
7
8
0
5
10
15
20
25

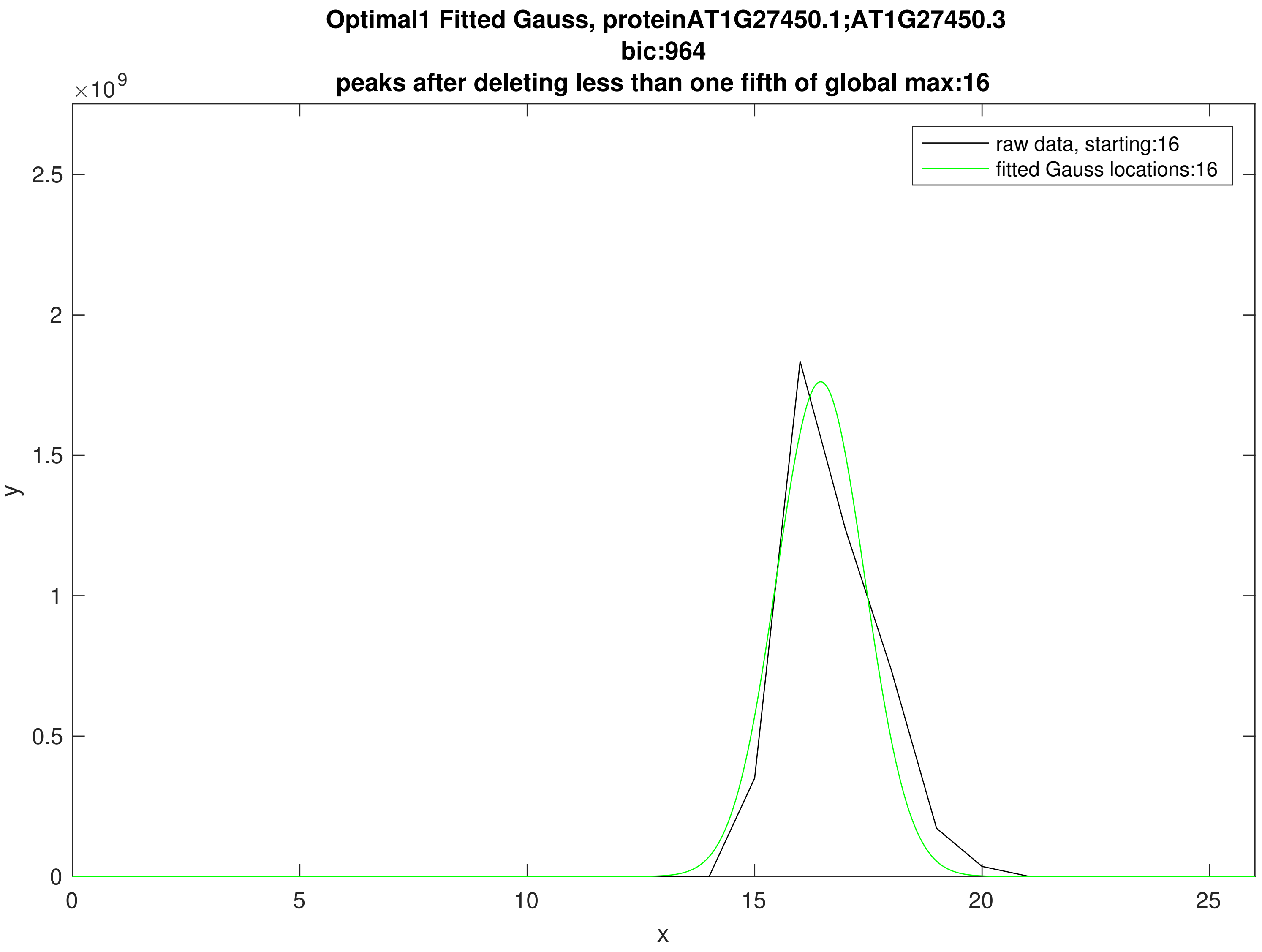

Optimal1 Fitted Gauss, proteinAT1G27450.1;AT1G27450.3
bic:964
peaks after deleting less than one fifth of global max:16
×10^9
raw data, starting:16
fitted Gauss locations:16
y
x
2.5
2
1.5
1
0.5
0
0
5
10
15
20
25

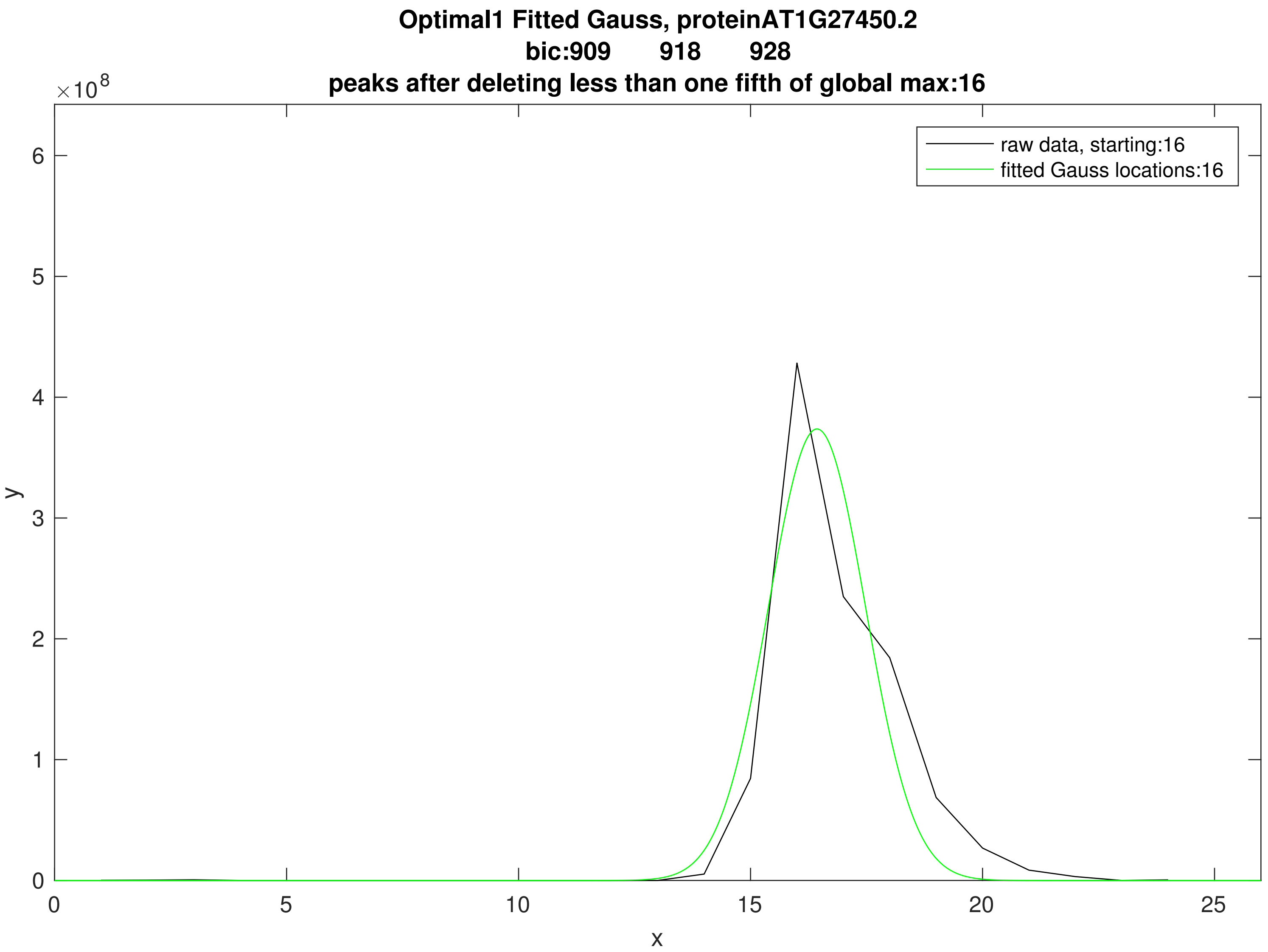

Optimal1 Fitted Gauss, proteinAT1G27450.2
bic:909 918 928
peaks after deleting less than one fifth of global max:16
×10^8
raw data, starting:16
fitted Gauss locations:16
y
x
0
1
2
3
4
5
6
0
5
10
15
20
25

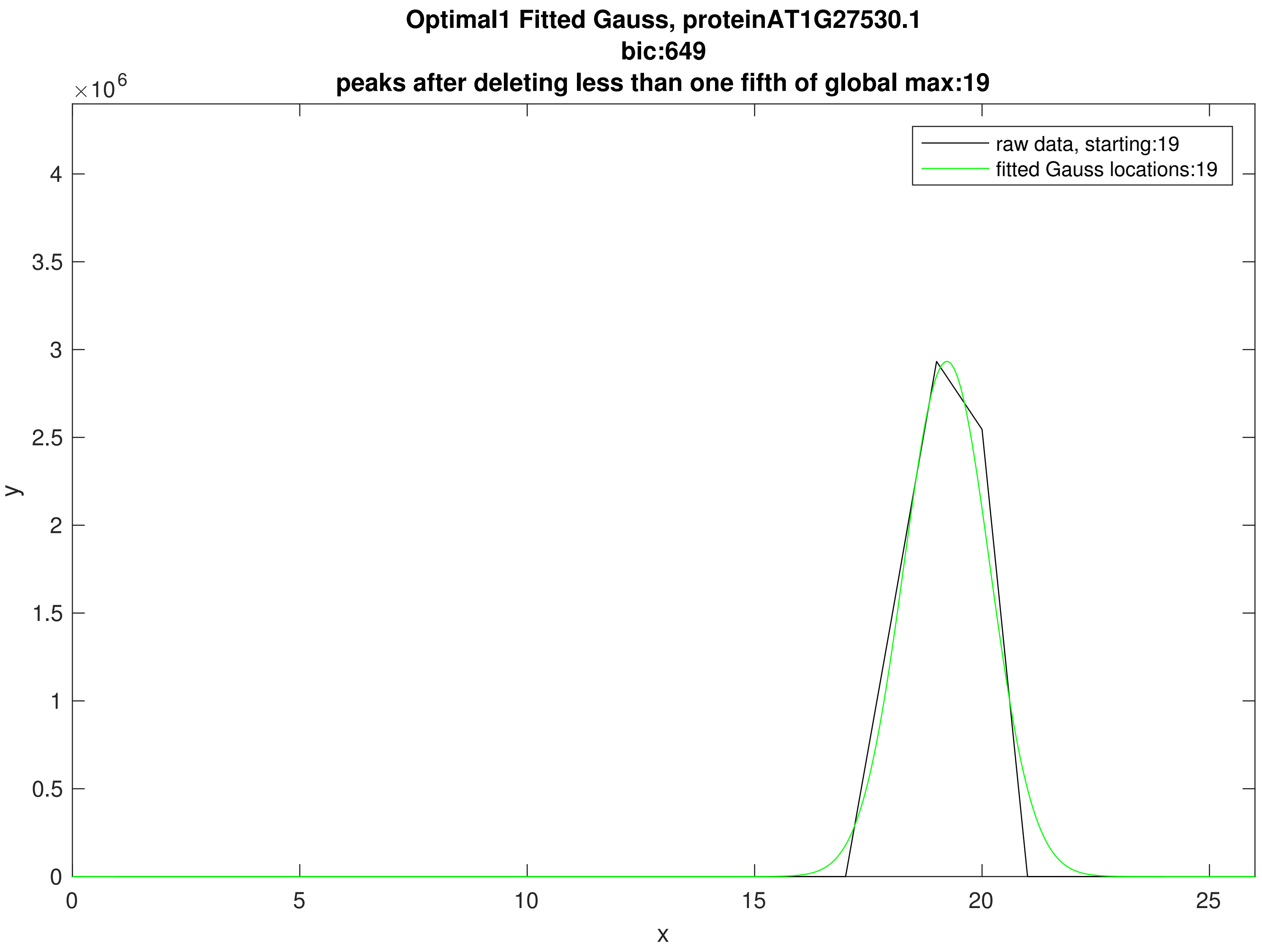

Optimal1 Fitted Gauss, proteinAT1G27530.1
bic:649
peaks after deleting less than one fifth of global max:19
×10^6
raw data, starting:19
fitted Gauss locations:19
y
x
0
0.5
1
1.5
2
2.5
3
3.5
4
0
5
10
15
20
25

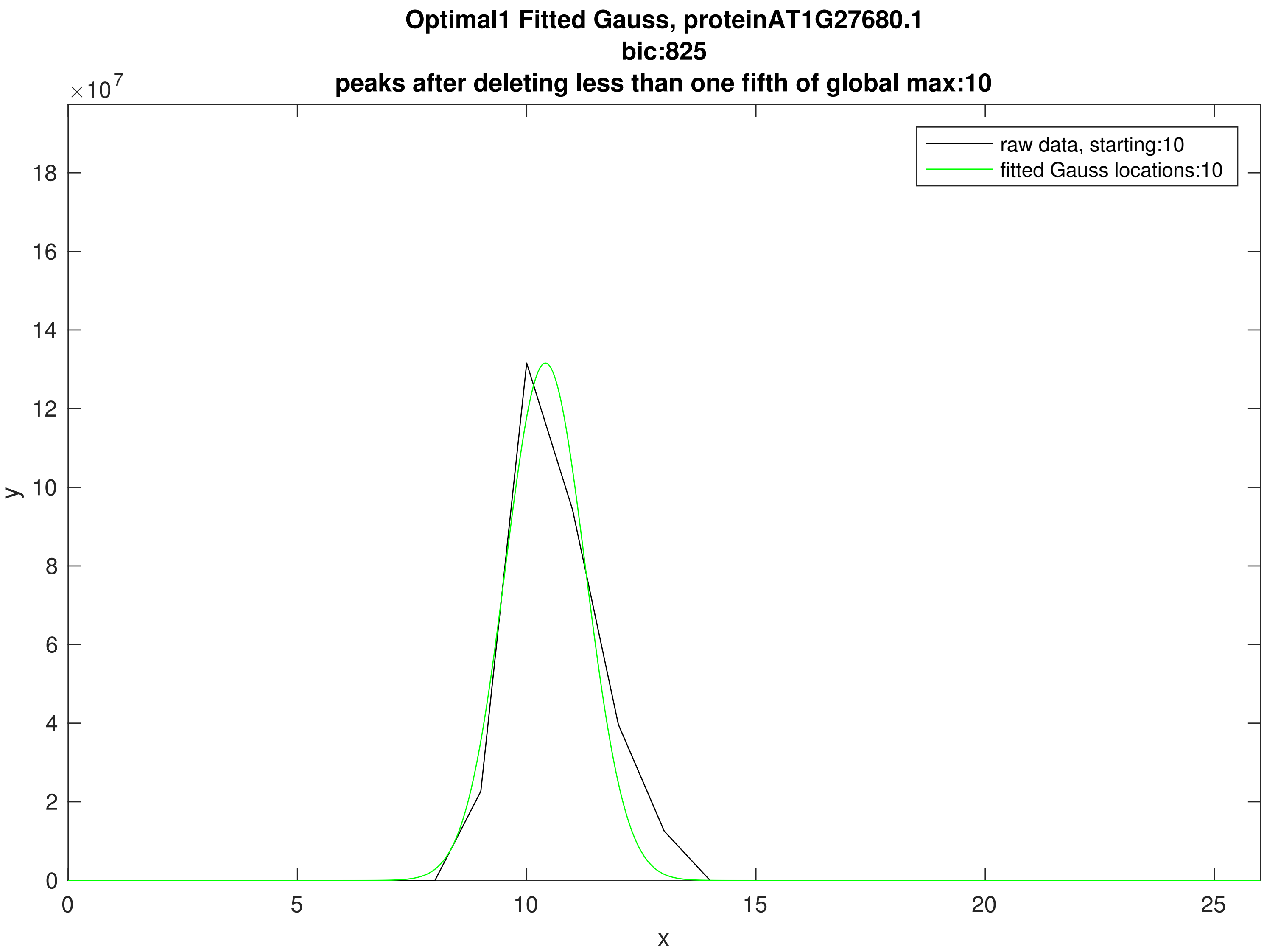
Optimal1 Fitted Gauss, proteinAT1G27680.1
bic:825
peaks after deleting less than one fifth of global max:10
×10^7
raw data, starting:10
fitted Gauss locations:10
y
x
0
2
4
6
8
10
12
14
16
18
0
5
10
15
20
25

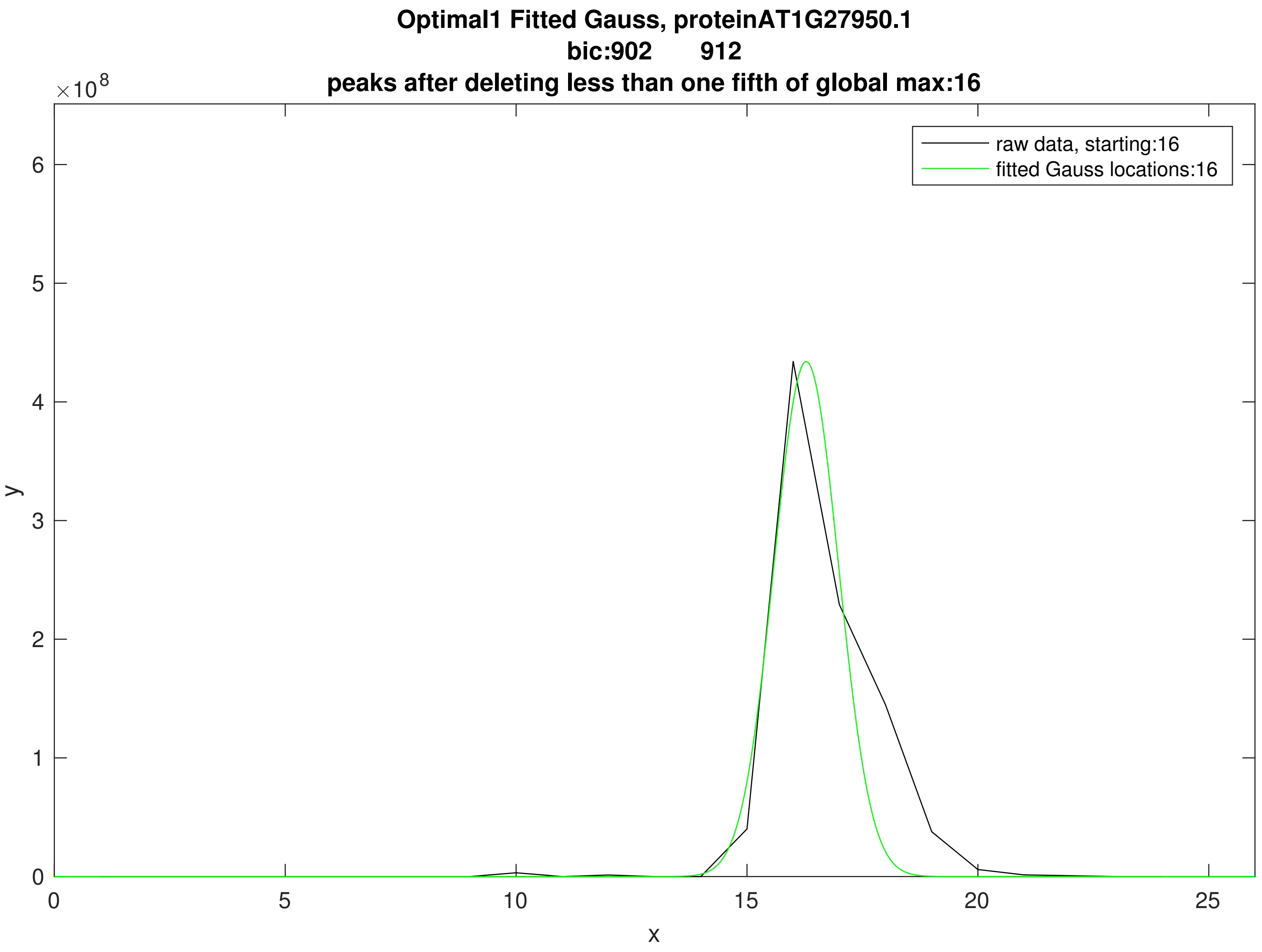

Optimal1 Fitted Gauss, proteinAT1G27950.1
bic:902 912
peaks after deleting less than one fifth of global max:16
×10^8
raw data, starting:16
fitted Gauss locations:16
y
x
0
1
2
3
4
5
6
10
15
20
25

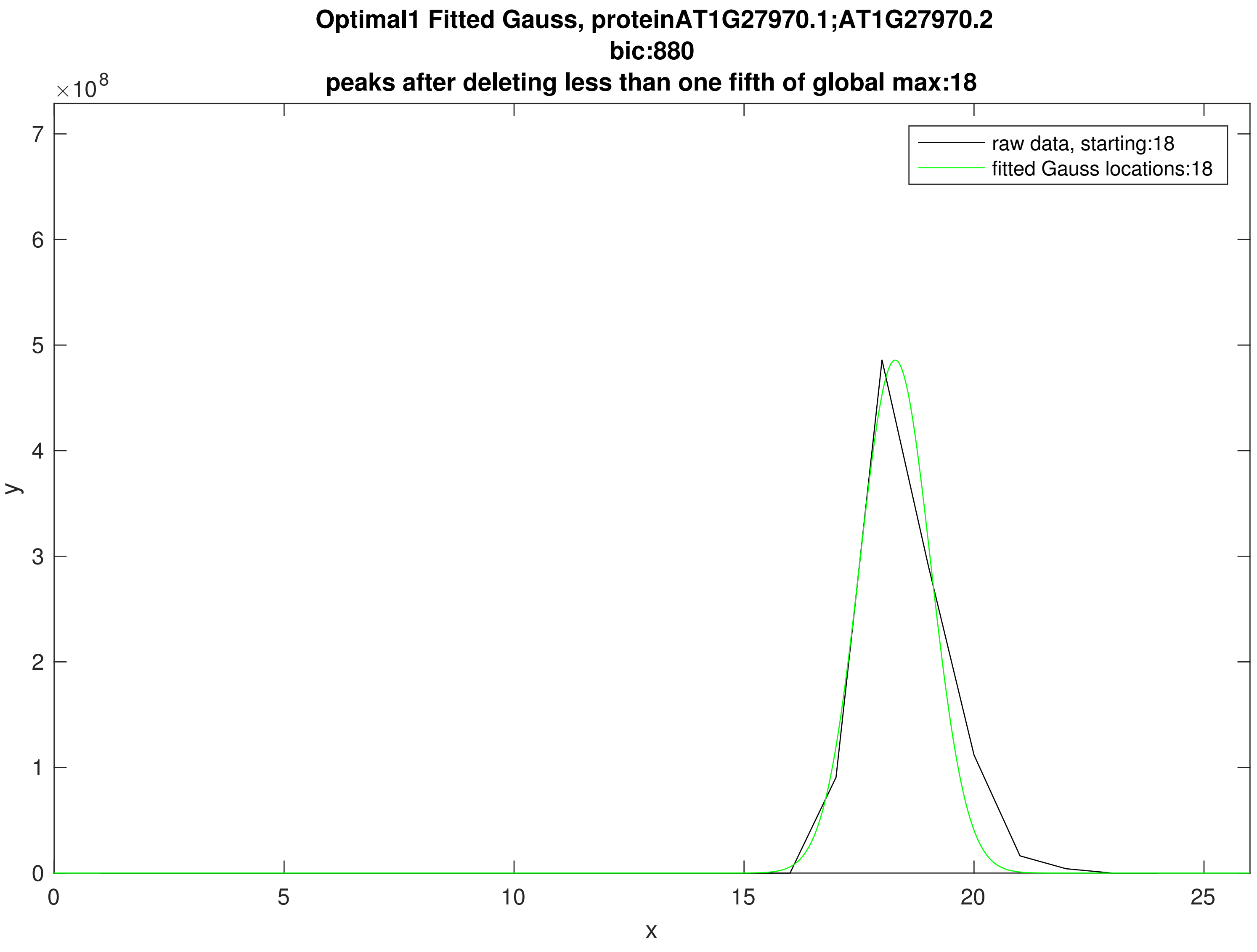
Optimal1 Fitted Gauss, proteinAT1G27970.1;AT1G27970.2
bic:880
peaks after deleting less than one fifth of global max:18
×10^8
raw data, starting:18
fitted Gauss locations:18
y
x
0
1
2
3
4
5
6
7
0
5
10
15
20
25

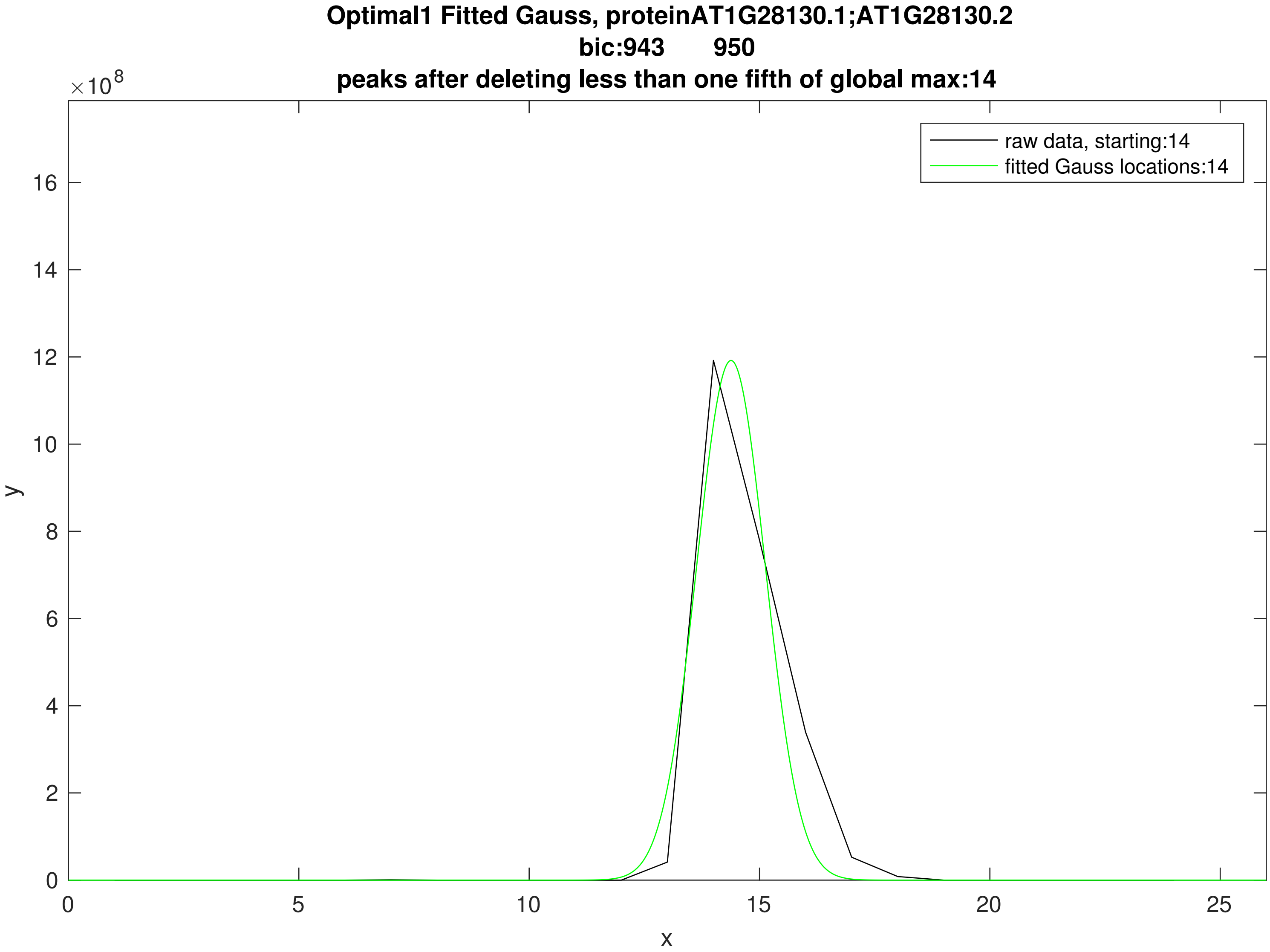
Optimal1 Fitted Gauss, proteinAT1G28130.1;AT1G28130.2
bic:943 950
peaks after deleting less than one fifth of global max:14
×10^8
16
14
12
10
8
6
4
2
0
y
0
5
10
15
20
25
x
raw data, starting:14
fitted Gauss locations:14

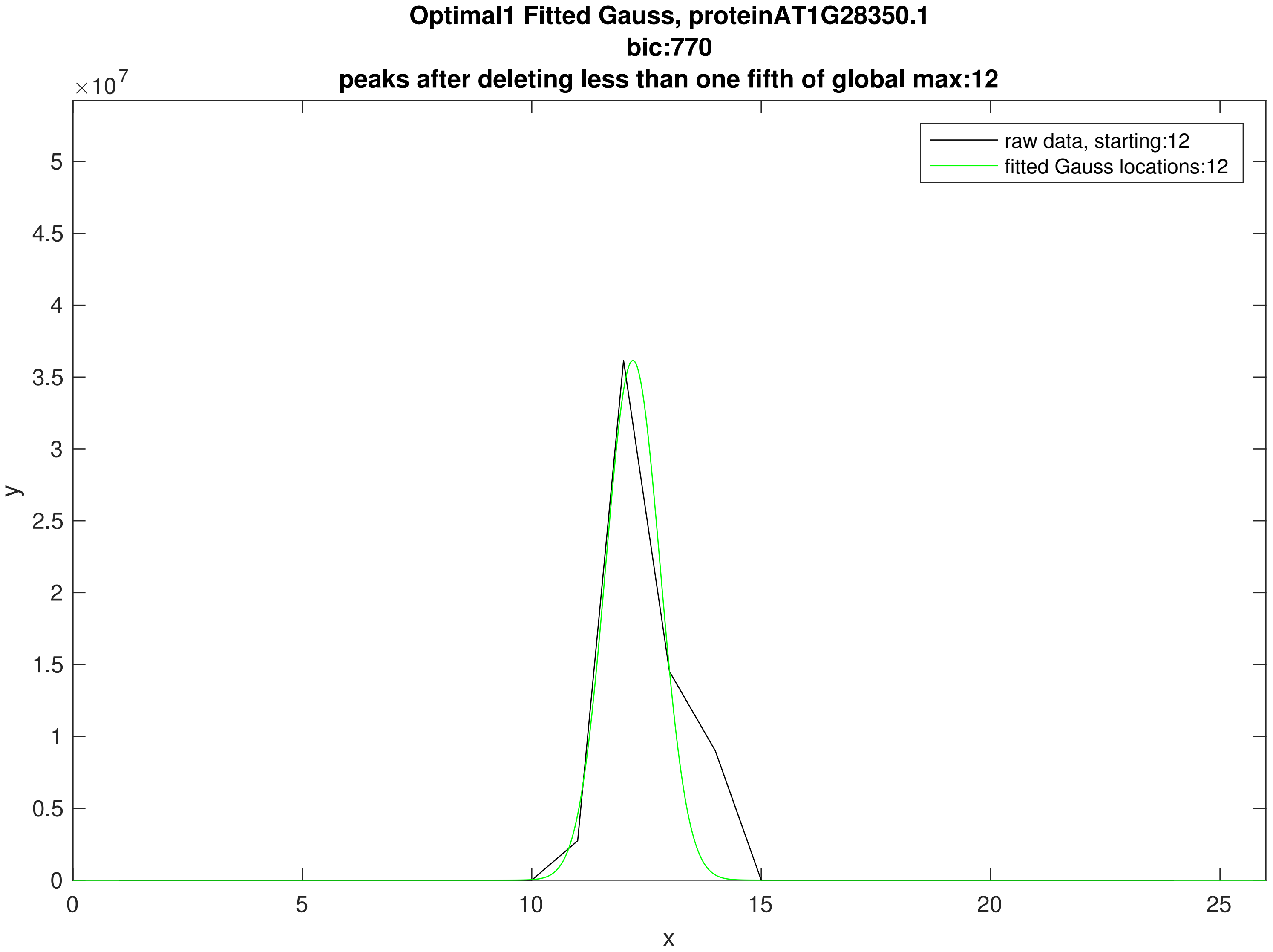
Optimal1 Fitted Gauss, proteinAT1G28350.1
bic:770
peaks after deleting less than one fifth of global max:12
×10^7
5
4.5
4
3.5
3
2.5
2
1.5
1
0.5
0
y
0
5
10
15
20
25
x
raw data, starting:12
fitted Gauss locations:12

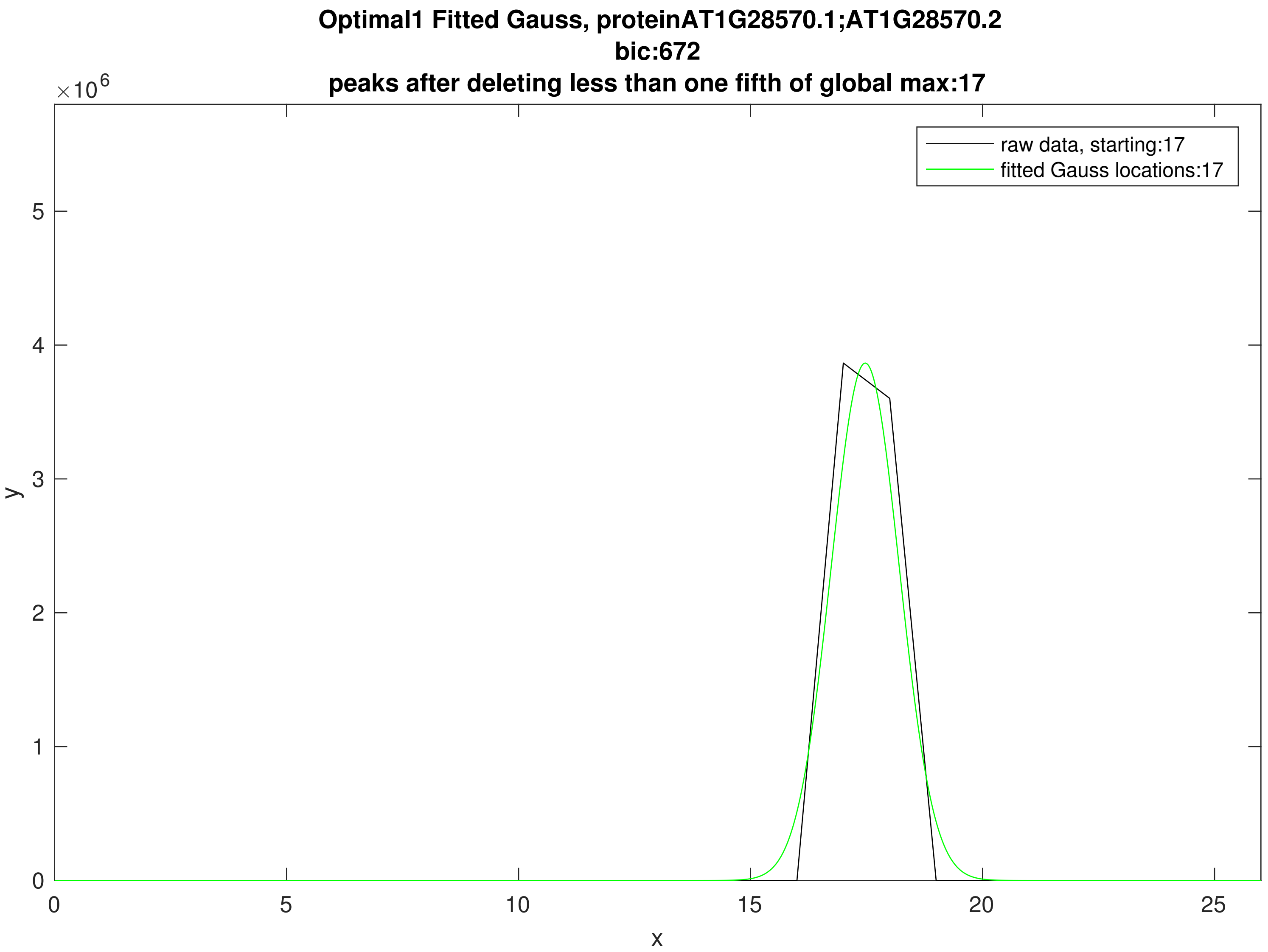
Optimal1 Fitted Gauss, proteinAT1G28570.1;AT1G28570.2
bic:672
peaks after deleting less than one fifth of global max:17
×10^6
raw data, starting:17
fitted Gauss locations:17
y
x
0
1
2
3
4
5
0
5
10
15
20
25

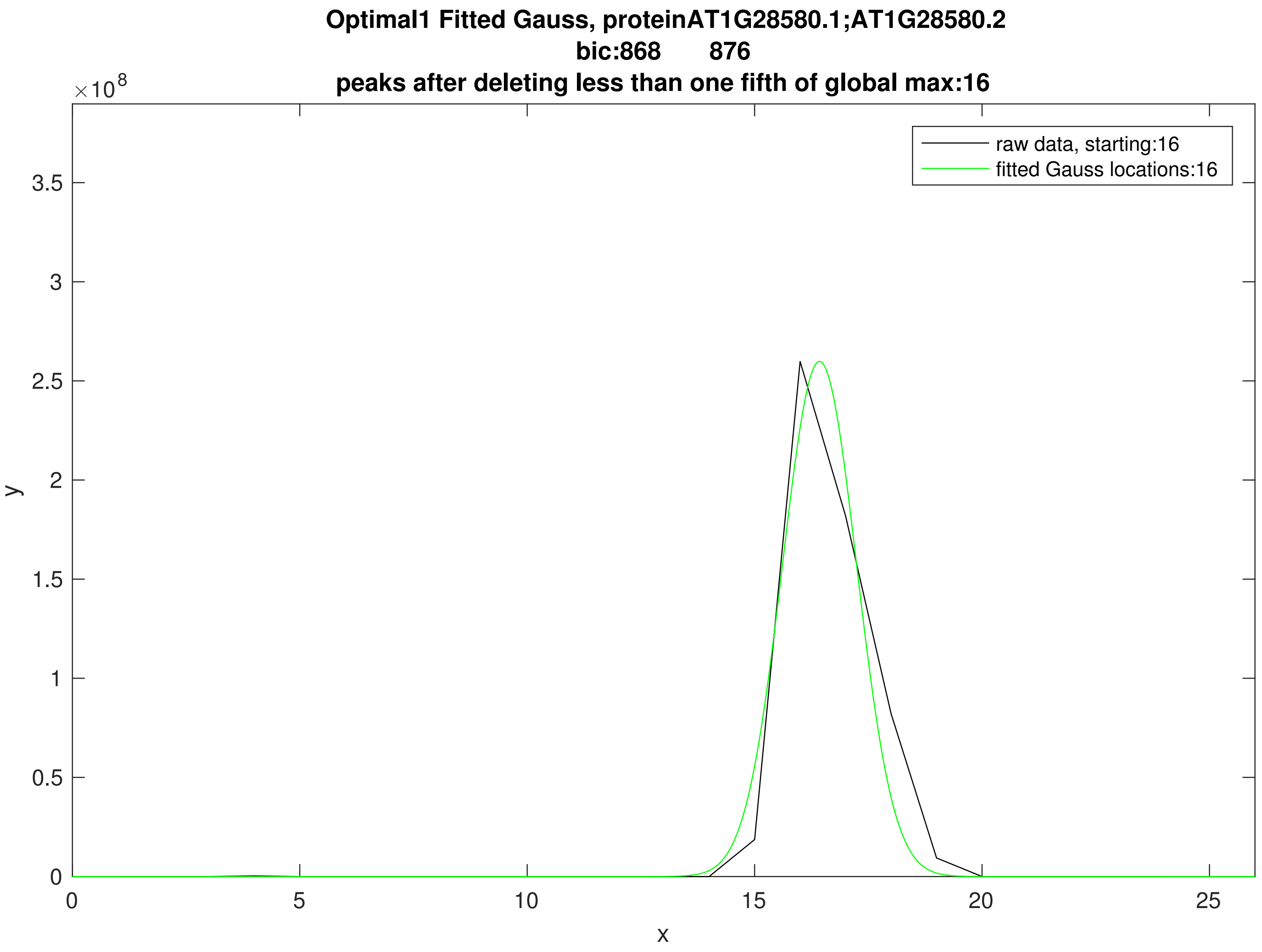

Optimal1 Fitted Gauss, proteinAT1G28580.1;AT1G28580.2
bic:868 876
peaks after deleting less than one fifth of global max:16
×10^8
3.5
3
2.5
2
1.5
1
0.5
0
y
0
5
10
15
20
25
x
raw data, starting:16
fitted Gauss locations:16

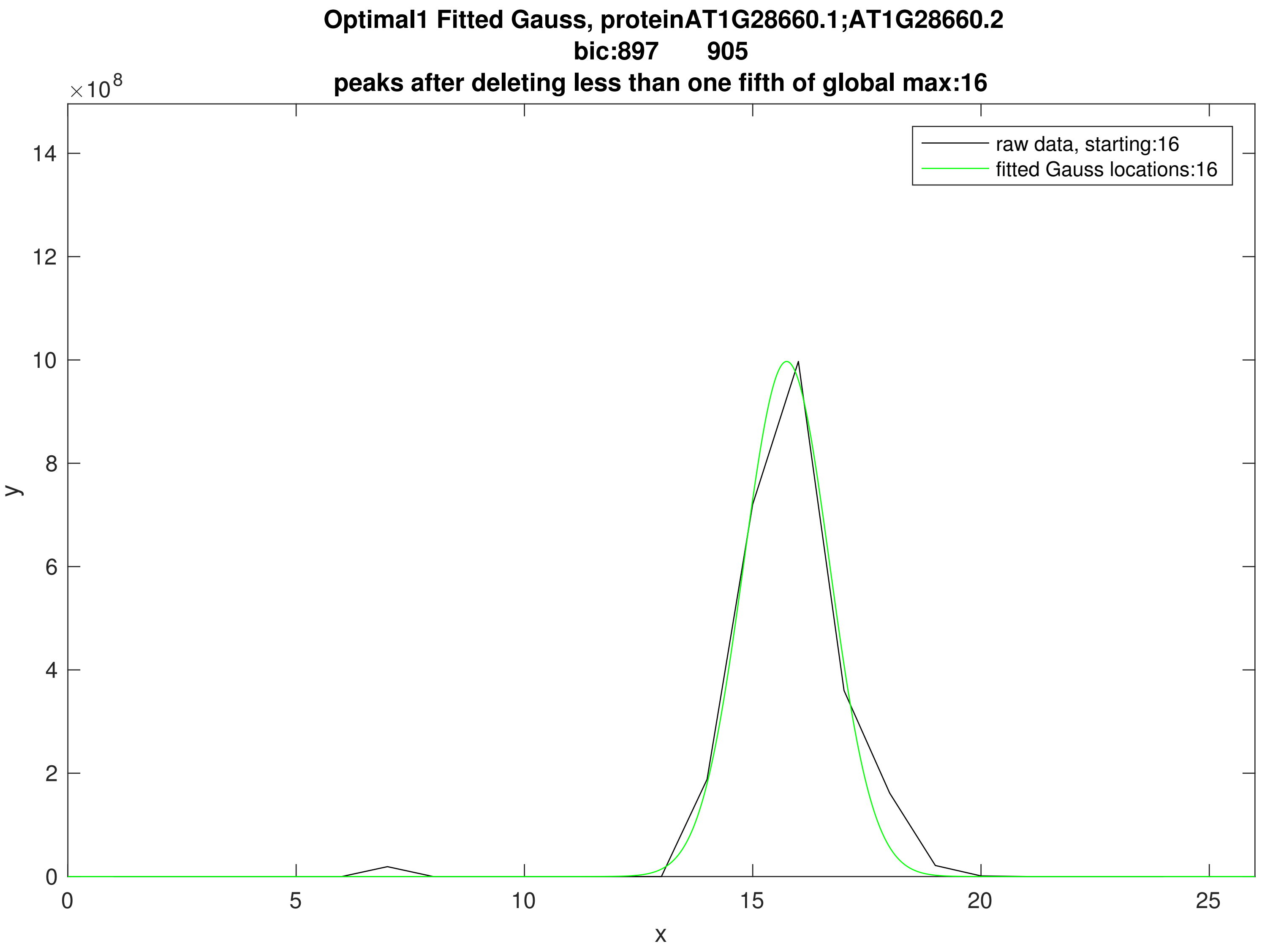

Optimal1 Fitted Gauss, proteinAT1G28660.1;AT1G28660.2
bic:897 905
peaks after deleting less than one fifth of global max:16
×10^8
raw data, starting:16
fitted Gauss locations:16
y
x
0
2
4
6
8
10
12
14
0
5
10
15
20
25

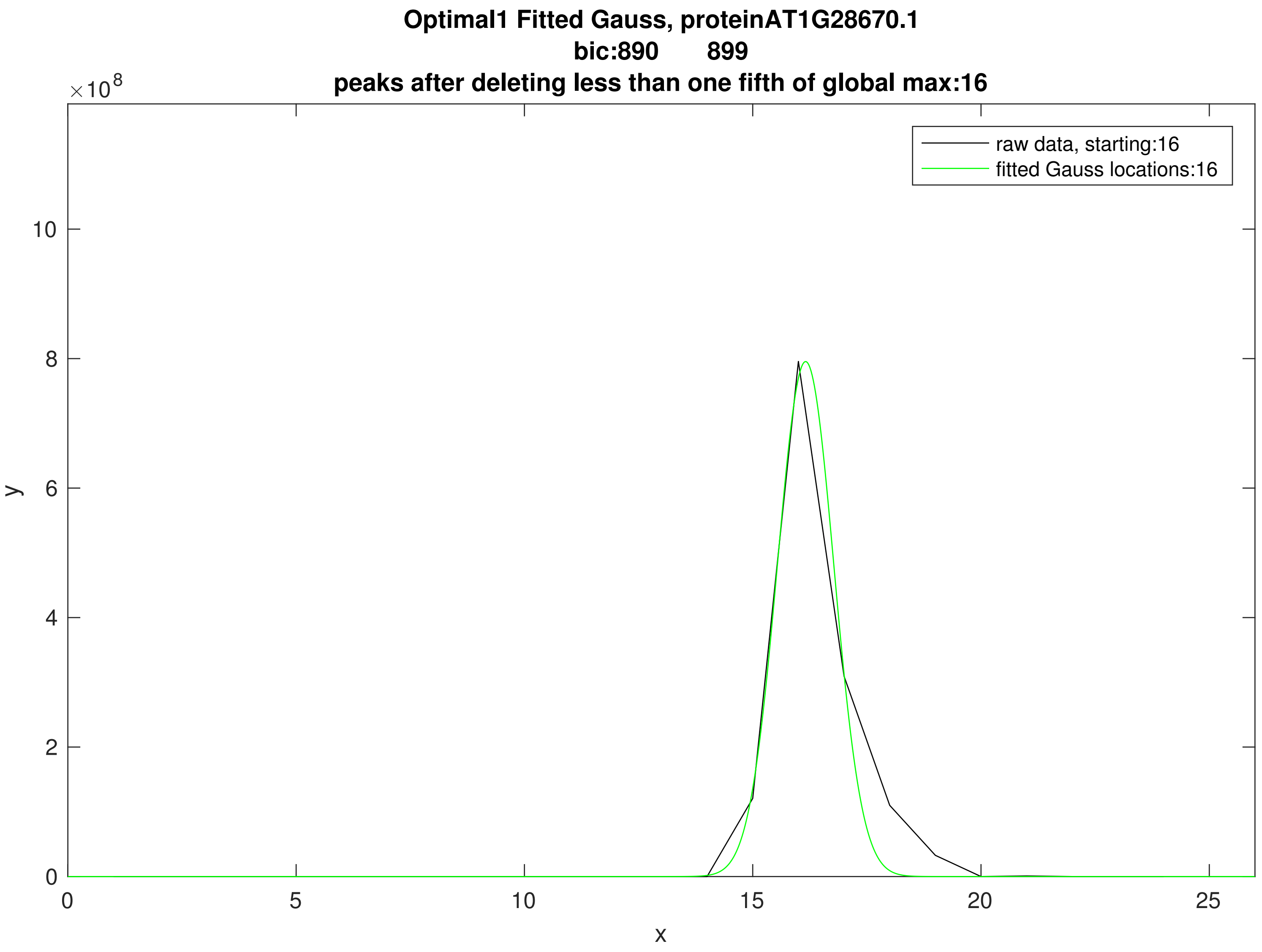

Optimal1 Fitted Gauss, proteinAT1G28670.1
bic:890 899
peaks after deleting less than one fifth of global max:16
×10^8
raw data, starting:16
fitted Gauss locations:16
y
x
0
2
4
6
8
10
0
5
10
15
20
25

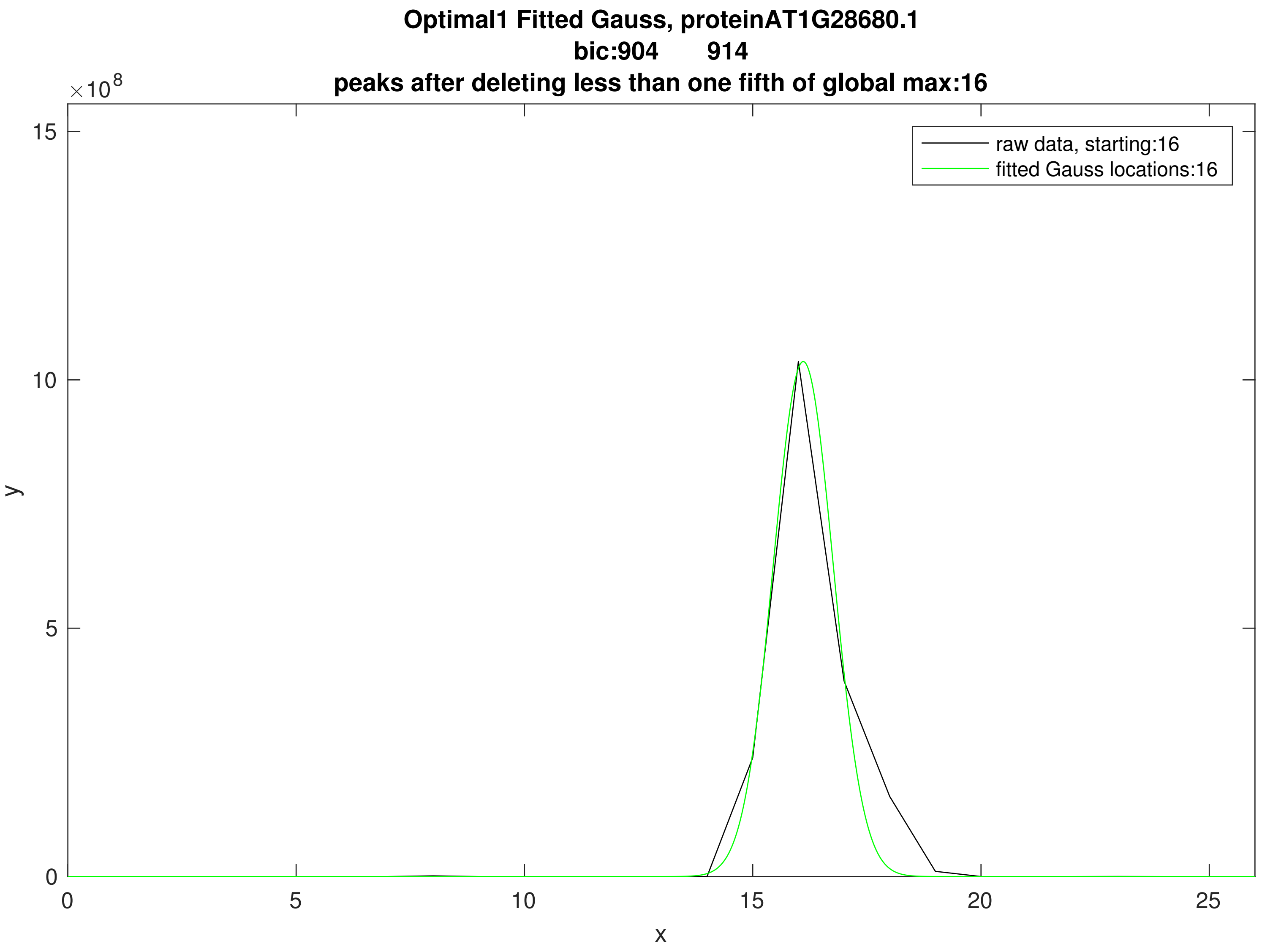
Optimal1 Fitted Gauss, proteinAT1G28680.1
bic:904 914
peaks after deleting less than one fifth of global max:16
×10^8
15
10
5
0
y
0
5
10
15
20
25
x
raw data, starting:16
fitted Gauss locations:16

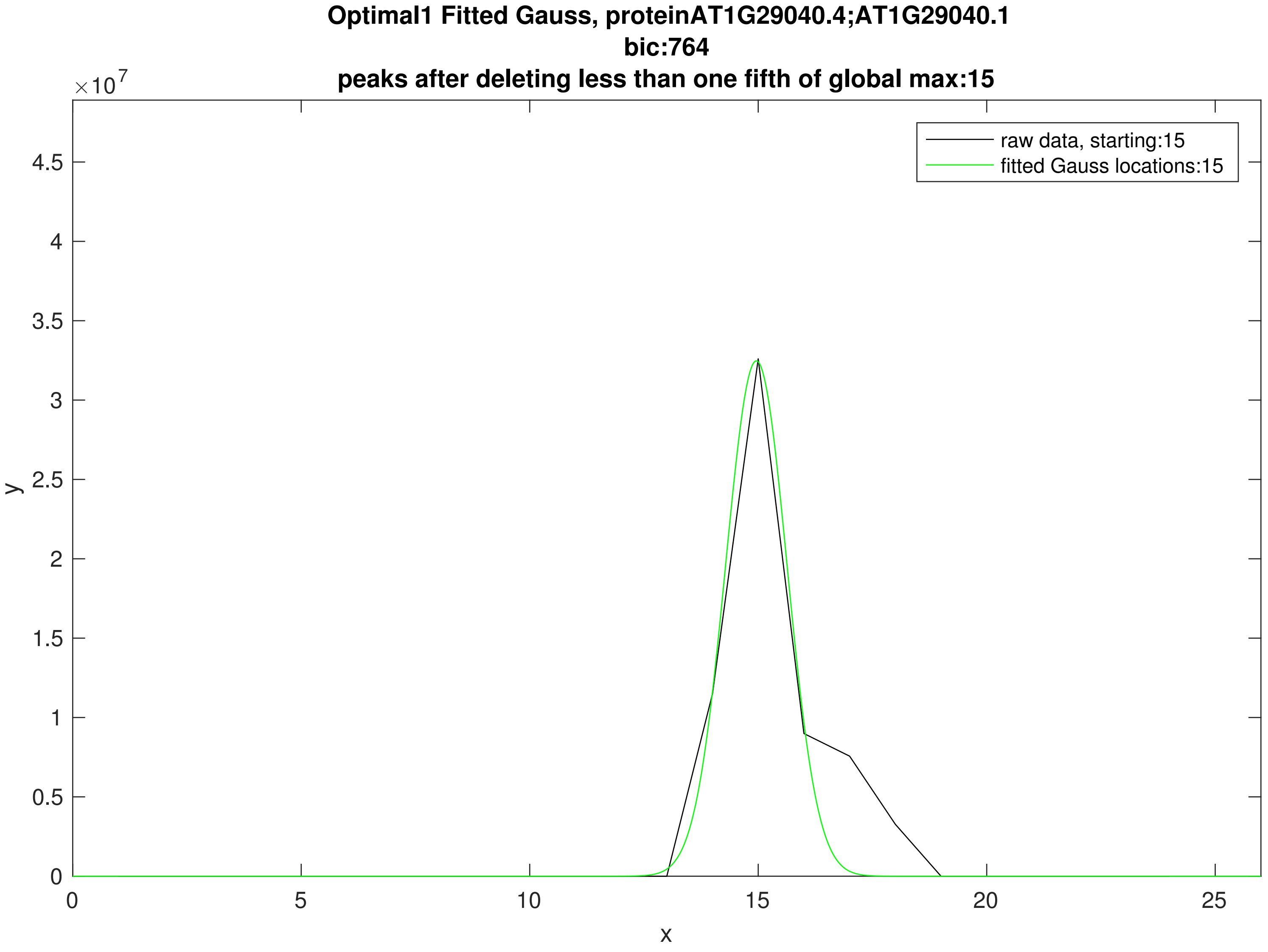
Optimal1 Fitted Gauss, proteinAT1G29040.4;AT1G29040.1
bic:764
peaks after deleting less than one fifth of global max:15
×10^7
raw data, starting:15
fitted Gauss locations:15
y
x
0
0.5
1
1.5
2
2.5
3
3.5
4
4.5
5
10
15
20
25

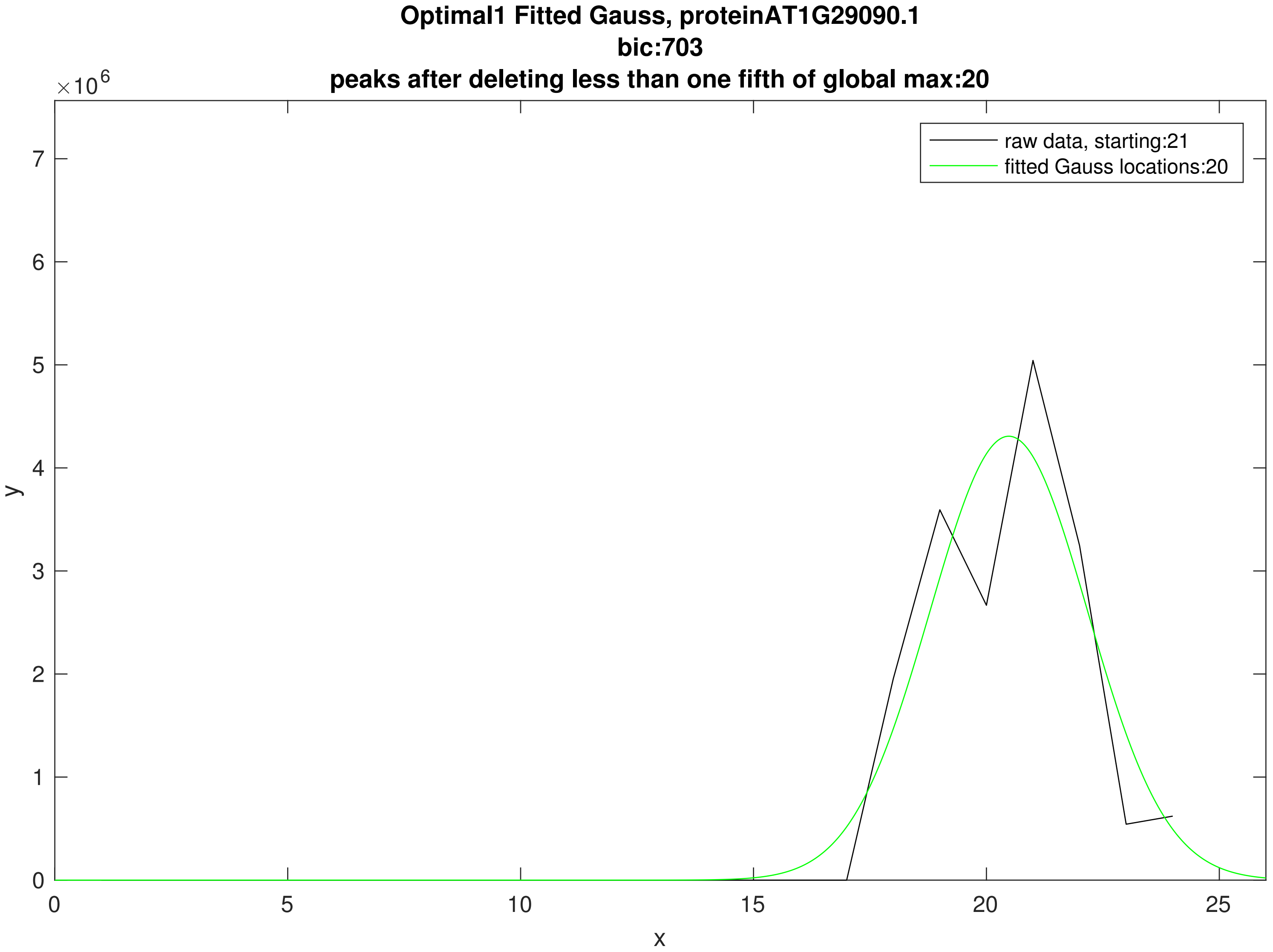
Optimal1 Fitted Gauss, proteinAT1G29090.1
bic:703
peaks after deleting less than one fifth of global max:20
×10^6
raw data, starting:21
fitted Gauss locations:20
y
x
0
1
2
3
4
5
6
7
0
5
10
15
20
25

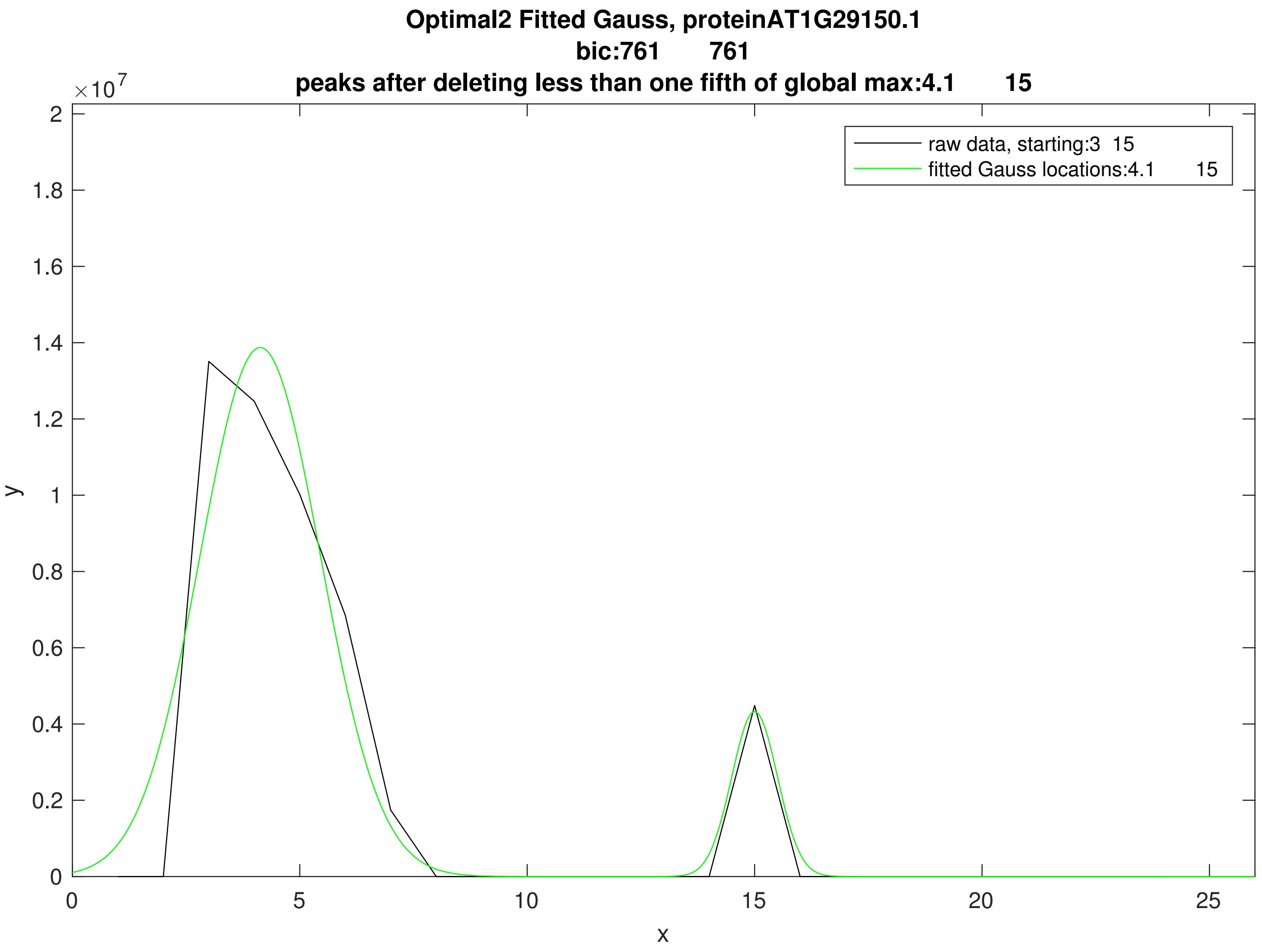

Optimal2 Fitted Gauss, proteinAT1G29150.1
bic:761 761
peaks after deleting less than one fifth of global max:4.1 15
×10^7
raw data, starting:3 15
fitted Gauss locations:4.1 15
y
x
0
0.2
0.4
0.6
0.8
1
1.2
1.4
1.6
1.8
2
5
10
15
20
25

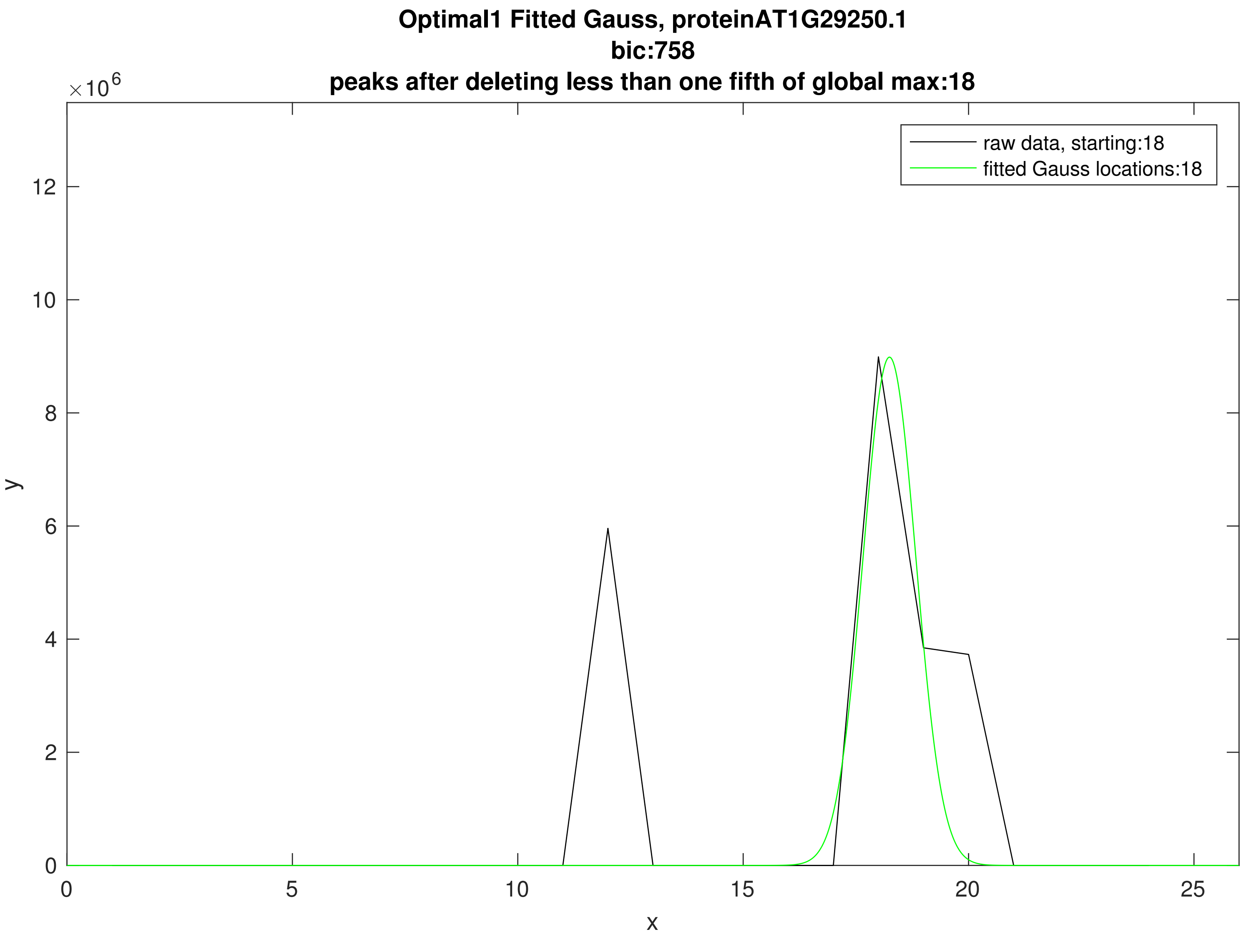
Optimal1 Fitted Gauss, proteinAT1G29250.1
bic:758
peaks after deleting less than one fifth of global max:18
×10^6
raw data, starting:18
fitted Gauss locations:18
12
10
8
6
4
2
0
y
0
5
10
15
20
25
x

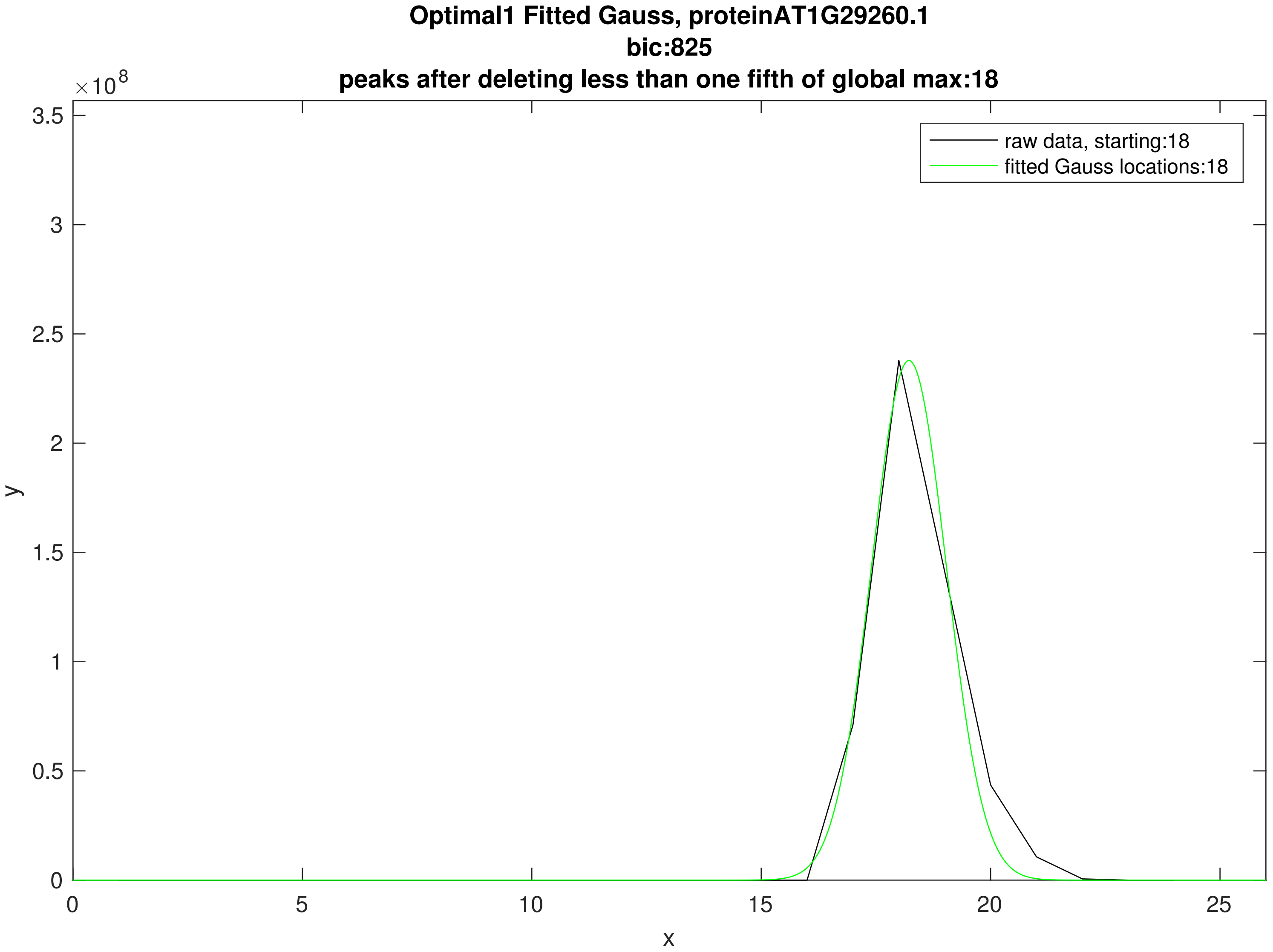

Optimal1 Fitted Gauss, proteinAT1G29260.1
bic:825
peaks after deleting less than one fifth of global max:18
×10^8
3.5
3
2.5
2
1.5
1
0.5
0
y
0
5
10
15
20
25
x
raw data, starting:18
fitted Gauss locations:18

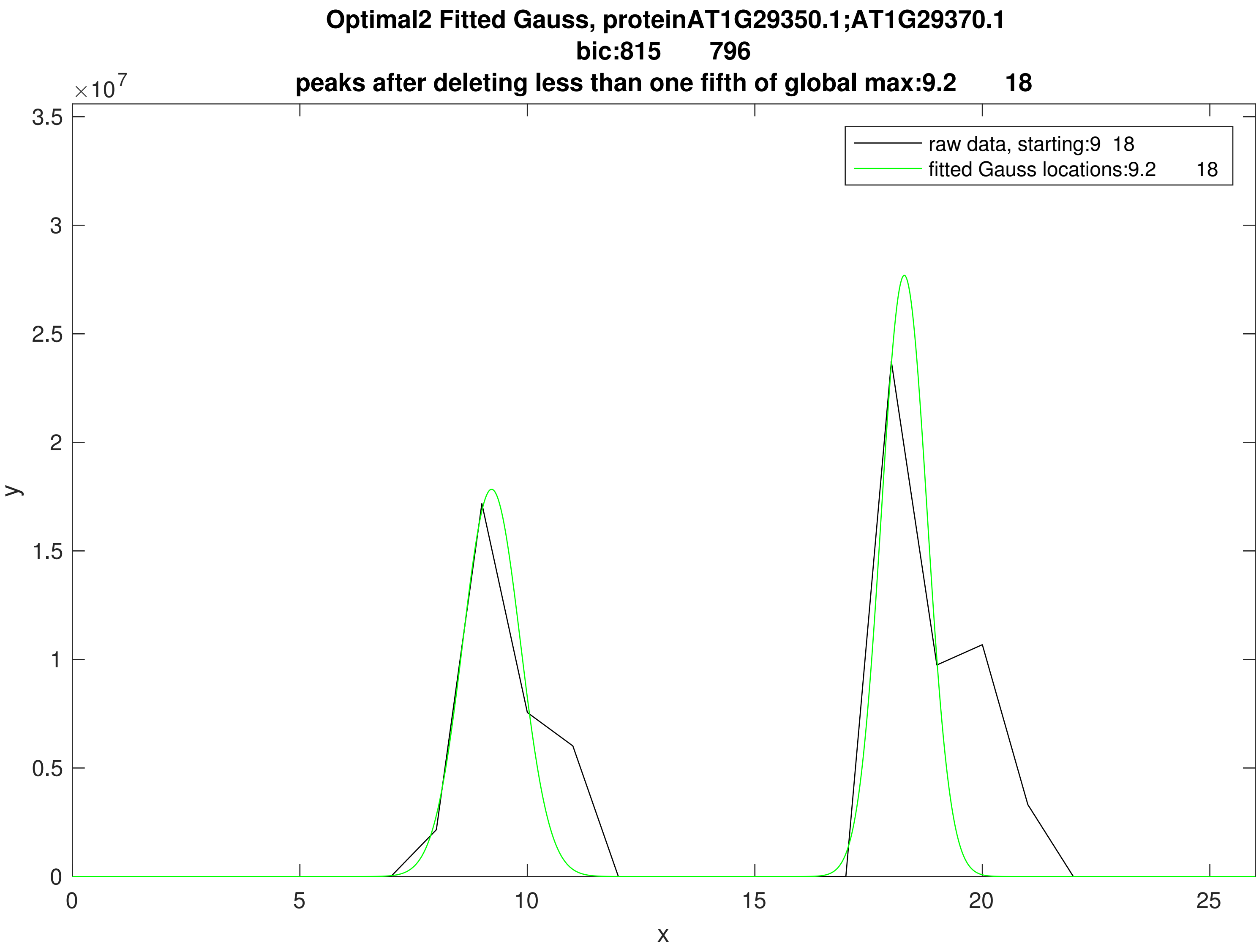

Optimal2 Fitted Gauss, proteinAT1G29350.1;AT1G29370.1
bic:815 796
peaks after deleting less than one fifth of global max:9.2 18
×10^7
3.5
3
2.5
2
1.5
1
0.5
0
y
0
5
10
15
20
25
x
raw data, starting:9 18
fitted Gauss locations:9.2 18

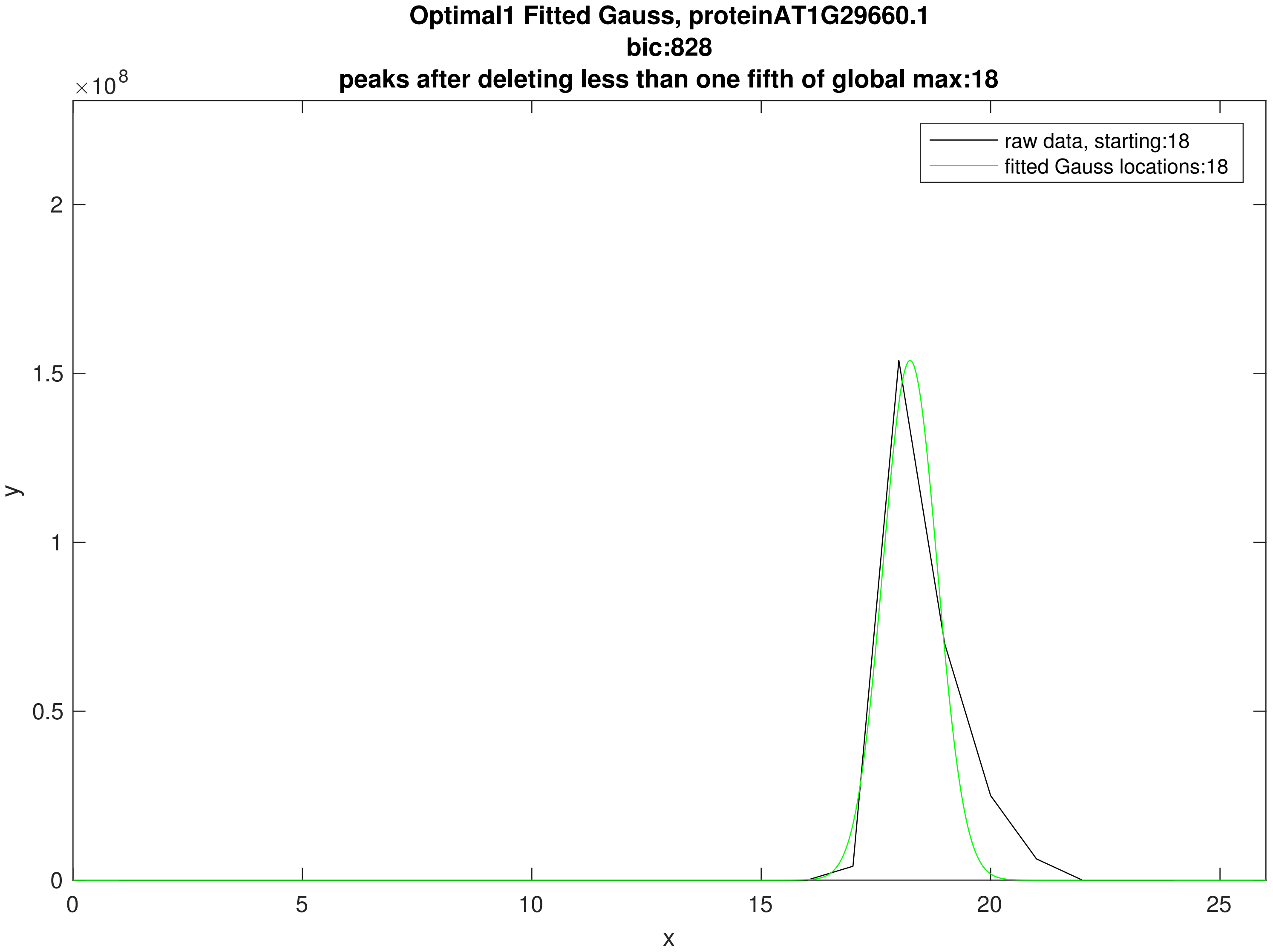

Optimal1 Fitted Gauss, proteinAT1G29660.1
bic:828
peaks after deleting less than one fifth of global max:18
×10^8
raw data, starting:18
fitted Gauss locations:18
y
x
0
0.5
1
1.5
2
0
5
10
15
20
25

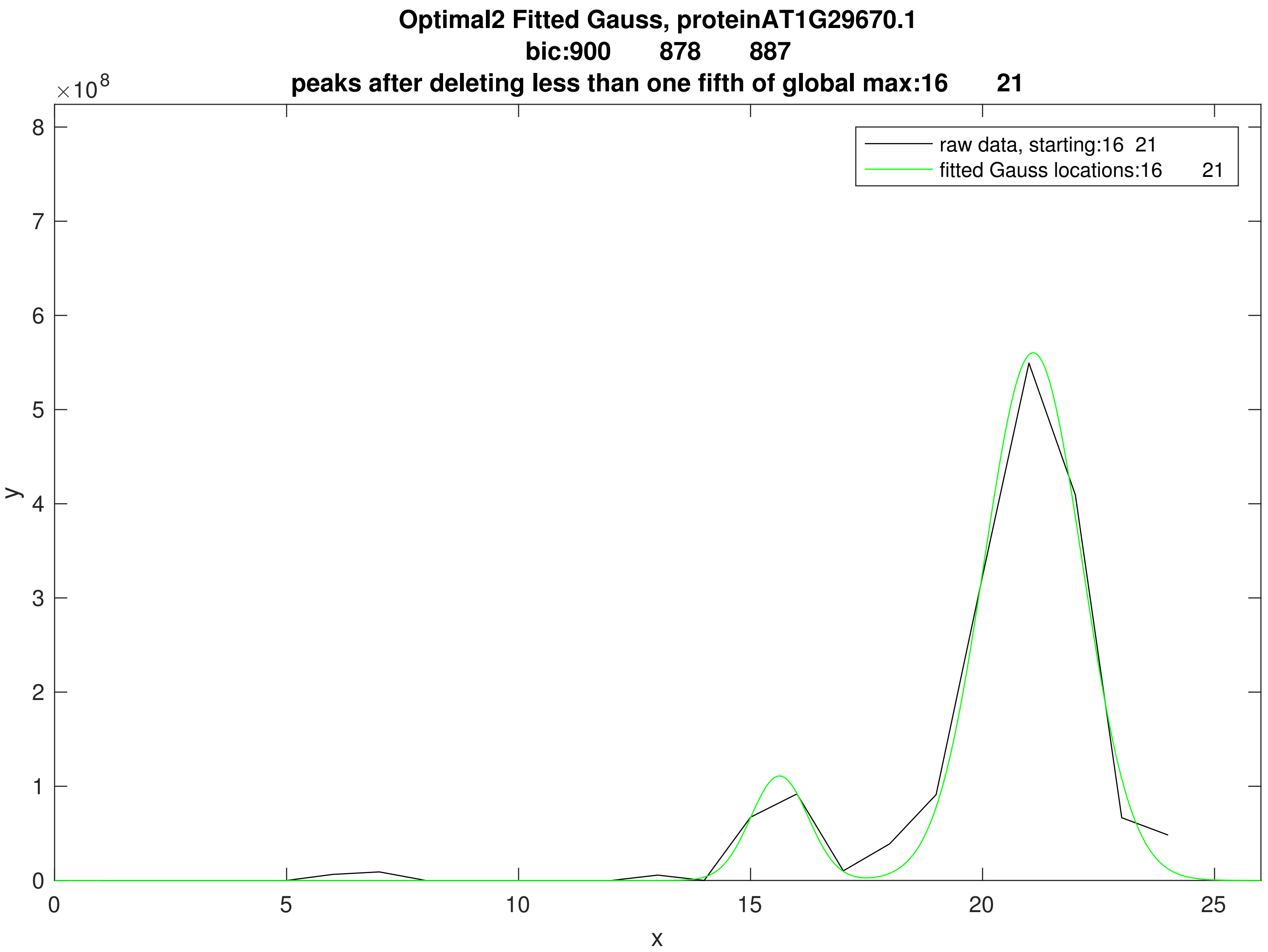
Optimal2 Fitted Gauss, proteinAT1G29670.1
bic:900 878 887
peaks after deleting less than one fifth of global max:16 21
×10^8
raw data, starting:16 21
fitted Gauss locations:16 21
y
x
0
1
2
3
4
5
6
7
8
0
5
10
15
20
25

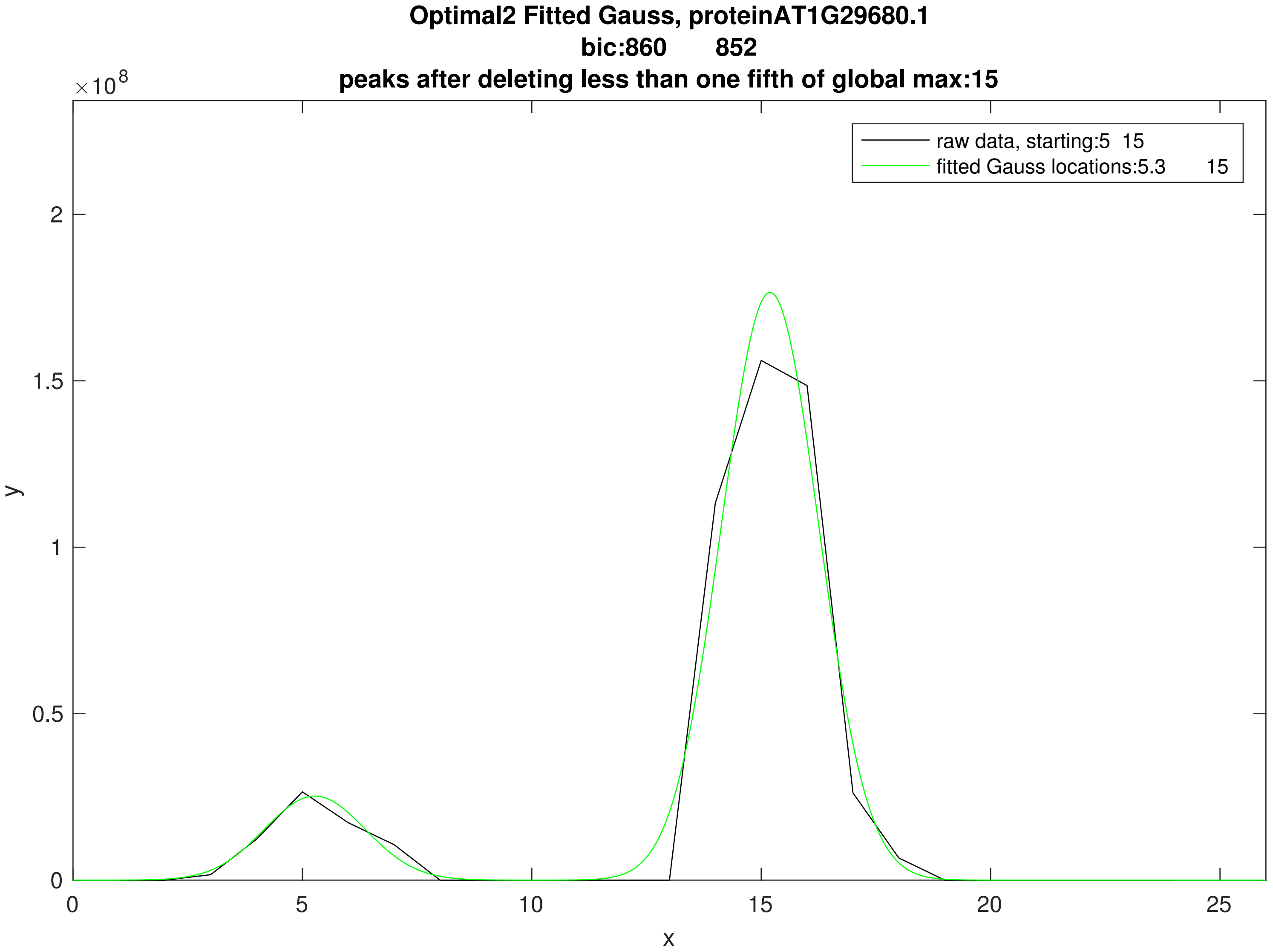

Optimal2 Fitted Gauss, proteinAT1G29680.1
bic:860 852
peaks after deleting less than one fifth of global max:15
×10^8
raw data, starting:5 15
fitted Gauss locations:5.3 15
y
x
0
0.5
1
1.5
2
0
5
10
15
20
25

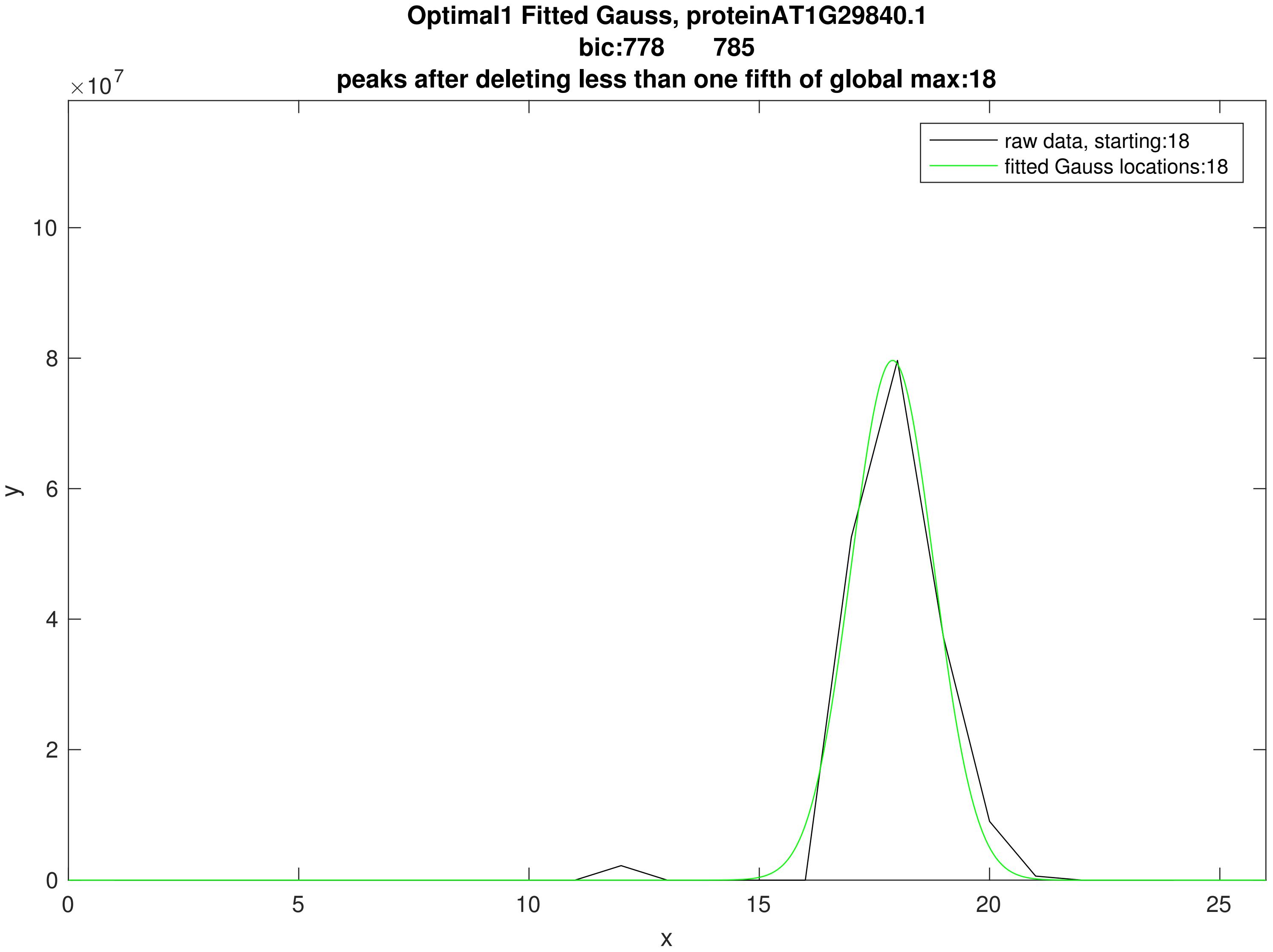
Optimal1 Fitted Gauss, proteinAT1G29840.1
bic:778 785
peaks after deleting less than one fifth of global max:18
×10^7
raw data, starting:18
fitted Gauss locations:18
10
8
6
4
2
0
y
0
5
10
15
20
25
x

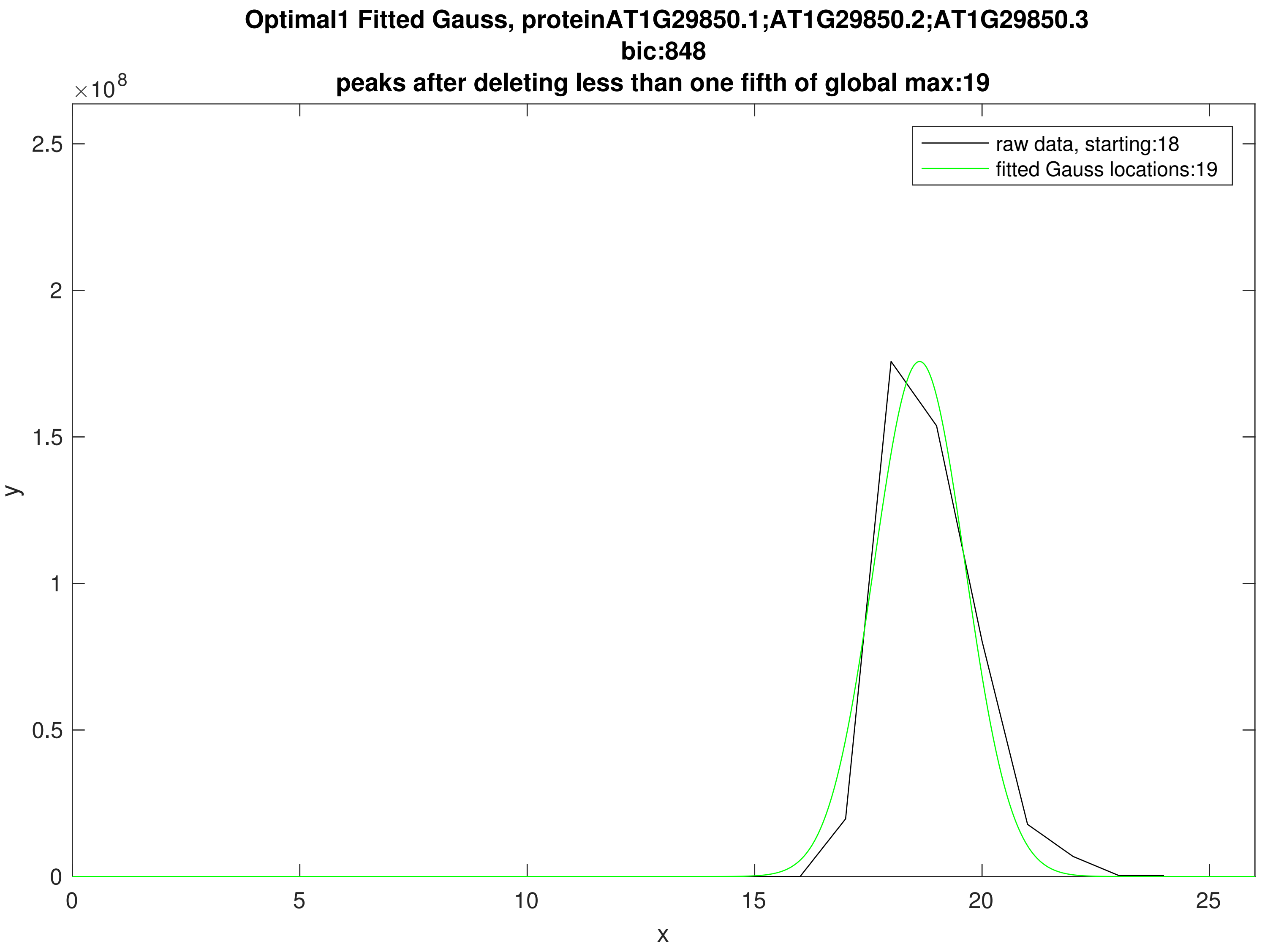

Optimal1 Fitted Gauss, proteinAT1G29850.1;AT1G29850.2;AT1G29850.3
bic:848
peaks after deleting less than one fifth of global max:19
×10^8
2.5
2
1.5
1
0.5
0
y
0
5
10
15
20
25
x
raw data, starting:18
fitted Gauss locations:19

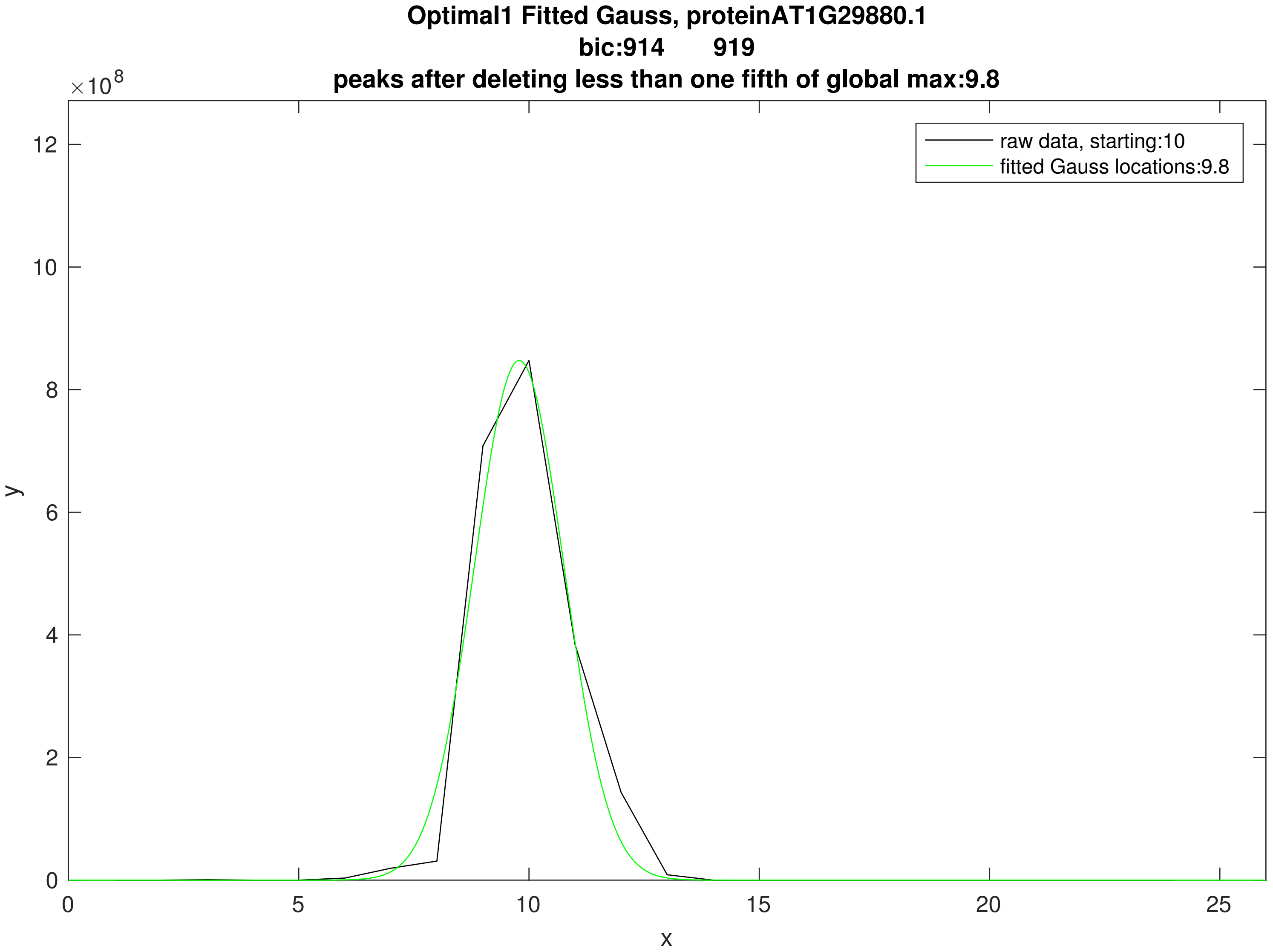
Optimal1 Fitted Gauss, proteinAT1G29880.1
bic:914 919
peaks after deleting less than one fifth of global max:9.8
×10^8
12
10
8
6
4
2
0
y
0
5
10
15
20
25
x
raw data, starting:10
fitted Gauss locations:9.8

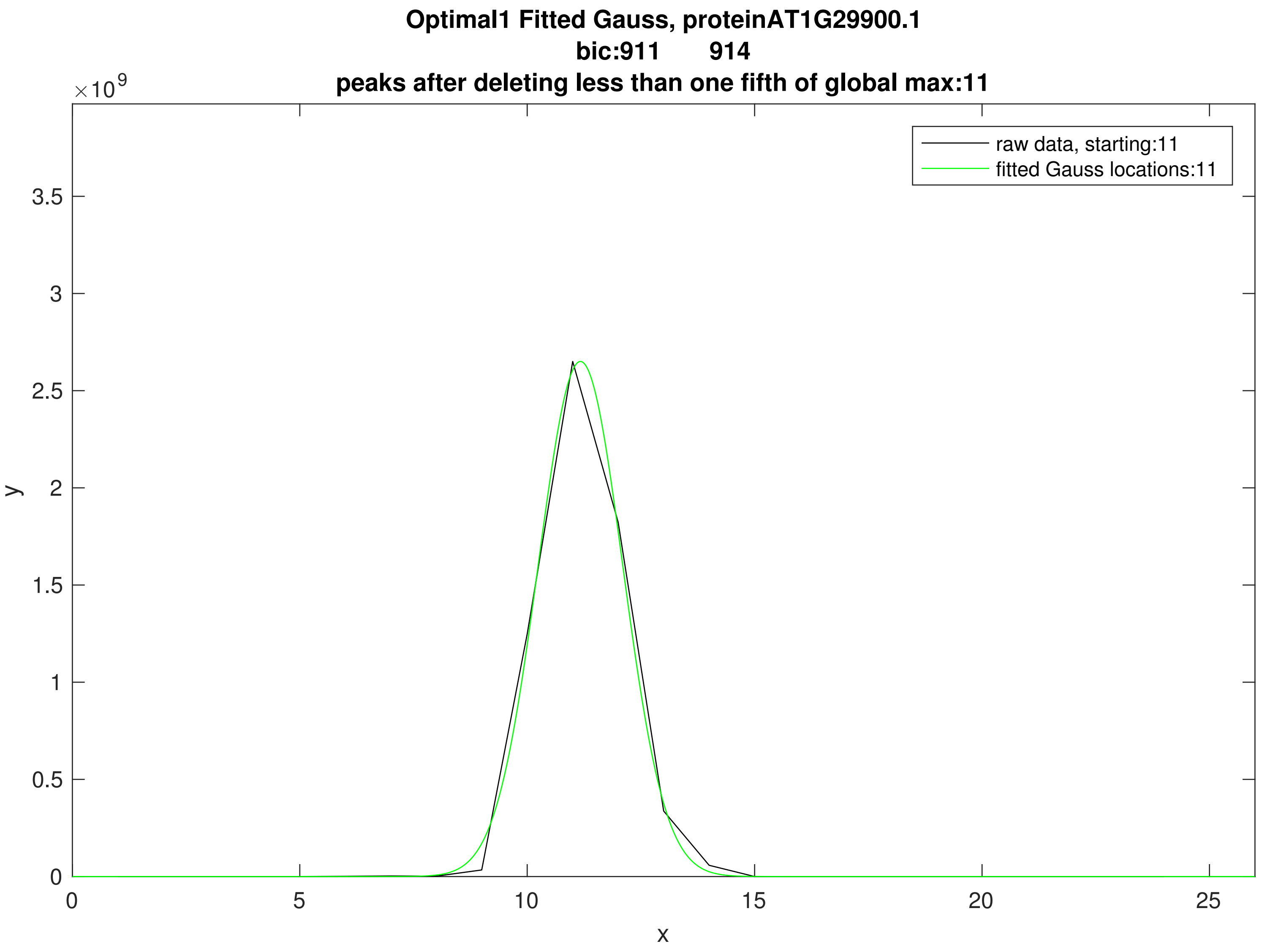
Optimal1 Fitted Gauss, proteinAT1G29900.1
bic:911 914
peaks after deleting less than one fifth of global max:11
×10^9
raw data, starting:11
fitted Gauss locations:11
3.5
3
2.5
2
1.5
1
0.5
0
y
0
5
10
15
20
25
x

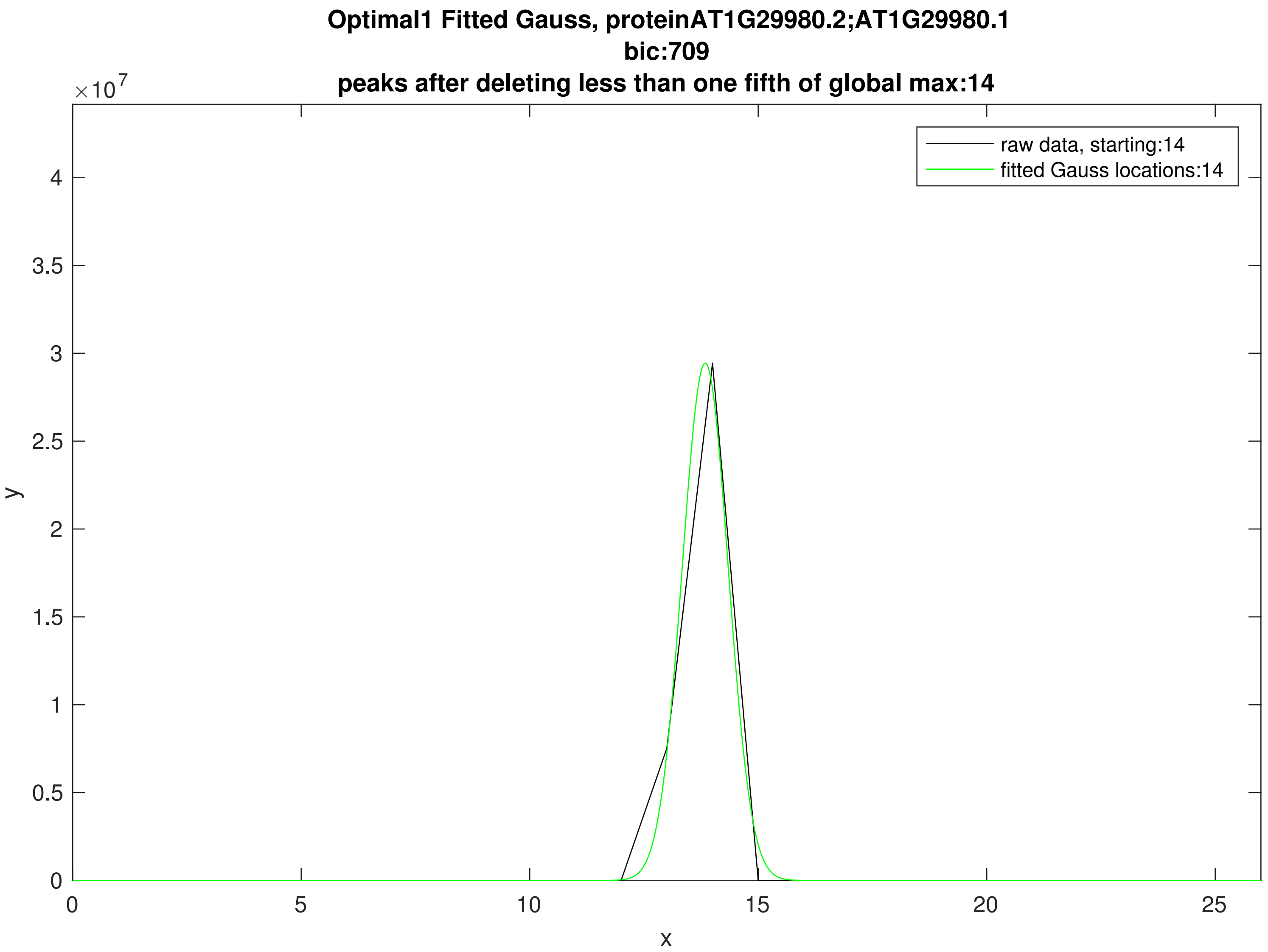
Optimal1 Fitted Gauss, proteinAT1G29980.2;AT1G29980.1
bic:709
peaks after deleting less than one fifth of global max:14
×10^7
raw data, starting:14
fitted Gauss locations:14
y
x
0
0.5
1
1.5
2
2.5
3
3.5
4
5
10
15
20
25

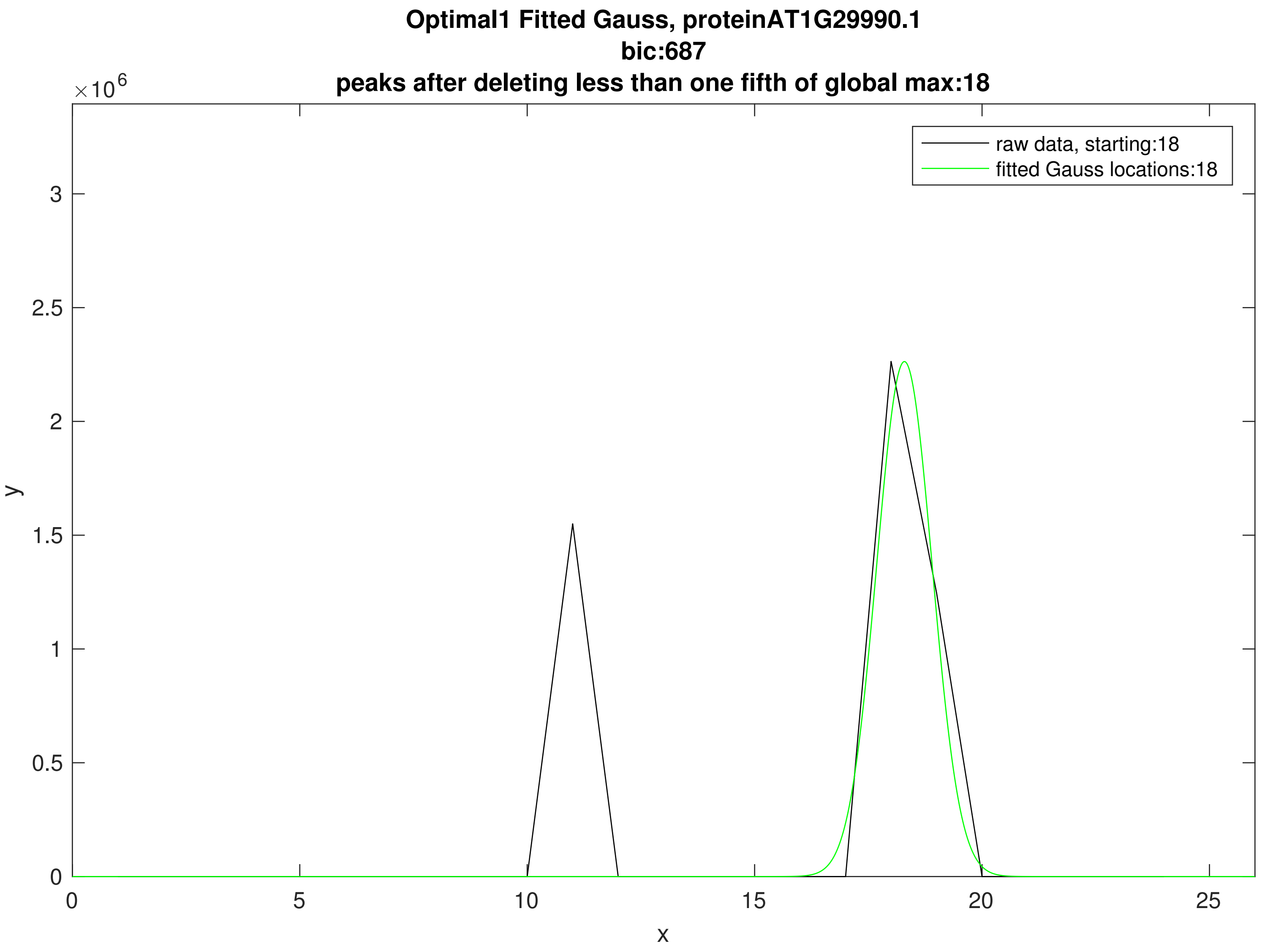
Optimal1 Fitted Gauss, proteinAT1G29990.1
bic:687
peaks after deleting less than one fifth of global max:18
×10^6
raw data, starting:18
fitted Gauss locations:18
y
x
3
2.5
2
1.5
1
0.5
0
5
10
15
20
25

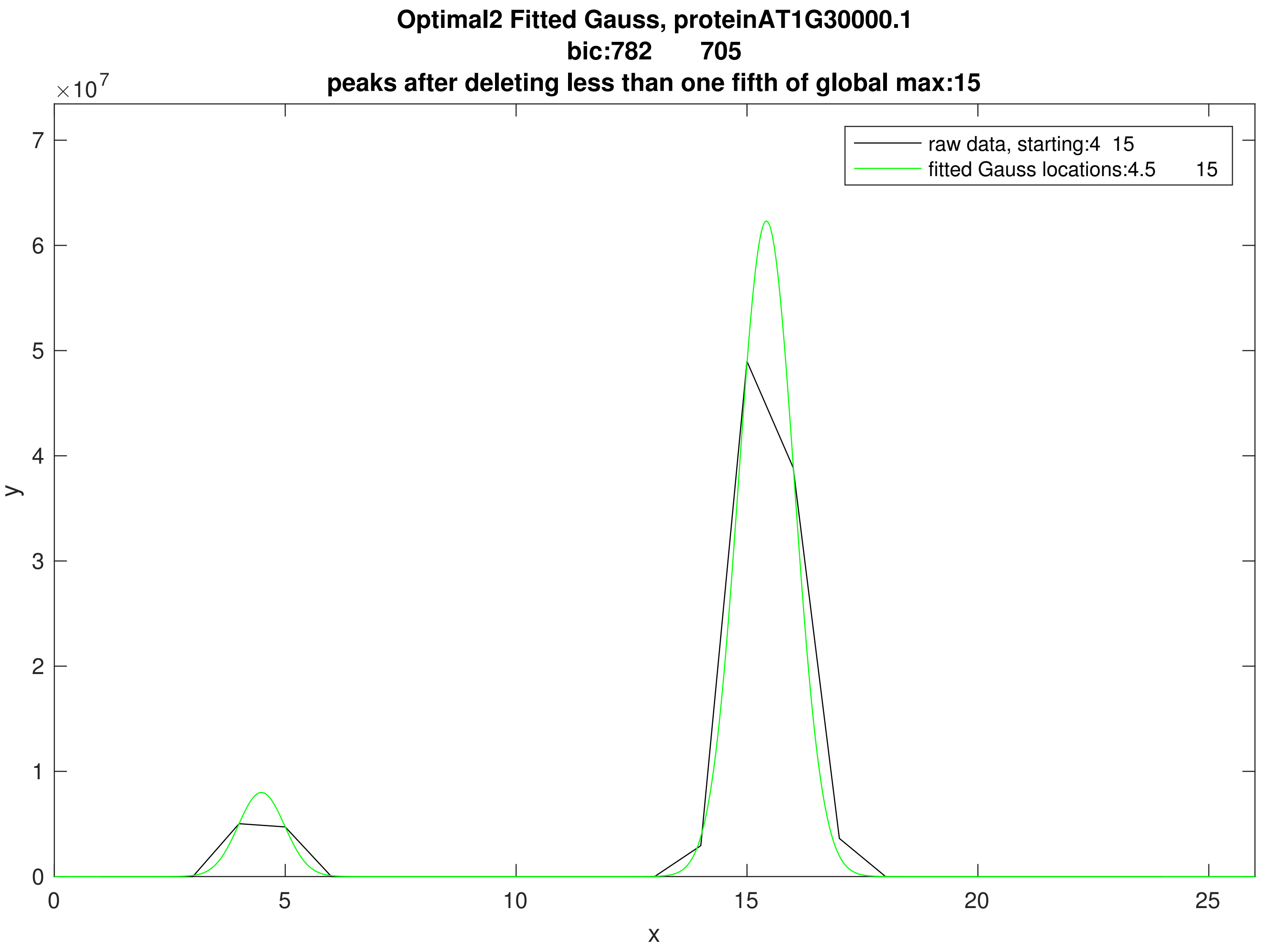
Optimal2 Fitted Gauss, proteinAT1G30000.1
bic:782 705
peaks after deleting less than one fifth of global max:15
×10^7
raw data, starting:4 15
fitted Gauss locations:4.5 15
y
x
0
1
2
3
4
5
6
7
10
15
20
25

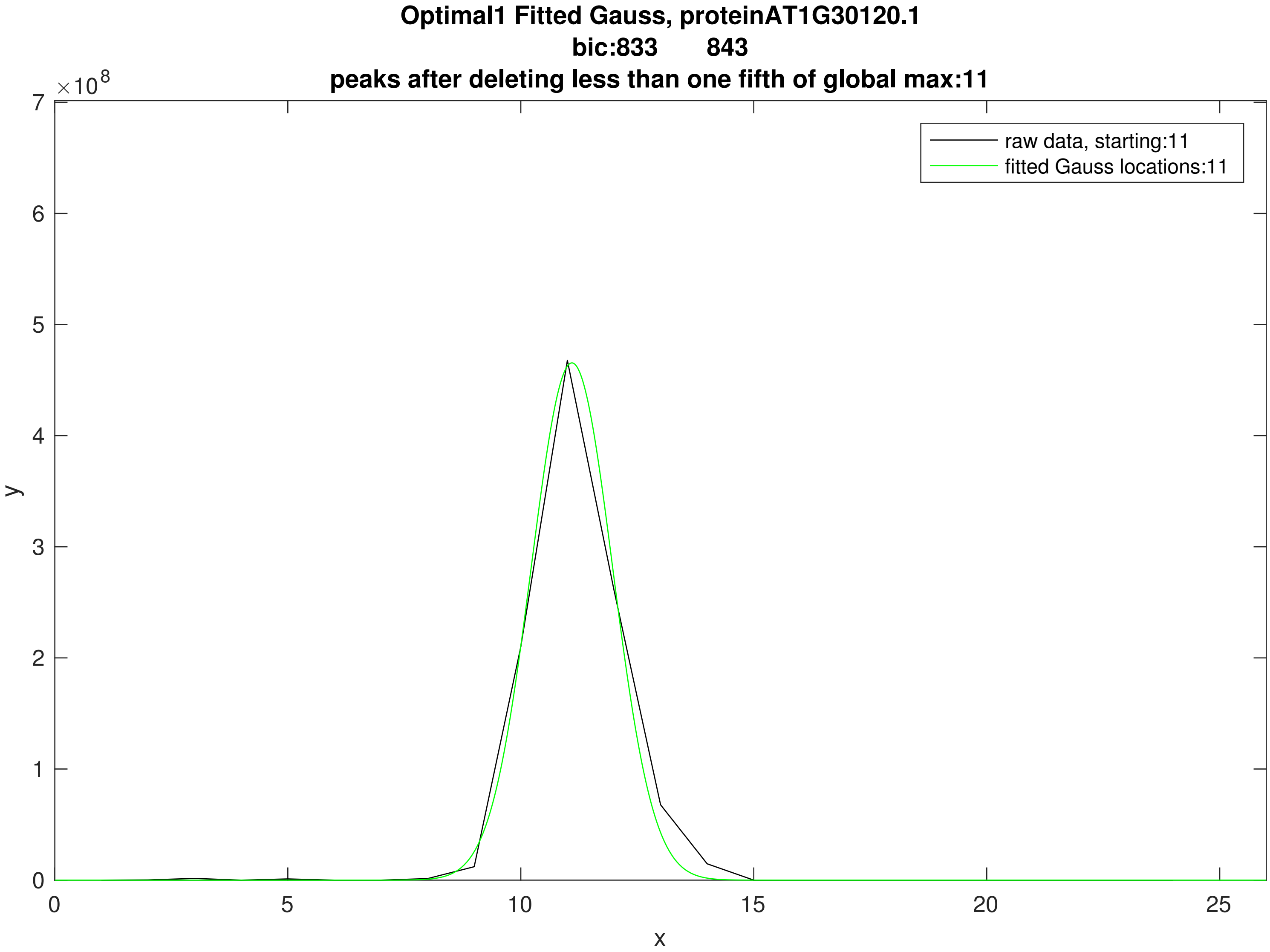

Optimal1 Fitted Gauss, proteinAT1G30120.1
bic:833 843
peaks after deleting less than one fifth of global max:11
×10^8
raw data, starting:11
fitted Gauss locations:11
y
x
0
1
2
3
4
5
6
7
0
5
10
15
20
25

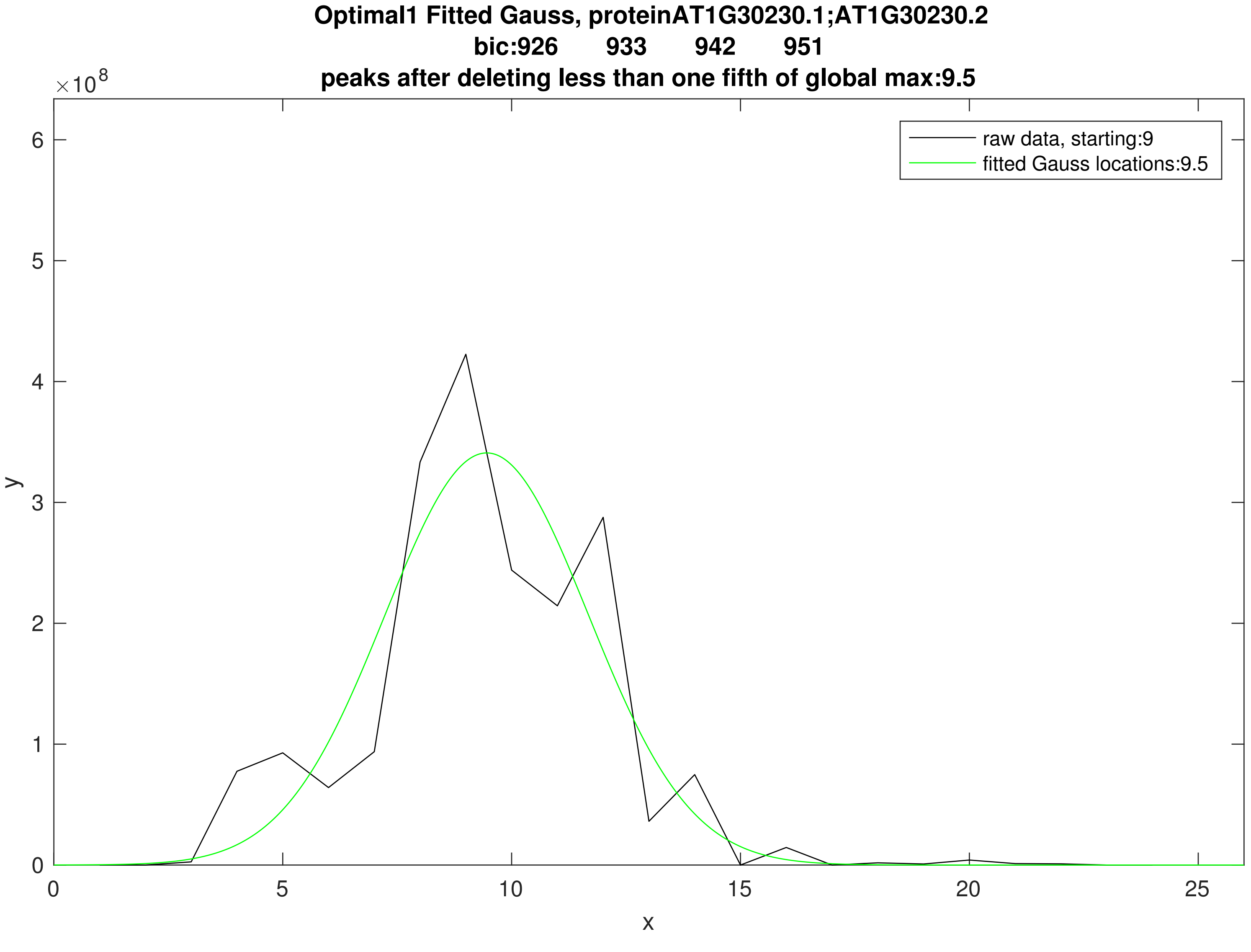
Optimal1 Fitted Gauss, proteinAT1G30230.1;AT1G30230.2
bic:926 933 942 951
peaks after deleting less than one fifth of global max:9.5
×10^8
y
x
raw data, starting:9
fitted Gauss locations:9.5
0
1
2
3
4
5
6
0
5
10
15
20
25

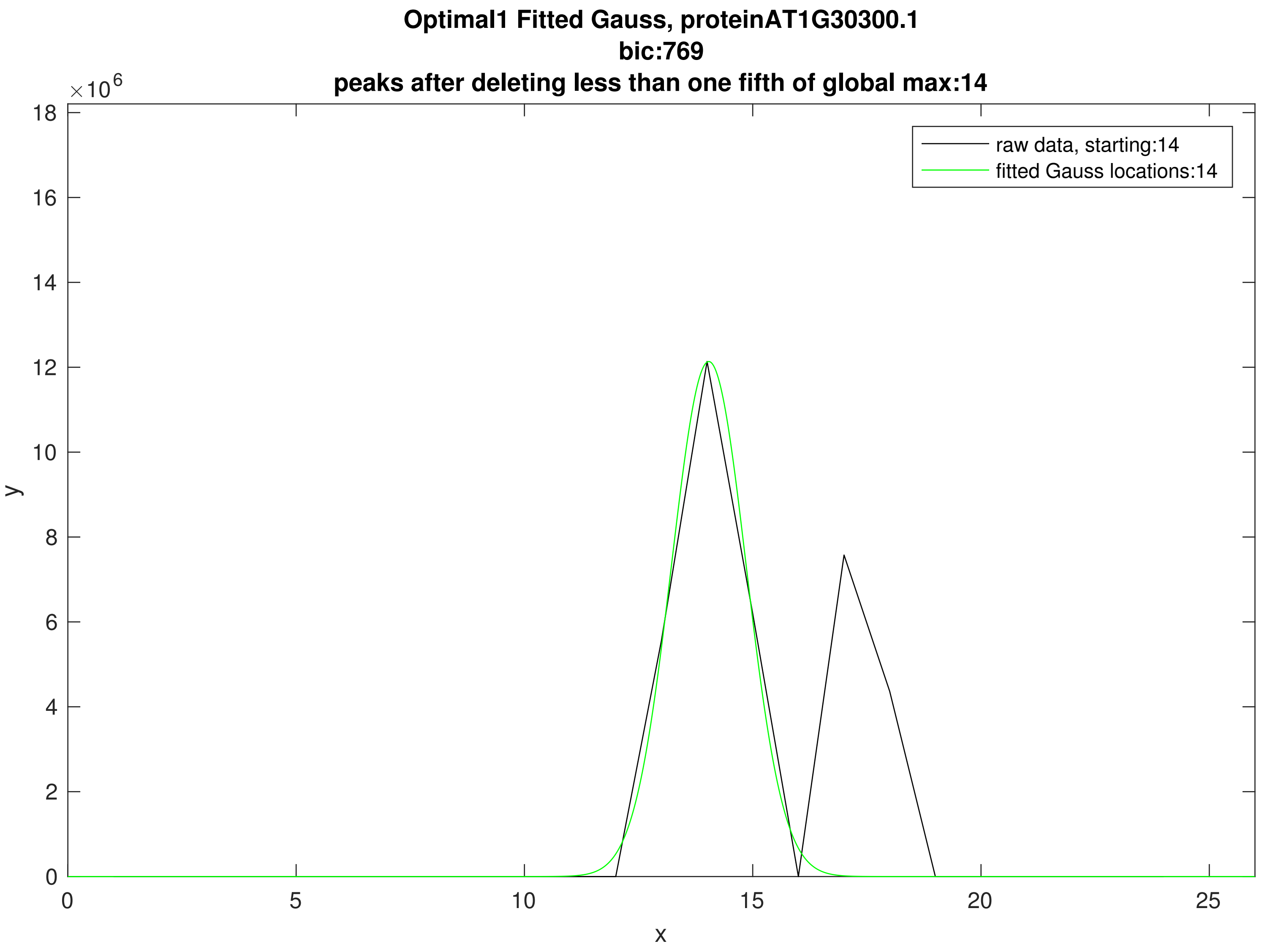

Optimal1 Fitted Gauss, proteinAT1G30300.1
bic:769
peaks after deleting less than one fifth of global max:14
×10^6
18
16
14
12
10
8
6
4
2
0
y
0
5
10
15
20
25
x
raw data, starting:14
fitted Gauss locations:14

#### Optimal1 Fitted Gauss, proteinAT1G30470.3;AT1G30470.2;AT1G30470.1
#### bic:775
#### peaks after deleting less than one fifth of global max:10

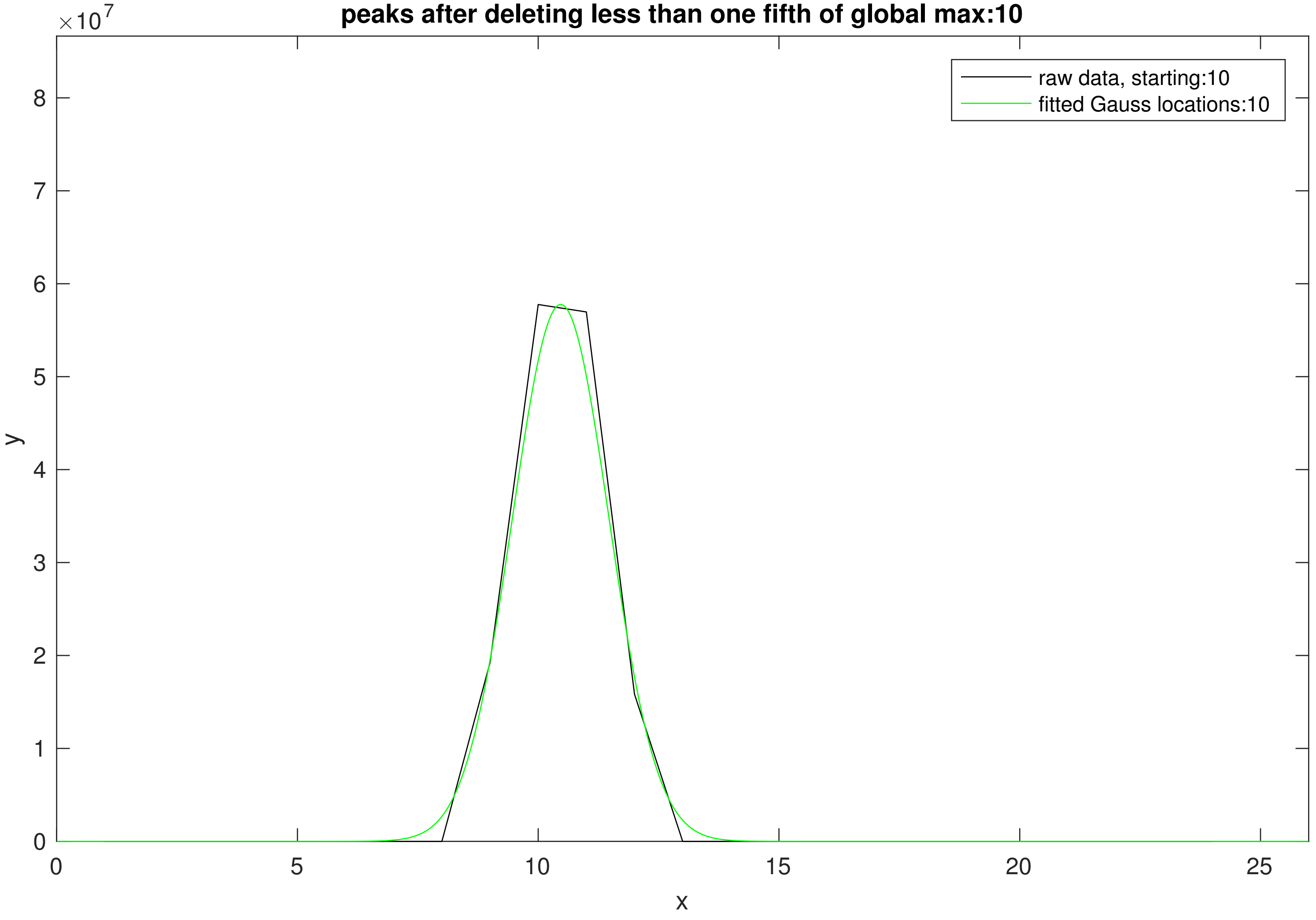

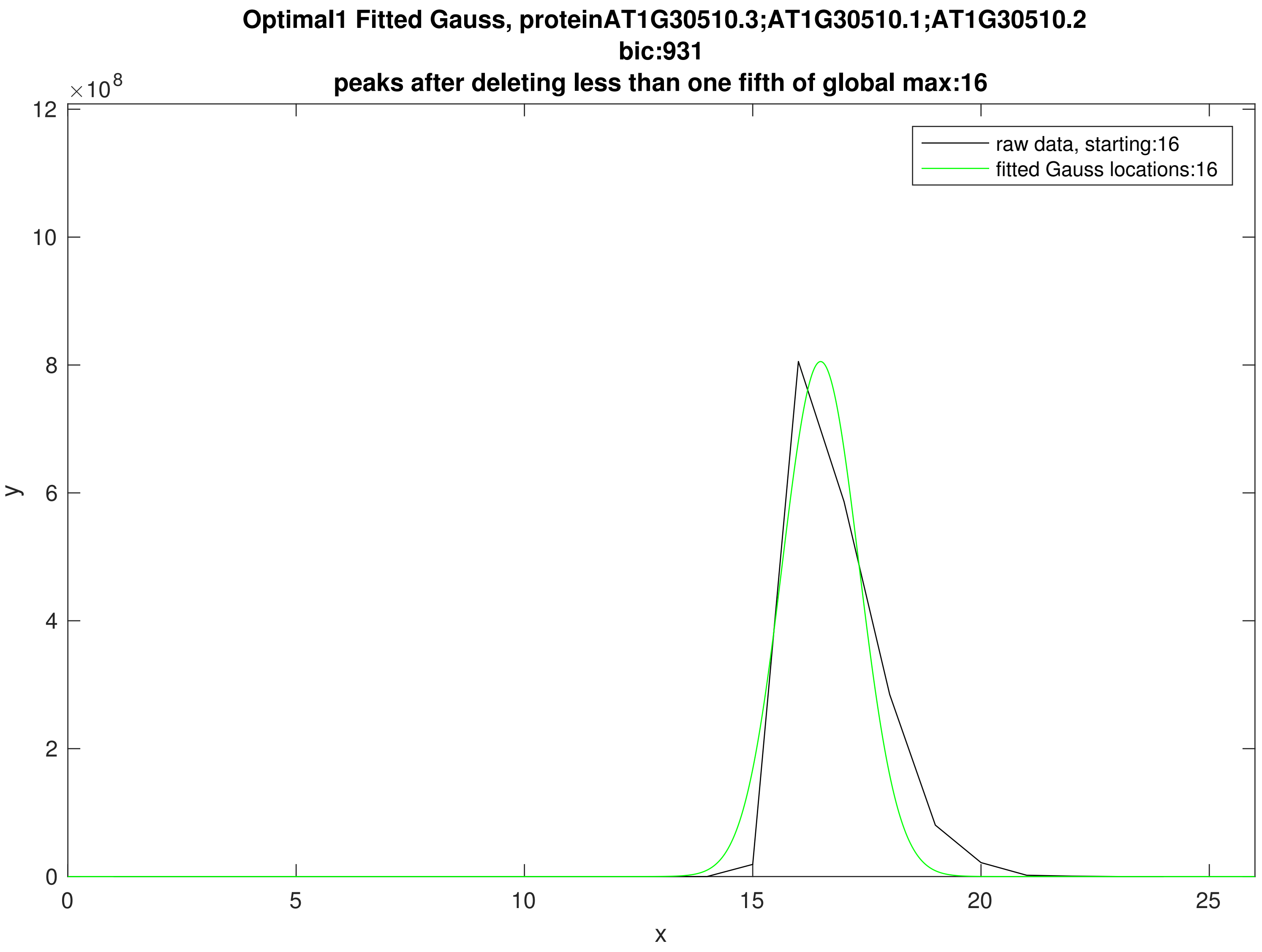

Optimal1 Fitted Gauss, proteinAT1G30510.3;AT1G30510.1;AT1G30510.2
bic:931
peaks after deleting less than one fifth of global max:16
×10^8
12
10
8
6
4
2
0
y
0
5
10
15
20
25
x
raw data, starting:16
fitted Gauss locations:16

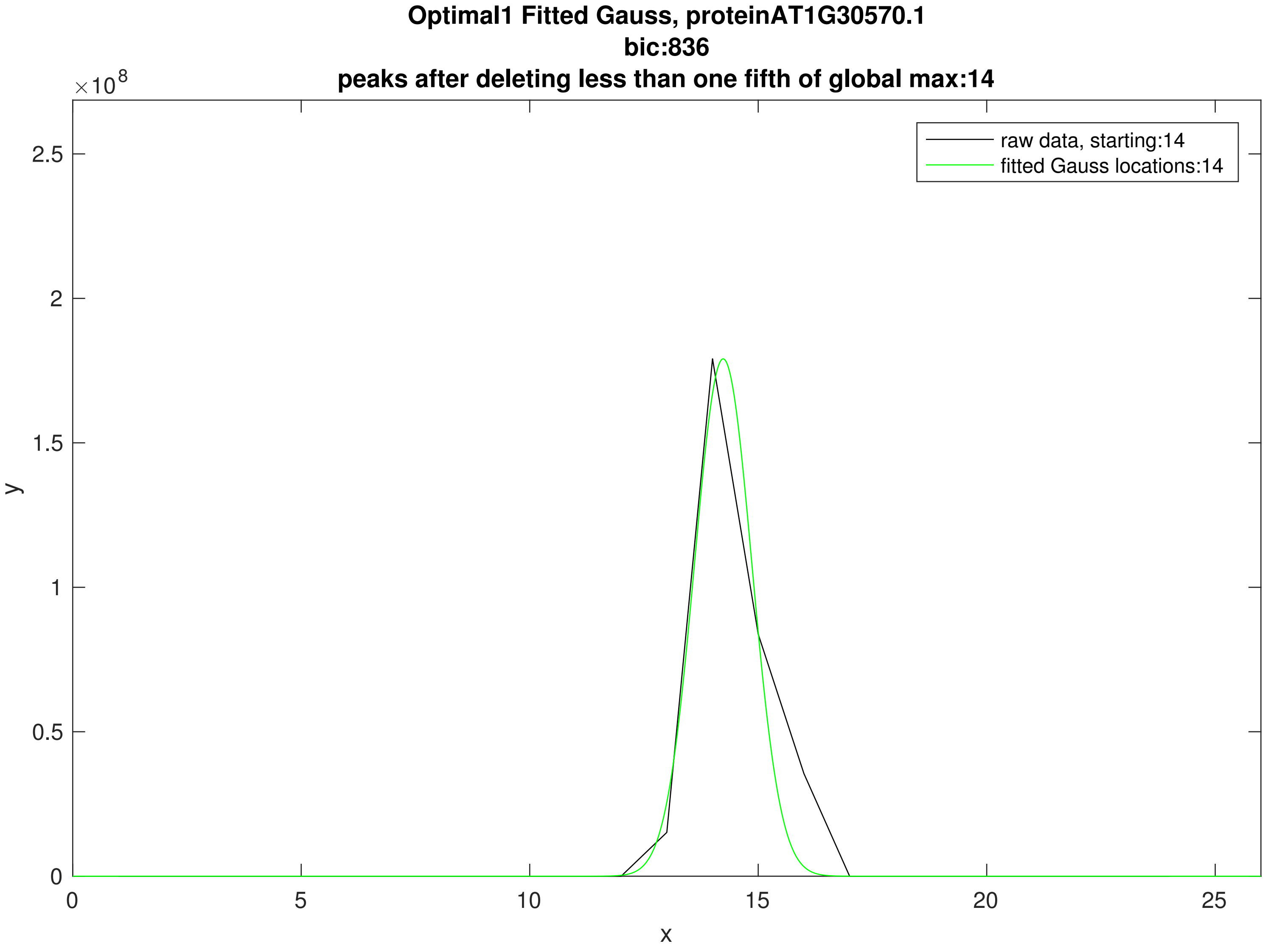
Optimal1 Fitted Gauss, proteinAT1G30570.1
bic:836
peaks after deleting less than one fifth of global max:14
×10^8
raw data, starting:14
fitted Gauss locations:14
y
x
0
0.5
1
1.5
2
2.5
0
5
10
15
20
25

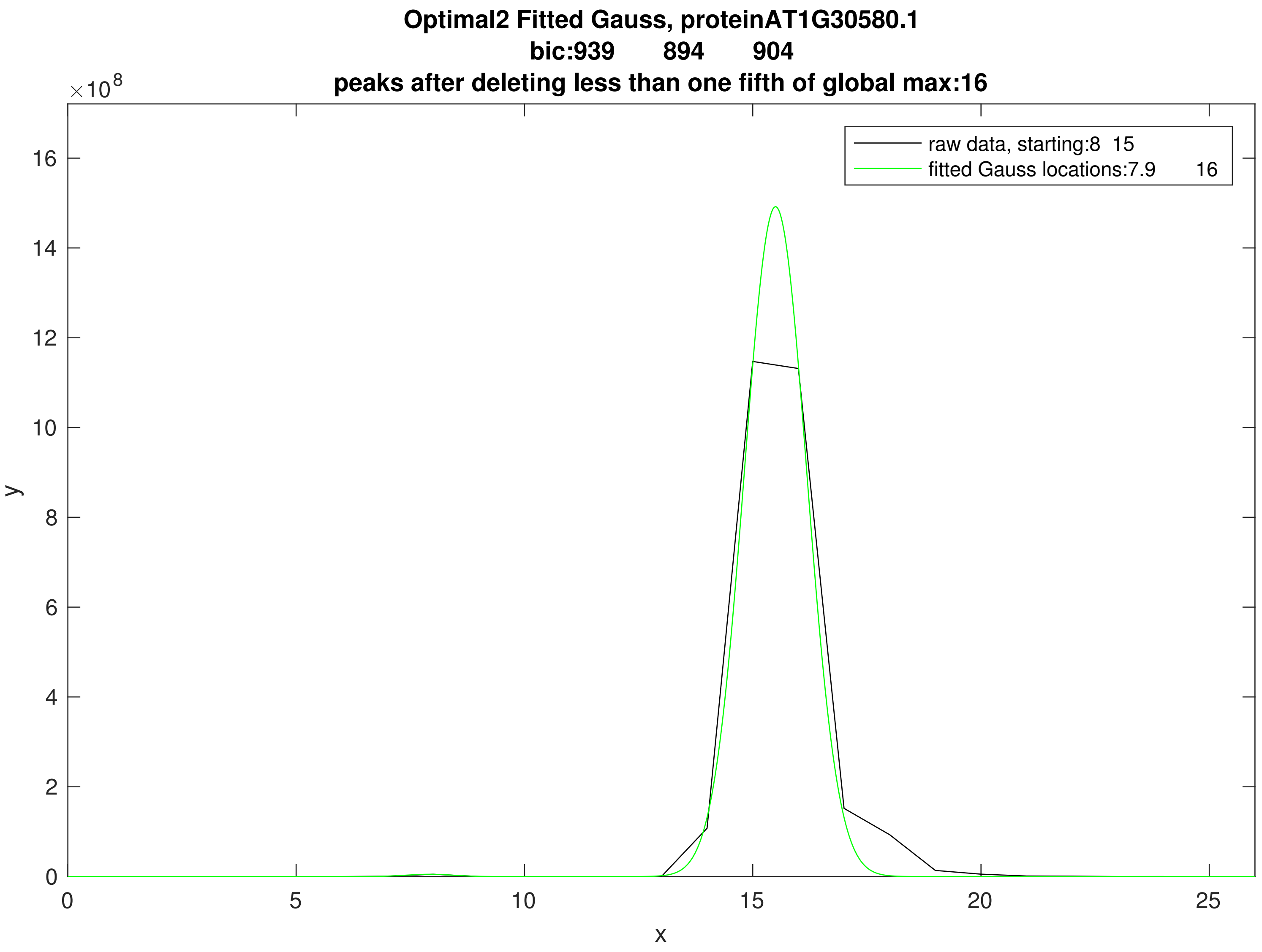

Optimal2 Fitted Gauss, proteinAT1G30580.1
bic:939 894 904
peaks after deleting less than one fifth of global max:16
×10^8
raw data, starting:8 15
fitted Gauss locations:7.9 16
y
x
0
2
4
6
8
10
12
14
16
5
10
15
20
25

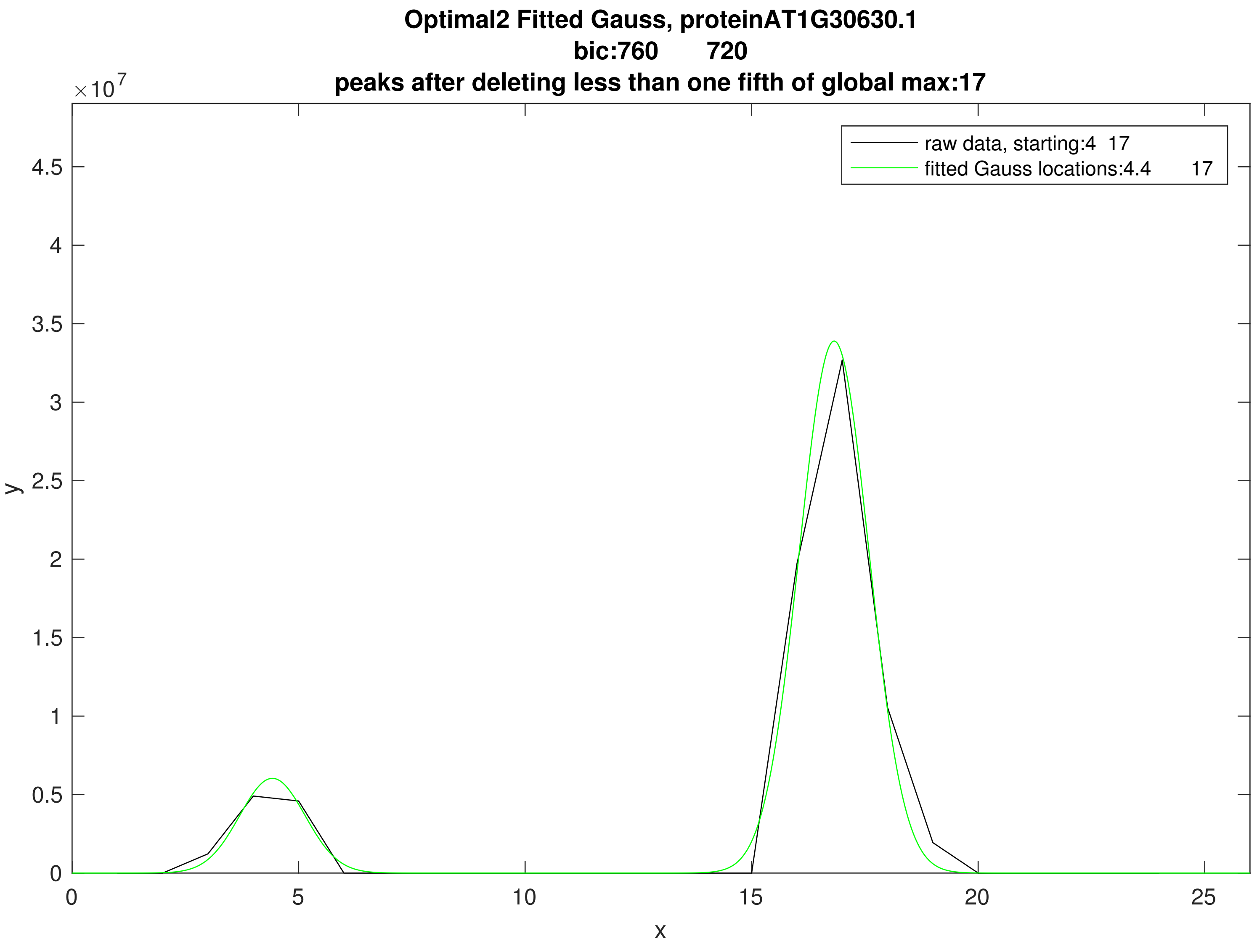
Optimal2 Fitted Gauss, proteinAT1G30630.1
bic:760 720
peaks after deleting less than one fifth of global max:17
×10^7
raw data, starting:4 17
fitted Gauss locations:4.4 17
y
x
0
0.5
1
1.5
2
2.5
3
3.5
4
4.5
0
5
10
15
20
25

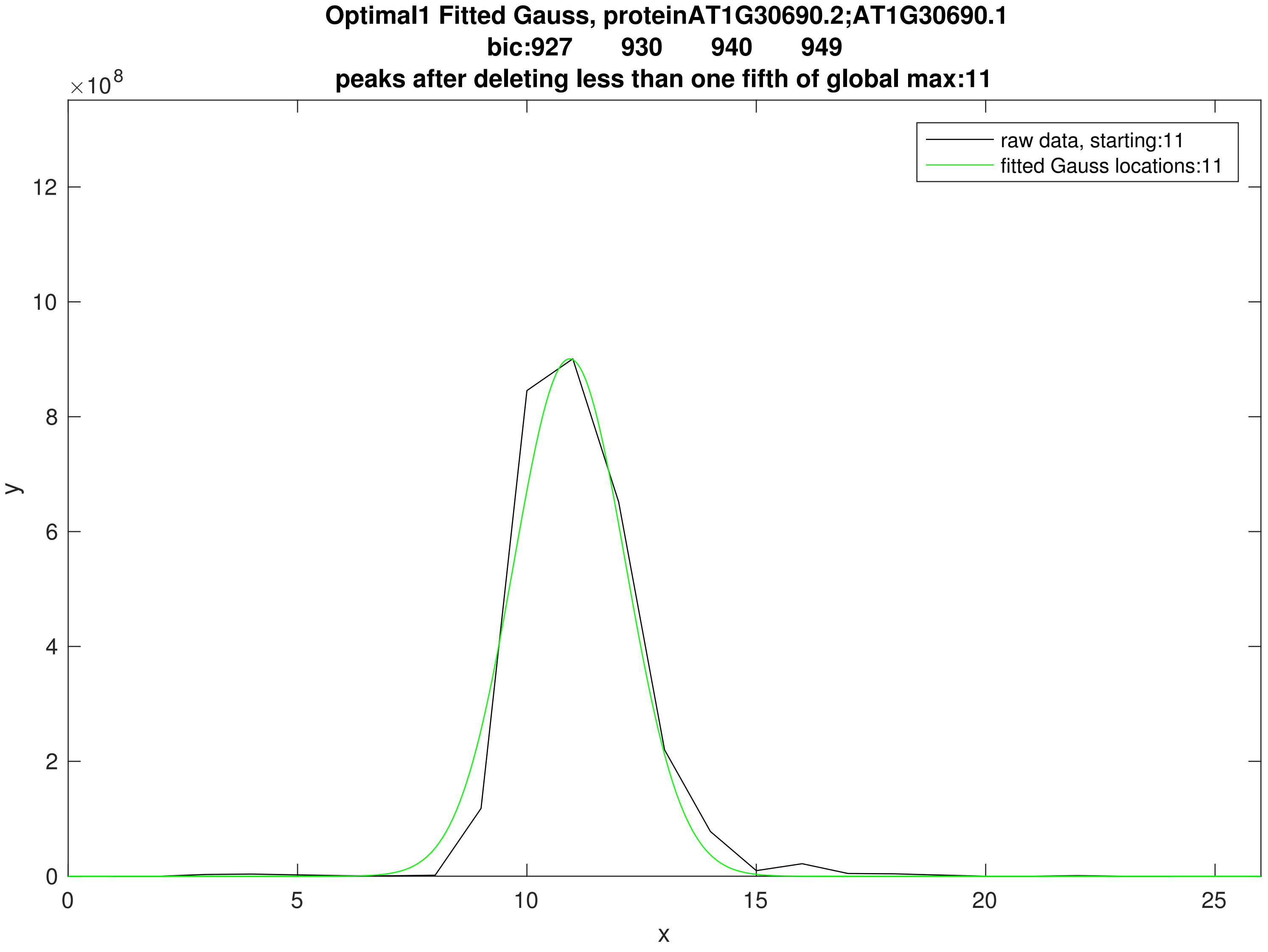
Optimal1 Fitted Gauss, proteinAT1G30690.2;AT1G30690.1
bic:927 930 940 949
peaks after deleting less than one fifth of global max:11
×10^8
raw data, starting:11
fitted Gauss locations:11
y
x
0
2
4
6
8
10
12
0
5
10
15
20
25

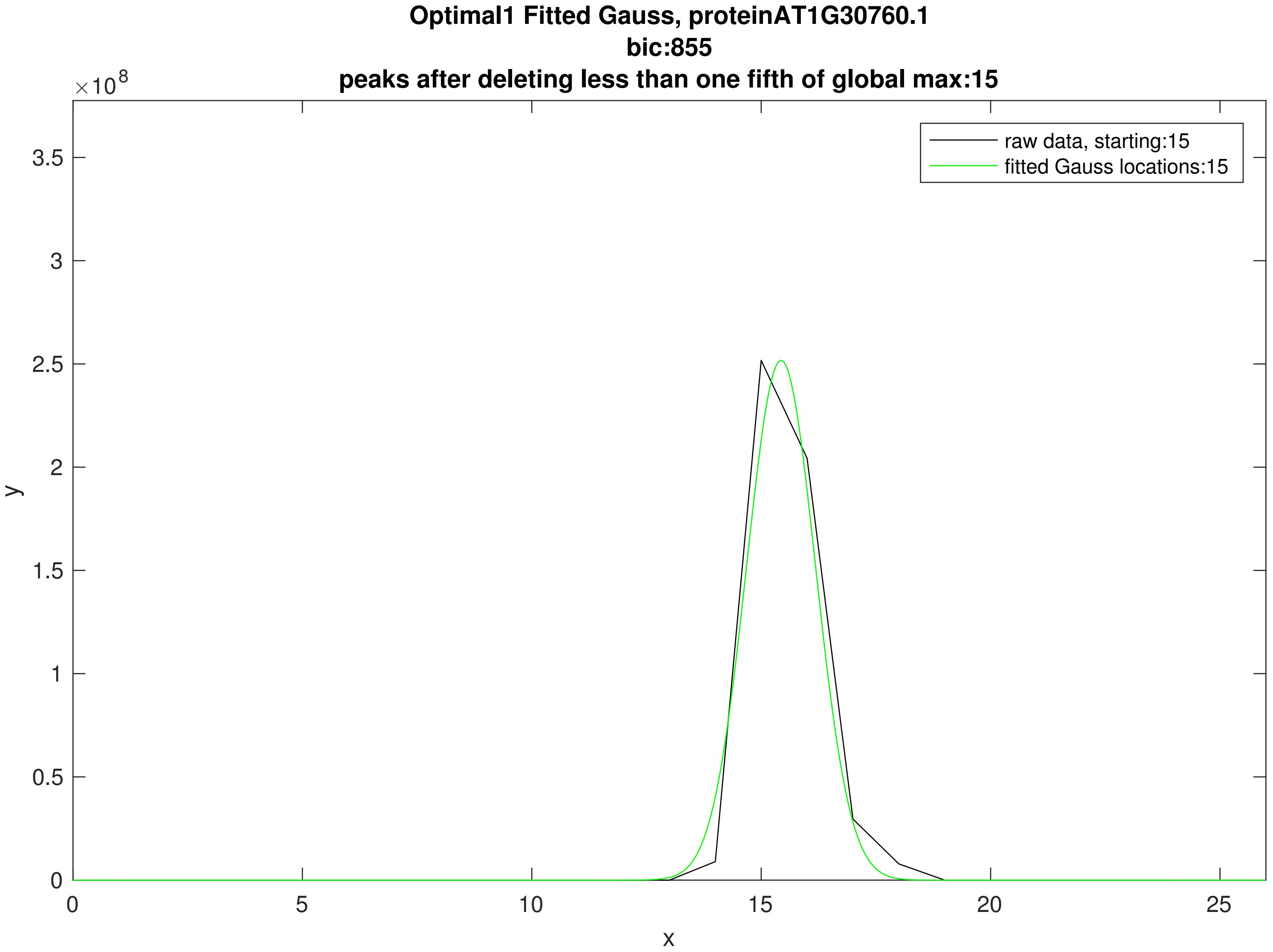
Optimal1 Fitted Gauss, proteinAT1G30760.1
bic:855
peaks after deleting less than one fifth of global max:15
×10^8
raw data, starting:15
fitted Gauss locations:15
y
x
0
0.5
1
1.5
2
2.5
3
3.5
0
5
10
15
20
25

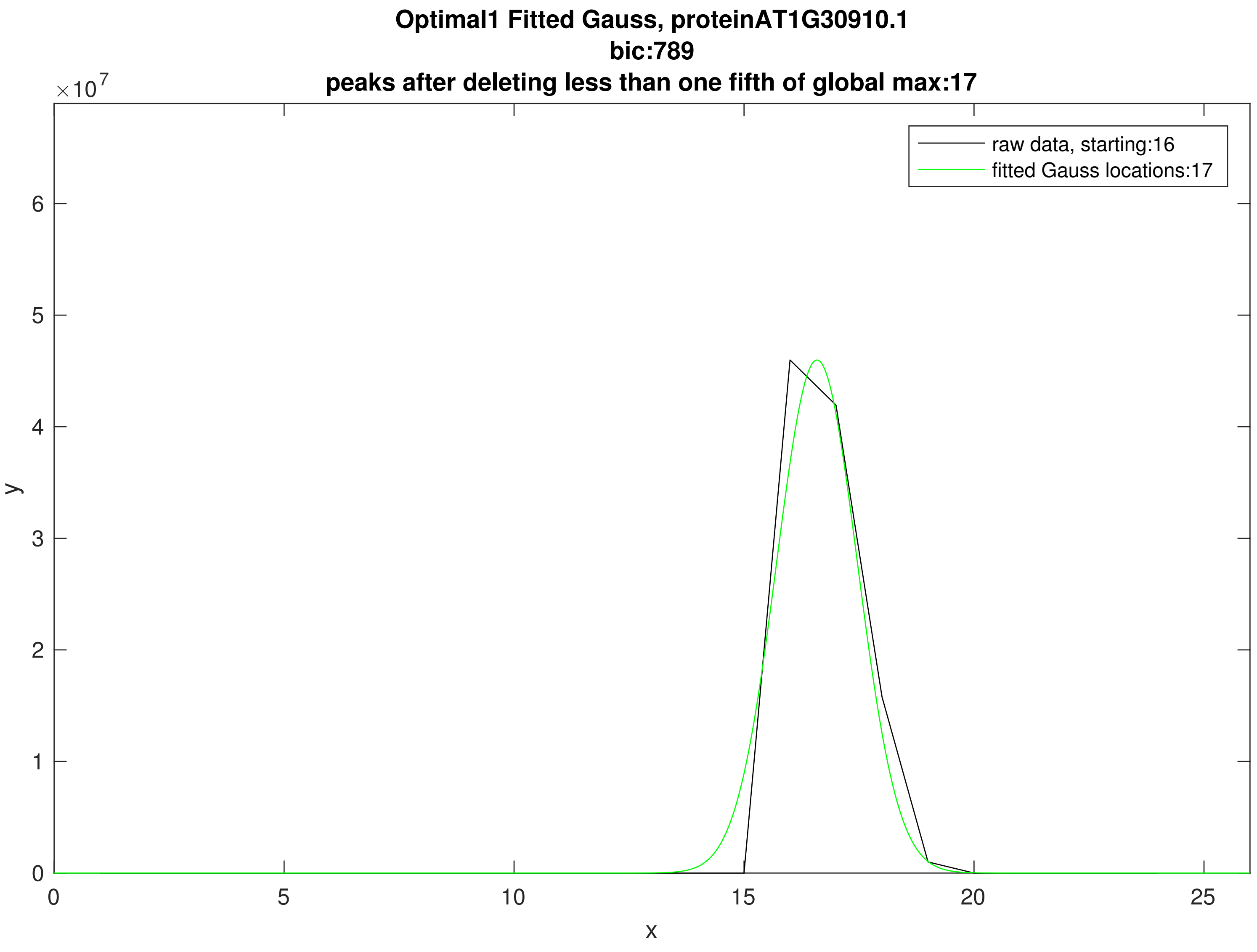

Optimal1 Fitted Gauss, proteinAT1G30910.1
bic:789
peaks after deleting less than one fifth of global max:17
×10^7
raw data, starting:16
fitted Gauss locations:17
y
x
0
1
2
3
4
5
6
0
5
10
15
20
25

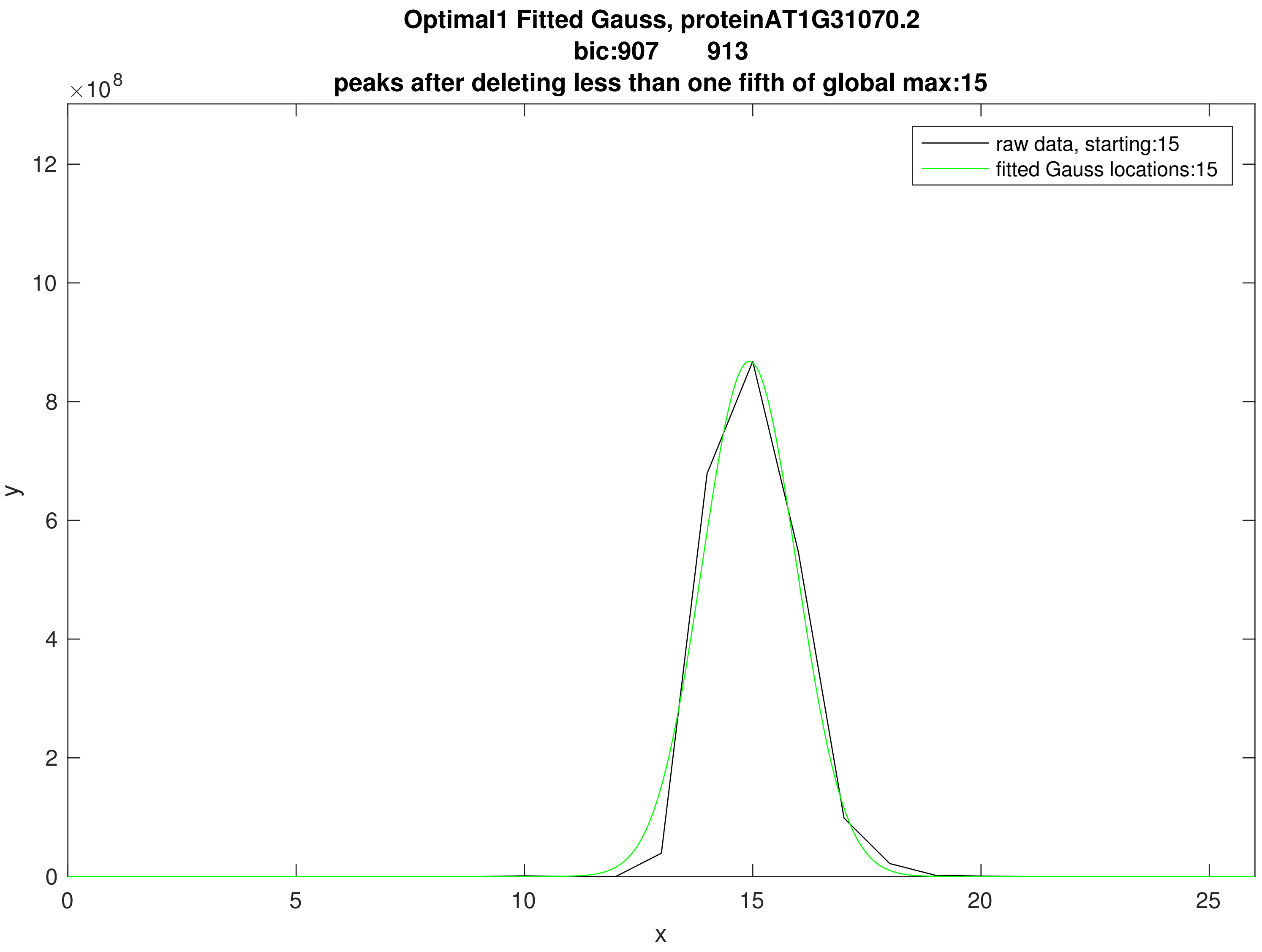
Optimal1 Fitted Gauss, proteinAT1G31070.2
bic:907 913
peaks after deleting less than one fifth of global max:15
×10^8
12
10
8
6
4
2
0
y
0
5
10
15
20
25
x
raw data, starting:15
fitted Gauss locations:15

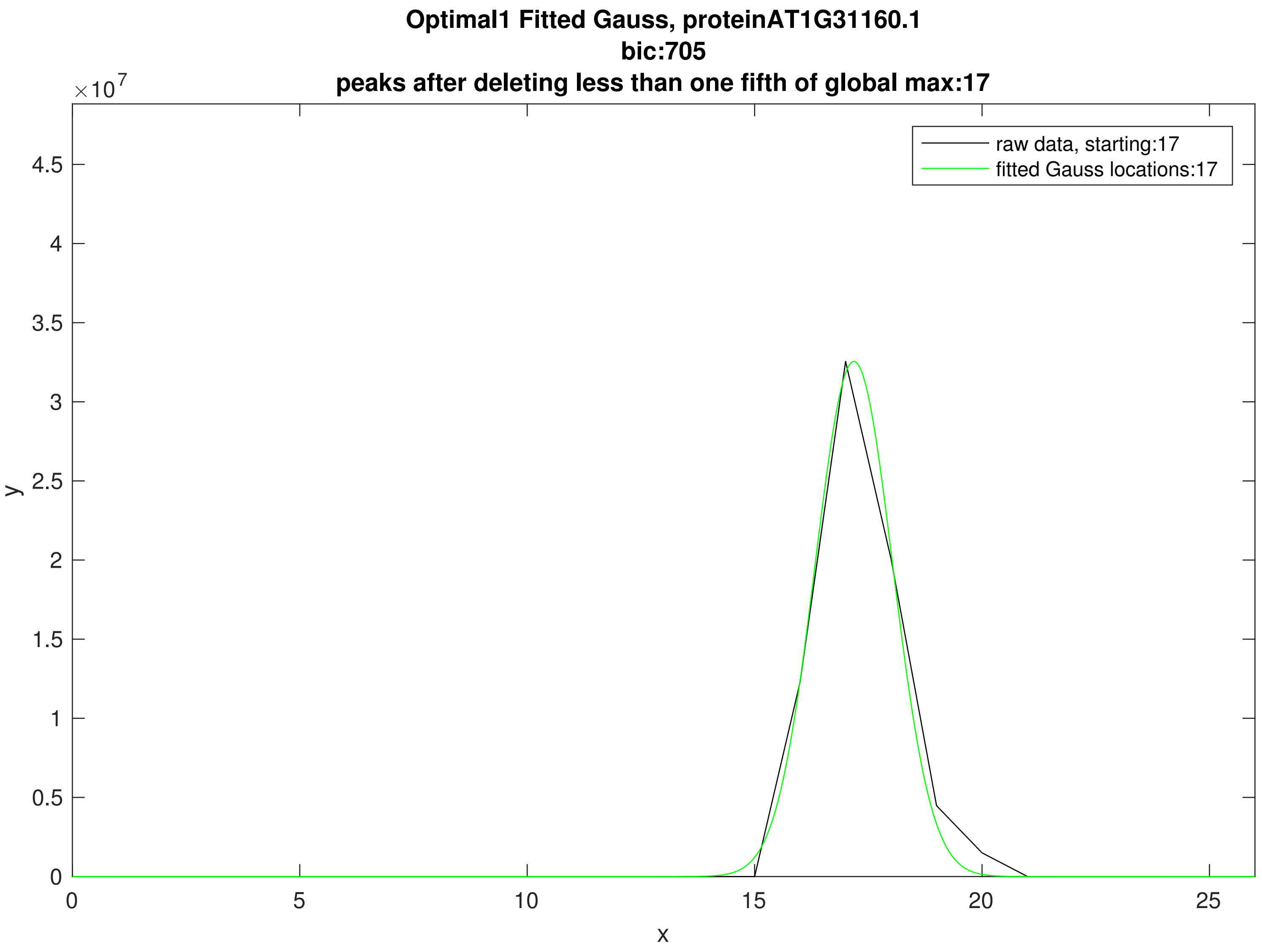

Optimal1 Fitted Gauss, proteinAT1G31160.1
bic:705
peaks after deleting less than one fifth of global max:17
×10^7
raw data, starting:17
fitted Gauss locations:17
y
x
0
0.5
1
1.5
2
2.5
3
3.5
4
4.5
5
10
15
20
25

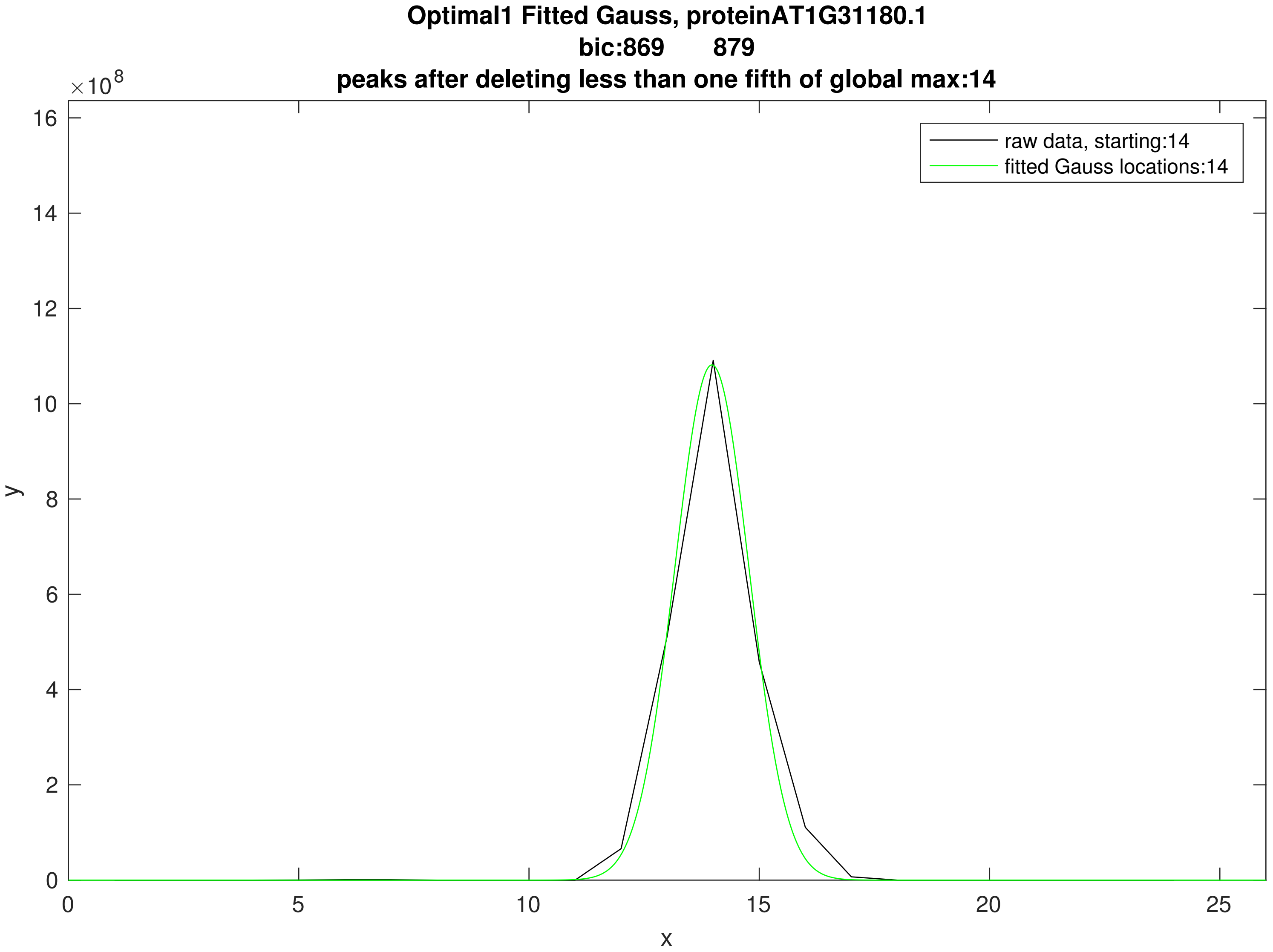
Optimal1 Fitted Gauss, proteinAT1G31180.1
bic:869 879
peaks after deleting less than one fifth of global max:14
×10^8
16
14
12
10
8
6
4
2
0
y
0
5
10
15
20
25
x
raw data, starting:14
fitted Gauss locations:14

Optimal1 Fitted Gauss, proteinAT1G31230.1
bic:867 877
peaks after deleting less than one fifth of global max:9.5

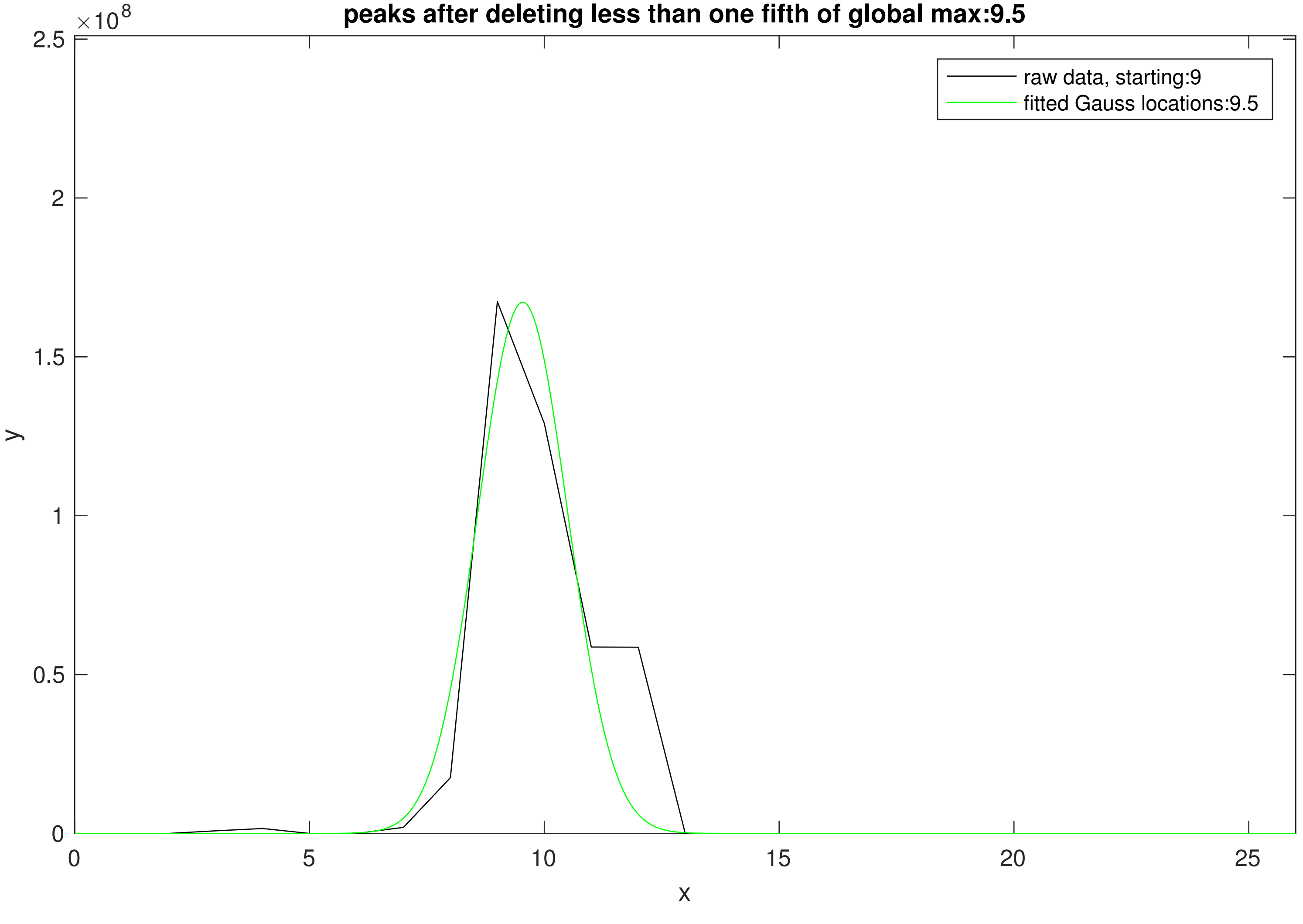

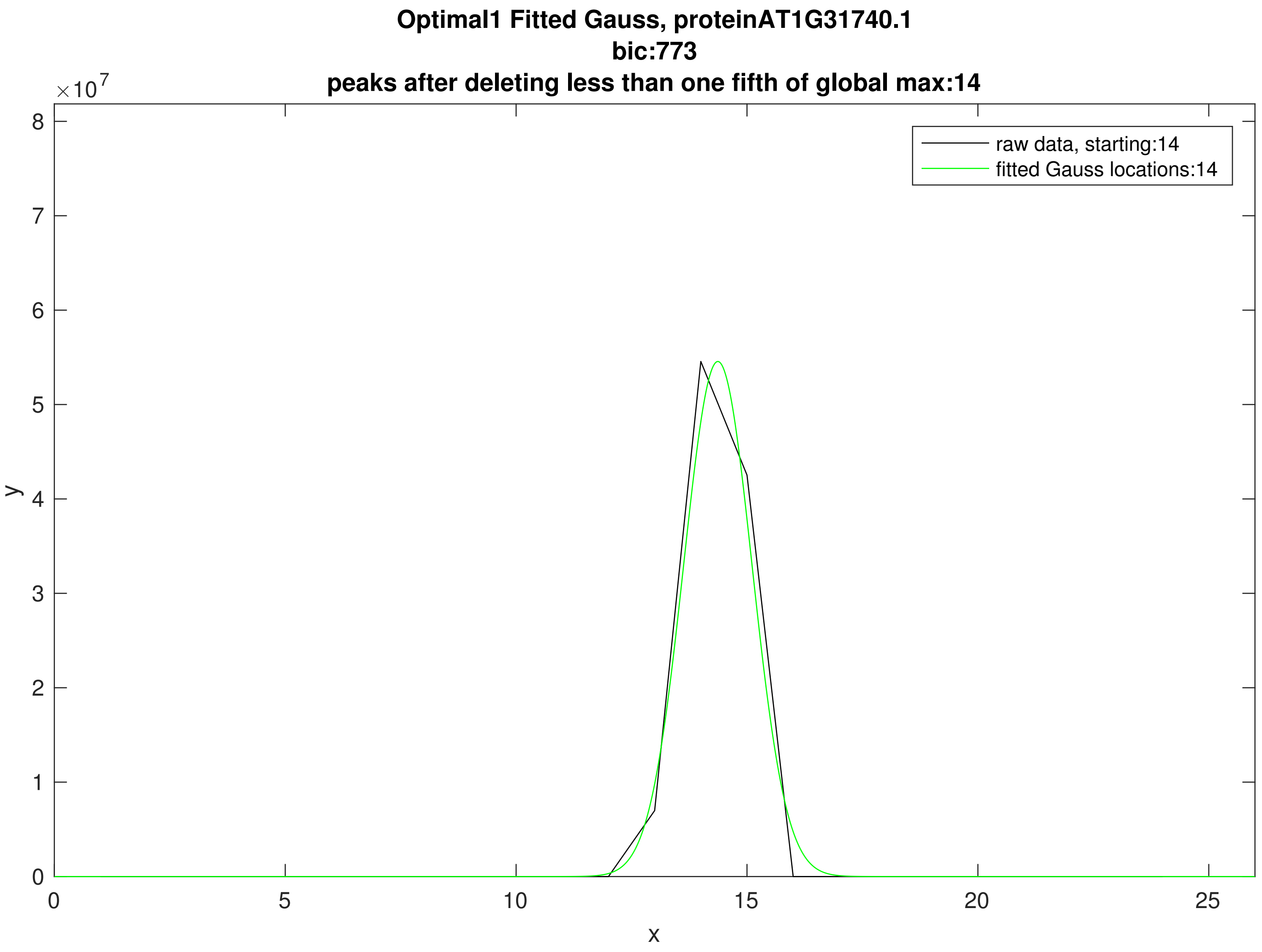
Optimal1 Fitted Gauss, proteinAT1G31740.1
bic:773
peaks after deleting less than one fifth of global max:14
×10^7
raw data, starting:14
fitted Gauss locations:14
y
x
0
1
2
3
4
5
6
7
8
0
5
10
15
20
25

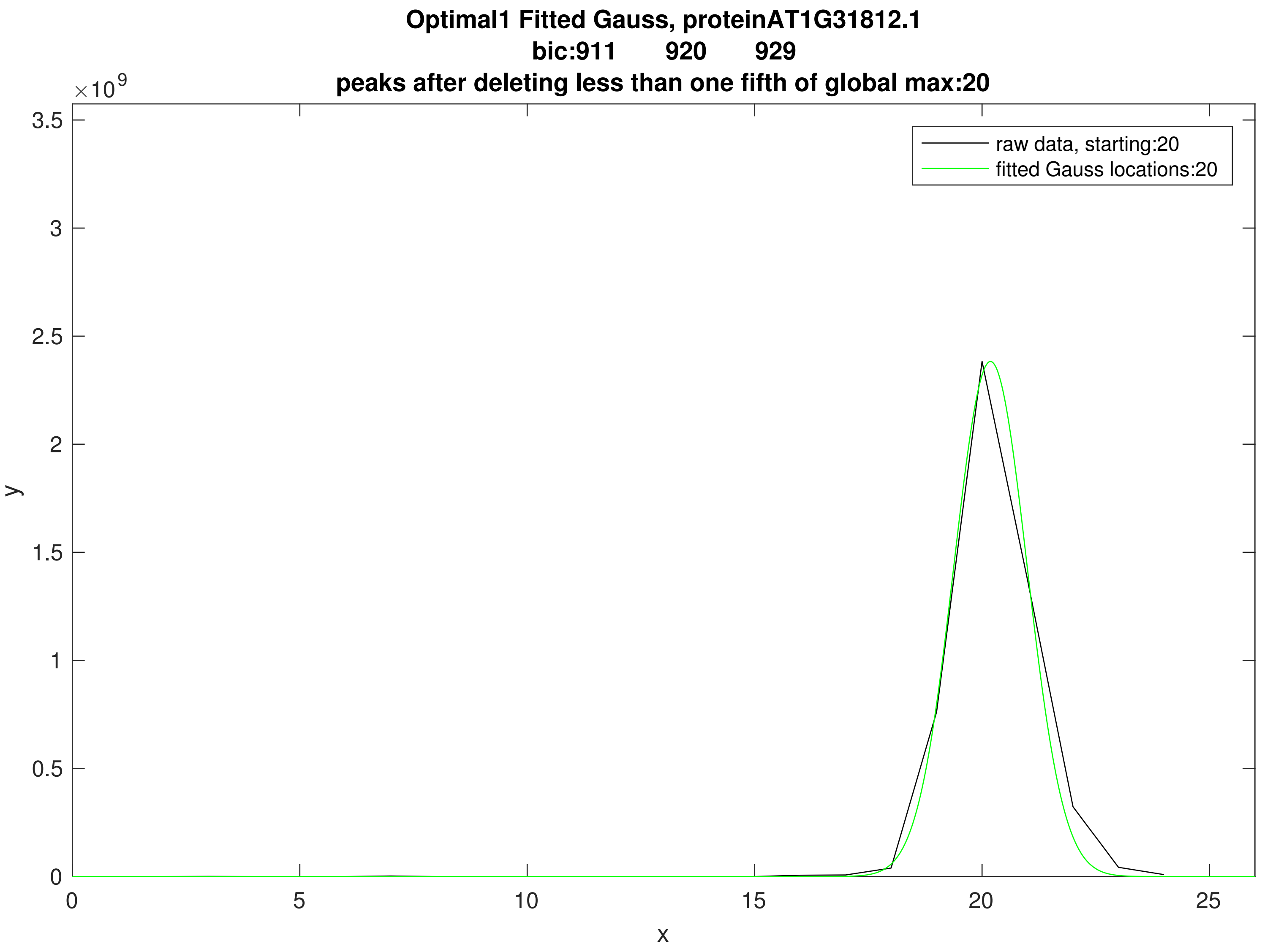

Optimal1 Fitted Gauss, proteinAT1G31812.1
bic:911 920 929
peaks after deleting less than one fifth of global max:20
×10^9
3.5
3
2.5
2
1.5
1
0.5
0
y
0
5
10
15
20
25
x
raw data, starting:20
fitted Gauss locations:20

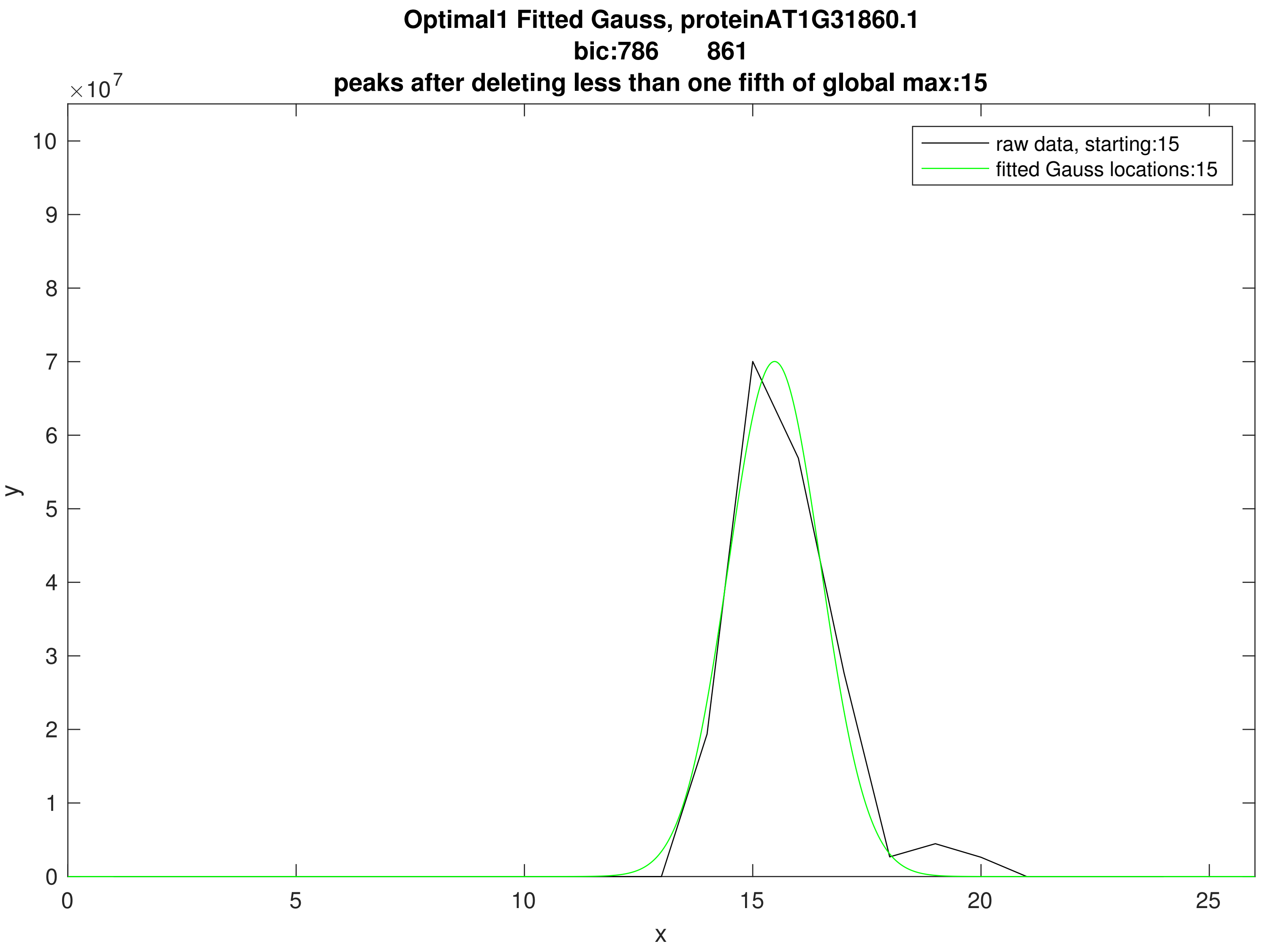

Optimal1 Fitted Gauss, proteinAT1G31860.1
bic:786 861
peaks after deleting less than one fifth of global max:15
×10^7
raw data, starting:15
fitted Gauss locations:15
y
x
0
1
2
3
4
5
6
7
8
9
10
0
5
10
15
20
25

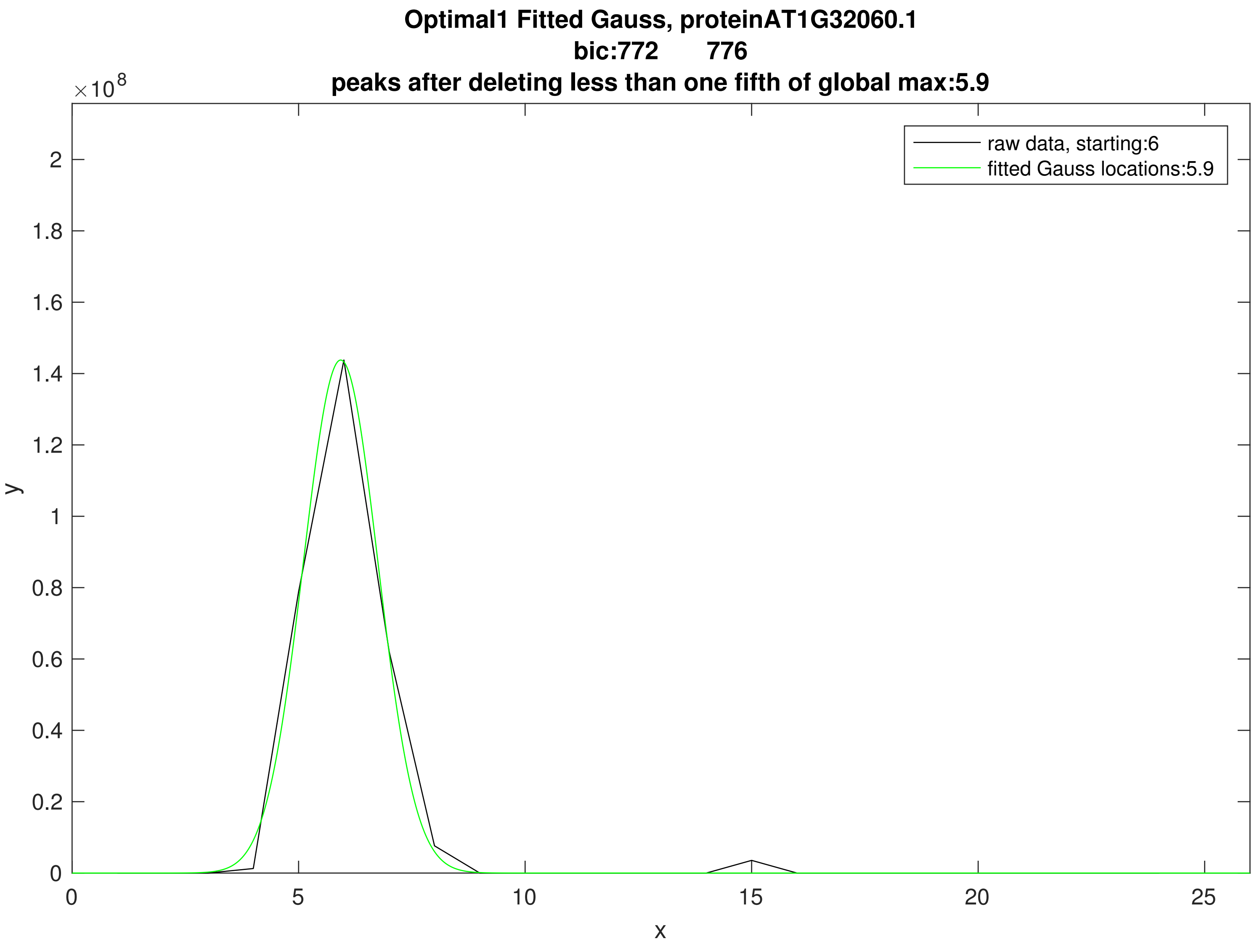
Optimal1 Fitted Gauss, proteinAT1G32060.1
bic:772 776
peaks after deleting less than one fifth of global max:5.9
×10^8
raw data, starting:6
fitted Gauss locations:5.9
0
0.2
0.4
0.6
0.8
1
1.2
1.4
1.6
1.8
2
y
0
5
10
15
20
25
x

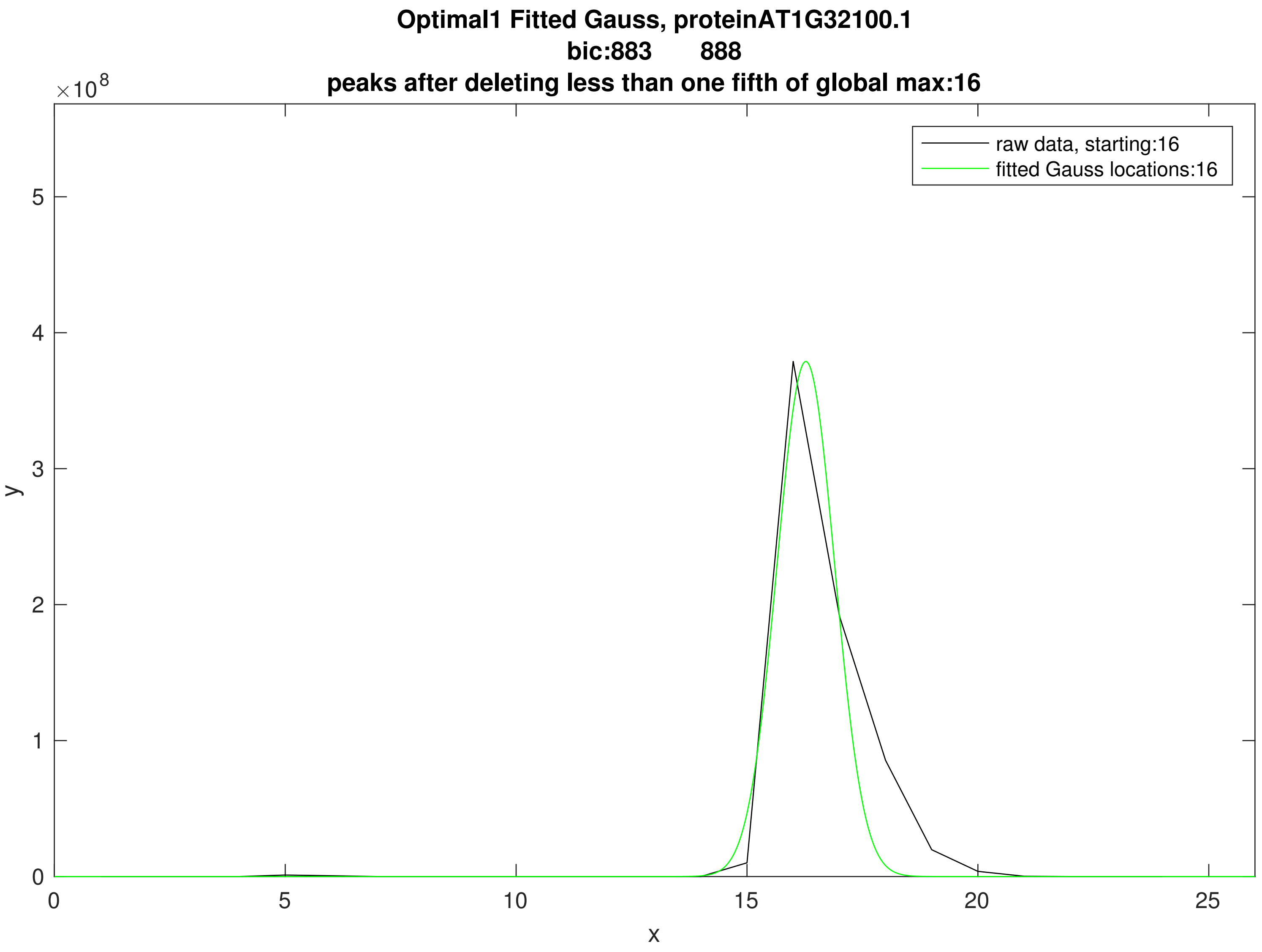
Optimal1 Fitted Gauss, proteinAT1G32100.1
bic:883 888
peaks after deleting less than one fifth of global max:16
×10^8
raw data, starting:16
fitted Gauss locations:16
y
x
0
1
2
3
4
5
0
5
10
15
20
25

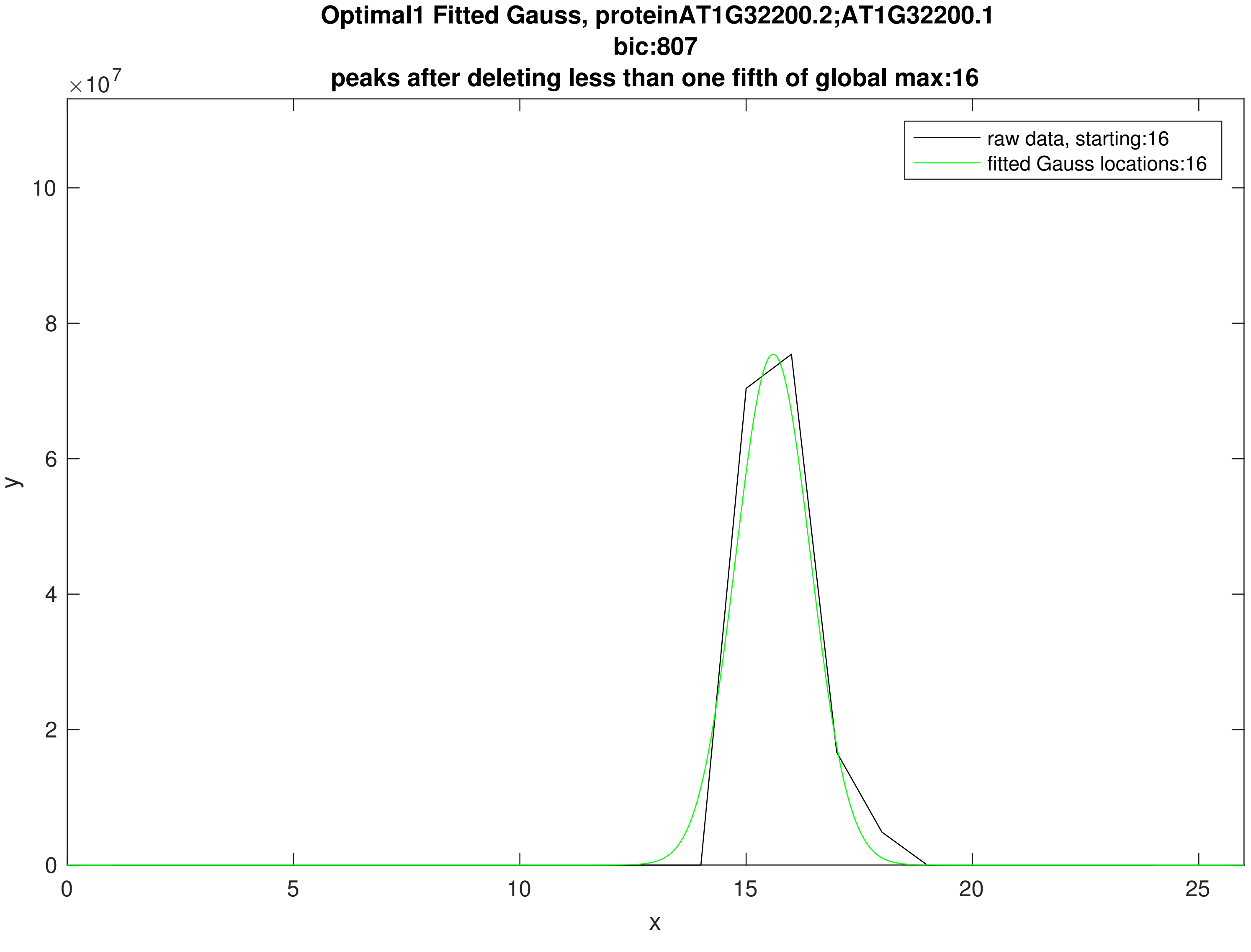
Optimal1 Fitted Gauss, proteinAT1G32200.2;AT1G32200.1
bic:807
peaks after deleting less than one fifth of global max:16
×10^7
raw data, starting:16
fitted Gauss locations:16
y
x
0
2
4
6
8
10
0
5
10
15
20
25

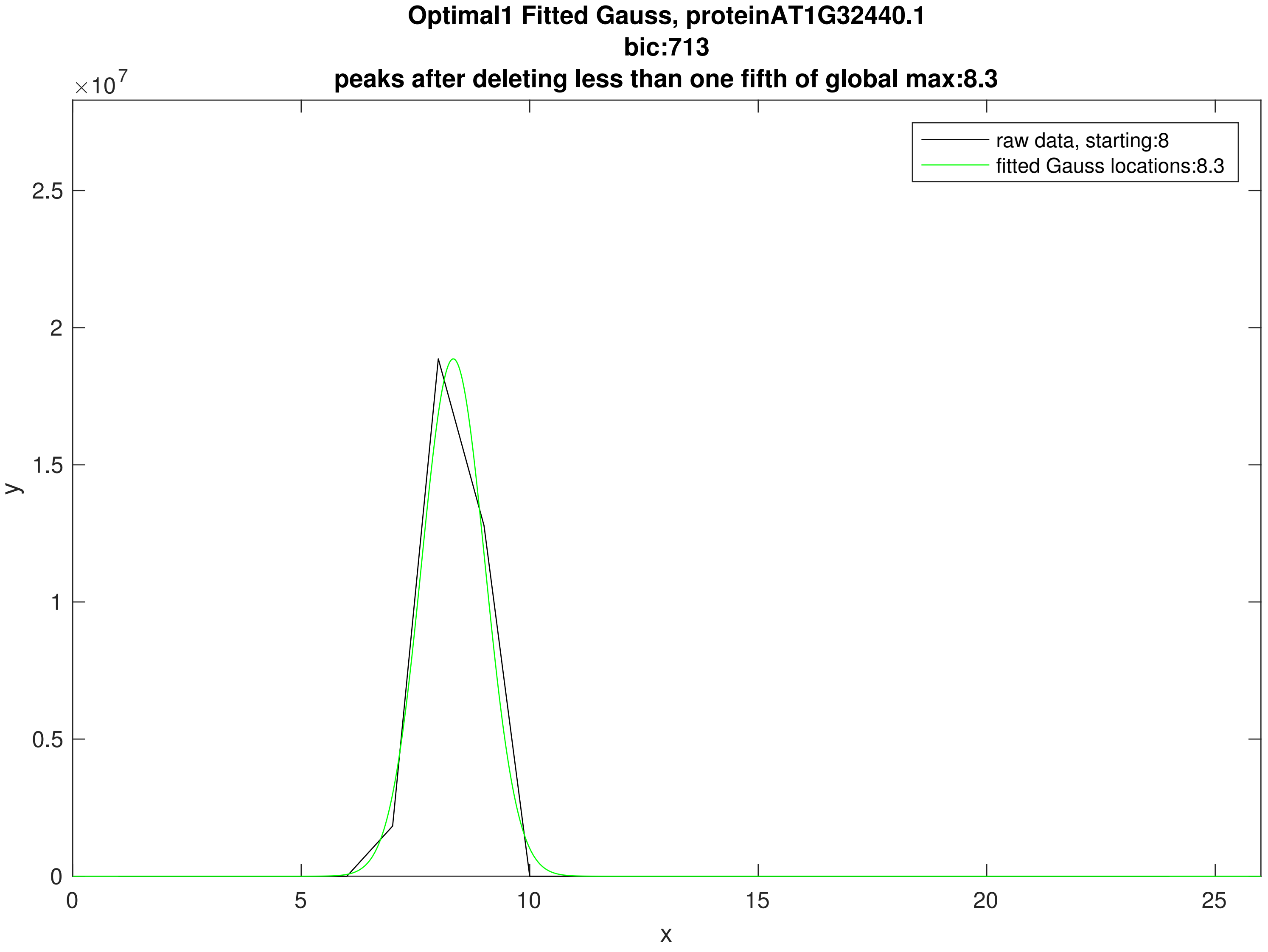
Optimal1 Fitted Gauss, proteinAT1G32440.1
bic:713
peaks after deleting less than one fifth of global max:8.3
×10^7
raw data, starting:8
fitted Gauss locations:8.3
2.5
2
1.5
1
0.5
0
y
0
5
10
15
20
25
x

#### Optimal1 Fitted Gauss, proteinAT1G32470.1
#### bic:773
#### peaks after deleting less than one fifth of global max:19

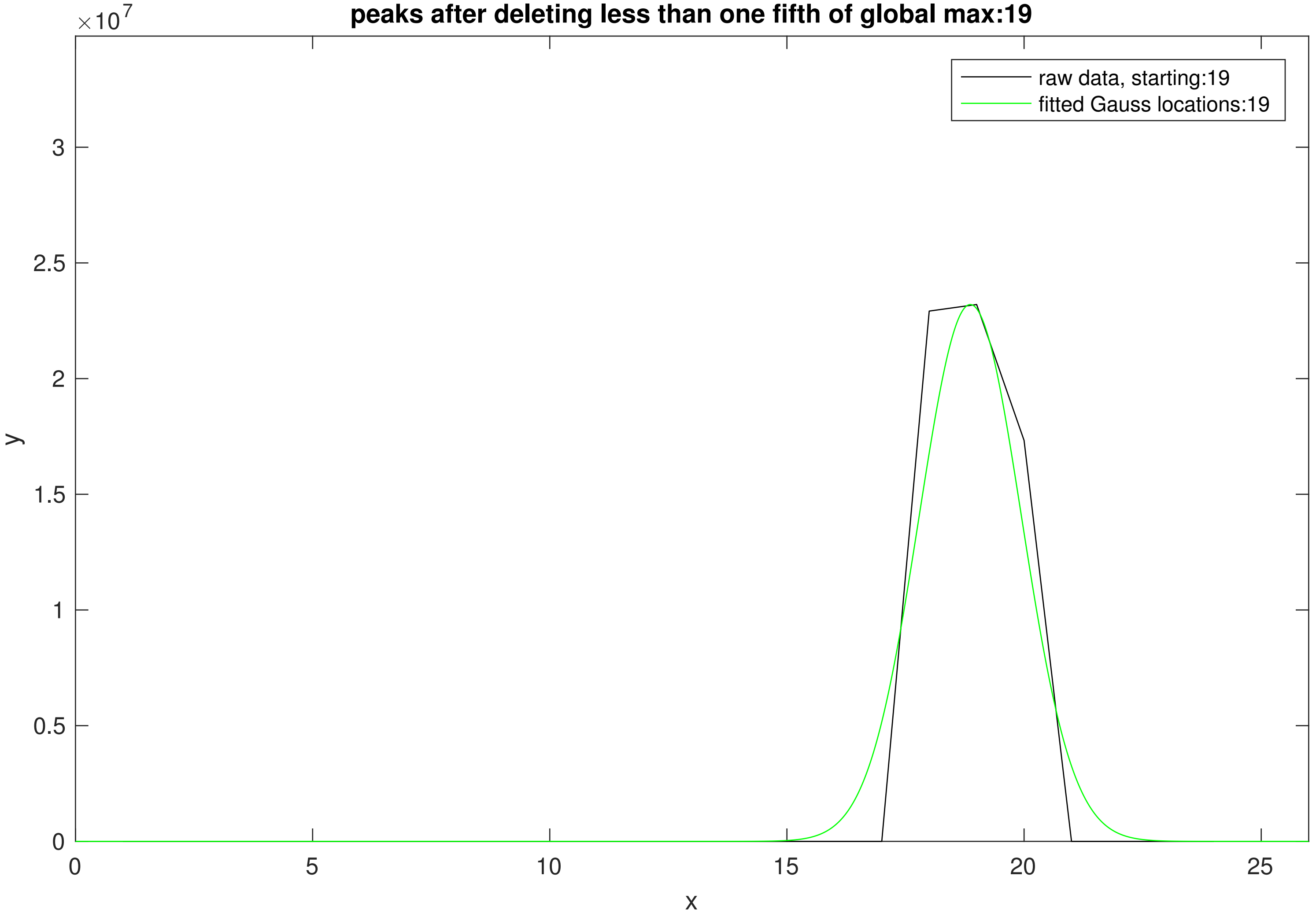

**Optimal1 Fitted Gauss, proteinAT1G32500.1**
**bic:717**
**peaks after deleting less than one fifth of global max:12**

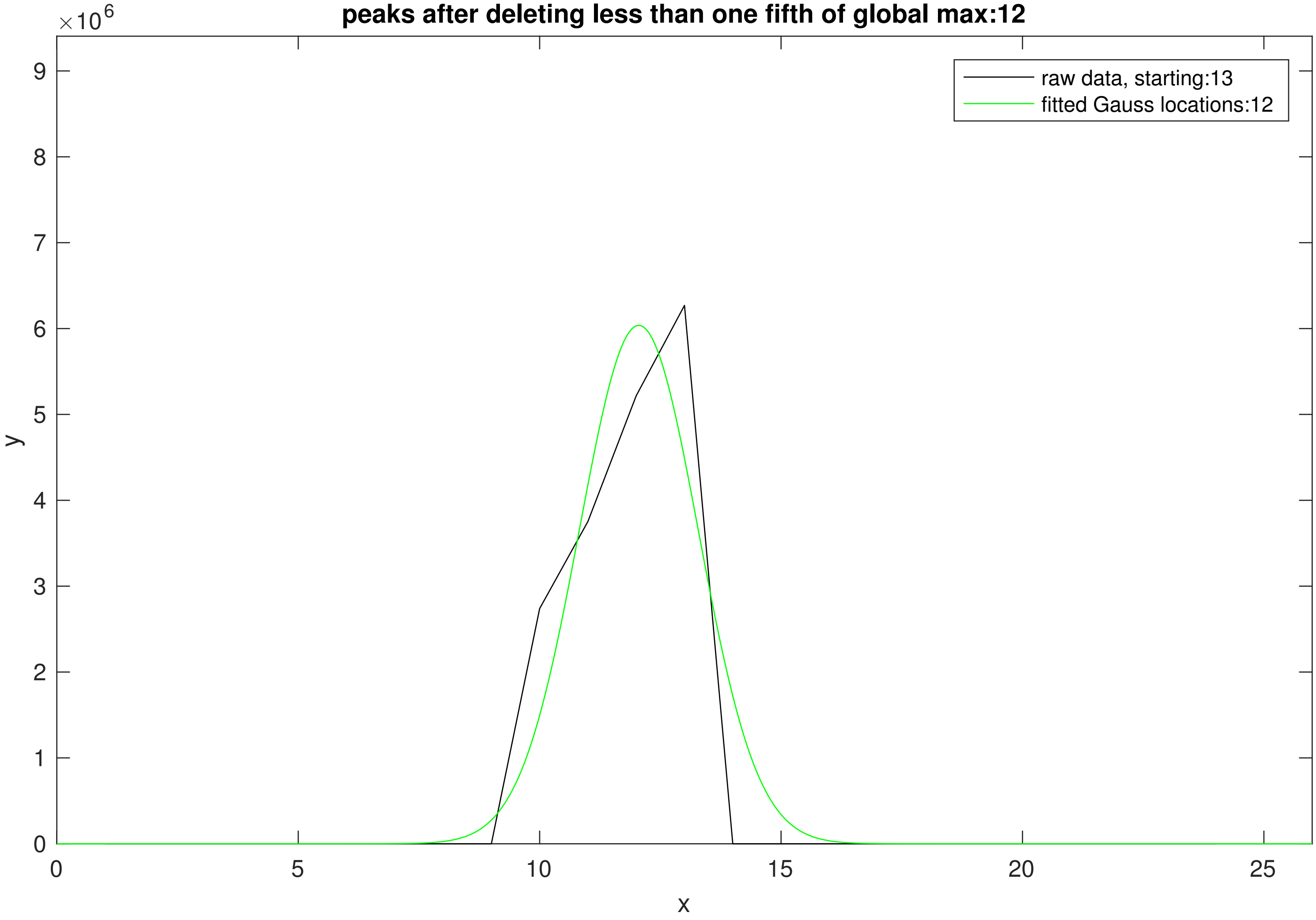

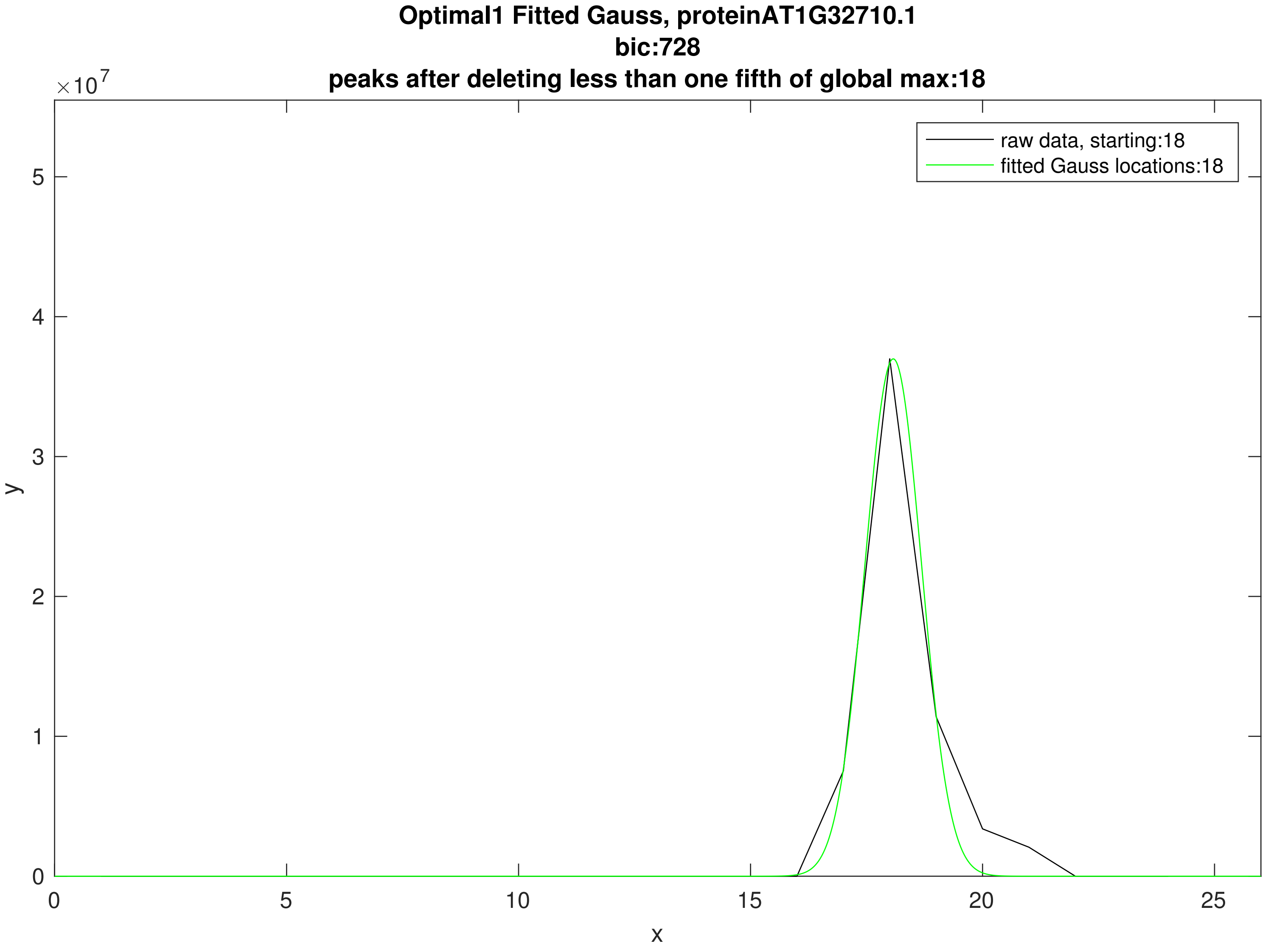
Optimal1 Fitted Gauss, proteinAT1G32710.1
bic:728
peaks after deleting less than one fifth of global max:18
×10^7
raw data, starting:18
fitted Gauss locations:18
y
x
0
1
2
3
4
5
0
5
10
15
20
25

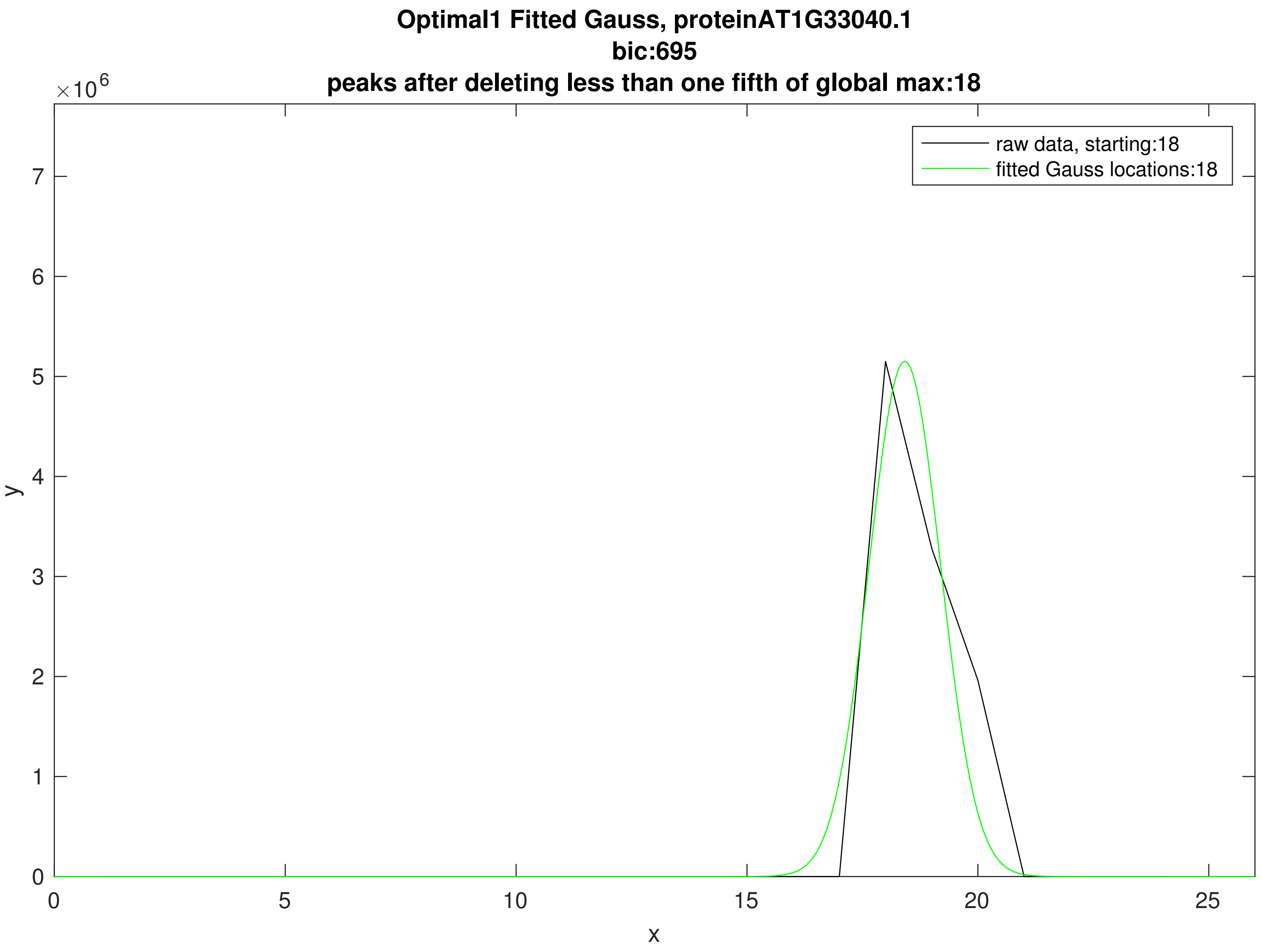

Optimal1 Fitted Gauss, proteinAT1G33040.1
bic:695
peaks after deleting less than one fifth of global max:18
×10^6
raw data, starting:18
fitted Gauss locations:18
y
x
0
1
2
3
4
5
6
7
0
5
10
15
20
25

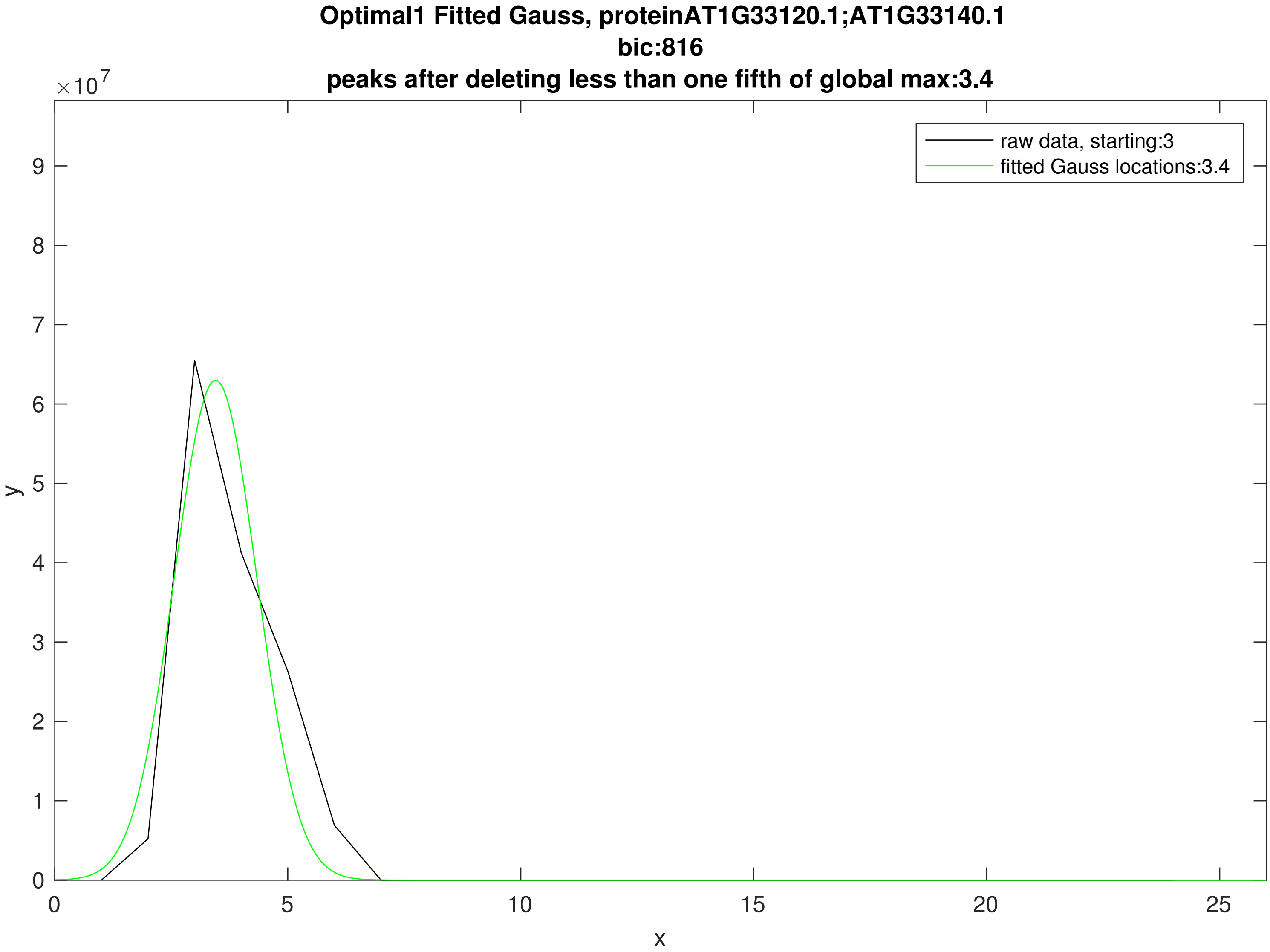

Optimal1 Fitted Gauss, proteinAT1G33120.1;AT1G33140.1
bic:816
peaks after deleting less than one fifth of global max:3.4
×10^7
raw data, starting:3
fitted Gauss locations:3.4
y
x
0
1
2
3
4
5
6
7
8
9
0
5
10
15
20
25

Optimal1 Fitted Gauss, proteinAT1G33790.2;AT1G33790.1
bic:797
peaks after deleting less than one fifth of global max:4.6

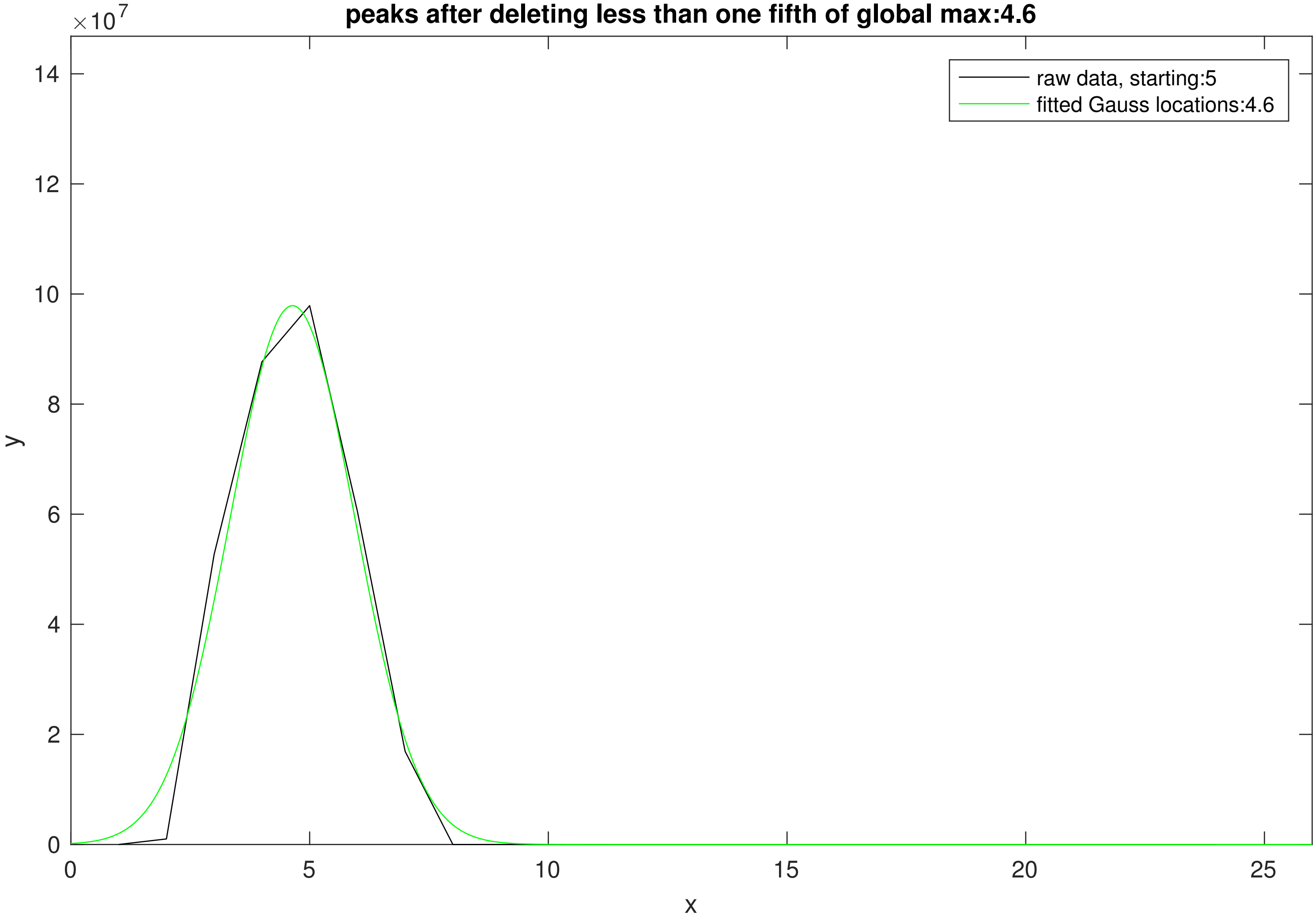

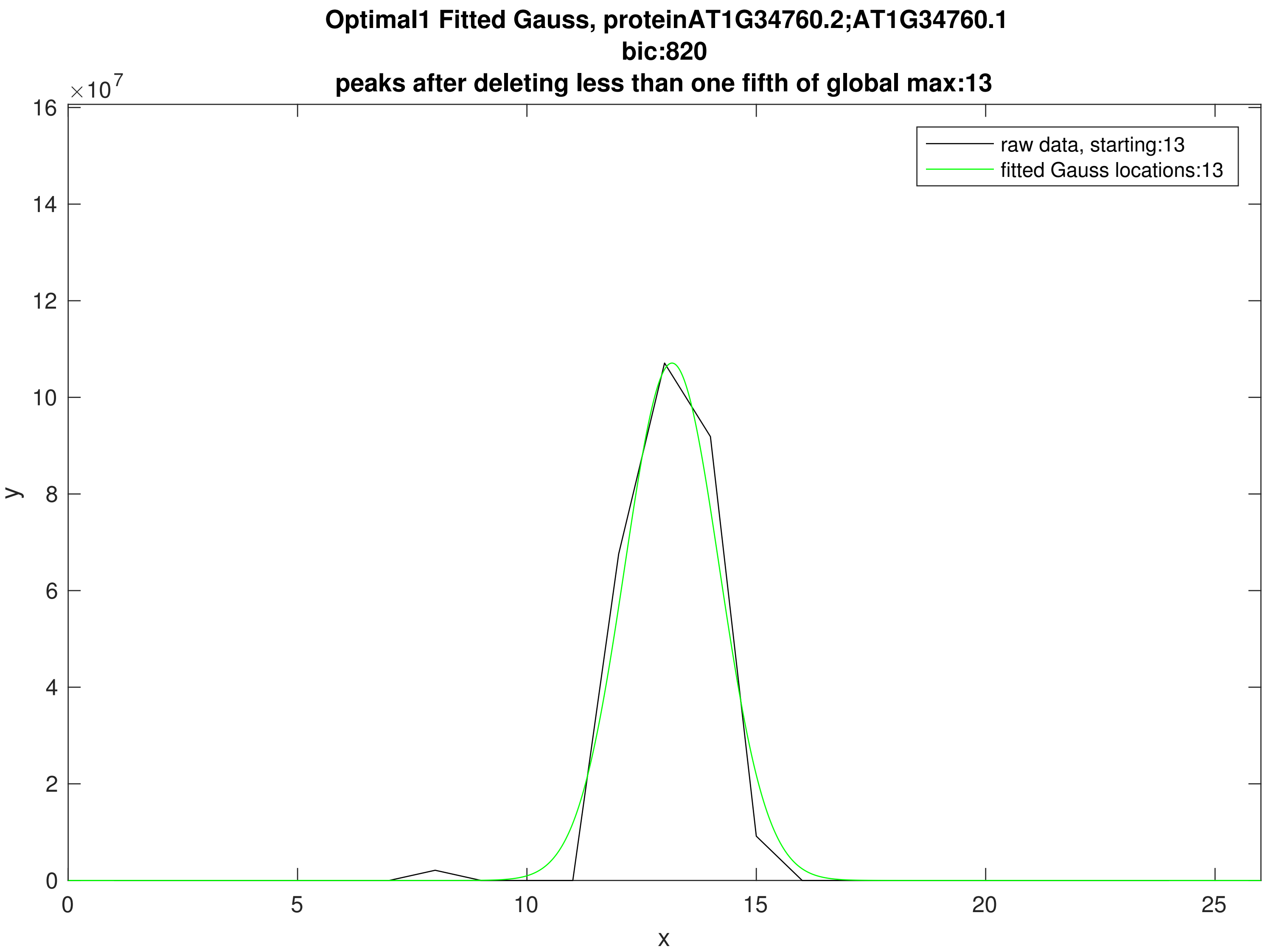
Optimal1 Fitted Gauss, proteinAT1G34760.2;AT1G34760.1
bic:820
peaks after deleting less than one fifth of global max:13
×10^7
16
14
12
10
8
6
4
2
0
y
0
5
10
15
20
25
x
raw data, starting:13
fitted Gauss locations:13

**Optimal1 Fitted Gauss, proteinAT1G35160.1;AT1G35160.2**
**bic:1.08e+03 1.09e+03 1.1e+03**
**peaks after deleting less than one fifth of global max:13**

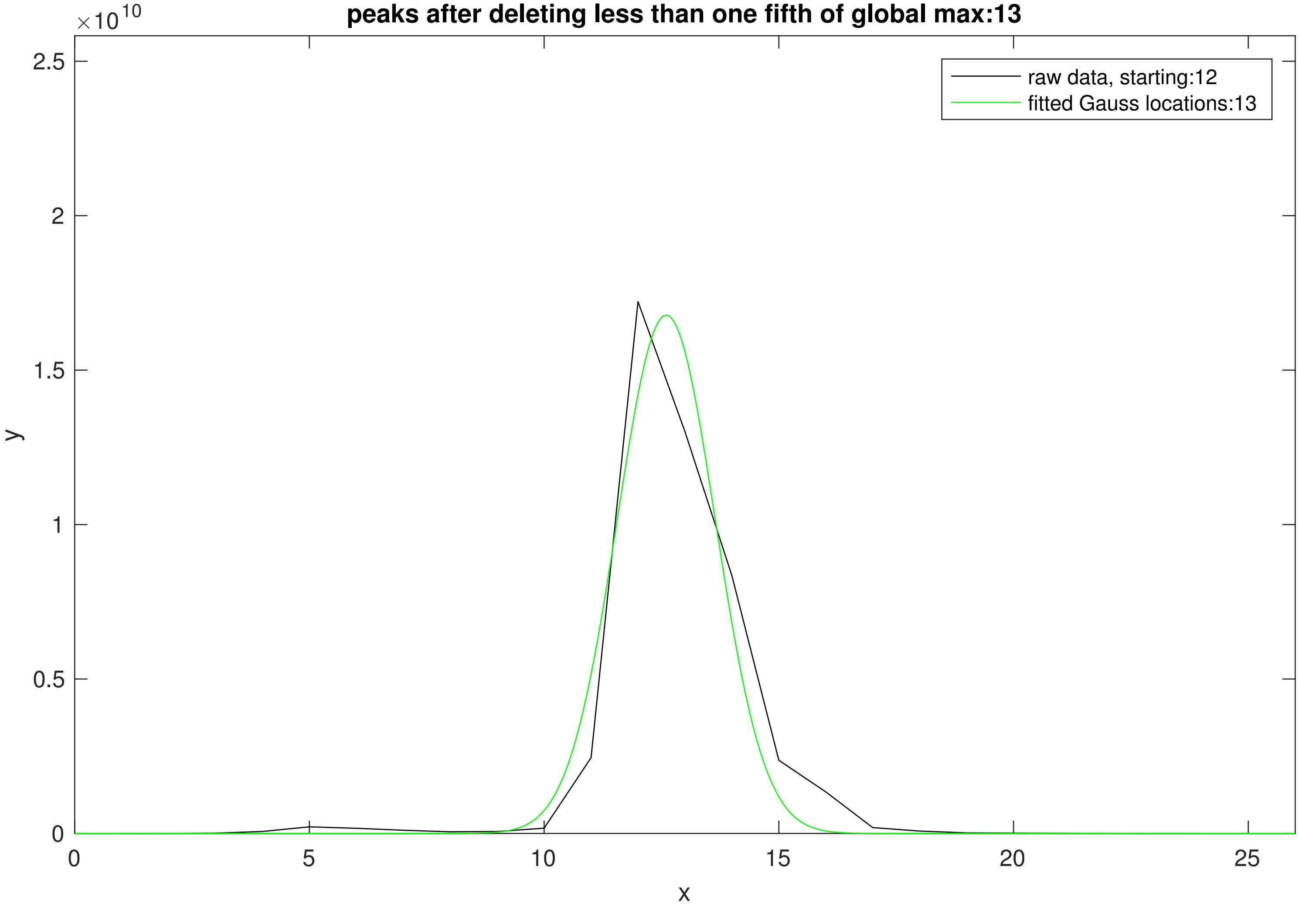

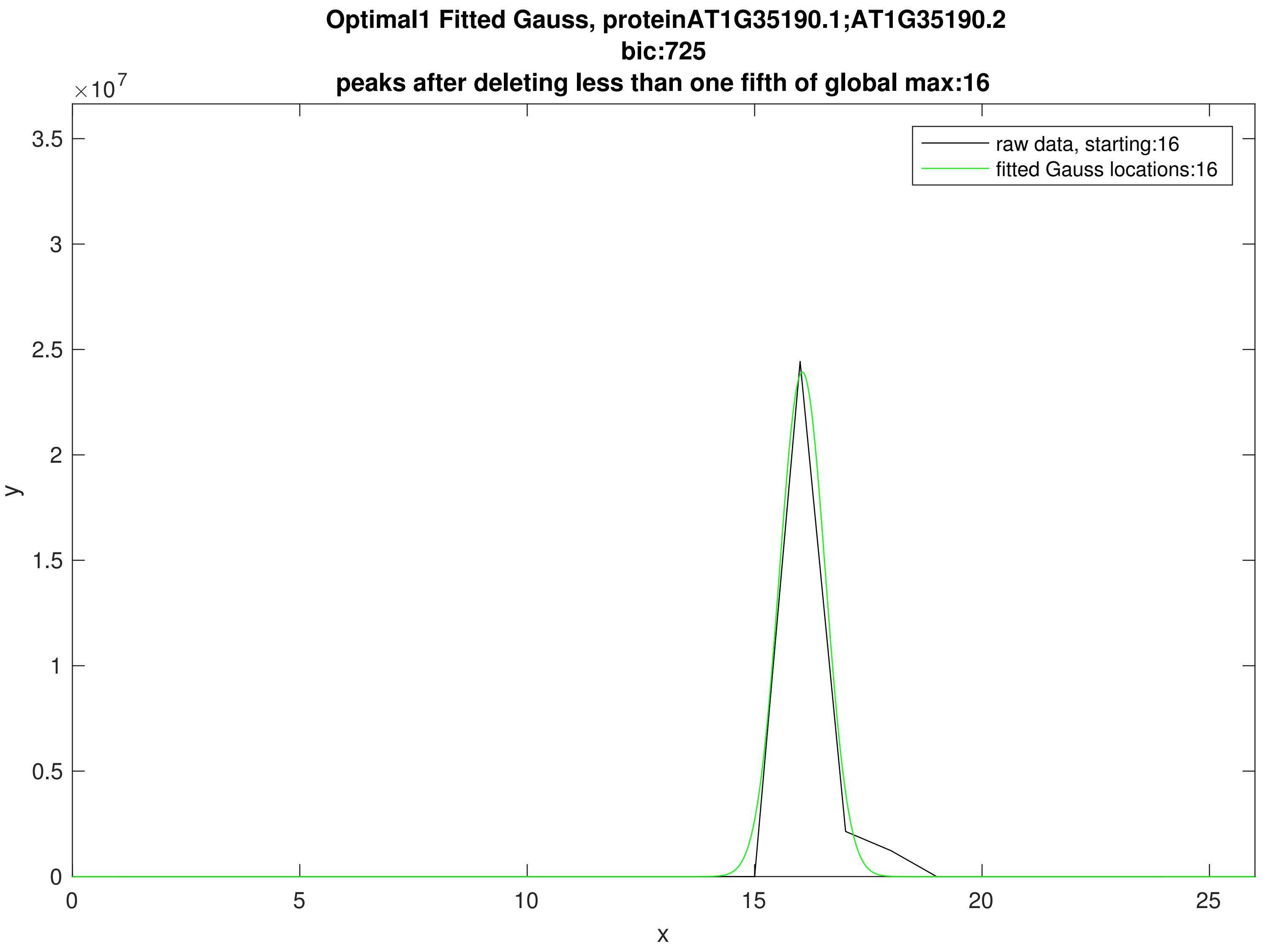
Optimal1 Fitted Gauss, proteinAT1G35190.1;AT1G35190.2
bic:725
peaks after deleting less than one fifth of global max:16
×10^7
3.5
3
2.5
2
1.5
1
0.5
0
y
0
5
10
15
20
25
x
raw data, starting:16
fitted Gauss locations:16

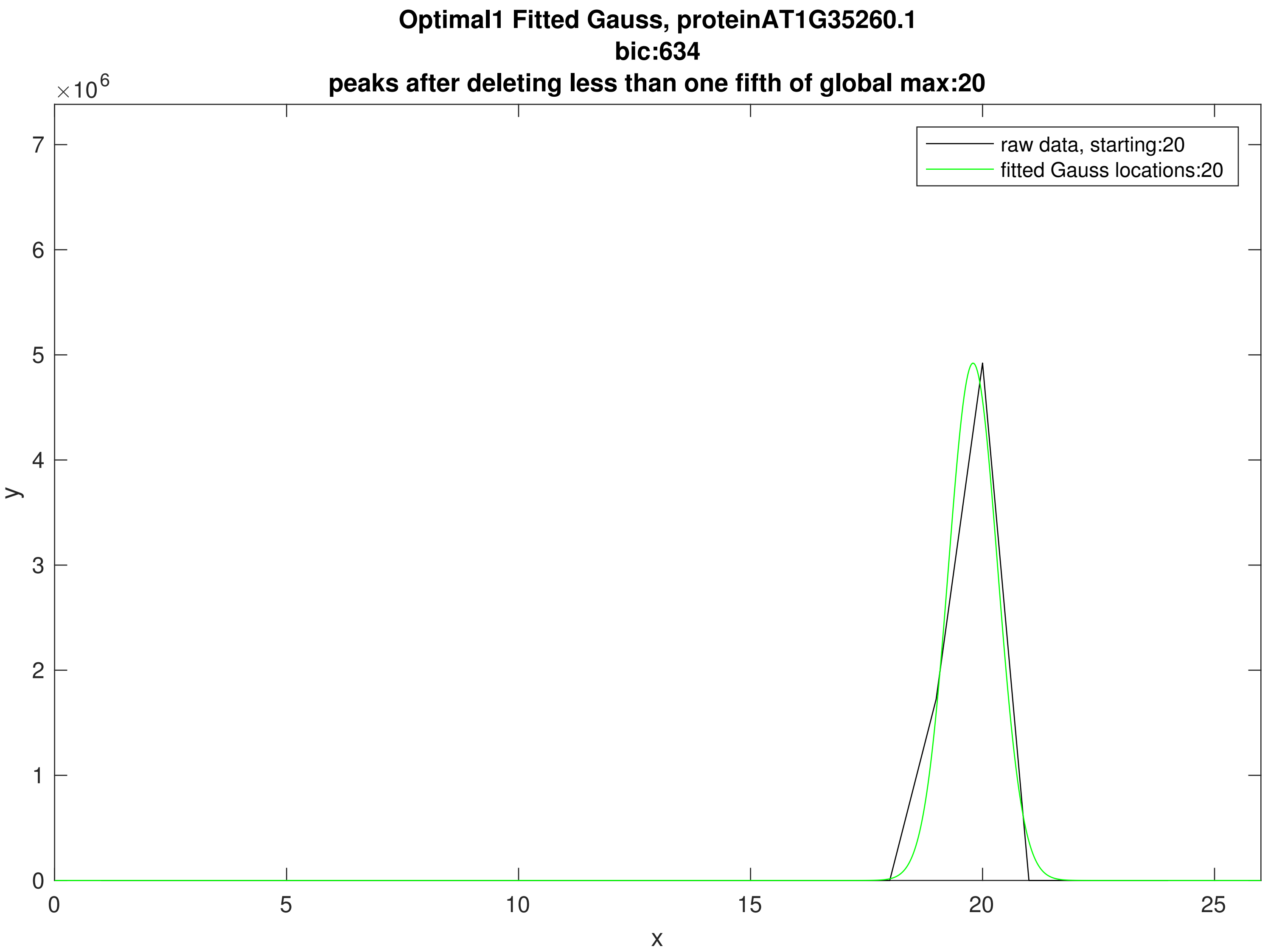
Optimal1 Fitted Gauss, proteinAT1G35260.1
bic:634
peaks after deleting less than one fifth of global max:20
×10^6
raw data, starting:20
fitted Gauss locations:20
y
x
0
1
2
3
4
5
6
7
0
5
10
15
20
25

#### Optimal1 Fitted Gauss, proteinAT1G35580.1;AT1G35580.2;AT1G35580.3
#### bic:896 899 908
#### peaks after deleting less than one fifth of global max:5.4

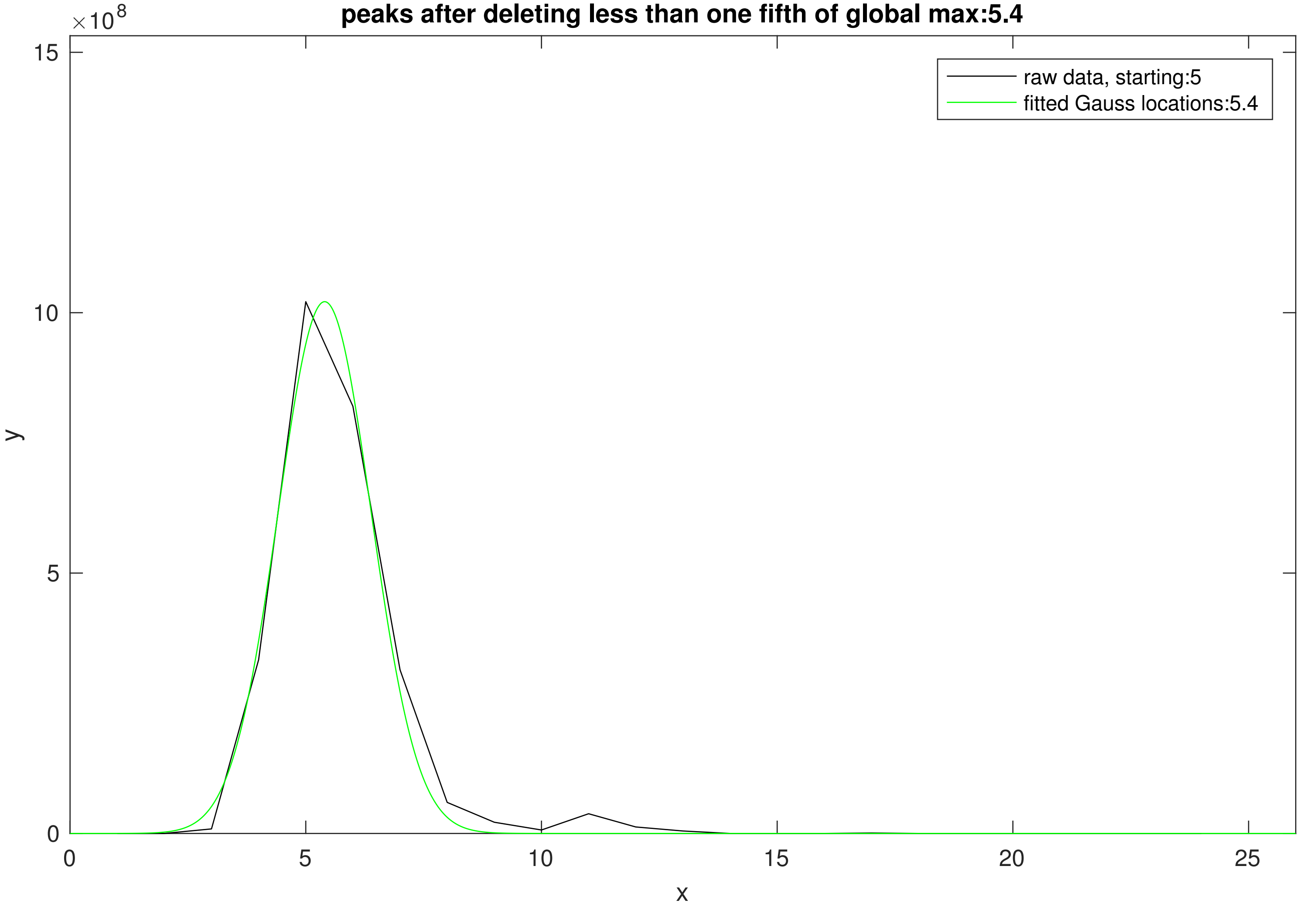

**Optimal1 Fitted Gauss, proteinAT1G35720.1**
**bic:1.09e+03   1.1e+03  1.11e+03  1.12e+03**
**peaks after deleting less than one fifth of global max:14**

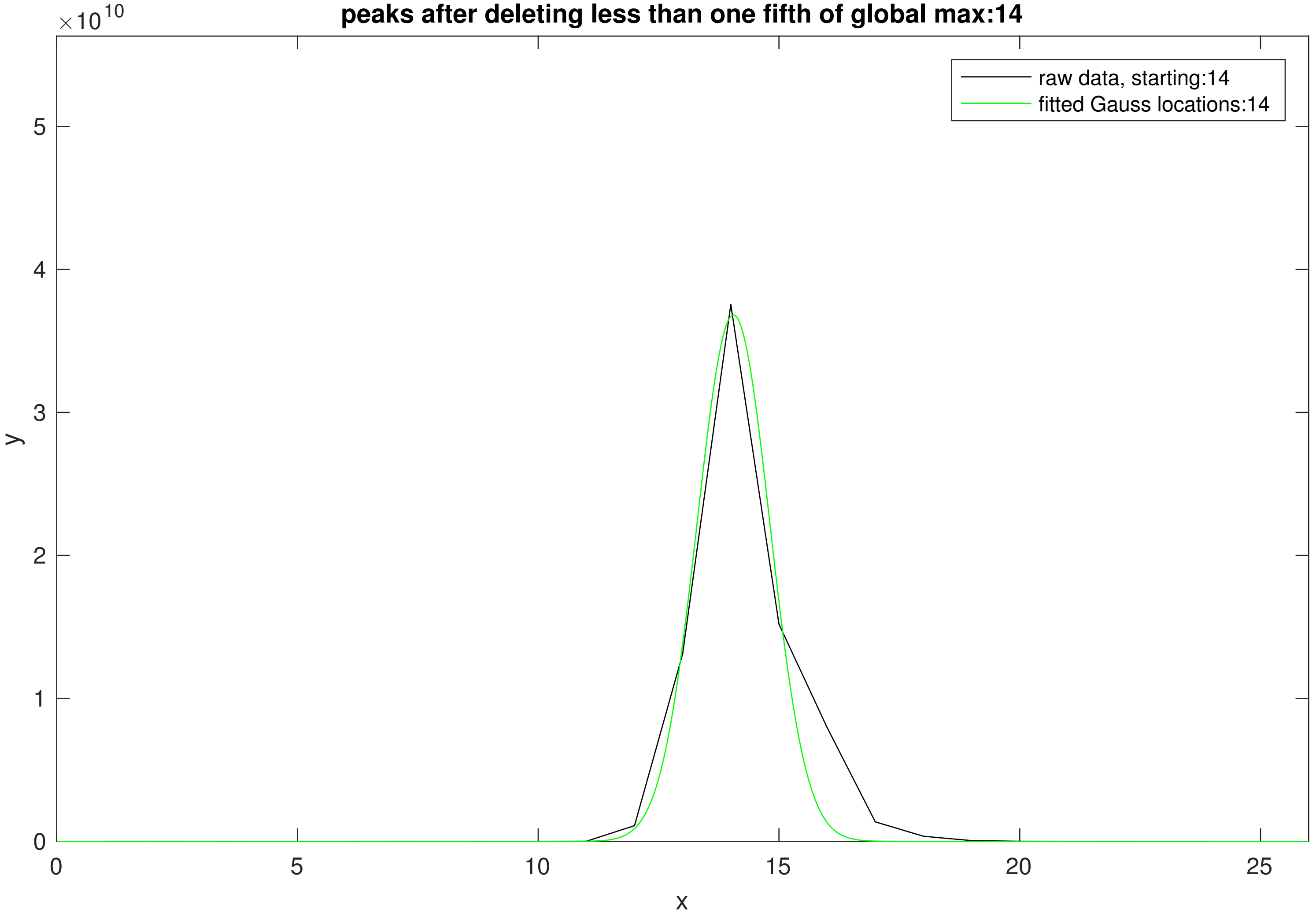

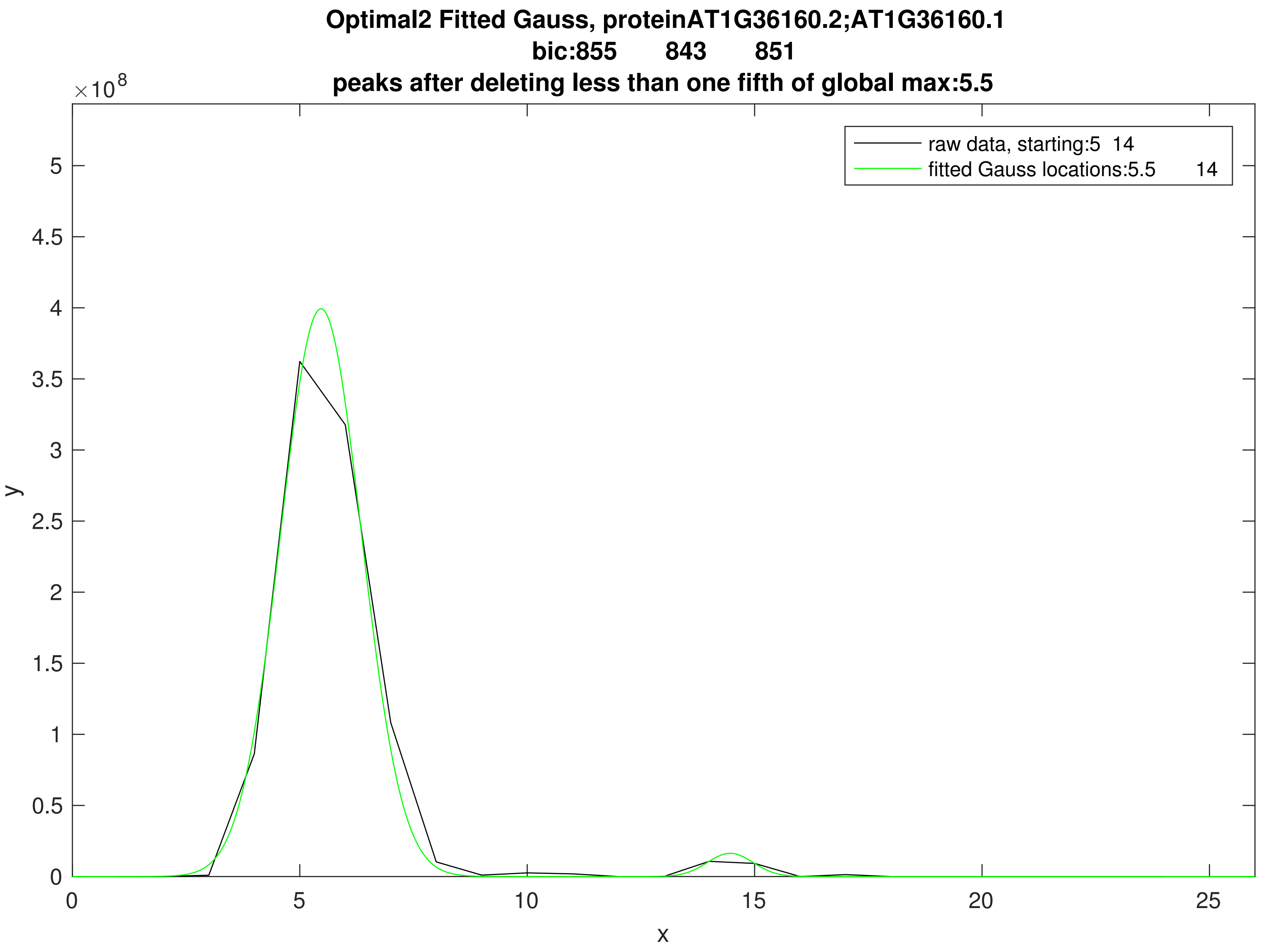
Optimal2 Fitted Gauss, proteinAT1G36160.2;AT1G36160.1
bic:855 843 851
peaks after deleting less than one fifth of global max:5.5
×10^8
raw data, starting:5 14
fitted Gauss locations:5.5 14
y
x
0
0.5
1
1.5
2
2.5
3
3.5
4
4.5
5
0
5
10
15
20
25

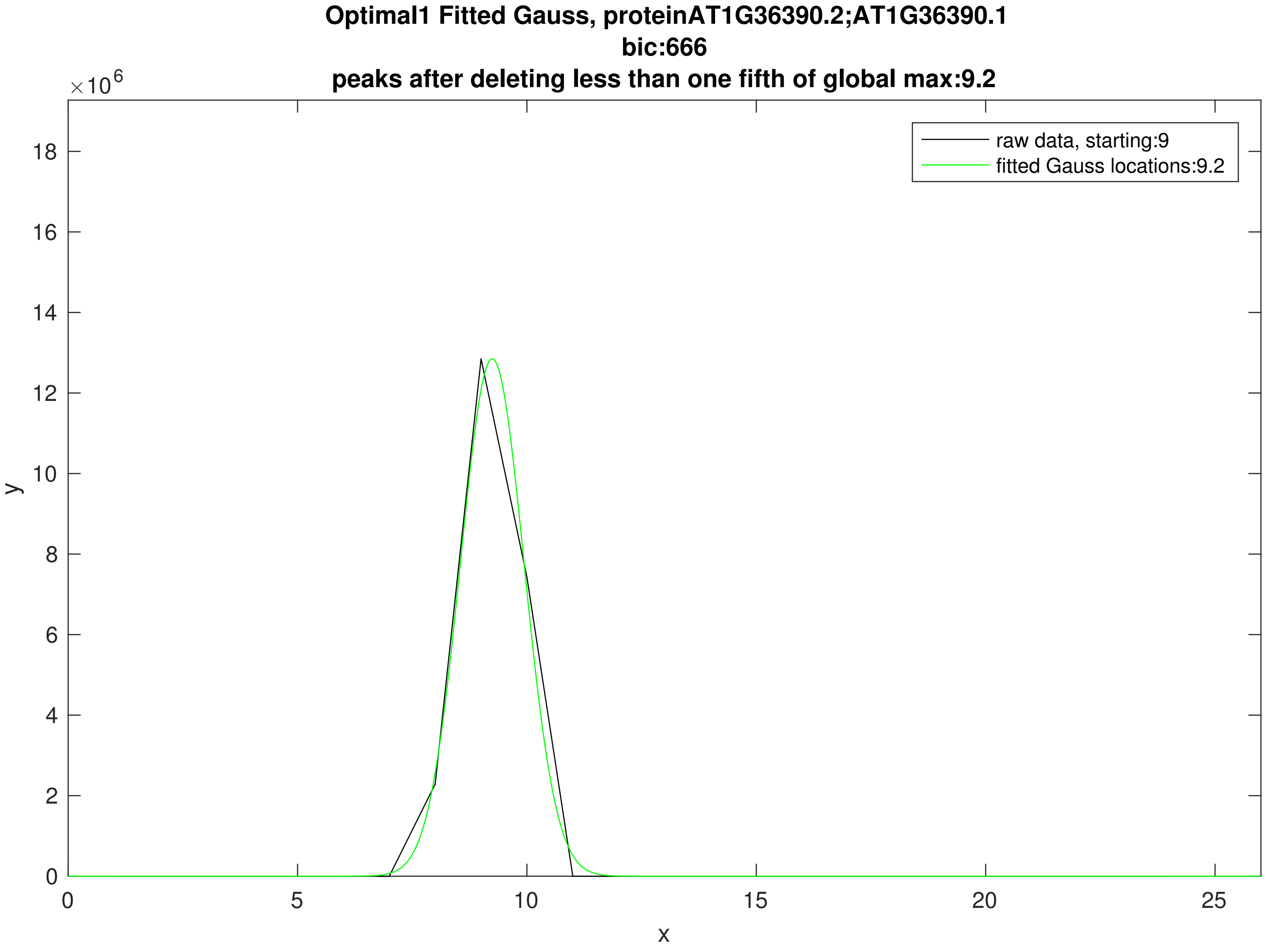
Optimal1 Fitted Gauss, proteinAT1G36390.2;AT1G36390.1
bic:666
peaks after deleting less than one fifth of global max:9.2
×10^6
raw data, starting:9
fitted Gauss locations:9.2
y
x
0
2
4
6
8
10
12
14
16
18
0
5
10
15
20
25

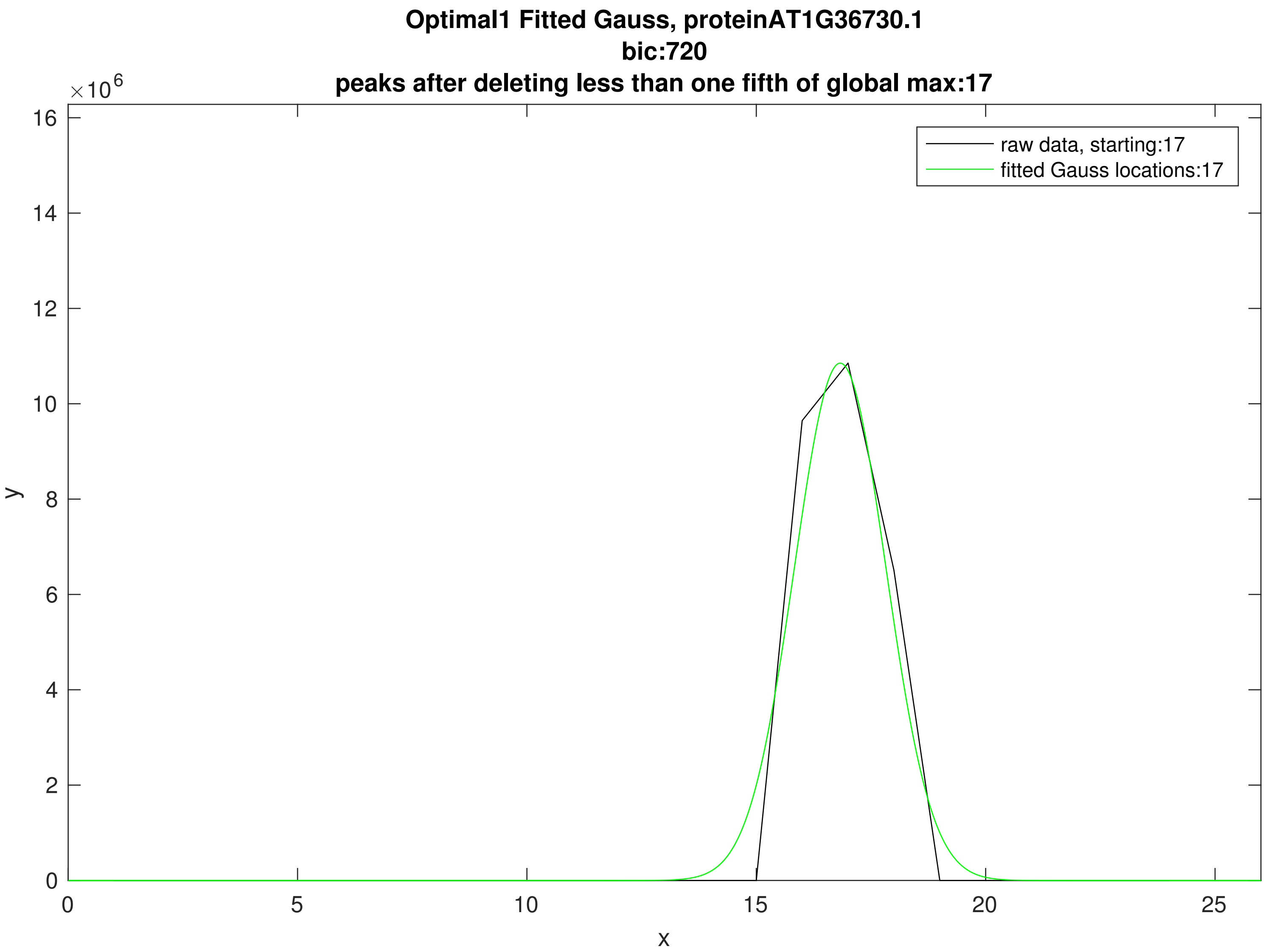

Optimal1 Fitted Gauss, proteinAT1G36730.1
bic:720
peaks after deleting less than one fifth of global max:17
×10^6
16
14
12
10
8
6
4
2
0
y
0
5
10
15
20
25
x
raw data, starting:17
fitted Gauss locations:17

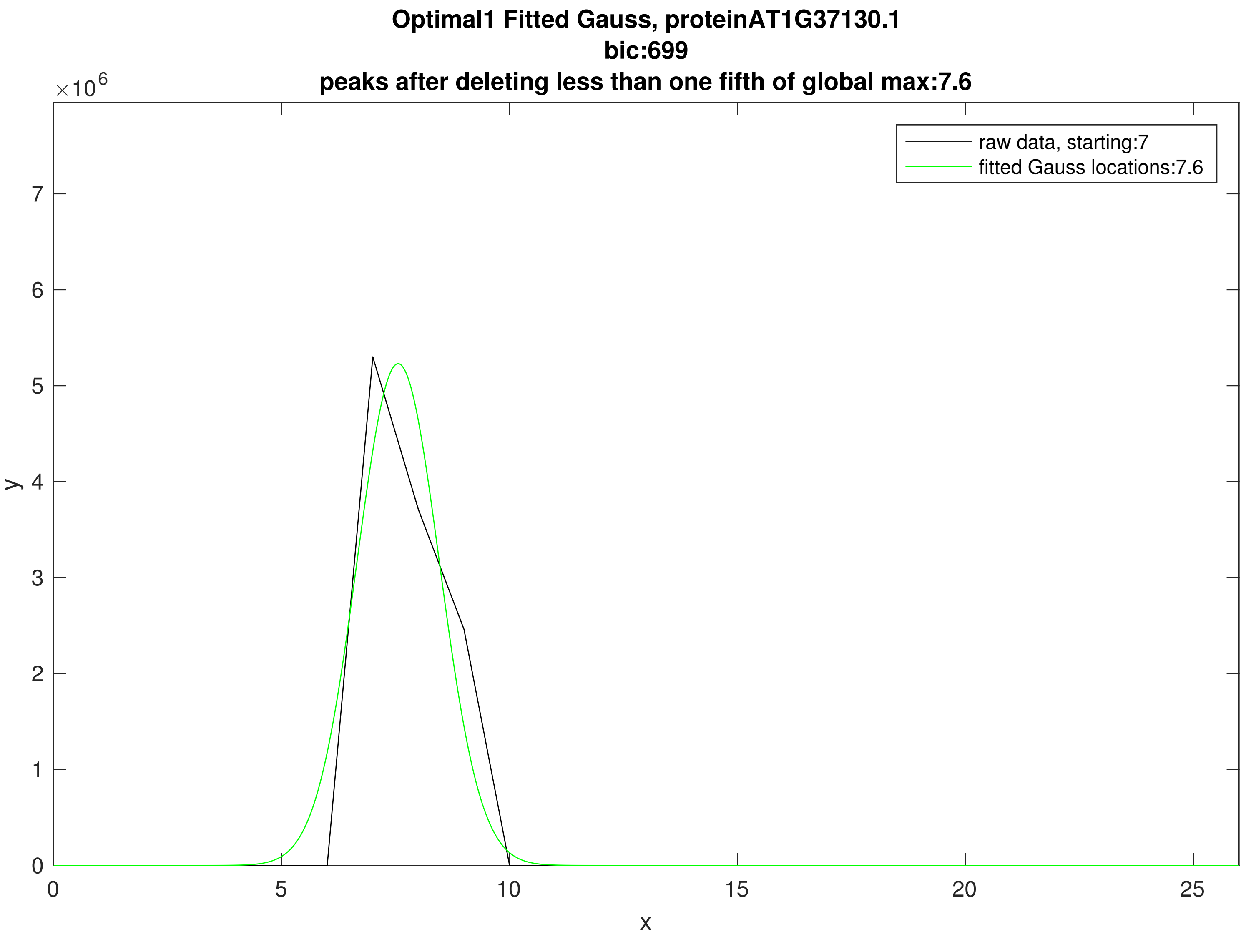
Optimal1 Fitted Gauss, proteinAT1G37130.1
bic:699
peaks after deleting less than one fifth of global max:7.6
×10^6
raw data, starting:7
fitted Gauss locations:7.6
y
x
0
1
2
3
4
5
6
7
0
5
10
15
20
25

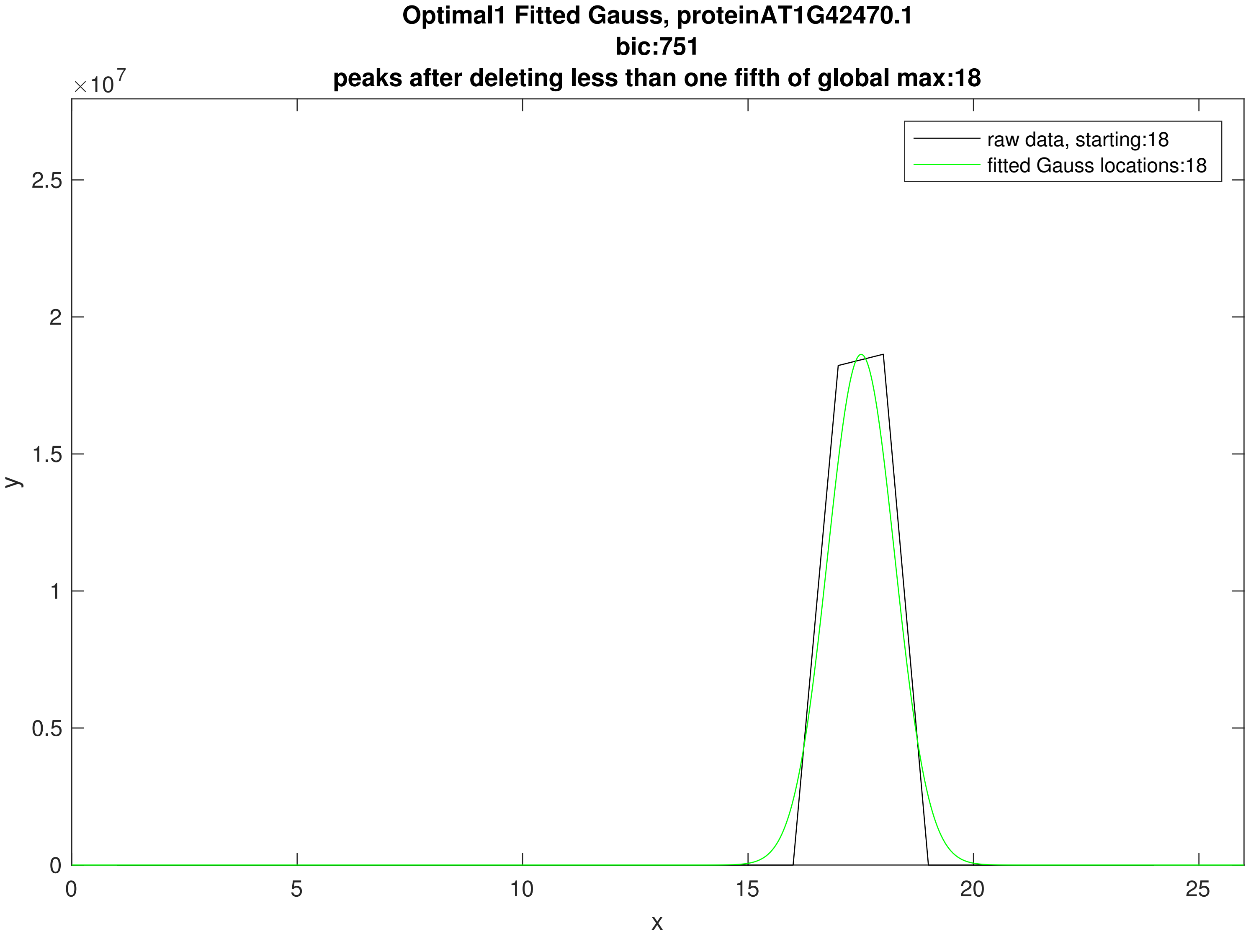
Optimal1 Fitted Gauss, proteinAT1G42470.1
bic:751
peaks after deleting less than one fifth of global max:18
×10^7
2.5
2
1.5
1
0.5
0
y
0
5
10
15
20
25
x
raw data, starting:18
fitted Gauss locations:18

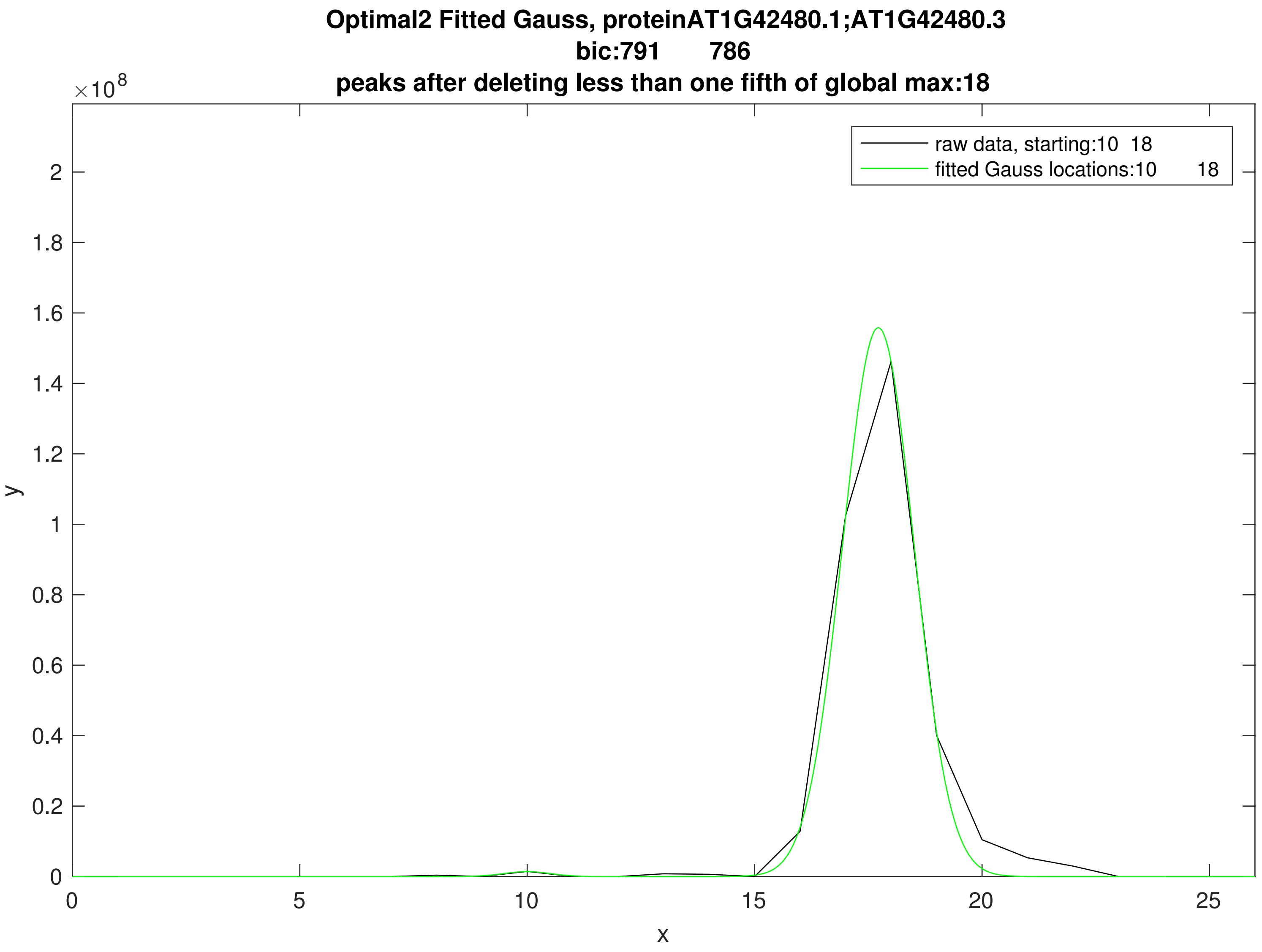
Optimal2 Fitted Gauss, proteinAT1G42480.1;AT1G42480.3
bic:791 786
peaks after deleting less than one fifth of global max:18
×10^8
raw data, starting:10 18
fitted Gauss locations:10 18
y
x
0
0.2
0.4
0.6
0.8
1
1.2
1.4
1.6
1.8
2
5
10
15
20
25

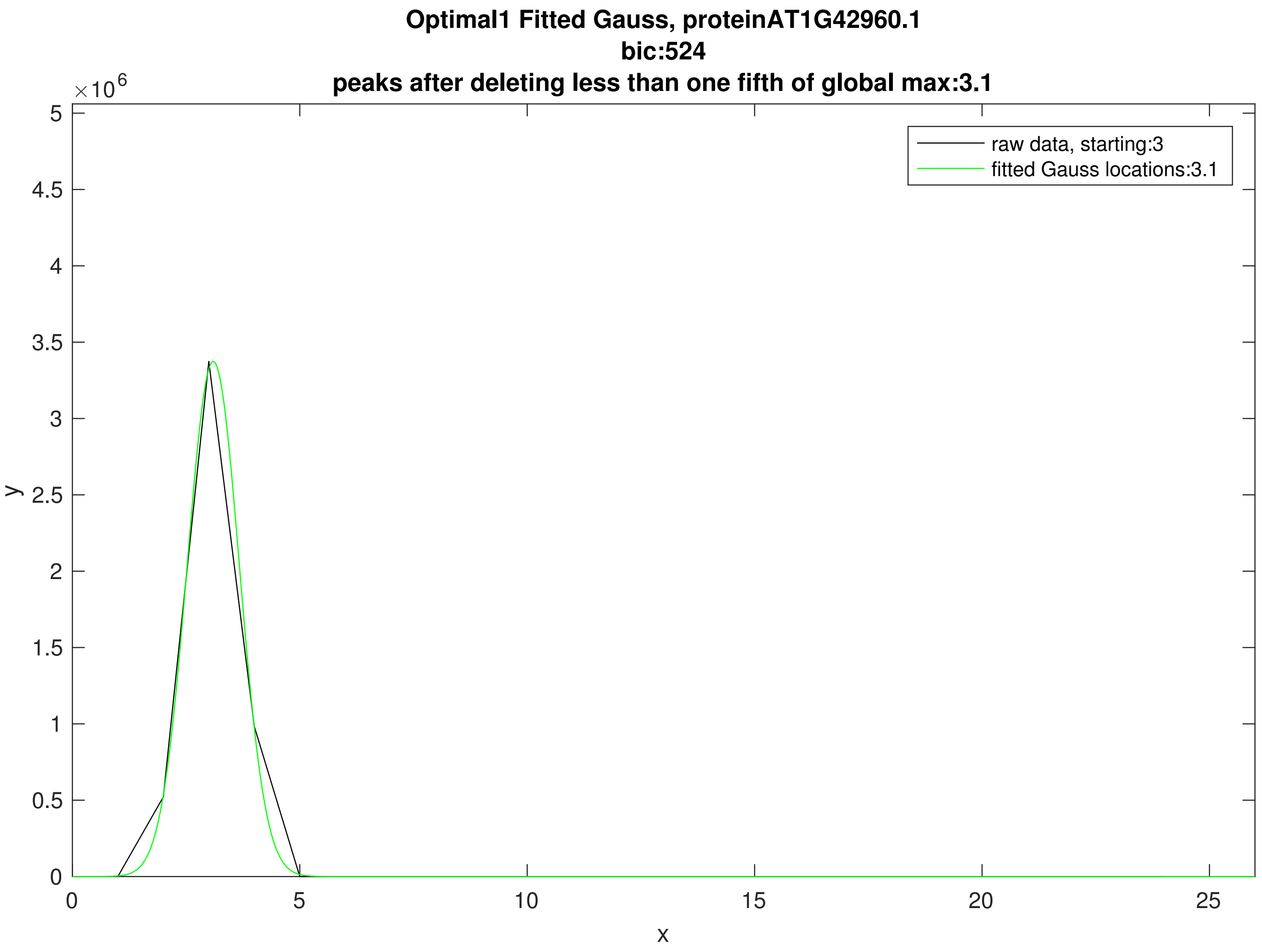

Optimal1 Fitted Gauss, proteinAT1G42960.1
bic:524
peaks after deleting less than one fifth of global max:3.1
×10^6
raw data, starting:3
fitted Gauss locations:3.1
y
x
0
0.5
1
1.5
2
2.5
3
3.5
4
4.5
5
0
5
10
15
20
25

**Optimal1 Fitted Gauss, proteinAT1G42970.1**
**bic:849**
**peaks after deleting less than one fifth of global max:5.8**

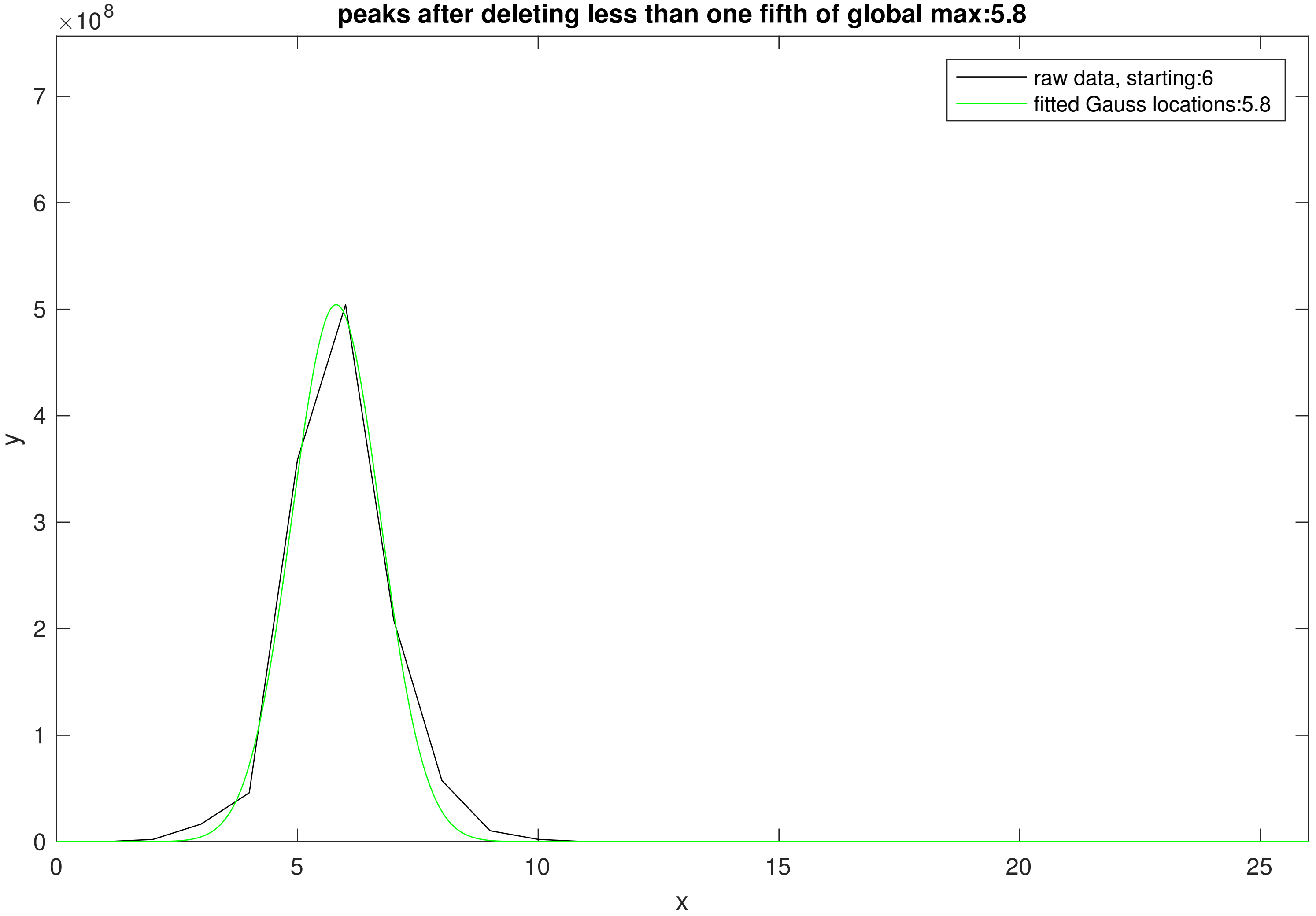

d Gauss, proteinAT1G43170.9;AT1G43170.8;AT1G43170.7;AT1G43170.6;AT1G43170.5;AT1G43170.3;AT1G43170.2;AT1

bic:872

peaks after deleting less than one fifth of global max:3.5

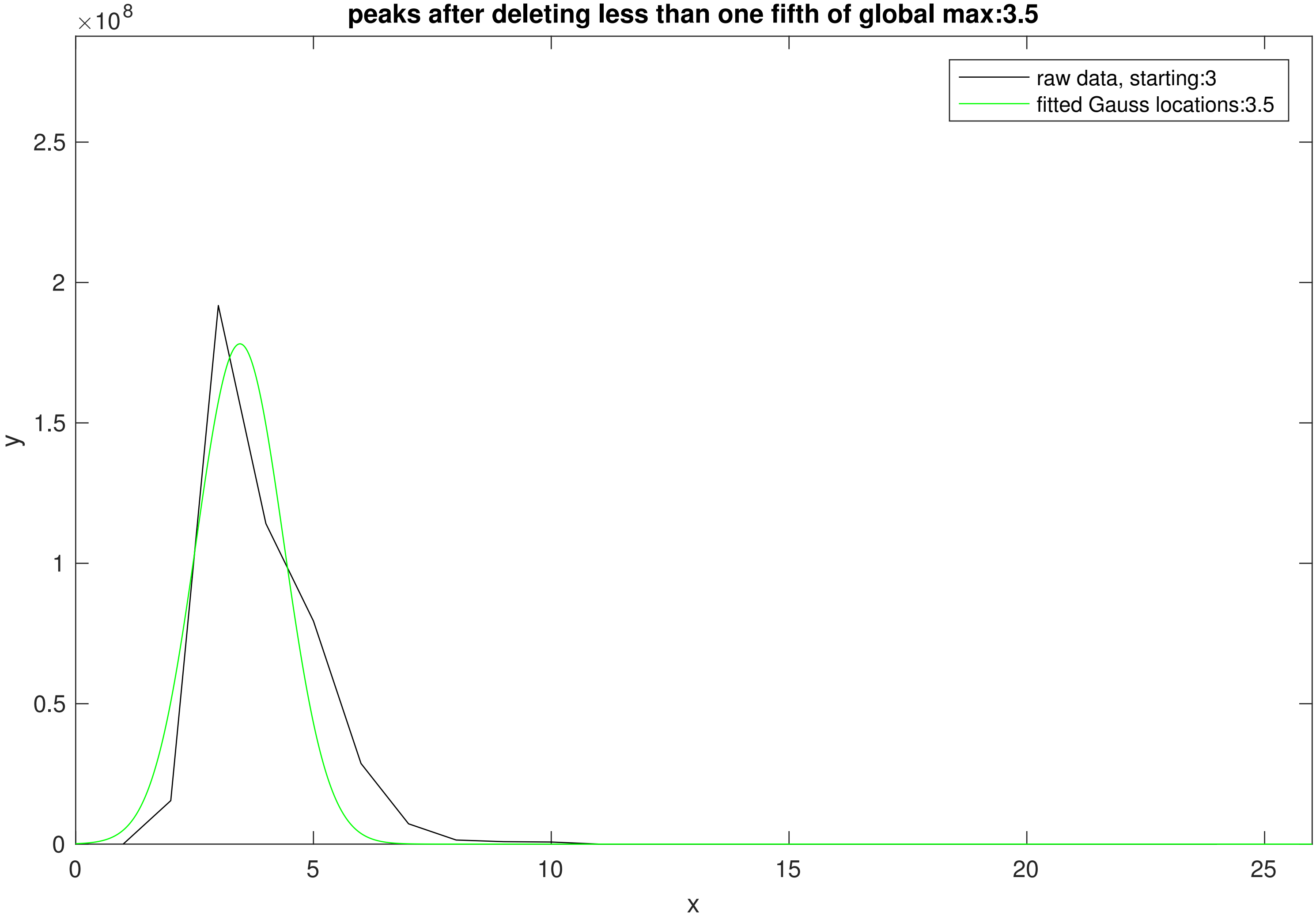

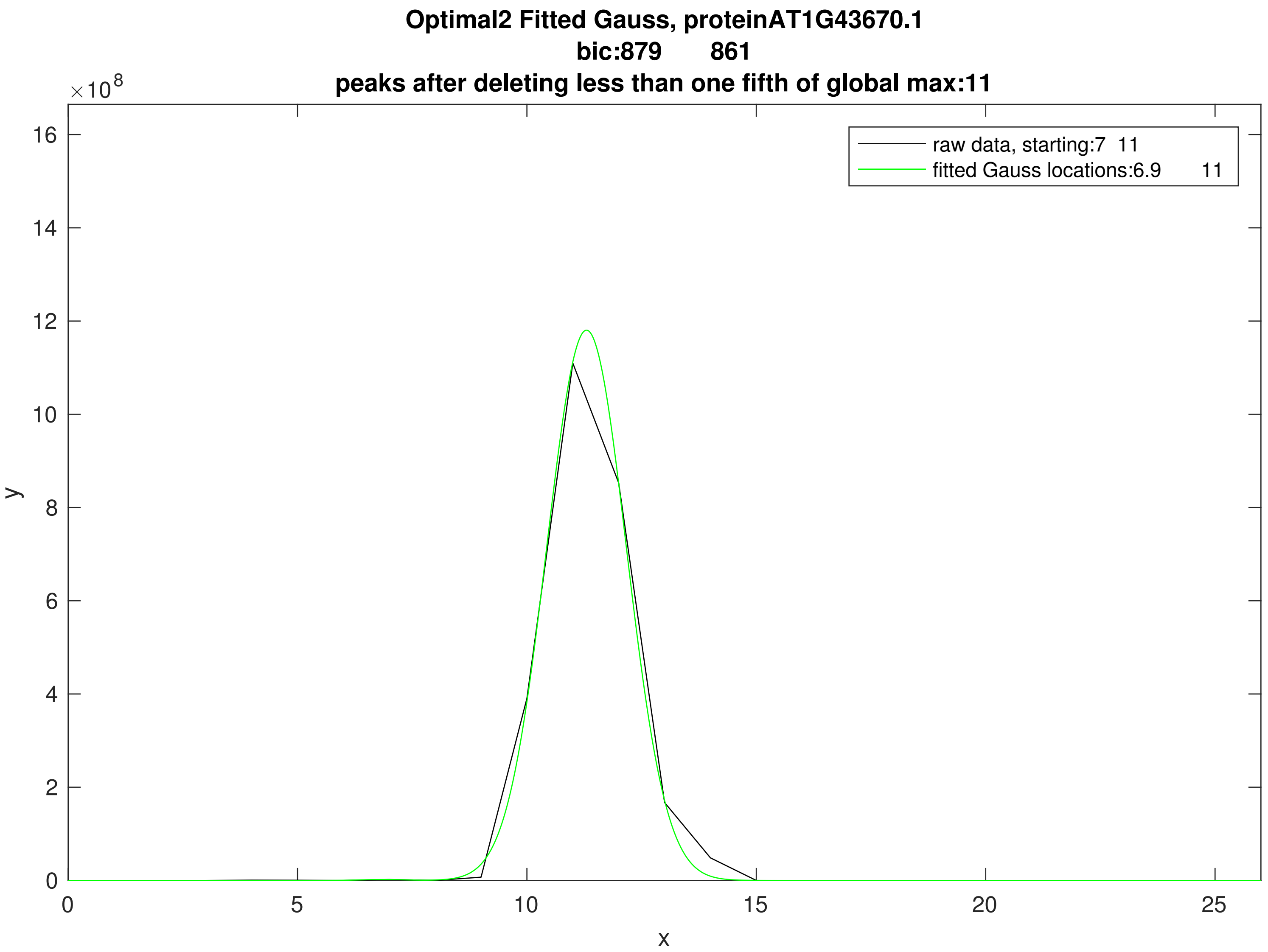

Optimal2 Fitted Gauss, proteinAT1G43670.1
bic:879 861
peaks after deleting less than one fifth of global max:11
×10^8
raw data, starting:7 11
fitted Gauss locations:6.9 11
y
x
0
2
4
6
8
10
12
14
16
0
5
10
15
20
25

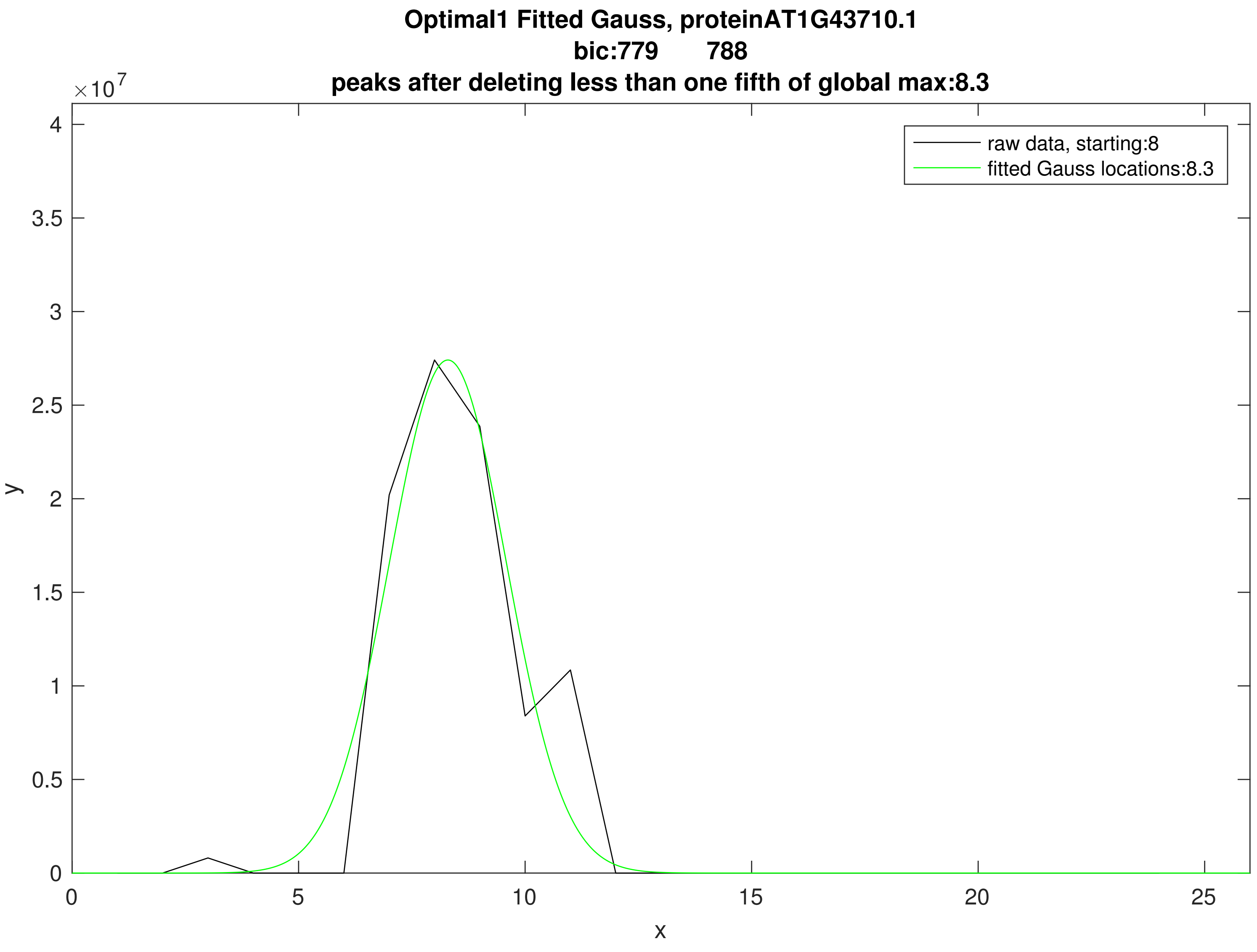
Optimal1 Fitted Gauss, proteinAT1G43710.1
bic:779 788
peaks after deleting less than one fifth of global max:8.3
×10^7
raw data, starting:8
fitted Gauss locations:8.3
y
x
0
0.5
1
1.5
2
2.5
3
3.5
4
0
5
10
15
20
25

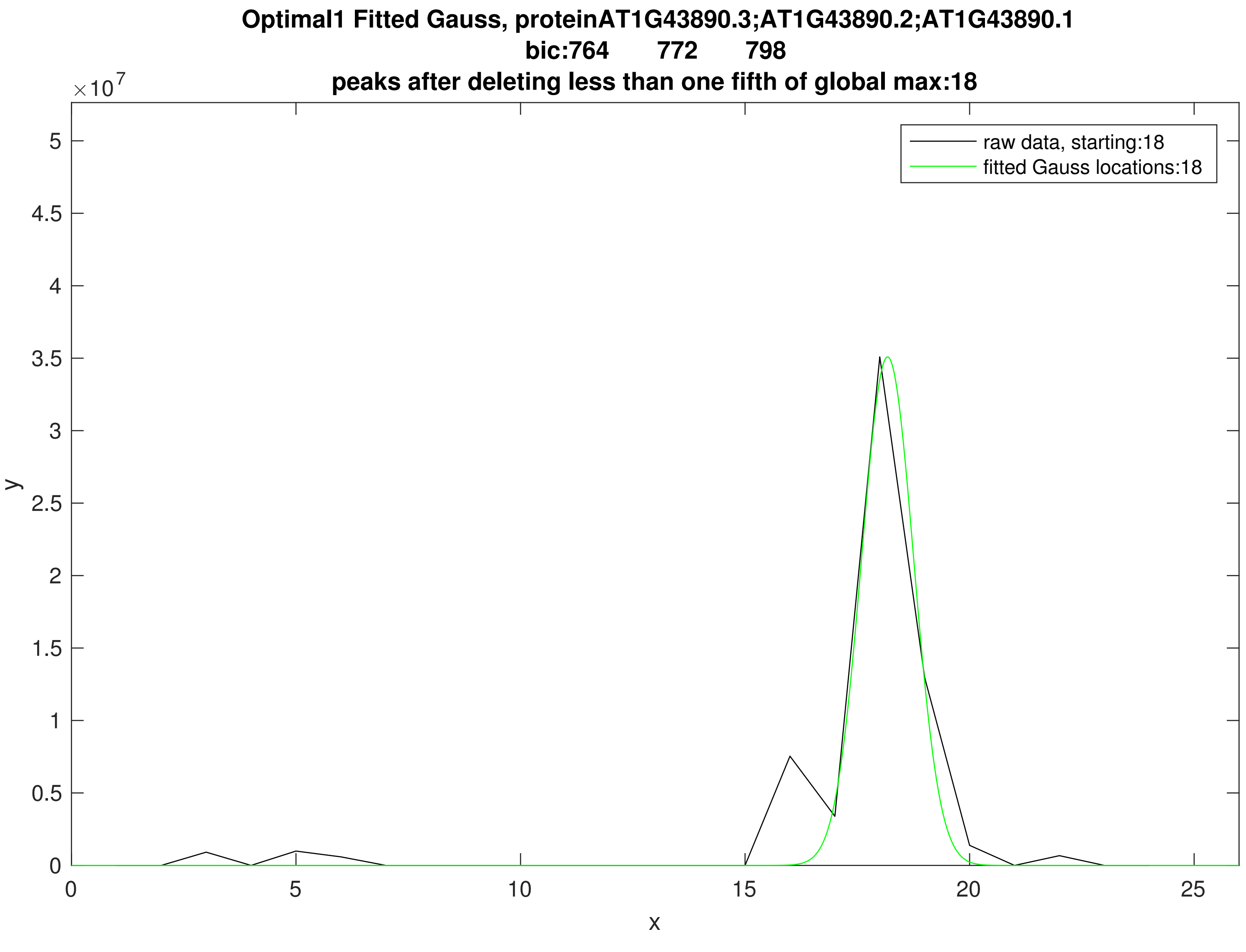

Optimal1 Fitted Gauss, proteinAT1G43890.3;AT1G43890.2;AT1G43890.1
bic:764 772 798
peaks after deleting less than one fifth of global max:18
×10^7
raw data, starting:18
fitted Gauss locations:18
y
x
0
0.5
1
1.5
2
2.5
3
3.5
4
4.5
5
0
5
10
15
20
25

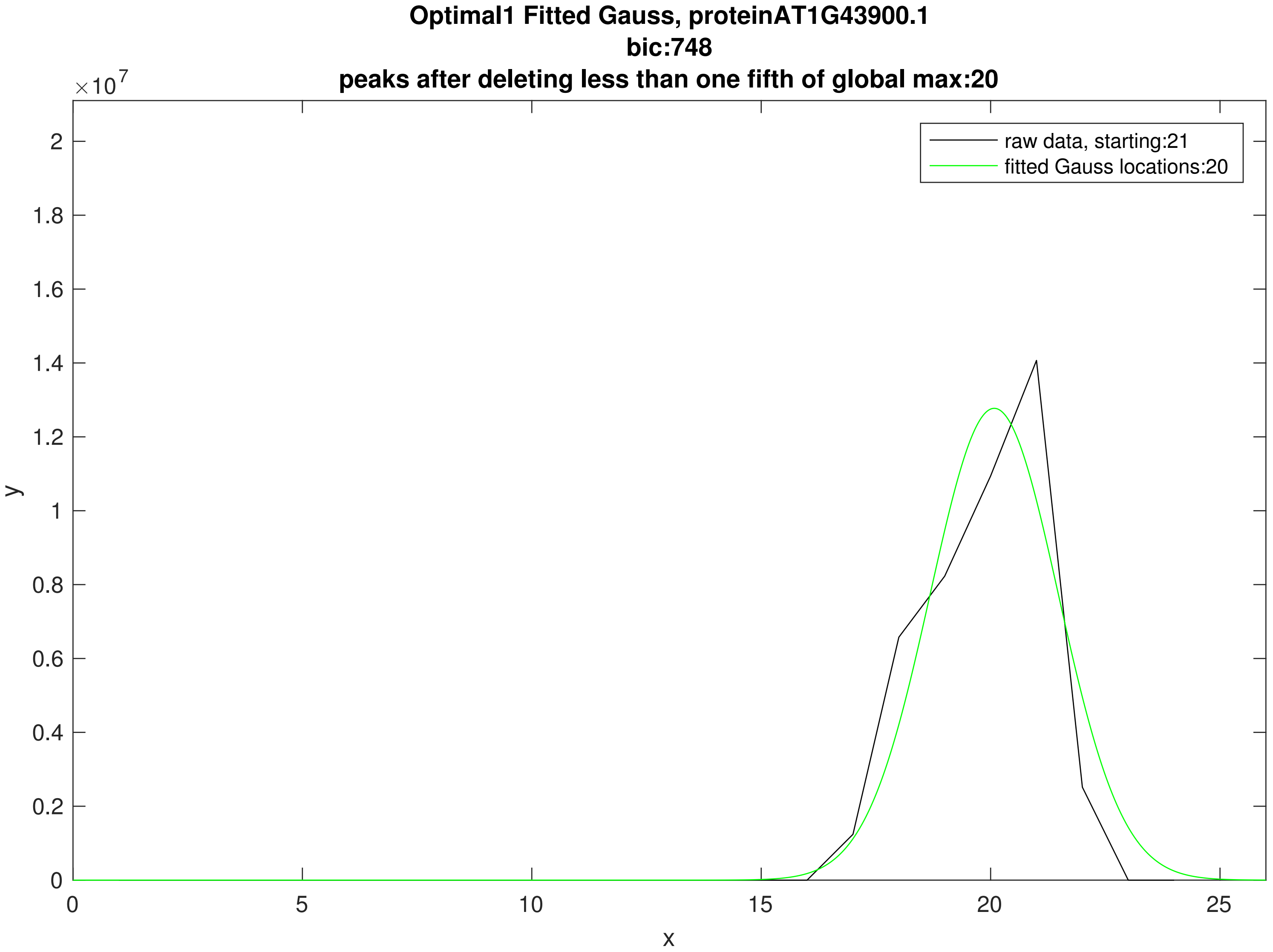

Optimal1 Fitted Gauss, proteinAT1G43900.1
bic:748
peaks after deleting less than one fifth of global max:20
×10^7
raw data, starting:21
fitted Gauss locations:20
y
x
0
0.2
0.4
0.6
0.8
1
1.2
1.4
1.6
1.8
2
0
5
10
15
20
25

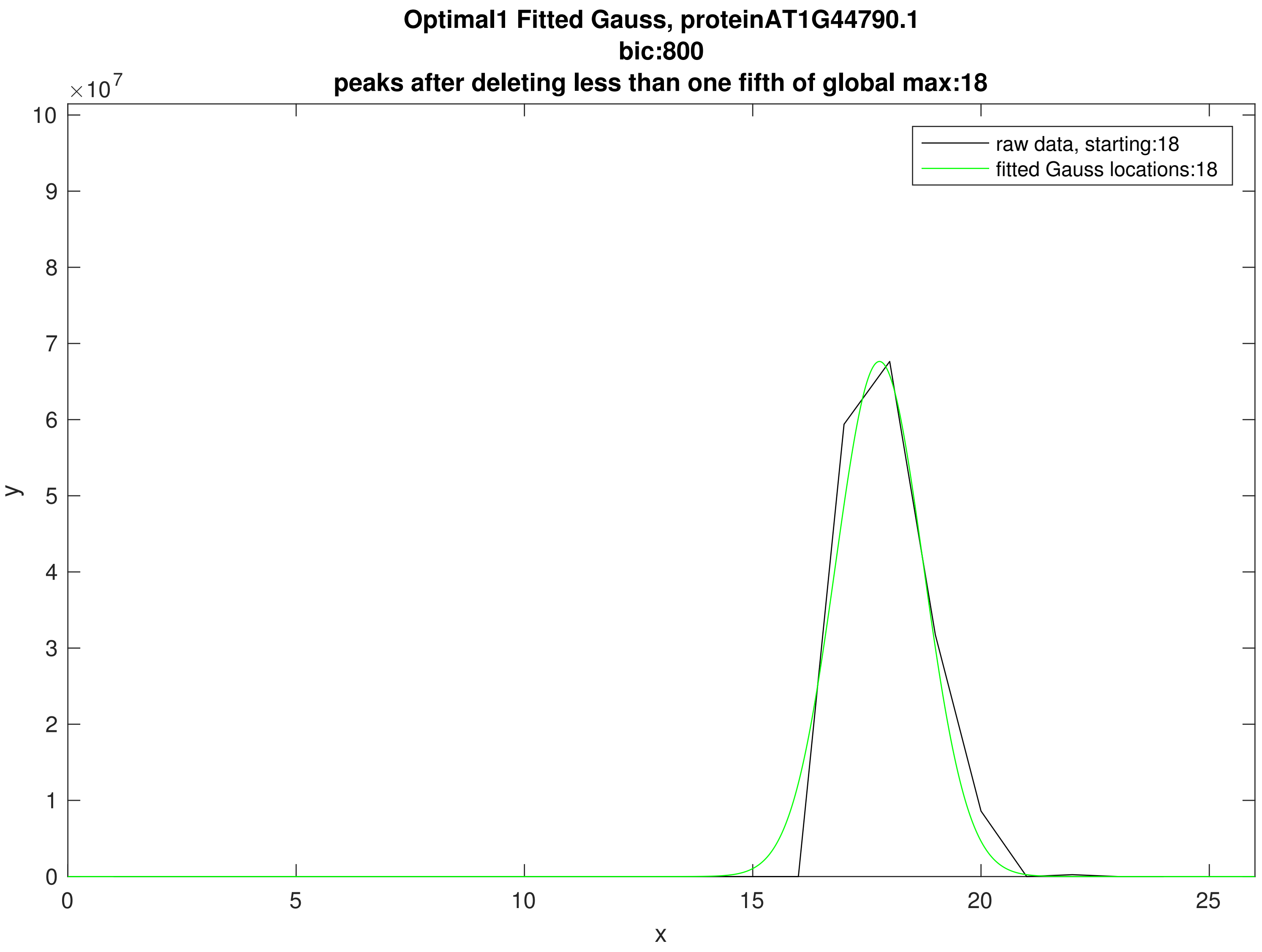
Optimal1 Fitted Gauss, proteinAT1G44790.1
bic:800
peaks after deleting less than one fifth of global max:18
raw data, starting:18
fitted Gauss locations:18
$\times10^7$
y
x

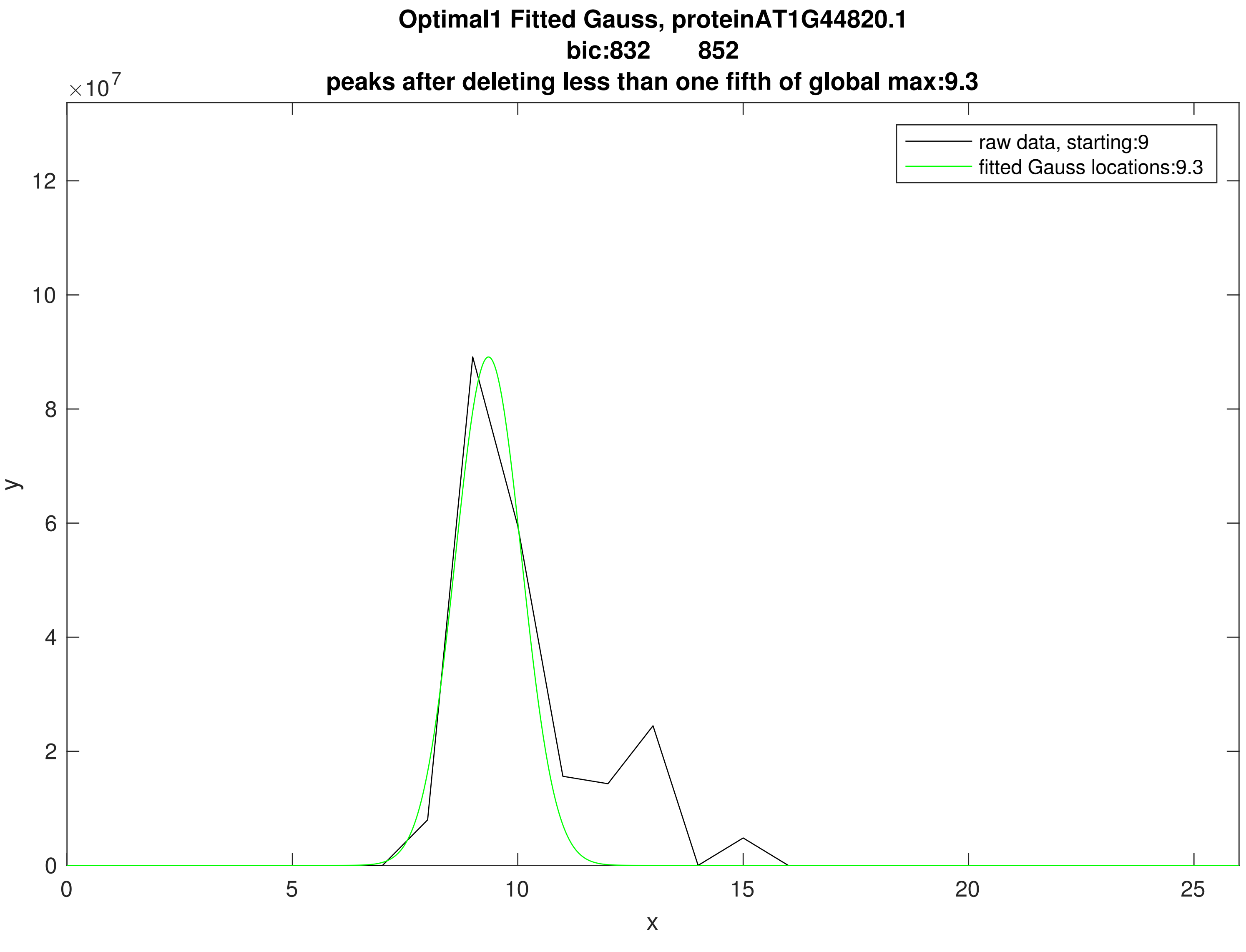
Optimal1 Fitted Gauss, proteinAT1G44820.1
bic:832 852
peaks after deleting less than one fifth of global max:9.3
×10^7
raw data, starting:9
fitted Gauss locations:9.3
y
x
0
2
4
6
8
10
12
0
5
10
15
20
25

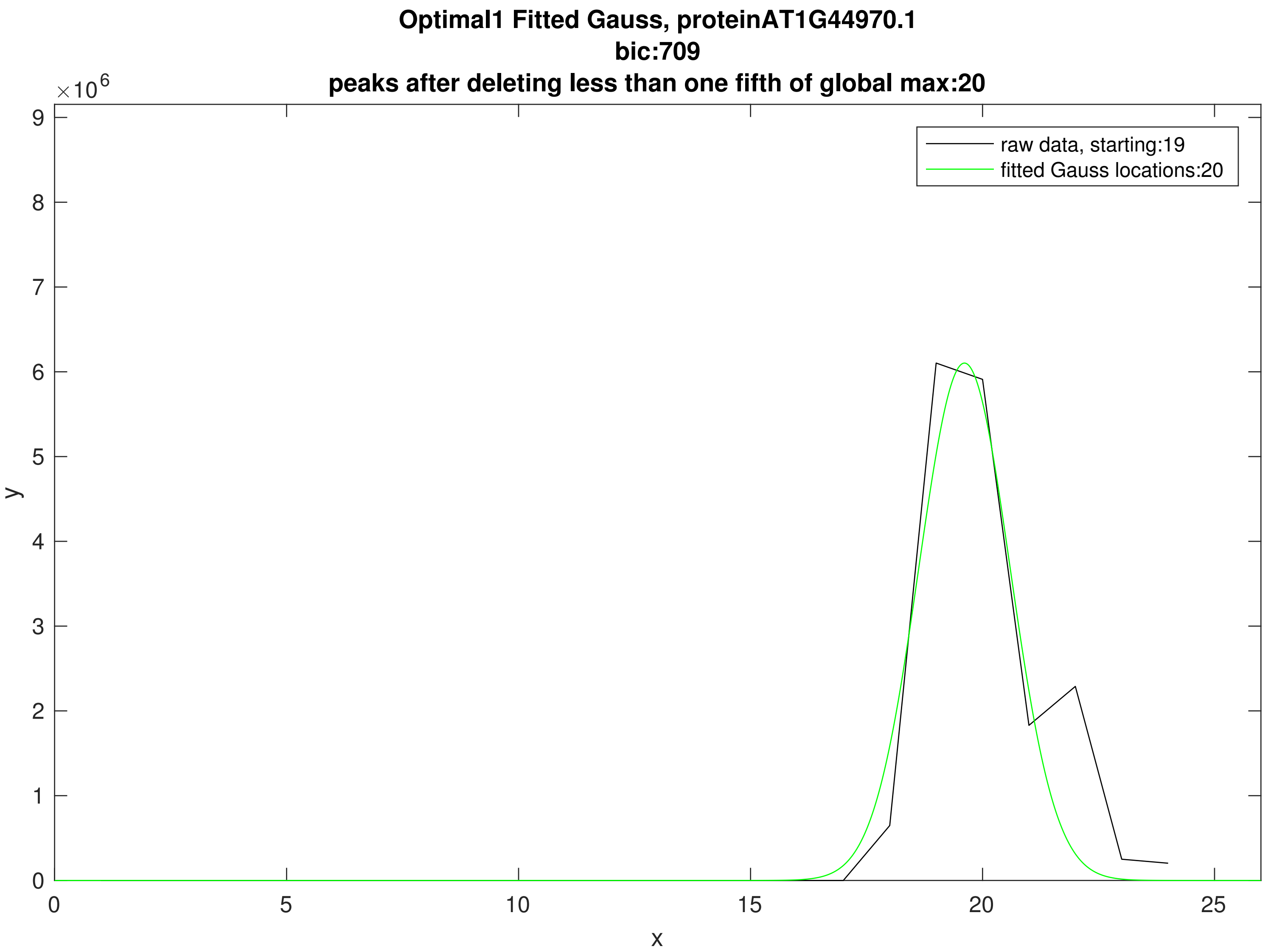

Optimal1 Fitted Gauss, proteinAT1G44970.1
bic:709
peaks after deleting less than one fifth of global max:20
×10^6
raw data, starting:19
fitted Gauss locations:20
y
x
0
1
2
3
4
5
6
7
8
9
0
5
10
15
20
25

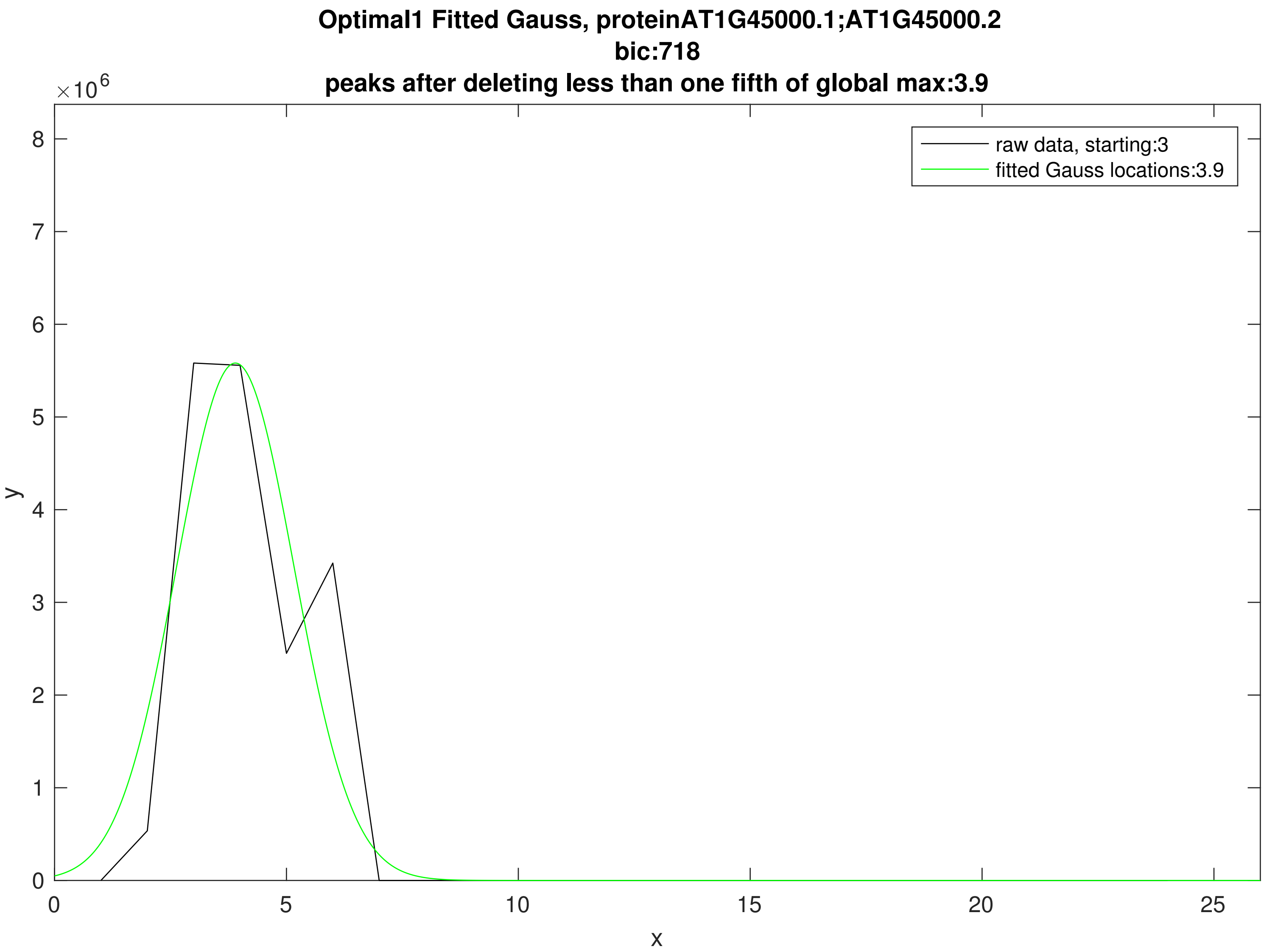
Optimal1 Fitted Gauss, proteinAT1G45000.1;AT1G45000.2
bic:718
peaks after deleting less than one fifth of global max:3.9
×10^6
y
x
raw data, starting:3
fitted Gauss locations:3.9
0
1
2
3
4
5
6
7
8
0
5
10
15
20
25

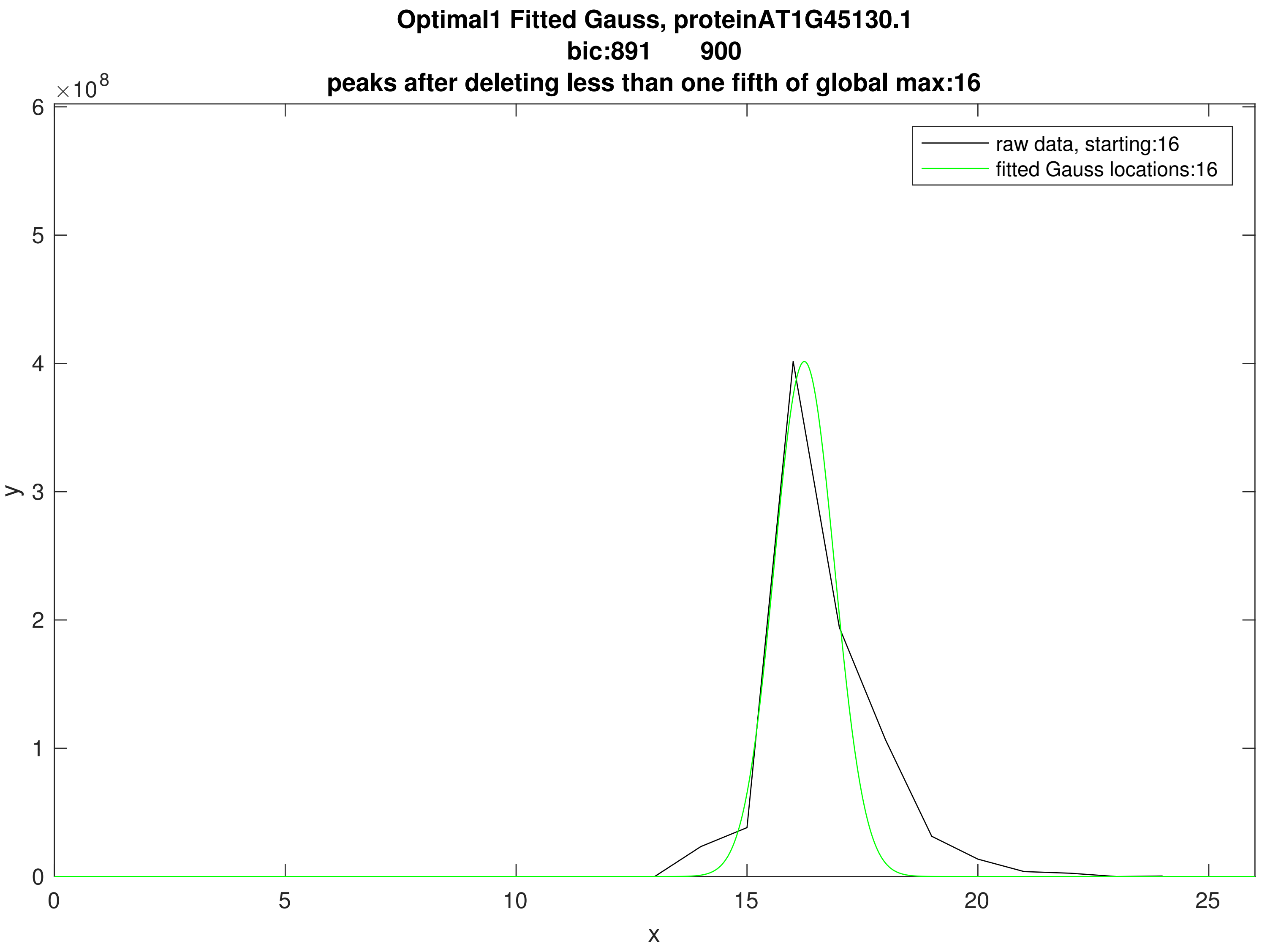

Optimal1 Fitted Gauss, proteinAT1G45130.1
bic:891 900
peaks after deleting less than one fifth of global max:16
×10^8
6
5
4
3
2
1
0
y
0
5
10
15
20
25
x
raw data, starting:16
fitted Gauss locations:16

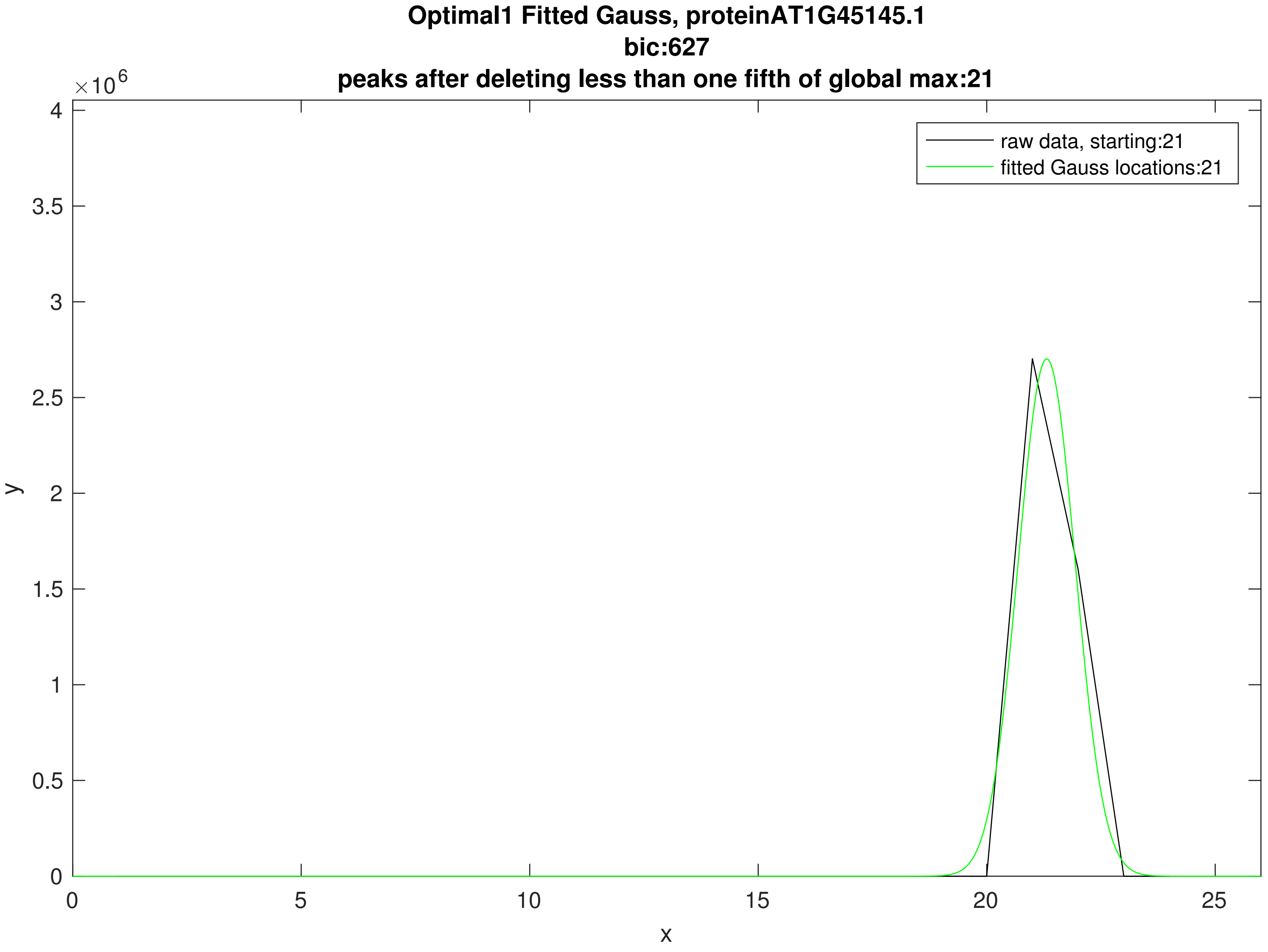
Optimal1 Fitted Gauss, proteinAT1G45145.1
bic:627
peaks after deleting less than one fifth of global max:21
×10^6
4
3.5
3
2.5
2
1.5
1
0.5
0
y
0
5
10
15
20
25
x
raw data, starting:21
fitted Gauss locations:21

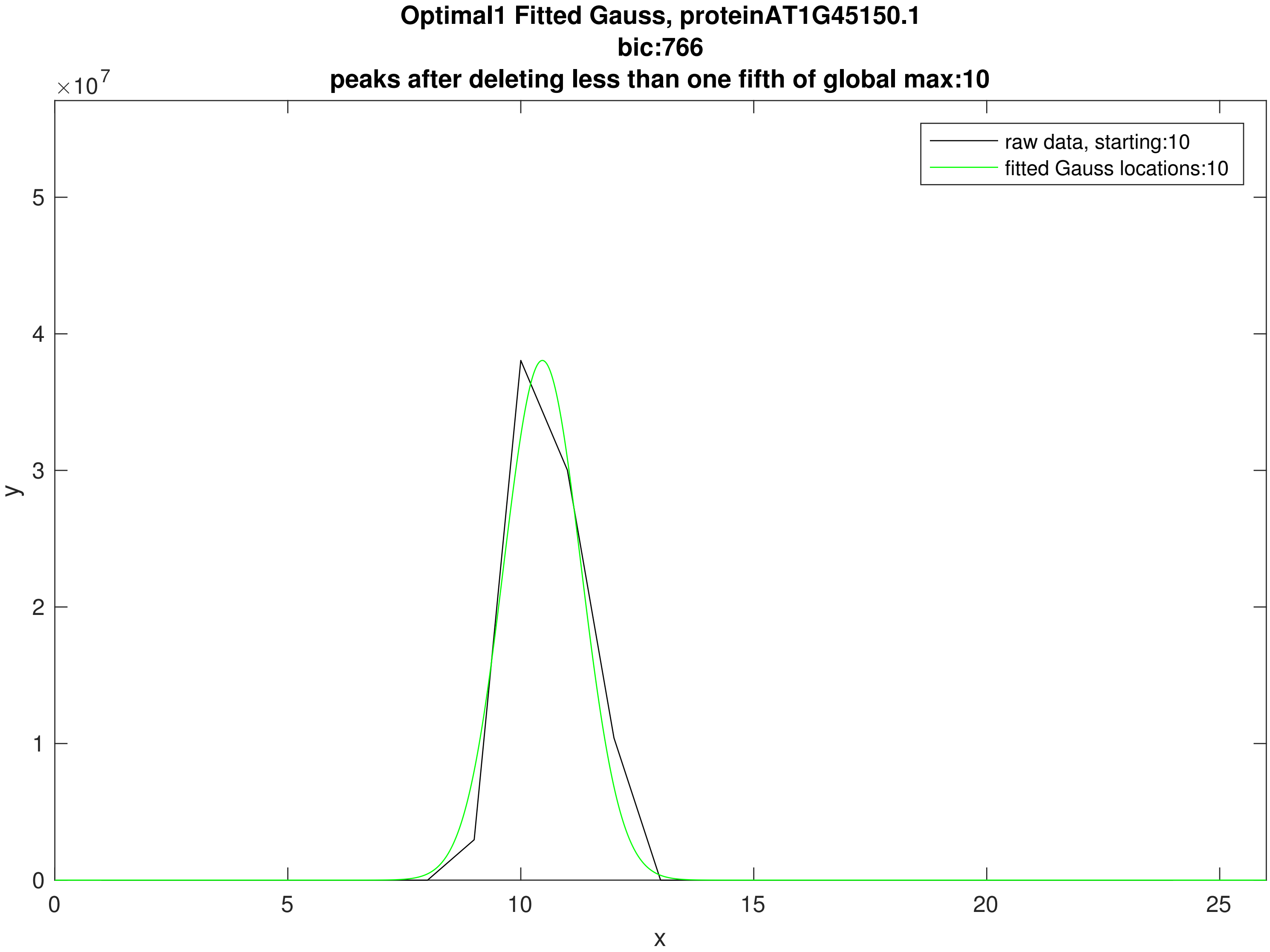
Optimal1 Fitted Gauss, proteinAT1G45150.1
bic:766
peaks after deleting less than one fifth of global max:10
×10^7
raw data, starting:10
fitted Gauss locations:10
y
x
0
1
2
3
4
5
0
5
10
15
20
25

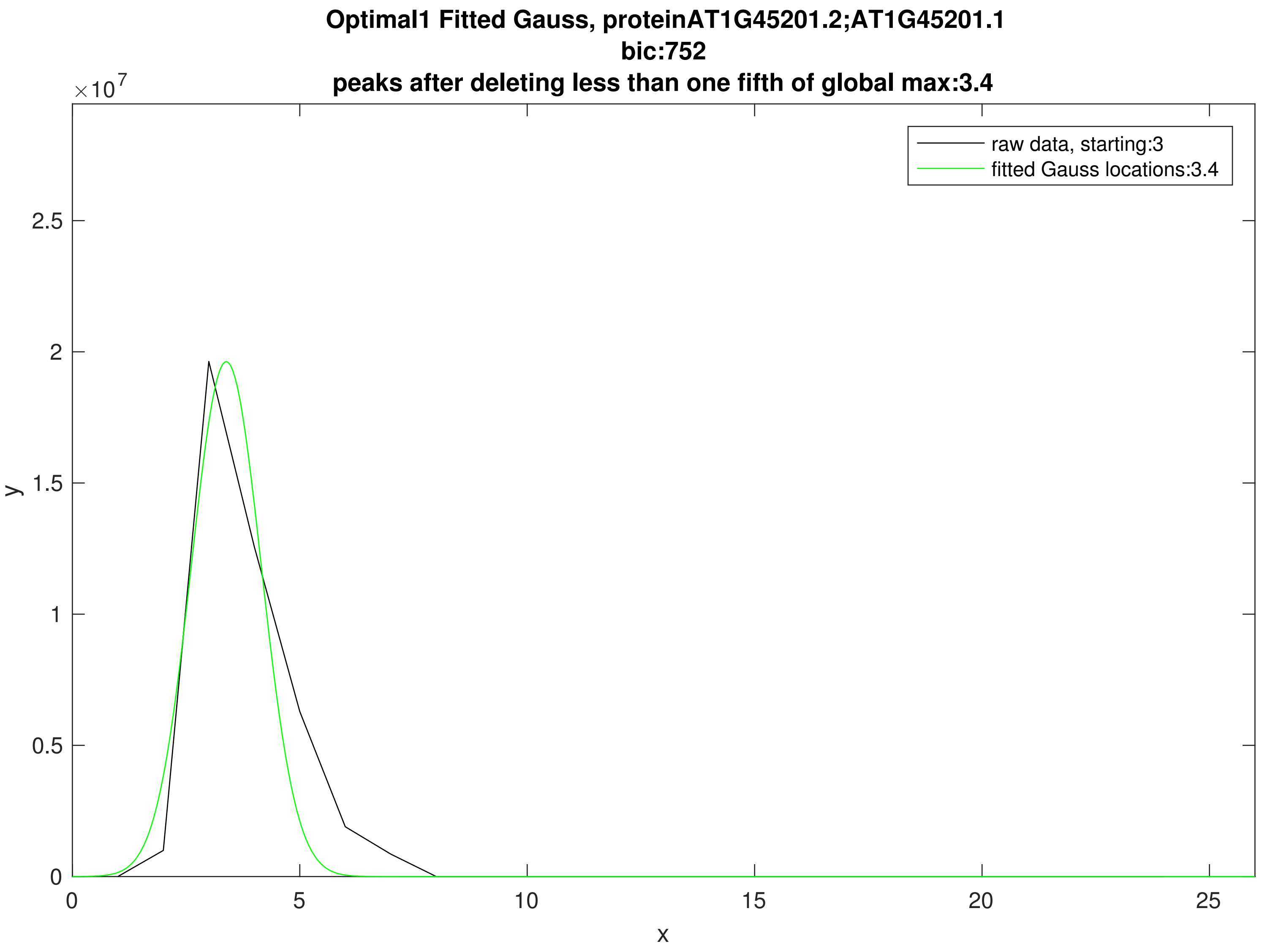

Optimal1 Fitted Gauss, proteinAT1G45201.2;AT1G45201.1
bic:752
peaks after deleting less than one fifth of global max:3.4
×10^7
raw data, starting:3
fitted Gauss locations:3.4
y
2.5
2
1.5
1
0.5
0
0
5
10
15
20
25
x

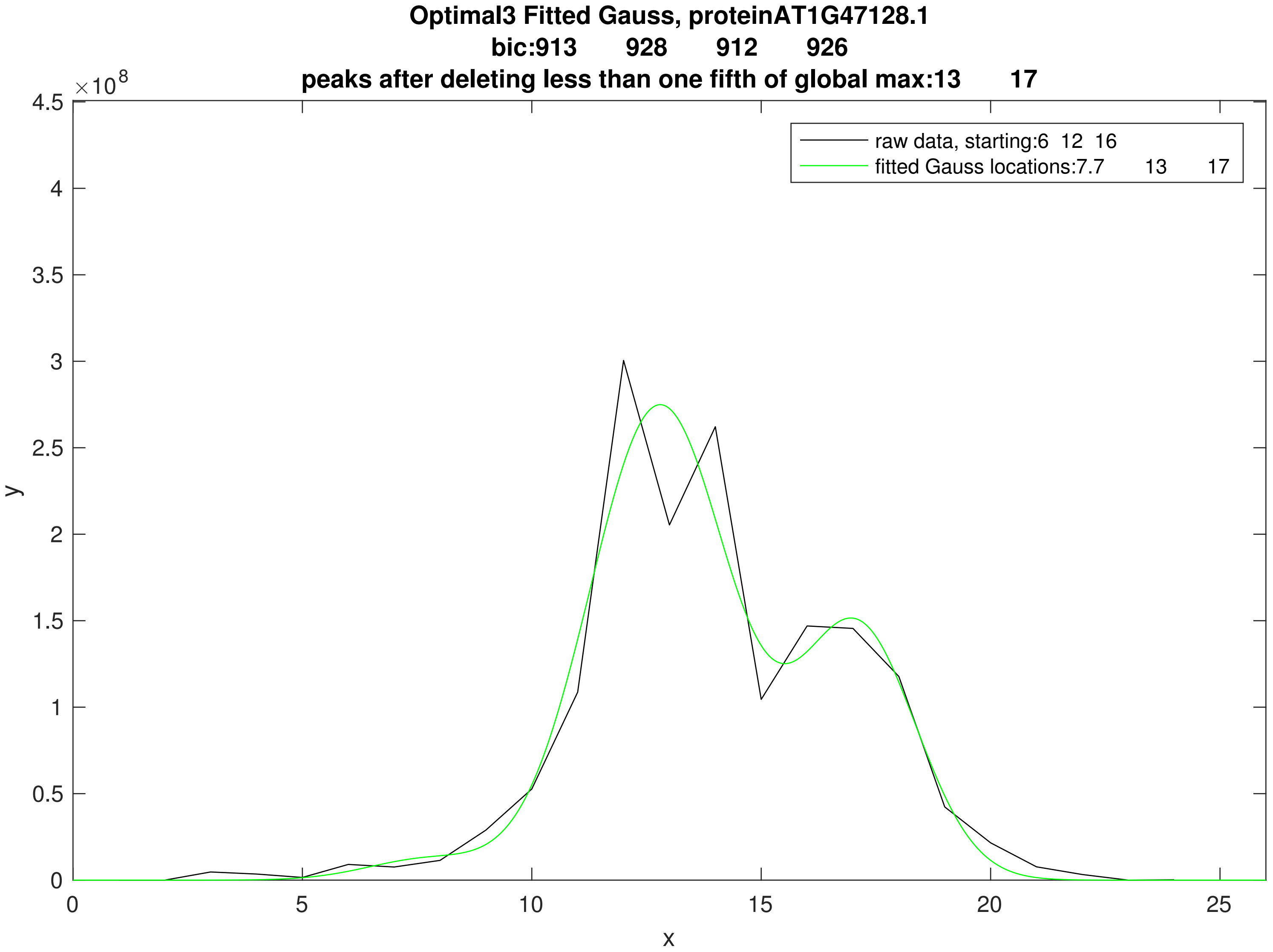
Optimal3 Fitted Gauss, proteinAT1G47128.1
bic:913 928 912 926
peaks after deleting less than one fifth of global max:13 17
raw data, starting:6 12 16
fitted Gauss locations:7.7 13 17
×10^8
4.5
4
3.5
3
2.5
2
1.5
1
0.5
0
y
0
5
10
15
20
25
x

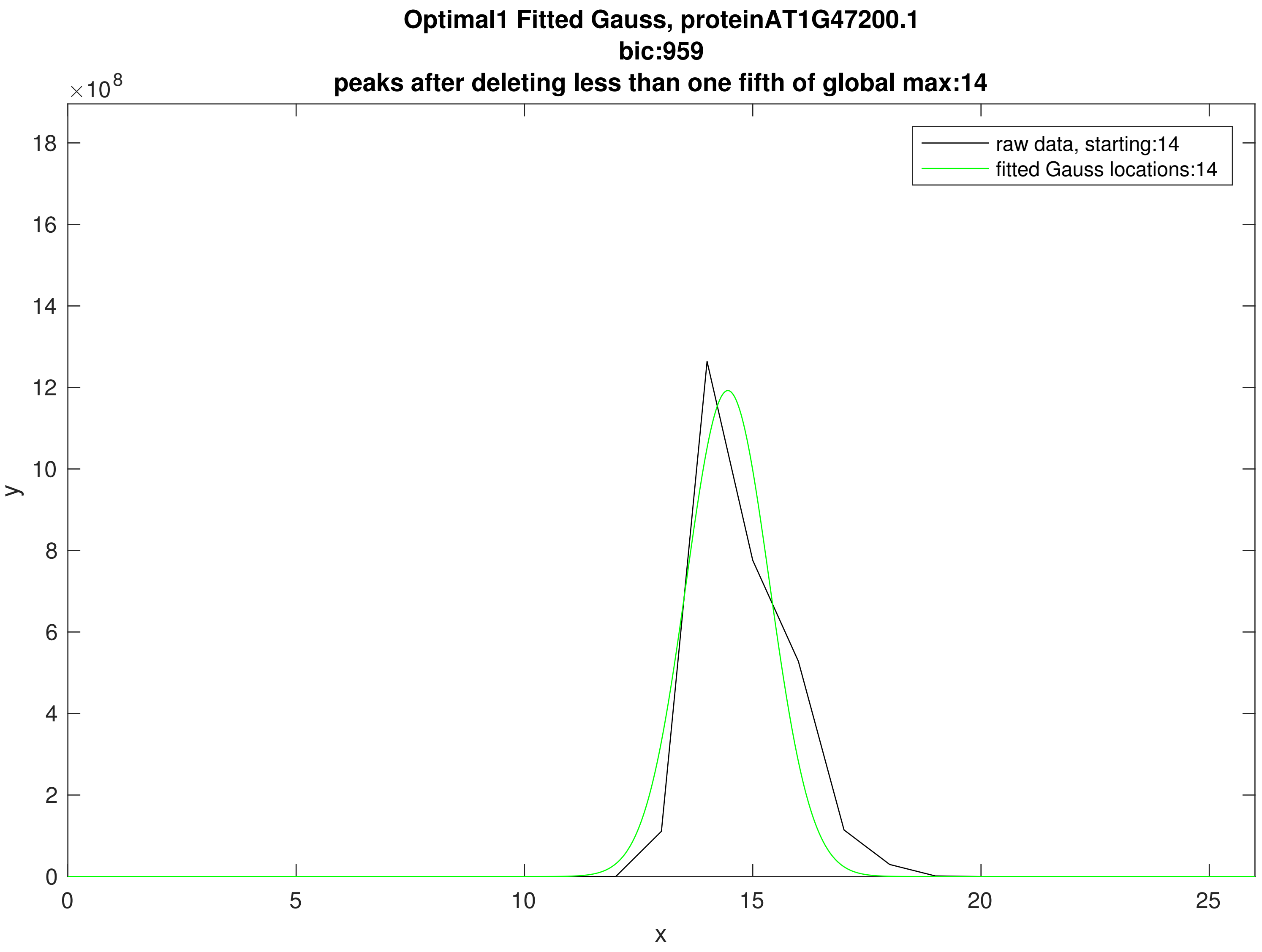

Optimal1 Fitted Gauss, proteinAT1G47200.1
bic:959
peaks after deleting less than one fifth of global max:14
$\times 10^8$
raw data, starting:14
fitted Gauss locations:14
y
x
0
2
4
6
8
10
12
14
16
18
0
5
10
15
20
25

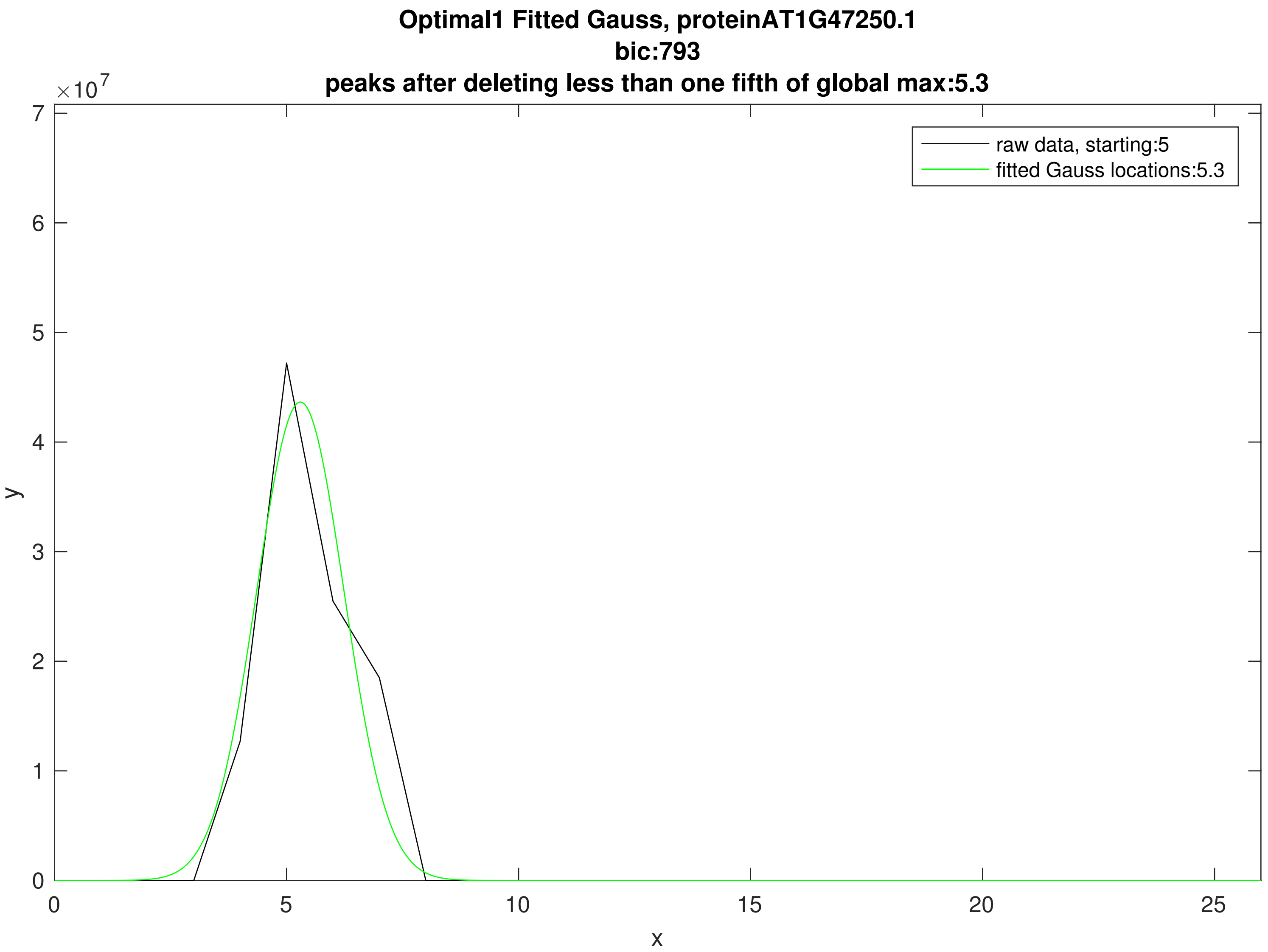
Optimal1 Fitted Gauss, proteinAT1G47250.1
bic:793
peaks after deleting less than one fifth of global max:5.3
×10^7
7
6
5
4
3
2
1
0
y
0
5
10
15
20
25
x
raw data, starting:5
fitted Gauss locations:5.3

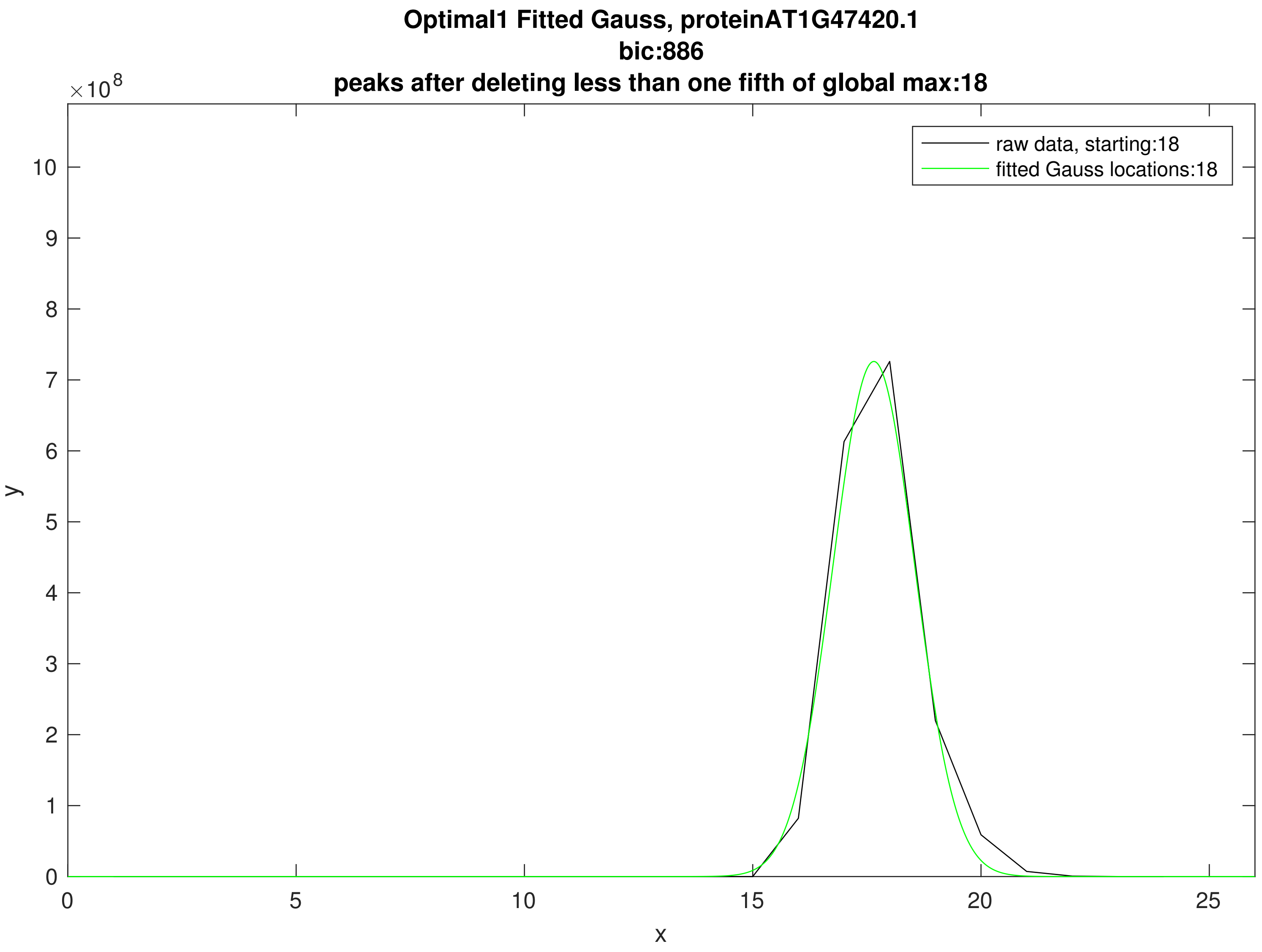

Optimal1 Fitted Gauss, proteinAT1G47420.1
bic:886
peaks after deleting less than one fifth of global max:18
×10^8
raw data, starting:18
fitted Gauss locations:18
y
x
0
1
2
3
4
5
6
7
8
9
10
15
20
25

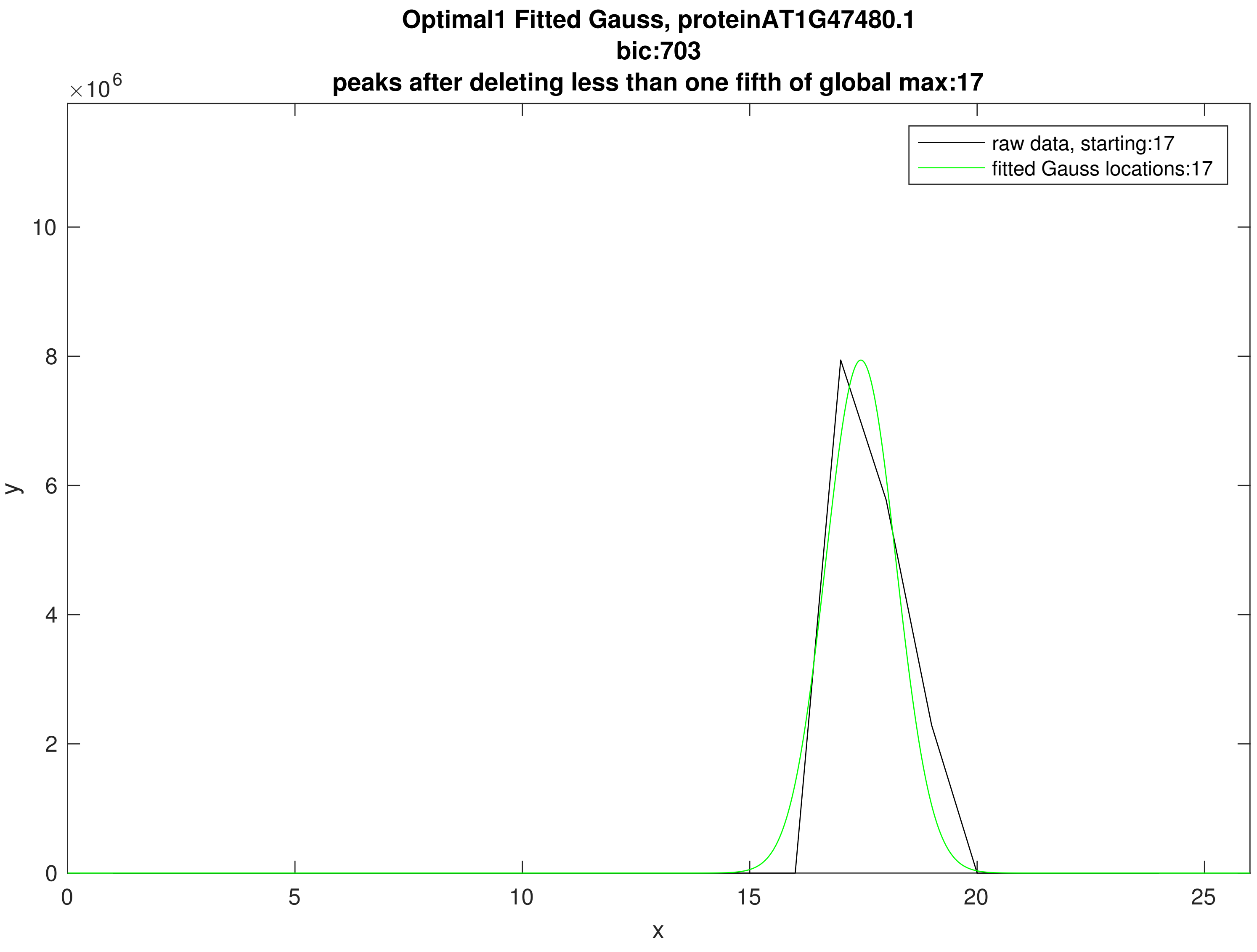
Optimal1 Fitted Gauss, proteinAT1G47480.1
bic:703
peaks after deleting less than one fifth of global max:17
×10^6
raw data, starting:17
fitted Gauss locations:17
y
x
0
2
4
6
8
10
0
5
10
15
20
25

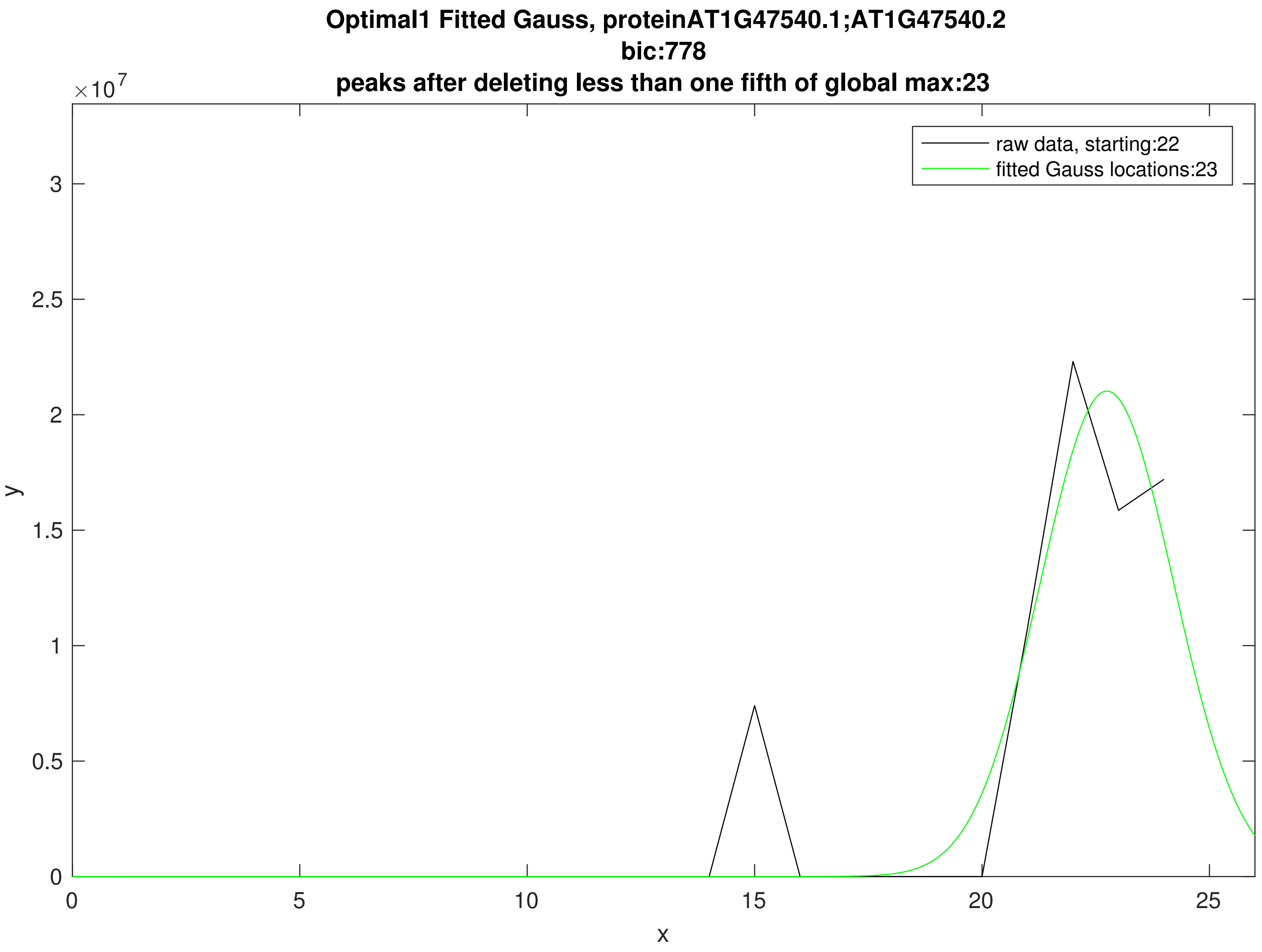

Optimal1 Fitted Gauss, proteinAT1G47540.1;AT1G47540.2
bic:778
peaks after deleting less than one fifth of global max:23
×10^7
raw data, starting:22
fitted Gauss locations:23
y
x
0
0.5
1
1.5
2
2.5
3
0
5
10
15
20
25

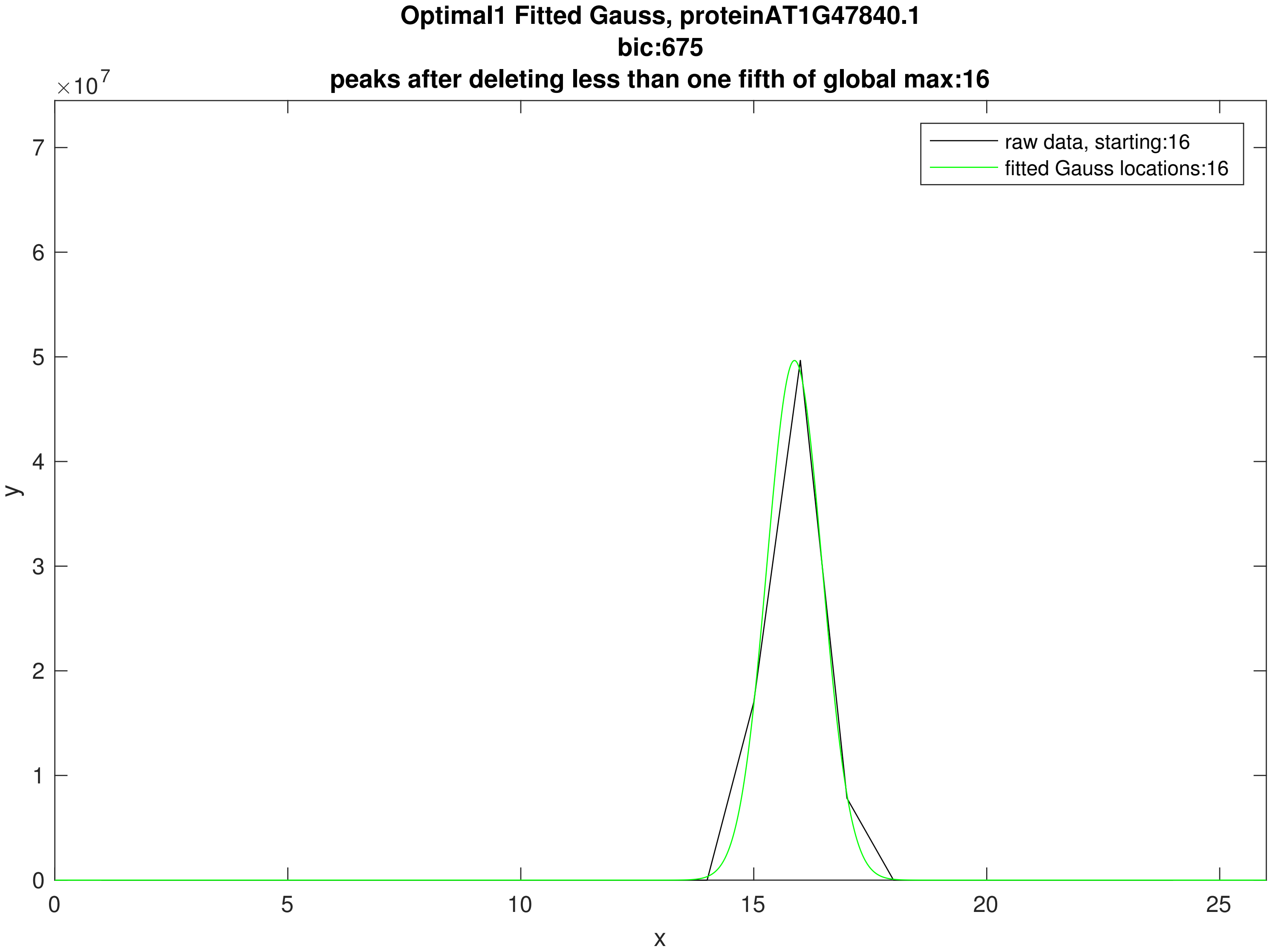

Optimal1 Fitted Gauss, proteinAT1G47840.1
bic:675
peaks after deleting less than one fifth of global max:16
×10^7
raw data, starting:16
fitted Gauss locations:16
y
x
0
1
2
3
4
5
6
7
0
5
10
15
20
25

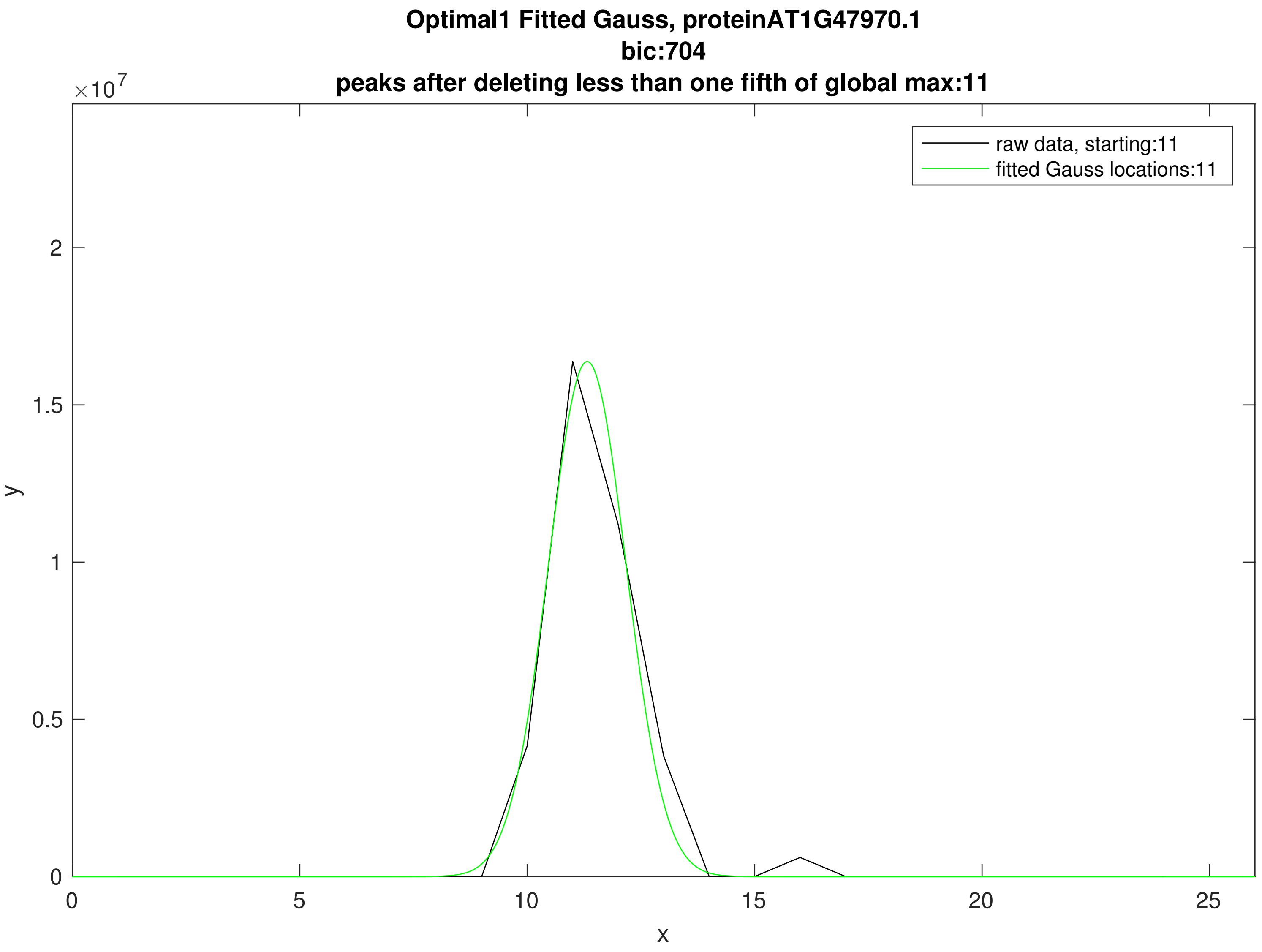
Optimal1 Fitted Gauss, proteinAT1G47970.1
bic:704
peaks after deleting less than one fifth of global max:11
×10^7
raw data, starting:11
fitted Gauss locations:11
y
x
0
0.5
1
1.5
2
0
5
10
15
20
25

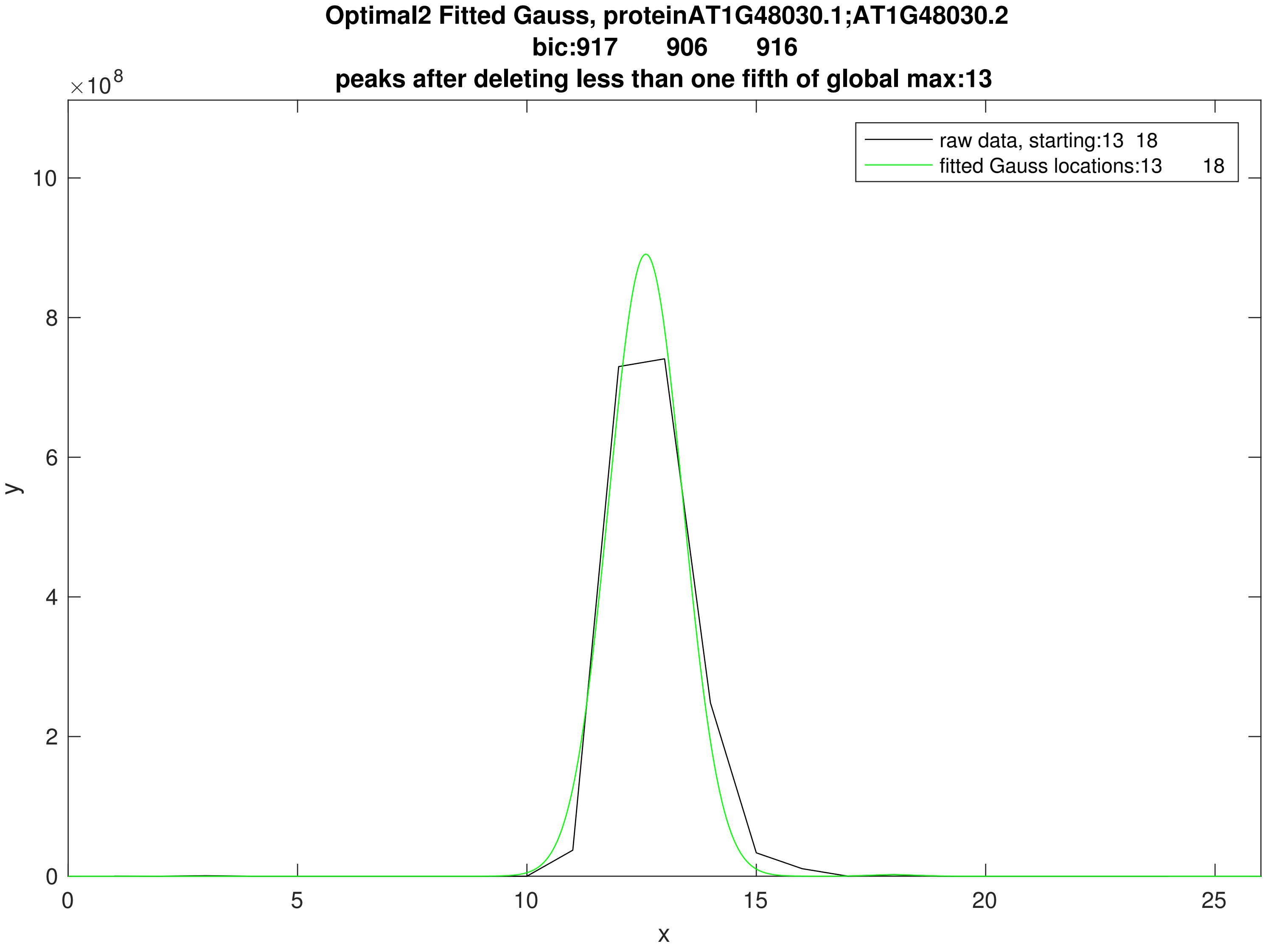
Optimal2 Fitted Gauss, proteinAT1G48030.1;AT1G48030.2
bic:917 906 916
peaks after deleting less than one fifth of global max:13
×10^8
raw data, starting:13 18
fitted Gauss locations:13 18
y
x
0
2
4
6
8
10
0
5
10
15
20
25

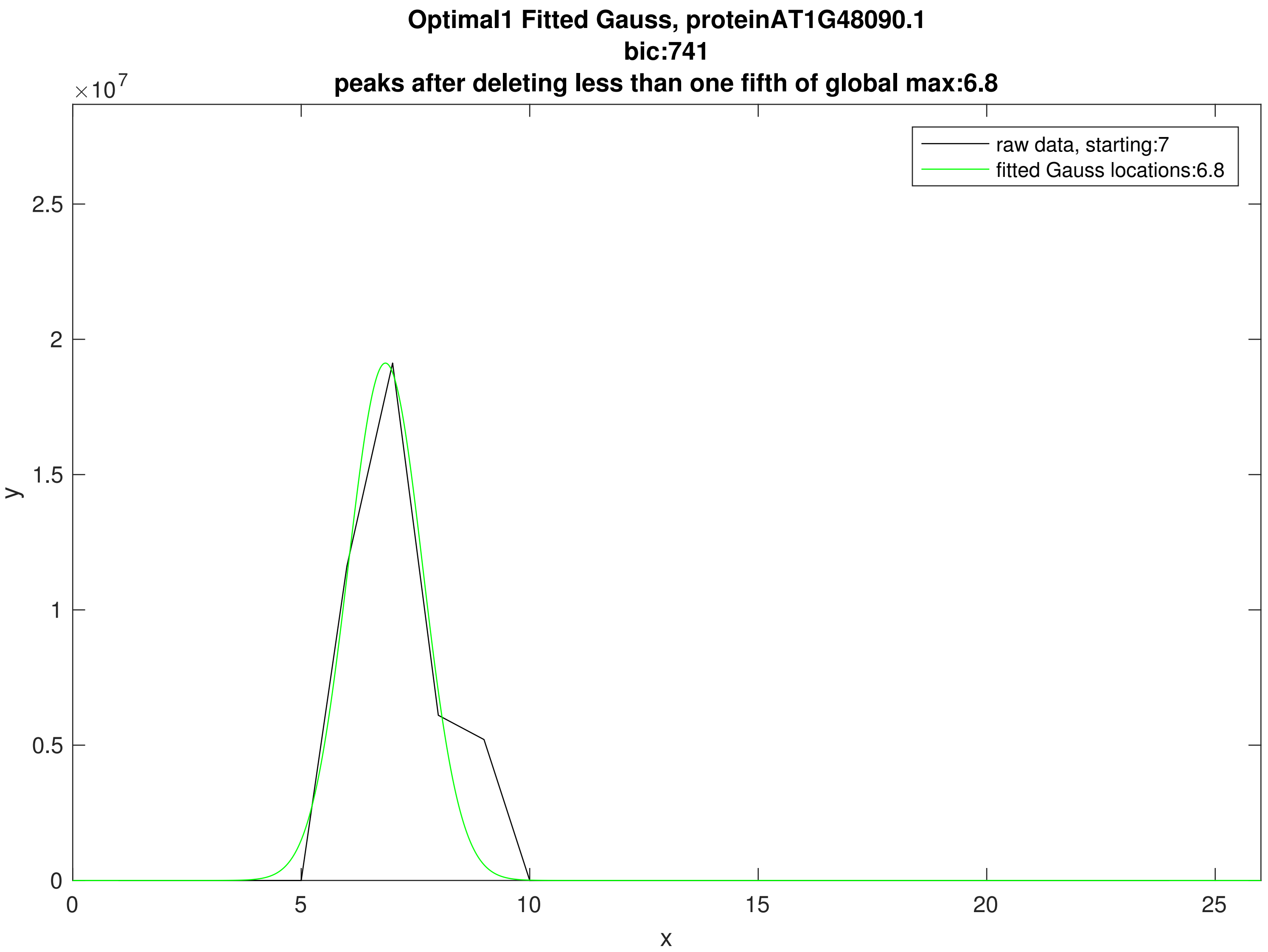

Optimal1 Fitted Gauss, proteinAT1G48090.1
bic:741
peaks after deleting less than one fifth of global max:6.8
×10^7
2.5
2
1.5
1
0.5
0
y
0
5
10
15
20
25
x
raw data, starting:7
fitted Gauss locations:6.8

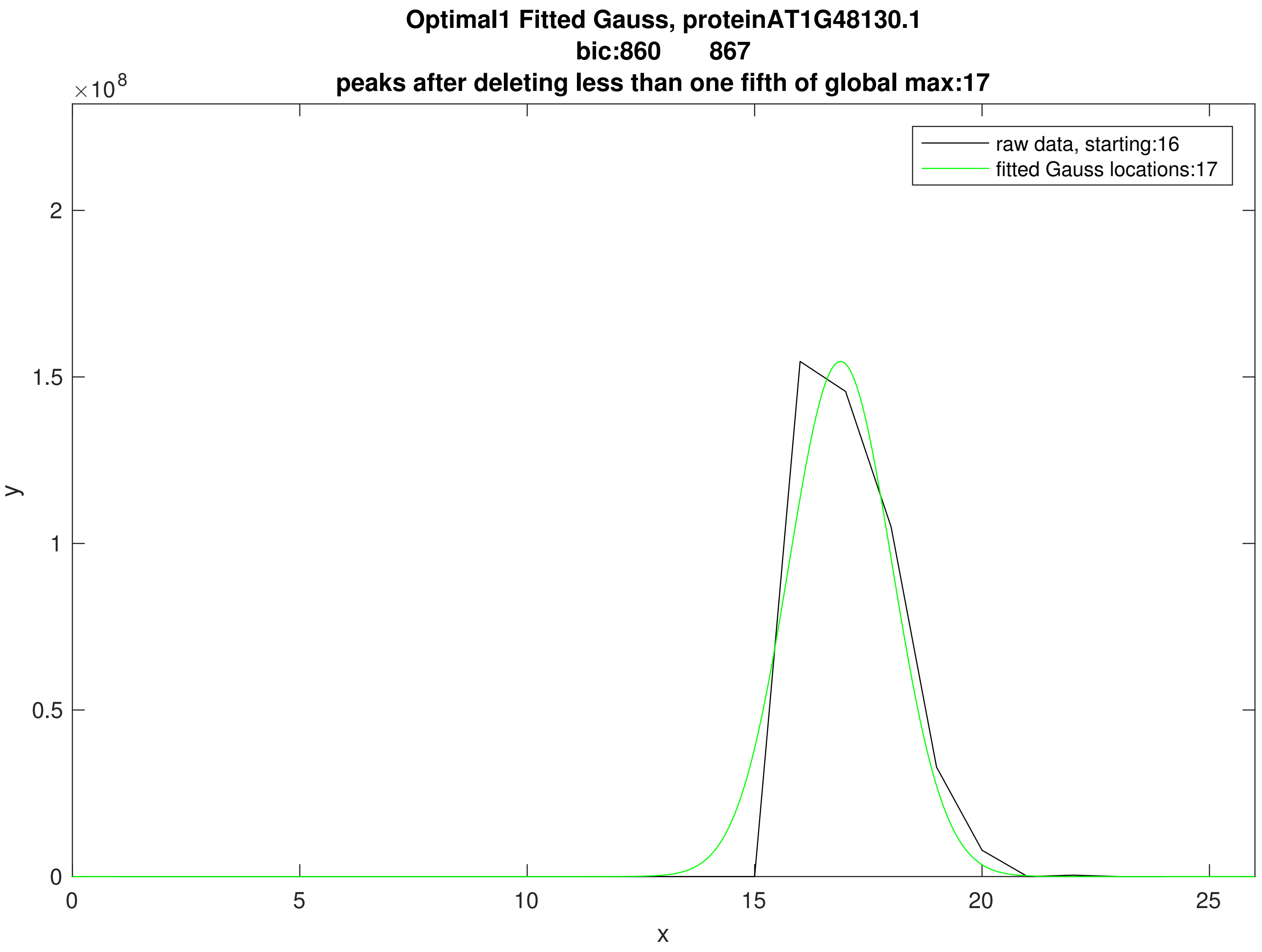

Optimal1 Fitted Gauss, proteinAT1G48130.1
bic:860 867
peaks after deleting less than one fifth of global max:17
×10^8
raw data, starting:16
fitted Gauss locations:17
y
x
0
0.5
1
1.5
2
0
5
10
15
20
25

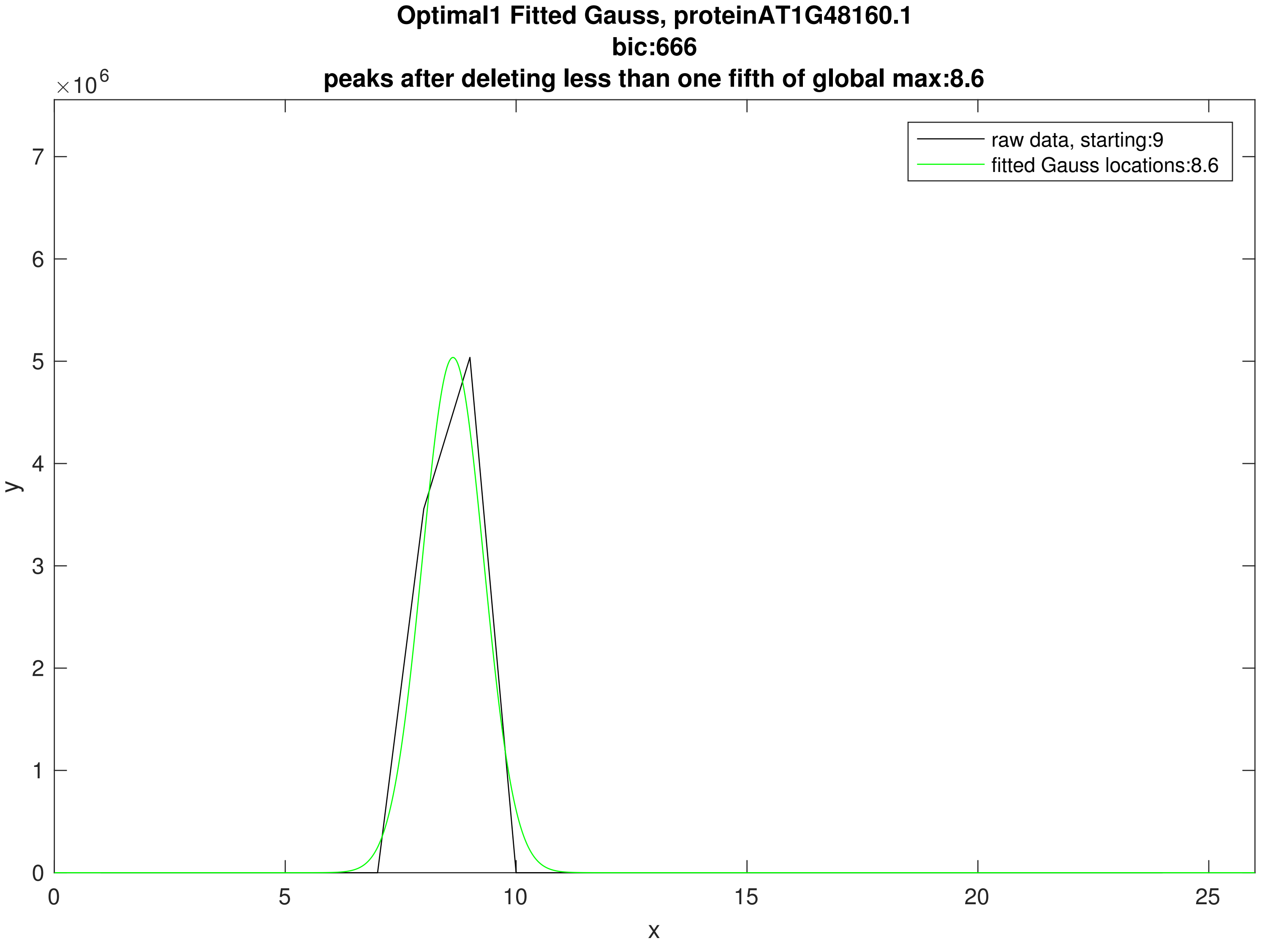
Optimal1 Fitted Gauss, proteinAT1G48160.1
bic:666
peaks after deleting less than one fifth of global max:8.6
×10^6
raw data, starting:9
fitted Gauss locations:8.6
y
x
0
1
2
3
4
5
6
7
0
5
10
15
20
25

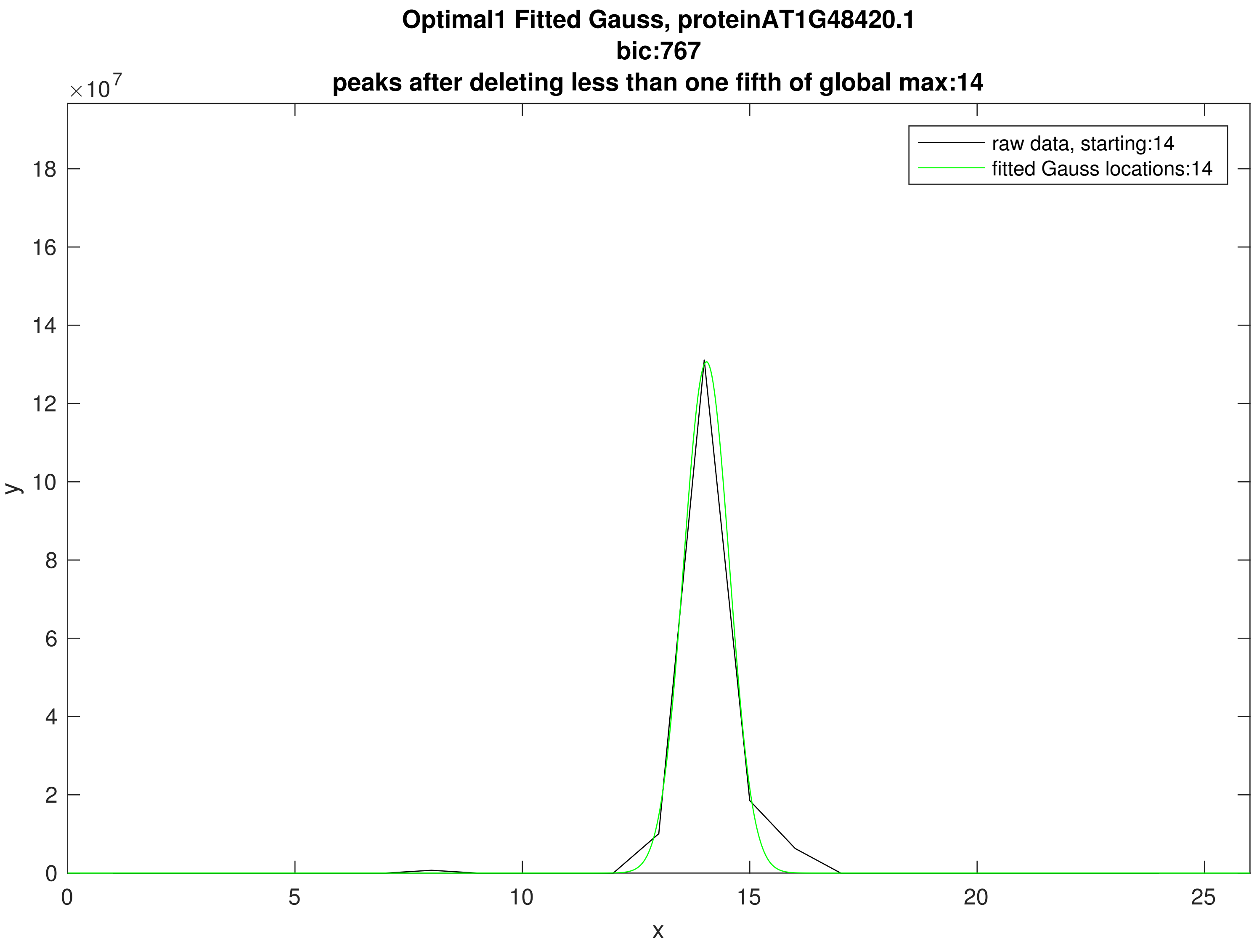

Optimal1 Fitted Gauss, proteinAT1G48420.1
bic:767
peaks after deleting less than one fifth of global max:14
×10^7
raw data, starting:14
fitted Gauss locations:14
y
x
0
2
4
6
8
10
12
14
16
18
0
5
10
15
20
25

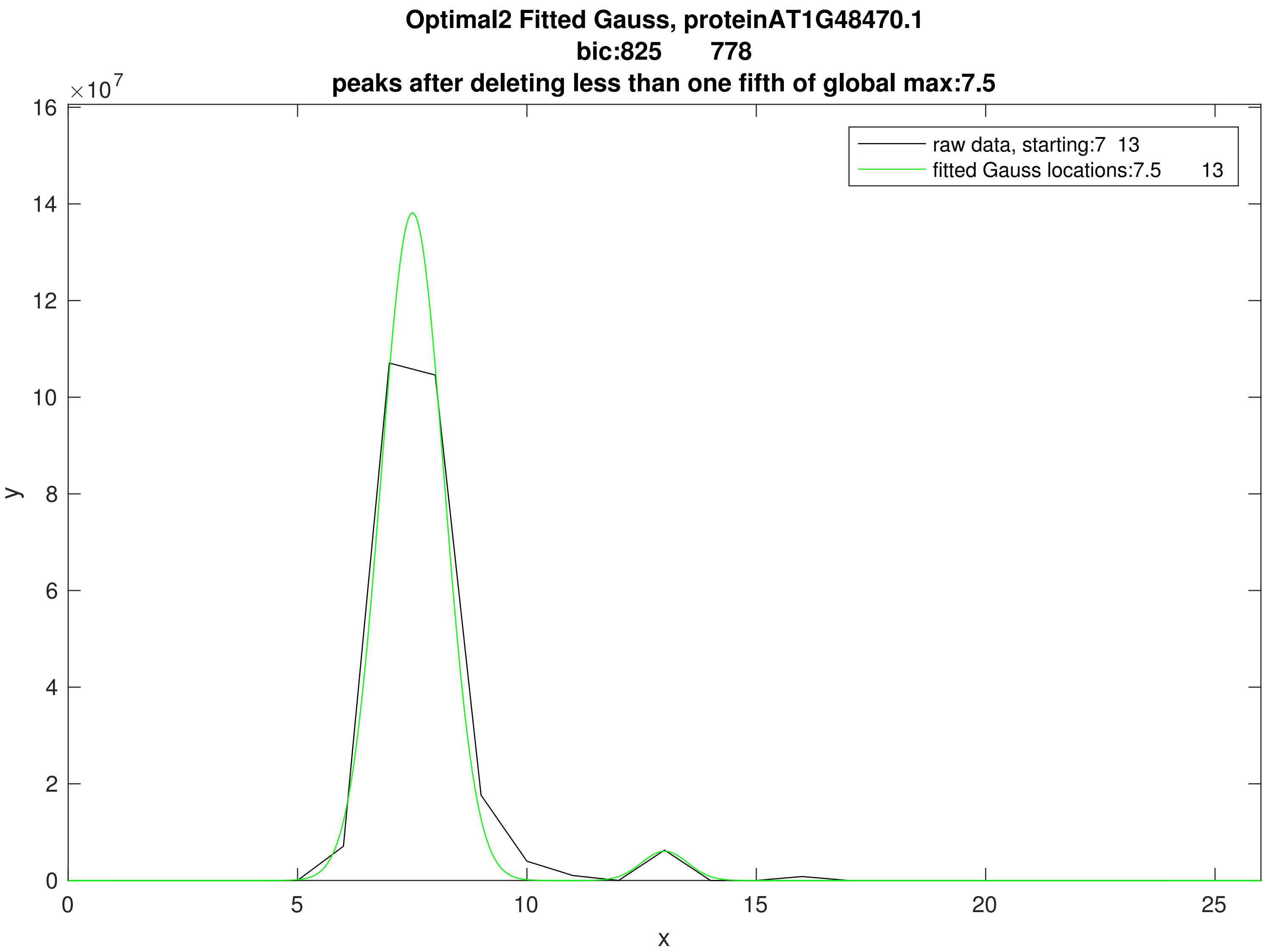
Optimal2 Fitted Gauss, proteinAT1G48470.1
bic:825 778
peaks after deleting less than one fifth of global max:7.5
×10^7
16
14
12
10
8
6
4
2
0
y
0
5
10
15
20
25
x
raw data, starting:7 13
fitted Gauss locations:7.5 13

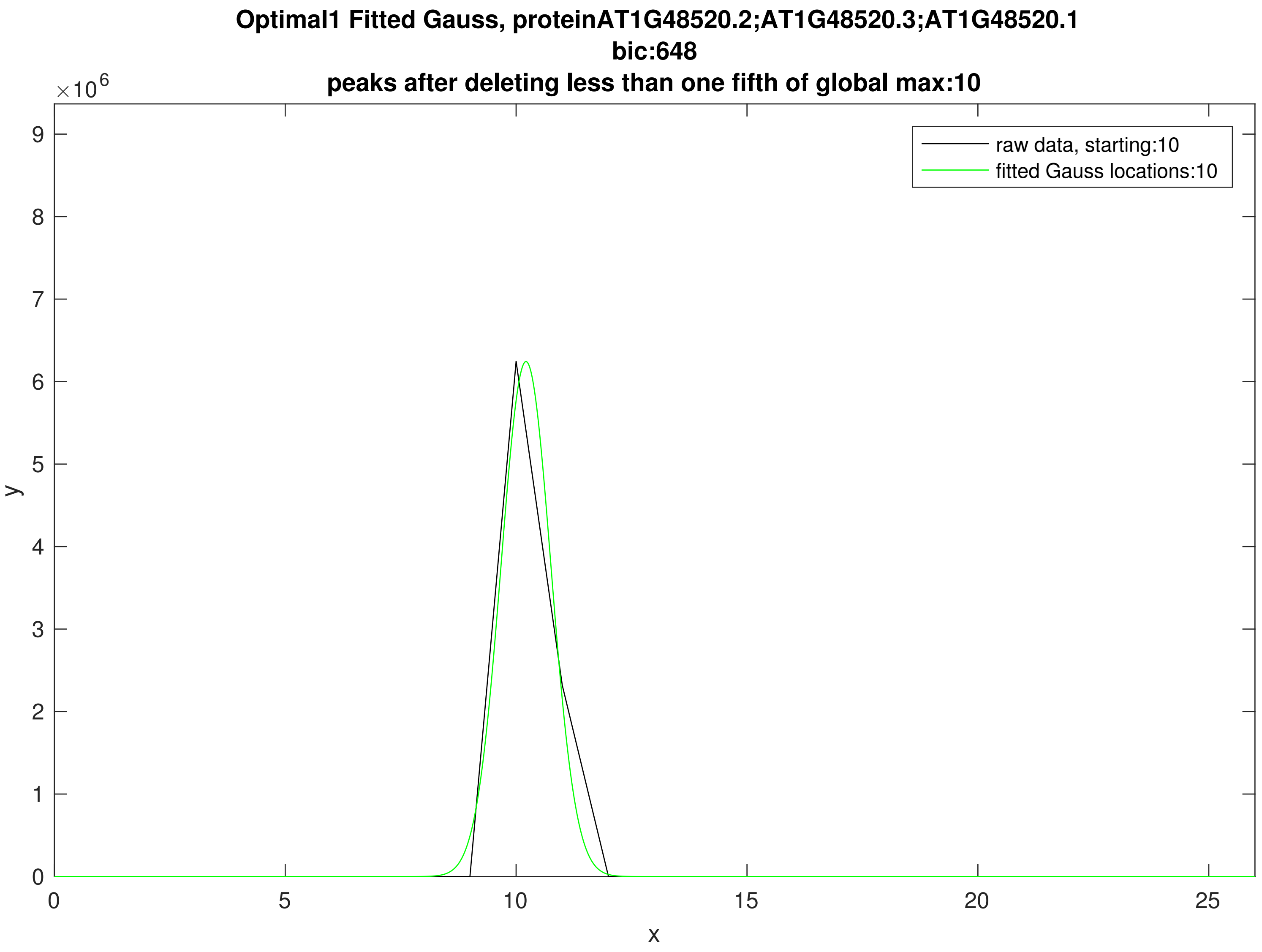
Optimal1 Fitted Gauss, proteinAT1G48520.2;AT1G48520.3;AT1G48520.1
bic:648
peaks after deleting less than one fifth of global max:10
×10^6
raw data, starting:10
fitted Gauss locations:10
y
x
0
1
2
3
4
5
6
7
8
9
0
5
10
15
20
25

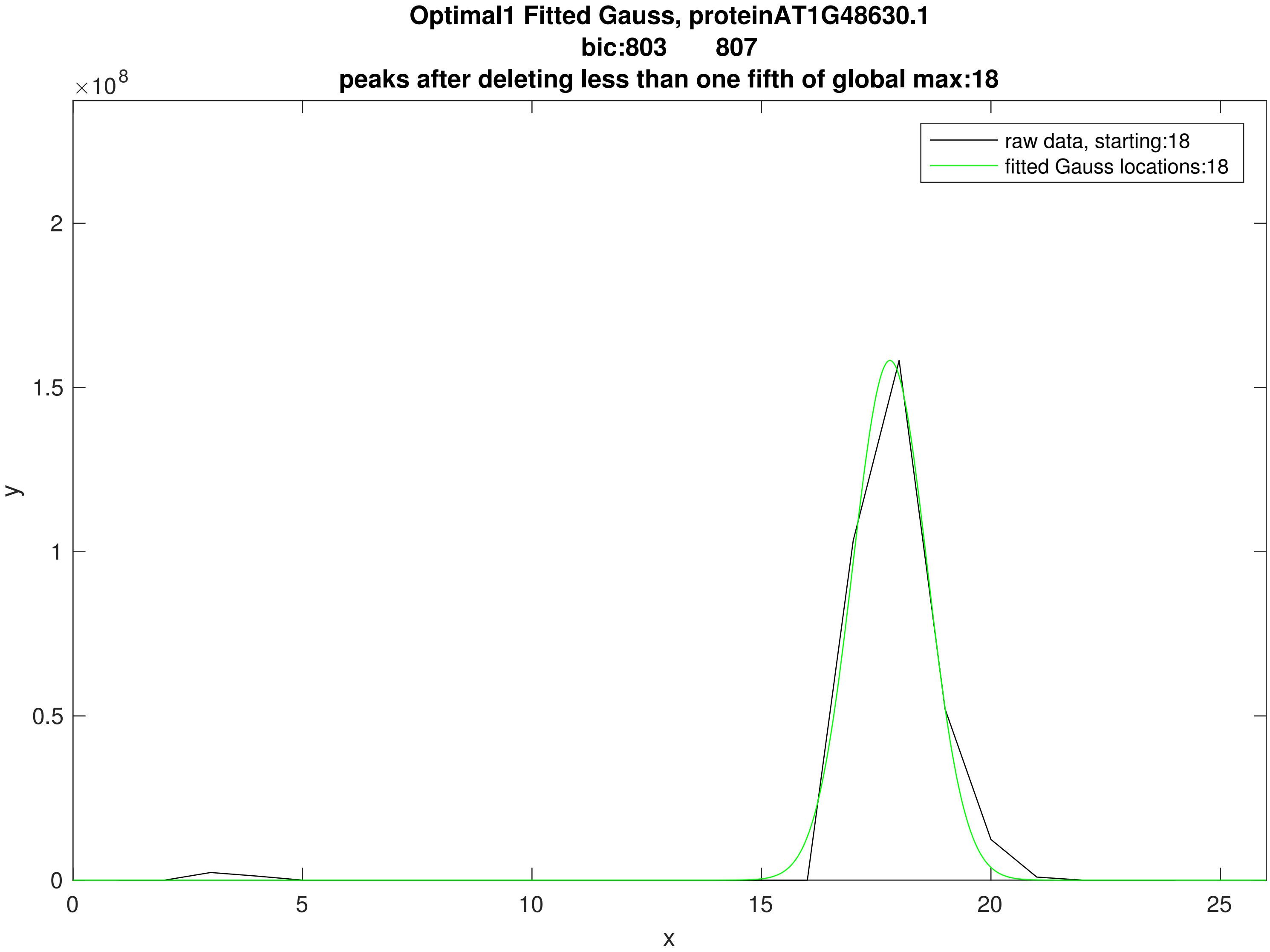

Optimal1 Fitted Gauss, proteinAT1G48630.1
bic:803 807
peaks after deleting less than one fifth of global max:18
×10^8
raw data, starting:18
fitted Gauss locations:18
y
x
0
0.5
1
1.5
2
0
5
10
15
20
25

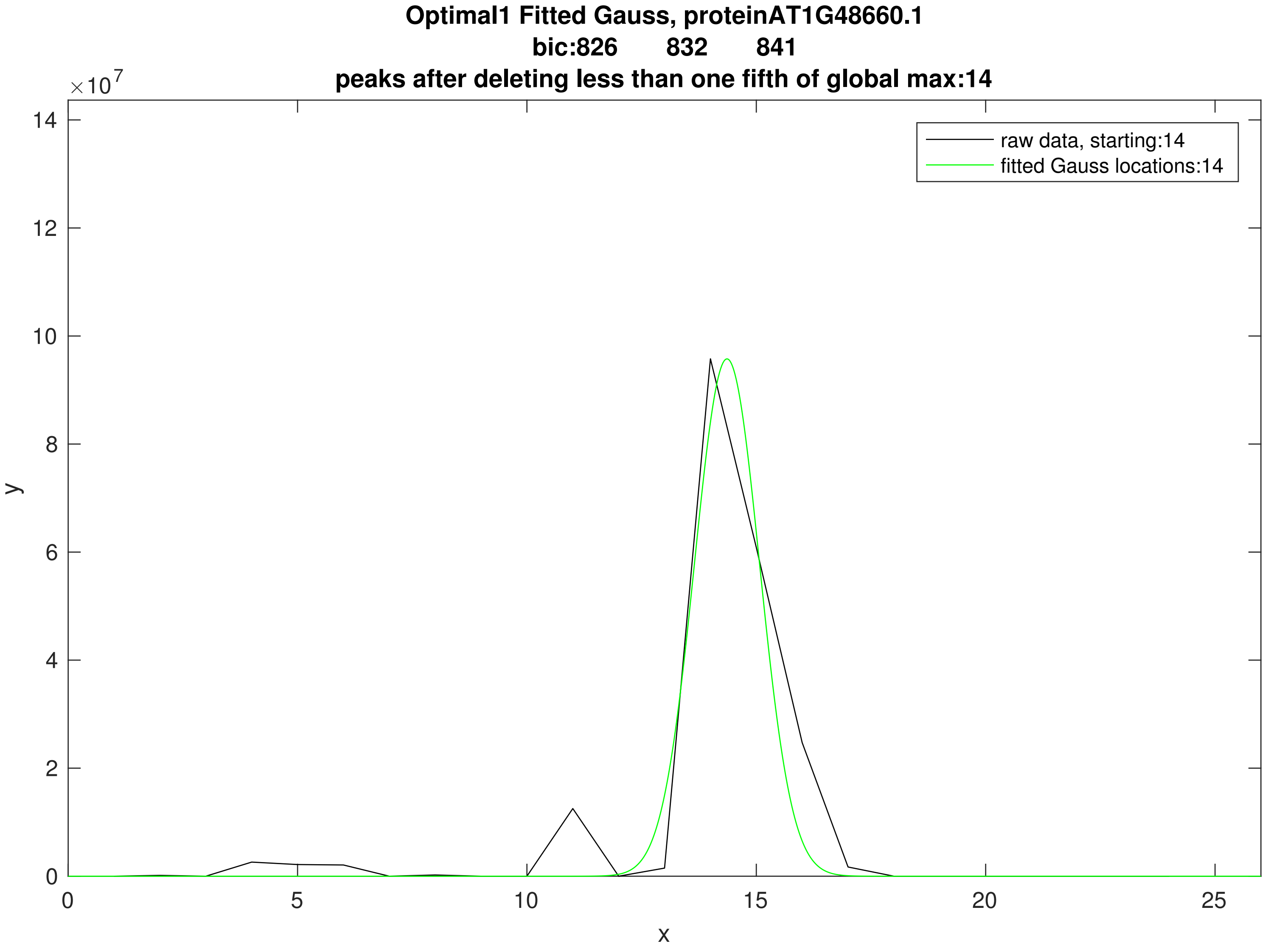
Optimal1 Fitted Gauss, proteinAT1G48660.1
bic:826 832 841
peaks after deleting less than one fifth of global max:14
×10^7
14
12
10
8
6
4
2
0
y
0
5
10
15
20
25
x
raw data, starting:14
fitted Gauss locations:14

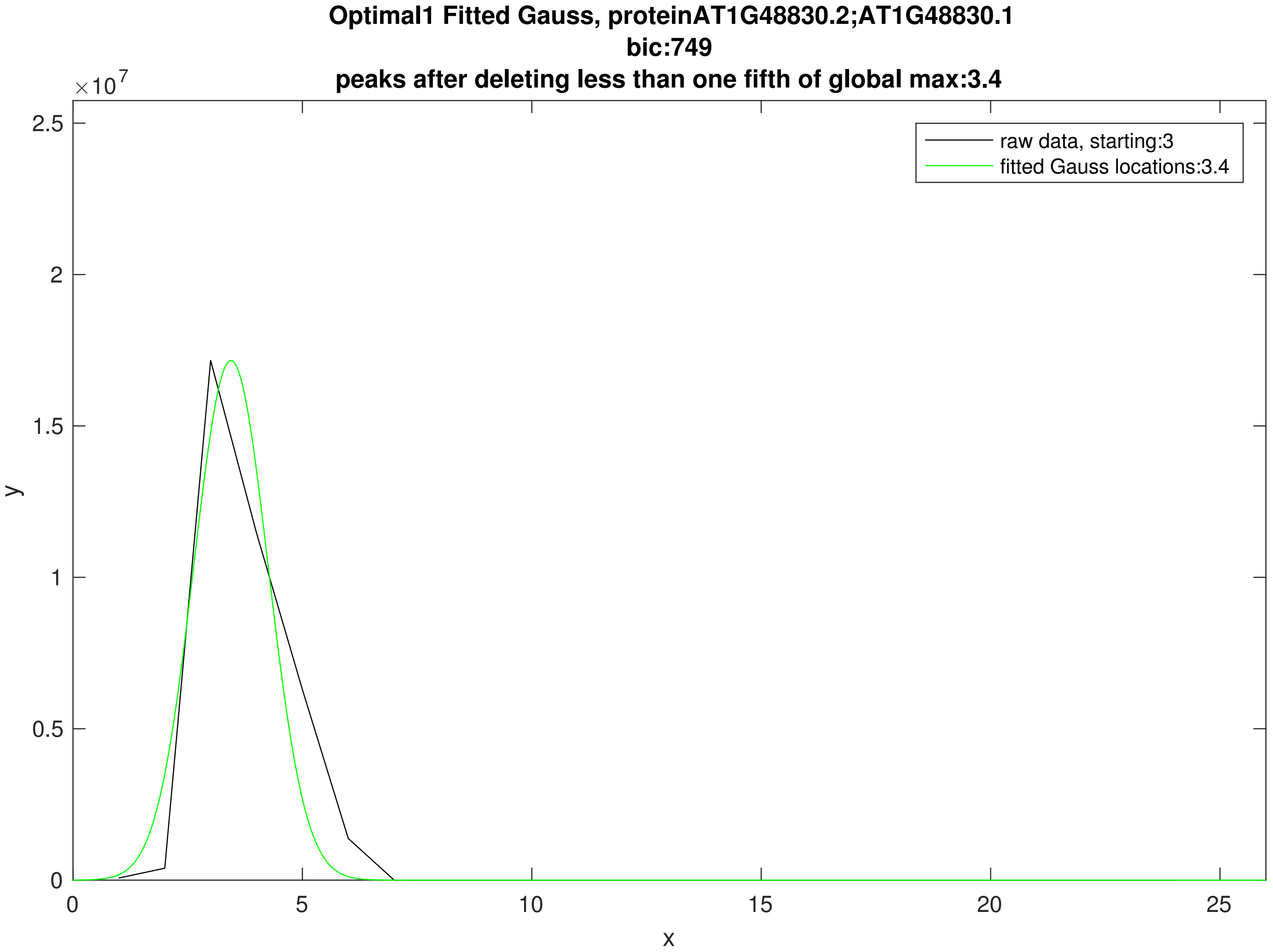

Optimal1 Fitted Gauss, proteinAT1G48830.2;AT1G48830.1
bic:749
peaks after deleting less than one fifth of global max:3.4
×10^7
2.5
2
1.5
1
0.5
0
y
0
5
10
15
20
25
x
raw data, starting:3
fitted Gauss locations:3.4

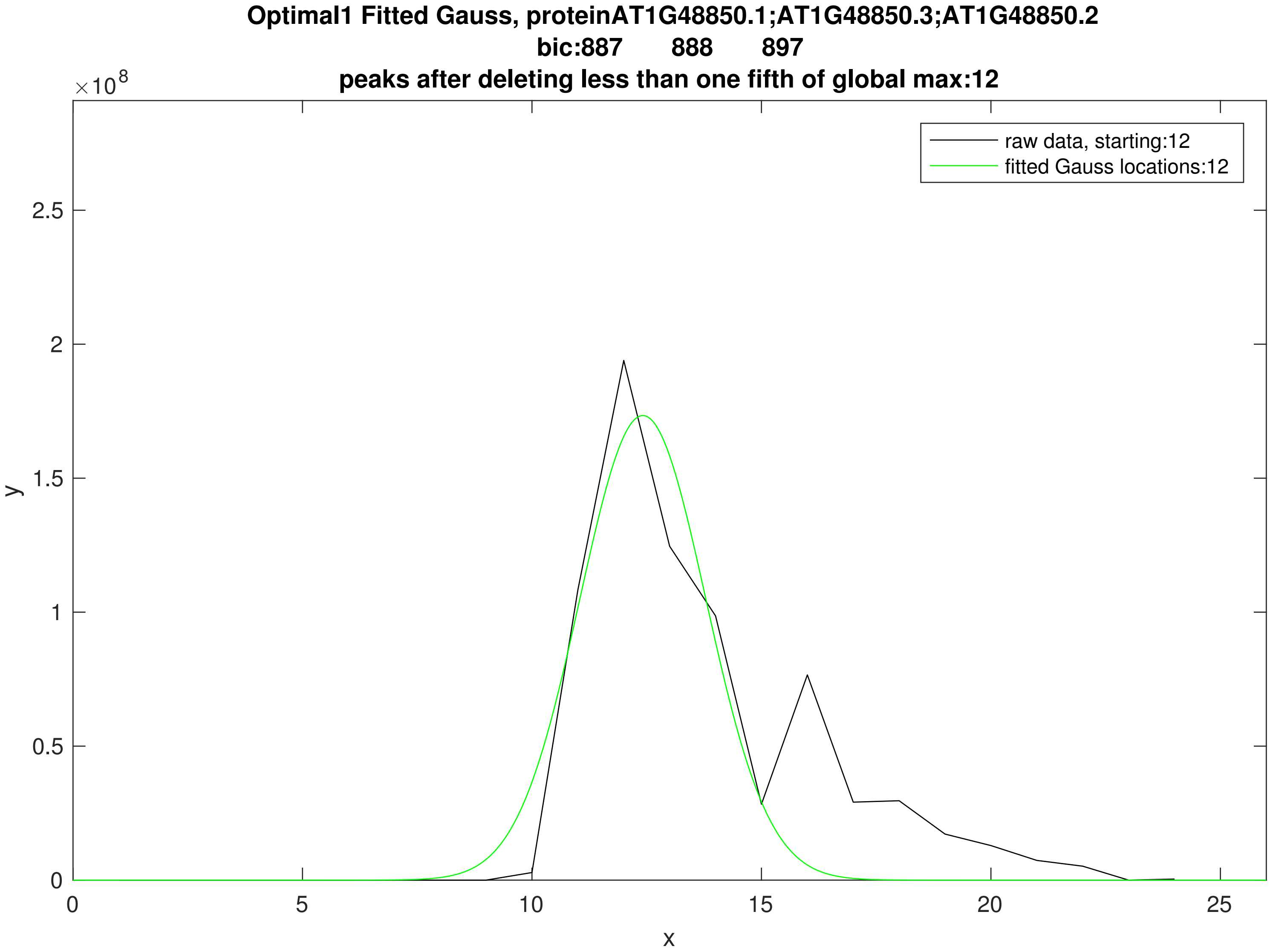

Optimal1 Fitted Gauss, proteinAT1G48850.1;AT1G48850.3;AT1G48850.2
bic:887 888 897
peaks after deleting less than one fifth of global max:12
×10^8
raw data, starting:12
fitted Gauss locations:12
y
x
0
0.5
1
1.5
2
2.5
5
10
15
20
25

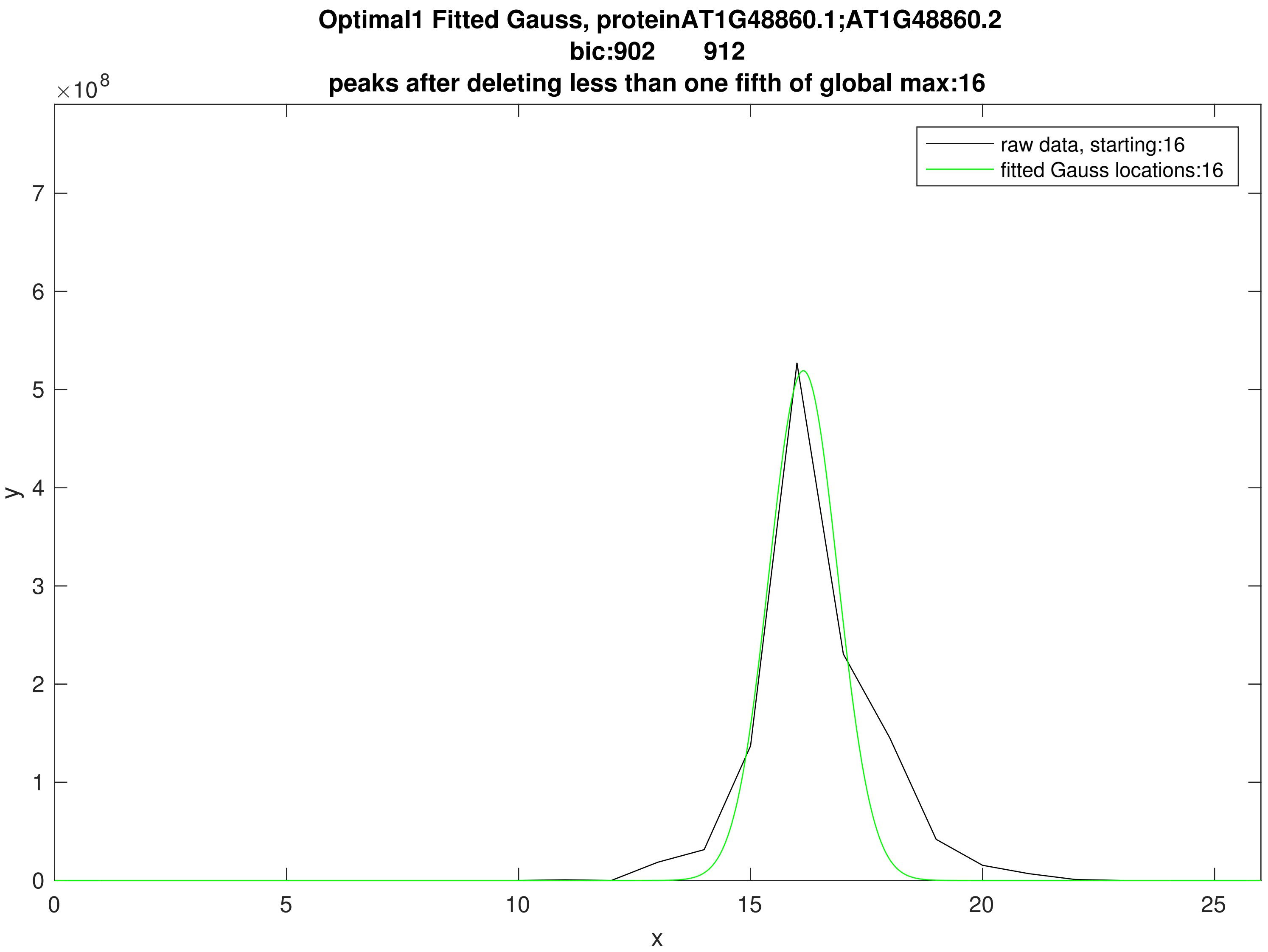
Optimal1 Fitted Gauss, proteinAT1G48860.1;AT1G48860.2
bic:902 912
peaks after deleting less than one fifth of global max:16
×10^8
raw data, starting:16
fitted Gauss locations:16
y
x
0
1
2
3
4
5
6
7
0
5
10
15
20
25

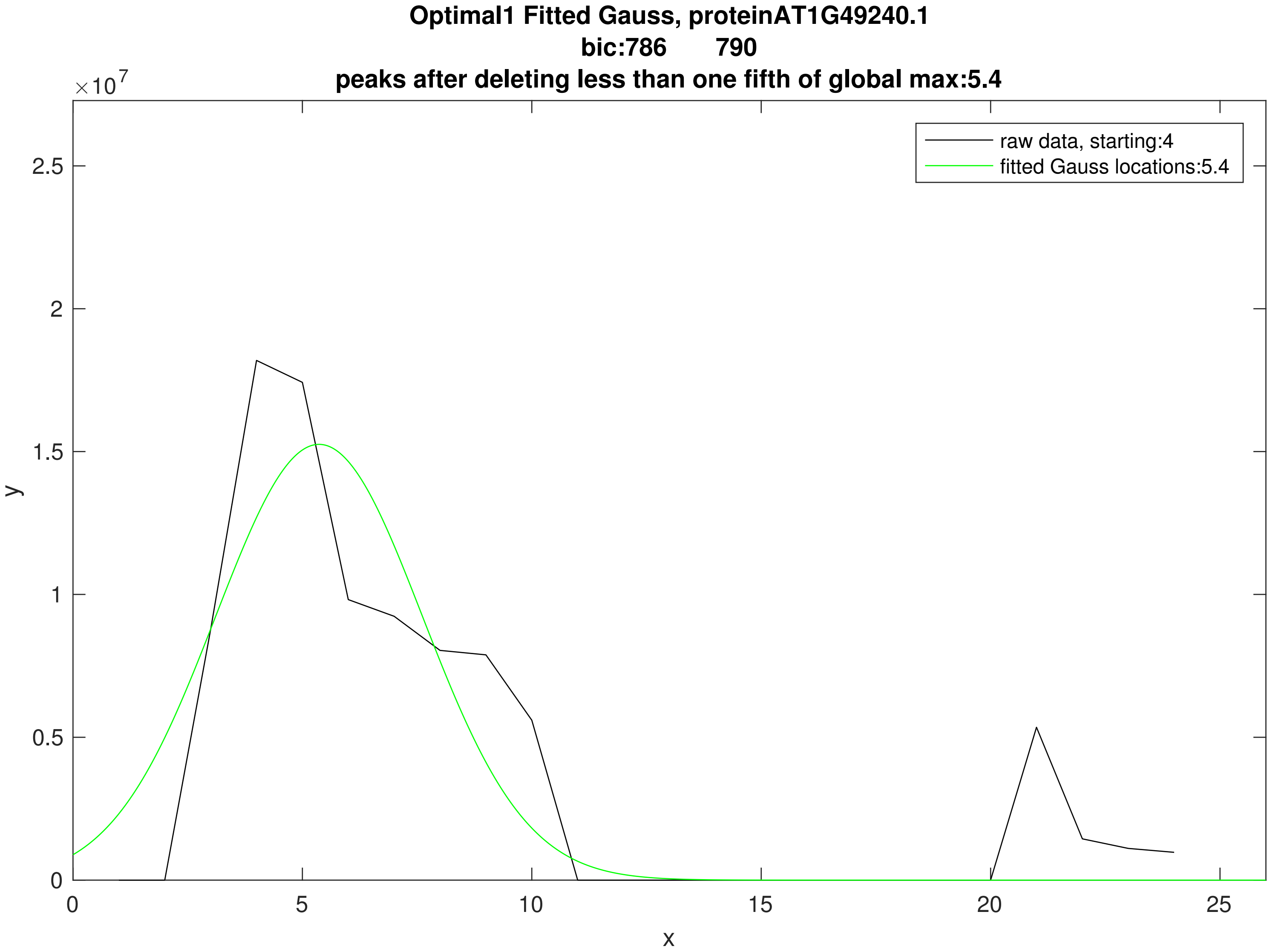

Optimal1 Fitted Gauss, proteinAT1G49240.1
bic:786 790
peaks after deleting less than one fifth of global max:5.4
×10^7
2.5
2
1.5
1
0.5
0
y
0
5
10
15
20
25
x
raw data, starting:4
fitted Gauss locations:5.4

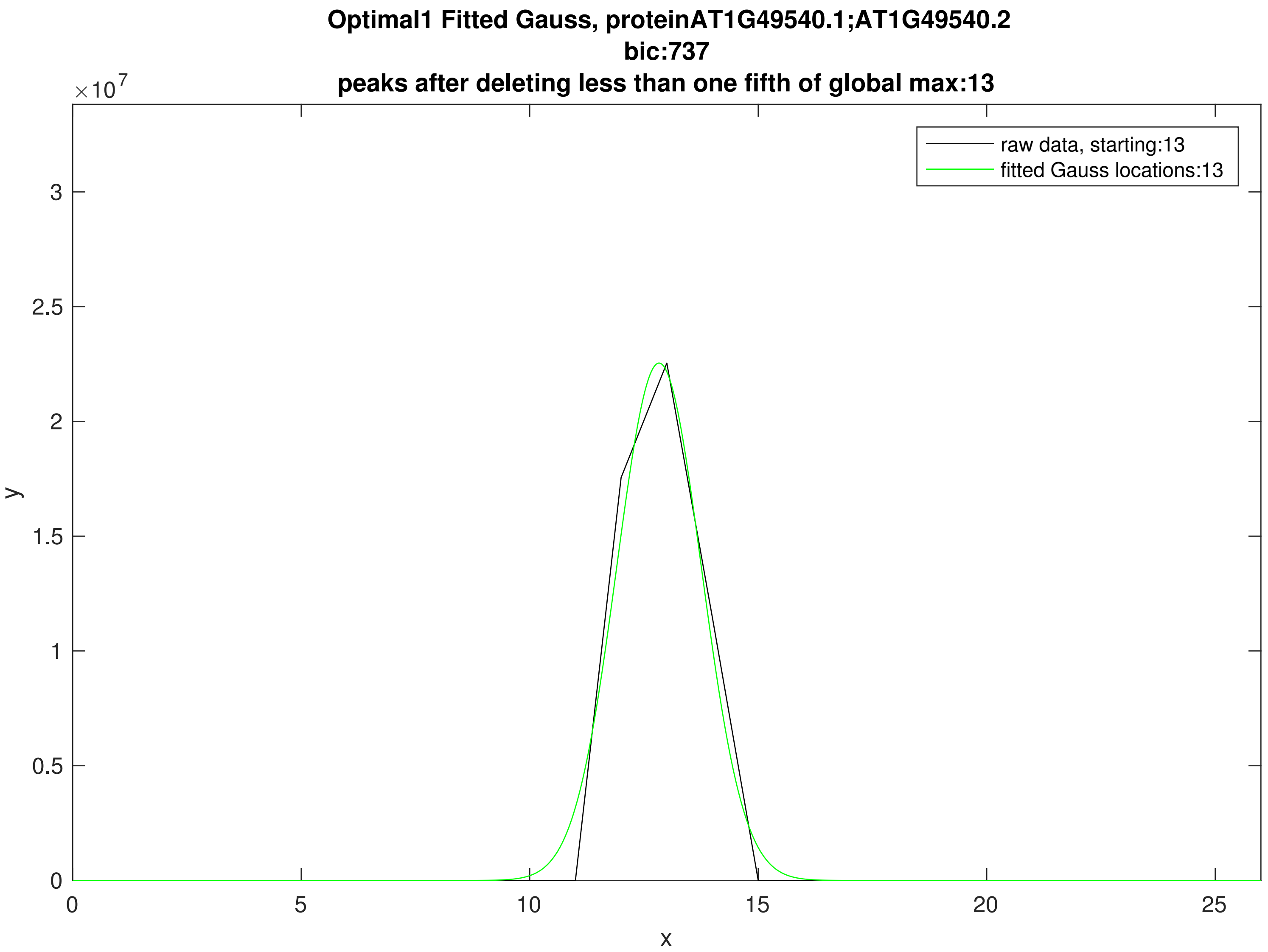
Optimal1 Fitted Gauss, proteinAT1G49540.1;AT1G49540.2
bic:737
peaks after deleting less than one fifth of global max:13
×10^7
raw data, starting:13
fitted Gauss locations:13
y
x
0
0.5
1
1.5
2
2.5
3
0
5
10
15
20
25

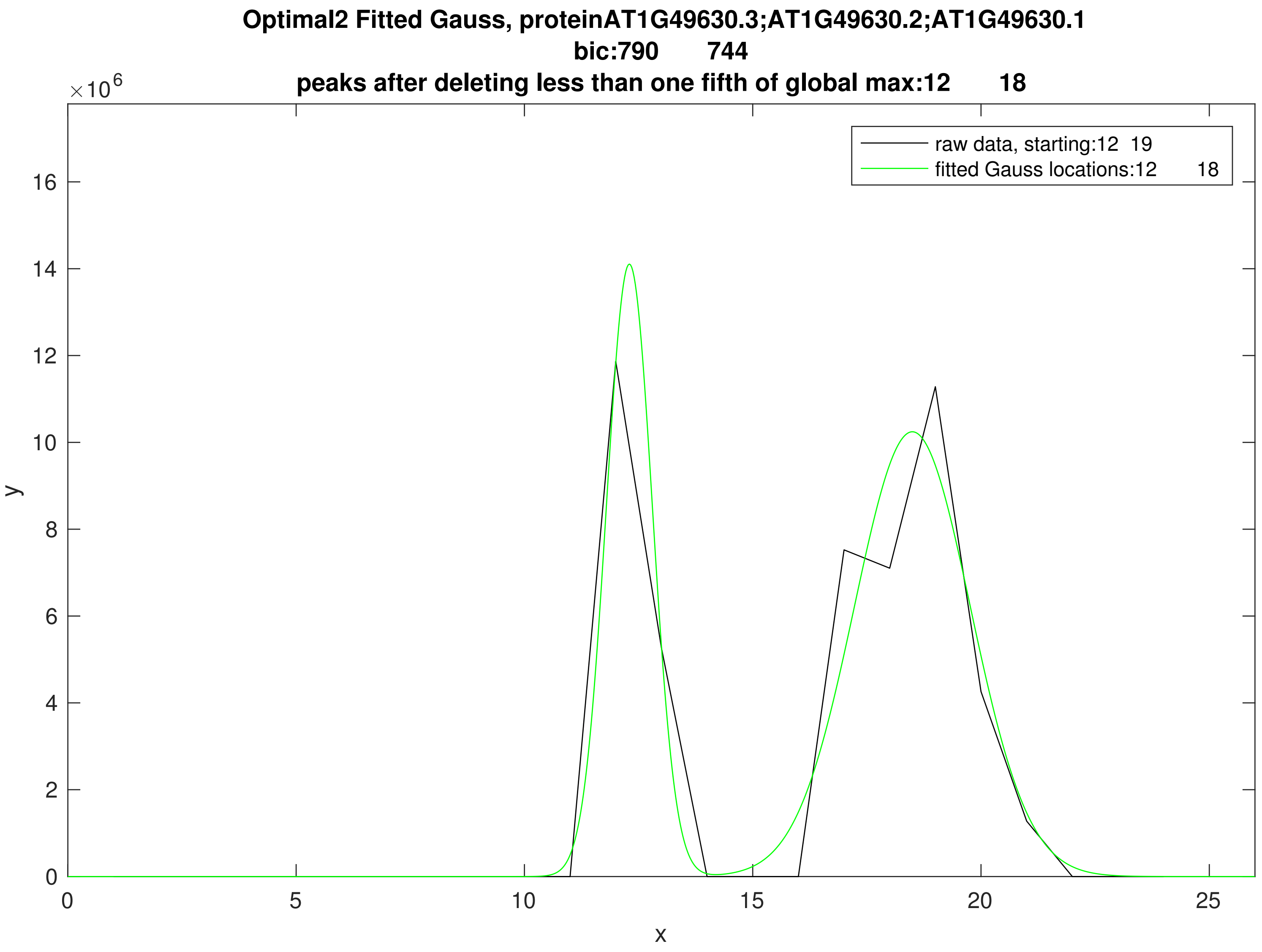

Optimal2 Fitted Gauss, proteinAT1G49630.3;AT1G49630.2;AT1G49630.1
bic:790 744
peaks after deleting less than one fifth of global max:12 18
×10^6
raw data, starting:12 19
fitted Gauss locations:12 18
16
14
12
10
8
6
4
2
0
y
0
5
10
15
20
25
x

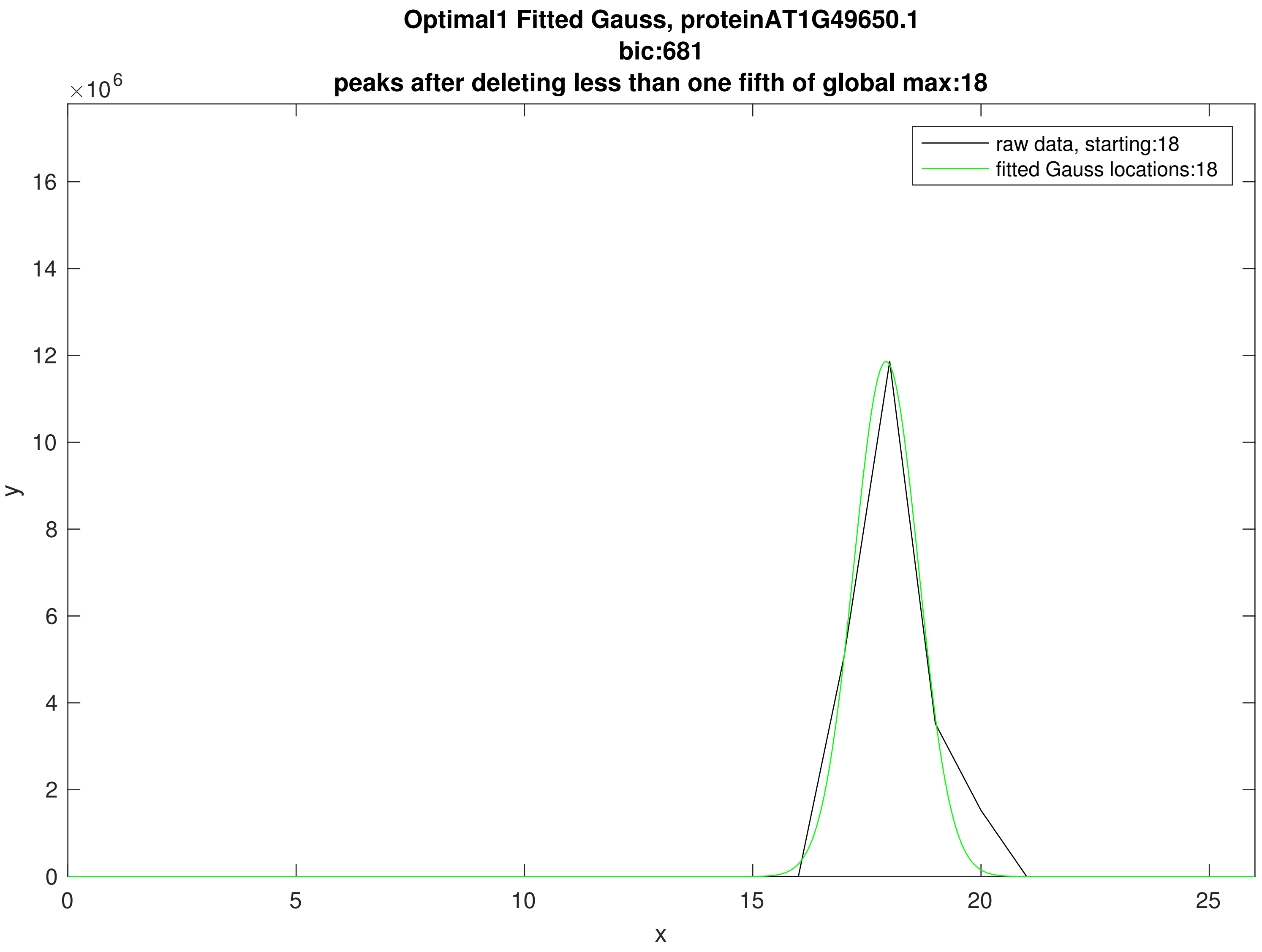

Optimal1 Fitted Gauss, proteinAT1G49650.1
bic:681
peaks after deleting less than one fifth of global max:18
×10^6
raw data, starting:18
fitted Gauss locations:18
y
x
0
2
4
6
8
10
12
14
16
0
5
10
15
20
25

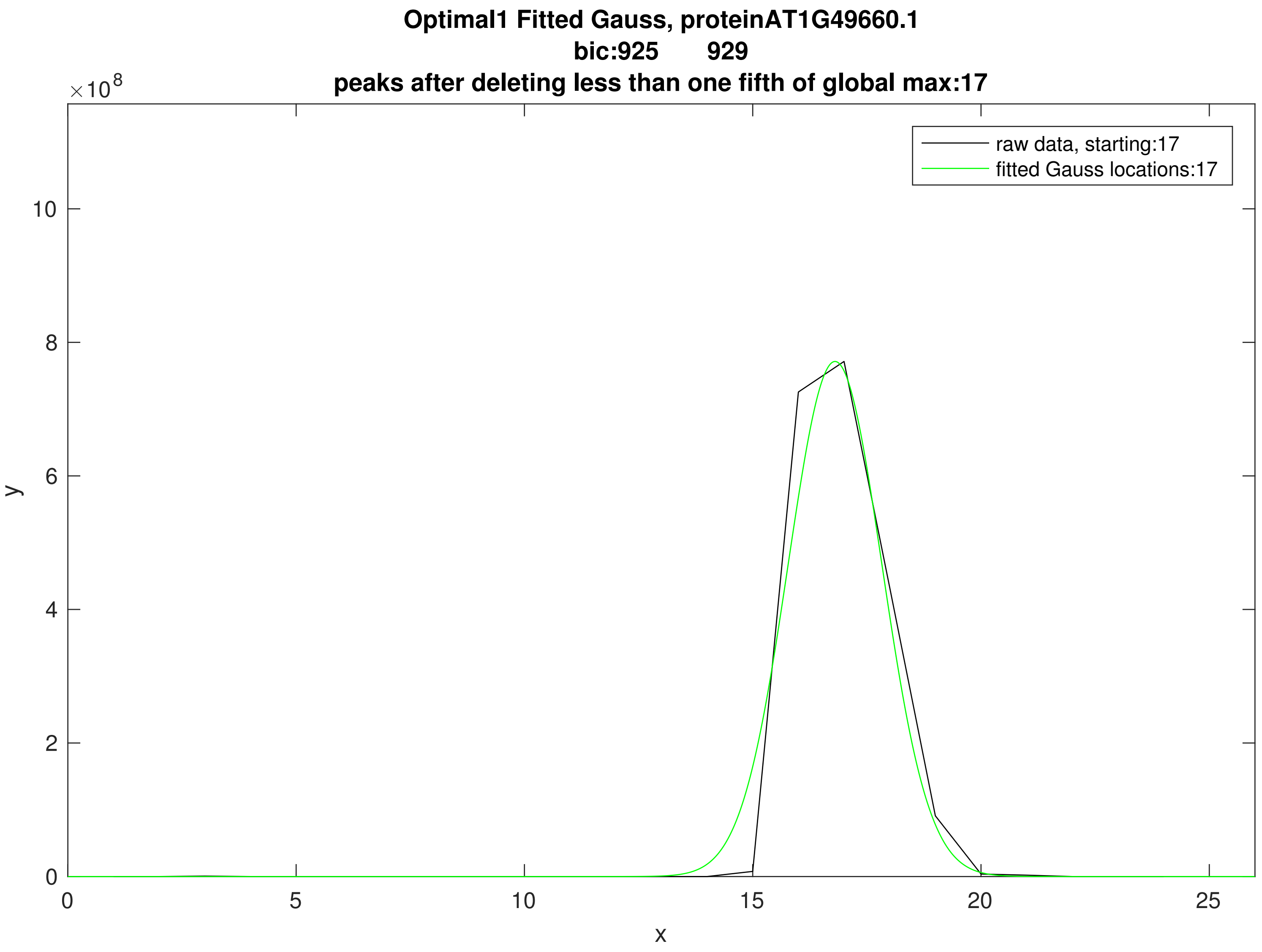
Optimal1 Fitted Gauss, proteinAT1G49660.1
bic:925 929
peaks after deleting less than one fifth of global max:17
×10^8
raw data, starting:17
fitted Gauss locations:17
y
x
0
2
4
6
8
10
0
5
10
15
20
25

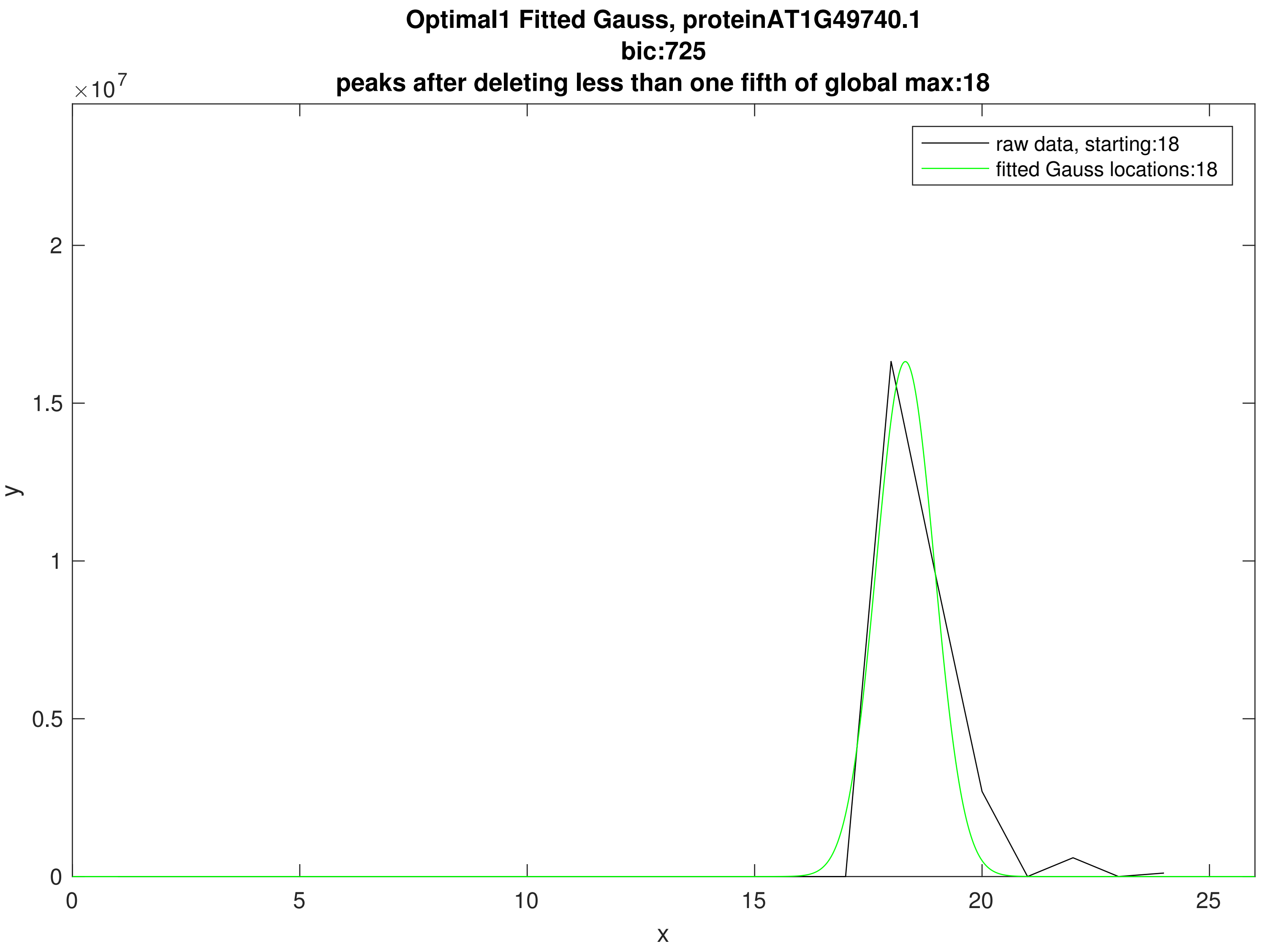
Optimal1 Fitted Gauss, proteinAT1G49740.1
bic:725
peaks after deleting less than one fifth of global max:18
×10^7
raw data, starting:18
fitted Gauss locations:18
2
1.5
1
0.5
0
y
0
5
10
15
20
25
x

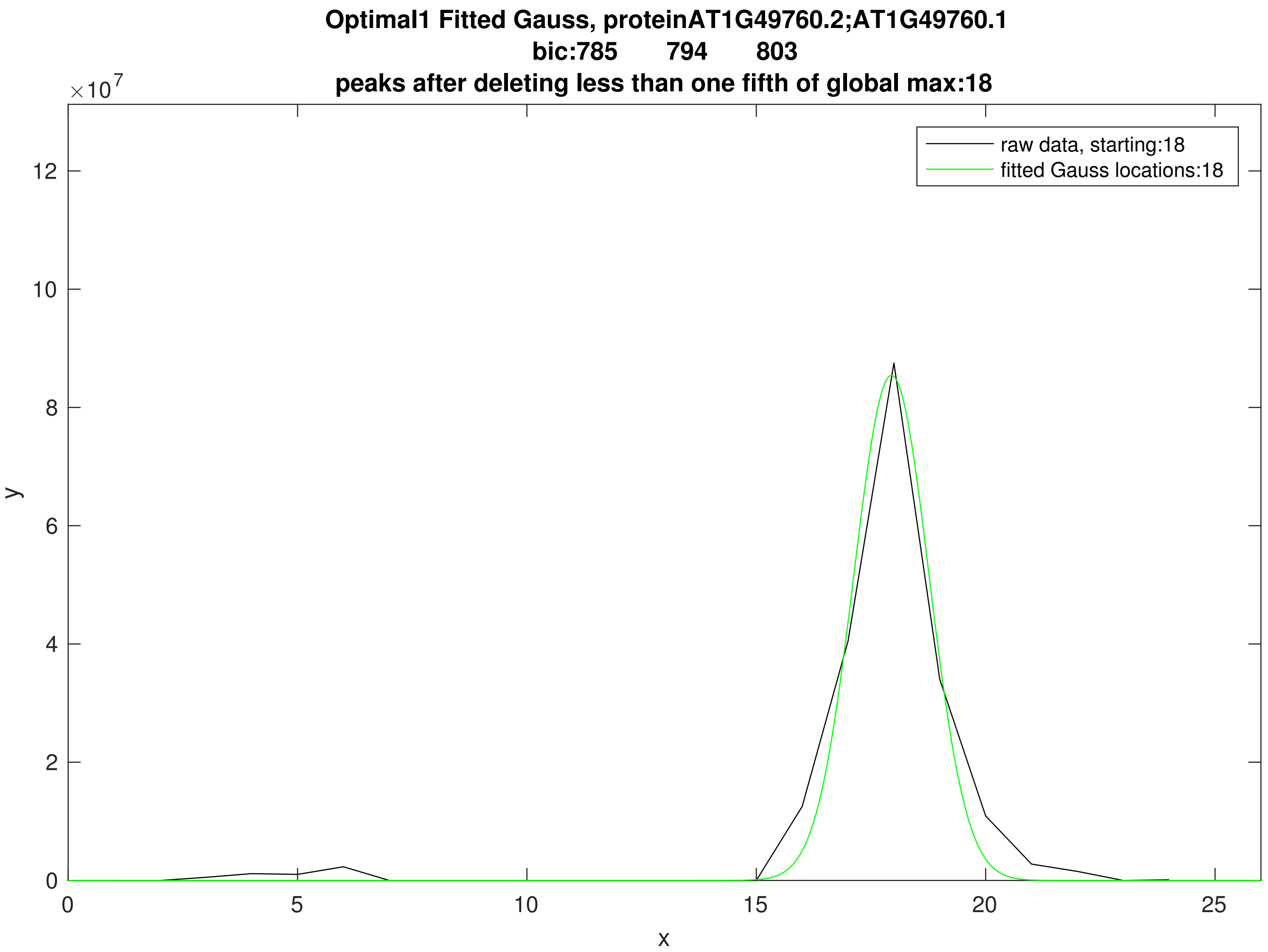
Optimal1 Fitted Gauss, proteinAT1G49760.2;AT1G49760.1
bic:785 794 803
peaks after deleting less than one fifth of global max:18
×10^7
raw data, starting:18
fitted Gauss locations:18
y
x
0
2
4
6
8
10
12
0
5
10
15
20
25

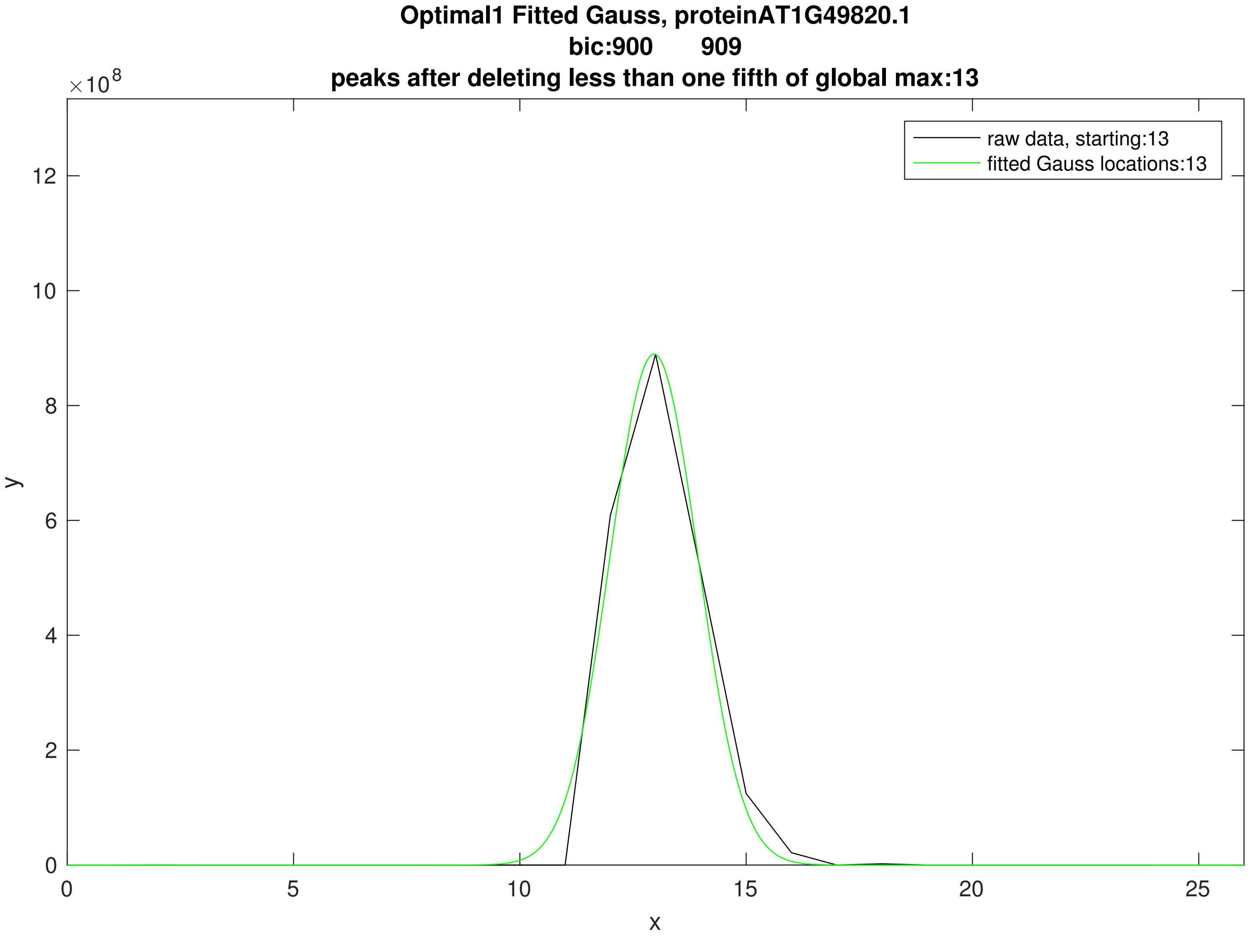
Optimal1 Fitted Gauss, proteinAT1G49820.1
bic:900 909
peaks after deleting less than one fifth of global max:13
×10^8
raw data, starting:13
fitted Gauss locations:13
y
x
0
2
4
6
8
10
12
5
15
20
25

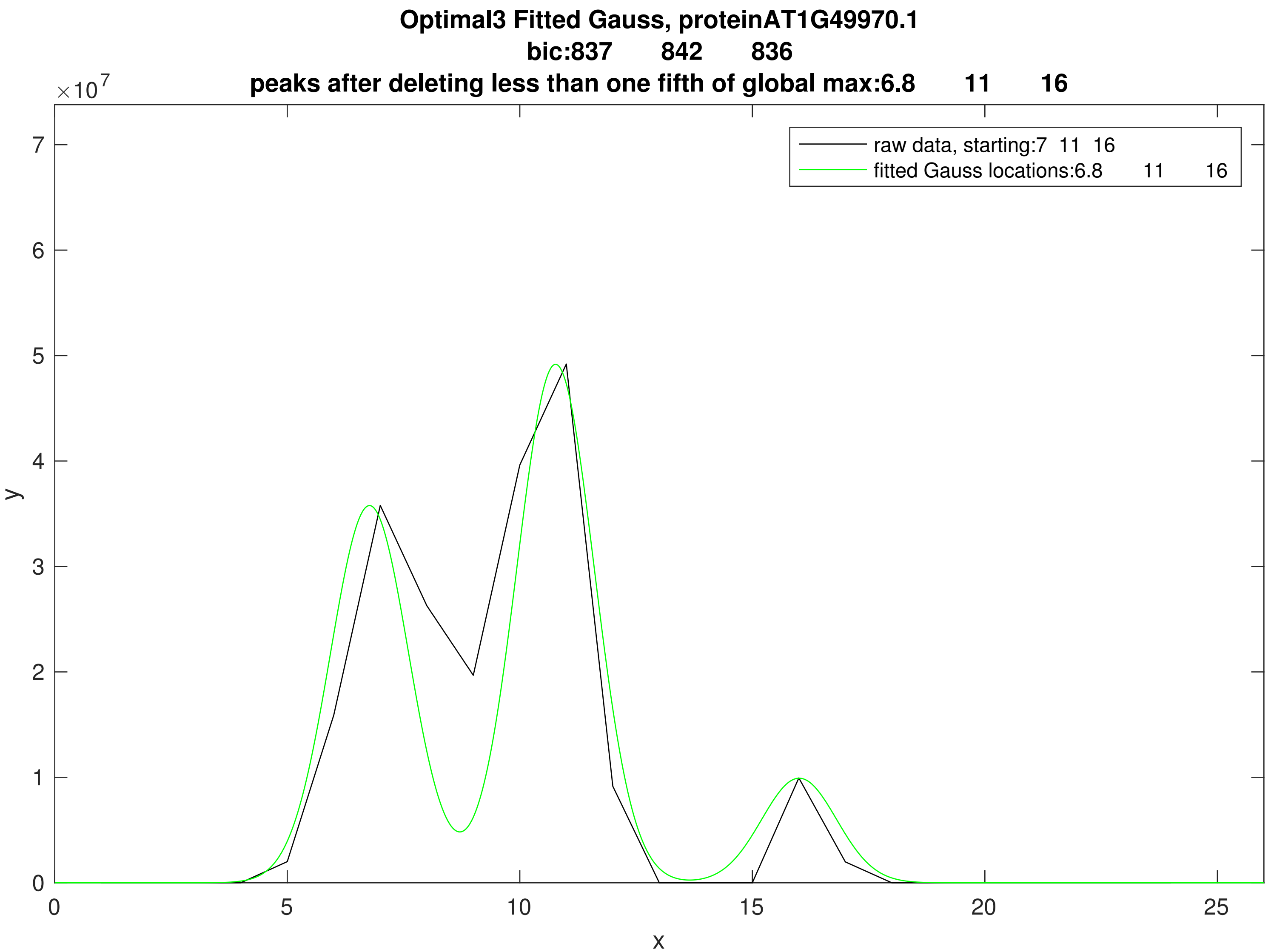
Optimal3 Fitted Gauss, proteinAT1G49970.1
bic:837 842 836
peaks after deleting less than one fifth of global max:6.8 11 16
×10^7
raw data, starting:7 11 16
fitted Gauss locations:6.8 11 16
y
x
0
1
2
3
4
5
6
7
0
5
10
15
20
25

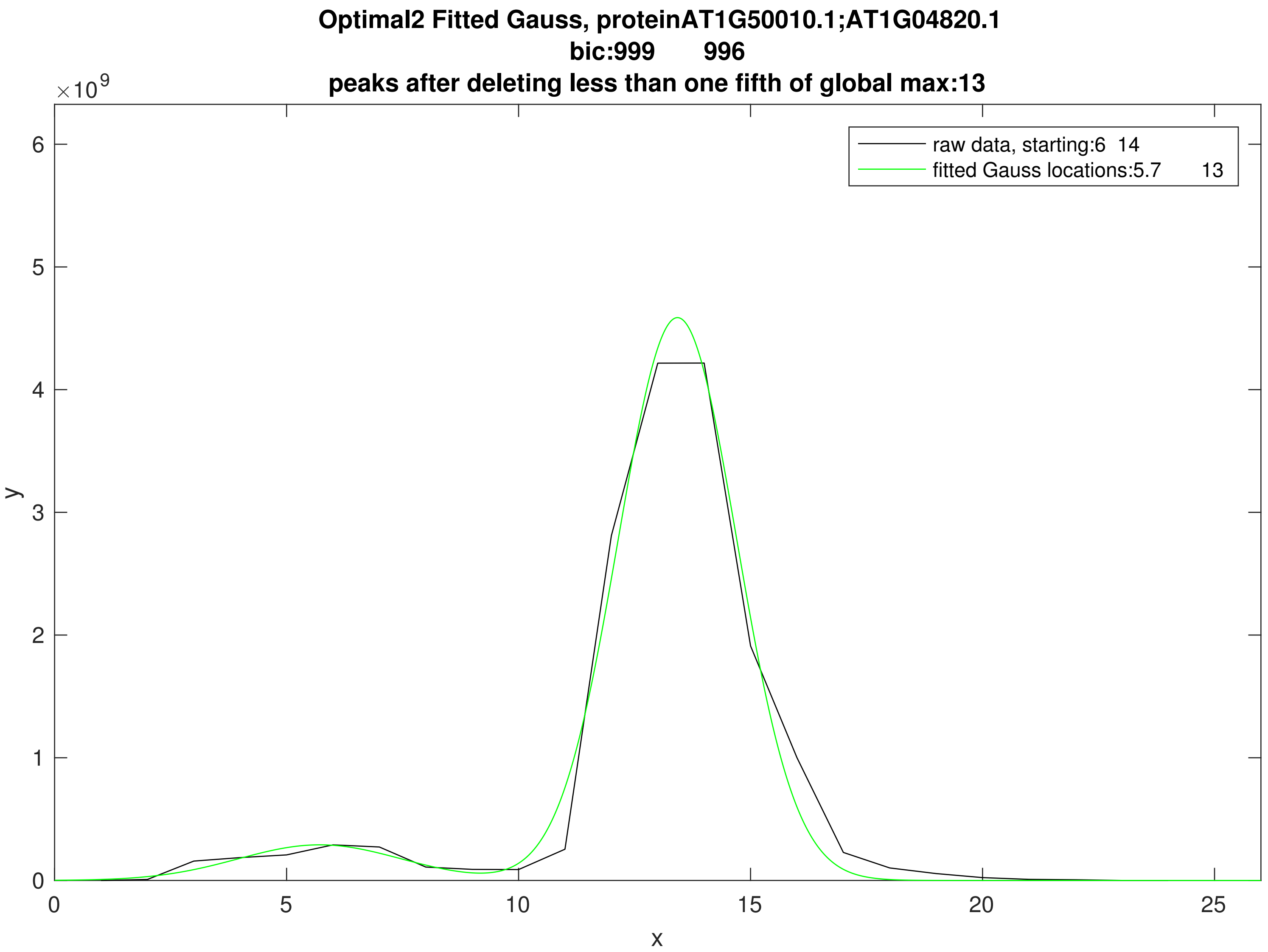
Optimal2 Fitted Gauss, proteinAT1G50010.1;AT1G04820.1
bic:999 996
peaks after deleting less than one fifth of global max:13
×10^9
raw data, starting:6 14
fitted Gauss locations:5.7 13
y
x

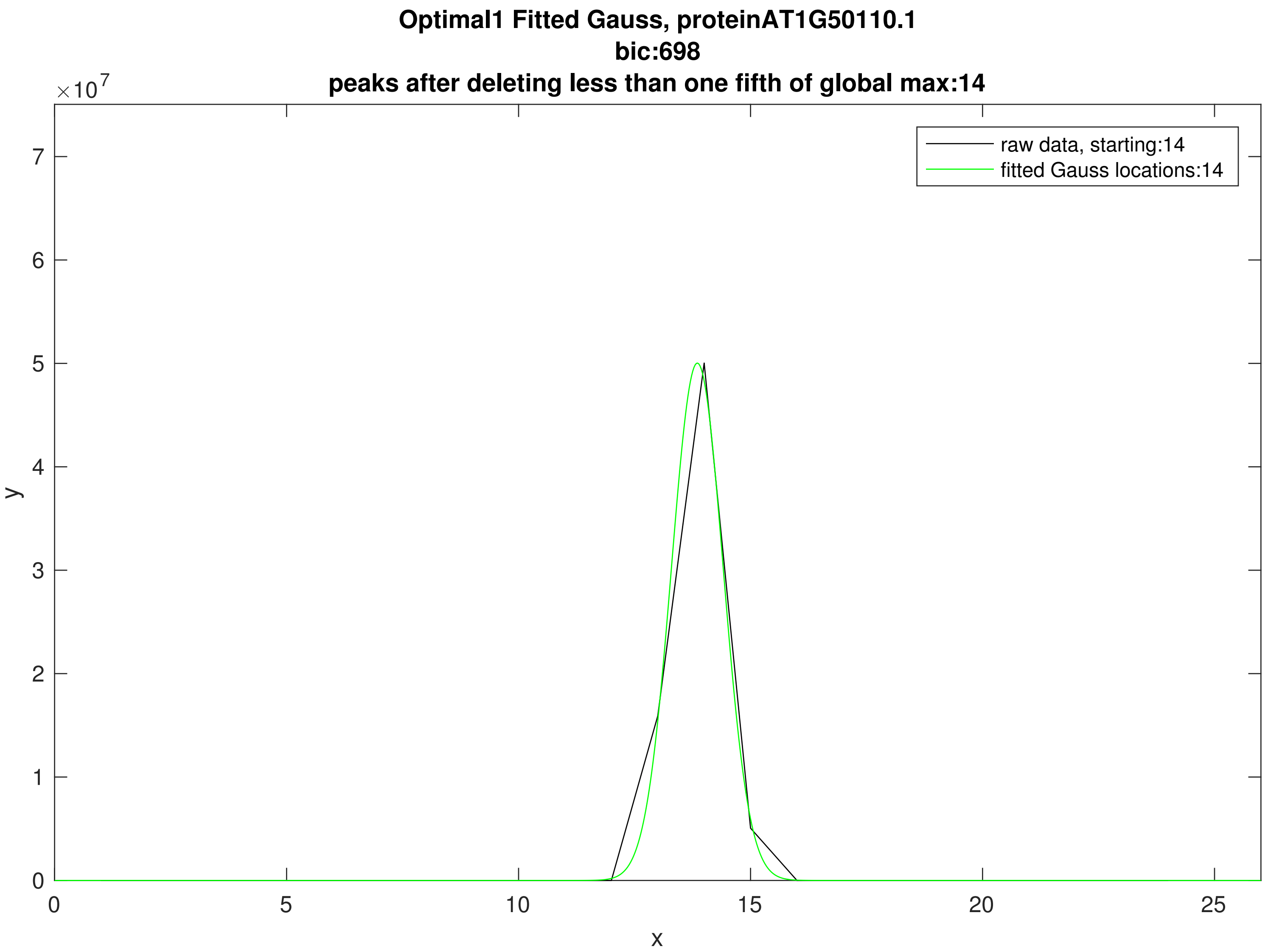
Optimal1 Fitted Gauss, proteinAT1G50110.1
bic:698
peaks after deleting less than one fifth of global max:14
×10^7
raw data, starting:14
fitted Gauss locations:14
y
x
0
1
2
3
4
5
6
7
0
5
10
15
20
25

**Optimal1 Fitted Gauss, proteinAT1G50200.1**
**bic:913 913 922 932**
**peaks after deleting less than one fifth of global max:11**

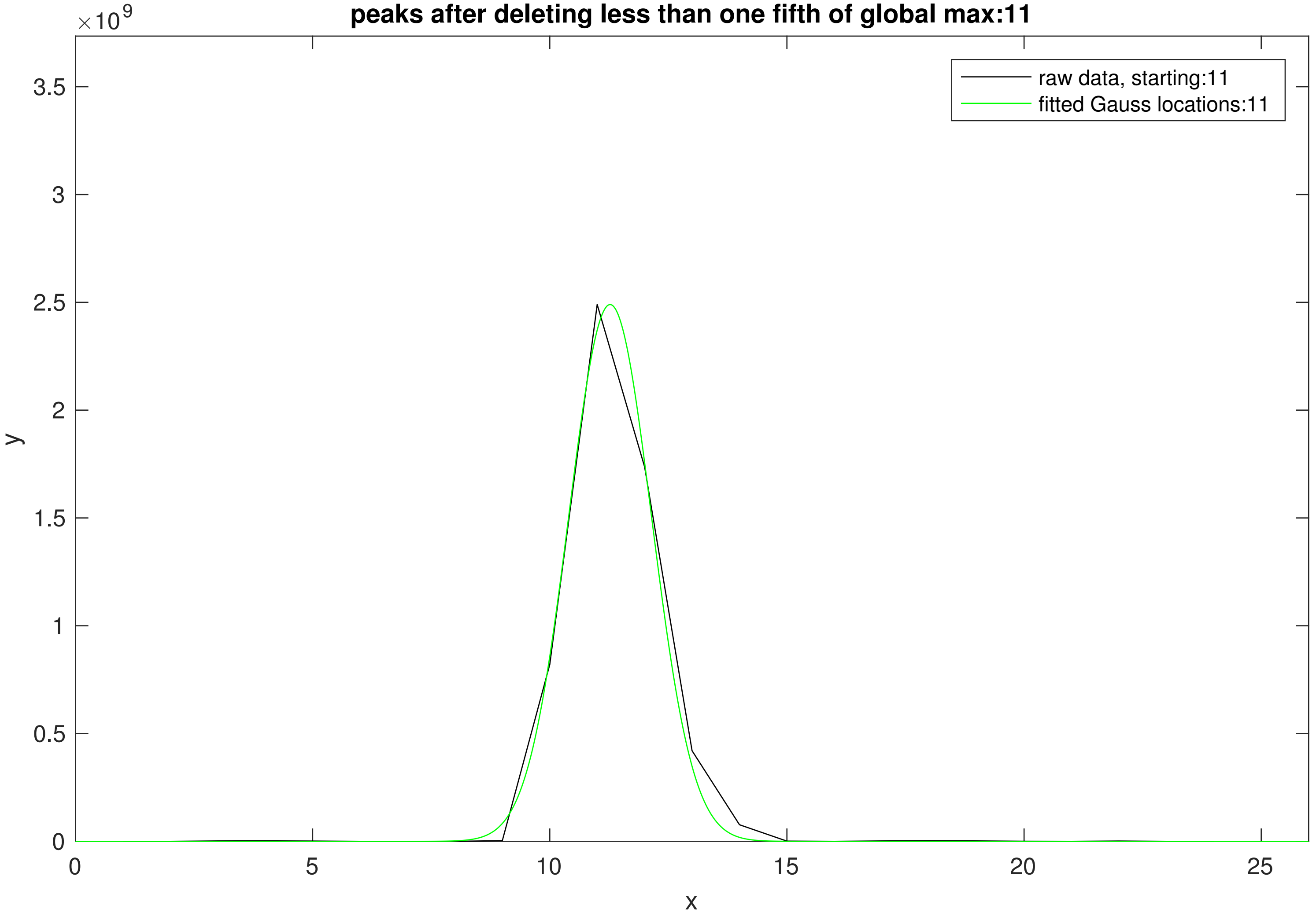

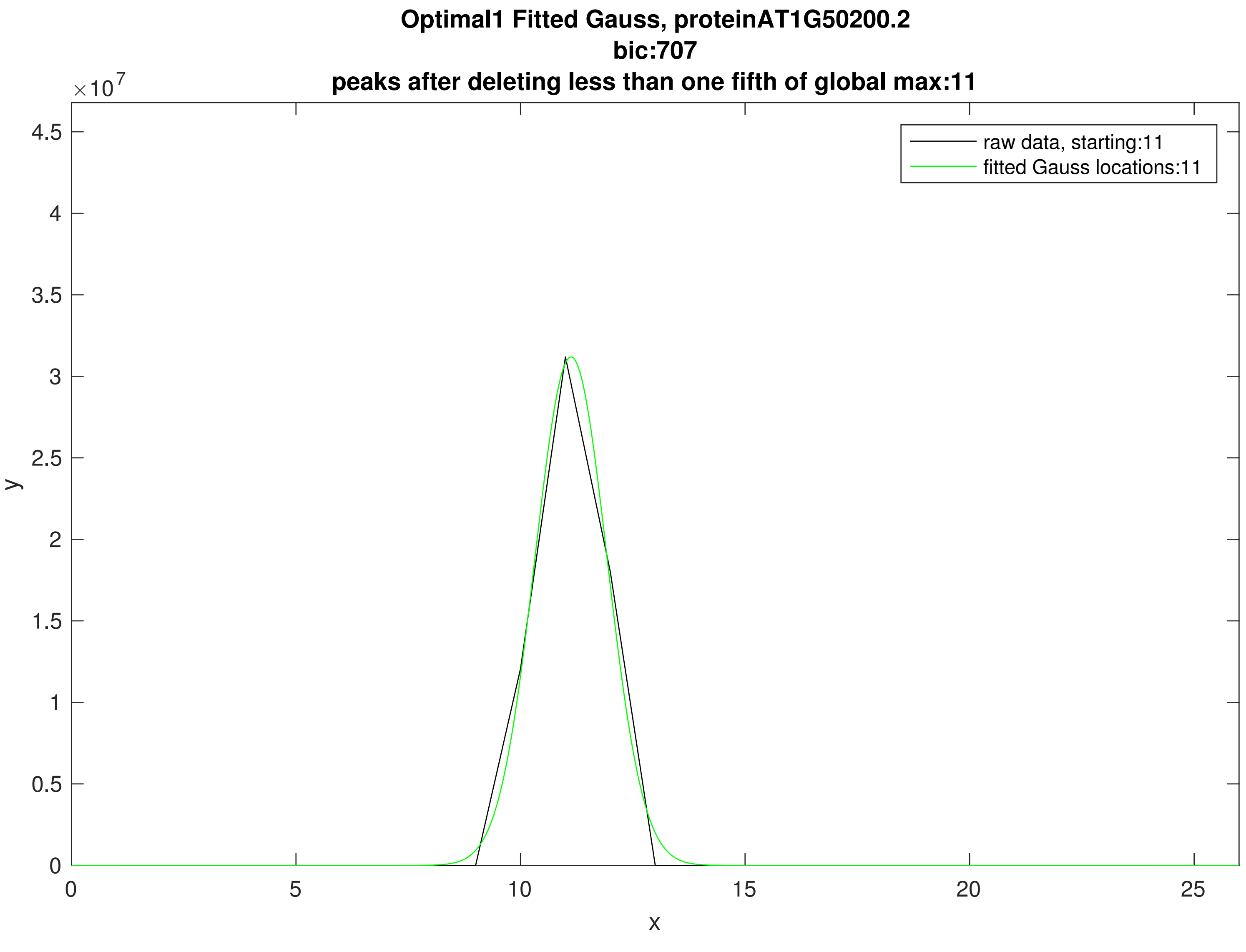

Optimal1 Fitted Gauss, proteinAT1G50200.2
bic:707
peaks after deleting less than one fifth of global max:11
×10^7
4.5
4
3.5
3
2.5
2
1.5
1
0.5
0
y
0
5
10
15
20
25
x
raw data, starting:11
fitted Gauss locations:11

**Optimal1 Fitted Gauss, proteinAT1G50370.1**
**bic:840 851**
**peaks after deleting less than one fifth of global max:18**

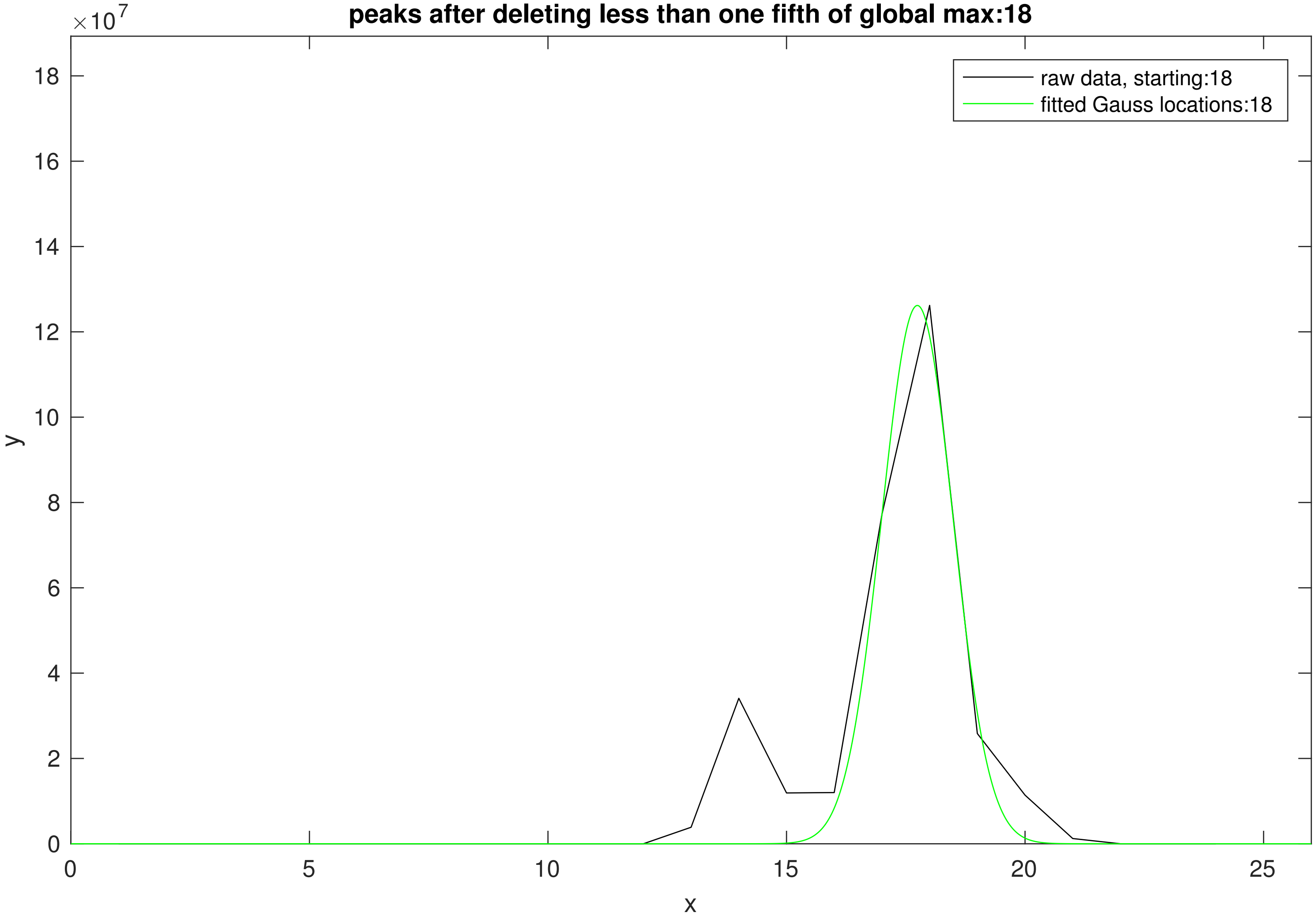

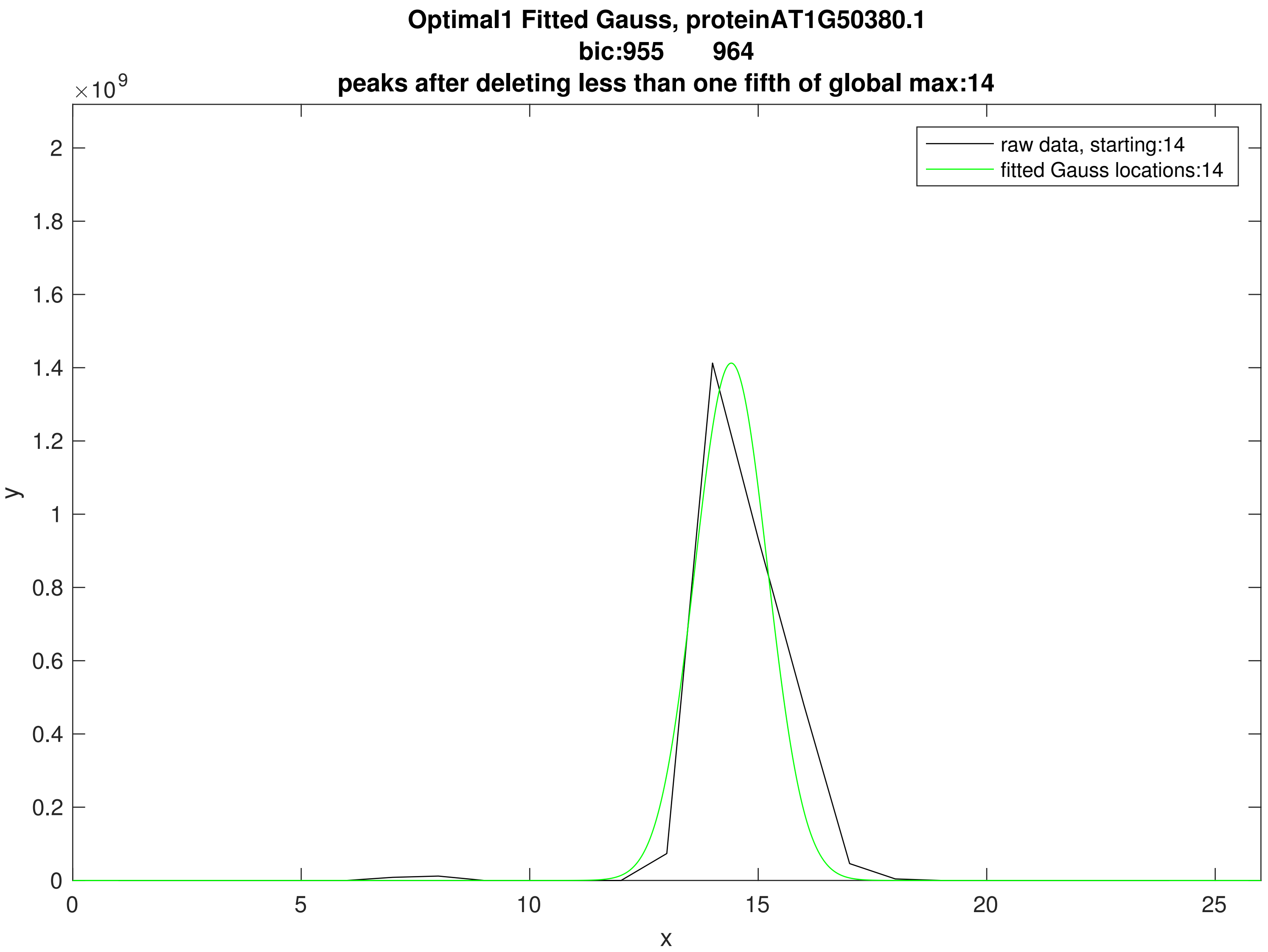
Optimal1 Fitted Gauss, proteinAT1G50380.1
bic:955 964
peaks after deleting less than one fifth of global max:14
×10^9
raw data, starting:14
fitted Gauss locations:14
y
x
0
0.2
0.4
0.6
0.8
1
1.2
1.4
1.6
1.8
2
0
5
10
15
20
25

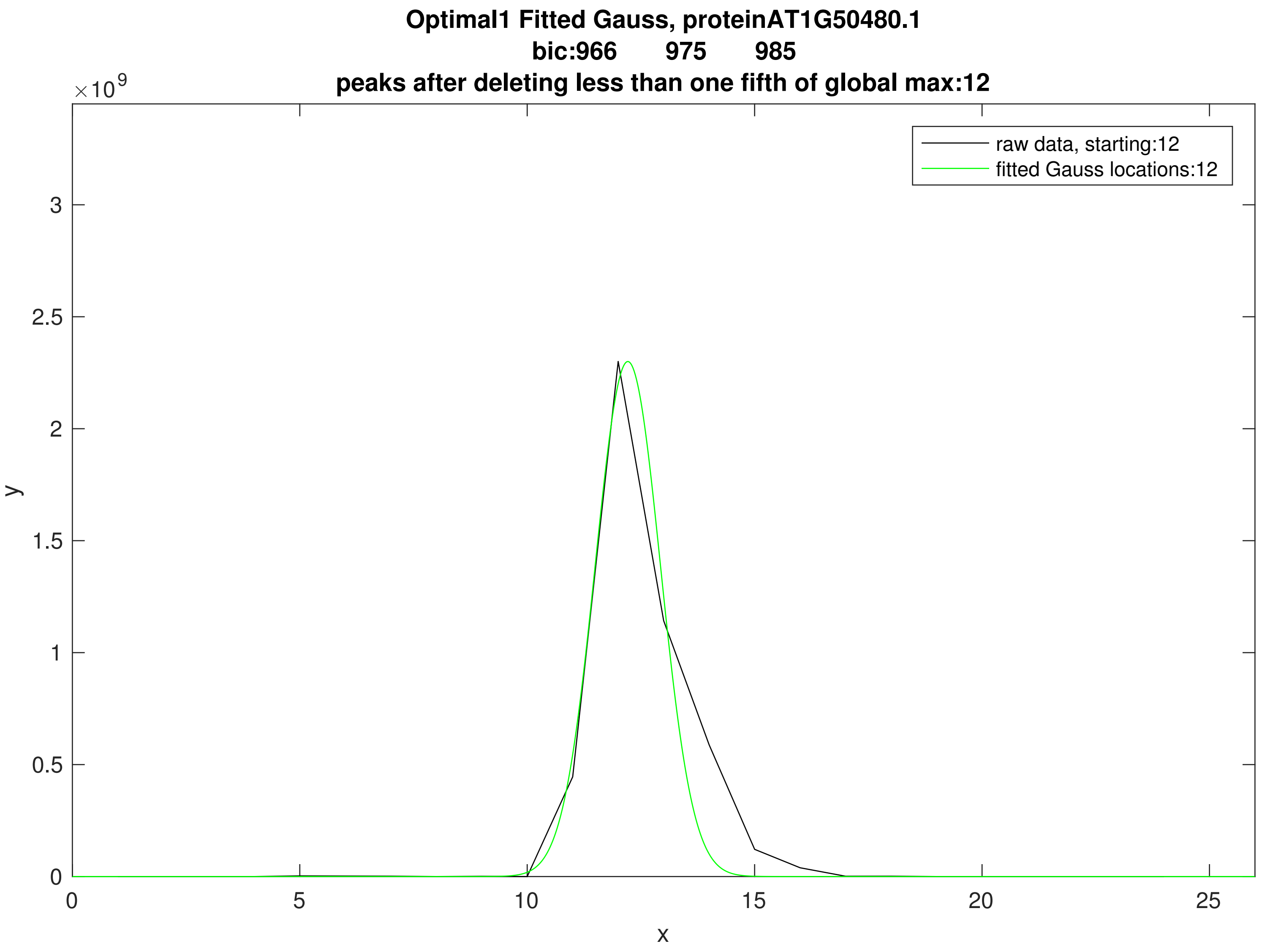
Optimal1 Fitted Gauss, proteinAT1G50480.1
bic:966 975 985
peaks after deleting less than one fifth of global max:12
×10^9
3
2.5
2
1.5
1
0.5
0
y
0
5
10
15
20
25
x
raw data, starting:12
fitted Gauss locations:12

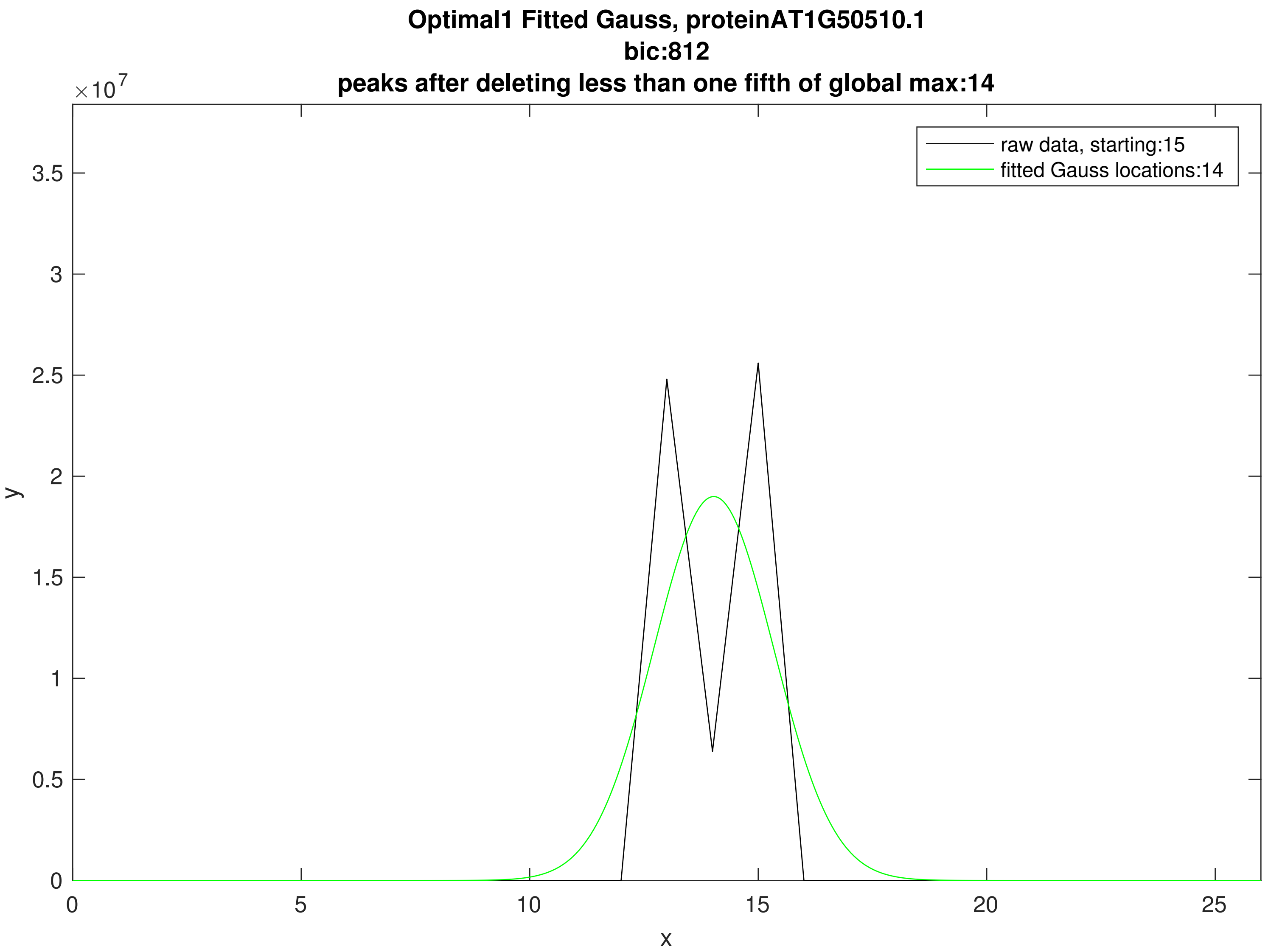

Optimal1 Fitted Gauss, proteinAT1G50510.1
bic:812
peaks after deleting less than one fifth of global max:14
×10^7
raw data, starting:15
fitted Gauss locations:14
y
x
0
0.5
1
1.5
2
2.5
3
3.5
0
5
10
15
20
25

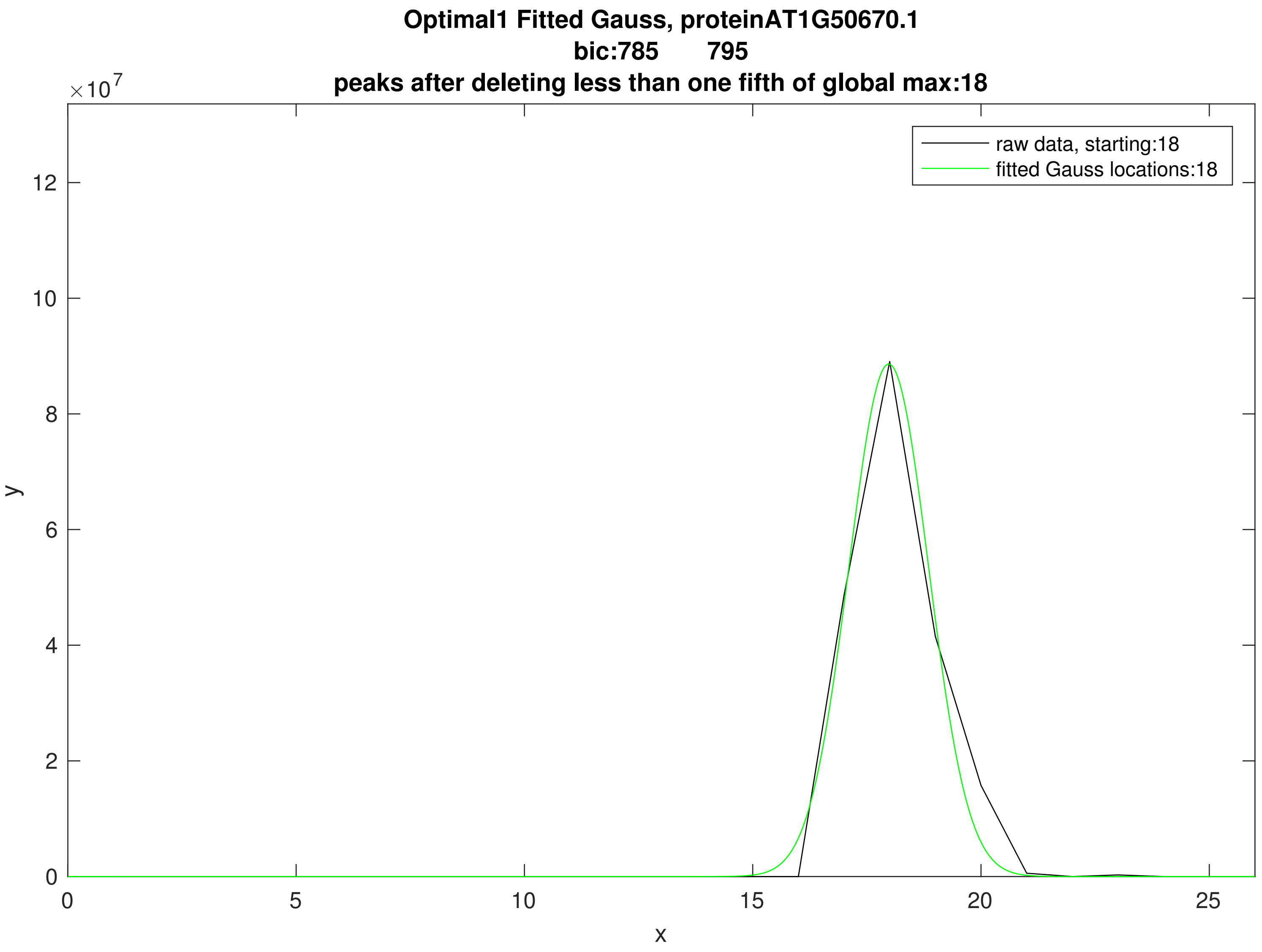
Optimal1 Fitted Gauss, proteinAT1G50670.1
bic:785 795
peaks after deleting less than one fifth of global max:18
×10^7
12
10
8
6
4
2
0
y
0
5
10
15
20
25
x
raw data, starting:18
fitted Gauss locations:18

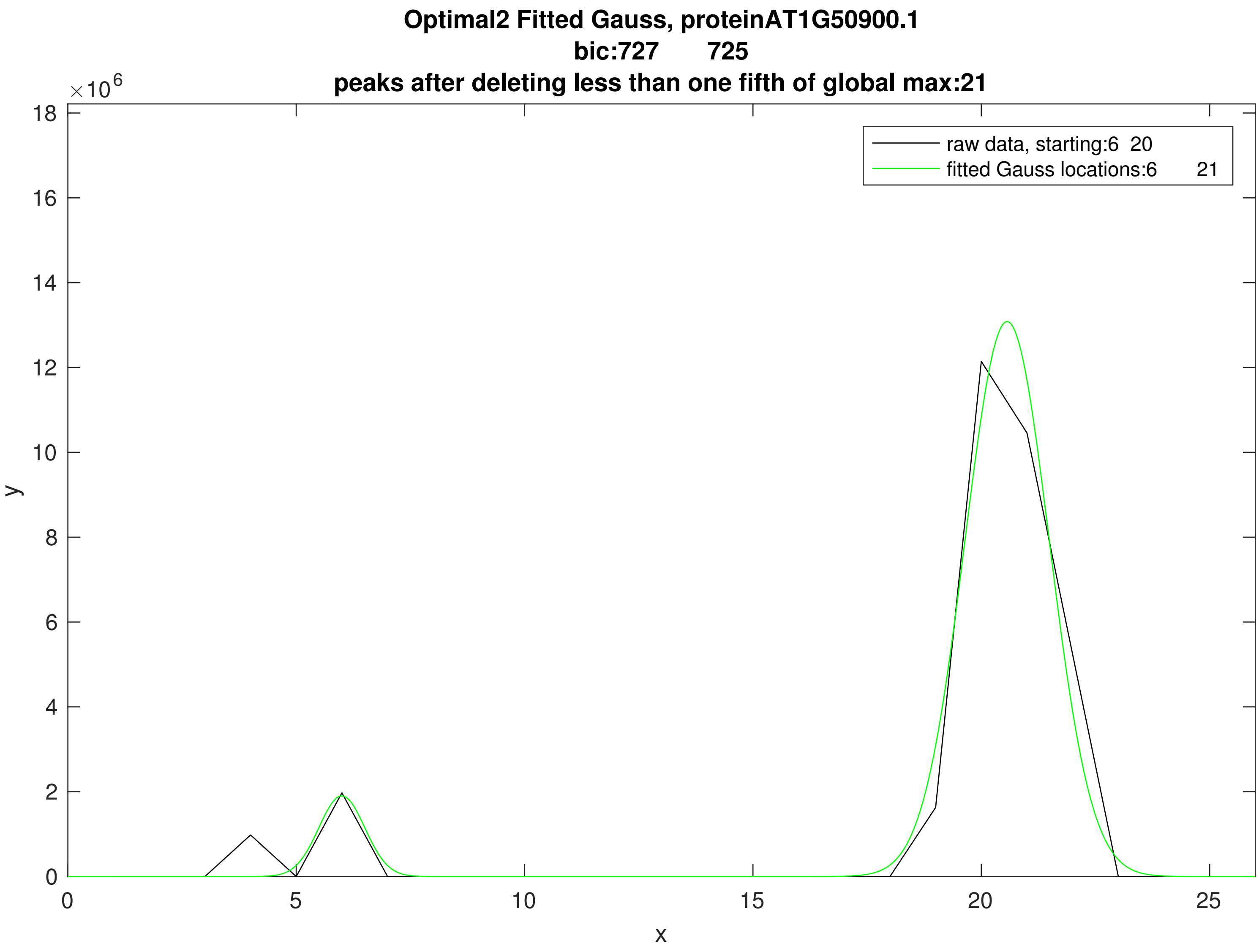
Optimal2 Fitted Gauss, proteinAT1G50900.1
bic:727 725
peaks after deleting less than one fifth of global max:21
×10^6
raw data, starting:6 20
fitted Gauss locations:6 21
y
x
0
2
4
6
8
10
12
14
16
18
0
5
10
15
20
25

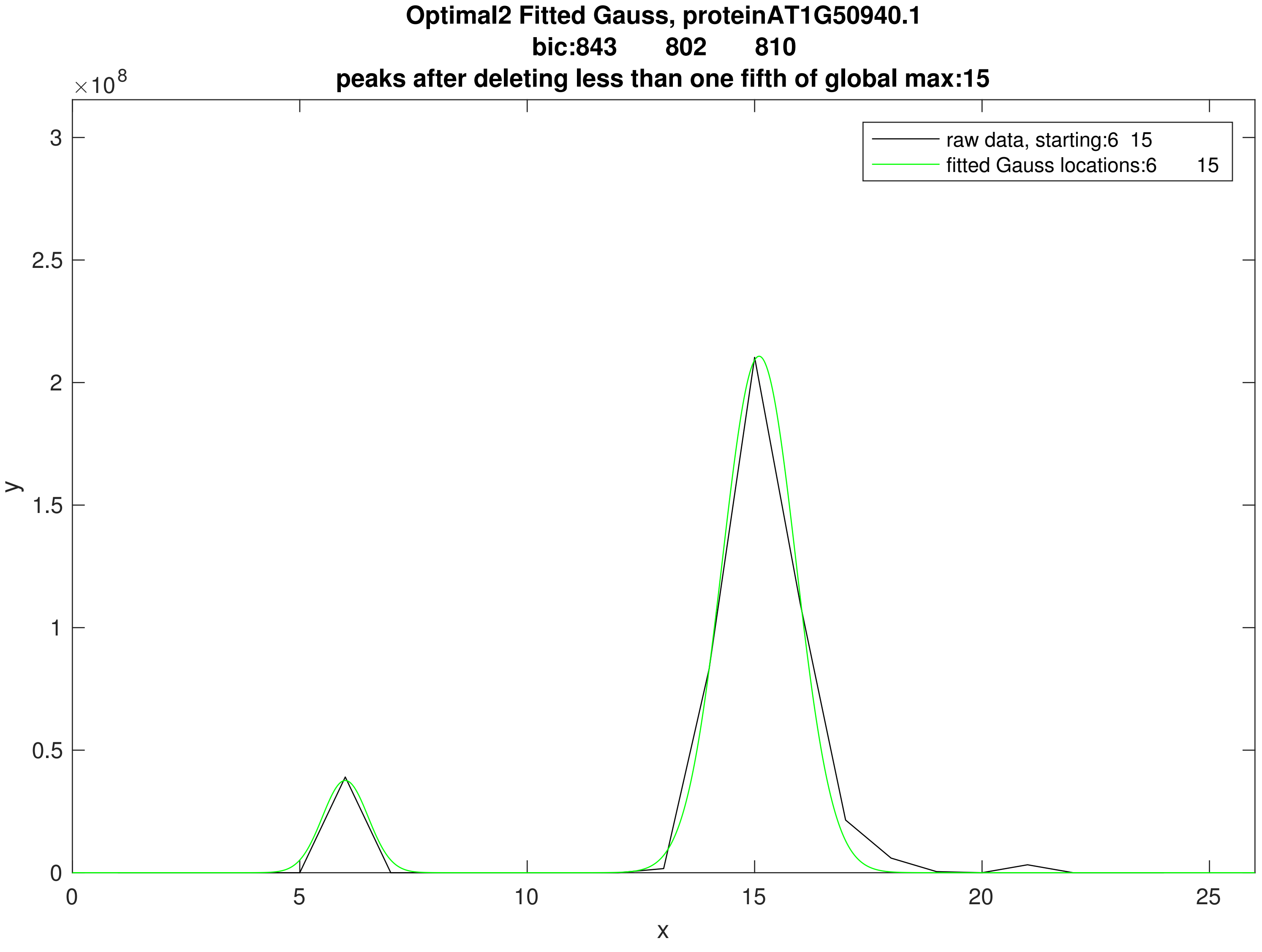

Optimal2 Fitted Gauss, proteinAT1G50940.1
bic:843 802 810
peaks after deleting less than one fifth of global max:15
×10^8
raw data, starting:6 15
fitted Gauss locations:6 15
y
x
0
0.5
1
1.5
2
2.5
3
0
5
10
15
20
25

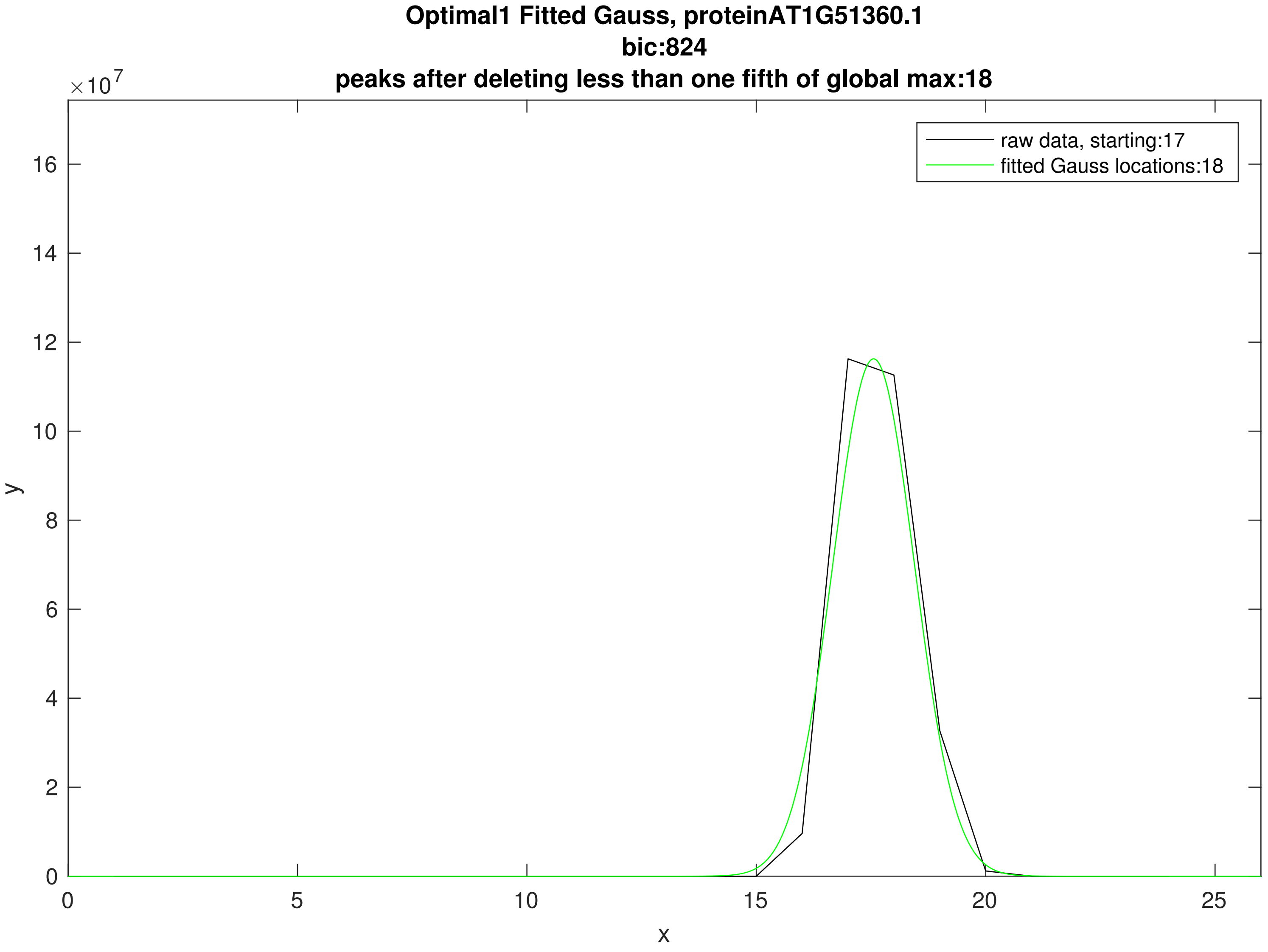

Optimal1 Fitted Gauss, proteinAT1G51360.1
bic:824
peaks after deleting less than one fifth of global max:18
×10^7
raw data, starting:17
fitted Gauss locations:18
16
14
12
10
8
6
4
2
0
y
0
5
10
15
20
25
x

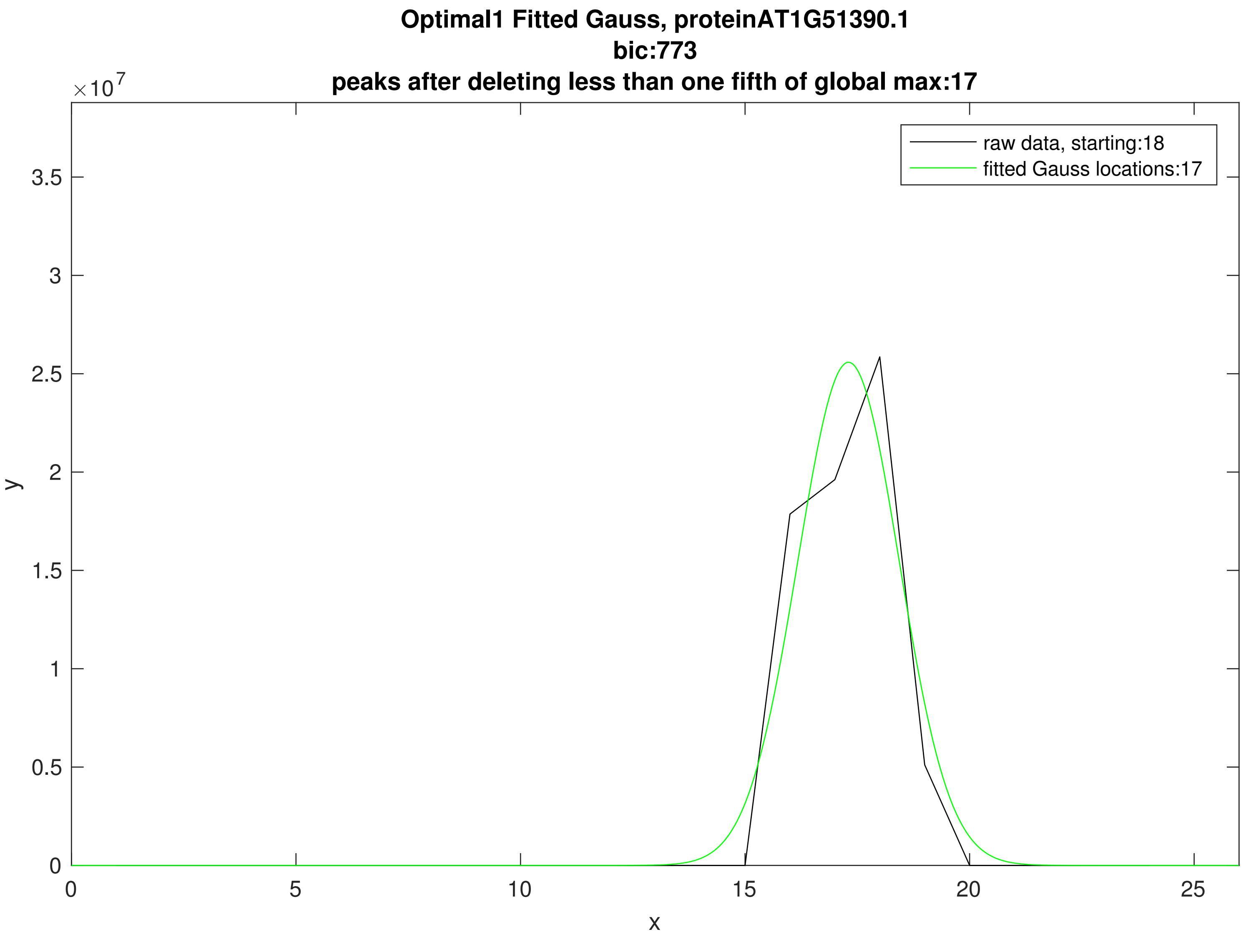
Optimal1 Fitted Gauss, proteinAT1G51390.1
bic:773
peaks after deleting less than one fifth of global max:17
×10^7
raw data, starting:18
fitted Gauss locations:17
y
x
0
0.5
1
1.5
2
2.5
3
3.5
5
10
15
20
25

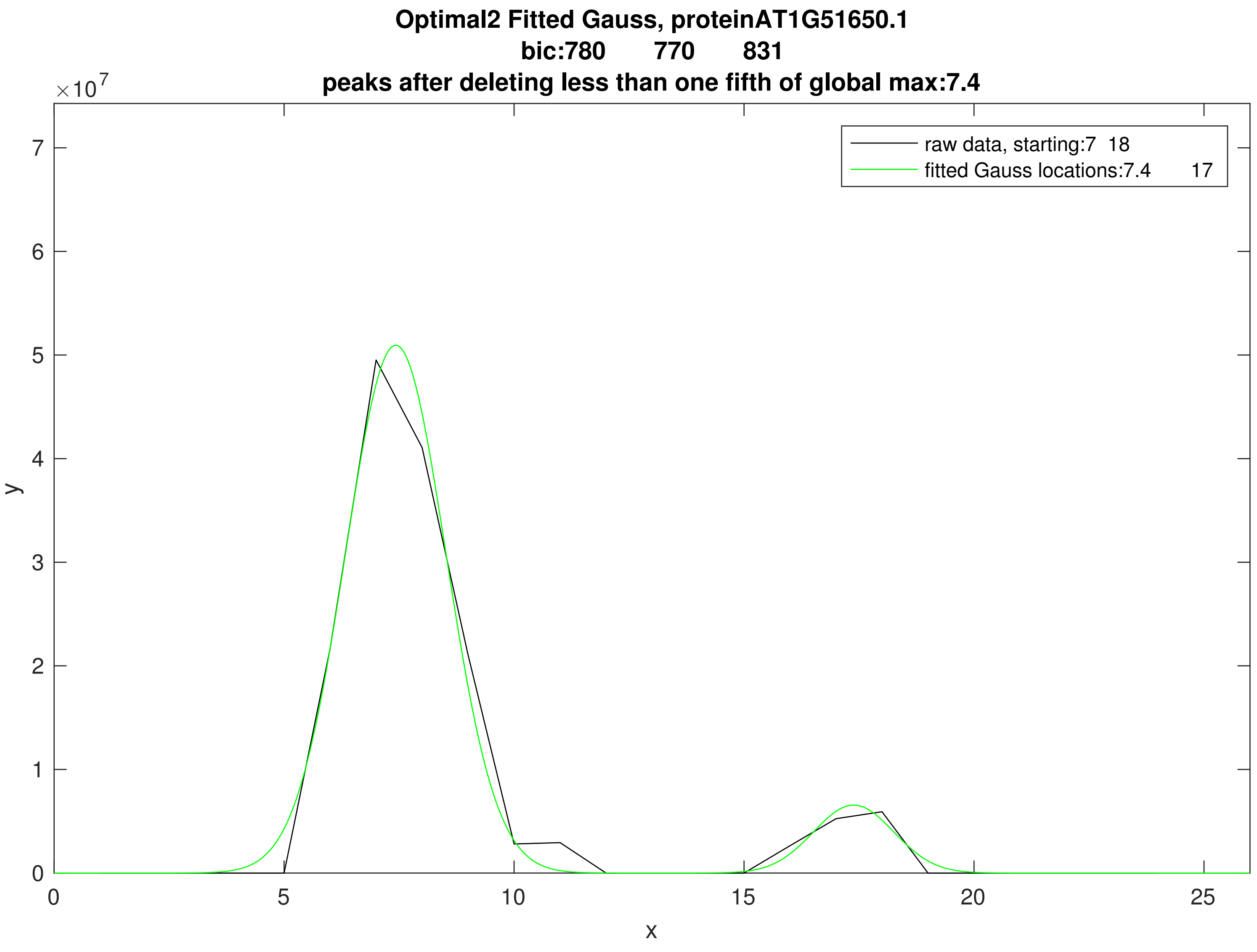

Optimal2 Fitted Gauss, proteinAT1G51650.1
bic:780 770 831
peaks after deleting less than one fifth of global max:7.4
×10^7
raw data, starting:7 18
fitted Gauss locations:7.4 17
y
x
0
1
2
3
4
5
6
7
0
5
10
15
20
25

**Optimal1 Fitted Gauss, proteinAT1G51680.1;AT1G51680.3;AT1G51680.2**
**bic:917 926**
**peaks after deleting less than one fifth of global max:14**

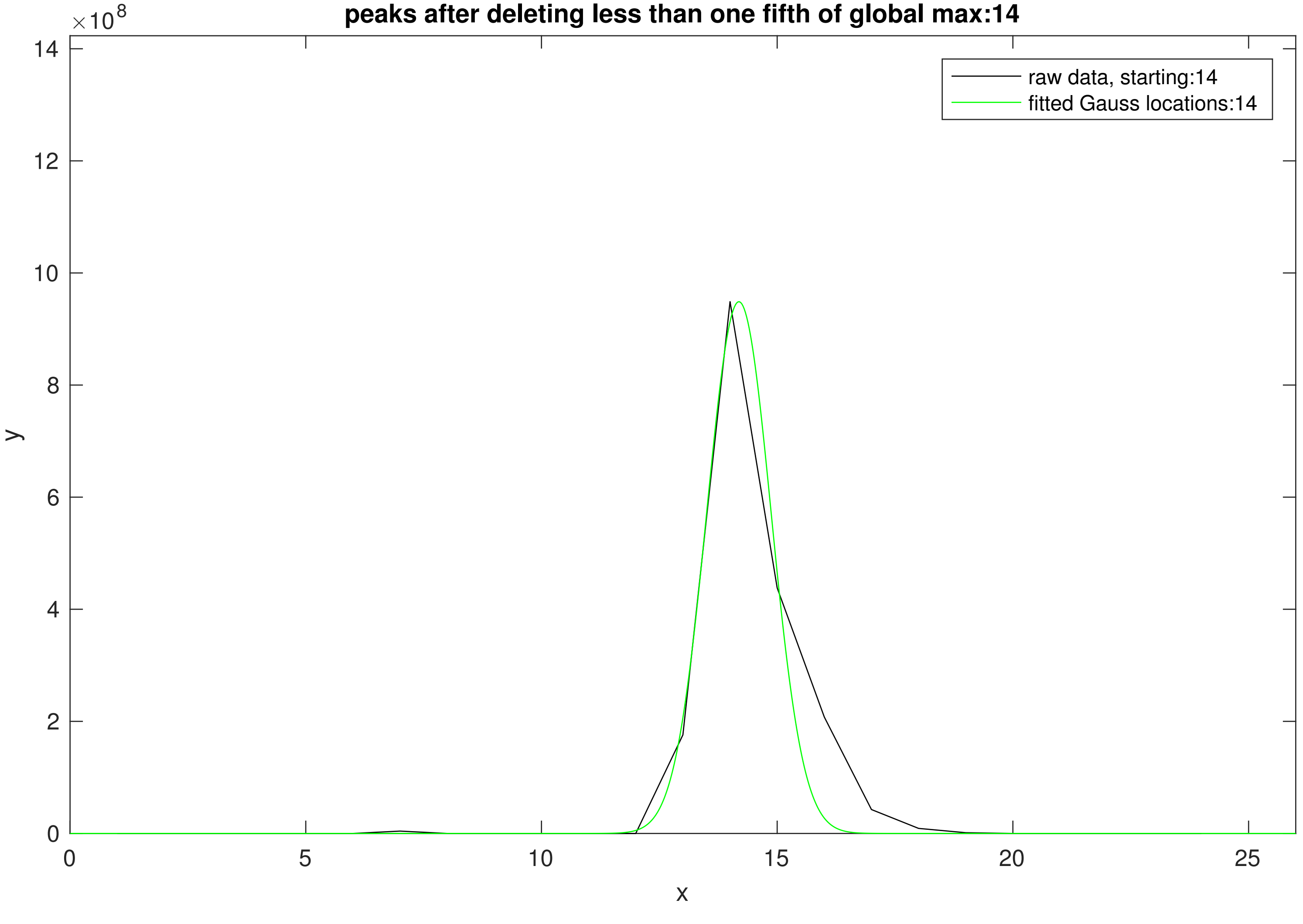

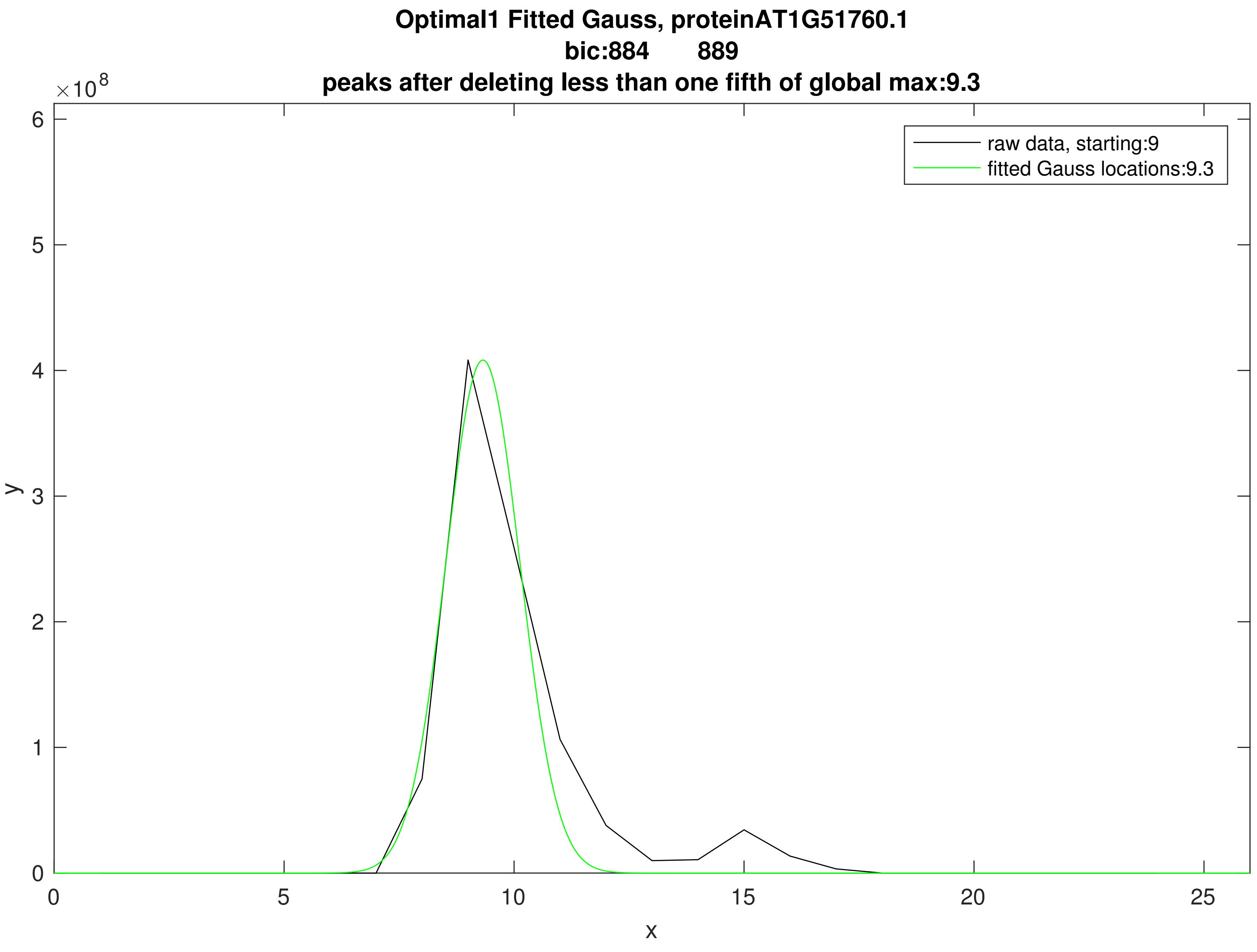
Optimal1 Fitted Gauss, proteinAT1G51760.1
bic:884 889
peaks after deleting less than one fifth of global max:9.3
×10^8
6
5
4
3
2
1
0
y
raw data, starting:9
fitted Gauss locations:9.3
0
5
10
15
20
25
x

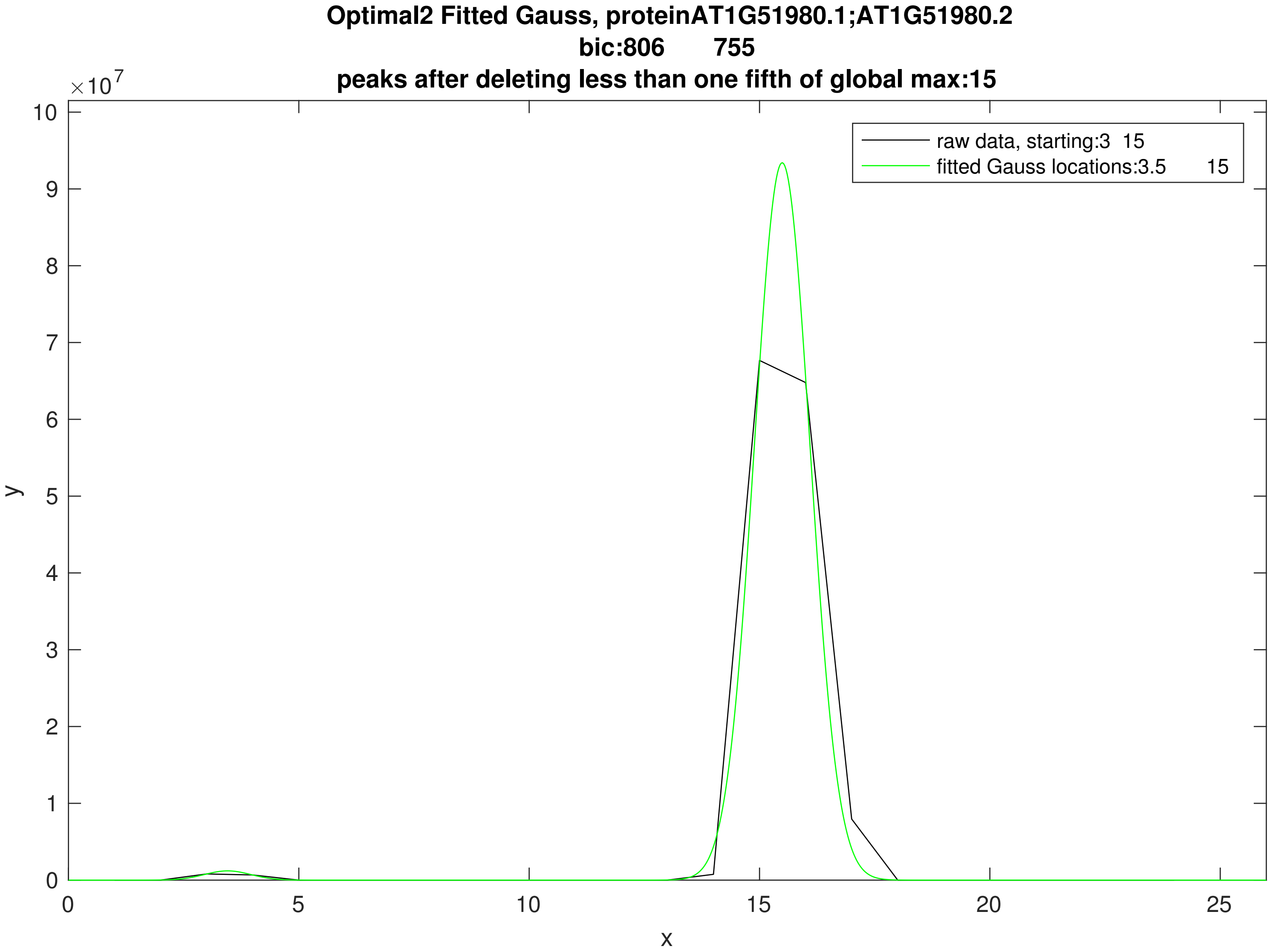

Optimal2 Fitted Gauss, proteinAT1G51980.1;AT1G51980.2
bic:806 755
peaks after deleting less than one fifth of global max:15
×10^7
raw data, starting:3 15
fitted Gauss locations:3.5 15
y
x
0
1
2
3
4
5
6
7
8
9
10
0
5
10
15
20
25

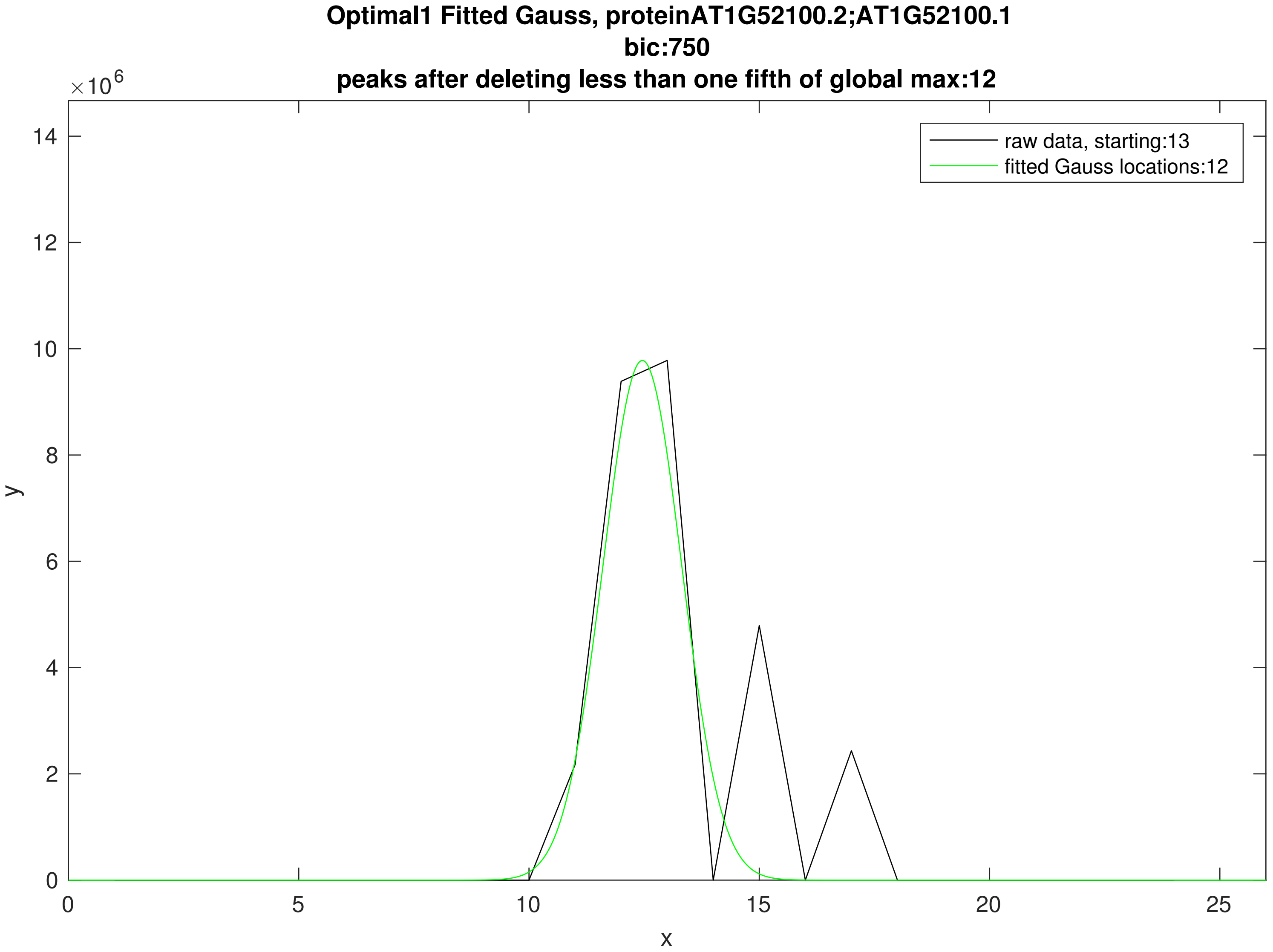
Optimal1 Fitted Gauss, proteinAT1G52100.2;AT1G52100.1
bic:750
peaks after deleting less than one fifth of global max:12
$\times 10^6$
raw data, starting:13
fitted Gauss locations:12
14
12
10
8
6
4
2
0
y
0
5
10
15
20
25
x

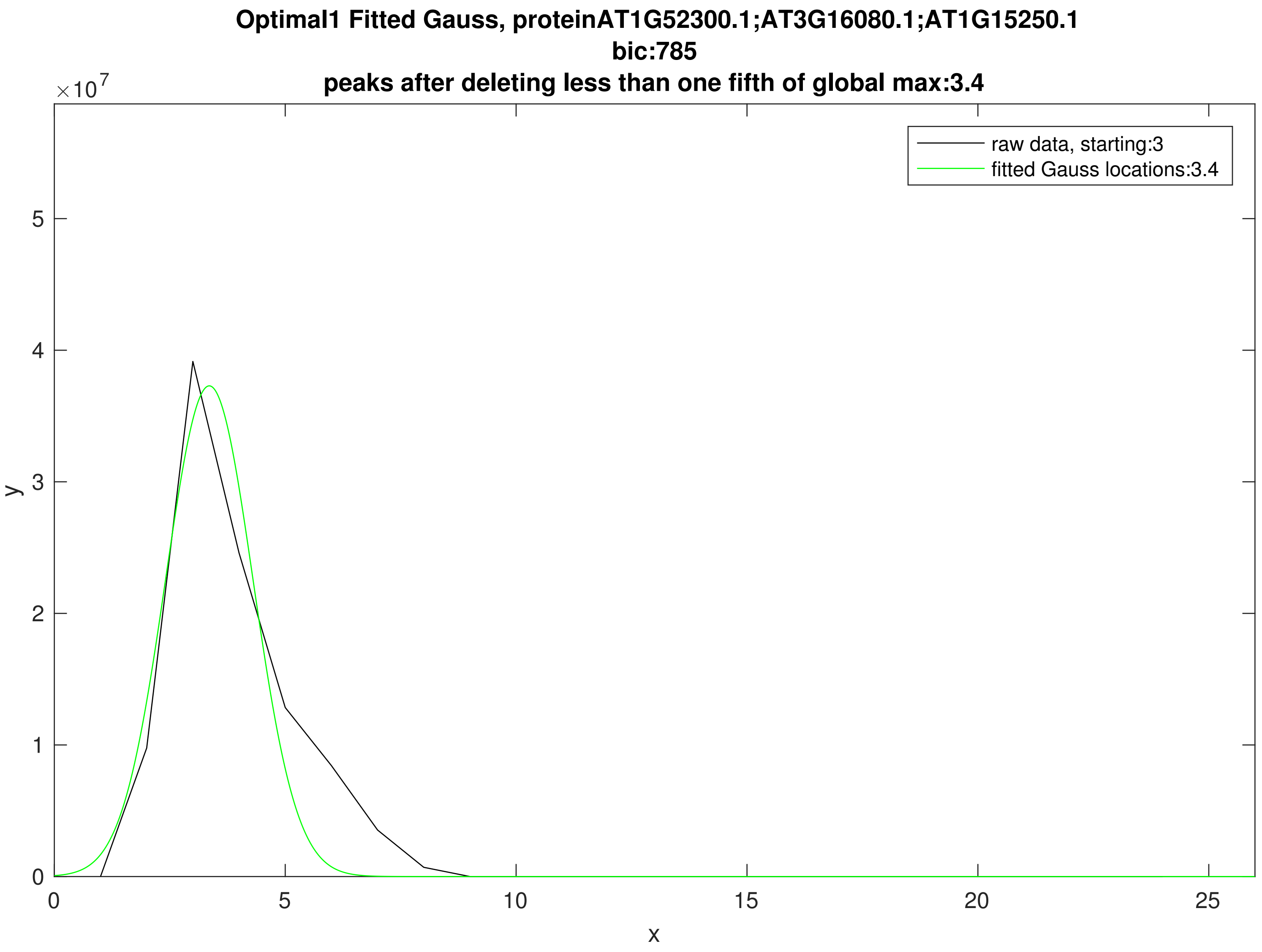
Optimal1 Fitted Gauss, proteinAT1G52300.1;AT3G16080.1;AT1G15250.1
bic:785
peaks after deleting less than one fifth of global max:3.4
×10^7
raw data, starting:3
fitted Gauss locations:3.4
y
x
0
1
2
3
4
5
0
5
10
15
20
25

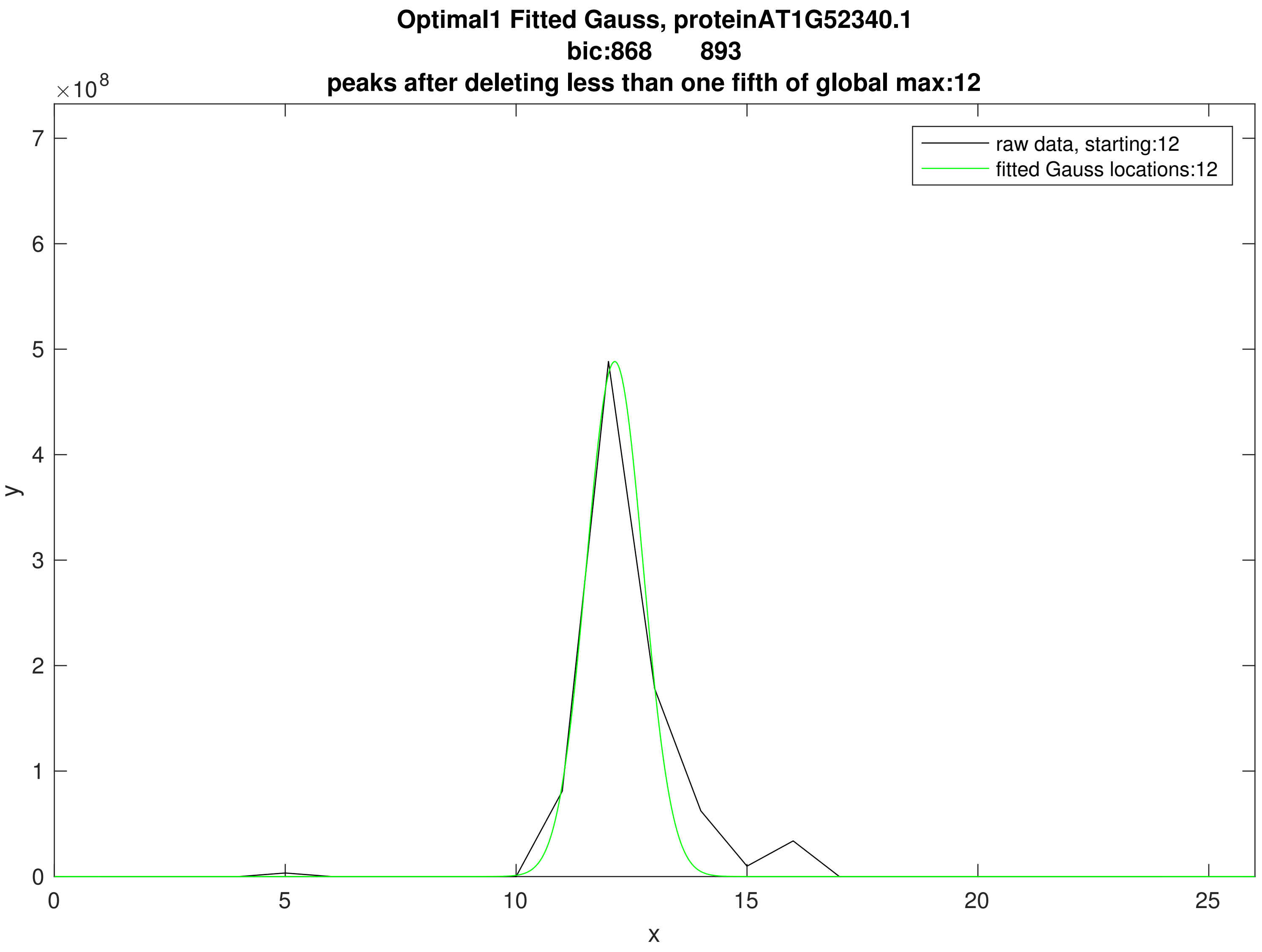

Optimal1 Fitted Gauss, proteinAT1G52340.1
bic:868 893
peaks after deleting less than one fifth of global max:12
×10^8
raw data, starting:12
fitted Gauss locations:12
y
x
0
1
2
3
4
5
6
7
10
15
20
25

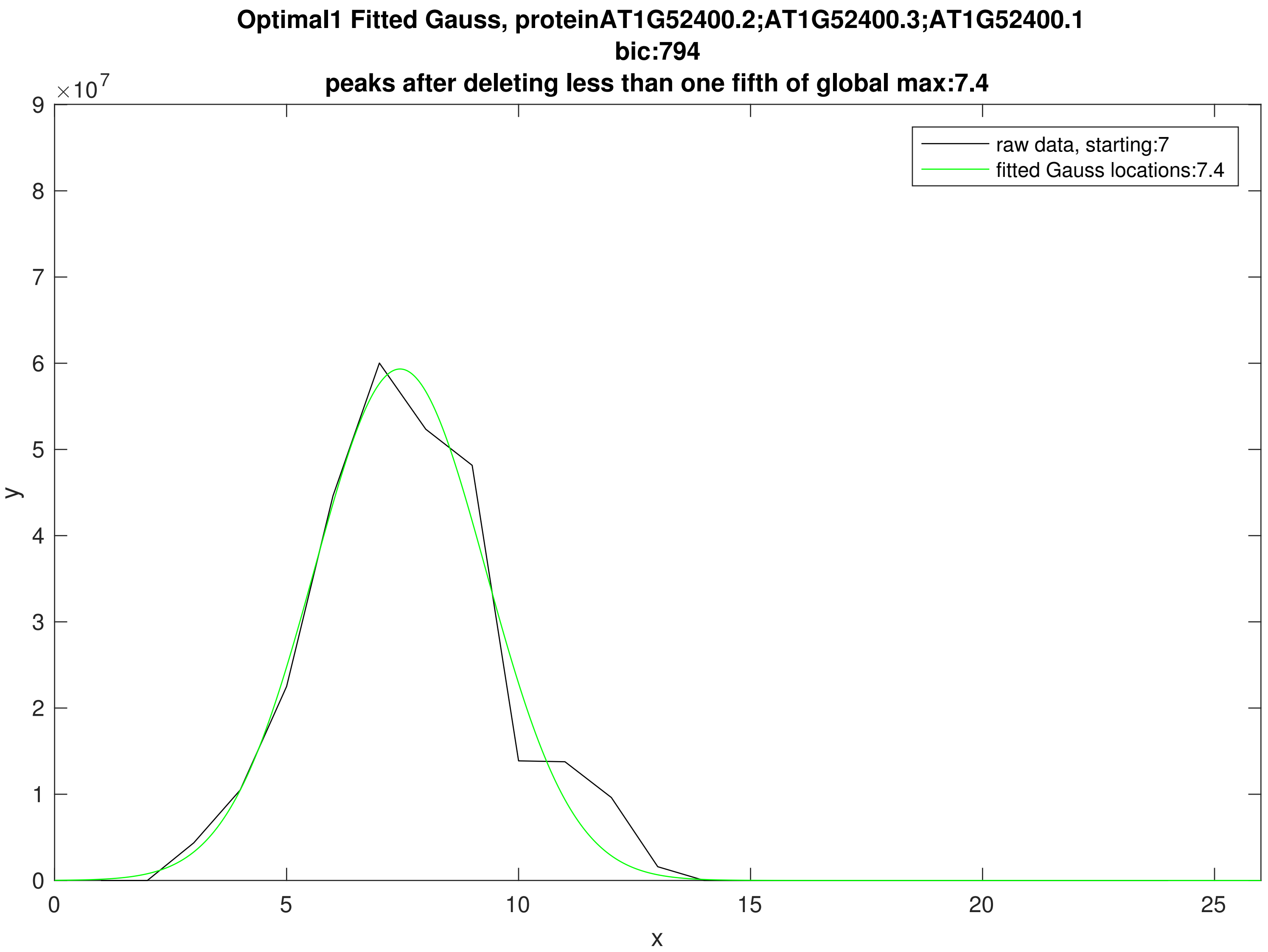
Optimal1 Fitted Gauss, proteinAT1G52400.2;AT1G52400.3;AT1G52400.1
bic:794
peaks after deleting less than one fifth of global max:7.4
×10^7
raw data, starting:7
fitted Gauss locations:7.4
y
x
0
1
2
3
4
5
6
7
8
9
0
5
10
15
20
25

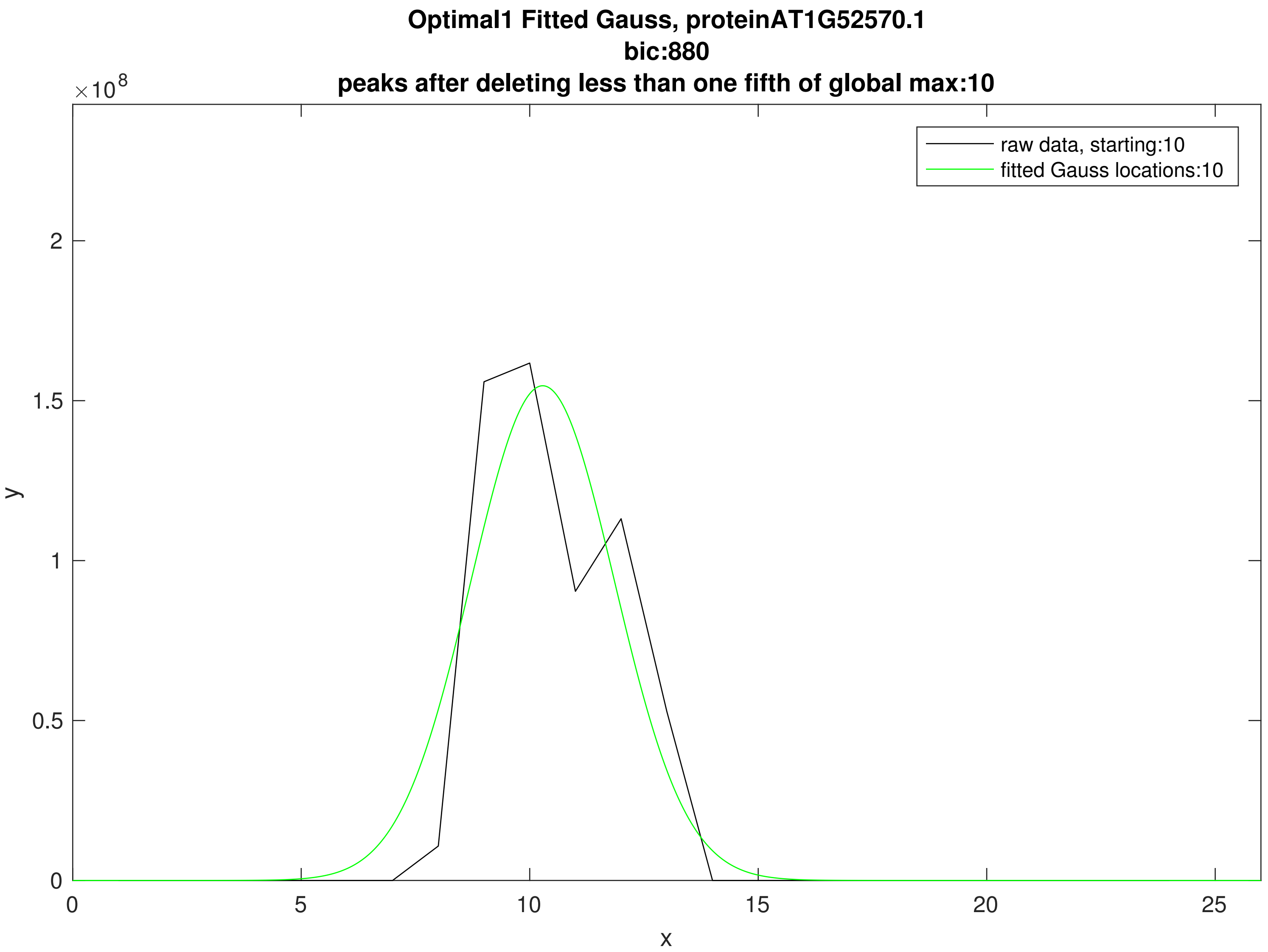
Optimal1 Fitted Gauss, proteinAT1G52570.1
bic:880
peaks after deleting less than one fifth of global max:10
×10^8
raw data, starting:10
fitted Gauss locations:10
2
1.5
1
0.5
0
y
0
5
10
15
20
25
x

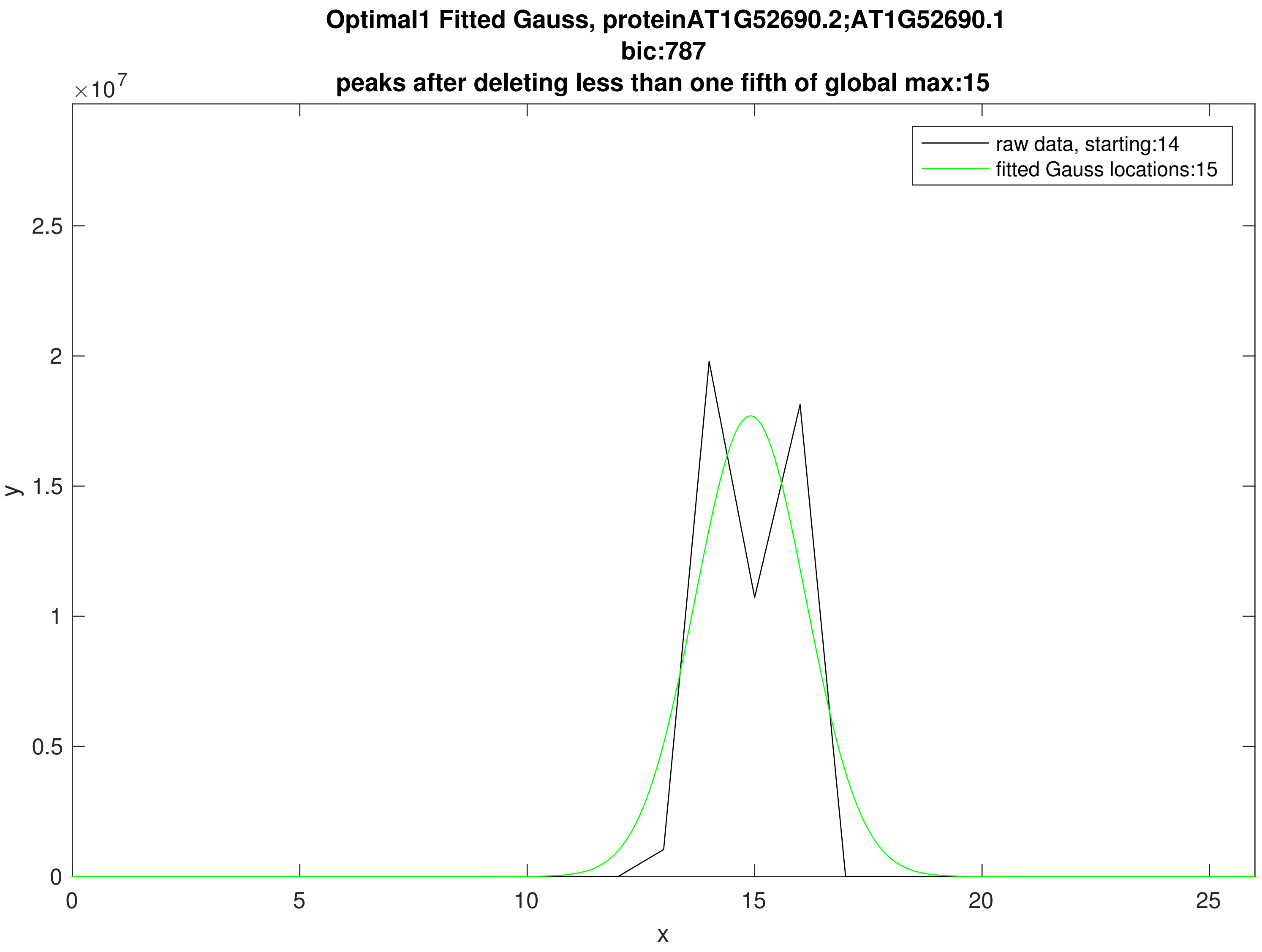

Optimal1 Fitted Gauss, proteinAT1G52690.2;AT1G52690.1
bic:787
peaks after deleting less than one fifth of global max:15
×10^7
raw data, starting:14
fitted Gauss locations:15
2.5
2
1.5
1
0.5
0
y
0
5
10
15
20
25
x

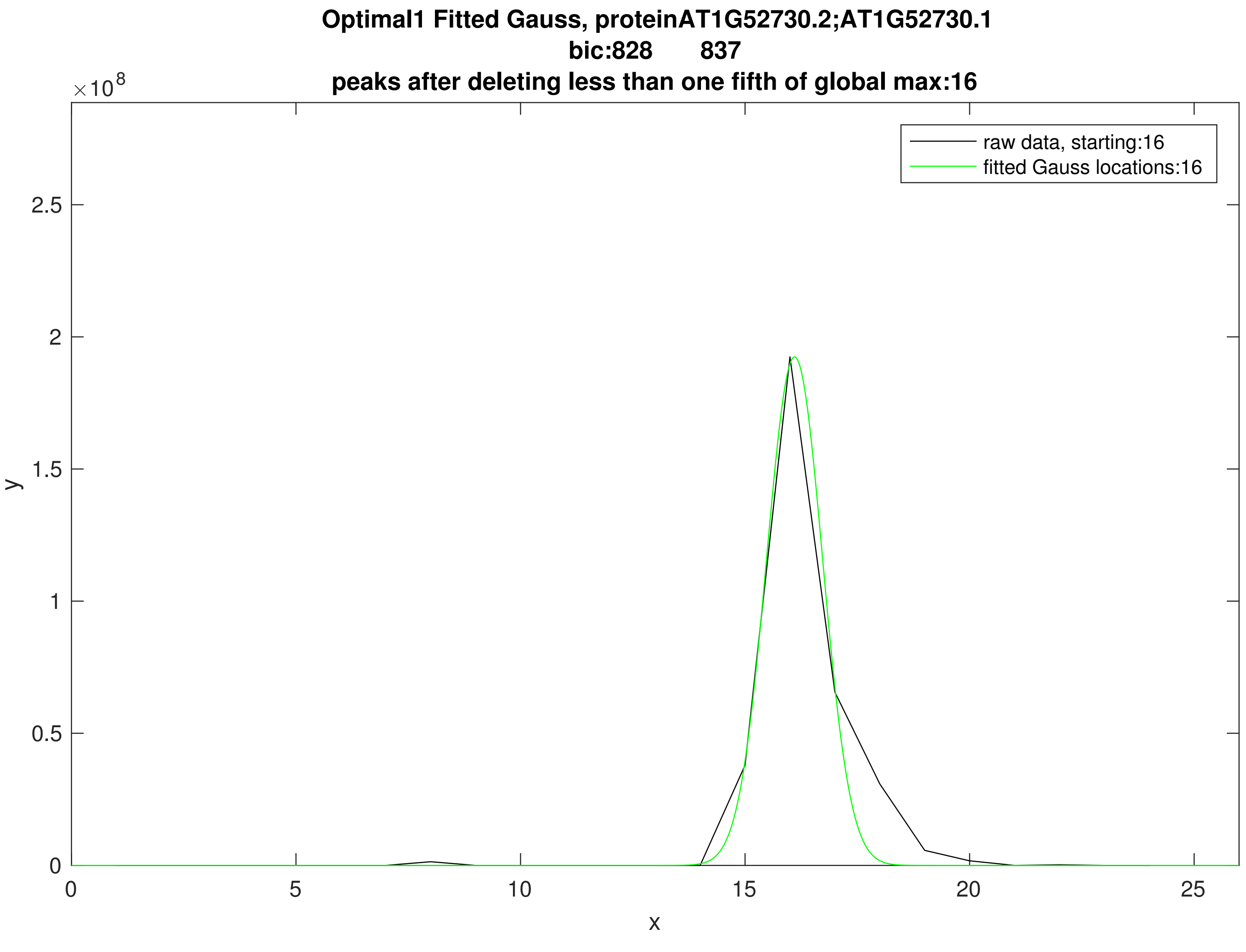
Optimal1 Fitted Gauss, proteinAT1G52730.2;AT1G52730.1
bic:828 837
peaks after deleting less than one fifth of global max:16
×10^8
raw data, starting:16
fitted Gauss locations:16
y
x
0
0.5
1
1.5
2
2.5
0
5
10
15
20
25

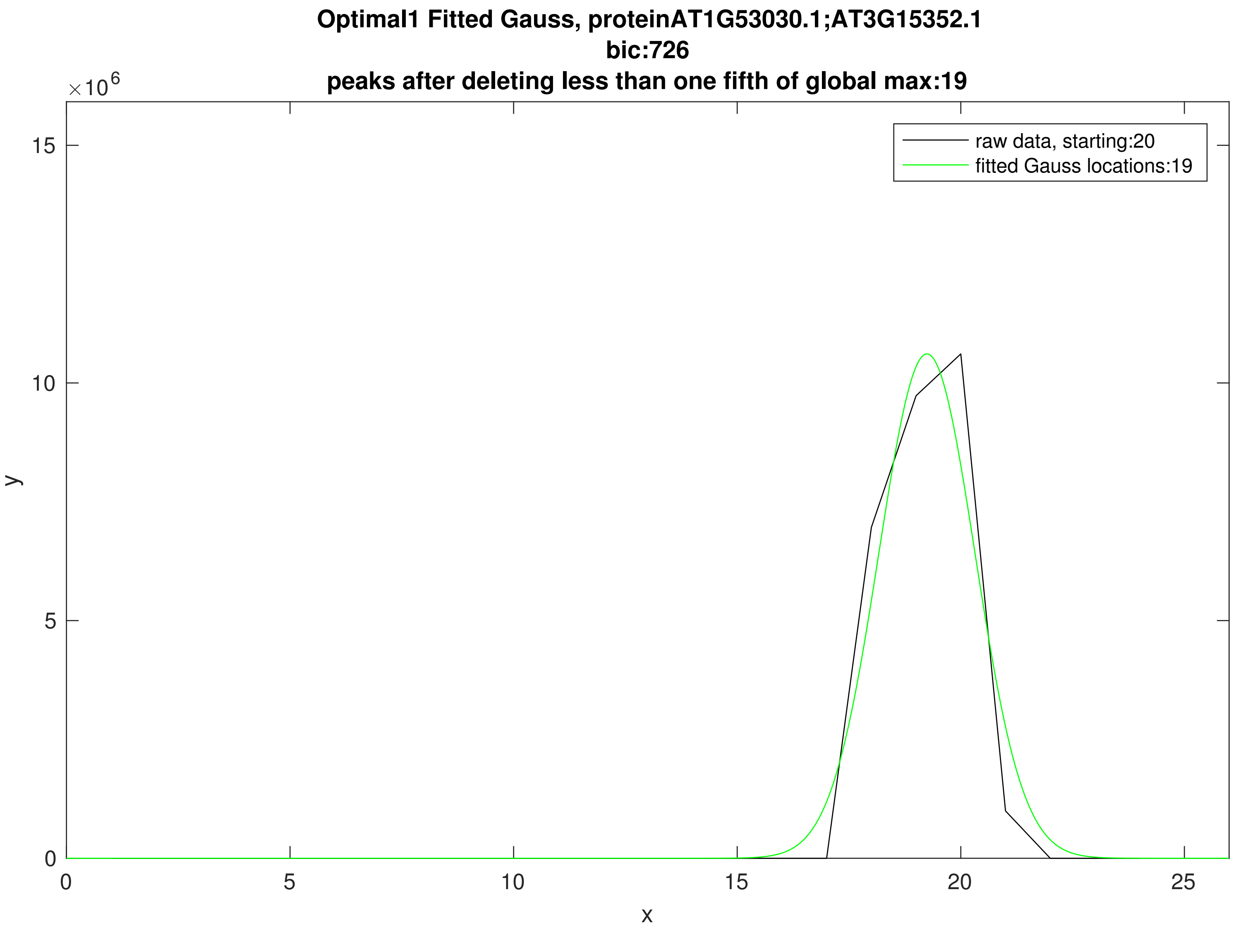
Optimal1 Fitted Gauss, proteinAT1G53030.1;AT3G15352.1
bic:726
peaks after deleting less than one fifth of global max:19
×10^6
raw data, starting:20
fitted Gauss locations:19
15
10
5
0
0
5
10
15
20
25
x
y

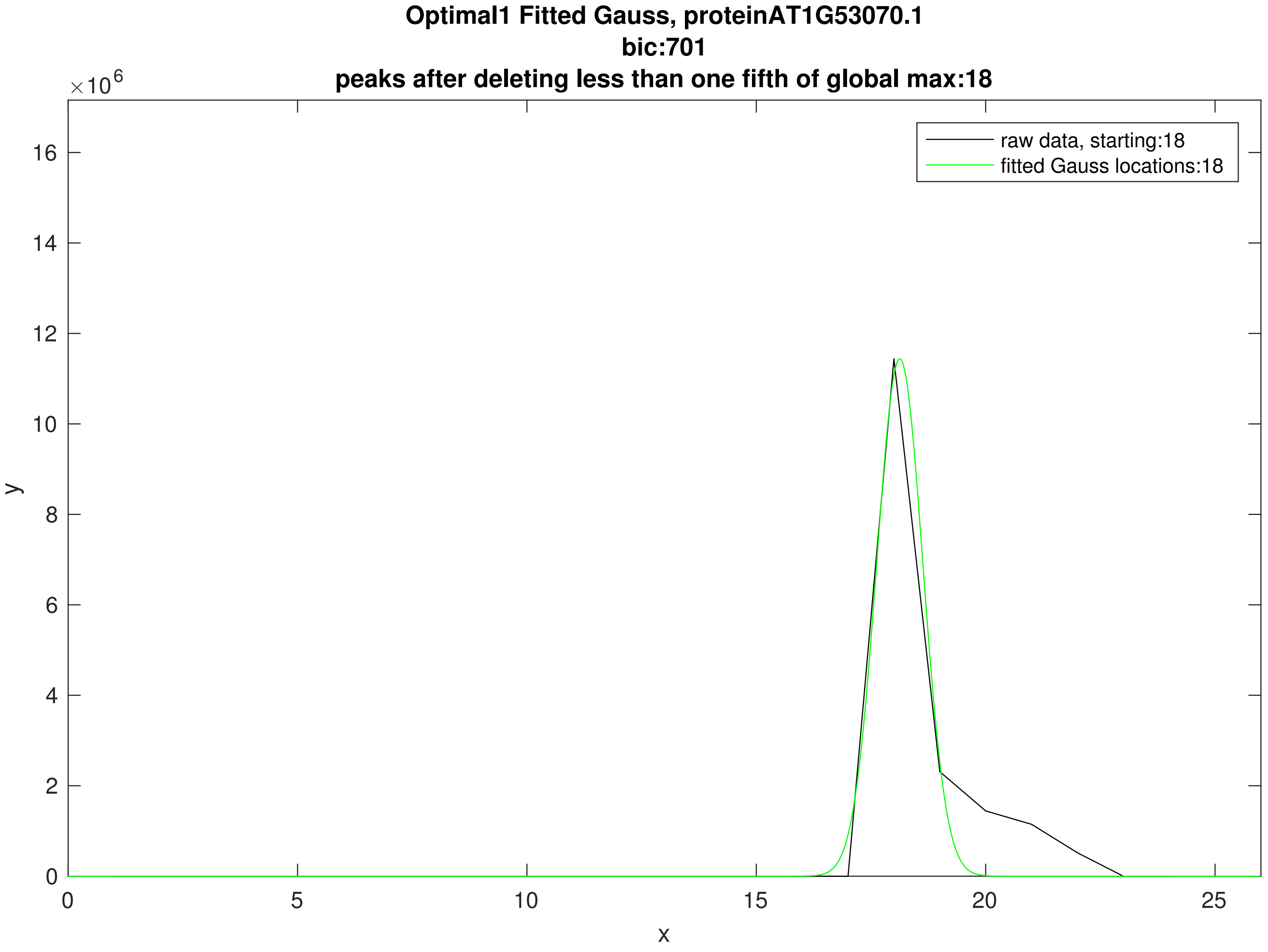
Optimal1 Fitted Gauss, proteinAT1G53070.1
bic:701
peaks after deleting less than one fifth of global max:18
×10^6
raw data, starting:18
fitted Gauss locations:18
y
x
0
2
4
6
8
10
12
14
16
0
5
10
15
20
25

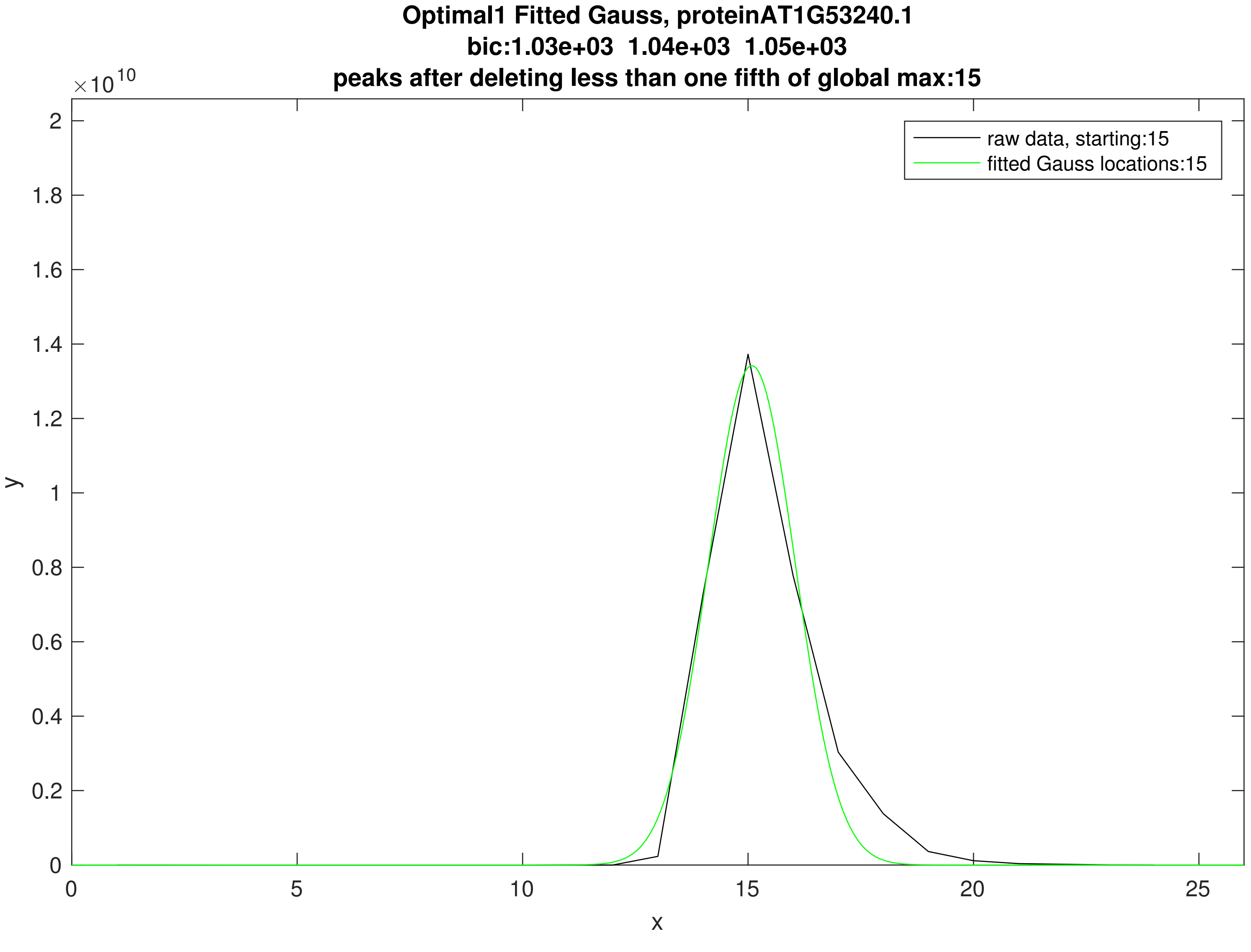

Optimal1 Fitted Gauss, proteinAT1G53240.1
bic:1.03e+03 1.04e+03 1.05e+03
peaks after deleting less than one fifth of global max:15
×10^10
2
1.8
1.6
1.4
1.2
1
0.8
0.6
0.4
0.2
0
y
0
5
10
15
20
25
x
raw data, starting:15
fitted Gauss locations:15

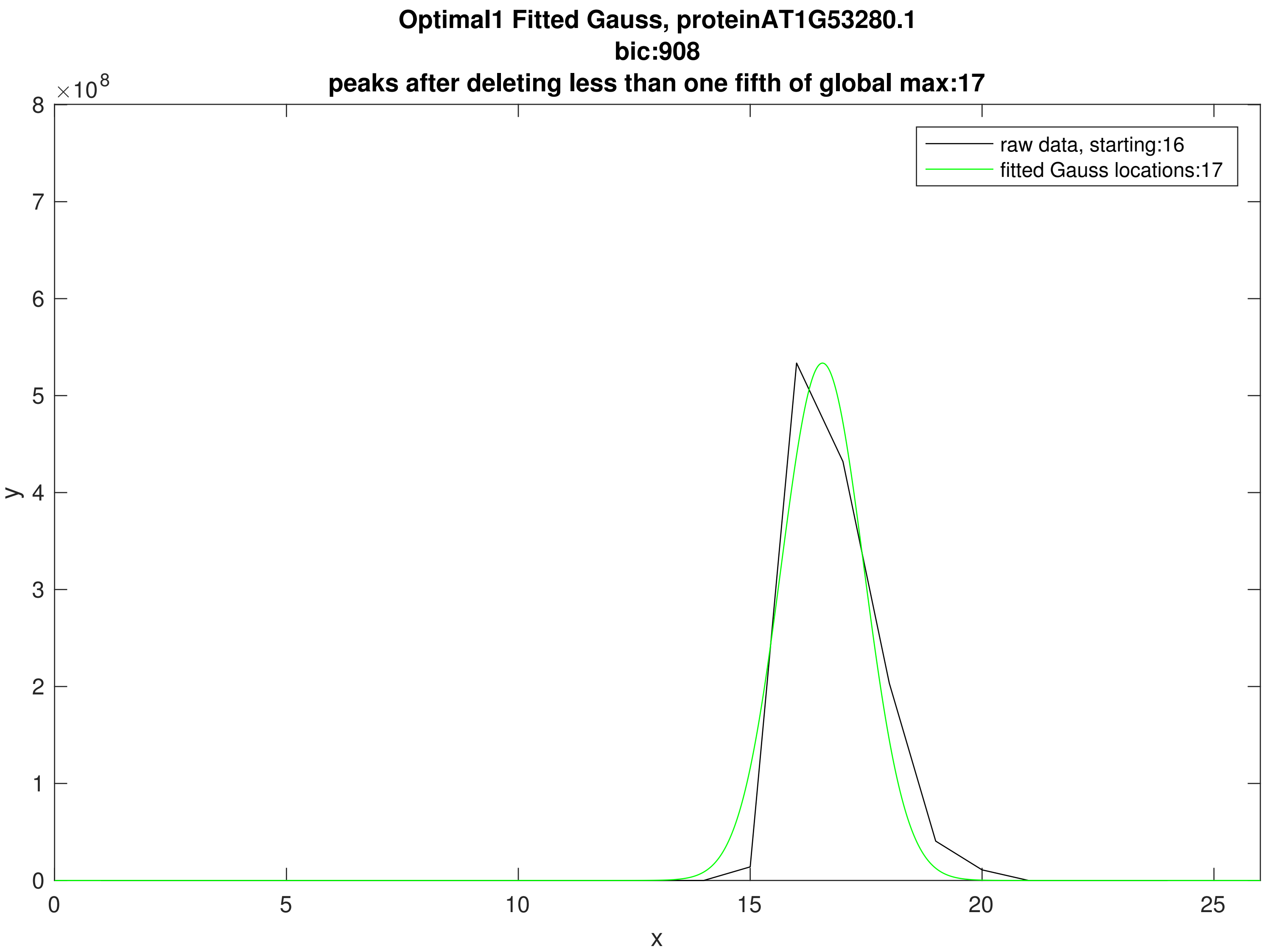

Optimal1 Fitted Gauss, proteinAT1G53280.1
bic:908
peaks after deleting less than one fifth of global max:17
×10^8
raw data, starting:16
fitted Gauss locations:17
y
x
0
1
2
3
4
5
6
7
8
0
5
10
15
20
25

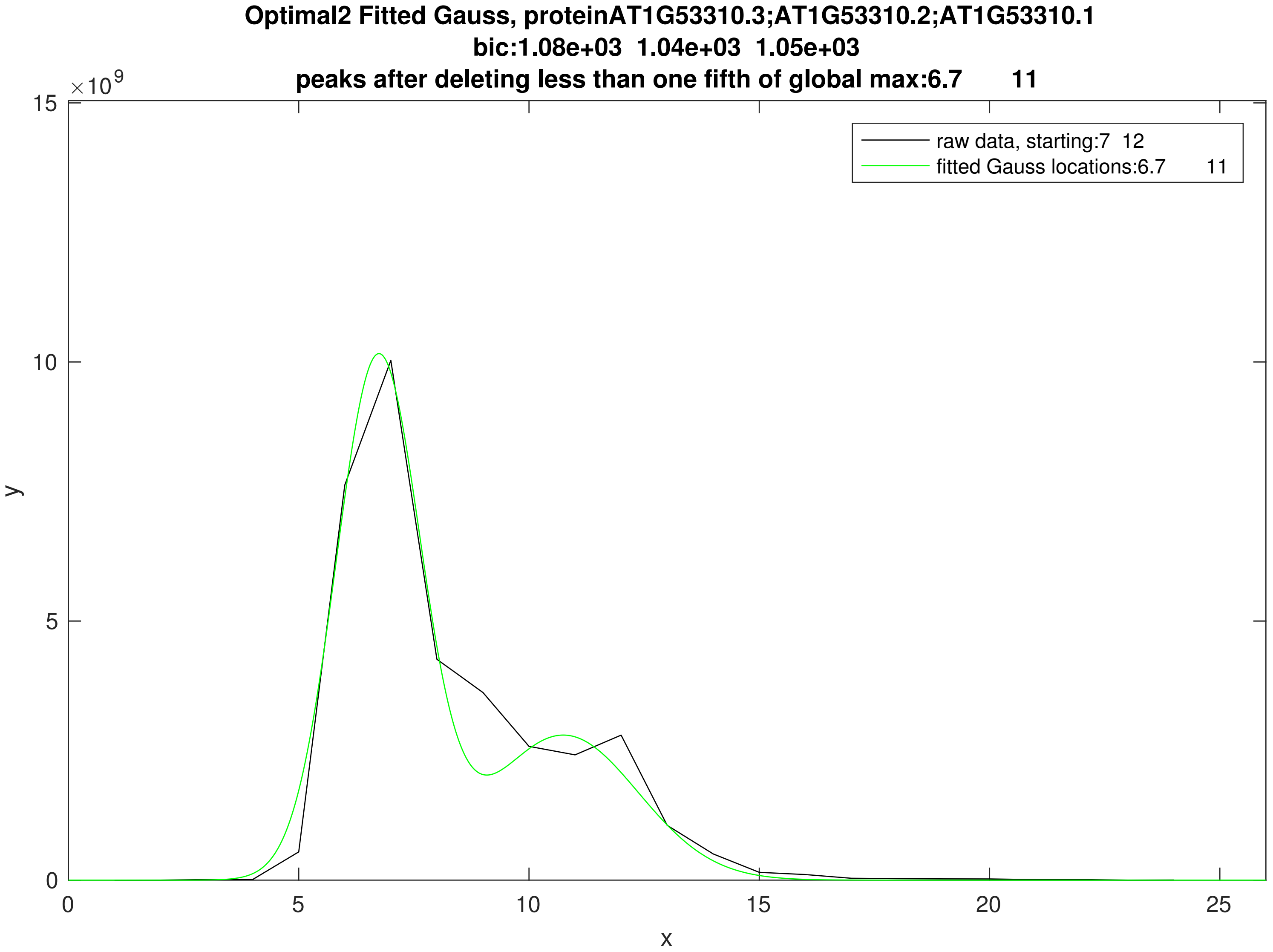

Optimal2 Fitted Gauss, proteinAT1G53310.3;AT1G53310.2;AT1G53310.1
bic:1.08e+03  1.04e+03  1.05e+03
peaks after deleting less than one fifth of global max:6.7       11
×10^9
15
10
5
0
y
0
5
10
15
20
25
x
raw data, starting:7  12
fitted Gauss locations:6.7       11

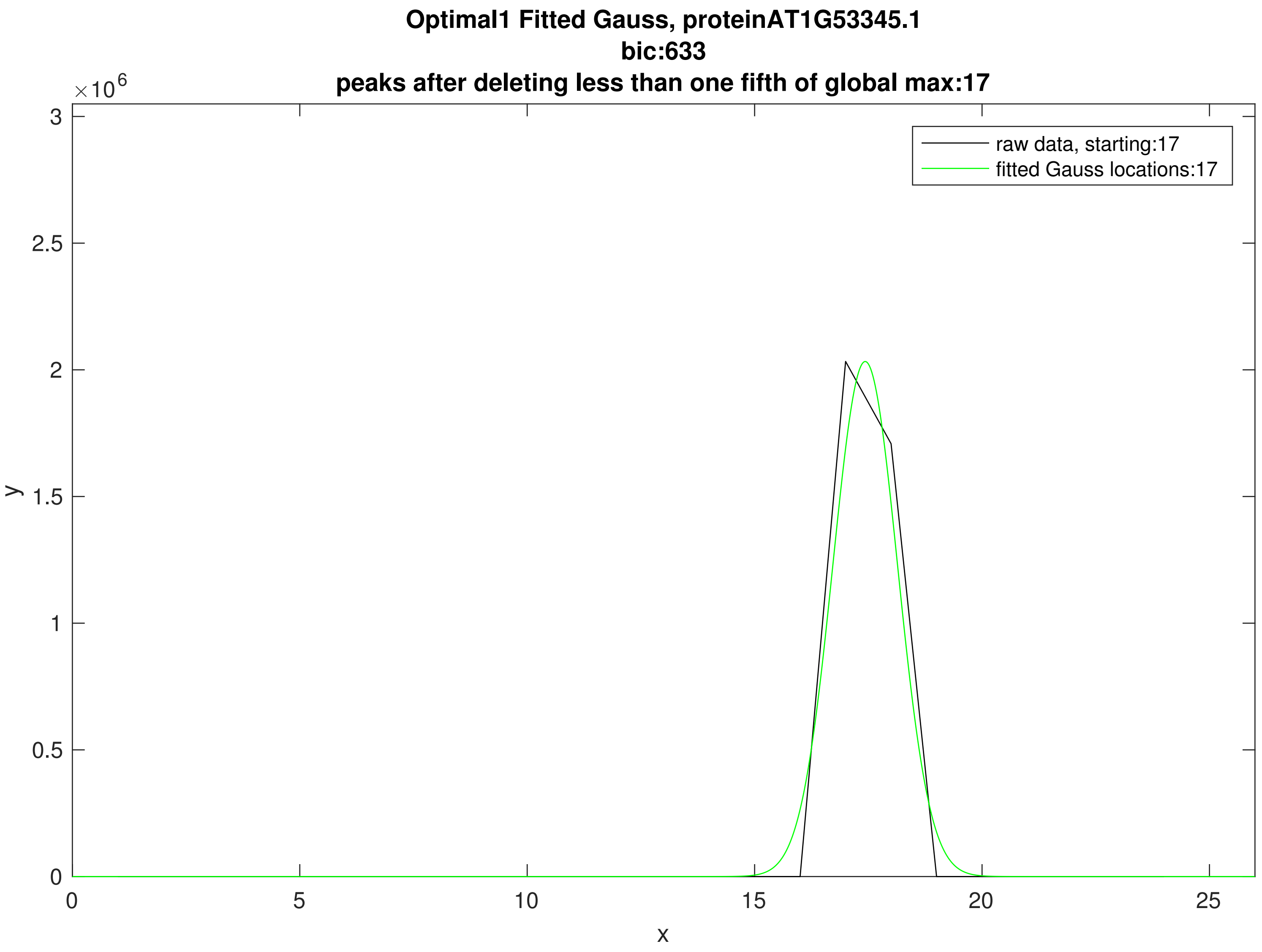
Optimal1 Fitted Gauss, proteinAT1G53345.1
bic:633
peaks after deleting less than one fifth of global max:17
×10^6
raw data, starting:17
fitted Gauss locations:17
y
x
3
2.5
2
1.5
1
0.5
0
0
5
10
15
20
25

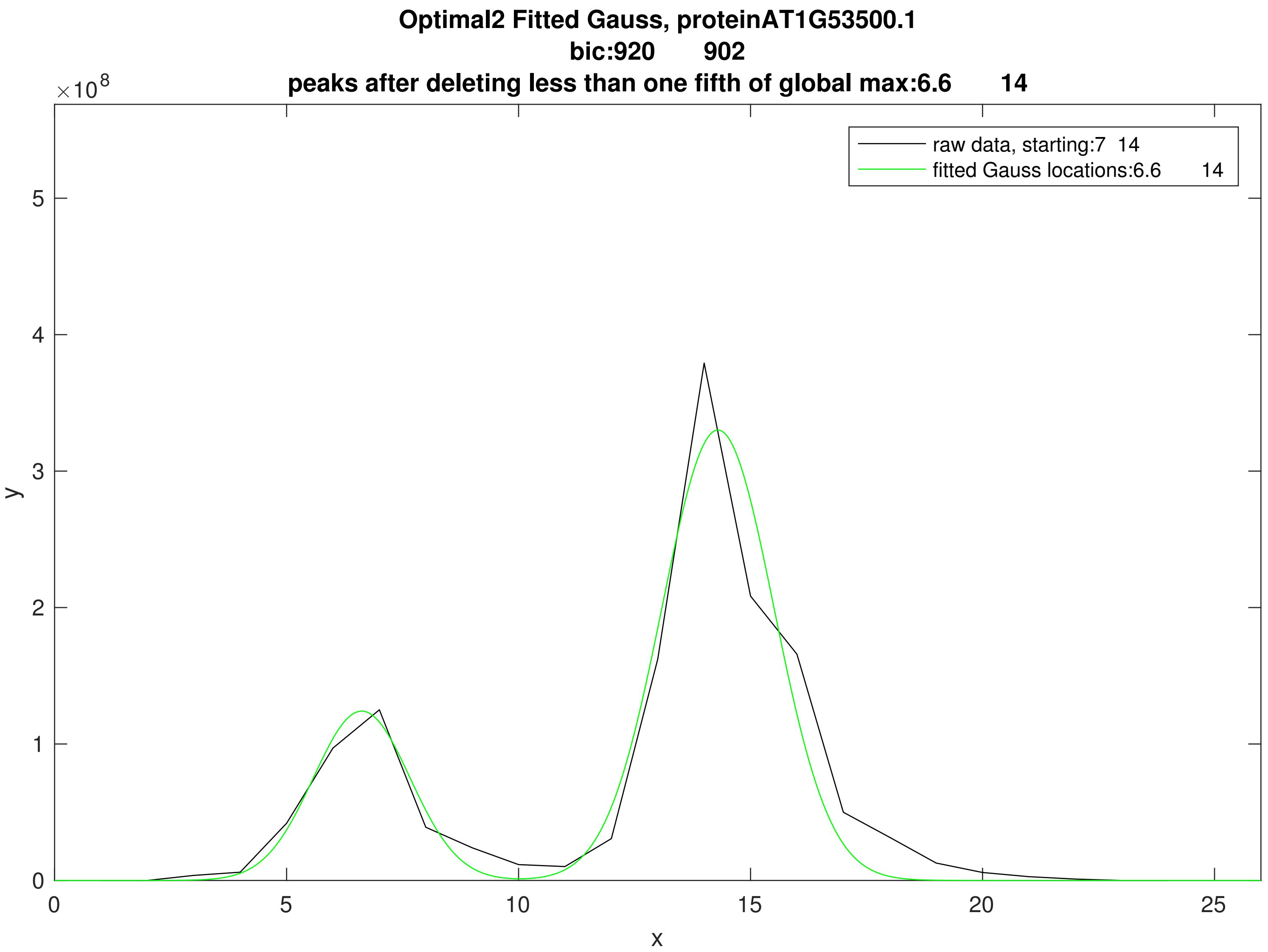

Optimal2 Fitted Gauss, proteinAT1G53500.1
bic:920 902
peaks after deleting less than one fifth of global max:6.6 14
×10^8
raw data, starting:7 14
fitted Gauss locations:6.6 14
y
x
0
1
2
3
4
5
0
5
10
15
20
25

**Optimal1 Fitted Gauss, proteinAT1G53580.1;AT1G53580.2**
**bic:923**
**peaks after deleting less than one fifth of global max:15**

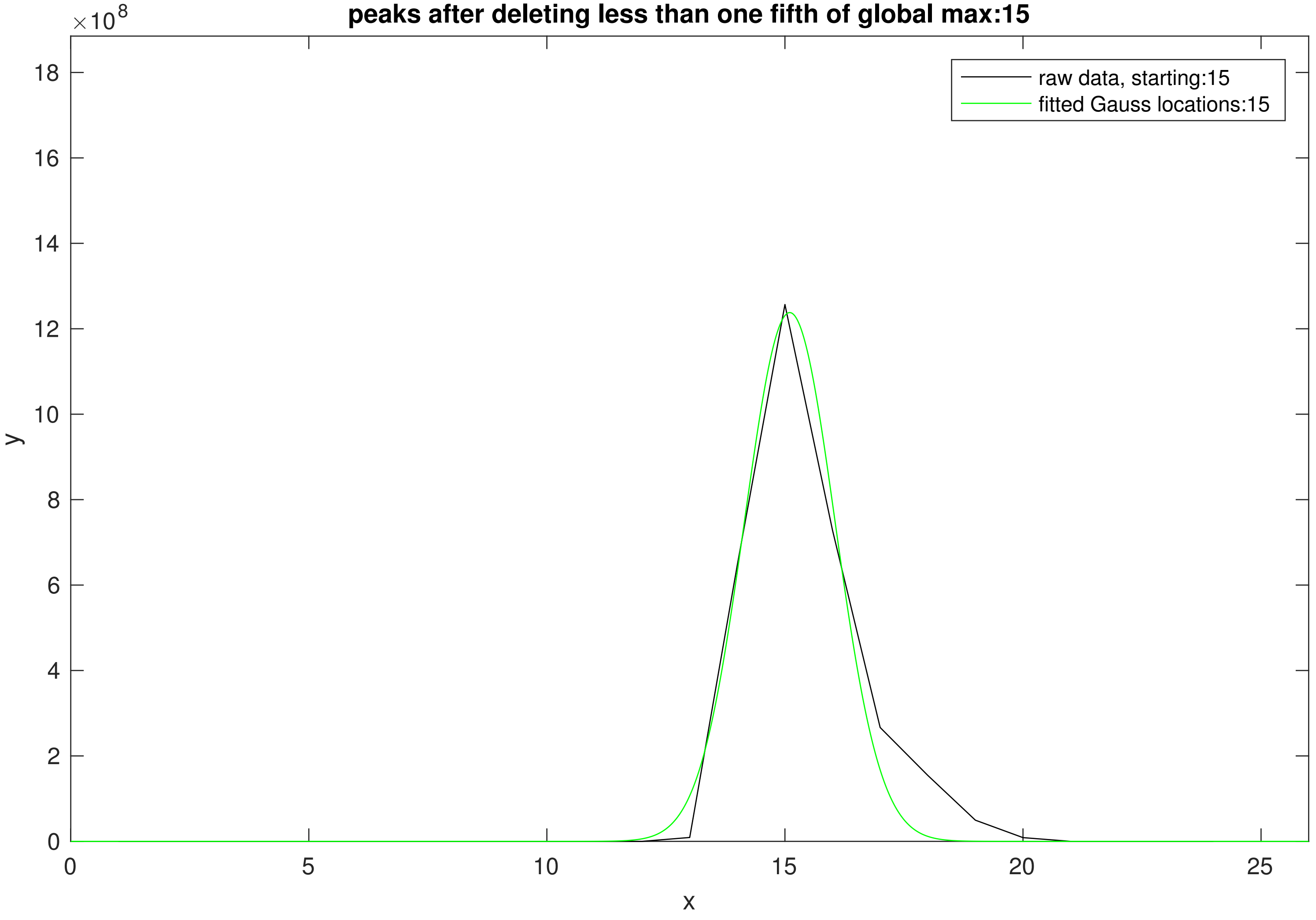

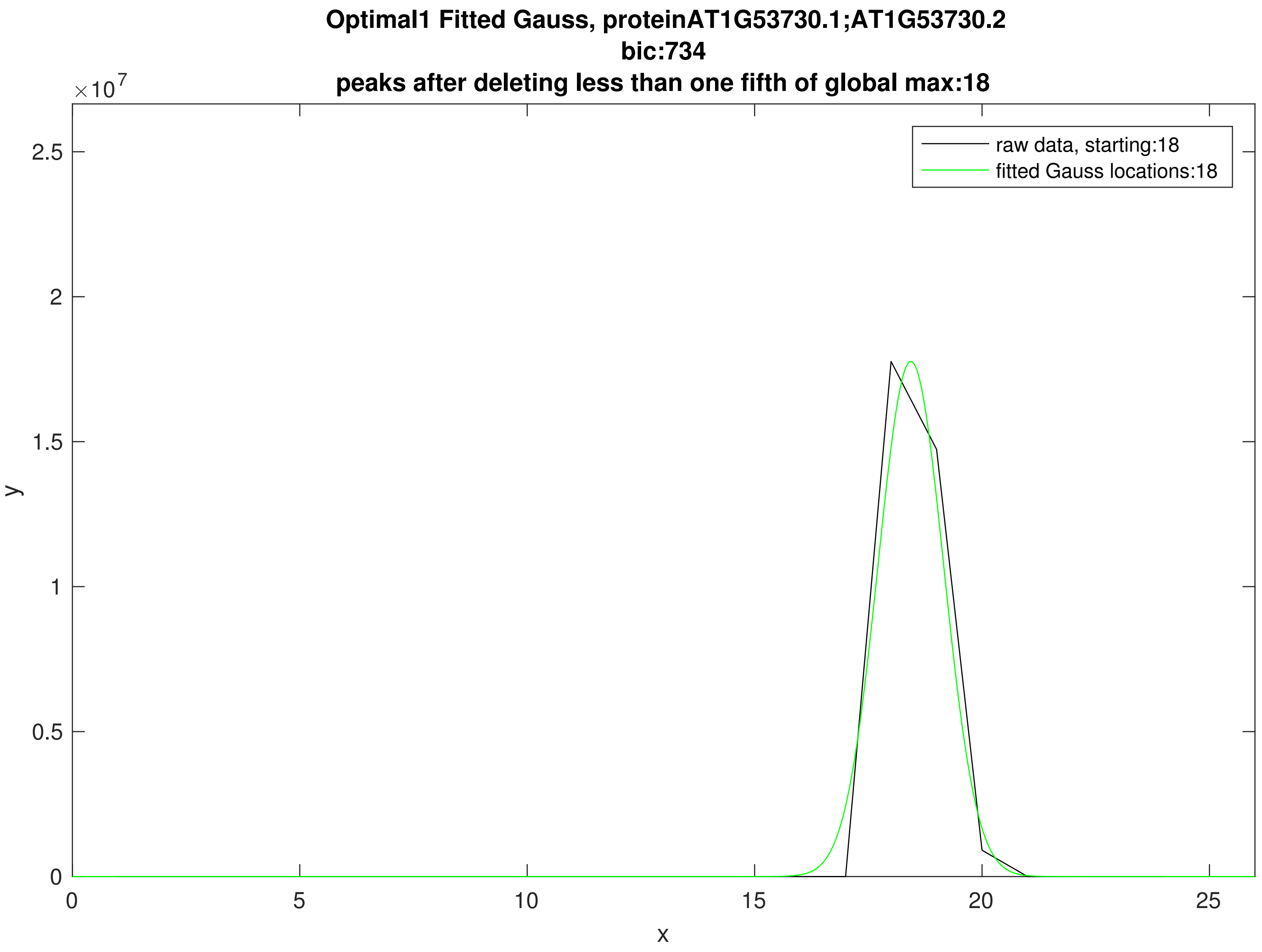

Optimal1 Fitted Gauss, proteinAT1G53730.1;AT1G53730.2
bic:734
peaks after deleting less than one fifth of global max:18
×10^7
2.5
2
1.5
1
0.5
0
y
0
5
10
15
20
25
x
raw data, starting:18
fitted Gauss locations:18

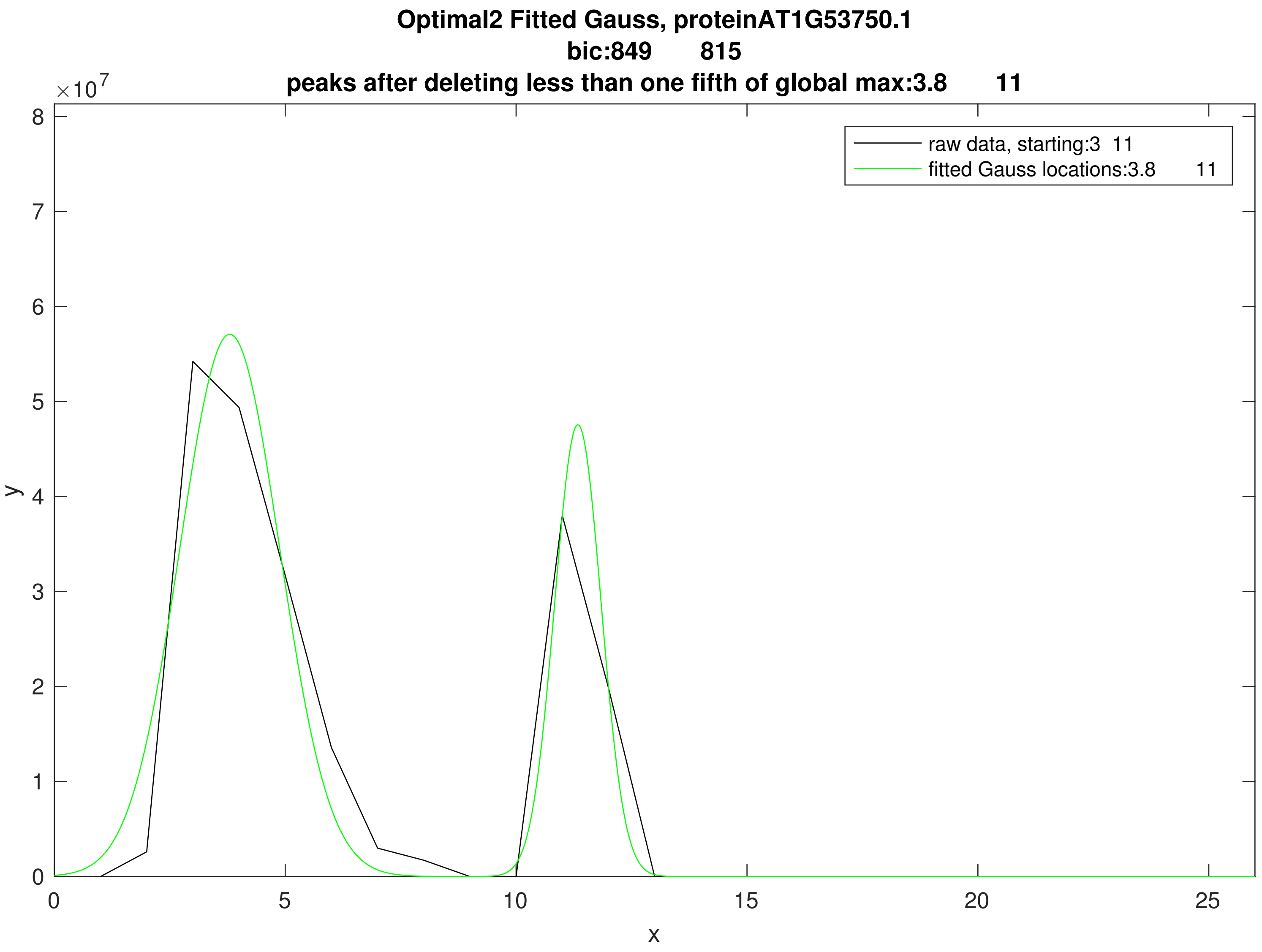
Optimal2 Fitted Gauss, proteinAT1G53750.1
bic:849 815
peaks after deleting less than one fifth of global max:3.8 11
×10^7
raw data, starting:3 11
fitted Gauss locations:3.8 11
y
x
0
1
2
3
4
5
6
7
8
0
5
10
15
20
25

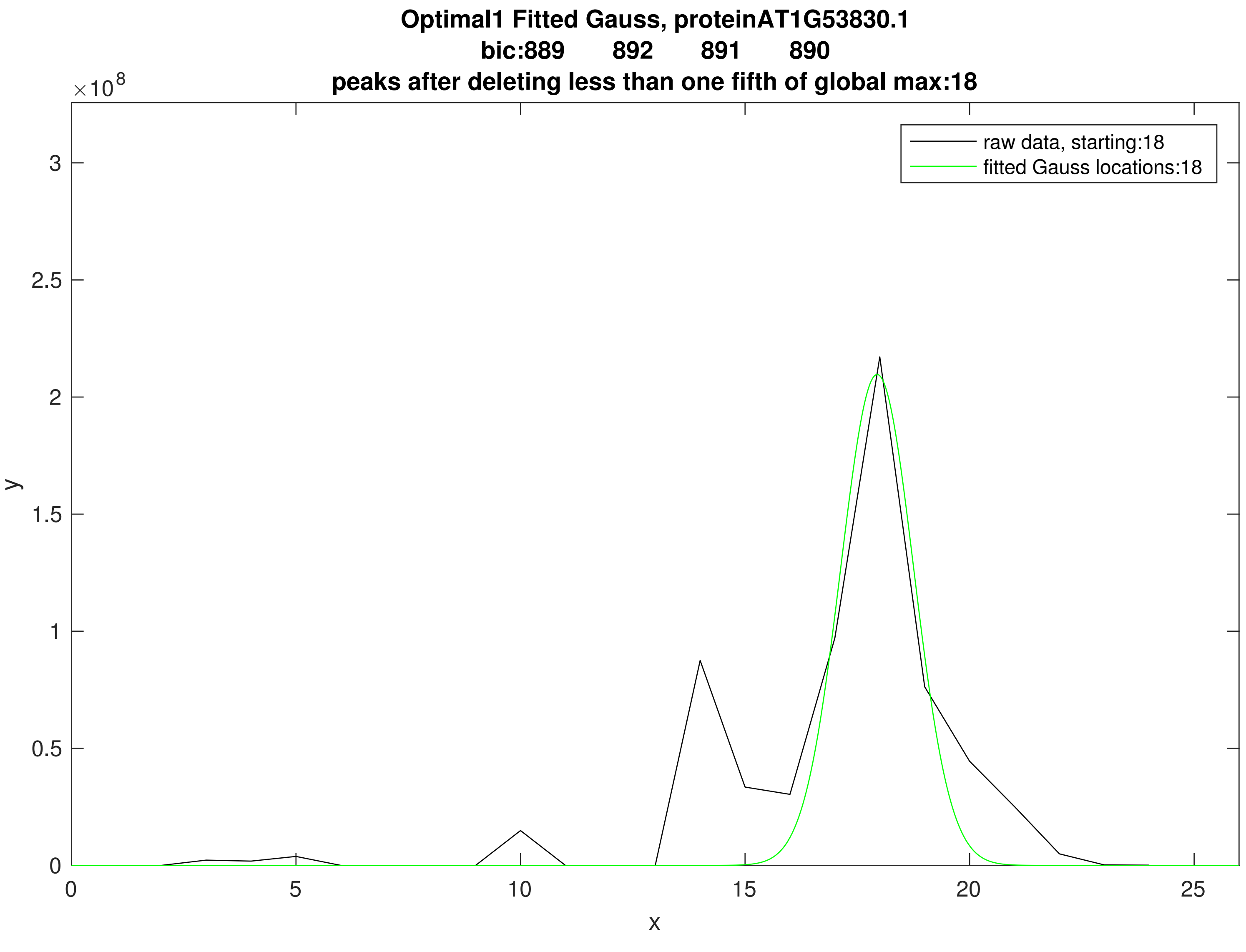
Optimal1 Fitted Gauss, proteinAT1G53830.1
bic:889 892 891 890
peaks after deleting less than one fifth of global max:18
×10^8
raw data, starting:18
fitted Gauss locations:18
y
x
0
0.5
1
1.5
2
2.5
3
5
10
15
20
25

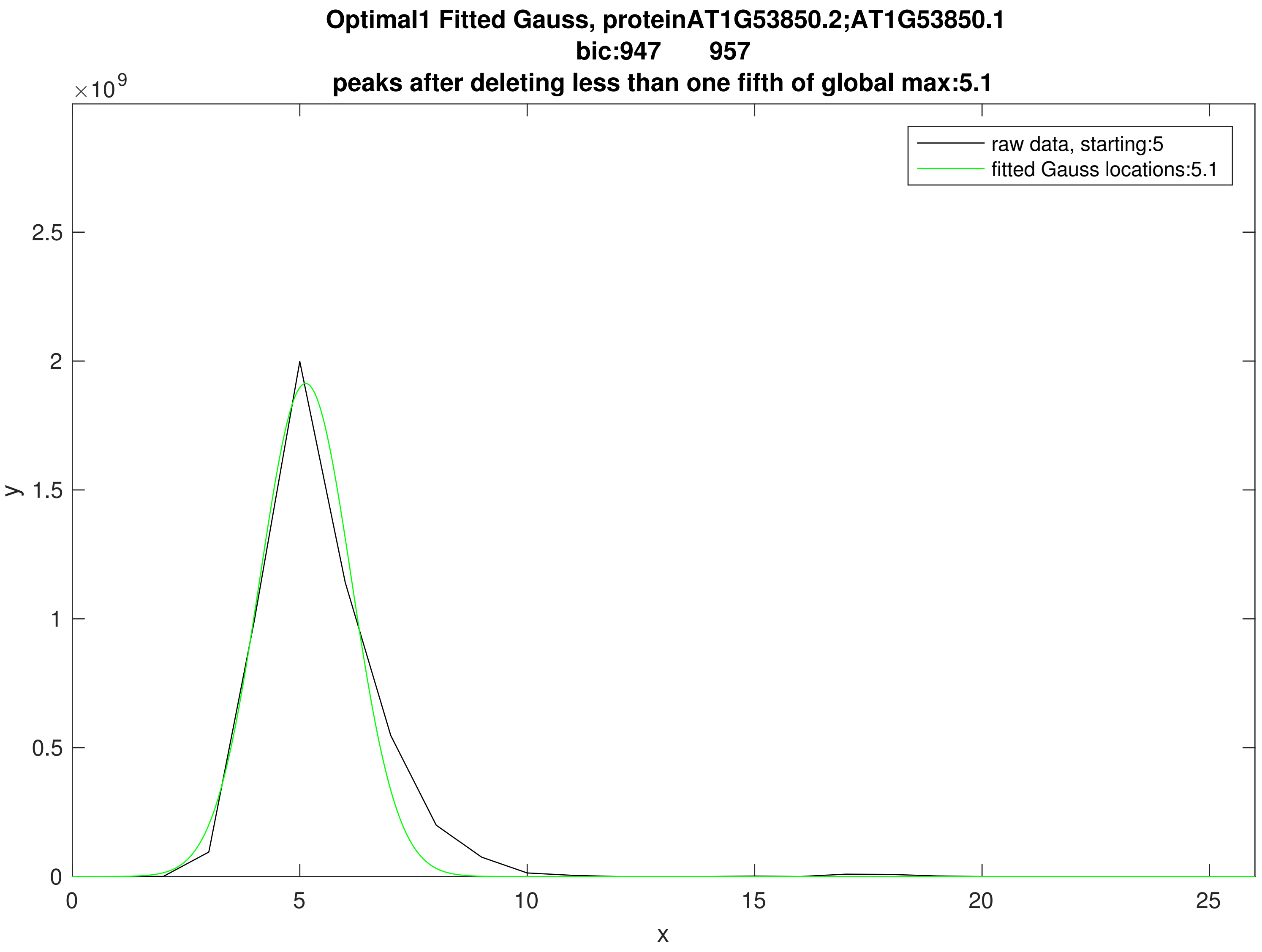

Optimal1 Fitted Gauss, proteinAT1G53850.2;AT1G53850.1
bic:947 957
peaks after deleting less than one fifth of global max:5.1
×10^9
raw data, starting:5
fitted Gauss locations:5.1
y
x
0
0.5
1
1.5
2
2.5
5
10
15
20
25

#### Optimal2 Fitted Gauss, proteinAT1G54100.2;AT1G54100.1
#### bic:870 804 813
#### peaks after deleting less than one fifth of global max:10

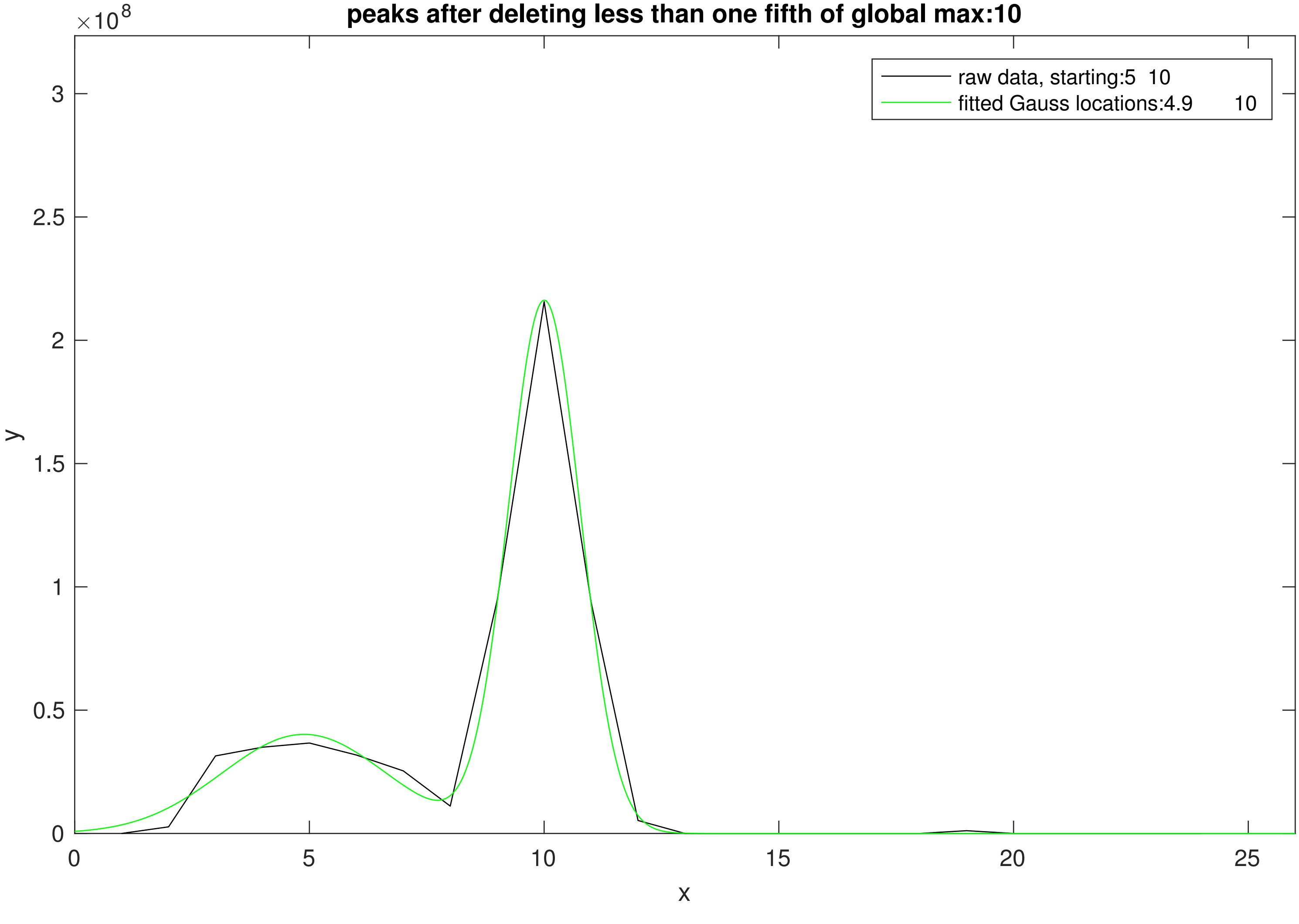

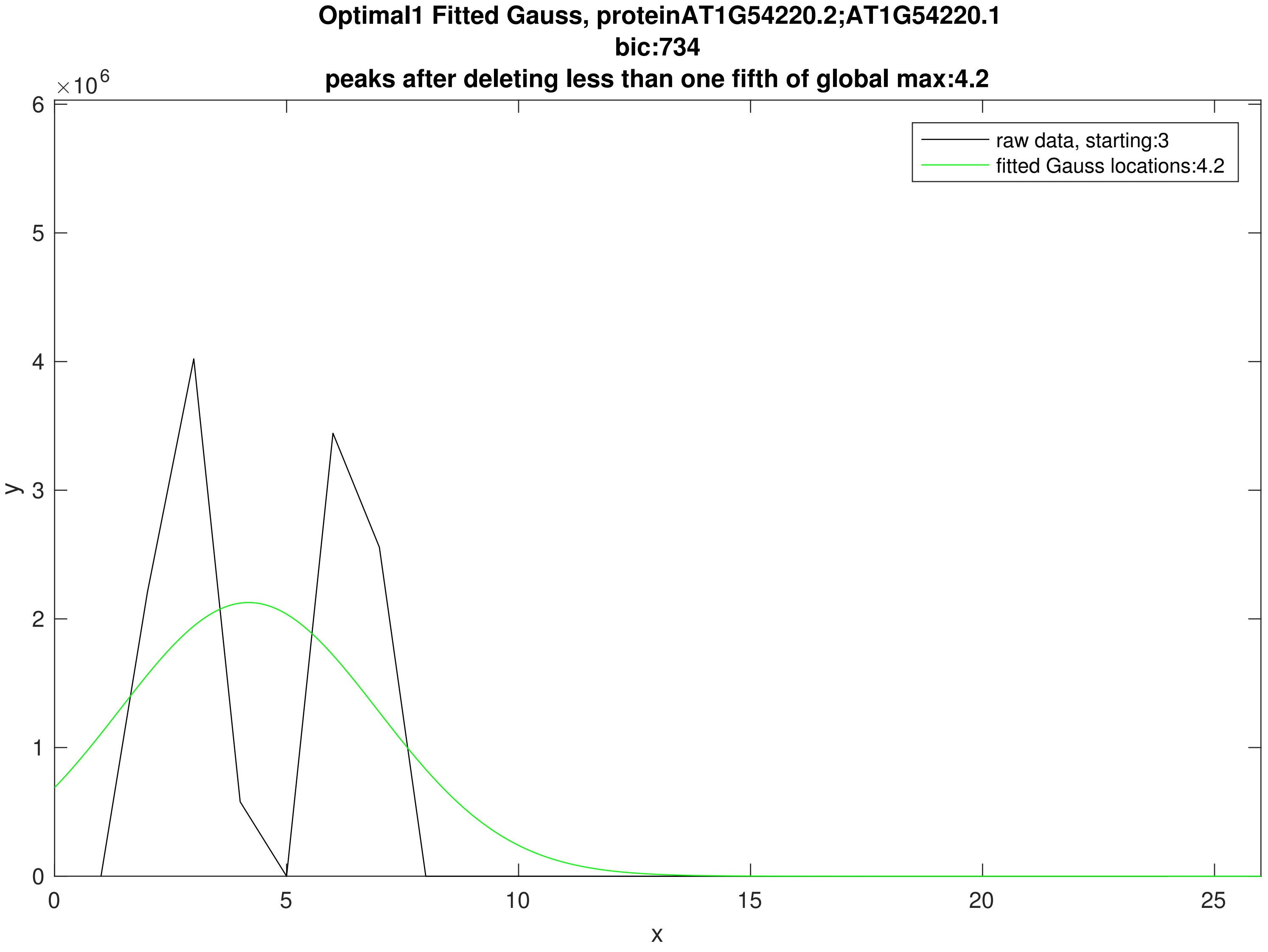

Optimal1 Fitted Gauss, proteinAT1G54220.2;AT1G54220.1
bic:734
peaks after deleting less than one fifth of global max:4.2
×10^6
raw data, starting:3
fitted Gauss locations:4.2
y
x
0
1
2
3
4
5
6
0
5
10
15
20
25

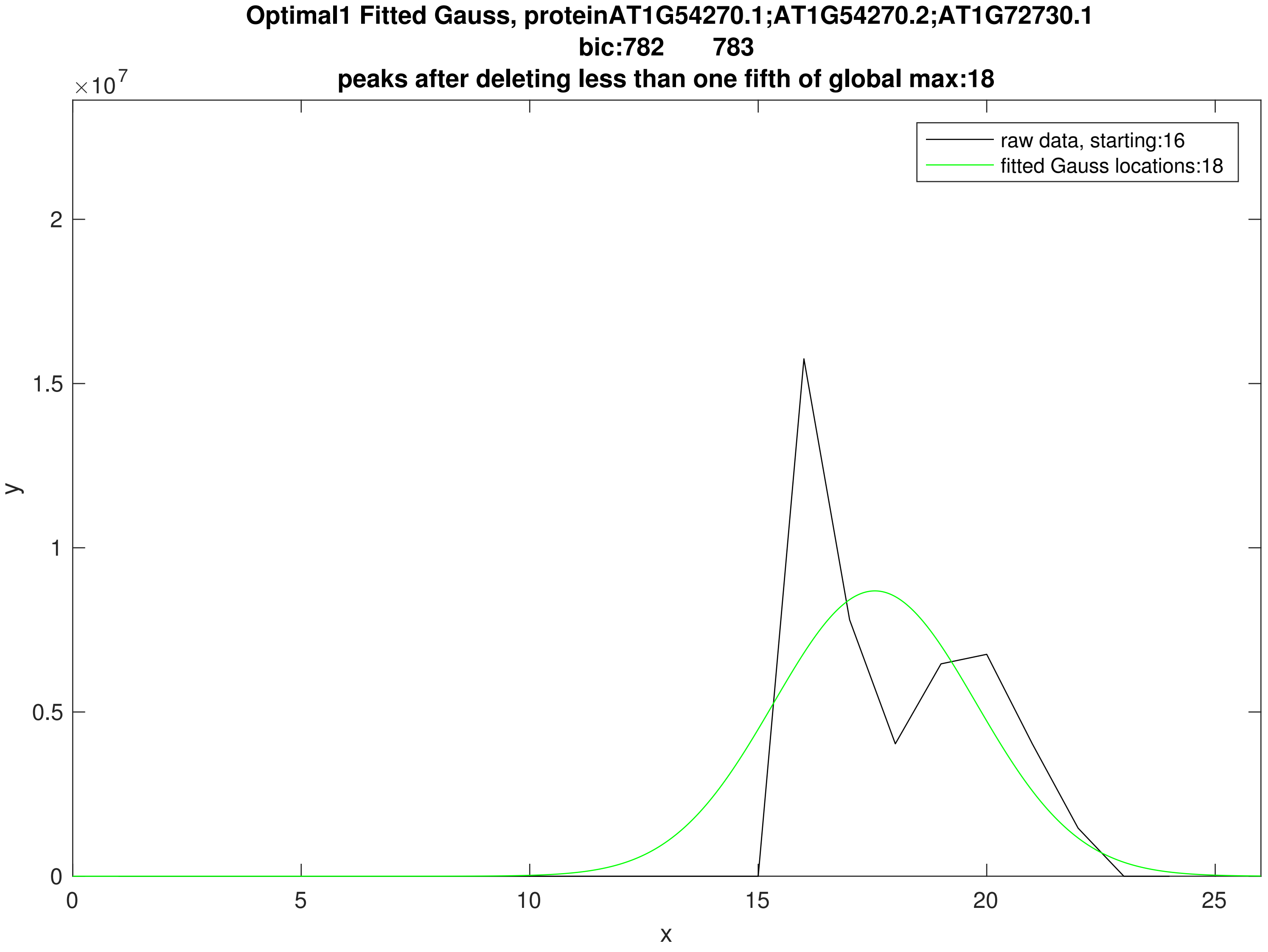

Optimal1 Fitted Gauss, proteinAT1G54270.1;AT1G54270.2;AT1G72730.1
bic:782 783
peaks after deleting less than one fifth of global max:18
×10^7
raw data, starting:16
fitted Gauss locations:18
2
1.5
1
0.5
0
y
0
5
10
15
20
25
x

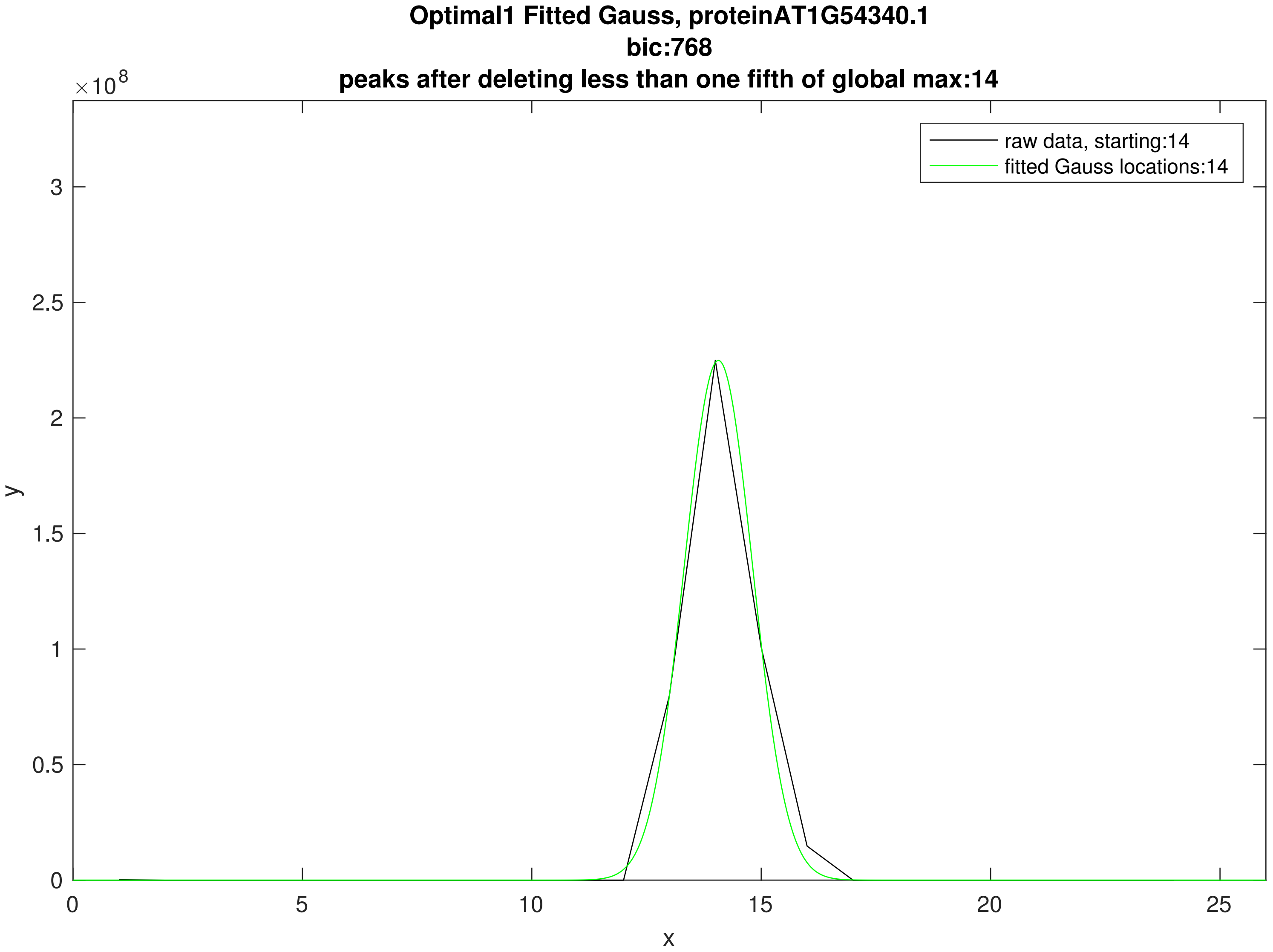

Optimal1 Fitted Gauss, proteinAT1G54340.1
bic:768
peaks after deleting less than one fifth of global max:14
×10^8
raw data, starting:14
fitted Gauss locations:14
y
x
0
0.5
1
1.5
2
2.5
3
0
5
10
15
20
25

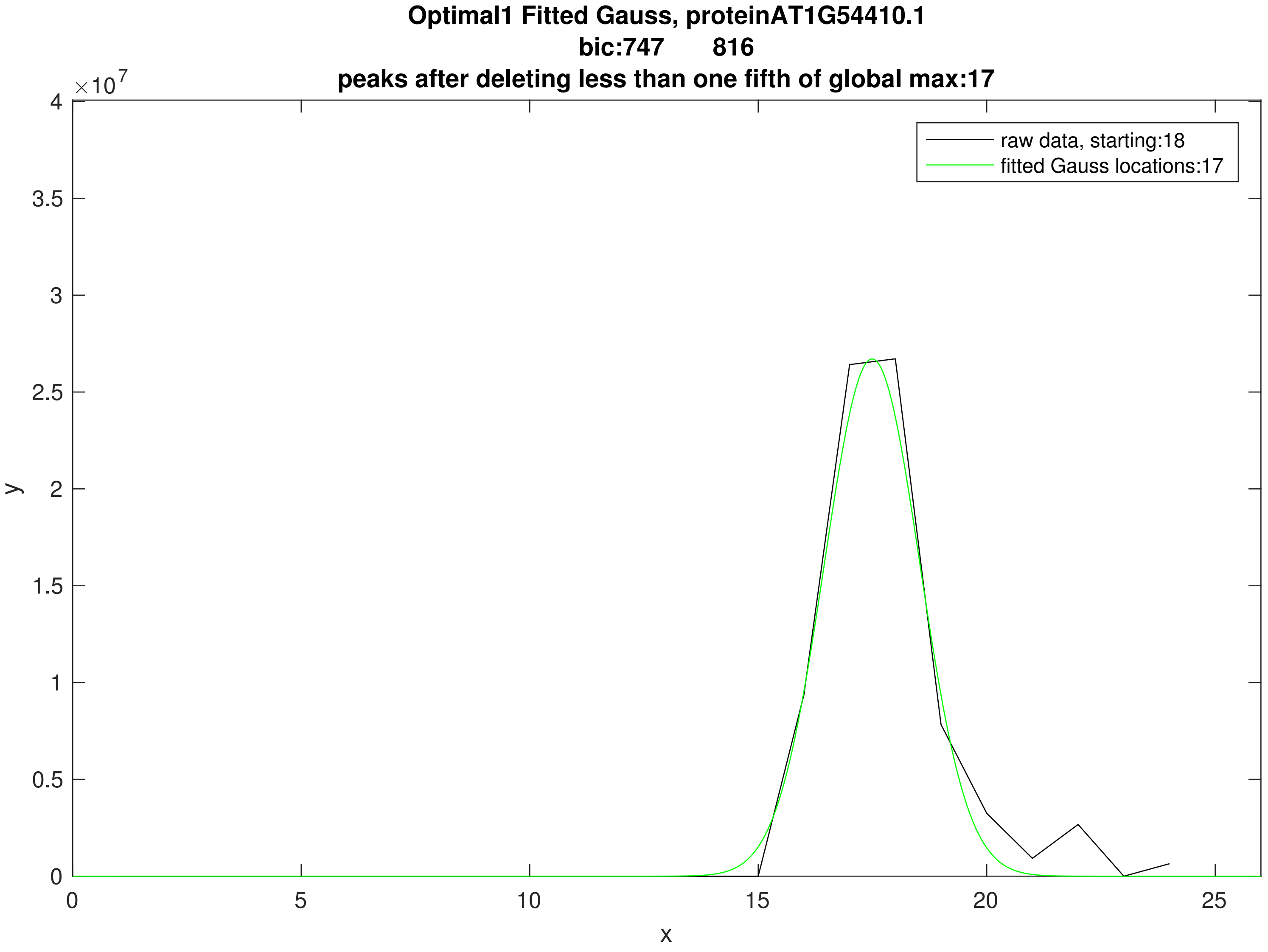
Optimal1 Fitted Gauss, proteinAT1G54410.1
bic:747 816
peaks after deleting less than one fifth of global max:17
×10^7
4
3.5
3
2.5
2
1.5
1
0.5
0
y
0
5
10
15
20
25
x
raw data, starting:18
fitted Gauss locations:17

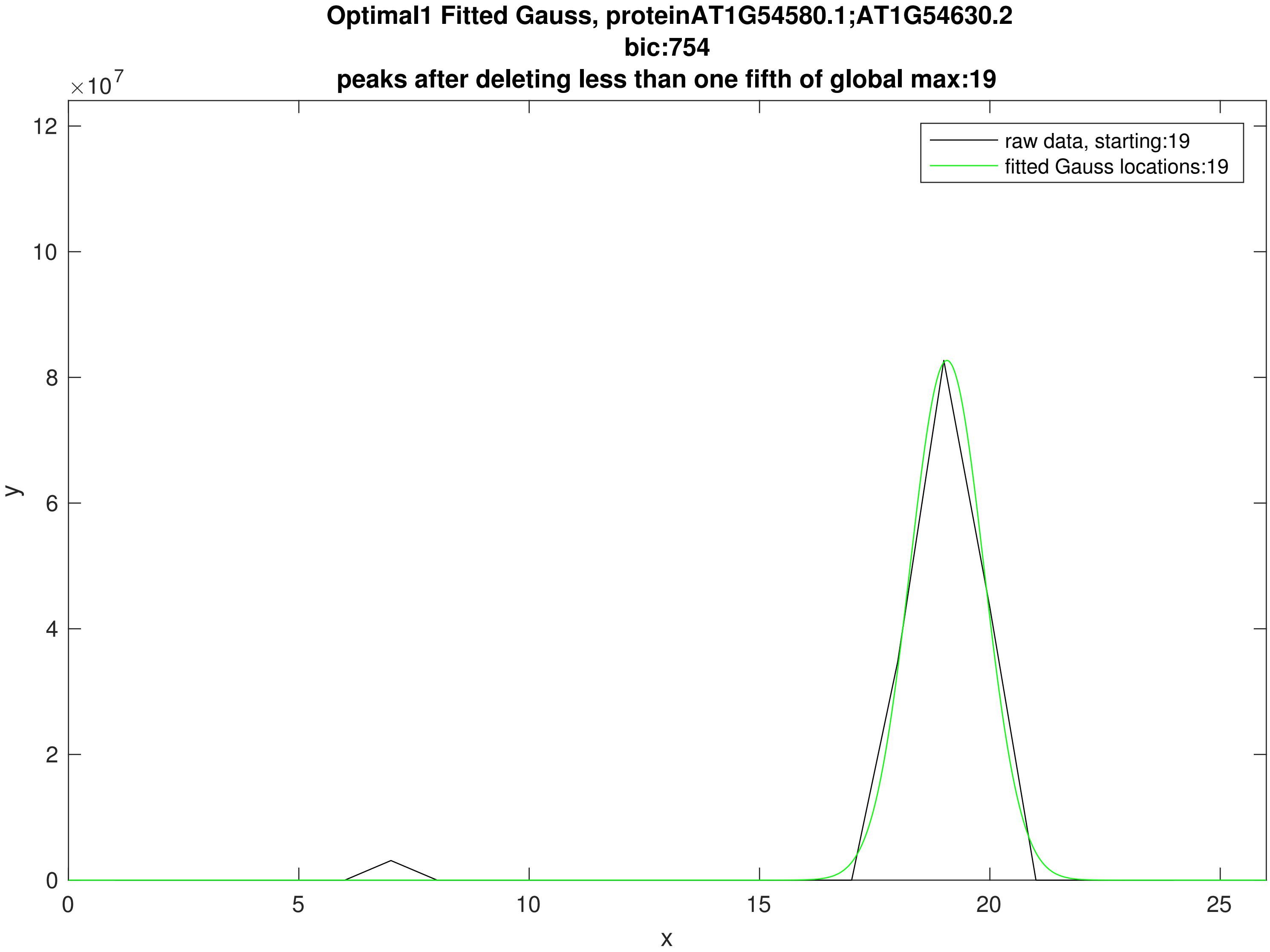
Optimal1 Fitted Gauss, proteinAT1G54580.1;AT1G54630.2
bic:754
peaks after deleting less than one fifth of global max:19
×10^7
12
10
8
6
4
2
0
y
0
5
10
15
20
25
x
raw data, starting:19
fitted Gauss locations:19

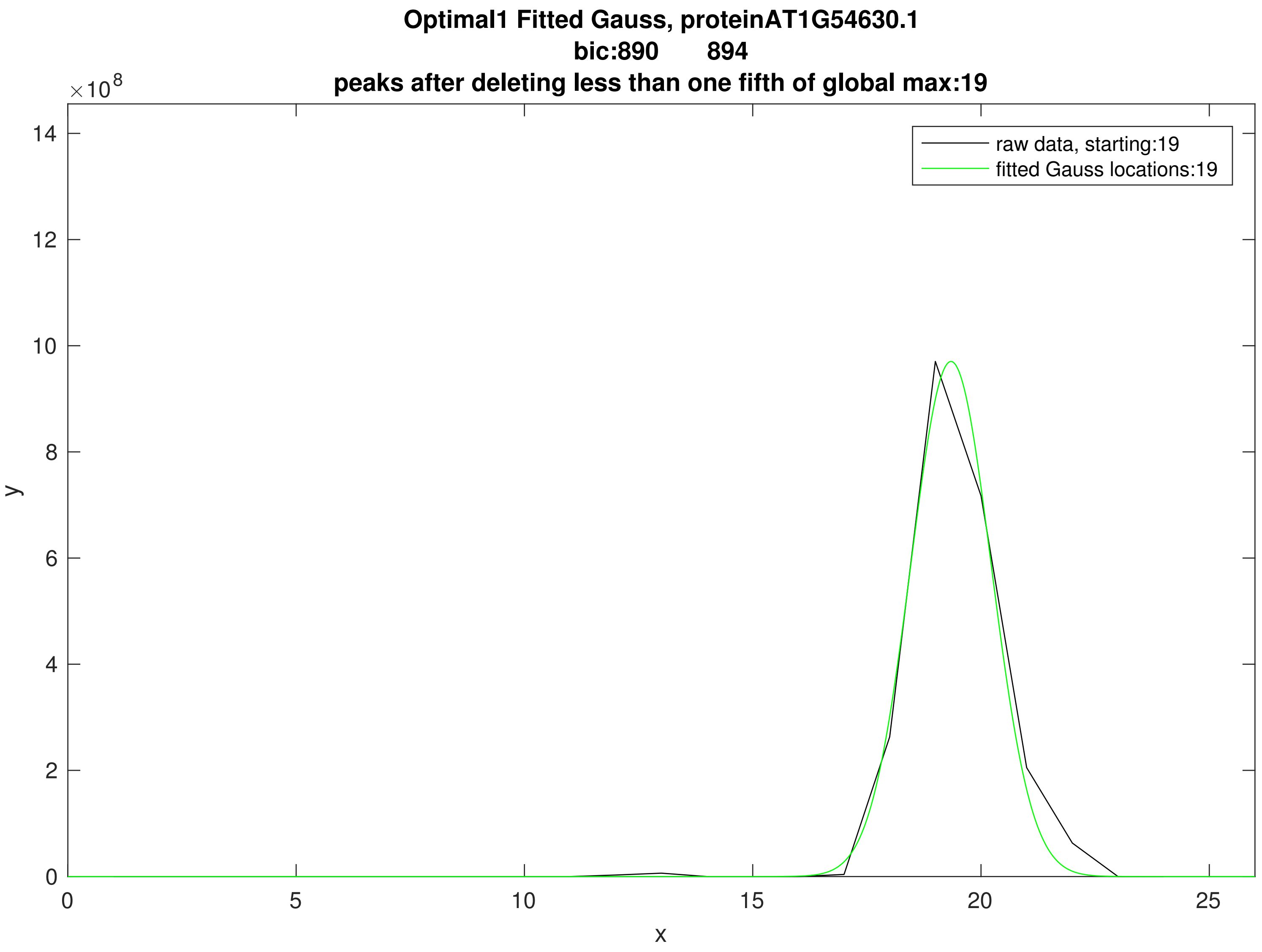
Optimal1 Fitted Gauss, proteinAT1G54630.1
bic:890 894
peaks after deleting less than one fifth of global max:19
×10^8
raw data, starting:19
fitted Gauss locations:19
14
12
10
8
6
4
2
0
y
0
5
10
15
20
25
x

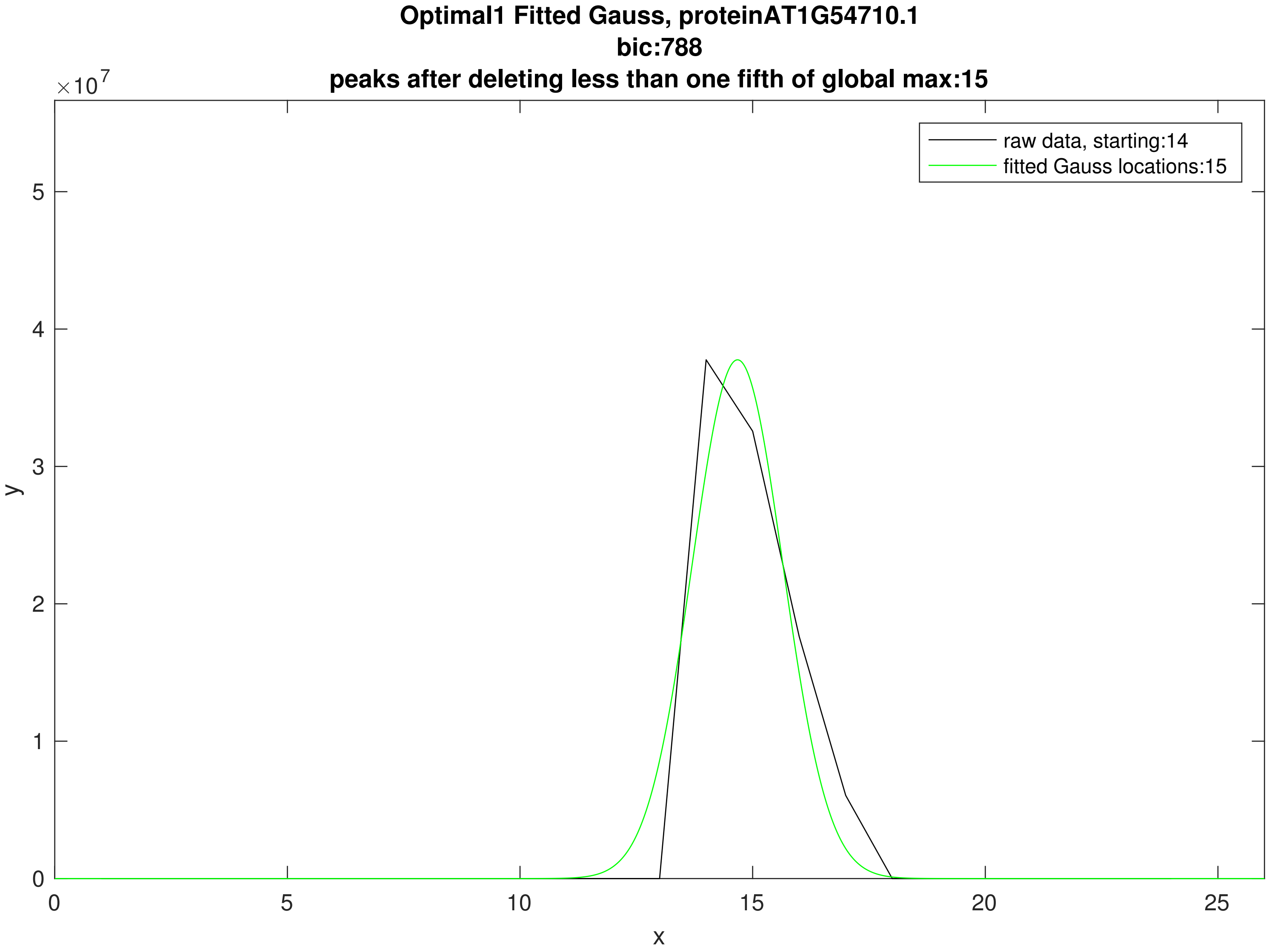
Optimal1 Fitted Gauss, proteinAT1G54710.1
bic:788
peaks after deleting less than one fifth of global max:15
×10^7
raw data, starting:14
fitted Gauss locations:15
0
1
2
3
4
5
y
0
5
10
15
20
25
x

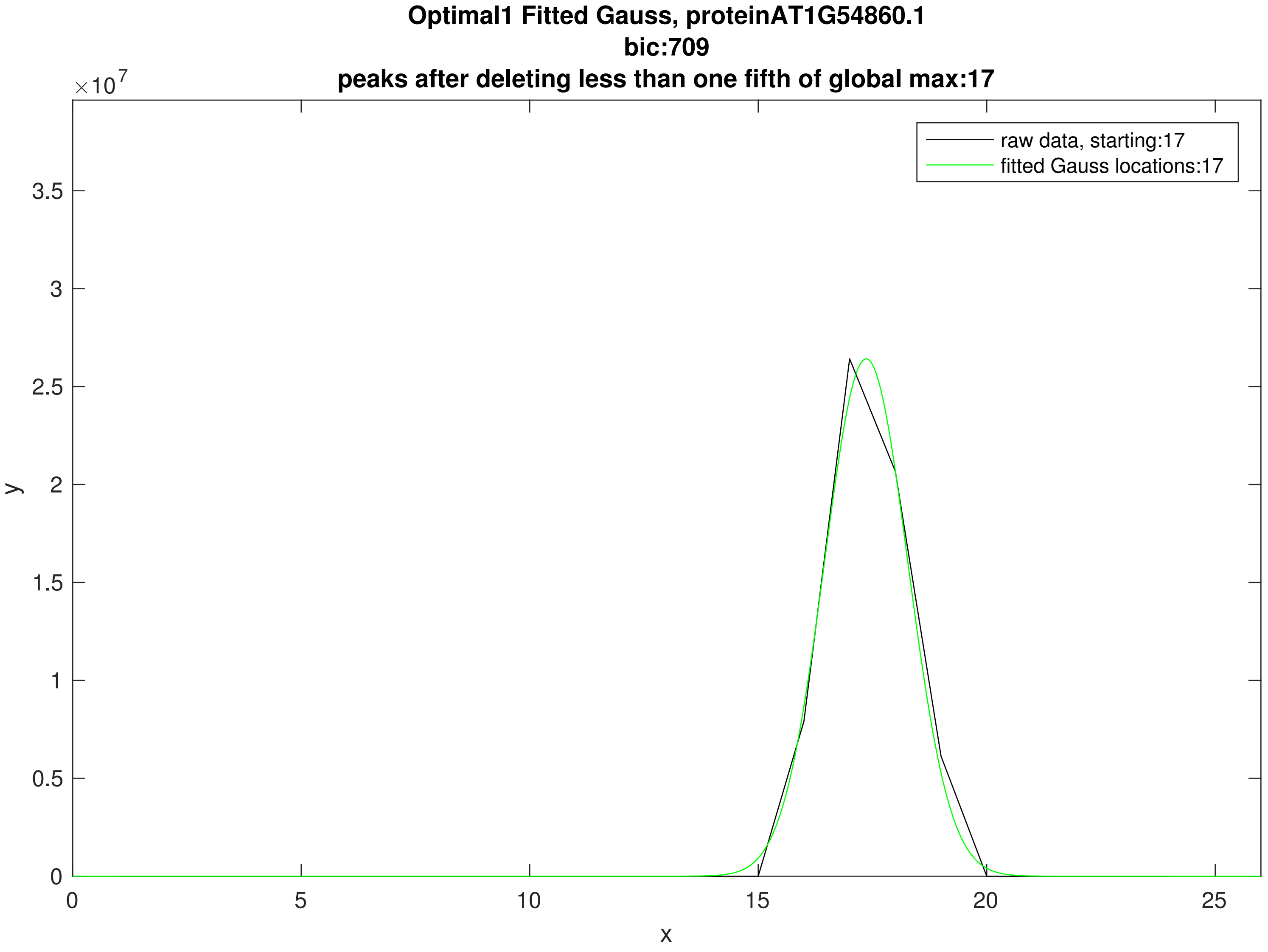

Optimal1 Fitted Gauss, proteinAT1G54860.1
bic:709
peaks after deleting less than one fifth of global max:17
×10^7
raw data, starting:17
fitted Gauss locations:17
3.5
3
2.5
2
1.5
1
0.5
0
y
0
5
10
15
20
25
x

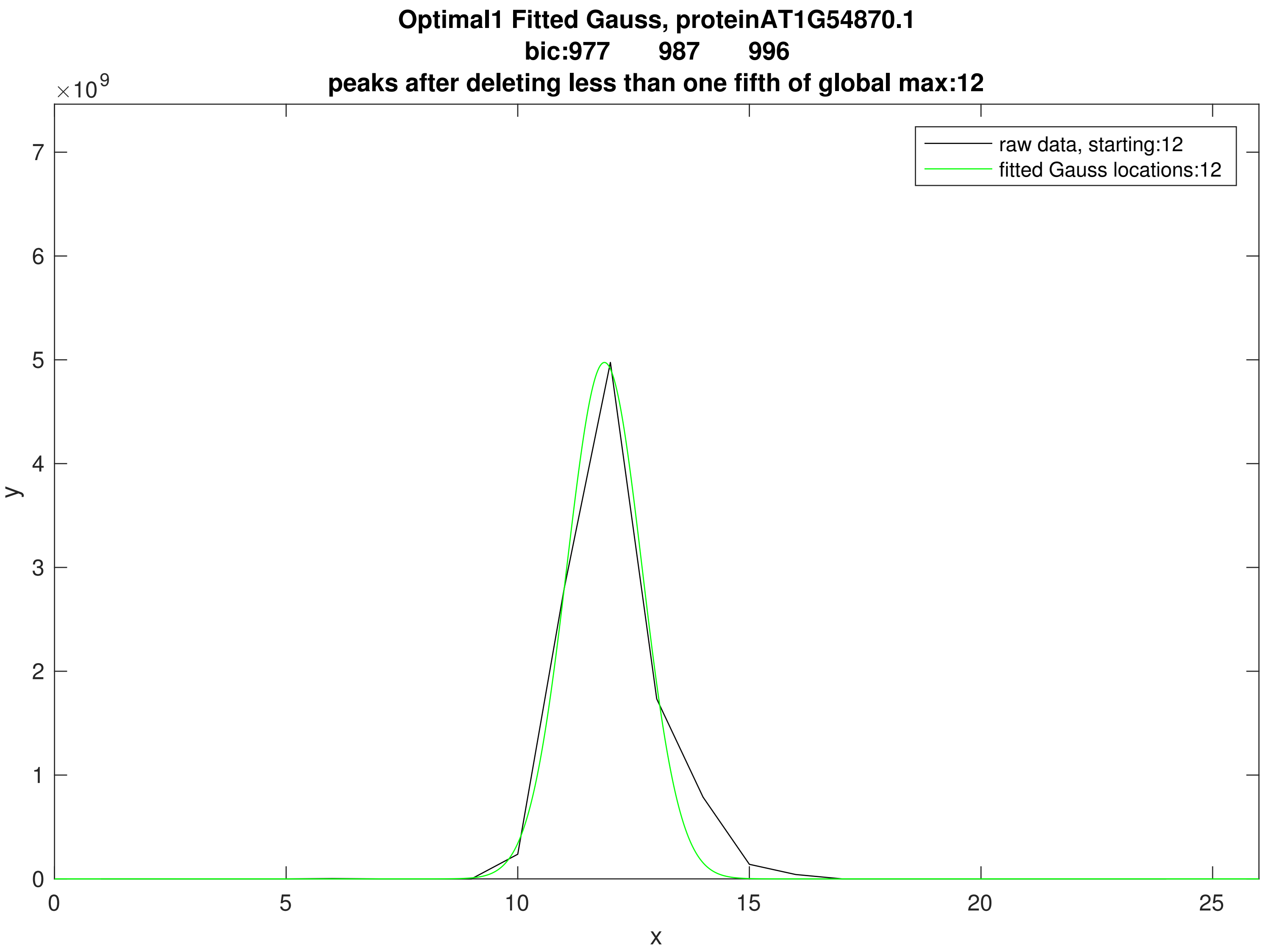
Optimal1 Fitted Gauss, proteinAT1G54870.1
bic:977 987 996
peaks after deleting less than one fifth of global max:12
×10^9
raw data, starting:12
fitted Gauss locations:12
y
x
0
1
2
3
4
5
6
7
0
5
10
15
20
25

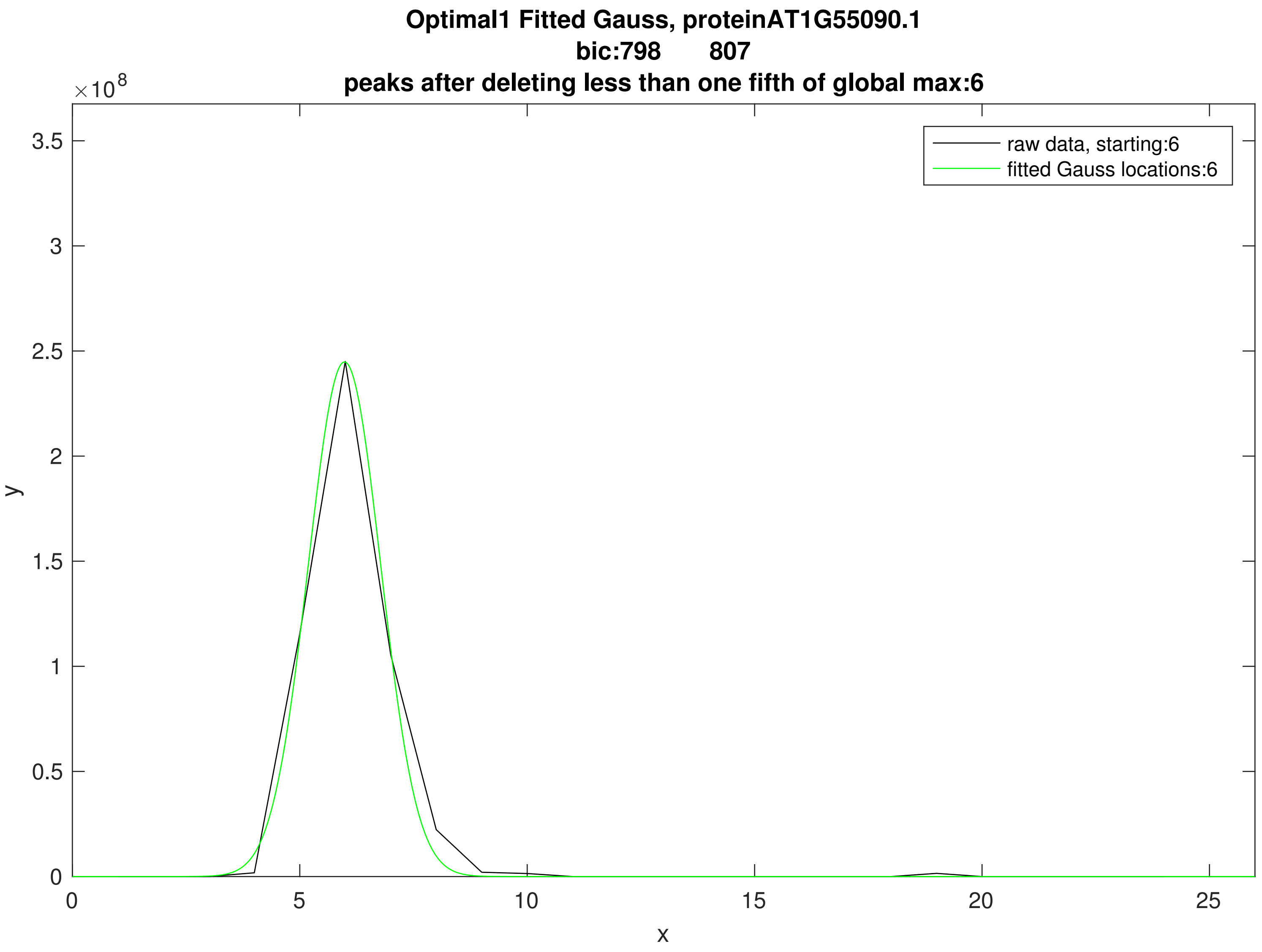
Optimal1 Fitted Gauss, proteinAT1G55090.1
bic:798 807
peaks after deleting less than one fifth of global max:6
×10^8
3.5
3
2.5
2
1.5
1
0.5
0
y
0
5
10
15
20
25
x
raw data, starting:6
fitted Gauss locations:6

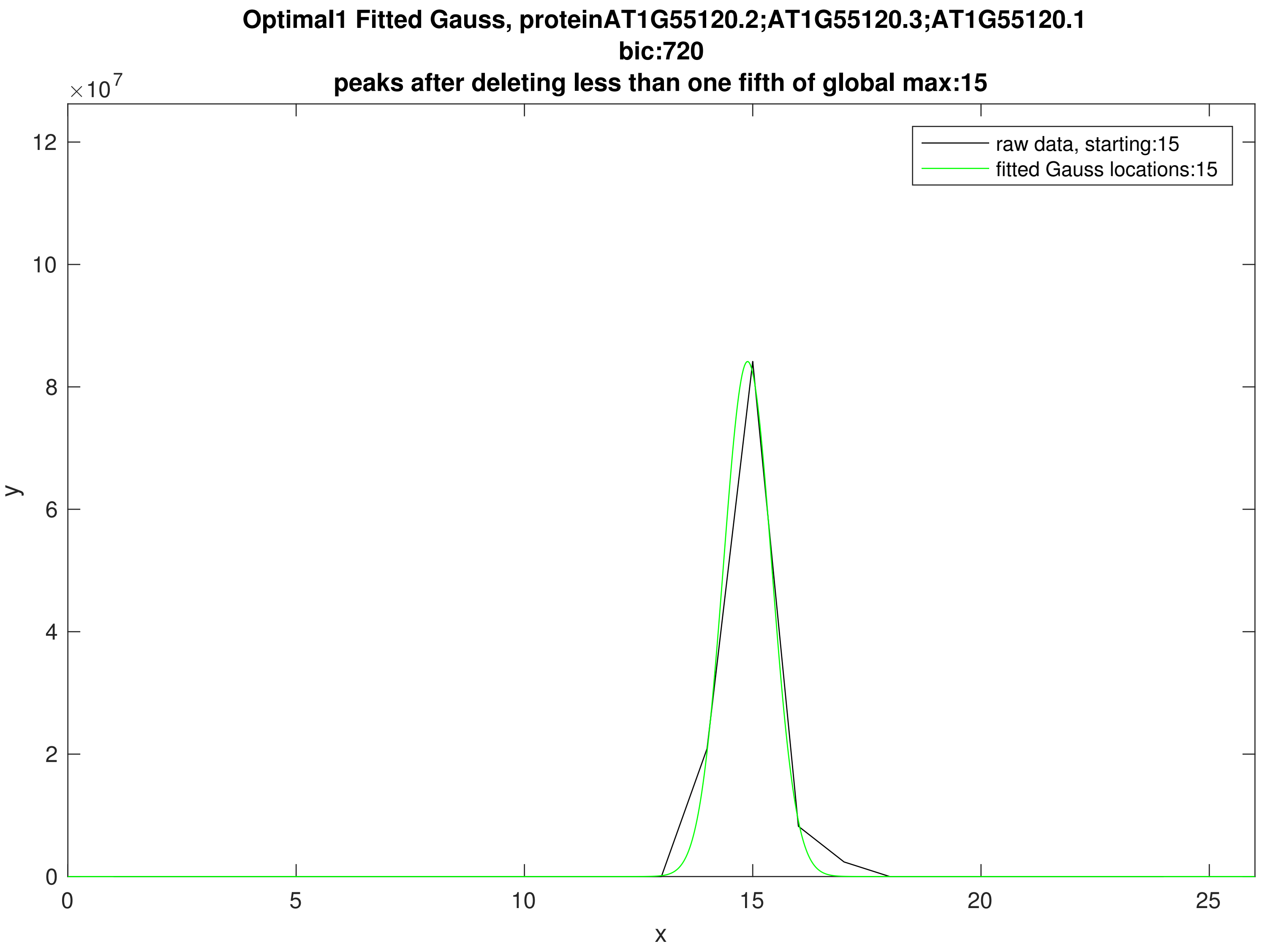
Optimal1 Fitted Gauss, proteinAT1G55120.2;AT1G55120.3;AT1G55120.1
bic:720
peaks after deleting less than one fifth of global max:15
×10^7
12
10
8
6
4
2
0
y
0
5
10
15
20
25
x
raw data, starting:15
fitted Gauss locations:15

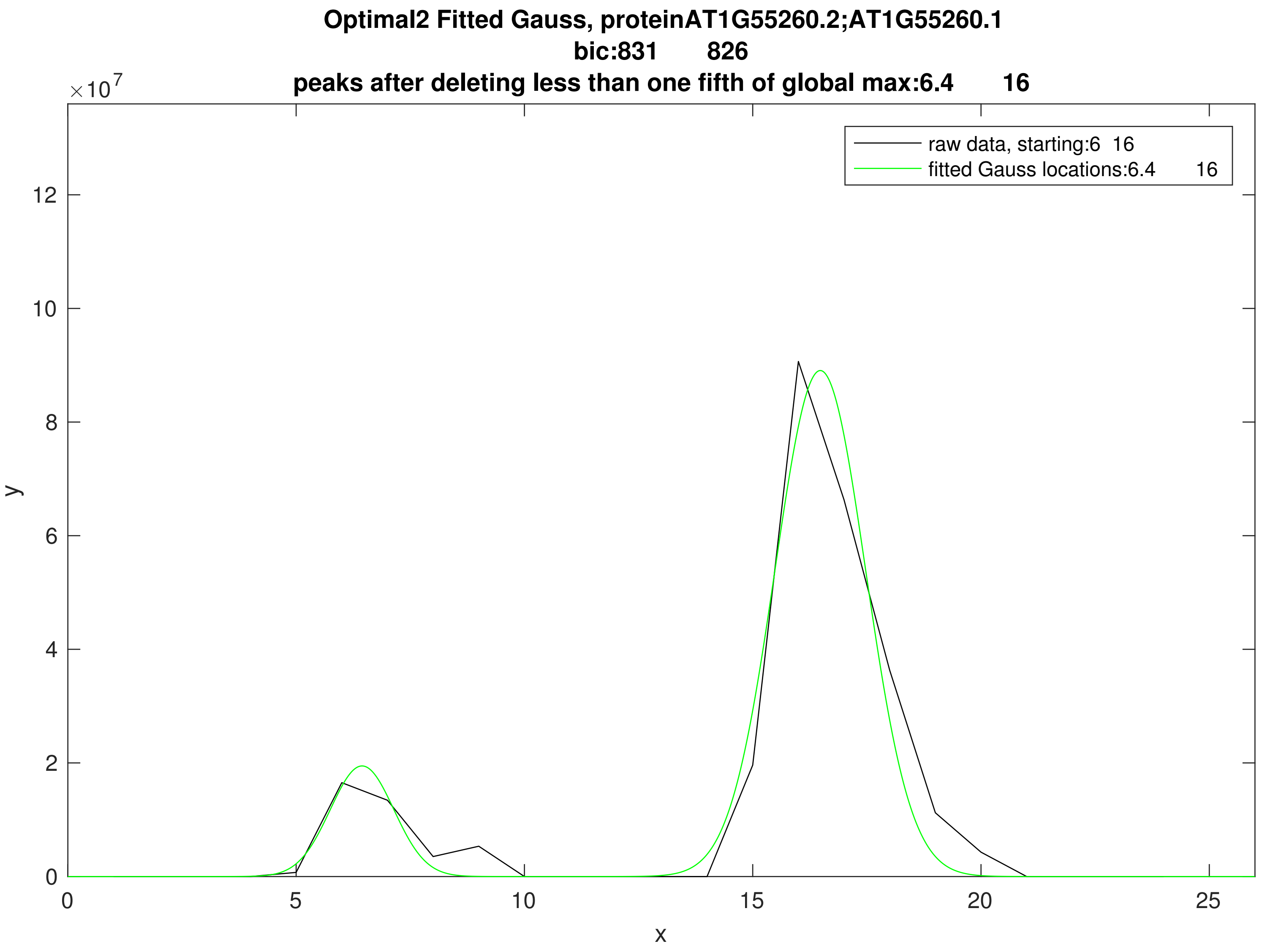
Optimal2 Fitted Gauss, proteinAT1G55260.2;AT1G55260.1
bic:831 826
peaks after deleting less than one fifth of global max:6.4 16
×10^7
raw data, starting:6 16
fitted Gauss locations:6.4 16
y
x
0
2
4
6
8
10
12
0
5
10
15
20
25

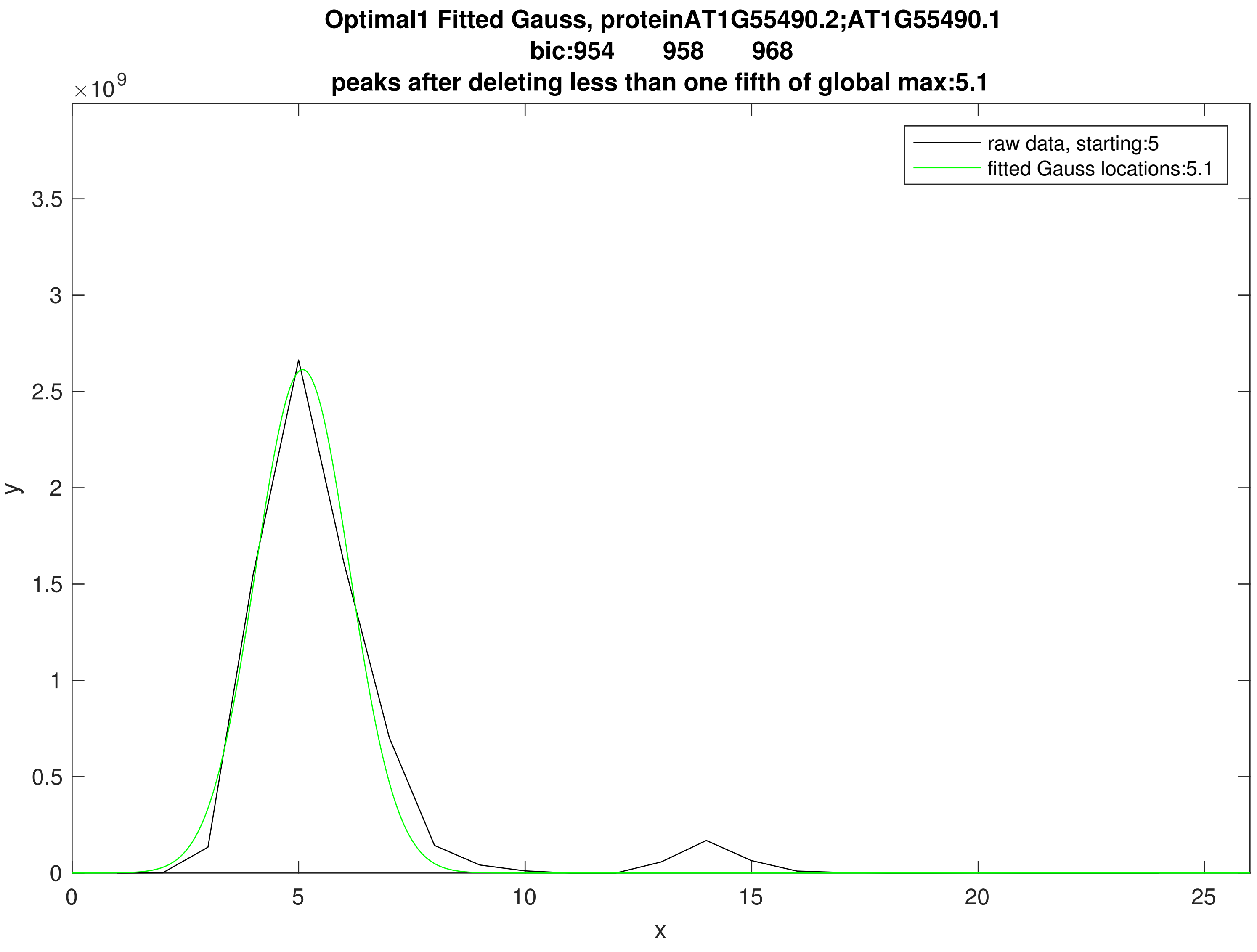
Optimal1 Fitted Gauss, proteinAT1G55490.2;AT1G55490.1
bic:954 958 968
peaks after deleting less than one fifth of global max:5.1
raw data, starting:5
fitted Gauss locations:5.1
×10^9
3.5
3
2.5
2
1.5
1
0.5
0
y
0
5
10
15
20
25
x

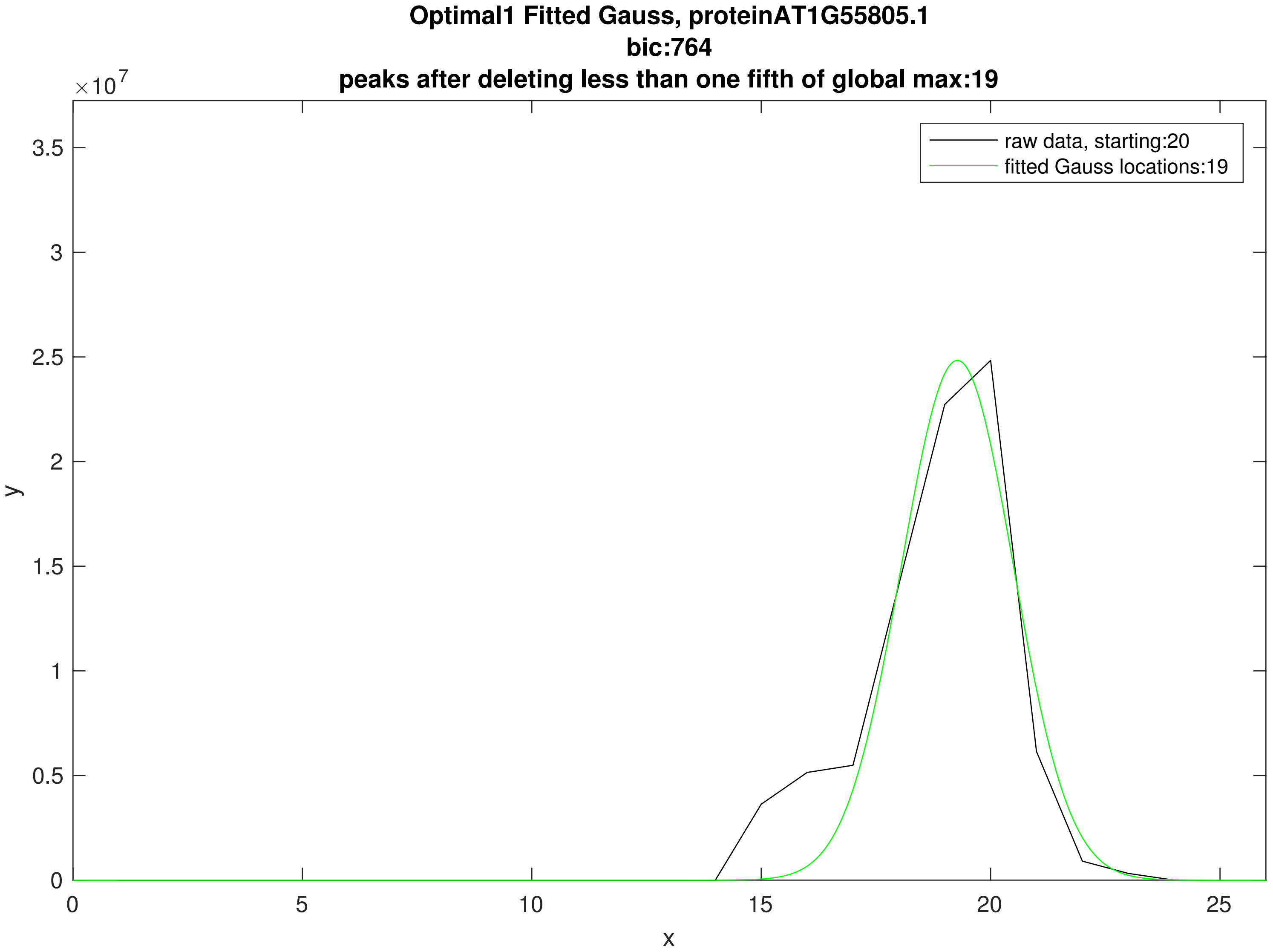

Optimal1 Fitted Gauss, proteinAT1G55805.1
bic:764
peaks after deleting less than one fifth of global max:19
×10^7
3.5
3
2.5
2
1.5
1
0.5
0
y
0
5
10
15
20
25
x
raw data, starting:20
fitted Gauss locations:19

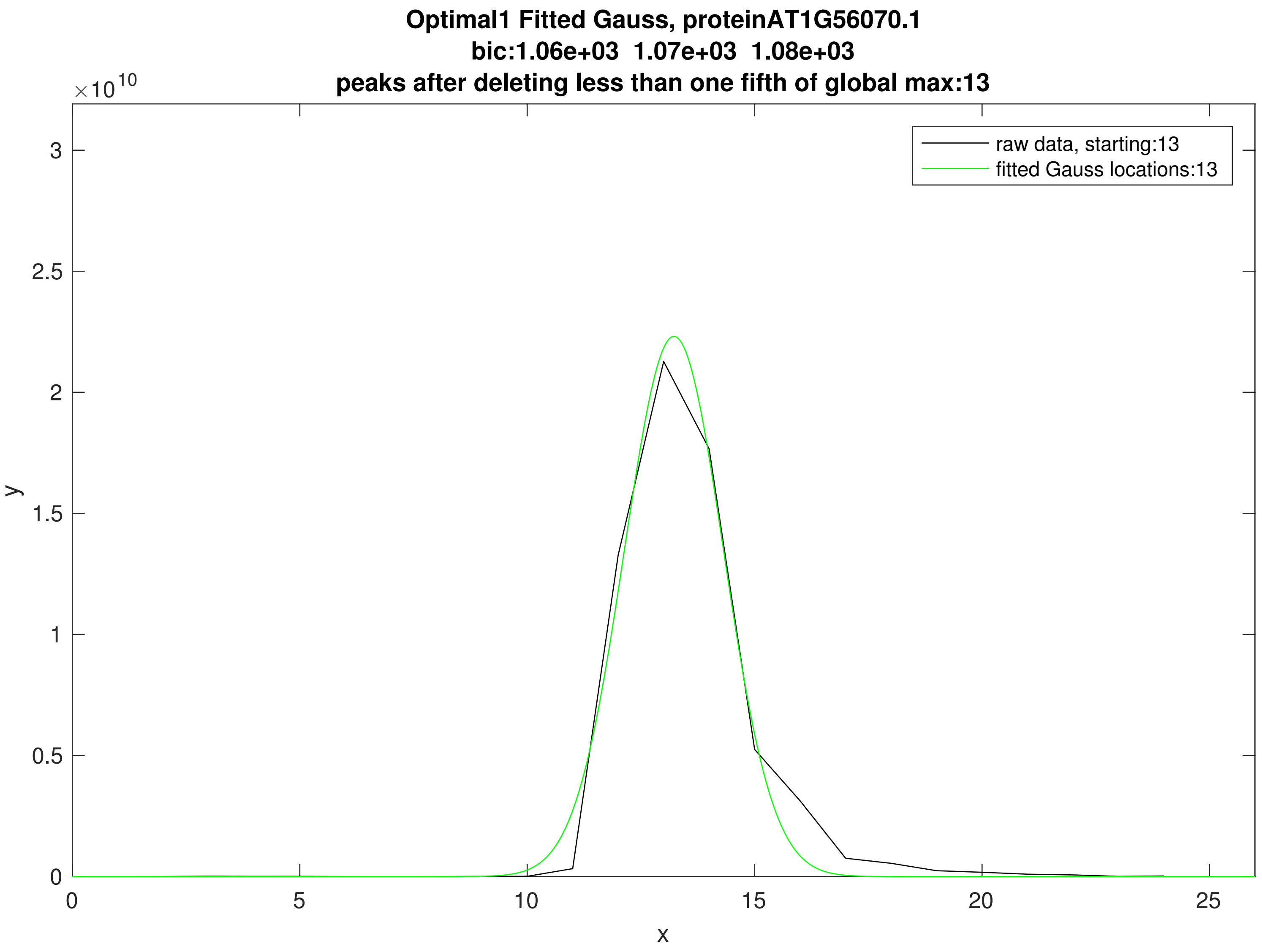
Optimal1 Fitted Gauss, proteinAT1G56070.1
bic:1.06e+03  1.07e+03  1.08e+03
peaks after deleting less than one fifth of global max:13
×10^10
raw data, starting:13
fitted Gauss locations:13
y
x
0
0.5
1
1.5
2
2.5
3
0
5
10
15
20
25

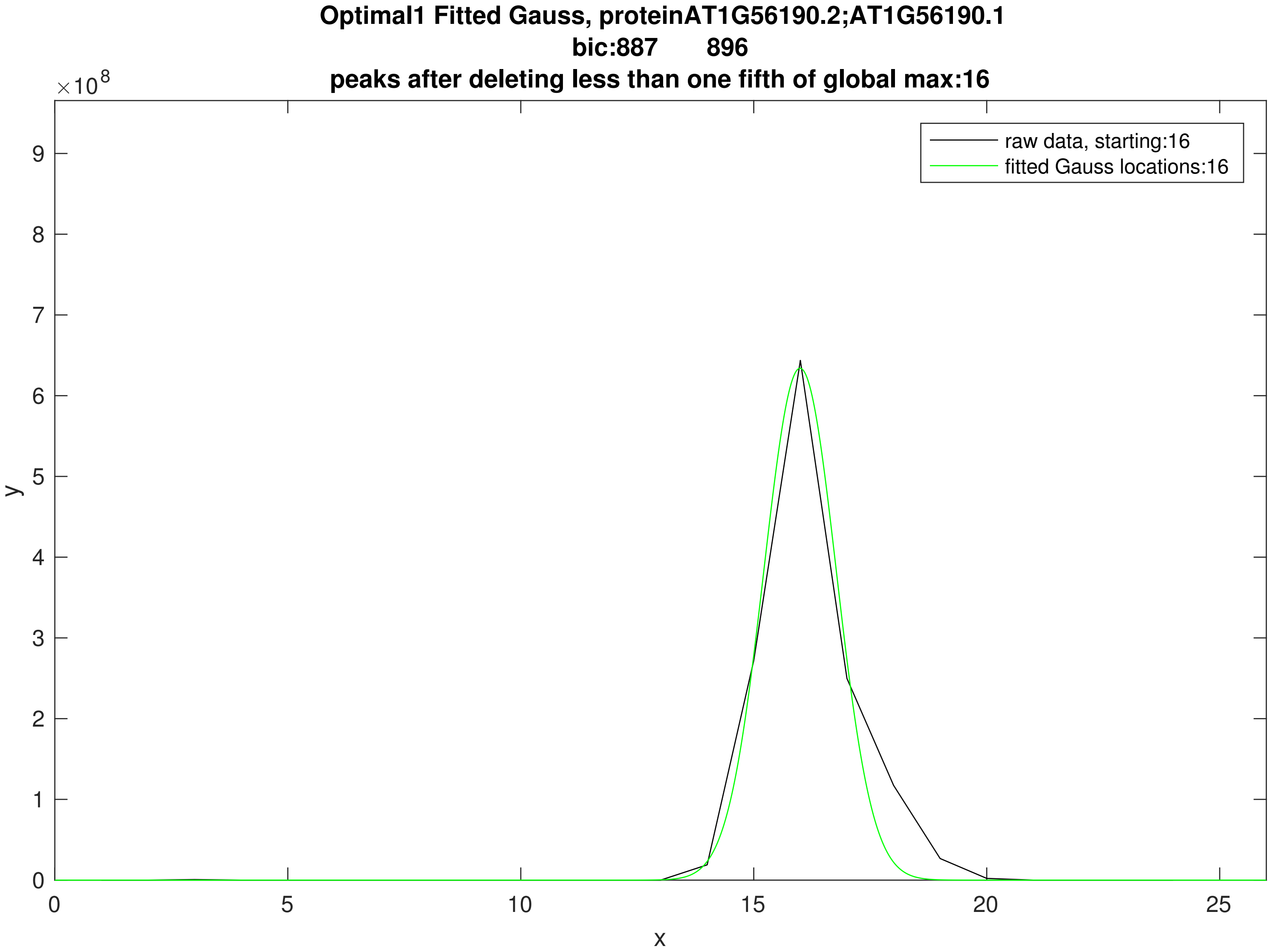
Optimal1 Fitted Gauss, proteinAT1G56190.2;AT1G56190.1
bic:887 896
peaks after deleting less than one fifth of global max:16
×10^8
raw data, starting:16
fitted Gauss locations:16
y
x
0
1
2
3
4
5
6
7
8
9
0
5
10
15
20
25

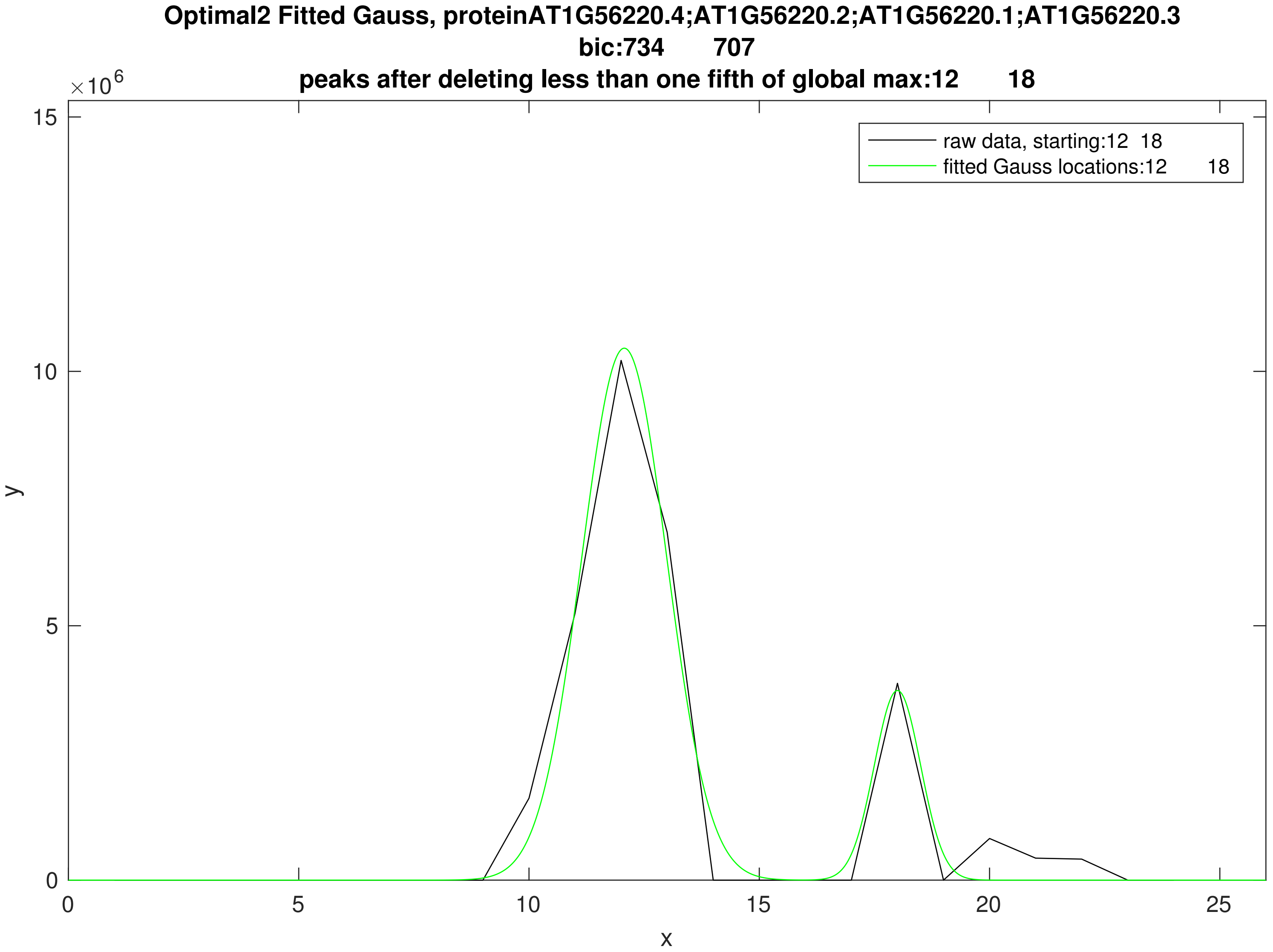

Optimal2 Fitted Gauss, proteinAT1G56220.4;AT1G56220.2;AT1G56220.1;AT1G56220.3
bic:734 707
peaks after deleting less than one fifth of global max:12 18
×10^6
raw data, starting:12 18
fitted Gauss locations:12 18
15
10
5
0
y
0
5
10
15
20
25
x

**Optimal2 Fitted Gauss, proteinAT1G56330.1;AT4G02080.1;AT3G62560.1**
**bic:819 749**
**peaks after deleting less than one fifth of global max:15**

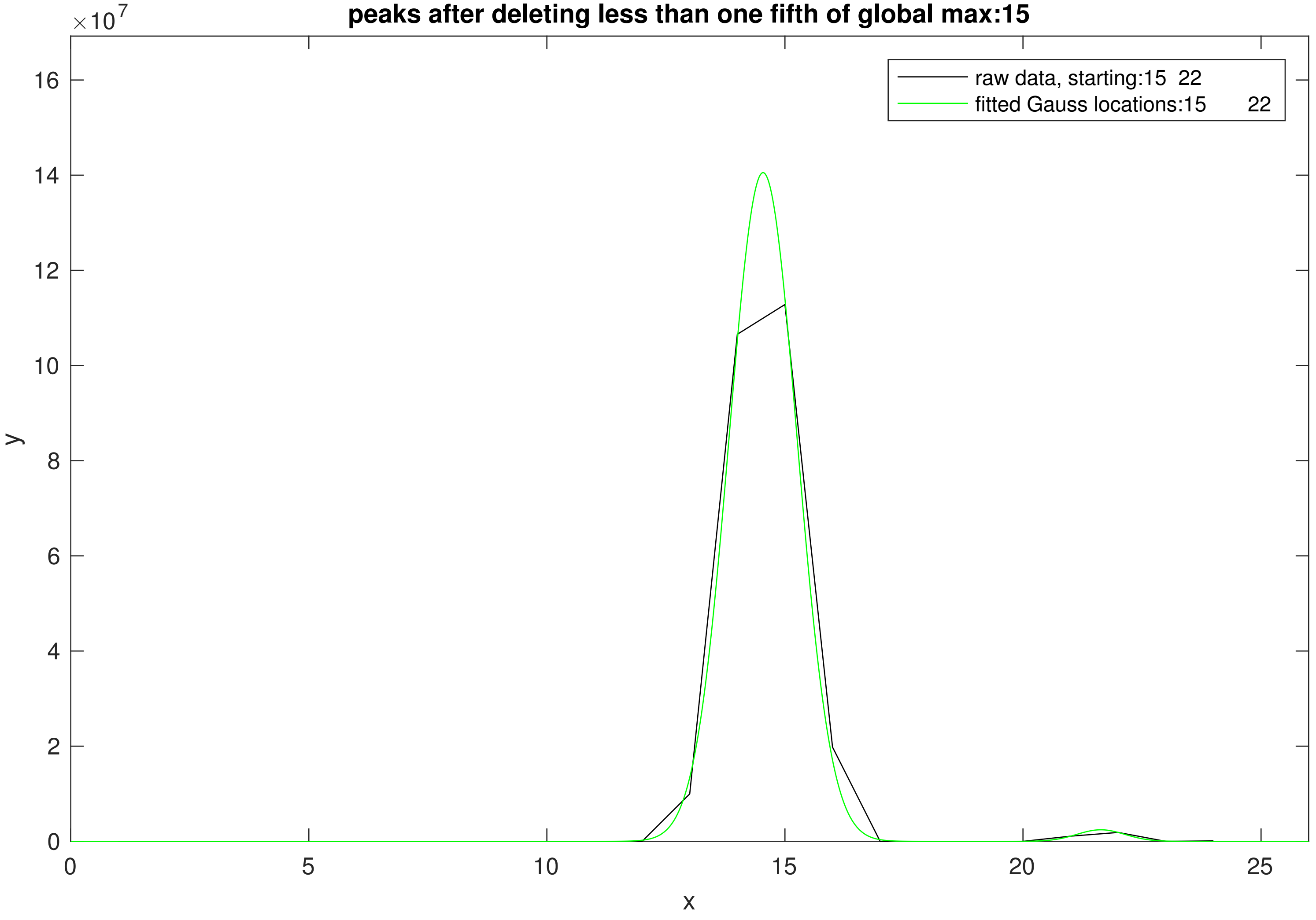

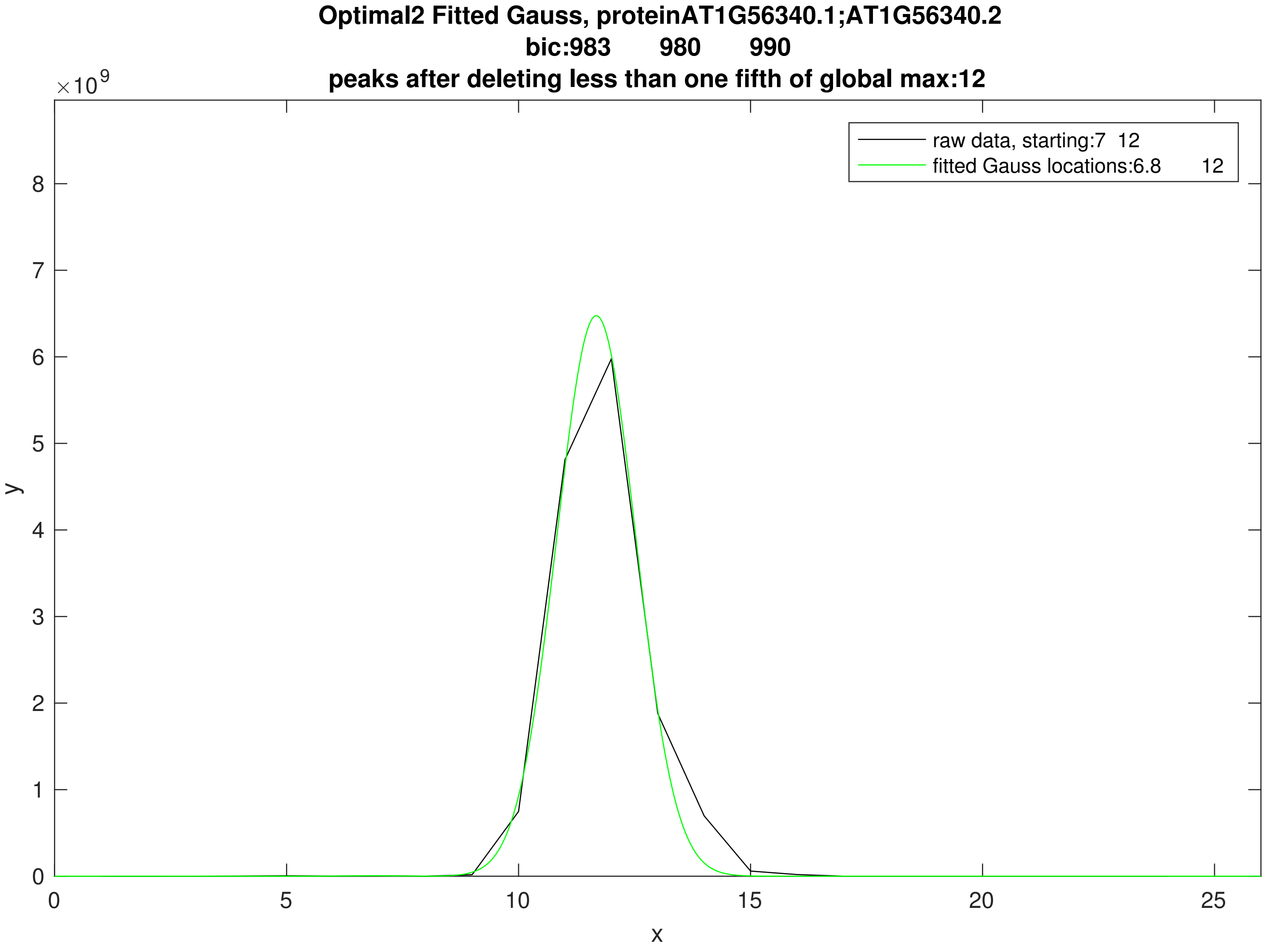

Optimal2 Fitted Gauss, proteinAT1G56340.1;AT1G56340.2
bic:983 980 990
peaks after deleting less than one fifth of global max:12
×10^9
raw data, starting:7 12
fitted Gauss locations:6.8 12
y
x
0
1
2
3
4
5
6
7
8
0
5
10
15
20
25

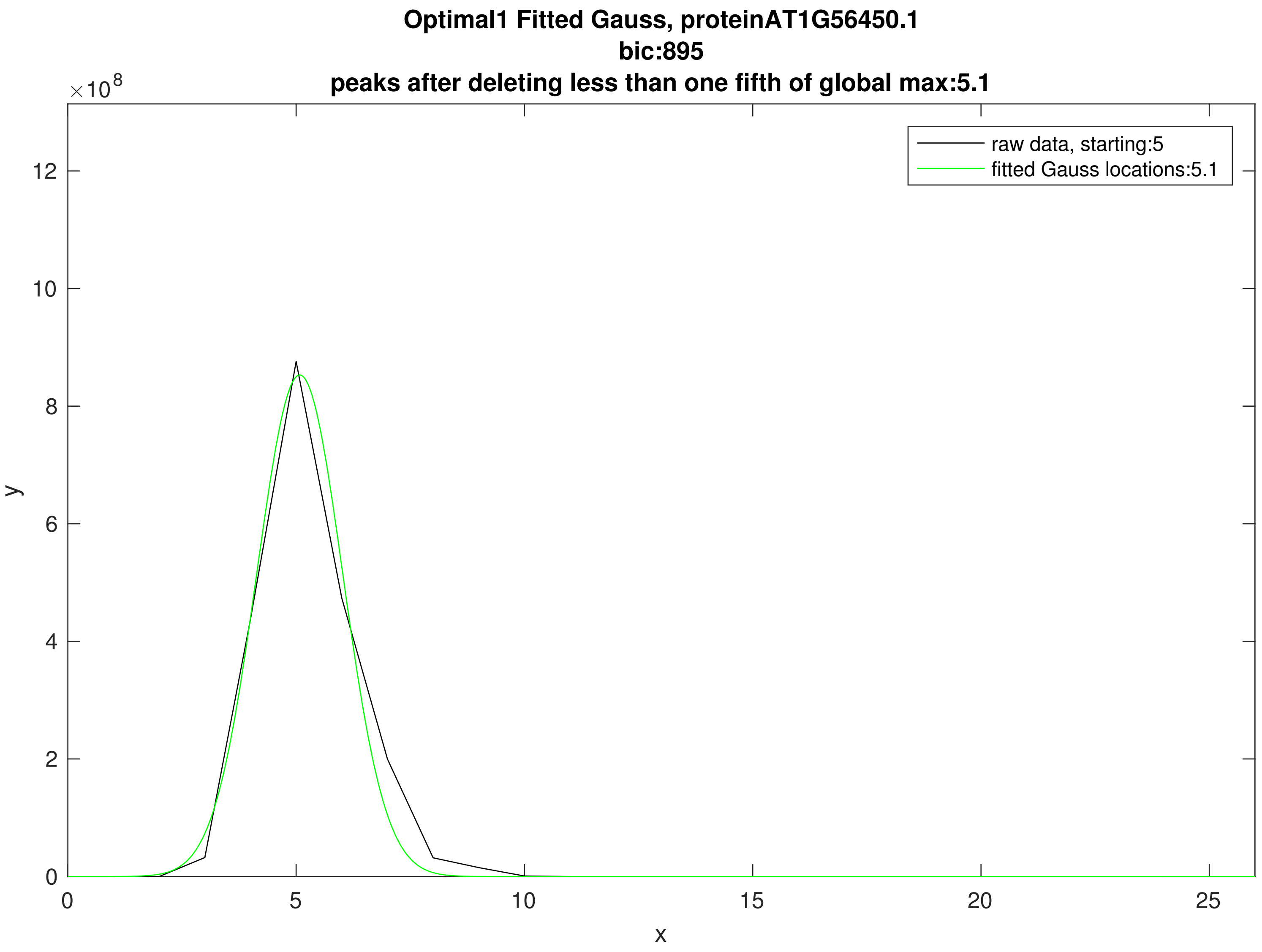

Optimal1 Fitted Gauss, proteinAT1G56450.1
bic:895
peaks after deleting less than one fifth of global max:5.1
×10^8
raw data, starting:5
fitted Gauss locations:5.1
y
x
0
2
4
6
8
10
12
0
5
10
15
20
25

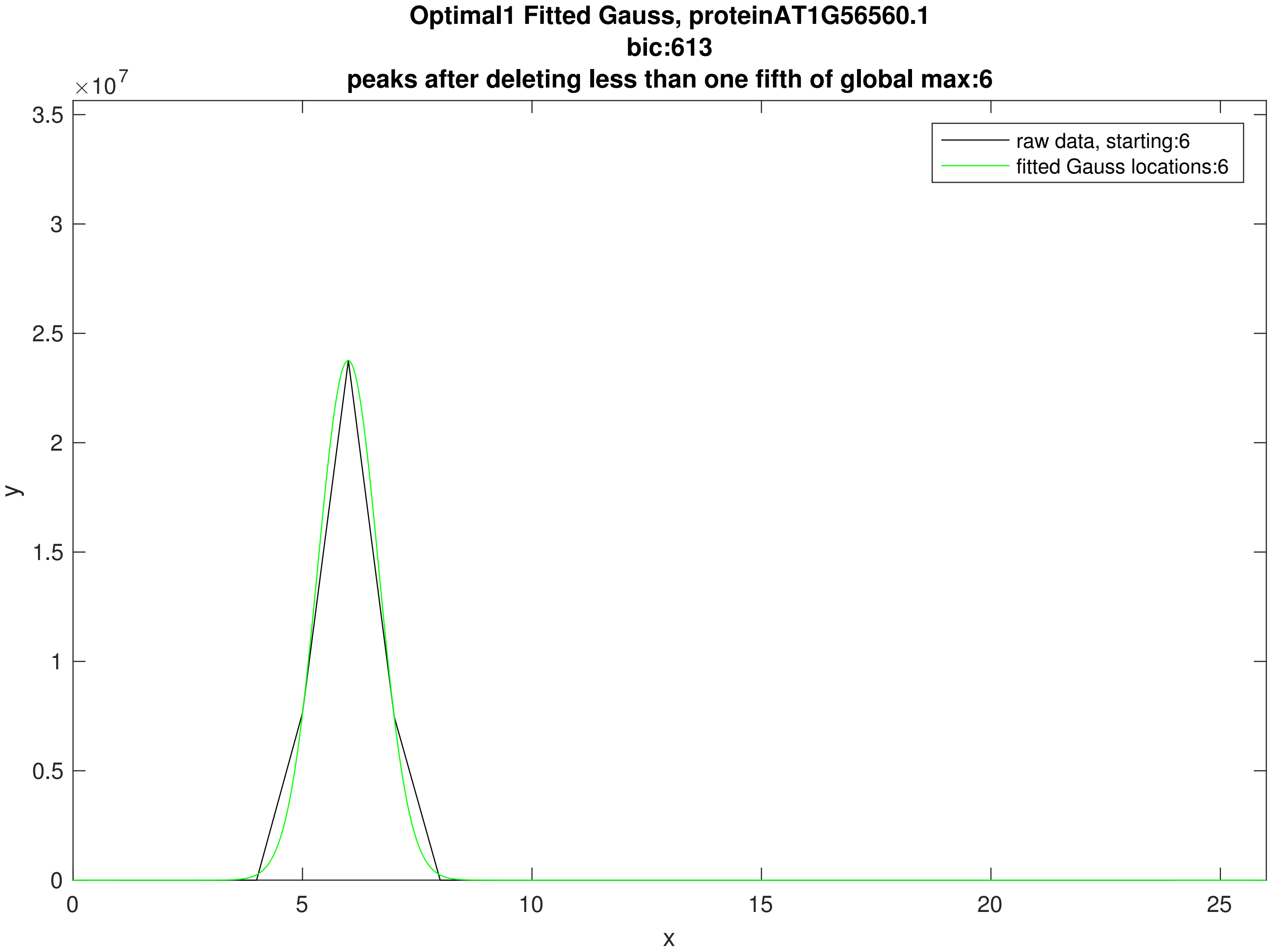

Optimal1 Fitted Gauss, proteinAT1G56560.1
bic:613
peaks after deleting less than one fifth of global max:6
×10^7
3.5
3
2.5
2
1.5
1
0.5
0
y
0
5
10
15
20
25
x
raw data, starting:6
fitted Gauss locations:6

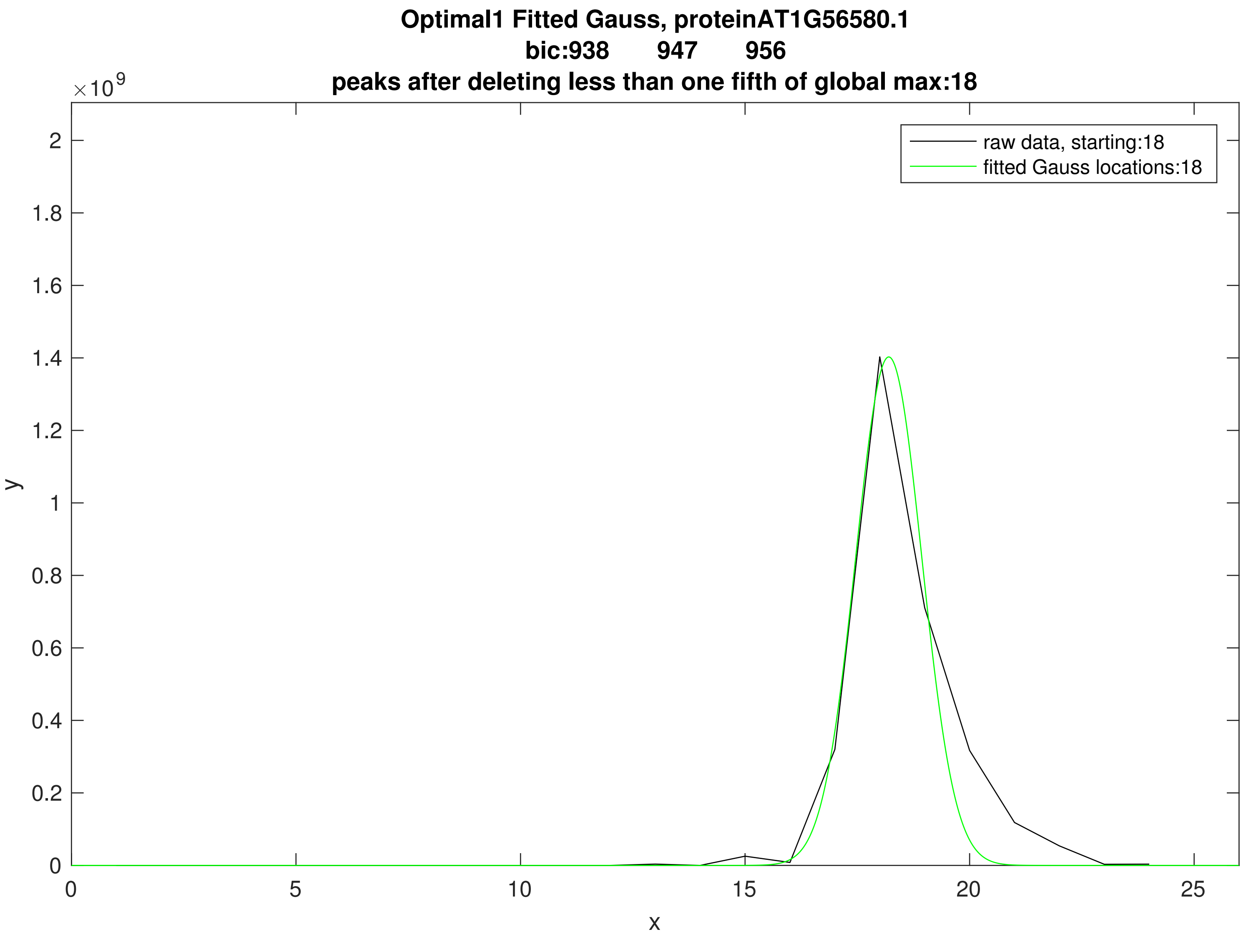
Optimal1 Fitted Gauss, proteinAT1G56580.1
bic:938 947 956
peaks after deleting less than one fifth of global max:18
×10^9
raw data, starting:18
fitted Gauss locations:18
y
x
2
1.8
1.6
1.4
1.2
1
0.8
0.6
0.4
0.2
0
5
10
15
20
25

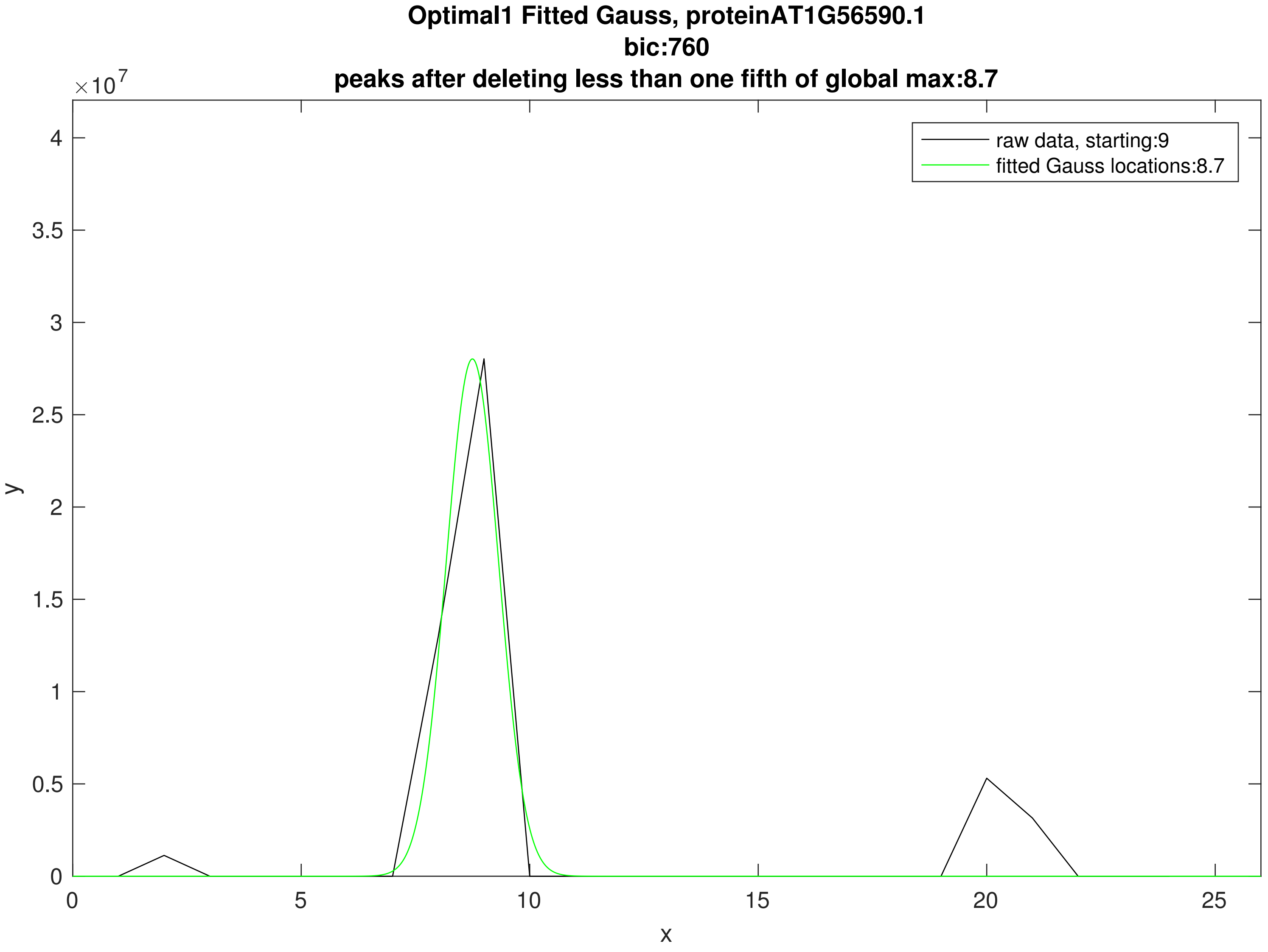

Optimal1 Fitted Gauss, proteinAT1G56590.1
bic:760
peaks after deleting less than one fifth of global max:8.7
×10^7
raw data, starting:9
fitted Gauss locations:8.7
y
x
0
0.5
1
1.5
2
2.5
3
3.5
4
5
10
15
20
25

Optimal1 Fitted Gauss, proteinAT1G56700.1;AT1G56700.2;AT1G56700.3
bic:752
peaks after deleting less than one fifth of global max:18

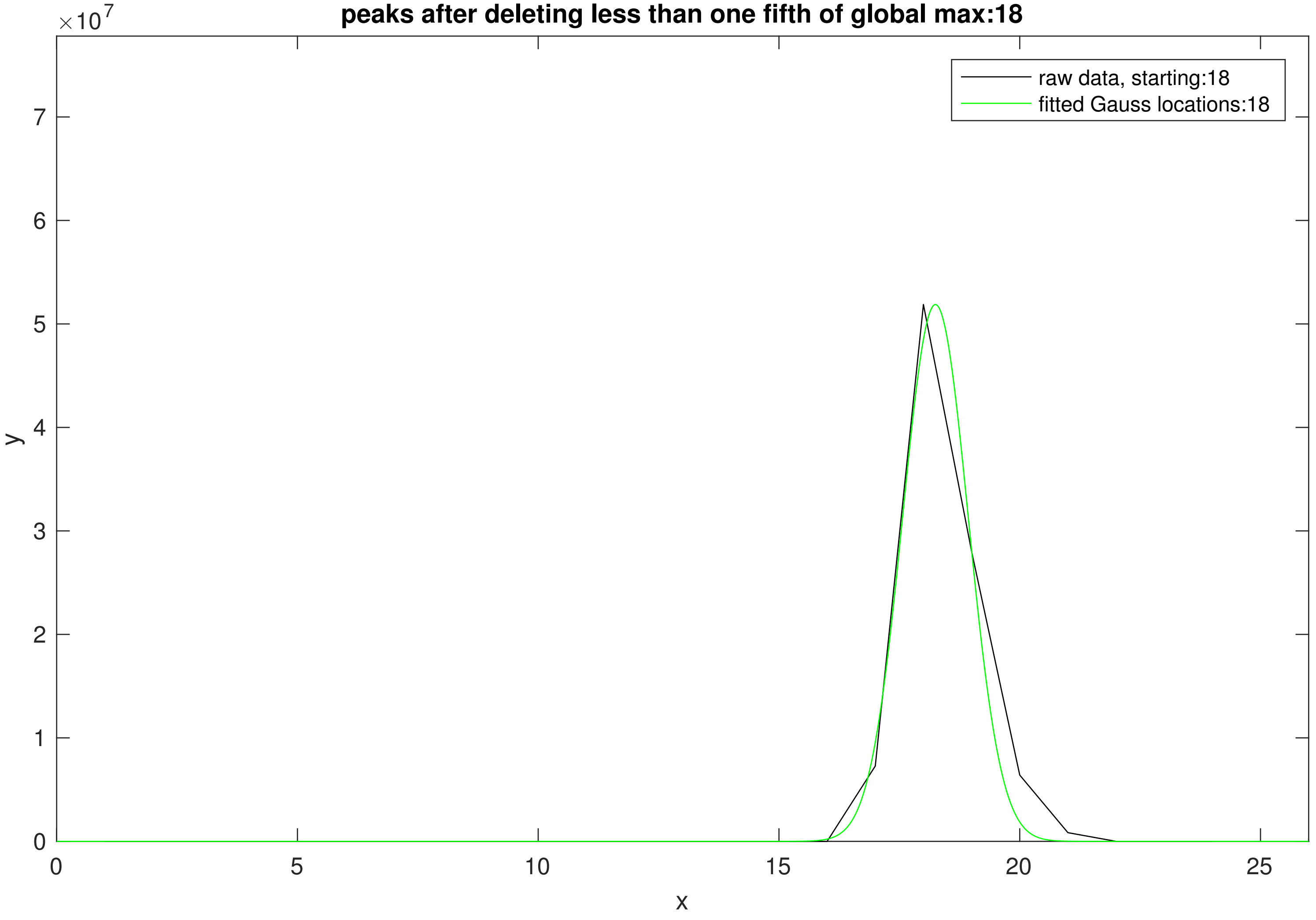

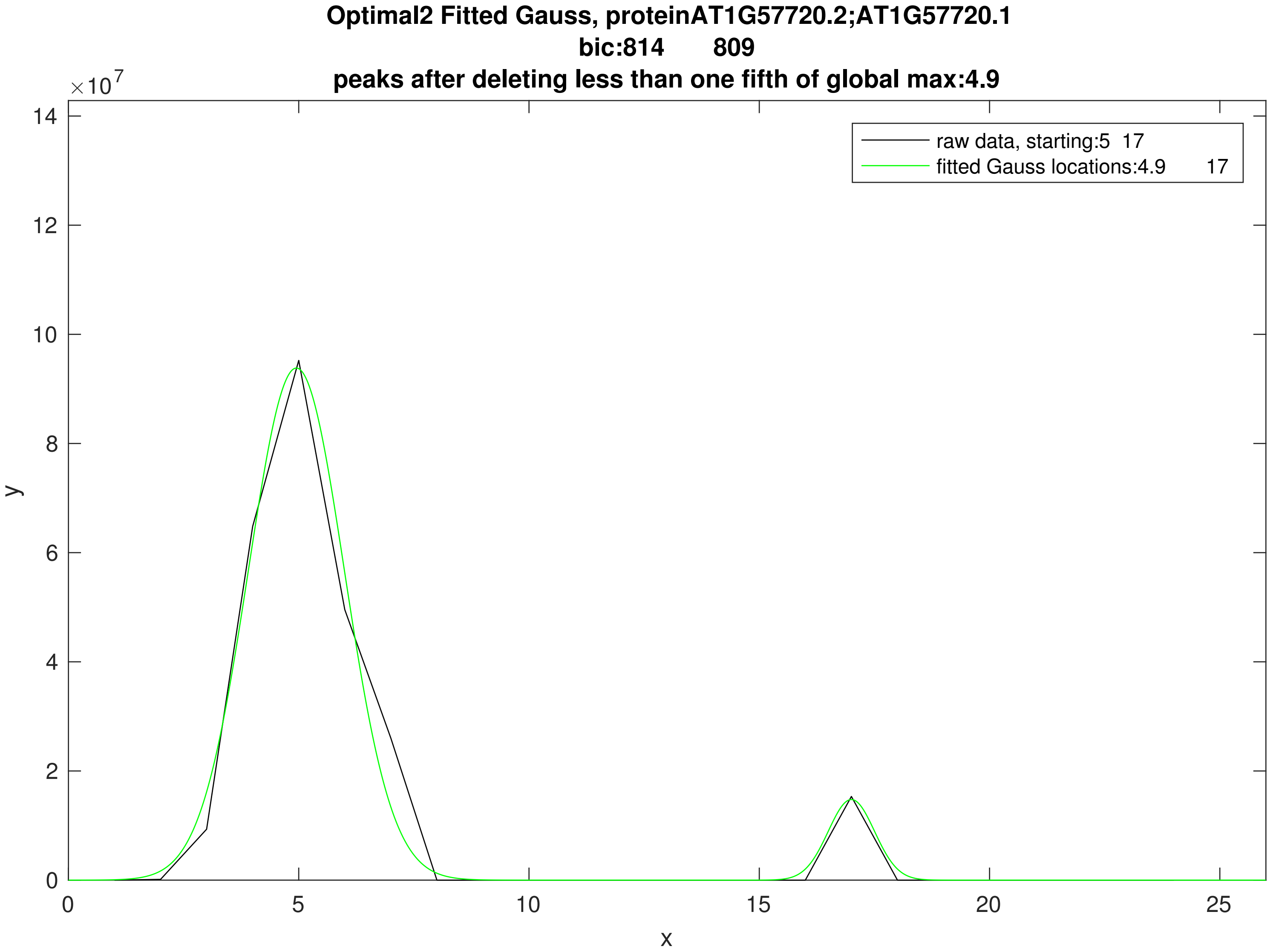

Optimal2 Fitted Gauss, proteinAT1G57720.2;AT1G57720.1
bic:814 809
peaks after deleting less than one fifth of global max:4.9
×10^7
raw data, starting:5 17
fitted Gauss locations:4.9 17
y
x
0
2
4
6
8
10
12
14
0
5
10
15
20
25

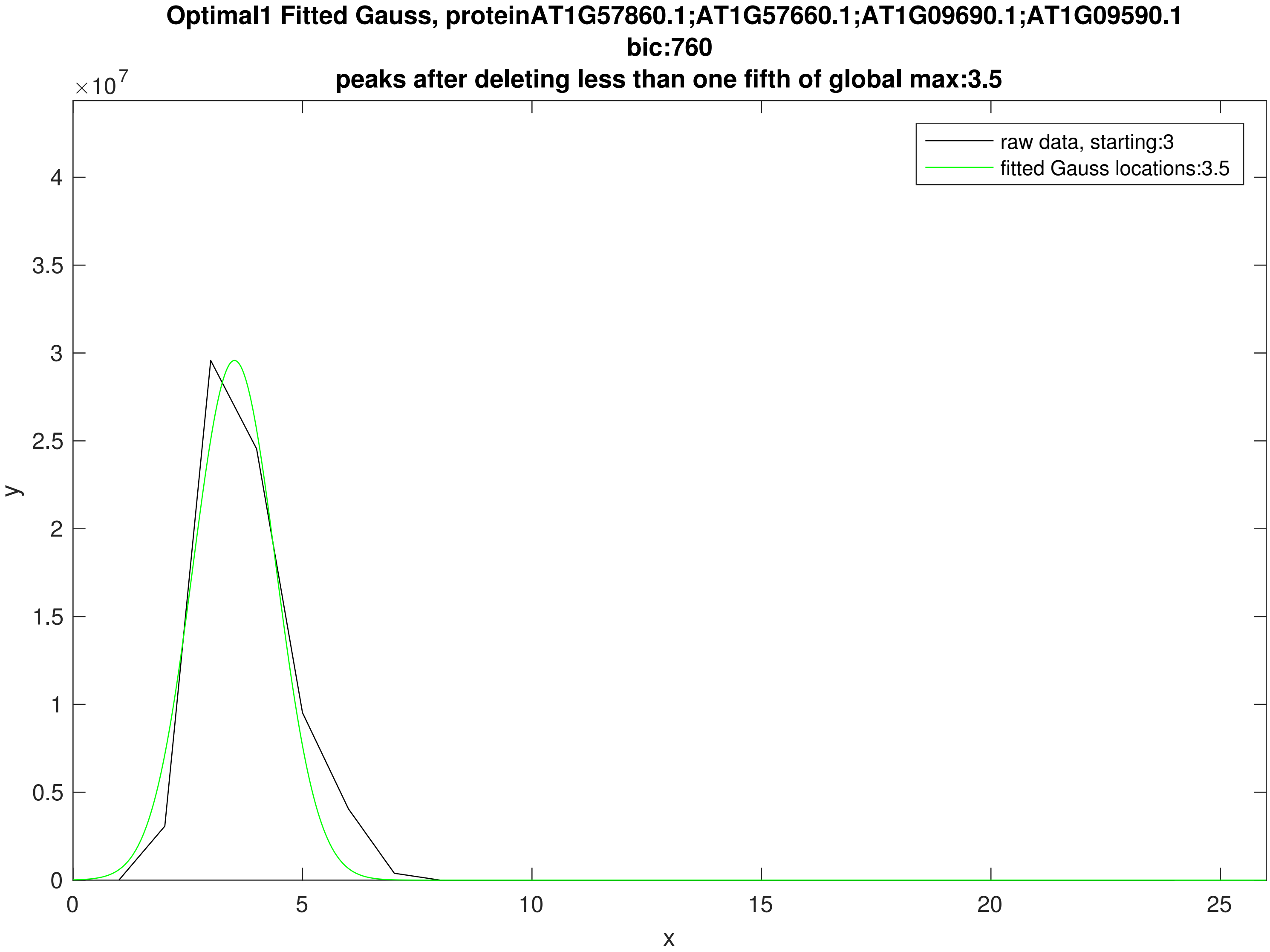
Optimal1 Fitted Gauss, proteinAT1G57860.1;AT1G57660.1;AT1G09690.1;AT1G09590.1
bic:760
peaks after deleting less than one fifth of global max:3.5
×10^7
raw data, starting:3
fitted Gauss locations:3.5
y
x
0
0.5
1
1.5
2
2.5
3
3.5
4
5
10
15
20
25

**Optimal1 Fitted Gauss, proteinAT1G58080.1**
**bic:796 797**
**peaks after deleting less than one fifth of global max:11**

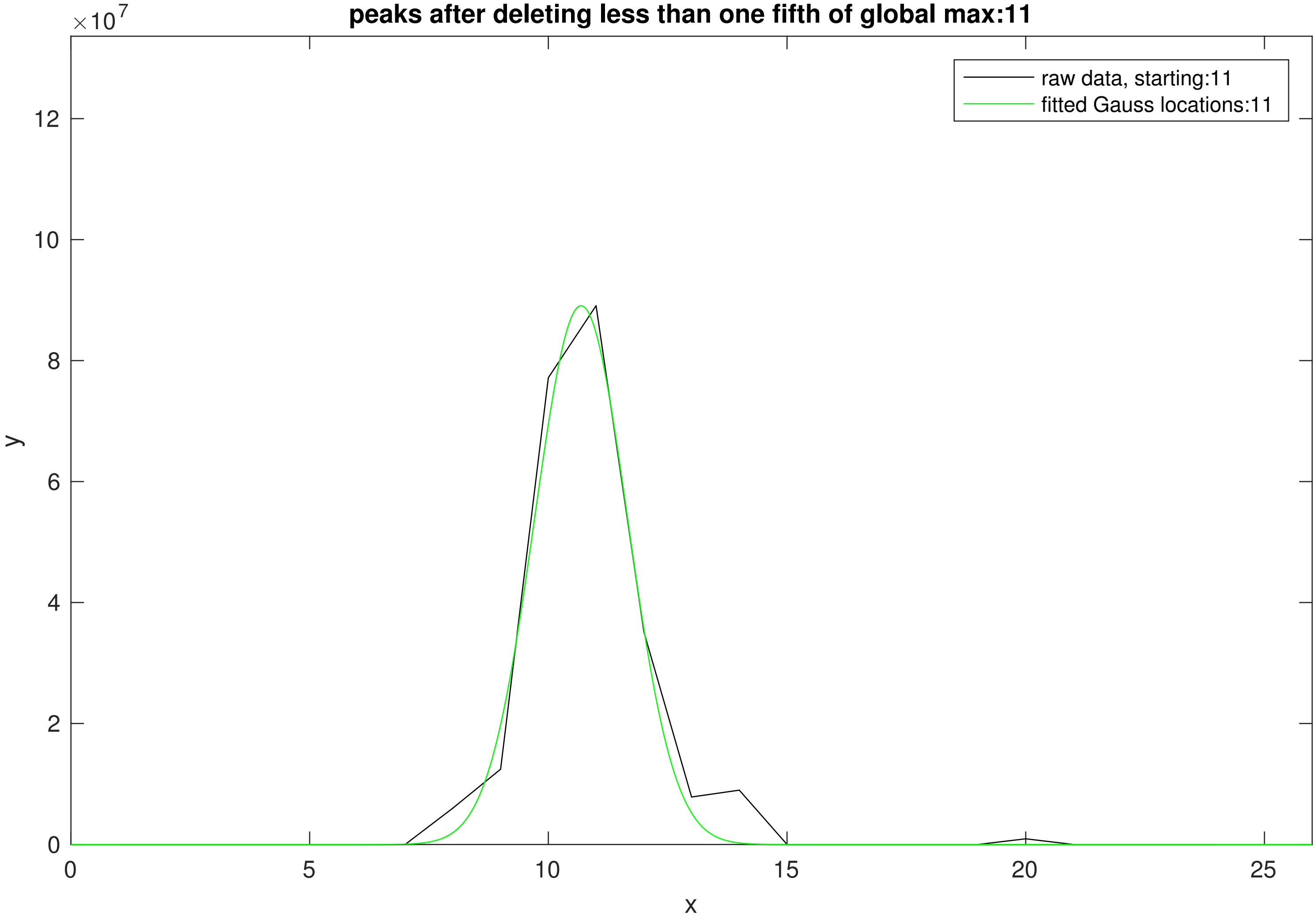

**Optimal1 Fitted Gauss, proteinAT1G59359.1;AT1G58983.1;AT1G58684.1;AT1G58380.1**
**bic:770 791**
**peaks after deleting less than one fifth of global max:3.7**

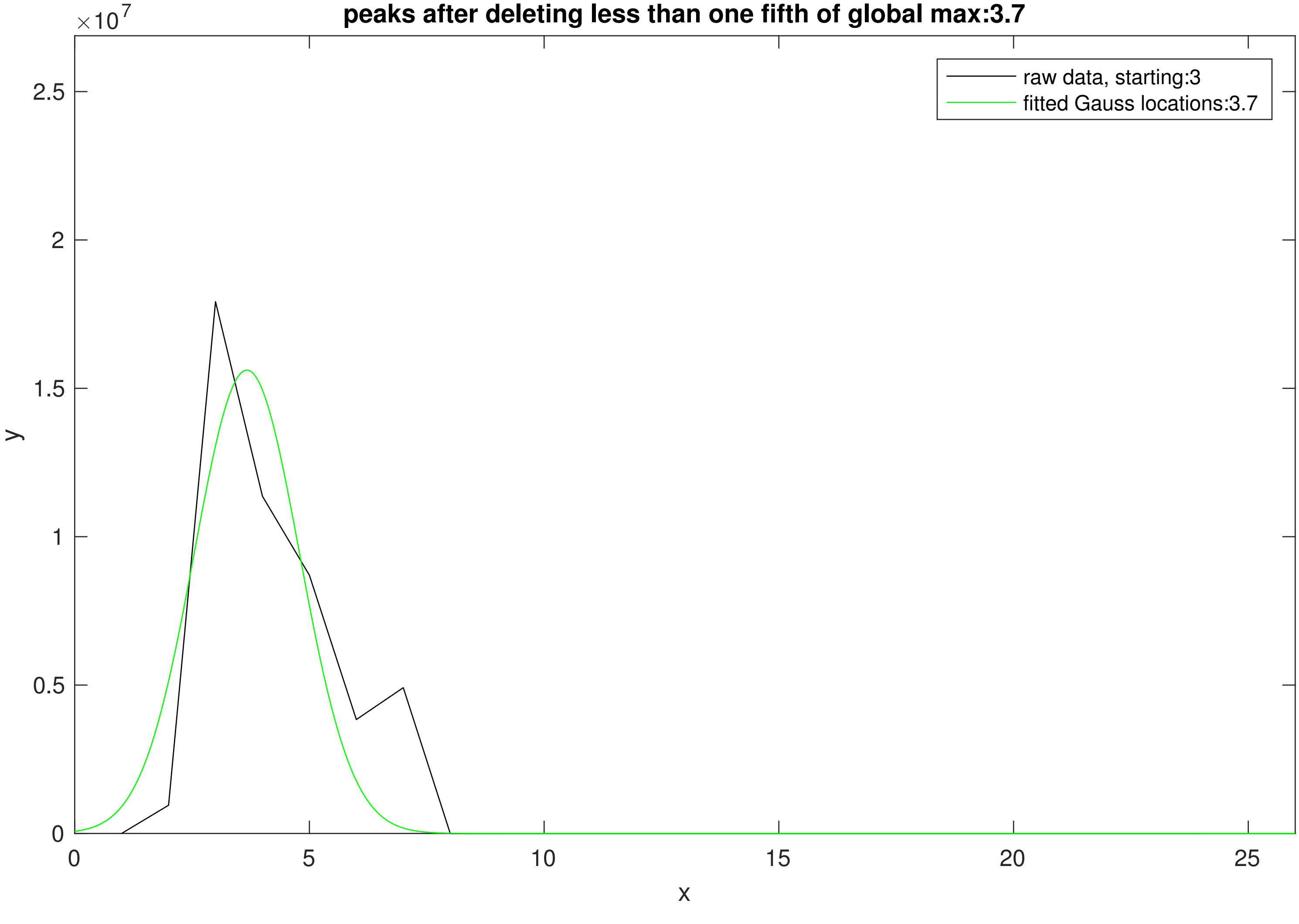

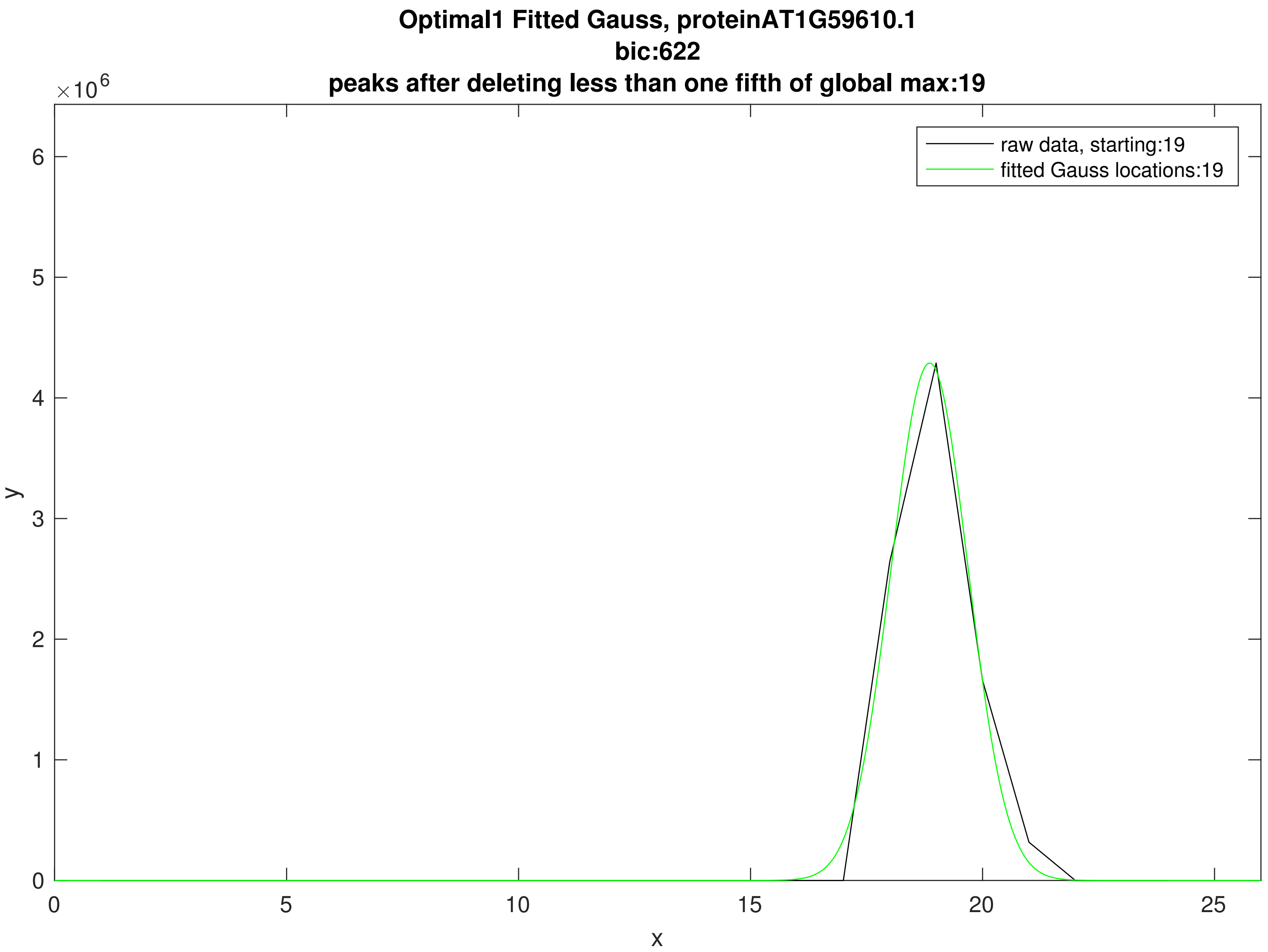

Optimal1 Fitted Gauss, proteinAT1G59610.1
bic:622
peaks after deleting less than one fifth of global max:19
×10^6
6
5
4
3
2
1
0
y
0
5
10
15
20
25
x
raw data, starting:19
fitted Gauss locations:19

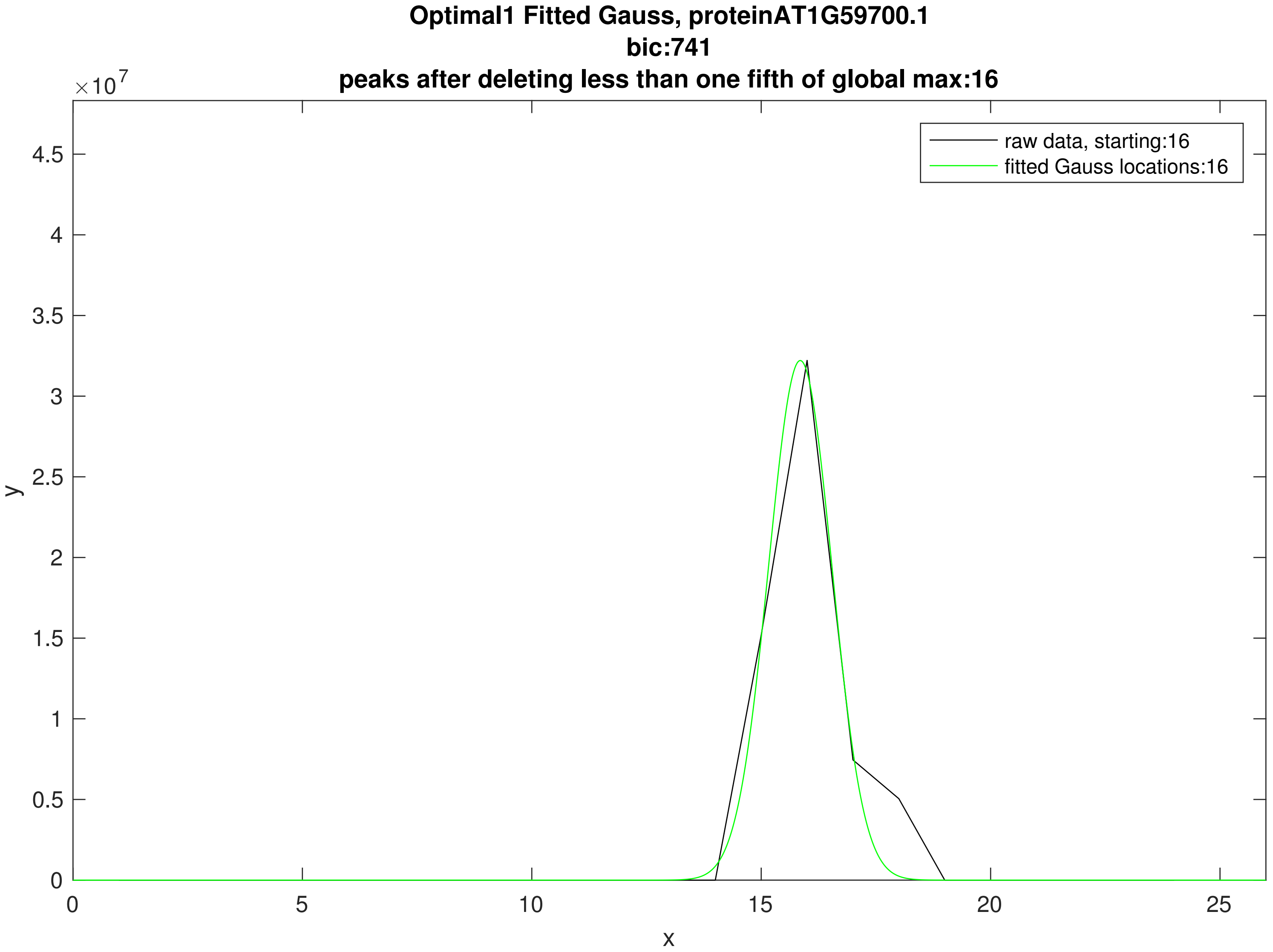
Optimal1 Fitted Gauss, proteinAT1G59700.1
bic:741
peaks after deleting less than one fifth of global max:16
×10^7
raw data, starting:16
fitted Gauss locations:16
y
x
0
0.5
1
1.5
2
2.5
3
3.5
4
4.5
0
5
10
15
20
25

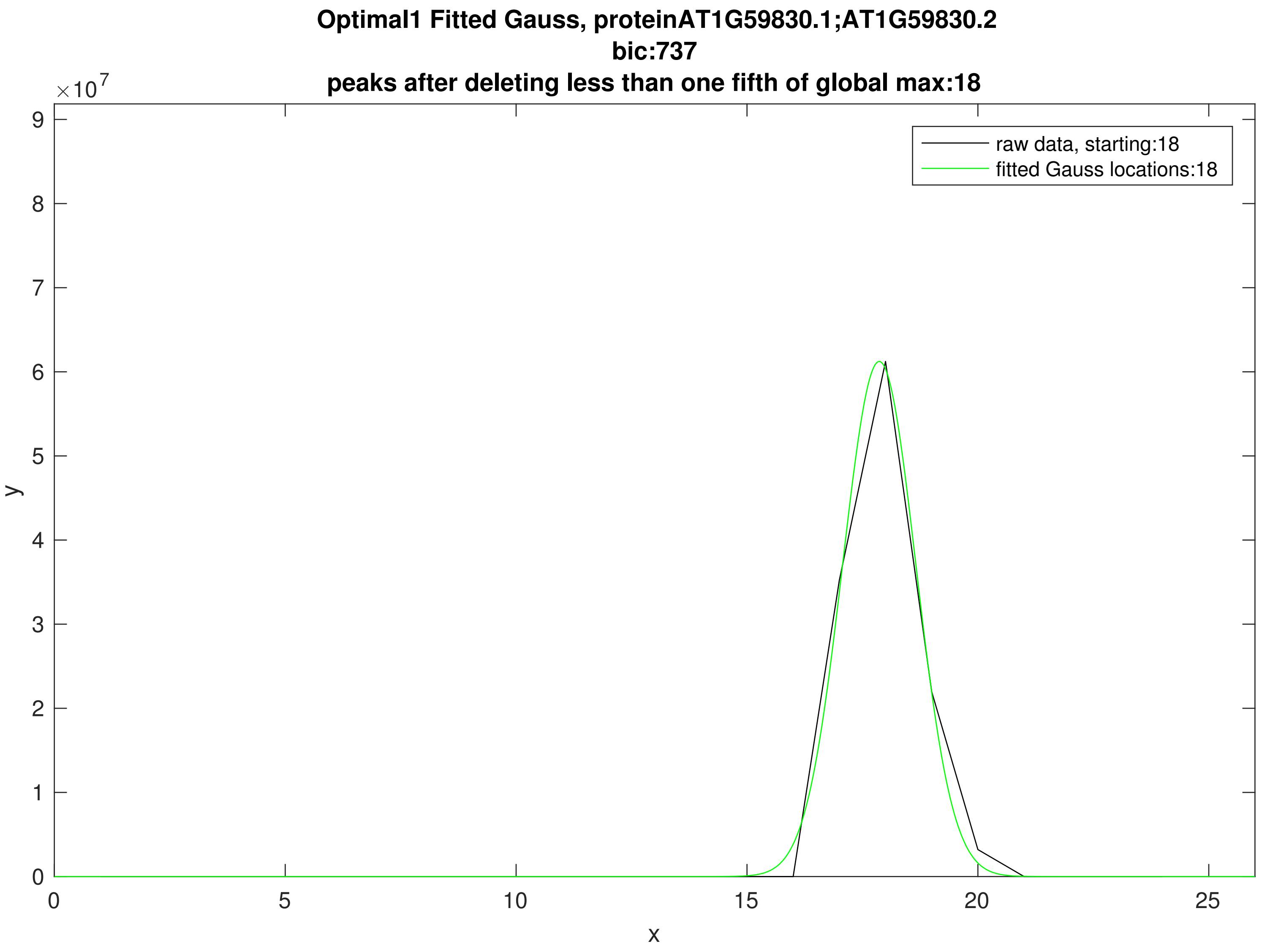
Optimal1 Fitted Gauss, proteinAT1G59830.1;AT1G59830.2
bic:737
peaks after deleting less than one fifth of global max:18
×10^7
raw data, starting:18
fitted Gauss locations:18
y
x
0
1
2
3
4
5
6
7
8
9
0
5
10
15
20
25

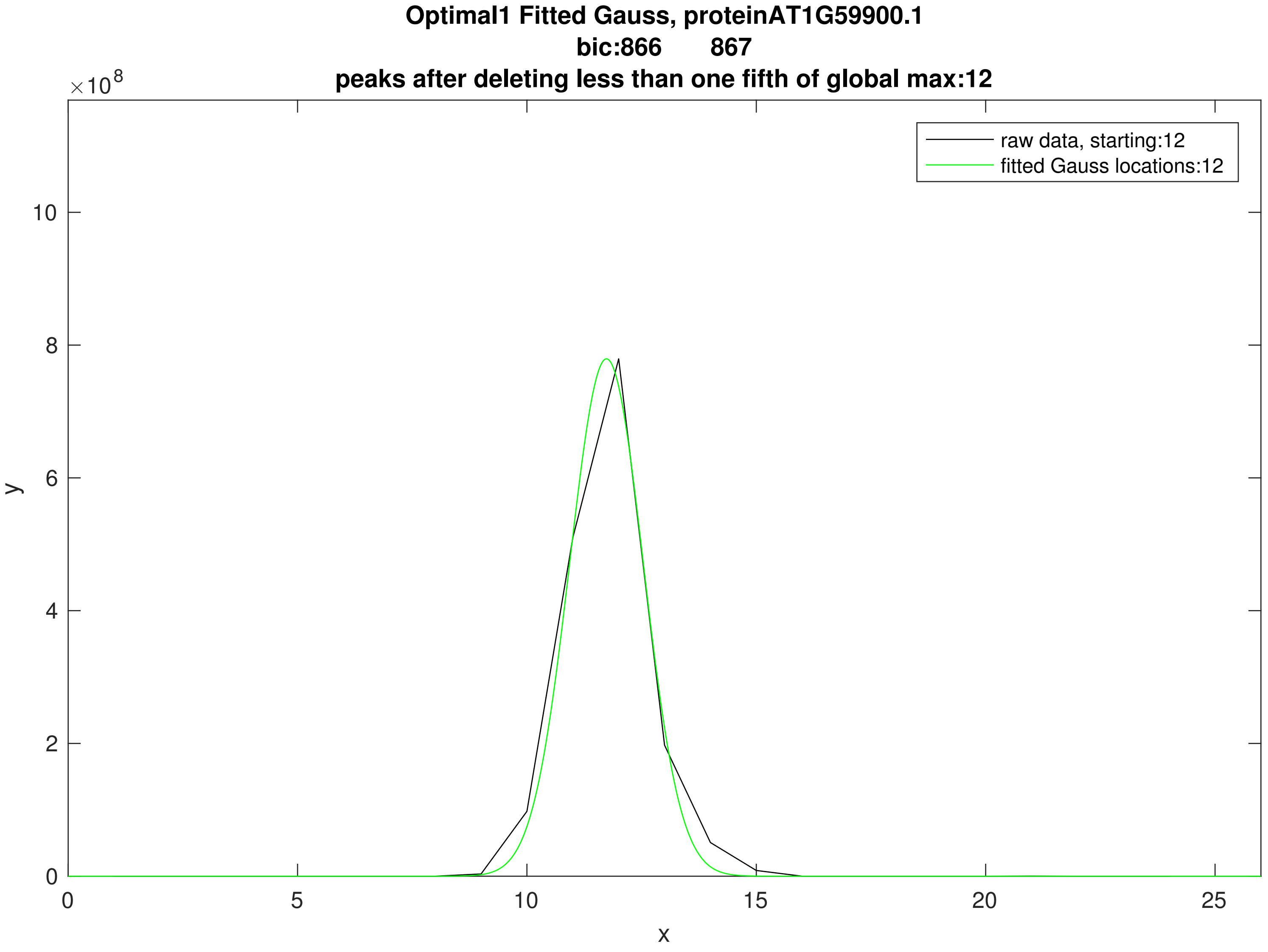

Optimal1 Fitted Gauss, proteinAT1G59900.1
bic:866 867
peaks after deleting less than one fifth of global max:12
×10^8
raw data, starting:12
fitted Gauss locations:12
10
8
6
4
2
0
y
0
5
10
15
20
25
x

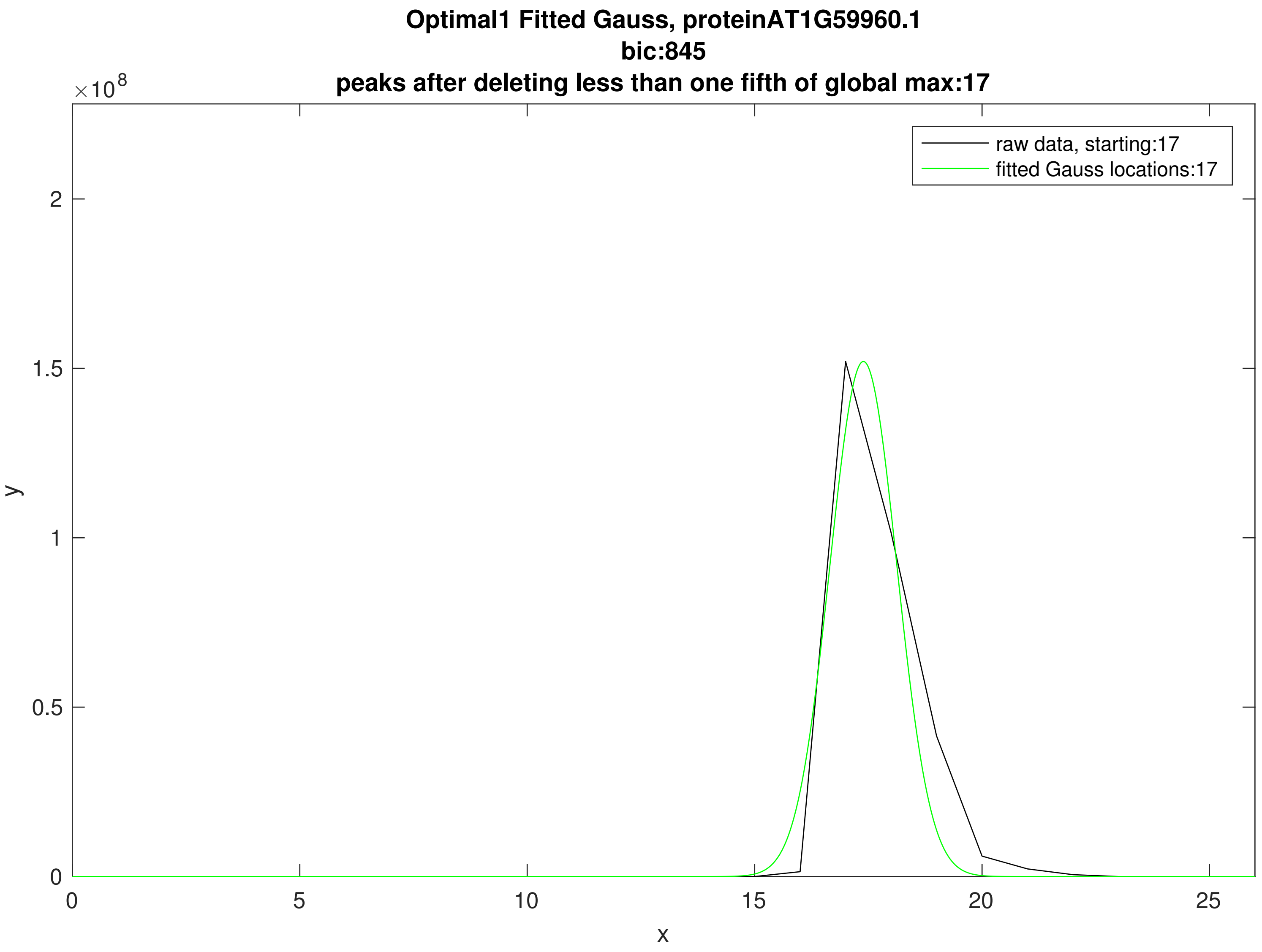
Optimal1 Fitted Gauss, proteinAT1G59960.1
bic:845
peaks after deleting less than one fifth of global max:17
×10^8
raw data, starting:17
fitted Gauss locations:17
y
x
0
0.5
1
1.5
2
0
5
10
15
20
25

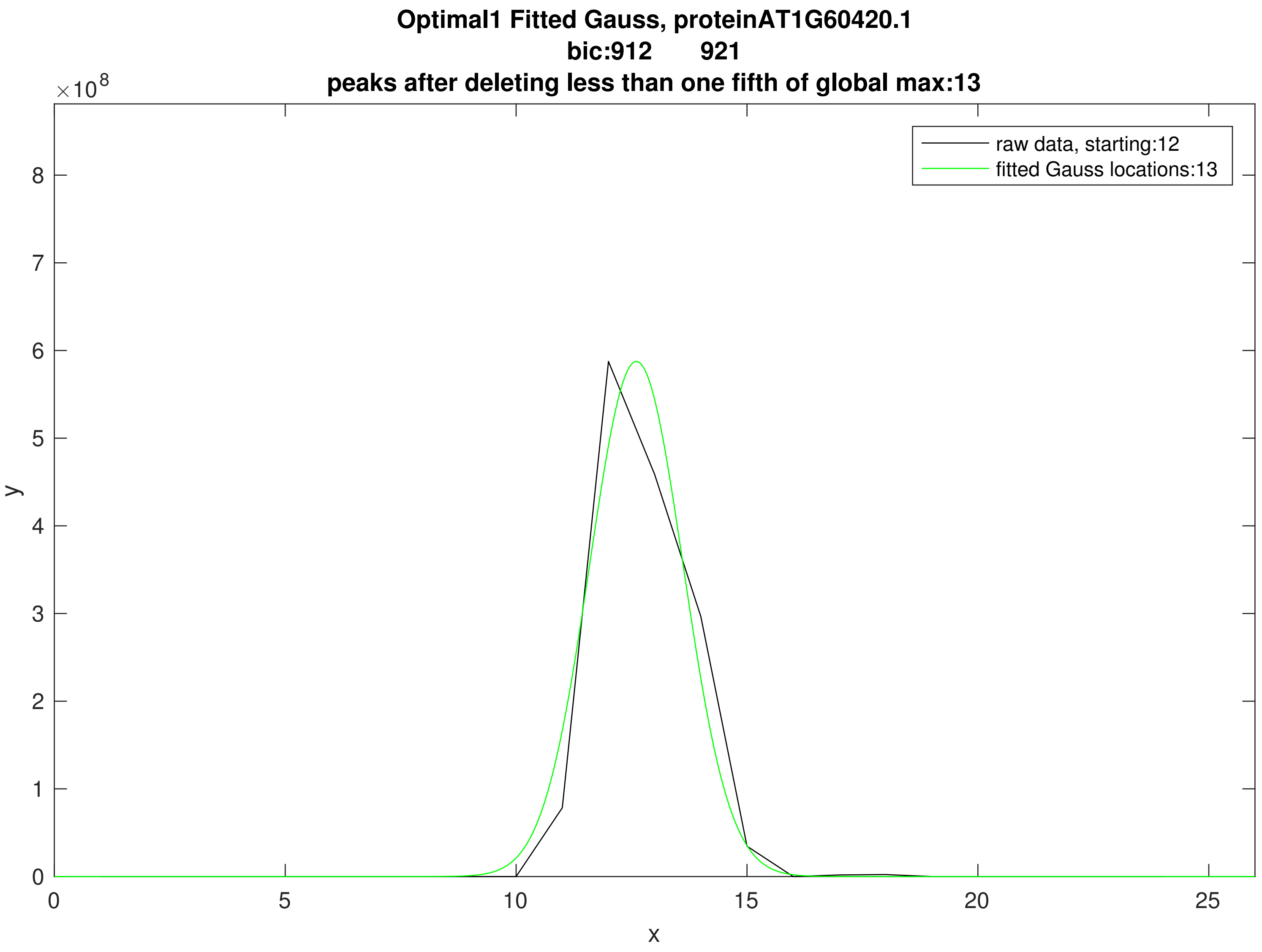
Optimal1 Fitted Gauss, proteinAT1G60420.1
bic:912 921
peaks after deleting less than one fifth of global max:13
×10^8
raw data, starting:12
fitted Gauss locations:13
y
x
0
1
2
3
4
5
6
7
8
0
5
10
15
20
25

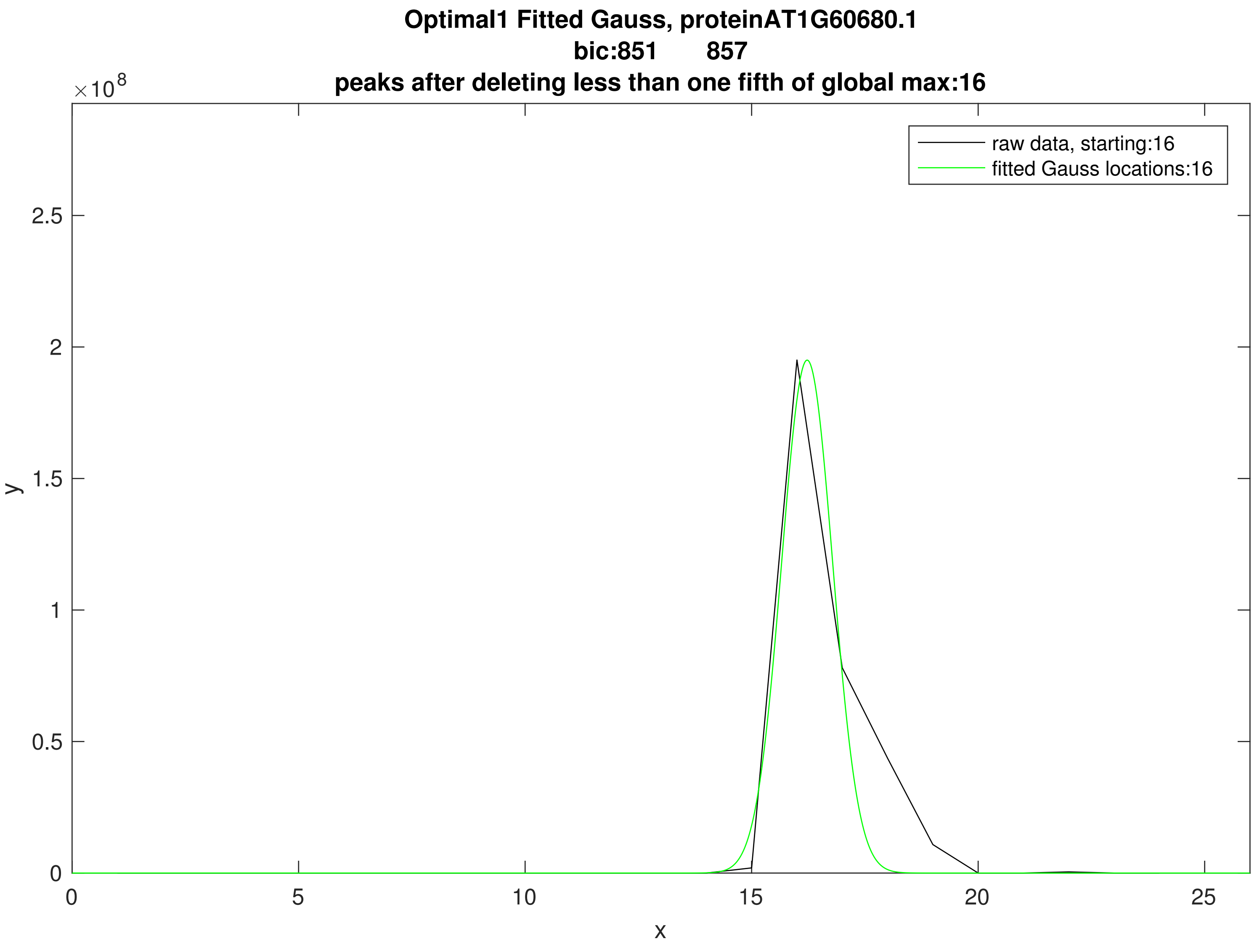

Optimal1 Fitted Gauss, proteinAT1G60680.1
bic:851 857
peaks after deleting less than one fifth of global max:16
×10^8
raw data, starting:16
fitted Gauss locations:16
y
x
0
0.5
1
1.5
2
2.5
0
5
10
15
20
25

**Optimal1 Fitted Gauss, proteinAT1G60710.1**
**bic:932 941**
**peaks after deleting less than one fifth of global max:16**

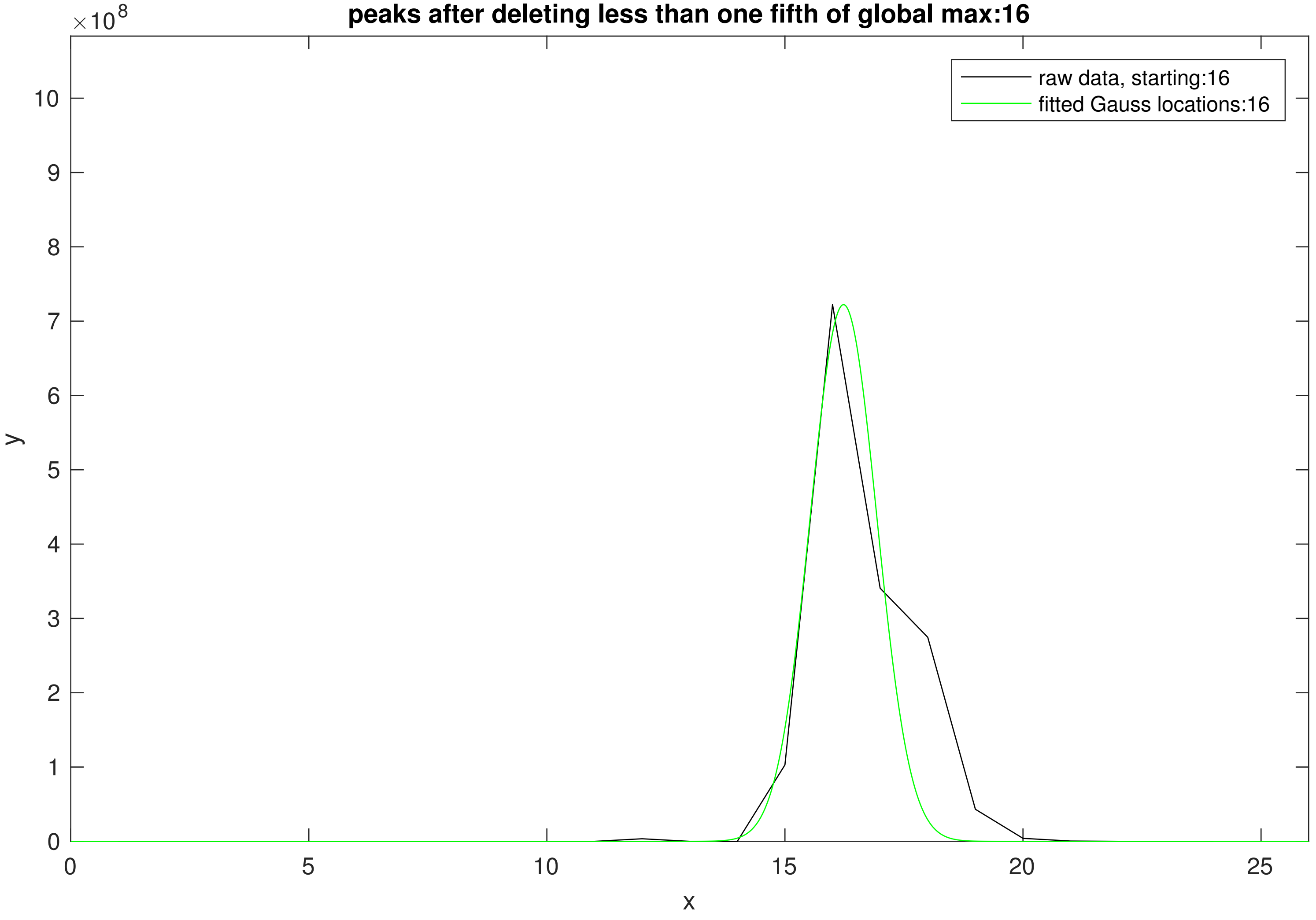

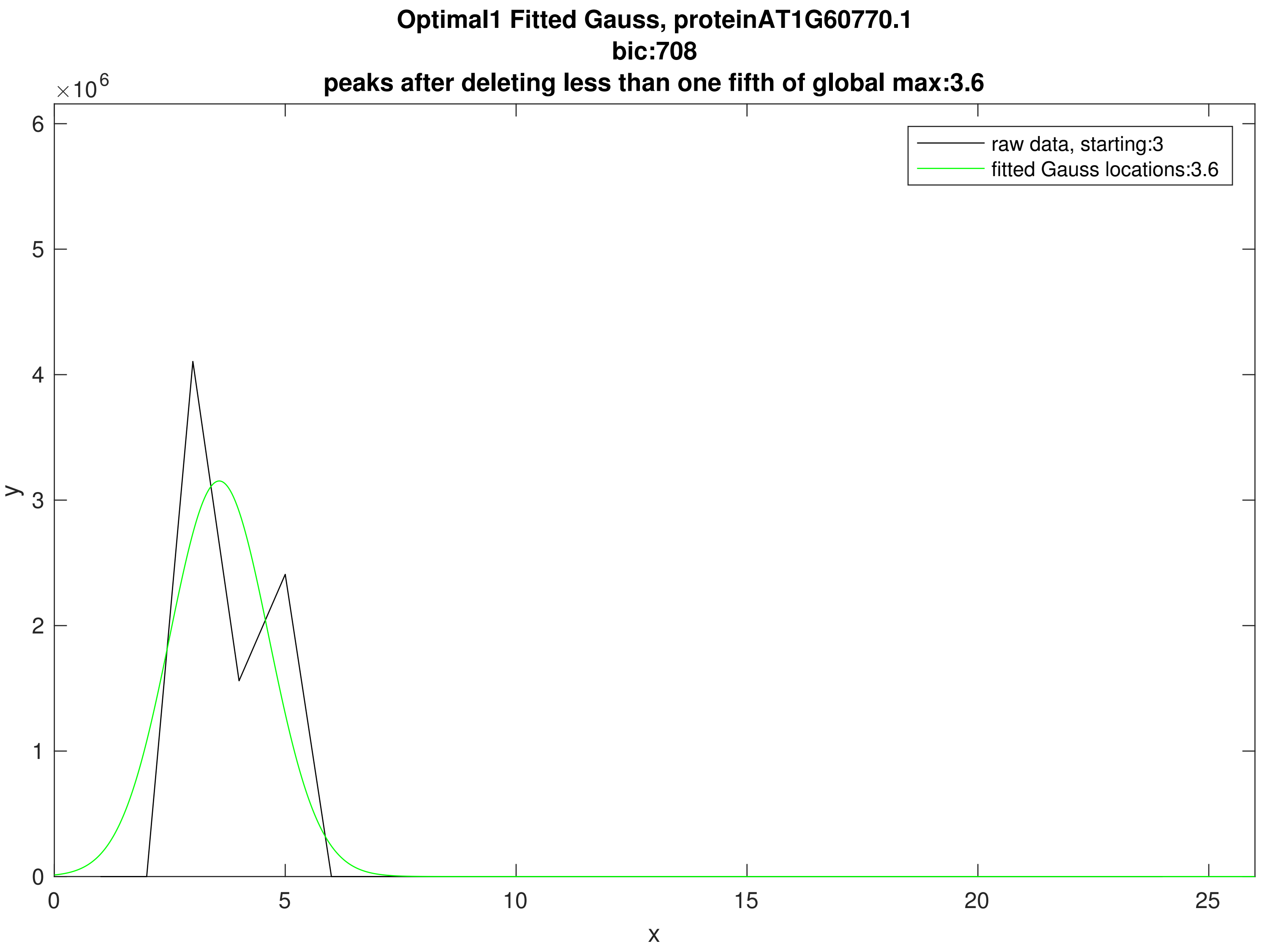
Optimal1 Fitted Gauss, proteinAT1G60770.1
bic:708
peaks after deleting less than one fifth of global max:3.6
×10^6
raw data, starting:3
fitted Gauss locations:3.6
y
x
0
1
2
3
4
5
6
0
5
10
15
20
25

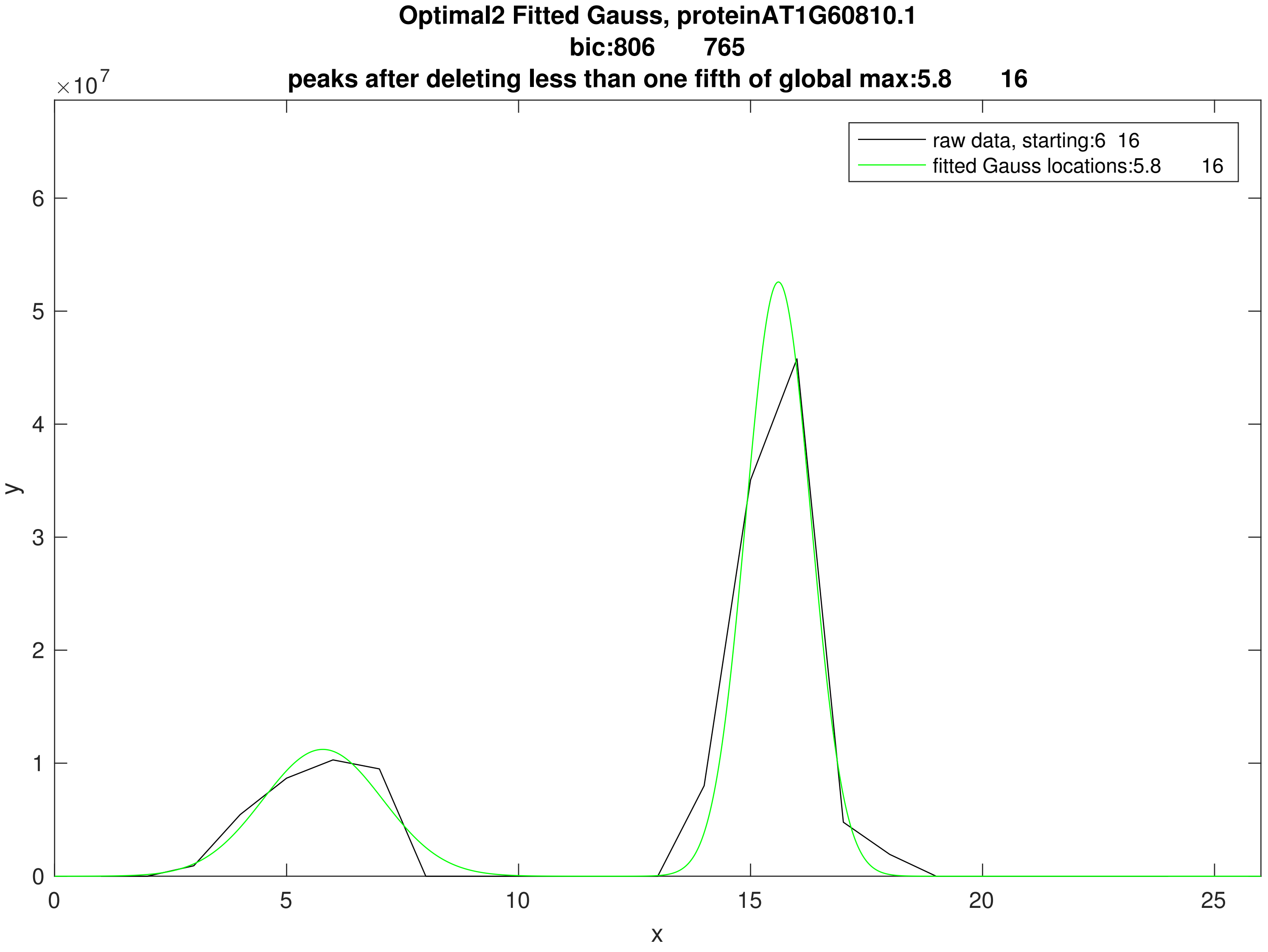
Optimal2 Fitted Gauss, proteinAT1G60810.1
bic:806 765
peaks after deleting less than one fifth of global max:5.8 16
×10^7
raw data, starting:6 16
fitted Gauss locations:5.8 16
y
x
0
1
2
3
4
5
6
0
5
10
15
20
25

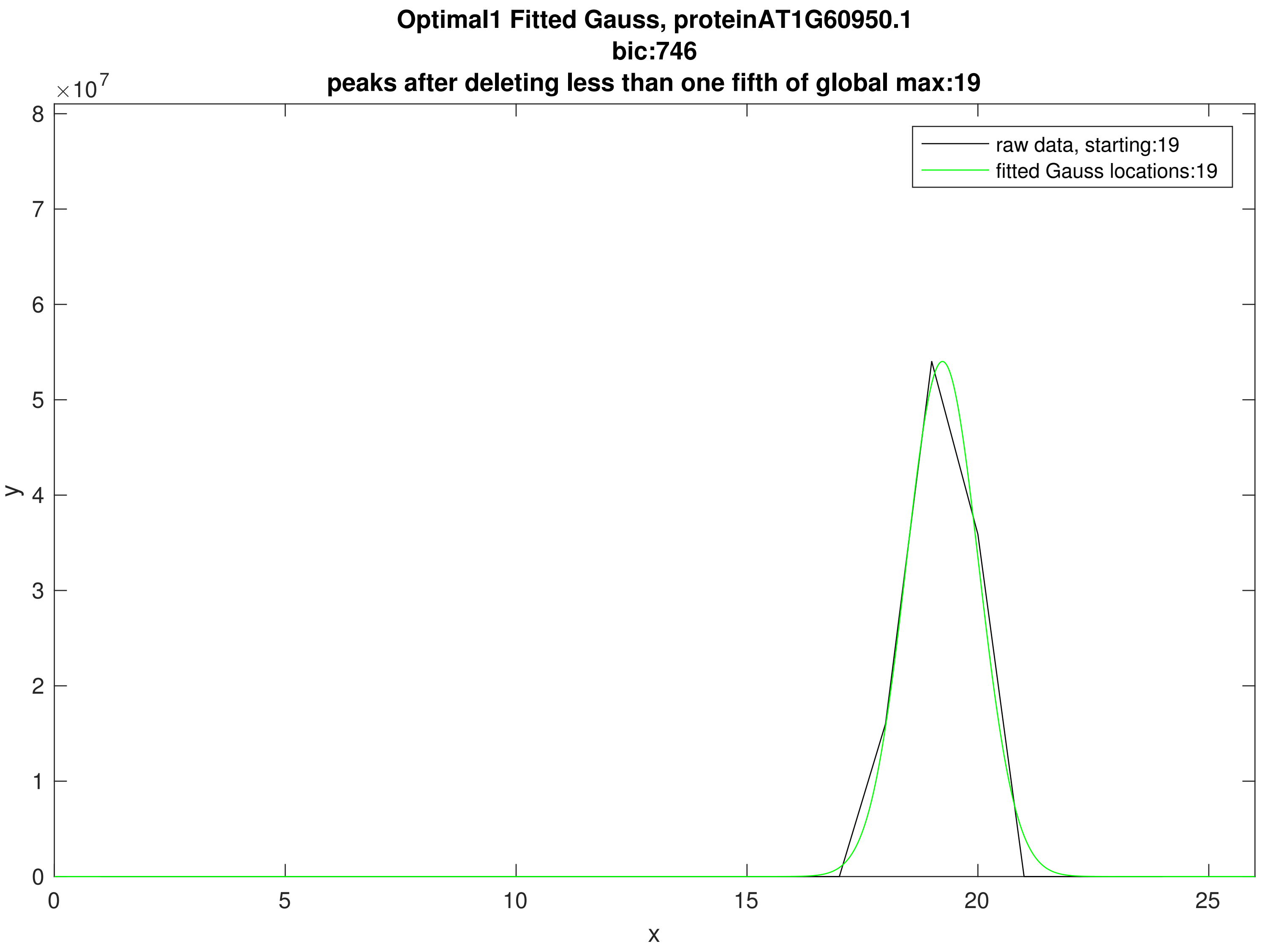
Optimal1 Fitted Gauss, proteinAT1G60950.1
bic:746
peaks after deleting less than one fifth of global max:19
×10^7
raw data, starting:19
fitted Gauss locations:19
y
x
0
1
2
3
4
5
6
7
8
0
5
10
15
20
25

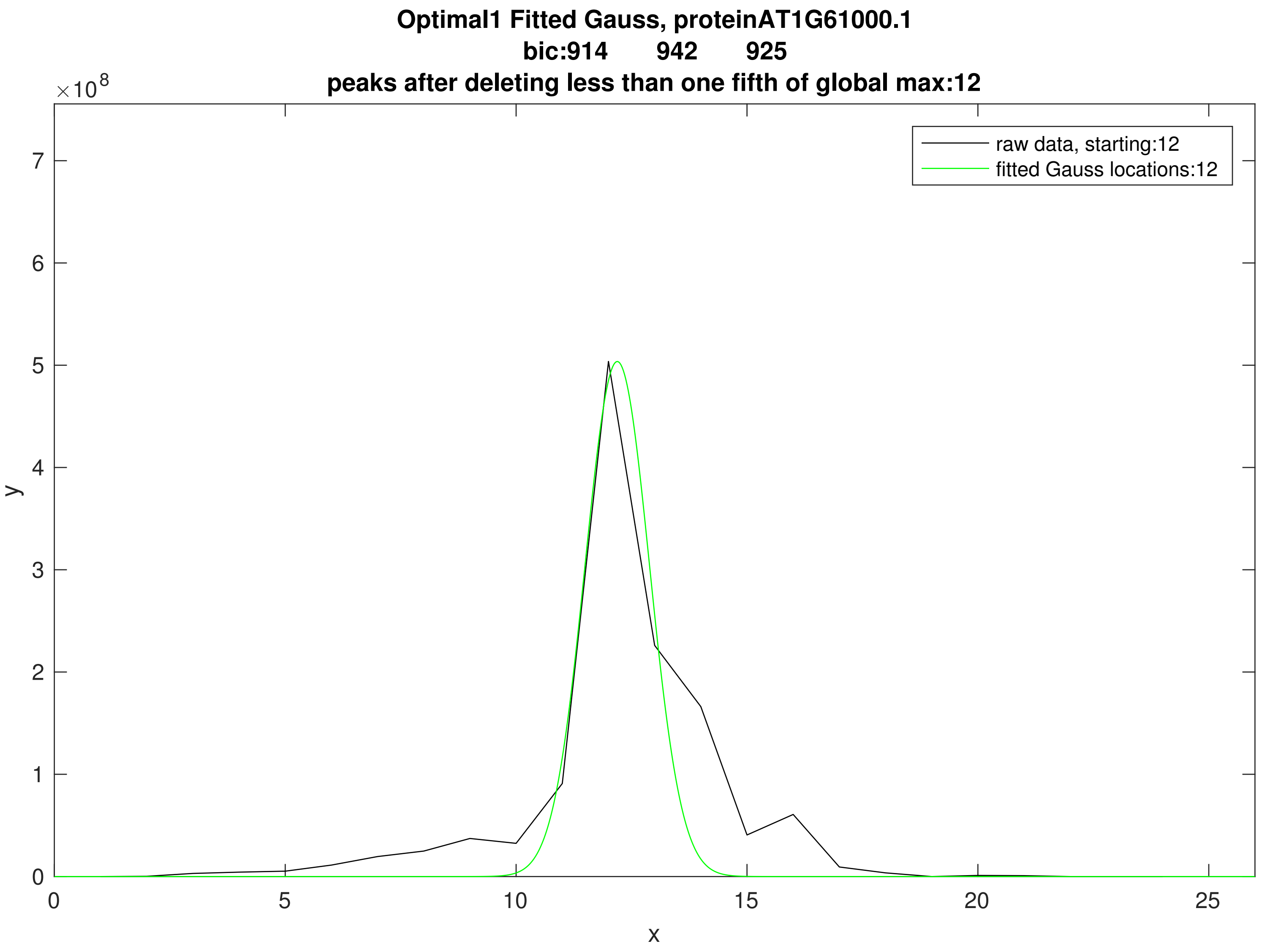

Optimal1 Fitted Gauss, proteinAT1G61000.1
bic:914 942 925
peaks after deleting less than one fifth of global max:12
×10^8
raw data, starting:12
fitted Gauss locations:12
y
x
0
1
2
3
4
5
6
7
0
5
10
15
20
25

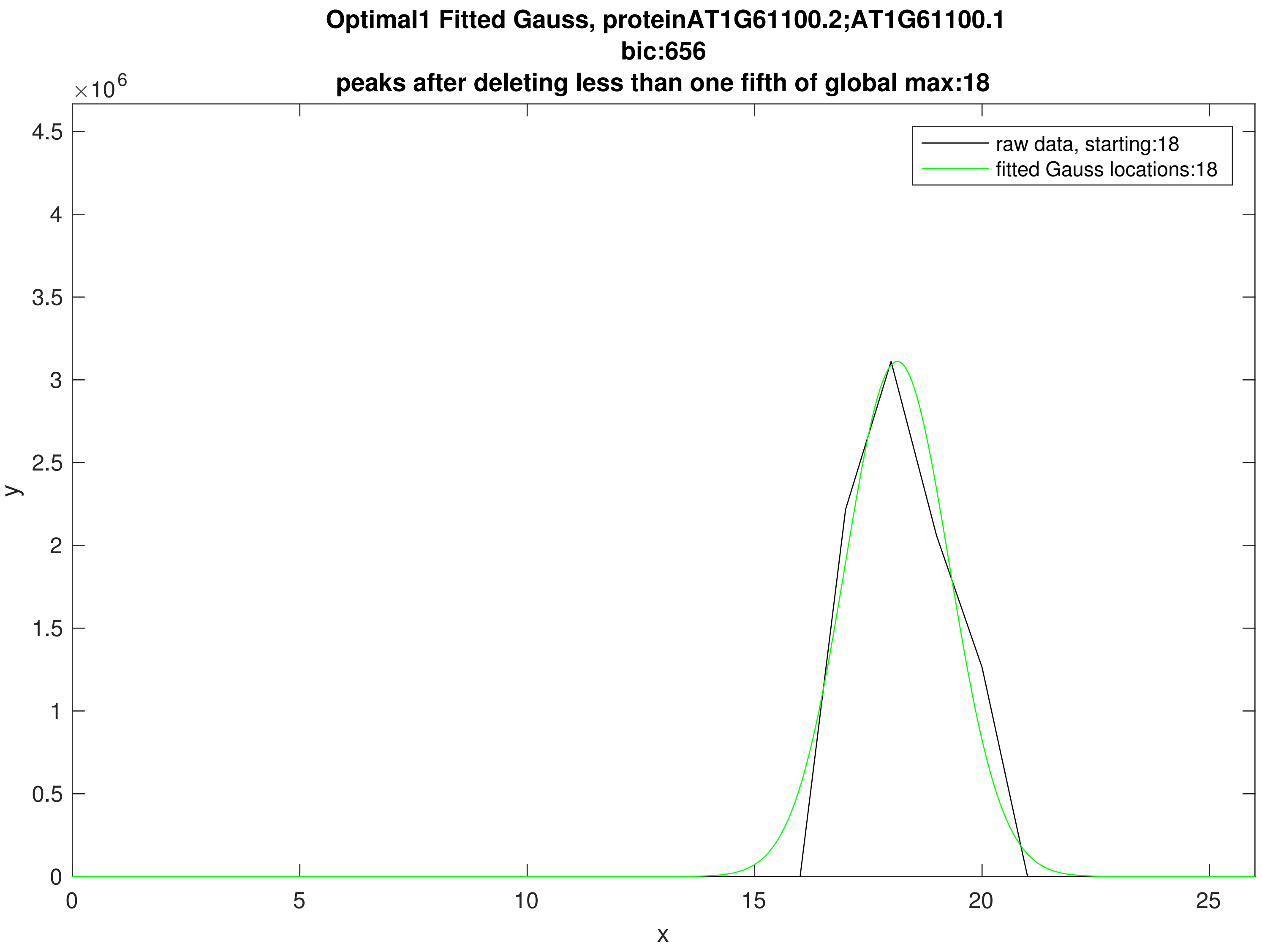

Optimal1 Fitted Gauss, proteinAT1G61100.2;AT1G61100.1
bic:656
peaks after deleting less than one fifth of global max:18
×10^6
raw data, starting:18
fitted Gauss locations:18
y
x
0
0.5
1
1.5
2
2.5
3
3.5
4
4.5
0
5
10
15
20
25

**Optimal1 Fitted Gauss, proteinAT1G61150.5;AT1G61150.6;AT1G61150.4;AT1G61150.2;AT1G61150.1;AT1G61150.7;AT1G**

**bic:813 821 830**

**peaks after deleting less than one fifth of global max:16**

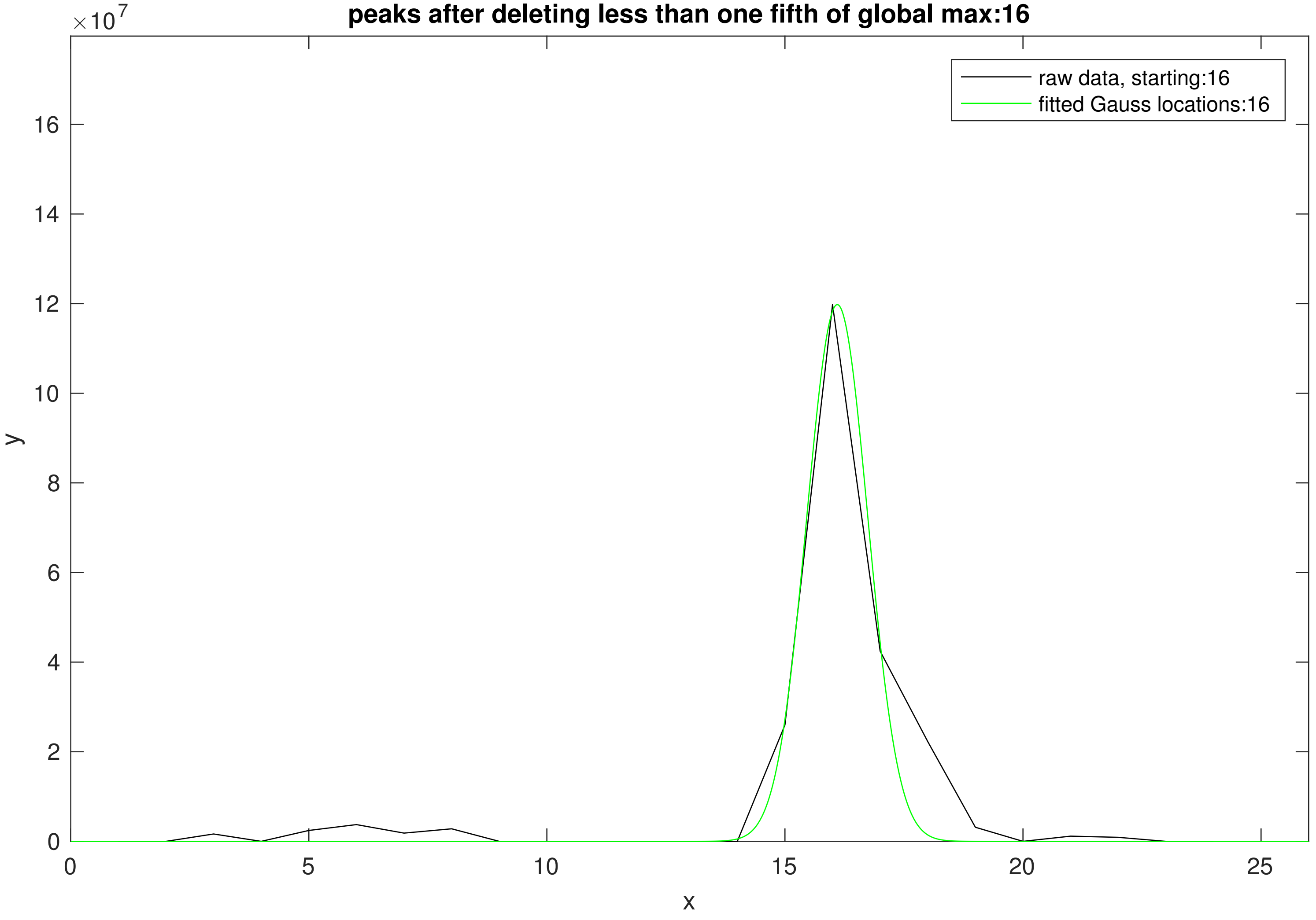

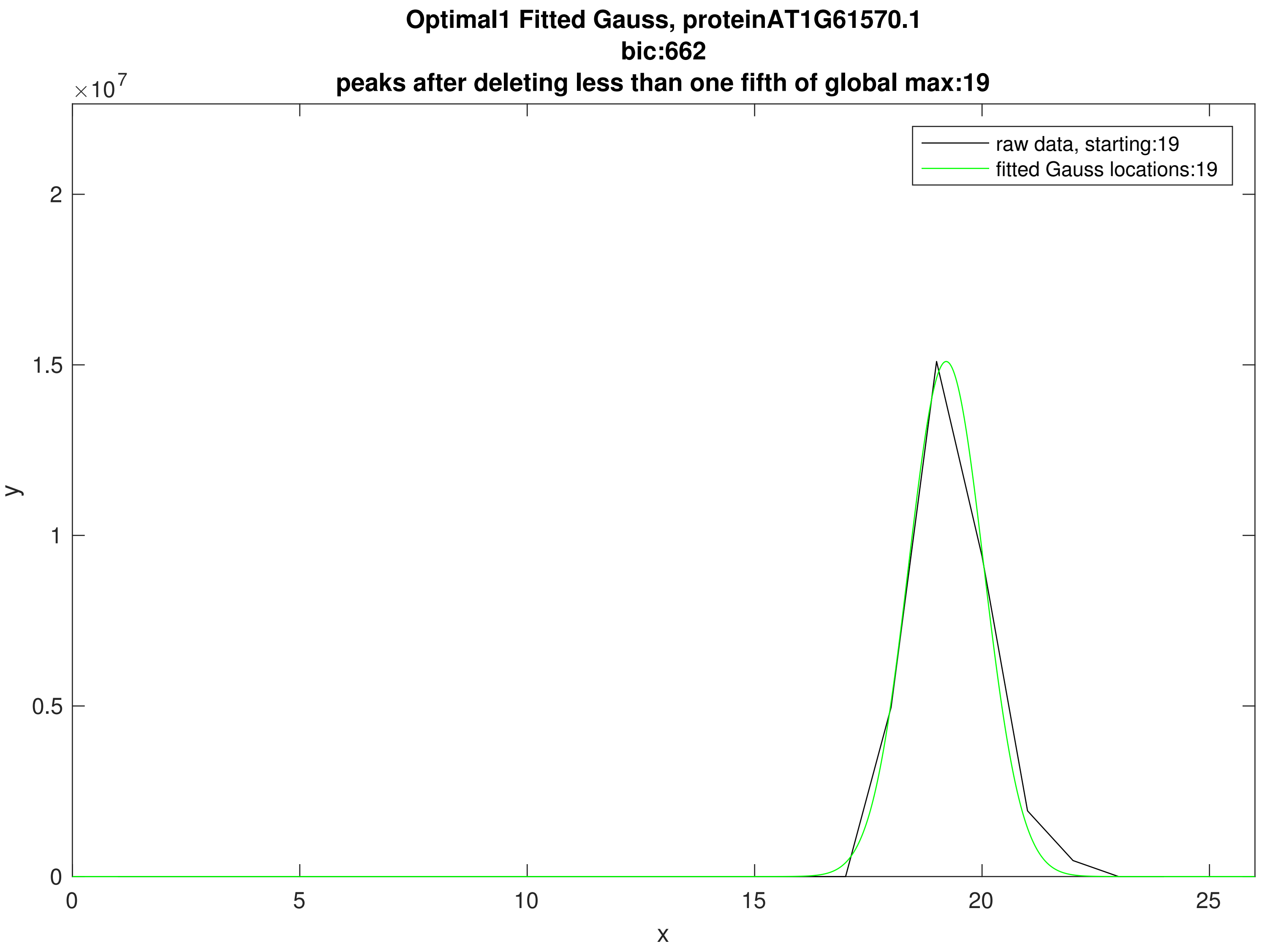
Optimal1 Fitted Gauss, proteinAT1G61570.1
bic:662
peaks after deleting less than one fifth of global max:19
×10^7
raw data, starting:19
fitted Gauss locations:19
y
x
0
0.5
1
1.5
2
0
5
10
15
20
25

**Optimal1 Fitted Gauss, proteinAT1G61820.1**
**bic:802**
**peaks after deleting less than one fifth of global max:16**

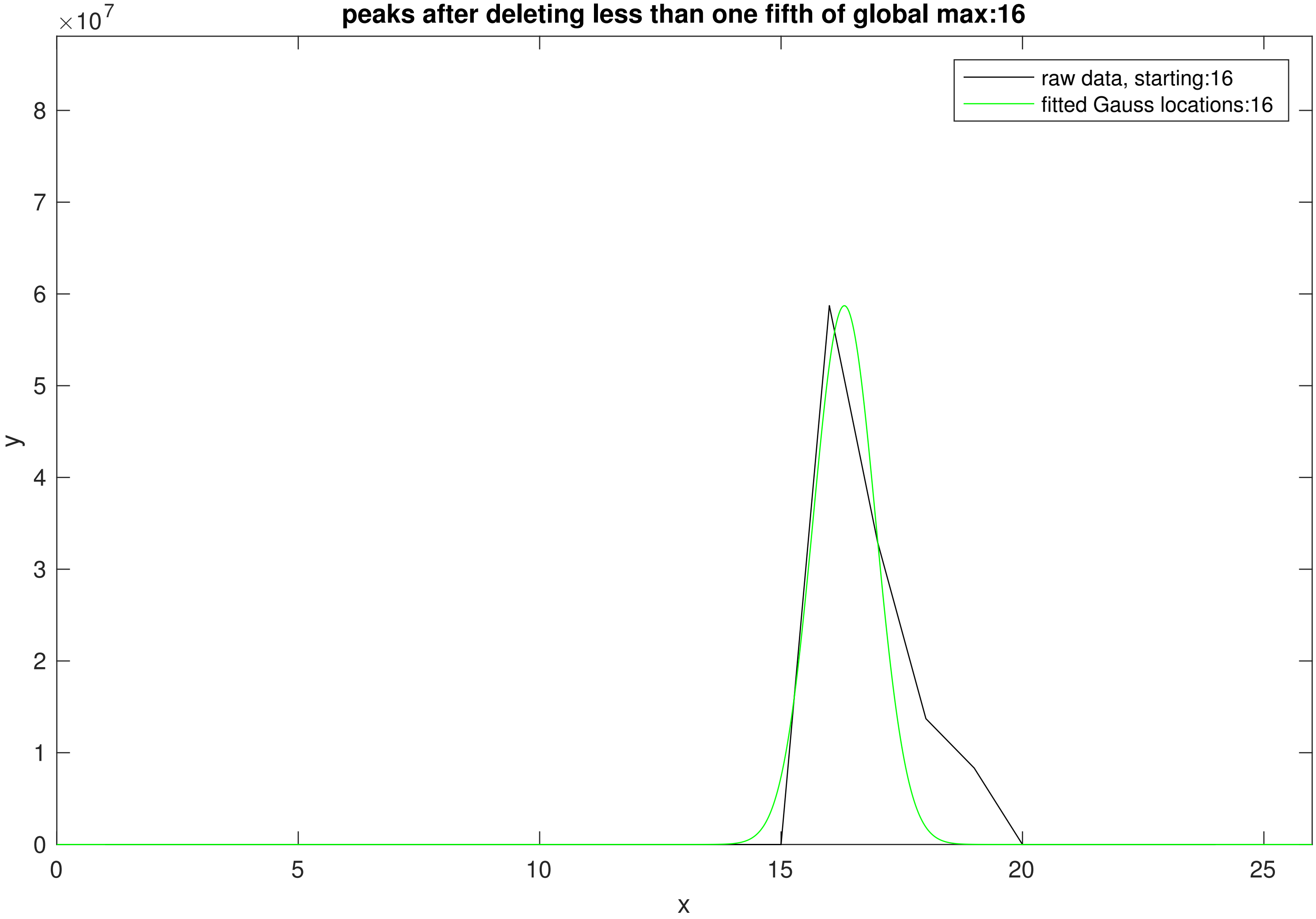

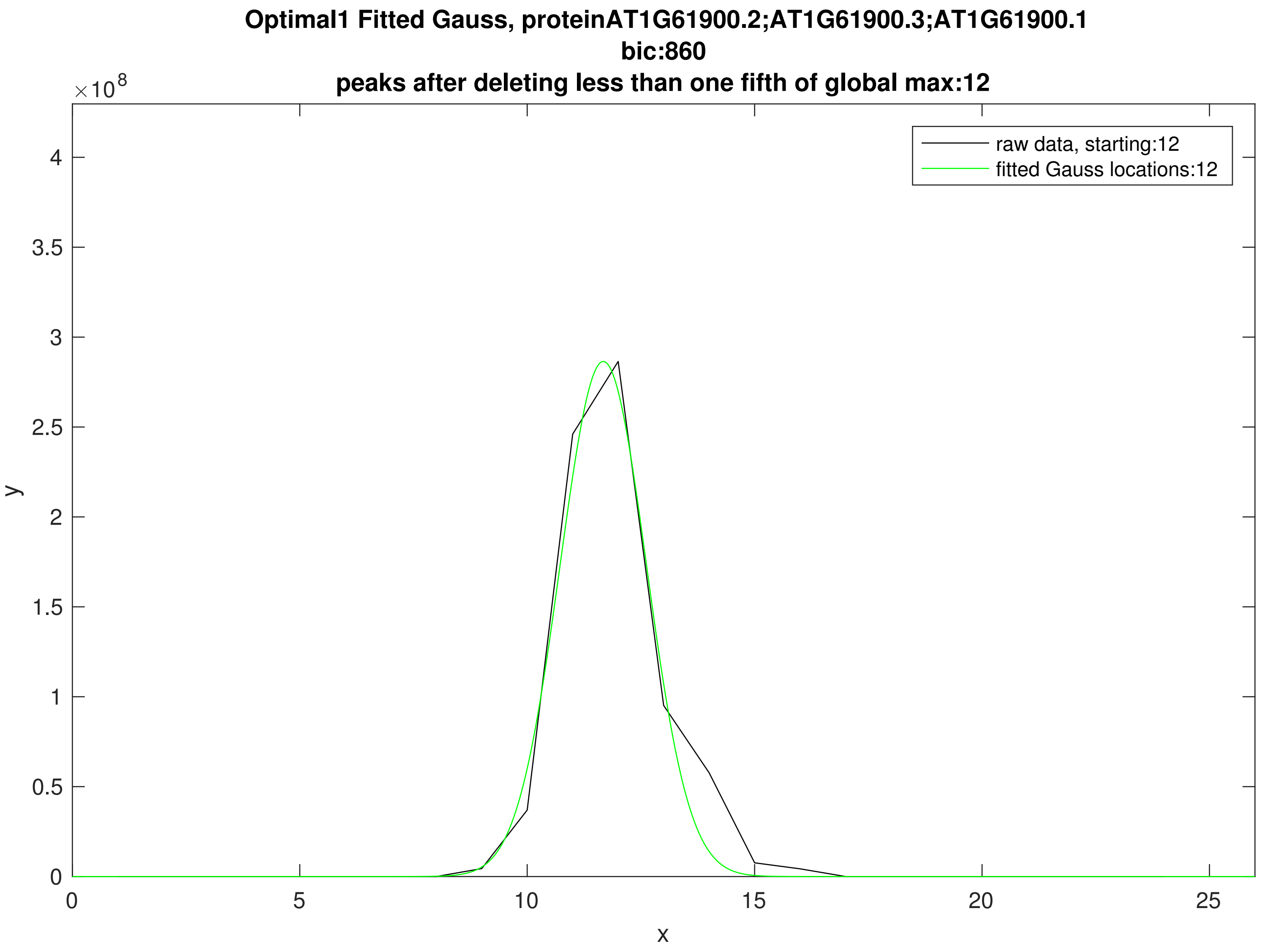
Optimal1 Fitted Gauss, proteinAT1G61900.2;AT1G61900.3;AT1G61900.1
bic:860
peaks after deleting less than one fifth of global max:12
×10^8
raw data, starting:12
fitted Gauss locations:12
y
x
0
0.5
1
1.5
2
2.5
3
3.5
4
5
10
15
20
25

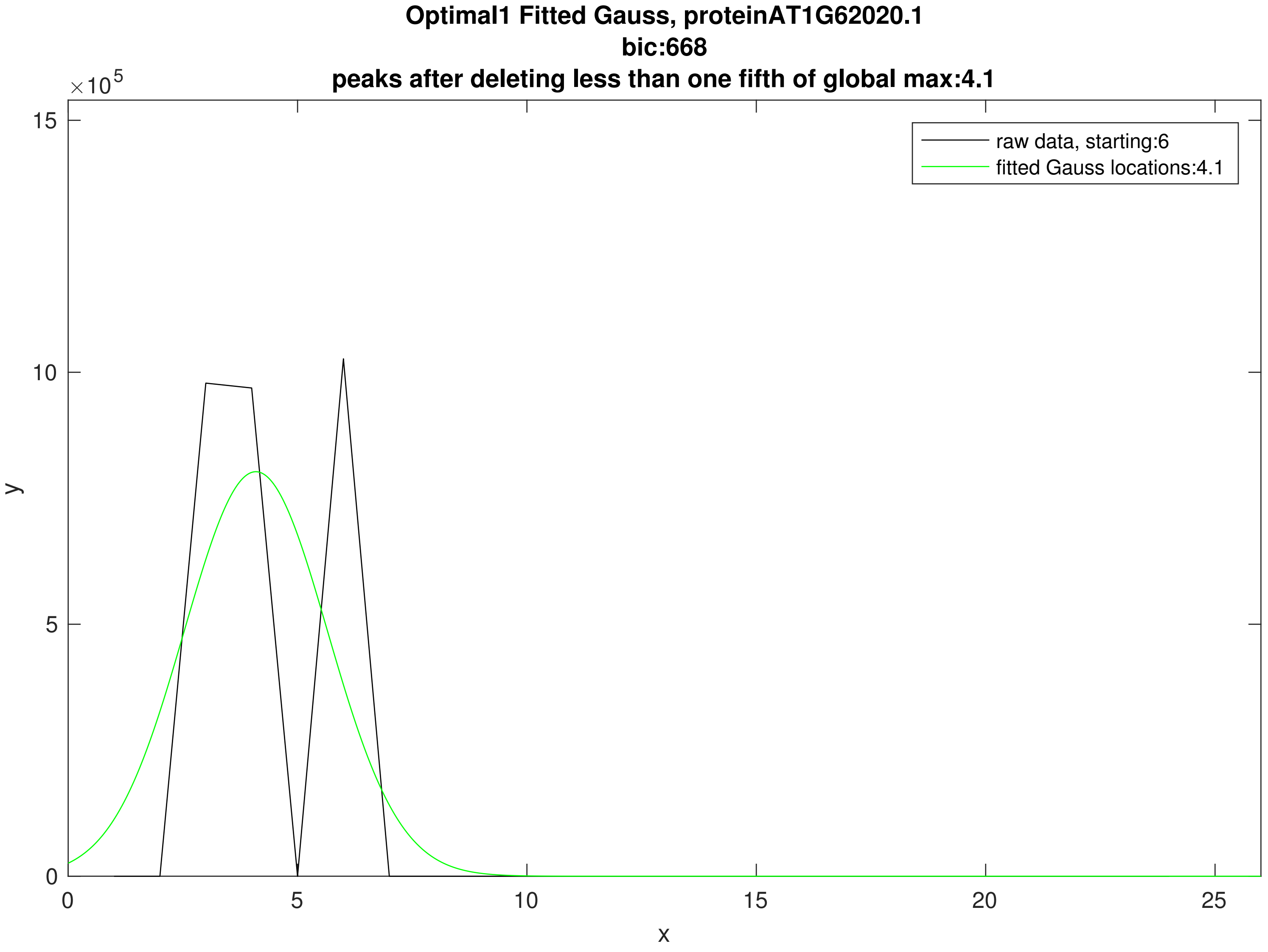

Optimal1 Fitted Gauss, proteinAT1G62020.1
bic:668
peaks after deleting less than one fifth of global max:4.1
×10^5
15
10
5
0
y
0
5
10
15
20
25
x
raw data, starting:6
fitted Gauss locations:4.1

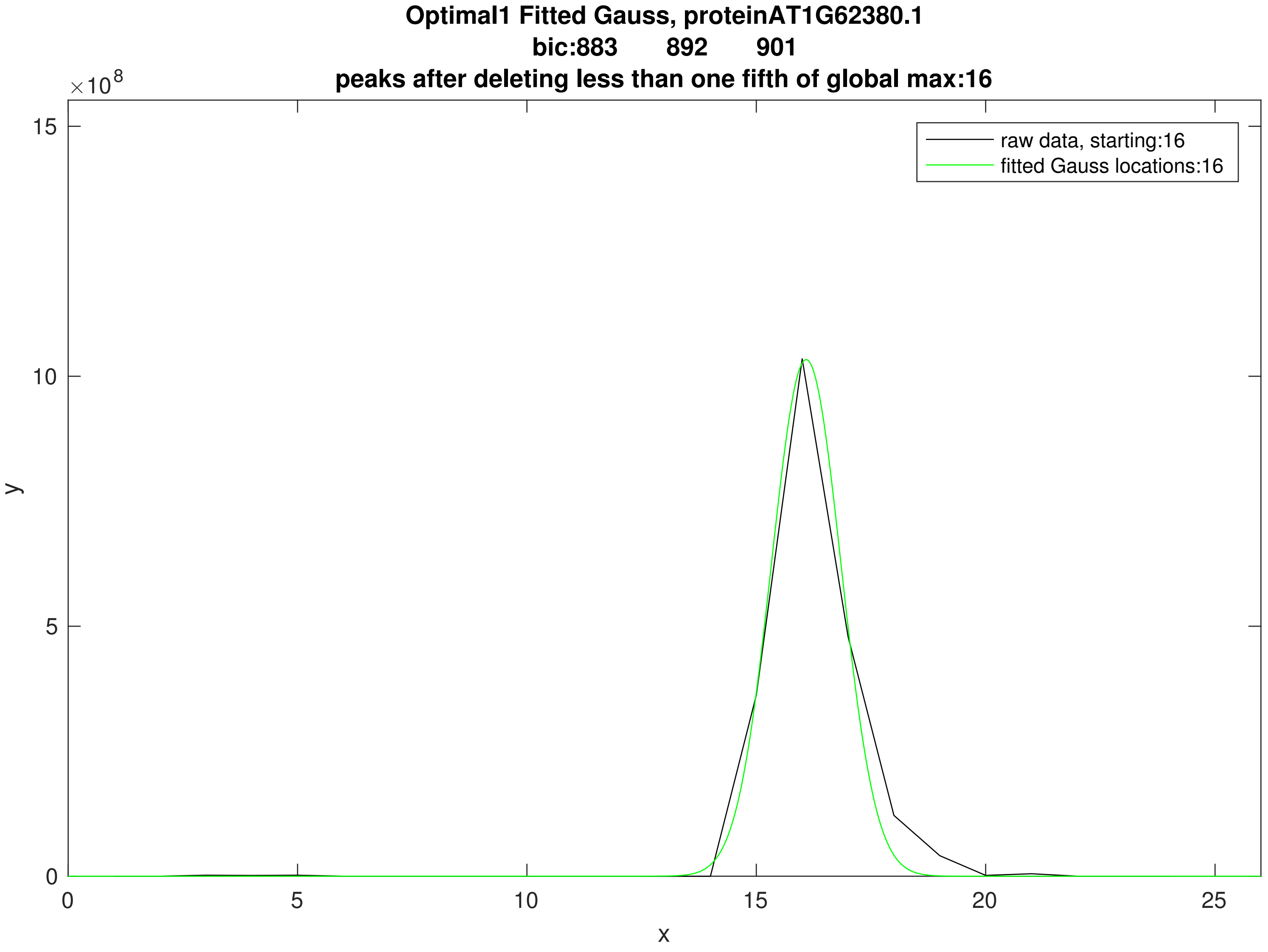

Optimal1 Fitted Gauss, proteinAT1G62380.1
bic:883 892 901
peaks after deleting less than one fifth of global max:16
×10^8
15
10
5
0
y
0
5
10
15
20
25
x
raw data, starting:16
fitted Gauss locations:16

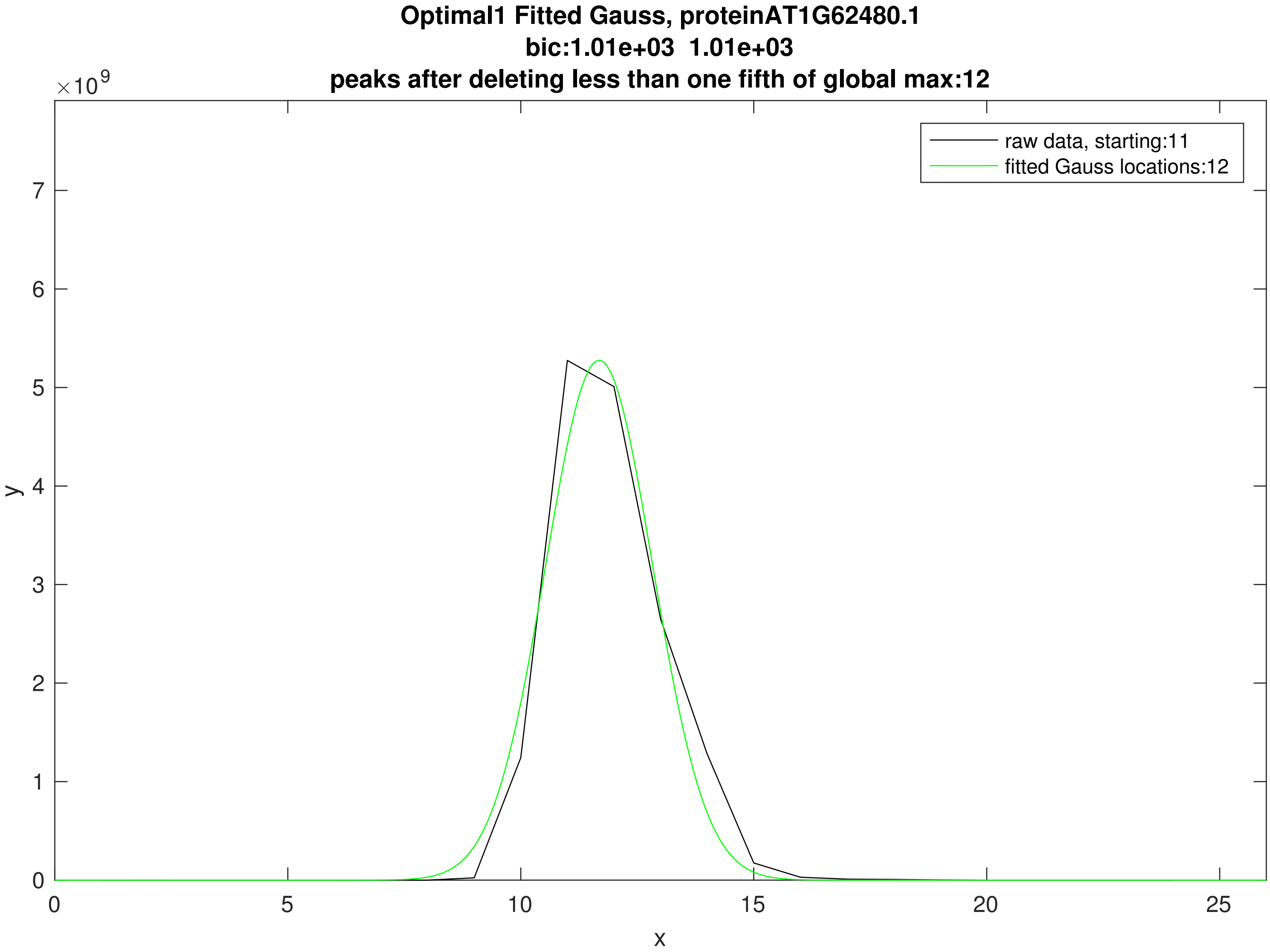
Optimal1 Fitted Gauss, proteinAT1G62480.1
bic:1.01e+03 1.01e+03
peaks after deleting less than one fifth of global max:12
×10^9
raw data, starting:11
fitted Gauss locations:12
y
x
0
1
2
3
4
5
6
7
0
5
10
15
20
25

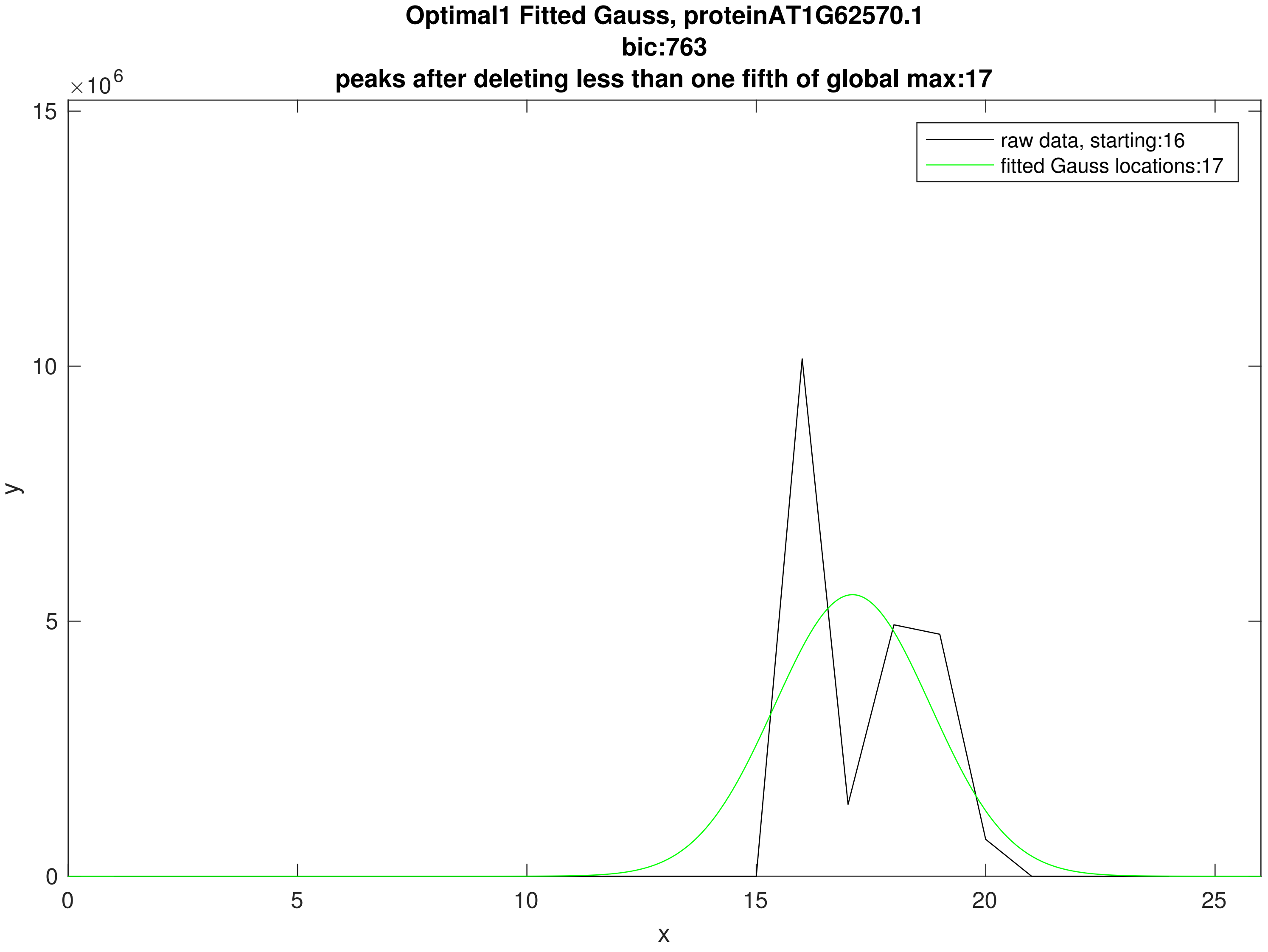
Optimal1 Fitted Gauss, proteinAT1G62570.1
bic:763
peaks after deleting less than one fifth of global max:17
×10^6
15
10
5
0
y
0
5
10
15
20
25
x
raw data, starting:16
fitted Gauss locations:17

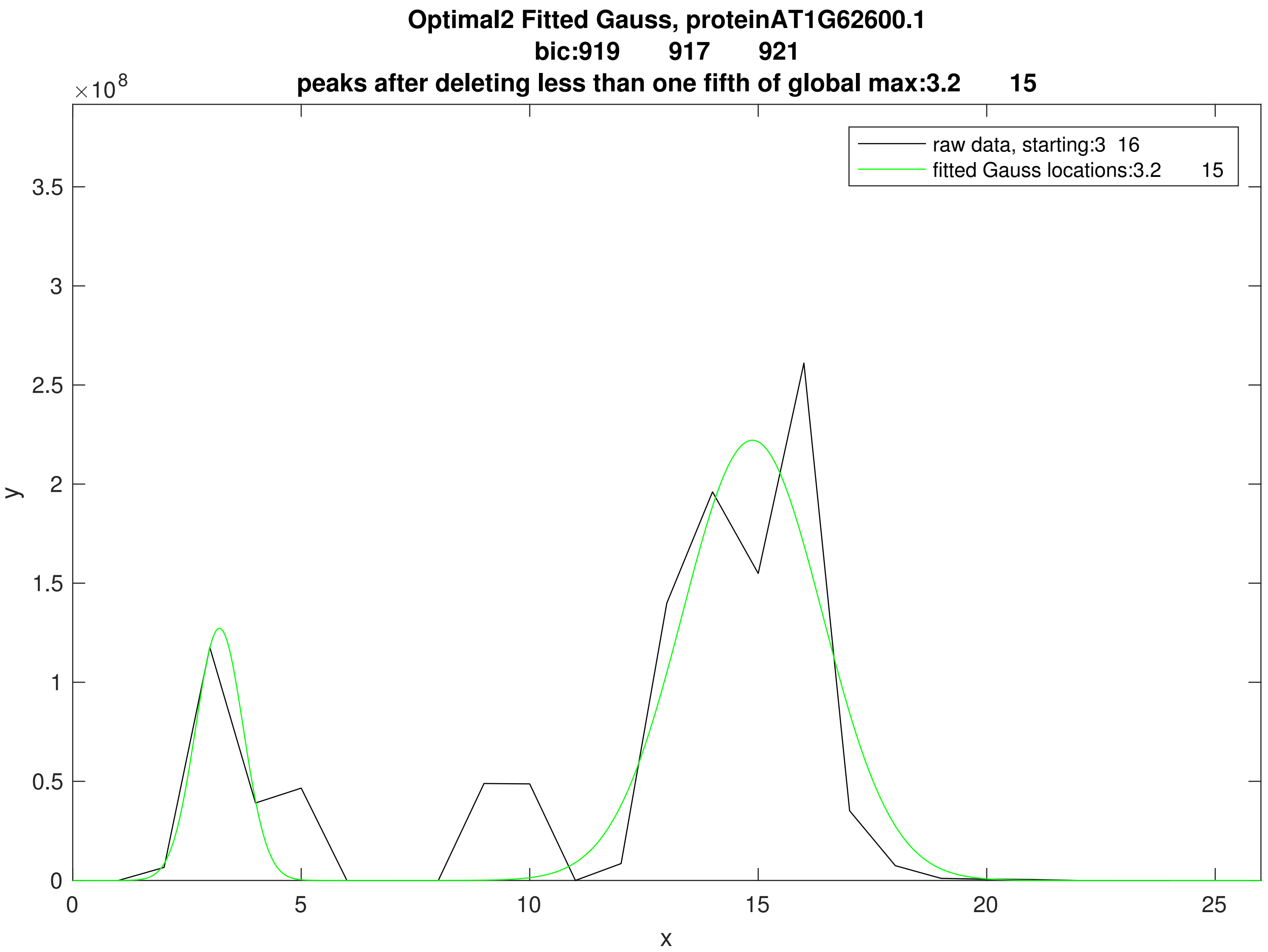

Optimal2 Fitted Gauss, proteinAT1G62600.1
bic:919 917 921
peaks after deleting less than one fifth of global max:3.2 15
×10^8
raw data, starting:3 16
fitted Gauss locations:3.2 15
3.5
3
2.5
2
1.5
1
0.5
0
y
0
5
10
15
20
25
x

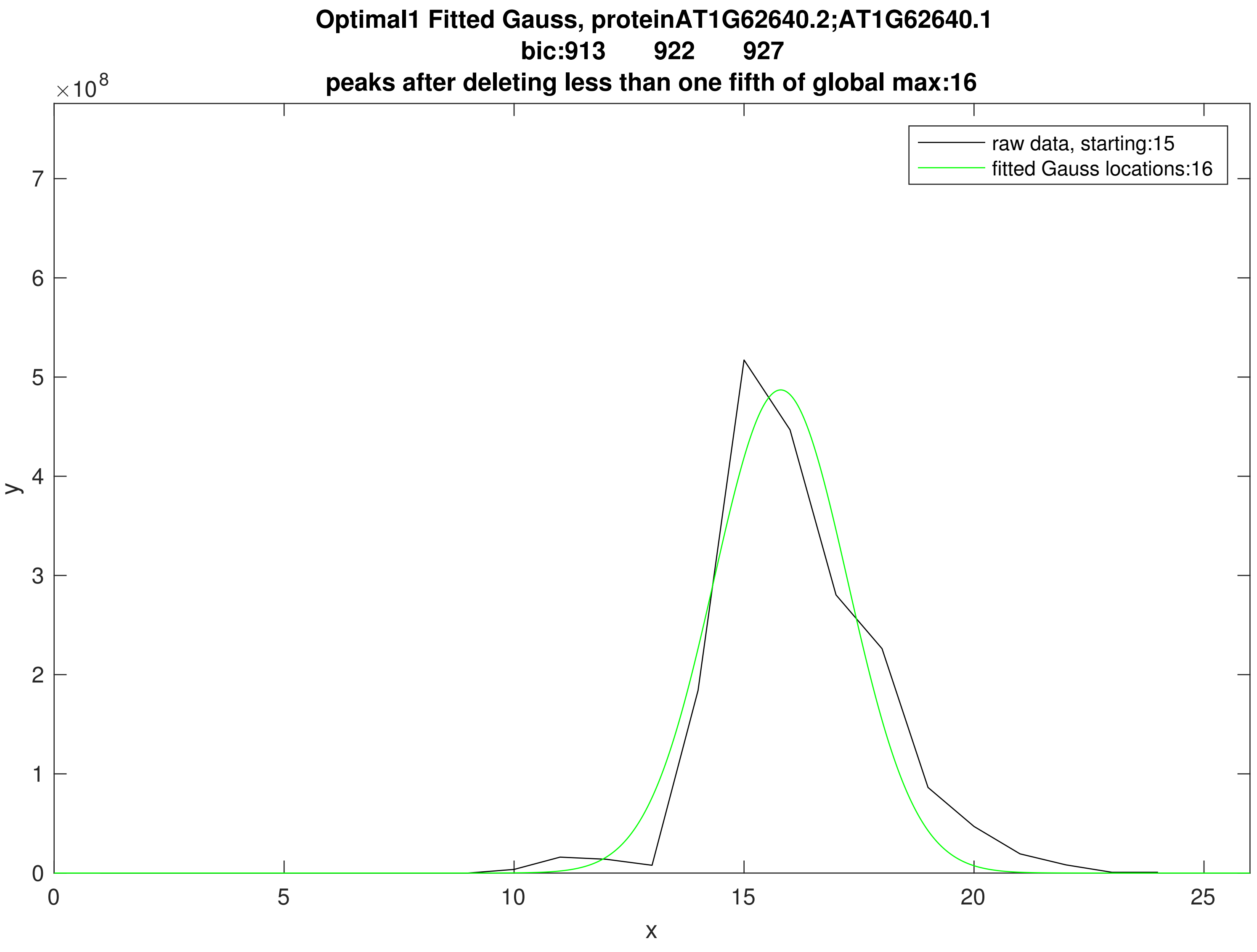
Optimal1 Fitted Gauss, proteinAT1G62640.2;AT1G62640.1
bic:913 922 927
peaks after deleting less than one fifth of global max:16
×10^8
raw data, starting:15
fitted Gauss locations:16
y
x
0
1
2
3
4
5
6
7
0
5
10
15
20
25

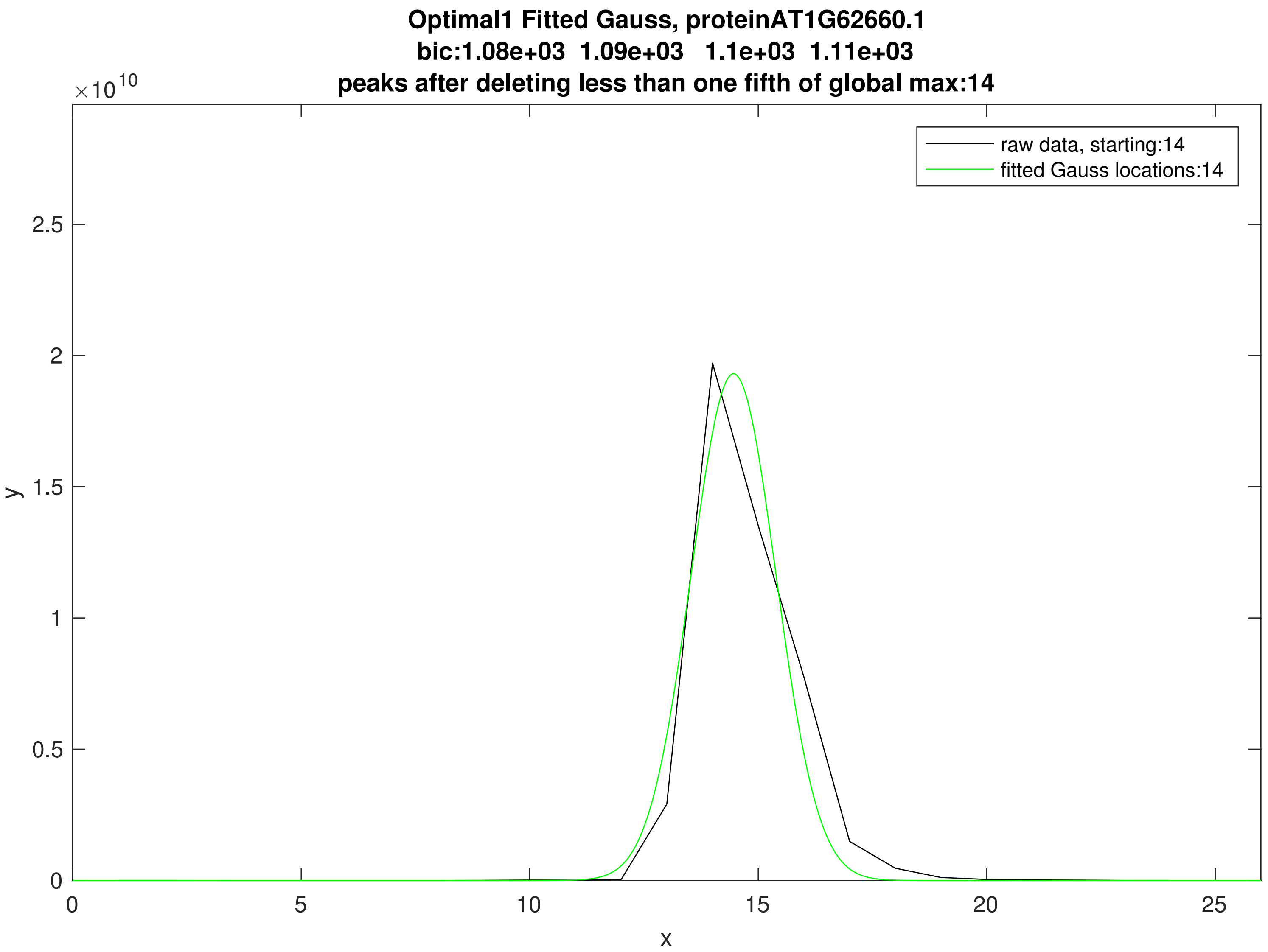

Optimal1 Fitted Gauss, proteinAT1G62660.1
bic:1.08e+03 1.09e+03 1.1e+03 1.11e+03
peaks after deleting less than one fifth of global max:14
×10^10
raw data, starting:14
fitted Gauss locations:14
2.5
2
1.5
1
0.5
0
y
0
5
10
15
20
25
x

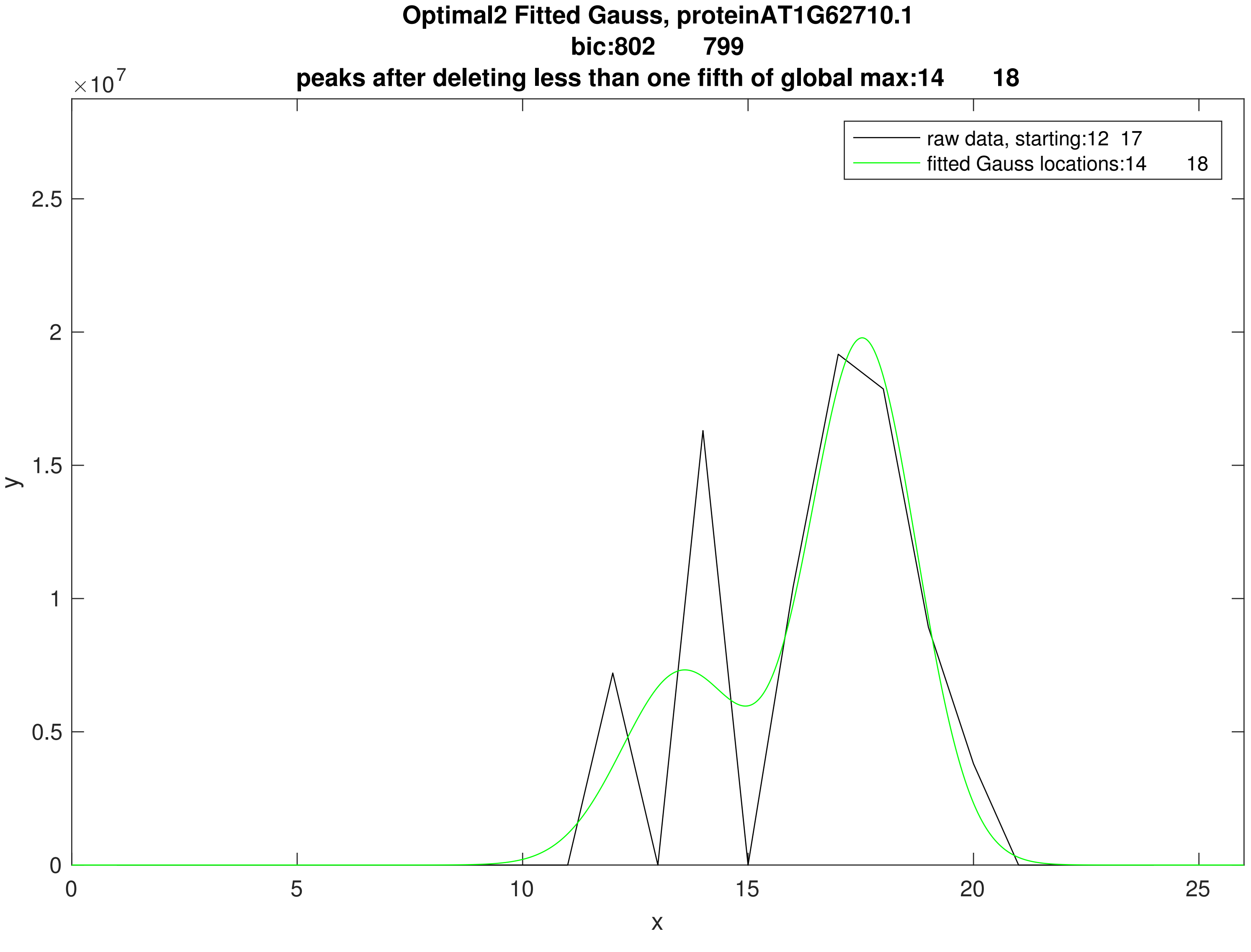

Optimal2 Fitted Gauss, proteinAT1G62710.1
bic:802 799
peaks after deleting less than one fifth of global max:14 18
×10^7
raw data, starting:12 17
fitted Gauss locations:14 18
2.5
2
1.5
1
0.5
0
y
0
5
10
15
20
25
x

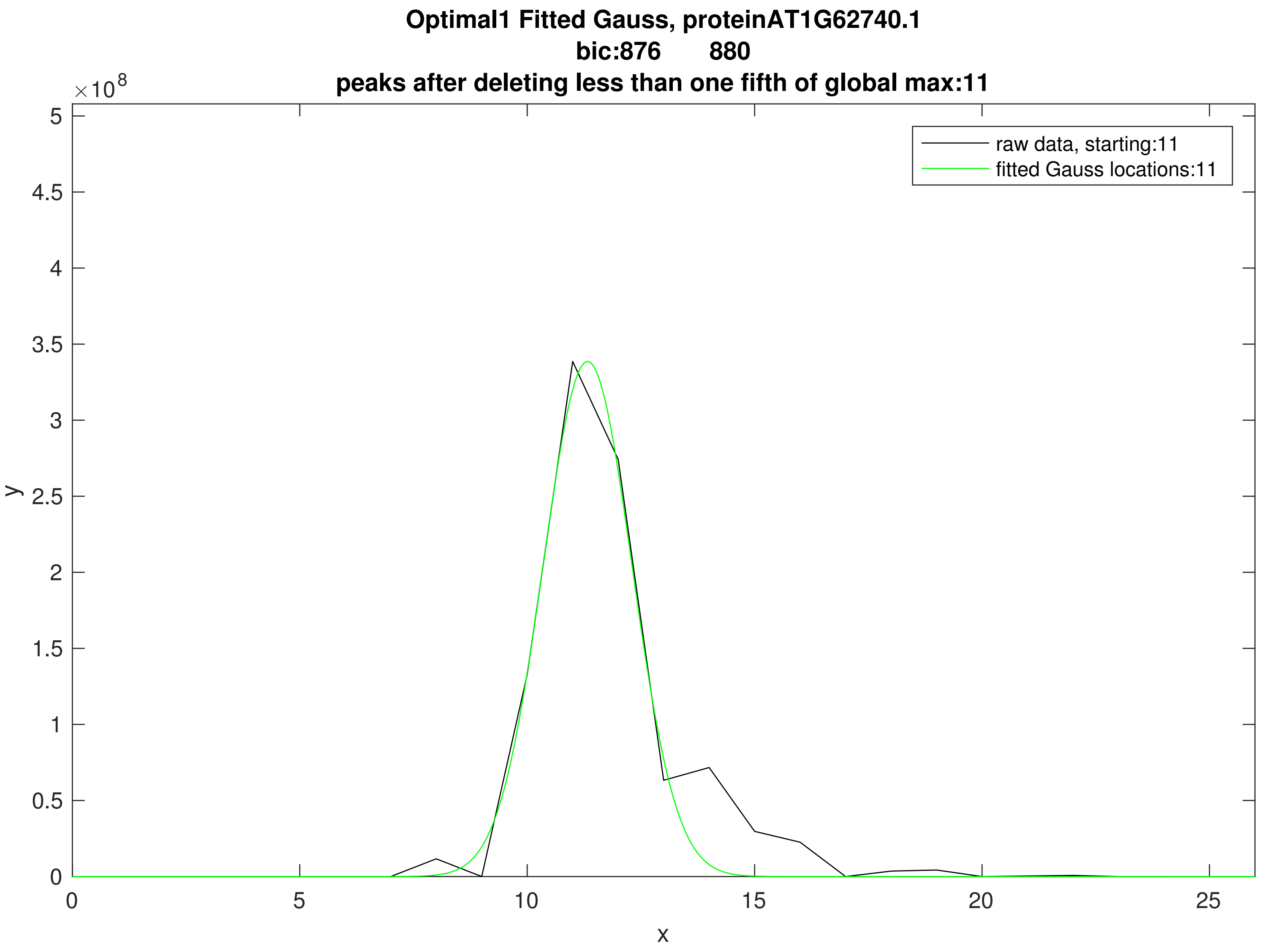

Optimal1 Fitted Gauss, proteinAT1G62740.1
bic:876 880
peaks after deleting less than one fifth of global max:11
×10^8
5
4.5
4
3.5
3
2.5
2
1.5
1
0.5
0
y
0
5
10
15
20
25
x
raw data, starting:11
fitted Gauss locations:11

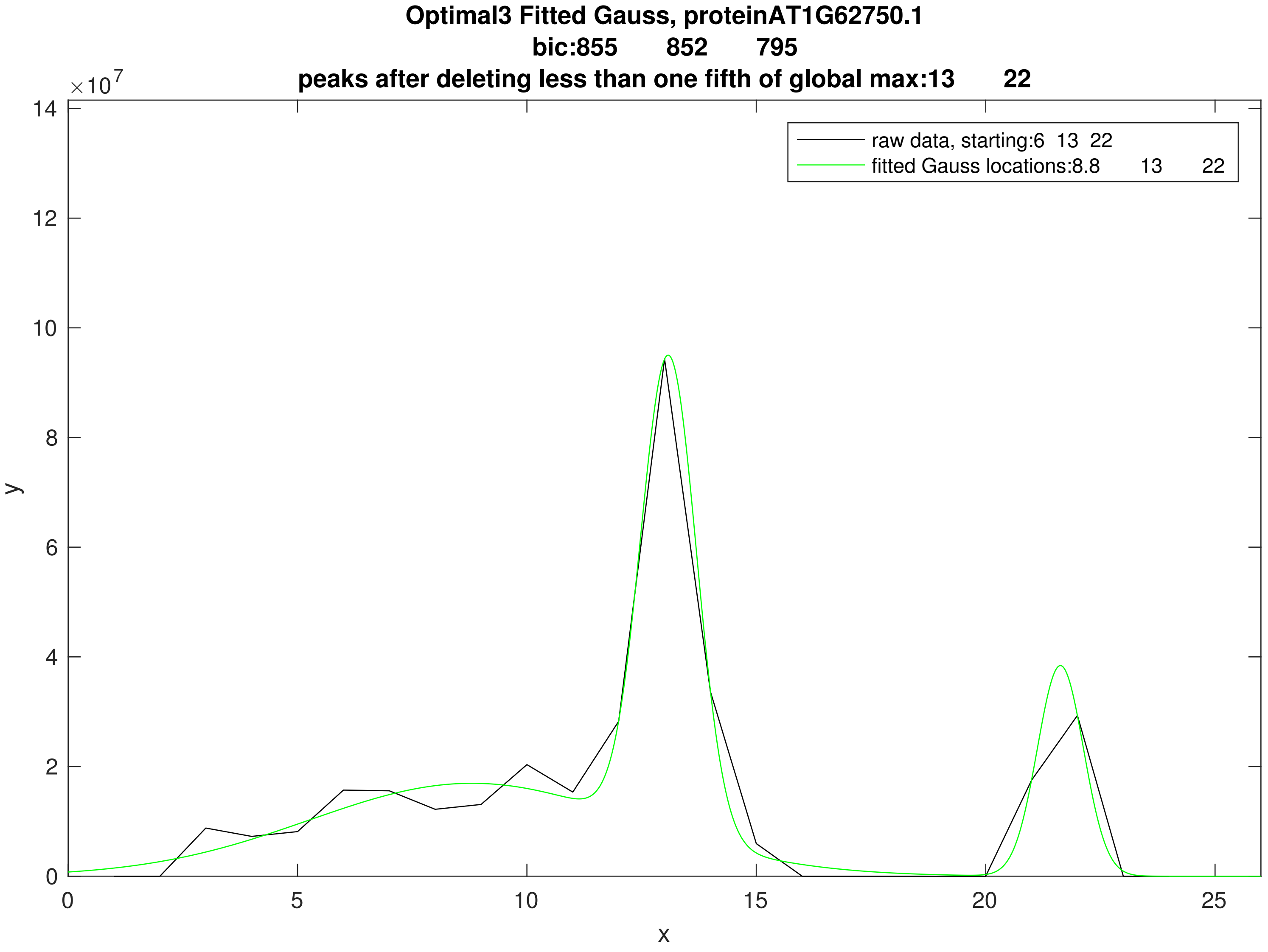
Optimal3 Fitted Gauss, proteinAT1G62750.1
bic:855 852 795
peaks after deleting less than one fifth of global max:13 22
×10^7
raw data, starting:6 13 22
fitted Gauss locations:8.8 13 22
y
x
0
2
4
6
8
10
12
14
5
15
20
25

Optimal1 Fitted Gauss, proteinAT1G62800.2;AT1G62800.1
bic:848
peaks after deleting less than one fifth of global max:14

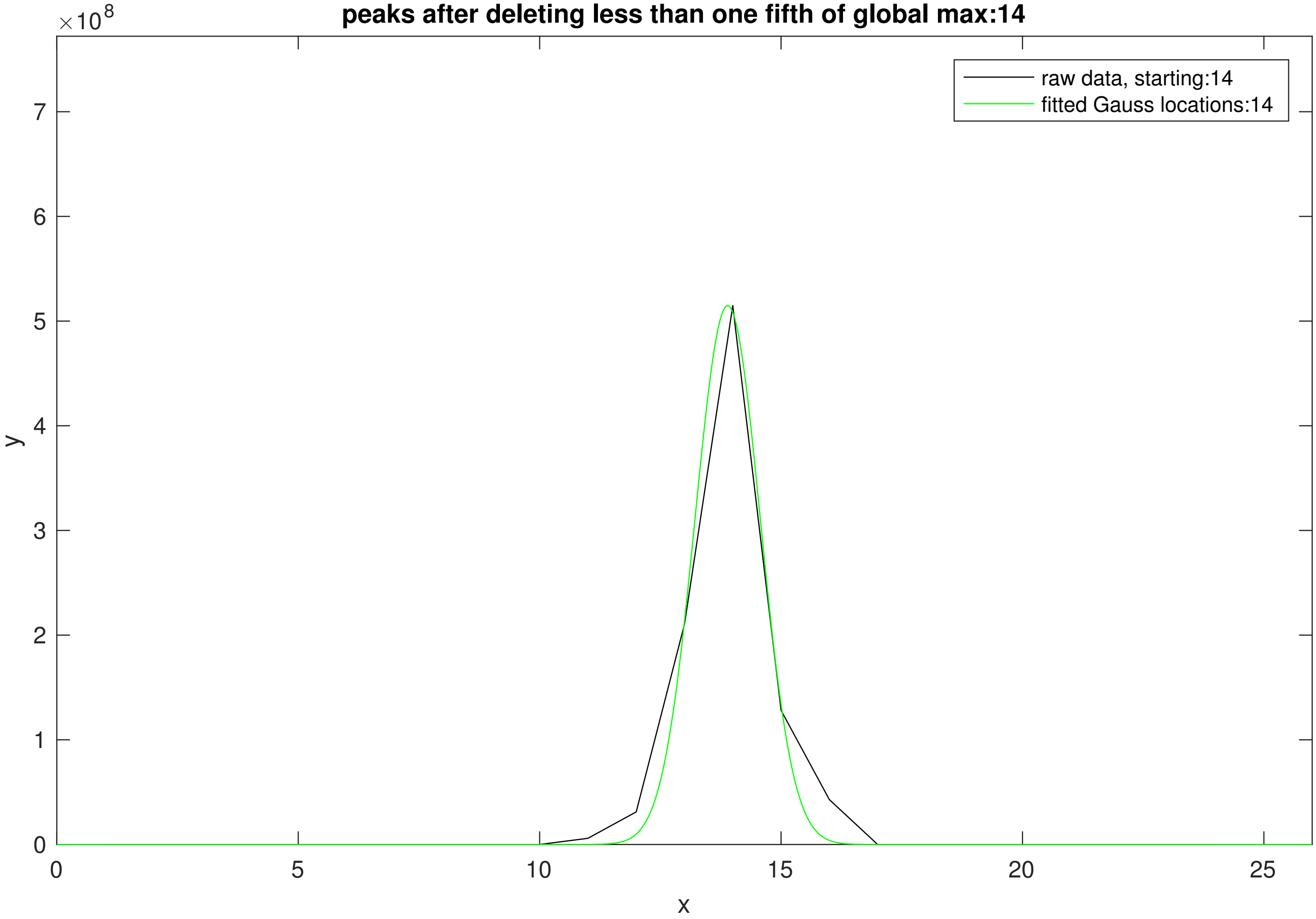

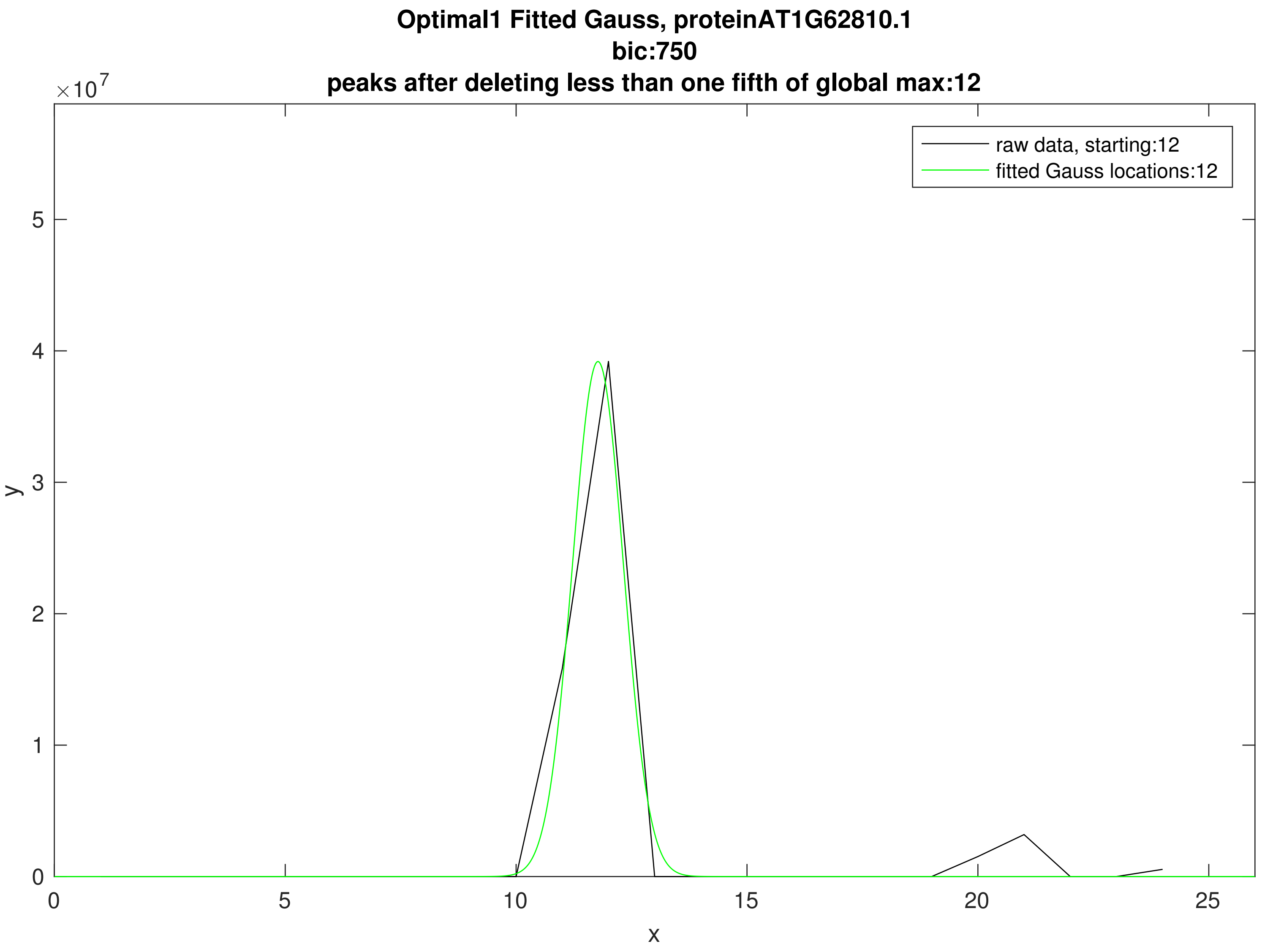
Optimal1 Fitted Gauss, proteinAT1G62810.1
bic:750
peaks after deleting less than one fifth of global max:12
×10^7
raw data, starting:12
fitted Gauss locations:12
y
x
0
1
2
3
4
5
0
5
10
15
20
25

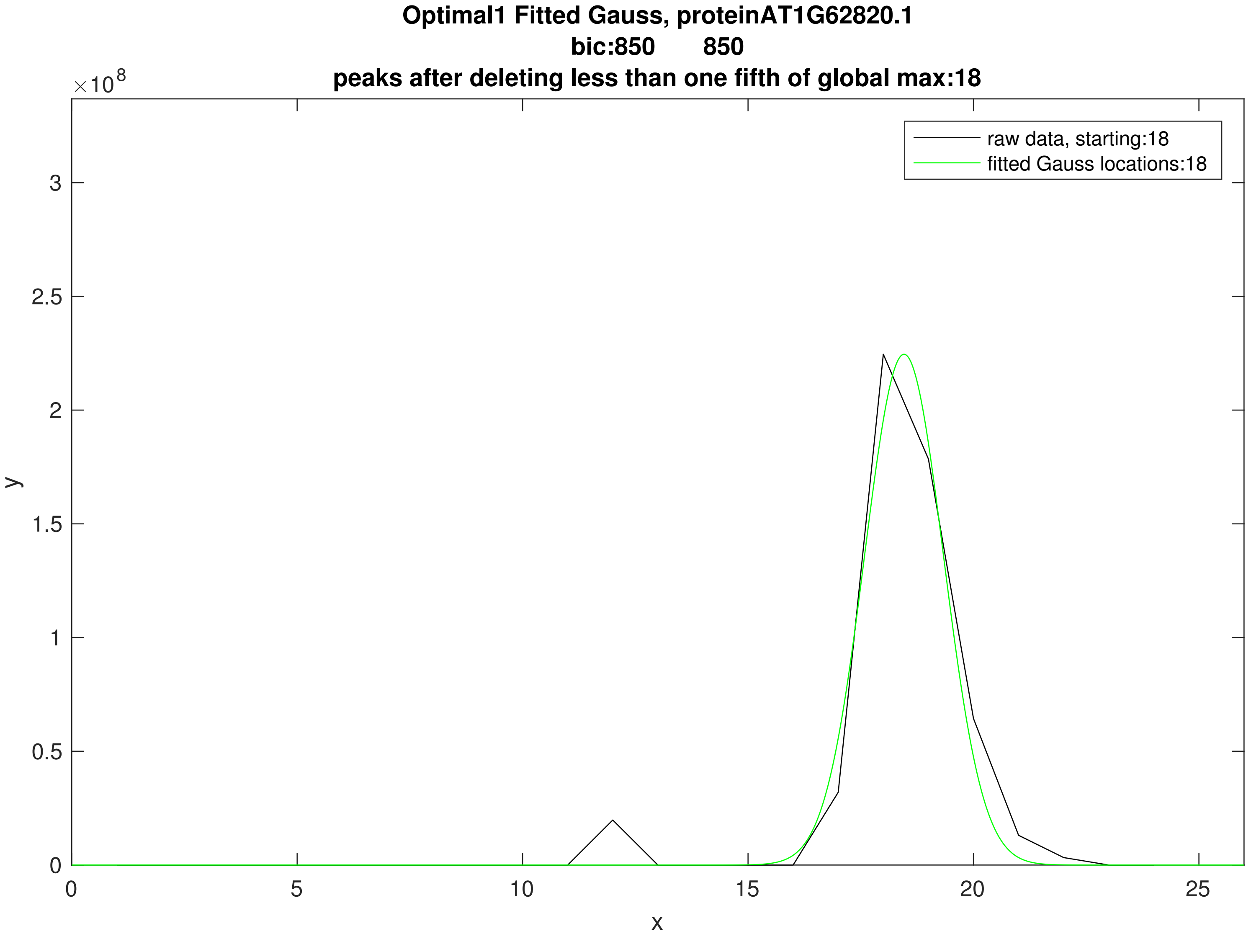
Optimal1 Fitted Gauss, proteinAT1G62820.1
bic:850 850
peaks after deleting less than one fifth of global max:18
×10^8
raw data, starting:18
fitted Gauss locations:18
y
x
0
0.5
1
1.5
2
2.5
3
0
5
10
15
20
25

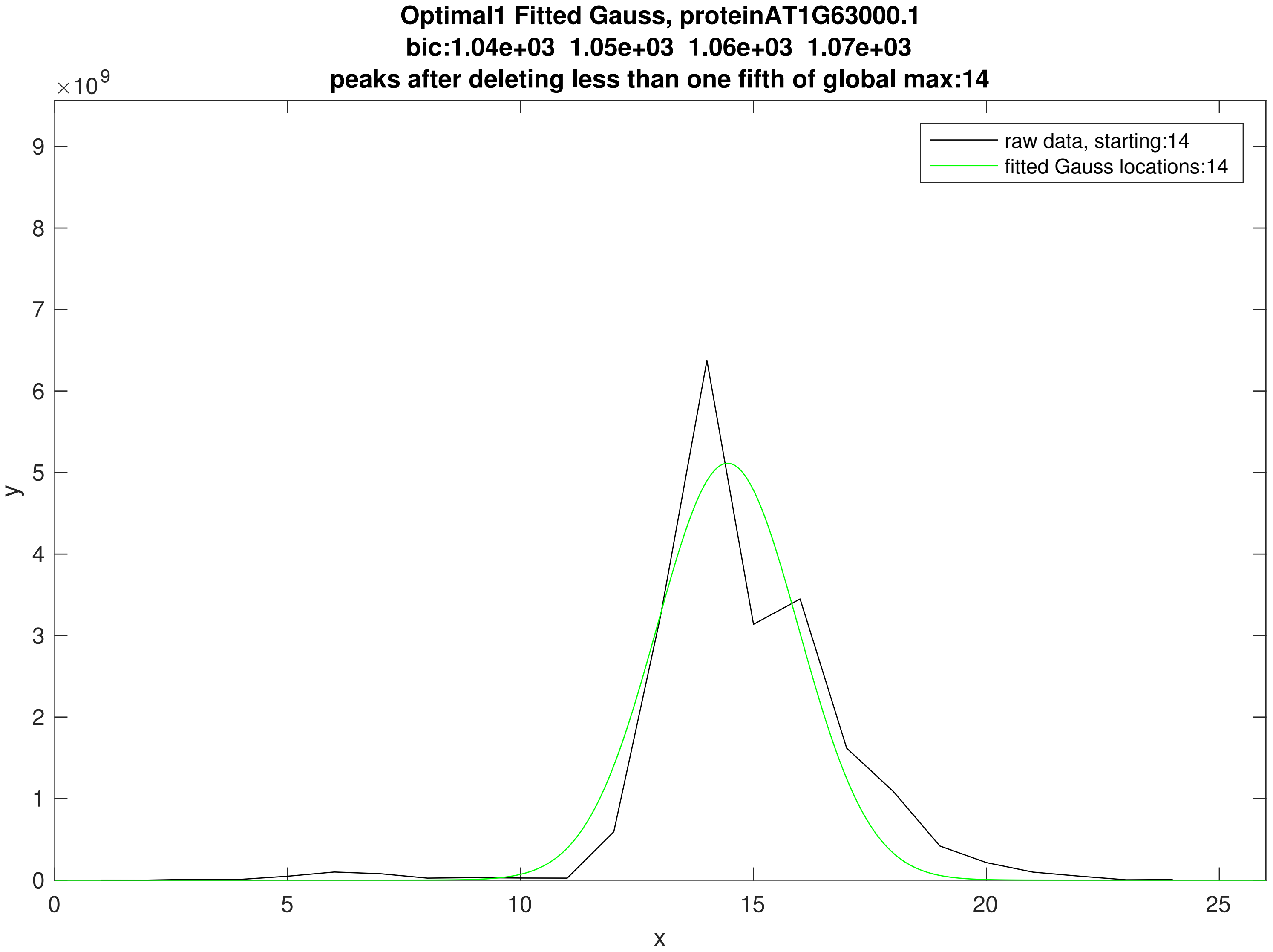
Optimal1 Fitted Gauss, proteinAT1G63000.1
bic:1.04e+03 1.05e+03 1.06e+03 1.07e+03
peaks after deleting less than one fifth of global max:14
×10^9
raw data, starting:14
fitted Gauss locations:14
y
x
0
1
2
3
4
5
6
7
8
9
0
5
10
15
20
25

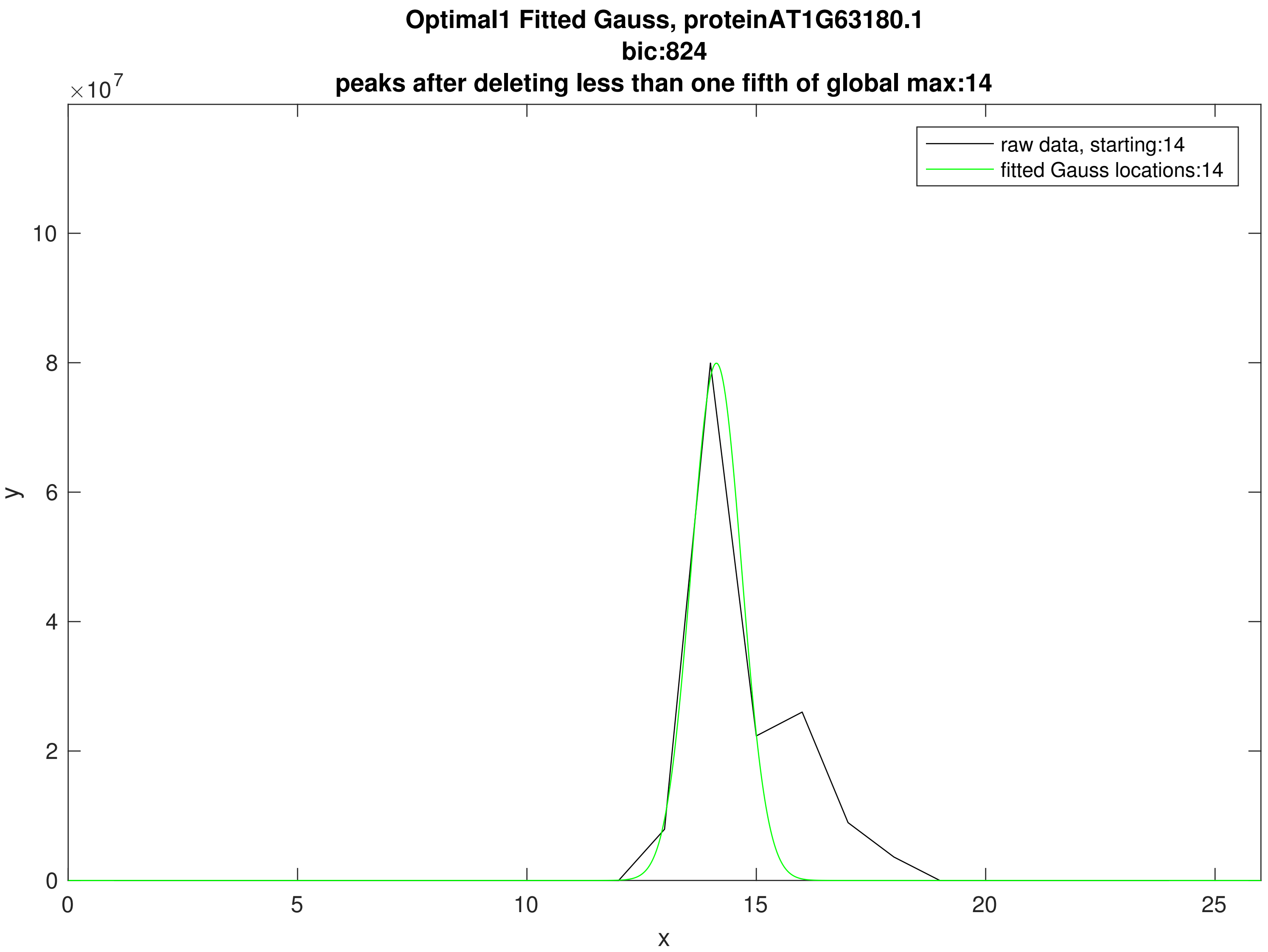

Optimal1 Fitted Gauss, proteinAT1G63180.1
bic:824
peaks after deleting less than one fifth of global max:14
×10^7
raw data, starting:14
fitted Gauss locations:14
y
x
0
2
4
6
8
10
0
5
10
15
20
25

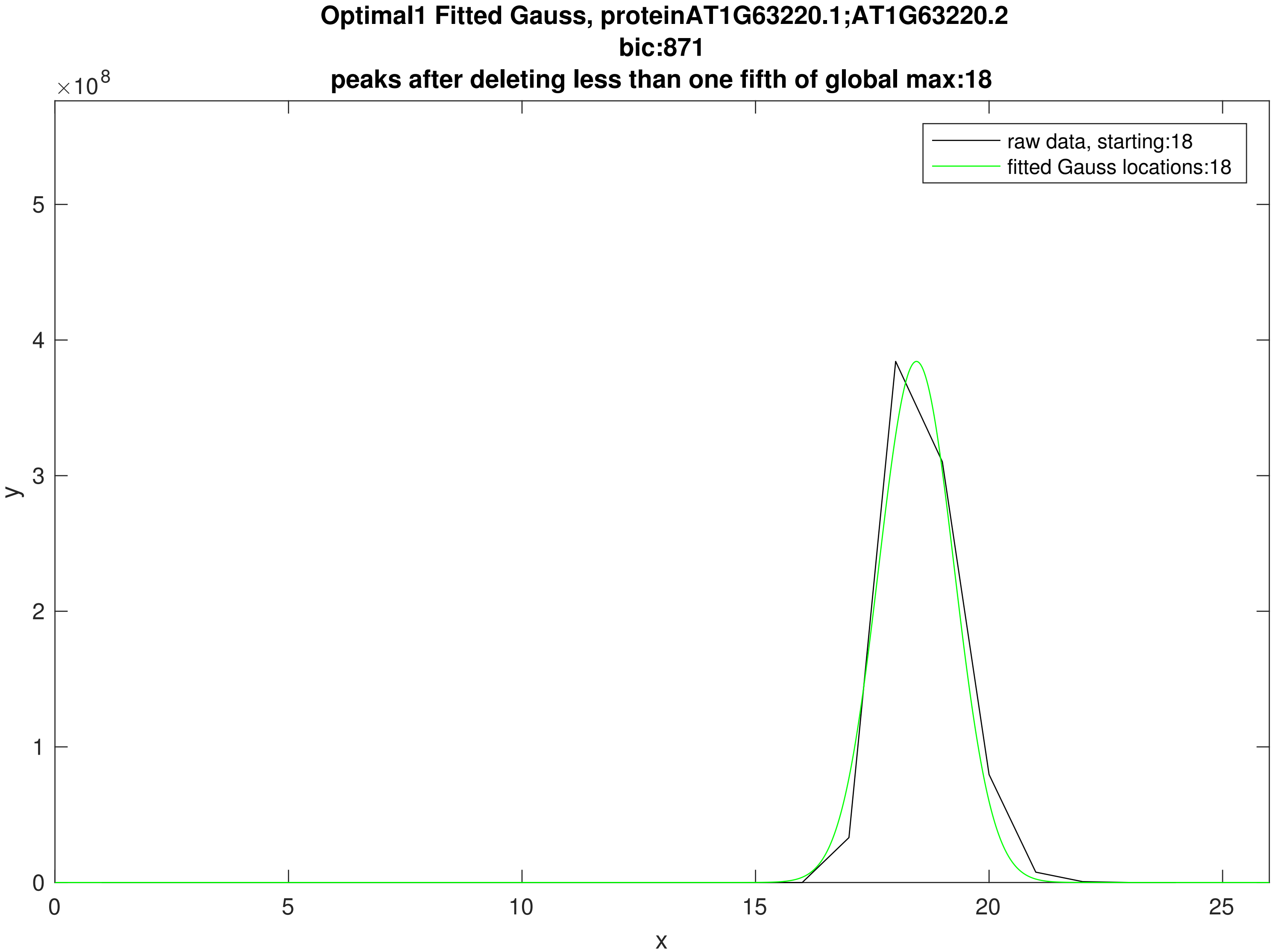

Optimal1 Fitted Gauss, proteinAT1G63220.1;AT1G63220.2
bic:871
peaks after deleting less than one fifth of global max:18
×10^8
raw data, starting:18
fitted Gauss locations:18
y
x
0
1
2
3
4
5
0
5
10
15
20
25

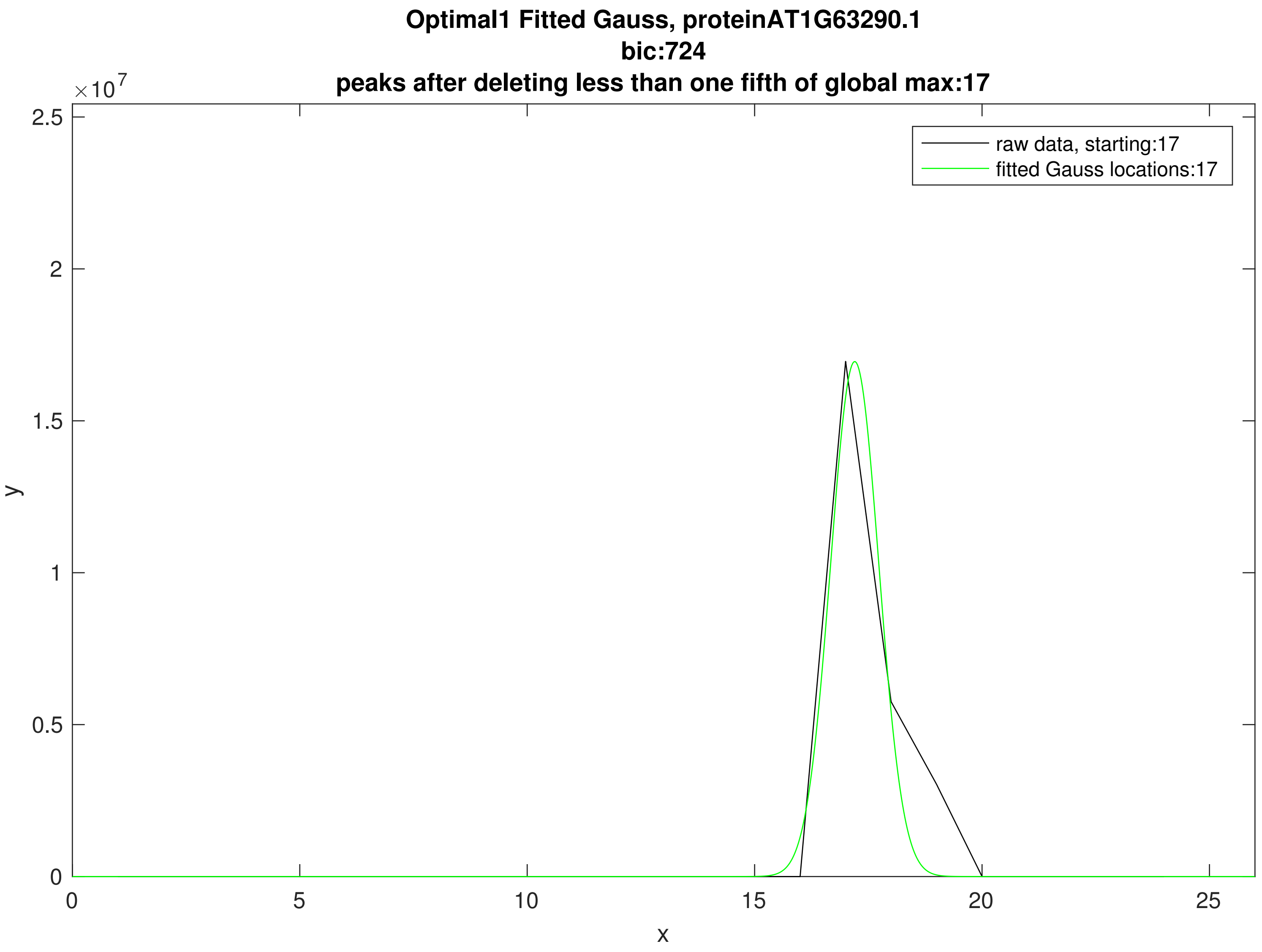
Optimal1 Fitted Gauss, proteinAT1G63290.1
bic:724
peaks after deleting less than one fifth of global max:17
×10^7
2.5
2
1.5
1
0.5
0
y
0
5
10
15
20
25
x
raw data, starting:17
fitted Gauss locations:17

**Optimal1 Fitted Gauss, proteinAT1G63460.1**
**bic:803 812**
**peaks after deleting less than one fifth of global max:18**

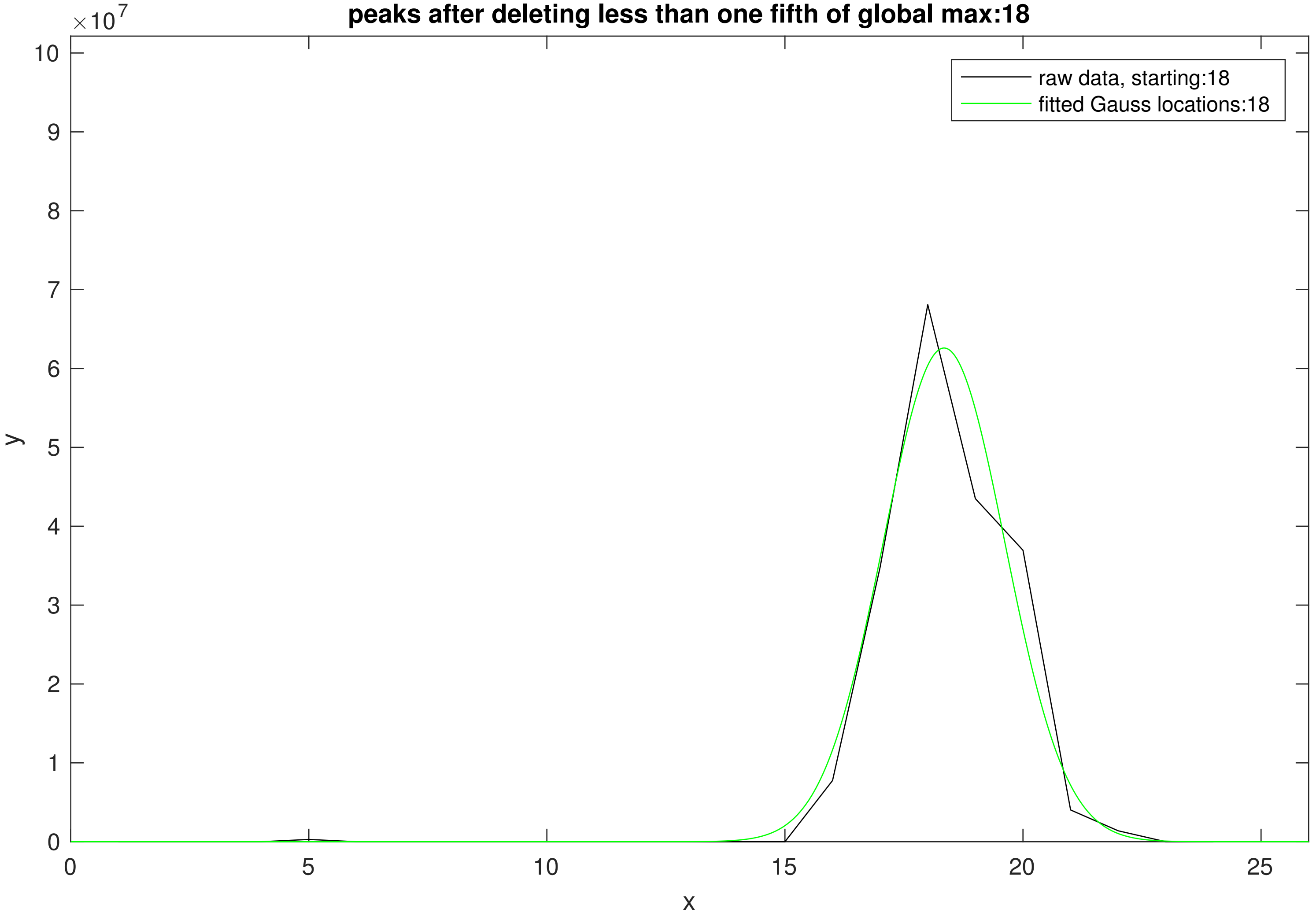

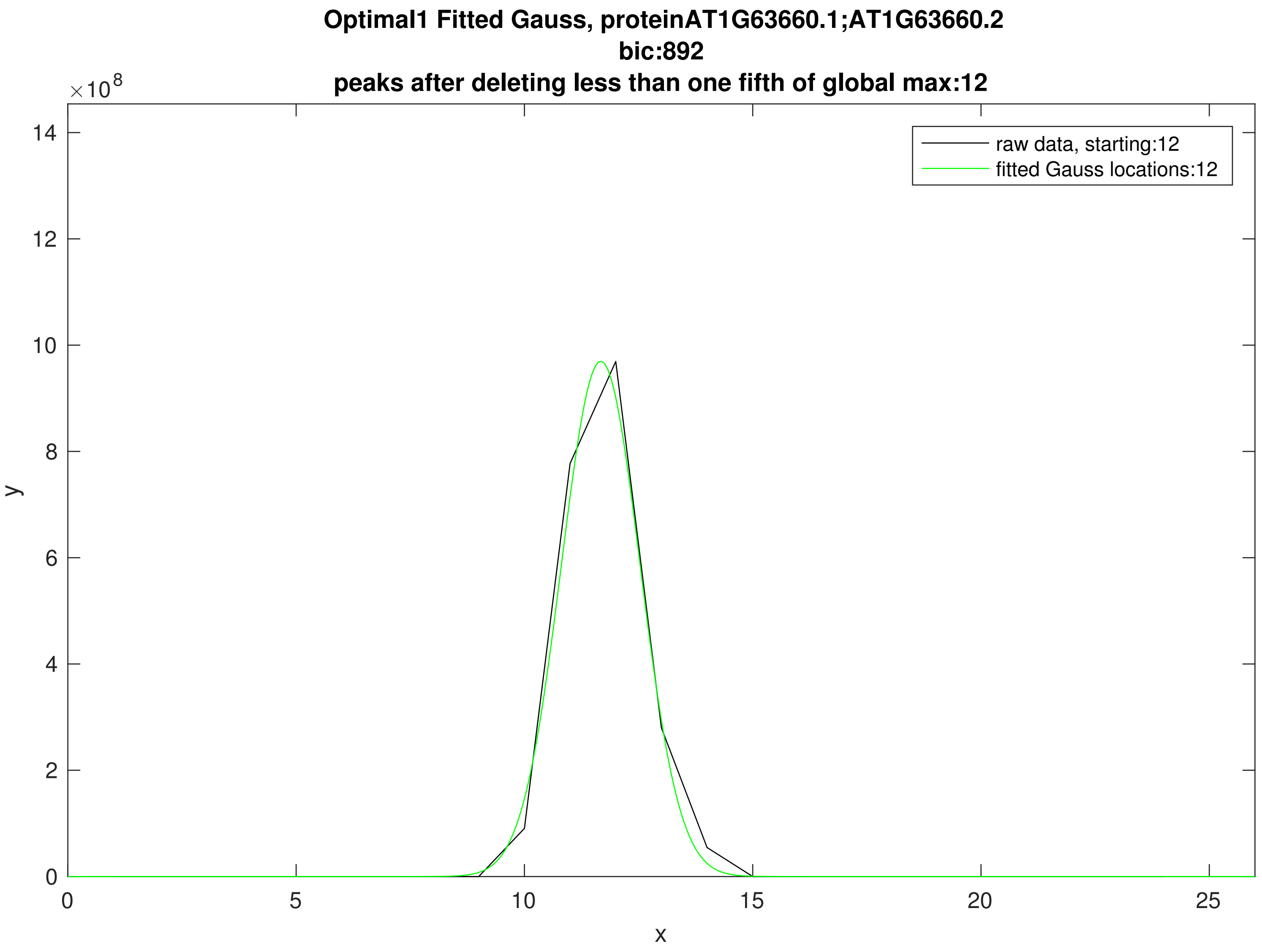

Optimal1 Fitted Gauss, proteinAT1G63660.1;AT1G63660.2
bic:892
peaks after deleting less than one fifth of global max:12
×10^8
14
12
10
8
6
4
2
0
y
0
5
10
15
20
25
x
raw data, starting:12
fitted Gauss locations:12

**Optimal2 Fitted Gauss, proteinAT1G63770.4;AT1G63770.3;AT1G63770.5;AT1G63770.2;AT1G63770.1**
**bic:1e+03 965**
**peaks after deleting less than one fifth of global max:13**

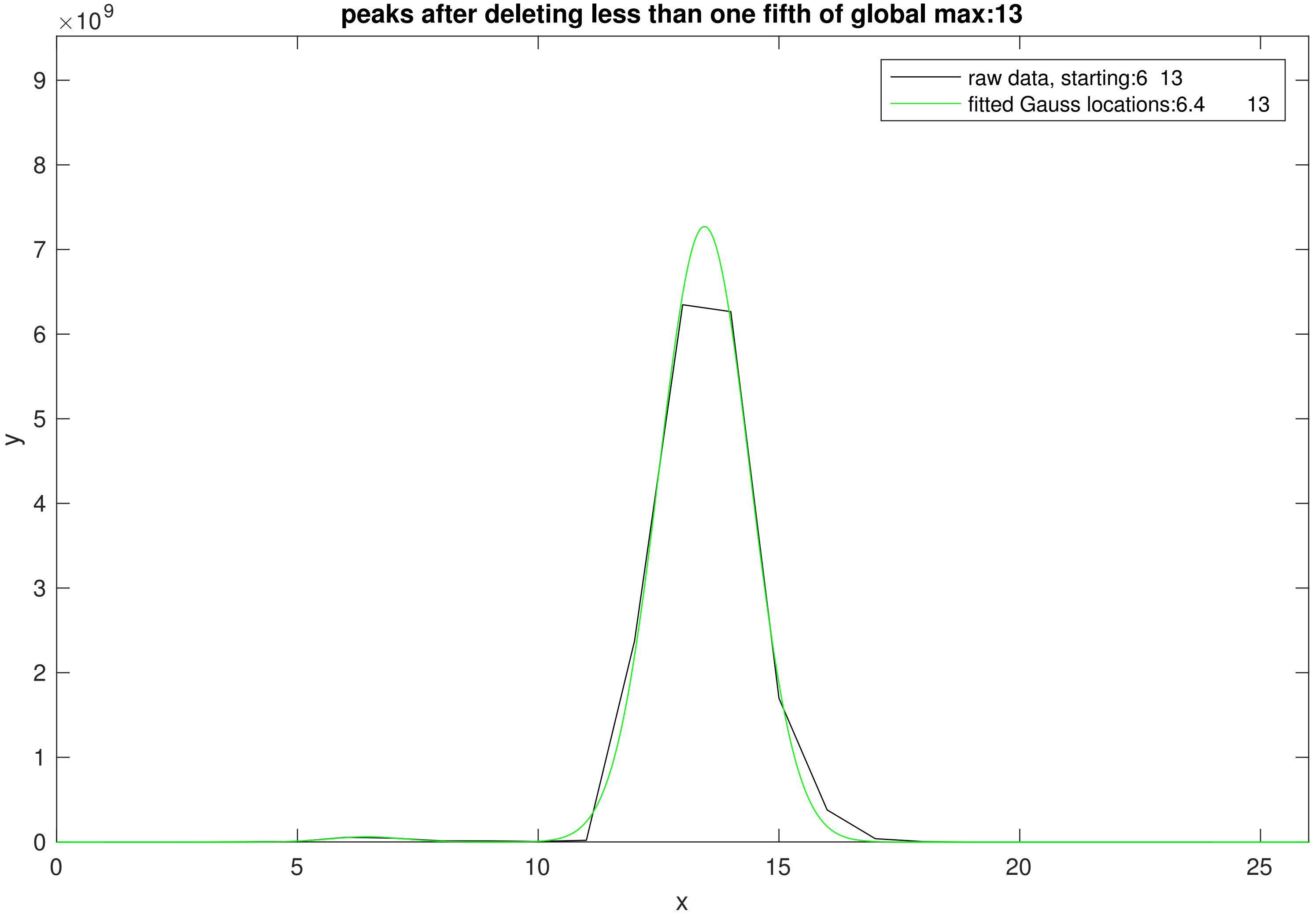

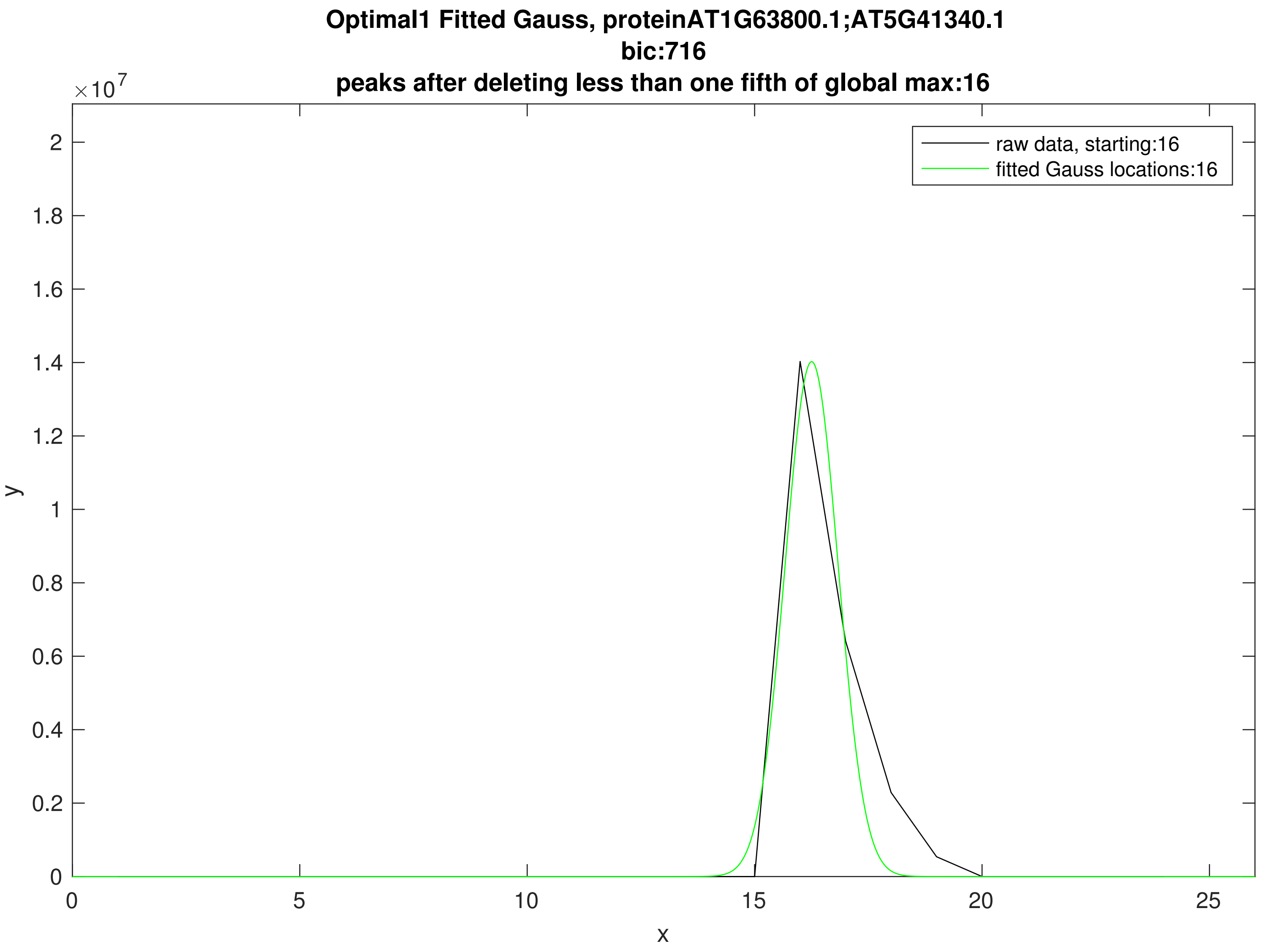
Optimal1 Fitted Gauss, proteinAT1G63800.1;AT5G41340.1
bic:716
peaks after deleting less than one fifth of global max:16
×10^7
2
1.8
1.6
1.4
1.2
1
0.8
0.6
0.4
0.2
0
y
0
5
10
15
20
25
x
raw data, starting:16
fitted Gauss locations:16

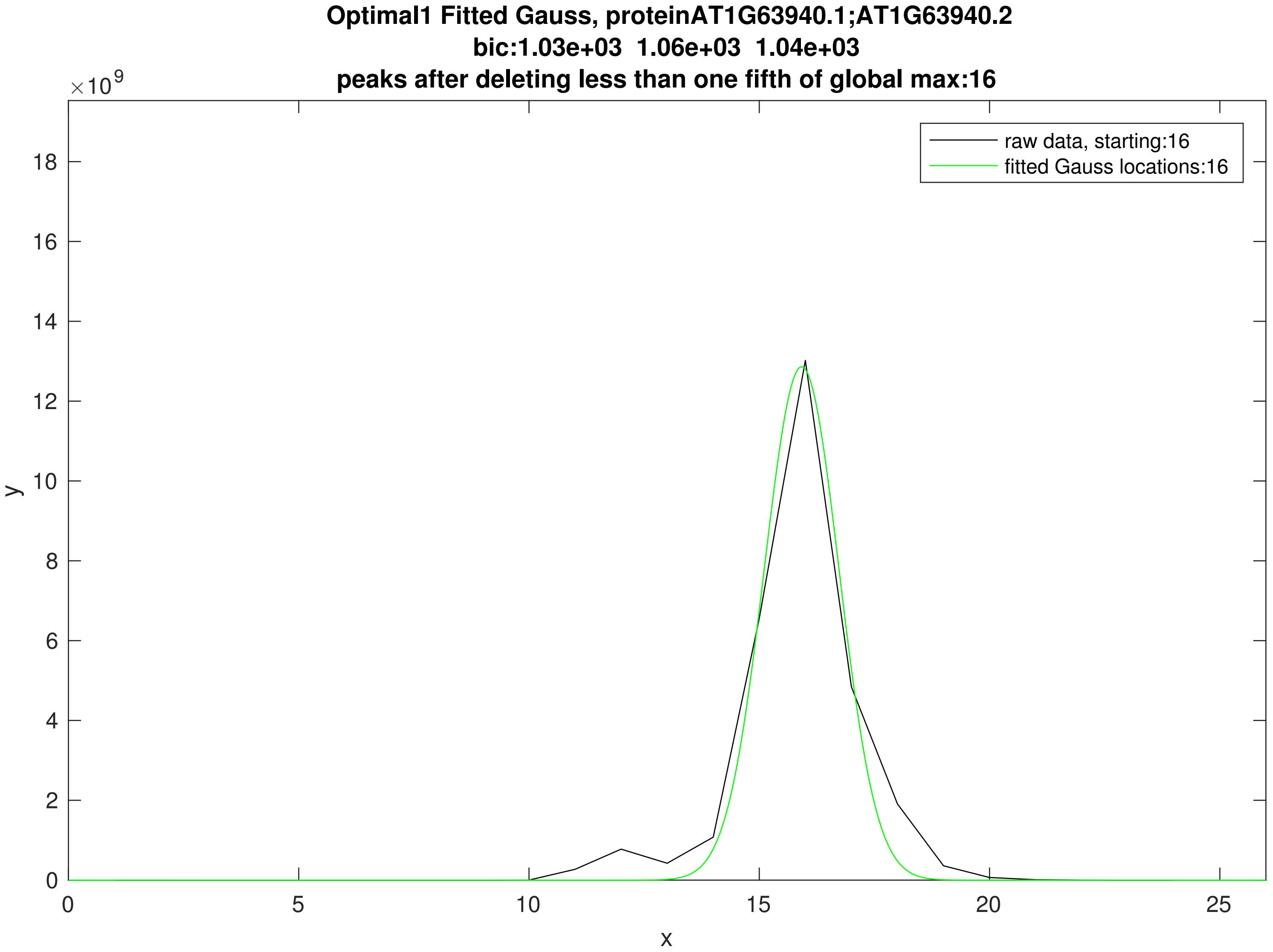

Optimal1 Fitted Gauss, proteinAT1G63940.1;AT1G63940.2
bic:1.03e+03 1.06e+03 1.04e+03
peaks after deleting less than one fifth of global max:16
×10^9
raw data, starting:16
fitted Gauss locations:16
y
x
0
2
4
6
8
10
12
14
16
18
0
5
10
15
20
25

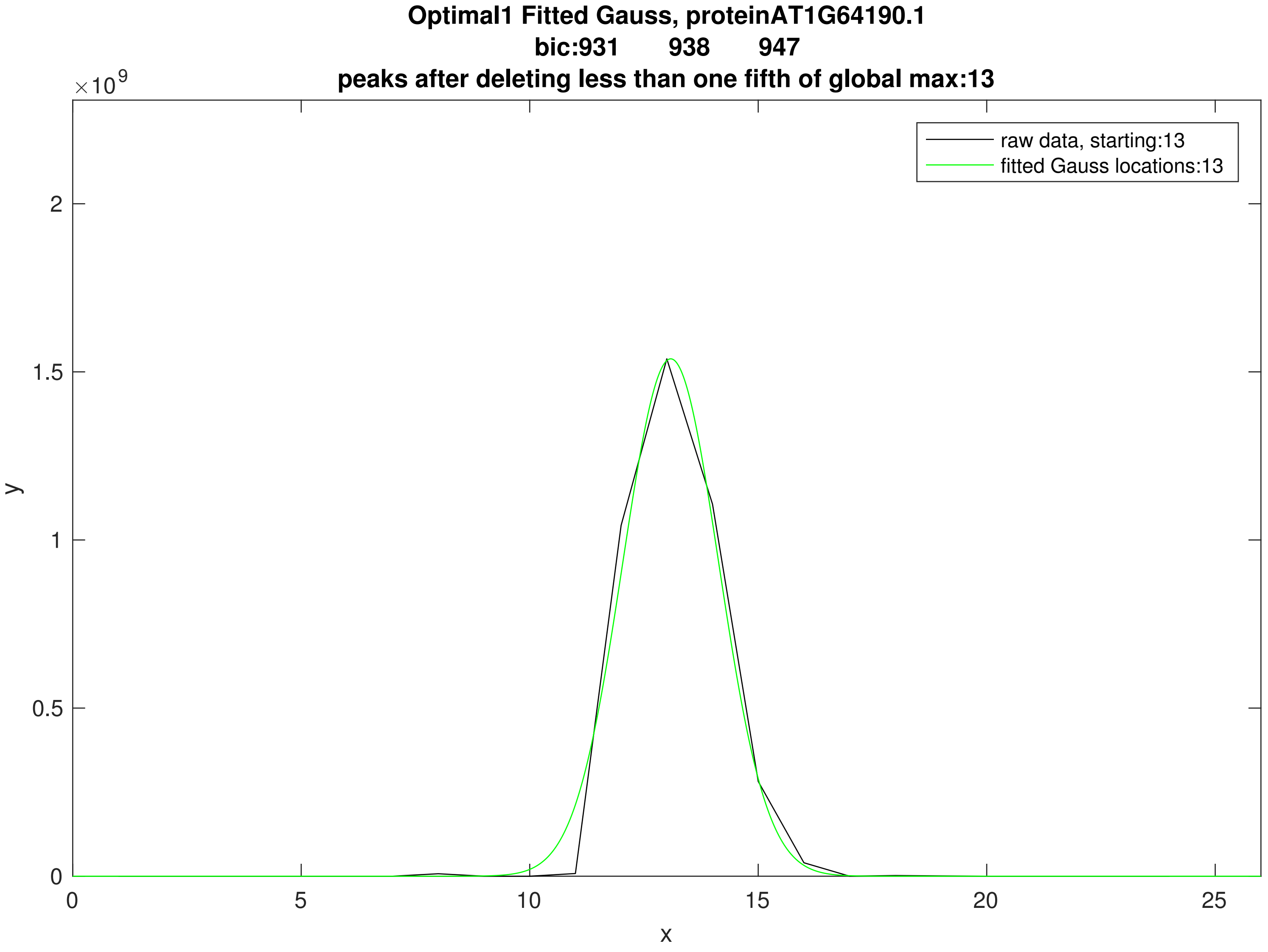
Optimal1 Fitted Gauss, proteinAT1G64190.1
bic:931 938 947
peaks after deleting less than one fifth of global max:13
×10^9
raw data, starting:13
fitted Gauss locations:13
y
x
0
0.5
1
1.5
2
0
5
10
15
20
25

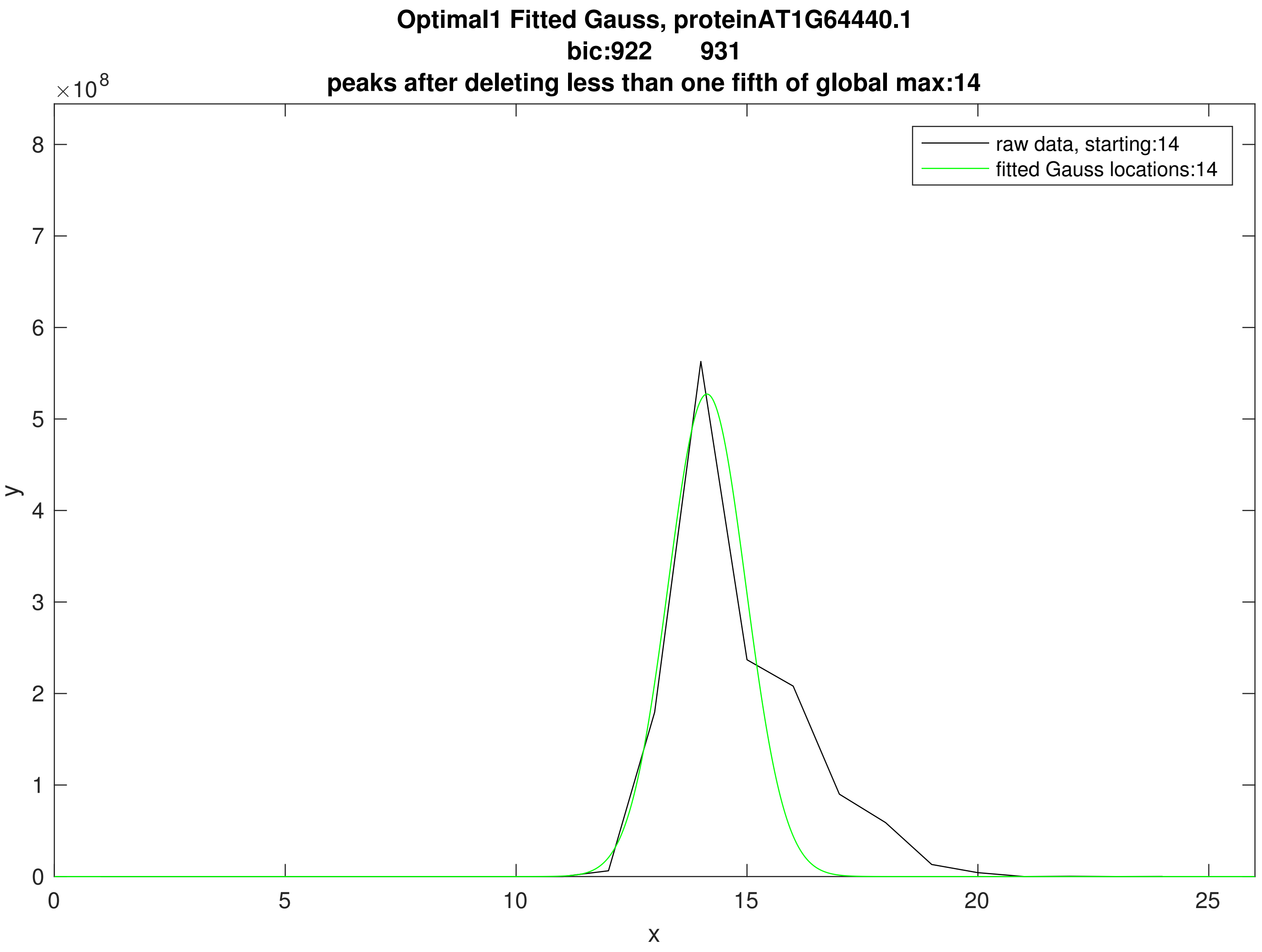
Optimal1 Fitted Gauss, proteinAT1G64440.1
bic:922 931
peaks after deleting less than one fifth of global max:14
×10^8
raw data, starting:14
fitted Gauss locations:14
y
x
0
1
2
3
4
5
6
7
8
0
5
10
15
20
25

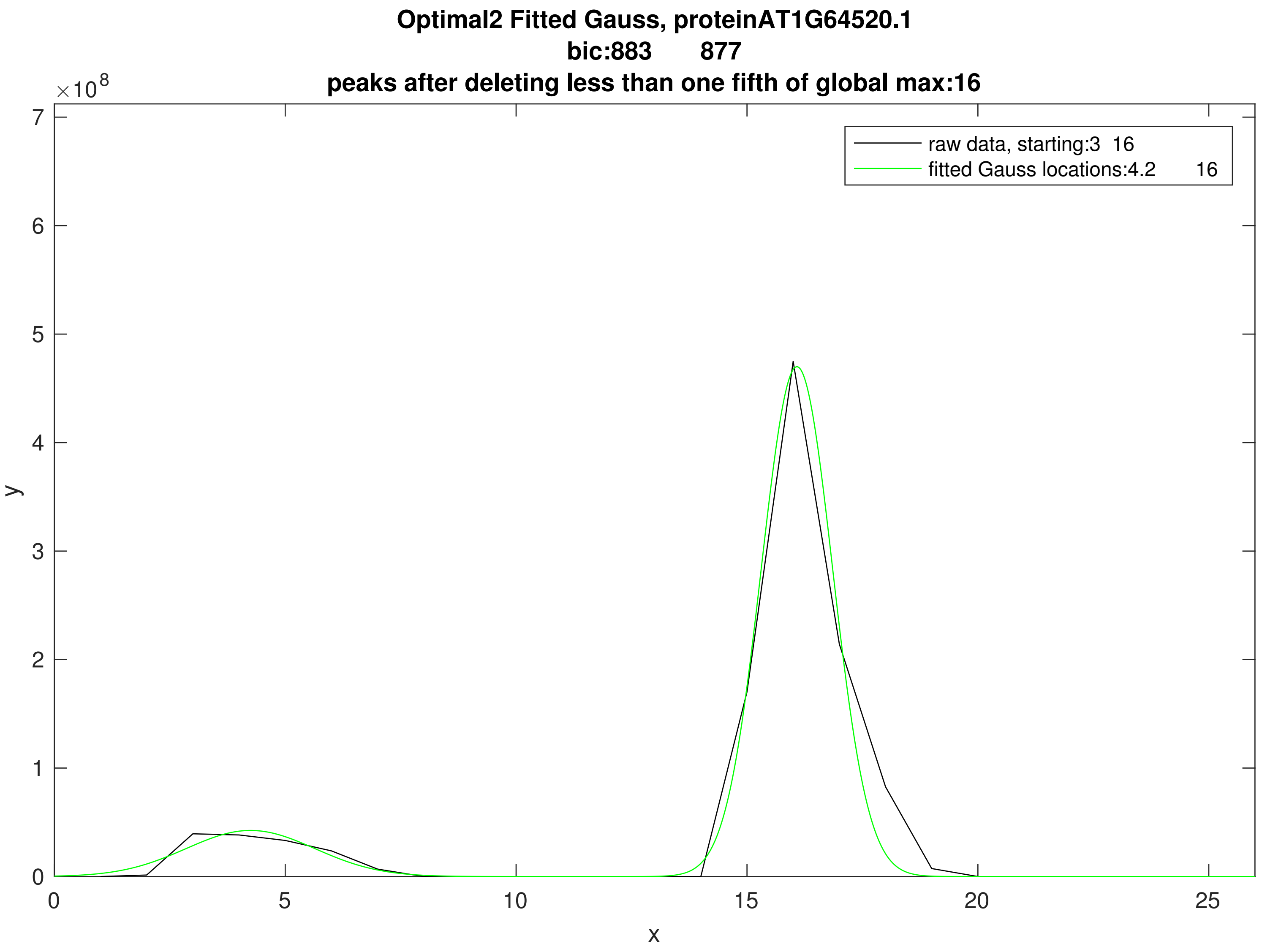
Optimal2 Fitted Gauss, proteinAT1G64520.1
bic:883 877
peaks after deleting less than one fifth of global max:16
×10^8
7
6
5
4
3
2
1
0
y
0
5
10
15
20
25
x
raw data, starting:3 16
fitted Gauss locations:4.2 16

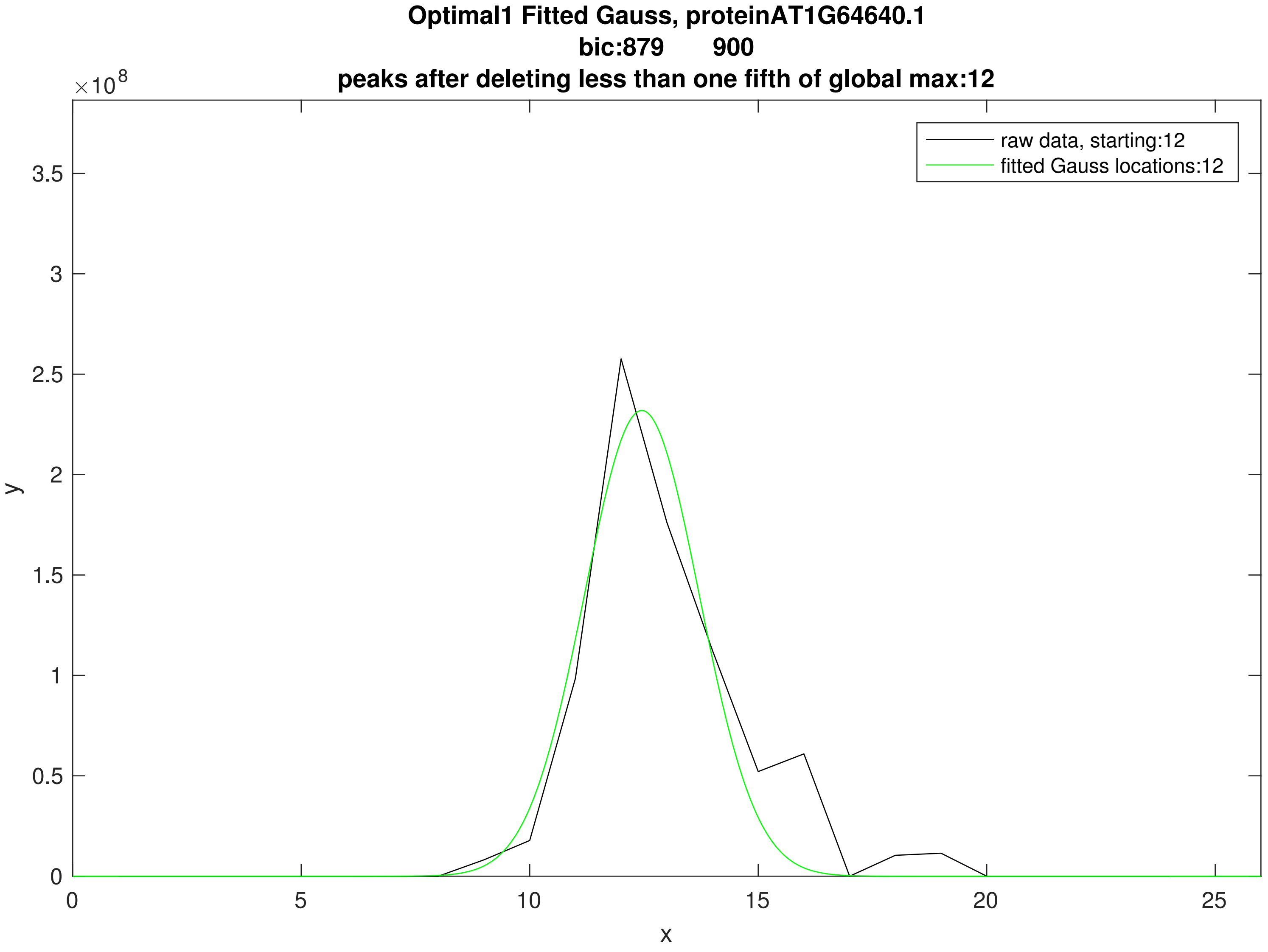
Optimal1 Fitted Gauss, proteinAT1G64640.1
bic:879 900
peaks after deleting less than one fifth of global max:12
×10^8
raw data, starting:12
fitted Gauss locations:12
y
x
0
0.5
1
1.5
2
2.5
3
3.5
5
10
15
20
25

**Optimal1 Fitted Gauss, proteinAT1G64660.1**
**bic:795 804**
**peaks after deleting less than one fifth of global max:8.4**

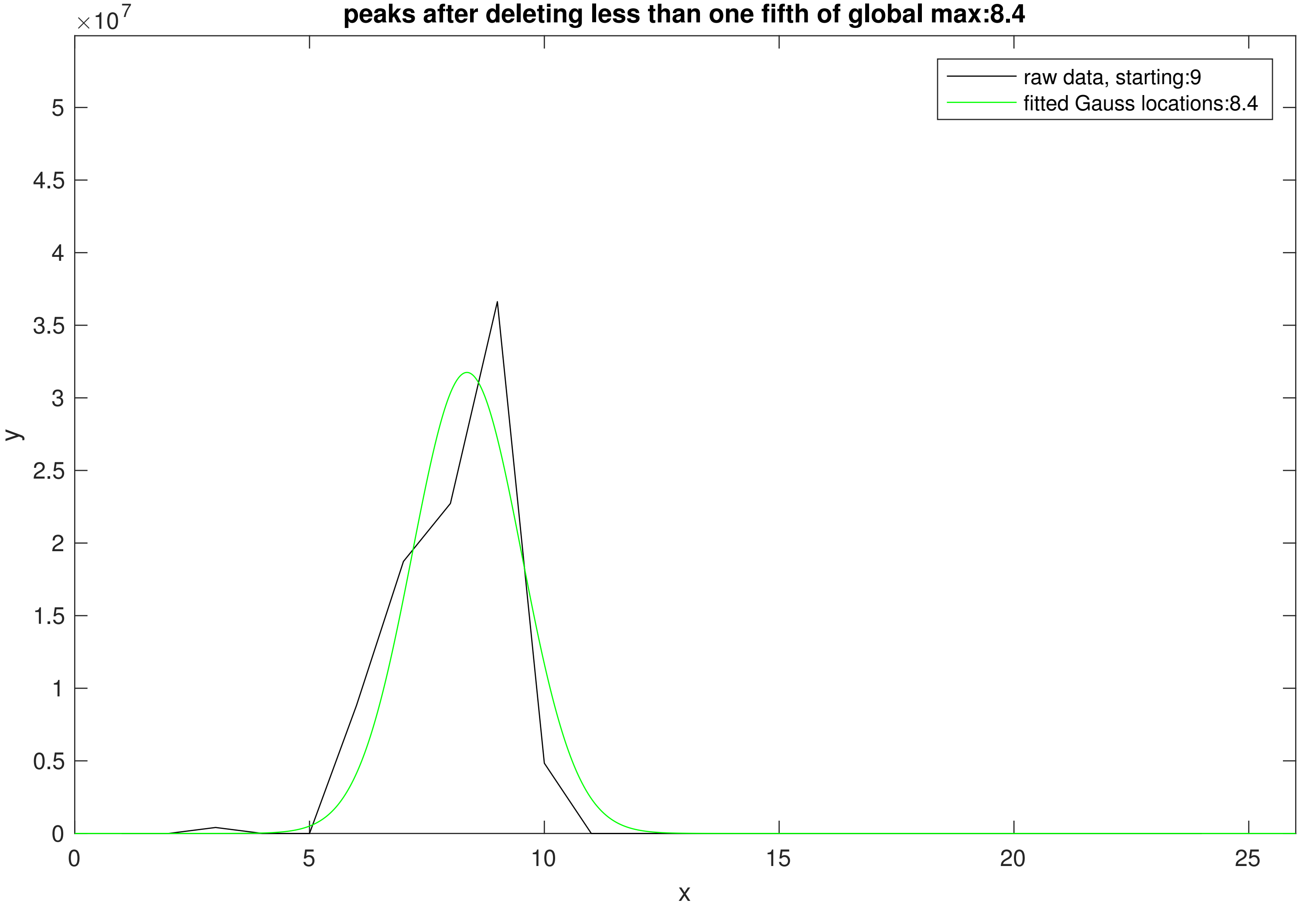

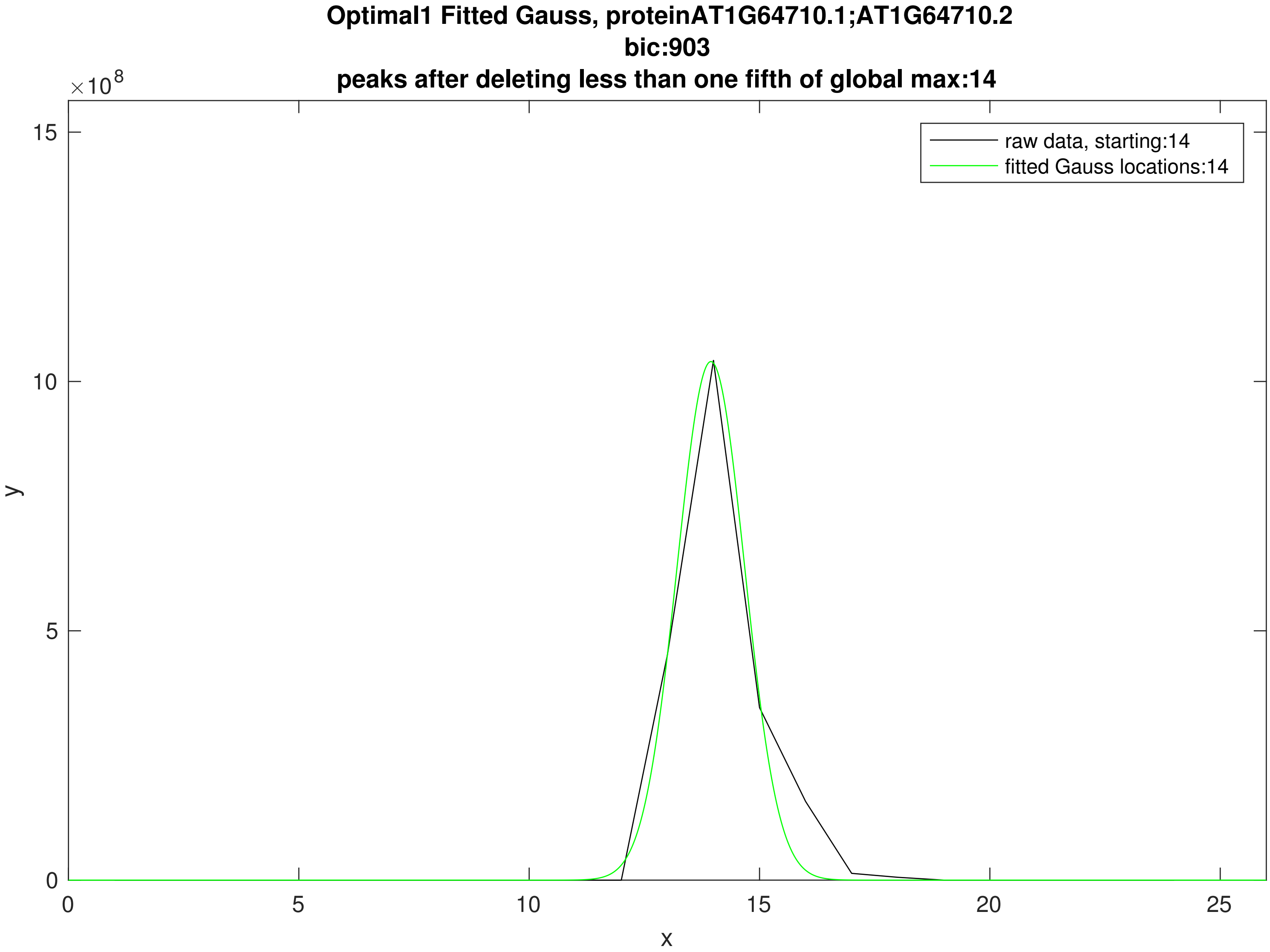
Optimal1 Fitted Gauss, proteinAT1G64710.1;AT1G64710.2
bic:903
peaks after deleting less than one fifth of global max:14
$\times 10^8$
15
10
5
0
y
0
5
10
15
20
25
x
raw data, starting:14
fitted Gauss locations:14

**Optimal1 Fitted Gauss, proteinAT1G64760.1;AT1G64760.2**
**bic:763**
**peaks after deleting less than one fifth of global max:16**

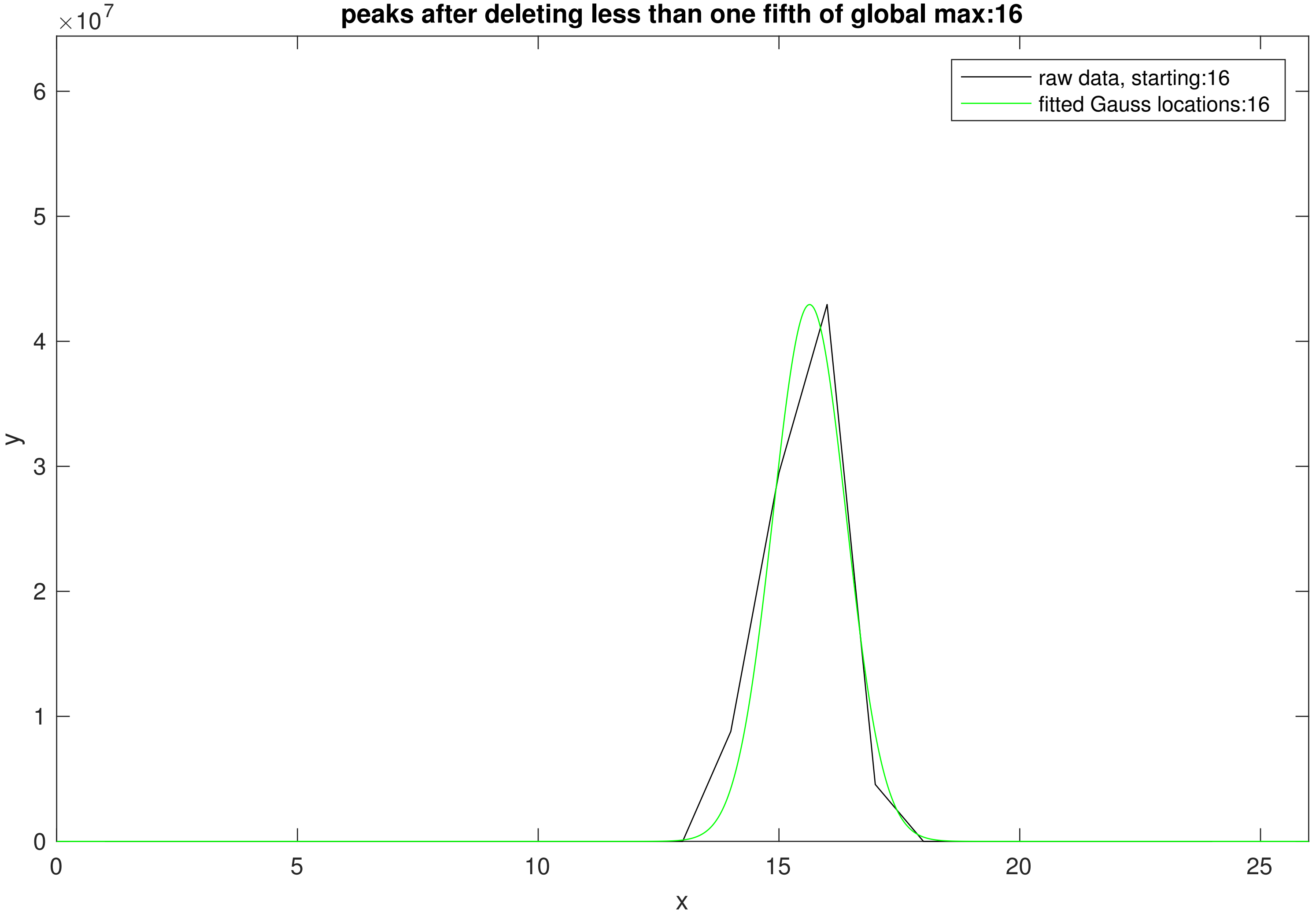

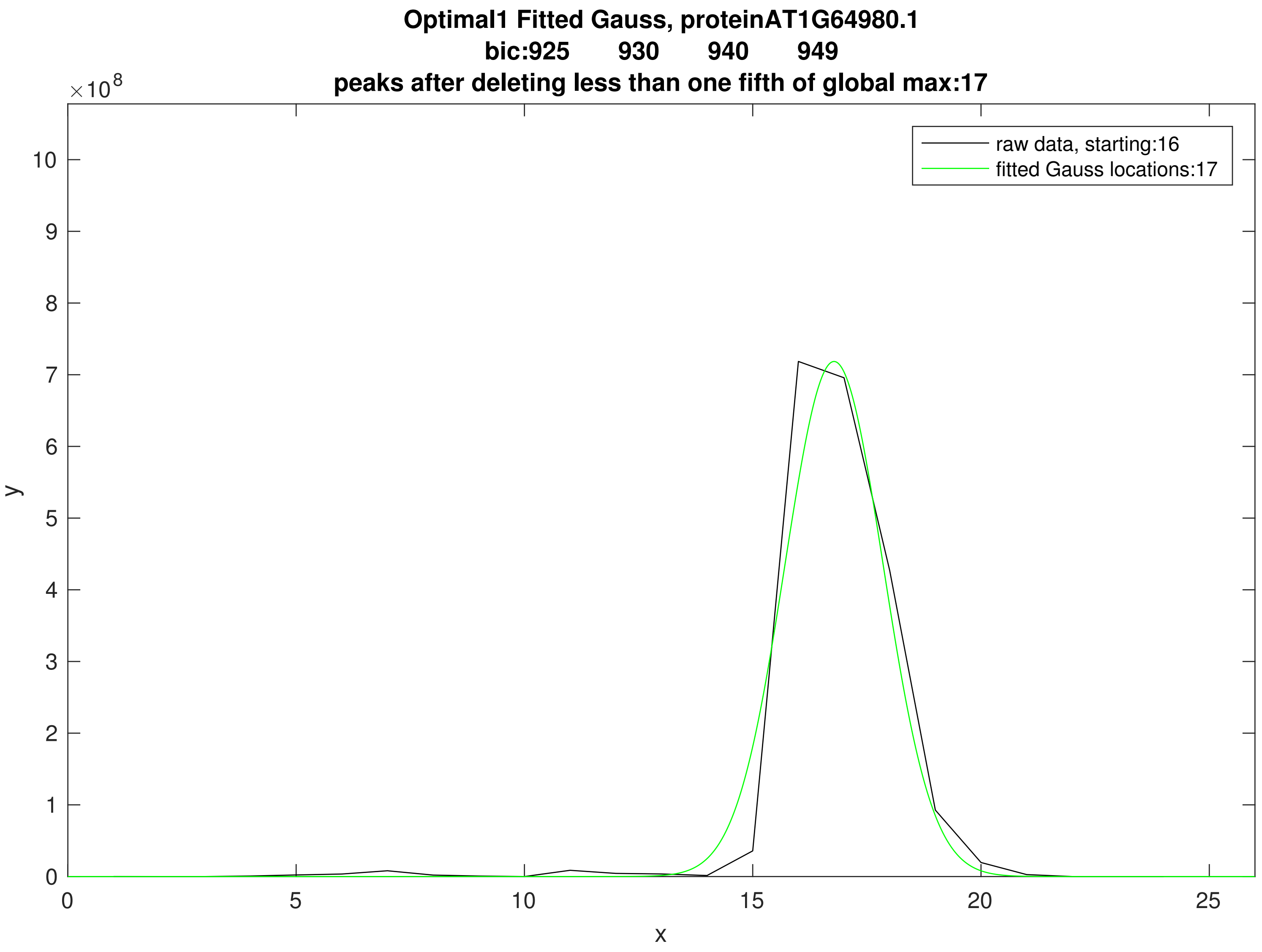
Optimal1 Fitted Gauss, proteinAT1G64980.1
bic:925 930 940 949
peaks after deleting less than one fifth of global max:17
×10^8
raw data, starting:16
fitted Gauss locations:17
y
x
0
1
2
3
4
5
6
7
8
9
10
0
5
10
15
20
25

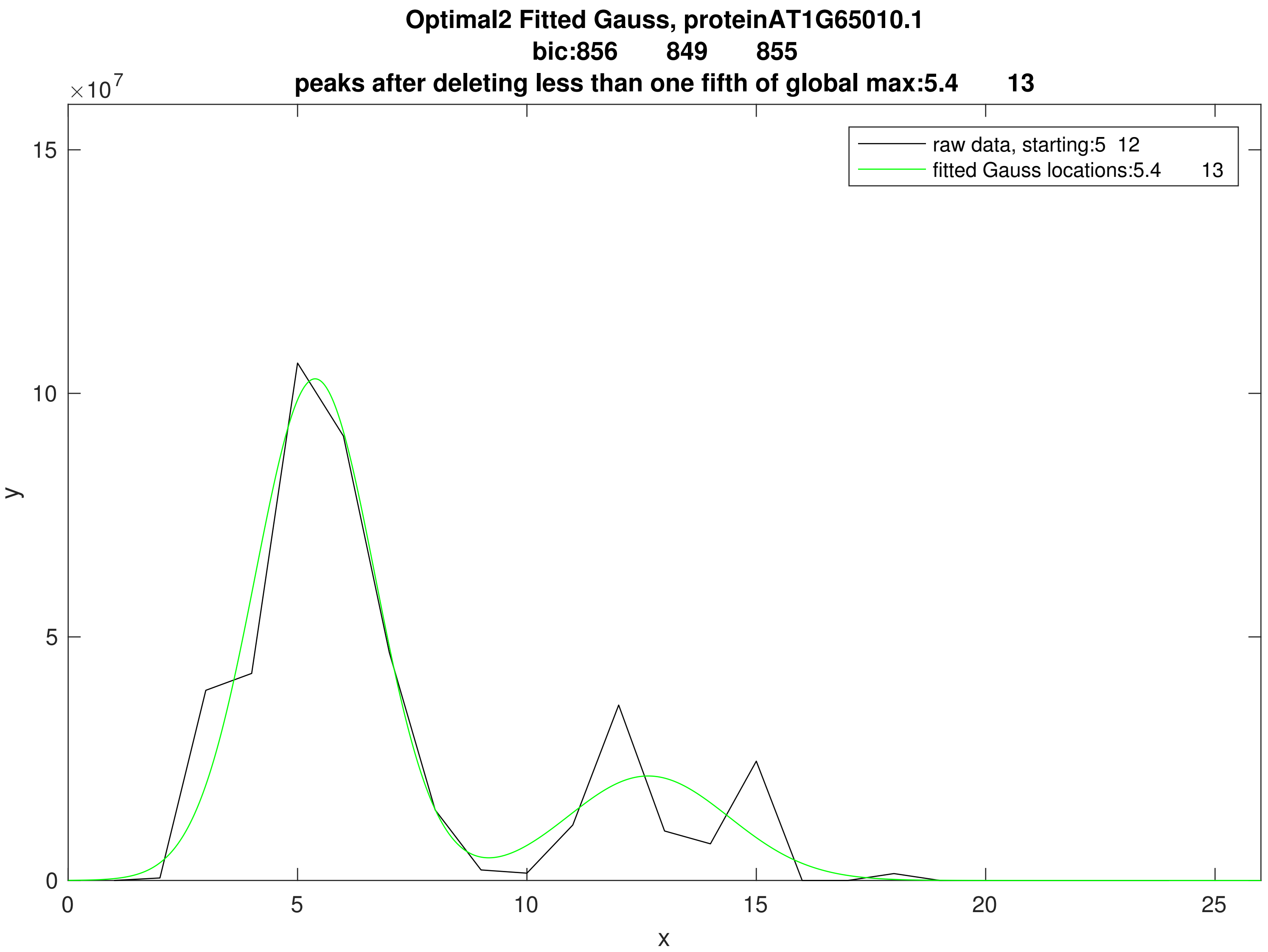

Optimal2 Fitted Gauss, proteinAT1G65010.1
bic:856 849 855
peaks after deleting less than one fifth of global max:5.4 13
×10^7
raw data, starting:5 12
fitted Gauss locations:5.4 13
y
x
0
5
10
15
20
25

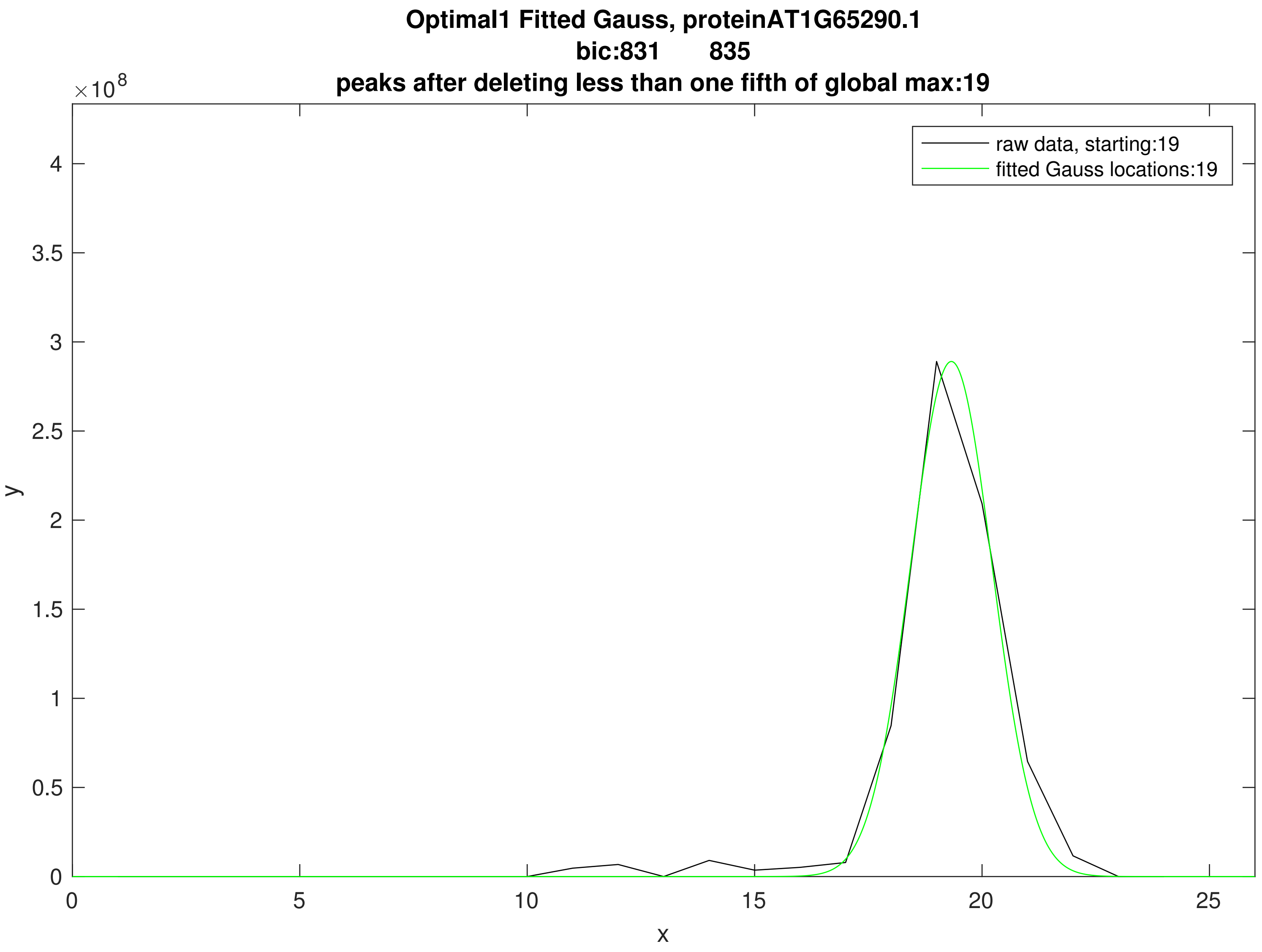
Optimal1 Fitted Gauss, proteinAT1G65290.1
bic:831 835
peaks after deleting less than one fifth of global max:19
×10^8
raw data, starting:19
fitted Gauss locations:19
y
x
0
0.5
1
1.5
2
2.5
3
3.5
4
0
5
10
15
20
25

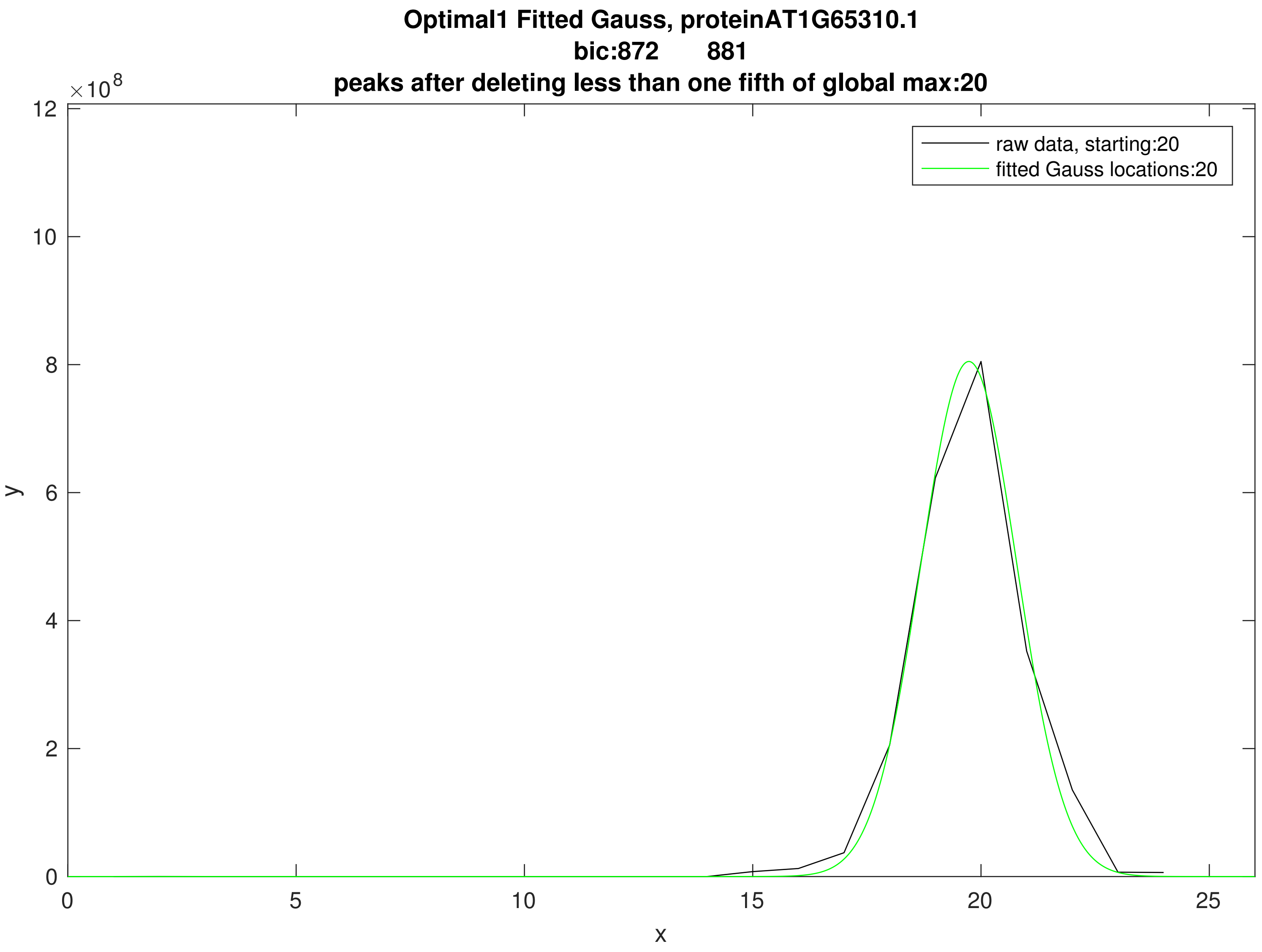
Optimal1 Fitted Gauss, proteinAT1G65310.1
bic:872 881
peaks after deleting less than one fifth of global max:20
raw data, starting:20
fitted Gauss locations:20
y
x
×10^8
0
2
4
6
8
10
12
0
5
10
15
20
25

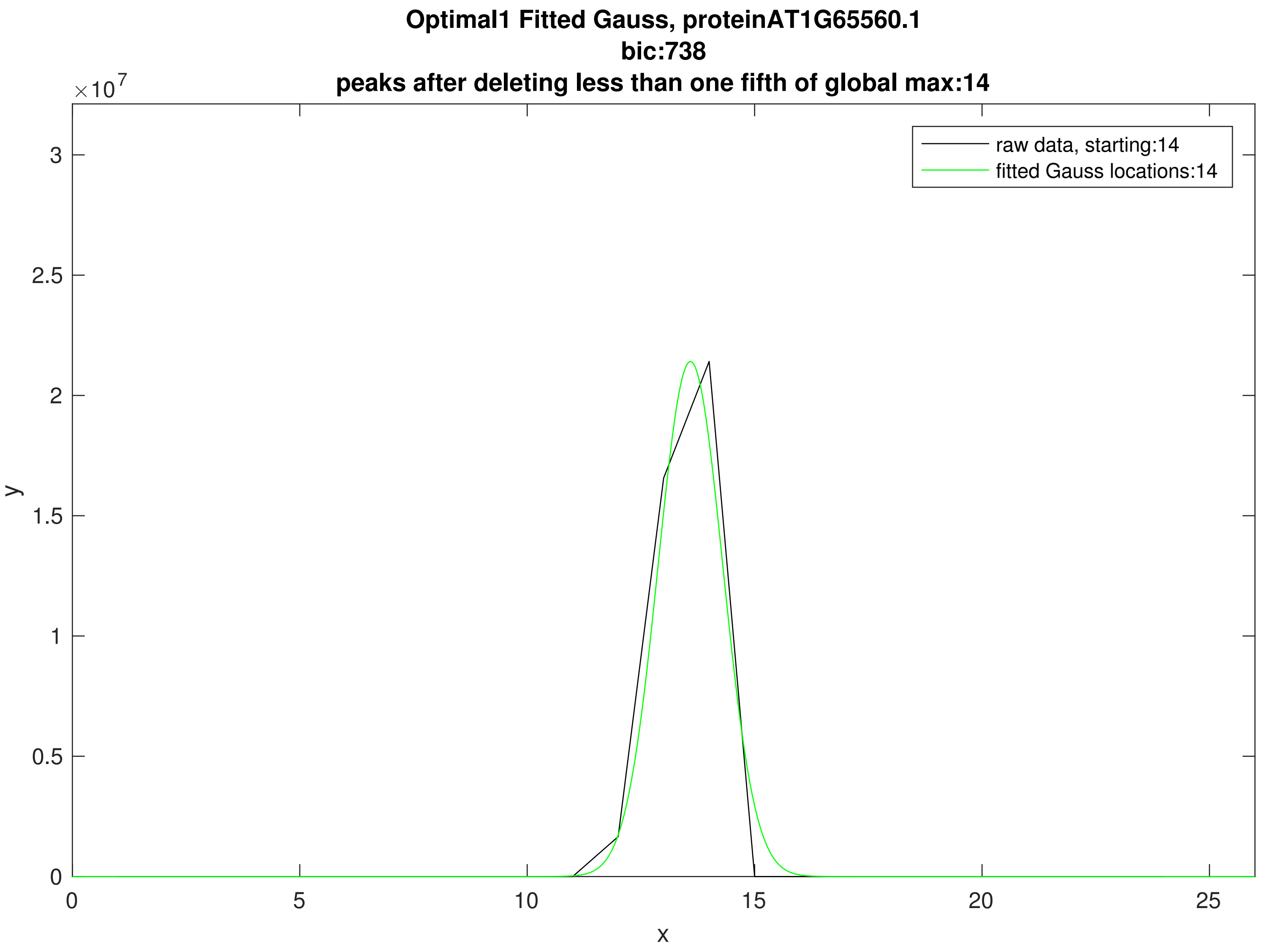

Optimal1 Fitted Gauss, proteinAT1G65560.1
bic:738
peaks after deleting less than one fifth of global max:14
×10^7
3
2.5
2
1.5
1
0.5
0
y
0
5
10
15
20
25
x
raw data, starting:14
fitted Gauss locations:14

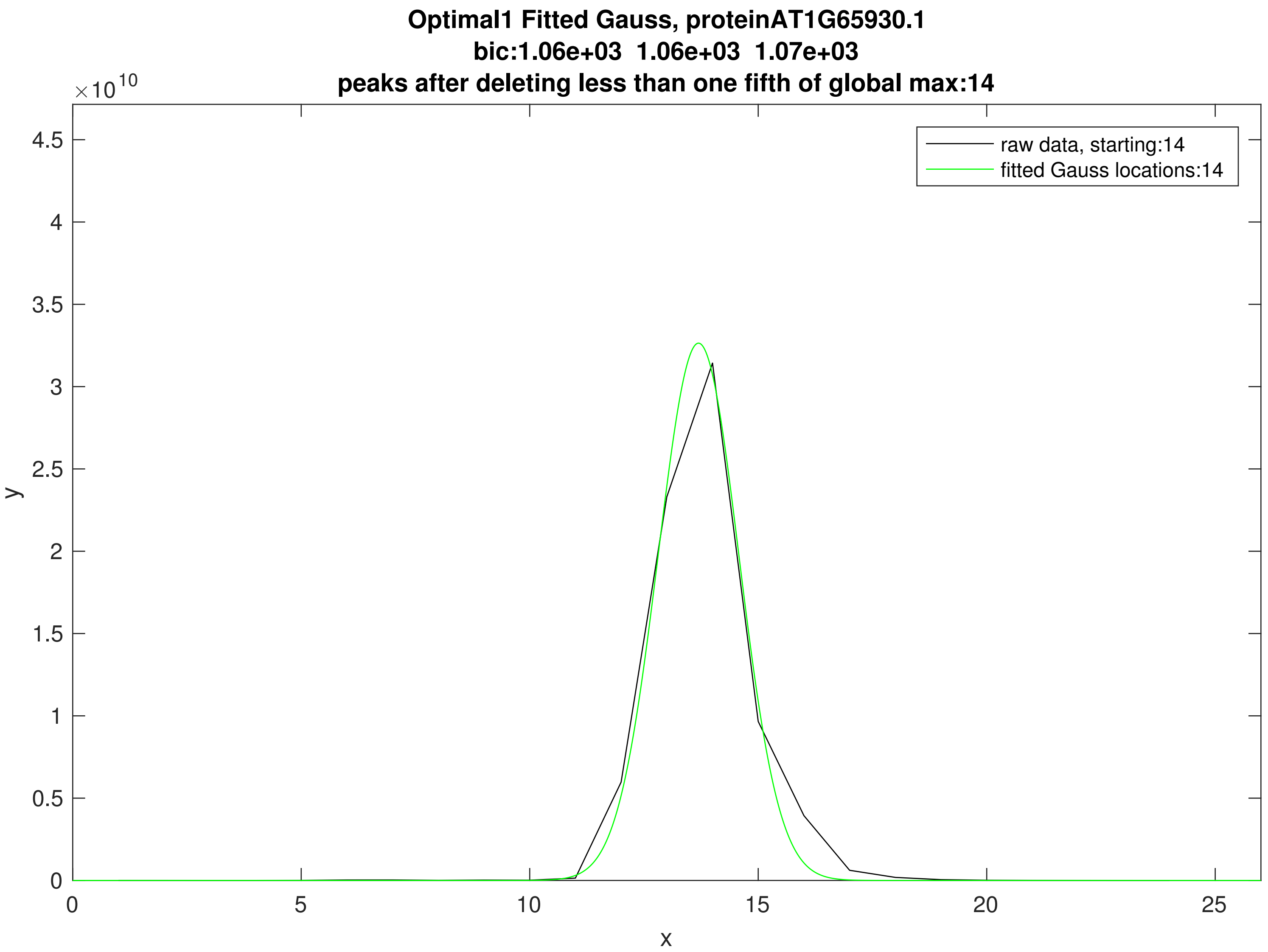
Optimal1 Fitted Gauss, proteinAT1G65930.1
bic:1.06e+03 1.06e+03 1.07e+03
peaks after deleting less than one fifth of global max:14
×10^10
raw data, starting:14
fitted Gauss locations:14
4.5
4
3.5
3
2.5
2
1.5
1
0.5
0
y
0
5
10
15
20
25
x

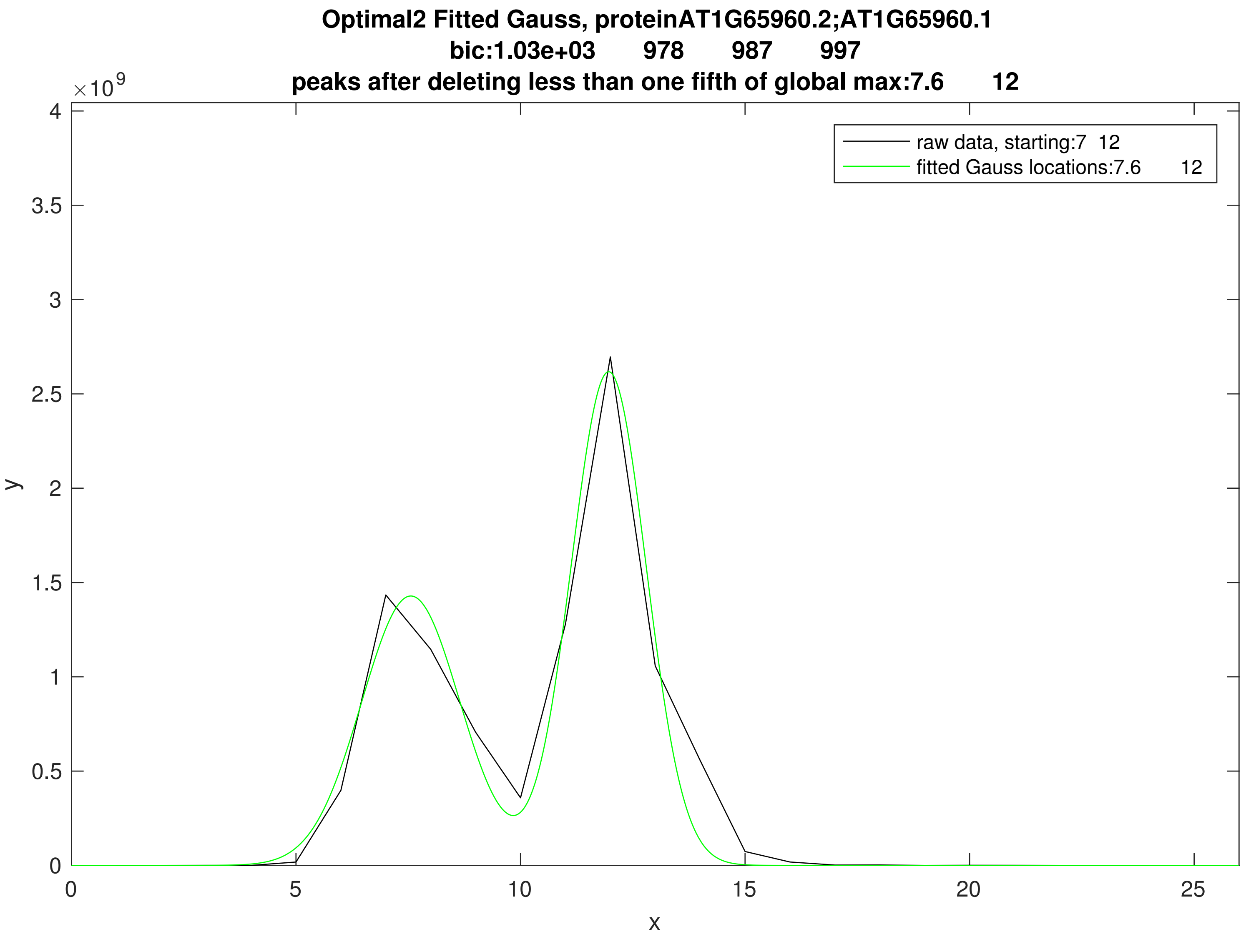
Optimal2 Fitted Gauss, proteinAT1G65960.2;AT1G65960.1
bic:1.03e+03 978 987 997
peaks after deleting less than one fifth of global max:7.6 12
raw data, starting:7 12
fitted Gauss locations:7.6 12
$\times 10^9$
y
x

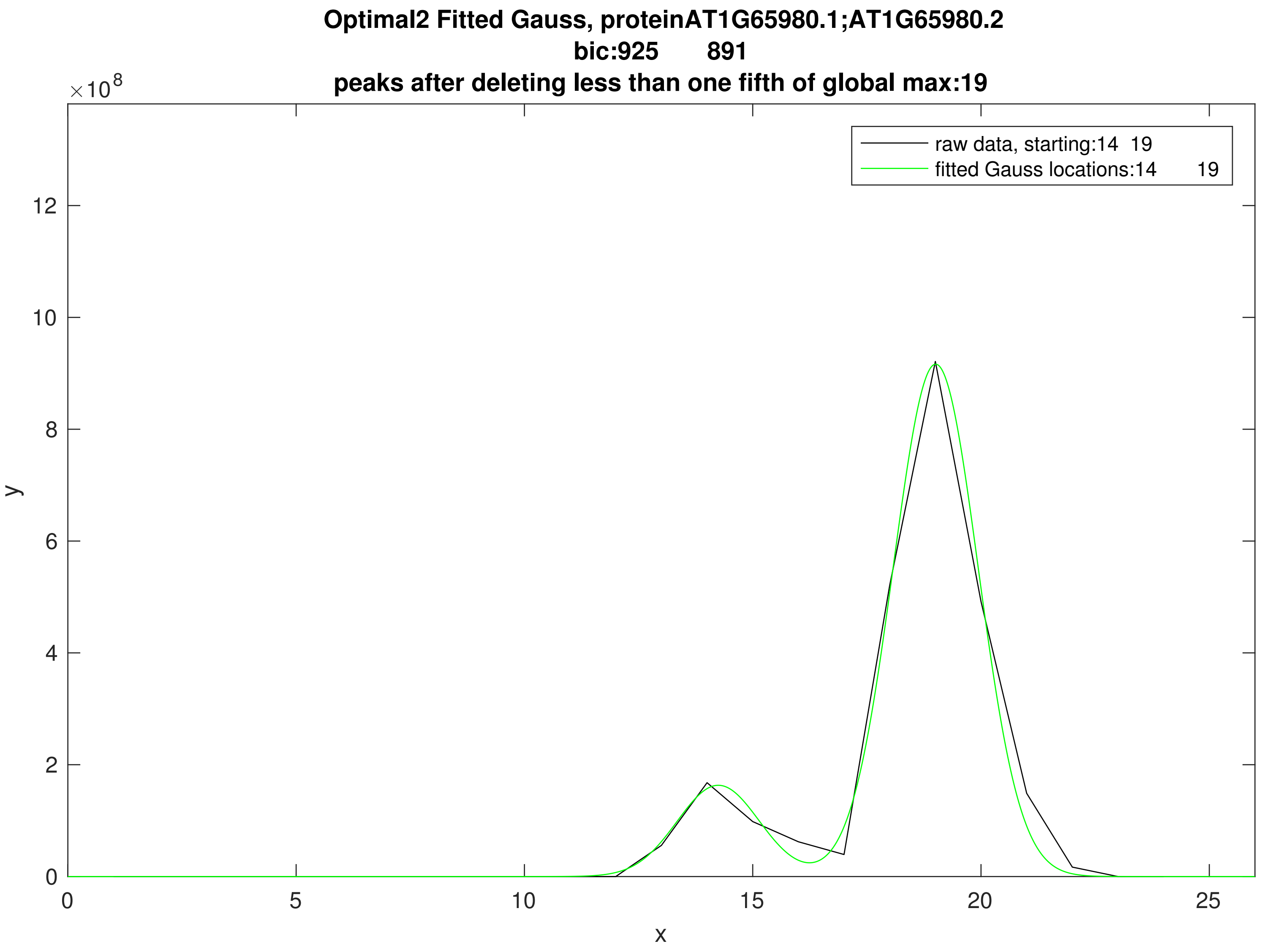
Optimal2 Fitted Gauss, proteinAT1G65980.1;AT1G65980.2
bic:925 891
peaks after deleting less than one fifth of global max:19
×10^8
raw data, starting:14 19
fitted Gauss locations:14 19
y
x
0
2
4
6
8
10
12
0
5
10
15
20
25

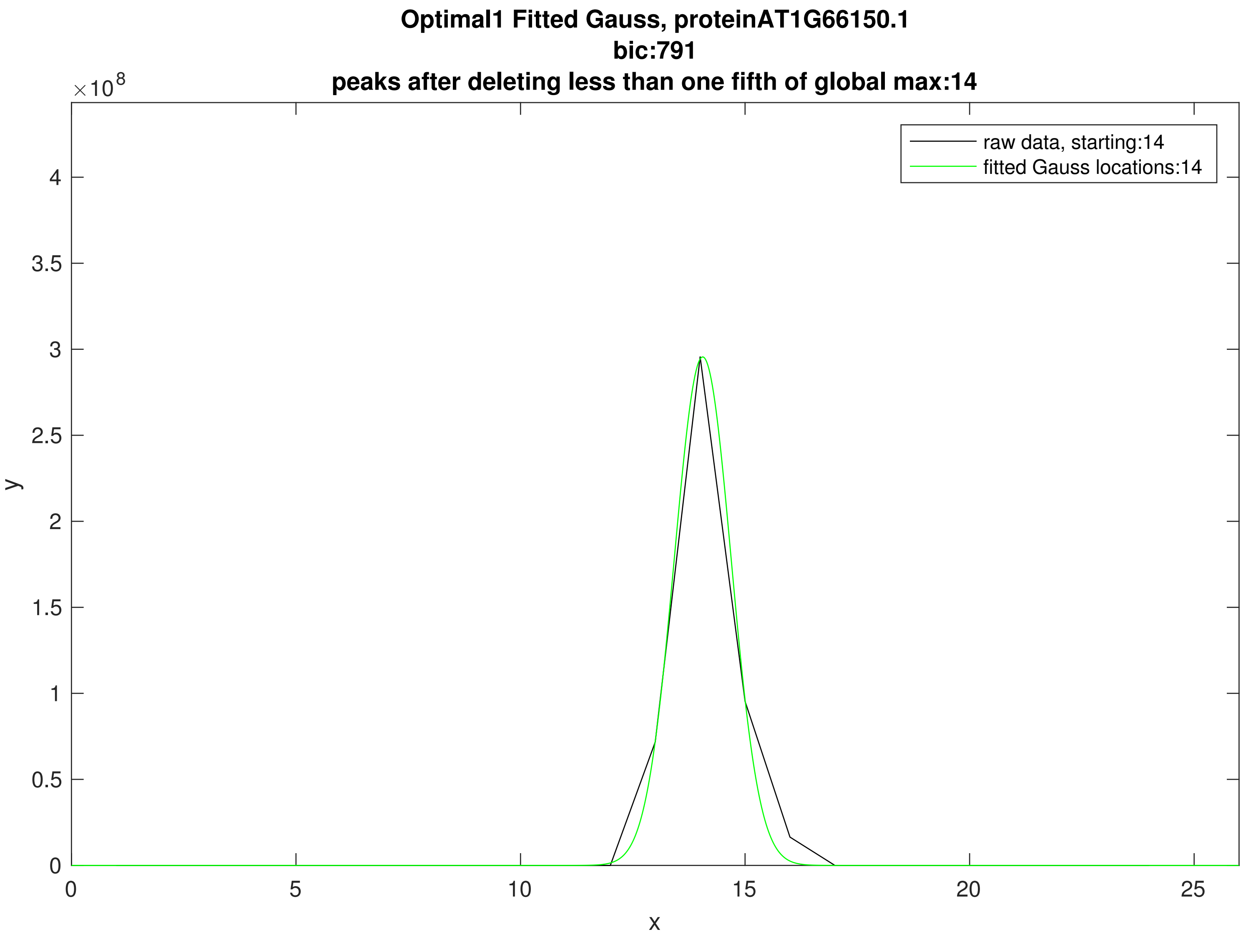
Optimal1 Fitted Gauss, proteinAT1G66150.1
bic:791
peaks after deleting less than one fifth of global max:14
×10^8
4
3.5
3
2.5
2
1.5
1
0.5
0
y
0
5
10
15
20
25
x
raw data, starting:14
fitted Gauss locations:14

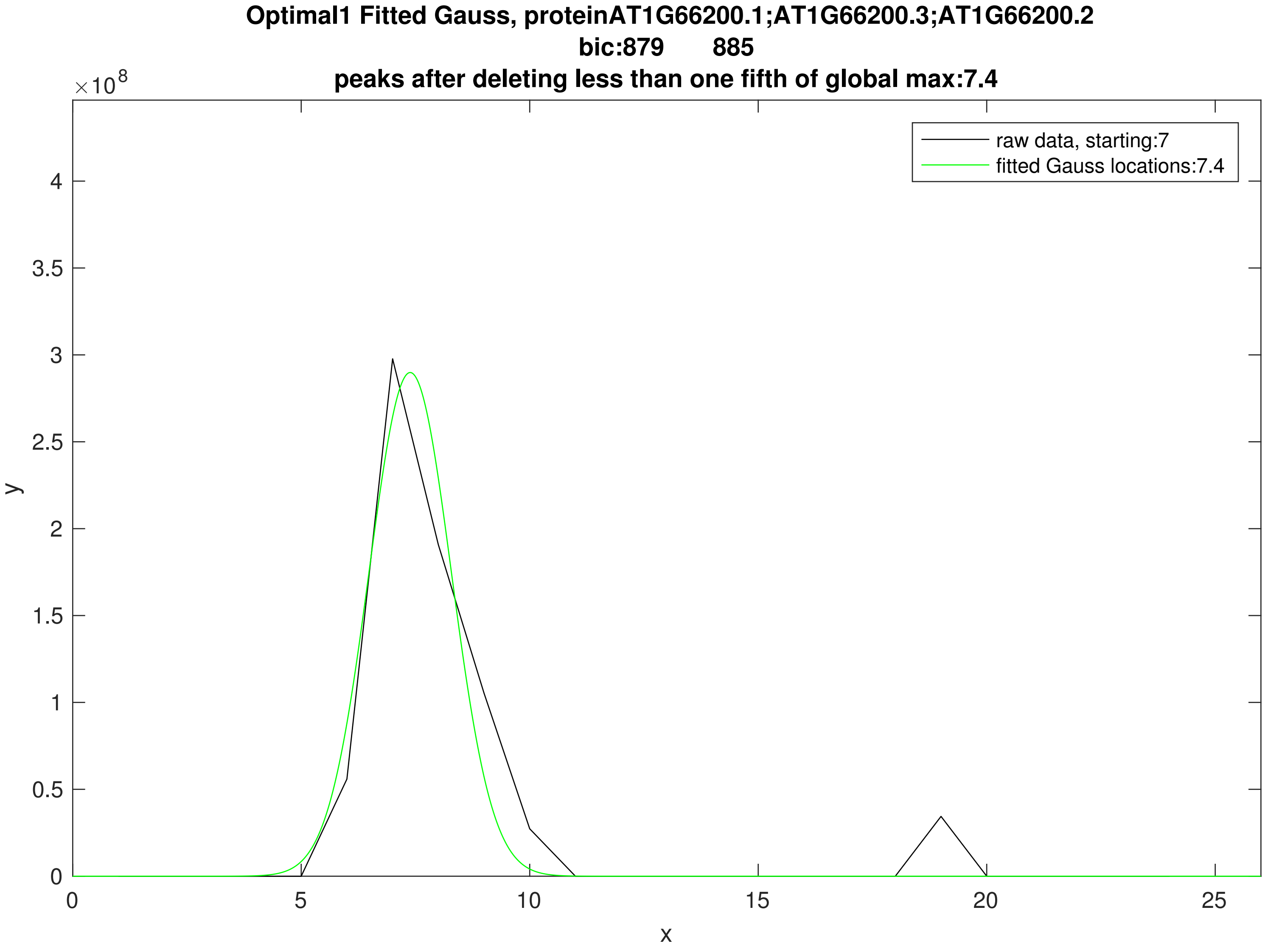
Optimal1 Fitted Gauss, proteinAT1G66200.1;AT1G66200.3;AT1G66200.2
bic:879 885
peaks after deleting less than one fifth of global max:7.4
×10^8
raw data, starting:7
fitted Gauss locations:7.4
y
x
0
0.5
1
1.5
2
2.5
3
3.5
4
0
5
10
15
20
25

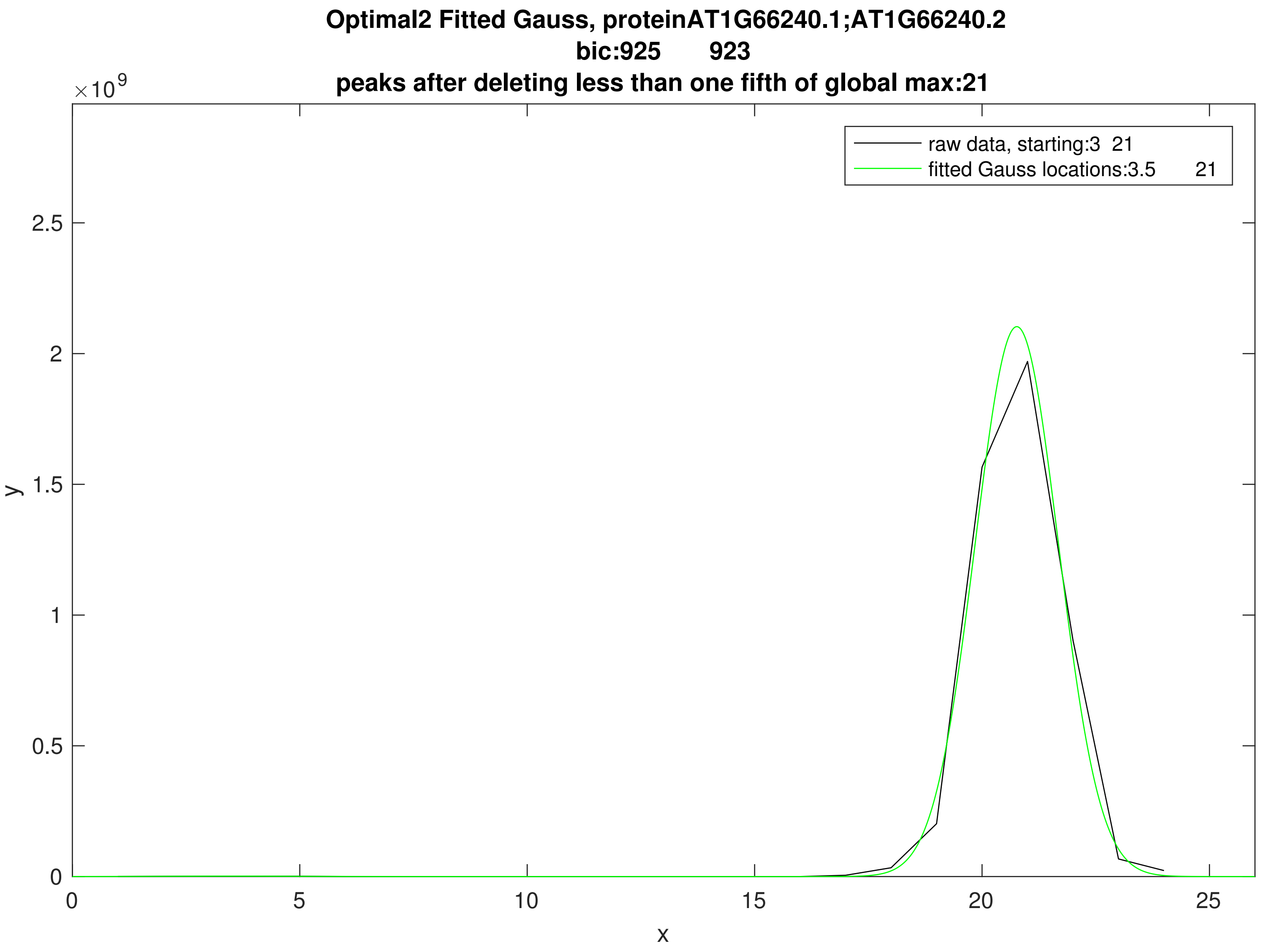
Optimal2 Fitted Gauss, proteinAT1G66240.1;AT1G66240.2
bic:925 923
peaks after deleting less than one fifth of global max:21
×10^9
raw data, starting:3 21
fitted Gauss locations:3.5 21
2.5
2
y 1.5
1
0.5
0
0
5
10
15
20
25
x

**Optimal1 Fitted Gauss, proteinAT1G66280.1**
**bic:812**
**peaks after deleting less than one fifth of global max:6.5**

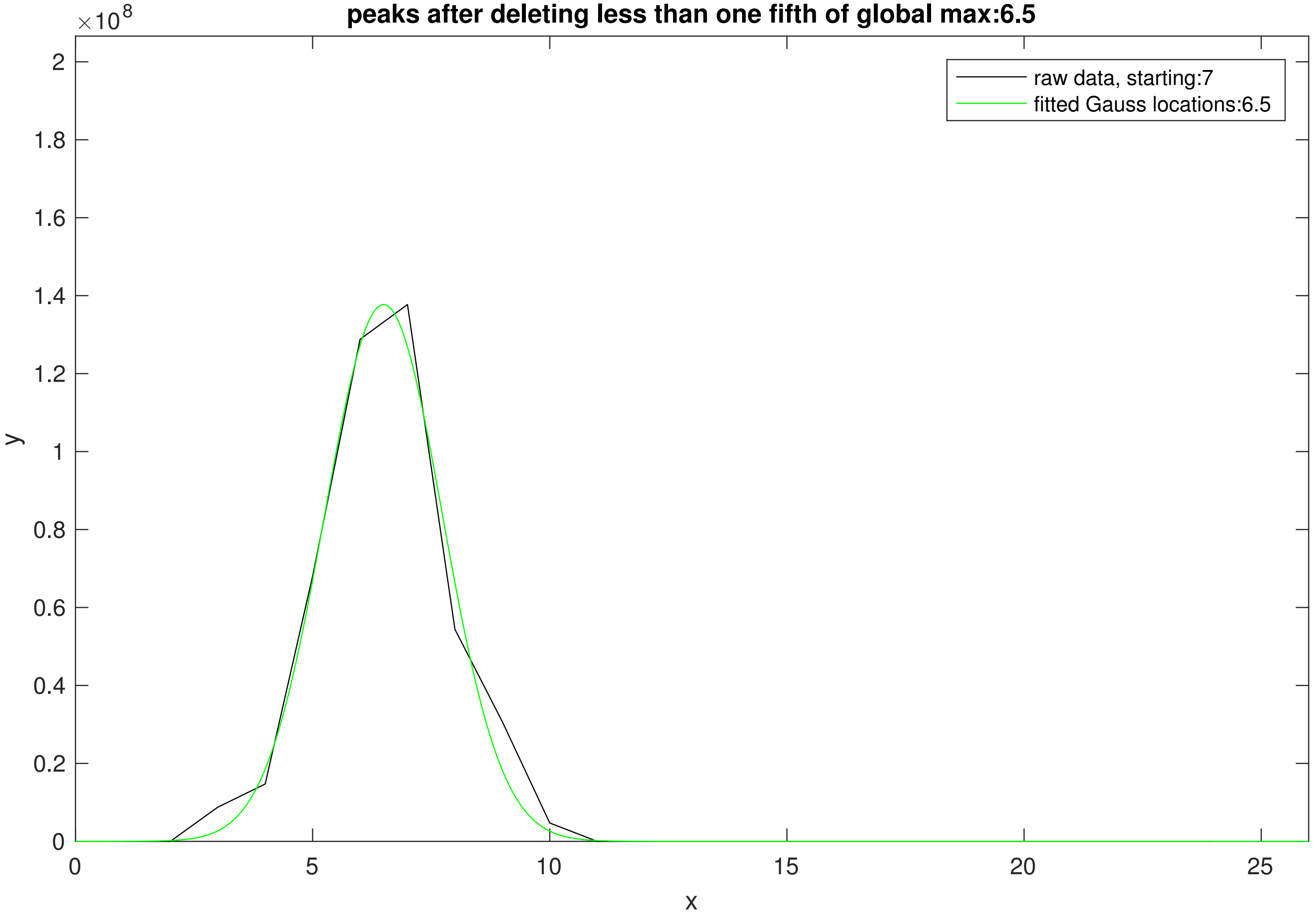

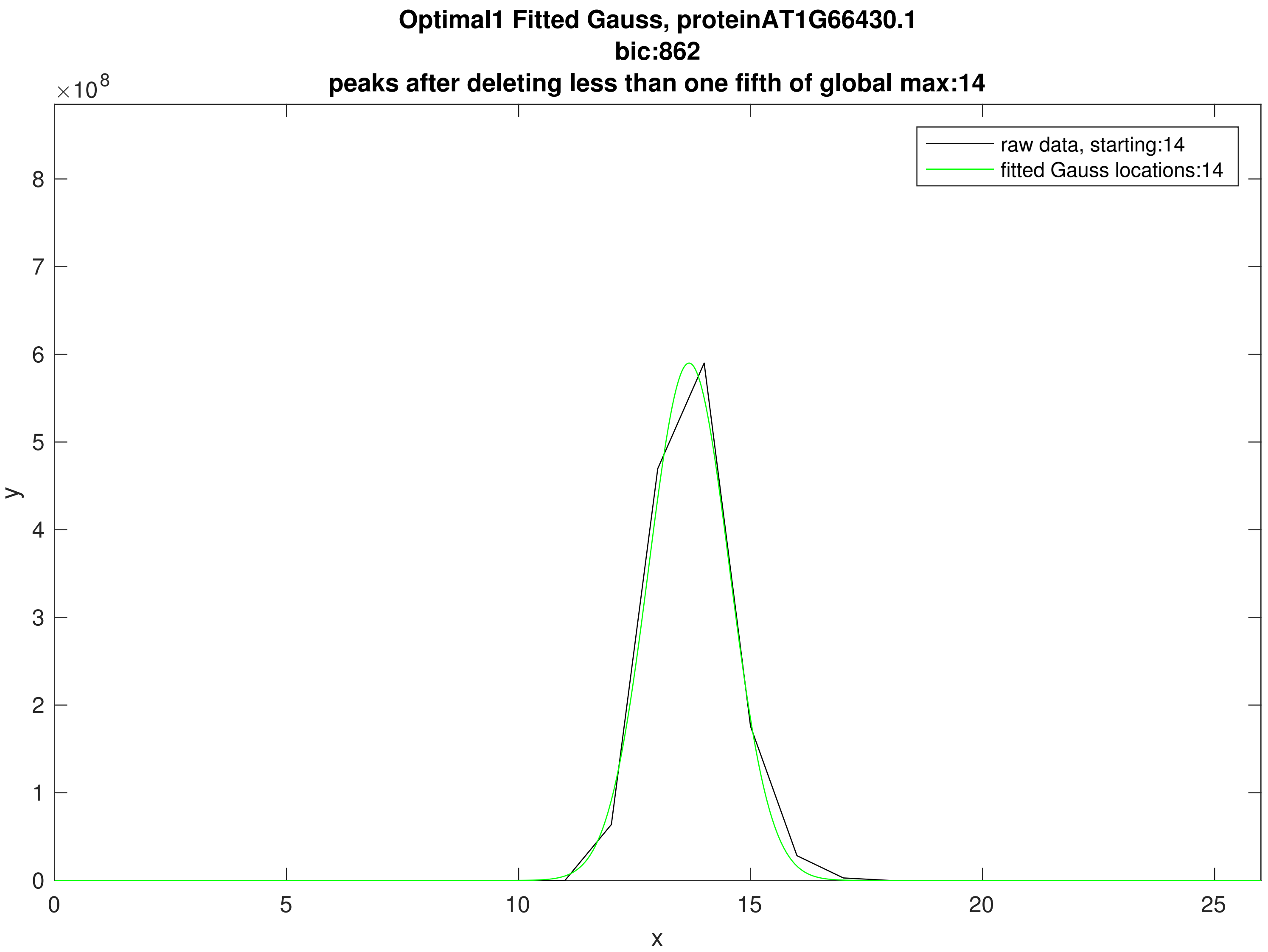
Optimal1 Fitted Gauss, proteinAT1G66430.1
bic:862
peaks after deleting less than one fifth of global max:14
×10^8
raw data, starting:14
fitted Gauss locations:14
y
x
0
1
2
3
4
5
6
7
8
0
5
10
15
20
25

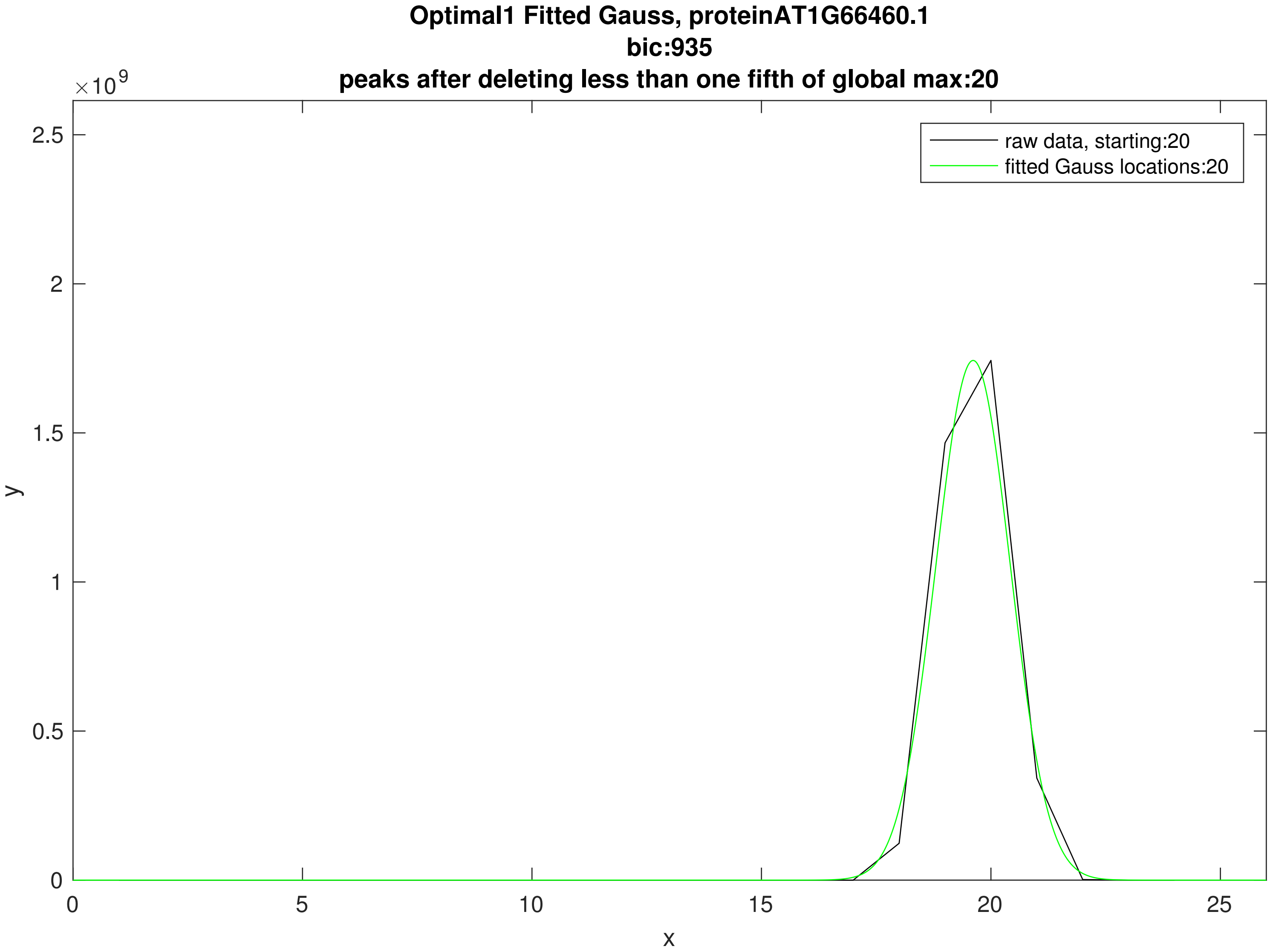
Optimal1 Fitted Gauss, proteinAT1G66460.1
bic:935
peaks after deleting less than one fifth of global max:20
×10^9
2.5
2
1.5
1
0.5
0
y
0
5
10
15
20
25
x
raw data, starting:20
fitted Gauss locations:20

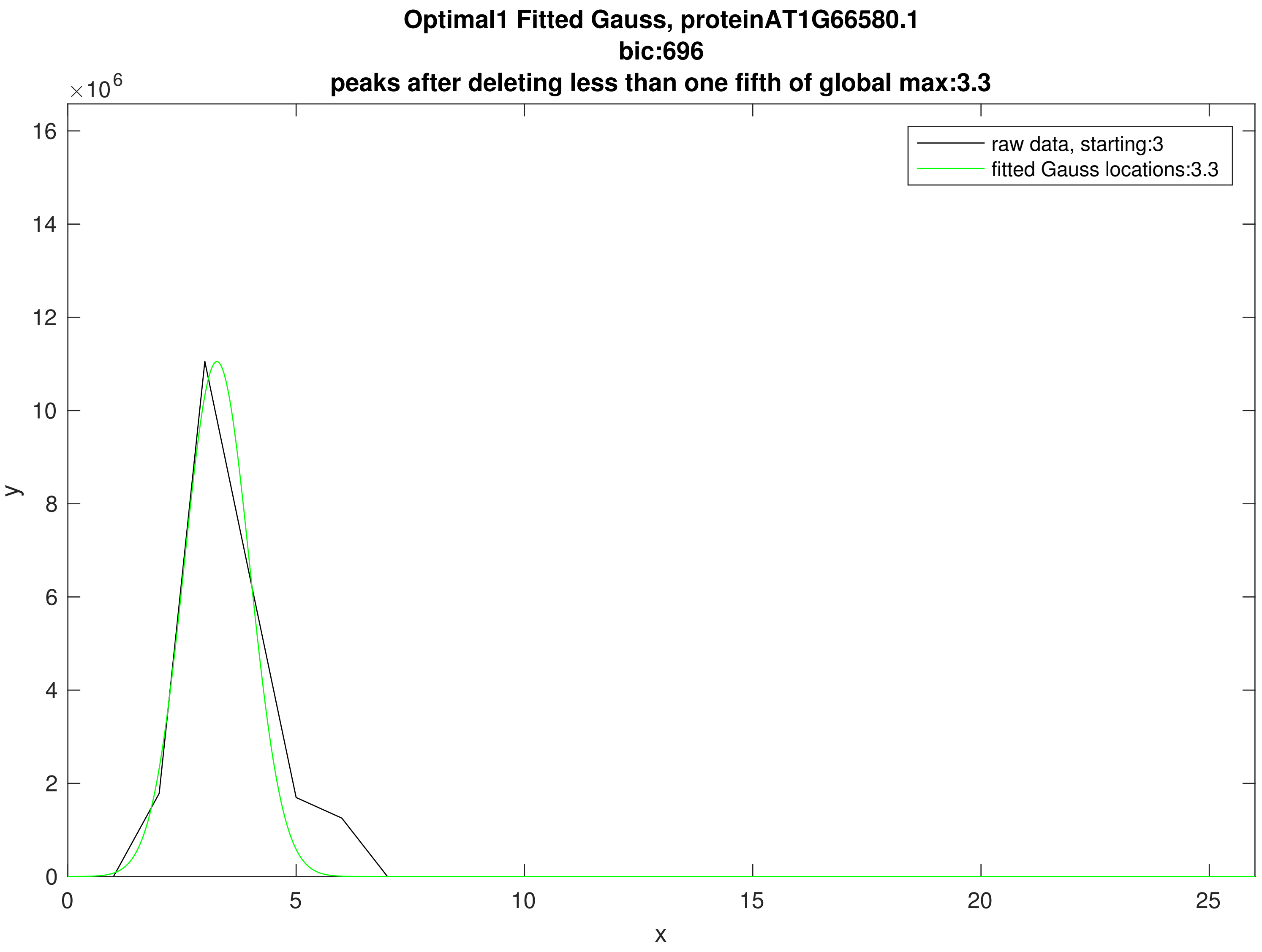

Optimal1 Fitted Gauss, proteinAT1G66580.1
bic:696
peaks after deleting less than one fifth of global max:3.3
×10^6
raw data, starting:3
fitted Gauss locations:3.3
y
x
0
2
4
6
8
10
12
14
16
0
5
10
15
20
25

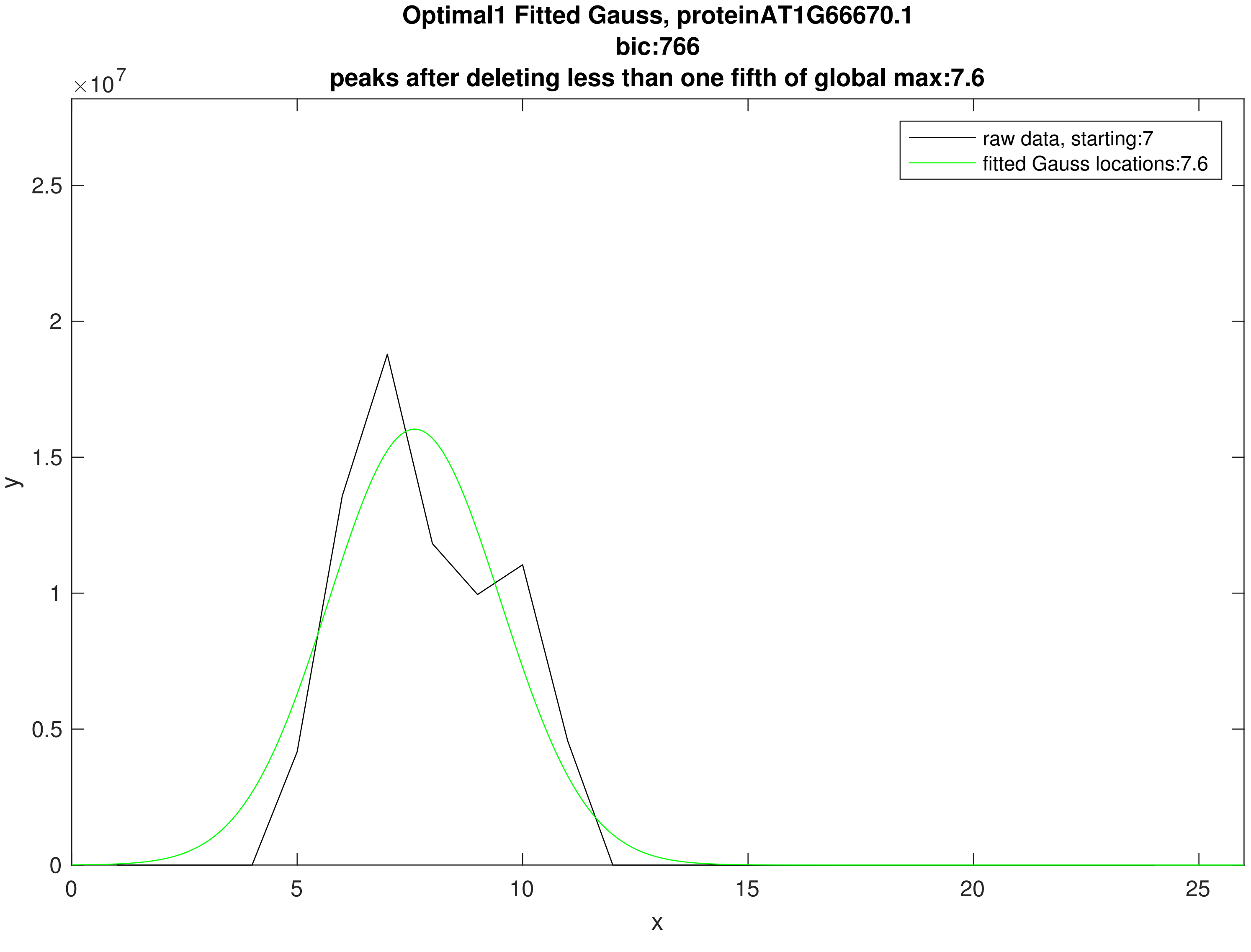
Optimal1 Fitted Gauss, proteinAT1G66670.1
bic:766
peaks after deleting less than one fifth of global max:7.6
×10^7
raw data, starting:7
fitted Gauss locations:7.6
y
x
0
0.5
1
1.5
2
2.5
0
5
10
15
20
25

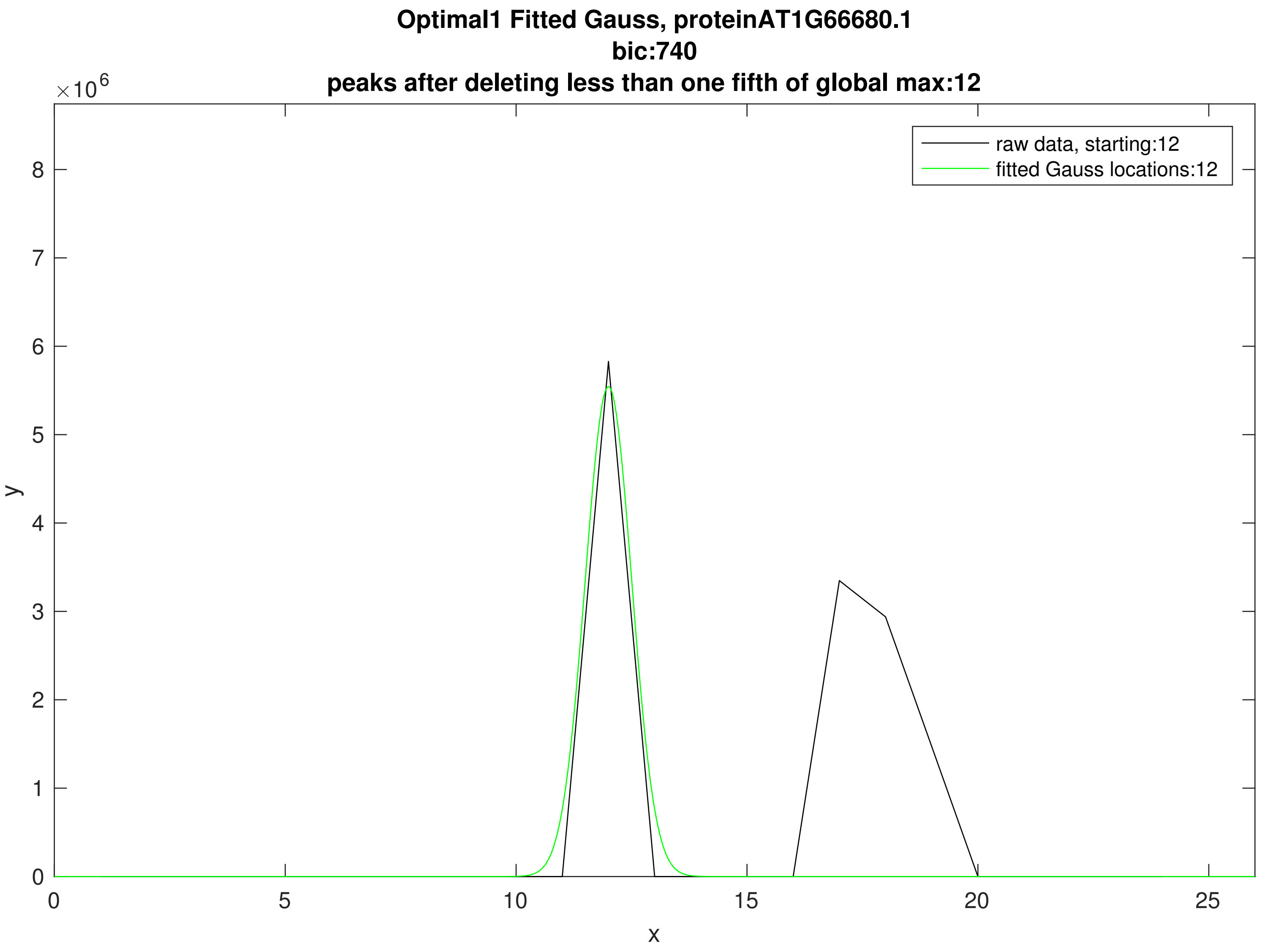
Optimal1 Fitted Gauss, proteinAT1G66680.1
bic:740
peaks after deleting less than one fifth of global max:12
×10^6
raw data, starting:12
fitted Gauss locations:12
y
x
0
1
2
3
4
5
6
7
8
0
5
10
15
20
25

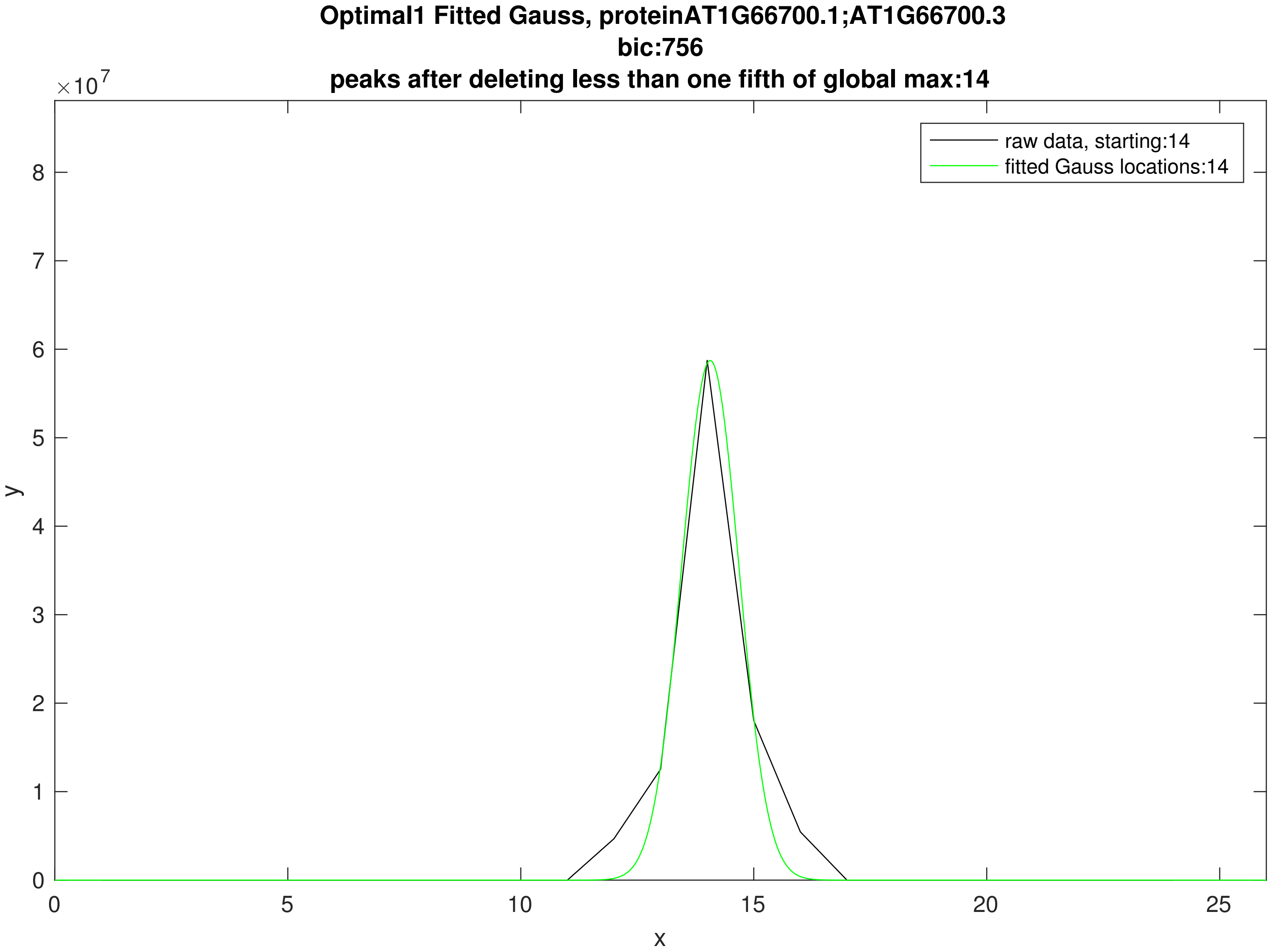

Optimal1 Fitted Gauss, proteinAT1G66700.1;AT1G66700.3
bic:756
peaks after deleting less than one fifth of global max:14
×10^7
raw data, starting:14
fitted Gauss locations:14
y
x
0
1
2
3
4
5
6
7
8
0
5
10
15
20
25

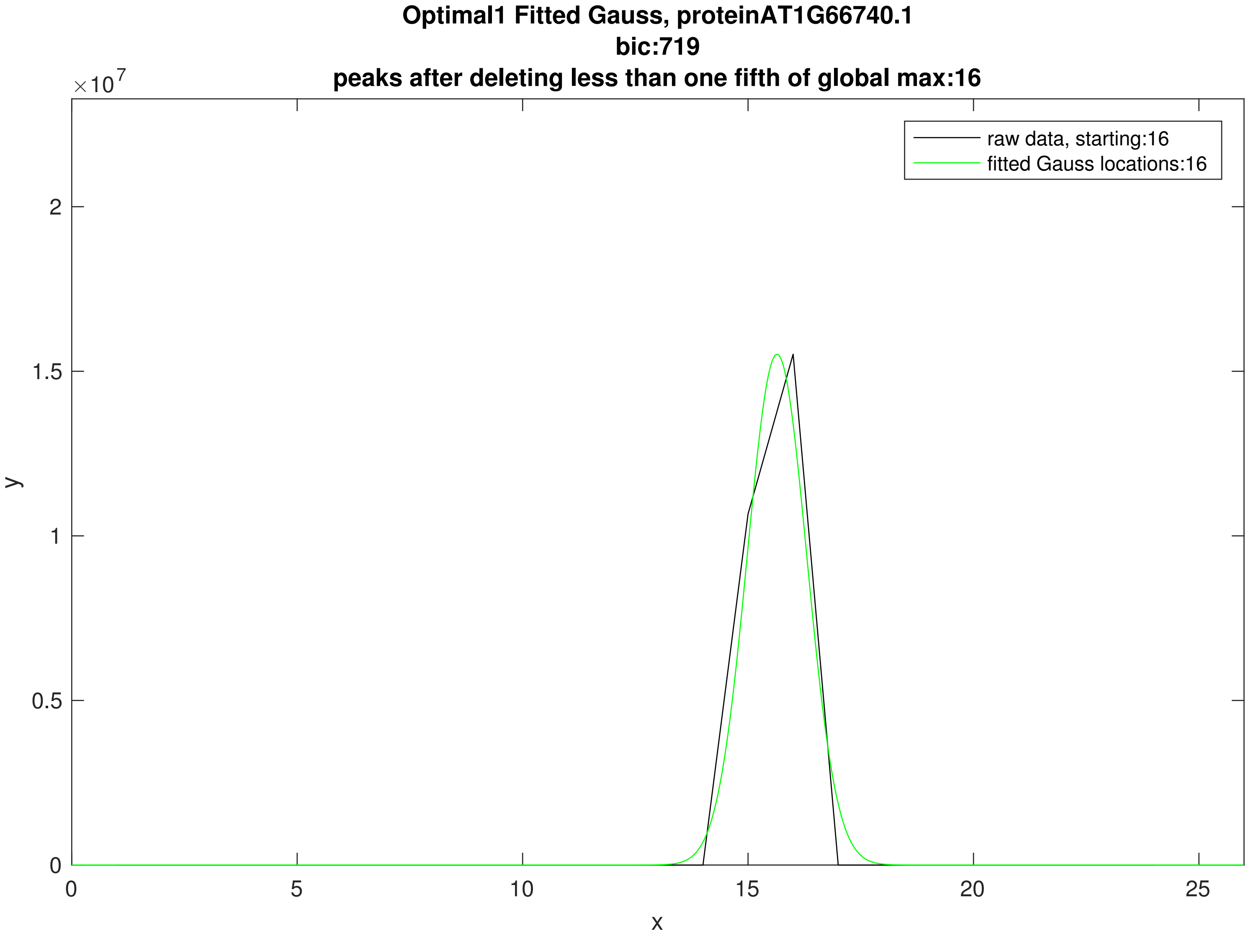

Optimal1 Fitted Gauss, proteinAT1G66740.1
bic:719
peaks after deleting less than one fifth of global max:16
×10^7
raw data, starting:16
fitted Gauss locations:16
y
x
0
0.5
1
1.5
2
0
5
10
15
20
25

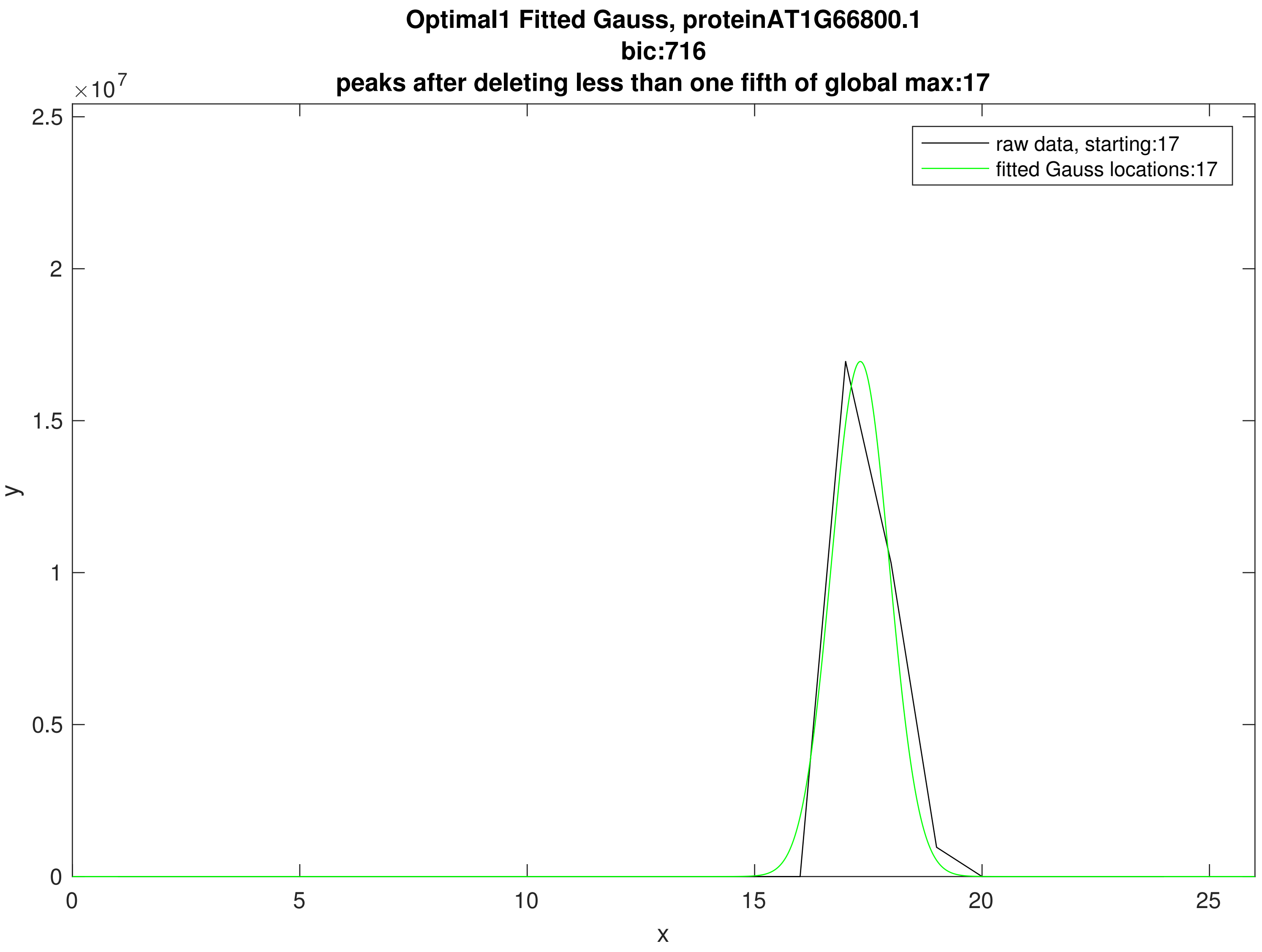
Optimal1 Fitted Gauss, proteinAT1G66800.1
bic:716
peaks after deleting less than one fifth of global max:17
×10^7
2.5
2
1.5
1
0.5
0
y
0
5
10
15
20
25
x
raw data, starting:17
fitted Gauss locations:17

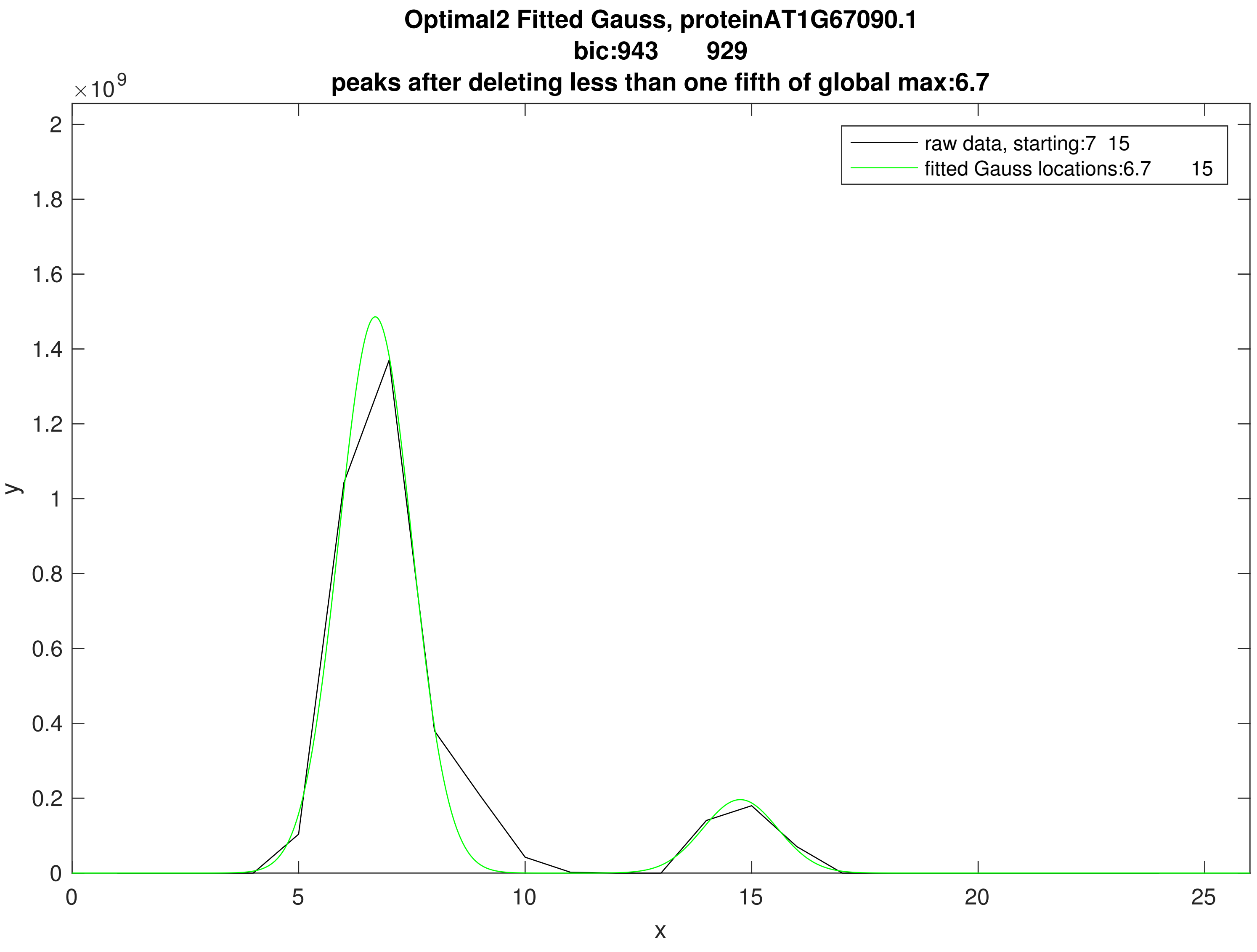
Optimal2 Fitted Gauss, proteinAT1G67090.1
bic:943 929
peaks after deleting less than one fifth of global max:6.7
×10^9
raw data, starting:7 15
fitted Gauss locations:6.7 15
y
x
0
0.2
0.4
0.6
0.8
1
1.2
1.4
1.6
1.8
2
0
5
10
15
20
25

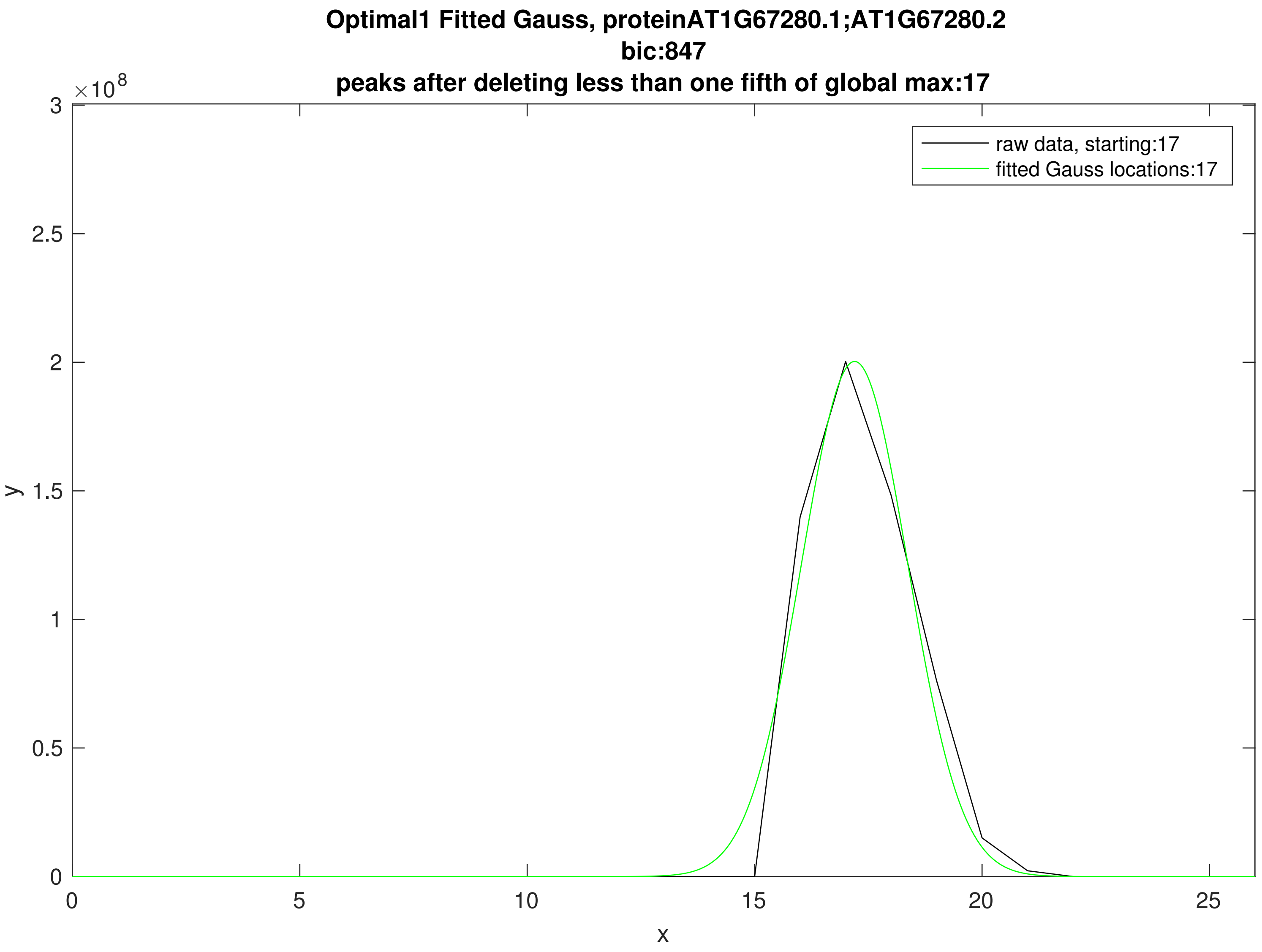

Optimal1 Fitted Gauss, proteinAT1G67280.1;AT1G67280.2
bic:847
peaks after deleting less than one fifth of global max:17
×10^8
3
2.5
2
1.5
1
0.5
0
y
0
5
10
15
20
25
x
raw data, starting:17
fitted Gauss locations:17

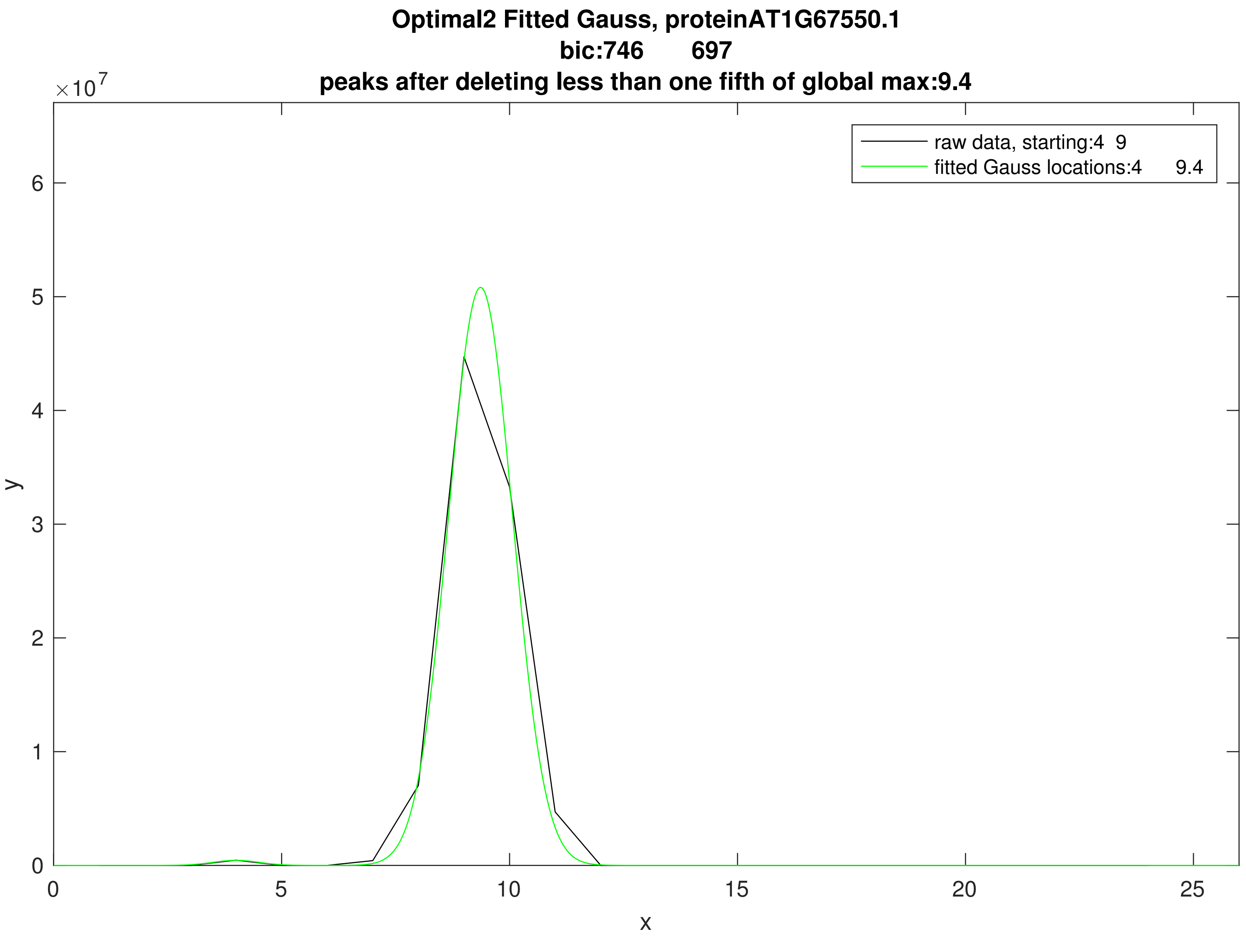
Optimal2 Fitted Gauss, proteinAT1G67550.1
bic:746 697
peaks after deleting less than one fifth of global max:9.4
×10^7
raw data, starting:4 9
fitted Gauss locations:4 9.4
y
x
0
1
2
3
4
5
6
0
5
10
15
20
25

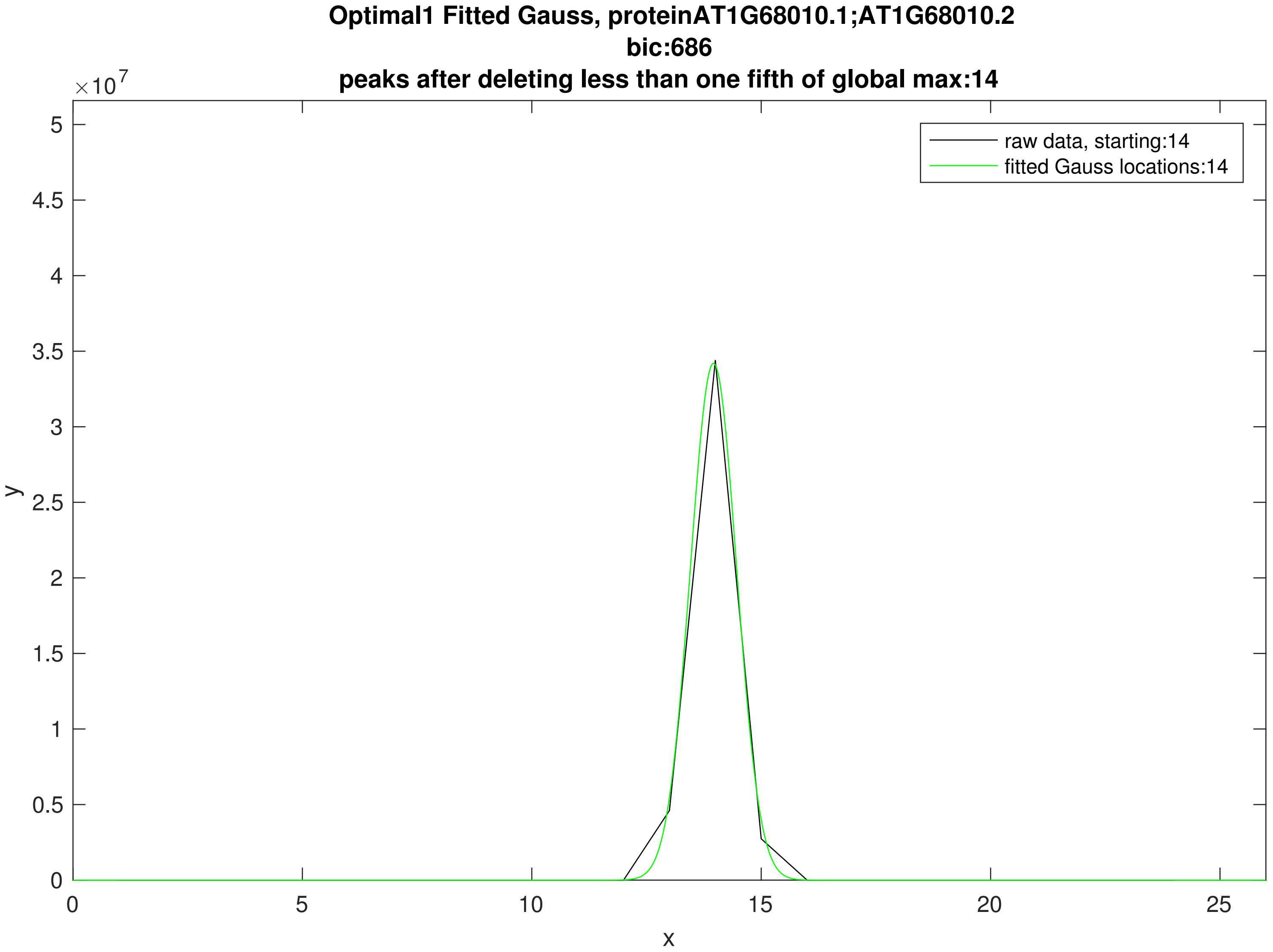
Optimal1 Fitted Gauss, proteinAT1G68010.1;AT1G68010.2
bic:686
peaks after deleting less than one fifth of global max:14
×10^7
raw data, starting:14
fitted Gauss locations:14
y
x
0
0.5
1
1.5
2
2.5
3
3.5
4
4.5
5
0
5
10
15
20
25

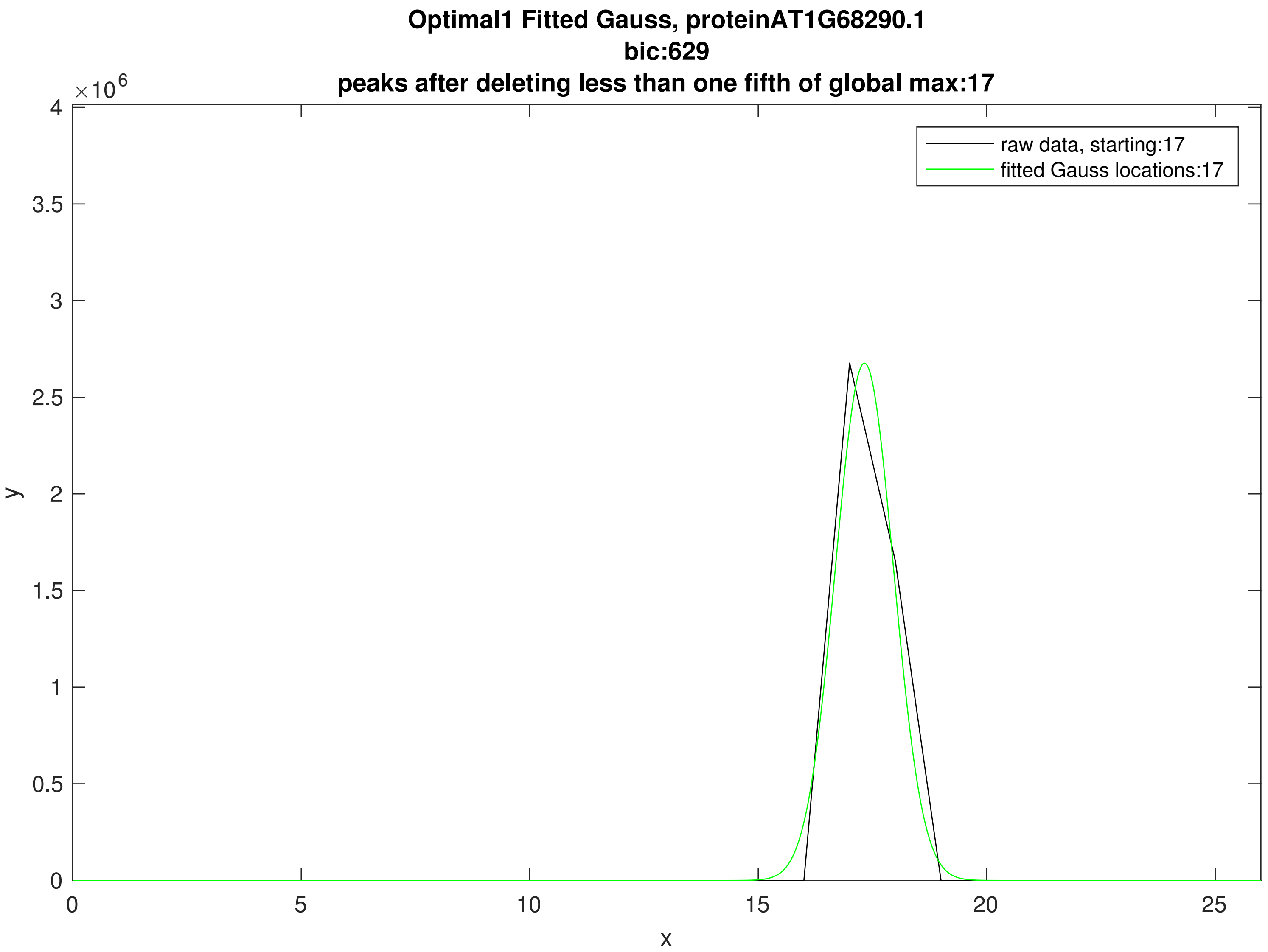

Optimal1 Fitted Gauss, proteinAT1G68290.1
bic:629
peaks after deleting less than one fifth of global max:17
×10^6
raw data, starting:17
fitted Gauss locations:17
y
x
0
0.5
1
1.5
2
2.5
3
3.5
4
0
5
10
15
20
25

Optimal1 Fitted Gauss, proteinAT1G68310.2;AT1G68310.1
bic:702
peaks after deleting less than one fifth of global max:14

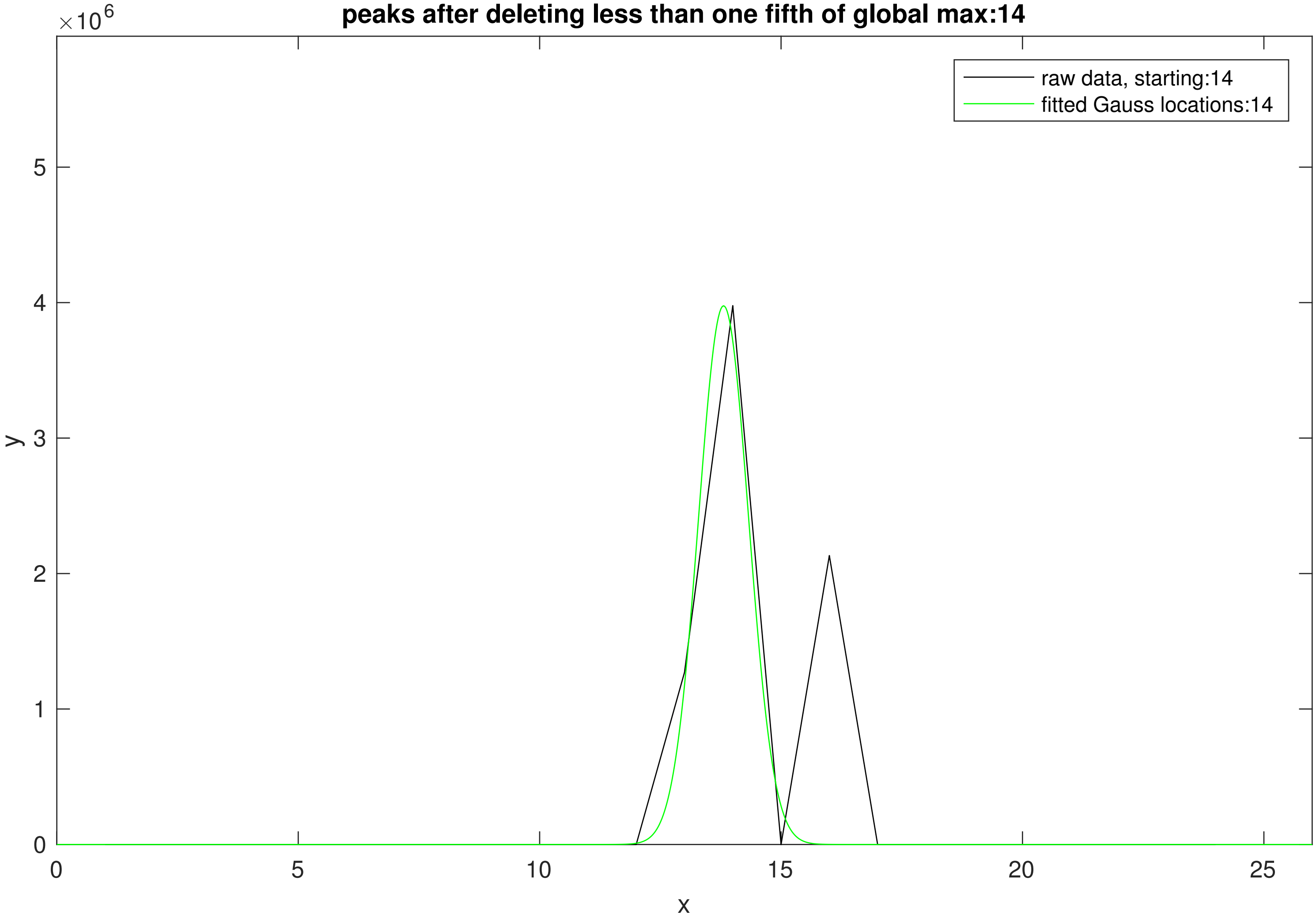

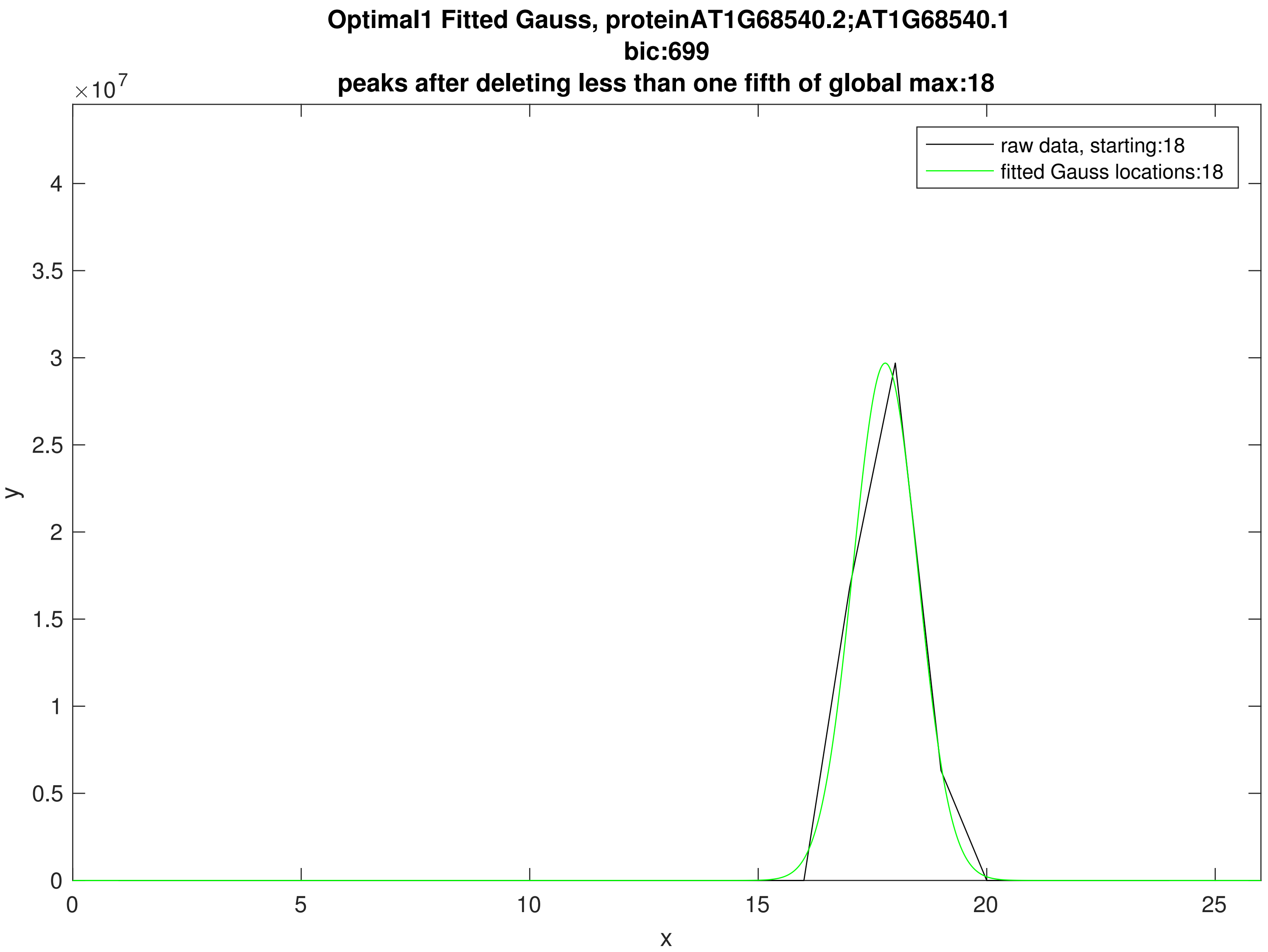
Optimal1 Fitted Gauss, proteinAT1G68540.2;AT1G68540.1
bic:699
peaks after deleting less than one fifth of global max:18
×10^7
raw data, starting:18
fitted Gauss locations:18
y
x
0
0.5
1
1.5
2
2.5
3
3.5
4
0
5
10
15
20
25

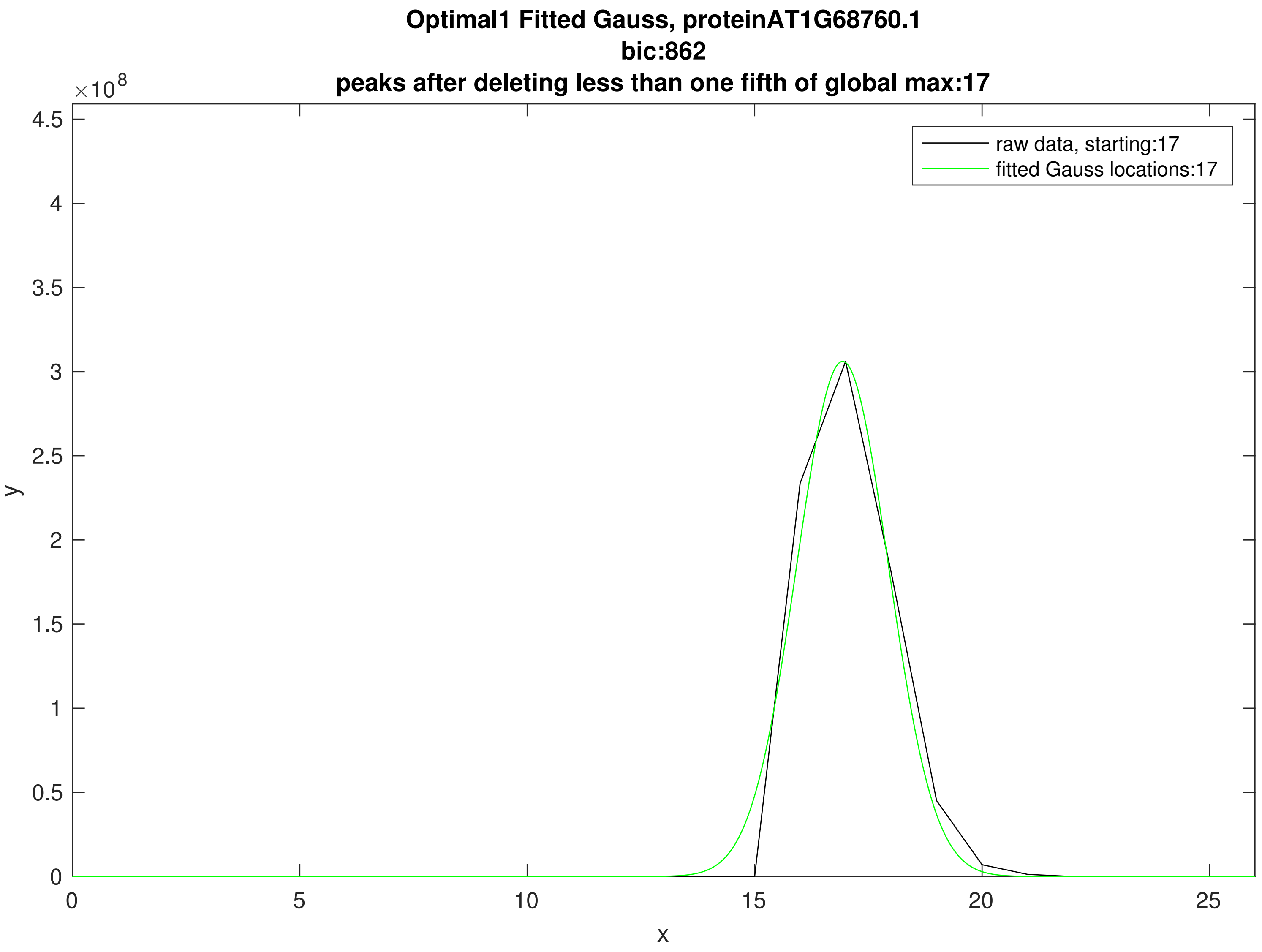
Optimal1 Fitted Gauss, proteinAT1G68760.1
bic:862
peaks after deleting less than one fifth of global max:17
×10^8
4.5
4
3.5
3
2.5
2
1.5
1
0.5
0
y
raw data, starting:17
fitted Gauss locations:17
0
5
10
15
20
25
x

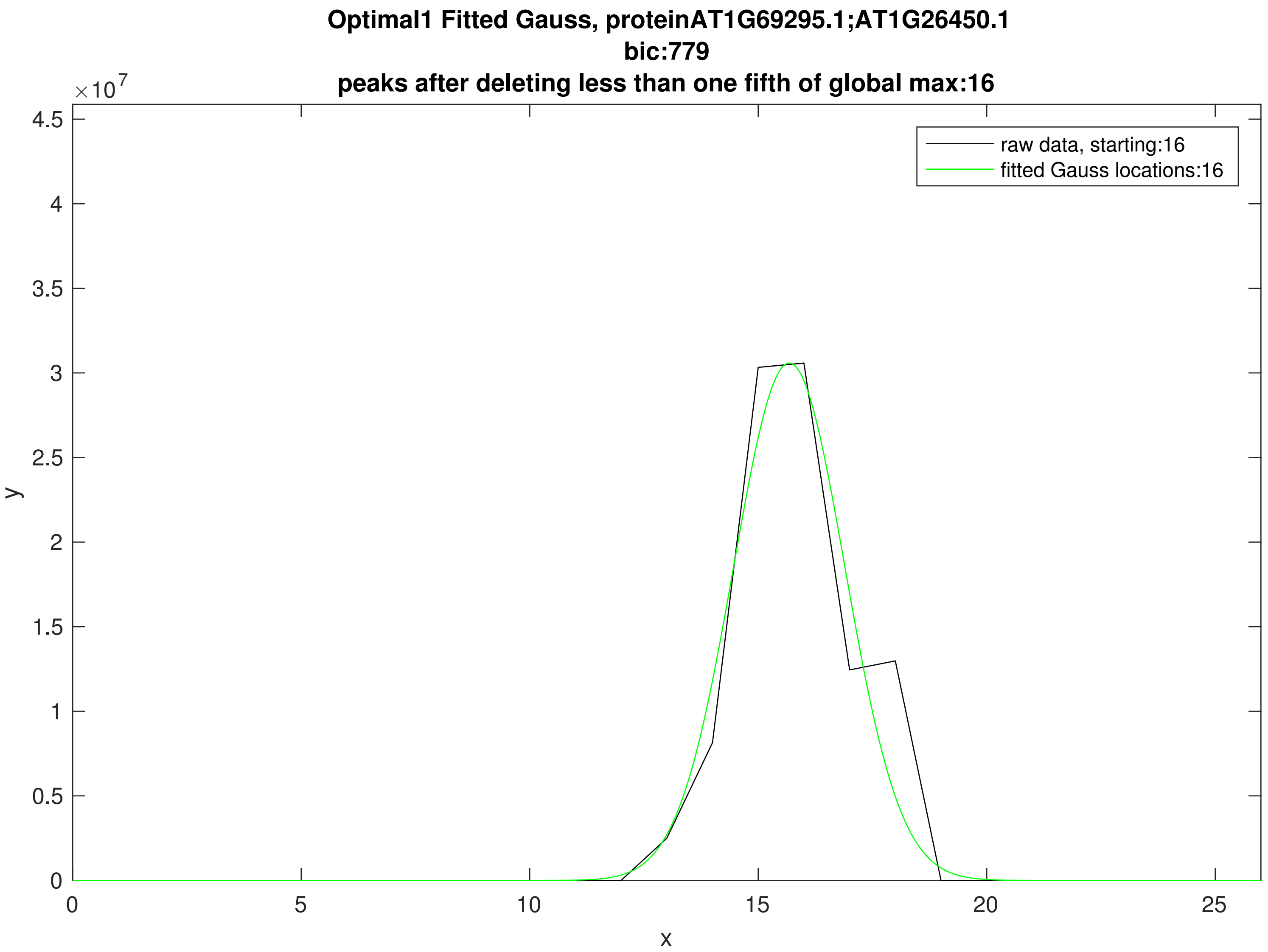
Optimal1 Fitted Gauss, proteinAT1G69295.1;AT1G26450.1
bic:779
peaks after deleting less than one fifth of global max:16
×10^7
4.5
4
3.5
3
2.5
2
1.5
1
0.5
0
y
0
5
10
15
20
25
x
raw data, starting:16
fitted Gauss locations:16

Optimal1 Fitted Gauss, proteinAT1G69340.1
bic:715
peaks after deleting less than one fifth of global max:16

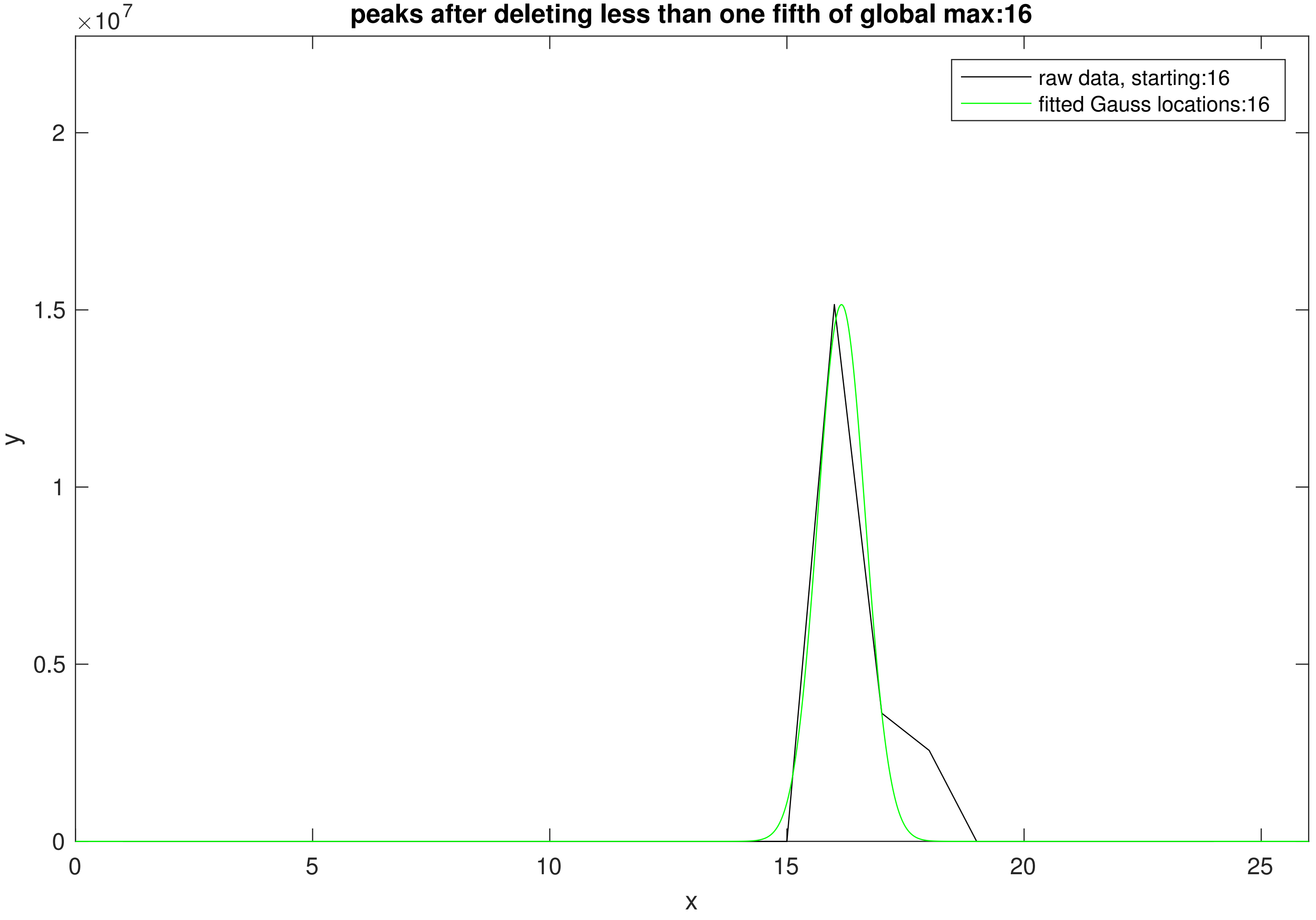

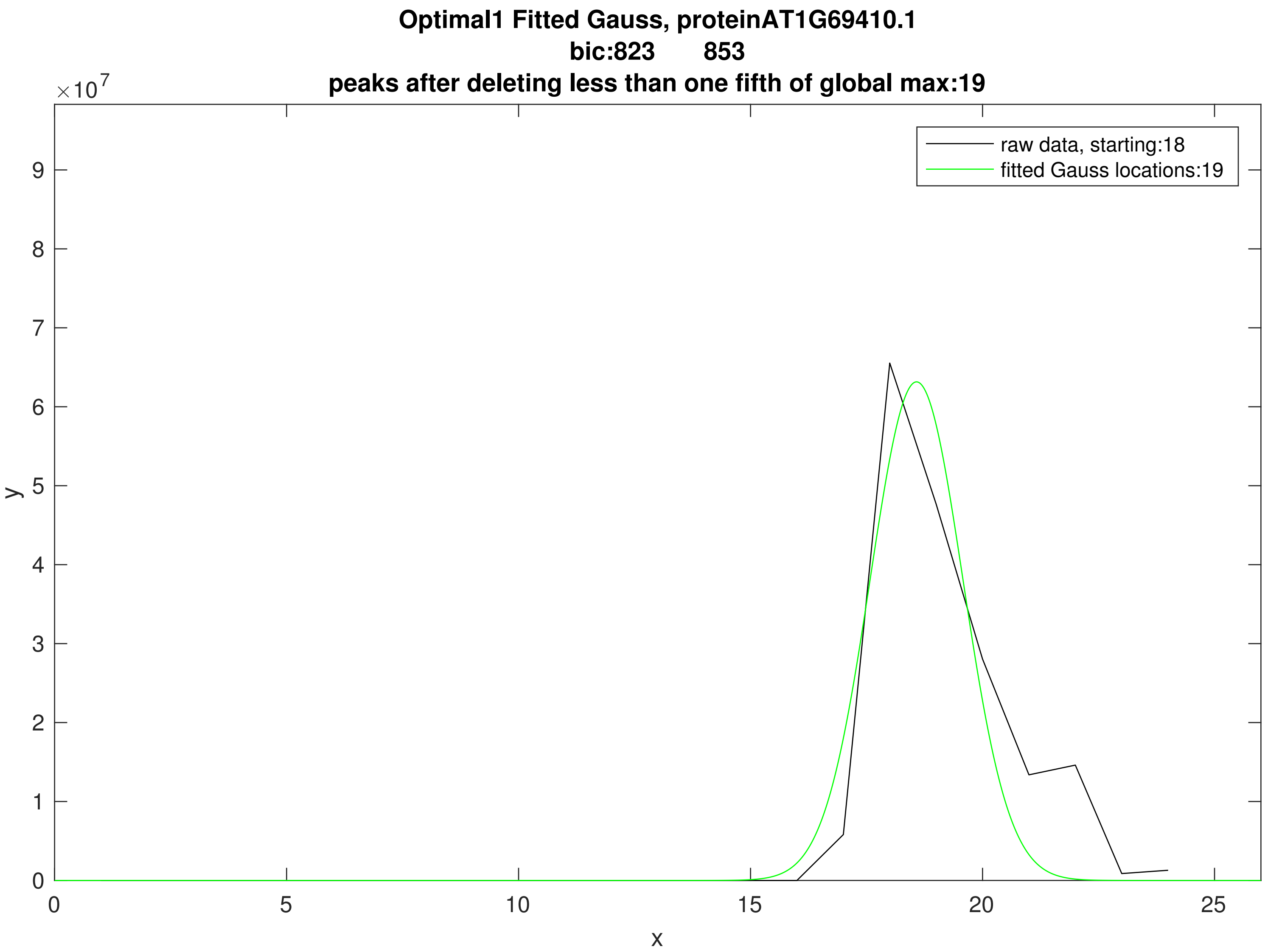
Optimal1 Fitted Gauss, proteinAT1G69410.1
bic:823 853
peaks after deleting less than one fifth of global max:19
×10^7
raw data, starting:18
fitted Gauss locations:19
y
x
0
1
2
3
4
5
6
7
8
9
0
5
10
15
20
25

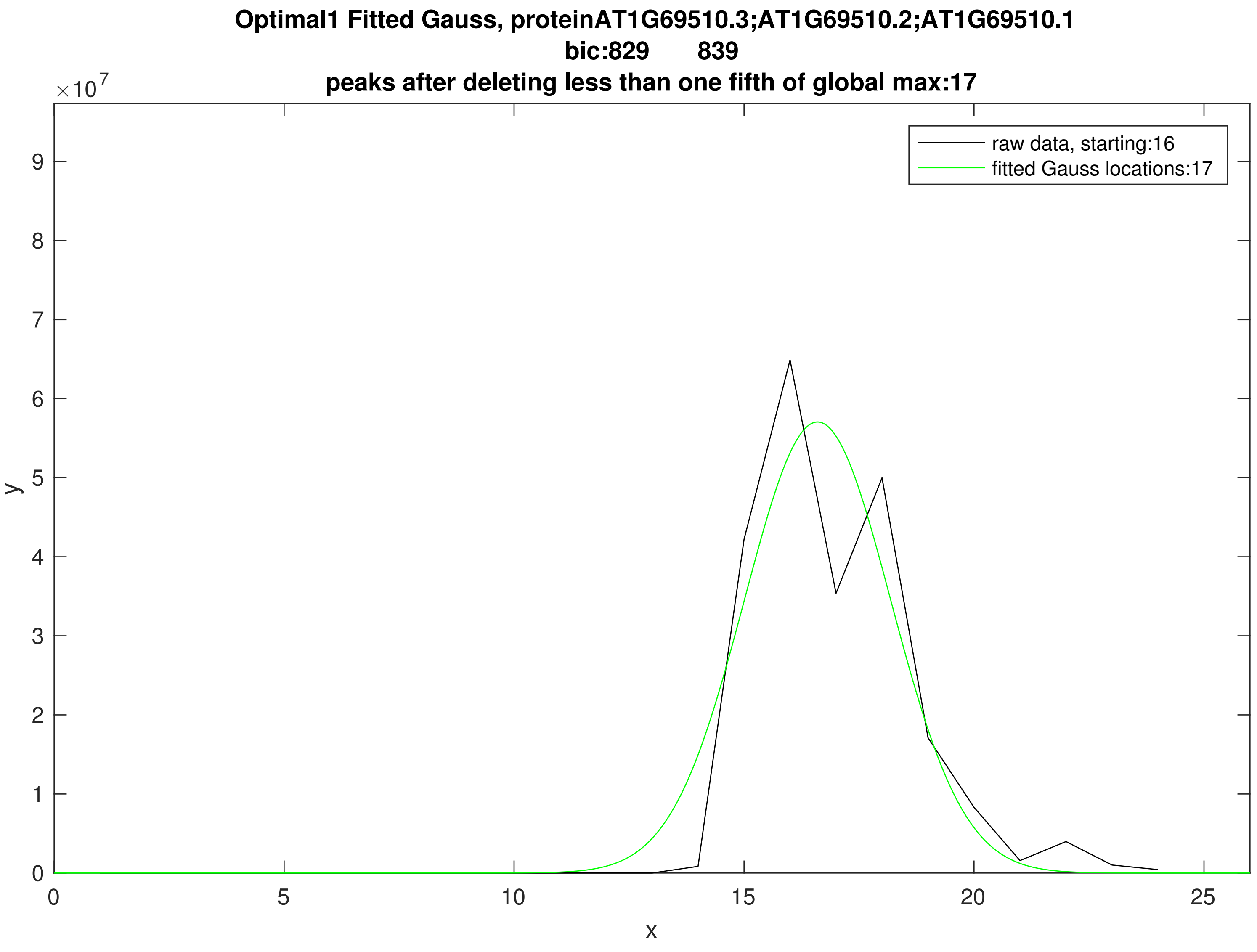

Optimal1 Fitted Gauss, proteinAT1G69510.3;AT1G69510.2;AT1G69510.1
bic:829 839
peaks after deleting less than one fifth of global max:17
×10^7
raw data, starting:16
fitted Gauss locations:17
y
x
0
1
2
3
4
5
6
7
8
9
0
5
10
15
20
25

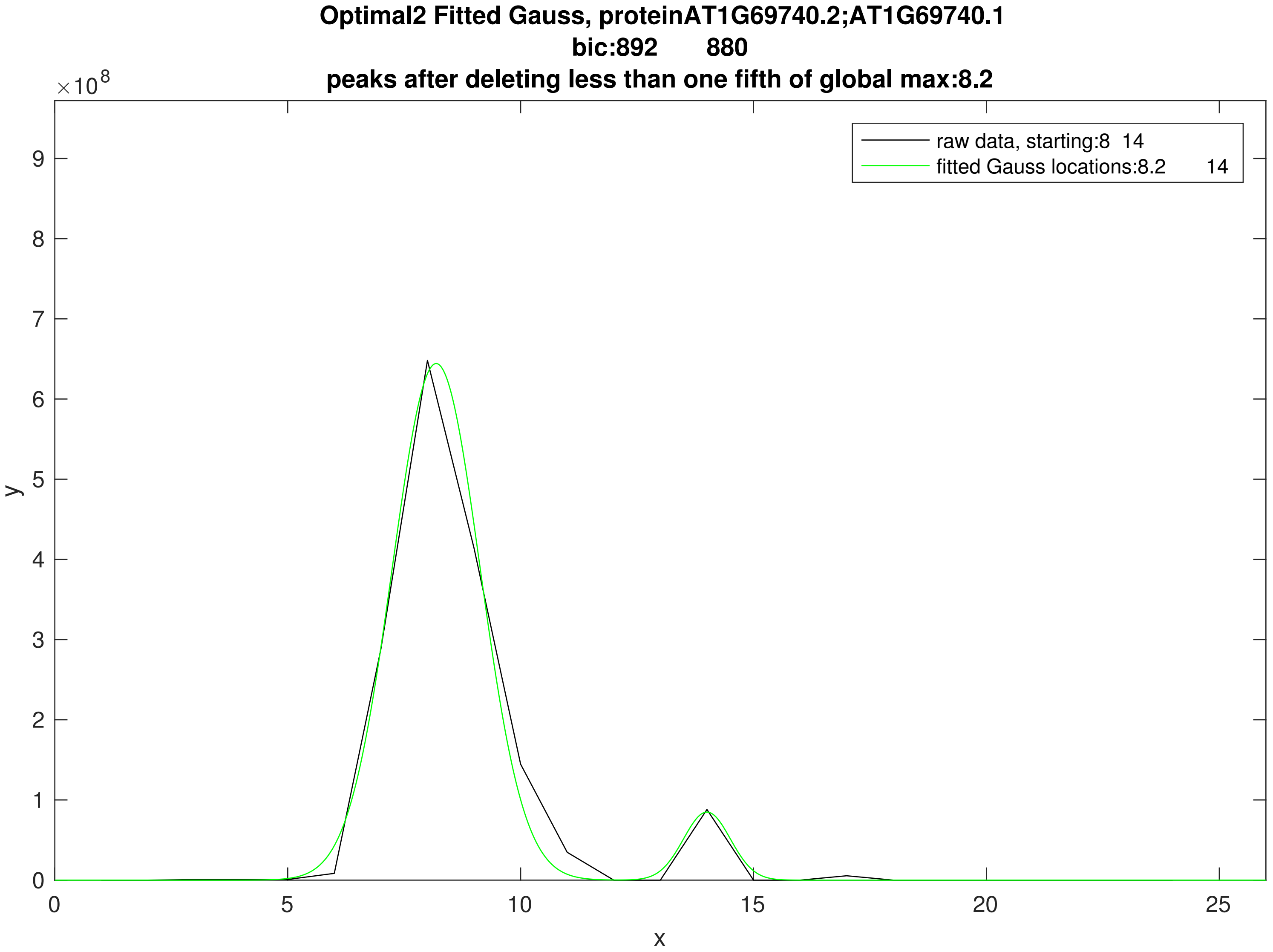
Optimal2 Fitted Gauss, proteinAT1G69740.2;AT1G69740.1
bic:892 880
peaks after deleting less than one fifth of global max:8.2
×10^8
raw data, starting:8 14
fitted Gauss locations:8.2 14
y
x
0
1
2
3
4
5
6
7
8
9
0
5
10
15
20
25

**Optimal1 Fitted Gauss, proteinAT1G69830.1**
**bic:687**
**peaks after deleting less than one fifth of global max:8.9**

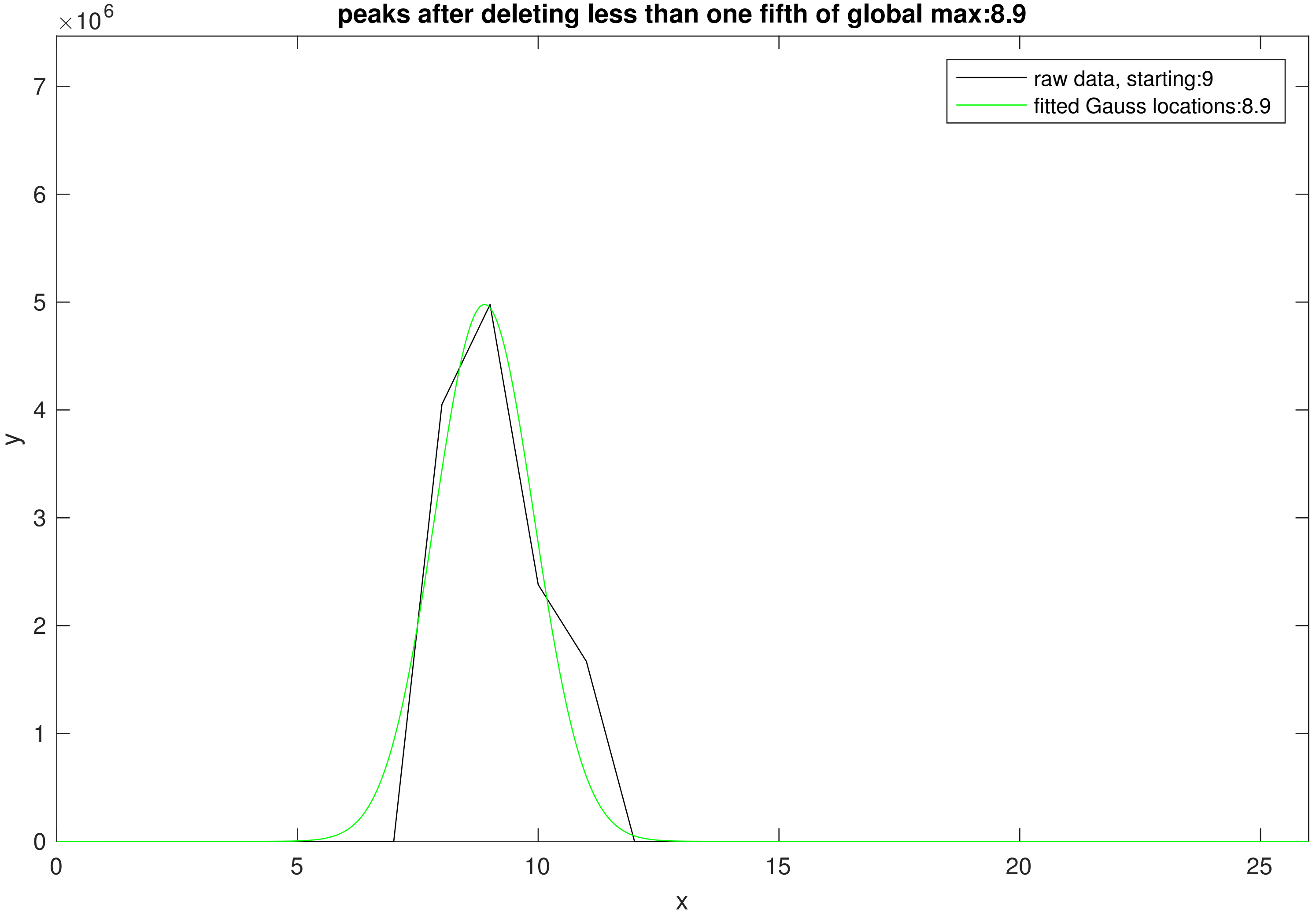

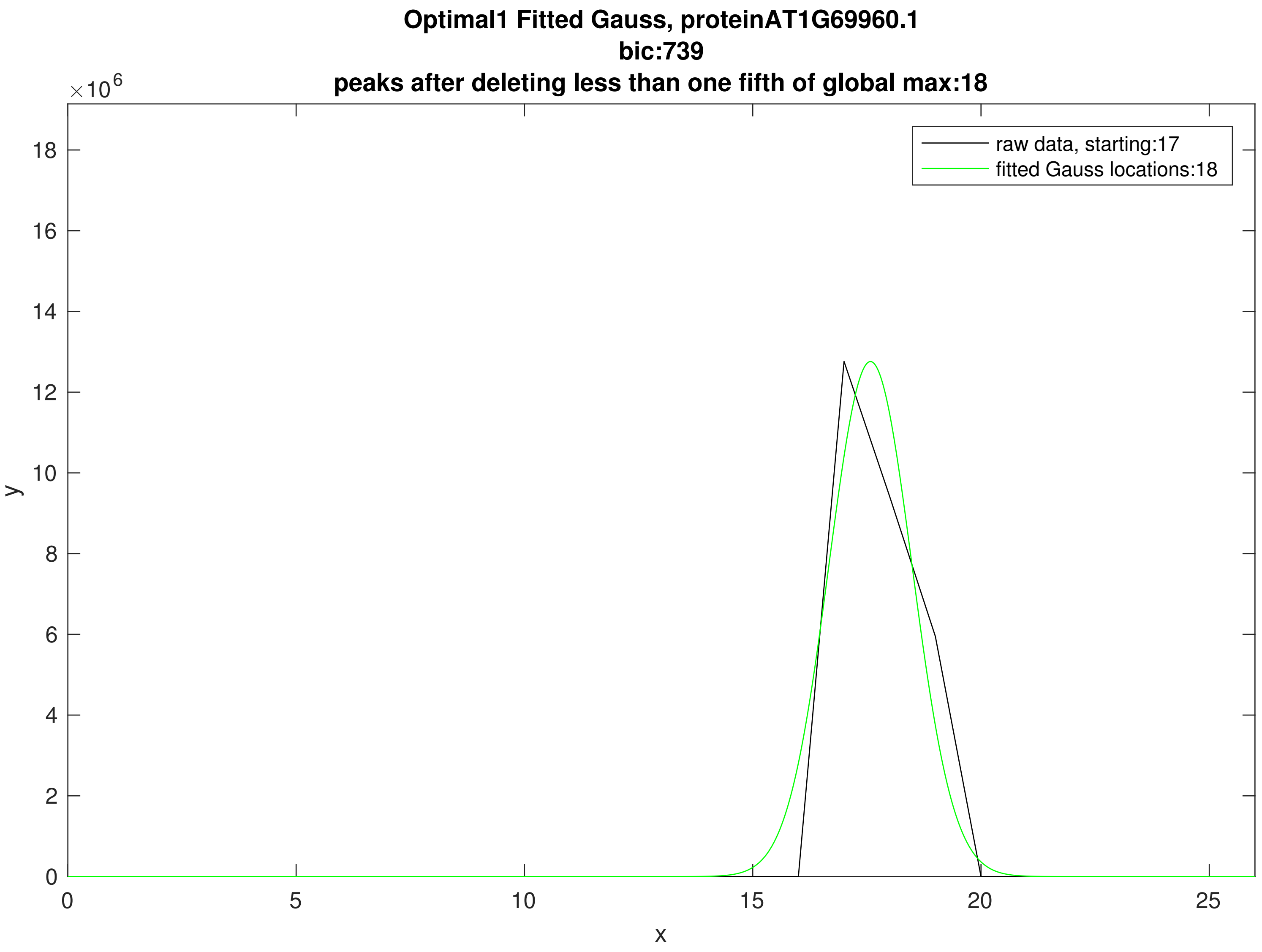
Optimal1 Fitted Gauss, proteinAT1G69960.1
bic:739
peaks after deleting less than one fifth of global max:18
×10^6
raw data, starting:17
fitted Gauss locations:18
y
x
0
2
4
6
8
10
12
14
16
18
0
5
10
15
20
25

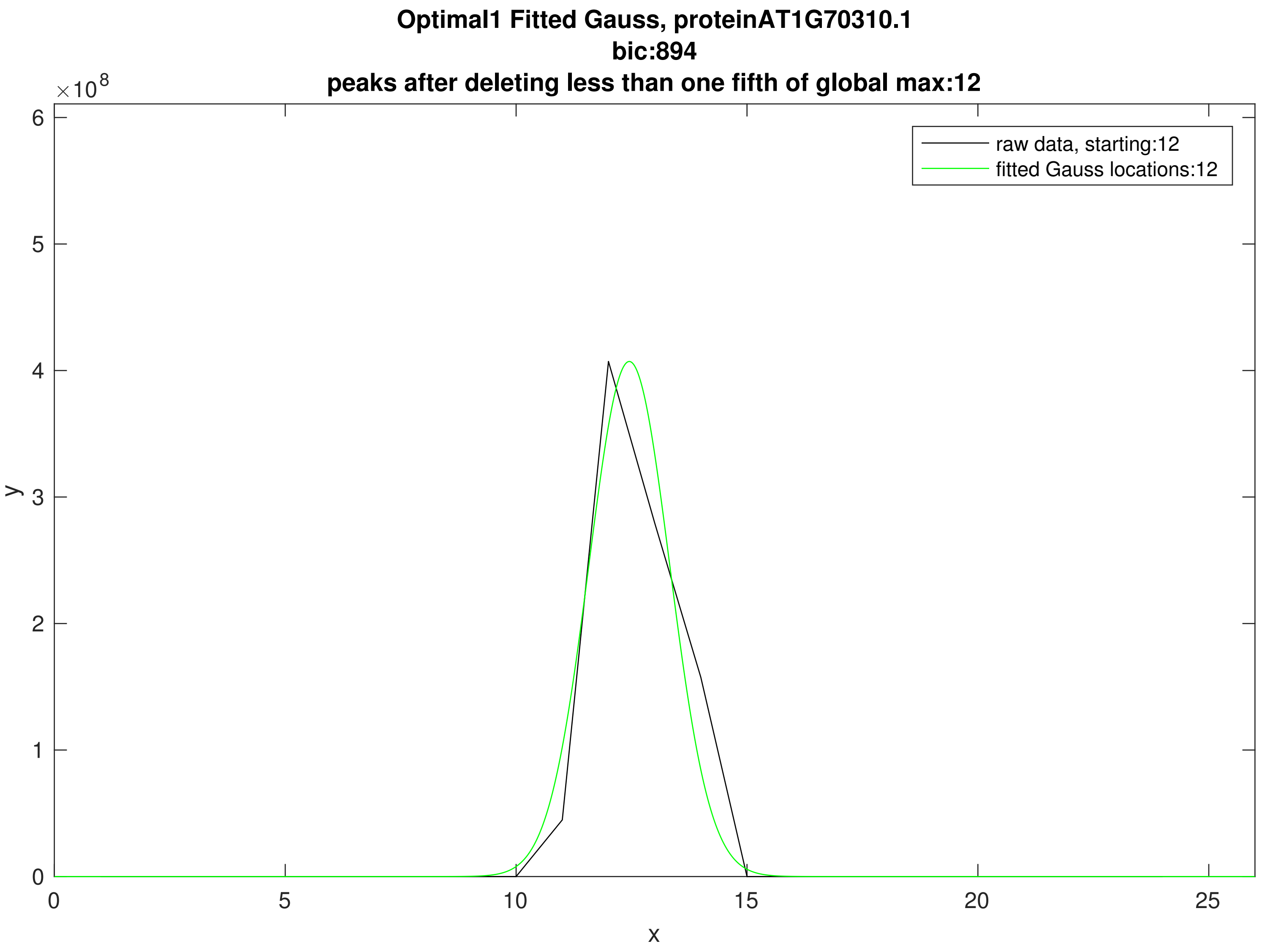
Optimal1 Fitted Gauss, proteinAT1G70310.1
bic:894
peaks after deleting less than one fifth of global max:12
$\times 10^8$
6
5
4
3
2
1
0
y
0
5
10
15
20
25
x
raw data, starting:12
fitted Gauss locations:12

Optimal1 Fitted Gauss, proteinAT1G70410.1;AT1G70410.3;AT1G70410.2
bic:975 985
peaks after deleting less than one fifth of global max:9.2

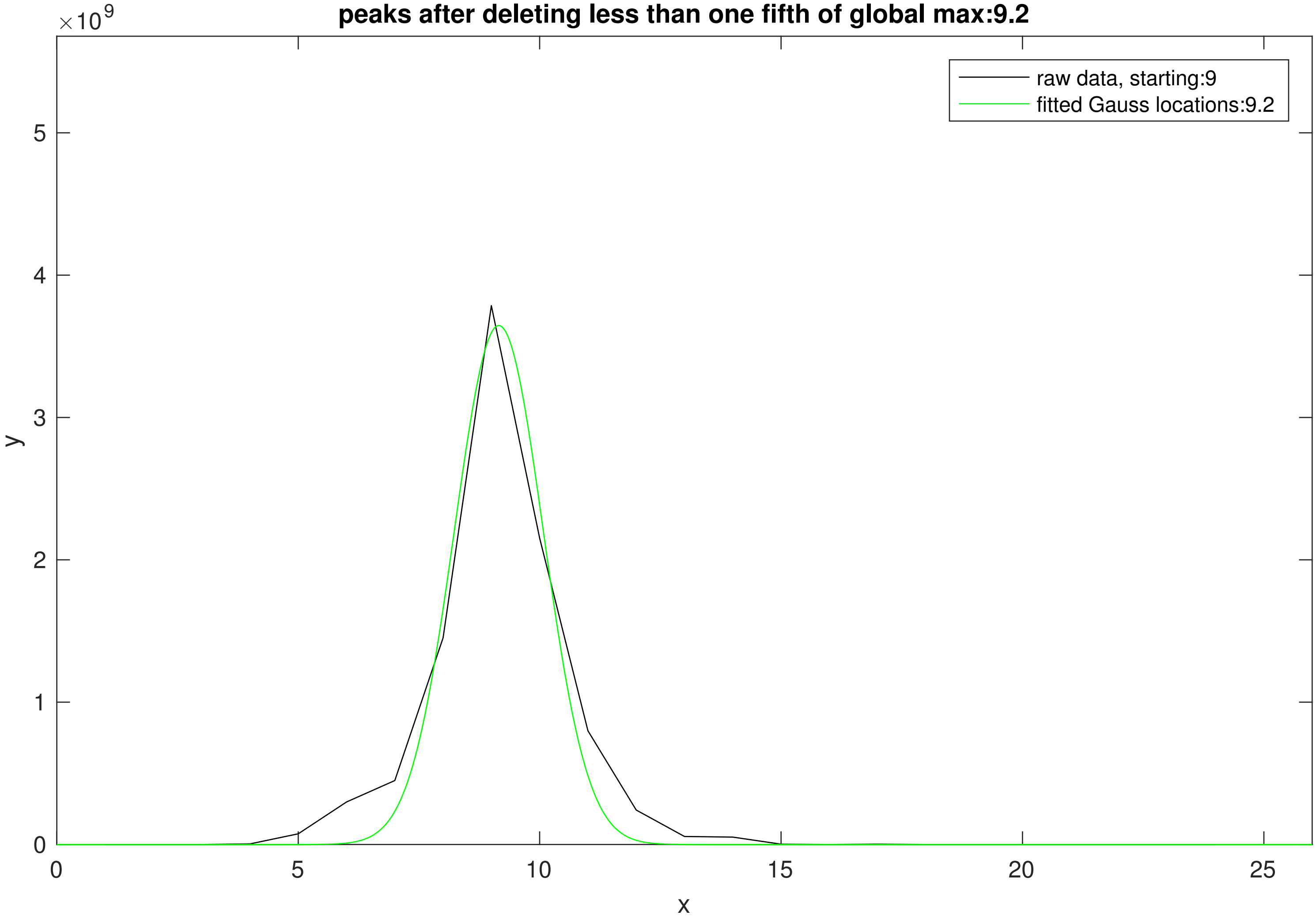

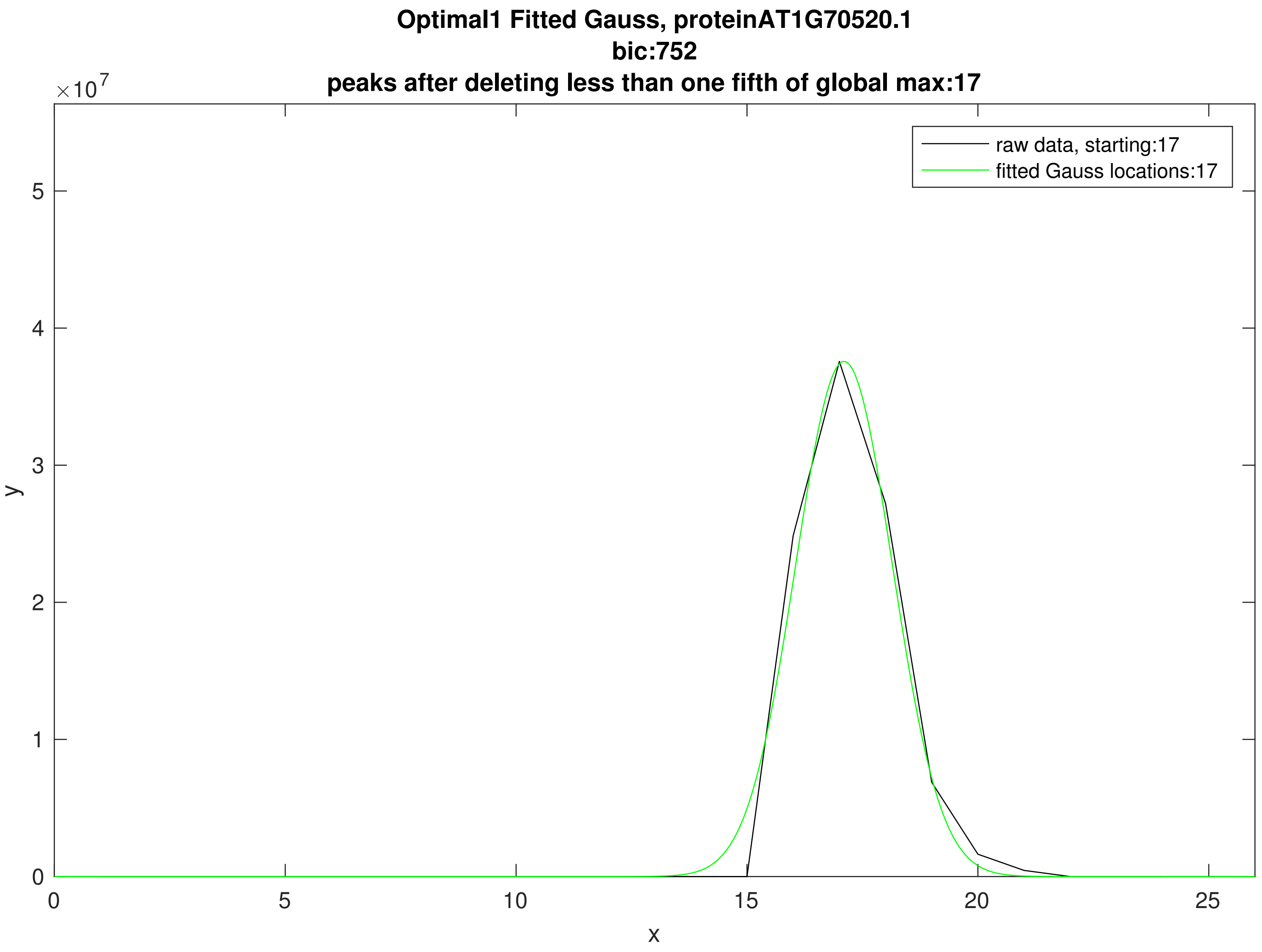
Optimal1 Fitted Gauss, proteinAT1G70520.1
bic:752
peaks after deleting less than one fifth of global max:17
×10^7
raw data, starting:17
fitted Gauss locations:17
5
4
3
2
1
0
y
0
5
10
15
20
25
x

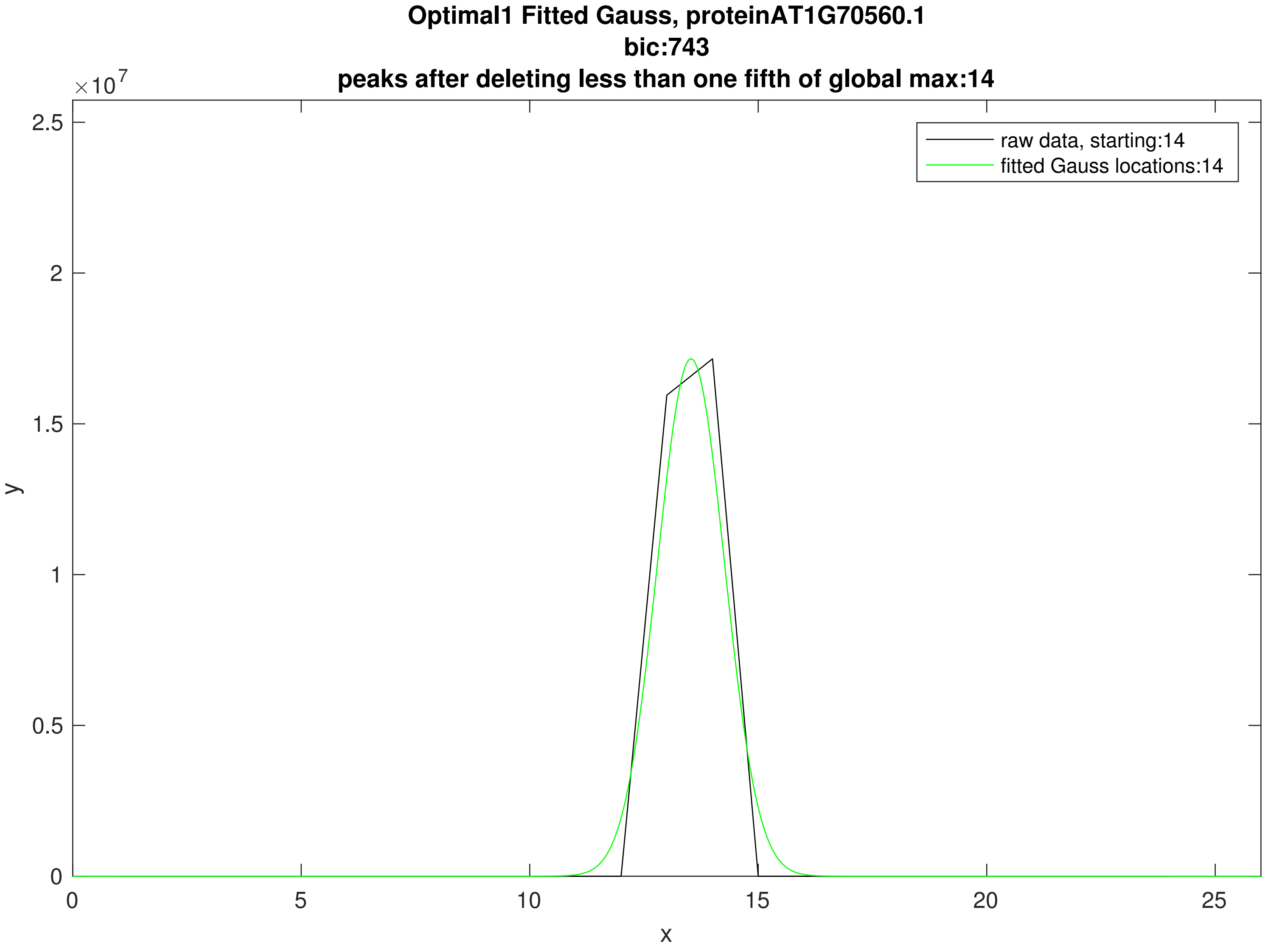
Optimal1 Fitted Gauss, proteinAT1G70560.1
bic:743
peaks after deleting less than one fifth of global max:14
×10^7
2.5
2
1.5
1
0.5
0
y
0
5
10
15
20
25
x
raw data, starting:14
fitted Gauss locations:14

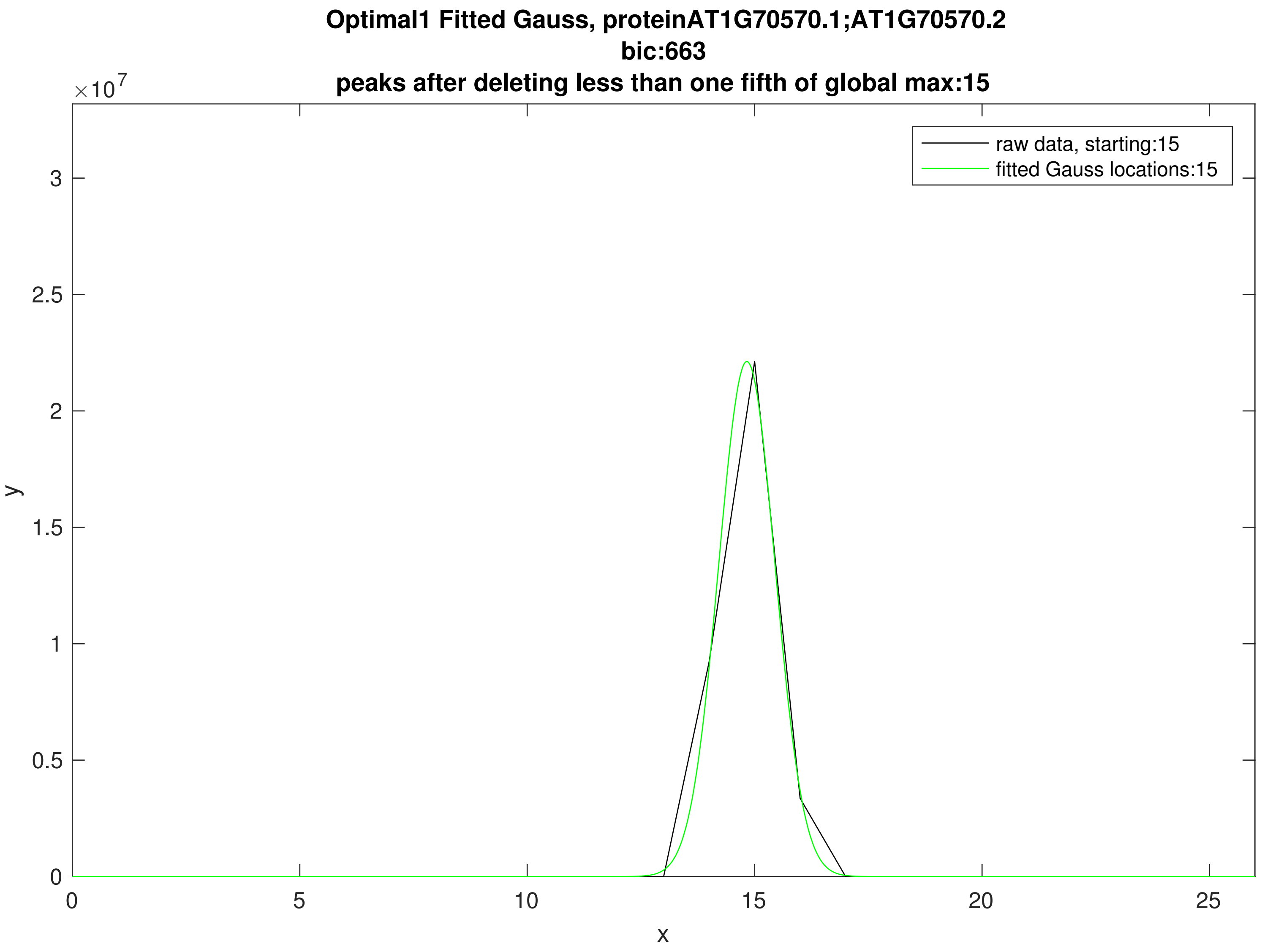

Optimal1 Fitted Gauss, proteinAT1G70570.1;AT1G70570.2
bic:663
peaks after deleting less than one fifth of global max:15
×10^7
raw data, starting:15
fitted Gauss locations:15
y
x
0
0.5
1
1.5
2
2.5
3
0
5
10
15
20
25

**Optimal1 Fitted Gauss, proteinAT1G70580.4;AT1G70580.3;AT1G70580.2;AT1G70580.1**
**bic:821 894**
**peaks after deleting less than one fifth of global max:14**

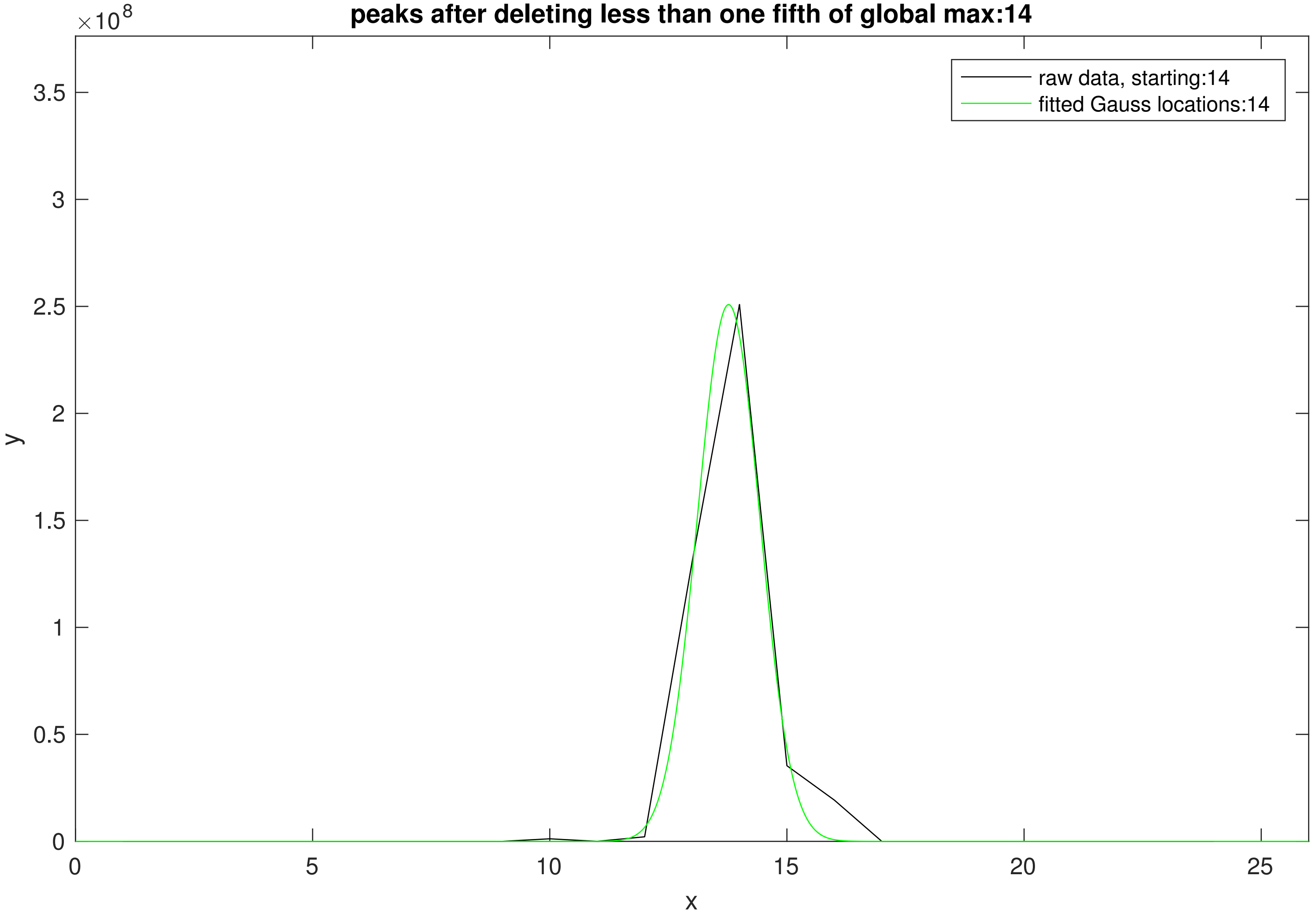

#### Optimal1 Fitted Gauss, proteinAT1G70600.1
## bic:794       802
#### peaks after deleting less than one fifth of global max:3.2

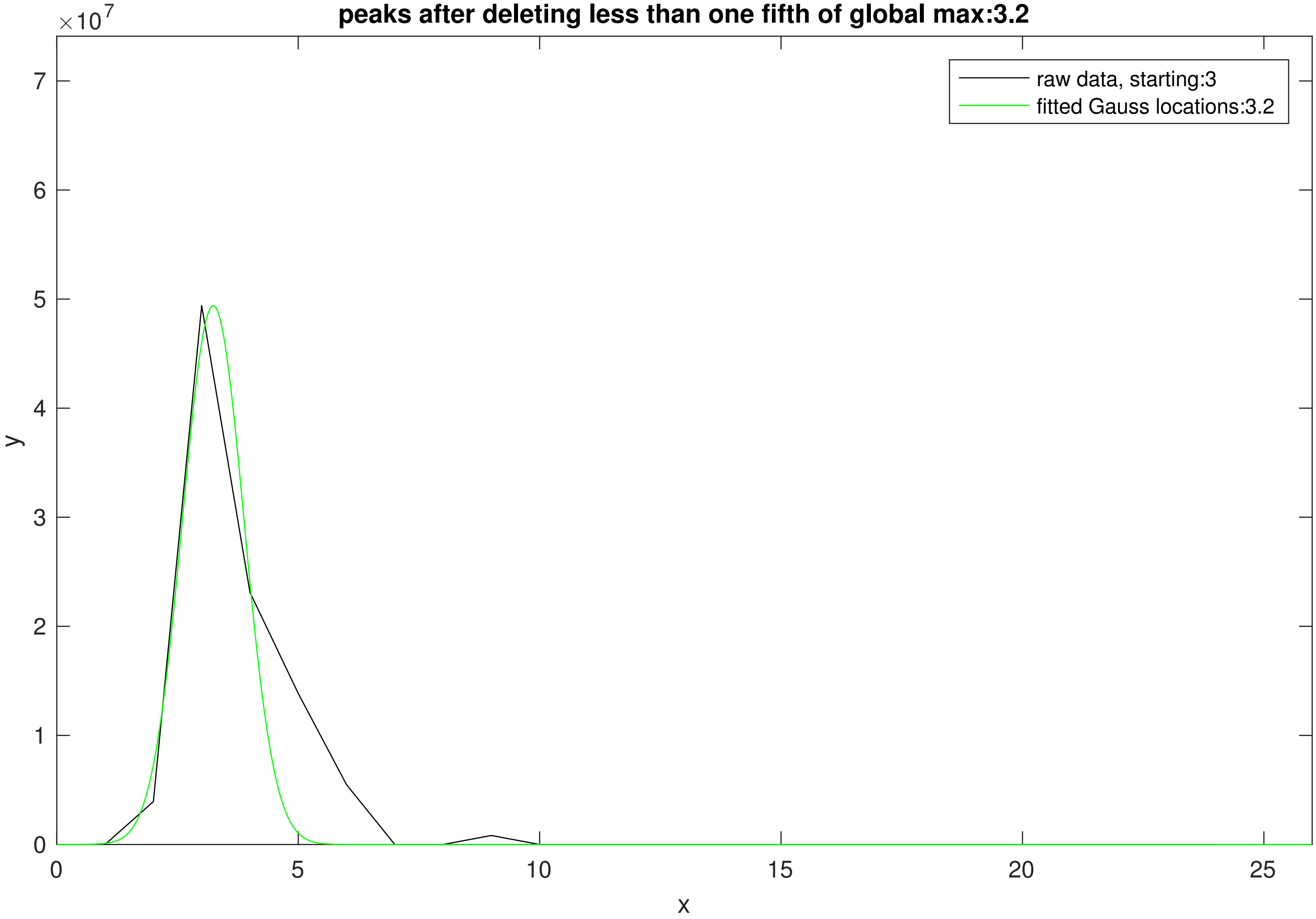

Optimal1 Fitted Gauss, proteinAT1G70730.1;AT1G70730.3;AT1G70730.2
bic:928 932
peaks after deleting less than one fifth of global max:15

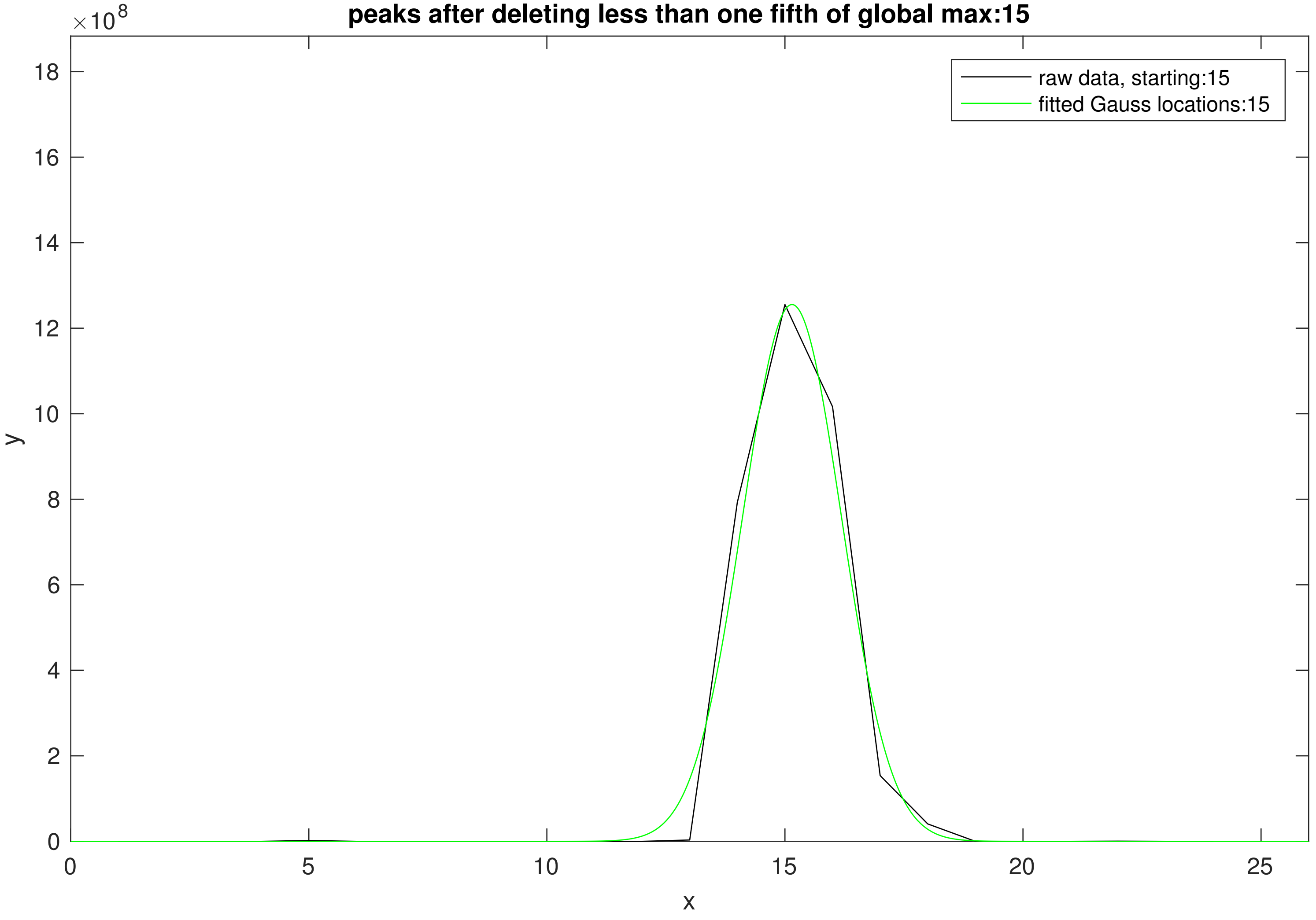

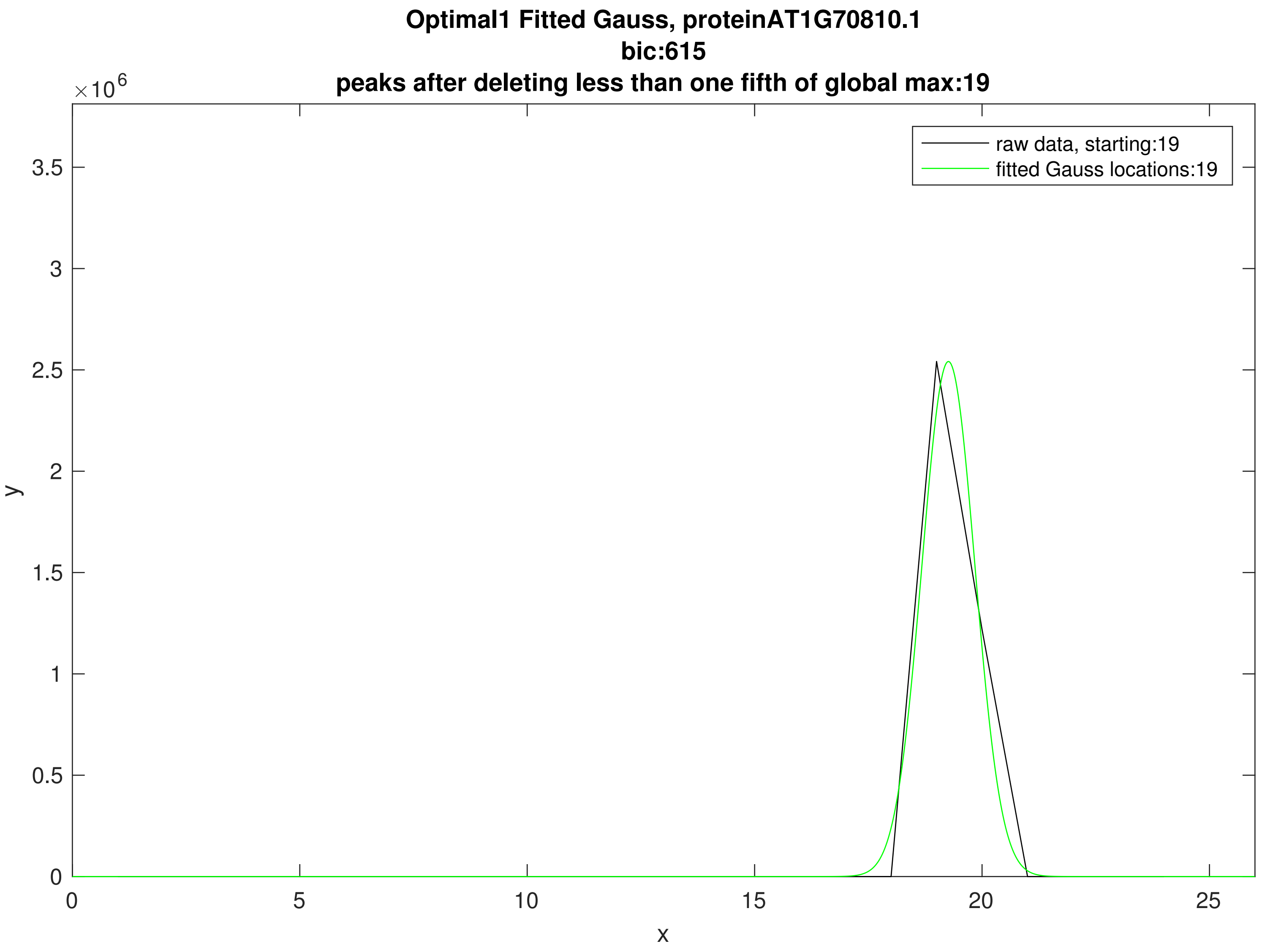

Optimal1 Fitted Gauss, proteinAT1G70810.1
bic:615
peaks after deleting less than one fifth of global max:19
×10^6
raw data, starting:19
fitted Gauss locations:19
y
x
0
0.5
1
1.5
2
2.5
3
3.5
0
5
10
15
20
25

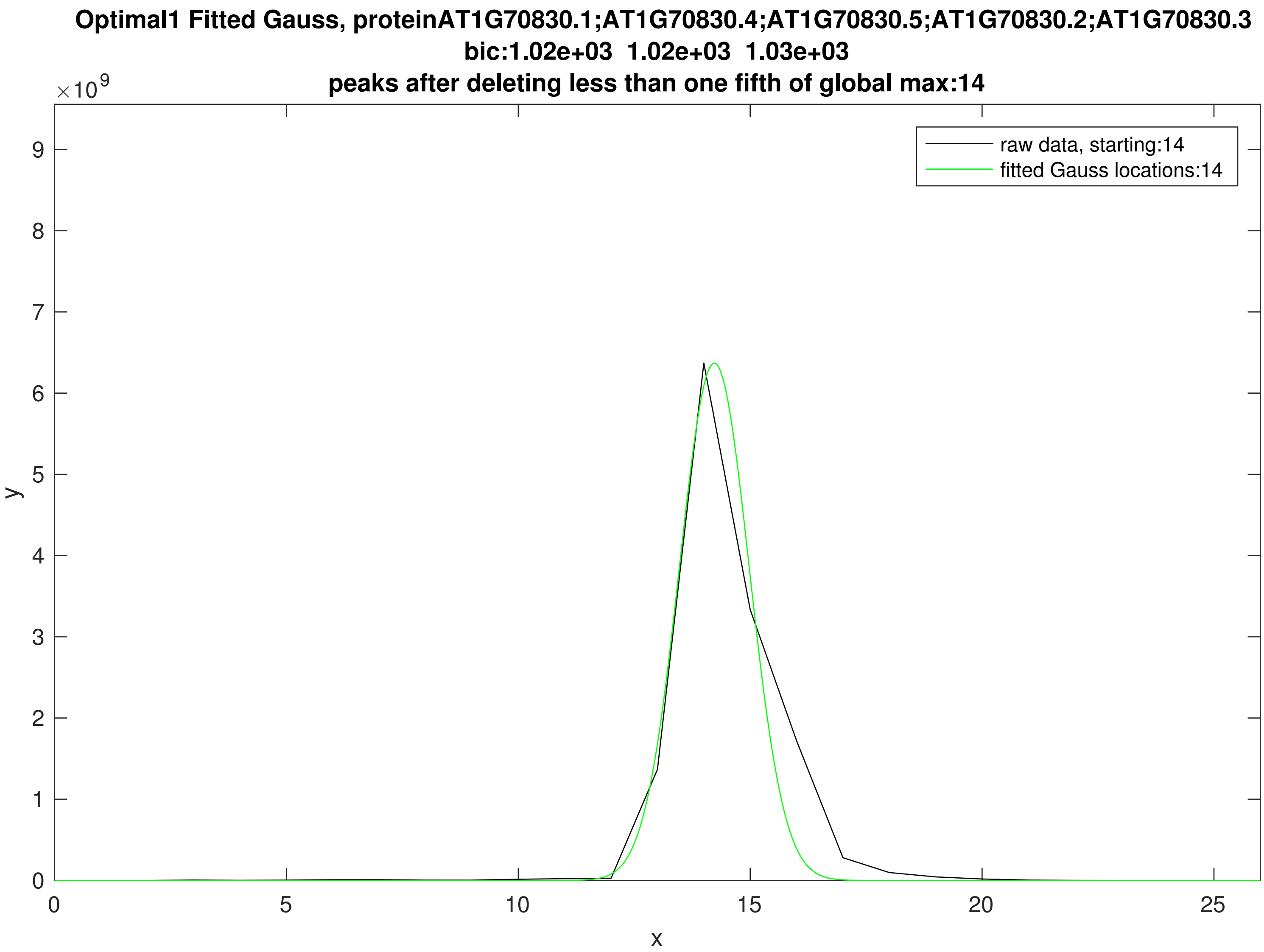

Optimal1 Fitted Gauss, proteinAT1G70830.1;AT1G70830.4;AT1G70830.5;AT1G70830.2;AT1G70830.3
bic:1.02e+03 1.02e+03 1.03e+03
peaks after deleting less than one fifth of global max:14
×10^9
raw data, starting:14
fitted Gauss locations:14
y
x
0
1
2
3
4
5
6
7
8
9
0
5
10
15
20
25

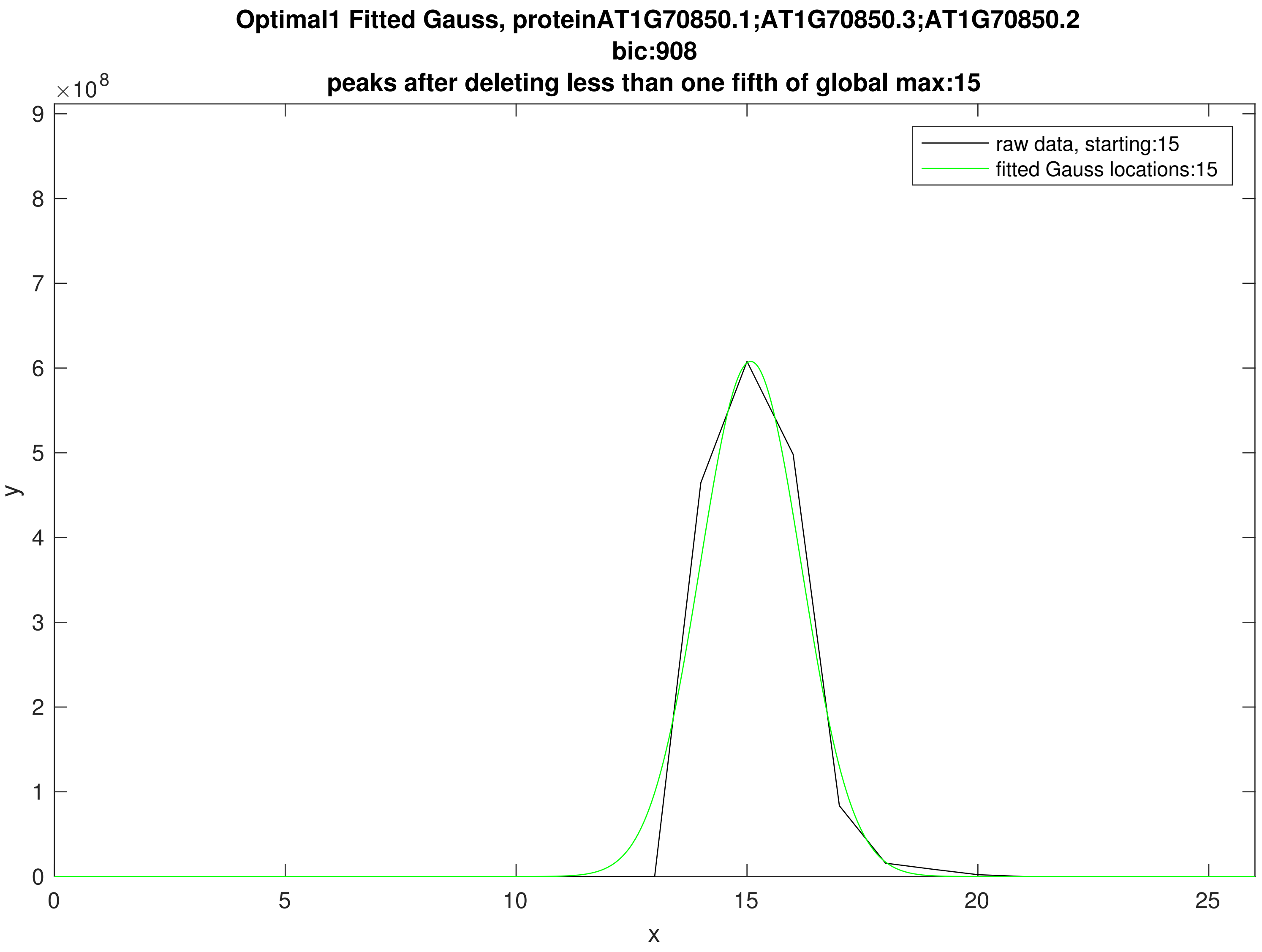

Optimal1 Fitted Gauss, proteinAT1G70850.1;AT1G70850.3;AT1G70850.2
bic:908
peaks after deleting less than one fifth of global max:15
×10^8
raw data, starting:15
fitted Gauss locations:15
y
x
0
1
2
3
4
5
6
7
8
9
0
5
10
15
20
25

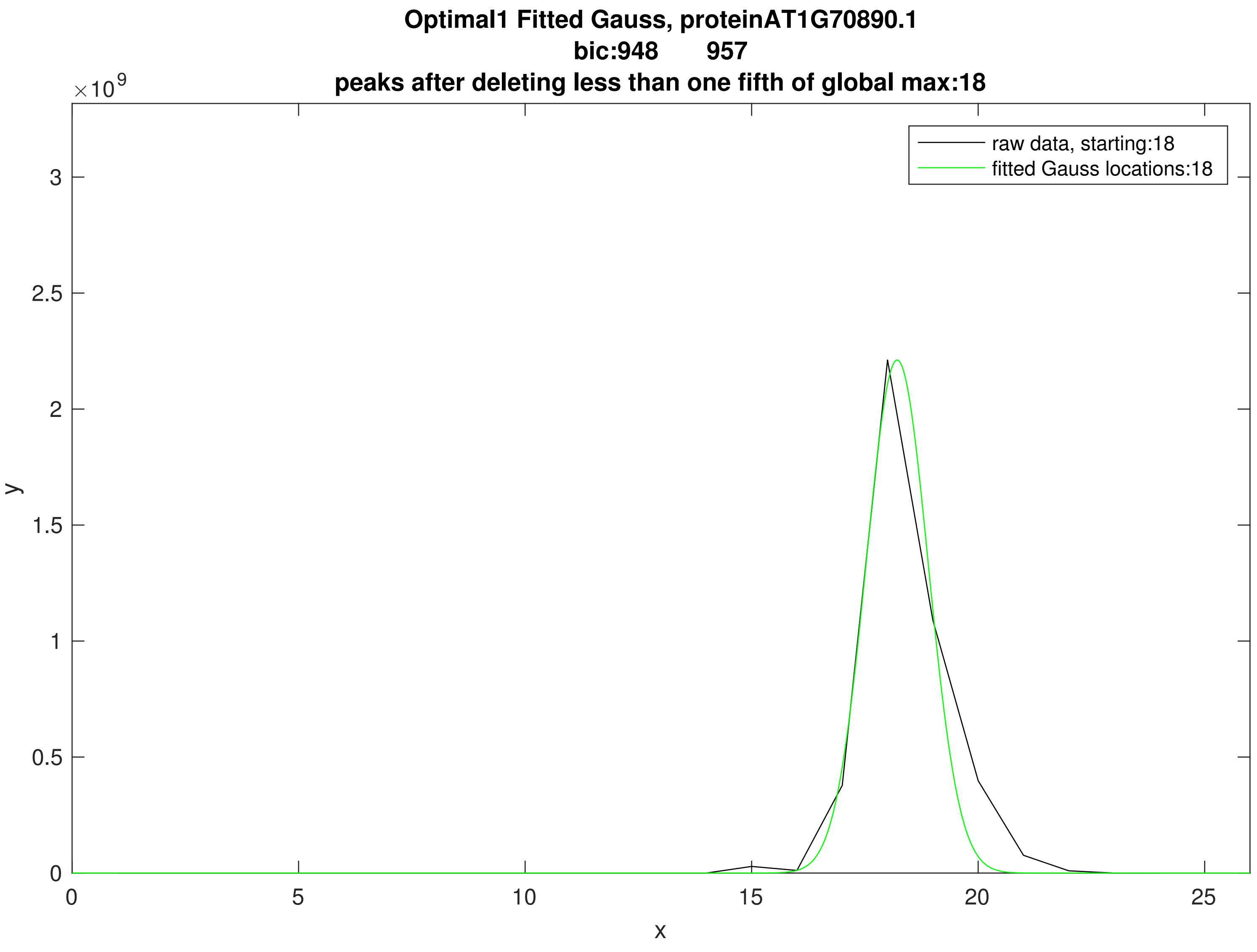
Optimal1 Fitted Gauss, proteinAT1G70890.1
bic:948 957
peaks after deleting less than one fifth of global max:18
×10^9
raw data, starting:18
fitted Gauss locations:18
3
2.5
2
1.5
1
0.5
0
y
0
5
10
15
20
25
x

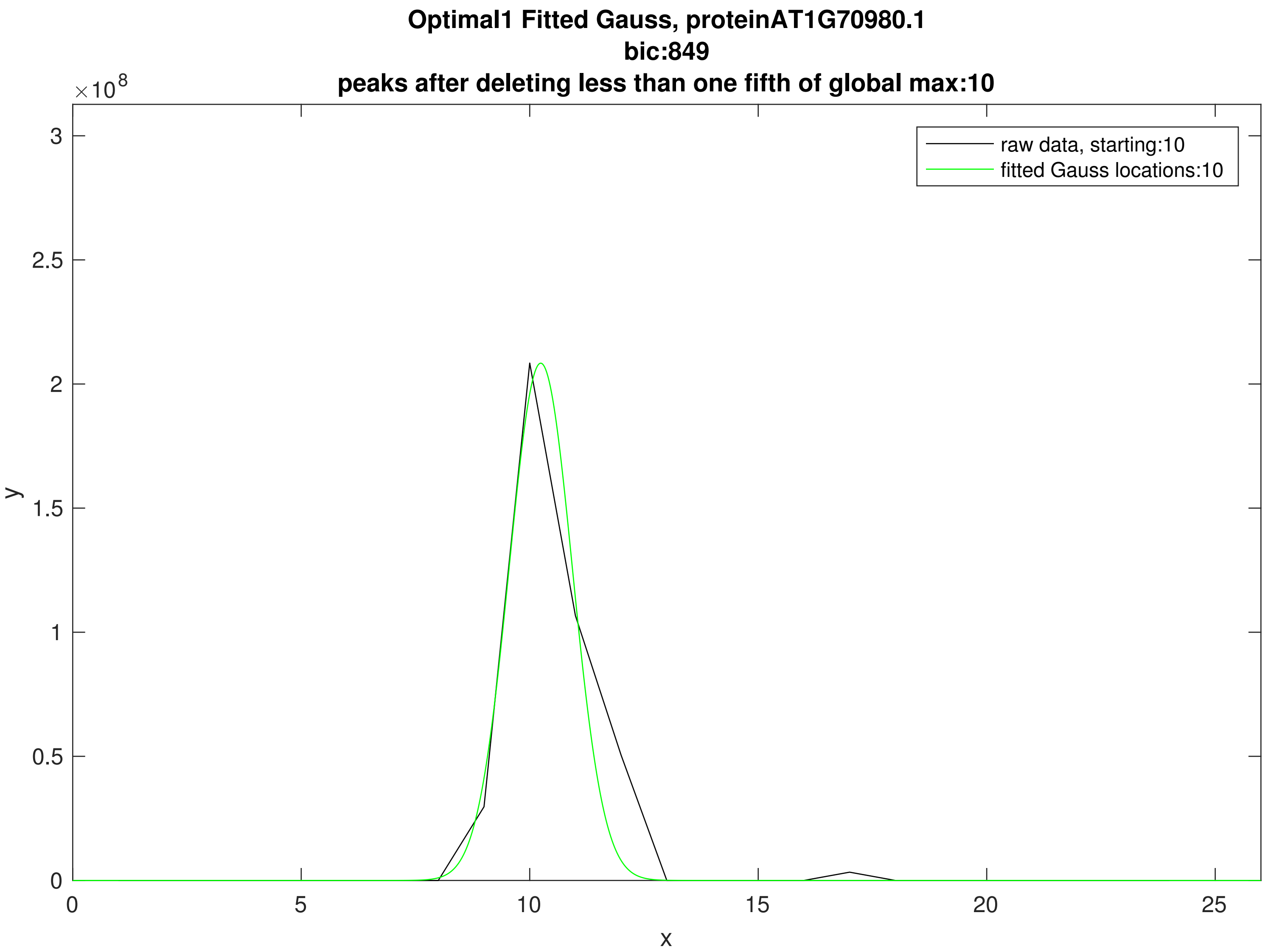
Optimal1 Fitted Gauss, proteinAT1G70980.1
bic:849
peaks after deleting less than one fifth of global max:10
×10^8
3
2.5
2
1.5
1
0.5
0
y
0
5
10
15
20
25
x
raw data, starting:10
fitted Gauss locations:10

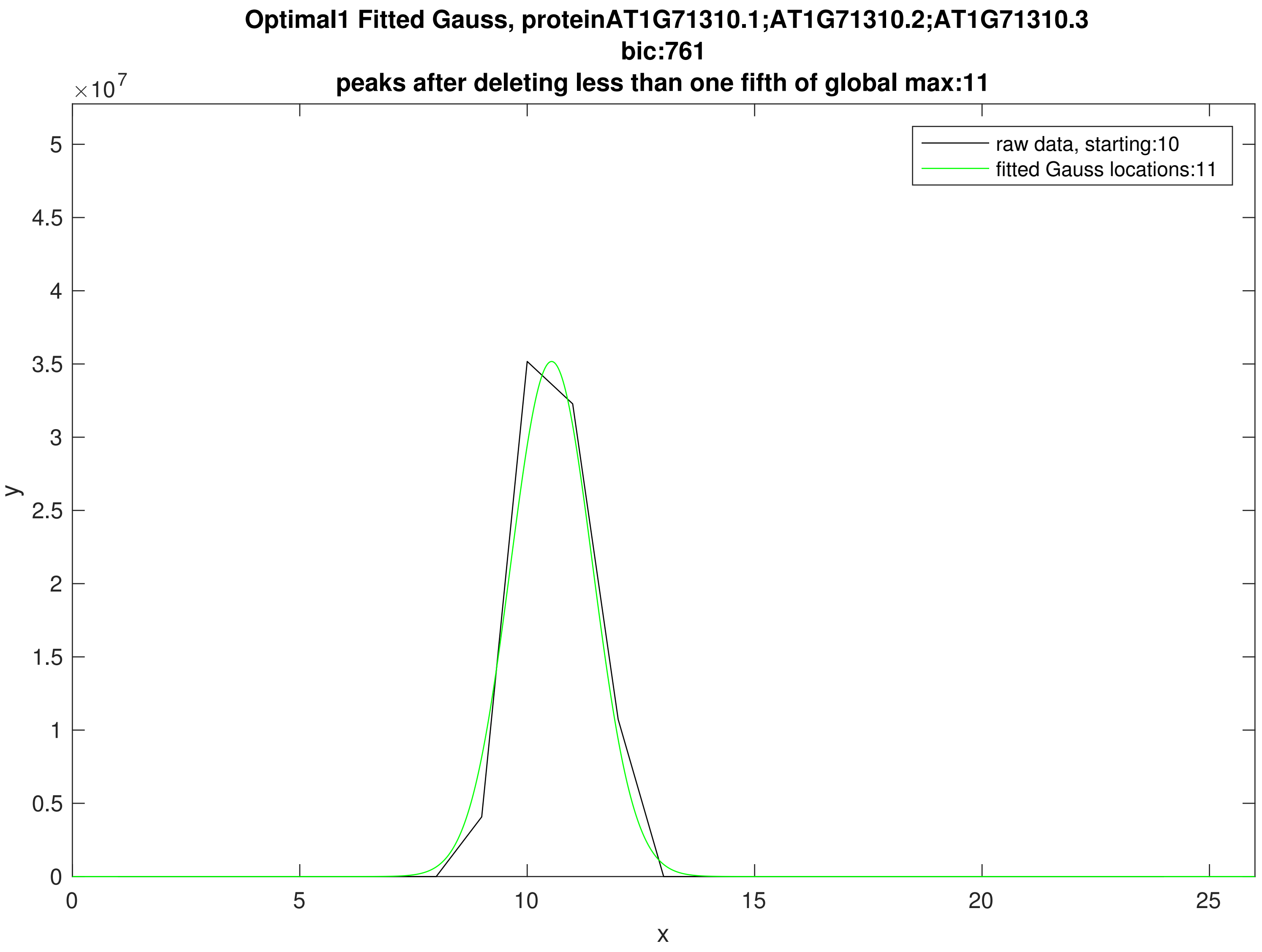
Optimal1 Fitted Gauss, proteinAT1G71310.1;AT1G71310.2;AT1G71310.3
bic:761
peaks after deleting less than one fifth of global max:11
×10^7
raw data, starting:10
fitted Gauss locations:11
y
x
0
0.5
1
1.5
2
2.5
3
3.5
4
4.5
5
0
5
10
15
20
25

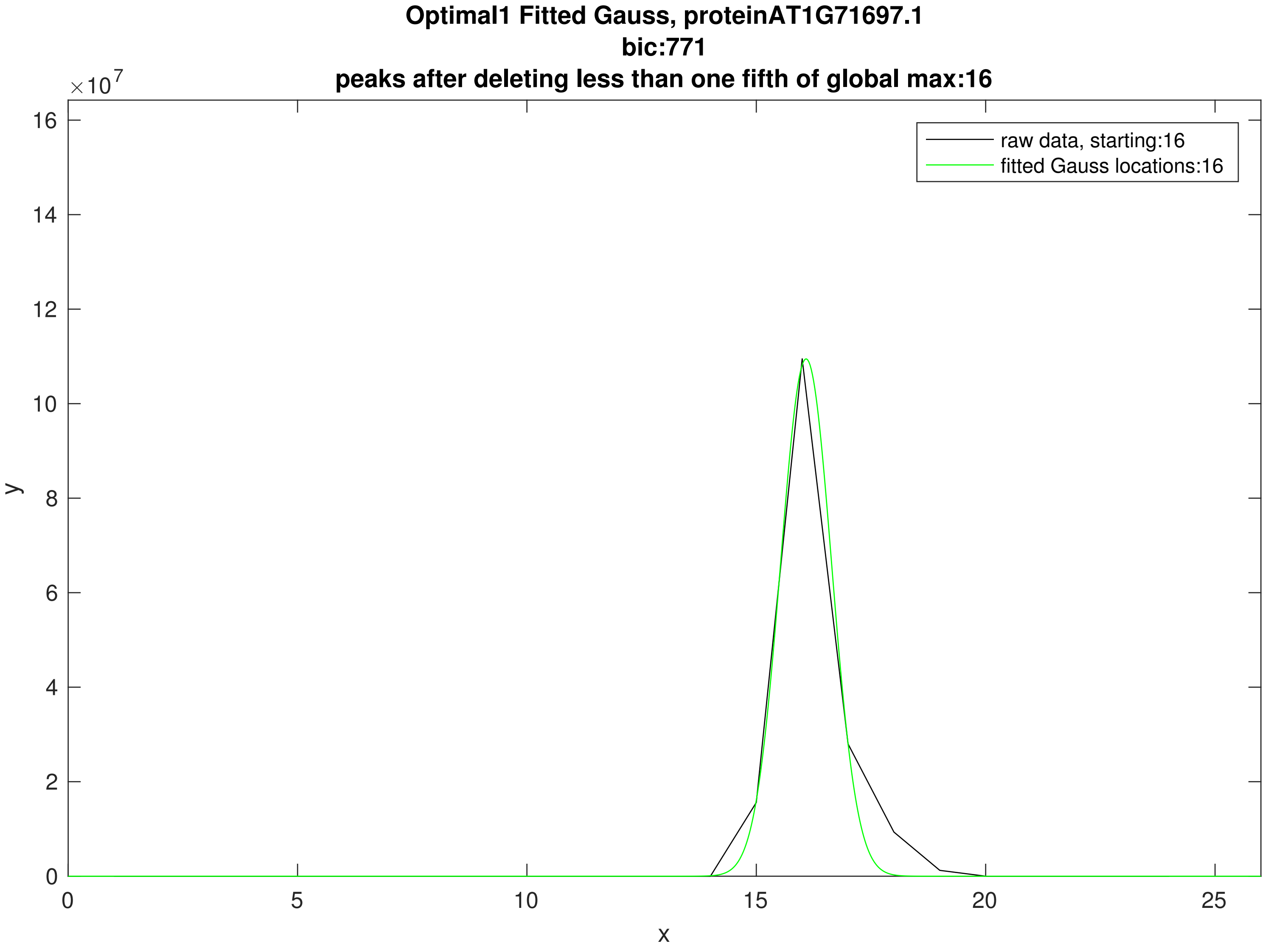
Optimal1 Fitted Gauss, proteinAT1G71697.1
bic:771
peaks after deleting less than one fifth of global max:16
×10^7
raw data, starting:16
fitted Gauss locations:16
y
x
0
2
4
6
8
10
12
14
16
0
5
10
15
20
25

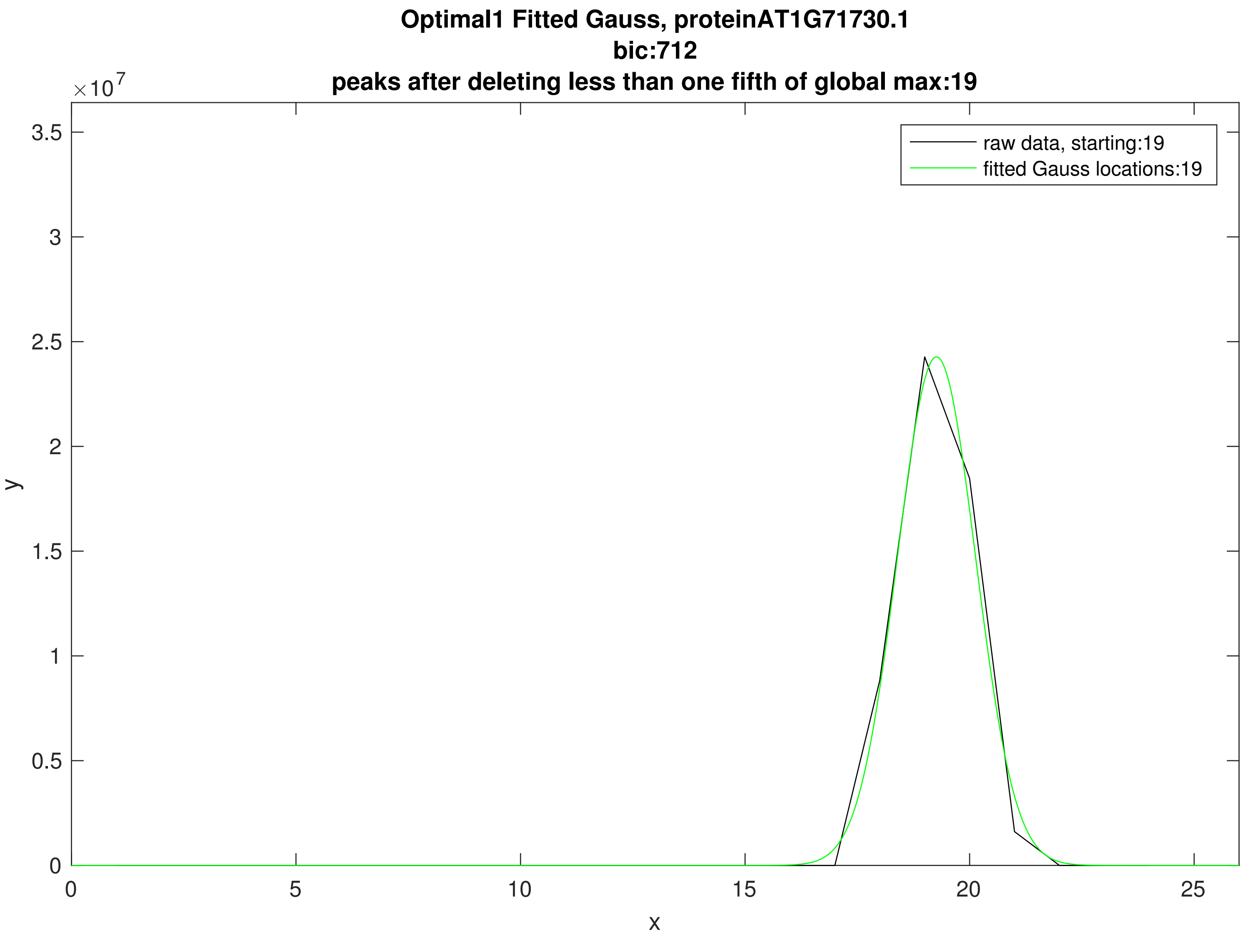

Optimal1 Fitted Gauss, proteinAT1G71730.1
bic:712
peaks after deleting less than one fifth of global max:19
×10^7
3.5
3
2.5
2
1.5
1
0.5
0
y
0
5
10
15
20
25
x
raw data, starting:19
fitted Gauss locations:19

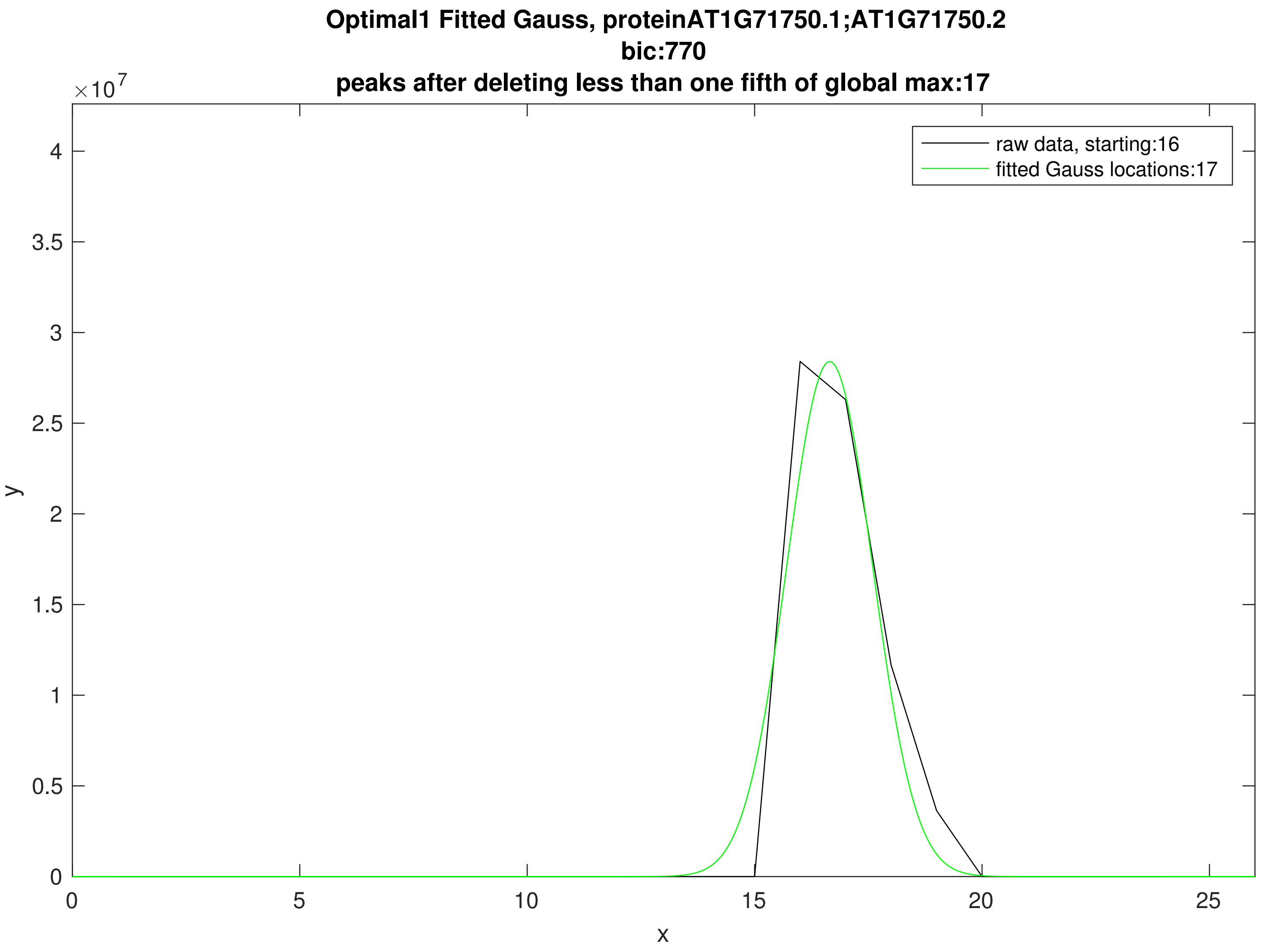
Optimal1 Fitted Gauss, proteinAT1G71750.1;AT1G71750.2
bic:770
peaks after deleting less than one fifth of global max:17
×10^7
raw data, starting:16
fitted Gauss locations:17
y
x
0
0.5
1
1.5
2
2.5
3
3.5
4
0
5
10
15
20
25

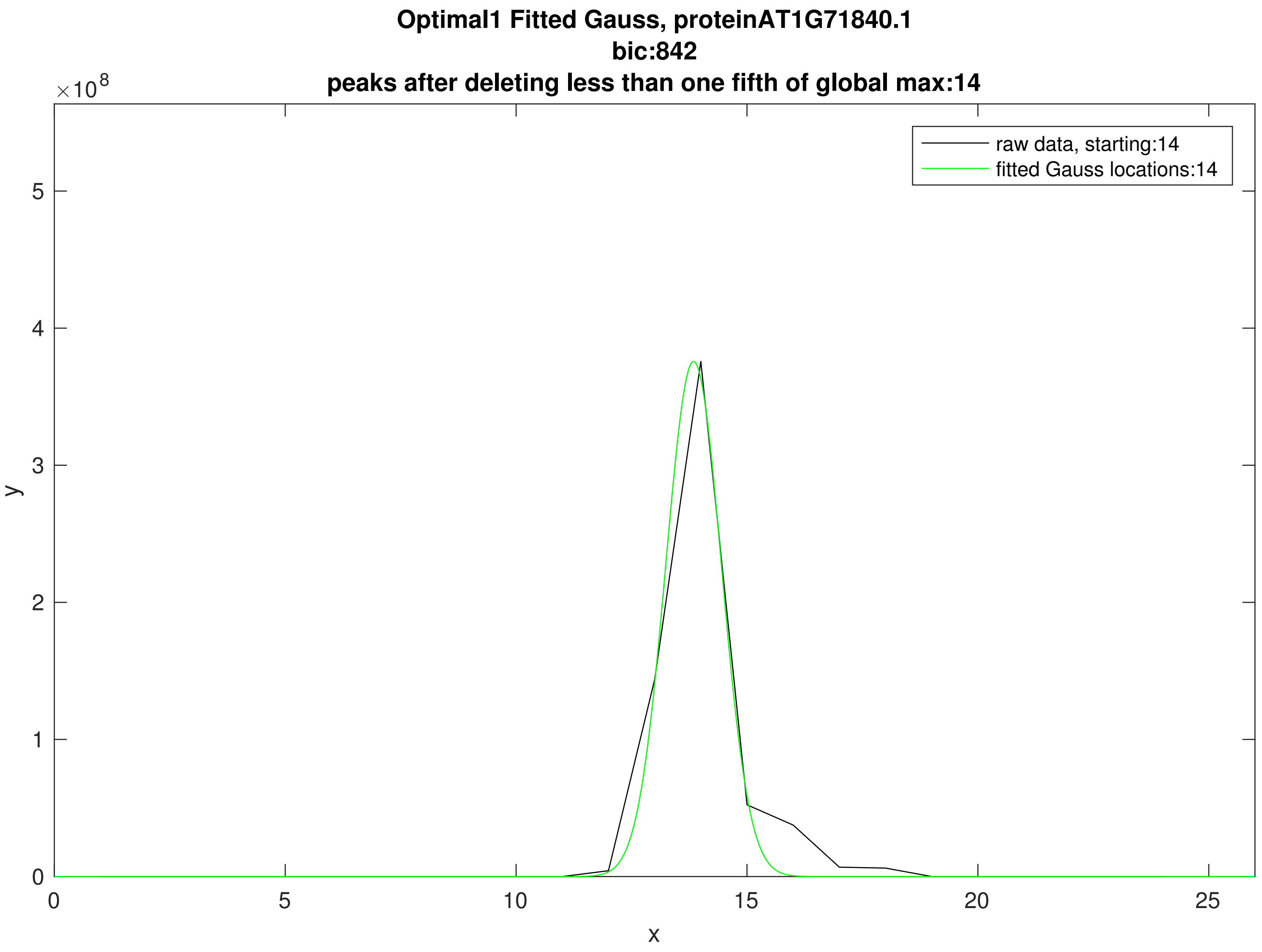

Optimal1 Fitted Gauss, proteinAT1G71840.1
bic:842
peaks after deleting less than one fifth of global max:14
×10^8
raw data, starting:14
fitted Gauss locations:14
y
x
0
1
2
3
4
5
0
5
10
15
20
25

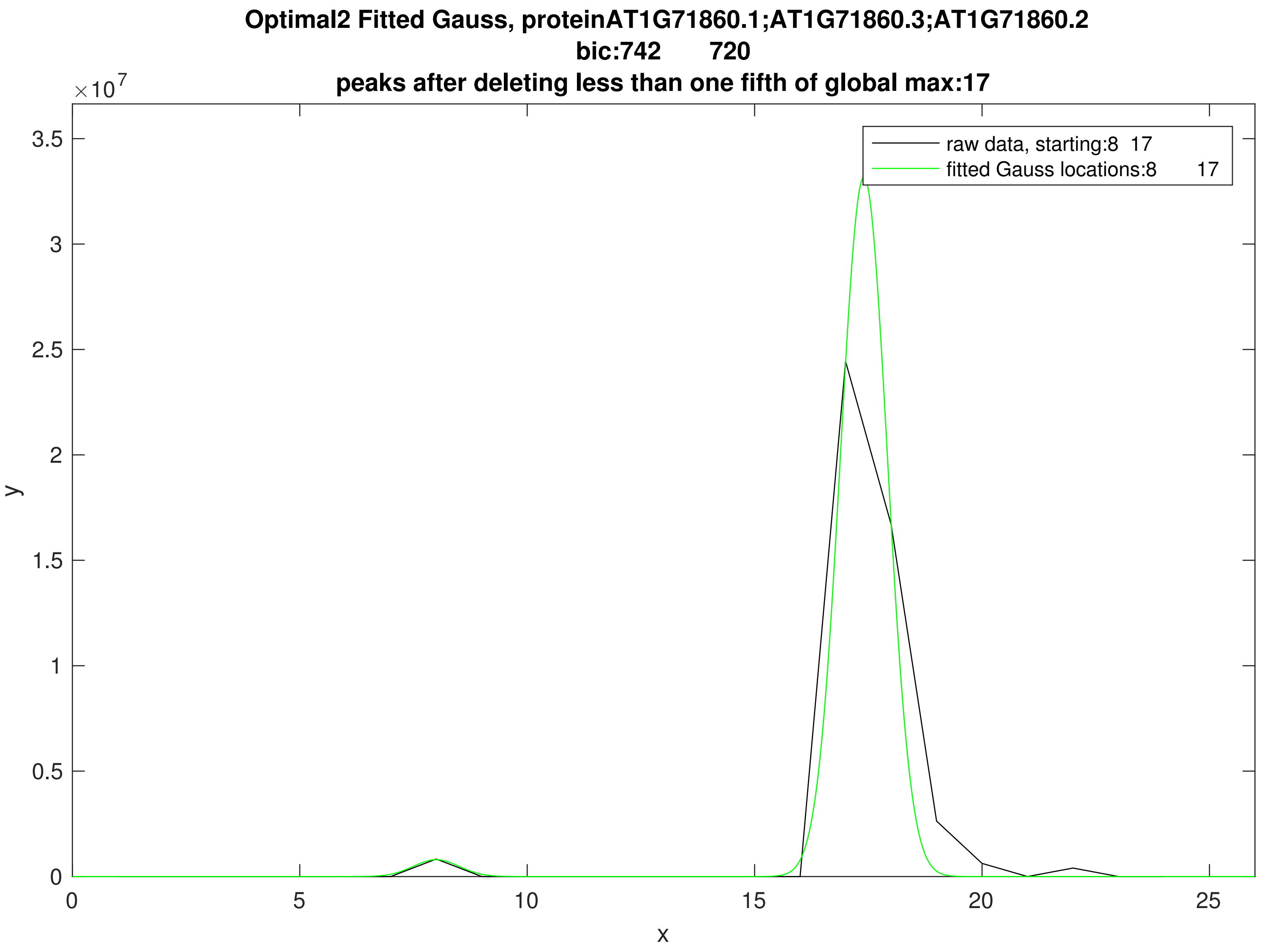
Optimal2 Fitted Gauss, proteinAT1G71860.1;AT1G71860.3;AT1G71860.2
bic:742 720
peaks after deleting less than one fifth of global max:17
×10^7
raw data, starting:8 17
fitted Gauss locations:8 17
3.5
3
2.5
2
1.5
1
0.5
0
y
0
5
10
15
20
25
x

**Optimal1 Fitted Gauss, proteinAT1G71950.1**
**bic:900**
**peaks after deleting less than one fifth of global max:13**

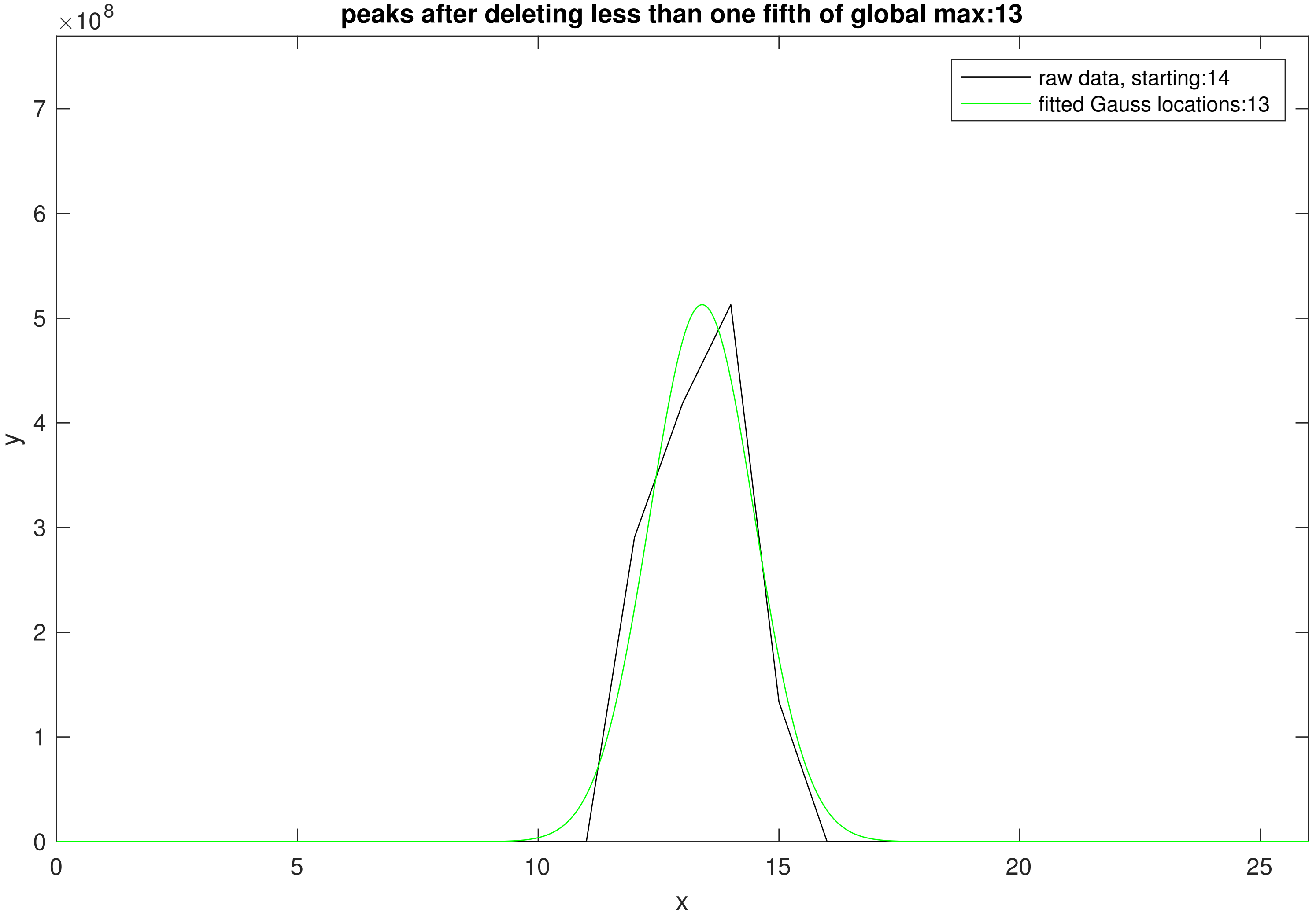

Optimal1 Fitted Gauss, proteinAT1G72150.1
bic:825 830 839
peaks after deleting less than one fifth of global max:9.7

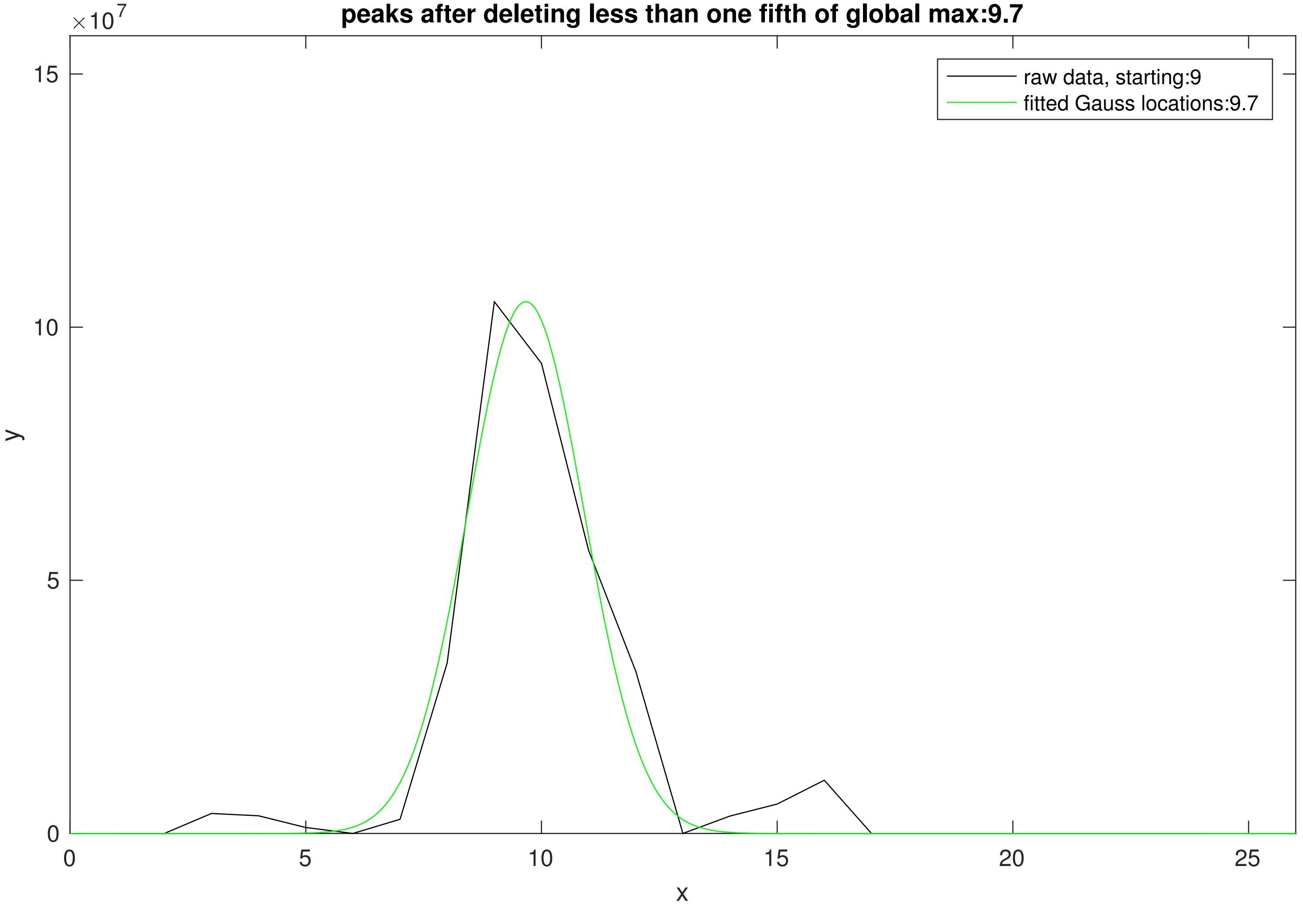

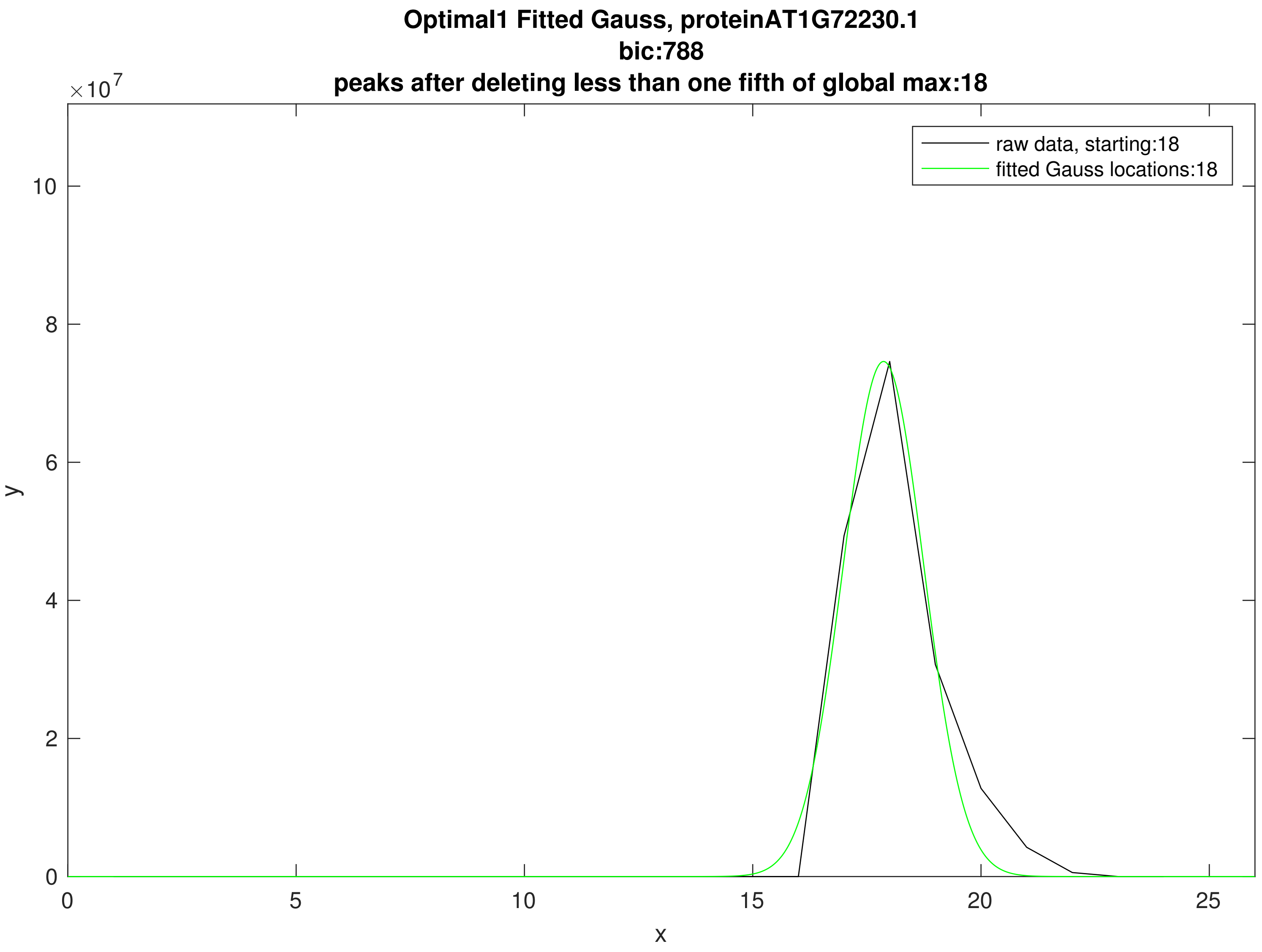
Optimal1 Fitted Gauss, proteinAT1G72230.1
bic:788
peaks after deleting less than one fifth of global max:18
×10^7
raw data, starting:18
fitted Gauss locations:18
y
x
0
2
4
6
8
10
0
5
10
15
20
25

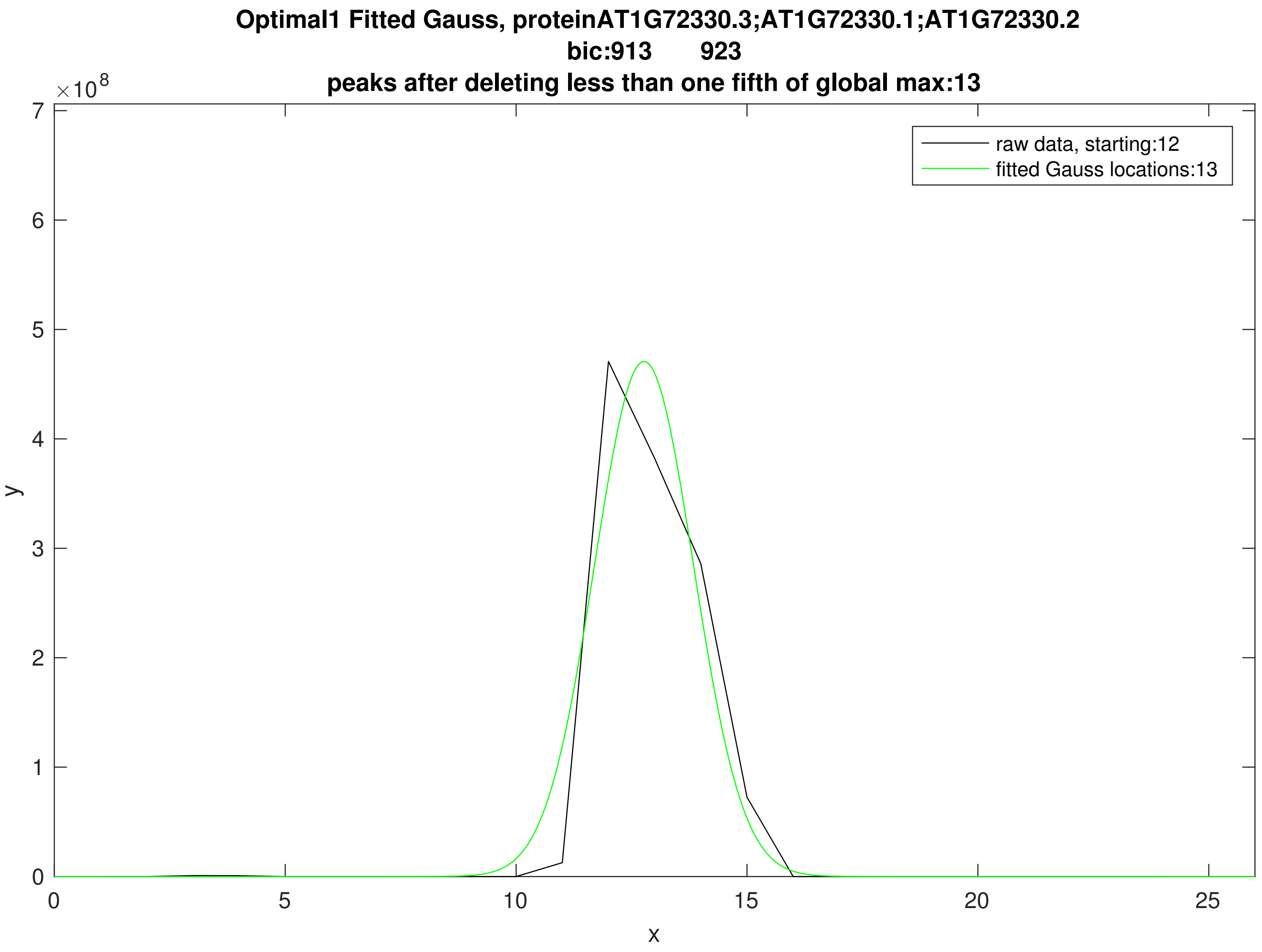
Optimal1 Fitted Gauss, proteinAT1G72330.3;AT1G72330.1;AT1G72330.2
bic:913 923
peaks after deleting less than one fifth of global max:13
×10^8
raw data, starting:12
fitted Gauss locations:13
y
x
0
1
2
3
4
5
6
7
0
5
10
15
20
25

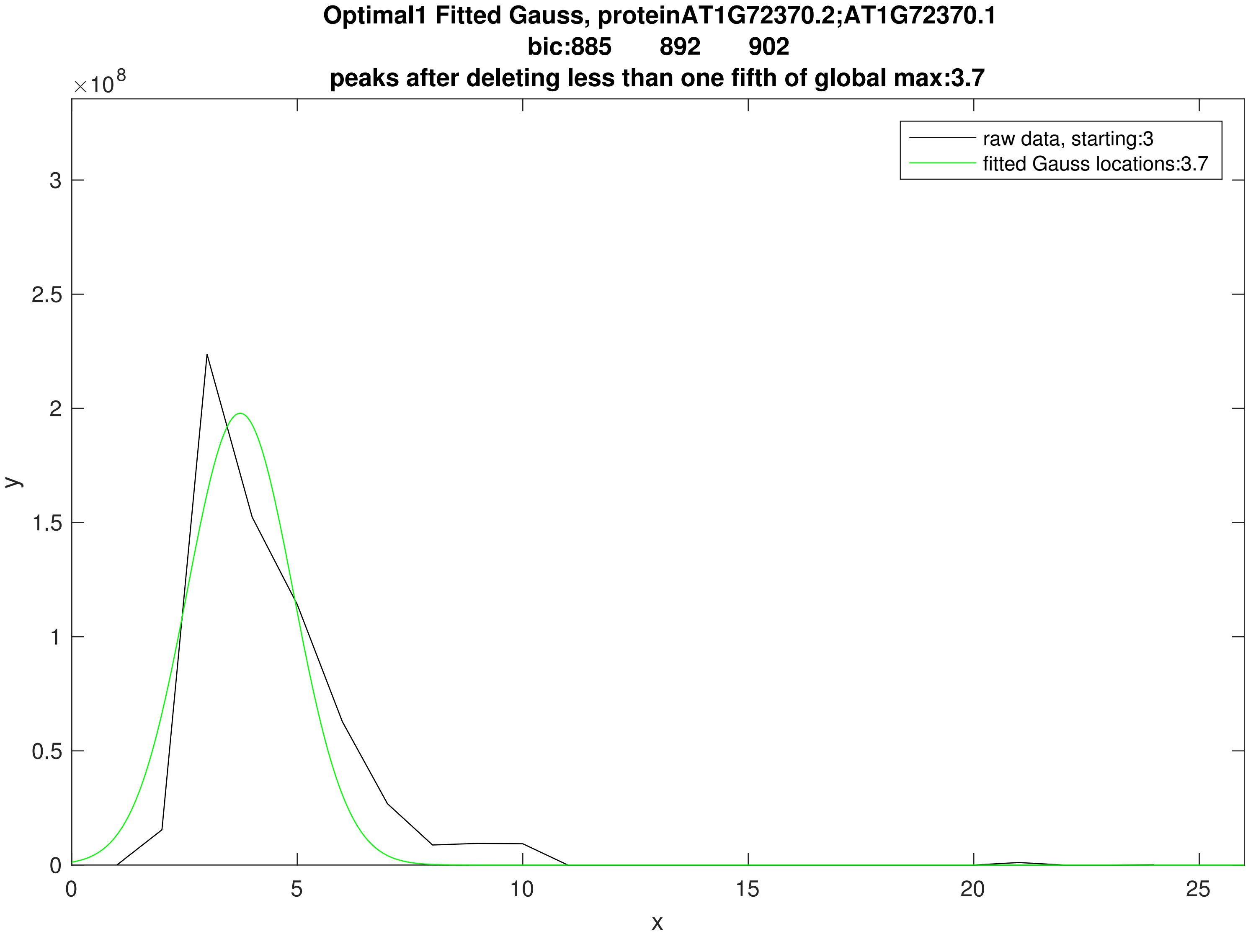
Optimal1 Fitted Gauss, proteinAT1G72370.2;AT1G72370.1
bic:885 892 902
peaks after deleting less than one fifth of global max:3.7
×10^8
raw data, starting:3
fitted Gauss locations:3.7
y
x
0
0.5
1
1.5
2
2.5
3
5
10
15
20
25

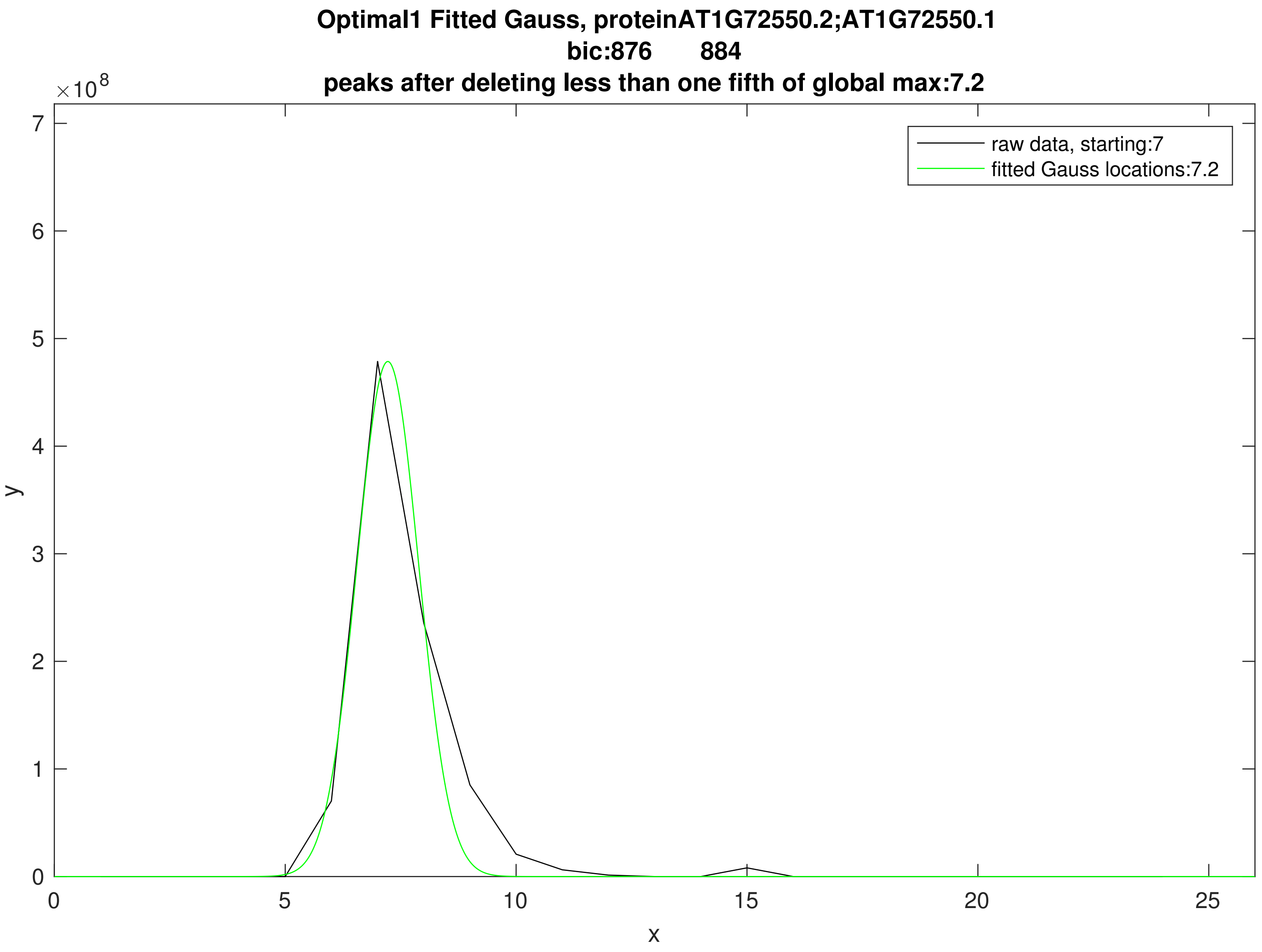
Optimal1 Fitted Gauss, proteinAT1G72550.2;AT1G72550.1
bic:876 884
peaks after deleting less than one fifth of global max:7.2
$\times 10^8$
raw data, starting:7
fitted Gauss locations:7.2
y
x
0
1
2
3
4
5
6
7
0
5
10
15
20
25

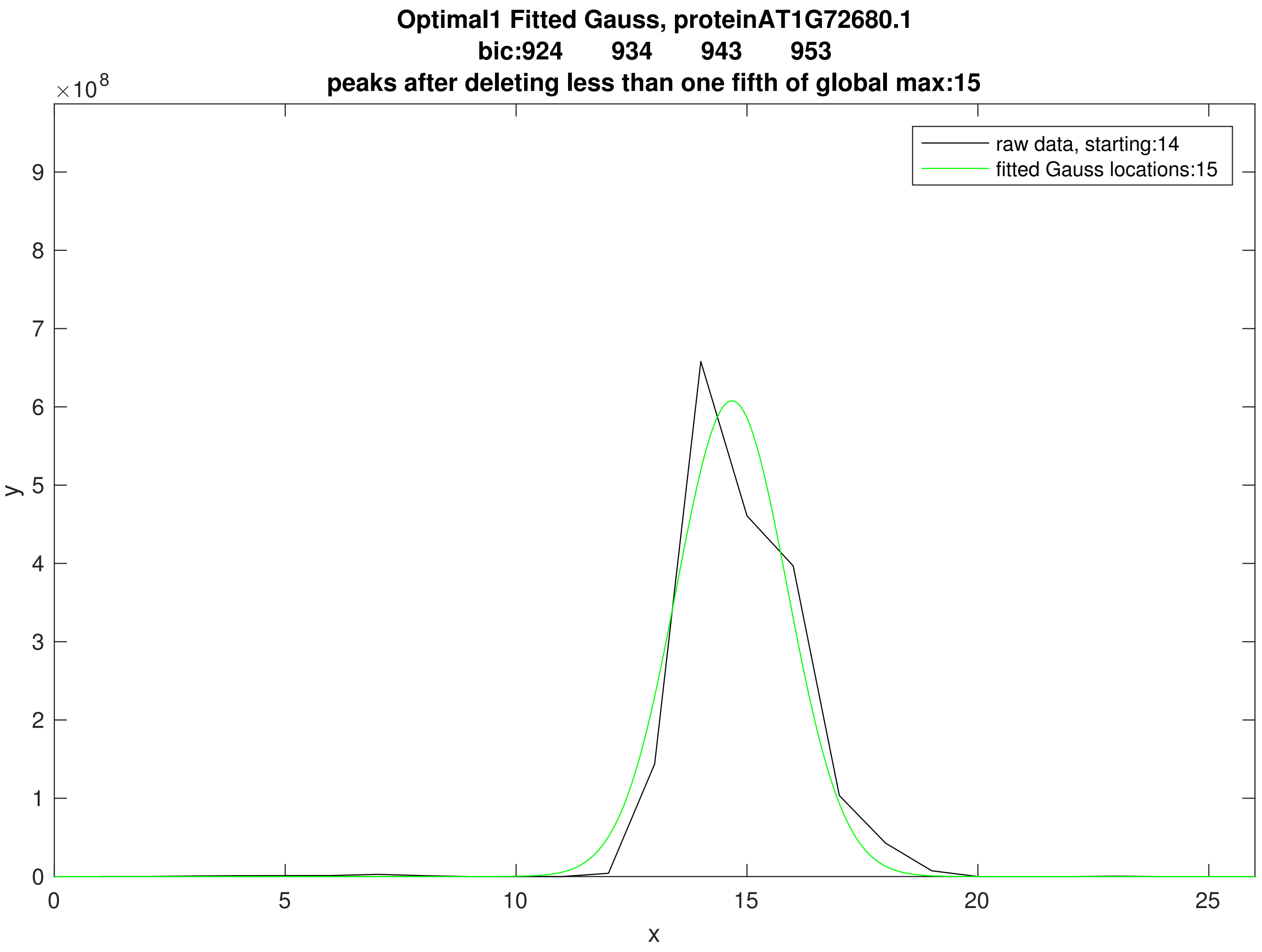
Optimal1 Fitted Gauss, proteinAT1G72680.1
bic:924 934 943 953
peaks after deleting less than one fifth of global max:15
×10^8
raw data, starting:14
fitted Gauss locations:15
y
x
0
1
2
3
4
5
6
7
8
9
0
5
10
15
20
25

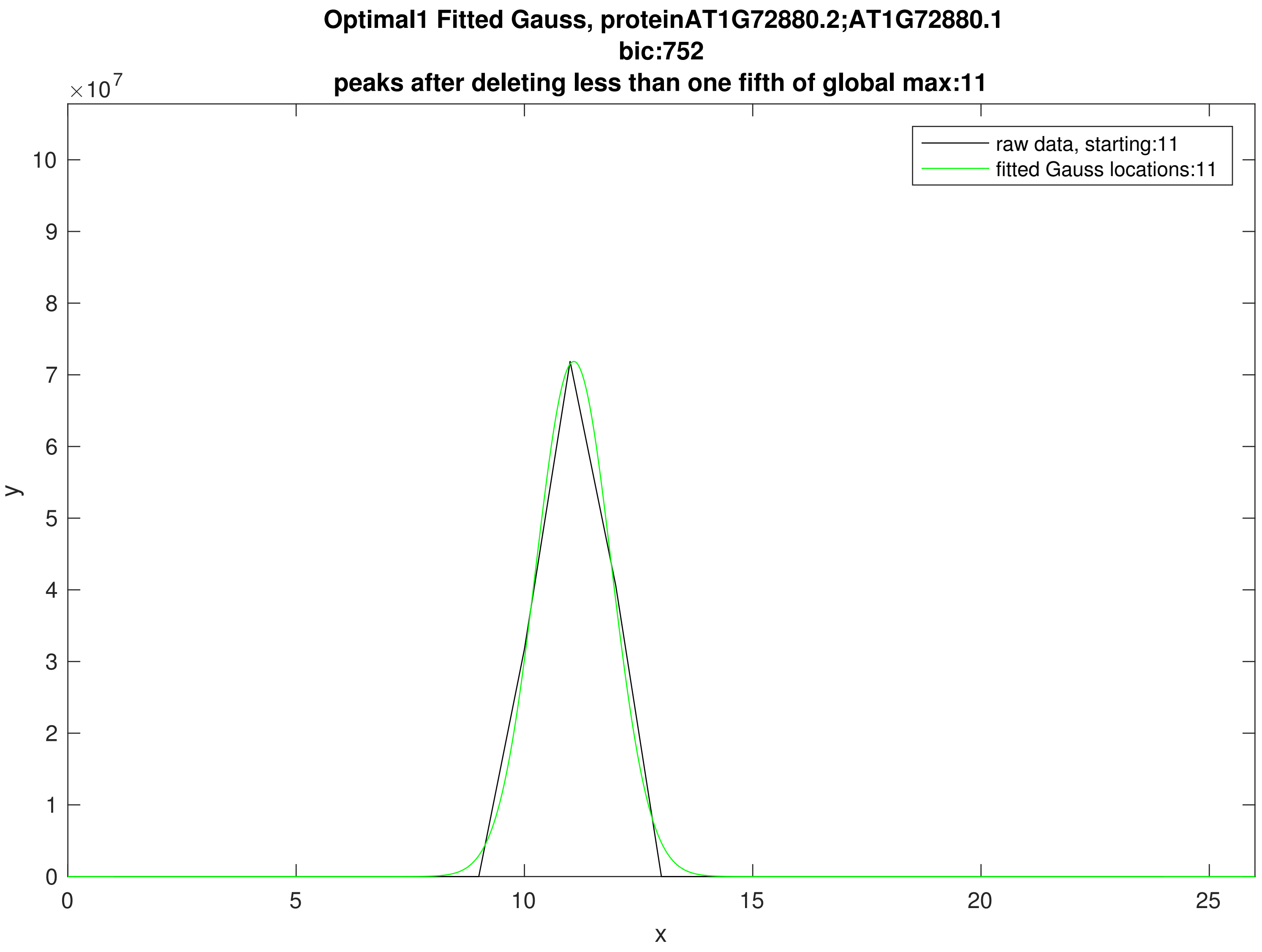

Optimal1 Fitted Gauss, proteinAT1G72880.2;AT1G72880.1
bic:752
peaks after deleting less than one fifth of global max:11
×10^7
raw data, starting:11
fitted Gauss locations:11
y
x
0
1
2
3
4
5
6
7
8
9
10
0
5
10
15
20
25

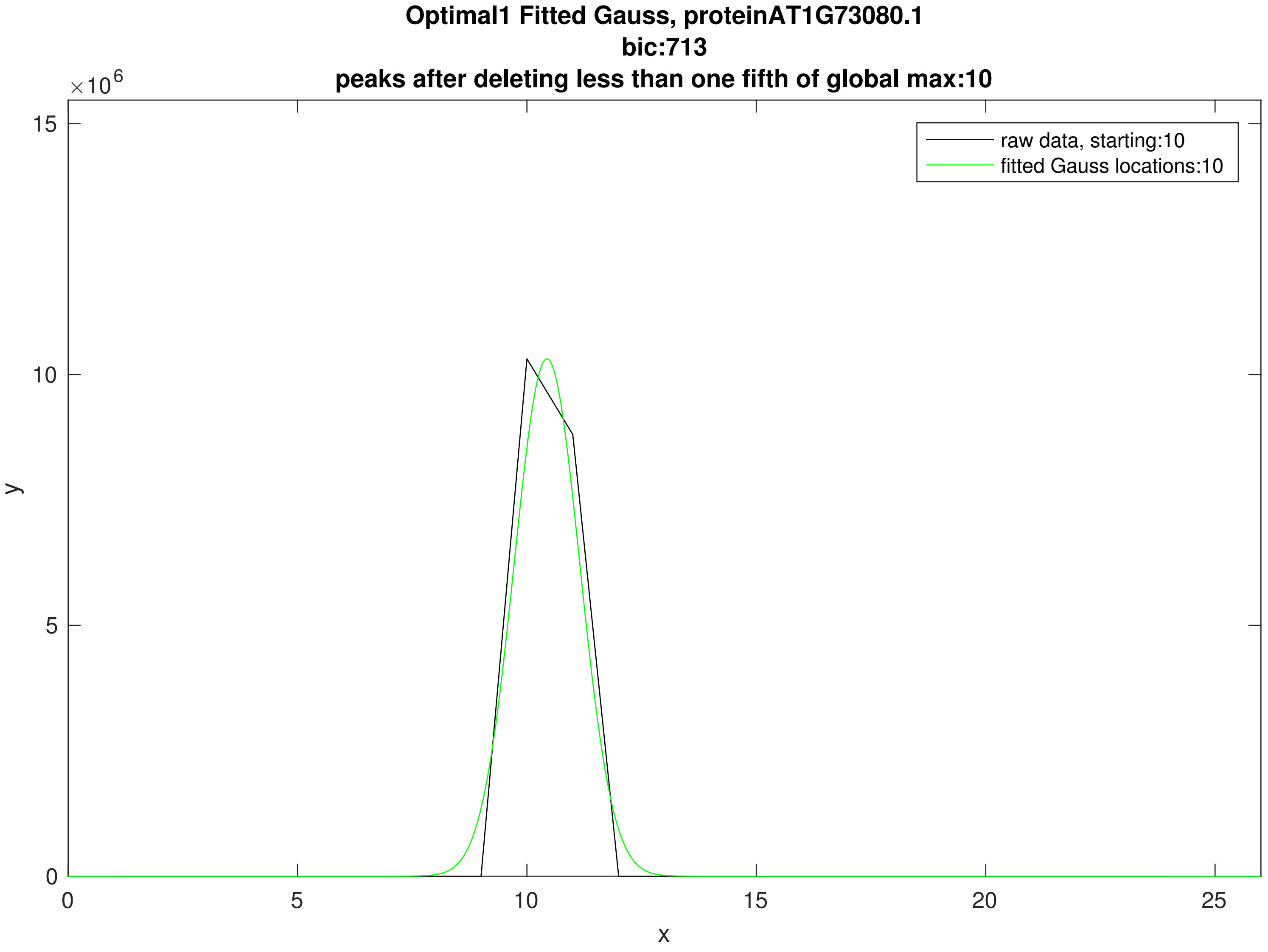
Optimal1 Fitted Gauss, proteinAT1G73080.1
bic:713
peaks after deleting less than one fifth of global max:10
×10^6
15
10
5
0
y
0
5
10
15
20
25
x
raw data, starting:10
fitted Gauss locations:10

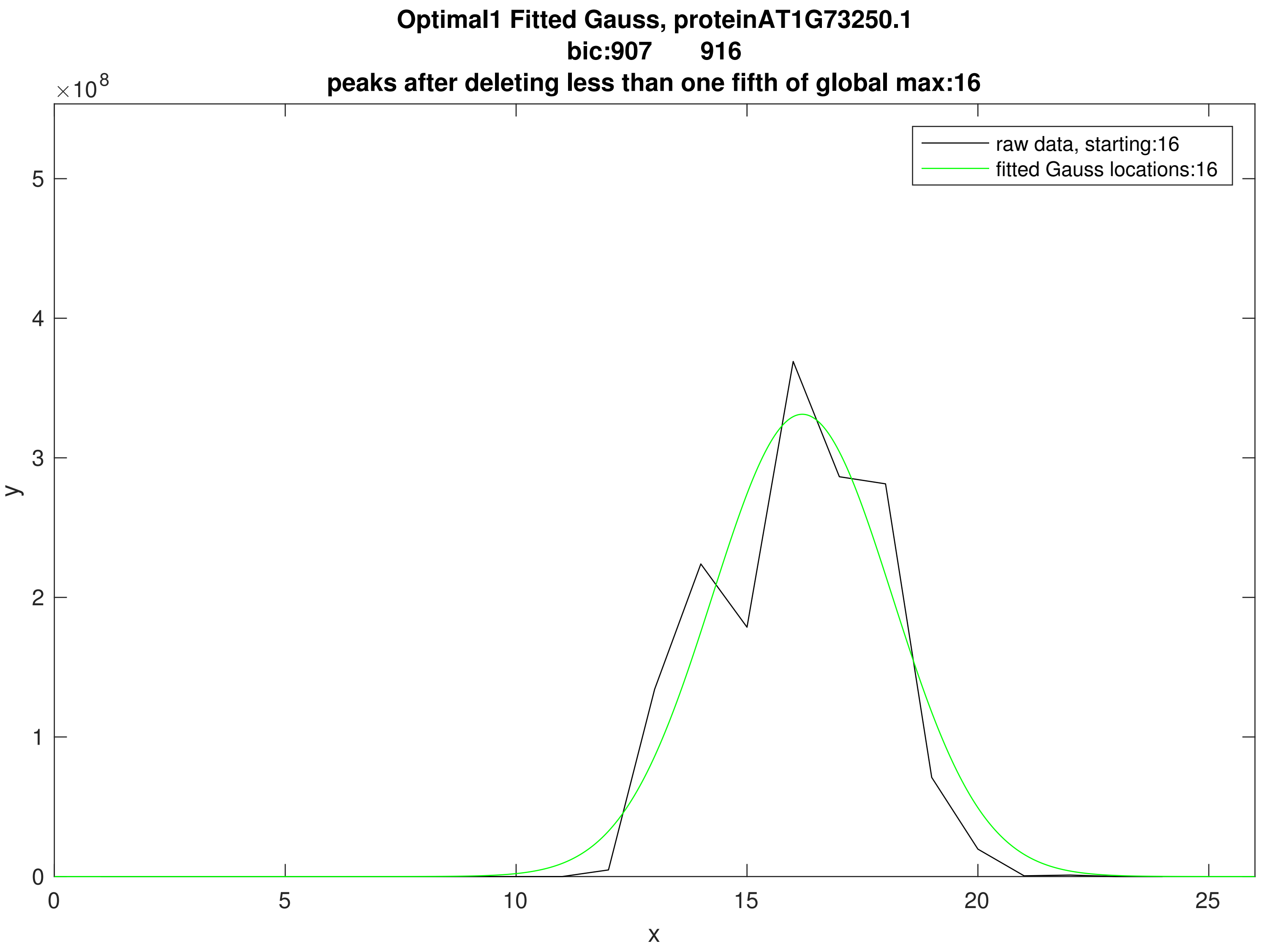
Optimal1 Fitted Gauss, proteinAT1G73250.1
bic:907 916
peaks after deleting less than one fifth of global max:16
×10^8
raw data, starting:16
fitted Gauss locations:16
y
x
0
1
2
3
4
5
0
5
10
15
20
25

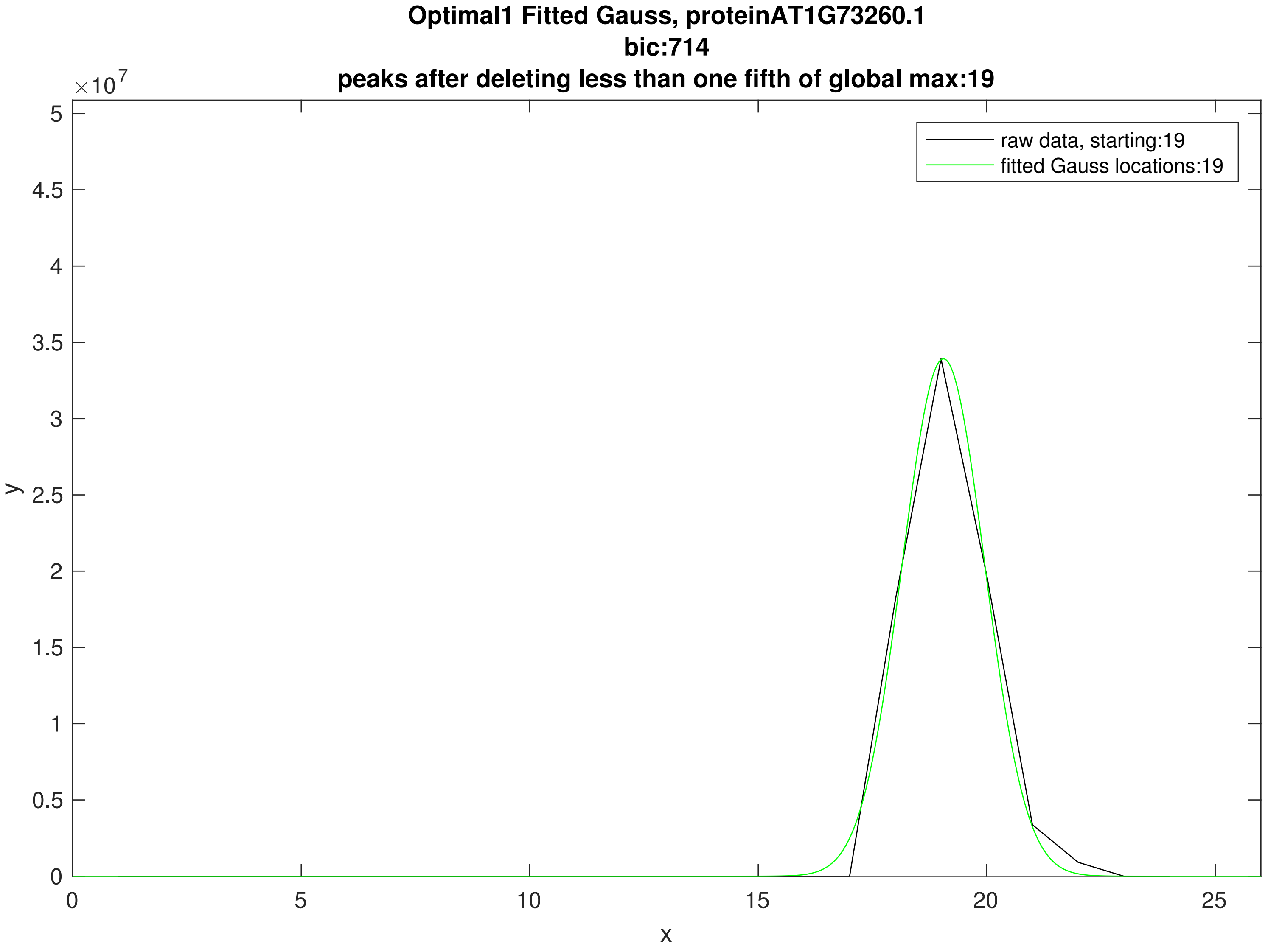
Optimal1 Fitted Gauss, proteinAT1G73260.1
bic:714
peaks after deleting less than one fifth of global max:19
×10^7
5
4.5
4
3.5
3
2.5
2
1.5
1
0.5
0
y
0
5
10
15
20
25
x
raw data, starting:19
fitted Gauss locations:19

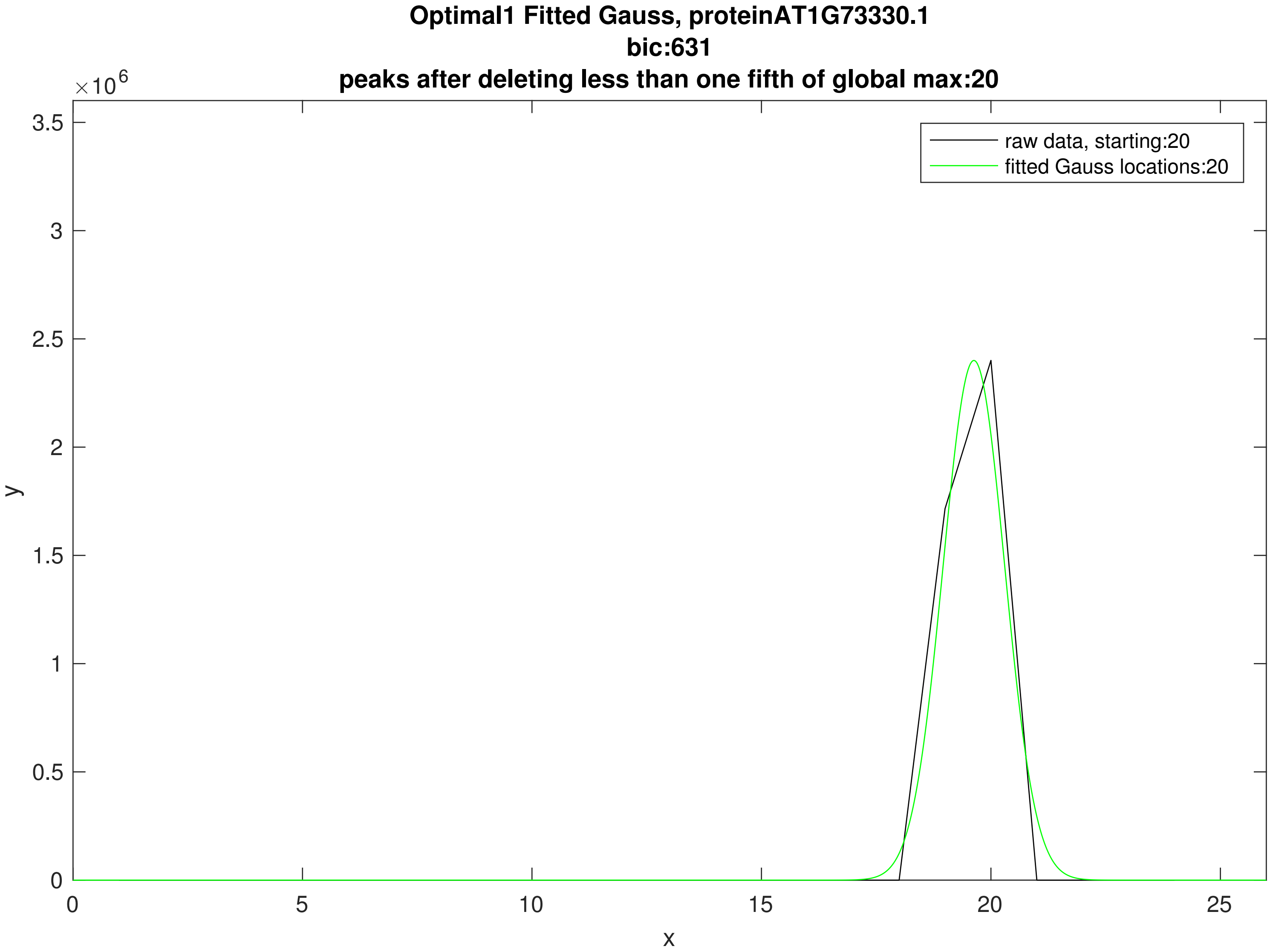

Optimal1 Fitted Gauss, proteinAT1G73330.1
bic:631
peaks after deleting less than one fifth of global max:20
×10^6
3.5
3
2.5
2
1.5
1
0.5
0
y
0
5
10
15
20
25
x
raw data, starting:20
fitted Gauss locations:20

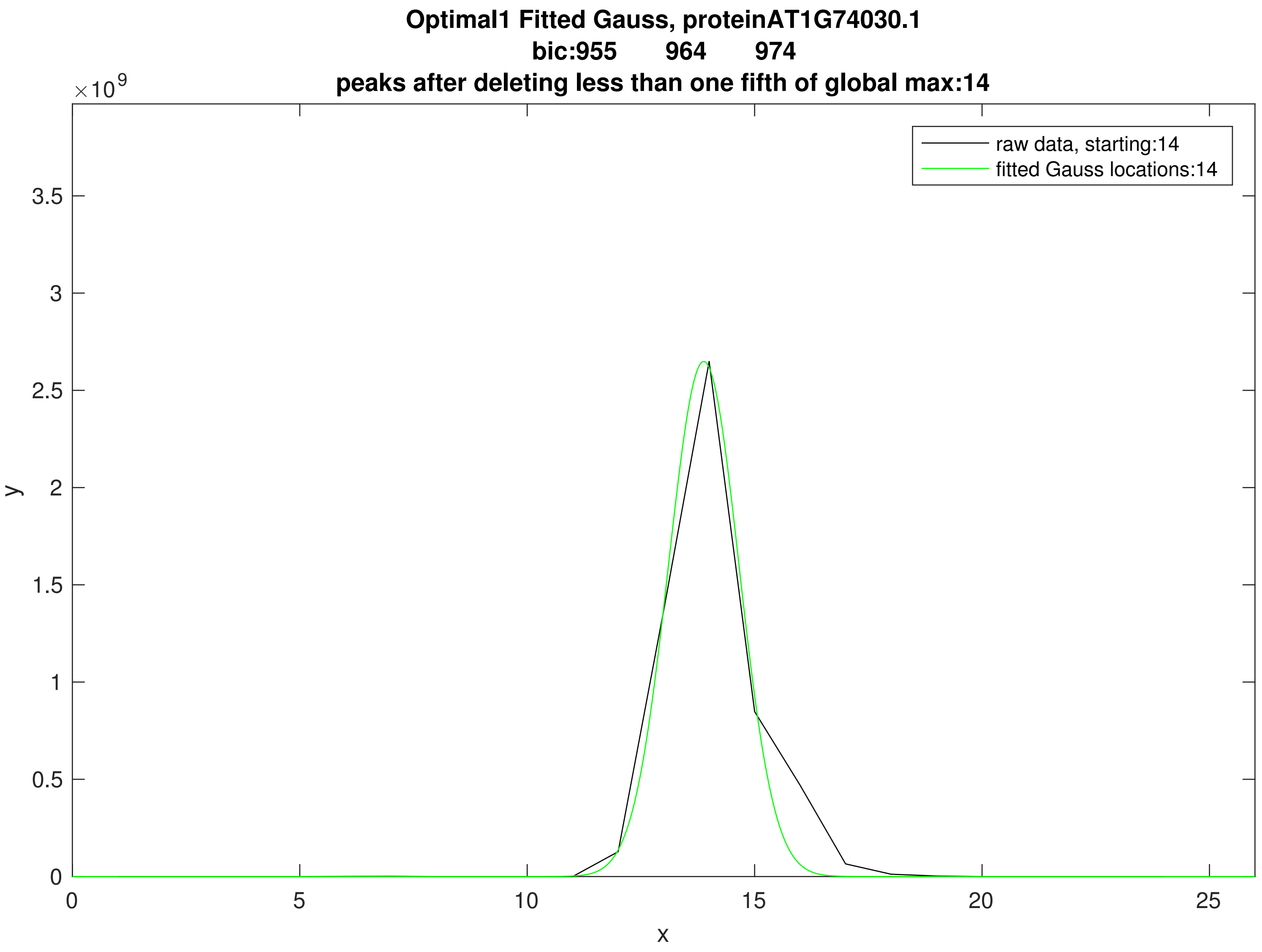
Optimal1 Fitted Gauss, proteinAT1G74030.1
bic:955 964 974
peaks after deleting less than one fifth of global max:14
×10^9
3.5
3
2.5
2
1.5
1
0.5
0
y
0
5
10
15
20
25
x
raw data, starting:14
fitted Gauss locations:14

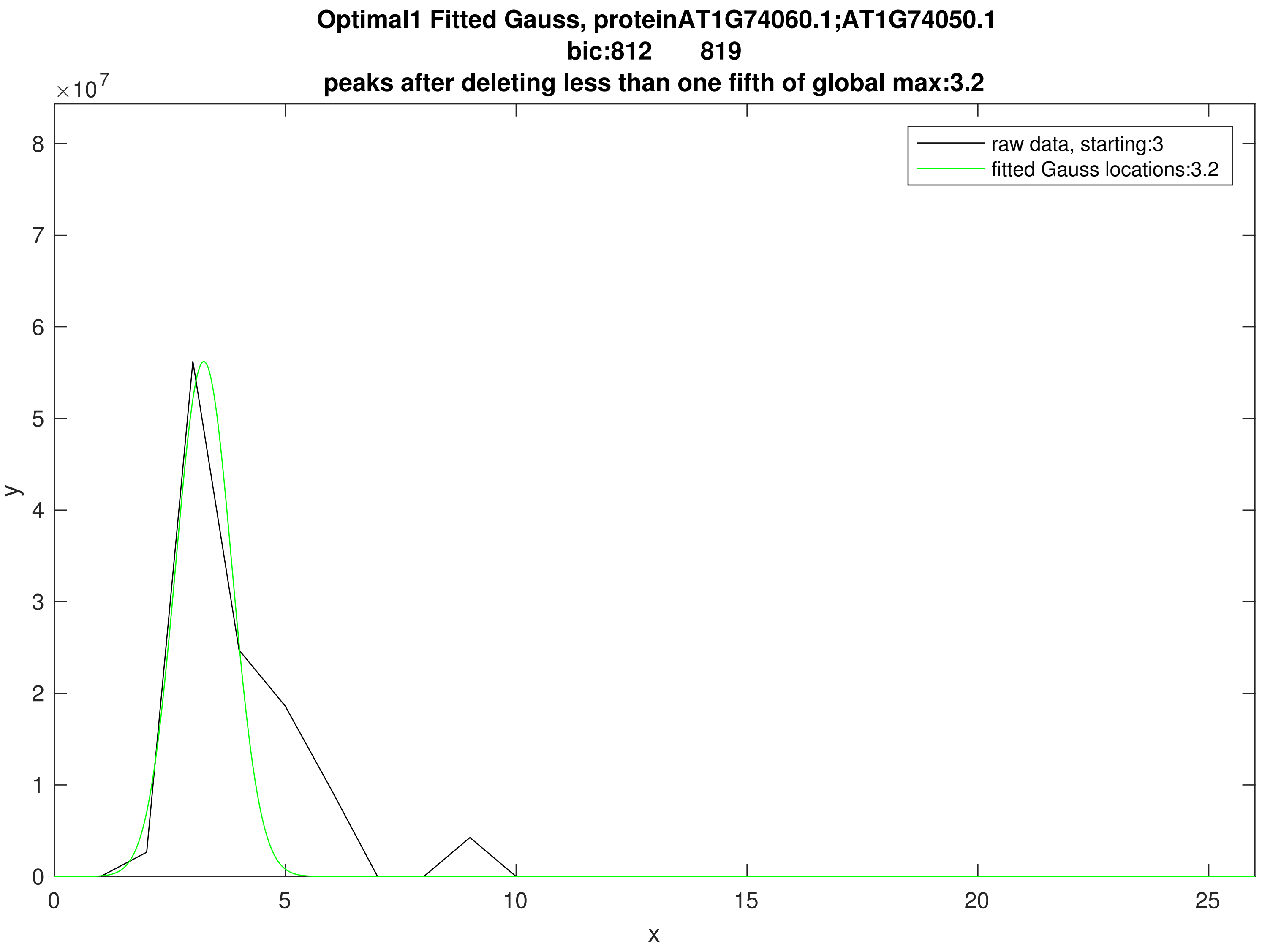
Optimal1 Fitted Gauss, proteinAT1G74060.1;AT1G74050.1
bic:812 819
peaks after deleting less than one fifth of global max:3.2
×10^7
raw data, starting:3
fitted Gauss locations:3.2
y
x
0
1
2
3
4
5
6
7
8
0
5
10
15
20
25

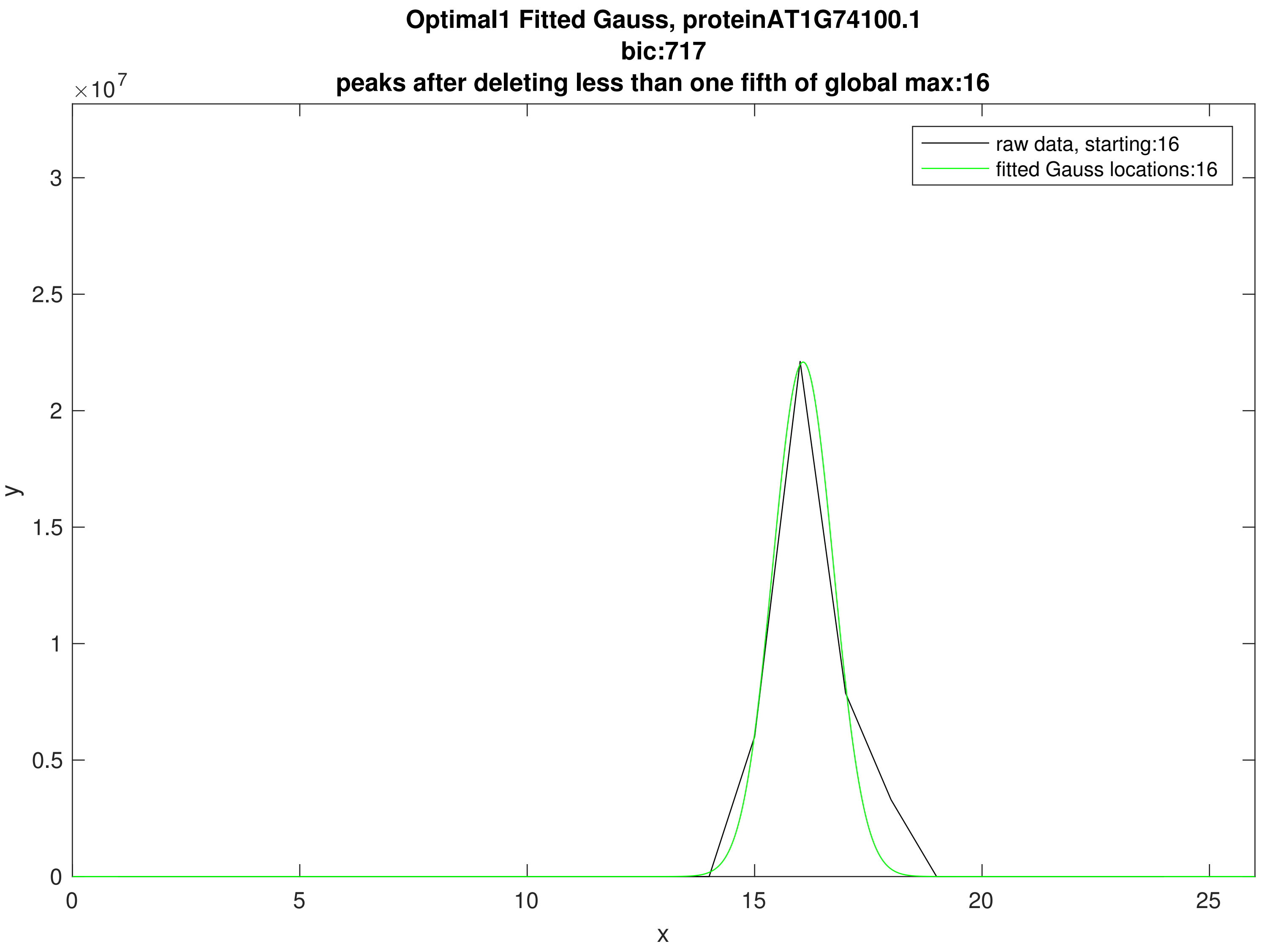
Optimal1 Fitted Gauss, proteinAT1G74100.1
bic:717
peaks after deleting less than one fifth of global max:16
×10^7
raw data, starting:16
fitted Gauss locations:16
y
x
0
0.5
1
1.5
2
2.5
3
0
5
10
15
20
25

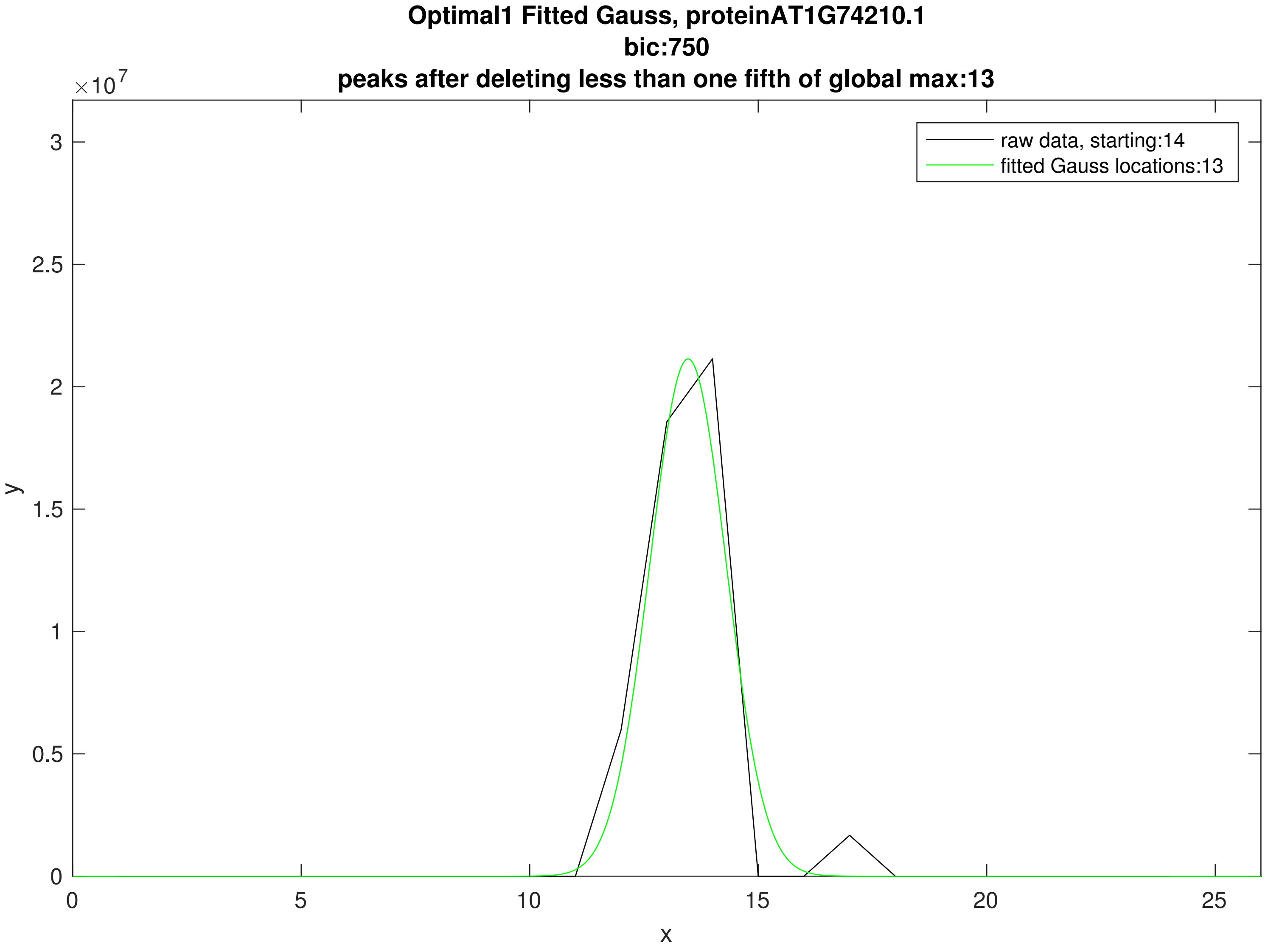

Optimal1 Fitted Gauss, proteinAT1G74210.1
bic:750
peaks after deleting less than one fifth of global max:13
×10^7
raw data, starting:14
fitted Gauss locations:13
y
x
0
0.5
1
1.5
2
2.5
3
0
5
10
15
20
25

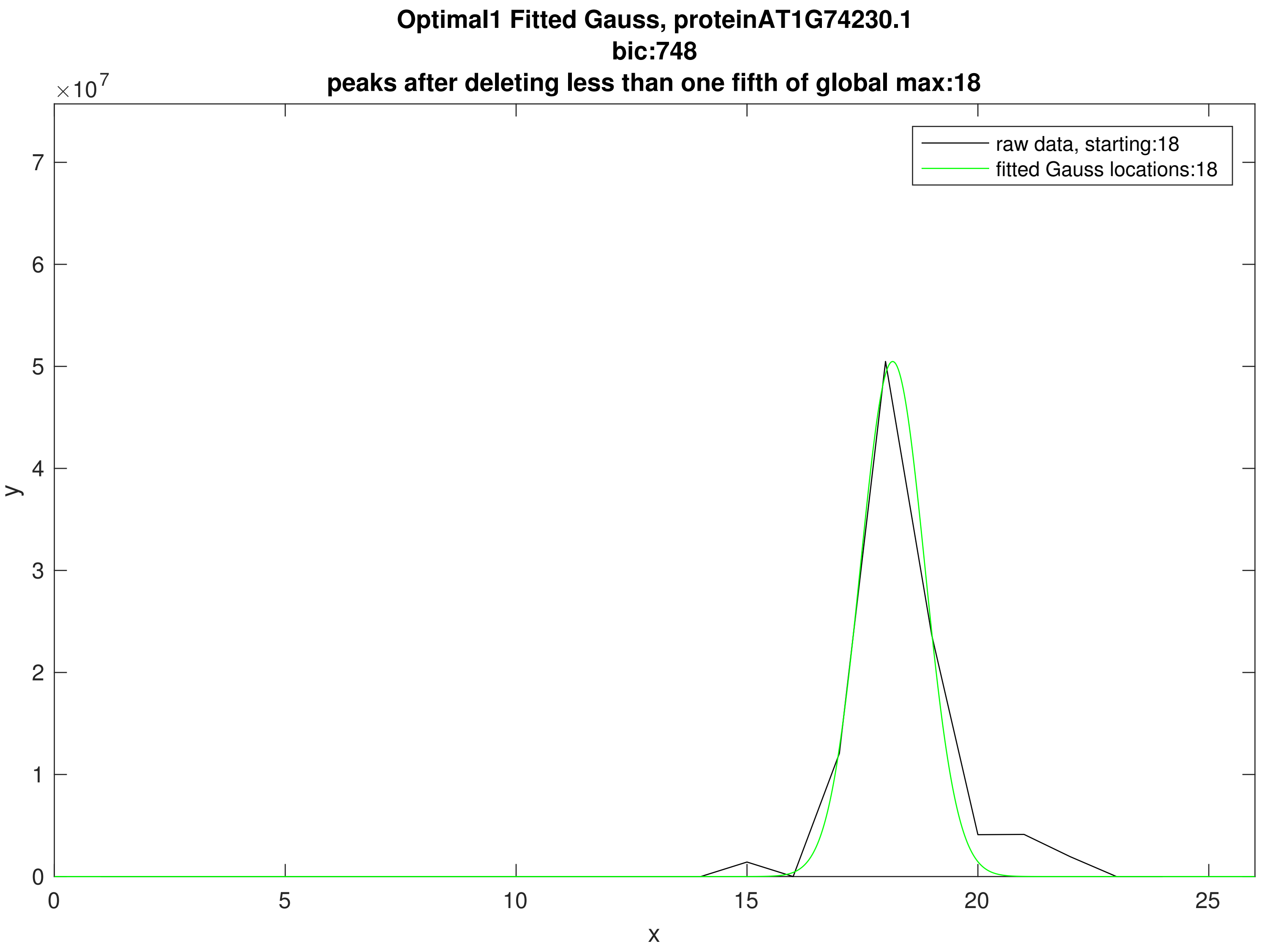
Optimal1 Fitted Gauss, proteinAT1G74230.1
bic:748
peaks after deleting less than one fifth of global max:18
×10^7
raw data, starting:18
fitted Gauss locations:18
y
x
0
1
2
3
4
5
6
7
0
5
10
15
20
25

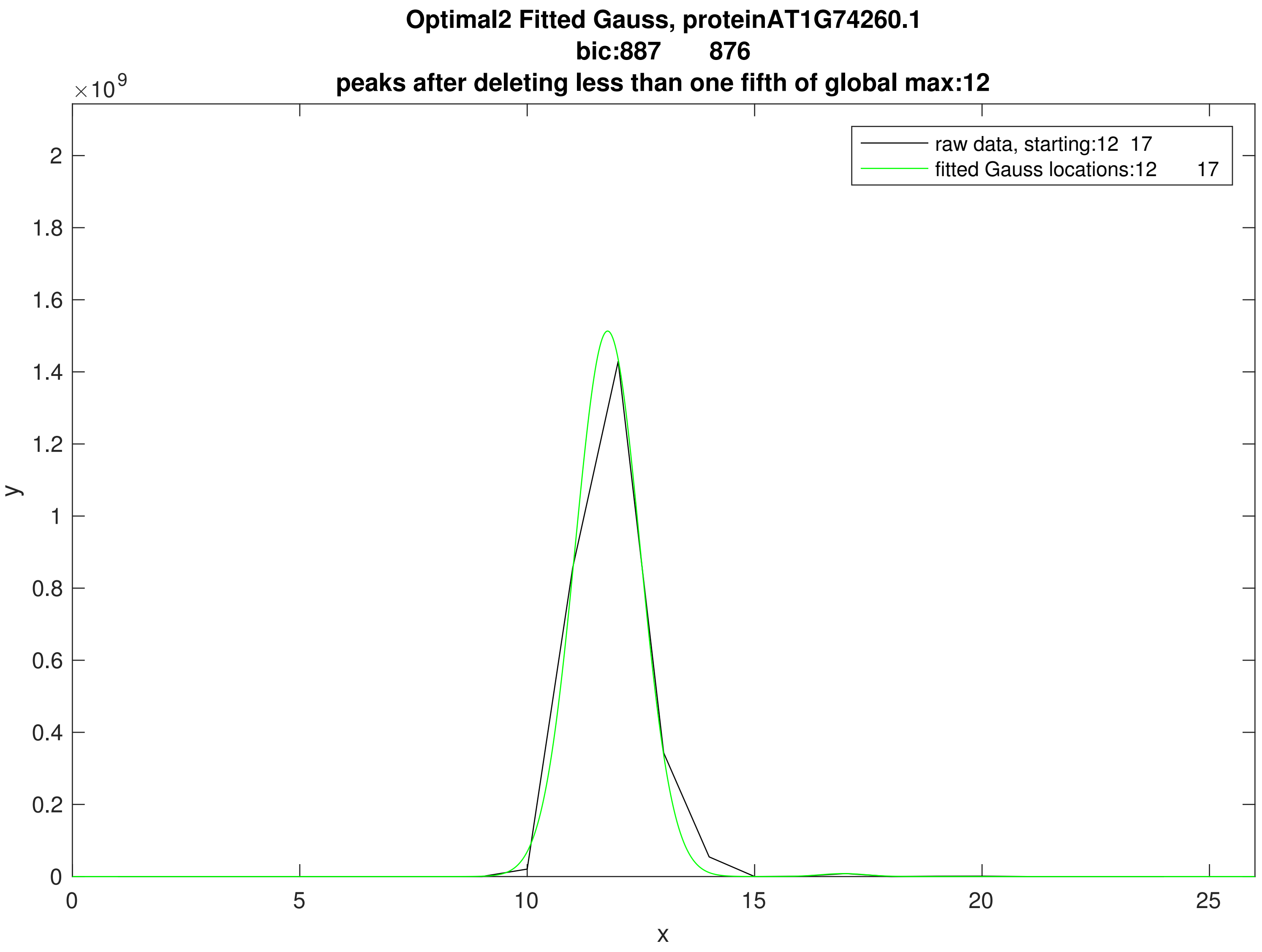
Optimal2 Fitted Gauss, proteinAT1G74260.1
bic:887 876
peaks after deleting less than one fifth of global max:12
×10^9
raw data, starting:12 17
fitted Gauss locations:12 17
y
x
0
0.2
0.4
0.6
0.8
1
1.2
1.4
1.6
1.8
2
0
5
10
15
20
25

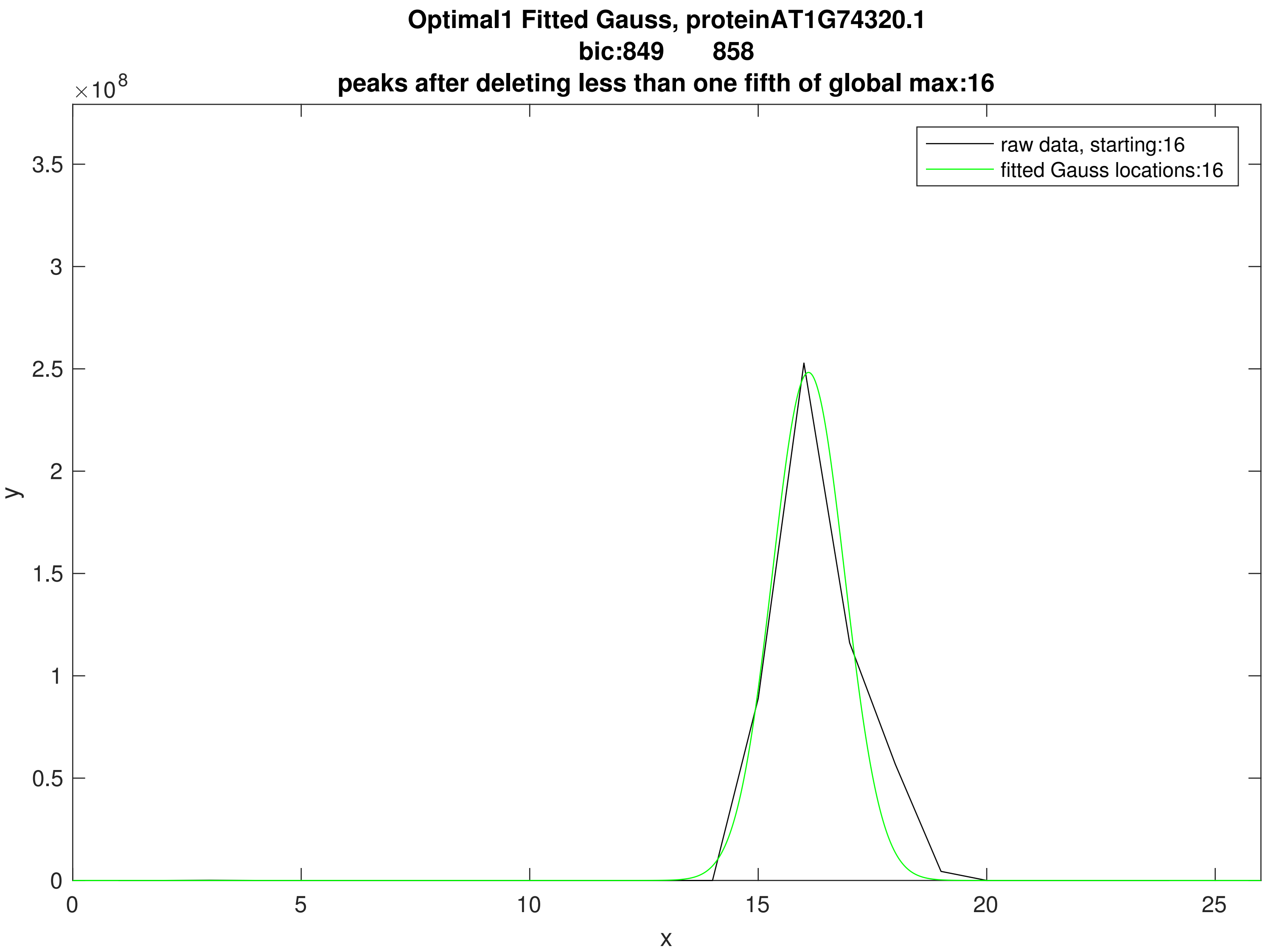
Optimal1 Fitted Gauss, proteinAT1G74320.1
bic:849 858
peaks after deleting less than one fifth of global max:16
×10^8
3.5
3
2.5
2
1.5
1
0.5
0
y
0
5
10
15
20
25
x
raw data, starting:16
fitted Gauss locations:16

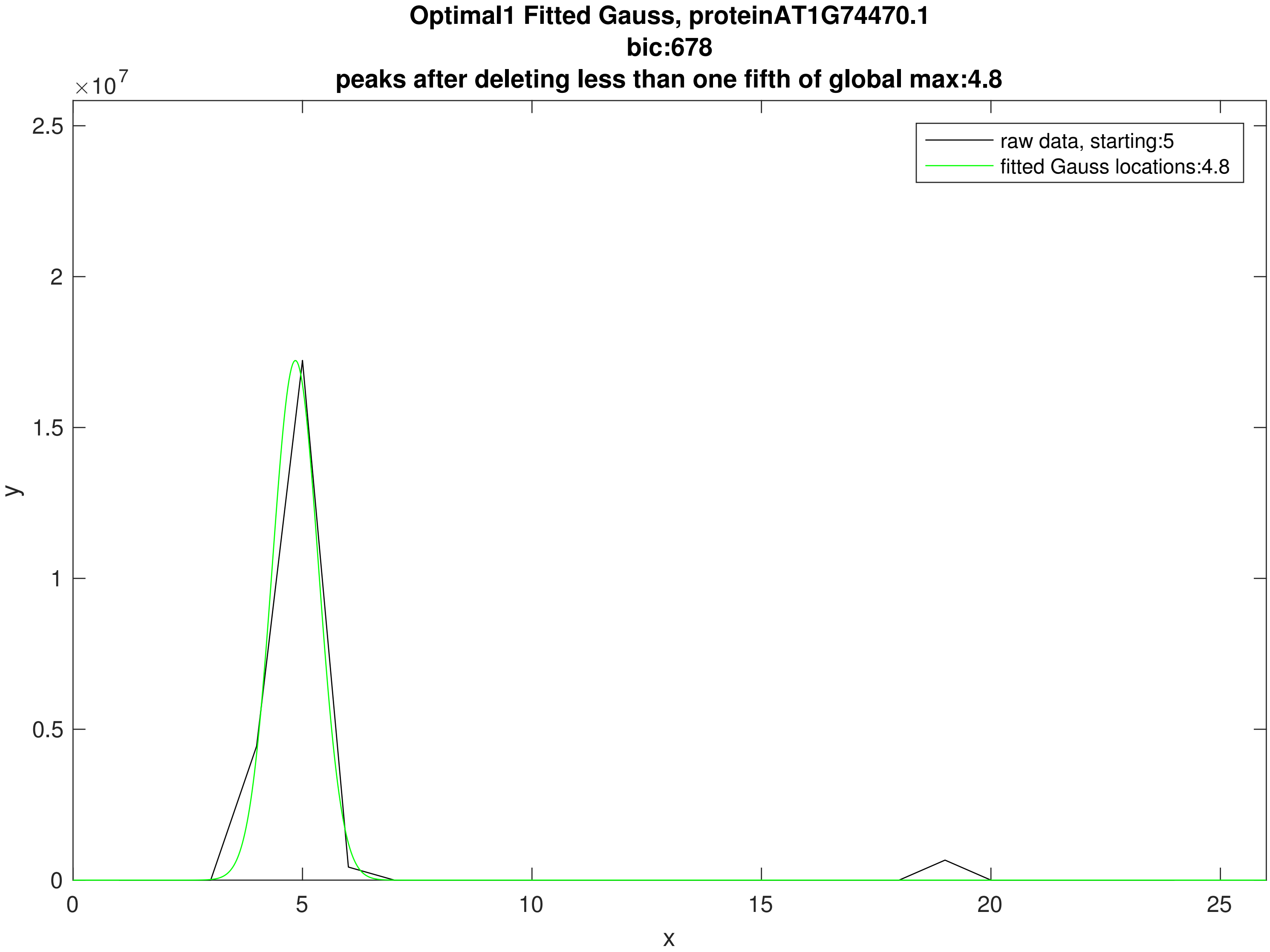
Optimal1 Fitted Gauss, proteinAT1G74470.1
bic:678
peaks after deleting less than one fifth of global max:4.8
×10^7
2.5
2
1.5
1
0.5
0
y
0
5
10
15
20
25
x
raw data, starting:5
fitted Gauss locations:4.8

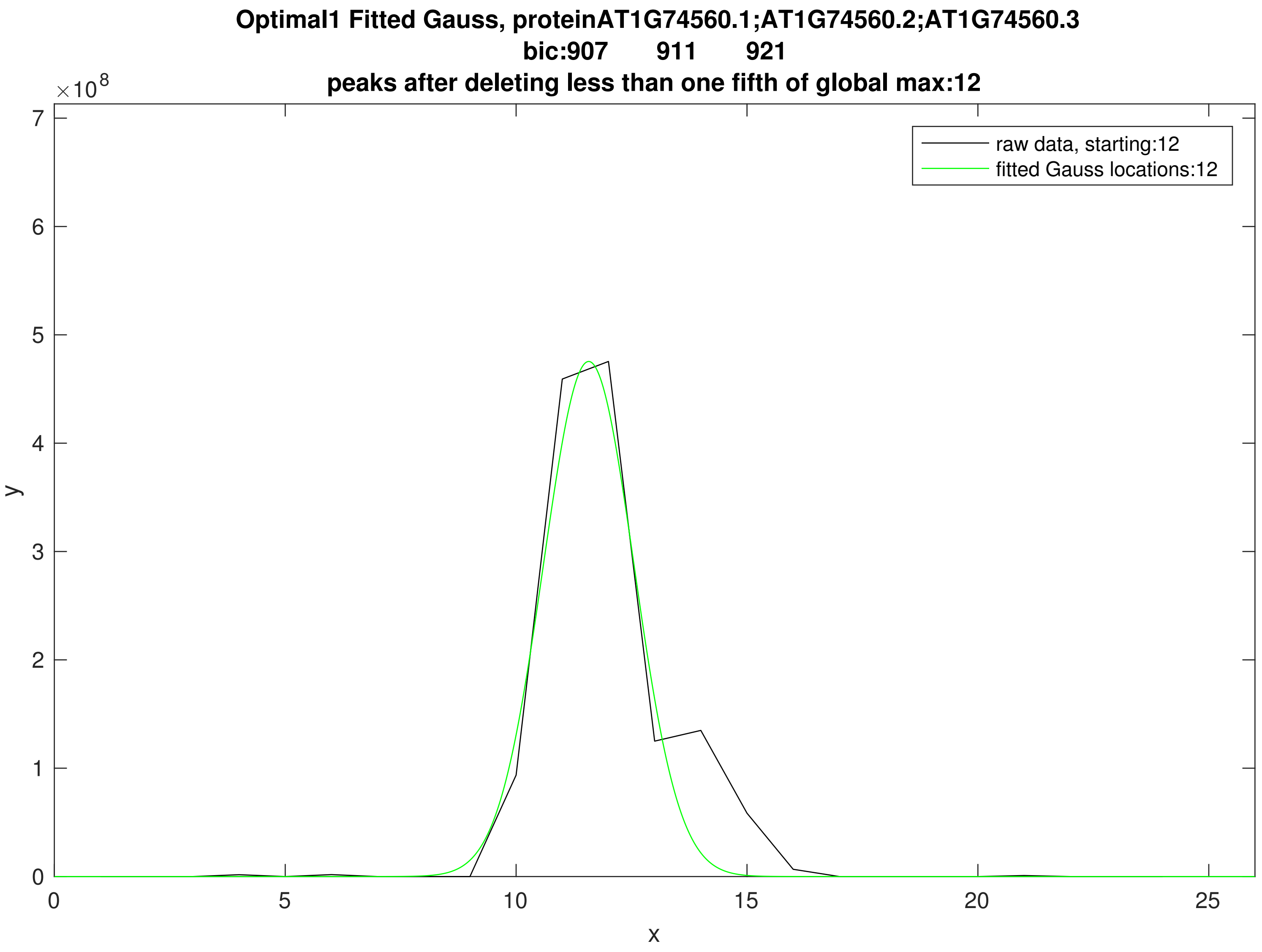

Optimal1 Fitted Gauss, proteinAT1G74560.1;AT1G74560.2;AT1G74560.3
bic:907 911 921
peaks after deleting less than one fifth of global max:12
raw data, starting:12
fitted Gauss locations:12
×10^8
y
x
0
1
2
3
4
5
6
7
0
5
10
15
20
25

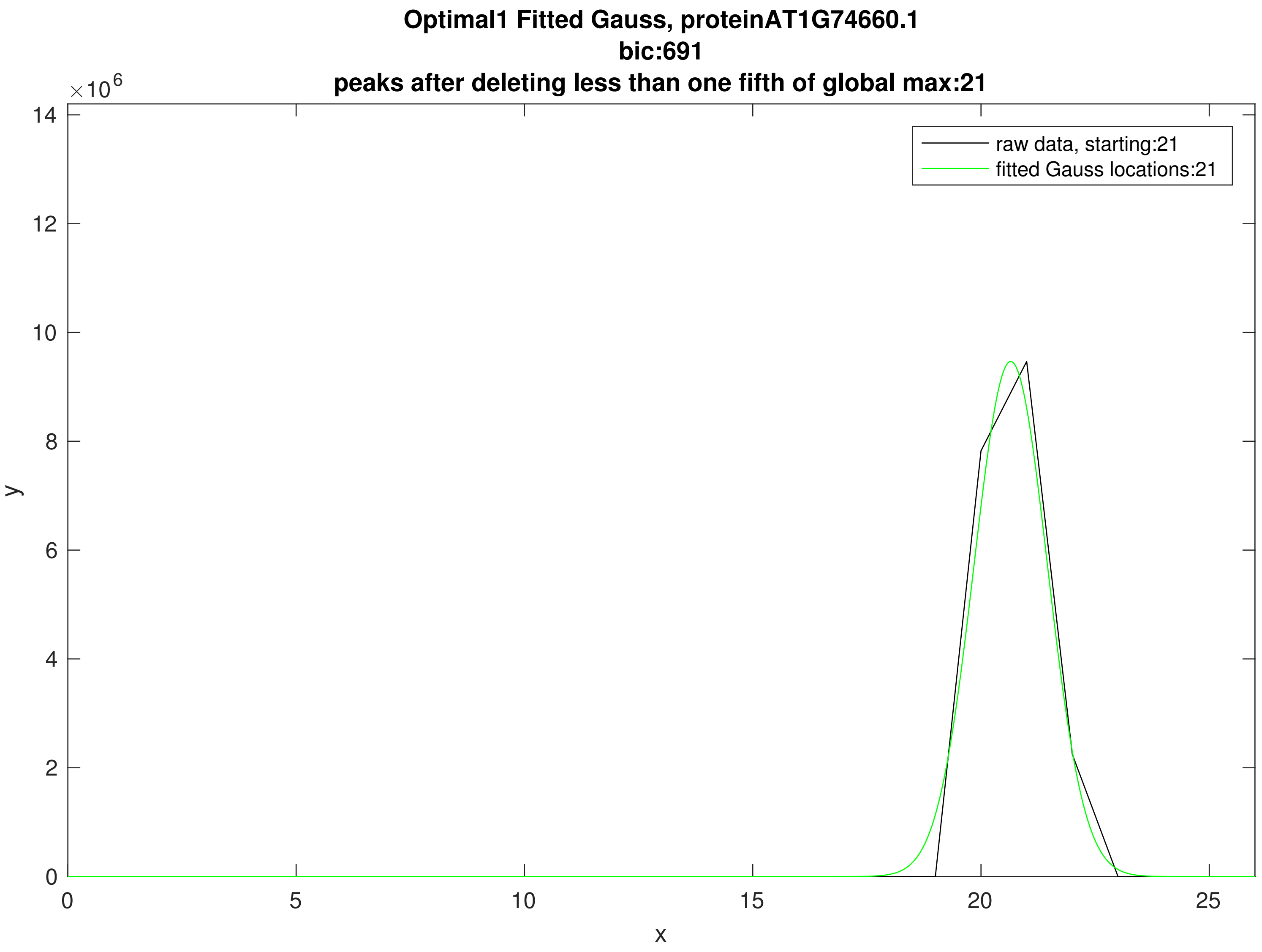
Optimal1 Fitted Gauss, proteinAT1G74660.1
bic:691
peaks after deleting less than one fifth of global max:21
×10^6
raw data, starting:21
fitted Gauss locations:21
y
x

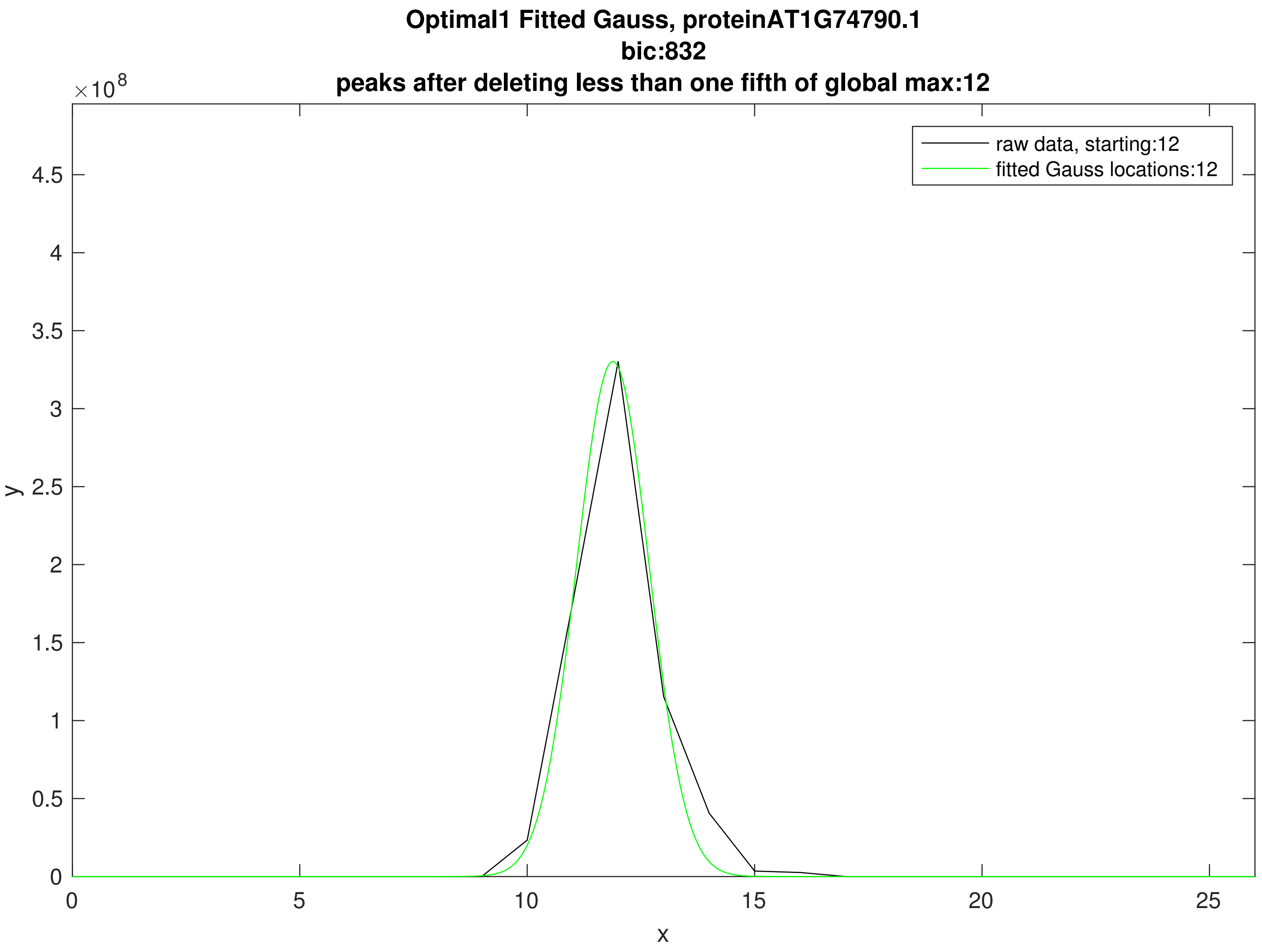
Optimal1 Fitted Gauss, proteinAT1G74790.1
bic:832
peaks after deleting less than one fifth of global max:12
×10^8
raw data, starting:12
fitted Gauss locations:12
y
x
0
0.5
1
1.5
2
2.5
3
3.5
4
4.5
5
10
15
20
25

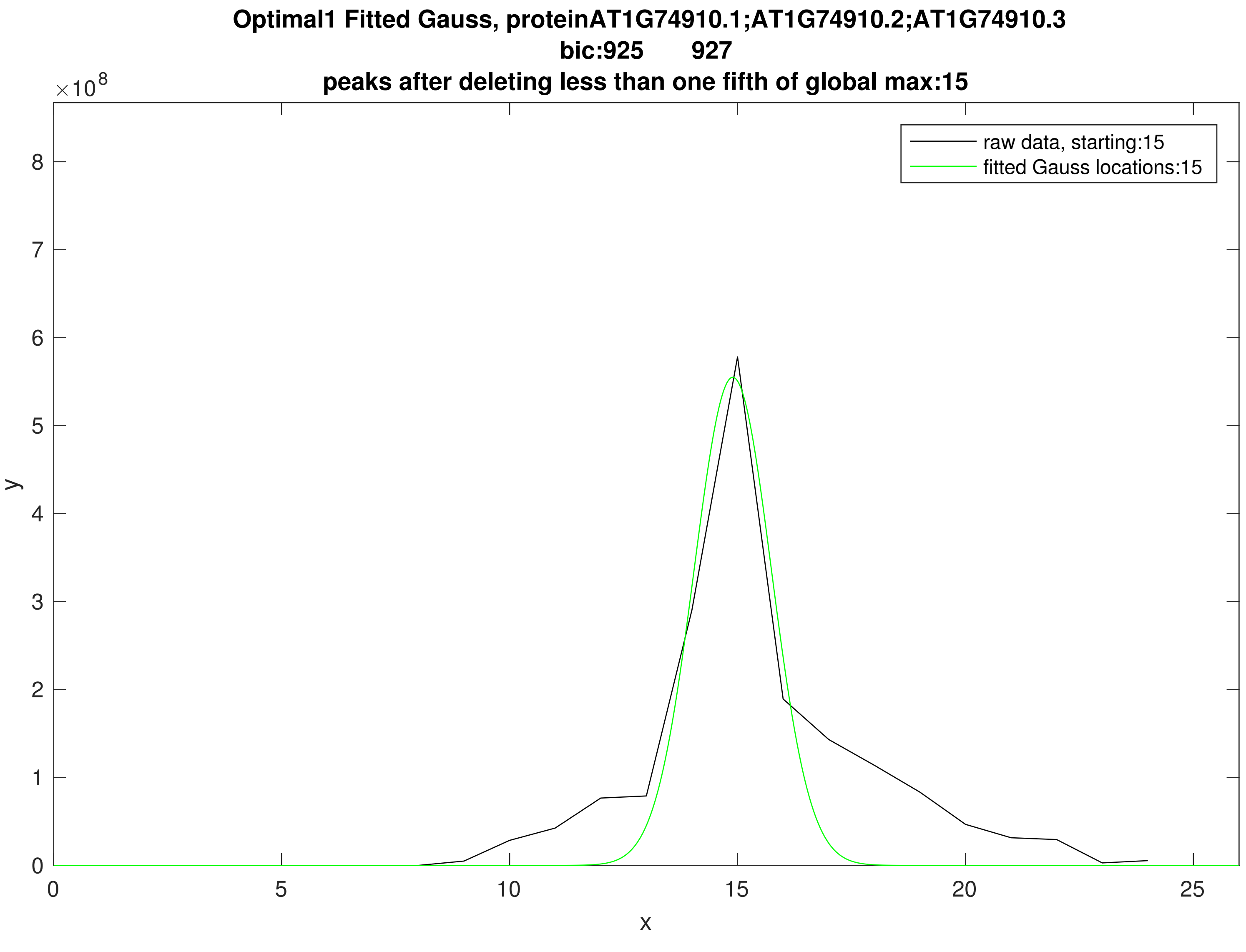

Optimal1 Fitted Gauss, proteinAT1G74910.1;AT1G74910.2;AT1G74910.3
bic:925 927
peaks after deleting less than one fifth of global max:15
×10^8
raw data, starting:15
fitted Gauss locations:15
y
x
0
1
2
3
4
5
6
7
8
0
5
10
15
20
25

**Optimal1 Fitted Gauss, proteinAT1G74920.1;AT1G74920.2**
**bic:964 972**
**peaks after deleting less than one fifth of global max:13**

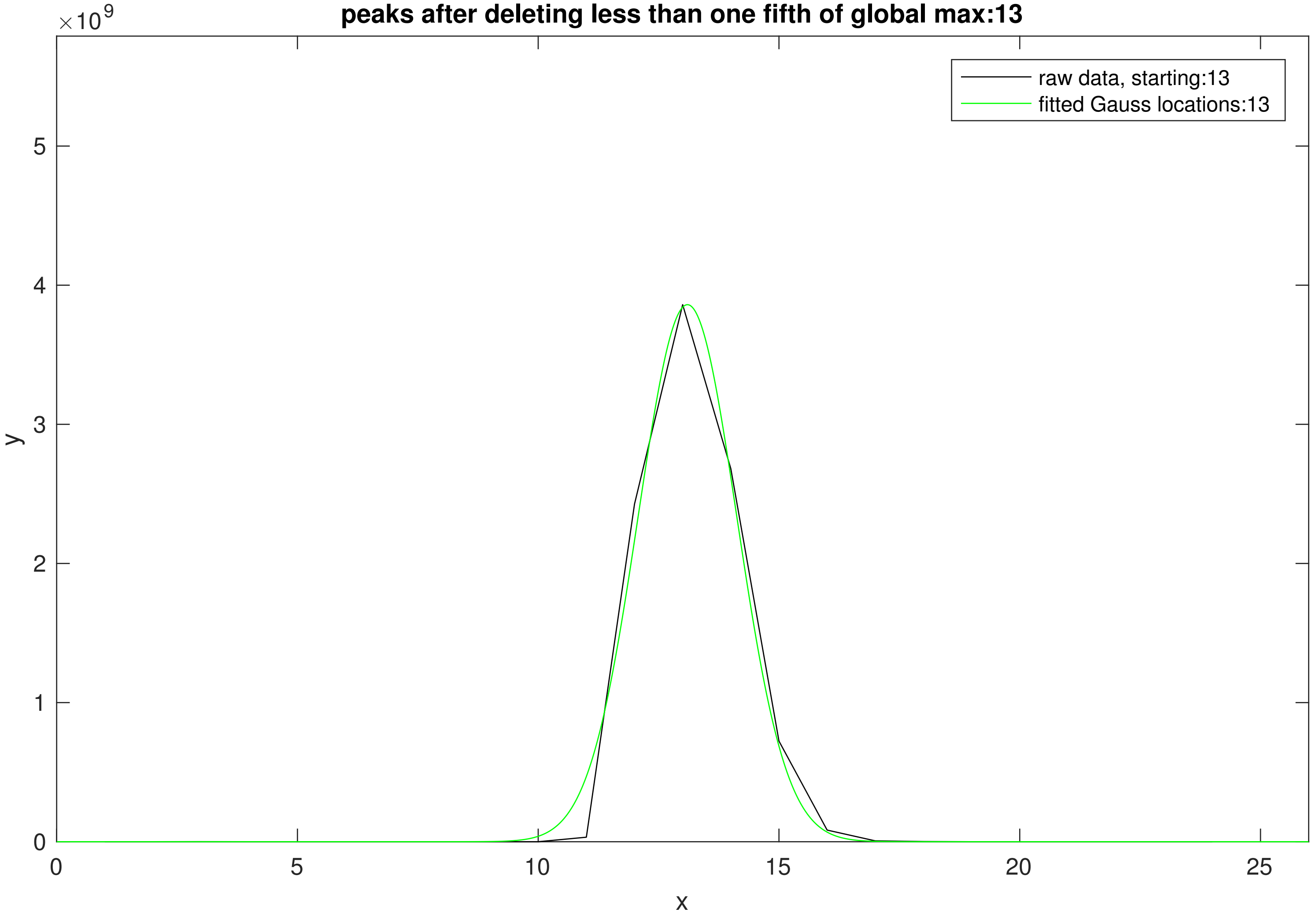

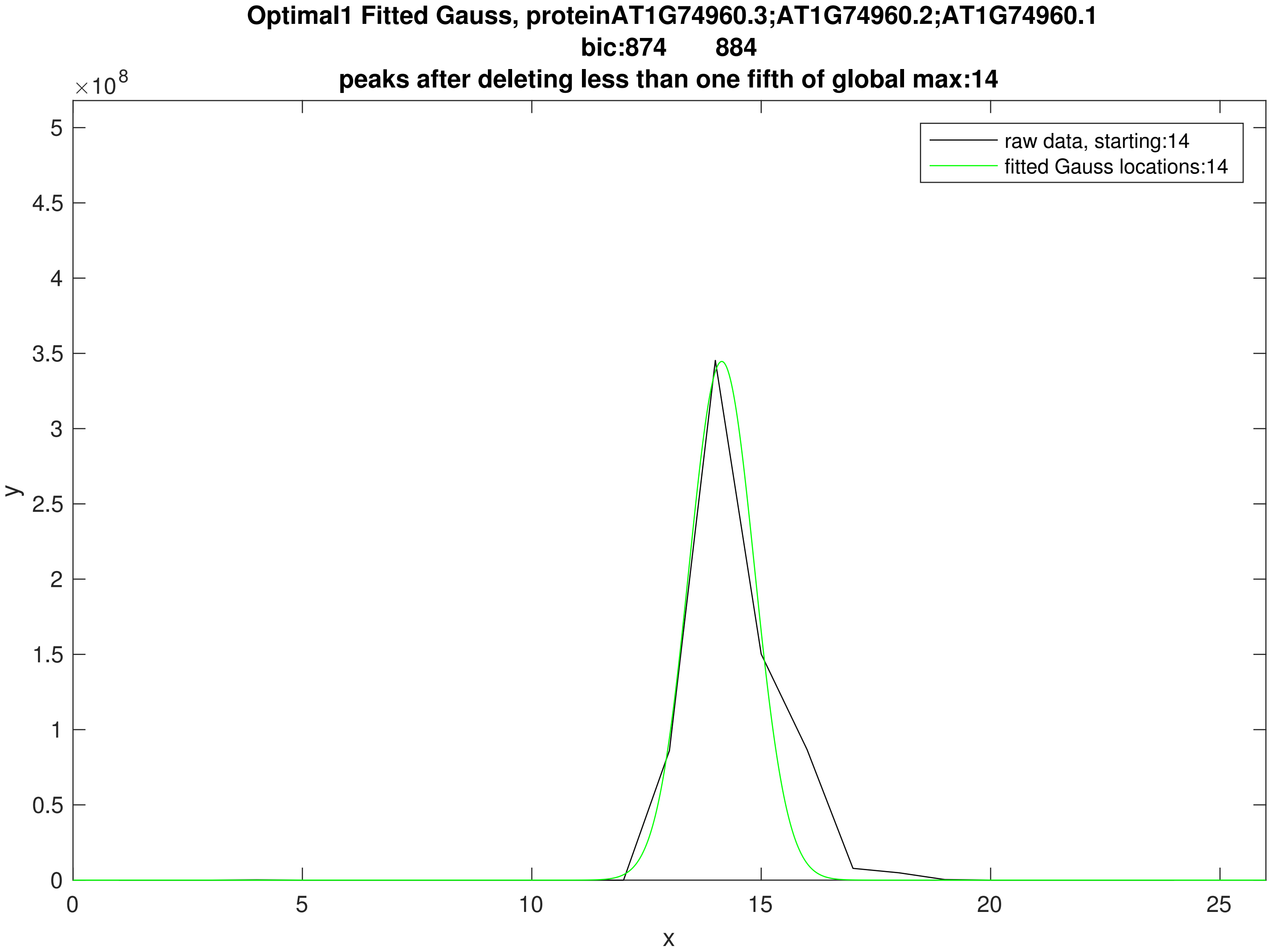

Optimal1 Fitted Gauss, proteinAT1G74960.3;AT1G74960.2;AT1G74960.1
bic:874 884
peaks after deleting less than one fifth of global max:14
×10^8
y
x
raw data, starting:14
fitted Gauss locations:14

**Optimal1 Fitted Gauss, proteinAT1G74970.1**
**bic:661**
**peaks after deleting less than one fifth of global max:5.3**

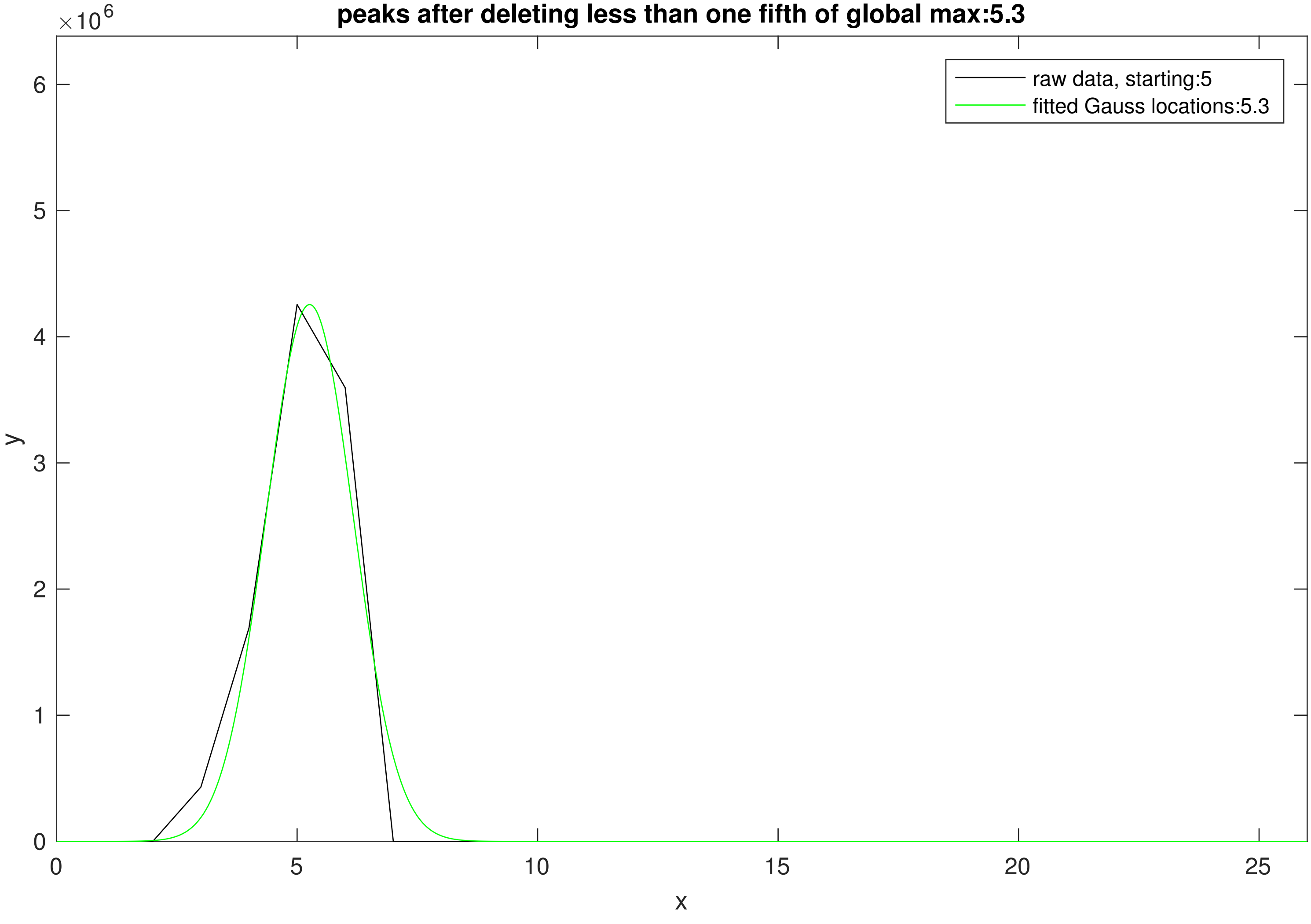

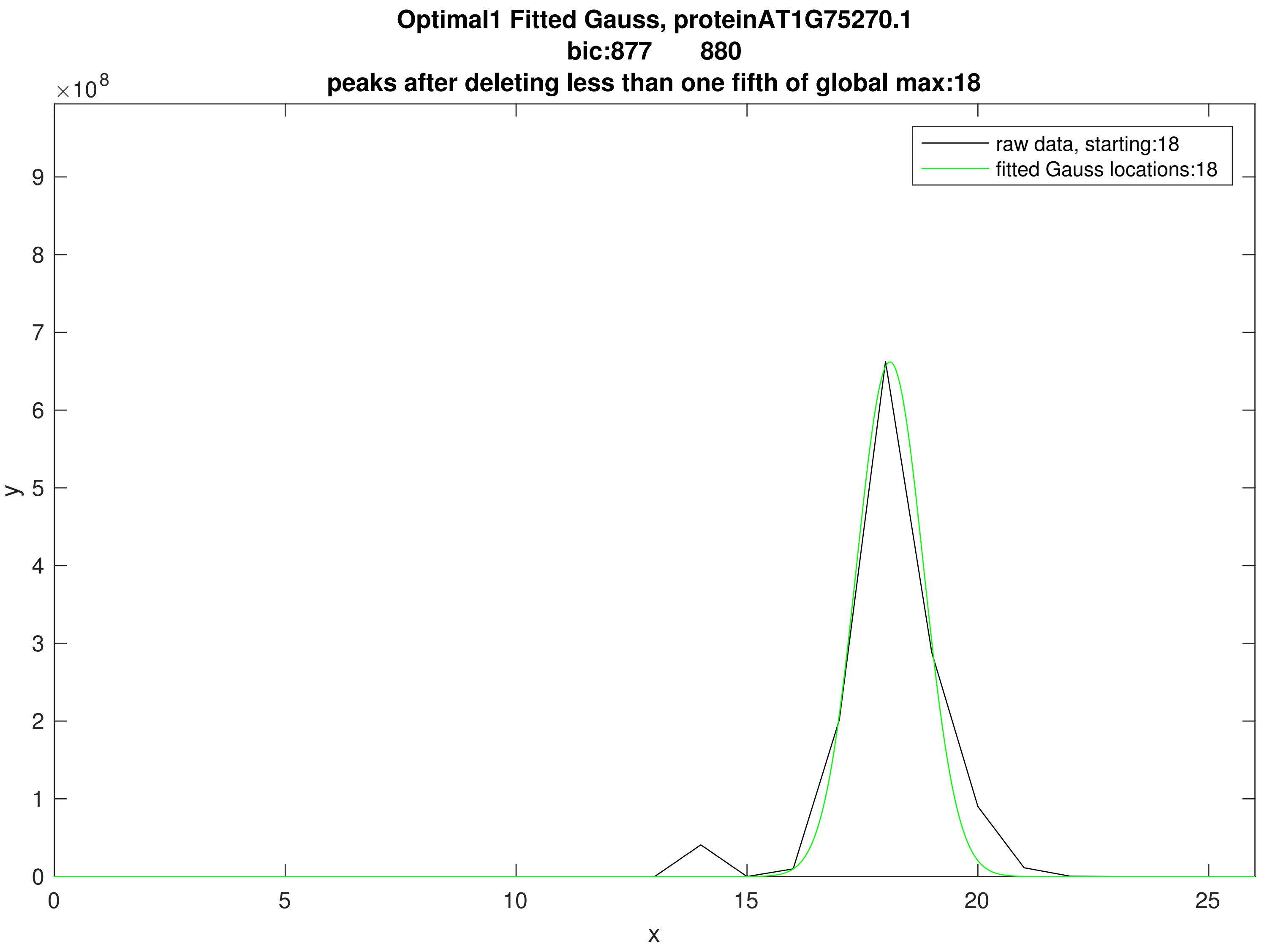

Optimal1 Fitted Gauss, proteinAT1G75270.1
bic:877 880
peaks after deleting less than one fifth of global max:18
×10^8
raw data, starting:18
fitted Gauss locations:18
y
x
0
1
2
3
4
5
6
7
8
9
0
5
10
15
20
25

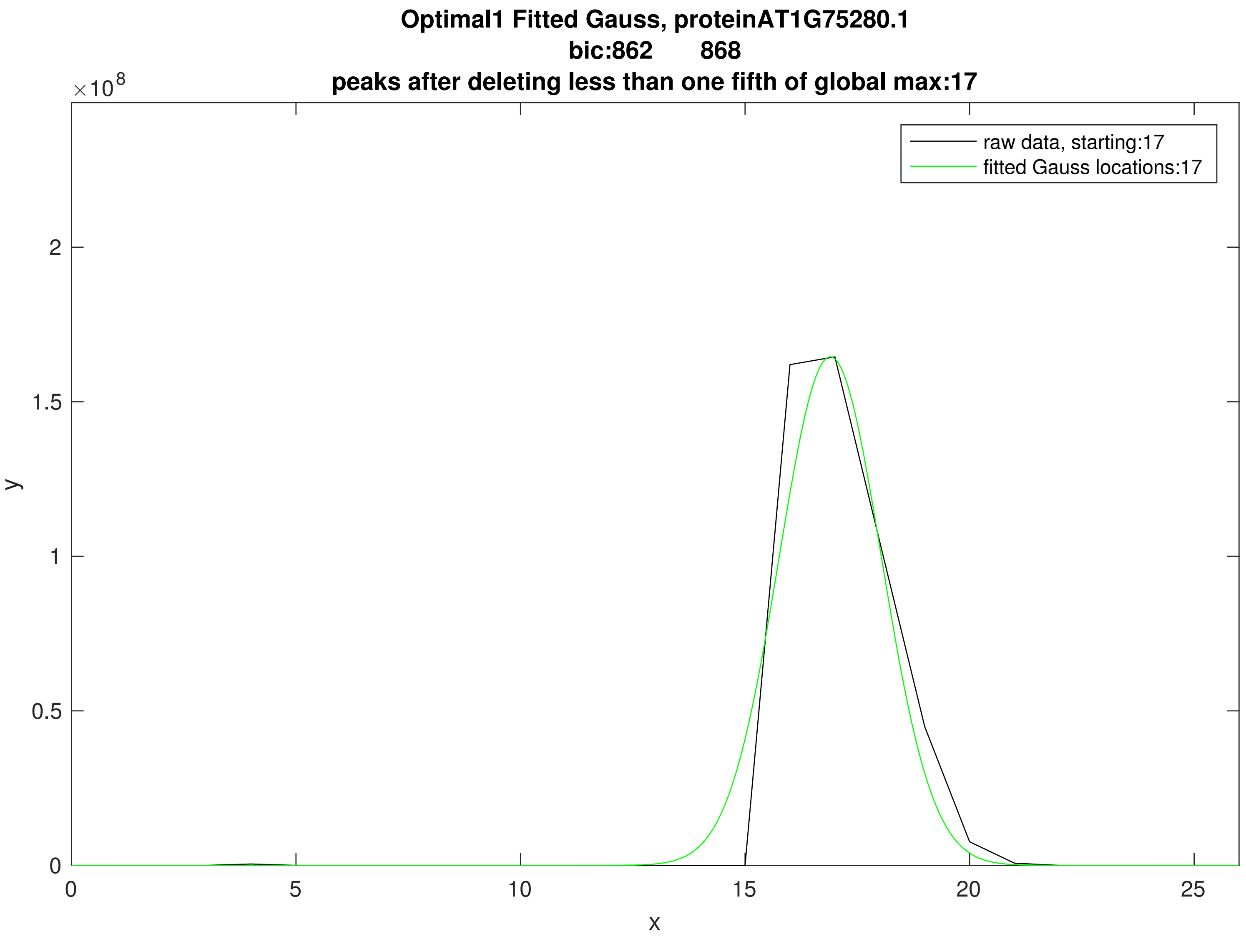

Optimal1 Fitted Gauss, proteinAT1G75280.1
bic:862 868
peaks after deleting less than one fifth of global max:17
×10^8
raw data, starting:17
fitted Gauss locations:17
2
1.5
1
0.5
0
y
0
5
10
15
20
25
x

**Optimal1 Fitted Gauss, proteinAT1G75330.1**
**bic:855**
**peaks after deleting less than one fifth of global max:12**

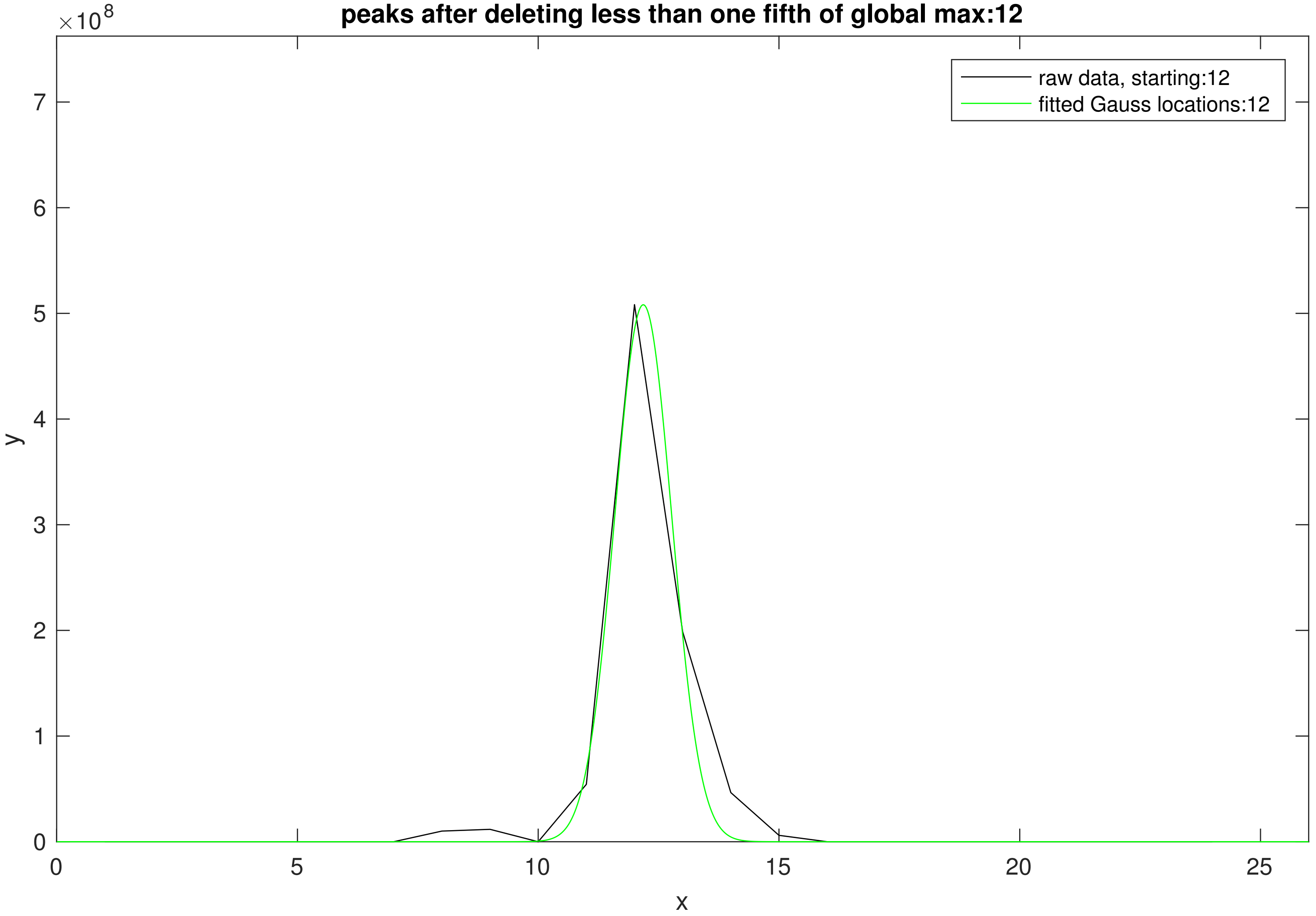

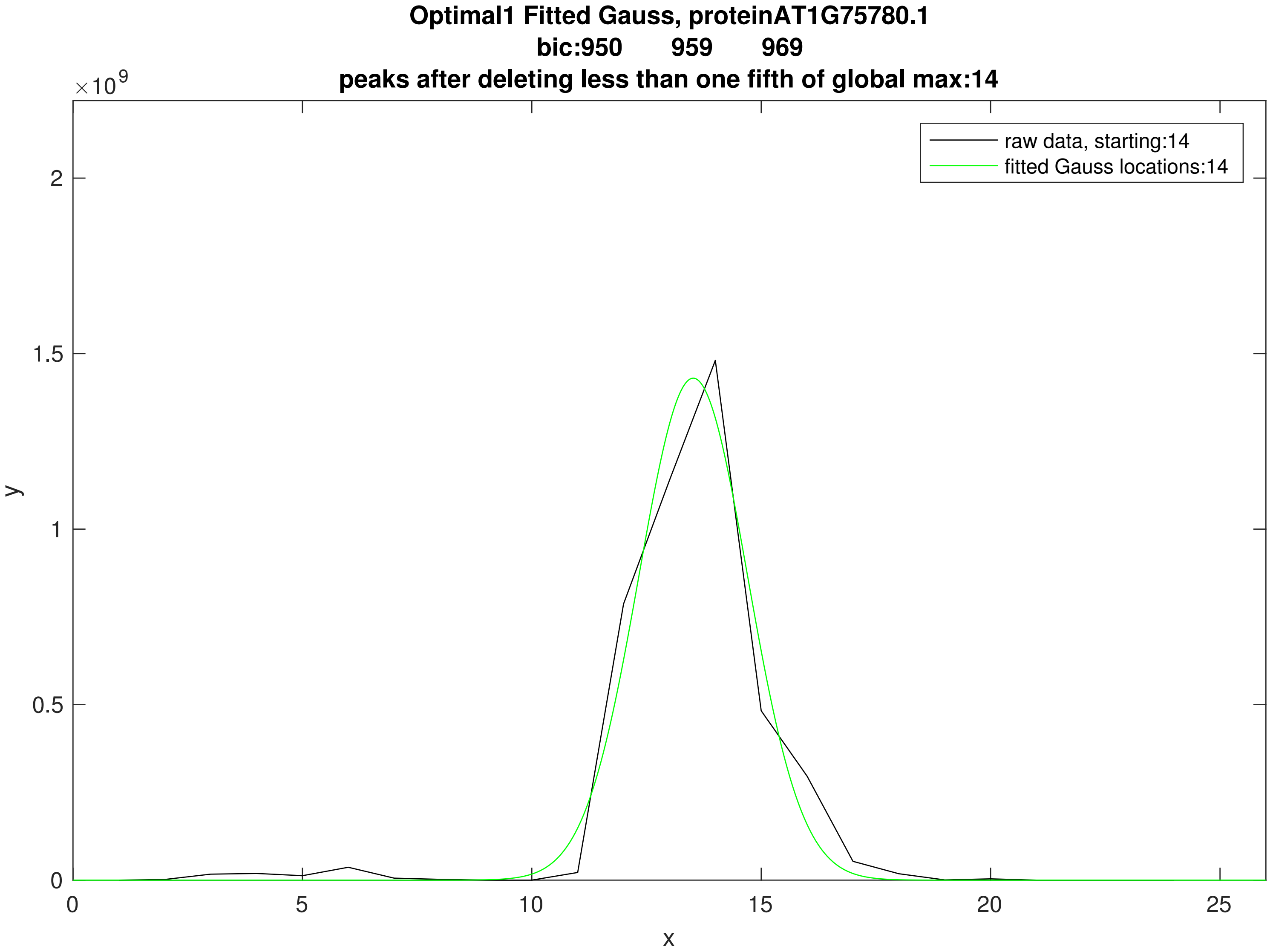
Optimal1 Fitted Gauss, proteinAT1G75780.1
bic:950 959 969
peaks after deleting less than one fifth of global max:14
×10^9
raw data, starting:14
fitted Gauss locations:14
2
1.5
1
0.5
0
y
0
5
10
15
20
25
x

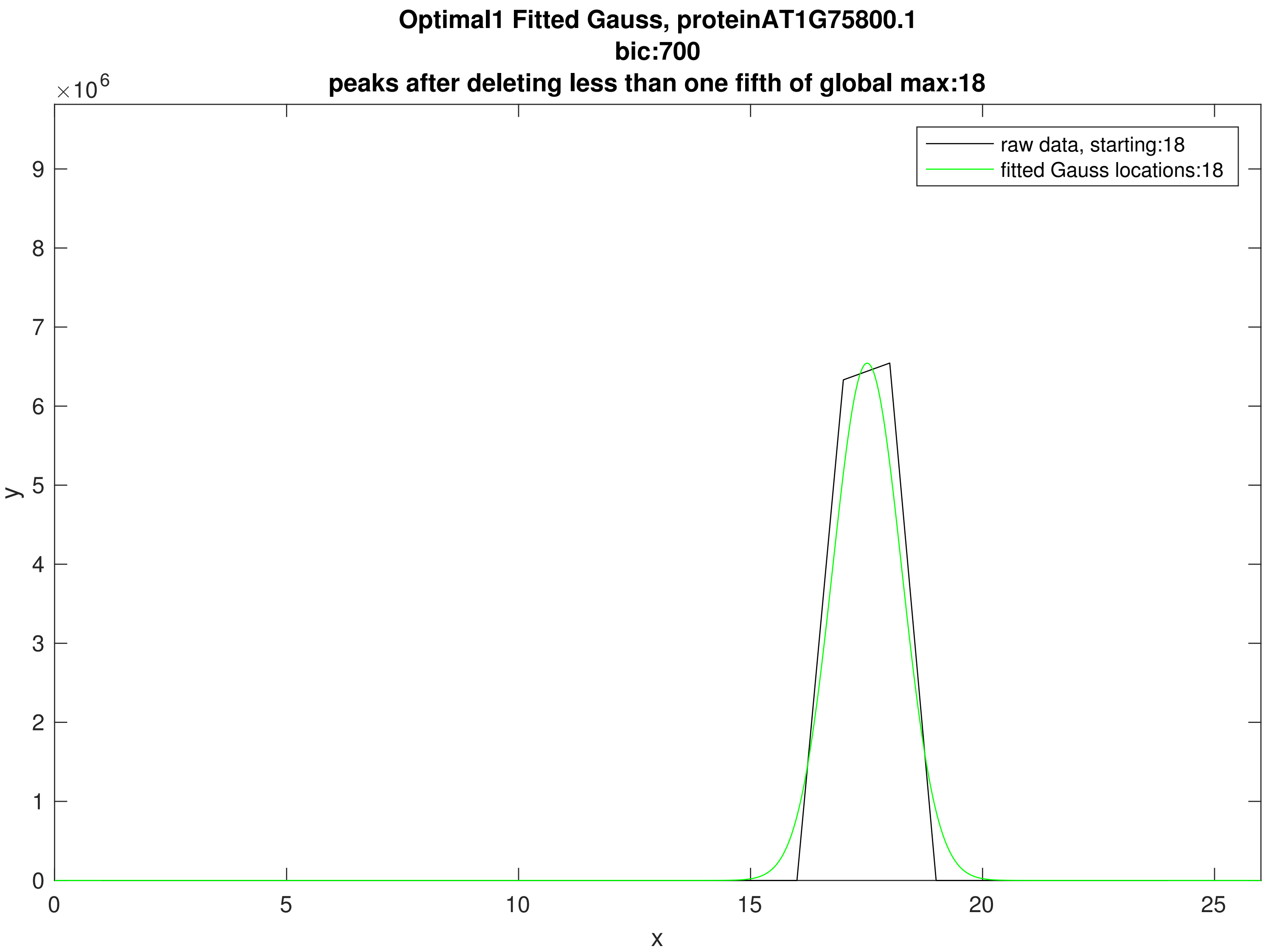

Optimal1 Fitted Gauss, proteinAT1G75800.1
bic:700
peaks after deleting less than one fifth of global max:18
×10^6
raw data, starting:18
fitted Gauss locations:18
y
x
0
1
2
3
4
5
6
7
8
9
0
5
10
15
20
25

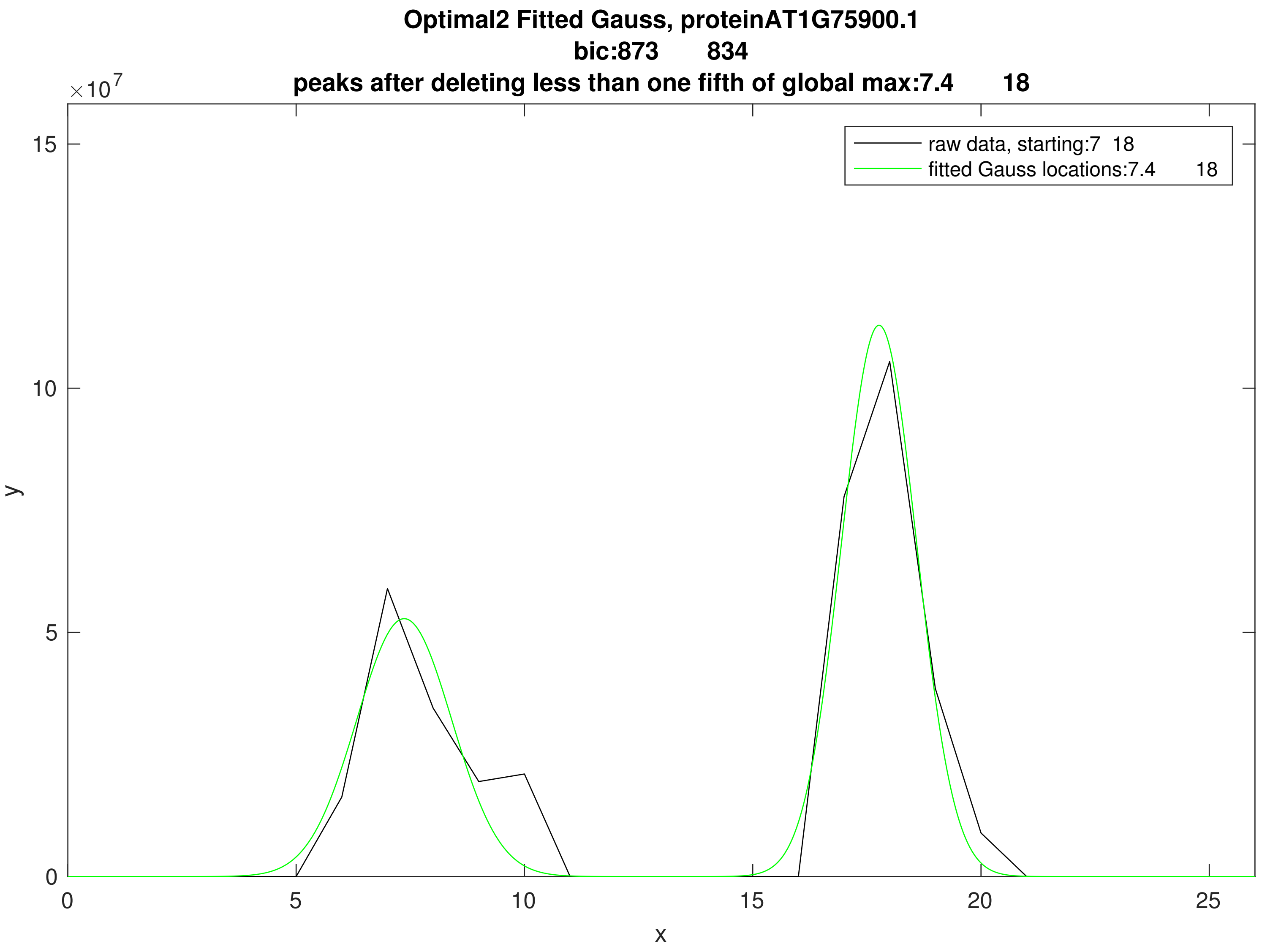
Optimal2 Fitted Gauss, proteinAT1G75900.1
bic:873 834
peaks after deleting less than one fifth of global max:7.4 18
×10^7
15
10
5
0
y
0
5
10
15
20
25
x
raw data, starting:7 18
fitted Gauss locations:7.4 18

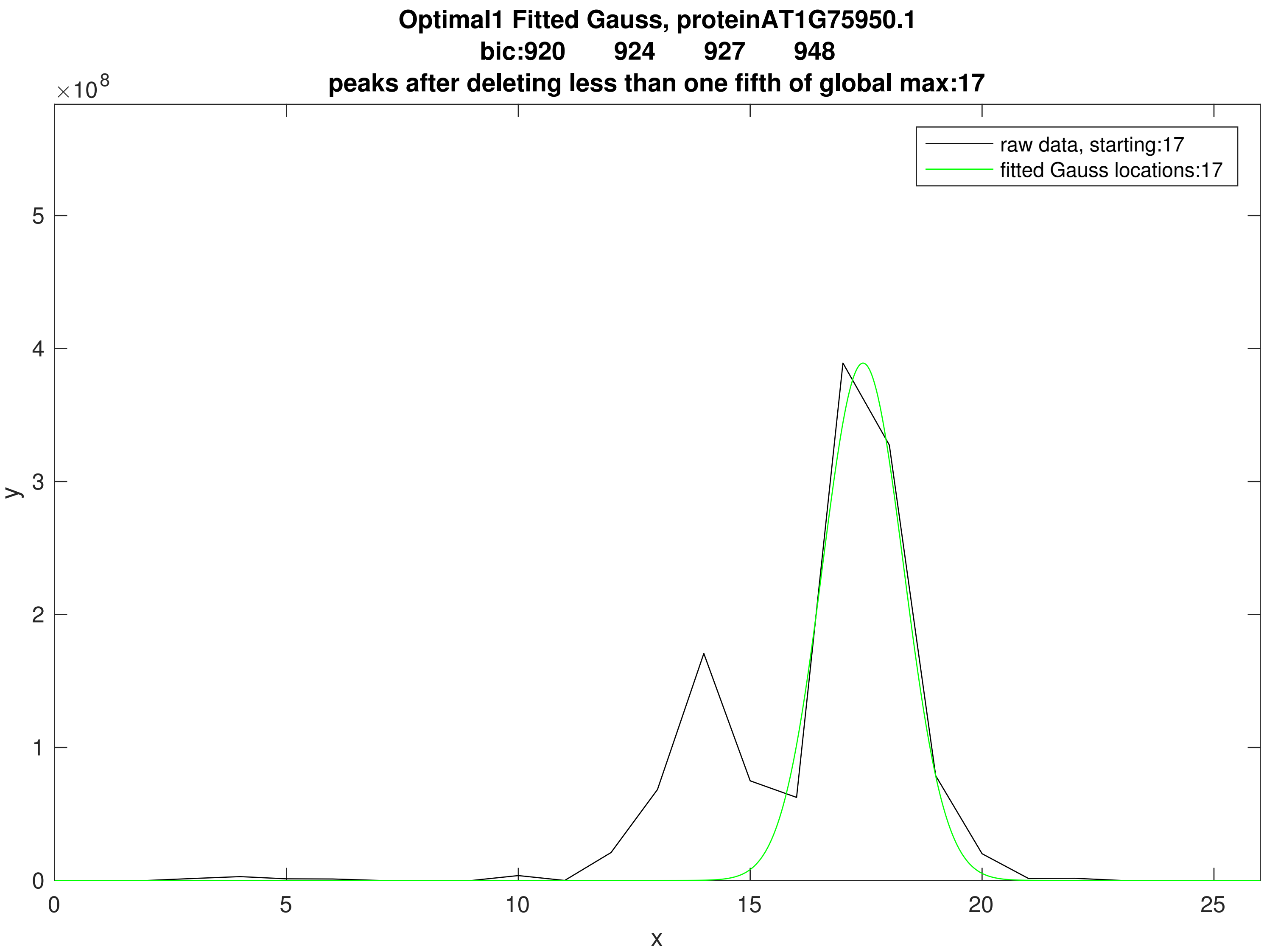
Optimal1 Fitted Gauss, proteinAT1G75950.1
bic:920 924 927 948
peaks after deleting less than one fifth of global max:17
×10^8
raw data, starting:17
fitted Gauss locations:17
y
x
0
1
2
3
4
5
0
5
10
15
20
25

#### Optimal1 Fitted Gauss, proteinAT1G75990.1
#### bic:726
#### peaks after deleting less than one fifth of global max:3.8

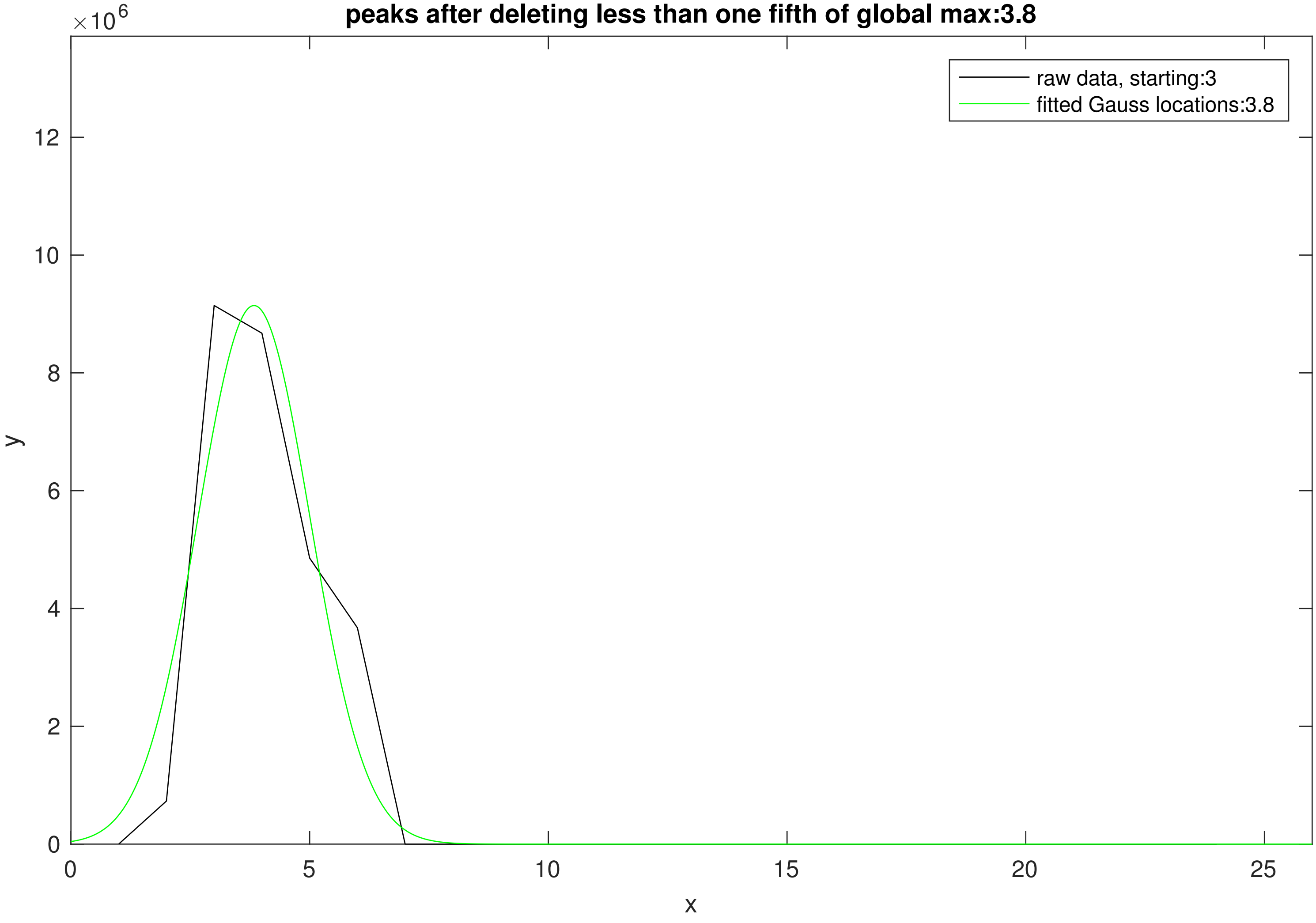

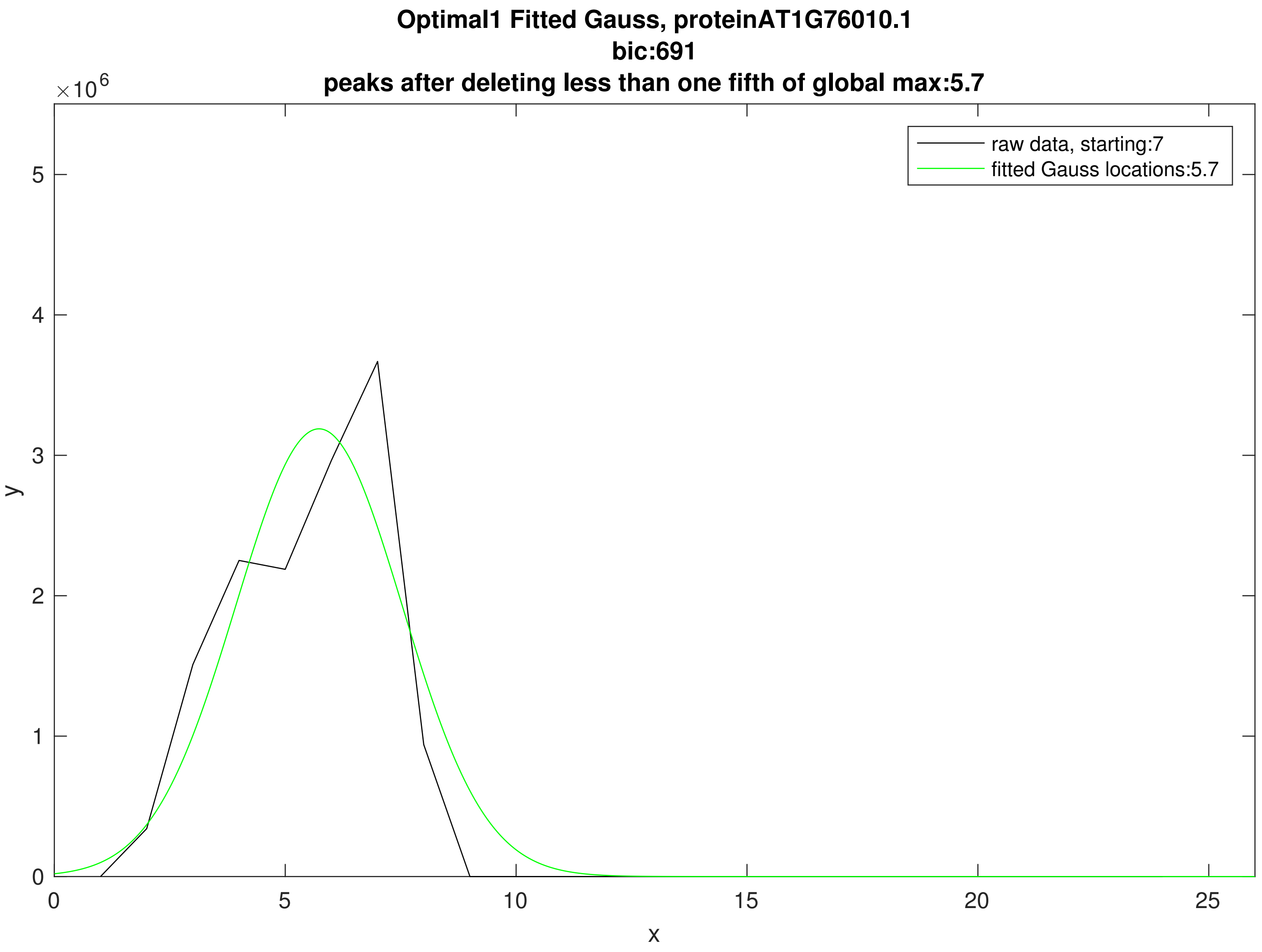

Optimal1 Fitted Gauss, proteinAT1G76010.1
bic:691
peaks after deleting less than one fifth of global max:5.7
×10^6
raw data, starting:7
fitted Gauss locations:5.7
y
x
0
1
2
3
4
5
0
5
10
15
20
25

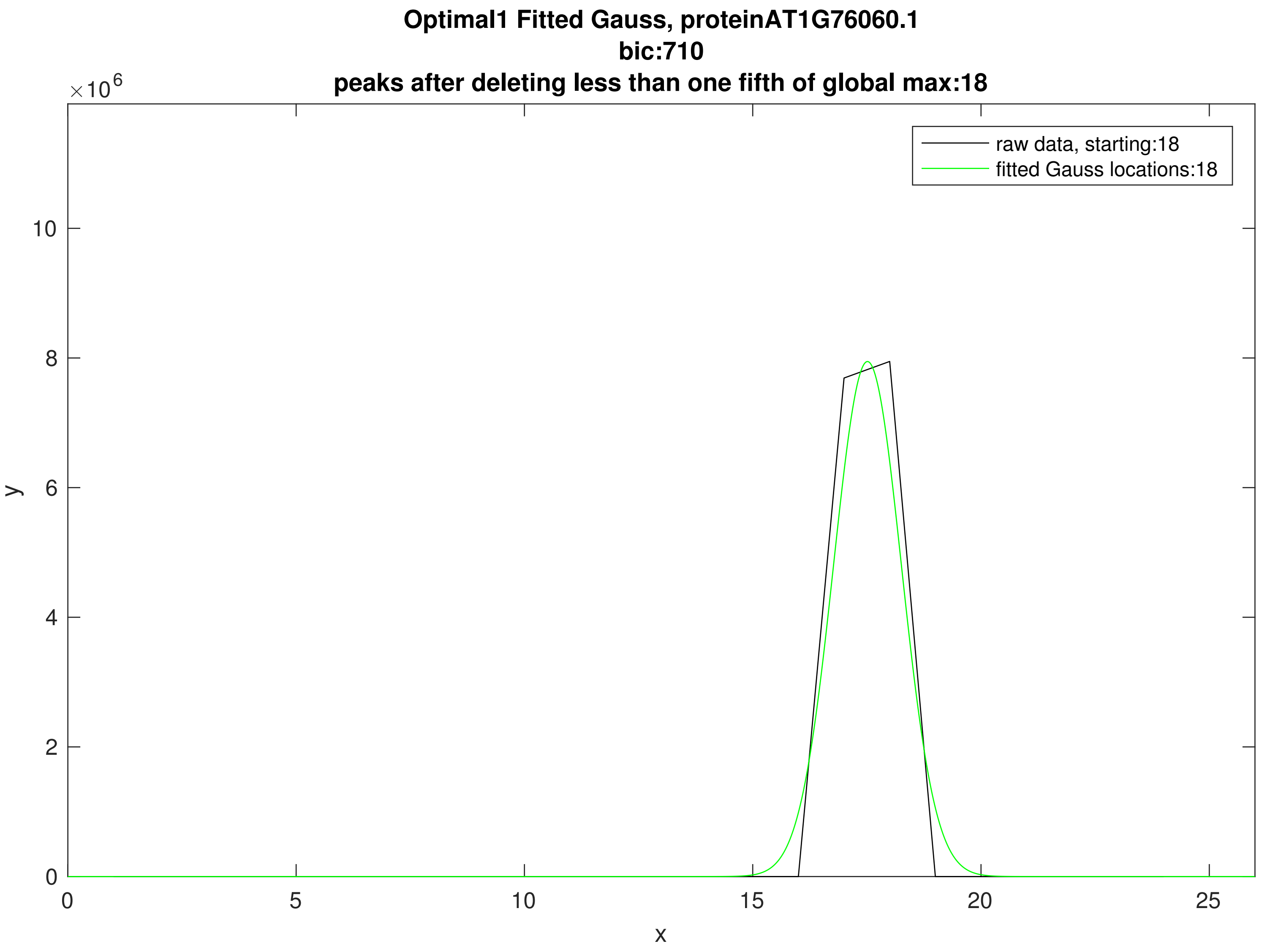
Optimal1 Fitted Gauss, proteinAT1G76060.1
bic:710
peaks after deleting less than one fifth of global max:18
×10^6
raw data, starting:18
fitted Gauss locations:18
y
x
0
2
4
6
8
10
0
5
10
15
20
25

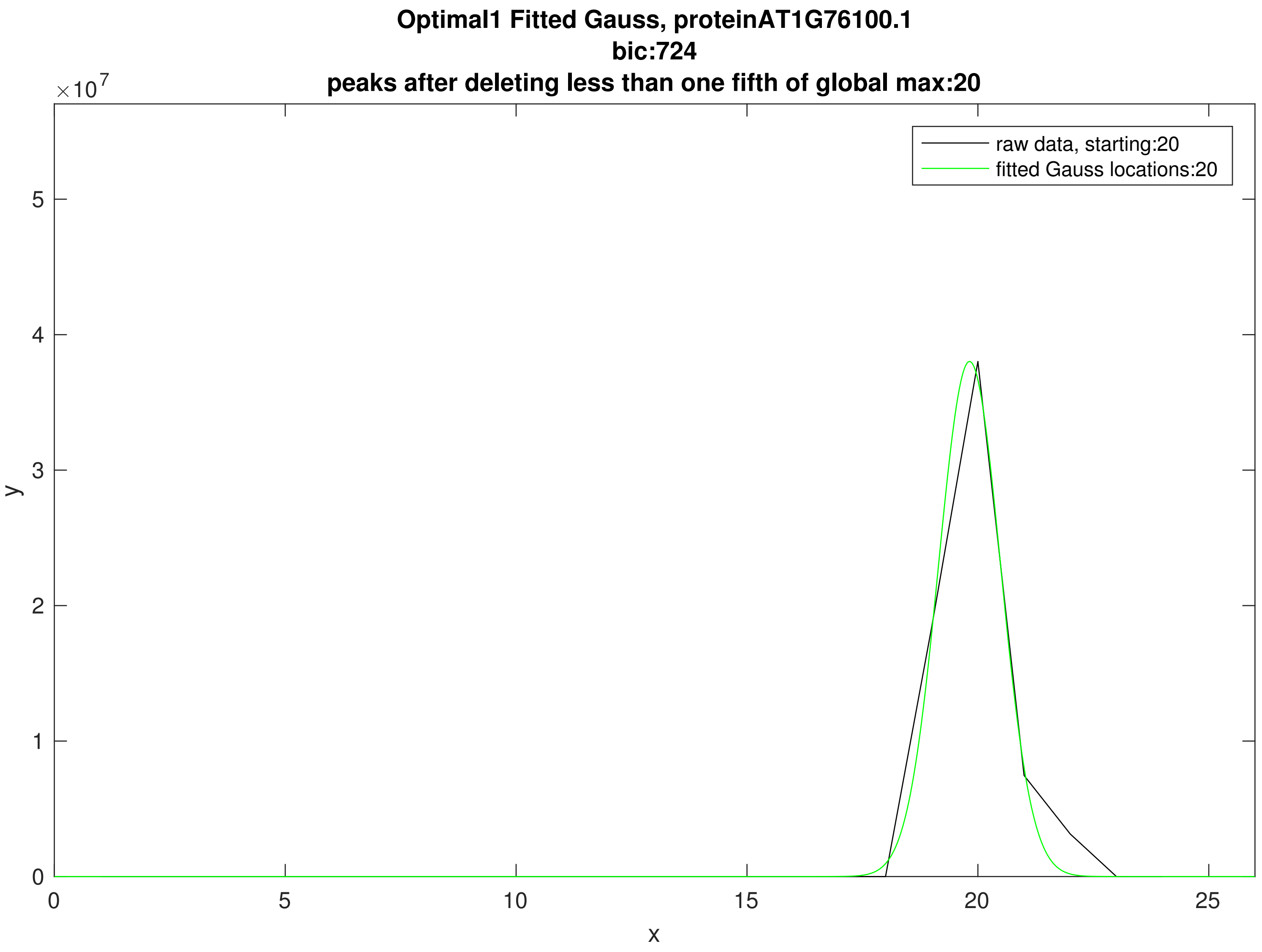

Optimal1 Fitted Gauss, proteinAT1G76100.1
bic:724
peaks after deleting less than one fifth of global max:20
×10^7
raw data, starting:20
fitted Gauss locations:20
y
x
0
1
2
3
4
5
0
5
10
15
20
25

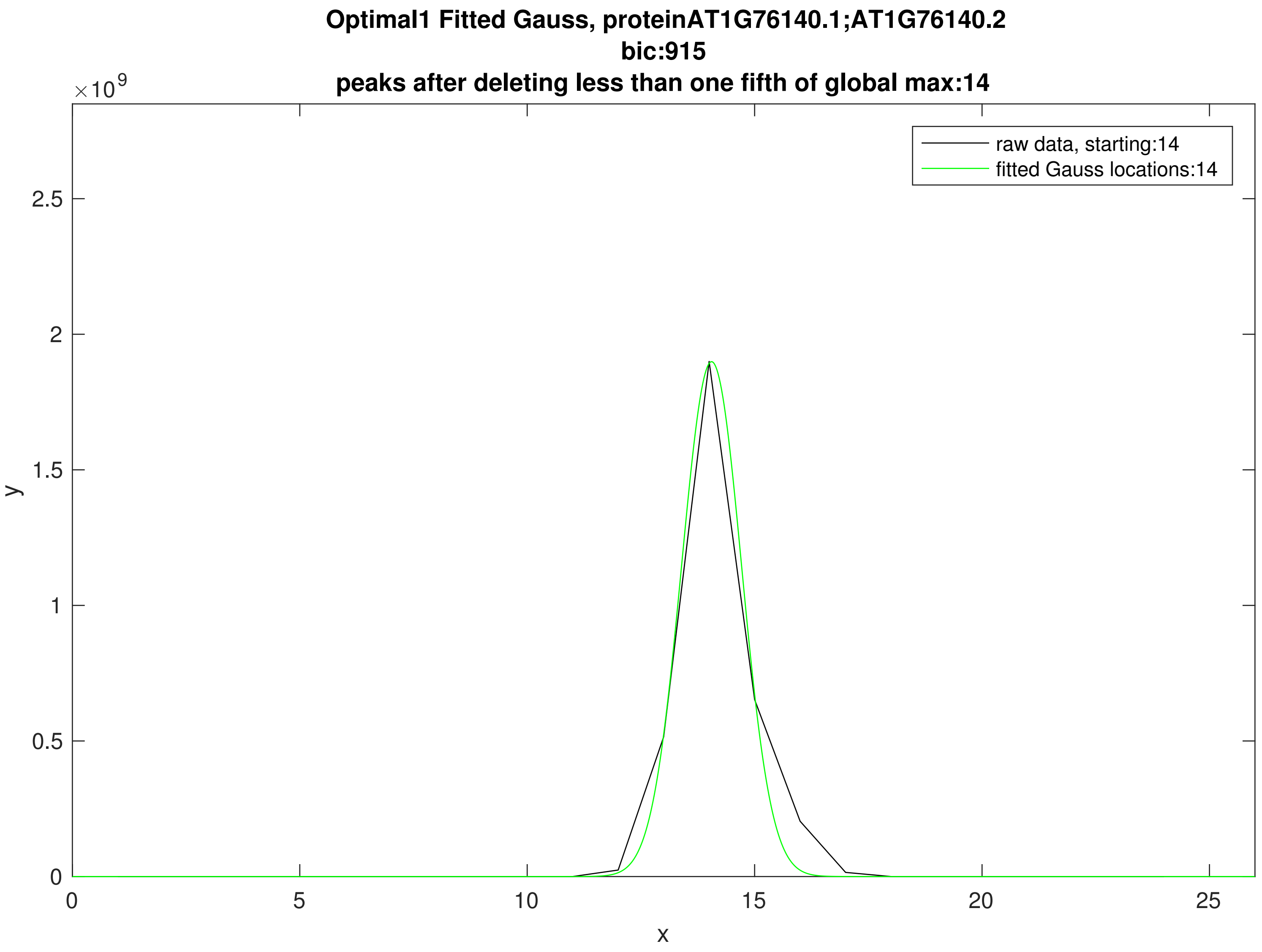
Optimal1 Fitted Gauss, proteinAT1G76140.1;AT1G76140.2
bic:915
peaks after deleting less than one fifth of global max:14
×10^9
raw data, starting:14
fitted Gauss locations:14
y
x
0
0.5
1
1.5
2
2.5
0
5
10
15
20
25

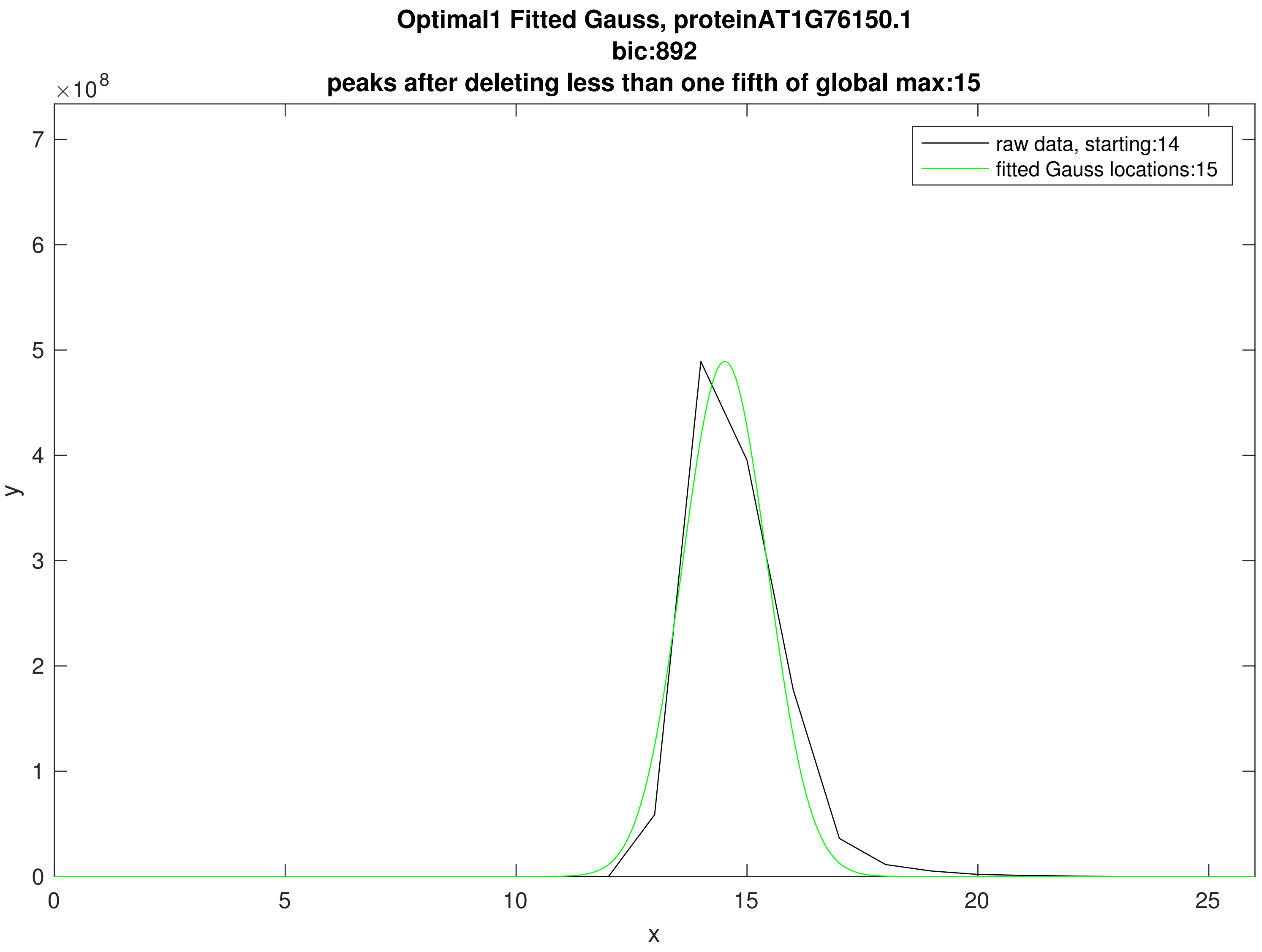
Optimal1 Fitted Gauss, proteinAT1G76150.1
bic:892
peaks after deleting less than one fifth of global max:15
×10^8
raw data, starting:14
fitted Gauss locations:15
y
x
0
1
2
3
4
5
6
7
0
5
10
15
20
25

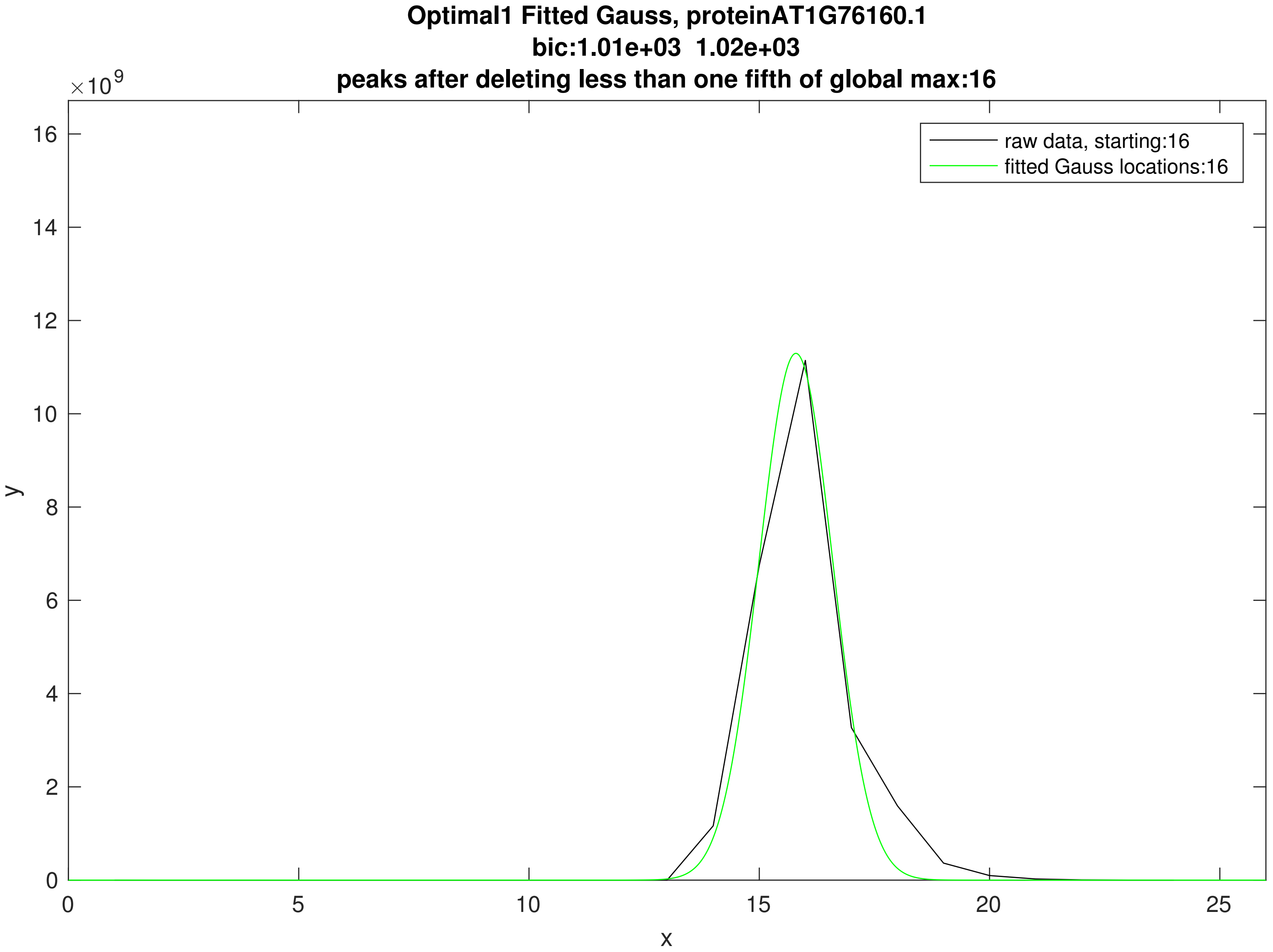

Optimal1 Fitted Gauss, proteinAT1G76160.1
bic:1.01e+03  1.02e+03
peaks after deleting less than one fifth of global max:16
×10^9
16
14
12
10
8
6
4
2
0
y
0
5
10
15
20
25
x
raw data, starting:16
fitted Gauss locations:16

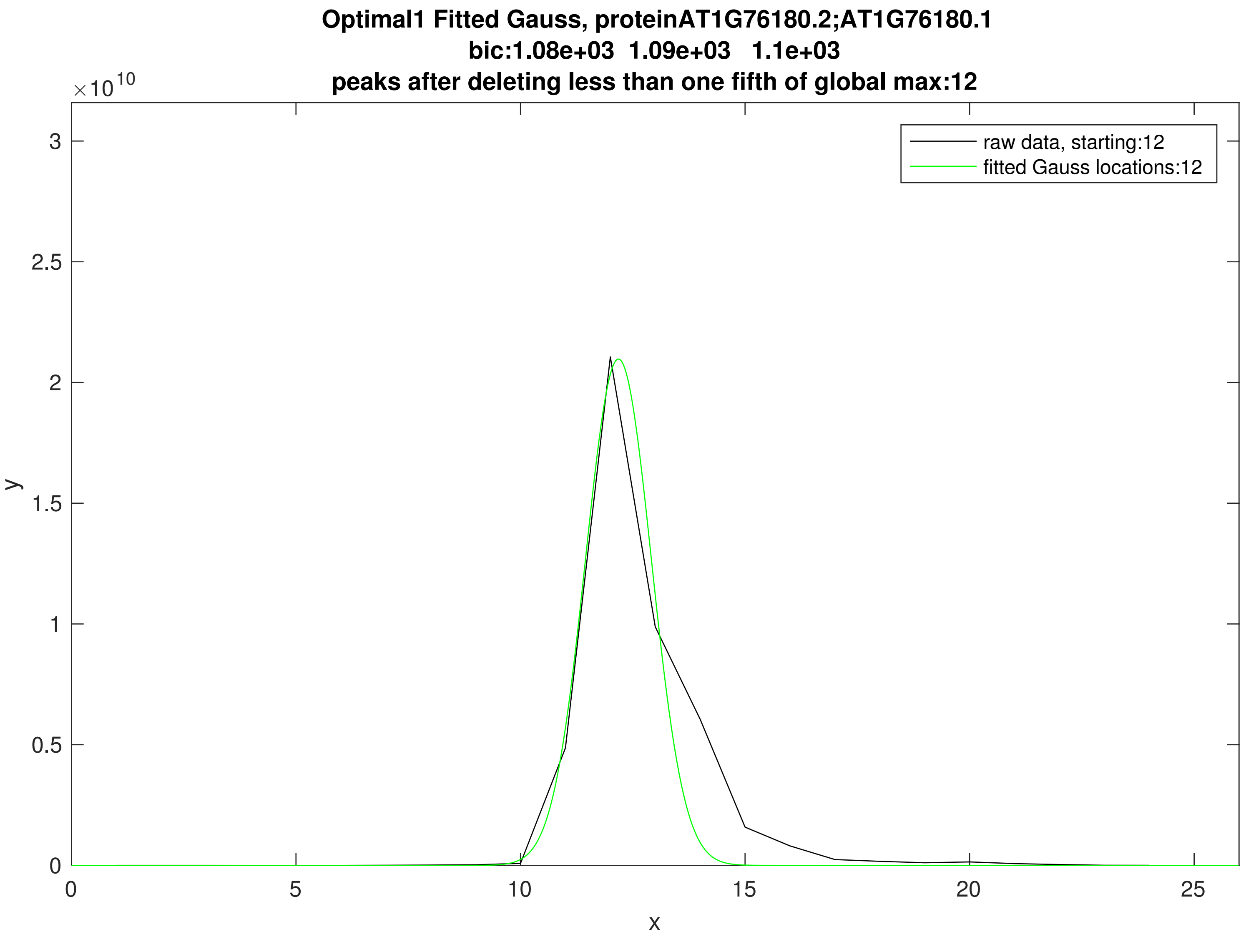
Optimal1 Fitted Gauss, proteinAT1G76180.2;AT1G76180.1
bic:1.08e+03 1.09e+03 1.1e+03
peaks after deleting less than one fifth of global max:12
×10^10
raw data, starting:12
fitted Gauss locations:12
y
x
0
0.5
1
1.5
2
2.5
3
0
5
10
15
20
25

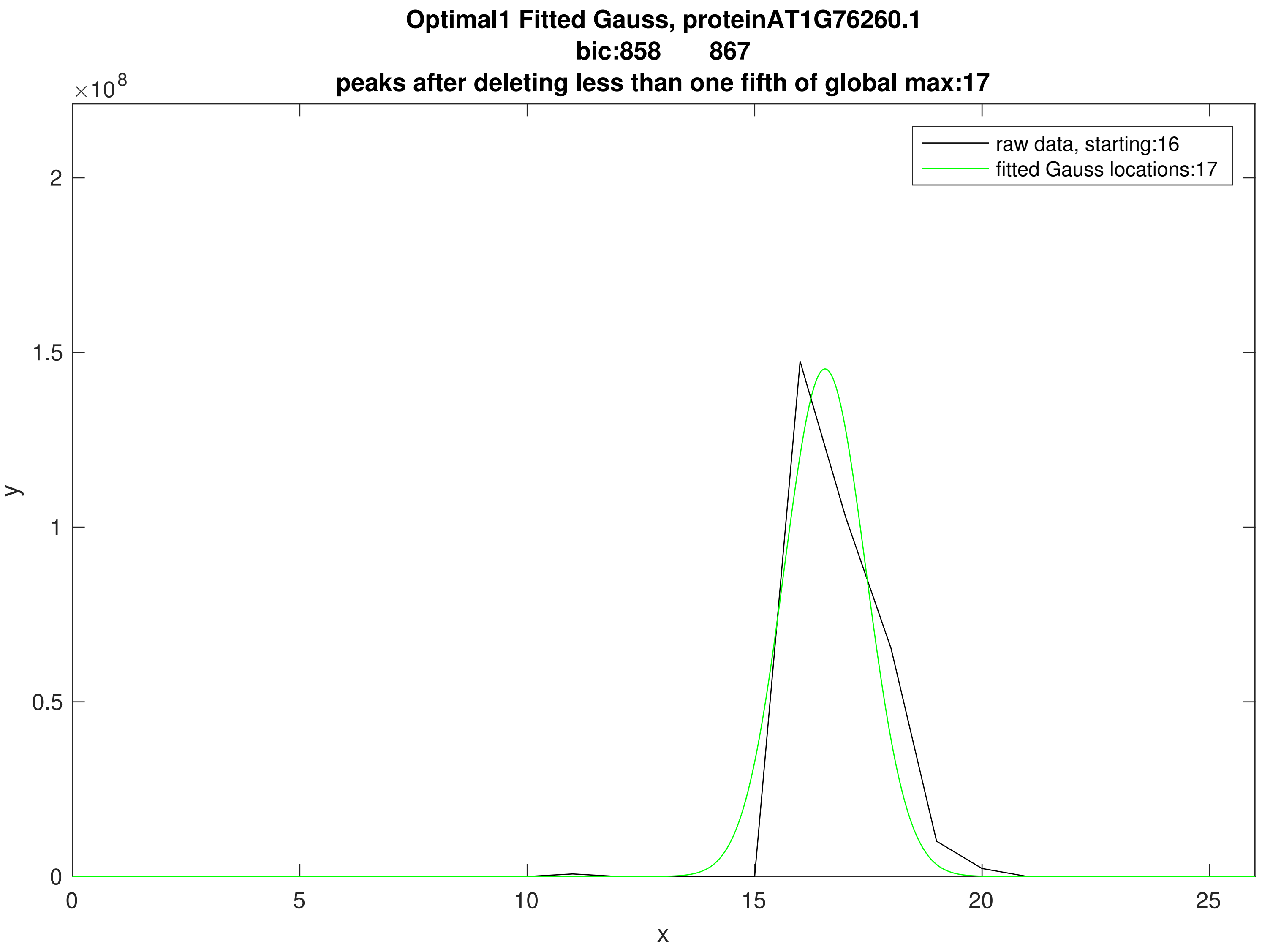
Optimal1 Fitted Gauss, proteinAT1G76260.1
bic:858 867
peaks after deleting less than one fifth of global max:17
×10^8
2
1.5
1
0.5
0
y
0
5
10
15
20
25
x
raw data, starting:16
fitted Gauss locations:17

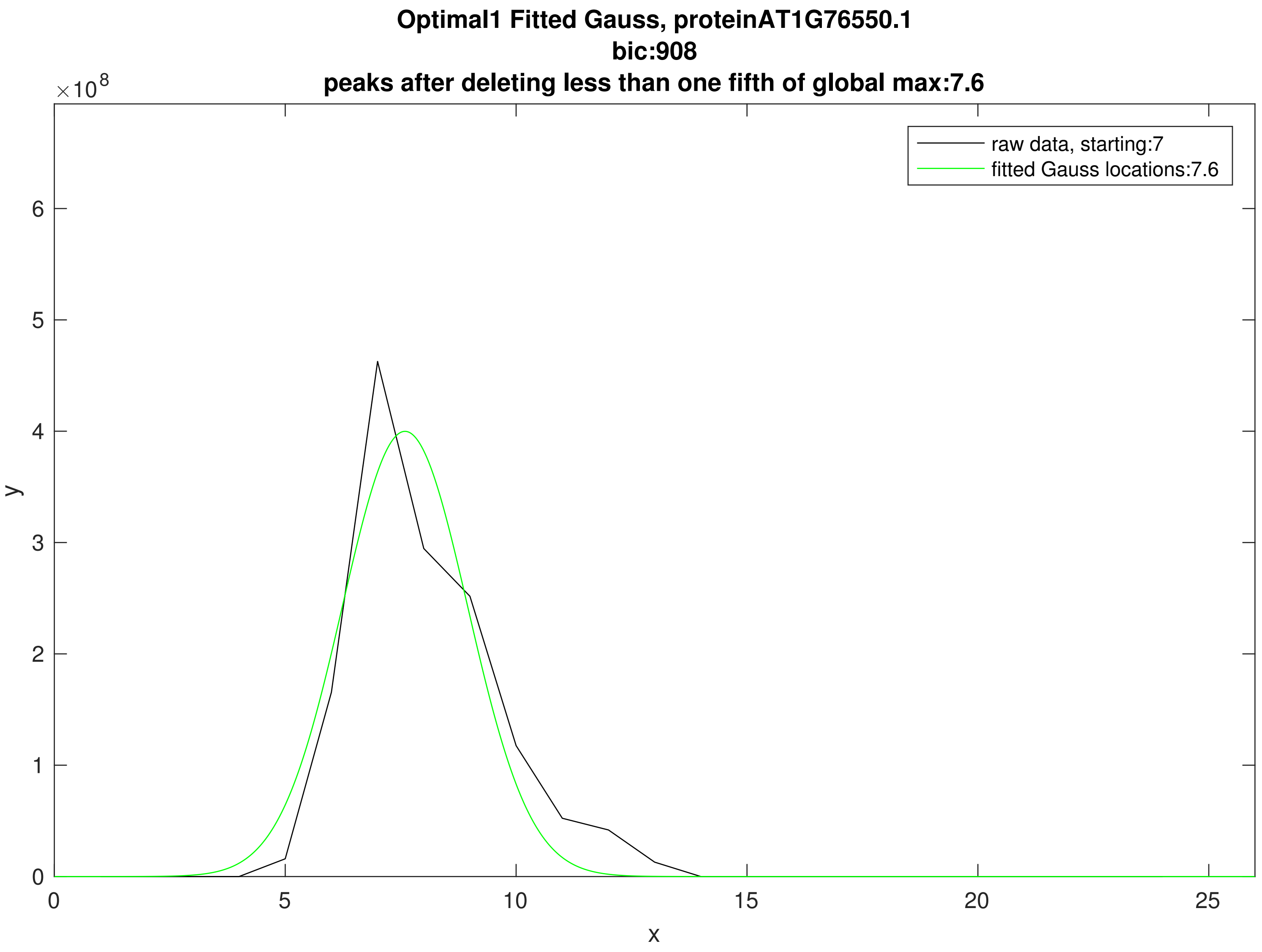

Optimal1 Fitted Gauss, proteinAT1G76550.1
bic:908
peaks after deleting less than one fifth of global max:7.6
×10^8
raw data, starting:7
fitted Gauss locations:7.6
y
x
0
1
2
3
4
5
6
0
5
10
15
20
25

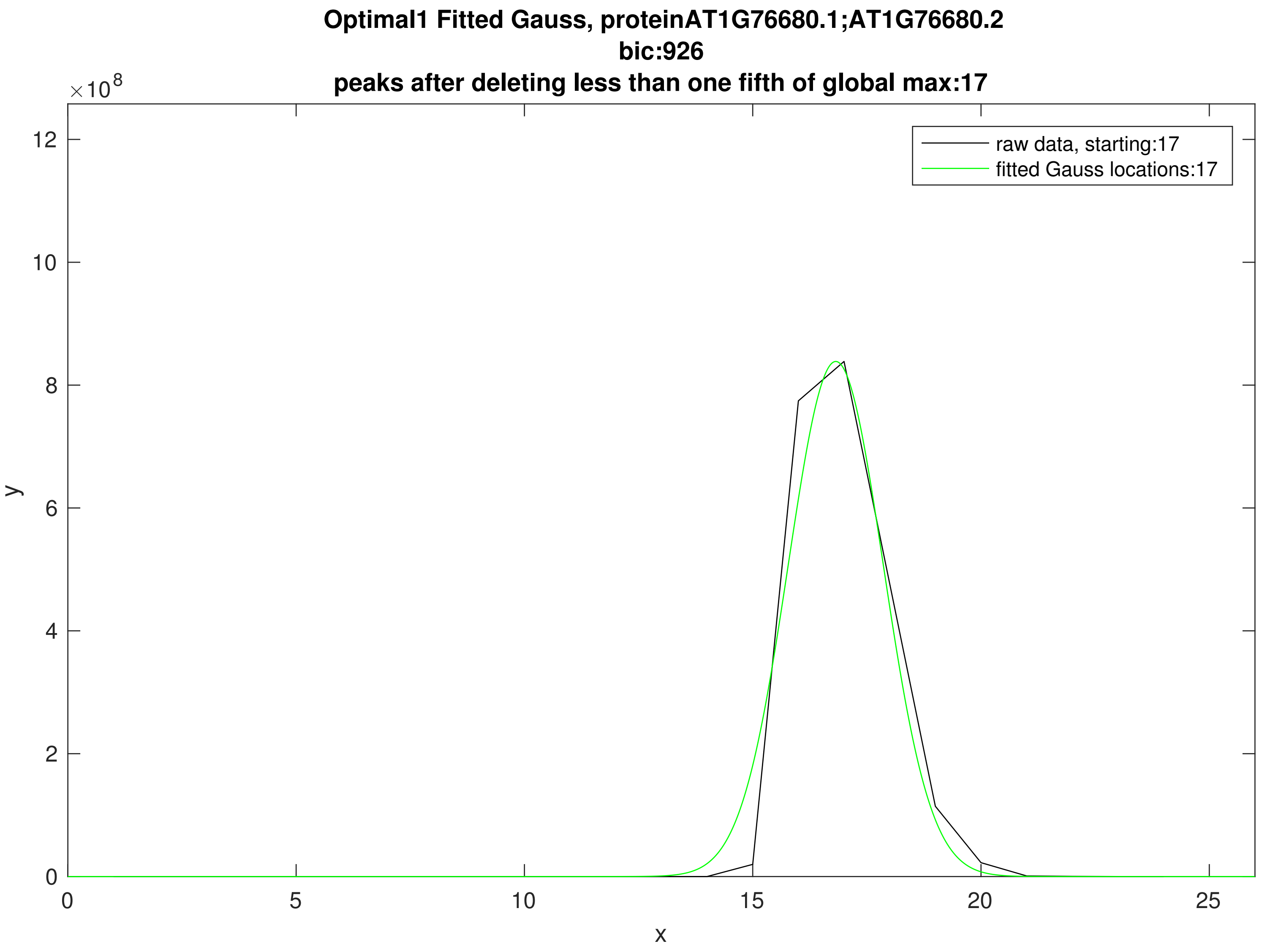

Optimal1 Fitted Gauss, proteinAT1G76680.1;AT1G76680.2
bic:926
peaks after deleting less than one fifth of global max:17
×10^8
raw data, starting:17
fitted Gauss locations:17
y
x
0
2
4
6
8
10
12
0
5
10
15
20
25

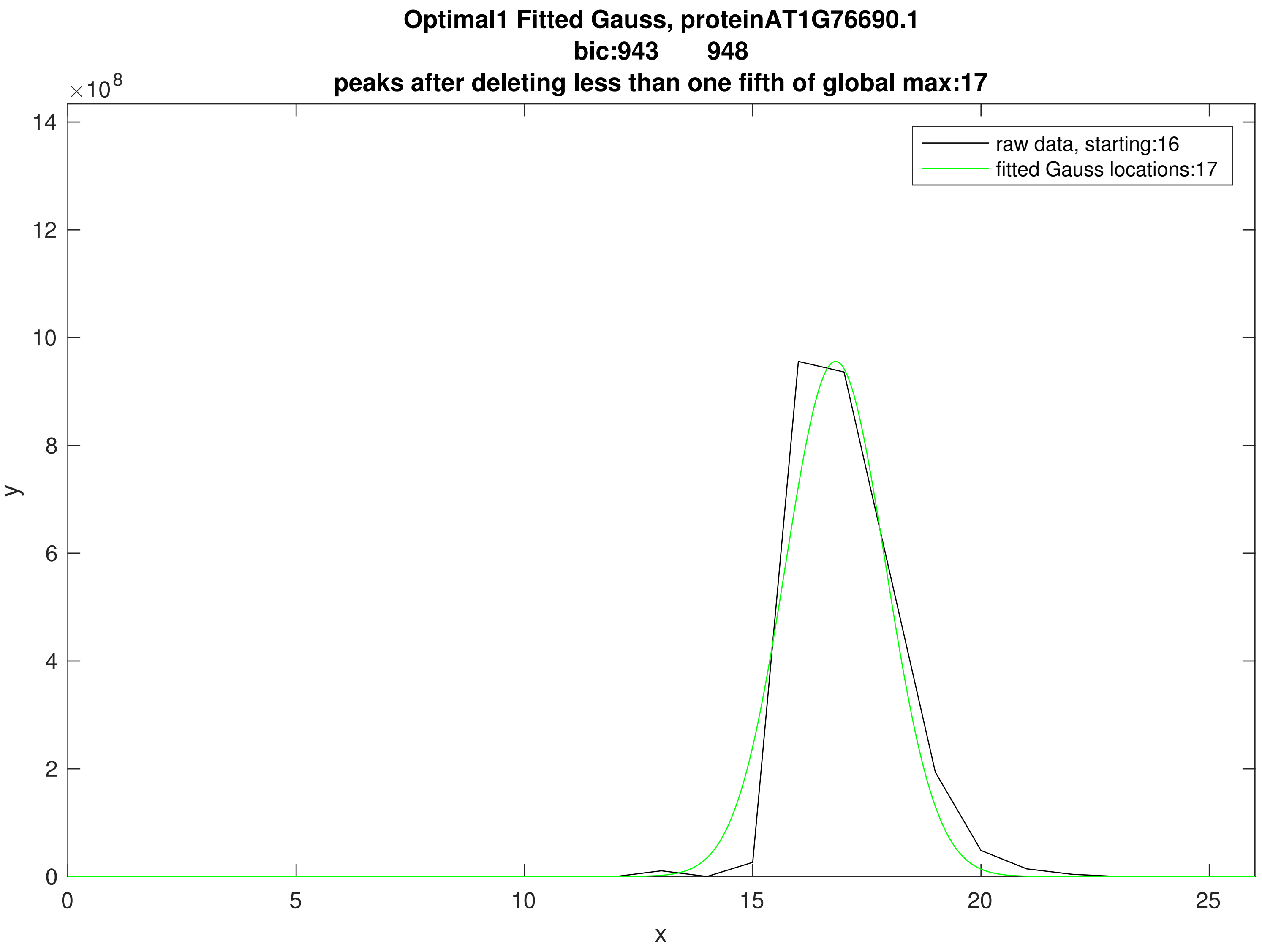
Optimal1 Fitted Gauss, proteinAT1G76690.1
bic:943 948
peaks after deleting less than one fifth of global max:17
×10^8
raw data, starting:16
fitted Gauss locations:17
0
2
4
6
8
10
12
14
y
0
5
10
15
20
25
x

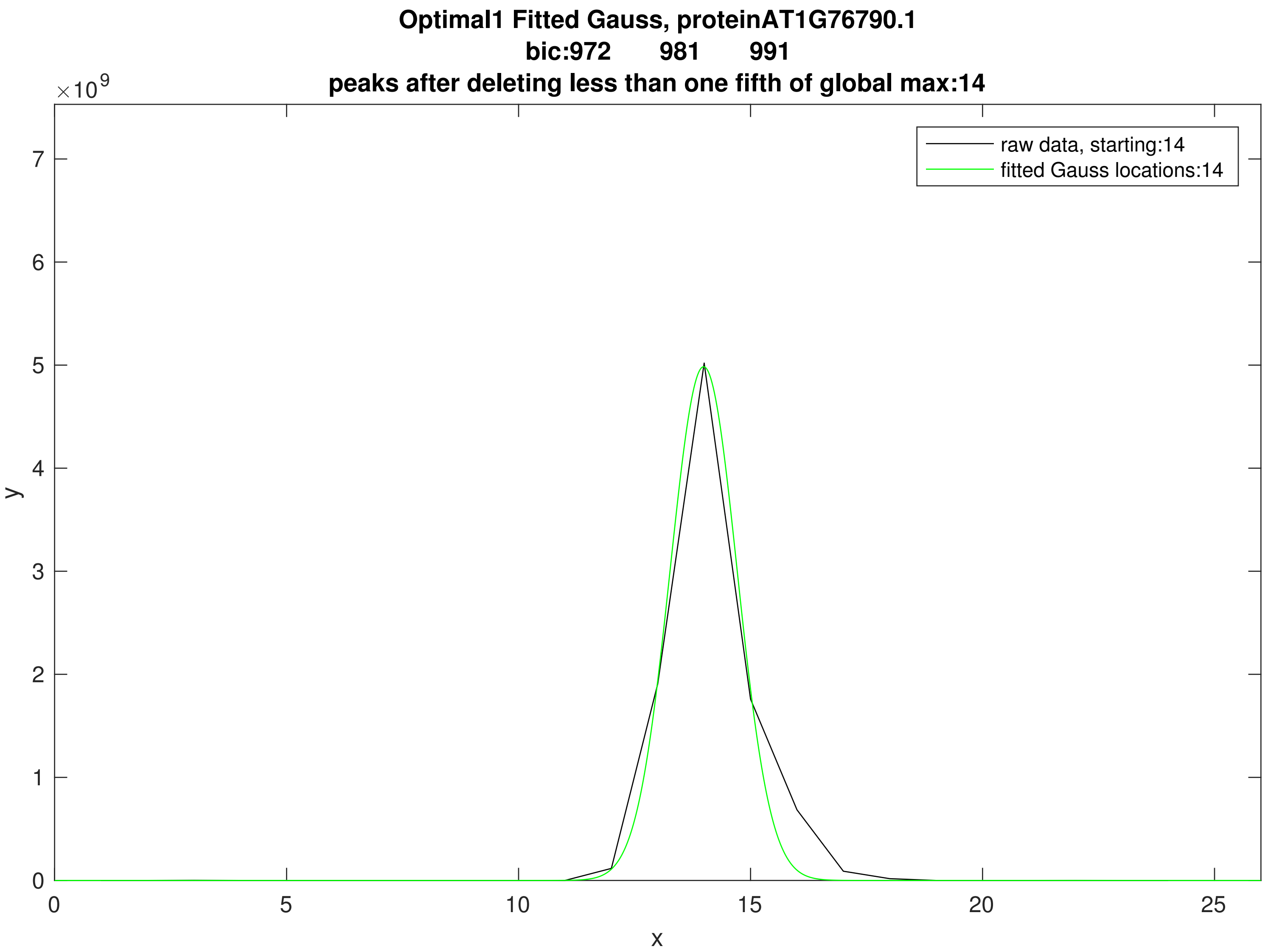
Optimal1 Fitted Gauss, proteinAT1G76790.1
bic:972 981 991
peaks after deleting less than one fifth of global max:14
×10^9
raw data, starting:14
fitted Gauss locations:14
y
x
0
1
2
3
4
5
6
7
0
5
10
15
20
25

#### Optimal1 Fitted Gauss, proteinAT1G76860.1;AT1G21190.1
#### bic:835 843
#### peaks after deleting less than one fifth of global max:14

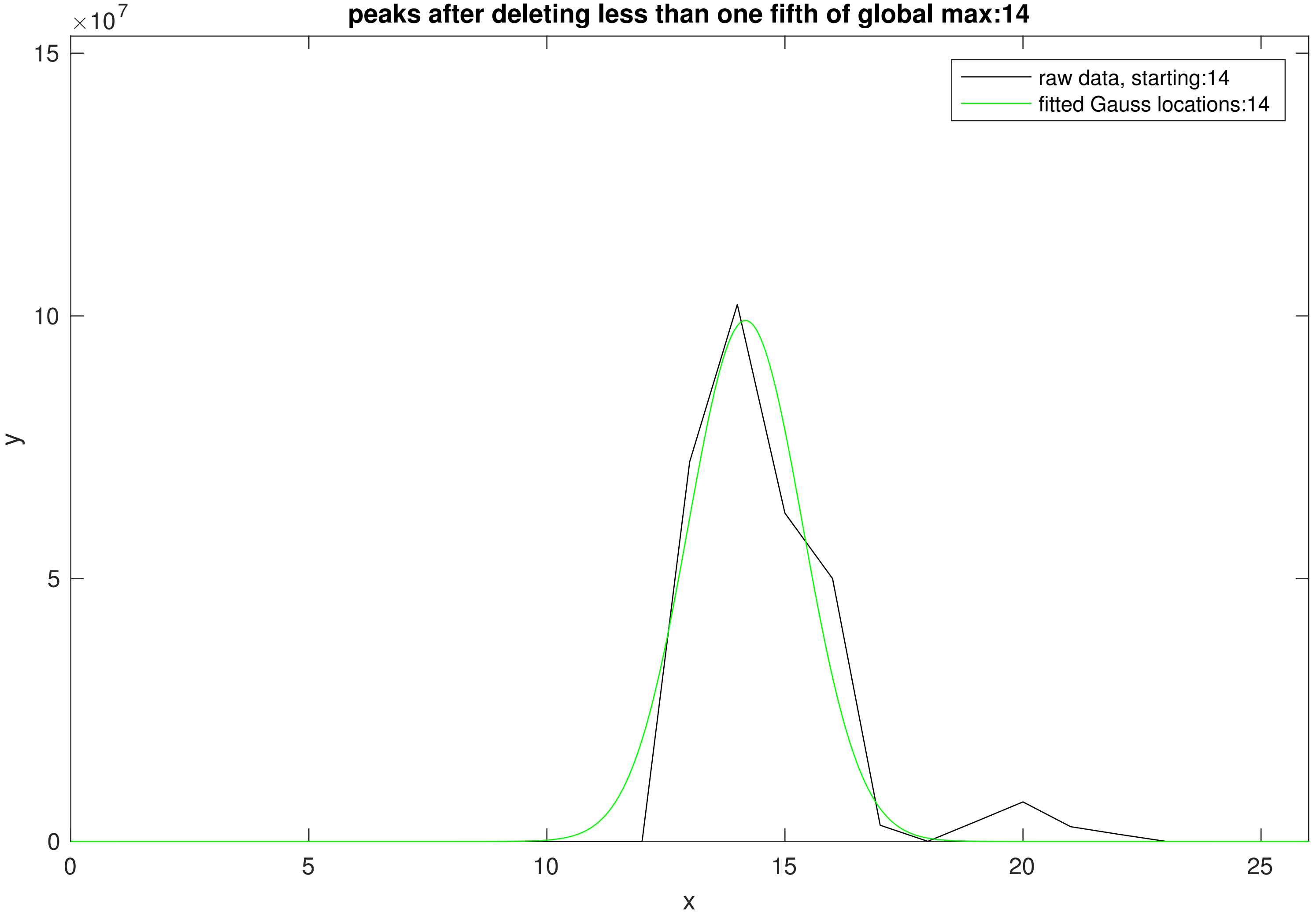

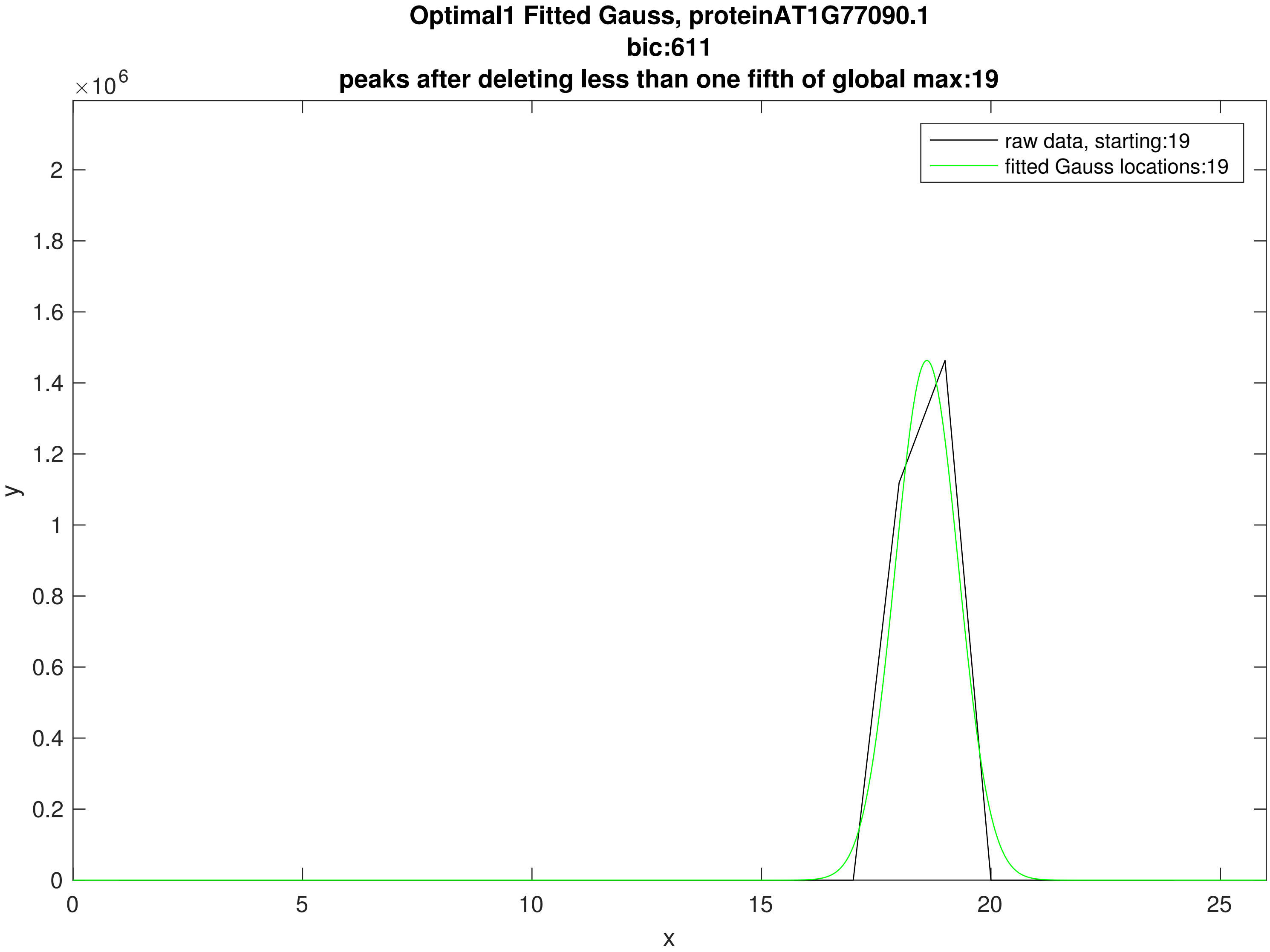
Optimal1 Fitted Gauss, proteinAT1G77090.1
bic:611
peaks after deleting less than one fifth of global max:19
×10^6
raw data, starting:19
fitted Gauss locations:19
2
1.8
1.6
1.4
1.2
1
0.8
0.6
0.4
0.2
0
y
0
5
10
15
20
25
x

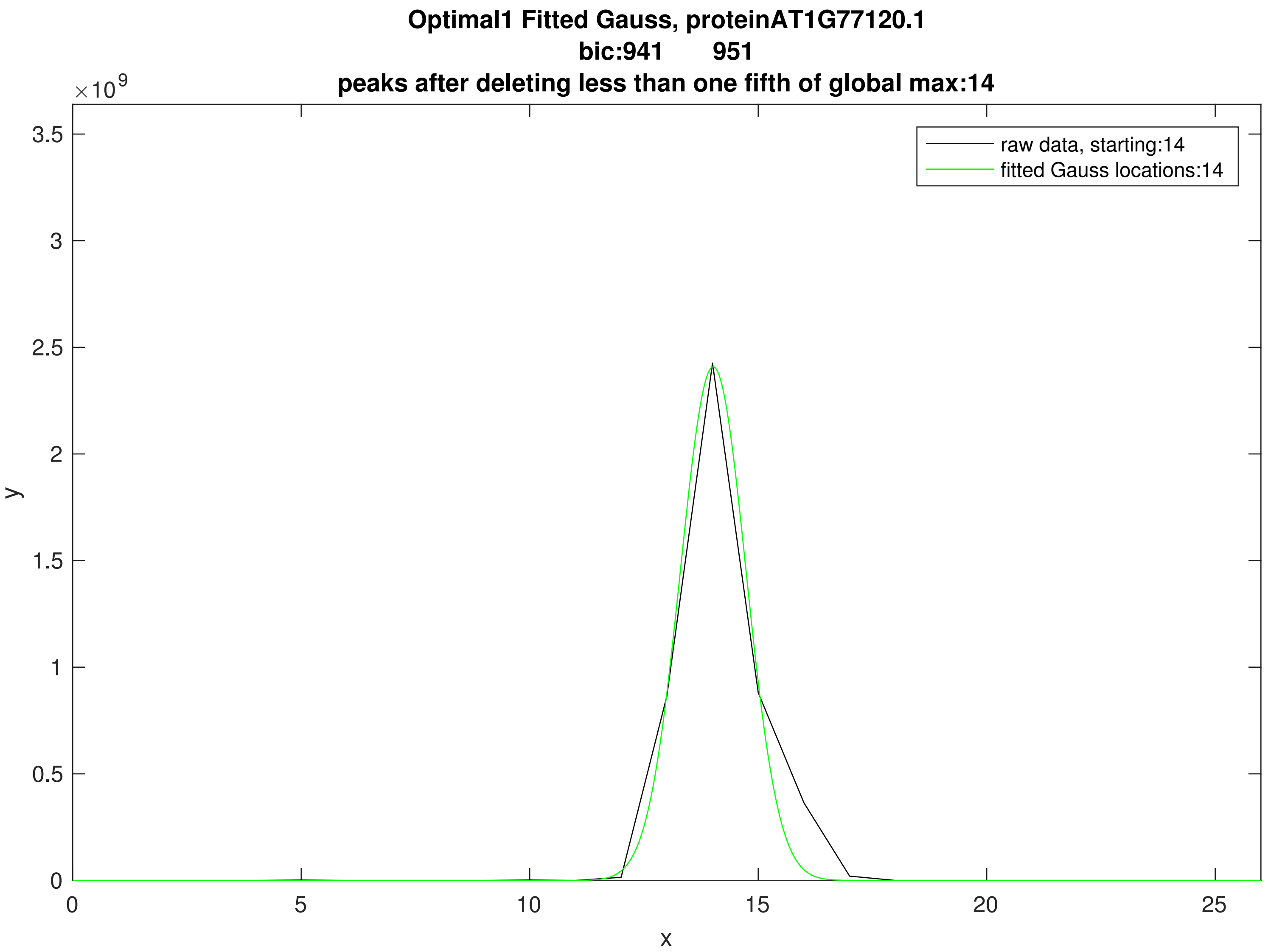

Optimal1 Fitted Gauss, proteinAT1G77120.1
bic:941 951
peaks after deleting less than one fifth of global max:14
×10^9
3.5
3
2.5
2
1.5
1
0.5
0
y
0
5
10
15
20
25
x
raw data, starting:14
fitted Gauss locations:14

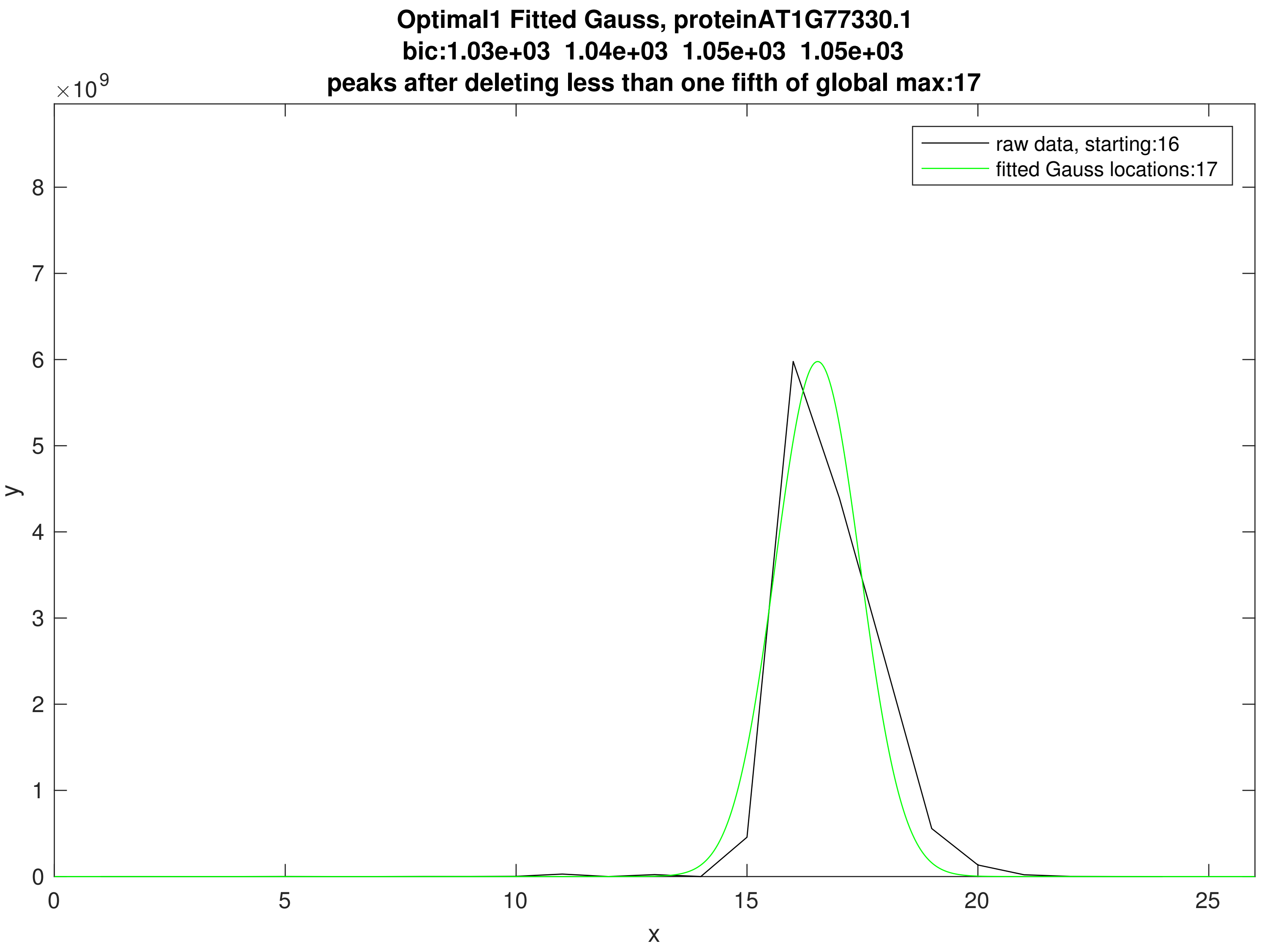

Optimal1 Fitted Gauss, proteinAT1G77330.1
bic:1.03e+03  1.04e+03  1.05e+03  1.05e+03
peaks after deleting less than one fifth of global max:17
×10^9
raw data, starting:16
fitted Gauss locations:17
y
x
0
1
2
3
4
5
6
7
8
0
5
10
15
20
25

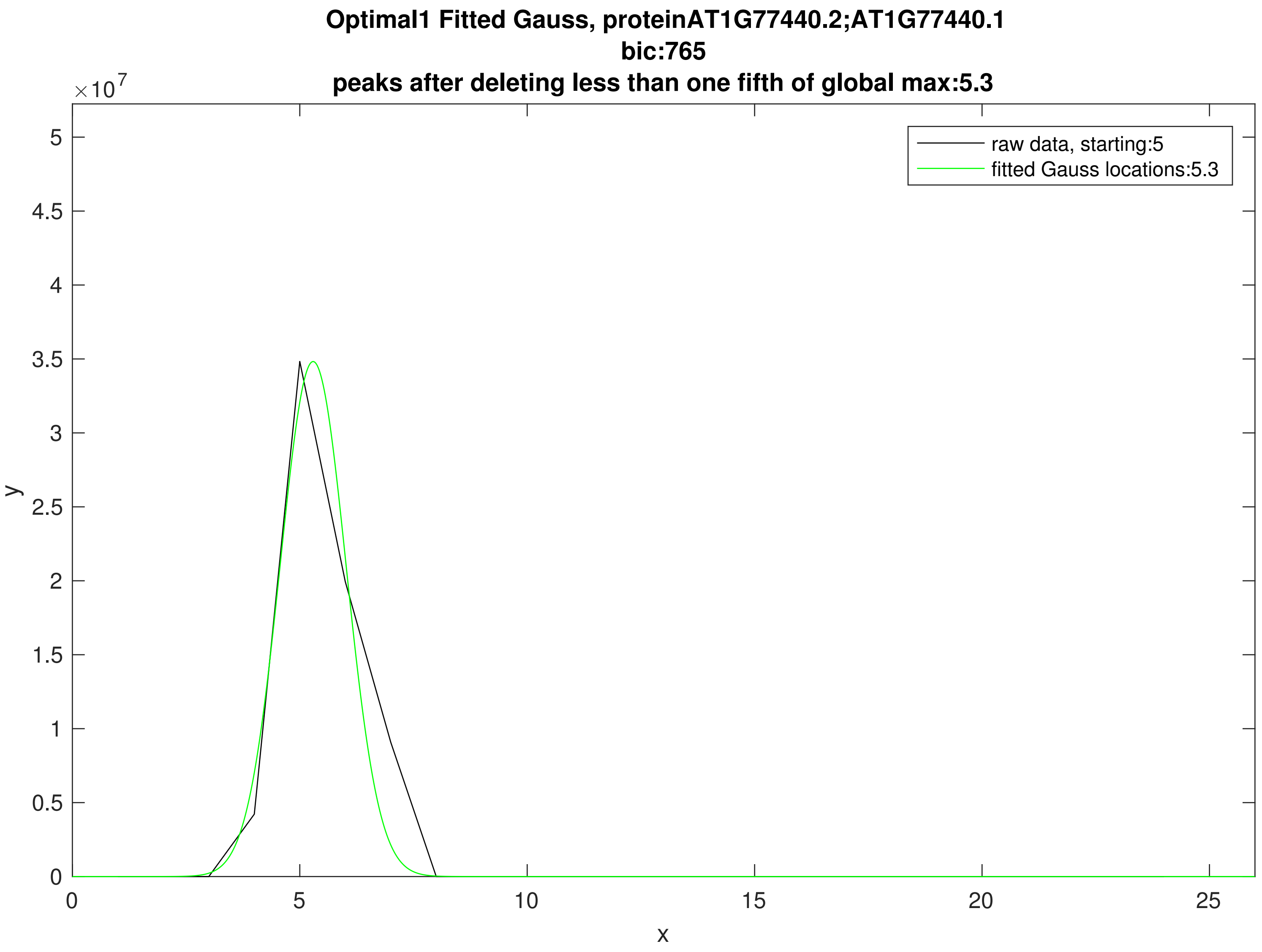

Optimal1 Fitted Gauss, proteinAT1G77440.2;AT1G77440.1
bic:765
peaks after deleting less than one fifth of global max:5.3
×10^7
raw data, starting:5
fitted Gauss locations:5.3
y
x
0
0.5
1
1.5
2
2.5
3
3.5
4
4.5
5
0
5
10
15
20
25

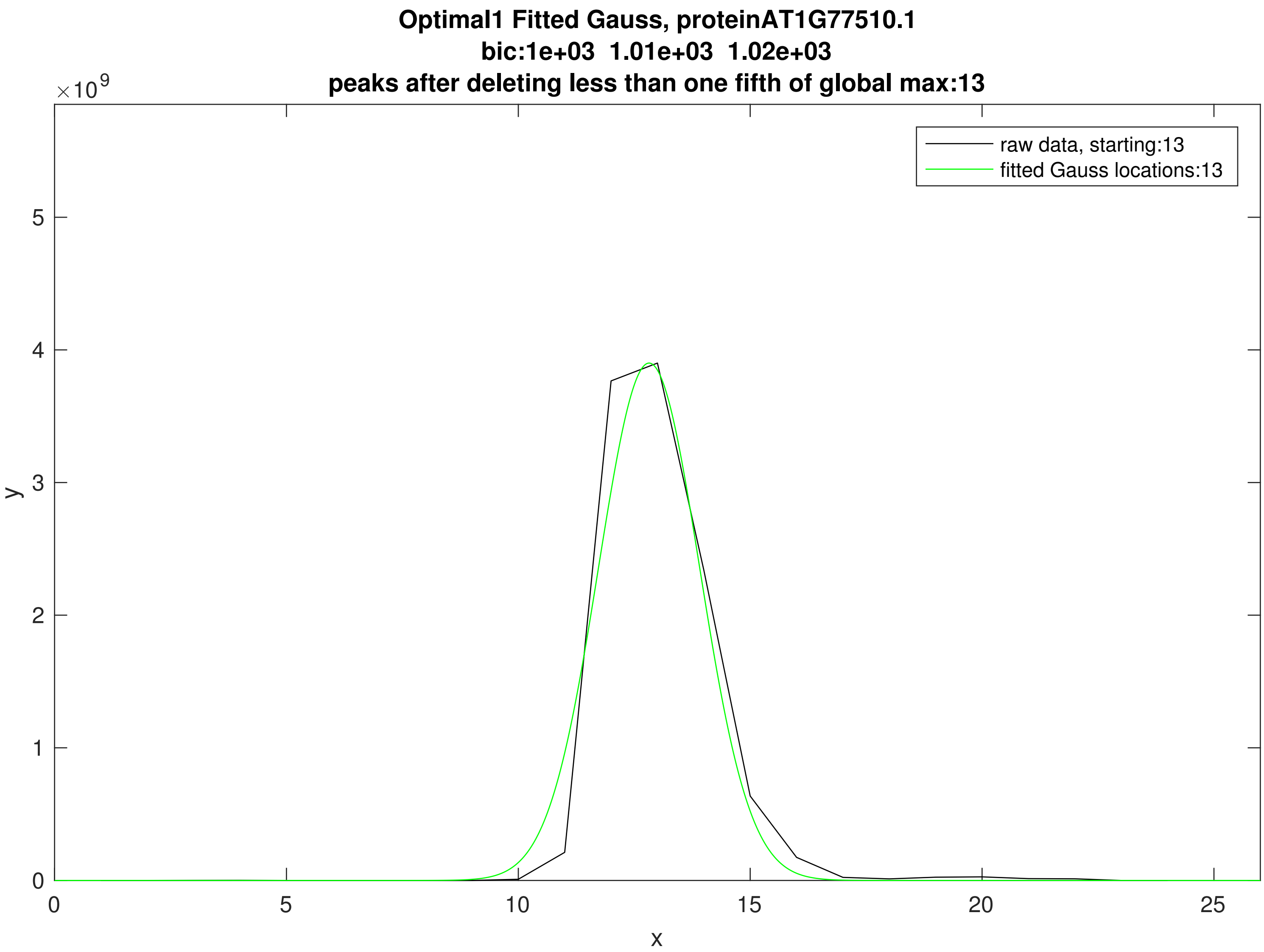
Optimal1 Fitted Gauss, proteinAT1G77510.1
bic:1e+03  1.01e+03  1.02e+03
peaks after deleting less than one fifth of global max:13
×10^9
raw data, starting:13
fitted Gauss locations:13
y
x
0
1
2
3
4
5
0
5
10
15
20
25

**Optimal1 Fitted Gauss, proteinAT1G77540.1**
**bic:734**
**peaks after deleting less than one fifth of global max:20**

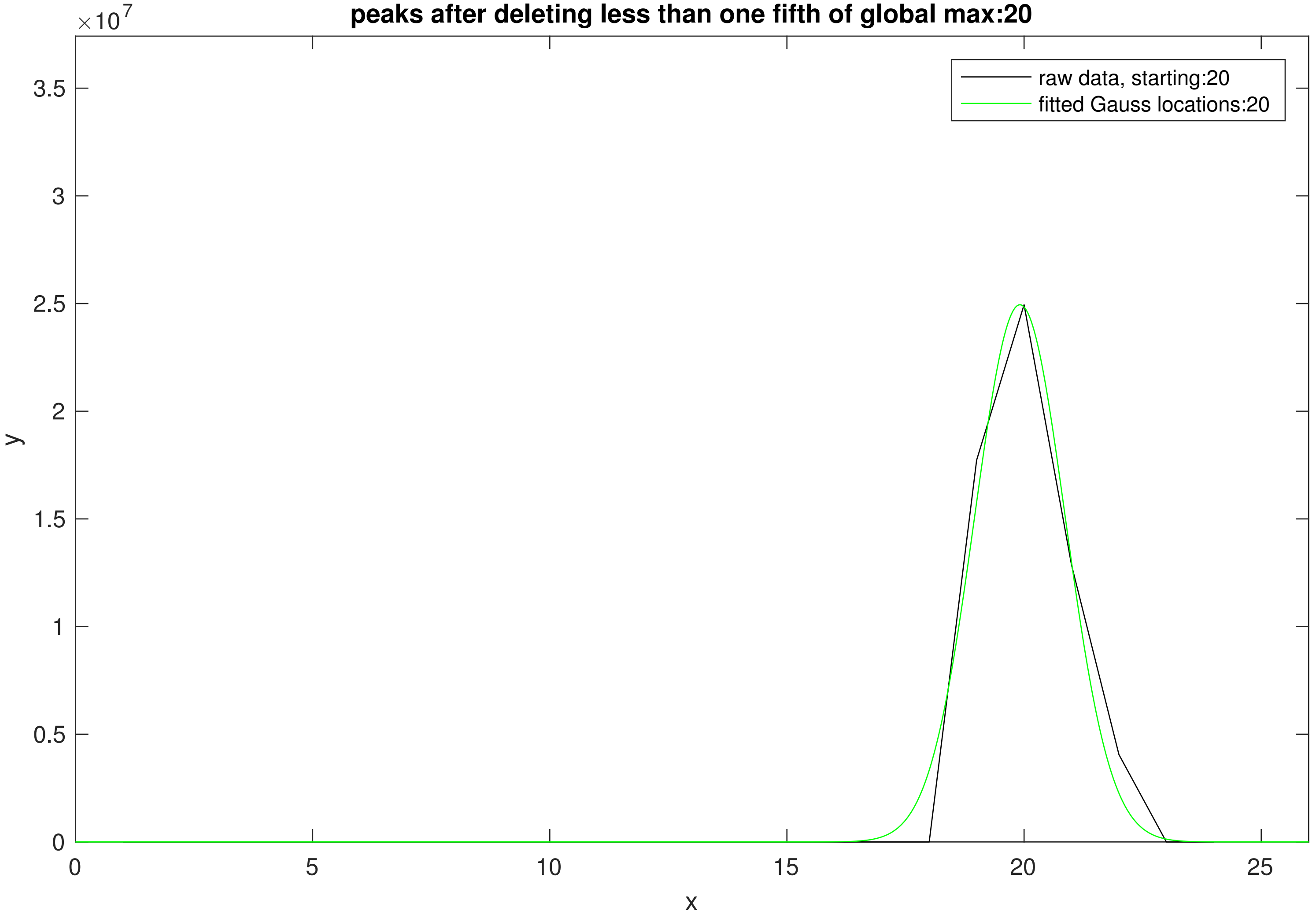

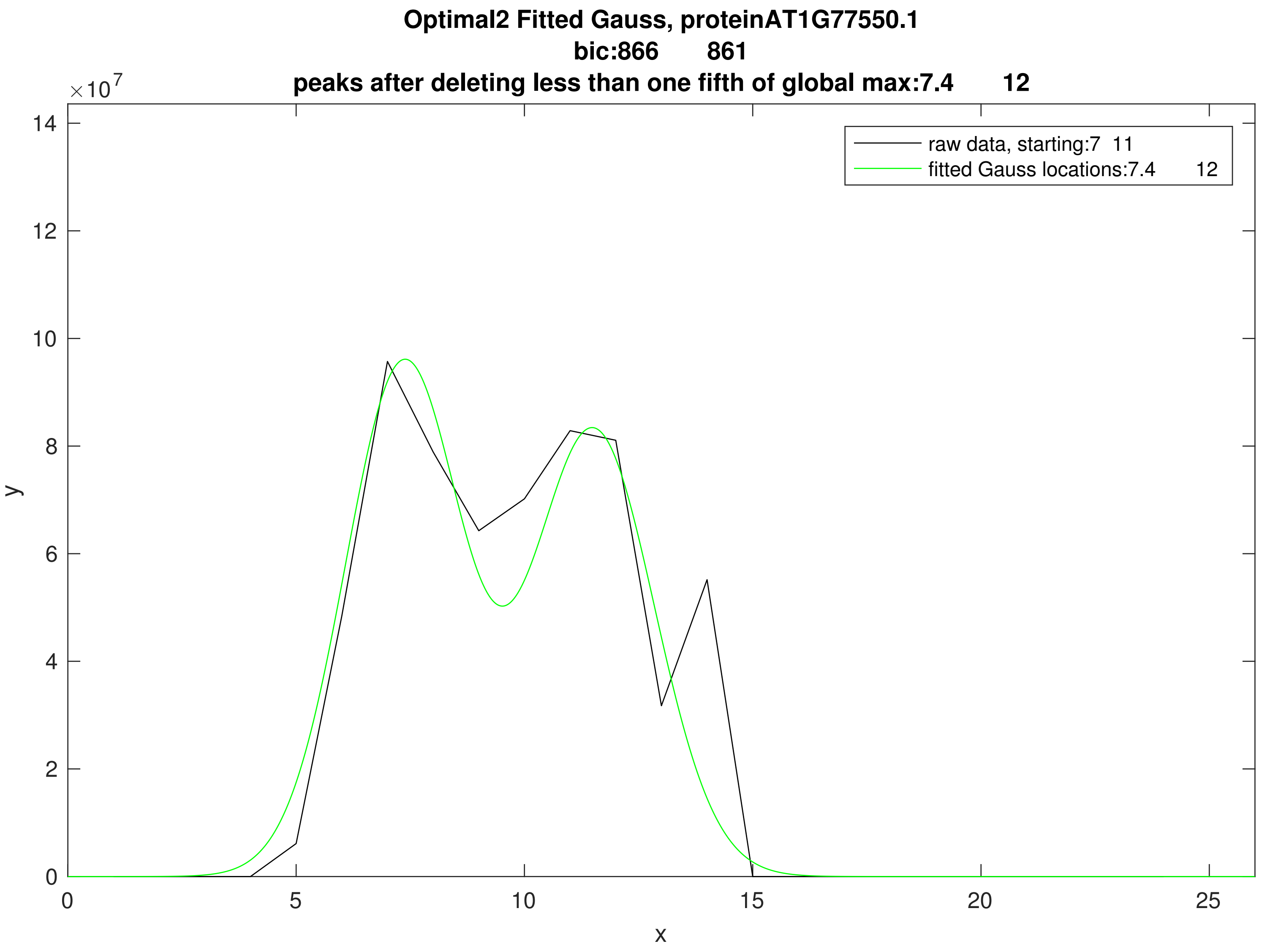
Optimal2 Fitted Gauss, proteinAT1G77550.1
bic:866 861
peaks after deleting less than one fifth of global max:7.4 12
×10^7
raw data, starting:7 11
fitted Gauss locations:7.4 12
y
x
0
2
4
6
8
10
12
14
0
5
10
15
20
25

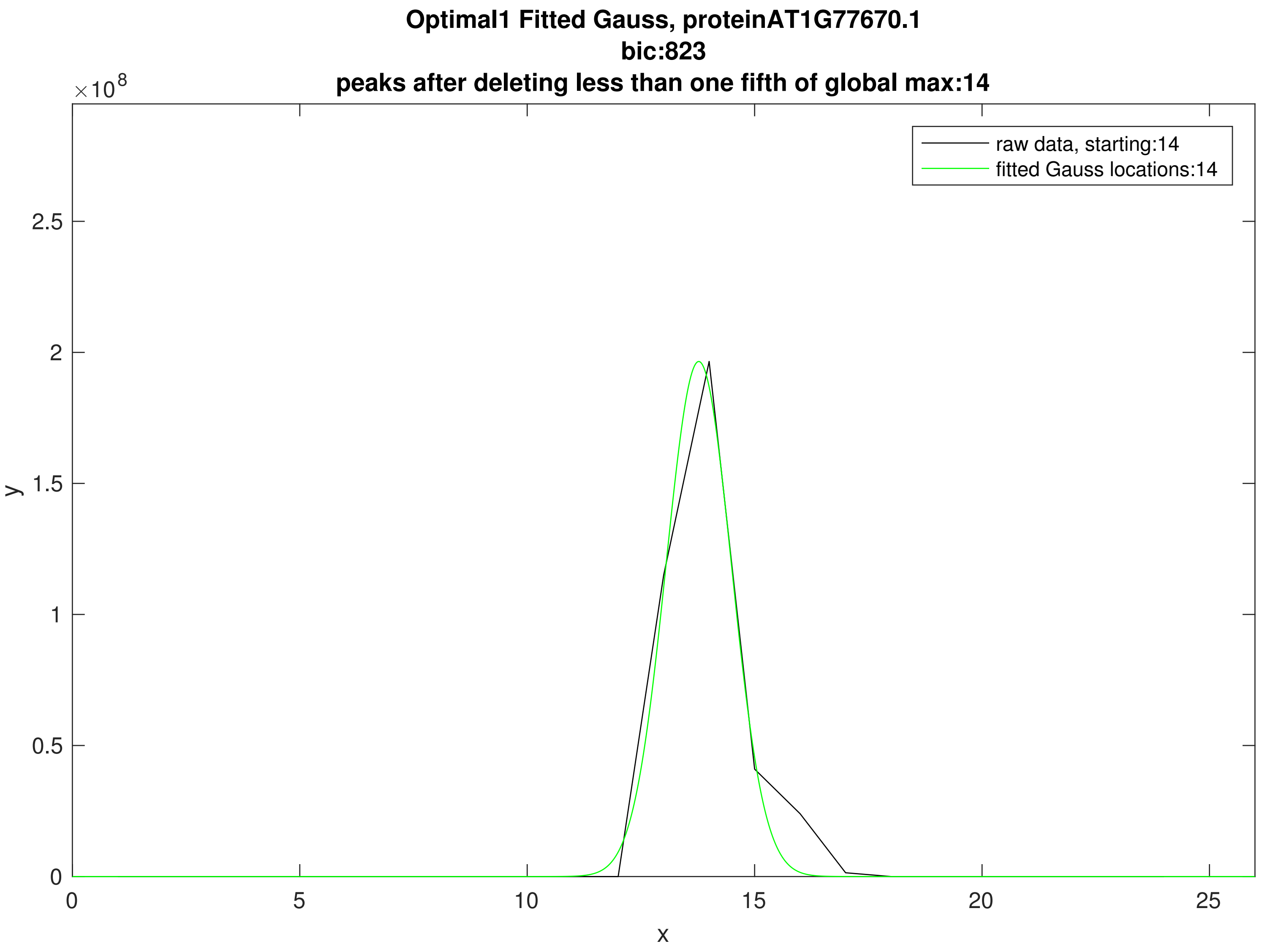

Optimal1 Fitted Gauss, proteinAT1G77670.1
bic:823
peaks after deleting less than one fifth of global max:14
×10^8
raw data, starting:14
fitted Gauss locations:14
2.5
2
1.5
1
0.5
0
y
0
5
10
15
20
25
x

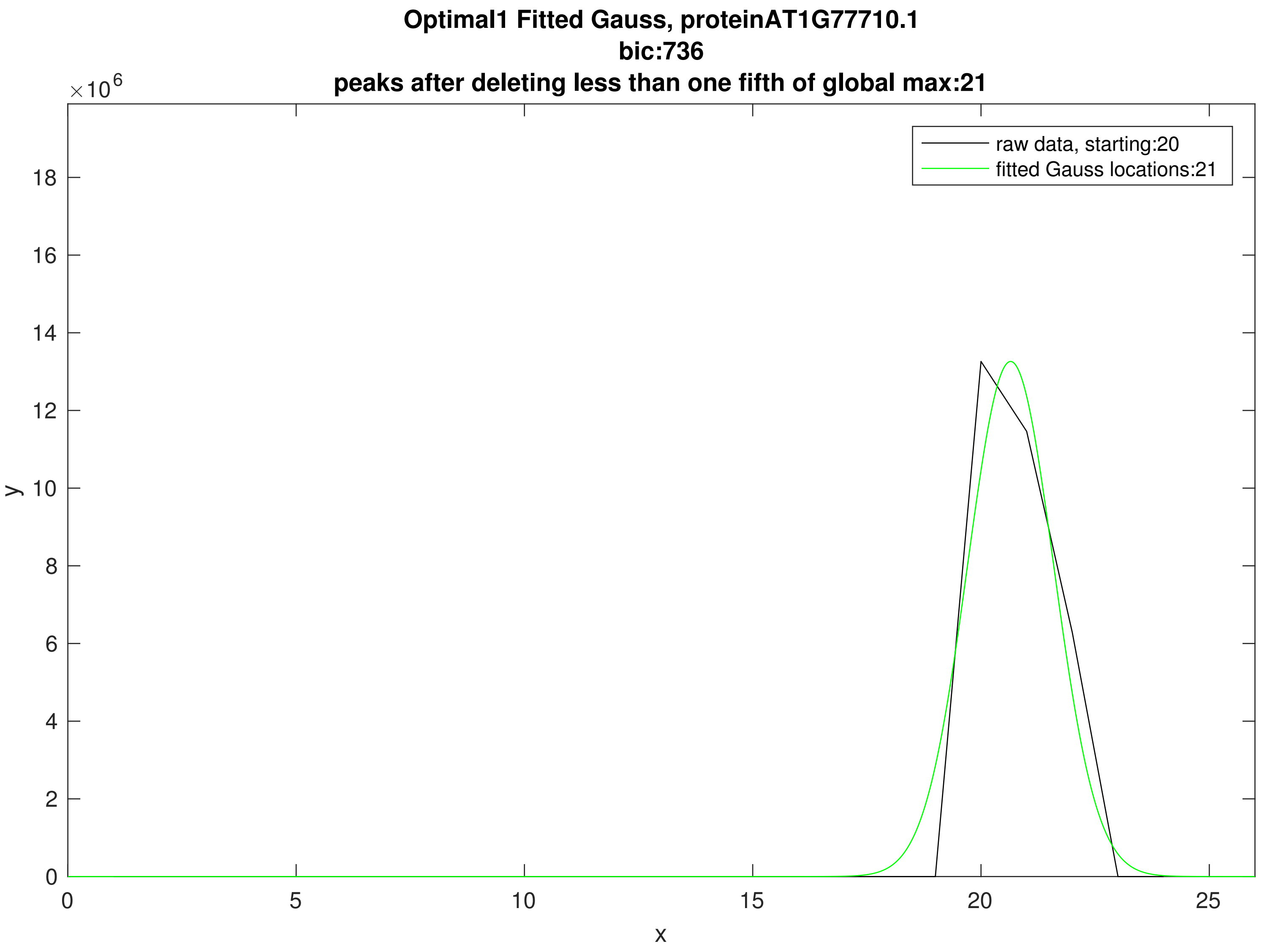

Optimal1 Fitted Gauss, proteinAT1G77710.1
bic:736
peaks after deleting less than one fifth of global max:21
×10^6
raw data, starting:20
fitted Gauss locations:21
y
x
0
2
4
6
8
10
12
14
16
18
0
5
10
15
20
25

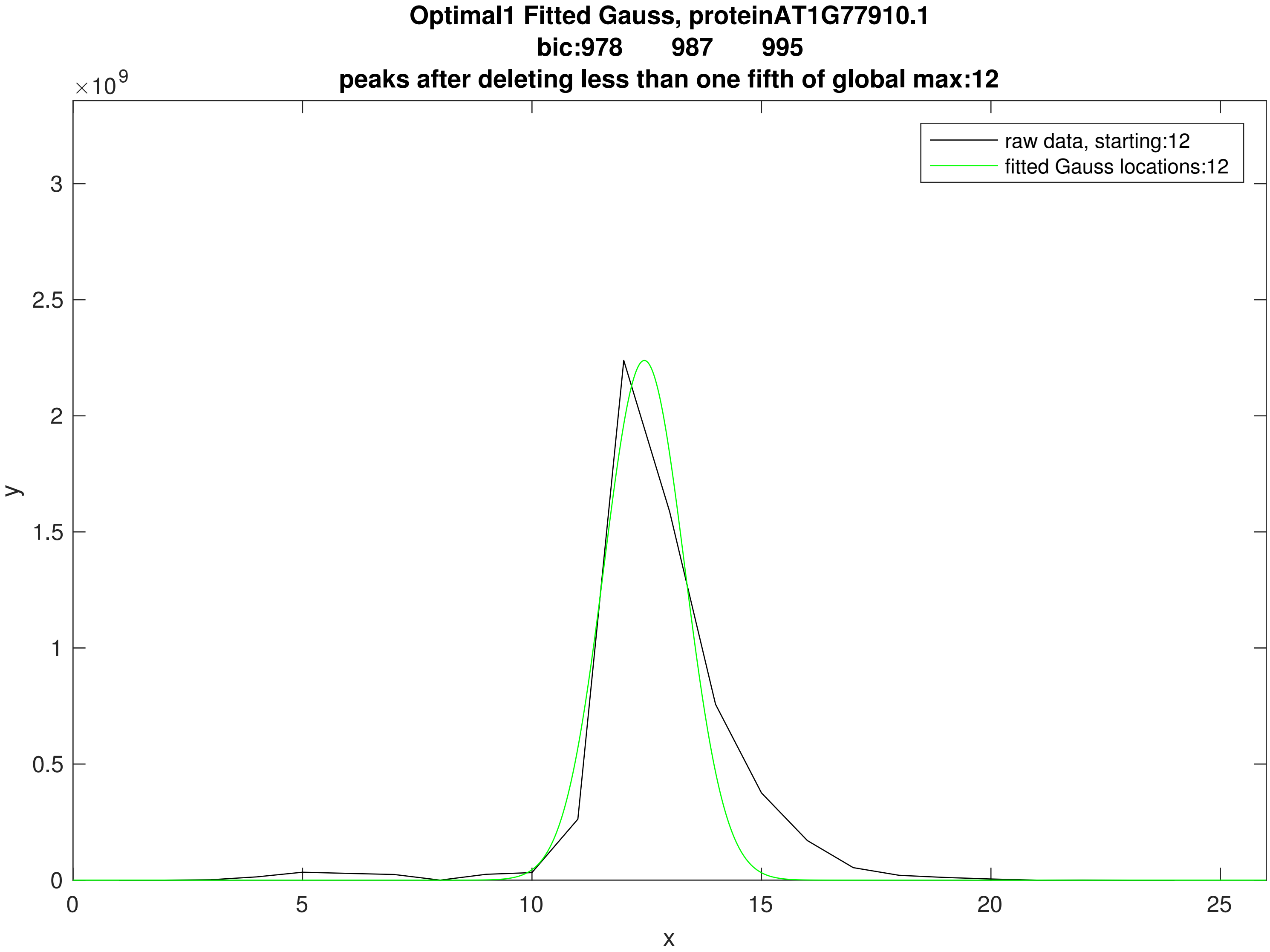

Optimal1 Fitted Gauss, proteinAT1G77910.1
bic:978 987 995
peaks after deleting less than one fifth of global max:12
×10^9
raw data, starting:12
fitted Gauss locations:12
y
x
0
0.5
1
1.5
2
2.5
3
5
10
15
20
25

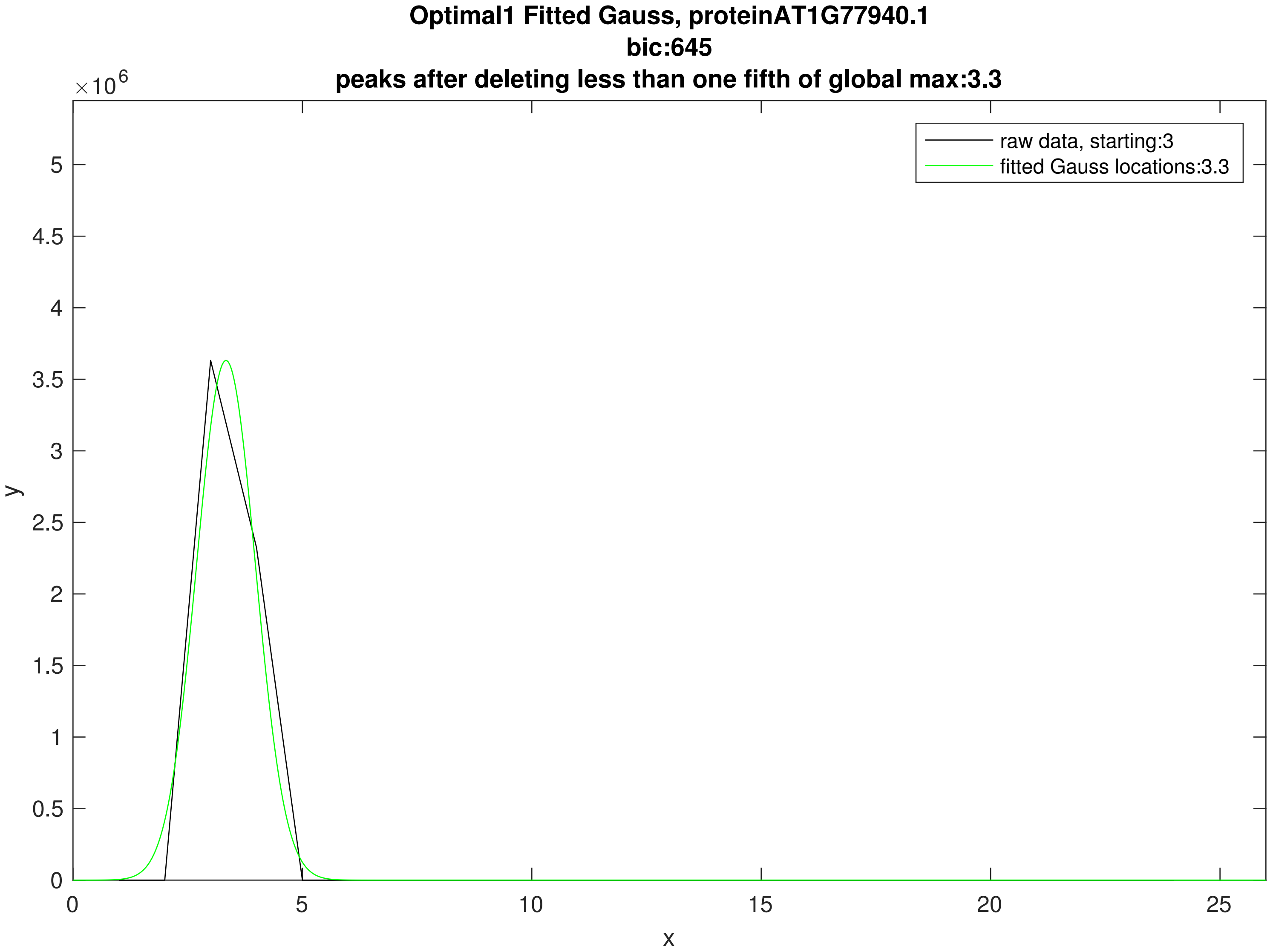

Optimal1 Fitted Gauss, proteinAT1G77940.1
bic:645
peaks after deleting less than one fifth of global max:3.3
×10^6
raw data, starting:3
fitted Gauss locations:3.3
5
4.5
4
3.5
3
2.5
2
1.5
1
0.5
0
y
0
5
10
15
20
25
x

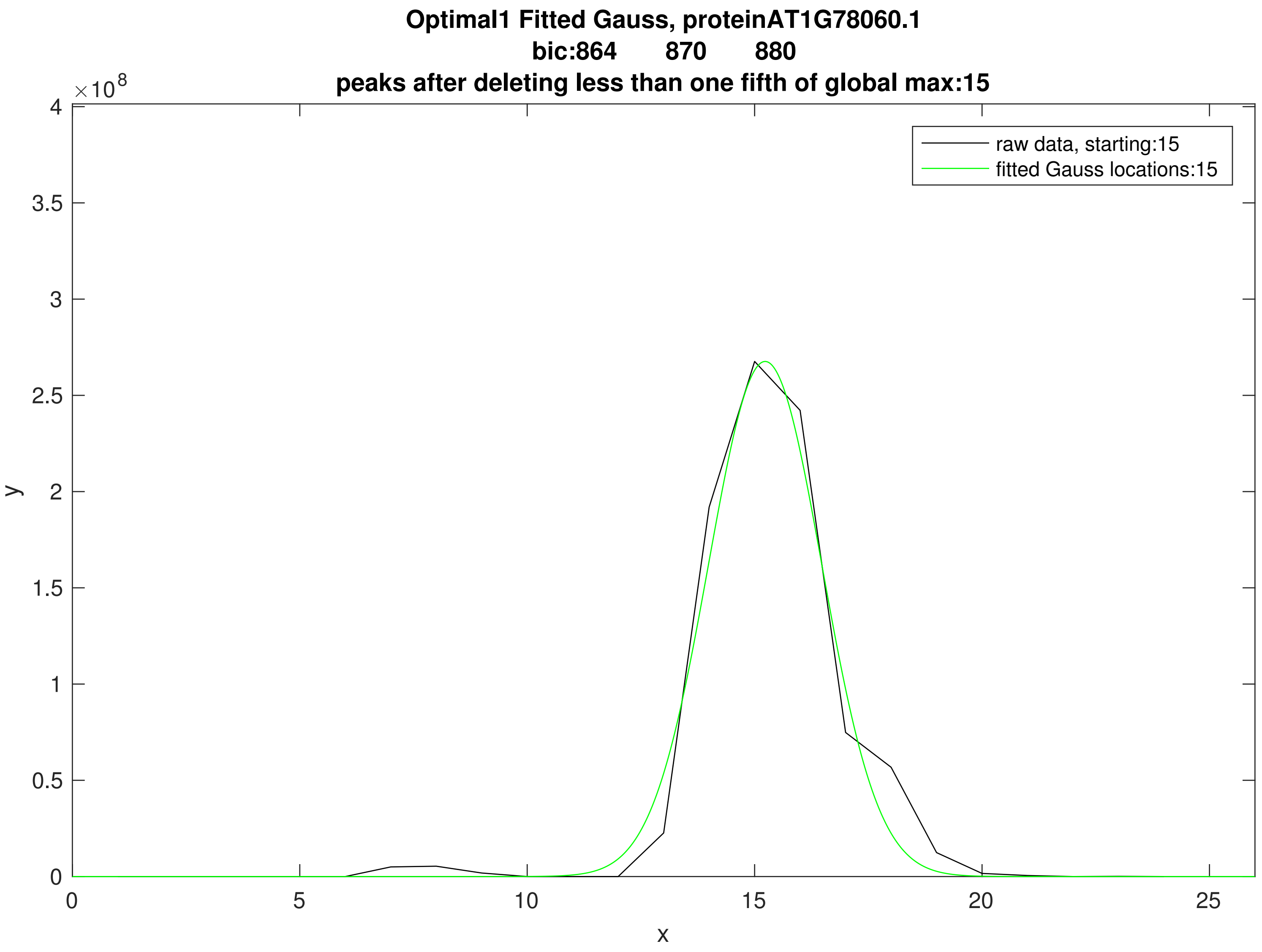
Optimal1 Fitted Gauss, proteinAT1G78060.1
bic:864 870 880
peaks after deleting less than one fifth of global max:15
×10^8
raw data, starting:15
fitted Gauss locations:15
y
x
0
0.5
1
1.5
2
2.5
3
3.5
4
0
5
10
15
20
25

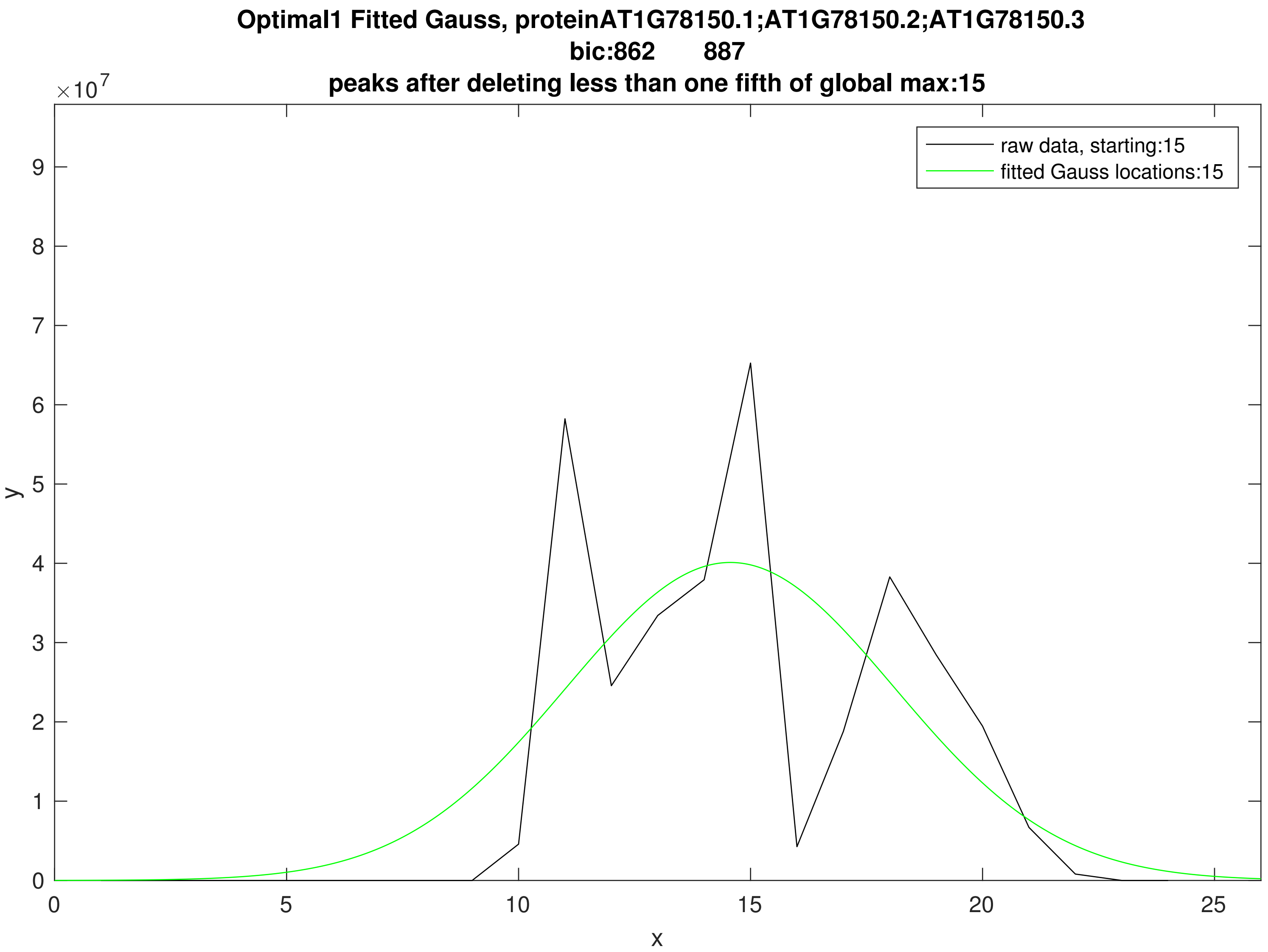

Optimal1 Fitted Gauss, proteinAT1G78150.1;AT1G78150.2;AT1G78150.3
bic:862 887
peaks after deleting less than one fifth of global max:15
×10^7
raw data, starting:15
fitted Gauss locations:15
y
x
0
1
2
3
4
5
6
7
8
9
0
5
10
15
20
25

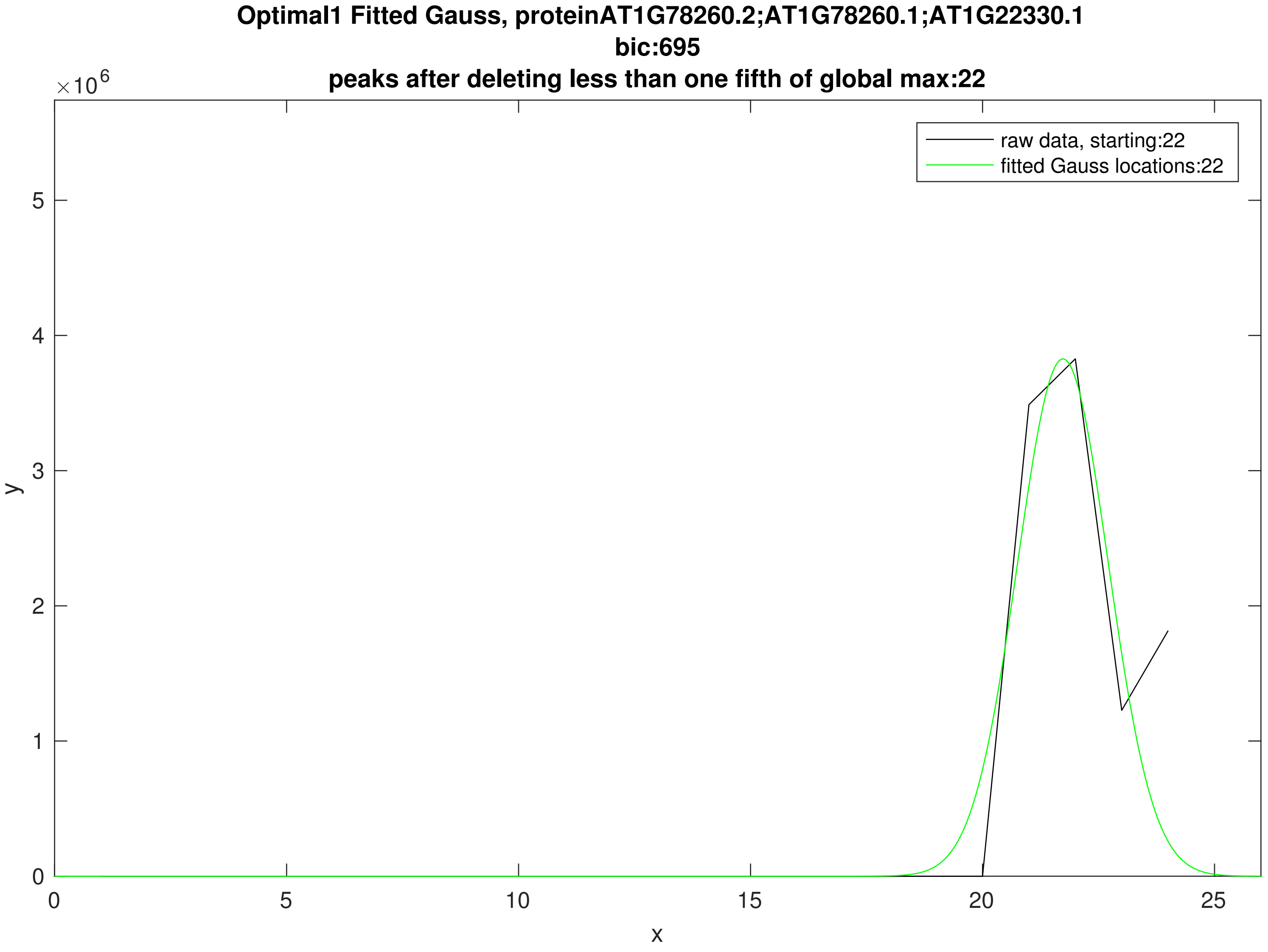

Optimal1 Fitted Gauss, proteinAT1G78260.2;AT1G78260.1;AT1G22330.1
bic:695
peaks after deleting less than one fifth of global max:22
×10^6
raw data, starting:22
fitted Gauss locations:22
y
x
0
1
2
3
4
5
0
5
10
15
20
25

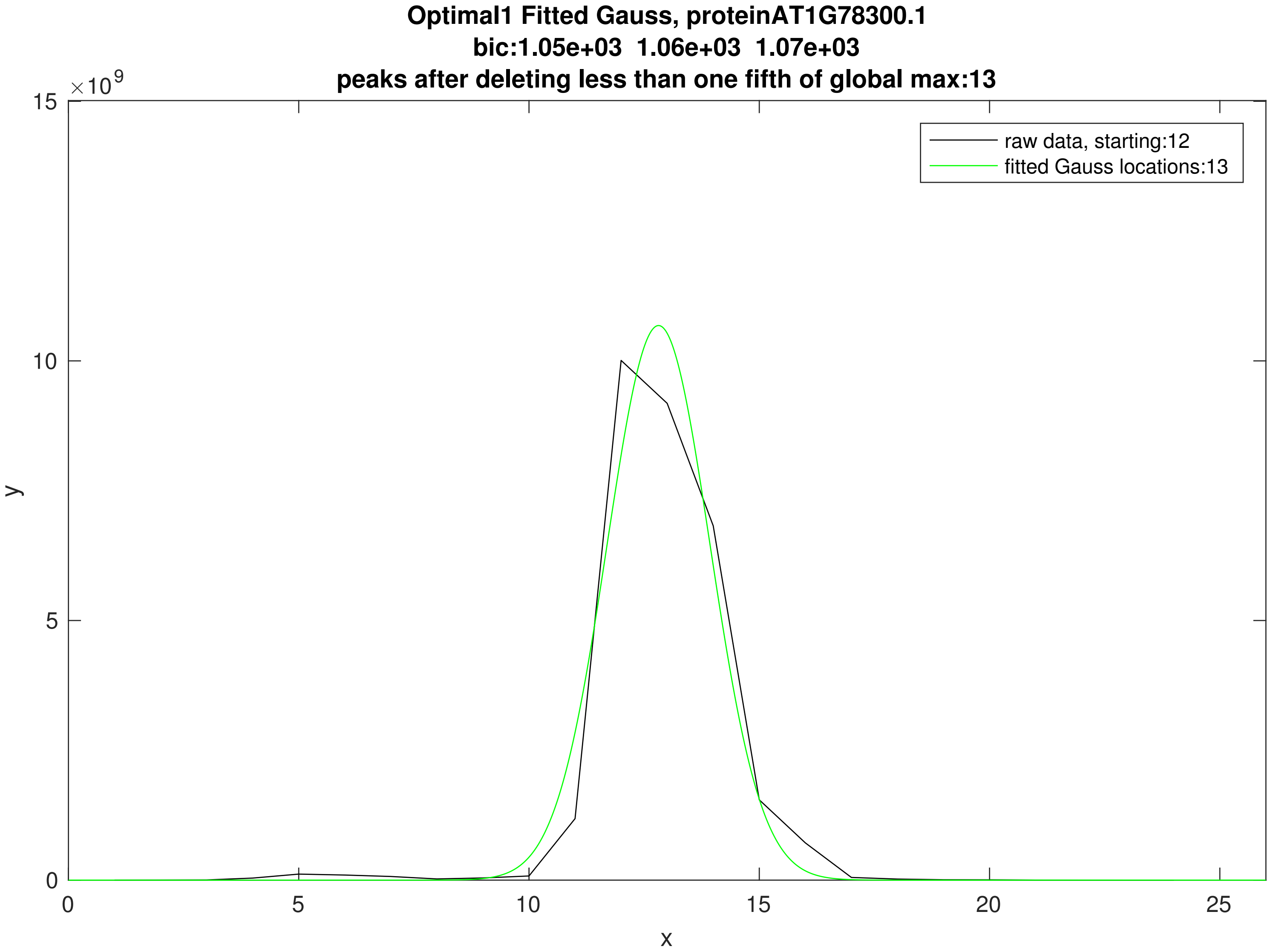
Optimal1 Fitted Gauss, proteinAT1G78300.1
bic:1.05e+03 1.06e+03 1.07e+03
peaks after deleting less than one fifth of global max:13
×10^9
15
10
5
0
y
0
5
10
15
20
25
x
raw data, starting:12
fitted Gauss locations:13

**Optimal1 Fitted Gauss, proteinAT1G78340.1**
**bic:681**
**peaks after deleting less than one fifth of global max:16**

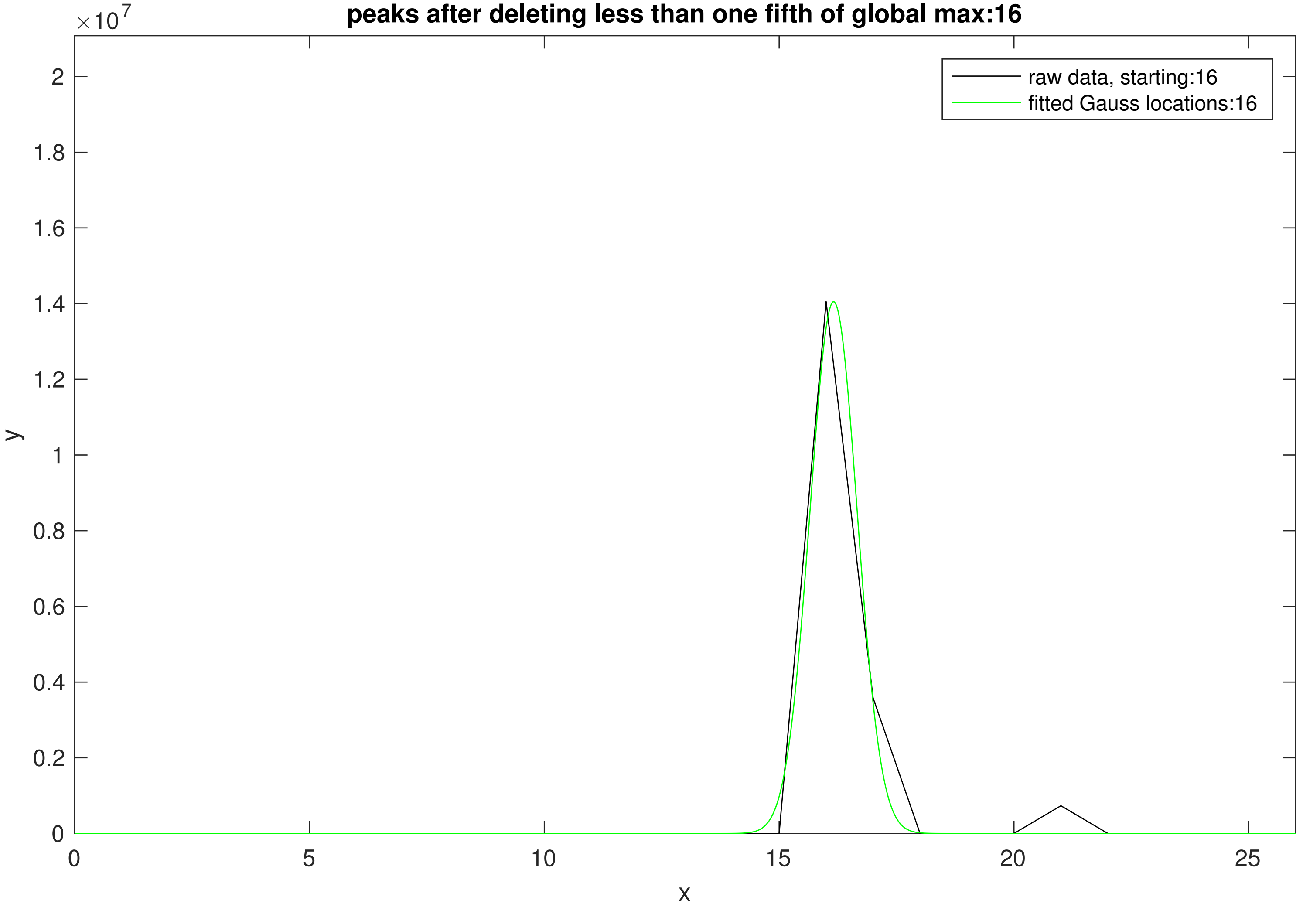

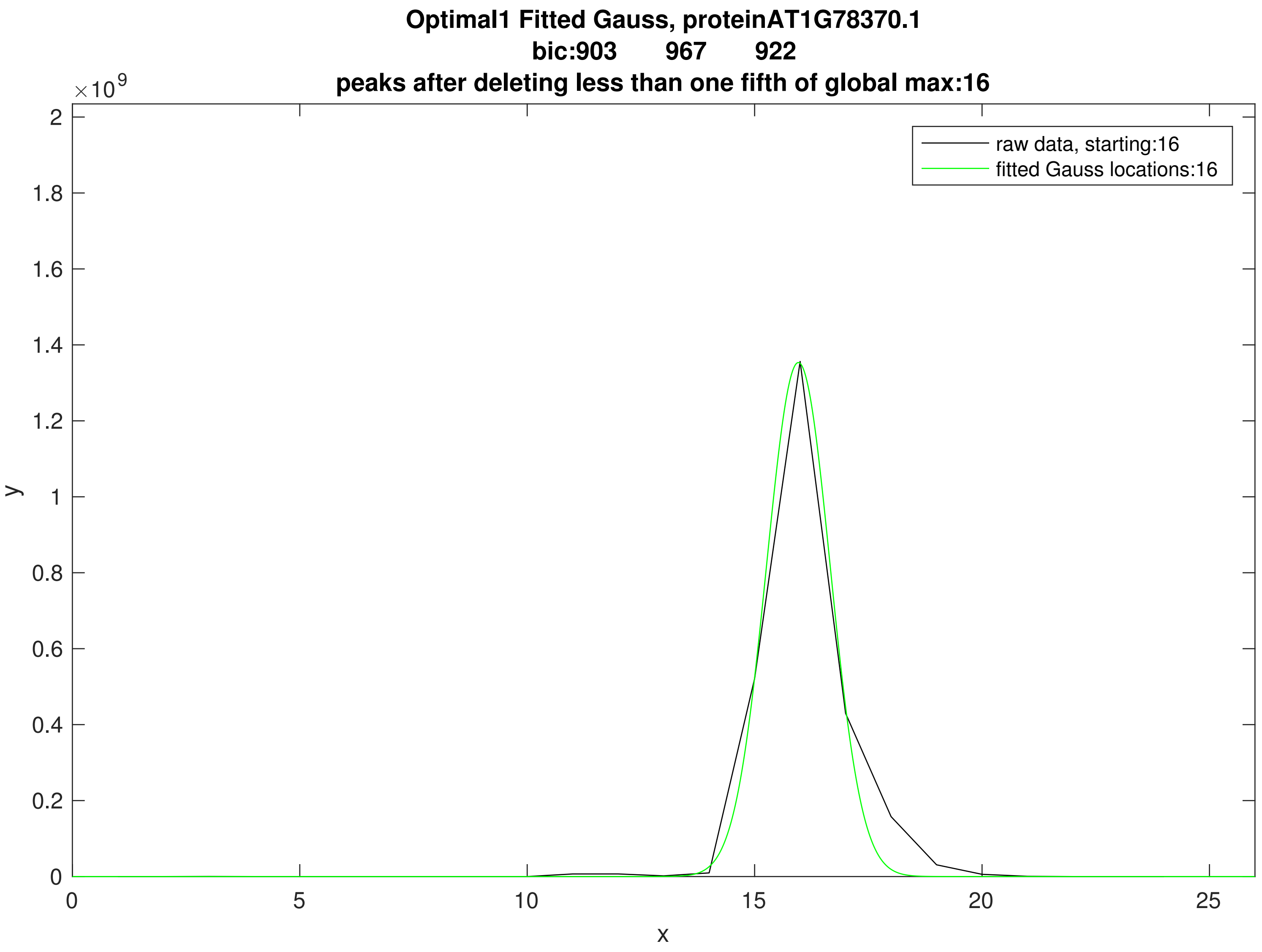
Optimal1 Fitted Gauss, proteinAT1G78370.1
bic:903 967 922
peaks after deleting less than one fifth of global max:16
×10^9
raw data, starting:16
fitted Gauss locations:16
y
x
0
0.2
0.4
0.6
0.8
1
1.2
1.4
1.6
1.8
2
0
5
10
15
20
25

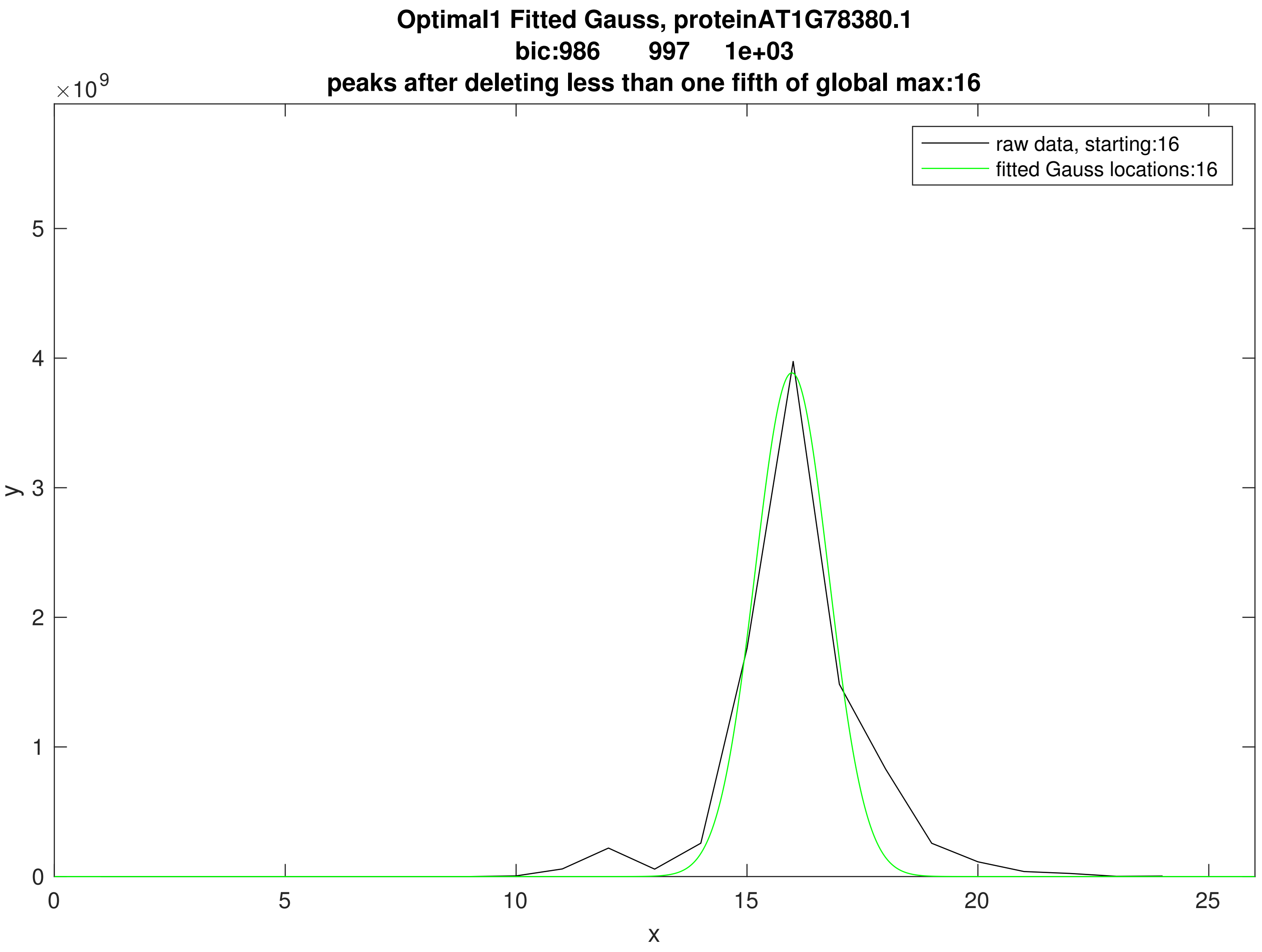
Optimal1 Fitted Gauss, proteinAT1G78380.1
bic:986 997 1e+03
peaks after deleting less than one fifth of global max:16
×10^9
raw data, starting:16
fitted Gauss locations:16
y
x
0
1
2
3
4
5
0
5
10
15
20
25

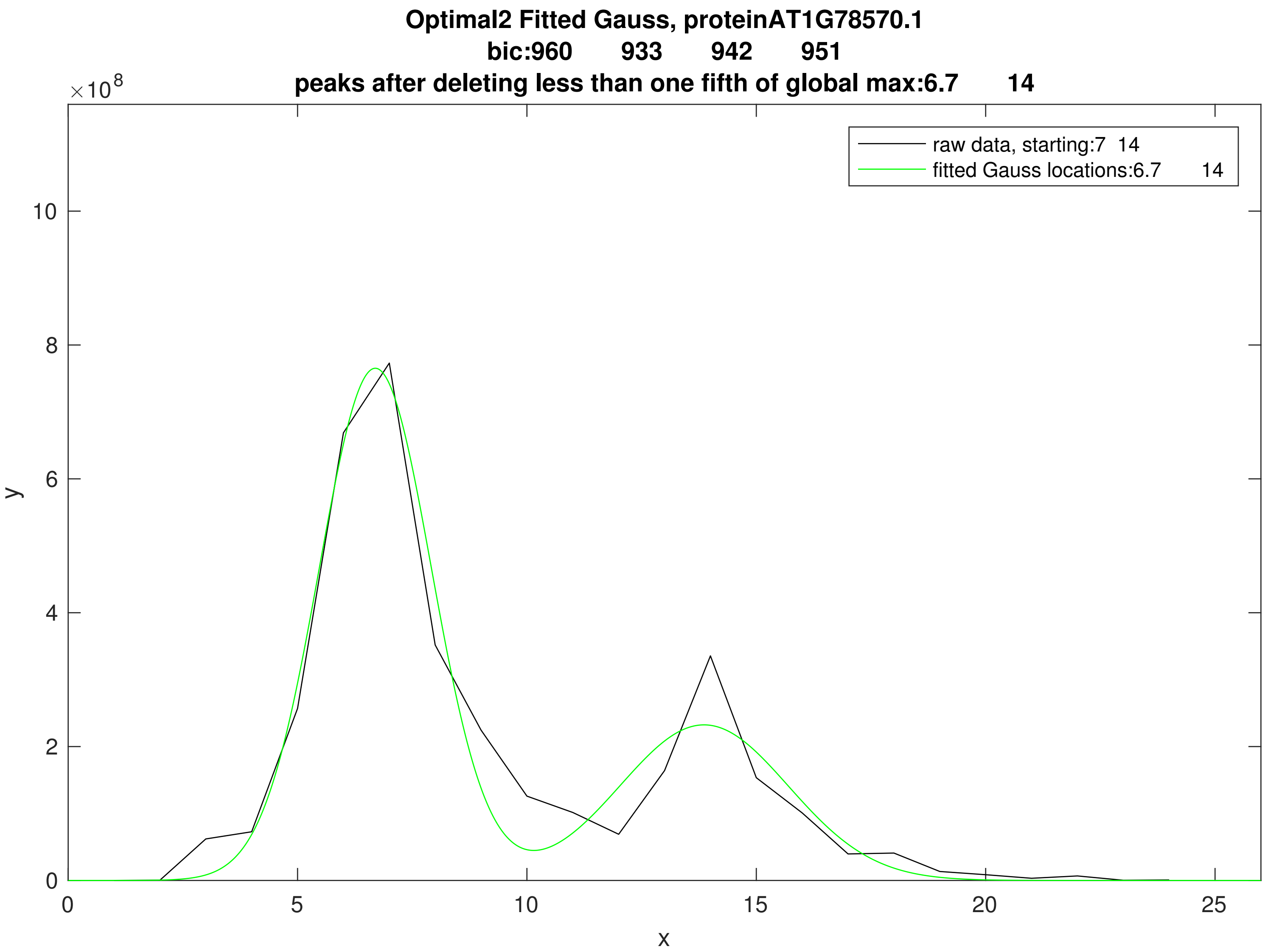

Optimal2 Fitted Gauss, proteinAT1G78570.1
bic:960 933 942 951
peaks after deleting less than one fifth of global max:6.7 14
×10^8
raw data, starting:7 14
fitted Gauss locations:6.7 14
y
x
0
2
4
6
8
10
5
15
20
25

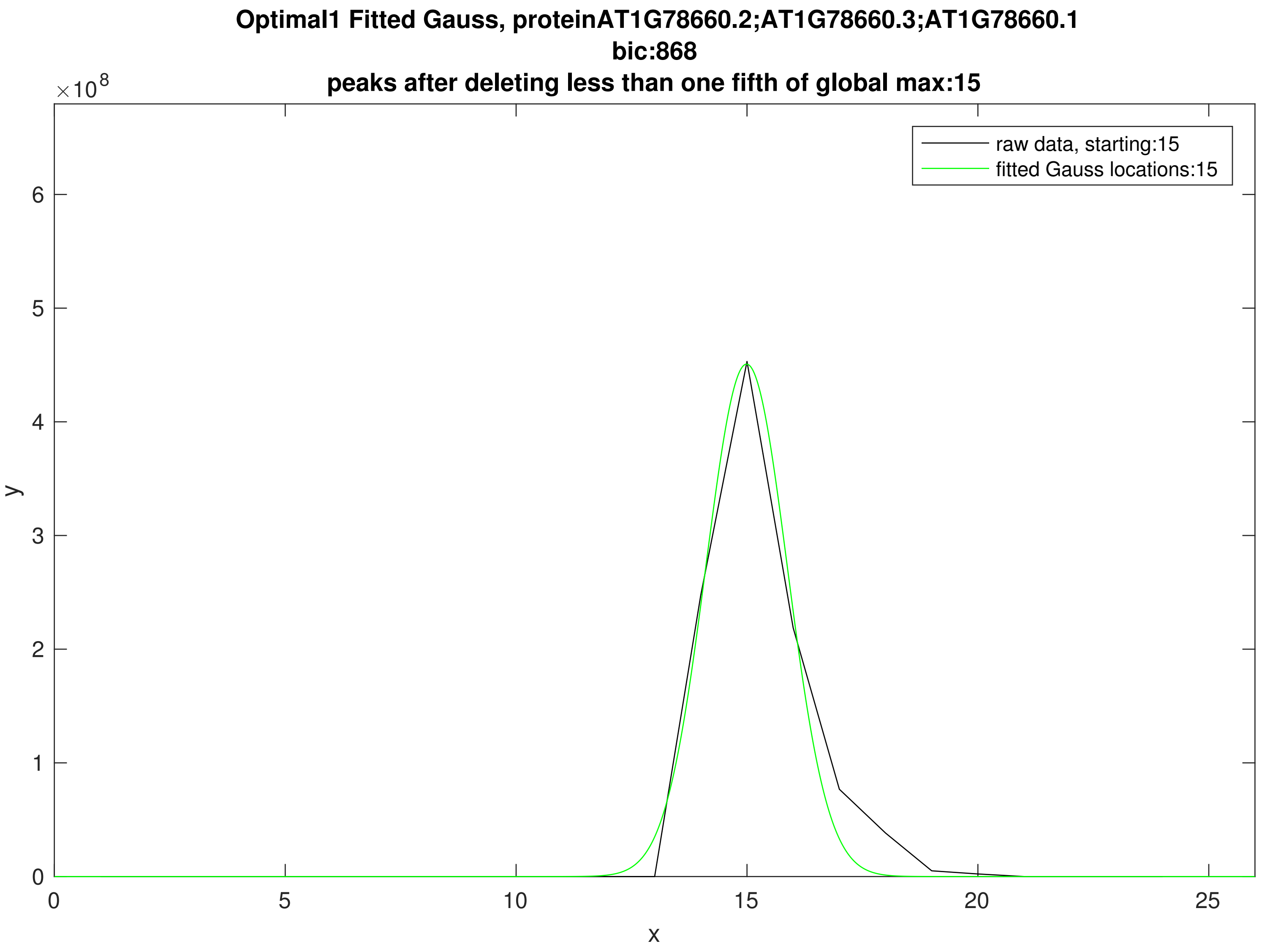
Optimal1 Fitted Gauss, proteinAT1G78660.2;AT1G78660.3;AT1G78660.1
bic:868
peaks after deleting less than one fifth of global max:15
×10^8
raw data, starting:15
fitted Gauss locations:15
y
x
0
1
2
3
4
5
6
0
5
10
15
20
25

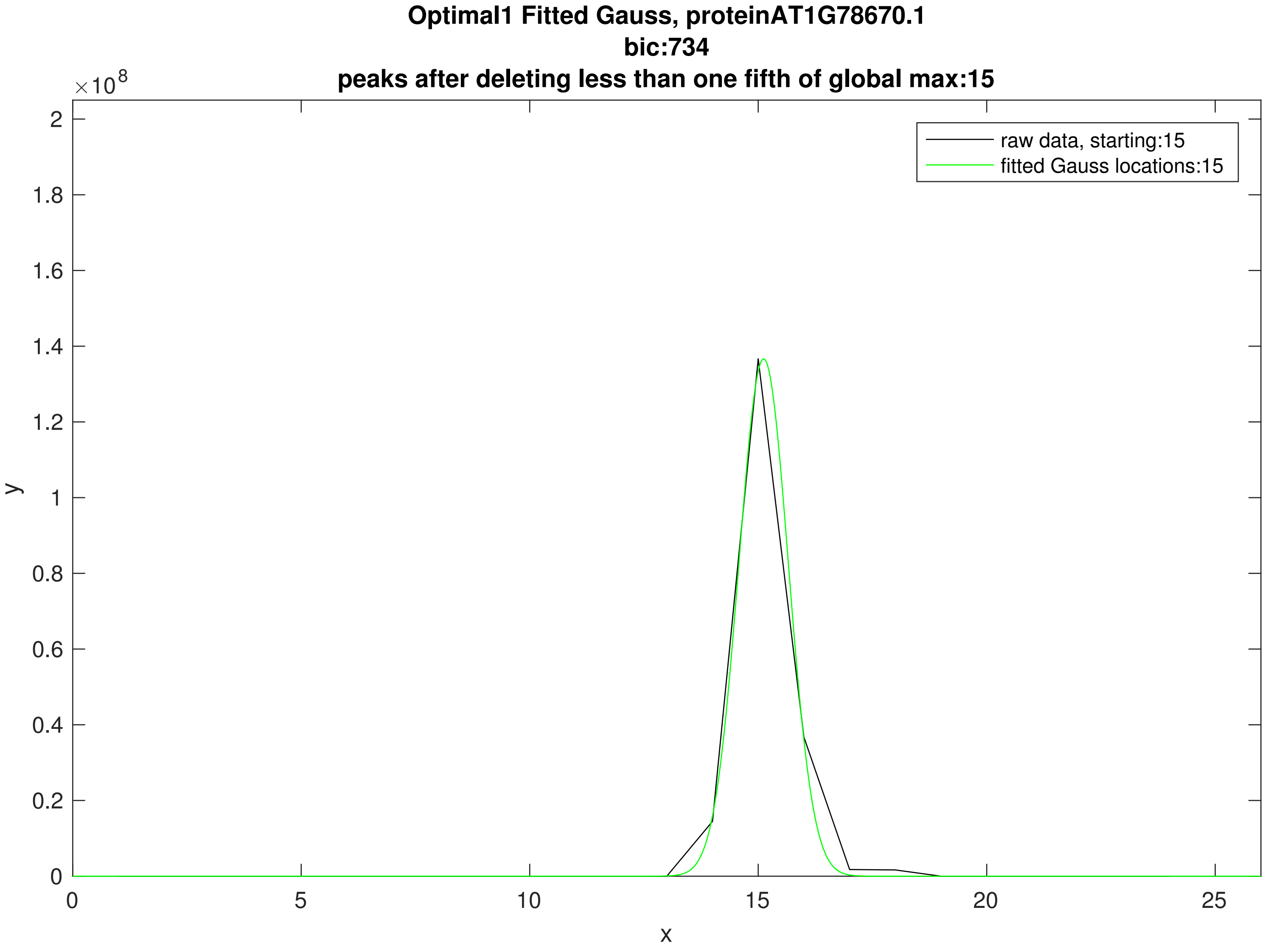

Optimal1 Fitted Gauss, proteinAT1G78670.1
bic:734
peaks after deleting less than one fifth of global max:15
×10^8
2
1.8
1.6
1.4
1.2
1
0.8
0.6
0.4
0.2
0
y
0
5
10
15
20
25
x
raw data, starting:15
fitted Gauss locations:15

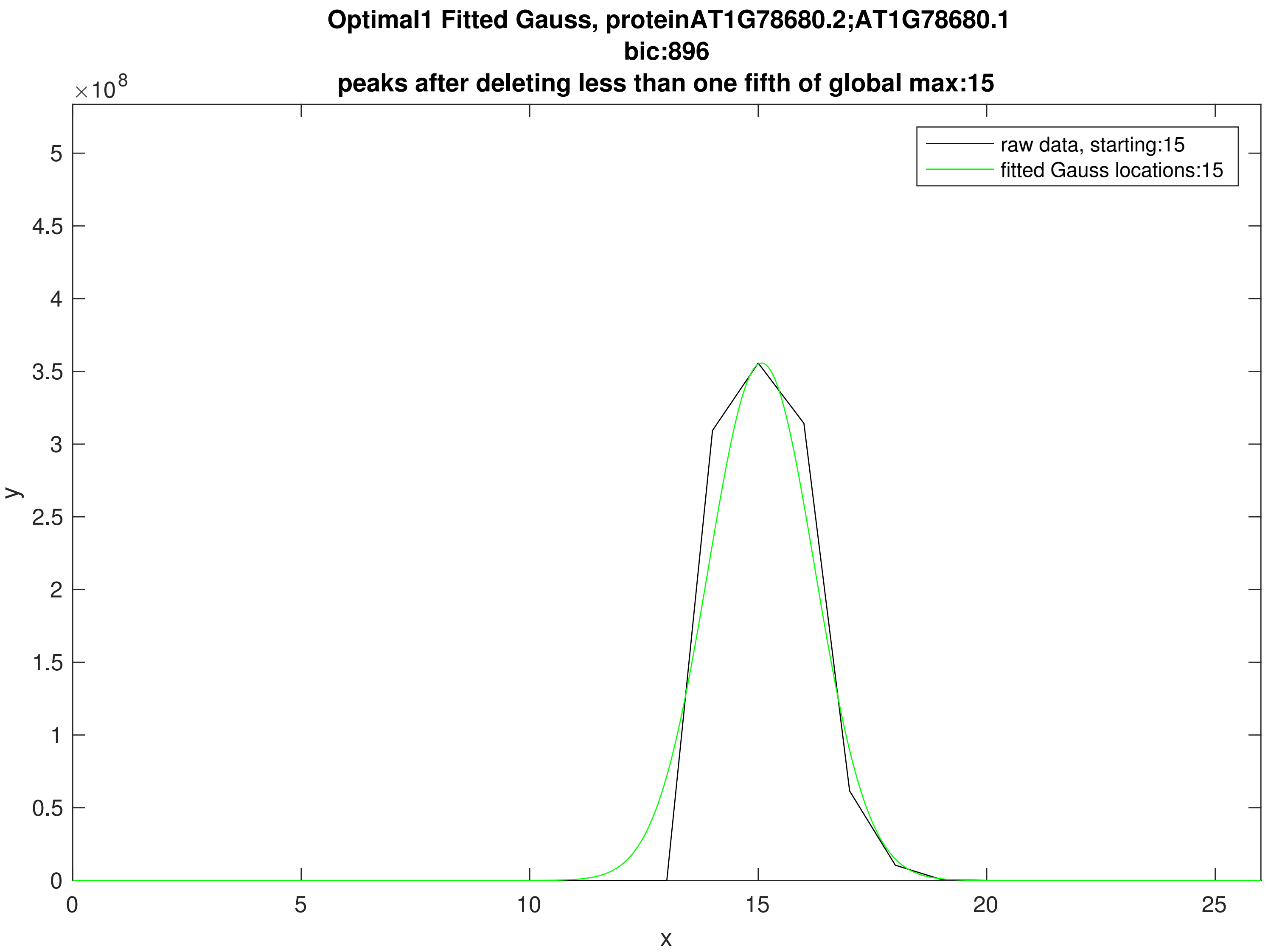
Optimal1 Fitted Gauss, proteinAT1G78680.2;AT1G78680.1
bic:896
peaks after deleting less than one fifth of global max:15
×10^8
raw data, starting:15
fitted Gauss locations:15
y
x
0
0.5
1
1.5
2
2.5
3
3.5
4
4.5
5
0
5
10
15
20
25

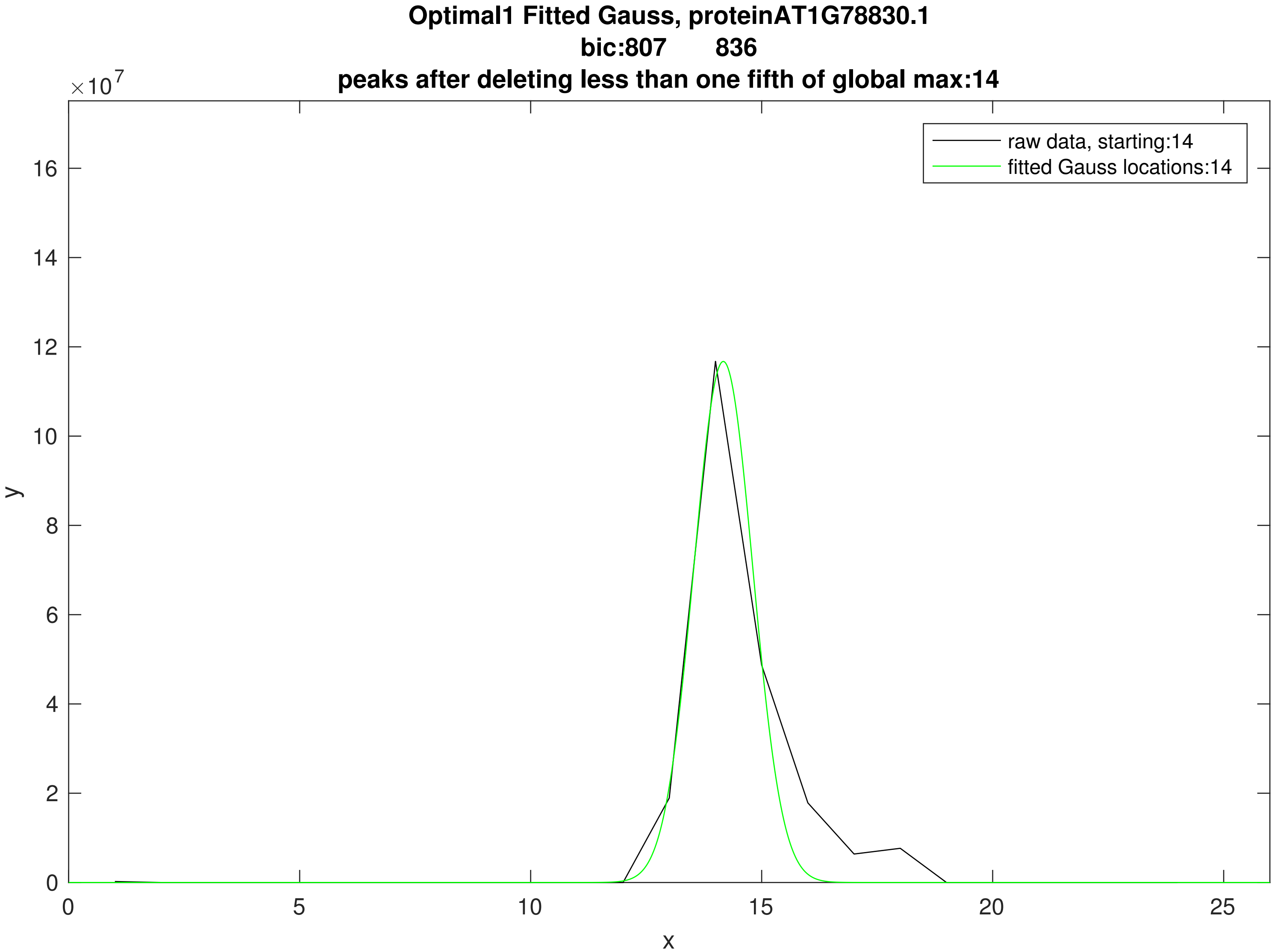
Optimal1 Fitted Gauss, proteinAT1G78830.1
bic:807 836
peaks after deleting less than one fifth of global max:14
×10^7
raw data, starting:14
fitted Gauss locations:14
y
x
0
2
4
6
8
10
12
14
16
0
5
10
15
20
25

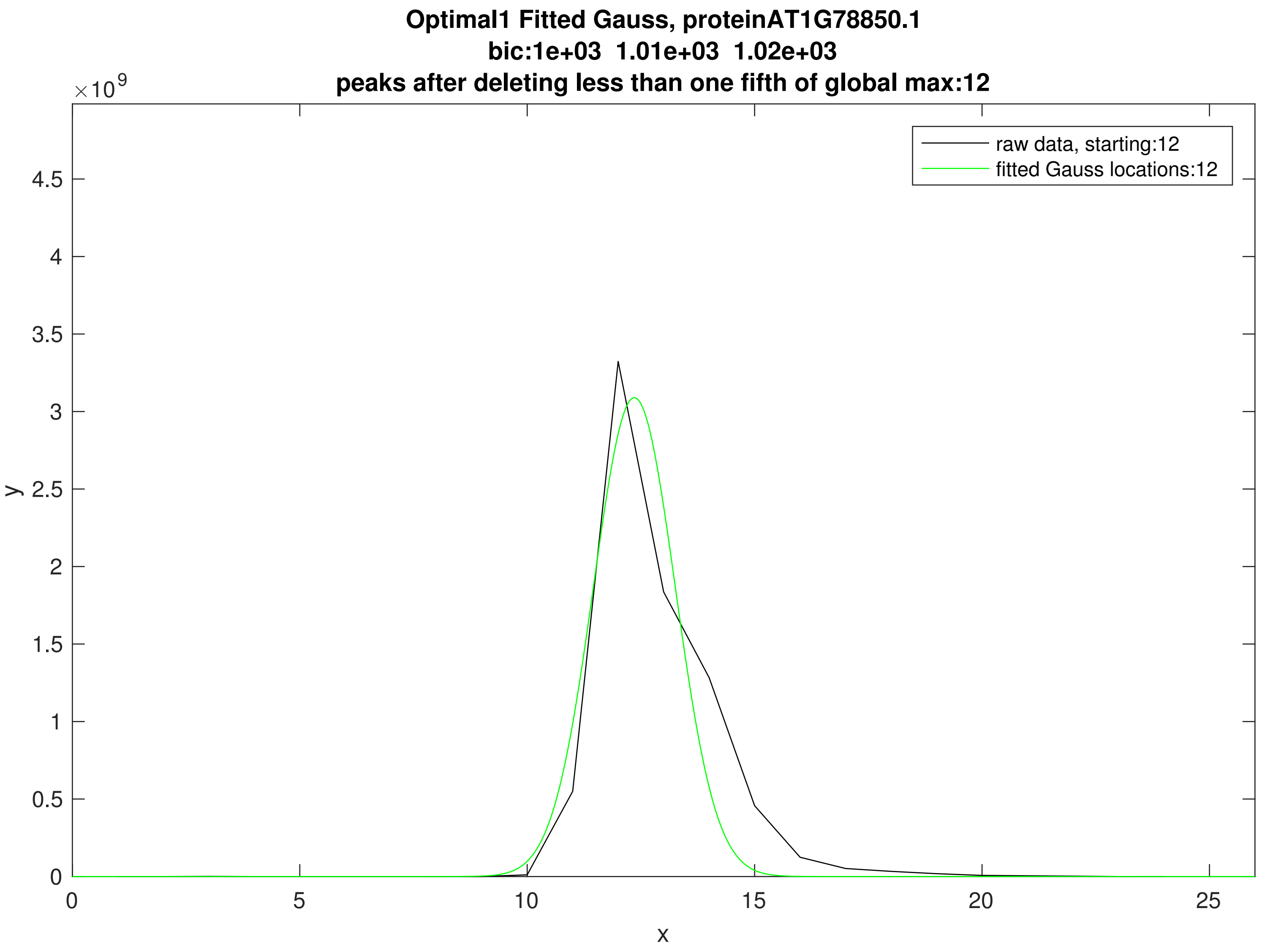
Optimal1 Fitted Gauss, proteinAT1G78850.1
bic:1e+03  1.01e+03  1.02e+03
peaks after deleting less than one fifth of global max:12
×10^9
raw data, starting:12
fitted Gauss locations:12
4.5
4
3.5
3
2.5
2
1.5
1
0.5
0
y
0
5
10
15
20
25
x

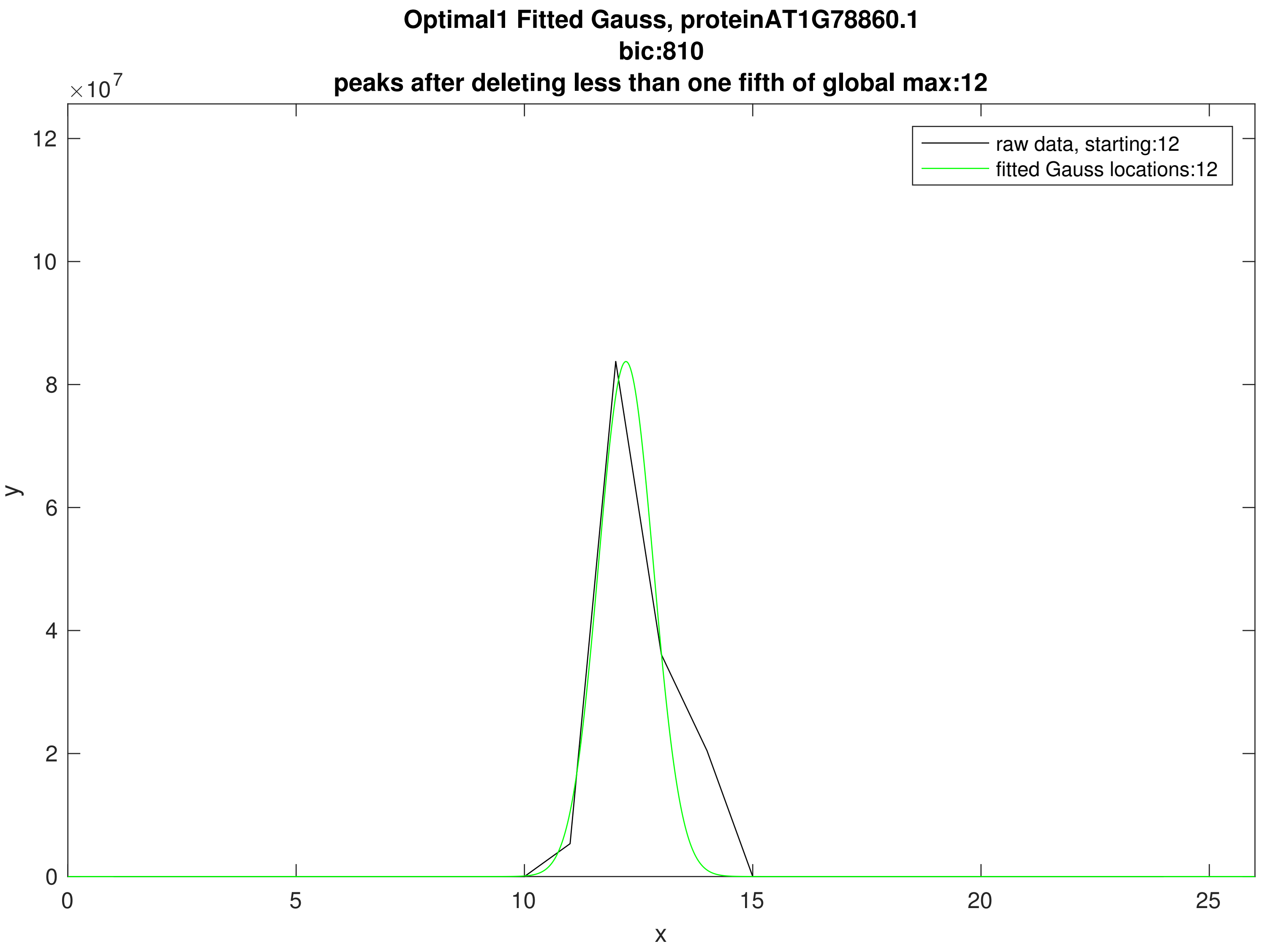
Optimal1 Fitted Gauss, proteinAT1G78860.1
bic:810
peaks after deleting less than one fifth of global max:12
×10^7
raw data, starting:12
fitted Gauss locations:12
y
x
0
2
4
6
8
10
12
0
5
10
15
20
25

**Optimal1 Fitted Gauss, proteinAT1G78870.1;AT1G78870.2;AT1G78870.3**
**bic:723**
**peaks after deleting less than one fifth of global max:19**

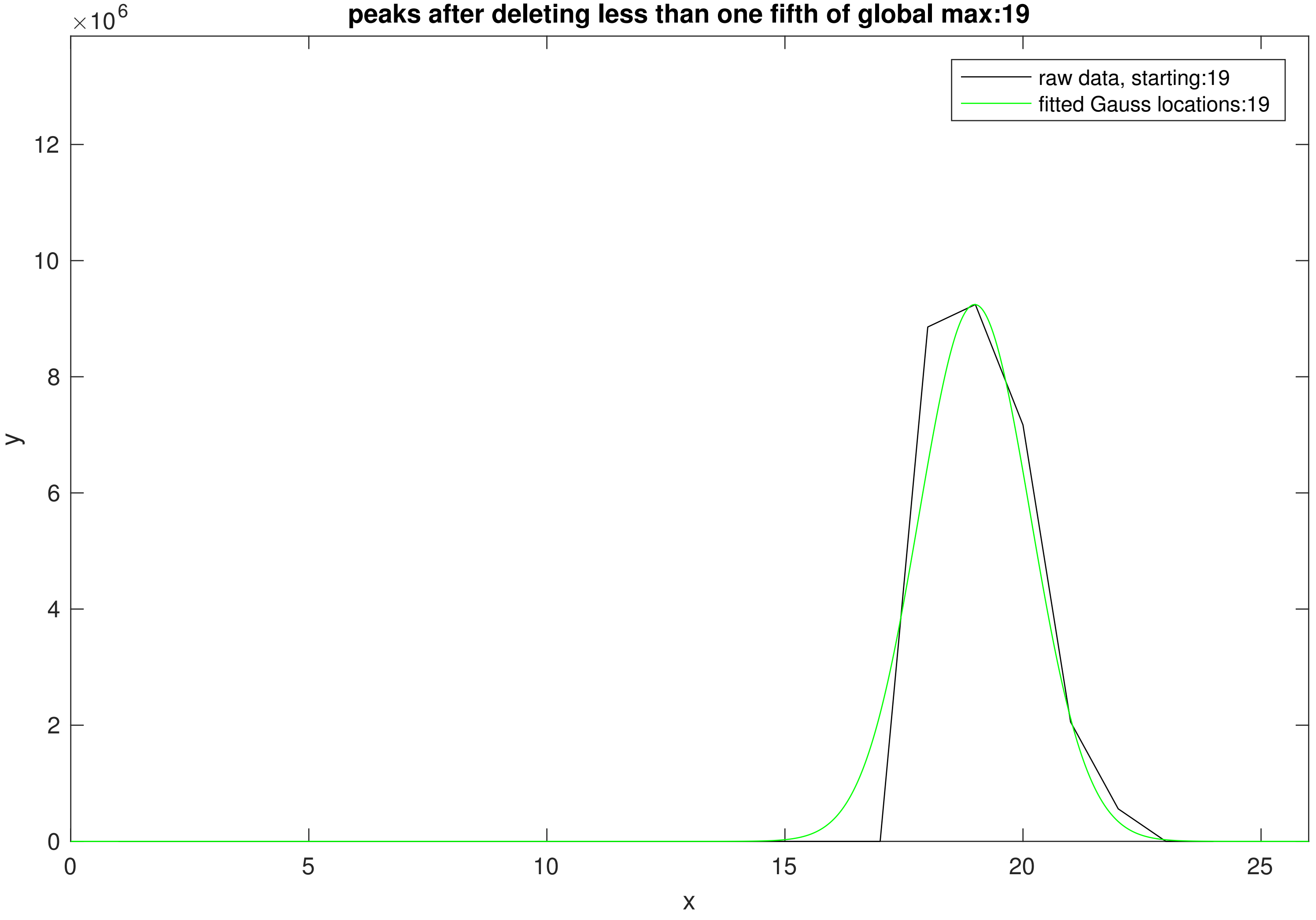

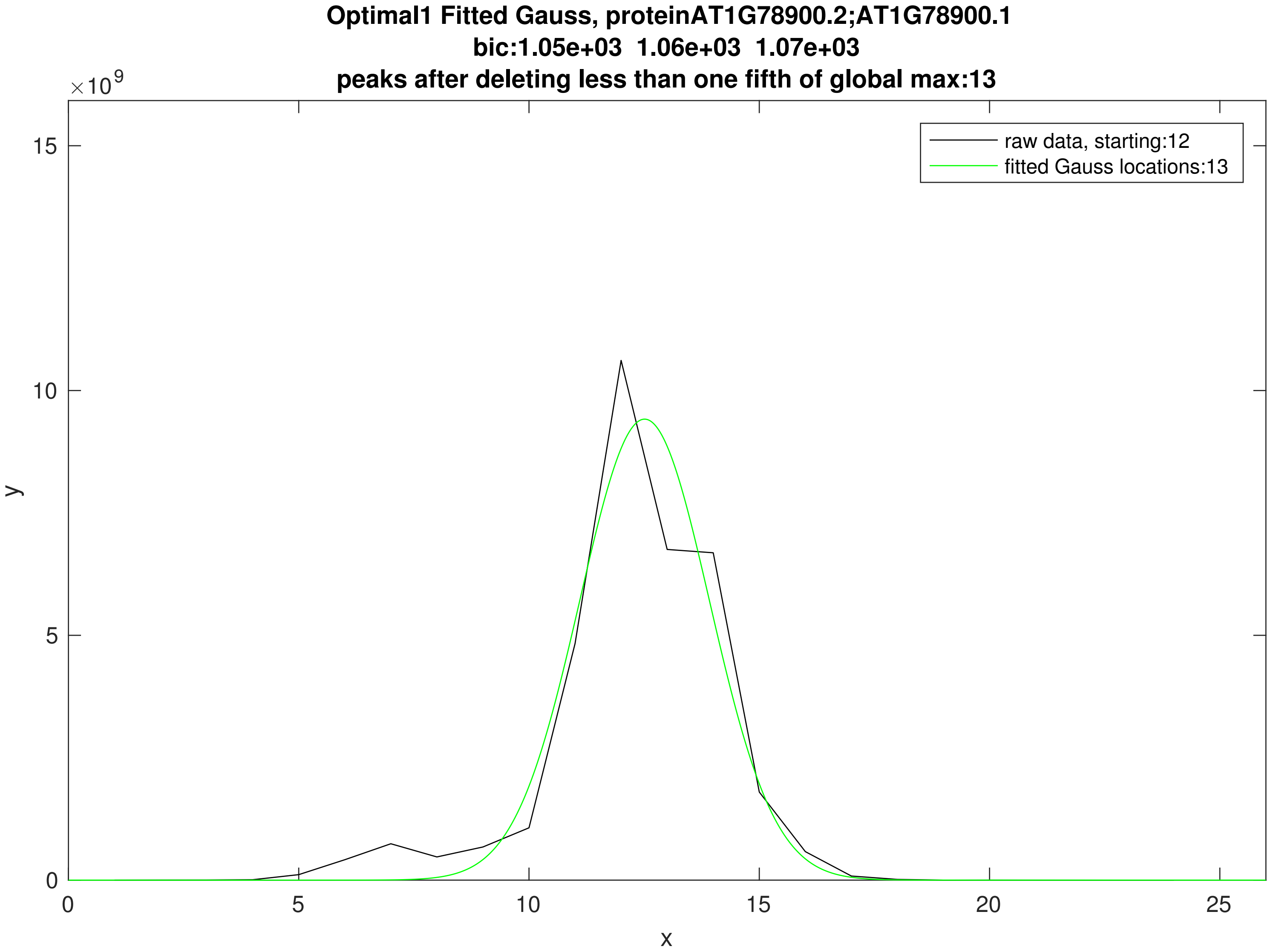

Optimal1 Fitted Gauss, proteinAT1G78900.2;AT1G78900.1
bic:1.05e+03 1.06e+03 1.07e+03
peaks after deleting less than one fifth of global max:13
×10^9
raw data, starting:12
fitted Gauss locations:13
15
10
5
0
y
0
5
10
15
20
25
x

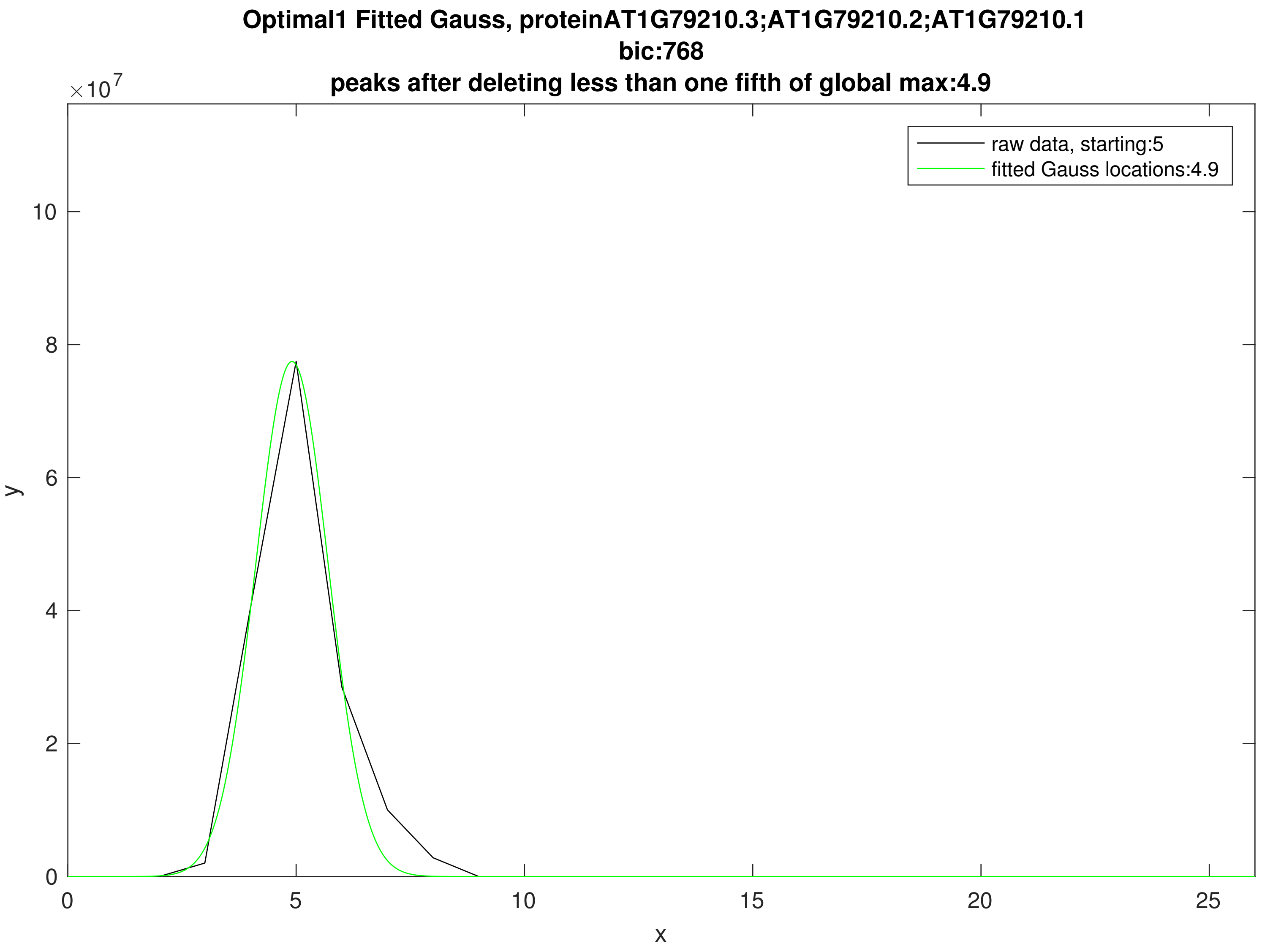
Optimal1 Fitted Gauss, proteinAT1G79210.3;AT1G79210.2;AT1G79210.1
bic:768
peaks after deleting less than one fifth of global max:4.9
×10^7
raw data, starting:5
fitted Gauss locations:4.9
y
x
0
2
4
6
8
10
0
5
10
15
20
25

**Optimal1 Fitted Gauss, proteinAT1G79230.1;AT1G79230.3;AT1G79230.2**
**bic:971 975**
**peaks after deleting less than one fifth of global max:17**

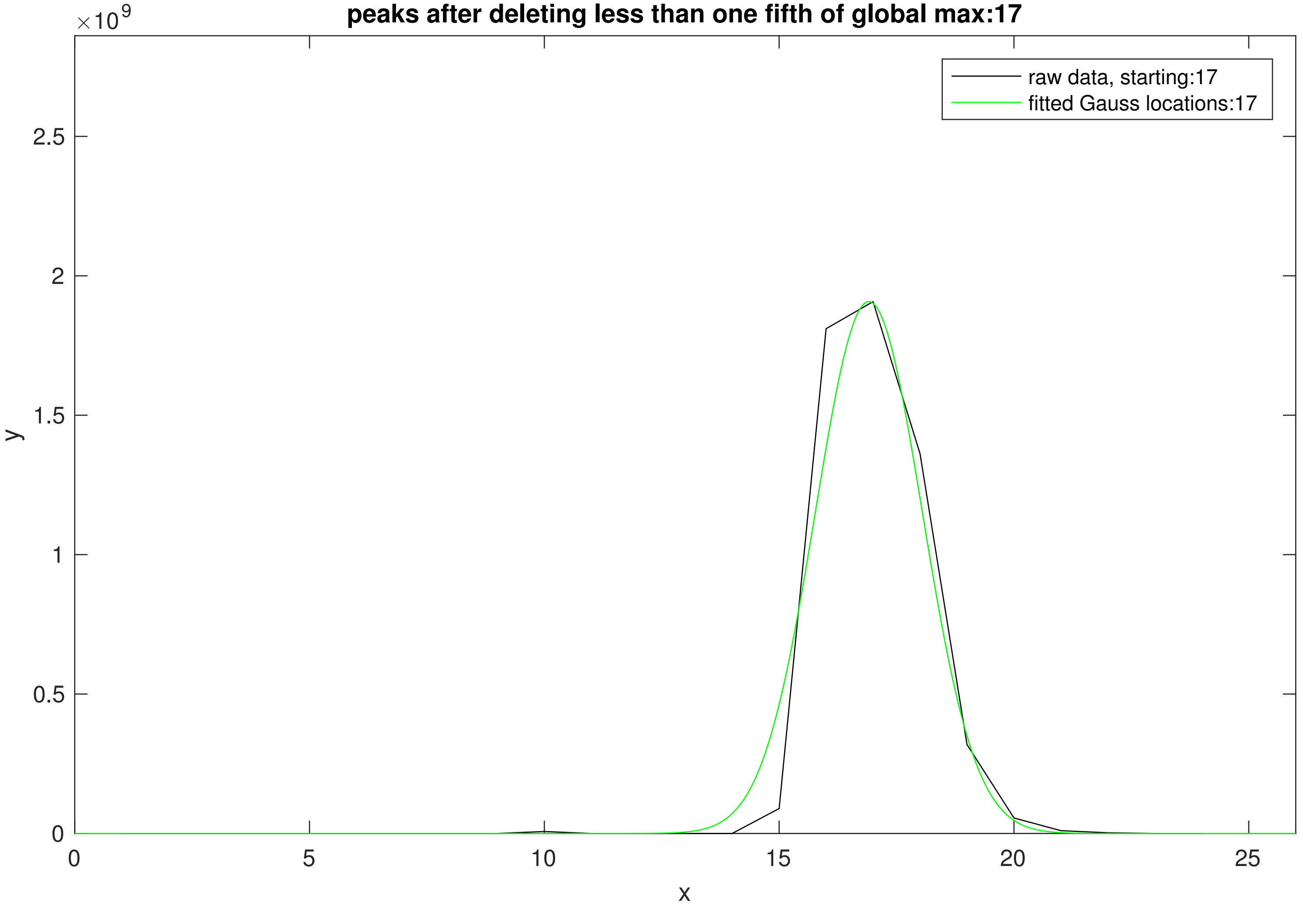

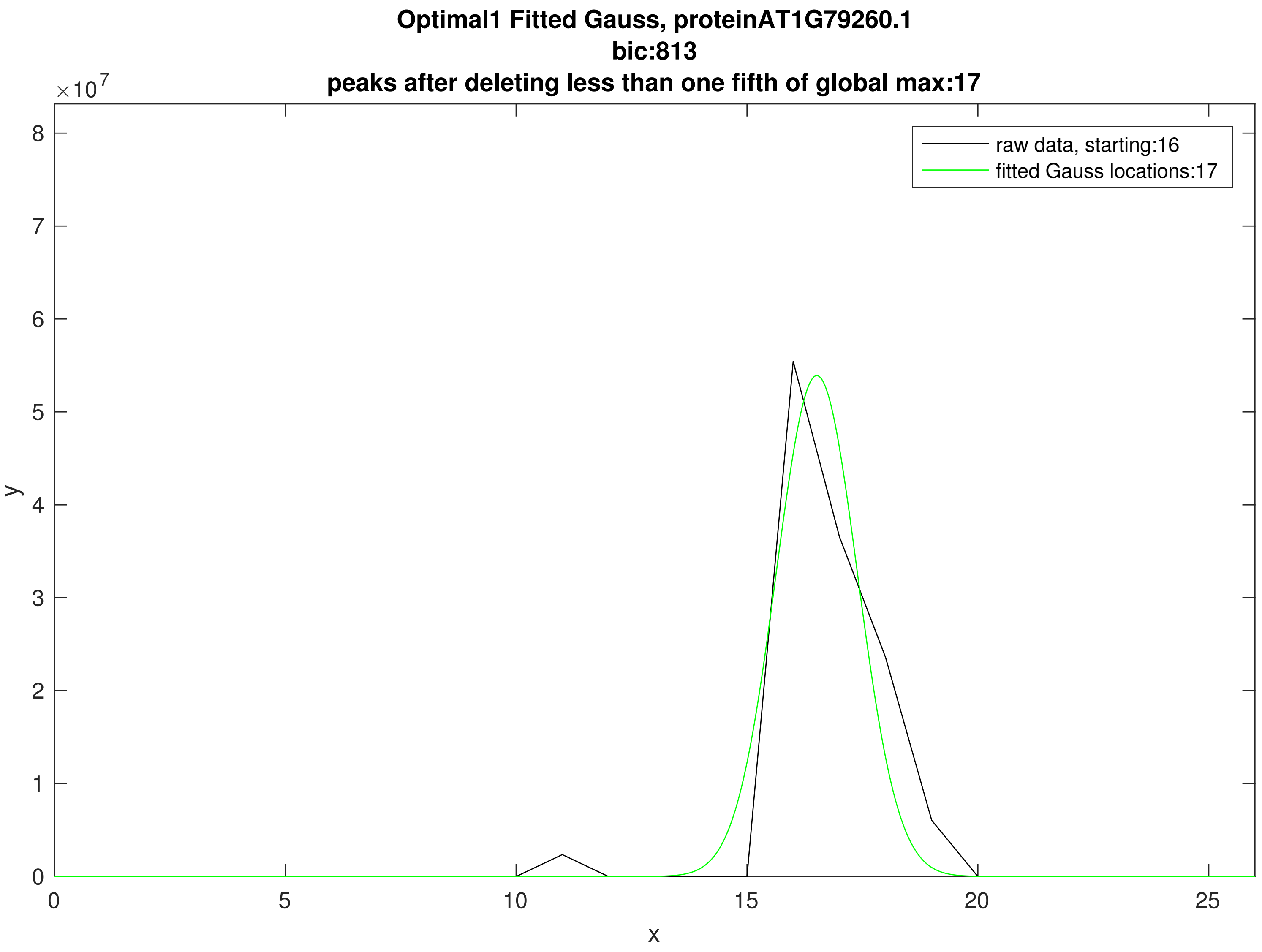

Optimal1 Fitted Gauss, proteinAT1G79260.1
bic:813
peaks after deleting less than one fifth of global max:17
×10^7
raw data, starting:16
fitted Gauss locations:17
y
x

**Optimal1 Fitted Gauss, proteinAT1G79340.1**
**bic:932 934 944**
**peaks after deleting less than one fifth of global max:15**

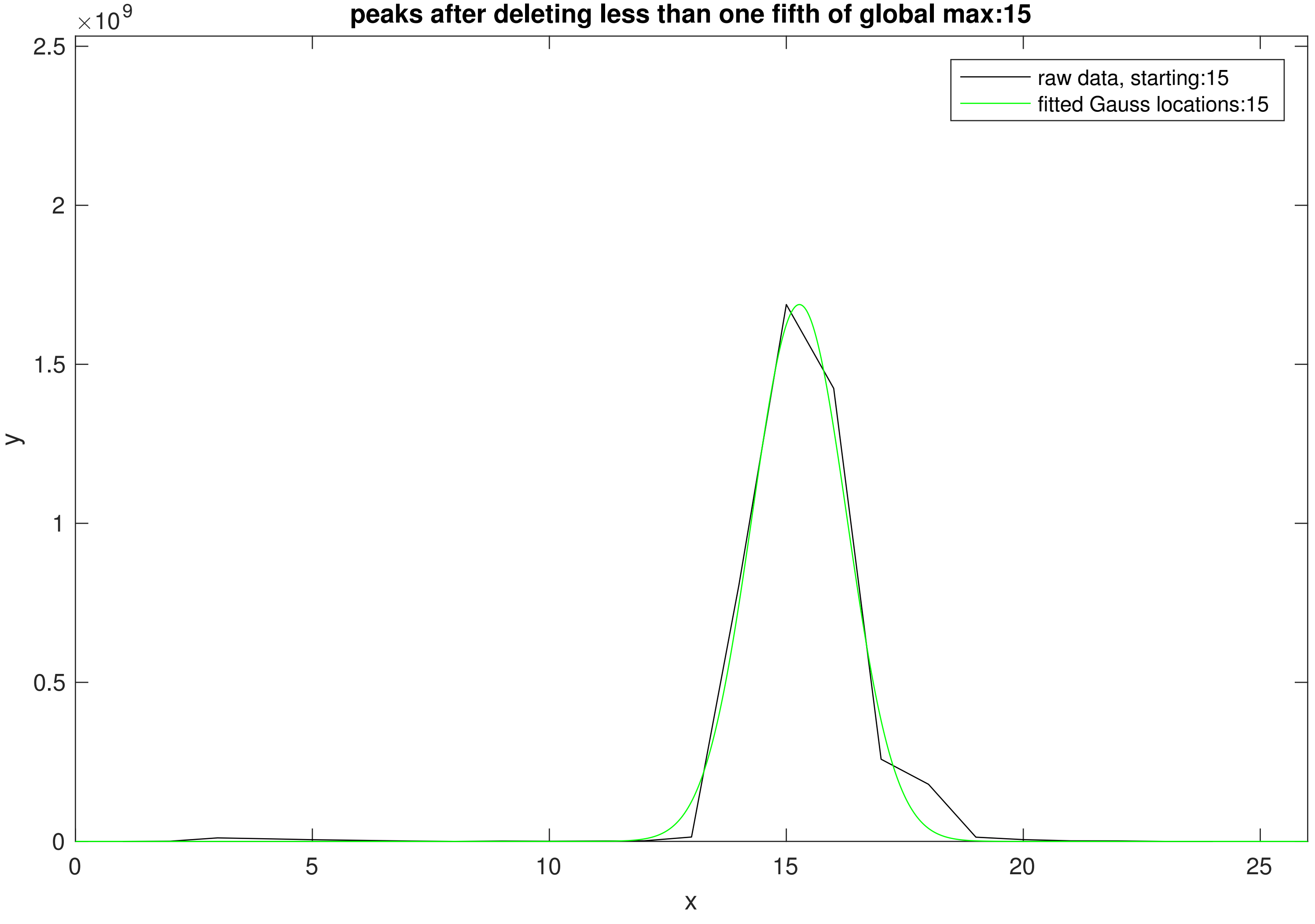

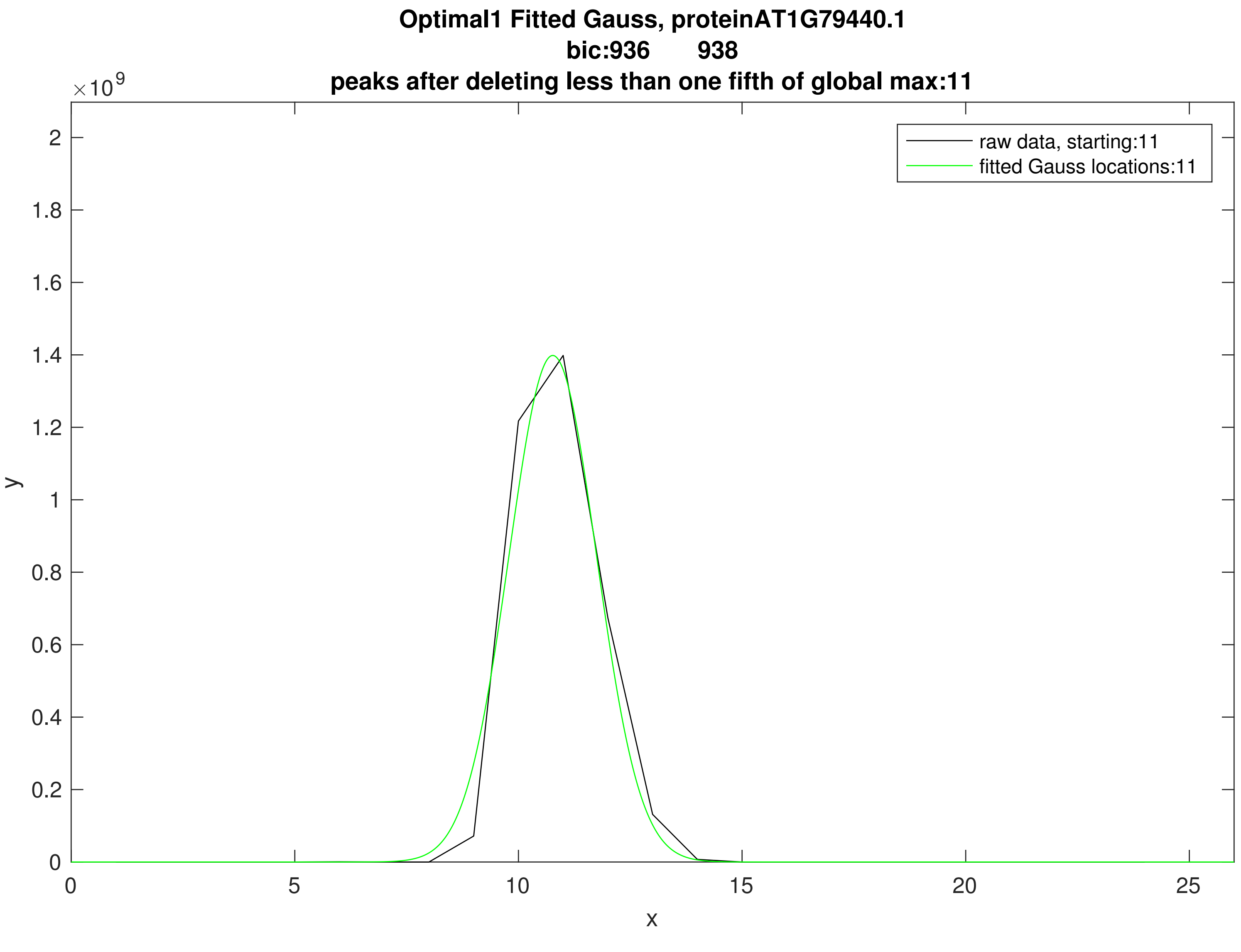

Optimal1 Fitted Gauss, proteinAT1G79440.1
bic:936 938
peaks after deleting less than one fifth of global max:11
×10^9
raw data, starting:11
fitted Gauss locations:11
y
x
0
0.2
0.4
0.6
0.8
1
1.2
1.4
1.6
1.8
2
5
10
15
20
25

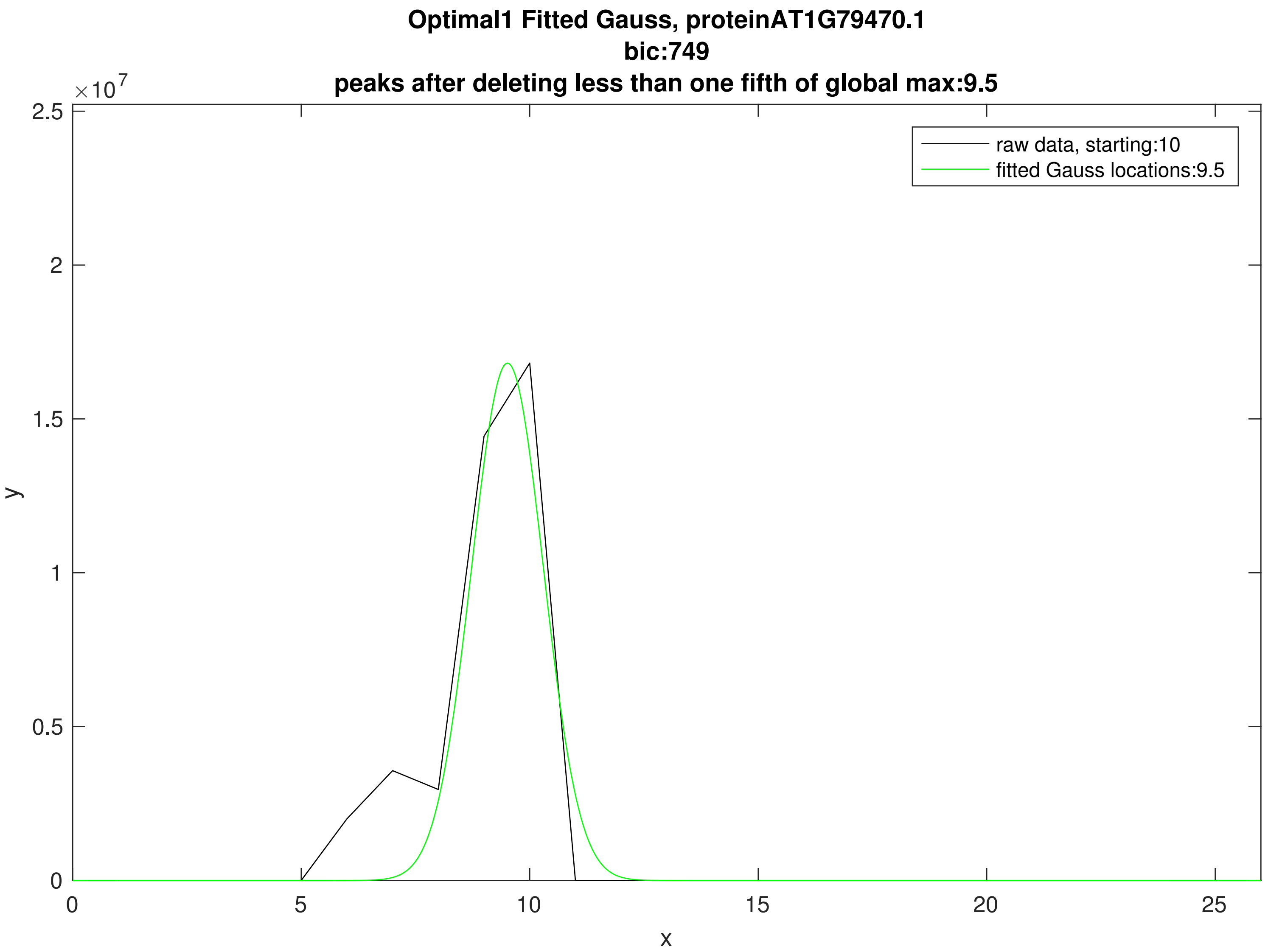
Optimal1 Fitted Gauss, proteinAT1G79470.1
bic:749
peaks after deleting less than one fifth of global max:9.5
×10^7
2.5
2
1.5
1
0.5
0
y
0
5
10
15
20
25
x
raw data, starting:10
fitted Gauss locations:9.5

#### Optimal1 Fitted Gauss, proteinAT1G79500.4;AT1G79500.3;AT1G79500.2;AT1G79500.1;AT1G79500.5
#### bic:931 940 949
#### peaks after deleting less than one fifth of global max:12

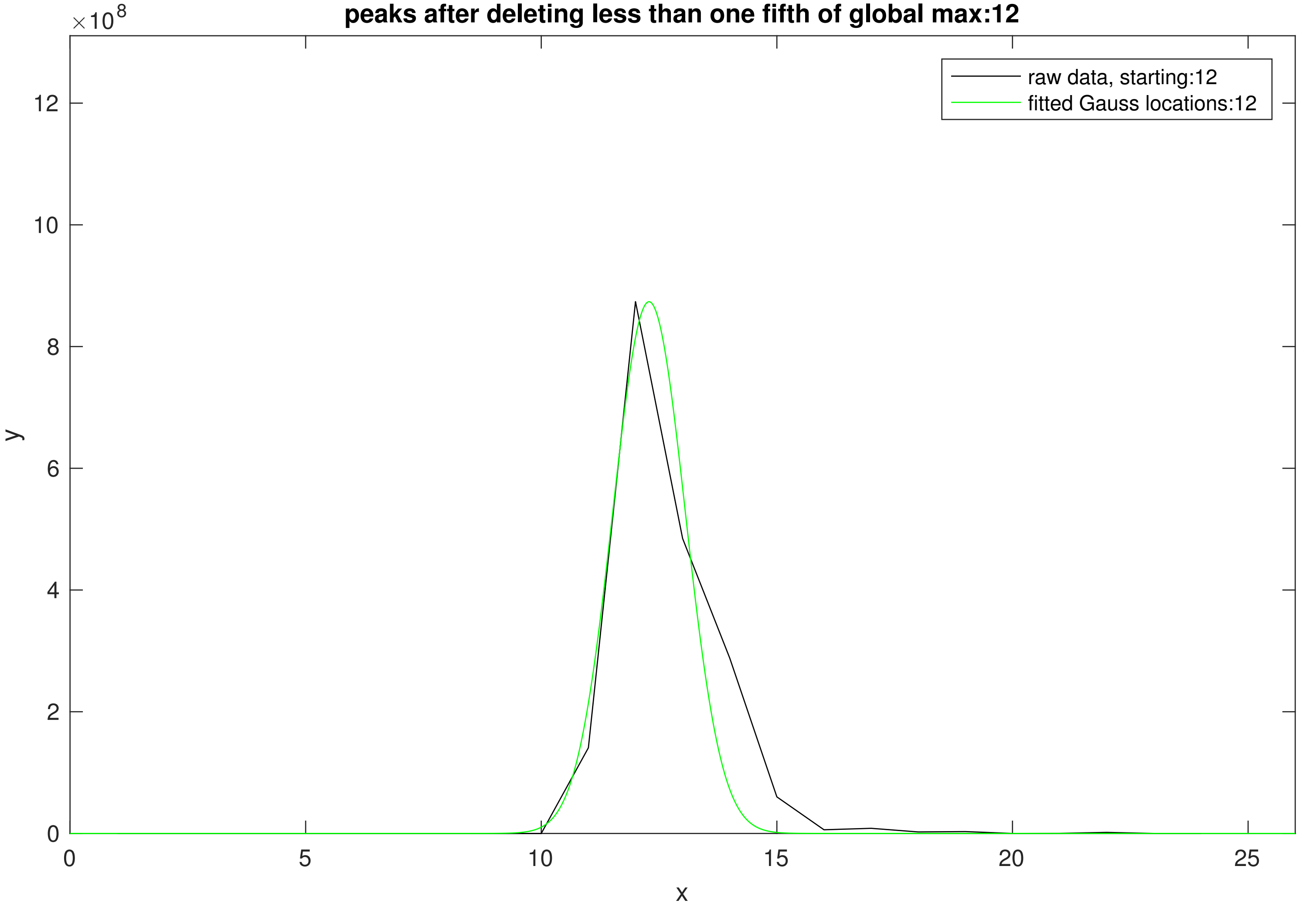

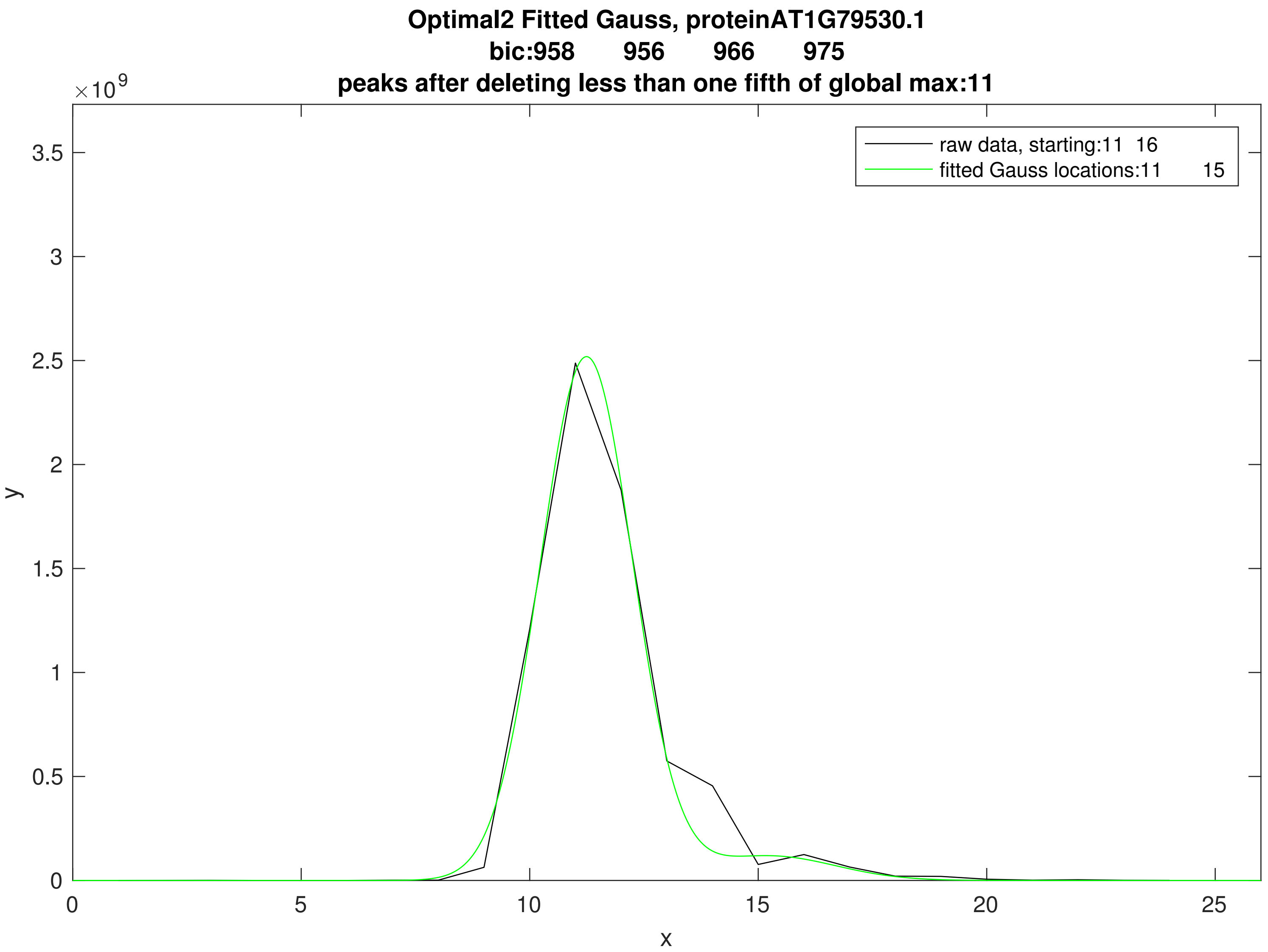

Optimal2 Fitted Gauss, proteinAT1G79530.1
bic:958 956 966 975
peaks after deleting less than one fifth of global max:11
×10^9
raw data, starting:11 16
fitted Gauss locations:11 15
y
x
0
0.5
1
1.5
2
2.5
3
3.5
0
5
10
15
20
25

#### Optimal1 Fitted Gauss, proteinAT1G79550.2;AT1G79550.1
## bic:1.07e+03  1.08e+03  1.09e+03   1.1e+03
#### peaks after deleting less than one fifth of global max:16

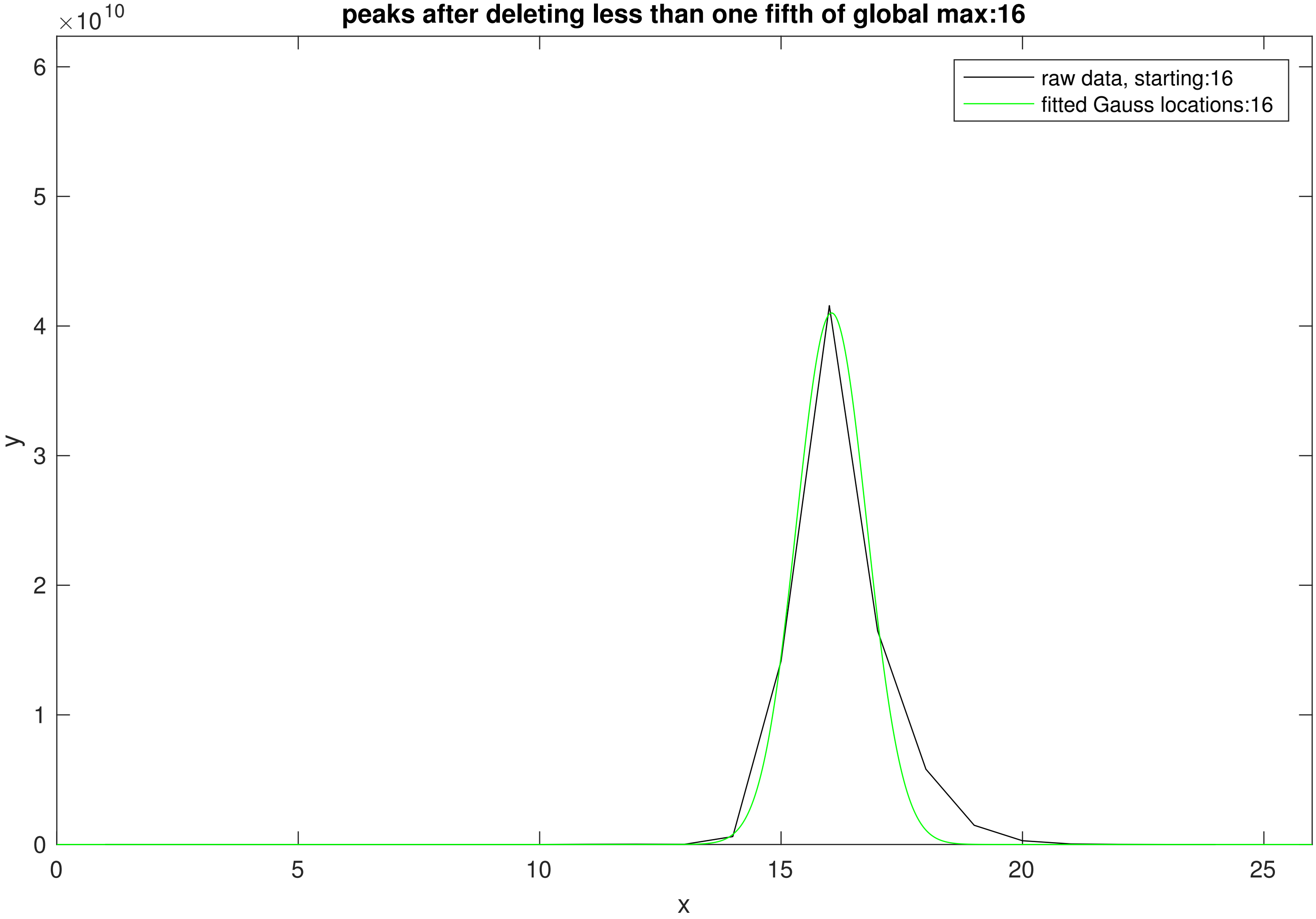

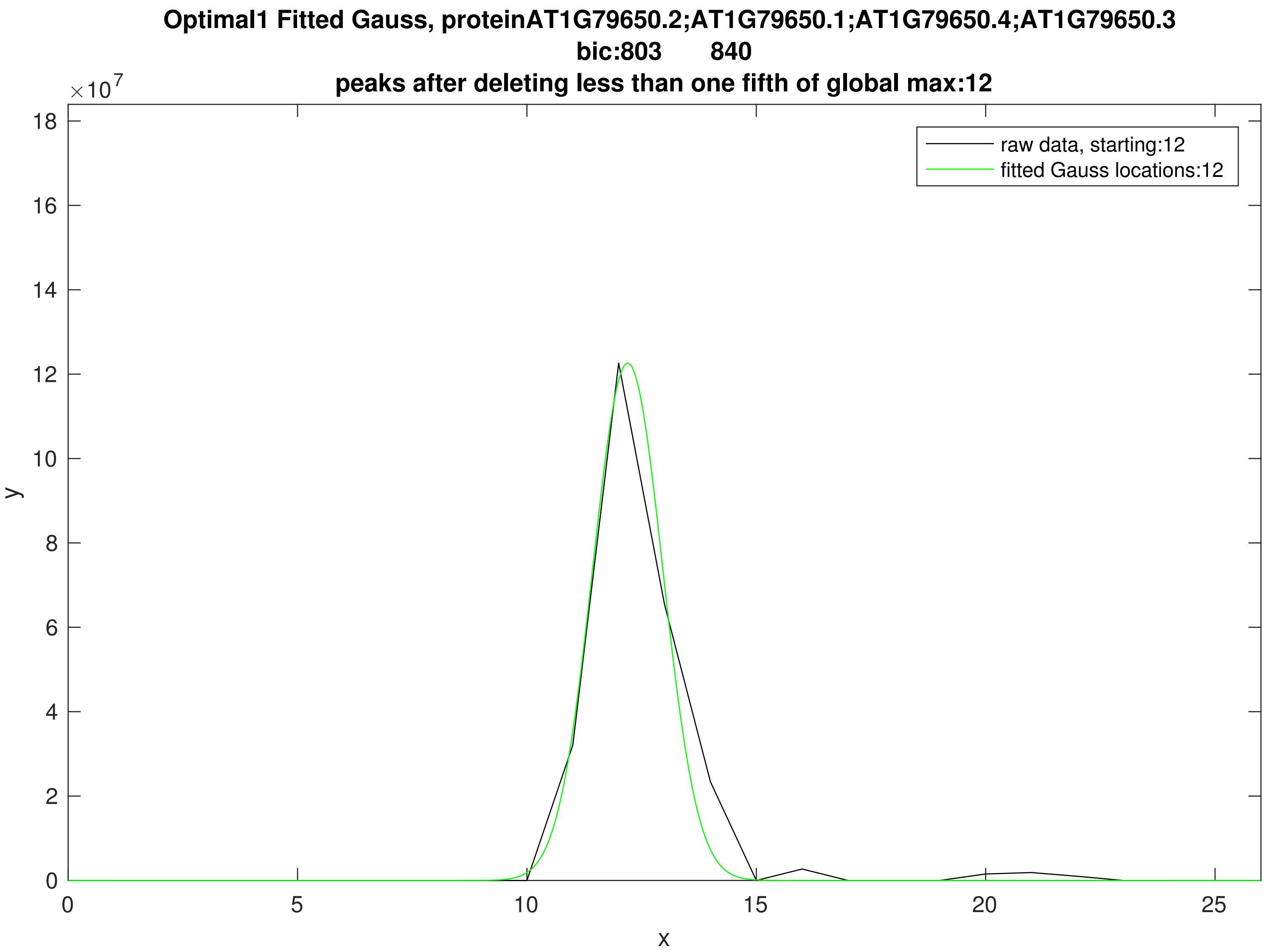
Optimal1 Fitted Gauss, proteinAT1G79650.2;AT1G79650.1;AT1G79650.4;AT1G79650.3
bic:803 840
peaks after deleting less than one fifth of global max:12
×10^7
raw data, starting:12
fitted Gauss locations:12
y
x
0
2
4
6
8
10
12
14
16
18
0
5
10
15
20
25

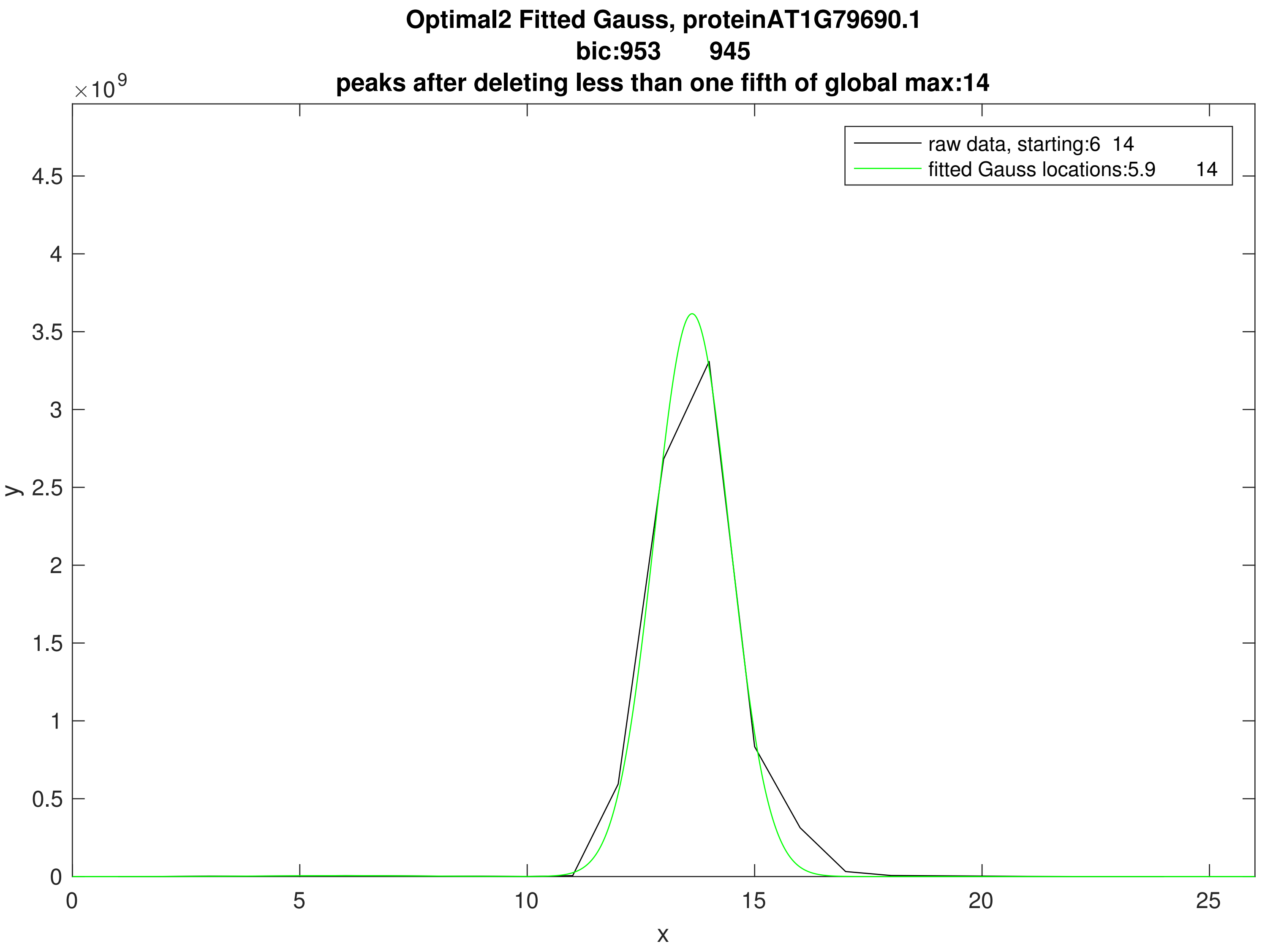

Optimal2 Fitted Gauss, proteinAT1G79690.1
bic:953 945
peaks after deleting less than one fifth of global max:14
×10^9
raw data, starting:6 14
fitted Gauss locations:5.9 14
4.5
4
3.5
3
2.5
2
1.5
1
0.5
0
y
0
5
10
15
20
25
x

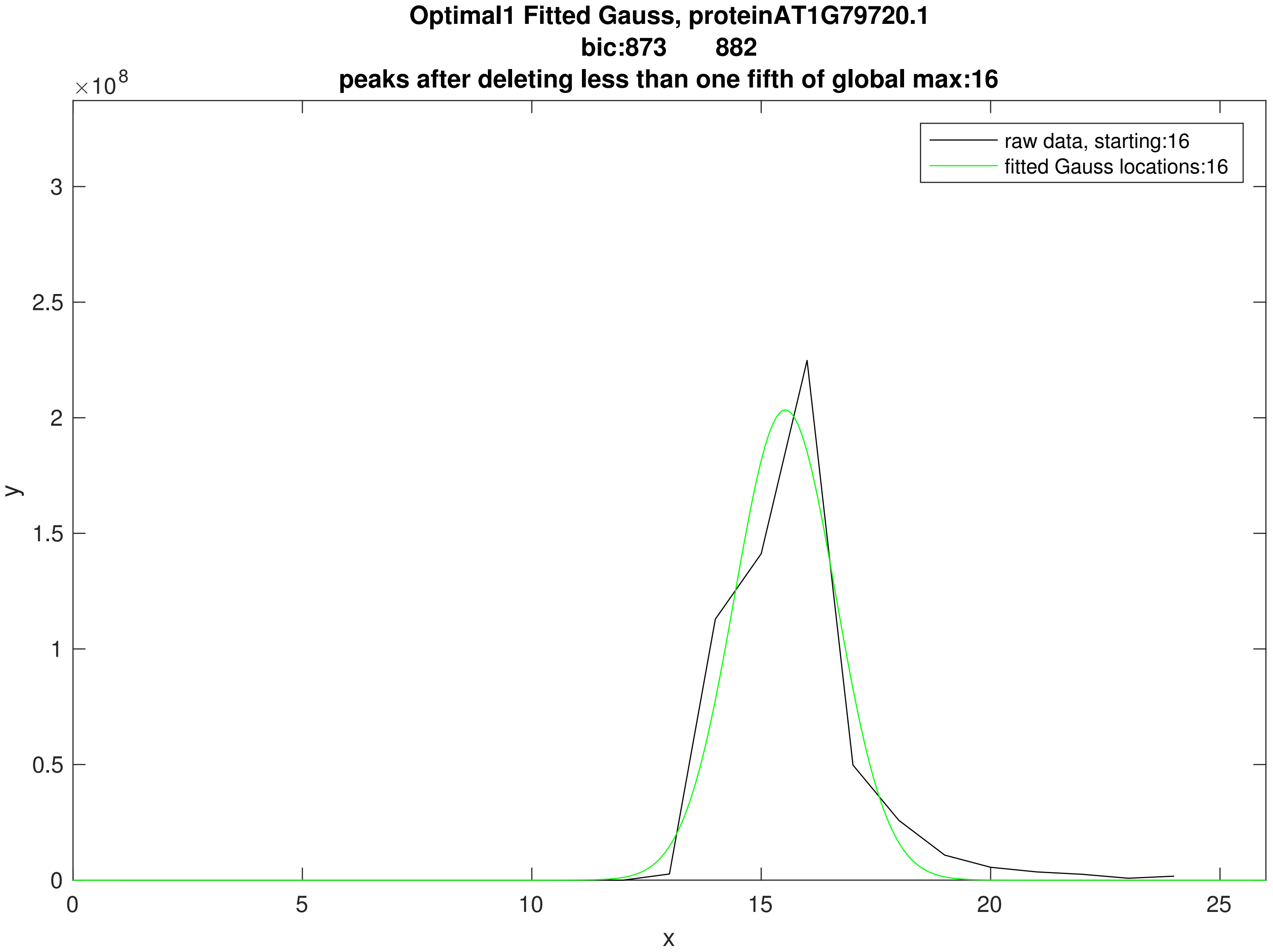
Optimal1 Fitted Gauss, proteinAT1G79720.1
bic:873 882
peaks after deleting less than one fifth of global max:16
×10^8
raw data, starting:16
fitted Gauss locations:16
y
x
0
0.5
1
1.5
2
2.5
3
5
10
15
20
25

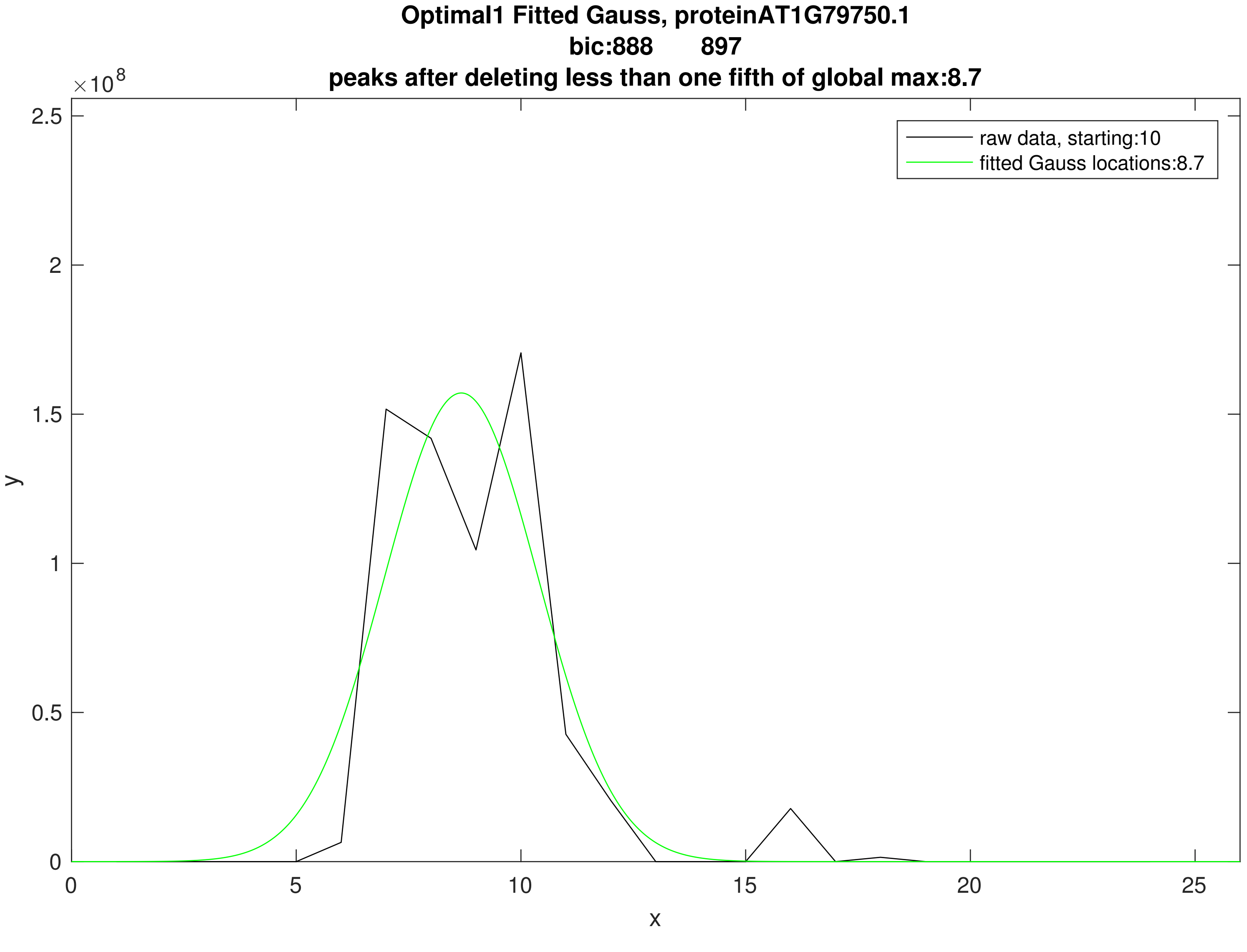
Optimal1 Fitted Gauss, proteinAT1G79750.1
bic:888 897
peaks after deleting less than one fifth of global max:8.7
×10^8
2.5
2
1.5
1
0.5
0
y
0
5
10
15
20
25
x
raw data, starting:10
fitted Gauss locations:8.7

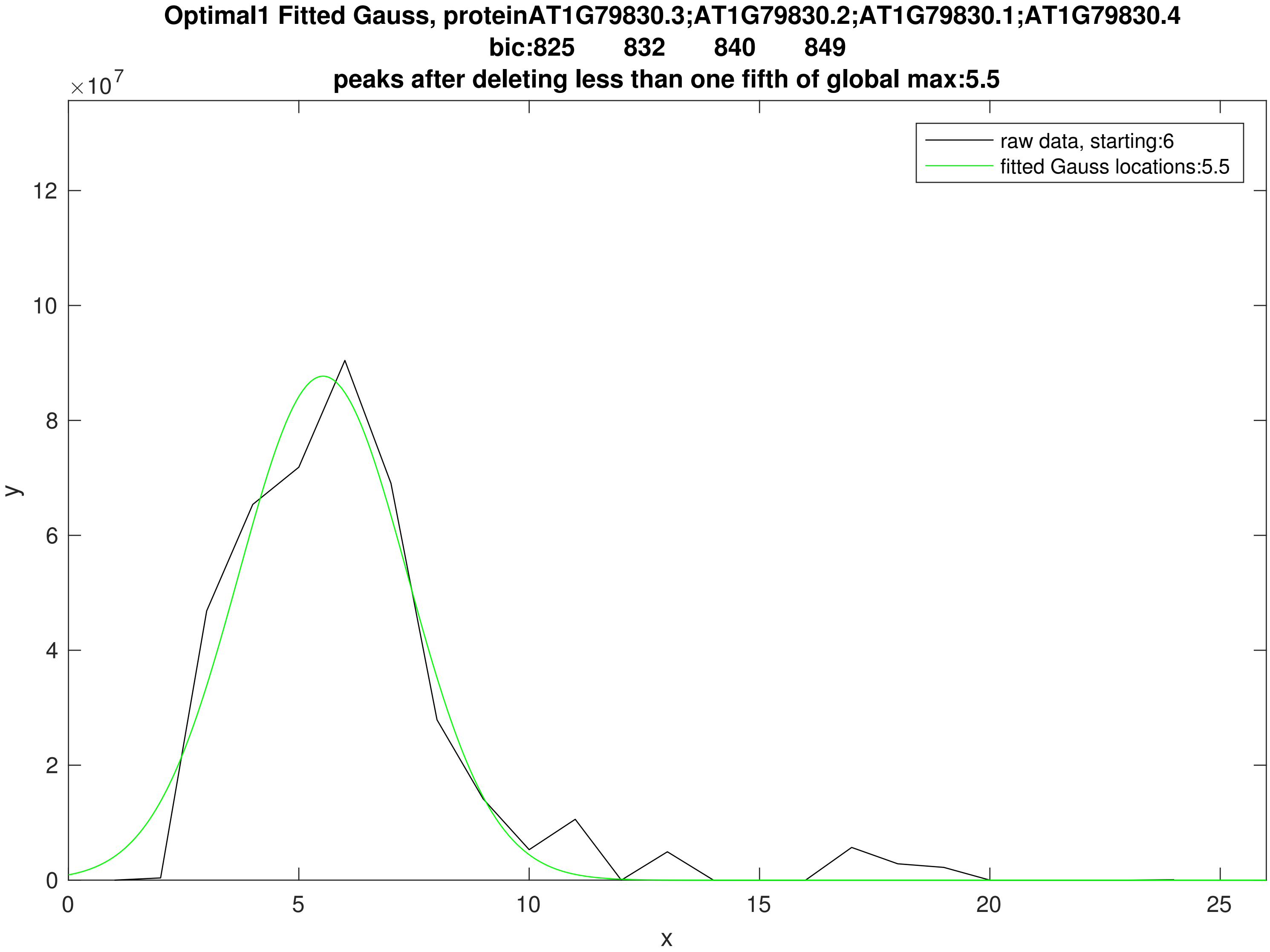

Optimal1 Fitted Gauss, proteinAT1G79830.3;AT1G79830.2;AT1G79830.1;AT1G79830.4
bic:825 832 840 849
peaks after deleting less than one fifth of global max:5.5
×10^7
raw data, starting:6
fitted Gauss locations:5.5
12
10
8
6
4
2
0
y
0
5
10
15
20
25
x

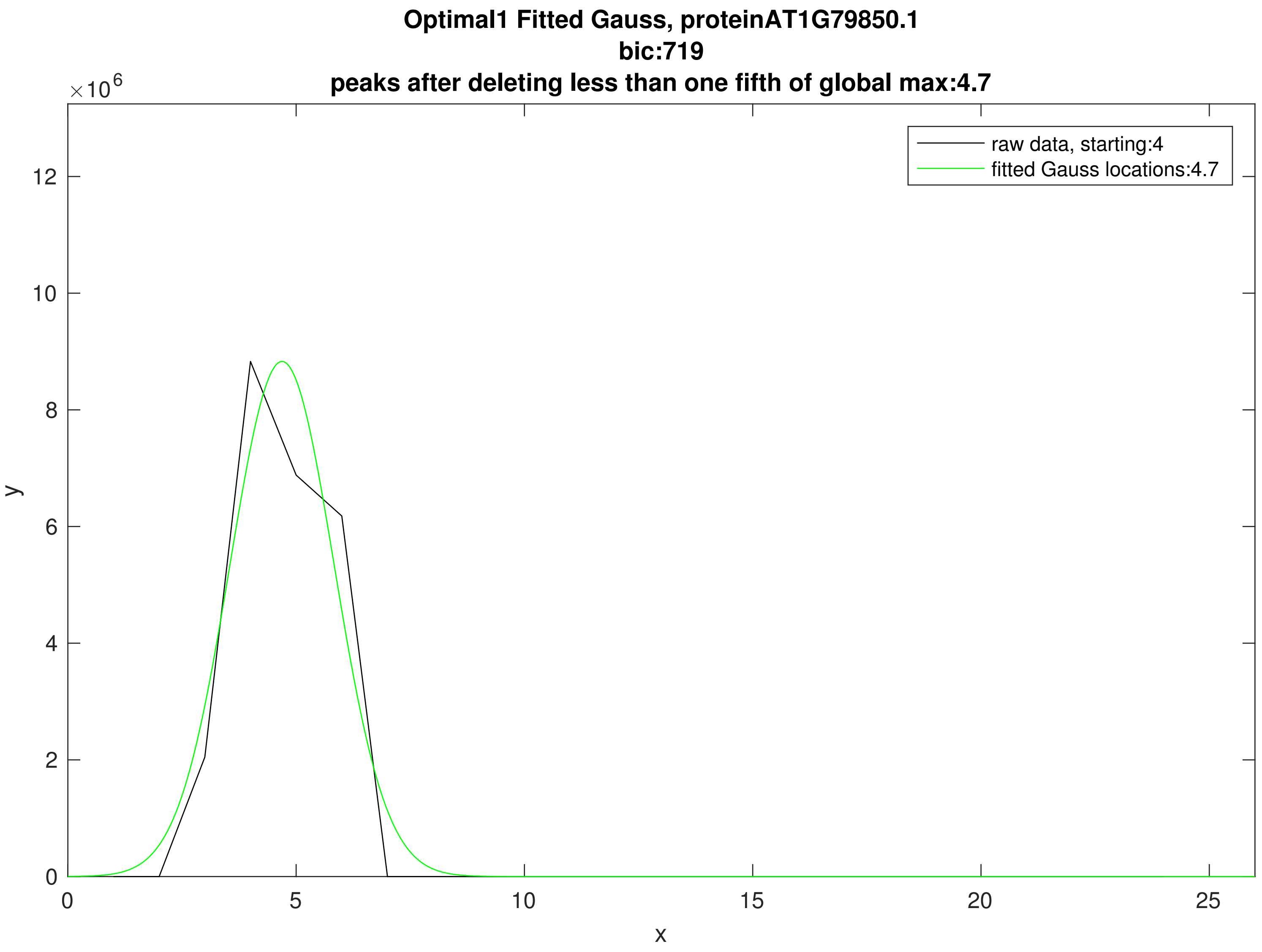

Optimal1 Fitted Gauss, proteinAT1G79850.1
bic:719
peaks after deleting less than one fifth of global max:4.7
×10^6
raw data, starting:4
fitted Gauss locations:4.7
y
x
0
2
4
6
8
10
12
0
5
10
15
20
25

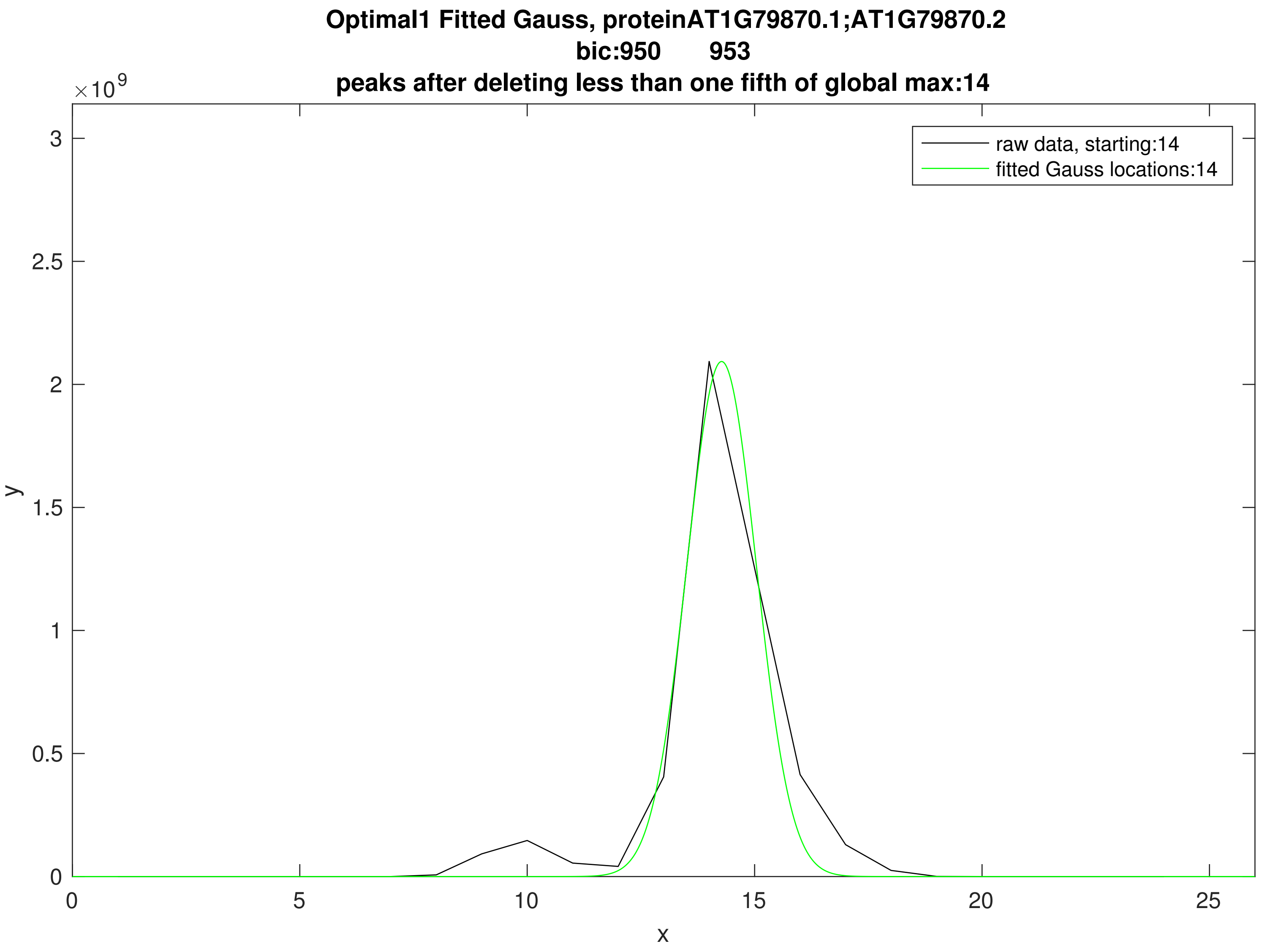
Optimal1 Fitted Gauss, proteinAT1G79870.1;AT1G79870.2
bic:950 953
peaks after deleting less than one fifth of global max:14
×10^9
3
2.5
2
1.5
1
0.5
0
y
0
5
10
15
20
25
x
raw data, starting:14
fitted Gauss locations:14

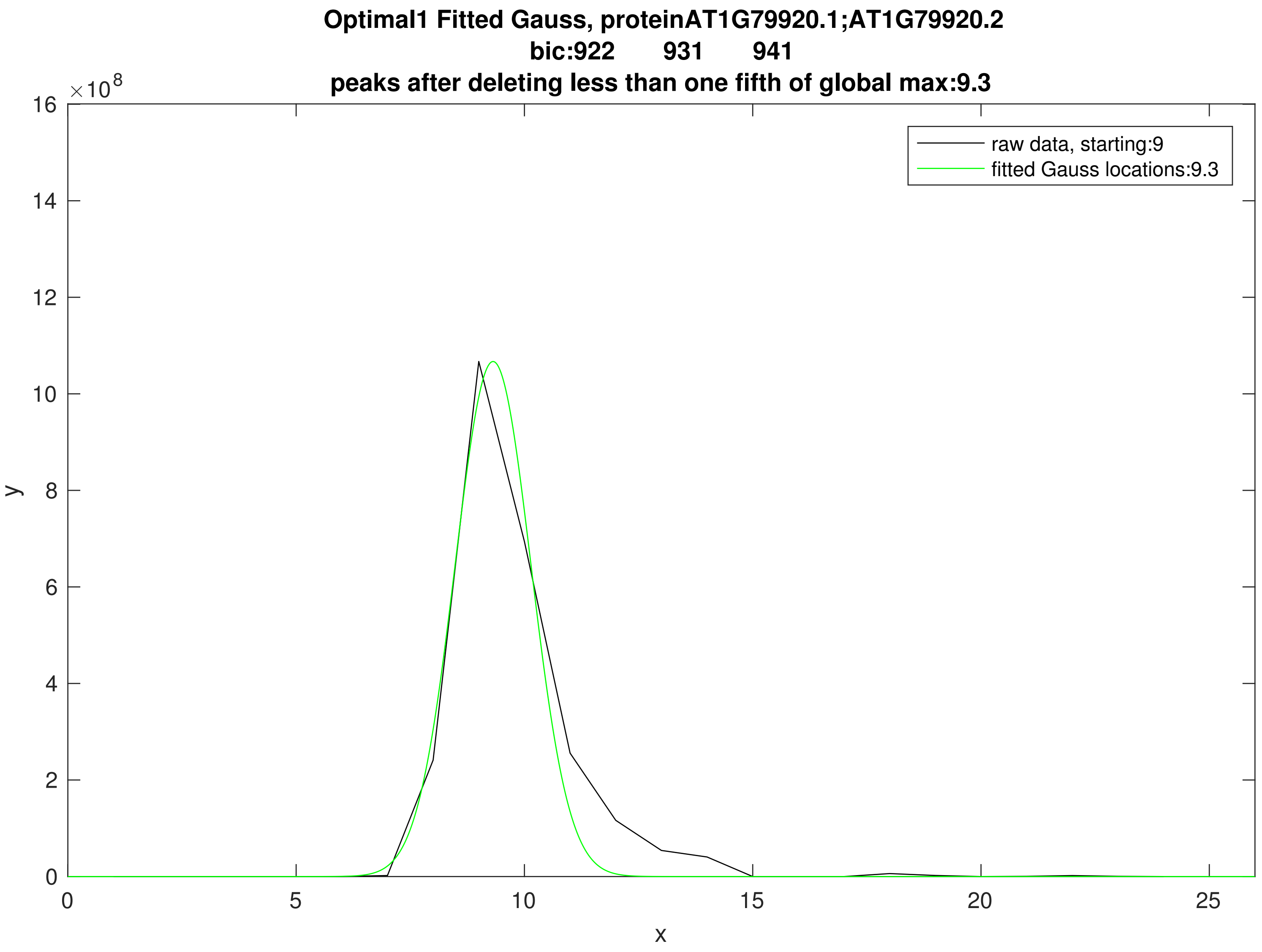
Optimal1 Fitted Gauss, proteinAT1G79920.1;AT1G79920.2
bic:922 931 941
peaks after deleting less than one fifth of global max:9.3
×10^8
16
14
12
10
8
6
4
2
0
y
0
5
10
15
20
25
x
raw data, starting:9
fitted Gauss locations:9.3

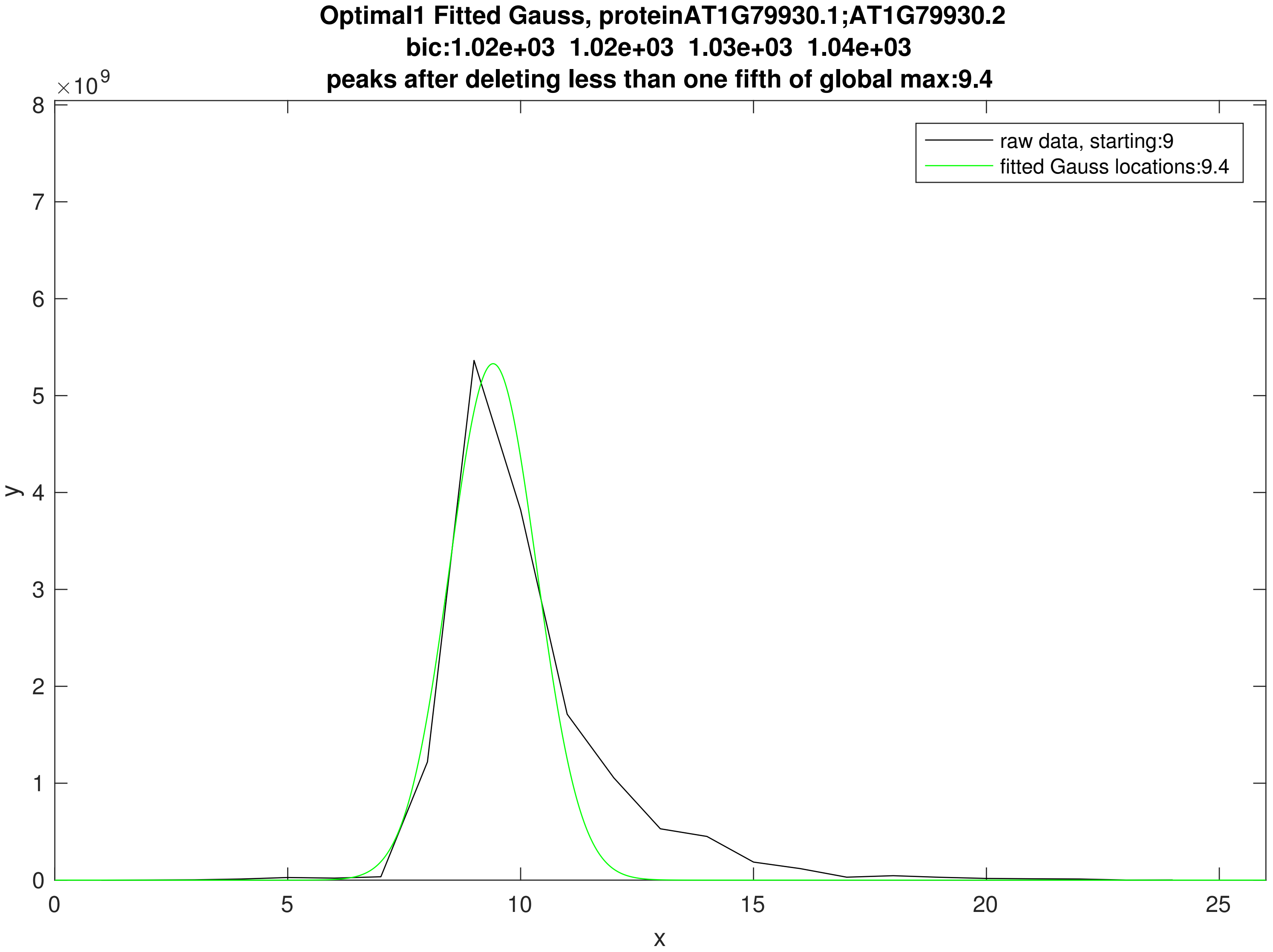
Optimal1 Fitted Gauss, proteinAT1G79930.1;AT1G79930.2
bic:1.02e+03 1.02e+03 1.03e+03 1.04e+03
peaks after deleting less than one fifth of global max:9.4
×10^9
raw data, starting:9
fitted Gauss locations:9.4
y
x
0
1
2
3
4
5
6
7
8
0
5
10
15
20
25

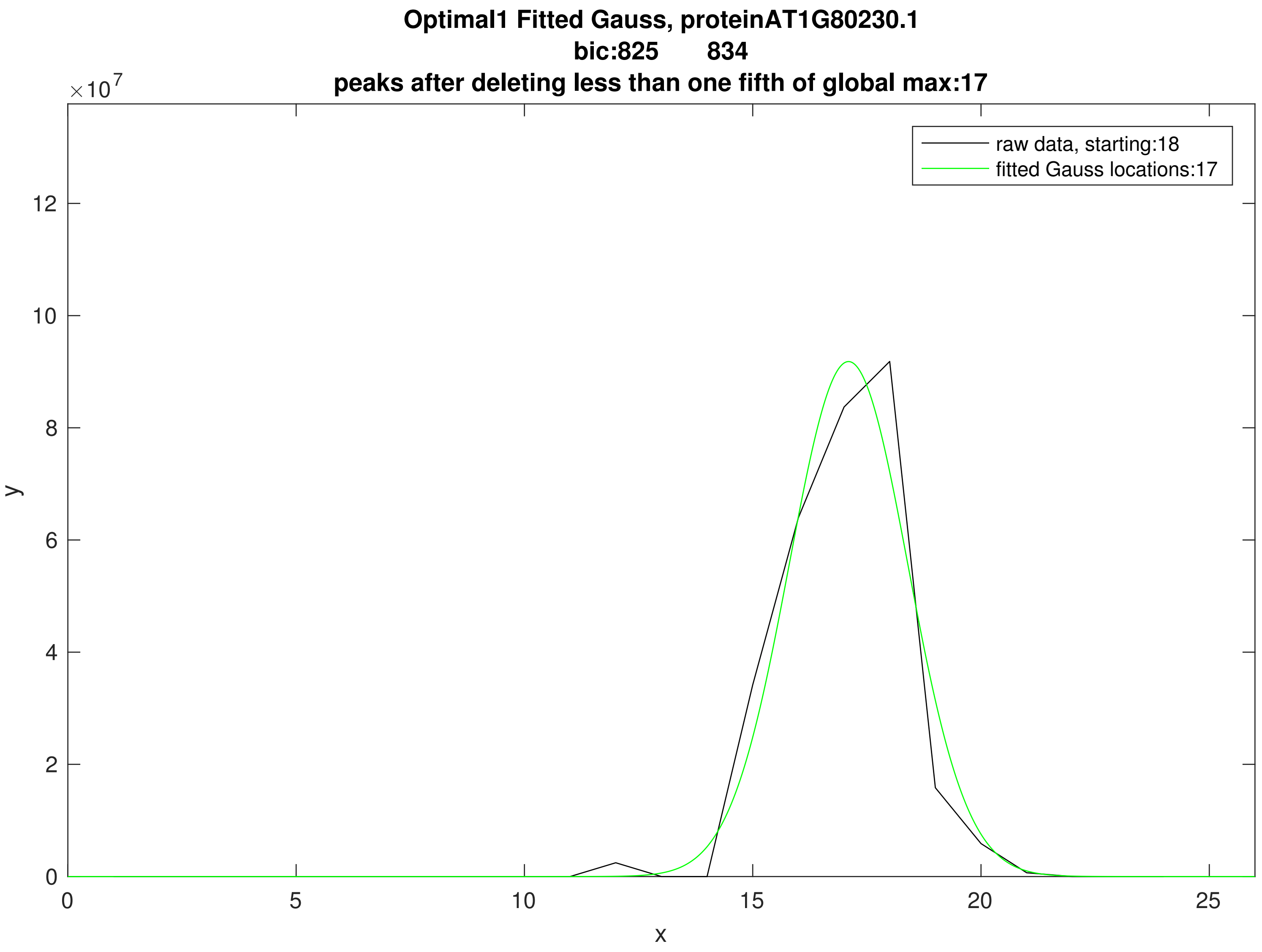

Optimal1 Fitted Gauss, proteinAT1G80230.1
bic:825 834
peaks after deleting less than one fifth of global max:17
×10^7
raw data, starting:18
fitted Gauss locations:17
y
x
0
2
4
6
8
10
12
0
5
10
15
20
25

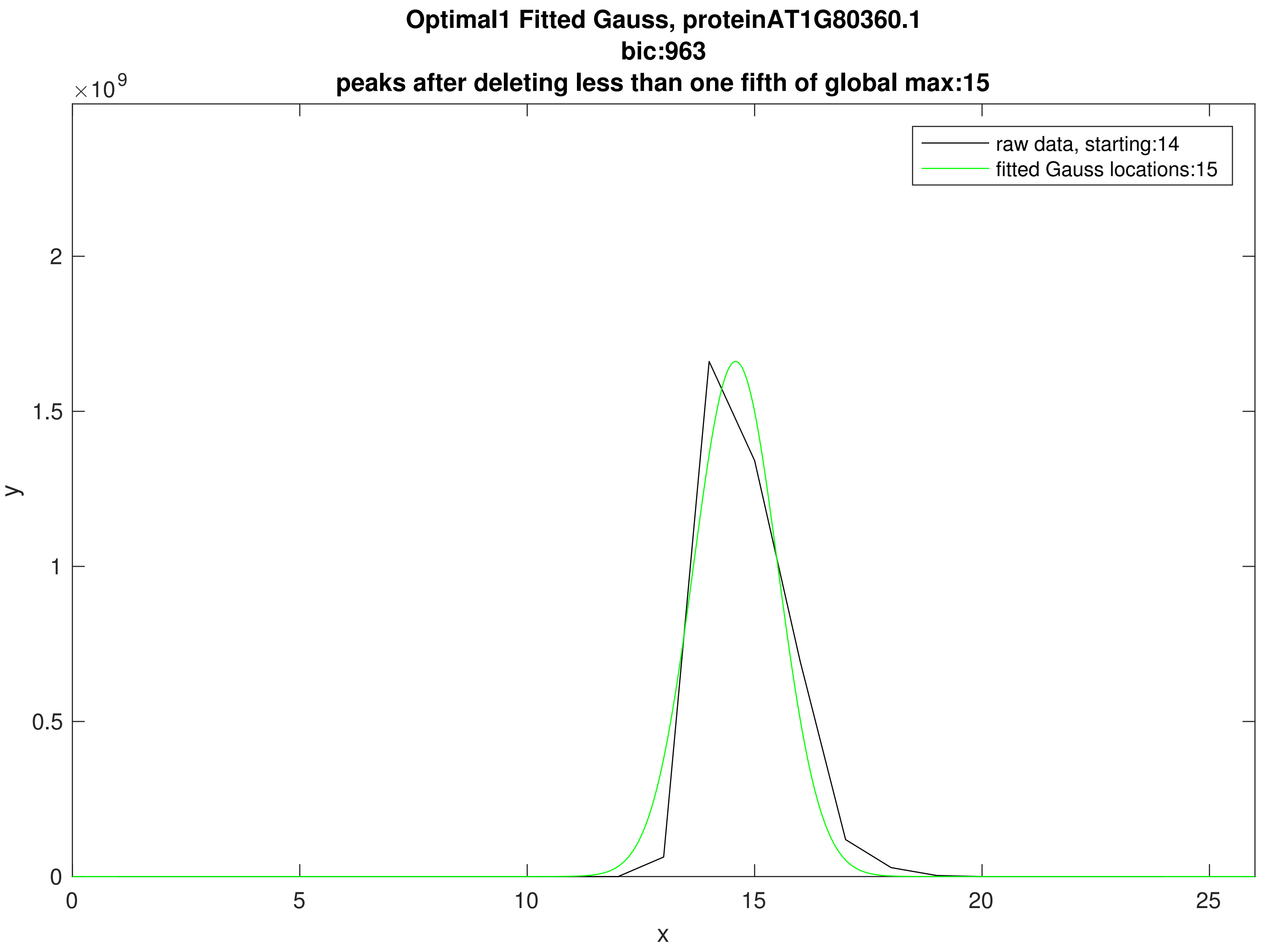

Optimal1 Fitted Gauss, proteinAT1G80360.1
bic:963
peaks after deleting less than one fifth of global max:15
×10^9
raw data, starting:14
fitted Gauss locations:15
y
x
0
0.5
1
1.5
2
5
10
15
20
25

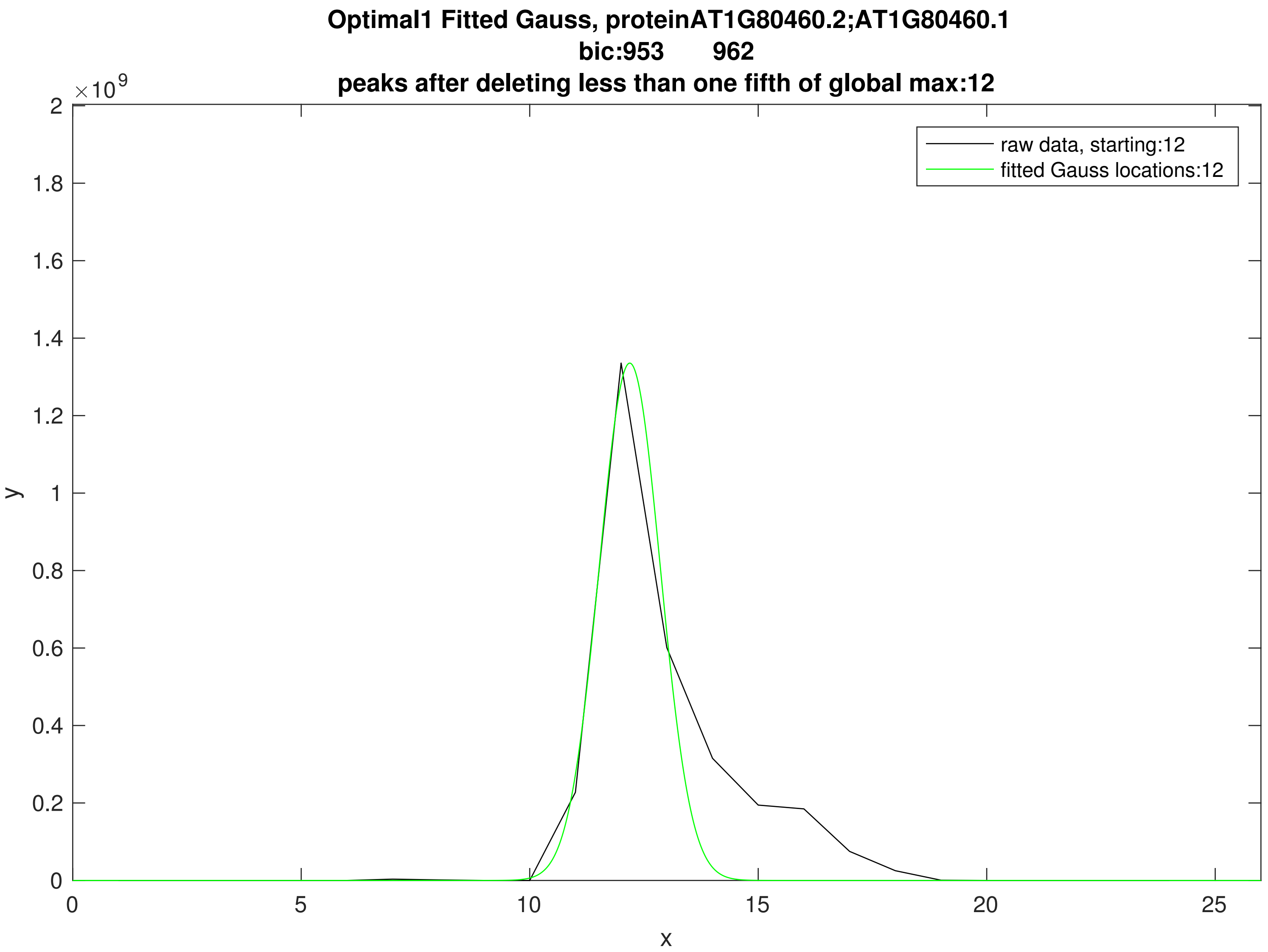
Optimal1 Fitted Gauss, proteinAT1G80460.2;AT1G80460.1
bic:953 962
peaks after deleting less than one fifth of global max:12
×10^9
raw data, starting:12
fitted Gauss locations:12
y
x
0
0.2
0.4
0.6
0.8
1
1.2
1.4
1.6
1.8
2
0
5
10
15
20
25

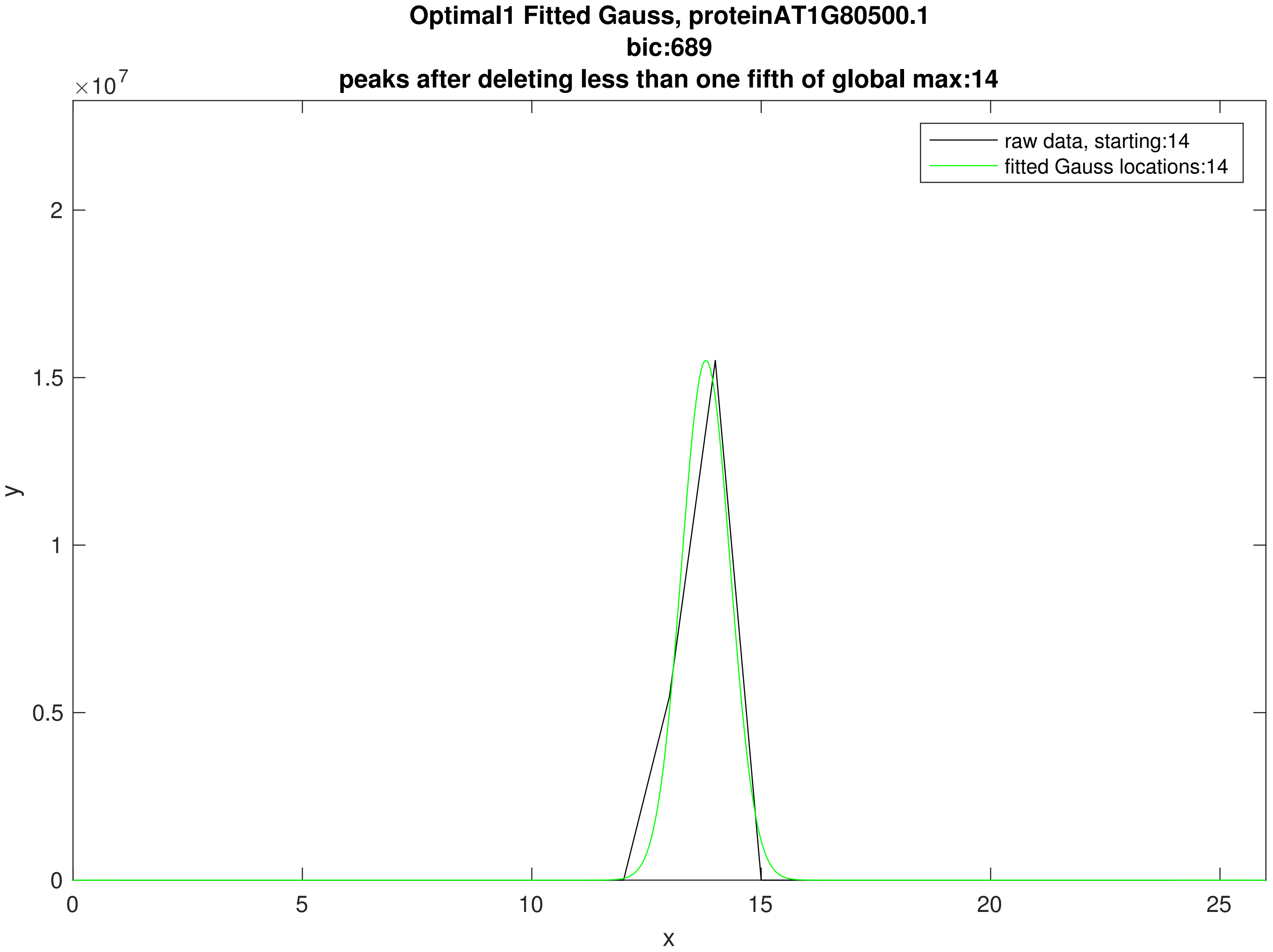
Optimal1 Fitted Gauss, proteinAT1G80500.1
bic:689
peaks after deleting less than one fifth of global max:14
×10^7
raw data, starting:14
fitted Gauss locations:14
y
x
0
0.5
1
1.5
2
0
5
10
15
20
25

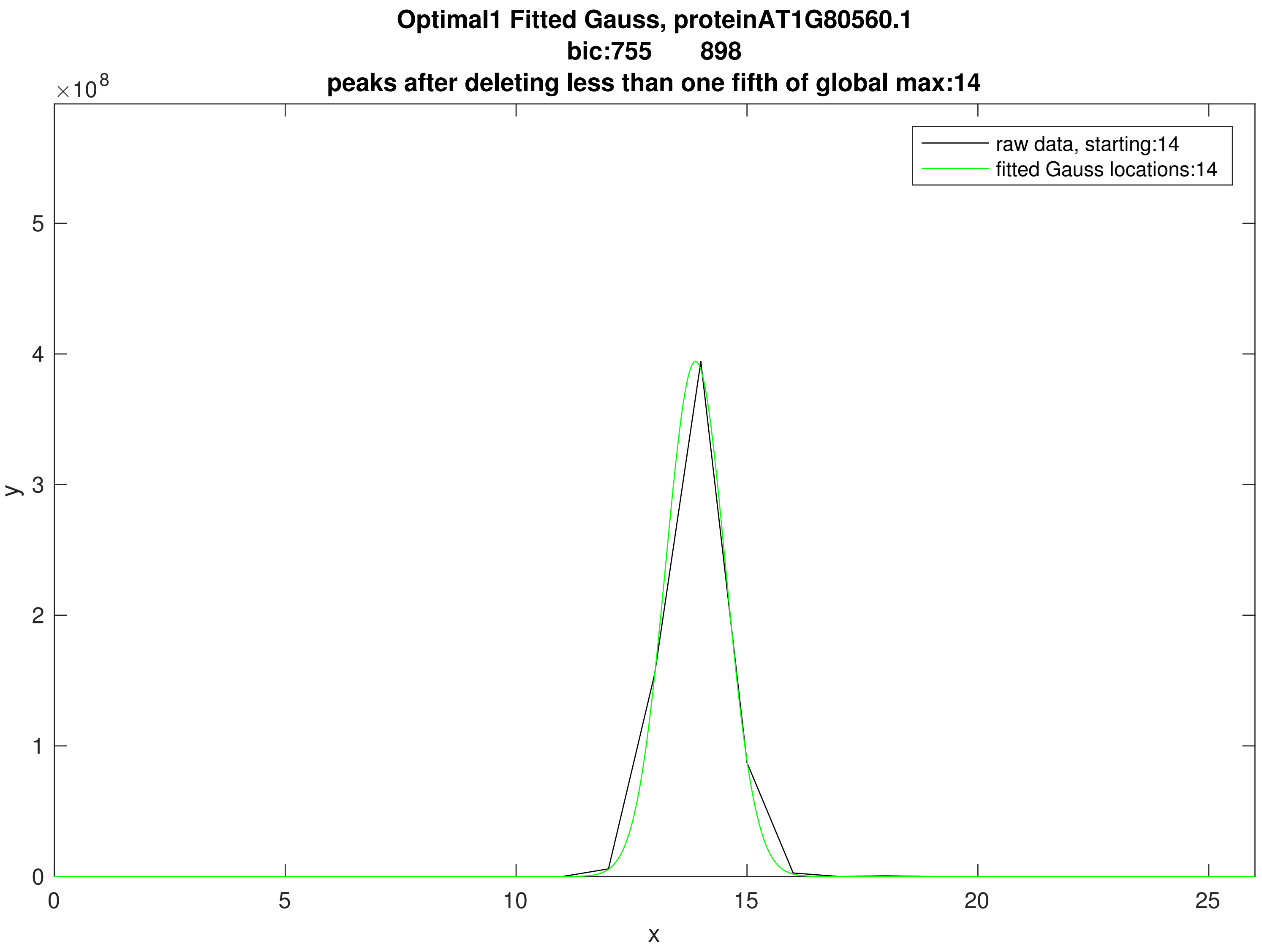

Optimal1 Fitted Gauss, proteinAT1G80560.1
bic:755 898
peaks after deleting less than one fifth of global max:14
×10^8
raw data, starting:14
fitted Gauss locations:14
y
x
0
1
2
3
4
5
0
5
10
15
20
25

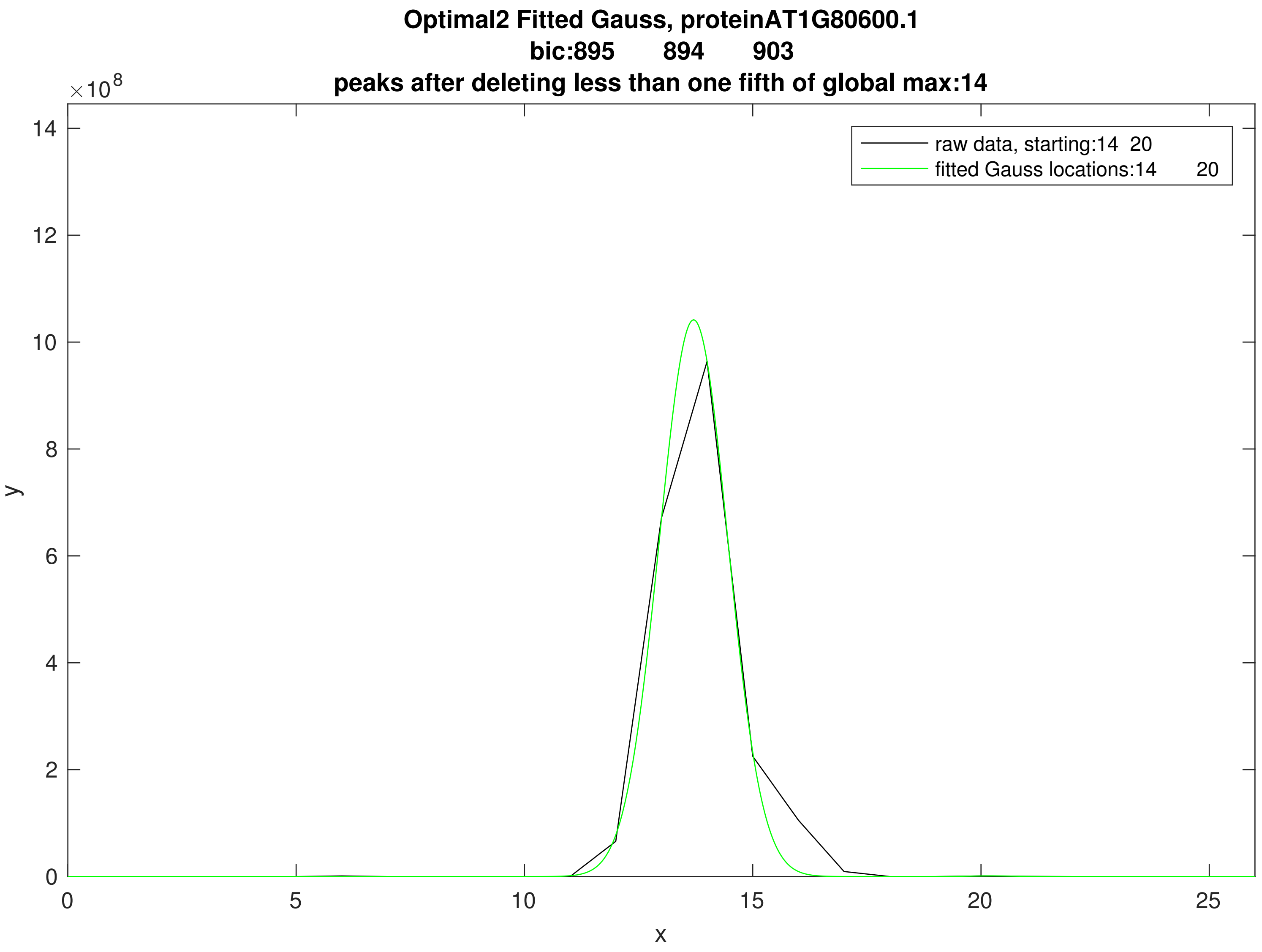
Optimal2 Fitted Gauss, proteinAT1G80600.1
bic:895 894 903
peaks after deleting less than one fifth of global max:14
×10^8
14
12
10
8
6
4
2
0
y
0
5
10
15
20
25
x
raw data, starting:14 20
fitted Gauss locations:14 20

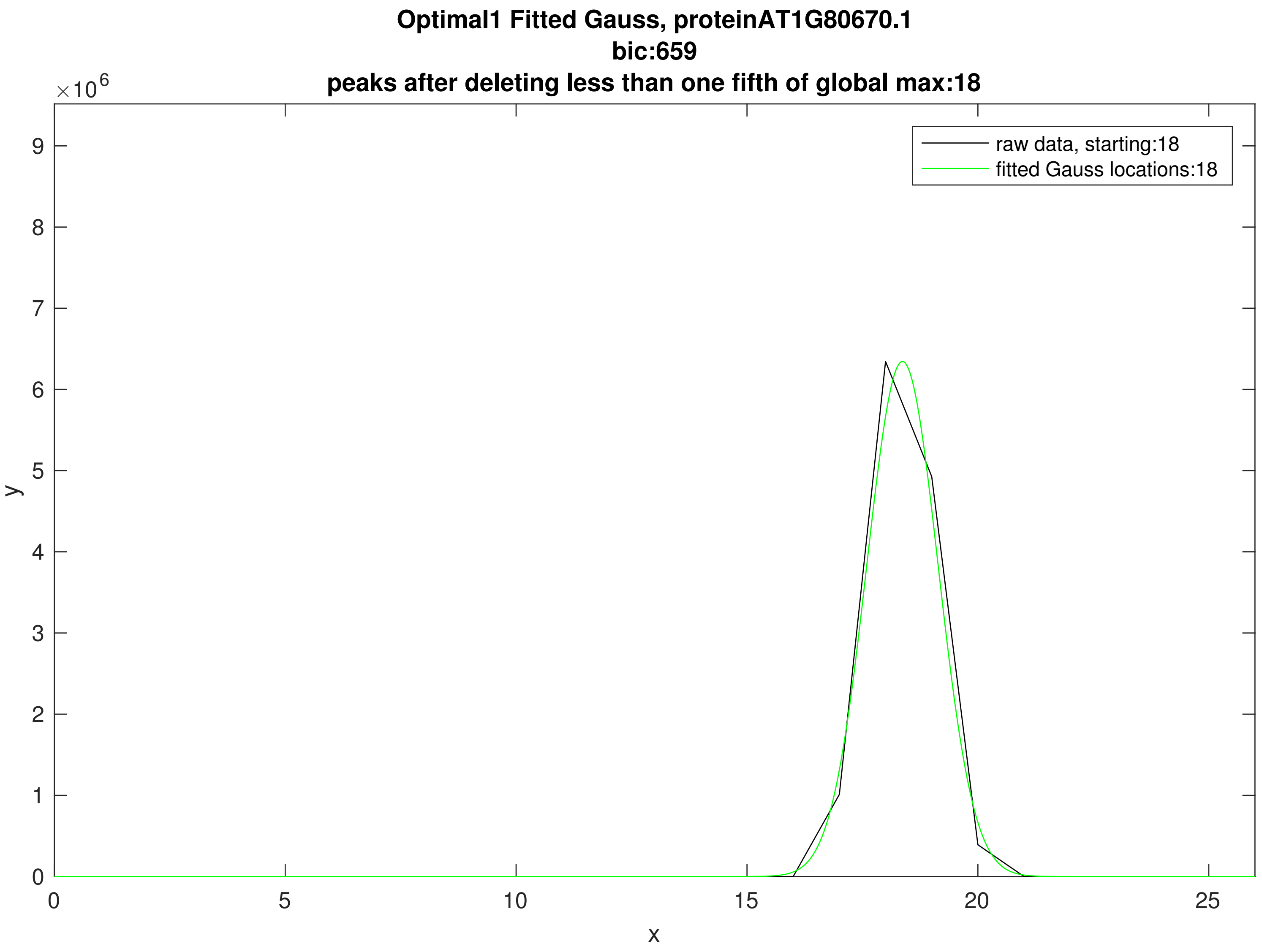
Optimal1 Fitted Gauss, proteinAT1G80670.1
bic:659
peaks after deleting less than one fifth of global max:18
×10^6
raw data, starting:18
fitted Gauss locations:18
y
x
0
1
2
3
4
5
6
7
8
9
0
5
10
15
20
25

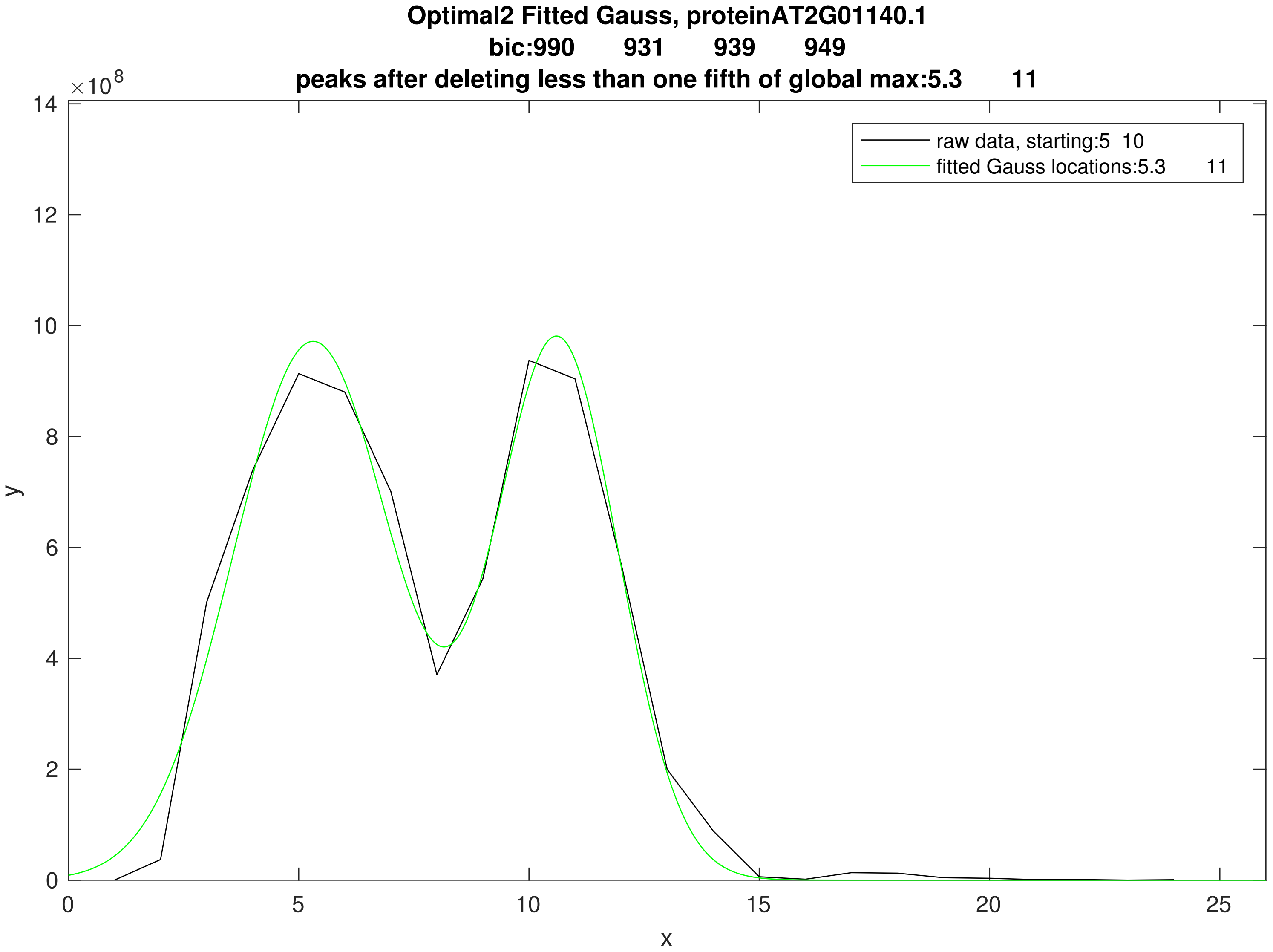

Optimal2 Fitted Gauss, proteinAT2G01140.1
bic:990 931 939 949
peaks after deleting less than one fifth of global max:5.3 11
×10^8
14
12
10
8
6
4
2
0
y
0
5
10
15
20
25
x
raw data, starting:5 10
fitted Gauss locations:5.3 11

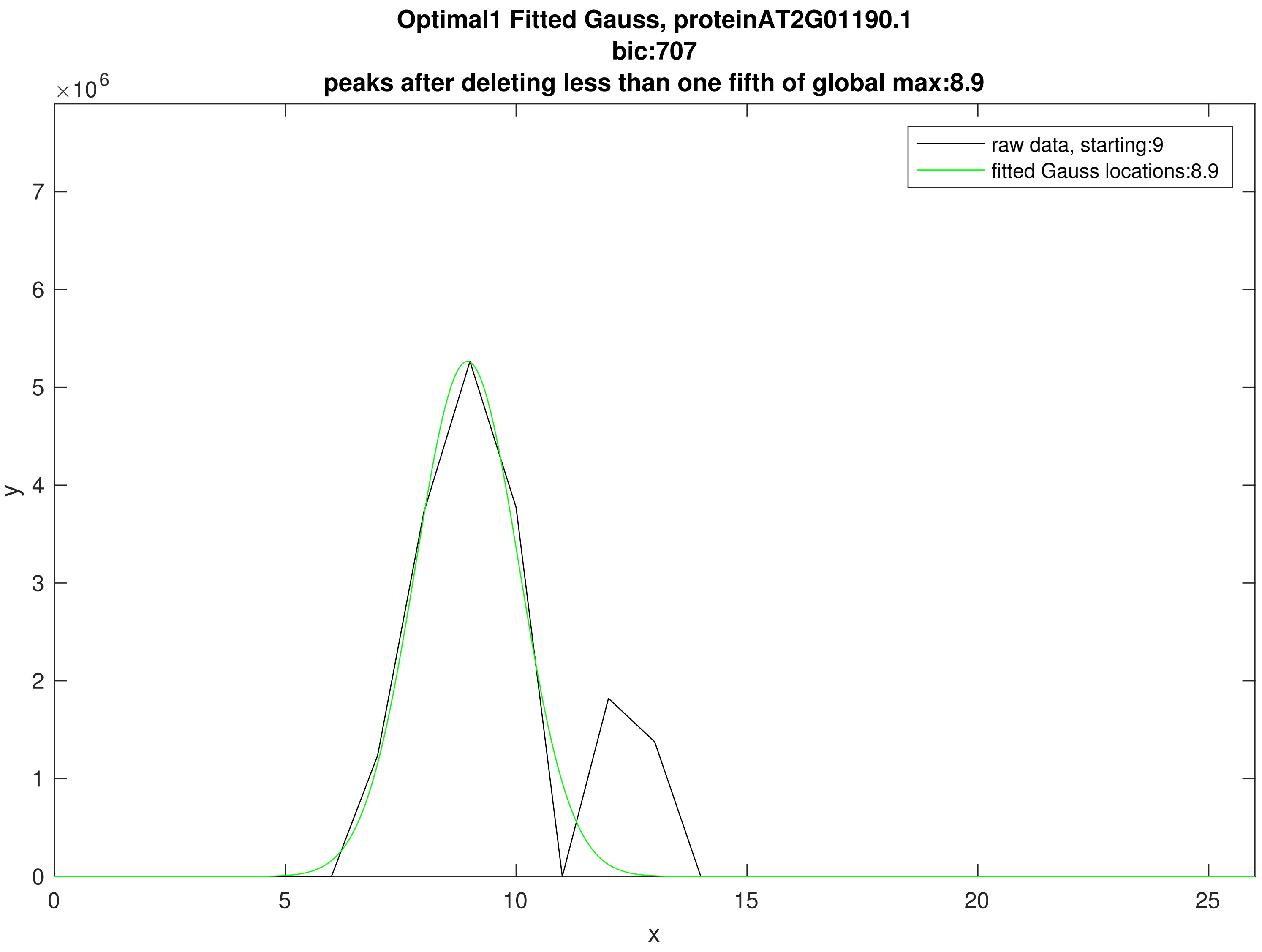
Optimal1 Fitted Gauss, proteinAT2G01190.1
bic:707
peaks after deleting less than one fifth of global max:8.9
×10^6
raw data, starting:9
fitted Gauss locations:8.9
y
x
0
1
2
3
4
5
6
7
0
5
10
15
20
25

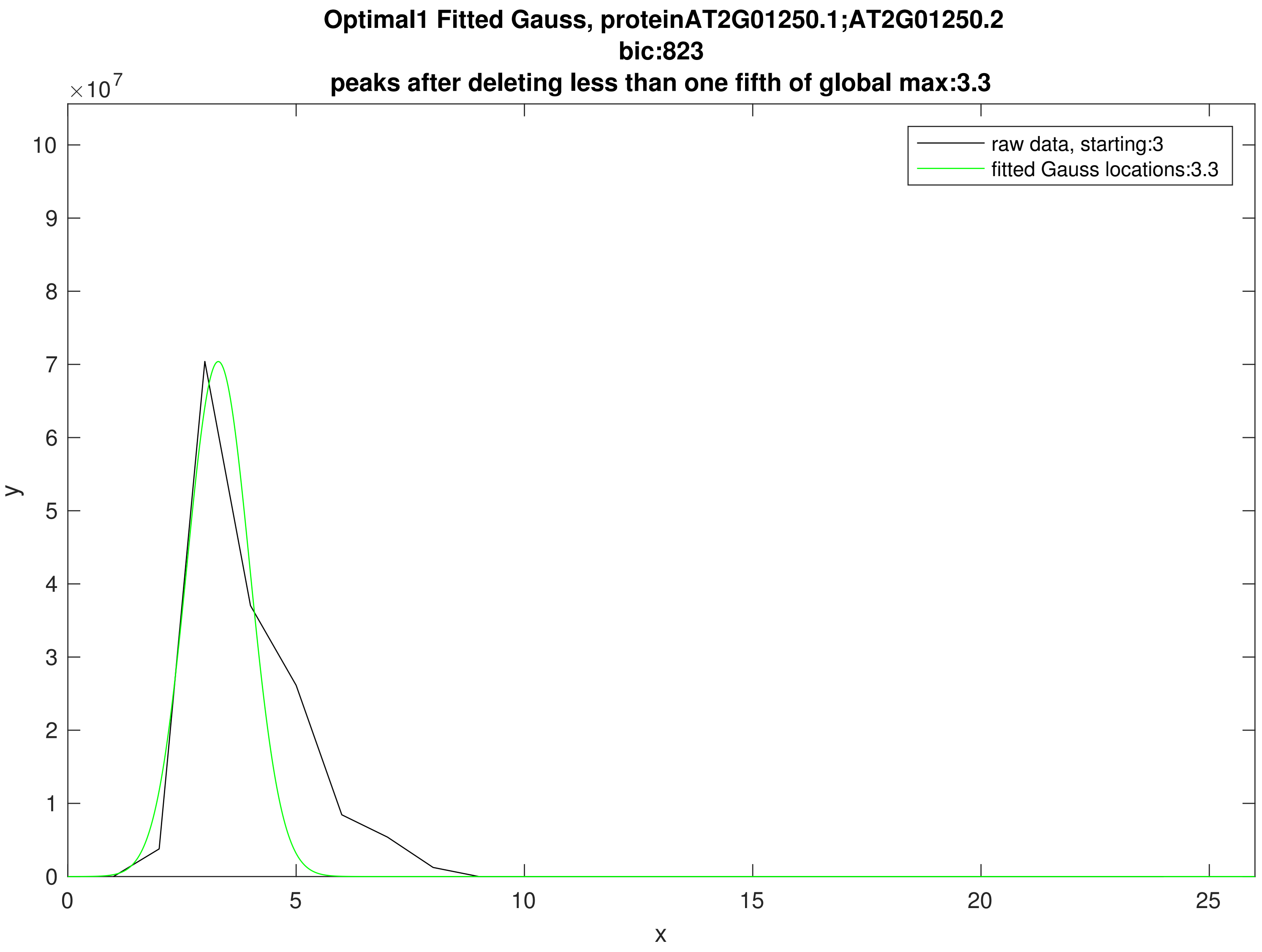
Optimal1 Fitted Gauss, proteinAT2G01250.1;AT2G01250.2
bic:823
peaks after deleting less than one fifth of global max:3.3
×10^7
raw data, starting:3
fitted Gauss locations:3.3
y
x
0
1
2
3
4
5
6
7
8
9
10
0
5
10
15
20
25

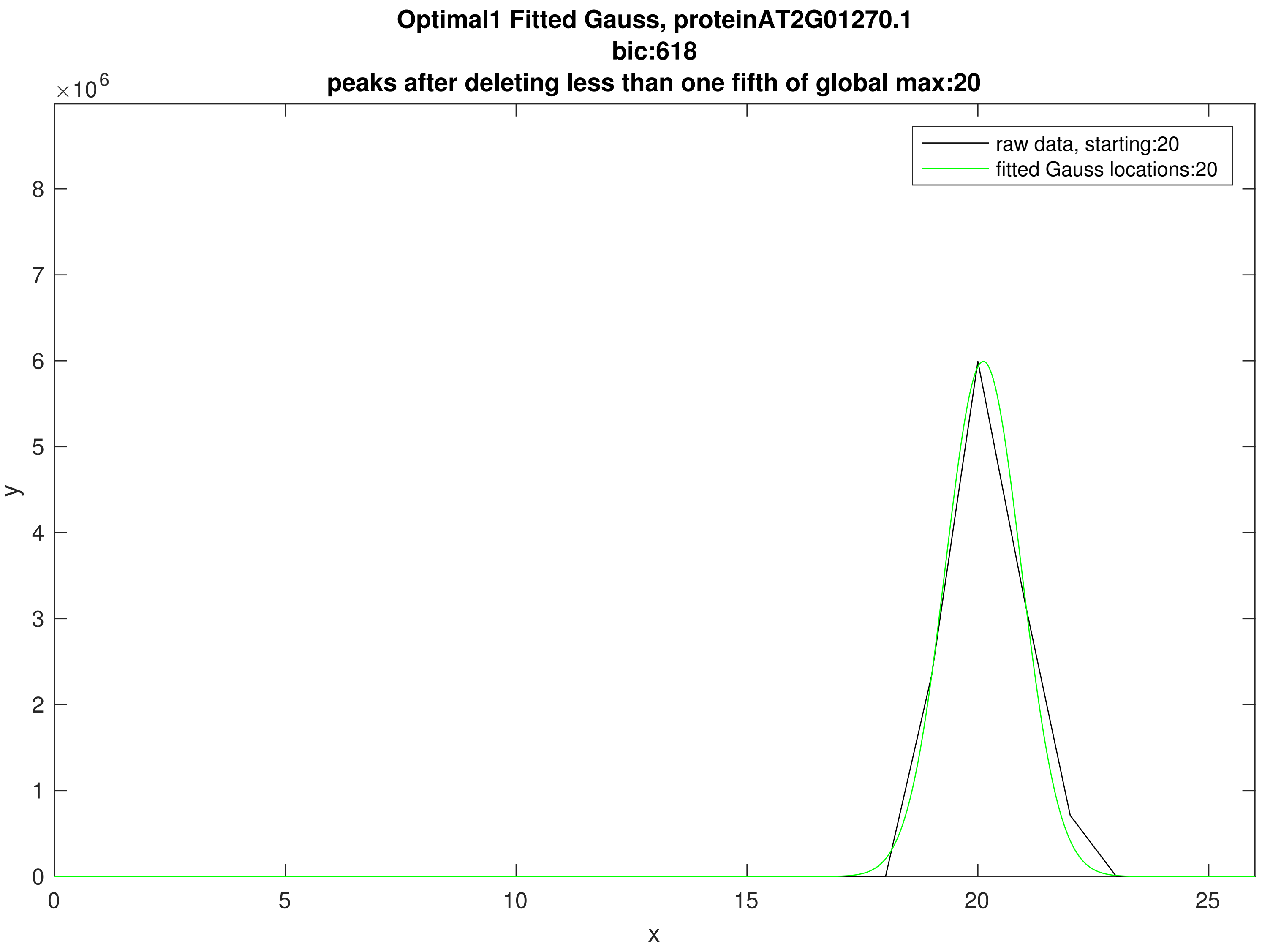
Optimal1 Fitted Gauss, proteinAT2G01270.1
bic:618
peaks after deleting less than one fifth of global max:20
×10^6
raw data, starting:20
fitted Gauss locations:20
y
x
0
1
2
3
4
5
6
7
8
0
5
10
15
20
25

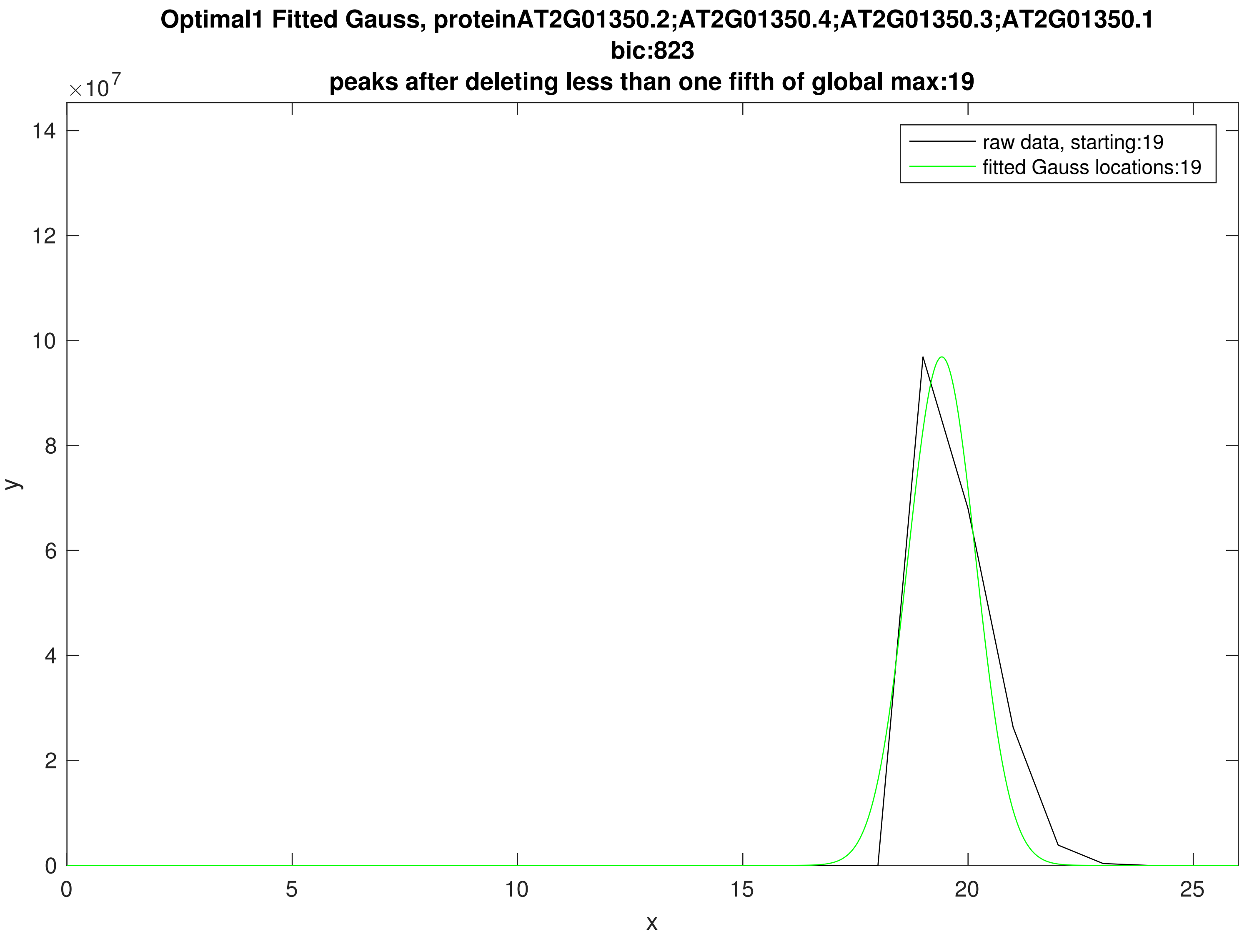
Optimal1 Fitted Gauss, proteinAT2G01350.2;AT2G01350.4;AT2G01350.3;AT2G01350.1
bic:823
peaks after deleting less than one fifth of global max:19
×10^7
raw data, starting:19
fitted Gauss locations:19
y
x
0
2
4
6
8
10
12
14
5
15
20
25

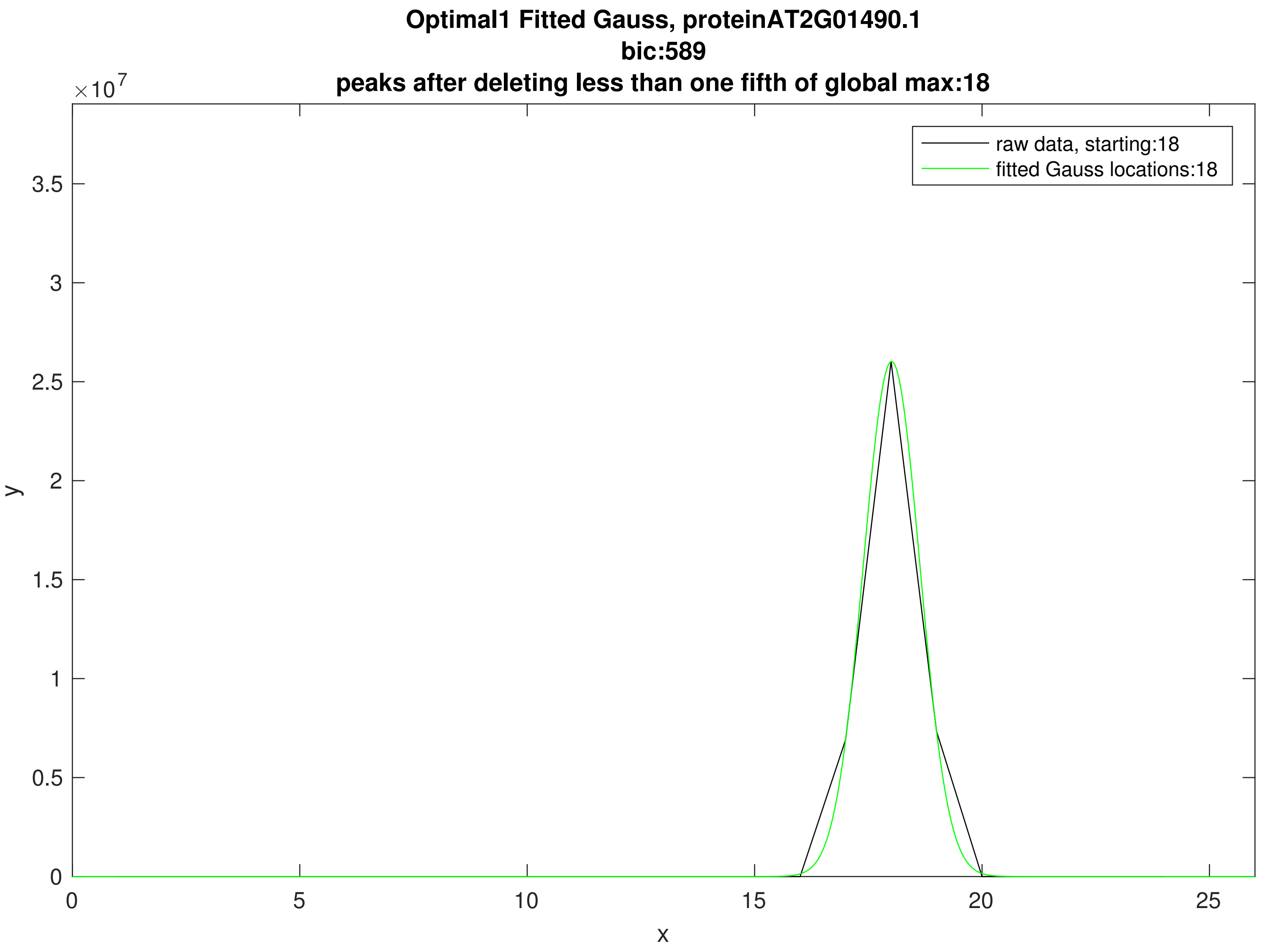
Optimal1 Fitted Gauss, proteinAT2G01490.1
bic:589
peaks after deleting less than one fifth of global max:18
×10^7
raw data, starting:18
fitted Gauss locations:18
y
x
0
0.5
1
1.5
2
2.5
3
3.5
0
5
10
15
20
25

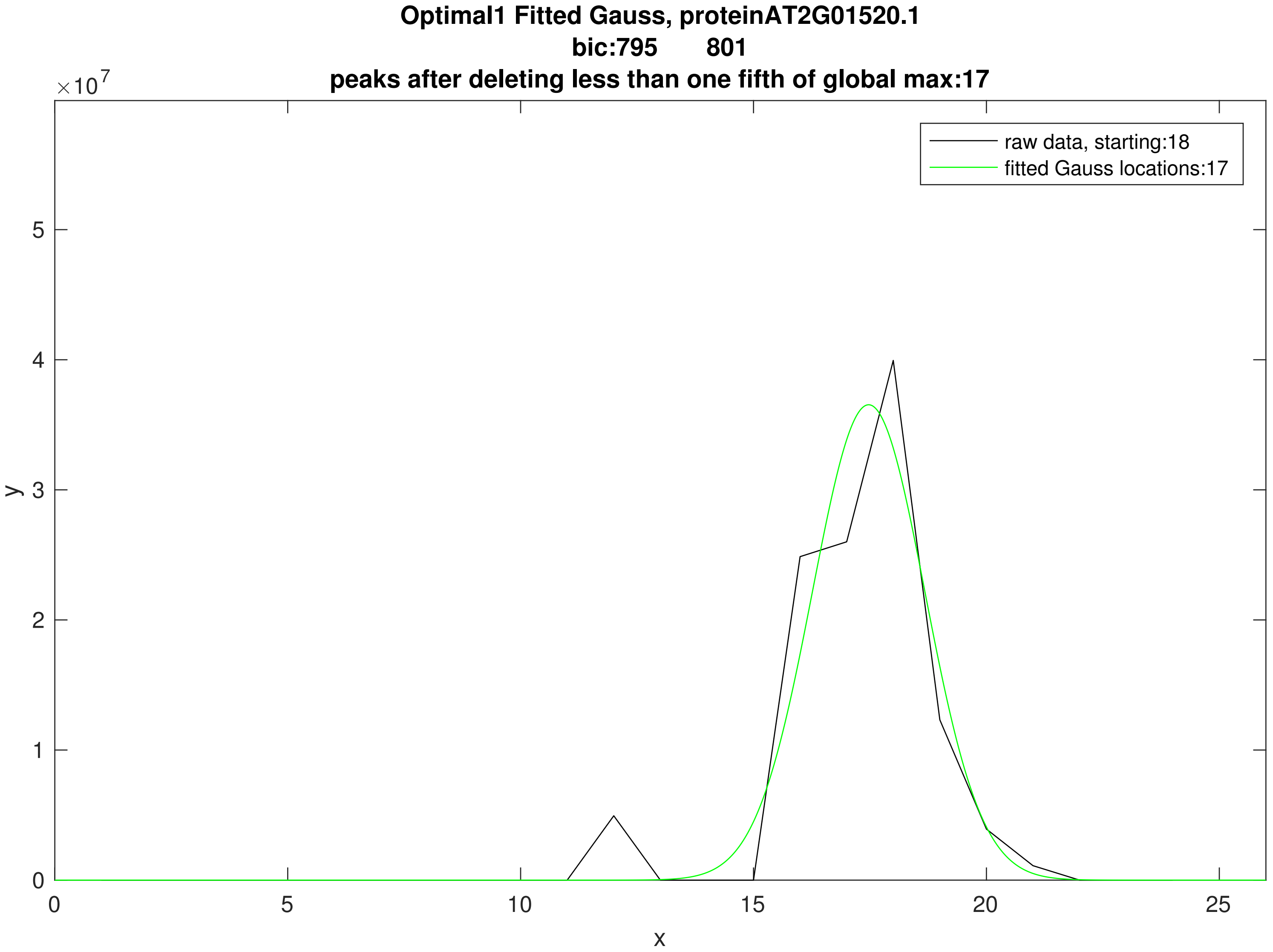
Optimal1 Fitted Gauss, proteinAT2G01520.1
bic:795 801
peaks after deleting less than one fifth of global max:17
×10^7
raw data, starting:18
fitted Gauss locations:17
y
x
0
1
2
3
4
5
0
5
10
15
20
25

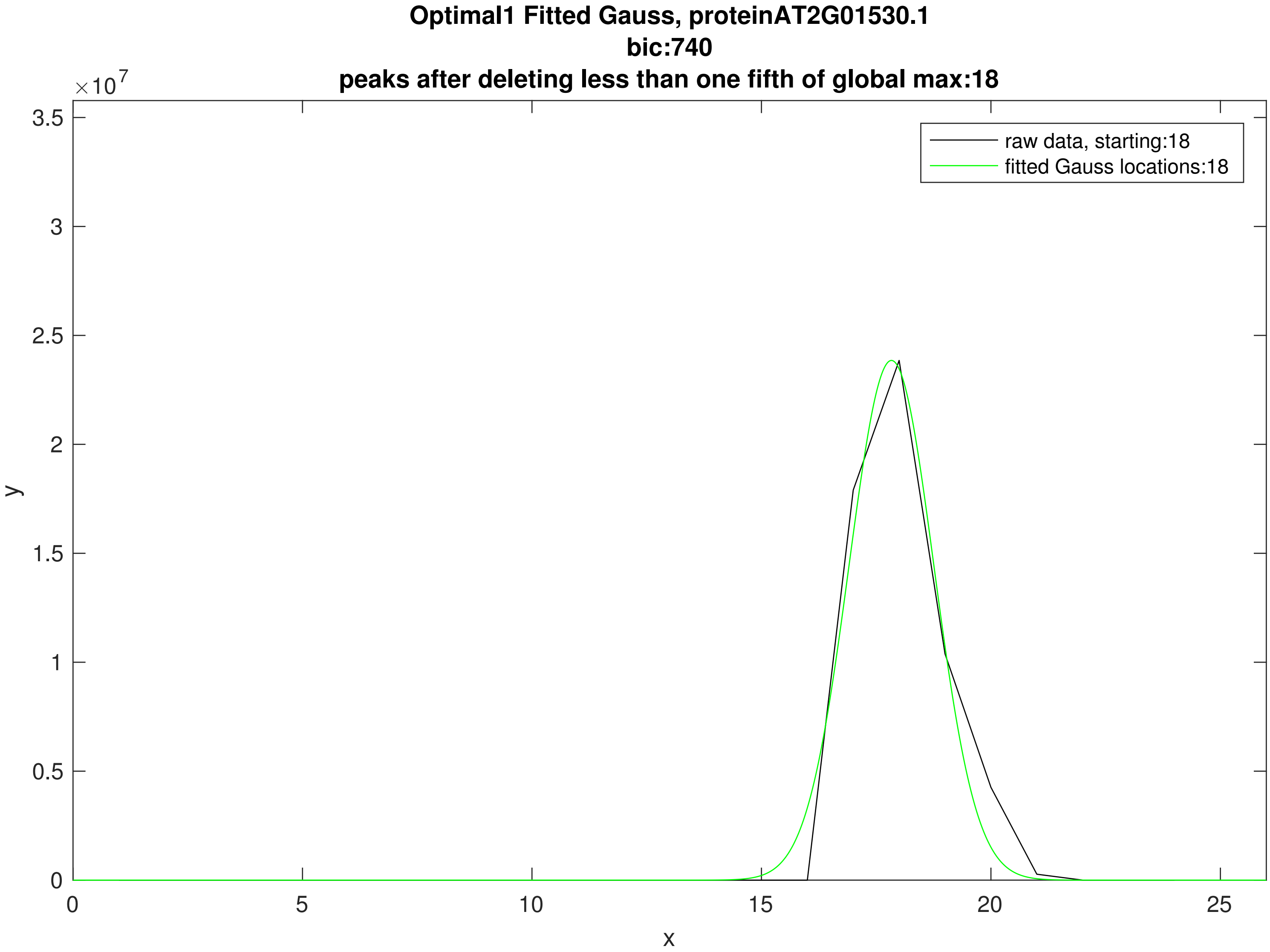
Optimal1 Fitted Gauss, proteinAT2G01530.1
bic:740
peaks after deleting less than one fifth of global max:18
×10^7
3.5
3
2.5
2
1.5
1
0.5
0
y
0
5
10
15
20
25
x
raw data, starting:18
fitted Gauss locations:18

#### Optimal1 Fitted Gauss, proteinAT2G01820.1
#### bic:853
#### peaks after deleting less than one fifth of global max:14

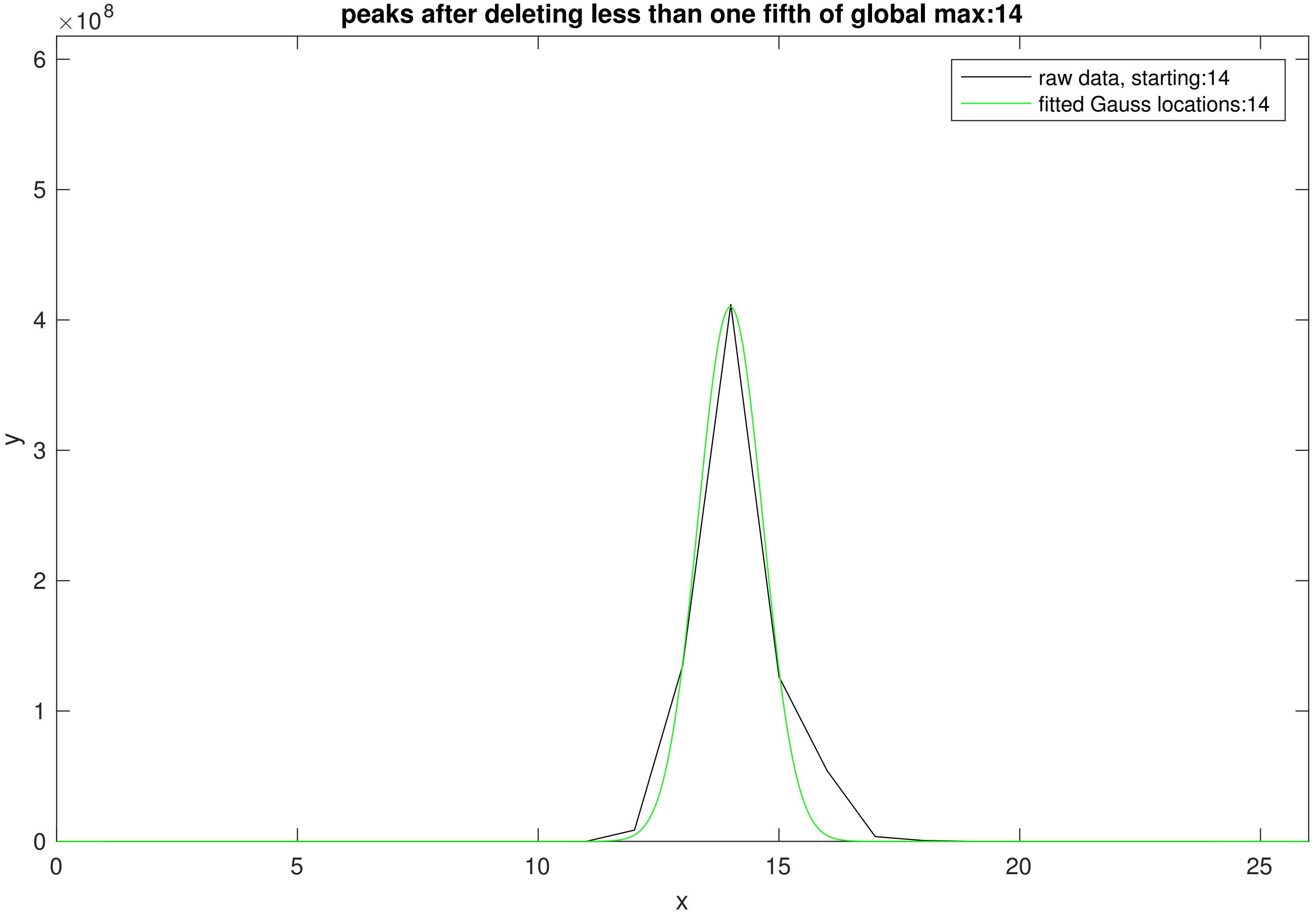

#### Optimal1 Fitted Gauss, proteinAT2G01890.1;AT2G01890.2
#### bic:759
#### peaks after deleting less than one fifth of global max:18

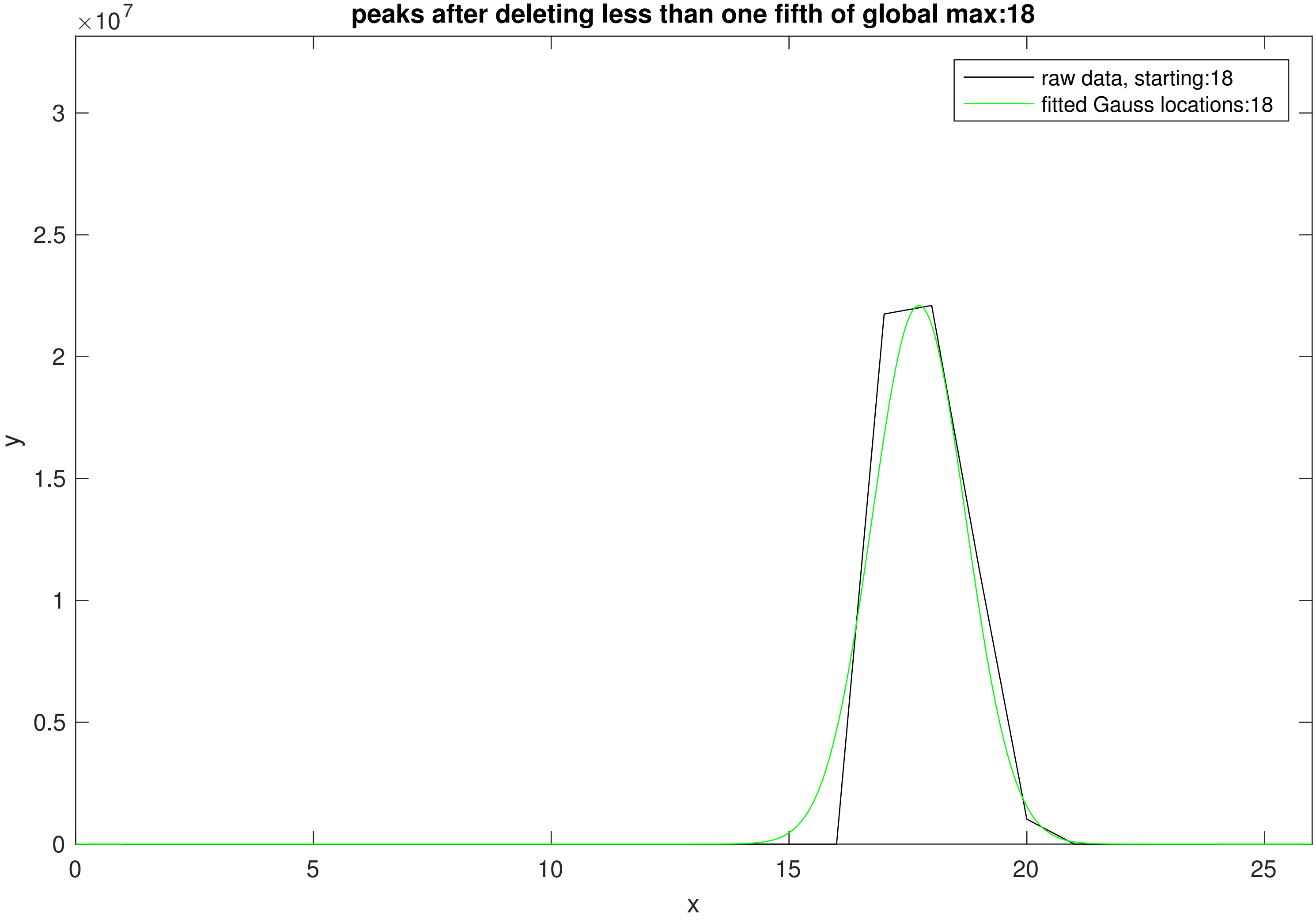

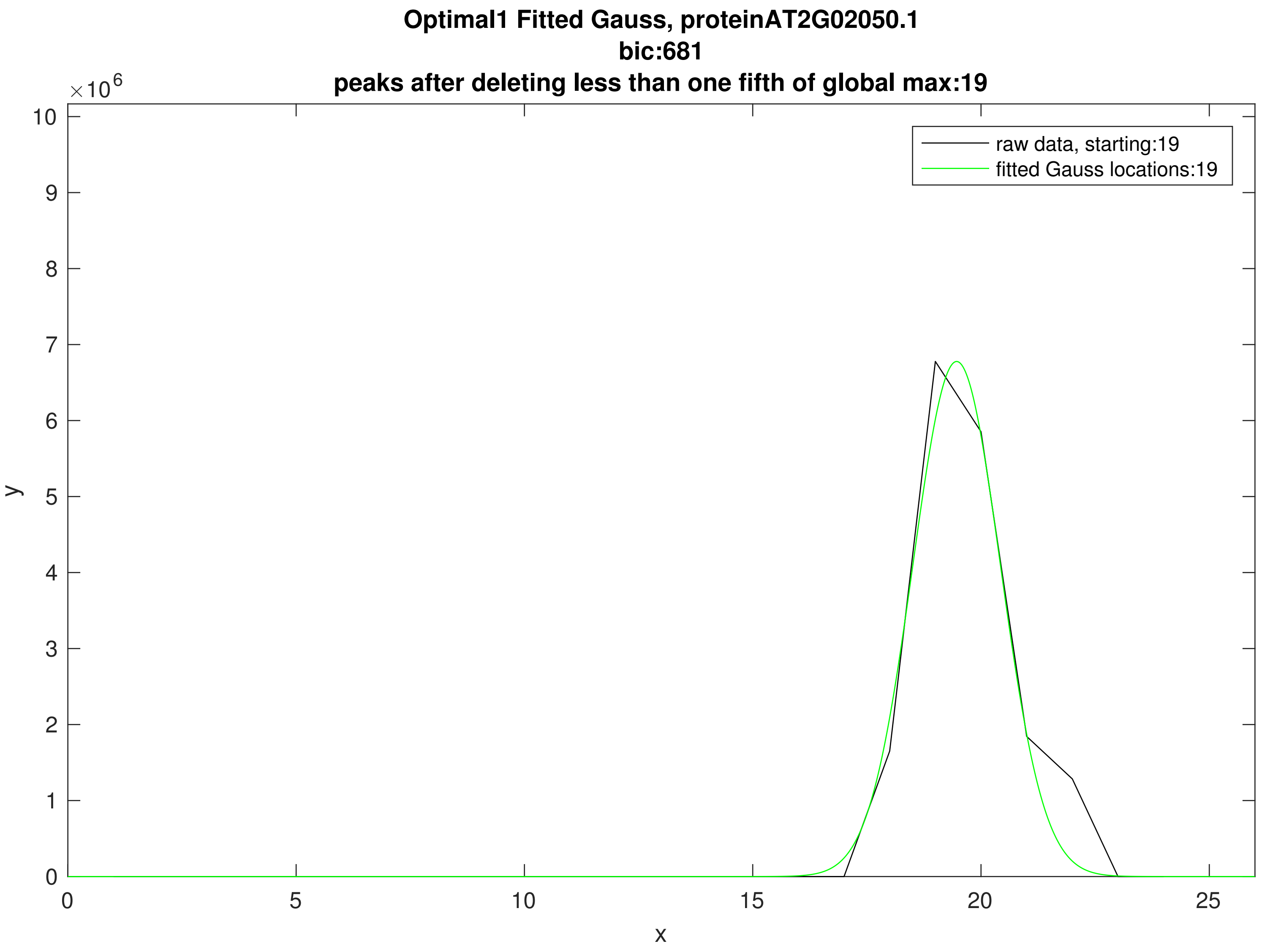
Optimal1 Fitted Gauss, proteinAT2G02050.1
bic:681
peaks after deleting less than one fifth of global max:19
×10^6
raw data, starting:19
fitted Gauss locations:19
y
x

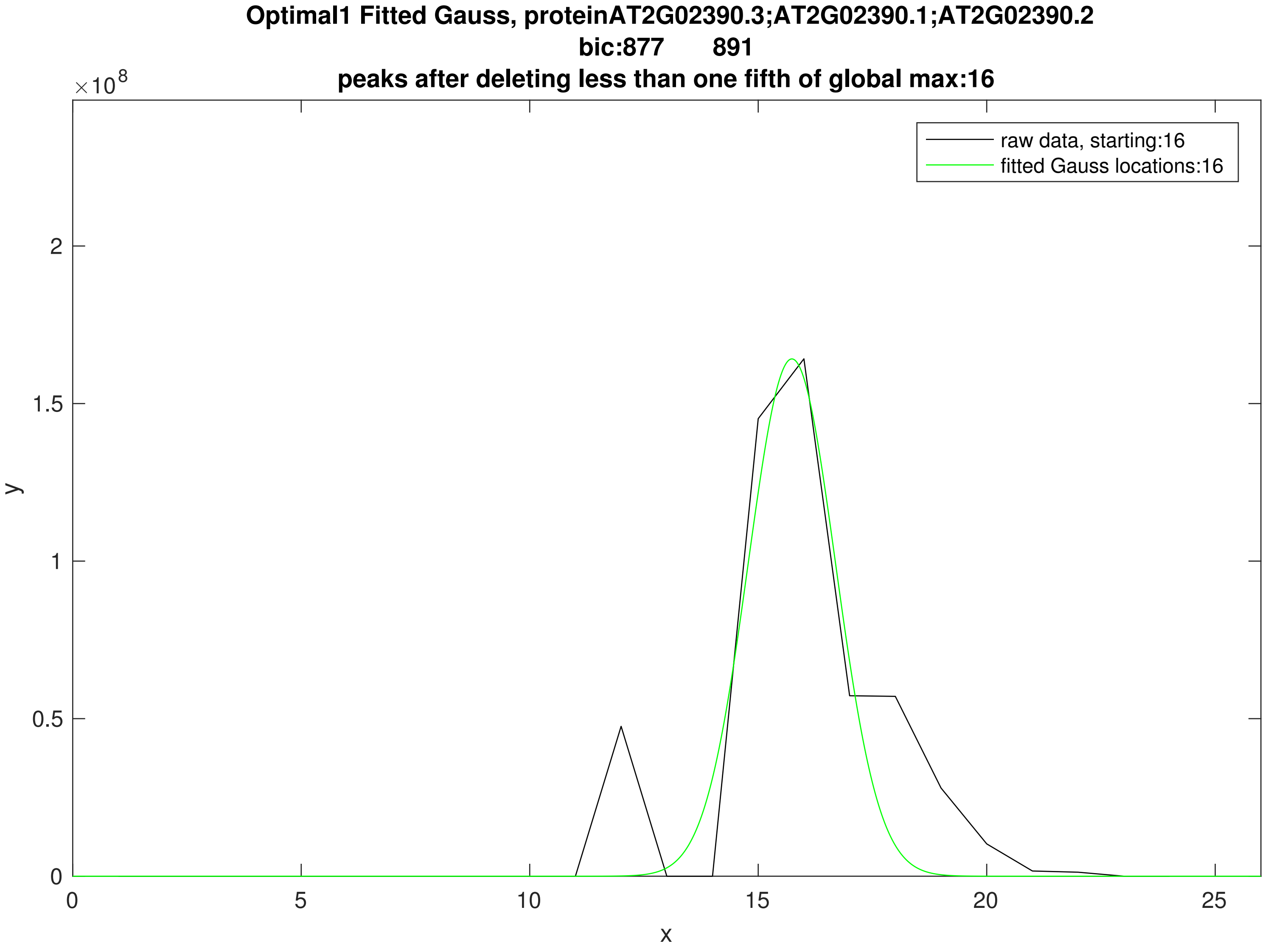
Optimal1 Fitted Gauss, proteinAT2G02390.3;AT2G02390.1;AT2G02390.2
bic:877 891
peaks after deleting less than one fifth of global max:16
×10^8
raw data, starting:16
fitted Gauss locations:16
y
x
0
0.5
1
1.5
2
0
5
10
15
20
25

#### Optimal1 Fitted Gauss, proteinAT2G02400.1
#### bic:791
#### peaks after deleting less than one fifth of global max:18

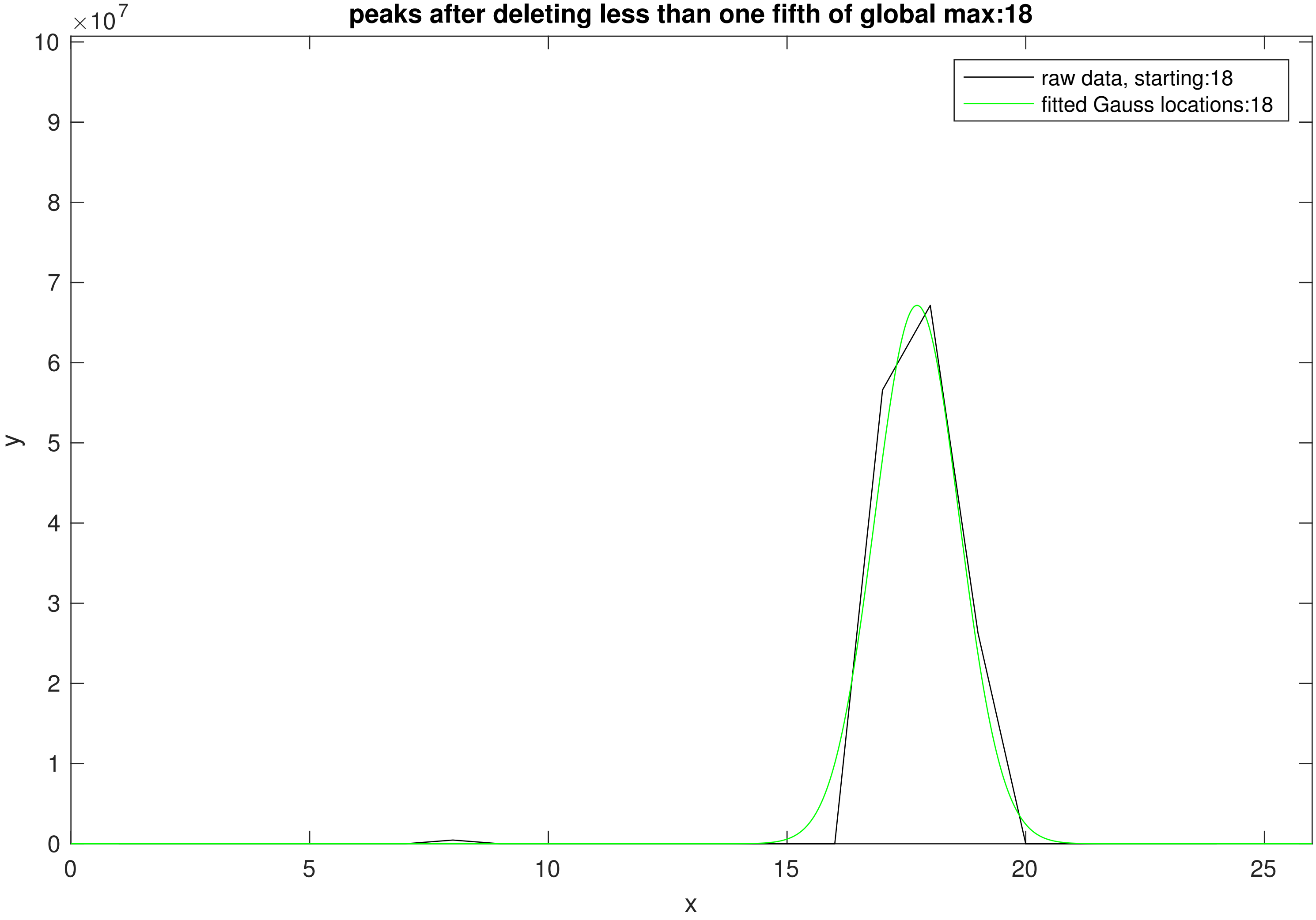

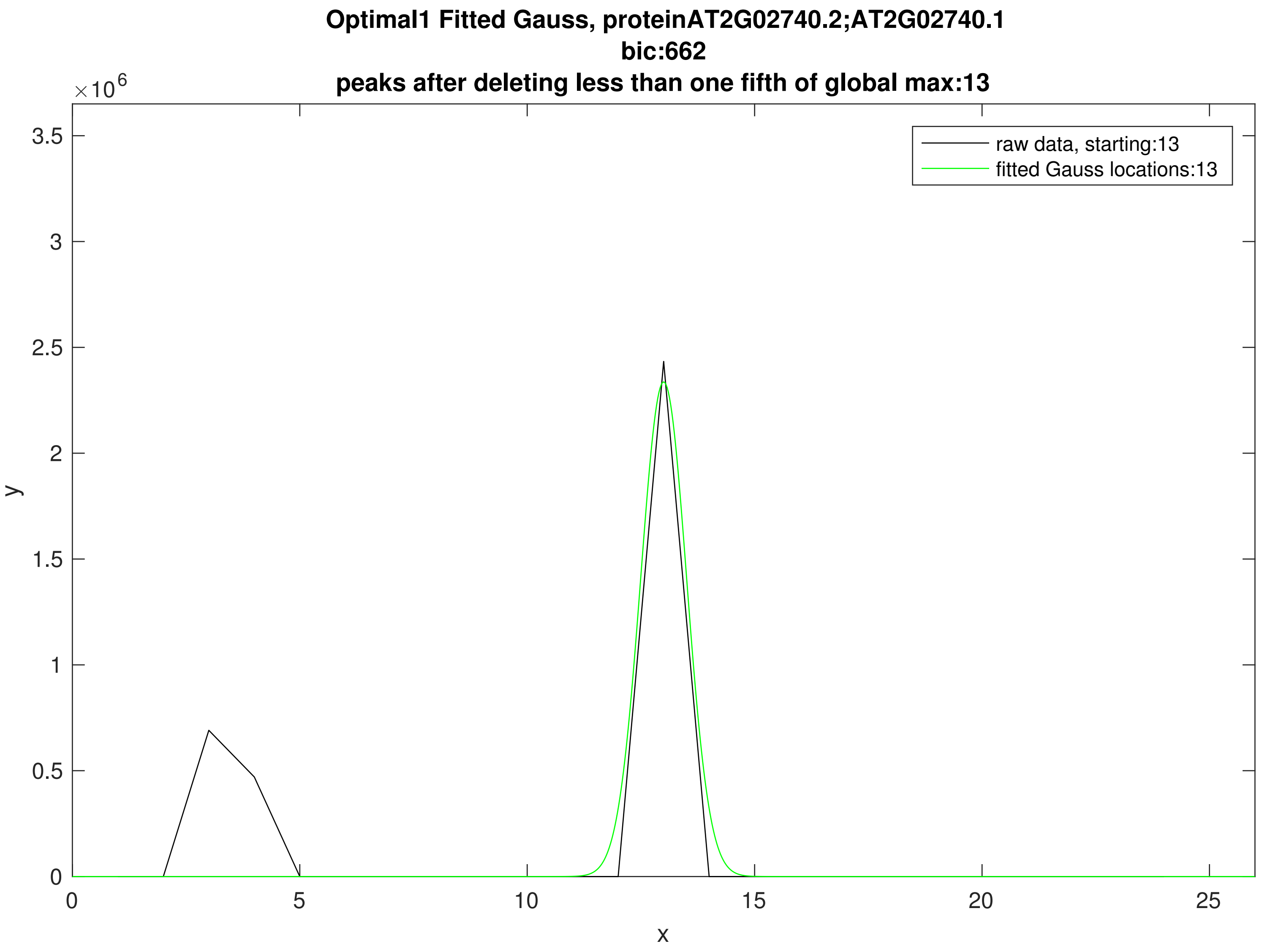
Optimal1 Fitted Gauss, proteinAT2G02740.2;AT2G02740.1
bic:662
peaks after deleting less than one fifth of global max:13
×10^6
3.5
3
2.5
2
1.5
1
0.5
0
y
0
5
10
15
20
25
x
raw data, starting:13
fitted Gauss locations:13

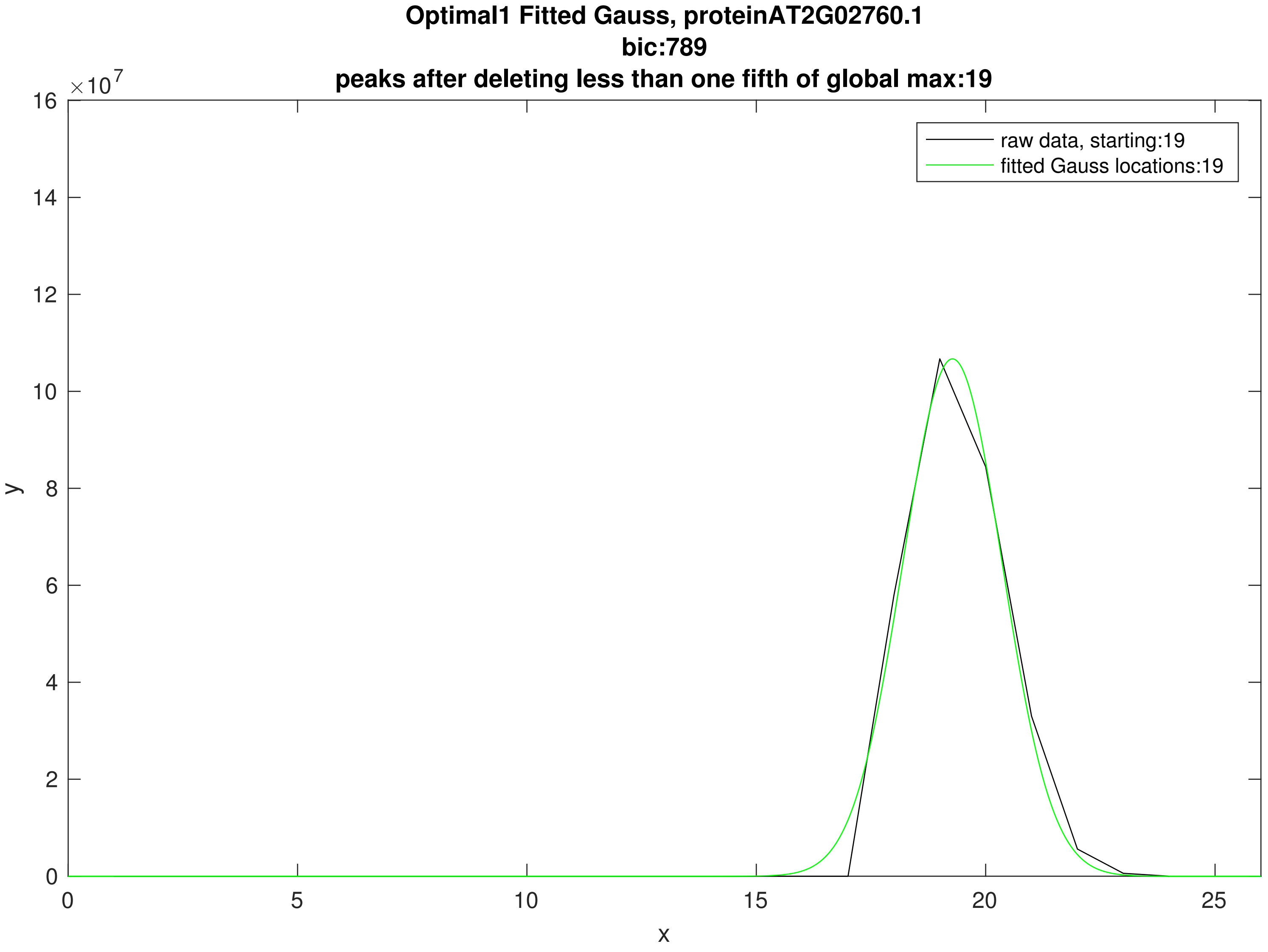

Optimal1 Fitted Gauss, proteinAT2G02760.1
bic:789
peaks after deleting less than one fifth of global max:19
×10^7
16
14
12
10
8
6
4
2
0
y
0
5
10
15
20
25
x
raw data, starting:19
fitted Gauss locations:19

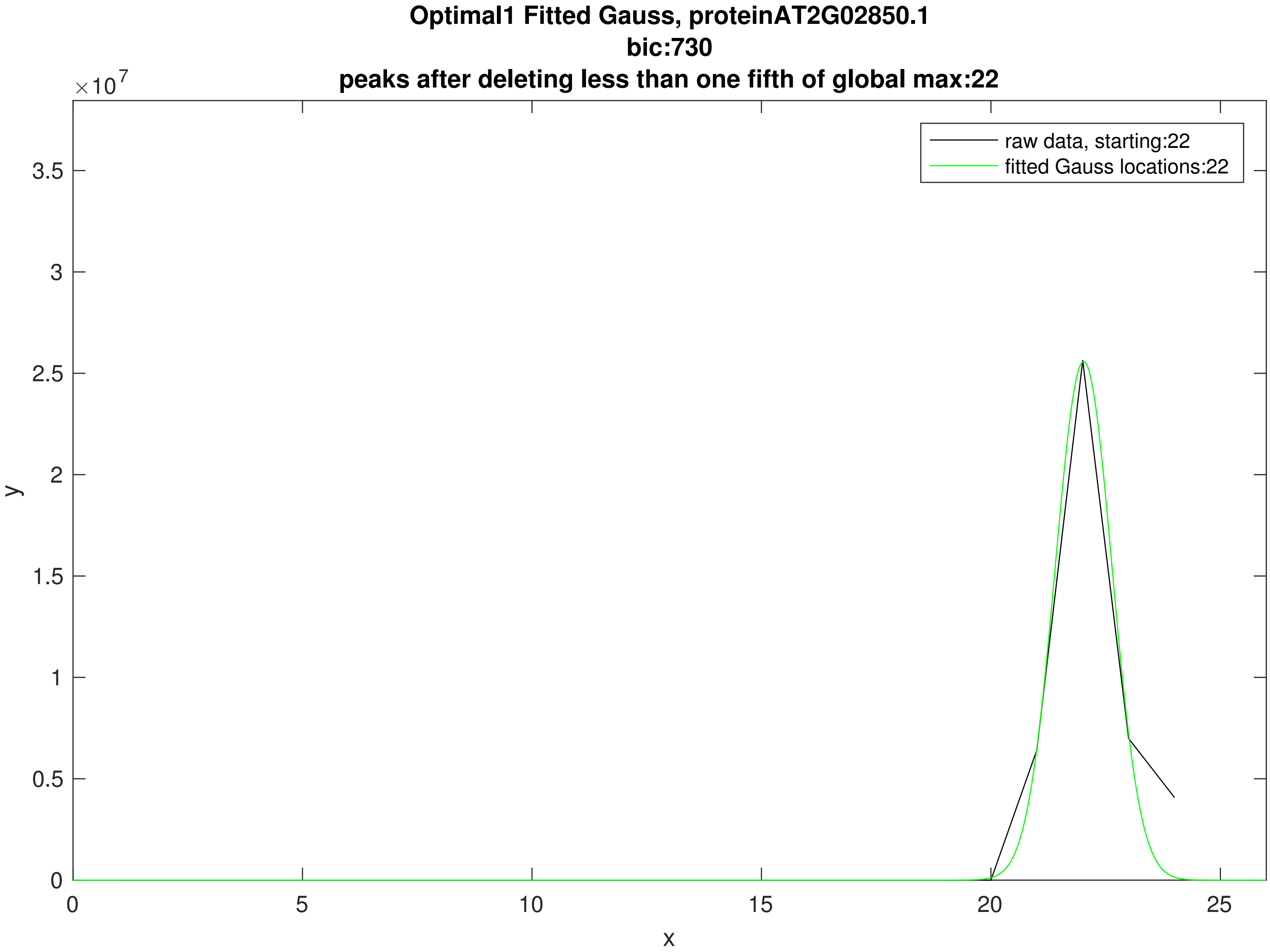
Optimal1 Fitted Gauss, proteinAT2G02850.1
bic:730
peaks after deleting less than one fifth of global max:22
×10^7
raw data, starting:22
fitted Gauss locations:22
3.5
3
2.5
2
1.5
1
0.5
0
y
0
5
10
15
20
25
x

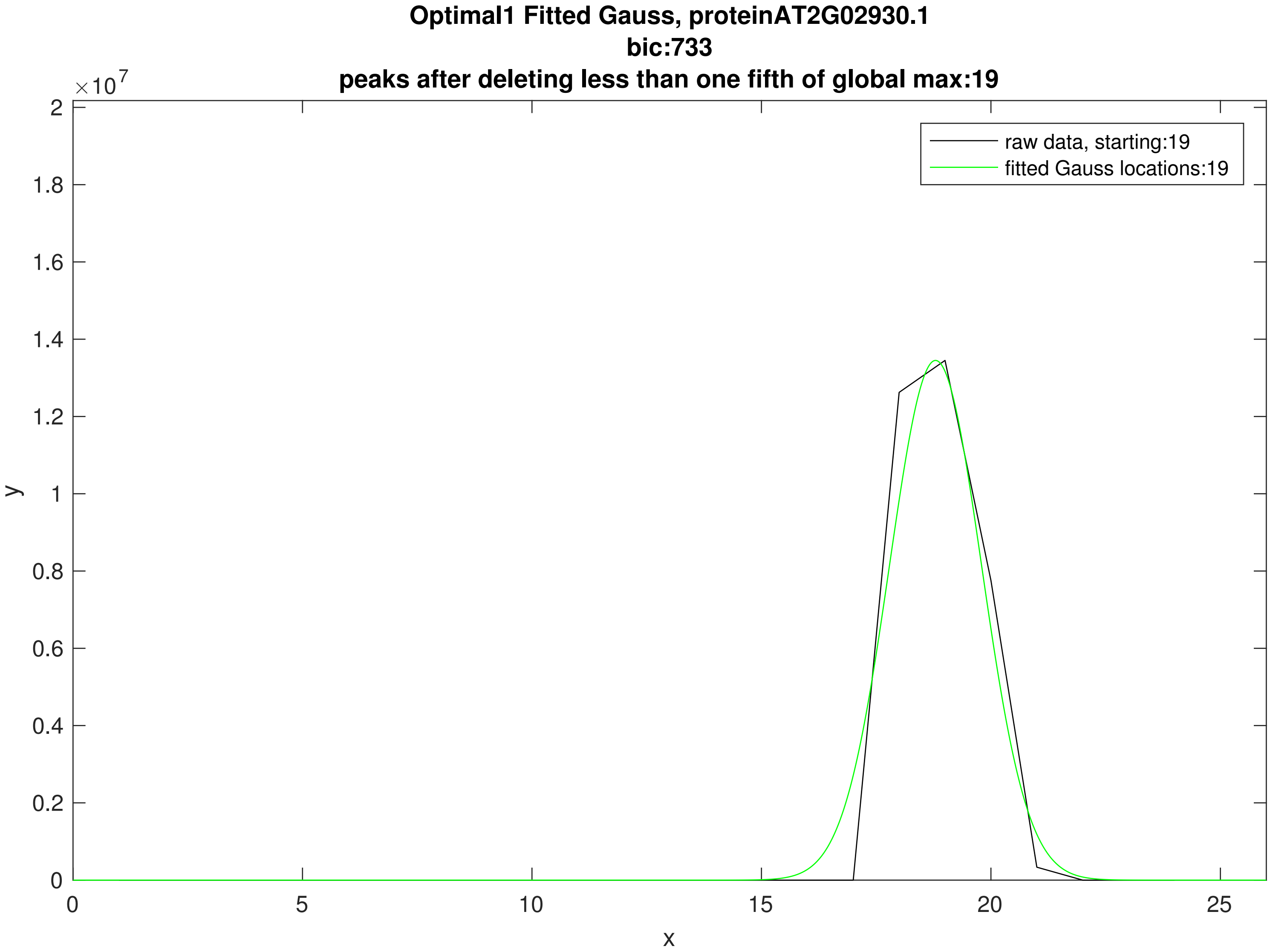
Optimal1 Fitted Gauss, proteinAT2G02930.1
bic:733
peaks after deleting less than one fifth of global max:19
×10^7
raw data, starting:19
fitted Gauss locations:19
y
x
0
0.2
0.4
0.6
0.8
1
1.2
1.4
1.6
1.8
2
5
10
15
20
25

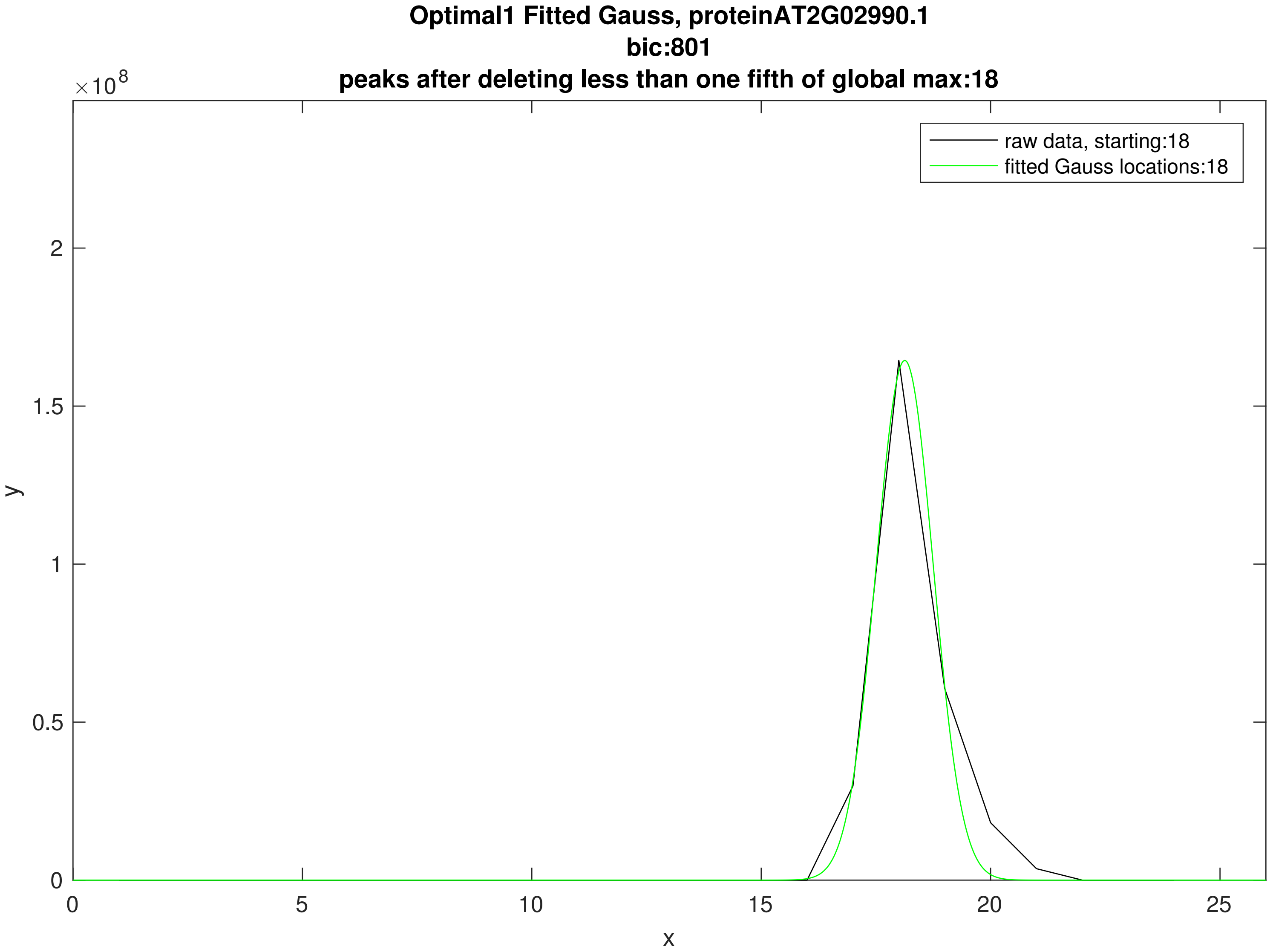

Optimal1 Fitted Gauss, proteinAT2G02990.1
bic:801
peaks after deleting less than one fifth of global max:18
×10^8
raw data, starting:18
fitted Gauss locations:18
y
x
0
0.5
1
1.5
2
0
5
10
15
20
25

Optimal1 Fitted Gauss, proteinAT2G03430.1
bic:701
peaks after deleting less than one fifth of global max:17

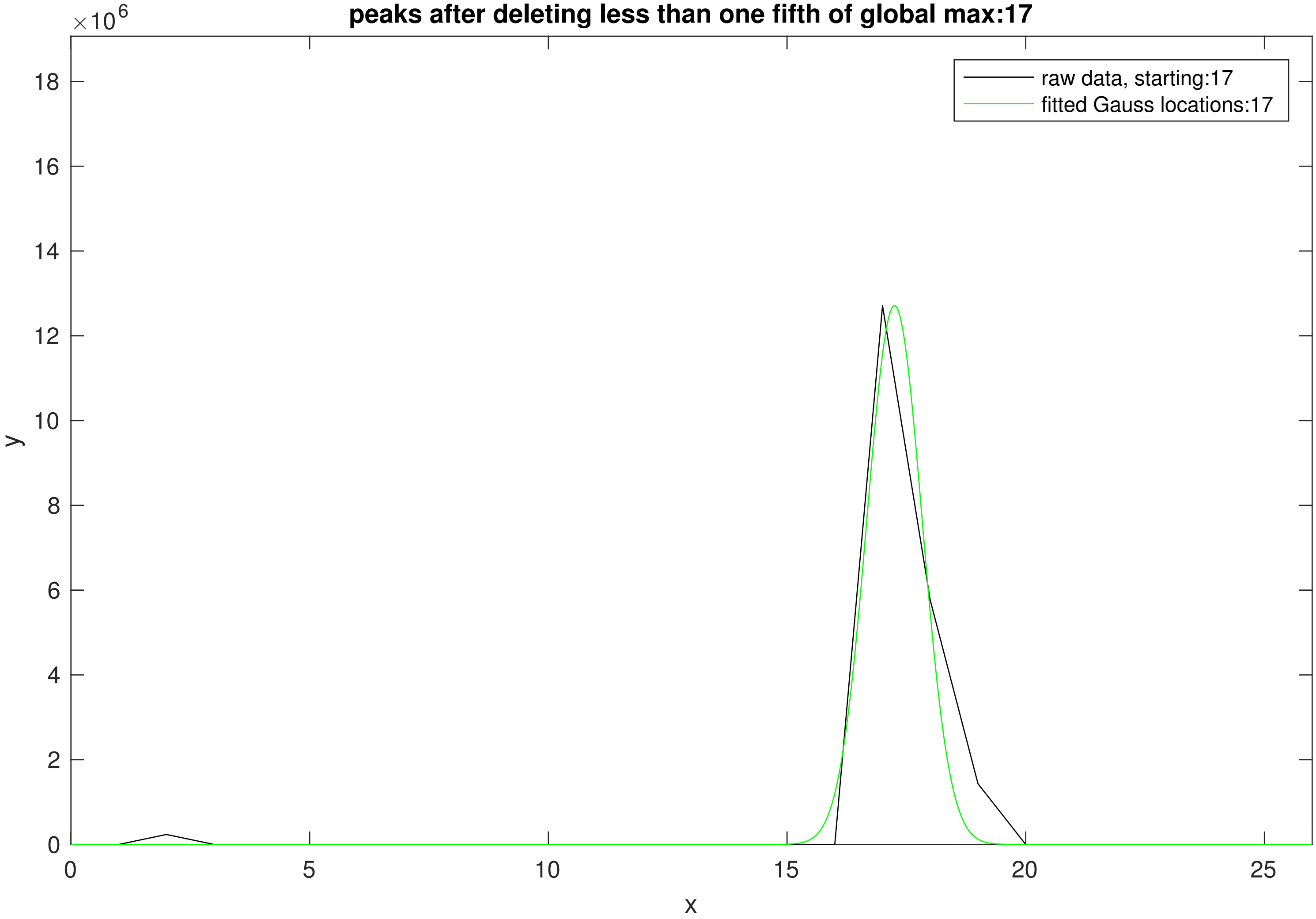

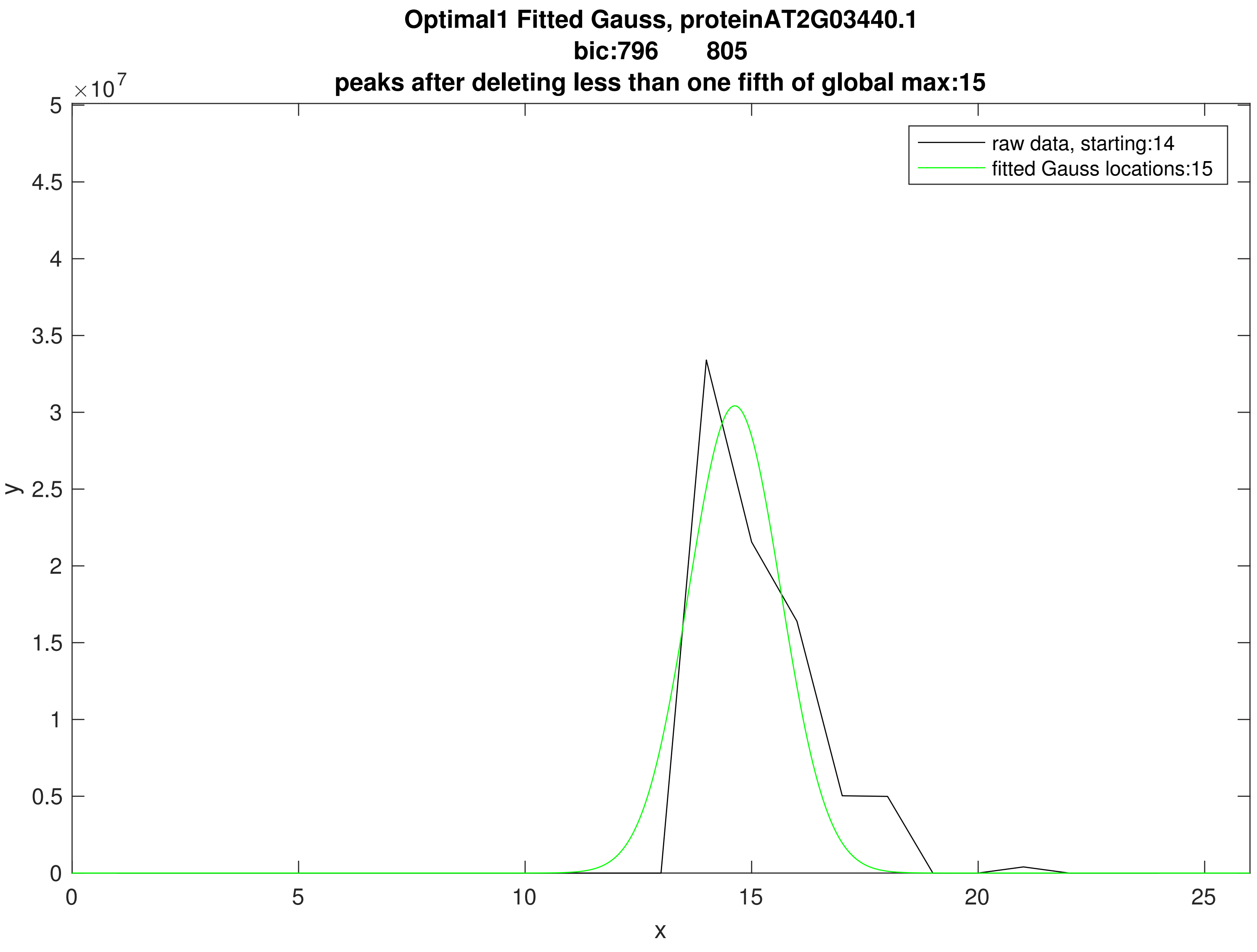
Optimal1 Fitted Gauss, proteinAT2G03440.1
bic:796 805
peaks after deleting less than one fifth of global max:15
×10^7
5
4.5
4
3.5
3
2.5
2
1.5
1
0.5
0
y
0
5
10
15
20
25
x
raw data, starting:14
fitted Gauss locations:15

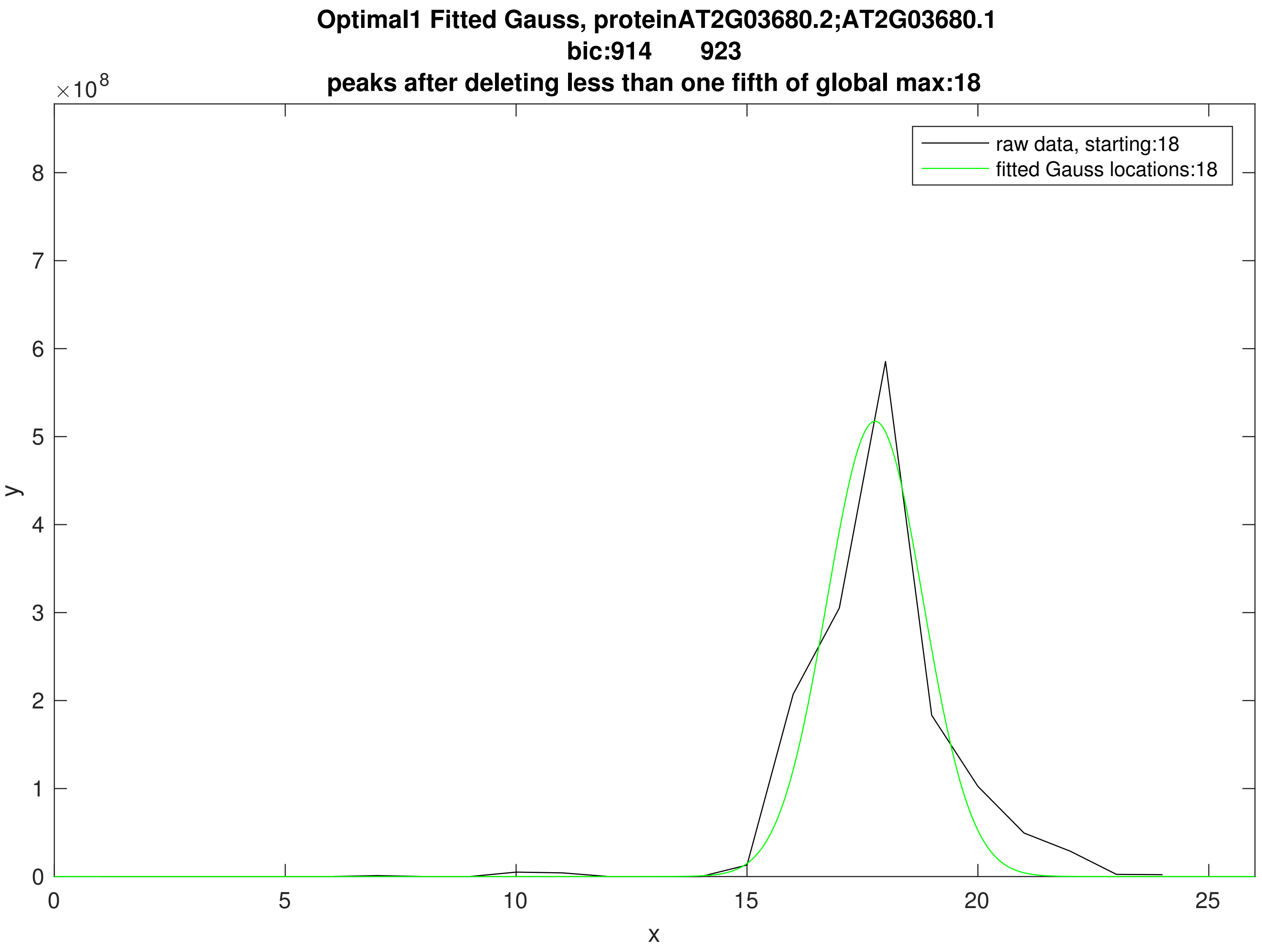
Optimal1 Fitted Gauss, proteinAT2G03680.2;AT2G03680.1
bic:914 923
peaks after deleting less than one fifth of global max:18
×10^8
8
7
6
5
4
3
2
1
0
y
0
5
10
15
20
25
x
raw data, starting:18
fitted Gauss locations:18

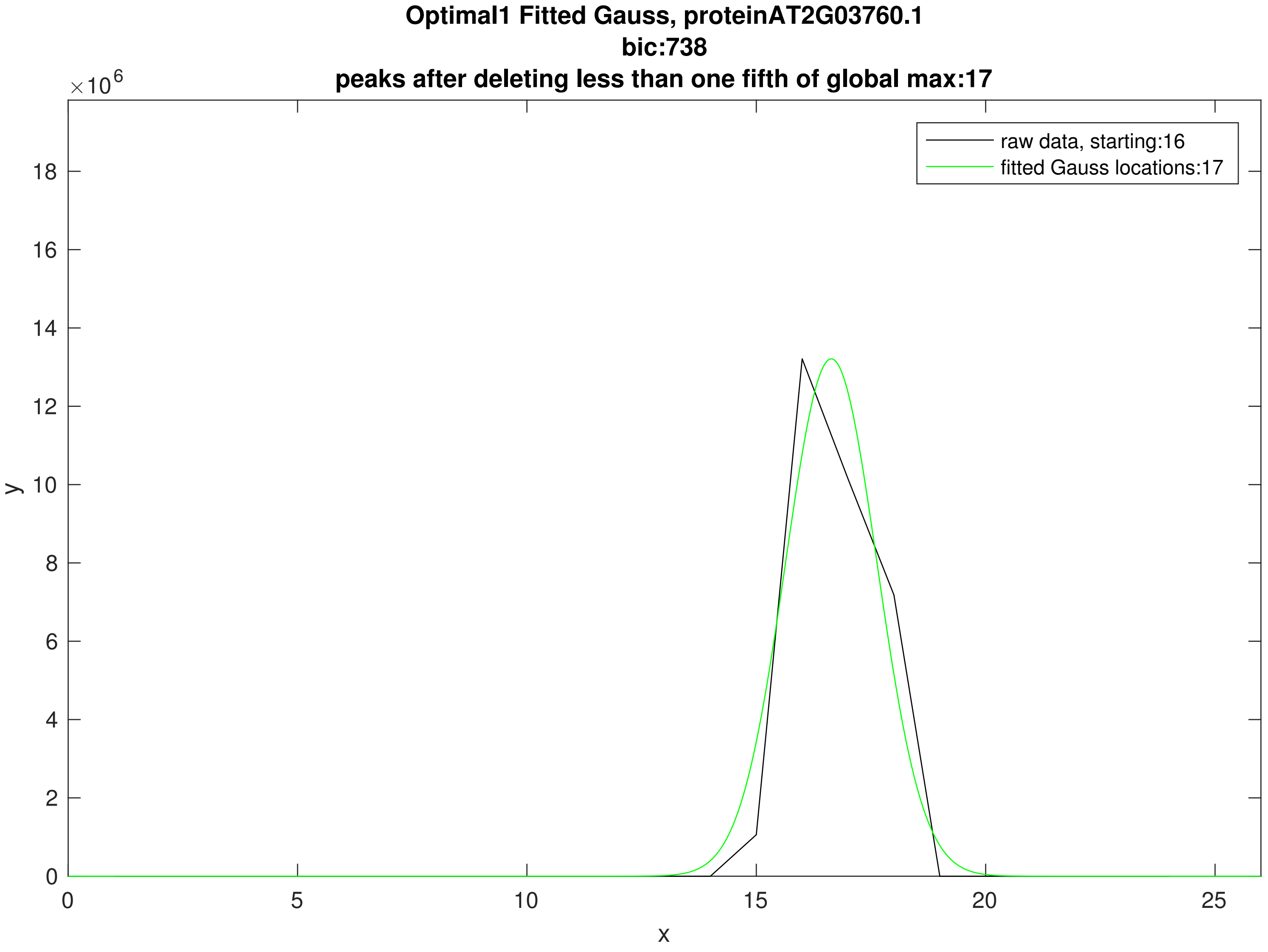
Optimal1 Fitted Gauss, proteinAT2G03760.1
bic:738
peaks after deleting less than one fifth of global max:17
×10^6
raw data, starting:16
fitted Gauss locations:17
y
x
0
2
4
6
8
10
12
14
16
18
0
5
10
15
20
25

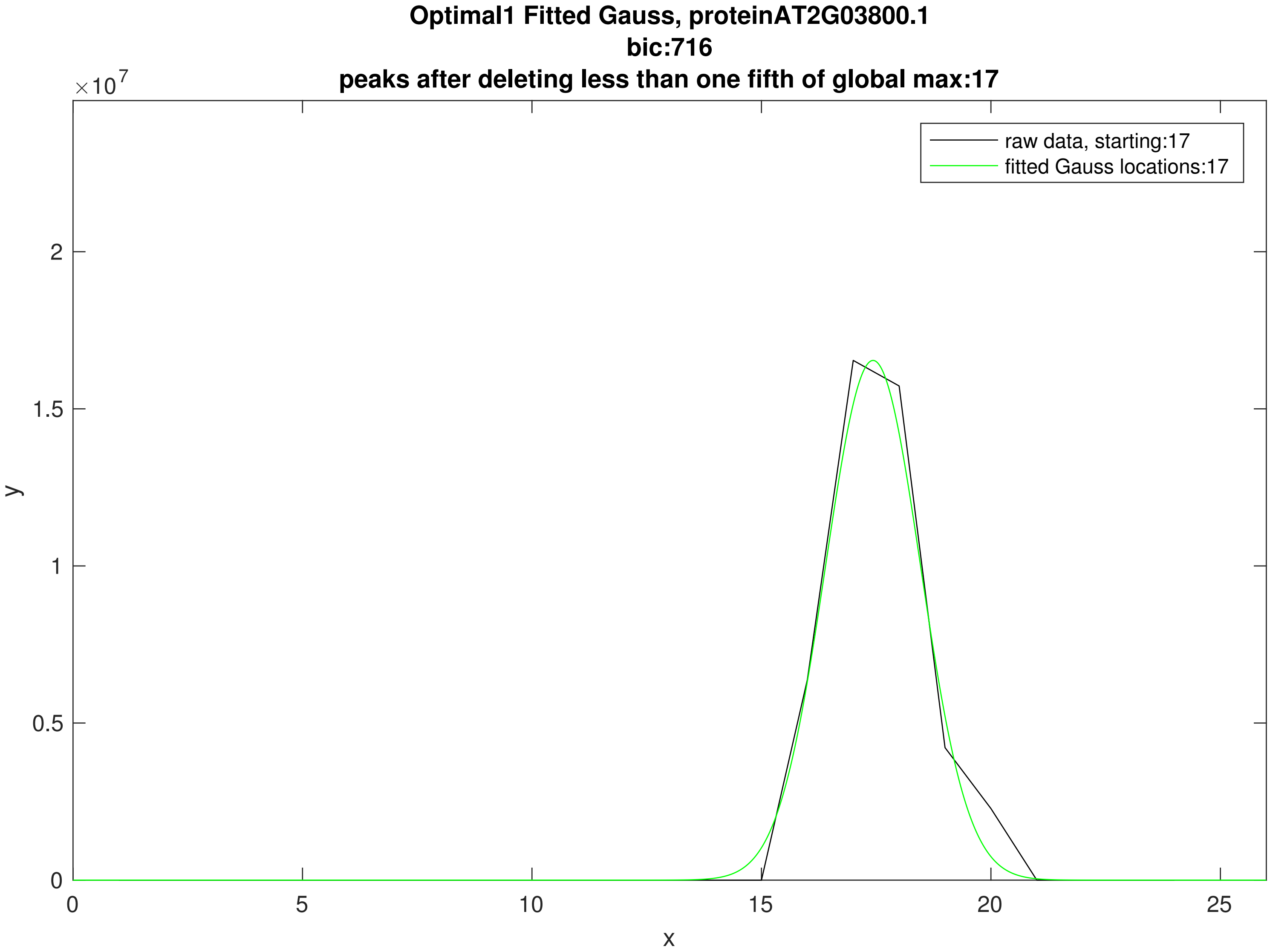
Optimal1 Fitted Gauss, proteinAT2G03800.1
bic:716
peaks after deleting less than one fifth of global max:17
×10^7
raw data, starting:17
fitted Gauss locations:17
y
0
0.5
1
1.5
2
x
0
5
10
15
20
25

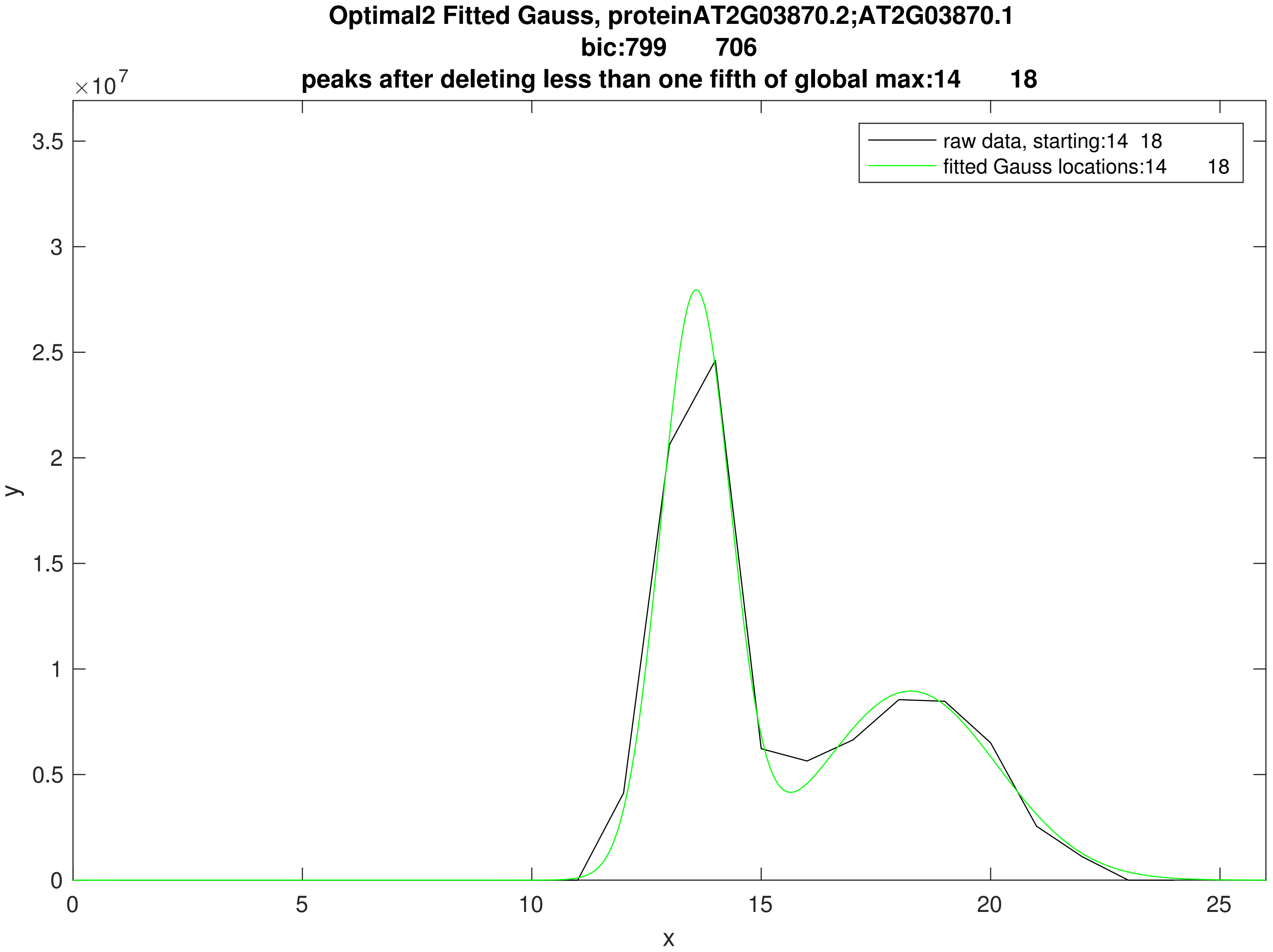
Optimal2 Fitted Gauss, proteinAT2G03870.2;AT2G03870.1
bic:799 706
peaks after deleting less than one fifth of global max:14 18
×10^7
raw data, starting:14 18
fitted Gauss locations:14 18
y
x
0
0.5
1
1.5
2
2.5
3
3.5
0
5
10
15
20
25

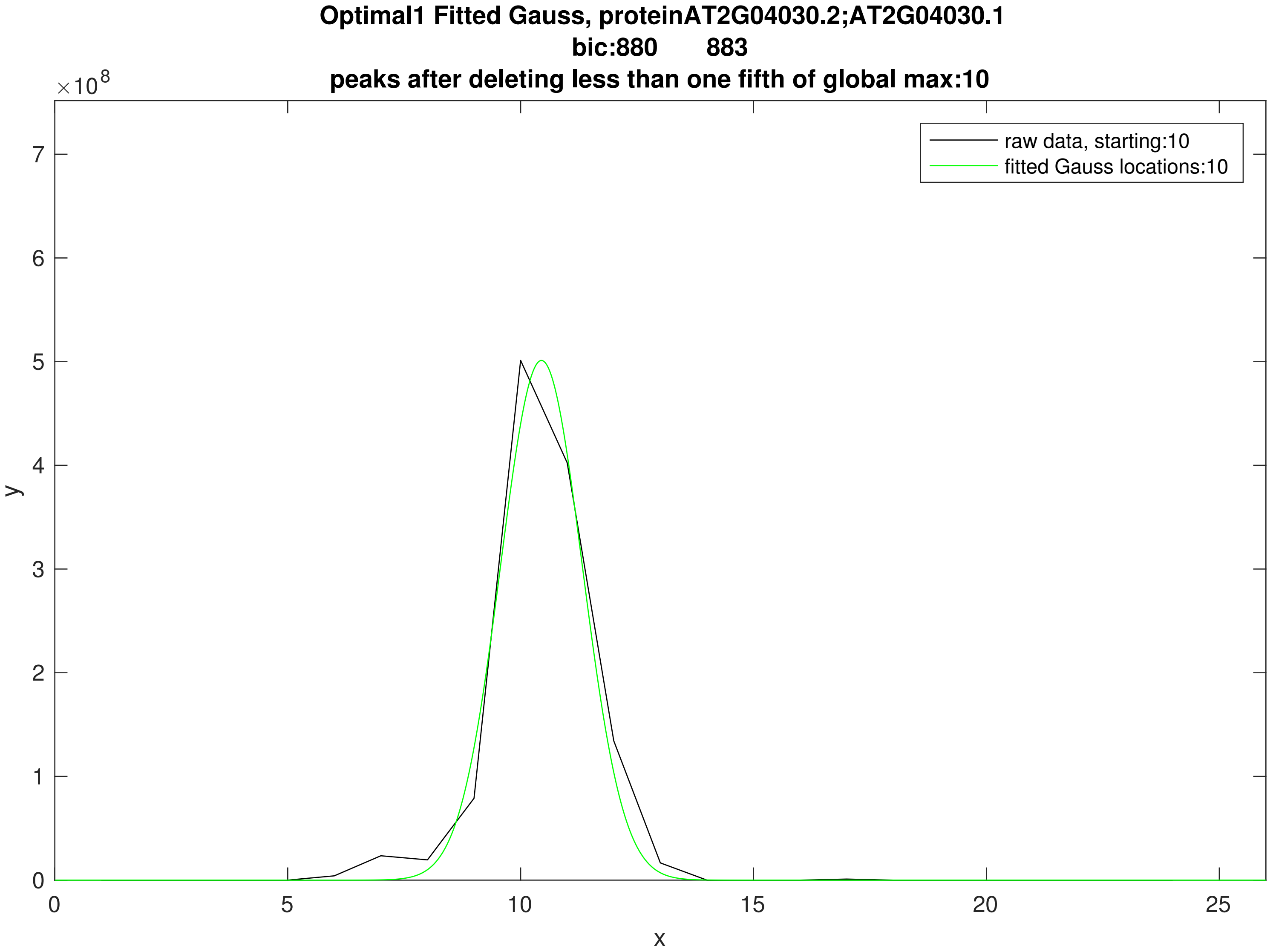
Optimal1 Fitted Gauss, proteinAT2G04030.2;AT2G04030.1
bic:880 883
peaks after deleting less than one fifth of global max:10
×10^8
raw data, starting:10
fitted Gauss locations:10
y
x
0
1
2
3
4
5
6
7
10
15
20
25

Optimal2 Fitted Gauss, proteinAT2G04570.1
bic:817 816
peaks after deleting less than one fifth of global max:17

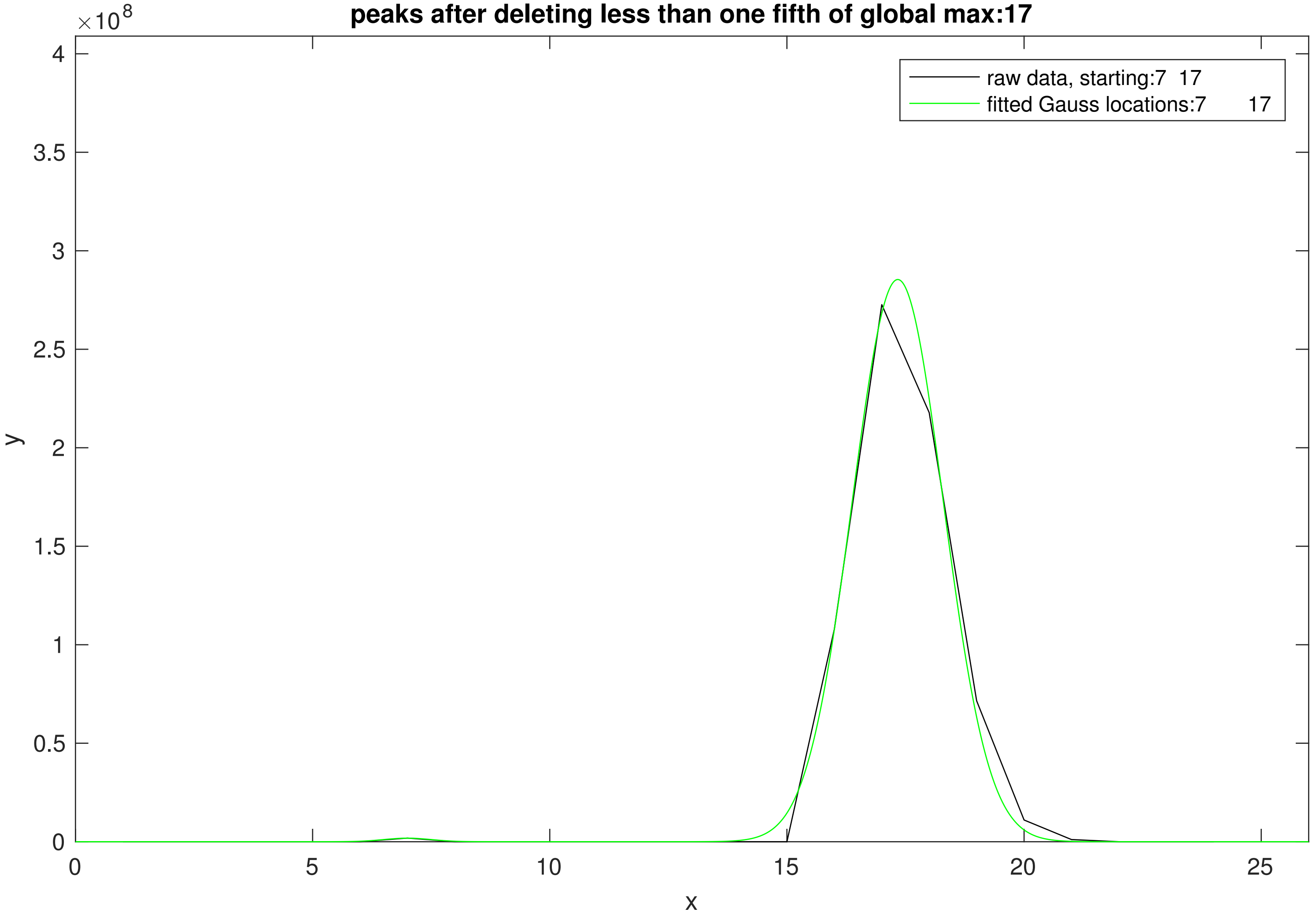

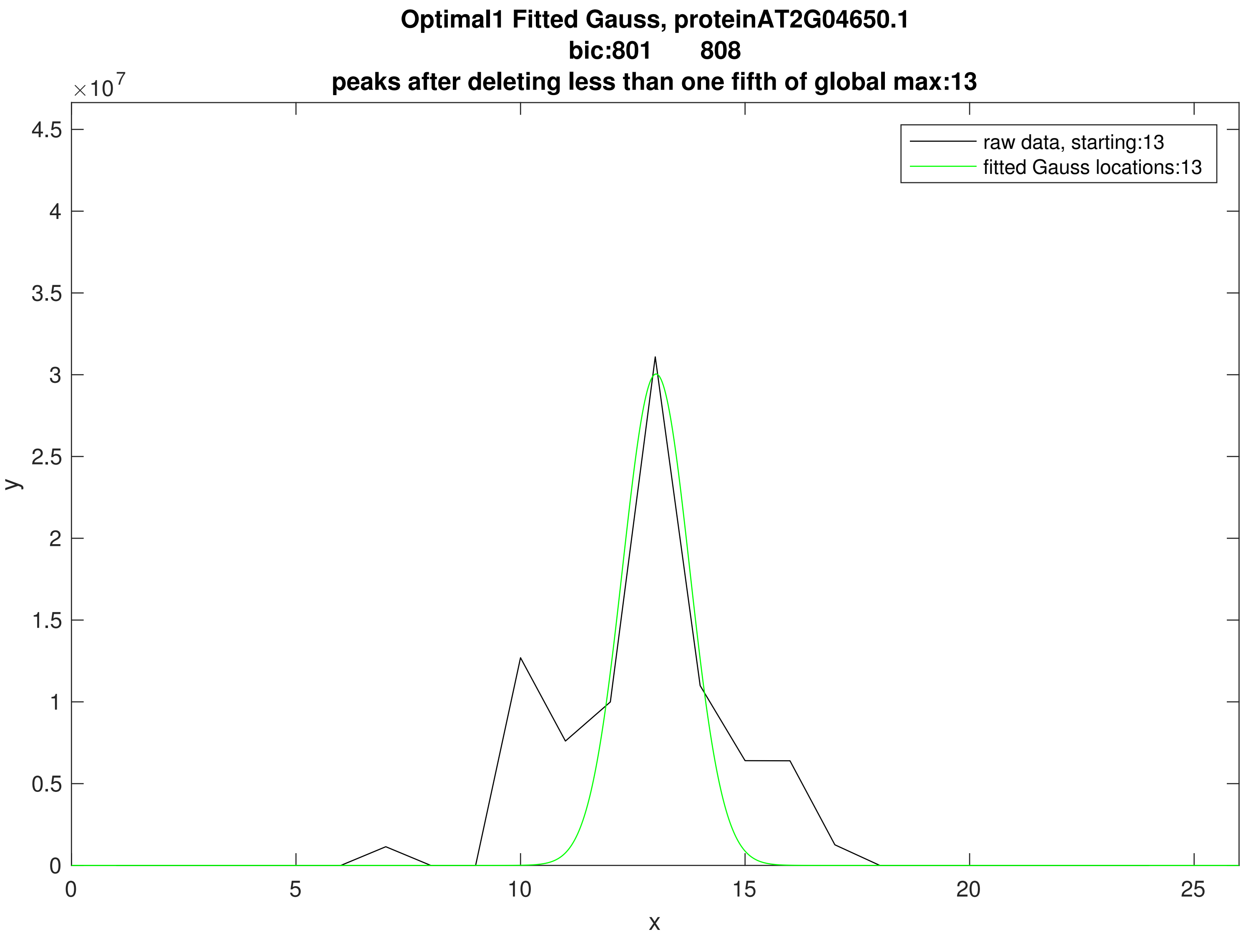
Optimal1 Fitted Gauss, proteinAT2G04650.1
bic:801 808
peaks after deleting less than one fifth of global max:13
×10^7
4.5
4
3.5
3
2.5
2
1.5
1
0.5
0
y
0
5
10
15
20
25
x
raw data, starting:13
fitted Gauss locations:13

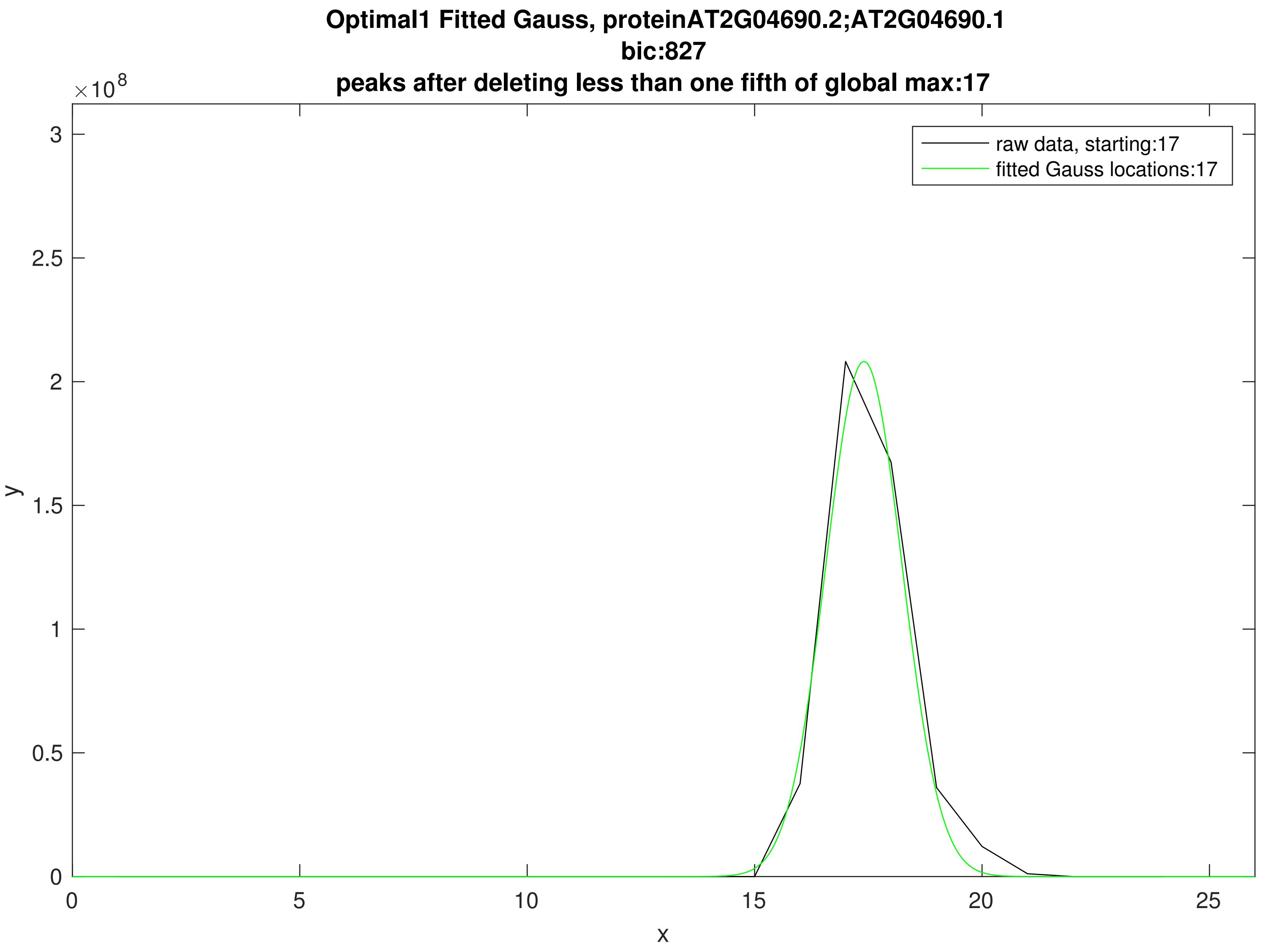

Optimal1 Fitted Gauss, proteinAT2G04690.2;AT2G04690.1
bic:827
peaks after deleting less than one fifth of global max:17
×10^8
3
2.5
2
1.5
1
0.5
0
y
0
5
10
15
20
25
x
raw data, starting:17
fitted Gauss locations:17

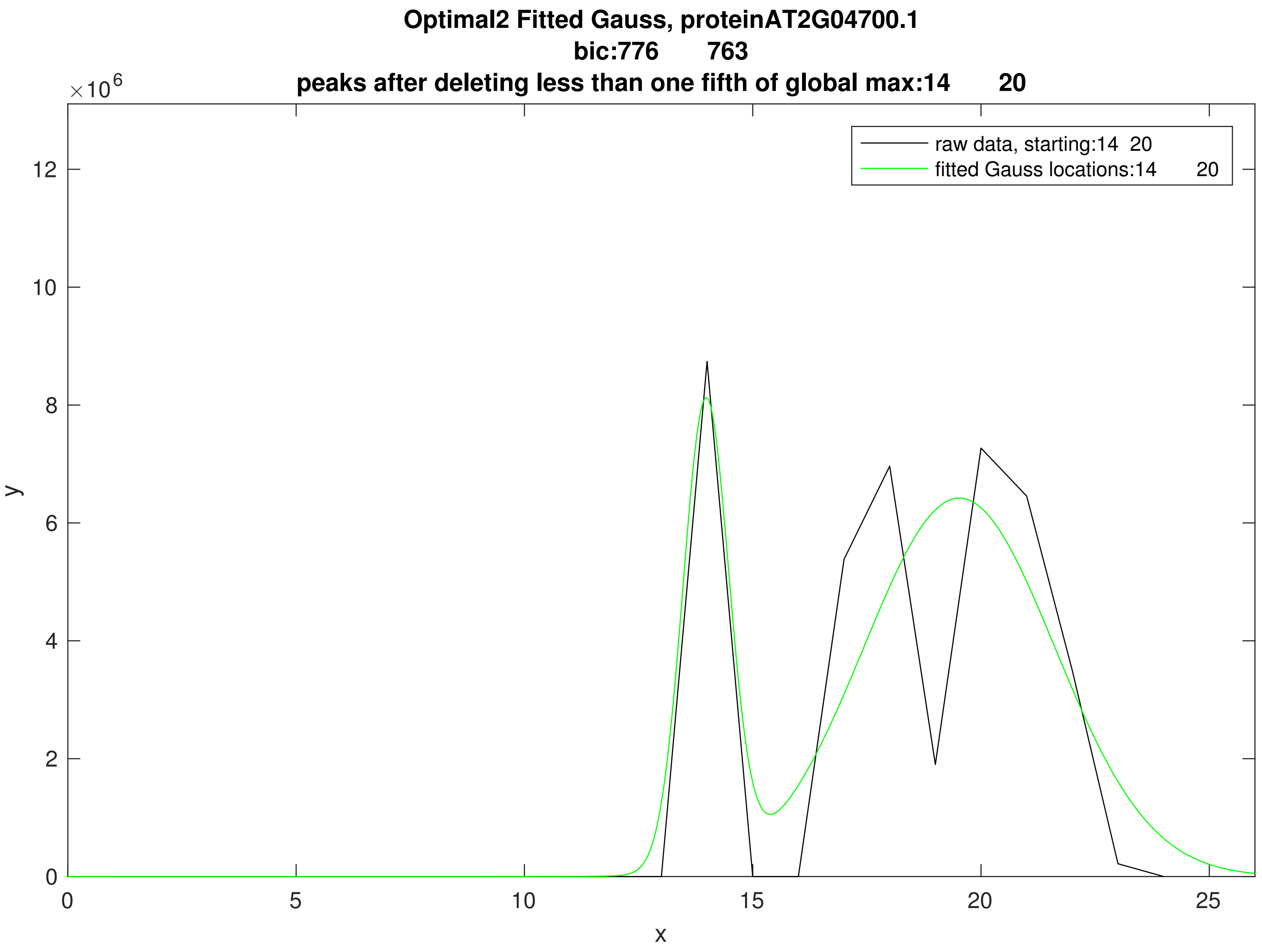

Optimal2 Fitted Gauss, proteinAT2G04700.1
bic:776 763
peaks after deleting less than one fifth of global max:14 20
×10^6
raw data, starting:14 20
fitted Gauss locations:14 20
y
x
0
2
4
6
8
10
12
0
5
10
15
20
25

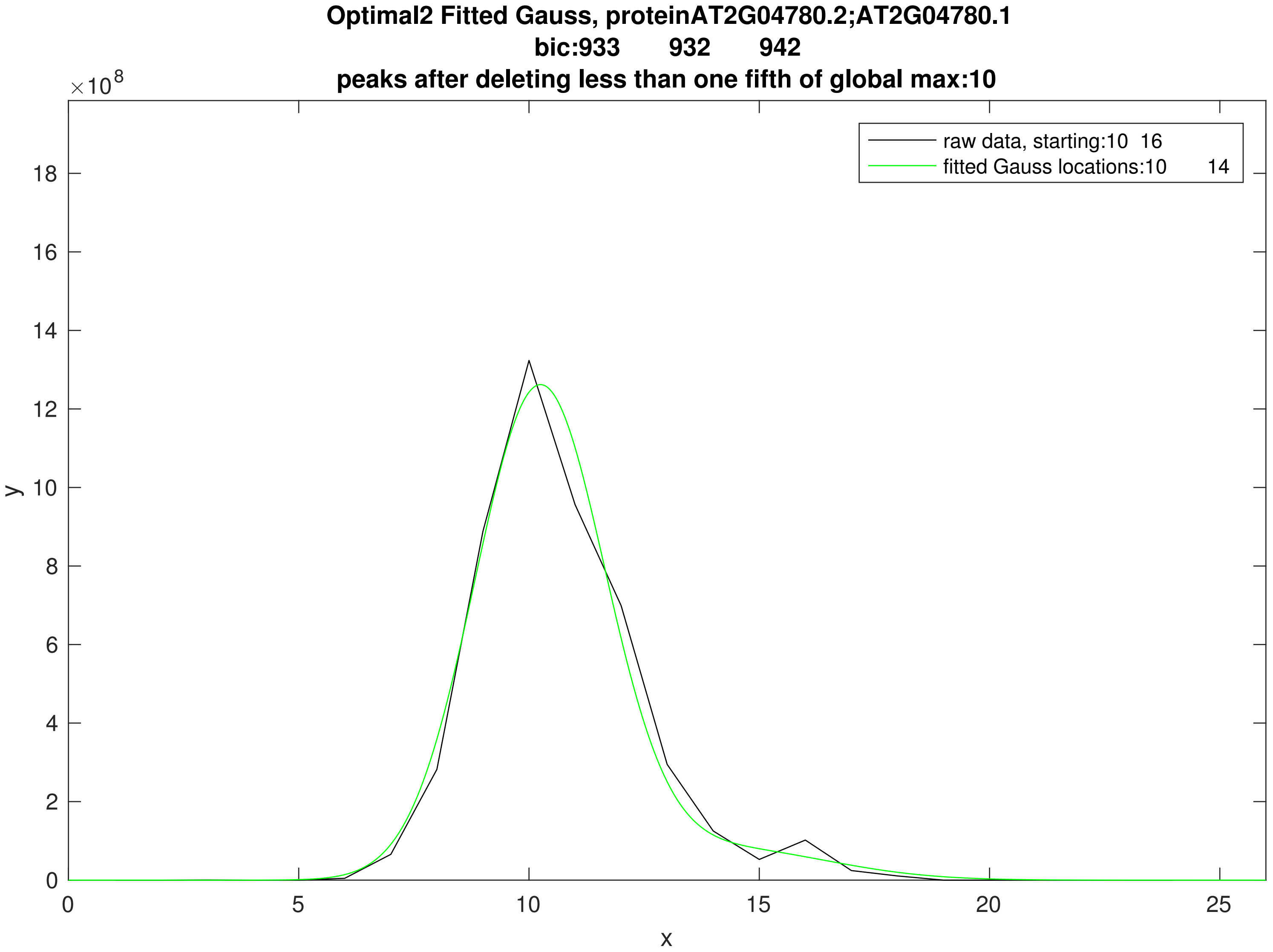

Optimal2 Fitted Gauss, proteinAT2G04780.2;AT2G04780.1
bic:933 932 942
peaks after deleting less than one fifth of global max:10
×10^8
raw data, starting:10 16
fitted Gauss locations:10 14
y
x
0
2
4
6
8
10
12
14
16
18
0
5
10
15
20
25

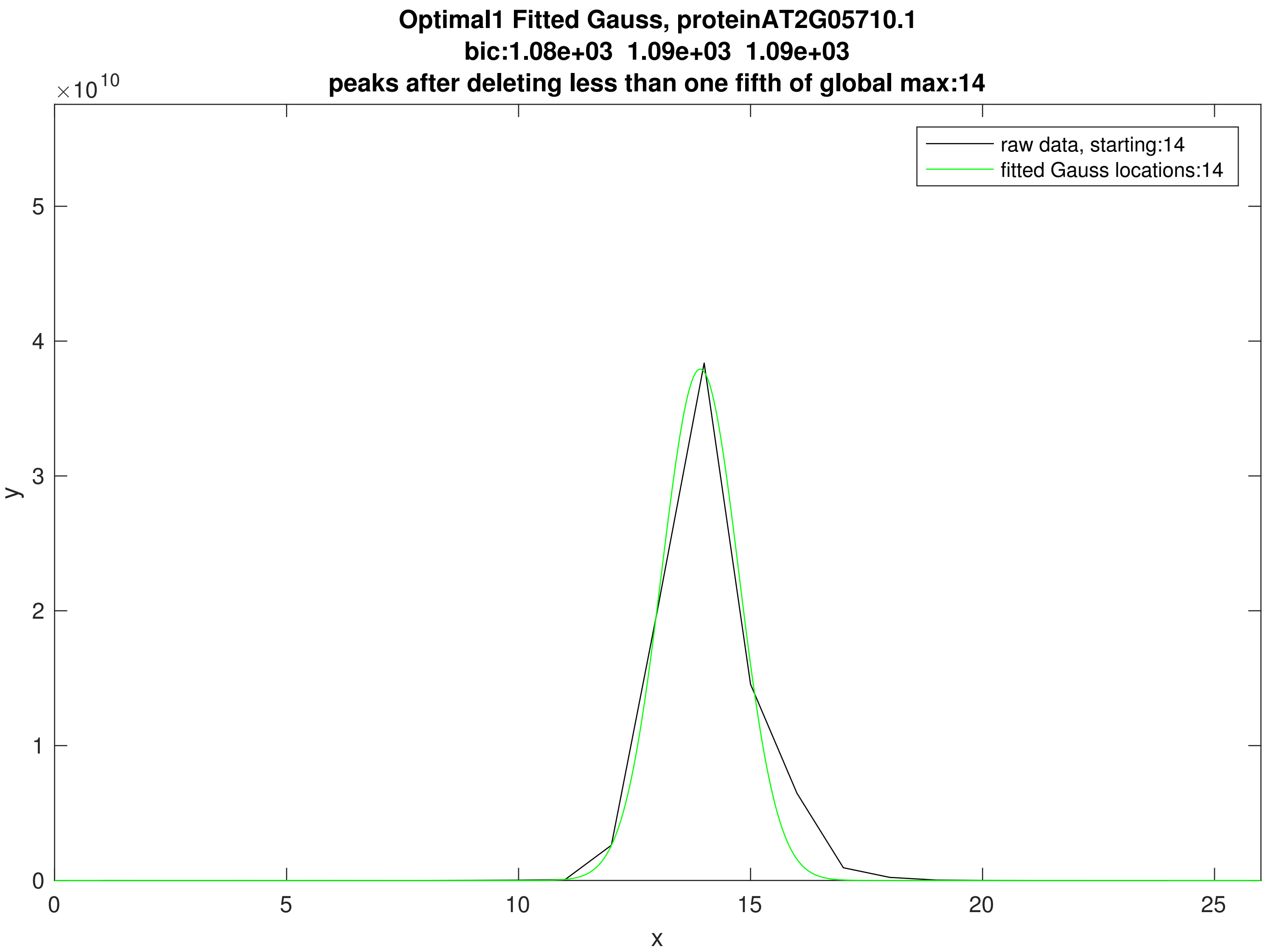
Optimal1 Fitted Gauss, proteinAT2G05710.1
bic:1.08e+03  1.09e+03  1.09e+03
peaks after deleting less than one fifth of global max:14
×10^10
raw data, starting:14
fitted Gauss locations:14
y
x
0
1
2
3
4
5
10
15
20
25

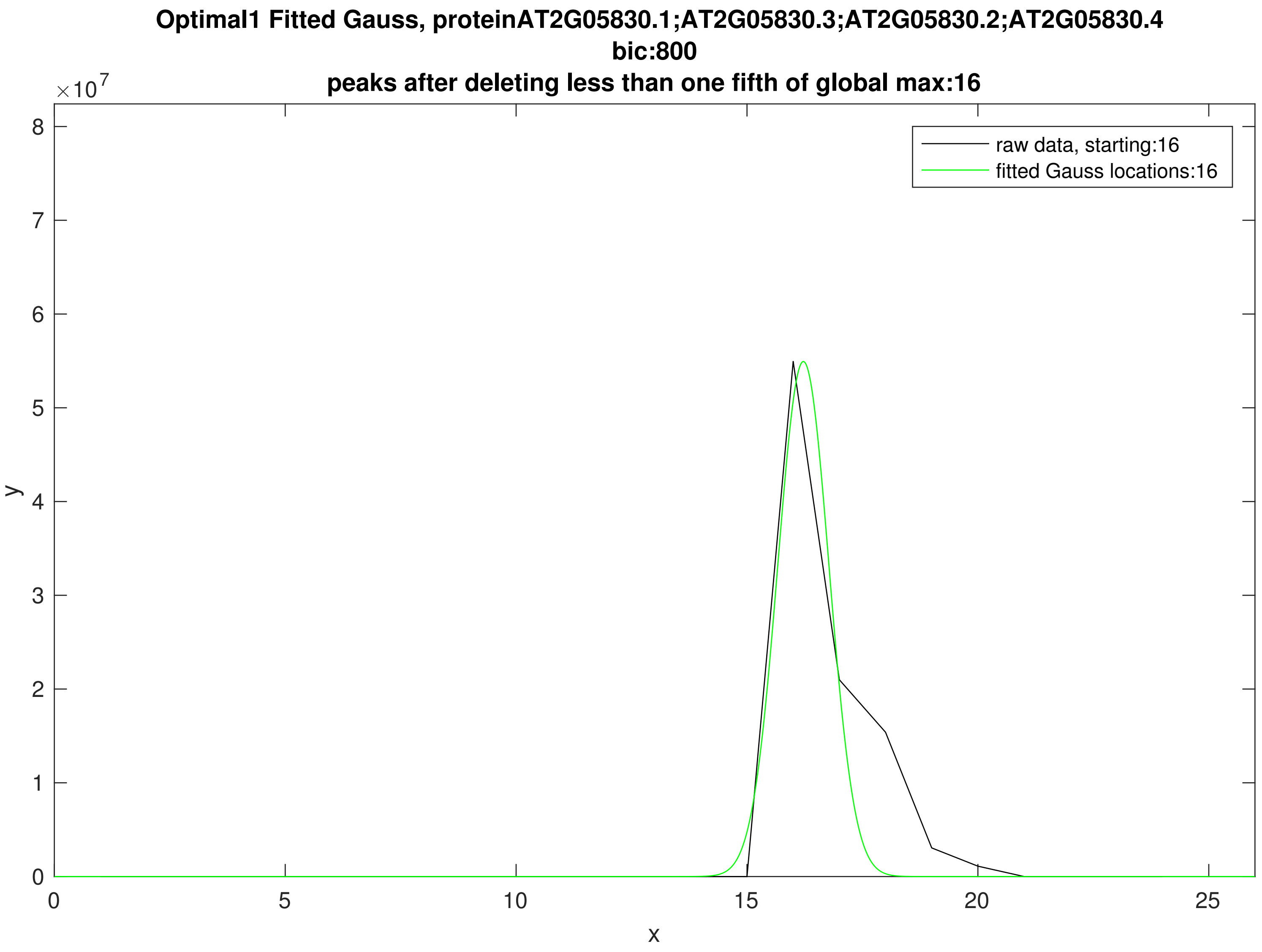
Optimal1 Fitted Gauss, proteinAT2G05830.1;AT2G05830.3;AT2G05830.2;AT2G05830.4
bic:800
peaks after deleting less than one fifth of global max:16
×10^7
raw data, starting:16
fitted Gauss locations:16
y
x
0
1
2
3
4
5
6
7
8
0
5
10
15
20
25

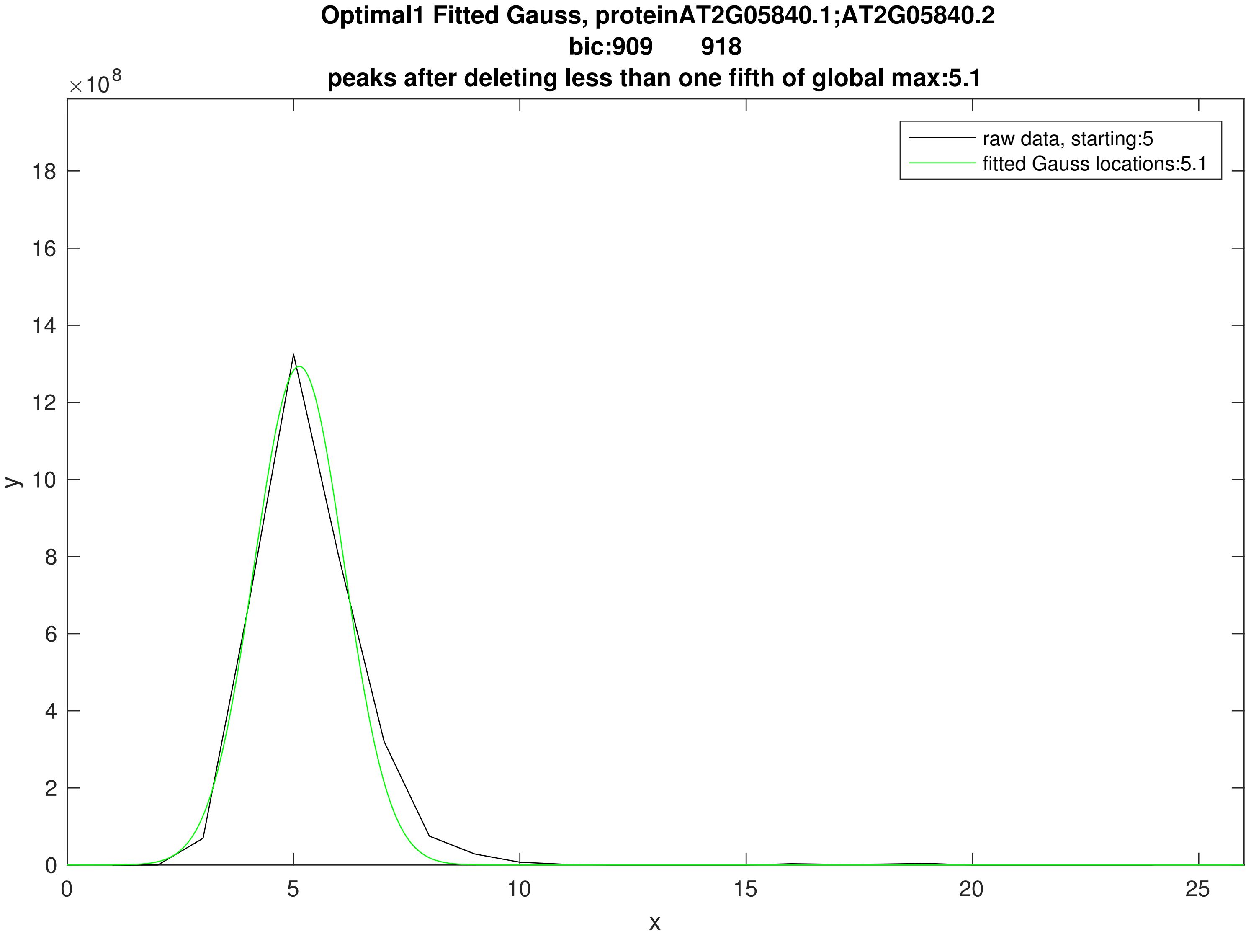

Optimal1 Fitted Gauss, proteinAT2G05840.1;AT2G05840.2
bic:909 918
peaks after deleting less than one fifth of global max:5.1
×10^8
raw data, starting:5
fitted Gauss locations:5.1
y
x
0
2
4
6
8
10
12
14
16
18
0
5
10
15
20
25

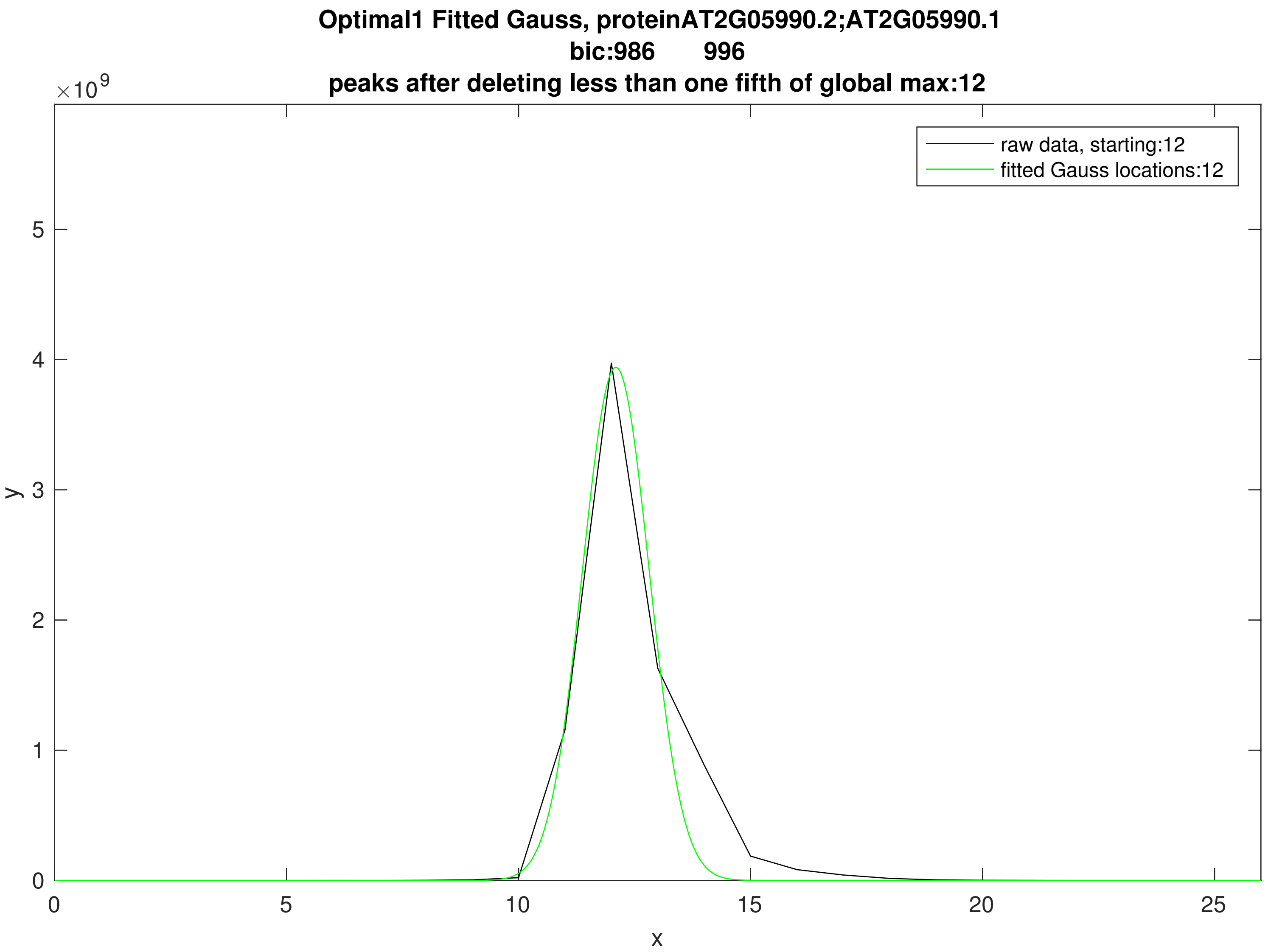
Optimal1 Fitted Gauss, proteinAT2G05990.2;AT2G05990.1
bic:986 996
peaks after deleting less than one fifth of global max:12
×10^9
raw data, starting:12
fitted Gauss locations:12
y
x
0
1
2
3
4
5
0
5
10
15
20
25

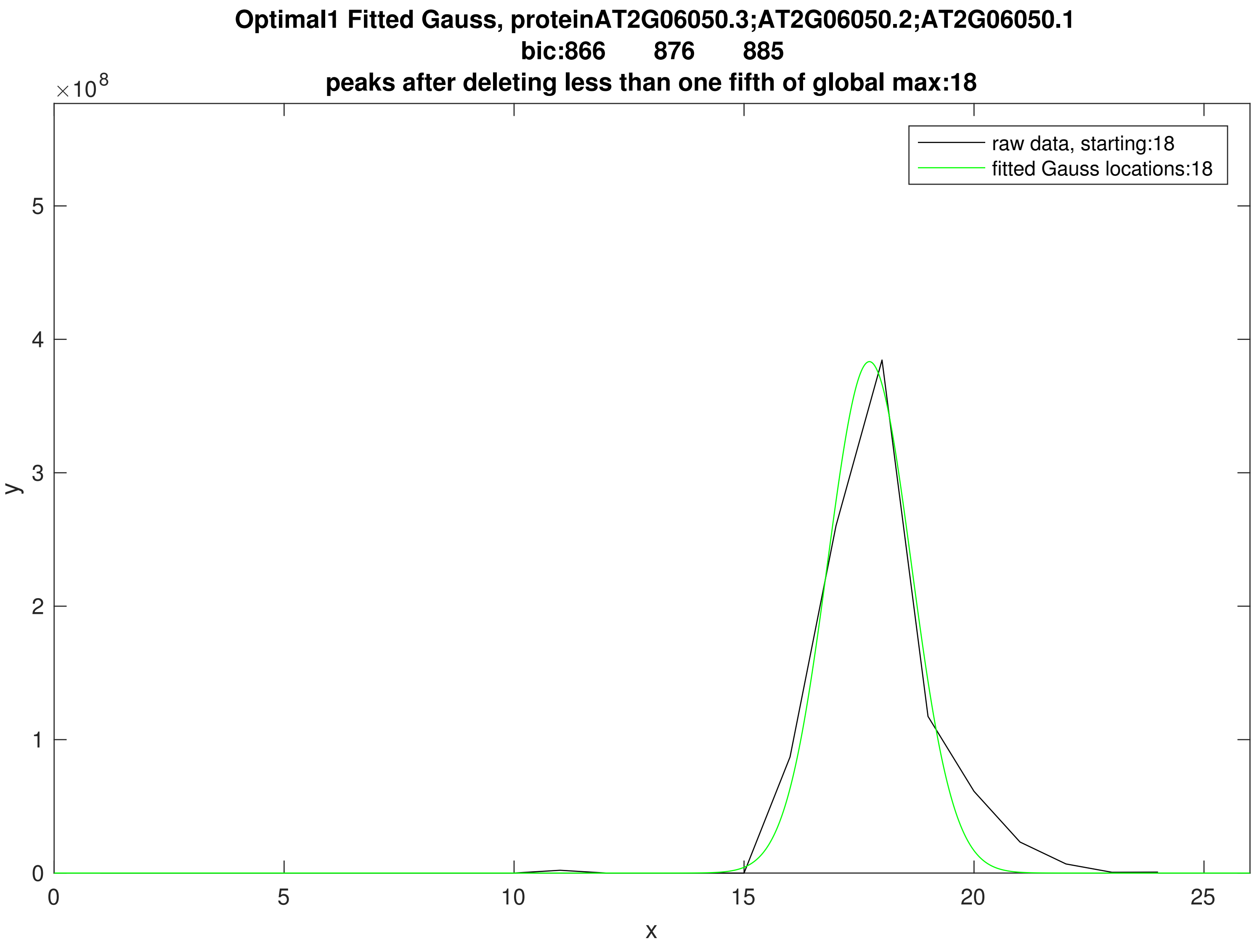
Optimal1 Fitted Gauss, proteinAT2G06050.3;AT2G06050.2;AT2G06050.1
bic:866 876 885
peaks after deleting less than one fifth of global max:18
×10^8
raw data, starting:18
fitted Gauss locations:18
y
x
0
1
2
3
4
5
0
5
10
15
20
25

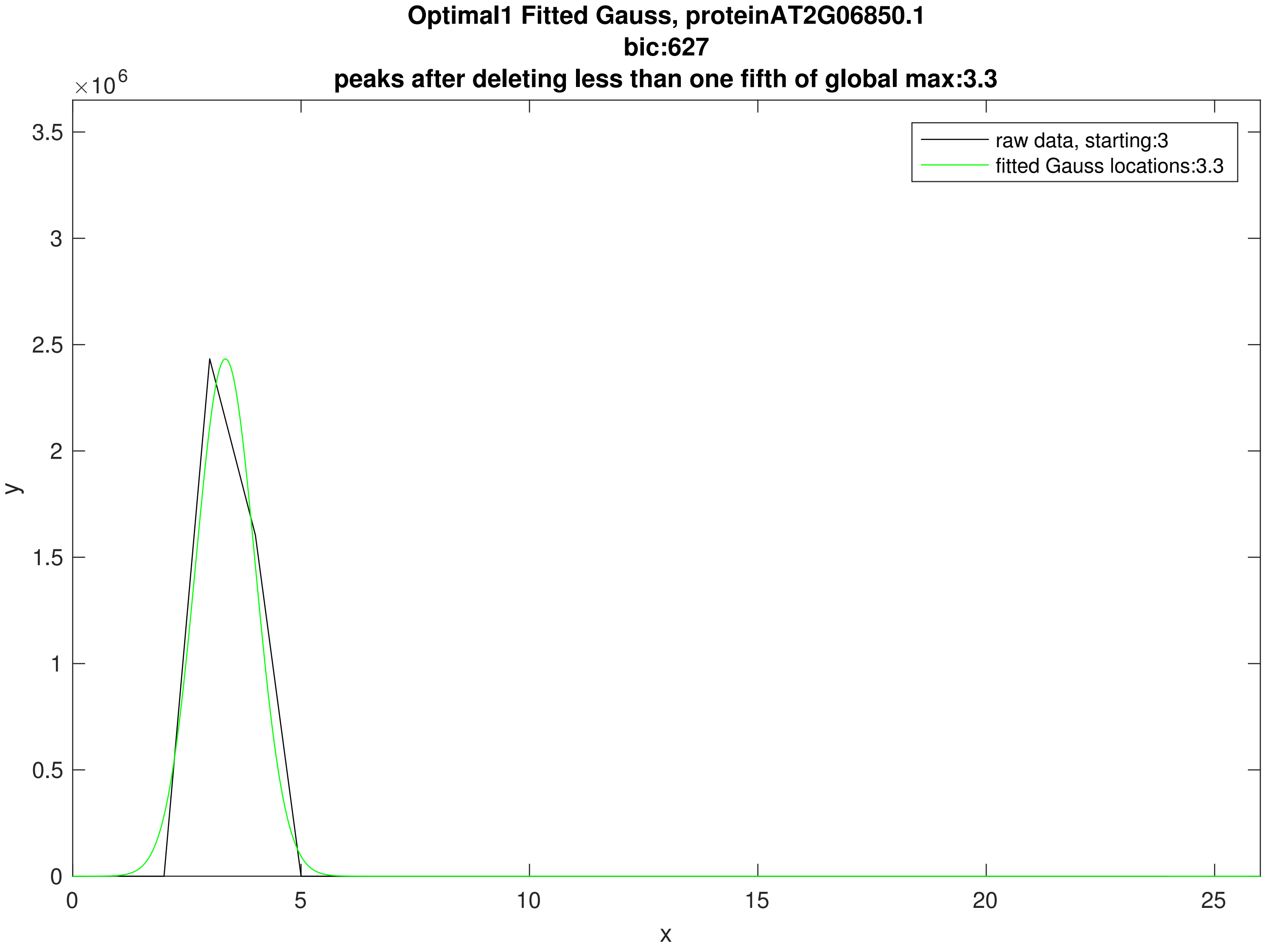
Optimal1 Fitted Gauss, proteinAT2G06850.1
bic:627
peaks after deleting less than one fifth of global max:3.3
×10^6
3.5
3
2.5
2
1.5
1
0.5
0
y
0
5
10
15
20
25
x
raw data, starting:3
fitted Gauss locations:3.3

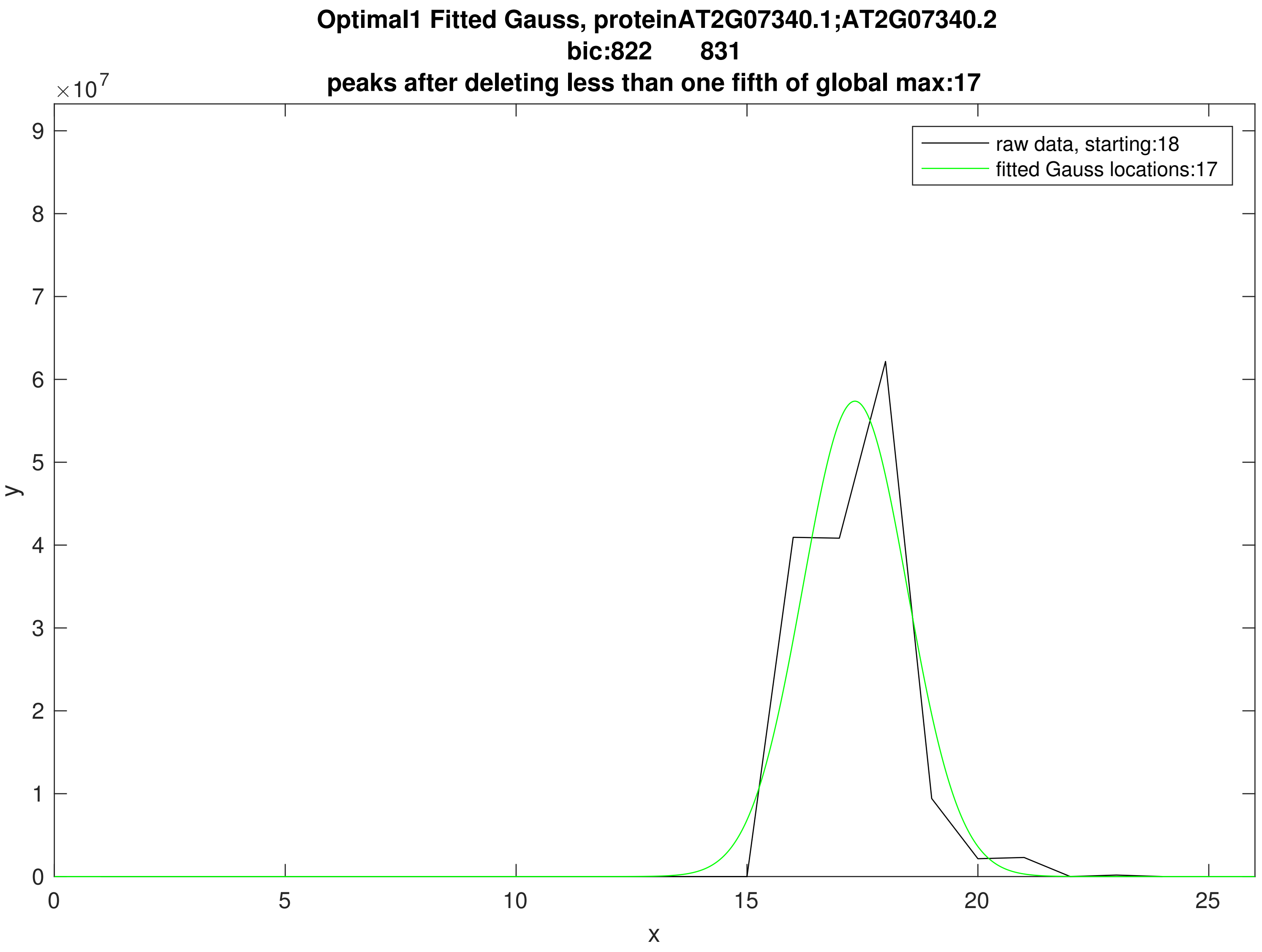
Optimal1 Fitted Gauss, proteinAT2G07340.1;AT2G07340.2
bic:822 831
peaks after deleting less than one fifth of global max:17
×10^7
raw data, starting:18
fitted Gauss locations:17
y
x
0
5
10
15
20
25
1
2
3
4
5
6
7
8
9

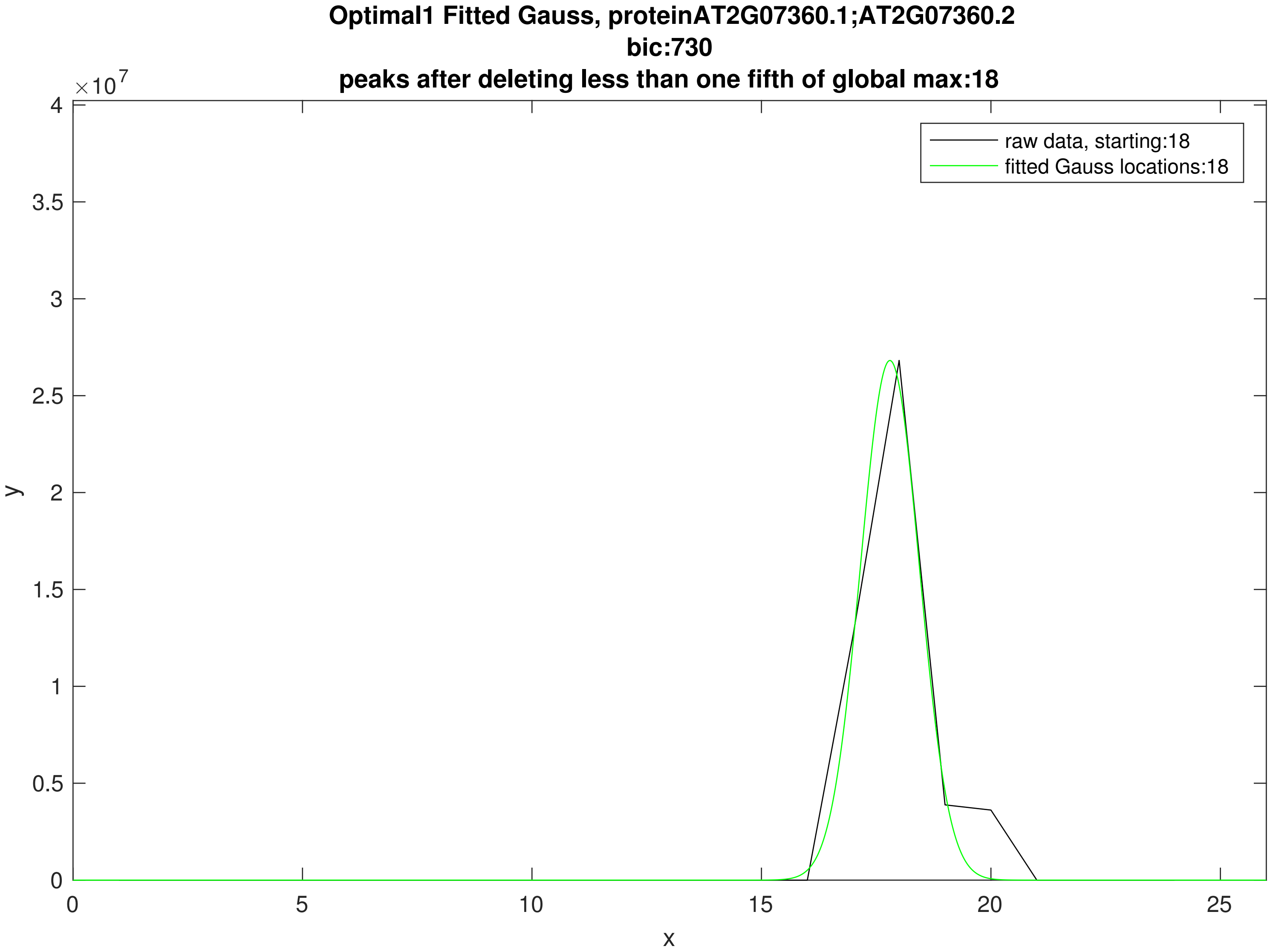

Optimal1 Fitted Gauss, proteinAT2G07360.1;AT2G07360.2
bic:730
peaks after deleting less than one fifth of global max:18
×10^7
raw data, starting:18
fitted Gauss locations:18
y
x
0
0.5
1
1.5
2
2.5
3
3.5
4
0
5
10
15
20
25

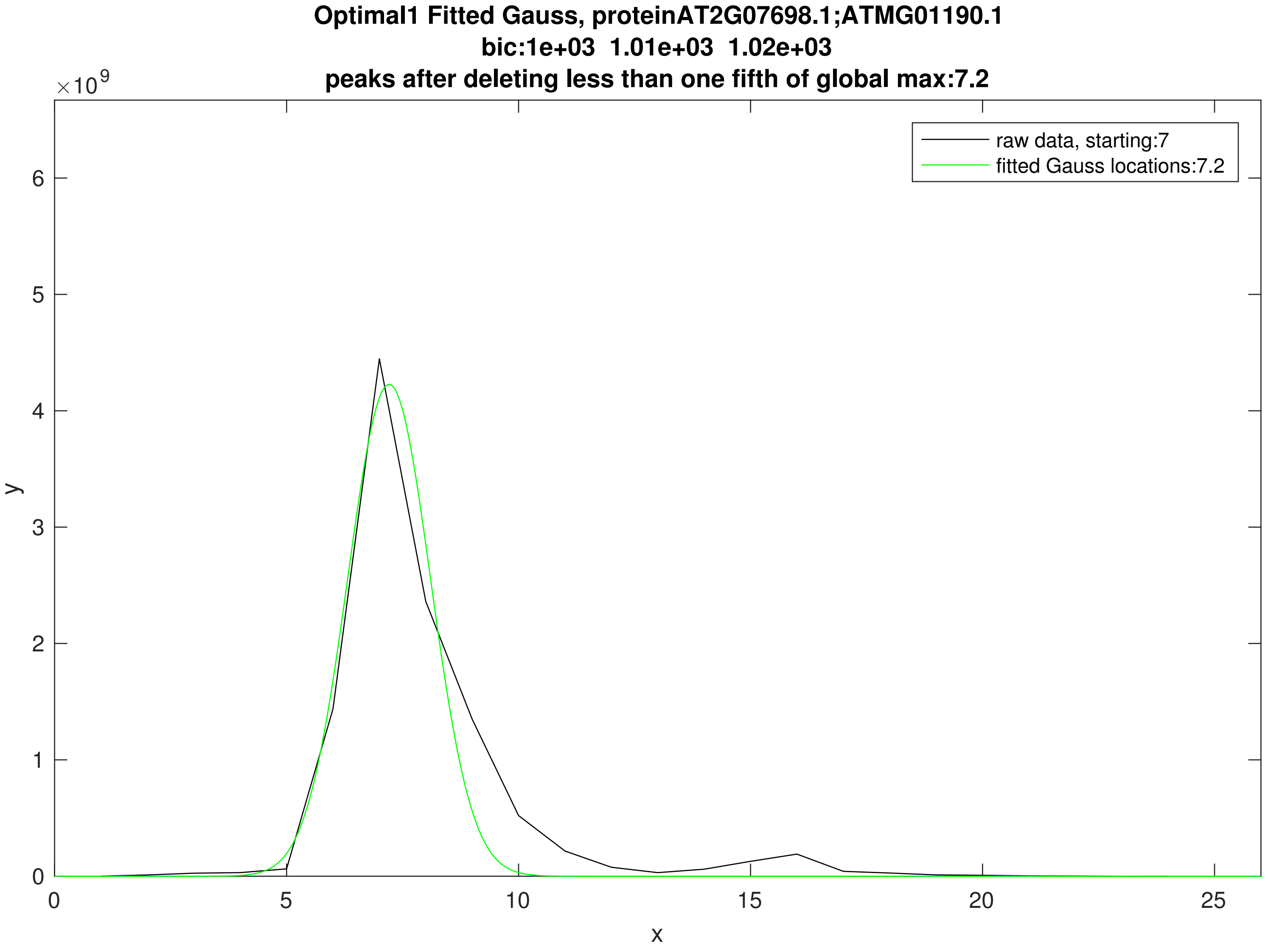
Optimal1 Fitted Gauss, proteinAT2G07698.1;ATMG01190.1
bic:1e+03  1.01e+03  1.02e+03
peaks after deleting less than one fifth of global max:7.2
×10^9
raw data, starting:7
fitted Gauss locations:7.2
y
x
0
1
2
3
4
5
6
0
5
10
15
20
25

**Optimal1 Fitted Gauss, proteinAT2G13360.2;AT2G13360.1**
**bic:685**
**peaks after deleting less than one fifth of global max:12**

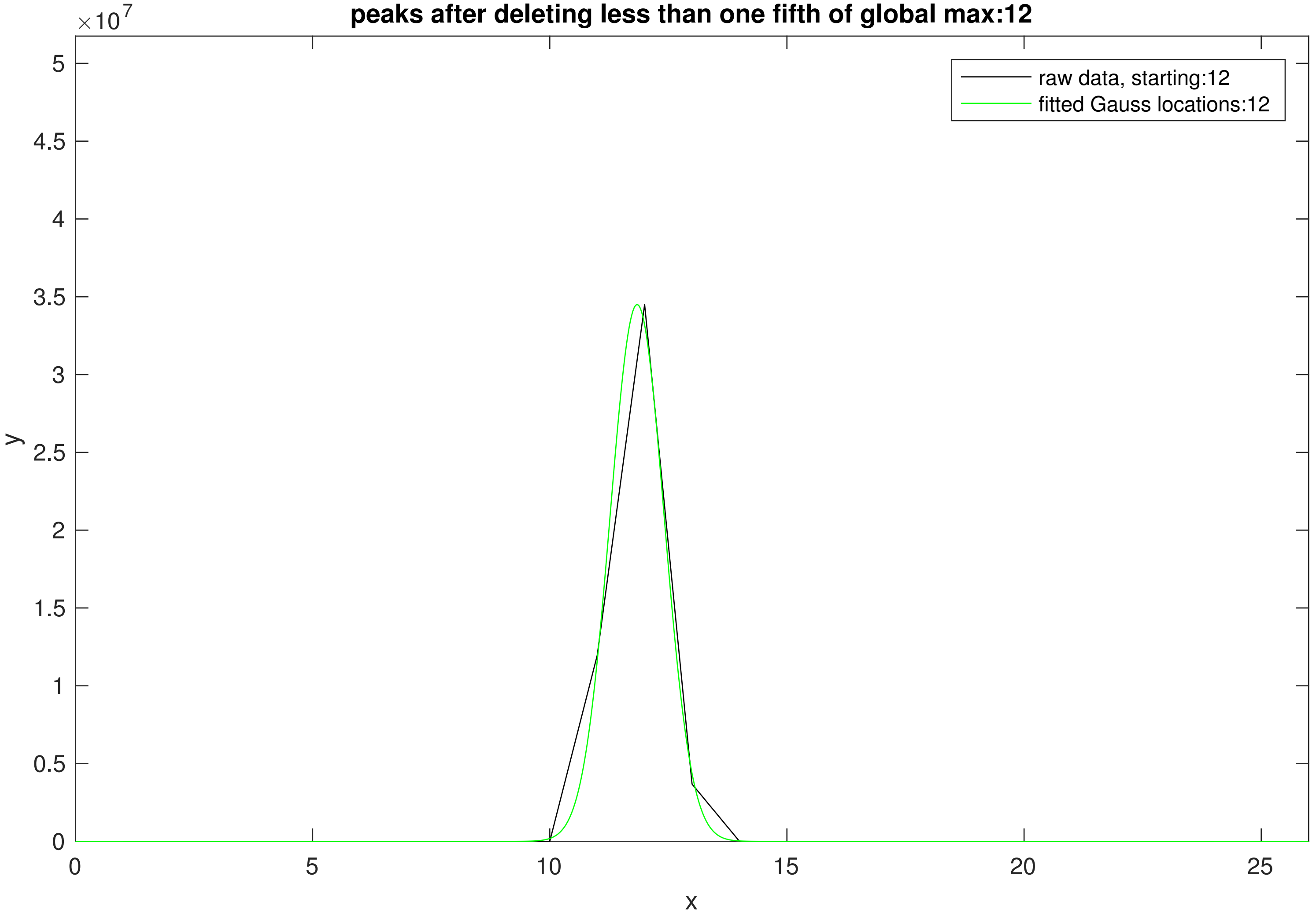

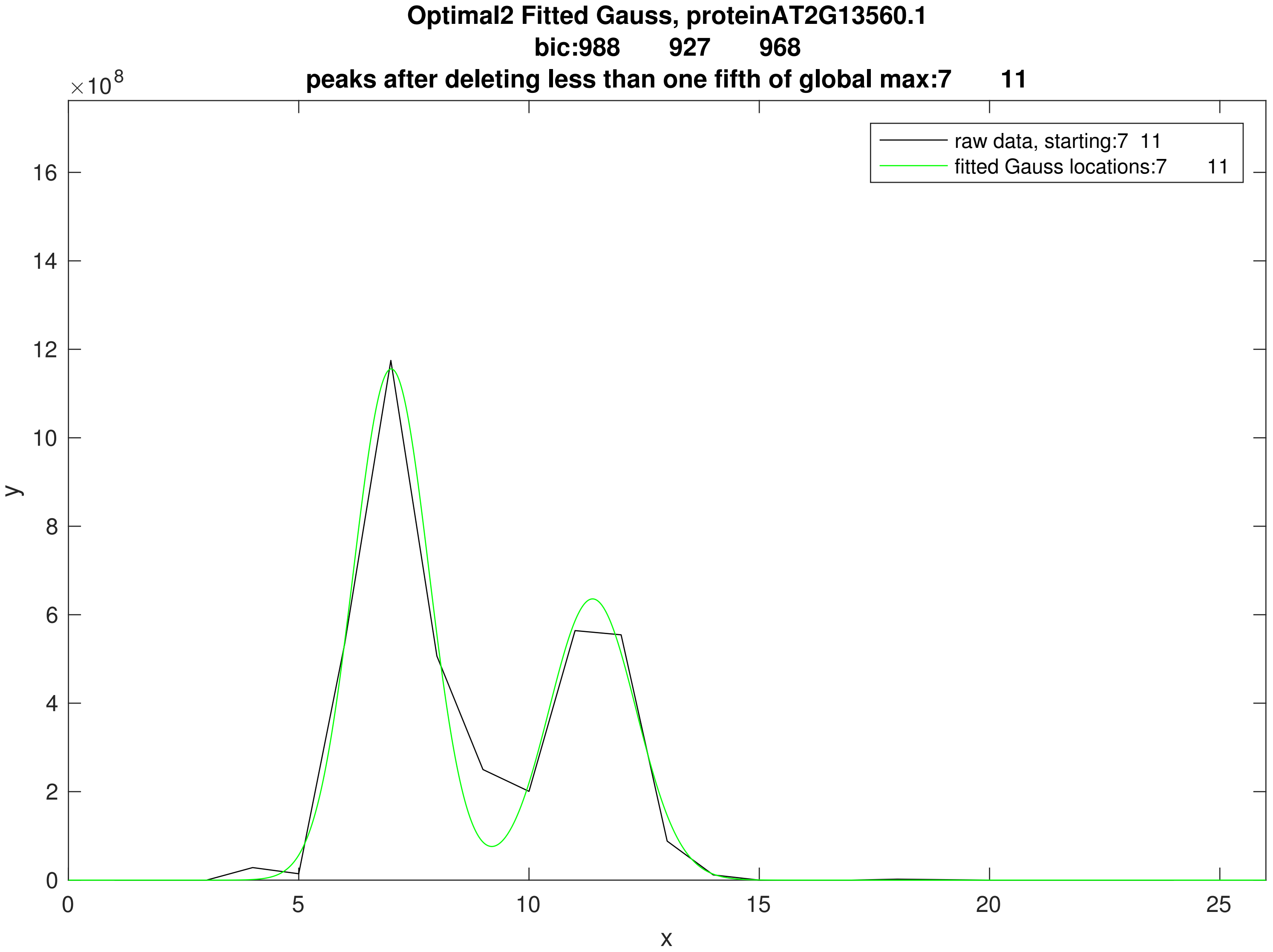

Optimal2 Fitted Gauss, proteinAT2G13560.1
bic:988 927 968
peaks after deleting less than one fifth of global max:7 11
×10^8
raw data, starting:7 11
fitted Gauss locations:7 11
y
x
0
2
4
6
8
10
12
14
16
0
5
10
15
20
25

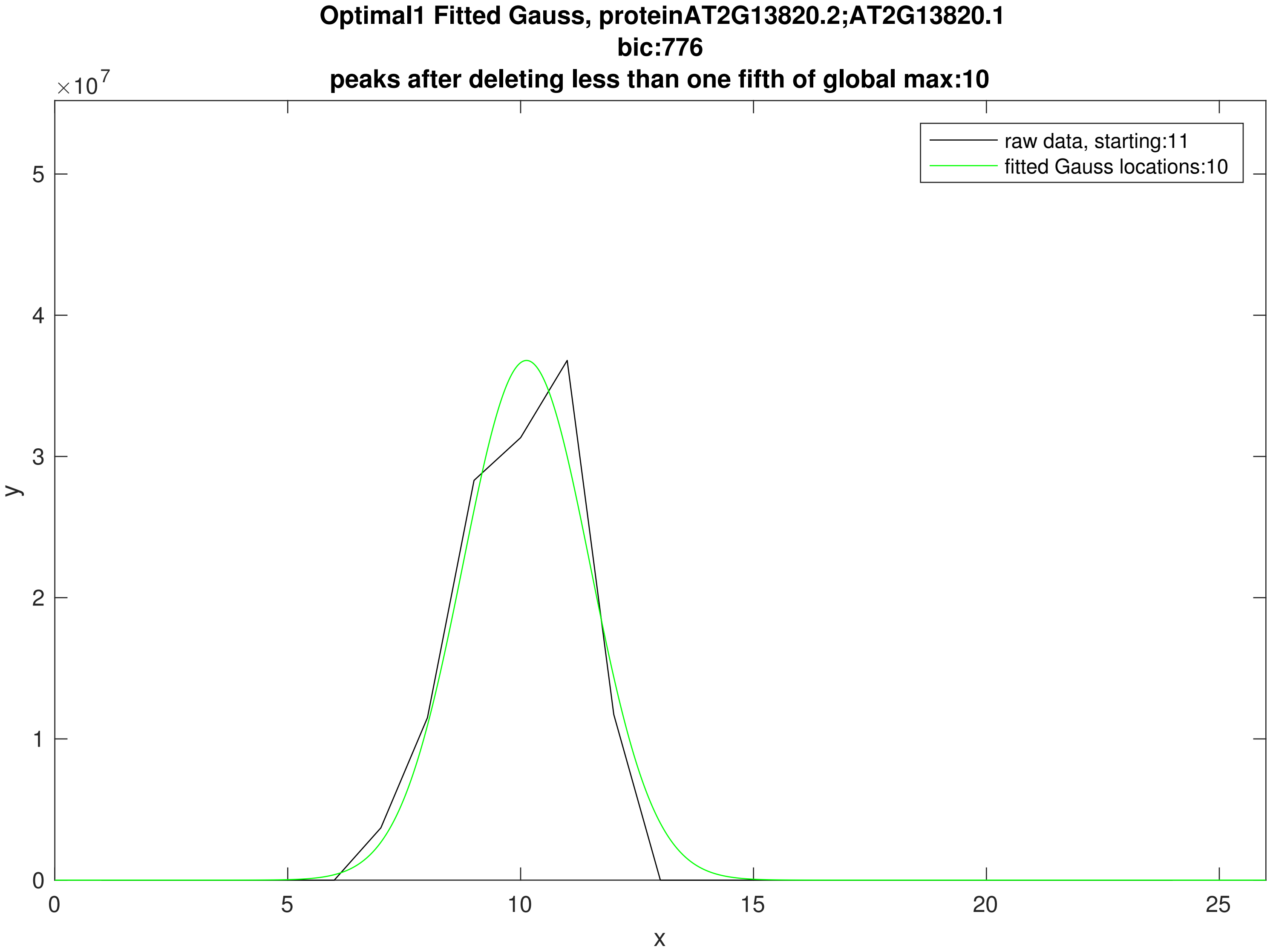
Optimal1 Fitted Gauss, proteinAT2G13820.2;AT2G13820.1
bic:776
peaks after deleting less than one fifth of global max:10
×10^7
raw data, starting:11
fitted Gauss locations:10
y
x
0
1
2
3
4
5
0
5
10
15
20
25

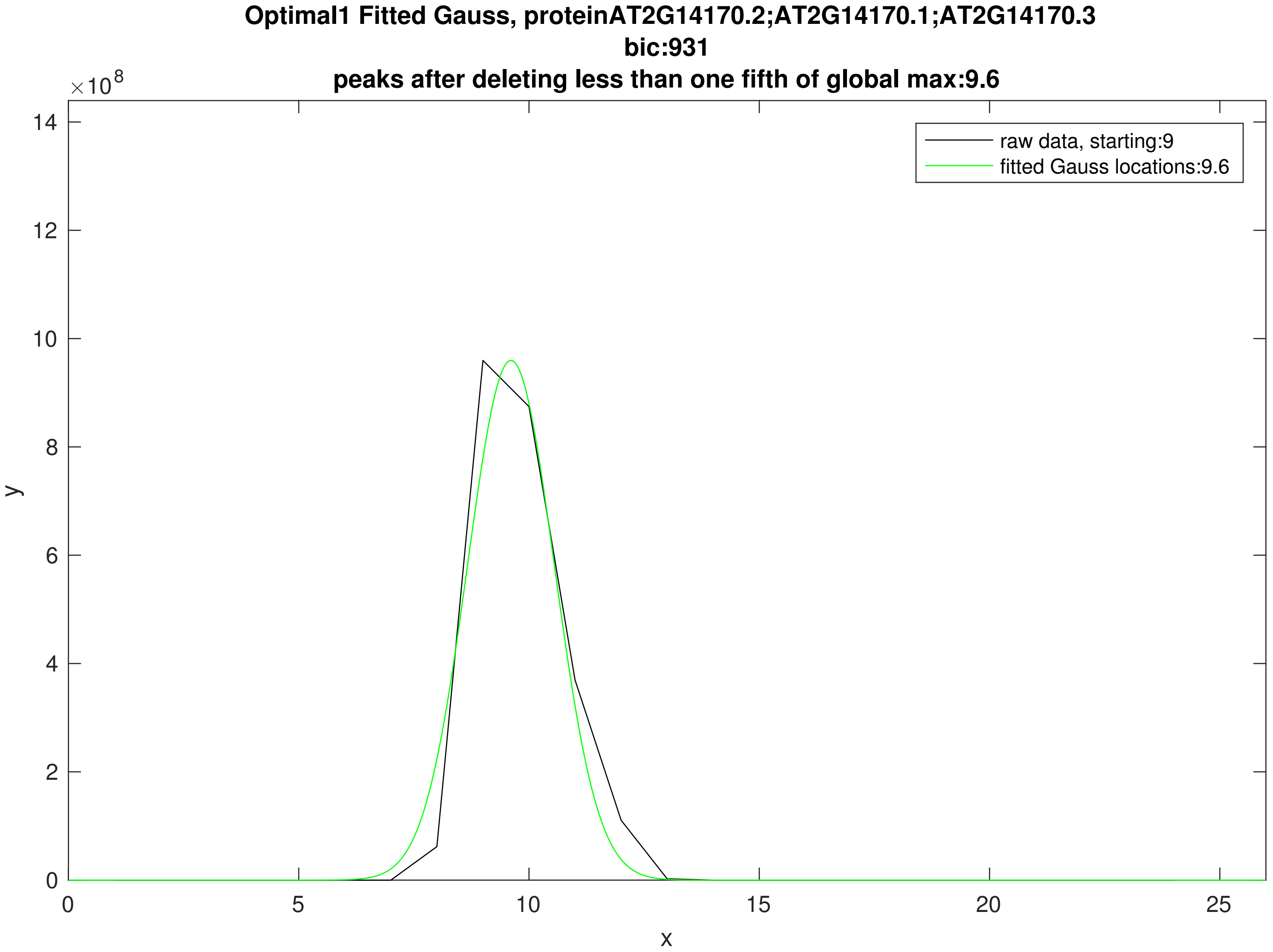
Optimal1 Fitted Gauss, proteinAT2G14170.2;AT2G14170.1;AT2G14170.3
bic:931
peaks after deleting less than one fifth of global max:9.6
×10^8
y
x
raw data, starting:9
fitted Gauss locations:9.6
0
2
4
6
8
10
12
14
0
5
10
15
20
25

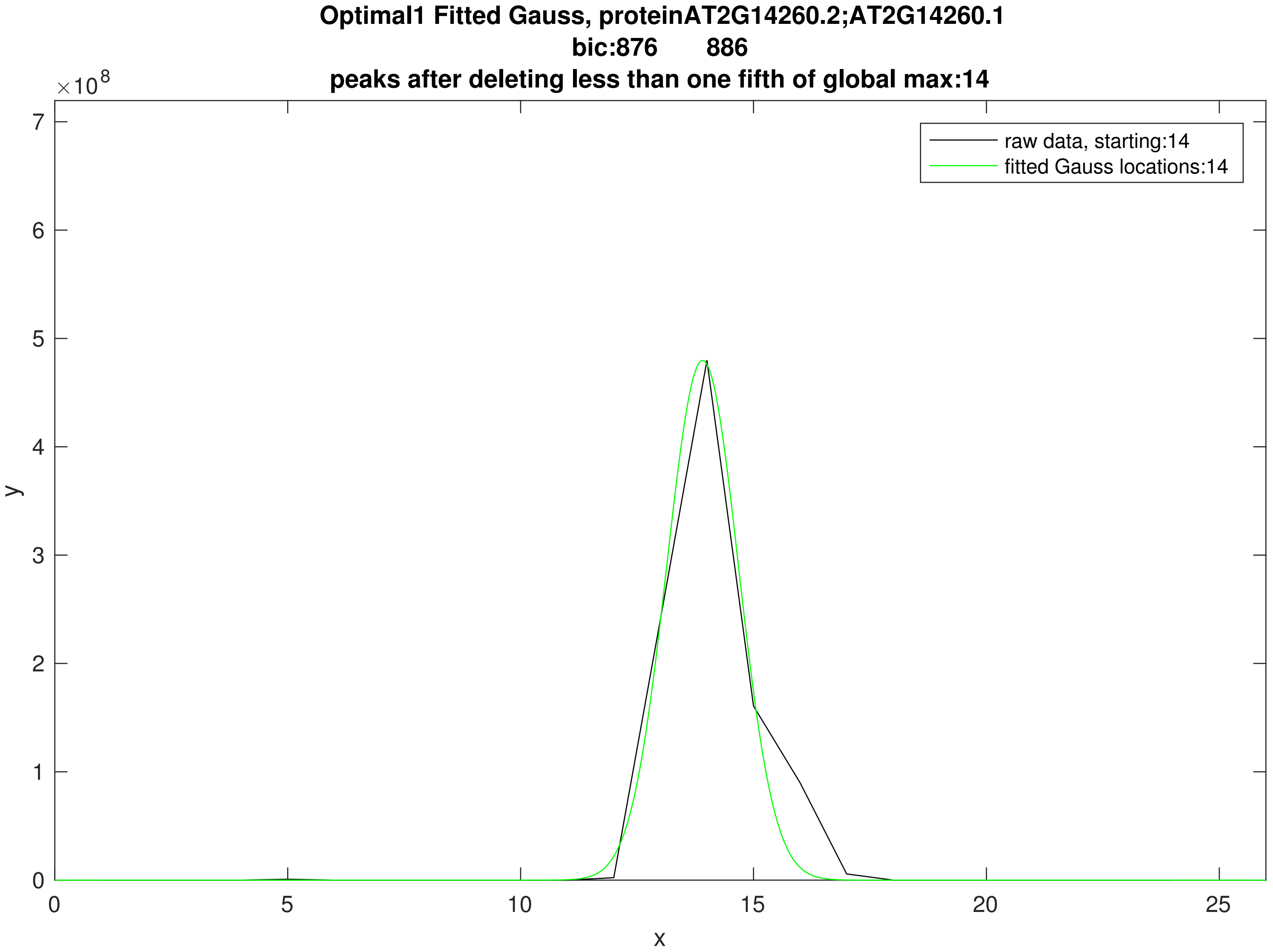

Optimal1 Fitted Gauss, proteinAT2G14260.2;AT2G14260.1
bic:876 886
peaks after deleting less than one fifth of global max:14
×10^8
raw data, starting:14
fitted Gauss locations:14
y
x
0
1
2
3
4
5
6
7
0
5
10
15
20
25

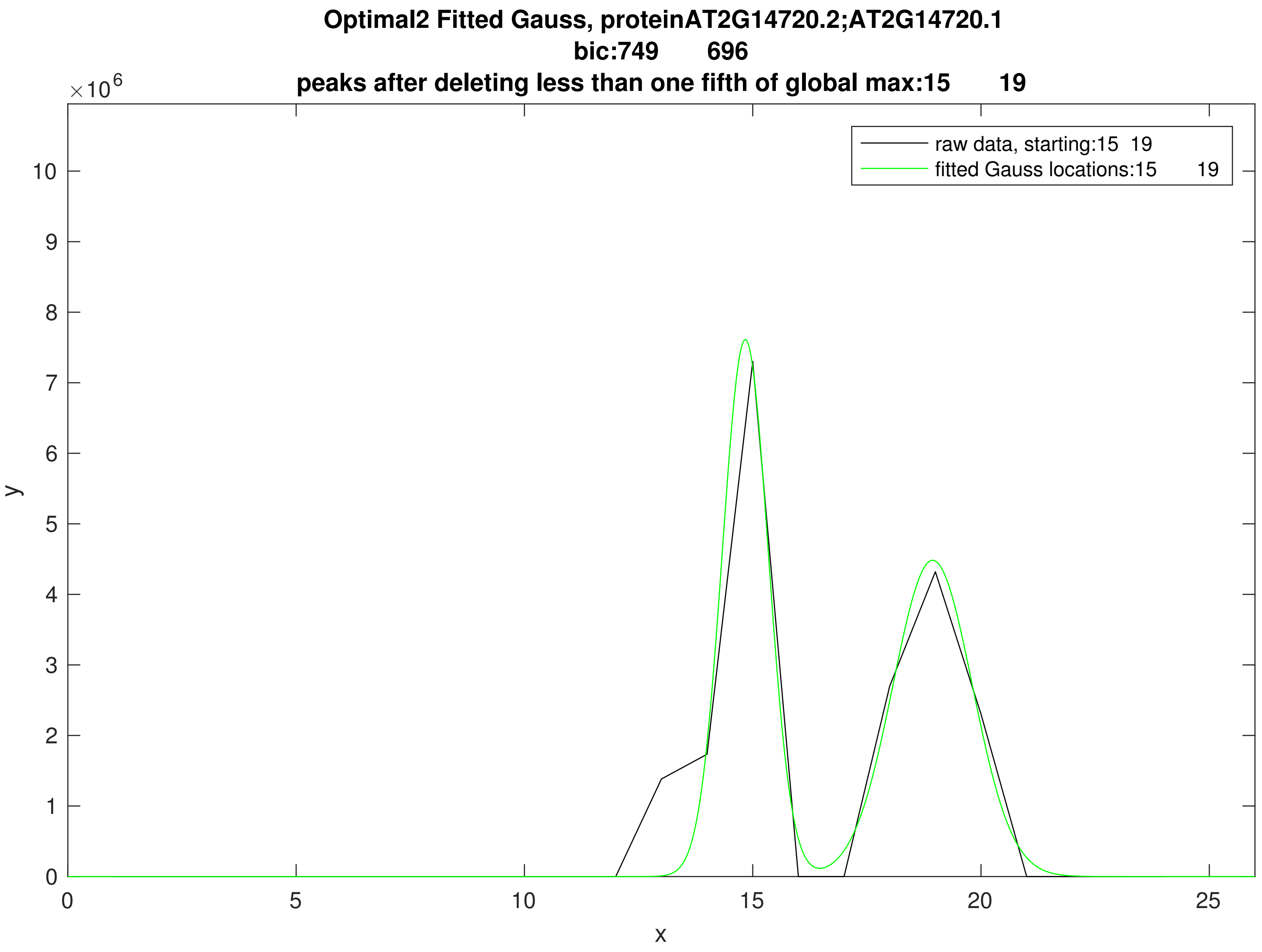
Optimal2 Fitted Gauss, proteinAT2G14720.2;AT2G14720.1
bic:749 696
peaks after deleting less than one fifth of global max:15 19
×10^6
raw data, starting:15 19
fitted Gauss locations:15 19
y
x
0
1
2
3
4
5
6
7
8
9
10
0
5
10
15
20
25

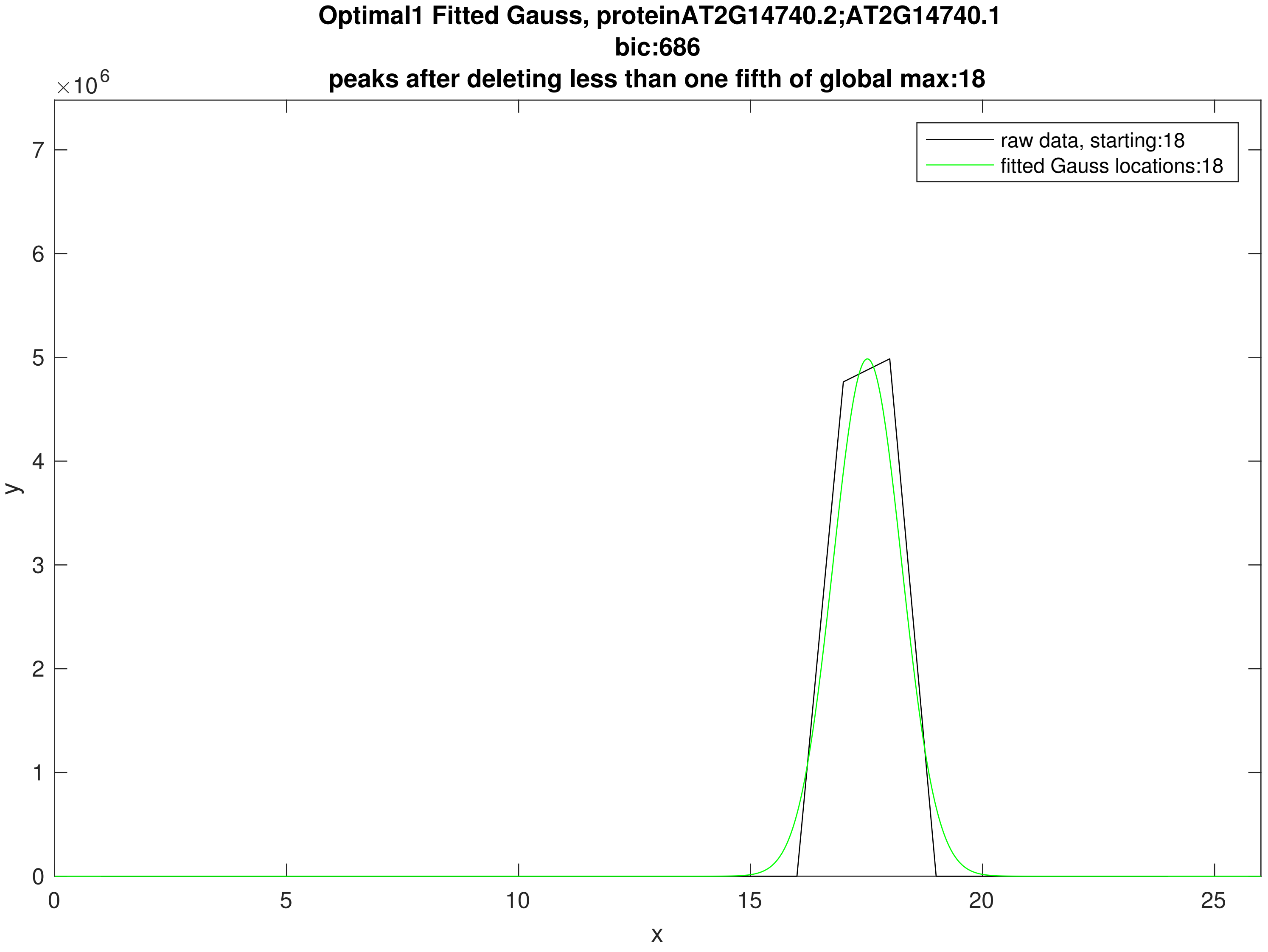
Optimal1 Fitted Gauss, proteinAT2G14740.2;AT2G14740.1
bic:686
peaks after deleting less than one fifth of global max:18
×10^6
raw data, starting:18
fitted Gauss locations:18
y
x
0
1
2
3
4
5
6
7
0
5
10
15
20
25

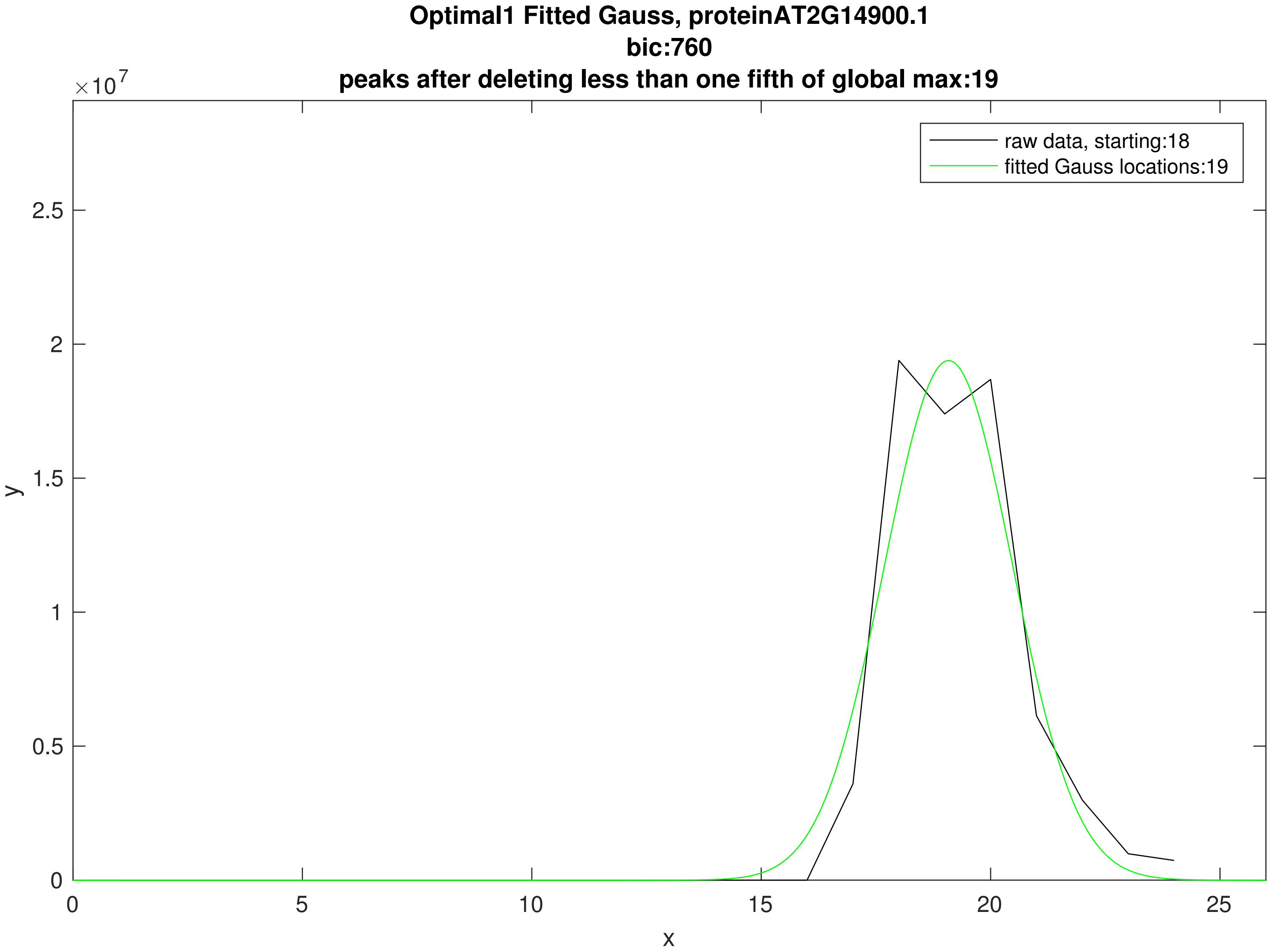

Optimal1 Fitted Gauss, proteinAT2G14900.1
bic:760
peaks after deleting less than one fifth of global max:19
×10^7
raw data, starting:18
fitted Gauss locations:19
y
x
0
0.5
1
1.5
2
2.5
0
5
10
15
20
25

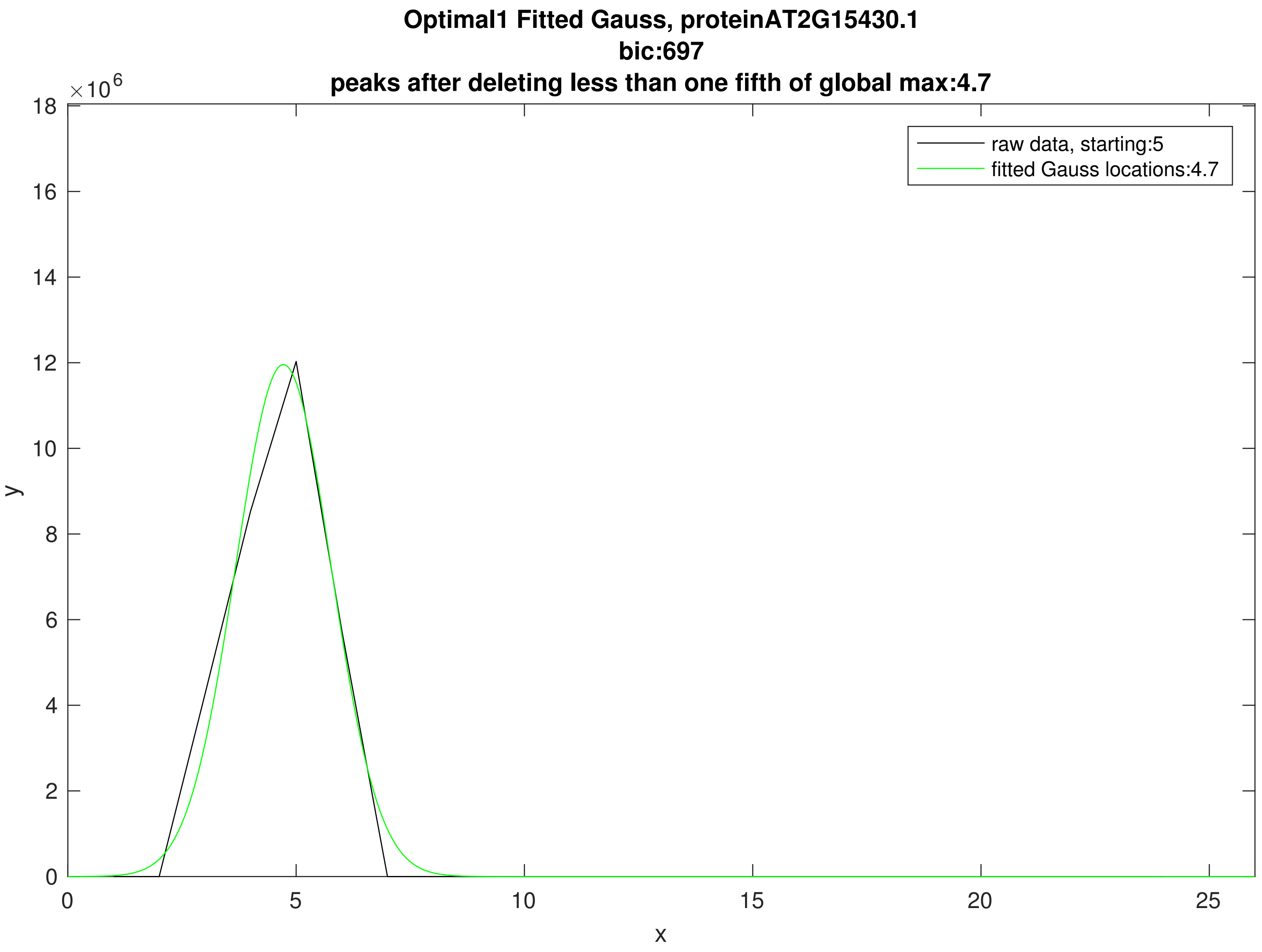
Optimal1 Fitted Gauss, proteinAT2G15430.1
bic:697
peaks after deleting less than one fifth of global max:4.7
×10^6
raw data, starting:5
fitted Gauss locations:4.7
y
x
0
2
4
6
8
10
12
14
16
18
0
5
10
15
20
25

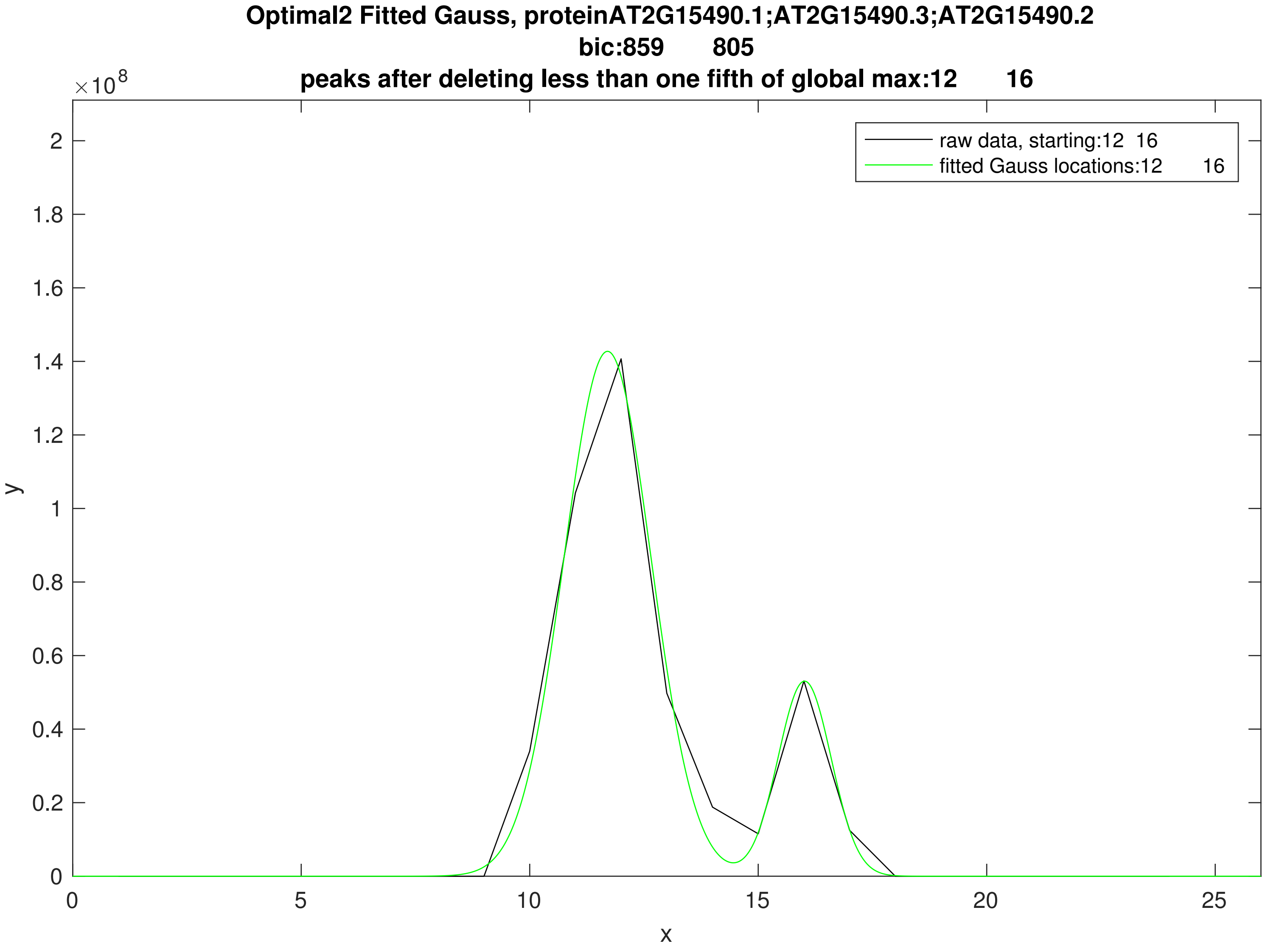
Optimal2 Fitted Gauss, proteinAT2G15490.1;AT2G15490.3;AT2G15490.2
bic:859 805
peaks after deleting less than one fifth of global max:12 16
×10^8
raw data, starting:12 16
fitted Gauss locations:12 16
y
x
0
0.2
0.4
0.6
0.8
1
1.2
1.4
1.6
1.8
2
0
5
10
15
20
25

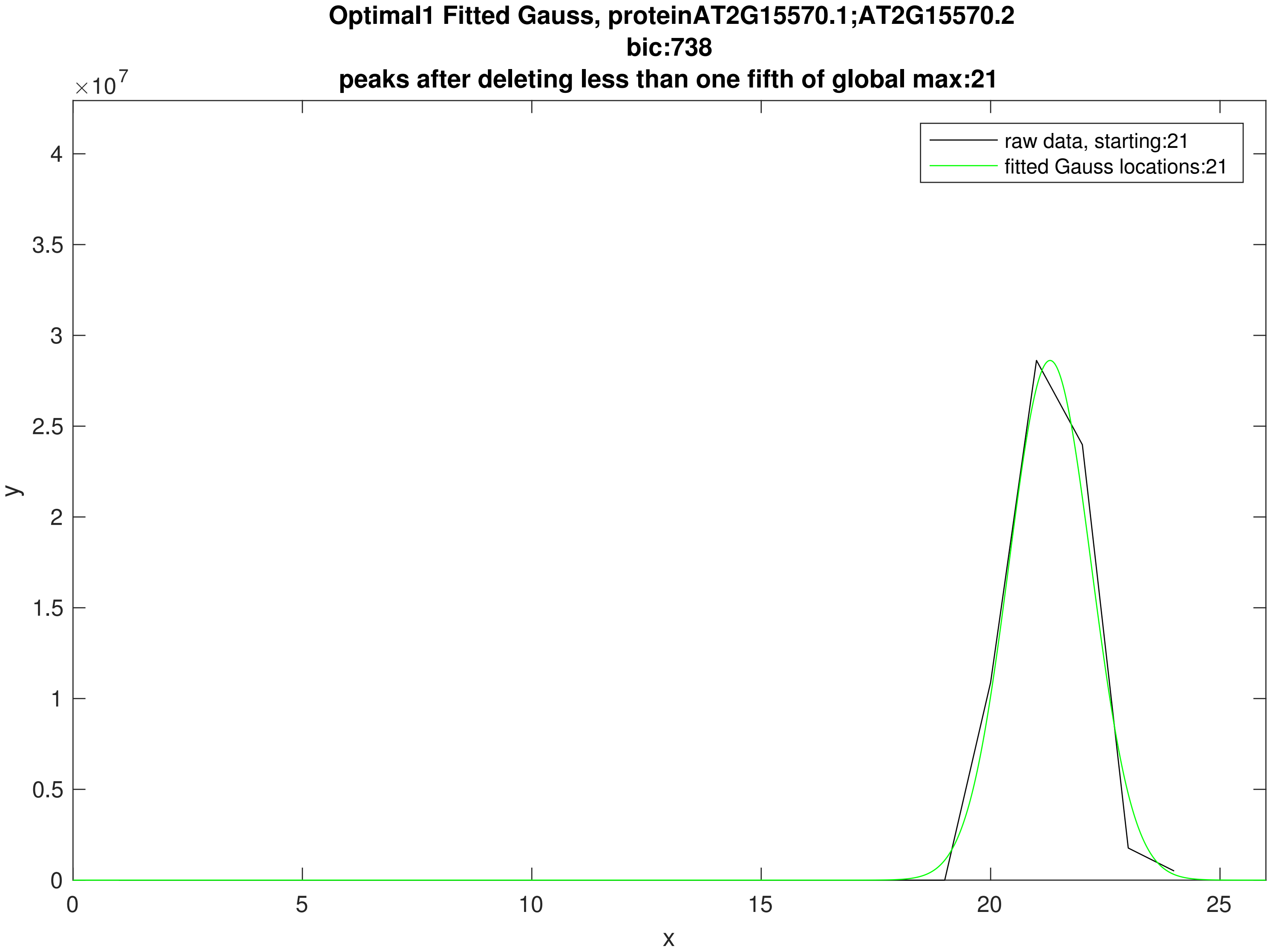
Optimal1 Fitted Gauss, proteinAT2G15570.1;AT2G15570.2
bic:738
peaks after deleting less than one fifth of global max:21
×10^7
raw data, starting:21
fitted Gauss locations:21
y
x
0
0.5
1
1.5
2
2.5
3
3.5
4
5
10
15
20
25

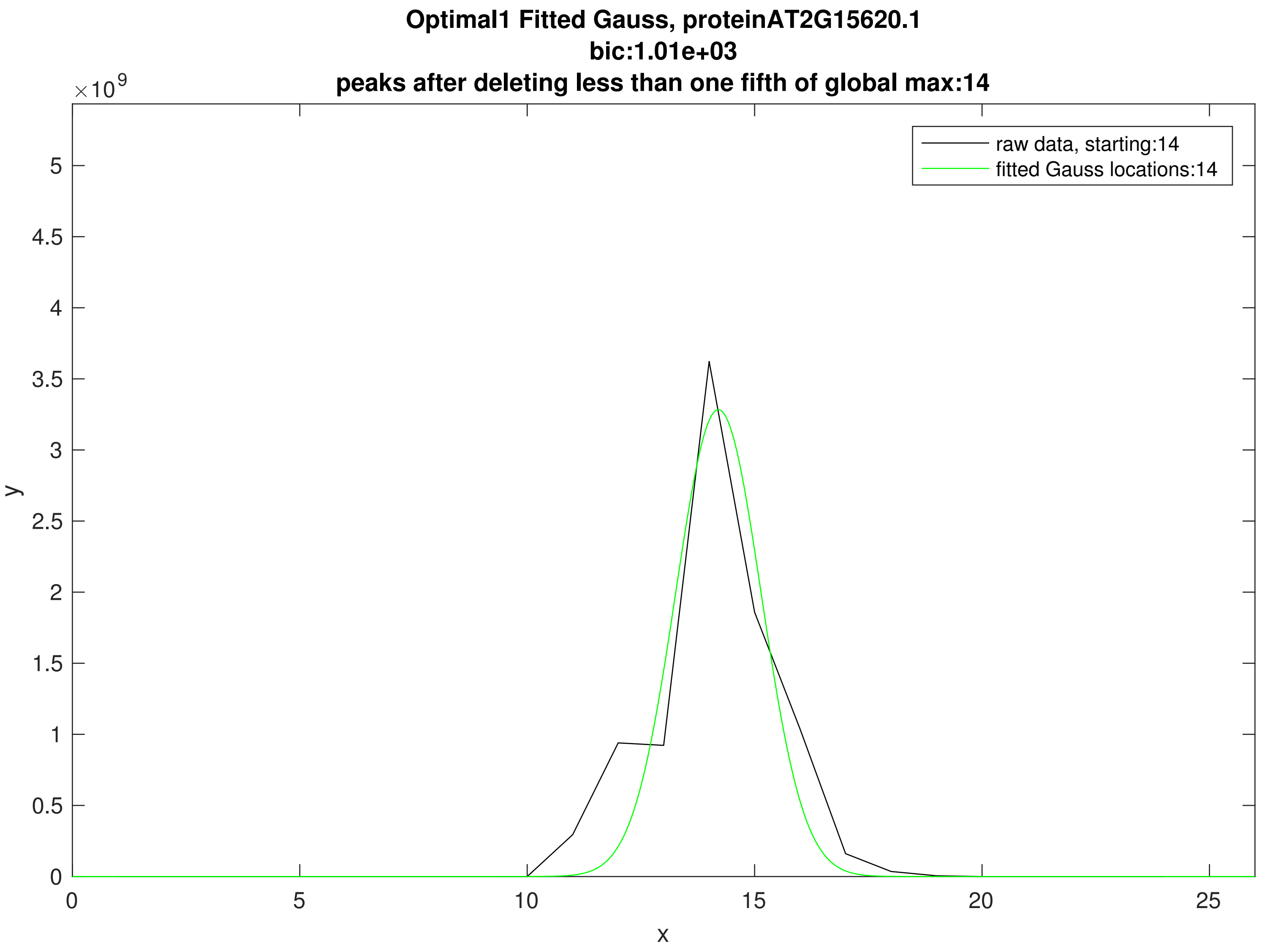
Optimal1 Fitted Gauss, proteinAT2G15620.1
bic:1.01e+03
peaks after deleting less than one fifth of global max:14
×10^9
raw data, starting:14
fitted Gauss locations:14
y
x
0
0.5
1
1.5
2
2.5
3
3.5
4
4.5
5
0
5
10
15
20
25

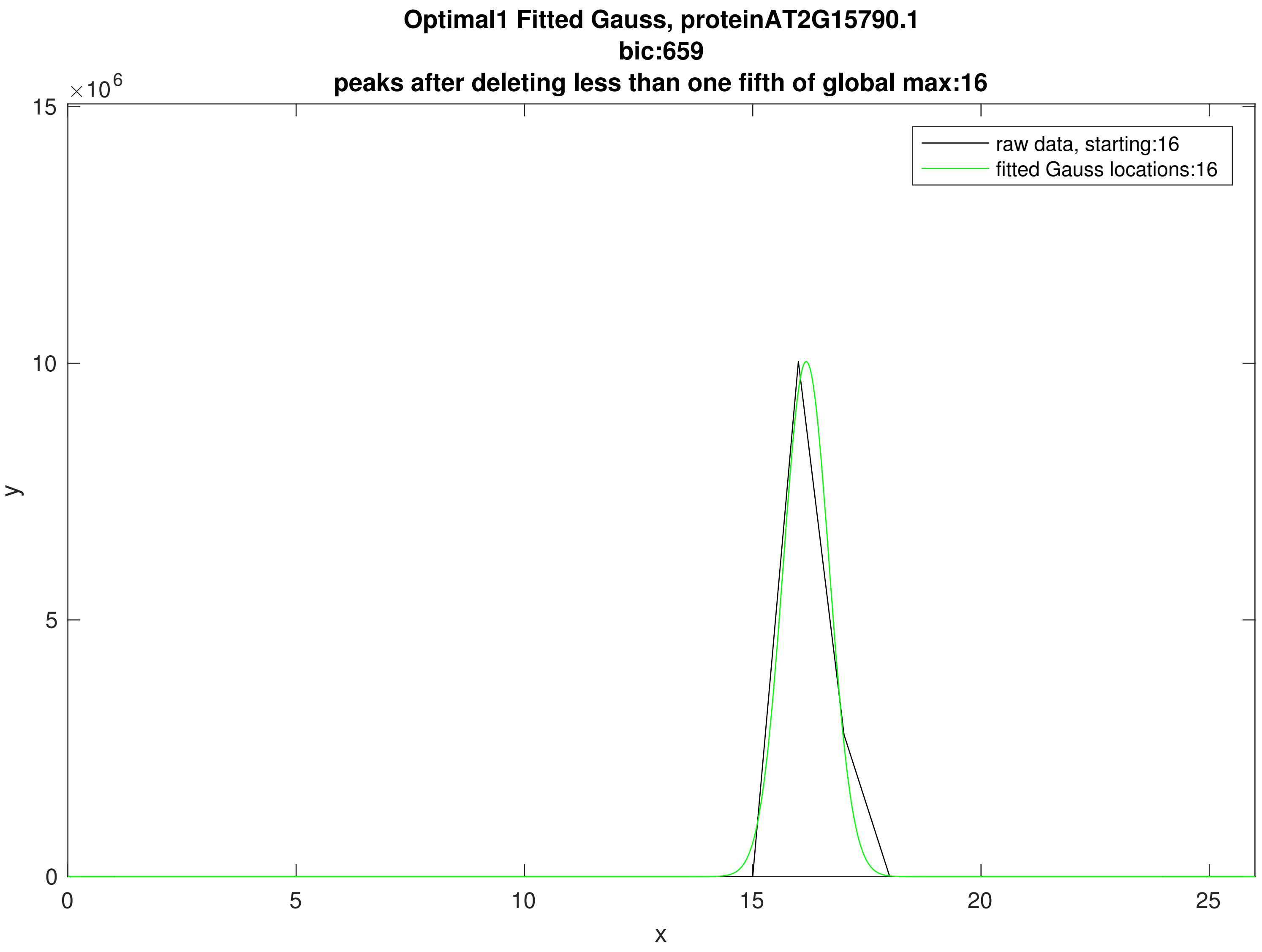
Optimal1 Fitted Gauss, proteinAT2G15790.1
bic:659
peaks after deleting less than one fifth of global max:16
×10^6
15
10
5
0
y
0
5
10
15
20
25
x
raw data, starting:16
fitted Gauss locations:16

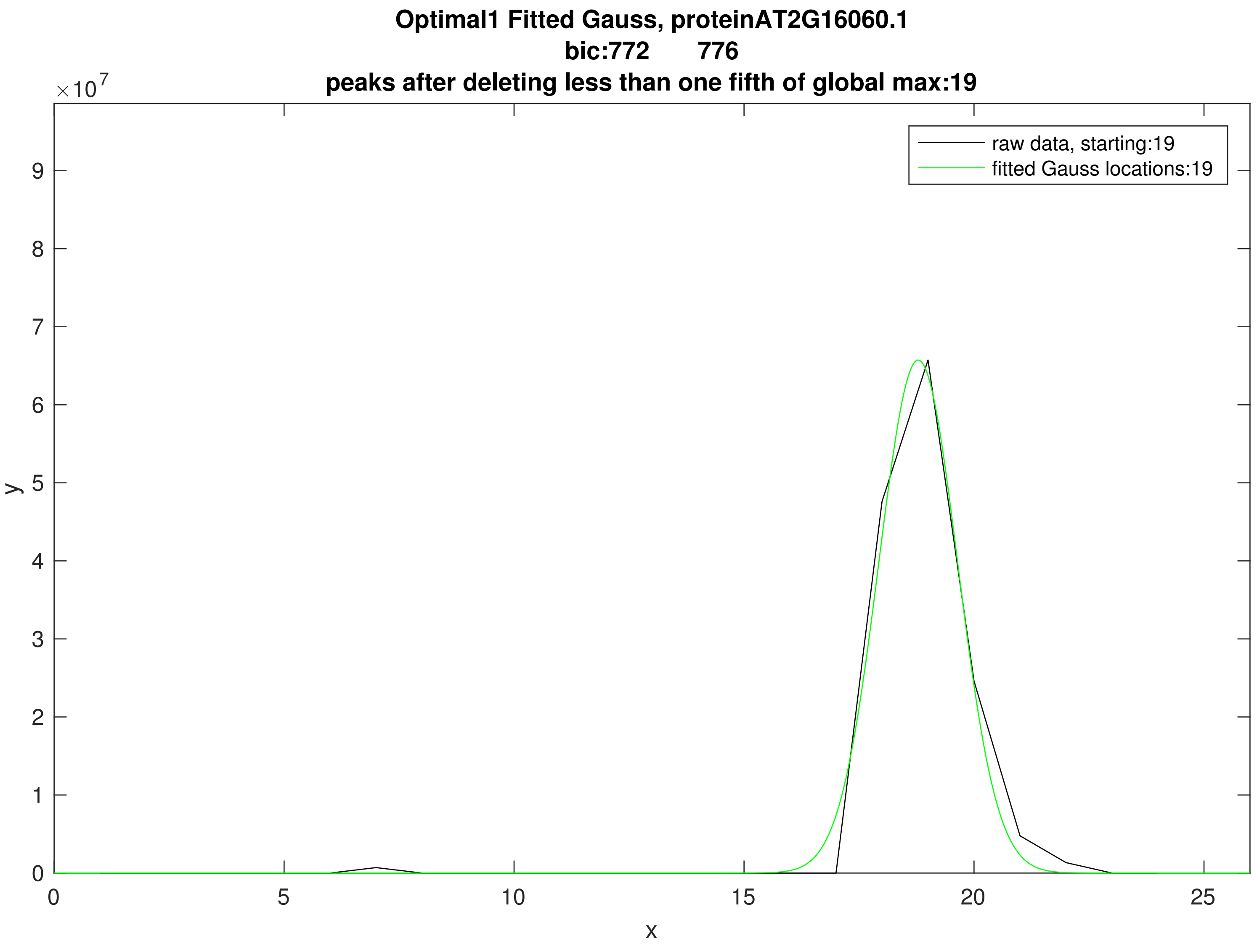
Optimal1 Fitted Gauss, proteinAT2G16060.1
bic:772 776
peaks after deleting less than one fifth of global max:19
×10^7
raw data, starting:19
fitted Gauss locations:19
y
x
0
1
2
3
4
5
6
7
8
9
0
5
10
15
20
25

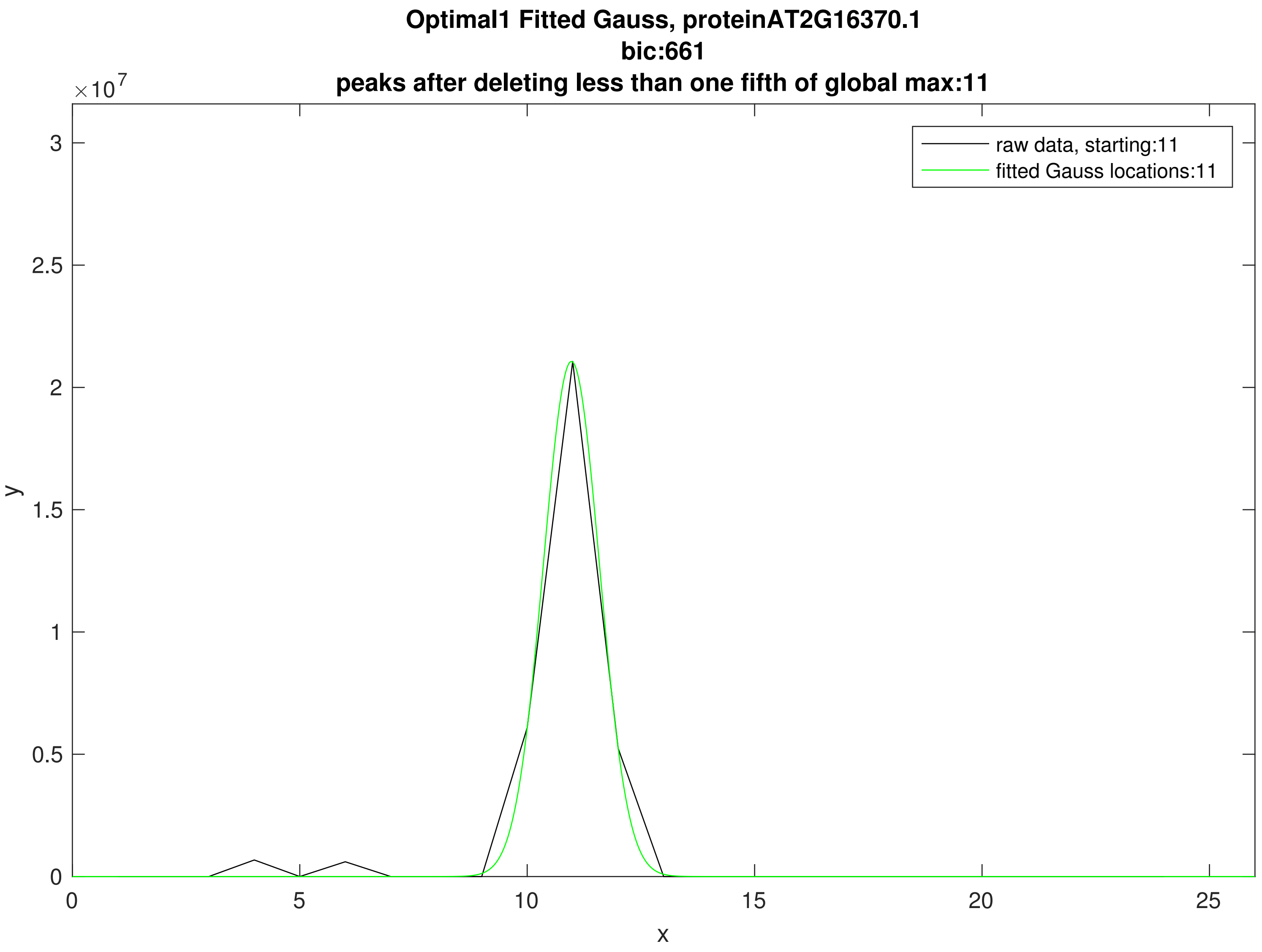
Optimal1 Fitted Gauss, proteinAT2G16370.1
bic:661
peaks after deleting less than one fifth of global max:11
×10^7
raw data, starting:11
fitted Gauss locations:11
y
x
0
0.5
1
1.5
2
2.5
3
0
5
10
15
20
25

**Optimal1 Fitted Gauss, proteinAT2G16570.1**
**bic:690**
**peaks after deleting less than one fifth of global max:16**

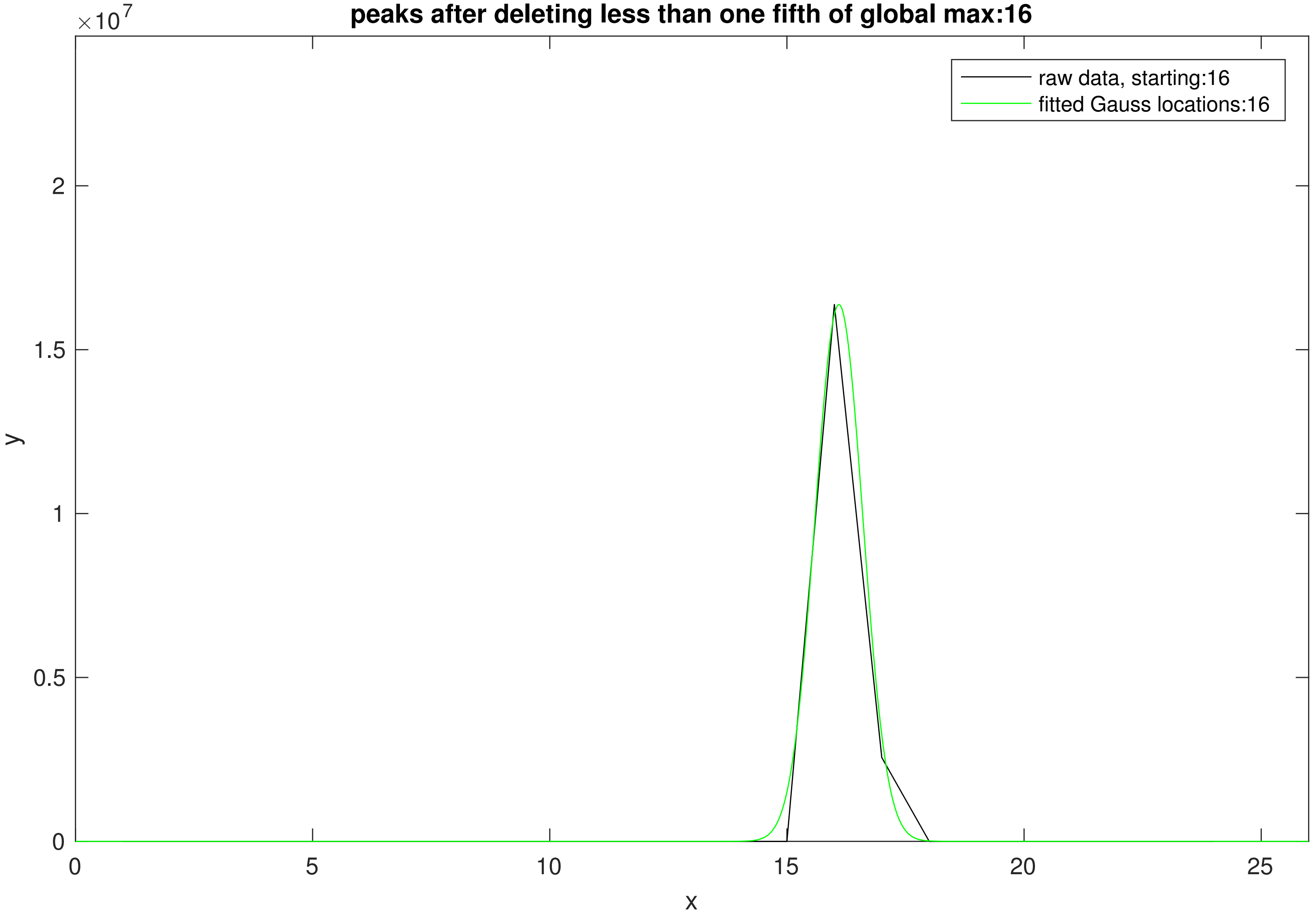

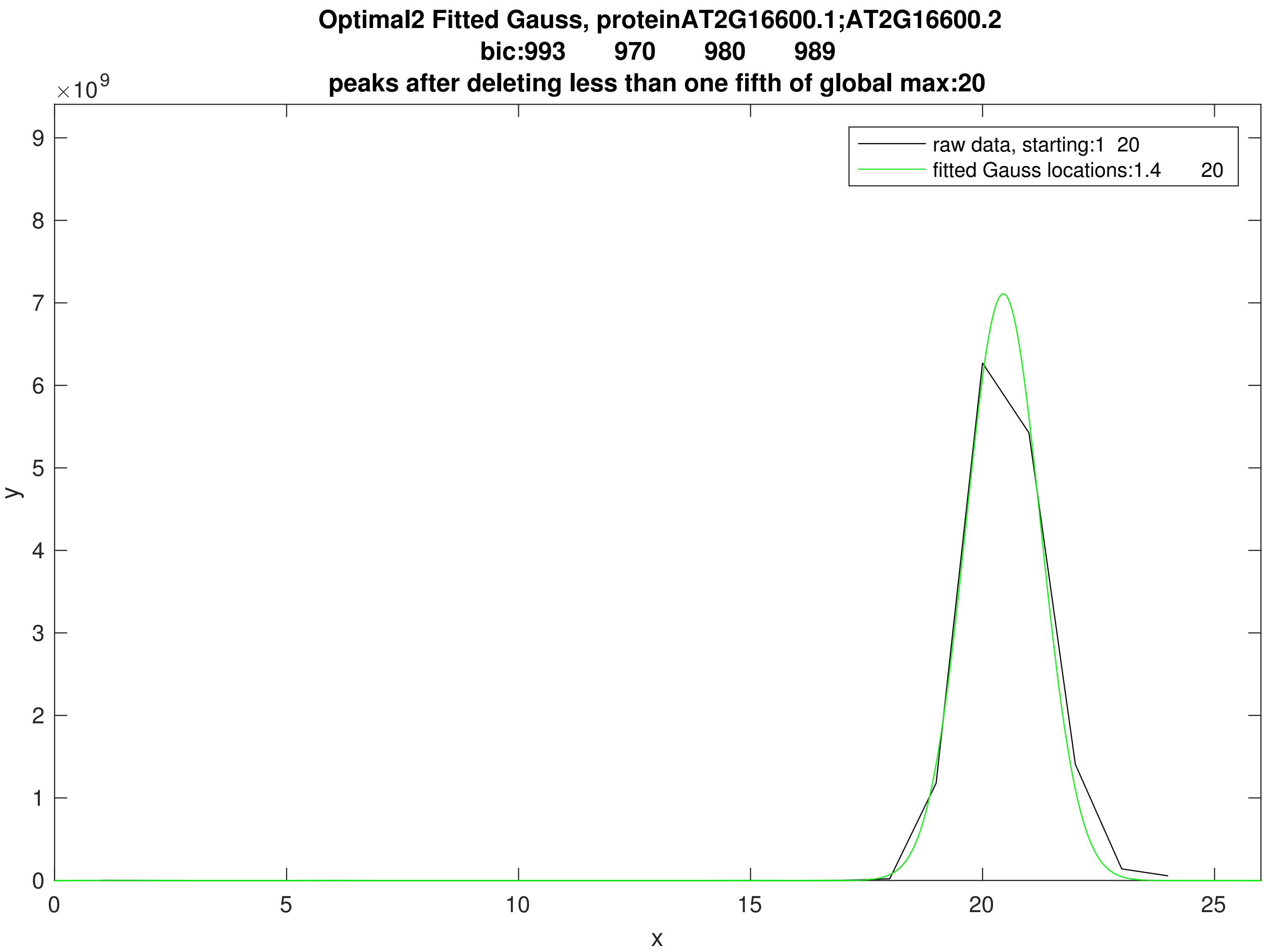

Optimal2 Fitted Gauss, proteinAT2G16600.1;AT2G16600.2
bic:993 970 980 989
peaks after deleting less than one fifth of global max:20
×10^9
raw data, starting:1 20
fitted Gauss locations:1.4 20
y
x
0
1
2
3
4
5
6
7
8
9
0
5
10
15
20
25

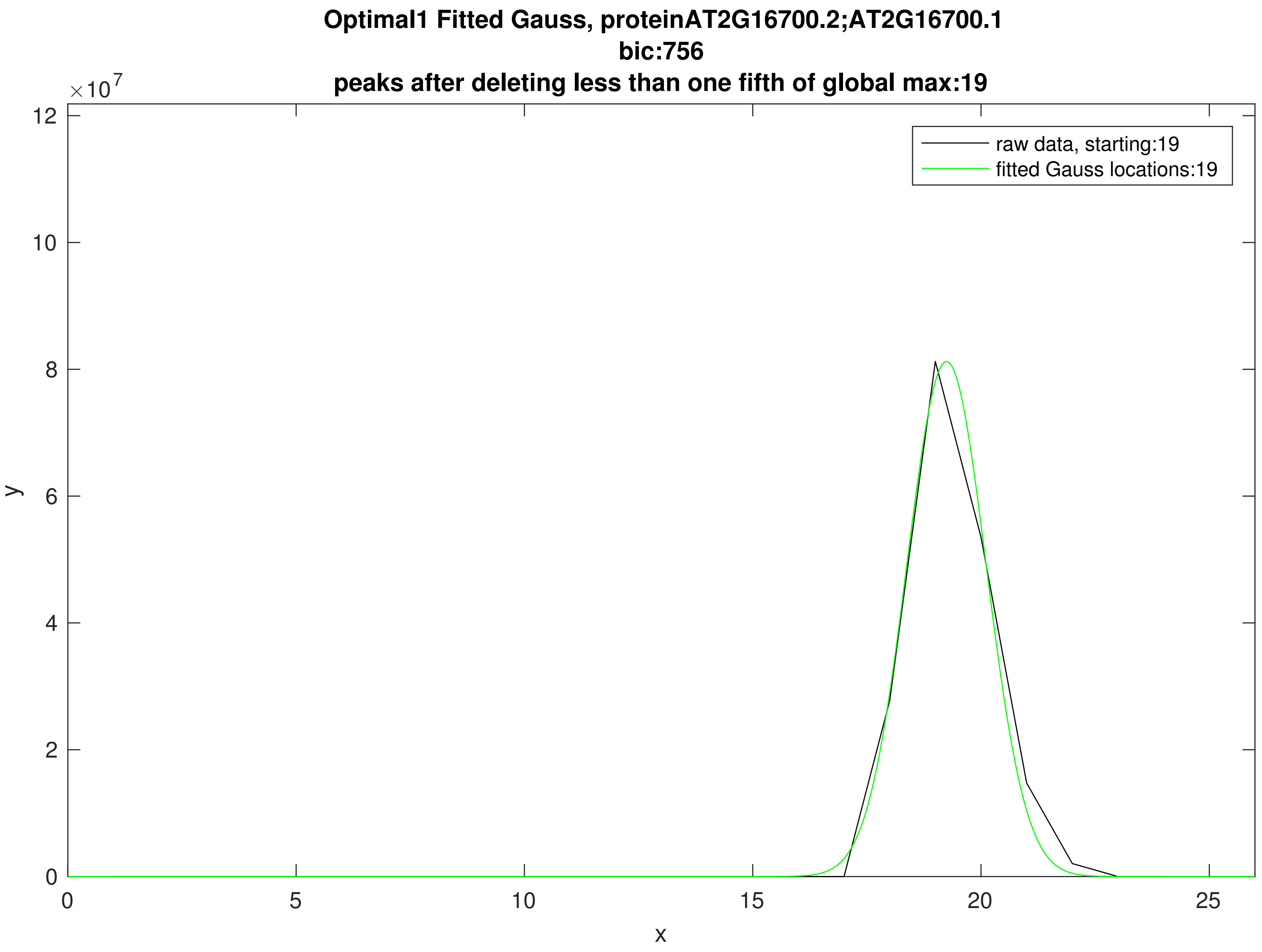
Optimal1 Fitted Gauss, proteinAT2G16700.2;AT2G16700.1
bic:756
peaks after deleting less than one fifth of global max:19
×10^7
raw data, starting:19
fitted Gauss locations:19
y
x
0
2
4
6
8
10
12
0
5
10
15
20
25

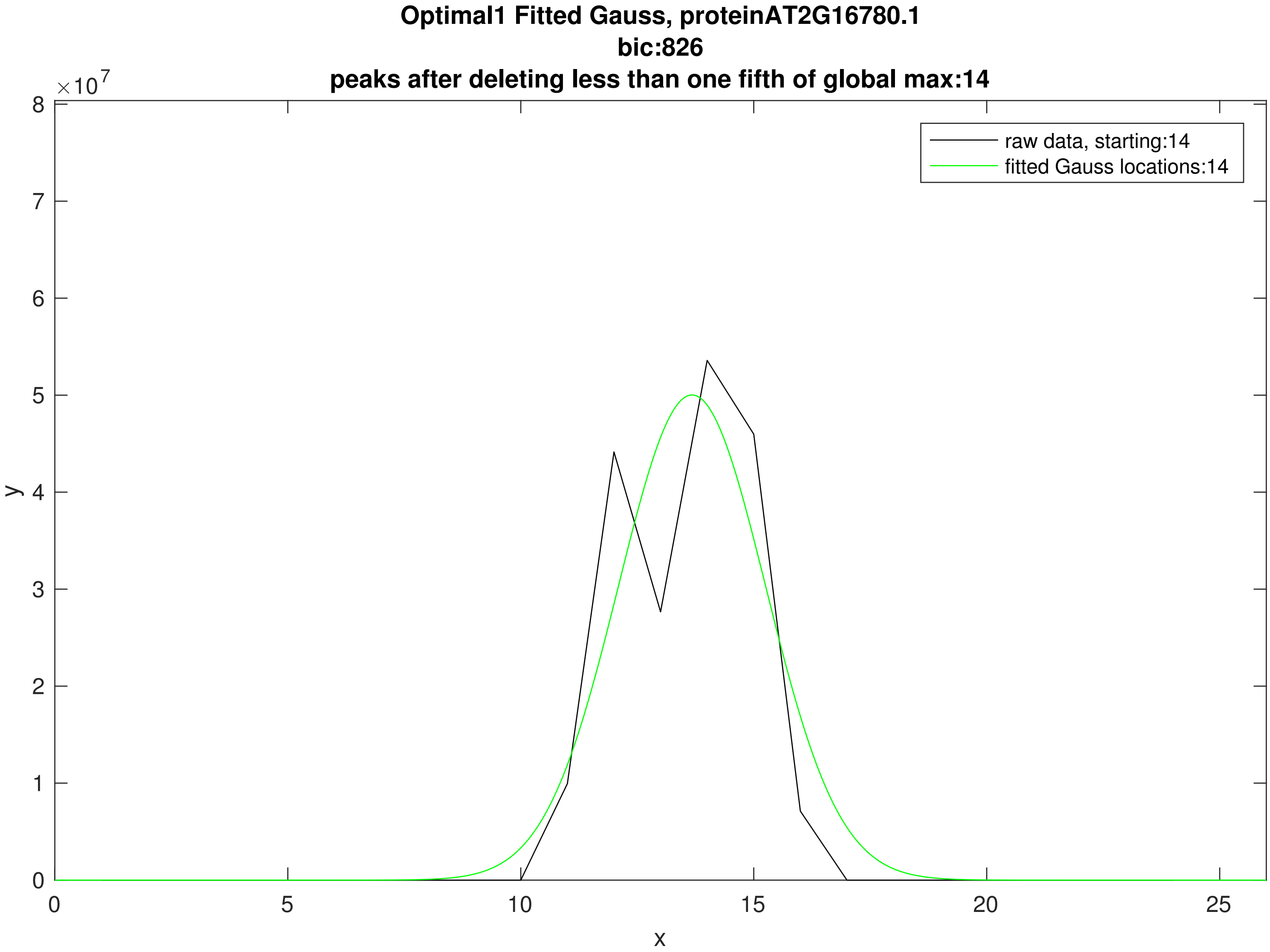
Optimal1 Fitted Gauss, proteinAT2G16780.1
bic:826
peaks after deleting less than one fifth of global max:14
×10^7
raw data, starting:14
fitted Gauss locations:14
y
x
0
1
2
3
4
5
6
7
8
0
5
10
15
20
25

**Optimal1 Fitted Gauss, proteinAT2G16950.2;AT2G16950.1**
**bic:629**
**peaks after deleting less than one fifth of global max:12**

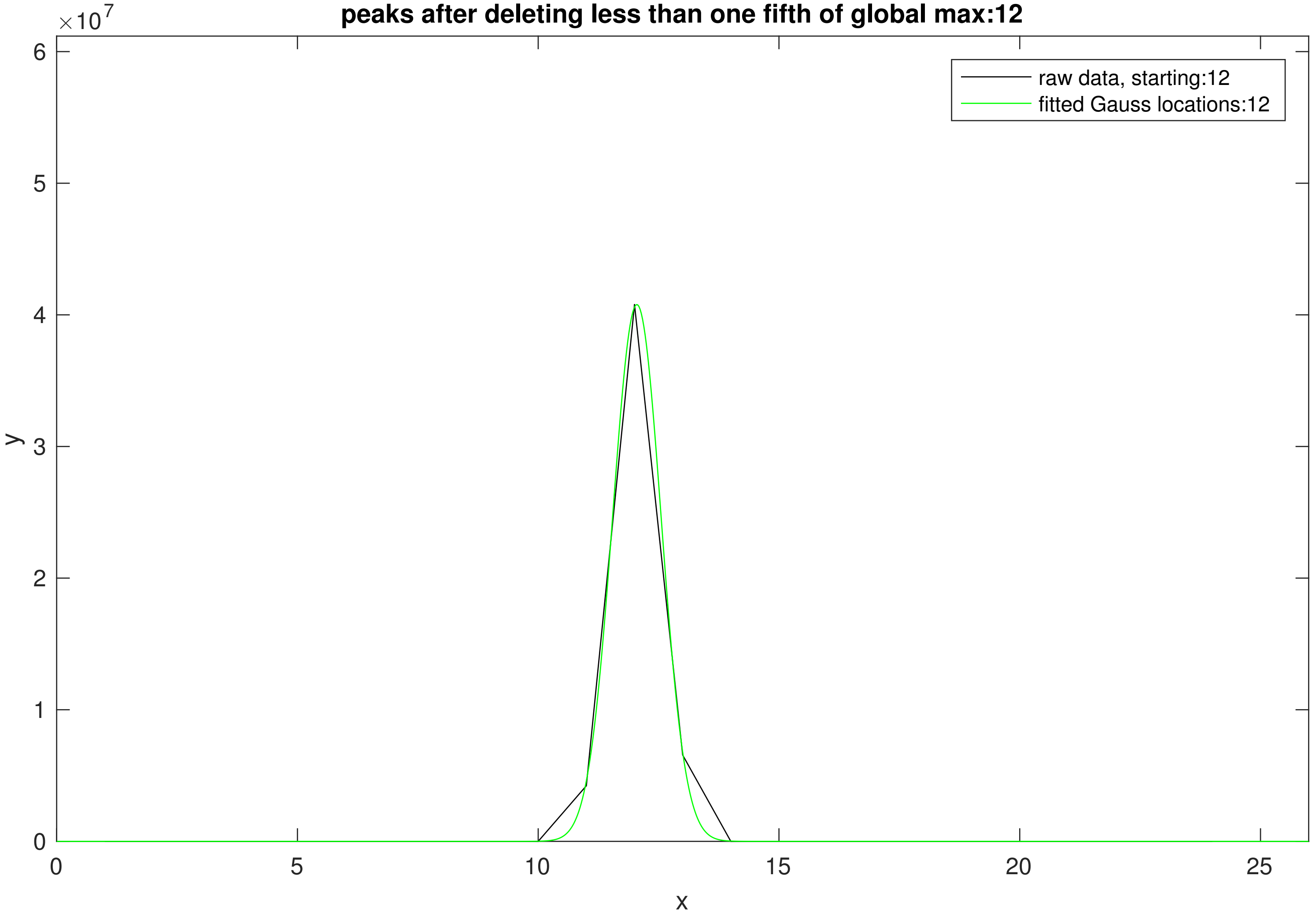

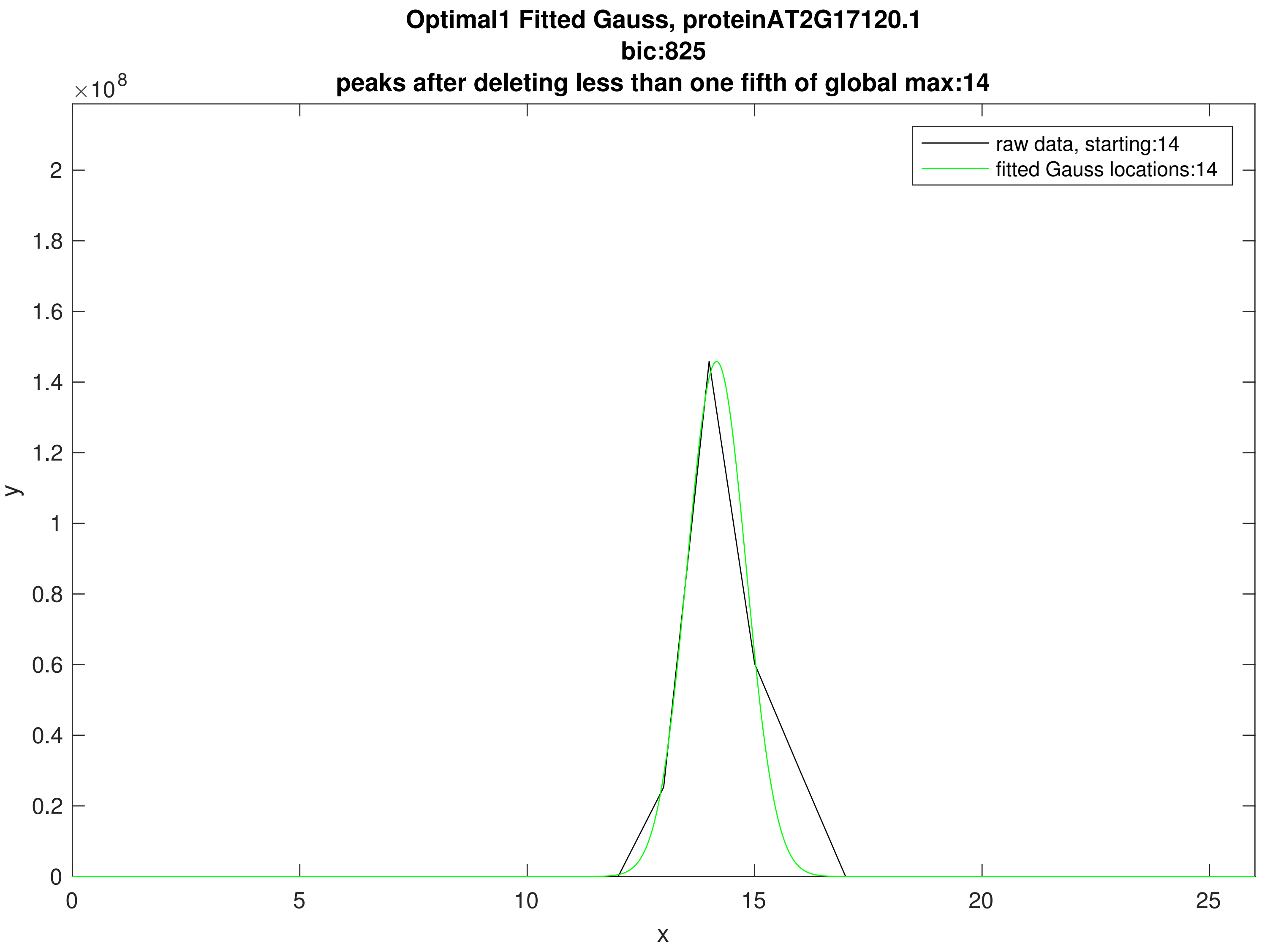
Optimal1 Fitted Gauss, proteinAT2G17120.1
bic:825
peaks after deleting less than one fifth of global max:14
×10^8
raw data, starting:14
fitted Gauss locations:14
y
x
0
0.2
0.4
0.6
0.8
1
1.2
1.4
1.6
1.8
2
0
5
10
15
20
25

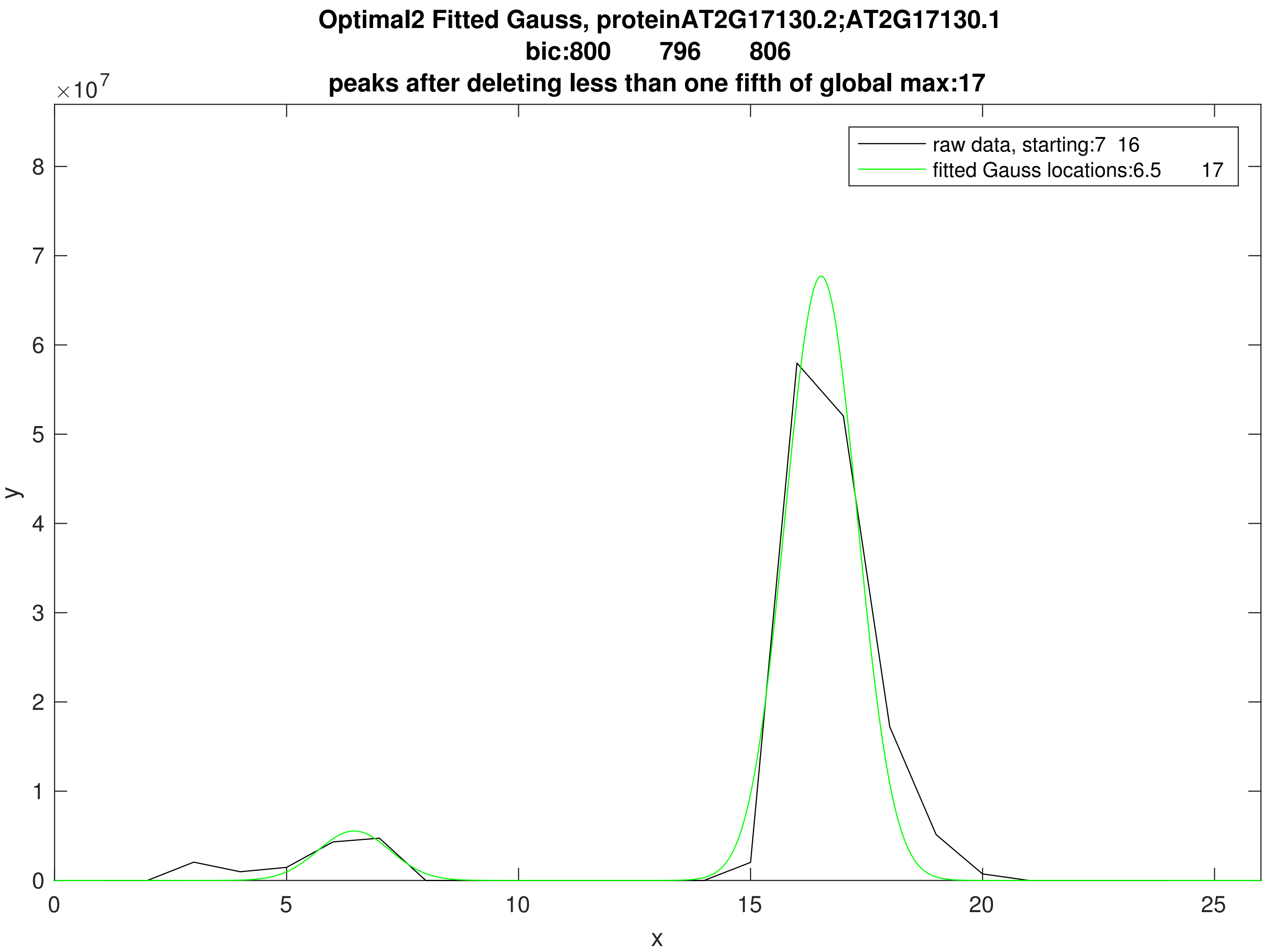

Optimal2 Fitted Gauss, proteinAT2G17130.2;AT2G17130.1
bic:800 796 806
peaks after deleting less than one fifth of global max:17
×10^7
raw data, starting:7 16
fitted Gauss locations:6.5 17
y
x
0
1
2
3
4
5
6
7
8
0
5
10
15
20
25

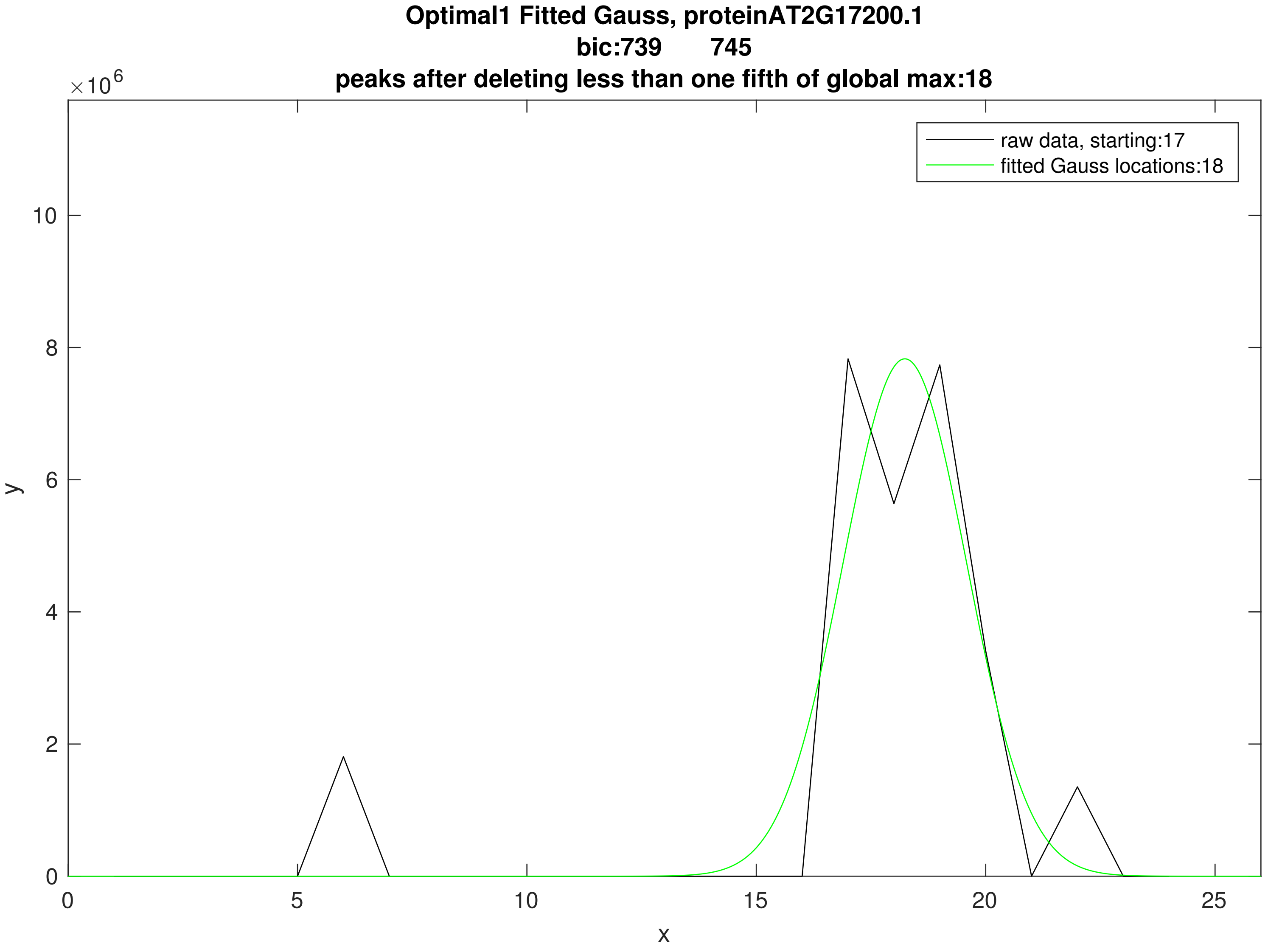
Optimal1 Fitted Gauss, proteinAT2G17200.1
bic:739 745
peaks after deleting less than one fifth of global max:18
×10^6
raw data, starting:17
fitted Gauss locations:18
y
x
0
2
4
6
8
10
0
5
10
15
20
25

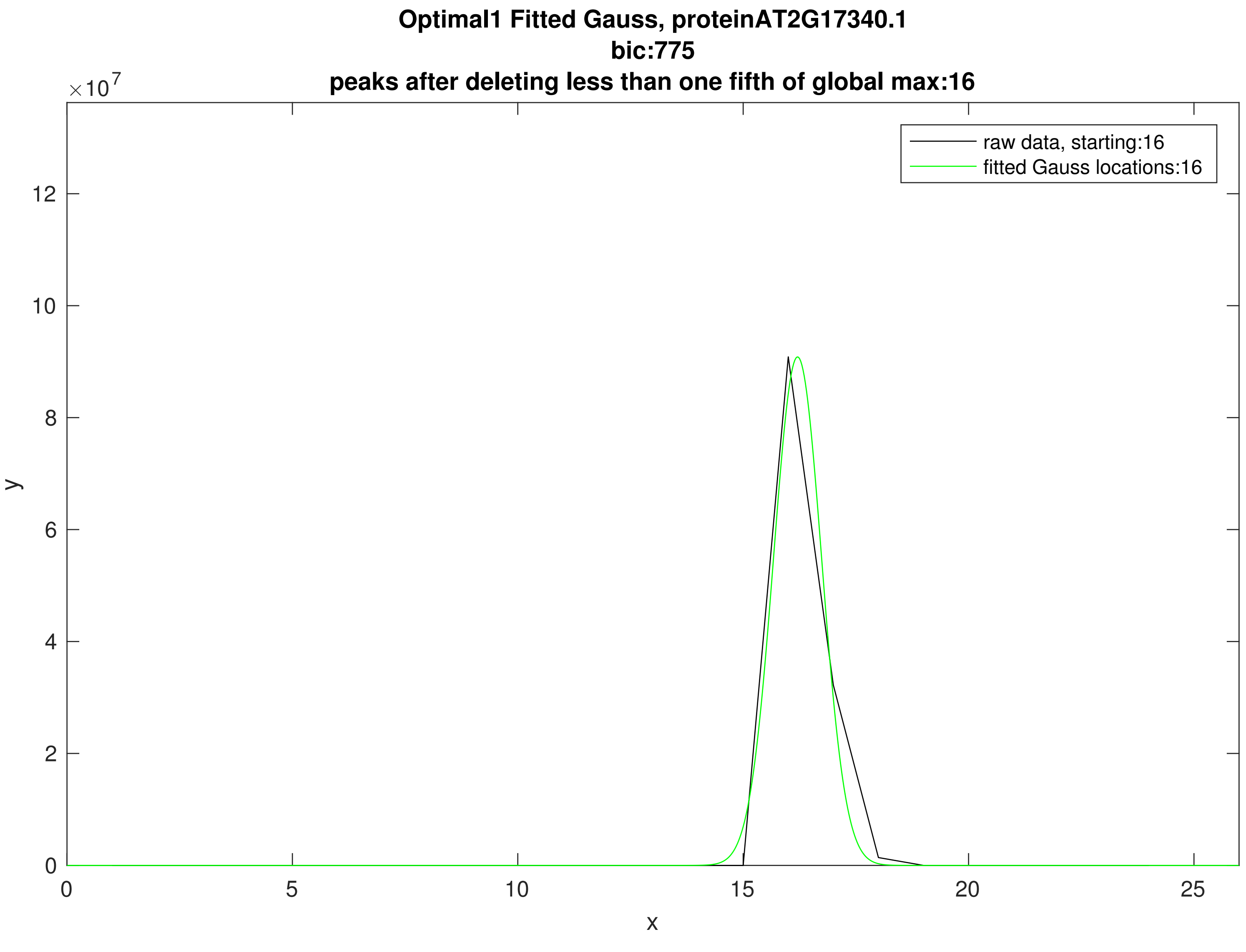

Optimal1 Fitted Gauss, proteinAT2G17340.1
bic:775
peaks after deleting less than one fifth of global max:16
×10^7
raw data, starting:16
fitted Gauss locations:16
12
10
8
6
4
2
0
y
0
5
10
15
20
25
x

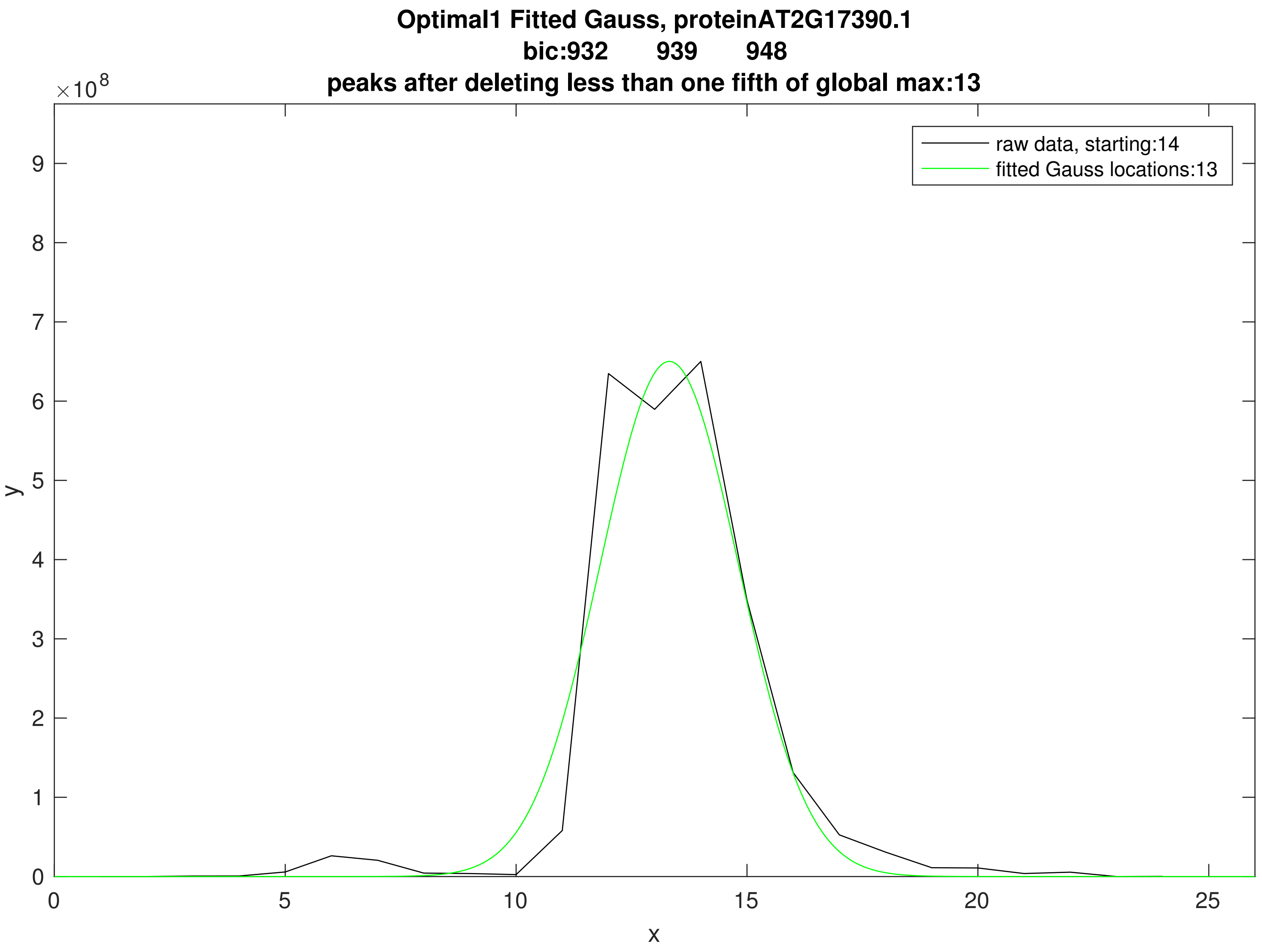
Optimal1 Fitted Gauss, proteinAT2G17390.1
bic:932 939 948
peaks after deleting less than one fifth of global max:13
×10^8
raw data, starting:14
fitted Gauss locations:13
y
x
0
1
2
3
4
5
6
7
8
9
0
5
10
15
20
25

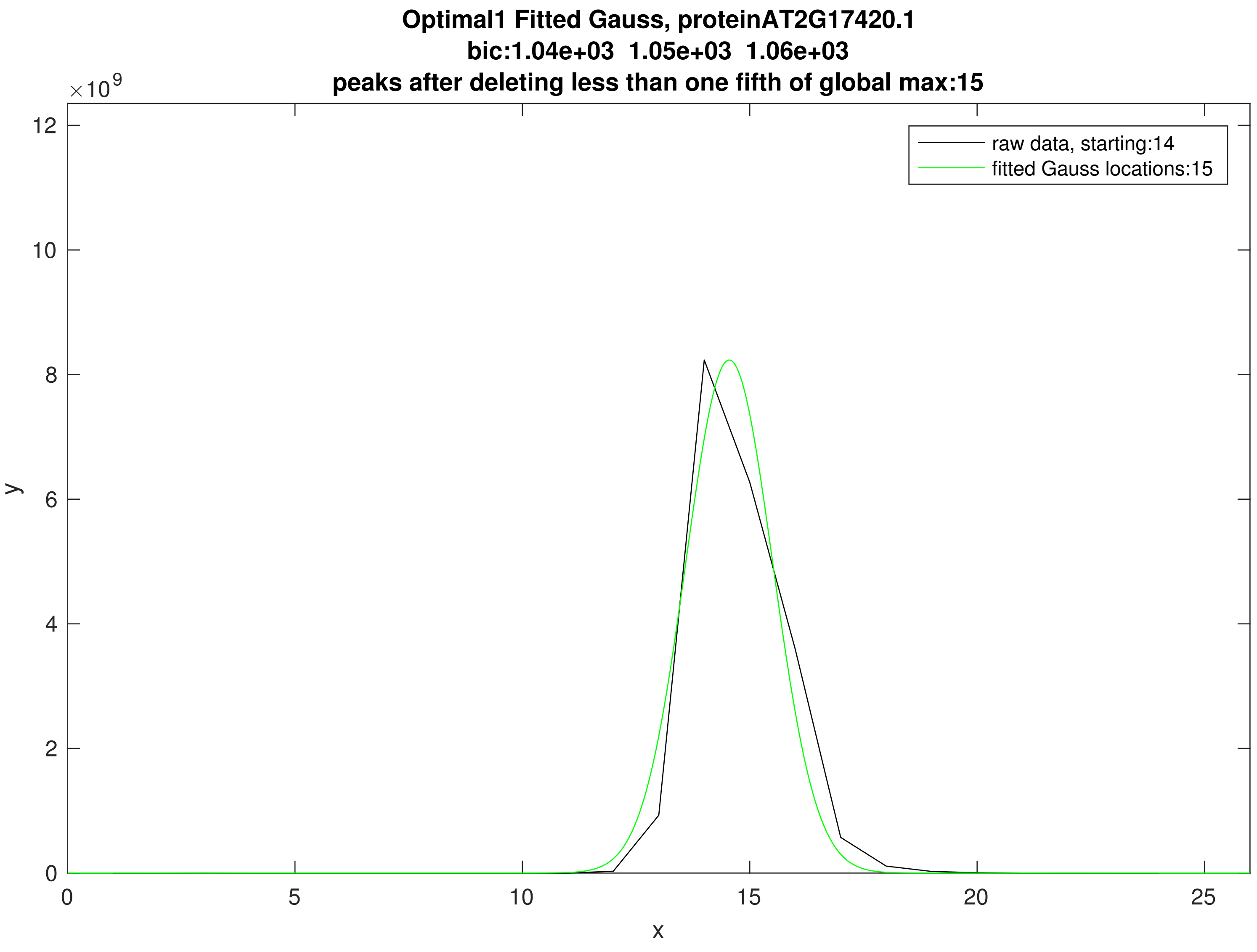

Optimal1 Fitted Gauss, proteinAT2G17420.1
bic:1.04e+03  1.05e+03  1.06e+03
peaks after deleting less than one fifth of global max:15
×10^9
raw data, starting:14
fitted Gauss locations:15
y
x

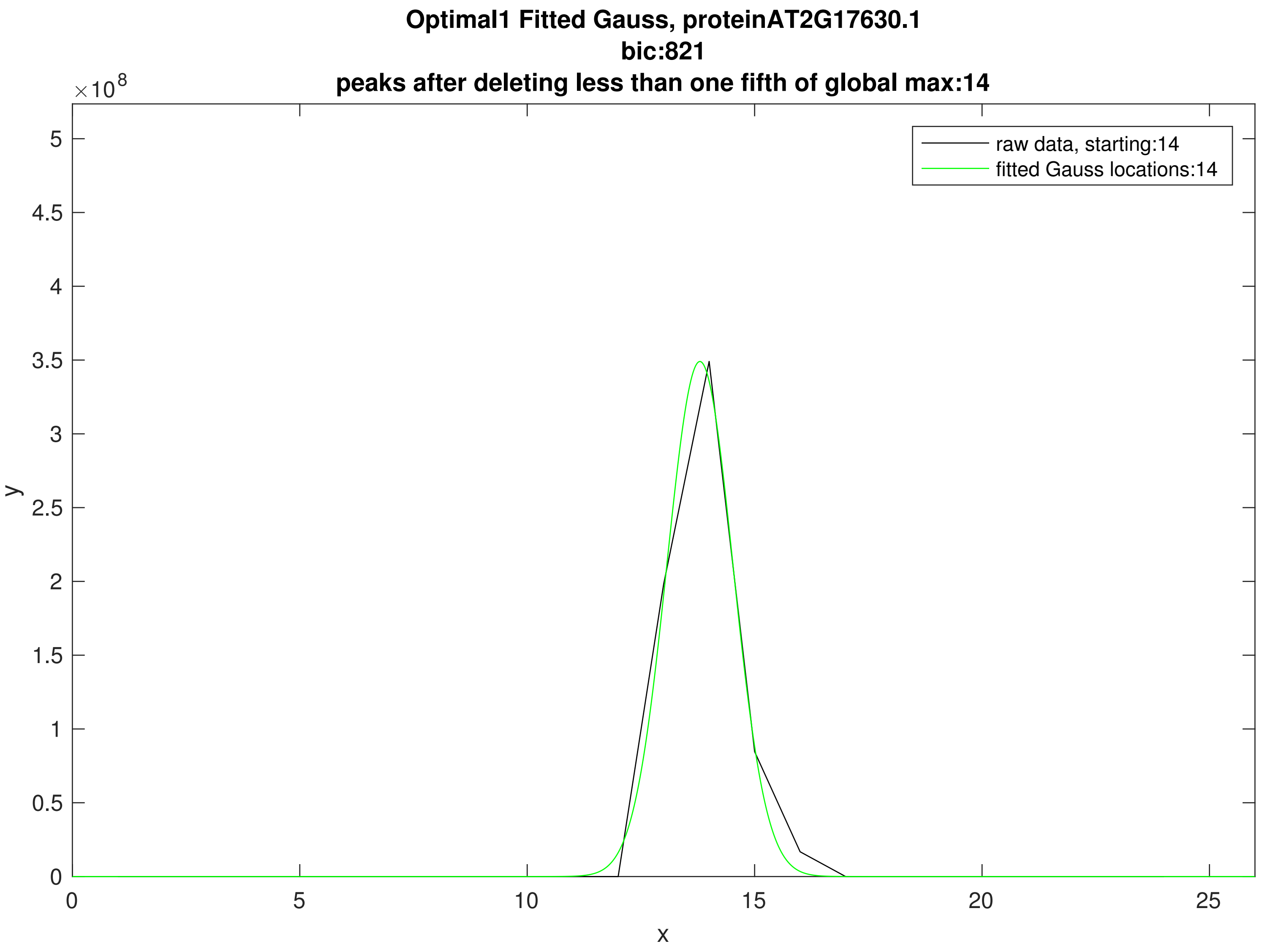

Optimal1 Fitted Gauss, proteinAT2G17630.1
bic:821
peaks after deleting less than one fifth of global max:14
×10^8
raw data, starting:14
fitted Gauss locations:14
y
x
0
0.5
1
1.5
2
2.5
3
3.5
4
4.5
5
0
5
10
15
20
25

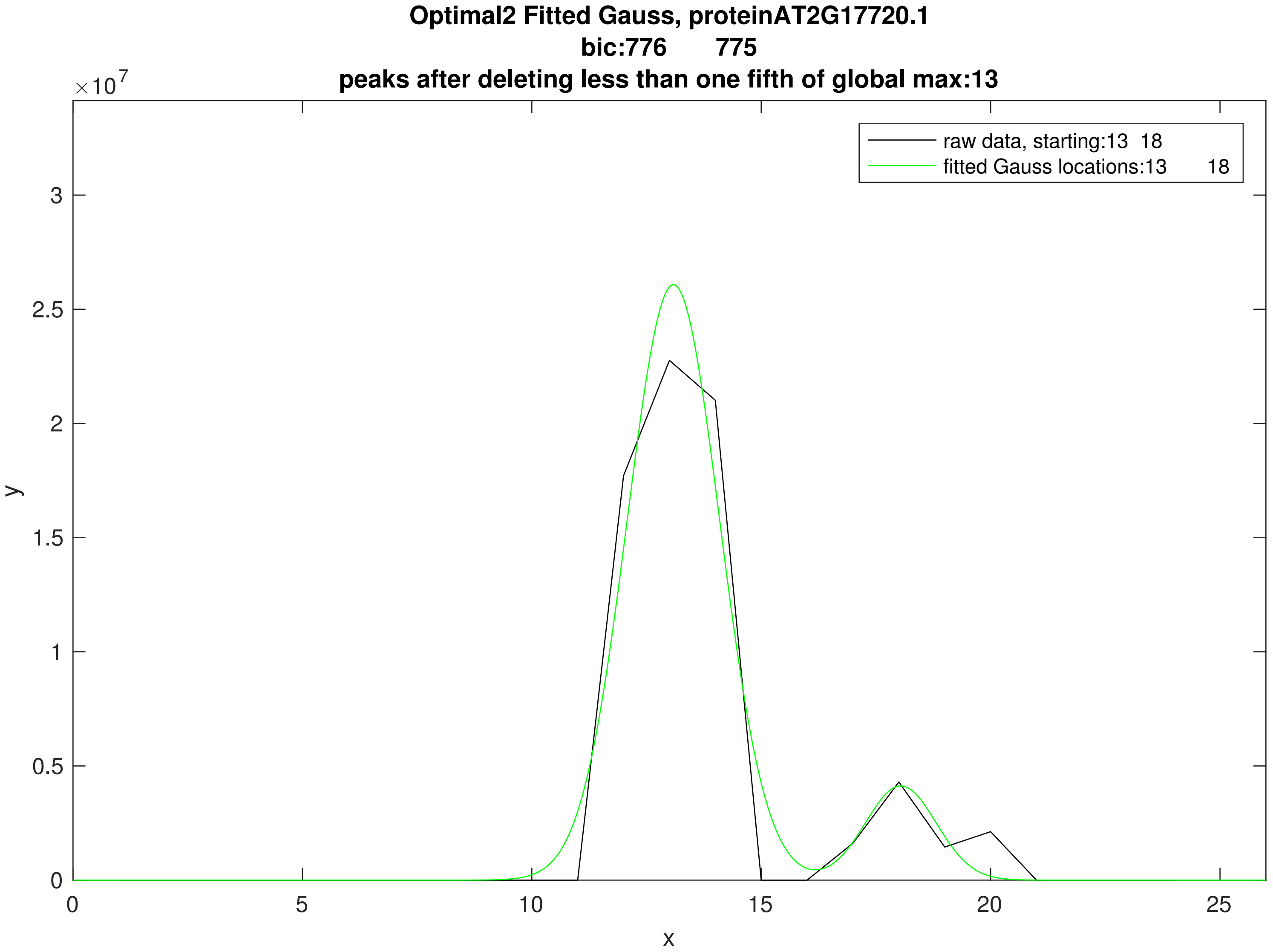
Optimal2 Fitted Gauss, proteinAT2G17720.1
bic:776 775
peaks after deleting less than one fifth of global max:13
×10^7
raw data, starting:13 18
fitted Gauss locations:13 18
y
x
0
0.5
1
1.5
2
2.5
3
5
10
15
20
25

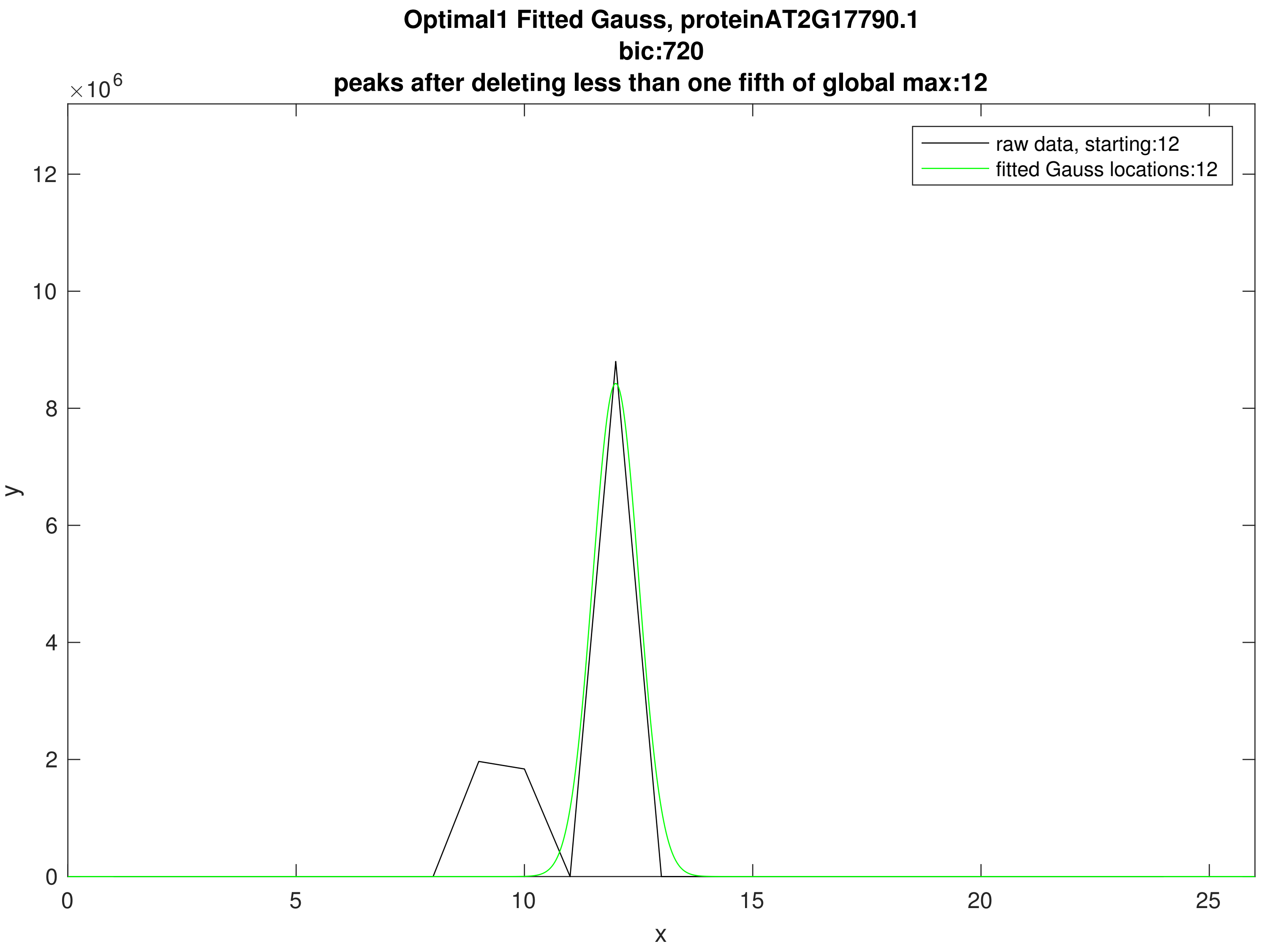
Optimal1 Fitted Gauss, proteinAT2G17790.1
bic:720
peaks after deleting less than one fifth of global max:12
×10^6
raw data, starting:12
fitted Gauss locations:12
y
x
0
2
4
6
8
10
12
0
5
10
15
20
25

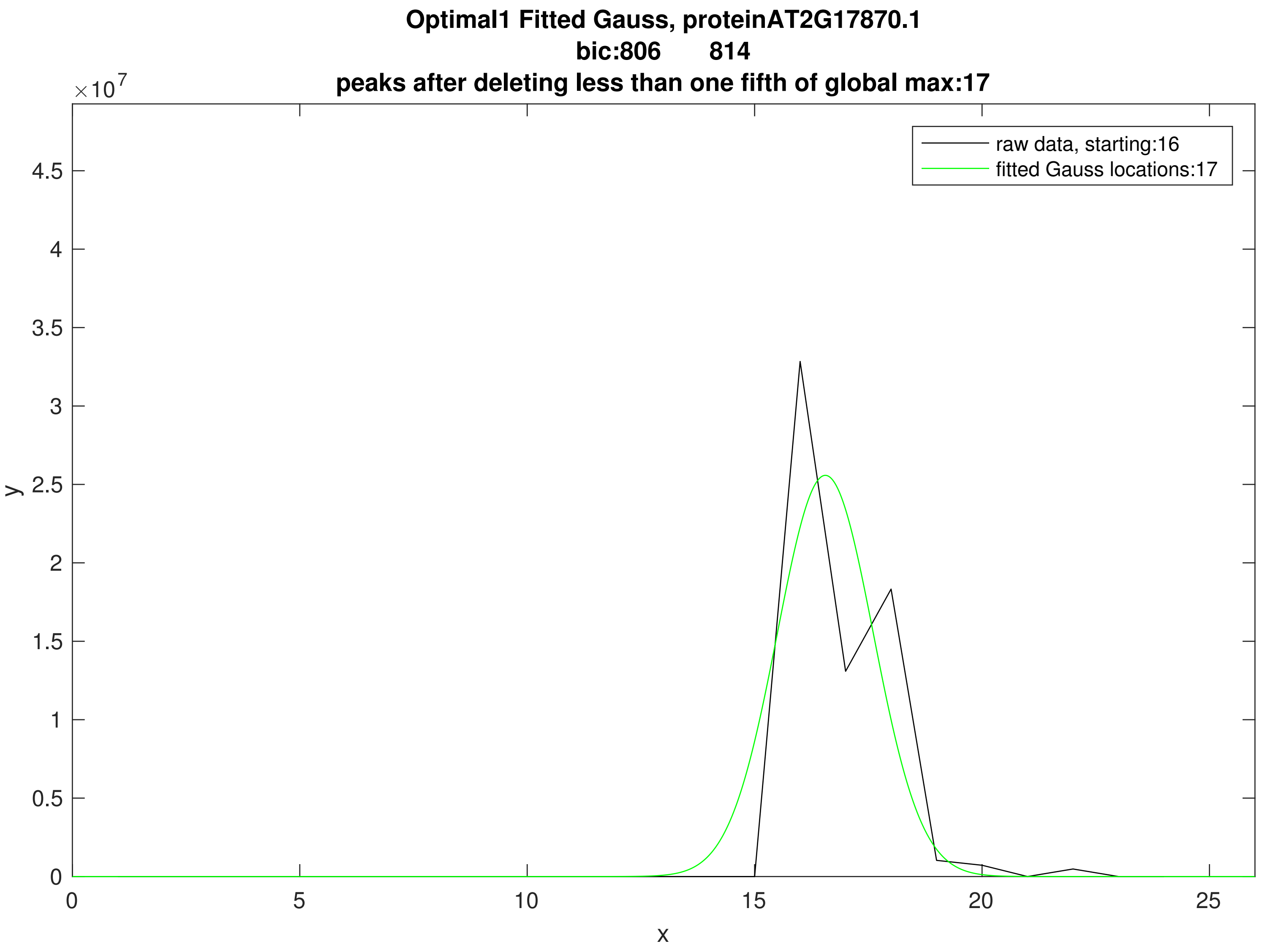
Optimal1 Fitted Gauss, proteinAT2G17870.1
bic:806 814
peaks after deleting less than one fifth of global max:17
×10^7
raw data, starting:16
fitted Gauss locations:17
4.5
4
3.5
3
2.5
2
1.5
1
0.5
0
y
0
5
10
15
20
25
x

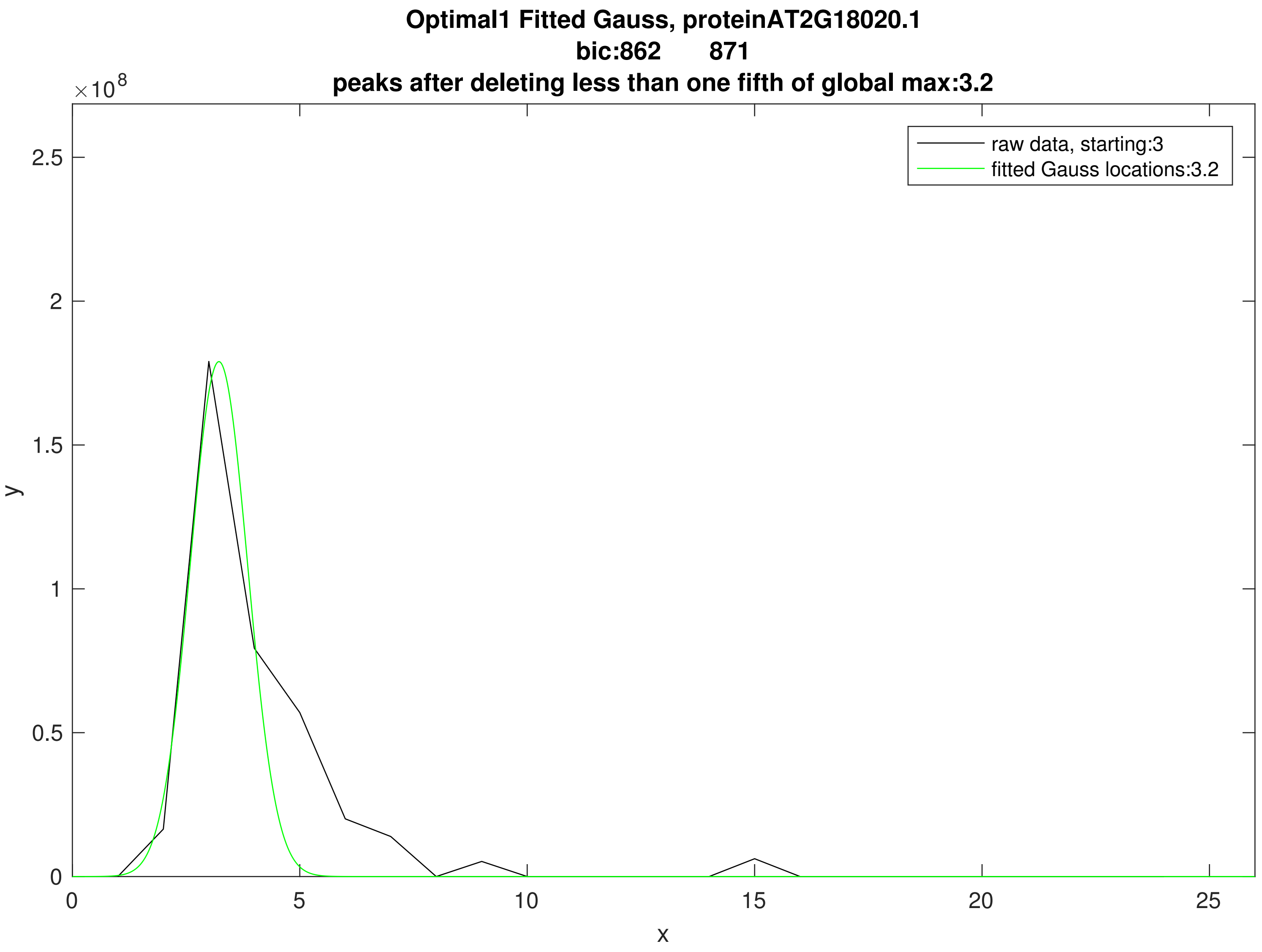

Optimal1 Fitted Gauss, proteinAT2G18020.1
bic:862 871
peaks after deleting less than one fifth of global max:3.2
×10^8
2.5
2
1.5
1
0.5
0
y
0
5
10
15
20
25
x
raw data, starting:3
fitted Gauss locations:3.2

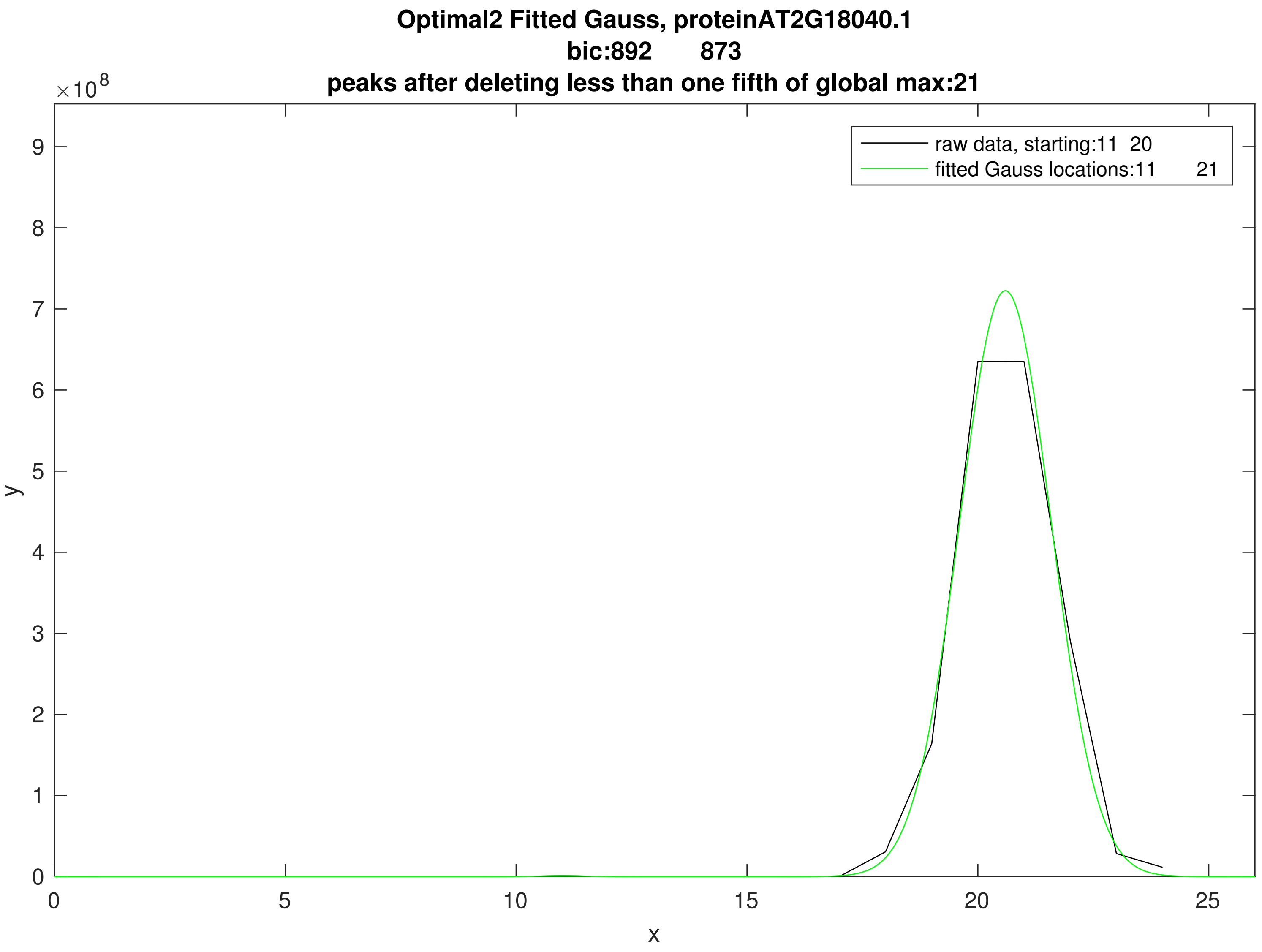
Optimal2 Fitted Gauss, proteinAT2G18040.1
bic:892 873
peaks after deleting less than one fifth of global max:21
×10^8
raw data, starting:11 20
fitted Gauss locations:11 21
y
x
0
1
2
3
4
5
6
7
8
9
0
5
10
15
20
25

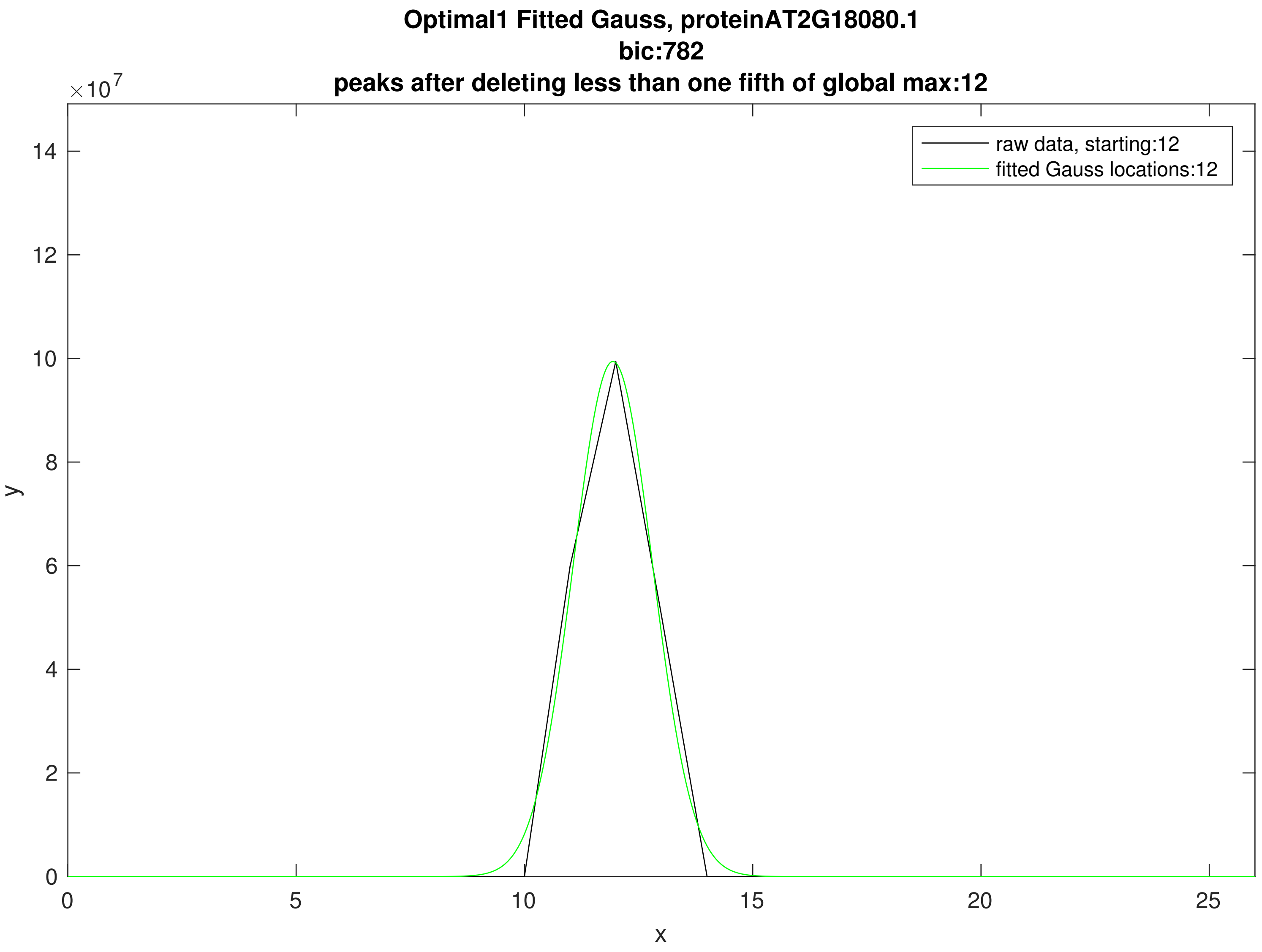
Optimal1 Fitted Gauss, proteinAT2G18080.1
bic:782
peaks after deleting less than one fifth of global max:12
×10^7
raw data, starting:12
fitted Gauss locations:12
y
x
0
2
4
6
8
10
12
14
0
5
10
15
20
25

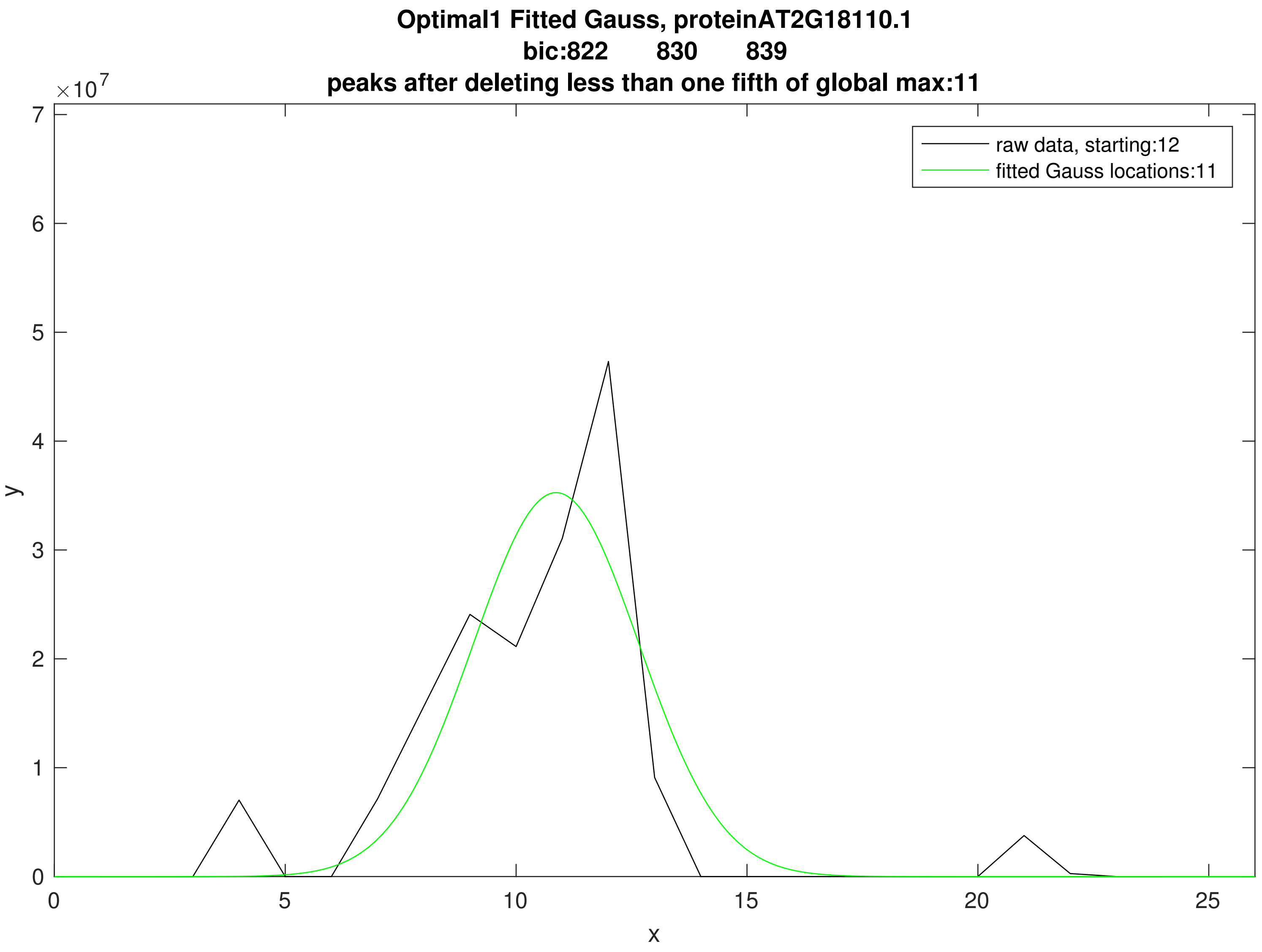
Optimal1 Fitted Gauss, proteinAT2G18110.1
bic:822 830 839
peaks after deleting less than one fifth of global max:11
×10^7
raw data, starting:12
fitted Gauss locations:11
y
x
0
1
2
3
4
5
6
7
0
5
10
15
20
25

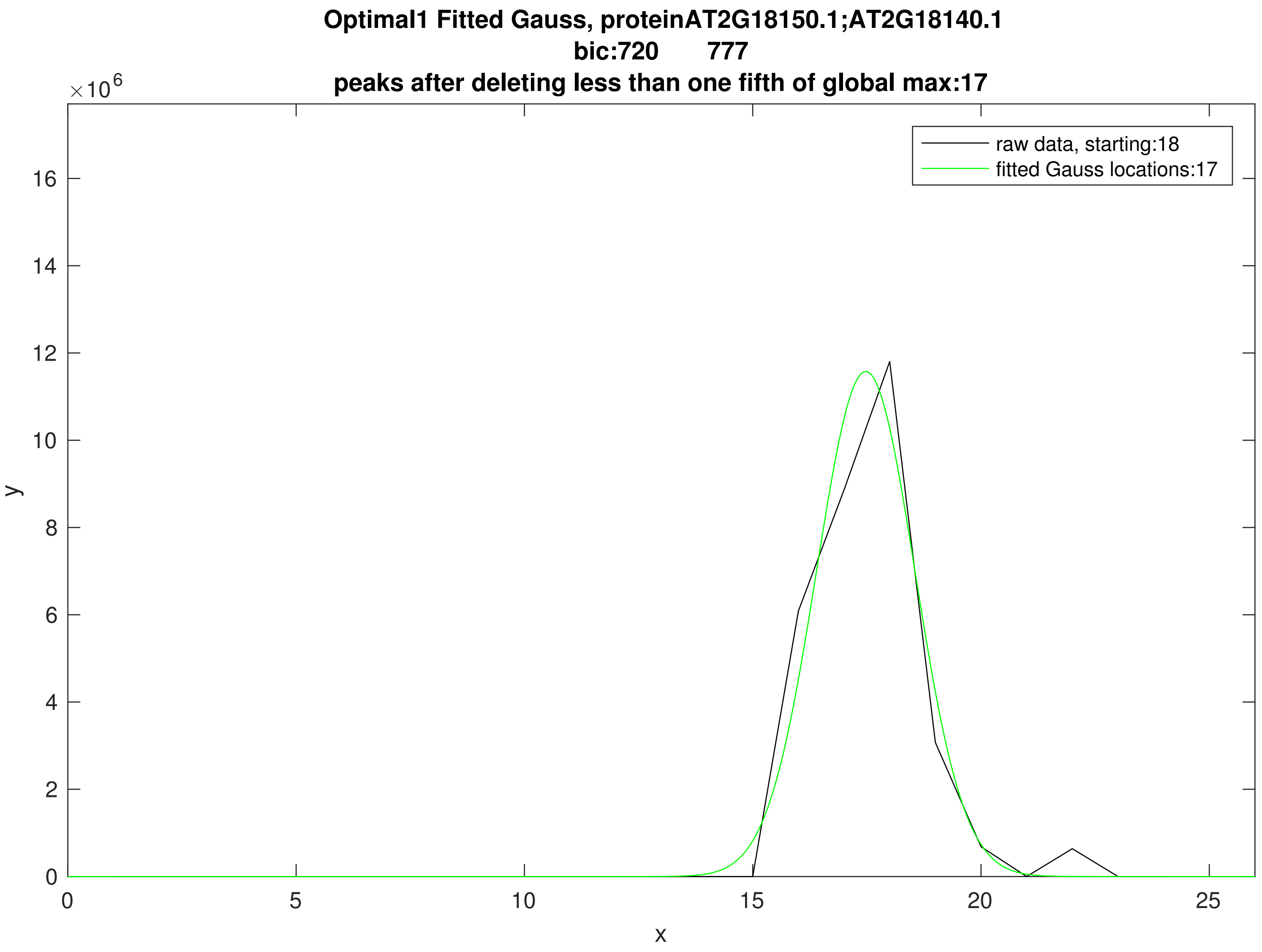
Optimal1 Fitted Gauss, proteinAT2G18150.1;AT2G18140.1
bic:720 777
peaks after deleting less than one fifth of global max:17
×10^6
raw data, starting:18
fitted Gauss locations:17
16
14
12
10
8
6
4
2
0
y
0
5
10
15
20
25
x

**Optimal2 Fitted Gauss, proteinAT2G18230.1**
**bic:816 790**
**peaks after deleting less than one fifth of global max:12 18**

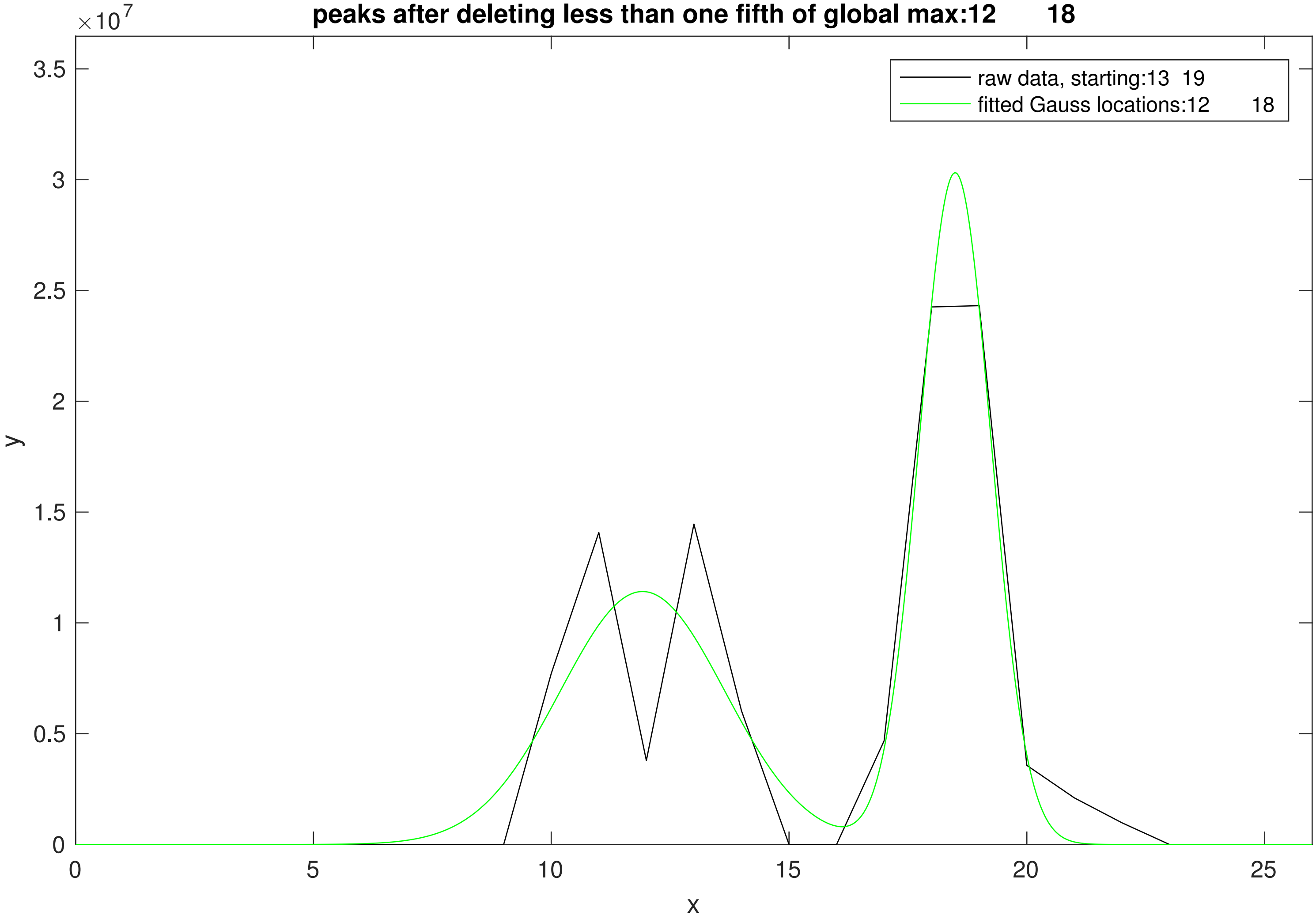

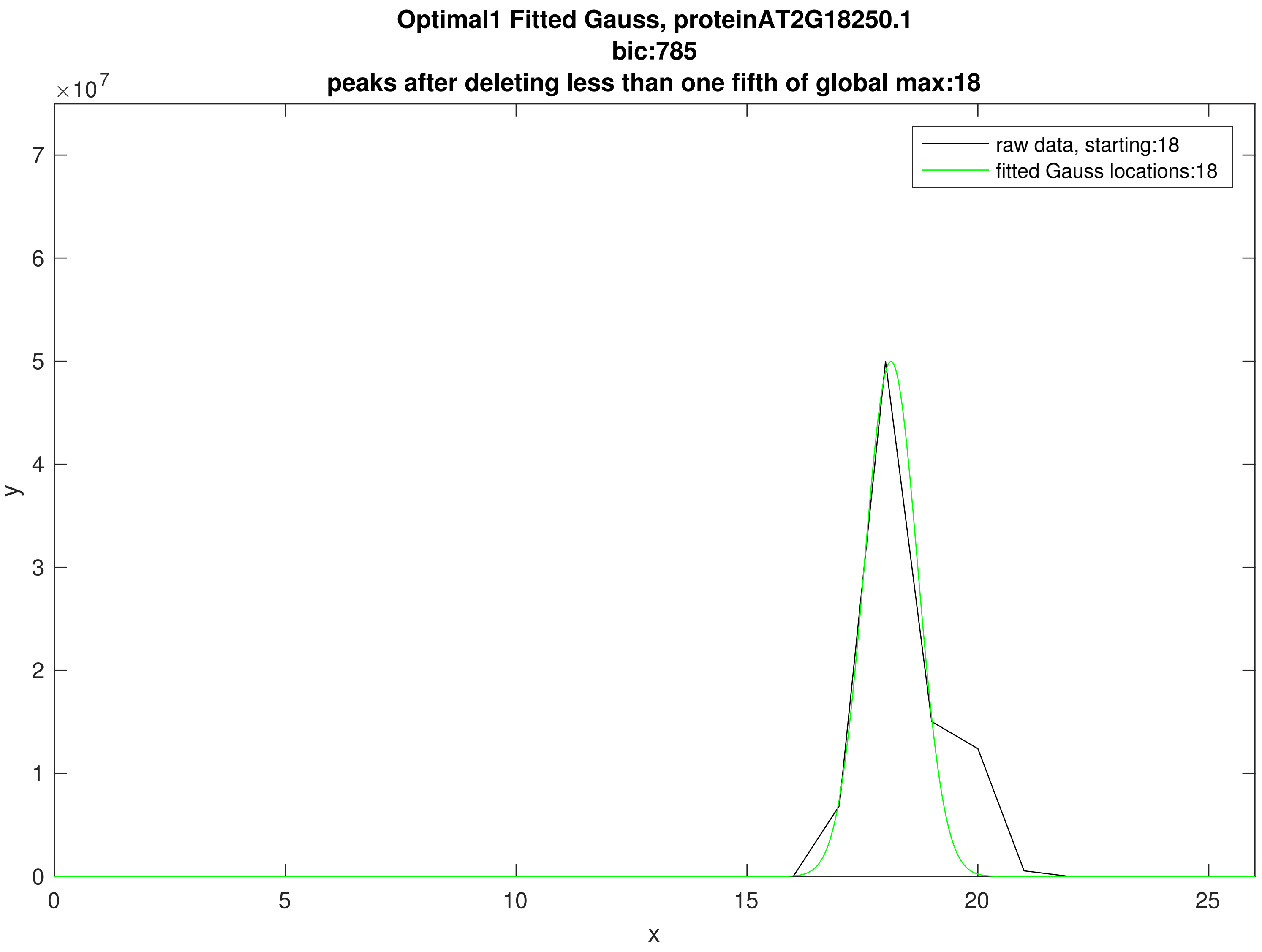
Optimal1 Fitted Gauss, proteinAT2G18250.1
bic:785
peaks after deleting less than one fifth of global max:18
×10^7
raw data, starting:18
fitted Gauss locations:18
y
x
0
1
2
3
4
5
6
7
0
5
10
15
20
25

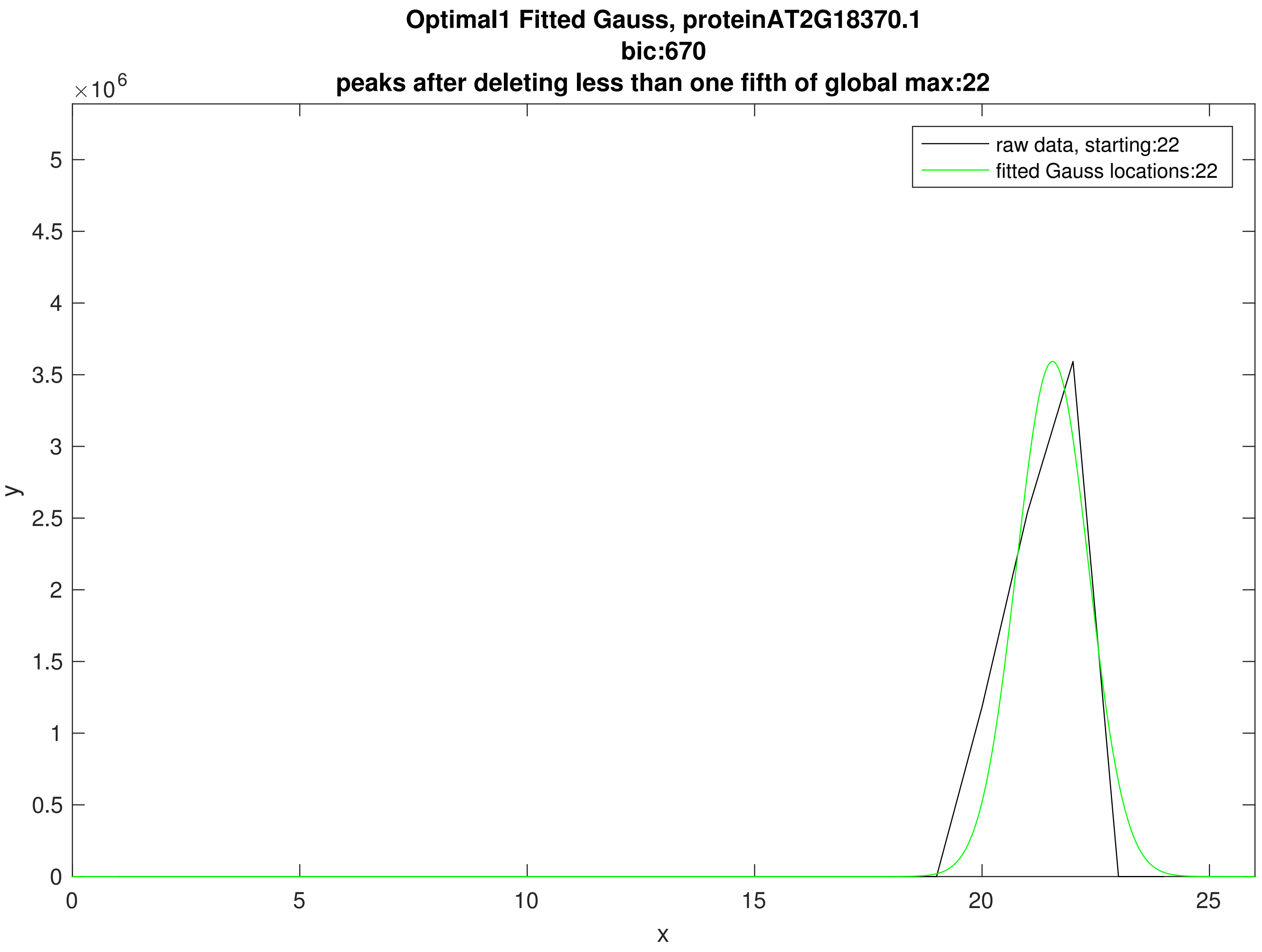
Optimal1 Fitted Gauss, proteinAT2G18370.1
bic:670
peaks after deleting less than one fifth of global max:22
×10^6
raw data, starting:22
fitted Gauss locations:22
y
x
0
0.5
1
1.5
2
2.5
3
3.5
4
4.5
5
0
5
10
15
20
25

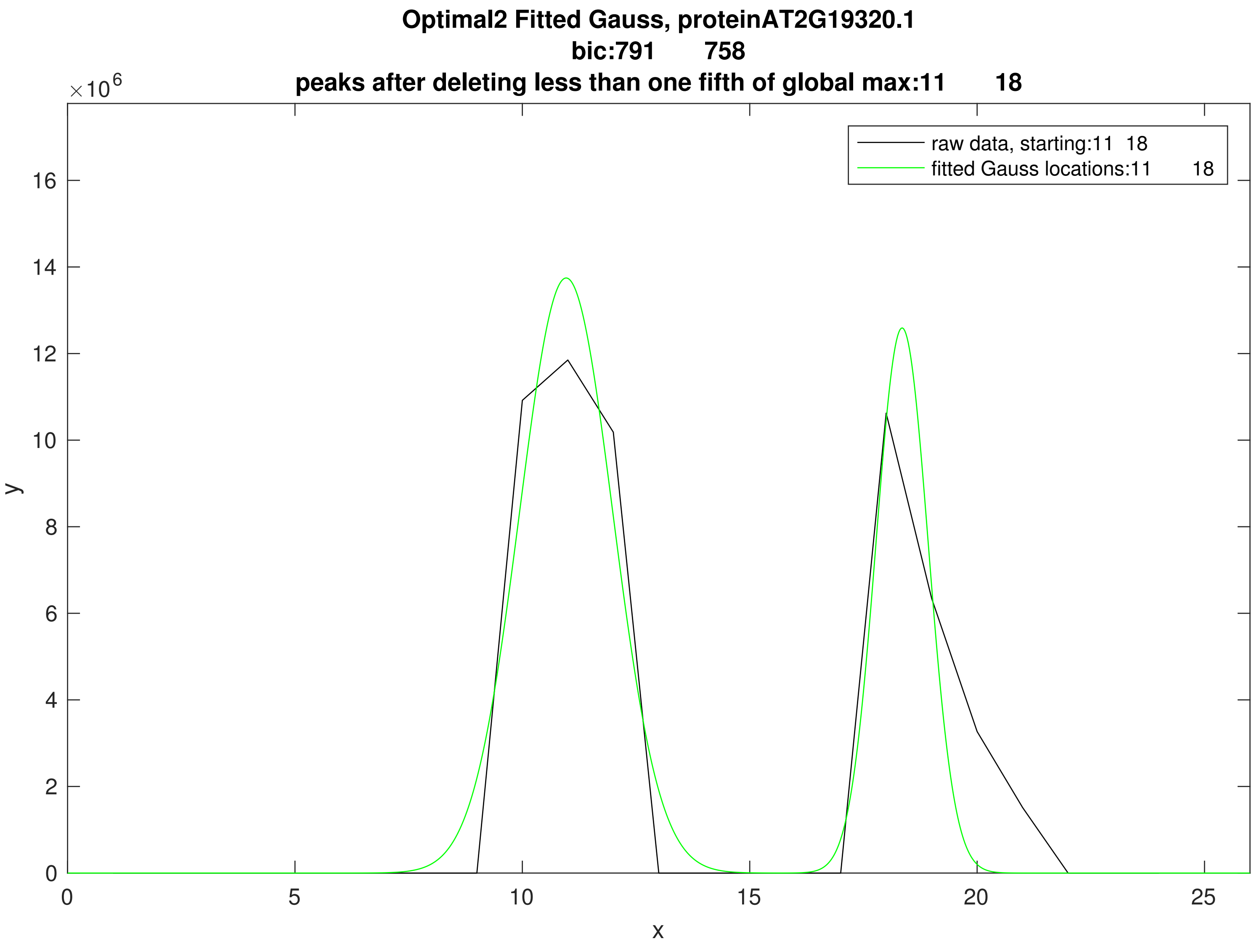
Optimal2 Fitted Gauss, proteinAT2G19320.1
bic:791 758
peaks after deleting less than one fifth of global max:11 18
×10^6
raw data, starting:11 18
fitted Gauss locations:11 18
y
x
0
2
4
6
8
10
12
14
16
0
5
10
15
20
25

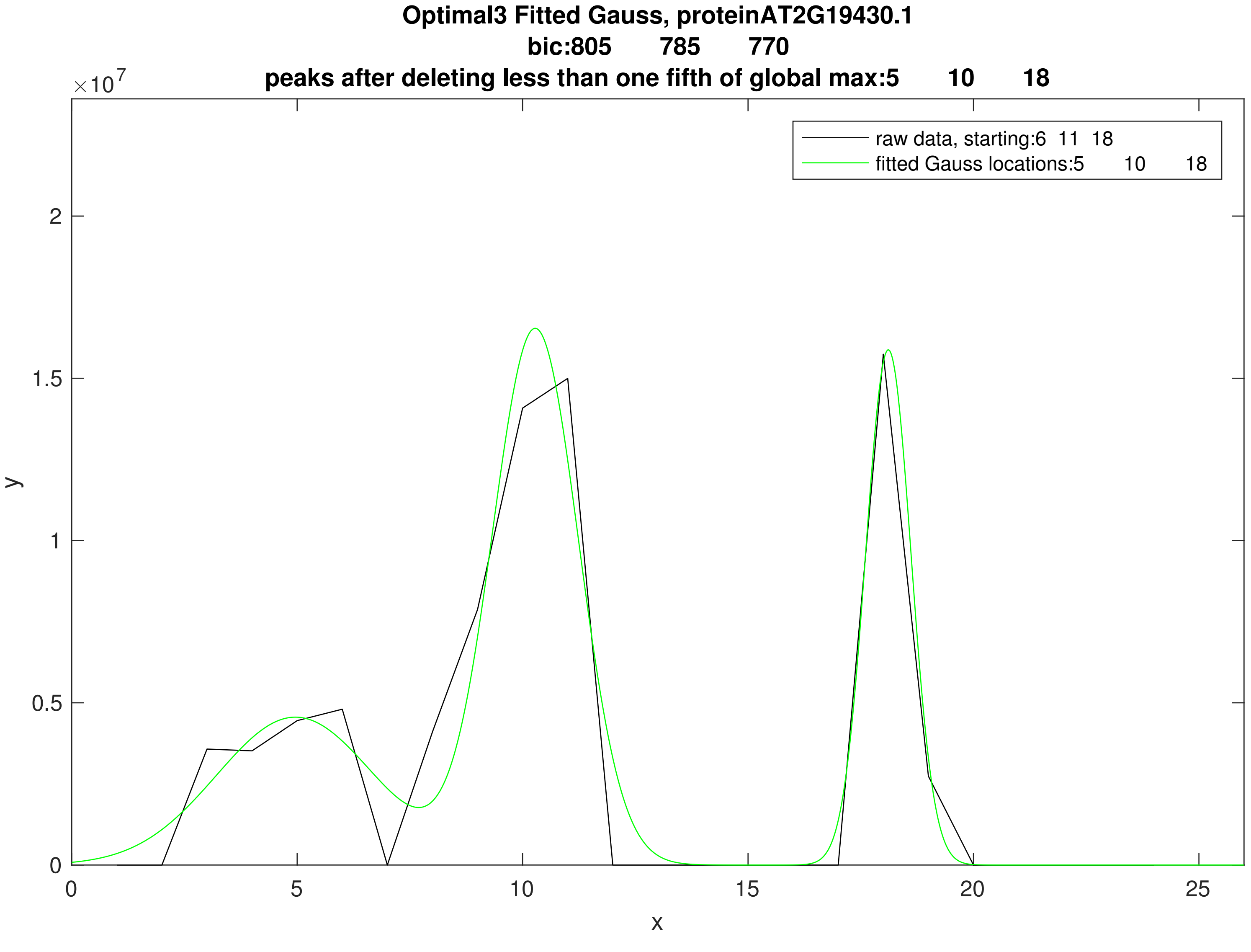

Optimal3 Fitted Gauss, proteinAT2G19430.1
bic:805 785 770
peaks after deleting less than one fifth of global max:5 10 18
×10^7
raw data, starting:6 11 18
fitted Gauss locations:5 10 18
y
x
0
0.5
1
1.5
2
0
5
10
15
20
25

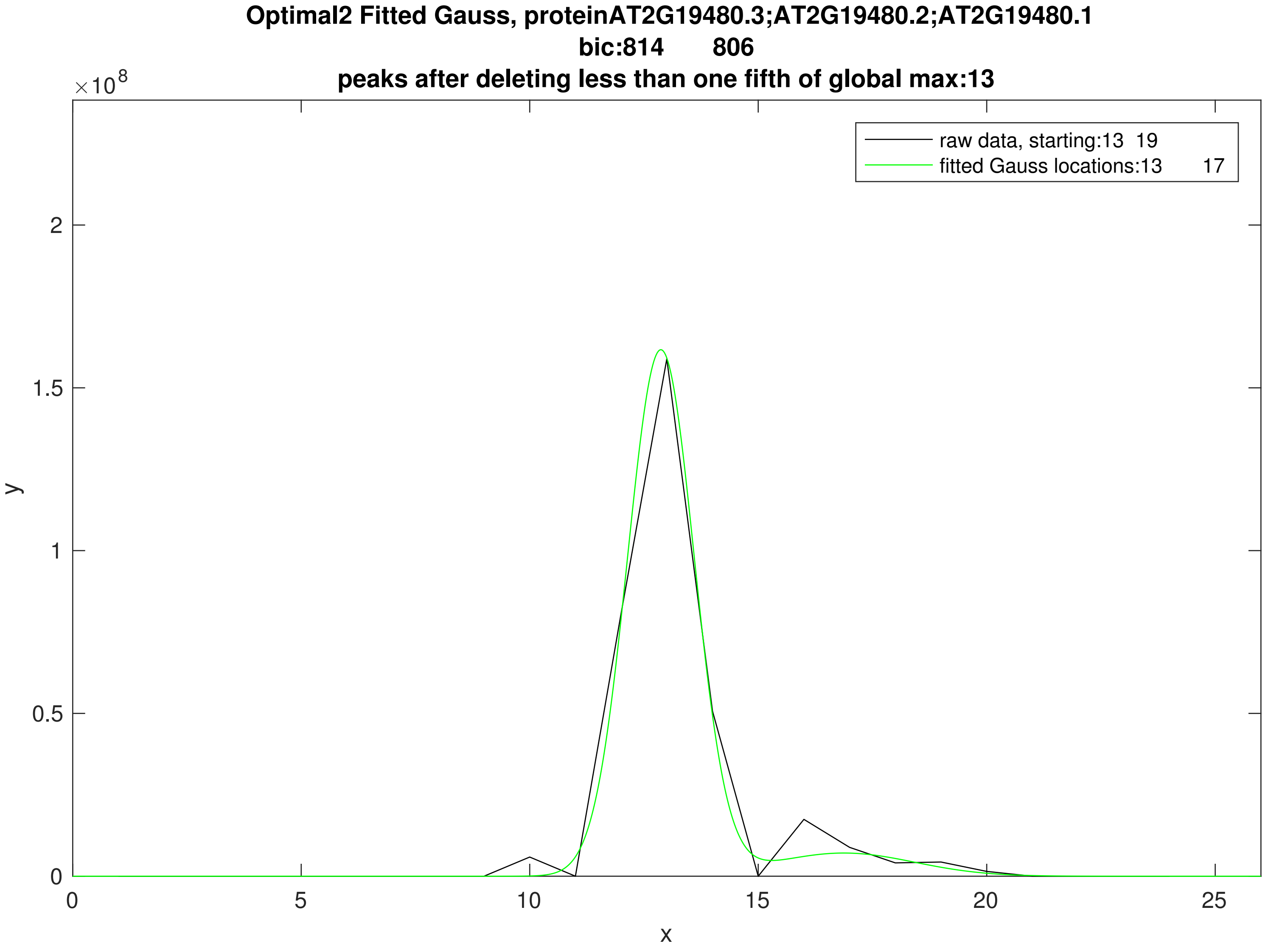

Optimal2 Fitted Gauss, proteinAT2G19480.3;AT2G19480.2;AT2G19480.1
bic:814 806
peaks after deleting less than one fifth of global max:13
×10^8
raw data, starting:13 19
fitted Gauss locations:13 17
y
x
0
0.5
1
1.5
2
0
5
10
15
20
25

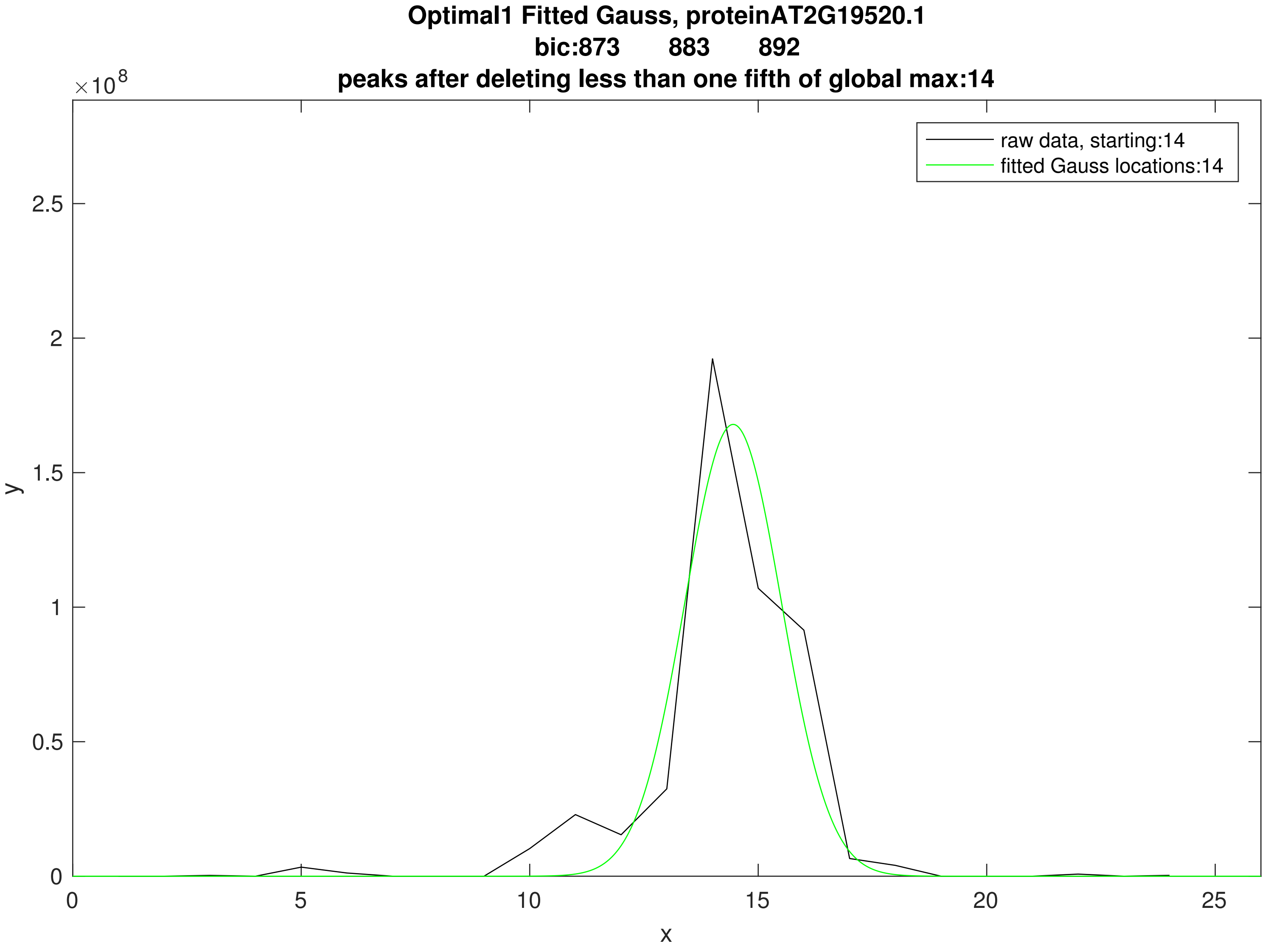
Optimal1 Fitted Gauss, proteinAT2G19520.1
bic:873 883 892
peaks after deleting less than one fifth of global max:14
×10^8
raw data, starting:14
fitted Gauss locations:14
2.5
2
1.5
1
0.5
0
y
0
5
10
15
20
25
x

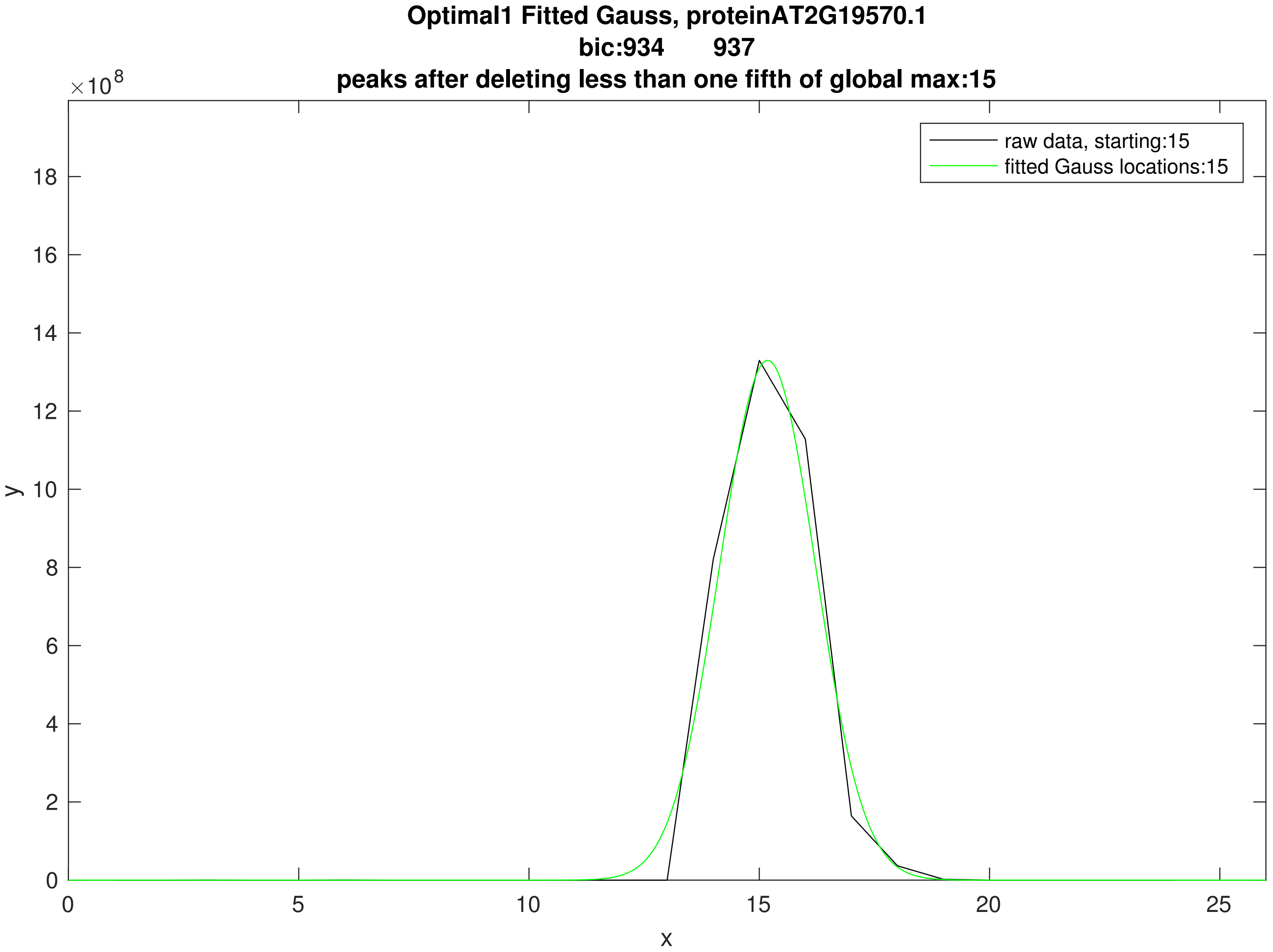
Optimal1 Fitted Gauss, proteinAT2G19570.1
bic:934 937
peaks after deleting less than one fifth of global max:15
×10^8
raw data, starting:15
fitted Gauss locations:15
y
x
0
2
4
6
8
10
12
14
16
18
0
5
10
15
20
25

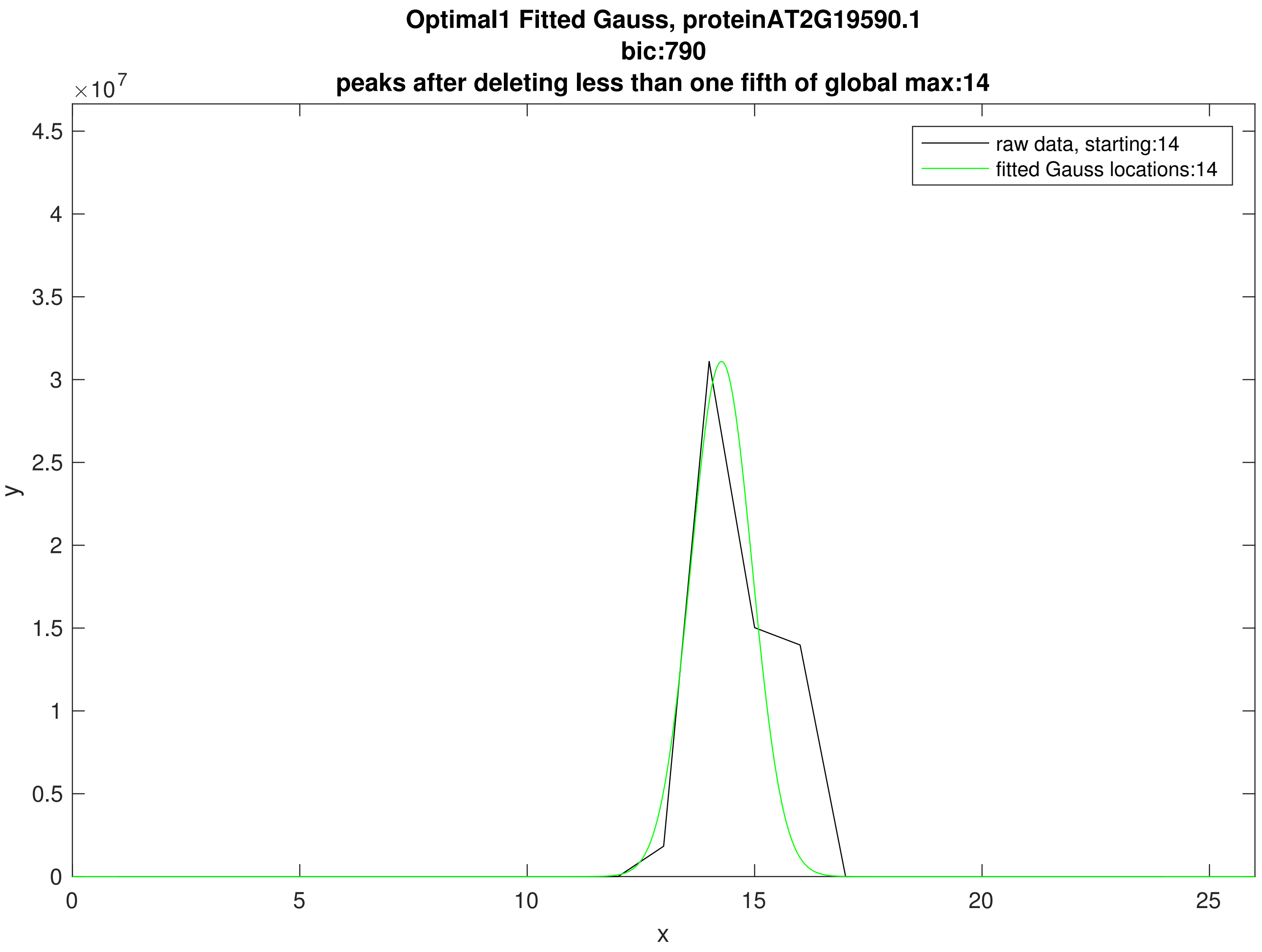

Optimal1 Fitted Gauss, proteinAT2G19590.1
bic:790
peaks after deleting less than one fifth of global max:14
×10^7
4.5
4
3.5
3
2.5
2
1.5
1
0.5
0
y
0
5
10
15
20
25
x
raw data, starting:14
fitted Gauss locations:14

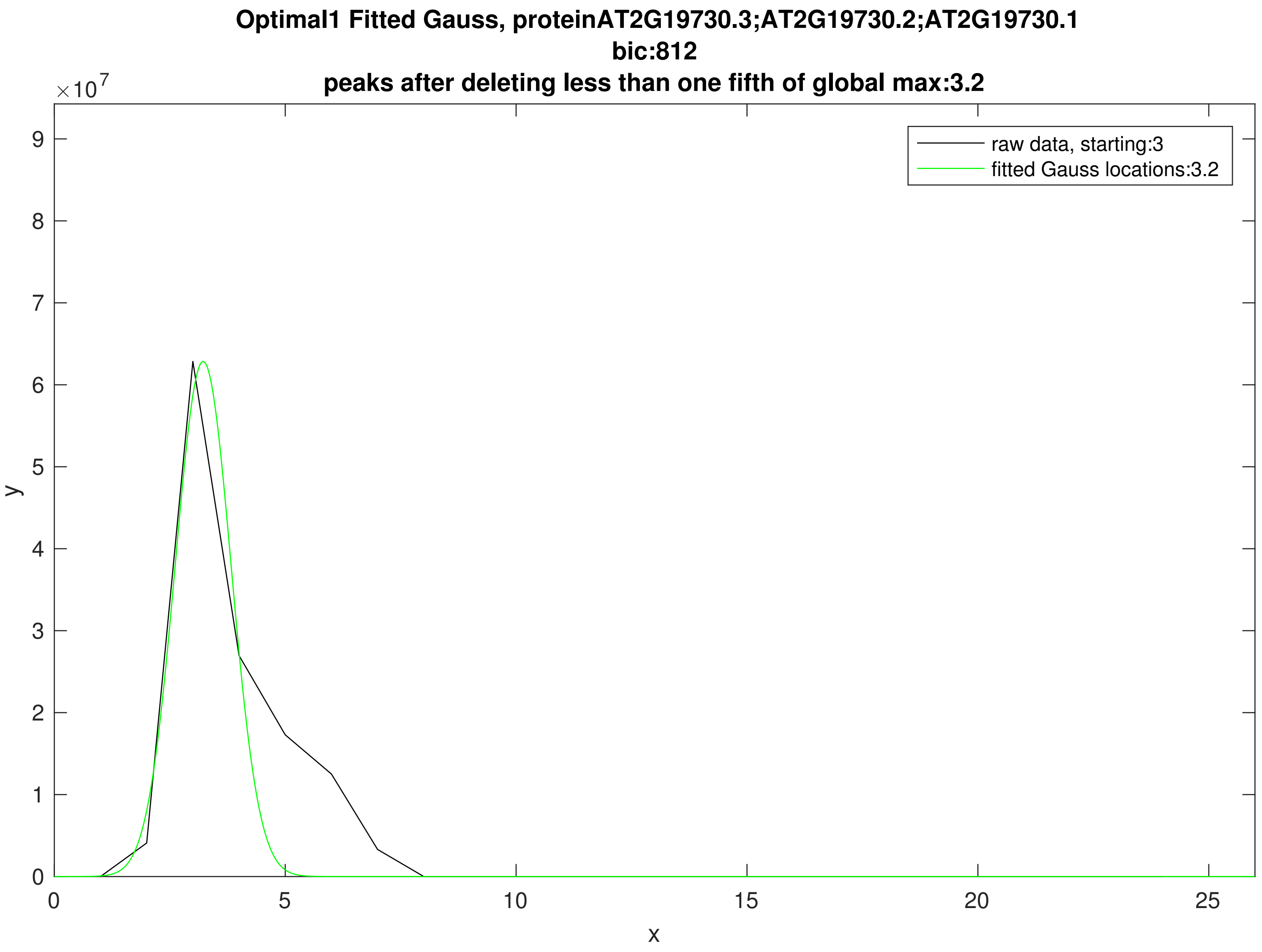
Optimal1 Fitted Gauss, proteinAT2G19730.3;AT2G19730.2;AT2G19730.1
bic:812
peaks after deleting less than one fifth of global max:3.2
×10^7
raw data, starting:3
fitted Gauss locations:3.2
y
x
0
1
2
3
4
5
6
7
8
9
0
5
10
15
20
25

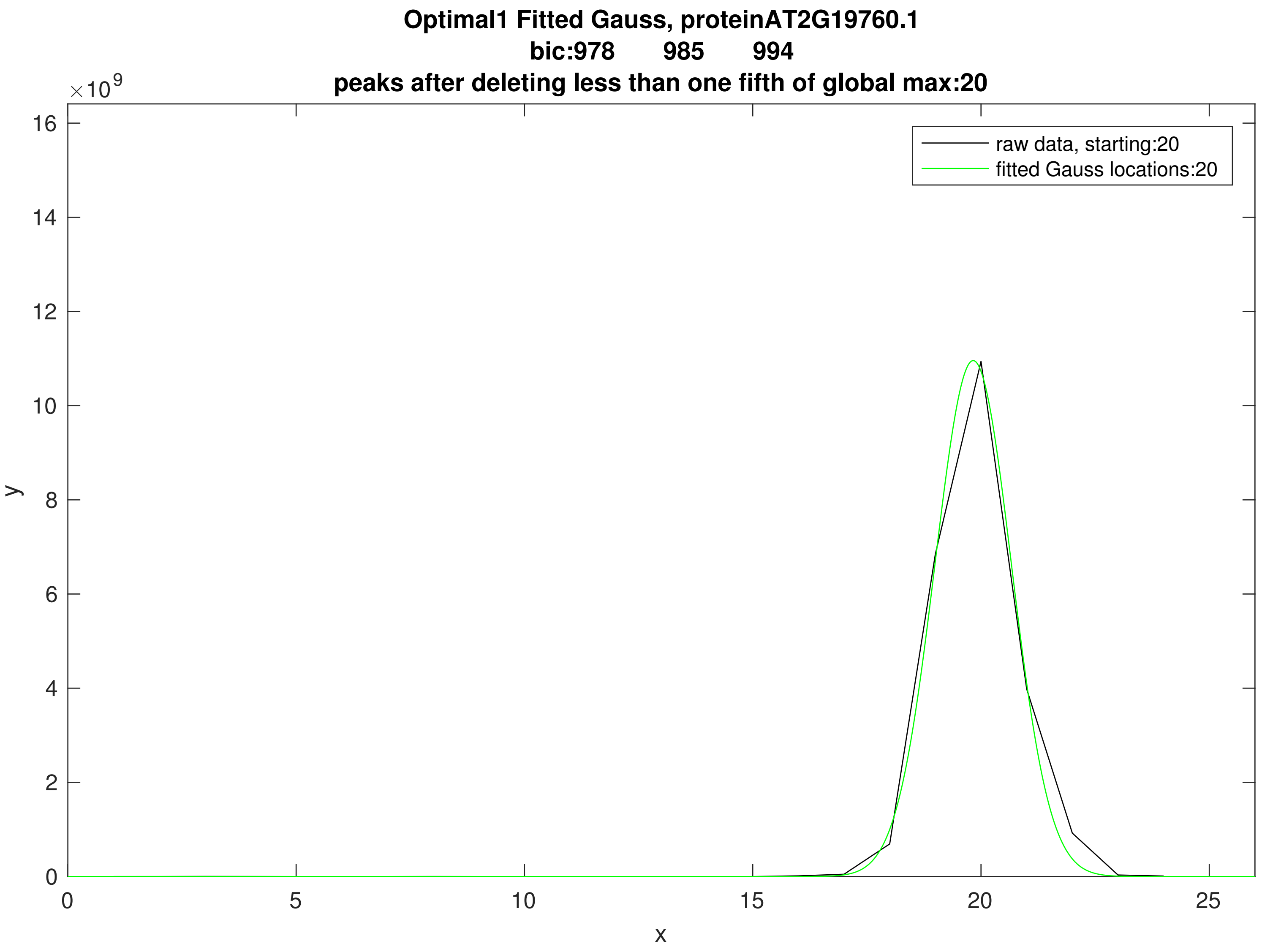
Optimal1 Fitted Gauss, proteinAT2G19760.1
bic:978 985 994
peaks after deleting less than one fifth of global max:20
×10^9
raw data, starting:20
fitted Gauss locations:20
y
x
0
2
4
6
8
10
12
14
16
0
5
10
15
20
25

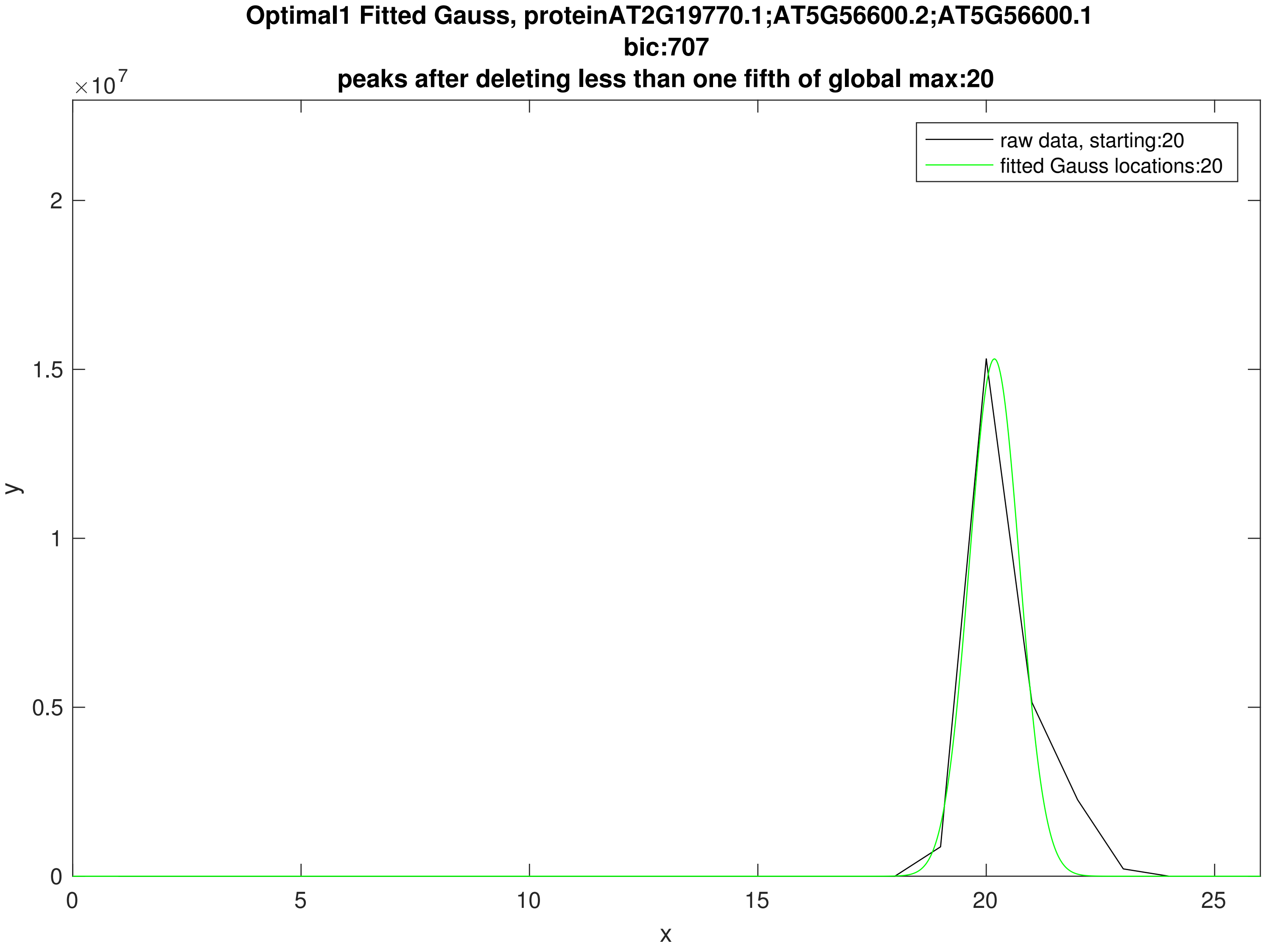

Optimal1 Fitted Gauss, proteinAT2G19770.1;AT5G56600.2;AT5G56600.1
bic:707
peaks after deleting less than one fifth of global max:20
×10^7
raw data, starting:20
fitted Gauss locations:20
2
1.5
1
0.5
0
y
0
5
10
15
20
25
x

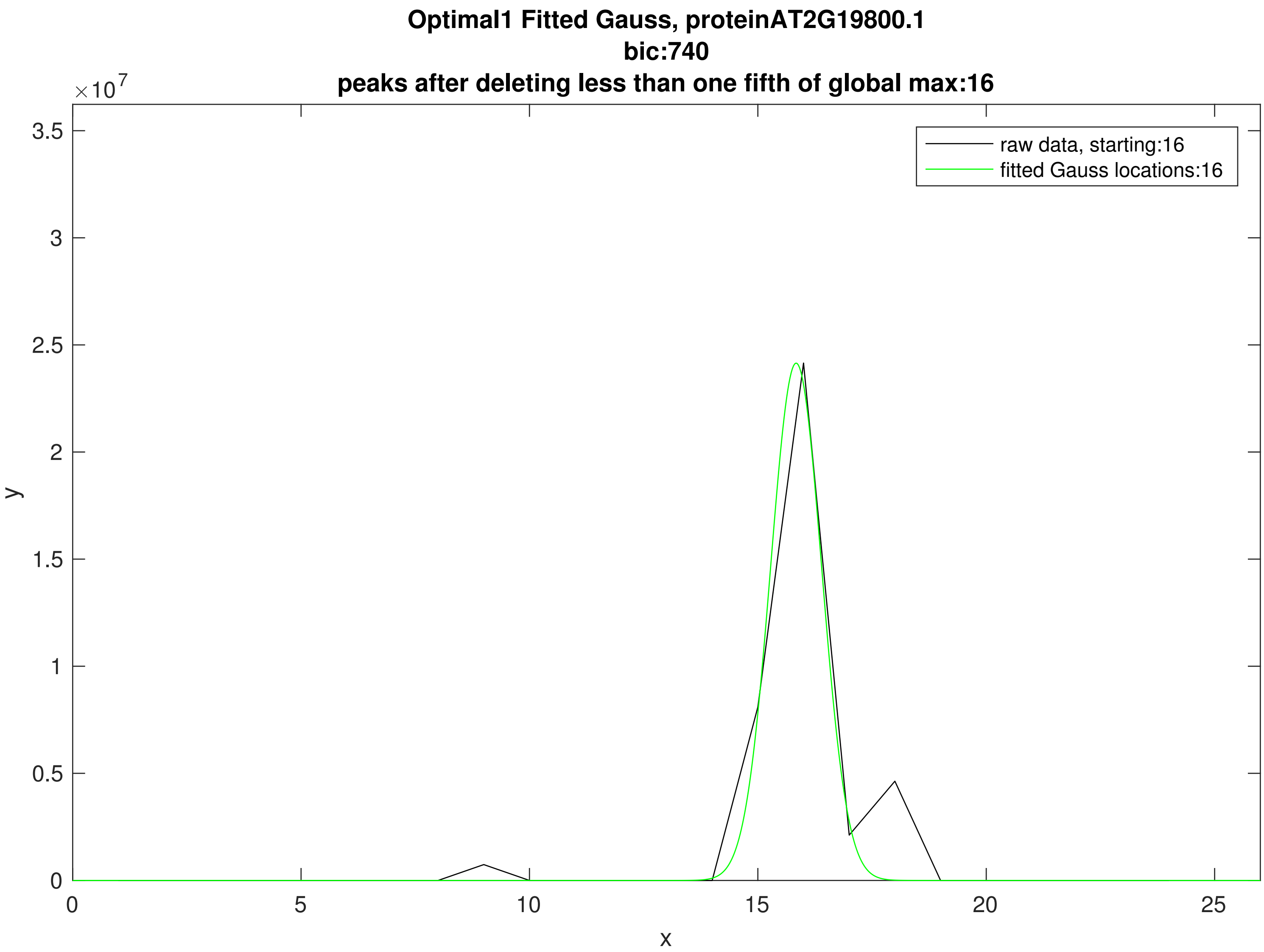
Optimal1 Fitted Gauss, proteinAT2G19800.1
bic:740
peaks after deleting less than one fifth of global max:16
×10^7
3.5
3
2.5
2
1.5
1
0.5
0
y
0
5
10
15
20
25
x
raw data, starting:16
fitted Gauss locations:16

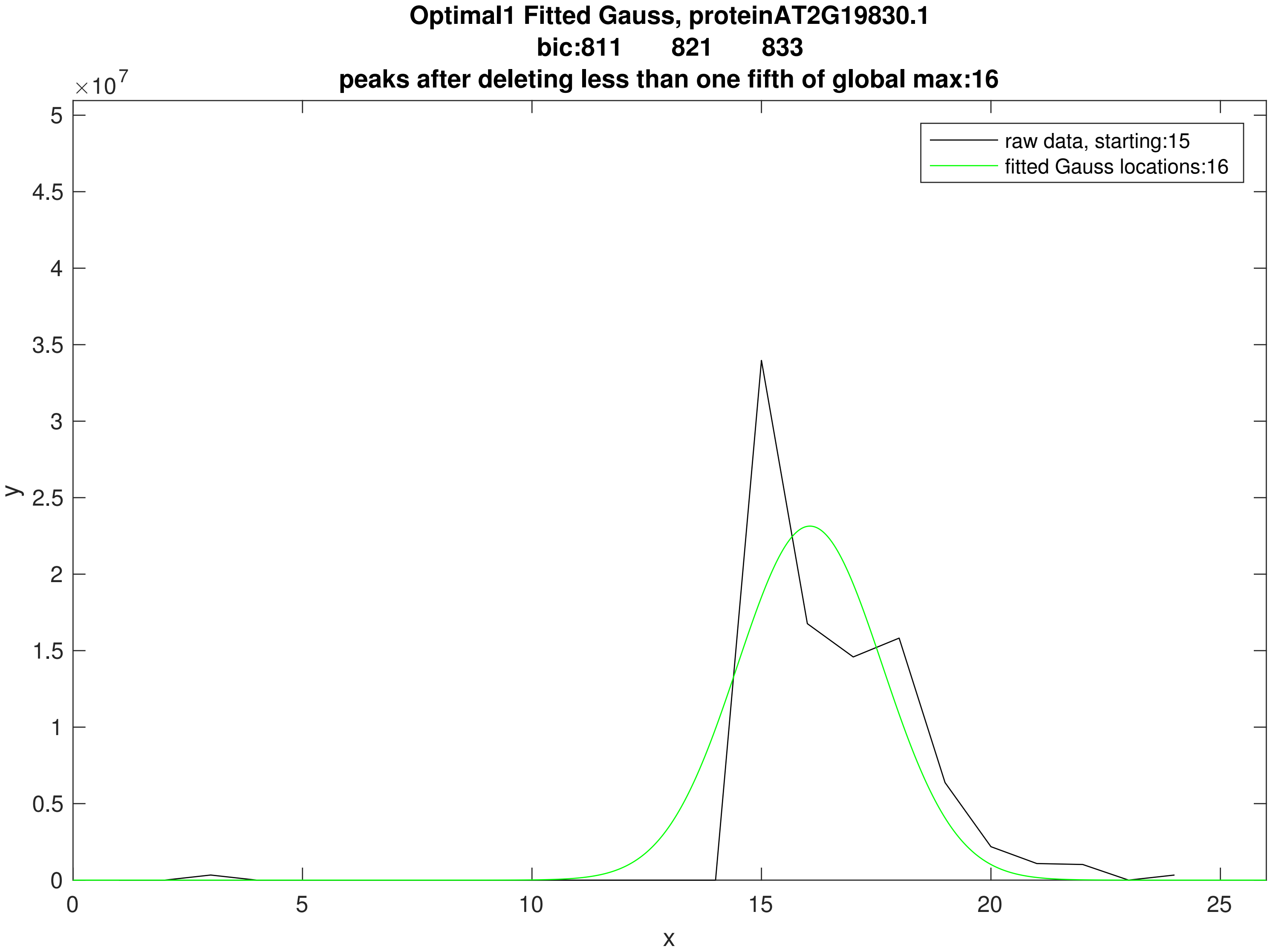

Optimal1 Fitted Gauss, proteinAT2G19830.1
bic:811 821 833
peaks after deleting less than one fifth of global max:16
×10^7
raw data, starting:15
fitted Gauss locations:16
y
x
0
0.5
1
1.5
2
2.5
3
3.5
4
4.5
5
0
5
10
15
20
25

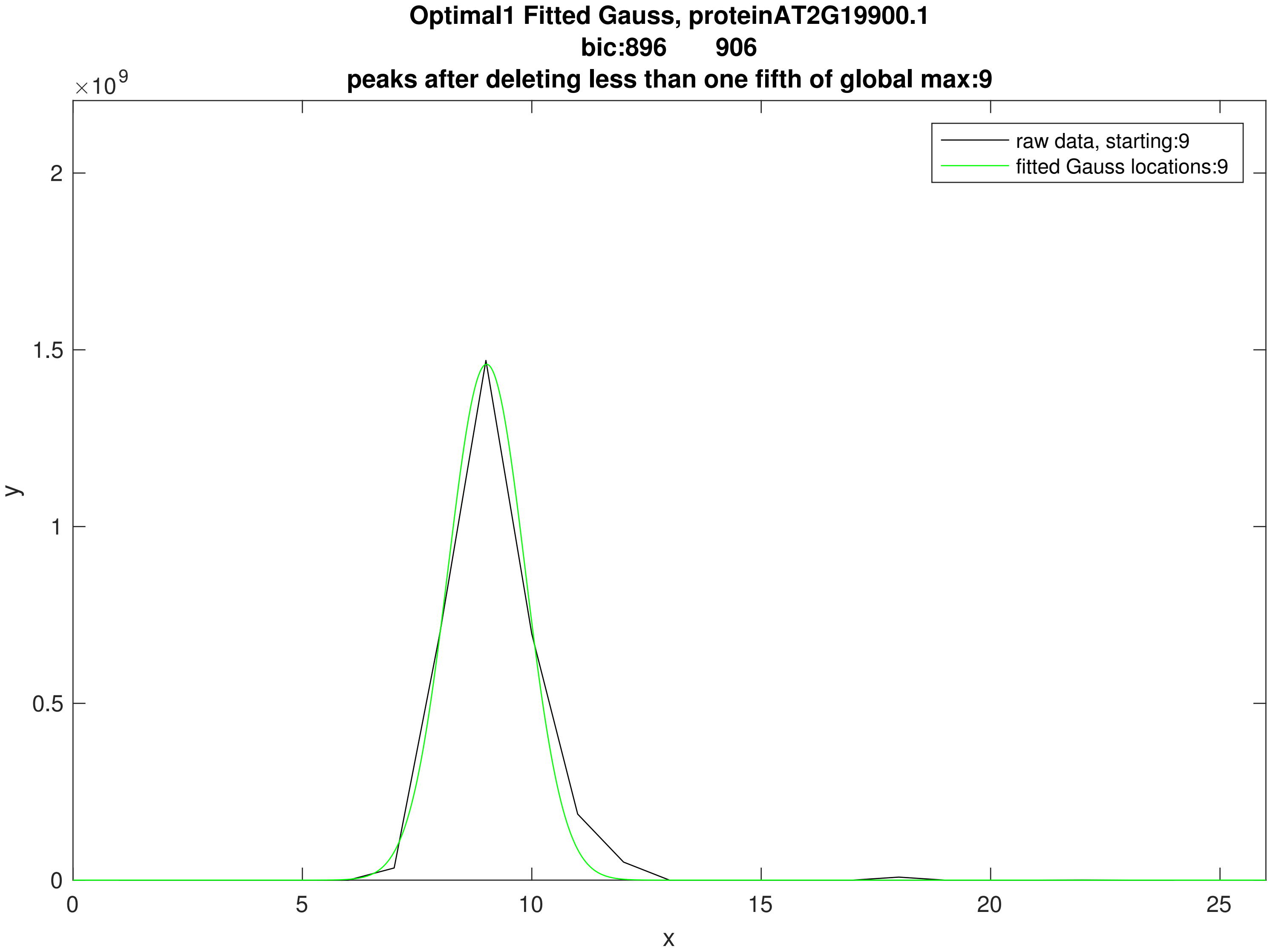
Optimal1 Fitted Gauss, proteinAT2G19900.1
bic:896 906
peaks after deleting less than one fifth of global max:9
×10^9
raw data, starting:9
fitted Gauss locations:9
y
x
0
0.5
1
1.5
2
5
10
15
20
25

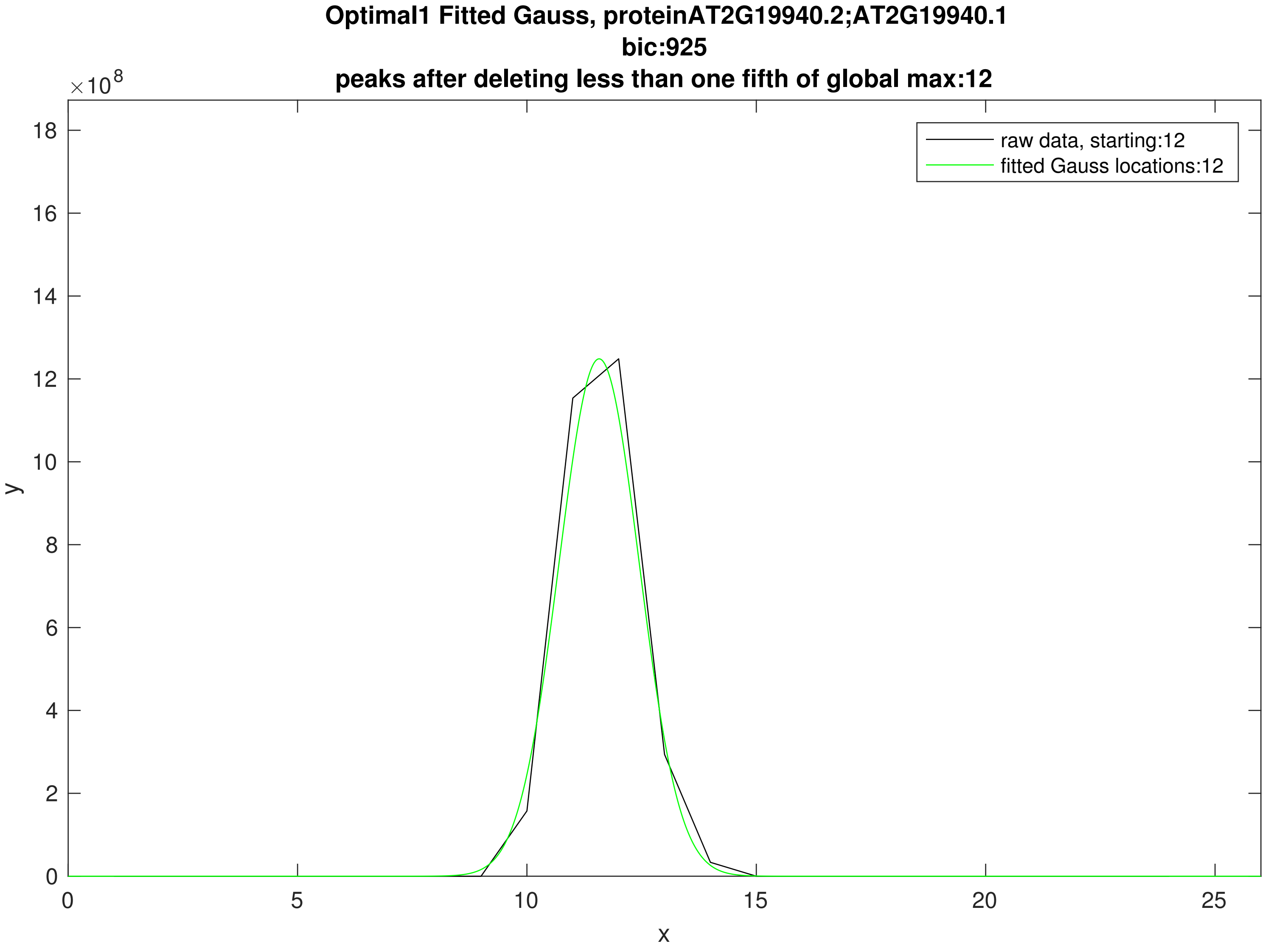
Optimal1 Fitted Gauss, proteinAT2G19940.2;AT2G19940.1
bic:925
peaks after deleting less than one fifth of global max:12
×10^8
18
16
14
12
10
8
6
4
2
0
y
0
5
10
15
20
25
x
raw data, starting:12
fitted Gauss locations:12

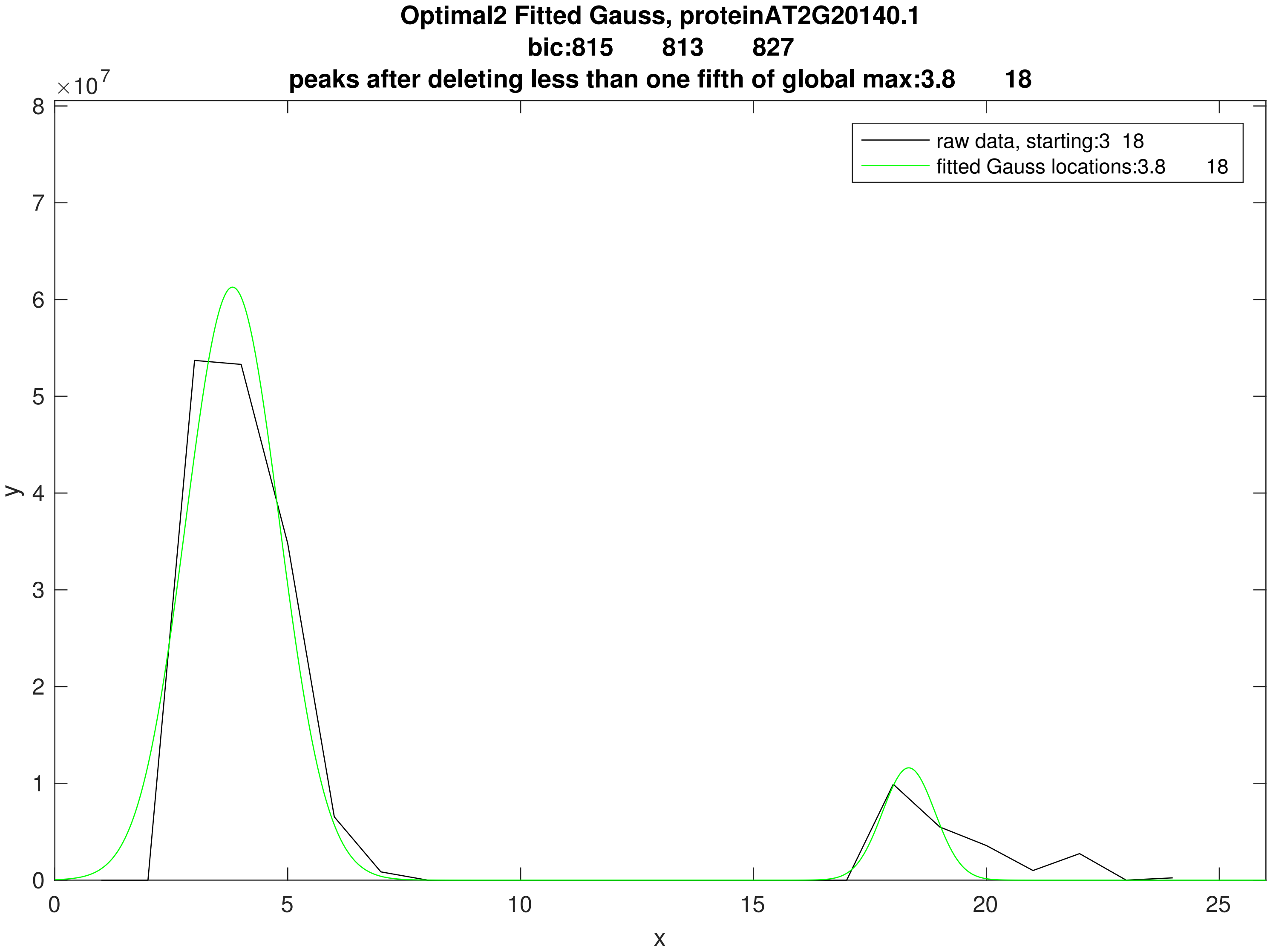

Optimal2 Fitted Gauss, proteinAT2G20140.1
bic:815 813 827
peaks after deleting less than one fifth of global max:3.8 18
×10^7
raw data, starting:3 18
fitted Gauss locations:3.8 18
y
x
0
1
2
3
4
5
6
7
8
0
5
10
15
20
25

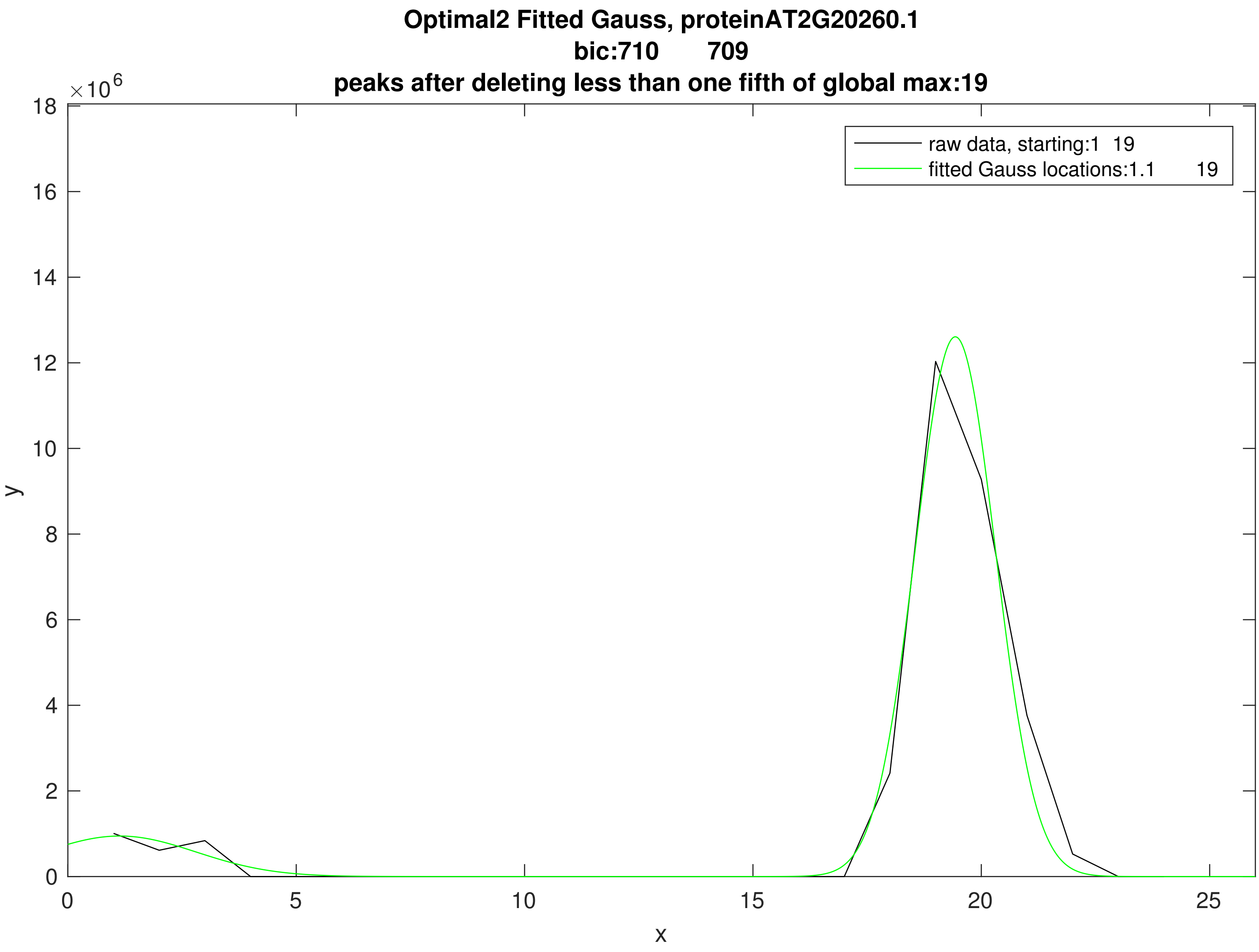
Optimal2 Fitted Gauss, proteinAT2G20260.1
bic:710 709
peaks after deleting less than one fifth of global max:19
×10^6
18
16
14
12
10
8
6
4
2
0
y
0
5
10
15
20
25
x
raw data, starting:1 19
fitted Gauss locations:1.1 19

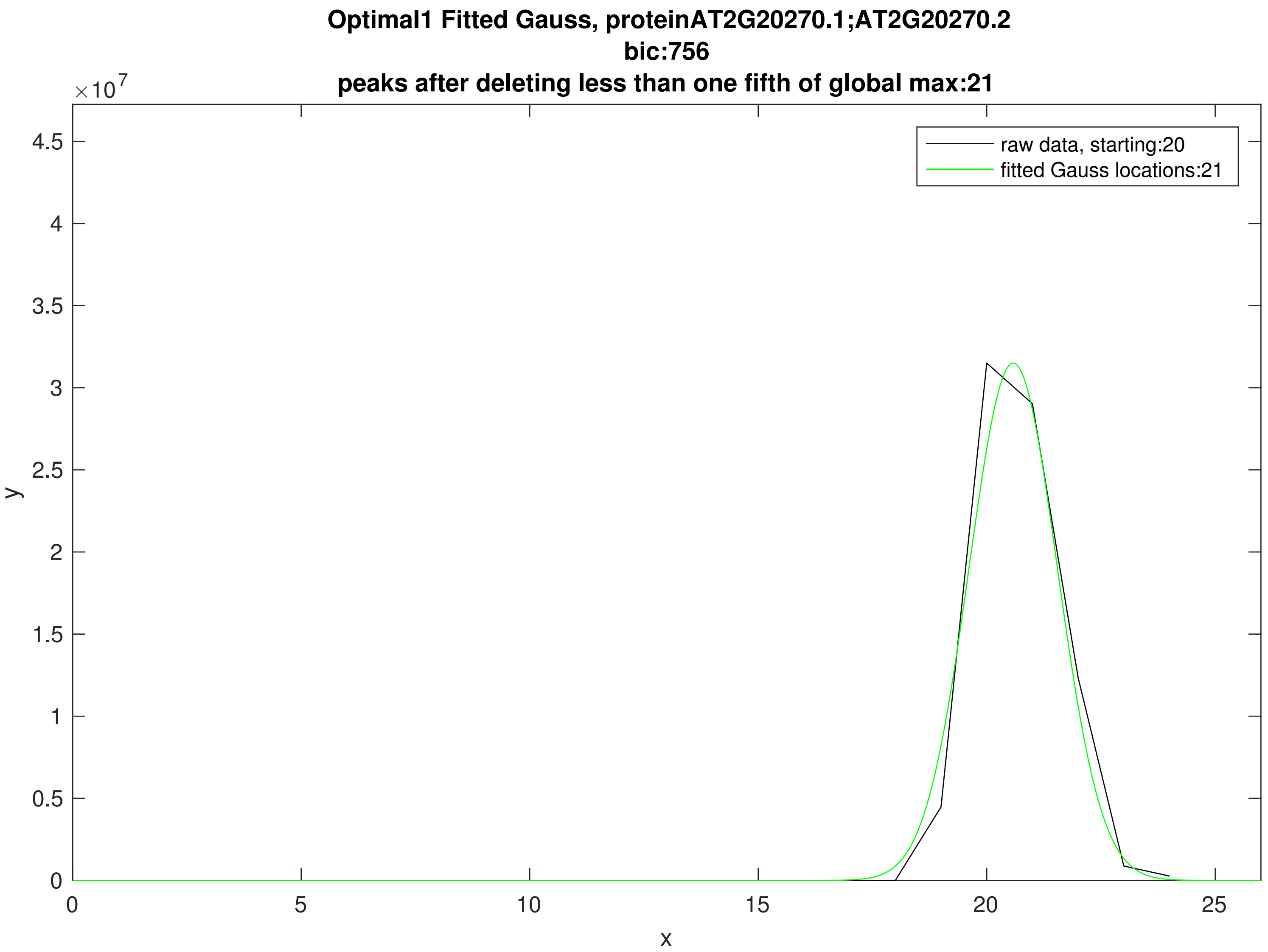
Optimal1 Fitted Gauss, proteinAT2G20270.1;AT2G20270.2
bic:756
peaks after deleting less than one fifth of global max:21
×10^7
4.5
4
3.5
3
2.5
2
1.5
1
0.5
0
y
0
5
10
15
20
25
x
raw data, starting:20
fitted Gauss locations:21

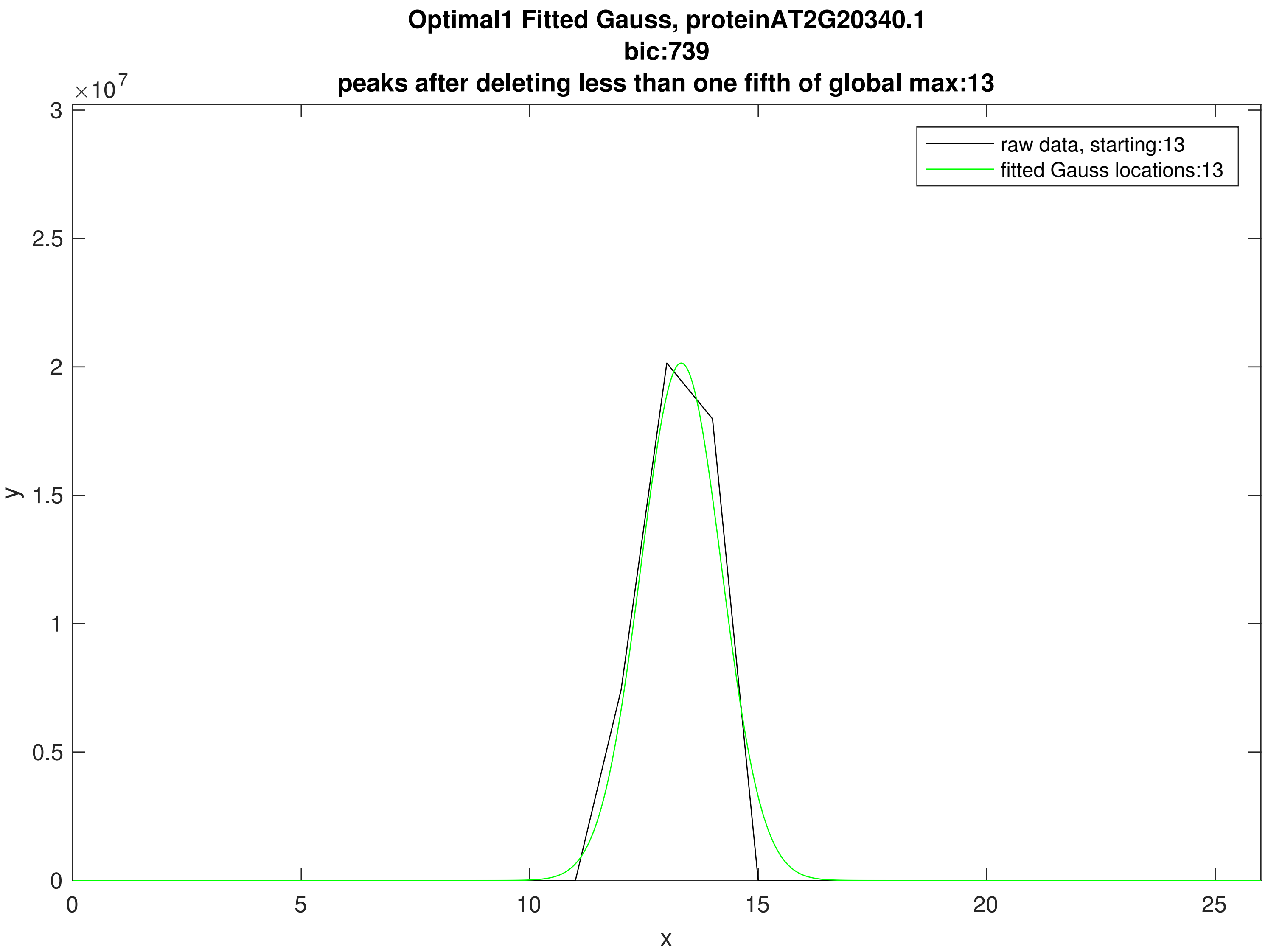

Optimal1 Fitted Gauss, proteinAT2G20340.1
bic:739
peaks after deleting less than one fifth of global max:13
×10^7
raw data, starting:13
fitted Gauss locations:13
y
x
0
0.5
1
1.5
2
2.5
3
0
5
10
15
20
25

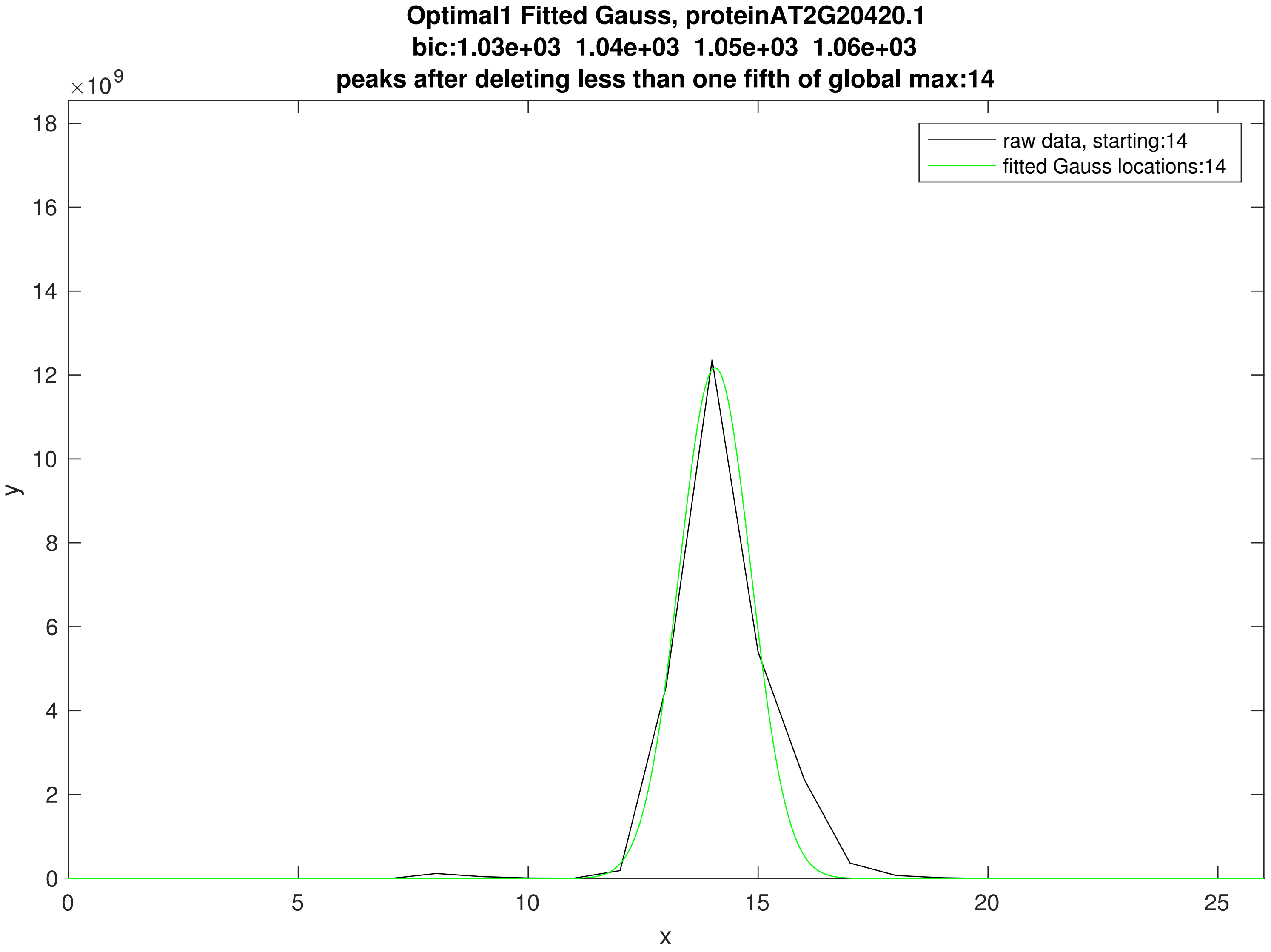
Optimal1 Fitted Gauss, proteinAT2G20420.1
bic:1.03e+03 1.04e+03 1.05e+03 1.06e+03
peaks after deleting less than one fifth of global max:14
raw data, starting:14
fitted Gauss locations:14
×10^9
y
x

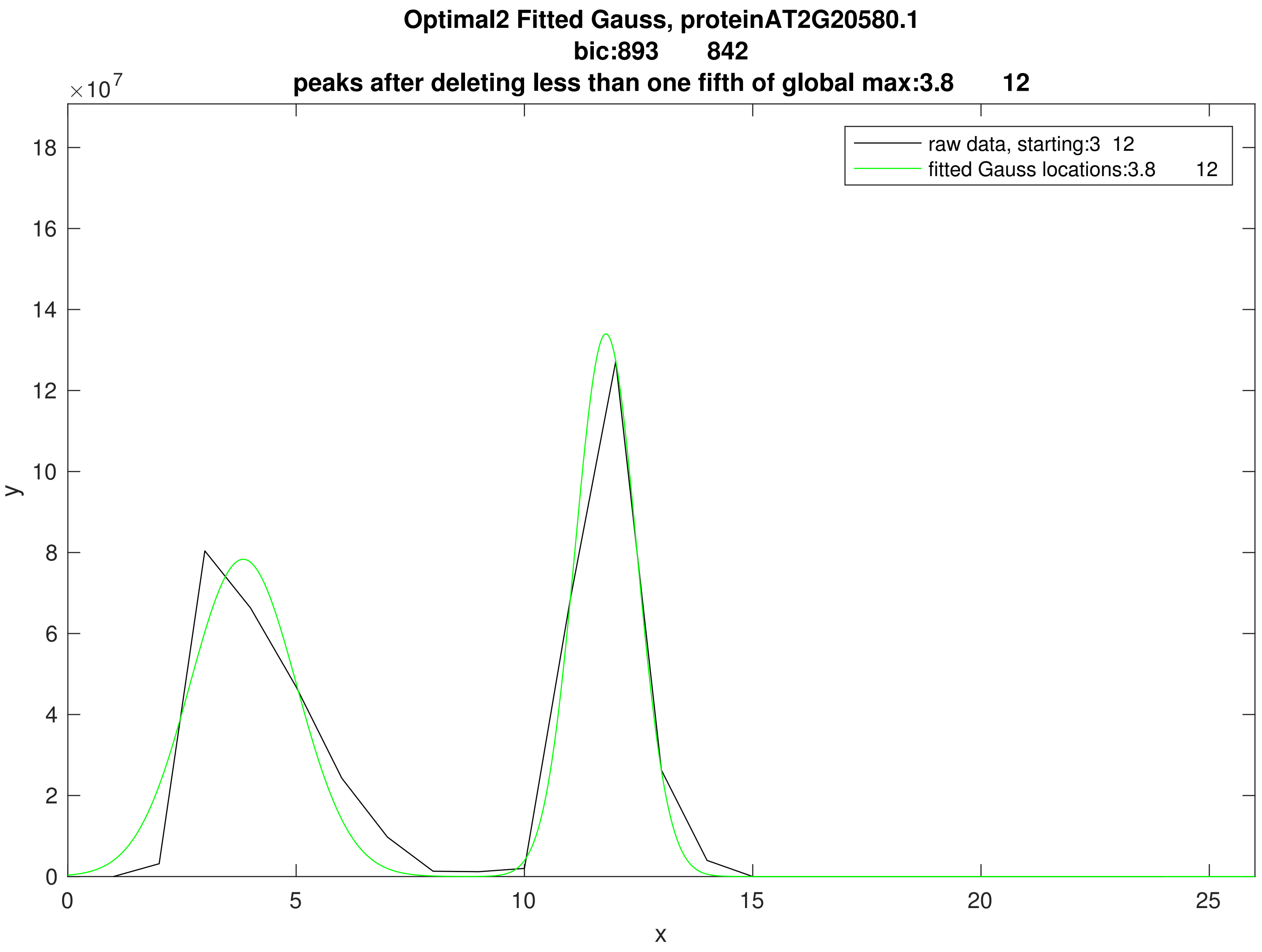

Optimal2 Fitted Gauss, proteinAT2G20580.1
bic:893 842
peaks after deleting less than one fifth of global max:3.8 12
×10^7
raw data, starting:3 12
fitted Gauss locations:3.8 12
y
x
0
2
4
6
8
10
12
14
16
18
0
5
10
15
20
25

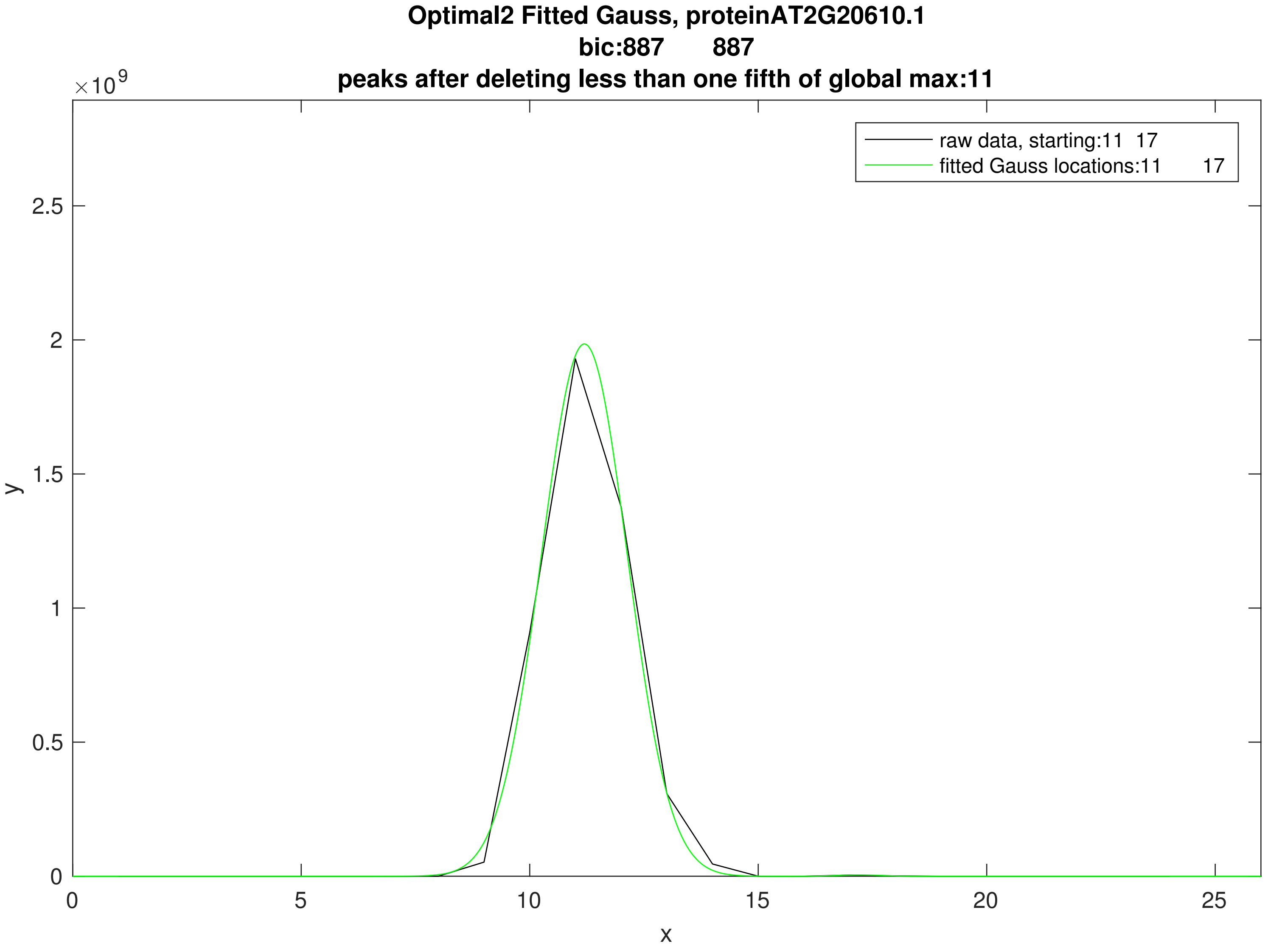
Optimal2 Fitted Gauss, proteinAT2G20610.1
bic:887 887
peaks after deleting less than one fifth of global max:11
×10^9
raw data, starting:11 17
fitted Gauss locations:11 17
2.5
2
1.5
1
0.5
0
y
0
5
10
15
20
25
x

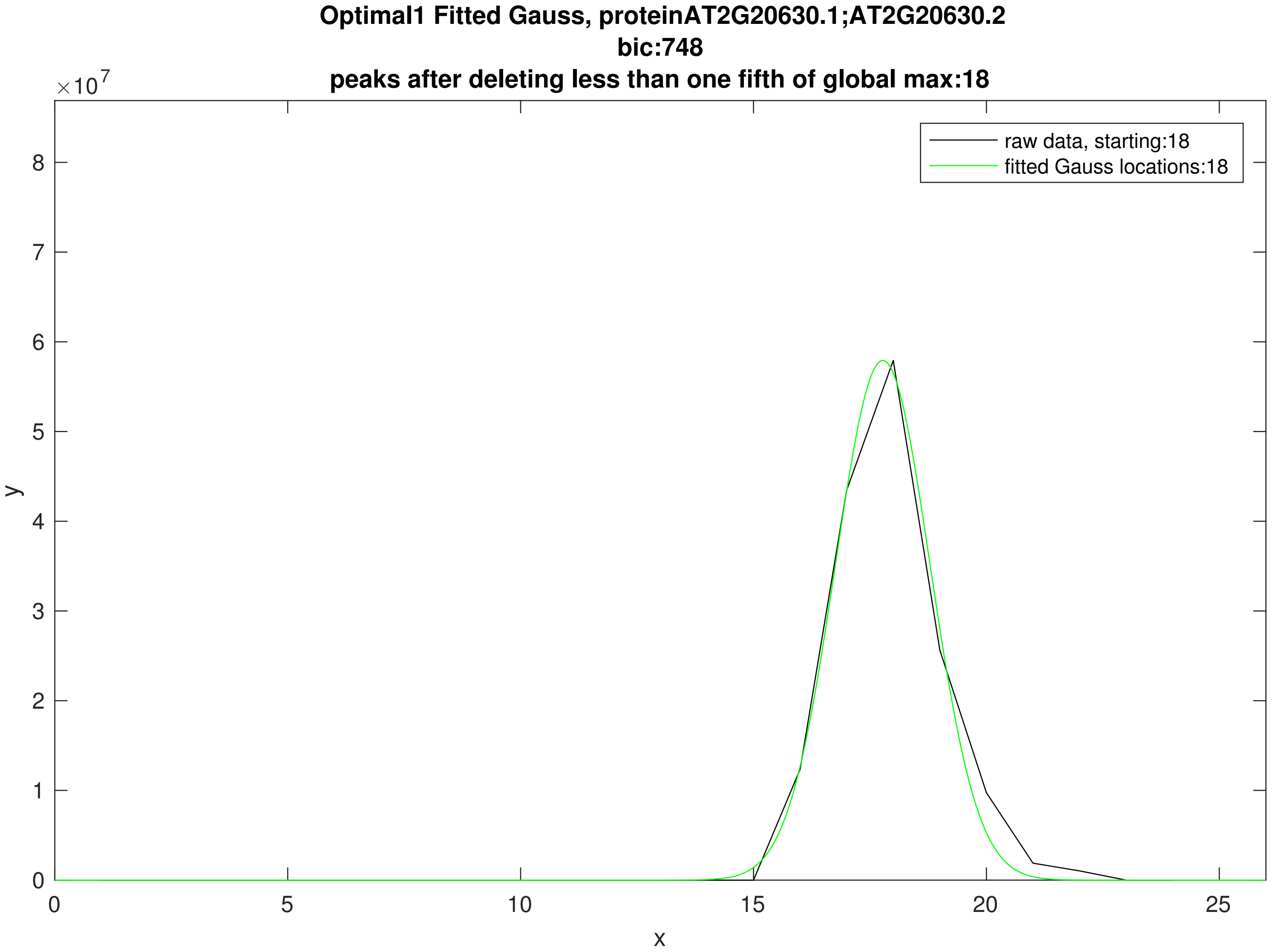

Optimal1 Fitted Gauss, proteinAT2G20630.1;AT2G20630.2
bic:748
peaks after deleting less than one fifth of global max:18
×10^7
raw data, starting:18
fitted Gauss locations:18
y
x
0
1
2
3
4
5
6
7
8
0
5
10
15
20
25

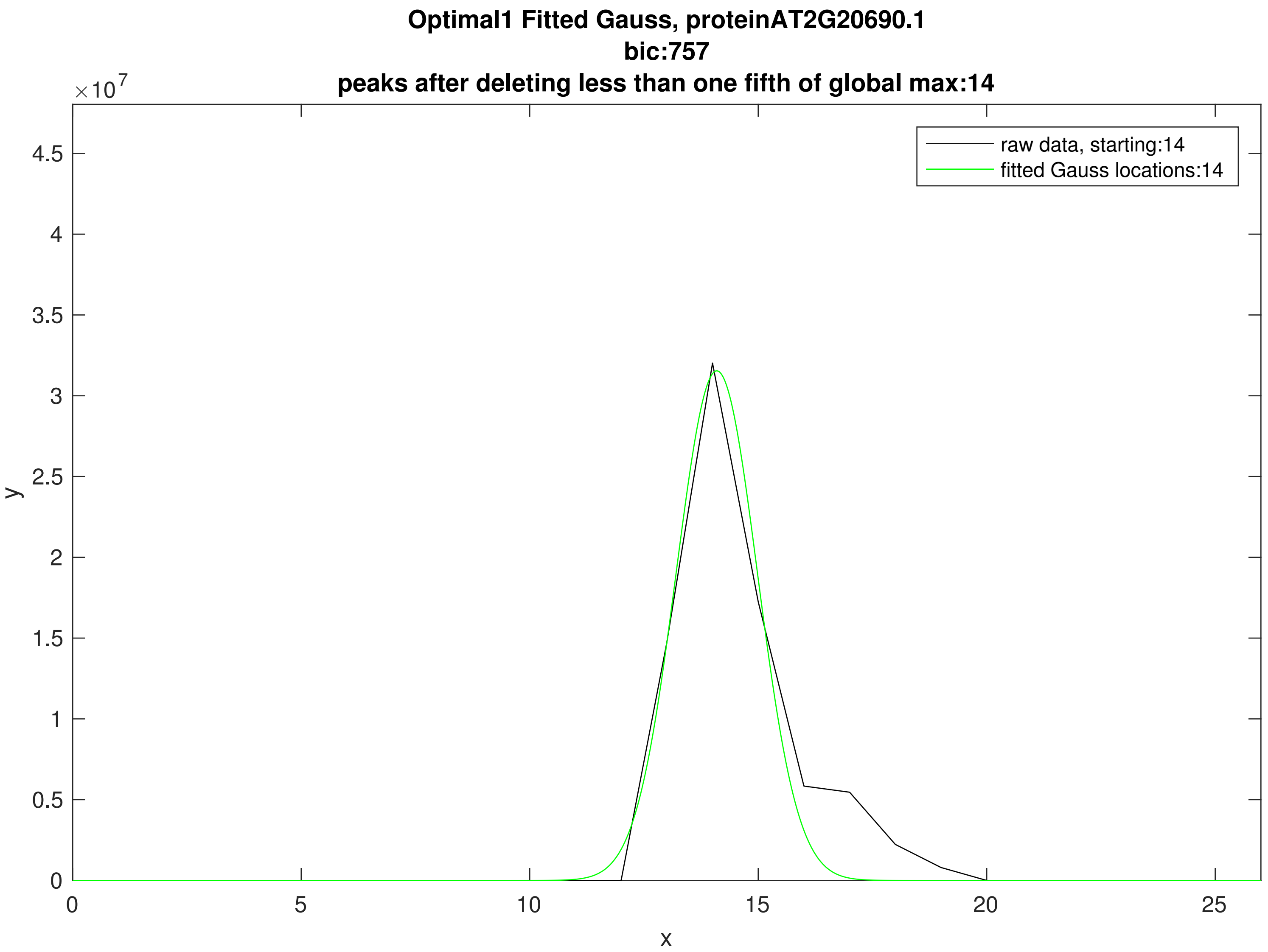
Optimal1 Fitted Gauss, proteinAT2G20690.1
bic:757
peaks after deleting less than one fifth of global max:14
×10^7
raw data, starting:14
fitted Gauss locations:14
y
x
0
0.5
1
1.5
2
2.5
3
3.5
4
4.5
0
5
10
15
20
25

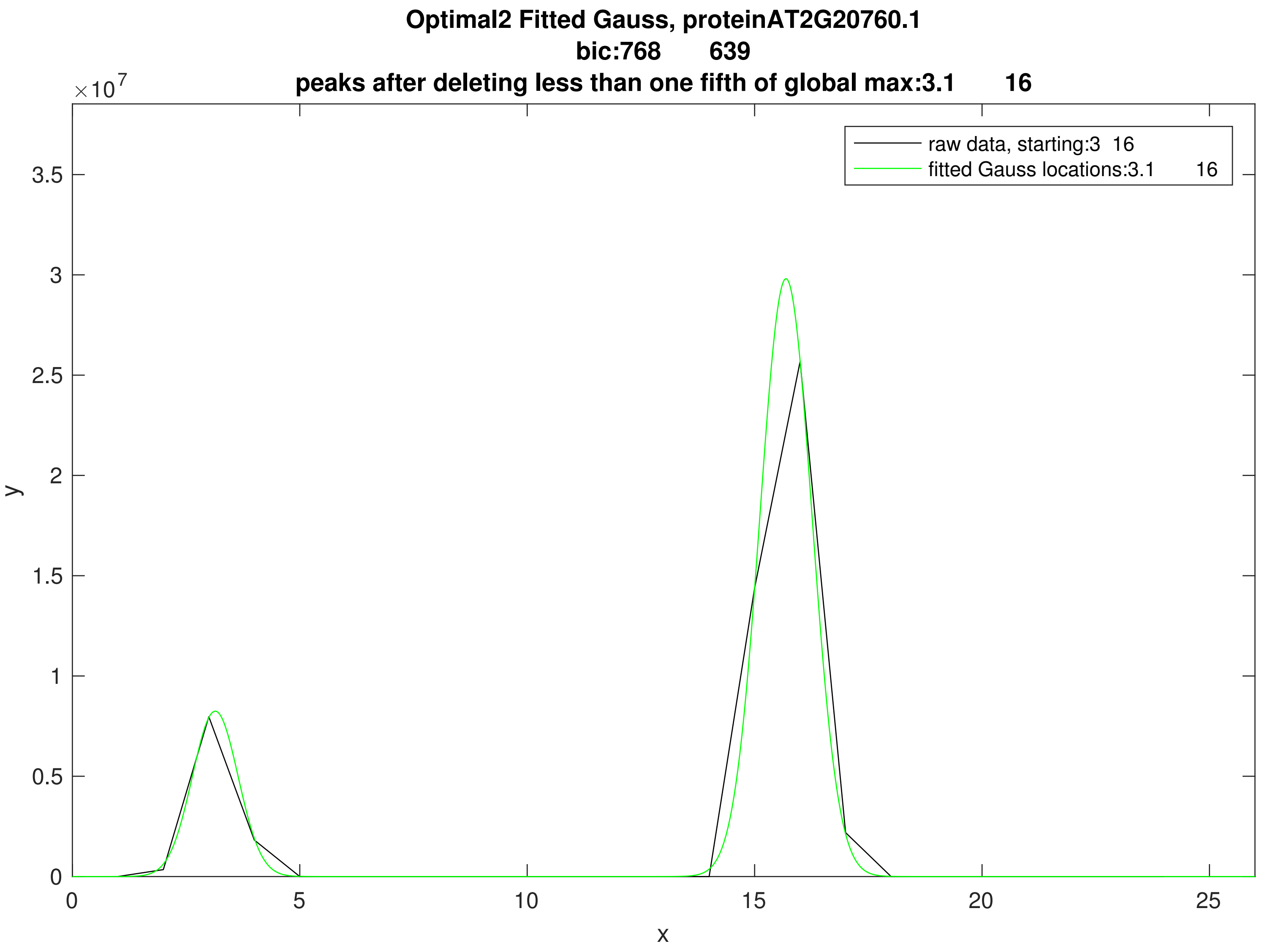
Optimal2 Fitted Gauss, proteinAT2G20760.1
bic:768 639
peaks after deleting less than one fifth of global max:3.1 16
×10^7
raw data, starting:3 16
fitted Gauss locations:3.1 16
y
x
0
0.5
1
1.5
2
2.5
3
3.5
0
5
10
15
20
25

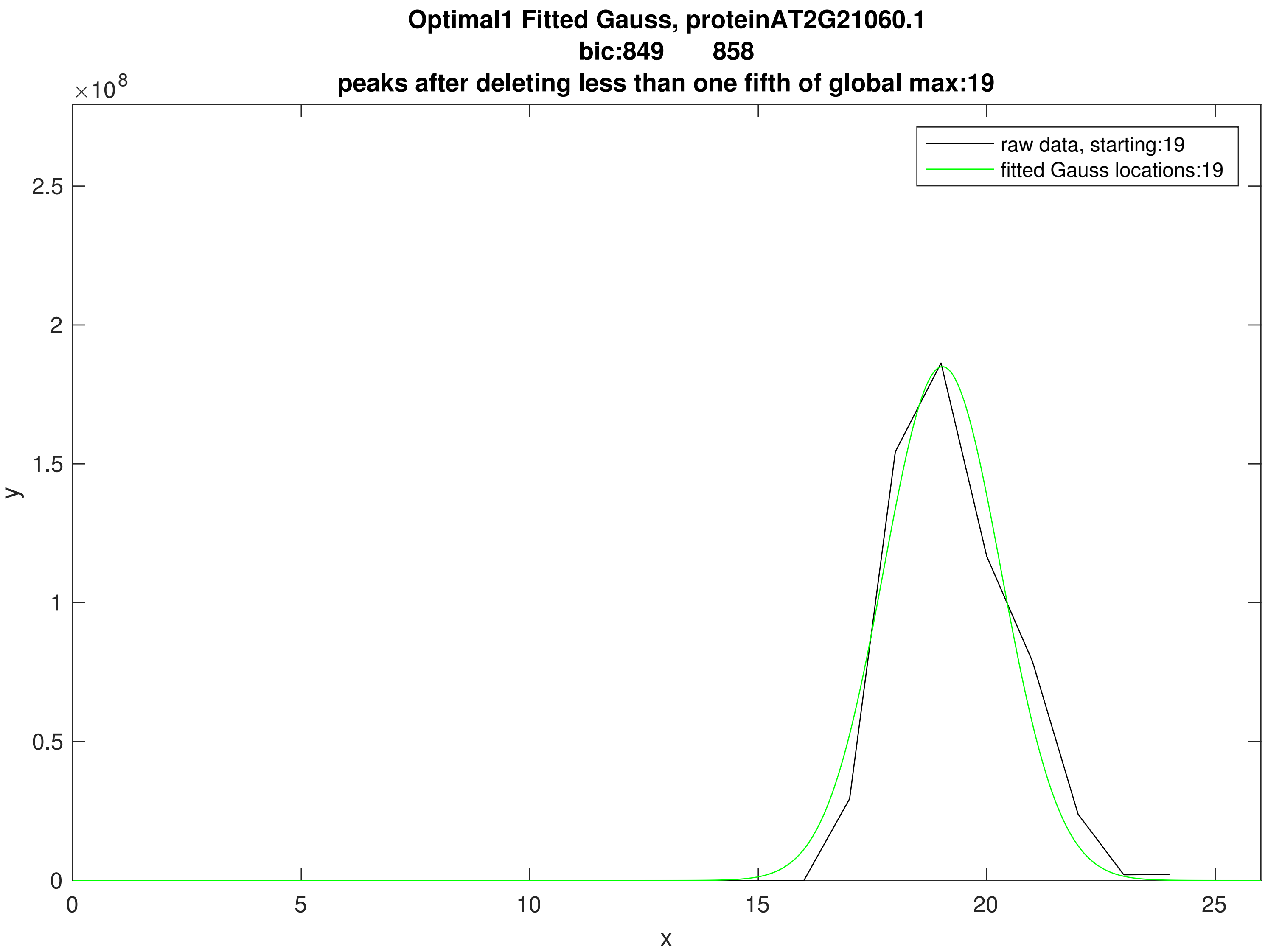
Optimal1 Fitted Gauss, proteinAT2G21060.1
bic:849 858
peaks after deleting less than one fifth of global max:19
raw data, starting:19
fitted Gauss locations:19
×10^8
y
x
0
0.5
1
1.5
2
2.5
5
10
15
20
25

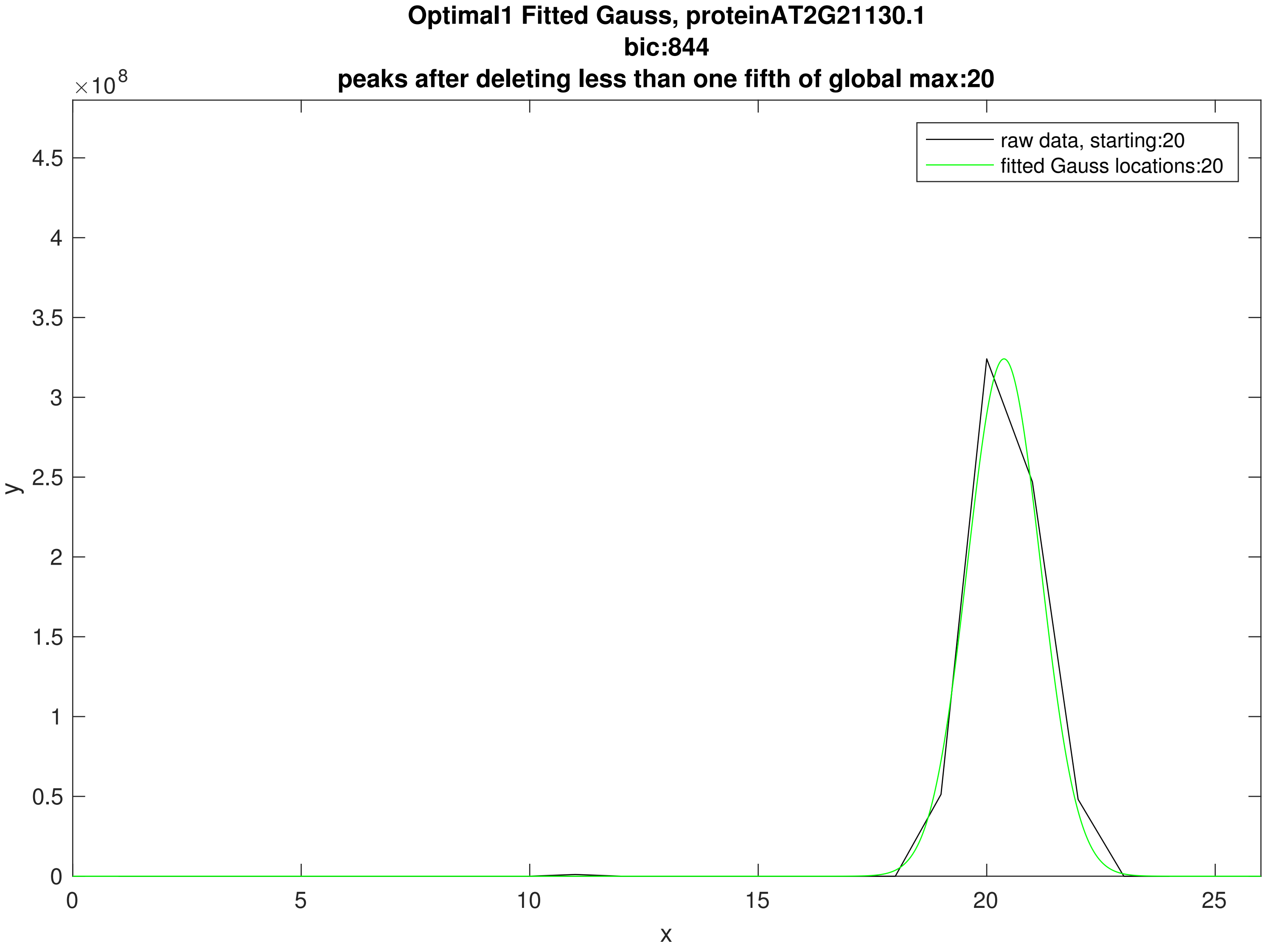

Optimal1 Fitted Gauss, proteinAT2G21130.1
bic:844
peaks after deleting less than one fifth of global max:20
×10^8
raw data, starting:20
fitted Gauss locations:20
4.5
4
3.5
3
2.5
2
1.5
1
0.5
0
y
0
5
10
15
20
25
x

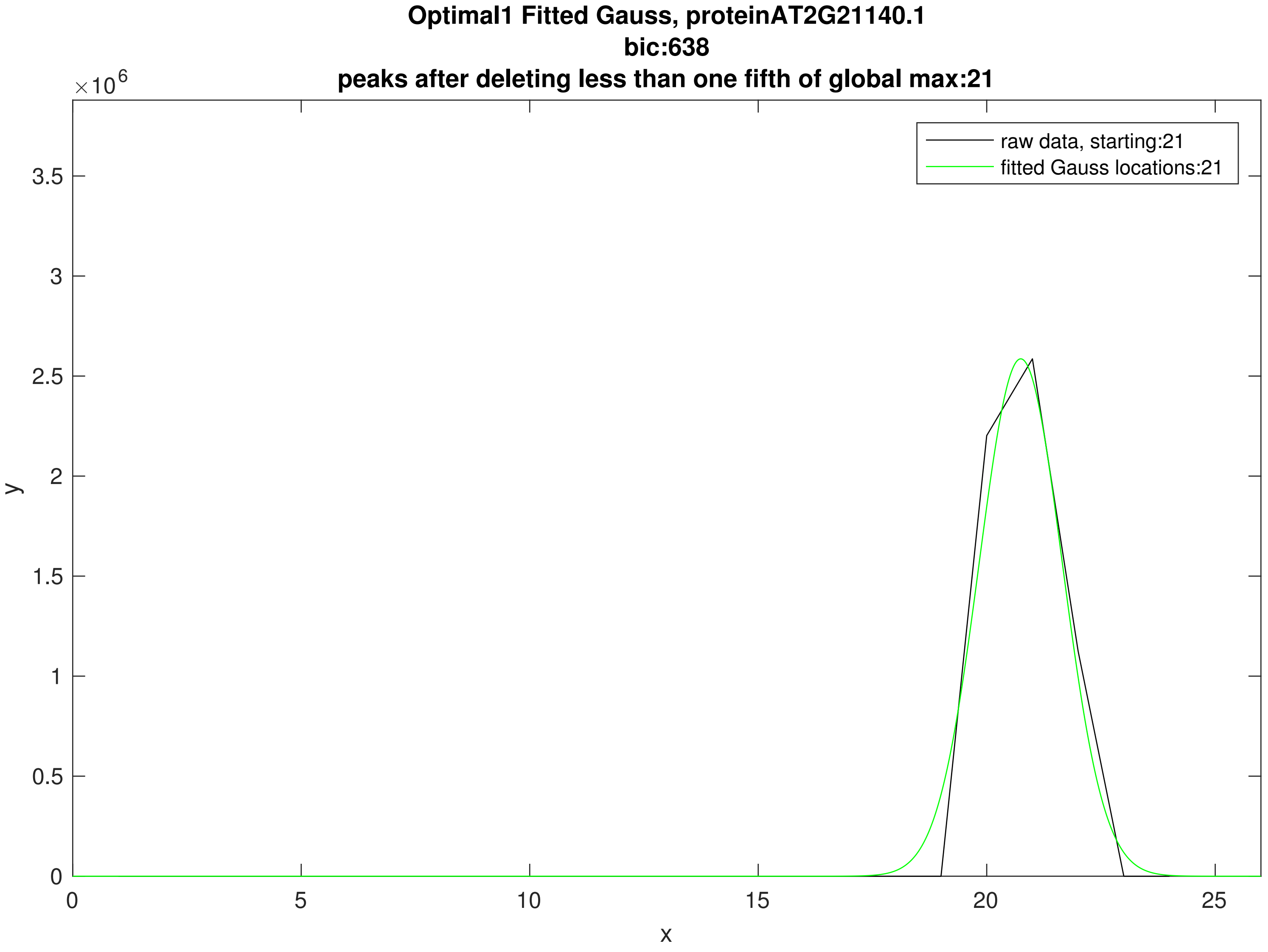

Optimal1 Fitted Gauss, proteinAT2G21140.1
bic:638
peaks after deleting less than one fifth of global max:21
×10^6
3.5
3
2.5
2
1.5
1
0.5
0
y
0
5
10
15
20
25
x
raw data, starting:21
fitted Gauss locations:21

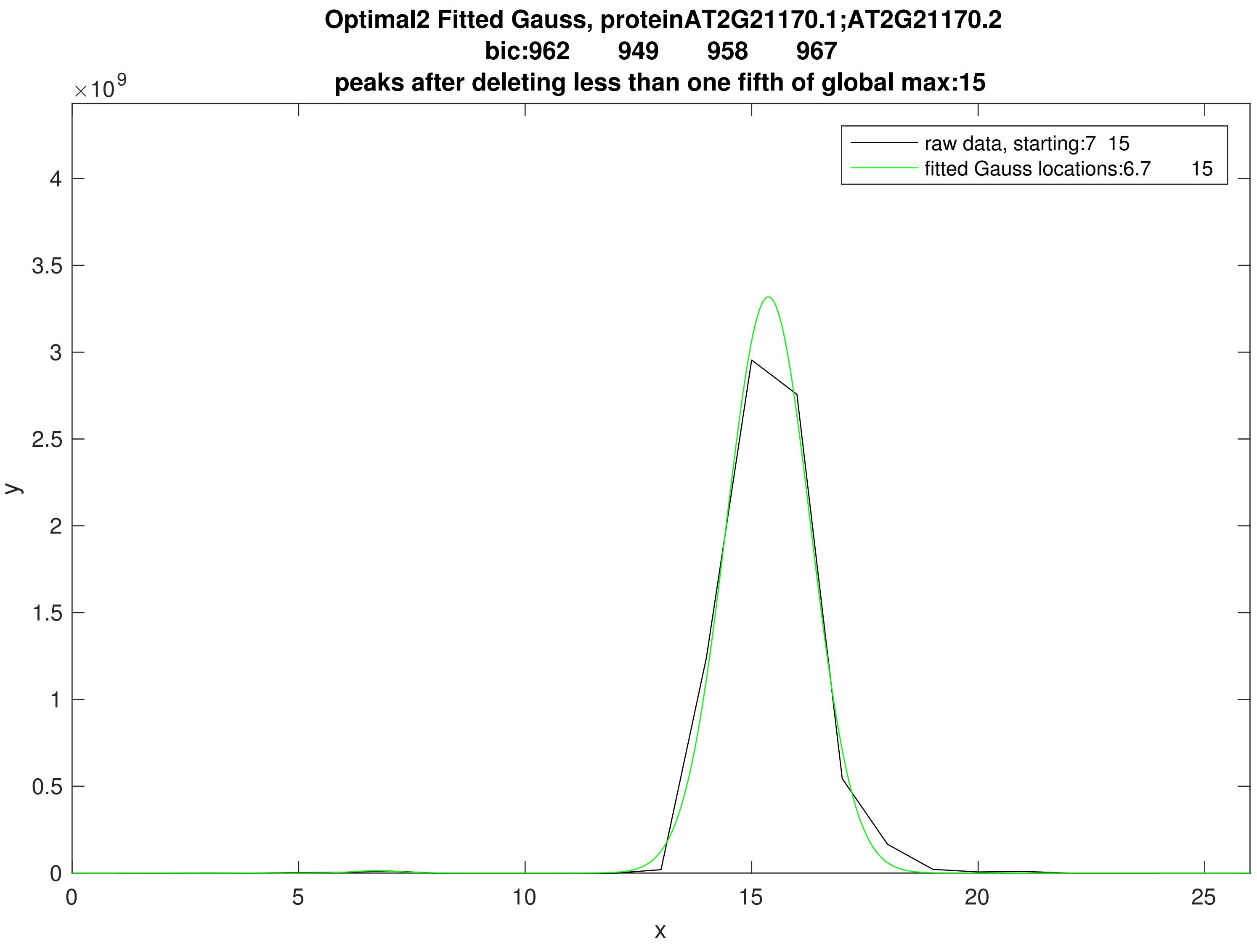

Optimal2 Fitted Gauss, proteinAT2G21170.1;AT2G21170.2
bic:962 949 958 967
peaks after deleting less than one fifth of global max:15
×10^9
raw data, starting:7 15
fitted Gauss locations:6.7 15
y
x
0
0.5
1
1.5
2
2.5
3
3.5
4
0
5
10
15
20
25

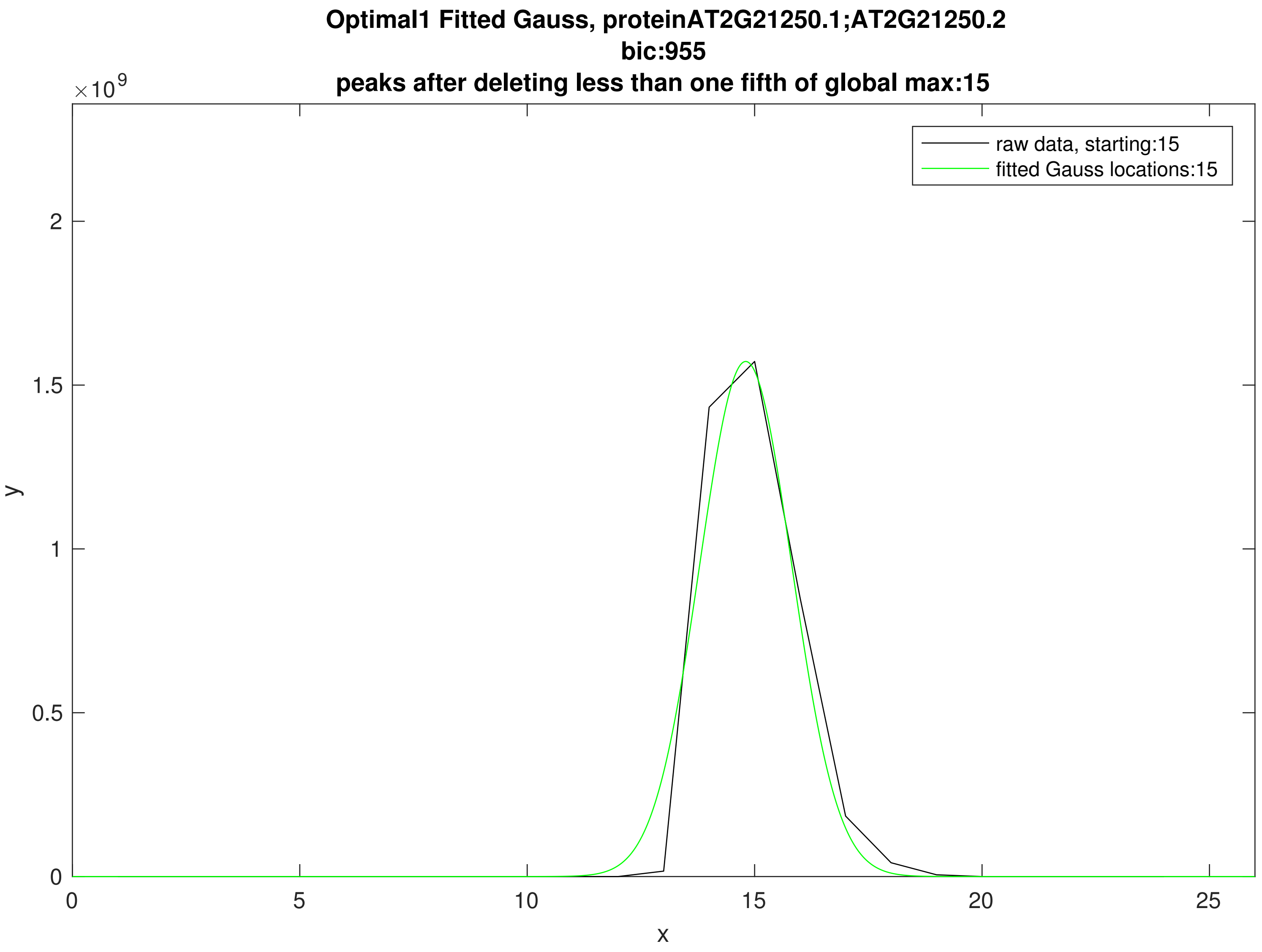
Optimal1 Fitted Gauss, proteinAT2G21250.1;AT2G21250.2
bic:955
peaks after deleting less than one fifth of global max:15
×10^9
raw data, starting:15
fitted Gauss locations:15
y
x
0
0.5
1
1.5
2
0
5
10
15
20
25

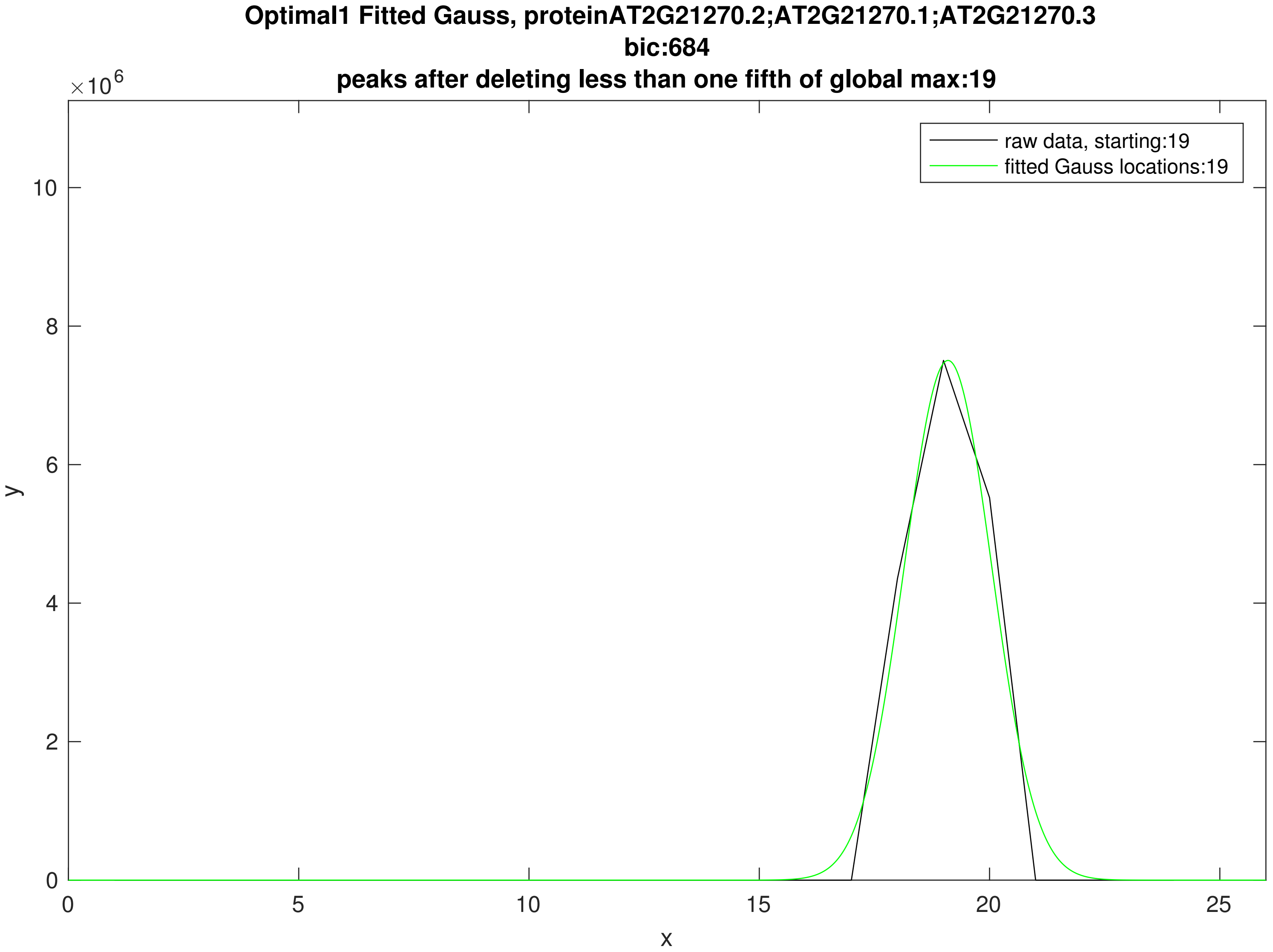

Optimal1 Fitted Gauss, proteinAT2G21270.2;AT2G21270.1;AT2G21270.3
bic:684
peaks after deleting less than one fifth of global max:19
×10^6
raw data, starting:19
fitted Gauss locations:19
10
8
6
4
2
0
y
0
5
10
15
20
25
x

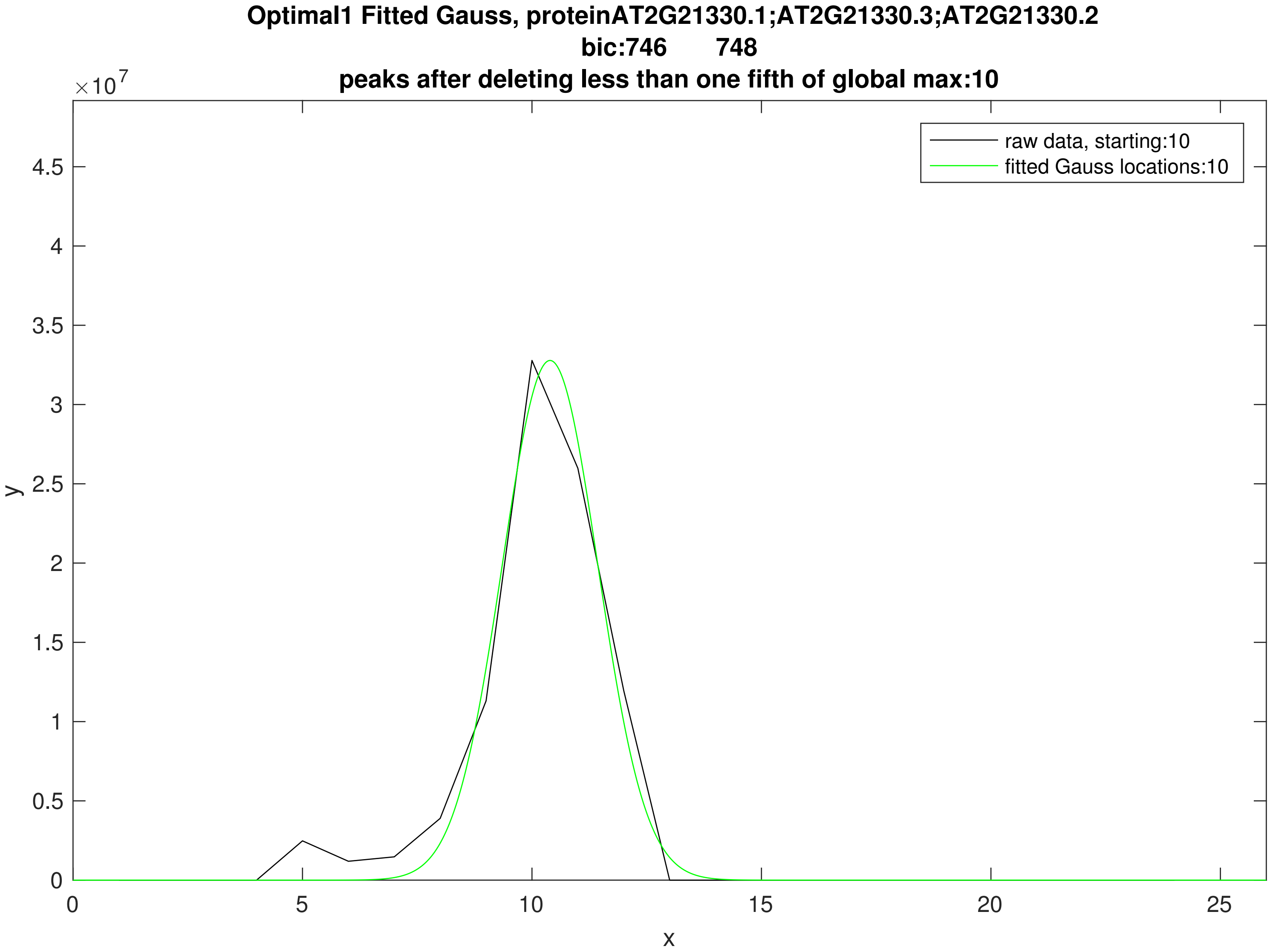

Optimal1 Fitted Gauss, proteinAT2G21330.1;AT2G21330.3;AT2G21330.2
bic:746 748
peaks after deleting less than one fifth of global max:10
×10^7
4.5
4
3.5
3
2.5
2
1.5
1
0.5
0
y
0
5
10
15
20
25
x
raw data, starting:10
fitted Gauss locations:10

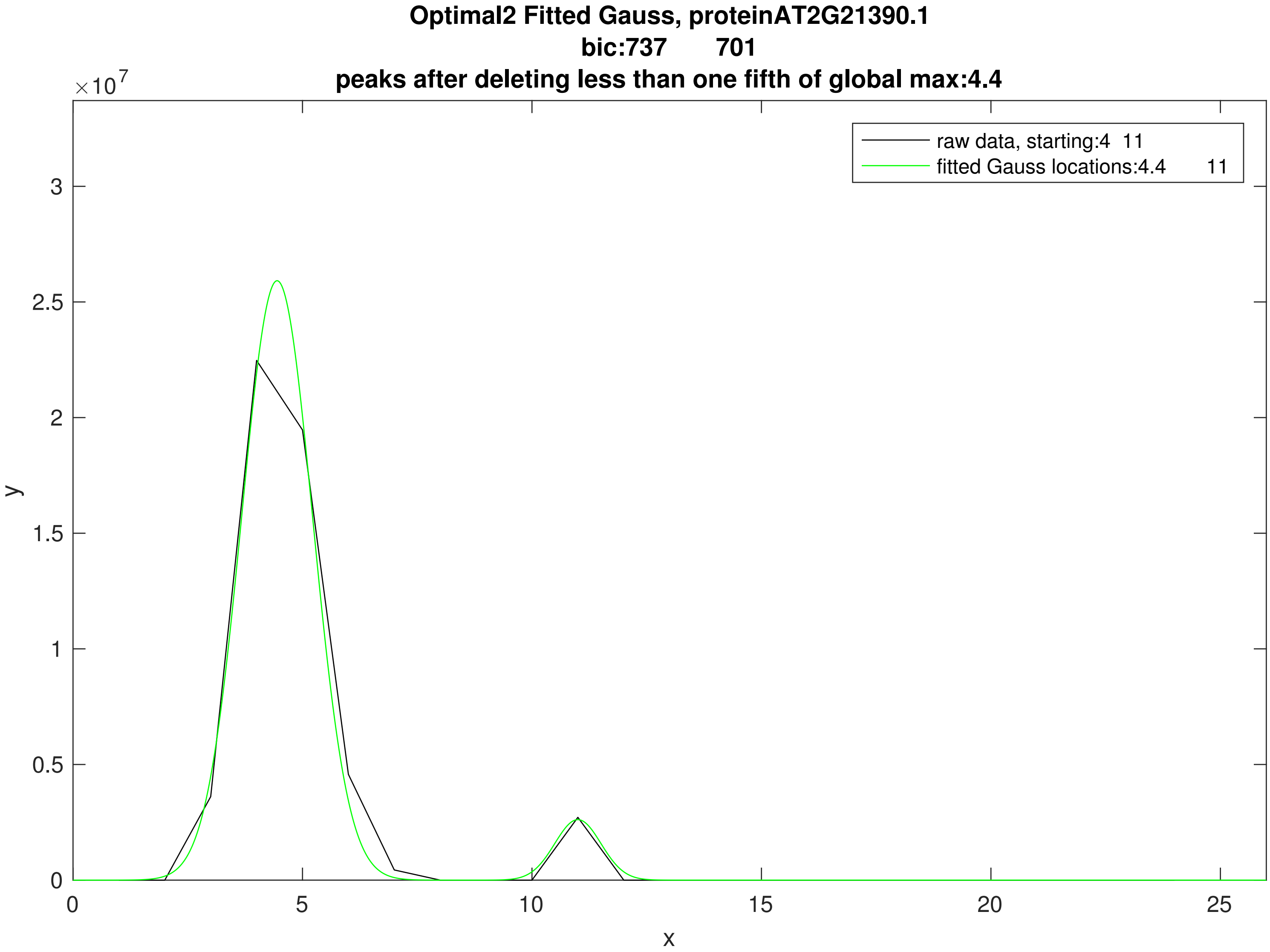
Optimal2 Fitted Gauss, proteinAT2G21390.1
bic:737 701
peaks after deleting less than one fifth of global max:4.4
×10^7
raw data, starting:4 11
fitted Gauss locations:4.4 11
y
x
0
0.5
1
1.5
2
2.5
3
0
5
10
15
20
25

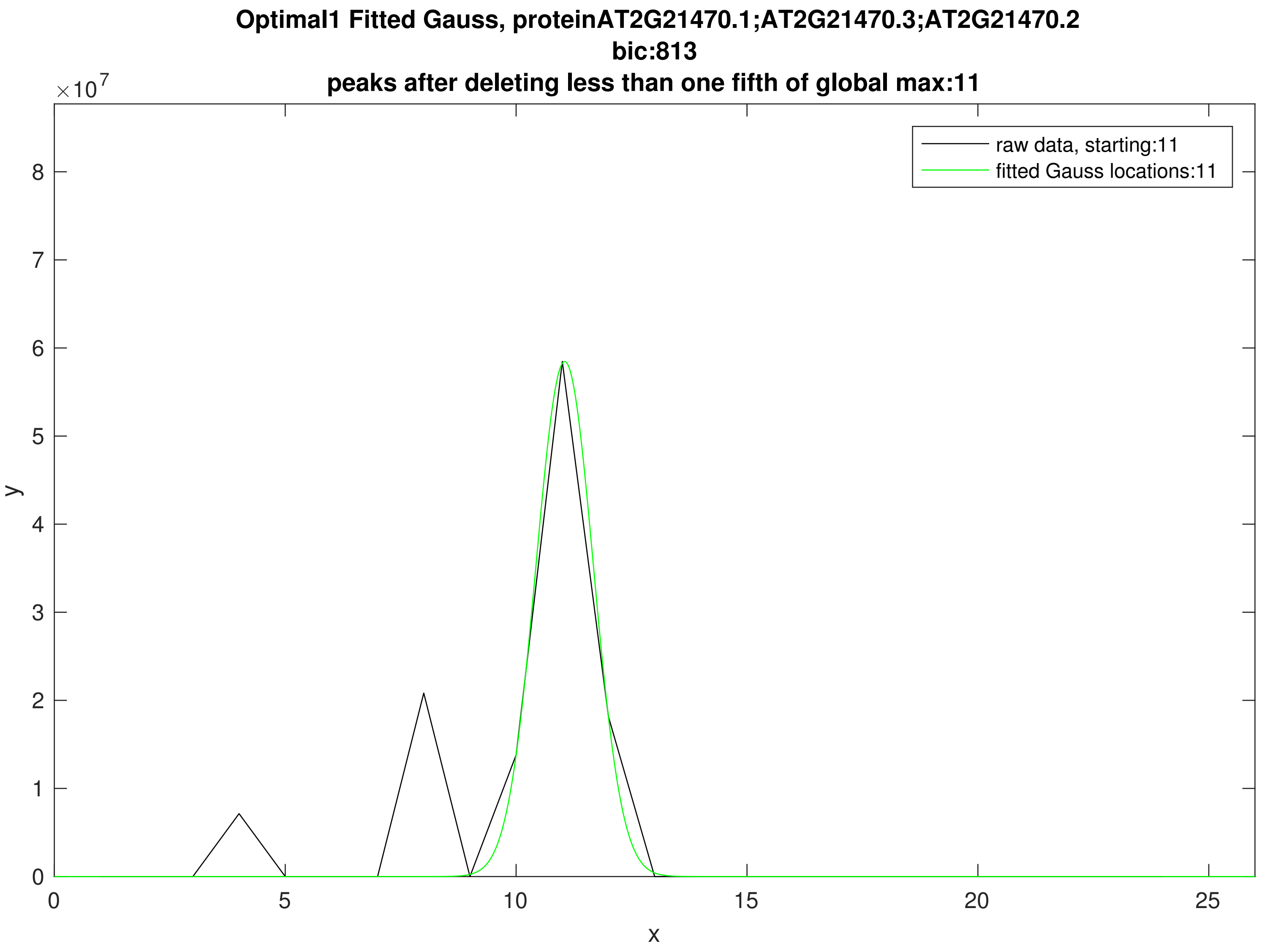
Optimal1 Fitted Gauss, proteinAT2G21470.1;AT2G21470.3;AT2G21470.2
bic:813
peaks after deleting less than one fifth of global max:11
×10^7
raw data, starting:11
fitted Gauss locations:11
y
x
0
1
2
3
4
5
6
7
8
0
5
10
15
20
25

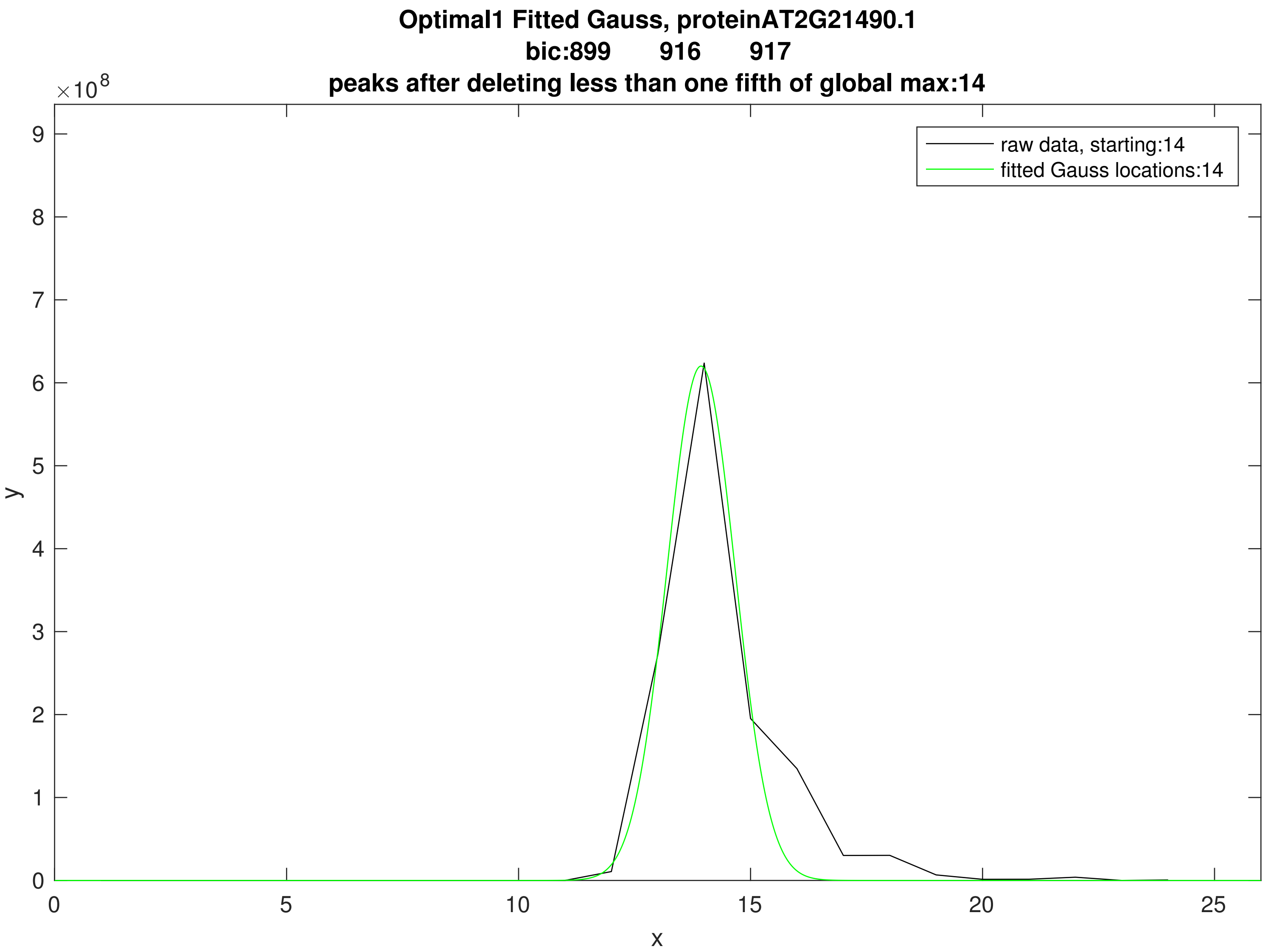
Optimal1 Fitted Gauss, proteinAT2G21490.1
bic:899 916 917
peaks after deleting less than one fifth of global max:14
×10^8
raw data, starting:14
fitted Gauss locations:14
y
x
0
1
2
3
4
5
6
7
8
9
0
5
10
15
20
25

**Optimal1 Fitted Gauss, proteinAT2G21580.2;AT2G21580.1**
**bic:746**
**peaks after deleting less than one fifth of global max:3.8**

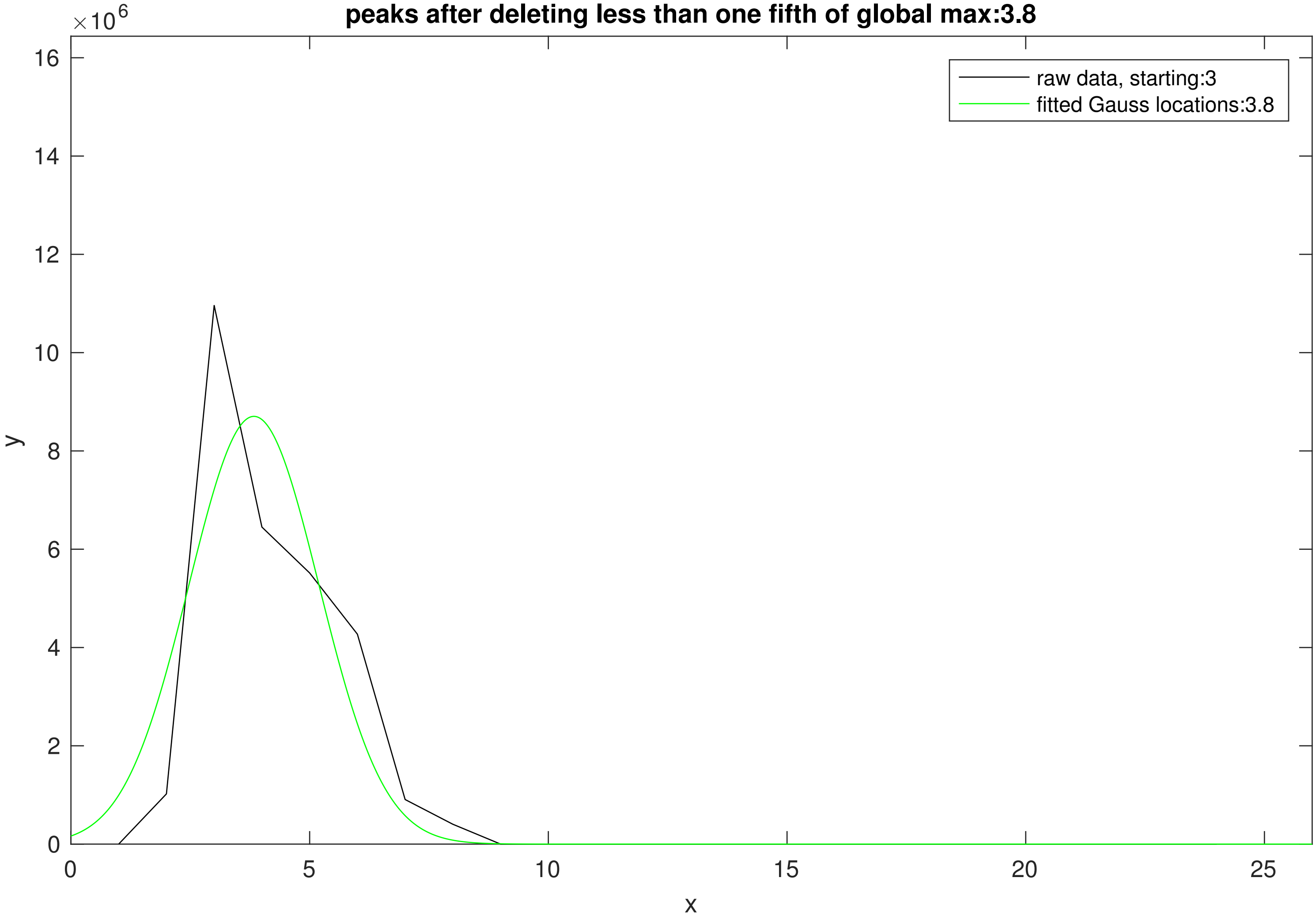

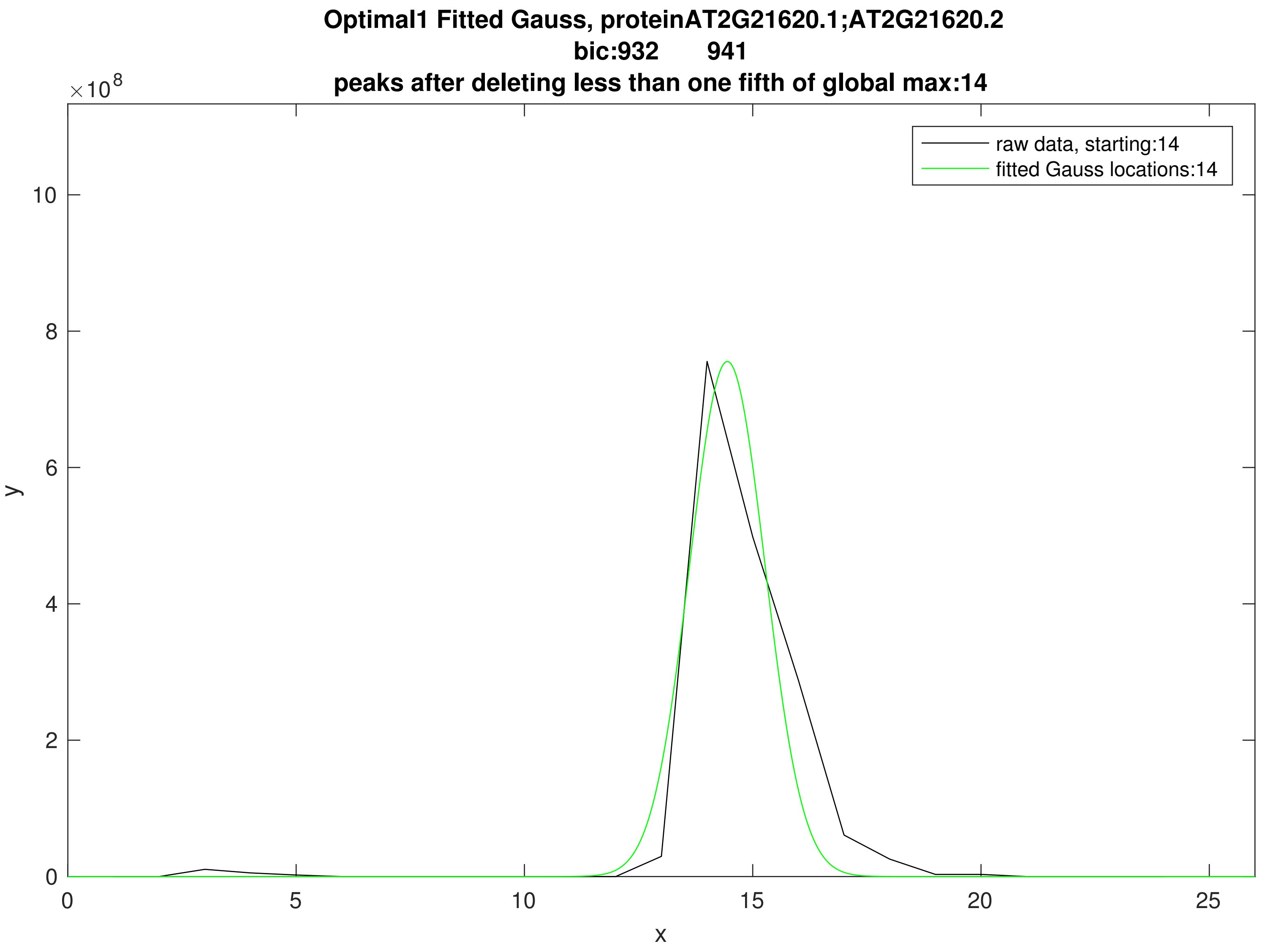
Optimal1 Fitted Gauss, proteinAT2G21620.1;AT2G21620.2
bic:932 941
peaks after deleting less than one fifth of global max:14
×10^8
raw data, starting:14
fitted Gauss locations:14
y
x
0
2
4
6
8
10
0
5
10
15
20
25

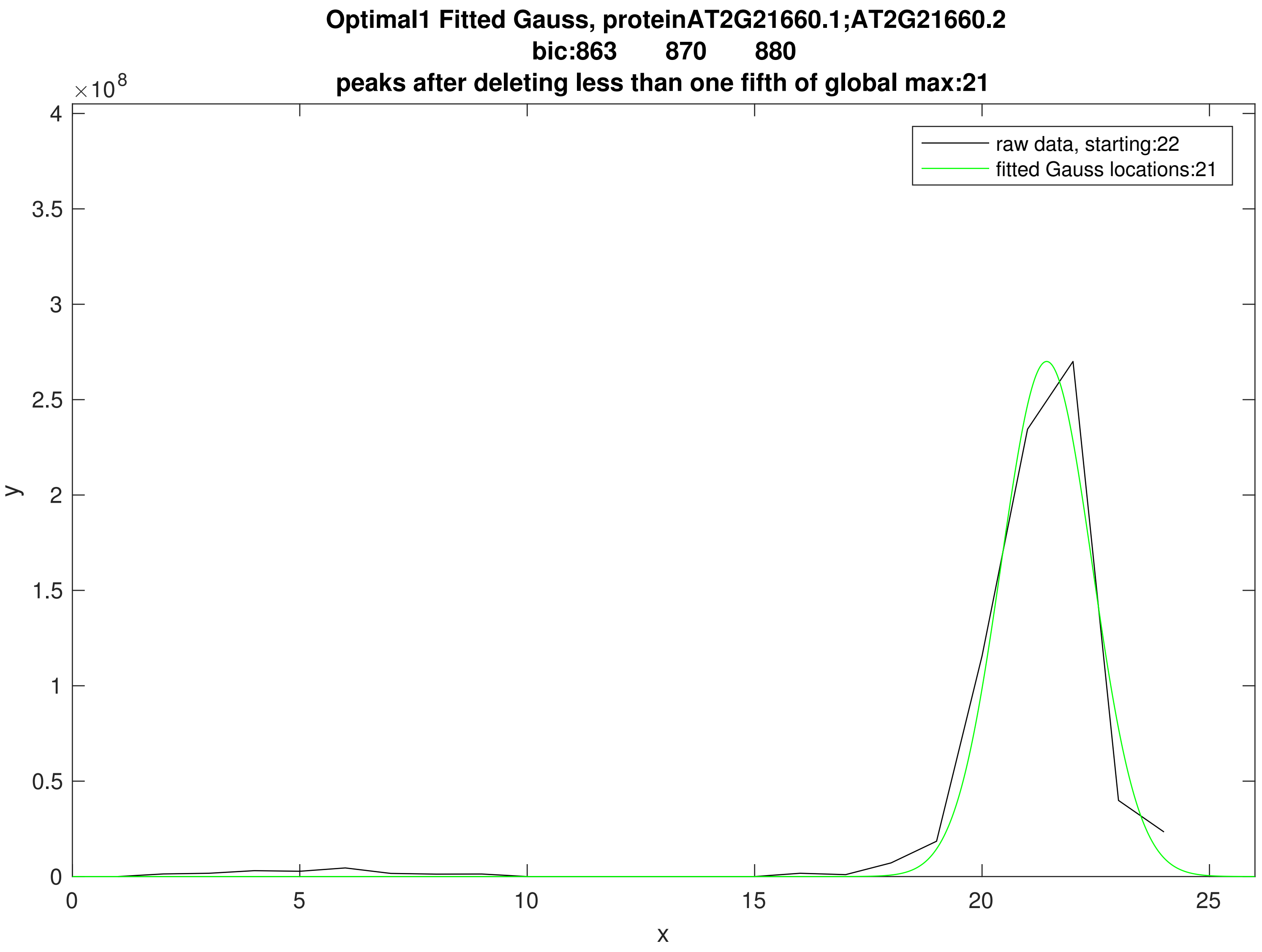
Optimal1 Fitted Gauss, proteinAT2G21660.1;AT2G21660.2
bic:863 870 880
peaks after deleting less than one fifth of global max:21
×10^8
4
3.5
3
2.5
2
1.5
1
0.5
0
y
0
5
10
15
20
25
x
raw data, starting:22
fitted Gauss locations:21

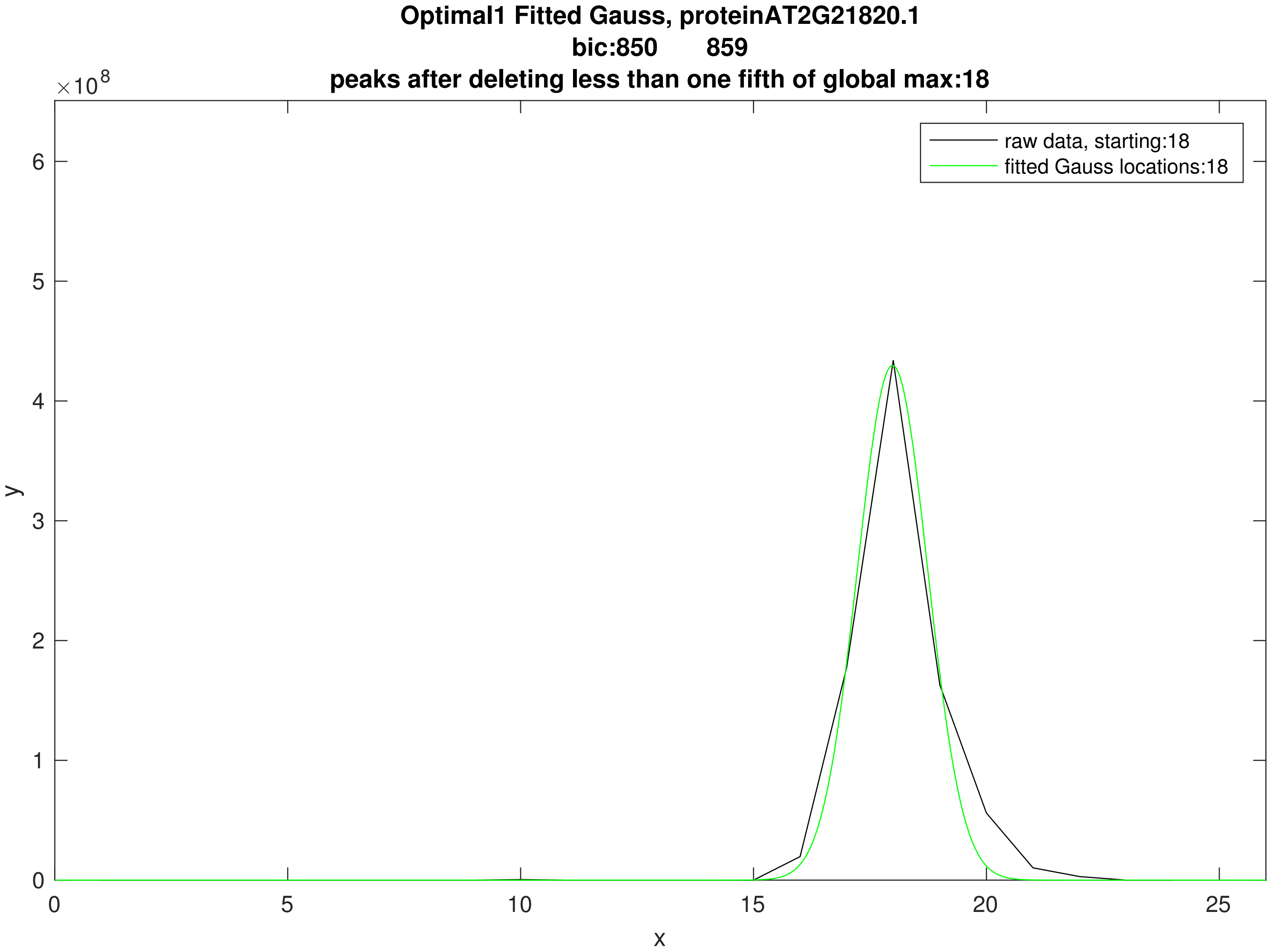

Optimal1 Fitted Gauss, proteinAT2G21820.1
bic:850 859
peaks after deleting less than one fifth of global max:18
×10^8
raw data, starting:18
fitted Gauss locations:18
y
x
0
1
2
3
4
5
6
0
5
10
15
20
25

Optimal1 Fitted Gauss, proteinAT2G21870.1;AT2G21870.2
bic:844 853
peaks after deleting less than one fifth of global max:16

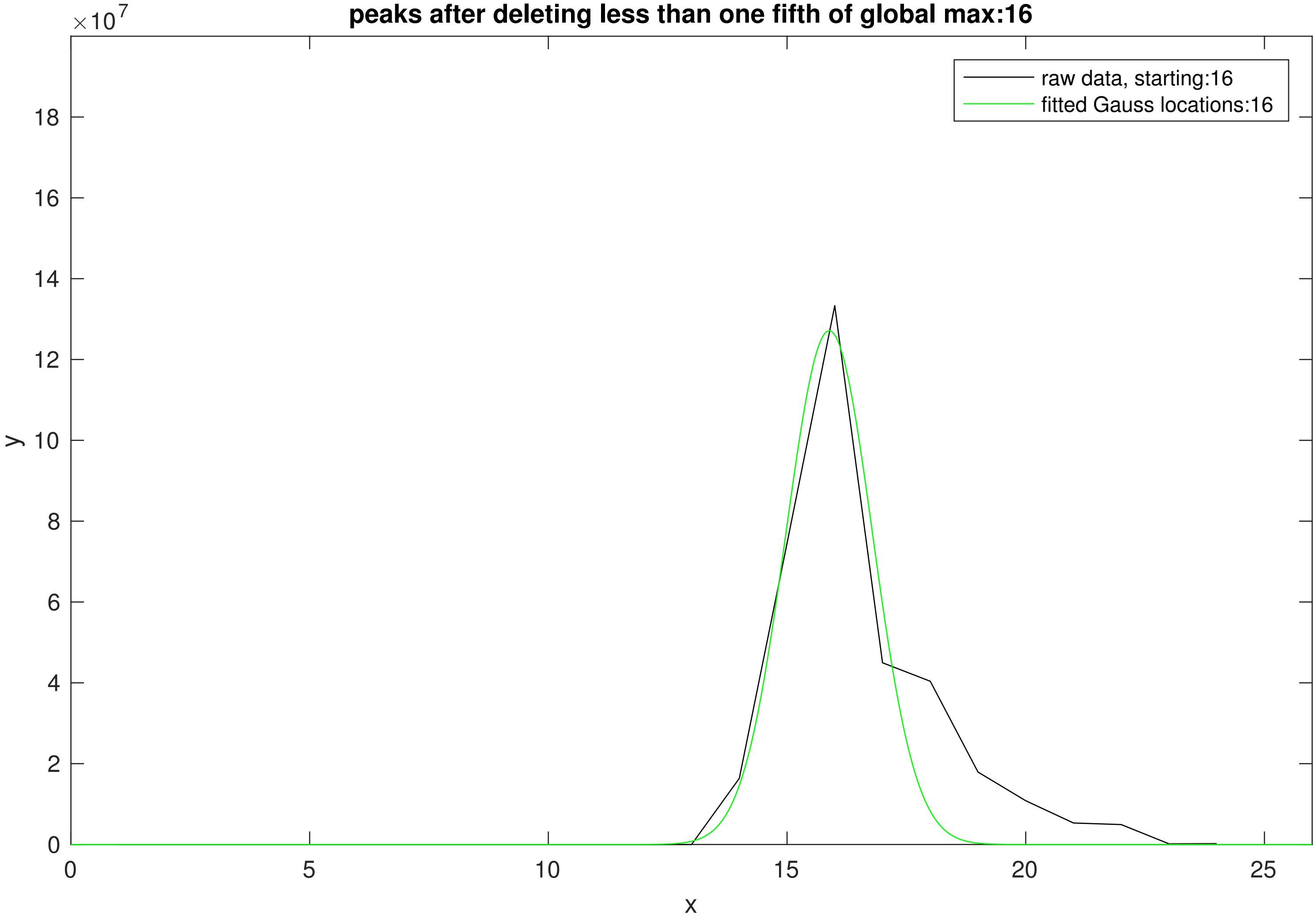

**Optimal1 Fitted Gauss, proteinAT2G22170.1**
**bic:939**
**peaks after deleting less than one fifth of global max:18**

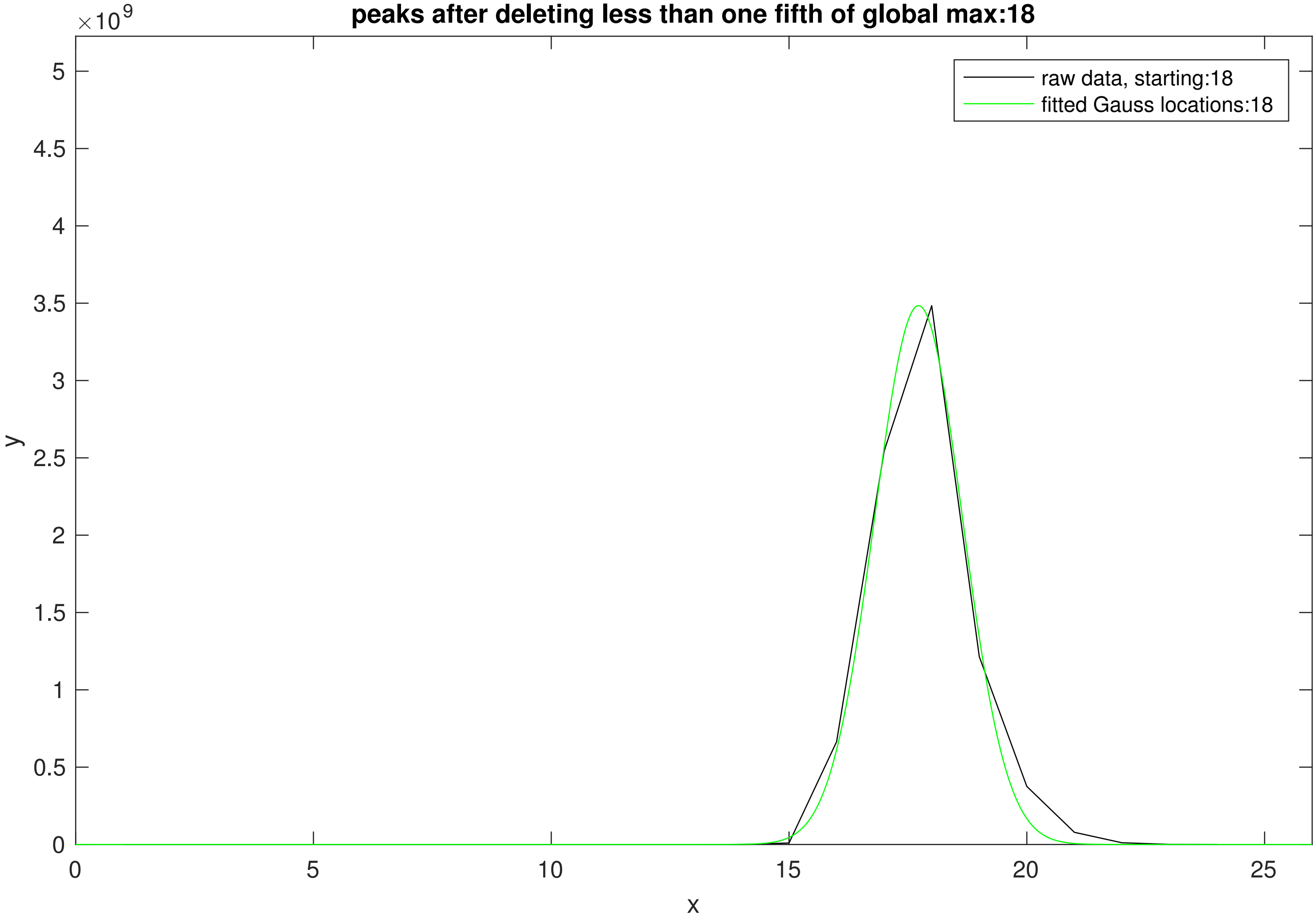

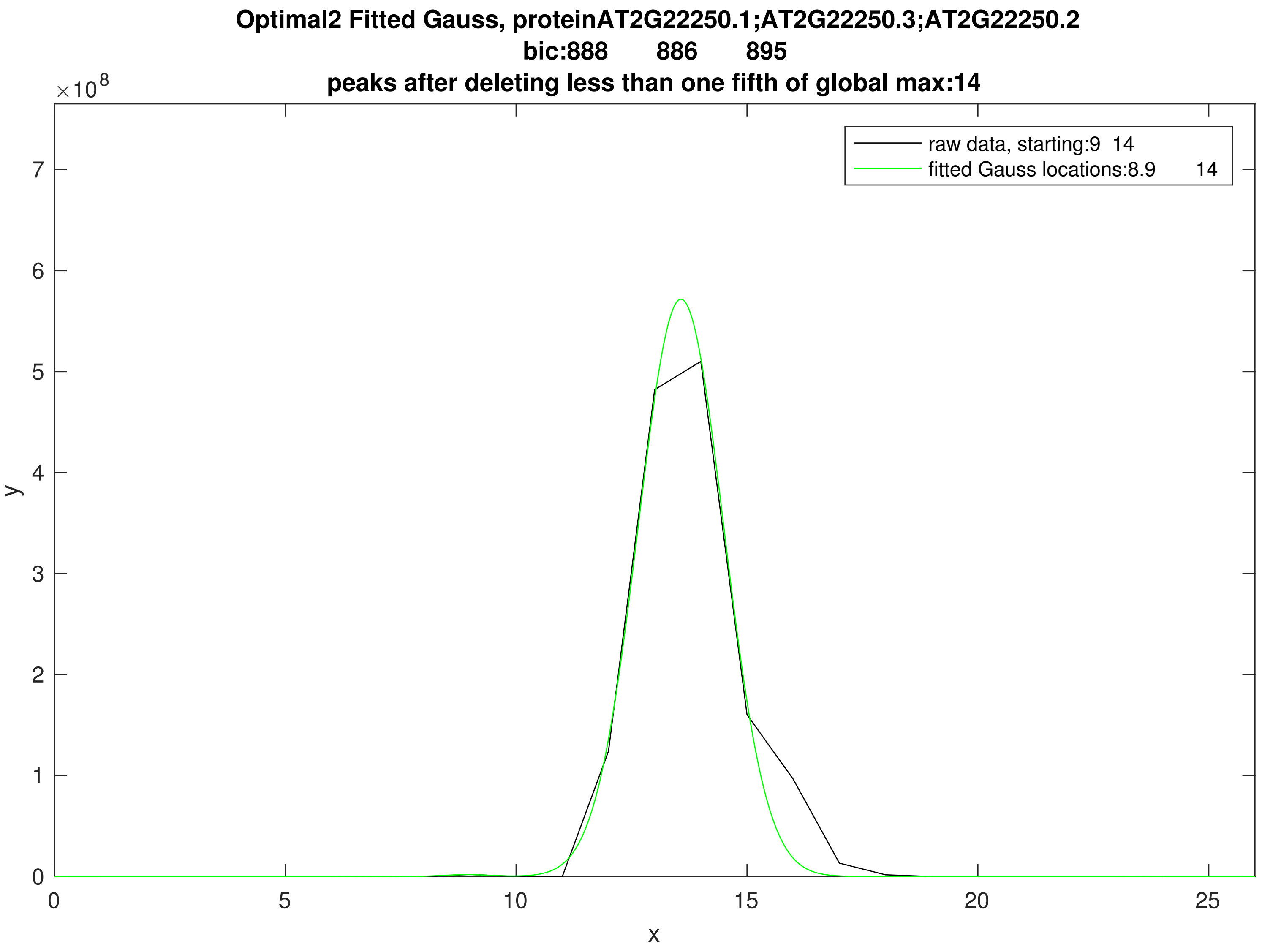
Optimal2 Fitted Gauss, proteinAT2G22250.1;AT2G22250.3;AT2G22250.2
bic:888 886 895
peaks after deleting less than one fifth of global max:14
×10^8
raw data, starting:9 14
fitted Gauss locations:8.9 14
y
x
0
1
2
3
4
5
6
7
0
5
10
15
20
25

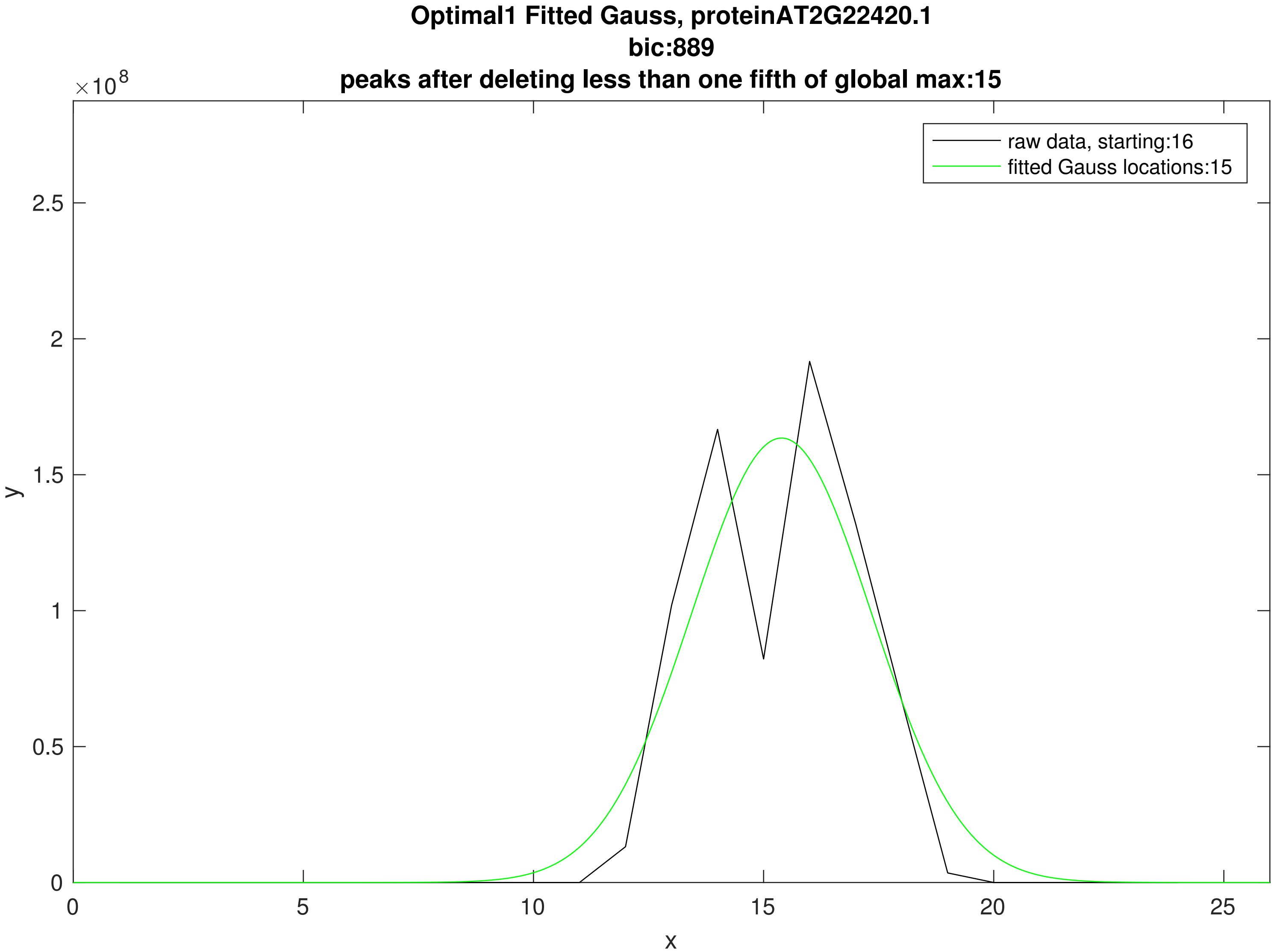
Optimal1 Fitted Gauss, proteinAT2G22420.1
bic:889
peaks after deleting less than one fifth of global max:15
×10^8
raw data, starting:16
fitted Gauss locations:15
y
x
0
0.5
1
1.5
2
2.5
0
5
10
15
20
25

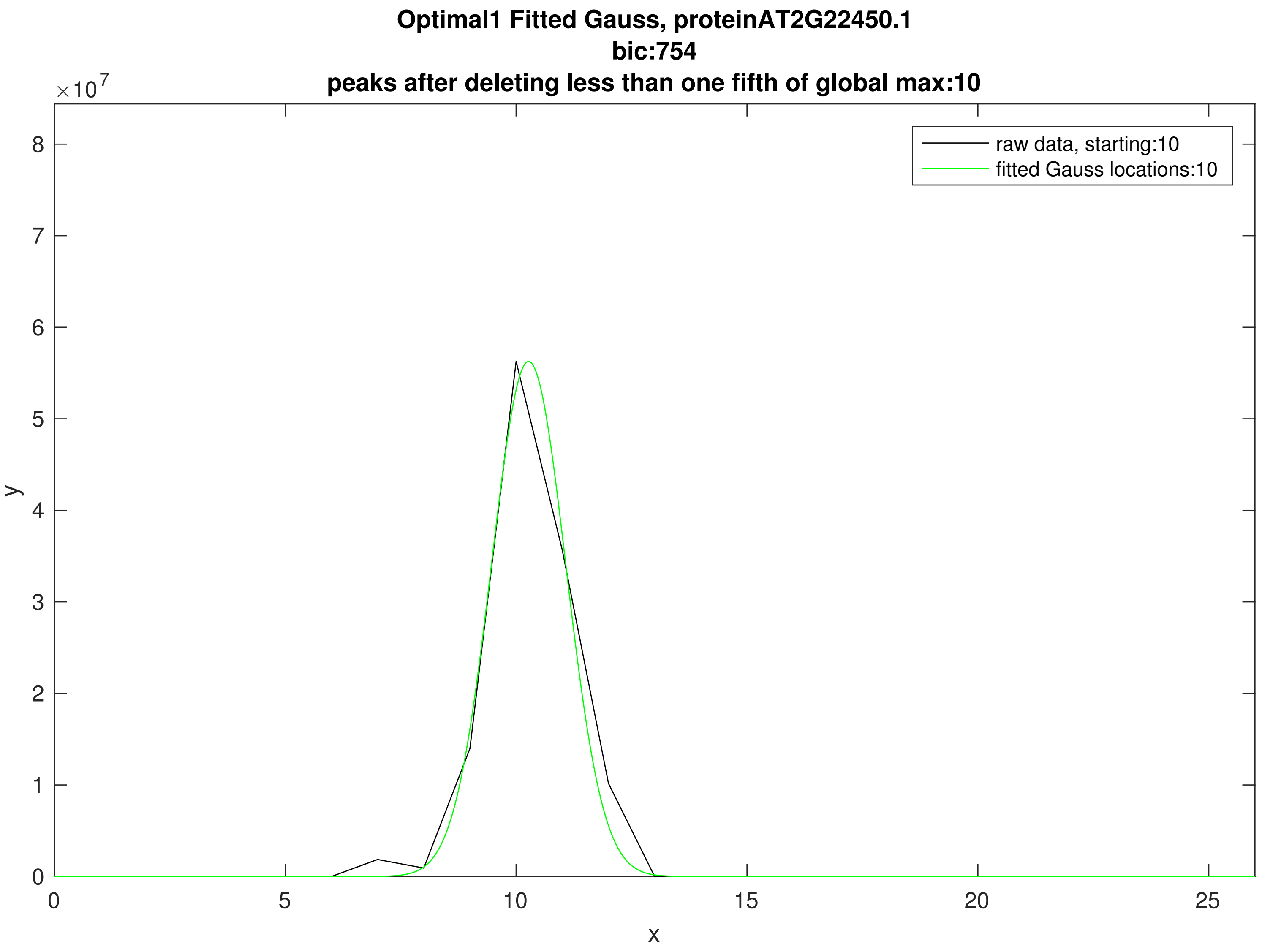

Optimal1 Fitted Gauss, proteinAT2G22450.1
bic:754
peaks after deleting less than one fifth of global max:10
×10^7
raw data, starting:10
fitted Gauss locations:10
y
x
0
1
2
3
4
5
6
7
8
0
5
10
15
20
25

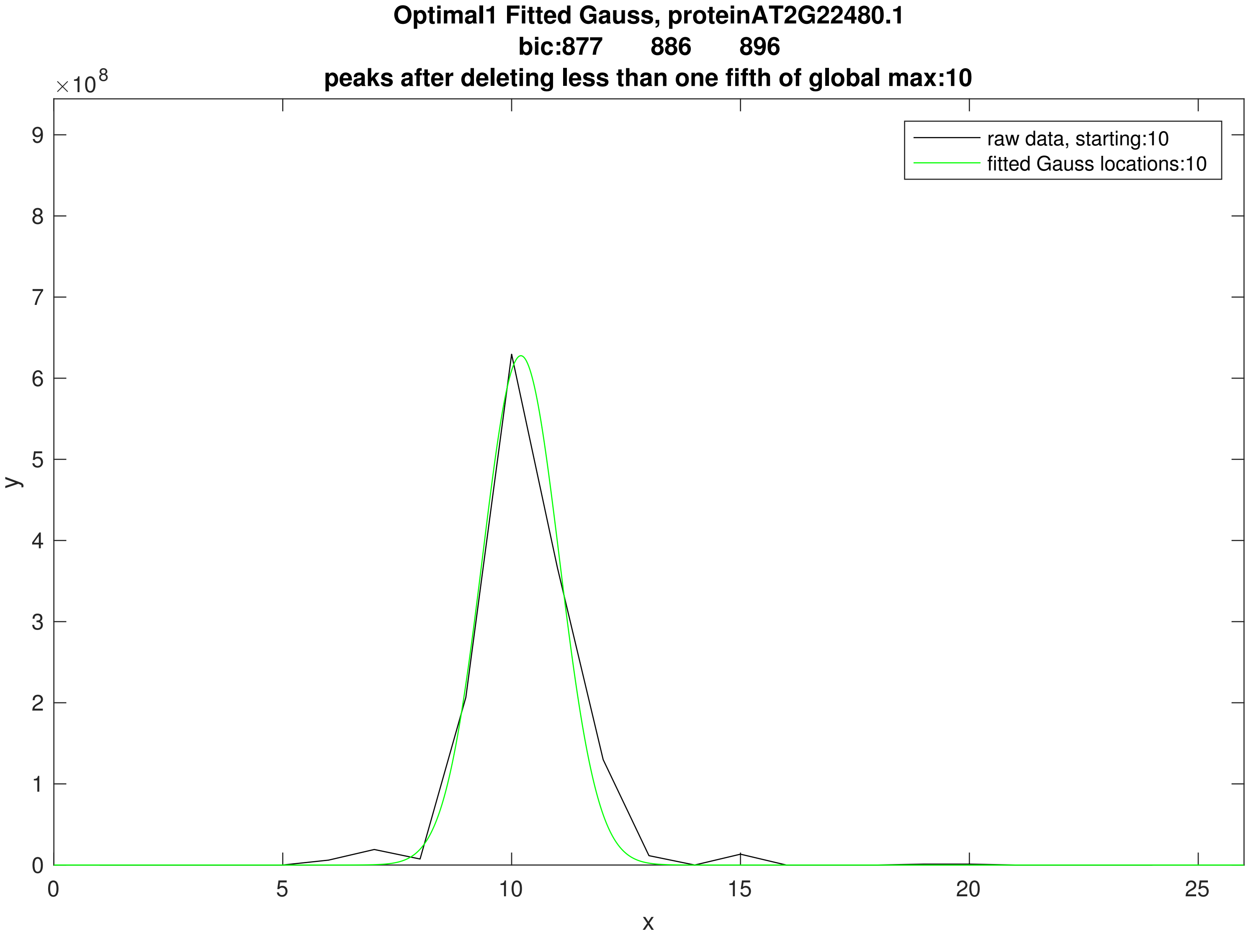

Optimal1 Fitted Gauss, proteinAT2G22480.1
bic:877 886 896
peaks after deleting less than one fifth of global max:10
×10^8
raw data, starting:10
fitted Gauss locations:10
y
x
0
1
2
3
4
5
6
7
8
9
0
5
10
15
20
25

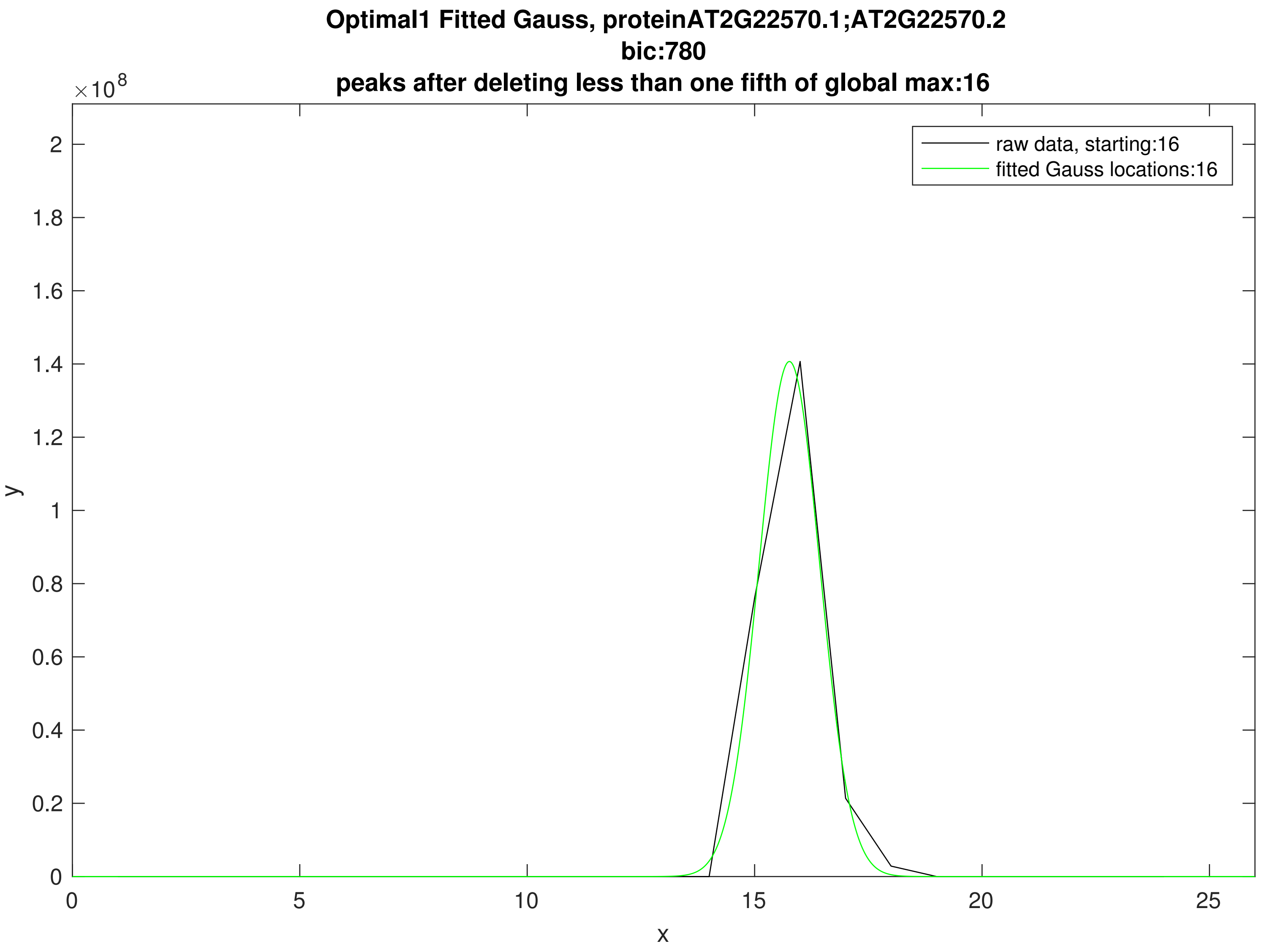
Optimal1 Fitted Gauss, proteinAT2G22570.1;AT2G22570.2
bic:780
peaks after deleting less than one fifth of global max:16
×10^8
raw data, starting:16
fitted Gauss locations:16
y
x
0
0.2
0.4
0.6
0.8
1
1.2
1.4
1.6
1.8
2
0
5
10
15
20
25

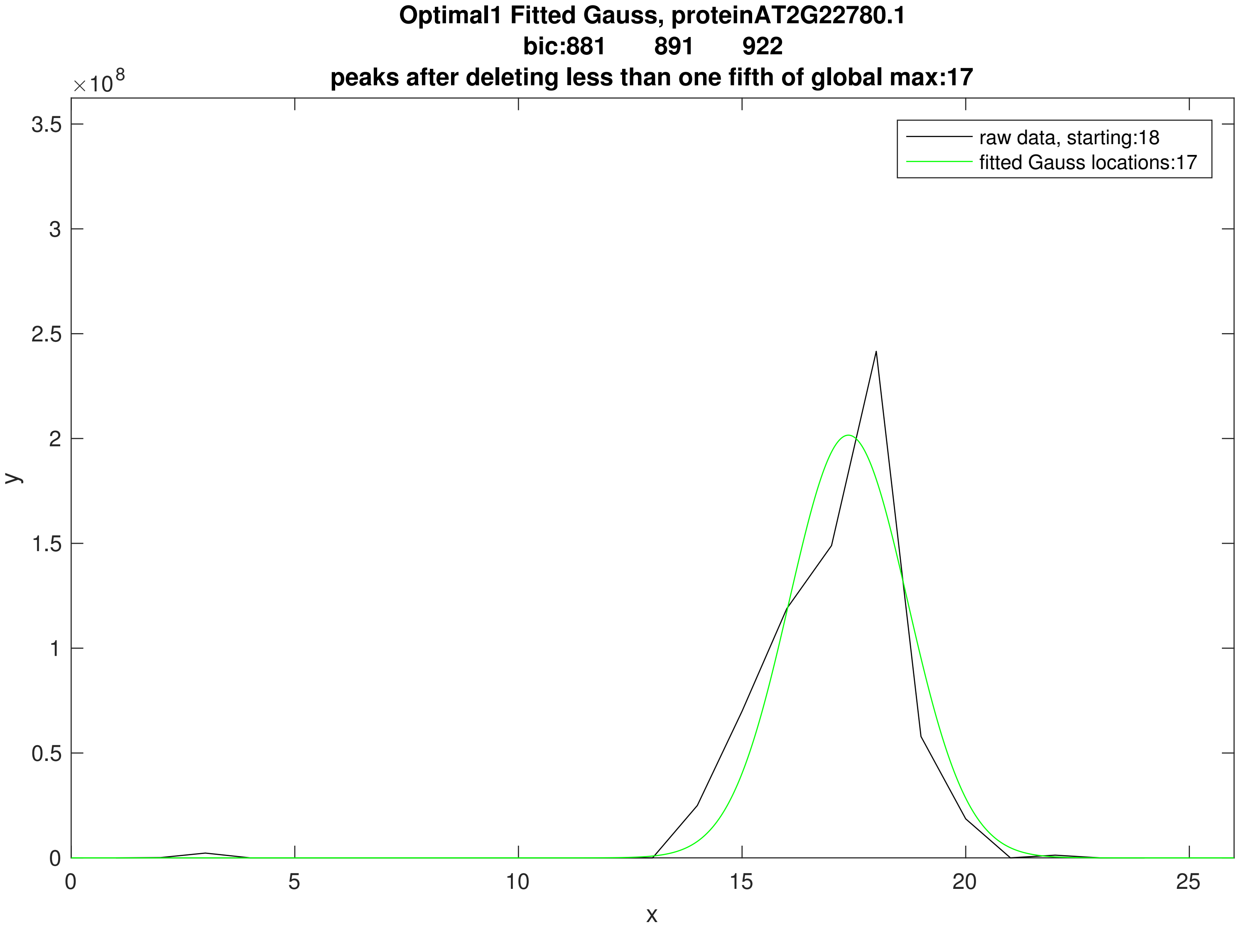
Optimal1 Fitted Gauss, proteinAT2G22780.1
bic:881 891 922
peaks after deleting less than one fifth of global max:17
×10^8
3.5
3
2.5
2
1.5
1
0.5
0
y
0
5
10
15
20
25
x
raw data, starting:18
fitted Gauss locations:17

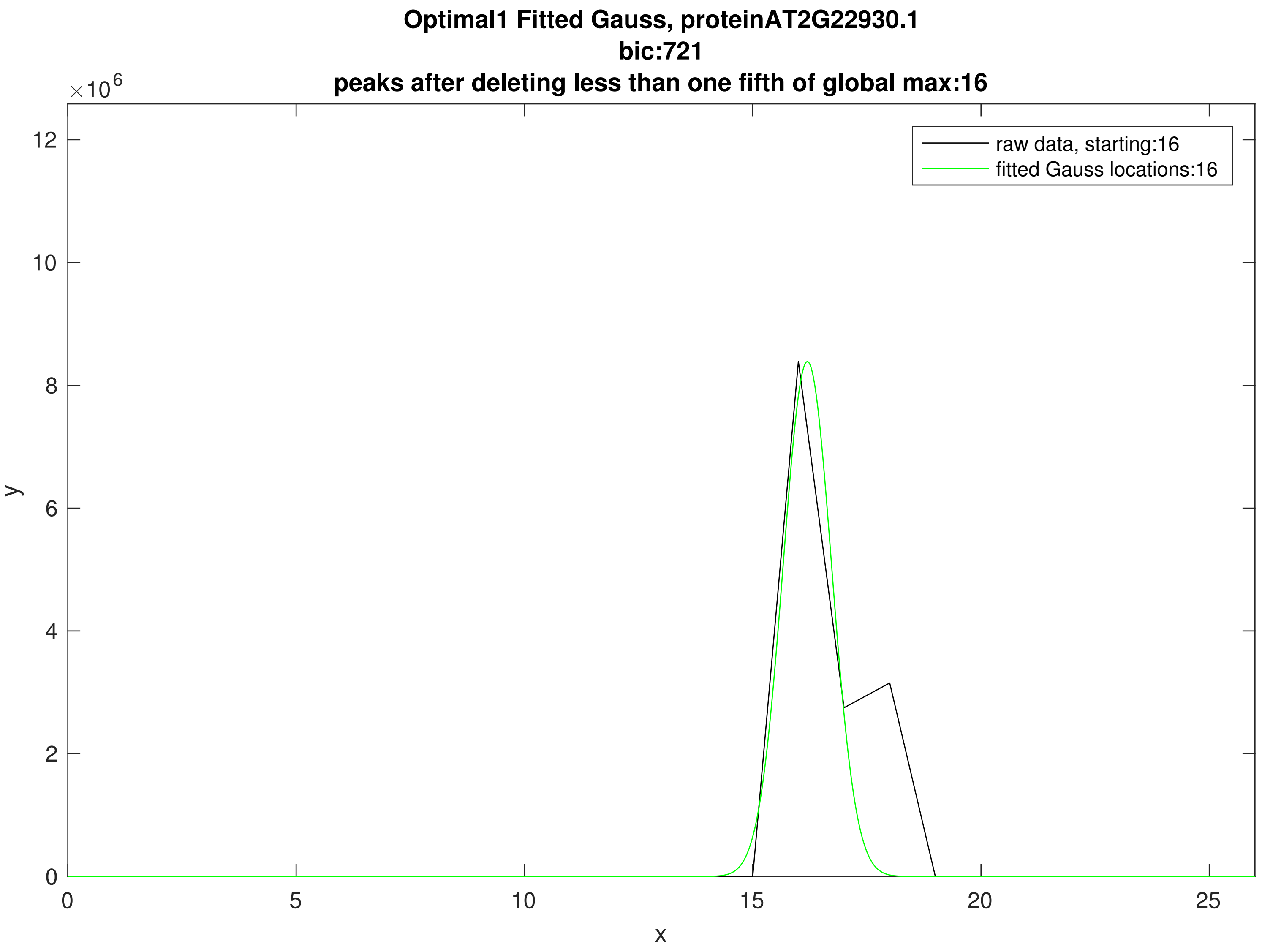
Optimal1 Fitted Gauss, proteinAT2G22930.1
bic:721
peaks after deleting less than one fifth of global max:16
$\times 10^6$
raw data, starting:16
fitted Gauss locations:16
y
x
0
2
4
6
8
10
12
0
5
10
15
20
25

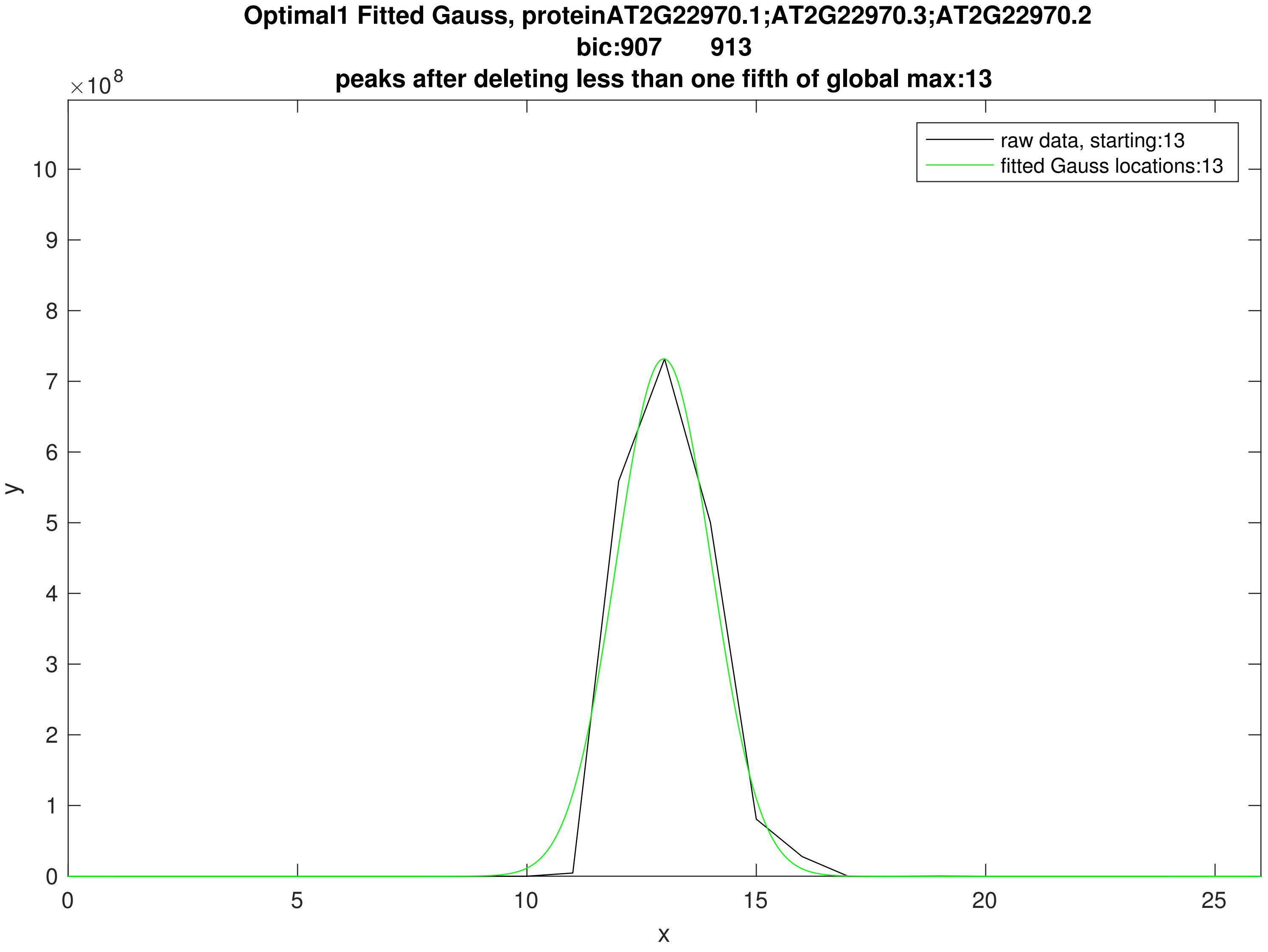

Optimal1 Fitted Gauss, proteinAT2G22970.1;AT2G22970.3;AT2G22970.2
bic:907 913
peaks after deleting less than one fifth of global max:13
×10^8
raw data, starting:13
fitted Gauss locations:13
y
x
0
1
2
3
4
5
6
7
8
9
10
0
5
10
15
20
25

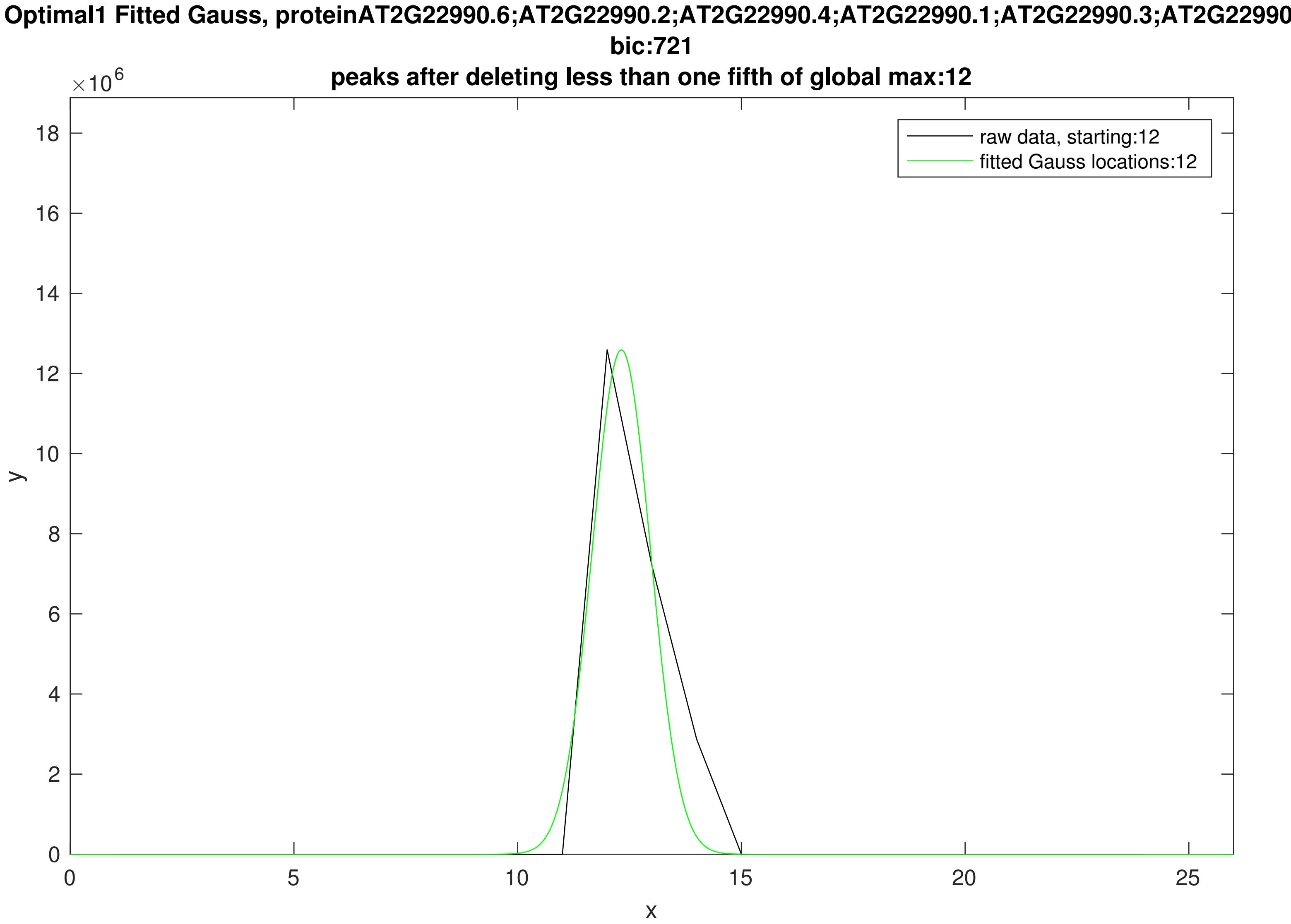
Optimal1 Fitted Gauss, proteinAT2G22990.6;AT2G22990.2;AT2G22990.4;AT2G22990.1;AT2G22990.3;AT2G22990
bic:721
peaks after deleting less than one fifth of global max:12
×10^6
raw data, starting:12
fitted Gauss locations:12
y
x
0
2
4
6
8
10
12
14
16
18
0
5
10
15
20
25

Optimal1 Fitted Gauss, proteinAT2G23010.1;AT2G23010.2
bic:720
peaks after deleting less than one fifth of global max:13

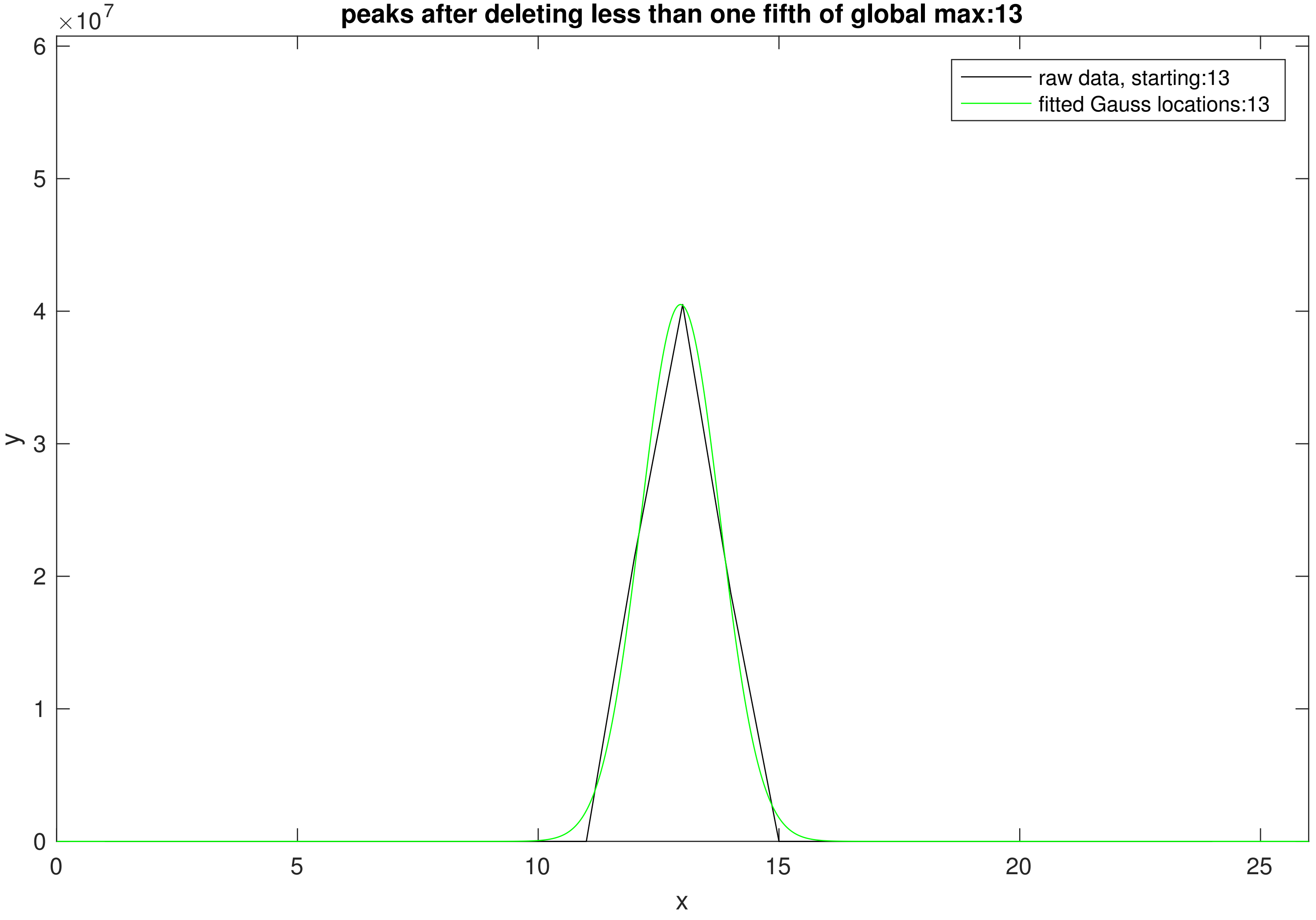

Optimal1 Fitted Gauss, proteinAT2G23110.2;AT2G23110.1
bic:694
peaks after deleting less than one fifth of global max:17

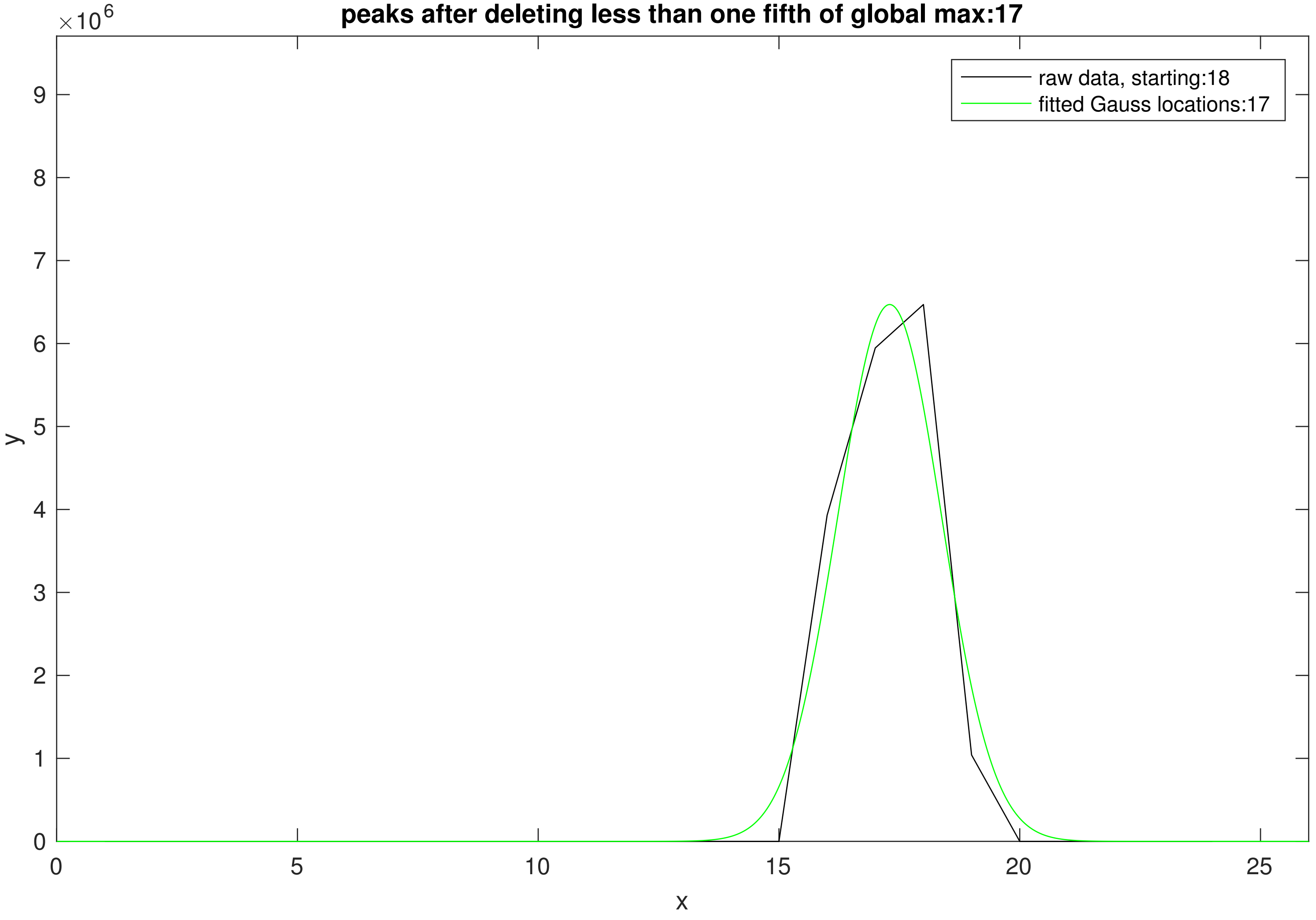

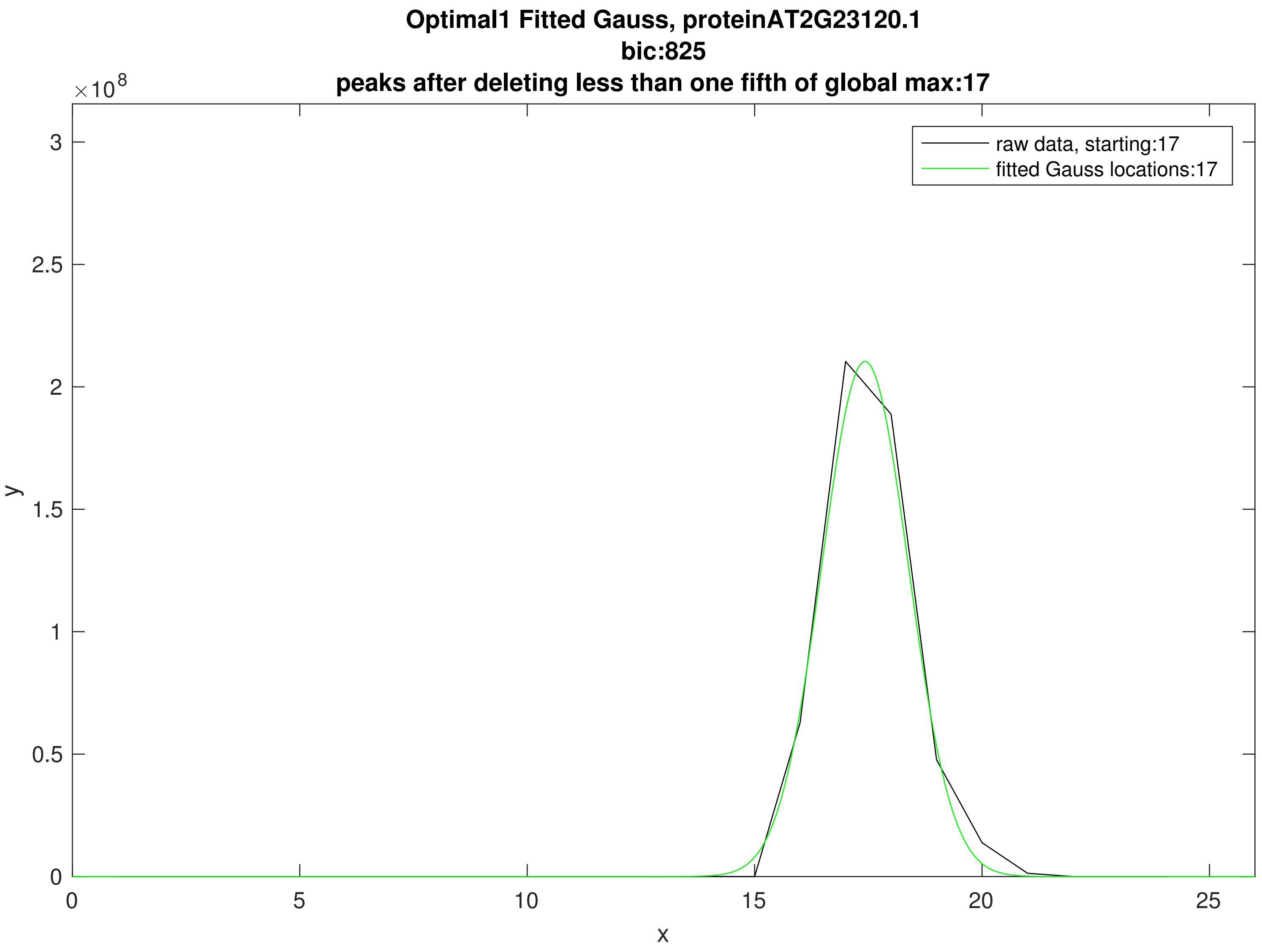
Optimal1 Fitted Gauss, proteinAT2G23120.1
bic:825
peaks after deleting less than one fifth of global max:17
×10^8
raw data, starting:17
fitted Gauss locations:17
y
x
0
0.5
1
1.5
2
2.5
3
0
5
10
15
20
25

Optimal1 Fitted Gauss, proteinAT2G23240.2;AT2G23240.1
bic:720
peaks after deleting less than one fifth of global max:20

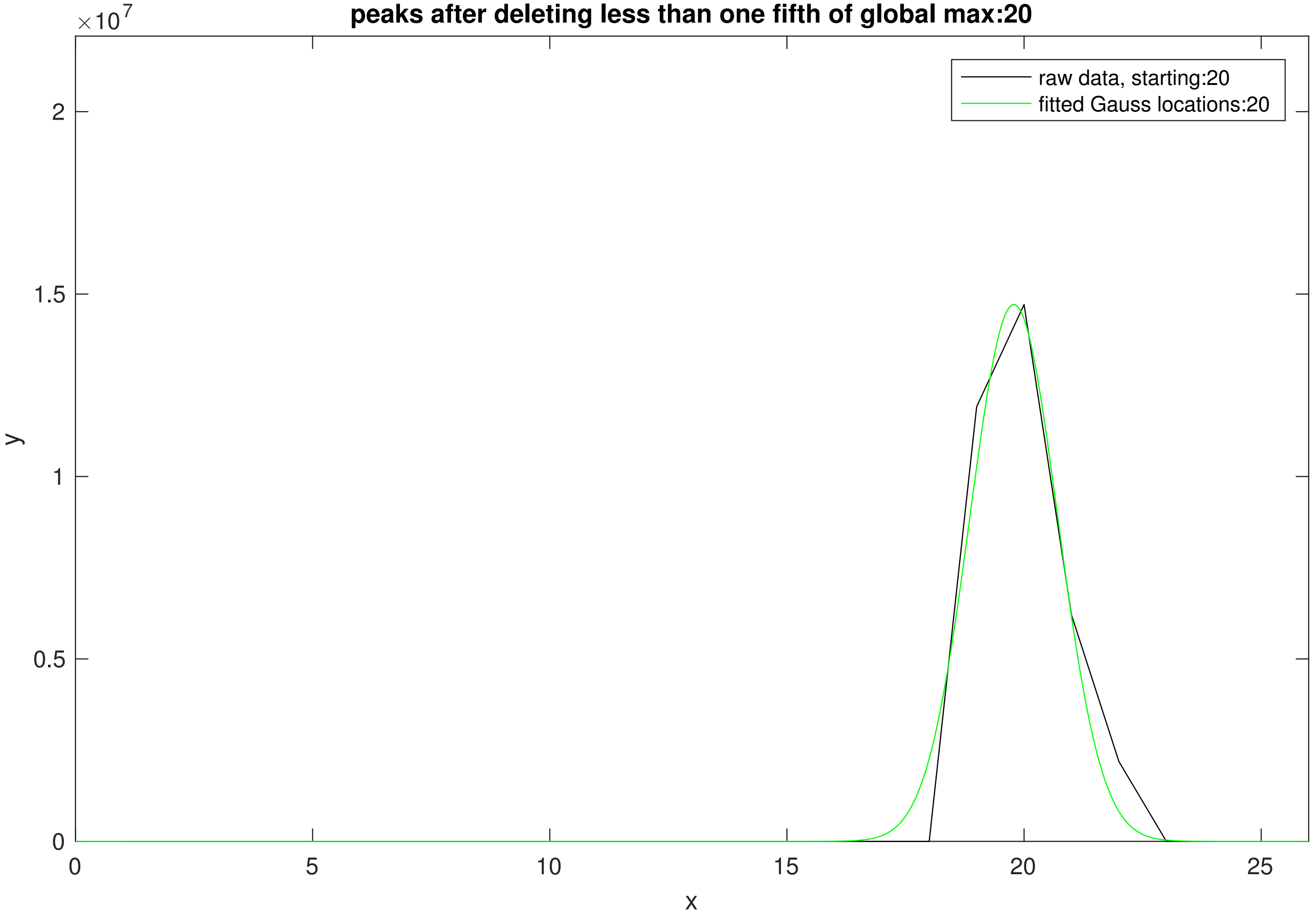

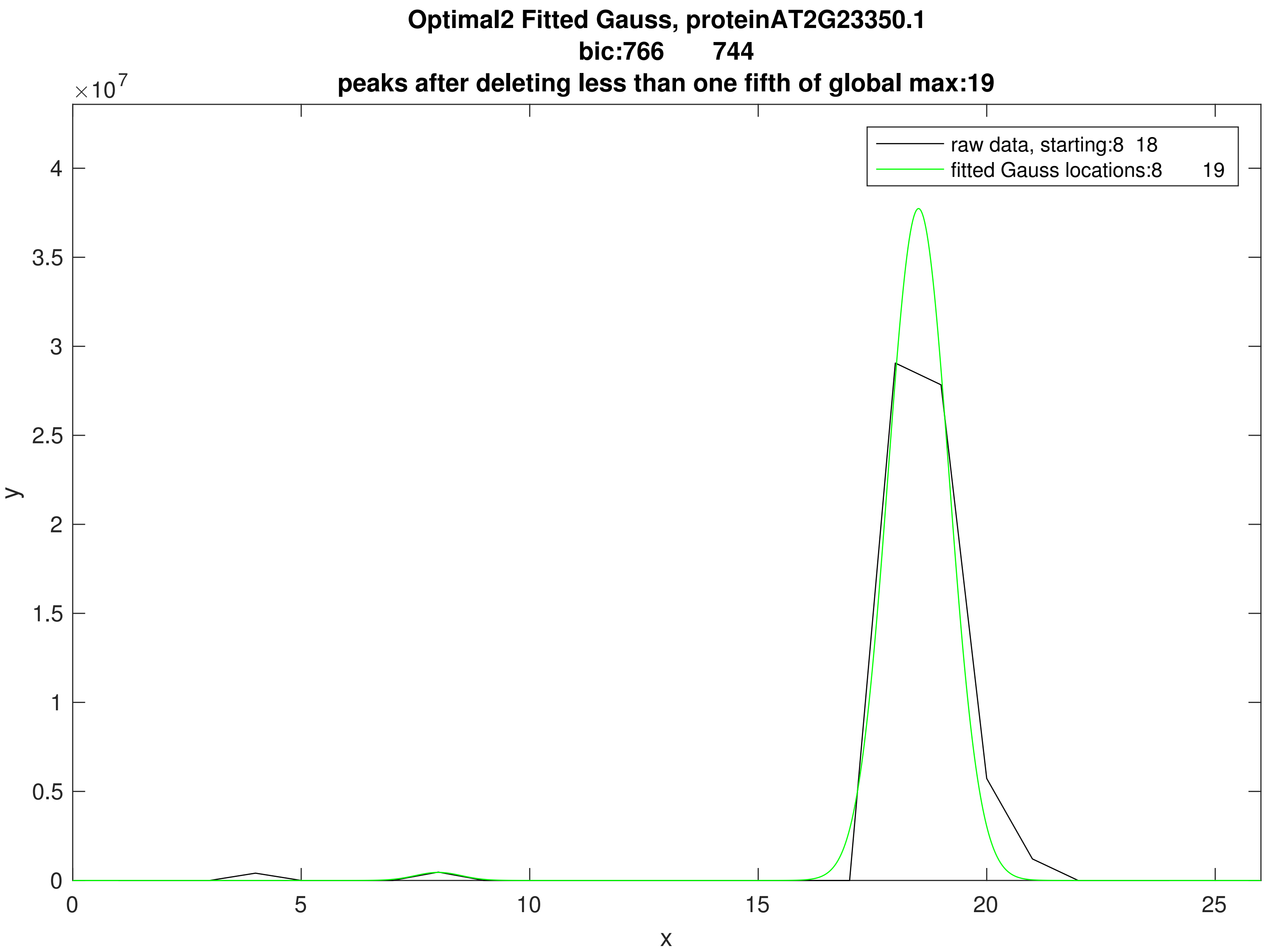
Optimal2 Fitted Gauss, proteinAT2G23350.1
bic:766 744
peaks after deleting less than one fifth of global max:19
×10^7
raw data, starting:8 18
fitted Gauss locations:8 19
y
x
0
0.5
1
1.5
2
2.5
3
3.5
4
0
5
10
15
20
25

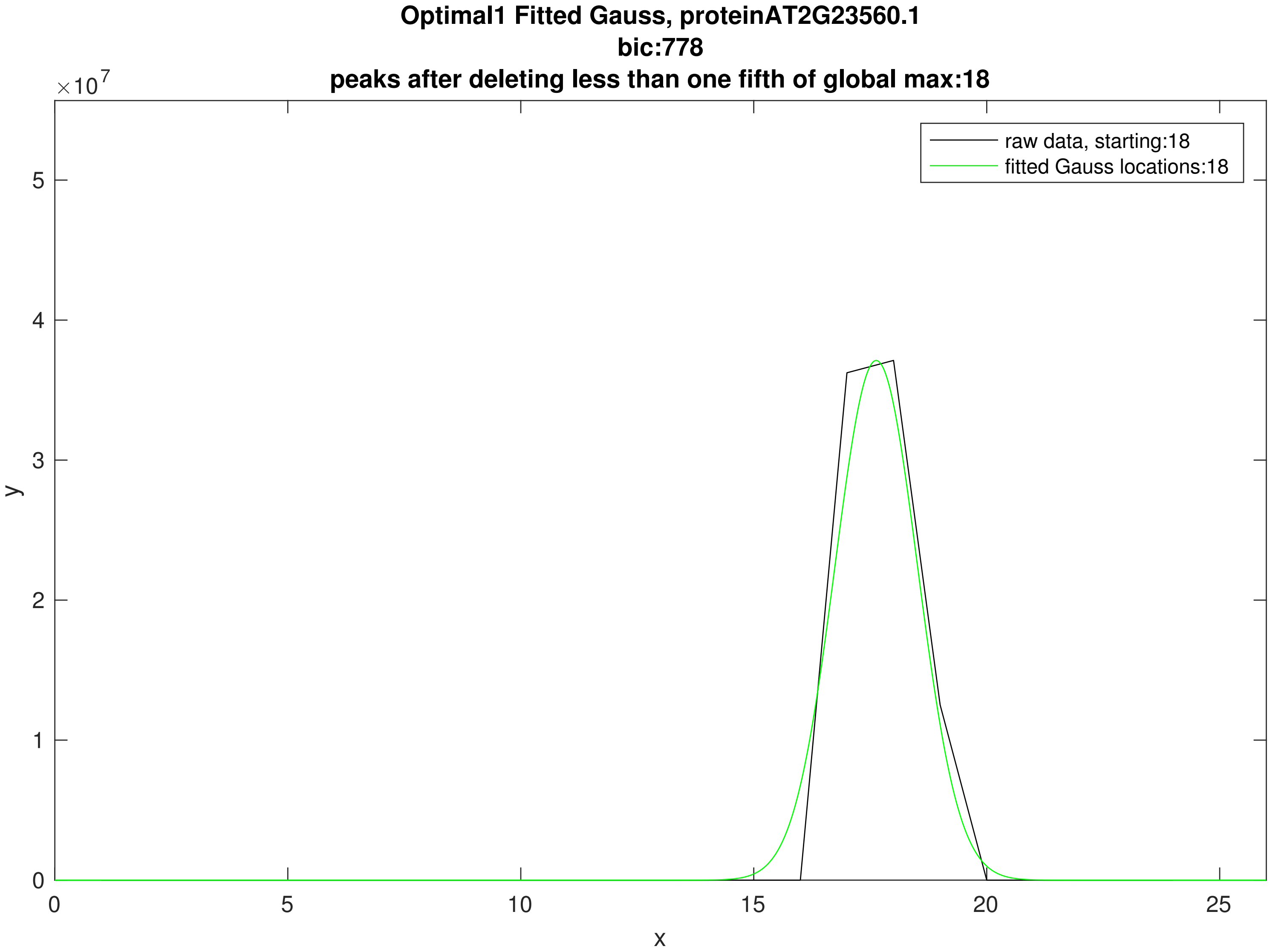

Optimal1 Fitted Gauss, proteinAT2G23560.1
bic:778
peaks after deleting less than one fifth of global max:18
×10^7
raw data, starting:18
fitted Gauss locations:18
y
x
0
1
2
3
4
5
0
5
10
15
20
25

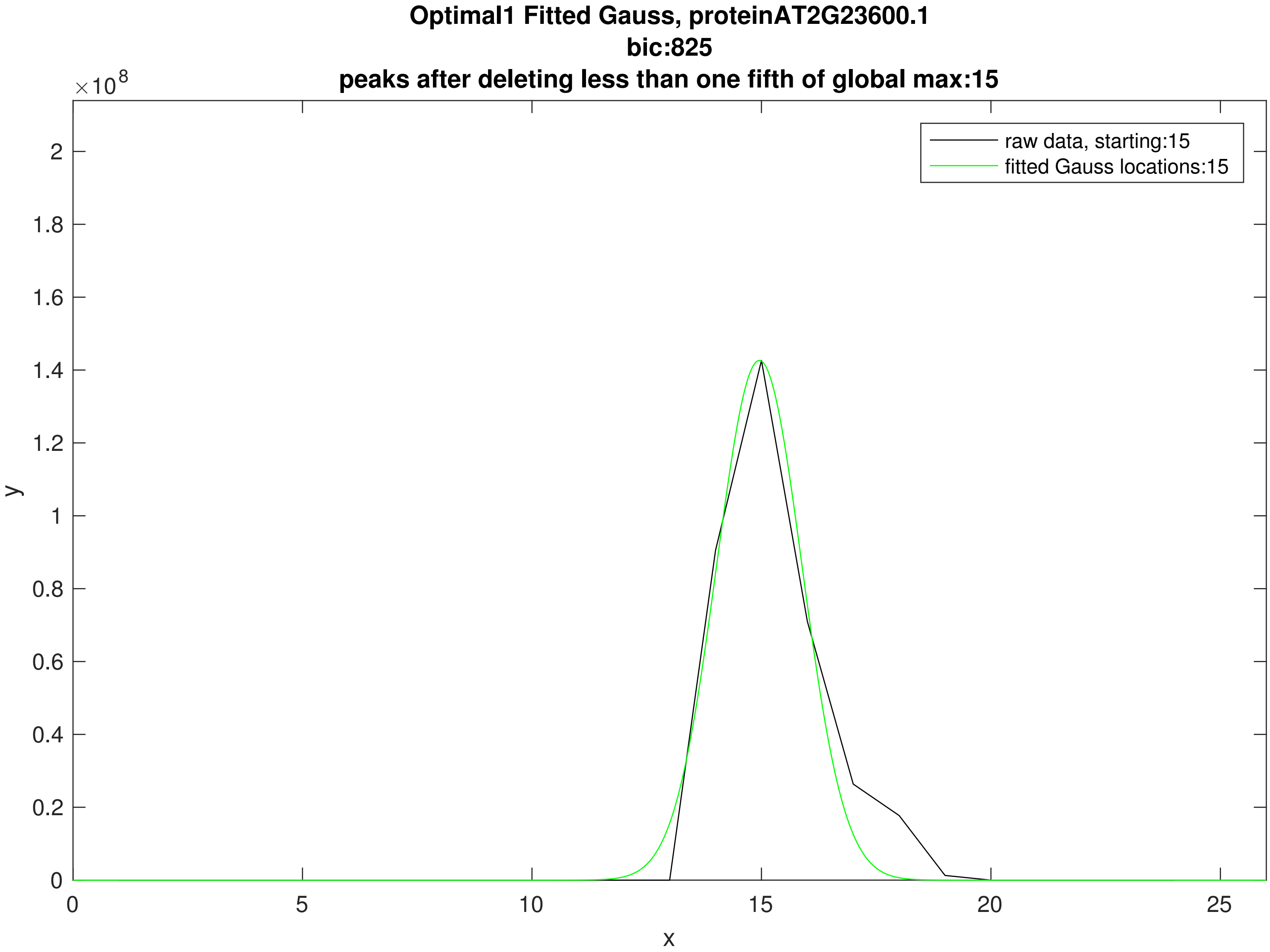
Optimal1 Fitted Gauss, proteinAT2G23600.1
bic:825
peaks after deleting less than one fifth of global max:15
×10^8
raw data, starting:15
fitted Gauss locations:15
y
x
0
0.2
0.4
0.6
0.8
1
1.2
1.4
1.6
1.8
2
0
5
10
15
20
25

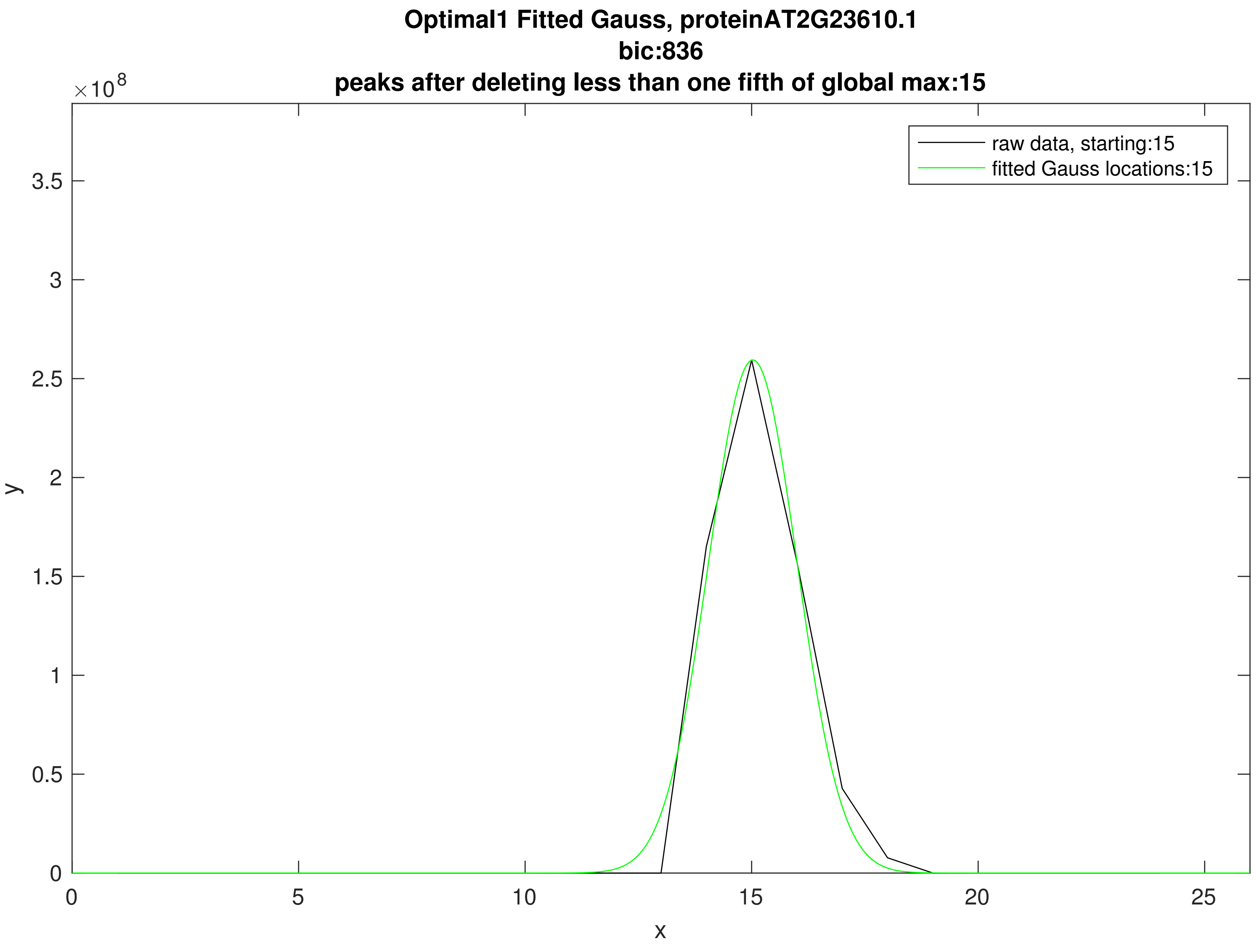
Optimal1 Fitted Gauss, proteinAT2G23610.1
bic:836
peaks after deleting less than one fifth of global max:15
×10^8
raw data, starting:15
fitted Gauss locations:15
y
x
0
0.5
1
1.5
2
2.5
3
3.5
0
5
10
15
20
25

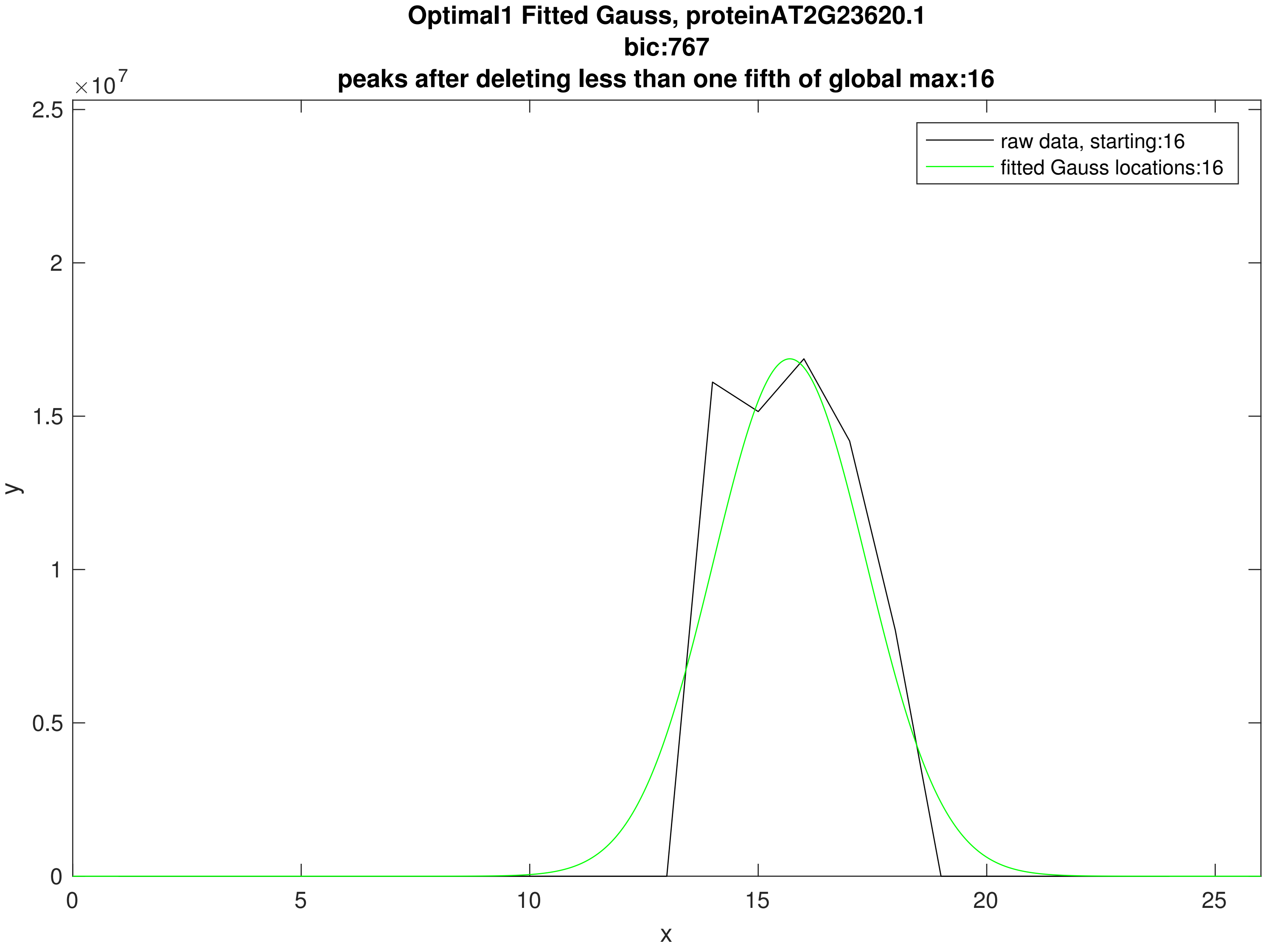

Optimal1 Fitted Gauss, proteinAT2G23620.1
bic:767
peaks after deleting less than one fifth of global max:16
×10^7
2.5
2
1.5
1
0.5
0
y
0
5
10
15
20
25
x
raw data, starting:16
fitted Gauss locations:16

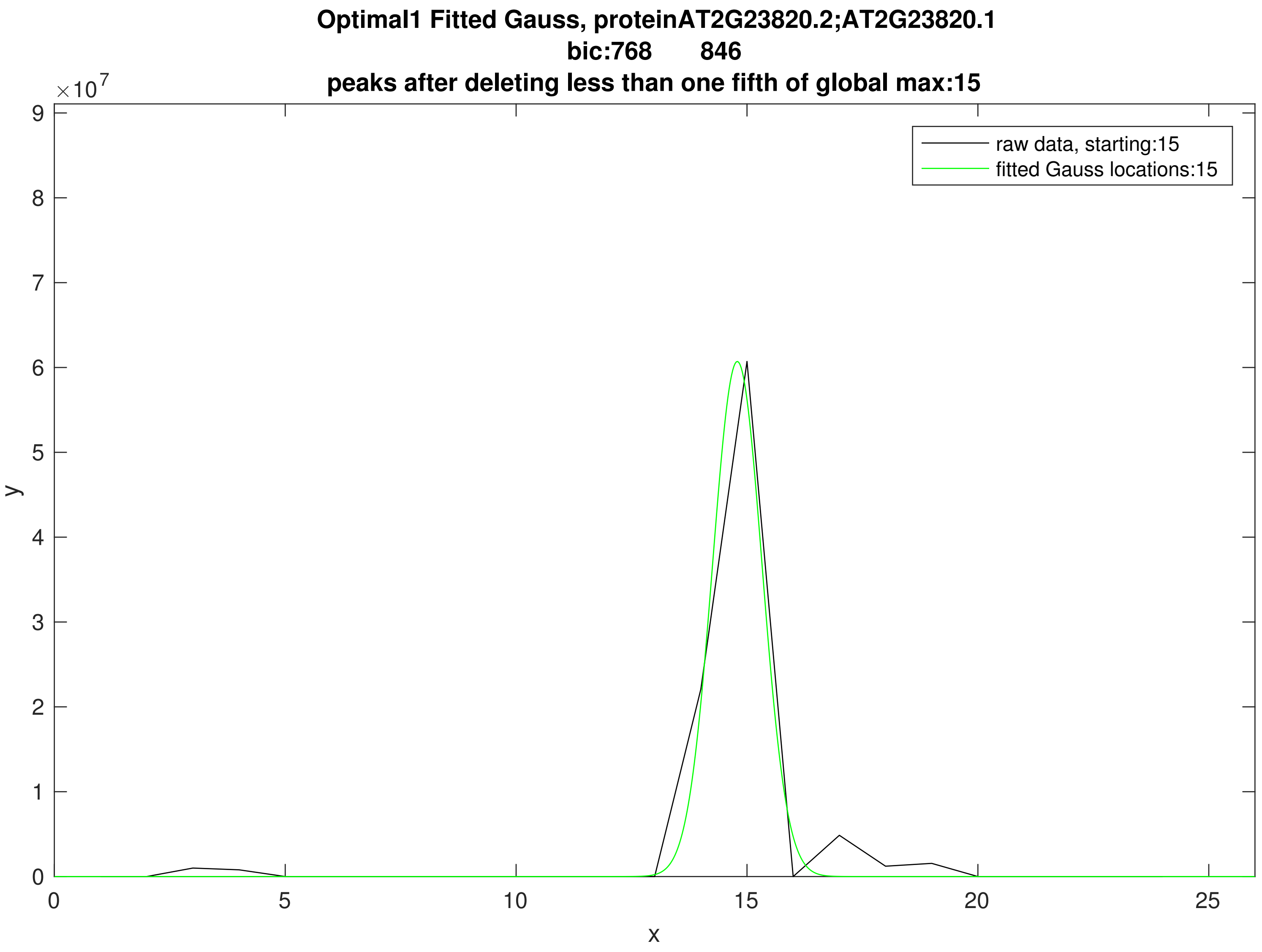
Optimal1 Fitted Gauss, proteinAT2G23820.2;AT2G23820.1
bic:768 846
peaks after deleting less than one fifth of global max:15
×10^7
raw data, starting:15
fitted Gauss locations:15
y
x
0
1
2
3
4
5
6
7
8
9
0
5
10
15
20
25

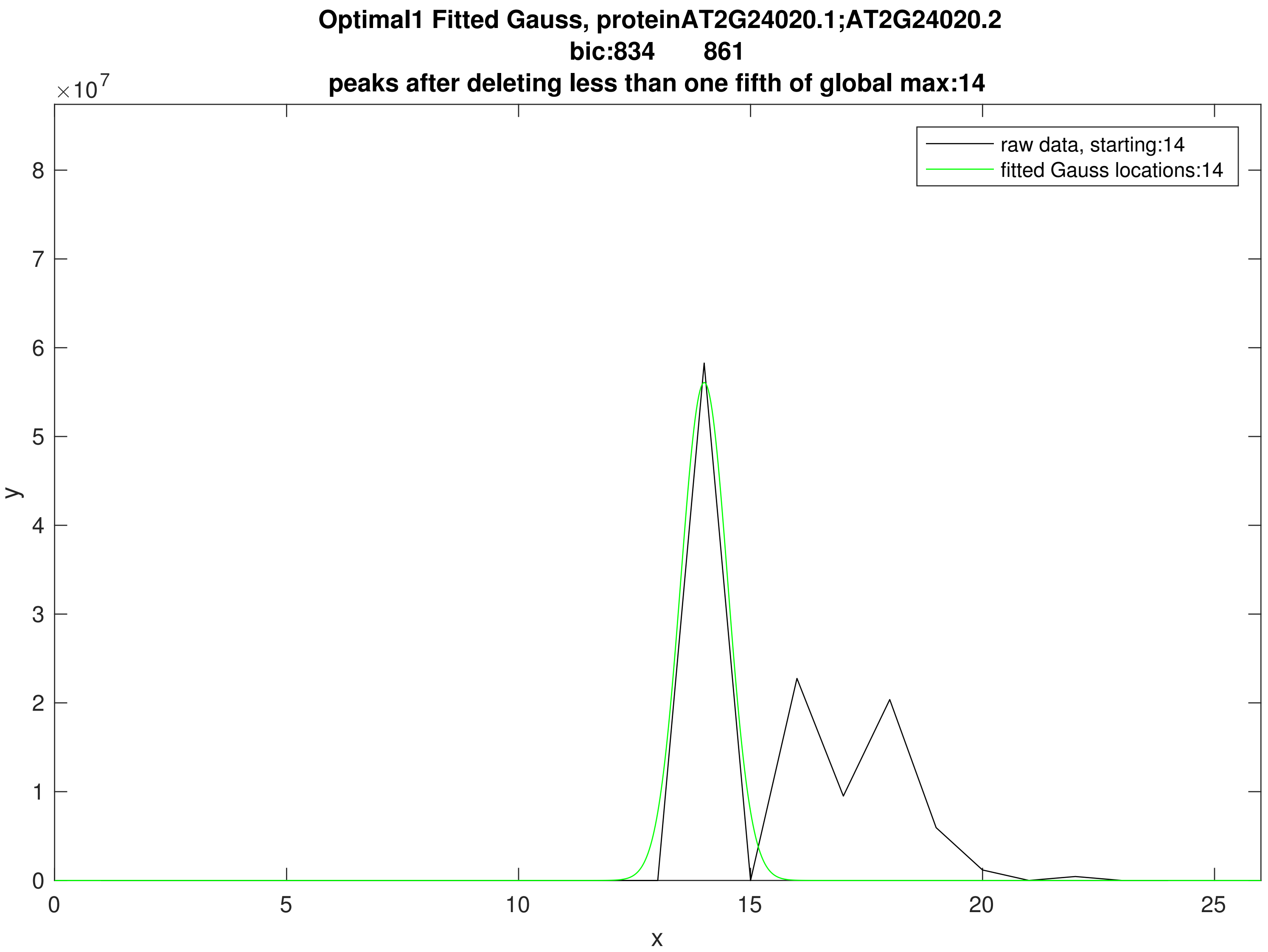

Optimal1 Fitted Gauss, proteinAT2G24020.1;AT2G24020.2
bic:834 861
peaks after deleting less than one fifth of global max:14
×10^7
8
7
6
5
4
3
2
1
0
y
0
5
10
15
20
25
x
raw data, starting:14
fitted Gauss locations:14

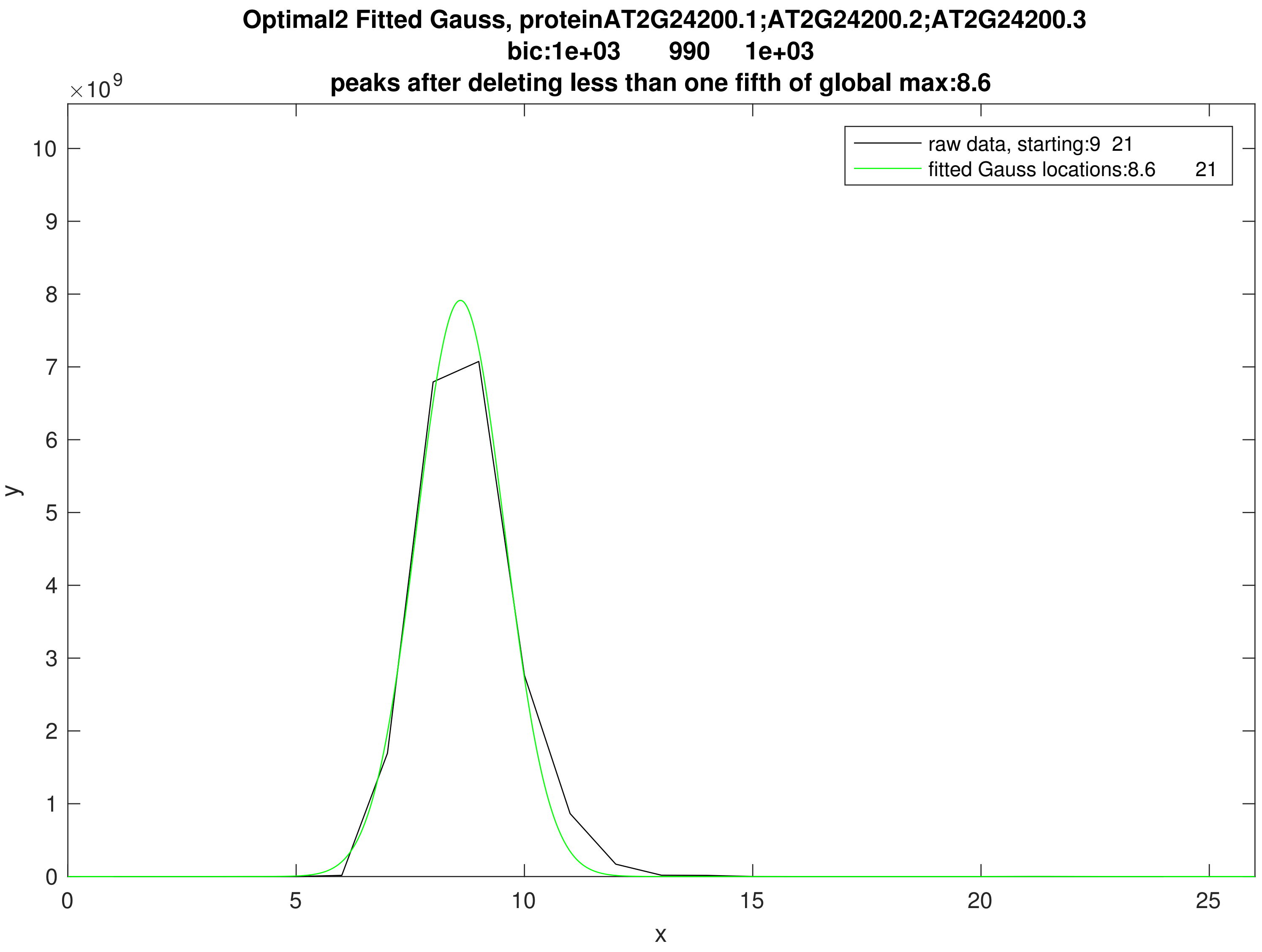
Optimal2 Fitted Gauss, proteinAT2G24200.1;AT2G24200.2;AT2G24200.3
bic:1e+03 990 1e+03
peaks after deleting less than one fifth of global max:8.6
×10^9
raw data, starting:9 21
fitted Gauss locations:8.6 21
y
x
0
1
2
3
4
5
6
7
8
9
10
0
5
10
15
20
25

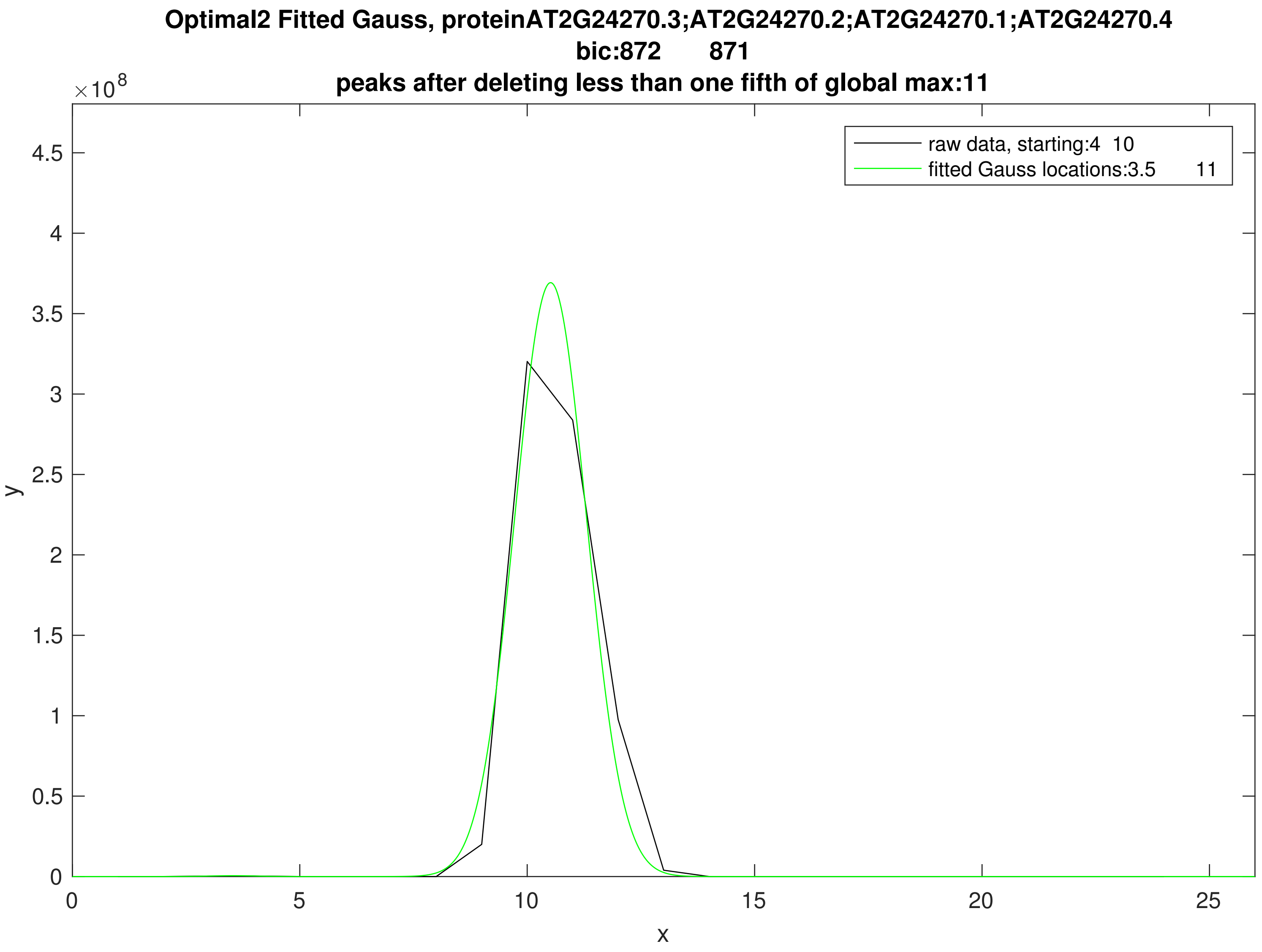

Optimal2 Fitted Gauss, proteinAT2G24270.3;AT2G24270.2;AT2G24270.1;AT2G24270.4
bic:872 871
peaks after deleting less than one fifth of global max:11
×10^8
raw data, starting:4 10
fitted Gauss locations:3.5 11
y
x
0
0.5
1
1.5
2
2.5
3
3.5
4
4.5
0
5
10
15
20
25

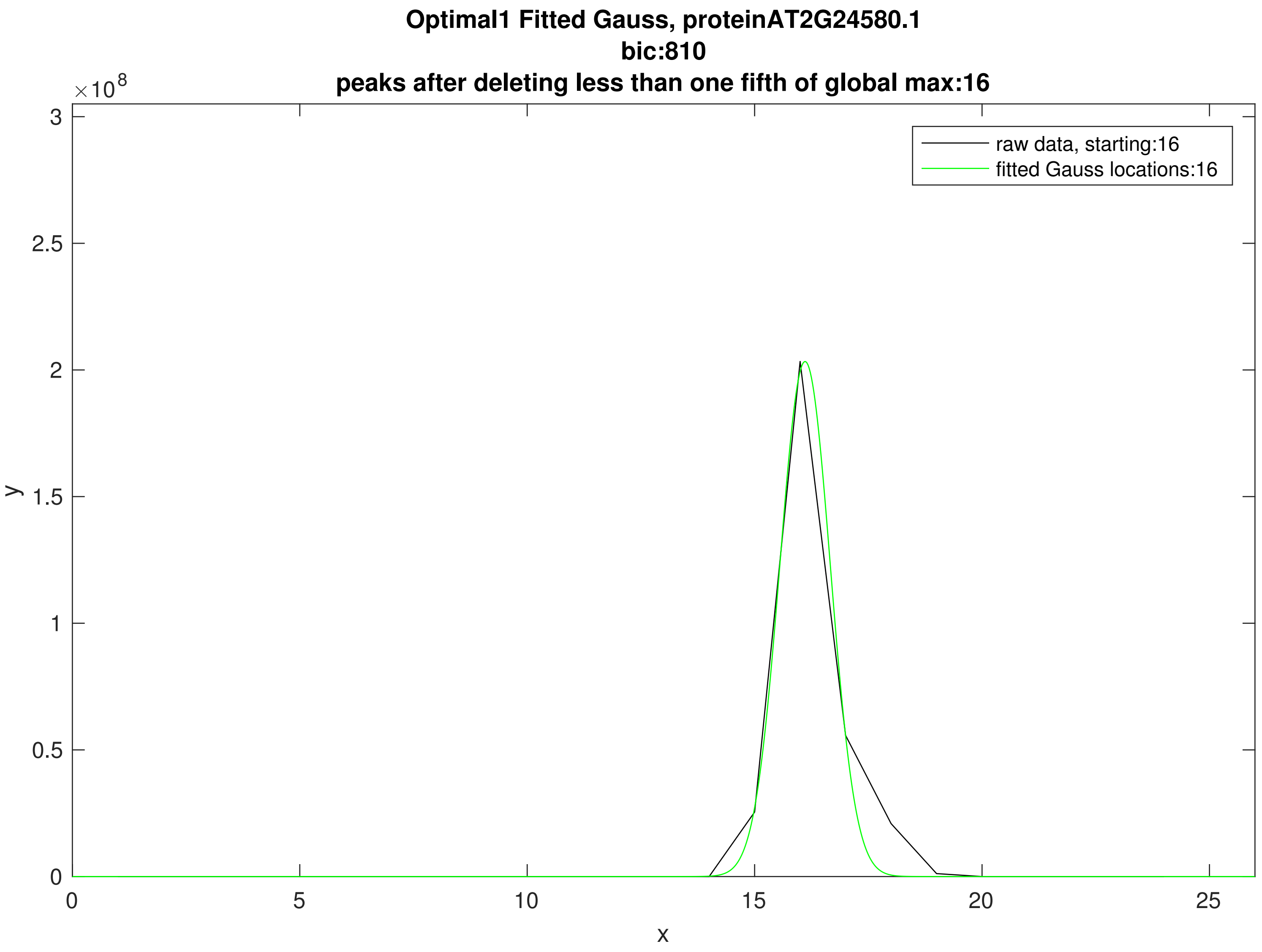
Optimal1 Fitted Gauss, proteinAT2G24580.1
bic:810
peaks after deleting less than one fifth of global max:16
×10^8
raw data, starting:16
fitted Gauss locations:16
y
x
0
0.5
1
1.5
2
2.5
3
0
5
10
15
20
25

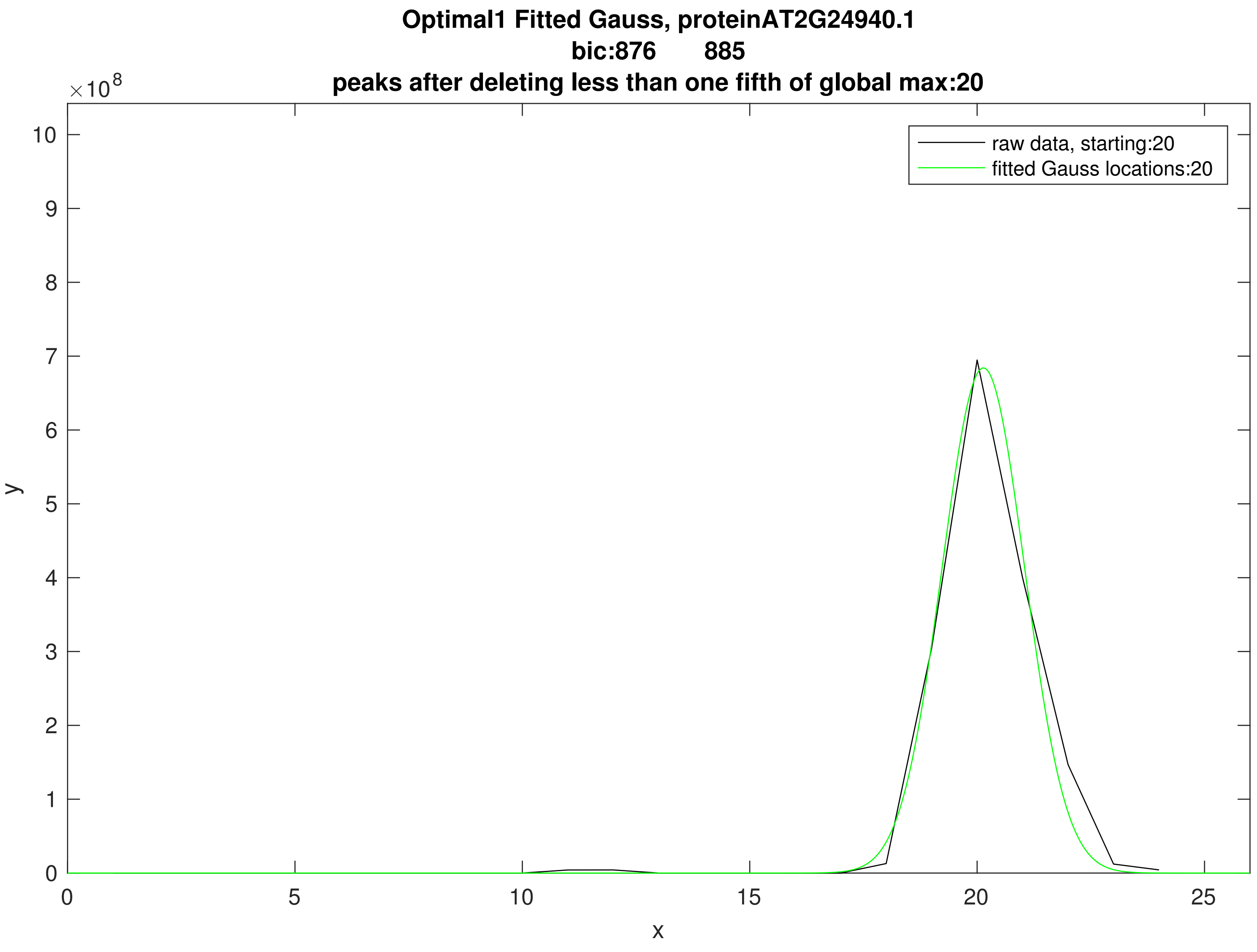

Optimal1 Fitted Gauss, proteinAT2G24940.1
bic:876 885
peaks after deleting less than one fifth of global max:20
×10^8
10
9
8
7
6
5
4
3
2
1
0
y
0
5
10
15
20
25
x
raw data, starting:20
fitted Gauss locations:20

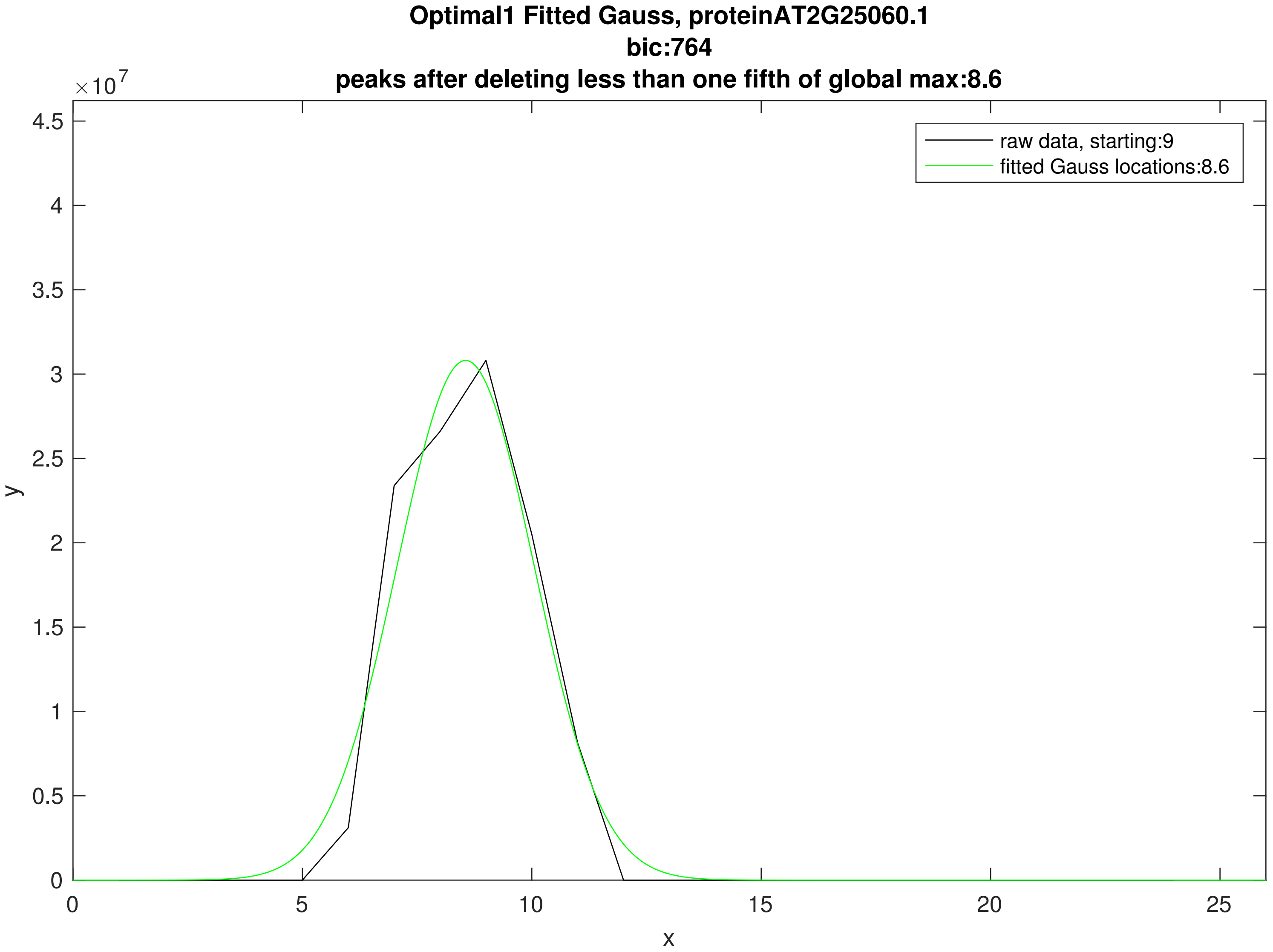

Optimal1 Fitted Gauss, proteinAT2G25060.1
bic:764
peaks after deleting less than one fifth of global max:8.6
×10^7
4.5
4
3.5
3
2.5
2
1.5
1
0.5
0
y
0
5
10
15
20
25
x
raw data, starting:9
fitted Gauss locations:8.6

**Optimal1 Fitted Gauss, proteinAT2G25070.1**
**bic:809 818**
**peaks after deleting less than one fifth of global max:16**

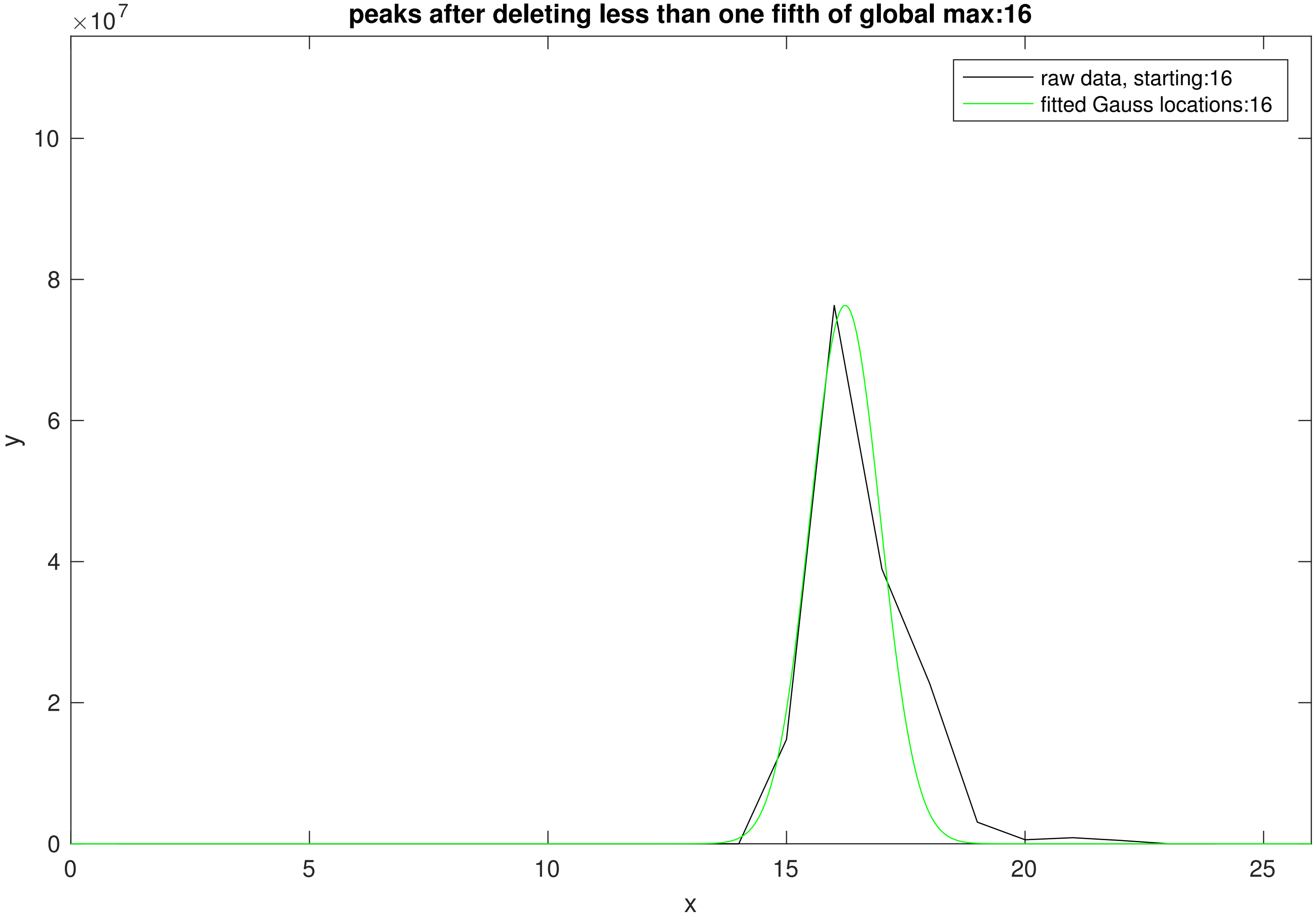

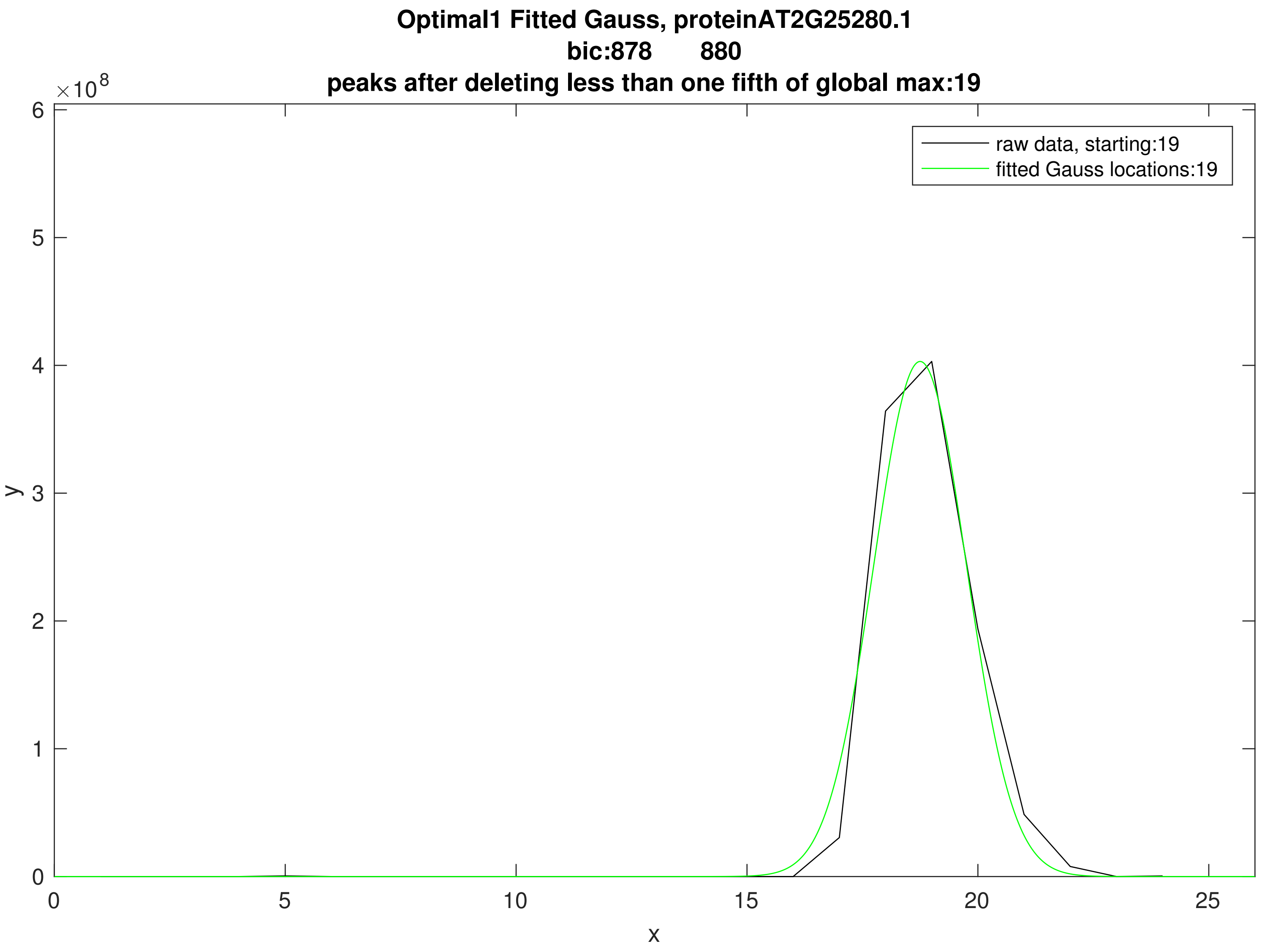

Optimal1 Fitted Gauss, proteinAT2G25280.1
bic:878 880
peaks after deleting less than one fifth of global max:19
×10^8
6
5
4
3
2
1
0
y
0
5
10
15
20
25
x
raw data, starting:19
fitted Gauss locations:19

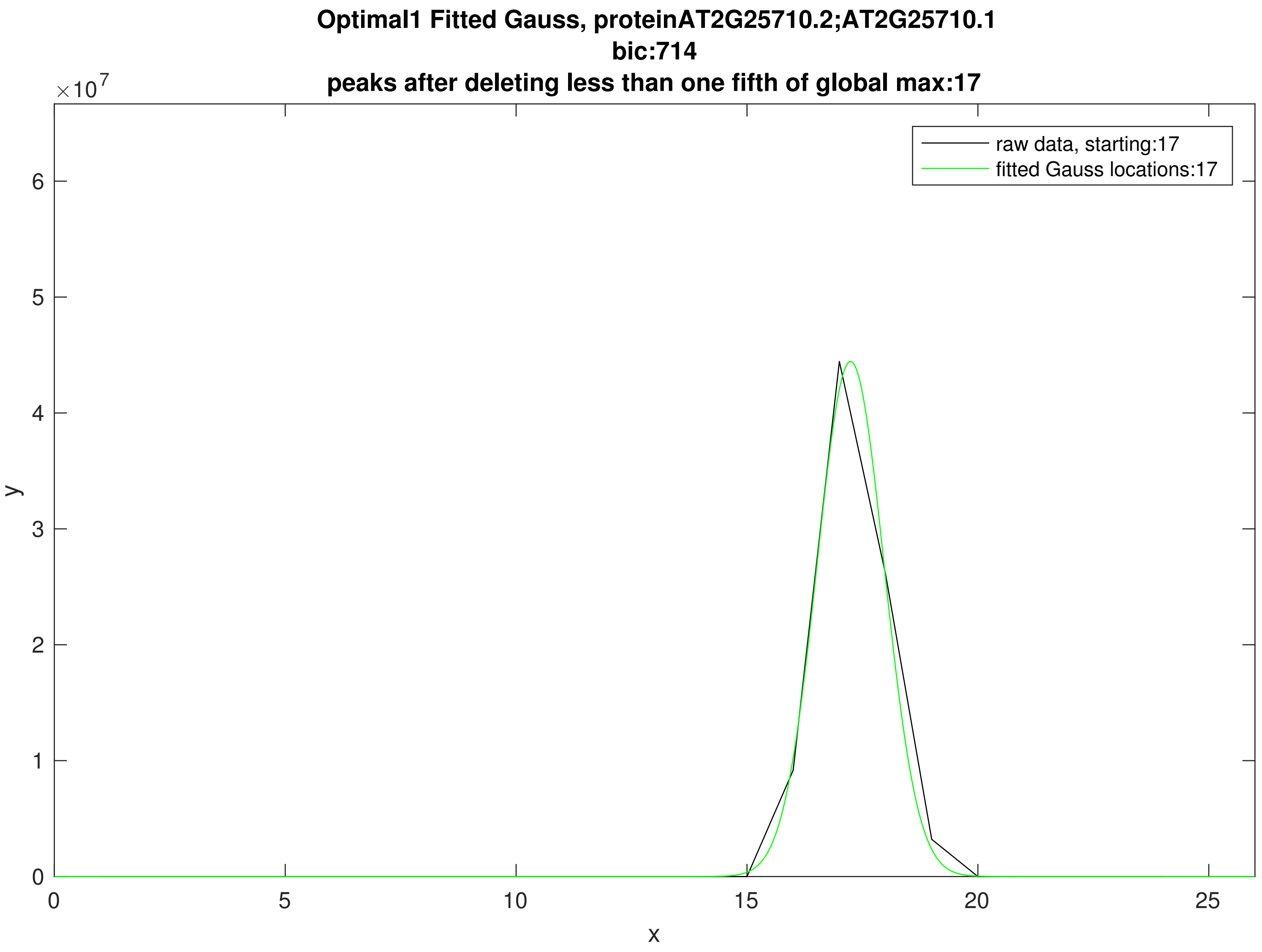
Optimal1 Fitted Gauss, proteinAT2G25710.2;AT2G25710.1
bic:714
peaks after deleting less than one fifth of global max:17
×10^7
raw data, starting:17
fitted Gauss locations:17
y
x
0
1
2
3
4
5
6
0
5
10
15
20
25

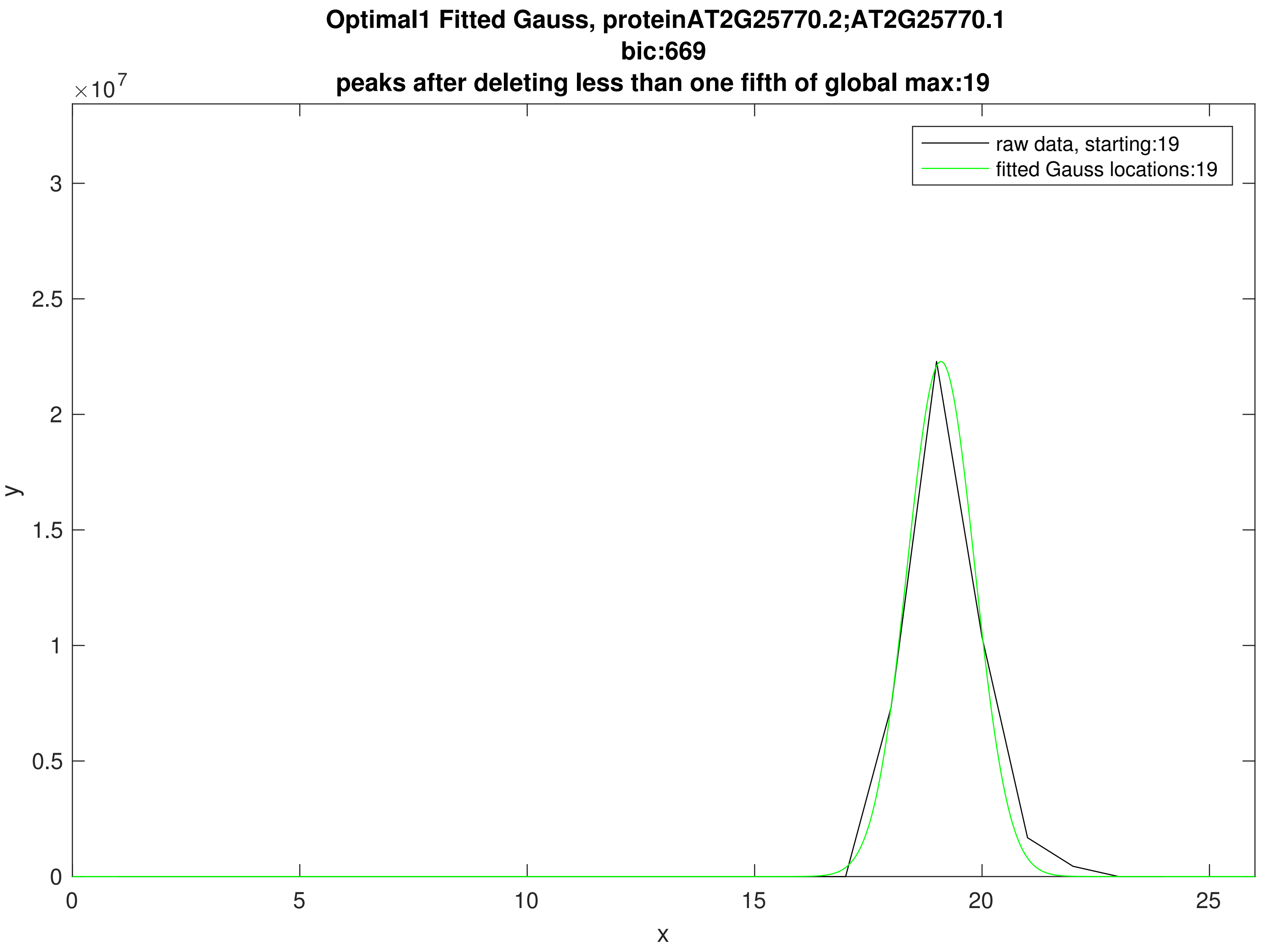
Optimal1 Fitted Gauss, proteinAT2G25770.2;AT2G25770.1
bic:669
peaks after deleting less than one fifth of global max:19
×10^7
raw data, starting:19
fitted Gauss locations:19
3
2.5
2
1.5
1
0.5
0
y
0
5
10
15
20
25
x

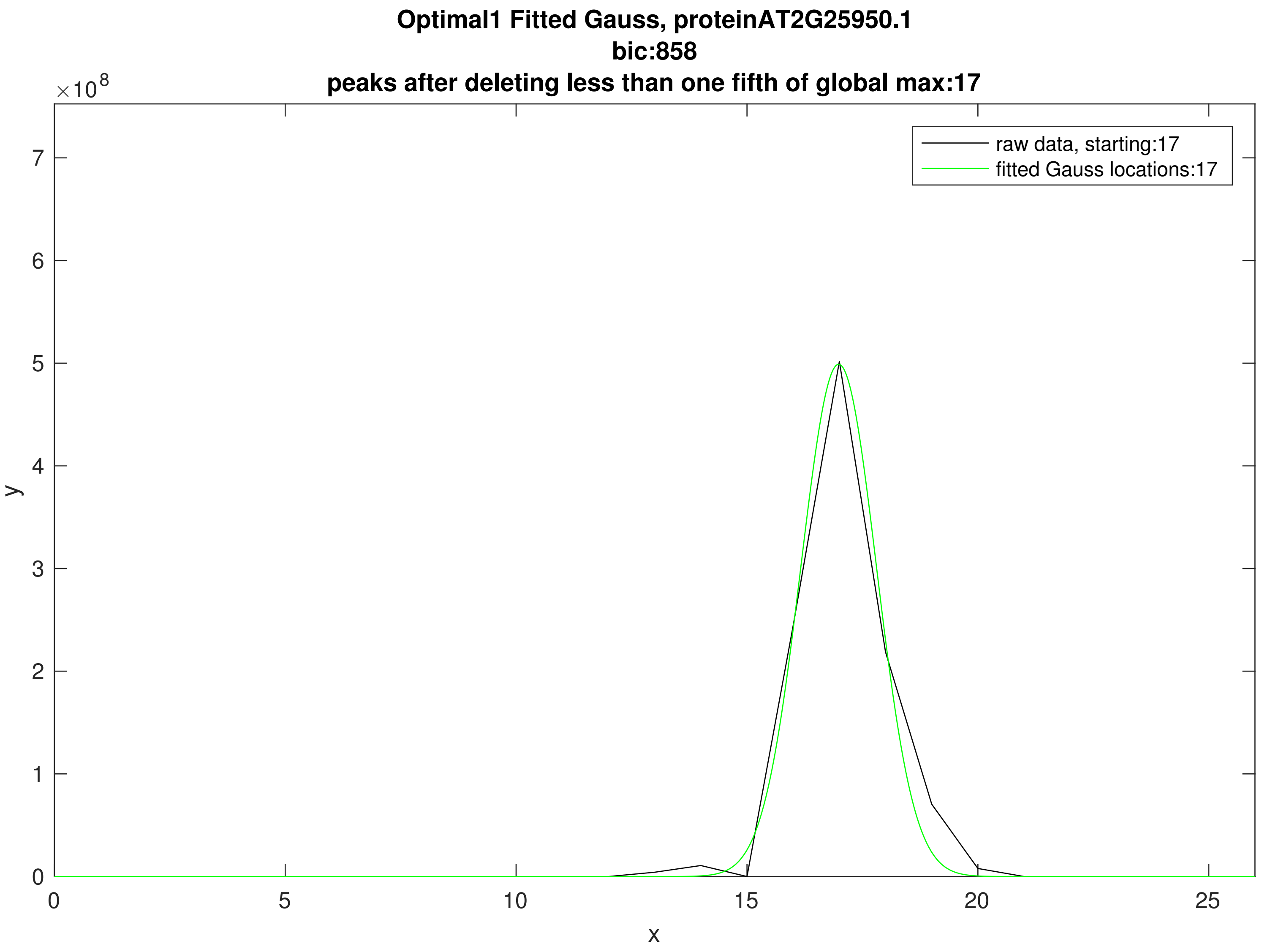
Optimal1 Fitted Gauss, proteinAT2G25950.1
bic:858
peaks after deleting less than one fifth of global max:17
×10^8
raw data, starting:17
fitted Gauss locations:17
y
x
0
1
2
3
4
5
6
7
0
5
10
15
20
25

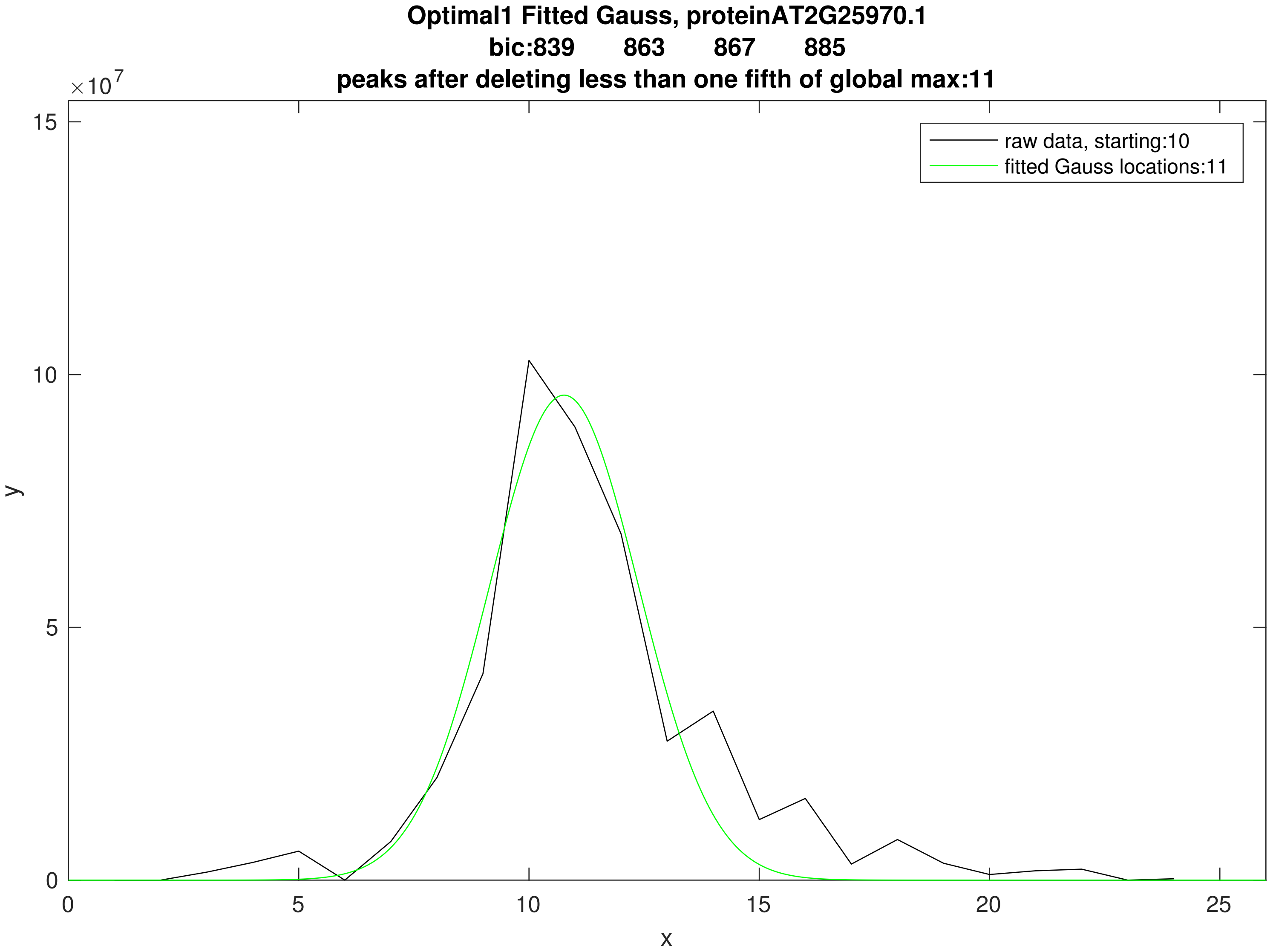

Optimal1 Fitted Gauss, proteinAT2G25970.1
bic:839 863 867 885
peaks after deleting less than one fifth of global max:11
×10^7
15
10
5
0
y
0
5
10
15
20
25
x
raw data, starting:10
fitted Gauss locations:11

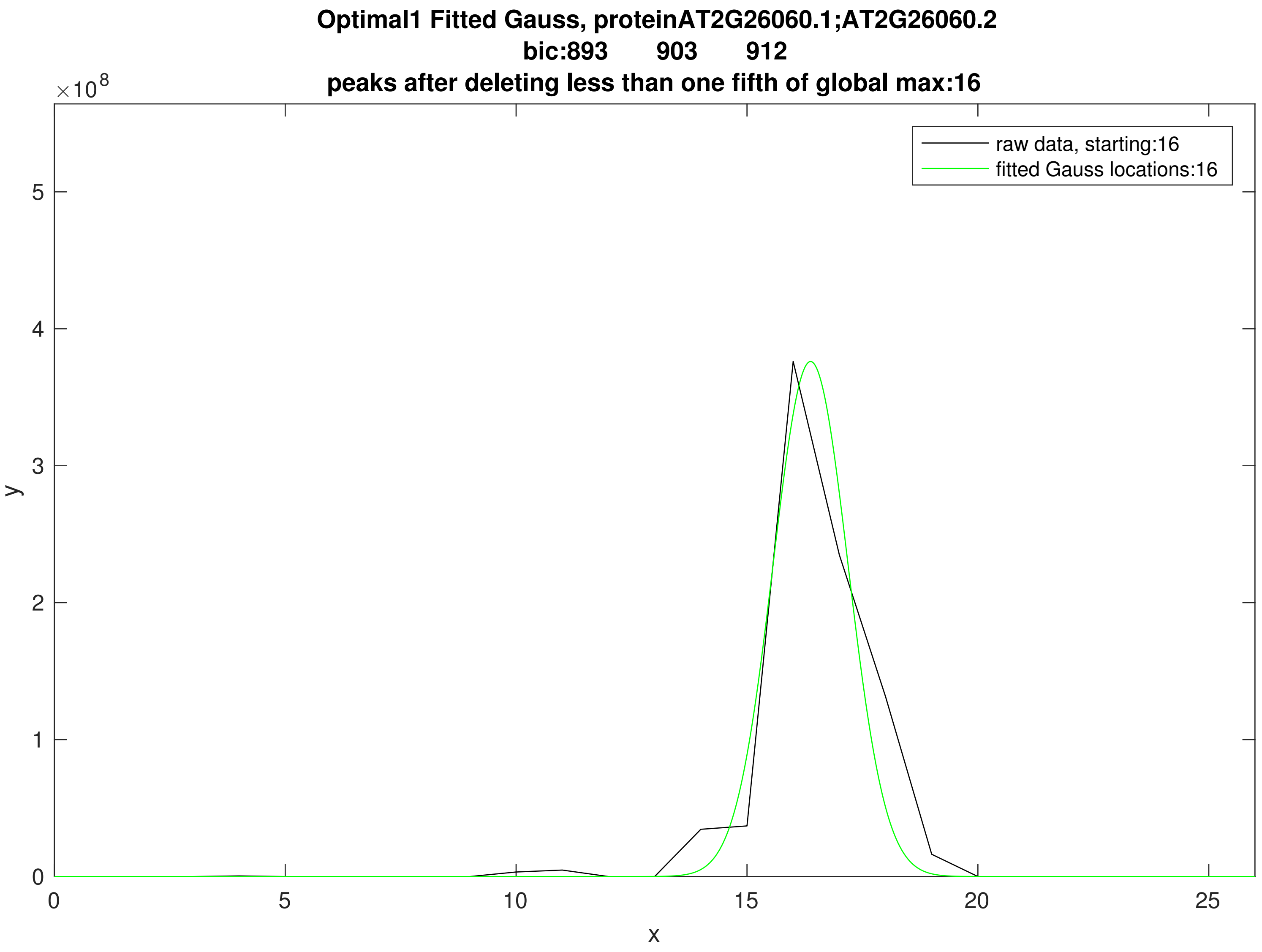
Optimal1 Fitted Gauss, proteinAT2G26060.1;AT2G26060.2
bic:893 903 912
peaks after deleting less than one fifth of global max:16
×10^8
raw data, starting:16
fitted Gauss locations:16
y
x
0
1
2
3
4
5
0
5
10
15
20
25

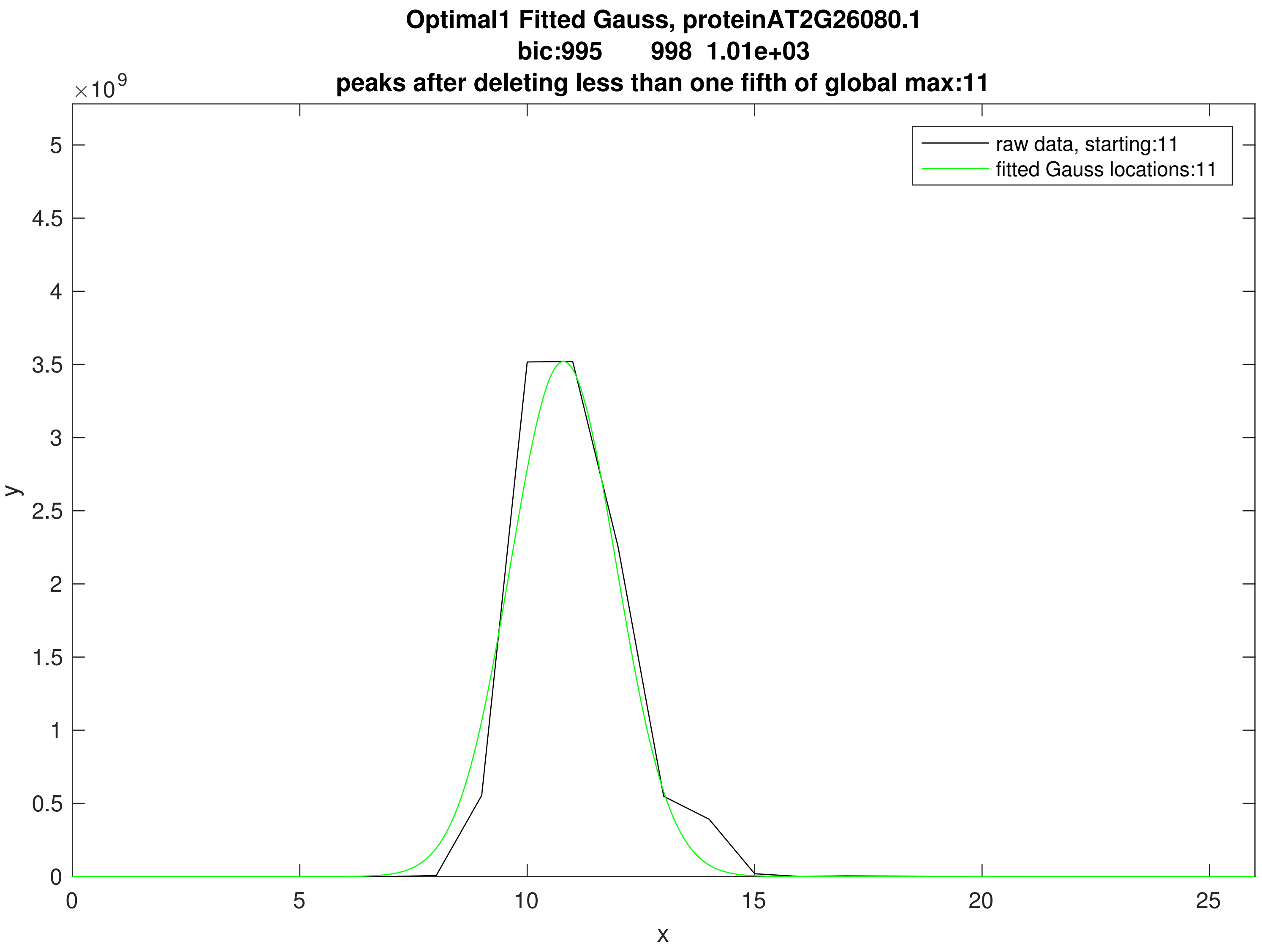
Optimal1 Fitted Gauss, proteinAT2G26080.1
bic:995 998 1.01e+03
peaks after deleting less than one fifth of global max:11
×10^9
raw data, starting:11
fitted Gauss locations:11
y
x
0
0.5
1
1.5
2
2.5
3
3.5
4
4.5
5
0
5
10
15
20
25

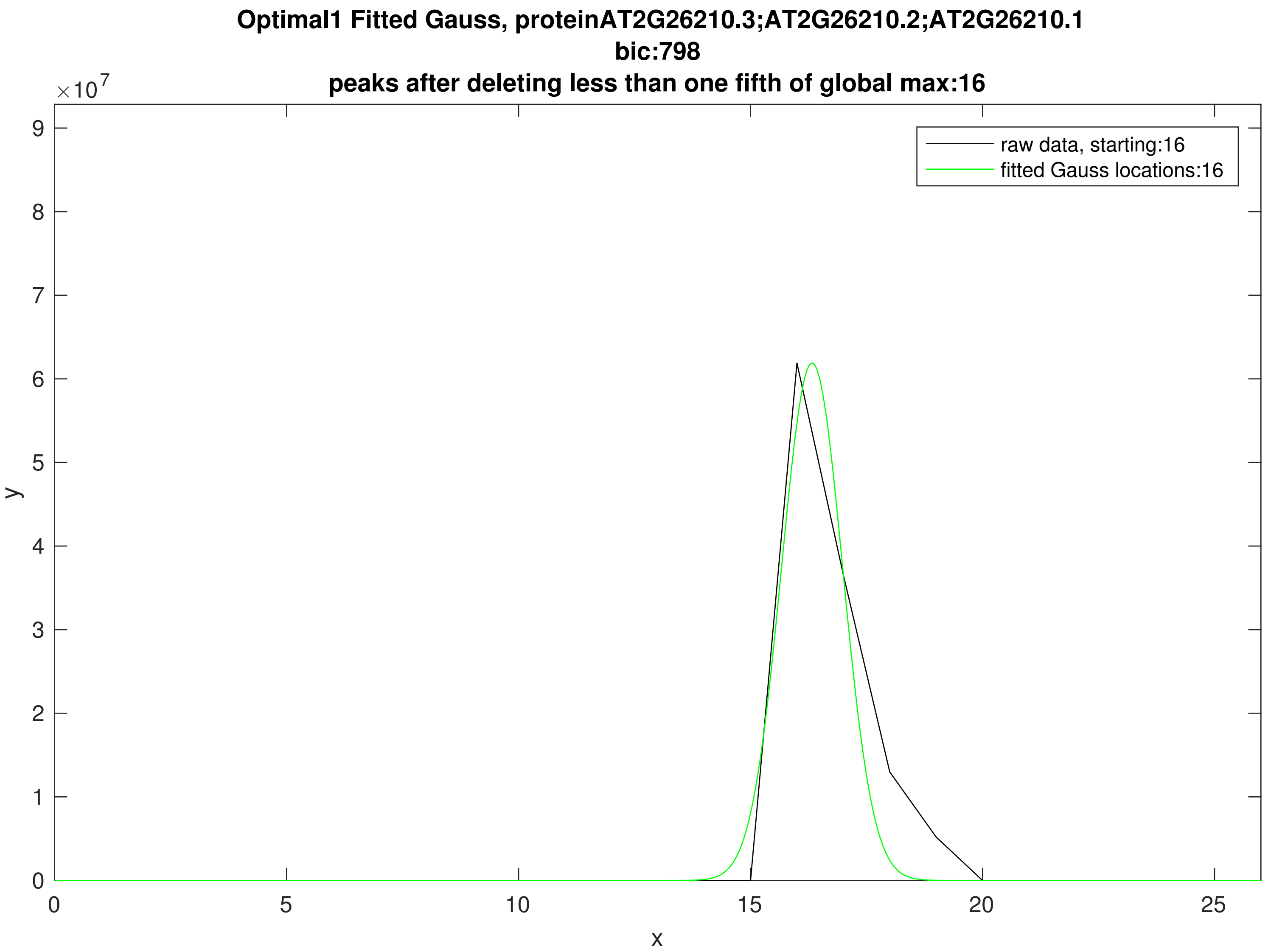
Optimal1 Fitted Gauss, proteinAT2G26210.3;AT2G26210.2;AT2G26210.1
bic:798
peaks after deleting less than one fifth of global max:16
×10^7
raw data, starting:16
fitted Gauss locations:16
y
x
0
1
2
3
4
5
6
7
8
9
0
5
10
15
20
25

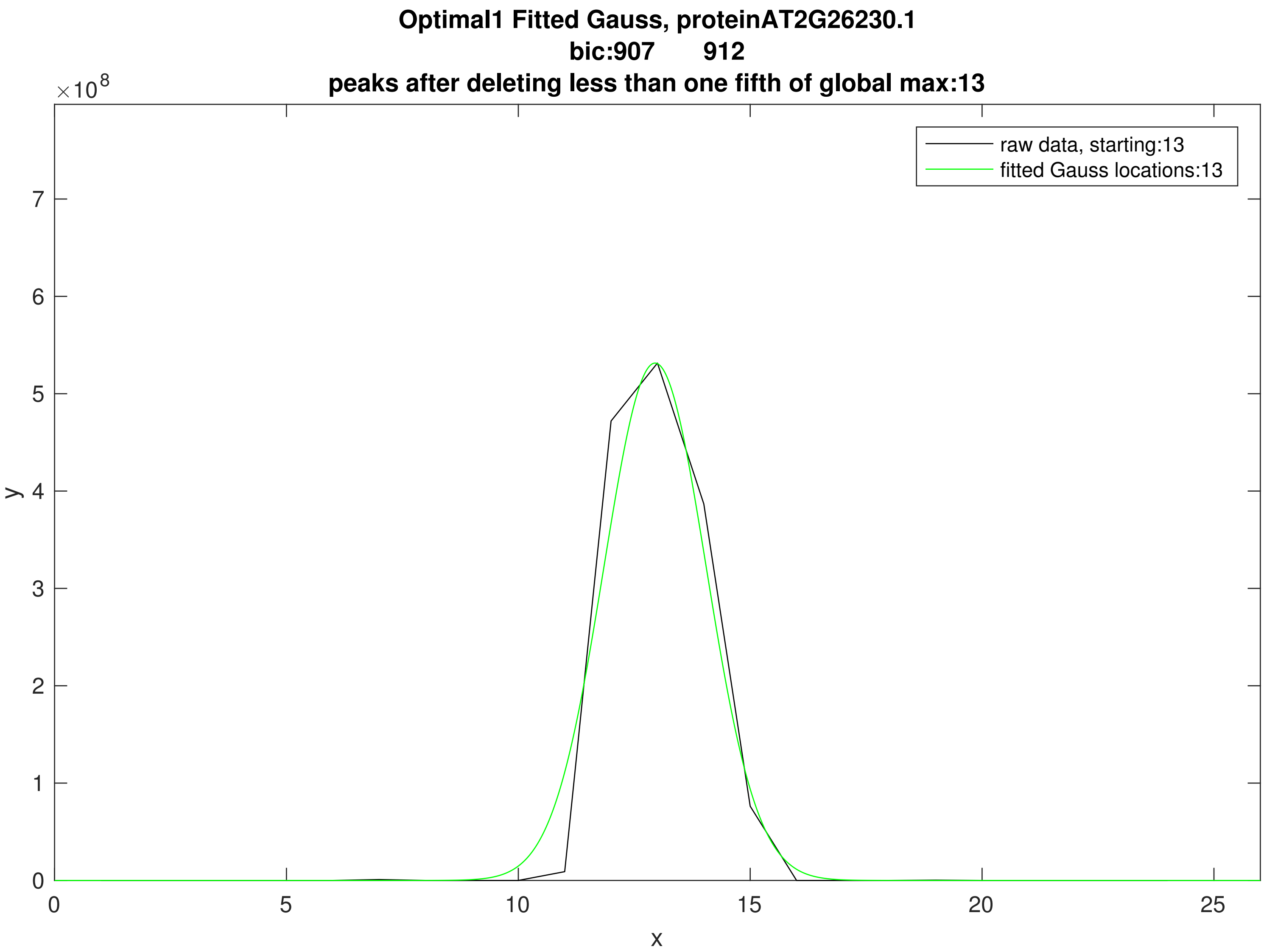
Optimal1 Fitted Gauss, proteinAT2G26230.1
bic:907 912
peaks after deleting less than one fifth of global max:13
×10^8
raw data, starting:13
fitted Gauss locations:13
y
x
0
1
2
3
4
5
6
7
0
5
10
15
20
25

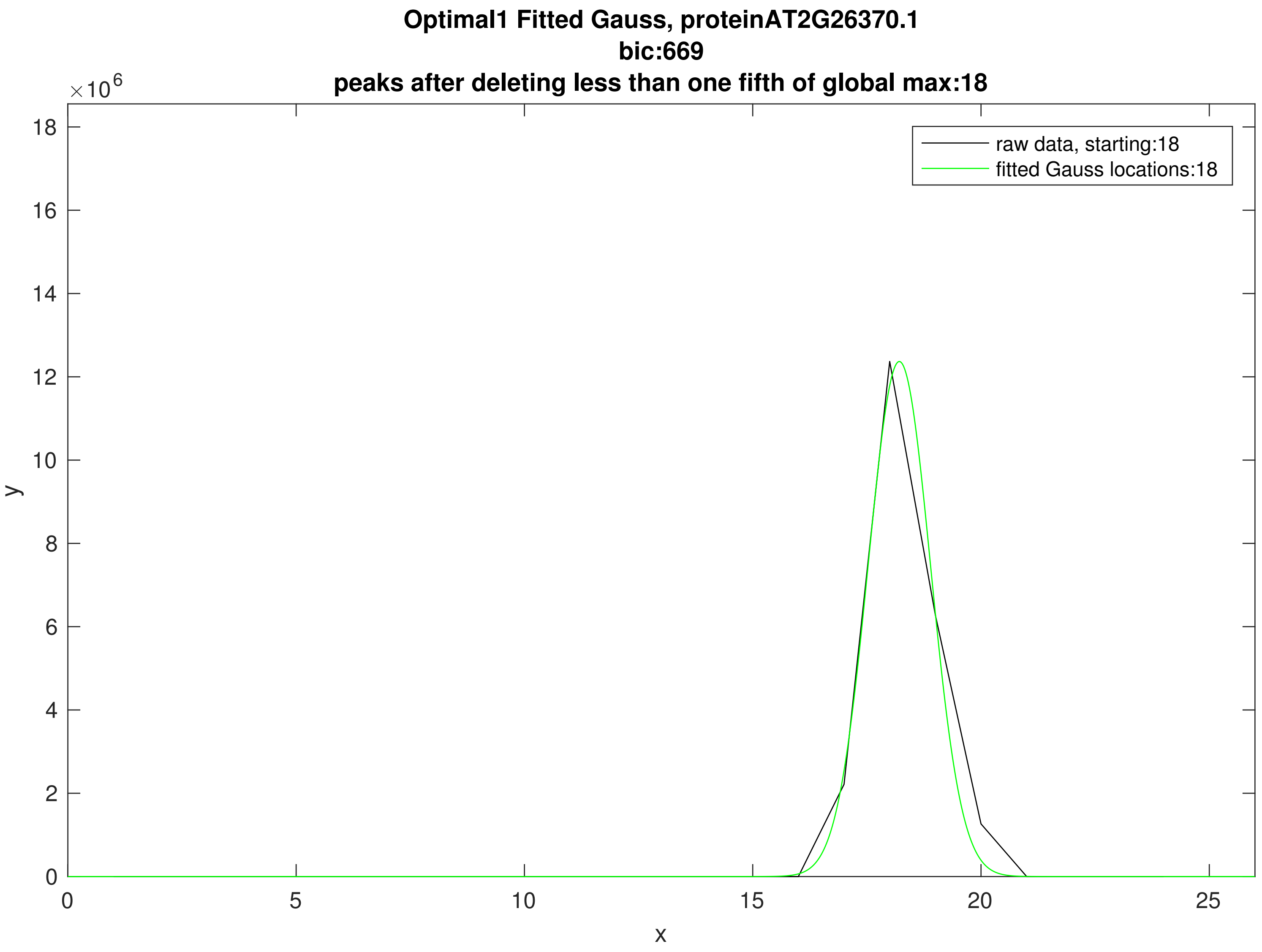
Optimal1 Fitted Gauss, proteinAT2G26370.1
bic:669
peaks after deleting less than one fifth of global max:18
×10^6
raw data, starting:18
fitted Gauss locations:18
y
x
0
2
4
6
8
10
12
14
16
18
0
5
10
15
20
25

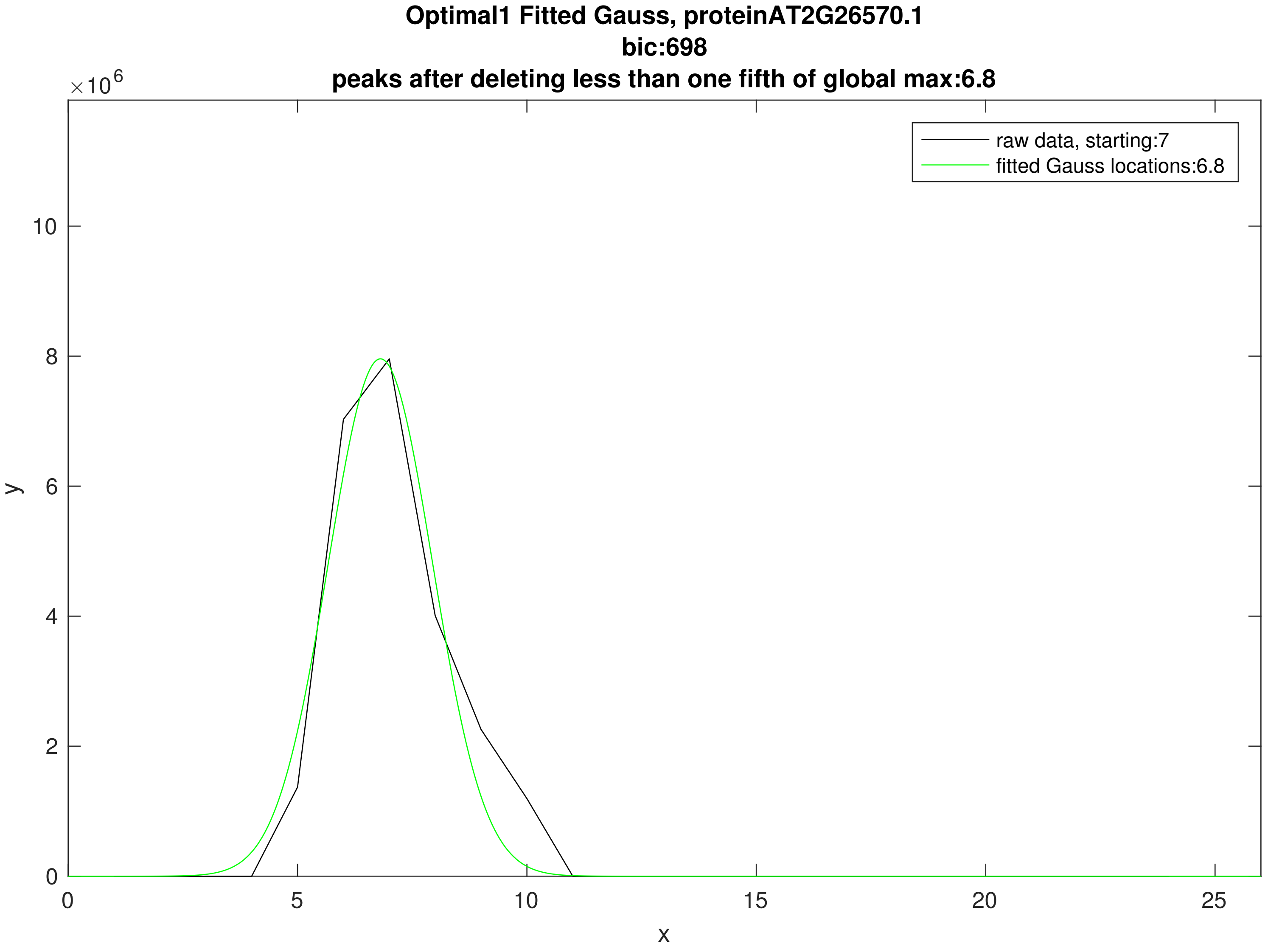
Optimal1 Fitted Gauss, proteinAT2G26570.1
bic:698
peaks after deleting less than one fifth of global max:6.8
×10^6
raw data, starting:7
fitted Gauss locations:6.8
y
x
0
2
4
6
8
10
0
5
10
15
20
25

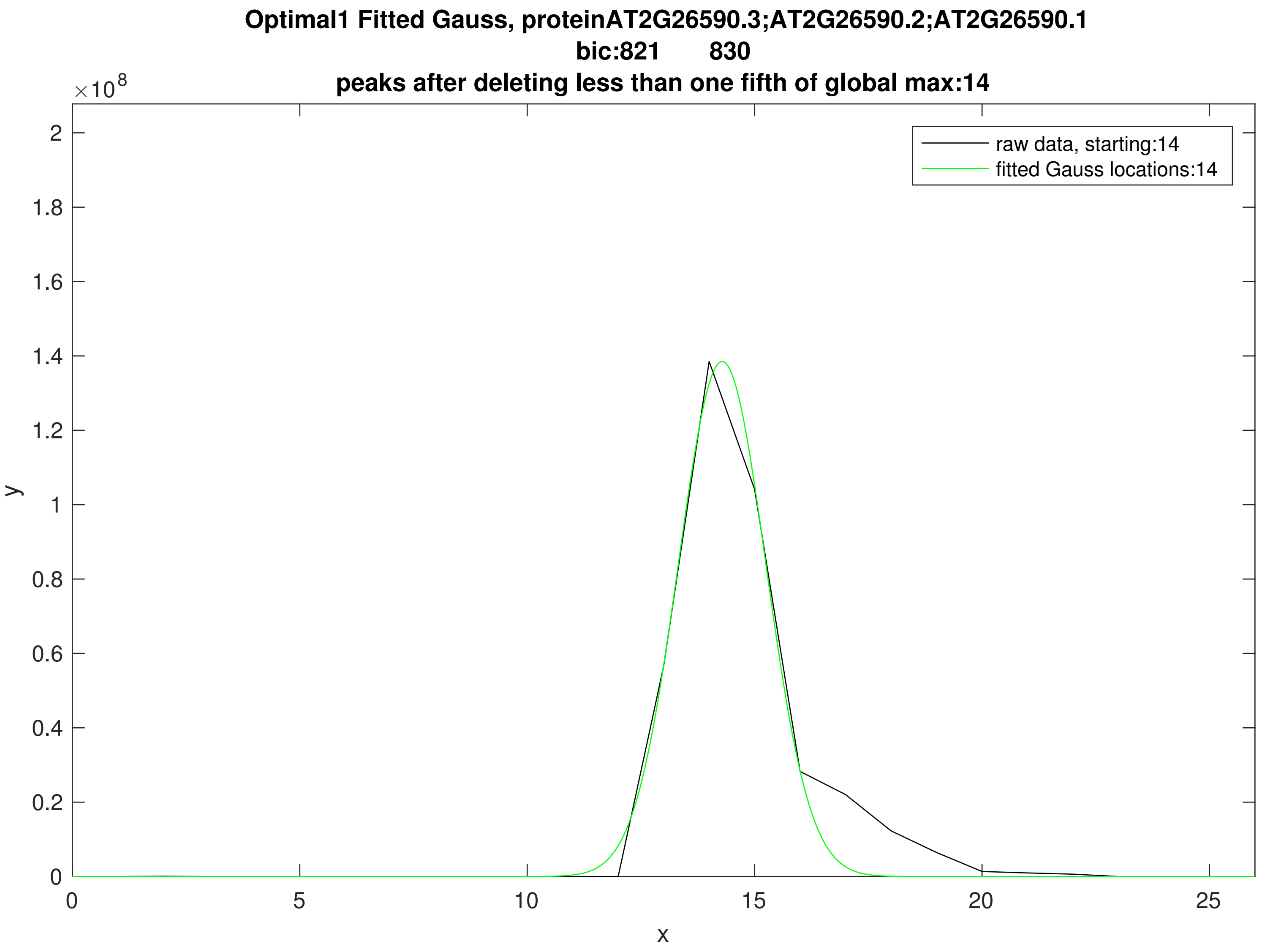
Optimal1 Fitted Gauss, proteinAT2G26590.3;AT2G26590.2;AT2G26590.1
bic:821 830
peaks after deleting less than one fifth of global max:14
×10^8
raw data, starting:14
fitted Gauss locations:14
y
x
0
0.2
0.4
0.6
0.8
1
1.2
1.4
1.6
1.8
2
0
5
10
15
20
25

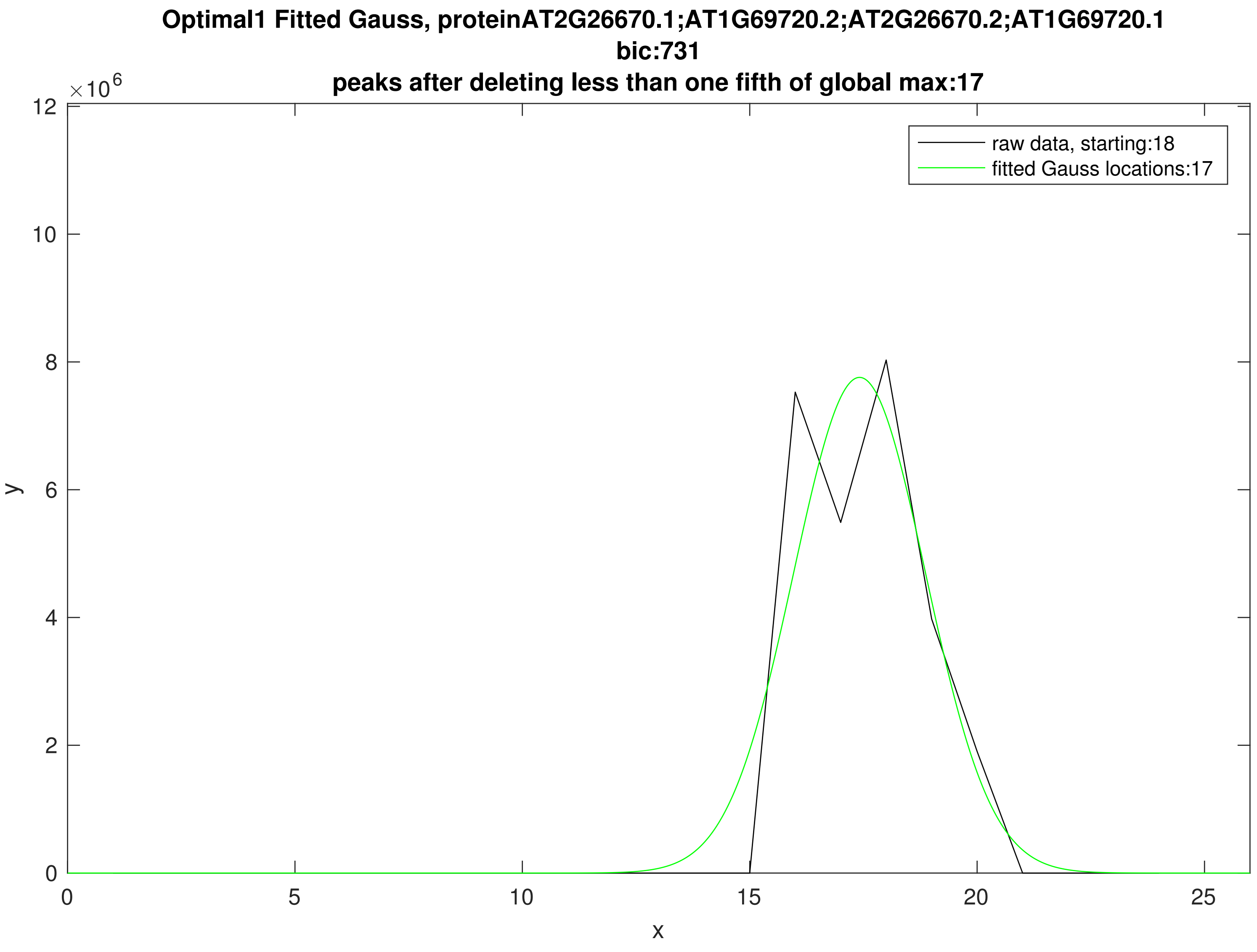
Optimal1 Fitted Gauss, proteinAT2G26670.1;AT1G69720.2;AT2G26670.2;AT1G69720.1
bic:731
peaks after deleting less than one fifth of global max:17
×10^6
12
10
8
6
4
2
0
y
0
5
10
15
20
25
x
raw data, starting:18
fitted Gauss locations:17

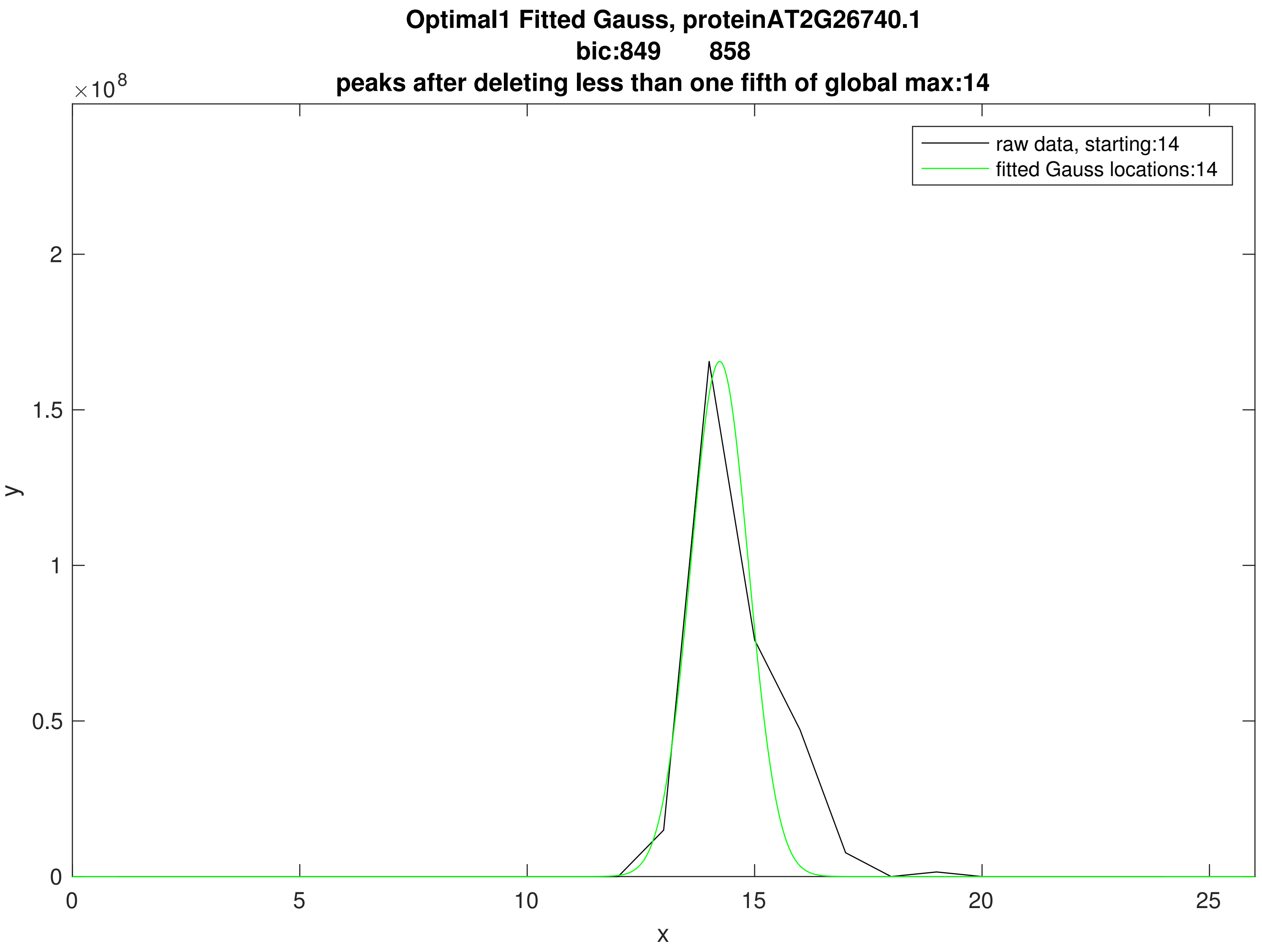

Optimal1 Fitted Gauss, proteinAT2G26740.1
bic:849 858
peaks after deleting less than one fifth of global max:14
×10^8
raw data, starting:14
fitted Gauss locations:14
y
x
0
0.5
1
1.5
2
0
5
10
15
20
25

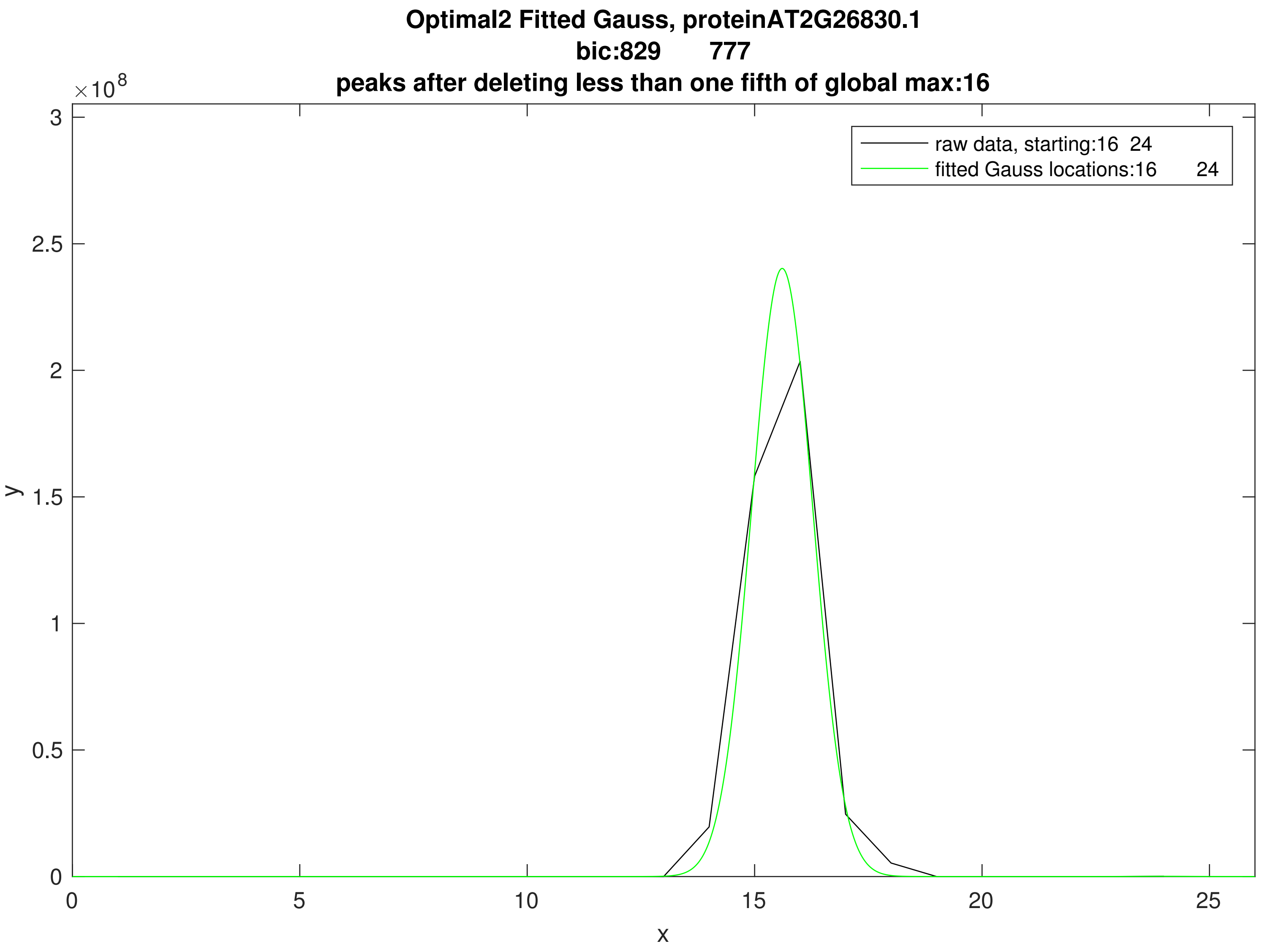
Optimal2 Fitted Gauss, proteinAT2G26830.1
bic:829 777
peaks after deleting less than one fifth of global max:16
raw data, starting:16 24
fitted Gauss locations:16 24
$\times10^8$
y
x
0
0.5
1
1.5
2
2.5
3
5
10
15
20
25

#### Optimal1 Fitted Gauss, proteinAT2G27020.1
#### bic:929 938
#### peaks after deleting less than one fifth of global max:5

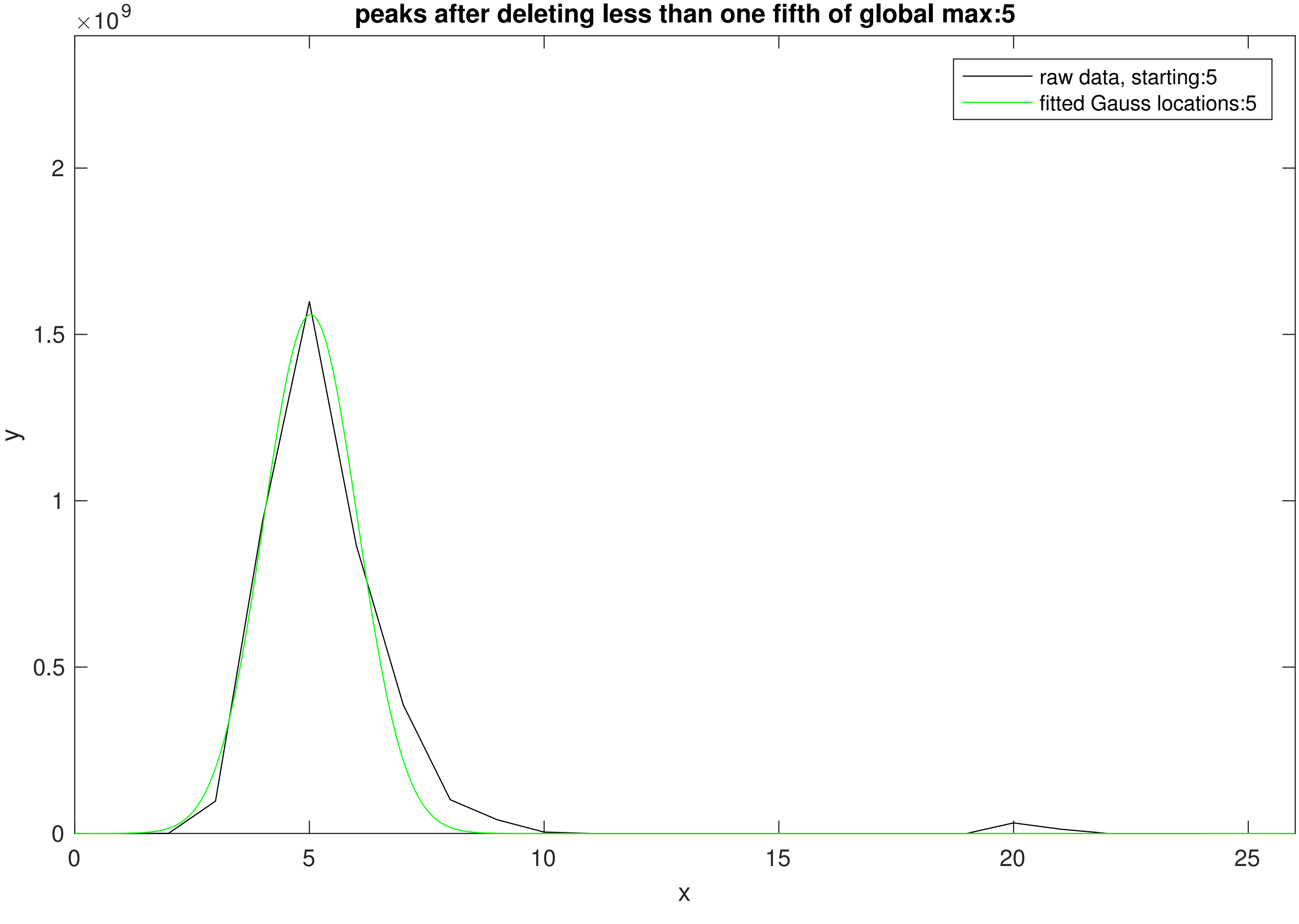

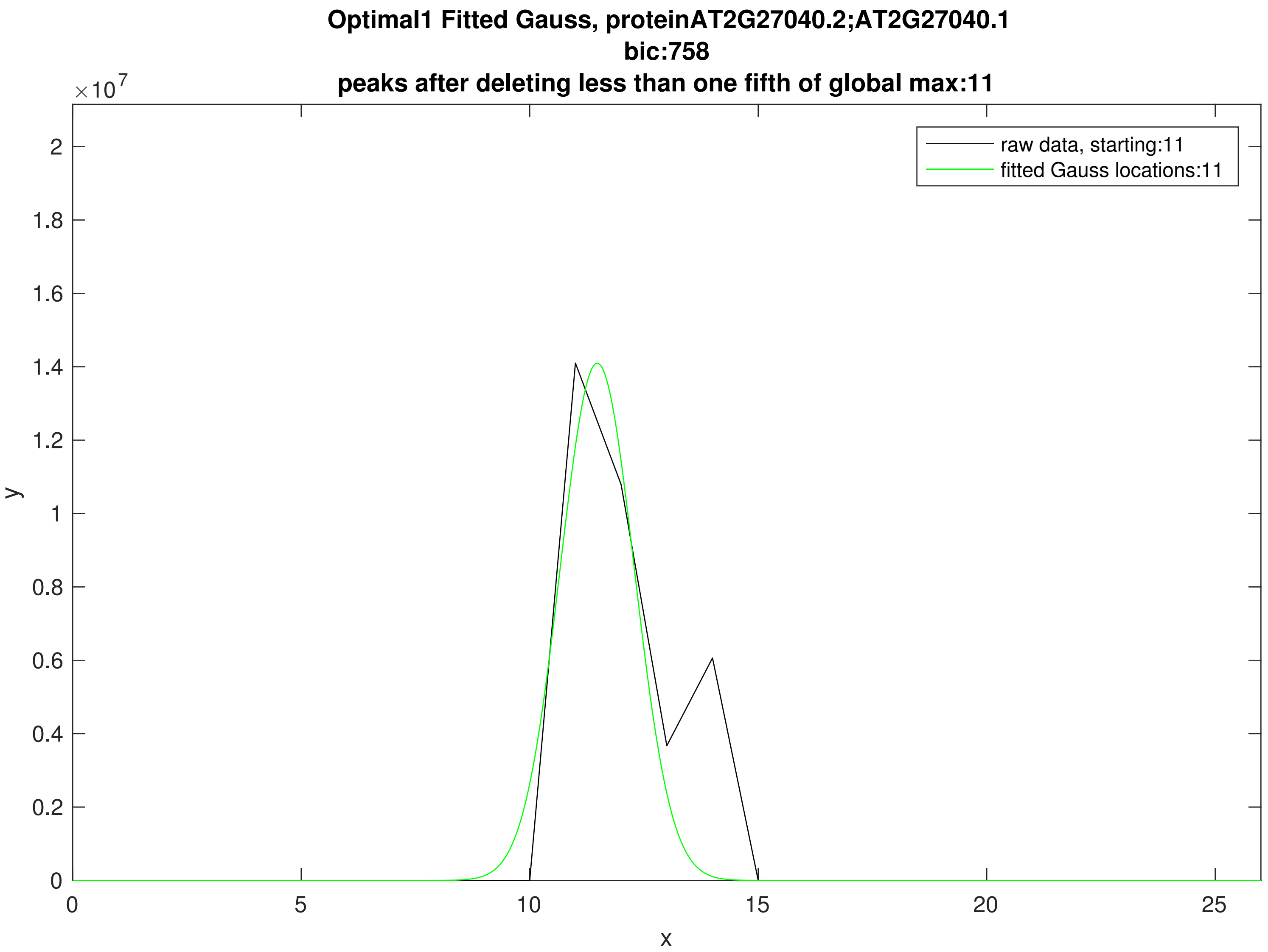
Optimal1 Fitted Gauss, proteinAT2G27040.2;AT2G27040.1
bic:758
peaks after deleting less than one fifth of global max:11
×10^7
2
1.8
1.6
1.4
1.2
1
0.8
0.6
0.4
0.2
0
y
0
5
10
15
20
25
x
raw data, starting:11
fitted Gauss locations:11

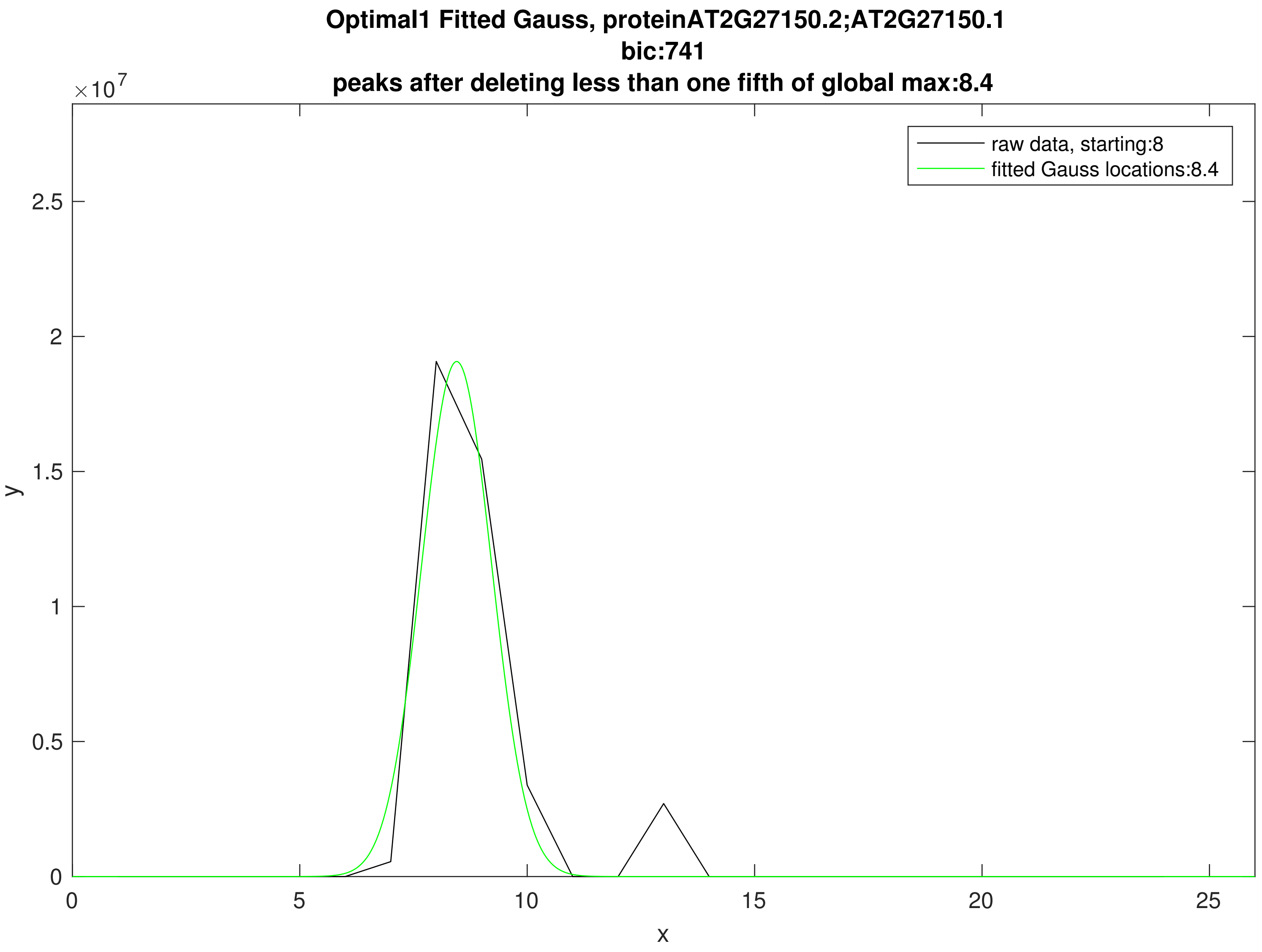
Optimal1 Fitted Gauss, proteinAT2G27150.2;AT2G27150.1
bic:741
peaks after deleting less than one fifth of global max:8.4
×10^7
raw data, starting:8
fitted Gauss locations:8.4
y
x
0
0.5
1
1.5
2
2.5
5
10
15
20
25

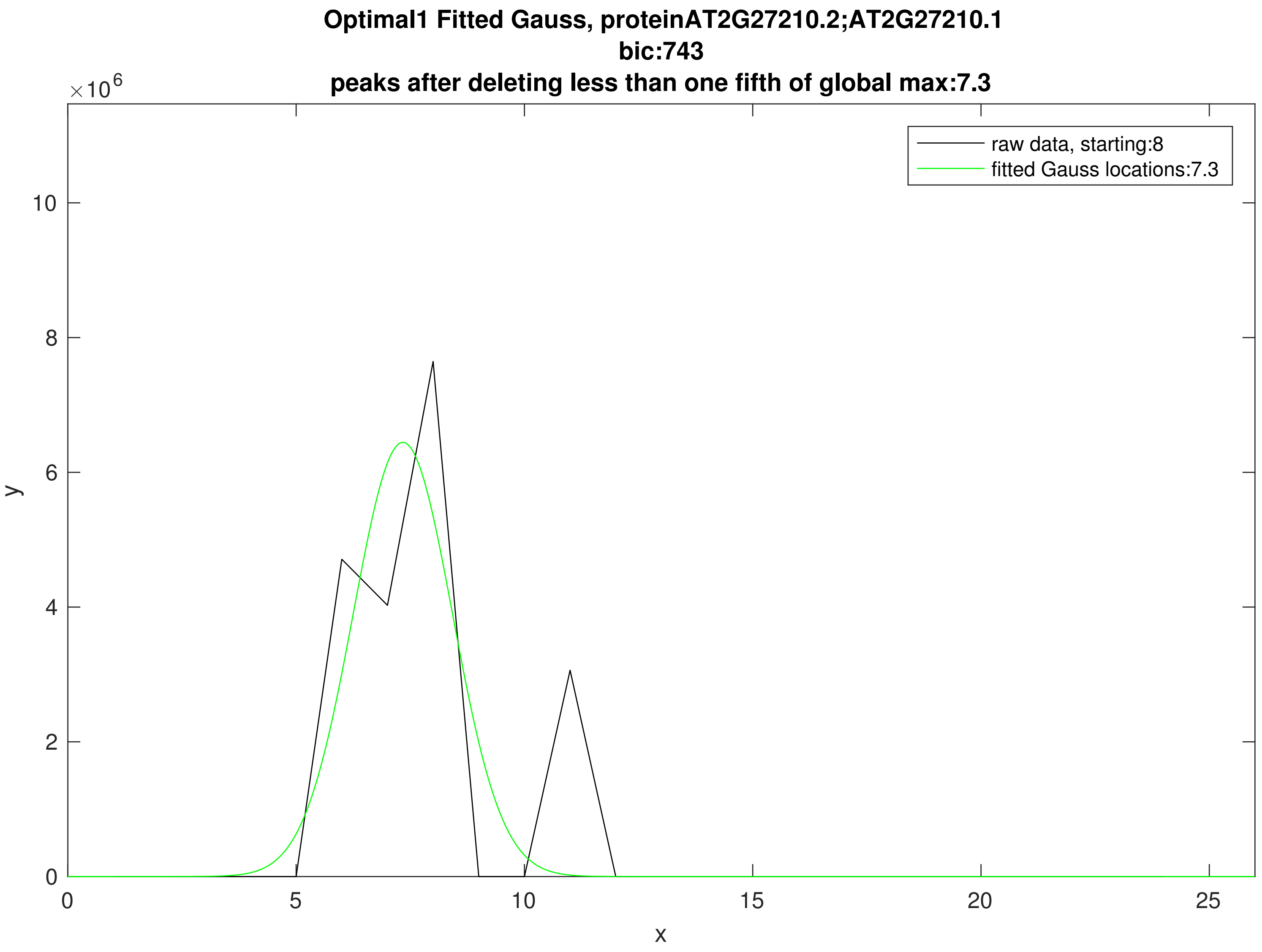

Optimal1 Fitted Gauss, proteinAT2G27210.2;AT2G27210.1
bic:743
peaks after deleting less than one fifth of global max:7.3
×10^6
raw data, starting:8
fitted Gauss locations:7.3
y
x
0
2
4
6
8
10
0
5
10
15
20
25

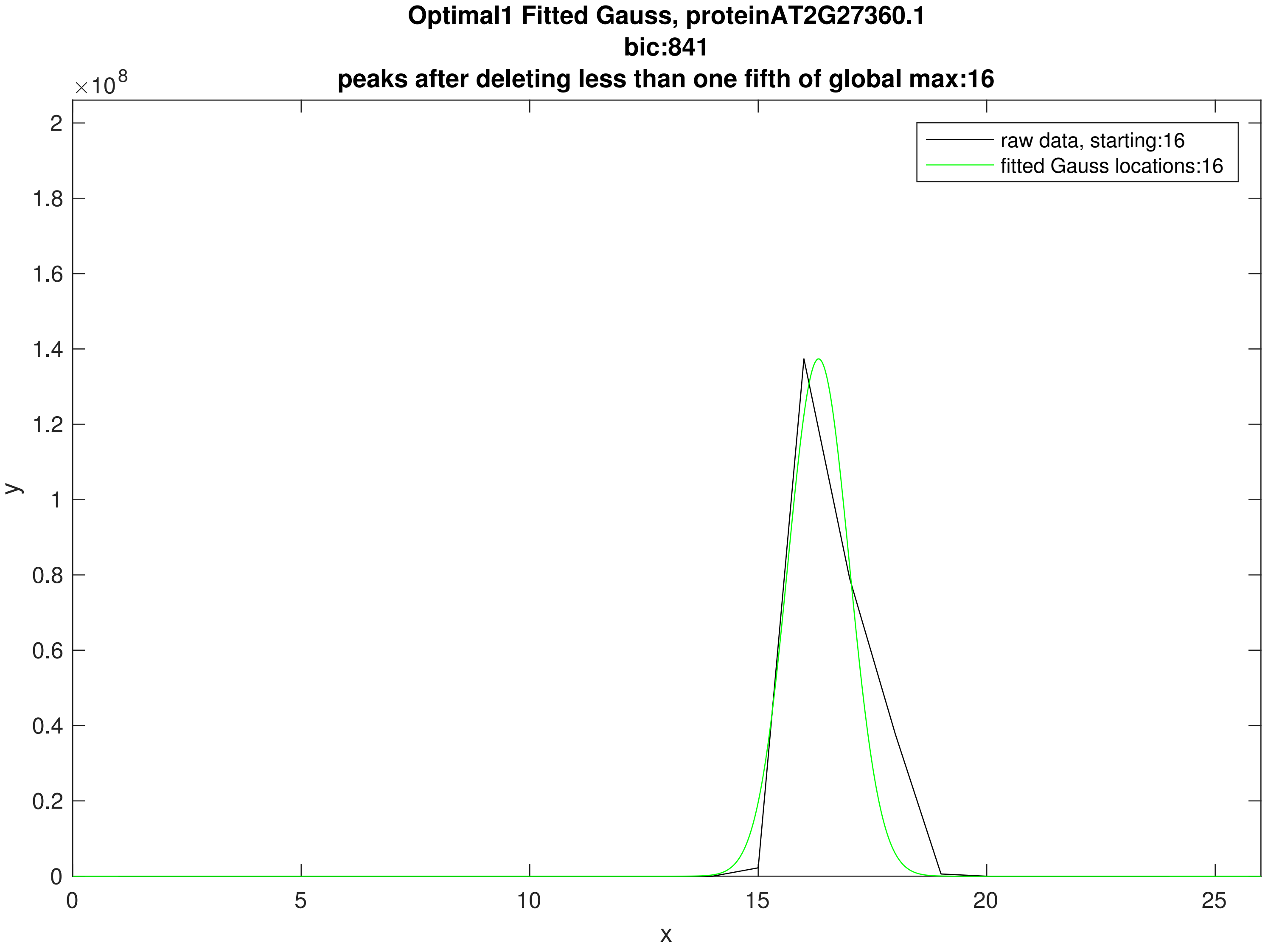
Optimal1 Fitted Gauss, proteinAT2G27360.1
bic:841
peaks after deleting less than one fifth of global max:16
×10^8
2
1.8
1.6
1.4
1.2
1
0.8
0.6
0.4
0.2
0
y
0
5
10
15
20
25
x
raw data, starting:16
fitted Gauss locations:16

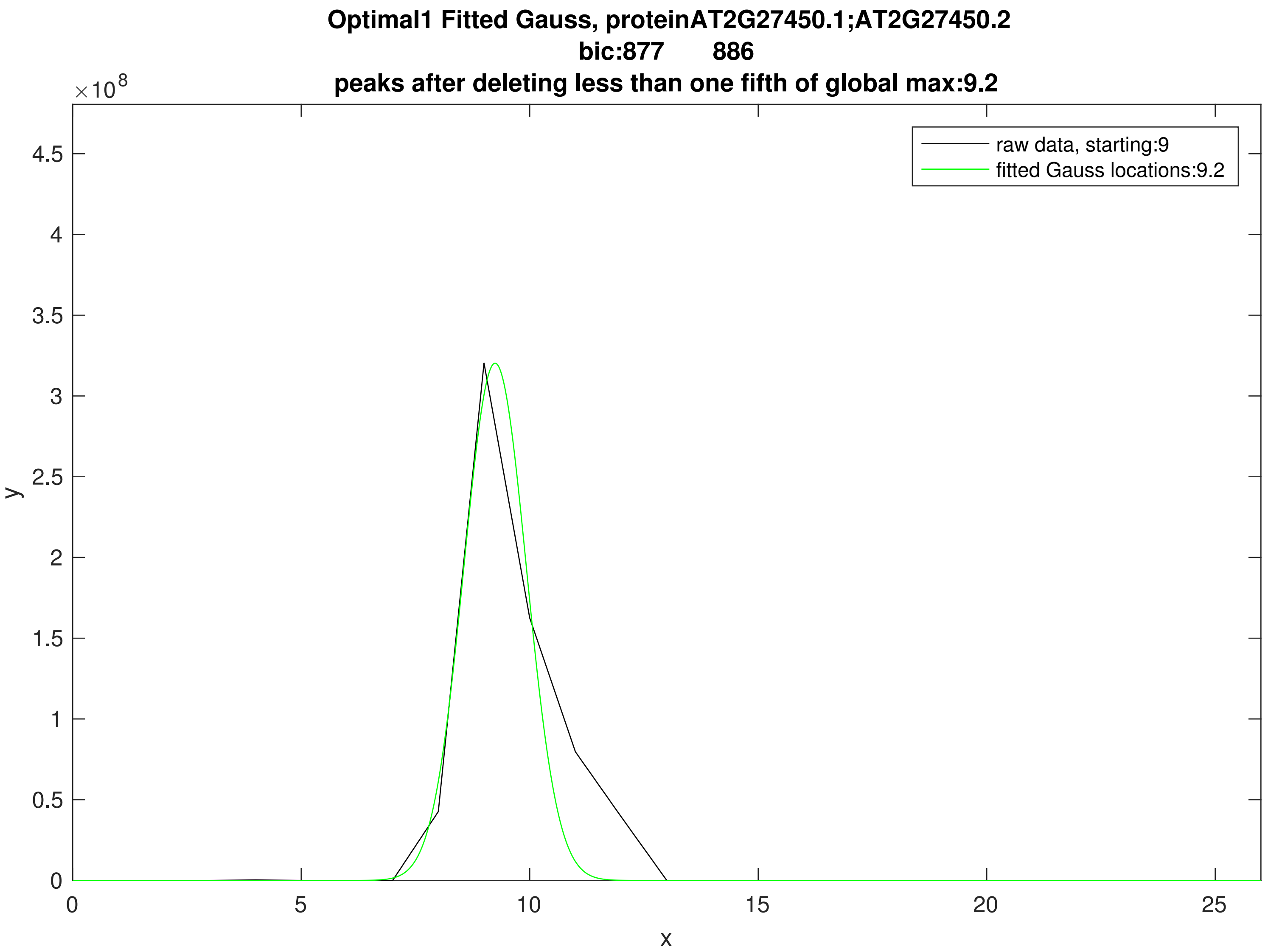

Optimal1 Fitted Gauss, proteinAT2G27450.1;AT2G27450.2
bic:877 886
peaks after deleting less than one fifth of global max:9.2
×10^8
4.5
4
3.5
3
2.5
2
1.5
1
0.5
0
y
0
5
10
15
20
25
x
raw data, starting:9
fitted Gauss locations:9.2

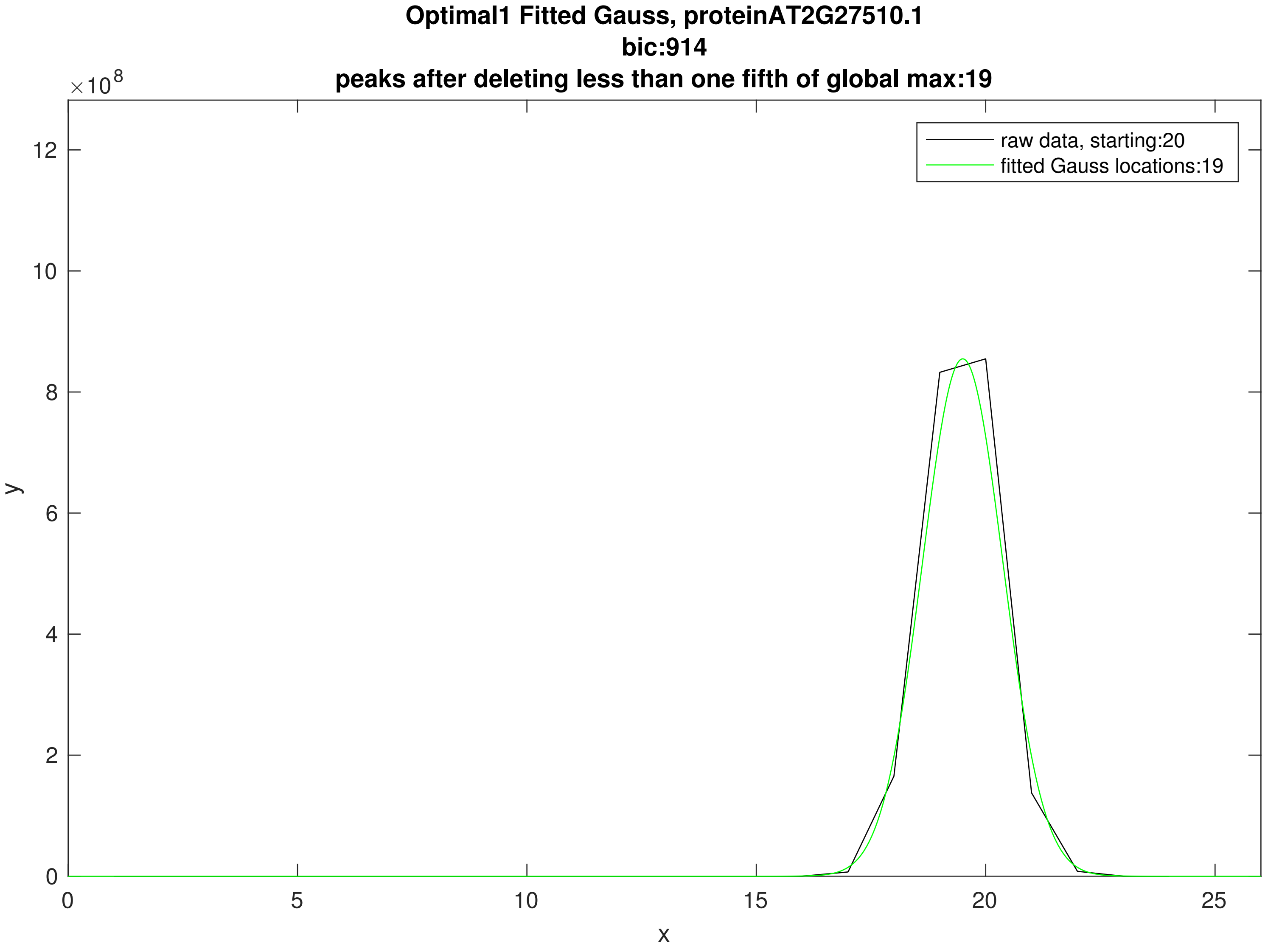

Optimal1 Fitted Gauss, proteinAT2G27510.1
bic:914
peaks after deleting less than one fifth of global max:19
×10^8
raw data, starting:20
fitted Gauss locations:19
y
x
0
2
4
6
8
10
12
0
5
10
15
20
25

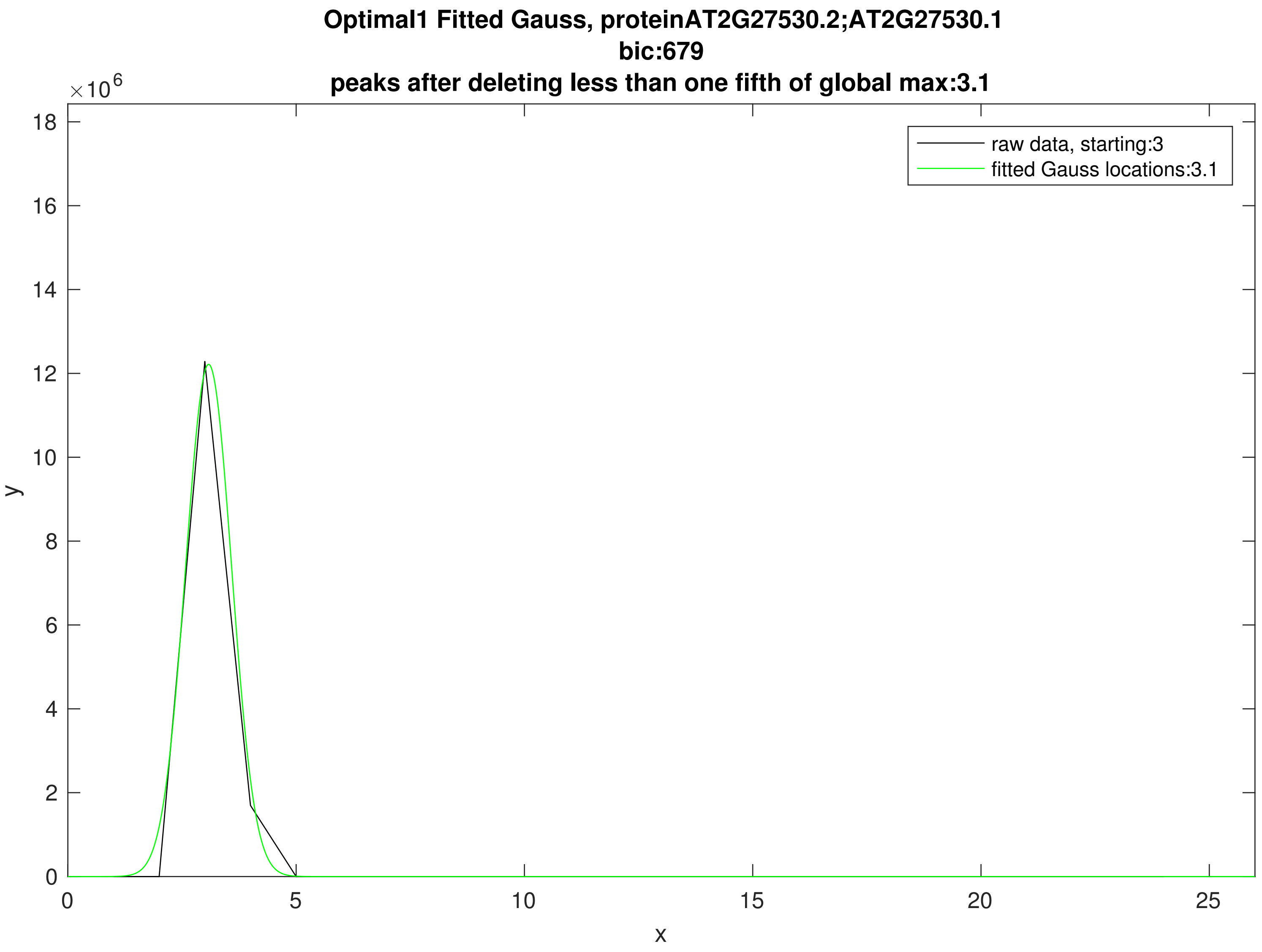
Optimal1 Fitted Gauss, proteinAT2G27530.2;AT2G27530.1
bic:679
peaks after deleting less than one fifth of global max:3.1
raw data, starting:3
fitted Gauss locations:3.1
y
x
×10^6
0
2
4
6
8
10
12
14
16
18
0
5
10
15
20
25

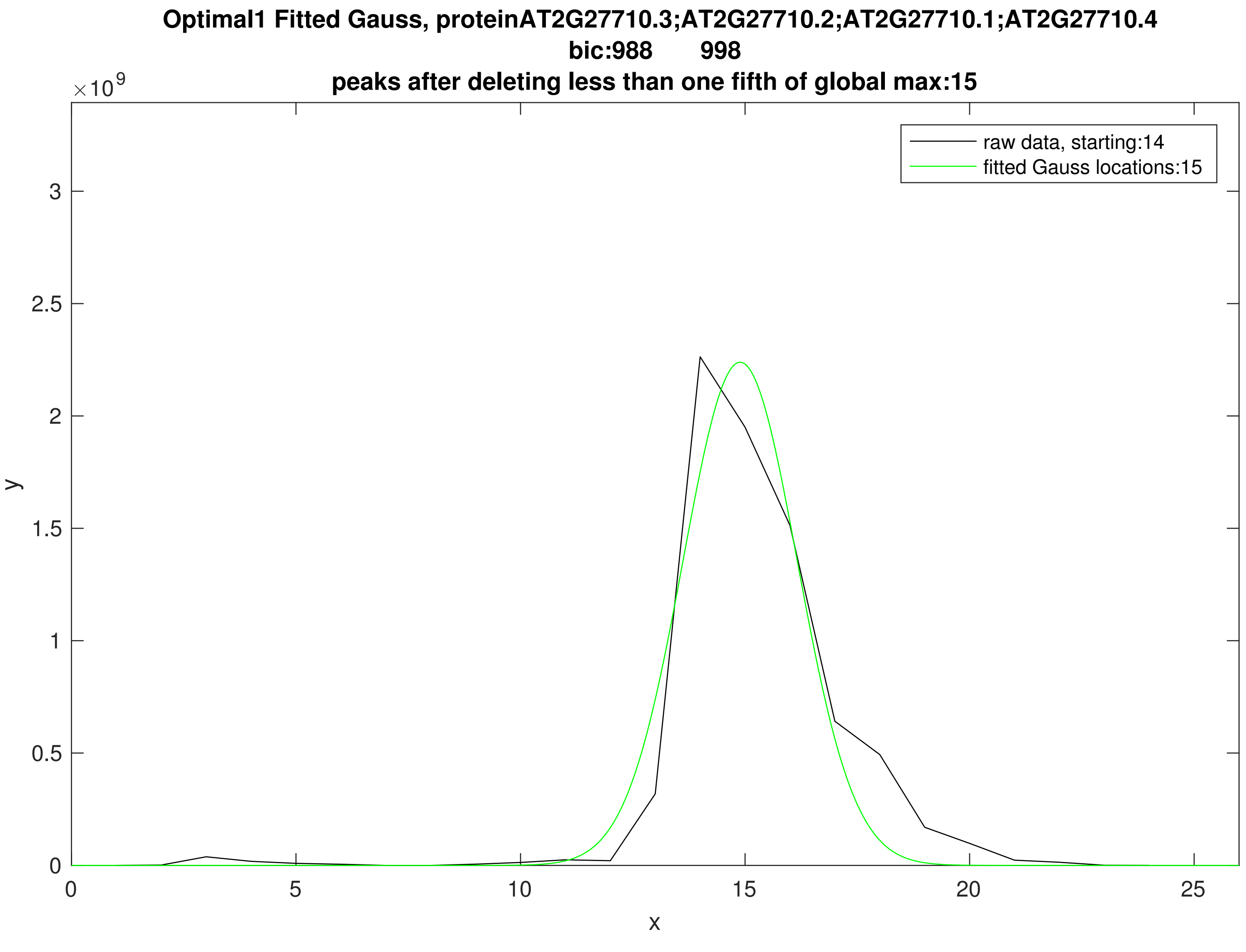
Optimal1 Fitted Gauss, proteinAT2G27710.3;AT2G27710.2;AT2G27710.1;AT2G27710.4
bic:988 998
peaks after deleting less than one fifth of global max:15
×10^9
raw data, starting:14
fitted Gauss locations:15
y
x
0
0.5
1
1.5
2
2.5
3
0
5
10
15
20
25

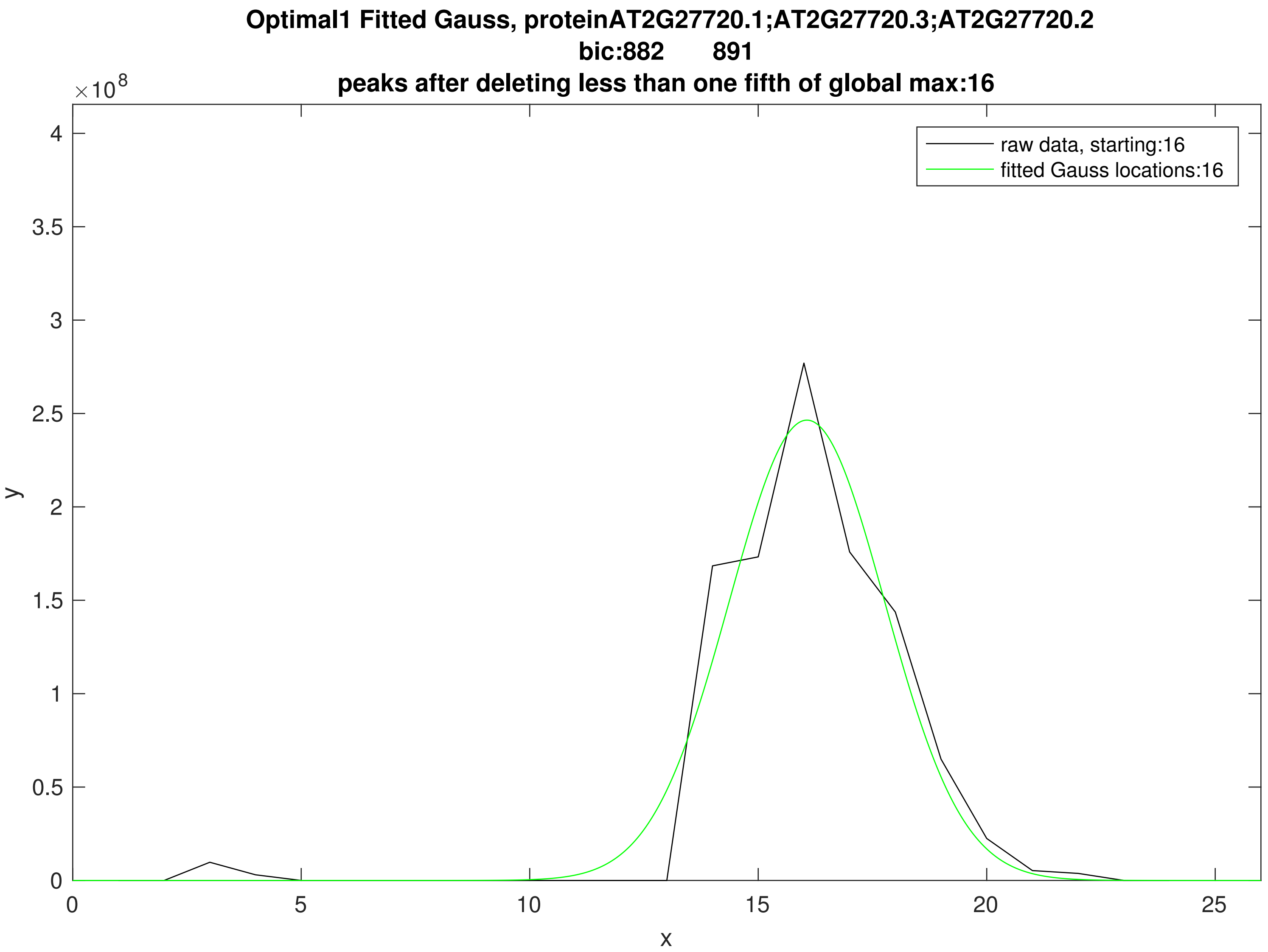

Optimal1 Fitted Gauss, proteinAT2G27720.1;AT2G27720.3;AT2G27720.2
bic:882 891
peaks after deleting less than one fifth of global max:16
×10^8
raw data, starting:16
fitted Gauss locations:16
y
x
0
0.5
1
1.5
2
2.5
3
3.5
4
0
5
10
15
20
25

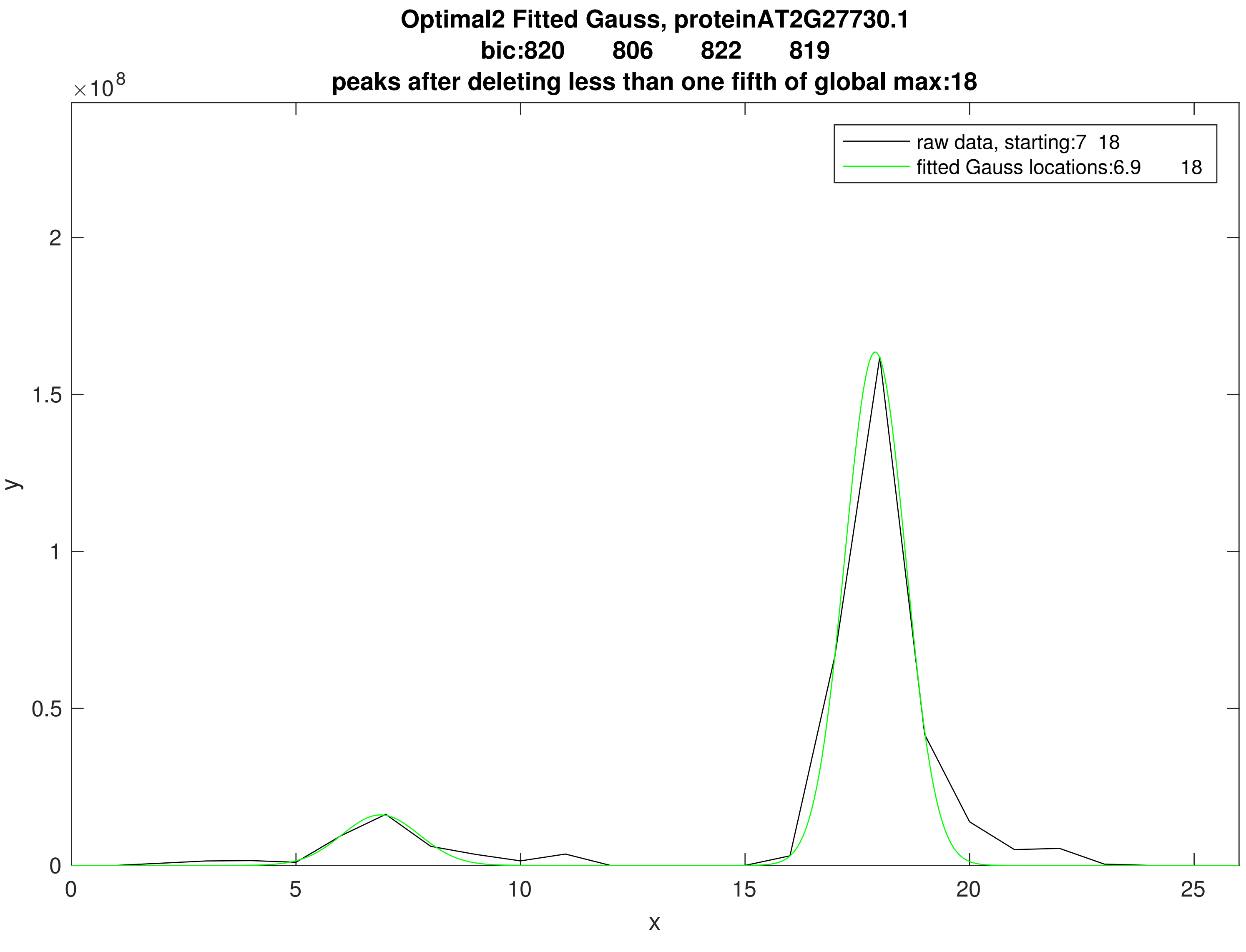
Optimal2 Fitted Gauss, proteinAT2G27730.1
bic:820 806 822 819
peaks after deleting less than one fifth of global max:18
×10^8
raw data, starting:7 18
fitted Gauss locations:6.9 18
y
0
0.5
1
1.5
2
x
0
5
10
15
20
25

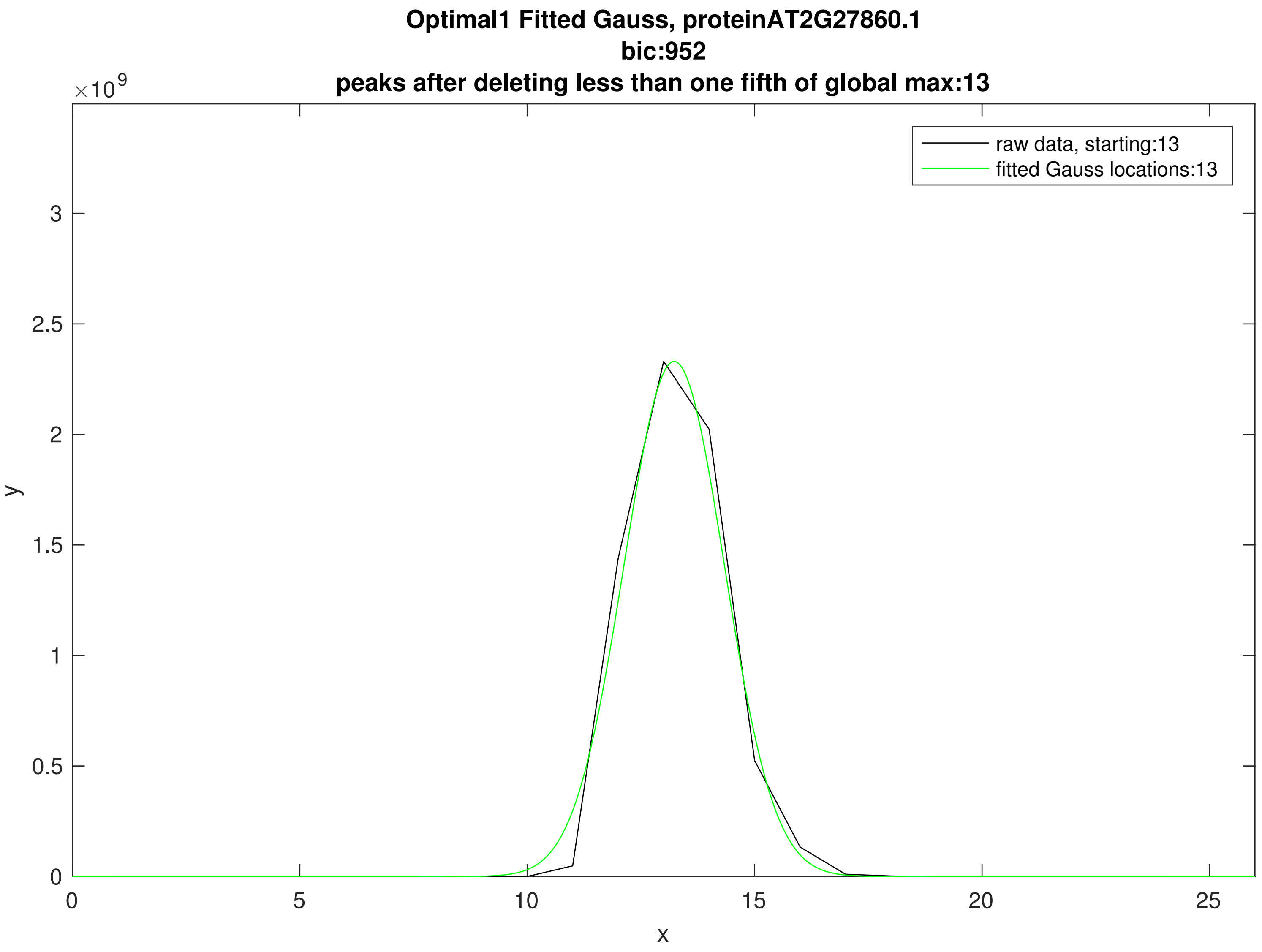
Optimal1 Fitted Gauss, proteinAT2G27860.1
bic:952
peaks after deleting less than one fifth of global max:13
×10^9
raw data, starting:13
fitted Gauss locations:13
y
x
0
0.5
1
1.5
2
2.5
3
0
5
10
15
20
25

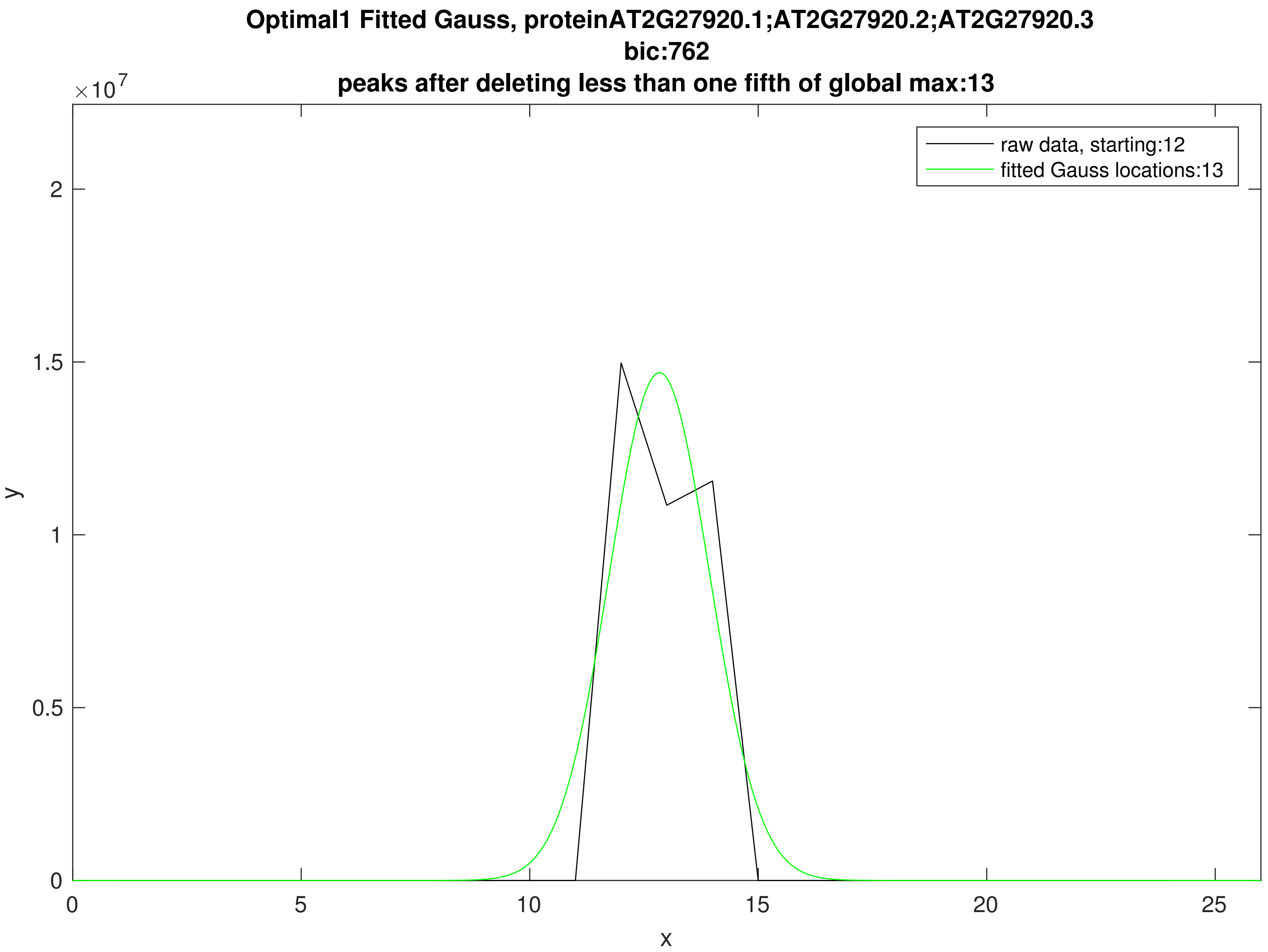

Optimal1 Fitted Gauss, proteinAT2G27920.1;AT2G27920.2;AT2G27920.3
bic:762
peaks after deleting less than one fifth of global max:13
×10^7
raw data, starting:12
fitted Gauss locations:13
2
1.5
1
0.5
0
y
0
5
10
15
20
25
x

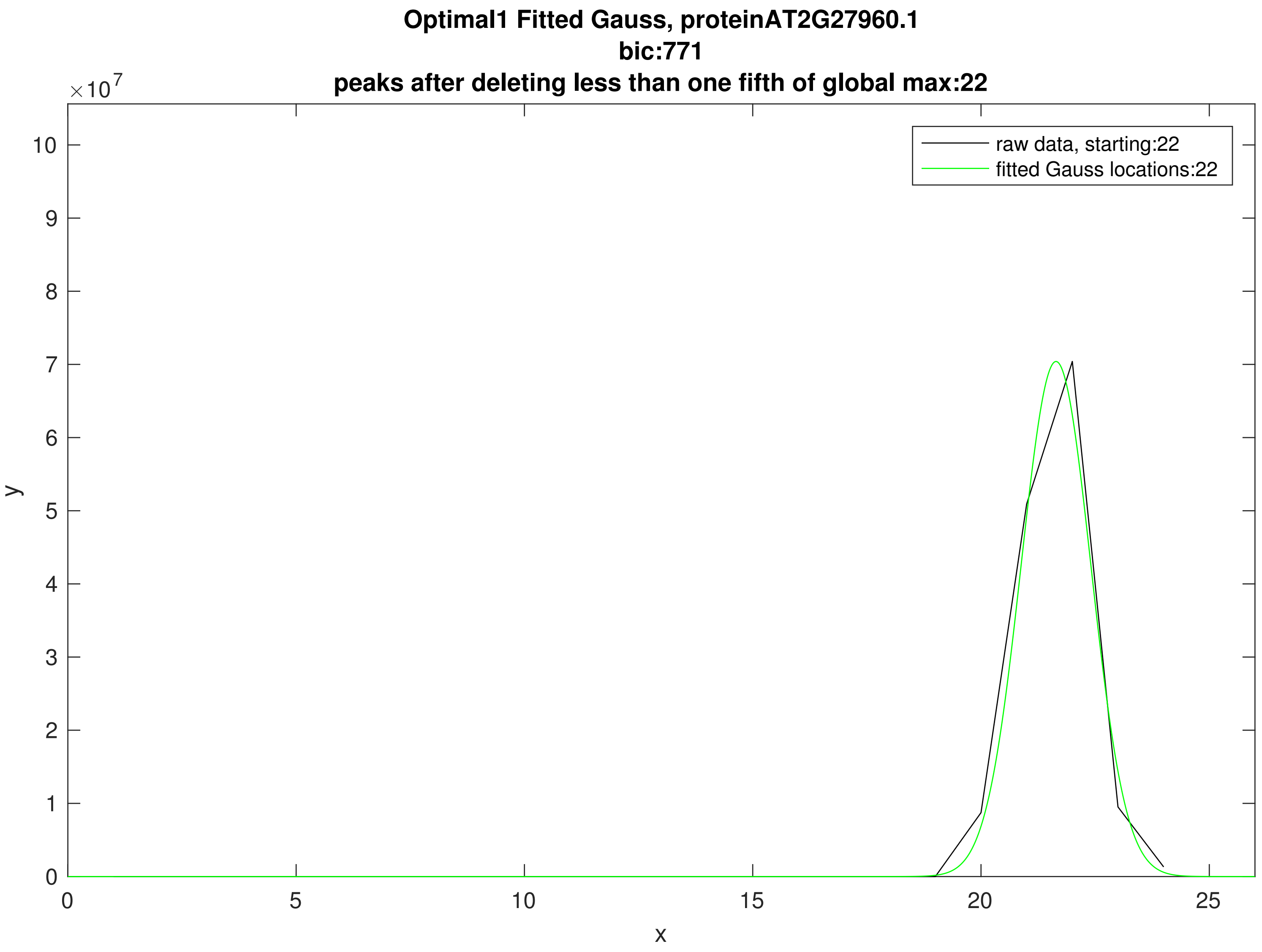
Optimal1 Fitted Gauss, proteinAT2G27960.1
bic:771
peaks after deleting less than one fifth of global max:22
×10^7
raw data, starting:22
fitted Gauss locations:22
y
x
0
1
2
3
4
5
6
7
8
9
10
0
5
10
15
20
25

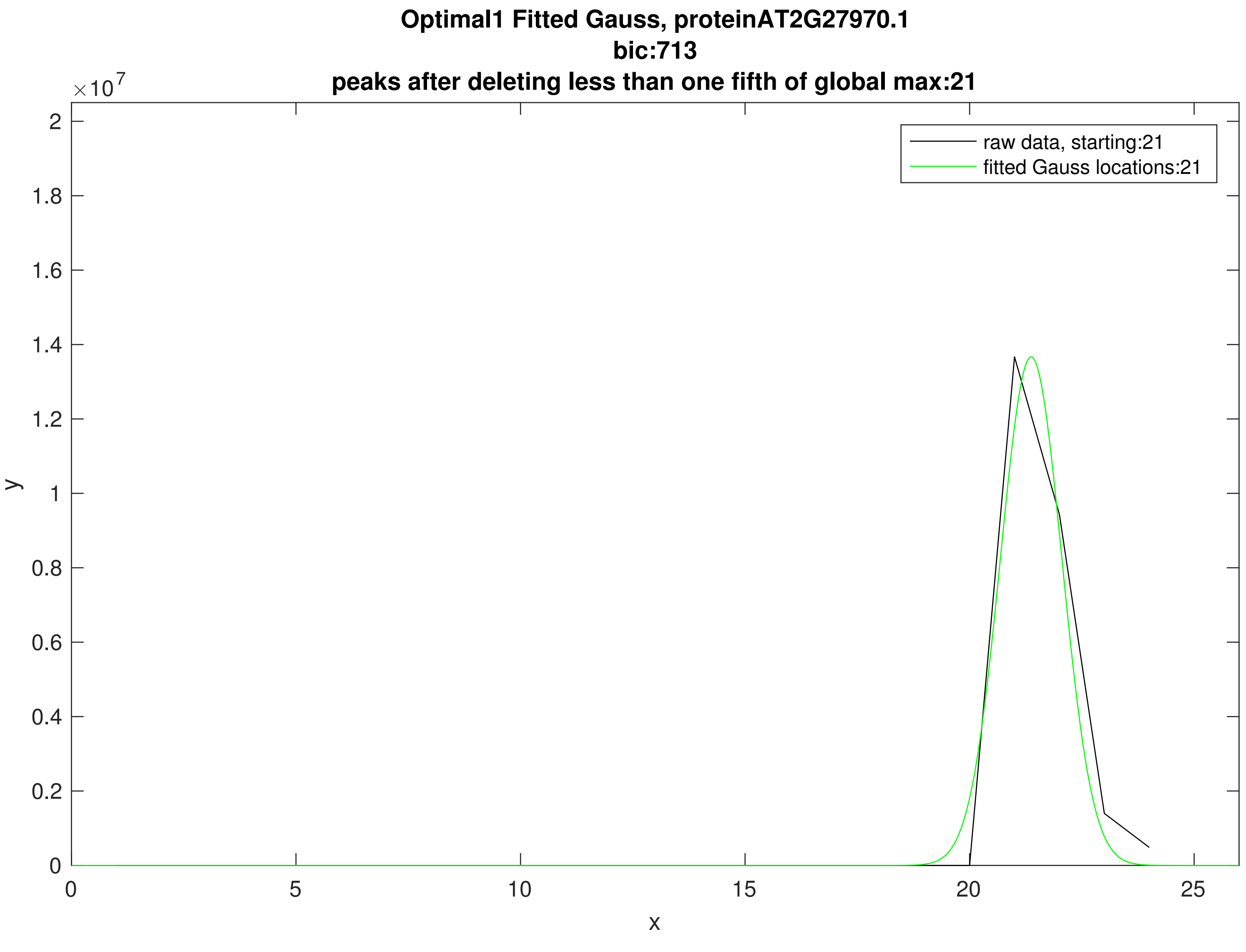
Optimal1 Fitted Gauss, proteinAT2G27970.1
bic:713
peaks after deleting less than one fifth of global max:21
×10^7
2
1.8
1.6
1.4
1.2
1
0.8
0.6
0.4
0.2
0
y
0
5
10
15
20
25
x
raw data, starting:21
fitted Gauss locations:21

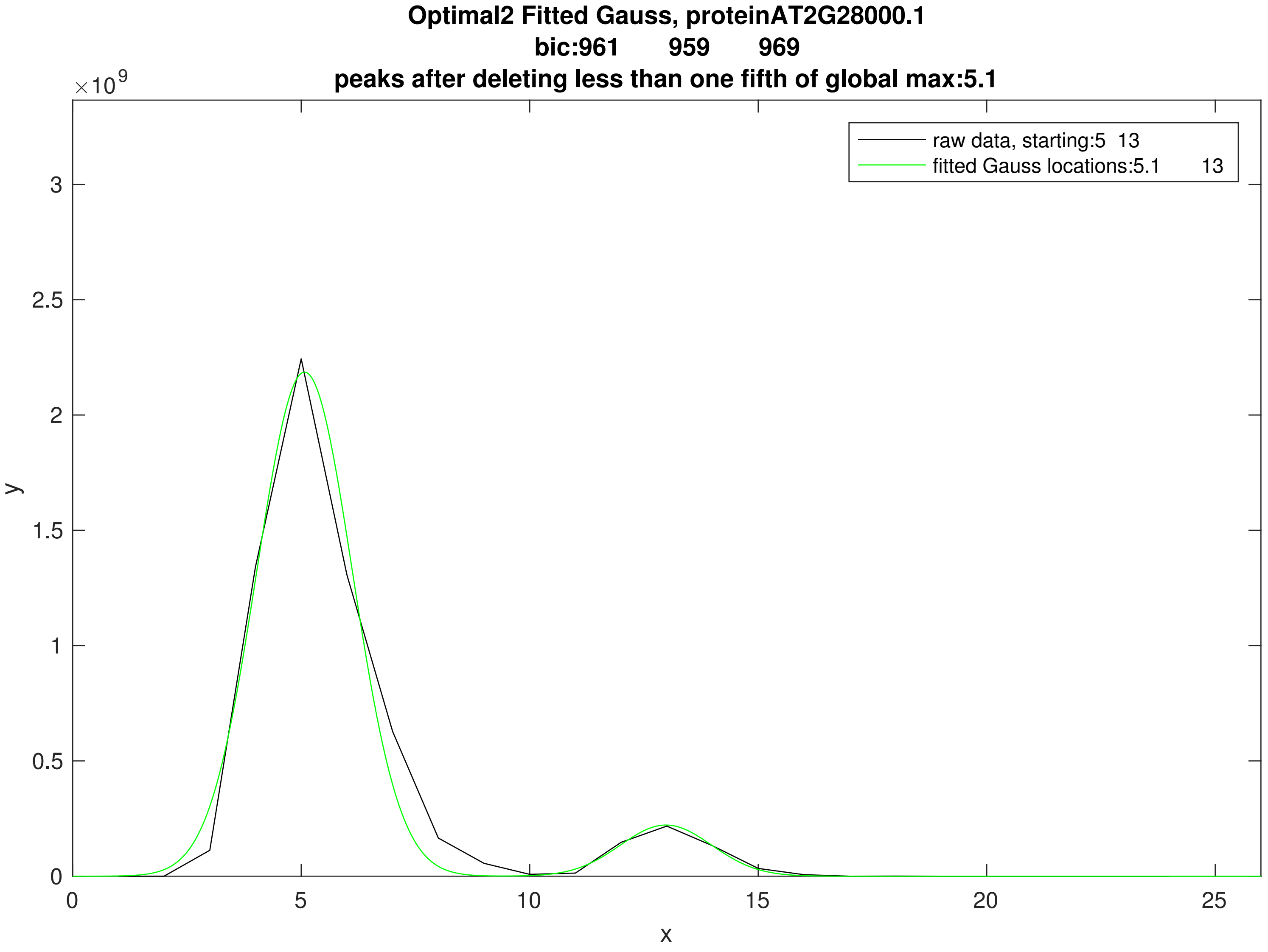

Optimal2 Fitted Gauss, proteinAT2G28000.1
bic:961 959 969
peaks after deleting less than one fifth of global max:5.1
×10^9
raw data, starting:5 13
fitted Gauss locations:5.1 13
y
x
0
0.5
1
1.5
2
2.5
3
0
5
10
15
20
25

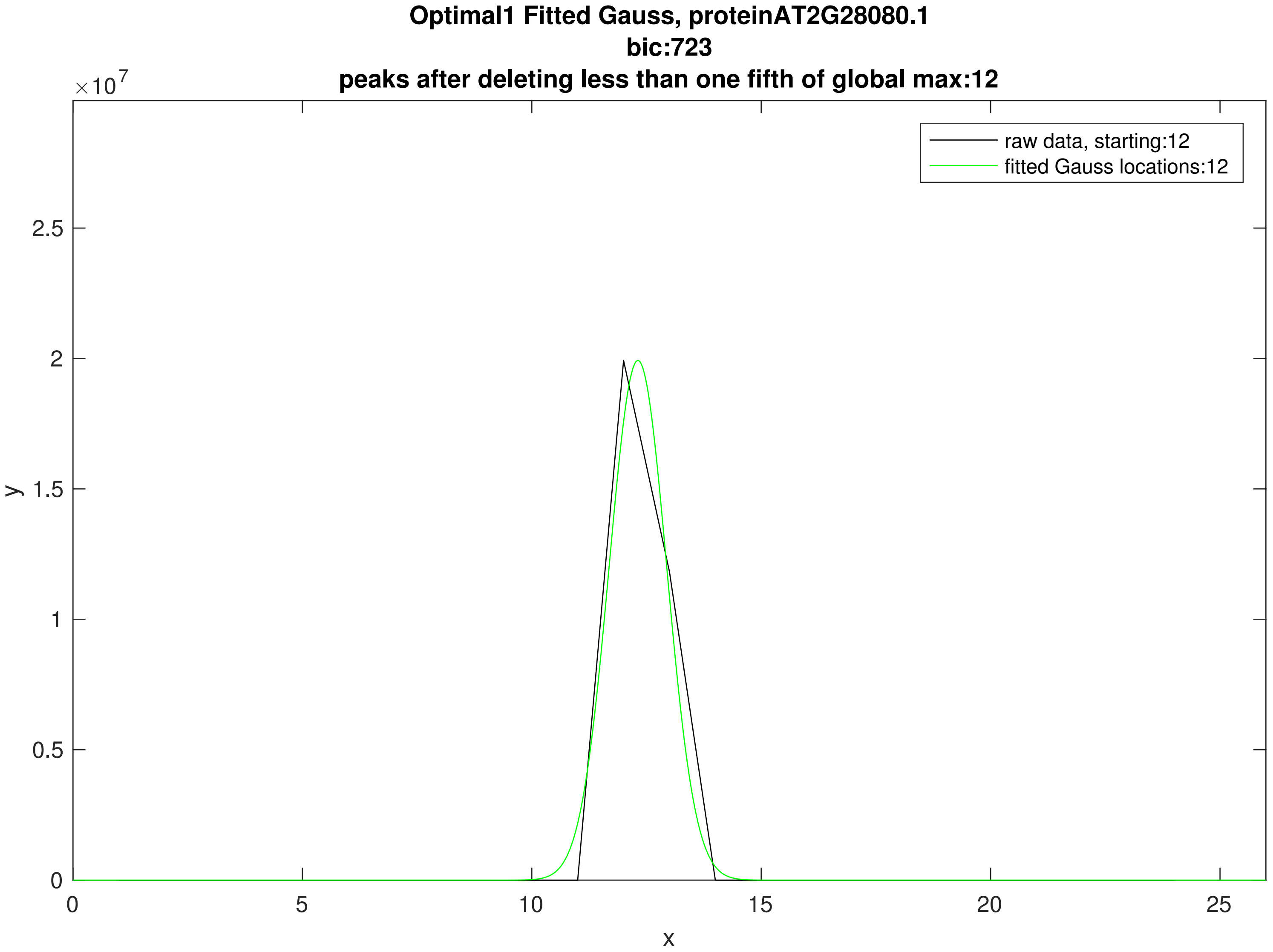

Optimal1 Fitted Gauss, proteinAT2G28080.1
bic:723
peaks after deleting less than one fifth of global max:12
×10^7
raw data, starting:12
fitted Gauss locations:12
y
x
0
0.5
1
1.5
2
2.5
0
5
10
15
20
25

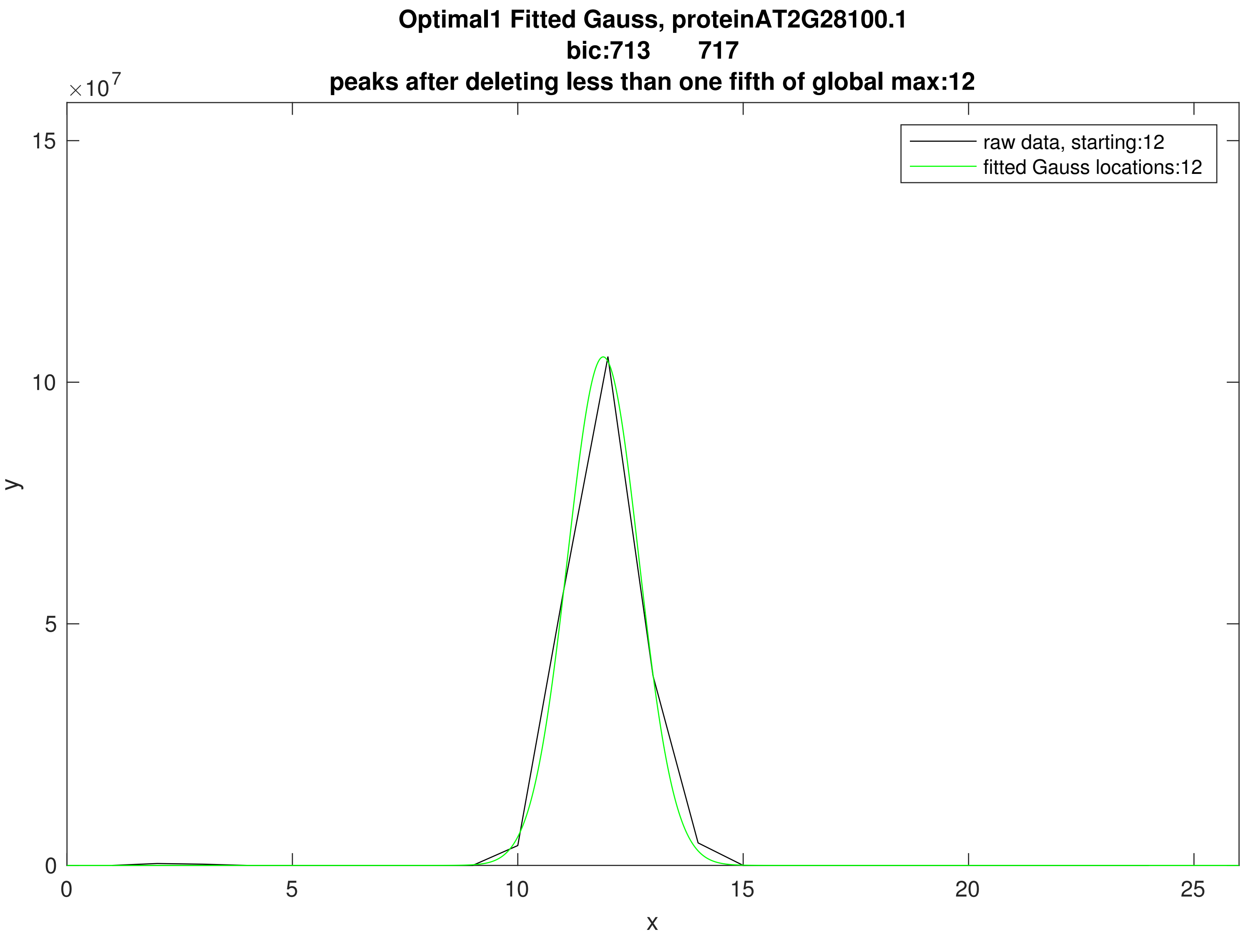
Optimal1 Fitted Gauss, proteinAT2G28100.1
bic:713 717
peaks after deleting less than one fifth of global max:12
×10^7
raw data, starting:12
fitted Gauss locations:12
y
x
0
5
10
15
20
25

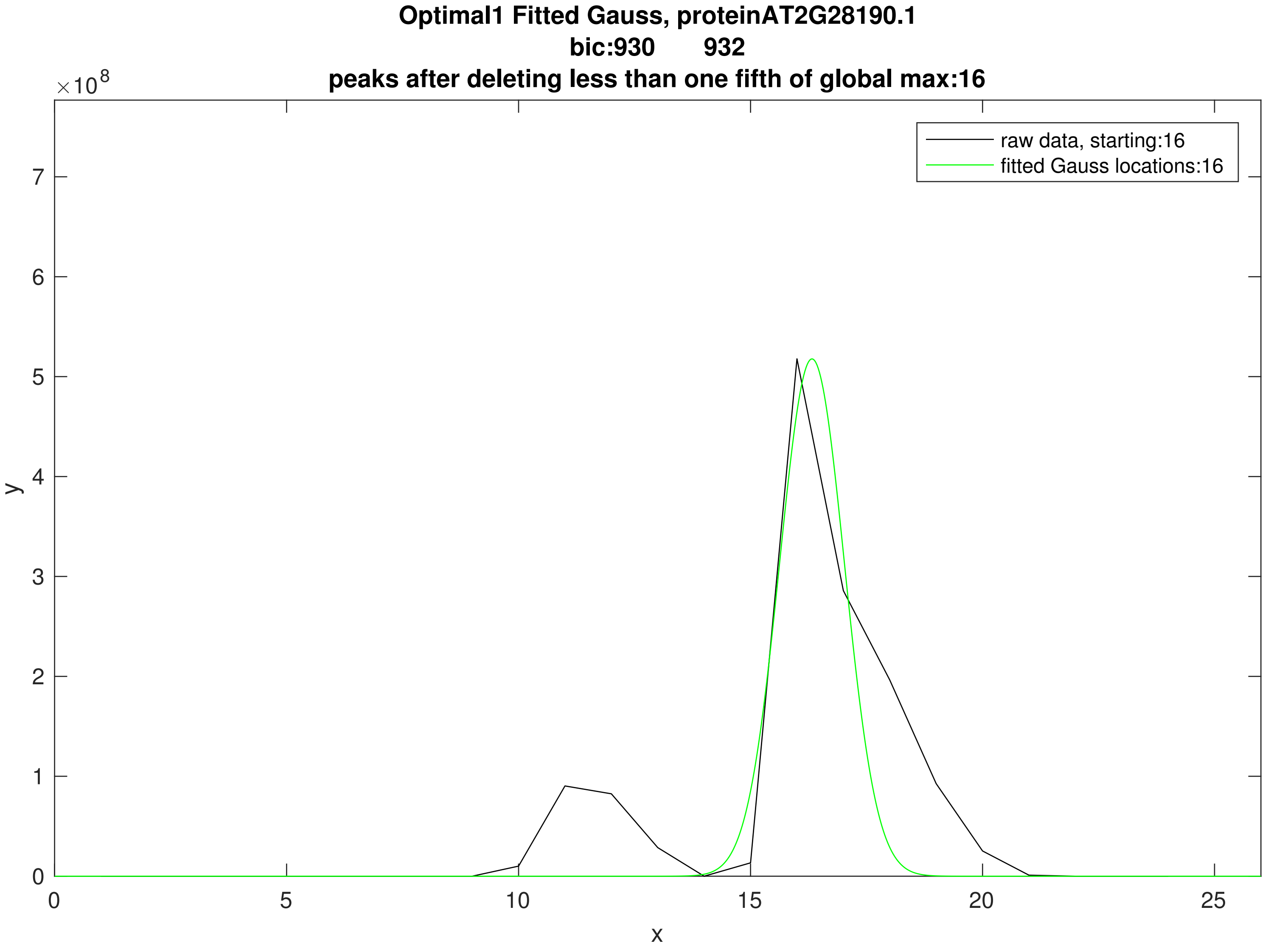
Optimal1 Fitted Gauss, proteinAT2G28190.1
bic:930 932
peaks after deleting less than one fifth of global max:16
×10^8
raw data, starting:16
fitted Gauss locations:16
y
x
0
1
2
3
4
5
6
7
0
5
10
15
20
25

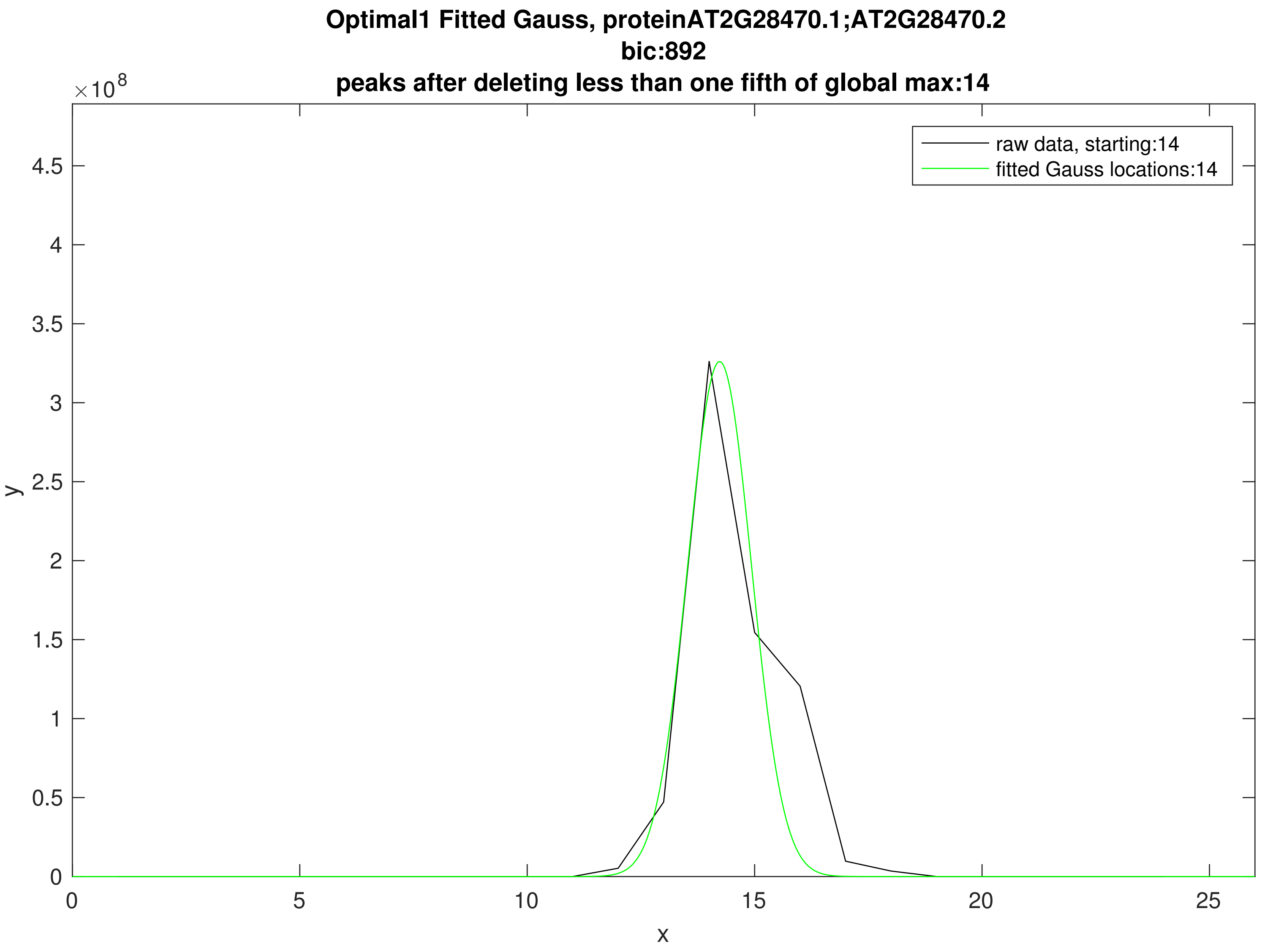
Optimal1 Fitted Gauss, proteinAT2G28470.1;AT2G28470.2
bic:892
peaks after deleting less than one fifth of global max:14
×10^8
4.5
4
3.5
3
2.5
2
1.5
1
0.5
0
y
0
5
10
15
20
25
x
raw data, starting:14
fitted Gauss locations:14

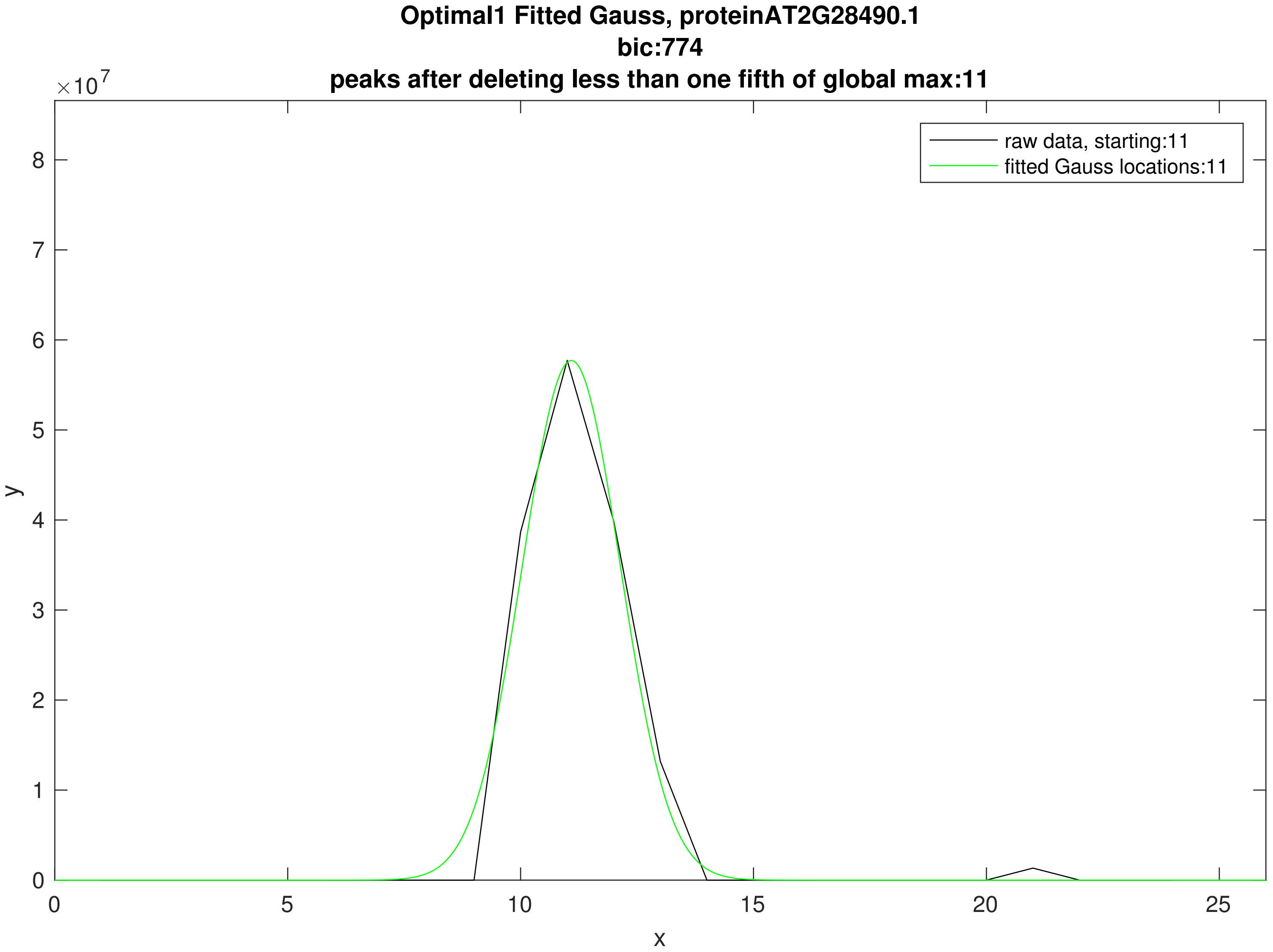
Optimal1 Fitted Gauss, proteinAT2G28490.1
bic:774
peaks after deleting less than one fifth of global max:11
×10^7
raw data, starting:11
fitted Gauss locations:11
y
x
0
1
2
3
4
5
6
7
8
0
5
10
15
20
25

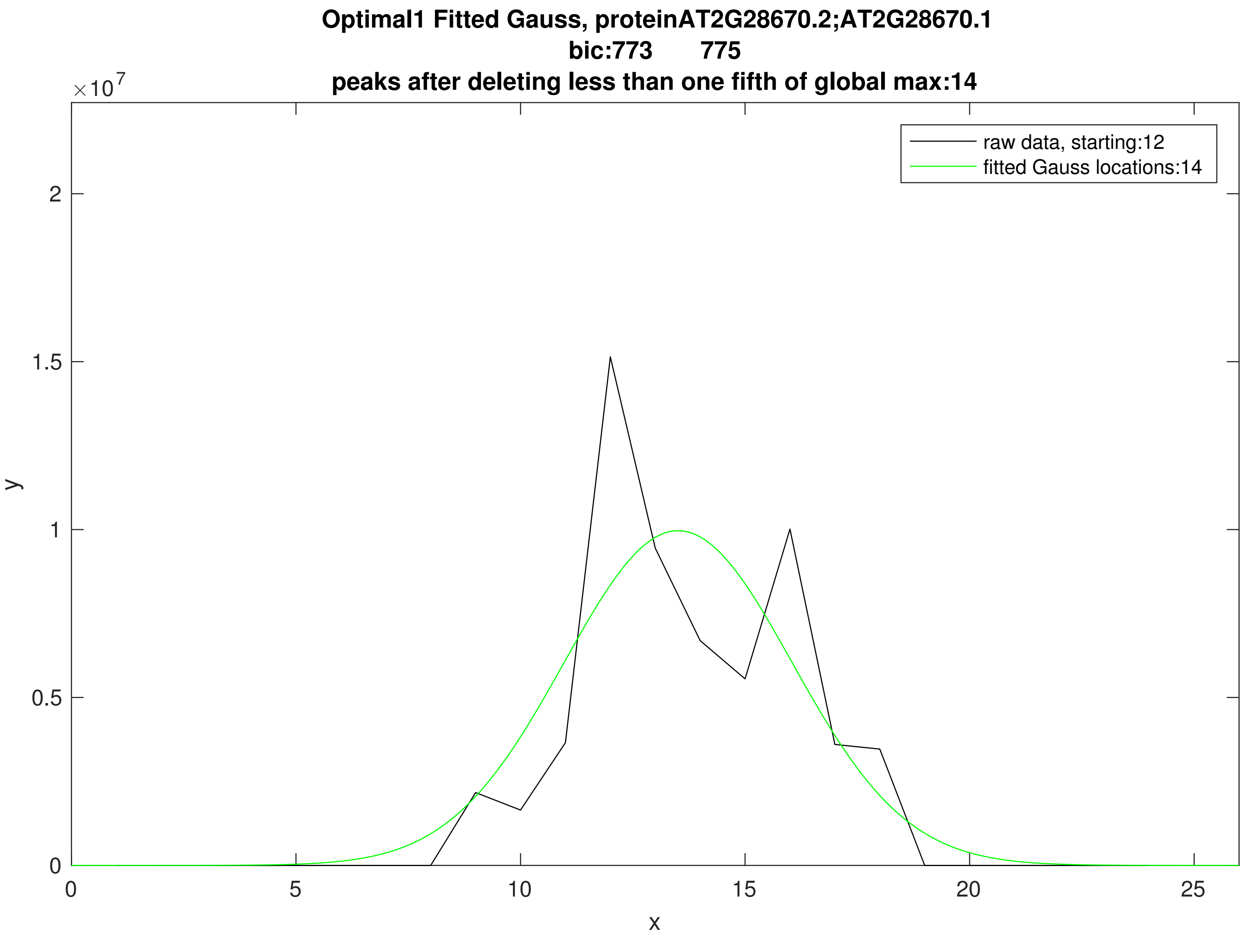
Optimal1 Fitted Gauss, proteinAT2G28670.2;AT2G28670.1
bic:773 775
peaks after deleting less than one fifth of global max:14
×10^7
raw data, starting:12
fitted Gauss locations:14
y
x
0
0.5
1
1.5
2
5
10
15
20
25

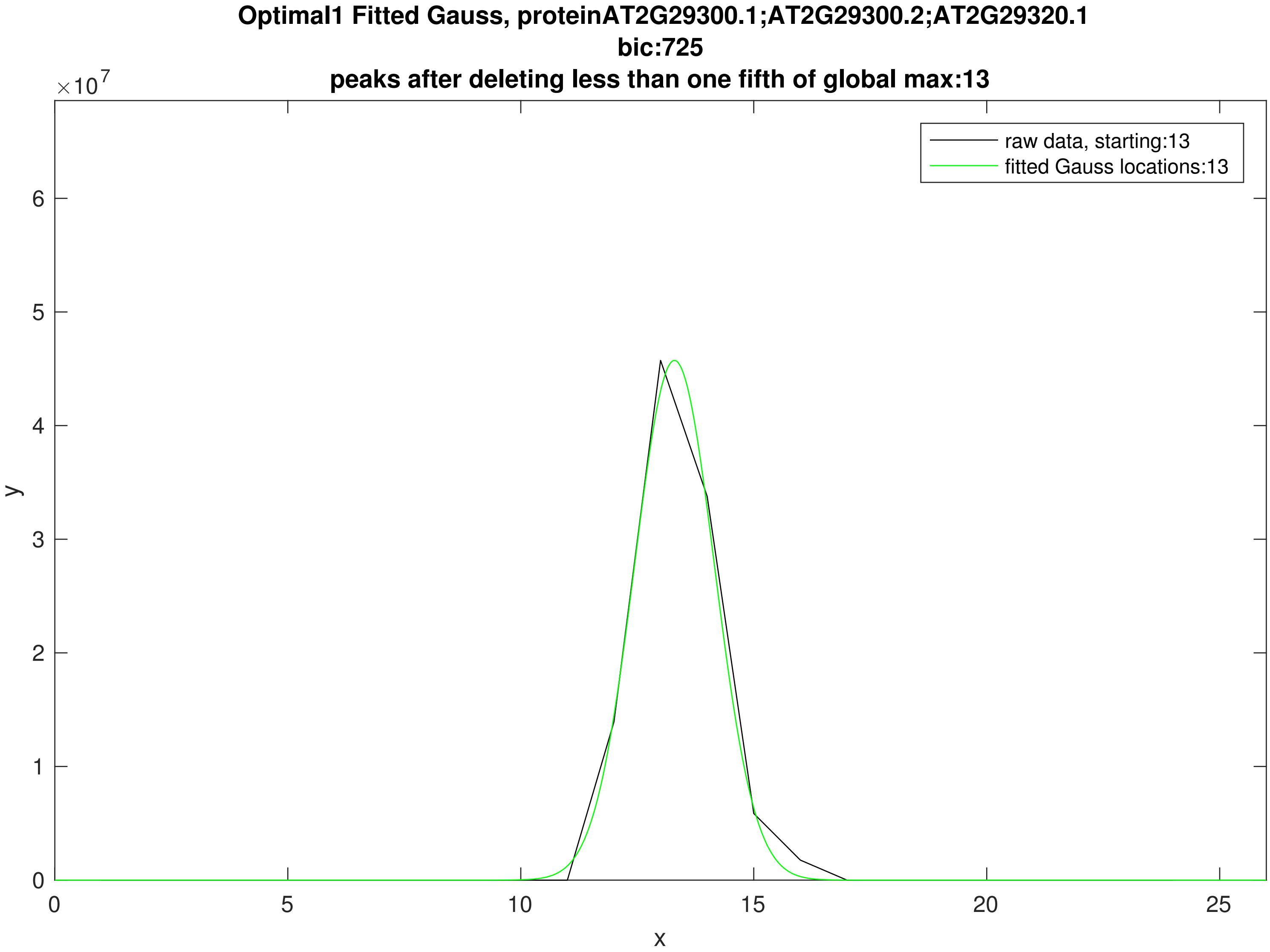
Optimal1 Fitted Gauss, proteinAT2G29300.1;AT2G29300.2;AT2G29320.1
bic:725
peaks after deleting less than one fifth of global max:13
×10^7
raw data, starting:13
fitted Gauss locations:13
y
x
0
1
2
3
4
5
6
0
5
10
15
20
25

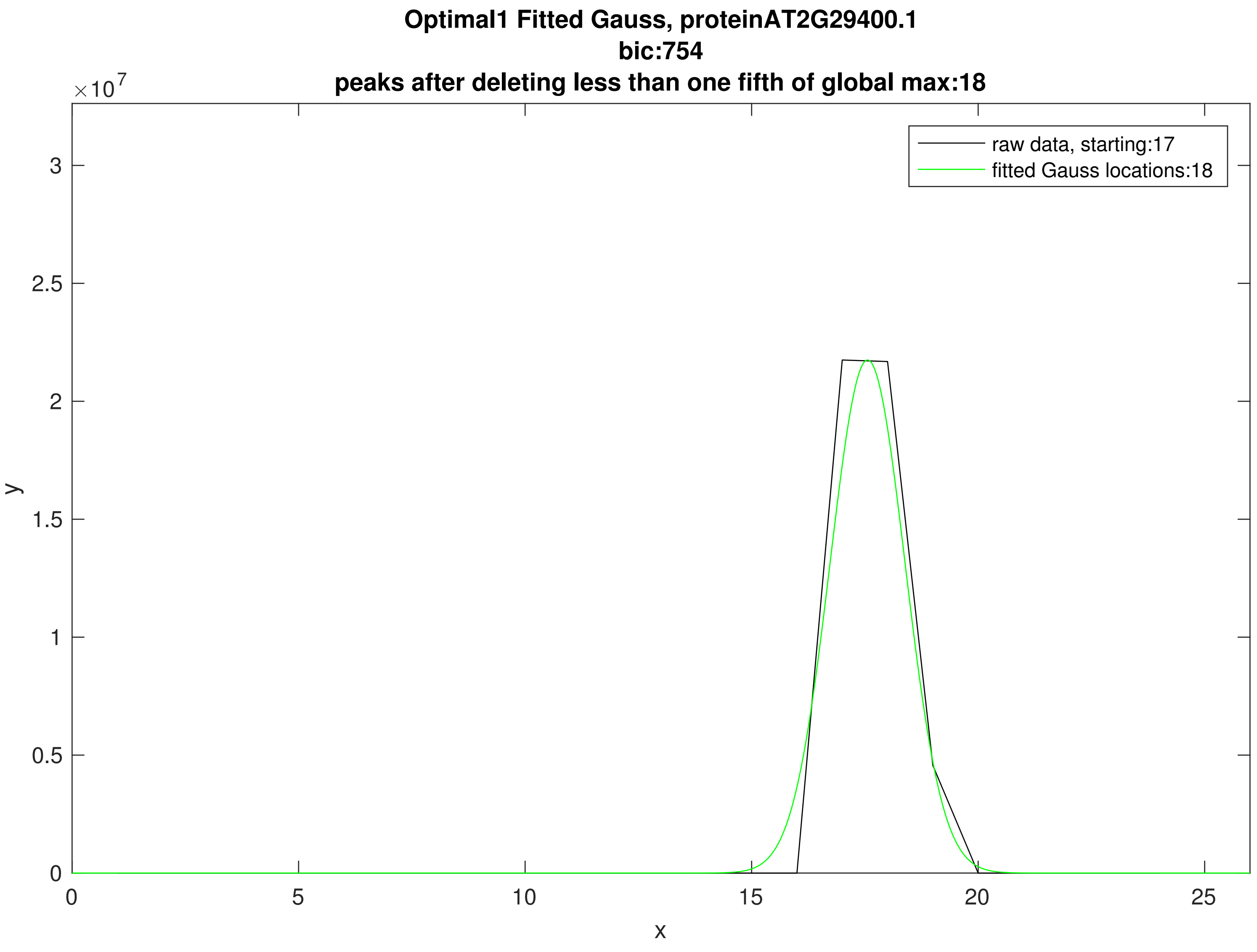
Optimal1 Fitted Gauss, proteinAT2G29400.1
bic:754
peaks after deleting less than one fifth of global max:18
×10^7
3
2.5
2
1.5
1
0.5
0
y
0
5
10
15
20
25
x
raw data, starting:17
fitted Gauss locations:18

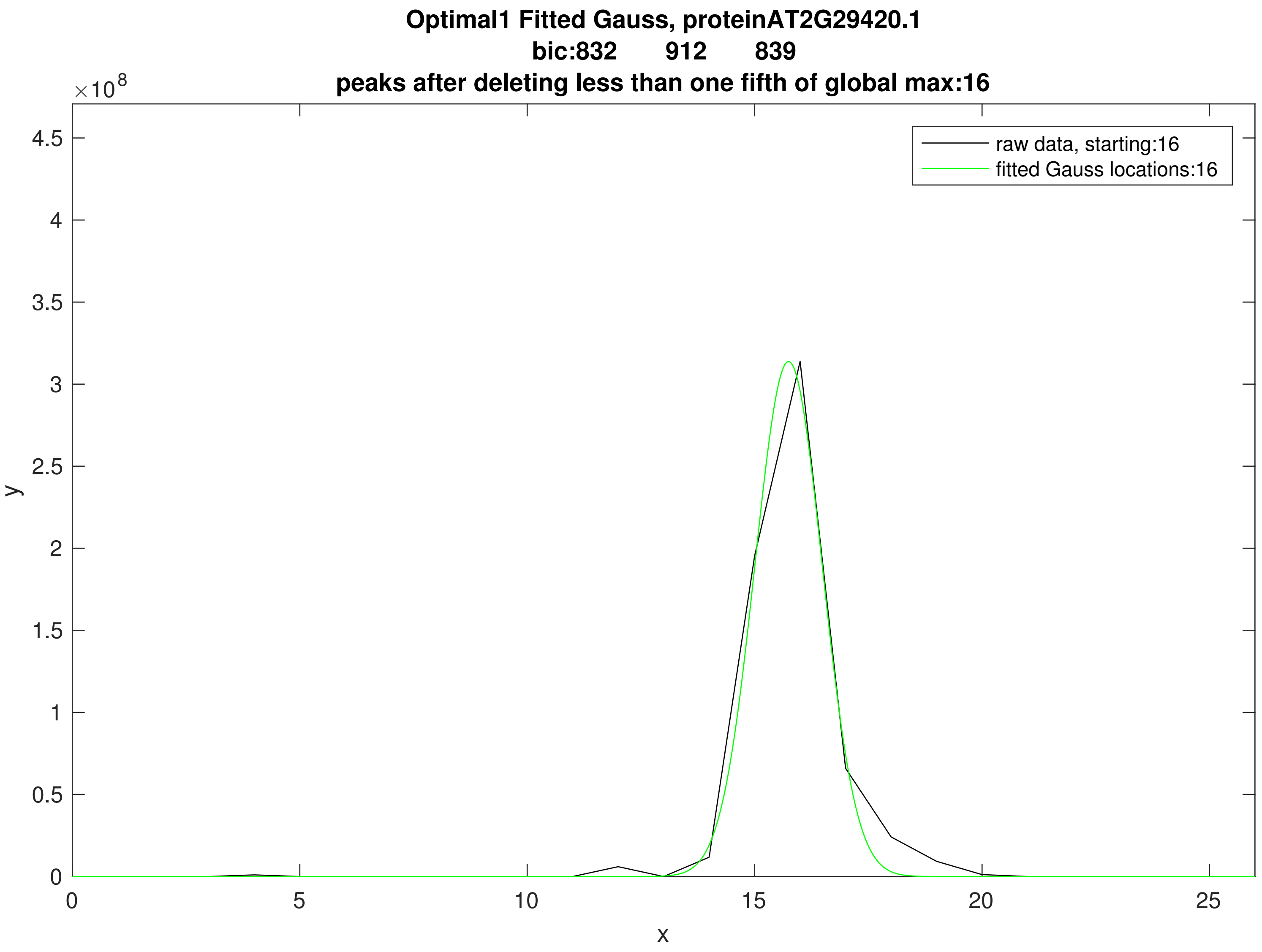
Optimal1 Fitted Gauss, proteinAT2G29420.1
bic:832 912 839
peaks after deleting less than one fifth of global max:16
×10^8
4.5
4
3.5
3
2.5
2
1.5
1
0.5
0
y
0
5
10
15
20
25
x
raw data, starting:16
fitted Gauss locations:16

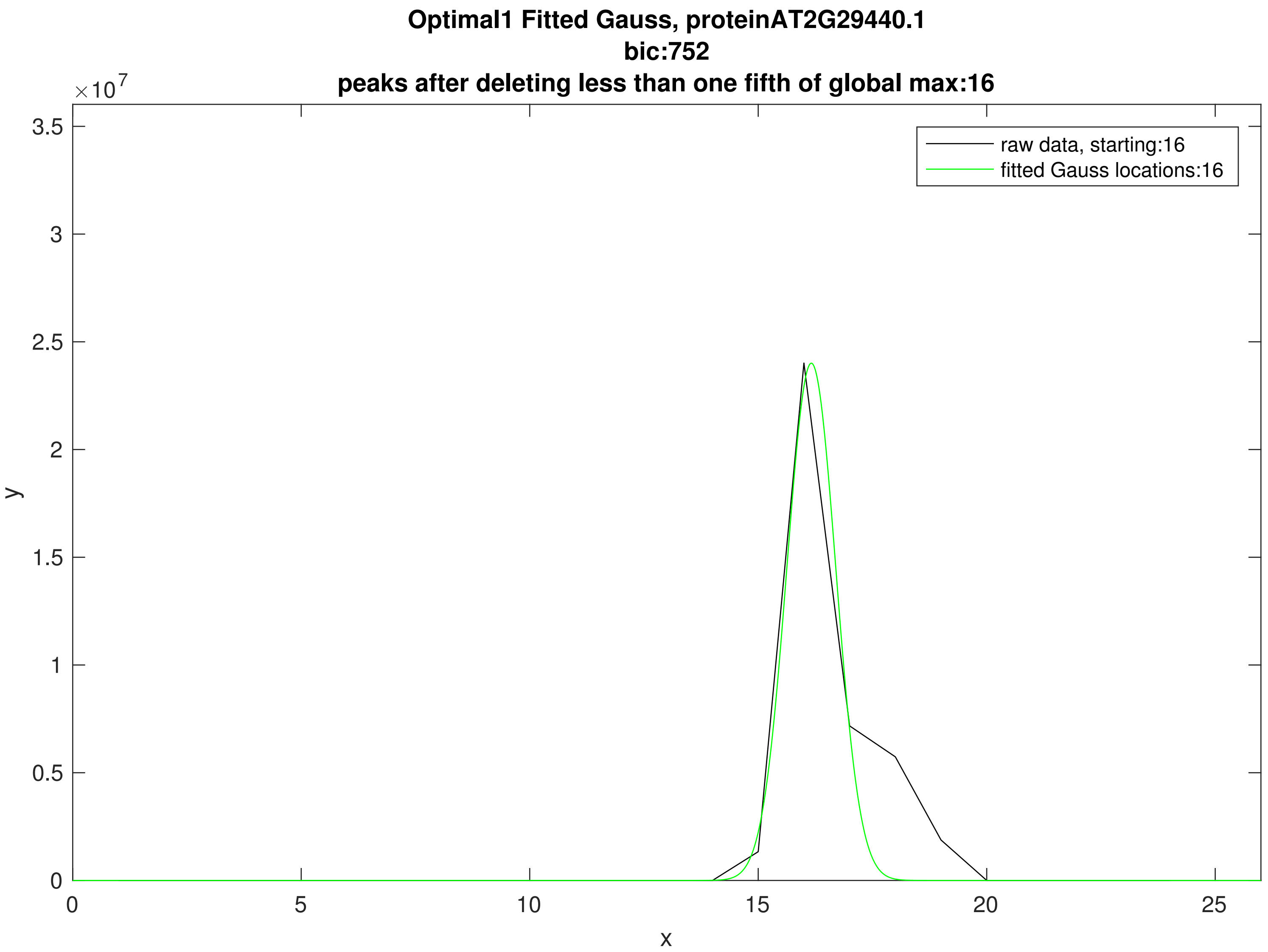
Optimal1 Fitted Gauss, proteinAT2G29440.1
bic:752
peaks after deleting less than one fifth of global max:16
×10^7
3.5
3
2.5
2
1.5
1
0.5
0
y
0
5
10
15
20
25
x
raw data, starting:16
fitted Gauss locations:16

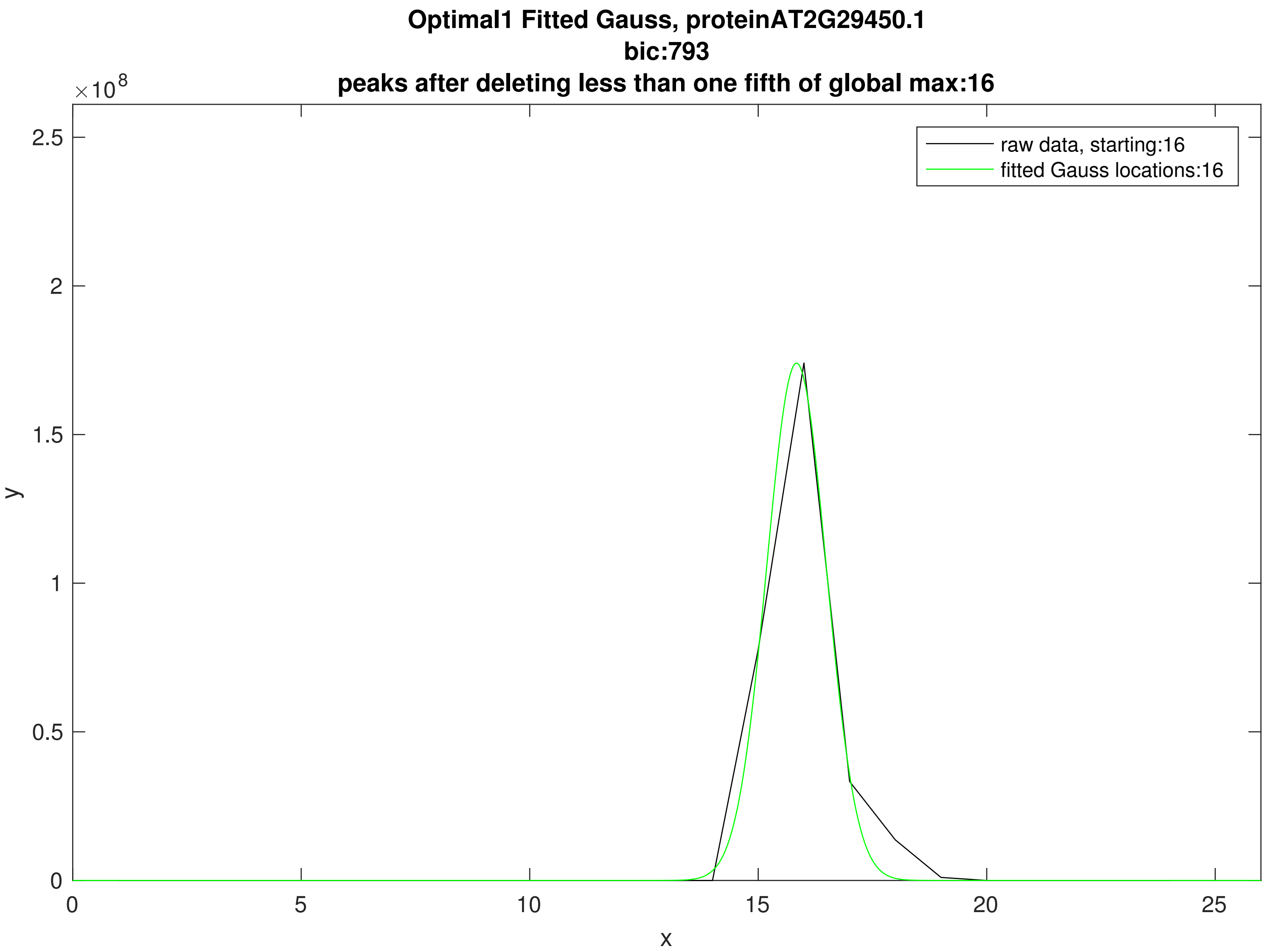

Optimal1 Fitted Gauss, proteinAT2G29450.1
bic:793
peaks after deleting less than one fifth of global max:16
×10^8
2.5
2
1.5
1
0.5
0
y
0
5
10
15
20
25
x
raw data, starting:16
fitted Gauss locations:16

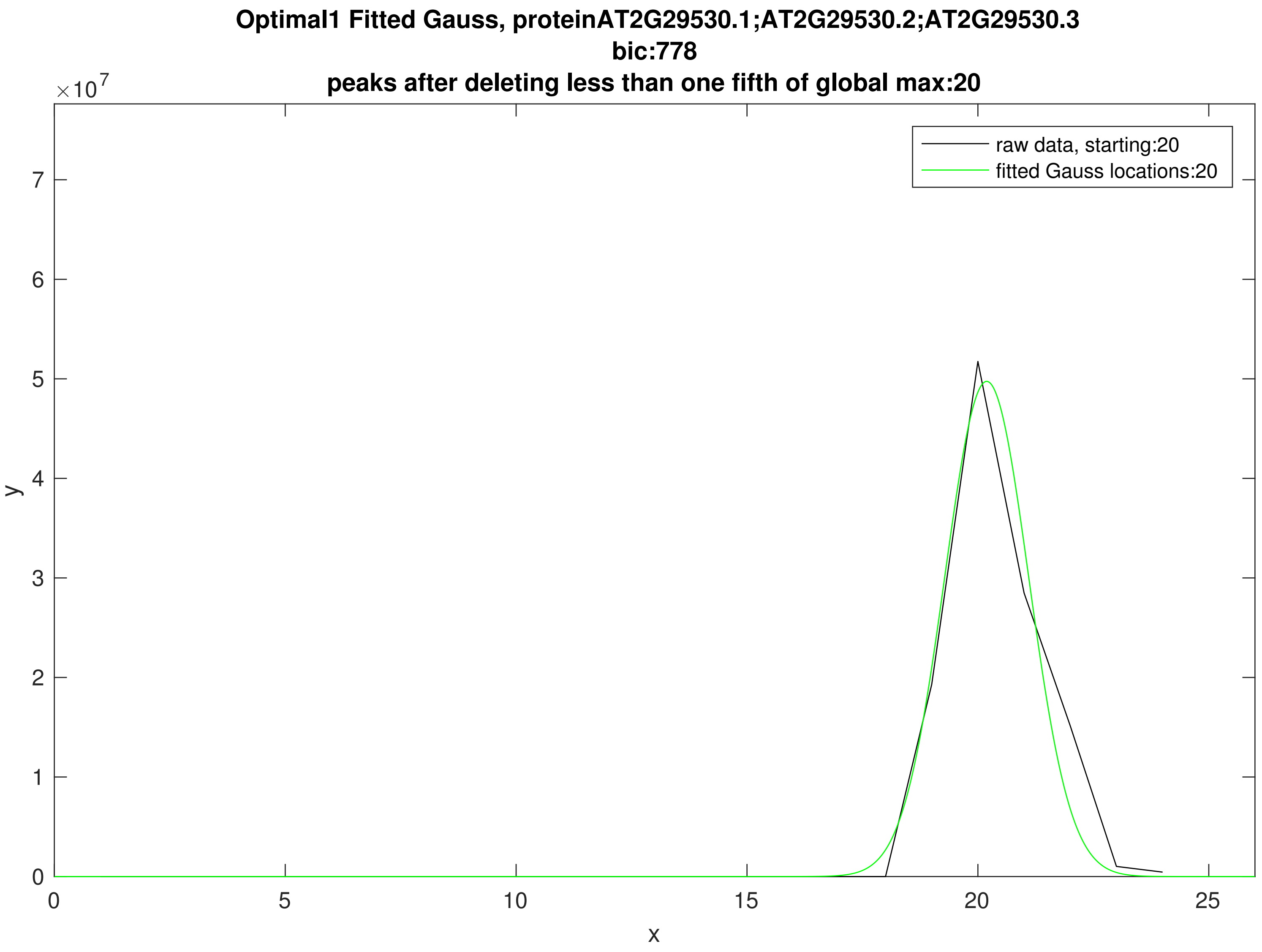
Optimal1 Fitted Gauss, proteinAT2G29530.1;AT2G29530.2;AT2G29530.3
bic:778
peaks after deleting less than one fifth of global max:20
×10^7
raw data, starting:20
fitted Gauss locations:20
y
x
0
1
2
3
4
5
6
7
0
5
10
15
20
25

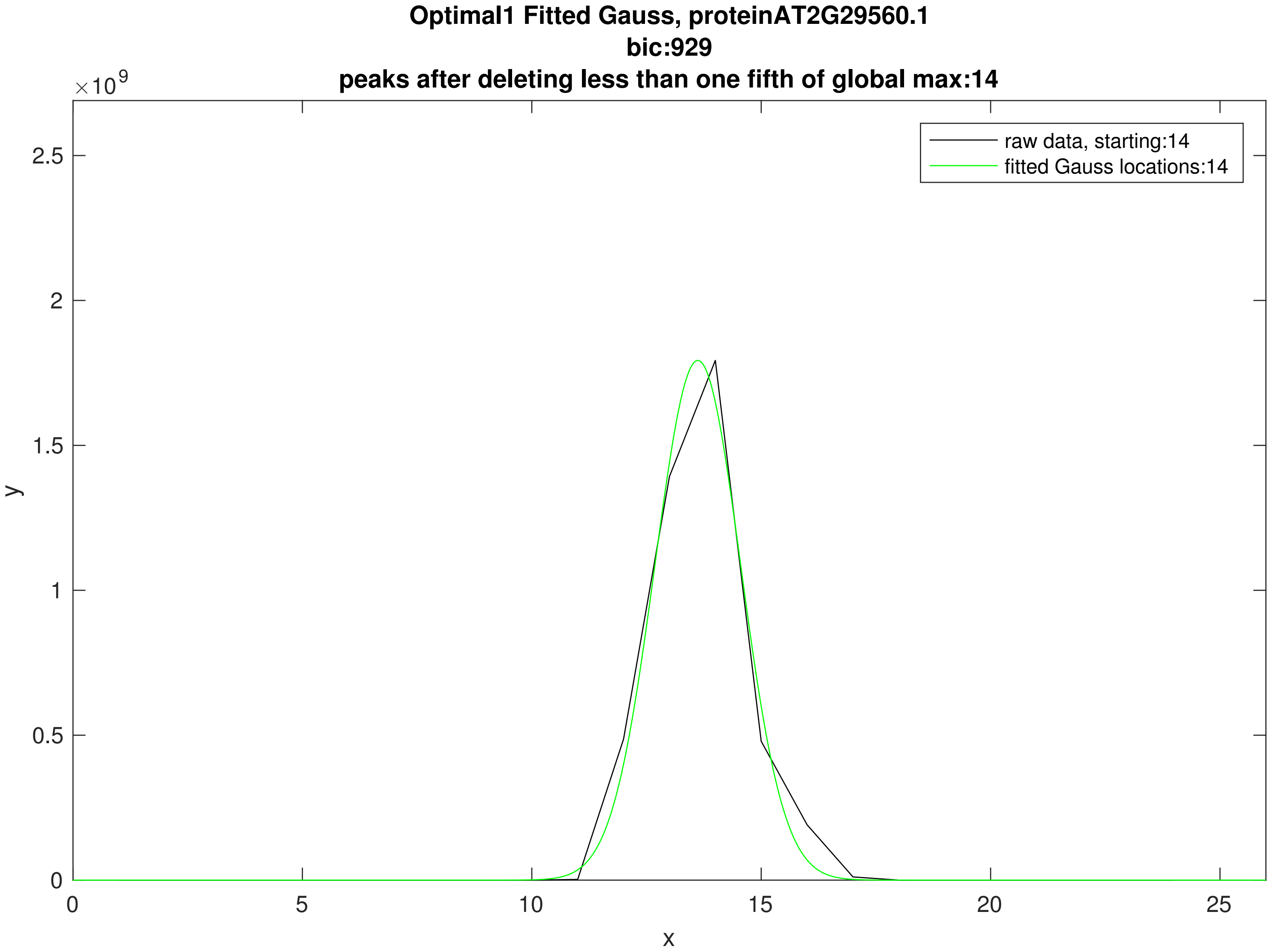
Optimal1 Fitted Gauss, proteinAT2G29560.1
bic:929
peaks after deleting less than one fifth of global max:14
×10^9
raw data, starting:14
fitted Gauss locations:14
y
x
0
0.5
1
1.5
2
2.5
0
5
10
15
20
25

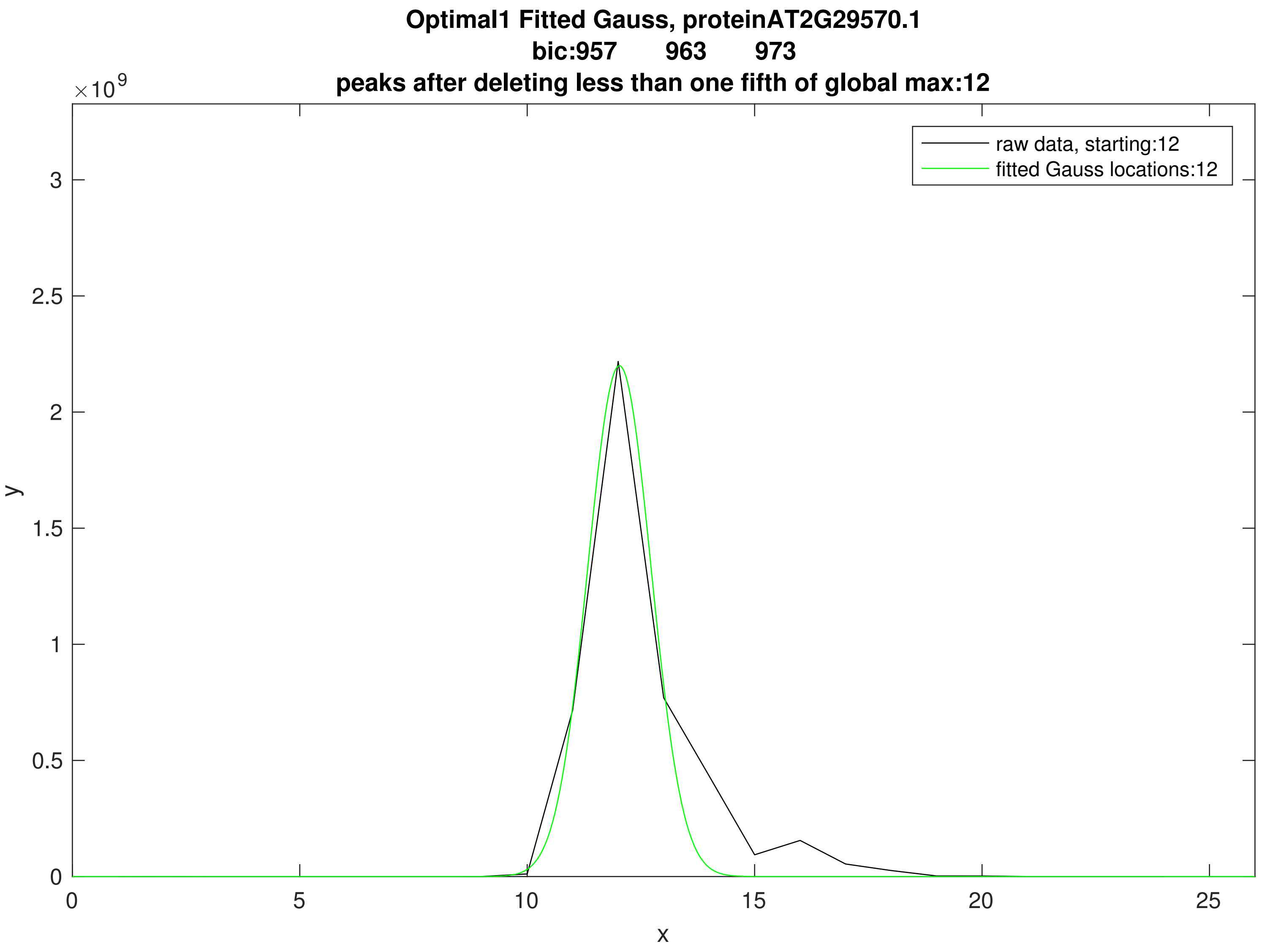
Optimal1 Fitted Gauss, proteinAT2G29570.1
bic:957 963 973
peaks after deleting less than one fifth of global max:12
×10^9
3
2.5
2
1.5
1
0.5
0
y
0
5
10
15
20
25
x
raw data, starting:12
fitted Gauss locations:12

**Optimal2 Fitted Gauss, proteinAT2G29690.1**
**bic:847 770**
**peaks after deleting less than one fifth of global max:6.1 15**

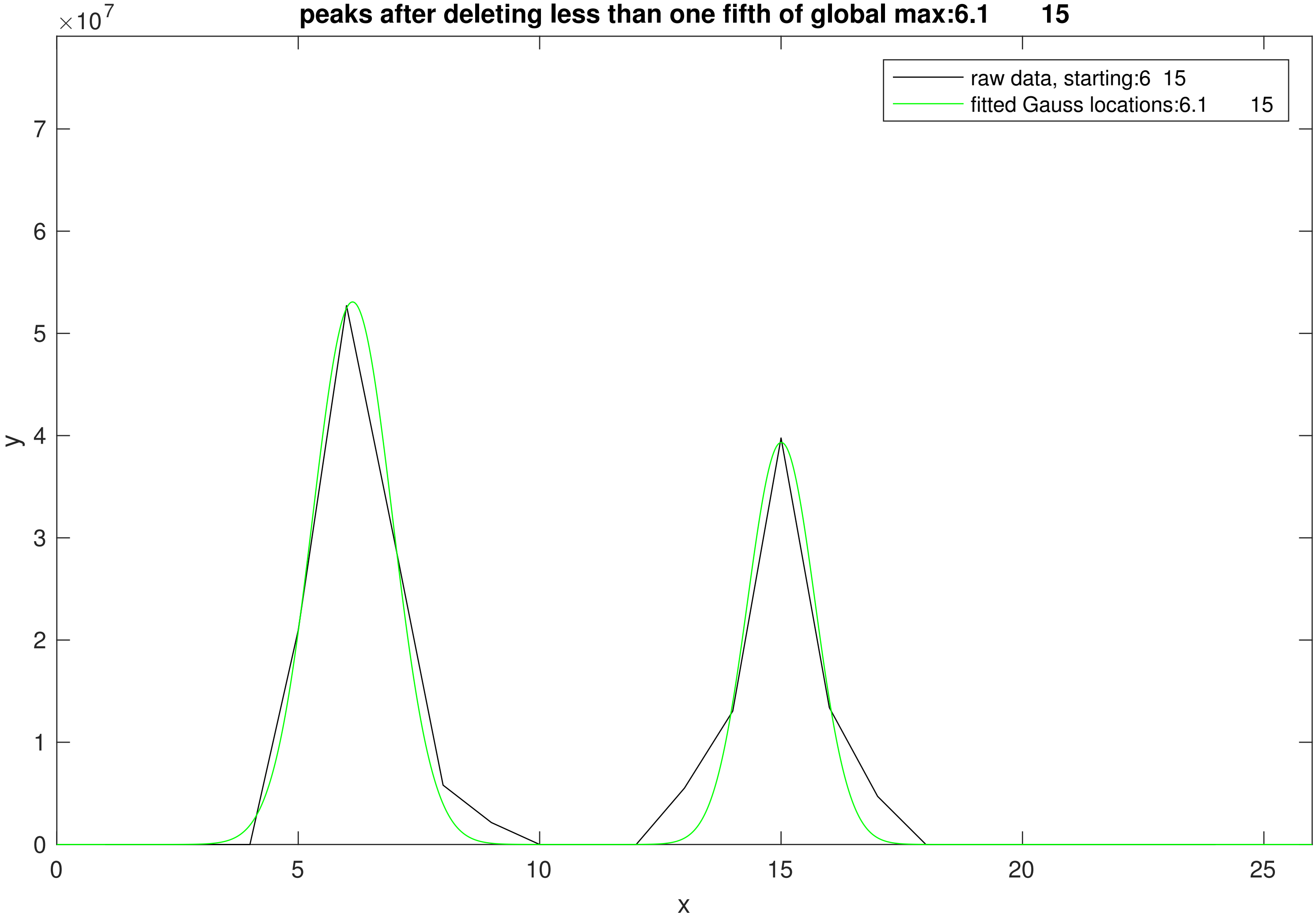

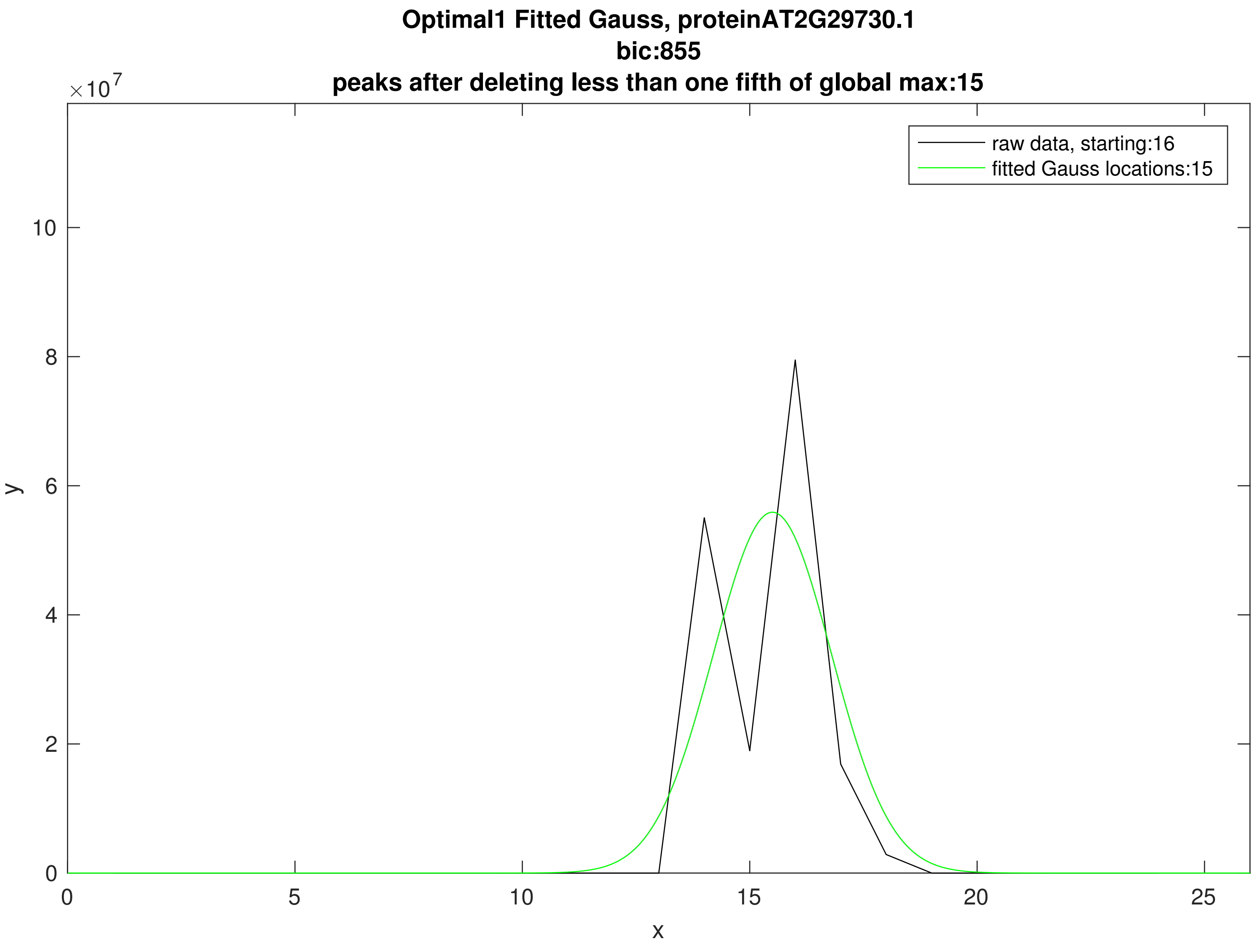
Optimal1 Fitted Gauss, proteinAT2G29730.1
bic:855
peaks after deleting less than one fifth of global max:15
×10^7
raw data, starting:16
fitted Gauss locations:15
y
x
0
2
4
6
8
10
0
5
10
15
20
25

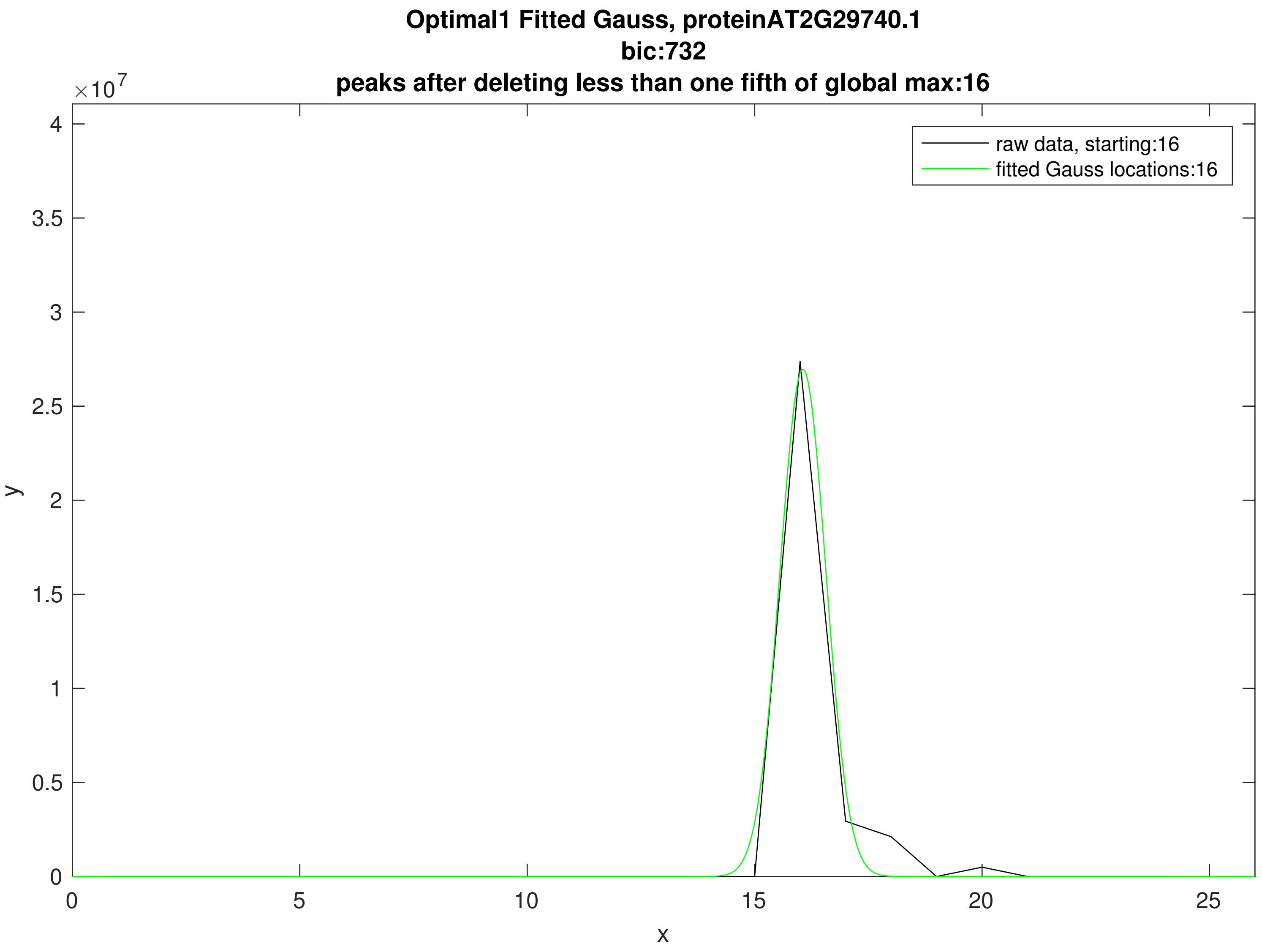

Optimal1 Fitted Gauss, proteinAT2G29740.1
bic:732
peaks after deleting less than one fifth of global max:16
×10^7
4
3.5
3
2.5
2
1.5
1
0.5
0
y
0
5
10
15
20
25
x
raw data, starting:16
fitted Gauss locations:16

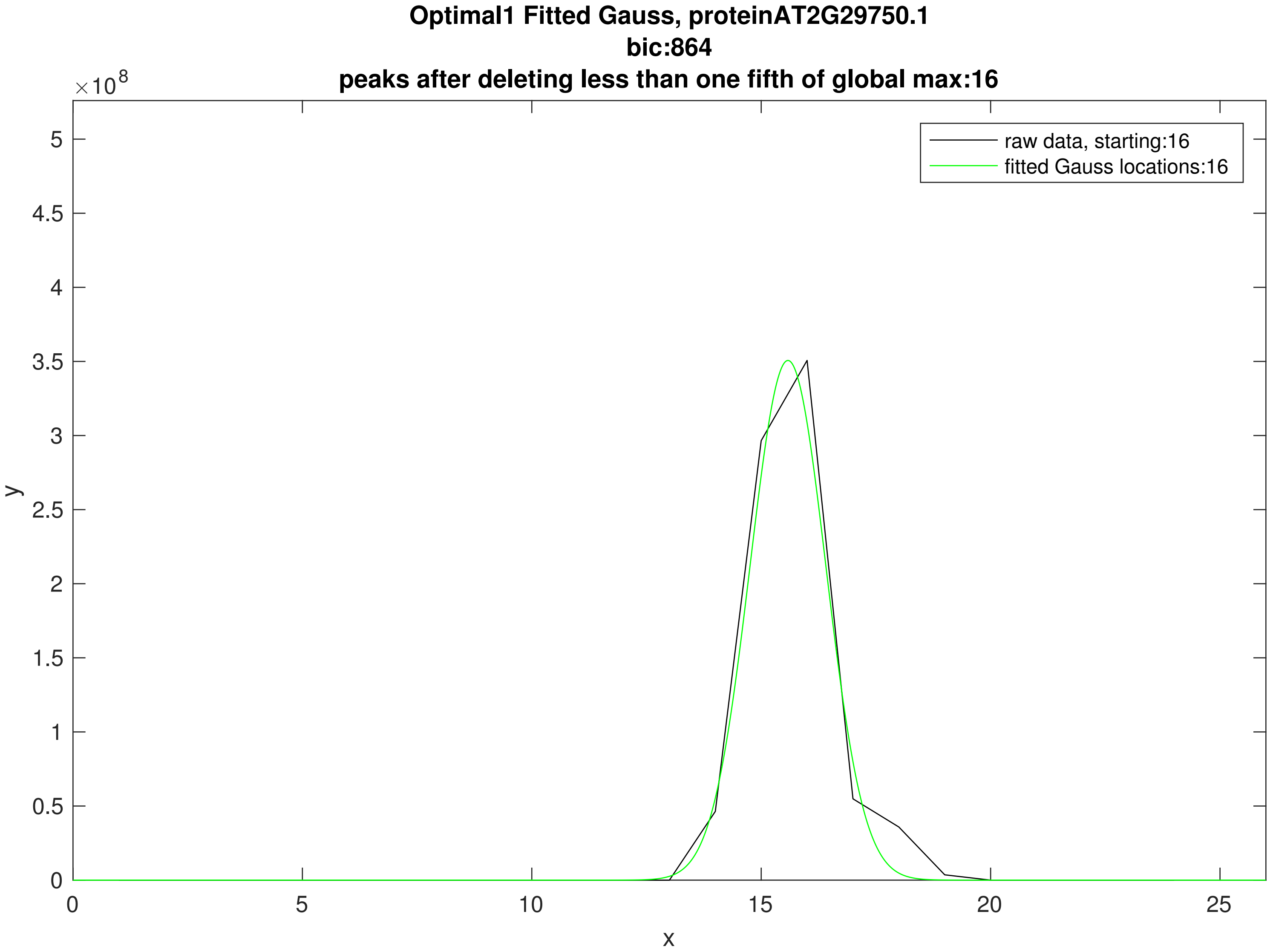

Optimal1 Fitted Gauss, proteinAT2G29750.1
bic:864
peaks after deleting less than one fifth of global max:16
×10^8
raw data, starting:16
fitted Gauss locations:16
y
x
0
0.5
1
1.5
2
2.5
3
3.5
4
4.5
5
0
5
10
15
20
25

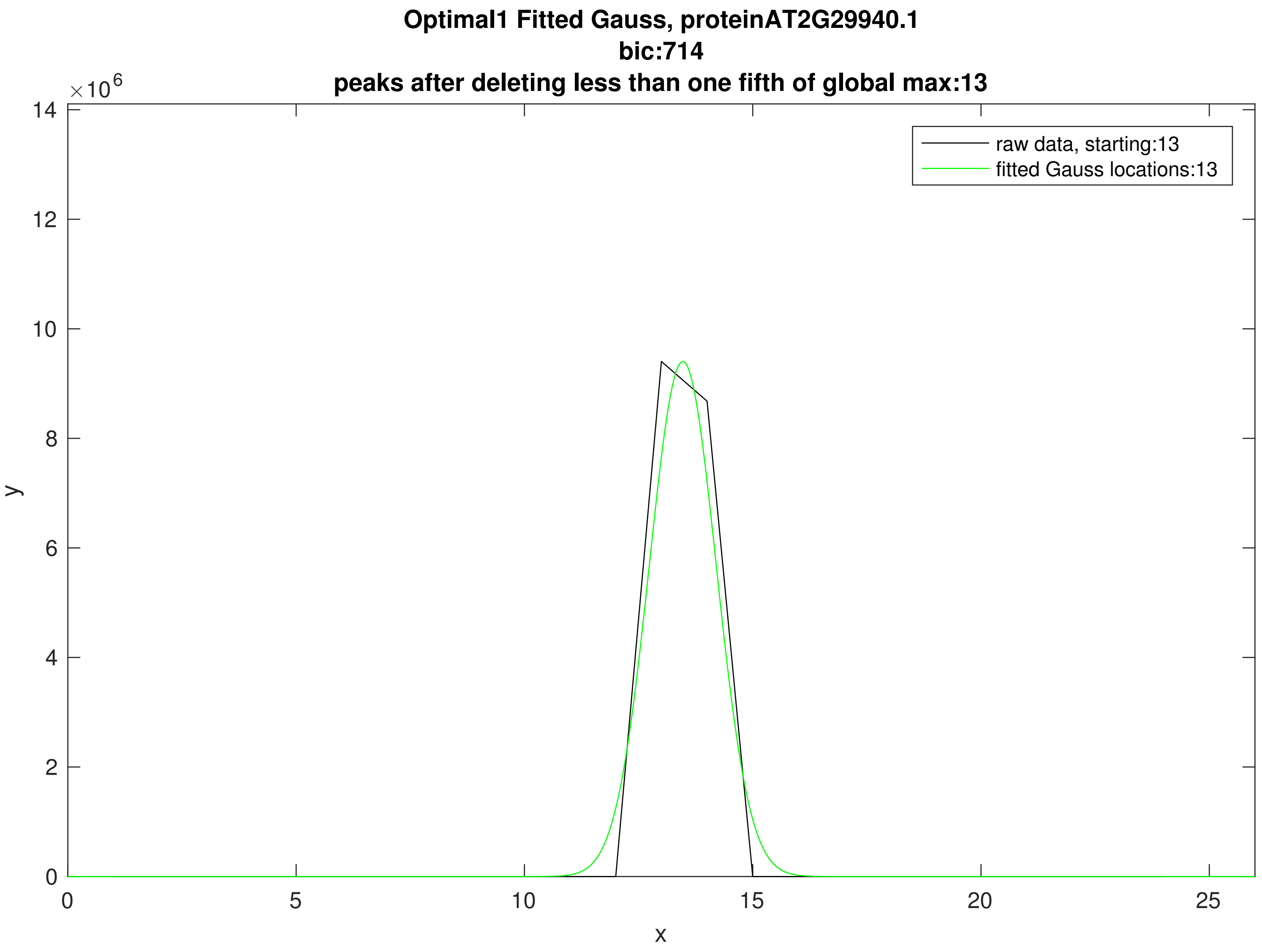
Optimal1 Fitted Gauss, proteinAT2G29940.1
bic:714
peaks after deleting less than one fifth of global max:13
×10^6
14
12
10
8
6
4
2
0
y
0
5
10
15
20
25
x
raw data, starting:13
fitted Gauss locations:13

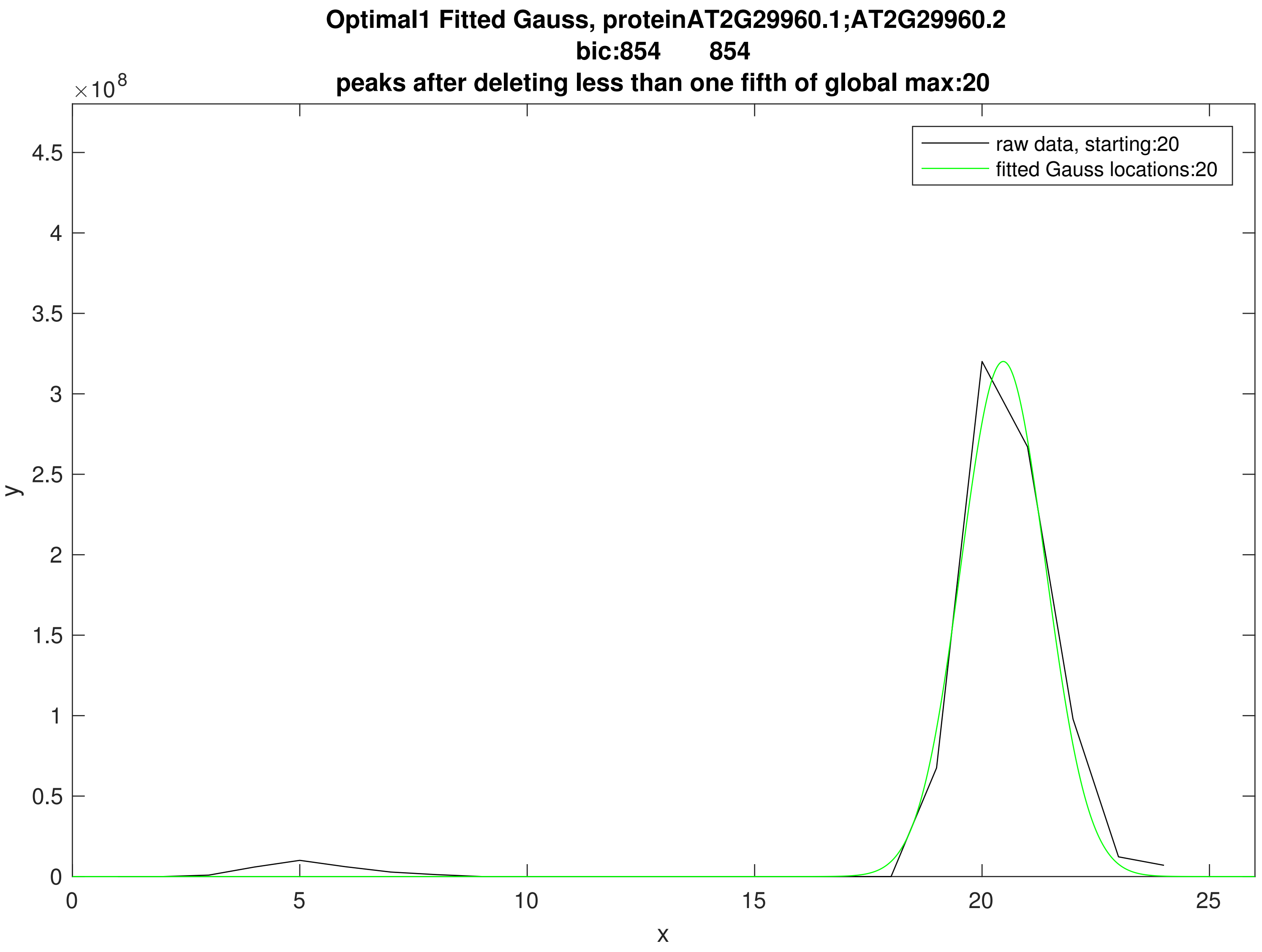

Optimal1 Fitted Gauss, proteinAT2G29960.1;AT2G29960.2
bic:854 854
peaks after deleting less than one fifth of global max:20
×10^8
4.5
4
3.5
3
2.5
2
1.5
1
0.5
0
y
0
5
10
15
20
25
x
raw data, starting:20
fitted Gauss locations:20

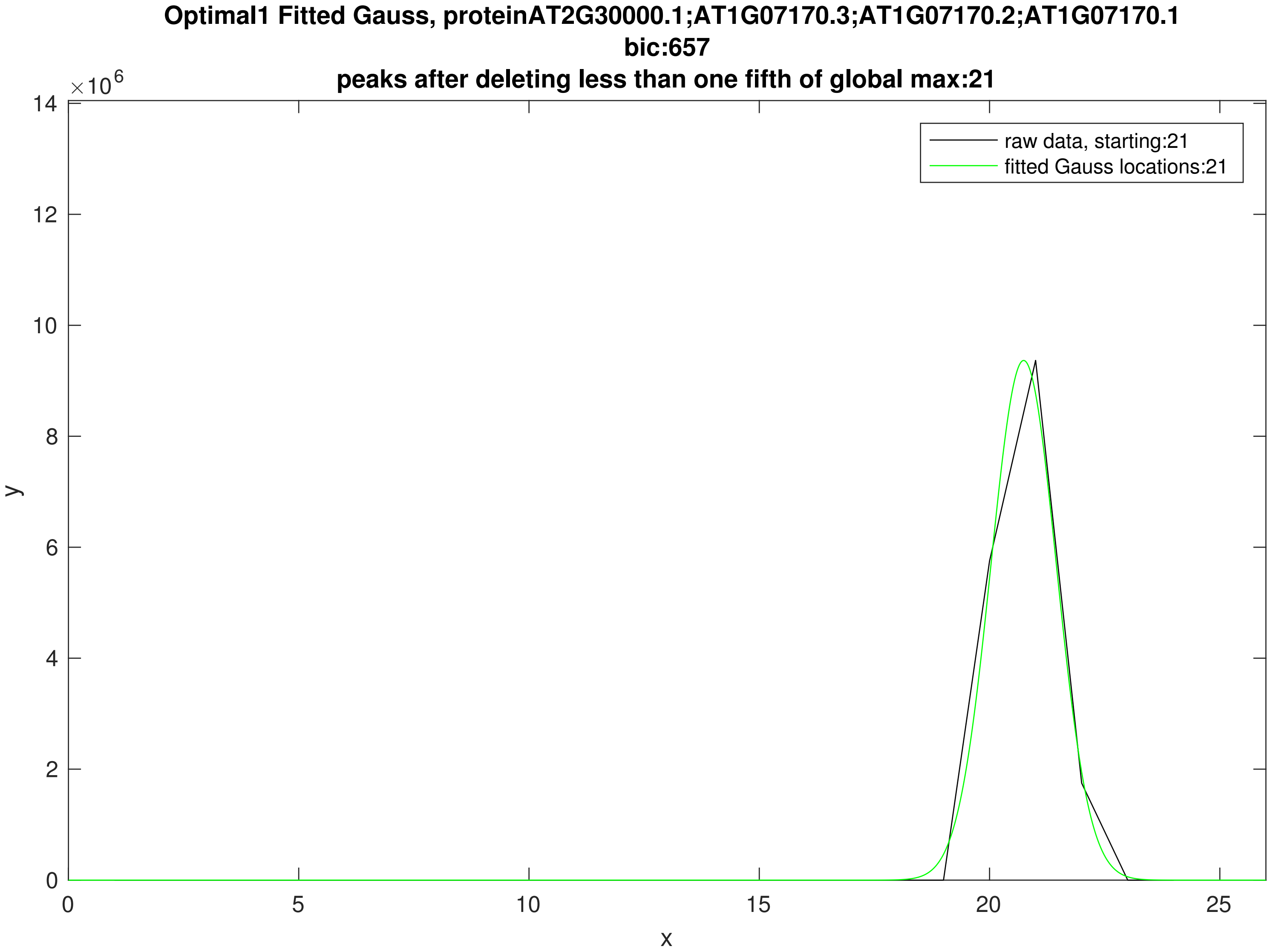
Optimal1 Fitted Gauss, proteinAT2G30000.1;AT1G07170.3;AT1G07170.2;AT1G07170.1
bic:657
peaks after deleting less than one fifth of global max:21
×10^6
14
12
10
8
6
4
2
0
y
0
5
10
15
20
25
x
raw data, starting:21
fitted Gauss locations:21

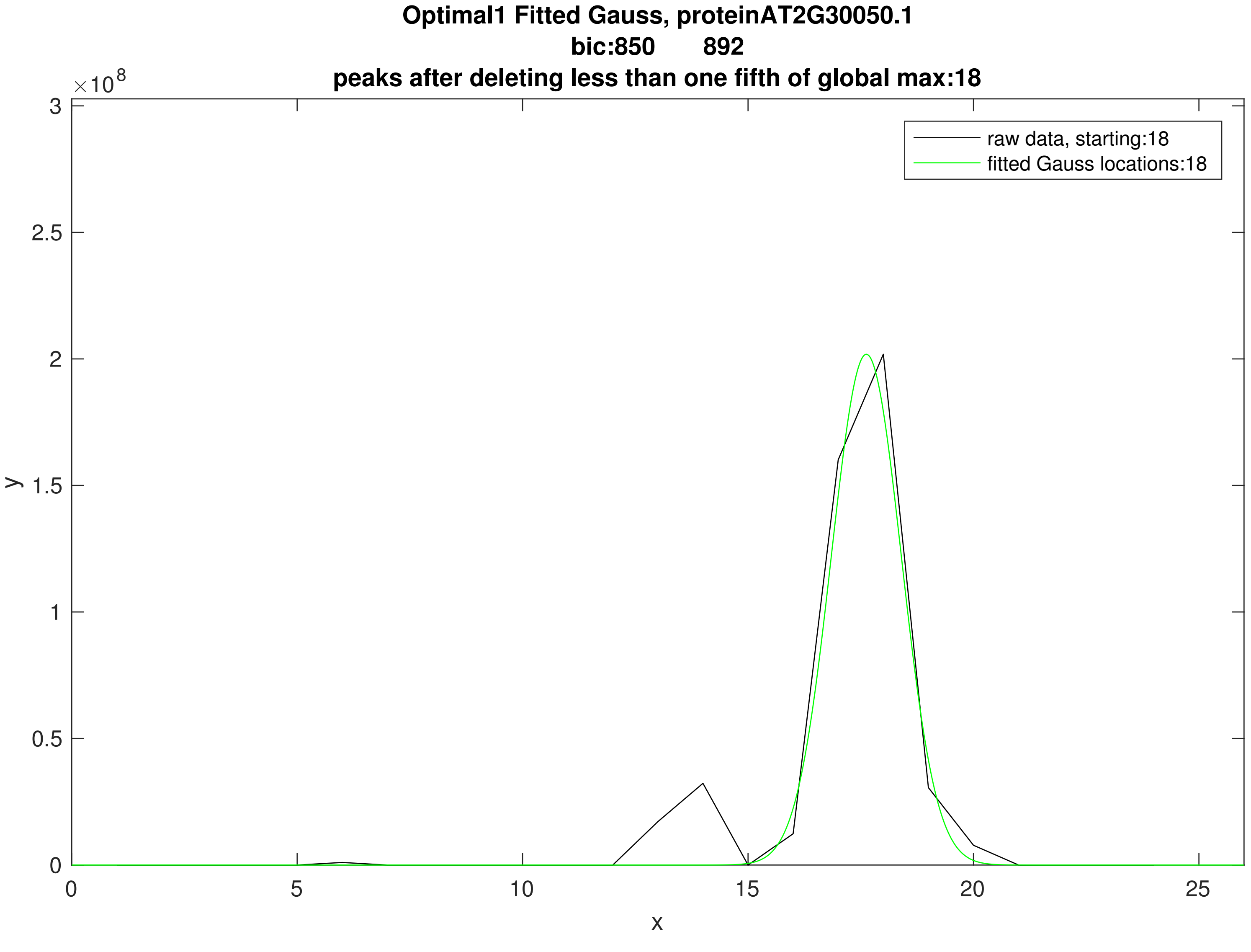
Optimal1 Fitted Gauss, proteinAT2G30050.1
bic:850 892
peaks after deleting less than one fifth of global max:18
raw data, starting:18
fitted Gauss locations:18
$\times 10^8$
y
x
0
0.5
1
1.5
2
2.5
3
5
10
15
20
25

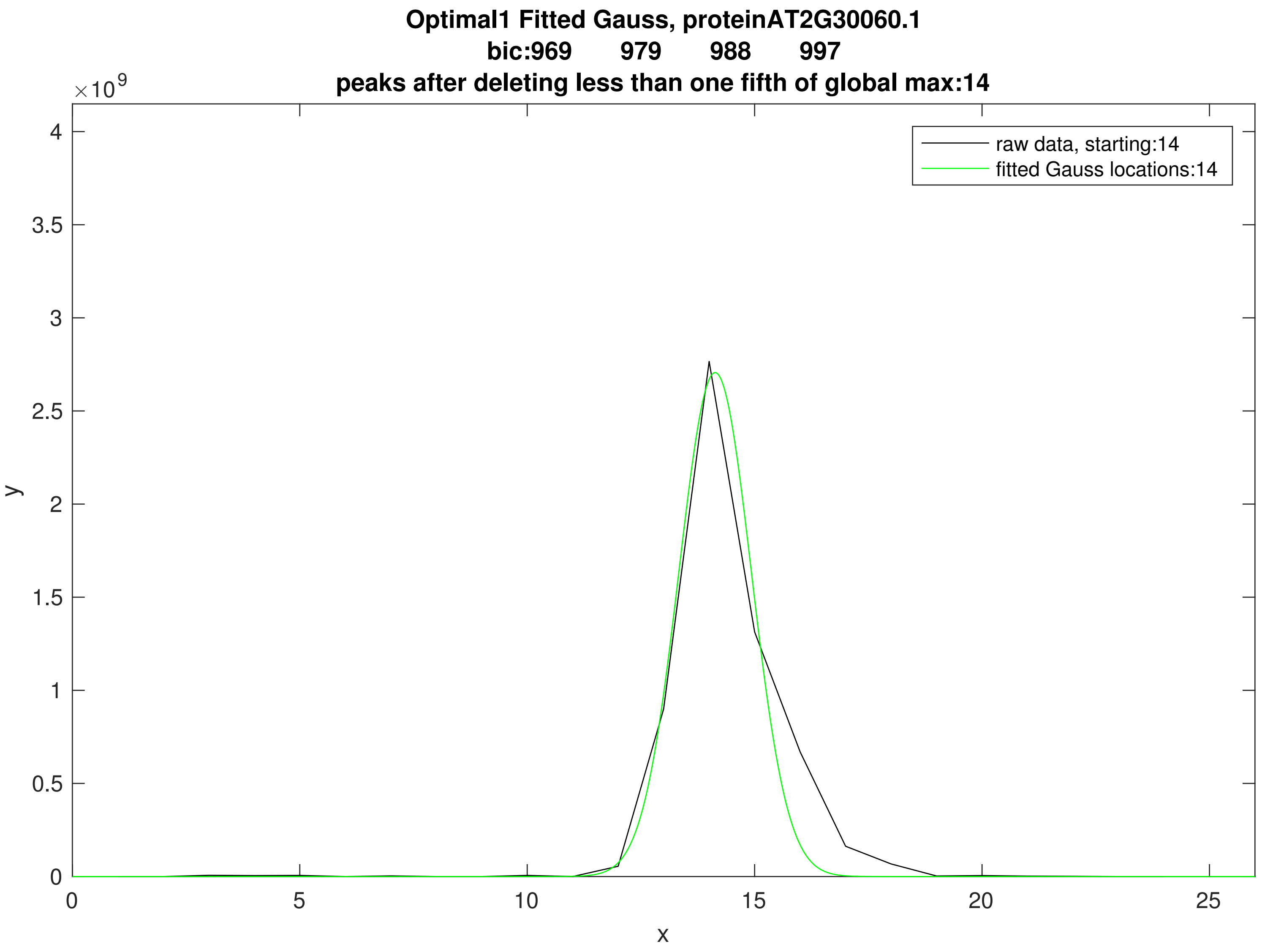

Optimal1 Fitted Gauss, proteinAT2G30060.1
bic:969 979 988 997
peaks after deleting less than one fifth of global max:14
×10^9
raw data, starting:14
fitted Gauss locations:14
y
x
0
0.5
1
1.5
2
2.5
3
3.5
4
5
10
15
20
25

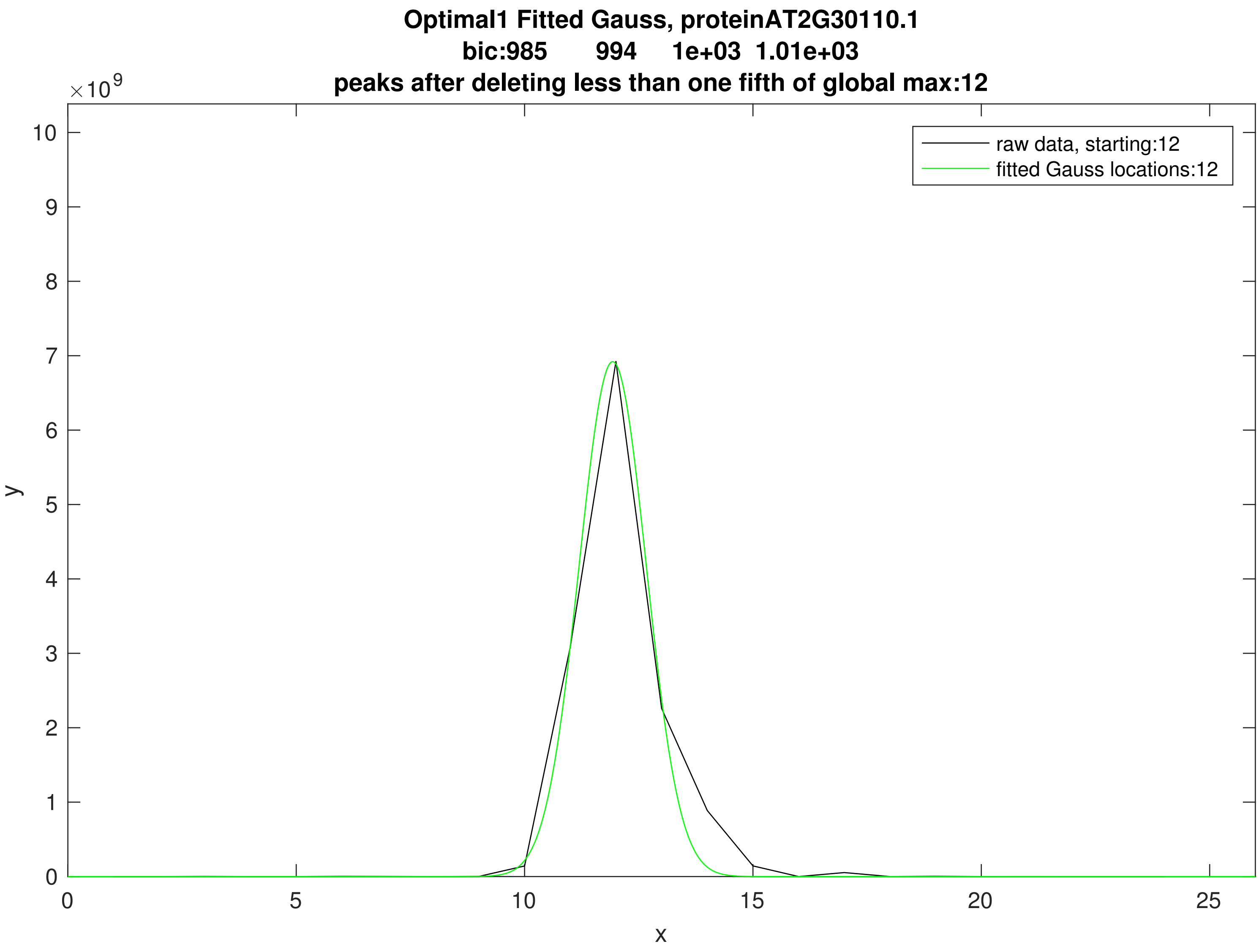
Optimal1 Fitted Gauss, proteinAT2G30110.1
bic:985 994 1e+03 1.01e+03
peaks after deleting less than one fifth of global max:12
×10^9
10
9
8
7
6
5
4
3
2
1
0
y
0
5
10
15
20
25
x
raw data, starting:12
fitted Gauss locations:12

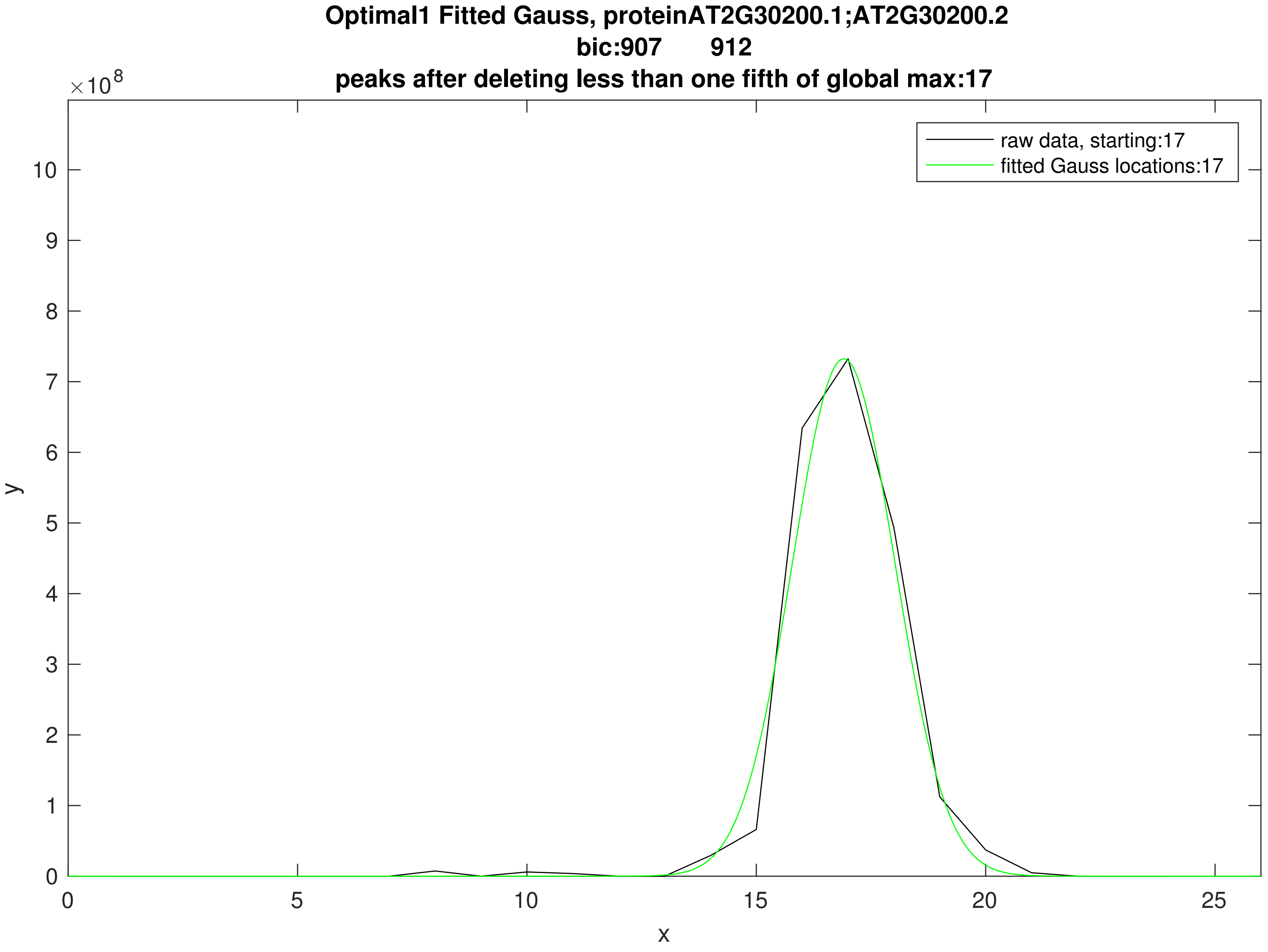
Optimal1 Fitted Gauss, proteinAT2G30200.1;AT2G30200.2
bic:907 912
peaks after deleting less than one fifth of global max:17
×10^8
raw data, starting:17
fitted Gauss locations:17
y
x
0
1
2
3
4
5
6
7
8
9
10
0
5
10
15
20
25

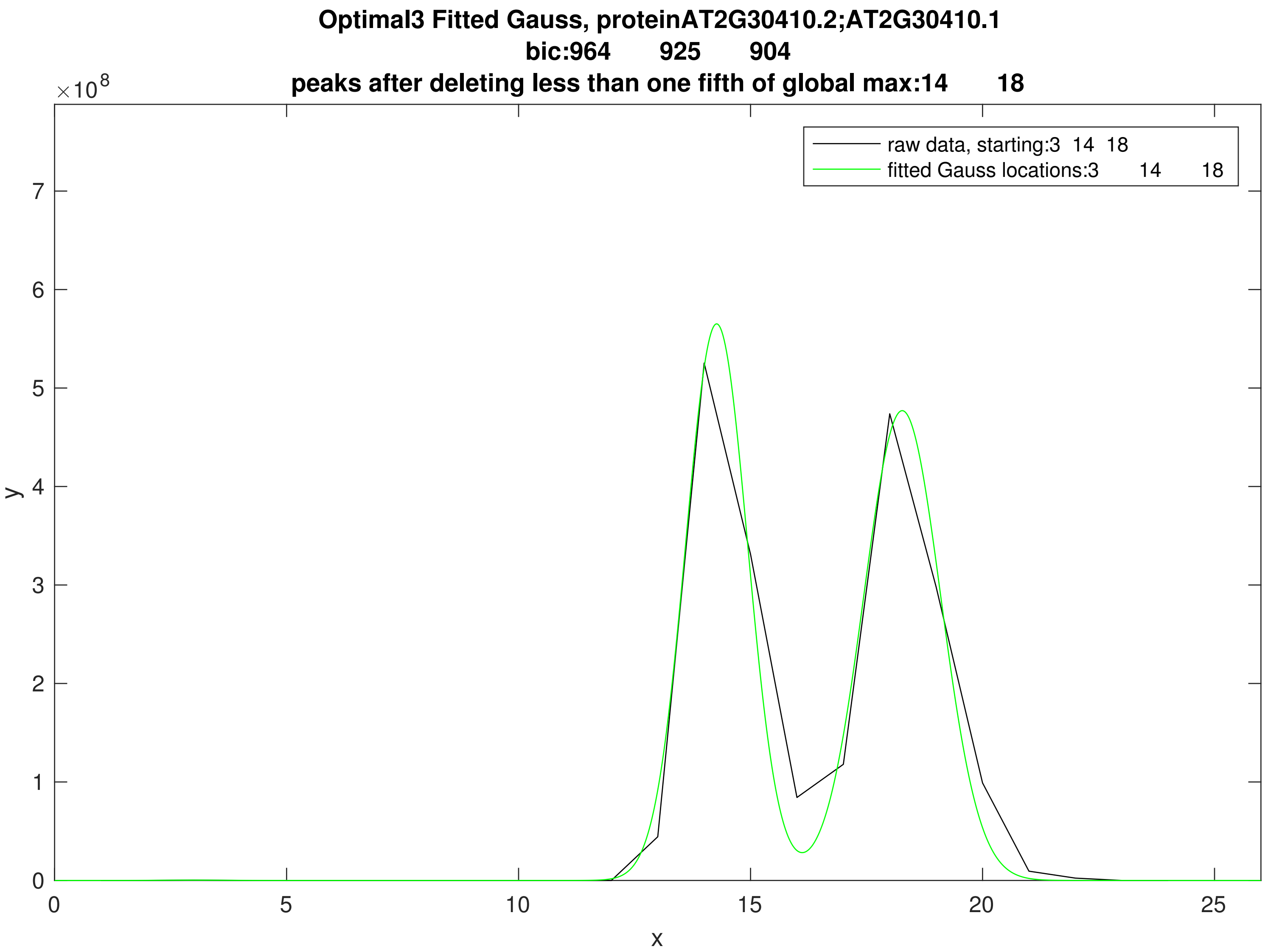
Optimal3 Fitted Gauss, proteinAT2G30410.2;AT2G30410.1
bic:964 925 904
peaks after deleting less than one fifth of global max:14 18
×10^8
raw data, starting:3 14 18
fitted Gauss locations:3 14 18
y
x
0
1
2
3
4
5
6
7
0
5
10
15
20
25

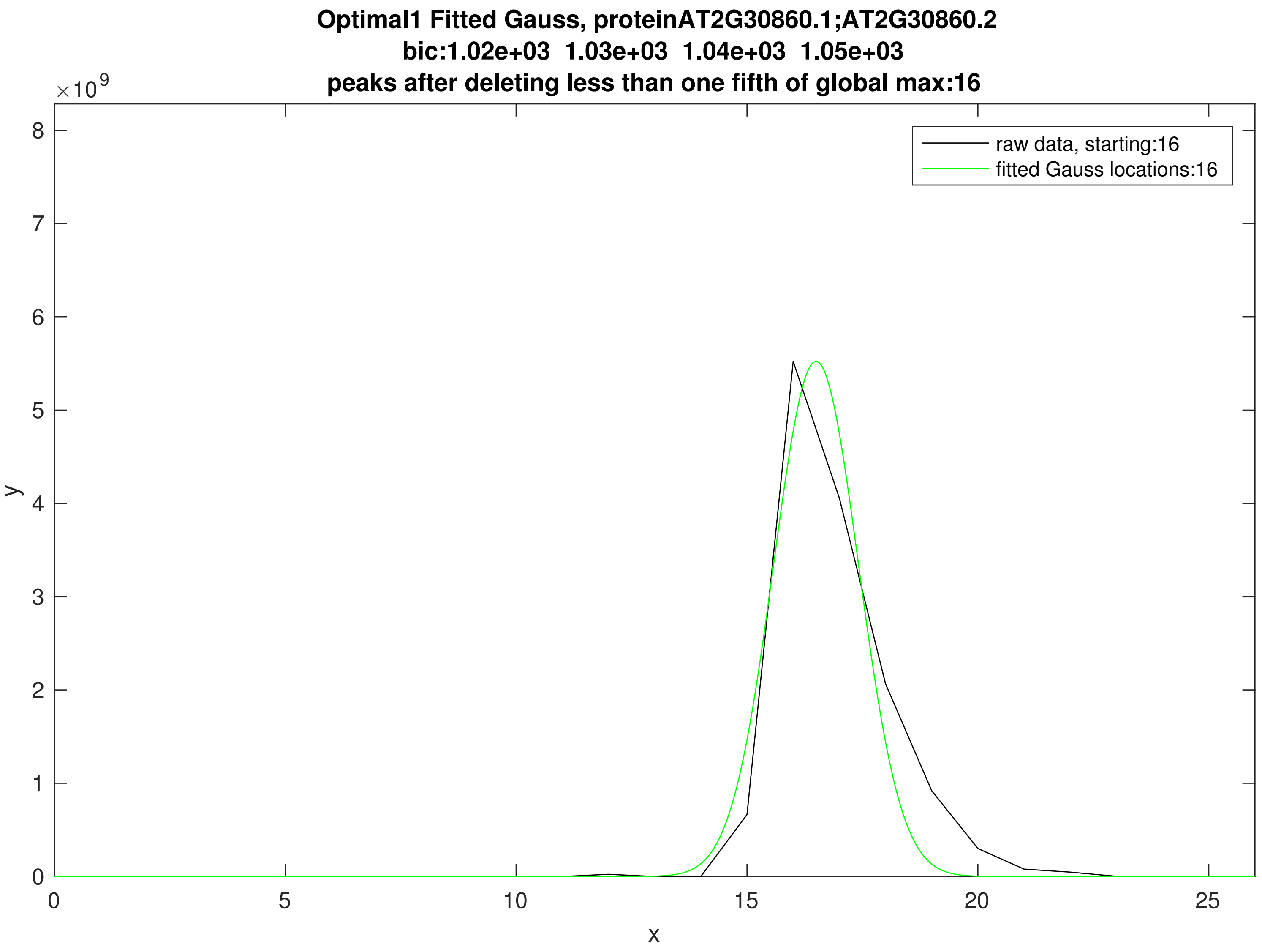

Optimal1 Fitted Gauss, proteinAT2G30860.1;AT2G30860.2
bic:1.02e+03 1.03e+03 1.04e+03 1.05e+03
peaks after deleting less than one fifth of global max:16
raw data, starting:16
fitted Gauss locations:16
×10^9
y
x

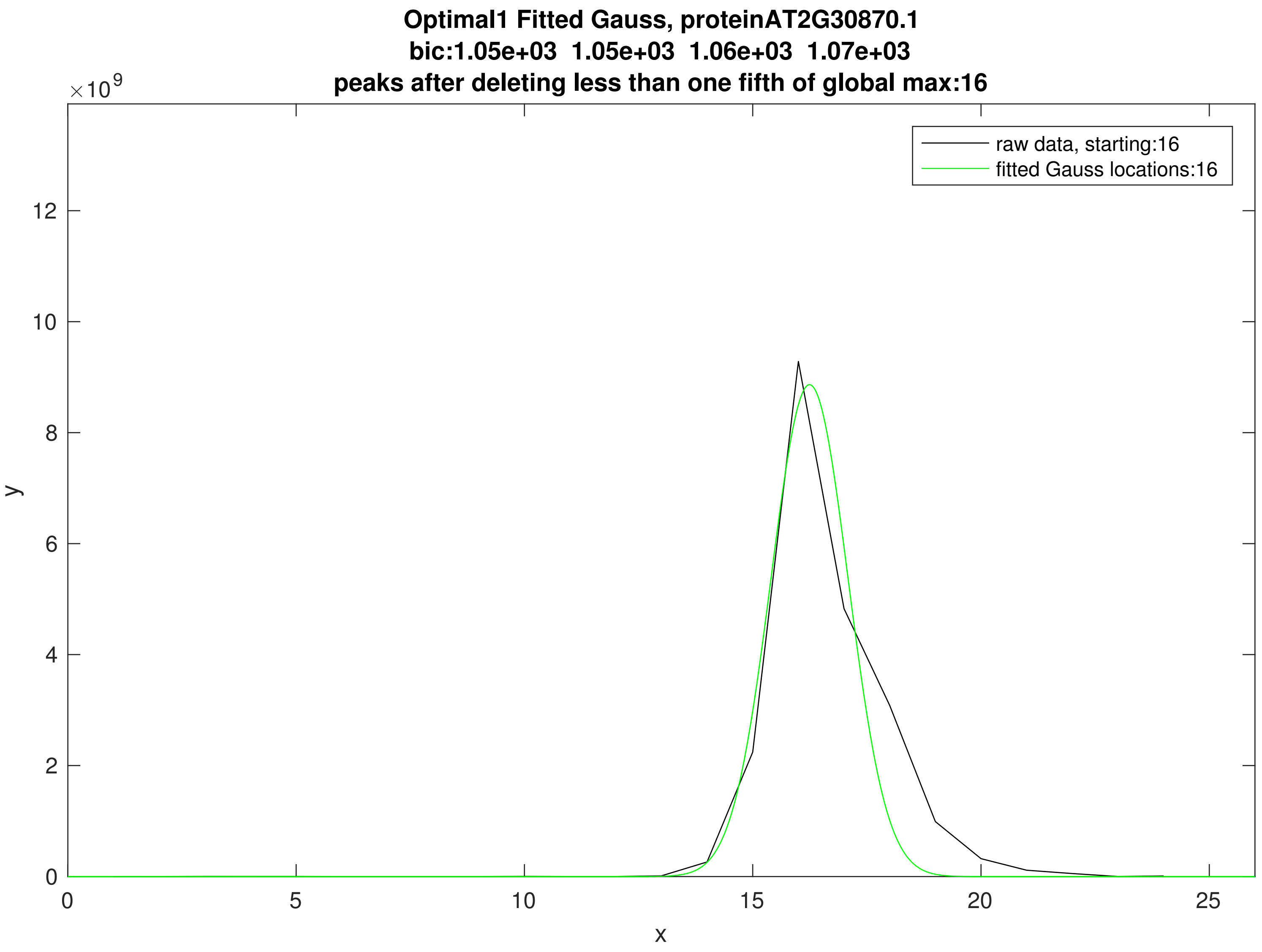

Optimal1 Fitted Gauss, proteinAT2G30870.1
bic:1.05e+03  1.05e+03  1.06e+03  1.07e+03
peaks after deleting less than one fifth of global max:16
×10^9
raw data, starting:16
fitted Gauss locations:16
y
x
0
2
4
6
8
10
12
0
5
10
15
20
25

Optimal1 Fitted Gauss, proteinAT2G30930.1
bic:872 881
peaks after deleting less than one fifth of global max:16

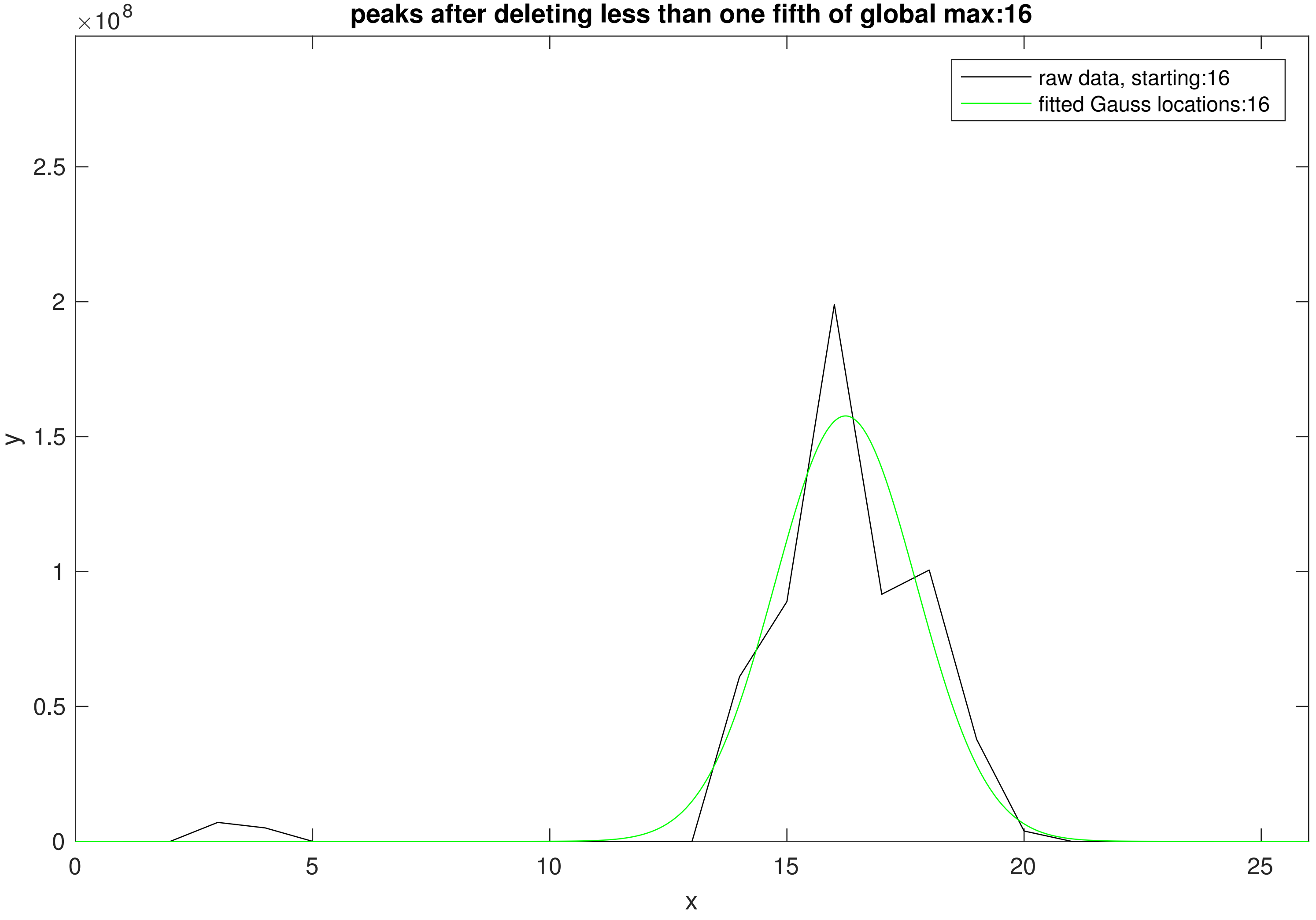

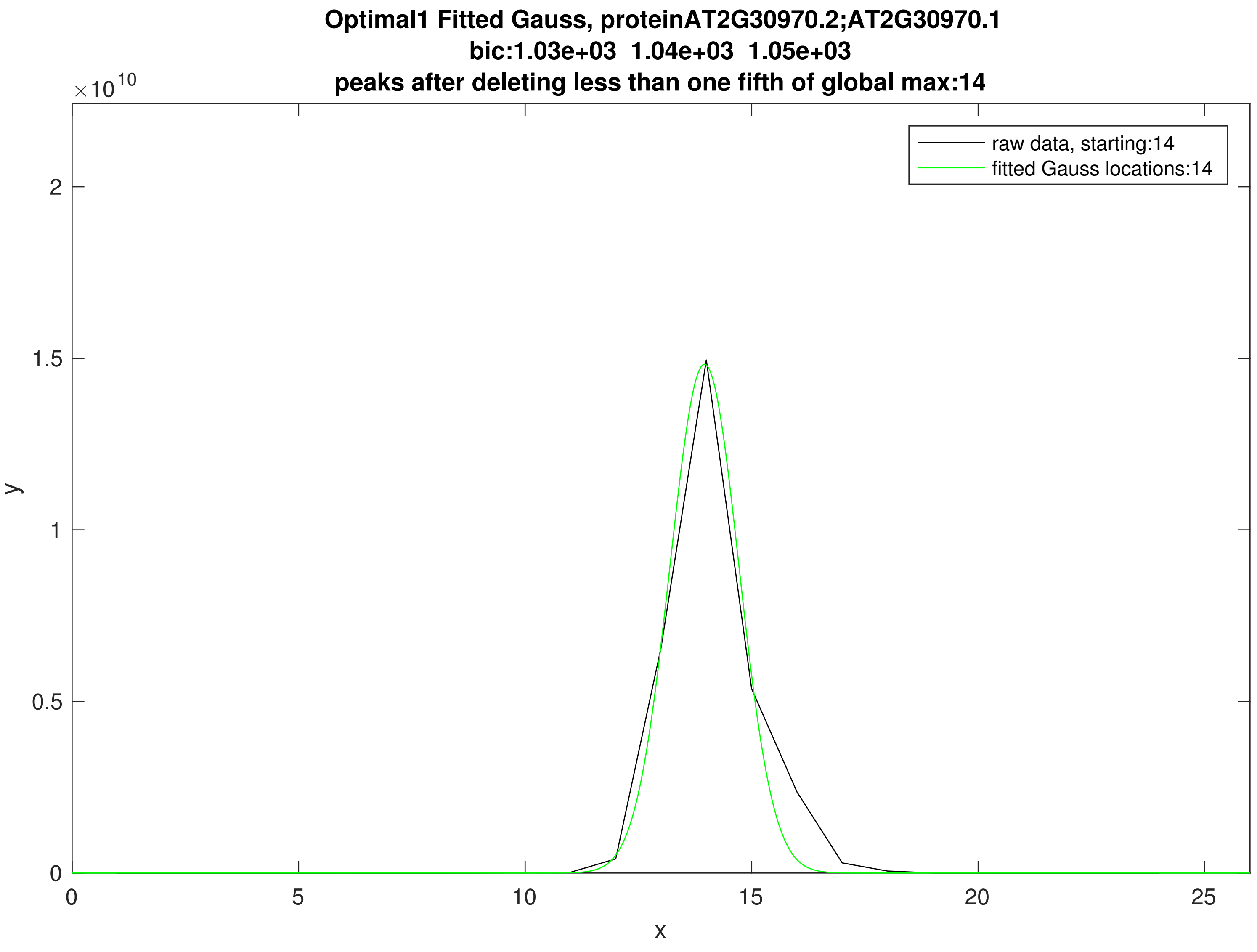
Optimal1 Fitted Gauss, proteinAT2G30970.2;AT2G30970.1
bic:1.03e+03 1.04e+03 1.05e+03
peaks after deleting less than one fifth of global max:14
×10^10
raw data, starting:14
fitted Gauss locations:14
y
x
0
0.5
1
1.5
2
0
5
10
15
20
25

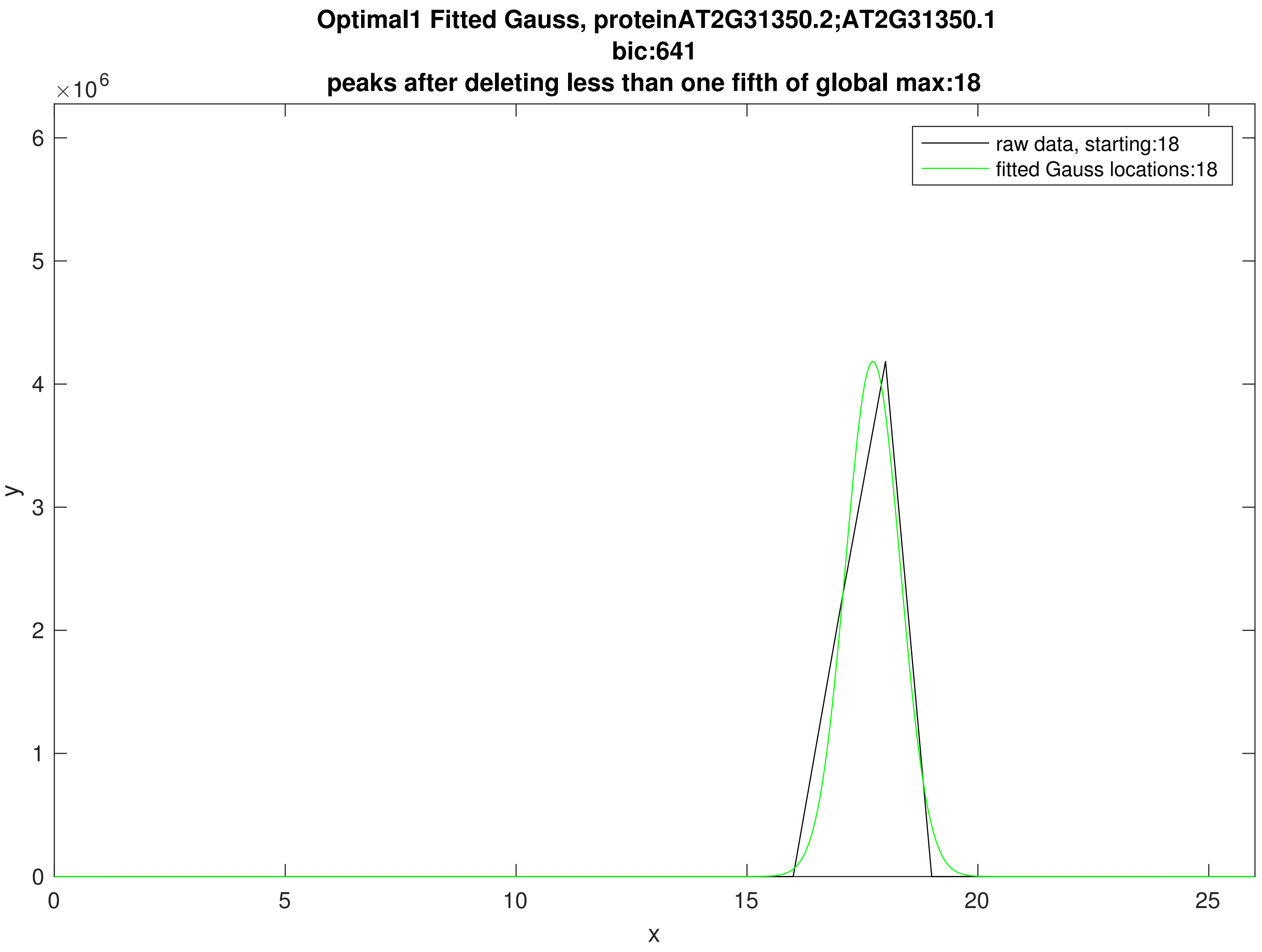
Optimal1 Fitted Gauss, proteinAT2G31350.2;AT2G31350.1
bic:641
peaks after deleting less than one fifth of global max:18
×10^6
raw data, starting:18
fitted Gauss locations:18
y
x
0
1
2
3
4
5
6
0
5
10
15
20
25

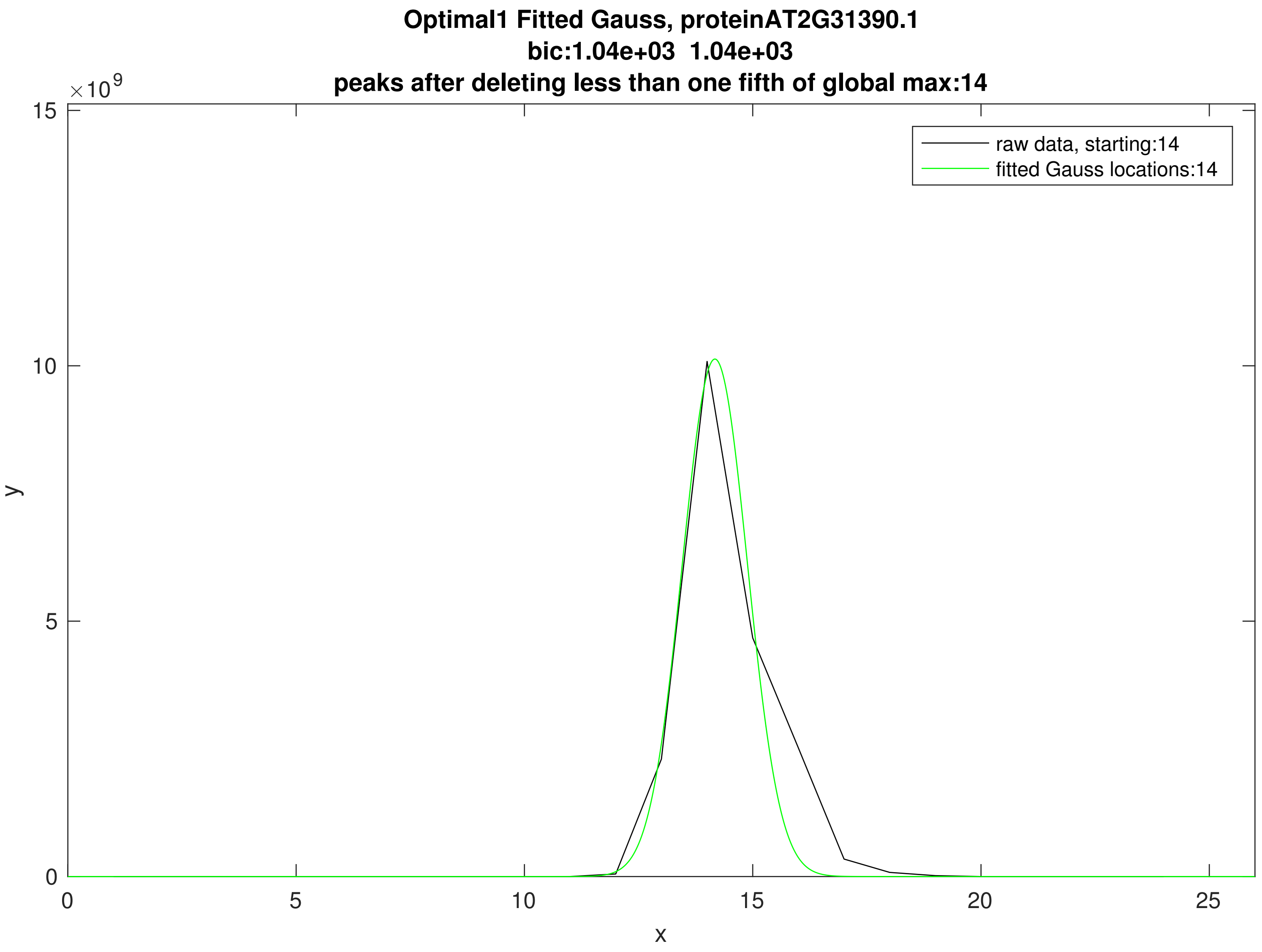
Optimal1 Fitted Gauss, proteinAT2G31390.1
bic:1.04e+03  1.04e+03
peaks after deleting less than one fifth of global max:14
×10^9
15
10
5
0
y
0
5
10
15
20
25
x
raw data, starting:14
fitted Gauss locations:14

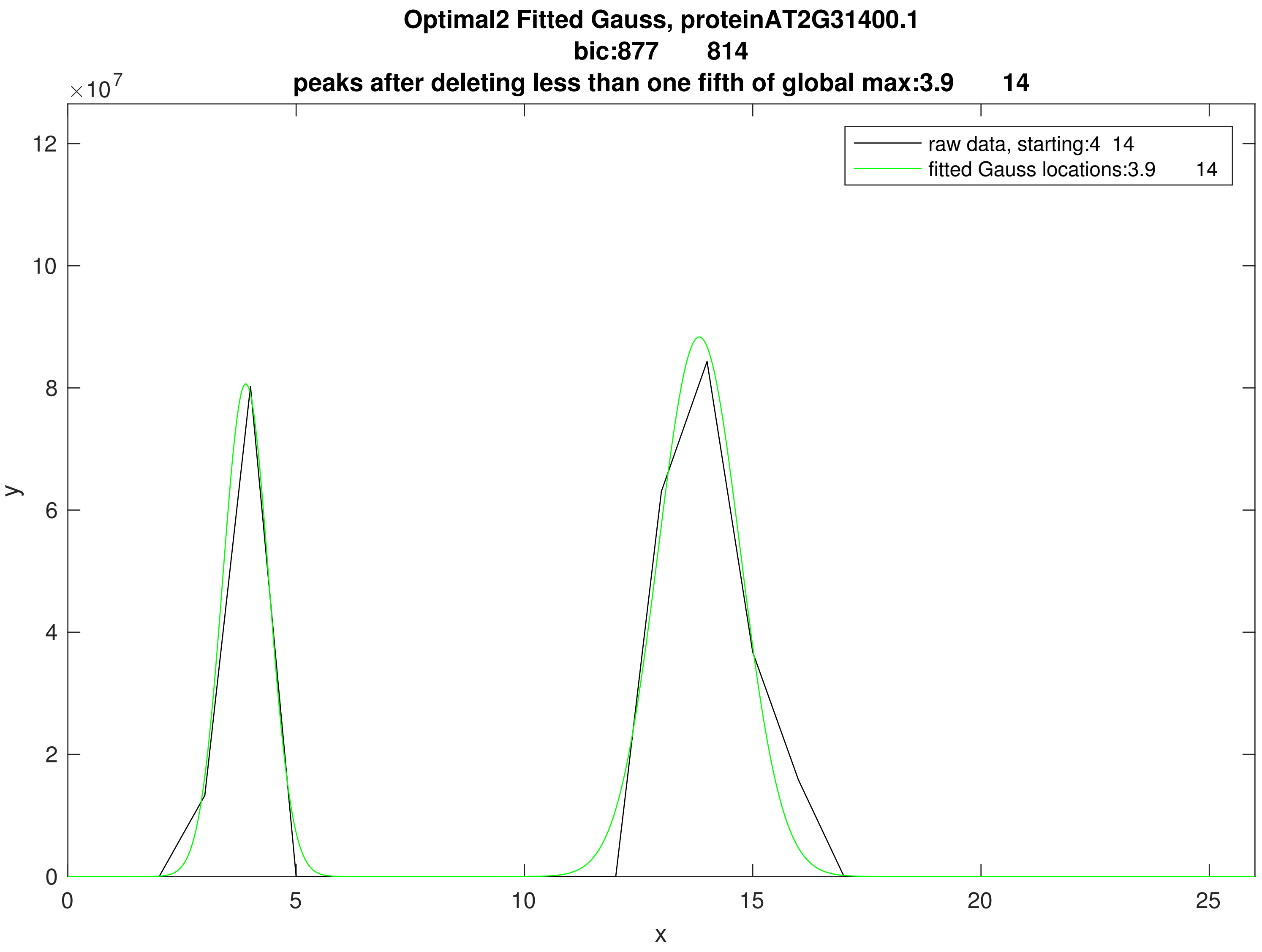
Optimal2 Fitted Gauss, proteinAT2G31400.1
bic:877 814
peaks after deleting less than one fifth of global max:3.9 14
×10^7
raw data, starting:4 14
fitted Gauss locations:3.9 14
y
x
0
2
4
6
8
10
12
5
15
20
25

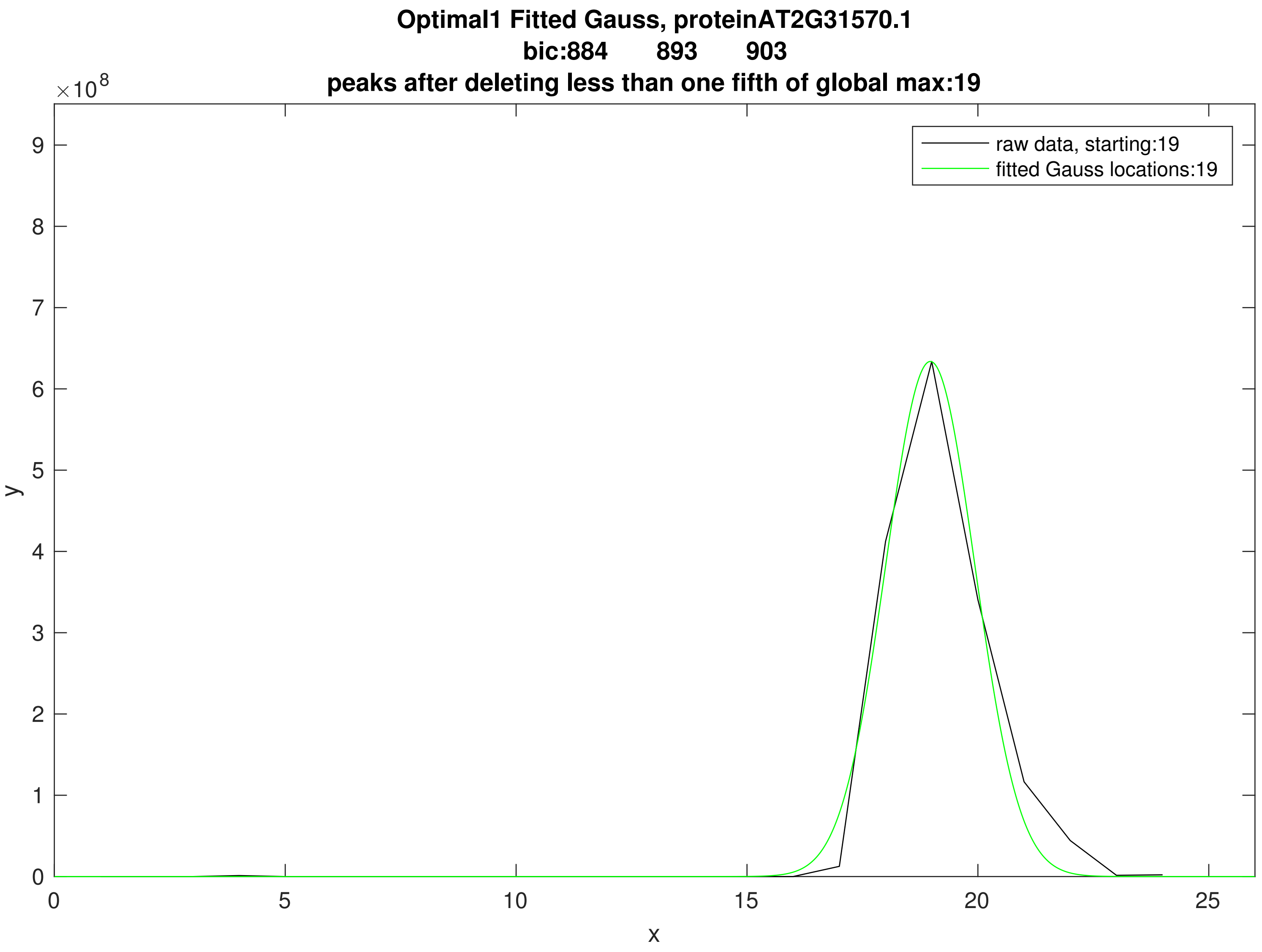

Optimal1 Fitted Gauss, proteinAT2G31570.1
bic:884 893 903
peaks after deleting less than one fifth of global max:19
×10^8
raw data, starting:19
fitted Gauss locations:19
y
x
0
1
2
3
4
5
6
7
8
9
0
5
10
15
20
25

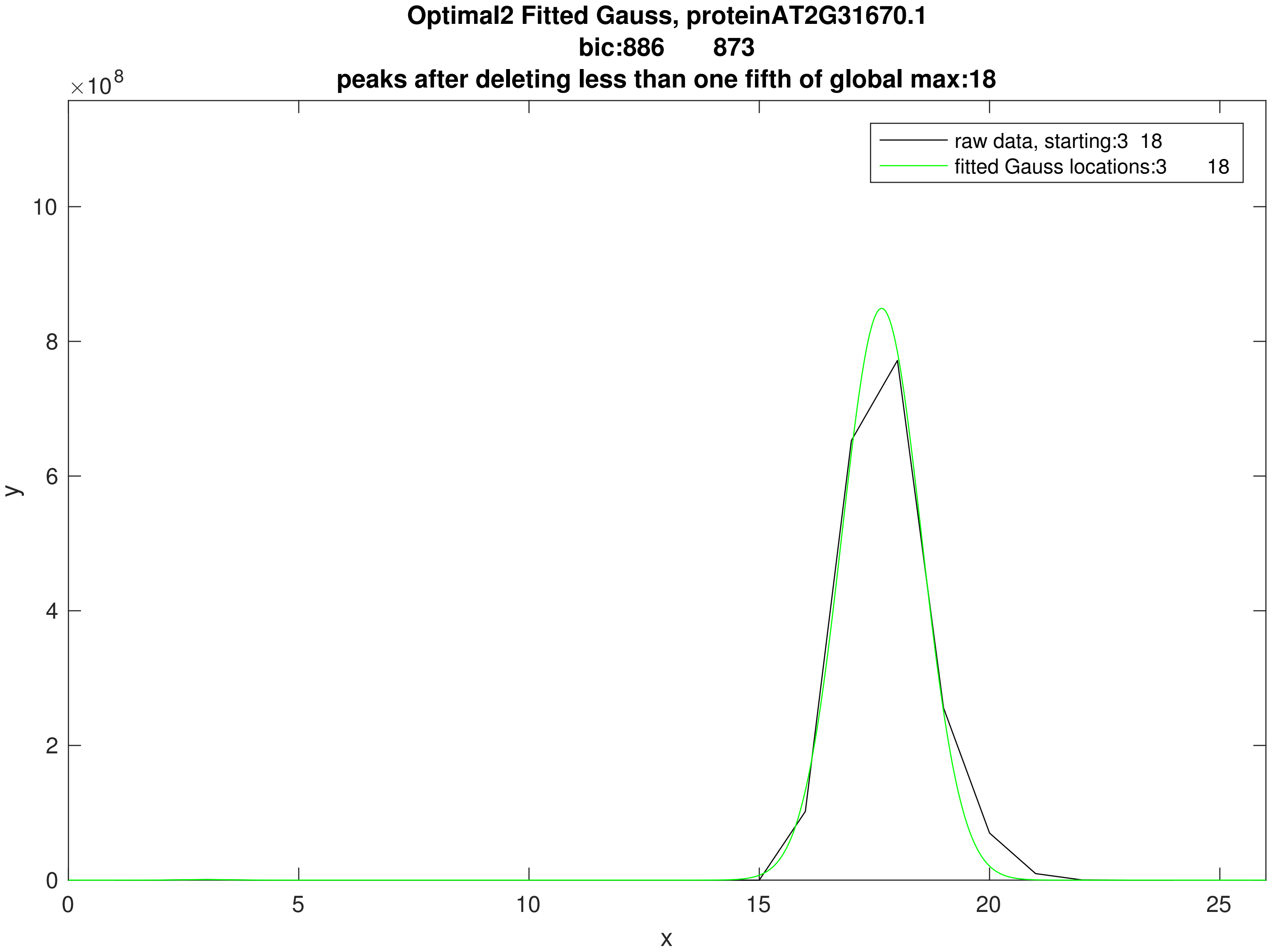
Optimal2 Fitted Gauss, proteinAT2G31670.1
bic:886 873
peaks after deleting less than one fifth of global max:18
×10^8
raw data, starting:3 18
fitted Gauss locations:3 18
10
8
6
4
2
0
y
0
5
10
15
20
25
x

**Optimal1 Fitted Gauss, proteinAT2G31725.1**
**bic:757**
**peaks after deleting less than one fifth of global max:15**

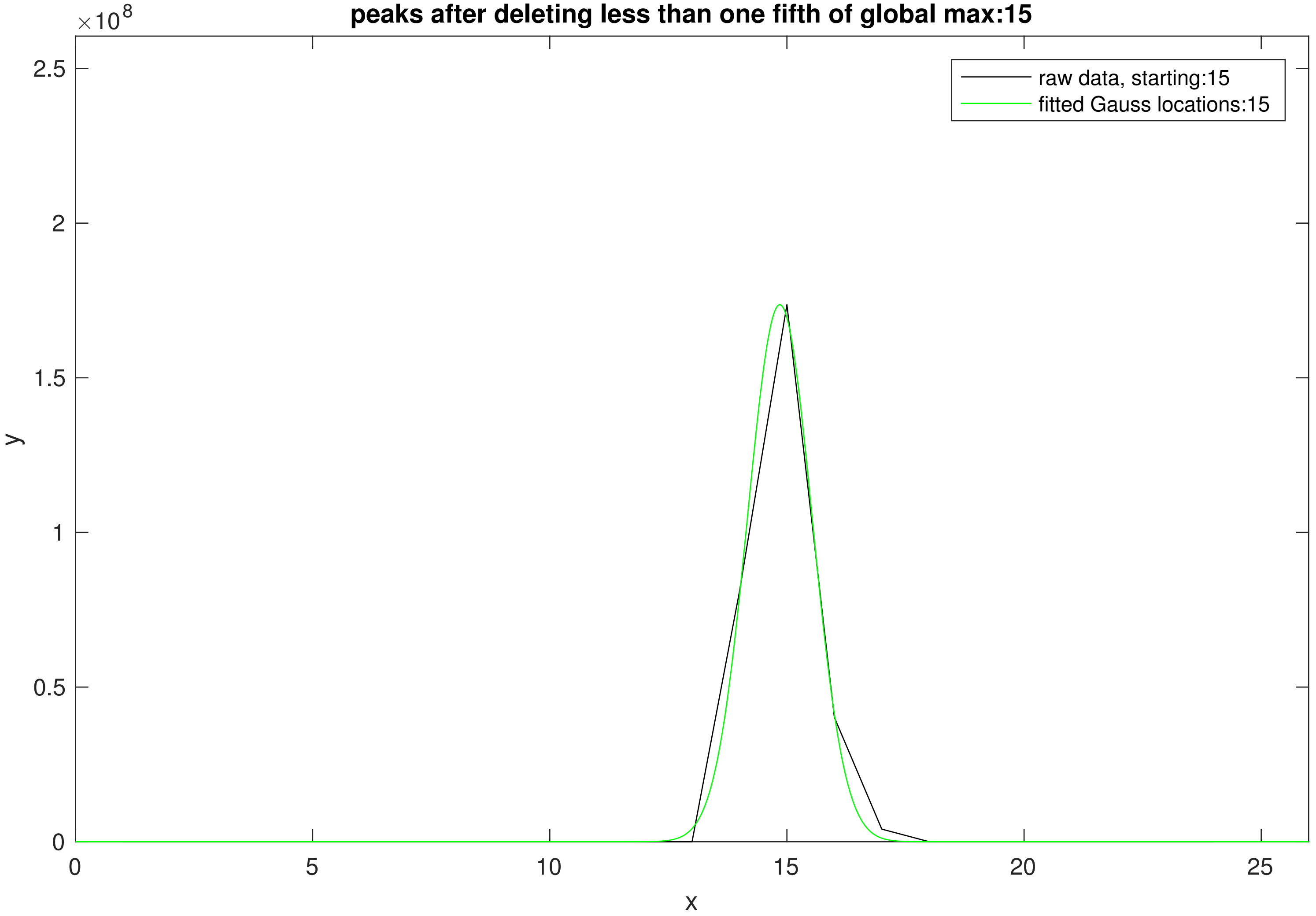

**Optimal1 Fitted Gauss, proteinAT2G31750.1**
**bic:800 802**
**peaks after deleting less than one fifth of global max:16**

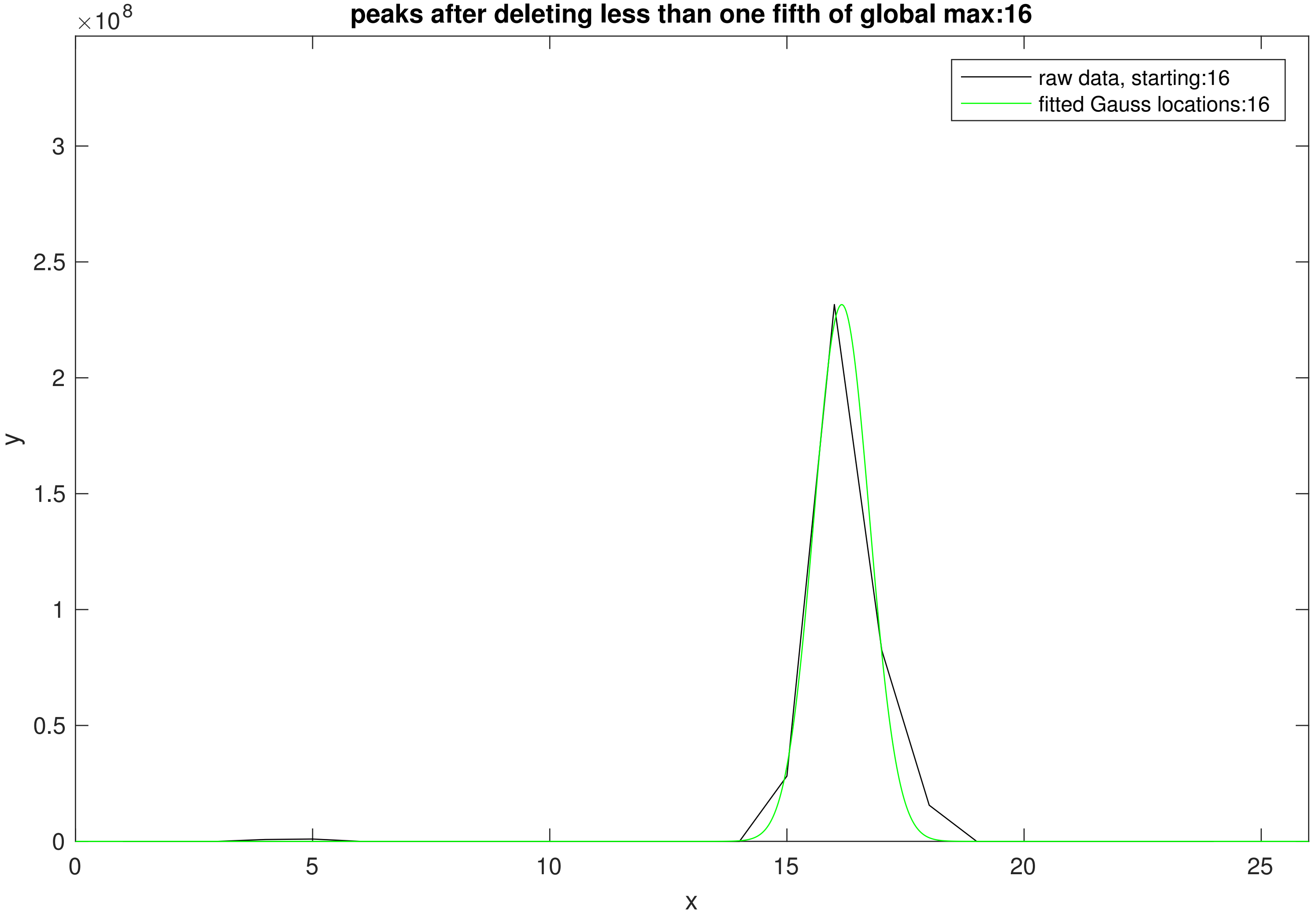

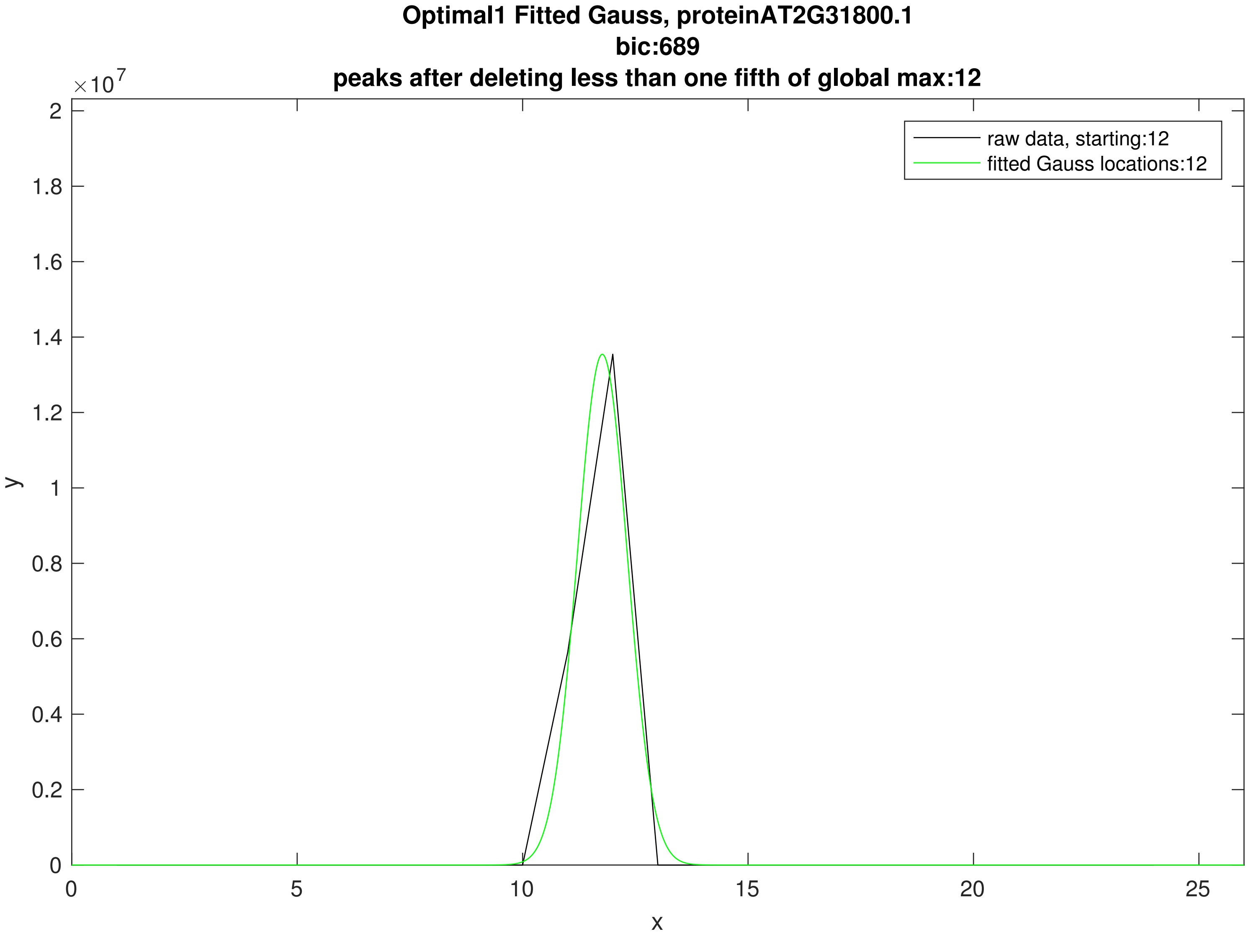

Optimal1 Fitted Gauss, proteinAT2G31800.1
bic:689
peaks after deleting less than one fifth of global max:12
×10^7
2
1.8
1.6
1.4
1.2
1
0.8
0.6
0.4
0.2
0
y
0
5
10
15
20
25
x
raw data, starting:12
fitted Gauss locations:12

**Optimal1 Fitted Gauss, proteinAT2G31985.1**
**bic:893**
**peaks after deleting less than one fifth of global max:13**

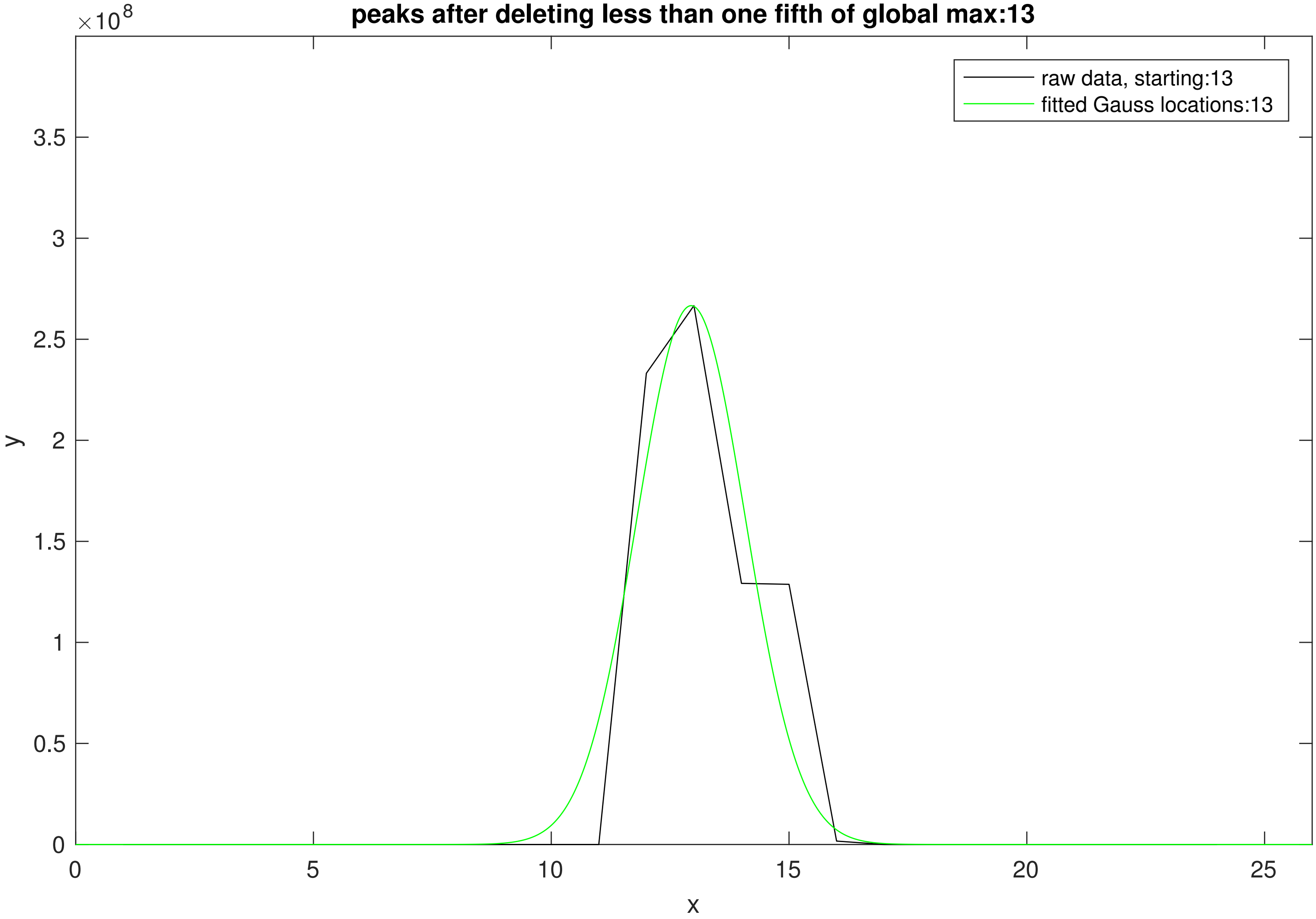

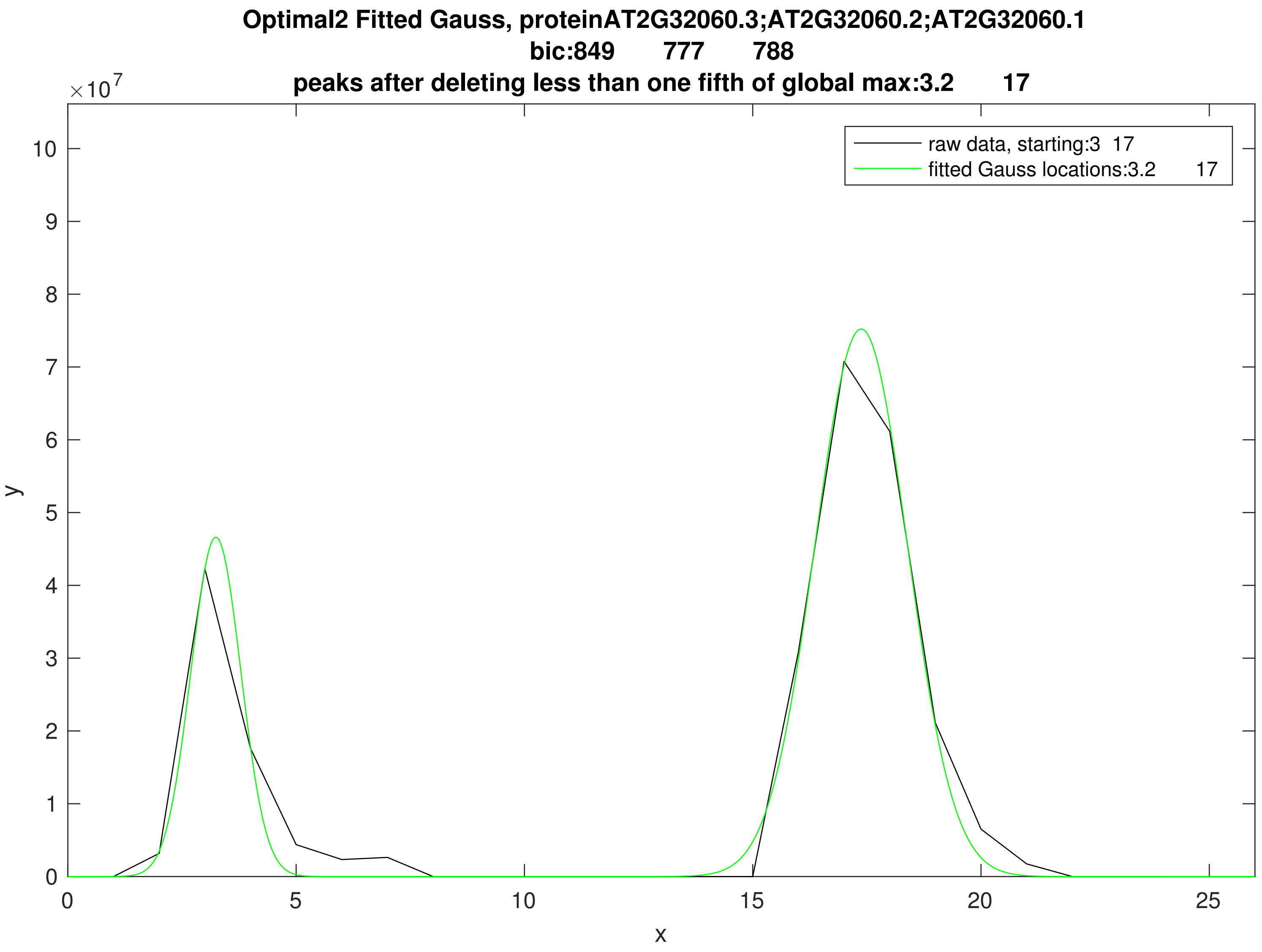

Optimal2 Fitted Gauss, proteinAT2G32060.3;AT2G32060.2;AT2G32060.1
bic:849 777 788
peaks after deleting less than one fifth of global max:3.2 17
×10^7
raw data, starting:3 17
fitted Gauss locations:3.2 17
y
x
0
1
2
3
4
5
6
7
8
9
10
0
5
10
15
20
25

Optimal1 Fitted Gauss, proteinAT2G32120.2;AT2G32120.1
bic:829
peaks after deleting less than one fifth of global max:12

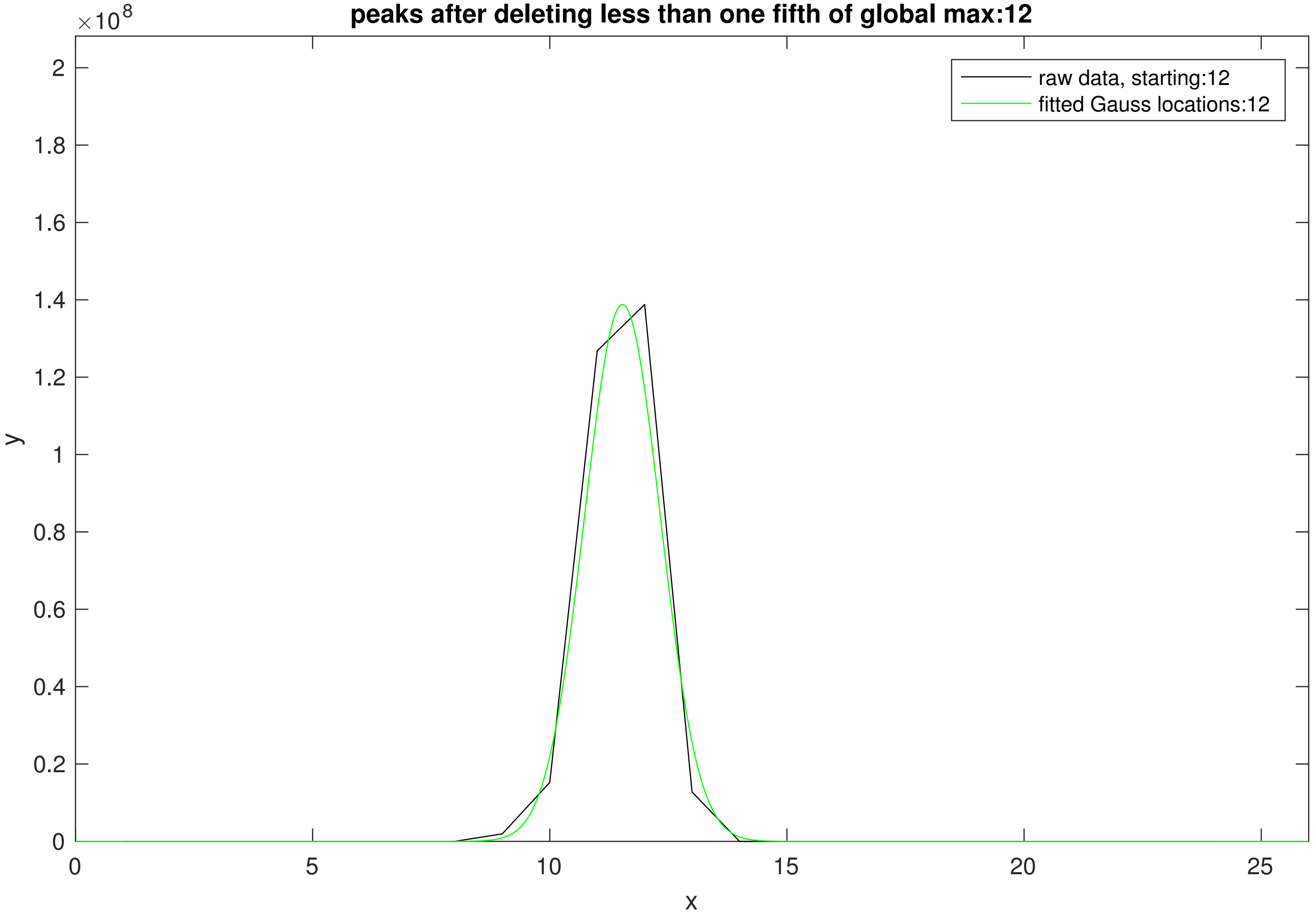

**Optimal1 Fitted Gauss, proteinAT2G32170.1;AT2G32170.2**
**bic:721**
**peaks after deleting less than one fifth of global max:10**

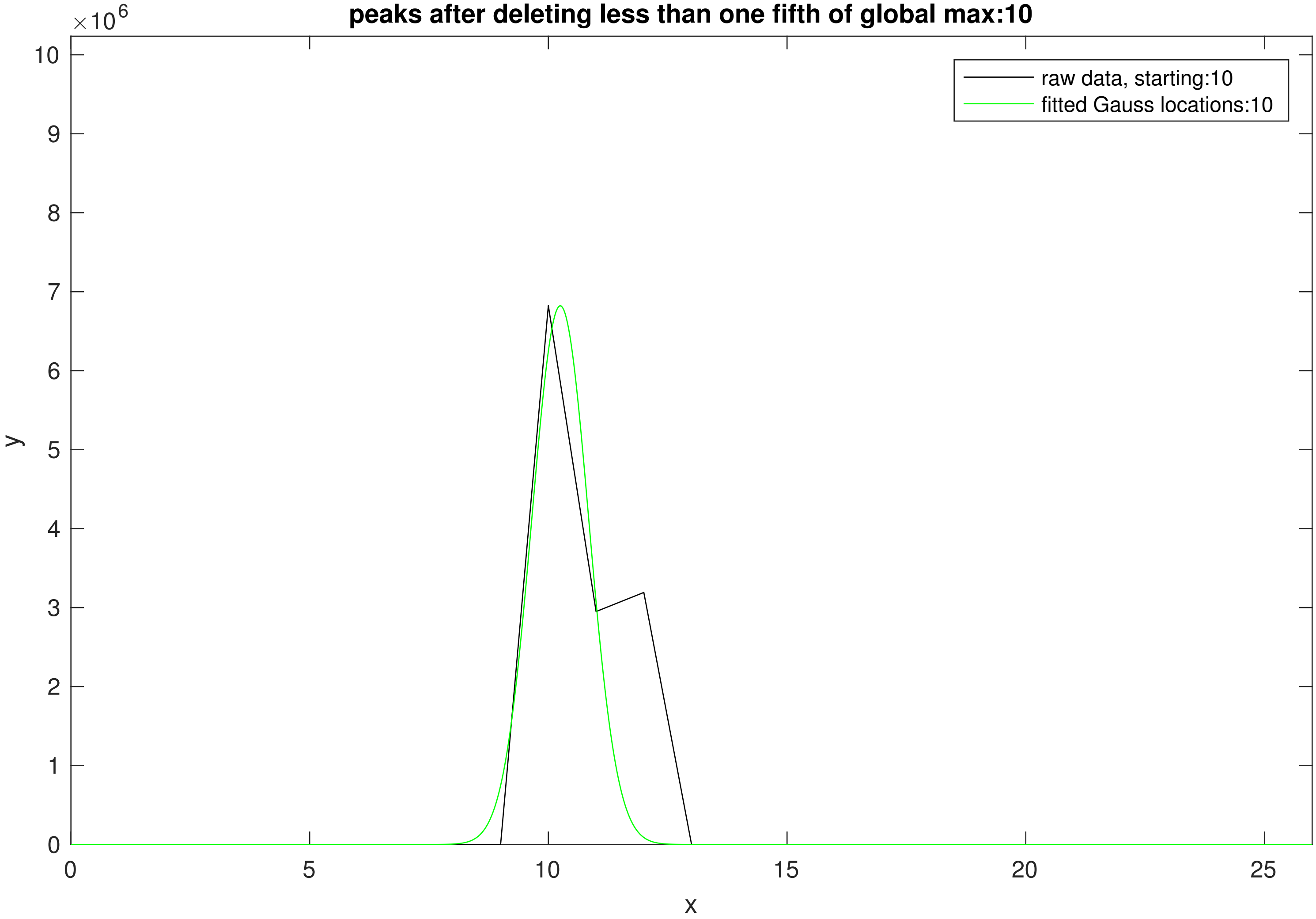

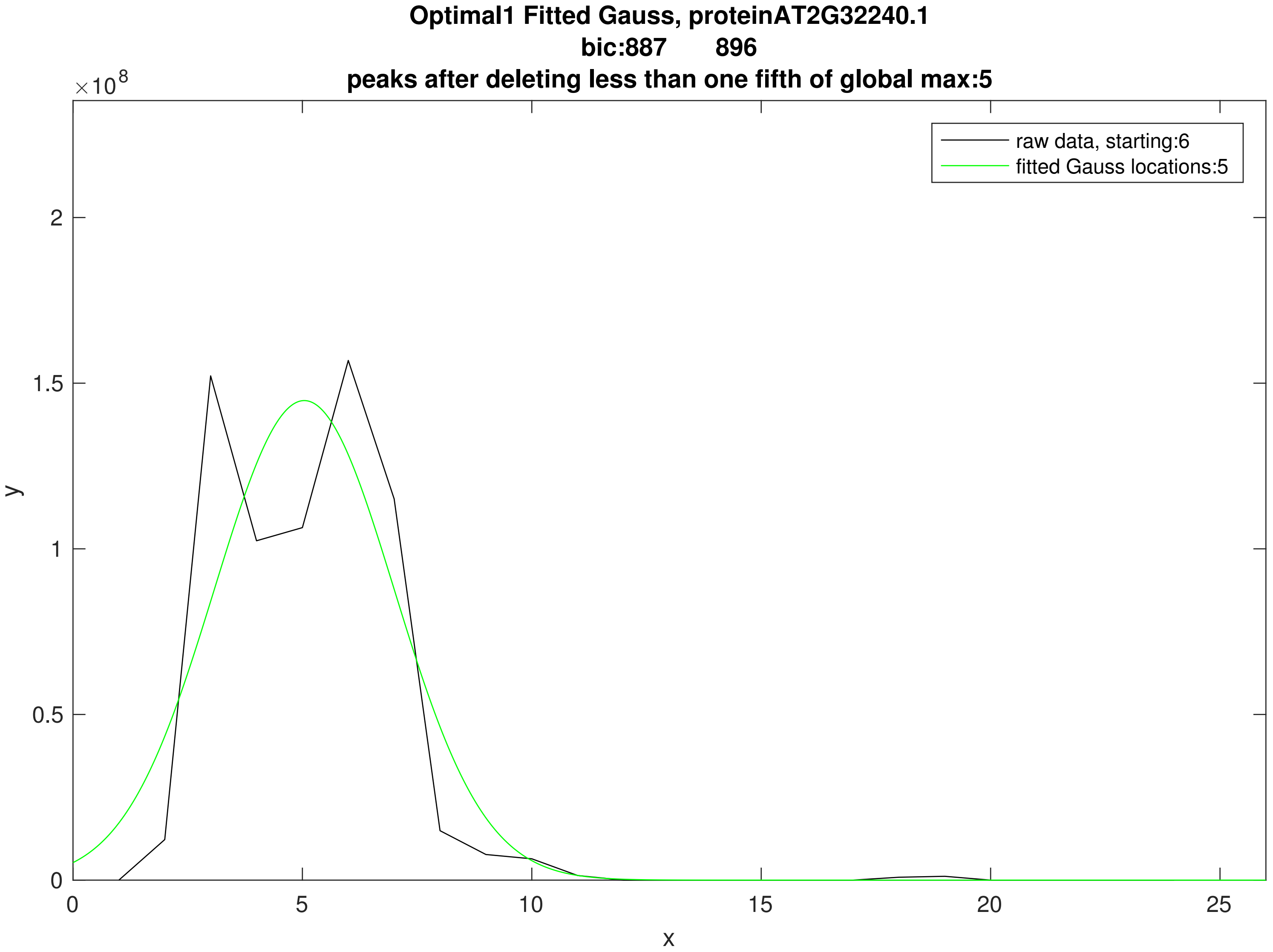
Optimal1 Fitted Gauss, proteinAT2G32240.1
bic:887 896
peaks after deleting less than one fifth of global max:5
×10^8
raw data, starting:6
fitted Gauss locations:5
y
x
0
0.5
1
1.5
2
0
5
10
15
20
25

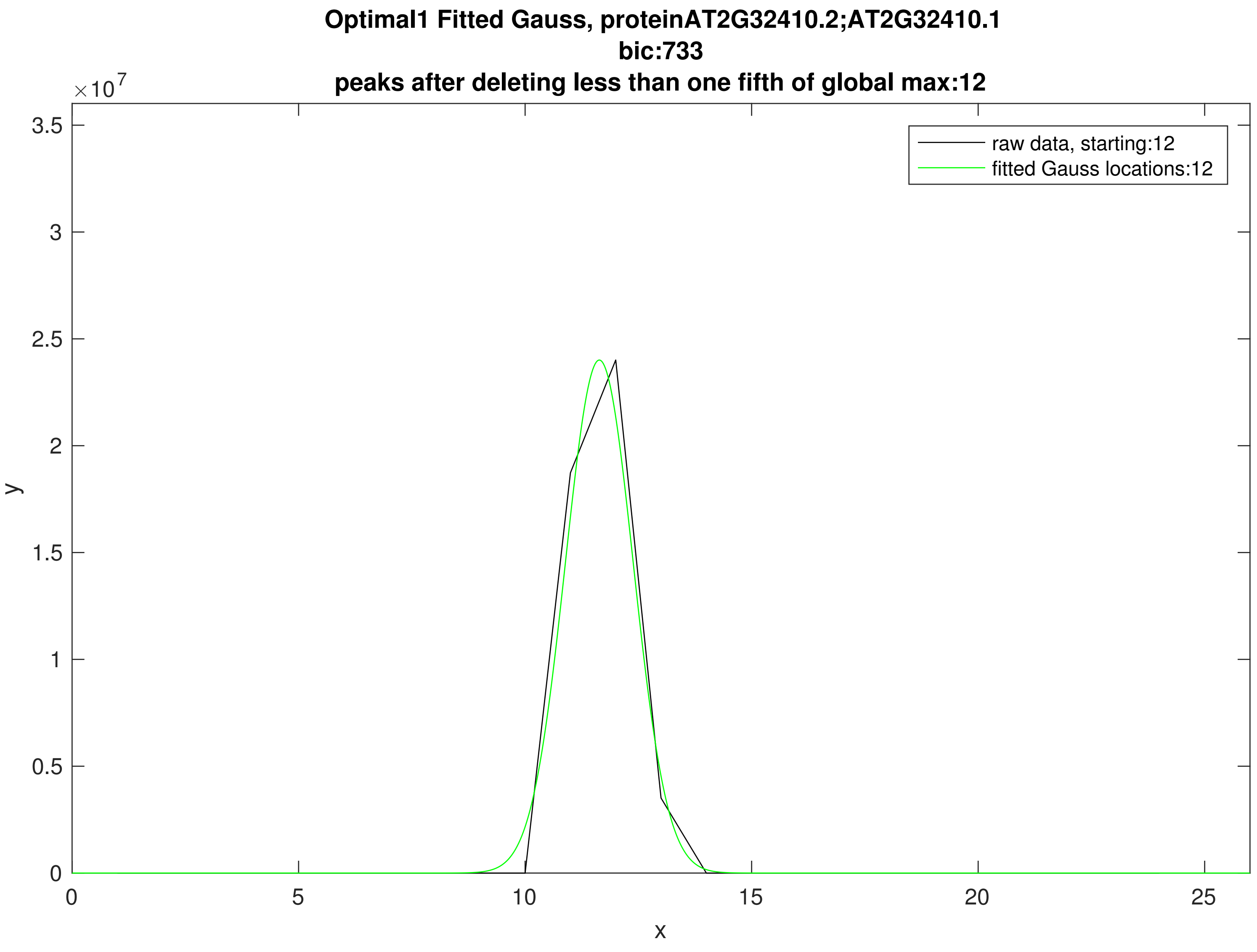
Optimal1 Fitted Gauss, proteinAT2G32410.2;AT2G32410.1
bic:733
peaks after deleting less than one fifth of global max:12
×10^7
raw data, starting:12
fitted Gauss locations:12
y
x
0
0.5
1
1.5
2
2.5
3
3.5
0
5
10
15
20
25

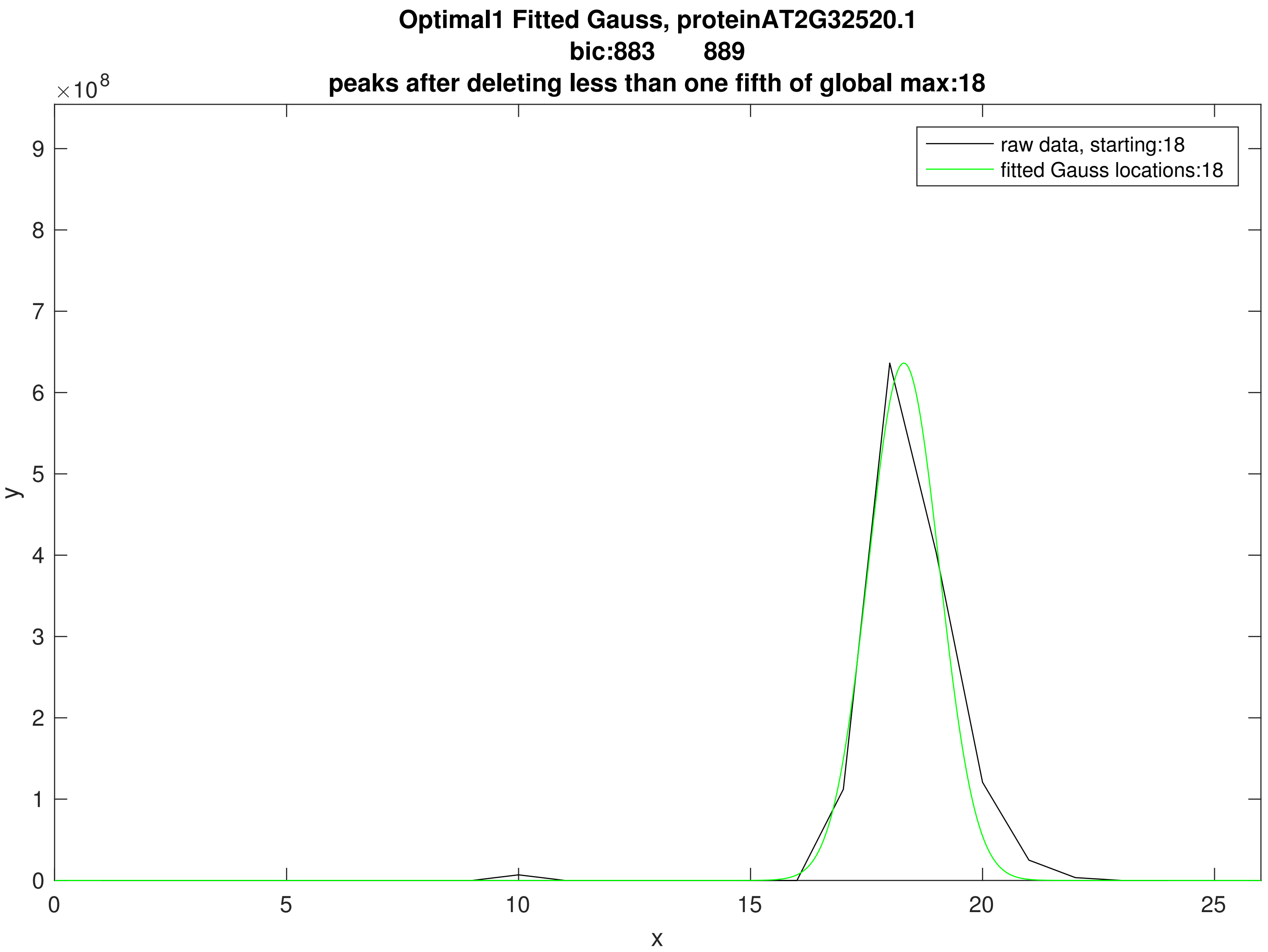
Optimal1 Fitted Gauss, proteinAT2G32520.1
bic:883 889
peaks after deleting less than one fifth of global max:18
×10^8
raw data, starting:18
fitted Gauss locations:18
y
x
0
1
2
3
4
5
6
7
8
9
0
5
10
15
20
25

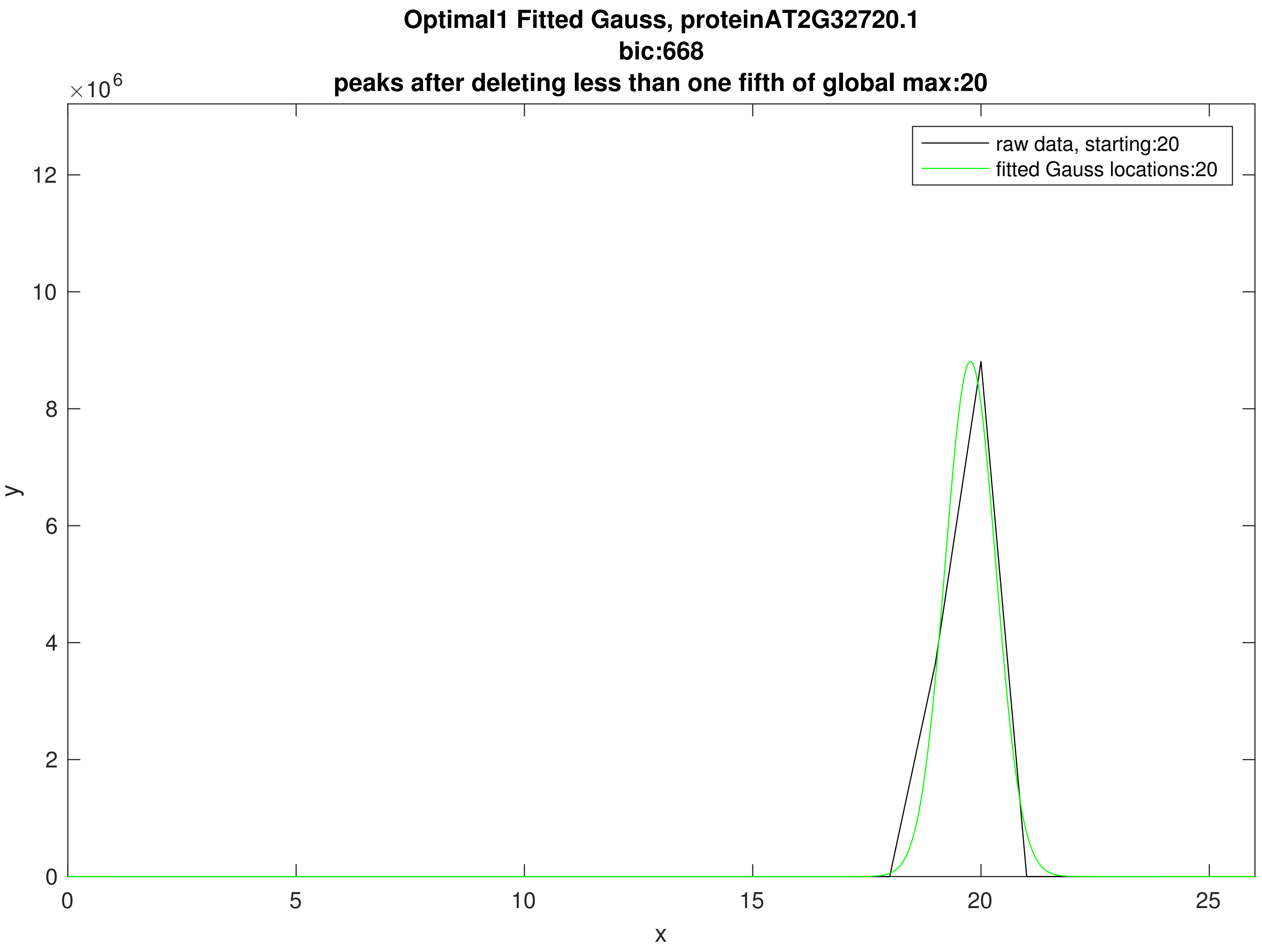

Optimal1 Fitted Gauss, proteinAT2G32720.1
bic:668
peaks after deleting less than one fifth of global max:20
×10^6
12
10
8
6
4
2
0
y
0
5
10
15
20
25
x
raw data, starting:20
fitted Gauss locations:20

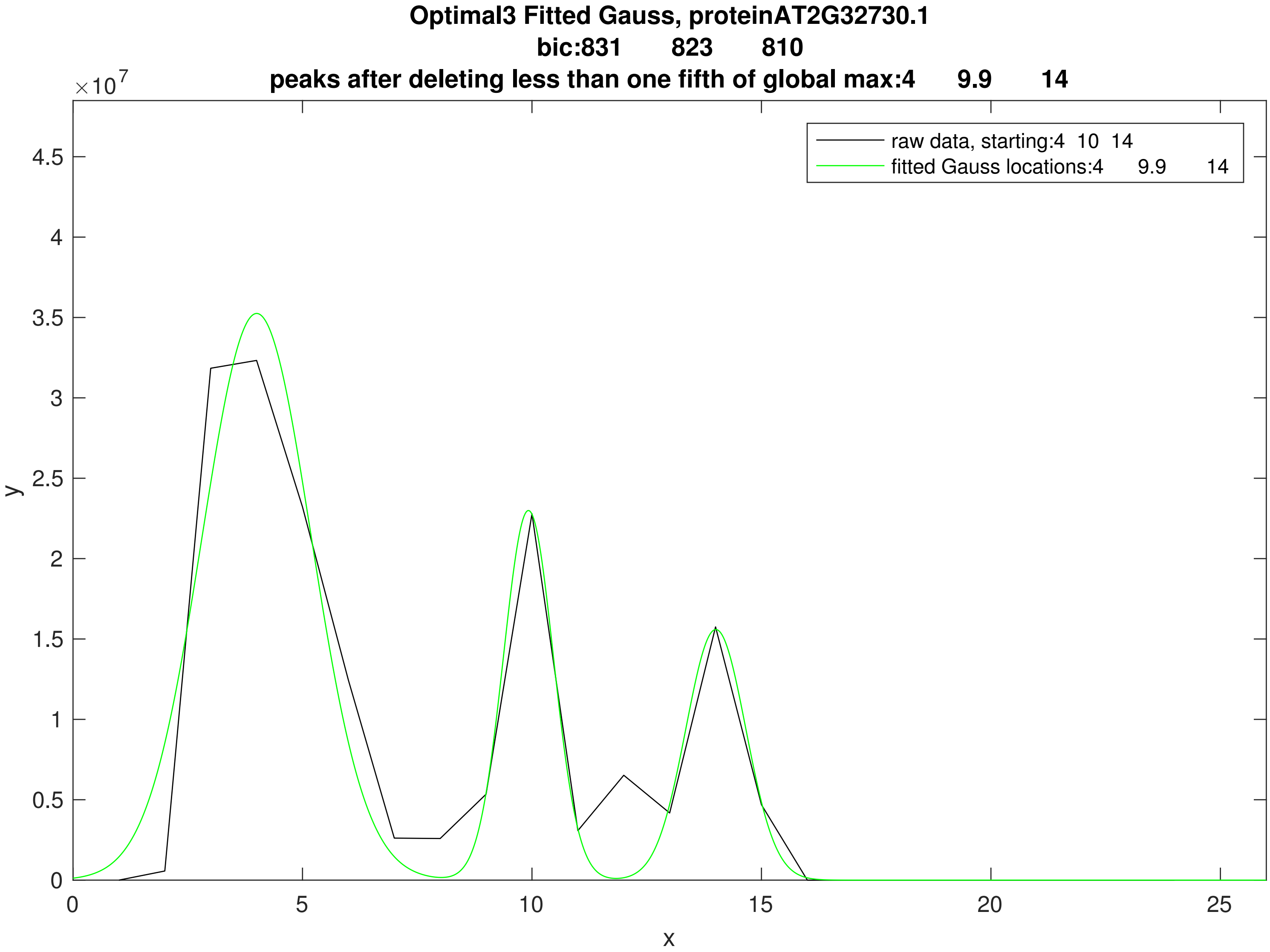
Optimal3 Fitted Gauss, proteinAT2G32730.1
bic:831 823 810
peaks after deleting less than one fifth of global max:4 9.9 14
×10^7
raw data, starting:4 10 14
fitted Gauss locations:4 9.9 14
y
x
0
0.5
1
1.5
2
2.5
3
3.5
4
4.5
5
10
15
20
25

**Optimal1 Fitted Gauss, proteinAT2G33040.1**
**bic:876 940**
**peaks after deleting less than one fifth of global max:7.2**

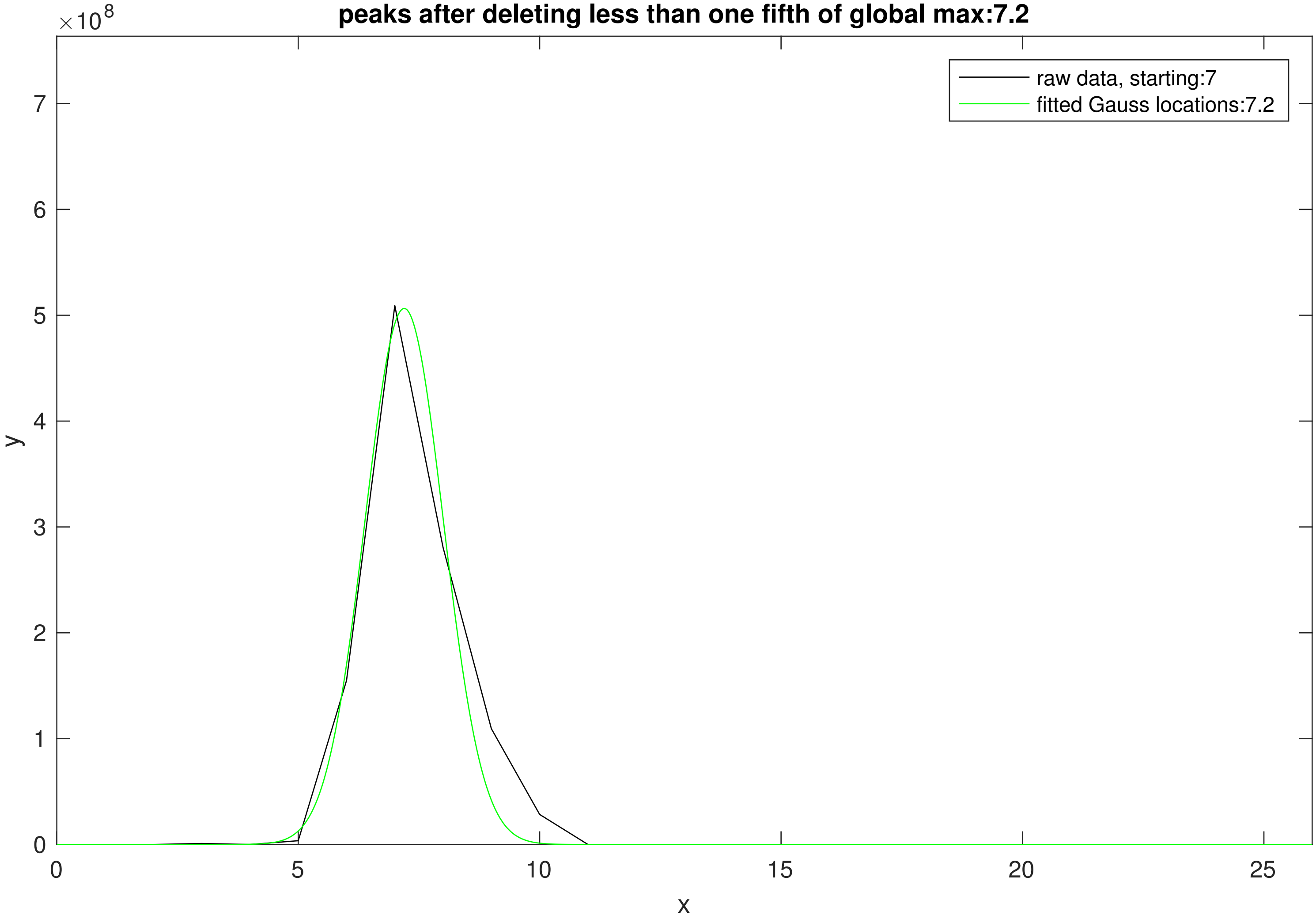

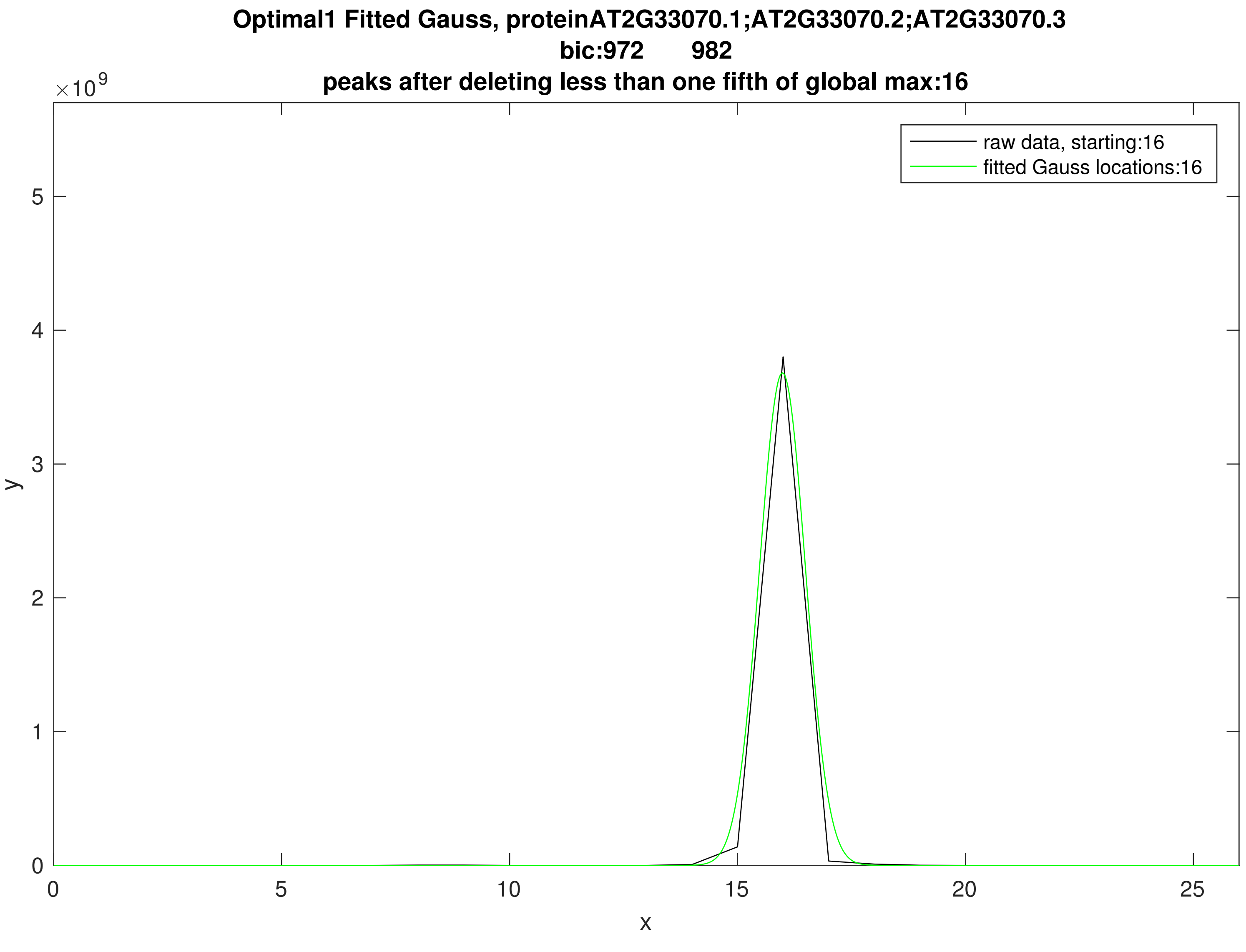
Optimal1 Fitted Gauss, proteinAT2G33070.1;AT2G33070.2;AT2G33070.3
bic:972 982
peaks after deleting less than one fifth of global max:16
×10^9
raw data, starting:16
fitted Gauss locations:16
y
x
0
1
2
3
4
5
0
5
10
15
20
25

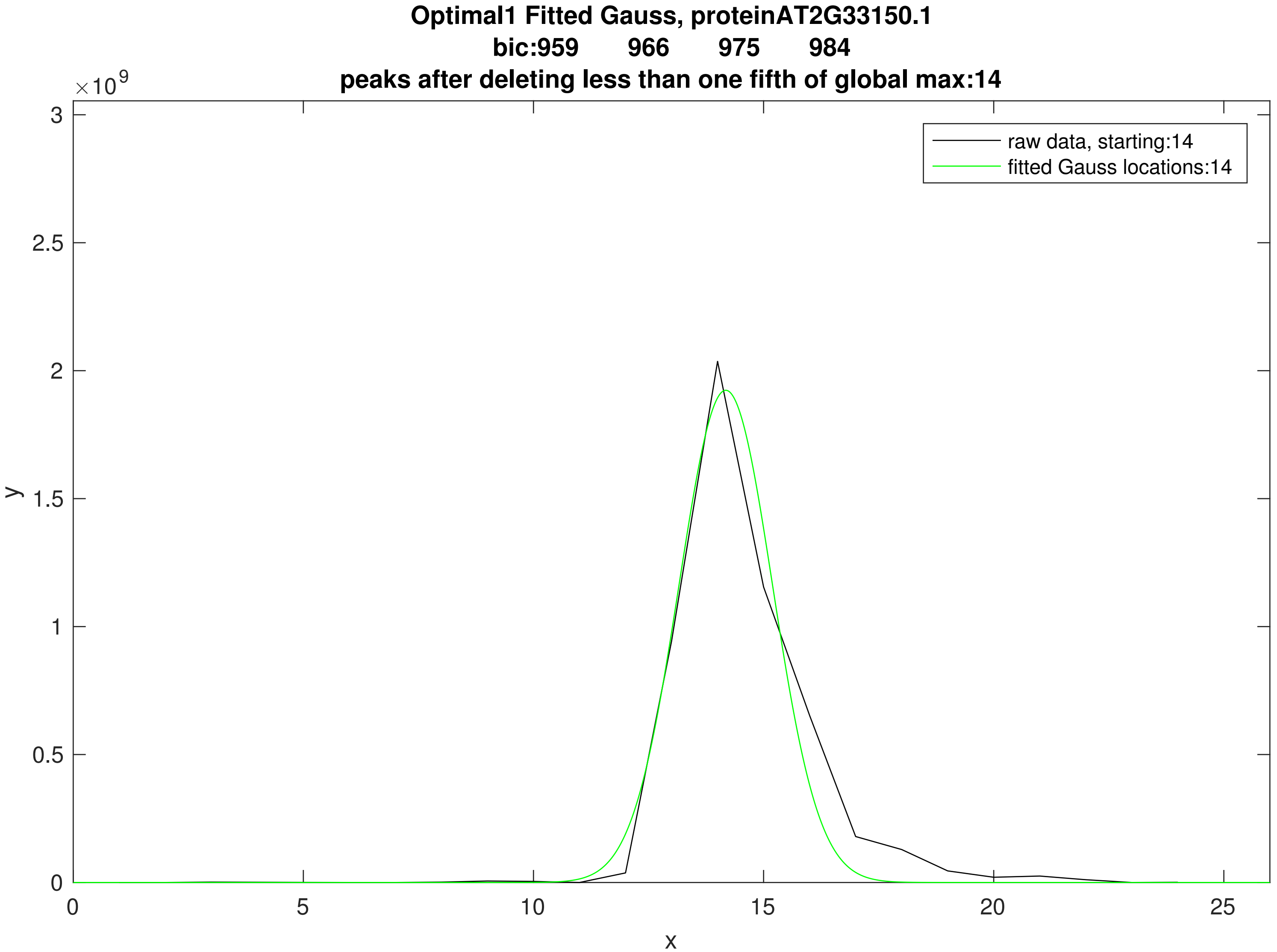
Optimal1 Fitted Gauss, proteinAT2G33150.1
bic:959 966 975 984
peaks after deleting less than one fifth of global max:14
×10^9
raw data, starting:14
fitted Gauss locations:14
y
x
0
0.5
1
1.5
2
2.5
3
0
5
10
15
20
25

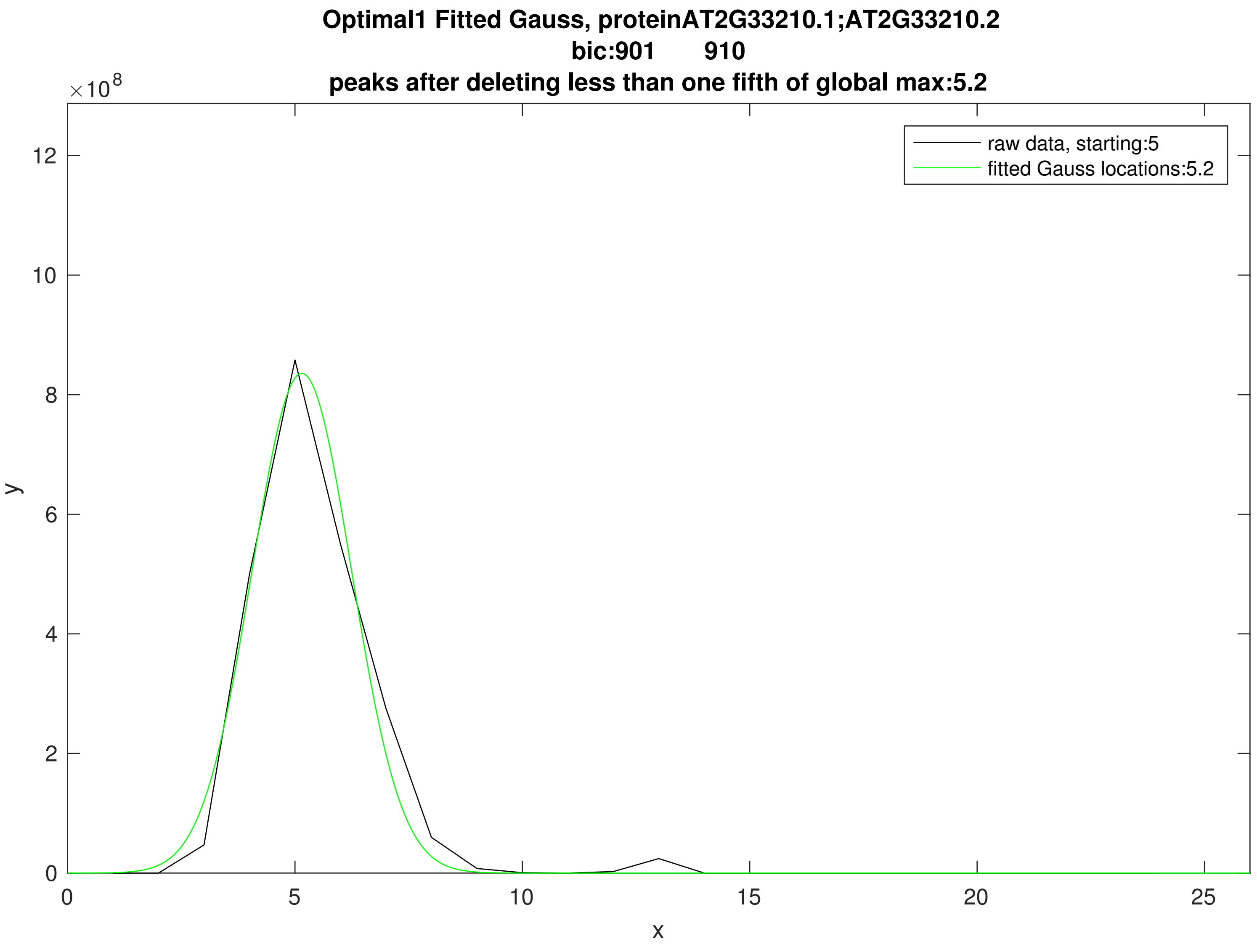

Optimal1 Fitted Gauss, proteinAT2G33210.1;AT2G33210.2
bic:901 910
peaks after deleting less than one fifth of global max:5.2
×10^8
12
10
8
6
4
2
0
y
0
5
10
15
20
25
x
raw data, starting:5
fitted Gauss locations:5.2

**Optimal1 Fitted Gauss, proteinAT2G33255.1**
**bic:683**
**peaks after deleting less than one fifth of global max:18**

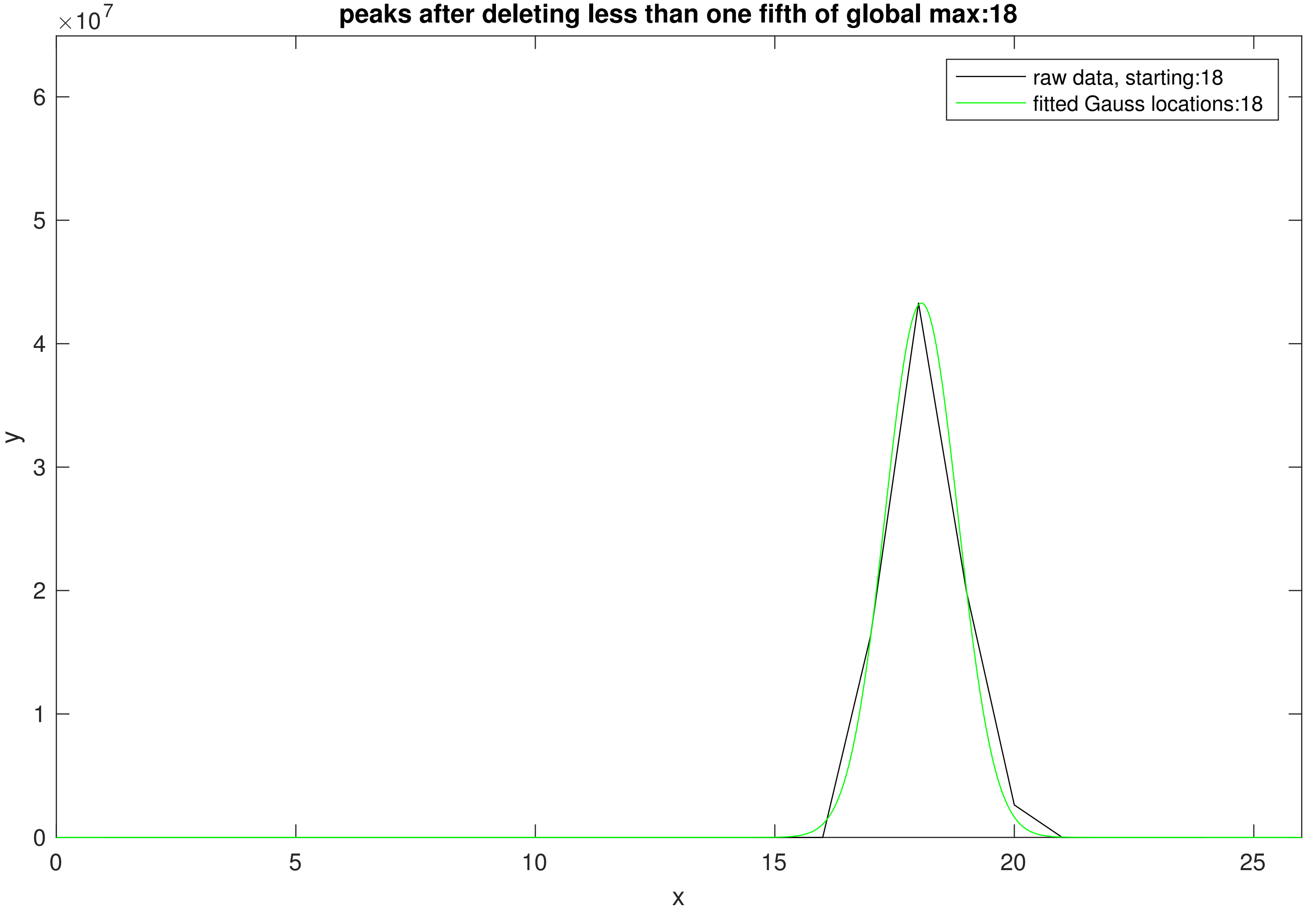

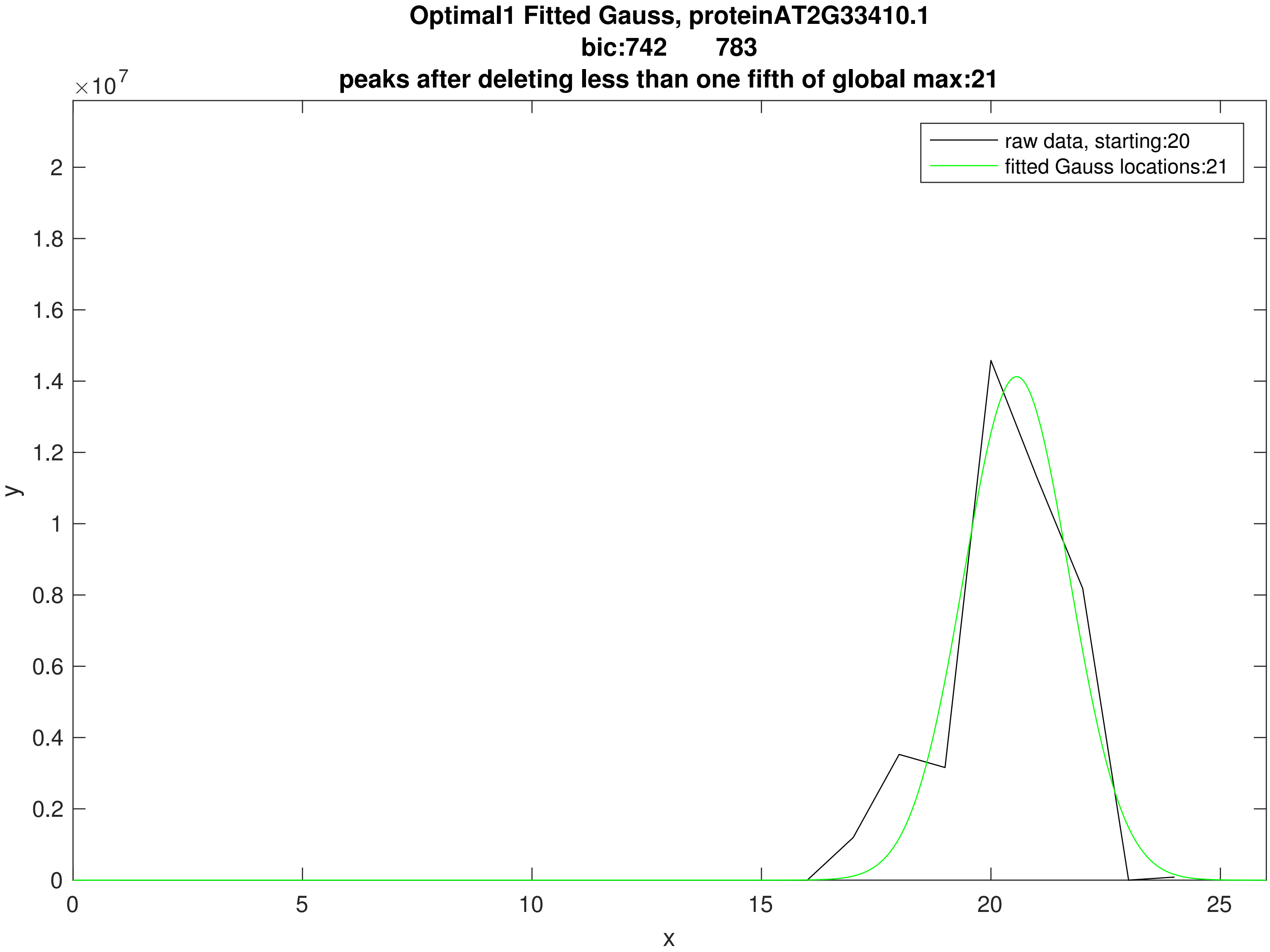
Optimal1 Fitted Gauss, proteinAT2G33410.1
bic:742 783
peaks after deleting less than one fifth of global max:21
×10^7
raw data, starting:20
fitted Gauss locations:21
y
x
0
0.2
0.4
0.6
0.8
1
1.2
1.4
1.6
1.8
2
0
5
10
15
20
25

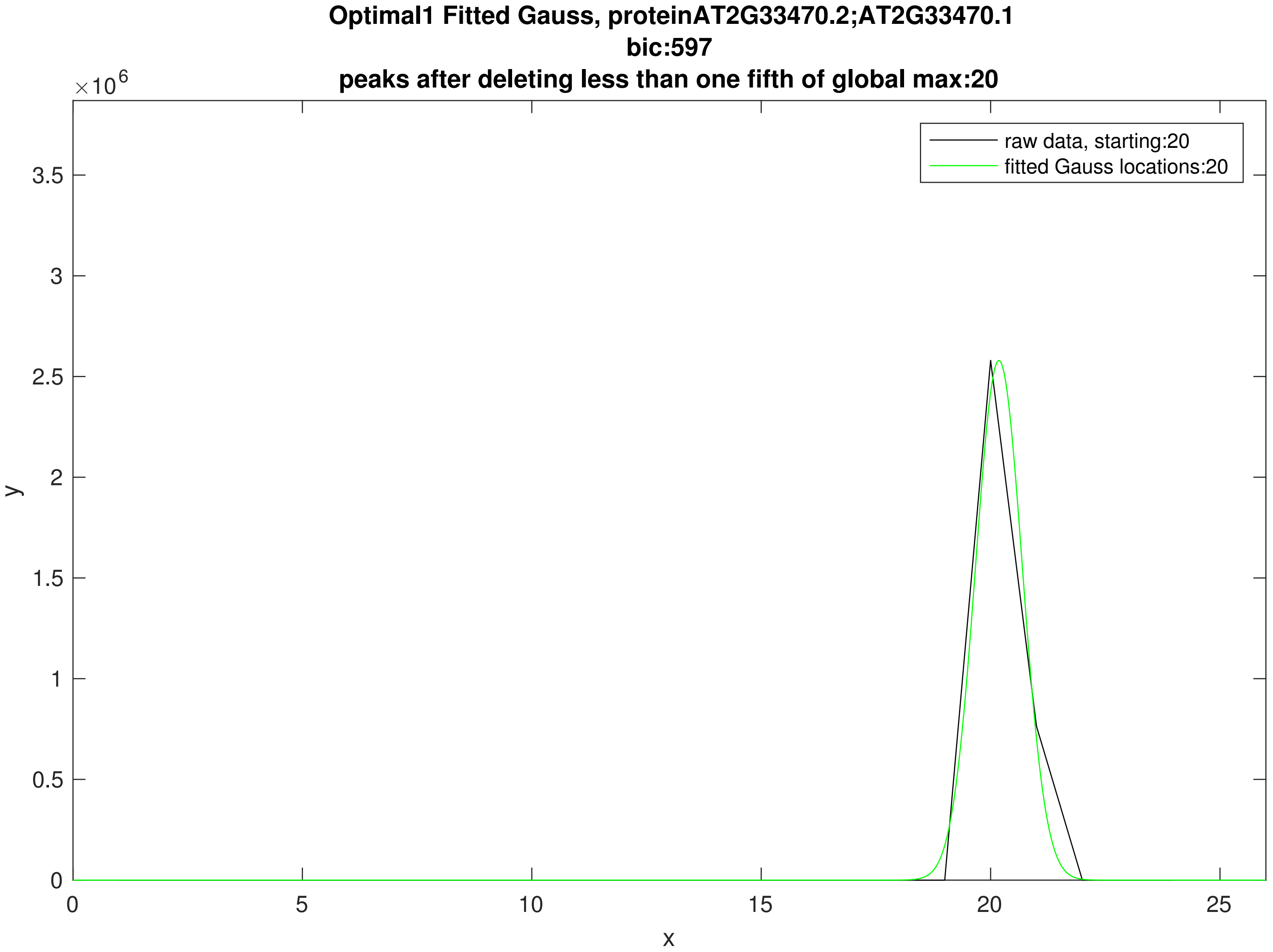
Optimal1 Fitted Gauss, proteinAT2G33470.2;AT2G33470.1
bic:597
peaks after deleting less than one fifth of global max:20
×10^6
3.5
3
2.5
2
1.5
1
0.5
0
y
0
5
10
15
20
25
x
raw data, starting:20
fitted Gauss locations:20

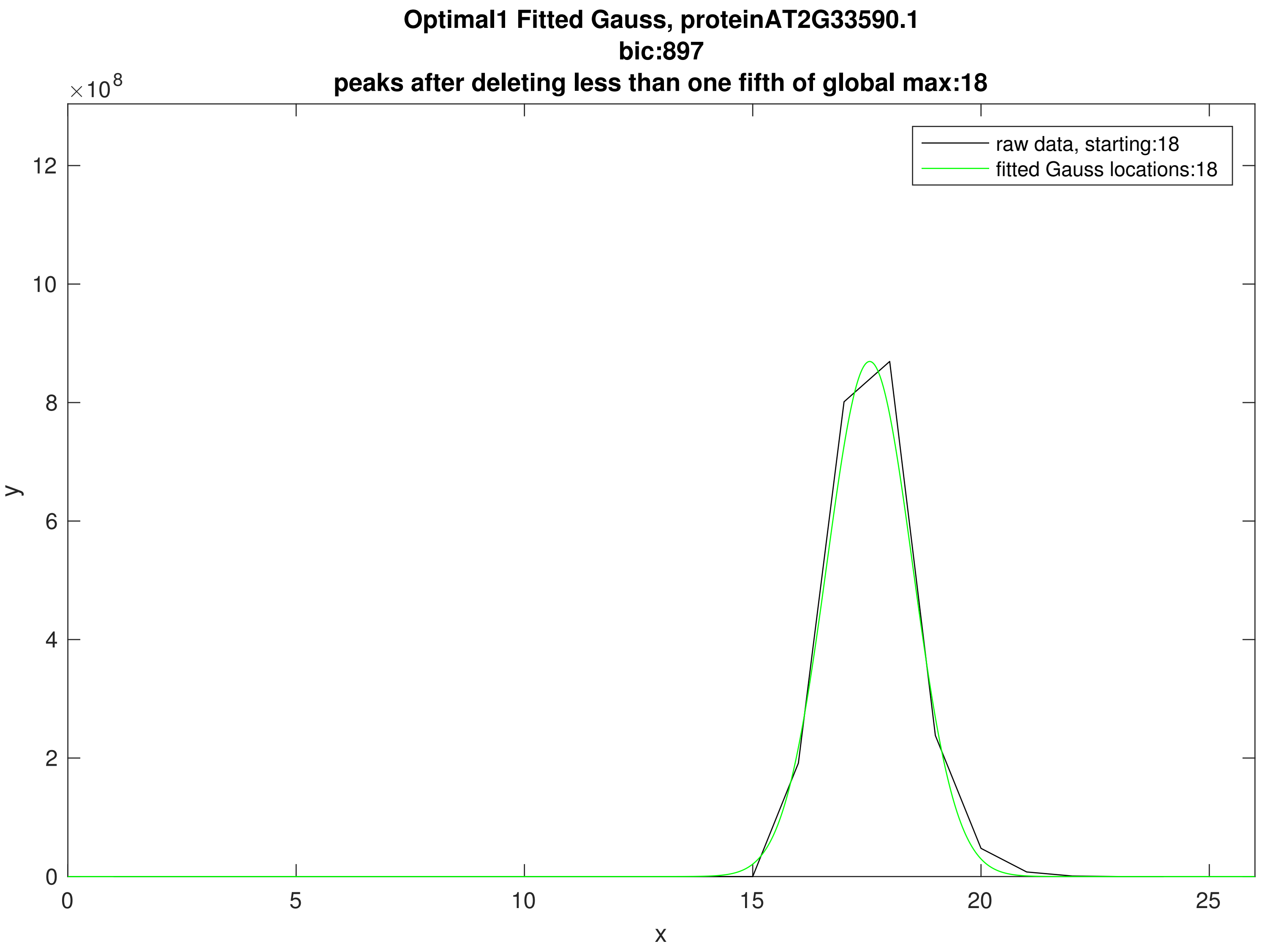

Optimal1 Fitted Gauss, proteinAT2G33590.1
bic:897
peaks after deleting less than one fifth of global max:18
×10^8
raw data, starting:18
fitted Gauss locations:18
y
x
0
2
4
6
8
10
12
0
5
10
15
20
25

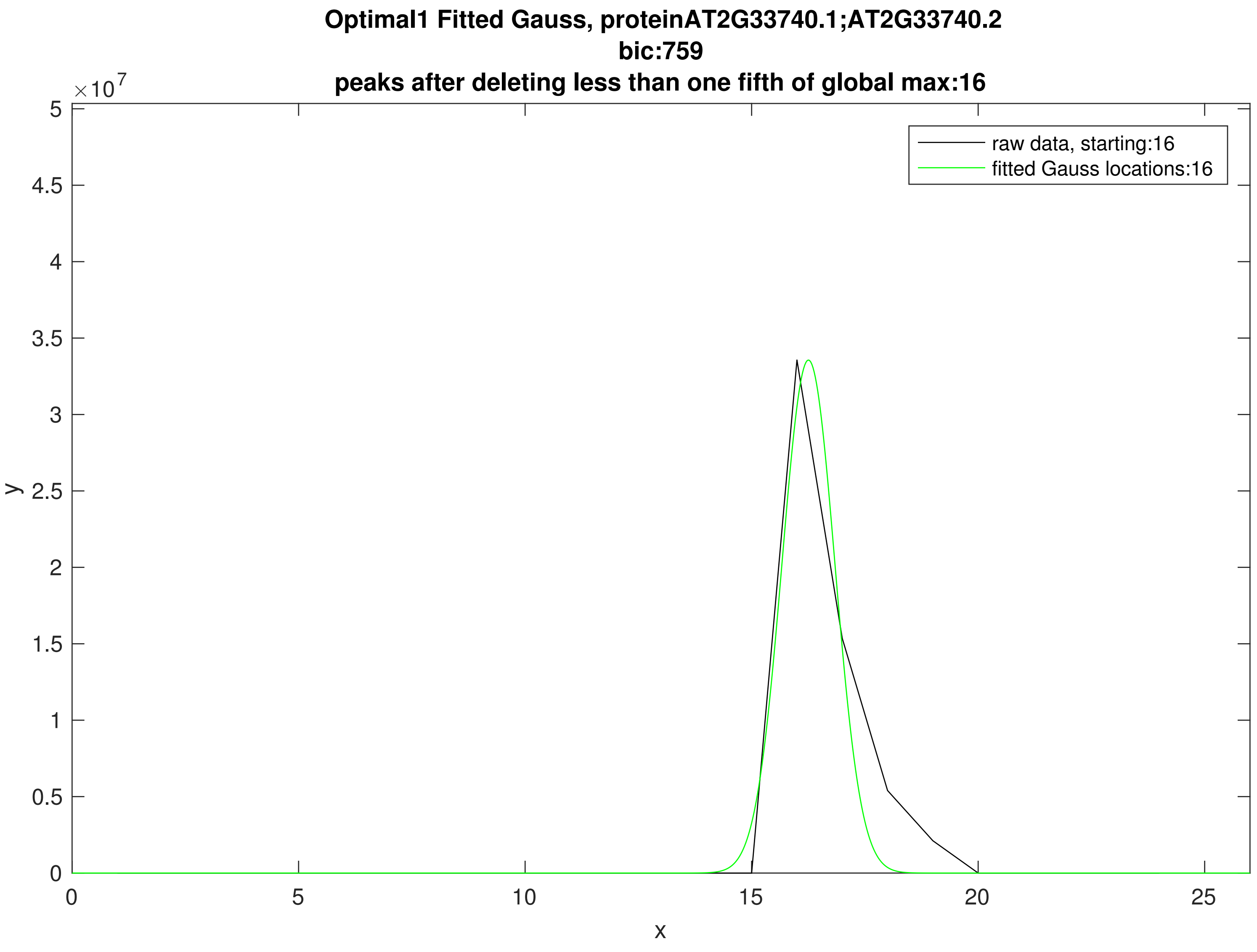

Optimal1 Fitted Gauss, proteinAT2G33740.1;AT2G33740.2
bic:759
peaks after deleting less than one fifth of global max:16
×10^7
raw data, starting:16
fitted Gauss locations:16
y
x
0
0.5
1
1.5
2
2.5
3
3.5
4
4.5
5
0
5
10
15
20
25

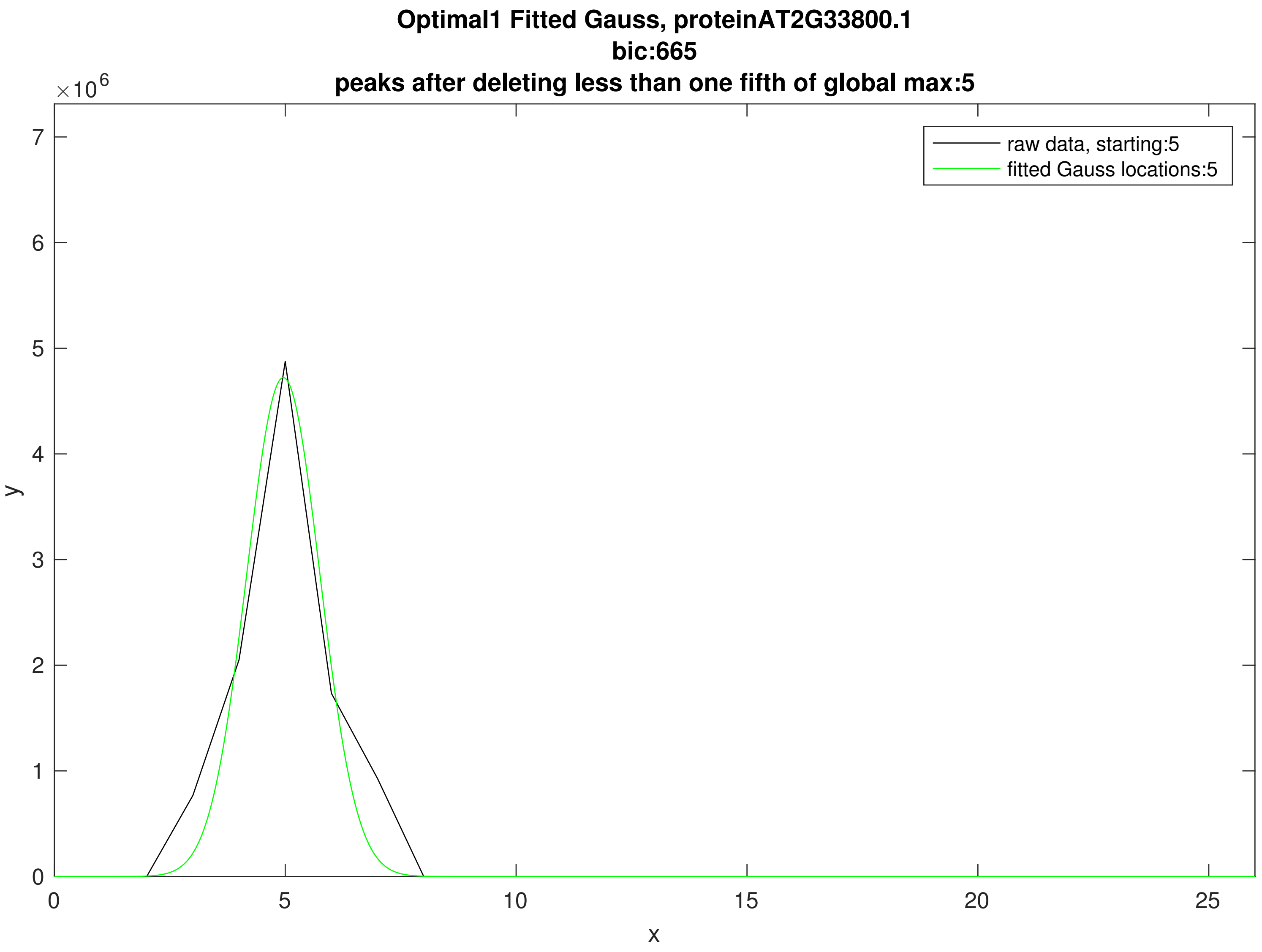
Optimal1 Fitted Gauss, proteinAT2G33800.1
bic:665
peaks after deleting less than one fifth of global max:5
×10^6
raw data, starting:5
fitted Gauss locations:5
y
x
0
1
2
3
4
5
6
7
0
5
10
15
20
25

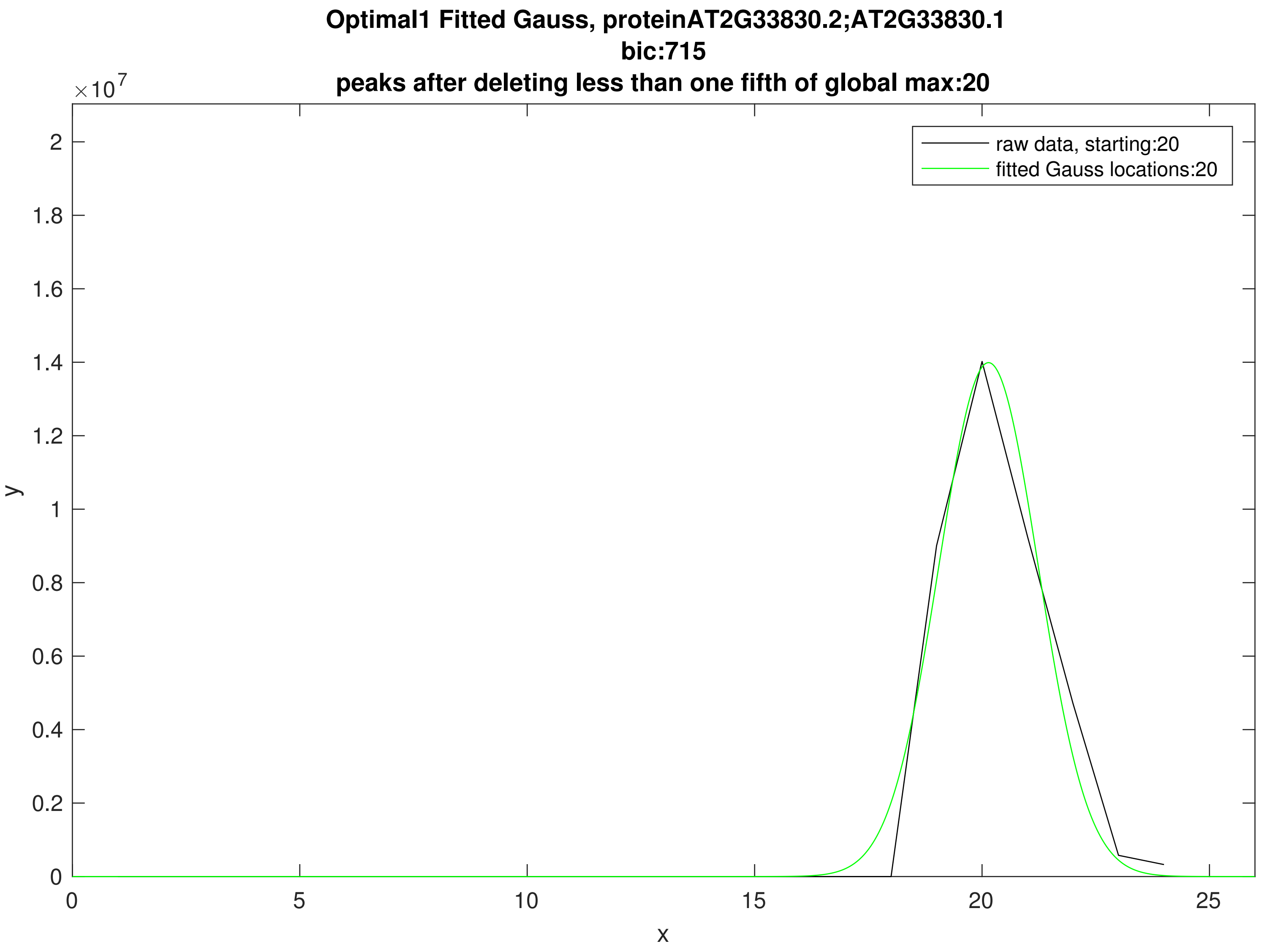
Optimal1 Fitted Gauss, proteinAT2G33830.2;AT2G33830.1
bic:715
peaks after deleting less than one fifth of global max:20
×10^7
raw data, starting:20
fitted Gauss locations:20
y
x
0
0.2
0.4
0.6
0.8
1
1.2
1.4
1.6
1.8
2
0
5
10
15
20
25

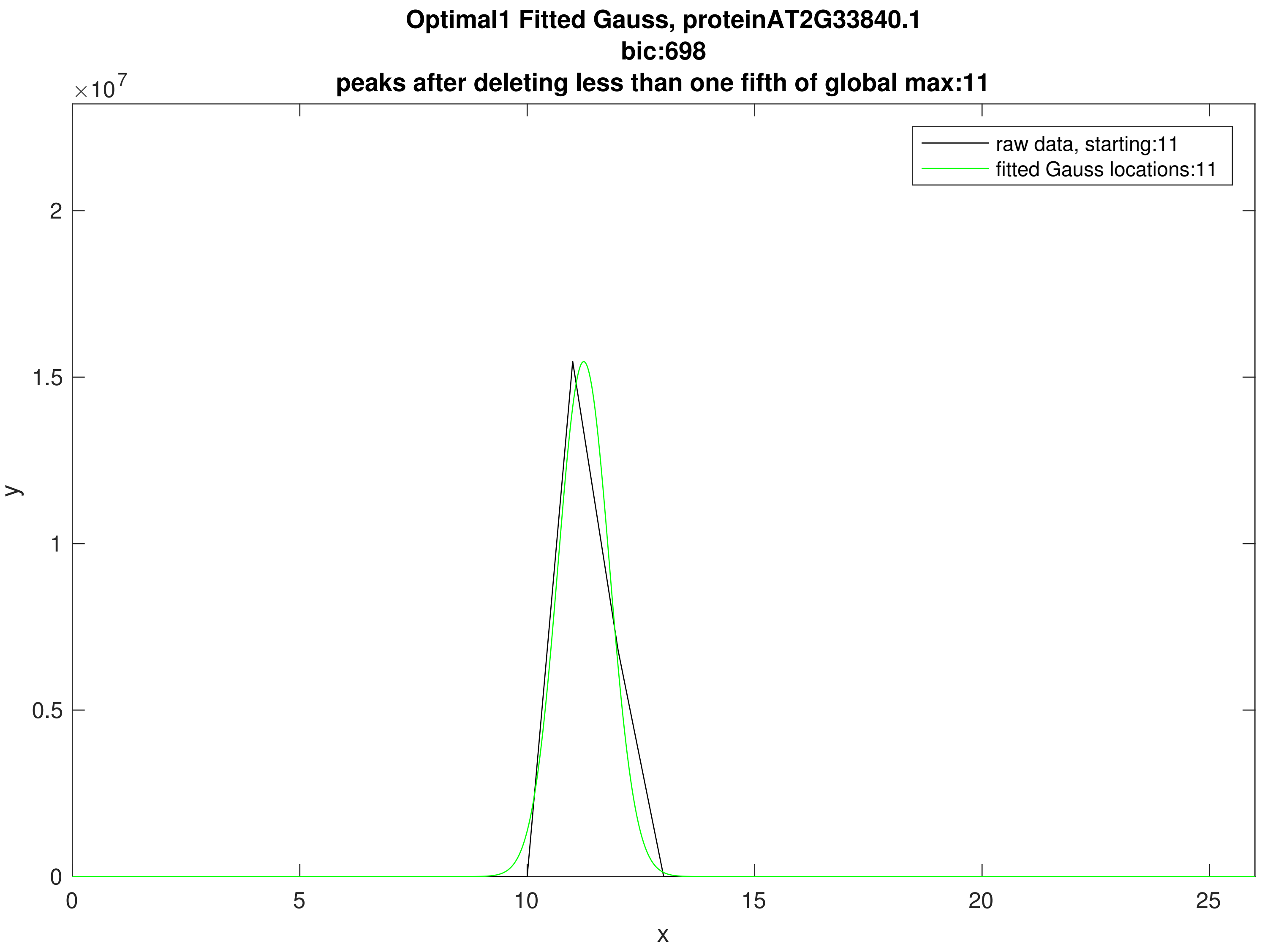
Optimal1 Fitted Gauss, proteinAT2G33840.1
bic:698
peaks after deleting less than one fifth of global max:11
×10^7
raw data, starting:11
fitted Gauss locations:11
y
x
0
0.5
1
1.5
2
0
5
10
15
20
25

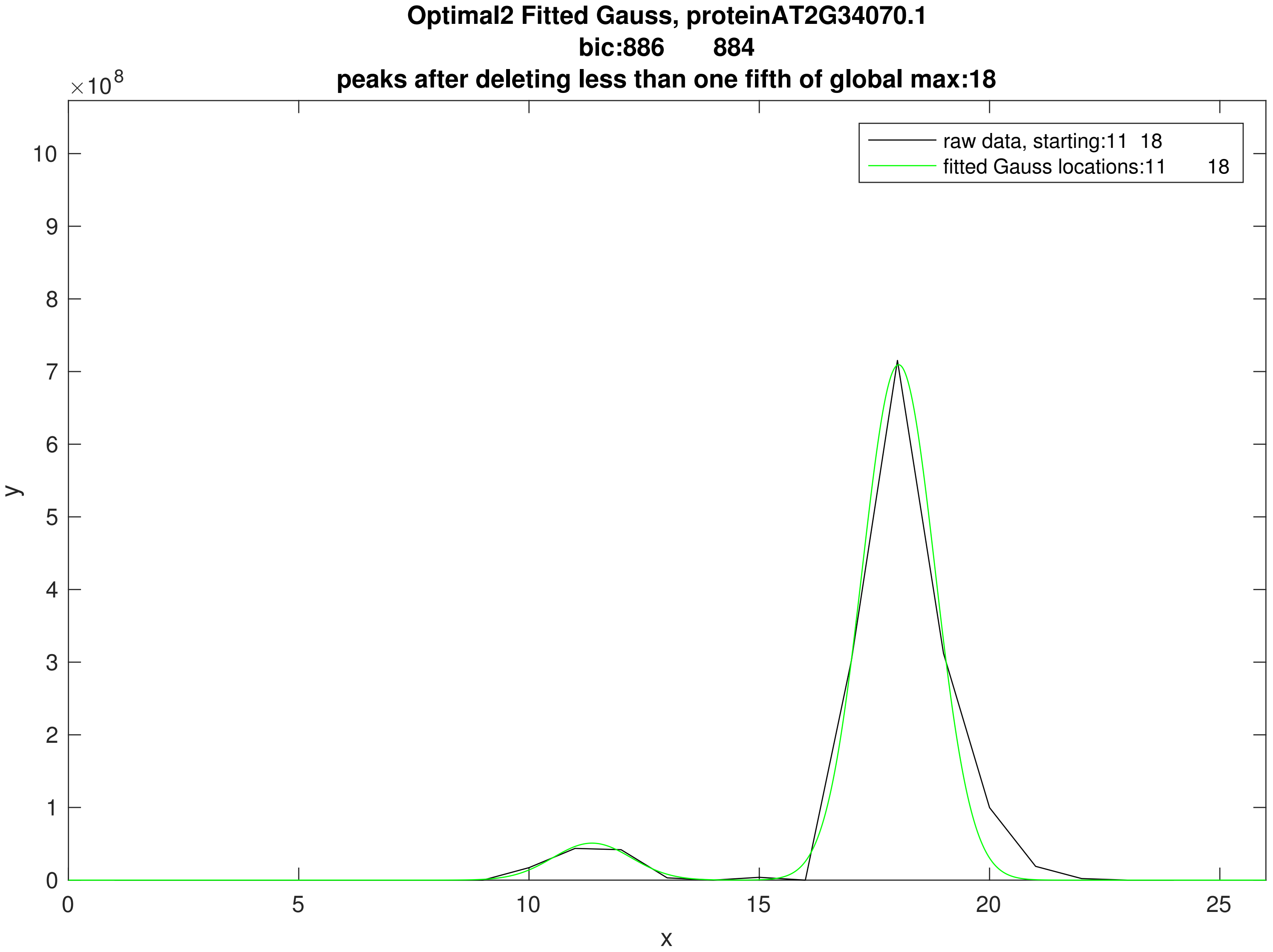
Optimal2 Fitted Gauss, proteinAT2G34070.1
bic:886 884
peaks after deleting less than one fifth of global max:18
×10^8
raw data, starting:11 18
fitted Gauss locations:11 18
y
x

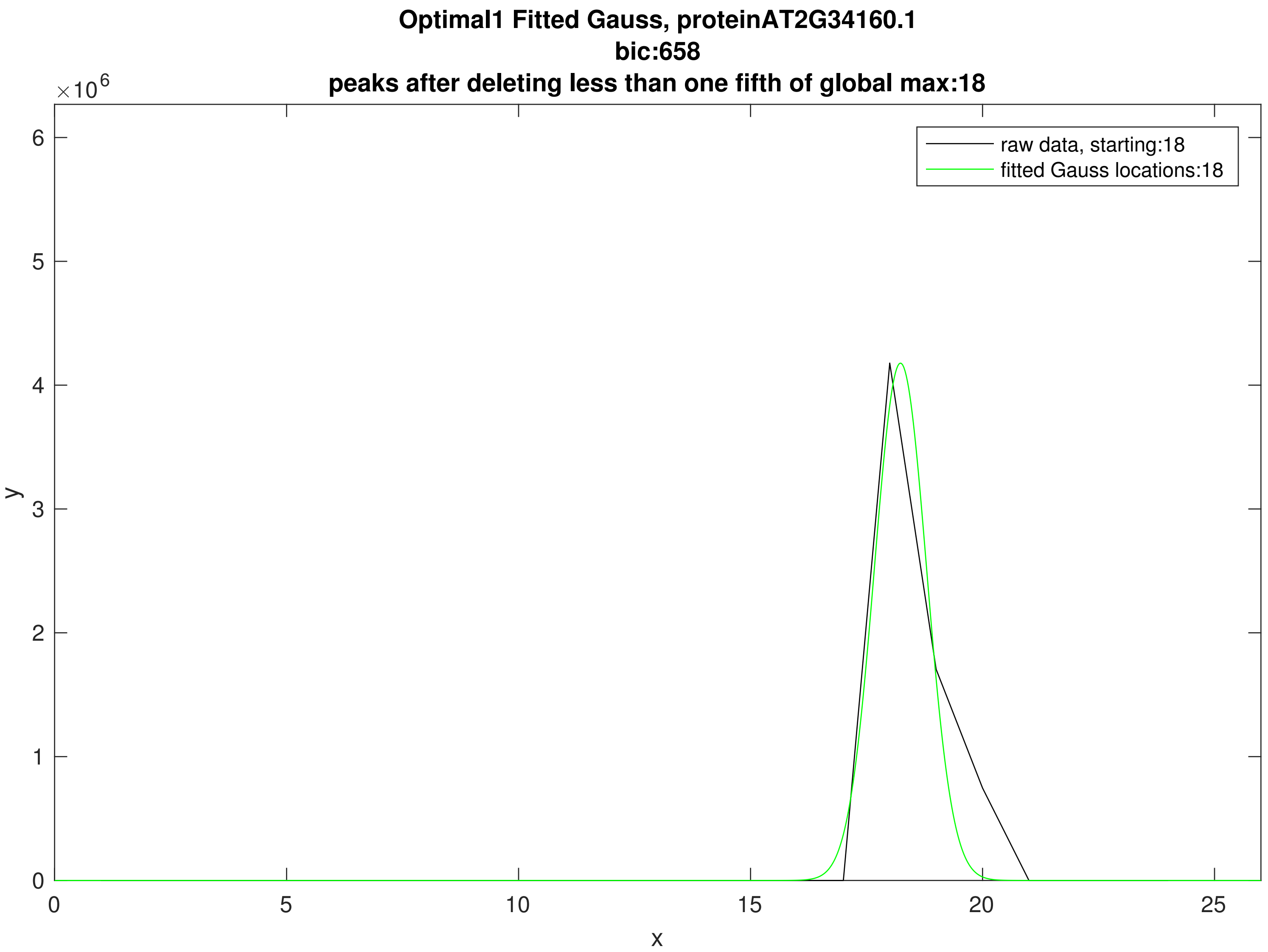

Optimal1 Fitted Gauss, proteinAT2G34160.1
bic:658
peaks after deleting less than one fifth of global max:18
×10^6
raw data, starting:18
fitted Gauss locations:18
y
x
0
1
2
3
4
5
6
0
5
10
15
20
25

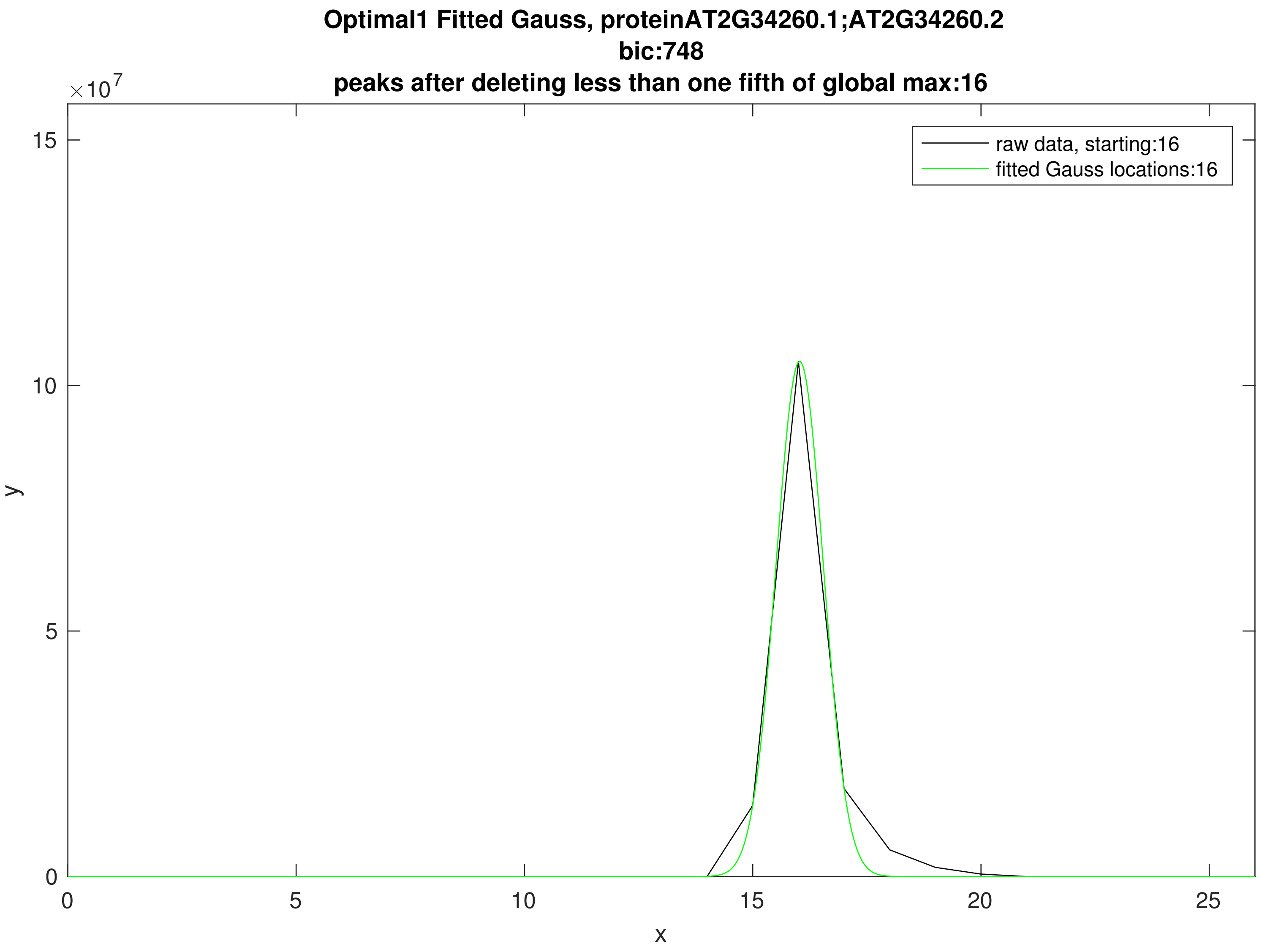
Optimal1 Fitted Gauss, proteinAT2G34260.1;AT2G34260.2
bic:748
peaks after deleting less than one fifth of global max:16
×10^7
15
10
5
0
y
0
5
10
15
20
25
x
raw data, starting:16
fitted Gauss locations:16

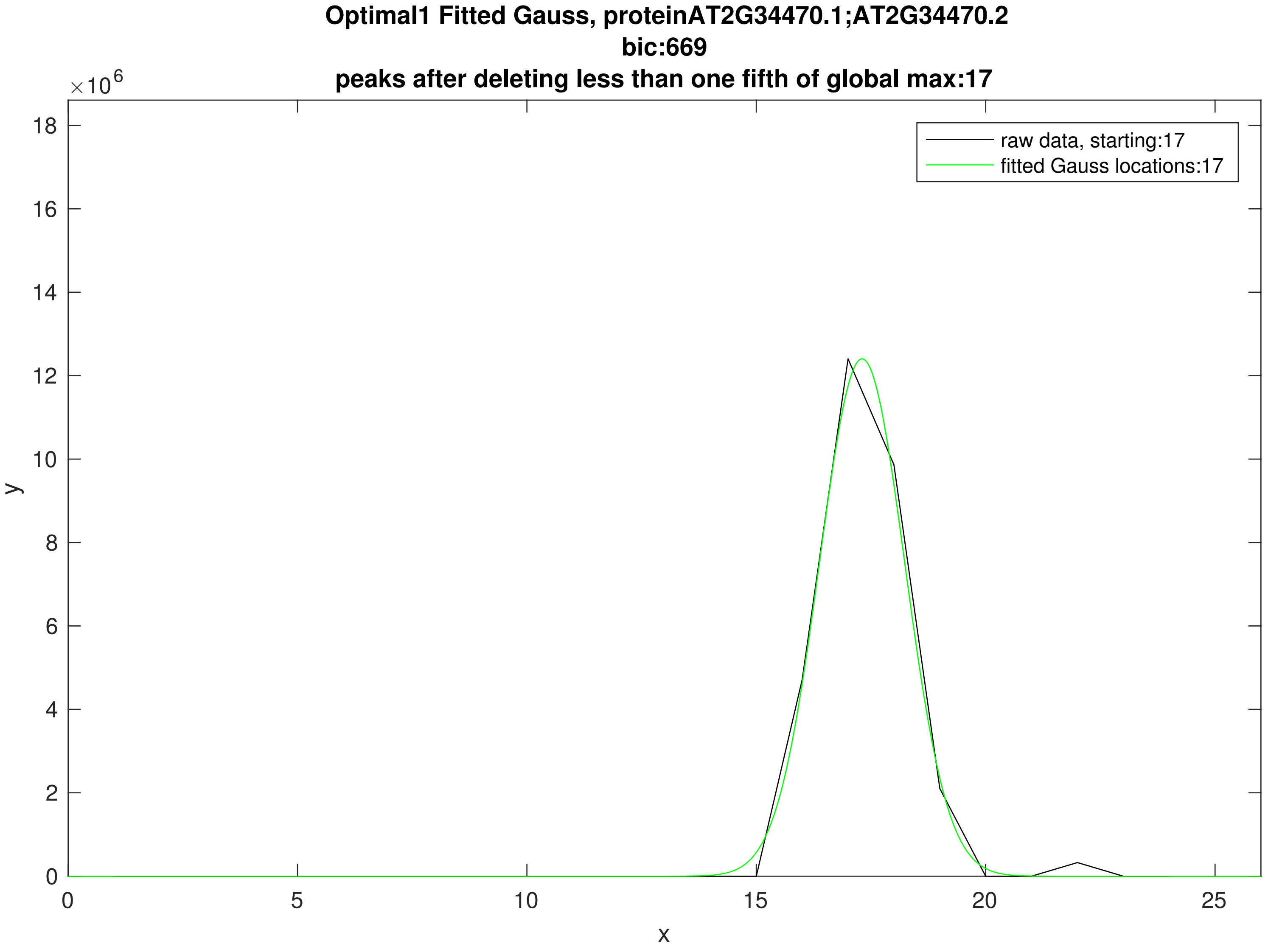
Optimal1 Fitted Gauss, proteinAT2G34470.1;AT2G34470.2
bic:669
peaks after deleting less than one fifth of global max:17
×10^6
raw data, starting:17
fitted Gauss locations:17
y
x
0
2
4
6
8
10
12
14
16
18
0
5
10
15
20
25

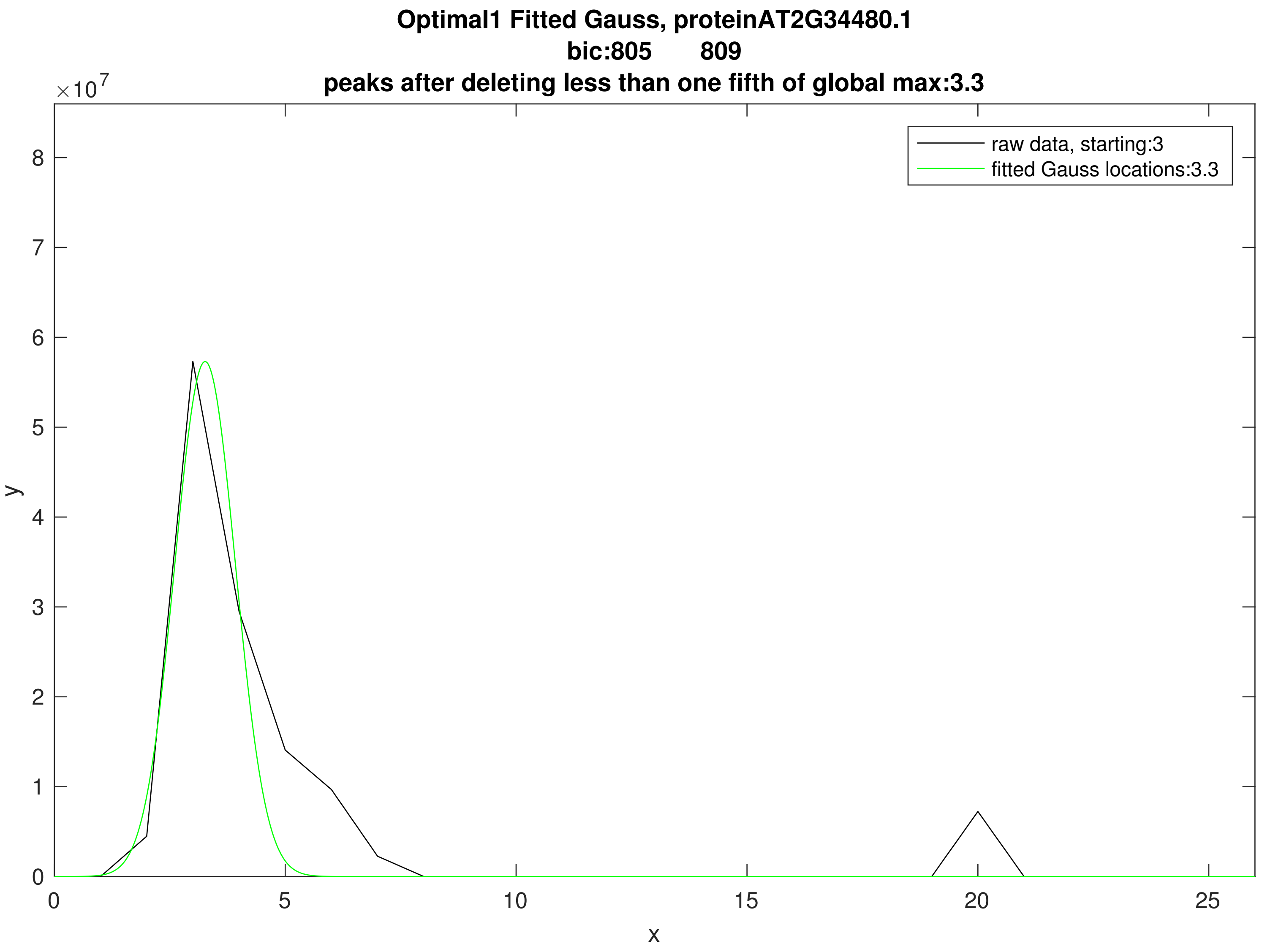
Optimal1 Fitted Gauss, proteinAT2G34480.1
bic:805 809
peaks after deleting less than one fifth of global max:3.3
×10^7
raw data, starting:3
fitted Gauss locations:3.3
y
x
0
1
2
3
4
5
6
7
8
0
5
10
15
20
25

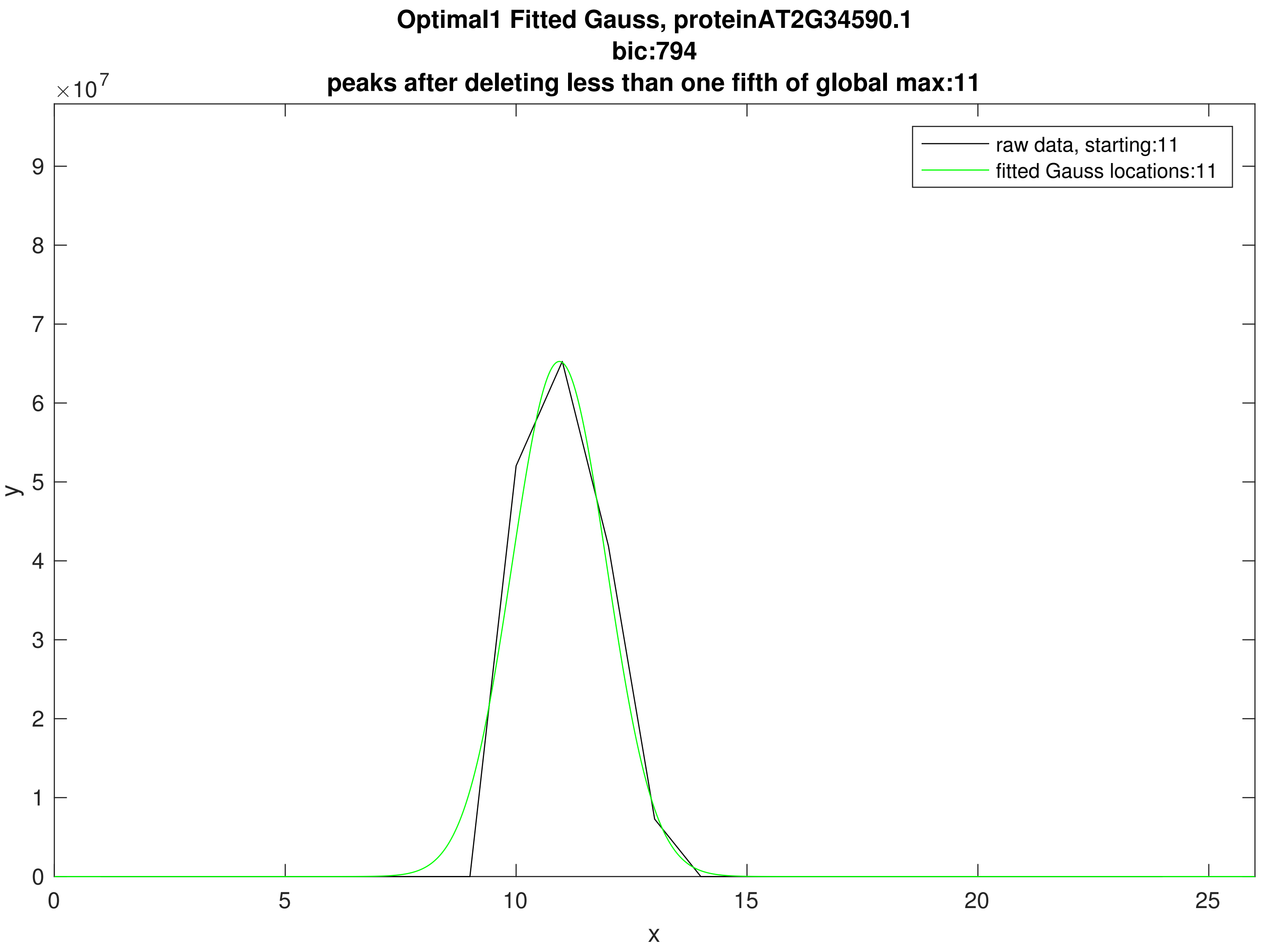

Optimal1 Fitted Gauss, proteinAT2G34590.1
bic:794
peaks after deleting less than one fifth of global max:11
×10^7
raw data, starting:11
fitted Gauss locations:11
y
x
0
1
2
3
4
5
6
7
8
9
0
5
10
15
20
25

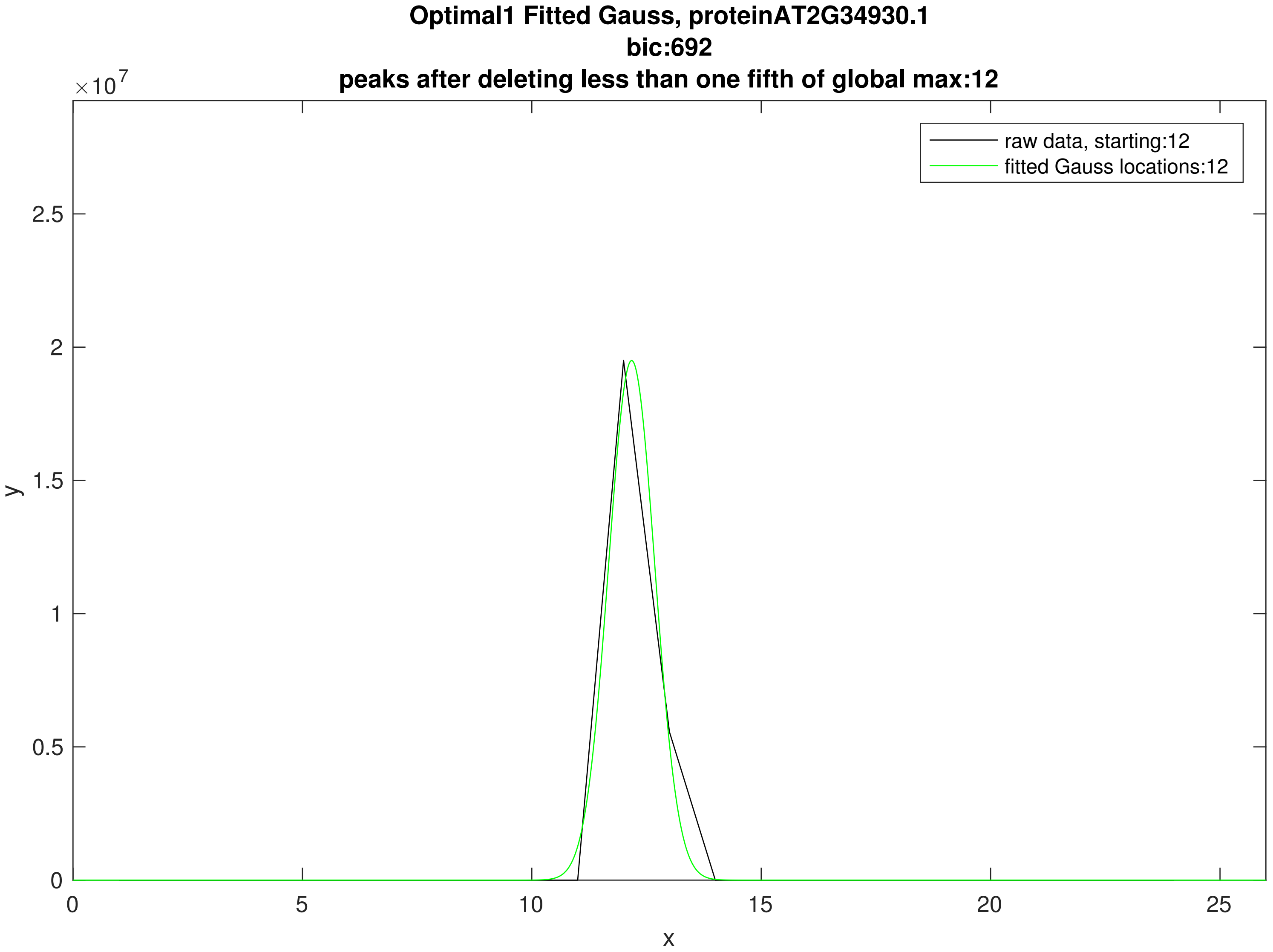

Optimal1 Fitted Gauss, proteinAT2G34930.1
bic:692
peaks after deleting less than one fifth of global max:12
×10^7
raw data, starting:12
fitted Gauss locations:12
2.5
2
1.5
1
0.5
0
y
0
5
10
15
20
25
x

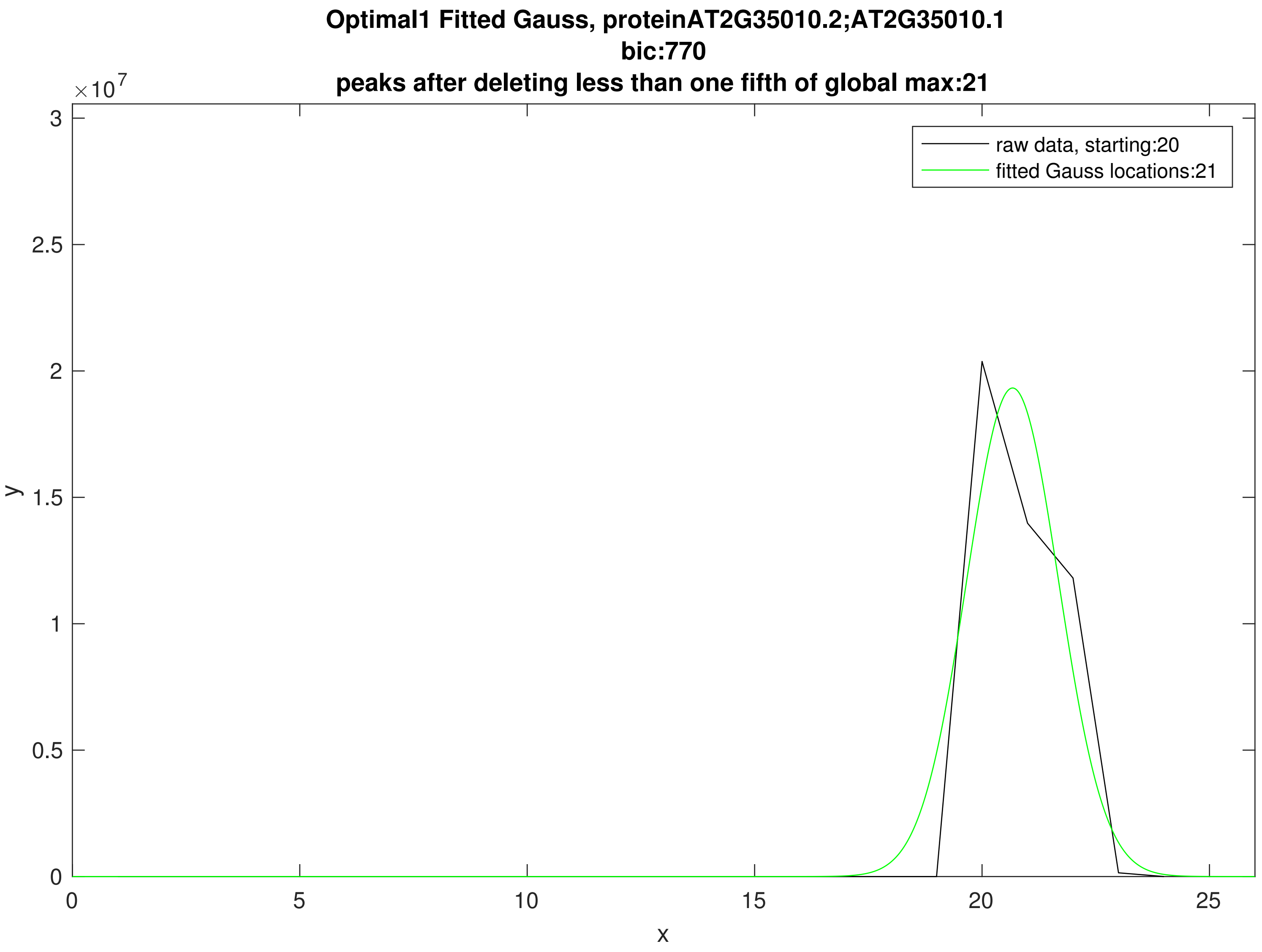

Optimal1 Fitted Gauss, proteinAT2G35010.2;AT2G35010.1
bic:770
peaks after deleting less than one fifth of global max:21
$\times 10^7$
3
2.5
2
1.5
1
0.5
0
y
0
5
10
15
20
25
x
raw data, starting:20
fitted Gauss locations:21

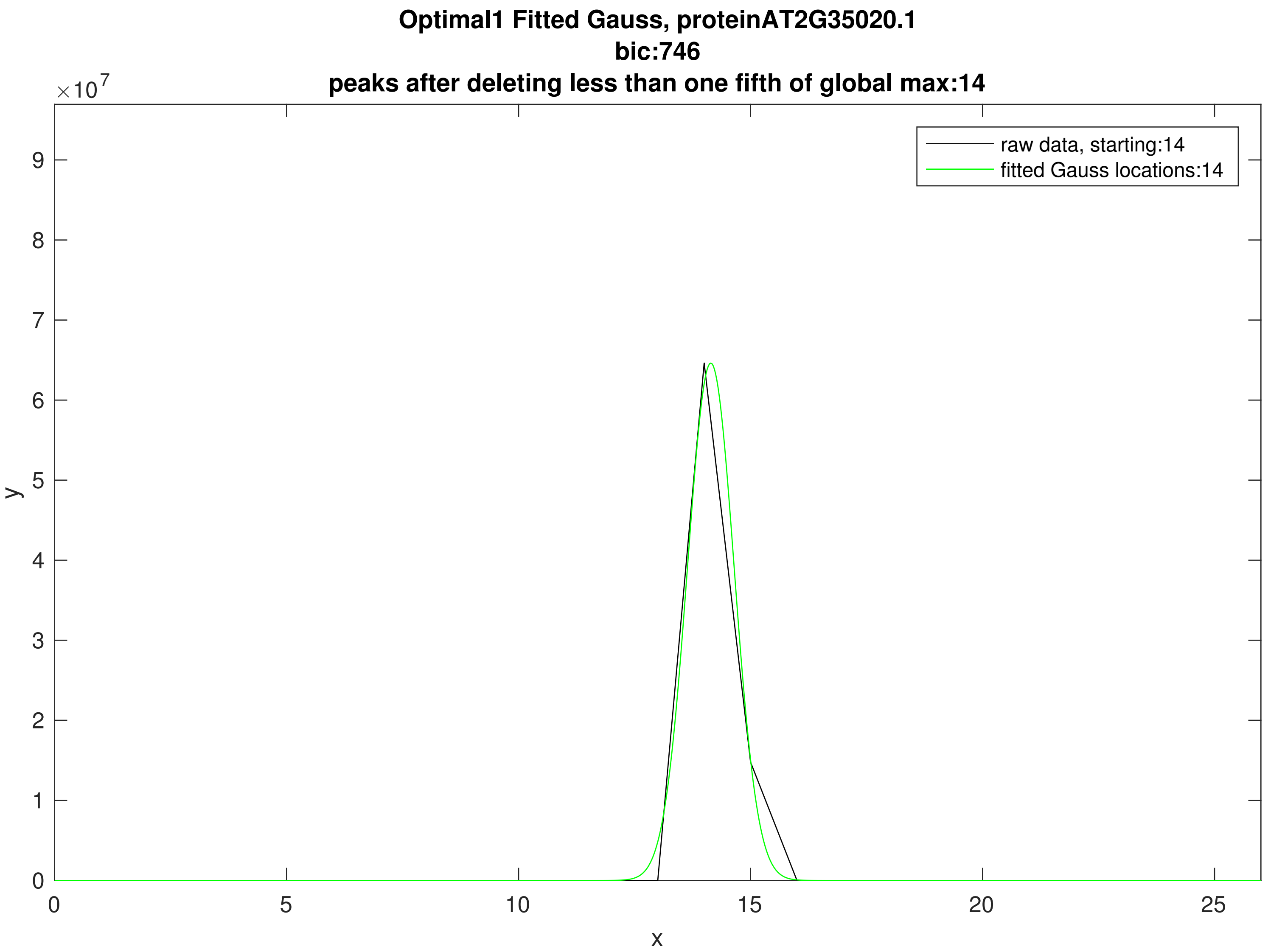
Optimal1 Fitted Gauss, proteinAT2G35020.1
bic:746
peaks after deleting less than one fifth of global max:14
×10^7
raw data, starting:14
fitted Gauss locations:14
9
8
7
6
5
4
3
2
1
0
y
0
5
10
15
20
25
x

Optimal1 Fitted Gauss, proteinAT2G35040.2;AT2G35040.1
bic:875
peaks after deleting less than one fifth of global max:12

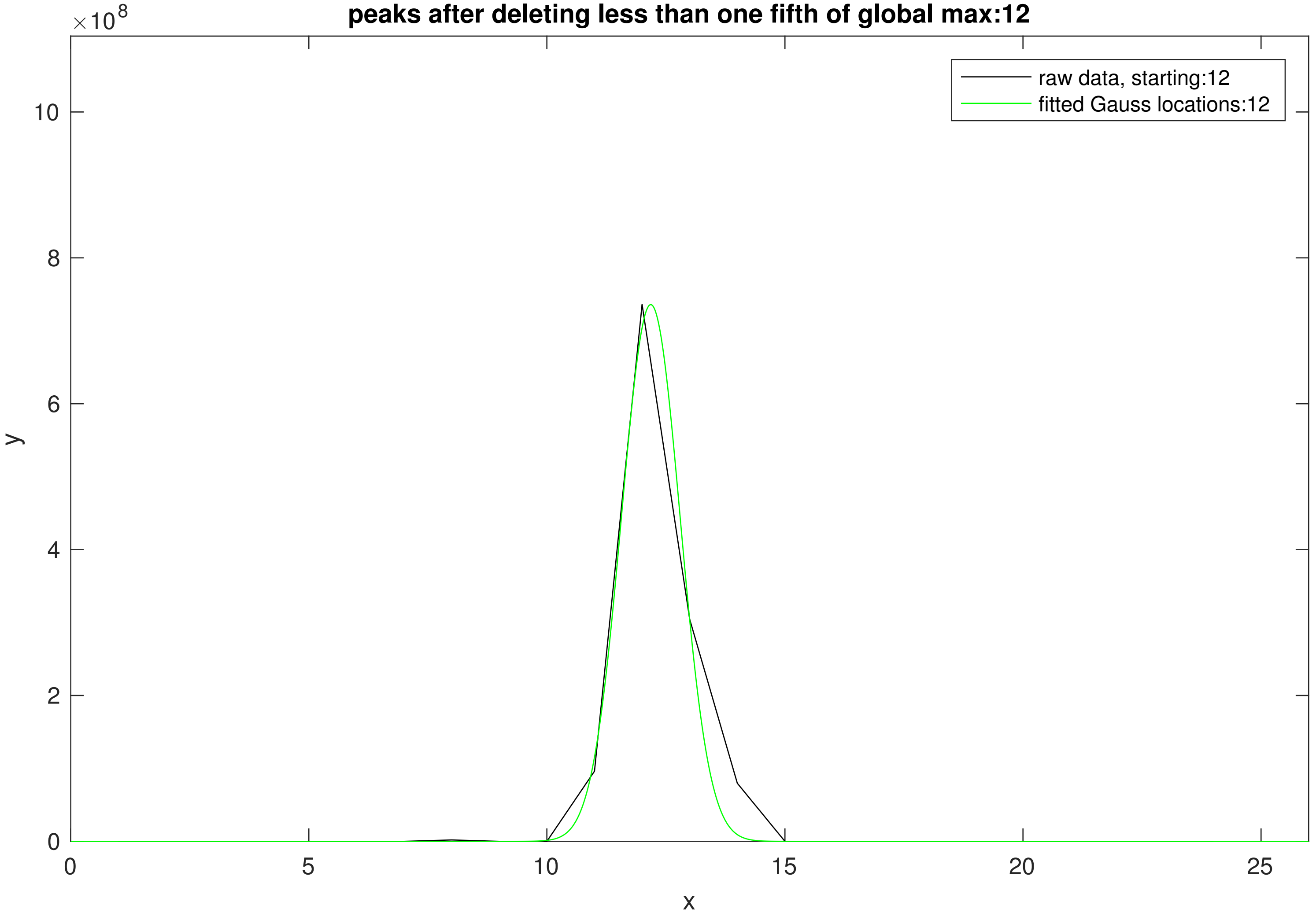

#### Optimal1 Fitted Gauss, proteinAT2G35050.1
#### bic:697
#### peaks after deleting less than one fifth of global max:19

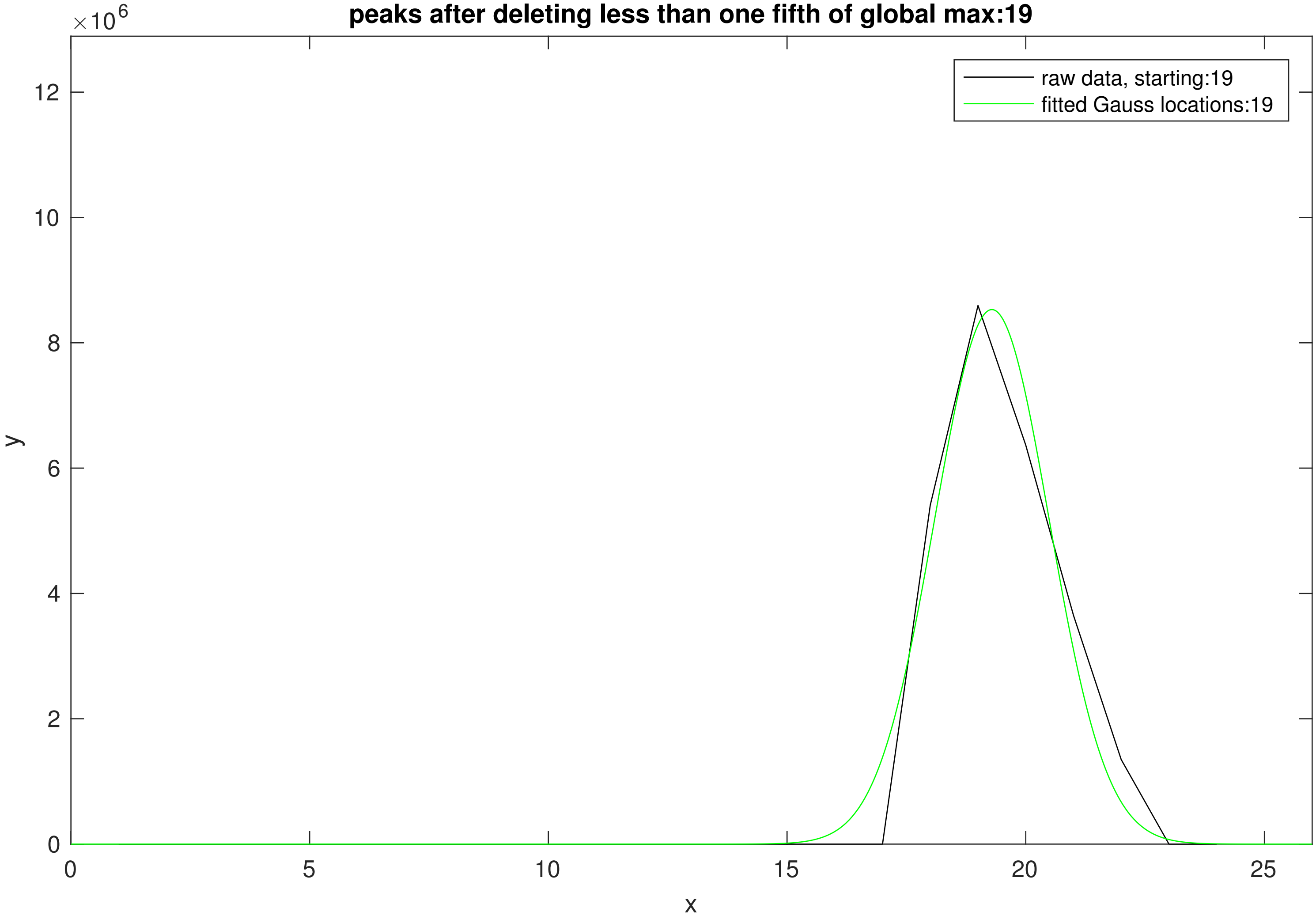

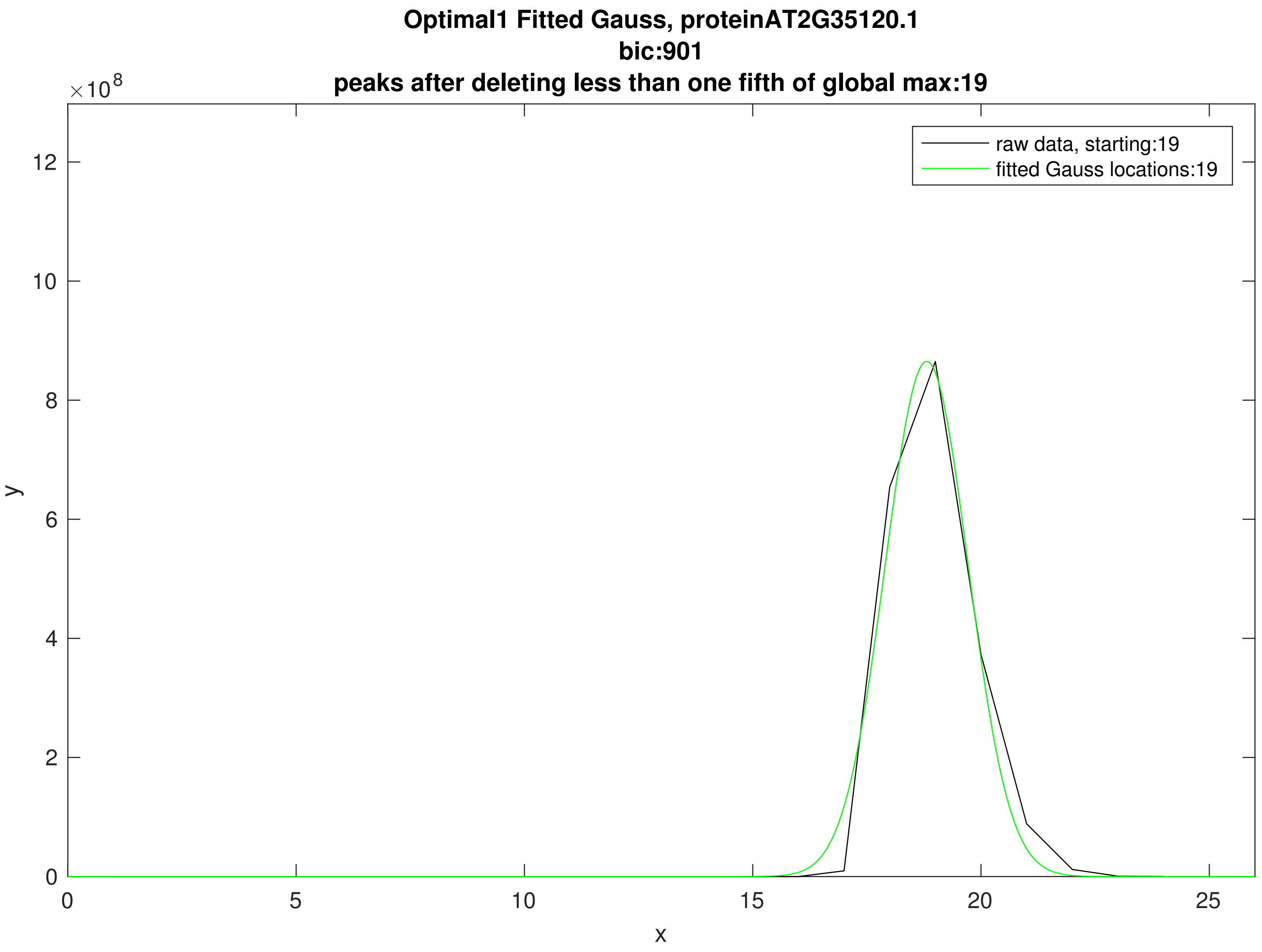
Optimal1 Fitted Gauss, proteinAT2G35120.1
bic:901
peaks after deleting less than one fifth of global max:19
×10^8
12
10
8
6
4
2
0
y
0
5
10
15
20
25
x
raw data, starting:19
fitted Gauss locations:19

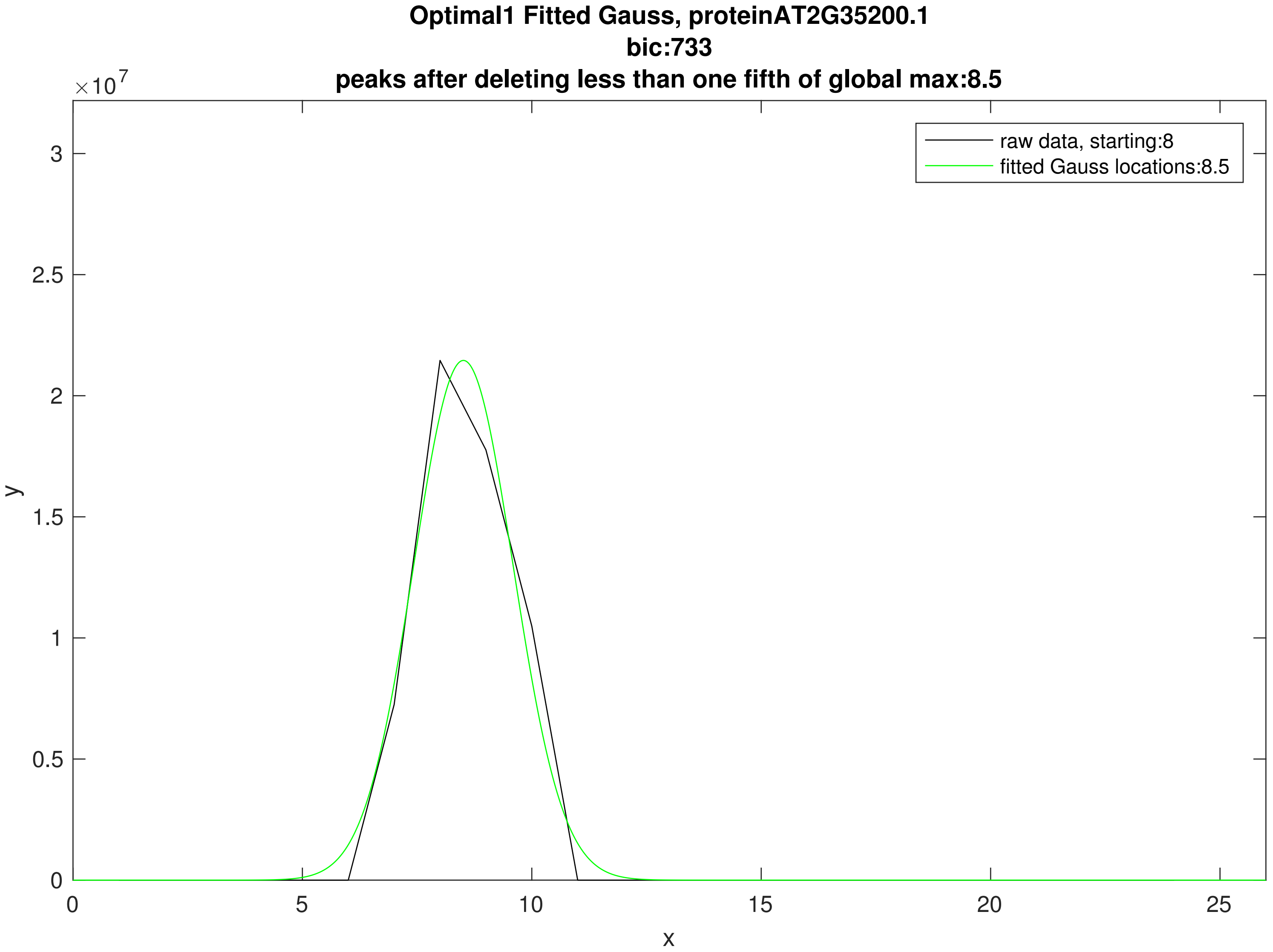
Optimal1 Fitted Gauss, proteinAT2G35200.1
bic:733
peaks after deleting less than one fifth of global max:8.5
×10^7
raw data, starting:8
fitted Gauss locations:8.5
3
2.5
2
1.5
1
0.5
0
y
0
5
10
15
20
25
x

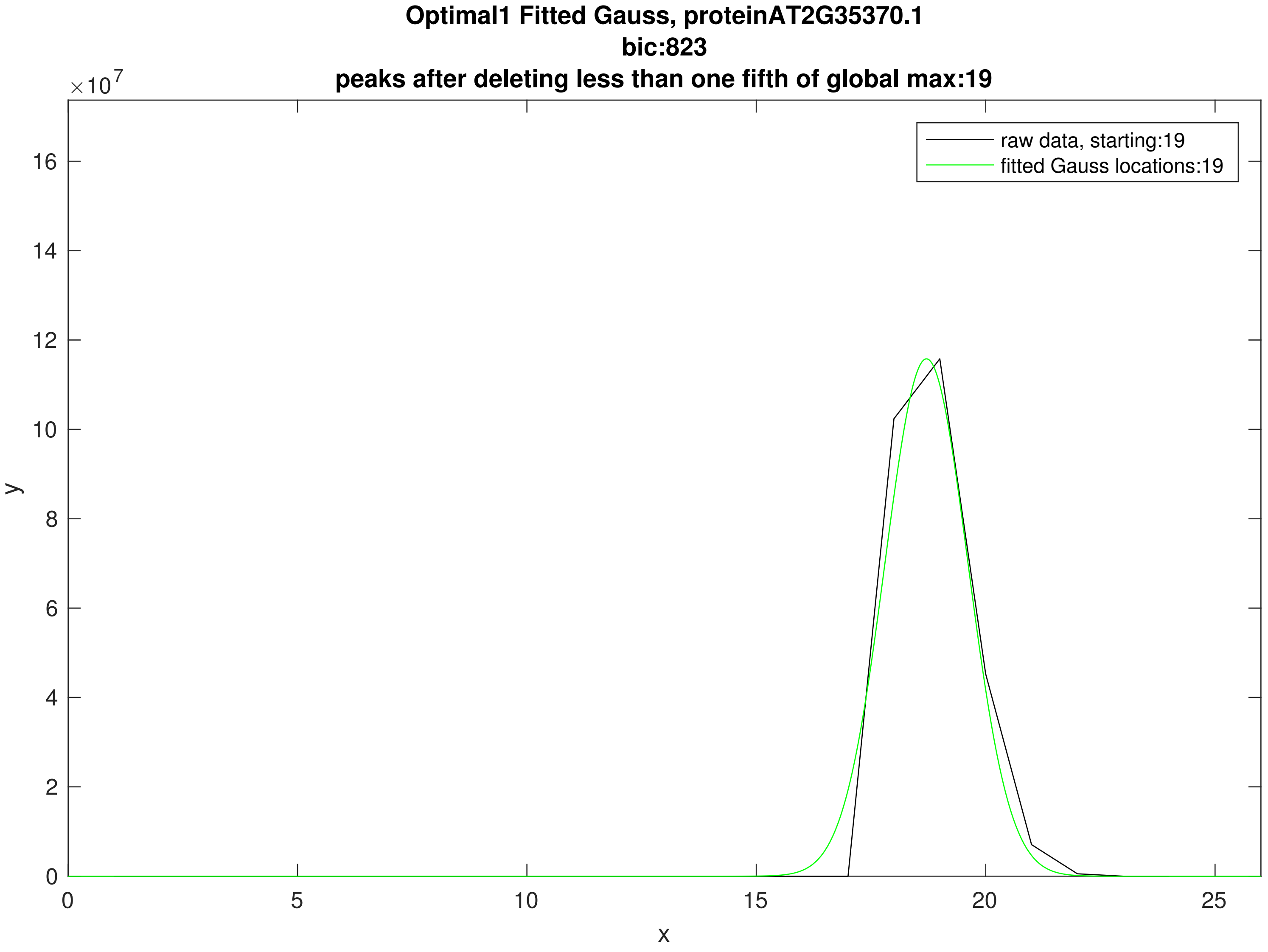

Optimal1 Fitted Gauss, proteinAT2G35370.1
bic:823
peaks after deleting less than one fifth of global max:19
×10^7
raw data, starting:19
fitted Gauss locations:19
y
x
0
2
4
6
8
10
12
14
16
0
5
10
15
20
25

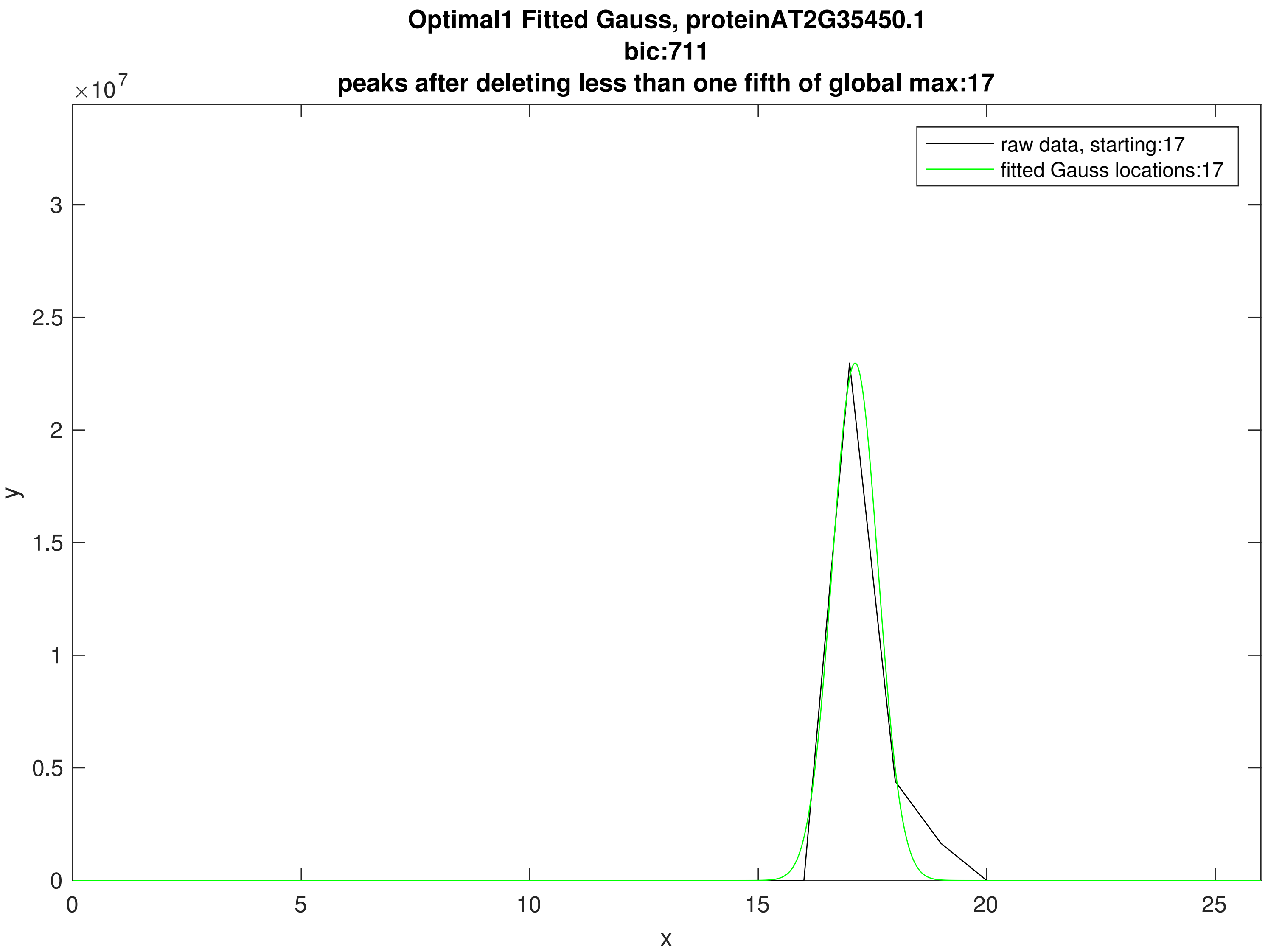

Optimal1 Fitted Gauss, proteinAT2G35450.1
bic:711
peaks after deleting less than one fifth of global max:17
×10^7
raw data, starting:17
fitted Gauss locations:17
y
x
0
0.5
1
1.5
2
2.5
3
0
5
10
15
20
25

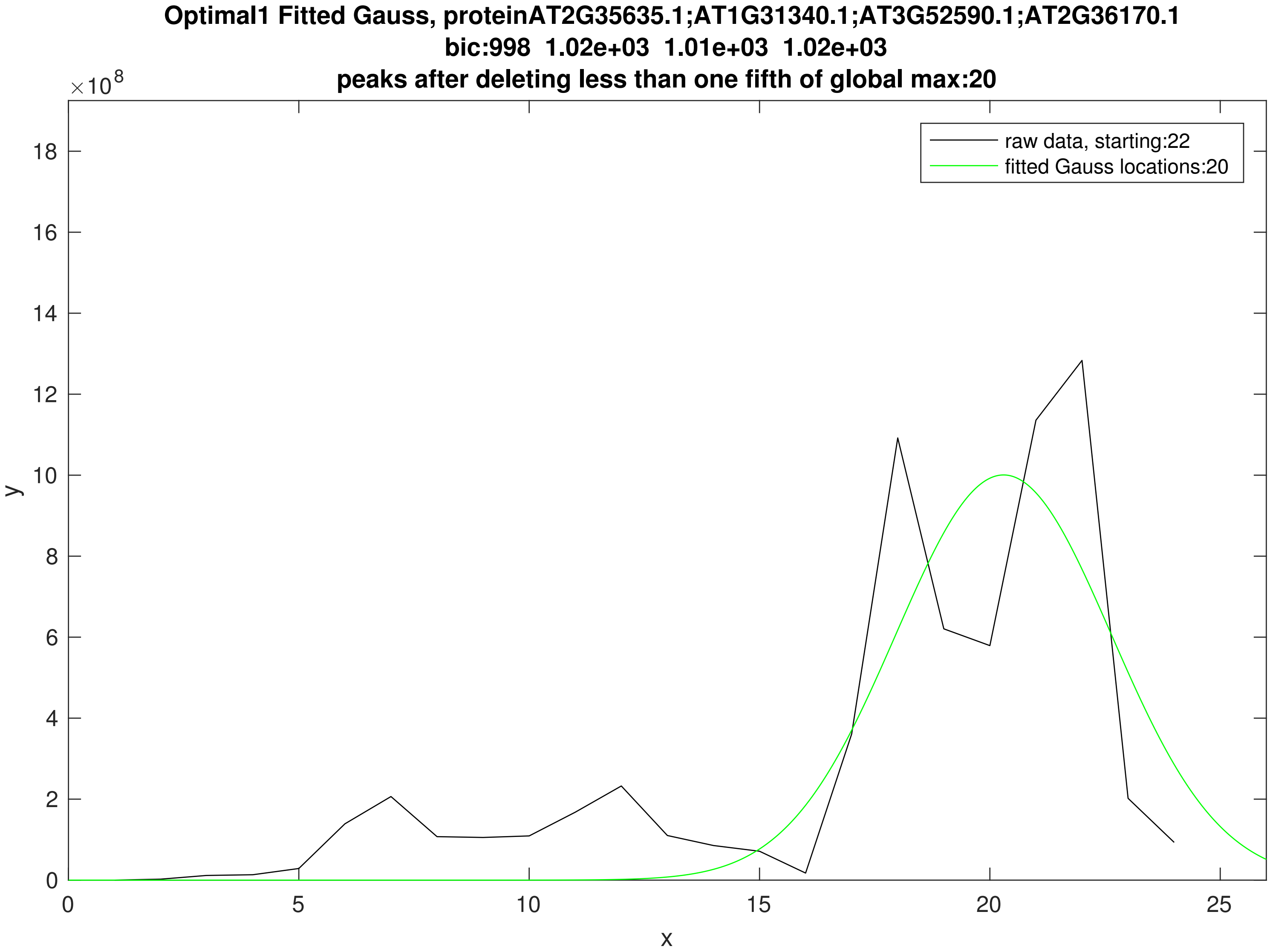
Optimal1 Fitted Gauss, proteinAT2G35635.1;AT1G31340.1;AT3G52590.1;AT2G36170.1
bic:998 1.02e+03 1.01e+03 1.02e+03
peaks after deleting less than one fifth of global max:20
×10^8
raw data, starting:22
fitted Gauss locations:20
y
x
0
2
4
6
8
10
12
14
16
18
0
5
10
15
20
25

**Optimal1 Fitted Gauss, proteinAT2G35690.1**
**bic:664**
**peaks after deleting less than one fifth of global max:13**

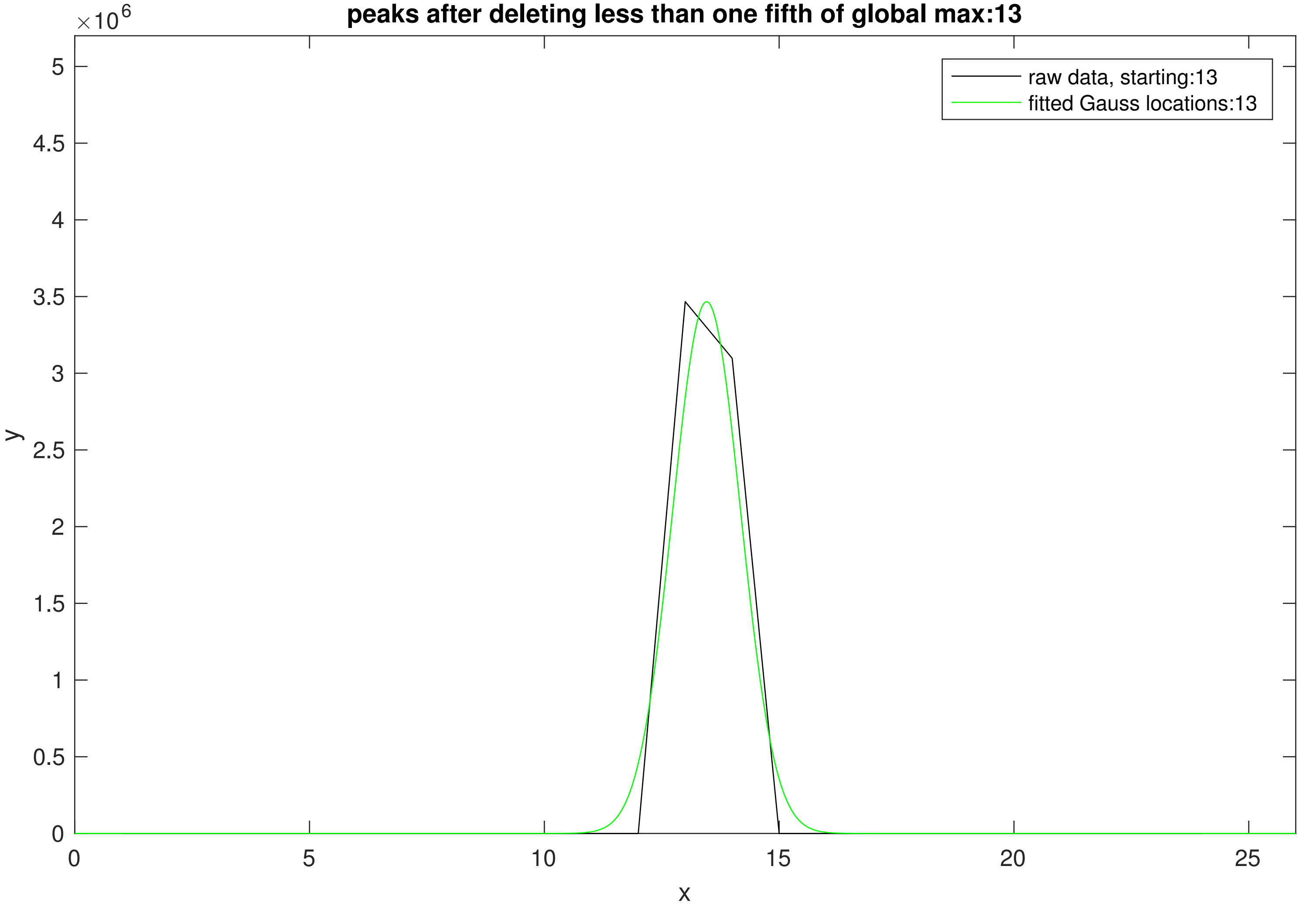

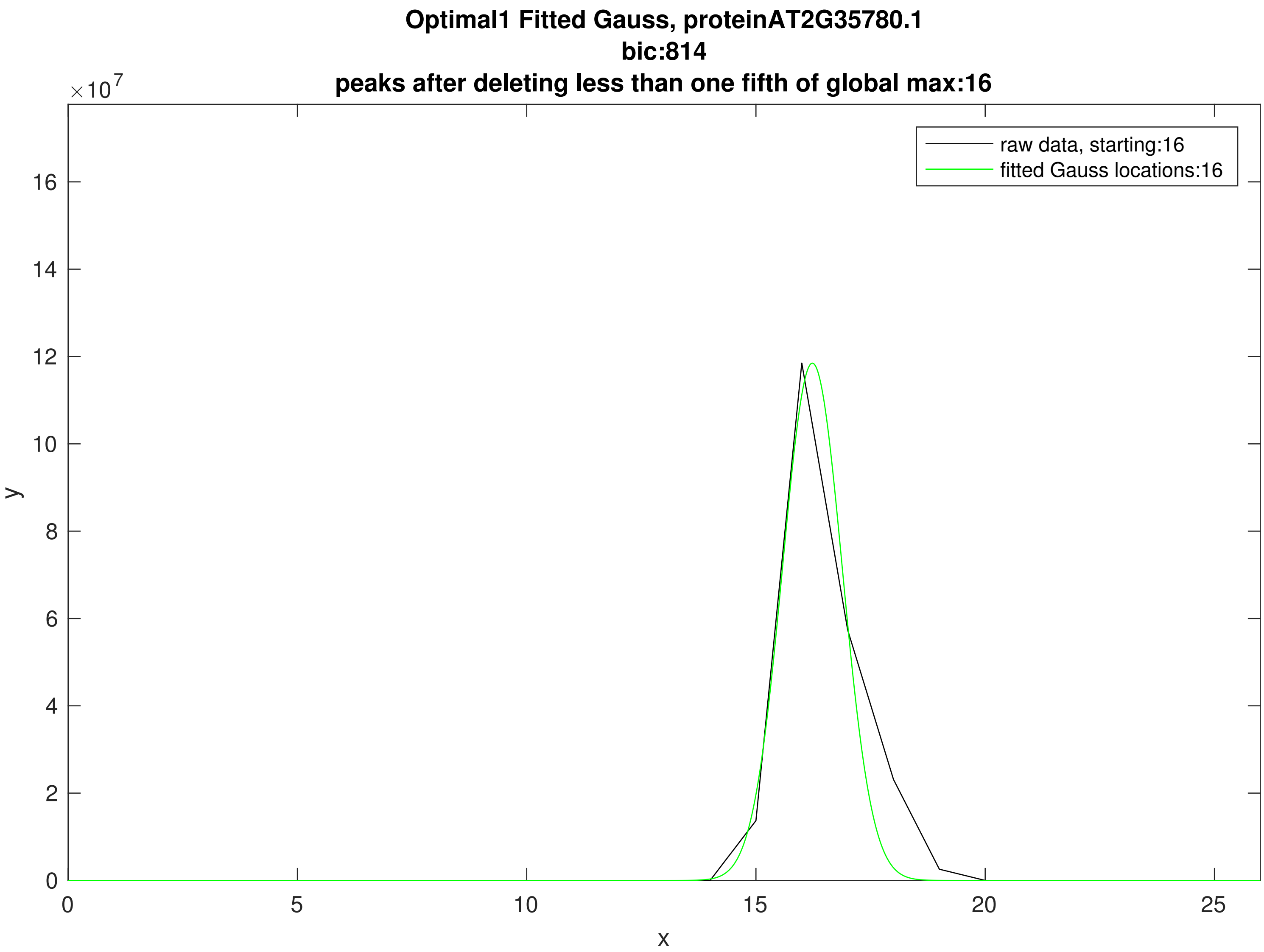
Optimal1 Fitted Gauss, proteinAT2G35780.1
bic:814
peaks after deleting less than one fifth of global max:16
×10^7
raw data, starting:16
fitted Gauss locations:16
16
14
12
10
8
6
4
2
0
y
0
5
10
15
20
25
x

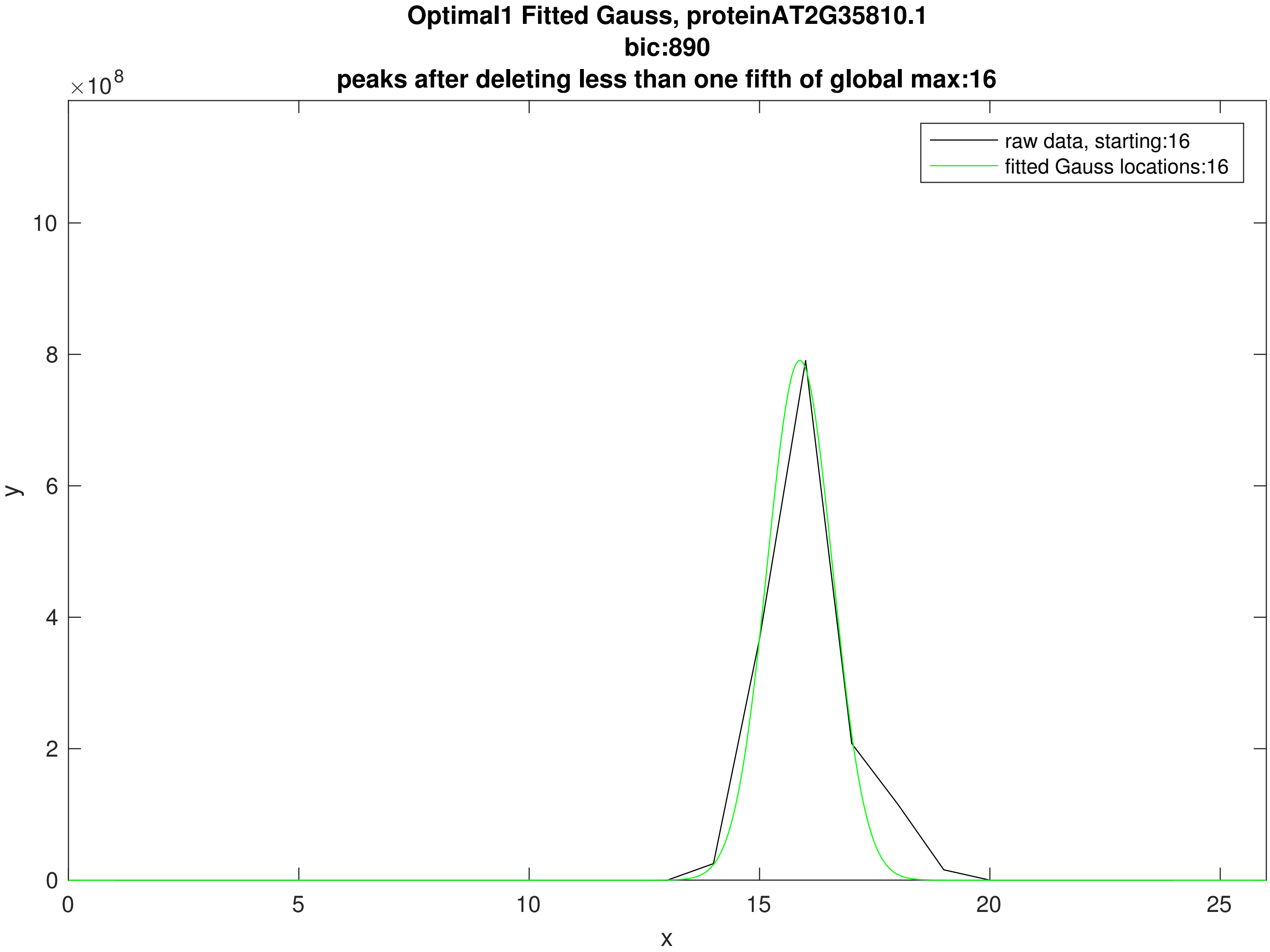

Optimal1 Fitted Gauss, proteinAT2G35810.1
bic:890
peaks after deleting less than one fifth of global max:16
×10^8
raw data, starting:16
fitted Gauss locations:16
10
8
6
4
2
0
y
0
5
10
15
20
25
x

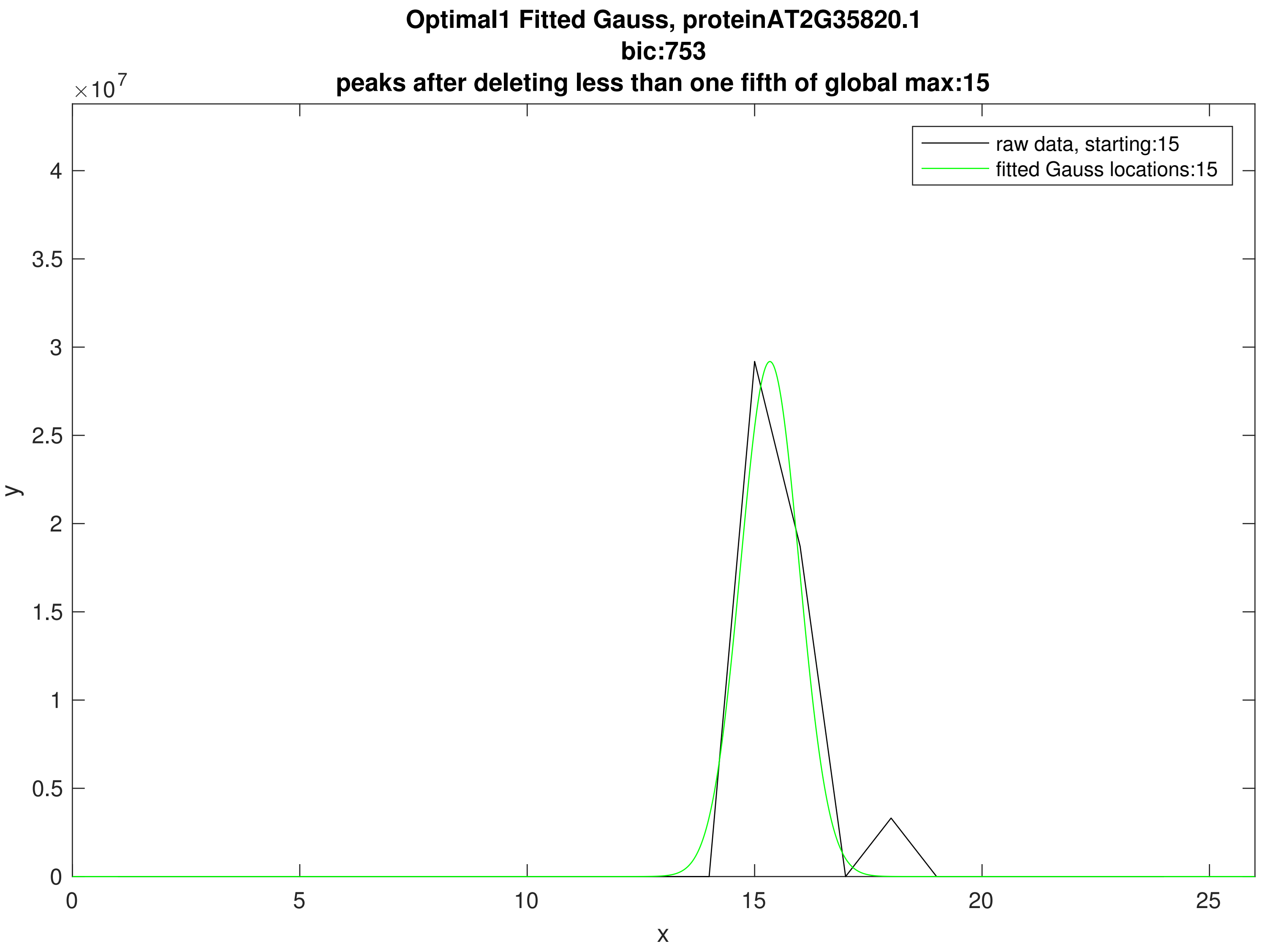

Optimal1 Fitted Gauss, proteinAT2G35820.1
bic:753
peaks after deleting less than one fifth of global max:15
×10^7
raw data, starting:15
fitted Gauss locations:15
y
x
0
0.5
1
1.5
2
2.5
3
3.5
4
0
5
10
15
20
25

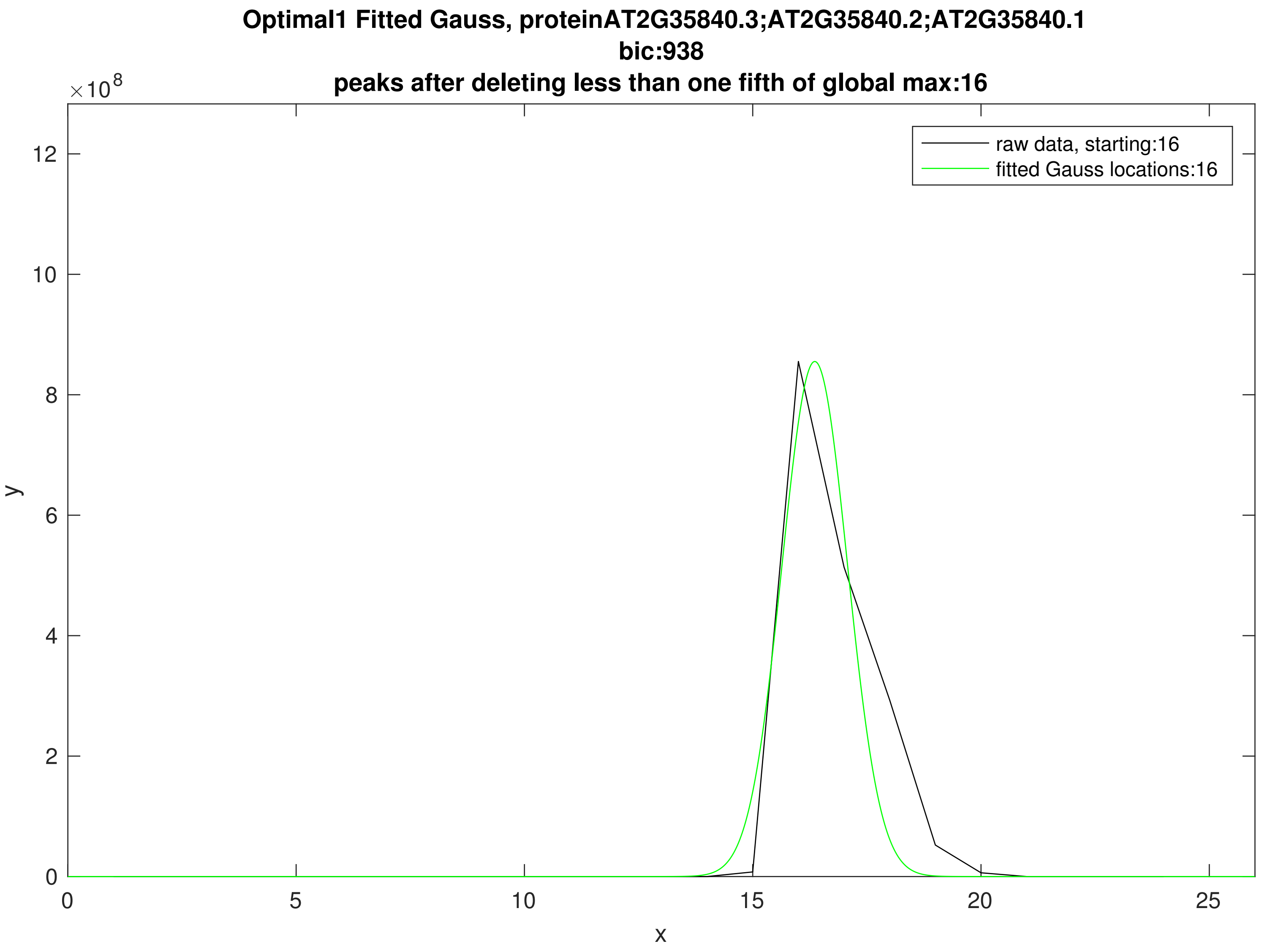

Optimal1 Fitted Gauss, proteinAT2G35840.3;AT2G35840.2;AT2G35840.1
bic:938
peaks after deleting less than one fifth of global max:16
×10^8
raw data, starting:16
fitted Gauss locations:16
y
x
0
2
4
6
8
10
12
0
5
10
15
20
25

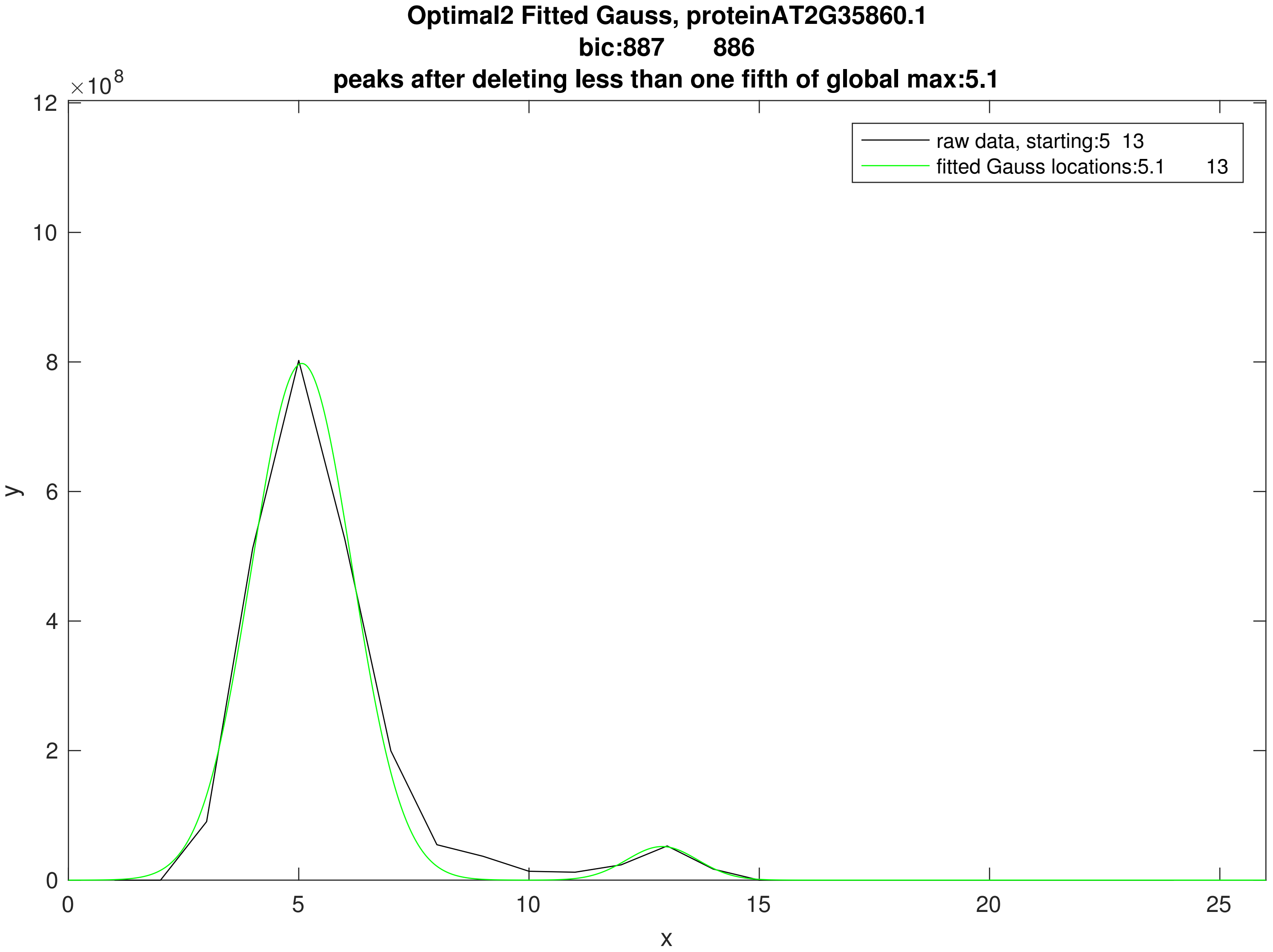
Optimal2 Fitted Gauss, proteinAT2G35860.1
bic:887 886
peaks after deleting less than one fifth of global max:5.1
×10^8
12
10
8
6
4
2
0
y
0
5
10
15
20
25
x
raw data, starting:5 13
fitted Gauss locations:5.1 13

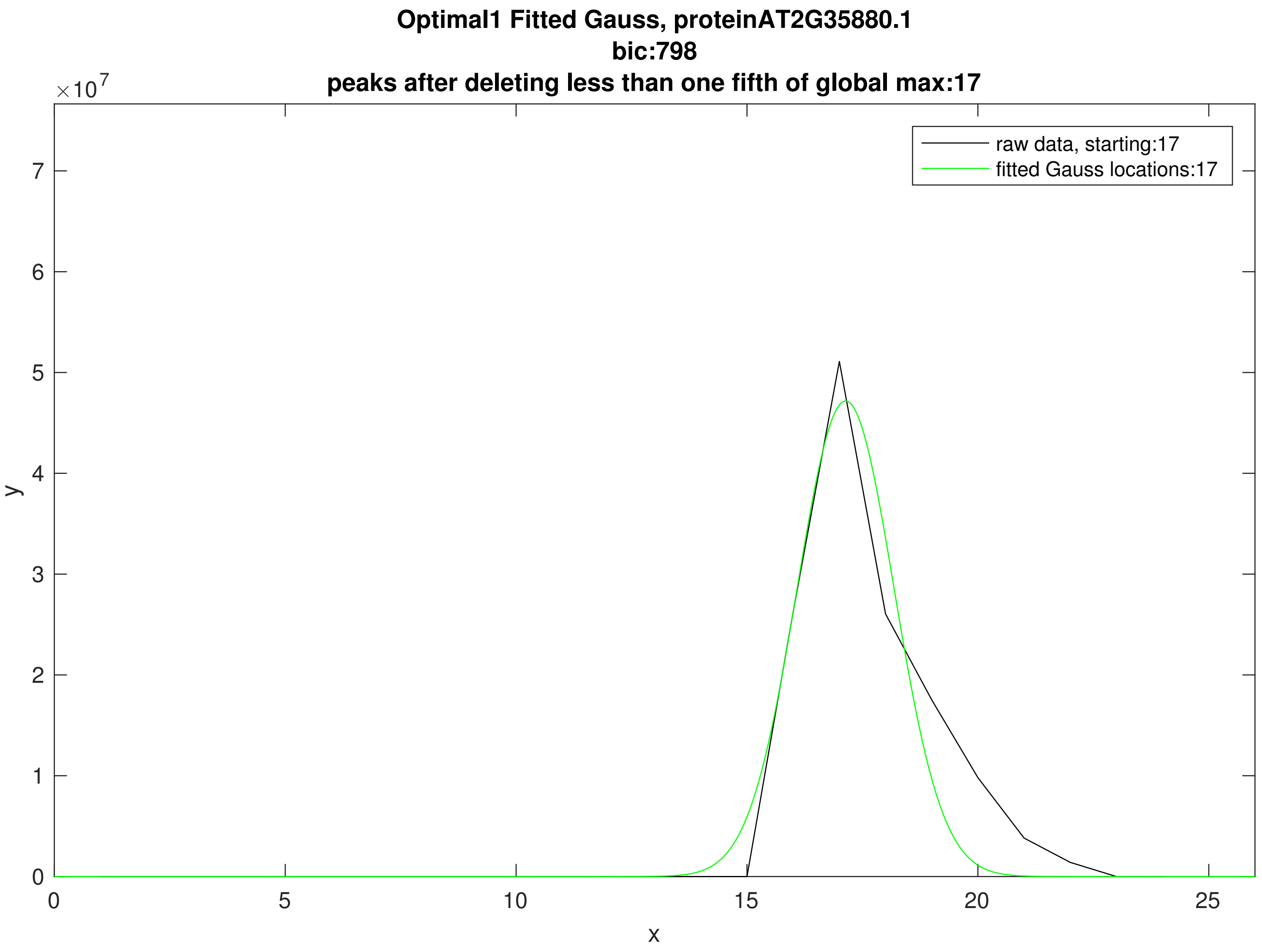
Optimal1 Fitted Gauss, proteinAT2G35880.1
bic:798
peaks after deleting less than one fifth of global max:17
×10^7
raw data, starting:17
fitted Gauss locations:17
y
x
0
1
2
3
4
5
6
7
0
5
10
15
20
25

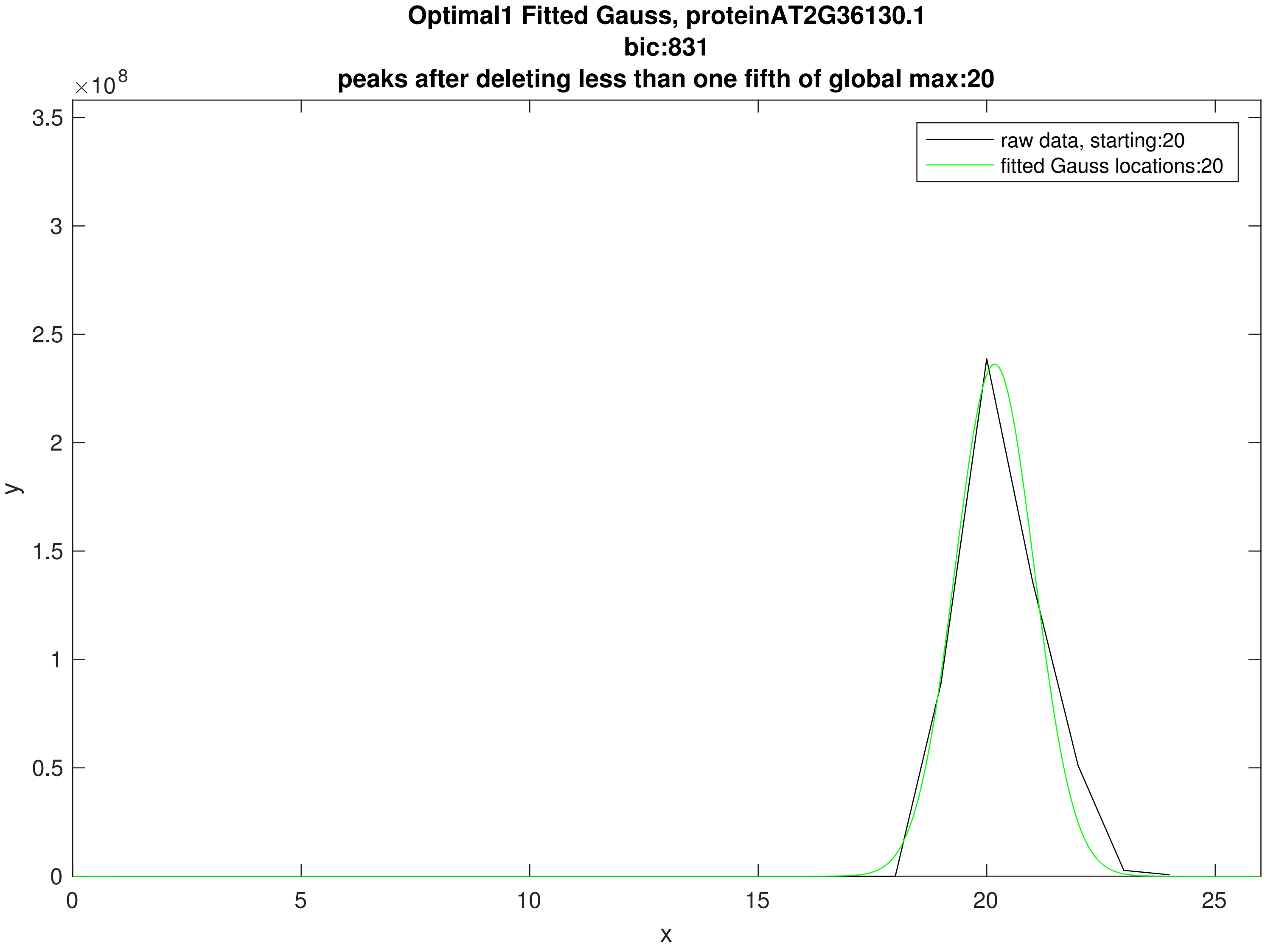
Optimal1 Fitted Gauss, proteinAT2G36130.1
bic:831
peaks after deleting less than one fifth of global max:20
×10^8
3.5
3
2.5
2
1.5
1
0.5
0
y
0
5
10
15
20
25
x
raw data, starting:20
fitted Gauss locations:20

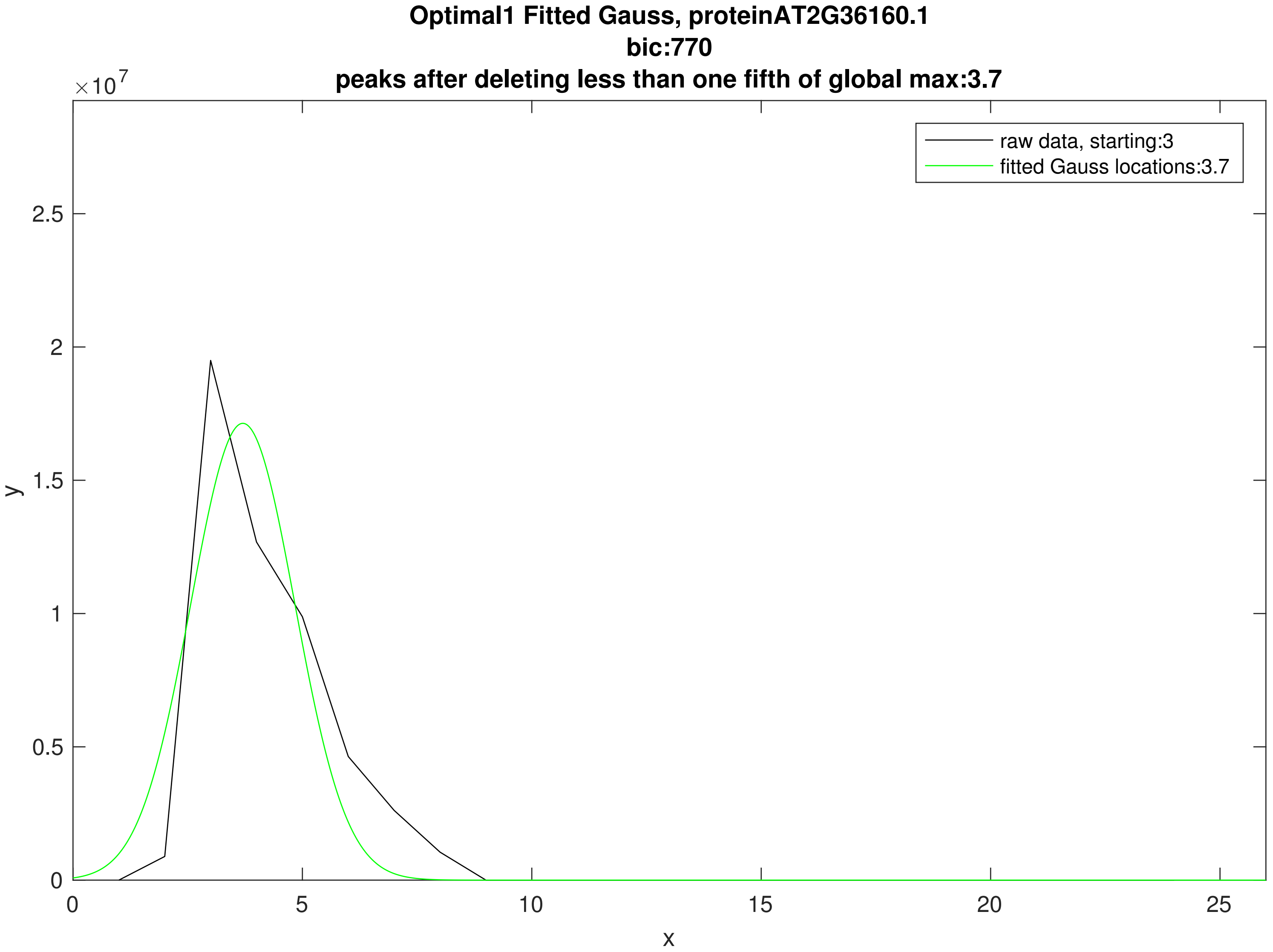

Optimal1 Fitted Gauss, proteinAT2G36160.1
bic:770
peaks after deleting less than one fifth of global max:3.7
×10^7
raw data, starting:3
fitted Gauss locations:3.7
y
0
0.5
1
1.5
2
2.5
x
0
5
10
15
20
25

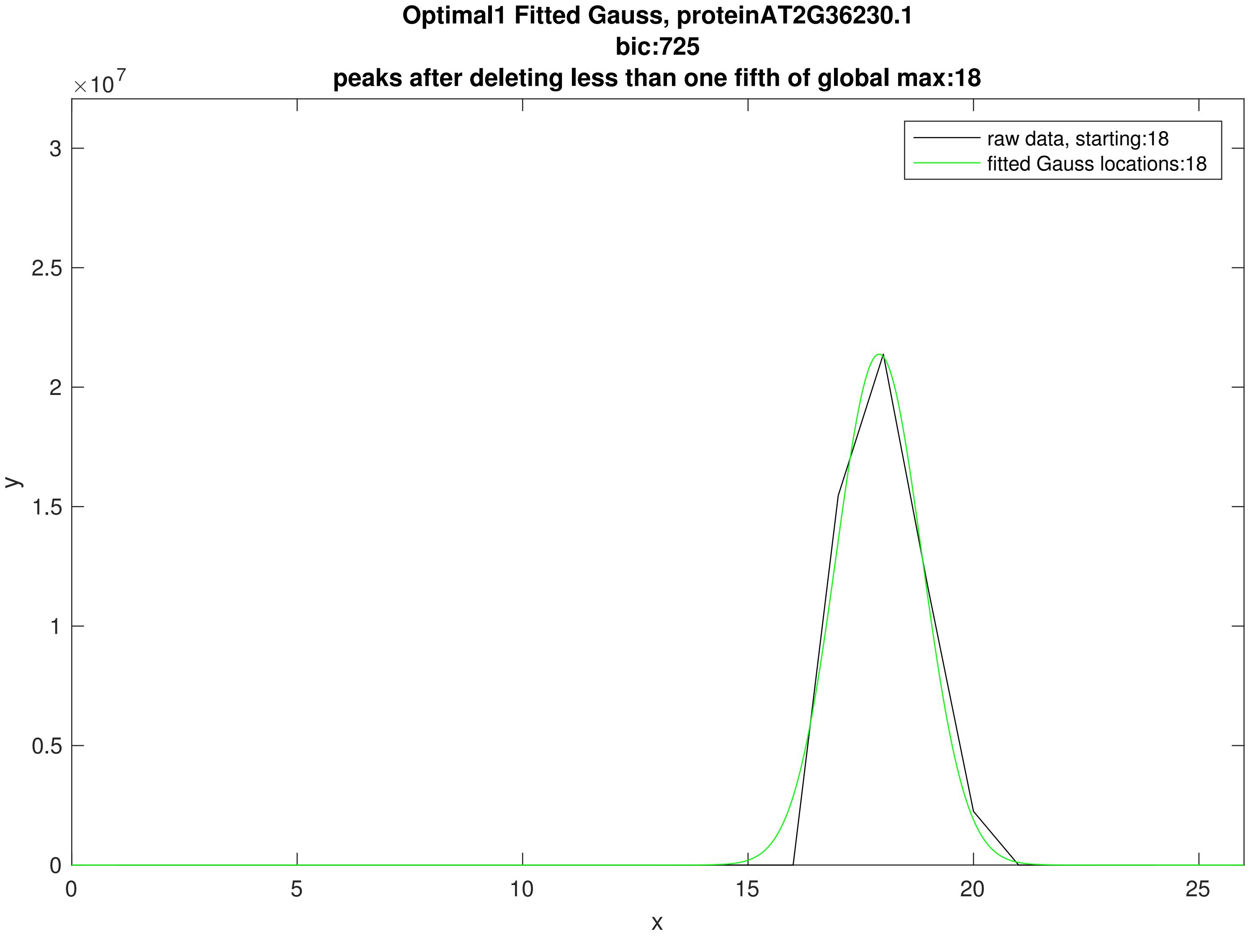

Optimal1 Fitted Gauss, proteinAT2G36230.1
bic:725
peaks after deleting less than one fifth of global max:18
×10^7
3
2.5
2
1.5
1
0.5
0
y
0
5
10
15
20
25
x
raw data, starting:18
fitted Gauss locations:18

**Optimal1 Fitted Gauss, proteinAT2G36310.1**
**bic:895**
**peaks after deleting less than one fifth of global max:14**

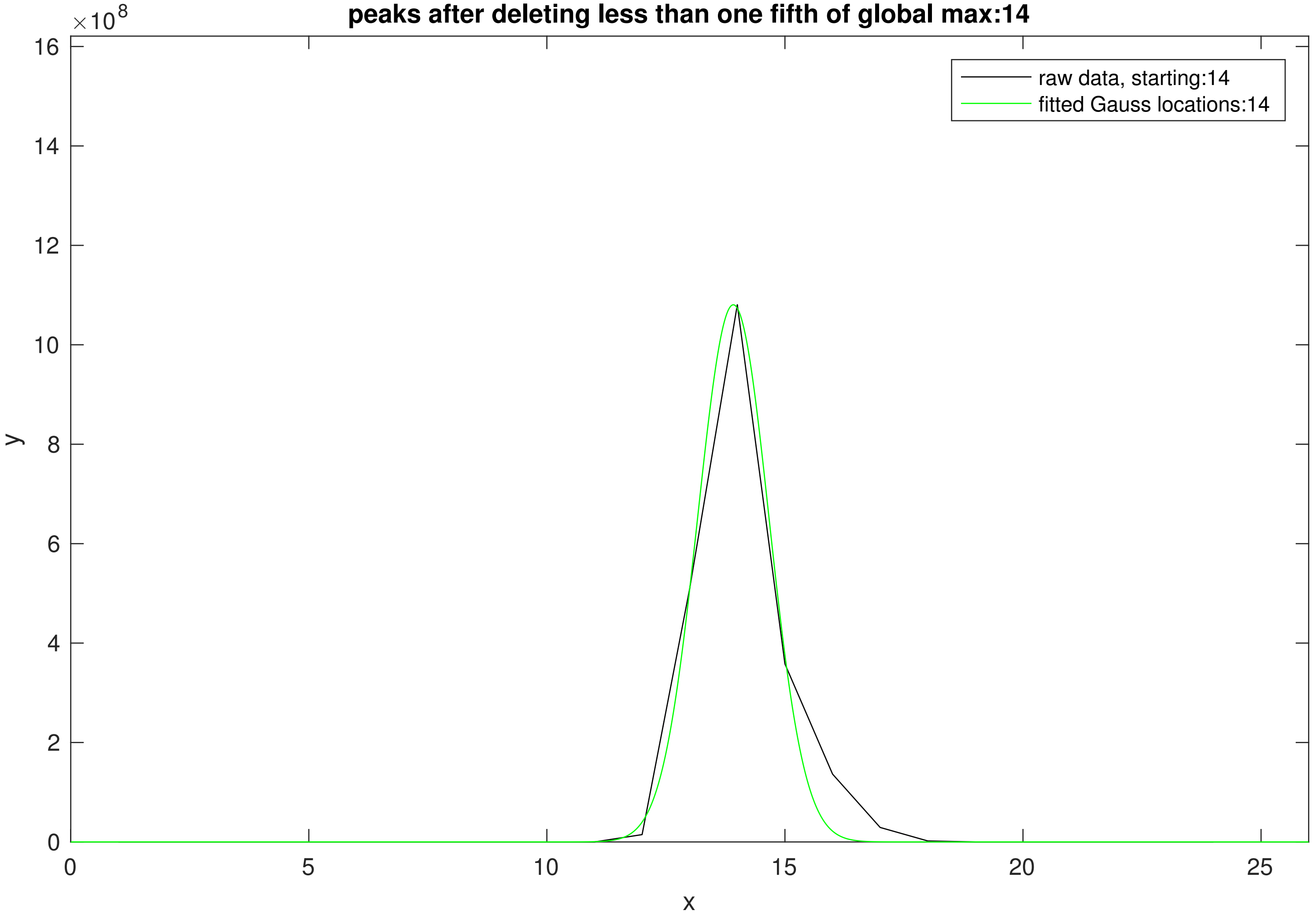

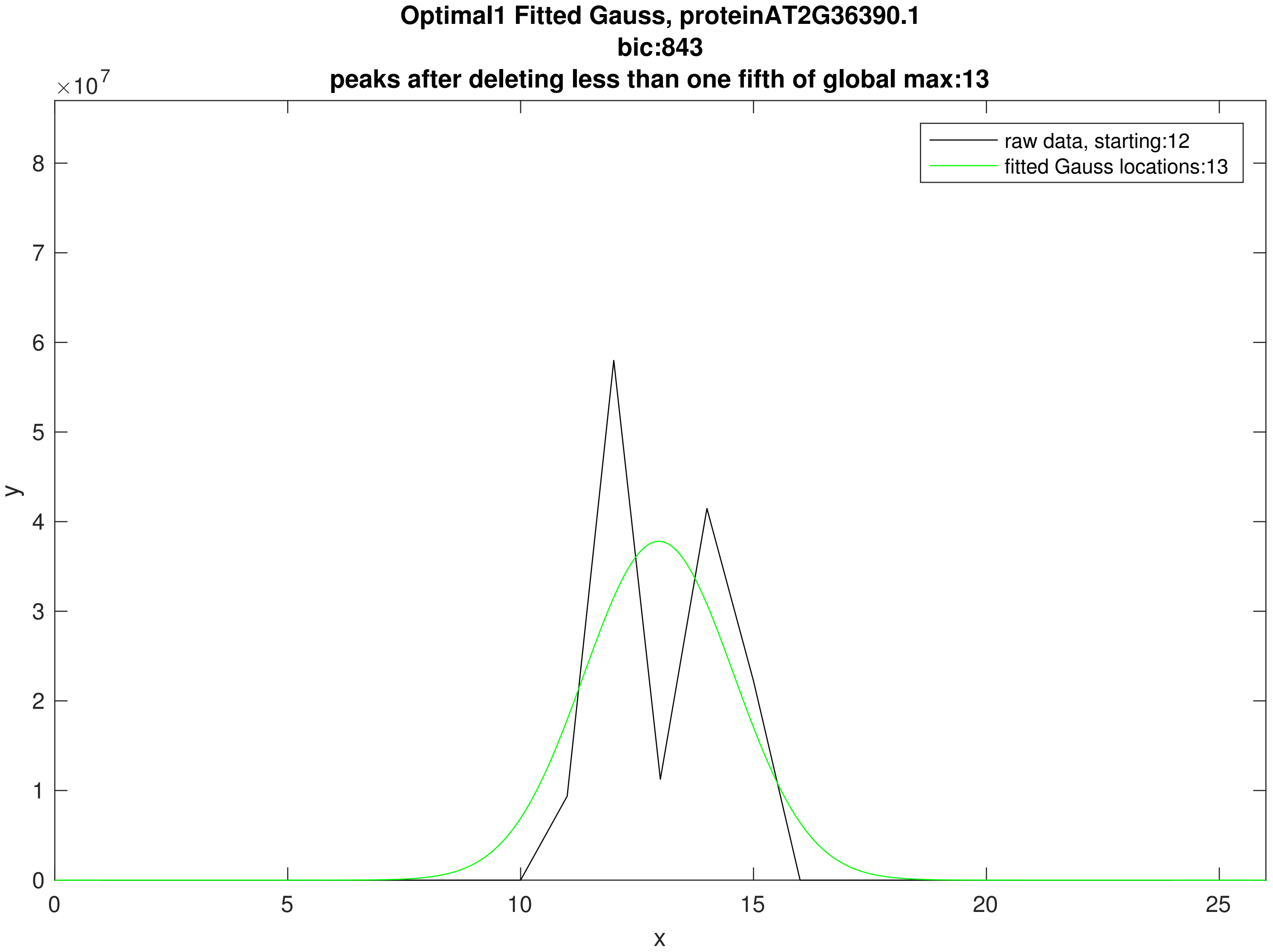

Optimal1 Fitted Gauss, proteinAT2G36390.1
bic:843
peaks after deleting less than one fifth of global max:13
×10^7
raw data, starting:12
fitted Gauss locations:13
y
x
0
1
2
3
4
5
6
7
8
0
5
10
15
20
25

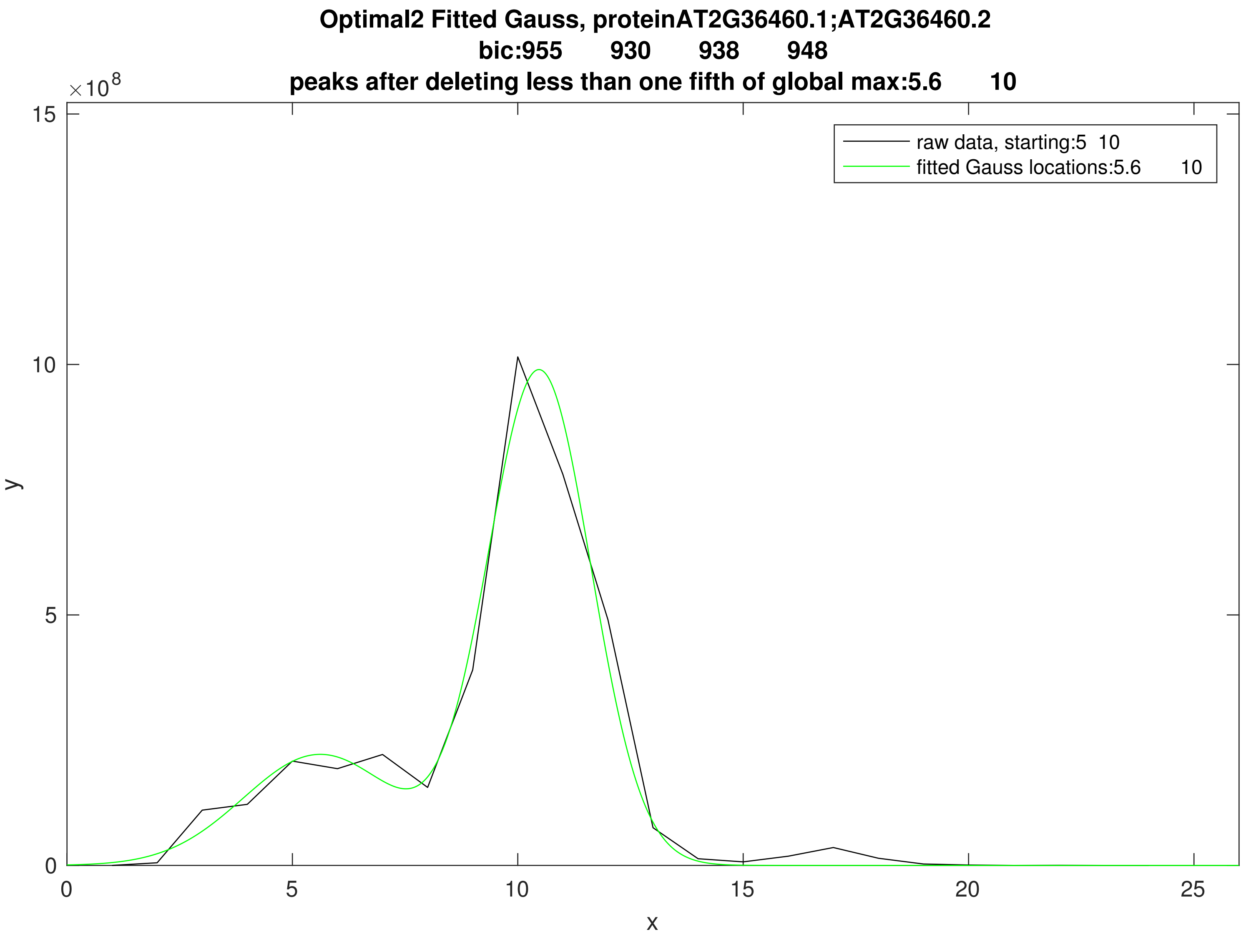
Optimal2 Fitted Gauss, proteinAT2G36460.1;AT2G36460.2
bic:955 930 938 948
peaks after deleting less than one fifth of global max:5.6 10
×10^8
raw data, starting:5 10
fitted Gauss locations:5.6 10
y
x
0
5
10
15
20
25

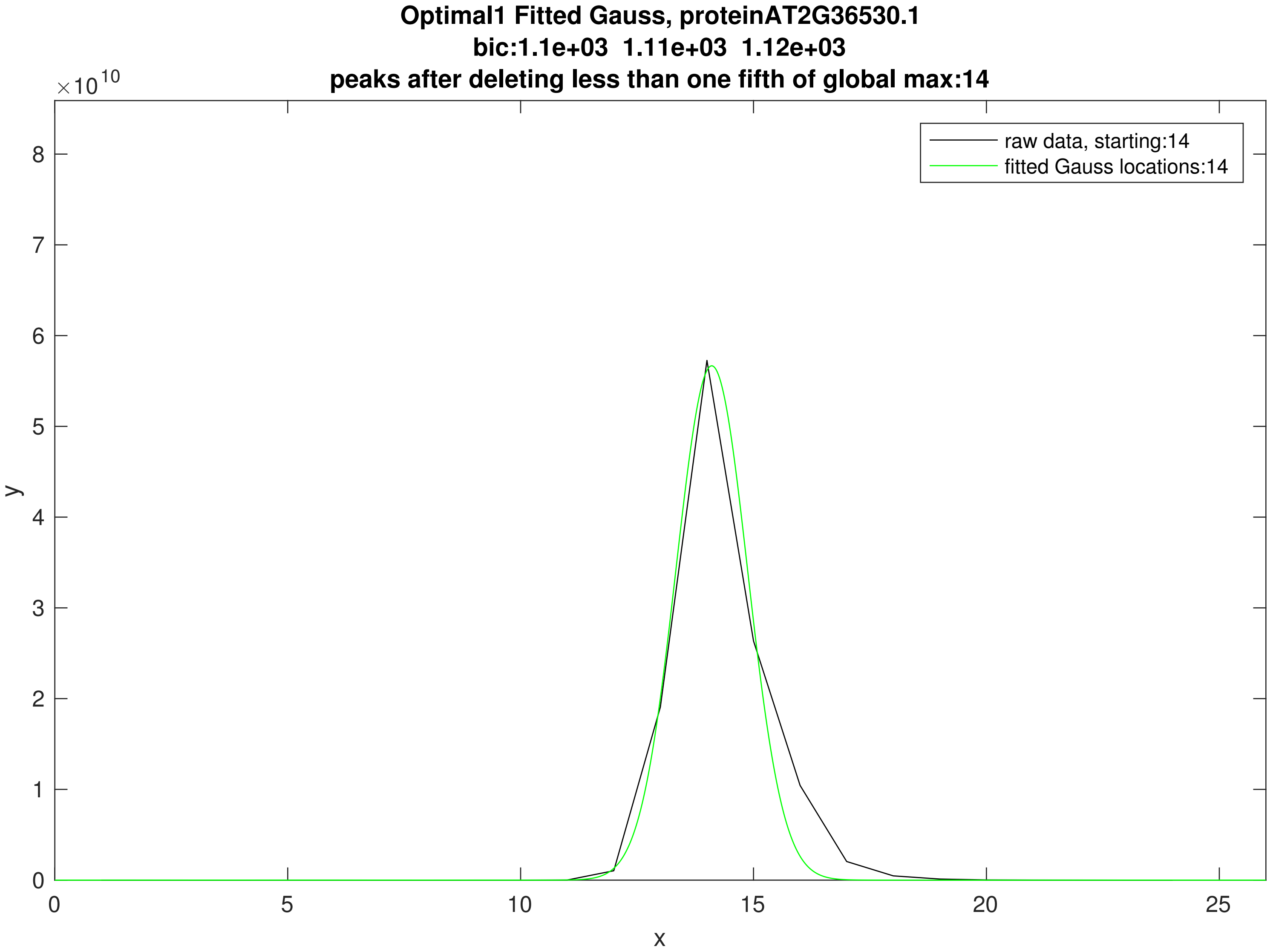

Optimal1 Fitted Gauss, proteinAT2G36530.1
bic:1.1e+03  1.11e+03  1.12e+03
peaks after deleting less than one fifth of global max:14
×10^10
raw data, starting:14
fitted Gauss locations:14
y
x
0
1
2
3
4
5
6
7
8
0
5
10
15
20
25

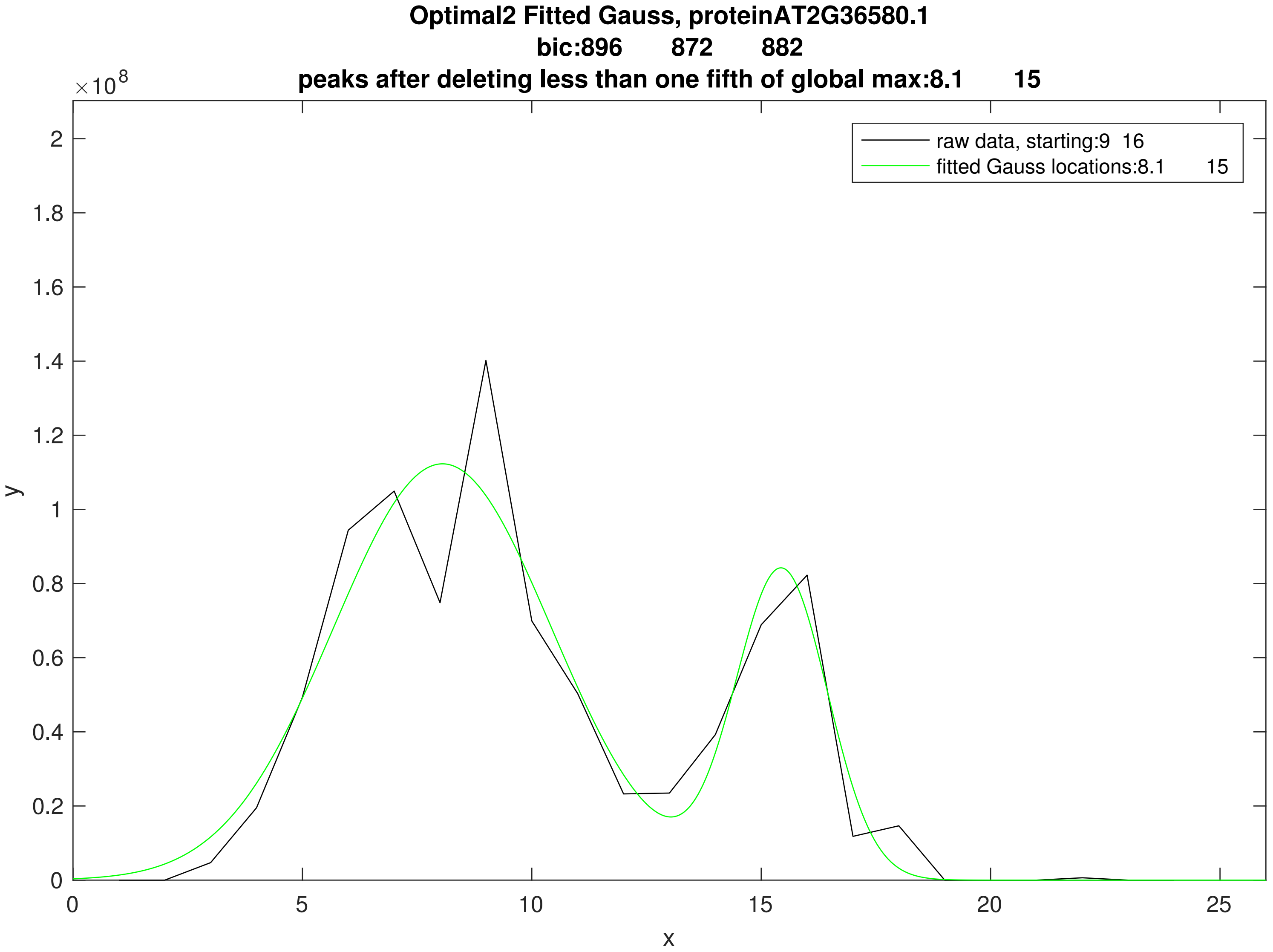

Optimal2 Fitted Gauss, proteinAT2G36580.1
bic:896 872 882
peaks after deleting less than one fifth of global max:8.1 15
×10^8
raw data, starting:9 16
fitted Gauss locations:8.1 15
y
x
0
0.2
0.4
0.6
0.8
1
1.2
1.4
1.6
1.8
2
0
5
10
15
20
25

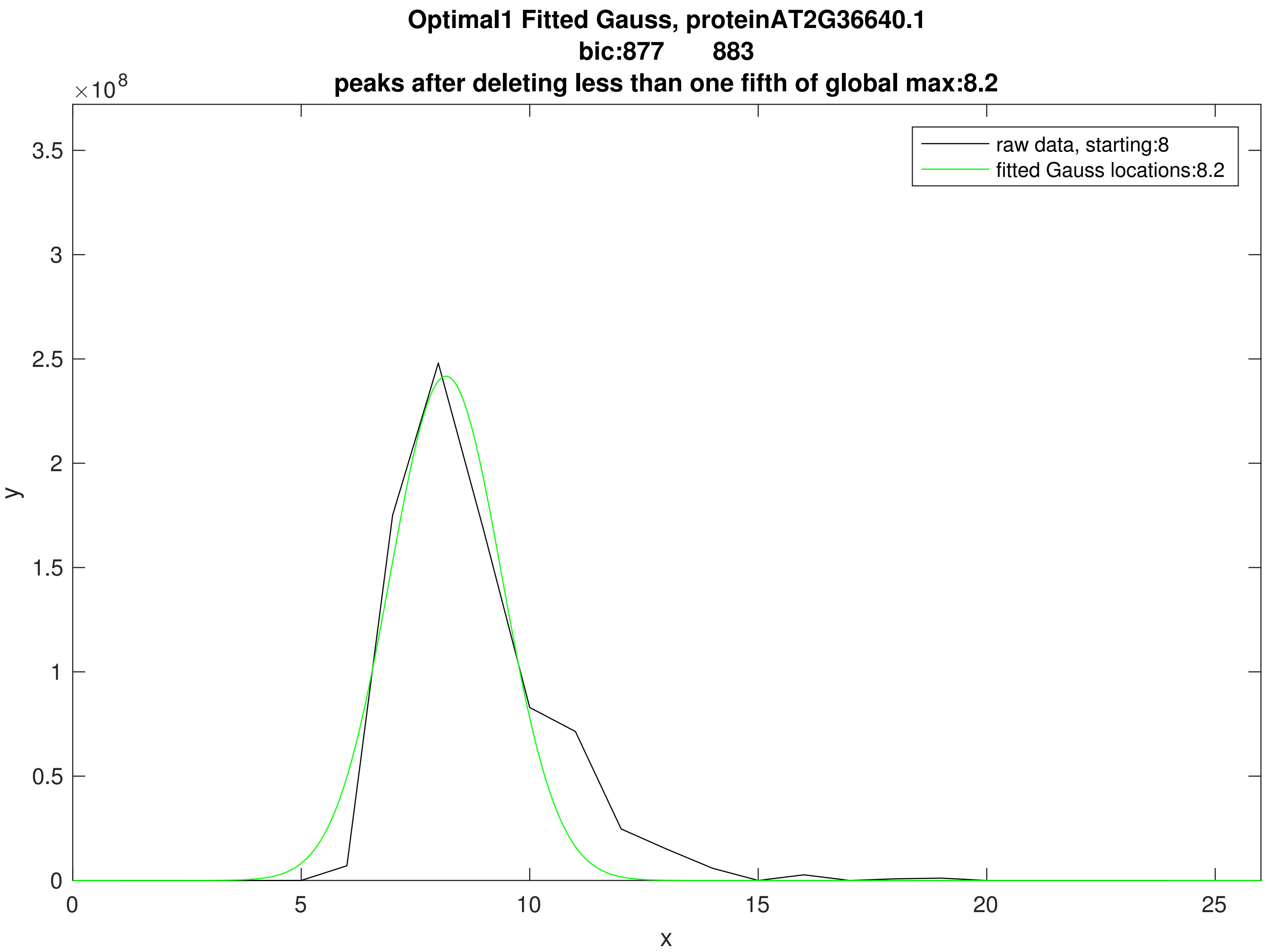

Optimal1 Fitted Gauss, proteinAT2G36640.1
bic:877 883
peaks after deleting less than one fifth of global max:8.2
×10^8
3.5
3
2.5
2
1.5
1
0.5
0
y
0
5
10
15
20
25
x
raw data, starting:8
fitted Gauss locations:8.2

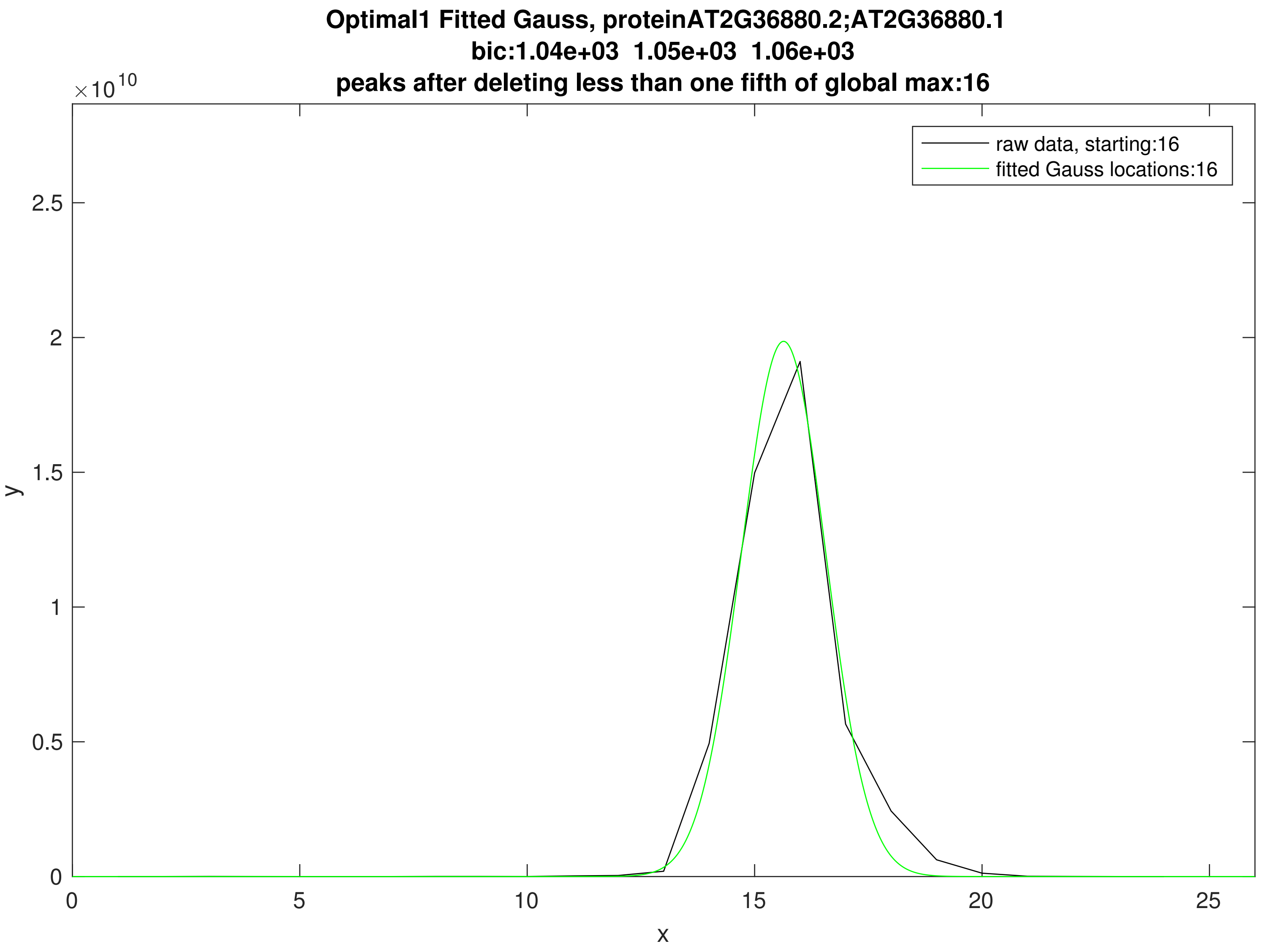
Optimal1 Fitted Gauss, proteinAT2G36880.2;AT2G36880.1
bic:1.04e+03  1.05e+03  1.06e+03
peaks after deleting less than one fifth of global max:16
×10^10
raw data, starting:16
fitted Gauss locations:16
y
x
0
0.5
1
1.5
2
2.5
0
5
10
15
20
25

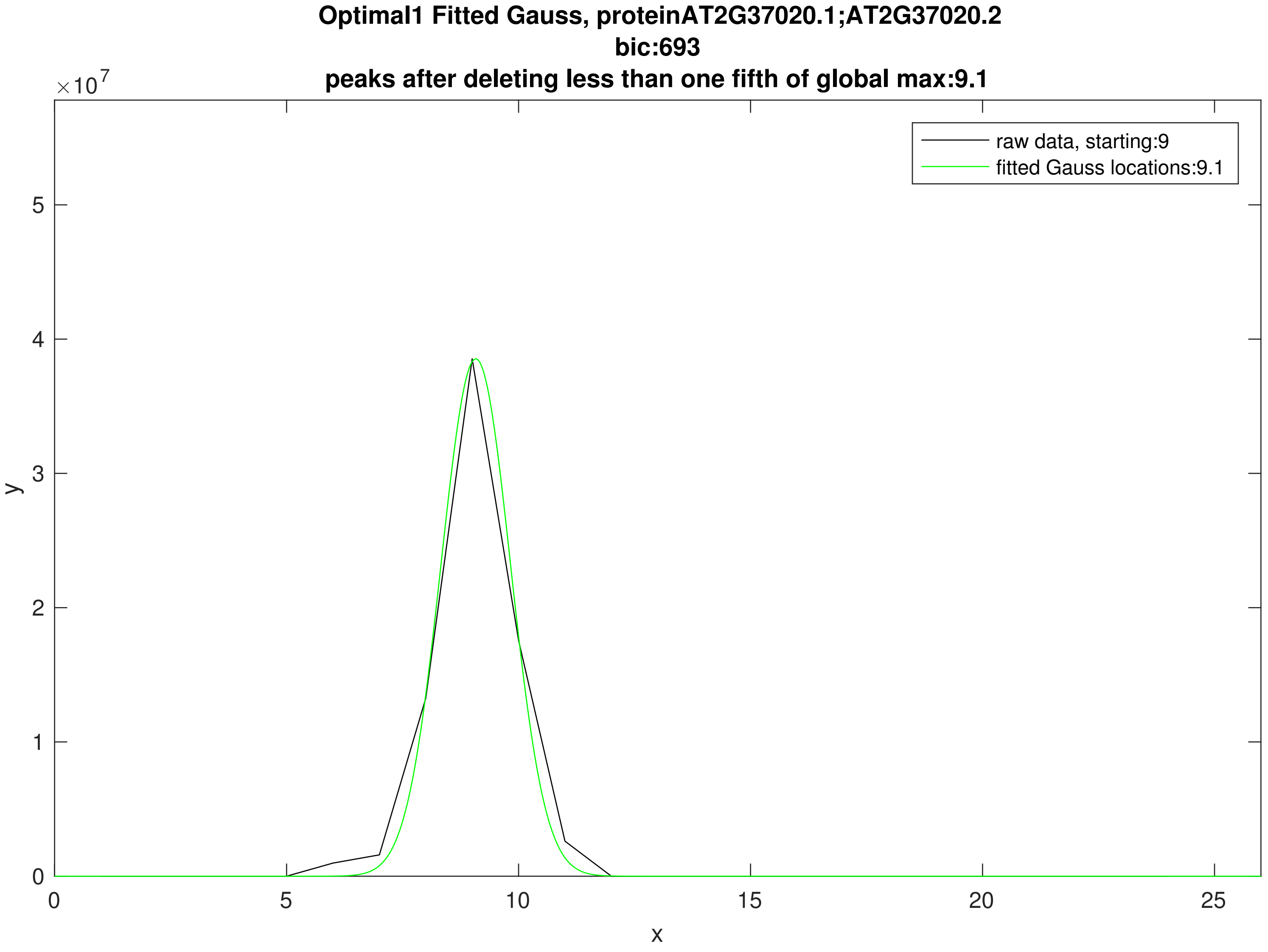
Optimal1 Fitted Gauss, proteinAT2G37020.1;AT2G37020.2
bic:693
peaks after deleting less than one fifth of global max:9.1
×10^7
raw data, starting:9
fitted Gauss locations:9.1
y
x
0
1
2
3
4
5
0
5
10
15
20
25

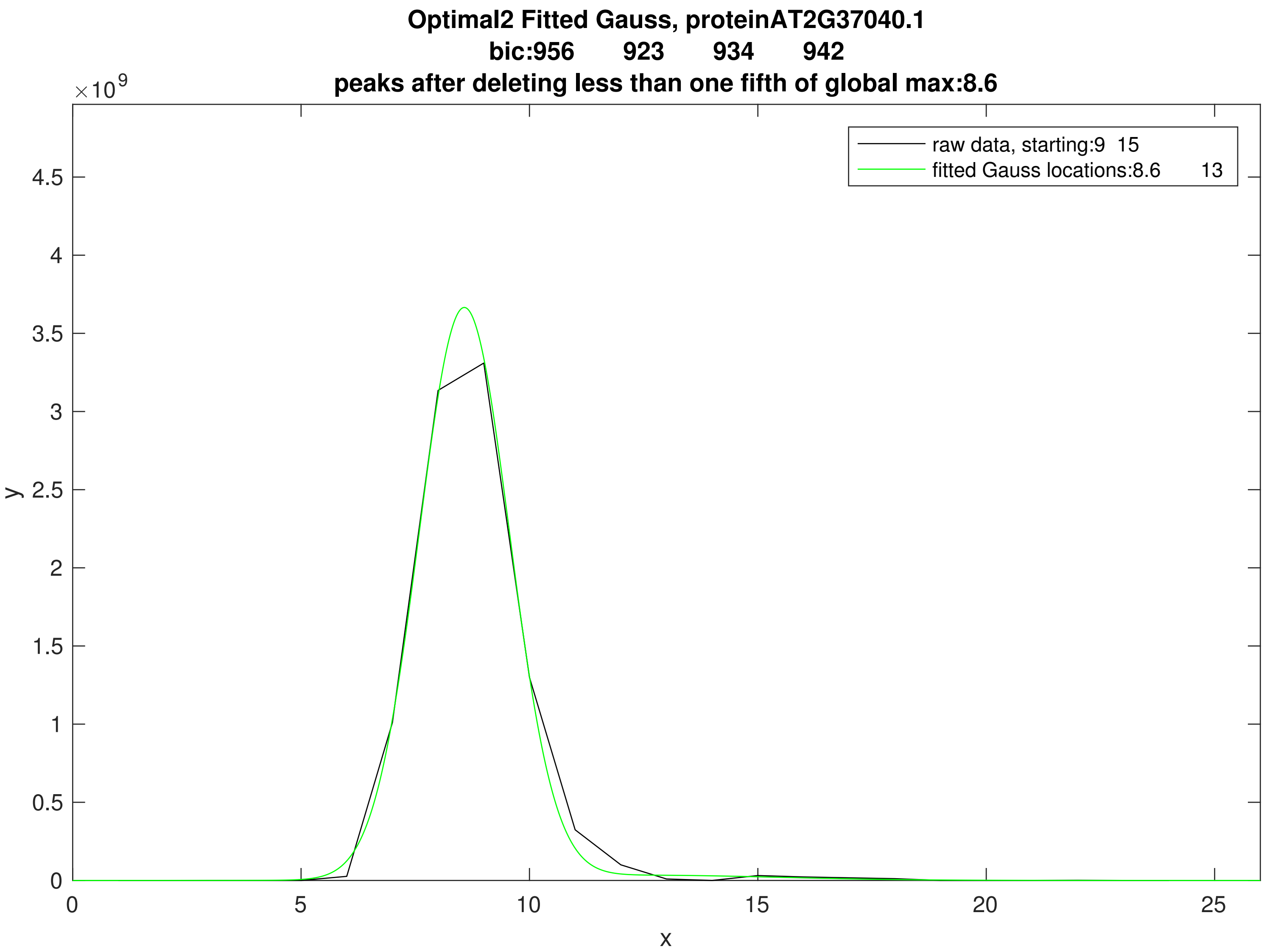
Optimal2 Fitted Gauss, proteinAT2G37040.1
bic:956 923 934 942
peaks after deleting less than one fifth of global max:8.6
×10^9
raw data, starting:9 15
fitted Gauss locations:8.6 13
y
x
0
0.5
1
1.5
2
2.5
3
3.5
4
4.5
5
10
15
20
25

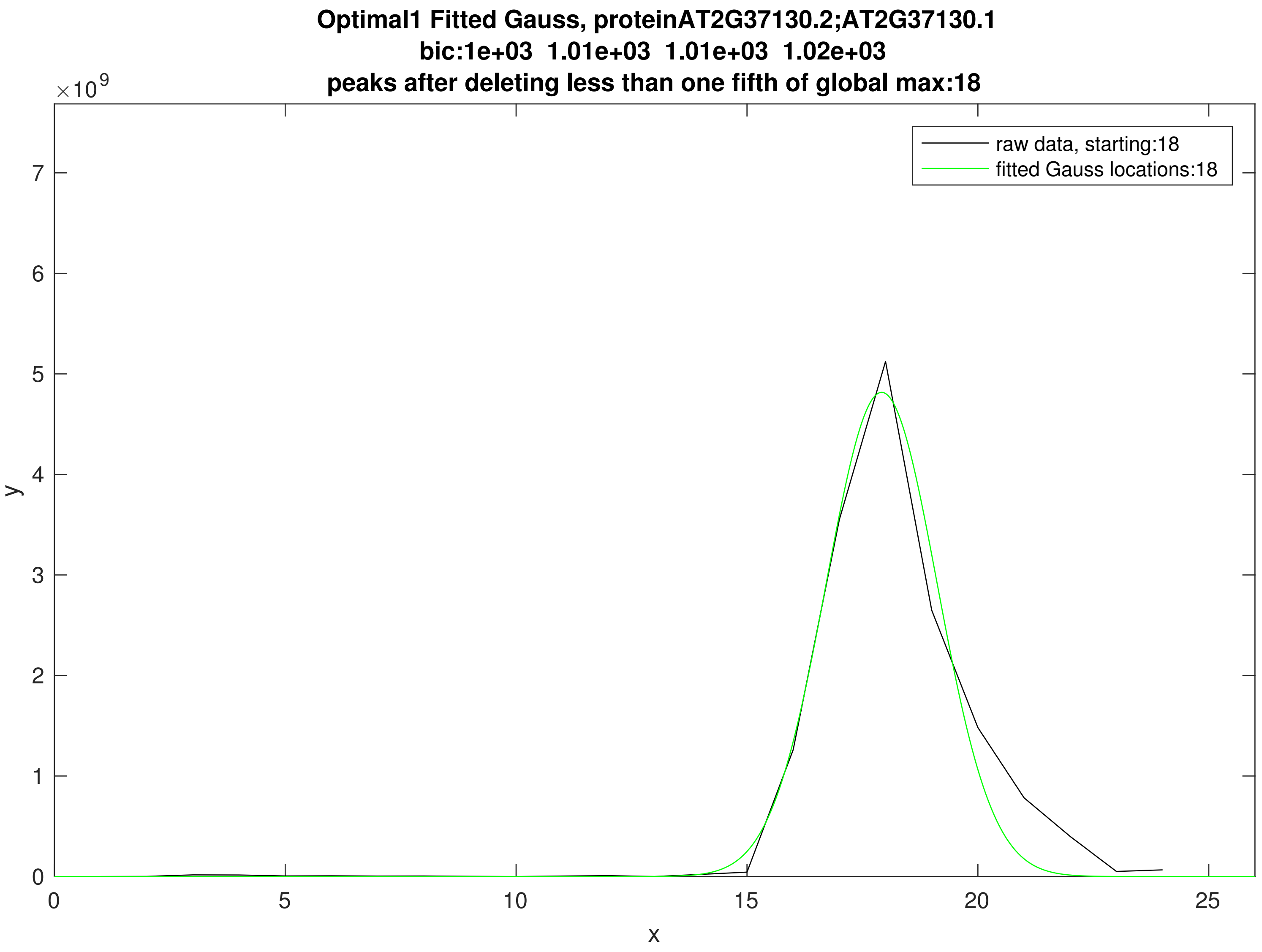
Optimal1 Fitted Gauss, proteinAT2G37130.2;AT2G37130.1
bic:1e+03  1.01e+03  1.01e+03  1.02e+03
peaks after deleting less than one fifth of global max:18
×10^9
raw data, starting:18
fitted Gauss locations:18
y
x
0
1
2
3
4
5
6
7
0
5
10
15
20
25

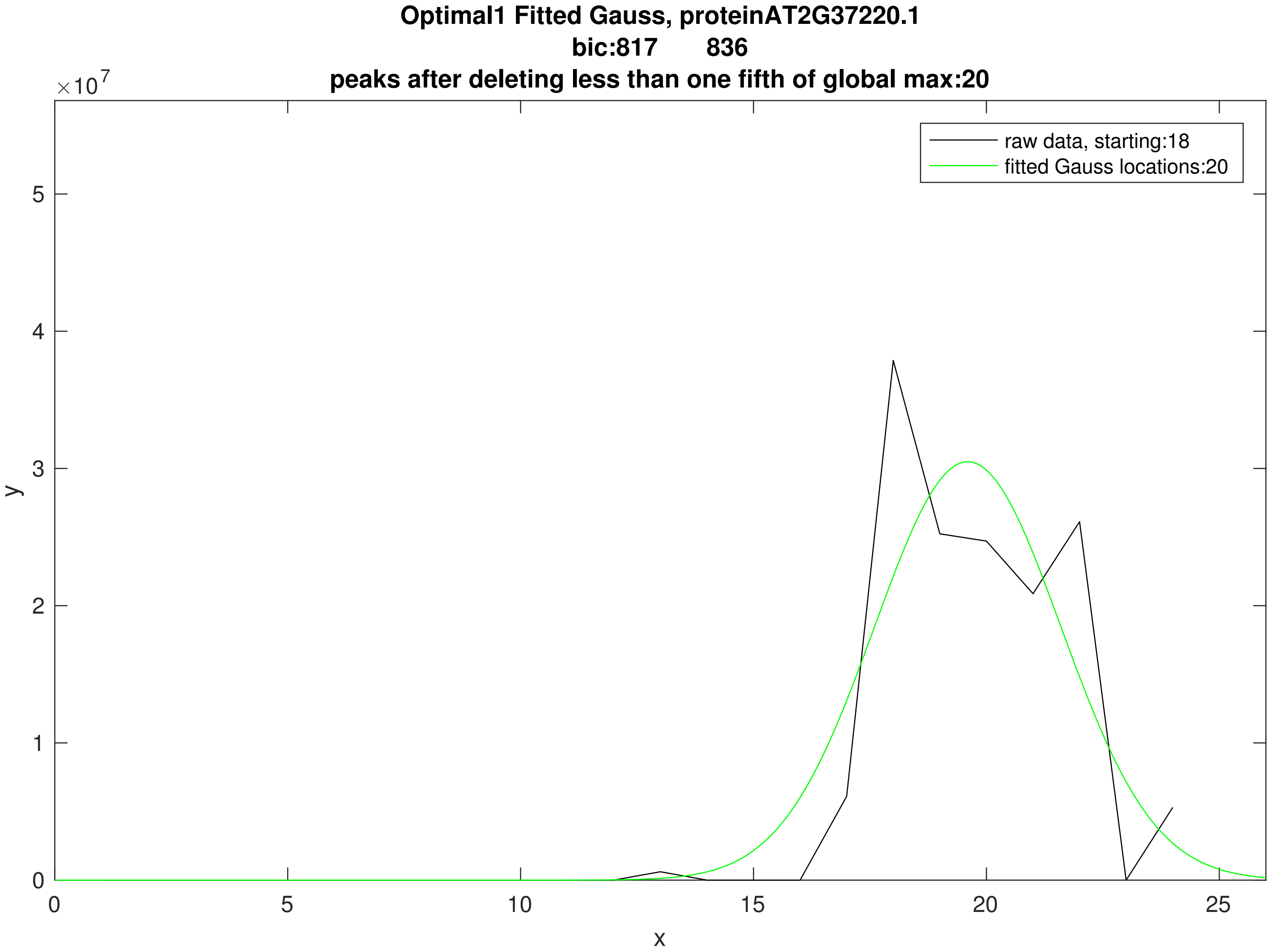

Optimal1 Fitted Gauss, proteinAT2G37220.1
bic:817 836
peaks after deleting less than one fifth of global max:20
×10^7
5
4
3
2
1
0
y
0
5
10
15
20
25
x
raw data, starting:18
fitted Gauss locations:20

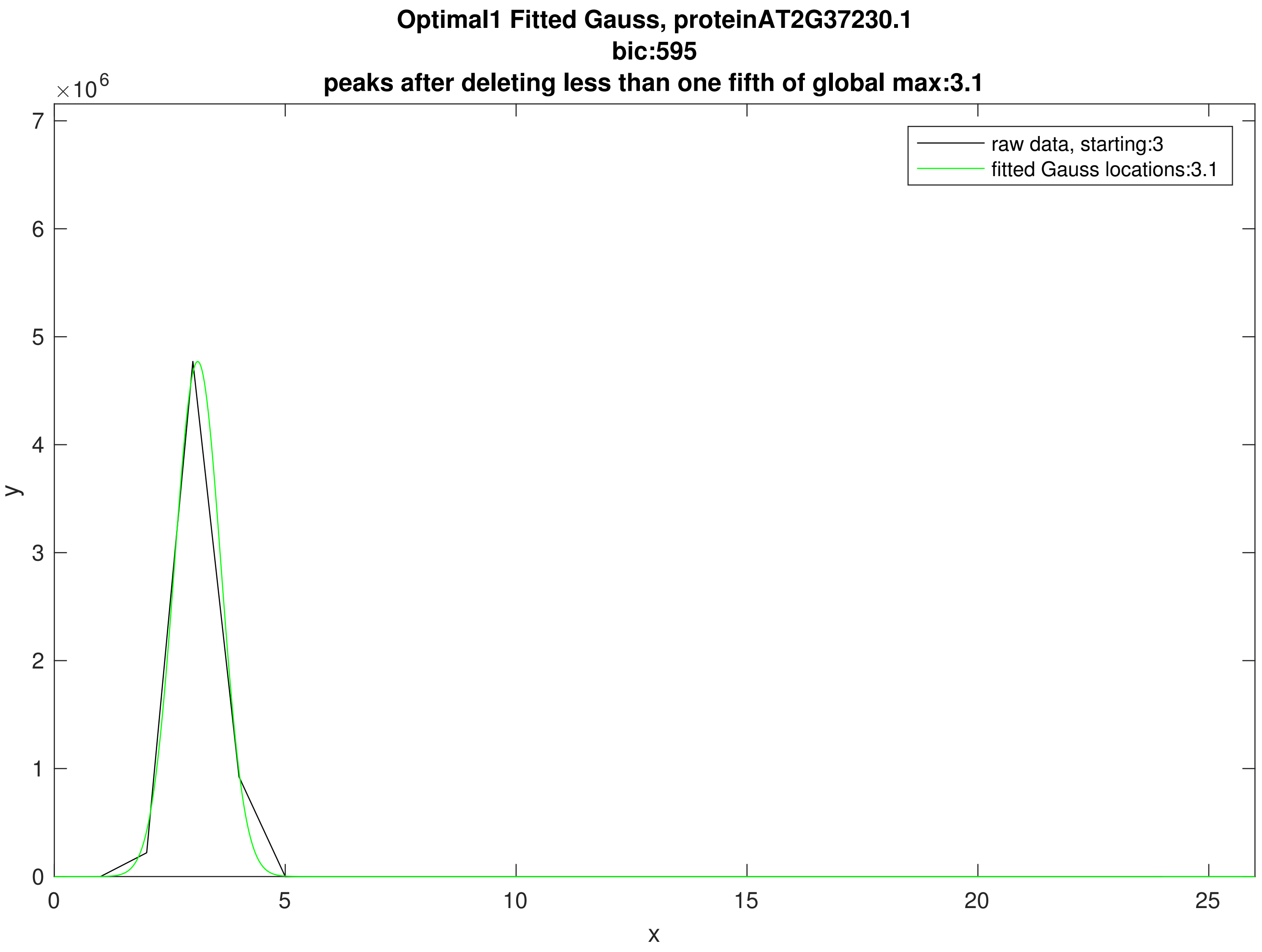
Optimal1 Fitted Gauss, proteinAT2G37230.1
bic:595
peaks after deleting less than one fifth of global max:3.1
×10^6
raw data, starting:3
fitted Gauss locations:3.1
y
x
0
1
2
3
4
5
6
7
0
5
10
15
20
25

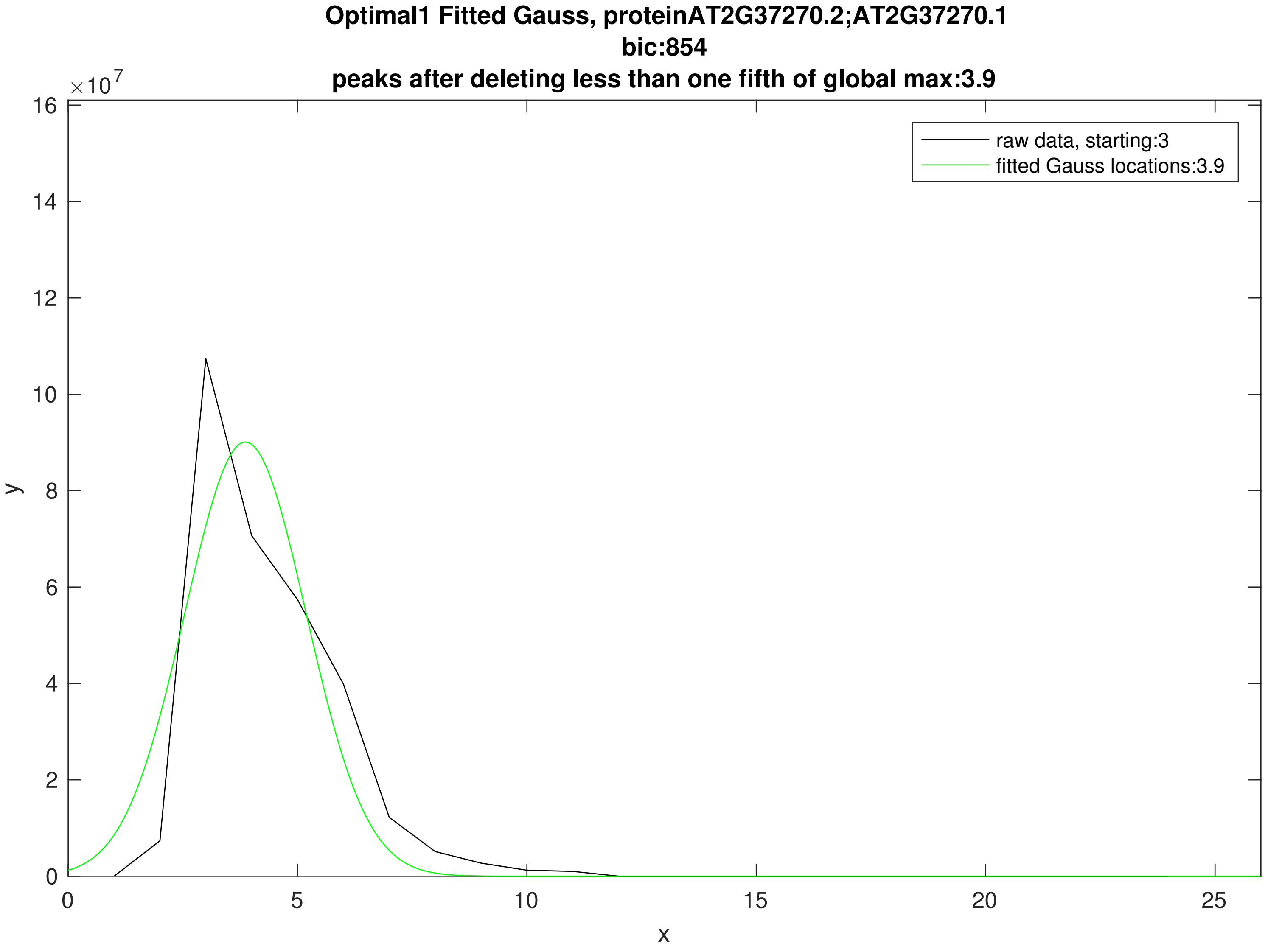
Optimal1 Fitted Gauss, proteinAT2G37270.2;AT2G37270.1
bic:854
peaks after deleting less than one fifth of global max:3.9
×10^7
raw data, starting:3
fitted Gauss locations:3.9
y
x
0
2
4
6
8
10
12
14
16
5
15
20
25

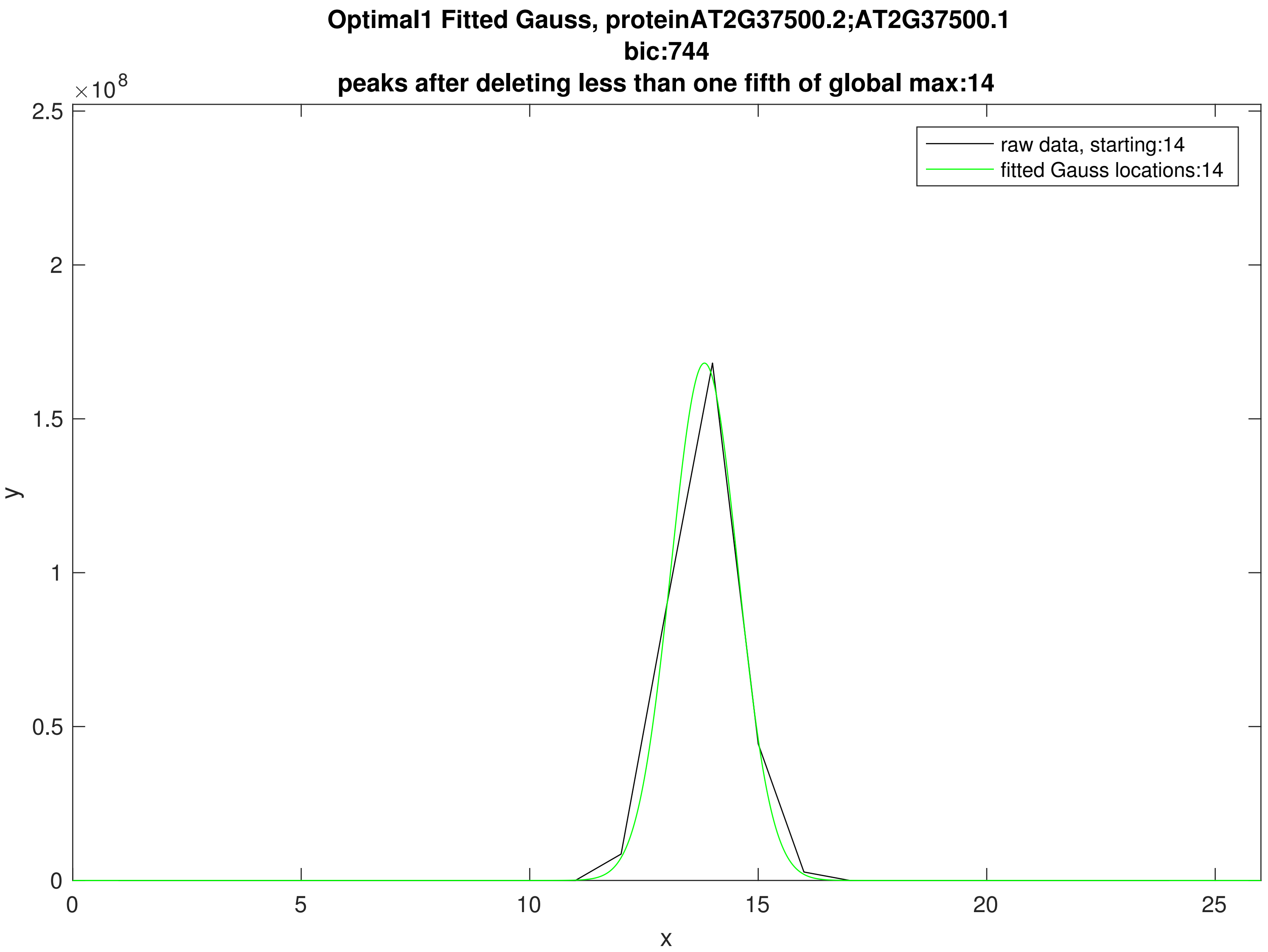
Optimal1 Fitted Gauss, proteinAT2G37500.2;AT2G37500.1
bic:744
peaks after deleting less than one fifth of global max:14
×10^8
2.5
2
1.5
1
0.5
0
y
0
5
10
15
20
25
x
raw data, starting:14
fitted Gauss locations:14

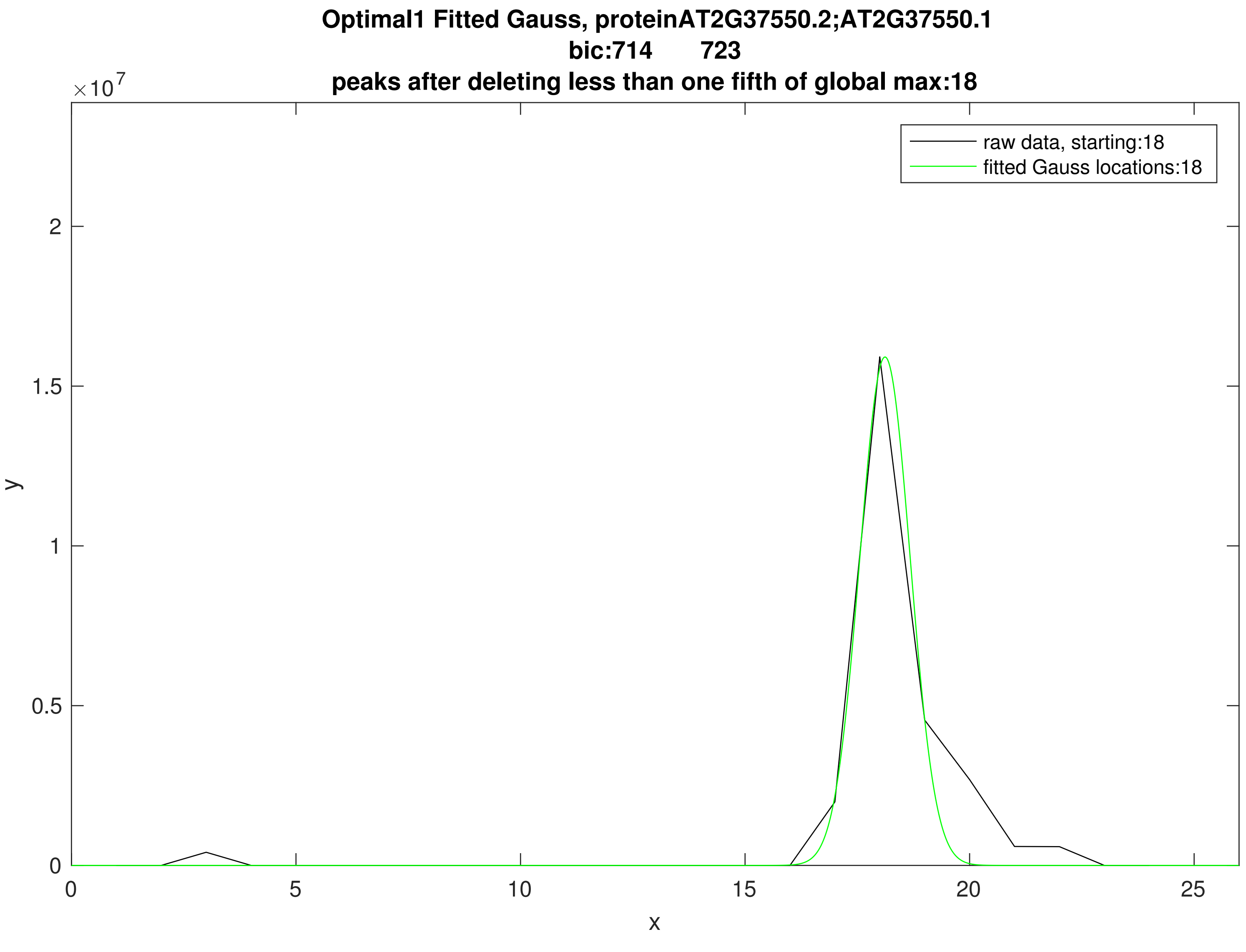

Optimal1 Fitted Gauss, proteinAT2G37550.2;AT2G37550.1
bic:714 723
peaks after deleting less than one fifth of global max:18
×10^7
raw data, starting:18
fitted Gauss locations:18
2
1.5
1
0.5
0
y
0
5
10
15
20
25
x

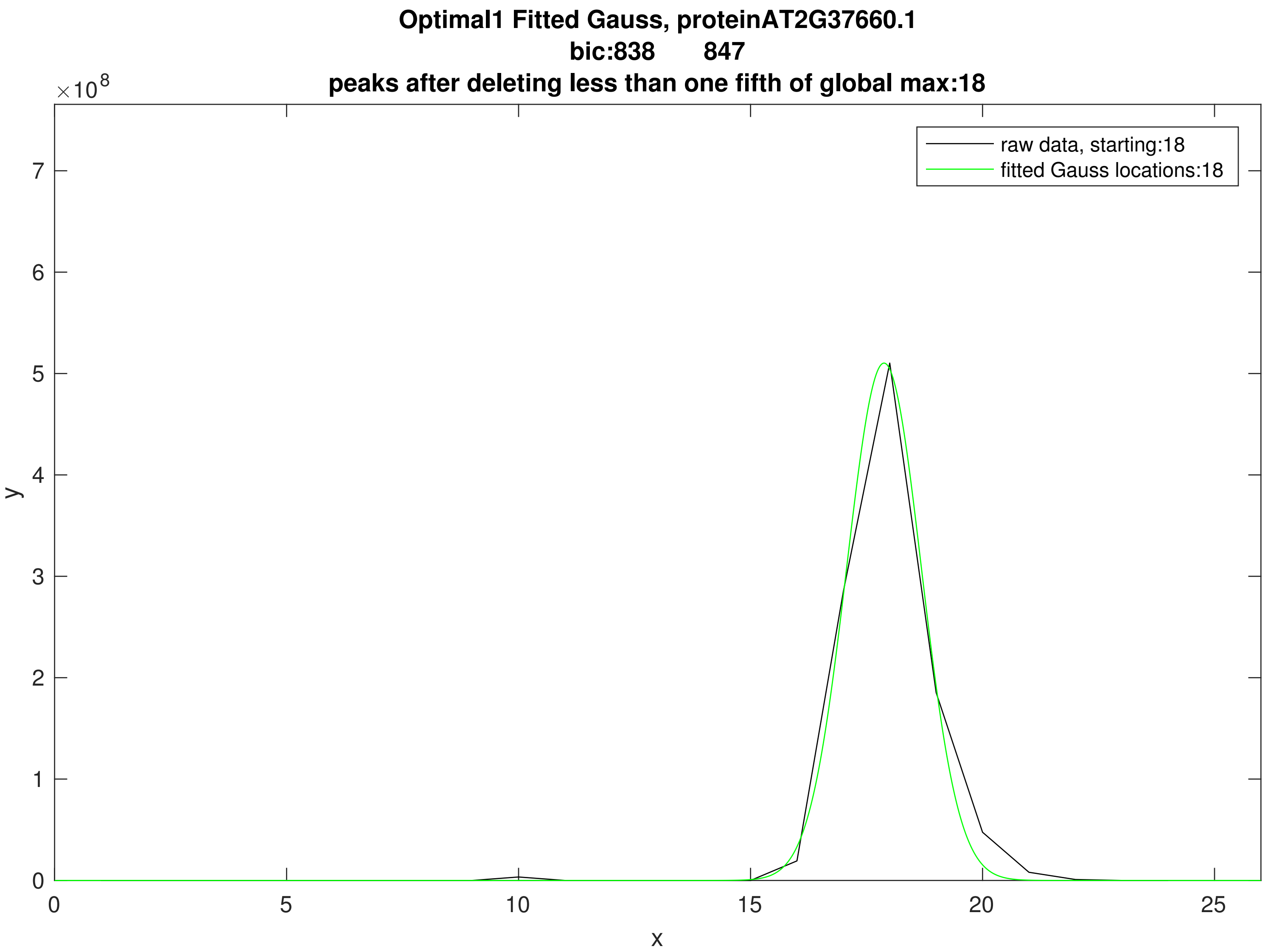
Optimal1 Fitted Gauss, proteinAT2G37660.1
bic:838 847
peaks after deleting less than one fifth of global max:18
×10^8
raw data, starting:18
fitted Gauss locations:18
y
x
0
1
2
3
4
5
6
7
0
5
10
15
20
25

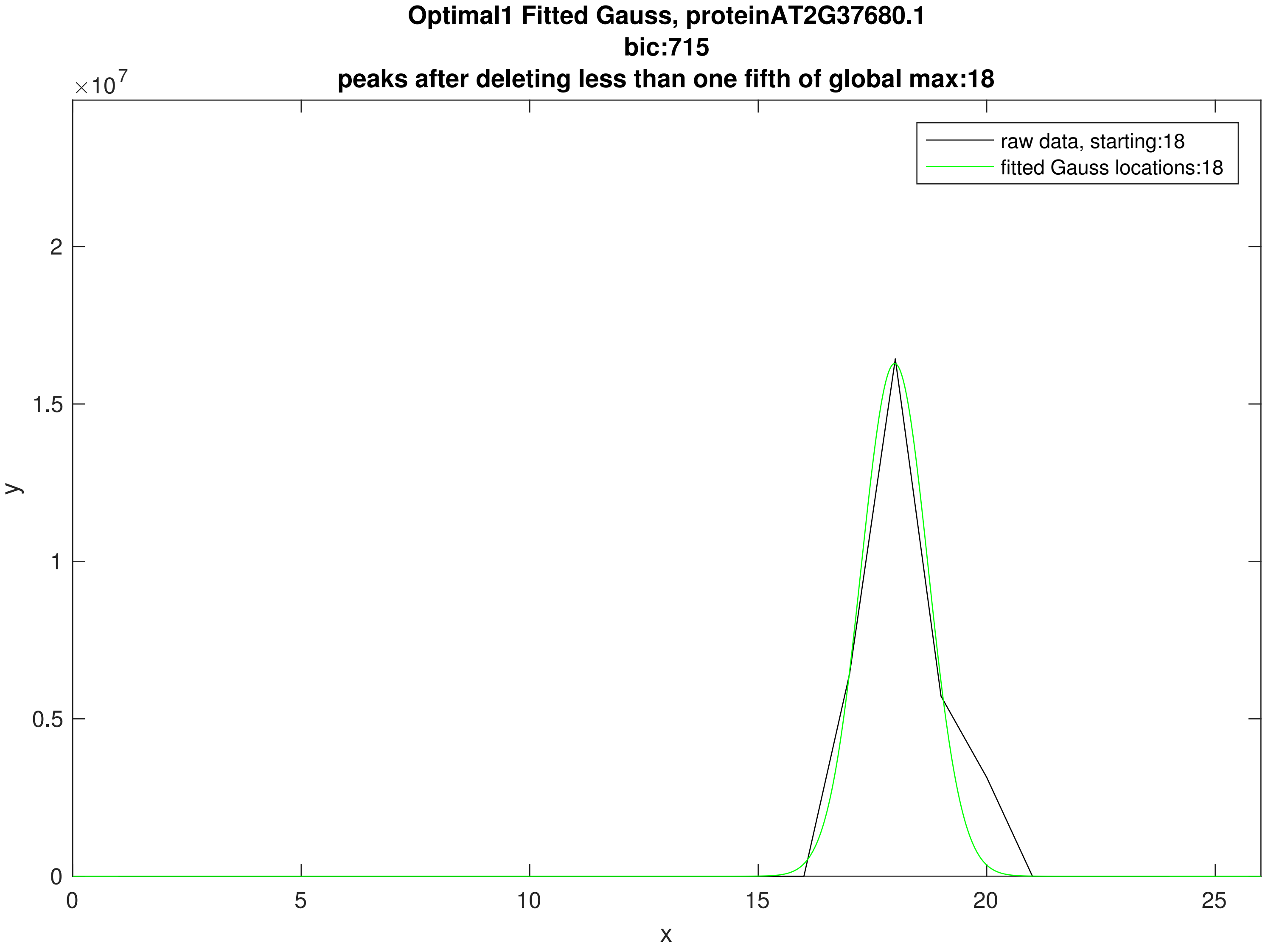

Optimal1 Fitted Gauss, proteinAT2G37680.1
bic:715
peaks after deleting less than one fifth of global max:18
×10^7
raw data, starting:18
fitted Gauss locations:18
y
x
0
0.5
1
1.5
2
0
5
10
15
20
25

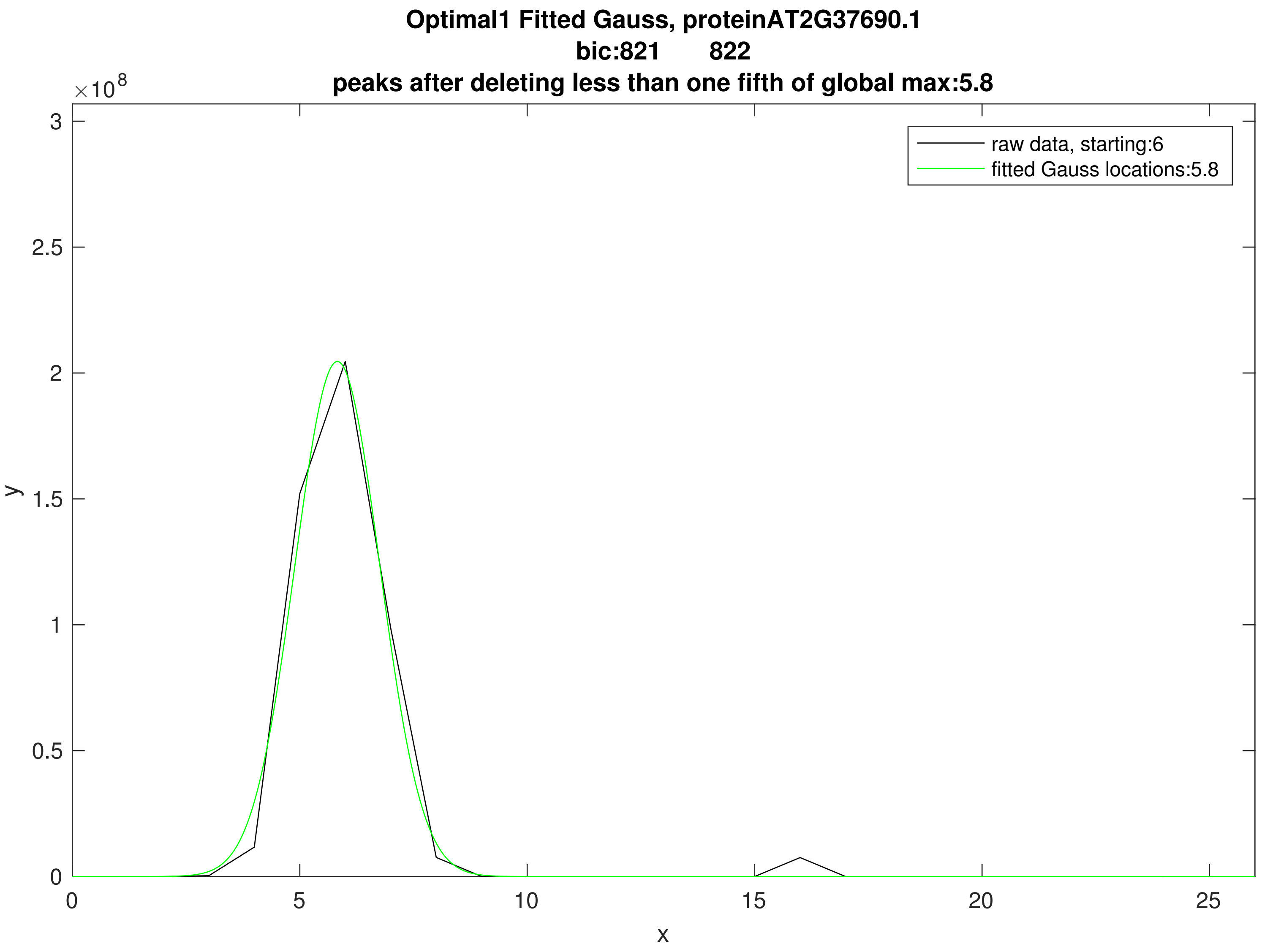
Optimal1 Fitted Gauss, proteinAT2G37690.1
bic:821 822
peaks after deleting less than one fifth of global max:5.8
×10^8
raw data, starting:6
fitted Gauss locations:5.8
y
x
0
0.5
1
1.5
2
2.5
3
0
5
10
15
20
25

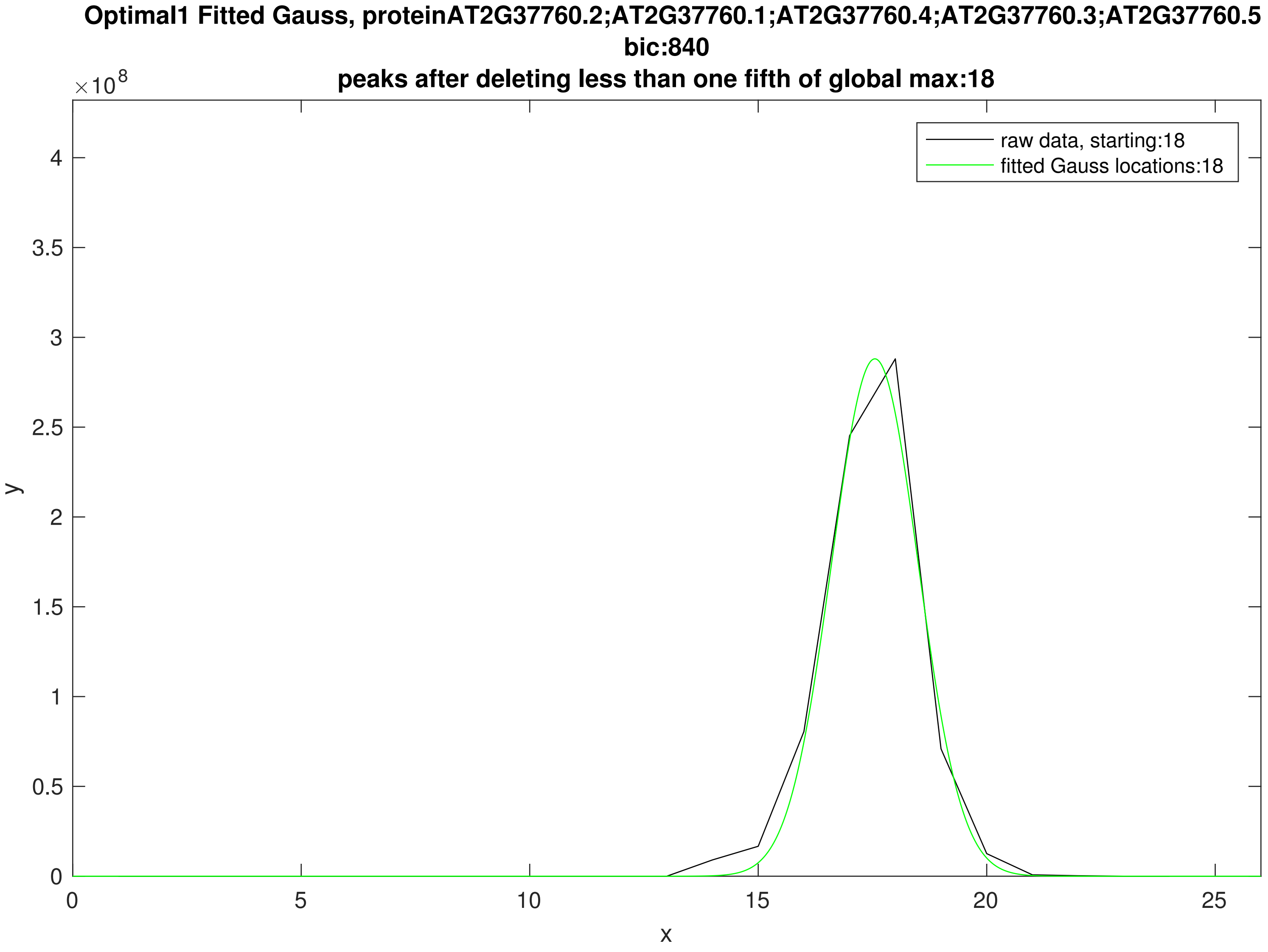

Optimal1 Fitted Gauss, proteinAT2G37760.2;AT2G37760.1;AT2G37760.4;AT2G37760.3;AT2G37760.5
bic:840
peaks after deleting less than one fifth of global max:18
×10^8
raw data, starting:18
fitted Gauss locations:18
y
x
0
0.5
1
1.5
2
2.5
3
3.5
4
5
10
15
20
25

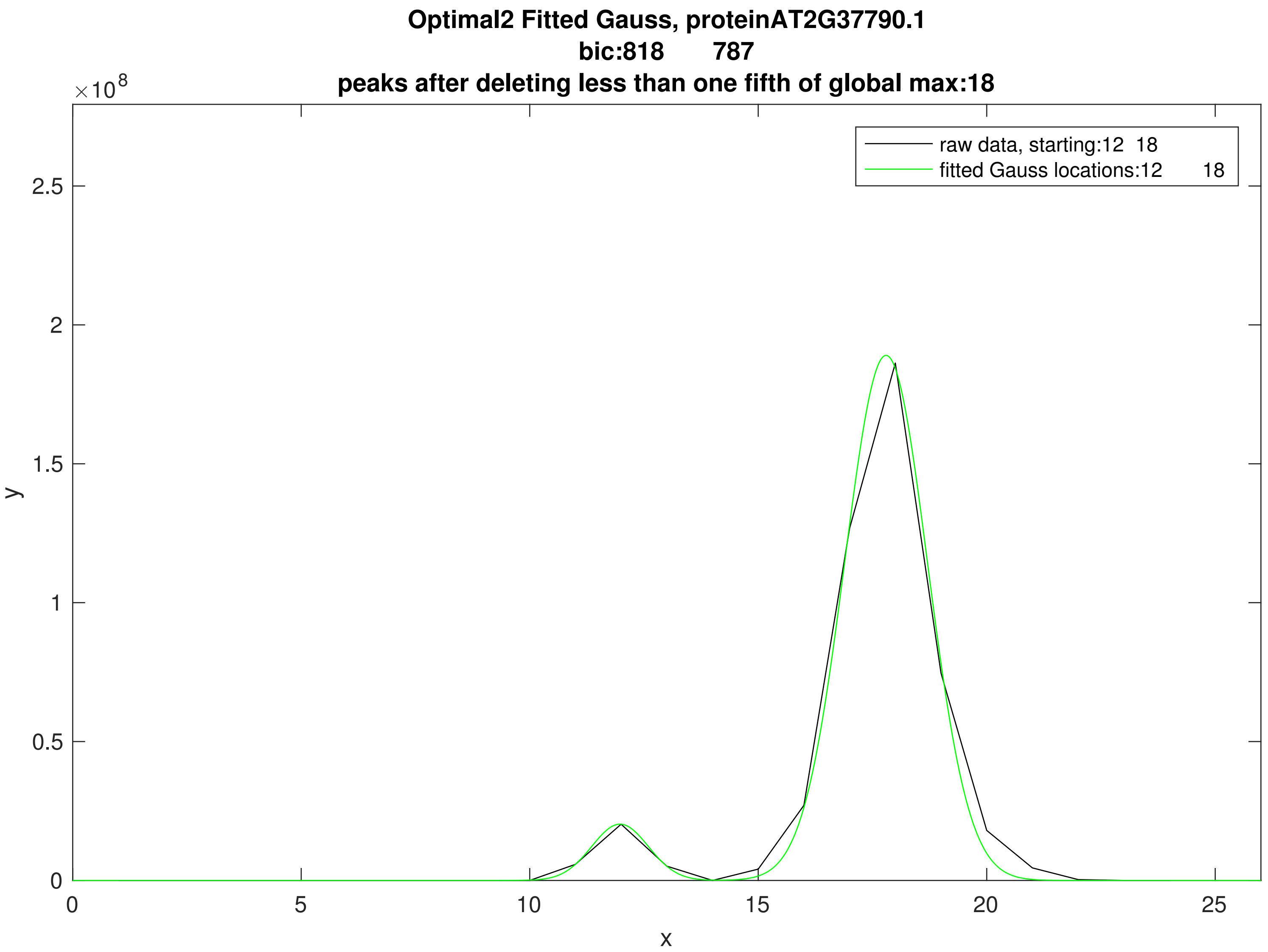

Optimal2 Fitted Gauss, proteinAT2G37790.1
bic:818 787
peaks after deleting less than one fifth of global max:18
×10^8
raw data, starting:12 18
fitted Gauss locations:12 18
2.5
2
1.5
1
0.5
0
y
0
5
10
15
20
25
x

Optimal2 Fitted Gauss, proteinAT2G38010.1;AT2G38010.2
bic:789 782
peaks after deleting less than one fifth of global max:9.4 15

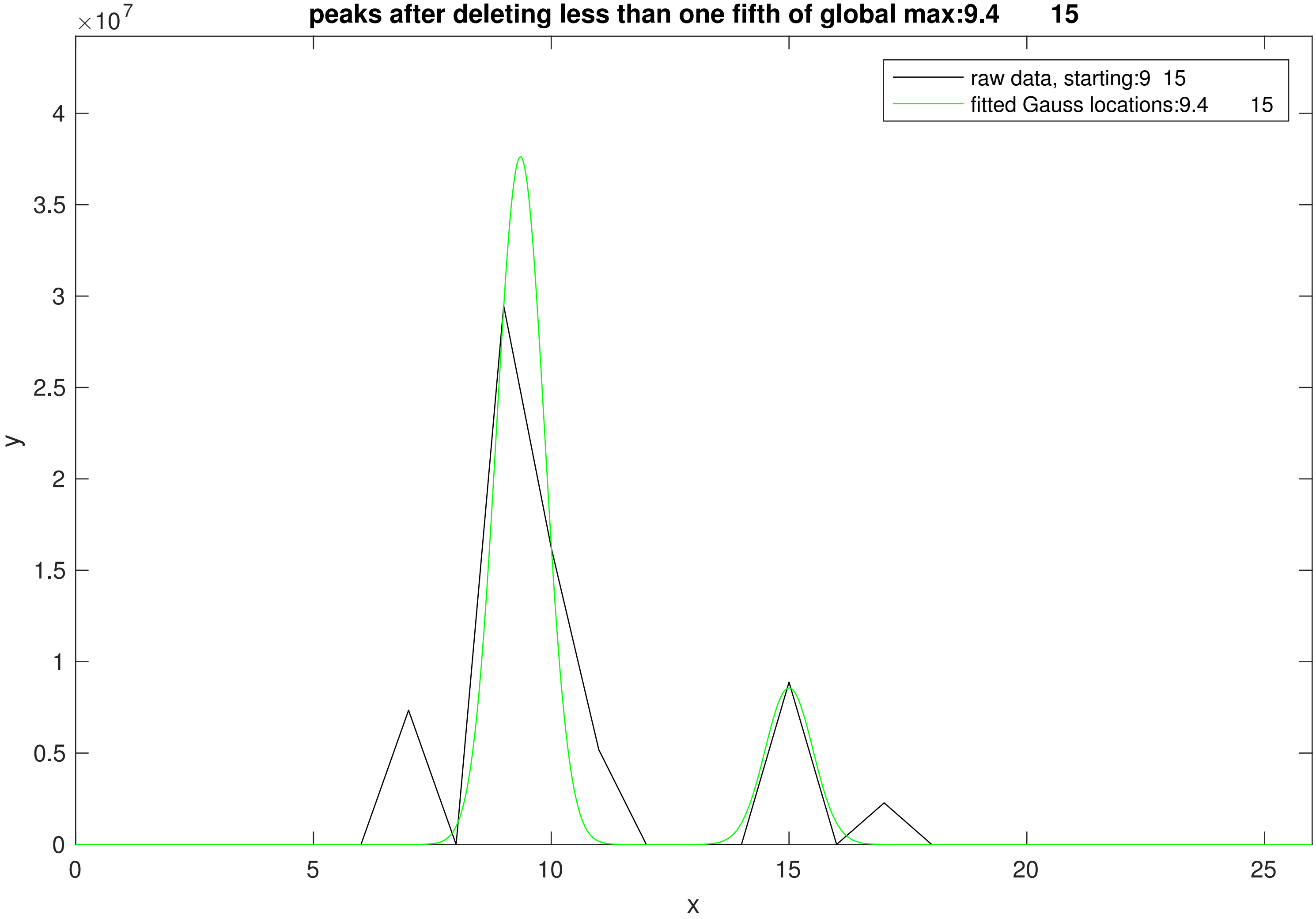

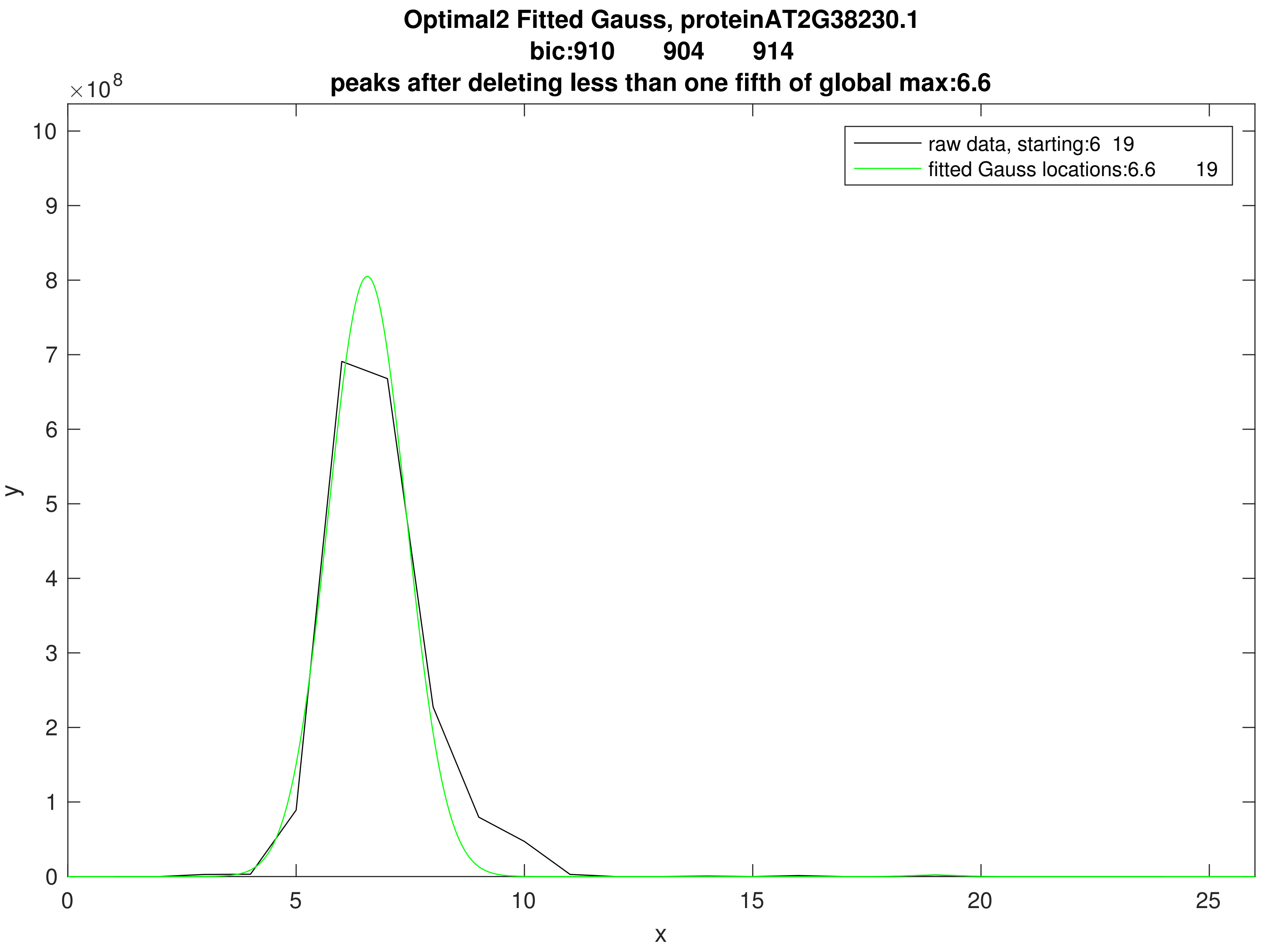

Optimal2 Fitted Gauss, proteinAT2G38230.1
bic:910 904 914
peaks after deleting less than one fifth of global max:6.6
×10^8
raw data, starting:6 19
fitted Gauss locations:6.6 19
y
x
0
1
2
3
4
5
6
7
8
9
10
15
20
25

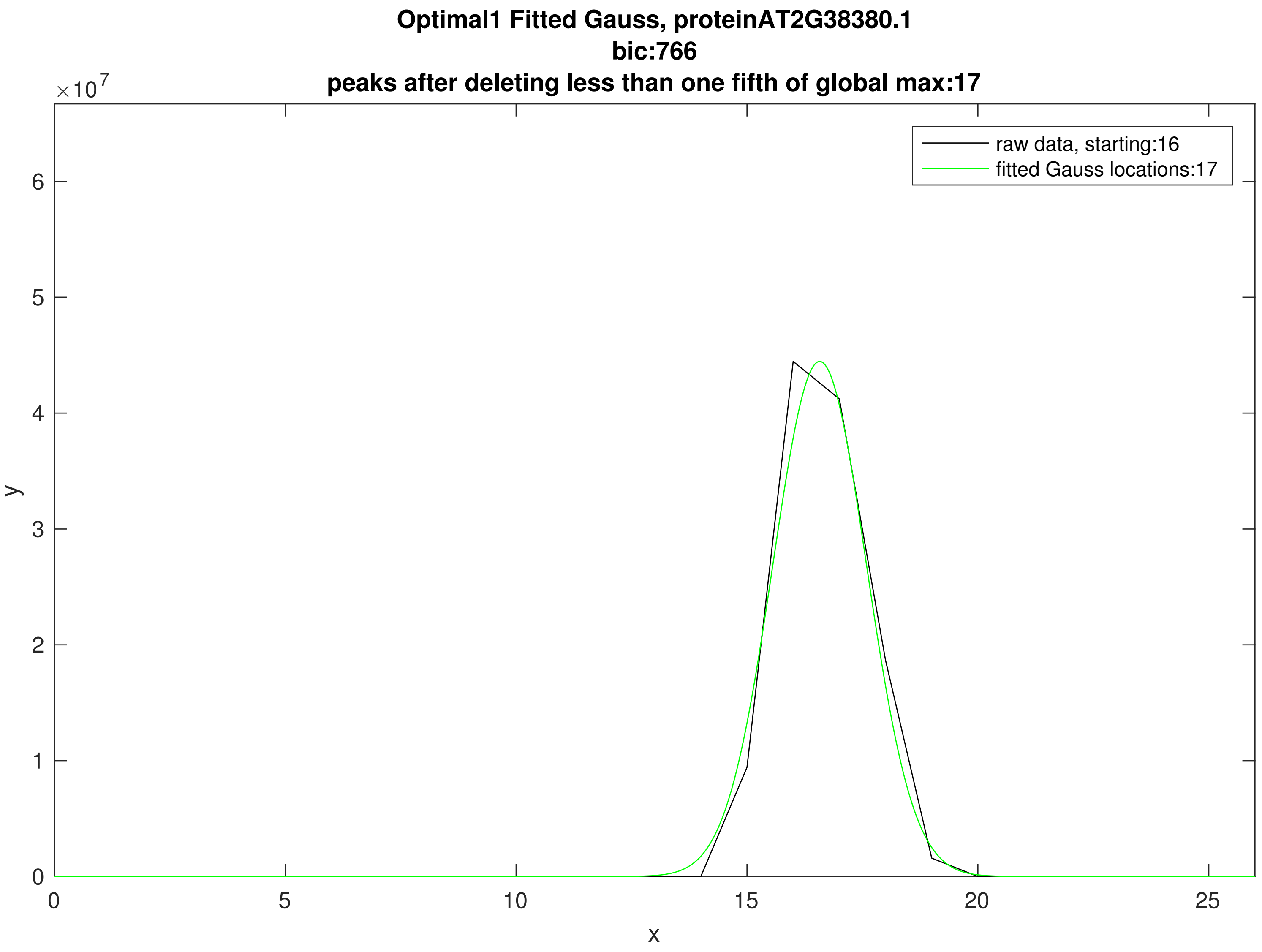
Optimal1 Fitted Gauss, proteinAT2G38380.1
bic:766
peaks after deleting less than one fifth of global max:17
×10^7
raw data, starting:16
fitted Gauss locations:17
y
x
0
1
2
3
4
5
6
0
5
10
15
20
25

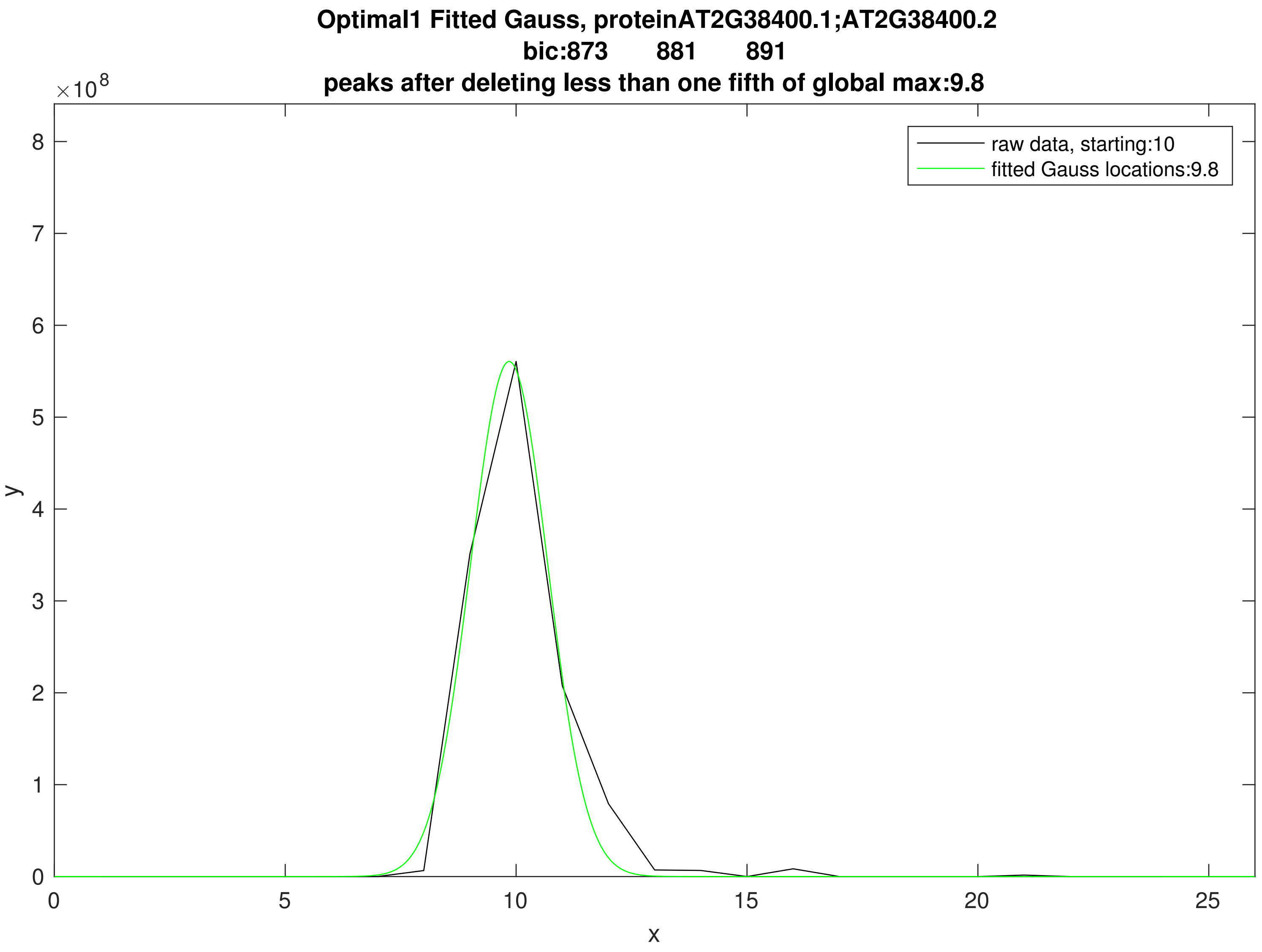
Optimal1 Fitted Gauss, proteinAT2G38400.1;AT2G38400.2
bic:873 881 891
peaks after deleting less than one fifth of global max:9.8
×10^8
raw data, starting:10
fitted Gauss locations:9.8
y
x
0
1
2
3
4
5
6
7
8
5
10
15
20
25

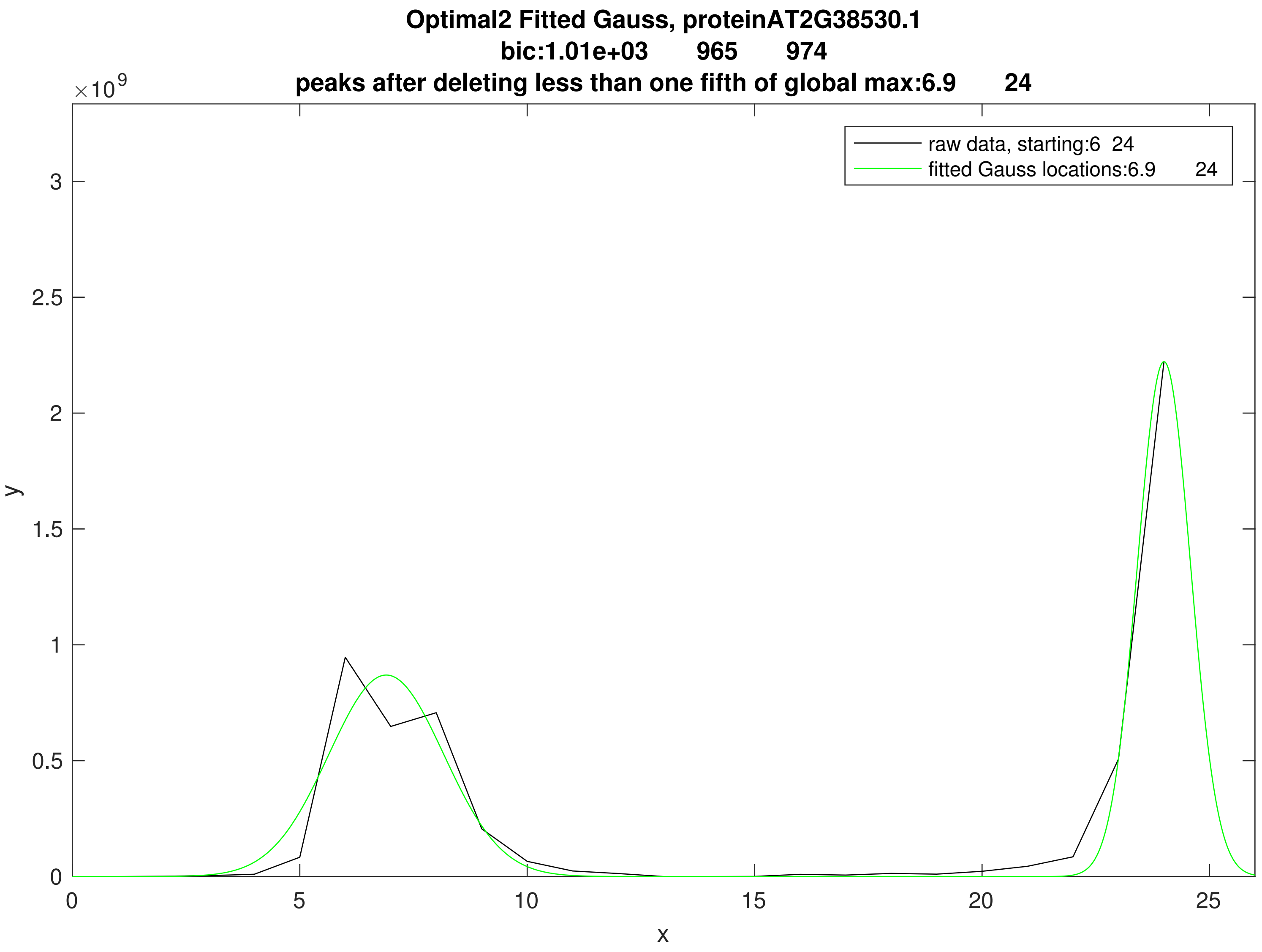

Optimal2 Fitted Gauss, proteinAT2G38530.1
bic:1.01e+03 965 974
peaks after deleting less than one fifth of global max:6.9 24
×10^9
raw data, starting:6 24
fitted Gauss locations:6.9 24
y
x
0
0.5
1
1.5
2
2.5
3
0
5
10
15
20
25

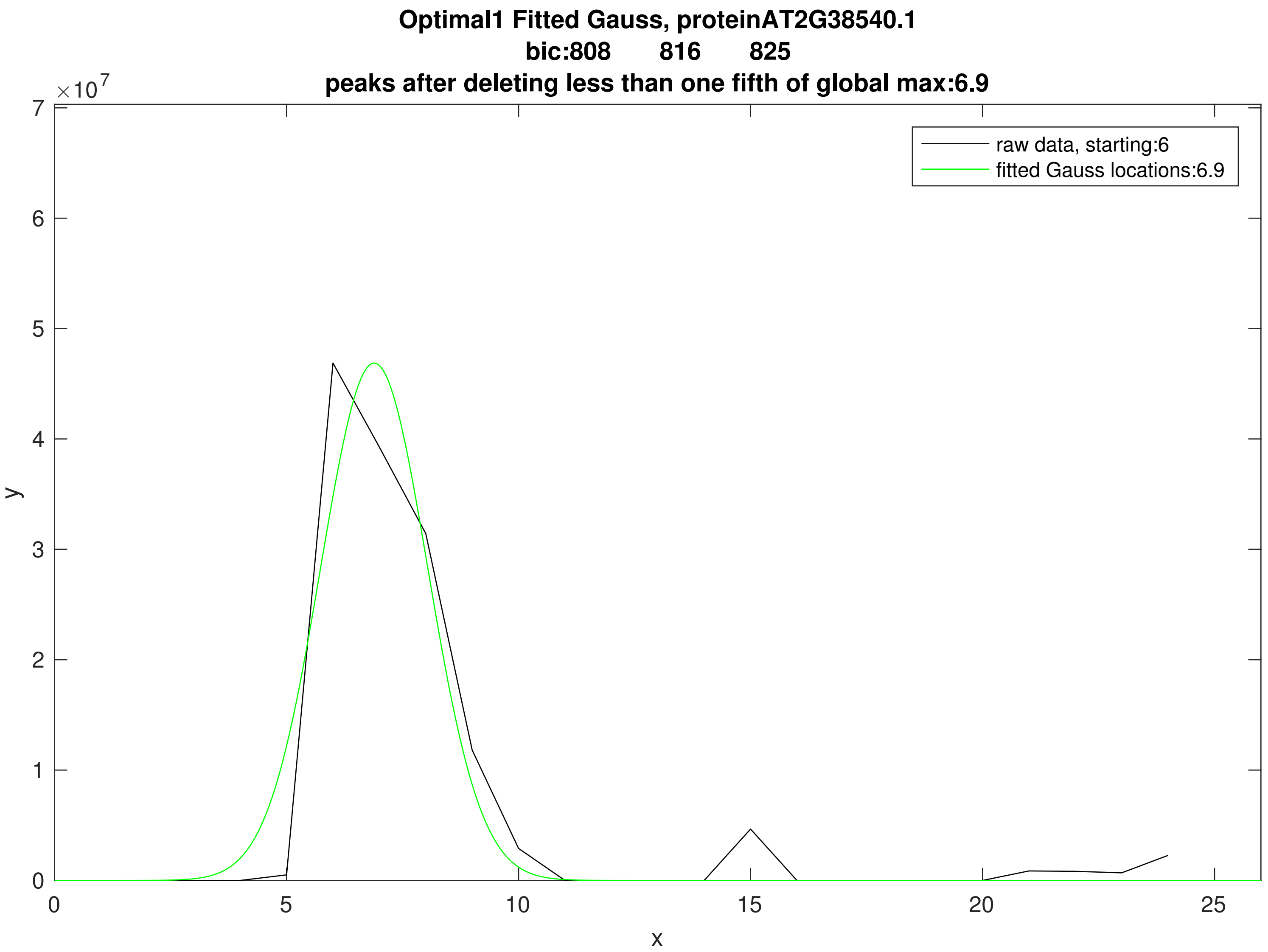
Optimal1 Fitted Gauss, proteinAT2G38540.1
bic:808 816 825
peaks after deleting less than one fifth of global max:6.9
×10^7
raw data, starting:6
fitted Gauss locations:6.9
y
x
0
1
2
3
4
5
6
7
0
5
10
15
20
25

**Optimal1 Fitted Gauss, proteinAT2G38700.1**
**bic:992**
**peaks after deleting less than one fifth of global max:13**

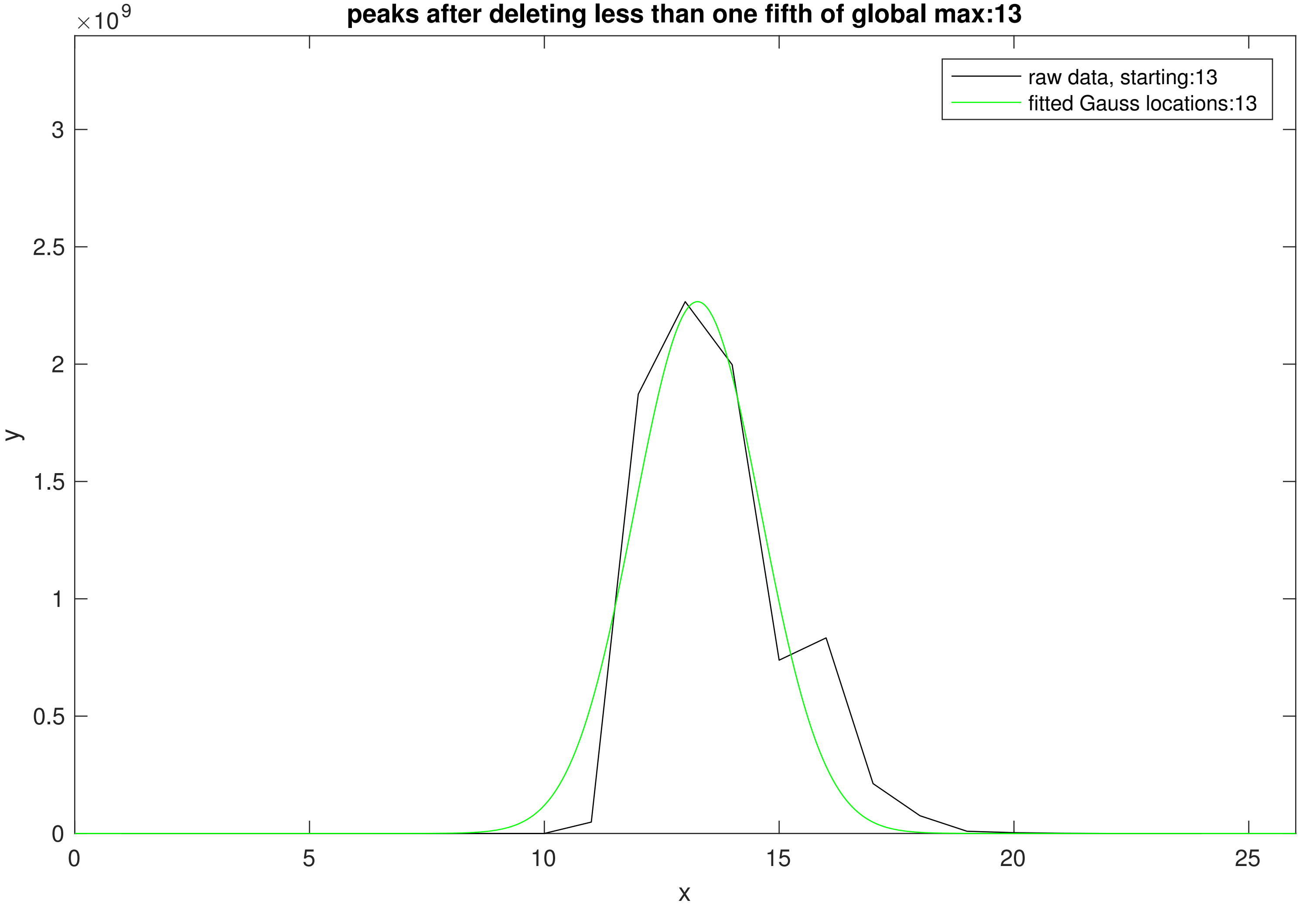

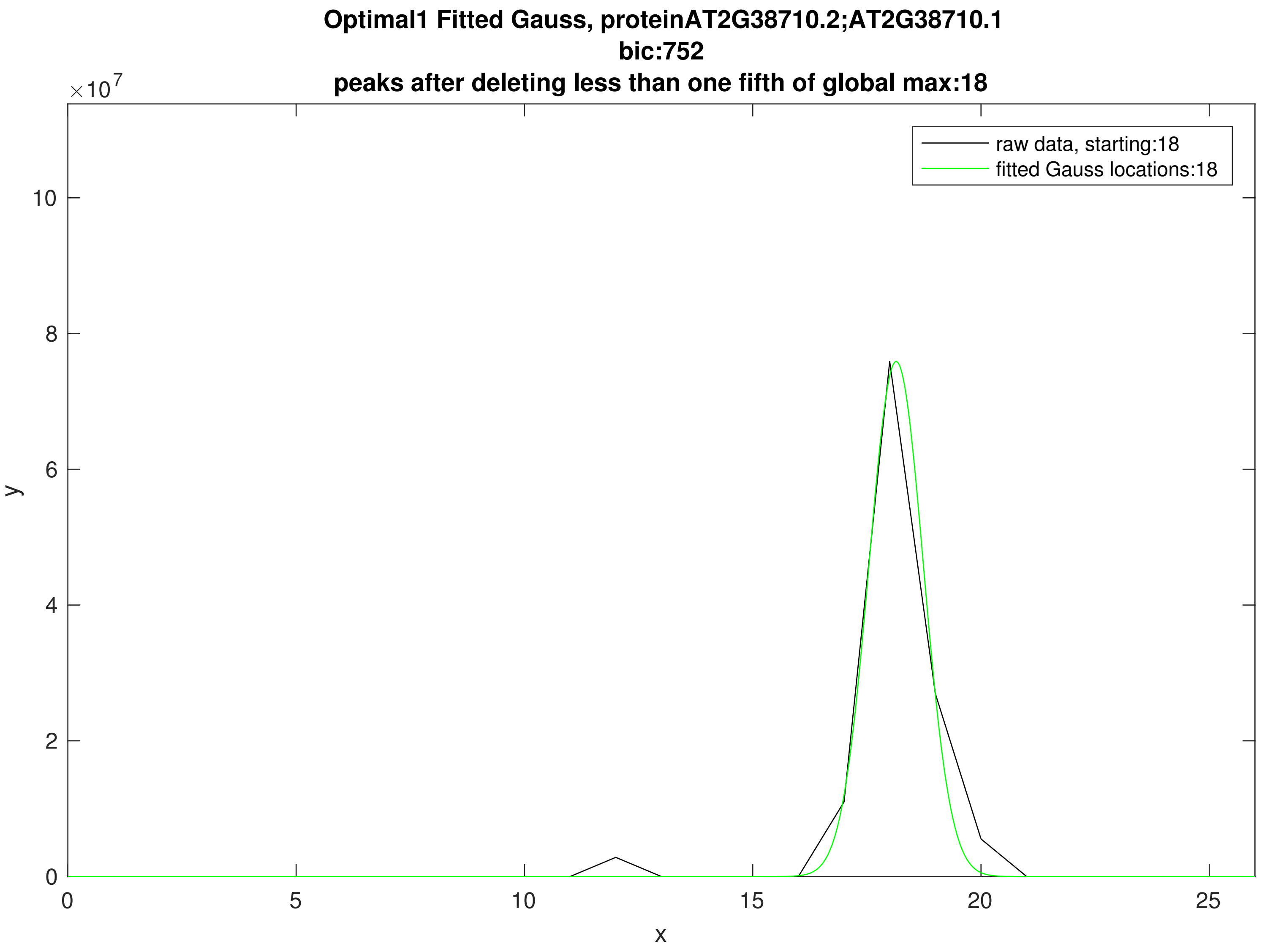

Optimal1 Fitted Gauss, proteinAT2G38710.2;AT2G38710.1
bic:752
peaks after deleting less than one fifth of global max:18
×10^7
raw data, starting:18
fitted Gauss locations:18
y
x
0
2
4
6
8
10
0
5
10
15
20
25

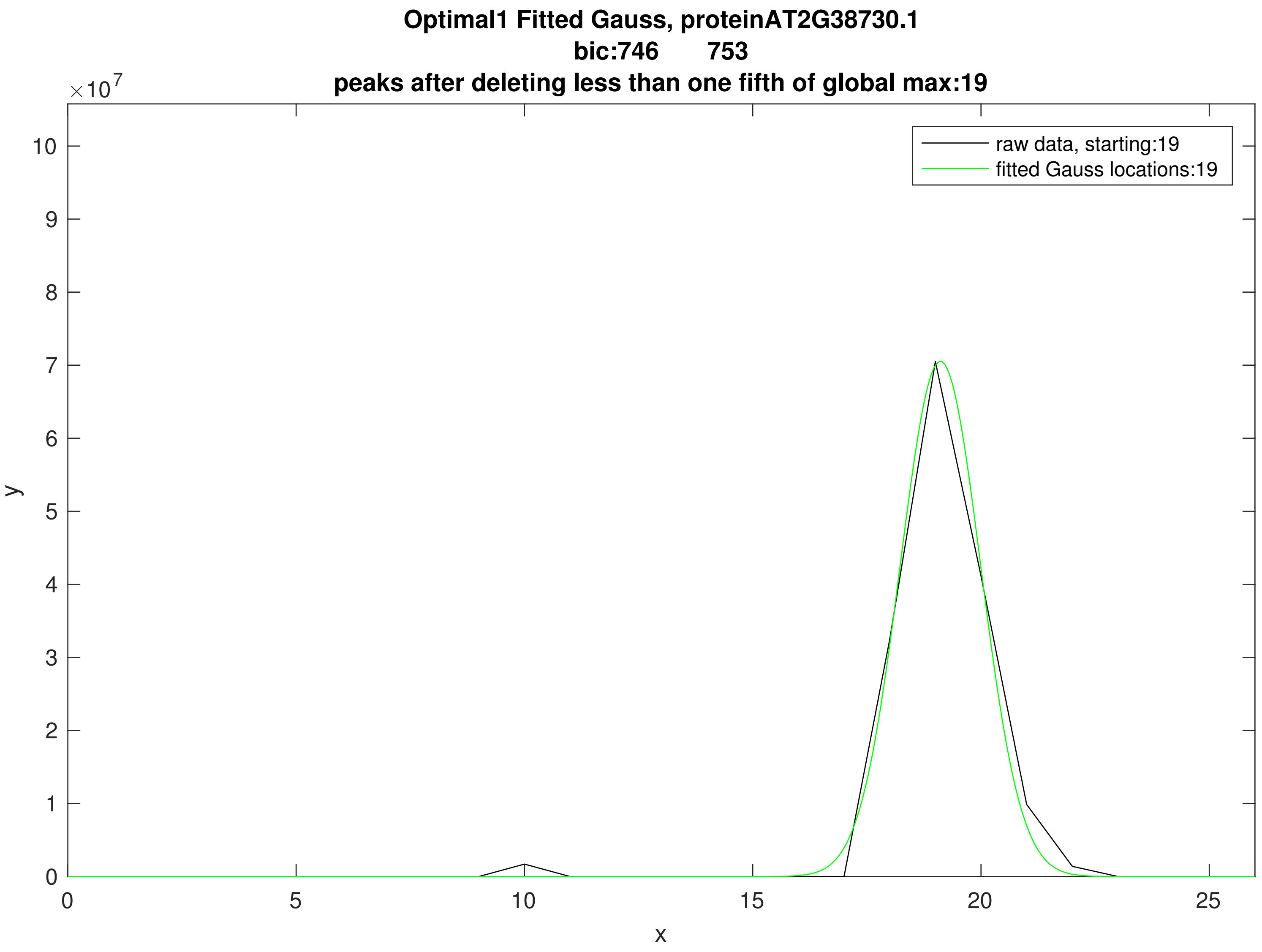

Optimal1 Fitted Gauss, proteinAT2G38730.1
bic:746 753
peaks after deleting less than one fifth of global max:19
×10^7
raw data, starting:19
fitted Gauss locations:19
y
x
0
1
2
3
4
5
6
7
8
9
10
0
5
10
15
20
25

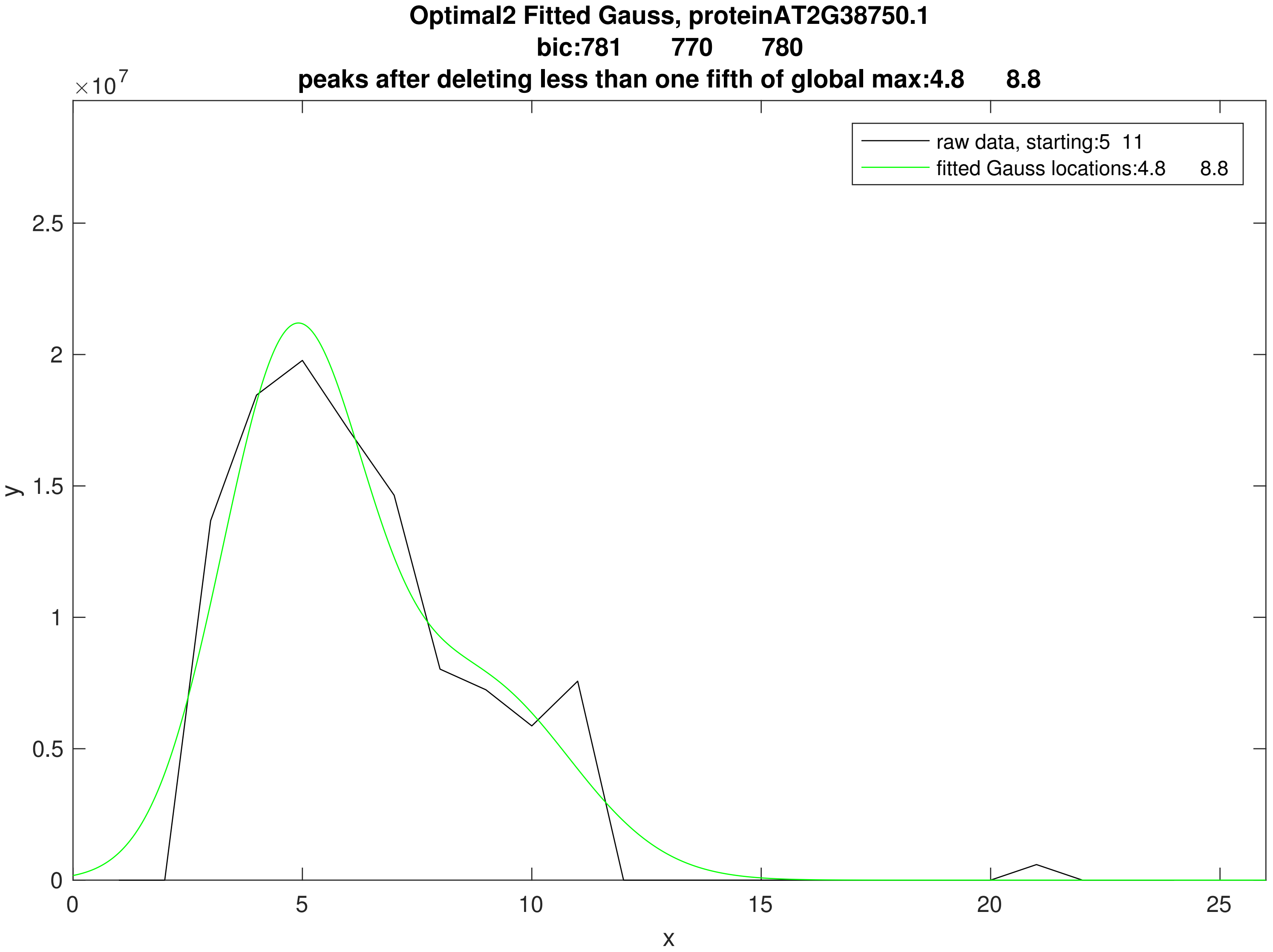
Optimal2 Fitted Gauss, proteinAT2G38750.1
bic:781 770 780
peaks after deleting less than one fifth of global max:4.8 8.8
×10^7
raw data, starting:5 11
fitted Gauss locations:4.8 8.8
y
x
0
0.5
1
1.5
2
2.5
5
10
15
20
25

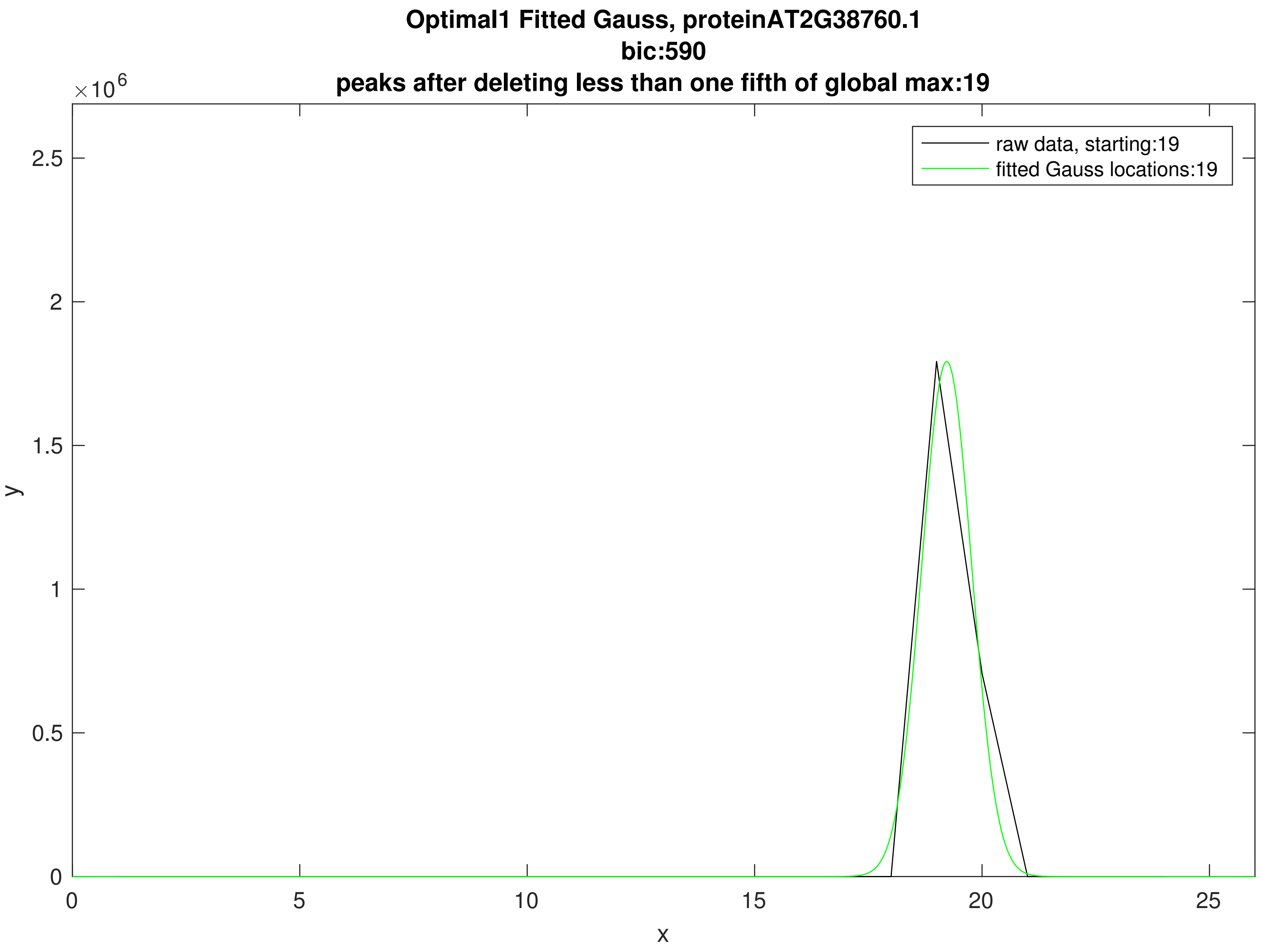

Optimal1 Fitted Gauss, proteinAT2G38760.1
bic:590
peaks after deleting less than one fifth of global max:19
×10^6
2.5
2
1.5
1
0.5
0
y
0
5
10
15
20
25
x
raw data, starting:19
fitted Gauss locations:19

**Optimal1 Fitted Gauss, proteinAT2G38860.2;AT2G38860.1**
**bic:682**
**peaks after deleting less than one fifth of global max:12**

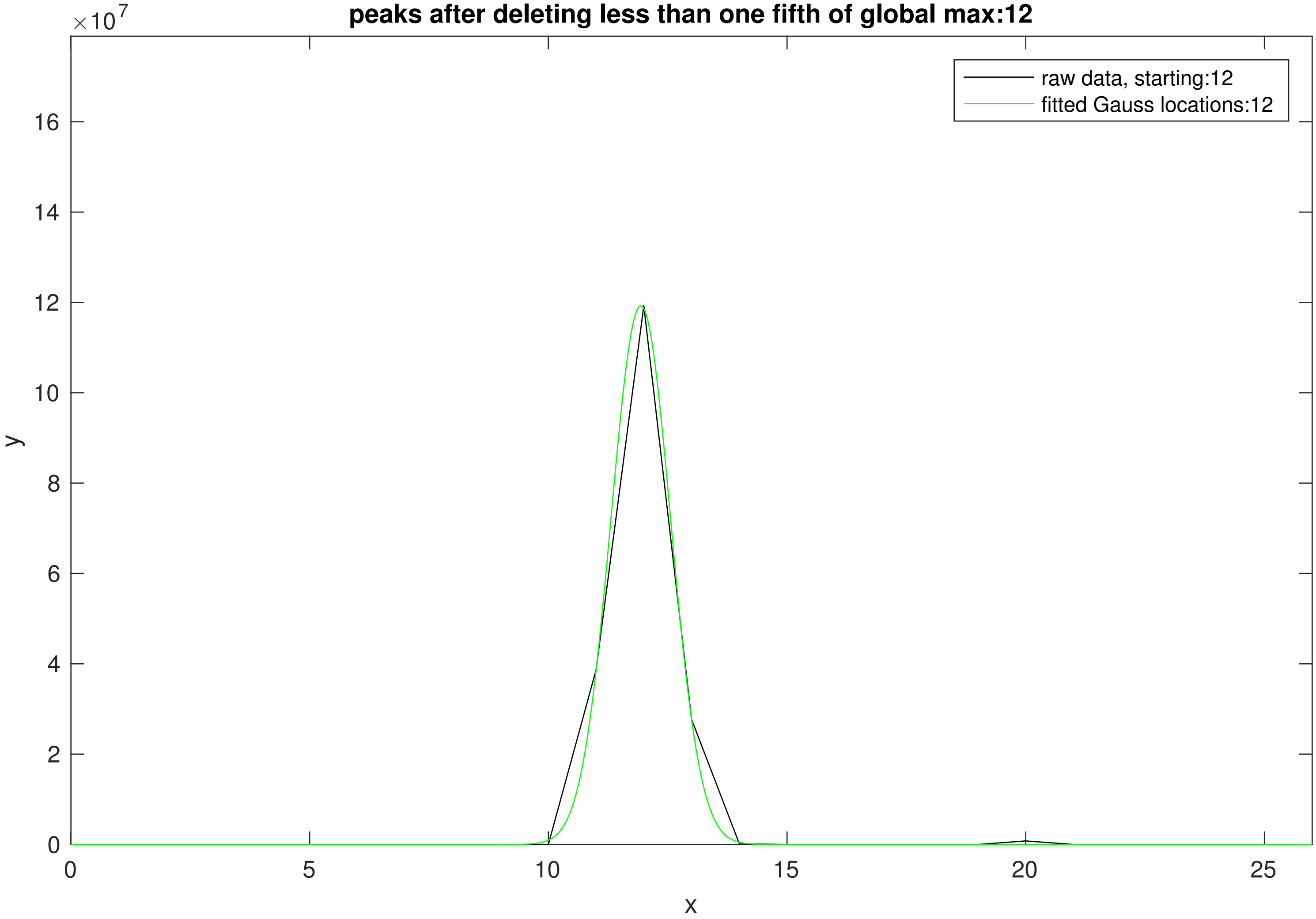

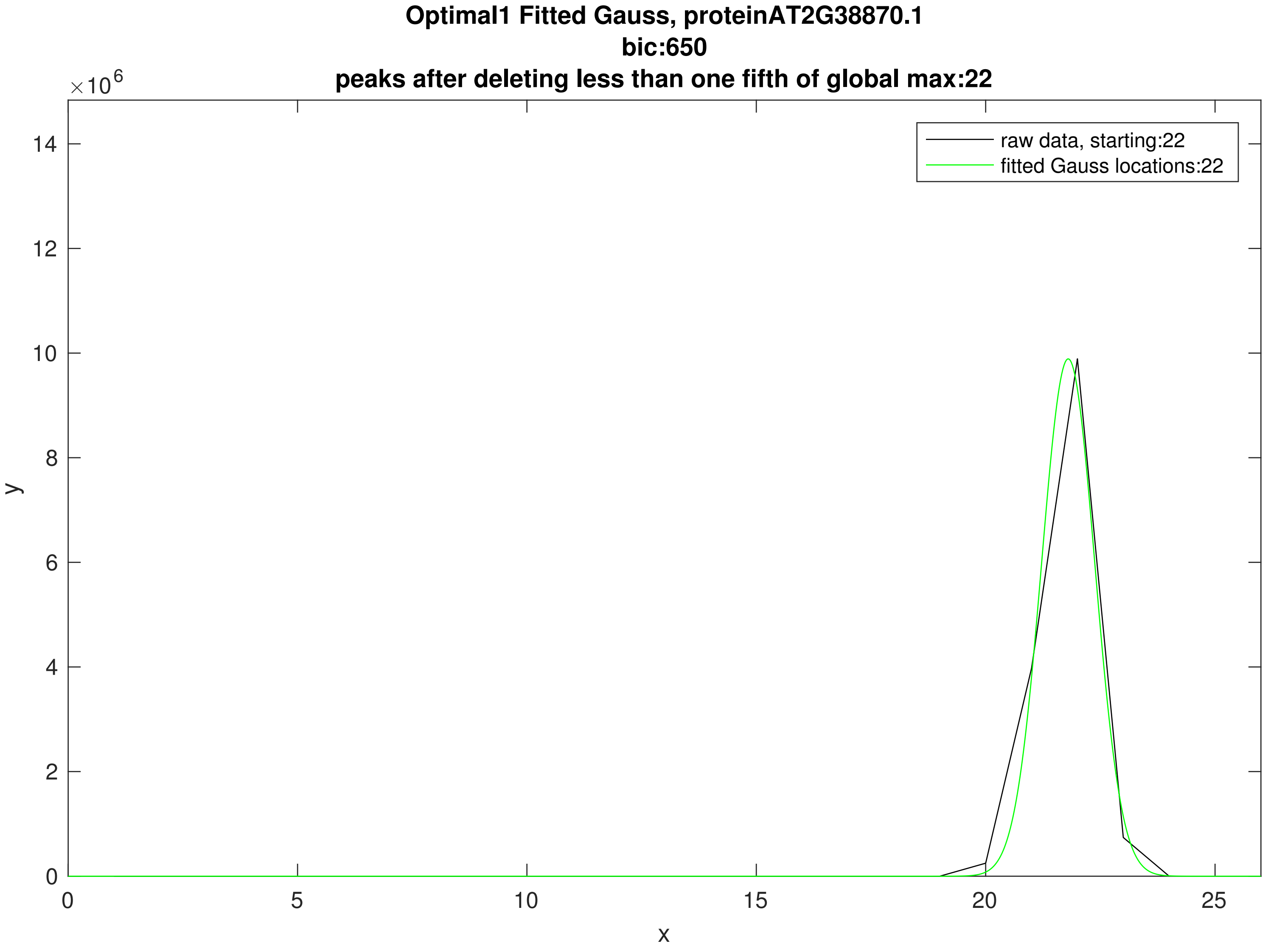
Optimal1 Fitted Gauss, proteinAT2G38870.1
bic:650
peaks after deleting less than one fifth of global max:22
×10^6
14
12
10
8
6
4
2
0
y
0
5
10
15
20
25
x
raw data, starting:22
fitted Gauss locations:22

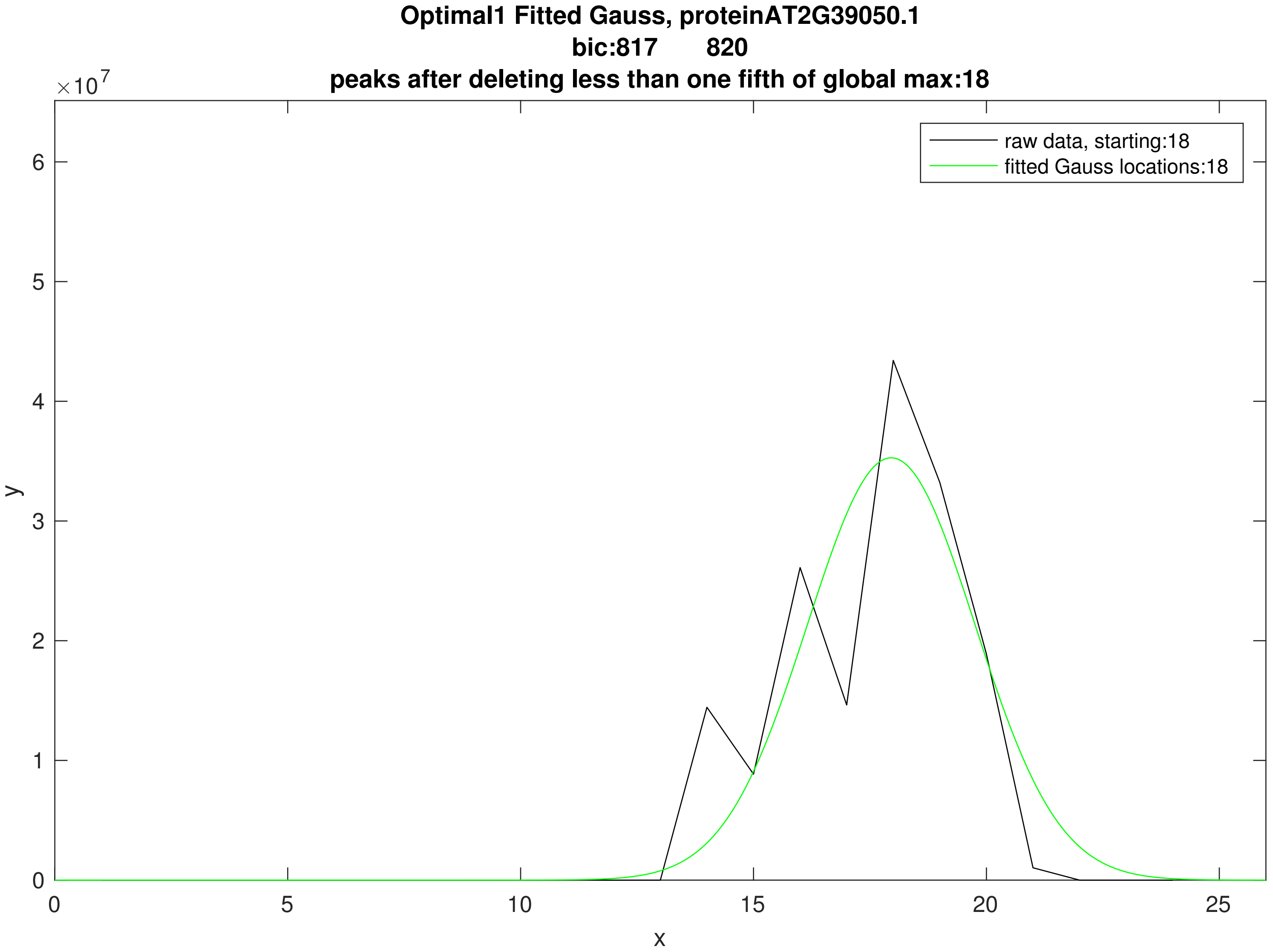
Optimal1 Fitted Gauss, proteinAT2G39050.1
bic:817 820
peaks after deleting less than one fifth of global max:18
×10^7
raw data, starting:18
fitted Gauss locations:18
y
x
0
1
2
3
4
5
6
0
5
10
15
20
25

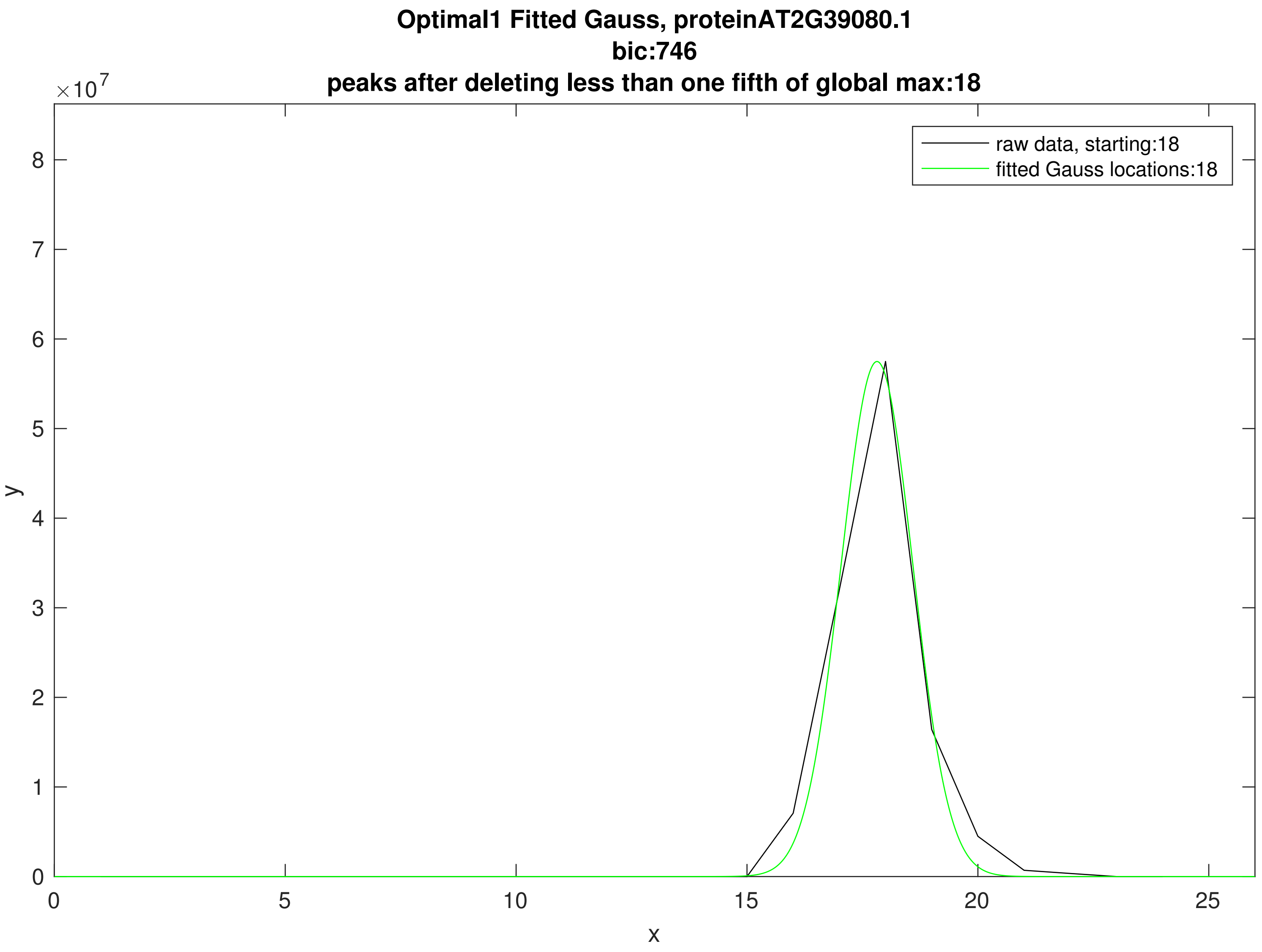
Optimal1 Fitted Gauss, proteinAT2G39080.1
bic:746
peaks after deleting less than one fifth of global max:18
×10^7
raw data, starting:18
fitted Gauss locations:18
y
x
0
1
2
3
4
5
6
7
8
0
5
10
15
20
25

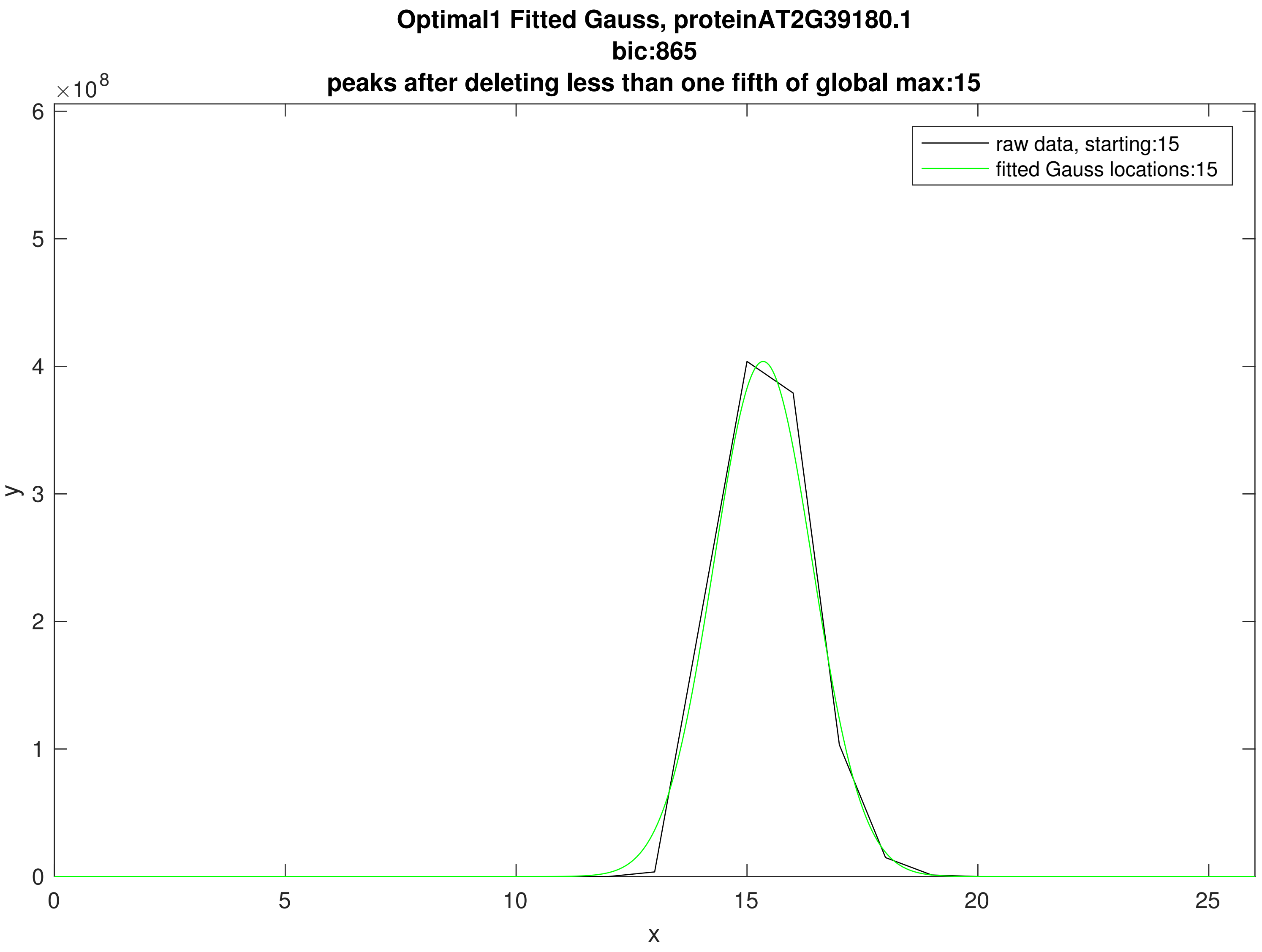
Optimal1 Fitted Gauss, proteinAT2G39180.1
bic:865
peaks after deleting less than one fifth of global max:15
×10^8
y
x
raw data, starting:15
fitted Gauss locations:15
0
1
2
3
4
5
6
0
5
10
15
20
25

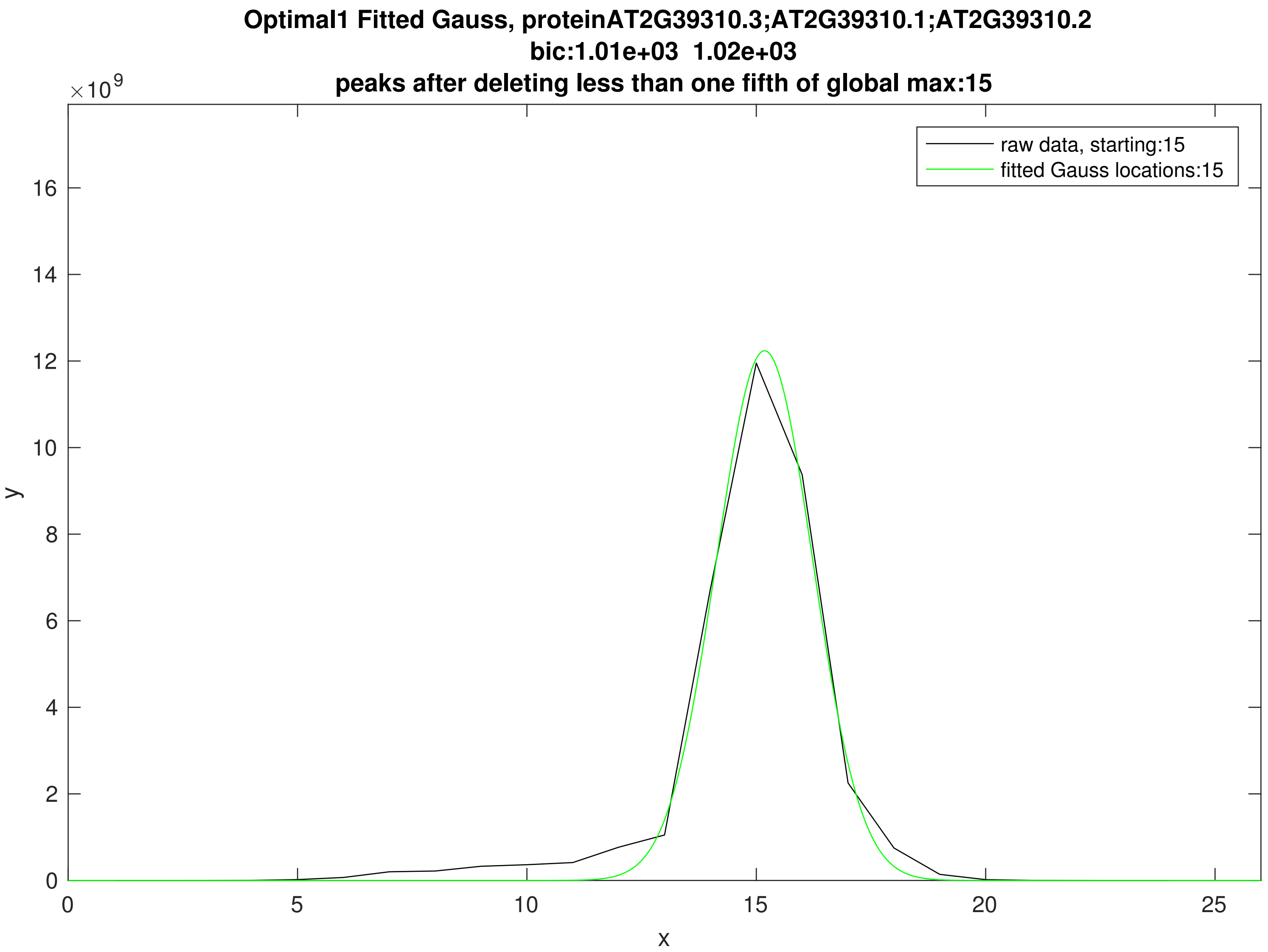
Optimal1 Fitted Gauss, proteinAT2G39310.3;AT2G39310.1;AT2G39310.2
bic:1.01e+03 1.02e+03
peaks after deleting less than one fifth of global max:15
×10^9
raw data, starting:15
fitted Gauss locations:15
y
x
0
2
4
6
8
10
12
14
16
0
5
10
15
20
25

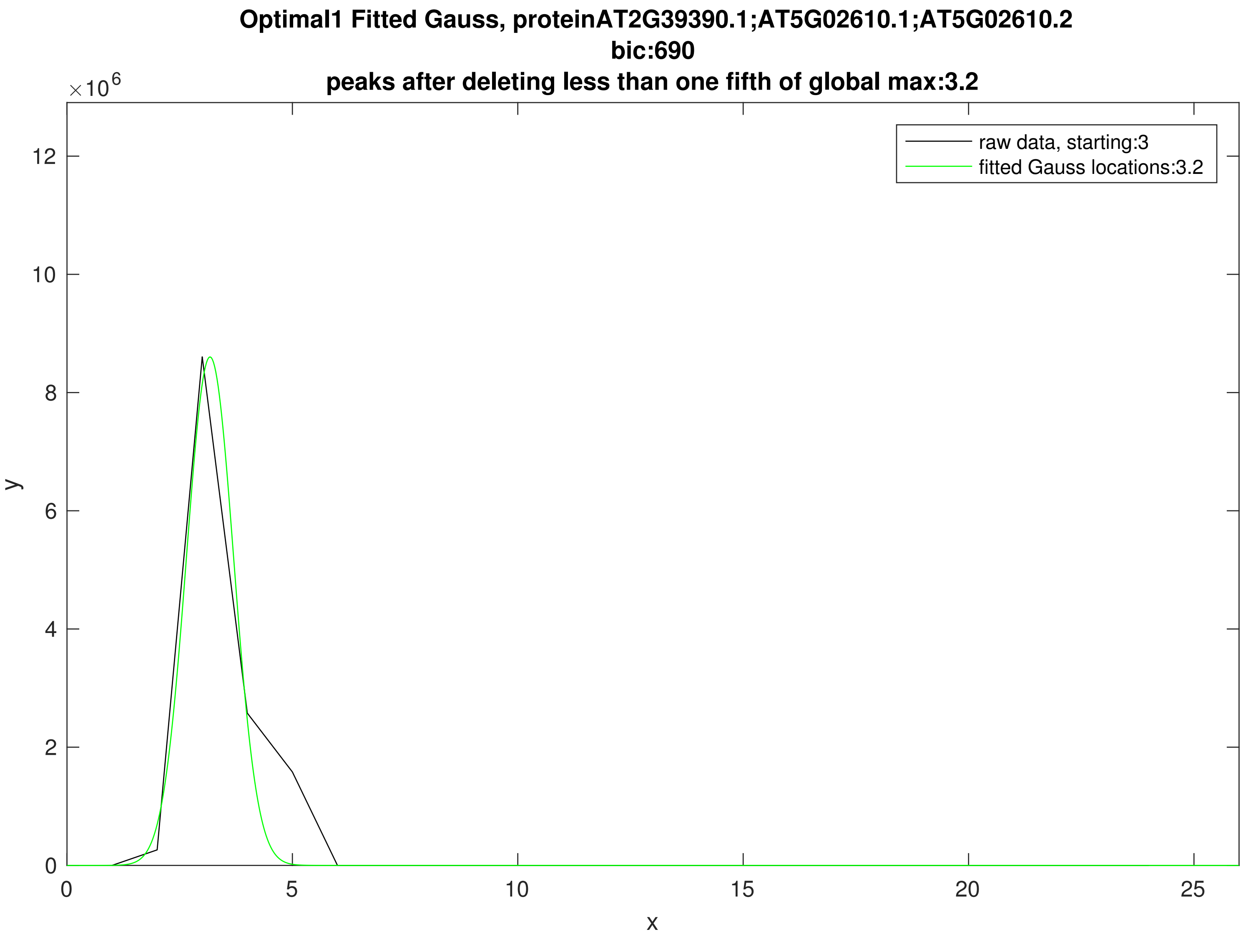
Optimal1 Fitted Gauss, proteinAT2G39390.1;AT5G02610.1;AT5G02610.2
bic:690
peaks after deleting less than one fifth of global max:3.2
×10^6
12
10
8
6
4
2
0
y
0
5
10
15
20
25
x
raw data, starting:3
fitted Gauss locations:3.2

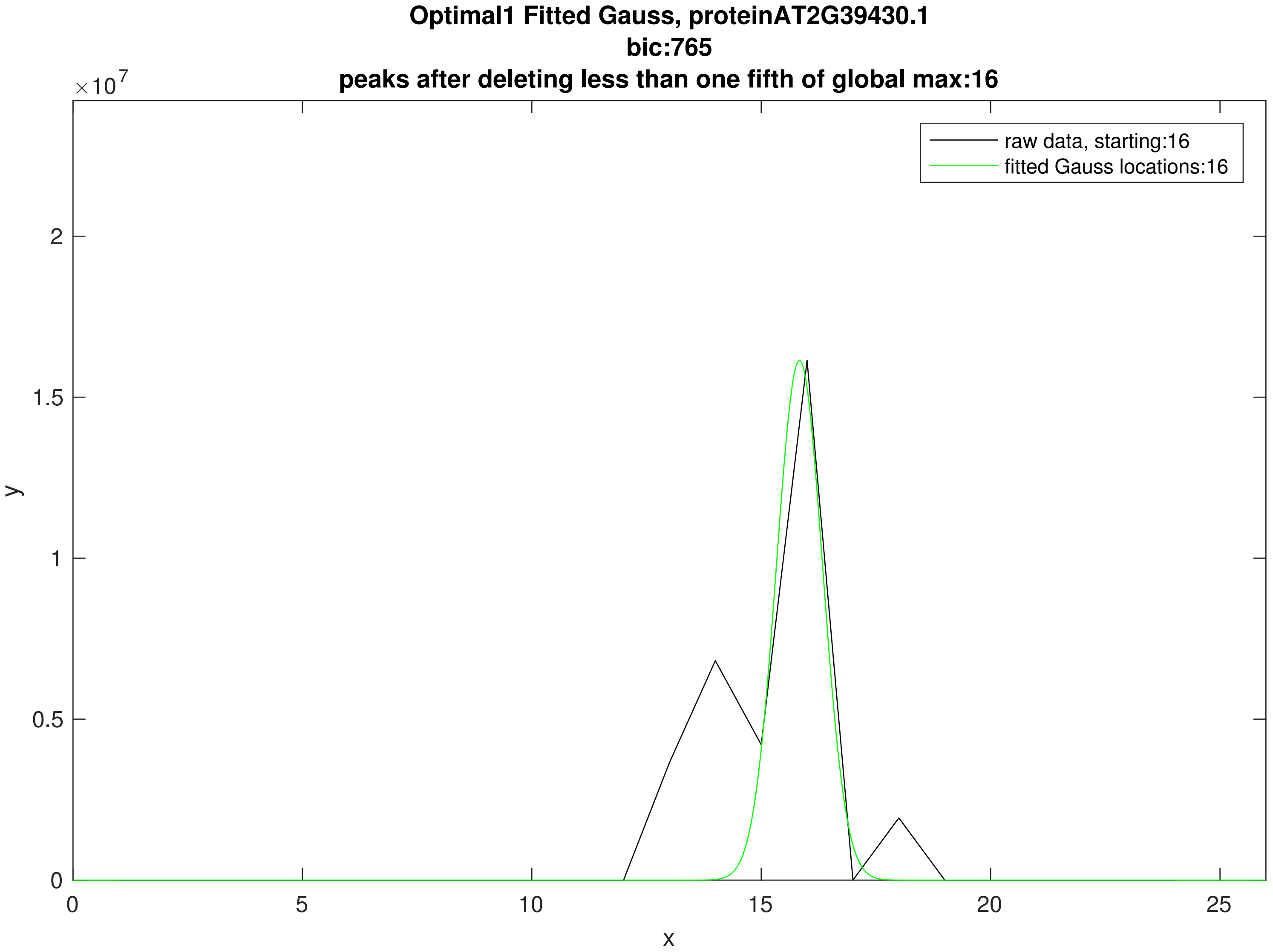

Optimal1 Fitted Gauss, proteinAT2G39430.1
bic:765
peaks after deleting less than one fifth of global max:16
×10^7
raw data, starting:16
fitted Gauss locations:16
y
x
0
0.5
1
1.5
2
0
5
10
15
20
25

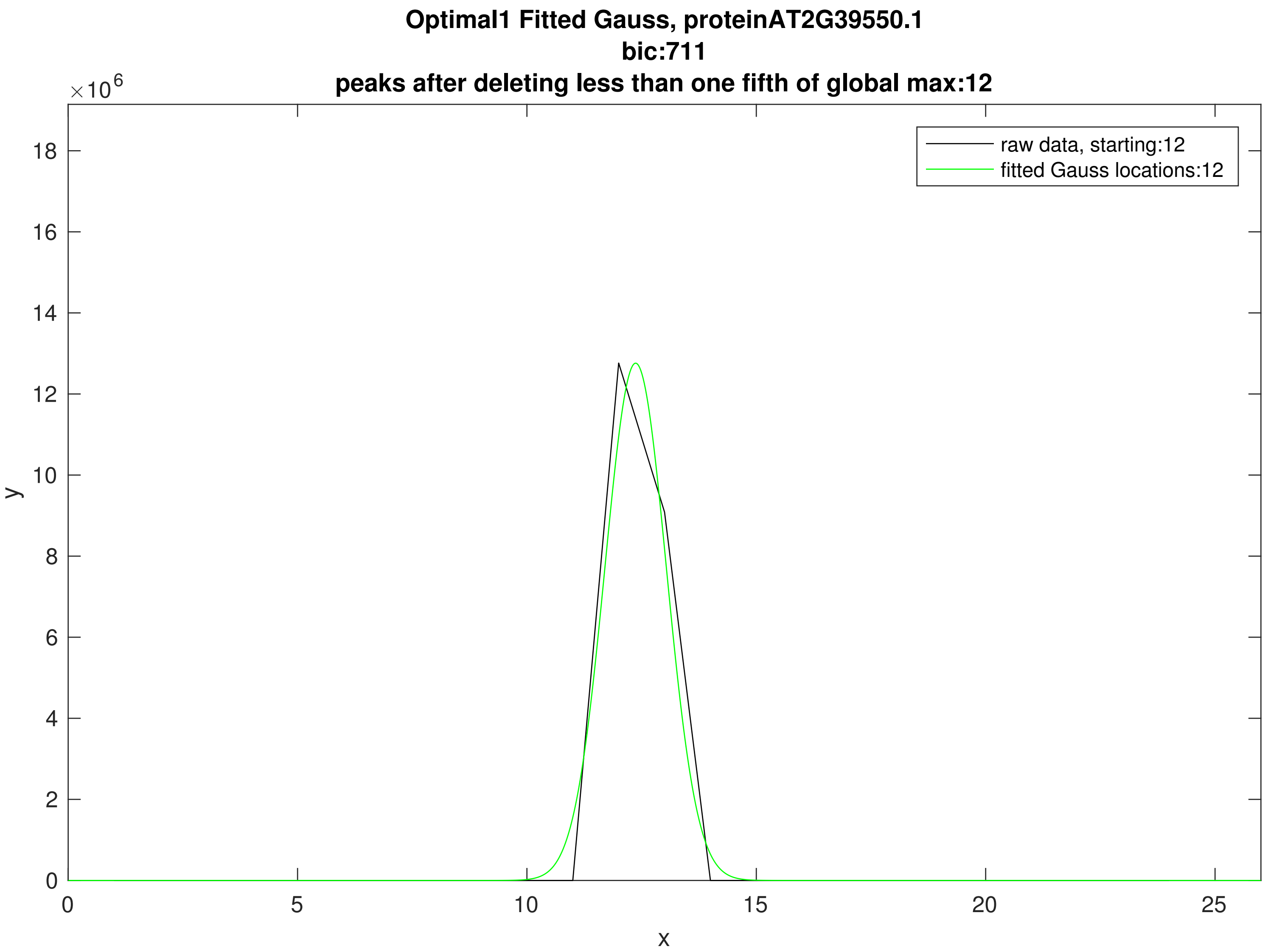

Optimal1 Fitted Gauss, proteinAT2G39550.1
bic:711
peaks after deleting less than one fifth of global max:12
×10^6
raw data, starting:12
fitted Gauss locations:12
y
x
0
2
4
6
8
10
12
14
16
18
0
5
10
15
20
25

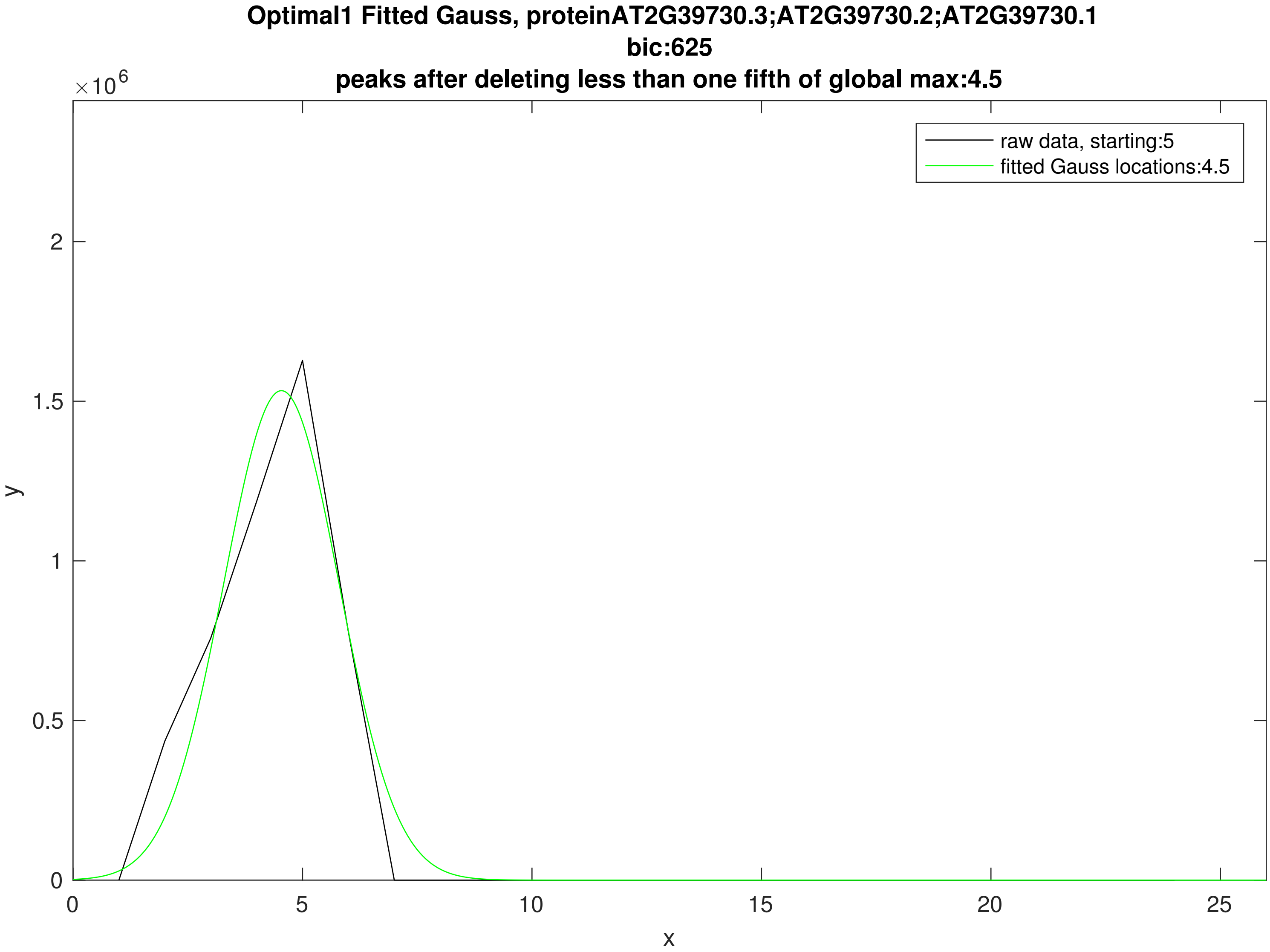
Optimal1 Fitted Gauss, proteinAT2G39730.3;AT2G39730.2;AT2G39730.1
bic:625
peaks after deleting less than one fifth of global max:4.5
×10^6
raw data, starting:5
fitted Gauss locations:4.5
y
x
0
0.5
1
1.5
2
5
10
15
20
25

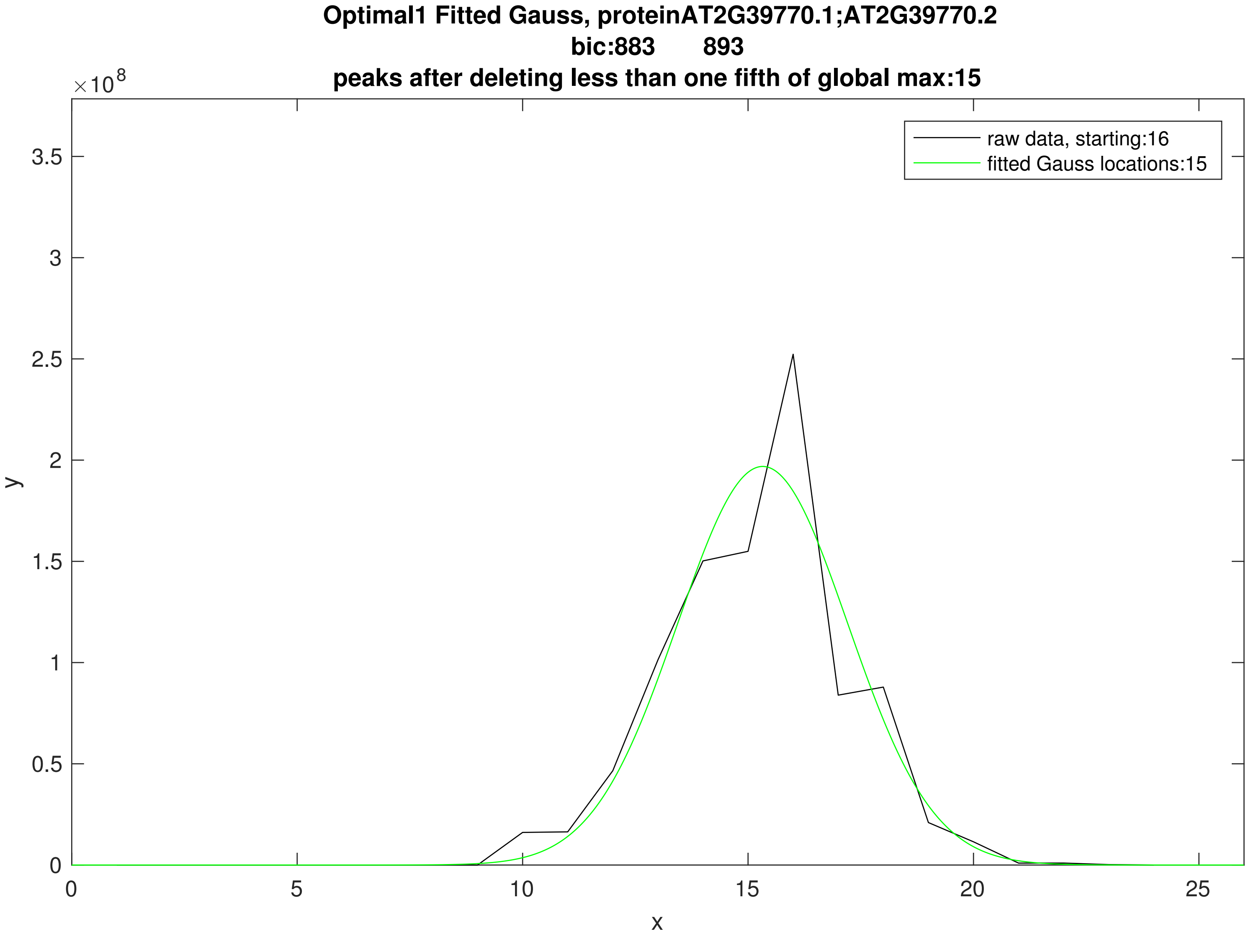
Optimal1 Fitted Gauss, proteinAT2G39770.1;AT2G39770.2
bic:883 893
peaks after deleting less than one fifth of global max:15
×10^8
raw data, starting:16
fitted Gauss locations:15
y
x
0
0.5
1
1.5
2
2.5
3
3.5
0
5
10
15
20
25

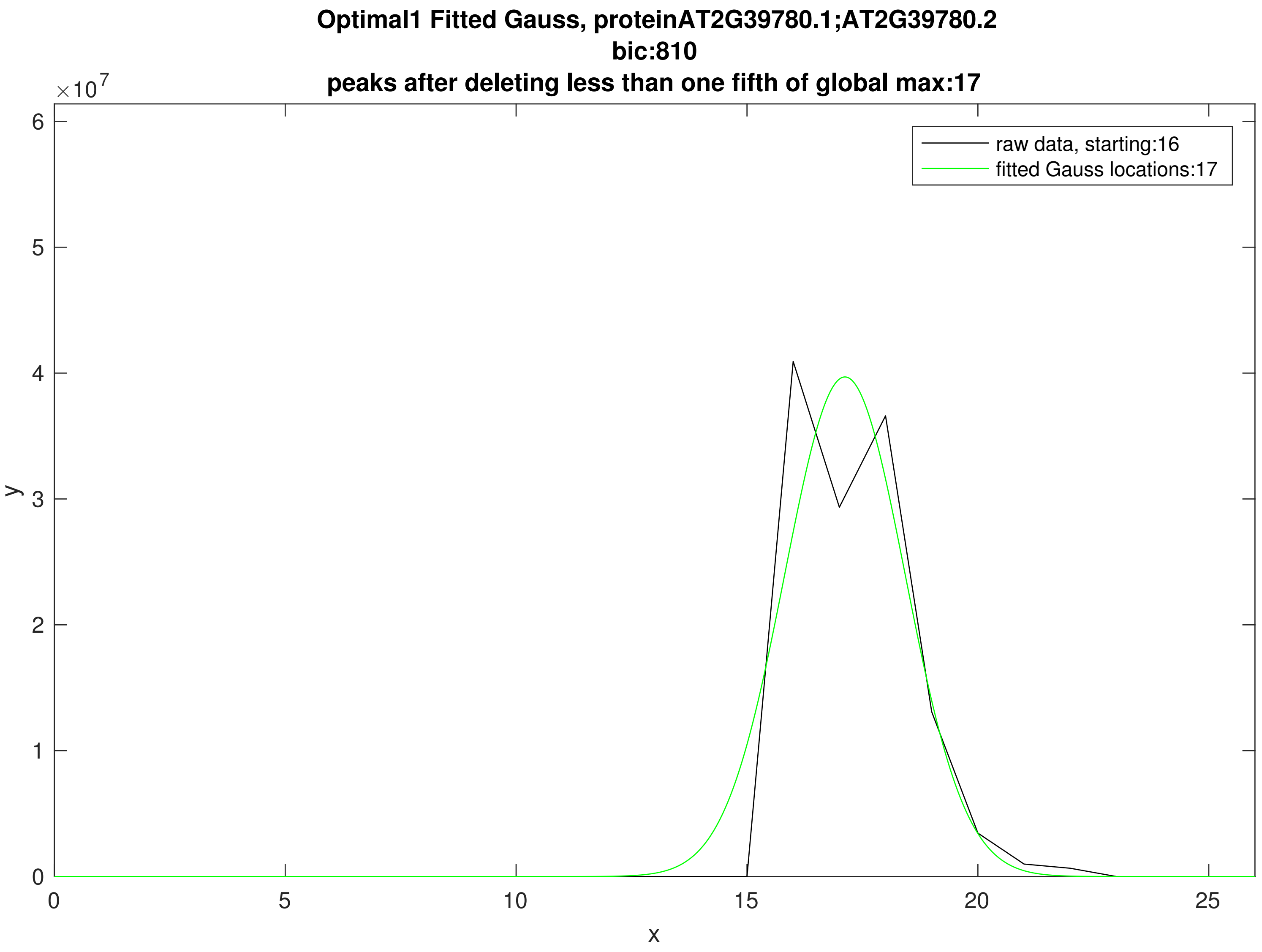
Optimal1 Fitted Gauss, proteinAT2G39780.1;AT2G39780.2
bic:810
peaks after deleting less than one fifth of global max:17
×10^7
raw data, starting:16
fitted Gauss locations:17
y
x
0
1
2
3
4
5
6
0
5
10
15
20
25

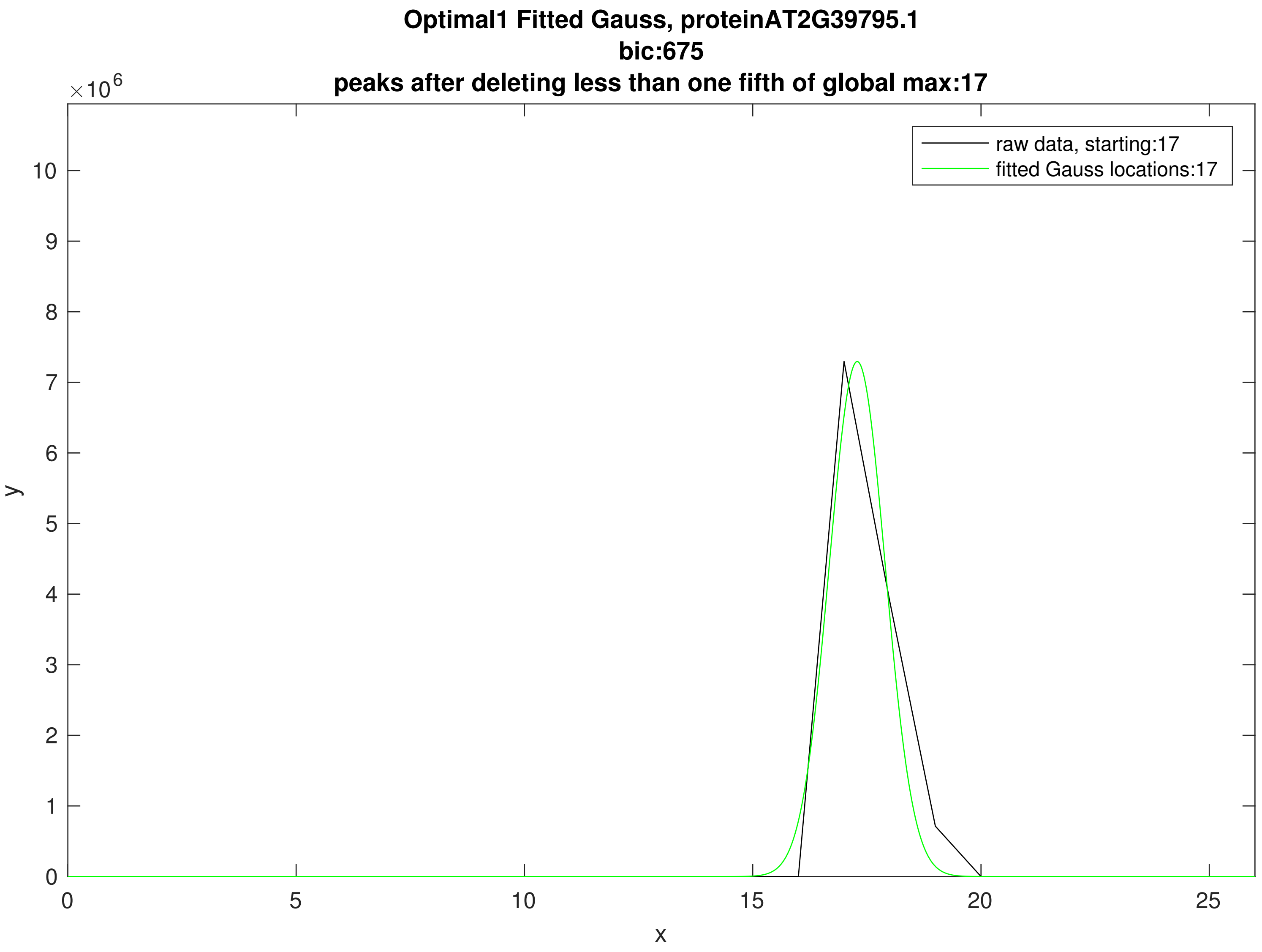

Optimal1 Fitted Gauss, proteinAT2G39795.1
bic:675
peaks after deleting less than one fifth of global max:17
×10^6
raw data, starting:17
fitted Gauss locations:17
10
9
8
7
6
5
4
3
2
1
0
y
0
5
10
15
20
25
x

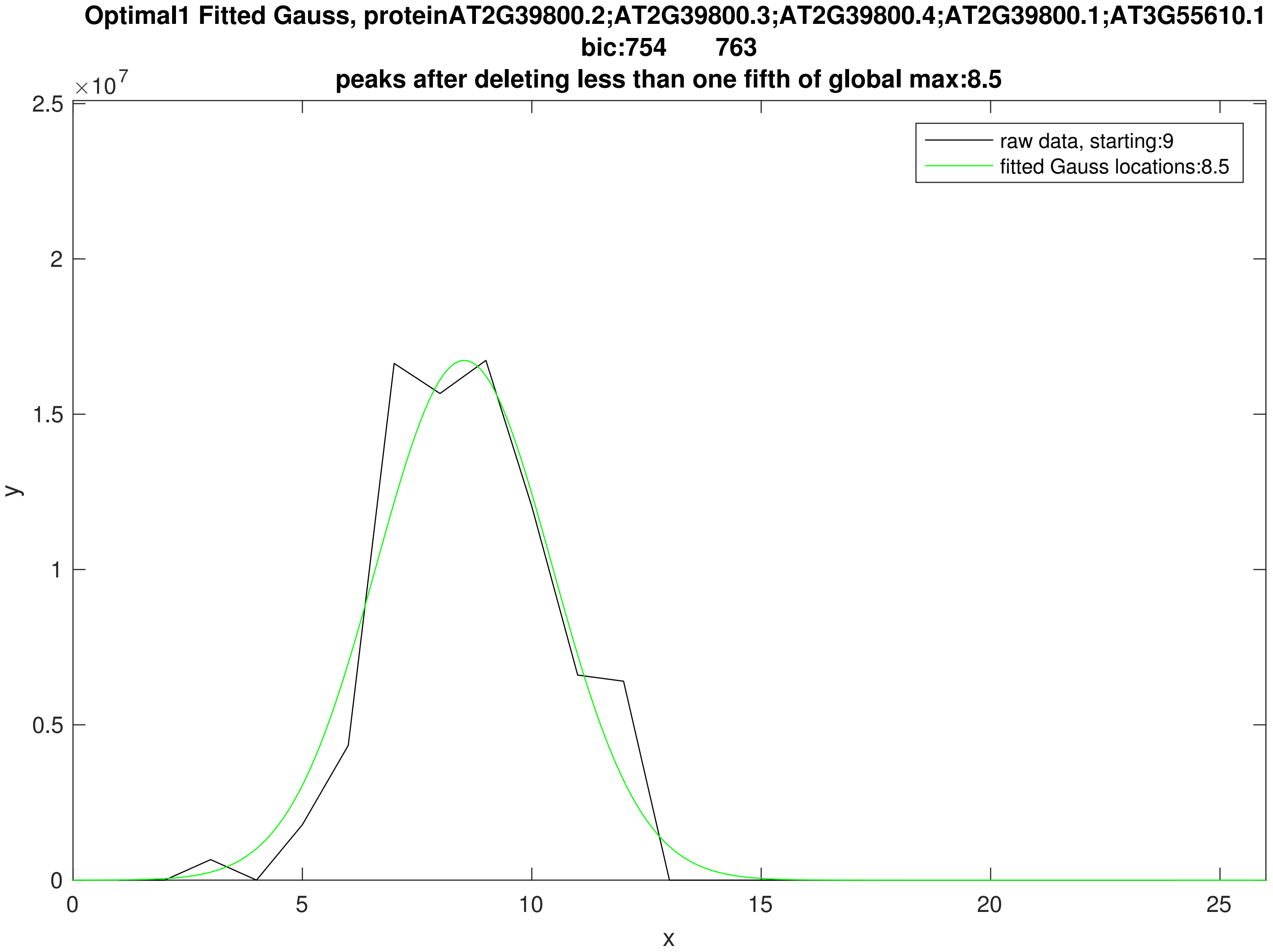
Optimal1 Fitted Gauss, proteinAT2G39800.2;AT2G39800.3;AT2G39800.4;AT2G39800.1;AT3G55610.1
bic:754 763
peaks after deleting less than one fifth of global max:8.5
×10^7
raw data, starting:9
fitted Gauss locations:8.5
y
x
0
0.5
1
1.5
2
2.5
5
10
15
20
25

**Optimal1 Fitted Gauss, proteinAT2G39930.1**
**bic:666**
**peaks after deleting less than one fifth of global max:6.5**

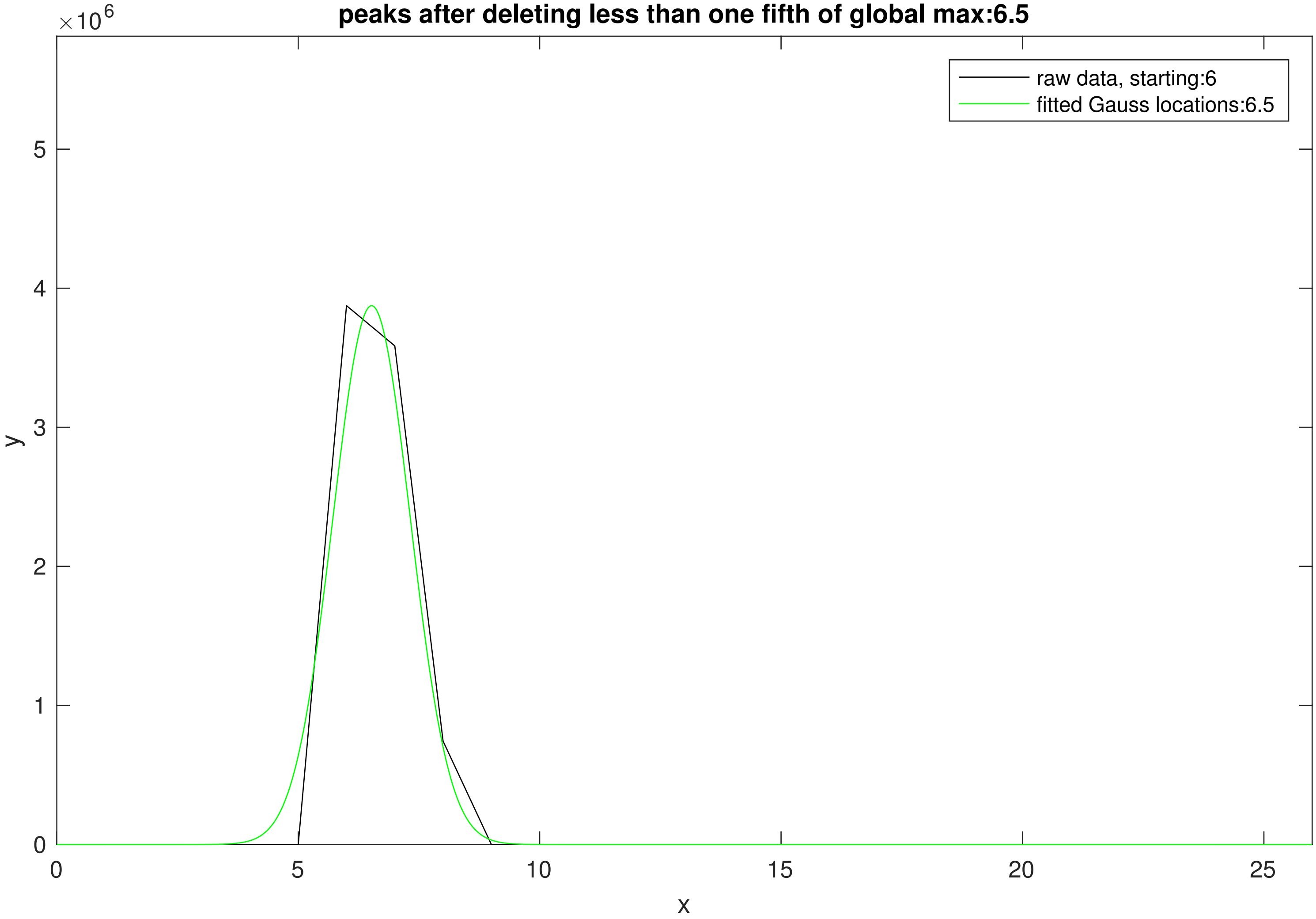

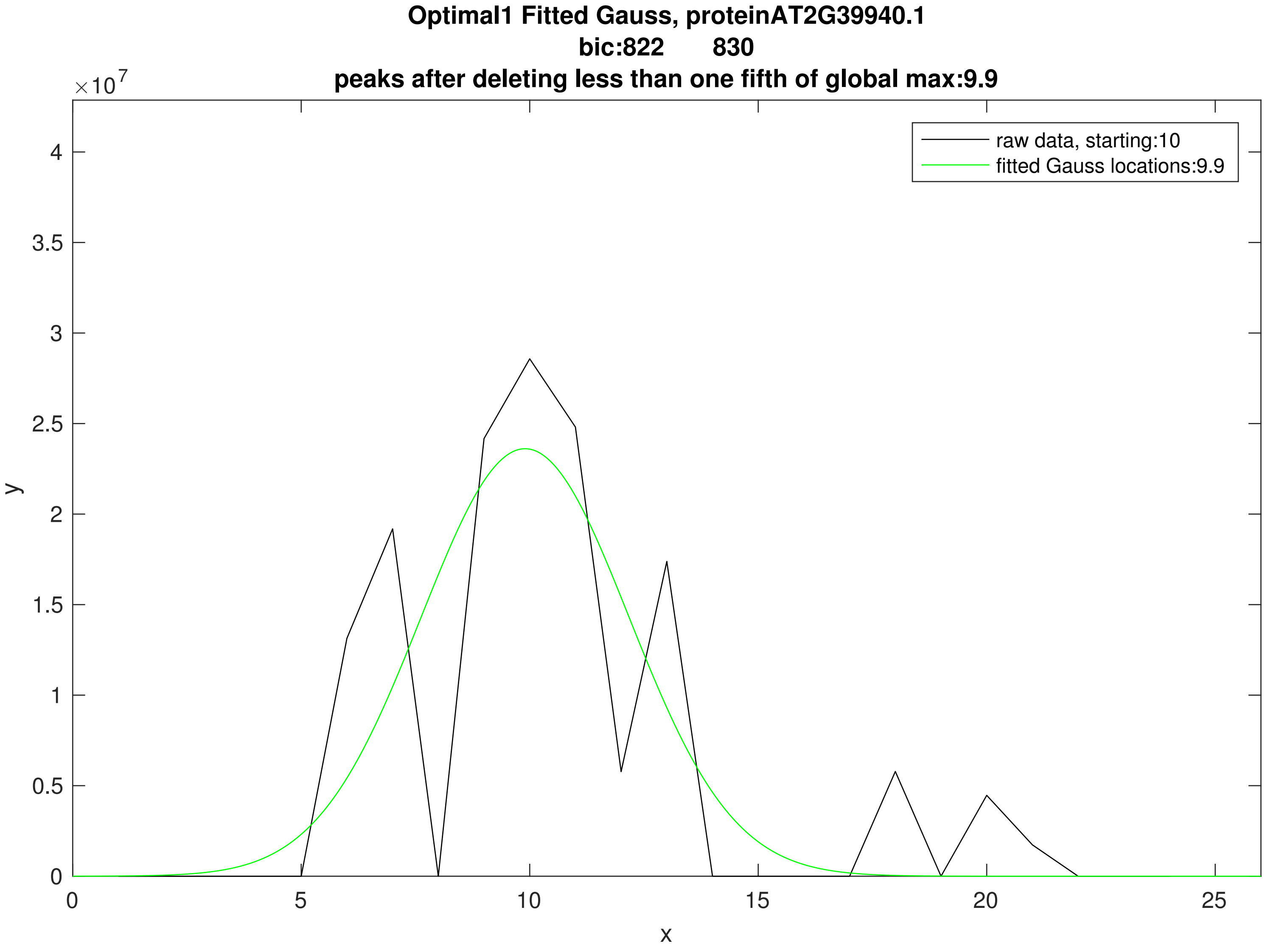
Optimal1 Fitted Gauss, proteinAT2G39940.1
bic:822 830
peaks after deleting less than one fifth of global max:9.9
×10^7
raw data, starting:10
fitted Gauss locations:9.9
y
x
0
0.5
1
1.5
2
2.5
3
3.5
4
0
5
10
15
20
25

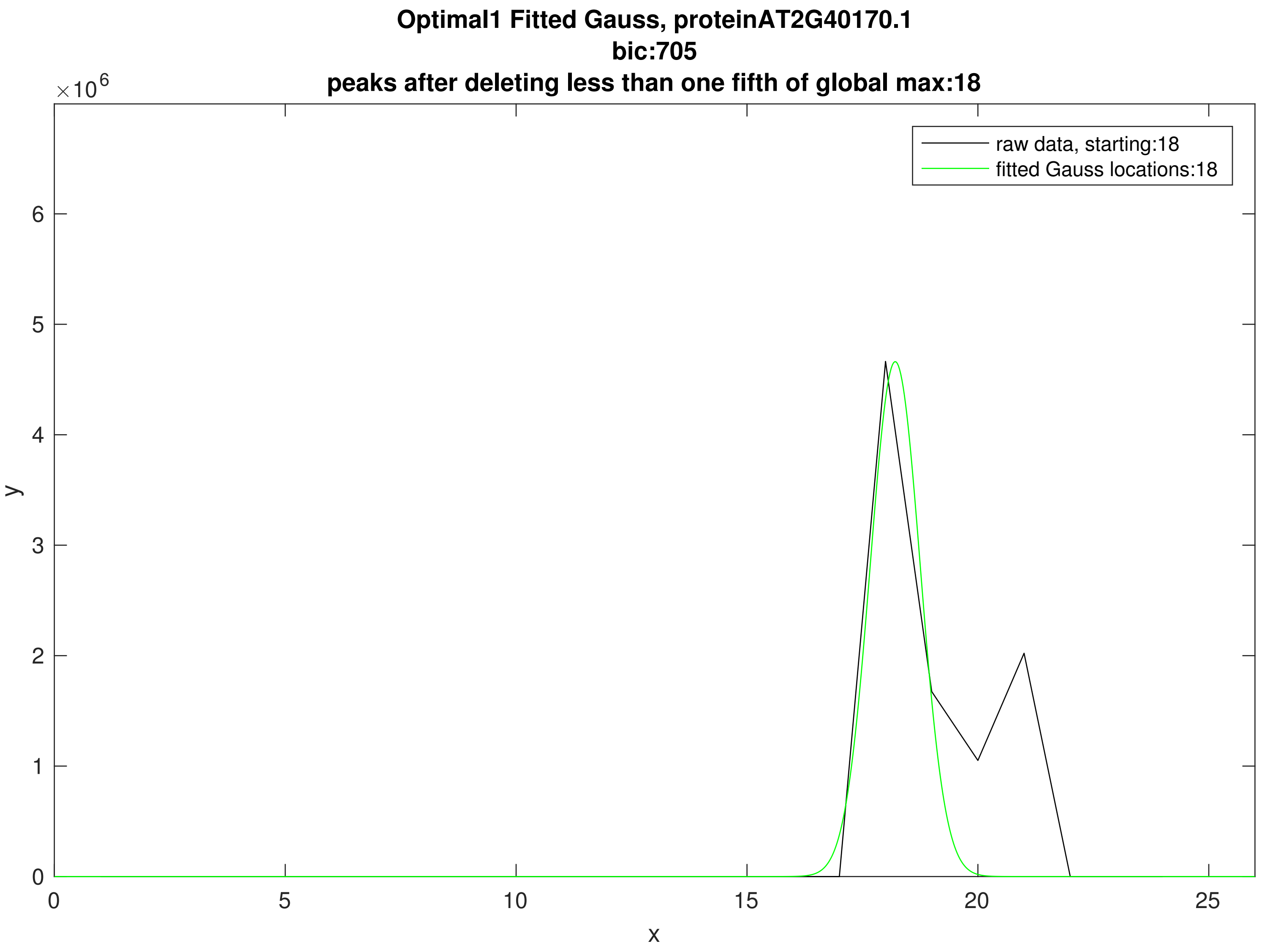
Optimal1 Fitted Gauss, proteinAT2G40170.1
bic:705
peaks after deleting less than one fifth of global max:18
×10^6
raw data, starting:18
fitted Gauss locations:18
0
1
2
3
4
5
6
y
0
5
10
15
20
25
x

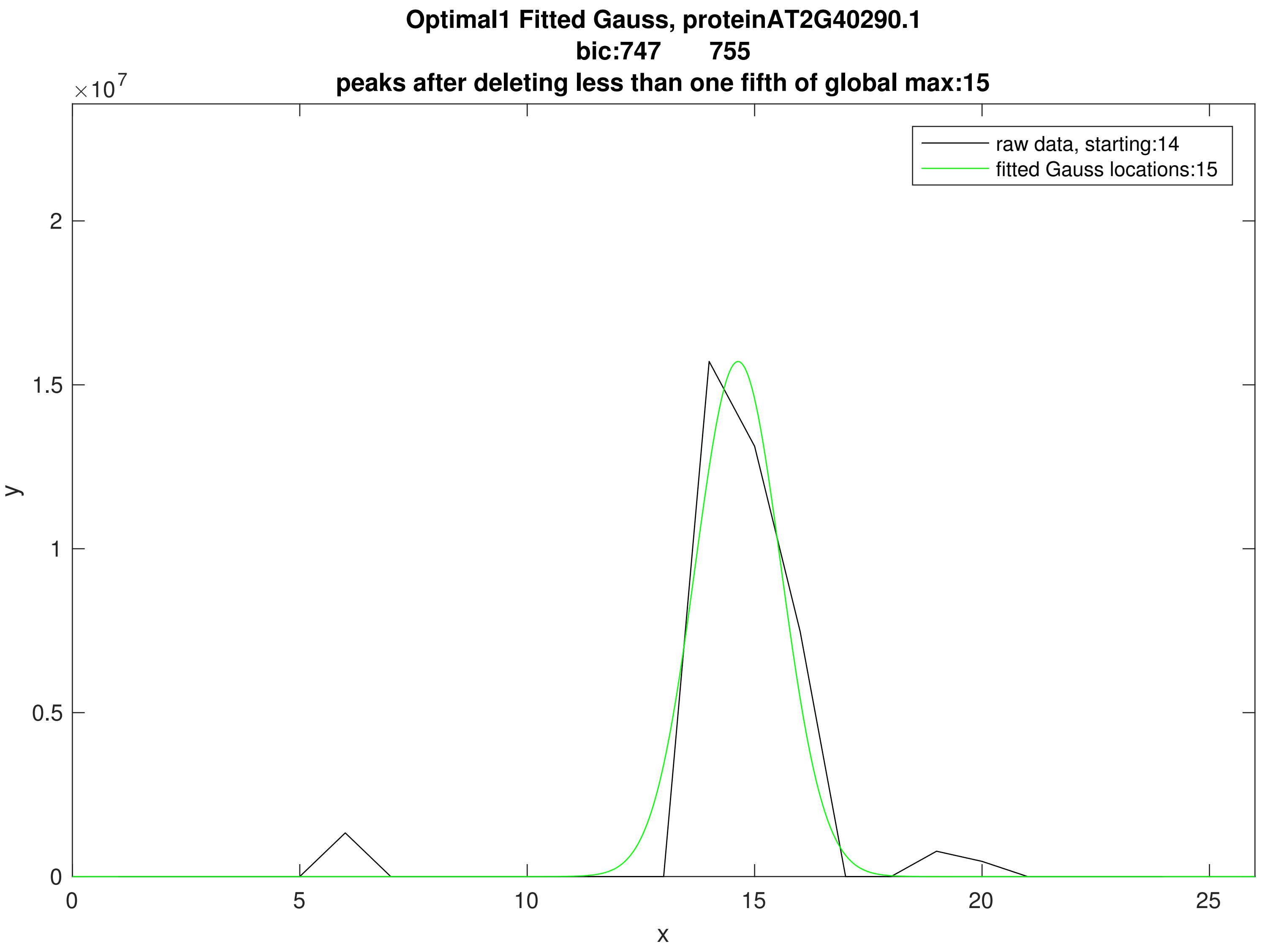

Optimal1 Fitted Gauss, proteinAT2G40290.1
bic:747 755
peaks after deleting less than one fifth of global max:15
×10^7
raw data, starting:14
fitted Gauss locations:15
y
x
0
0.5
1
1.5
2
0
5
10
15
20
25

#### Optimal1 Fitted Gauss, proteinAT2G40300.1
#### bic:1.01e+03  1.01e+03  1.02e+03
#### peaks after deleting less than one fifth of global max:12

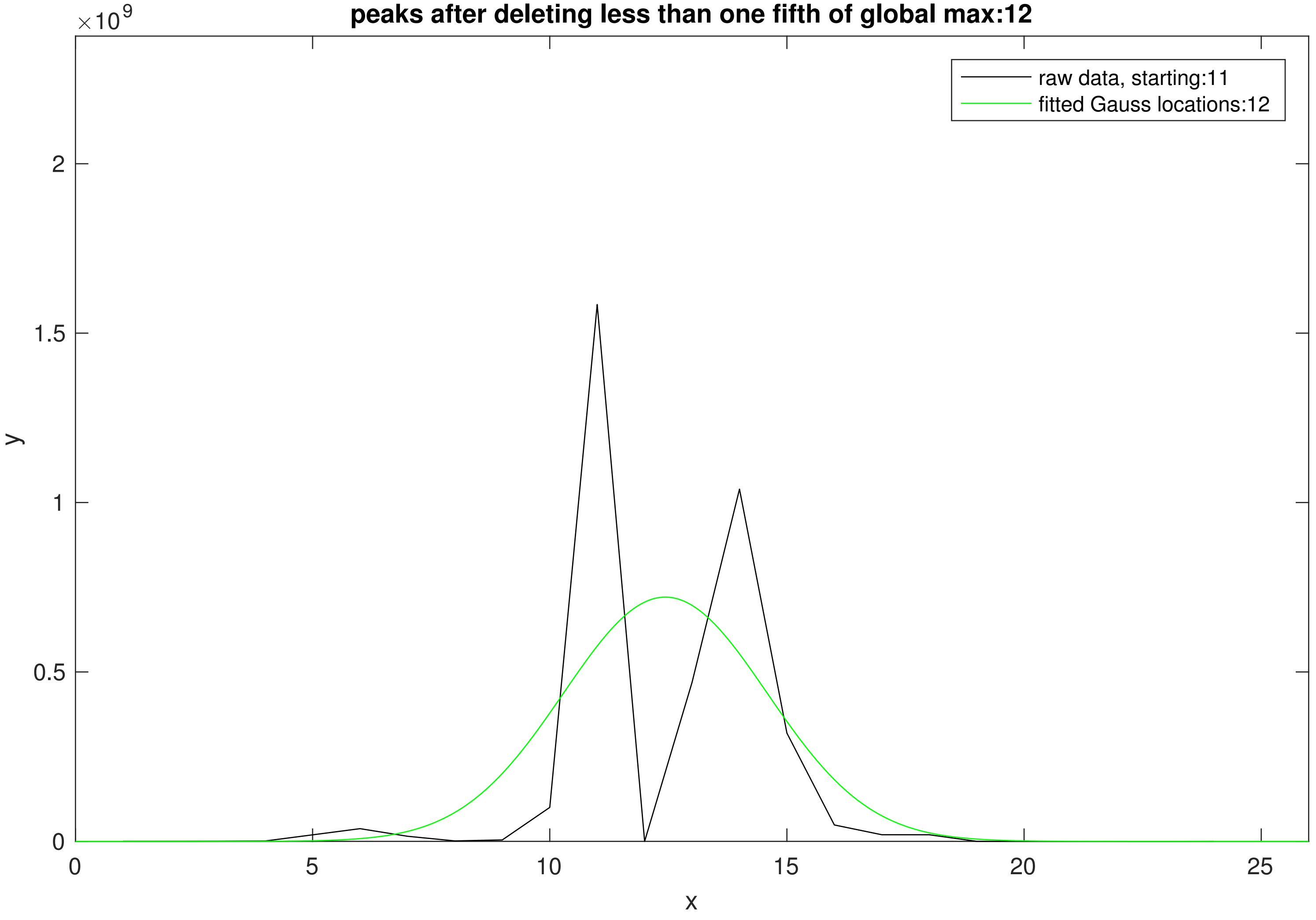

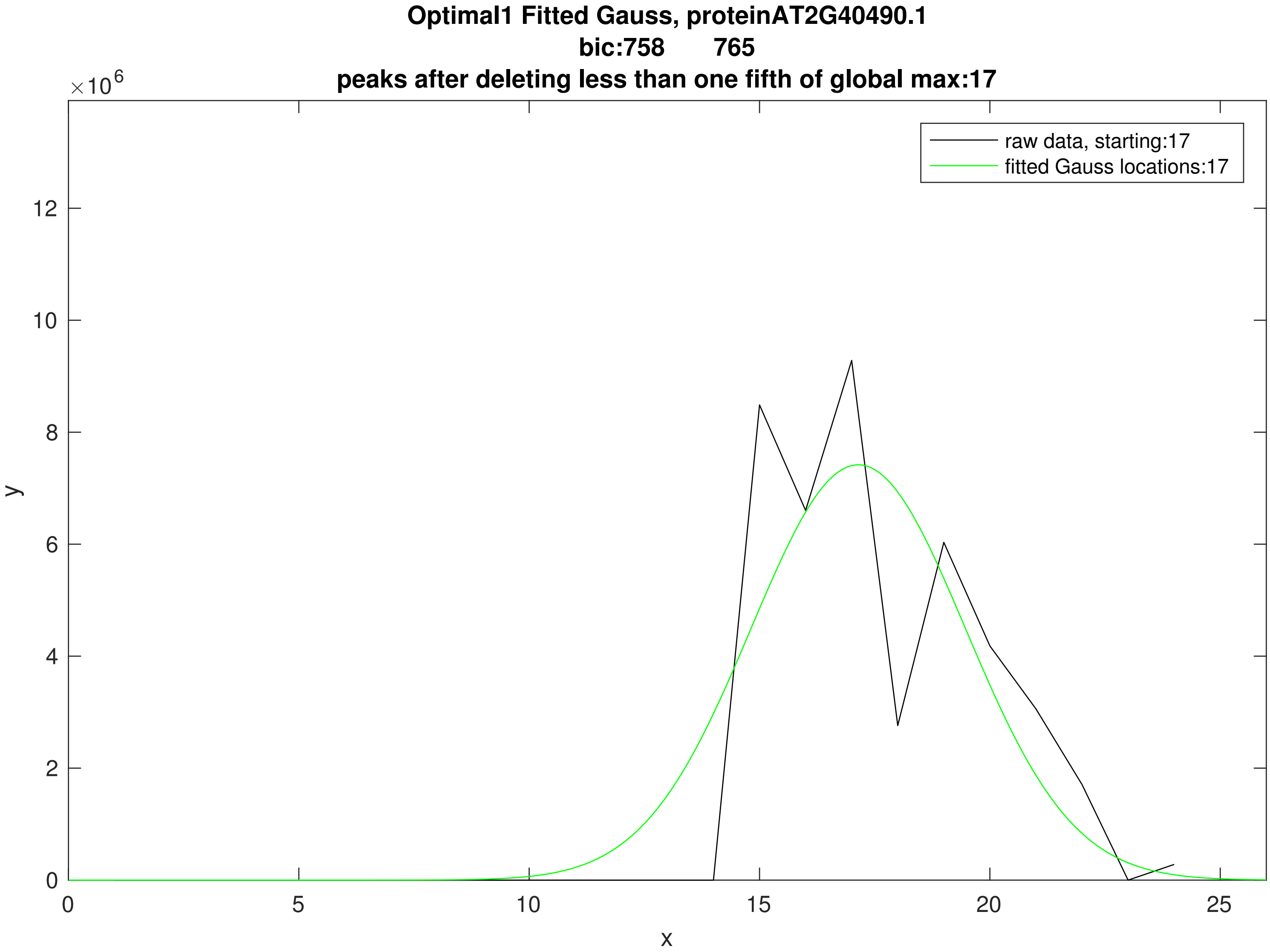
Optimal1 Fitted Gauss, proteinAT2G40490.1
bic:758 765
peaks after deleting less than one fifth of global max:17
×10^6
raw data, starting:17
fitted Gauss locations:17
y
x
0
2
4
6
8
10
12
0
5
10
15
20
25

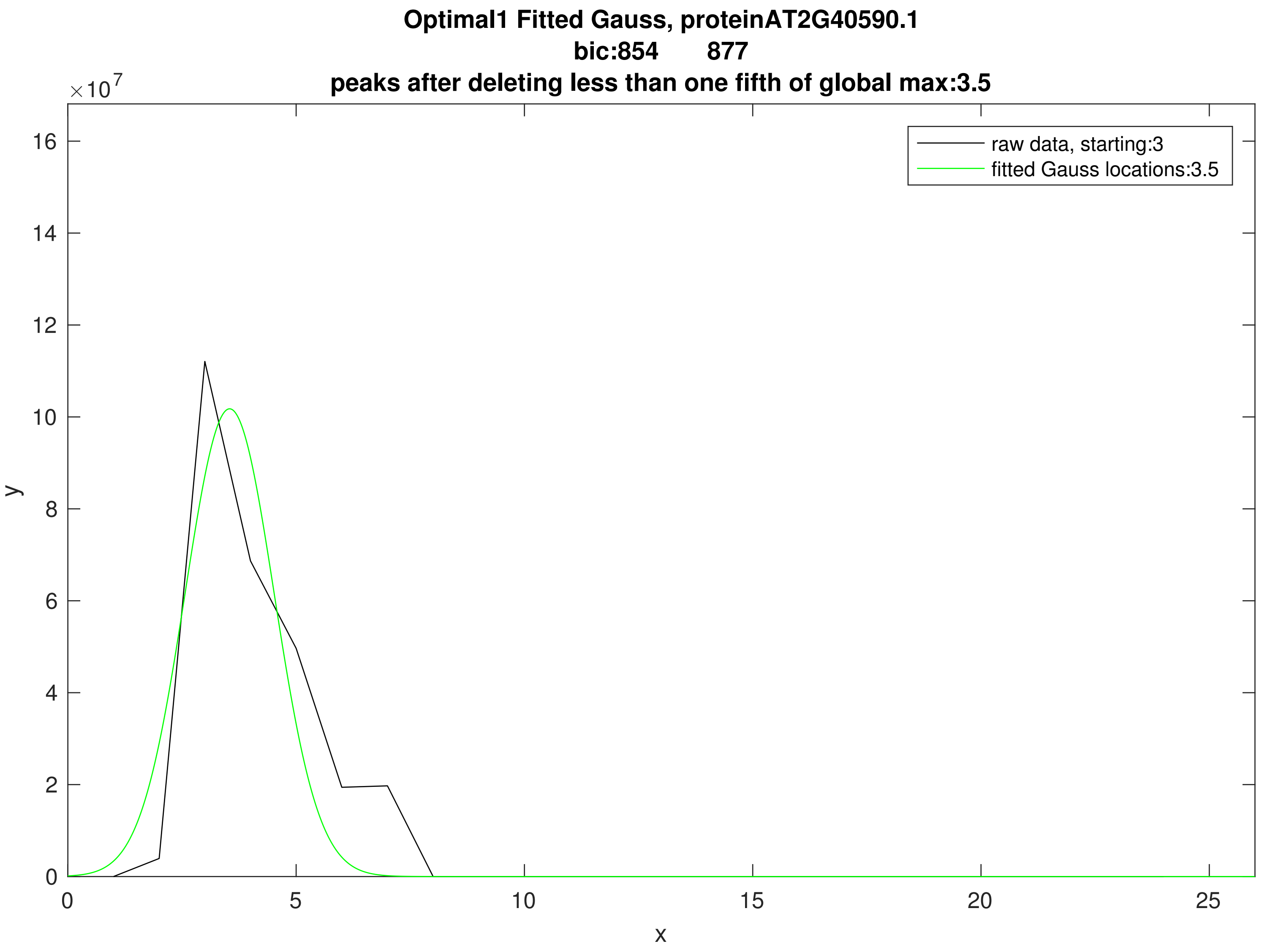
Optimal1 Fitted Gauss, proteinAT2G40590.1
bic:854 877
peaks after deleting less than one fifth of global max:3.5
×10^7
16
14
12
10
8
6
4
2
0
y
0
5
10
15
20
25
x
raw data, starting:3
fitted Gauss locations:3.5

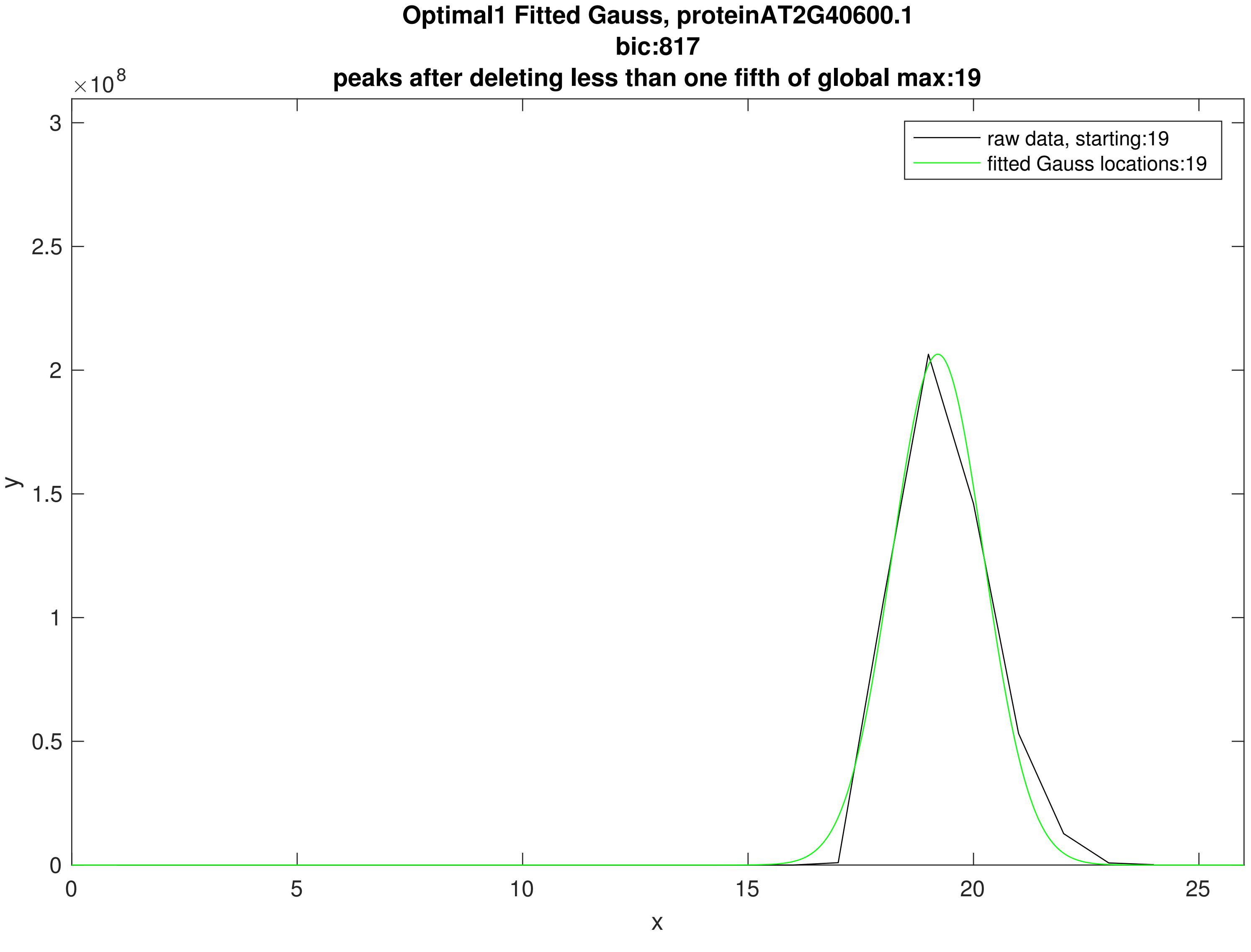
Optimal1 Fitted Gauss, proteinAT2G40600.1
bic:817
peaks after deleting less than one fifth of global max:19
×10^8
raw data, starting:19
fitted Gauss locations:19
y
x
0
0.5
1
1.5
2
2.5
3
0
5
10
15
20
25

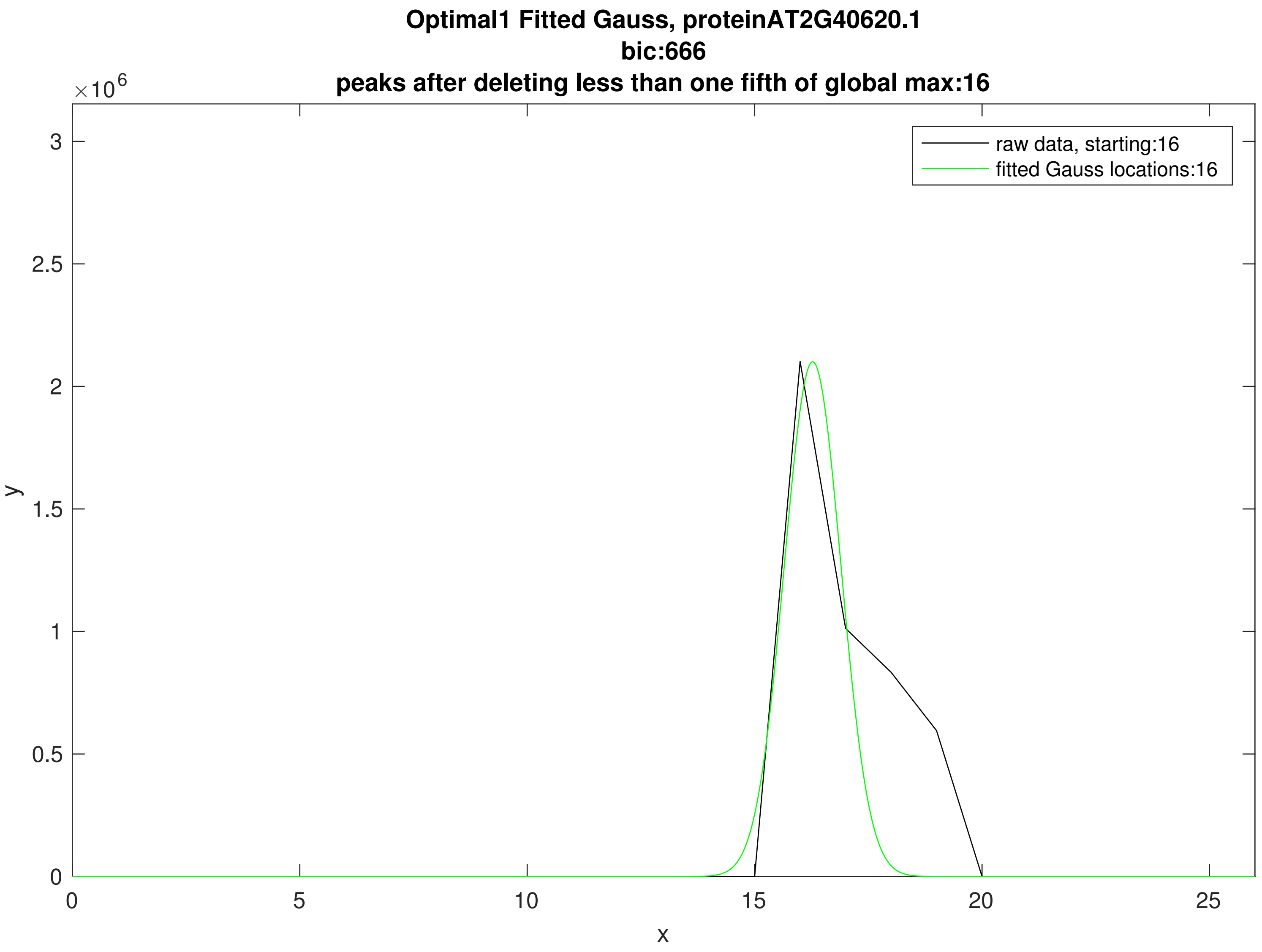
Optimal1 Fitted Gauss, proteinAT2G40620.1
bic:666
peaks after deleting less than one fifth of global max:16
×10^6
3
2.5
2
1.5
1
0.5
0
y
0
5
10
15
20
25
x
raw data, starting:16
fitted Gauss locations:16

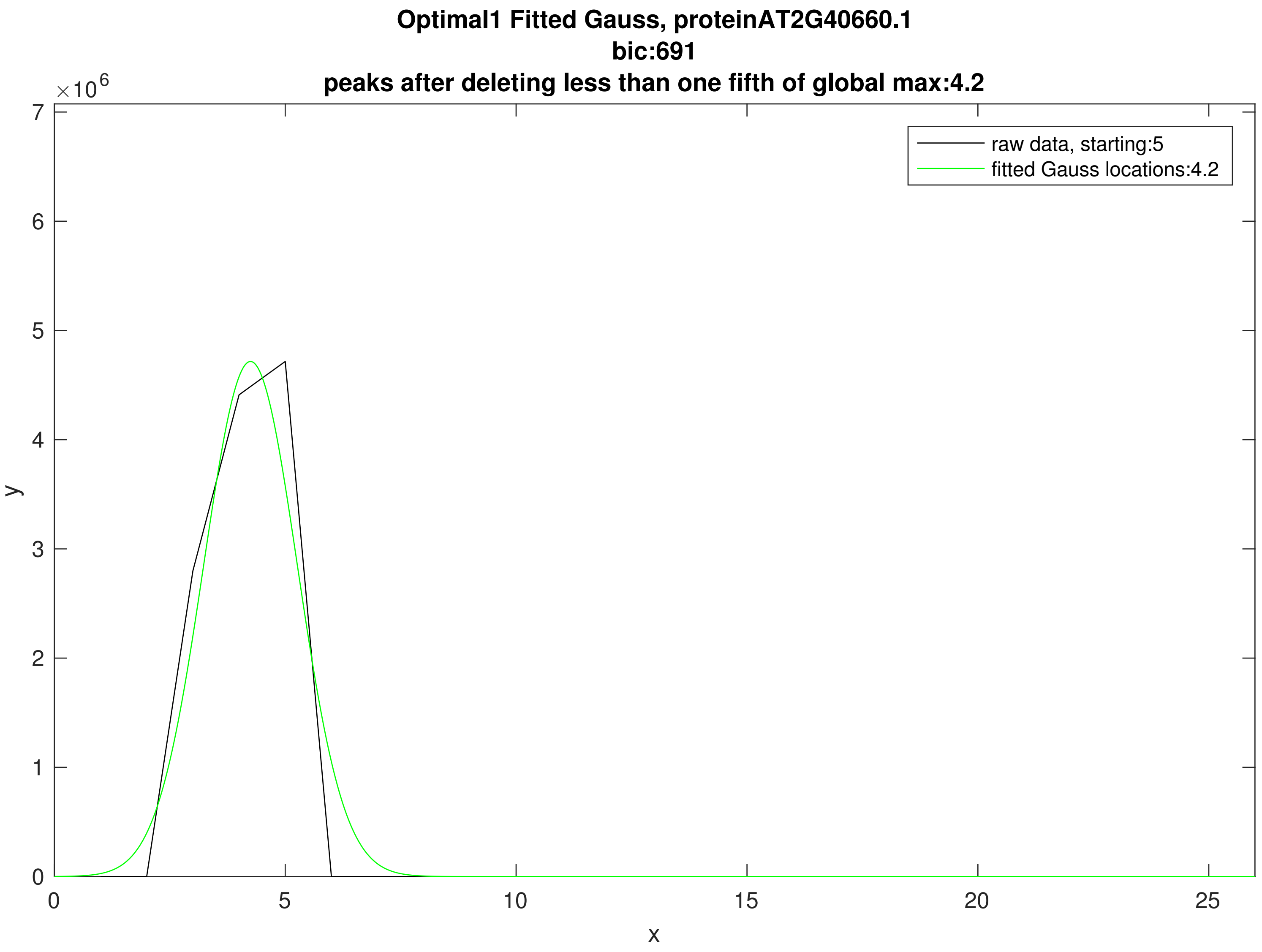

Optimal1 Fitted Gauss, proteinAT2G40660.1
bic:691
peaks after deleting less than one fifth of global max:4.2
$\times 10^6$
raw data, starting:5
fitted Gauss locations:4.2
y
x
0
1
2
3
4
5
6
7
0
5
10
15
20
25

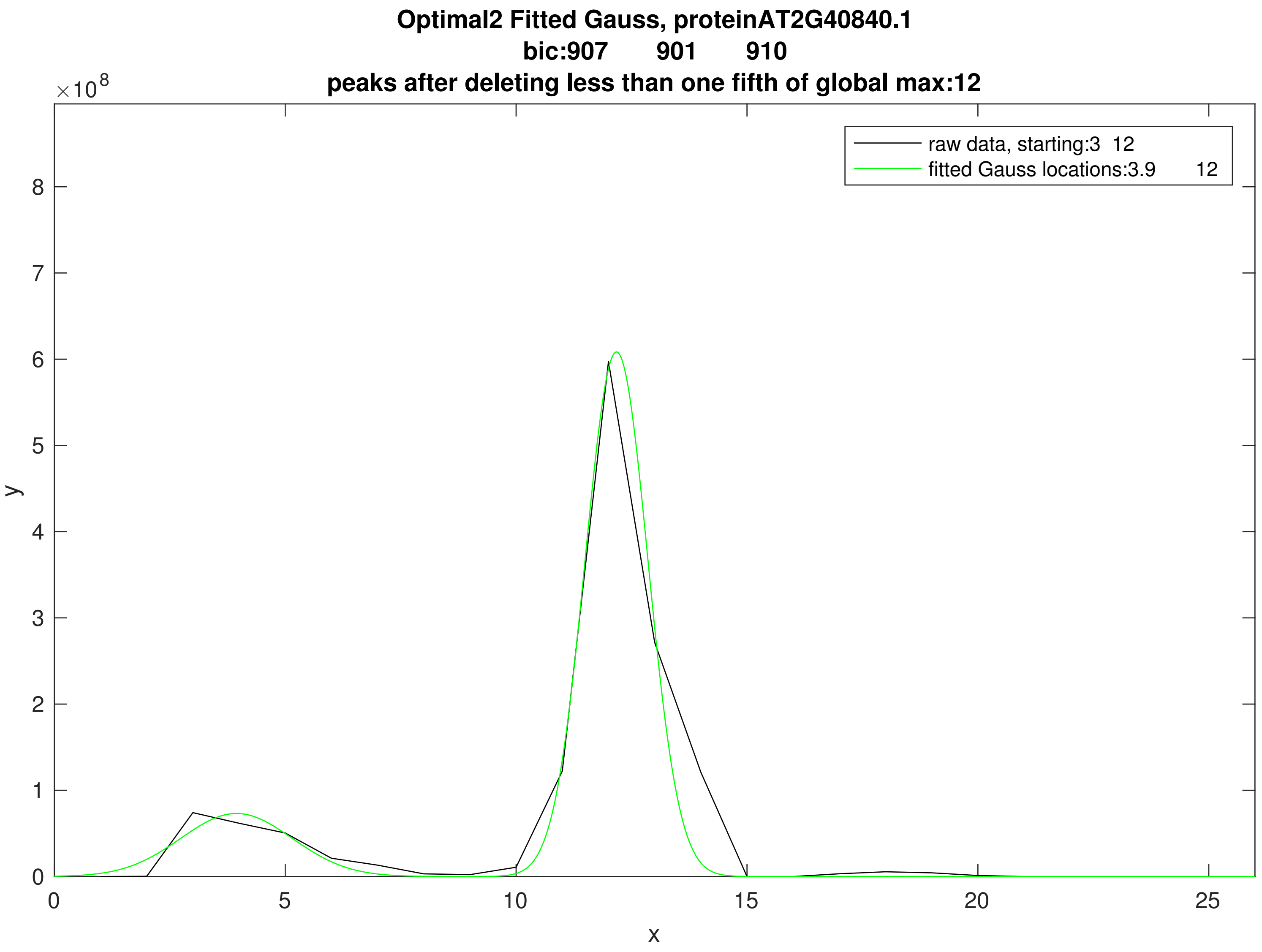

Optimal2 Fitted Gauss, proteinAT2G40840.1
bic:907 901 910
peaks after deleting less than one fifth of global max:12
×10^8
raw data, starting:3 12
fitted Gauss locations:3.9 12
y
x
0
1
2
3
4
5
6
7
8
0
5
10
15
20
25

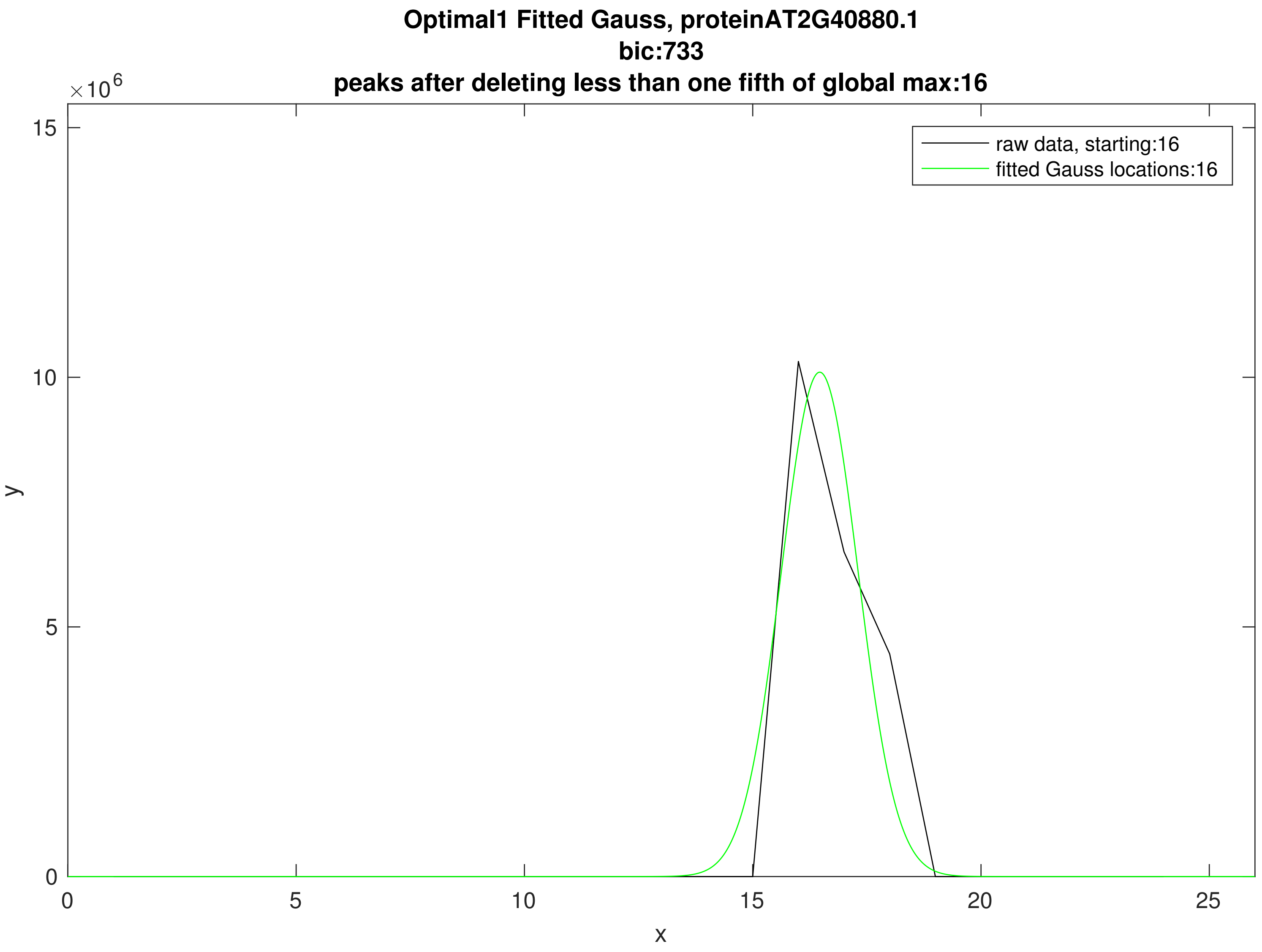
Optimal1 Fitted Gauss, proteinAT2G40880.1
bic:733
peaks after deleting less than one fifth of global max:16
×10^6
raw data, starting:16
fitted Gauss locations:16
y
x
0
5
10
15
20
25

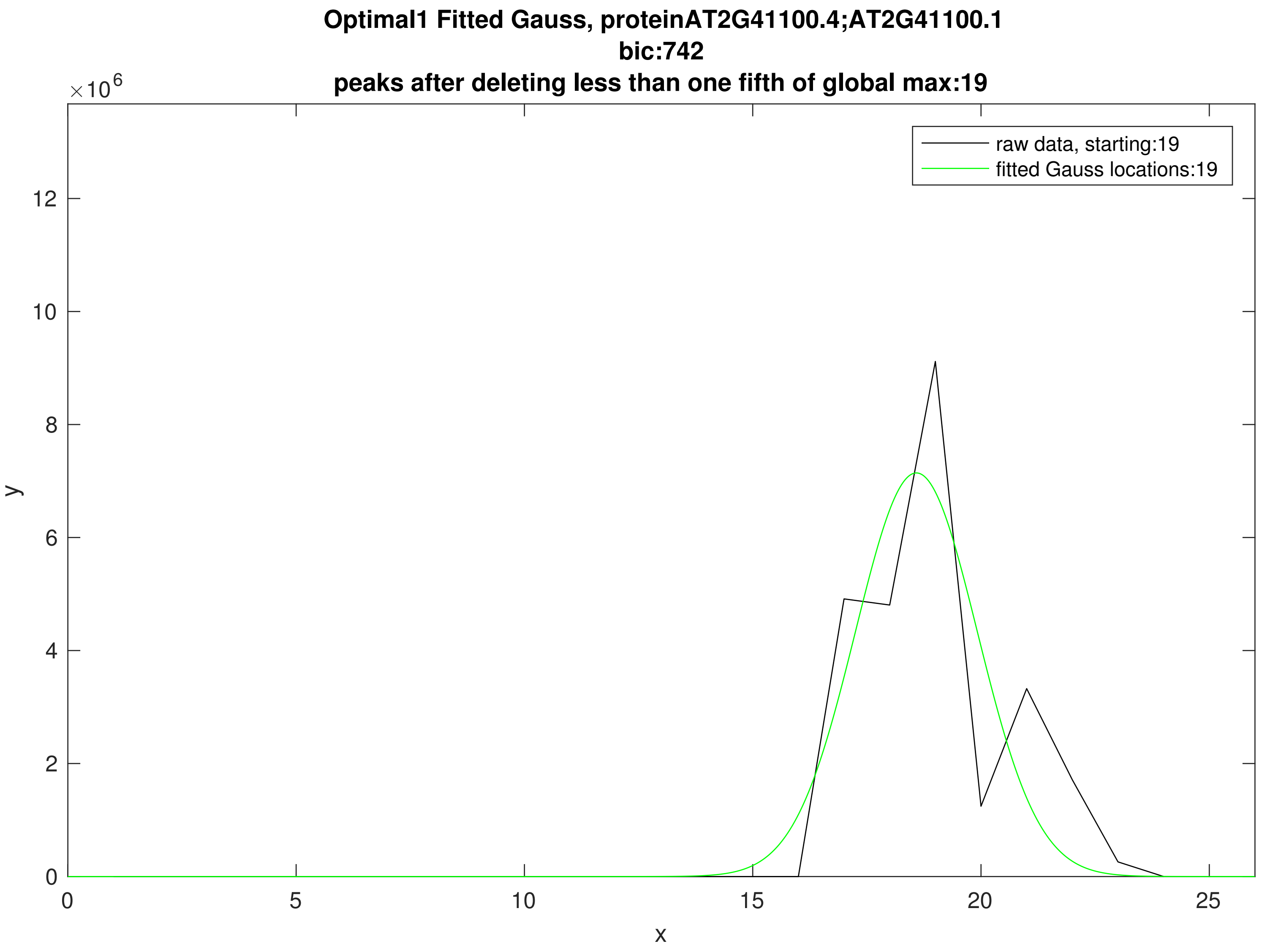
Optimal1 Fitted Gauss, proteinAT2G41100.4;AT2G41100.1
bic:742
peaks after deleting less than one fifth of global max:19
×10^6
raw data, starting:19
fitted Gauss locations:19
y
x
0
2
4
6
8
10
12
0
5
10
15
20
25

**Optimal1 Fitted Gauss, proteinAT2G41220.1**
**bic:935  1.01e+03       994**
**peaks after deleting less than one fifth of global max:12**

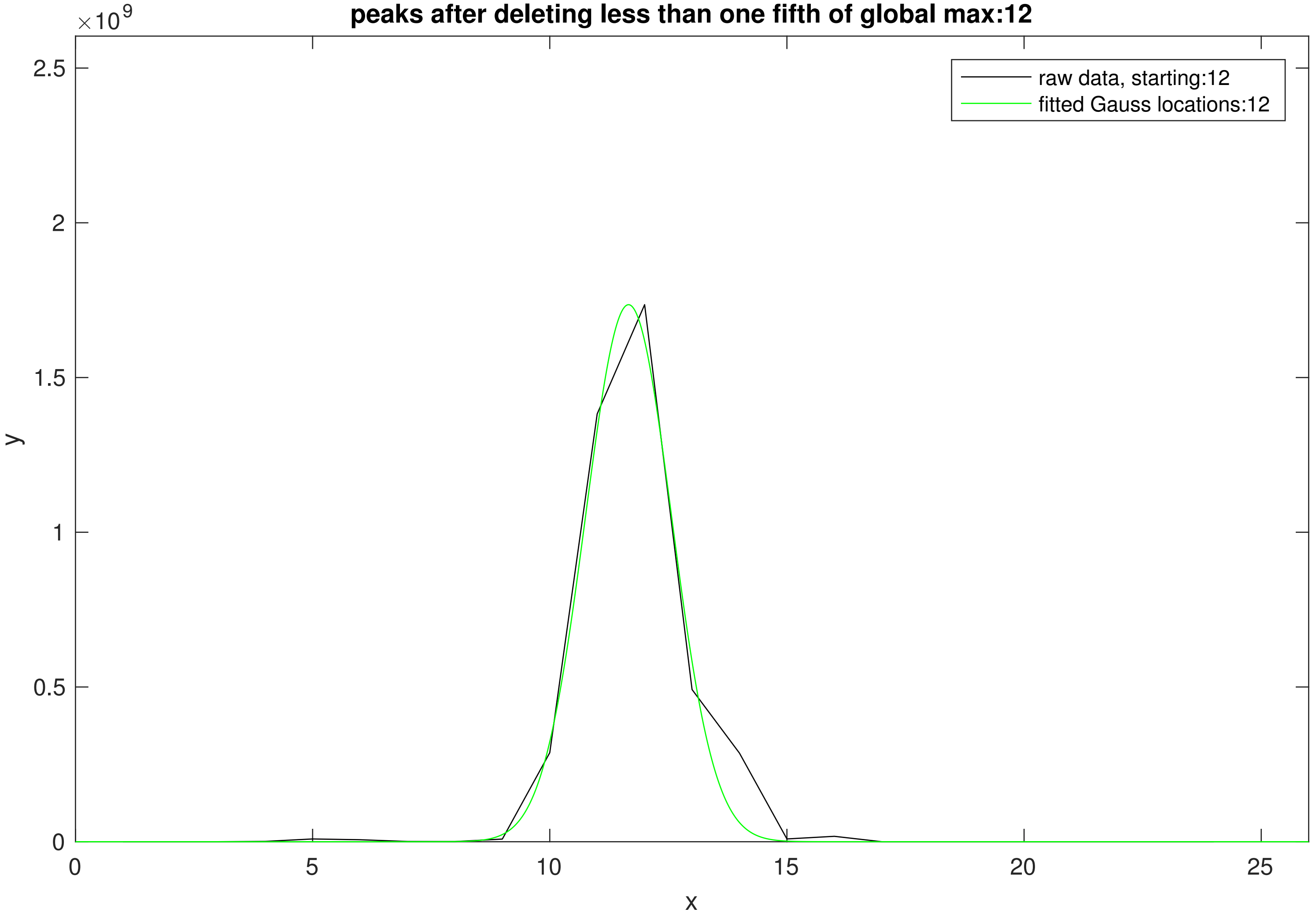

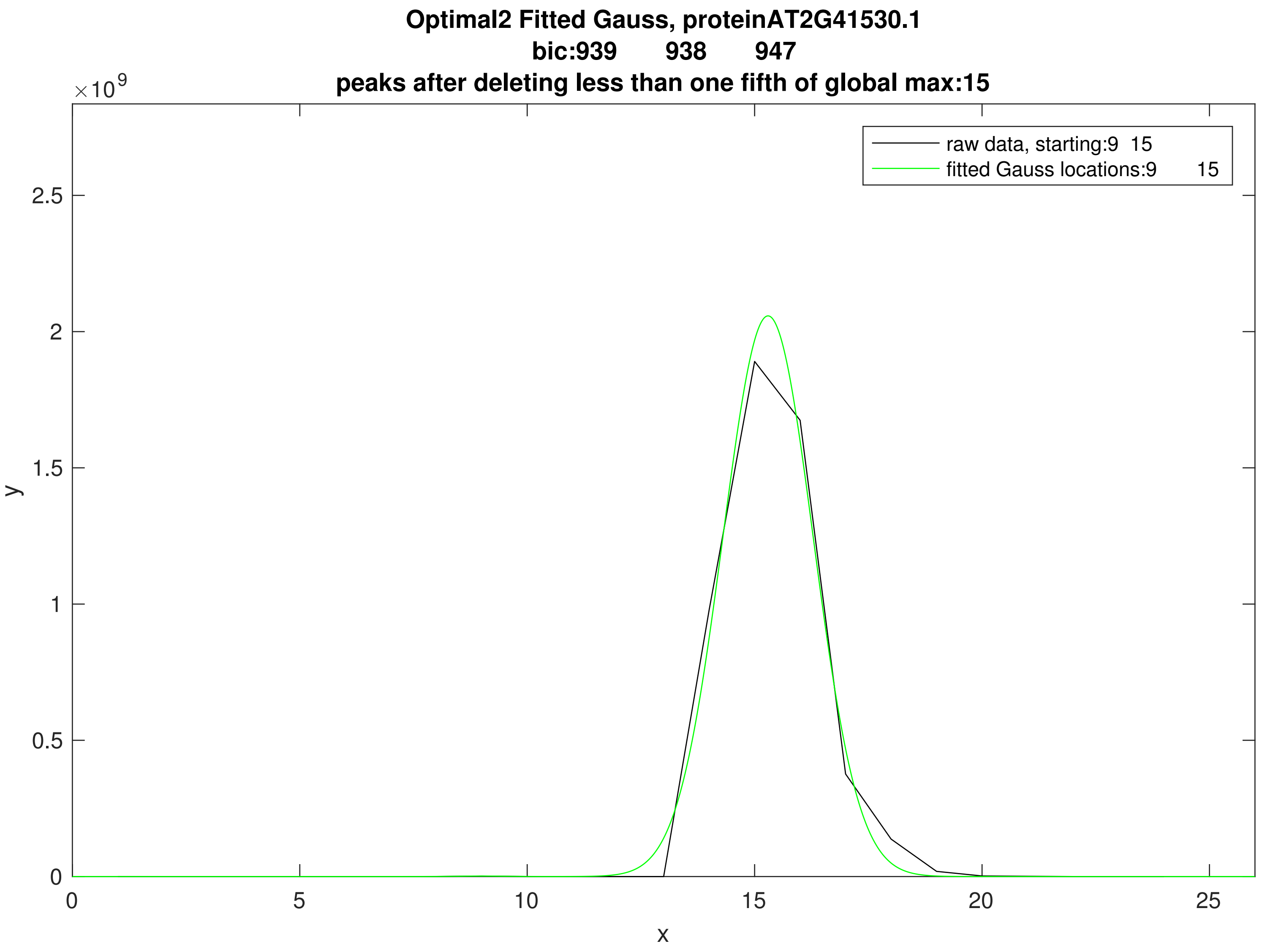
Optimal2 Fitted Gauss, proteinAT2G41530.1
bic:939 938 947
peaks after deleting less than one fifth of global max:15
×10^9
raw data, starting:9 15
fitted Gauss locations:9 15
y
x
0
0.5
1
1.5
2
2.5
0
5
10
15
20
25

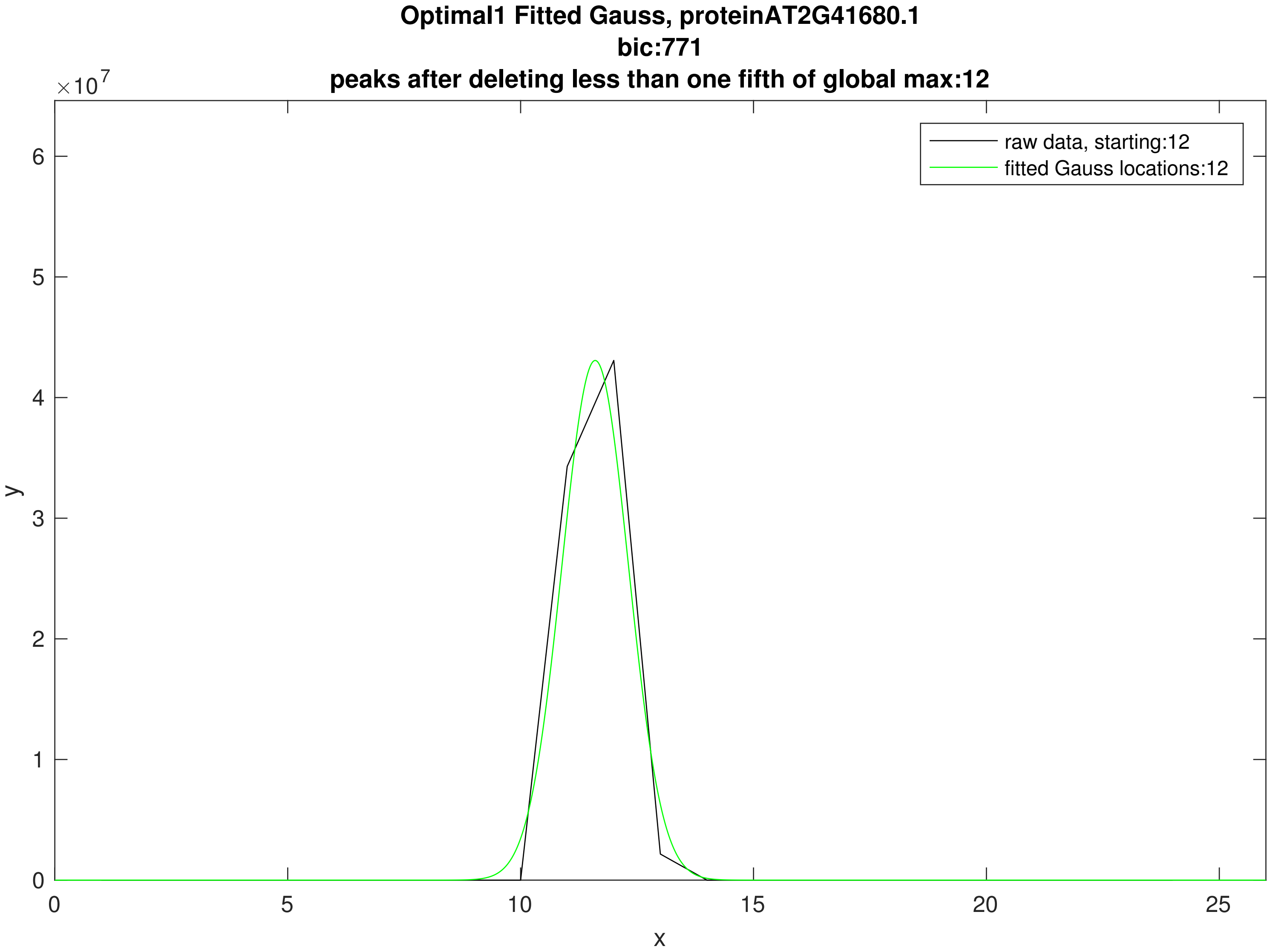
Optimal1 Fitted Gauss, proteinAT2G41680.1
bic:771
peaks after deleting less than one fifth of global max:12
×10^7
raw data, starting:12
fitted Gauss locations:12
y
x

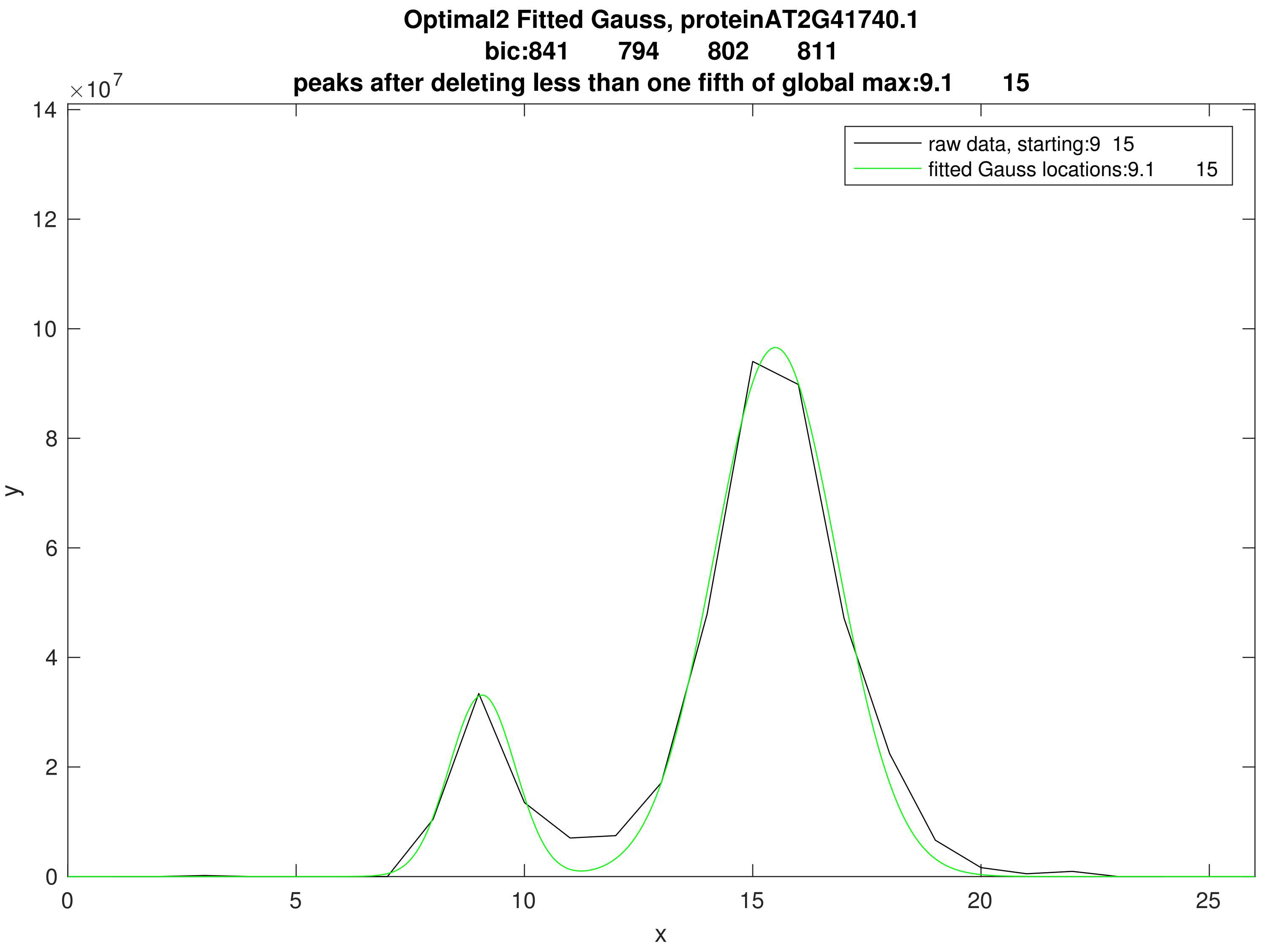
Optimal2 Fitted Gauss, proteinAT2G41740.1
bic:841 794 802 811
peaks after deleting less than one fifth of global max:9.1 15
×10^7
14
12
10
8
6
4
2
0
y
0
5
10
15
20
25
x
raw data, starting:9 15
fitted Gauss locations:9.1 15

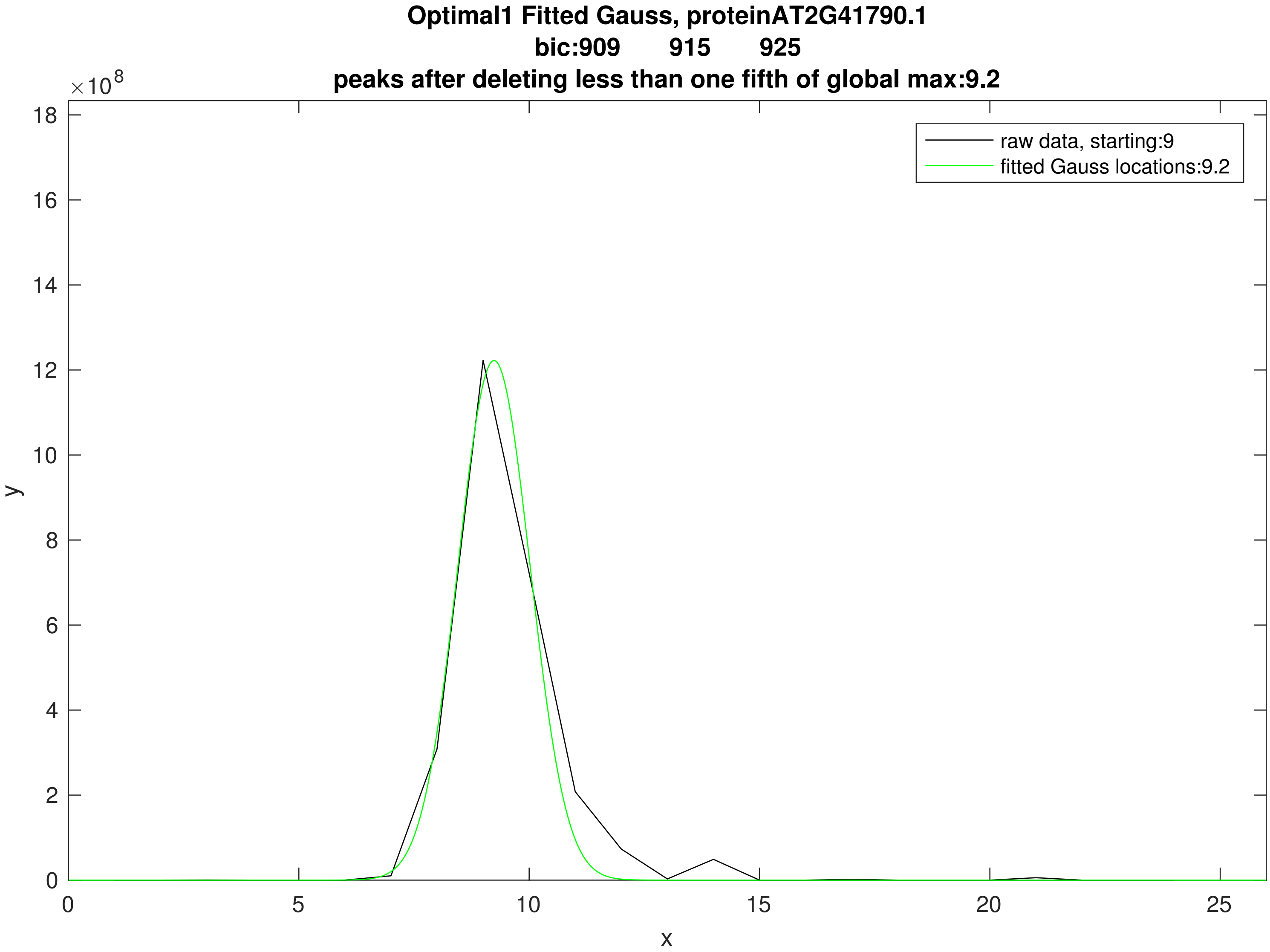

Optimal1 Fitted Gauss, proteinAT2G41790.1
bic:909 915 925
peaks after deleting less than one fifth of global max:9.2
×10^8
18
16
14
12
10
8
6
4
2
0
y
0
5
10
15
20
25
x
raw data, starting:9
fitted Gauss locations:9.2

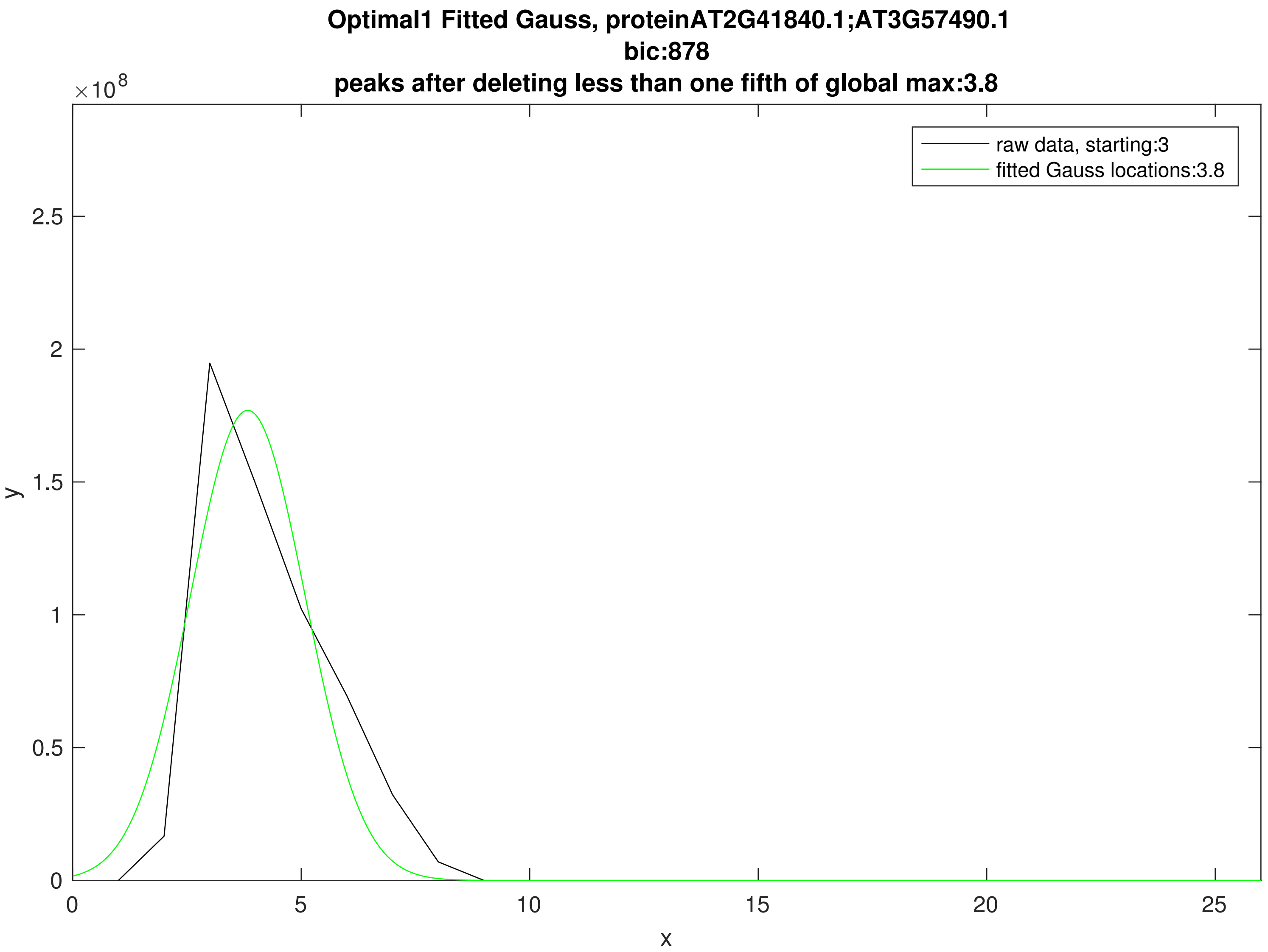
Optimal1 Fitted Gauss, proteinAT2G41840.1;AT3G57490.1
bic:878
peaks after deleting less than one fifth of global max:3.8
×10^8
raw data, starting:3
fitted Gauss locations:3.8
y
x
0
0.5
1
1.5
2
2.5
5
10
15
20
25

**Optimal1 Fitted Gauss, proteinAT2G42500.1;AT2G42500.2**
**bic:809 825**
**peaks after deleting less than one fifth of global max:16**

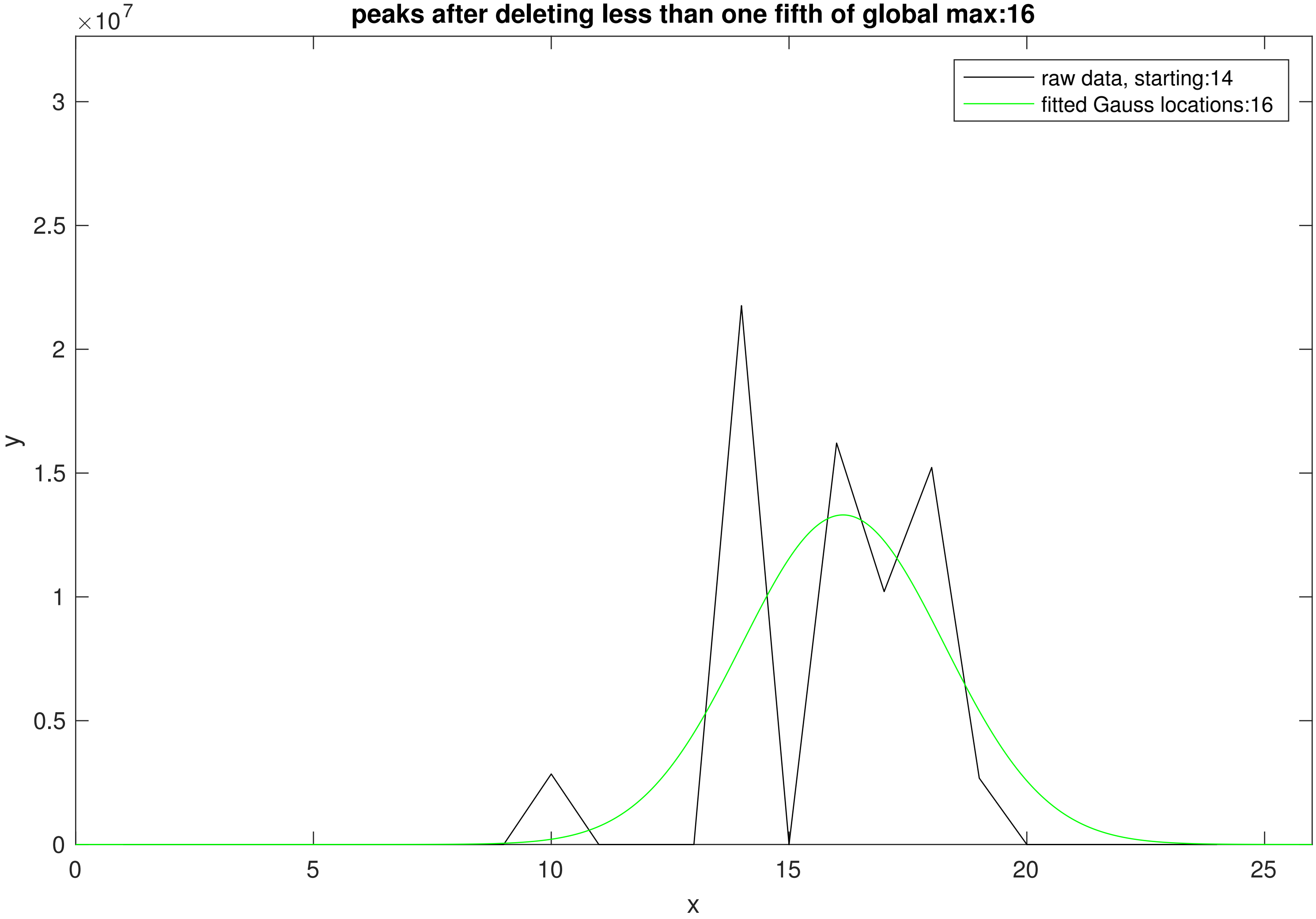

**Optimal1 Fitted Gauss, proteinAT2G42560.1**
**bic:751**
**peaks after deleting less than one fifth of global max:14**

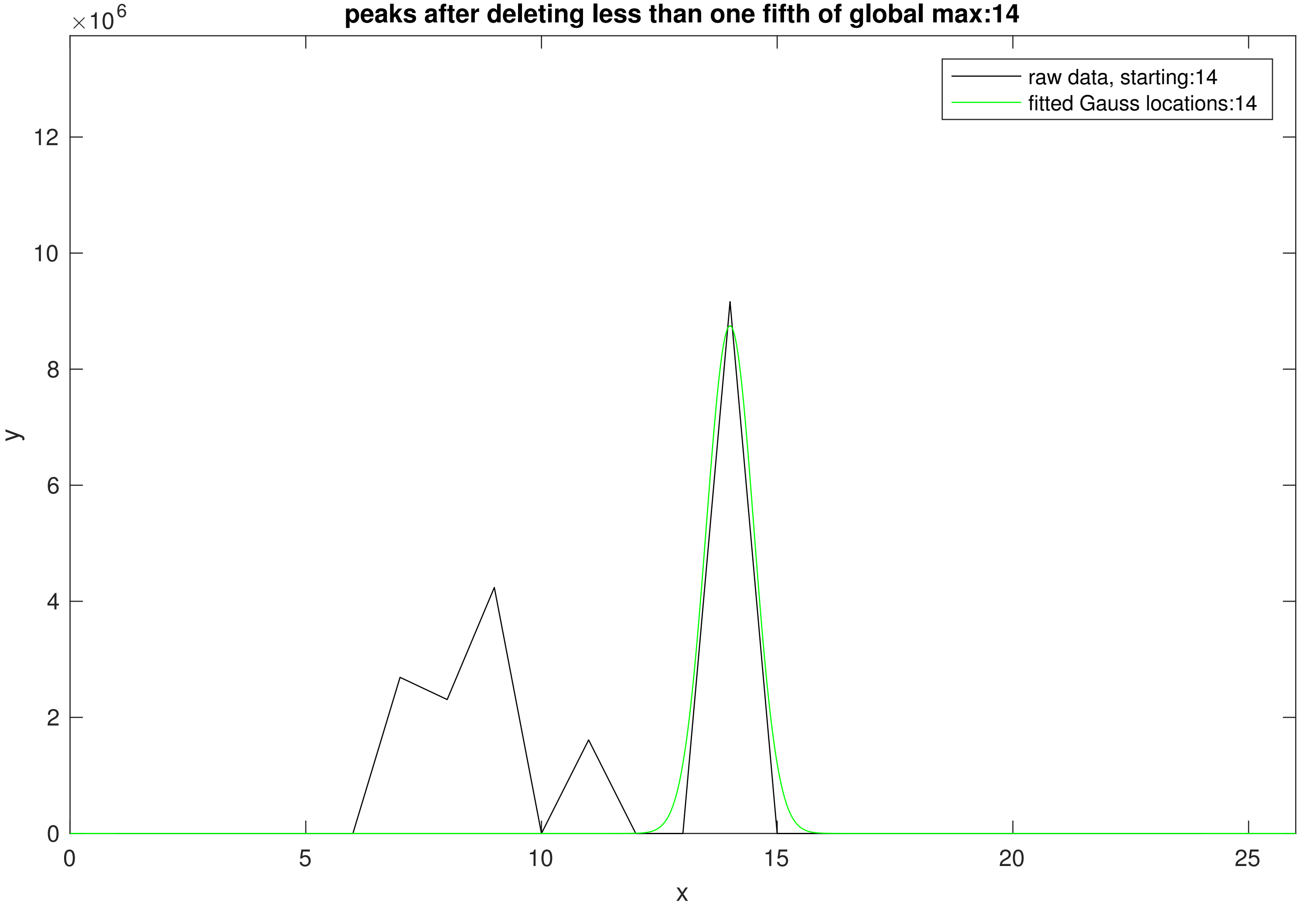

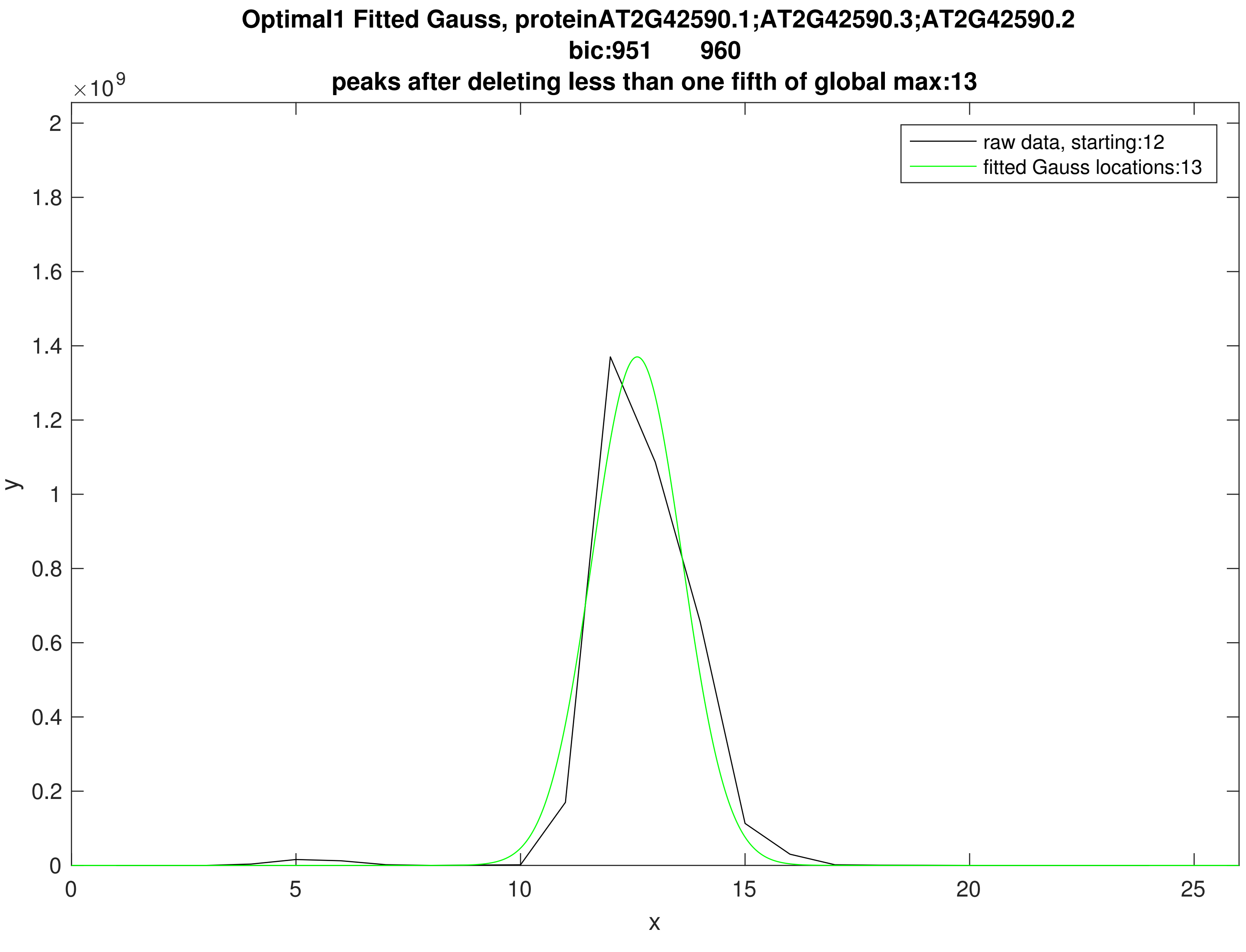
Optimal1 Fitted Gauss, proteinAT2G42590.1;AT2G42590.3;AT2G42590.2
bic:951 960
peaks after deleting less than one fifth of global max:13
×10^9
2
1.8
1.6
1.4
1.2
1
0.8
0.6
0.4
0.2
0
y
0
5
10
15
20
25
x
raw data, starting:12
fitted Gauss locations:13

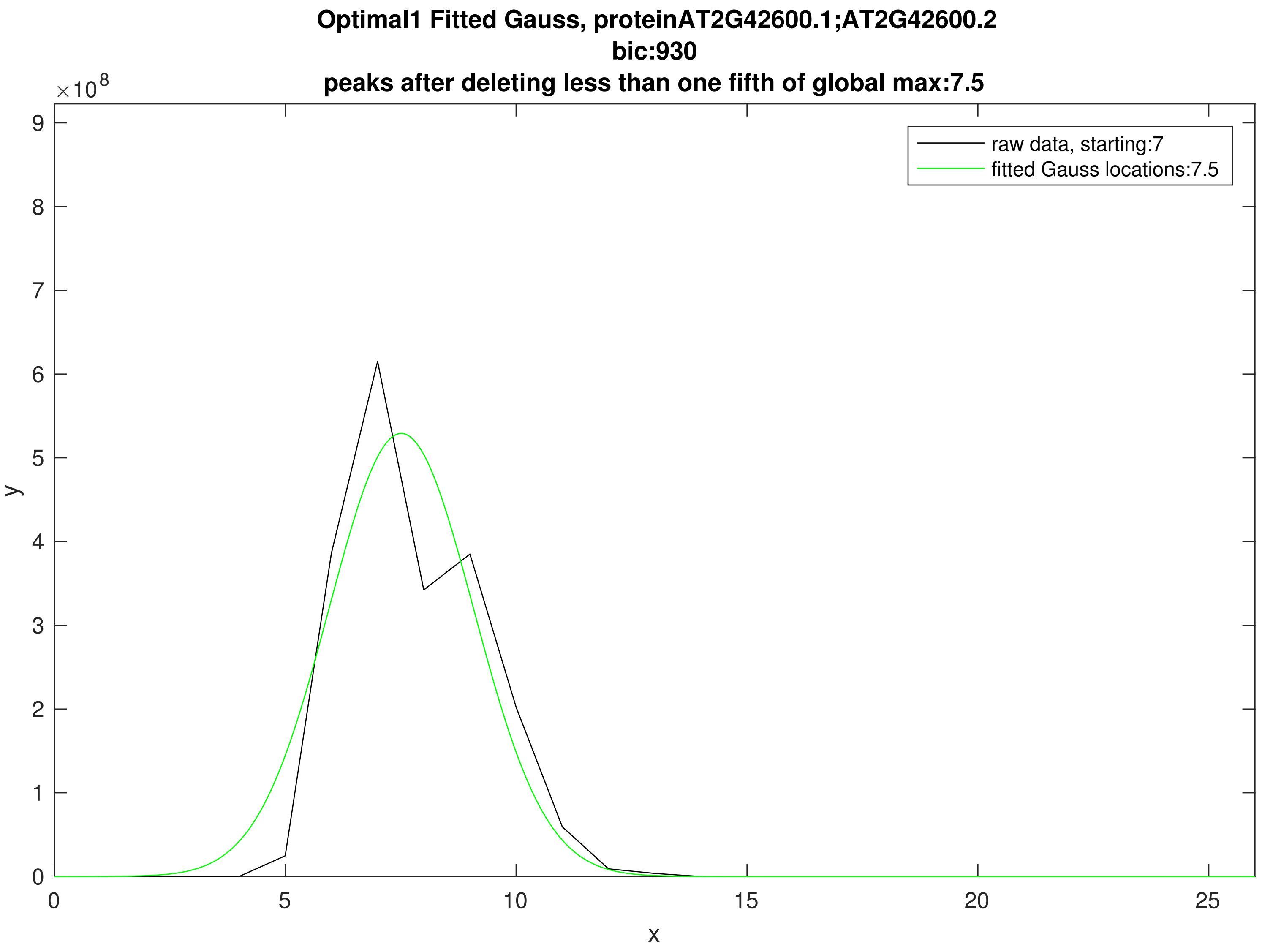
Optimal1 Fitted Gauss, proteinAT2G42600.1;AT2G42600.2
bic:930
peaks after deleting less than one fifth of global max:7.5
×10^8
raw data, starting:7
fitted Gauss locations:7.5
y
x
0
1
2
3
4
5
6
7
8
9
0
5
10
15
20
25

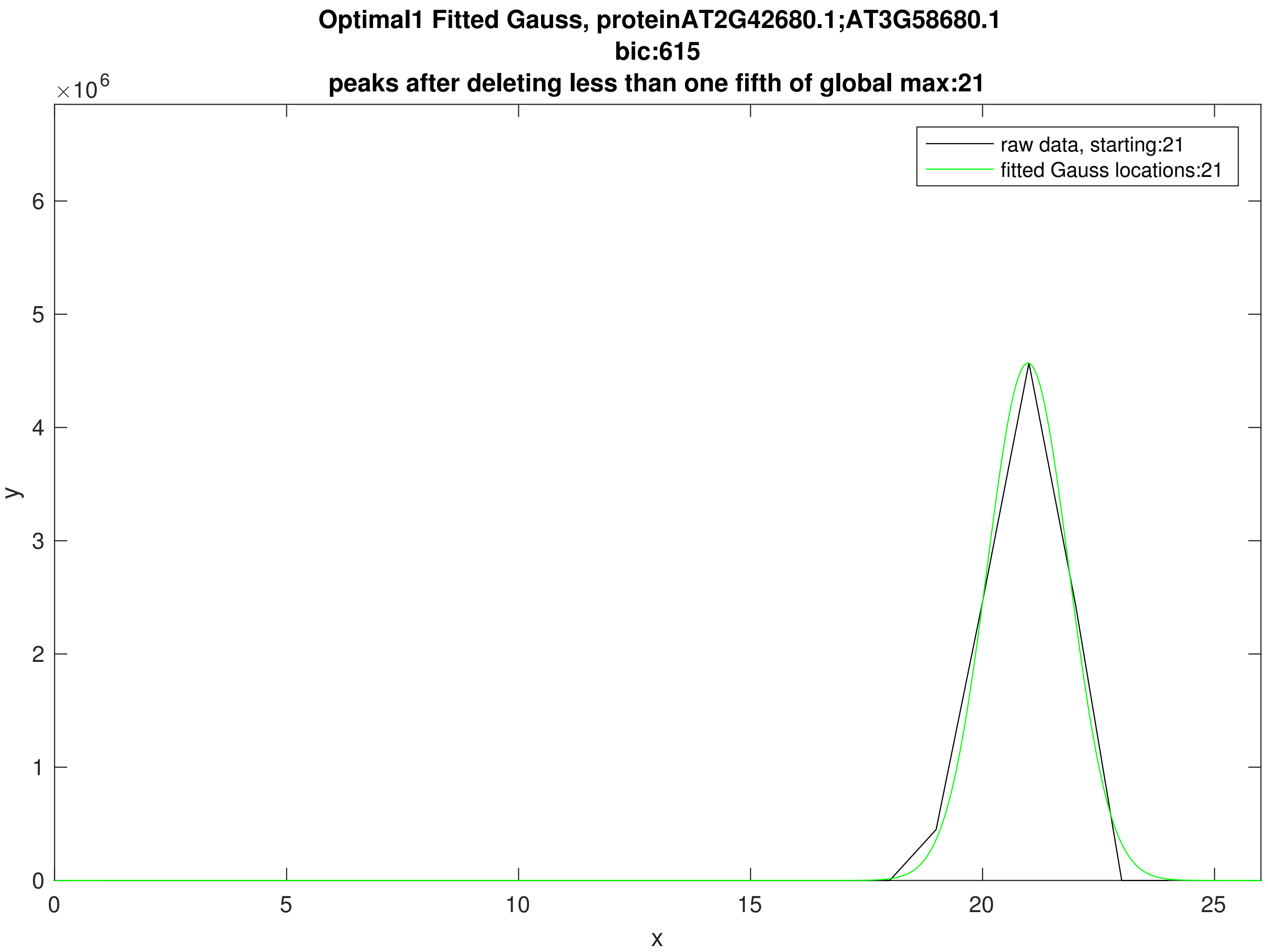
Optimal1 Fitted Gauss, proteinAT2G42680.1;AT3G58680.1
bic:615
peaks after deleting less than one fifth of global max:21
×10^6
raw data, starting:21
fitted Gauss locations:21
y
x

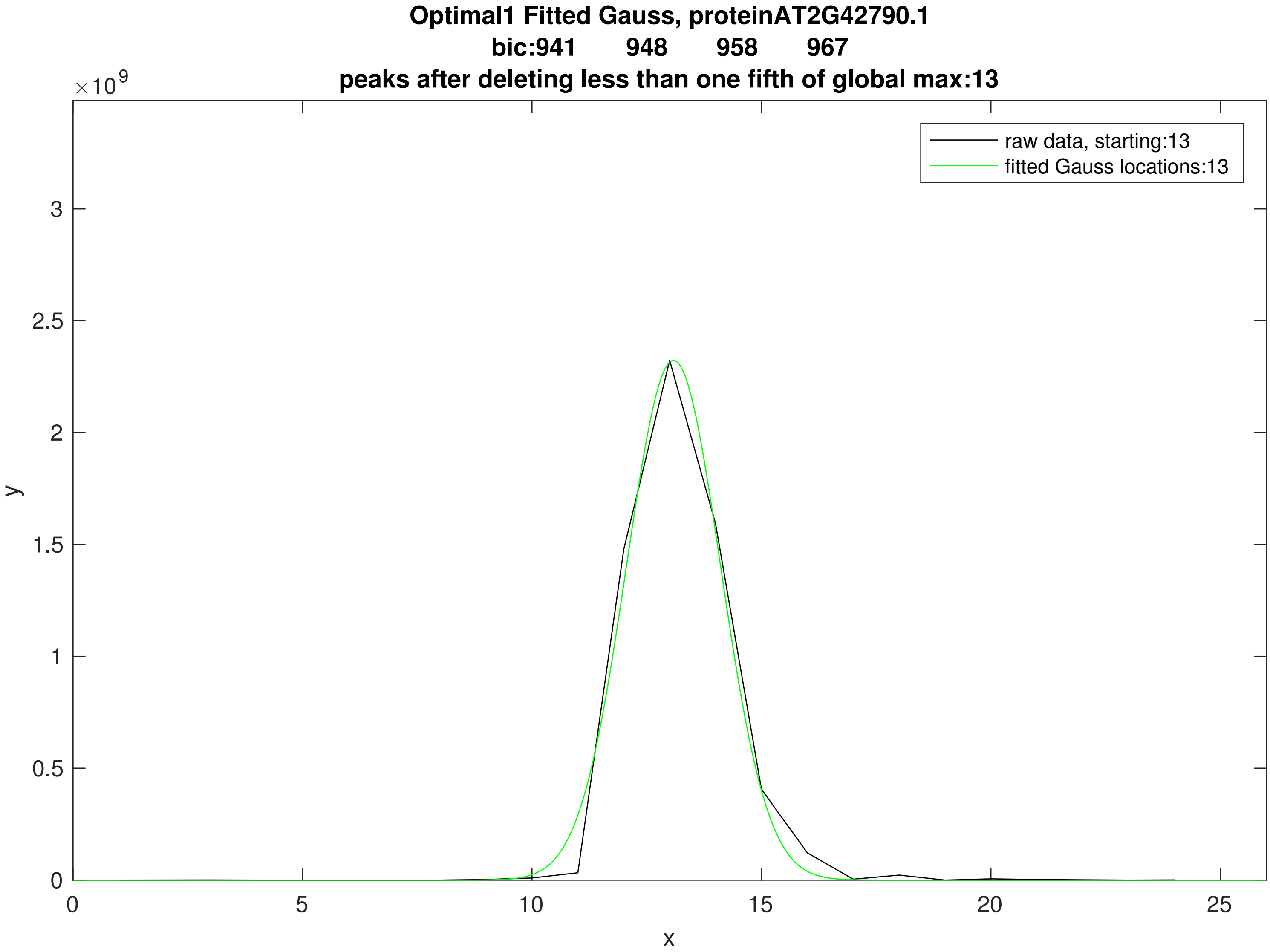
Optimal1 Fitted Gauss, proteinAT2G42790.1
bic:941 948 958 967
peaks after deleting less than one fifth of global max:13
×10^9
raw data, starting:13
fitted Gauss locations:13
y
x
0
0.5
1
1.5
2
2.5
3
0
5
10
15
20
25

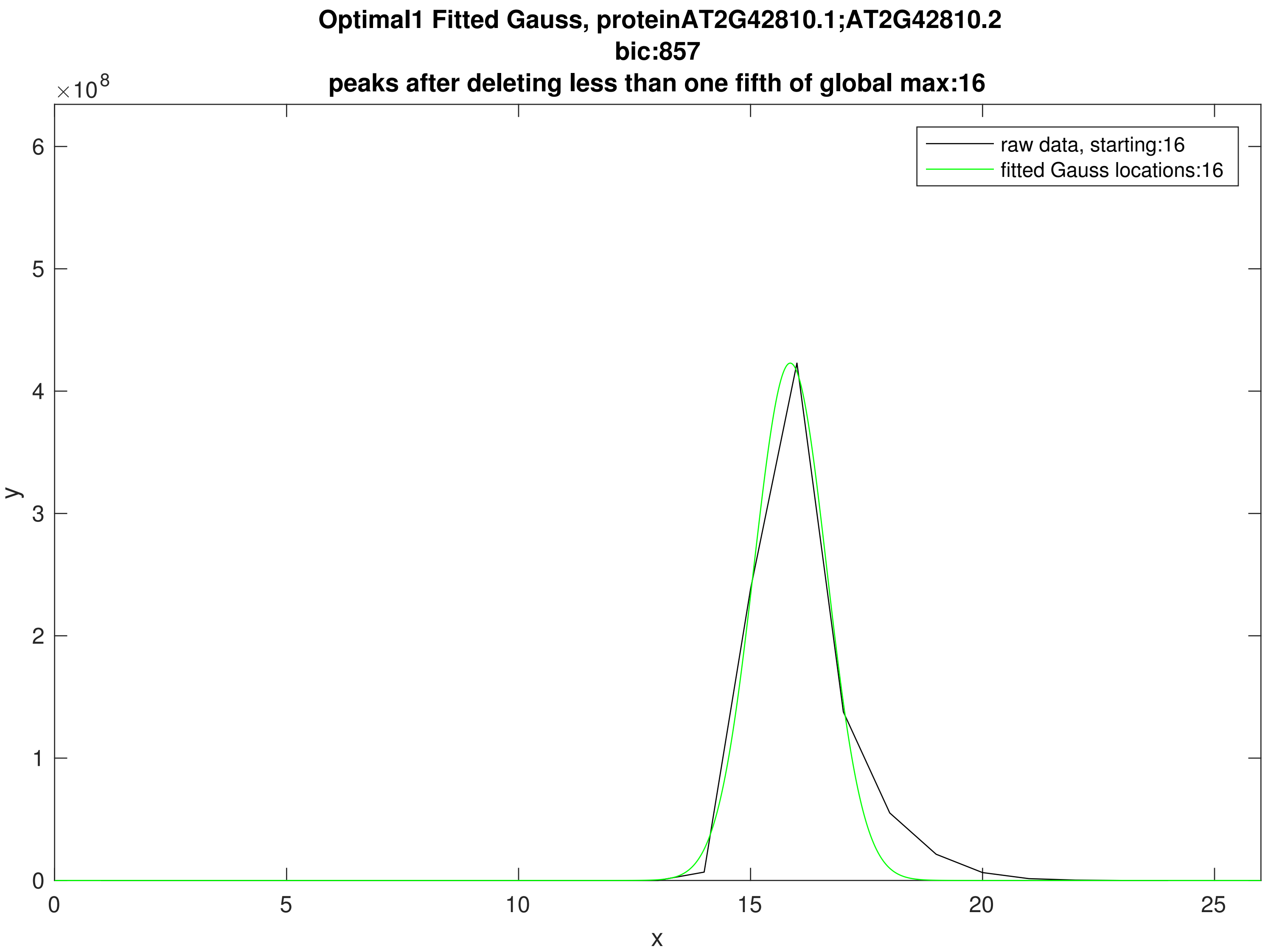

Optimal1 Fitted Gauss, proteinAT2G42810.1;AT2G42810.2
bic:857
peaks after deleting less than one fifth of global max:16
×10^8
raw data, starting:16
fitted Gauss locations:16
y
x
0
1
2
3
4
5
6
0
5
10
15
20
25

**Optimal1 Fitted Gauss, proteinAT2G42910.1**
**bic:830 838**
**peaks after deleting less than one fifth of global max:6.2**

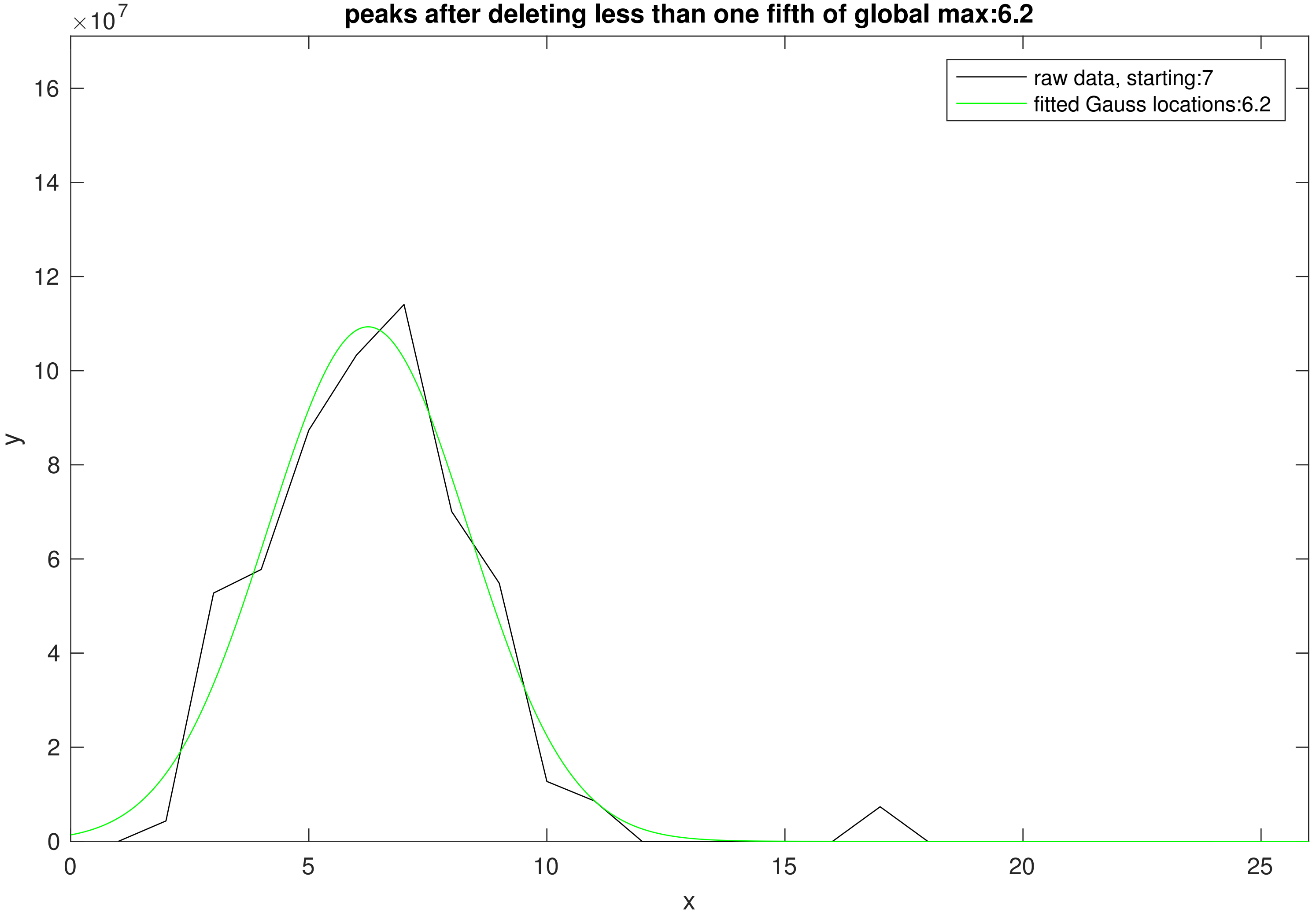

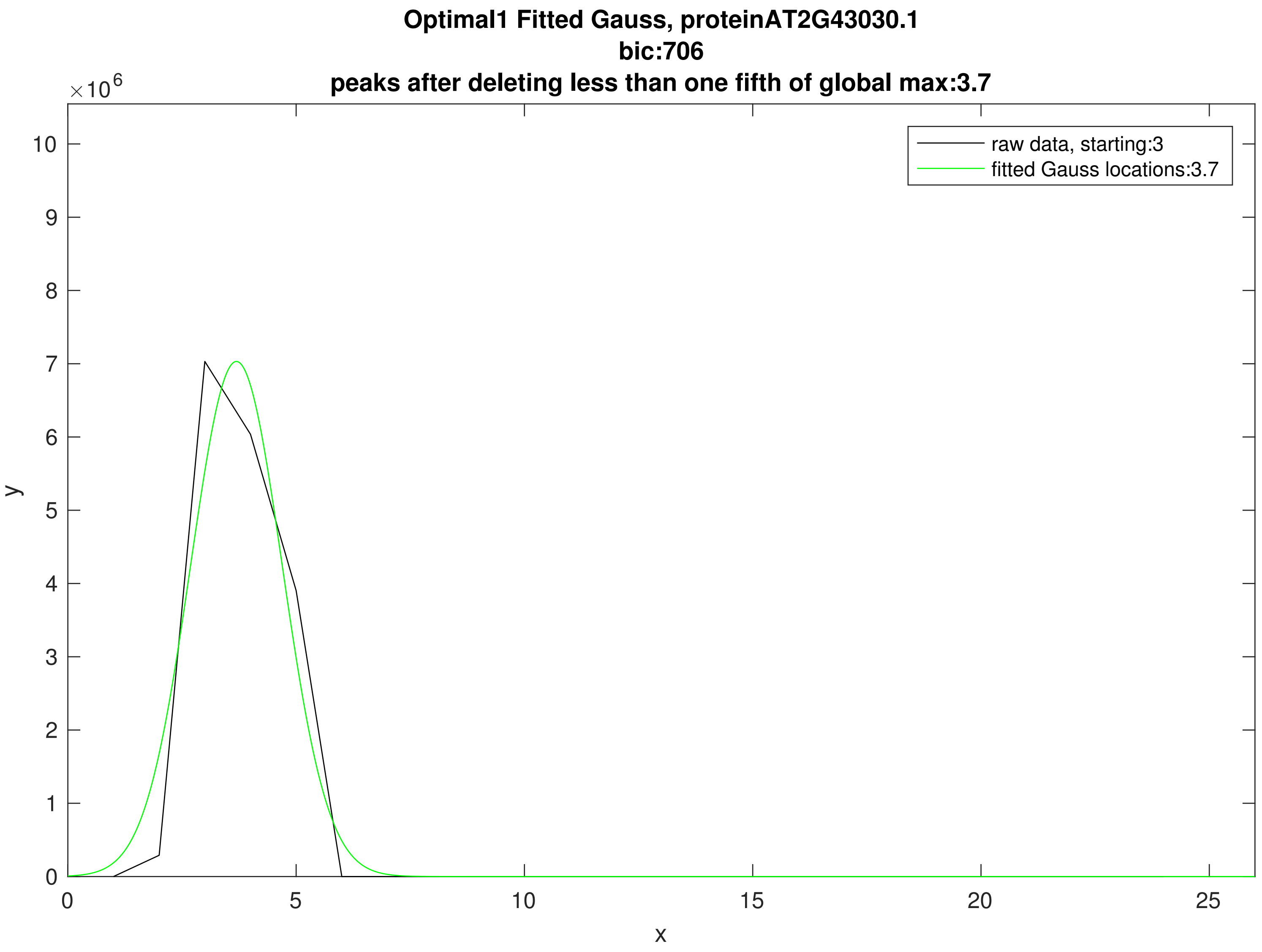
Optimal1 Fitted Gauss, proteinAT2G43030.1
bic:706
peaks after deleting less than one fifth of global max:3.7
×10^6
raw data, starting:3
fitted Gauss locations:3.7
y
x
0
1
2
3
4
5
6
7
8
9
10
0
5
10
15
20
25

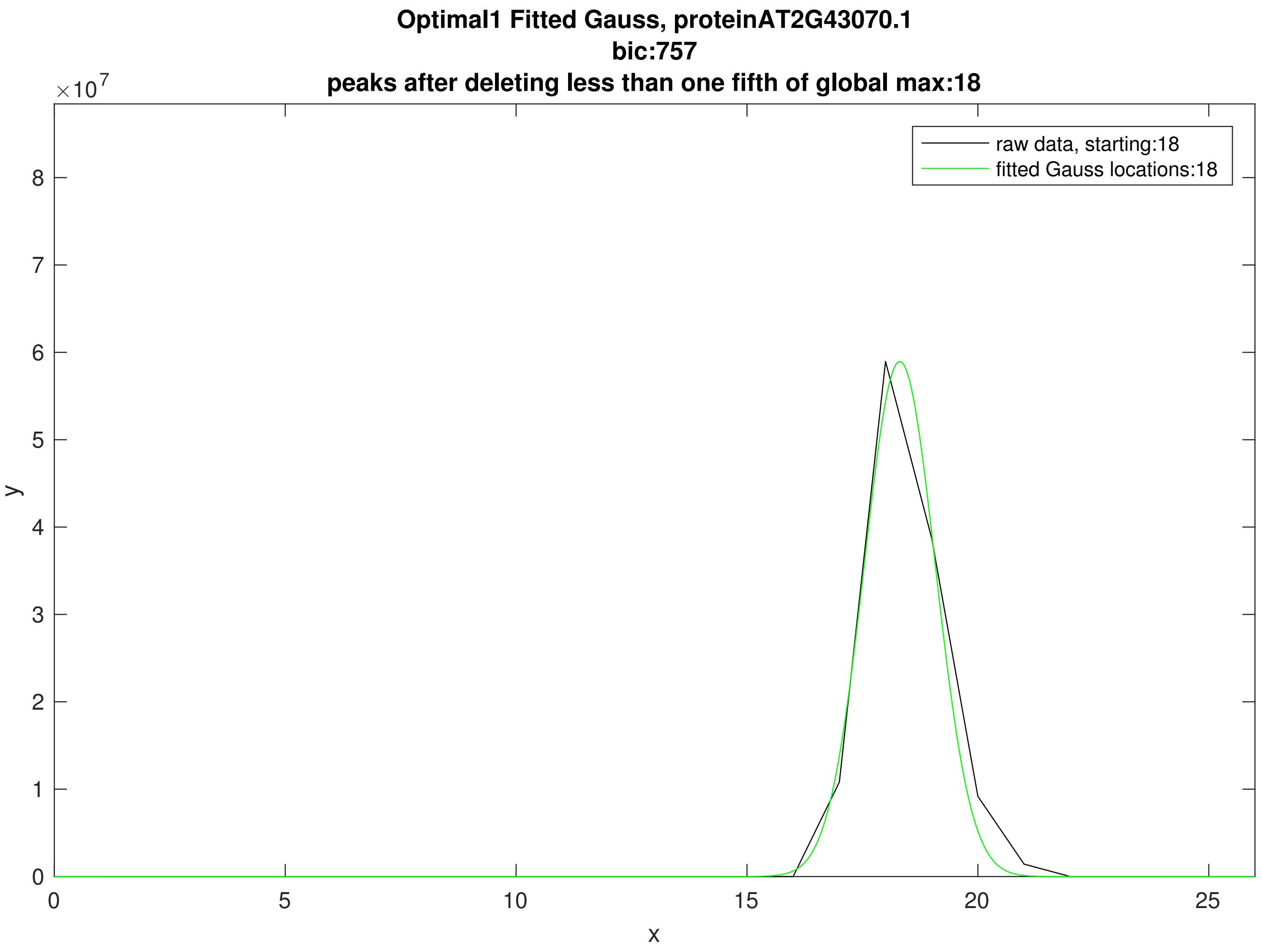
Optimal1 Fitted Gauss, proteinAT2G43070.1
bic:757
peaks after deleting less than one fifth of global max:18
×10^7
raw data, starting:18
fitted Gauss locations:18
y
x
0
1
2
3
4
5
6
7
8
0
5
10
15
20
25

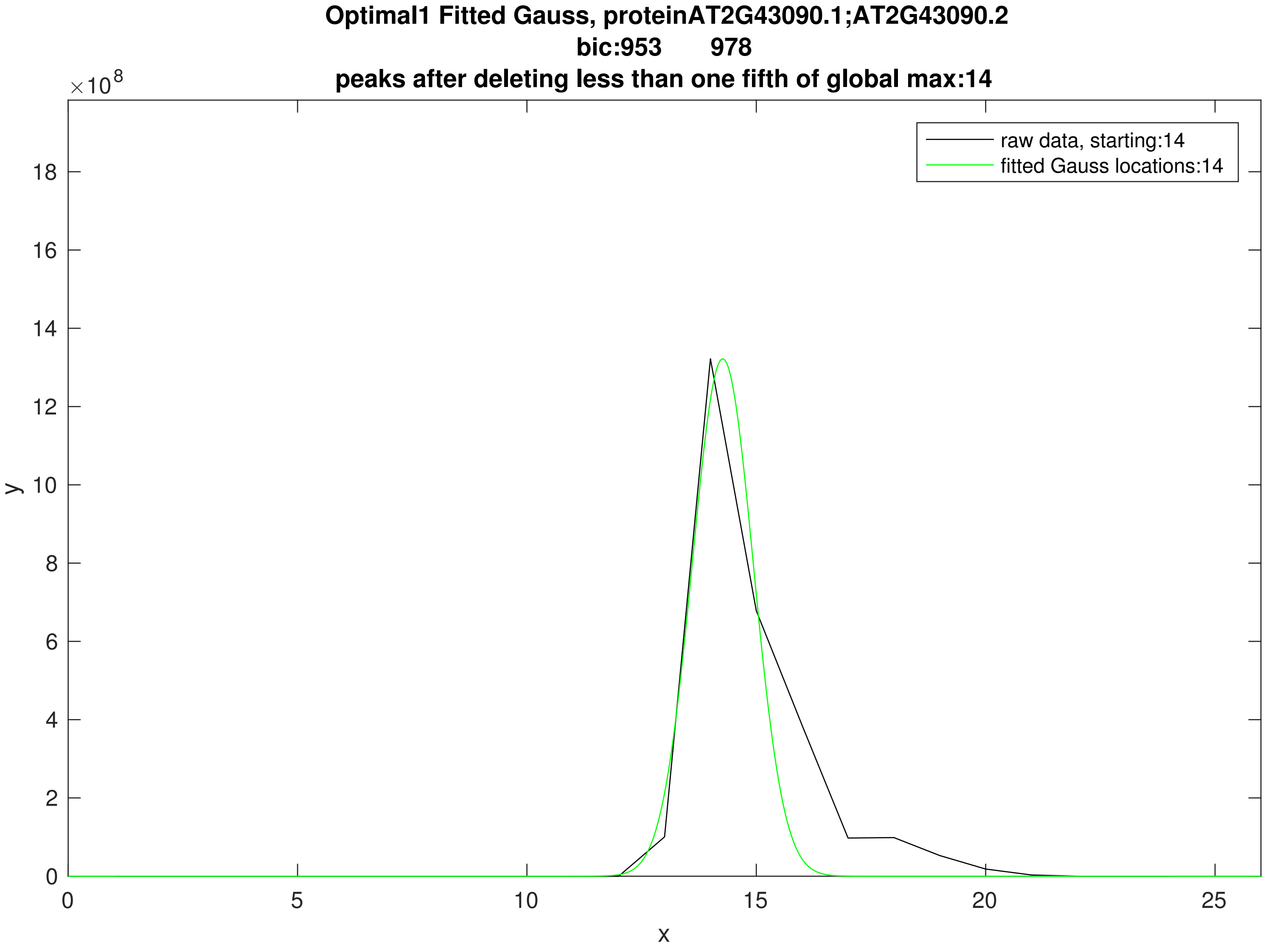
Optimal1 Fitted Gauss, proteinAT2G43090.1;AT2G43090.2
bic:953 978
peaks after deleting less than one fifth of global max:14
×10^8
raw data, starting:14
fitted Gauss locations:14
y
x
0
2
4
6
8
10
12
14
16
18
0
5
10
15
20
25

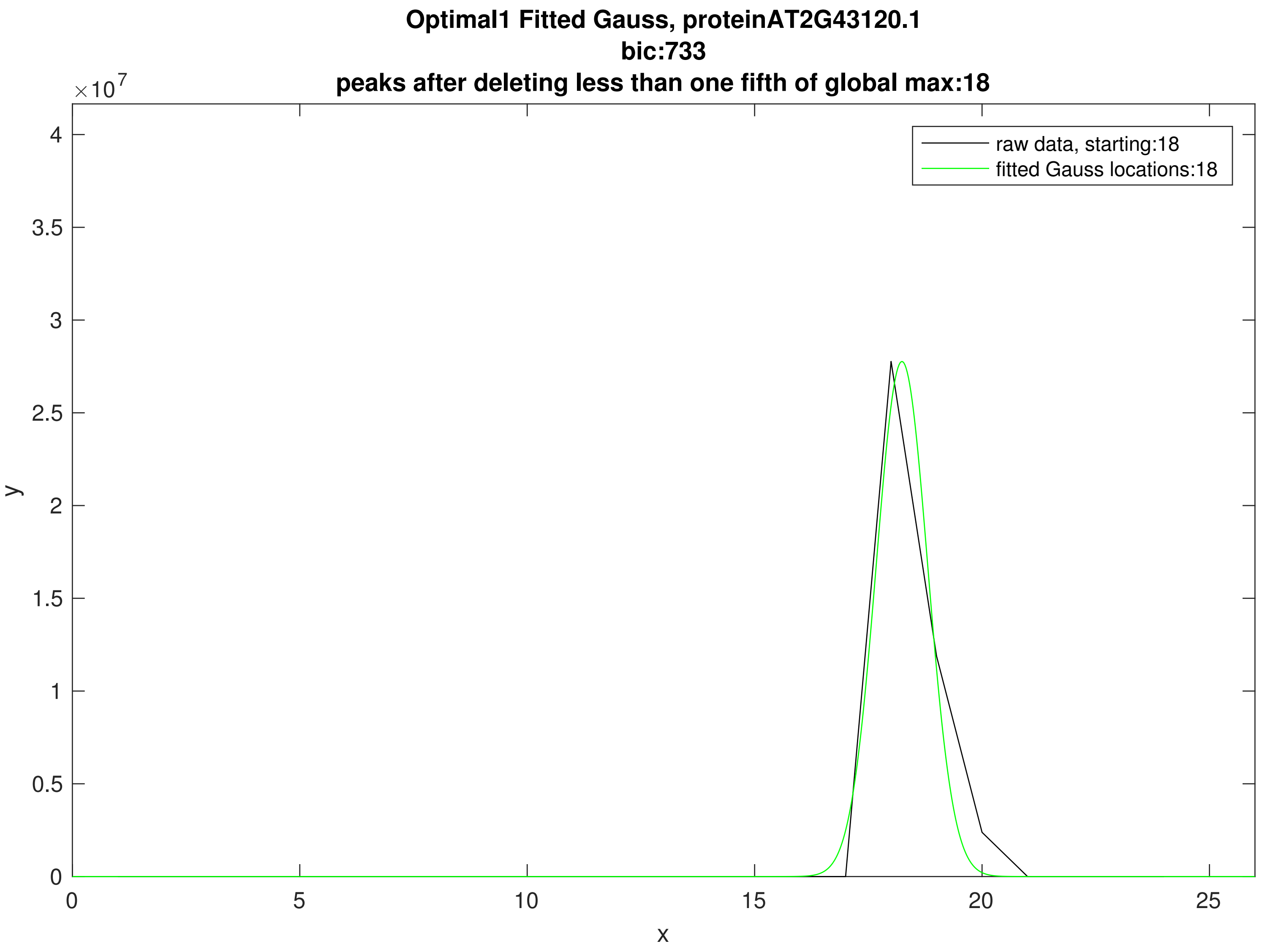

Optimal1 Fitted Gauss, proteinAT2G43120.1
bic:733
peaks after deleting less than one fifth of global max:18
×10^7
raw data, starting:18
fitted Gauss locations:18
y
x
0
0.5
1
1.5
2
2.5
3
3.5
4
0
5
10
15
20
25

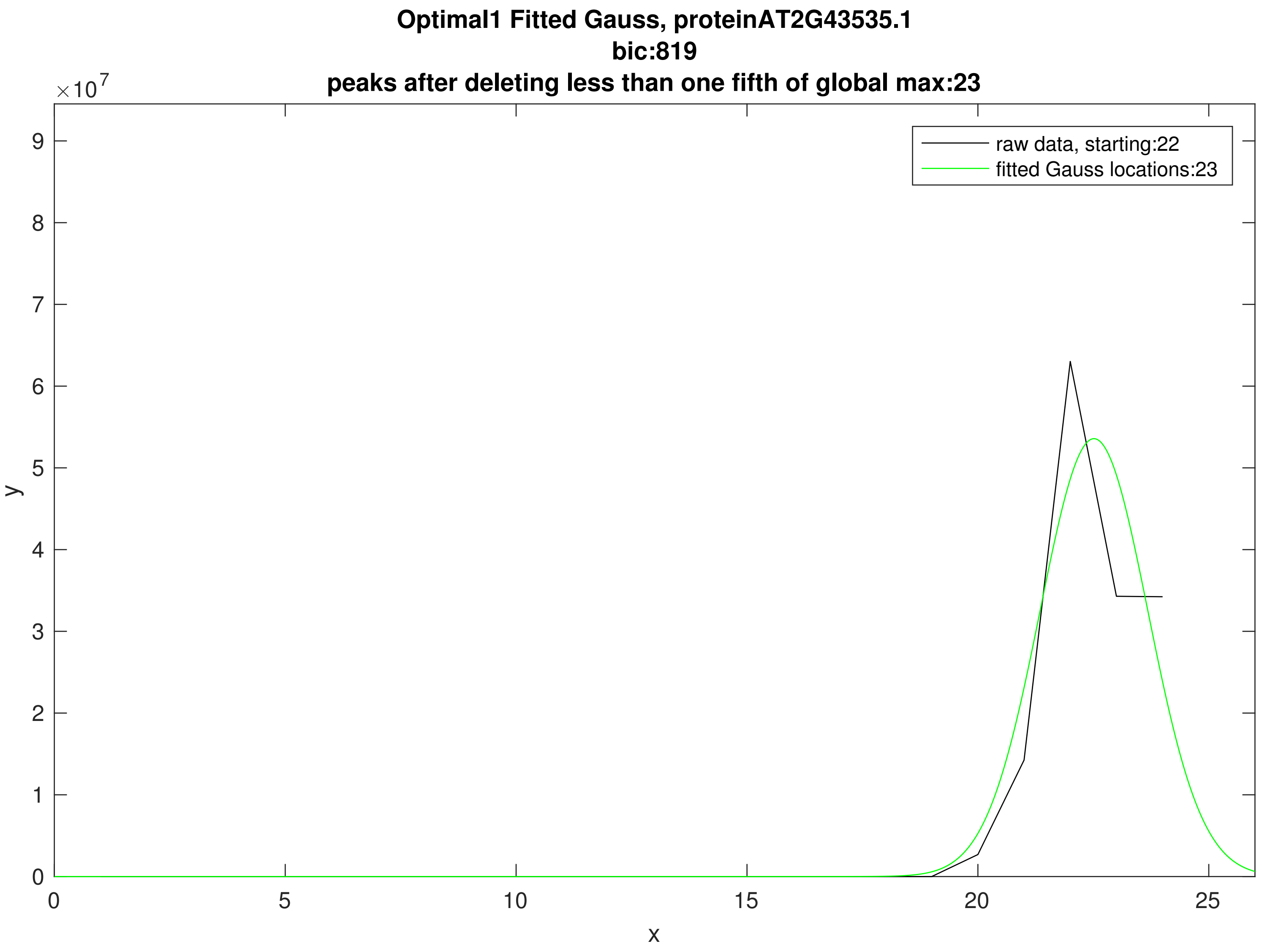

Optimal1 Fitted Gauss, proteinAT2G43535.1
bic:819
peaks after deleting less than one fifth of global max:23
×10^7
raw data, starting:22
fitted Gauss locations:23
y
x
0
1
2
3
4
5
6
7
8
9
0
5
10
15
20
25

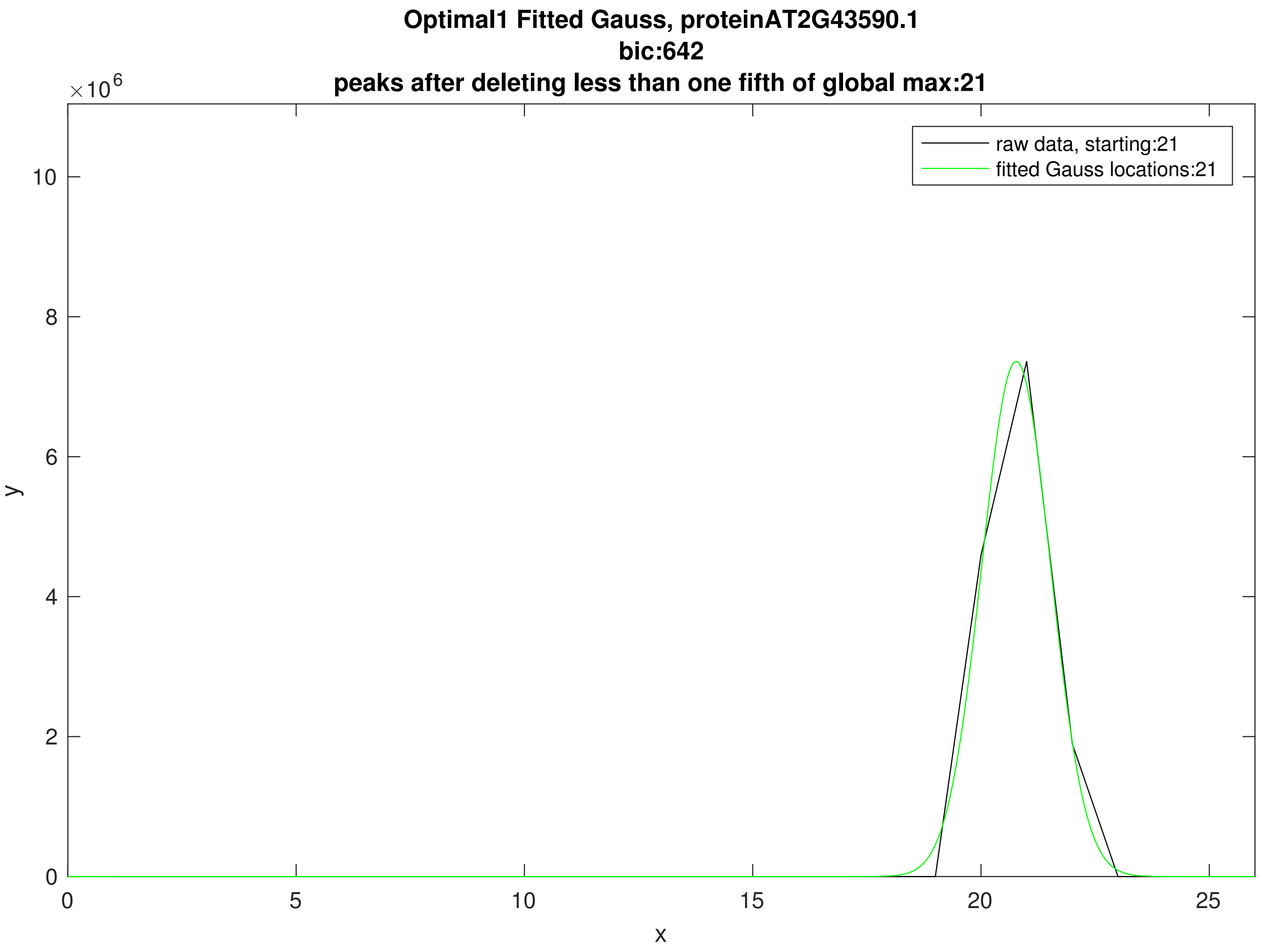

Optimal1 Fitted Gauss, proteinAT2G43590.1
bic:642
peaks after deleting less than one fifth of global max:21
×10^6
raw data, starting:21
fitted Gauss locations:21
y
x
0
2
4
6
8
10
0
5
10
15
20
25

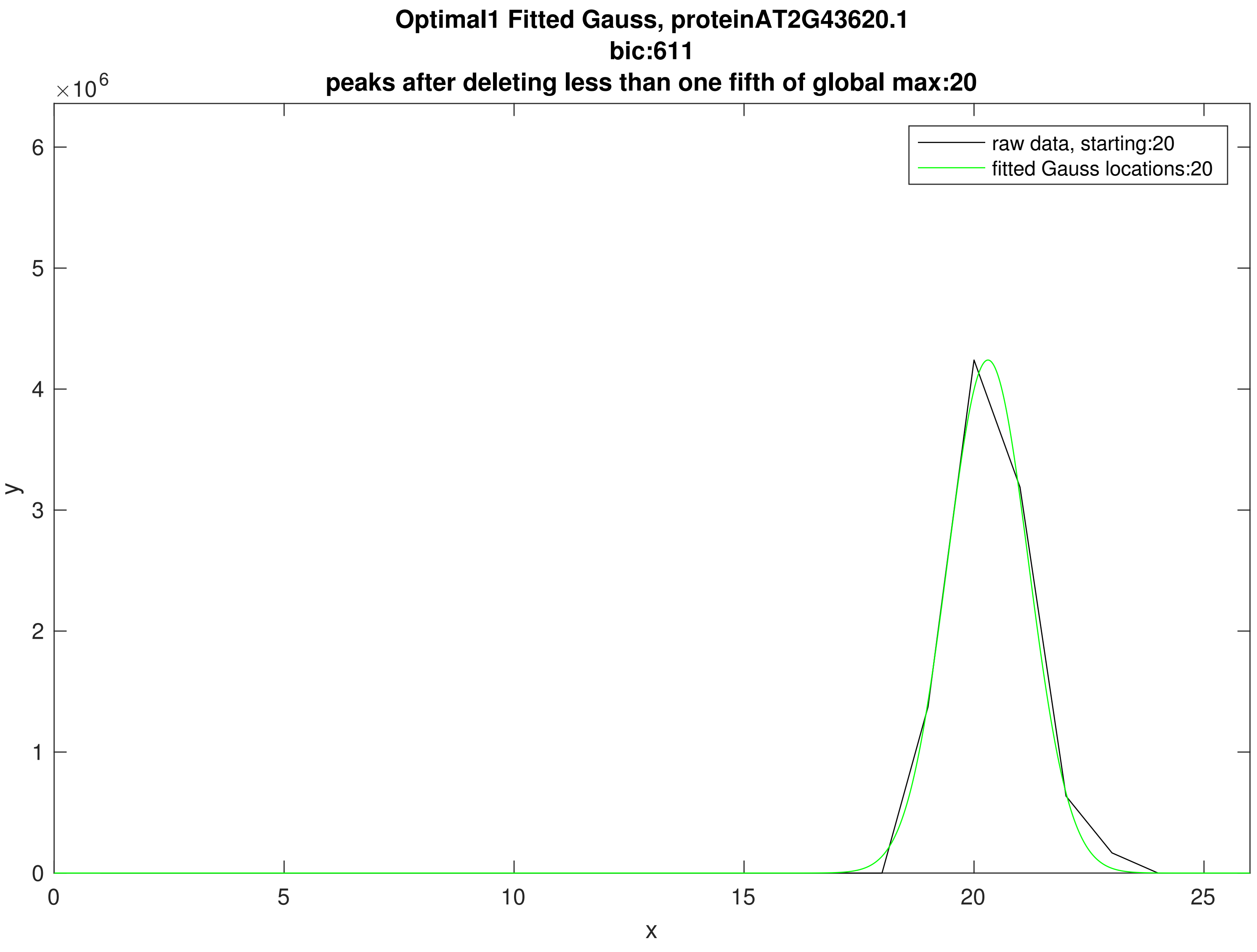

Optimal1 Fitted Gauss, proteinAT2G43620.1
bic:611
peaks after deleting less than one fifth of global max:20
×10^6
raw data, starting:20
fitted Gauss locations:20
y
x
0
1
2
3
4
5
6
0
5
10
15
20
25

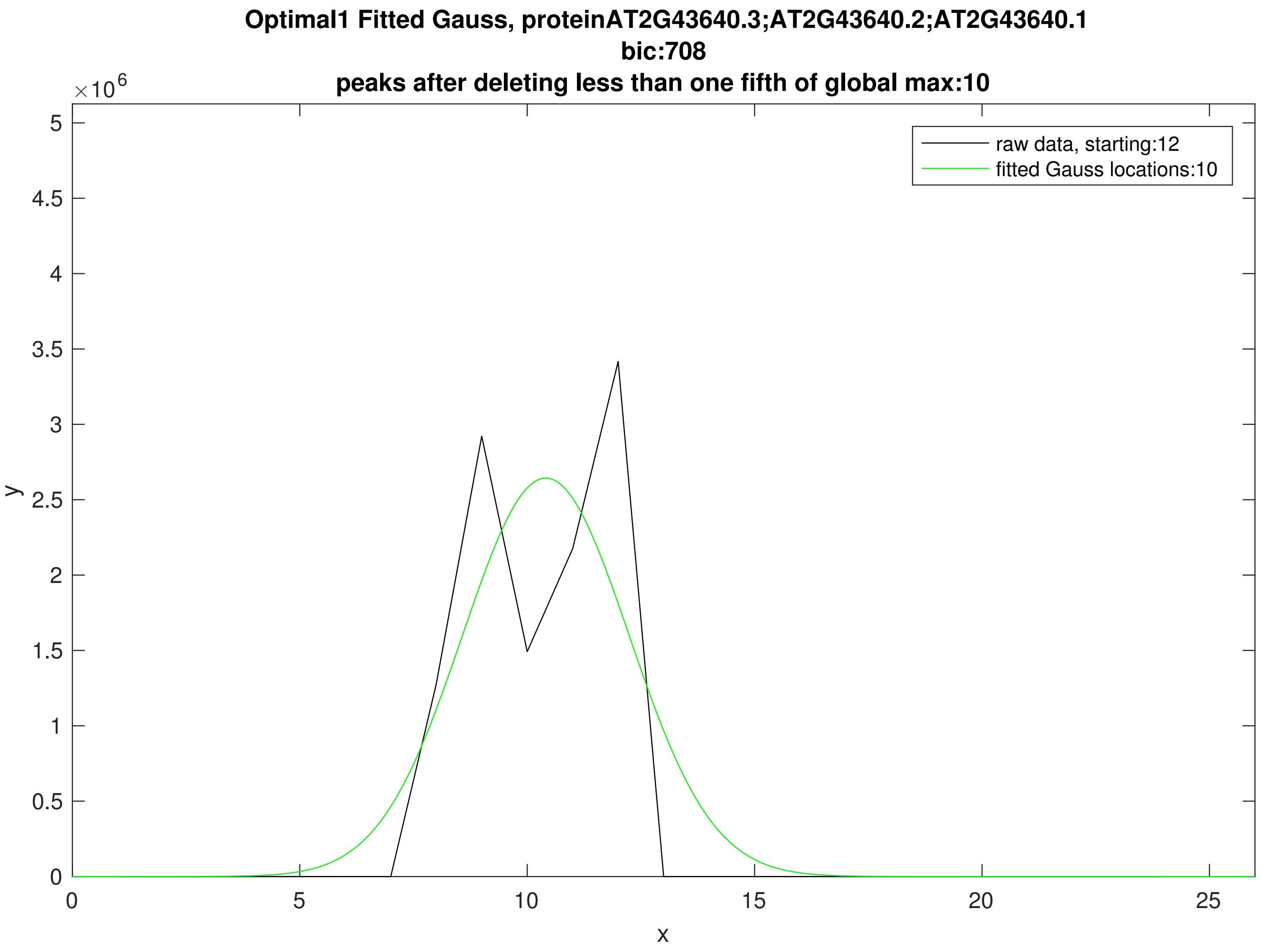
Optimal1 Fitted Gauss, proteinAT2G43640.3;AT2G43640.2;AT2G43640.1
bic:708
peaks after deleting less than one fifth of global max:10
×10^6
raw data, starting:12
fitted Gauss locations:10
y
x
0
0.5
1
1.5
2
2.5
3
3.5
4
4.5
5
0
5
10
15
20
25

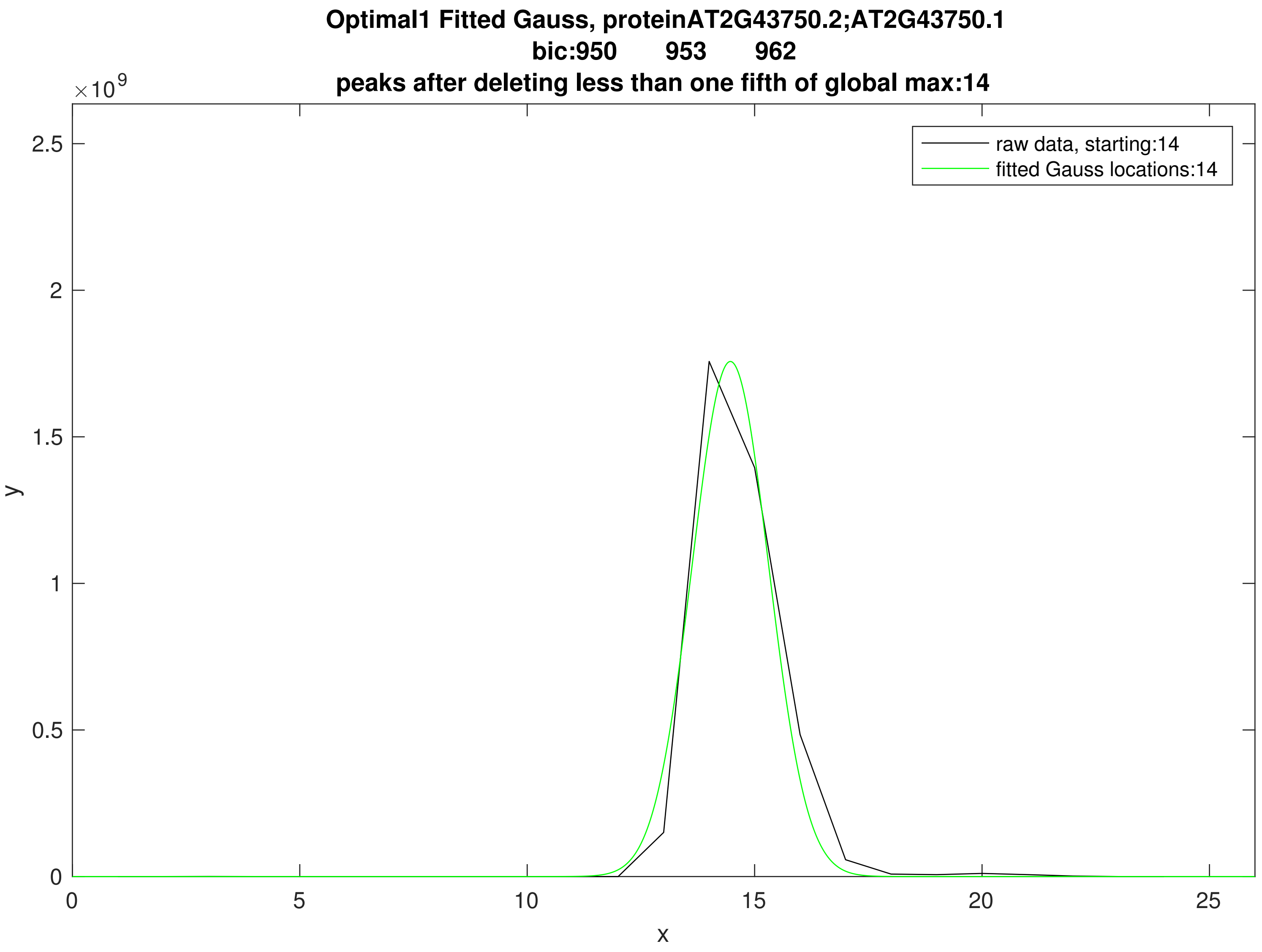

Optimal1 Fitted Gauss, proteinAT2G43750.2;AT2G43750.1
bic:950 953 962
peaks after deleting less than one fifth of global max:14
×10^9
2.5
2
1.5
1
0.5
0
y
0
5
10
15
20
25
x
raw data, starting:14
fitted Gauss locations:14

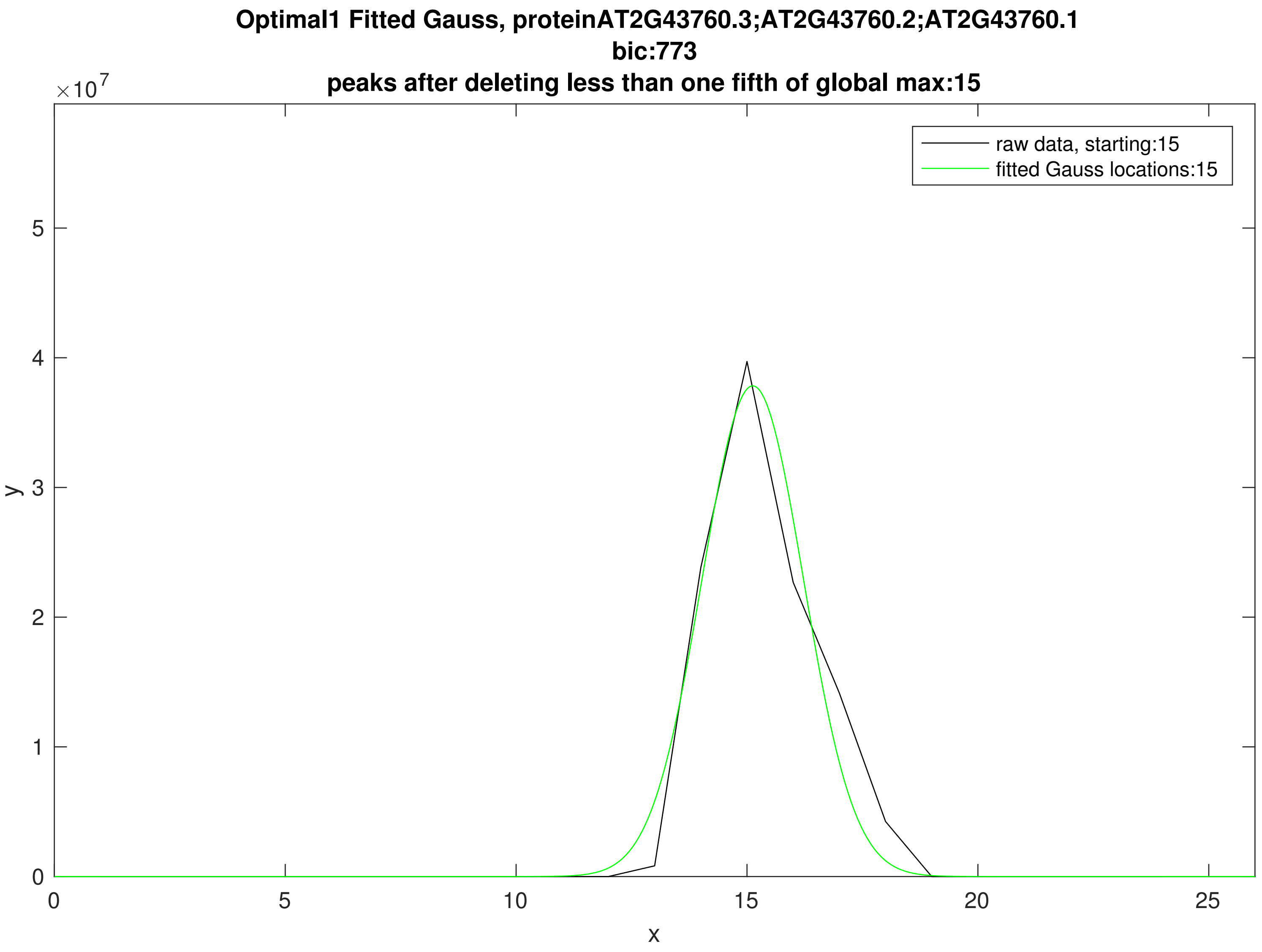

Optimal1 Fitted Gauss, proteinAT2G43760.3;AT2G43760.2;AT2G43760.1
bic:773
peaks after deleting less than one fifth of global max:15
×10^7
raw data, starting:15
fitted Gauss locations:15
y
x
0
1
2
3
4
5
0
5
10
15
20
25

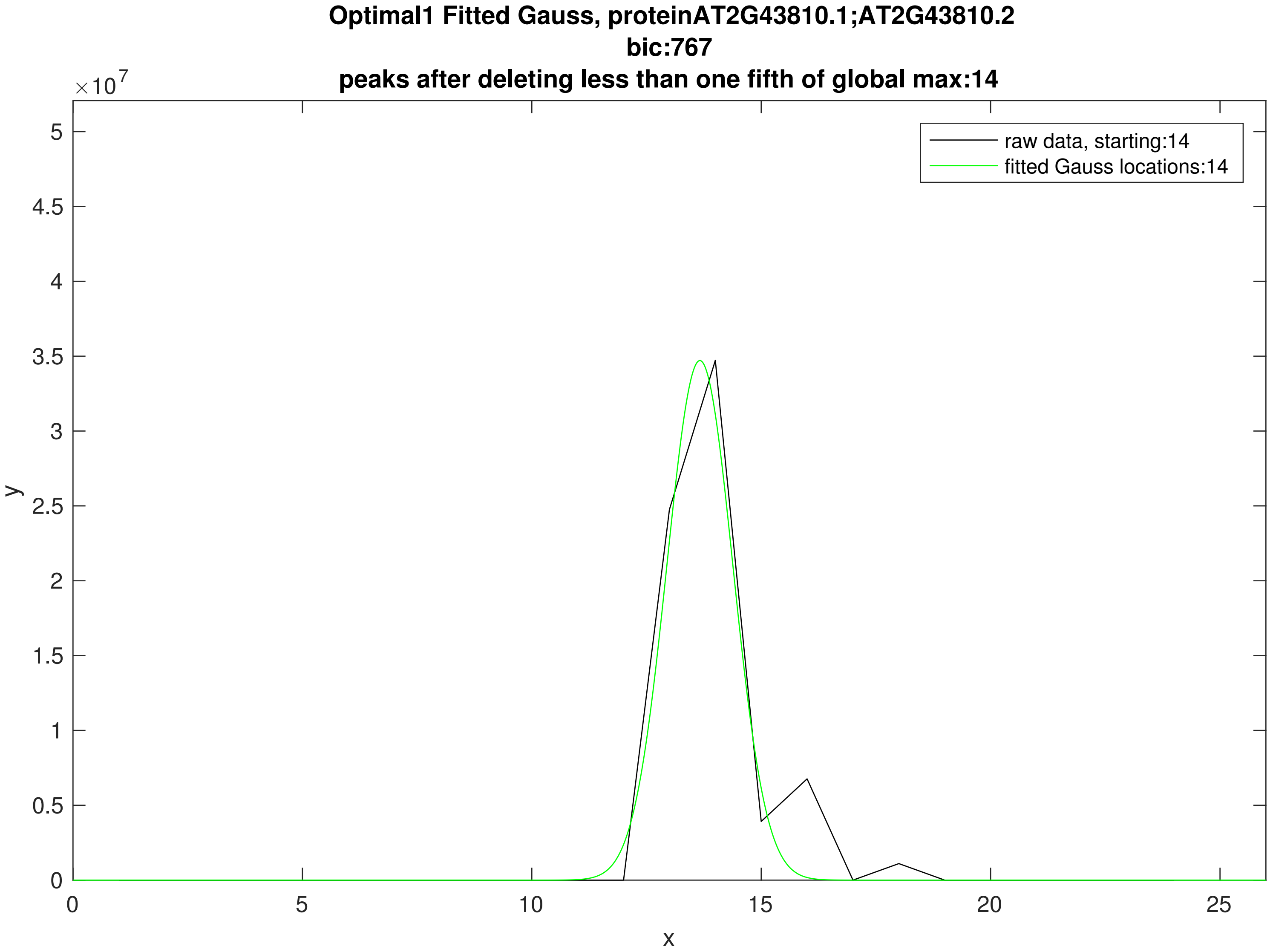
Optimal1 Fitted Gauss, proteinAT2G43810.1;AT2G43810.2
bic:767
peaks after deleting less than one fifth of global max:14
×10^7
raw data, starting:14
fitted Gauss locations:14
y
x
0
0.5
1
1.5
2
2.5
3
3.5
4
4.5
5
0
5
10
15
20
25

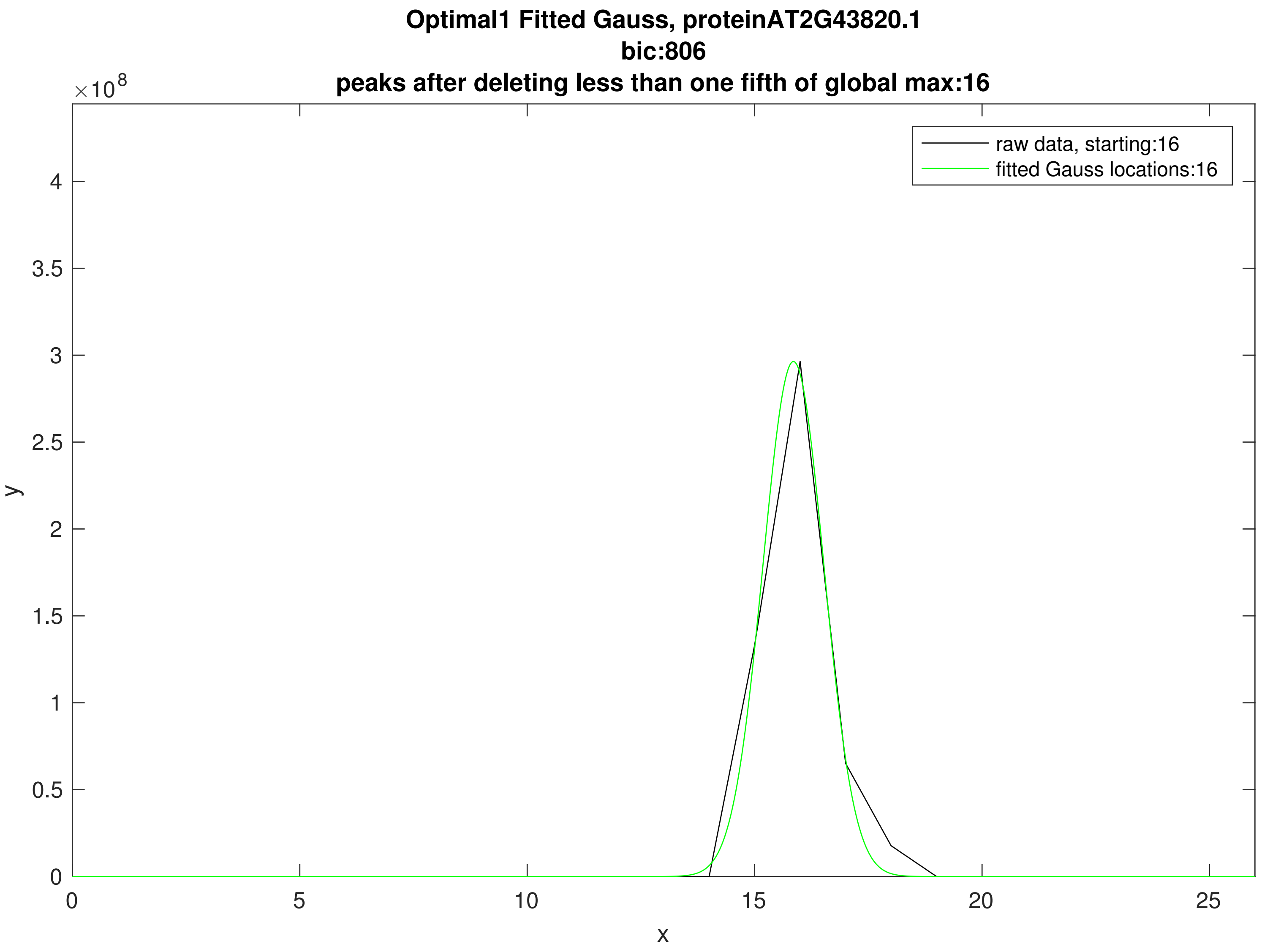
Optimal1 Fitted Gauss, proteinAT2G43820.1
bic:806
peaks after deleting less than one fifth of global max:16
×10^8
raw data, starting:16
fitted Gauss locations:16
4
3.5
3
2.5
2
1.5
1
0.5
0
y
0
5
10
15
20
25
x

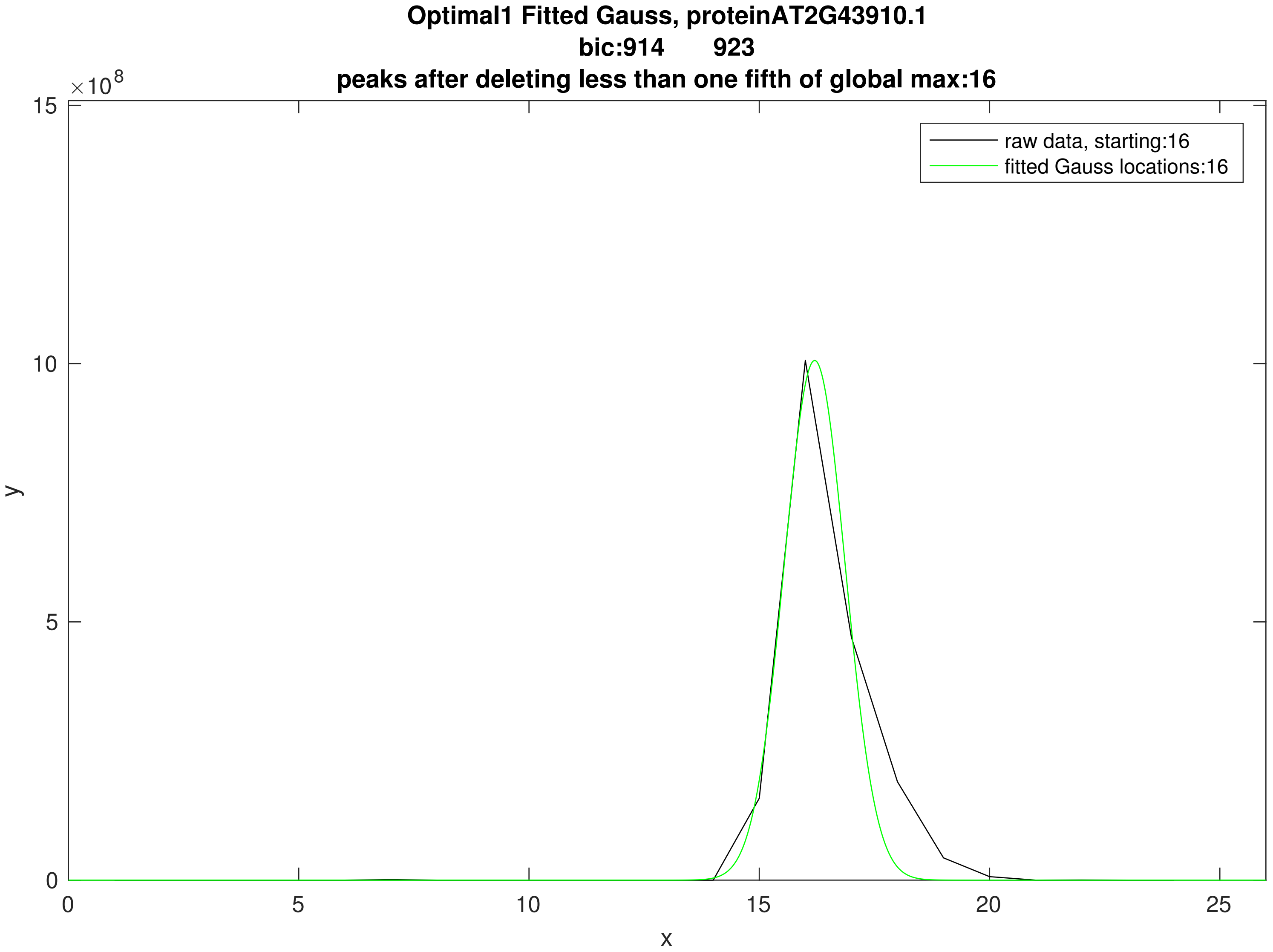
Optimal1 Fitted Gauss, proteinAT2G43910.1
bic:914 923
peaks after deleting less than one fifth of global max:16
×10^8
15
10
5
0
y
0
5
10
15
20
25
x
raw data, starting:16
fitted Gauss locations:16

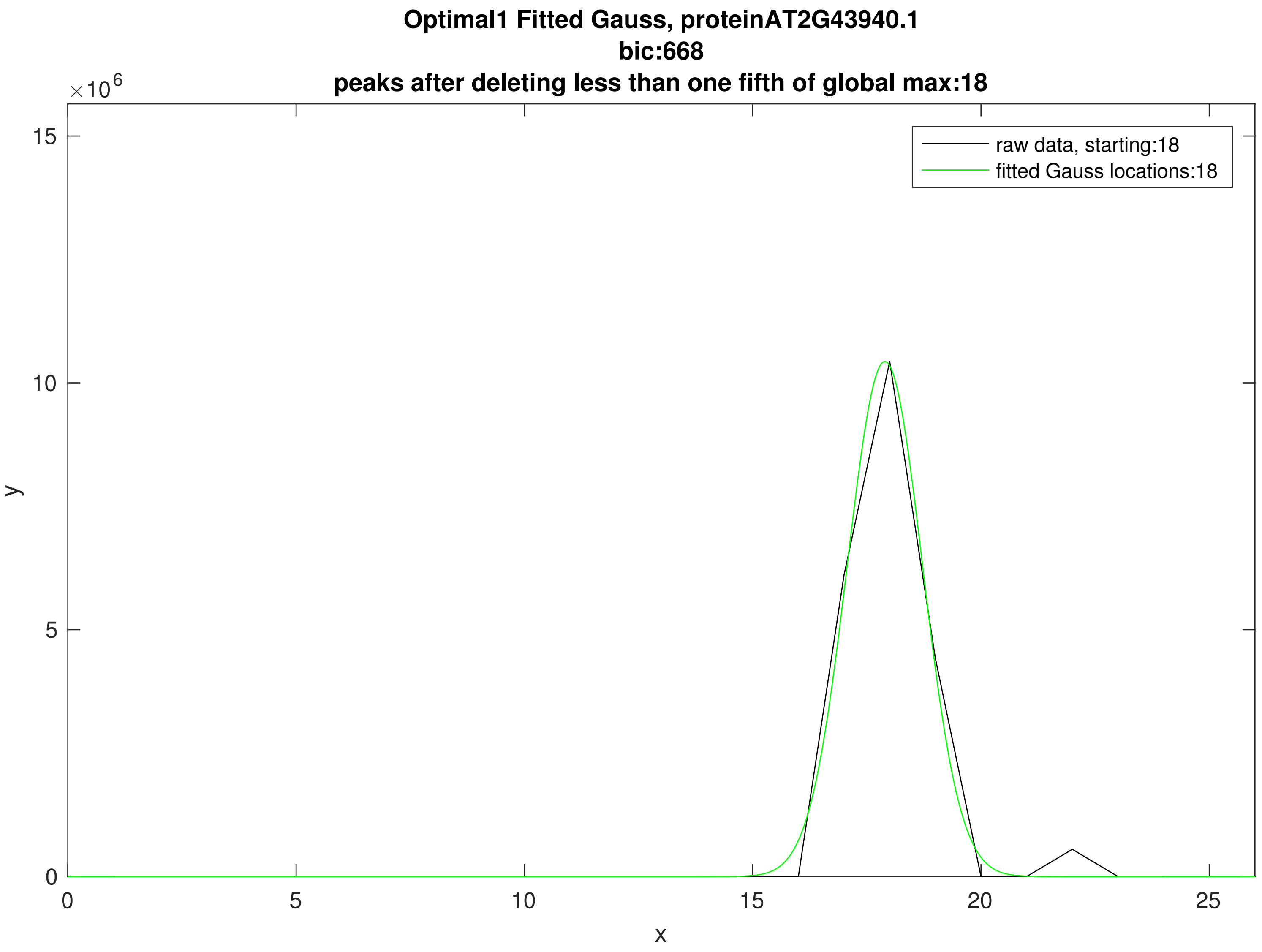

Optimal1 Fitted Gauss, proteinAT2G43940.1
bic:668
peaks after deleting less than one fifth of global max:18
×10^6
15
10
5
0
y
0
5
10
15
20
25
x
raw data, starting:18
fitted Gauss locations:18

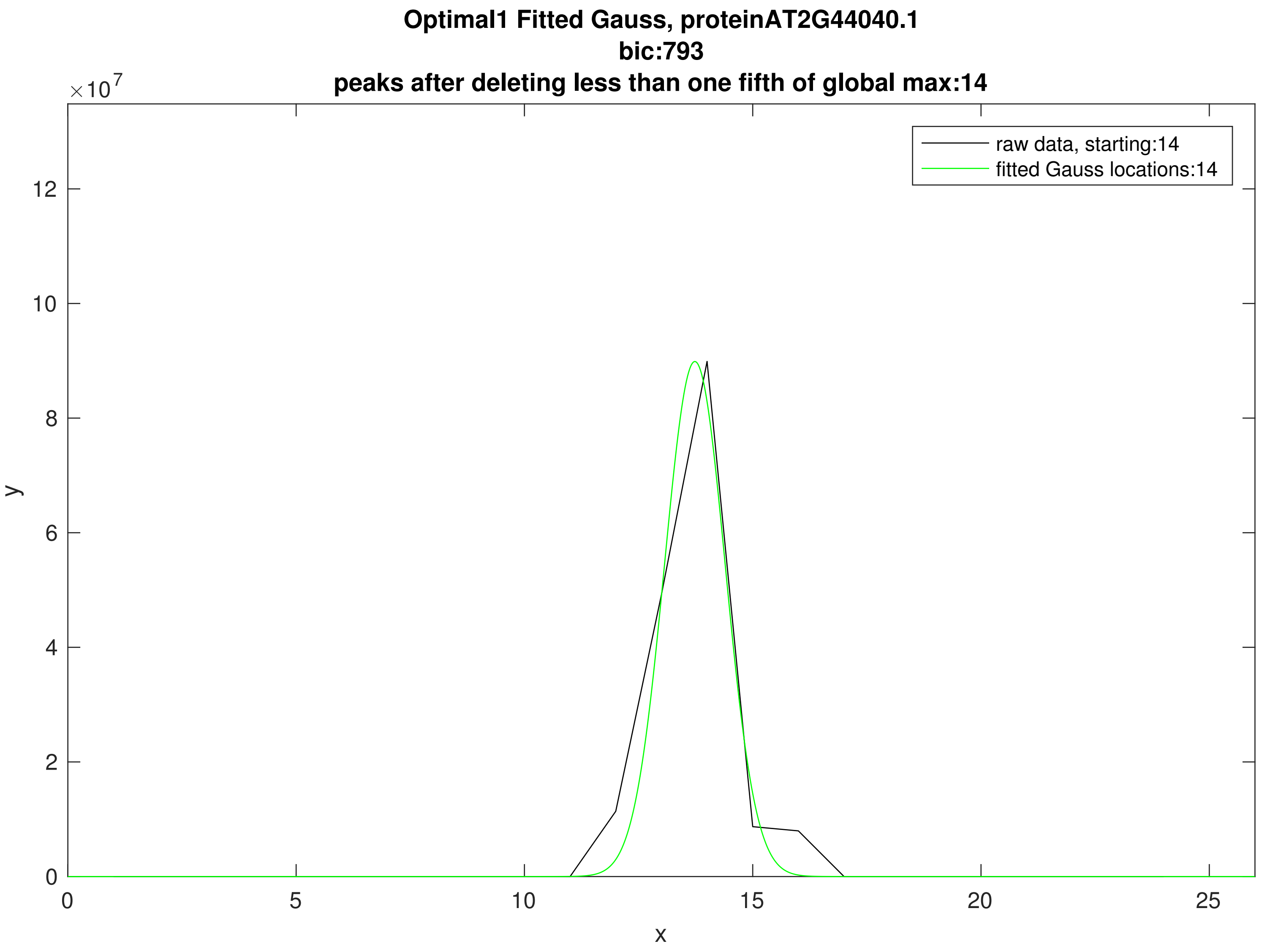

Optimal1 Fitted Gauss, proteinAT2G44040.1
bic:793
peaks after deleting less than one fifth of global max:14
×10^7
raw data, starting:14
fitted Gauss locations:14
y
x
0
2
4
6
8
10
12
0
5
10
15
20
25

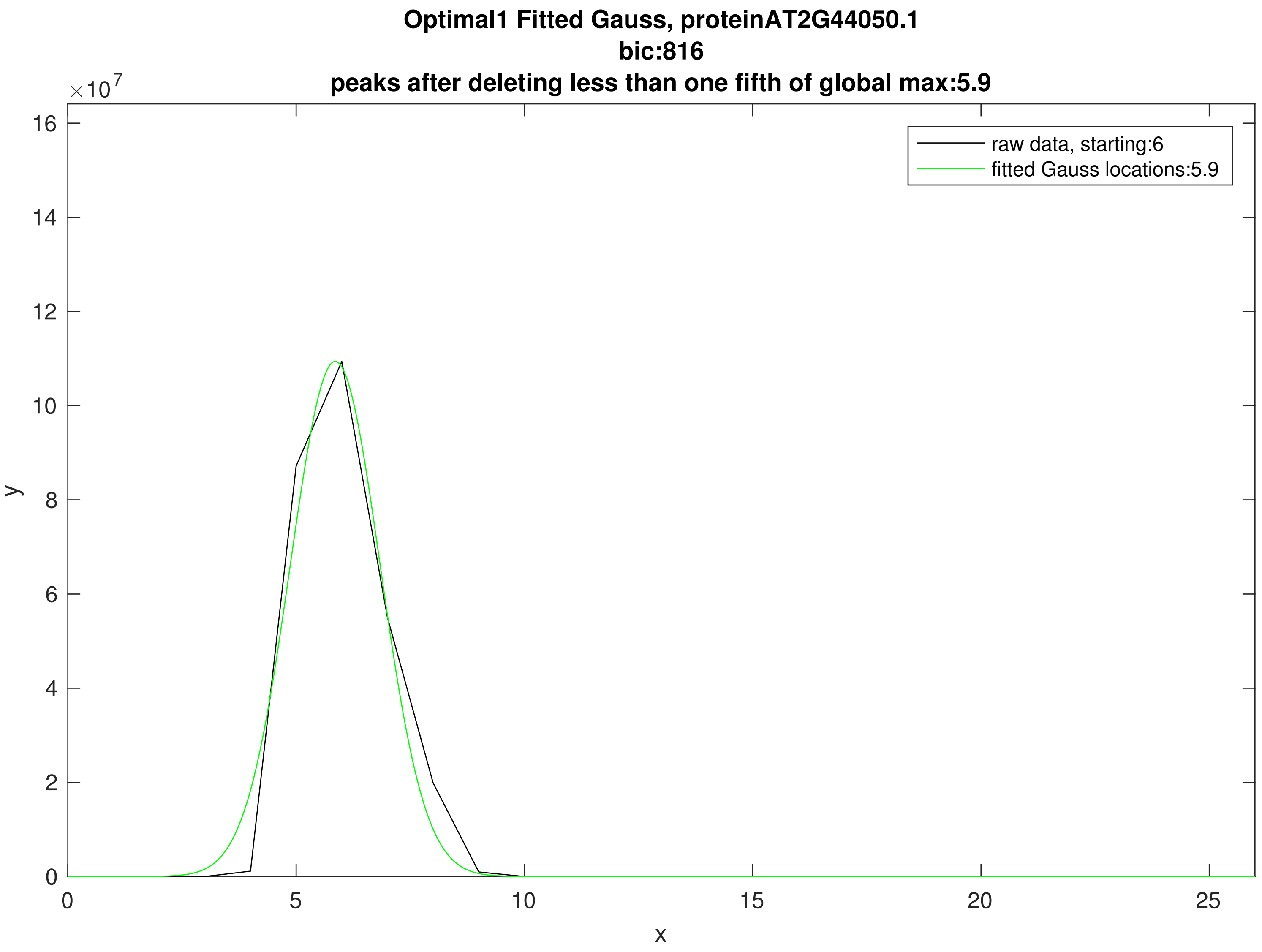
Optimal1 Fitted Gauss, proteinAT2G44050.1
bic:816
peaks after deleting less than one fifth of global max:5.9
×10^7
raw data, starting:6
fitted Gauss locations:5.9
y
x
0
2
4
6
8
10
12
14
16
0
5
10
15
20
25

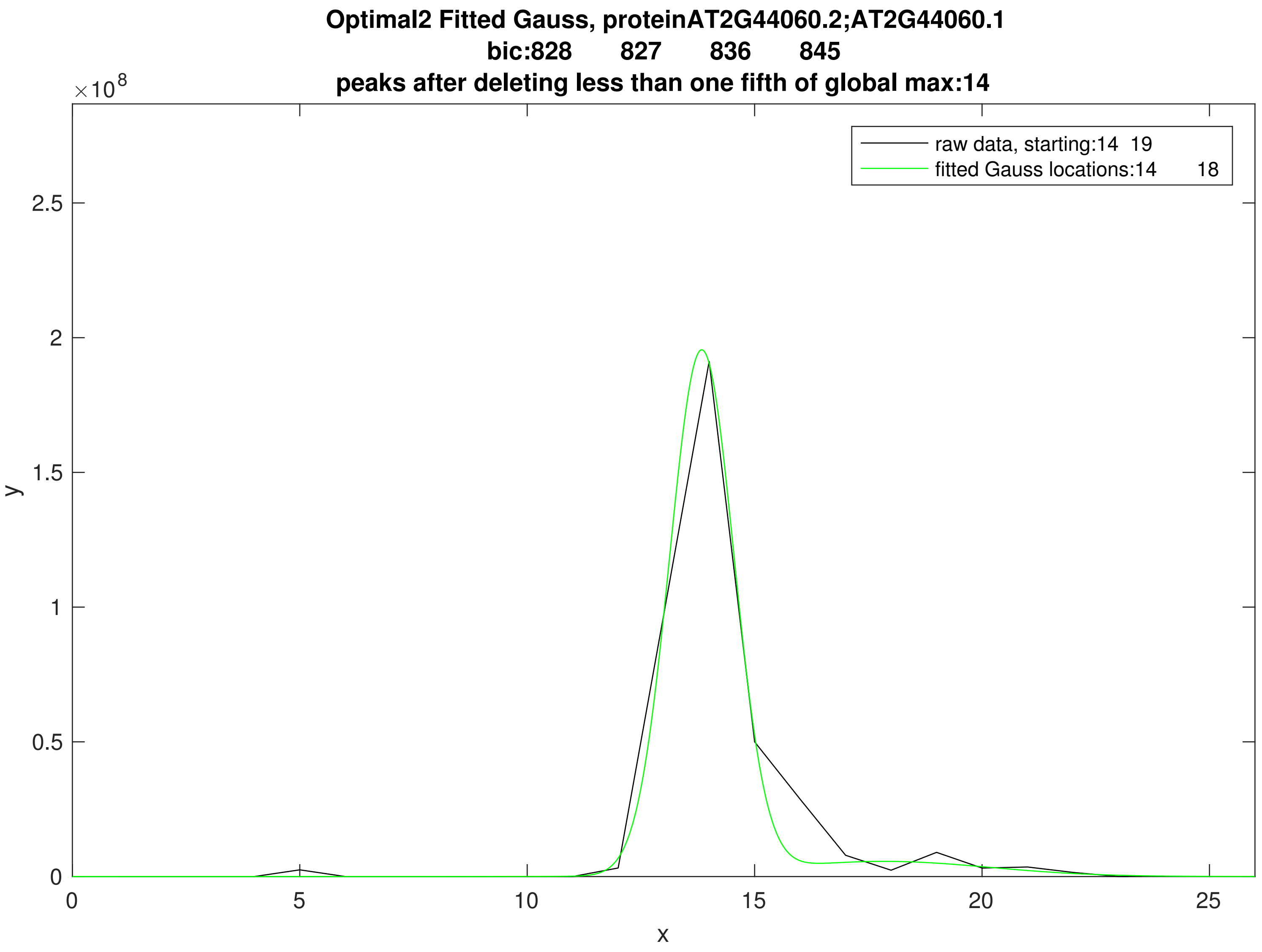
Optimal2 Fitted Gauss, proteinAT2G44060.2;AT2G44060.1
bic:828 827 836 845
peaks after deleting less than one fifth of global max:14
×10^8
raw data, starting:14 19
fitted Gauss locations:14 18
y
x
0
0.5
1
1.5
2
2.5
0
5
10
15
20
25

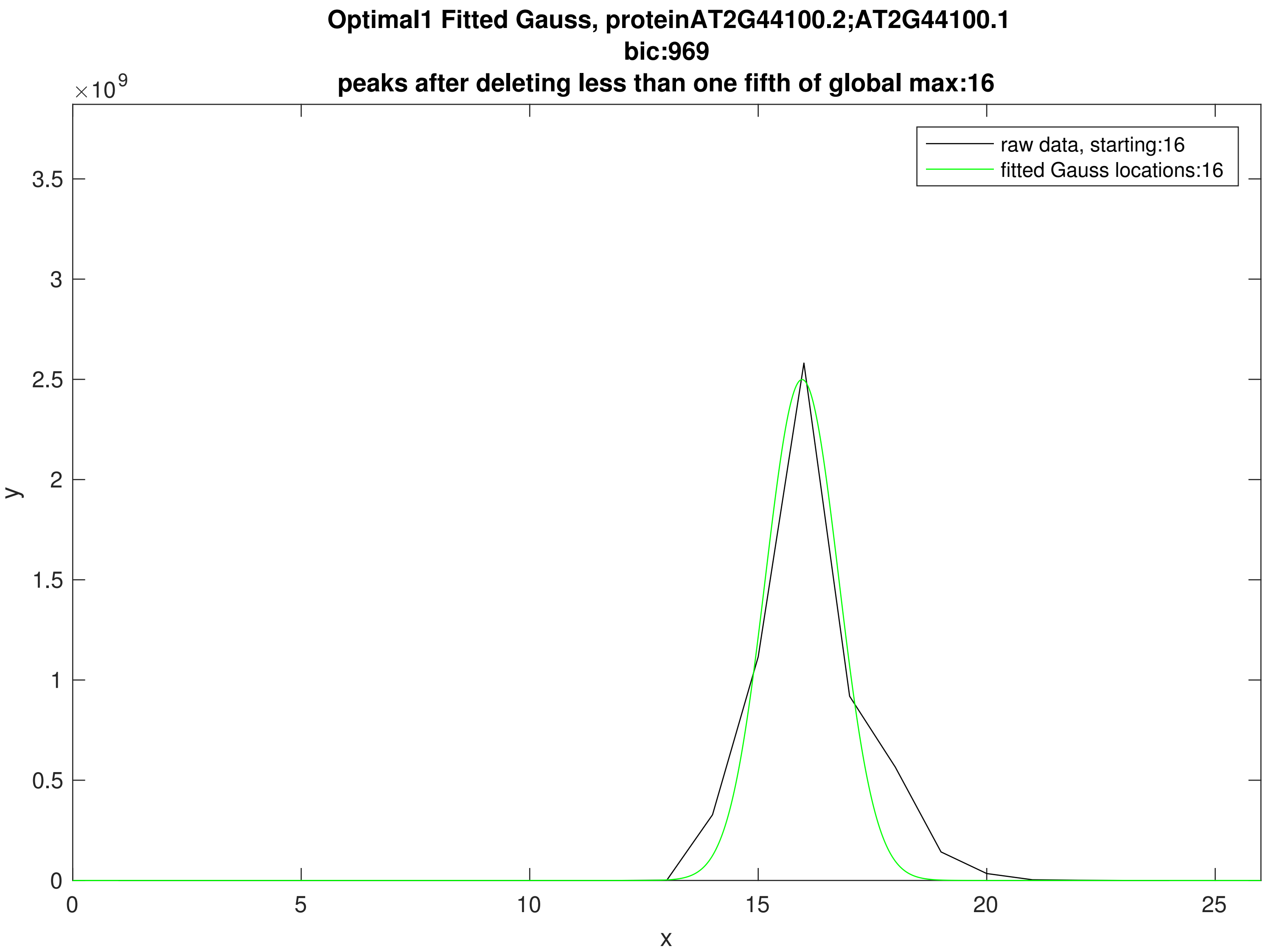

Optimal1 Fitted Gauss, proteinAT2G44100.2;AT2G44100.1
bic:969
peaks after deleting less than one fifth of global max:16
raw data, starting:16
fitted Gauss locations:16
y
x

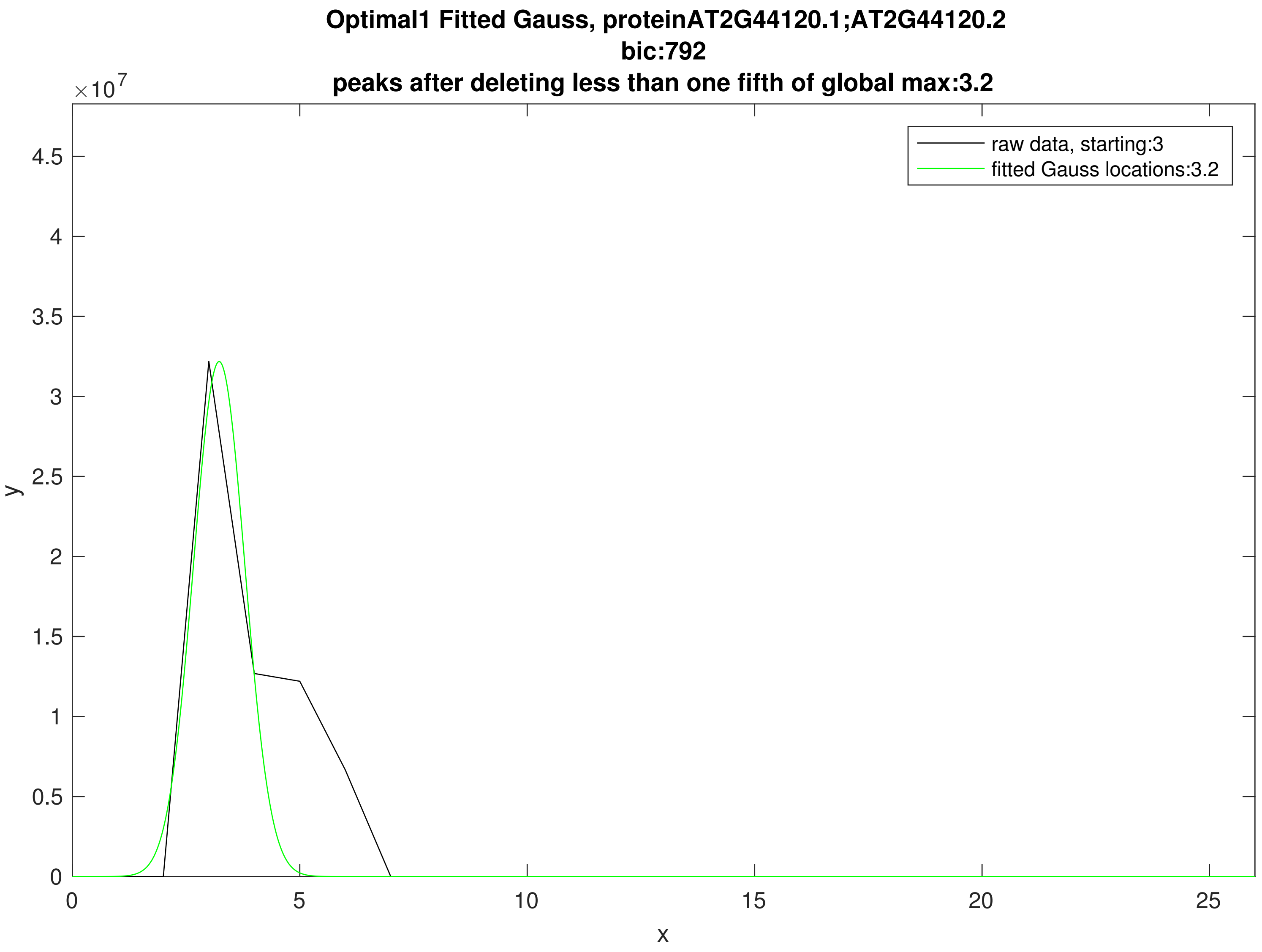

Optimal1 Fitted Gauss, proteinAT2G44120.1;AT2G44120.2
bic:792
peaks after deleting less than one fifth of global max:3.2
×10^7
4.5
4
3.5
3
2.5
2
1.5
1
0.5
0
y
0
5
10
15
20
25
x
raw data, starting:3
fitted Gauss locations:3.2

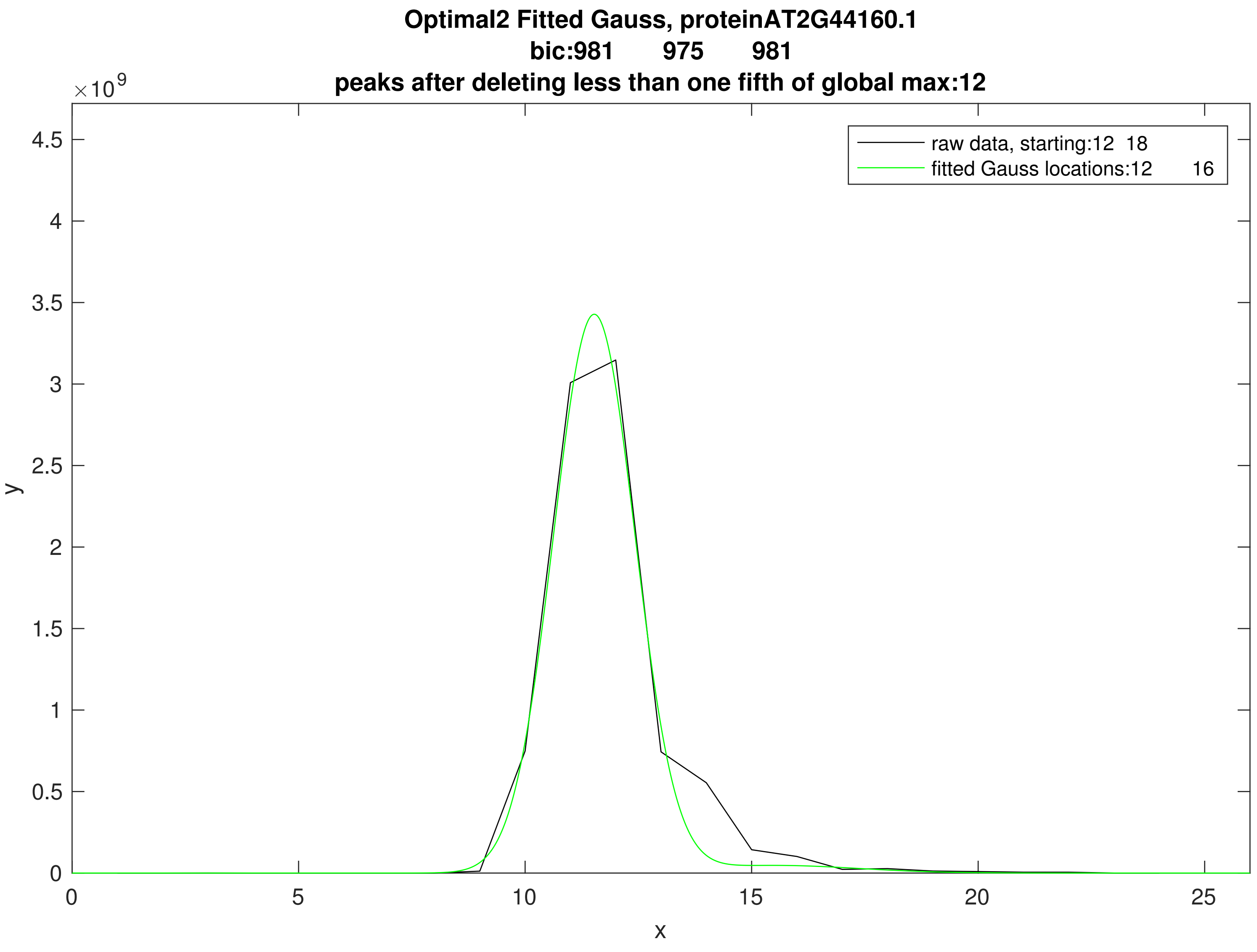
Optimal2 Fitted Gauss, proteinAT2G44160.1
bic:981 975 981
peaks after deleting less than one fifth of global max:12
raw data, starting:12 18
fitted Gauss locations:12 16
y
x

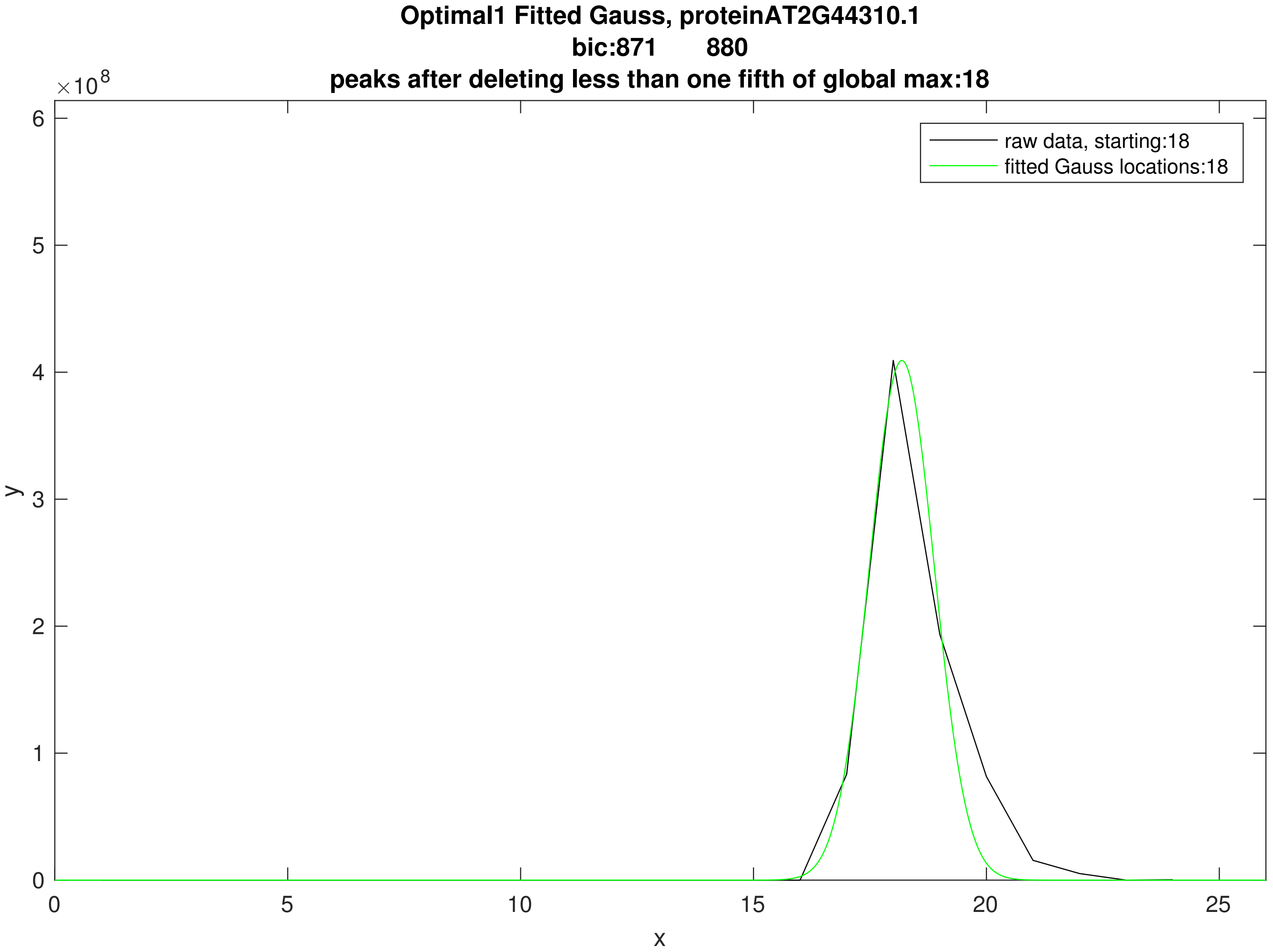
Optimal1 Fitted Gauss, proteinAT2G44310.1
bic:871 880
peaks after deleting less than one fifth of global max:18
raw data, starting:18
fitted Gauss locations:18
y
x

Optimal2 Fitted Gauss, proteinAT2G44350.1;AT2G44350.2
bic:978 978 987
peaks after deleting less than one fifth of global max:14

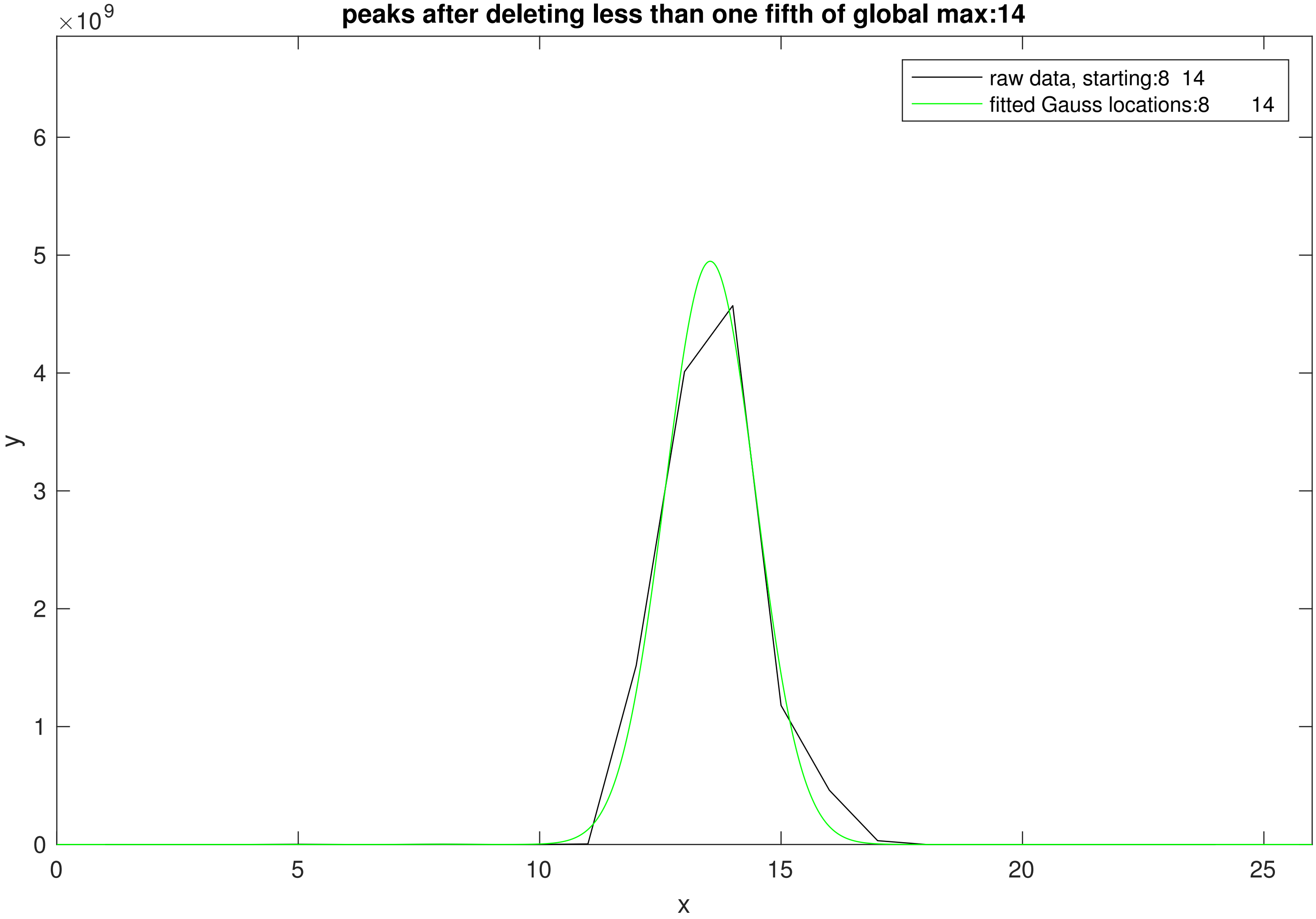

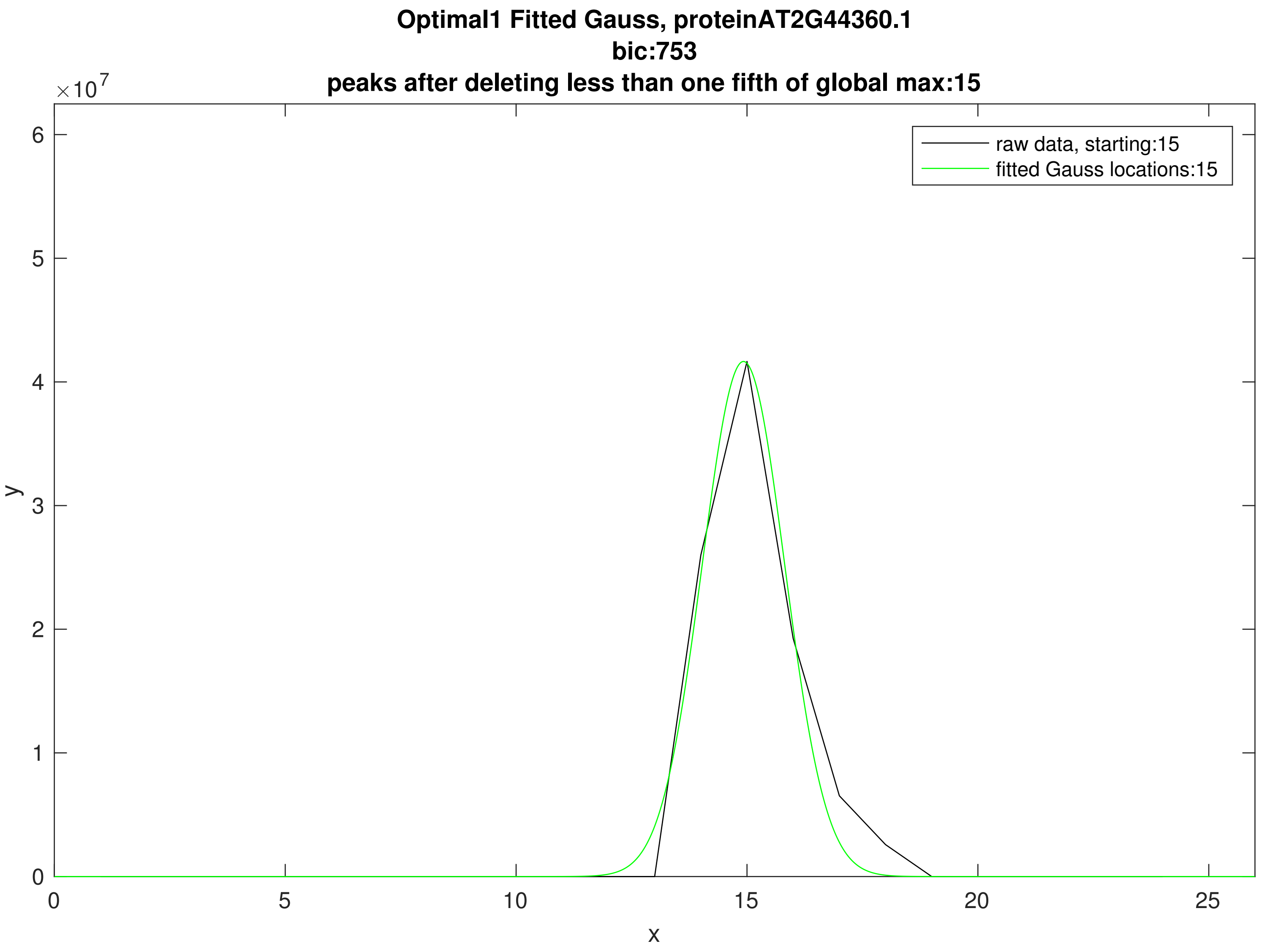
Optimal1 Fitted Gauss, proteinAT2G44360.1
bic:753
peaks after deleting less than one fifth of global max:15
×10^7
6
5
4
3
2
1
0
y
0
5
10
15
20
25
x
raw data, starting:15
fitted Gauss locations:15

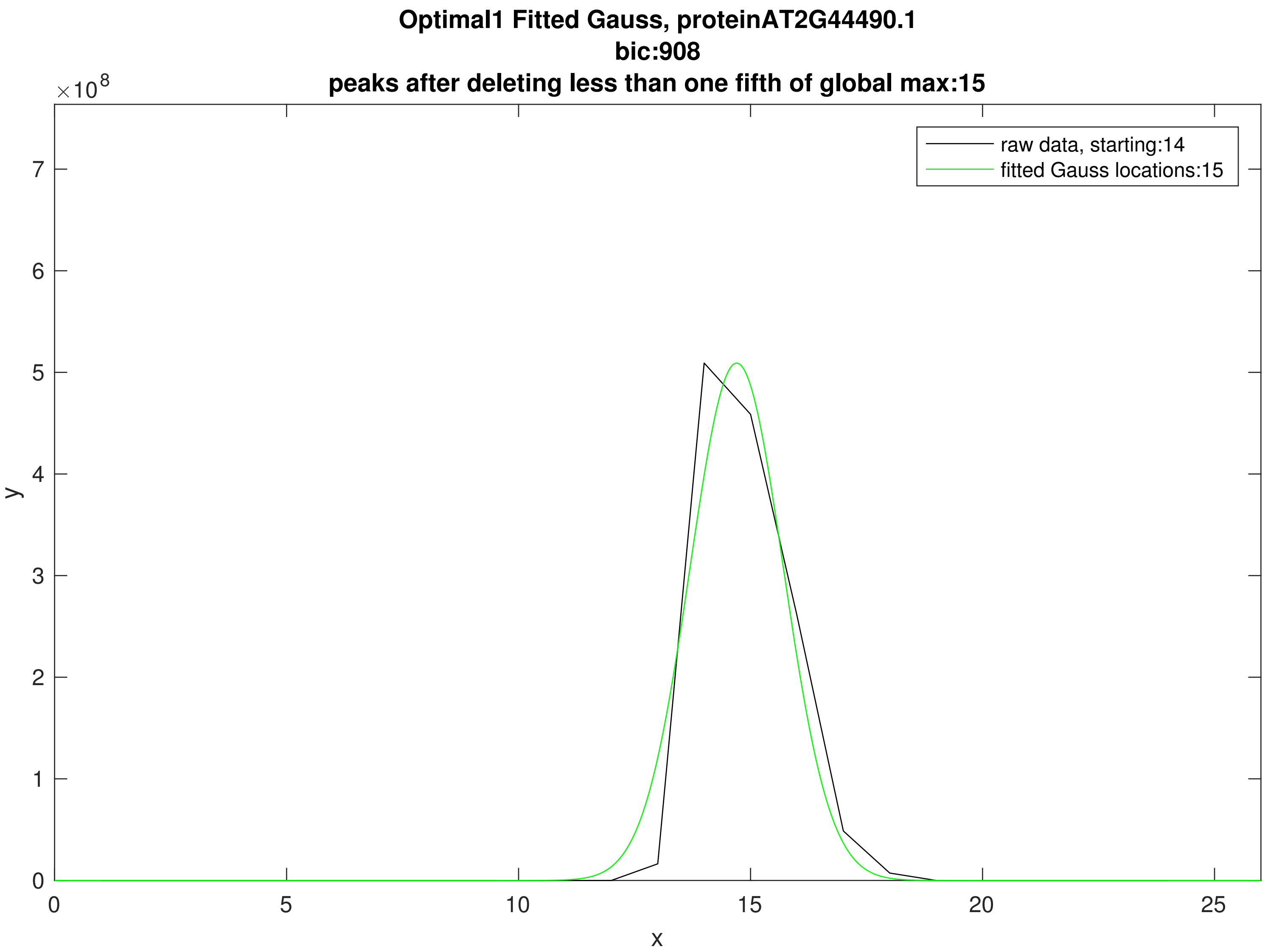

Optimal1 Fitted Gauss, proteinAT2G44490.1
bic:908
peaks after deleting less than one fifth of global max:15
×10^8
raw data, starting:14
fitted Gauss locations:15
y
x
0
1
2
3
4
5
6
7
0
5
10
15
20
25

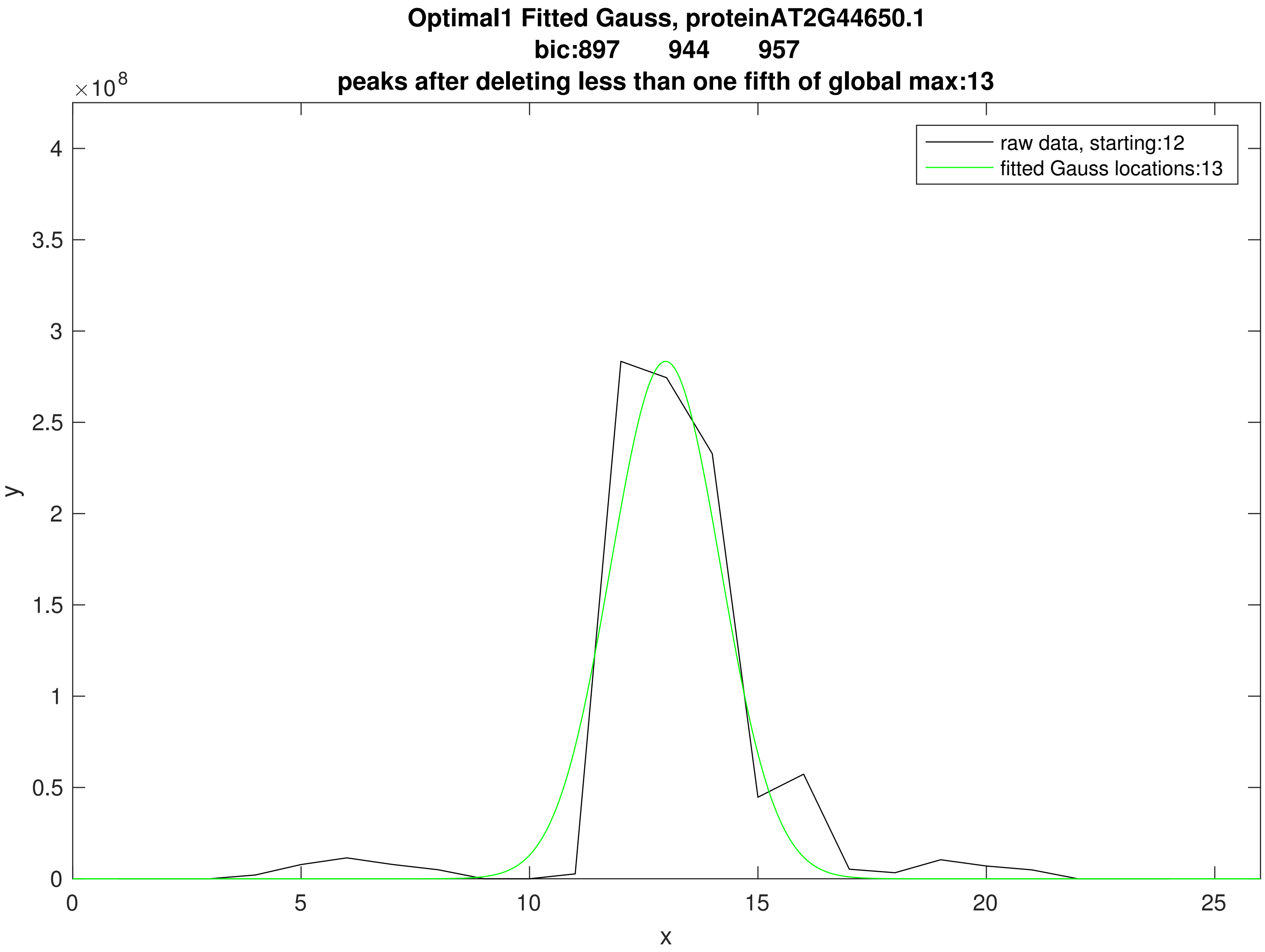
Optimal1 Fitted Gauss, proteinAT2G44650.1
bic:897 944 957
peaks after deleting less than one fifth of global max:13
×10^8
raw data, starting:12
fitted Gauss locations:13
y
x
0
0.5
1
1.5
2
2.5
3
3.5
4
0
5
10
15
20
25

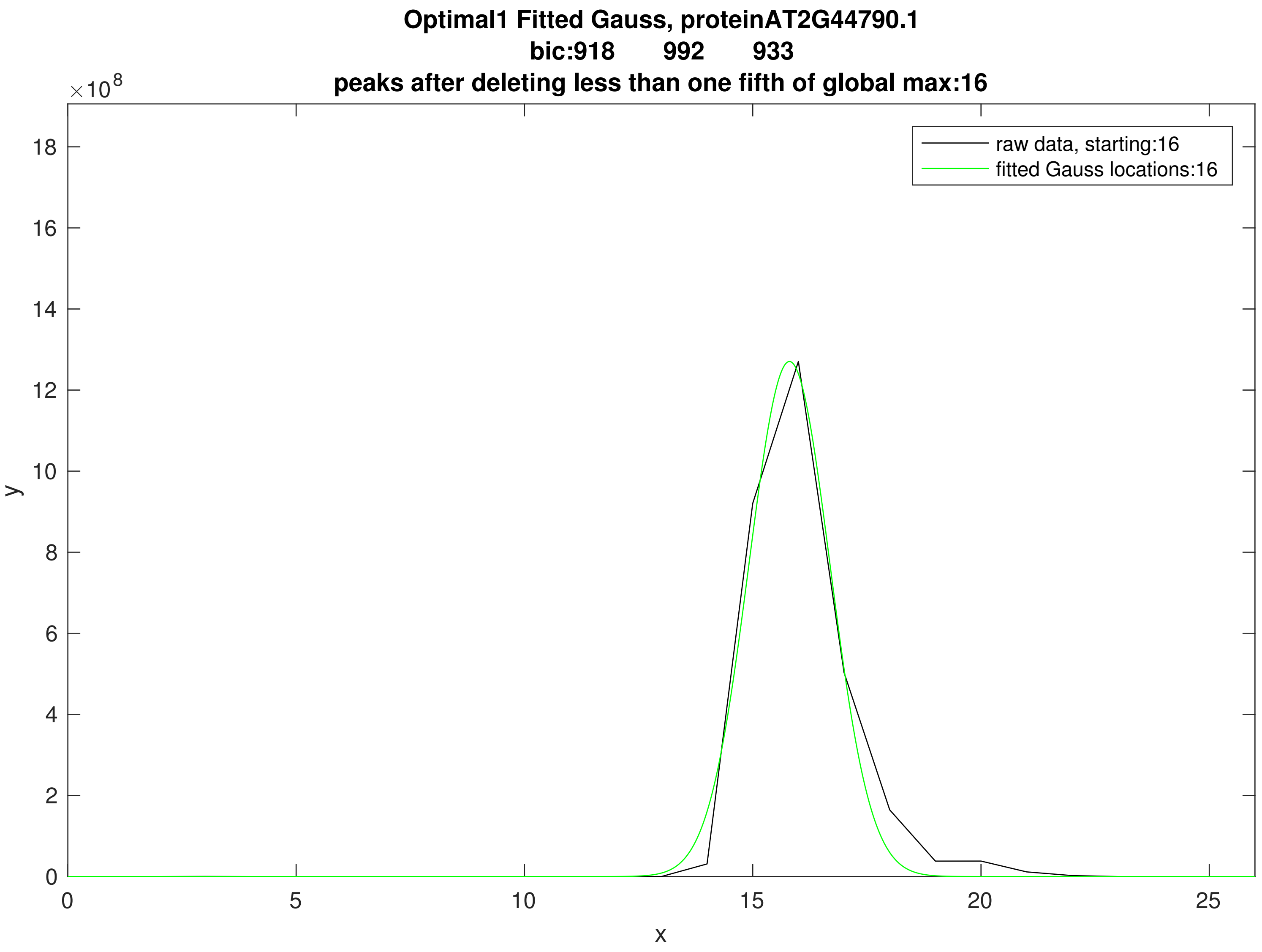
Optimal1 Fitted Gauss, proteinAT2G44790.1
bic:918 992 933
peaks after deleting less than one fifth of global max:16
×10^8
raw data, starting:16
fitted Gauss locations:16
y
x
0
2
4
6
8
10
12
14
16
18
0
5
10
15
20
25

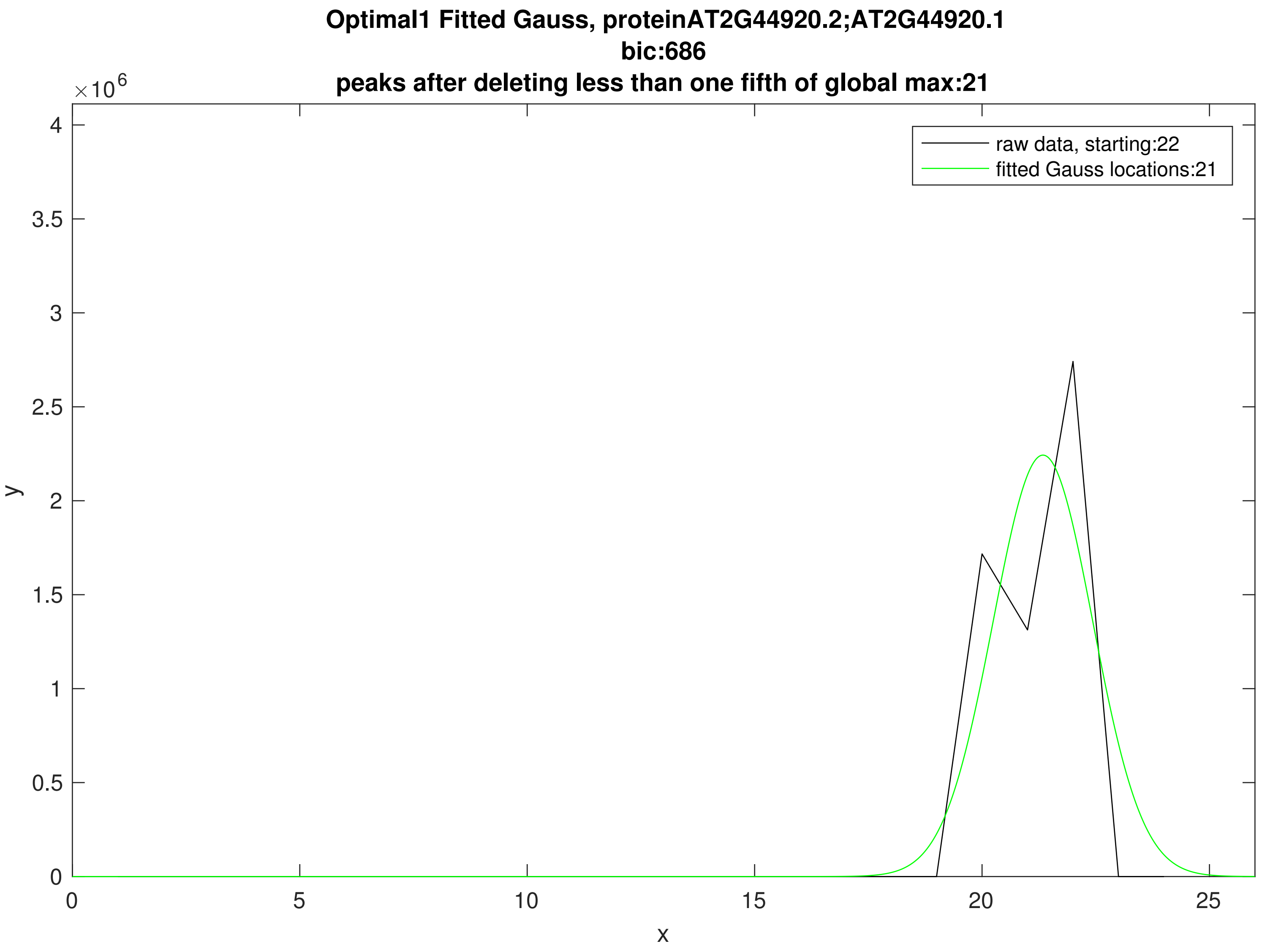

Optimal1 Fitted Gauss, proteinAT2G44920.2;AT2G44920.1
bic:686
peaks after deleting less than one fifth of global max:21
×10^6
raw data, starting:22
fitted Gauss locations:21
y
x
0
0.5
1
1.5
2
2.5
3
3.5
4
0
5
10
15
20
25

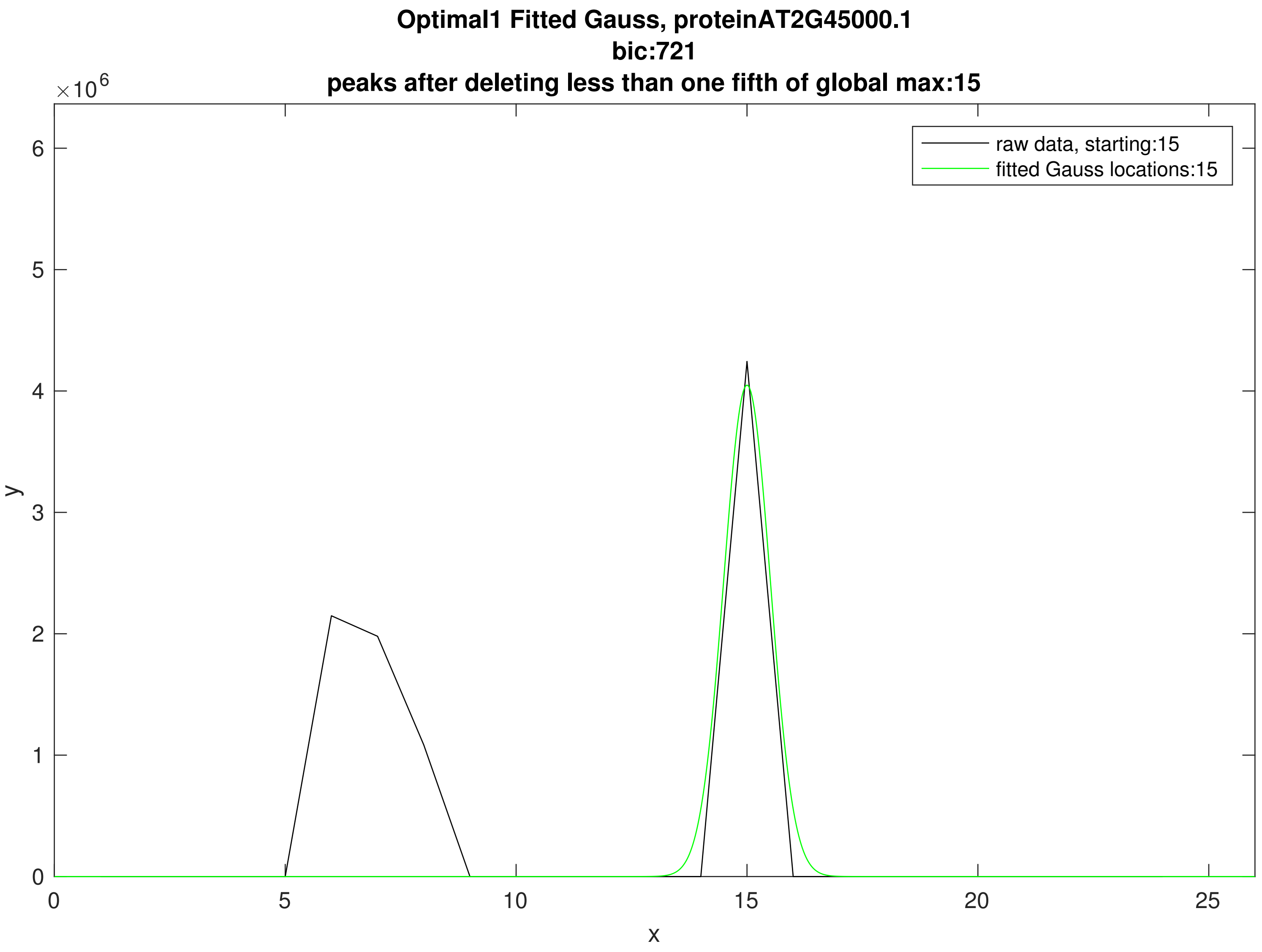
Optimal1 Fitted Gauss, proteinAT2G45000.1
bic:721
peaks after deleting less than one fifth of global max:15
×10^6
raw data, starting:15
fitted Gauss locations:15
y
x
0
1
2
3
4
5
6
0
5
10
15
20
25

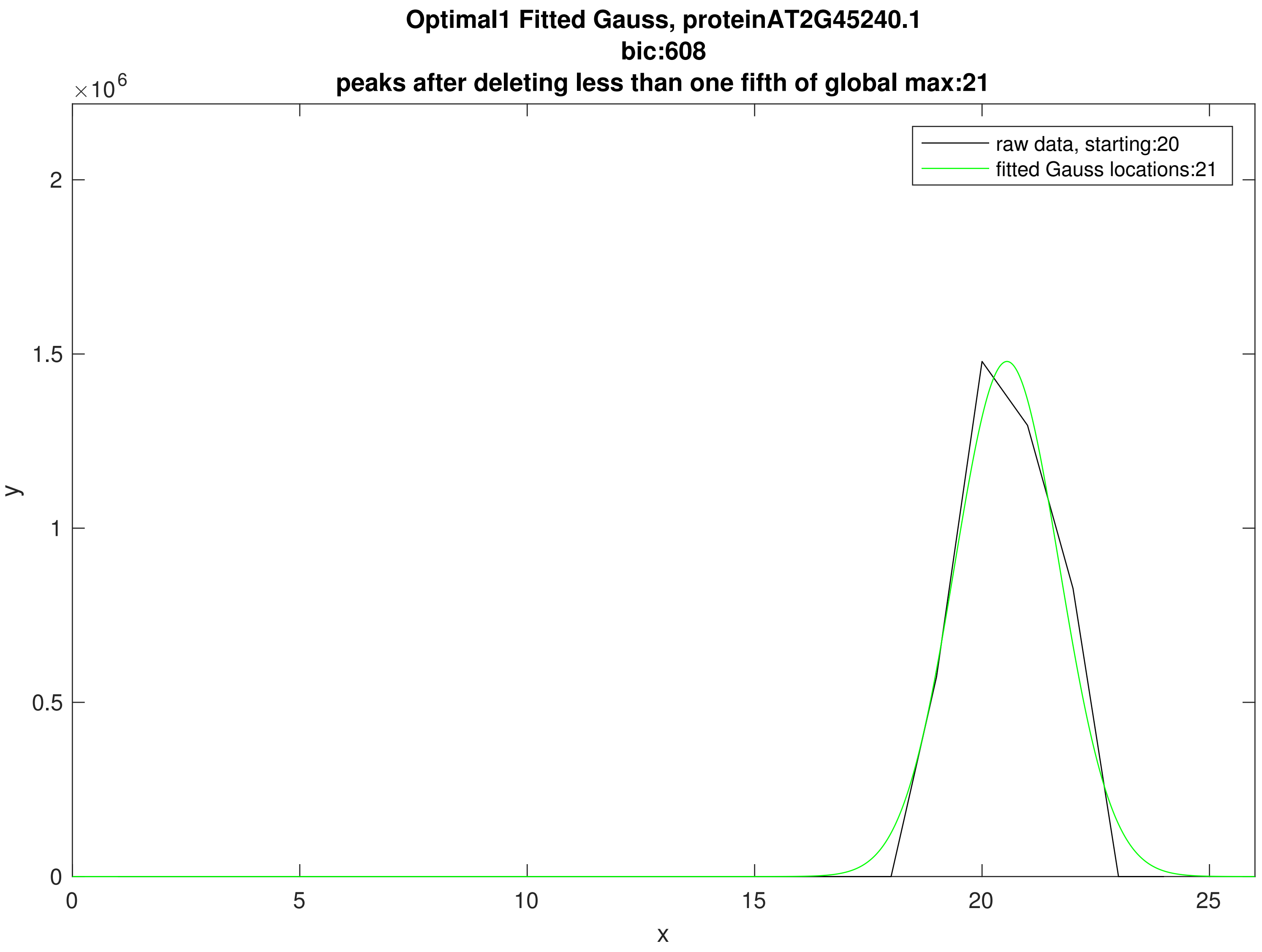
Optimal1 Fitted Gauss, proteinAT2G45240.1
bic:608
peaks after deleting less than one fifth of global max:21
×10^6
raw data, starting:20
fitted Gauss locations:21
y
x
0
0.5
1
1.5
2
0
5
10
15
20
25

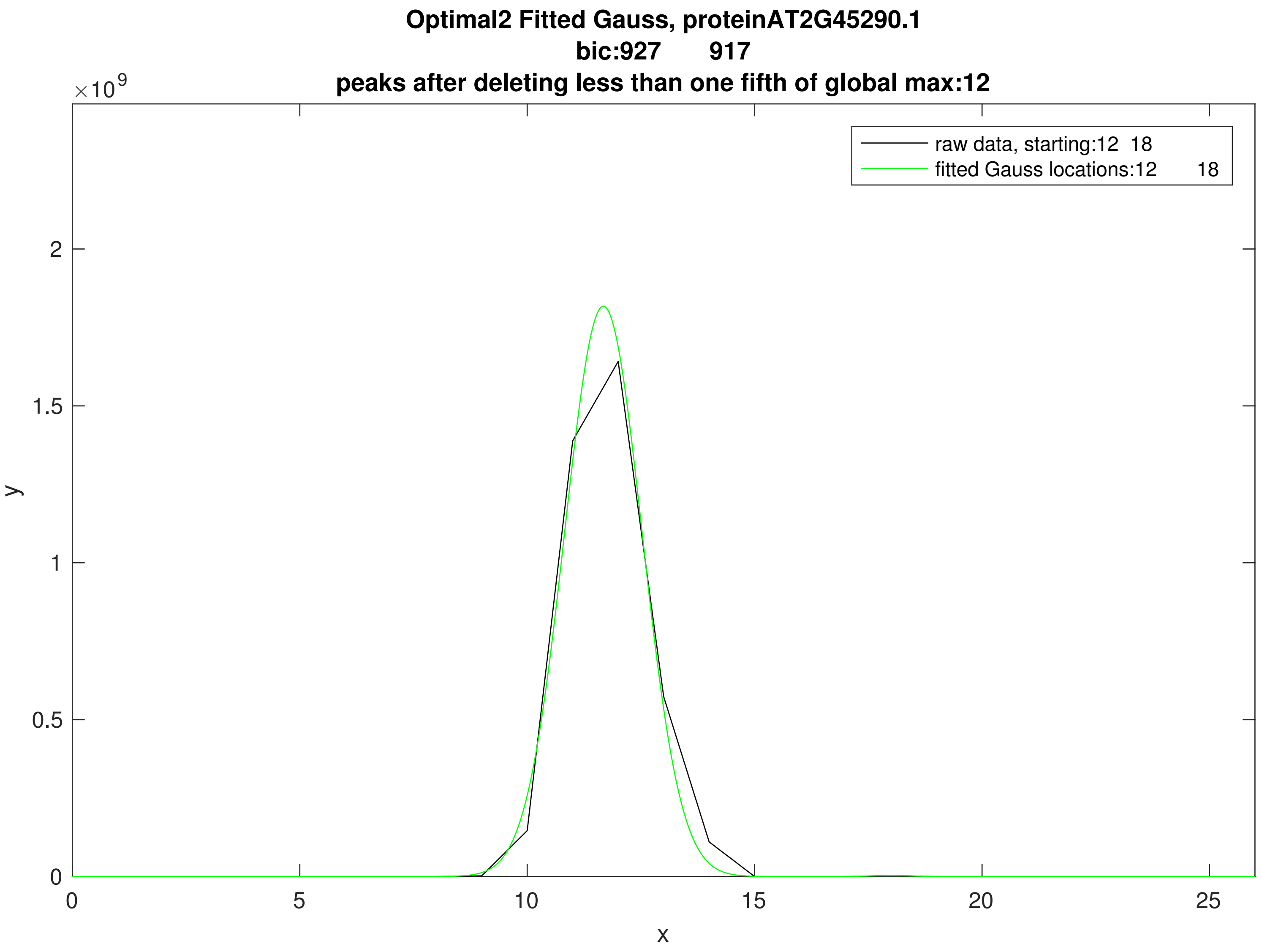
Optimal2 Fitted Gauss, proteinAT2G45290.1
bic:927 917
peaks after deleting less than one fifth of global max:12
×10^9
raw data, starting:12 18
fitted Gauss locations:12 18
y
x
0
0.5
1
1.5
2
5
10
15
20
25

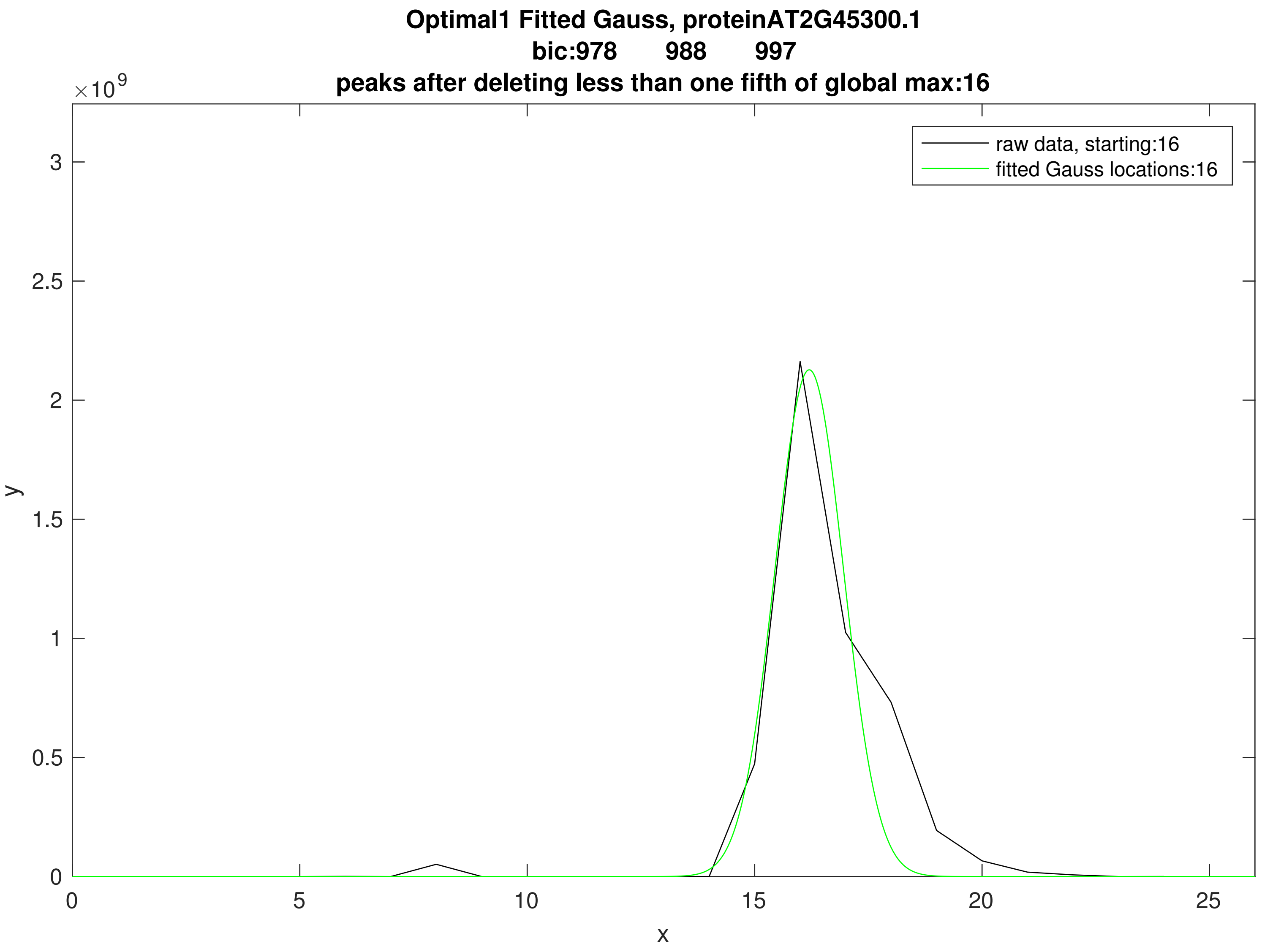
Optimal1 Fitted Gauss, proteinAT2G45300.1
bic:978 988 997
peaks after deleting less than one fifth of global max:16
×10^9
3
2.5
2
1.5
1
0.5
0
y
0
5
10
15
20
25
x
raw data, starting:16
fitted Gauss locations:16

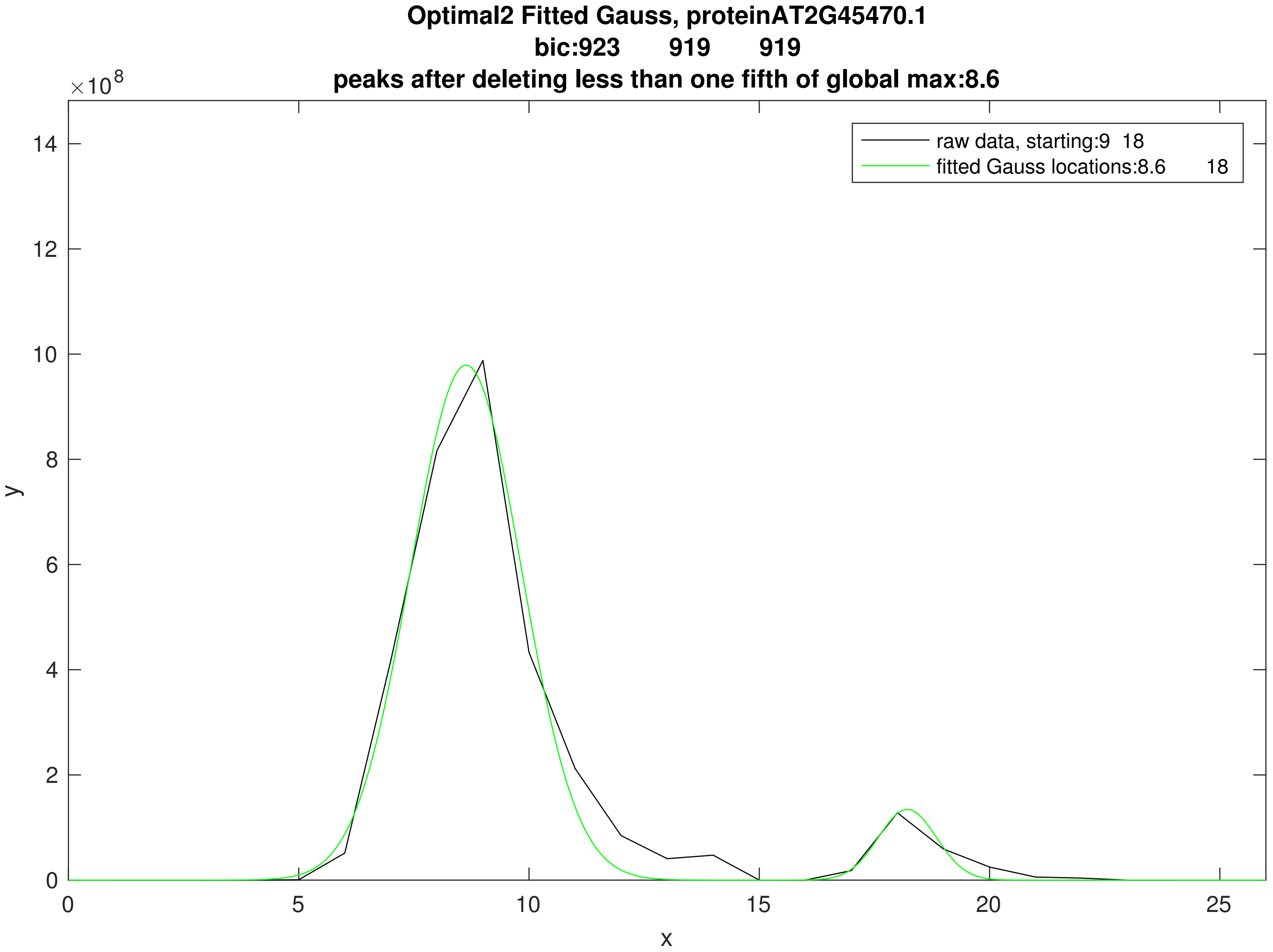
Optimal2 Fitted Gauss, proteinAT2G45470.1
bic:923 919 919
peaks after deleting less than one fifth of global max:8.6
×10^8
raw data, starting:9 18
fitted Gauss locations:8.6 18
y
x
0
2
4
6
8
10
12
14
5
15
20
25

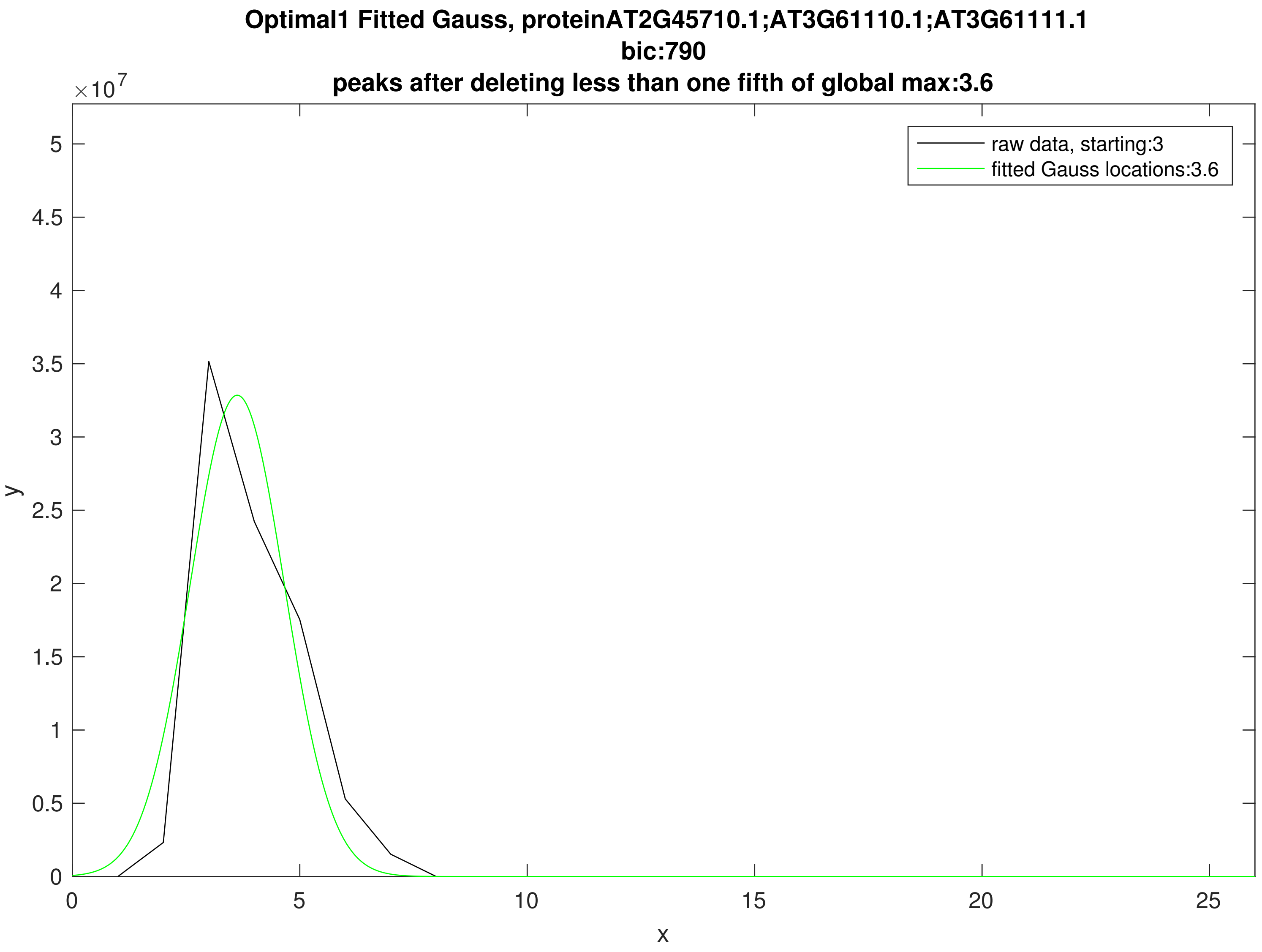

Optimal1 Fitted Gauss, proteinAT2G45710.1;AT3G61110.1;AT3G61111.1
bic:790
peaks after deleting less than one fifth of global max:3.6
×10^7
5
4.5
4
3.5
3
2.5
2
1.5
1
0.5
0
y
0
5
10
15
20
25
x
raw data, starting:3
fitted Gauss locations:3.6

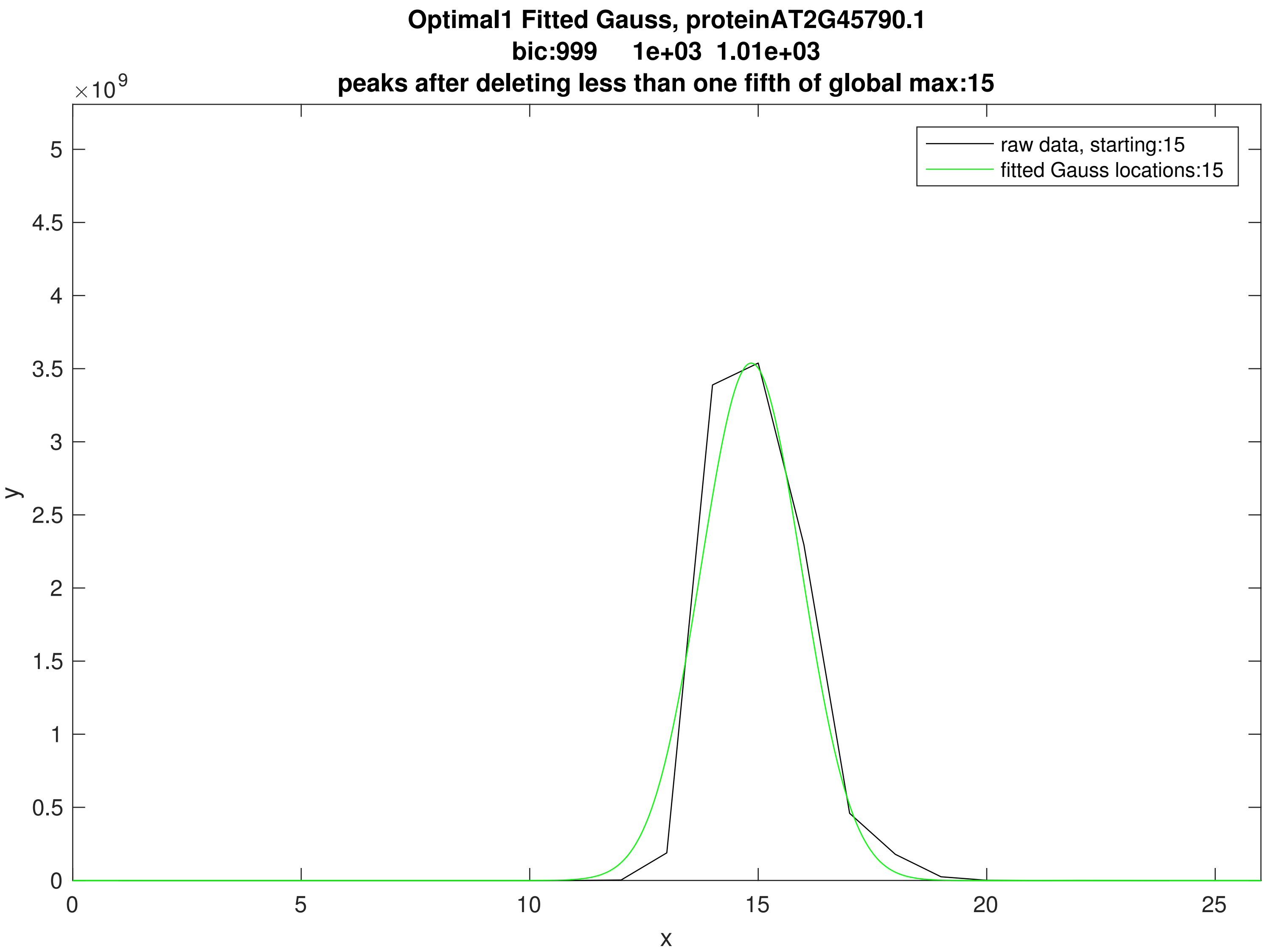
Optimal1 Fitted Gauss, proteinAT2G45790.1
bic:999 1e+03 1.01e+03
peaks after deleting less than one fifth of global max:15
×10^9
raw data, starting:15
fitted Gauss locations:15
y
x
0
0.5
1
1.5
2
2.5
3
3.5
4
4.5
5
0
5
10
15
20
25

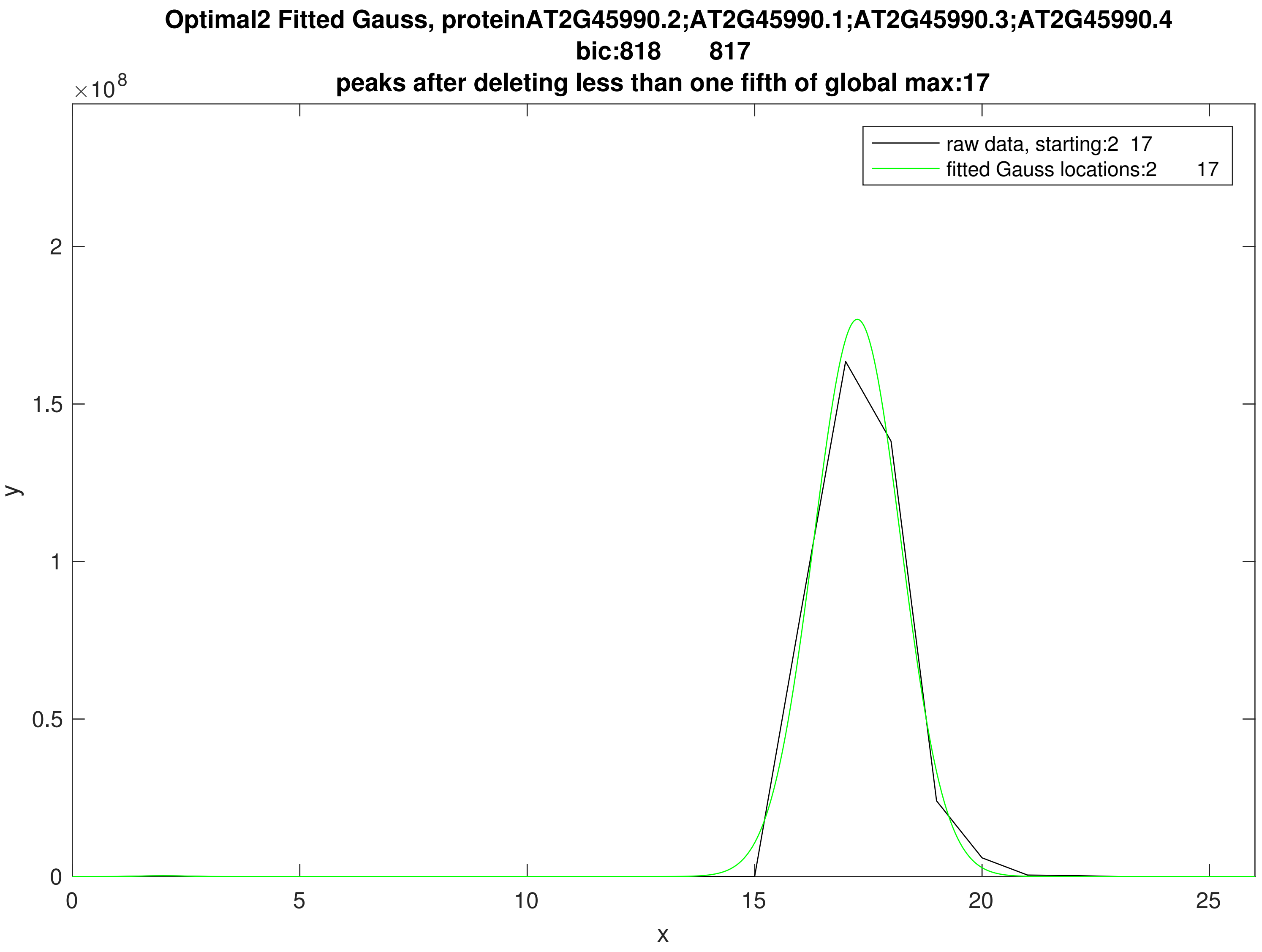

Optimal2 Fitted Gauss, proteinAT2G45990.2;AT2G45990.1;AT2G45990.3;AT2G45990.4
bic:818 817
peaks after deleting less than one fifth of global max:17
×10^8
raw data, starting:2 17
fitted Gauss locations:2 17
2
1.5
1
0.5
0
y
0
5
10
15
20
25
x

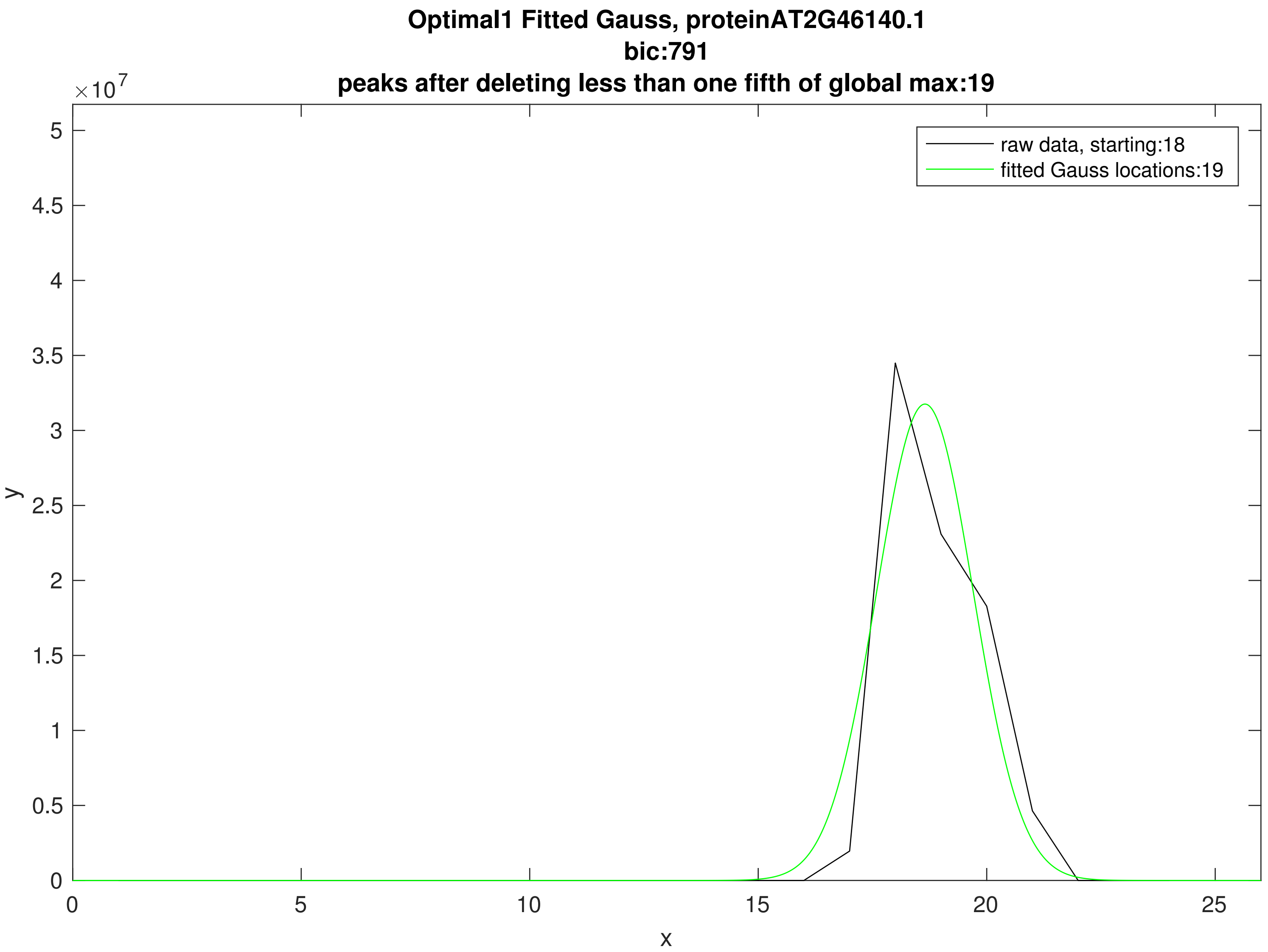
Optimal1 Fitted Gauss, proteinAT2G46140.1
bic:791
peaks after deleting less than one fifth of global max:19
×10^7
5
4.5
4
3.5
3
2.5
2
1.5
1
0.5
0
y
0
5
10
15
20
25
x
raw data, starting:18
fitted Gauss locations:19

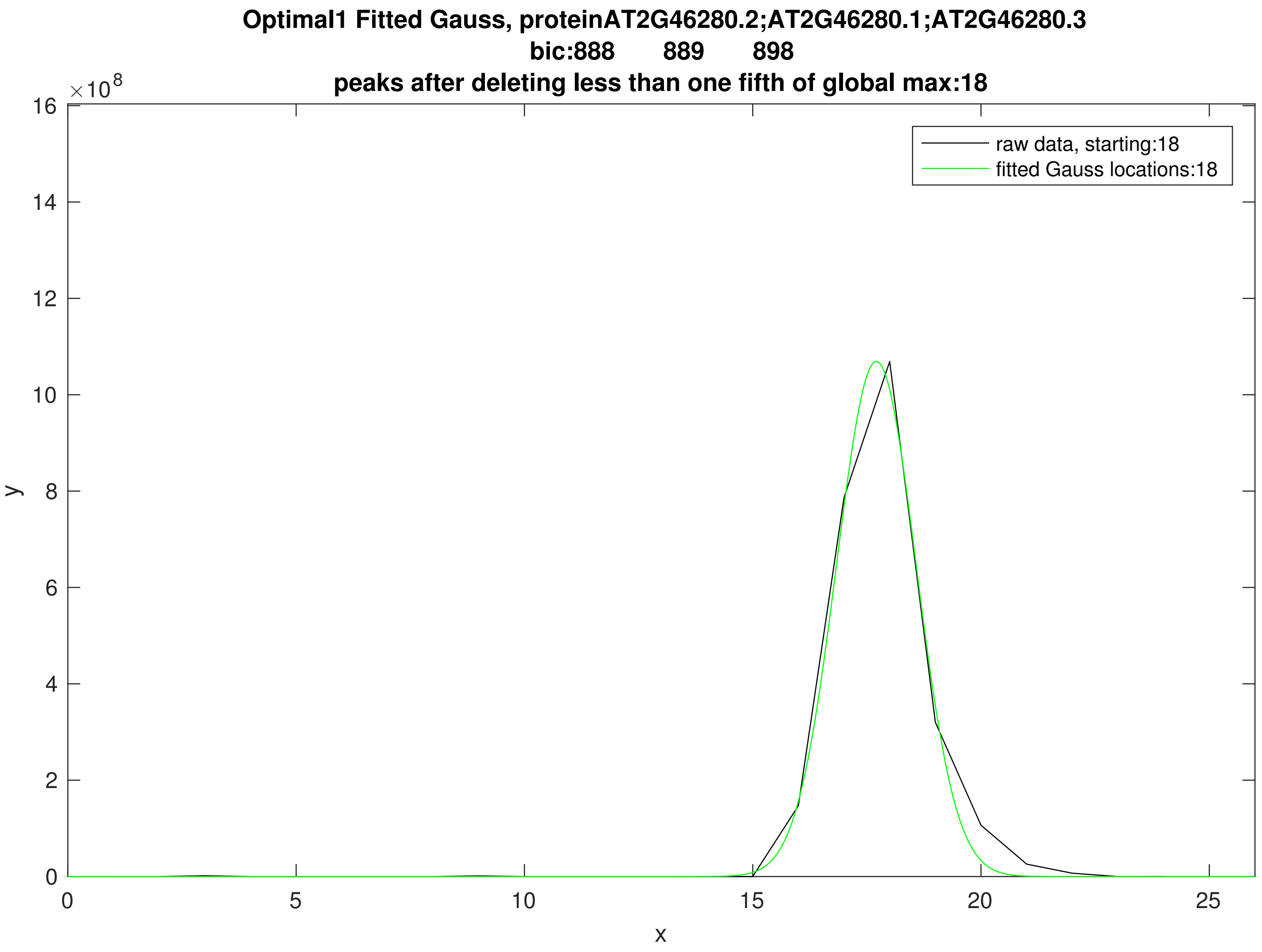
Optimal1 Fitted Gauss, proteinAT2G46280.2;AT2G46280.1;AT2G46280.3
bic:888 889 898
peaks after deleting less than one fifth of global max:18
×10^8
raw data, starting:18
fitted Gauss locations:18
y
x
0
2
4
6
8
10
12
14
16
0
5
10
15
20
25

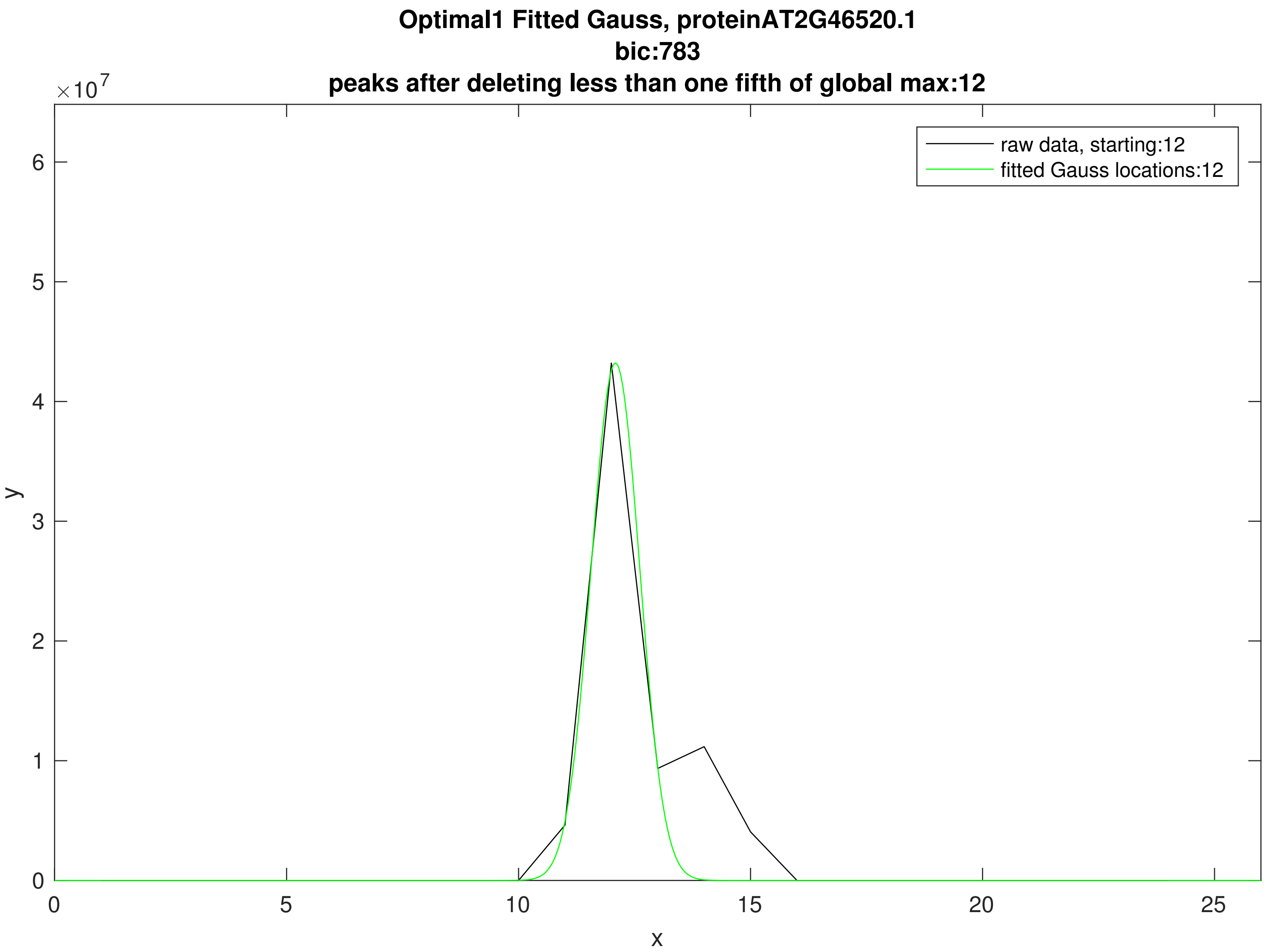
Optimal1 Fitted Gauss, proteinAT2G46520.1
bic:783
peaks after deleting less than one fifth of global max:12
×10^7
raw data, starting:12
fitted Gauss locations:12
y
x
0
1
2
3
4
5
6
0
5
10
15
20
25

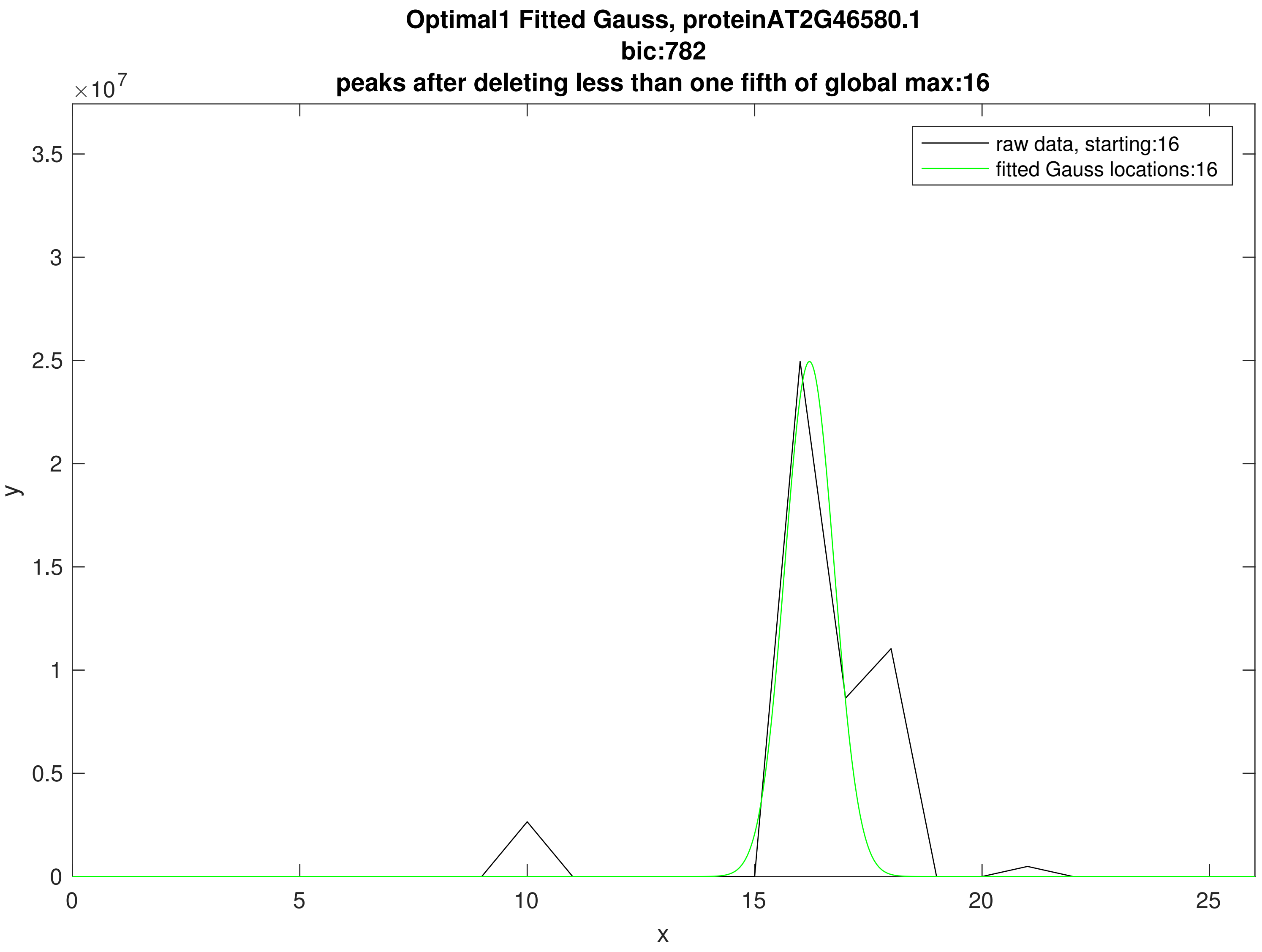

Optimal1 Fitted Gauss, proteinAT2G46580.1
bic:782
peaks after deleting less than one fifth of global max:16
×10^7
3.5
3
2.5
2
1.5
1
0.5
0
y
0
5
10
15
20
25
x
raw data, starting:16
fitted Gauss locations:16

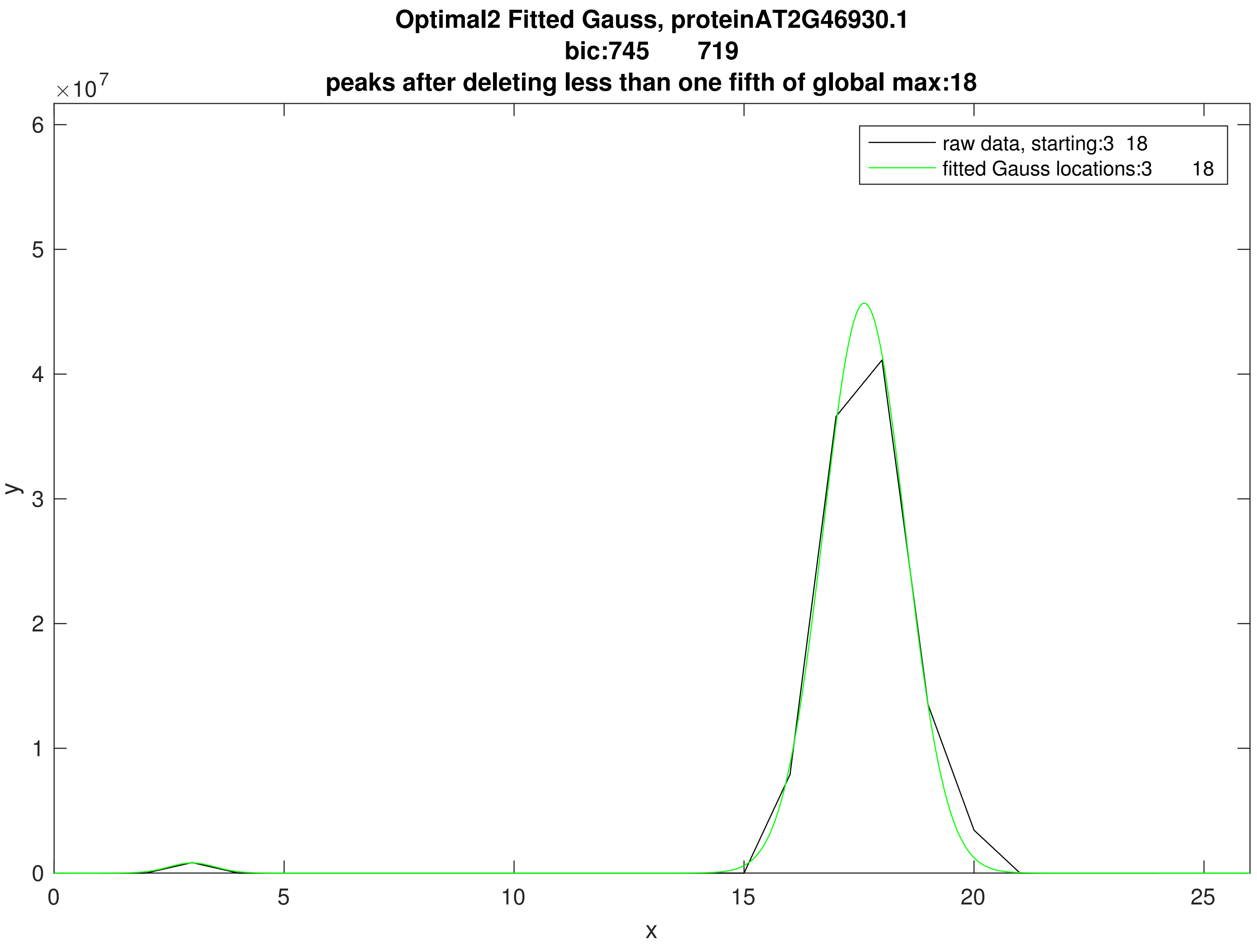
Optimal2 Fitted Gauss, proteinAT2G46930.1
bic:745 719
peaks after deleting less than one fifth of global max:18
×10^7
raw data, starting:3 18
fitted Gauss locations:3 18
y
x
0
1
2
3
4
5
6
0
5
10
15
20
25

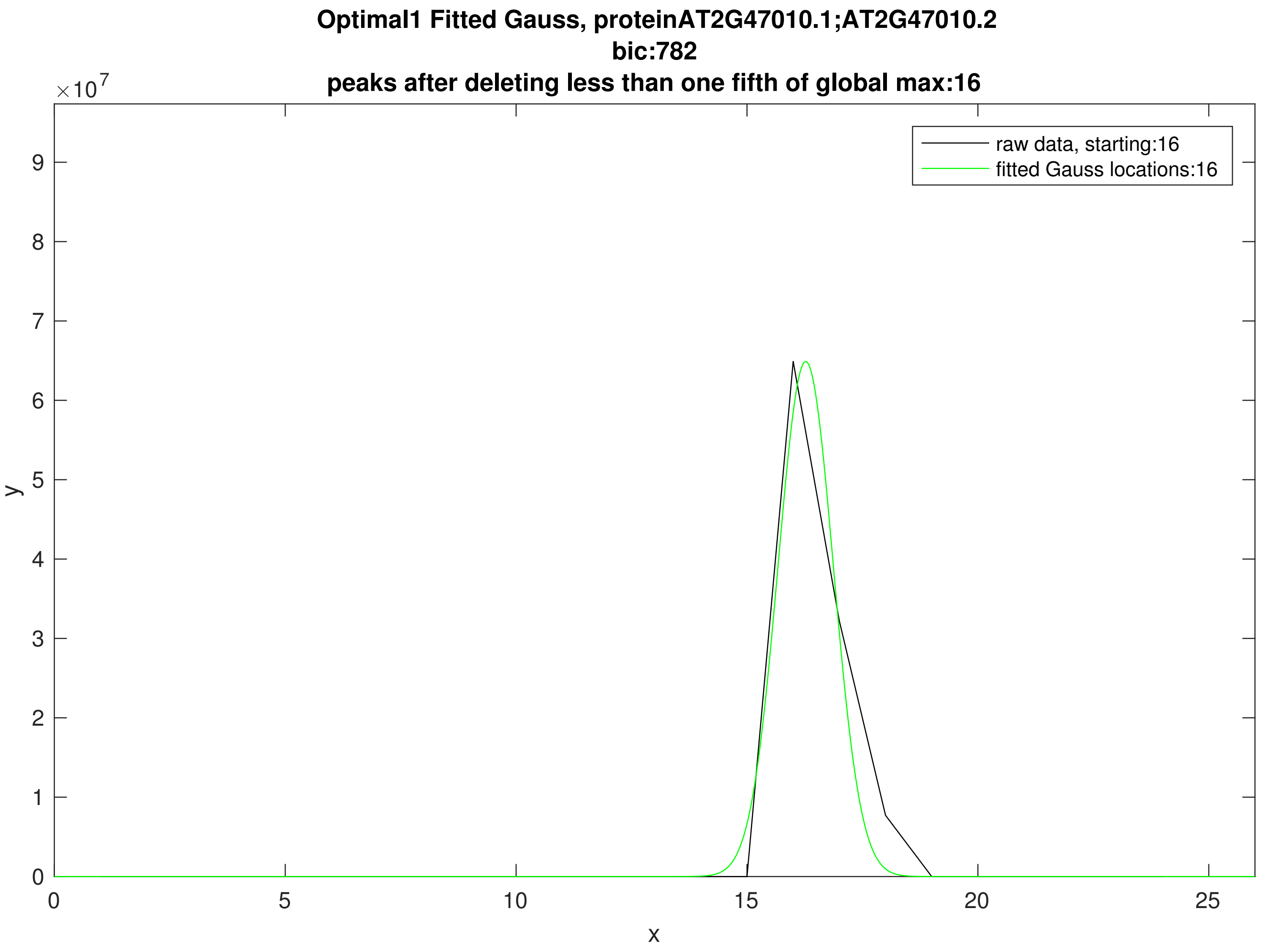
Optimal1 Fitted Gauss, proteinAT2G47010.1;AT2G47010.2
bic:782
peaks after deleting less than one fifth of global max:16
×10^7
raw data, starting:16
fitted Gauss locations:16
y
x
0
1
2
3
4
5
6
7
8
9
0
5
10
15
20
25

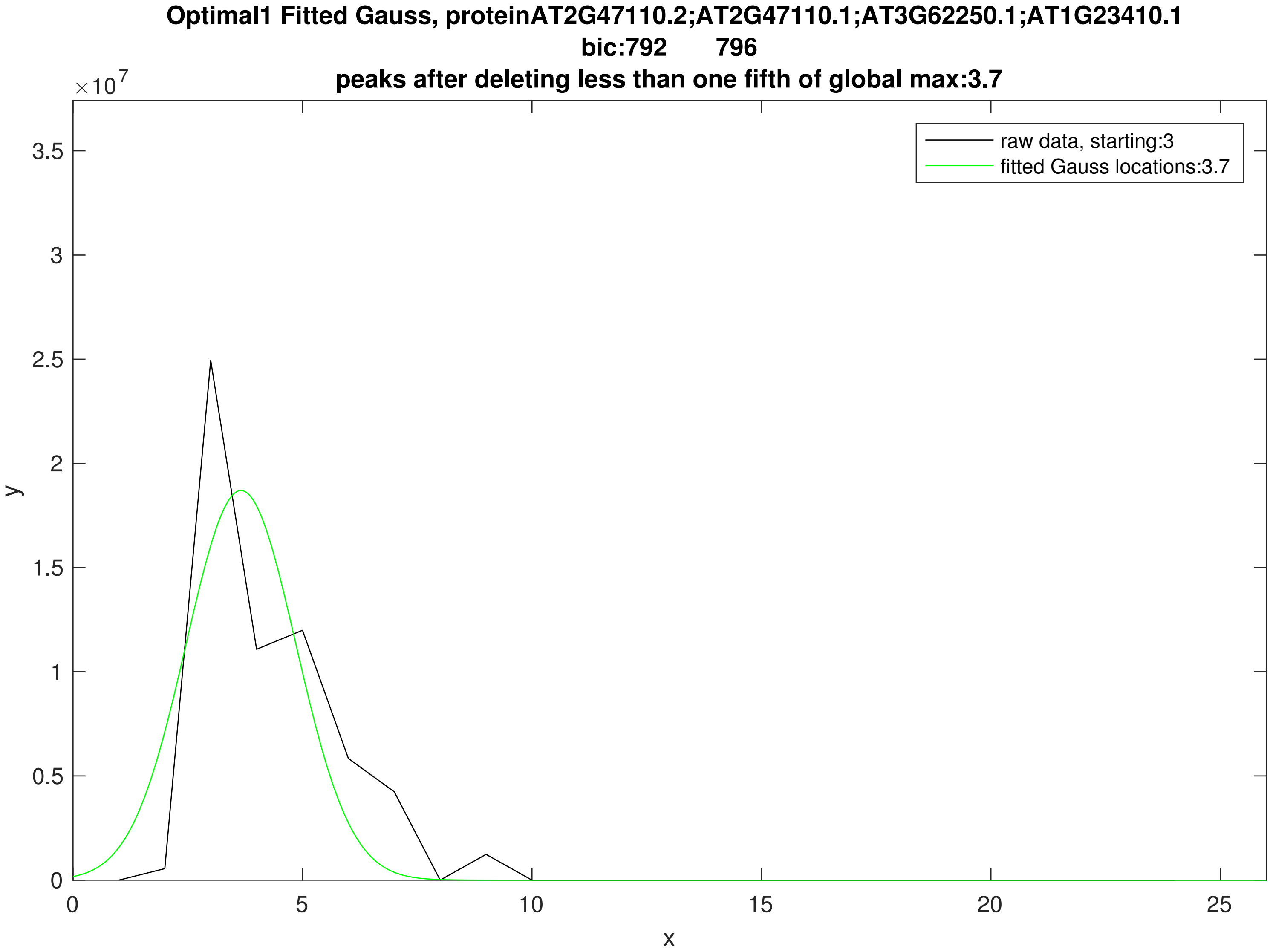

Optimal1 Fitted Gauss, proteinAT2G47110.2;AT2G47110.1;AT3G62250.1;AT1G23410.1
bic:792 796
peaks after deleting less than one fifth of global max:3.7
×10^7
raw data, starting:3
fitted Gauss locations:3.7
y
x
0
0.5
1
1.5
2
2.5
3
3.5
5
10
15
20
25

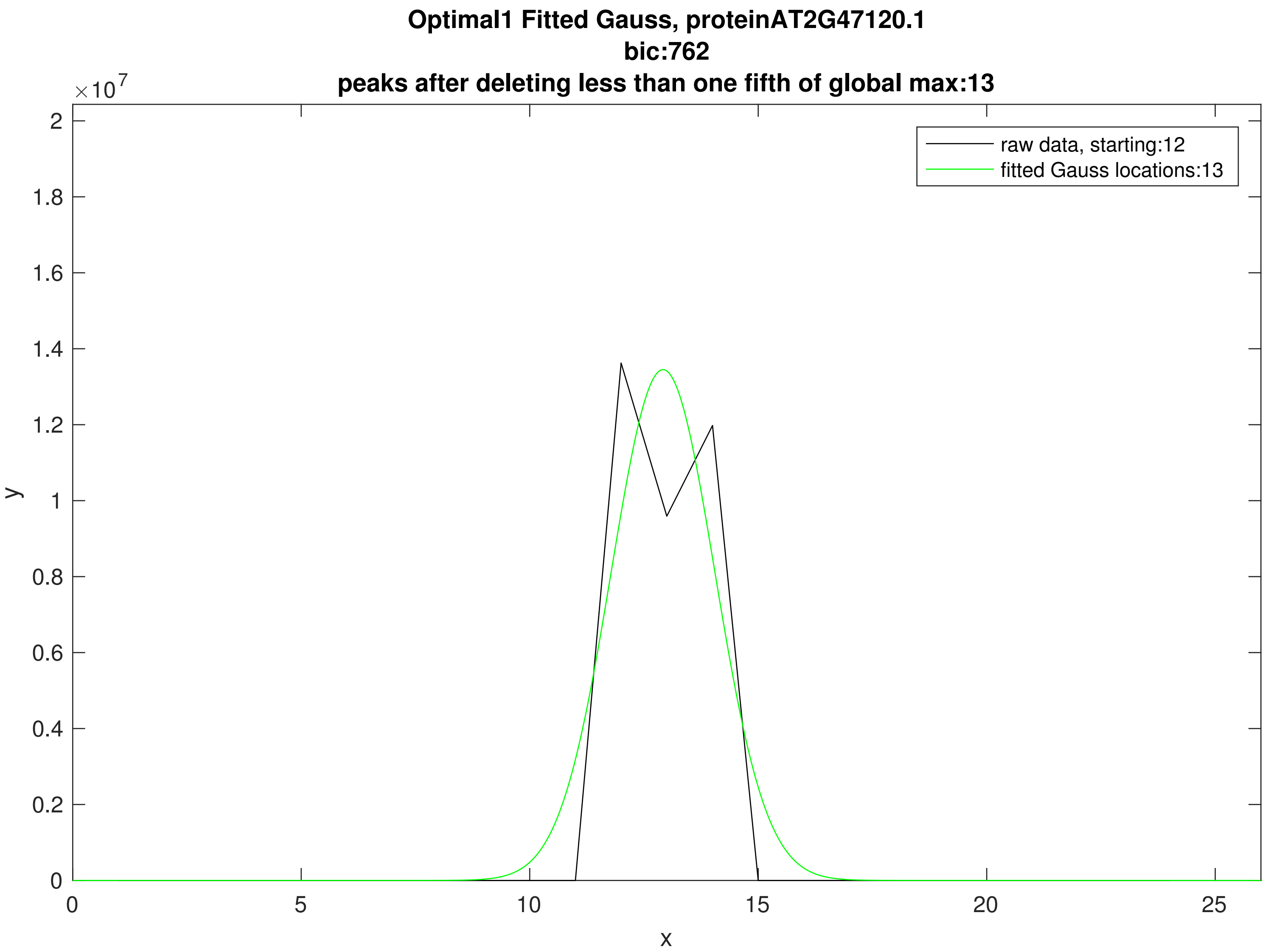
Optimal1 Fitted Gauss, proteinAT2G47120.1
bic:762
peaks after deleting less than one fifth of global max:13
×10^7
2
1.8
1.6
1.4
1.2
1
0.8
0.6
0.4
0.2
0
y
0
5
10
15
20
25
x
raw data, starting:12
fitted Gauss locations:13

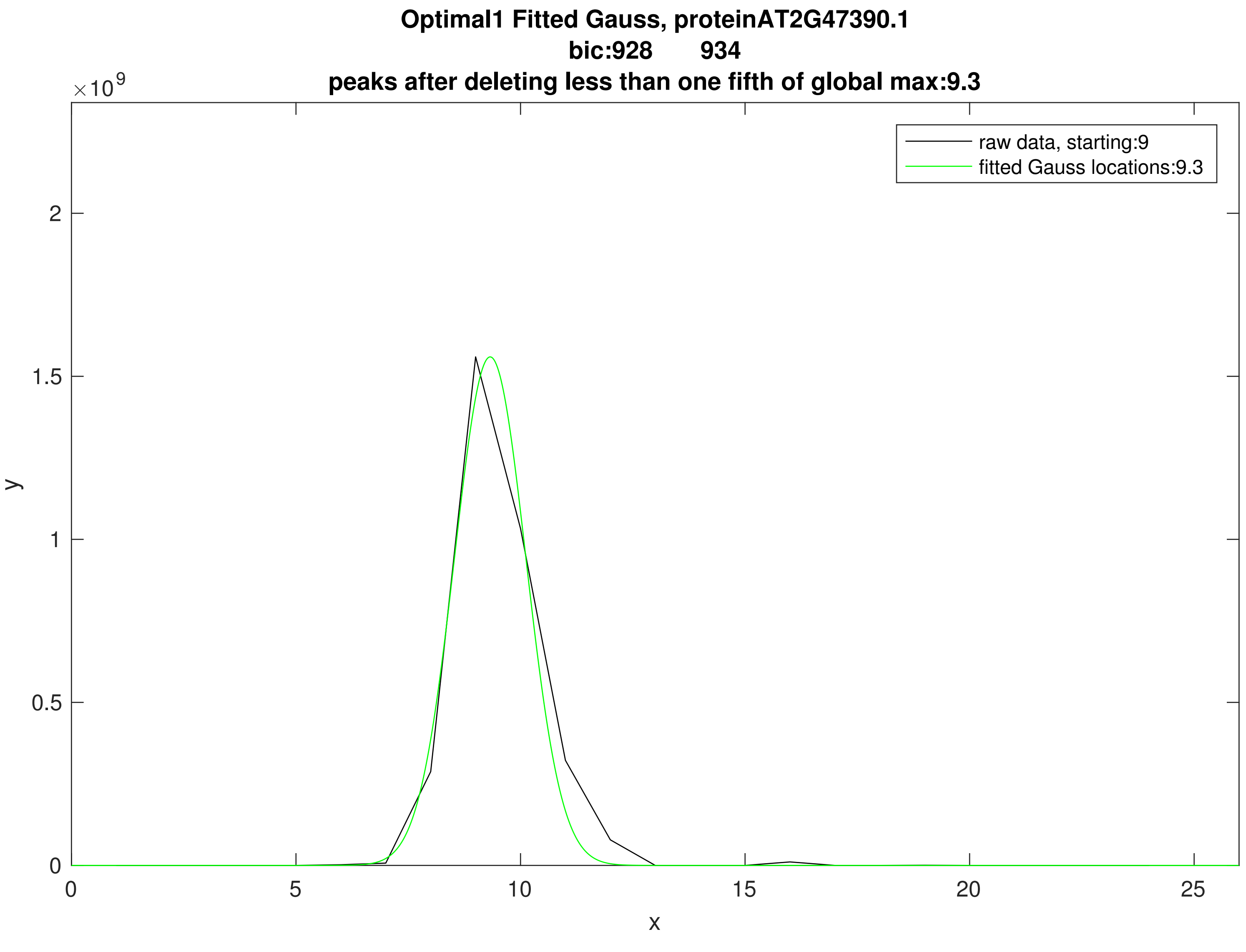
Optimal1 Fitted Gauss, proteinAT2G47390.1
bic:928 934
peaks after deleting less than one fifth of global max:9.3
×10^9
raw data, starting:9
fitted Gauss locations:9.3
y
x
0
0.5
1
1.5
2
0
5
10
15
20
25

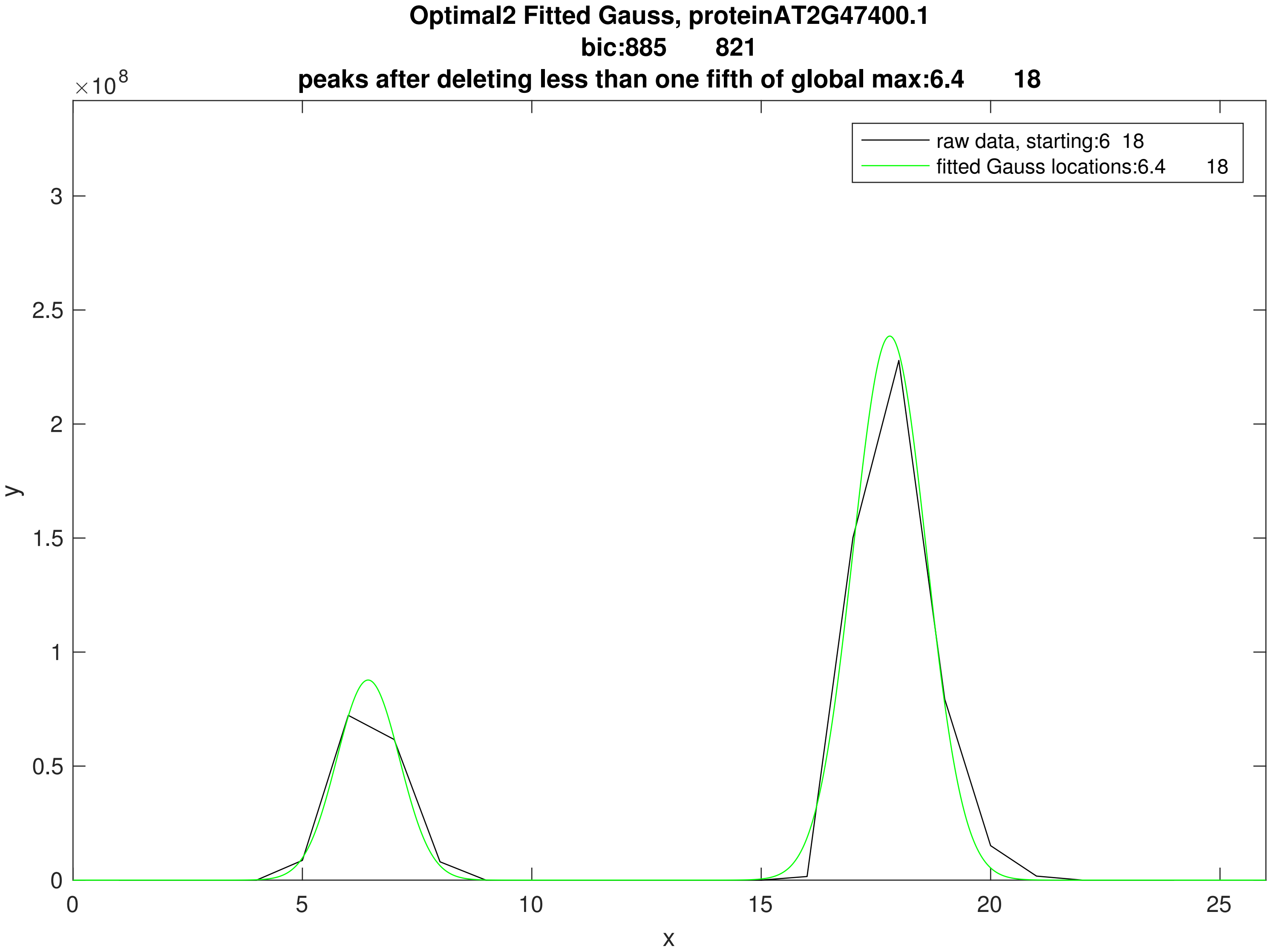

Optimal2 Fitted Gauss, proteinAT2G47400.1
bic:885 821
peaks after deleting less than one fifth of global max:6.4 18
×10^8
raw data, starting:6 18
fitted Gauss locations:6.4 18
y
x
0
0.5
1
1.5
2
2.5
3
0
5
10
15
20
25

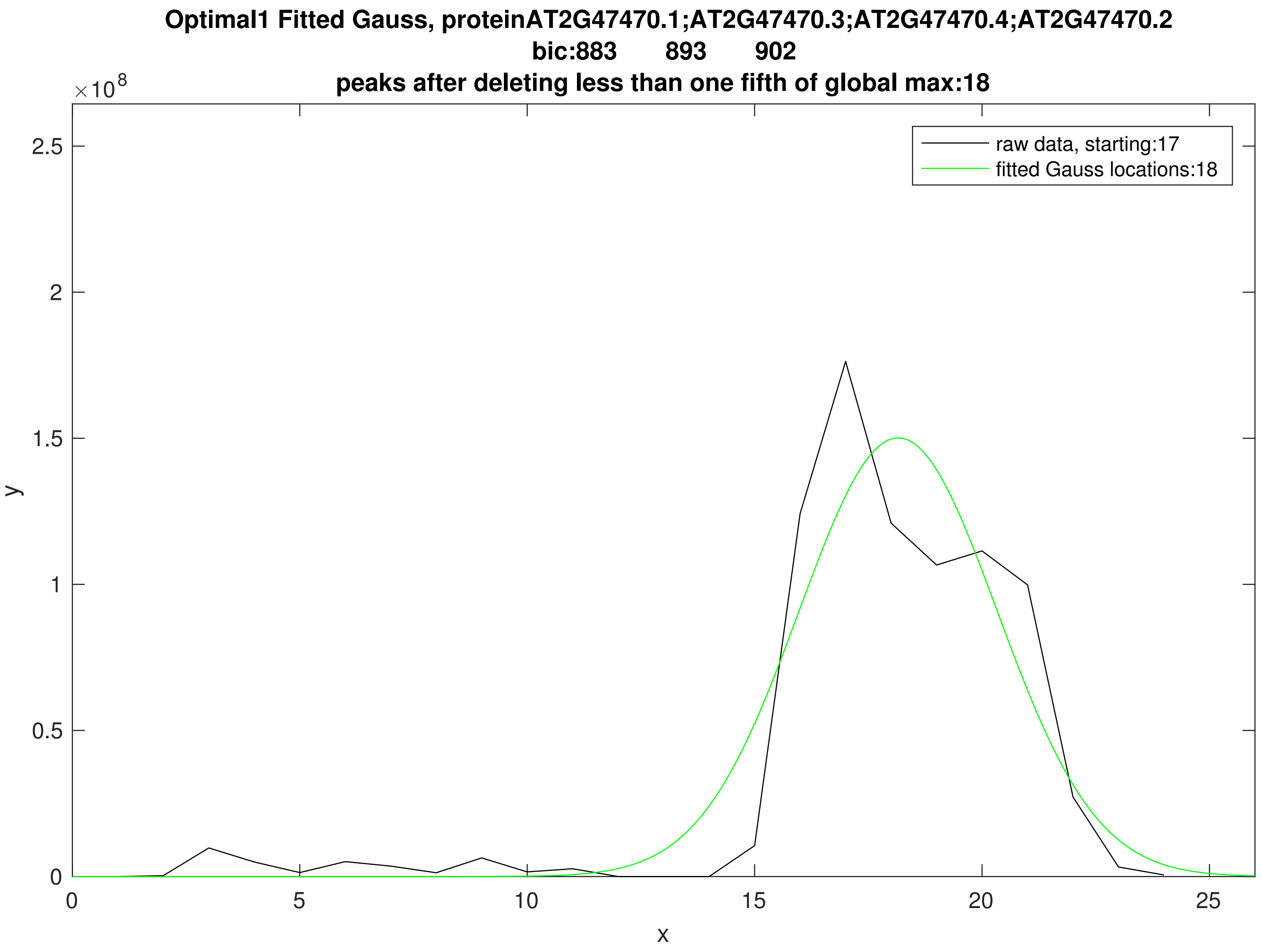
Optimal1 Fitted Gauss, proteinAT2G47470.1;AT2G47470.3;AT2G47470.4;AT2G47470.2
bic:883 893 902
peaks after deleting less than one fifth of global max:18
×10^8
2.5
2
1.5
1
0.5
0
y
0
5
10
15
20
25
x
raw data, starting:17
fitted Gauss locations:18

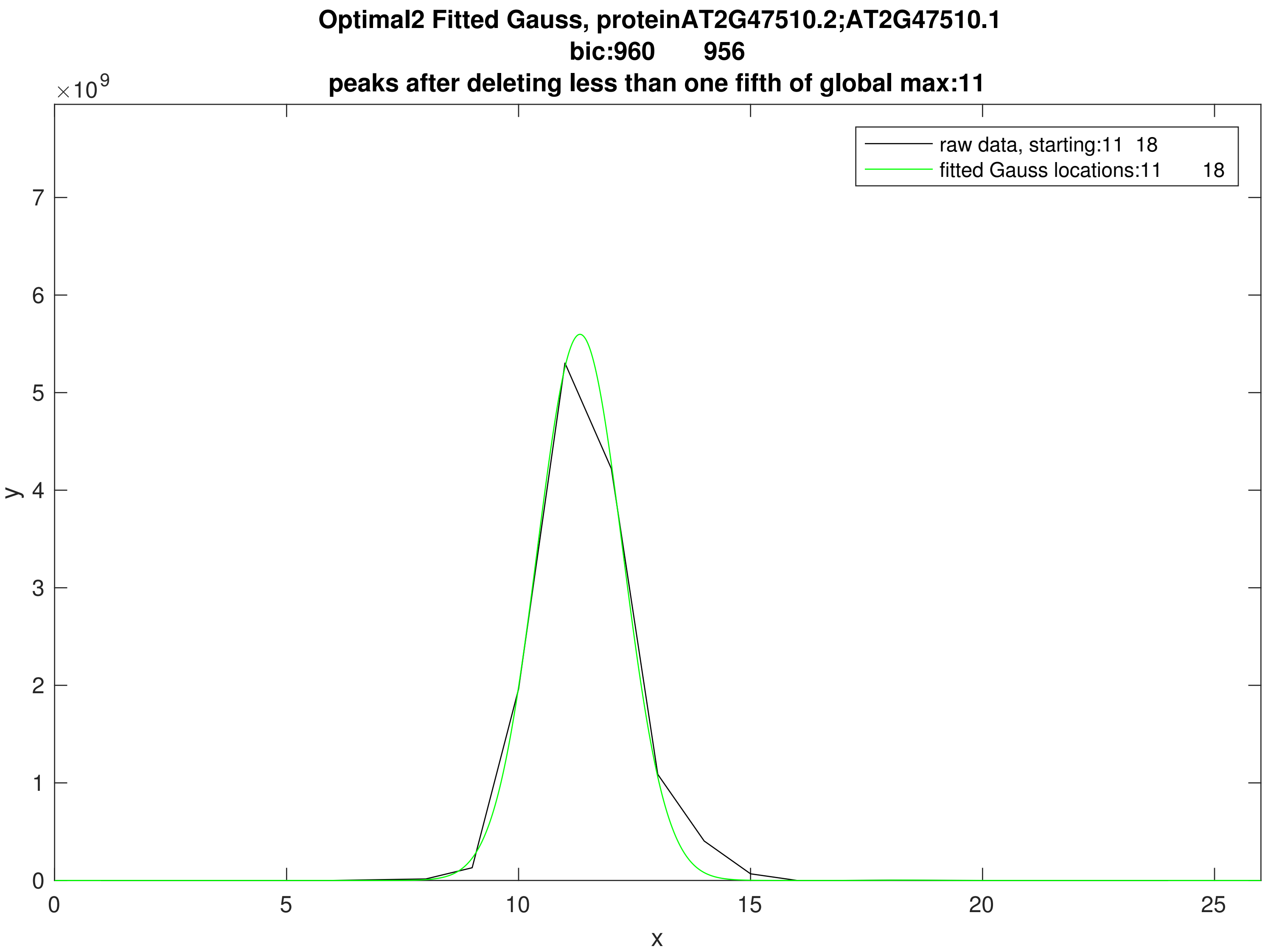
Optimal2 Fitted Gauss, proteinAT2G47510.2;AT2G47510.1
bic:960 956
peaks after deleting less than one fifth of global max:11
×10^9
raw data, starting:11 18
fitted Gauss locations:11 18
y
x
0
1
2
3
4
5
6
7
0
5
10
15
20
25

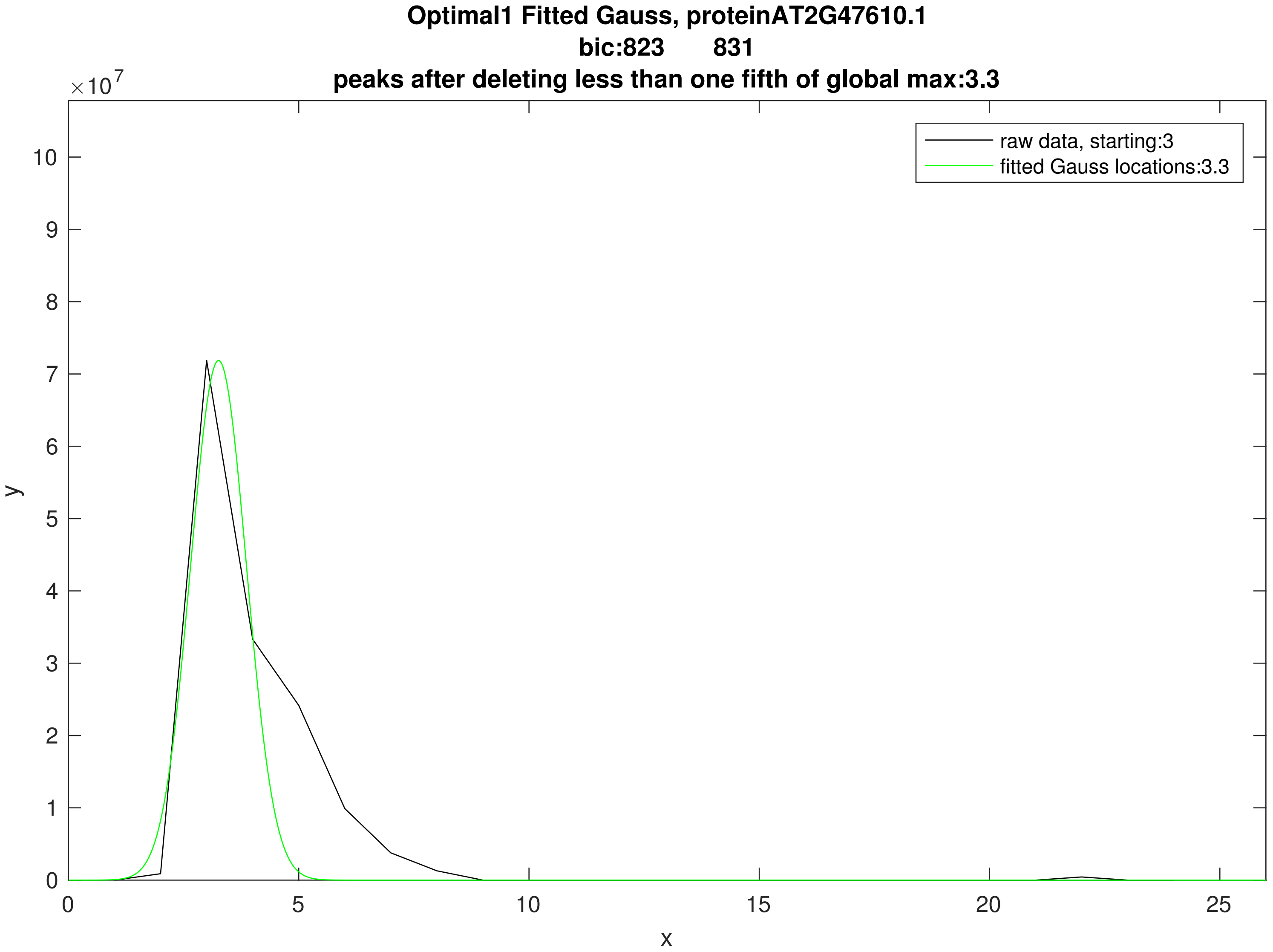

Optimal1 Fitted Gauss, proteinAT2G47610.1
bic:823 831
peaks after deleting less than one fifth of global max:3.3
×10^7
raw data, starting:3
fitted Gauss locations:3.3
y
x
0
1
2
3
4
5
6
7
8
9
10
0
5
10
15
20
25

**Optimal1 Fitted Gauss, proteinAT2G47650.1;AT3G62830.2;AT3G62830.1;AT2G47650.2**
**bic:703**
**peaks after deleting less than one fifth of global max:3.2**

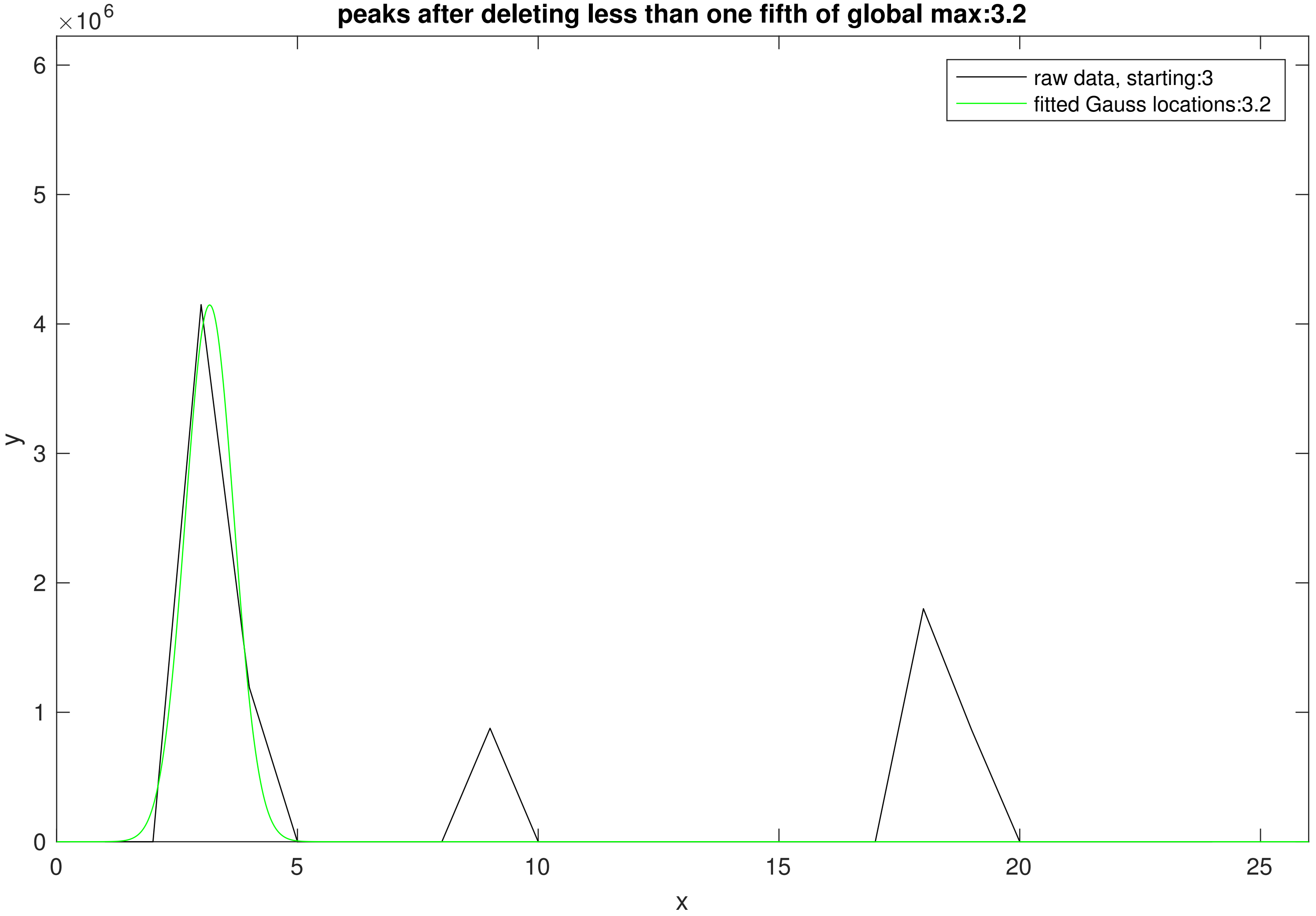

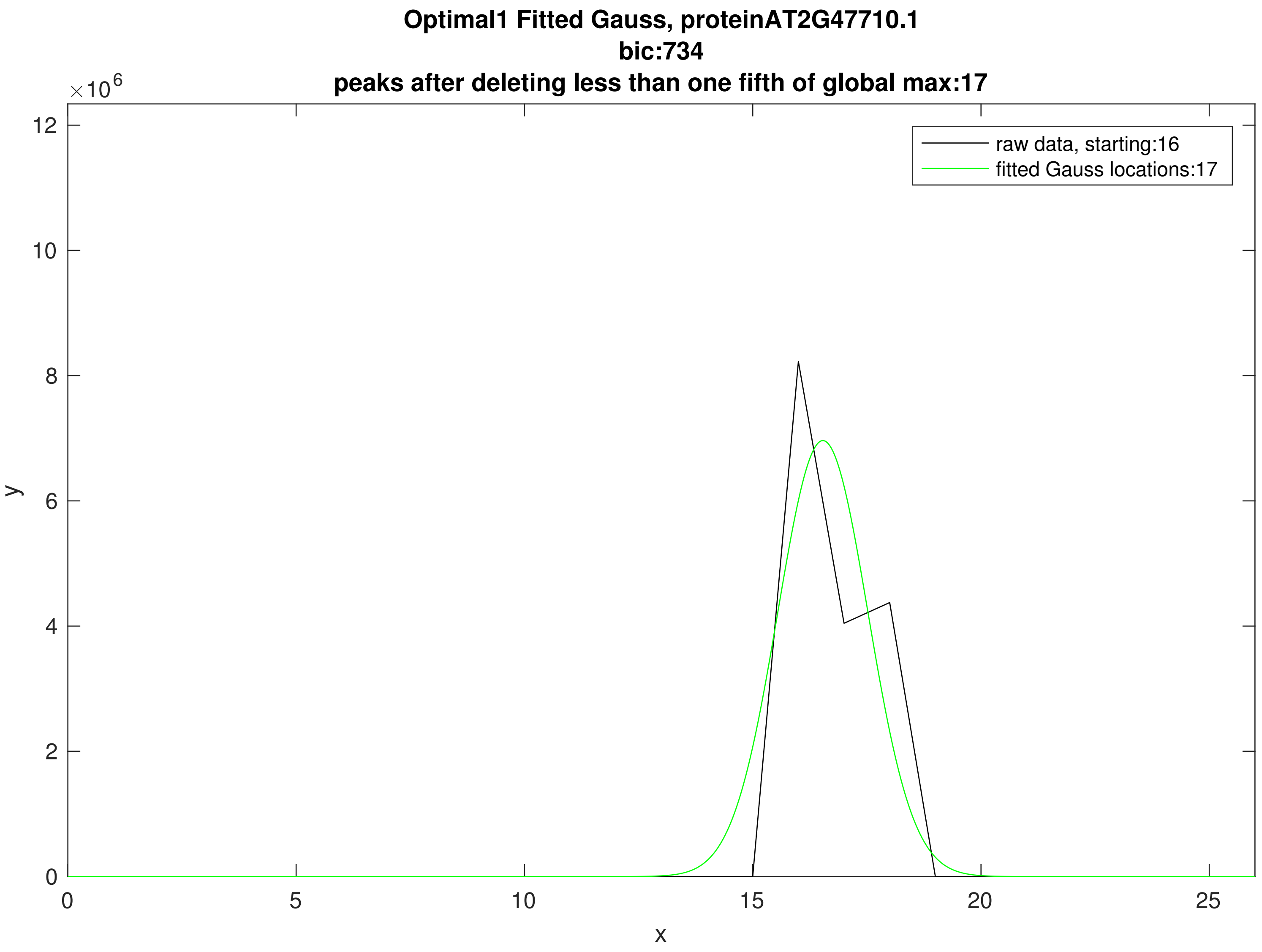
Optimal1 Fitted Gauss, proteinAT2G47710.1
bic:734
peaks after deleting less than one fifth of global max:17
×10^6
raw data, starting:16
fitted Gauss locations:17
y
x
0
2
4
6
8
10
12
0
5
10
15
20
25

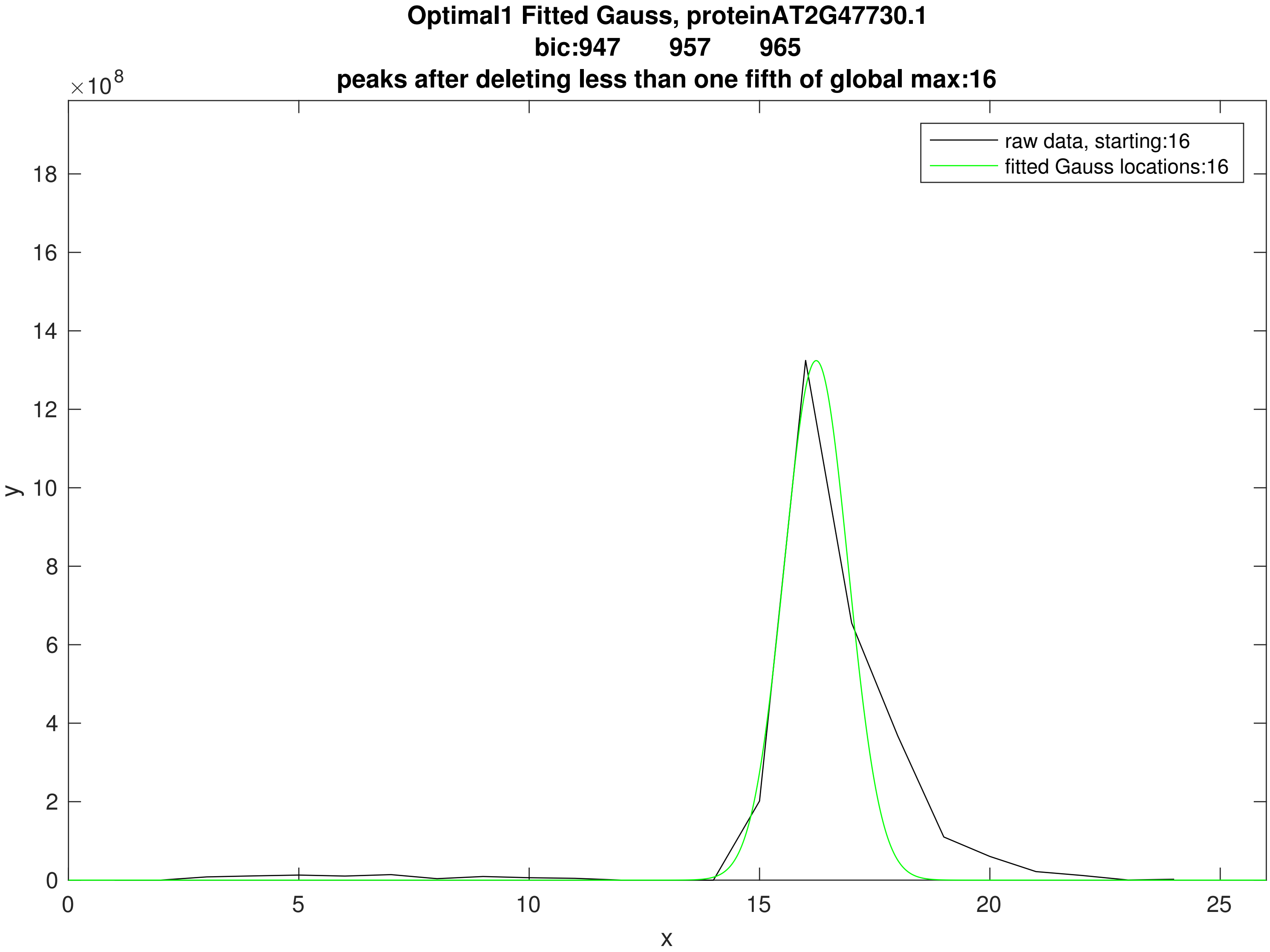
Optimal1 Fitted Gauss, proteinAT2G47730.1
bic:947 957 965
peaks after deleting less than one fifth of global max:16
×10^8
raw data, starting:16
fitted Gauss locations:16
y
x
0
2
4
6
8
10
12
14
16
18
0
5
10
15
20
25

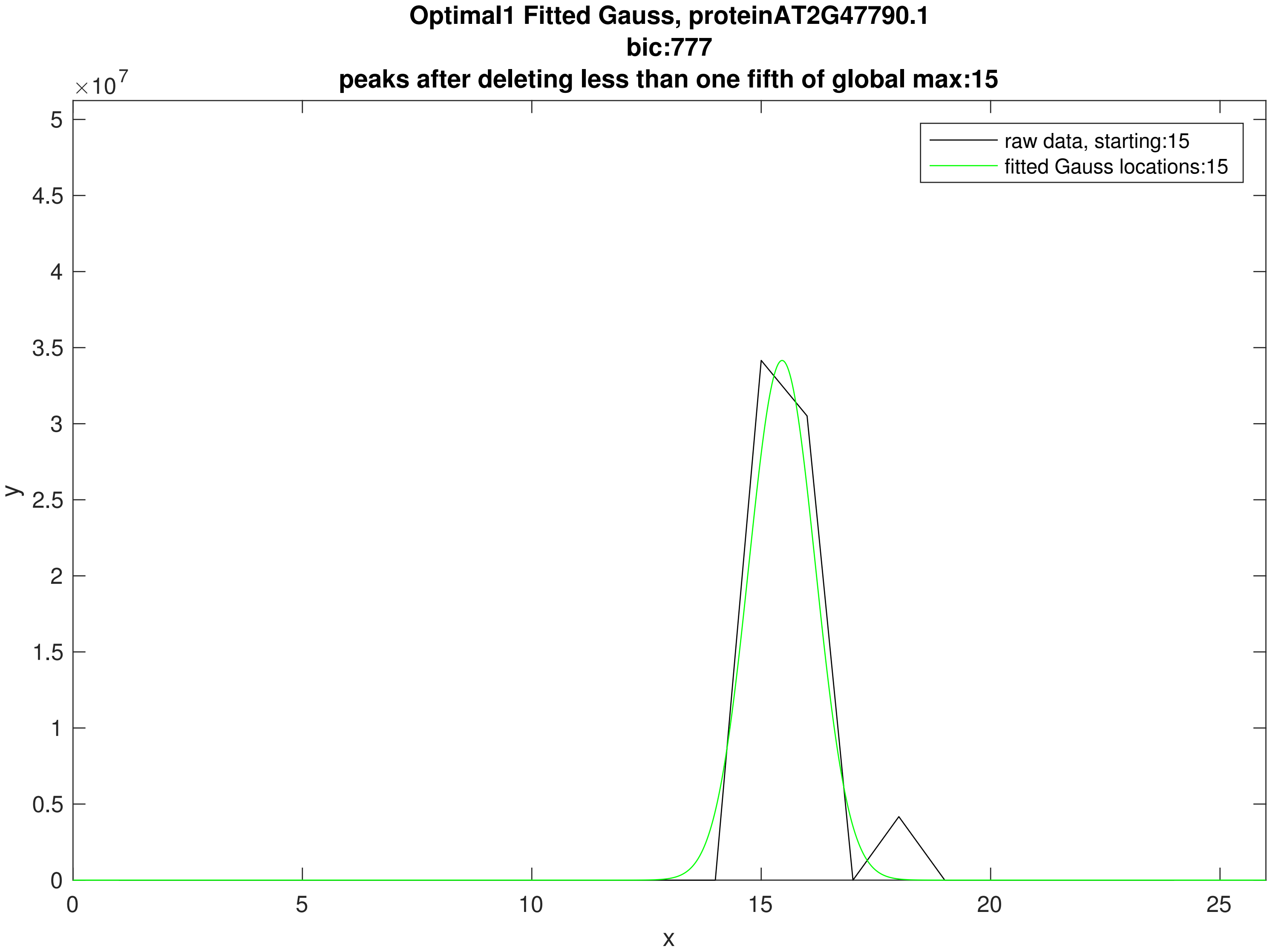
Optimal1 Fitted Gauss, proteinAT2G47790.1
bic:777
peaks after deleting less than one fifth of global max:15
×10^7
raw data, starting:15
fitted Gauss locations:15
y
x
0
0.5
1
1.5
2
2.5
3
3.5
4
4.5
5
0
5
10
15
20
25

Optimal1 Fitted Gauss, proteinAT2G47970.1;AT2G47970.2
bic:874
peaks after deleting less than one fifth of global max:15

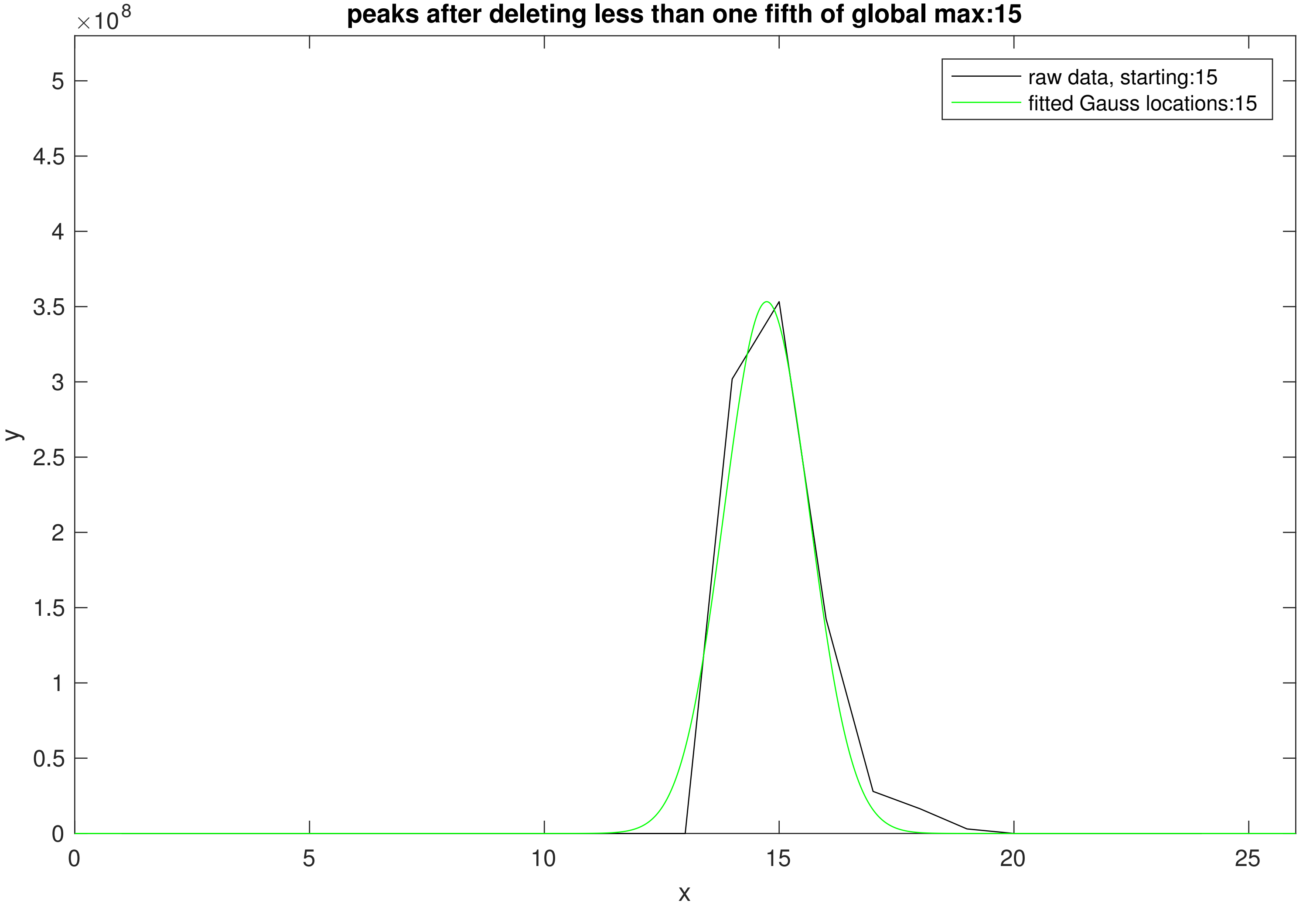

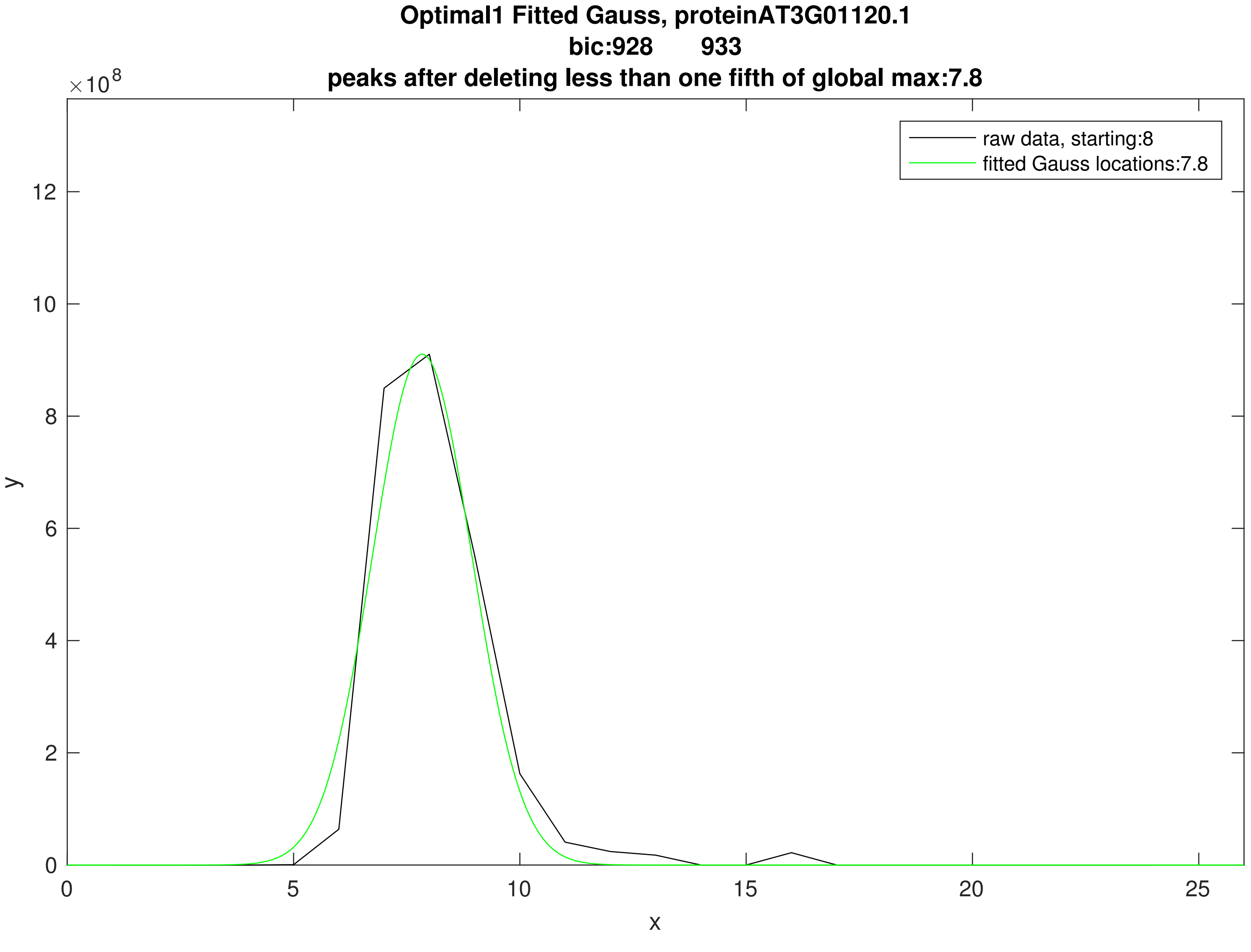

Optimal1 Fitted Gauss, proteinAT3G01120.1
bic:928 933
peaks after deleting less than one fifth of global max:7.8
×10^8
raw data, starting:8
fitted Gauss locations:7.8
y
x
0
2
4
6
8
10
12
0
5
10
15
20
25

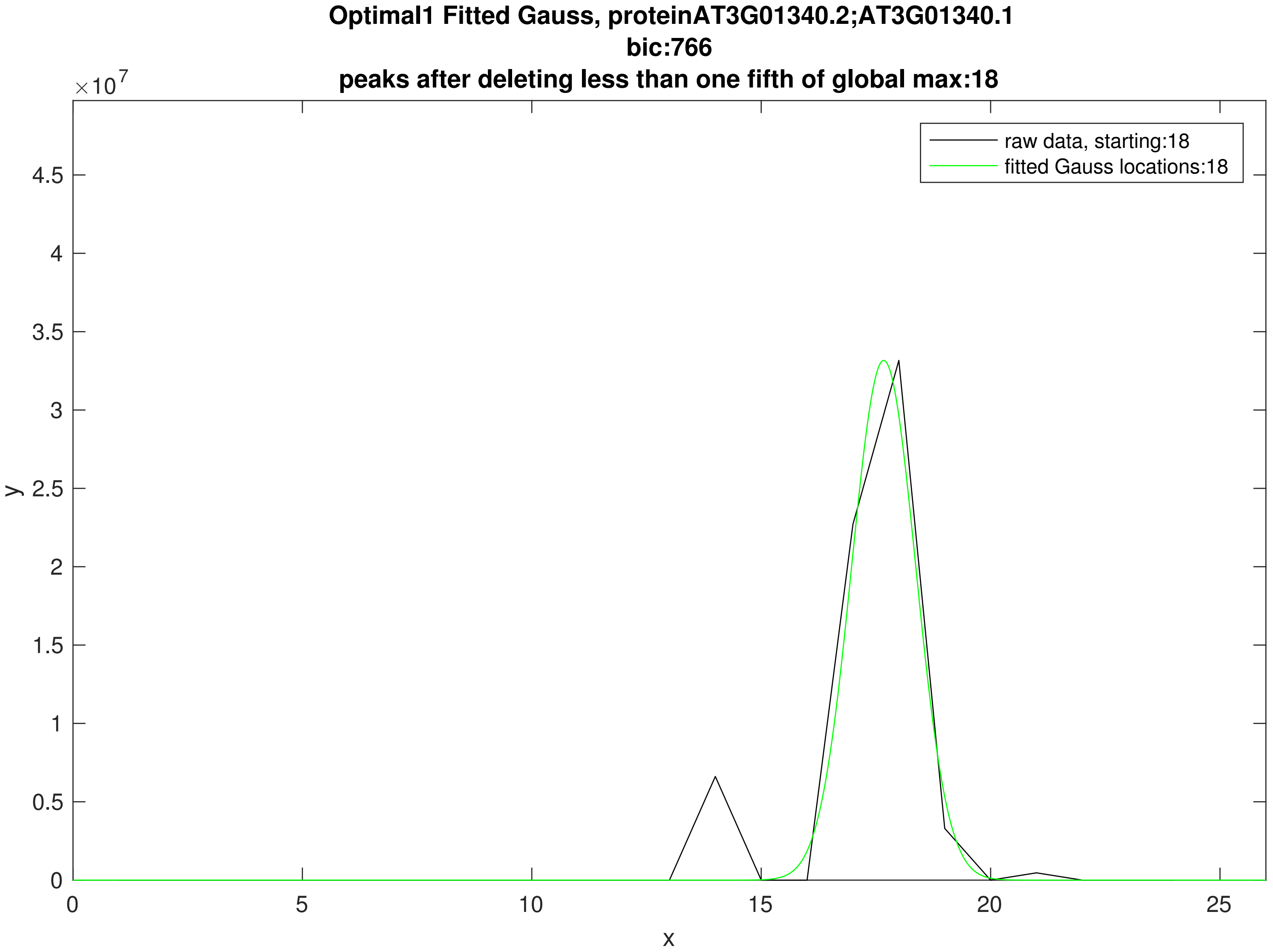

Optimal1 Fitted Gauss, proteinAT3G01340.2;AT3G01340.1
bic:766
peaks after deleting less than one fifth of global max:18
×10^7
raw data, starting:18
fitted Gauss locations:18
4.5
4
3.5
3
2.5
2
1.5
1
0.5
0
y
0
5
10
15
20
25
x

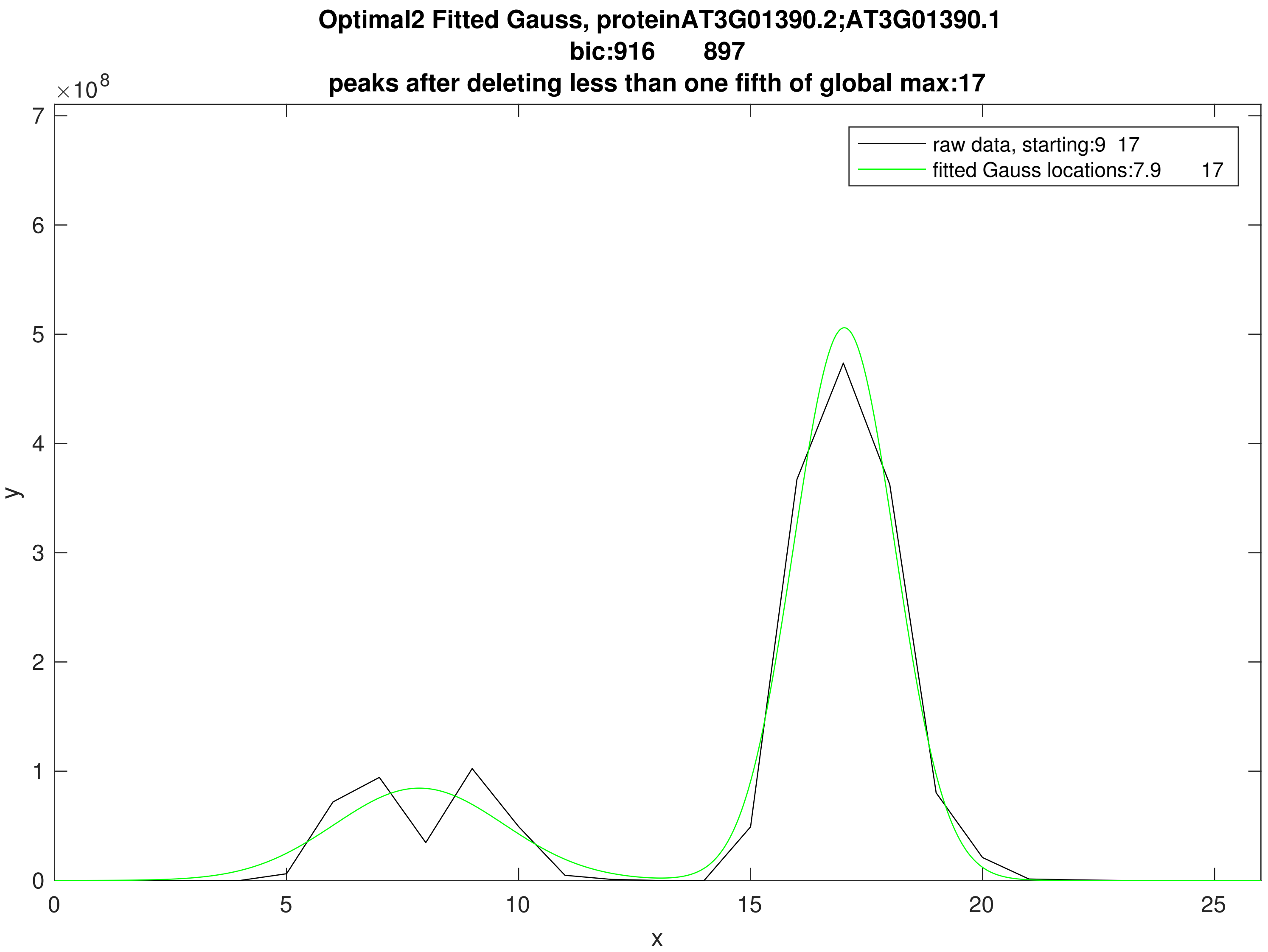
Optimal2 Fitted Gauss, proteinAT3G01390.2;AT3G01390.1
bic:916 897
peaks after deleting less than one fifth of global max:17
×10^8
raw data, starting:9 17
fitted Gauss locations:7.9 17
y
x
0
1
2
3
4
5
6
7
0
5
10
15
20
25

##### Optimal1 Fitted Gauss, proteinAT3G01420.1
##### bic:801
##### peaks after deleting less than one fifth of global max:6.6

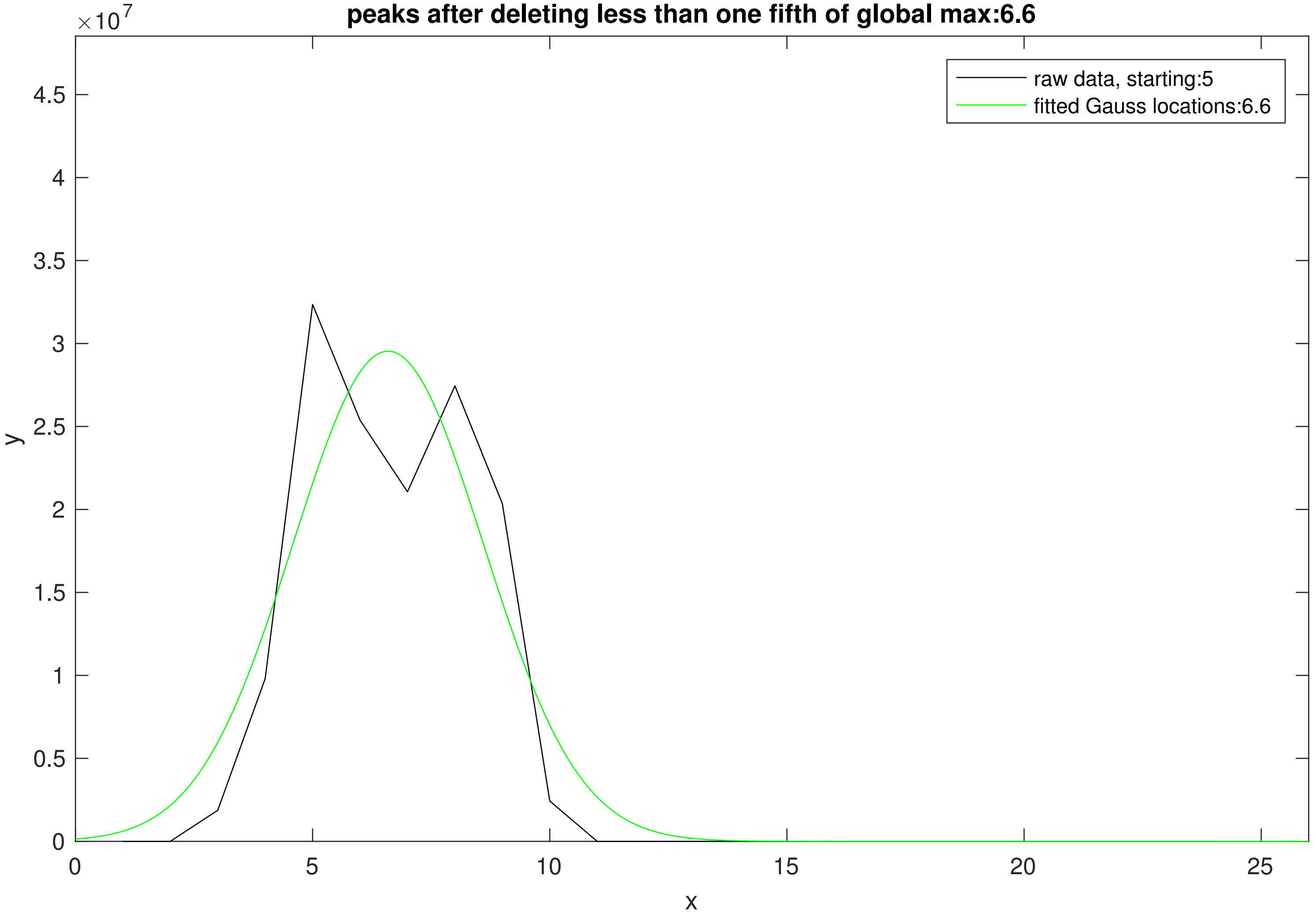

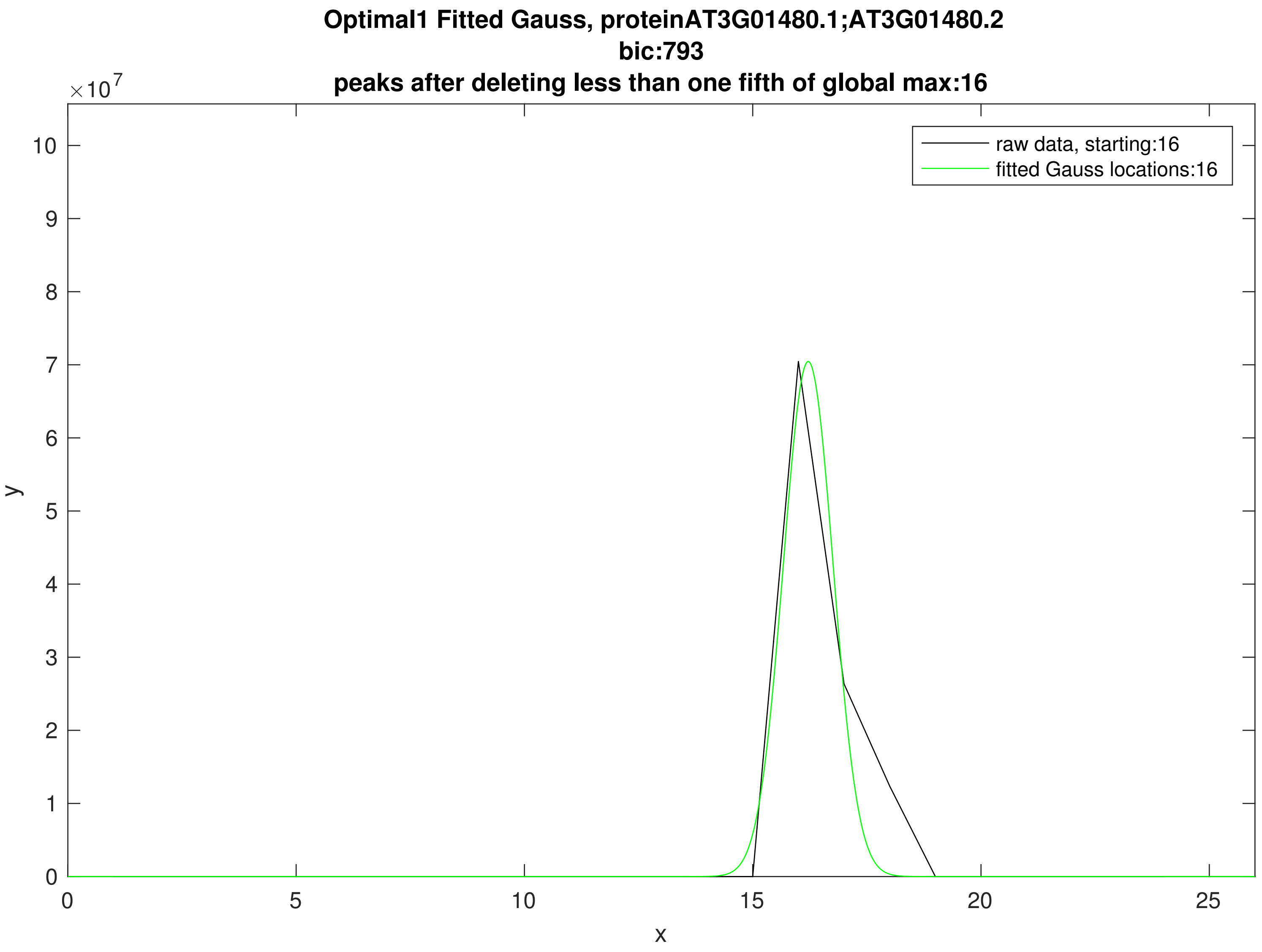
Optimal1 Fitted Gauss, proteinAT3G01480.1;AT3G01480.2
bic:793
peaks after deleting less than one fifth of global max:16
×10^7
raw data, starting:16
fitted Gauss locations:16
0
1
2
3
4
5
6
7
8
9
10
y
0
5
10
15
20
25
x

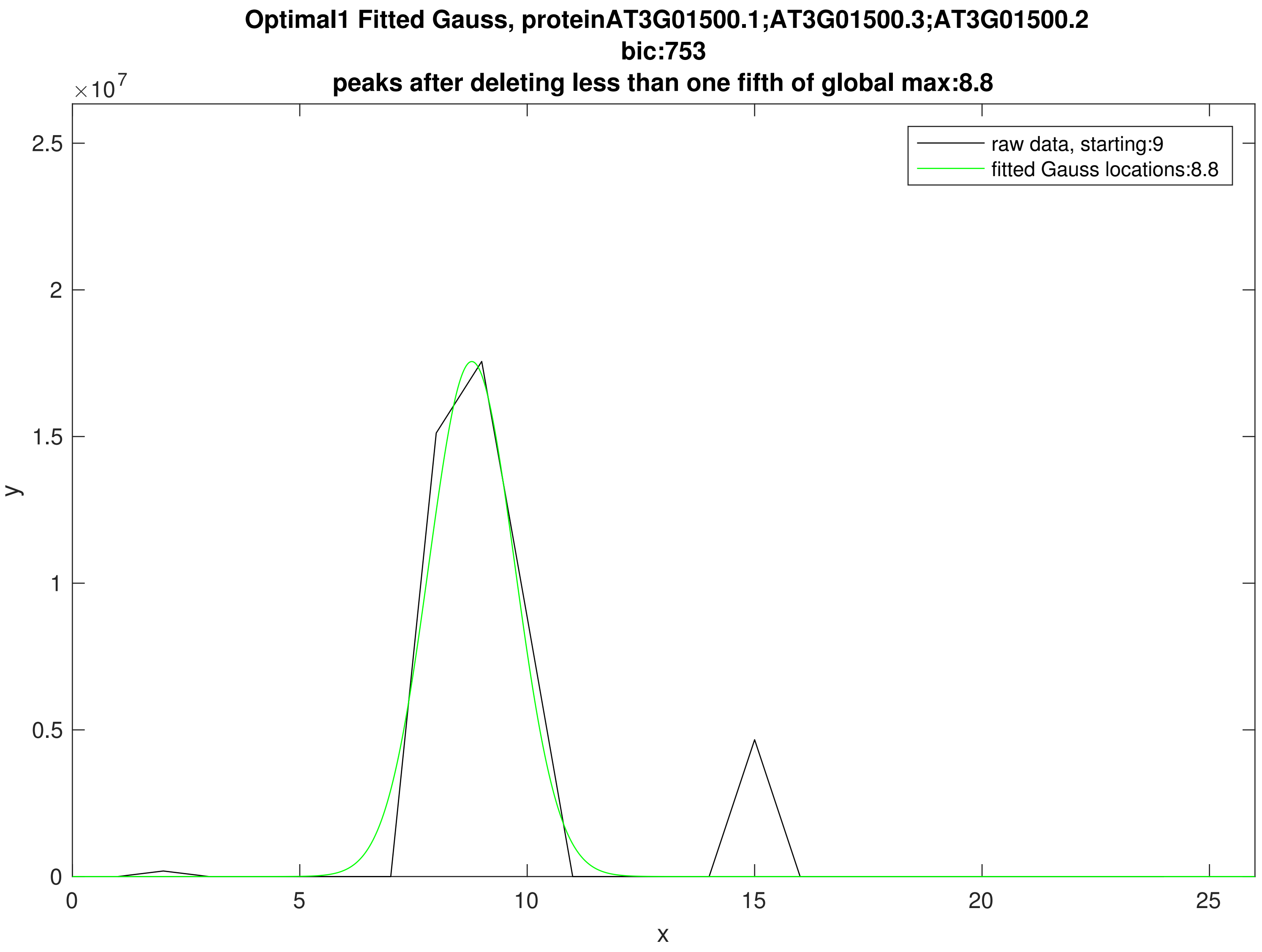
Optimal1 Fitted Gauss, proteinAT3G01500.1;AT3G01500.3;AT3G01500.2
bic:753
peaks after deleting less than one fifth of global max:8.8
raw data, starting:9
fitted Gauss locations:8.8
×10^7
2.5
2
1.5
1
0.5
0
y
0
5
10
15
20
25
x

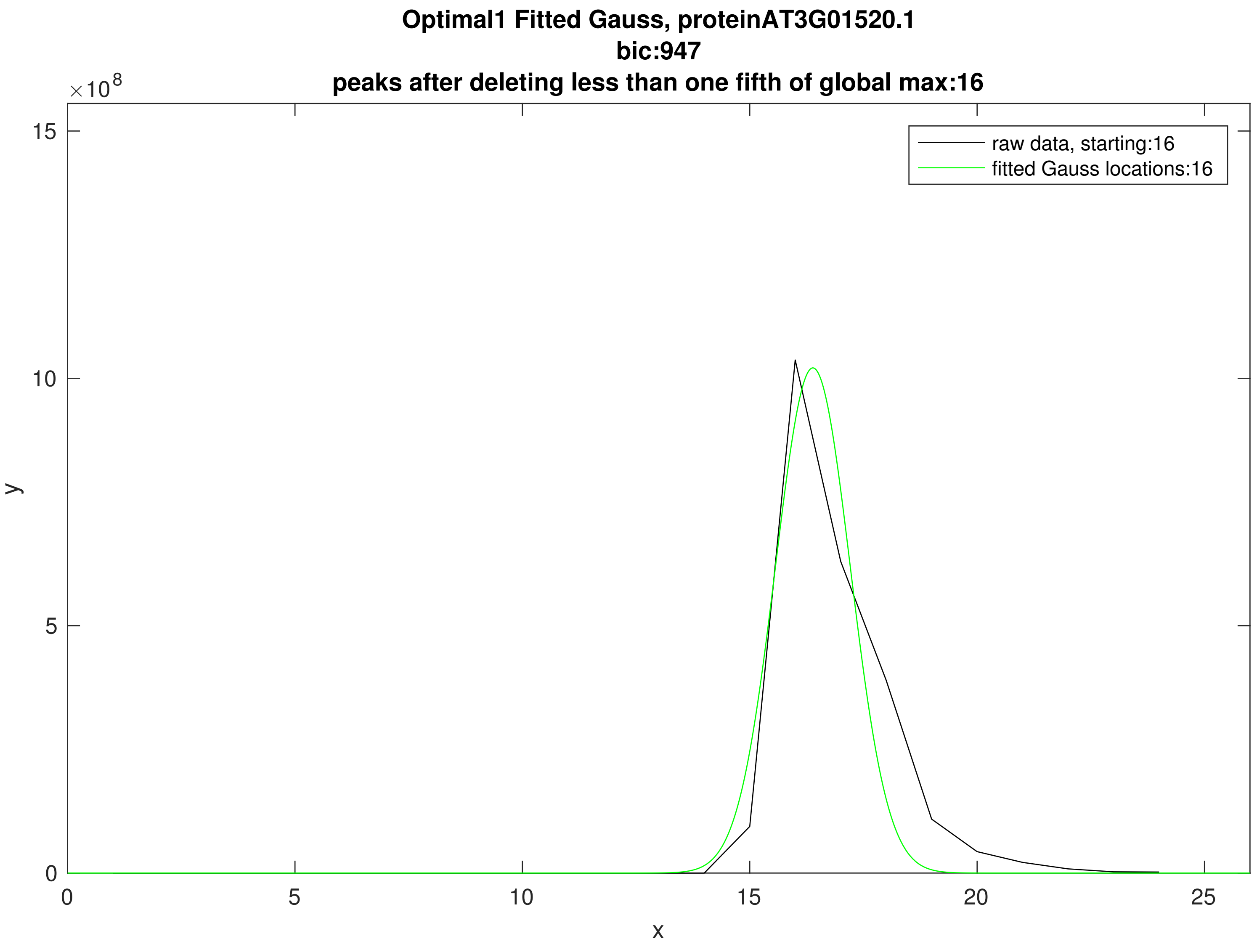

Optimal1 Fitted Gauss, proteinAT3G01520.1
bic:947
peaks after deleting less than one fifth of global max:16
×10^8
15
10
5
0
y
0
5
10
15
20
25
x
raw data, starting:16
fitted Gauss locations:16

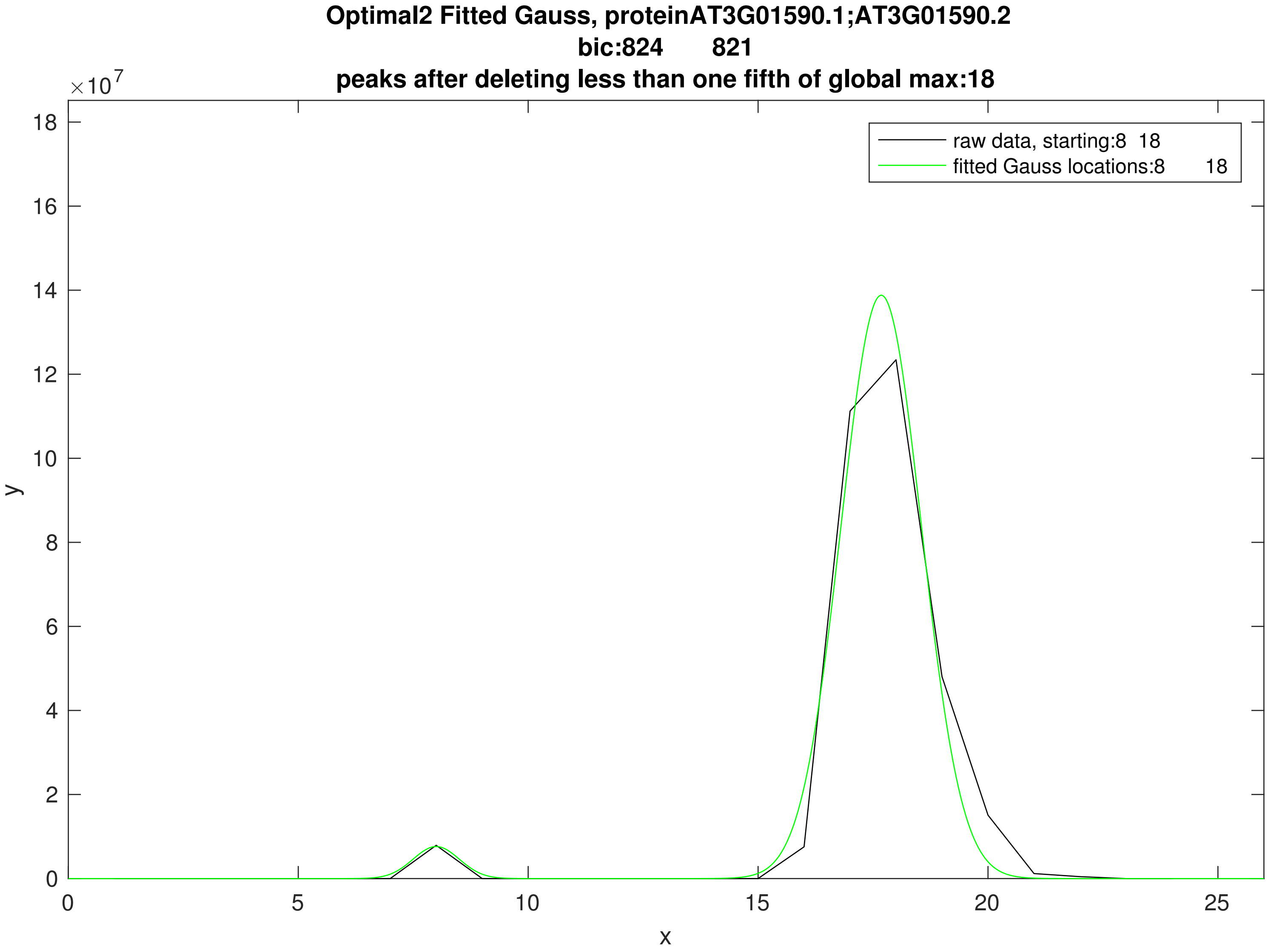

Optimal2 Fitted Gauss, proteinAT3G01590.1;AT3G01590.2
bic:824 821
peaks after deleting less than one fifth of global max:18
×10^7
raw data, starting:8 18
fitted Gauss locations:8 18
y
x
0
2
4
6
8
10
12
14
16
18
0
5
10
15
20
25

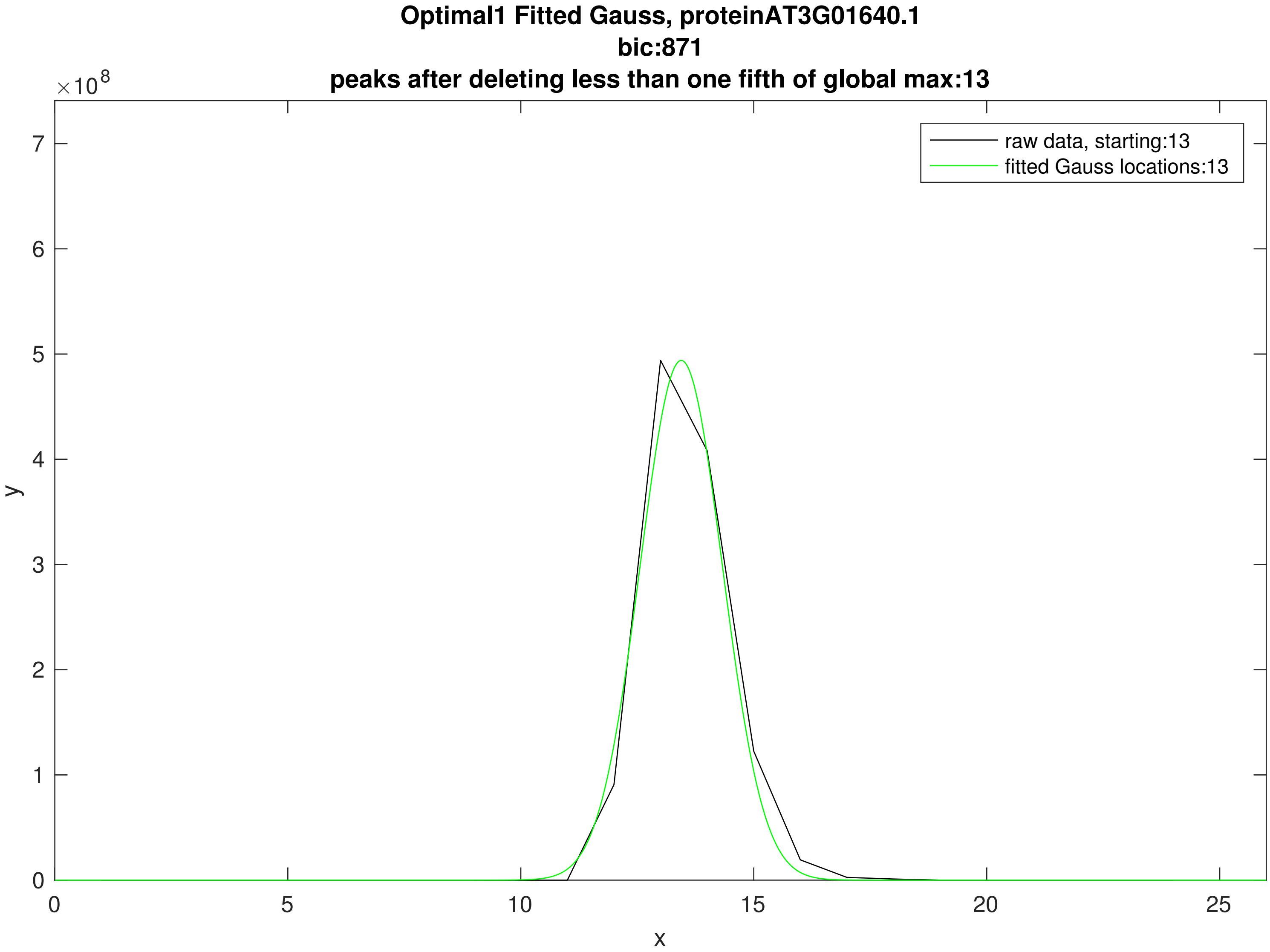
Optimal1 Fitted Gauss, proteinAT3G01640.1
bic:871
peaks after deleting less than one fifth of global max:13
×10^8
raw data, starting:13
fitted Gauss locations:13
y
x
0
1
2
3
4
5
6
7
0
5
10
15
20
25

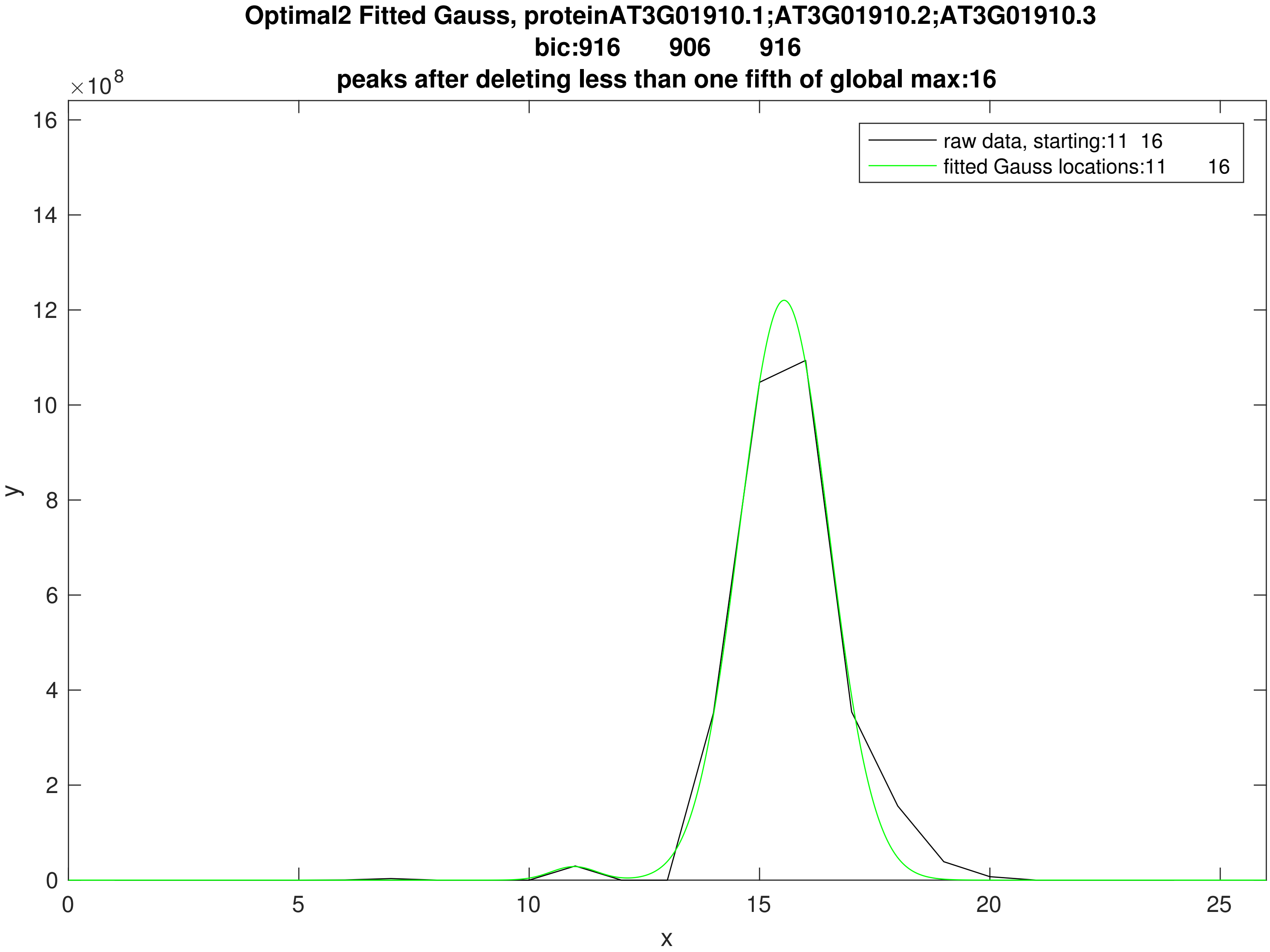

Optimal2 Fitted Gauss, proteinAT3G01910.1;AT3G01910.2;AT3G01910.3
bic:916 906 916
peaks after deleting less than one fifth of global max:16
×10^8
raw data, starting:11 16
fitted Gauss locations:11 16
y
x
0
2
4
6
8
10
12
14
16
0
5
10
15
20
25

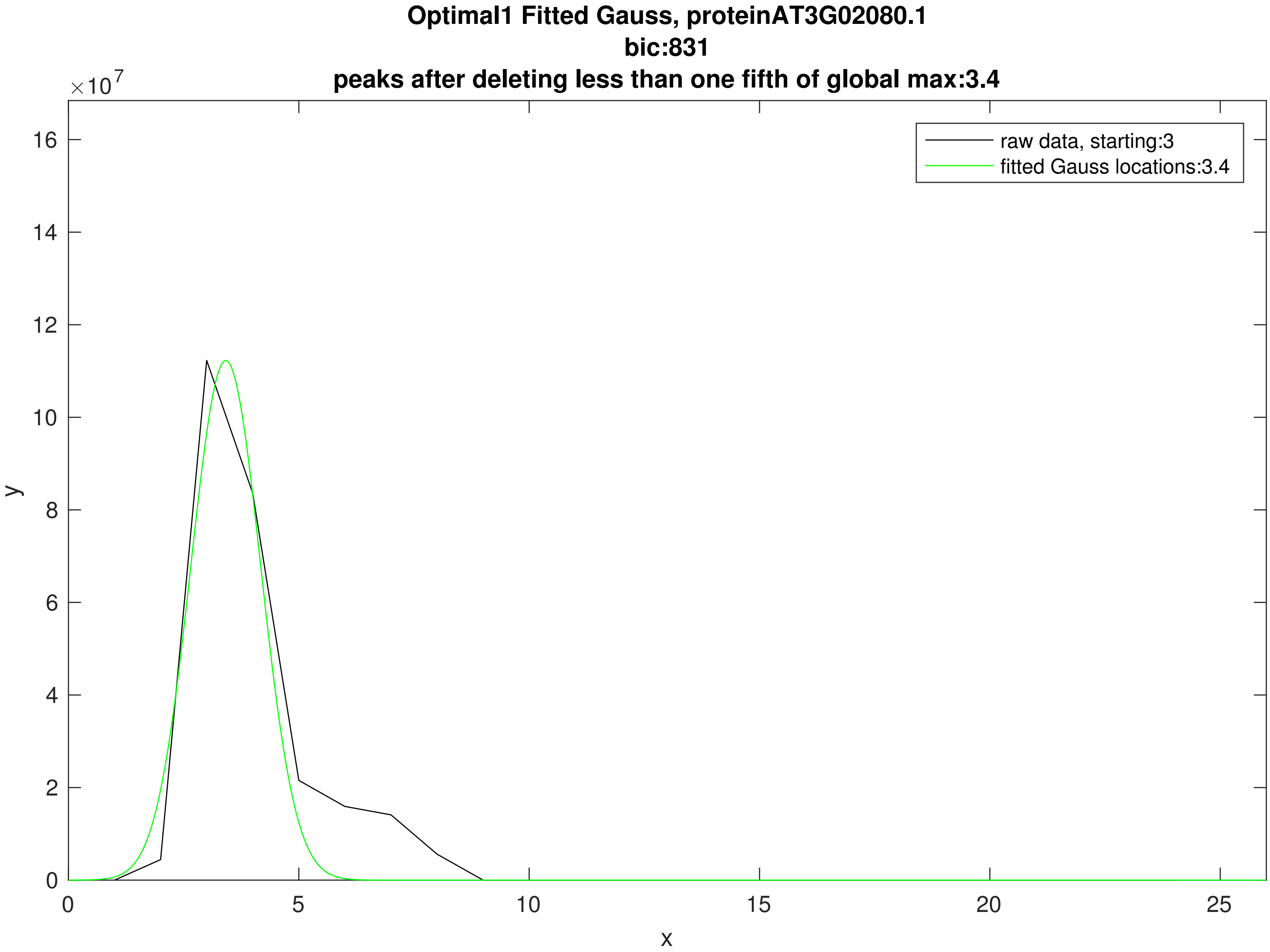
Optimal1 Fitted Gauss, proteinAT3G02080.1
bic:831
peaks after deleting less than one fifth of global max:3.4
×10^7
raw data, starting:3
fitted Gauss locations:3.4
y
x
0
2
4
6
8
10
12
14
16
0
5
10
15
20
25

**Optimal1 Fitted Gauss, proteinAT3G02090.1;AT3G02090.2**
**bic:824 830 839 849**
**peaks after deleting less than one fifth of global max:16**

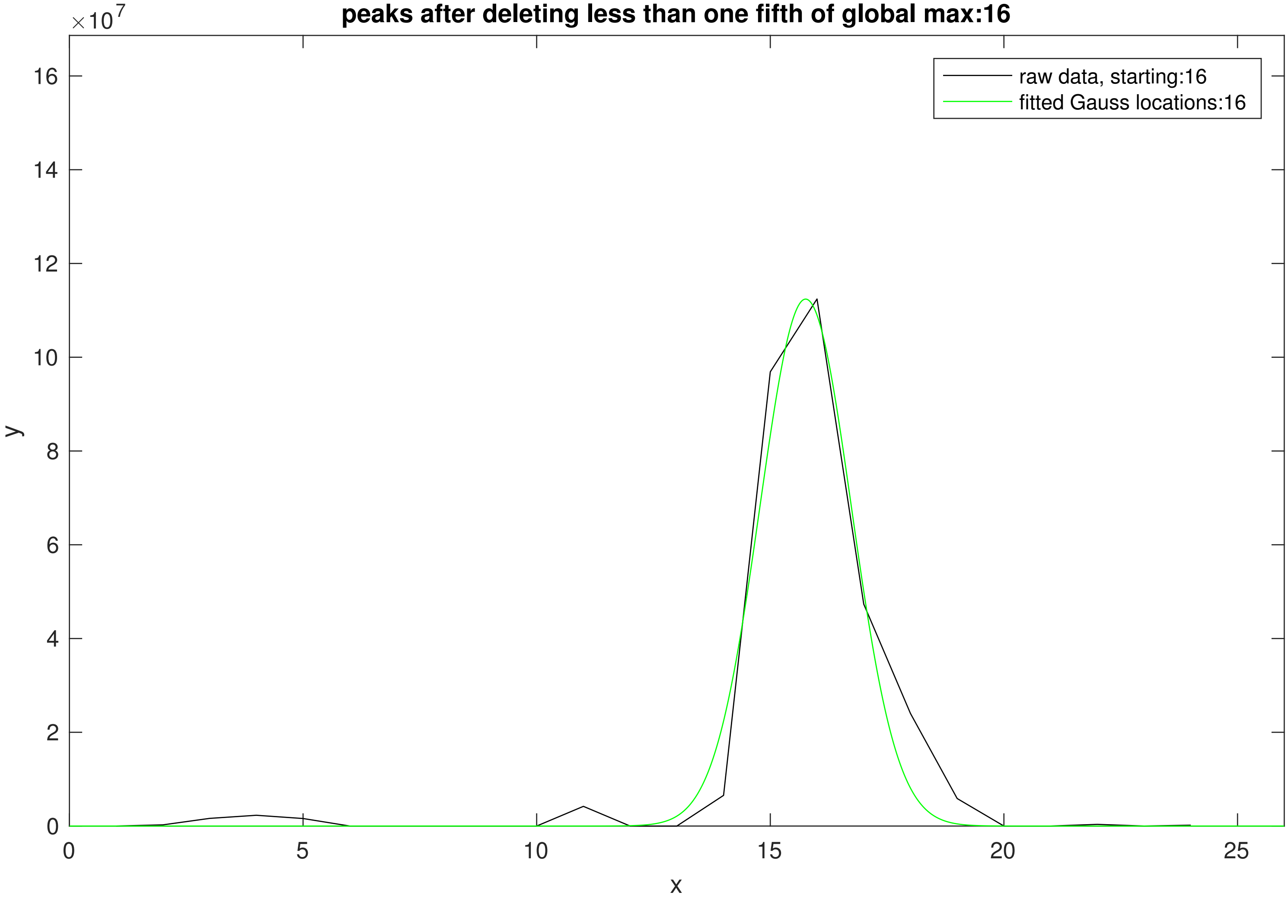

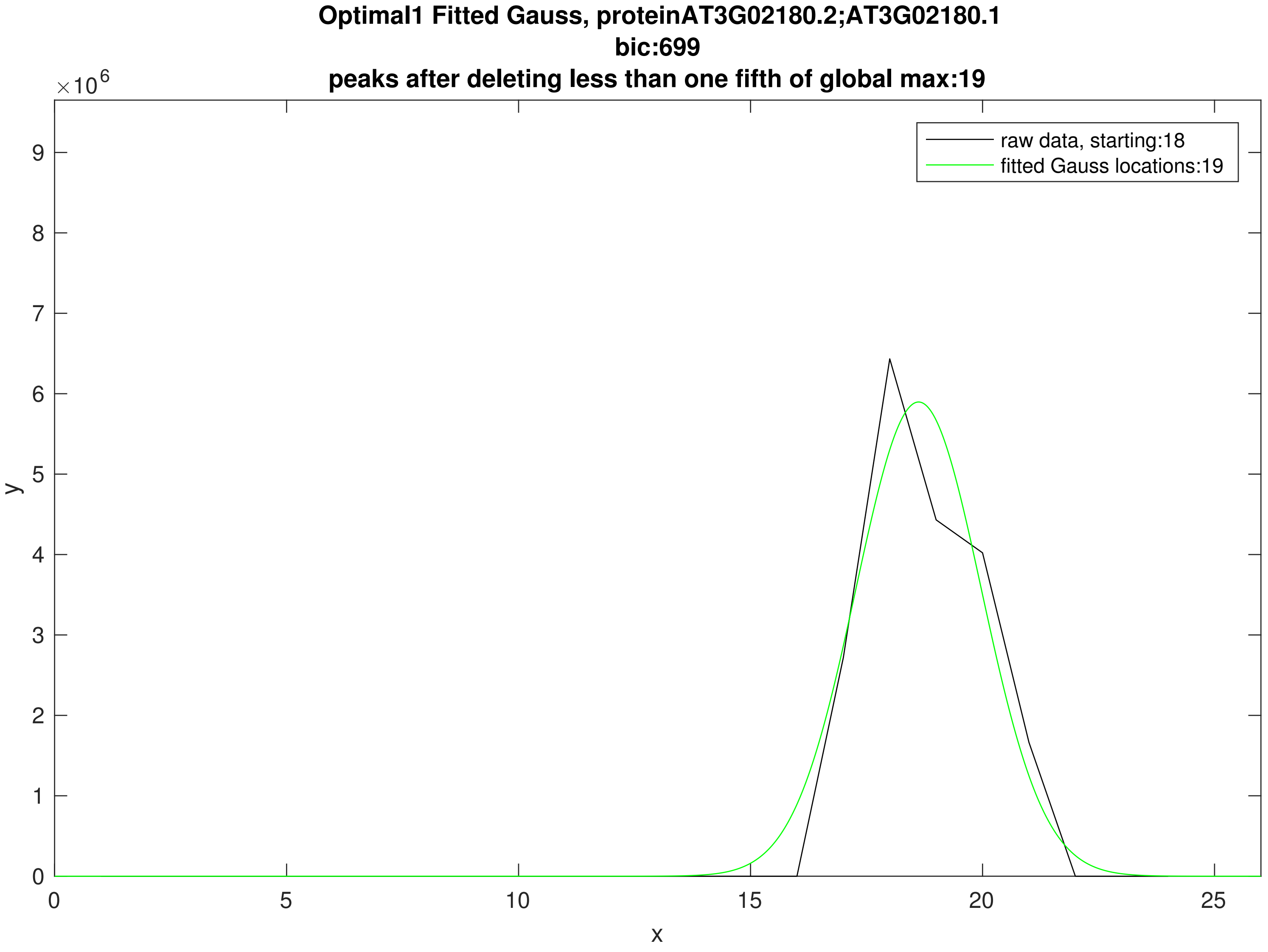
Optimal1 Fitted Gauss, proteinAT3G02180.2;AT3G02180.1
bic:699
peaks after deleting less than one fifth of global max:19
×10^6
raw data, starting:18
fitted Gauss locations:19
y
x
0
1
2
3
4
5
6
7
8
9
0
5
10
15
20
25

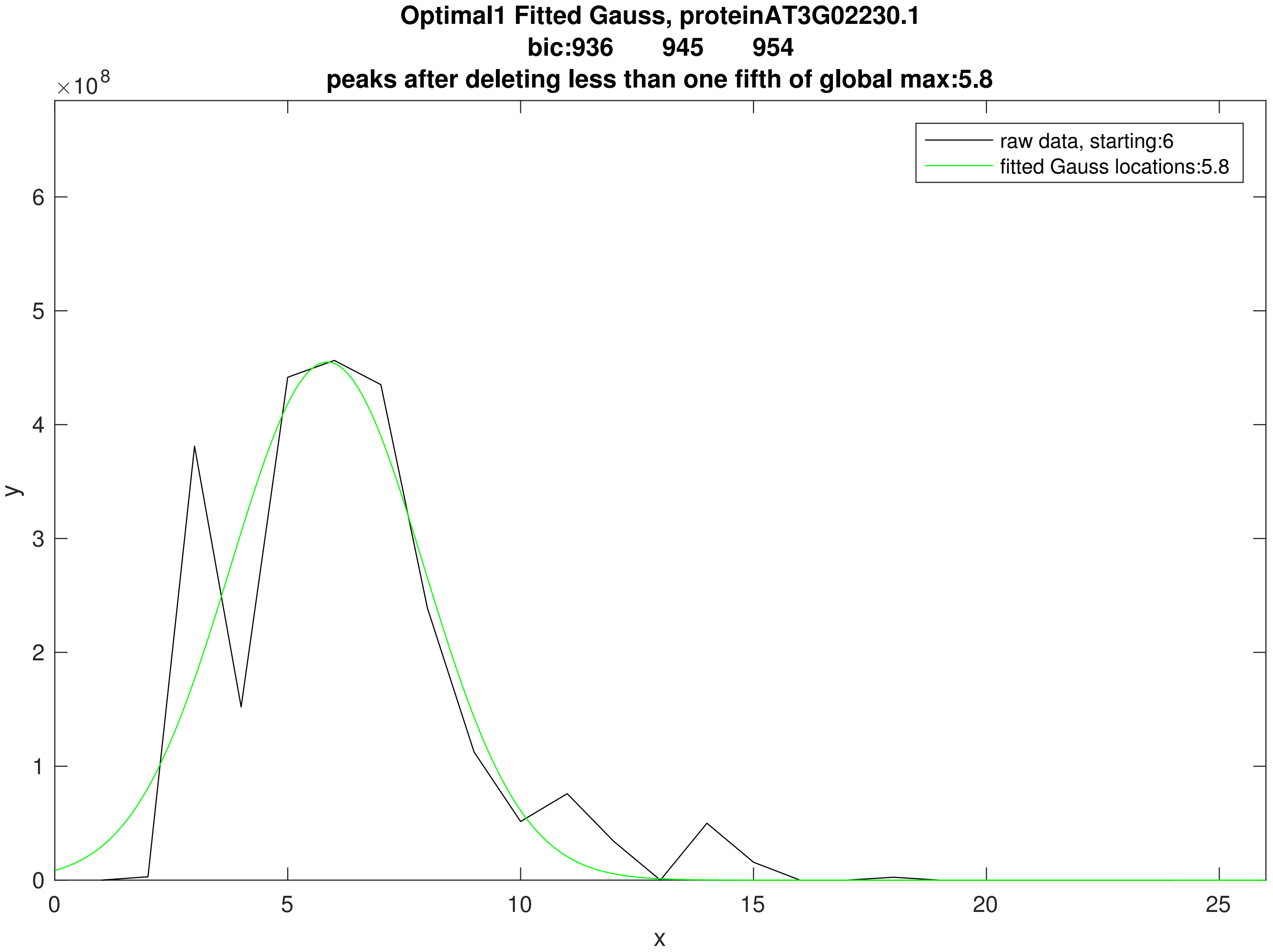
Optimal1 Fitted Gauss, proteinAT3G02230.1
bic:936 945 954
peaks after deleting less than one fifth of global max:5.8
×10^8
raw data, starting:6
fitted Gauss locations:5.8
y
x
0
1
2
3
4
5
6
0
5
10
15
20
25

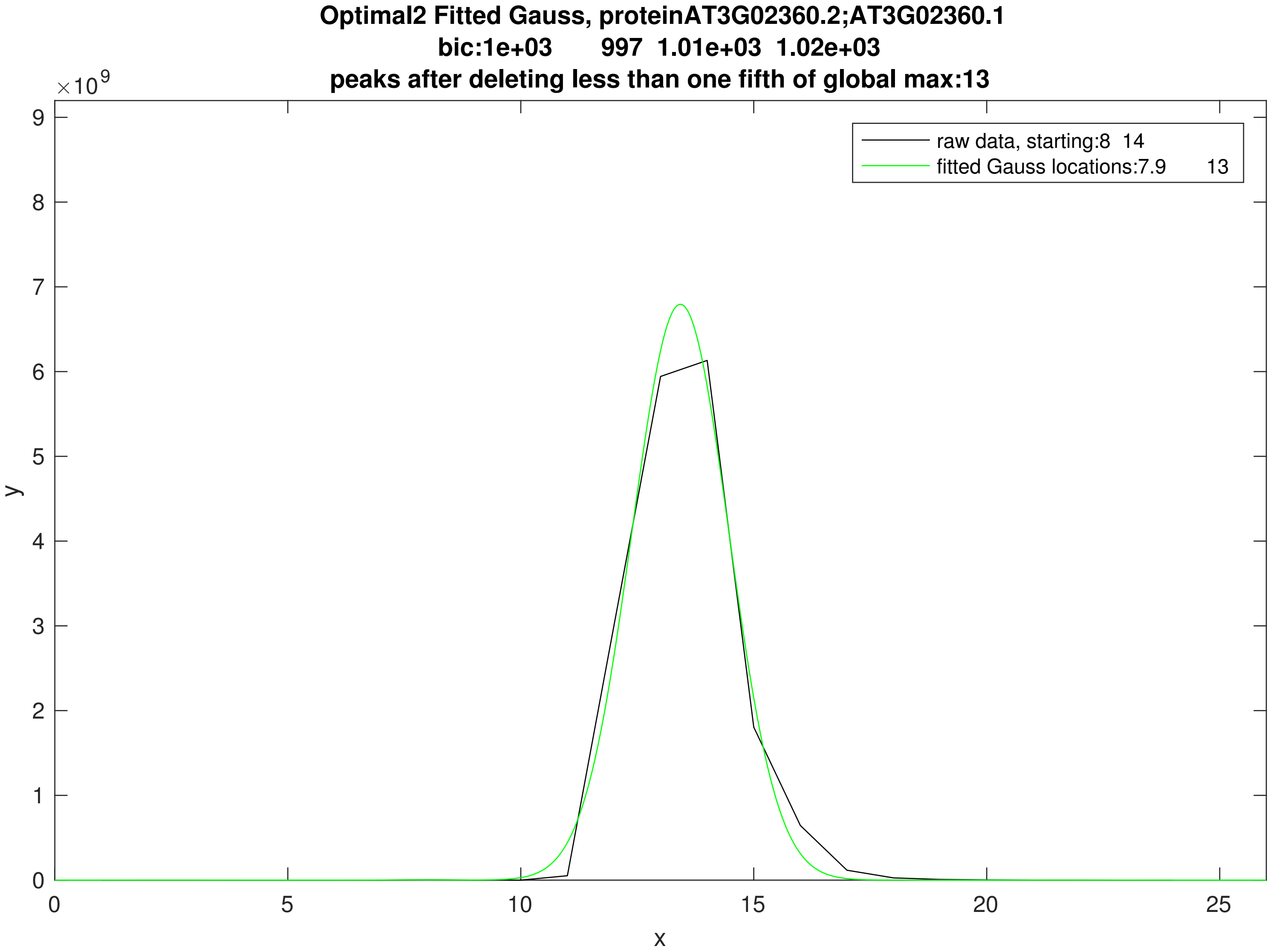

Optimal2 Fitted Gauss, proteinAT3G02360.2;AT3G02360.1
bic:1e+03 997 1.01e+03 1.02e+03
peaks after deleting less than one fifth of global max:13
×10^9
raw data, starting:8 14
fitted Gauss locations:7.9 13
y
x
0
1
2
3
4
5
6
7
8
9
0
5
10
15
20
25

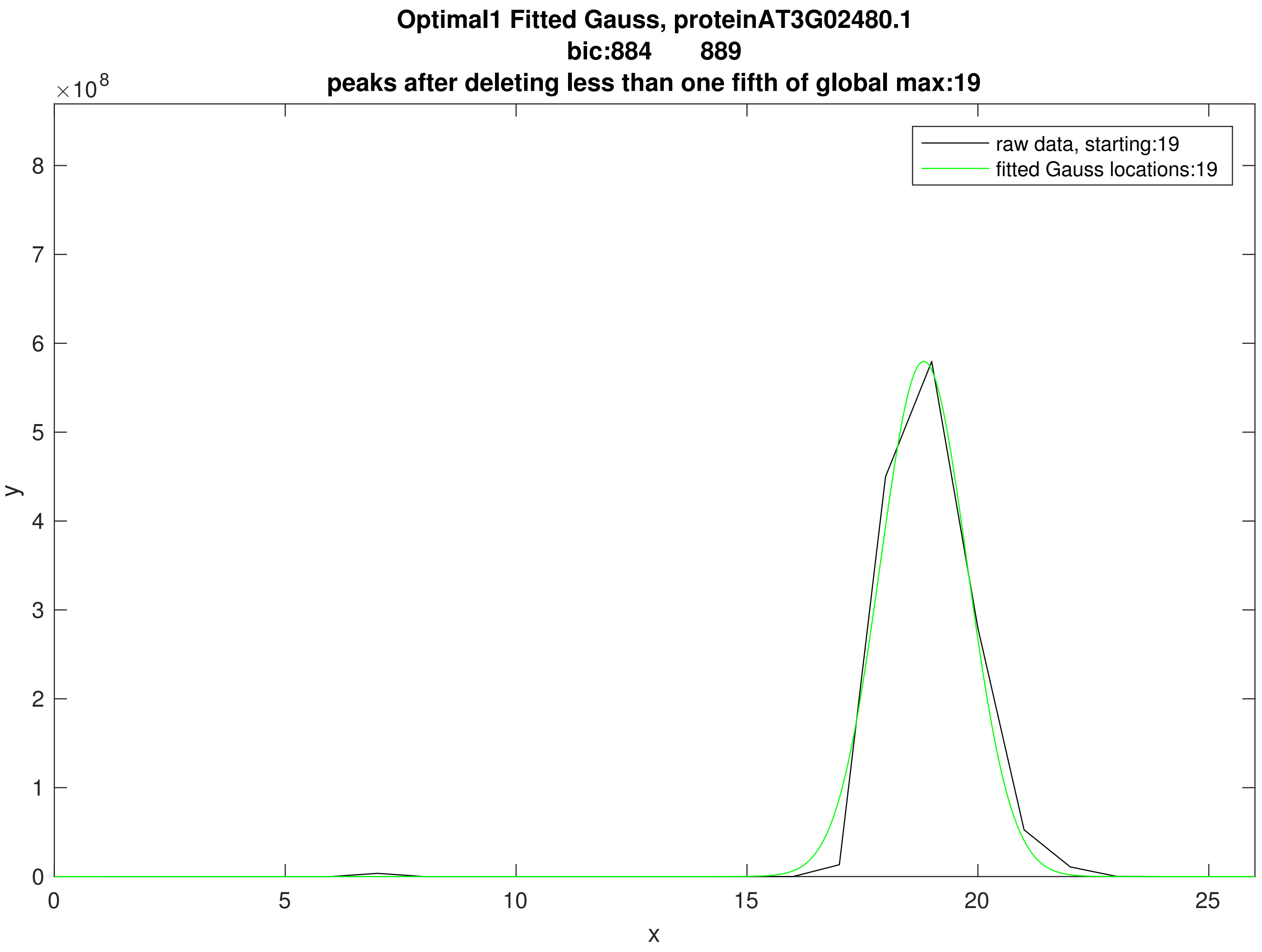

Optimal1 Fitted Gauss, proteinAT3G02480.1
bic:884 889
peaks after deleting less than one fifth of global max:19
×10^8
raw data, starting:19
fitted Gauss locations:19
y
x
0
1
2
3
4
5
6
7
8
0
5
10
15
20
25

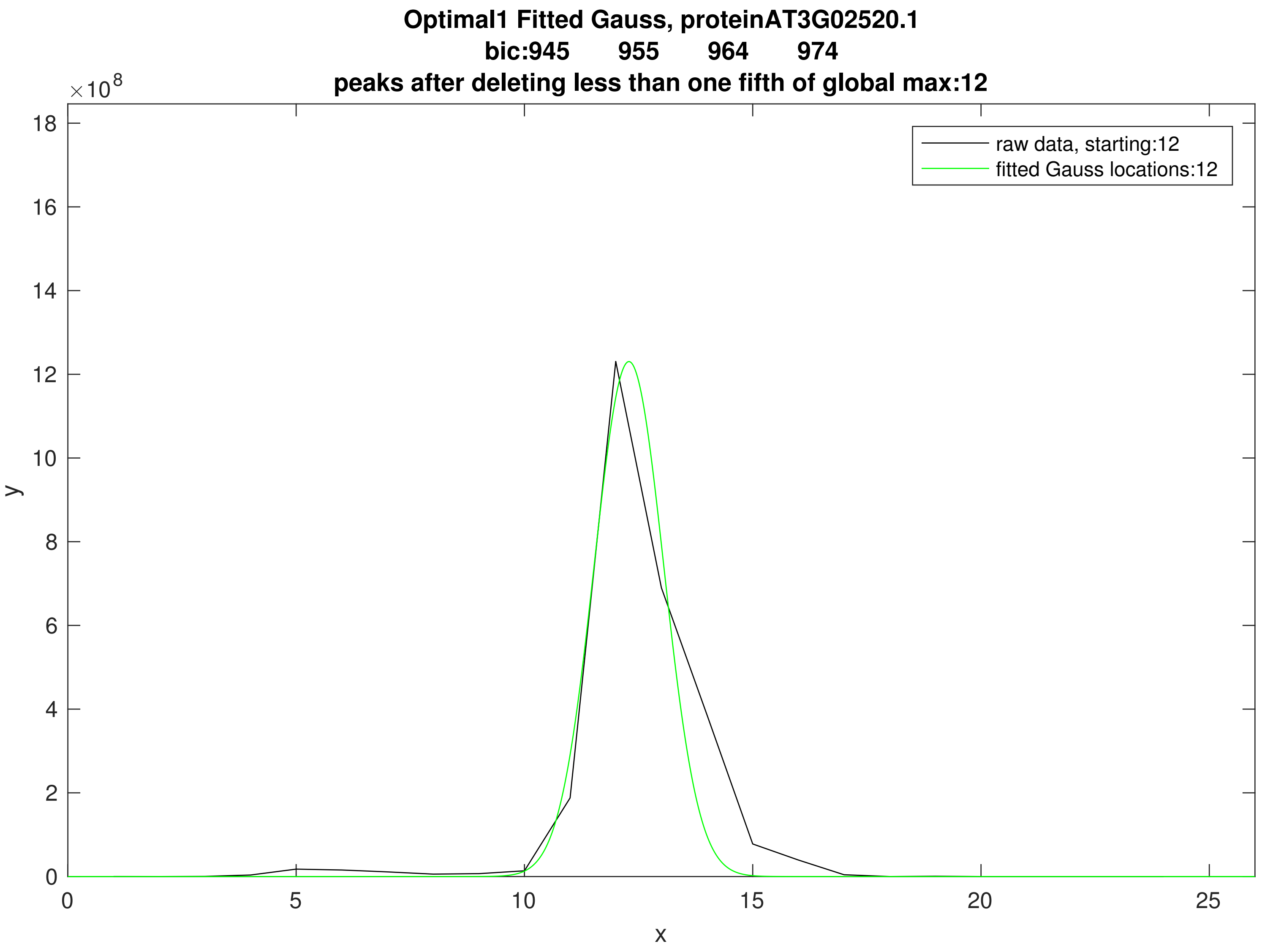

Optimal1 Fitted Gauss, proteinAT3G02520.1
bic:945 955 964 974
peaks after deleting less than one fifth of global max:12
×10^8
raw data, starting:12
fitted Gauss locations:12
y
x
0
2
4
6
8
10
12
14
16
18
0
5
10
15
20
25

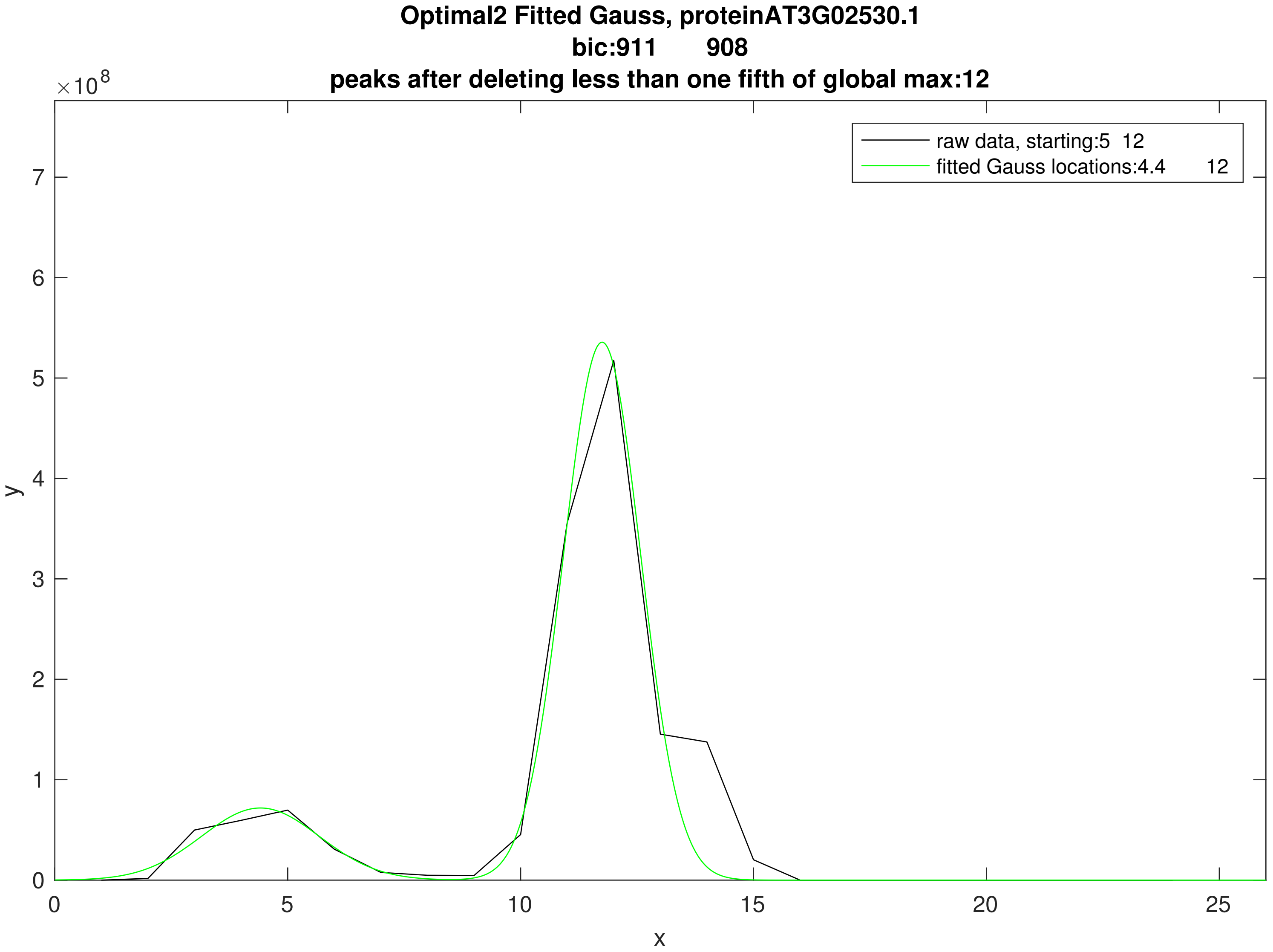
Optimal2 Fitted Gauss, proteinAT3G02530.1
bic:911 908
peaks after deleting less than one fifth of global max:12
×10^8
raw data, starting:5 12
fitted Gauss locations:4.4 12
y
x
0
1
2
3
4
5
6
7
0
5
10
15
20
25

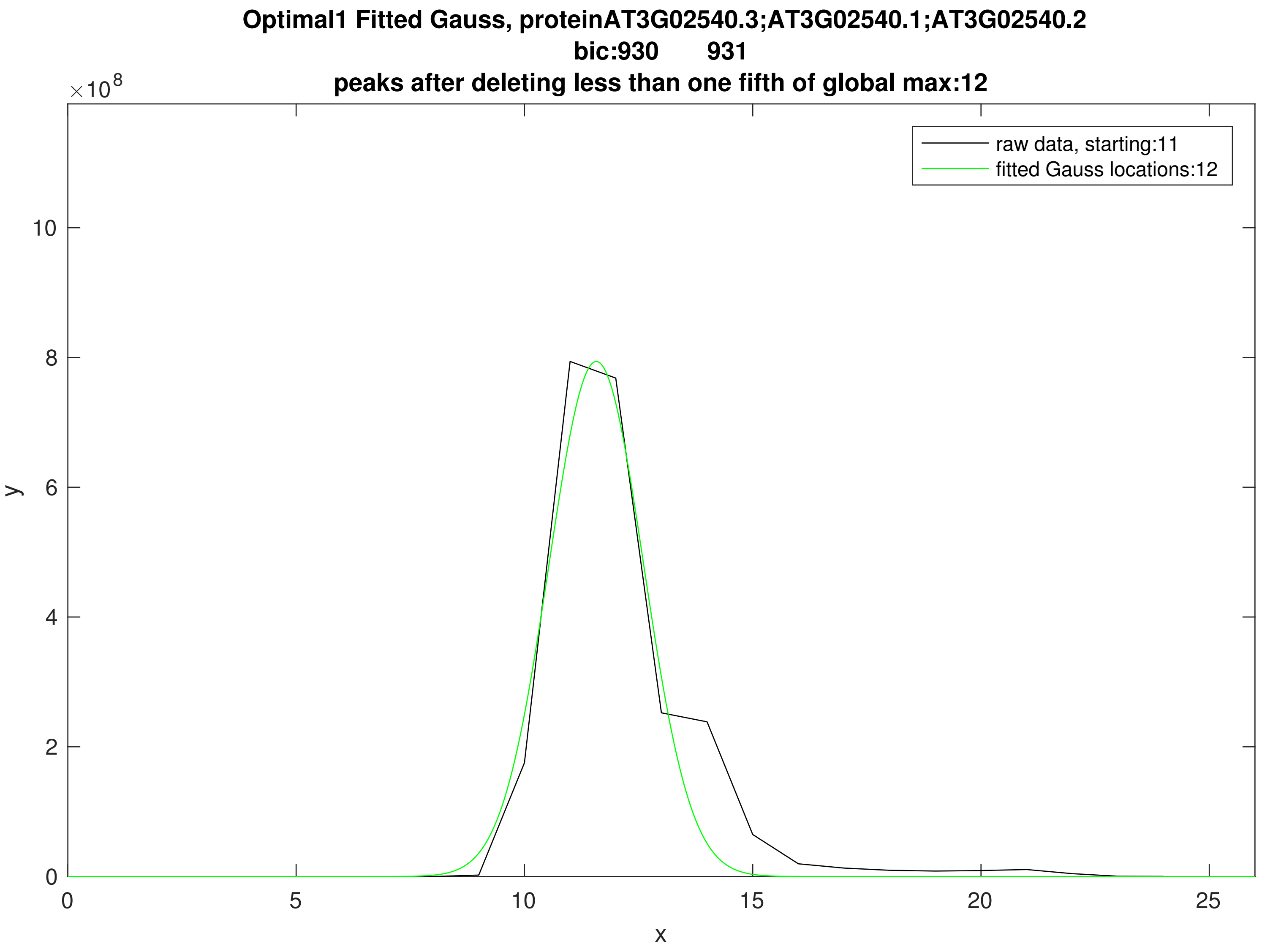
Optimal1 Fitted Gauss, proteinAT3G02540.3;AT3G02540.1;AT3G02540.2
bic:930 931
peaks after deleting less than one fifth of global max:12
×10^8
raw data, starting:11
fitted Gauss locations:12
10
8
6
4
2
0
y
0
5
10
15
20
25
x

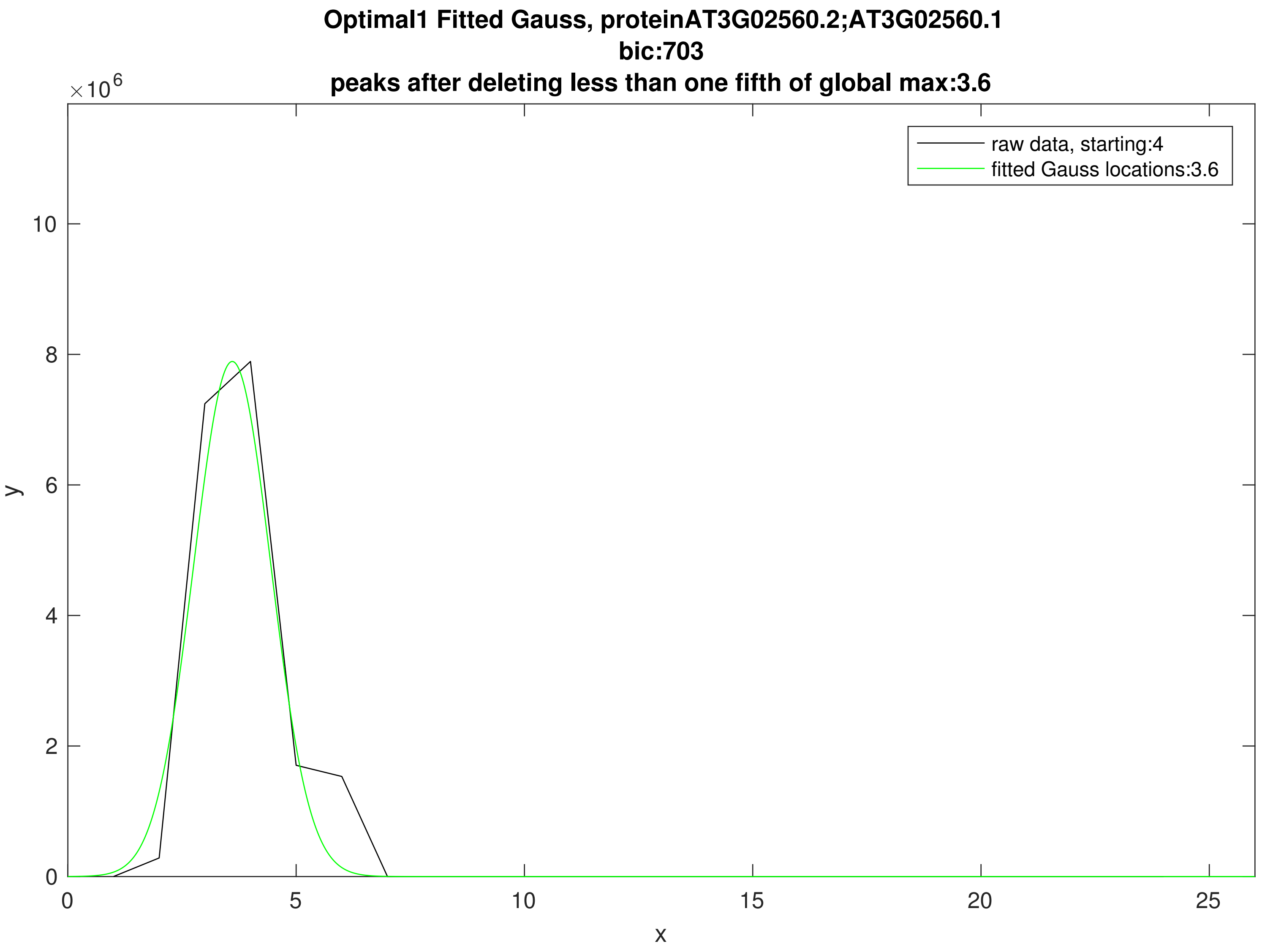
Optimal1 Fitted Gauss, proteinAT3G02560.2;AT3G02560.1
bic:703
peaks after deleting less than one fifth of global max:3.6
×10^6
raw data, starting:4
fitted Gauss locations:3.6
y
x
0
2
4
6
8
10
0
5
10
15
20
25

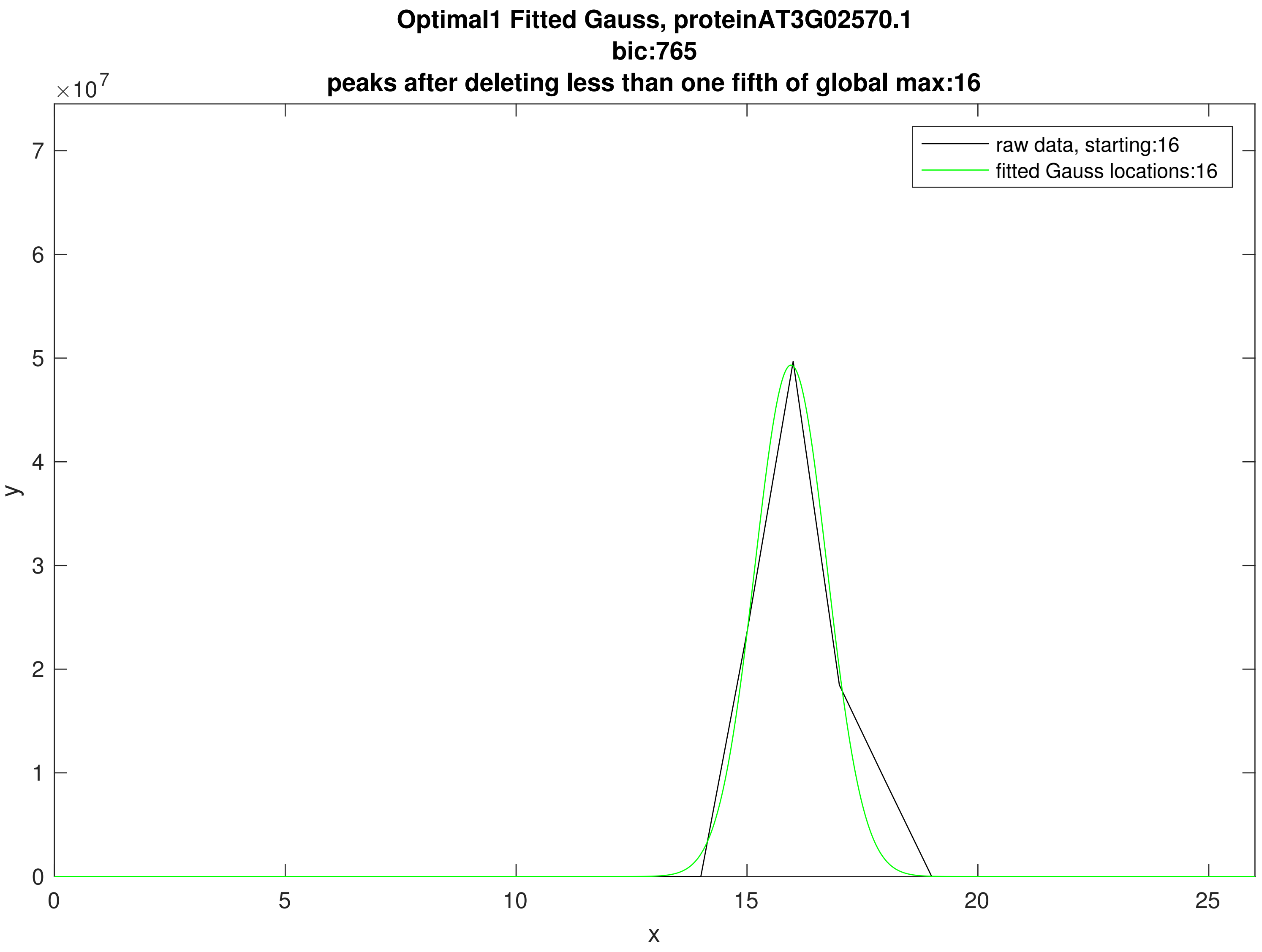

Optimal1 Fitted Gauss, proteinAT3G02570.1
bic:765
peaks after deleting less than one fifth of global max:16
×10^7
raw data, starting:16
fitted Gauss locations:16
y
x
0
1
2
3
4
5
6
7
0
5
10
15
20
25

**Optimal2 Fitted Gauss, proteinAT3G02660.1**
**bic:838 833**
**peaks after deleting less than one fifth of global max:15**

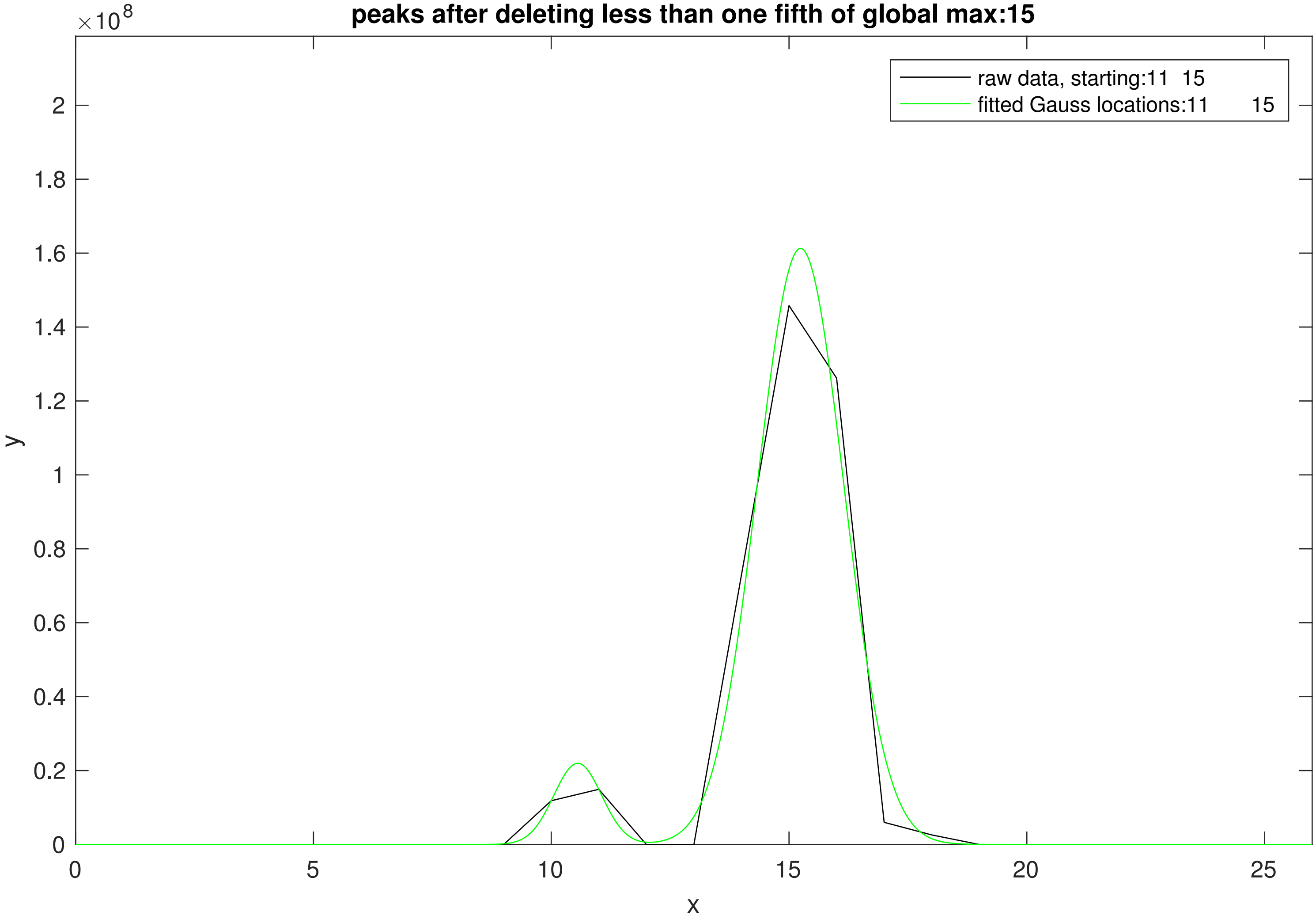

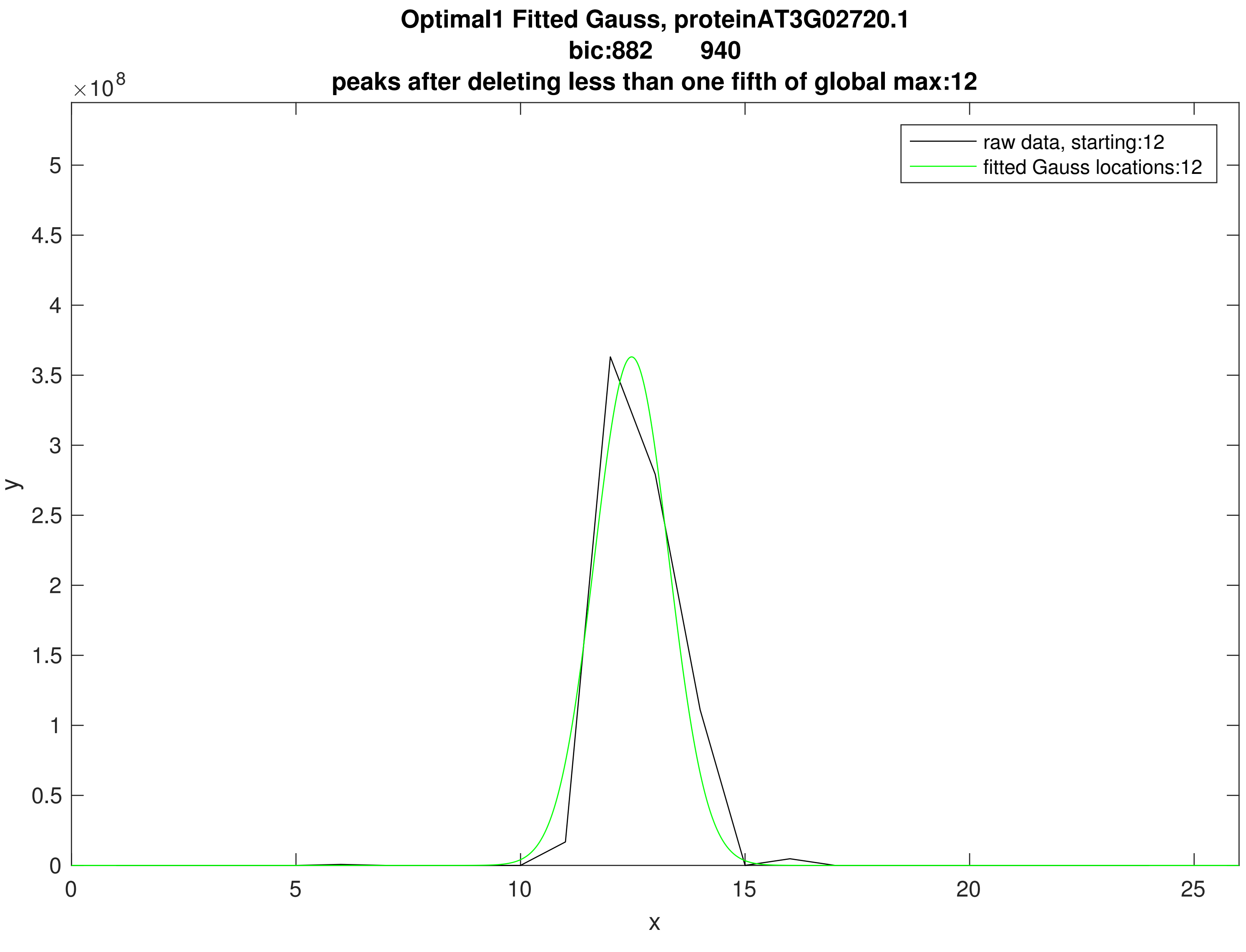
Optimal1 Fitted Gauss, proteinAT3G02720.1
bic:882 940
peaks after deleting less than one fifth of global max:12
×10^8
5
4.5
4
3.5
3
2.5
2
1.5
1
0.5
0
y
0
5
10
15
20
25
x
raw data, starting:12
fitted Gauss locations:12

**Optimal1 Fitted Gauss, proteinAT3G02740.1**
**bic:826**
**peaks after deleting less than one fifth of global max:12**

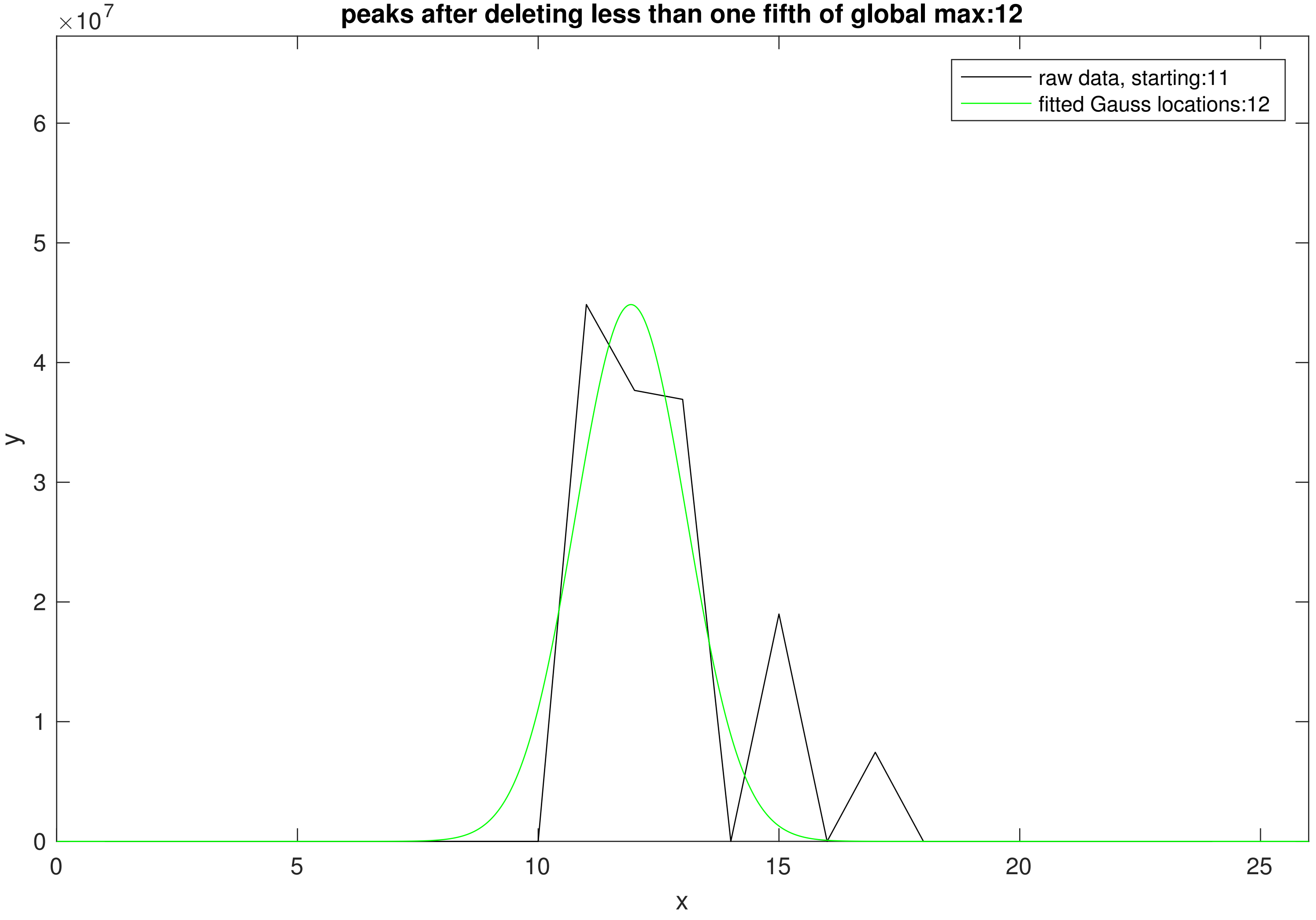

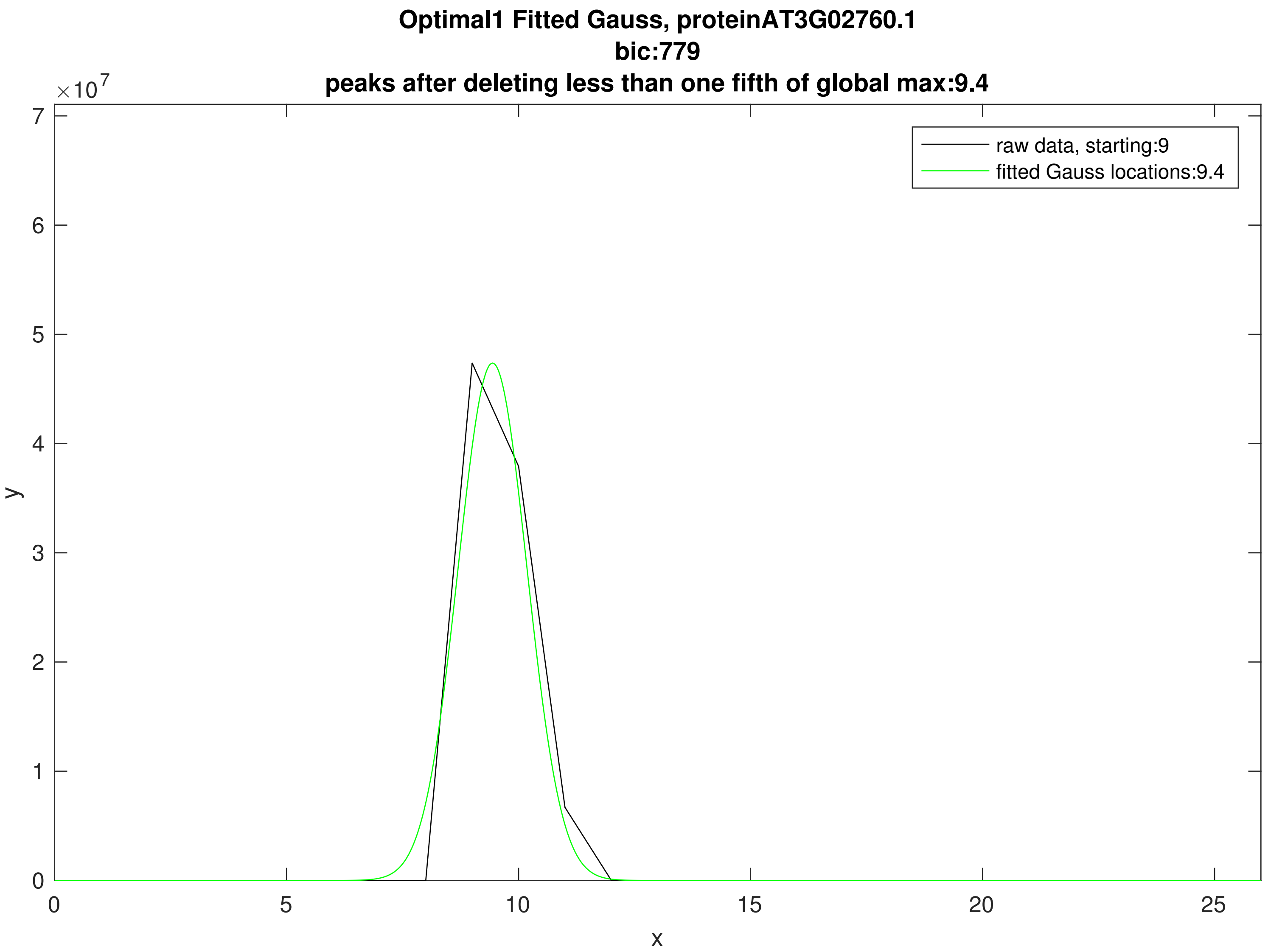
Optimal1 Fitted Gauss, proteinAT3G02760.1
bic:779
peaks after deleting less than one fifth of global max:9.4
×10^7
raw data, starting:9
fitted Gauss locations:9.4
y
x
0
1
2
3
4
5
6
7
0
5
10
15
20
25

**Optimal1 Fitted Gauss, proteinAT3G02770.1**
**bic:894 901**
**peaks after deleting less than one fifth of global max:14**

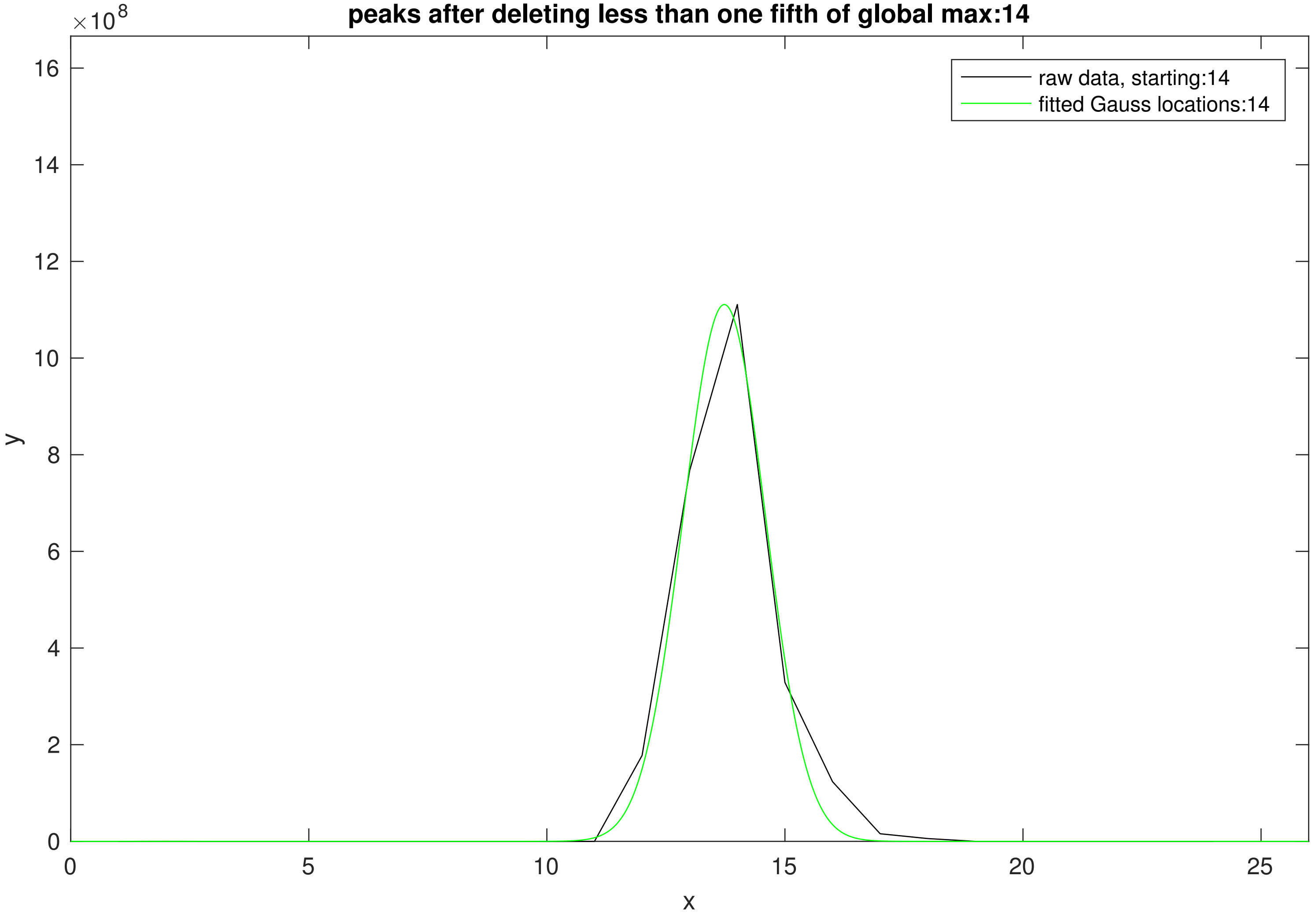

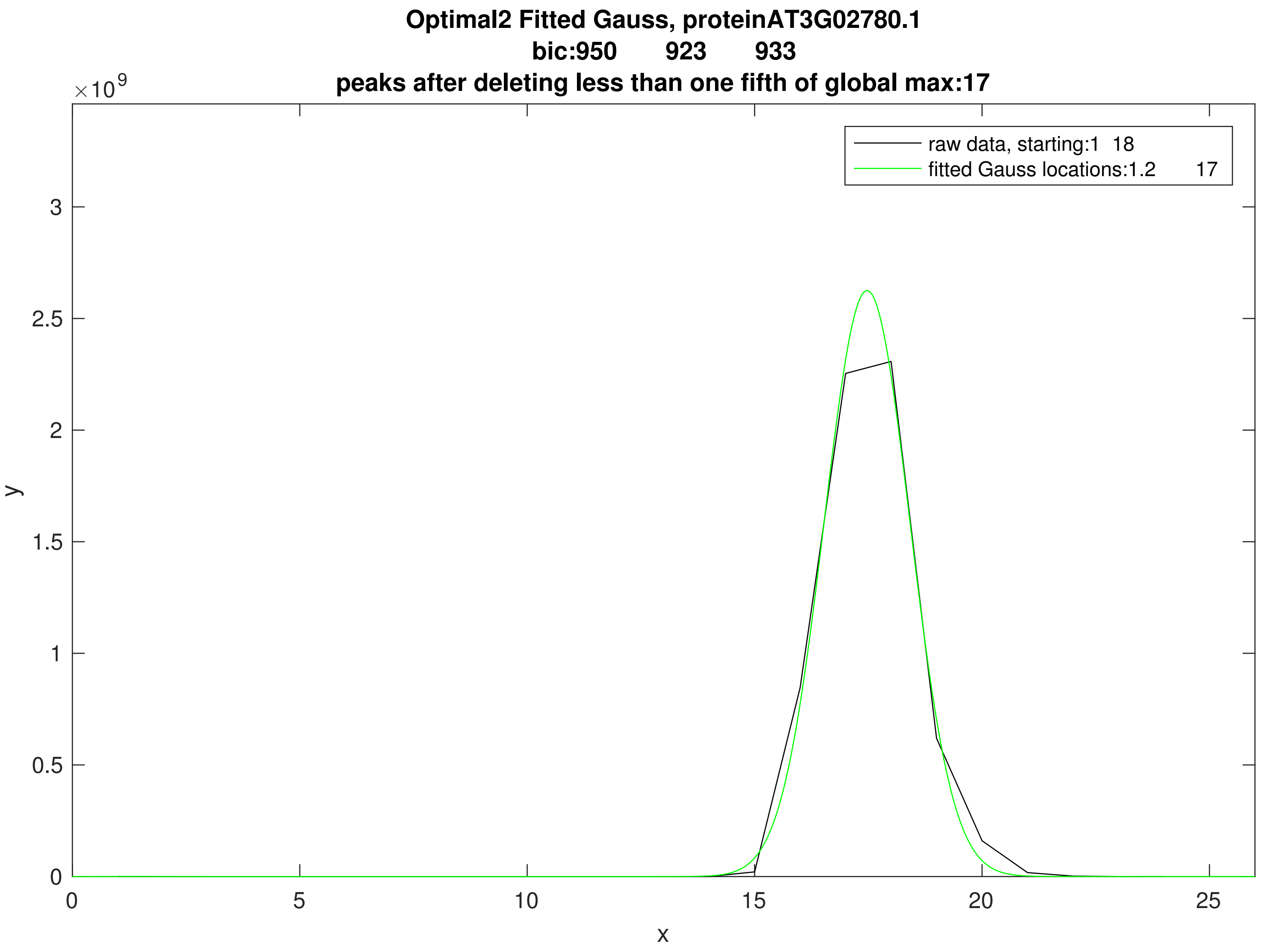

Optimal2 Fitted Gauss, proteinAT3G02780.1
bic:950 923 933
peaks after deleting less than one fifth of global max:17
×10^9
raw data, starting:1 18
fitted Gauss locations:1.2 17
0
0.5
1
1.5
2
2.5
3
y
0
5
10
15
20
25
x

Optimal1 Fitted Gauss, proteinAT3G02870.3;AT3G02870.1;AT3G02870.2
bic:850
peaks after deleting less than one fifth of global max:15

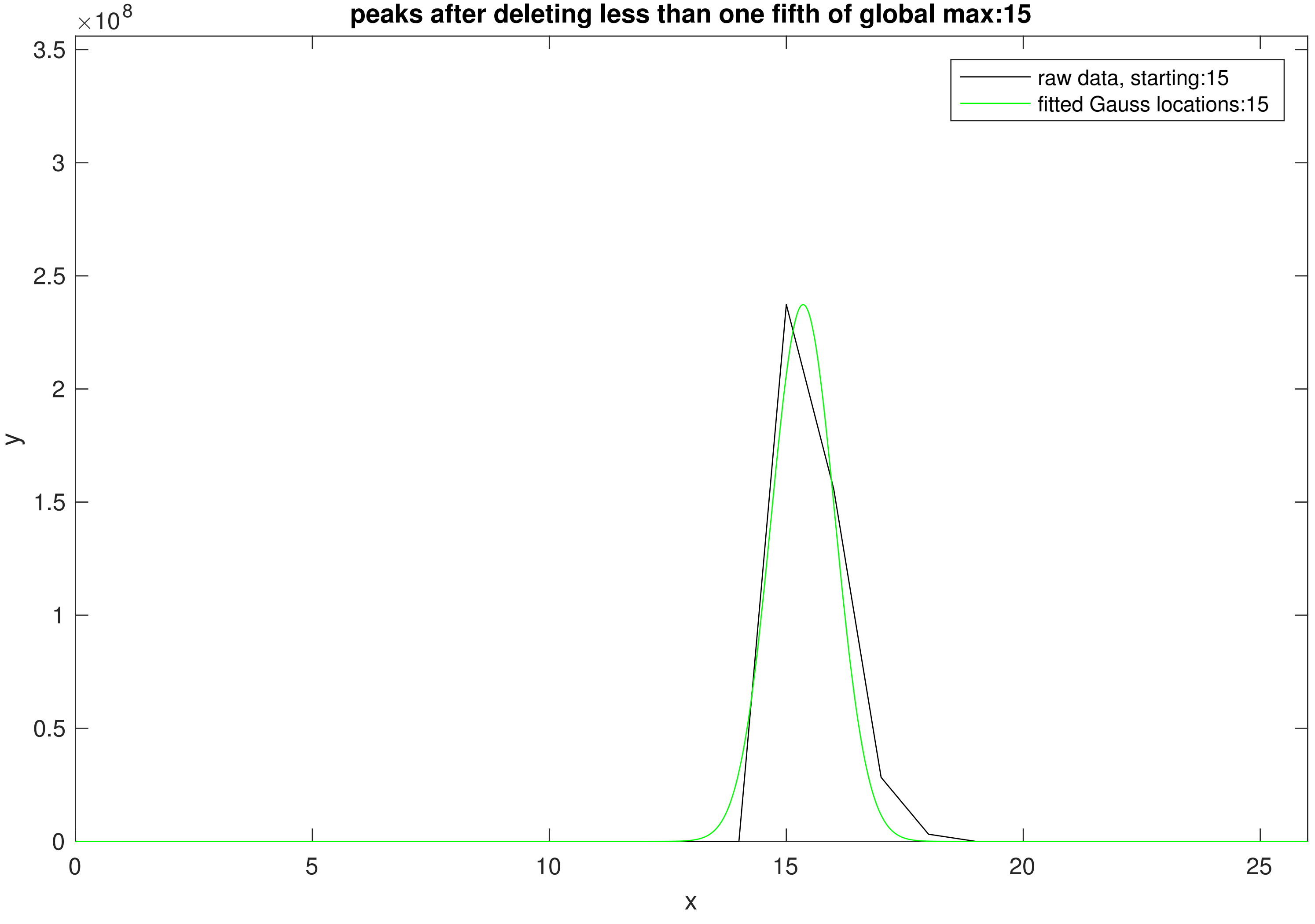

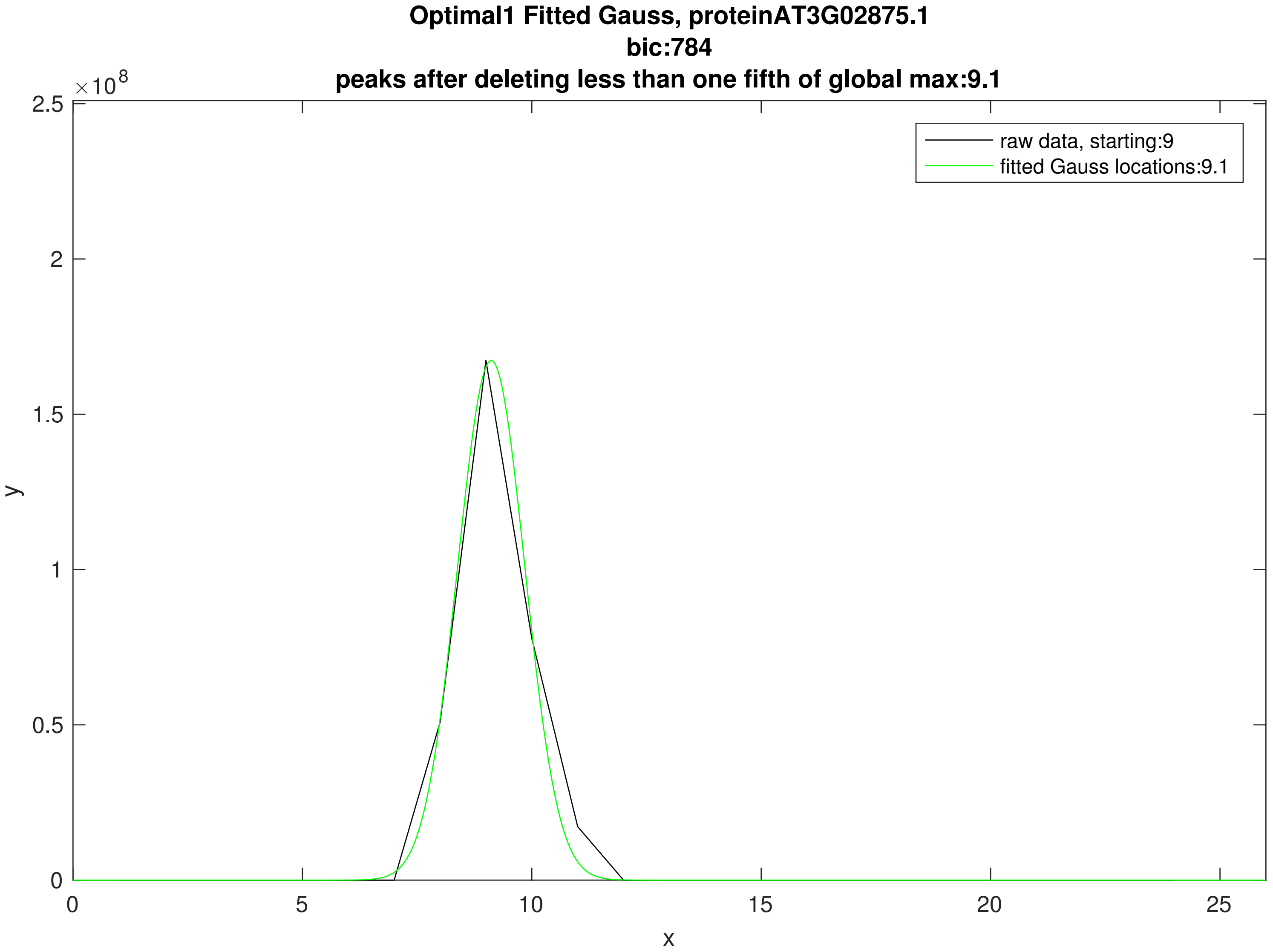

Optimal1 Fitted Gauss, proteinAT3G02875.1
bic:784
peaks after deleting less than one fifth of global max:9.1
×10^8
2.5
2
1.5
1
0.5
0
y
0
5
10
15
20
25
x
raw data, starting:9
fitted Gauss locations:9.1

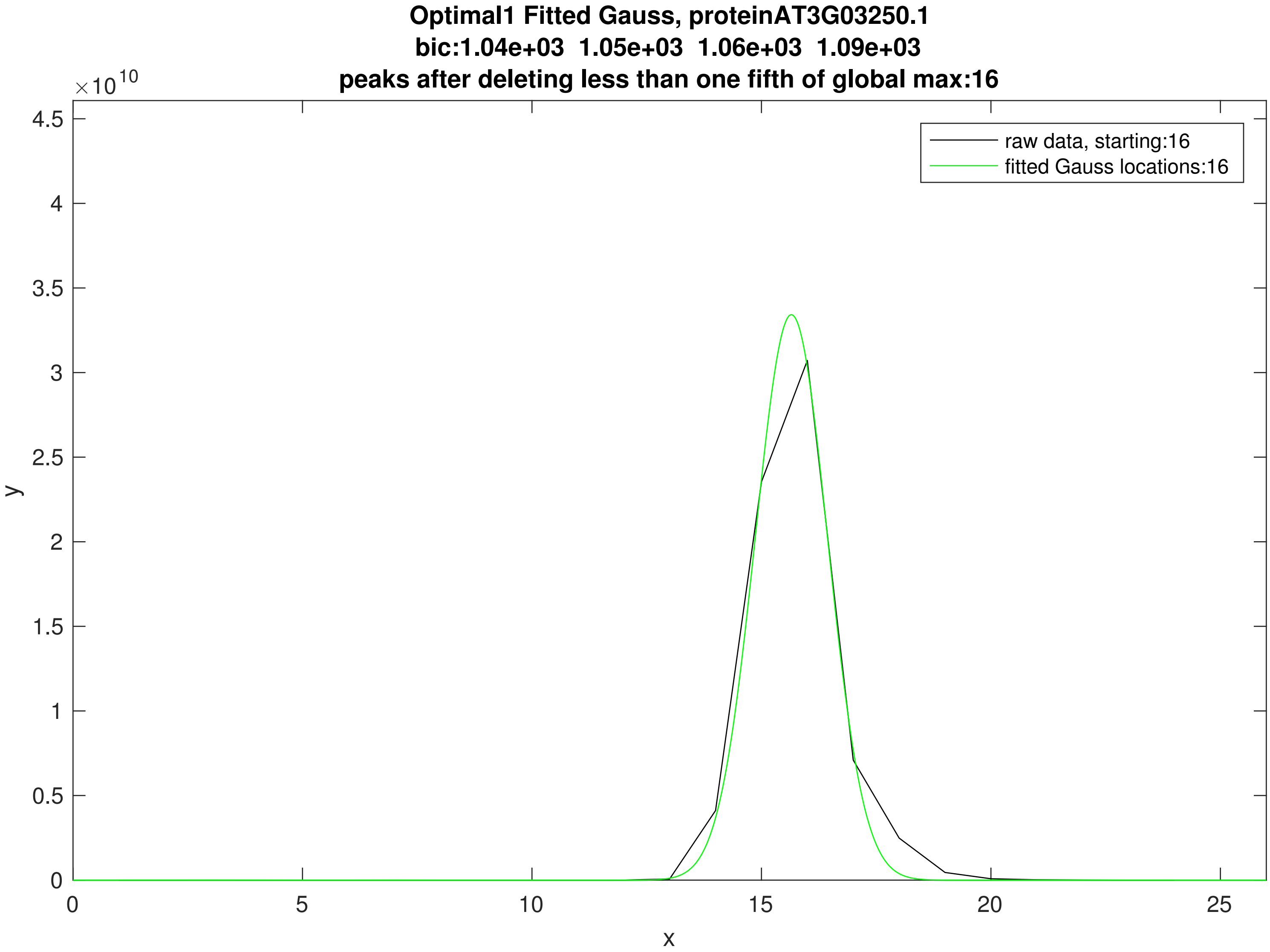

Optimal1 Fitted Gauss, proteinAT3G03250.1
bic:1.04e+03 1.05e+03 1.06e+03 1.09e+03
peaks after deleting less than one fifth of global max:16
×10^10
4.5
4
3.5
3
2.5
2
1.5
1
0.5
0
y
0
5
10
15
20
25
x
raw data, starting:16
fitted Gauss locations:16

**Optimal1 Fitted Gauss, proteinAT3G03320.1**
**bic:735**
**peaks after deleting less than one fifth of global max:17**

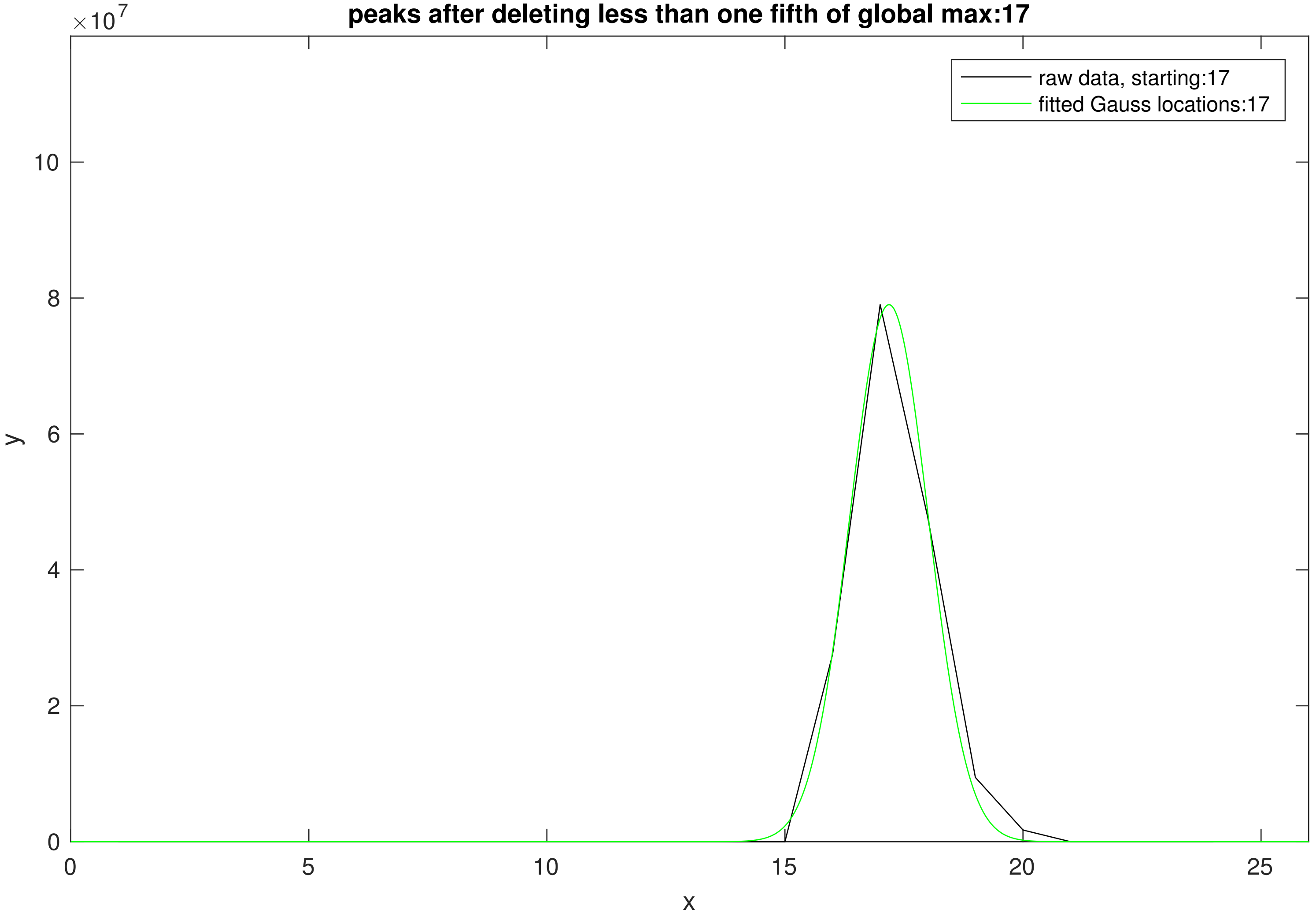

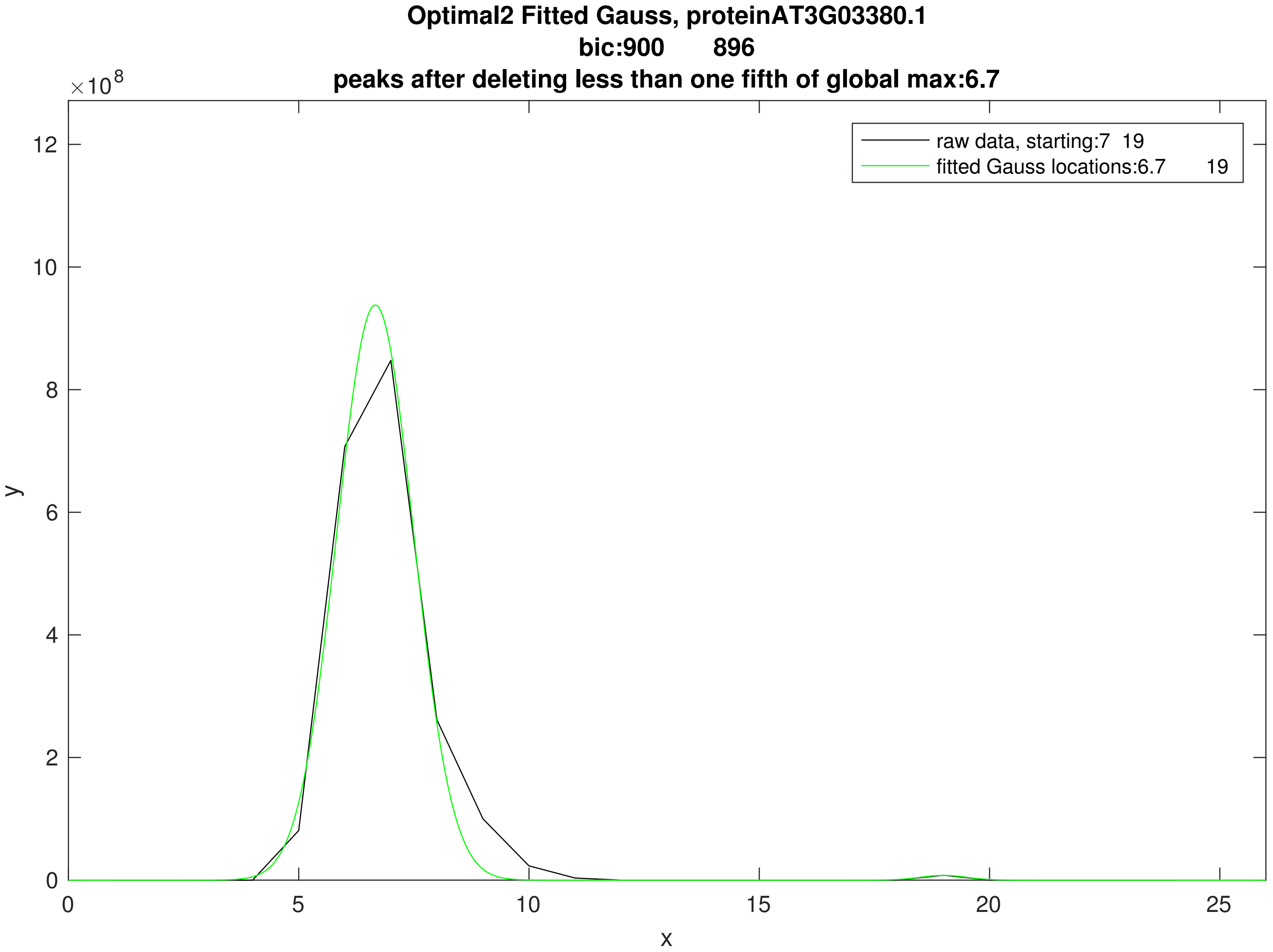

Optimal2 Fitted Gauss, proteinAT3G03380.1
bic:900 896
peaks after deleting less than one fifth of global max:6.7
×10^8
raw data, starting:7 19
fitted Gauss locations:6.7 19
y
x
0
2
4
6
8
10
12
0
5
10
15
20
25

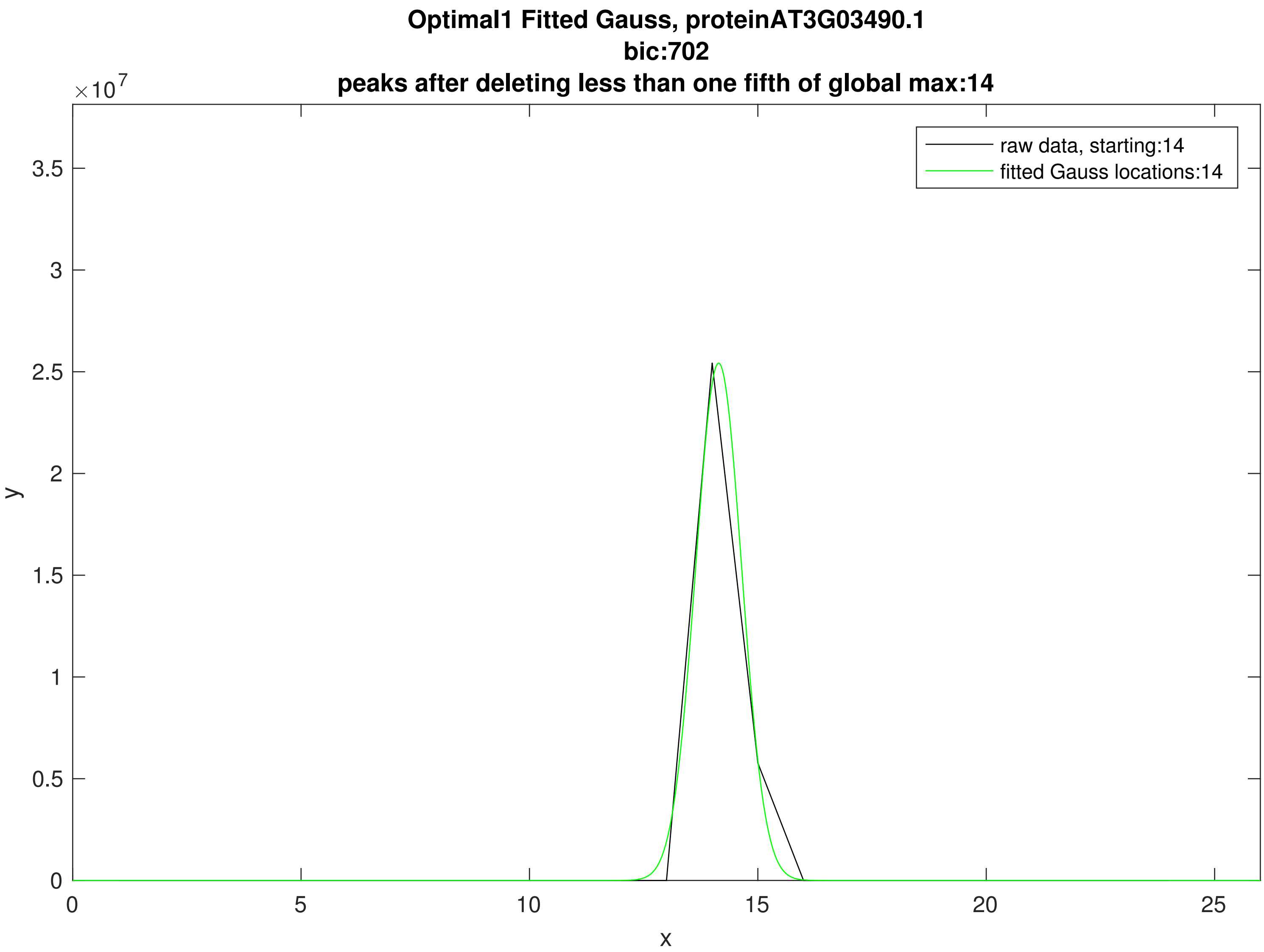
Optimal1 Fitted Gauss, proteinAT3G03490.1
bic:702
peaks after deleting less than one fifth of global max:14
×10^7
3.5
3
2.5
2
1.5
1
0.5
0
y
0
5
10
15
20
25
x
raw data, starting:14
fitted Gauss locations:14

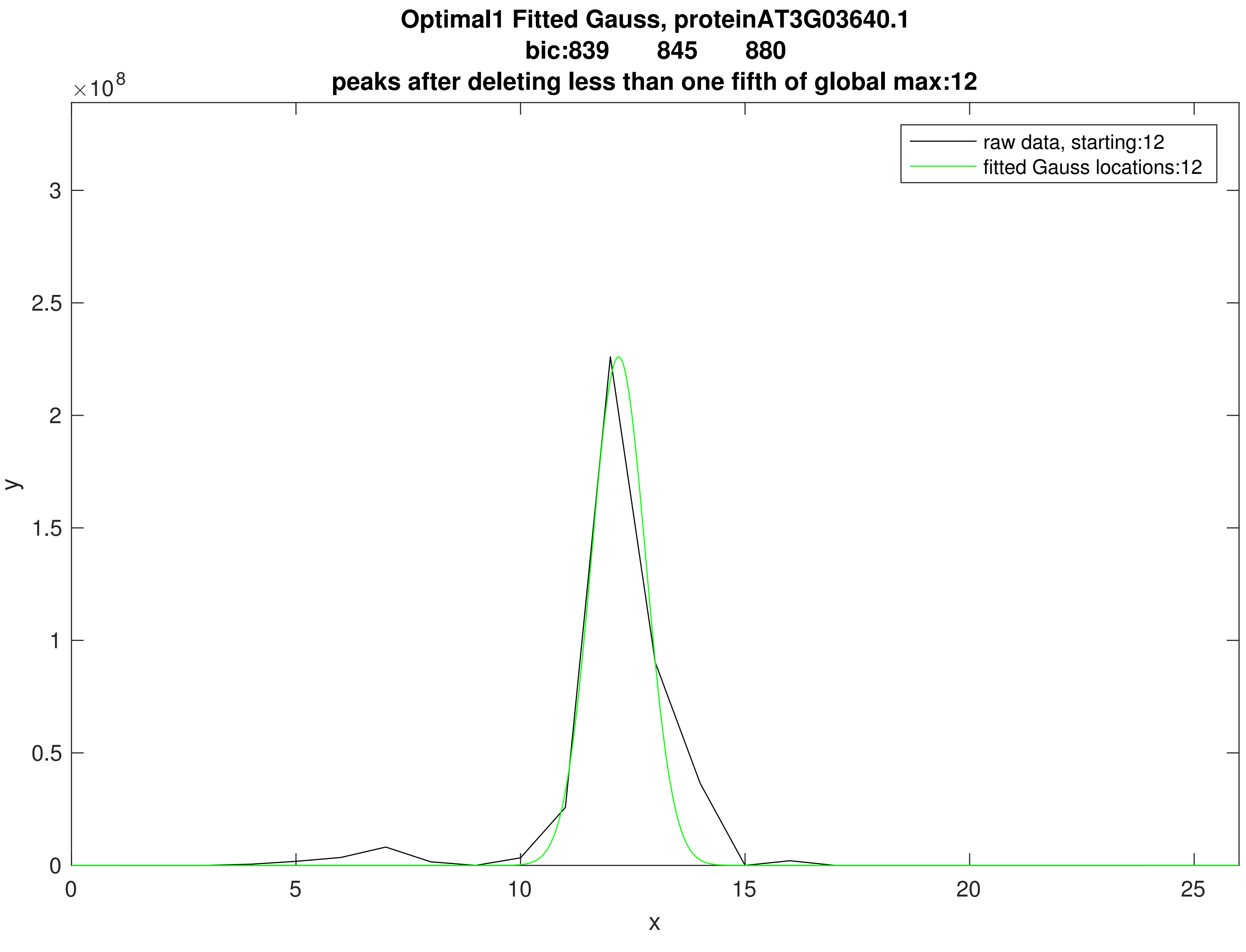

Optimal1 Fitted Gauss, proteinAT3G03640.1
bic:839 845 880
peaks after deleting less than one fifth of global max:12
×10^8
raw data, starting:12
fitted Gauss locations:12
3
2.5
2
1.5
1
0.5
0
y
0
5
10
15
20
25
x

Optimal1 Fitted Gauss, proteinAT3G03780.3;AT3G03780.2;AT3G03780.1
bic:1.07e+03  1.12e+03  1.09e+03   1.1e+03
peaks after deleting less than one fifth of global max:14

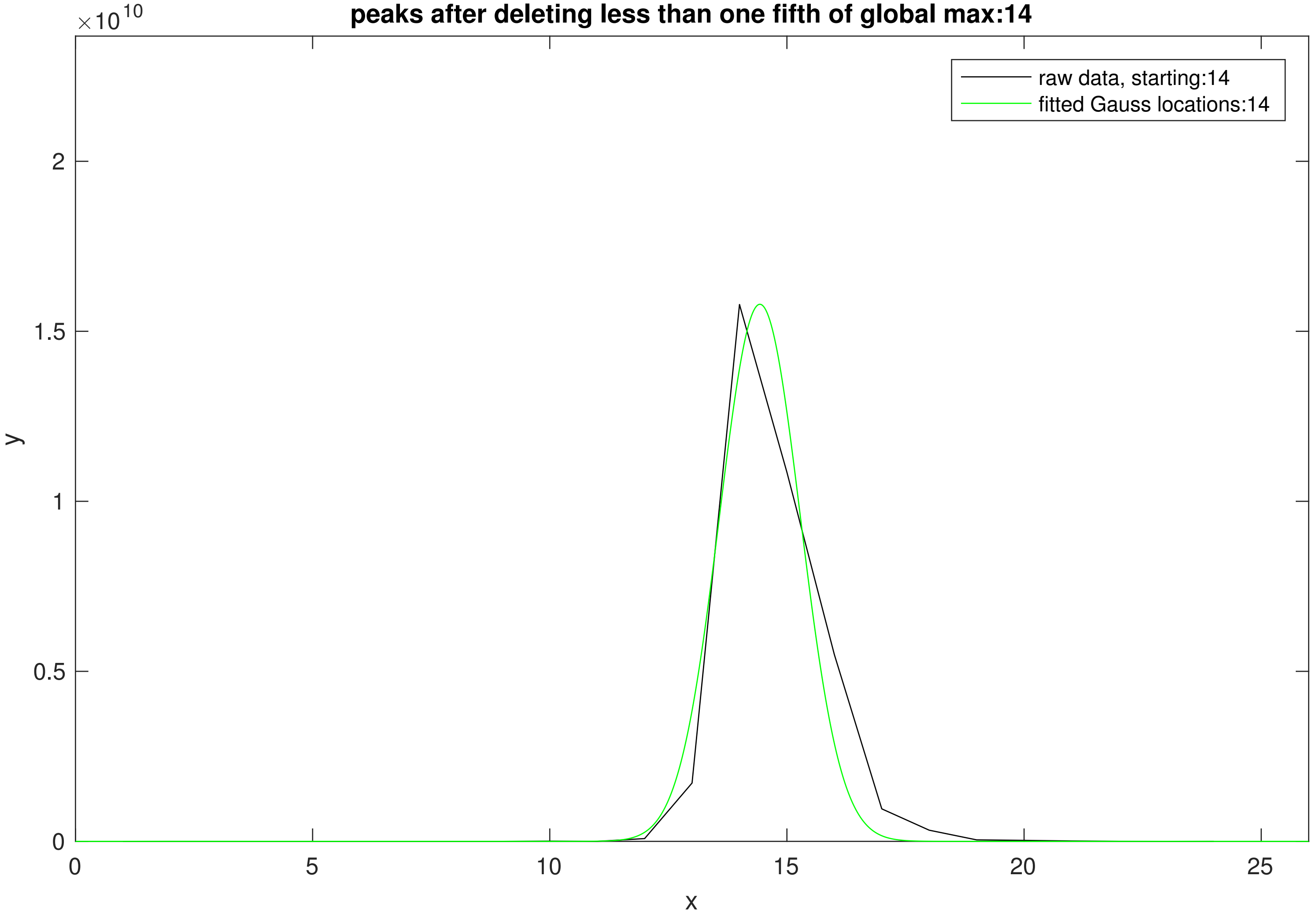

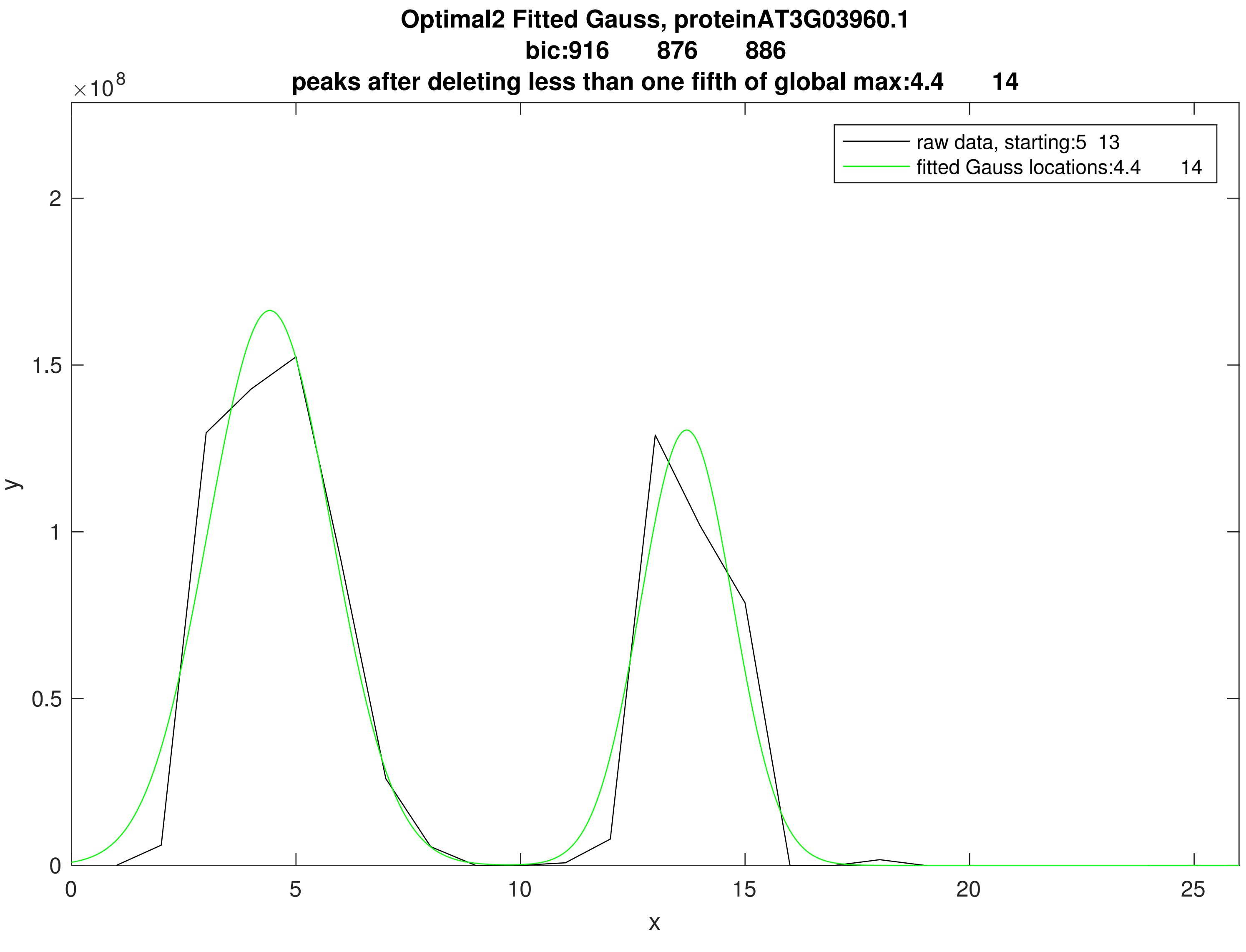
Optimal2 Fitted Gauss, proteinAT3G03960.1
bic:916 876 886
peaks after deleting less than one fifth of global max:4.4 14
×10^8
raw data, starting:5 13
fitted Gauss locations:4.4 14
y
x
0
0.5
1
1.5
2
5
10
15
20
25

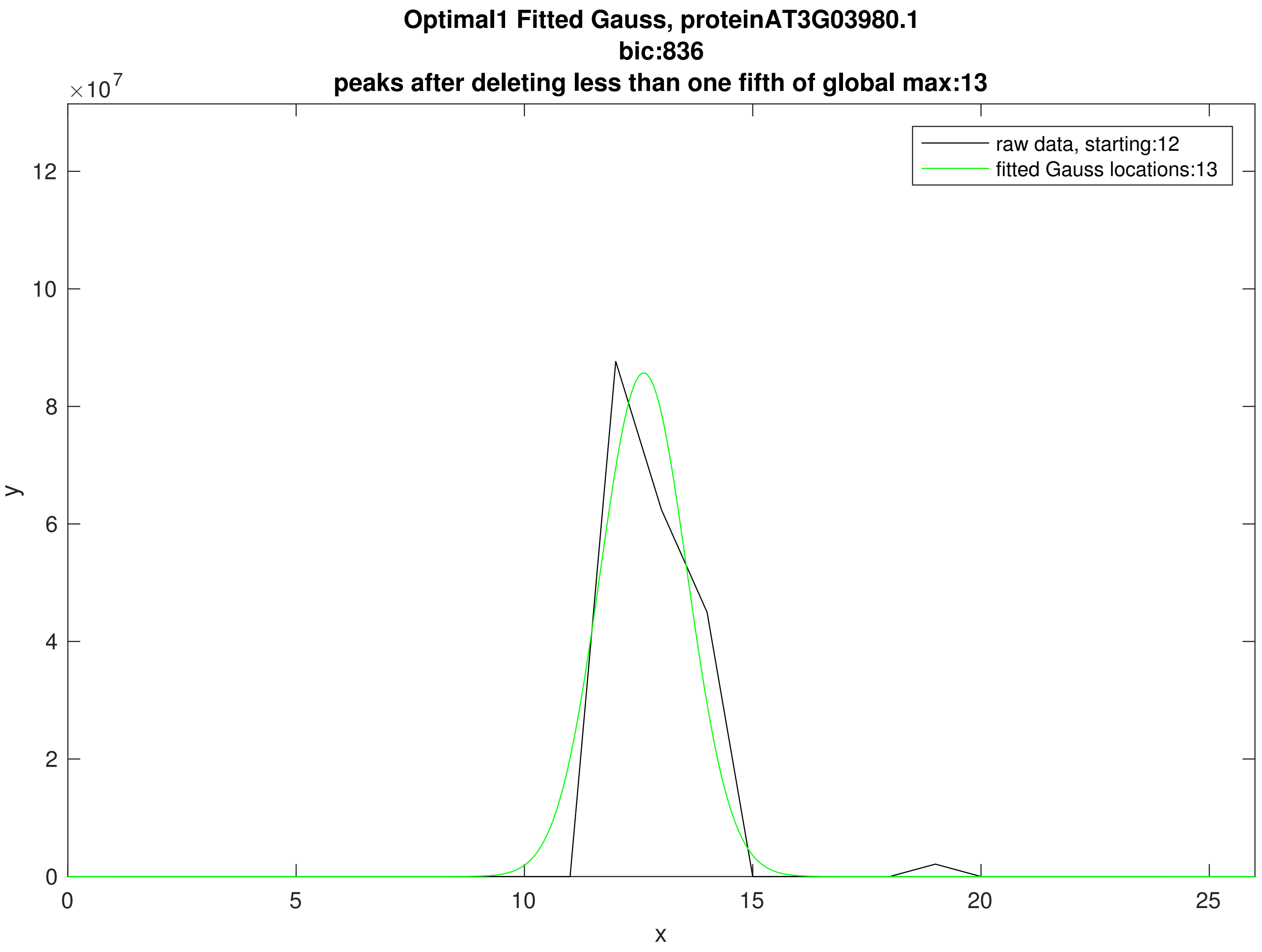
Optimal1 Fitted Gauss, proteinAT3G03980.1
bic:836
peaks after deleting less than one fifth of global max:13
$\times 10^7$
raw data, starting:12
fitted Gauss locations:13
y
x
0
2
4
6
8
10
12
0
5
10
15
20
25

**Optimal2 Fitted Gauss, proteinAT3G03990.1**
**bic:837 831**
**peaks after deleting less than one fifth of global max:18**

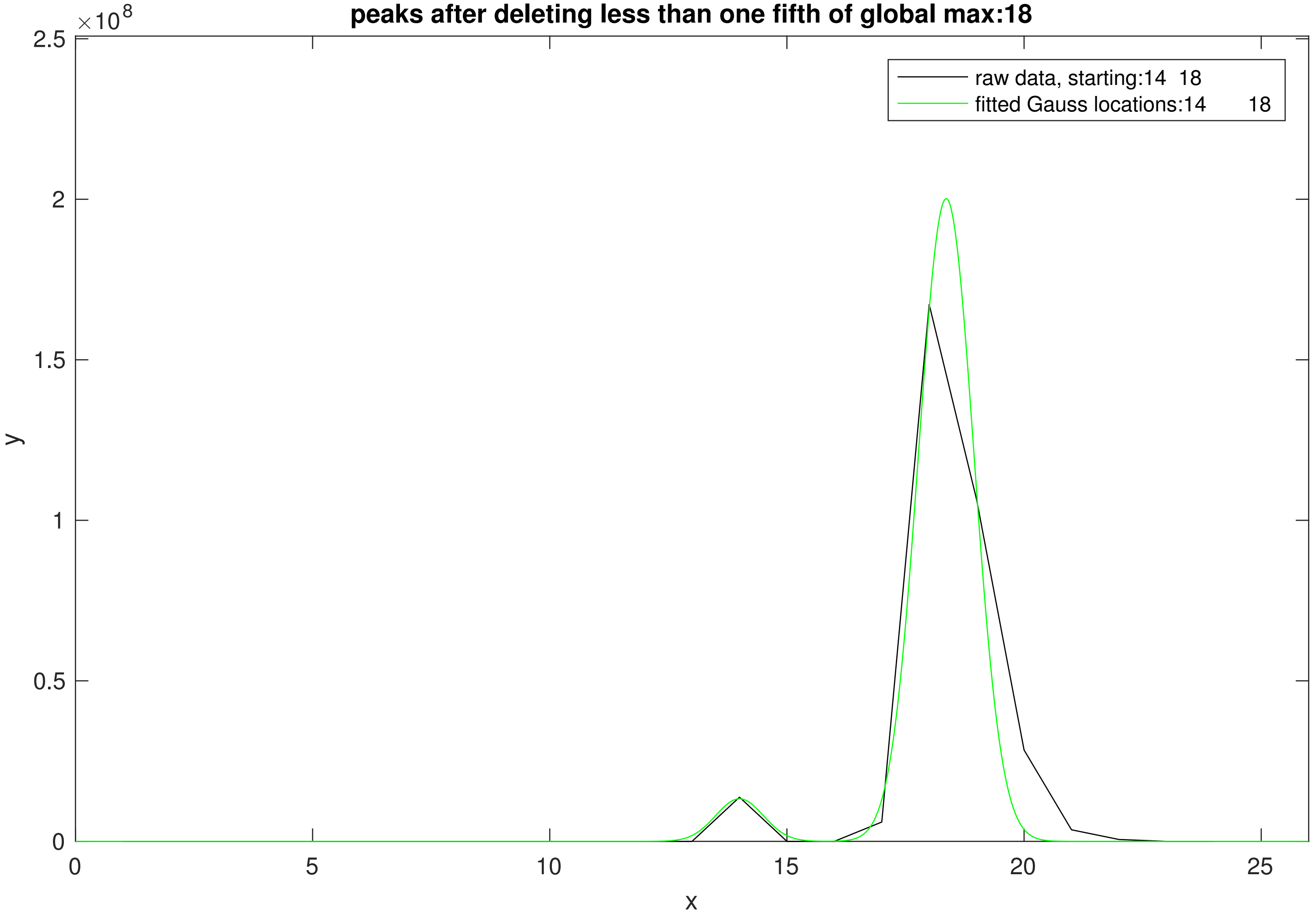

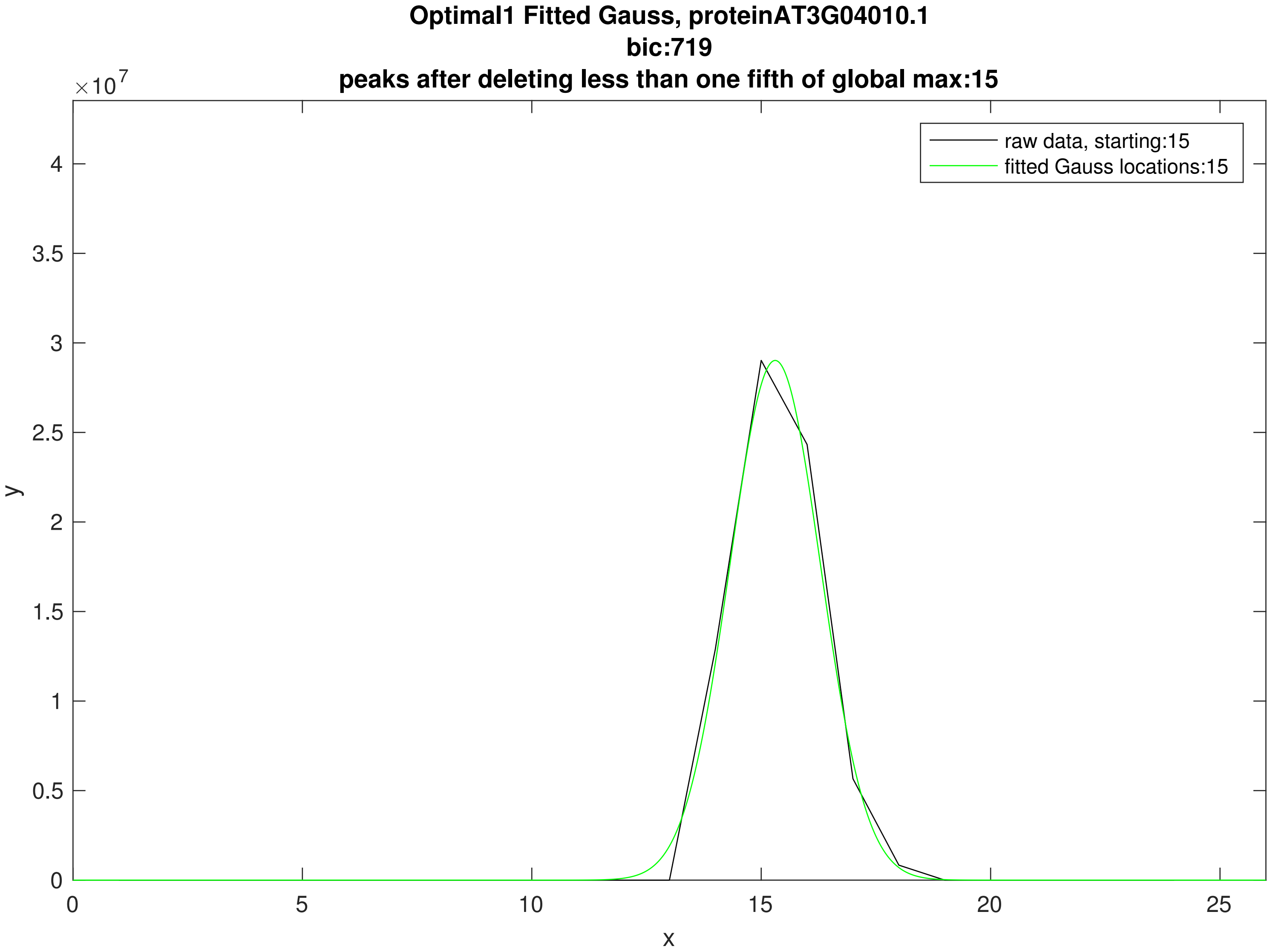
Optimal1 Fitted Gauss, proteinAT3G04010.1
bic:719
peaks after deleting less than one fifth of global max:15
×10^7
raw data, starting:15
fitted Gauss locations:15
y
x

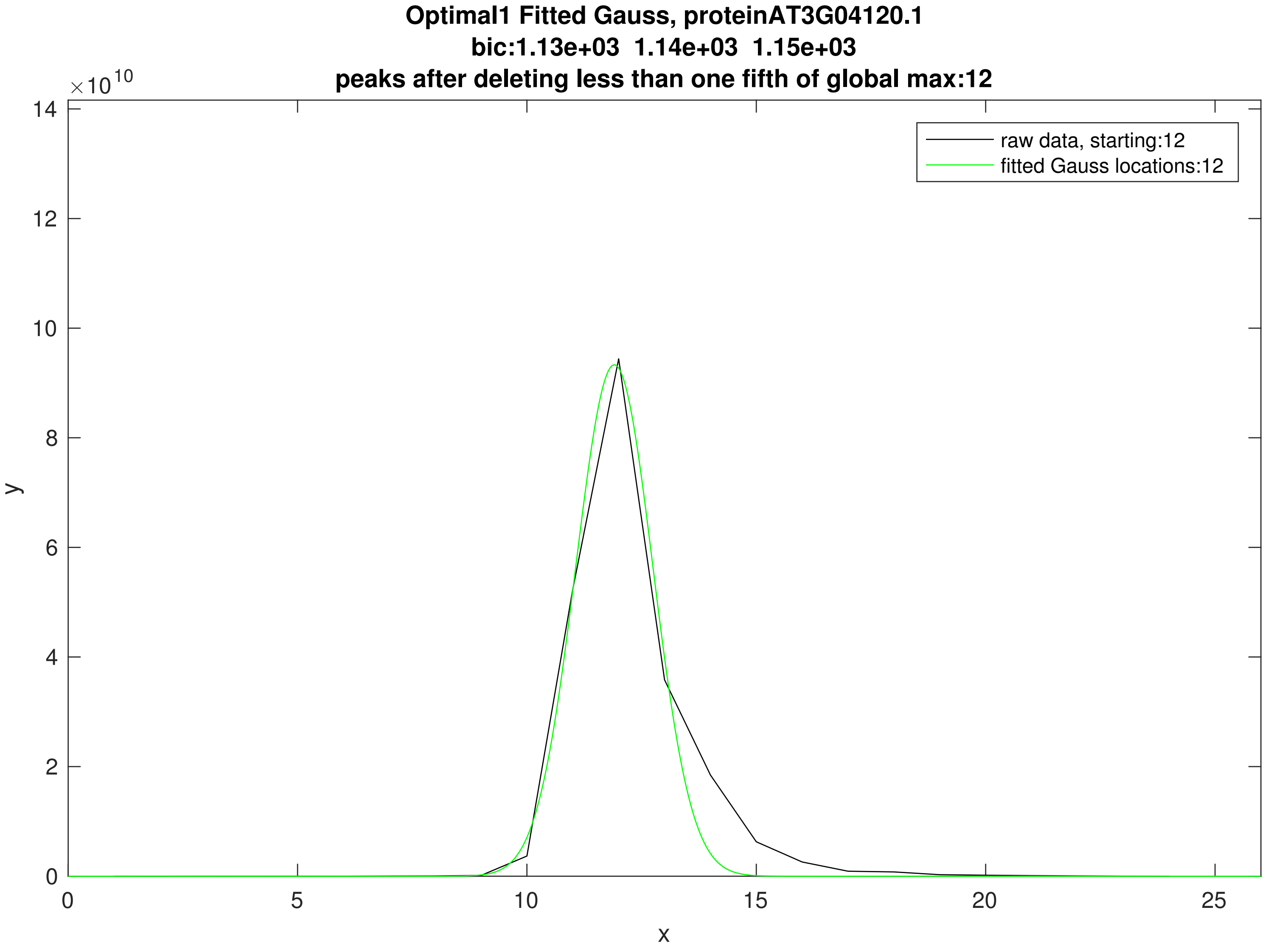

Optimal1 Fitted Gauss, proteinAT3G04120.1
bic:1.13e+03 1.14e+03 1.15e+03
peaks after deleting less than one fifth of global max:12
×10^10
14
12
10
8
6
4
2
0
y
0
5
10
15
20
25
x
raw data, starting:12
fitted Gauss locations:12

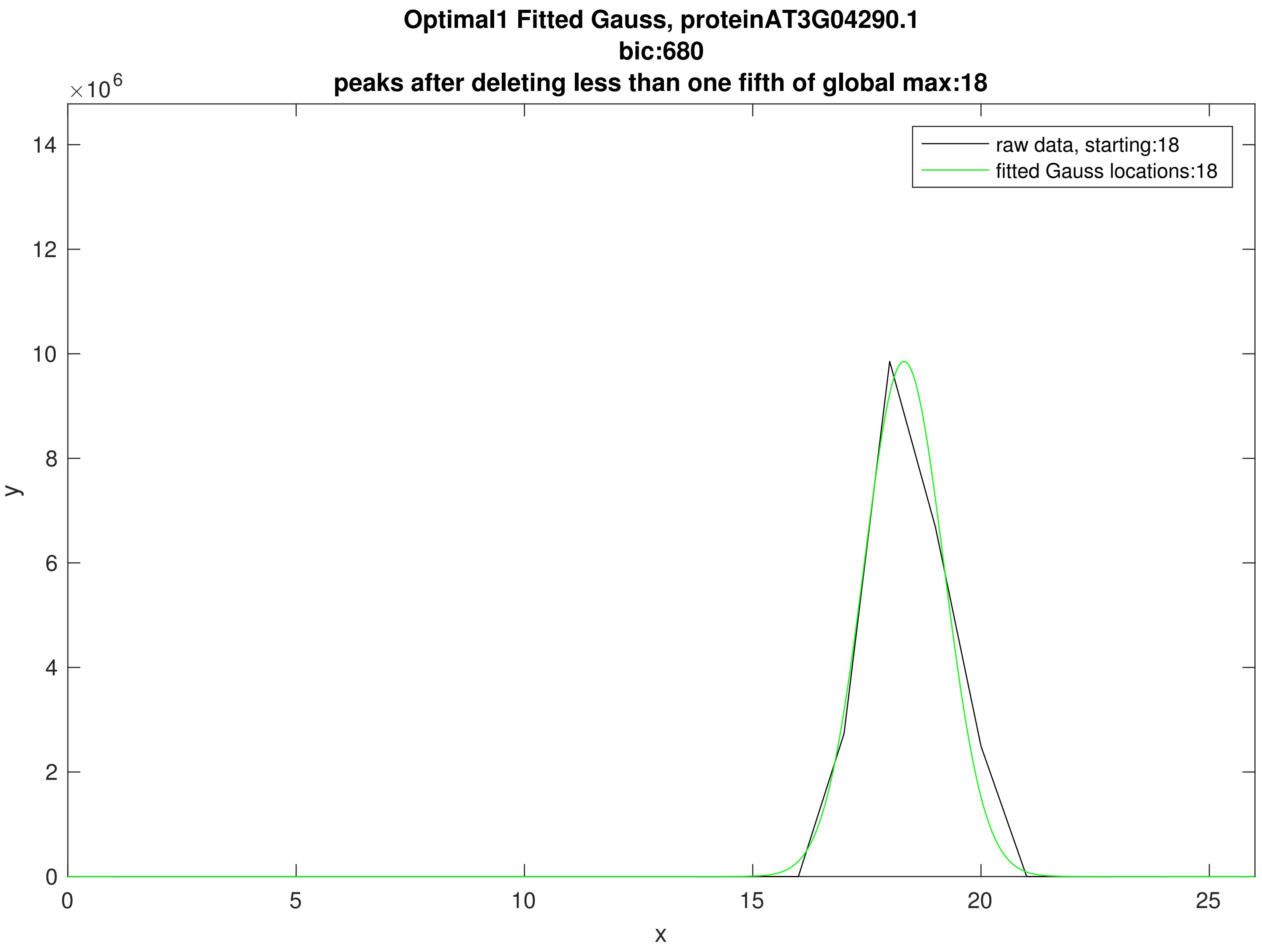

Optimal1 Fitted Gauss, proteinAT3G04290.1
bic:680
peaks after deleting less than one fifth of global max:18
×10^6
raw data, starting:18
fitted Gauss locations:18
y
x
0
2
4
6
8
10
12
14
0
5
10
15
20
25

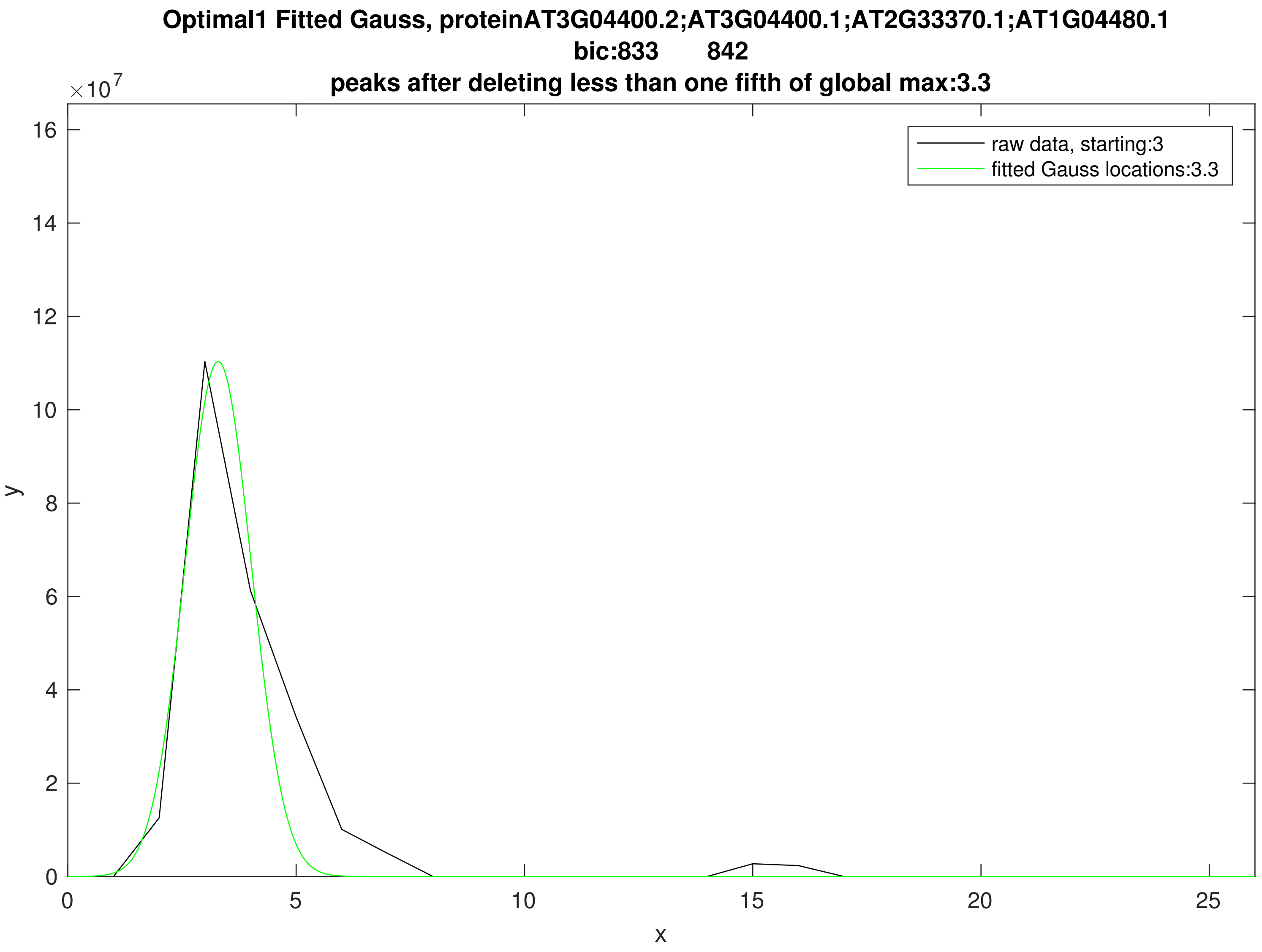
Optimal1 Fitted Gauss, proteinAT3G04400.2;AT3G04400.1;AT2G33370.1;AT1G04480.1
bic:833 842
peaks after deleting less than one fifth of global max:3.3
×10^7
raw data, starting:3
fitted Gauss locations:3.3
y
x
0
2
4
6
8
10
12
14
16
0
5
10
15
20
25

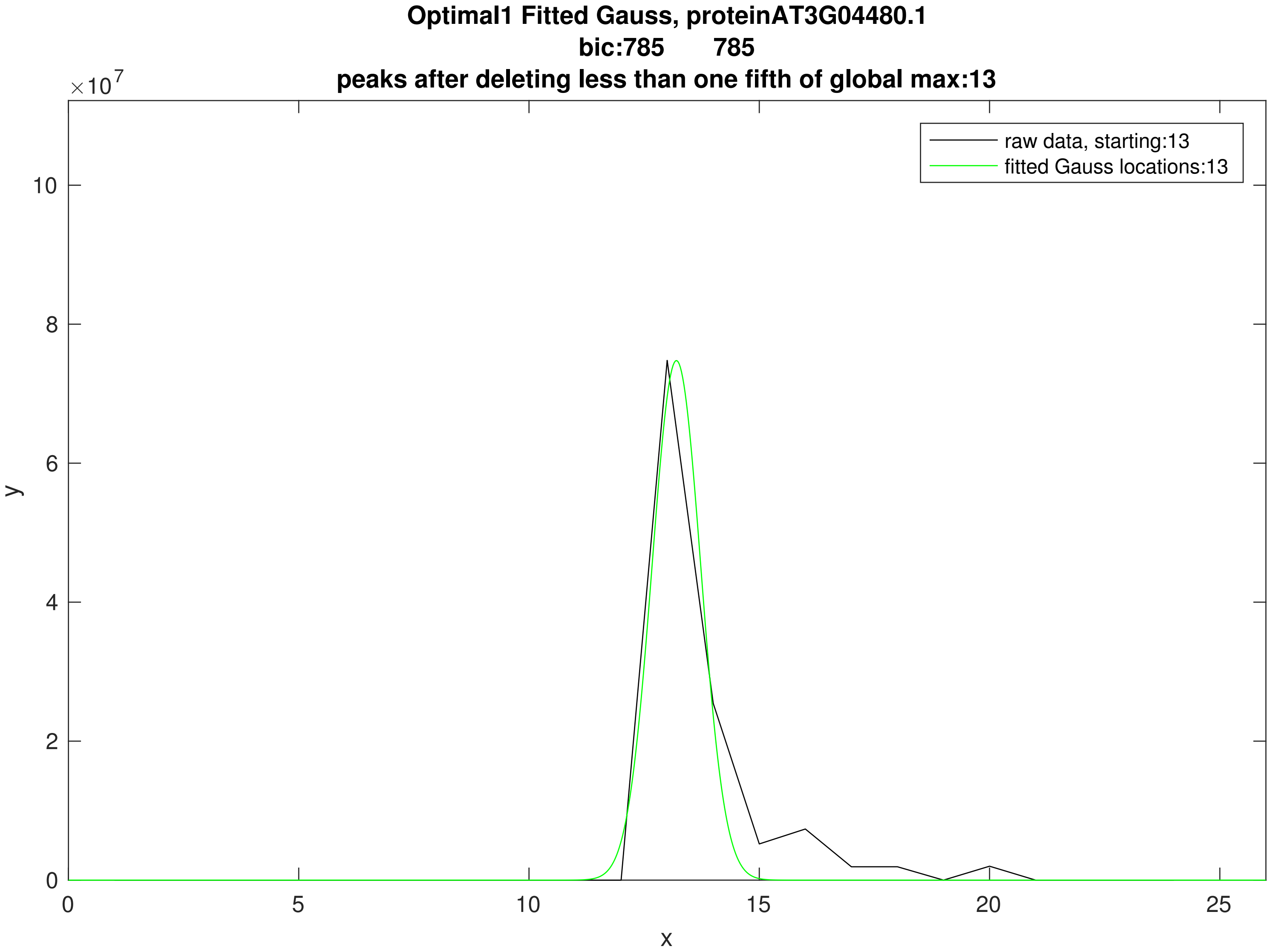

Optimal1 Fitted Gauss, proteinAT3G04480.1
bic:785 785
peaks after deleting less than one fifth of global max:13
×10^7
raw data, starting:13
fitted Gauss locations:13
10
8
6
4
2
0
y
0
5
10
15
20
25
x

Optimal1 Fitted Gauss, proteinAT3G04520.1;AT3G04520.2
bic:781
peaks after deleting less than one fifth of global max:11

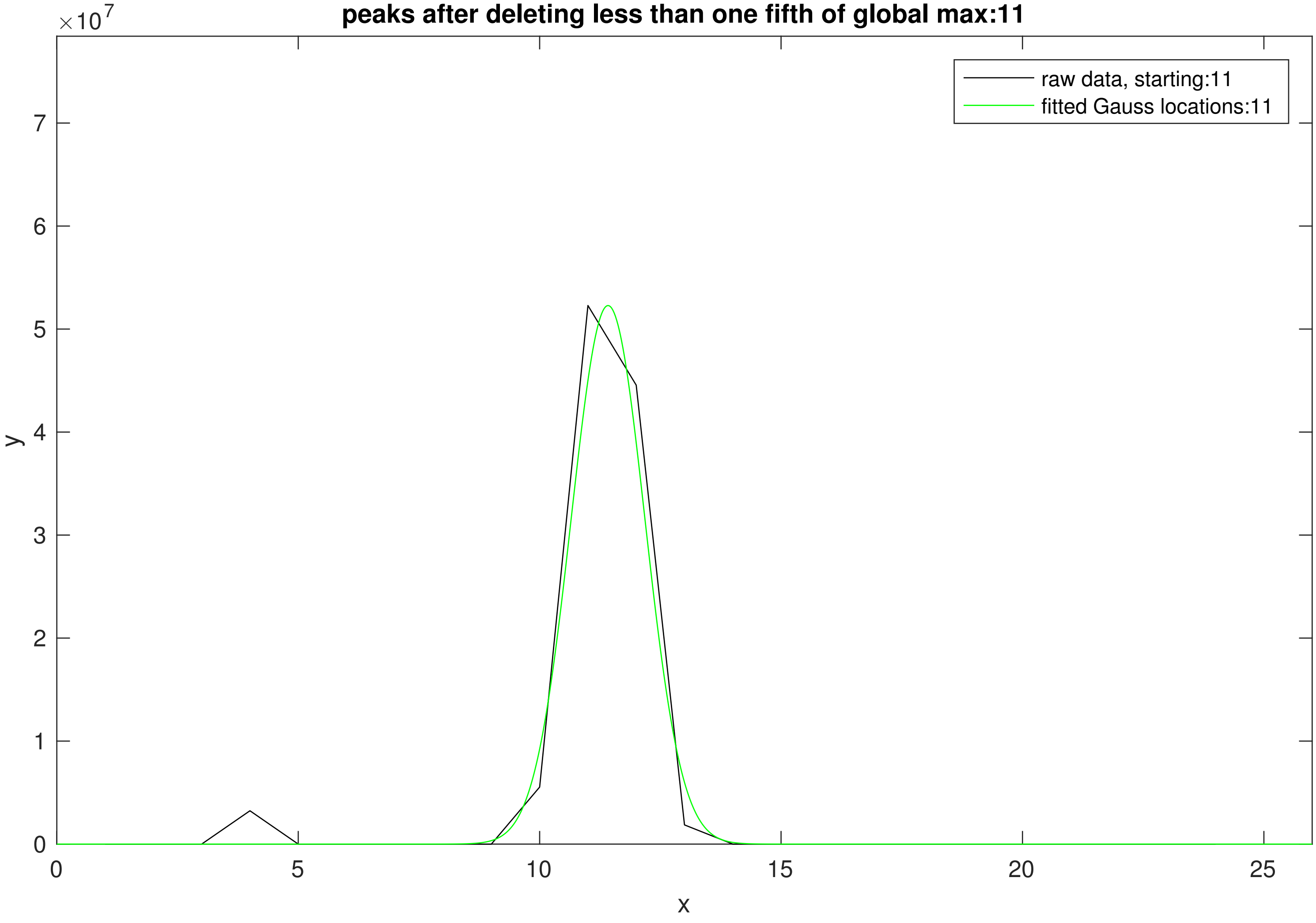

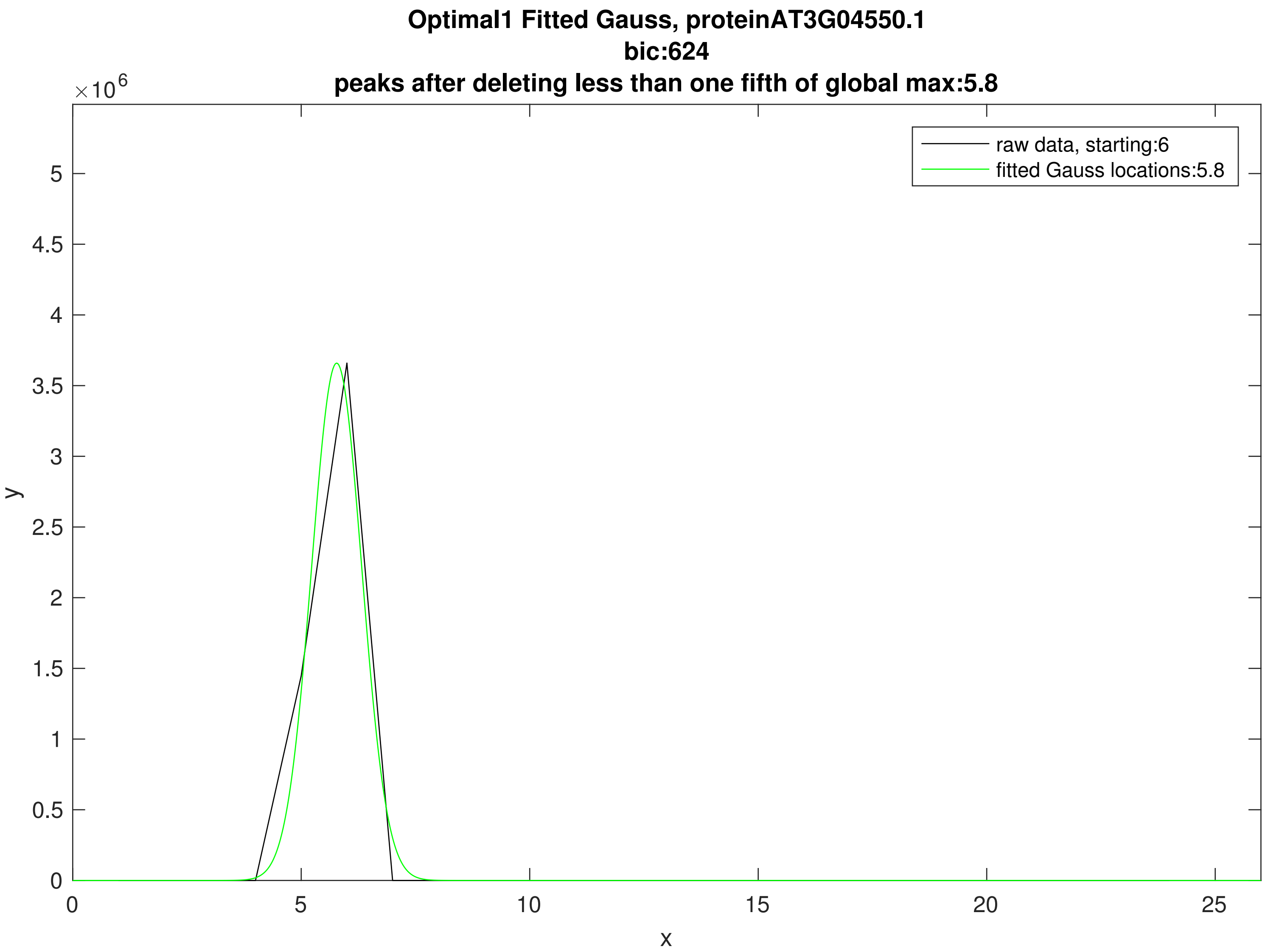
Optimal1 Fitted Gauss, proteinAT3G04550.1
bic:624
peaks after deleting less than one fifth of global max:5.8
×10^6
raw data, starting:6
fitted Gauss locations:5.8
y
x
0
0.5
1
1.5
2
2.5
3
3.5
4
4.5
5
0
5
10
15
20
25

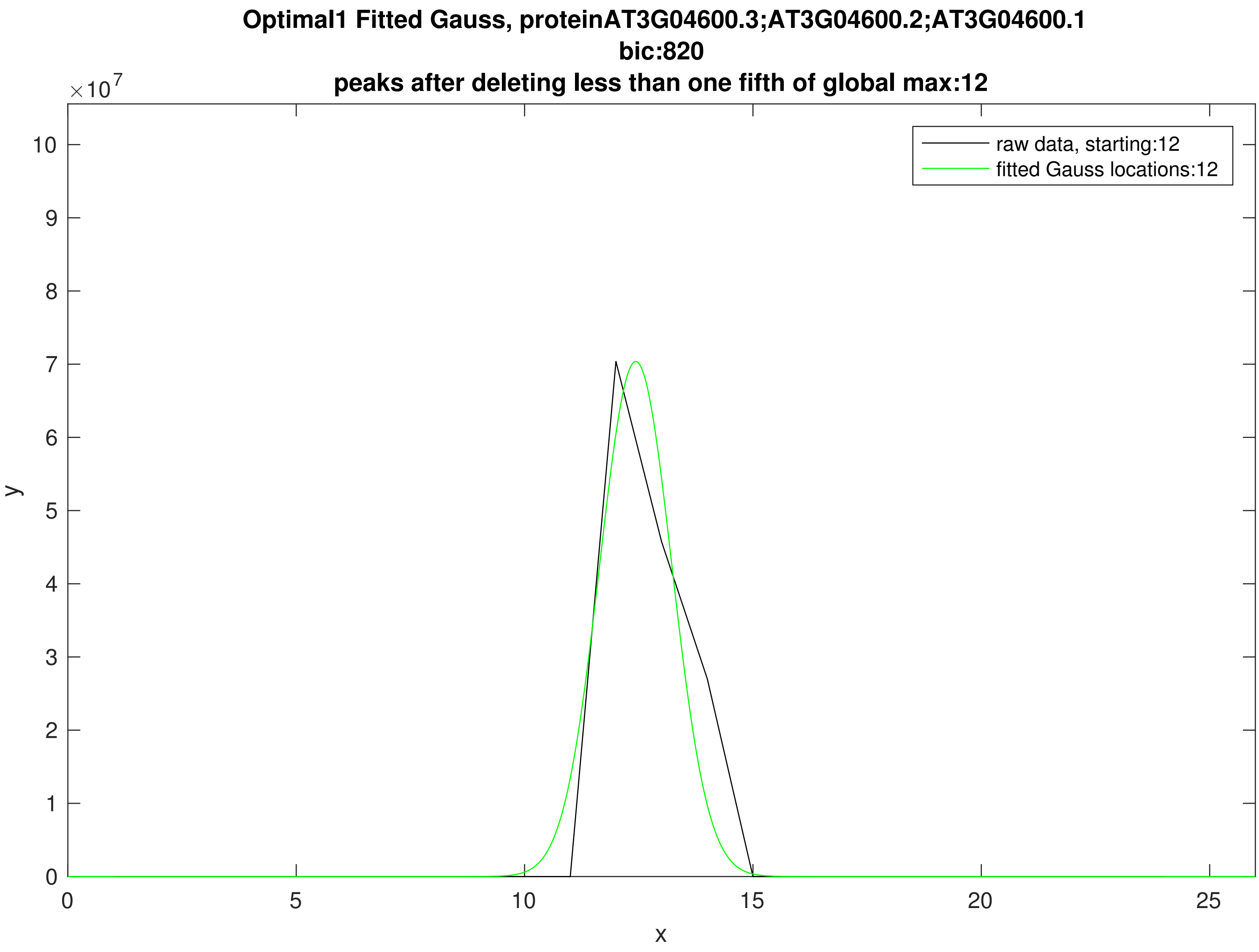

Optimal1 Fitted Gauss, proteinAT3G04600.3;AT3G04600.2;AT3G04600.1
bic:820
peaks after deleting less than one fifth of global max:12
×10^7
raw data, starting:12
fitted Gauss locations:12
y
x
0
1
2
3
4
5
6
7
8
9
10
0
5
10
15
20
25

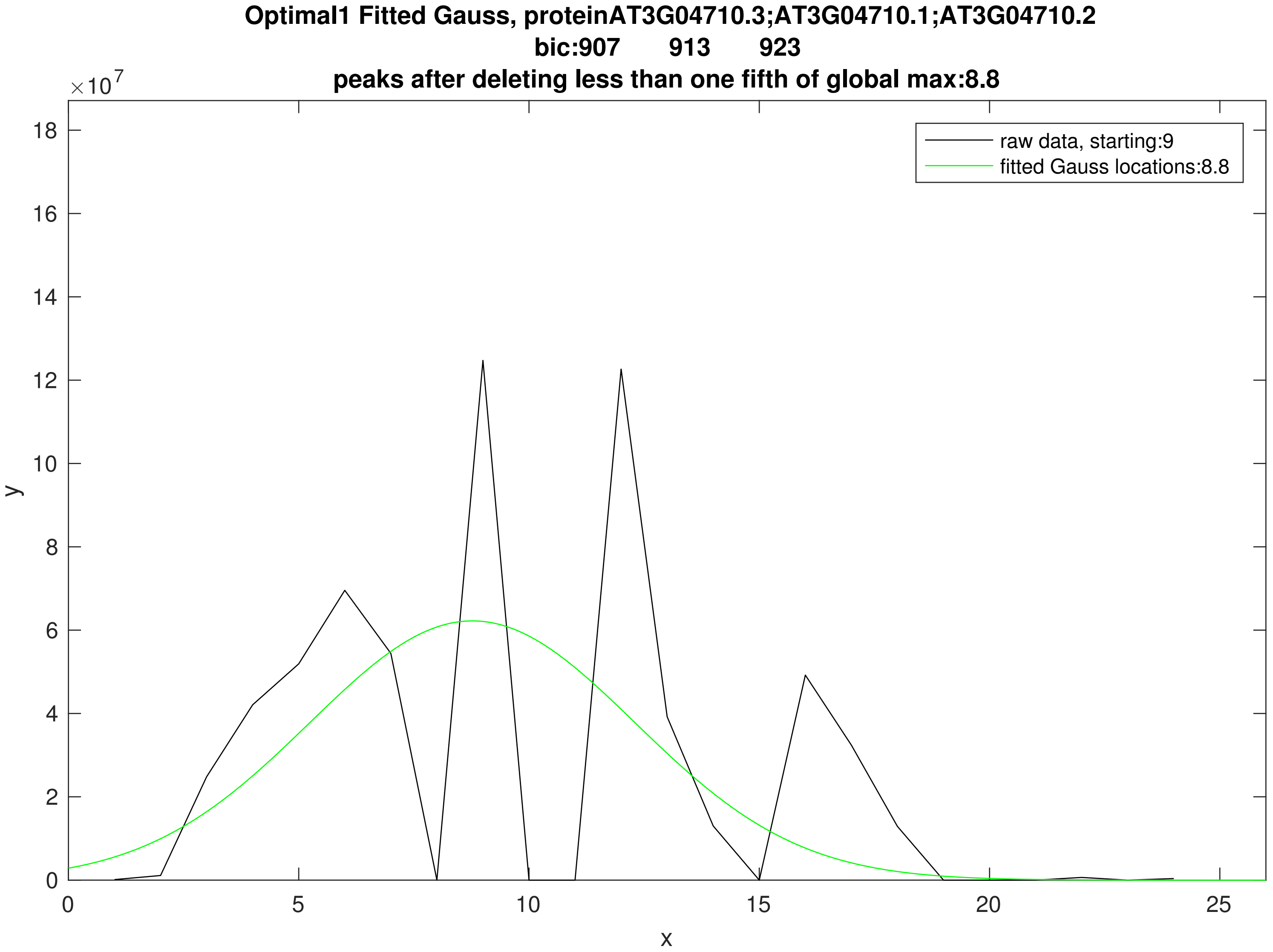

Optimal1 Fitted Gauss, proteinAT3G04710.3;AT3G04710.1;AT3G04710.2
bic:907 913 923
peaks after deleting less than one fifth of global max:8.8
×10^7
18
16
14
12
10
8
6
4
2
0
y
0
5
10
15
20
25
x
raw data, starting:9
fitted Gauss locations:8.8

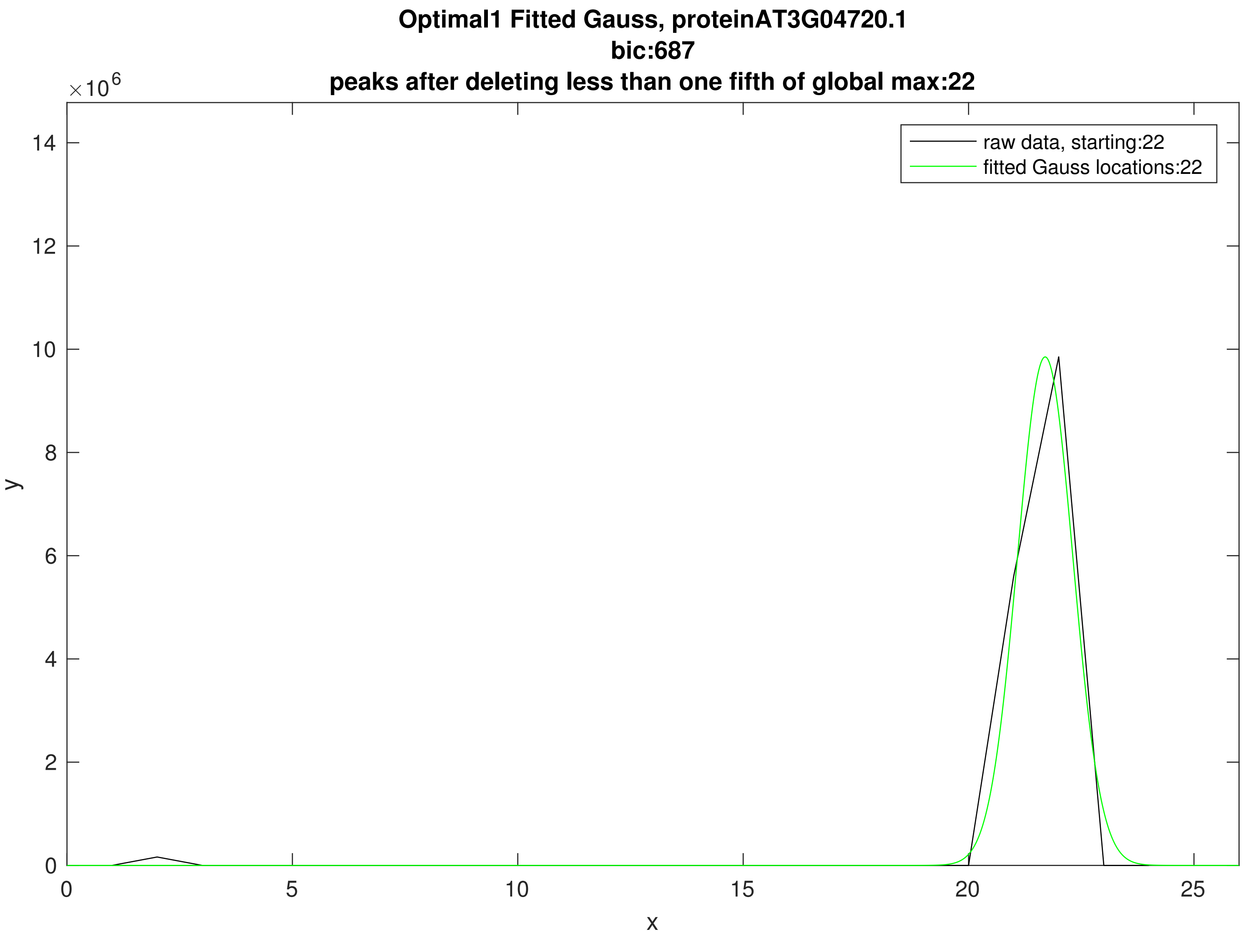

Optimal1 Fitted Gauss, proteinAT3G04720.1
bic:687
peaks after deleting less than one fifth of global max:22
$\times 10^6$
raw data, starting:22
fitted Gauss locations:22
14
12
10
8
6
4
2
0
y
0
5
10
15
20
25
x

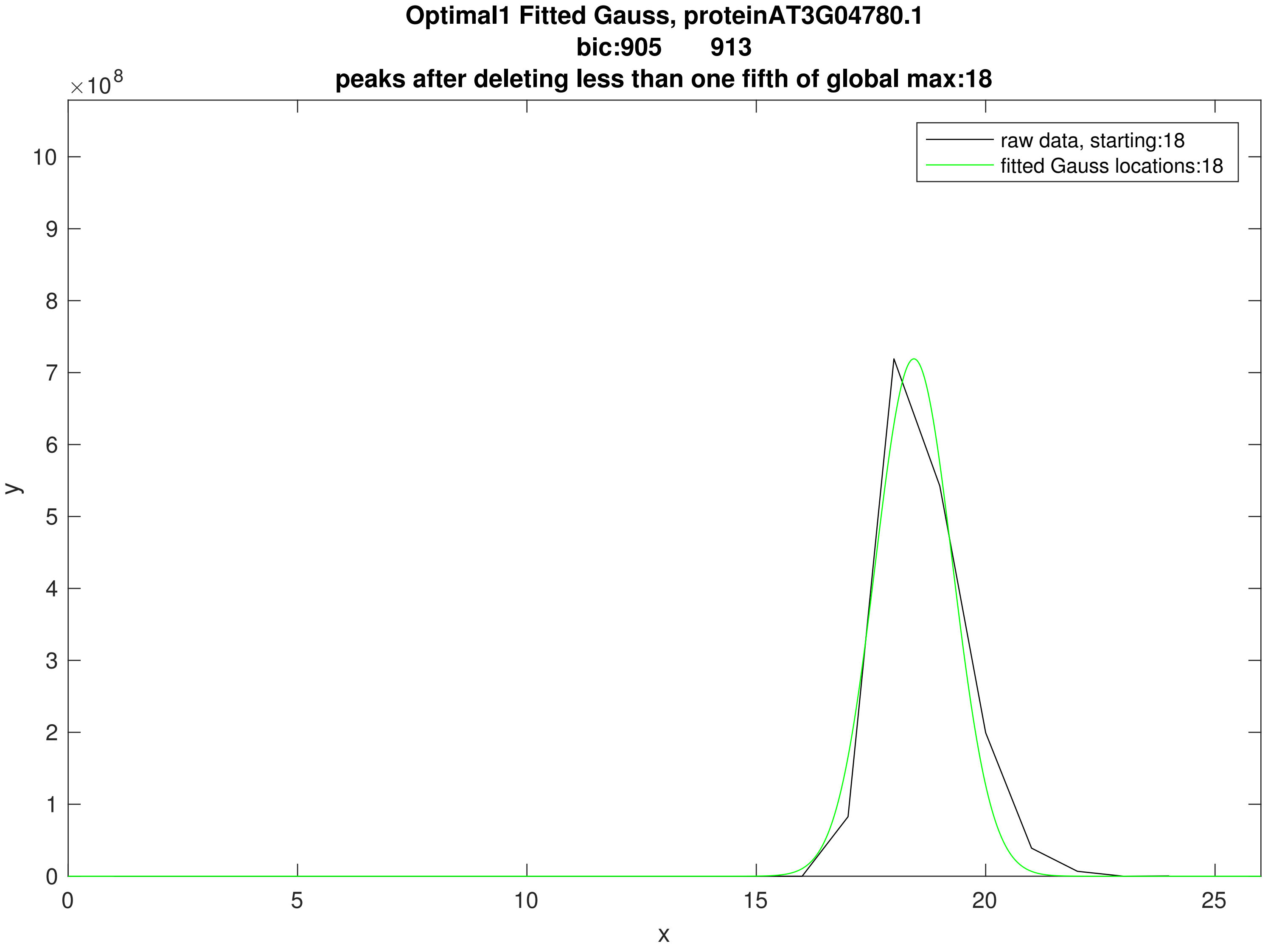
Optimal1 Fitted Gauss, proteinAT3G04780.1
bic:905 913
peaks after deleting less than one fifth of global max:18
×10^8
raw data, starting:18
fitted Gauss locations:18
y
x
0
1
2
3
4
5
6
7
8
9
10
0
5
10
15
20
25

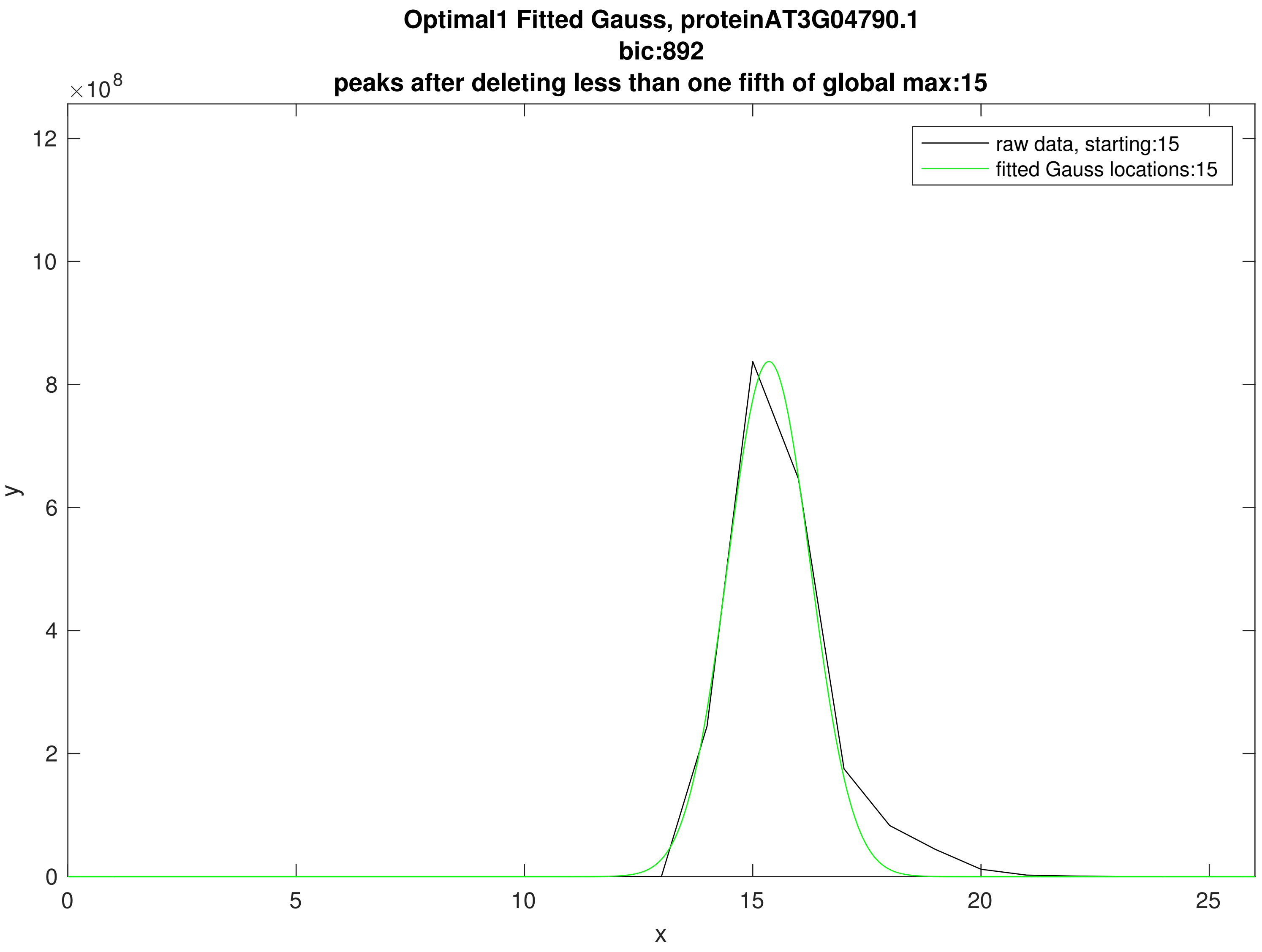

Optimal1 Fitted Gauss, proteinAT3G04790.1
bic:892
peaks after deleting less than one fifth of global max:15
×10^8
raw data, starting:15
fitted Gauss locations:15
y
x
0
2
4
6
8
10
12
0
5
10
15
20
25

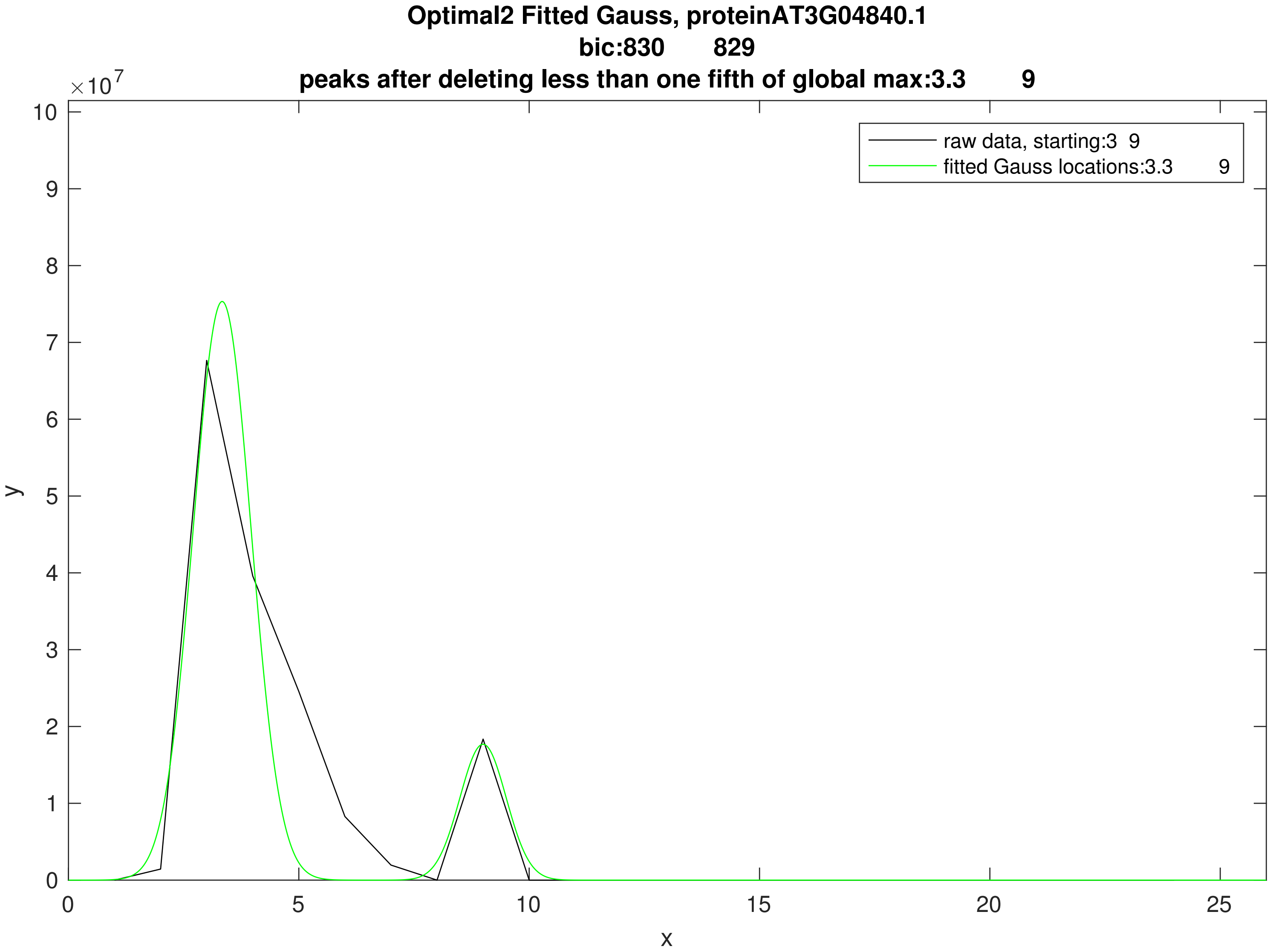

Optimal2 Fitted Gauss, proteinAT3G04840.1
bic:830 829
peaks after deleting less than one fifth of global max:3.3 9
×10^7
10
9
8
7
6
5
4
3
2
1
0
y
0
5
10
15
20
25
x
raw data, starting:3 9
fitted Gauss locations:3.3 9

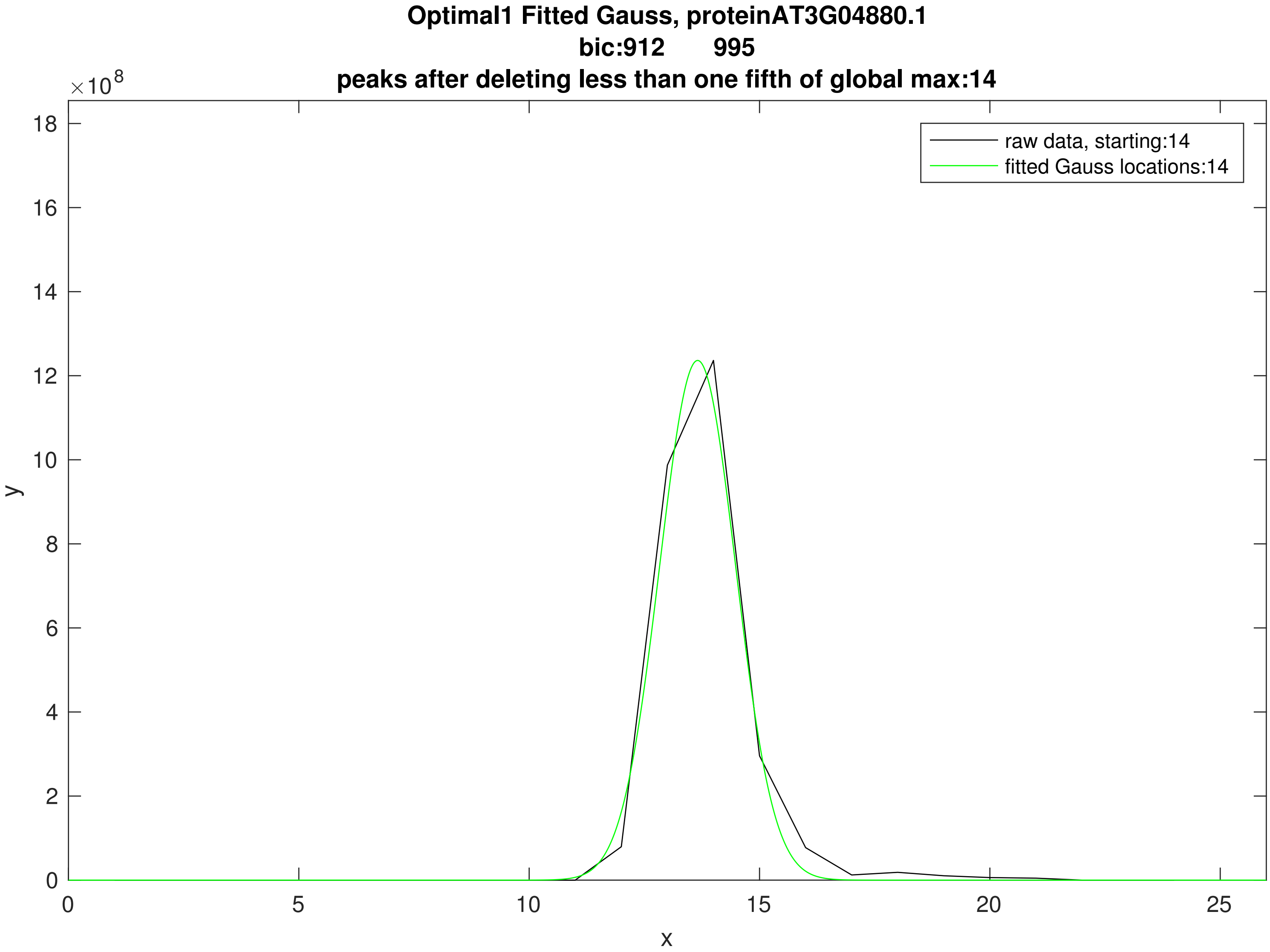
Optimal1 Fitted Gauss, proteinAT3G04880.1
bic:912 995
peaks after deleting less than one fifth of global max:14
×10^8
18
16
14
12
10
8
6
4
2
0
y
0
5
10
15
20
25
x
raw data, starting:14
fitted Gauss locations:14

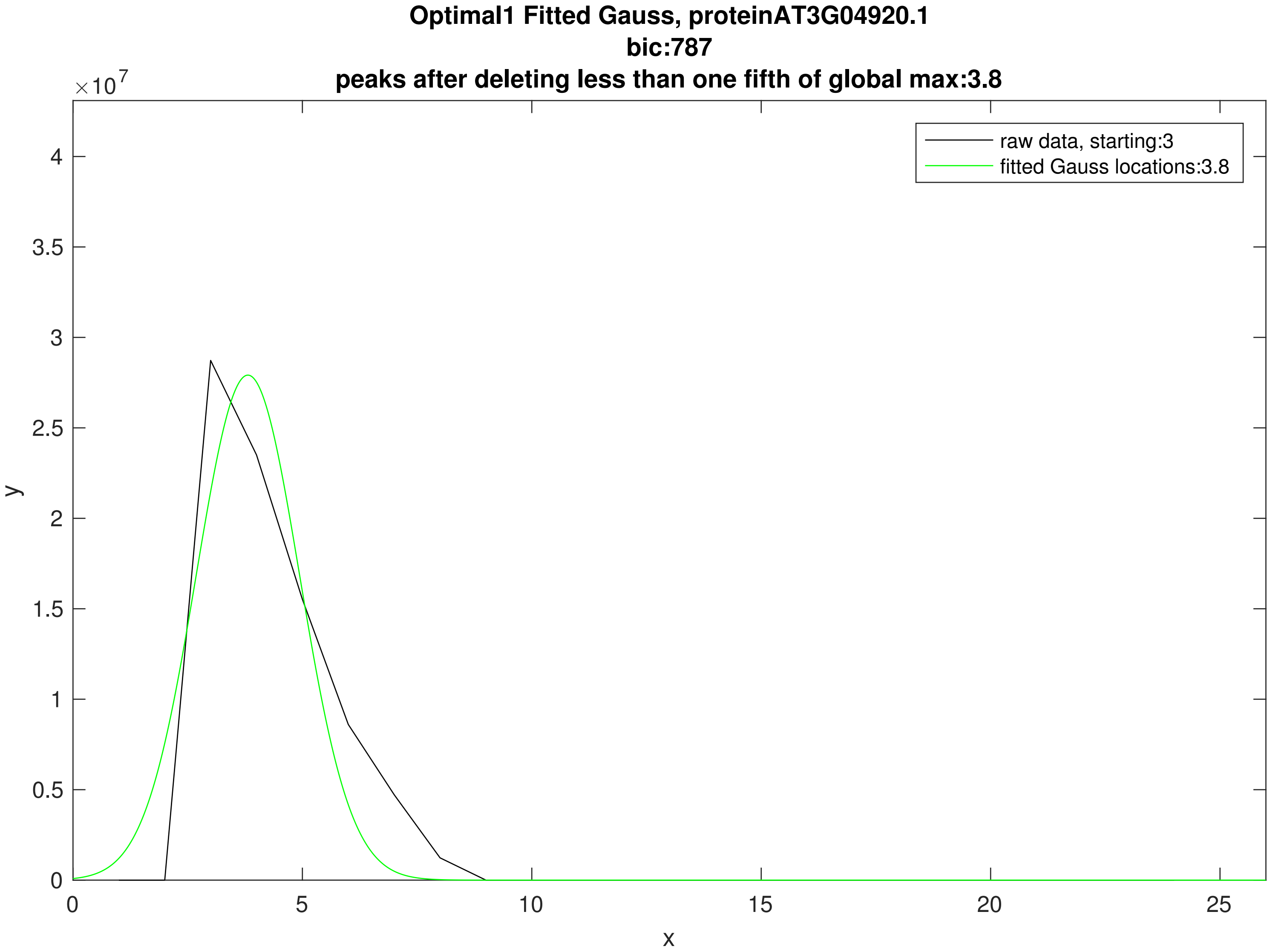

Optimal1 Fitted Gauss, proteinAT3G04920.1
bic:787
peaks after deleting less than one fifth of global max:3.8
×10^7
raw data, starting:3
fitted Gauss locations:3.8
y
x
0
0.5
1
1.5
2
2.5
3
3.5
4
5
10
15
20
25

Optimal1 Fitted Gauss, proteinAT3G04940.1
bic:818
peaks after deleting less than one fifth of global max:15

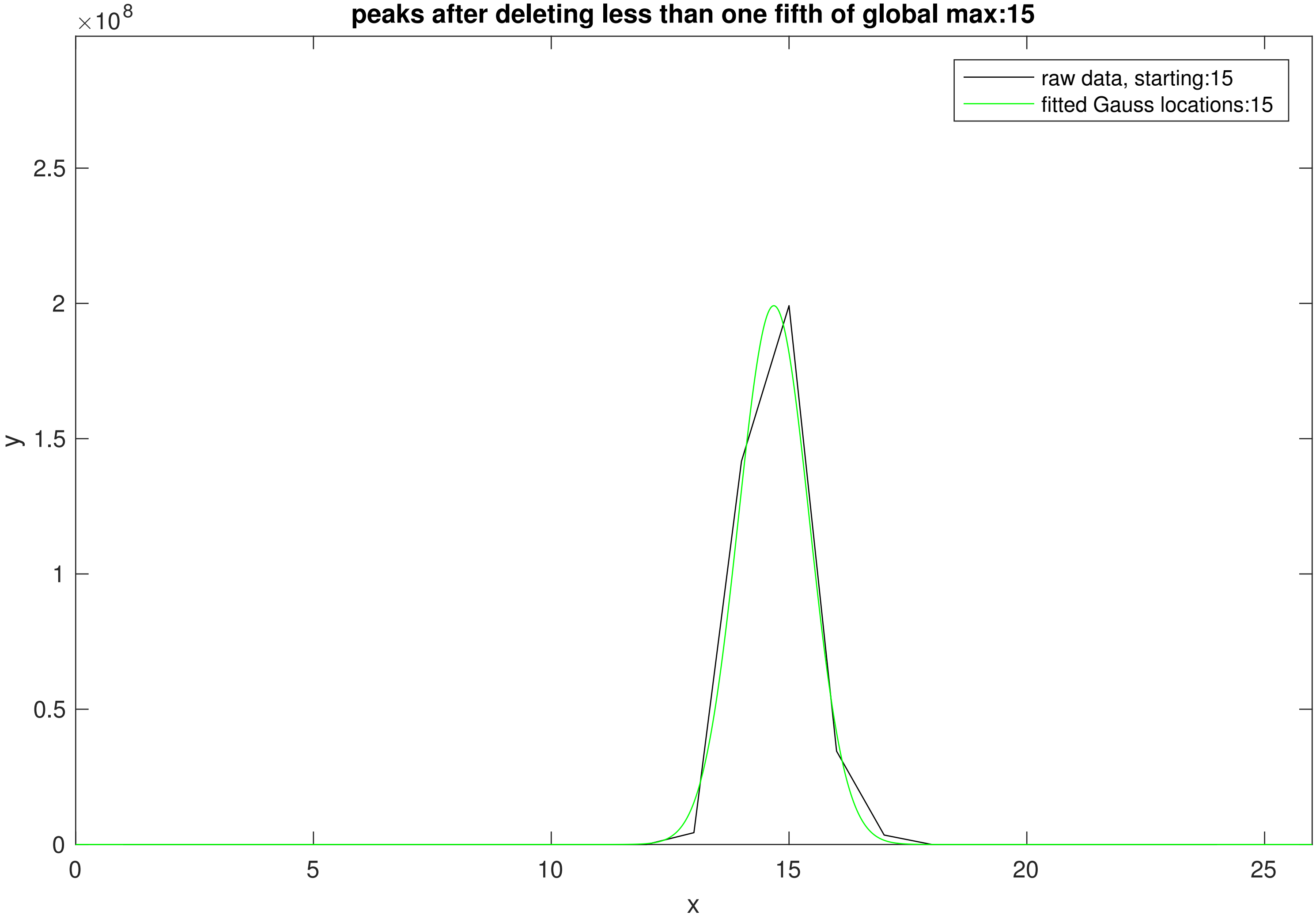

**Optimal1 Fitted Gauss, proteinAT3G05000.1**
**bic:760**
**peaks after deleting less than one fifth of global max:16**

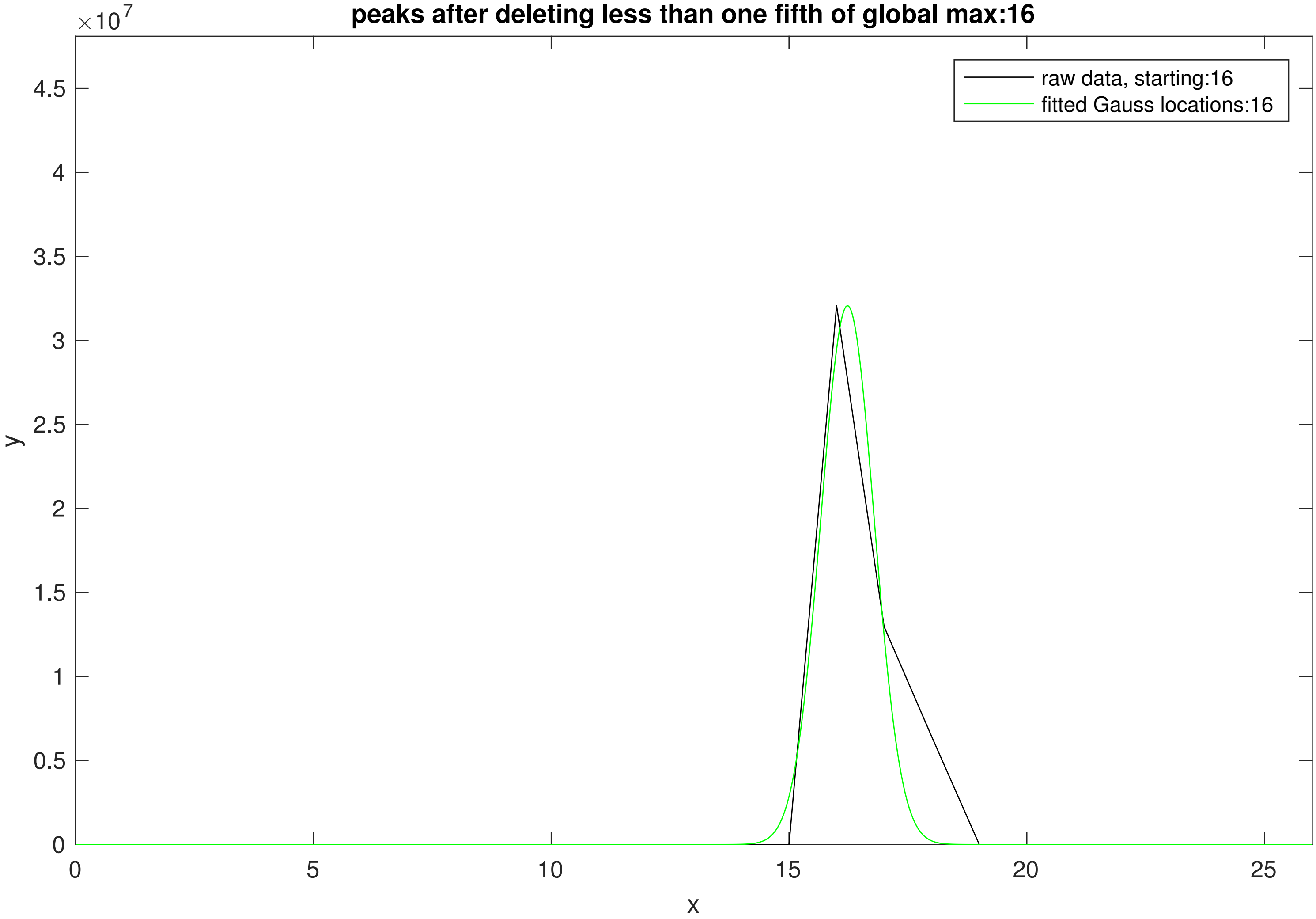

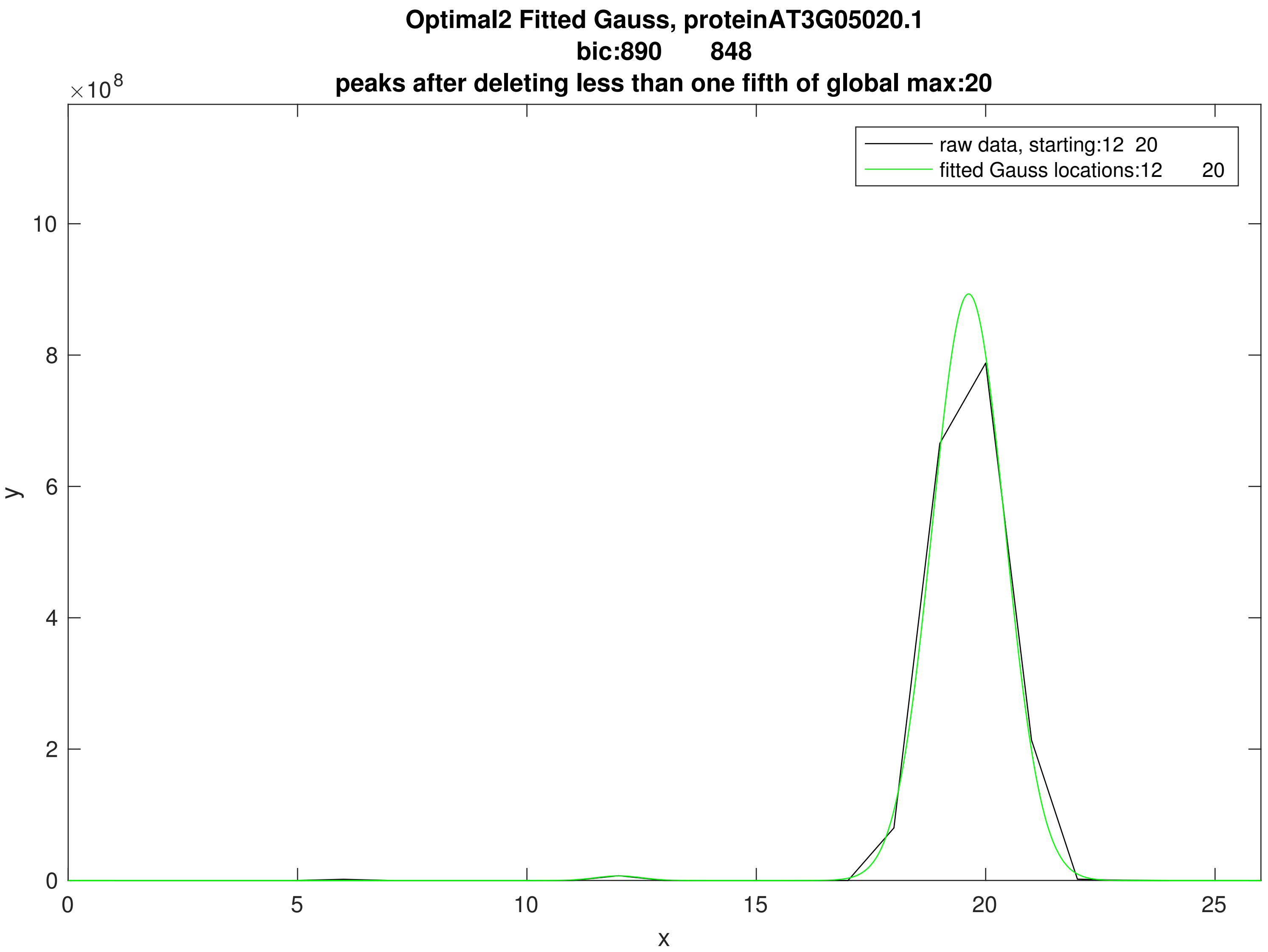
Optimal2 Fitted Gauss, proteinAT3G05020.1
bic:890 848
peaks after deleting less than one fifth of global max:20
×10^8
raw data, starting:12 20
fitted Gauss locations:12 20
y
x
0
2
4
6
8
10
0
5
10
15
20
25

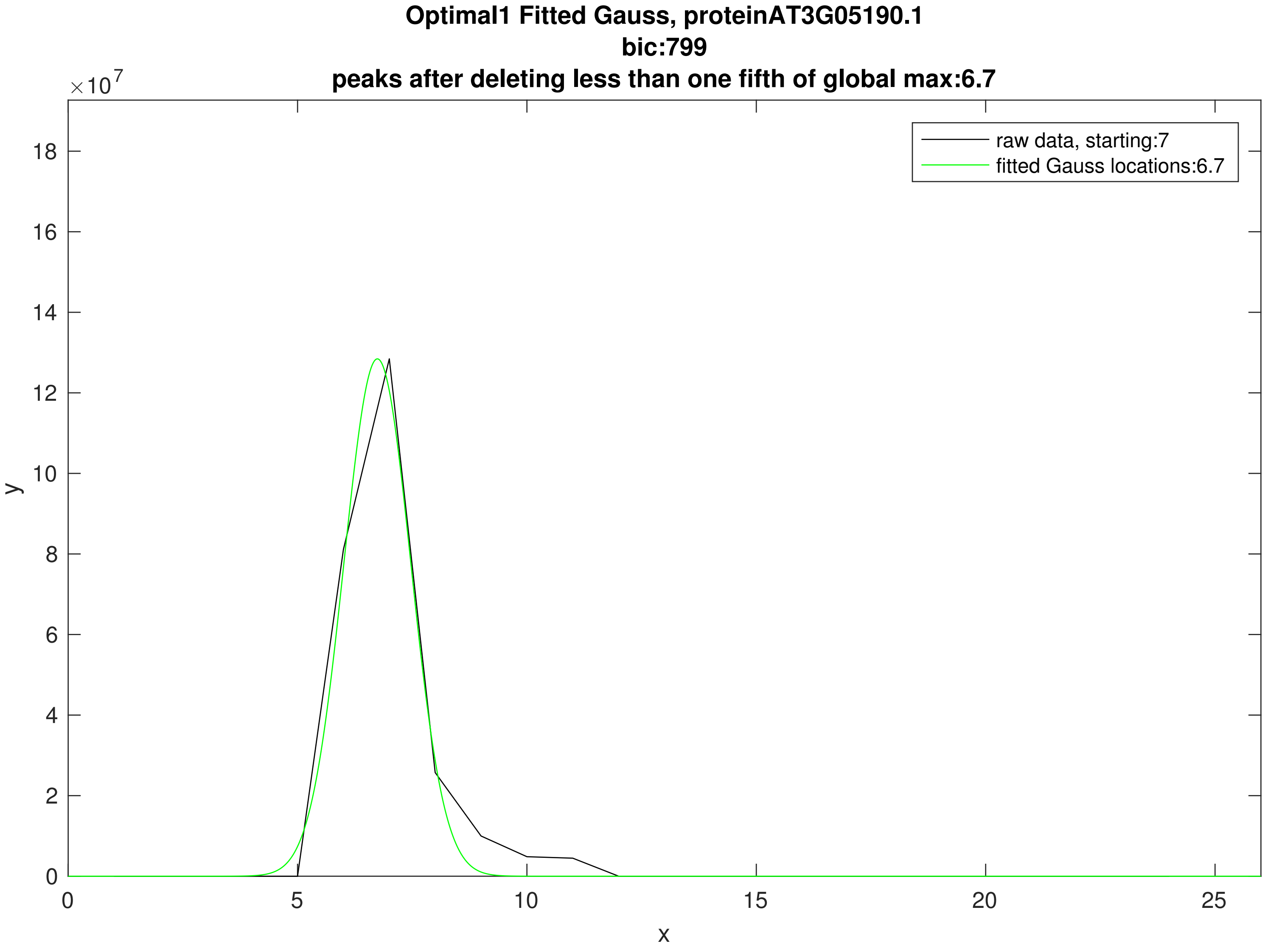
Optimal1 Fitted Gauss, proteinAT3G05190.1
bic:799
peaks after deleting less than one fifth of global max:6.7
$\times 10^7$
raw data, starting:7
fitted Gauss locations:6.7
y
x
0
2
4
6
8
10
12
14
16
18
0
5
10
15
20
25

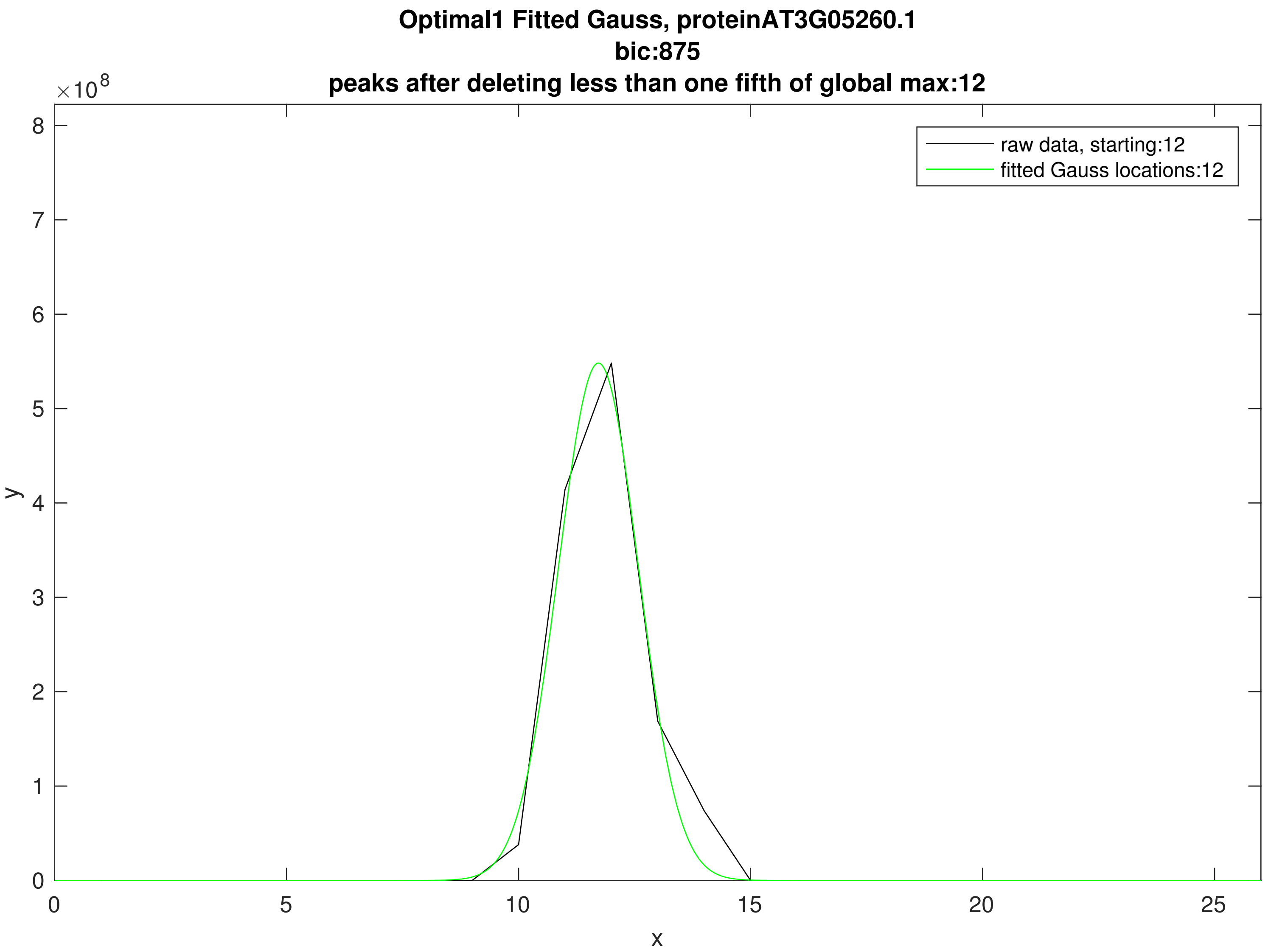
Optimal1 Fitted Gauss, proteinAT3G05260.1
bic:875
peaks after deleting less than one fifth of global max:12
×10^8
raw data, starting:12
fitted Gauss locations:12
y
x
0
1
2
3
4
5
6
7
8
0
5
10
15
20
25

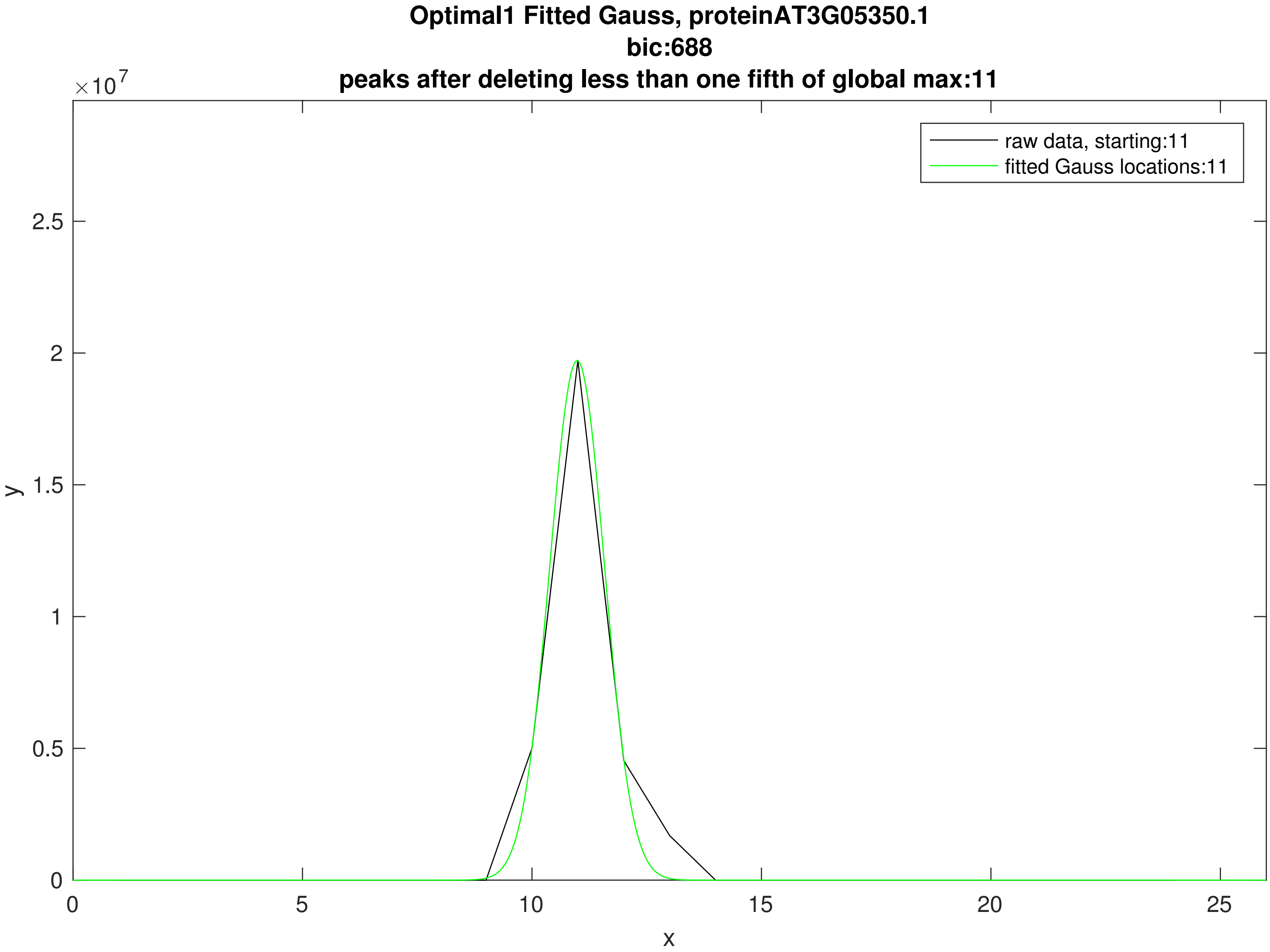

Optimal1 Fitted Gauss, proteinAT3G05350.1
bic:688
peaks after deleting less than one fifth of global max:11
×10^7
raw data, starting:11
fitted Gauss locations:11
y
x
0
0.5
1
1.5
2
2.5
5
10
15
20
25

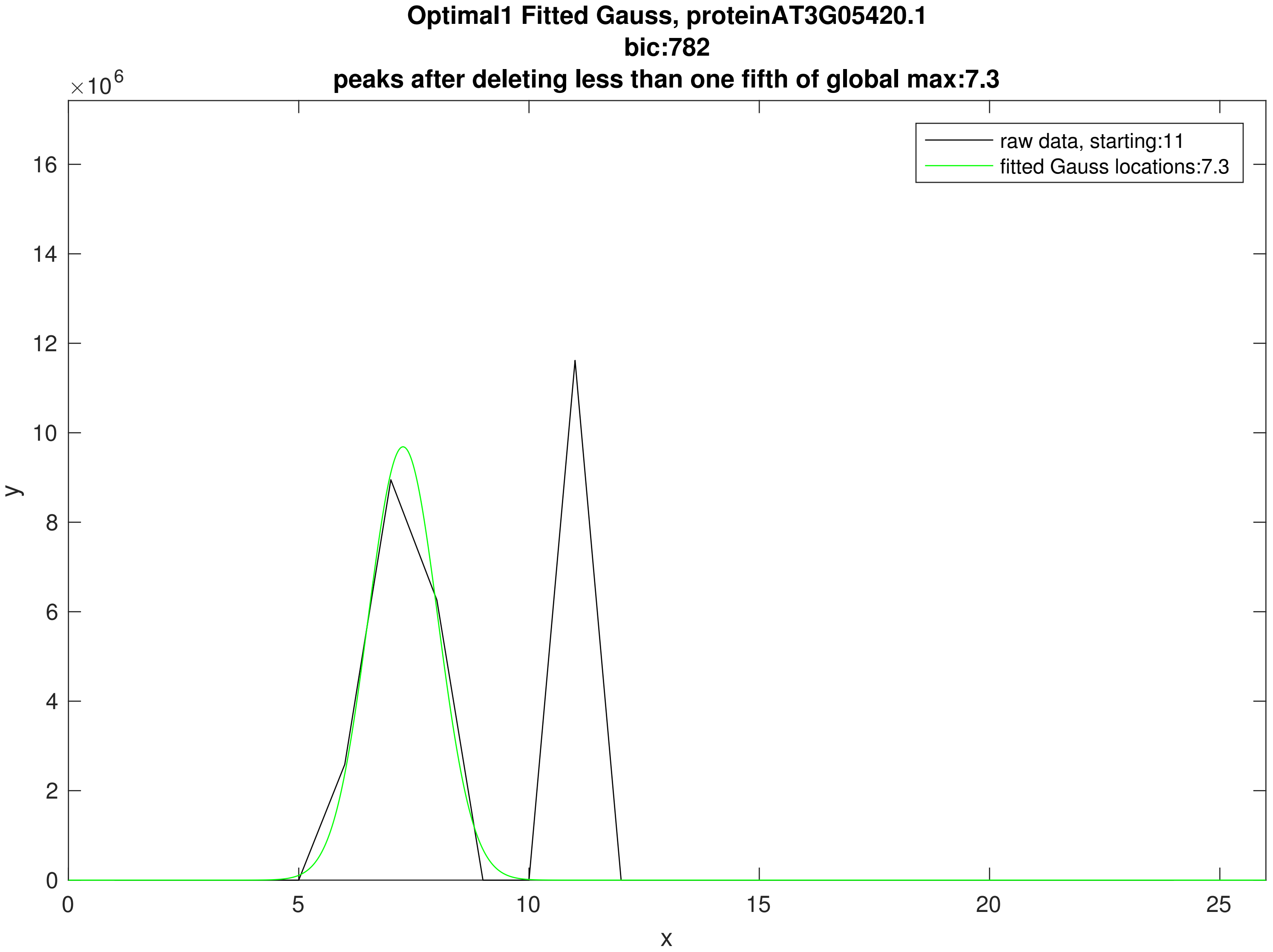

Optimal1 Fitted Gauss, proteinAT3G05420.1
bic:782
peaks after deleting less than one fifth of global max:7.3
×10^6
raw data, starting:11
fitted Gauss locations:7.3
16
14
12
10
8
6
4
2
0
y
0
5
10
15
20
25
x

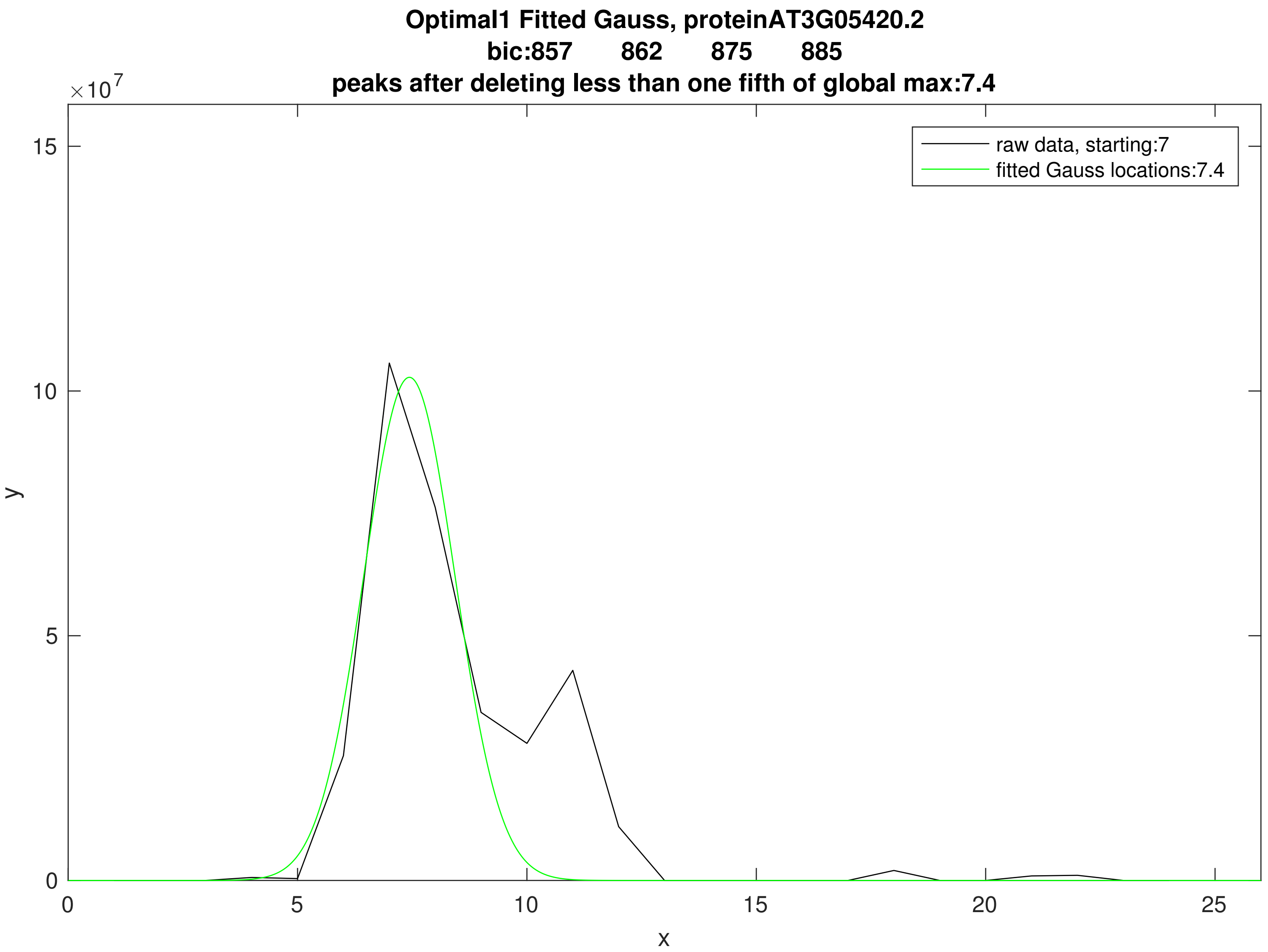
Optimal1 Fitted Gauss, proteinAT3G05420.2
bic:857 862 875 885
peaks after deleting less than one fifth of global max:7.4
×10^7
15
10
5
0
0
5
10
15
20
25
x
y
raw data, starting:7
fitted Gauss locations:7.4

**Optimal2 Fitted Gauss, proteinAT3G05530.1;AT1G09100.1**
**bic:843 841 850 860**
**peaks after deleting less than one fifth of global max:3.9 13**

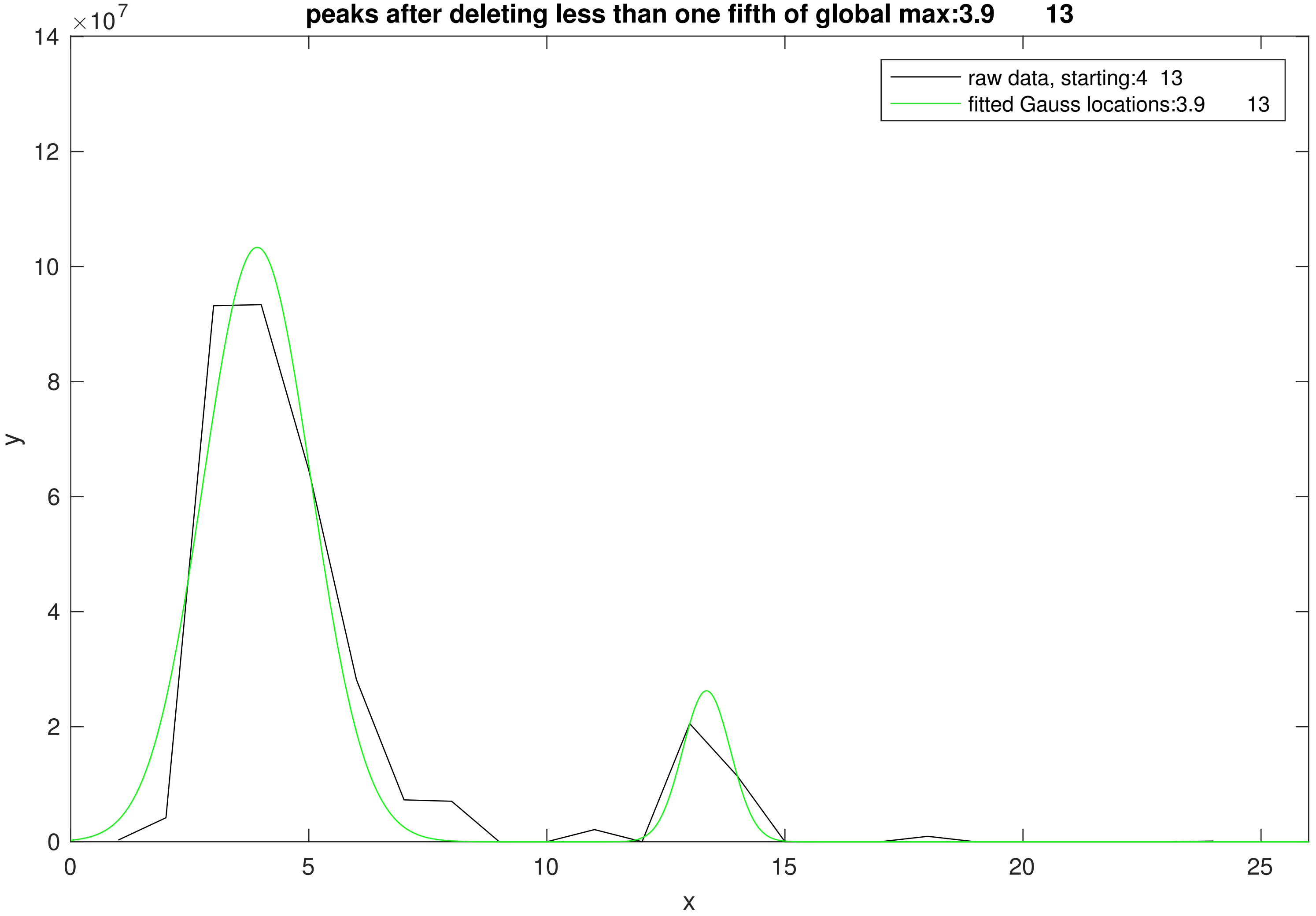

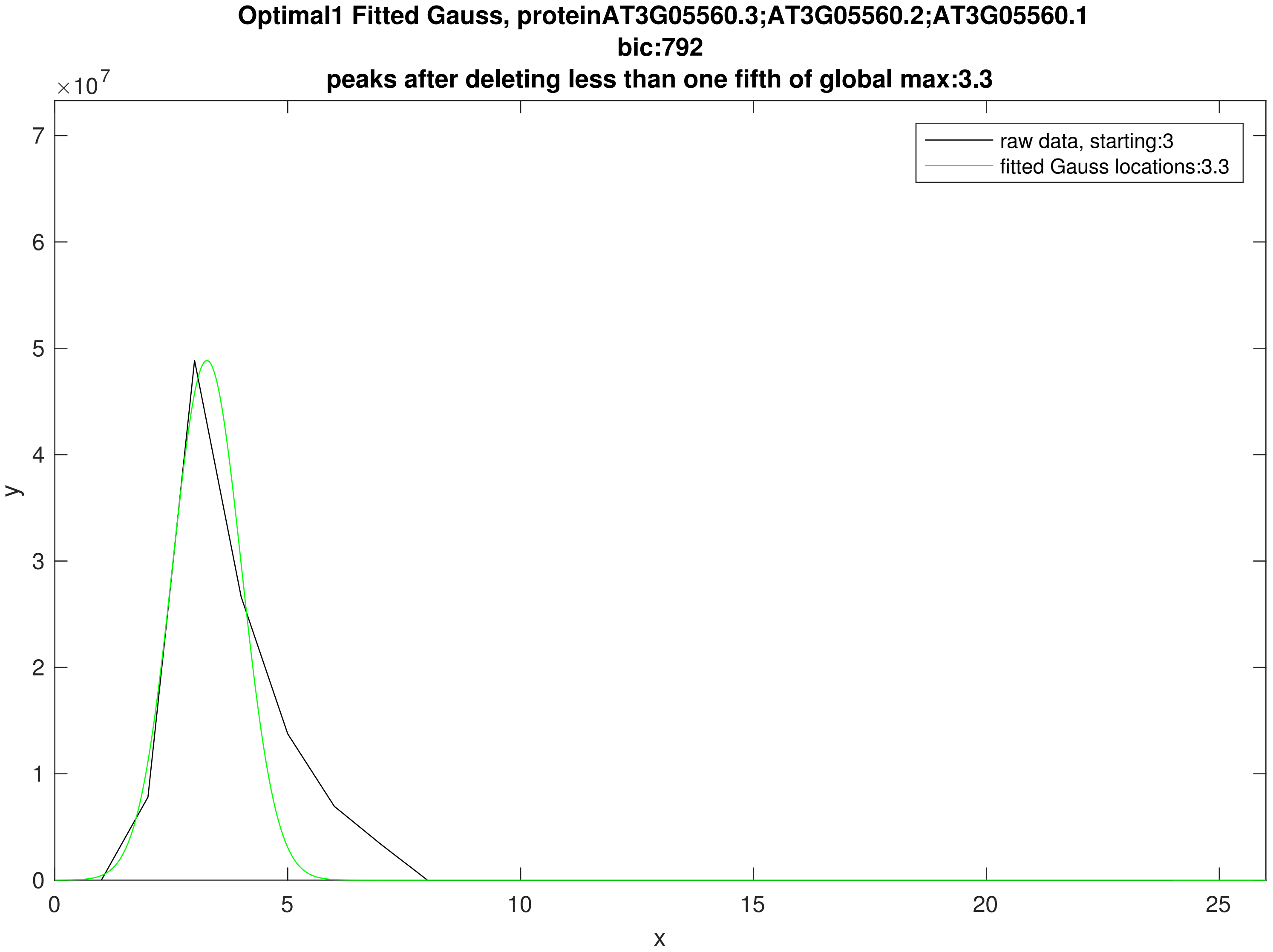
Optimal1 Fitted Gauss, proteinAT3G05560.3;AT3G05560.2;AT3G05560.1
bic:792
peaks after deleting less than one fifth of global max:3.3
×10^7
raw data, starting:3
fitted Gauss locations:3.3
0
1
2
3
4
5
6
7
y
0
5
10
15
20
25
x

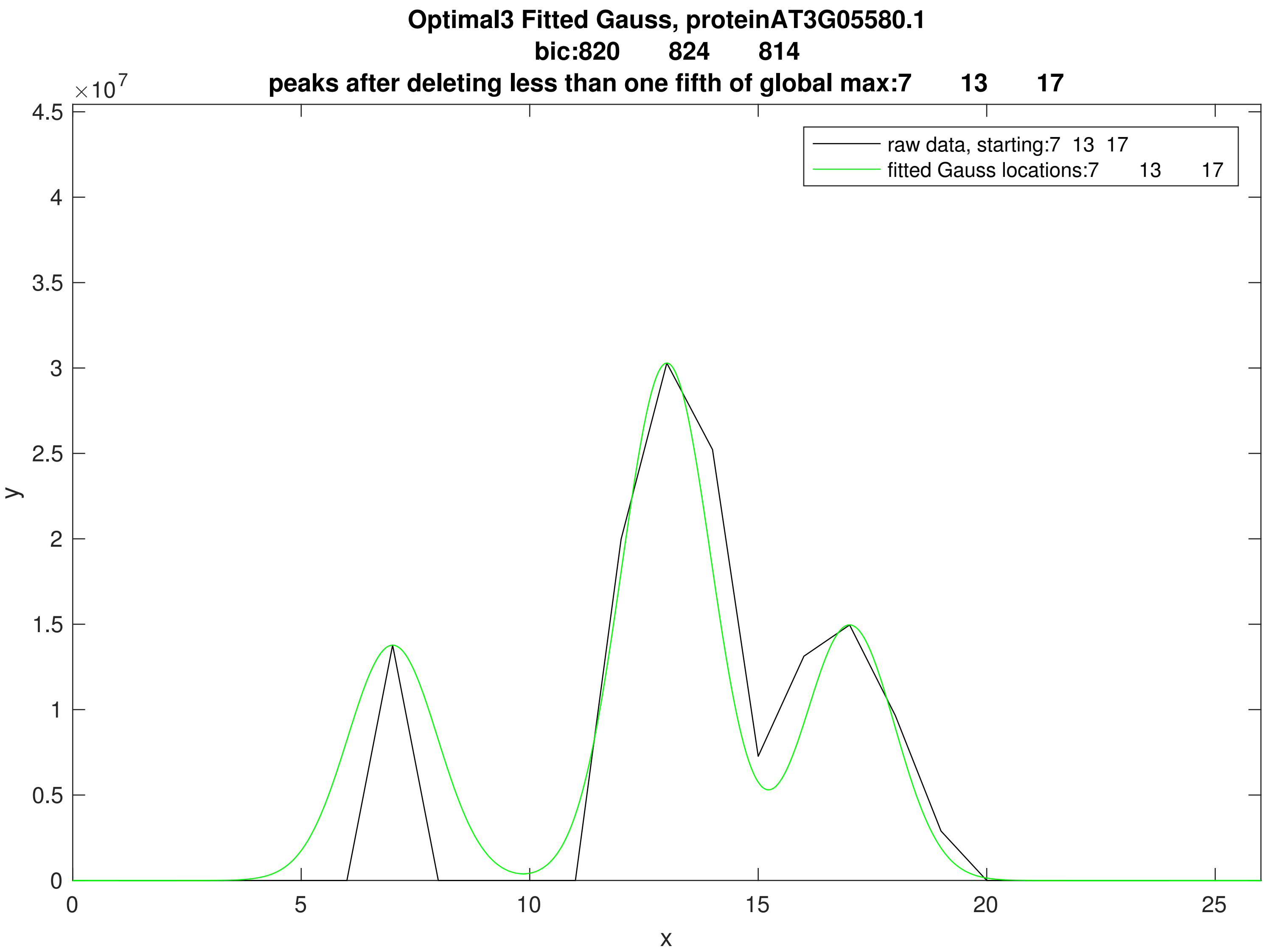
Optimal3 Fitted Gauss, proteinAT3G05580.1
bic:820 824 814
peaks after deleting less than one fifth of global max:7 13 17
×10^7
raw data, starting:7 13 17
fitted Gauss locations:7 13 17
y
x
0
0.5
1
1.5
2
2.5
3
3.5
4
4.5
5
10
15
20
25

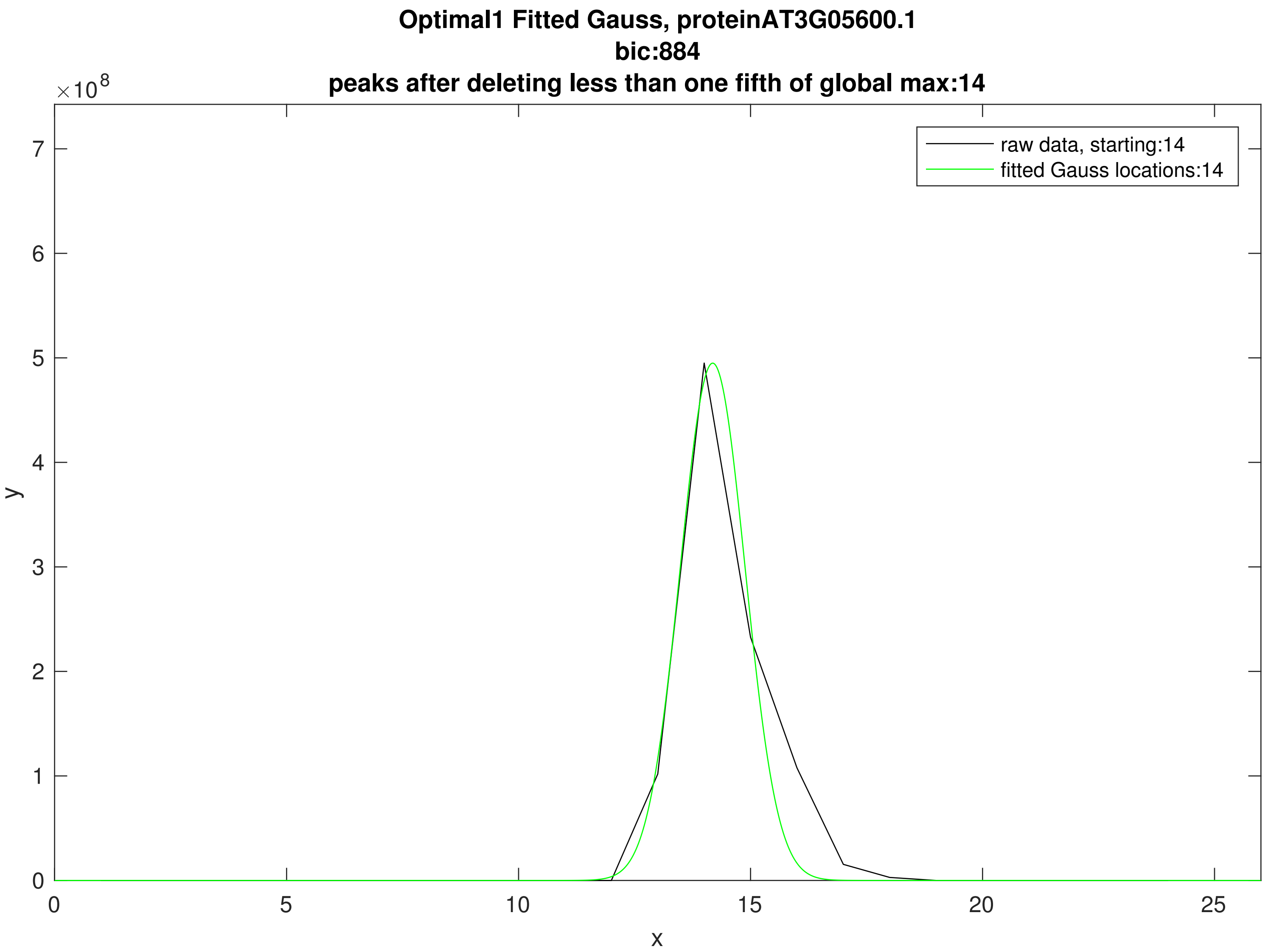

Optimal1 Fitted Gauss, proteinAT3G05600.1
bic:884
peaks after deleting less than one fifth of global max:14
×10^8
raw data, starting:14
fitted Gauss locations:14
0
1
2
3
4
5
6
7
y
0
5
10
15
20
25
x

**Optimal1 Fitted Gauss, proteinAT3G05625.1**
**bic:689**
**peaks after deleting less than one fifth of global max:19**

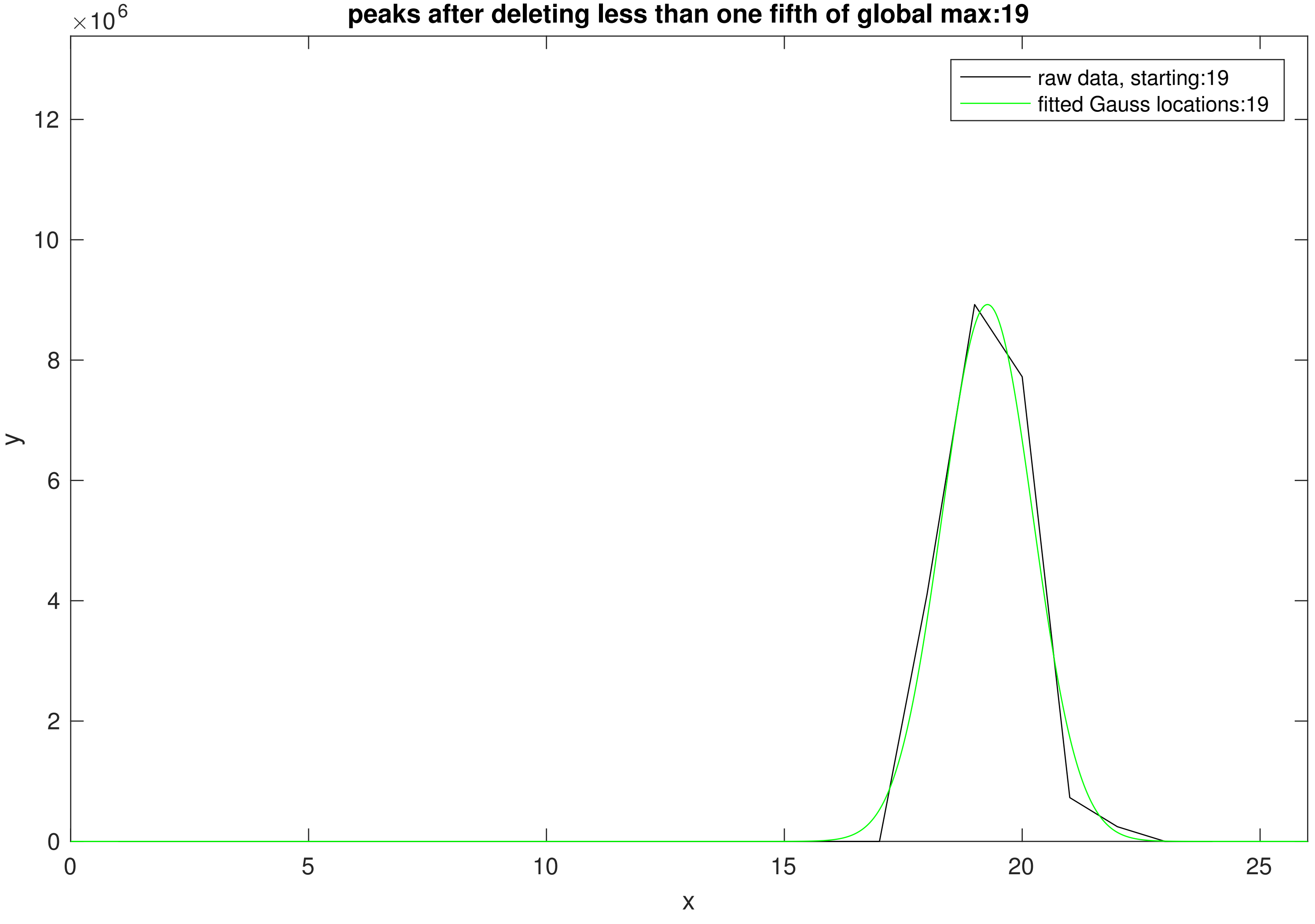

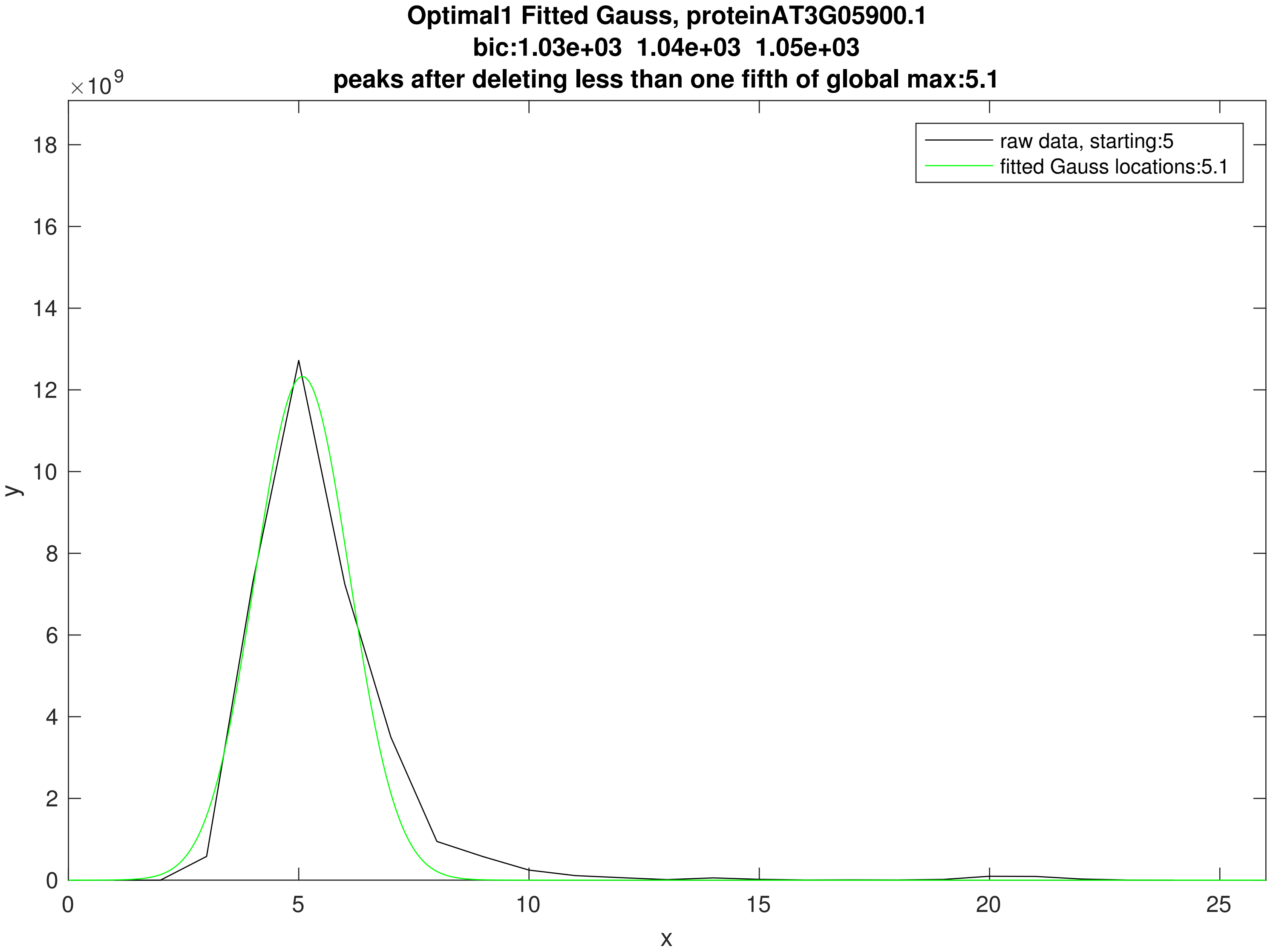

Optimal1 Fitted Gauss, proteinAT3G05900.1
bic:1.03e+03 1.04e+03 1.05e+03
peaks after deleting less than one fifth of global max:5.1
×10^9
18
16
14
12
10
8
6
4
2
0
y
0
5
10
15
20
25
x
raw data, starting:5
fitted Gauss locations:5.1

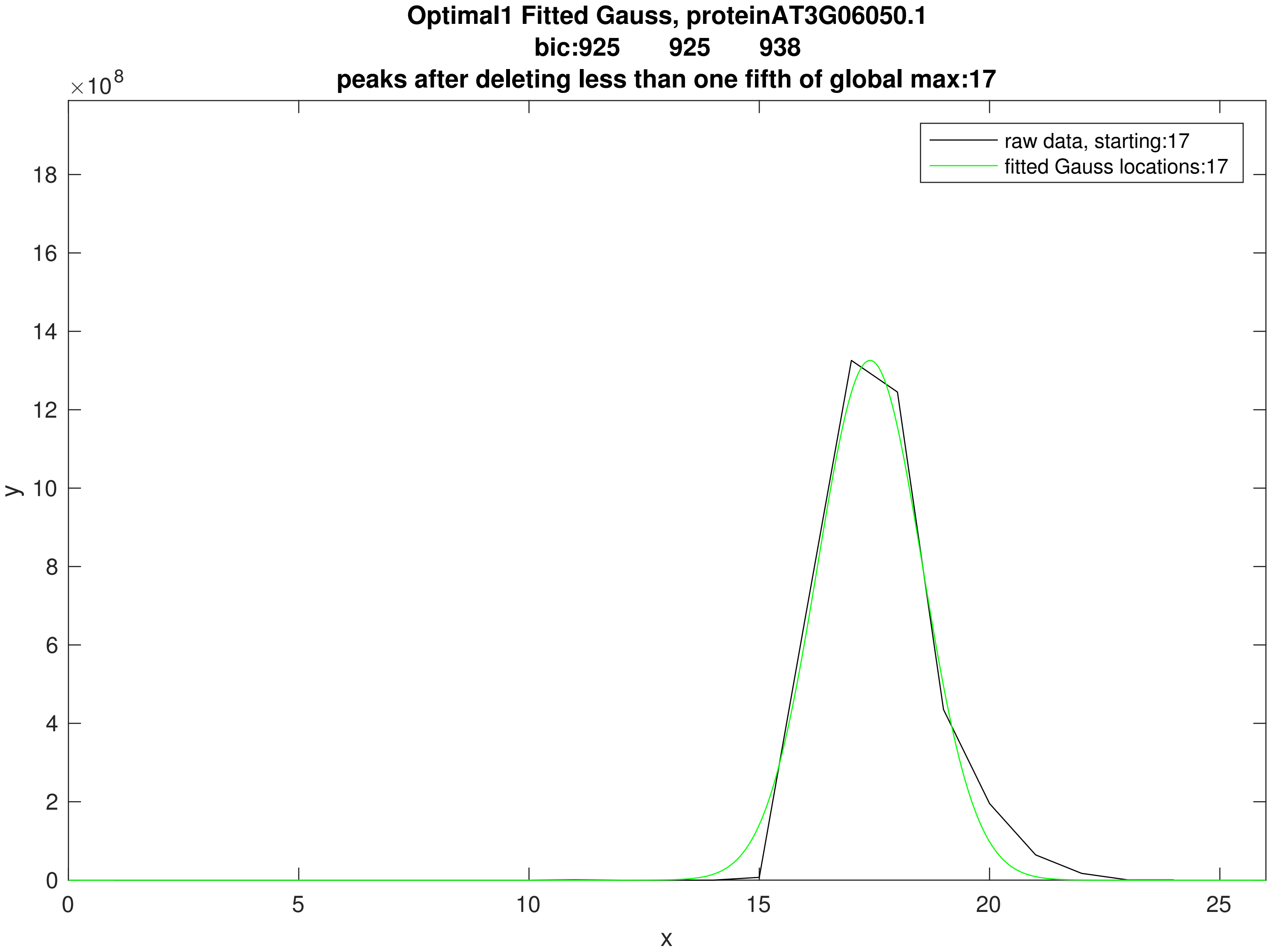
Optimal1 Fitted Gauss, proteinAT3G06050.1
bic:925 925 938
peaks after deleting less than one fifth of global max:17
×10^8
raw data, starting:17
fitted Gauss locations:17
y
x
0
2
4
6
8
10
12
14
16
18
0
5
10
15
20
25

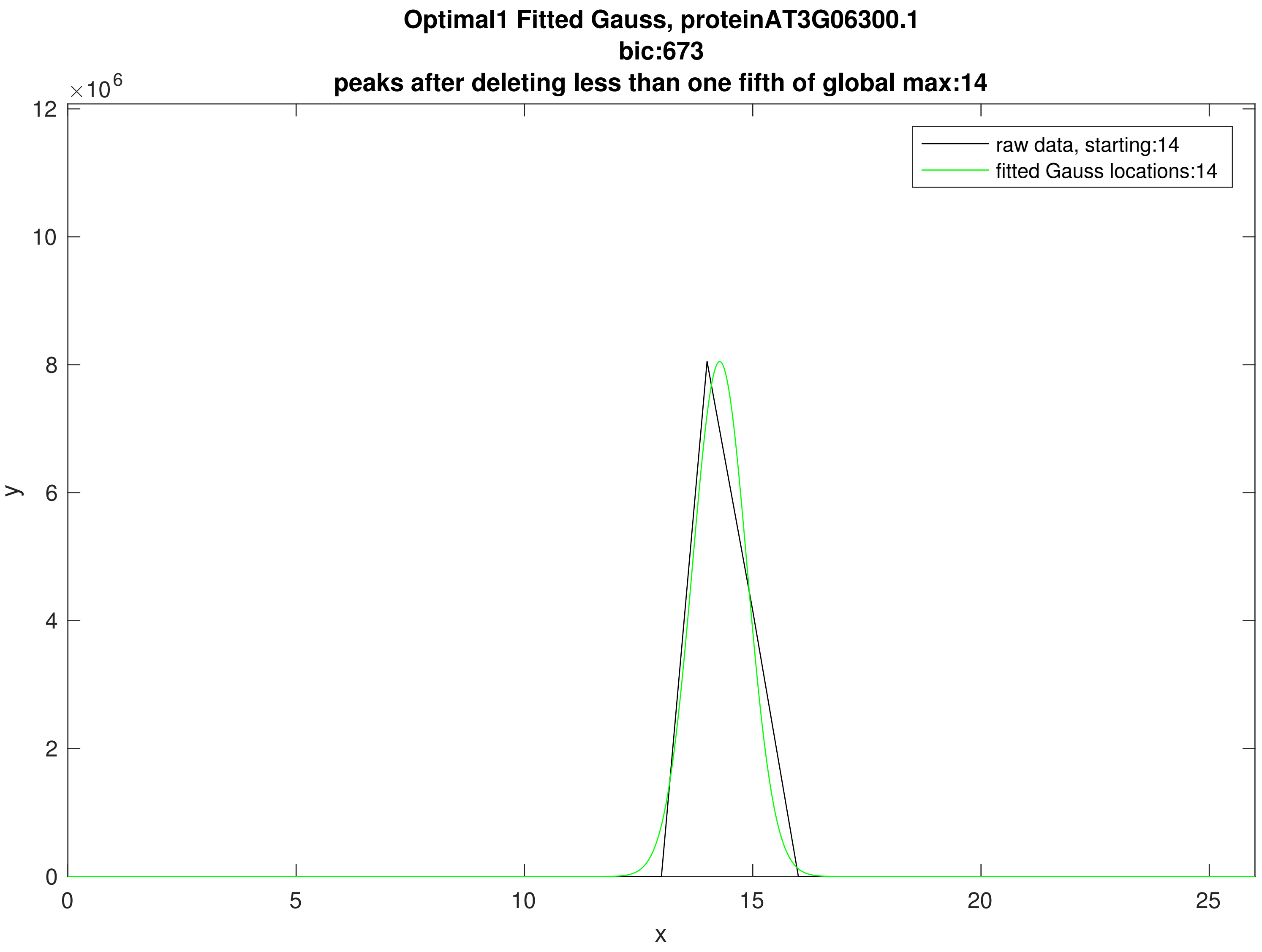

Optimal1 Fitted Gauss, proteinAT3G06300.1
bic:673
peaks after deleting less than one fifth of global max:14
×10^6
y
x
raw data, starting:14
fitted Gauss locations:14

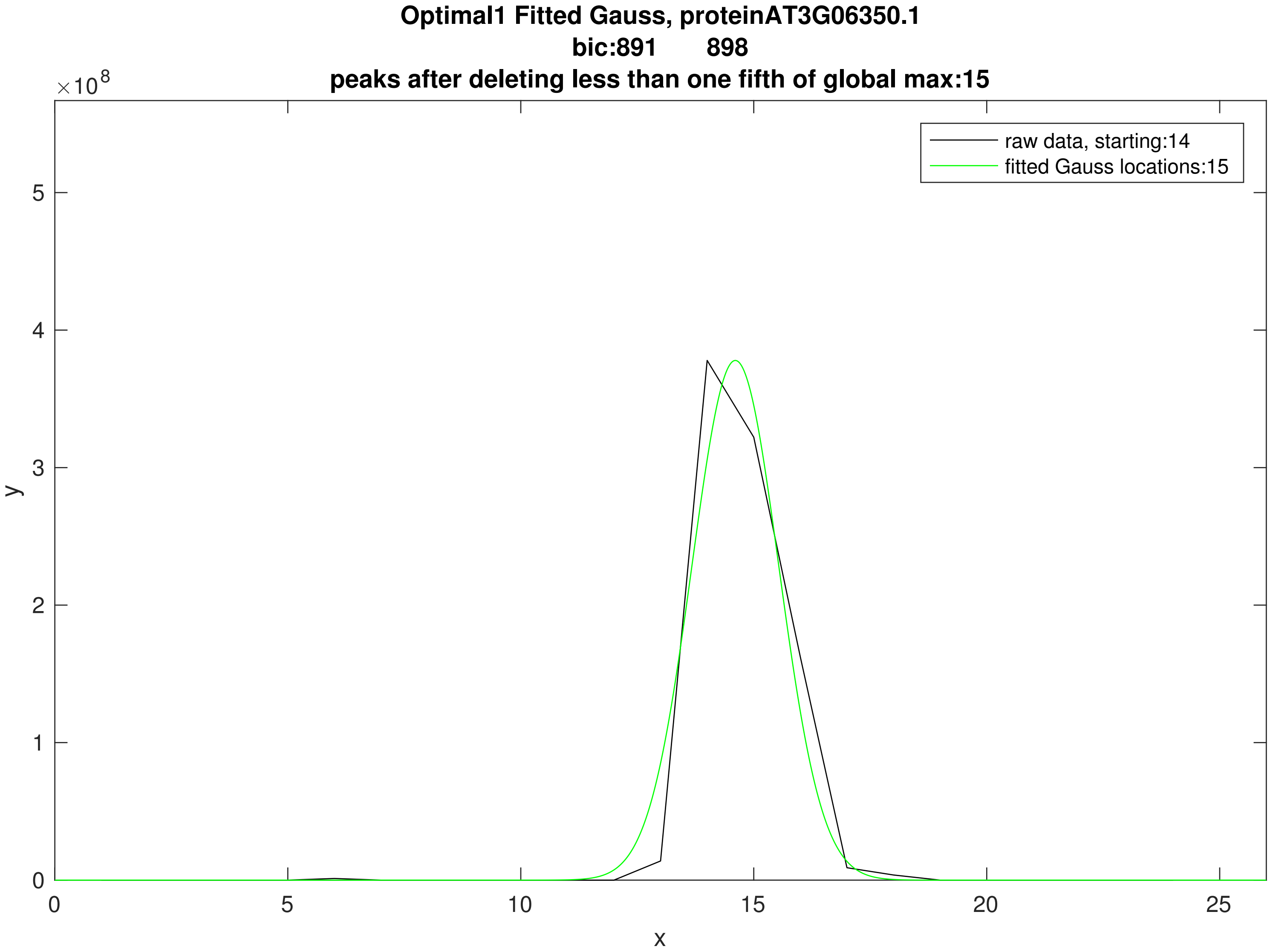

Optimal1 Fitted Gauss, proteinAT3G06350.1
bic:891 898
peaks after deleting less than one fifth of global max:15
×10^8
raw data, starting:14
fitted Gauss locations:15
y
x
0
1
2
3
4
5
0
5
10
15
20
25

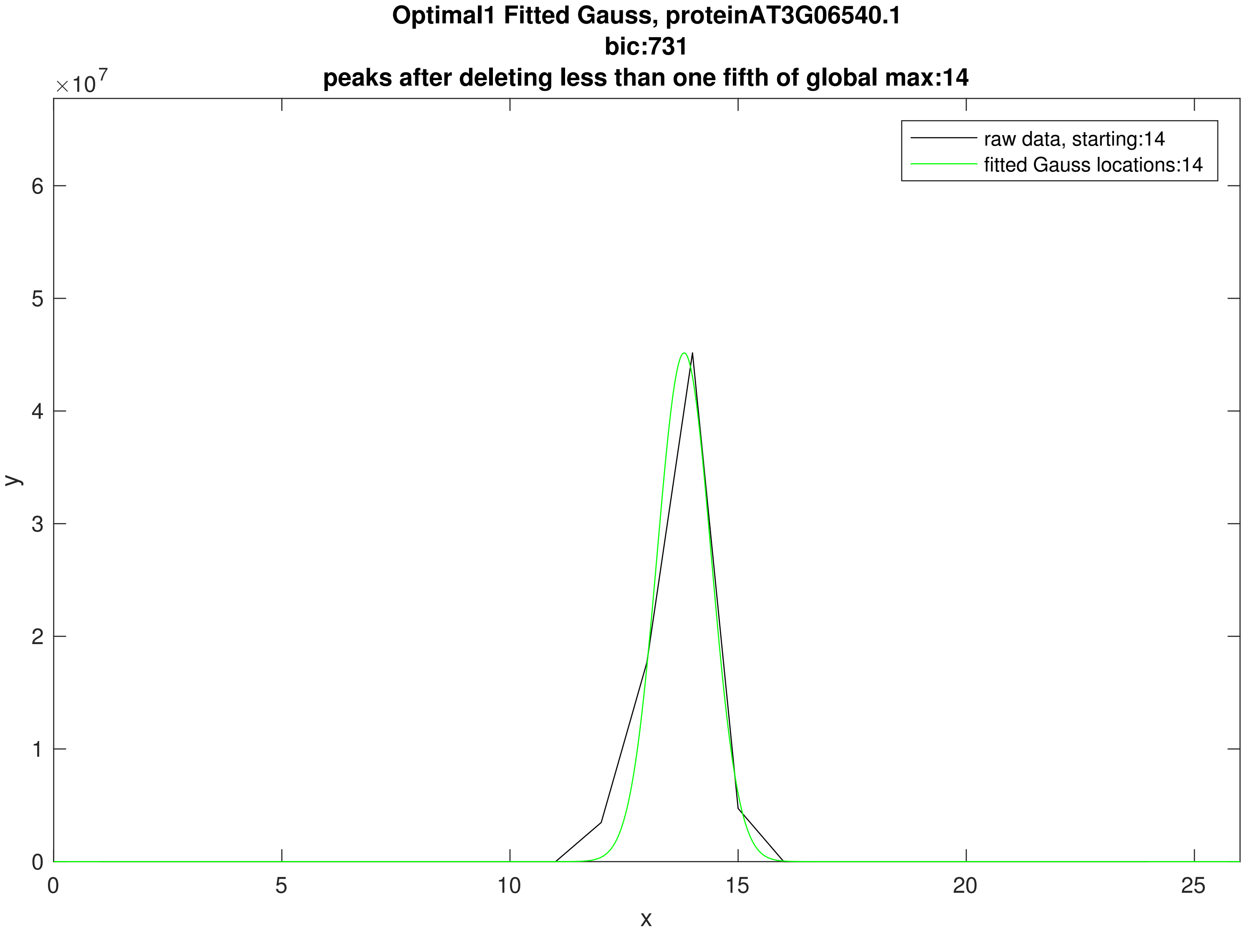

Optimal1 Fitted Gauss, proteinAT3G06540.1
bic:731
peaks after deleting less than one fifth of global max:14
×10^7
raw data, starting:14
fitted Gauss locations:14
y
x
0
1
2
3
4
5
6
0
5
10
15
20
25

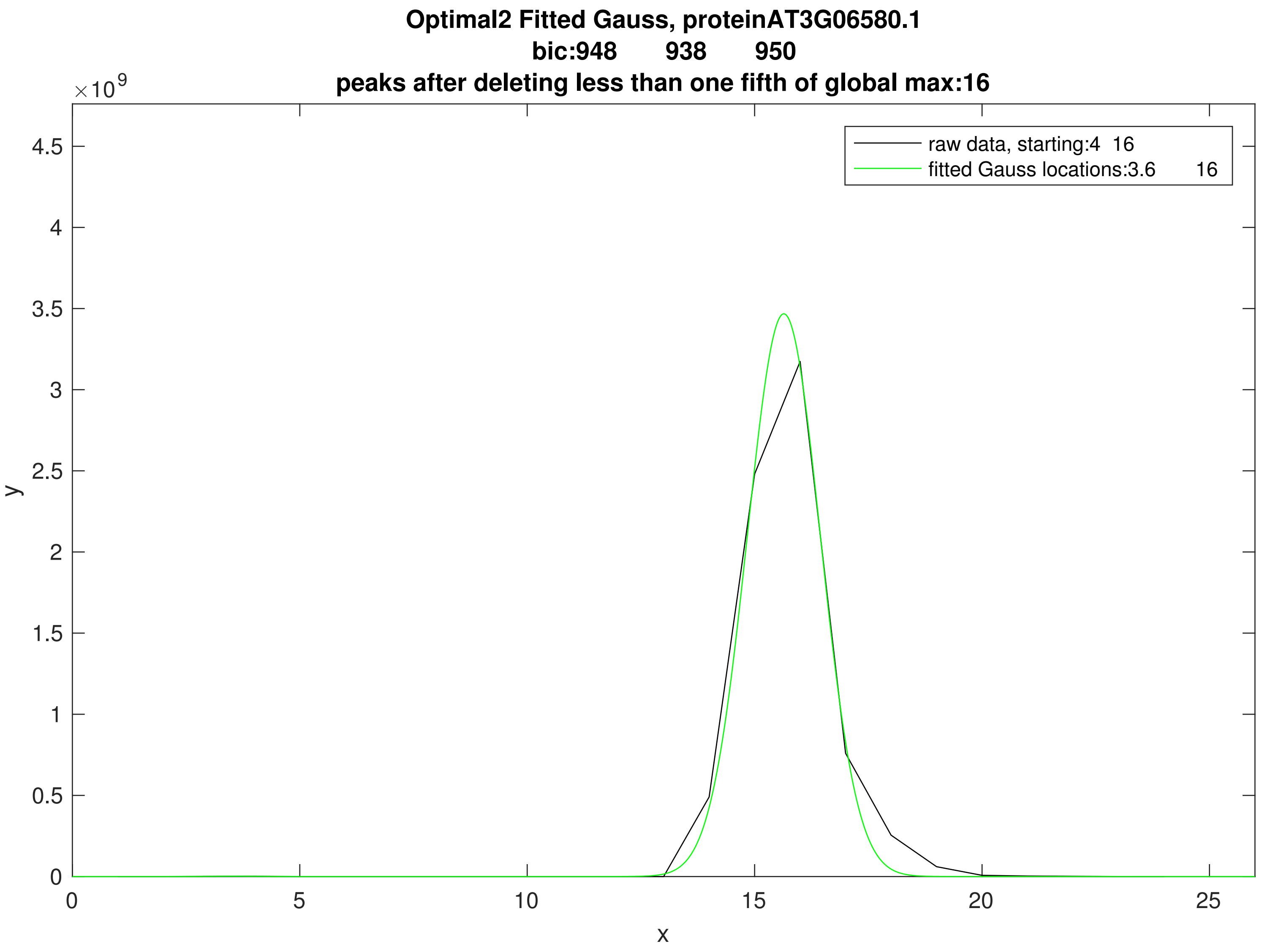
Optimal2 Fitted Gauss, proteinAT3G06580.1
bic:948 938 950
peaks after deleting less than one fifth of global max:16
×10^9
raw data, starting:4 16
fitted Gauss locations:3.6 16
y
x
0
0.5
1
1.5
2
2.5
3
3.5
4
4.5
5
10
15
20
25

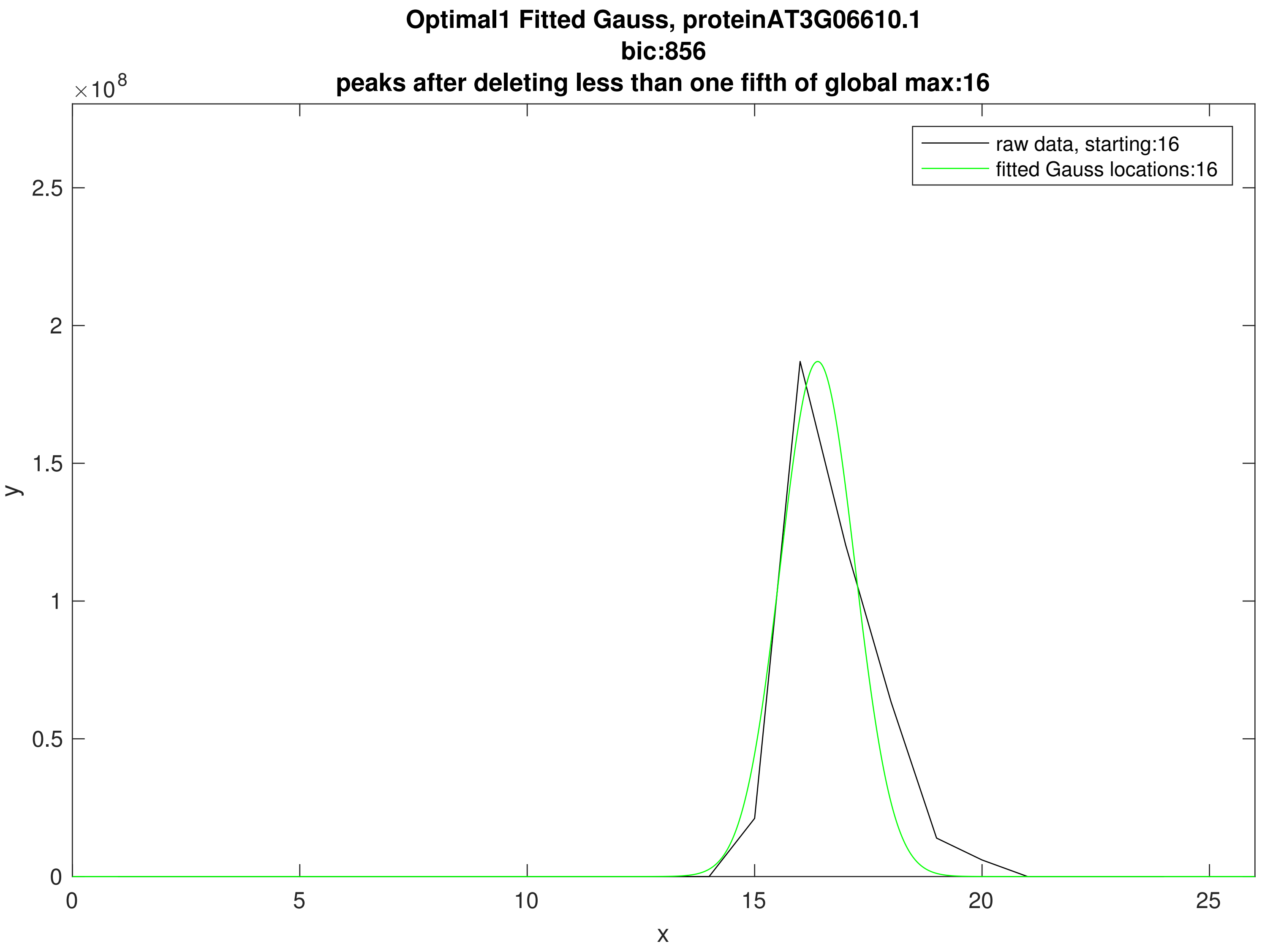
Optimal1 Fitted Gauss, proteinAT3G06610.1
bic:856
peaks after deleting less than one fifth of global max:16
×10^8
raw data, starting:16
fitted Gauss locations:16
2.5
2
1.5
1
0.5
0
y
0
5
10
15
20
25
x

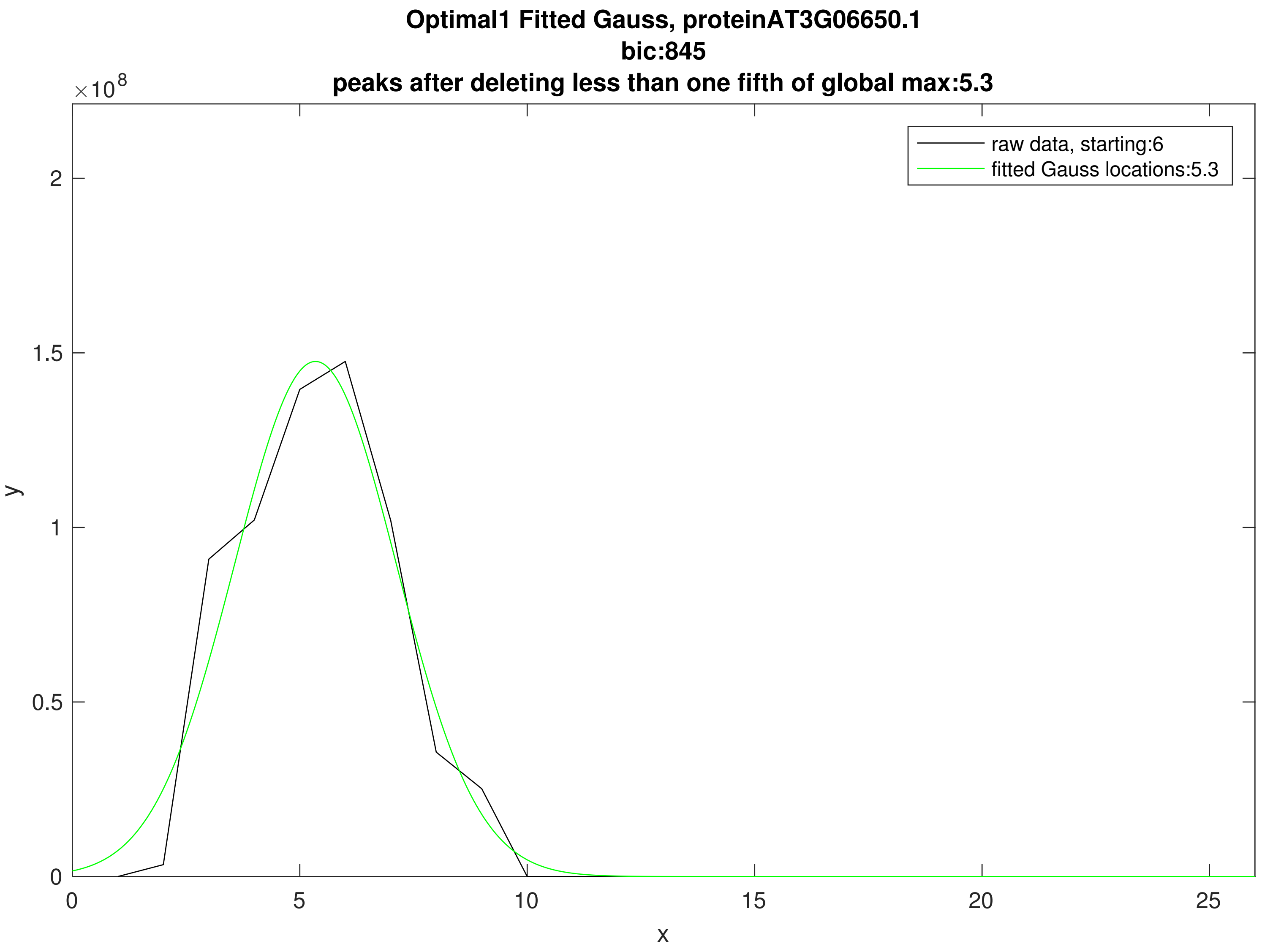
Optimal1 Fitted Gauss, proteinAT3G06650.1
bic:845
peaks after deleting less than one fifth of global max:5.3
×10^8
raw data, starting:6
fitted Gauss locations:5.3
y
x
0
0.5
1
1.5
2
0
5
10
15
20
25

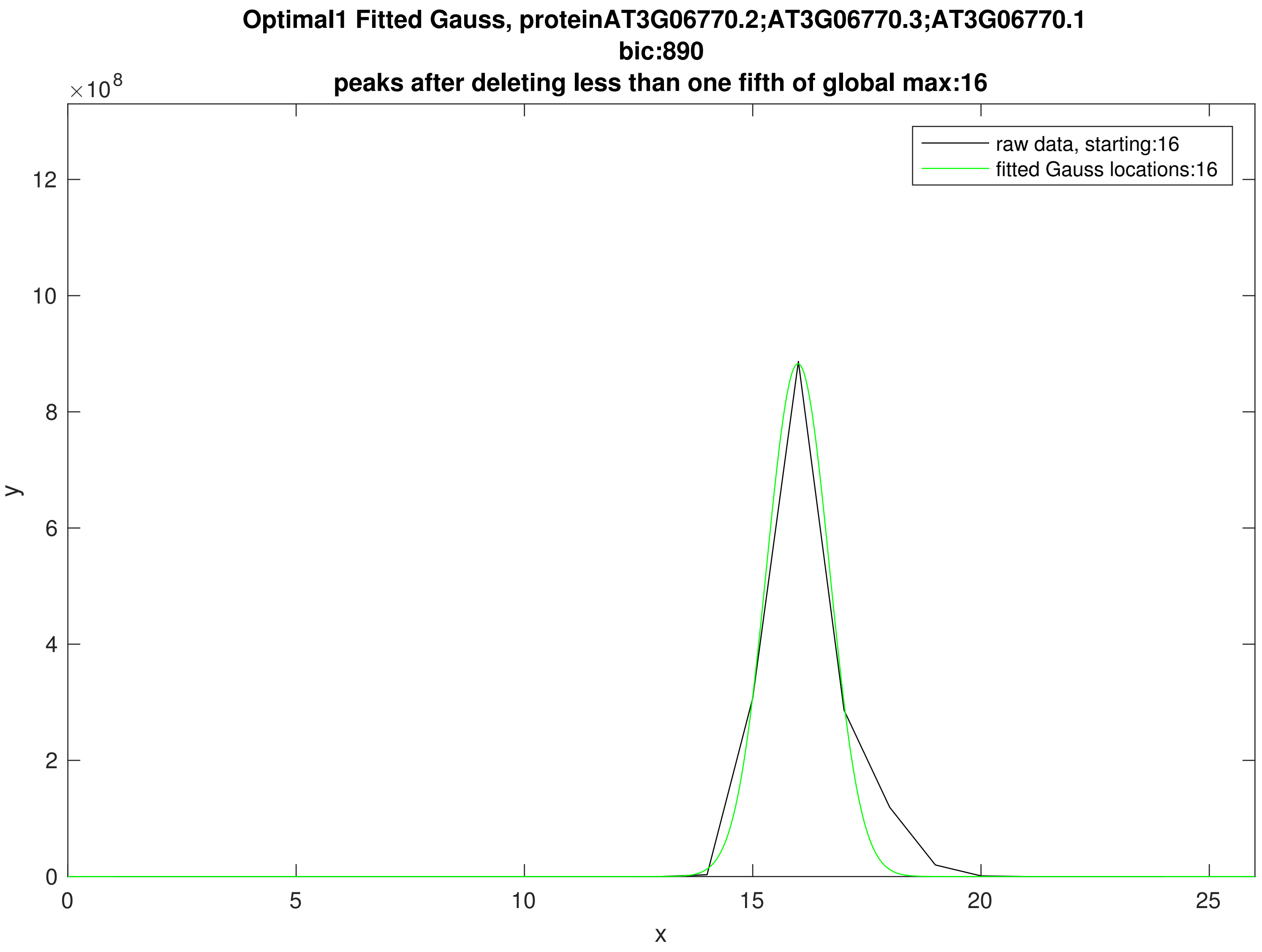

Optimal1 Fitted Gauss, proteinAT3G06770.2;AT3G06770.3;AT3G06770.1
bic:890
peaks after deleting less than one fifth of global max:16
×10^8
12
10
8
6
4
2
0
y
0
5
10
15
20
25
x
raw data, starting:16
fitted Gauss locations:16

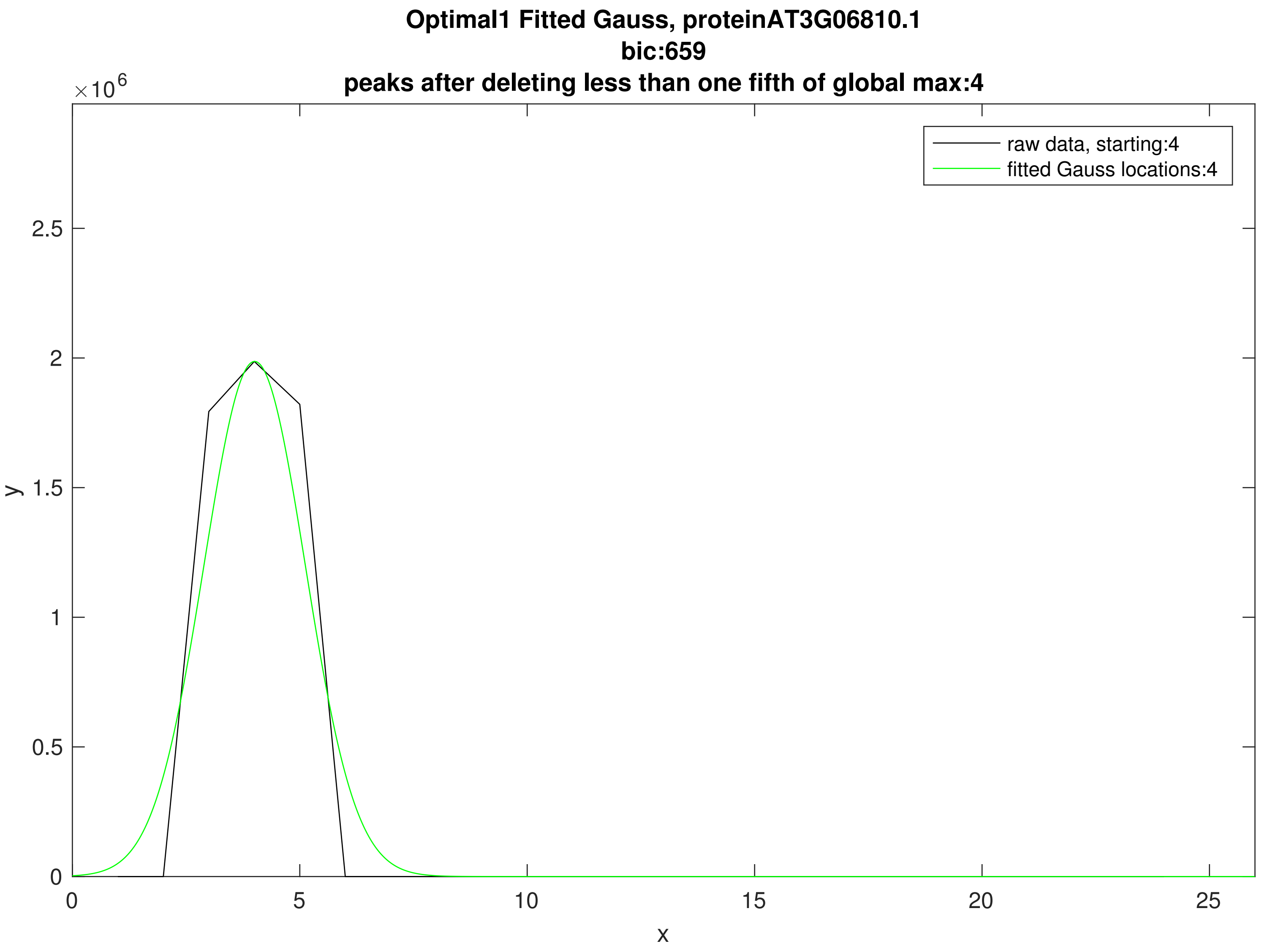
Optimal1 Fitted Gauss, proteinAT3G06810.1
bic:659
peaks after deleting less than one fifth of global max:4
×10^6
raw data, starting:4
fitted Gauss locations:4
y
0
0.5
1
1.5
2
2.5
x
0
5
10
15
20
25

#### Optimal1 Fitted Gauss, proteinAT3G06860.1
#### bic:871 905 919
#### peaks after deleting less than one fifth of global max:15

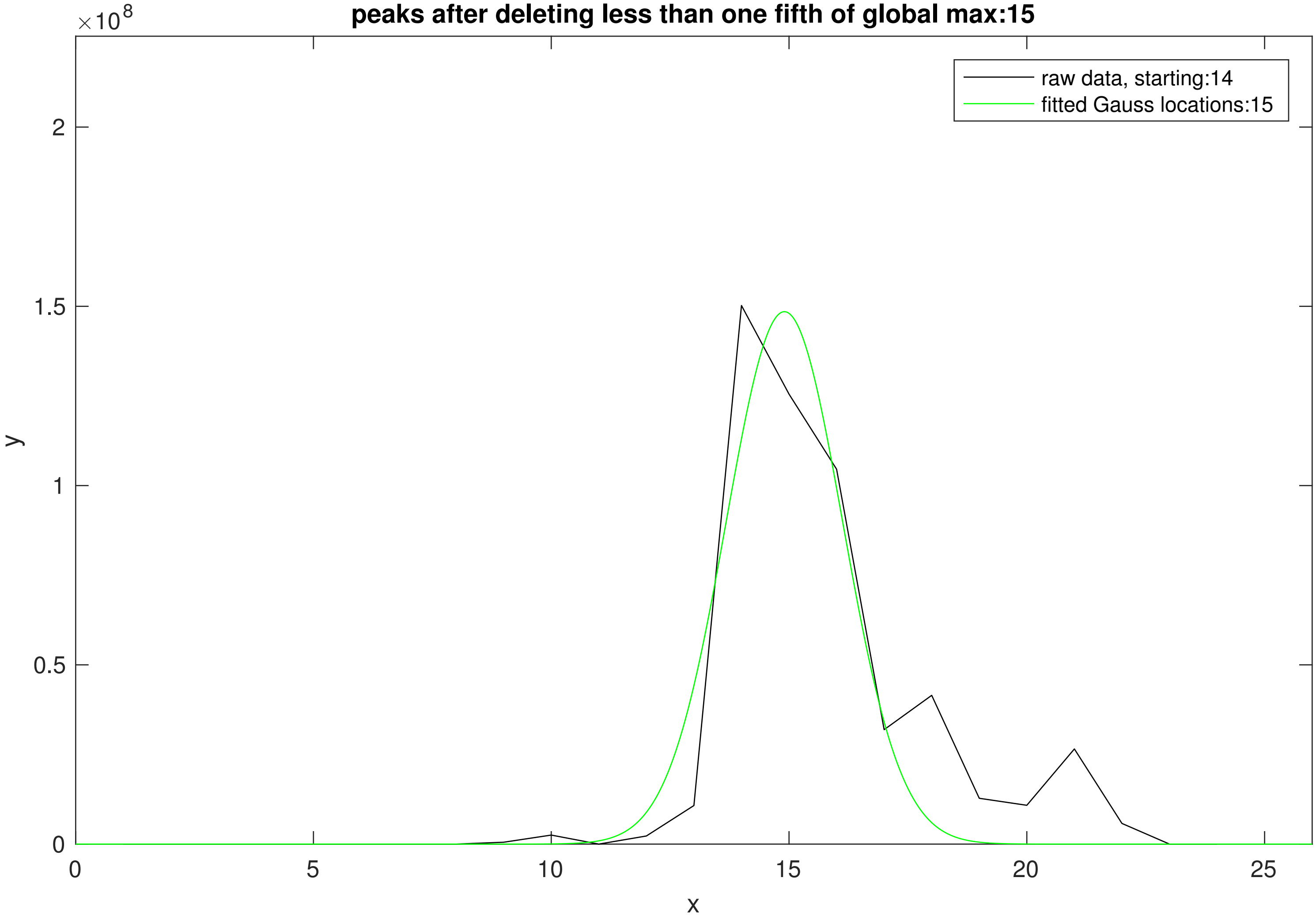

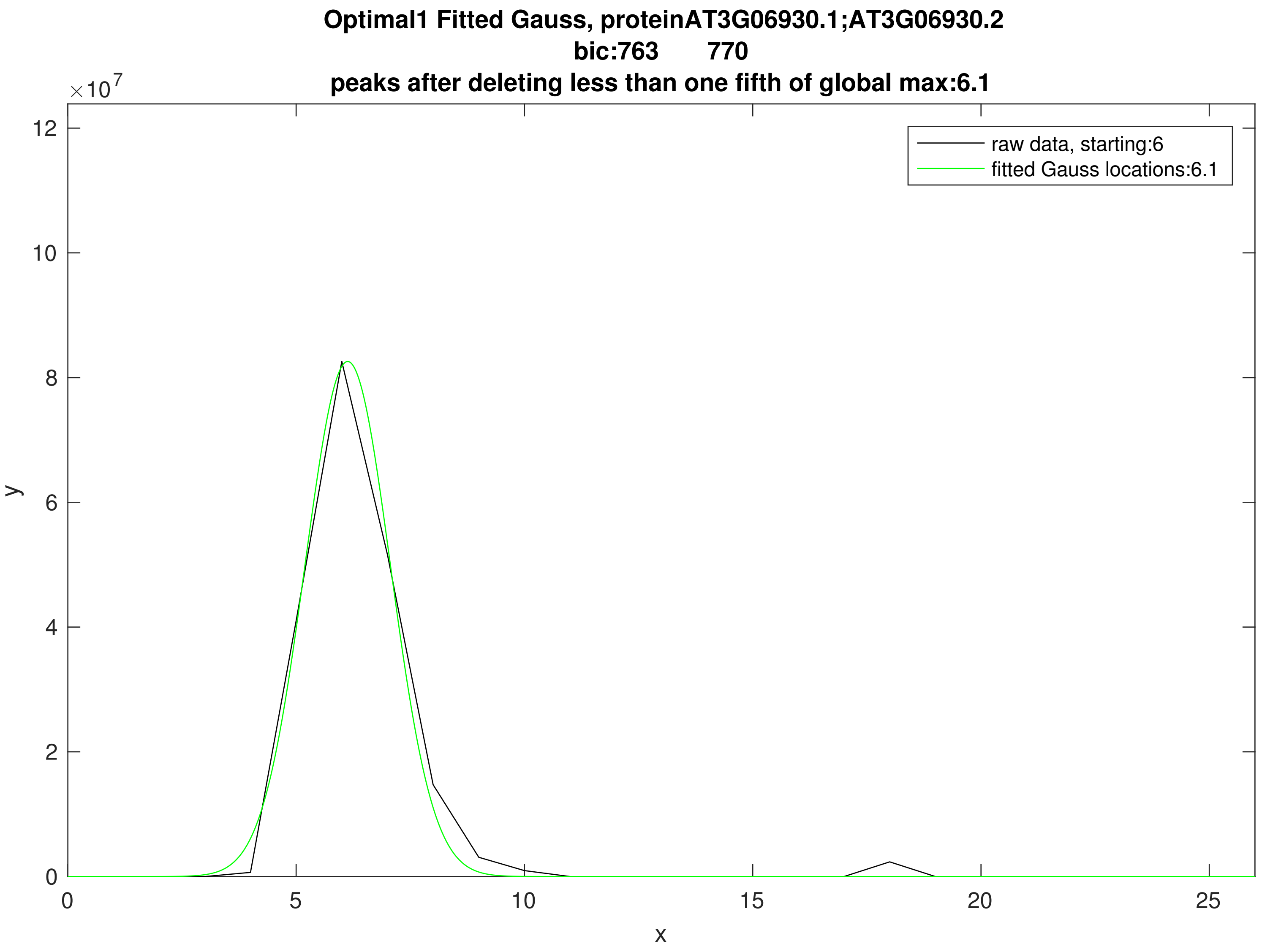
Optimal1 Fitted Gauss, proteinAT3G06930.1;AT3G06930.2
bic:763 770
peaks after deleting less than one fifth of global max:6.1
×10^7
raw data, starting:6
fitted Gauss locations:6.1
y
x
0
2
4
6
8
10
12
0
5
10
15
20
25

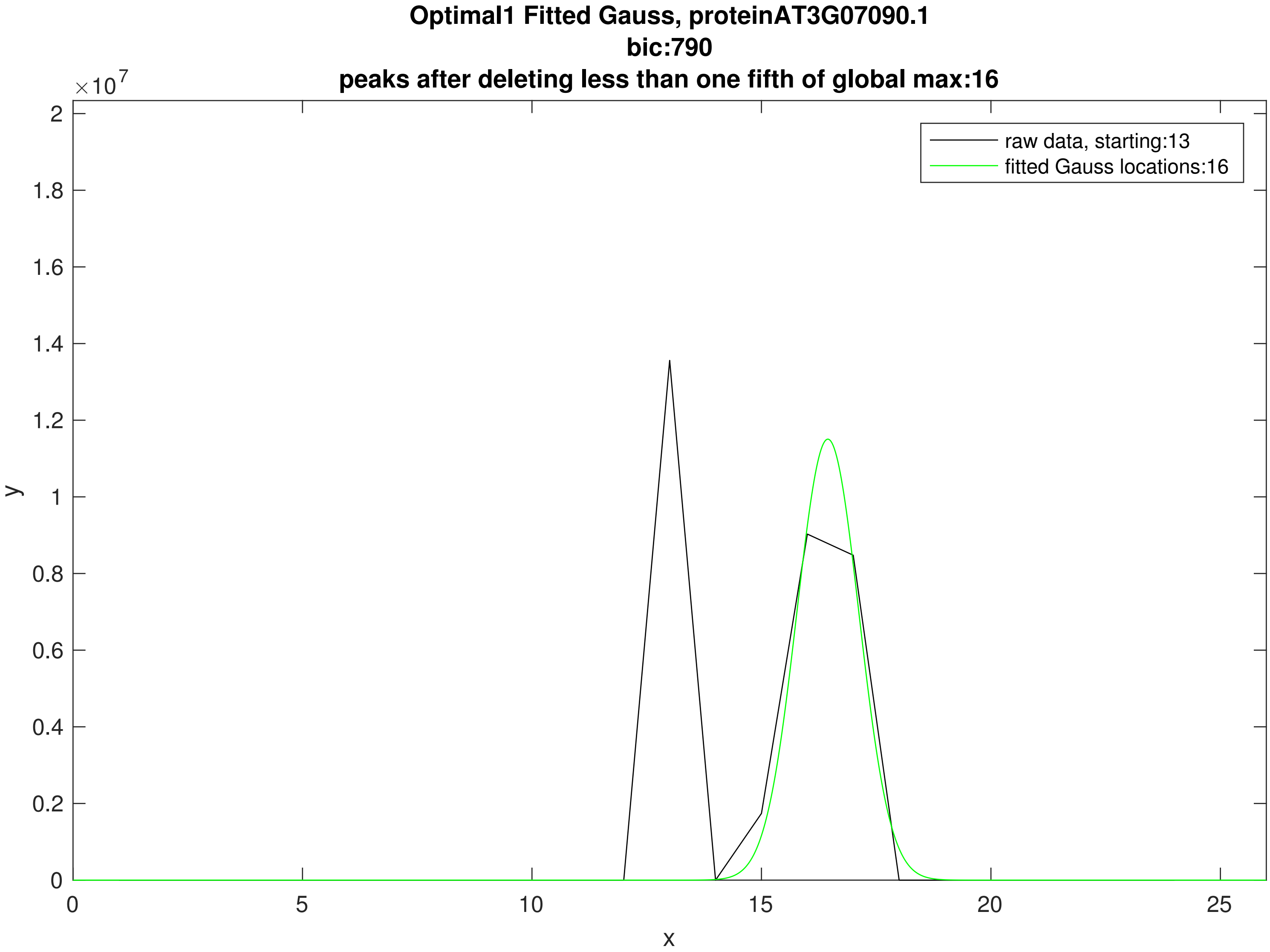
Optimal1 Fitted Gauss, proteinAT3G07090.1
bic:790
peaks after deleting less than one fifth of global max:16
×10^7
raw data, starting:13
fitted Gauss locations:16
y
x
0
0.2
0.4
0.6
0.8
1
1.2
1.4
1.6
1.8
2
0
5
10
15
20
25

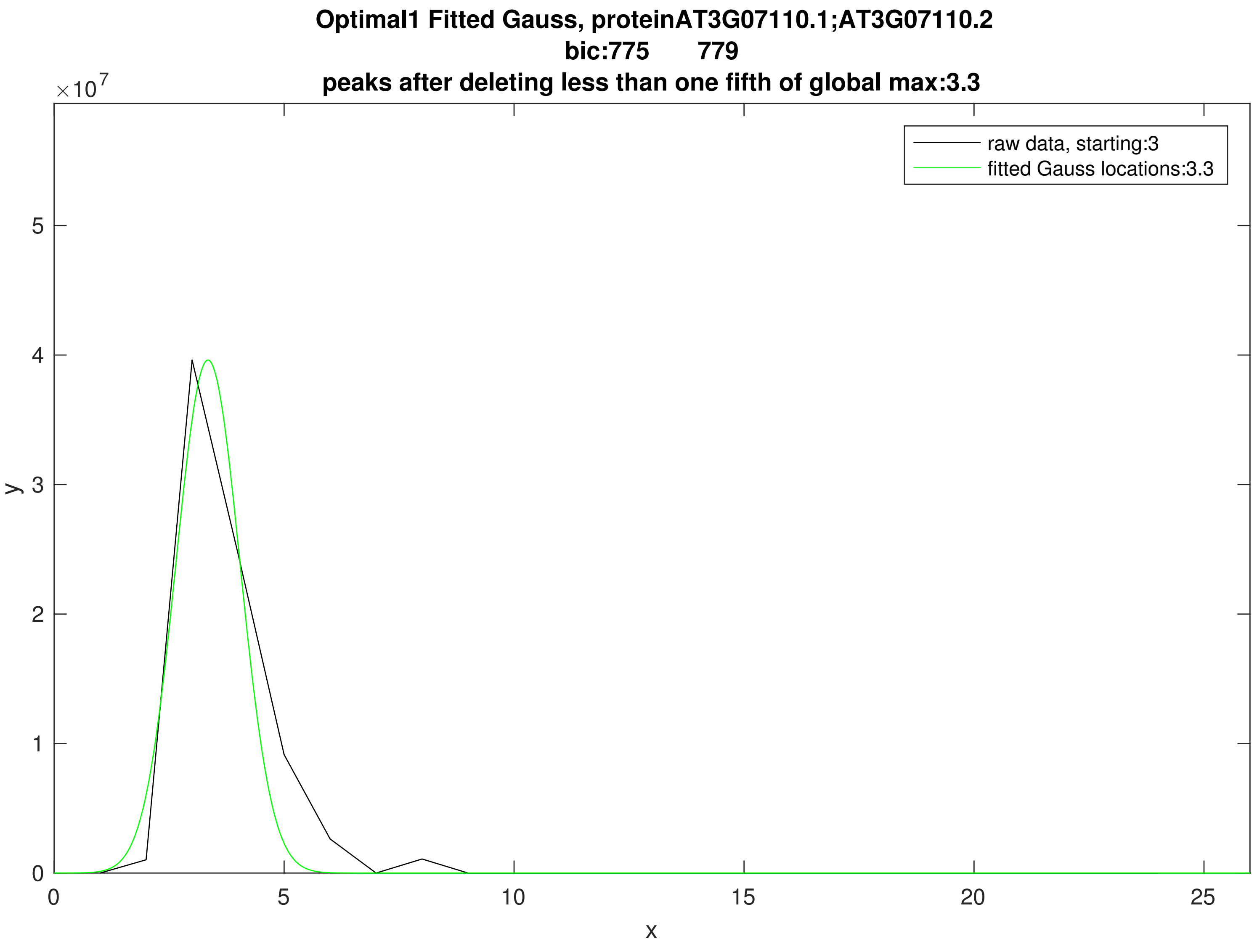
Optimal1 Fitted Gauss, proteinAT3G07110.1;AT3G07110.2
bic:775 779
peaks after deleting less than one fifth of global max:3.3
×10^7
raw data, starting:3
fitted Gauss locations:3.3
y
x
0
1
2
3
4
5
0
5
10
15
20
25

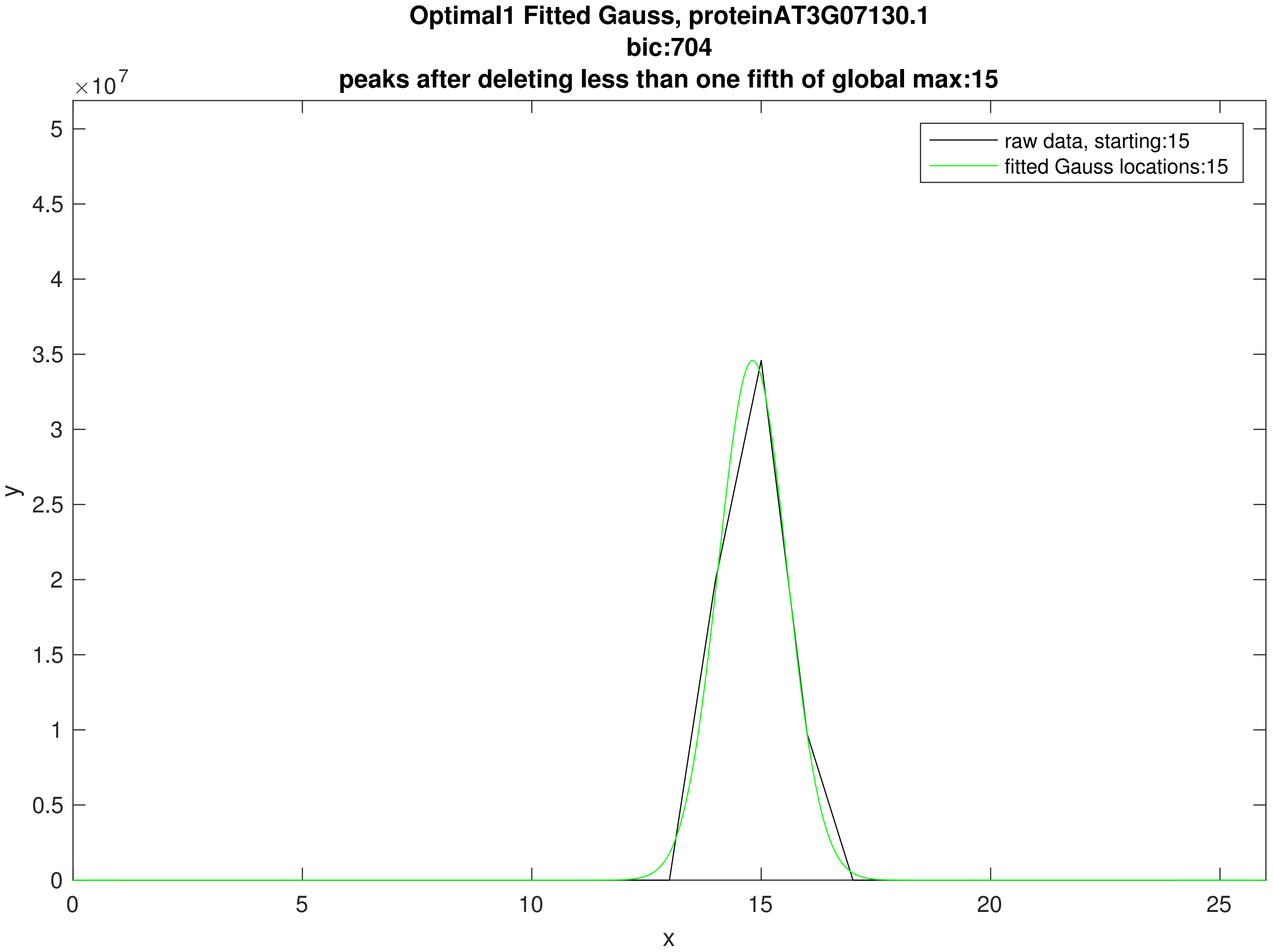

Optimal1 Fitted Gauss, proteinAT3G07130.1
bic:704
peaks after deleting less than one fifth of global max:15
×10^7
raw data, starting:15
fitted Gauss locations:15
y
x
0
0.5
1
1.5
2
2.5
3
3.5
4
4.5
5
0
5
10
15
20
25

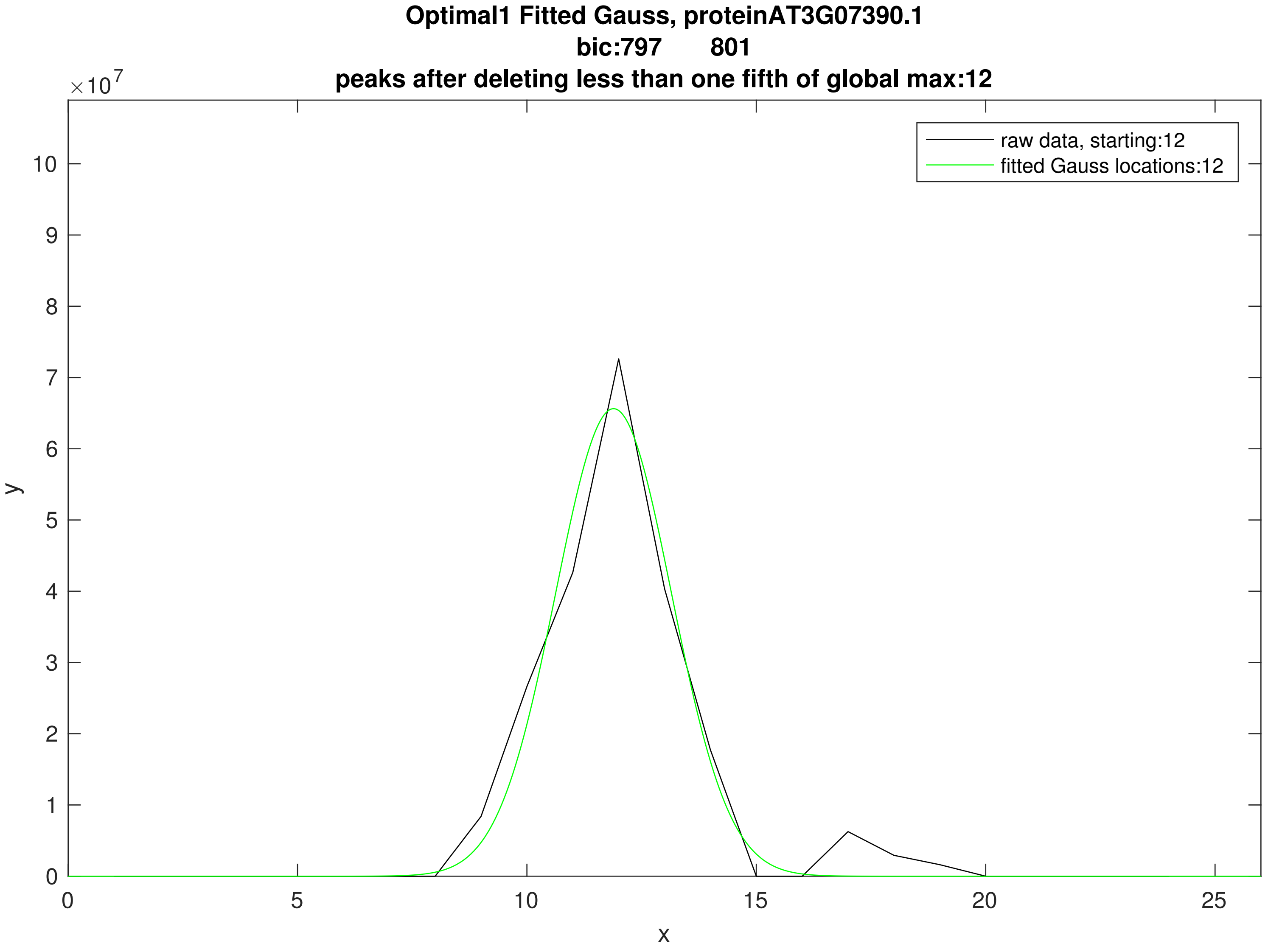
Optimal1 Fitted Gauss, proteinAT3G07390.1
bic:797 801
peaks after deleting less than one fifth of global max:12
×10^7
raw data, starting:12
fitted Gauss locations:12
y
x
0
1
2
3
4
5
6
7
8
9
10
0
5
10
15
20
25

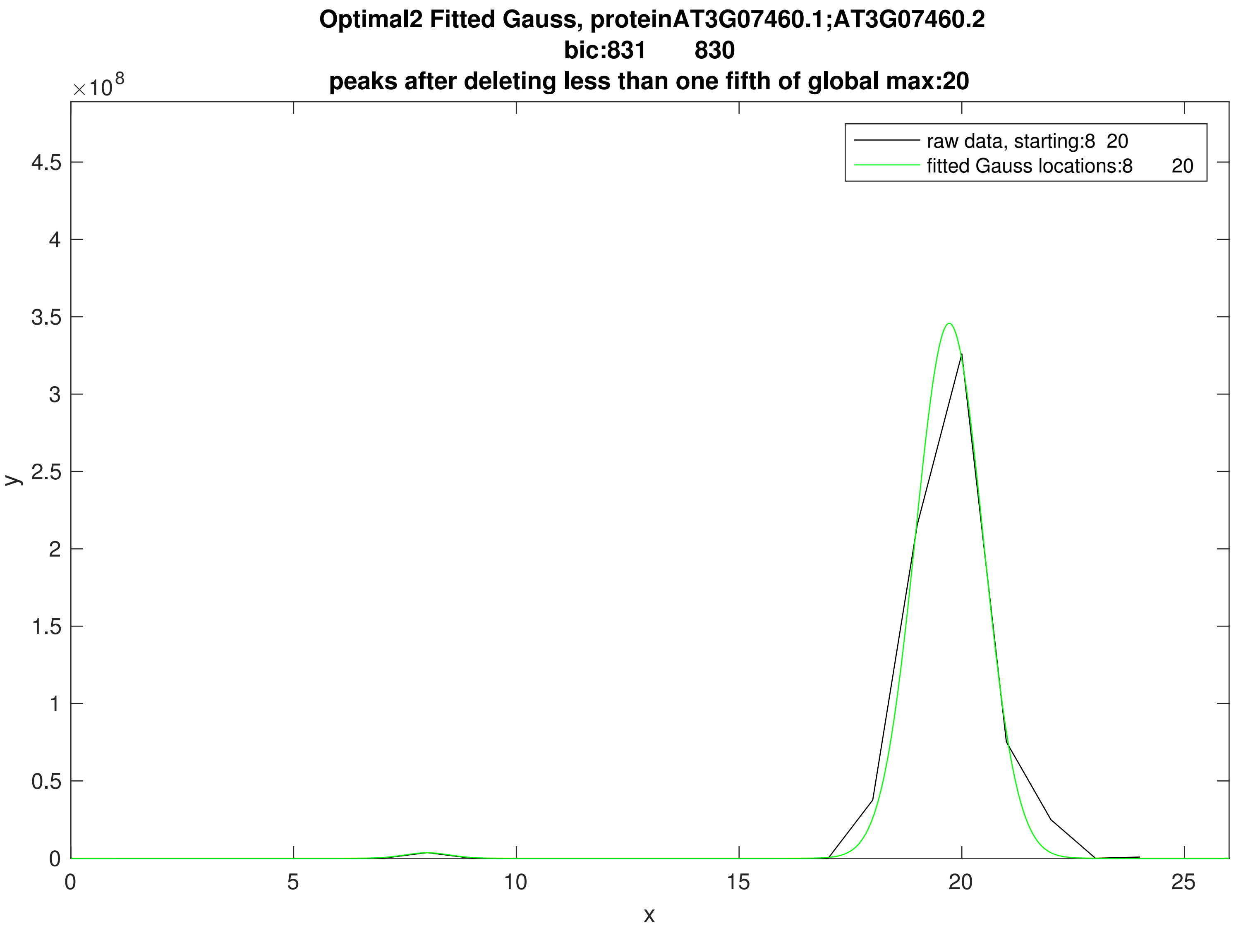
Optimal2 Fitted Gauss, proteinAT3G07460.1;AT3G07460.2
bic:831 830
peaks after deleting less than one fifth of global max:20
×10^8
raw data, starting:8 20
fitted Gauss locations:8 20
y
x
0
0.5
1
1.5
2
2.5
3
3.5
4
4.5
5
10
15
20
25

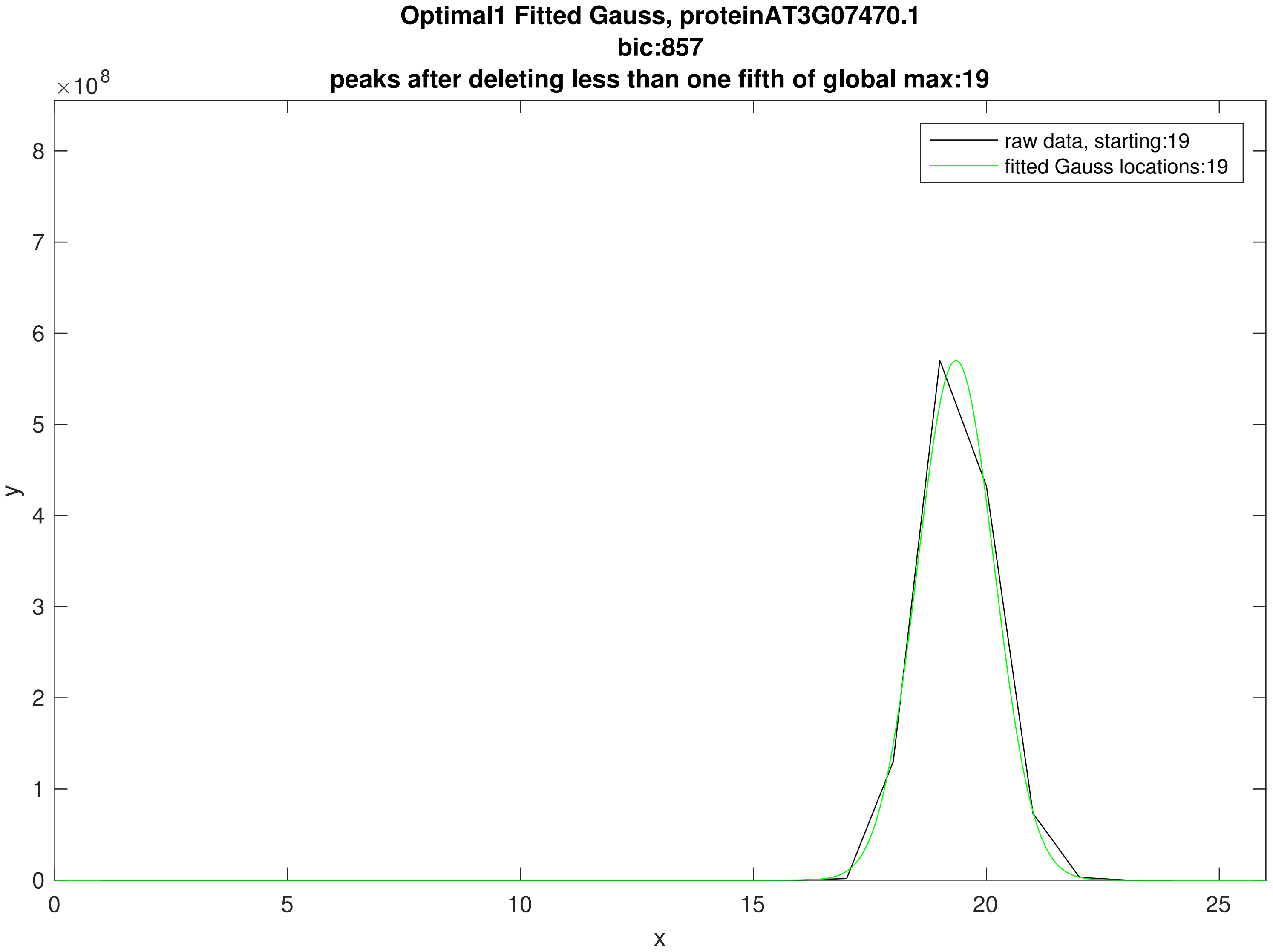
Optimal1 Fitted Gauss, proteinAT3G07470.1
bic:857
peaks after deleting less than one fifth of global max:19
×10^8
raw data, starting:19
fitted Gauss locations:19
y
x
0
1
2
3
4
5
6
7
8
0
5
10
15
20
25

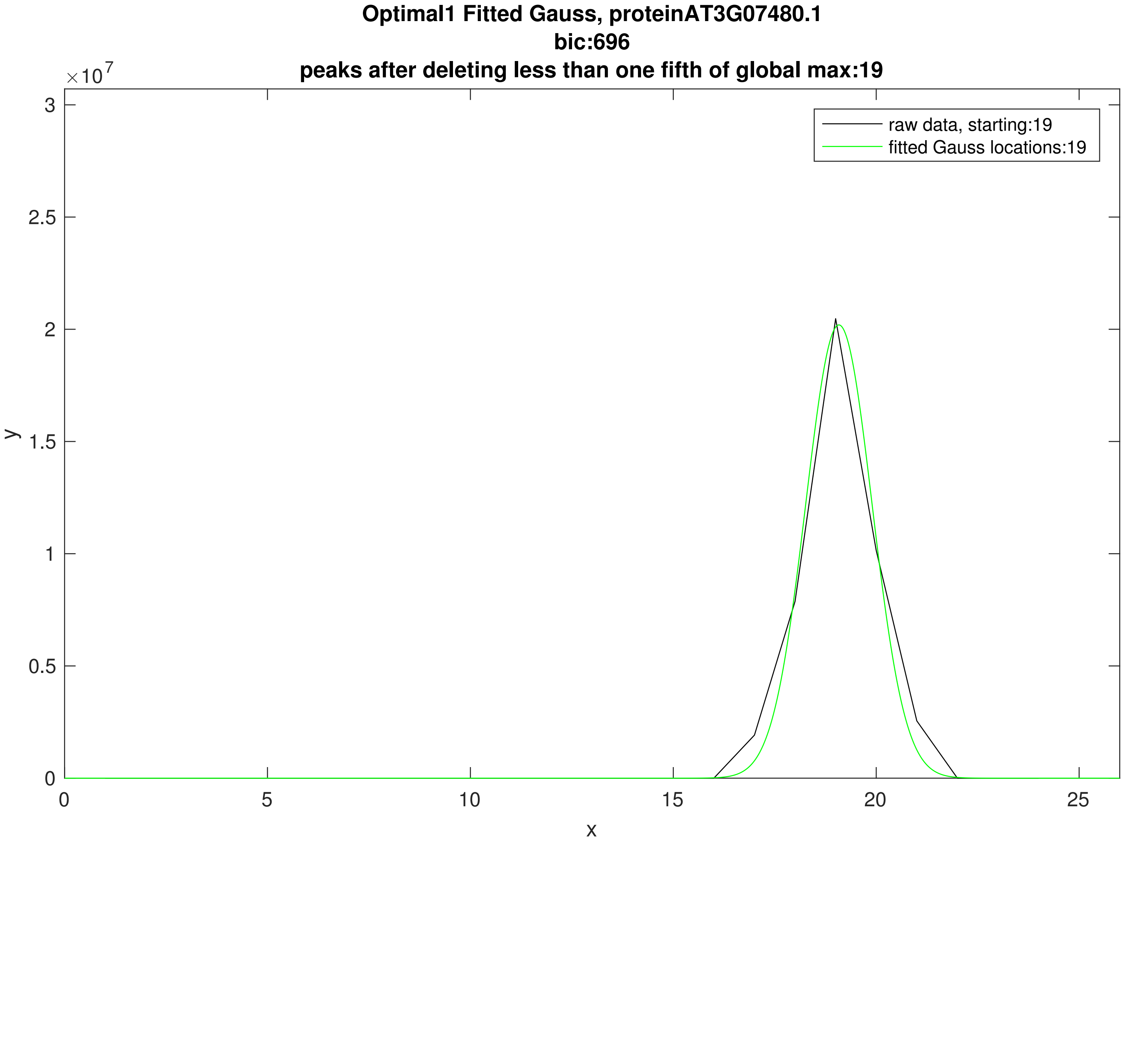
Optimal1 Fitted Gauss, proteinAT3G07480.1
bic:696
peaks after deleting less than one fifth of global max:19
×10^7
3
2.5
2
1.5
1
0.5
0
y
0
5
10
15
20
25
x
raw data, starting:19
fitted Gauss locations:19

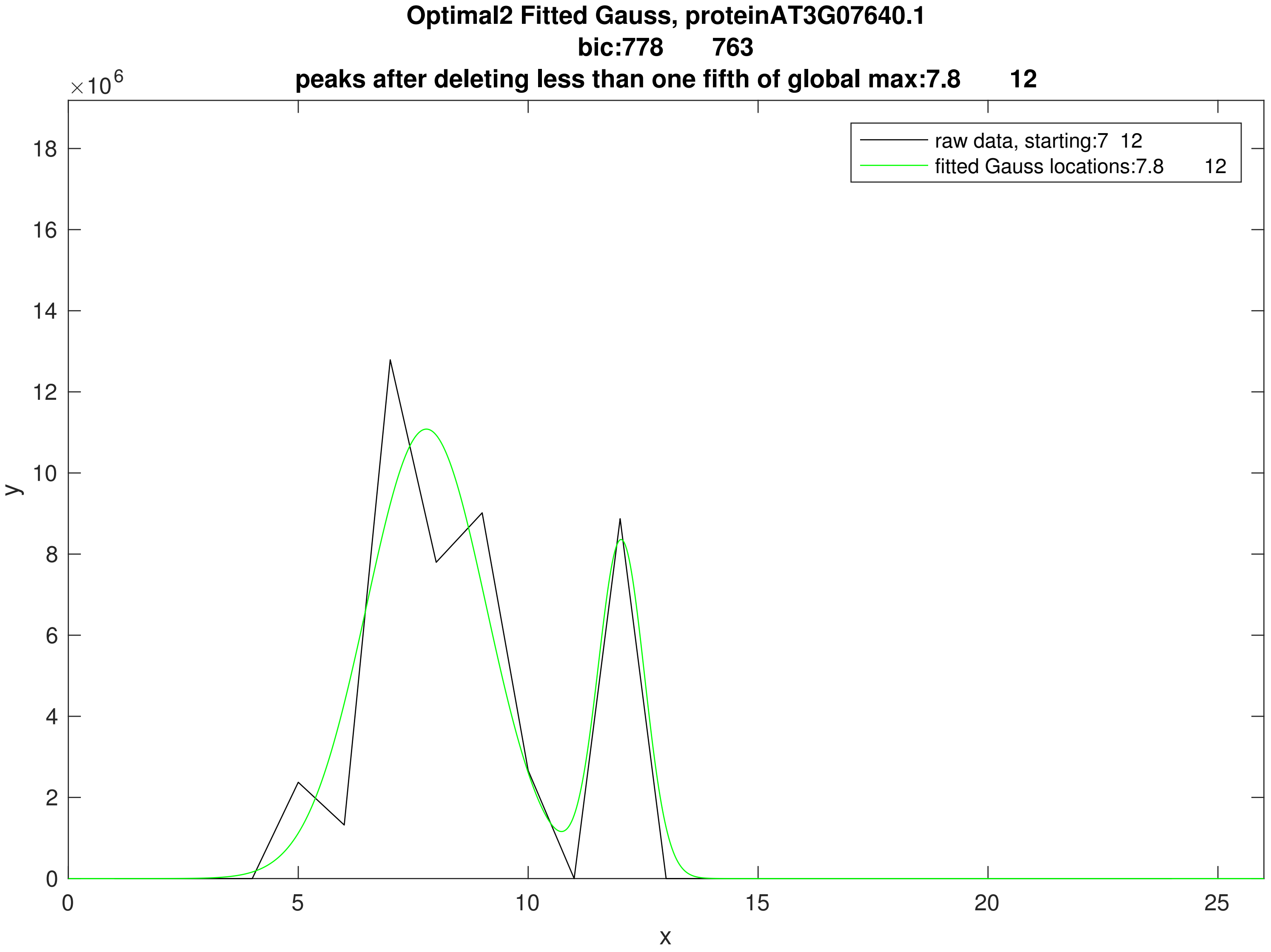

Optimal2 Fitted Gauss, proteinAT3G07640.1
bic:778 763
peaks after deleting less than one fifth of global max:7.8 12
×10^6
raw data, starting:7 12
fitted Gauss locations:7.8 12
y
x
0
2
4
6
8
10
12
14
16
18
5
15
20
25

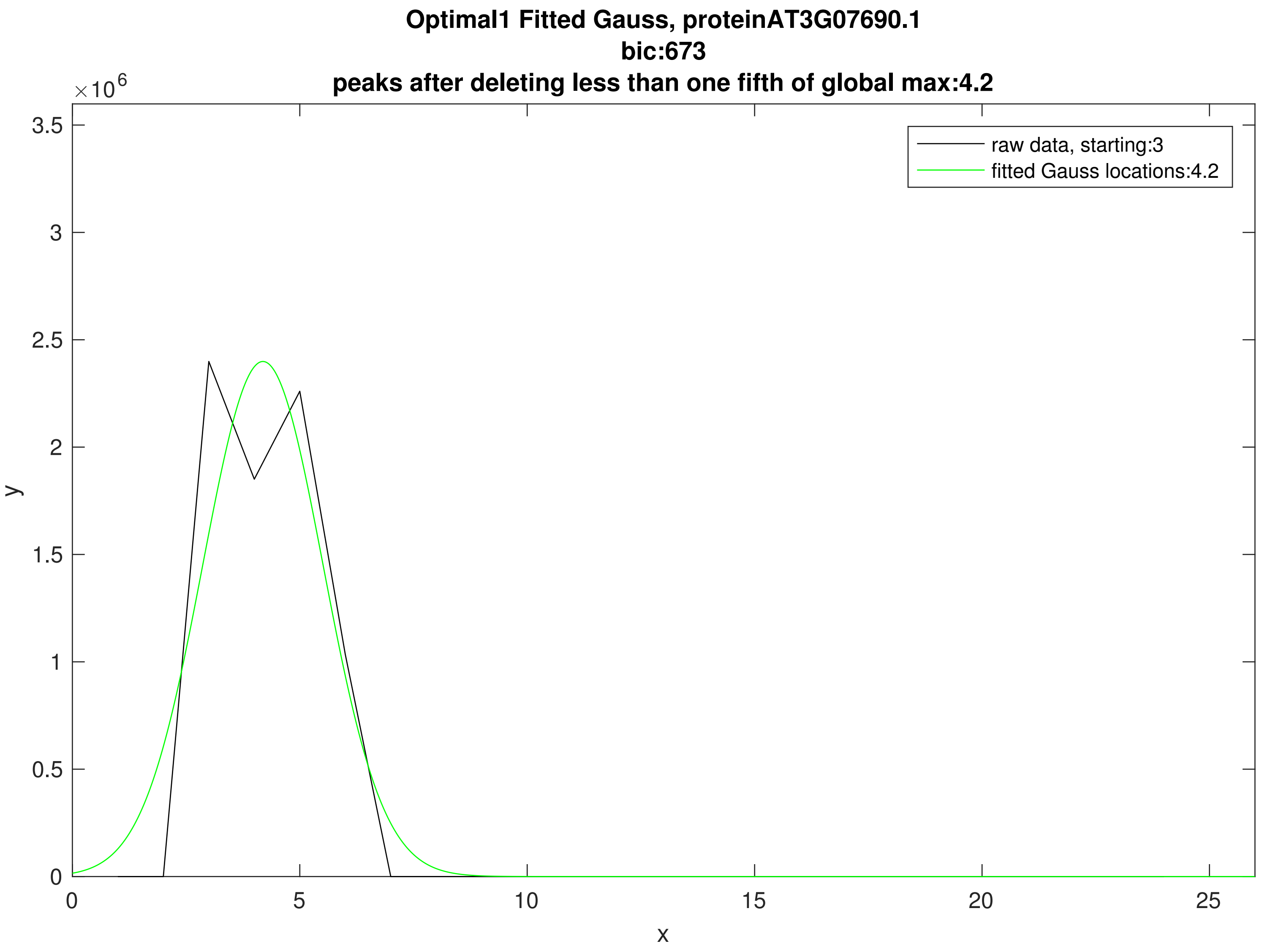

Optimal1 Fitted Gauss, proteinAT3G07690.1
bic:673
peaks after deleting less than one fifth of global max:4.2
×10^6
3.5
3
2.5
2
1.5
1
0.5
0
y
0
5
10
15
20
25
x
raw data, starting:3
fitted Gauss locations:4.2

**Optimal1 Fitted Gauss, proteinAT3G07720.1**
**bic:866**
**peaks after deleting less than one fifth of global max:17**

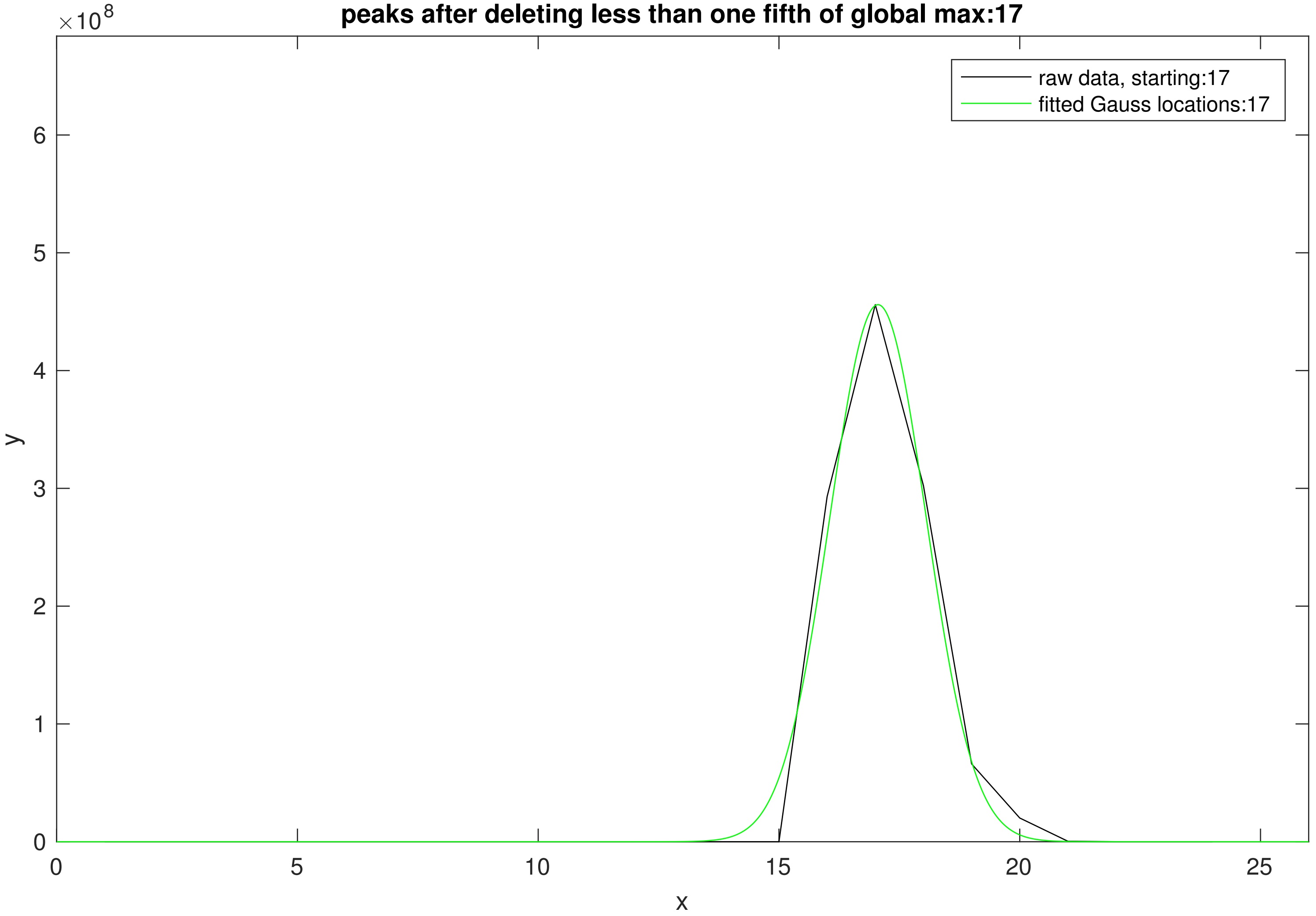

**Optimal1 Fitted Gauss, proteinAT3G07770.1**
**bic:777 836**
**peaks after deleting less than one fifth of global max:11**

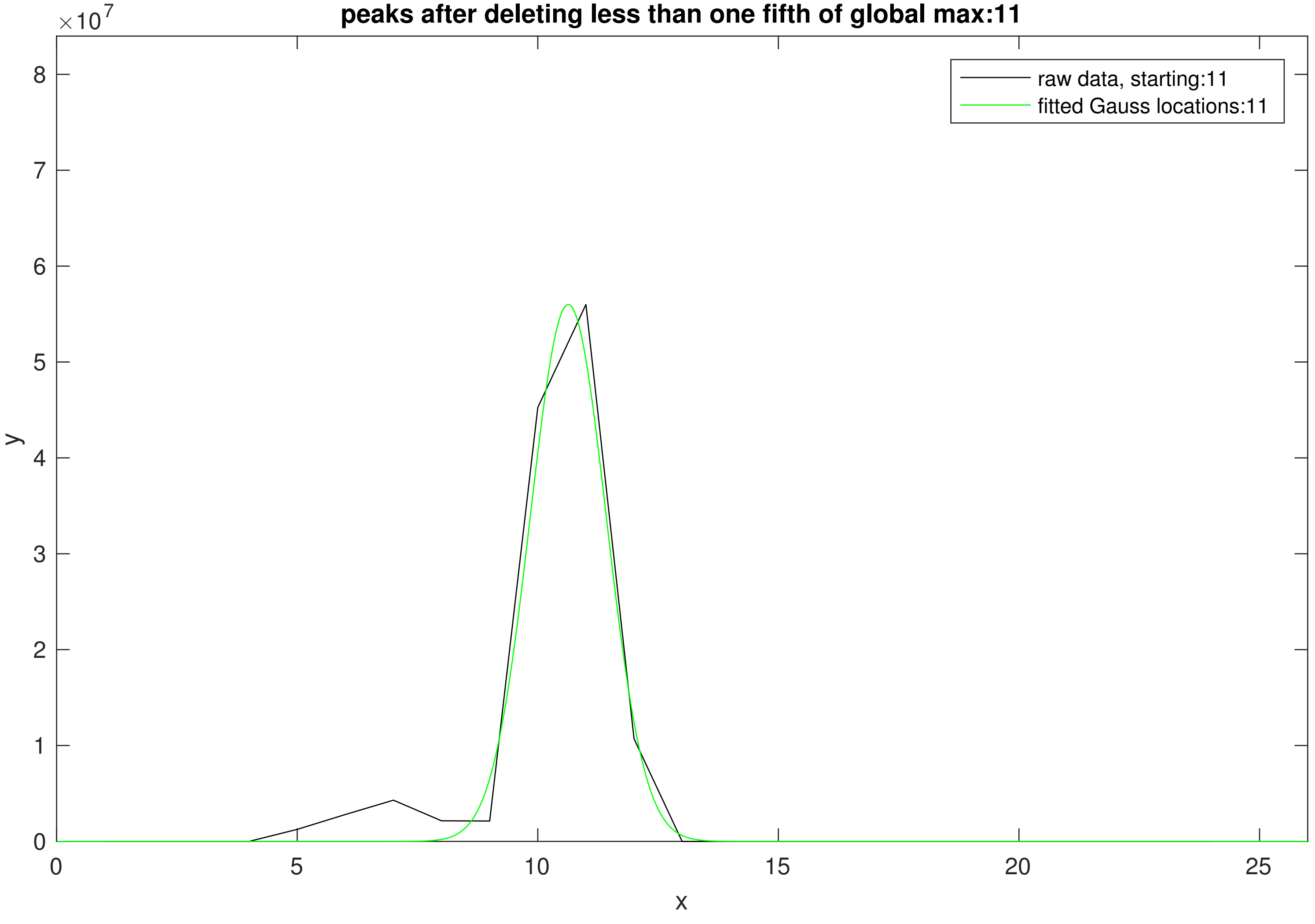

#### Optimal1 Fitted Gauss, proteinAT3G07810.1;AT3G07810.2
#### bic:736
#### peaks after deleting less than one fifth of global max:17

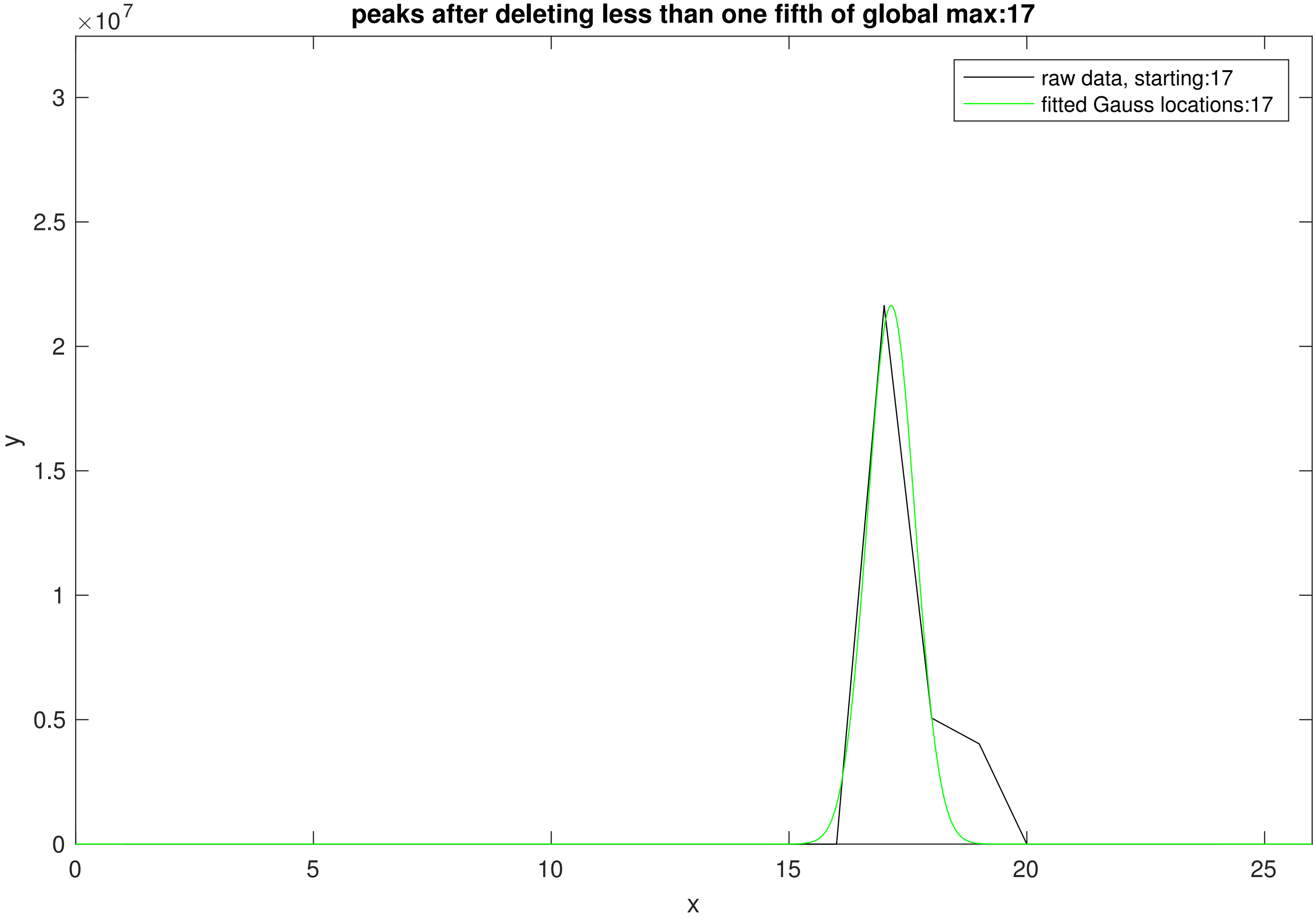

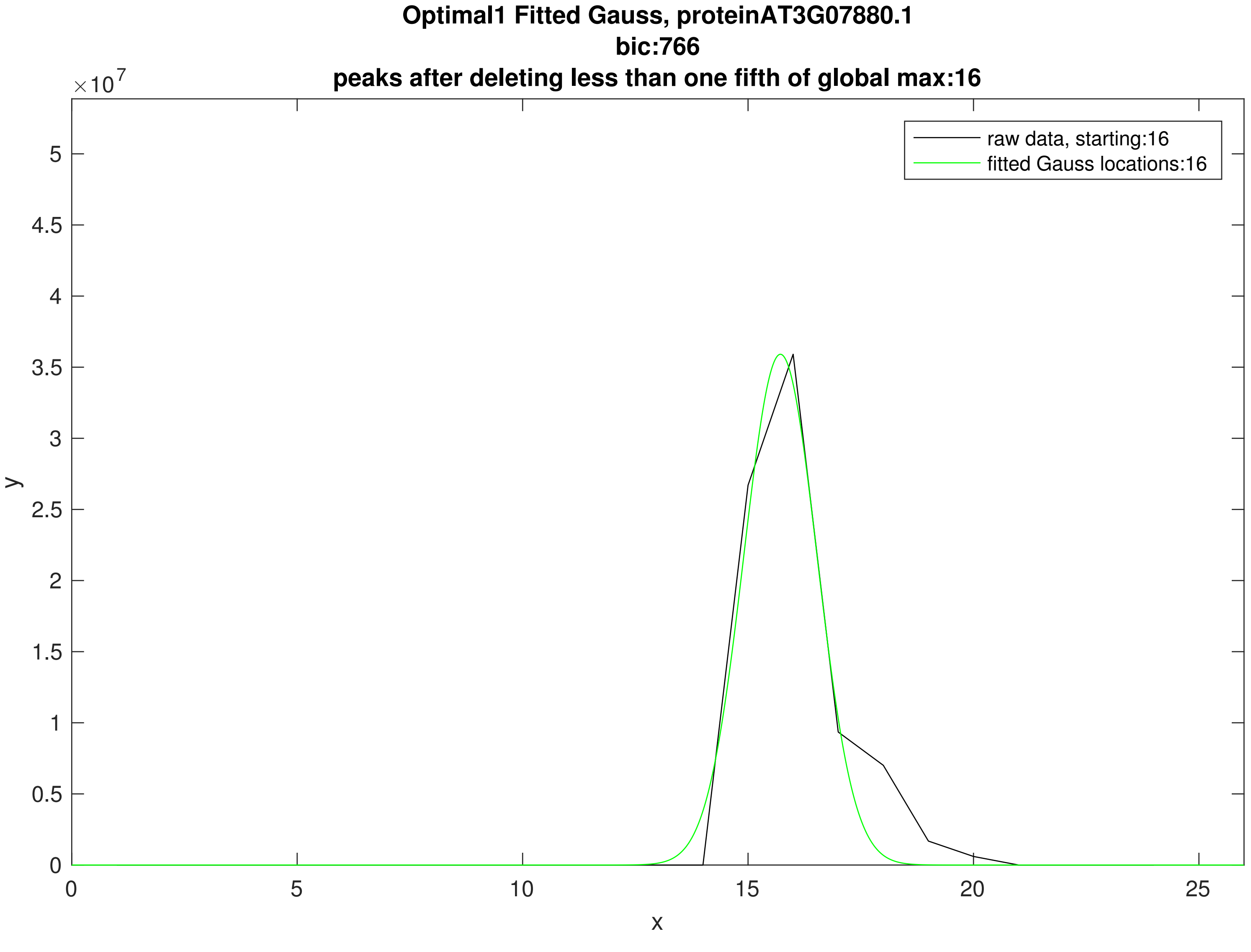
Optimal1 Fitted Gauss, proteinAT3G07880.1
bic:766
peaks after deleting less than one fifth of global max:16
×10^7
raw data, starting:16
fitted Gauss locations:16
y
x
0
0.5
1
1.5
2
2.5
3
3.5
4
4.5
5
0
5
10
15
20
25

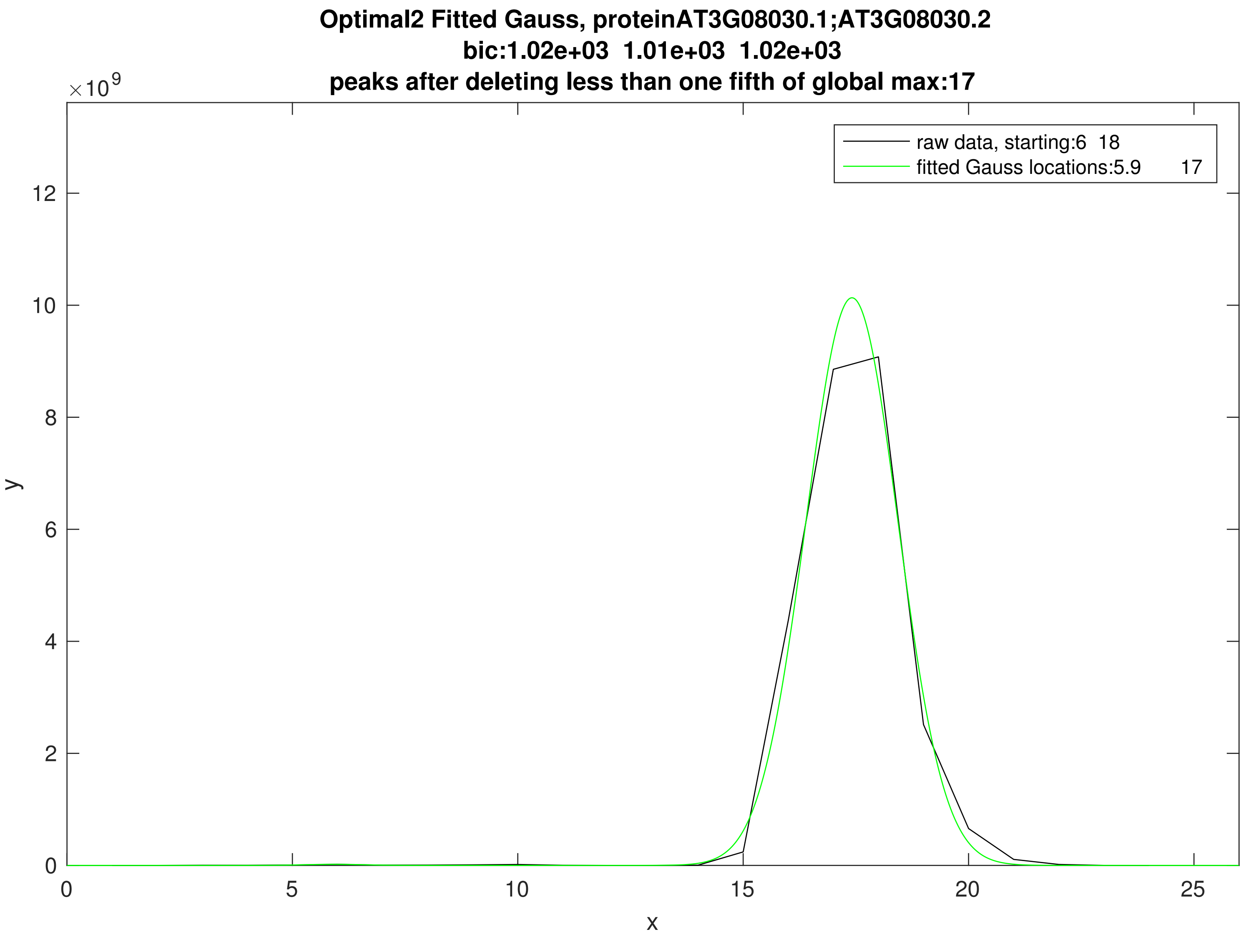
Optimal2 Fitted Gauss, proteinAT3G08030.1;AT3G08030.2
bic:1.02e+03 1.01e+03 1.02e+03
peaks after deleting less than one fifth of global max:17
×10^9
raw data, starting:6 18
fitted Gauss locations:5.9 17
12
10
8
6
4
2
0
y
0
5
10
15
20
25
x

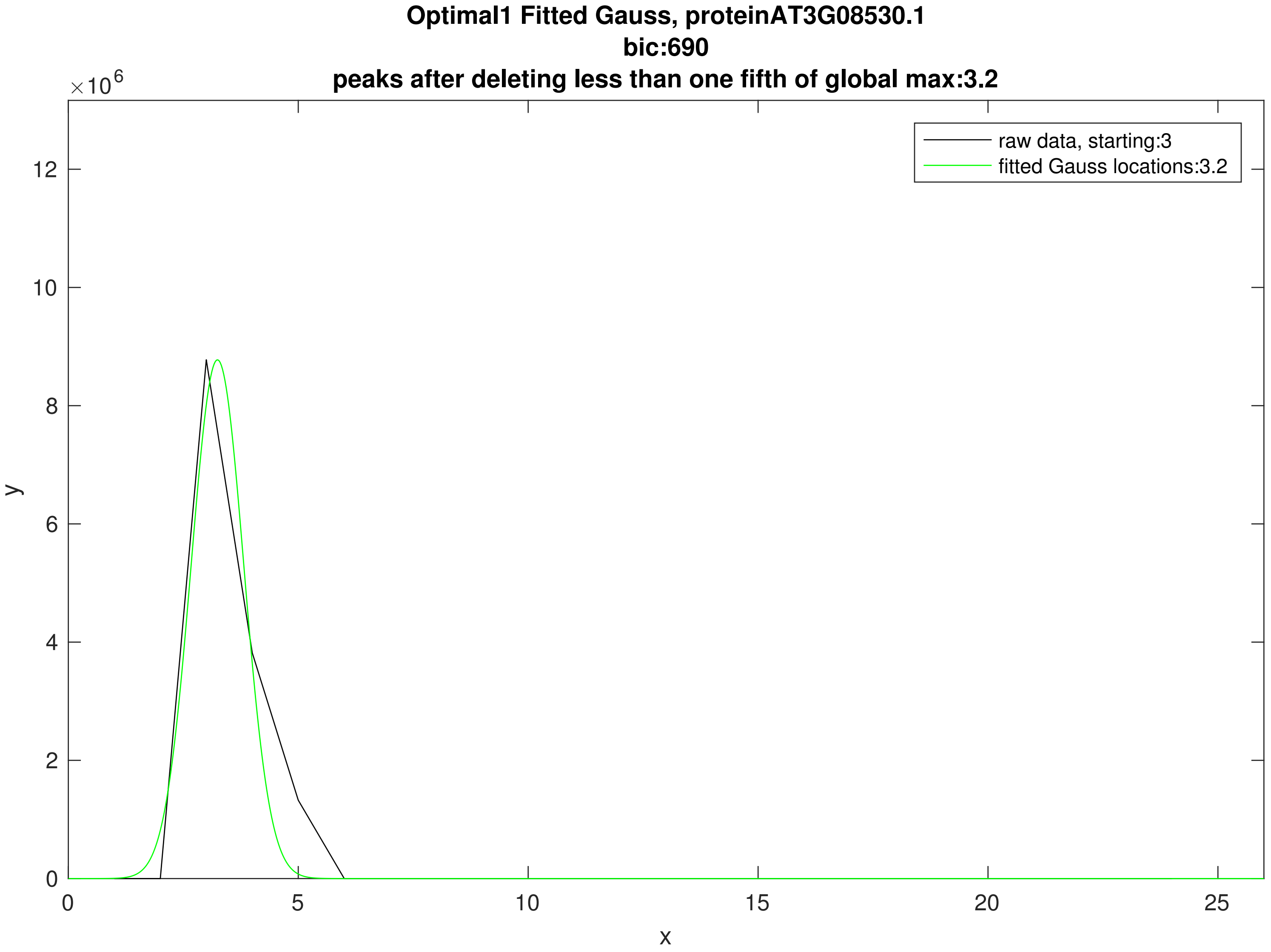

Optimal1 Fitted Gauss, proteinAT3G08530.1
bic:690
peaks after deleting less than one fifth of global max:3.2
×10^6
raw data, starting:3
fitted Gauss locations:3.2
y
x
0
2
4
6
8
10
12
0
5
10
15
20
25

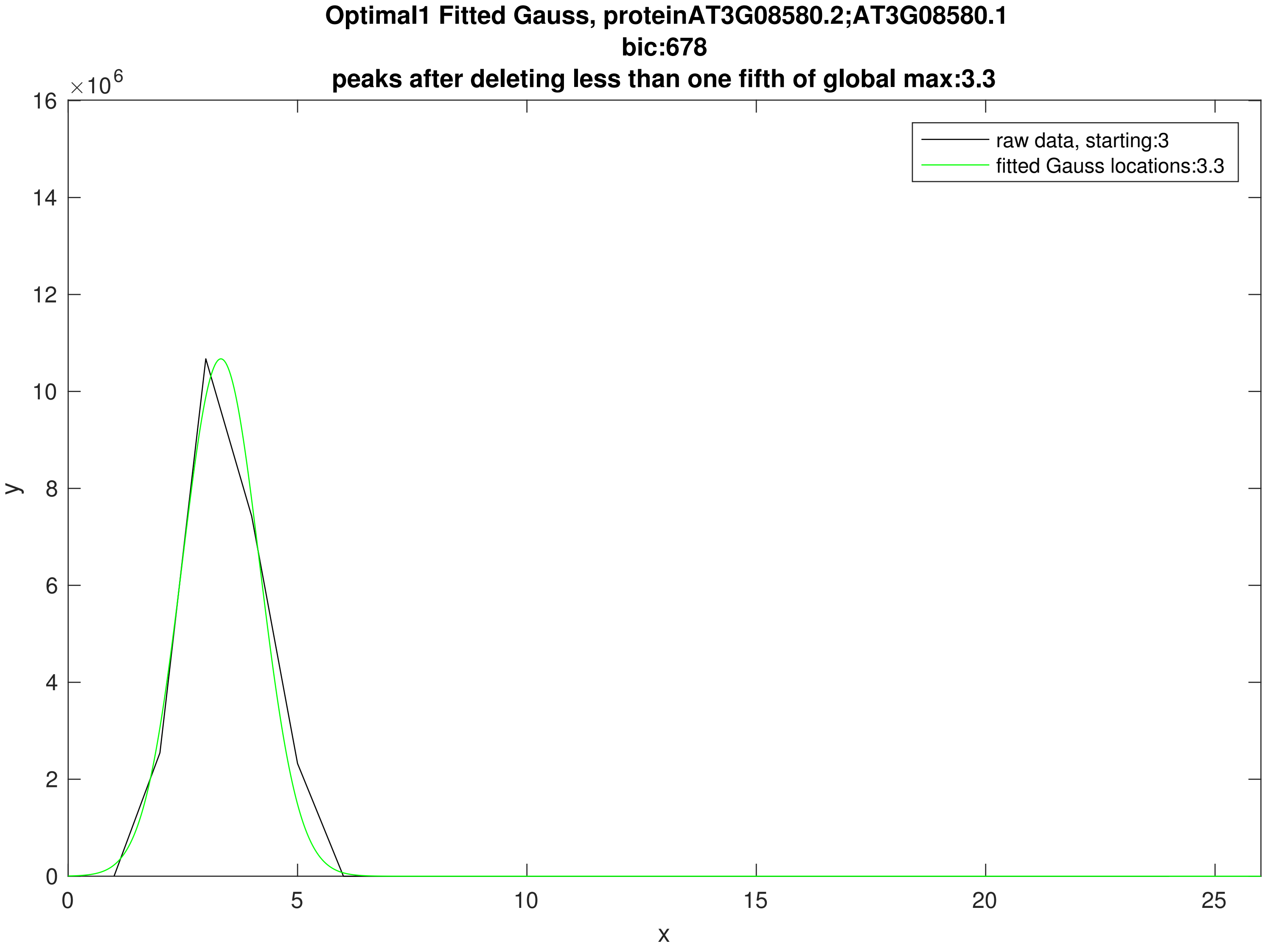
Optimal1 Fitted Gauss, proteinAT3G08580.2;AT3G08580.1
bic:678
peaks after deleting less than one fifth of global max:3.3
×10^6
16
14
12
10
8
6
4
2
0
y
0
5
10
15
20
25
x
raw data, starting:3
fitted Gauss locations:3.3

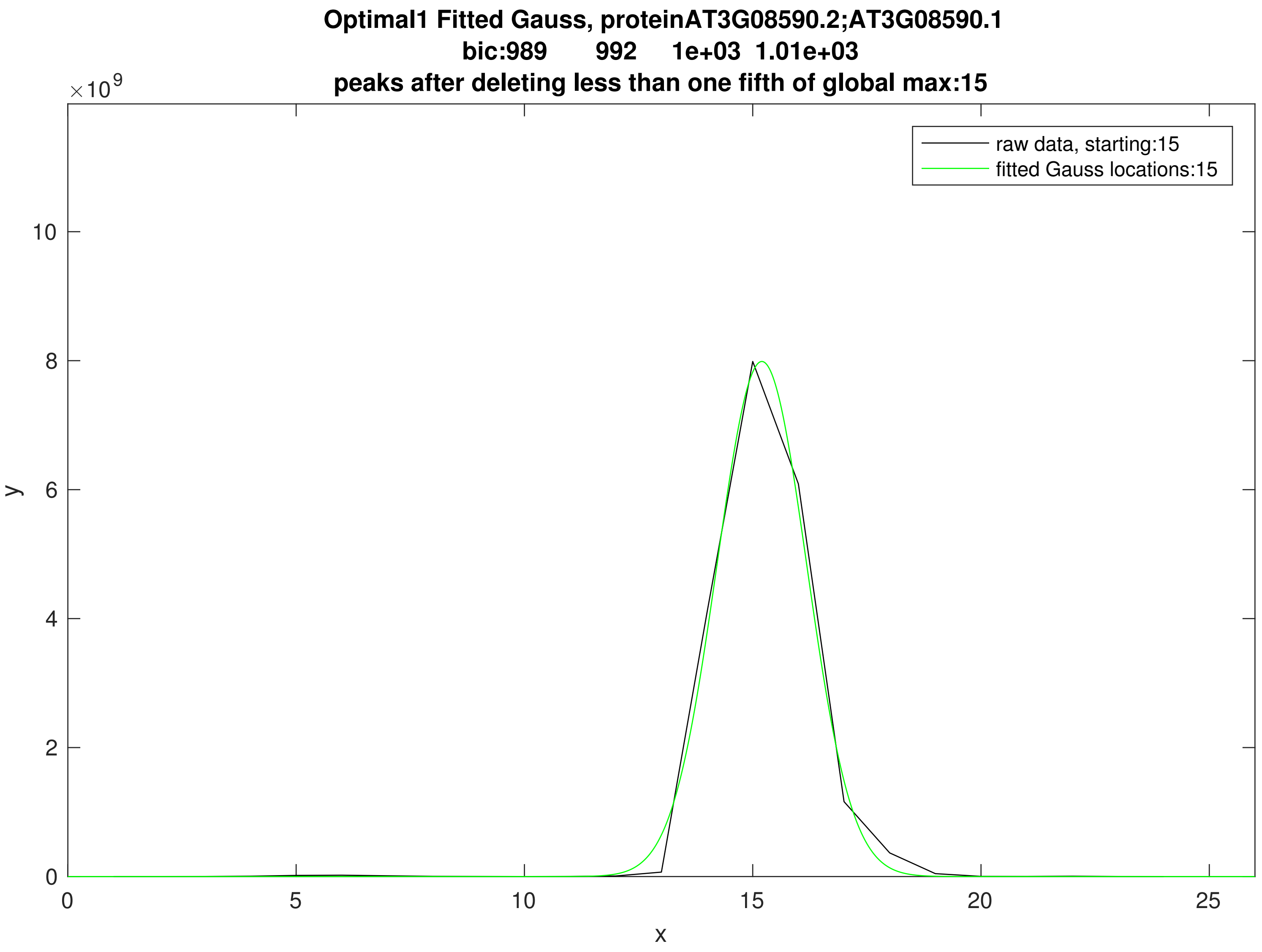
Optimal1 Fitted Gauss, proteinAT3G08590.2;AT3G08590.1
bic:989 992 1e+03 1.01e+03
peaks after deleting less than one fifth of global max:15
×10^9
raw data, starting:15
fitted Gauss locations:15
10
8
6
4
2
0
y
0
5
10
15
20
25
x

**Optimal1 Fitted Gauss, proteinAT3G08740.1**
**bic:728**
**peaks after deleting less than one fifth of global max:19**

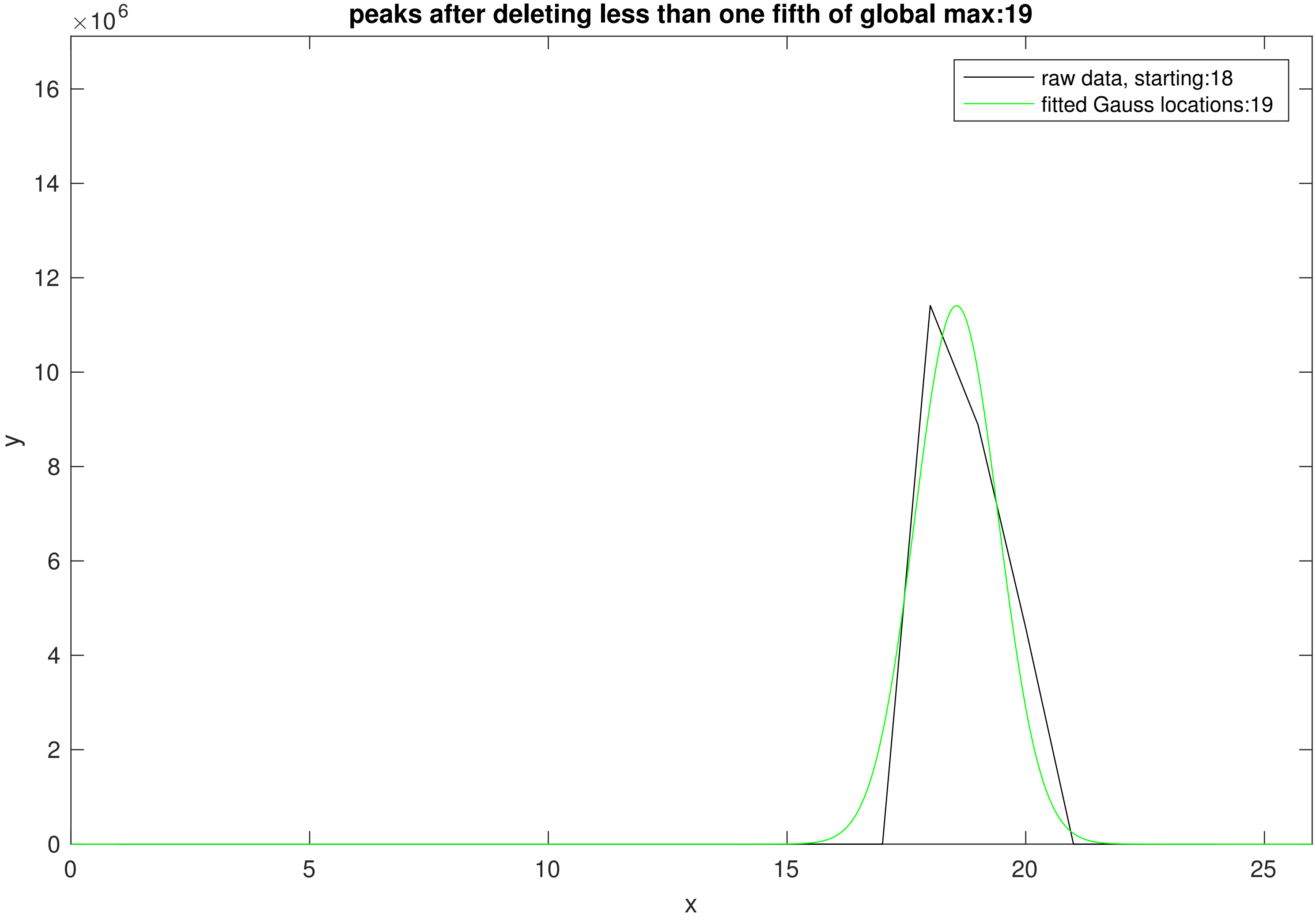

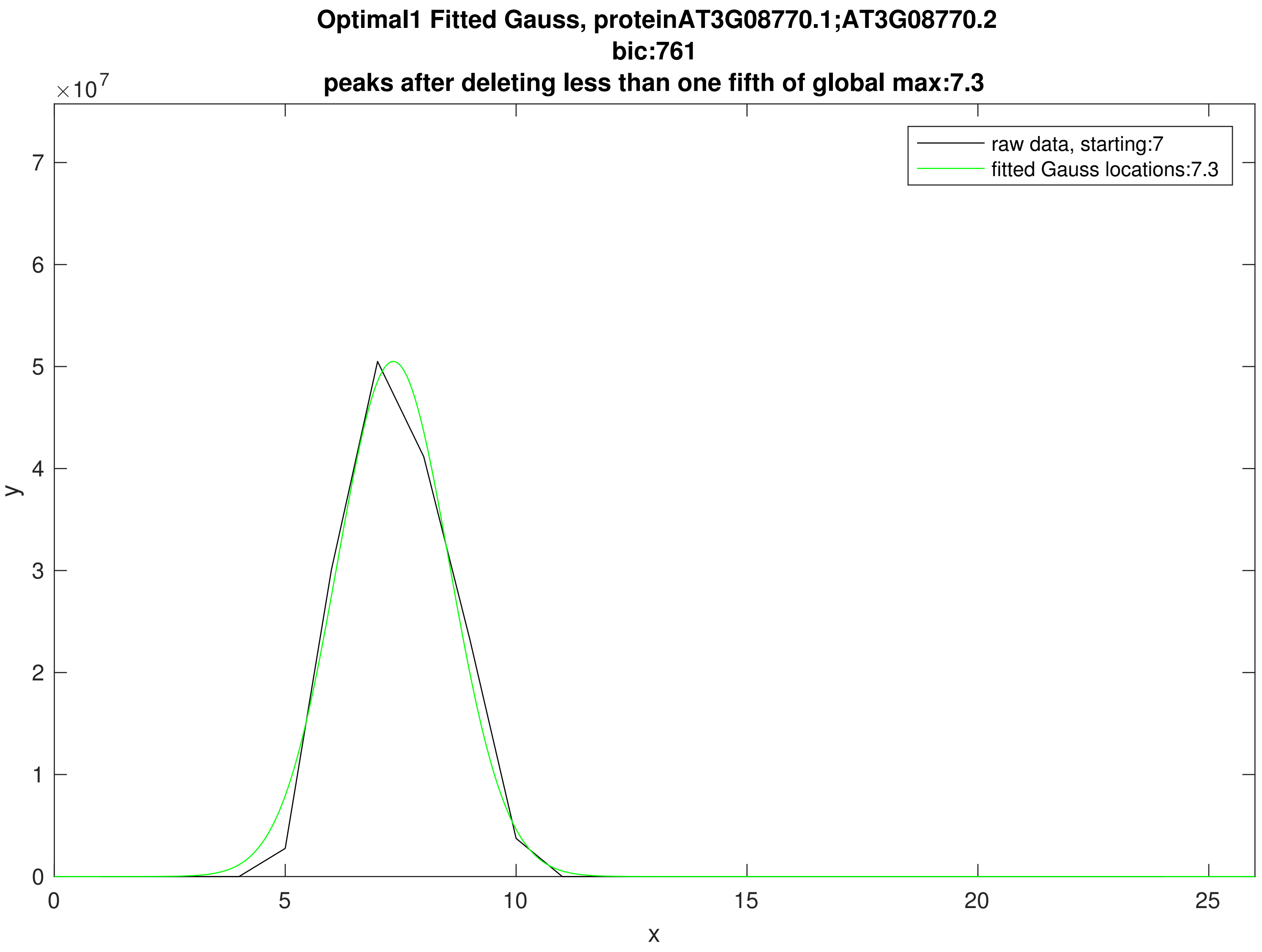
Optimal1 Fitted Gauss, proteinAT3G08770.1;AT3G08770.2
bic:761
peaks after deleting less than one fifth of global max:7.3
×10^7
raw data, starting:7
fitted Gauss locations:7.3
y
x
0
1
2
3
4
5
6
7
0
5
10
15
20
25

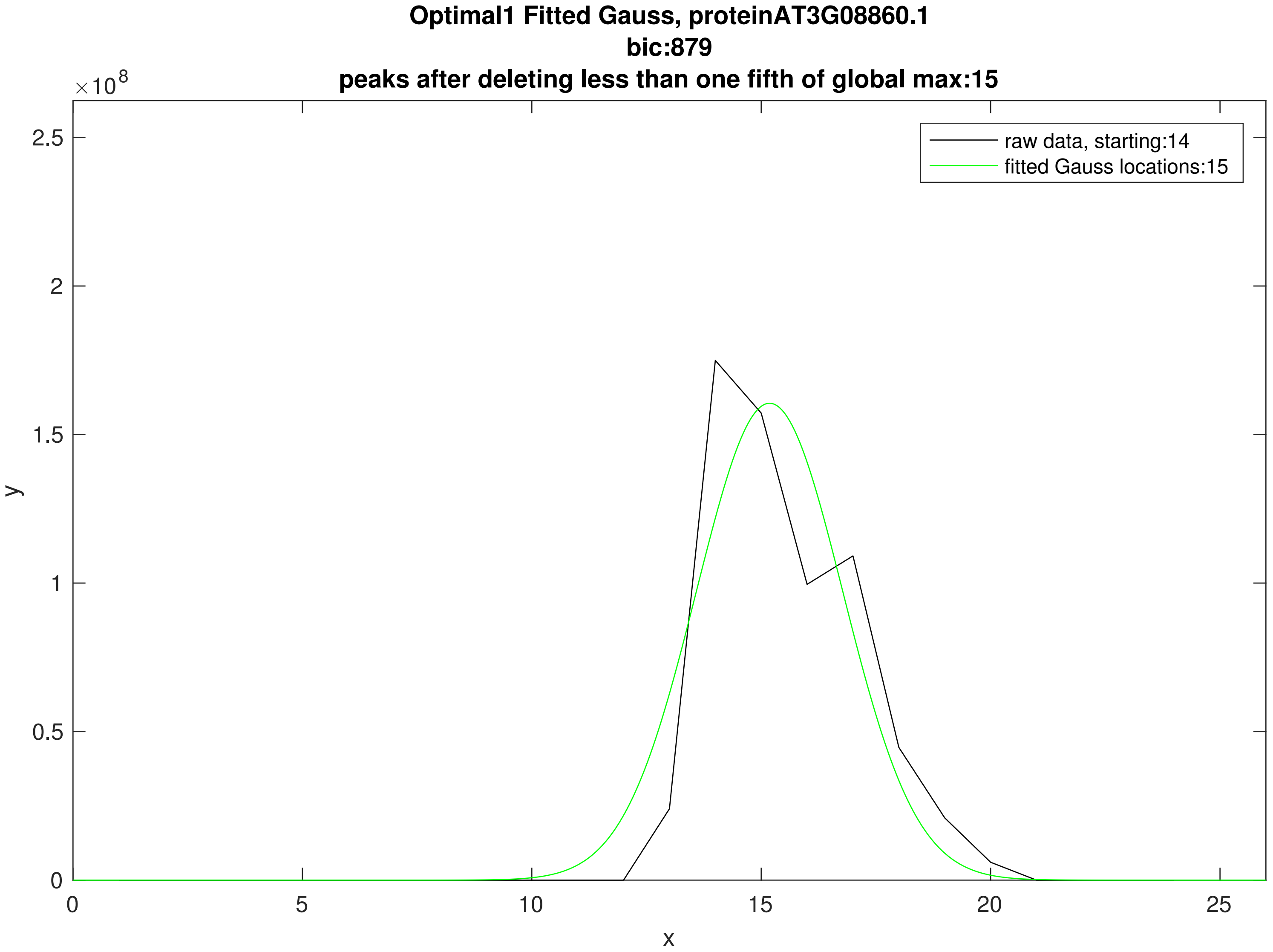
Optimal1 Fitted Gauss, proteinAT3G08860.1
bic:879
peaks after deleting less than one fifth of global max:15
×10^8
2.5
2
1.5
1
0.5
0
y
0
5
10
15
20
25
x
raw data, starting:14
fitted Gauss locations:15

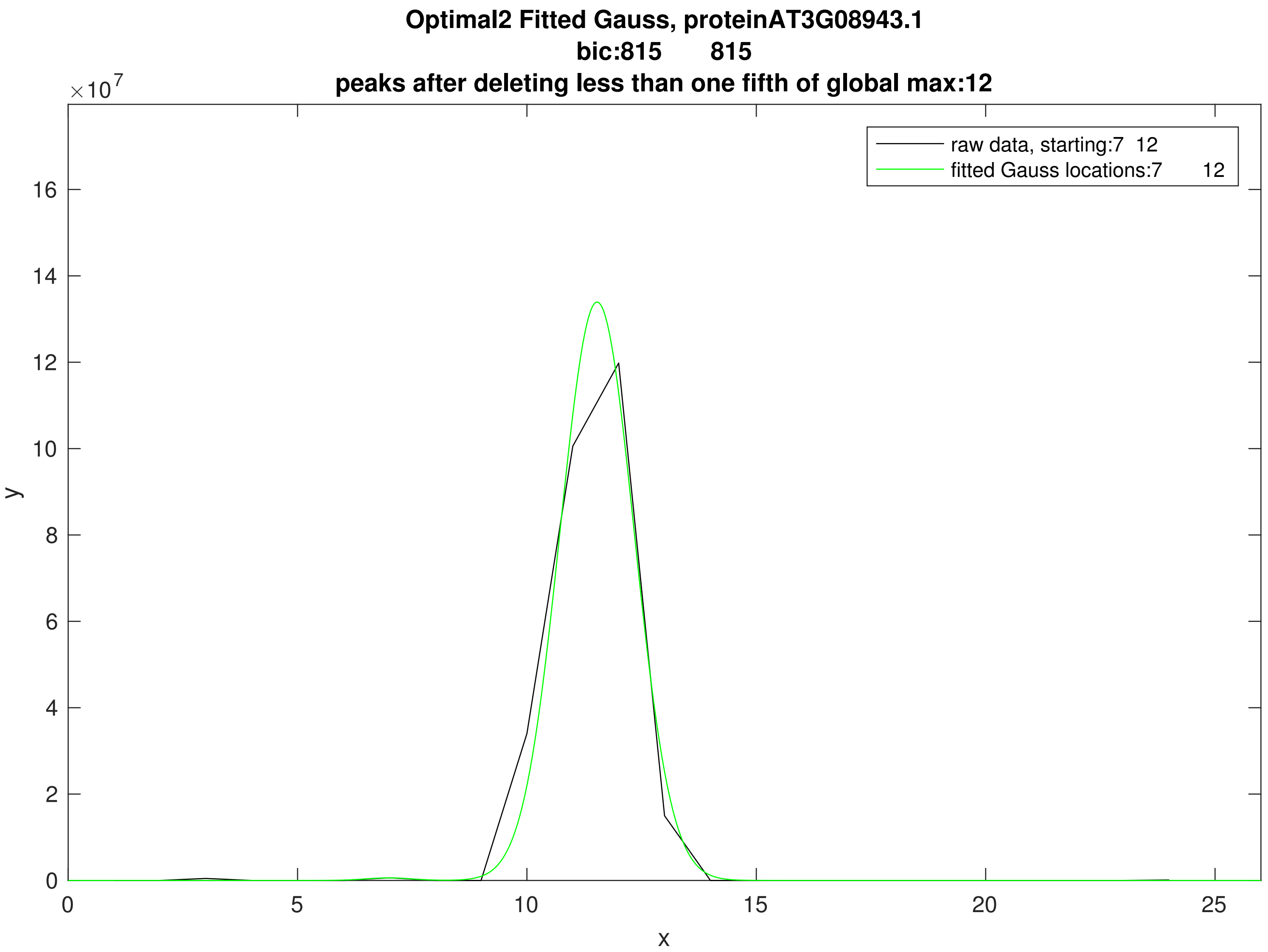
Optimal2 Fitted Gauss, proteinAT3G08943.1
bic:815 815
peaks after deleting less than one fifth of global max:12
×10^7
raw data, starting:7 12
fitted Gauss locations:7 12
0
2
4
6
8
10
12
14
16
y
0
5
10
15
20
25
x

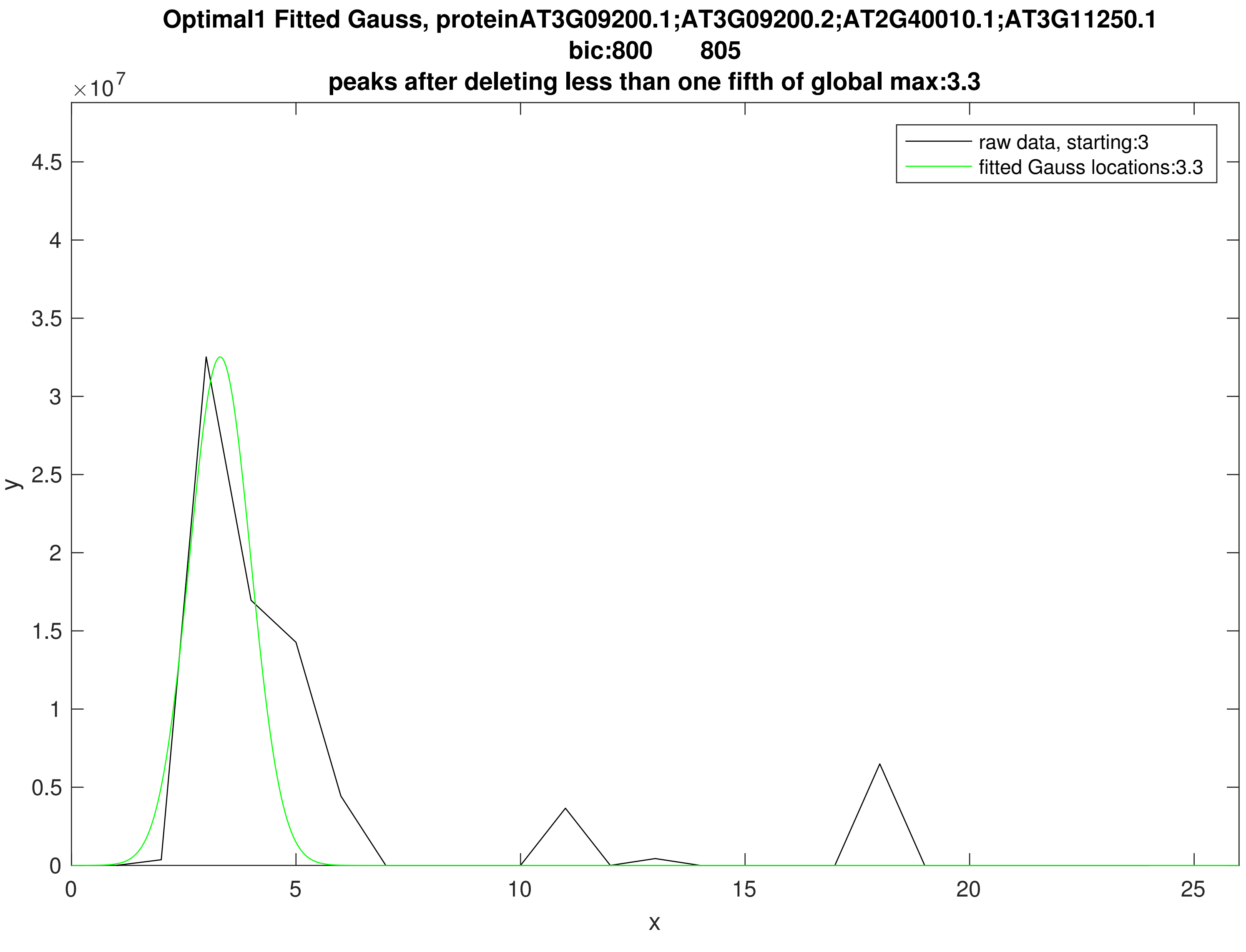
Optimal1 Fitted Gauss, proteinAT3G09200.1;AT3G09200.2;AT2G40010.1;AT3G11250.1
bic:800 805
peaks after deleting less than one fifth of global max:3.3
×10^7
4.5
4
3.5
3
2.5
2
1.5
1
0.5
0
y
0
5
10
15
20
25
x
raw data, starting:3
fitted Gauss locations:3.3

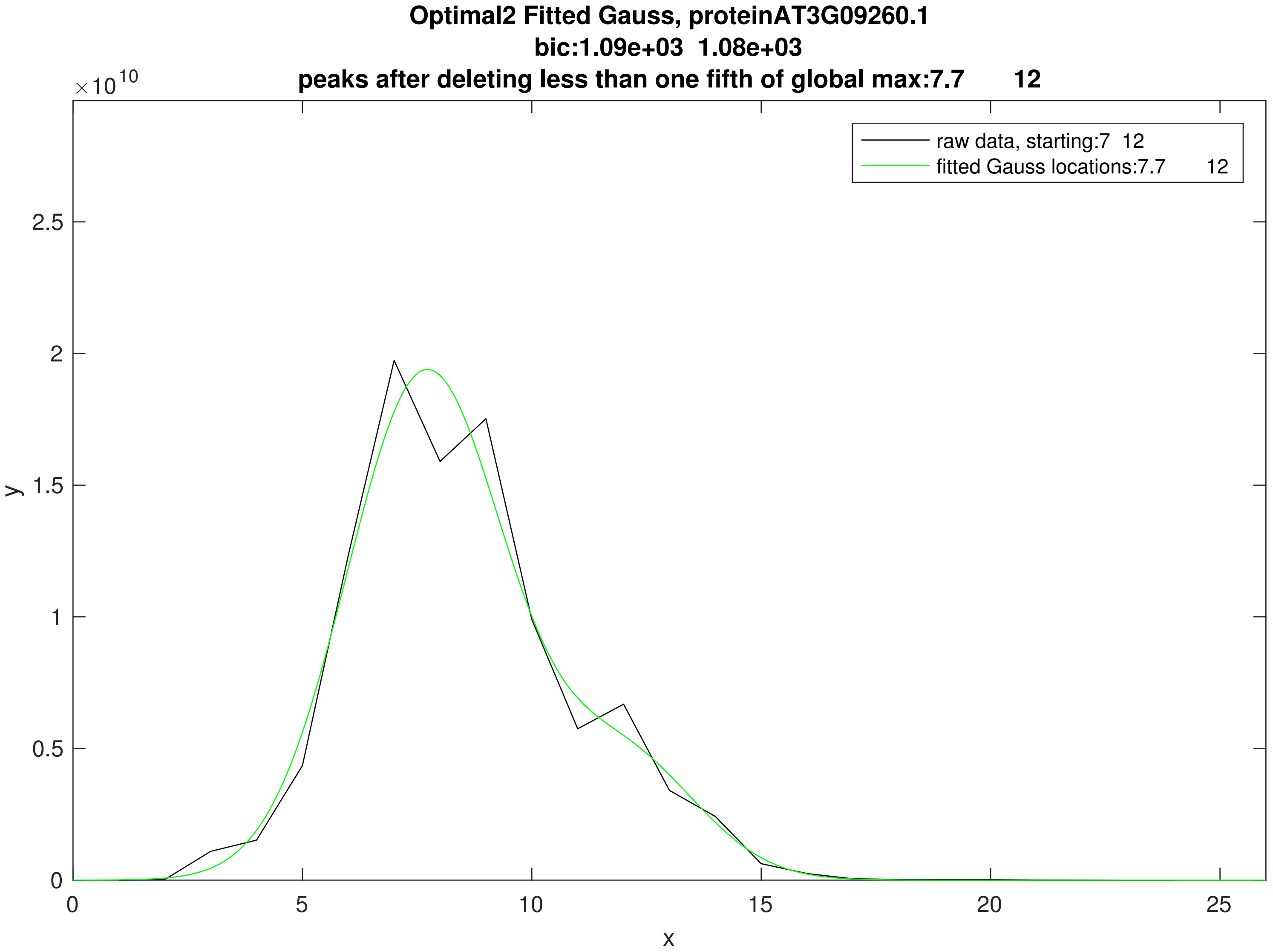
Optimal2 Fitted Gauss, proteinAT3G09260.1
bic:1.09e+03  1.08e+03
peaks after deleting less than one fifth of global max:7.7       12
×10^10
raw data, starting:7  12
fitted Gauss locations:7.7       12
y
x
0
0.5
1
1.5
2
2.5
5
10
15
20
25

**Optimal1 Fitted Gauss, proteinAT3G09270.1**
**bic:825       834**
**peaks after deleting less than one fifth of global max:16**

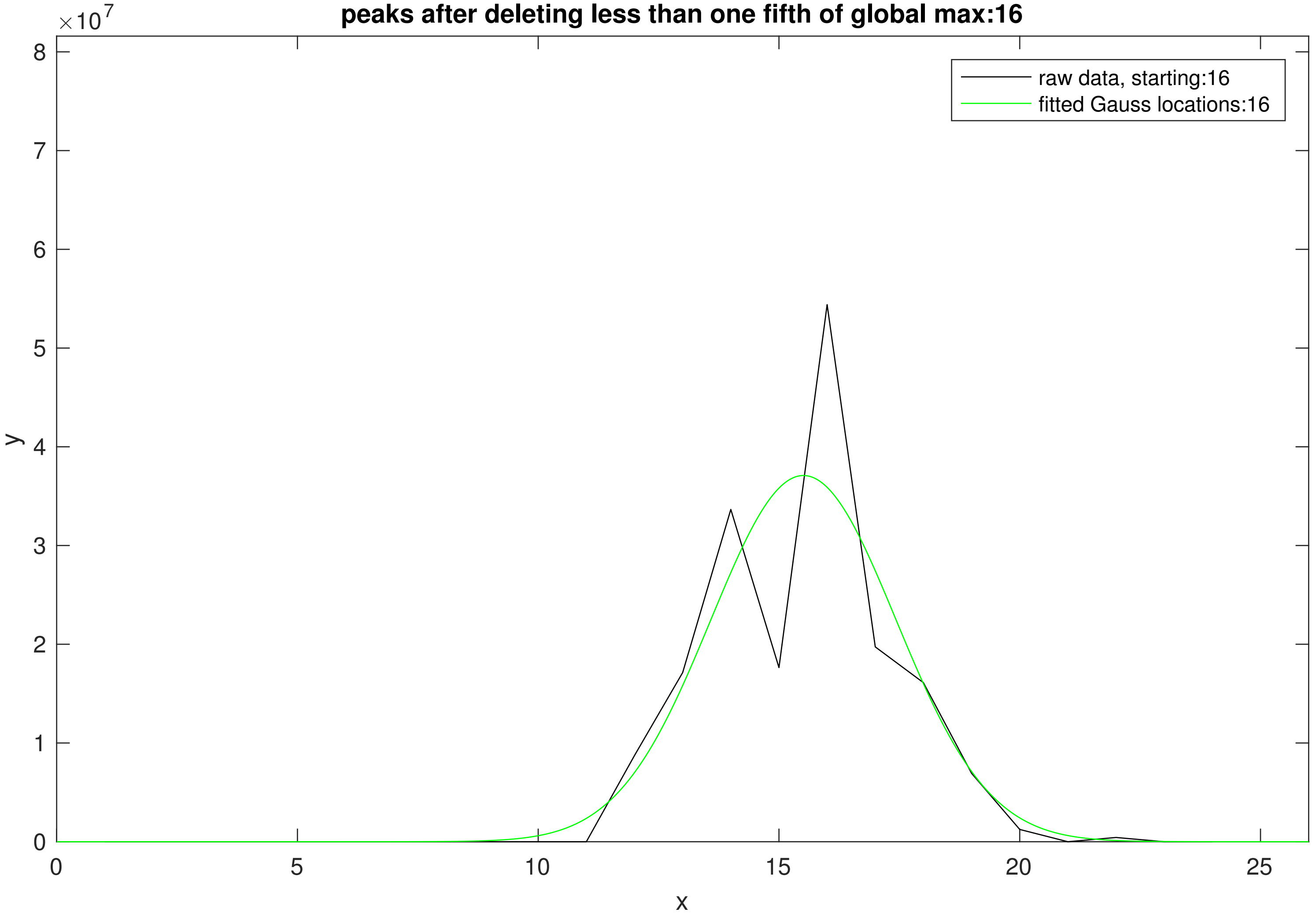

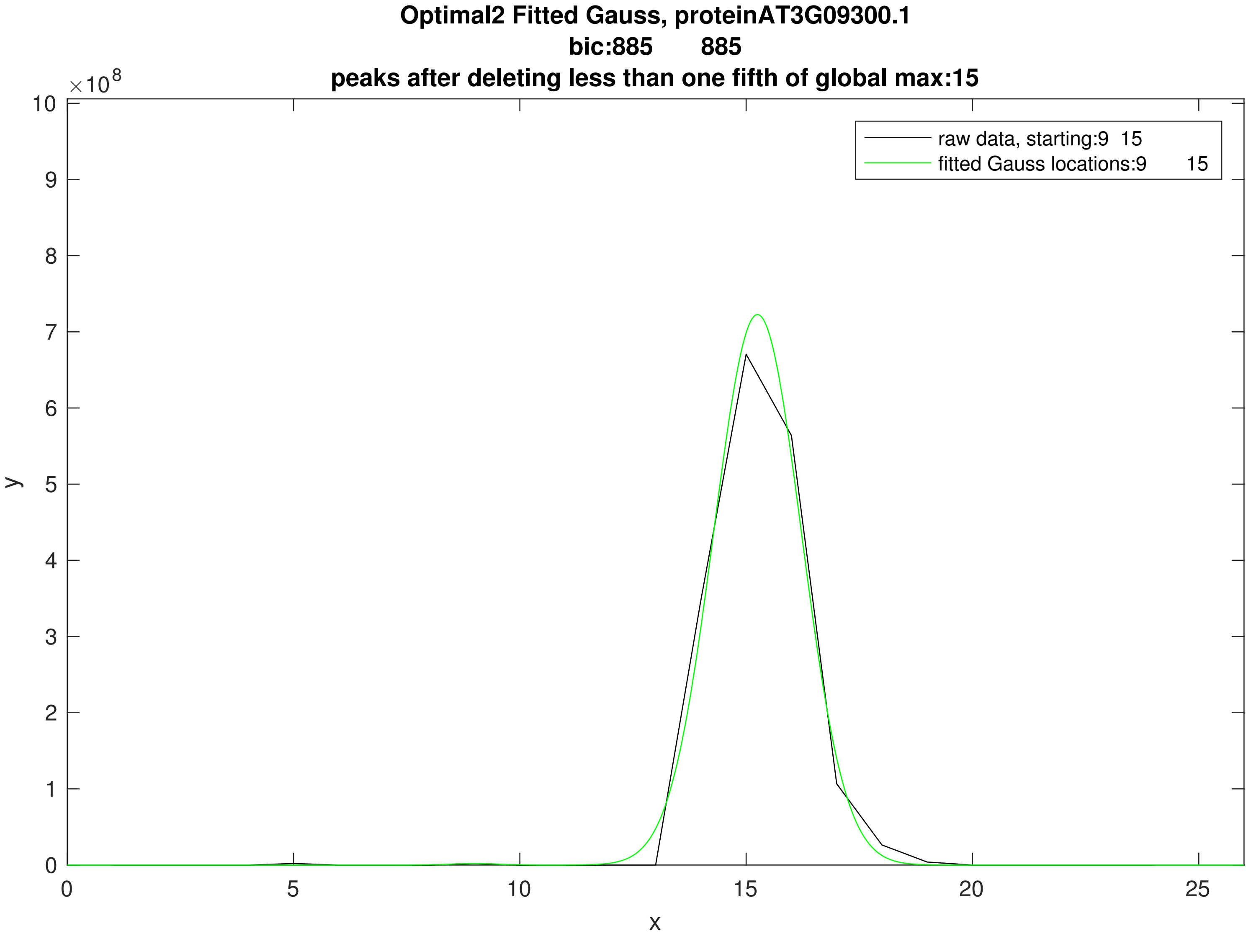

Optimal2 Fitted Gauss, proteinAT3G09300.1
bic:885 885
peaks after deleting less than one fifth of global max:15
×10^8
10
9
8
7
6
5
4
3
2
1
0
y
0
5
10
15
20
25
x
raw data, starting:9 15
fitted Gauss locations:9 15

**Optimal1 Fitted Gauss, proteinAT3G09350.1;AT3G09350.3;AT3G09350.2**
**bic:765**
**peaks after deleting less than one fifth of global max:11**

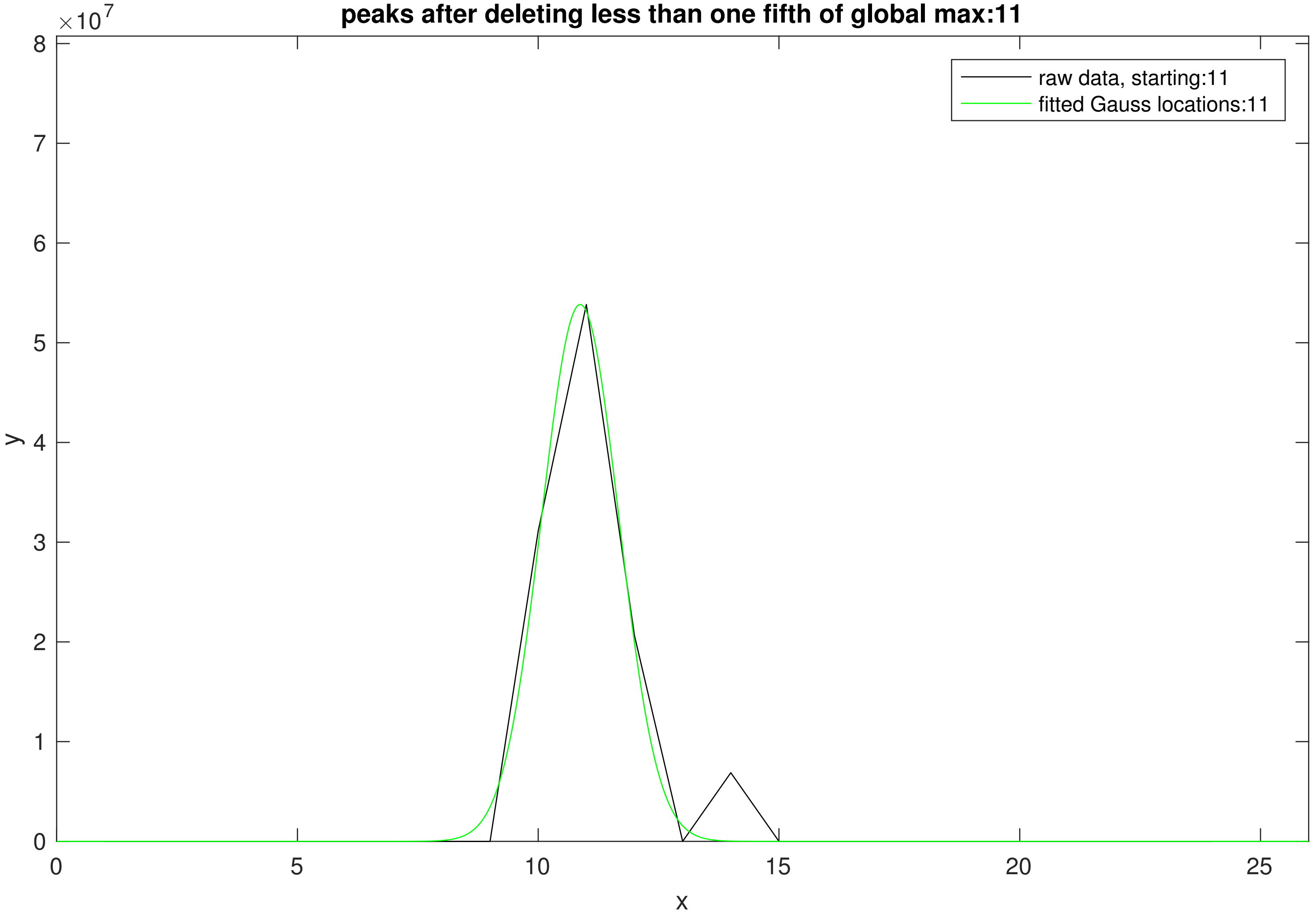

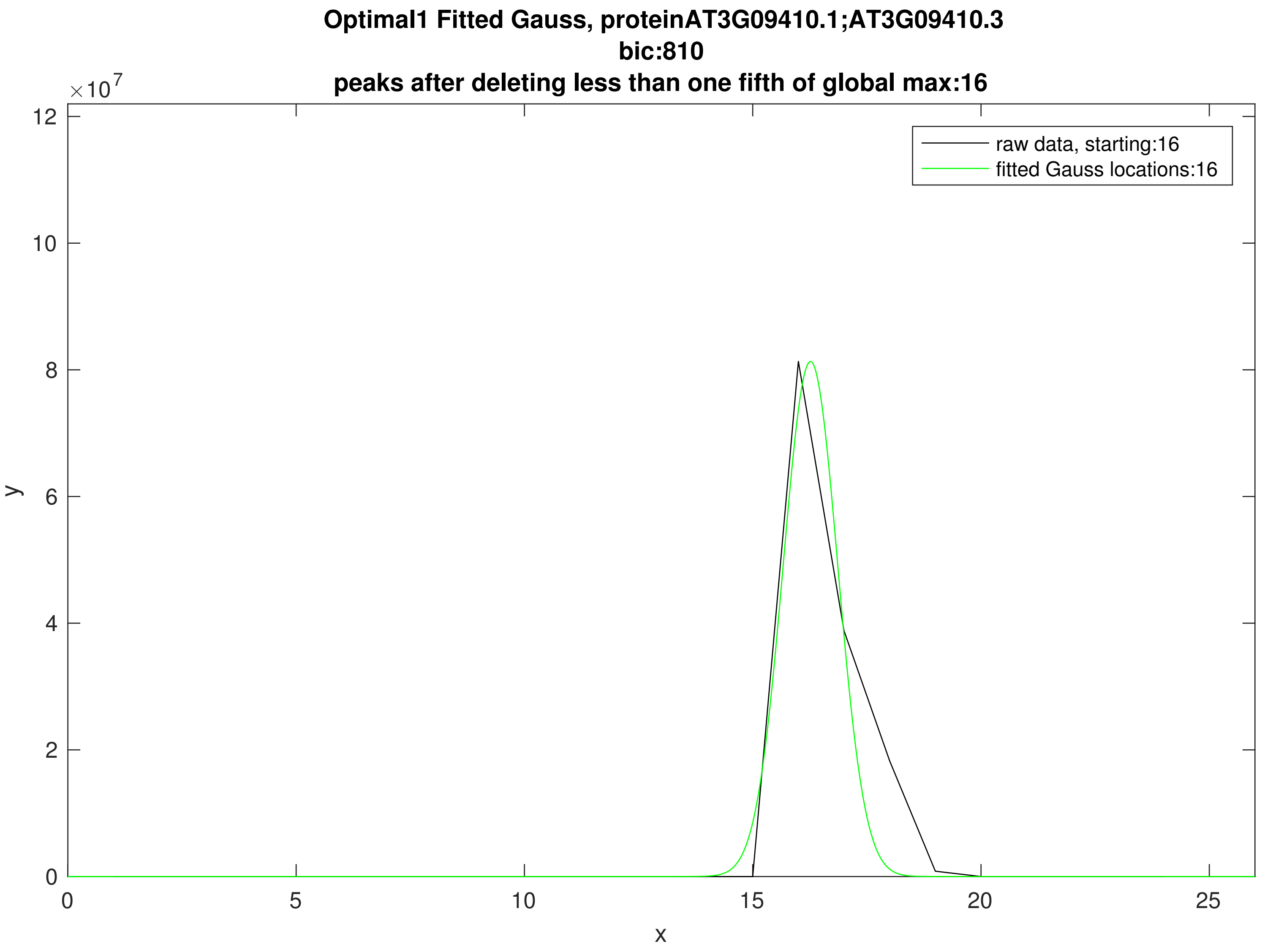
Optimal1 Fitted Gauss, proteinAT3G09410.1;AT3G09410.3
bic:810
peaks after deleting less than one fifth of global max:16
×10^7
12
10
8
6
4
2
0
y
0
5
10
15
20
25
x
raw data, starting:16
fitted Gauss locations:16

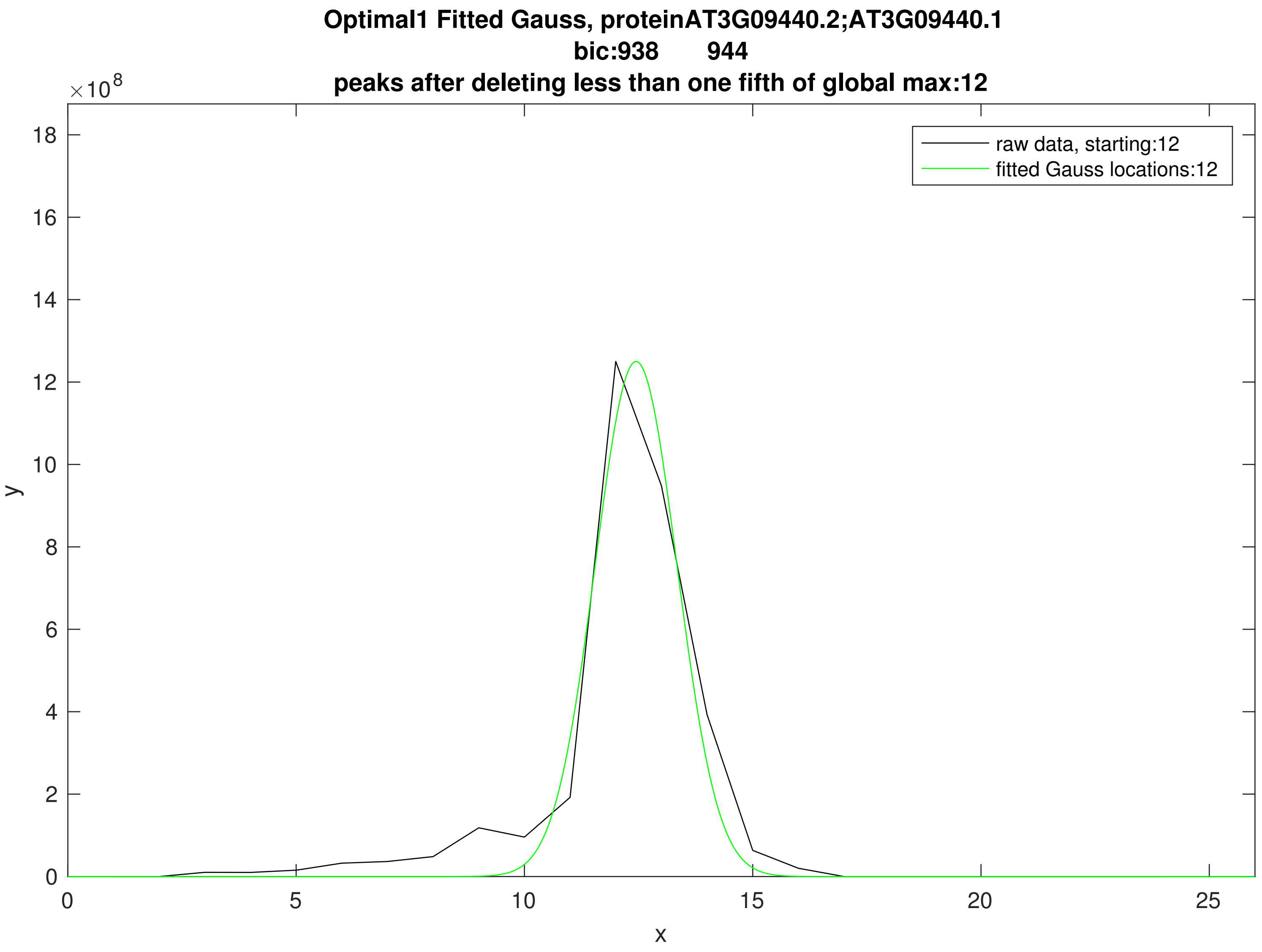

Optimal1 Fitted Gauss, proteinAT3G09440.2;AT3G09440.1
bic:938 944
peaks after deleting less than one fifth of global max:12
×10^8
18
16
14
12
10
8
6
4
2
0
y
0
5
10
15
20
25
x
raw data, starting:12
fitted Gauss locations:12

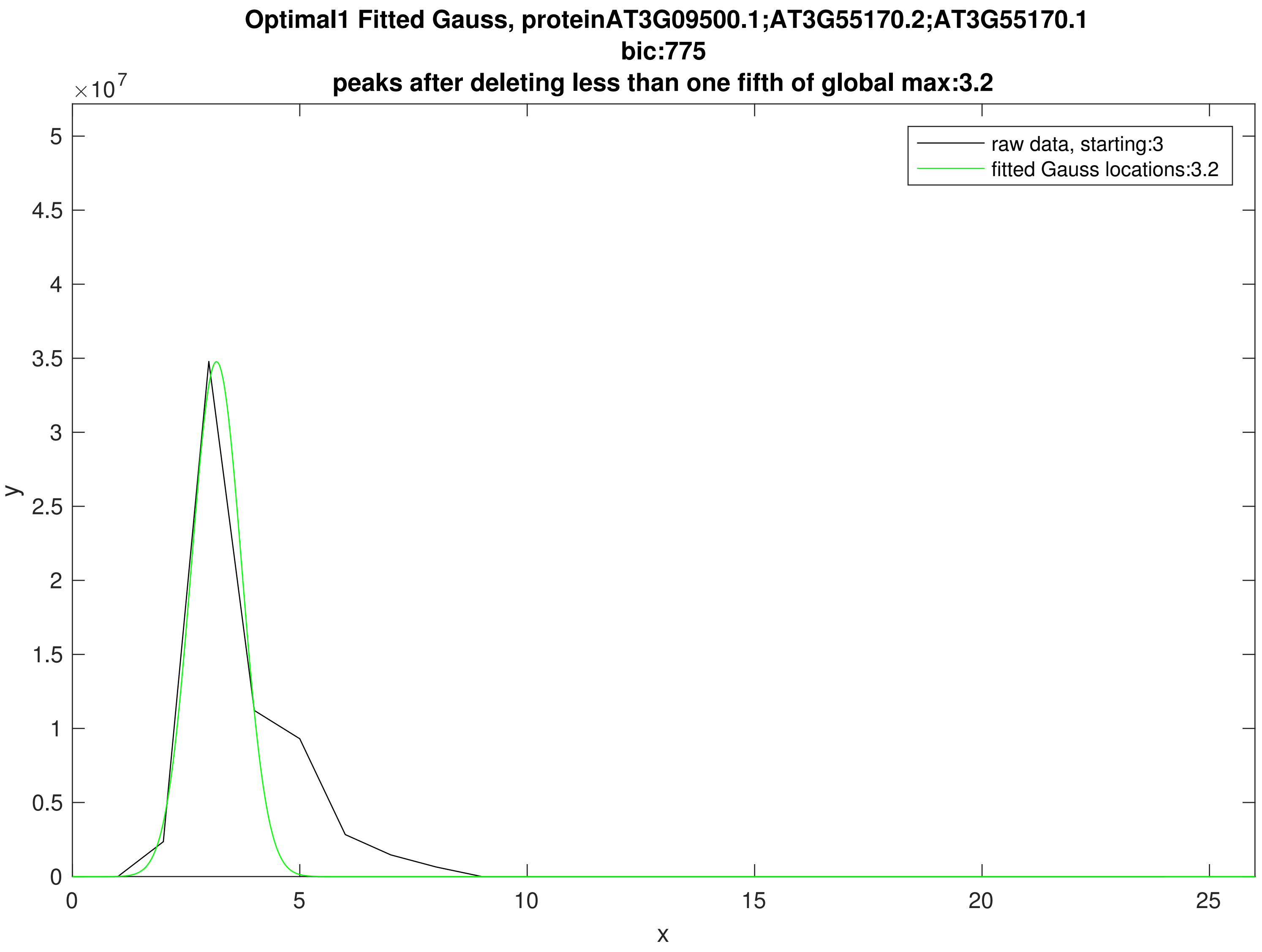

Optimal1 Fitted Gauss, proteinAT3G09500.1;AT3G55170.2;AT3G55170.1
bic:775
peaks after deleting less than one fifth of global max:3.2
×10^7
raw data, starting:3
fitted Gauss locations:3.2
y
x
0
0.5
1
1.5
2
2.5
3
3.5
4
4.5
5
0
5
10
15
20
25

Optimal1 Fitted Gauss, proteinAT3G09630.1;AT3G09630.2
bic:869 877 886
peaks after deleting less than one fifth of global max:3.3

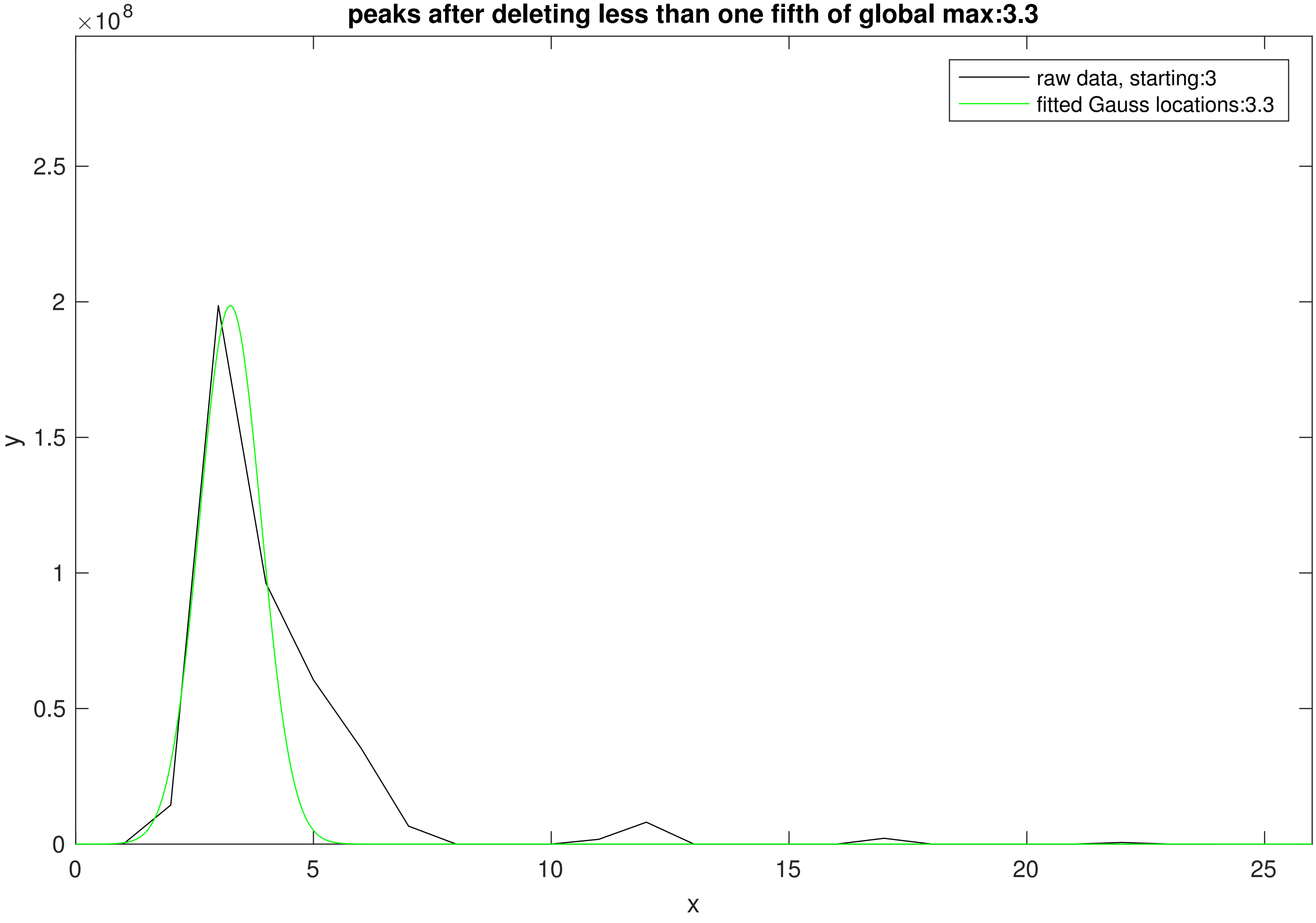

**Optimal2 Fitted Gauss, proteinAT3G09810.1**
**bic:886 803 811**
**peaks after deleting less than one fifth of global max:5.2 18**

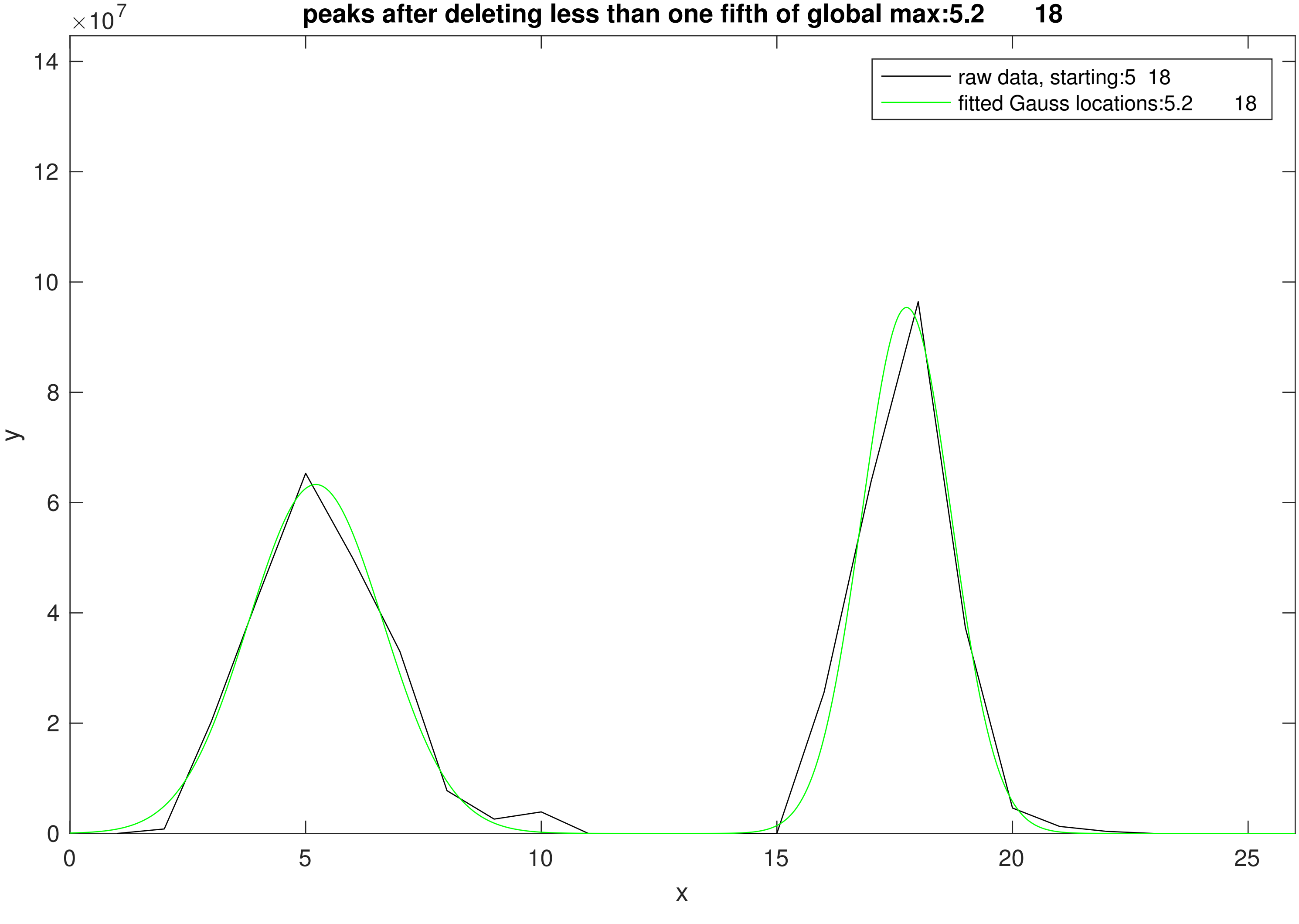

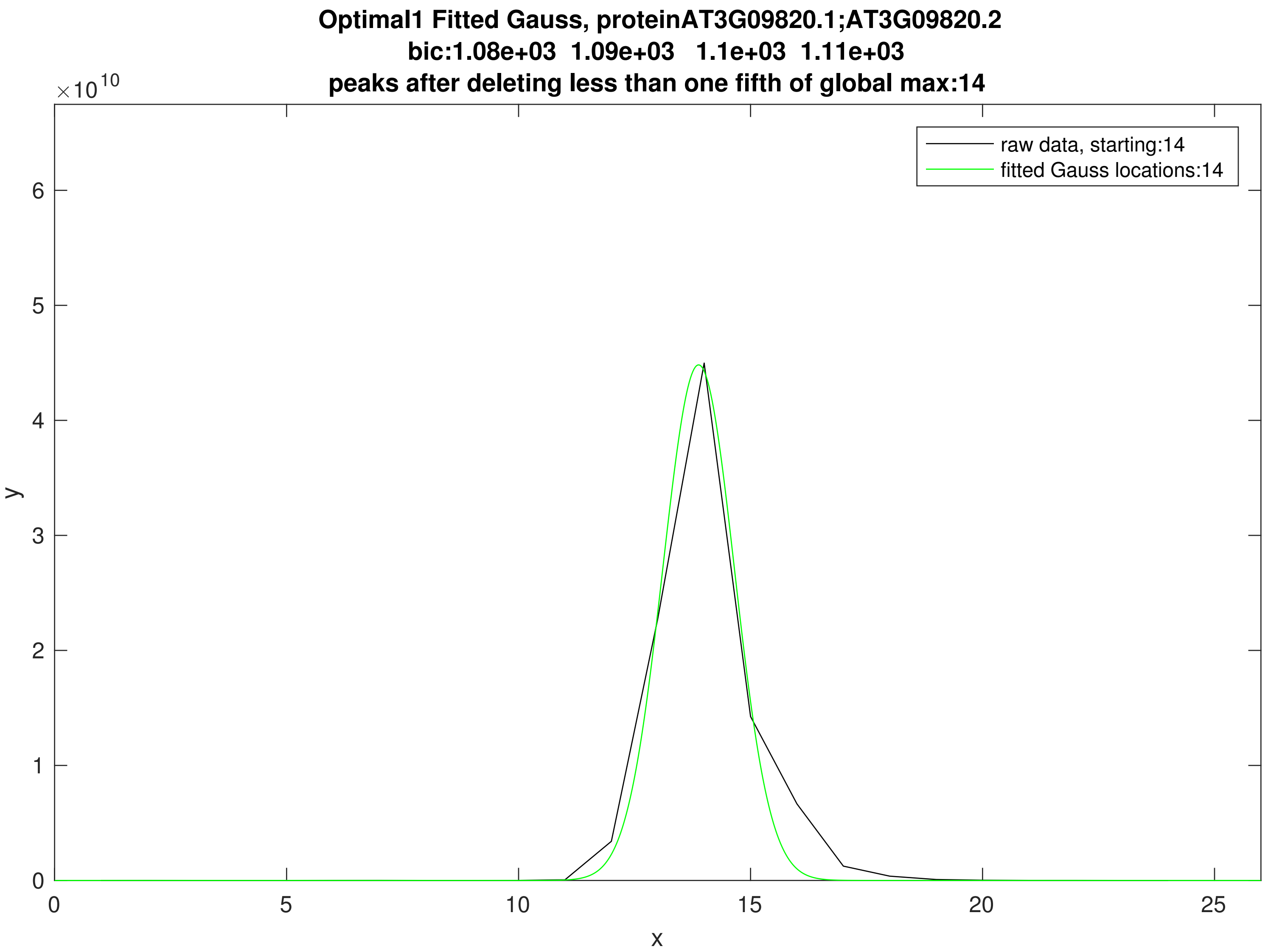
Optimal1 Fitted Gauss, proteinAT3G09820.1;AT3G09820.2
bic:1.08e+03 1.09e+03 1.1e+03 1.11e+03
peaks after deleting less than one fifth of global max:14
×10^10
raw data, starting:14
fitted Gauss locations:14
y
x

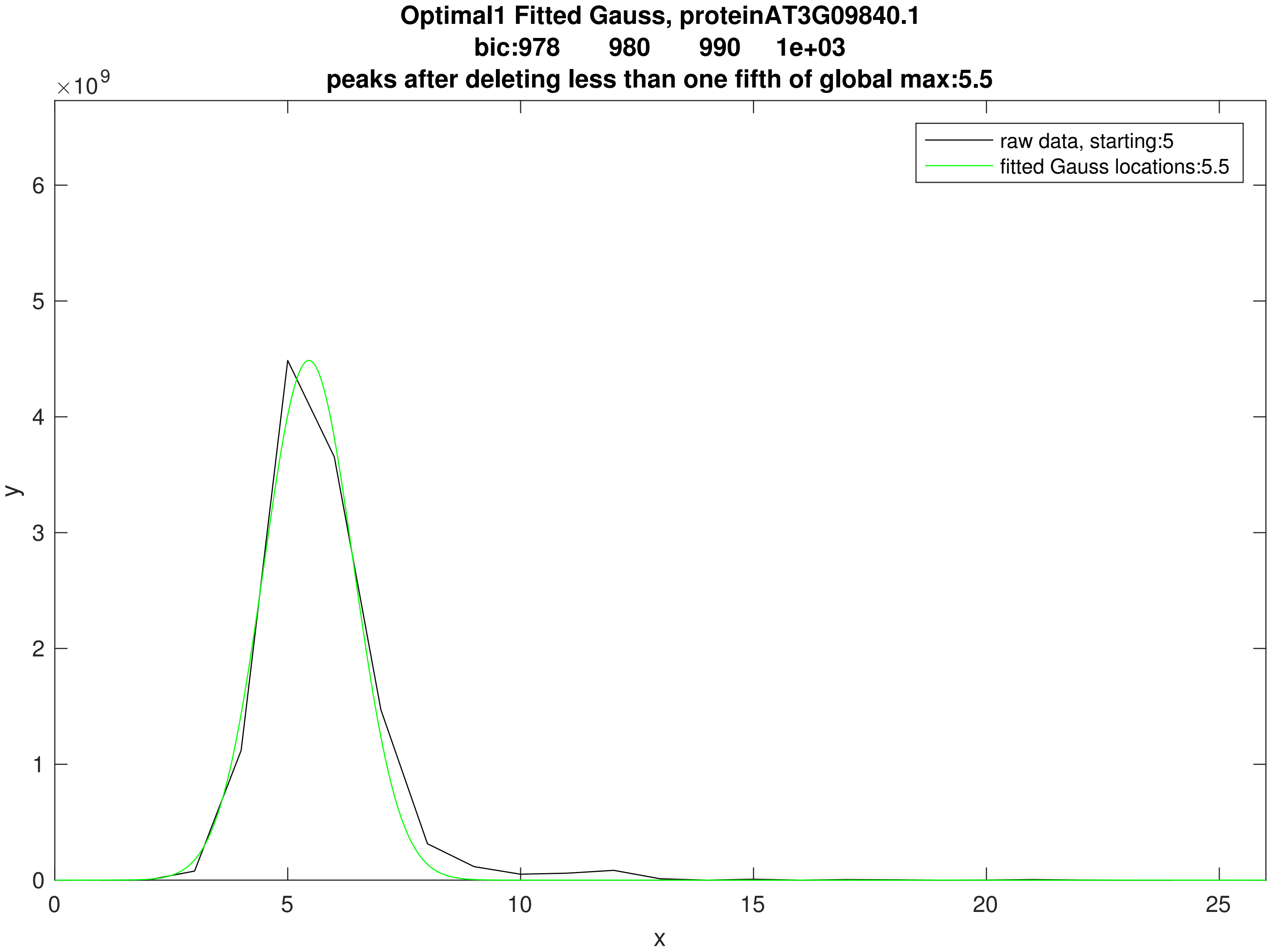
Optimal1 Fitted Gauss, proteinAT3G09840.1
bic:978 980 990 1e+03
peaks after deleting less than one fifth of global max:5.5
×10^9
raw data, starting:5
fitted Gauss locations:5.5
y
x
0
1
2
3
4
5
6
0
5
10
15
20
25

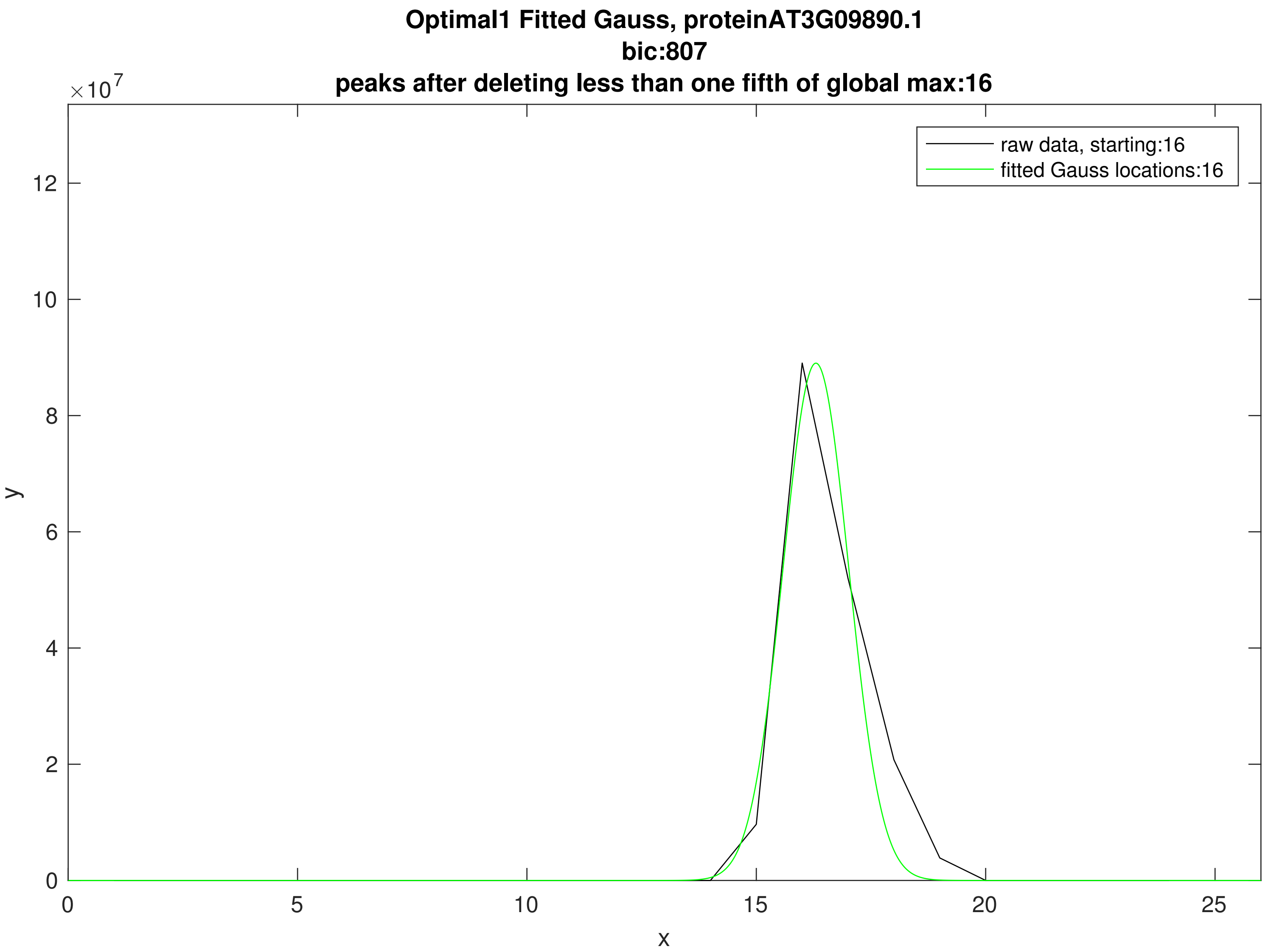

Optimal1 Fitted Gauss, proteinAT3G09890.1
bic:807
peaks after deleting less than one fifth of global max:16
×10^7
raw data, starting:16
fitted Gauss locations:16
y
x
0
2
4
6
8
10
12
0
5
10
15
20
25

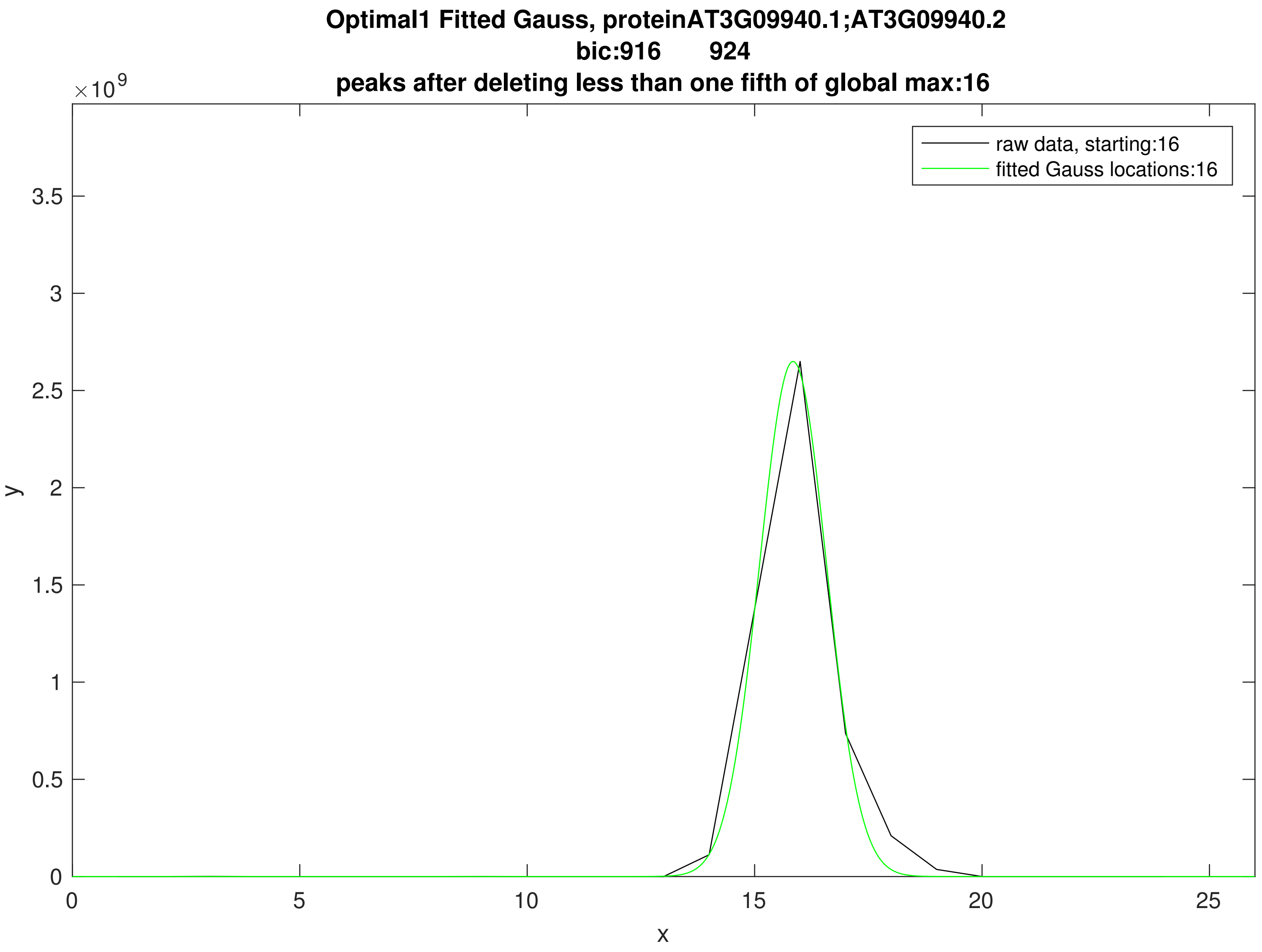

Optimal1 Fitted Gauss, proteinAT3G09940.1;AT3G09940.2
bic:916 924
peaks after deleting less than one fifth of global max:16
×10^9
raw data, starting:16
fitted Gauss locations:16
3.5
3
2.5
2
1.5
1
0.5
0
y
0
5
10
15
20
25
x

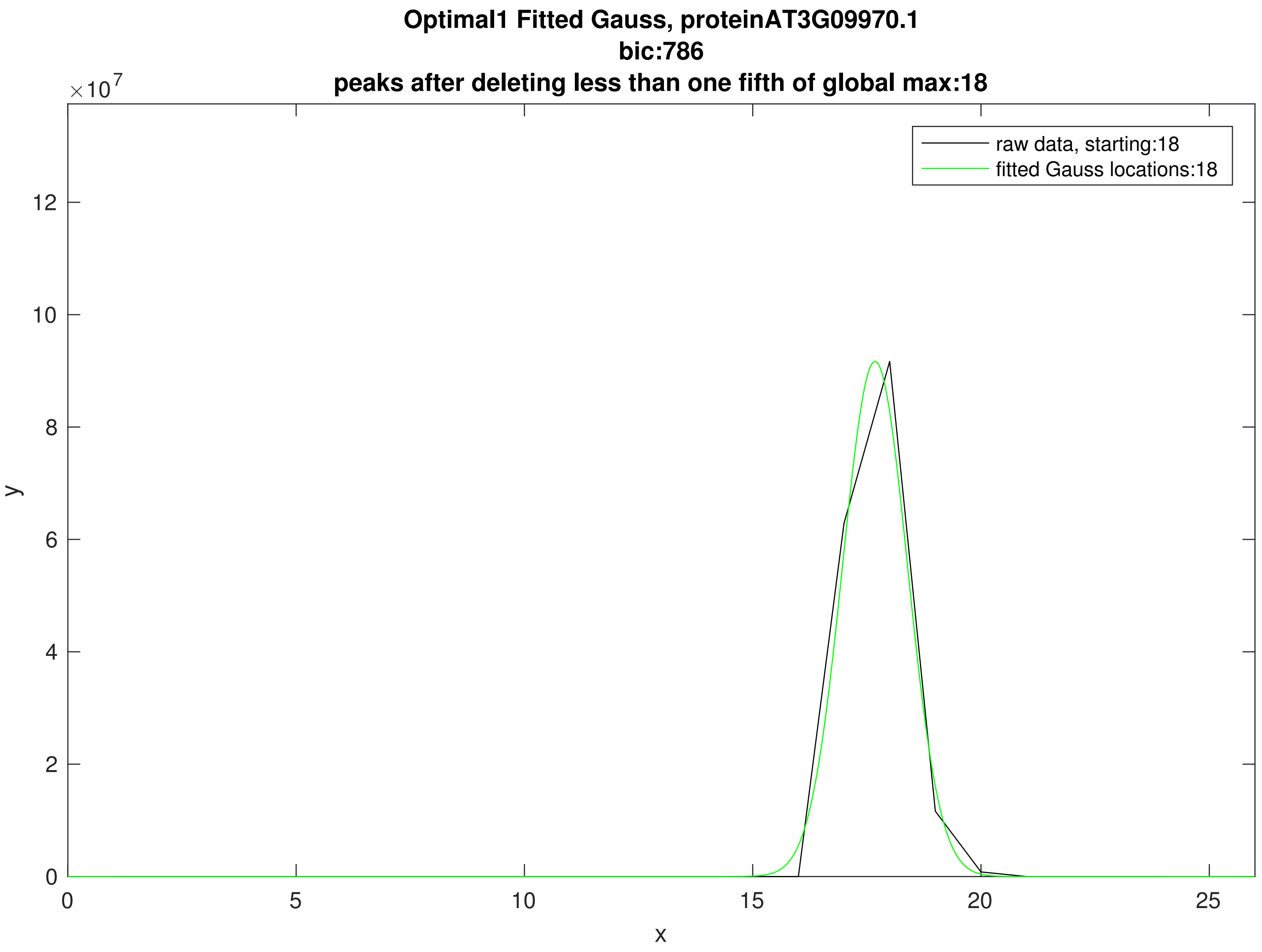
Optimal1 Fitted Gauss, proteinAT3G09970.1
bic:786
peaks after deleting less than one fifth of global max:18
×10^7
raw data, starting:18
fitted Gauss locations:18
y
x
0
2
4
6
8
10
12
0
5
10
15
20
25

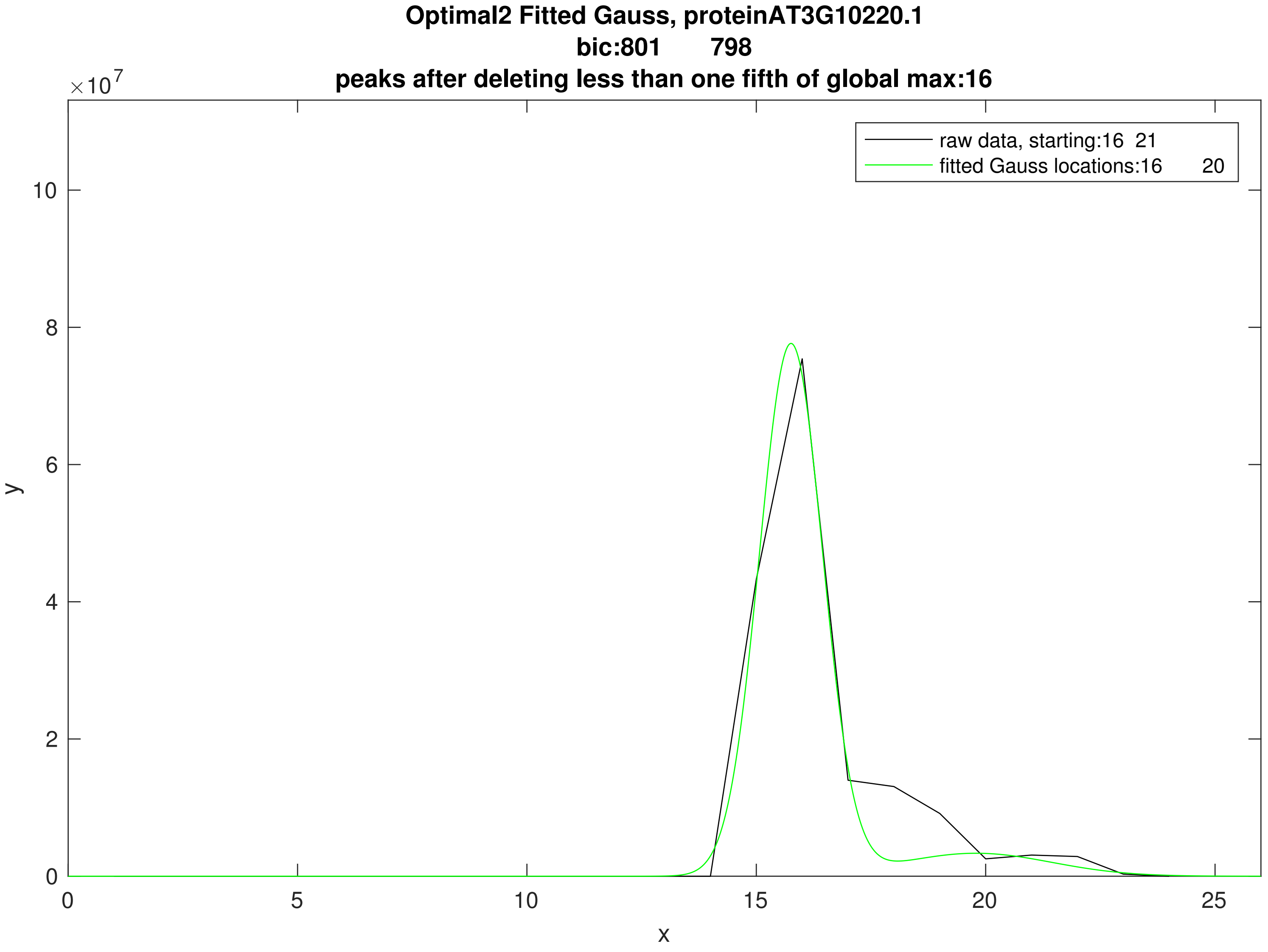
Optimal2 Fitted Gauss, proteinAT3G10220.1
bic:801 798
peaks after deleting less than one fifth of global max:16
×10^7
raw data, starting:16 21
fitted Gauss locations:16 20
y
x
0
2
4
6
8
10
0
5
10
15
20
25

**Optimal2 Fitted Gauss, proteinAT3G10340.1**
**bic:843 837 846**
**peaks after deleting less than one fifth of global max:8.7**

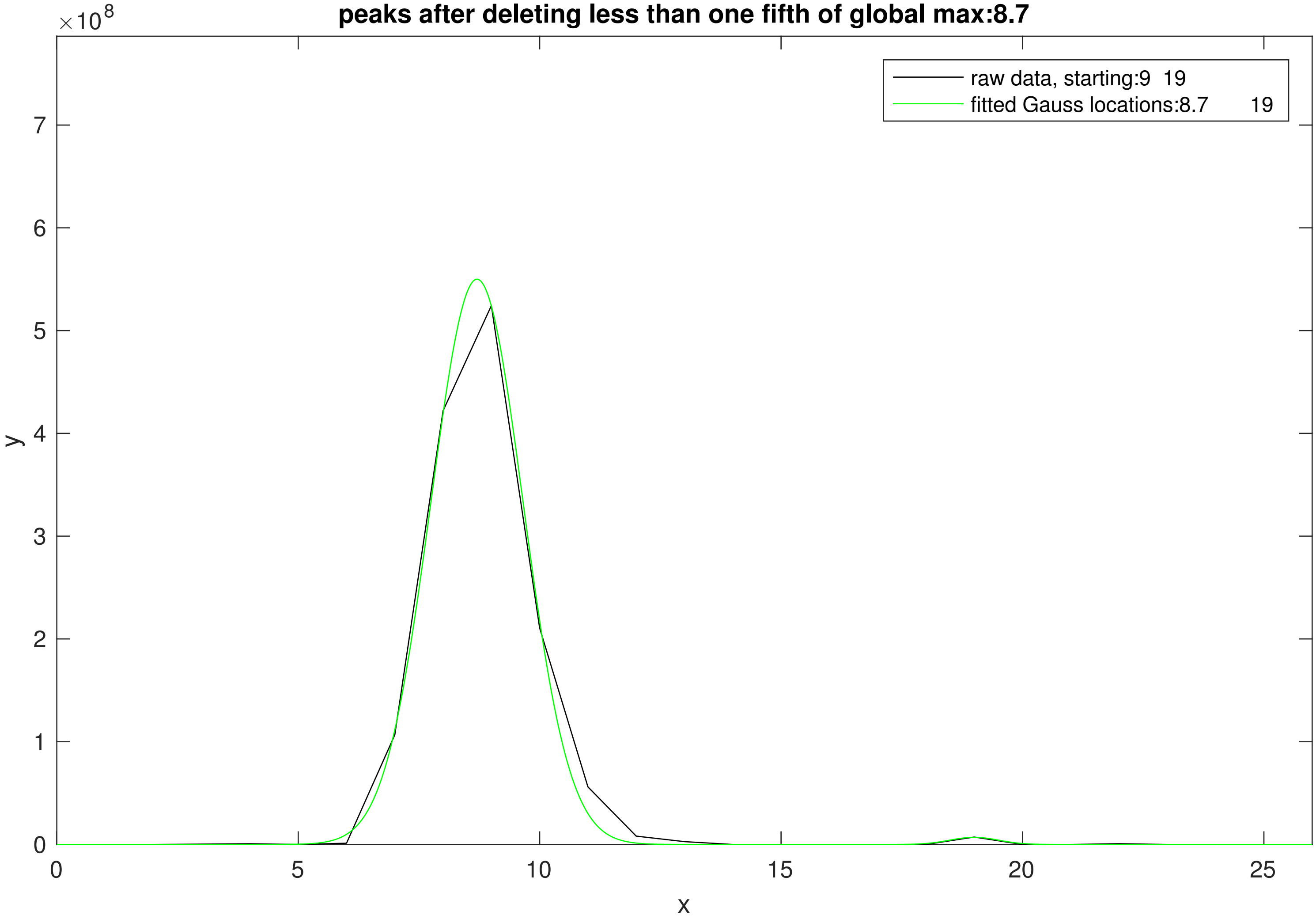

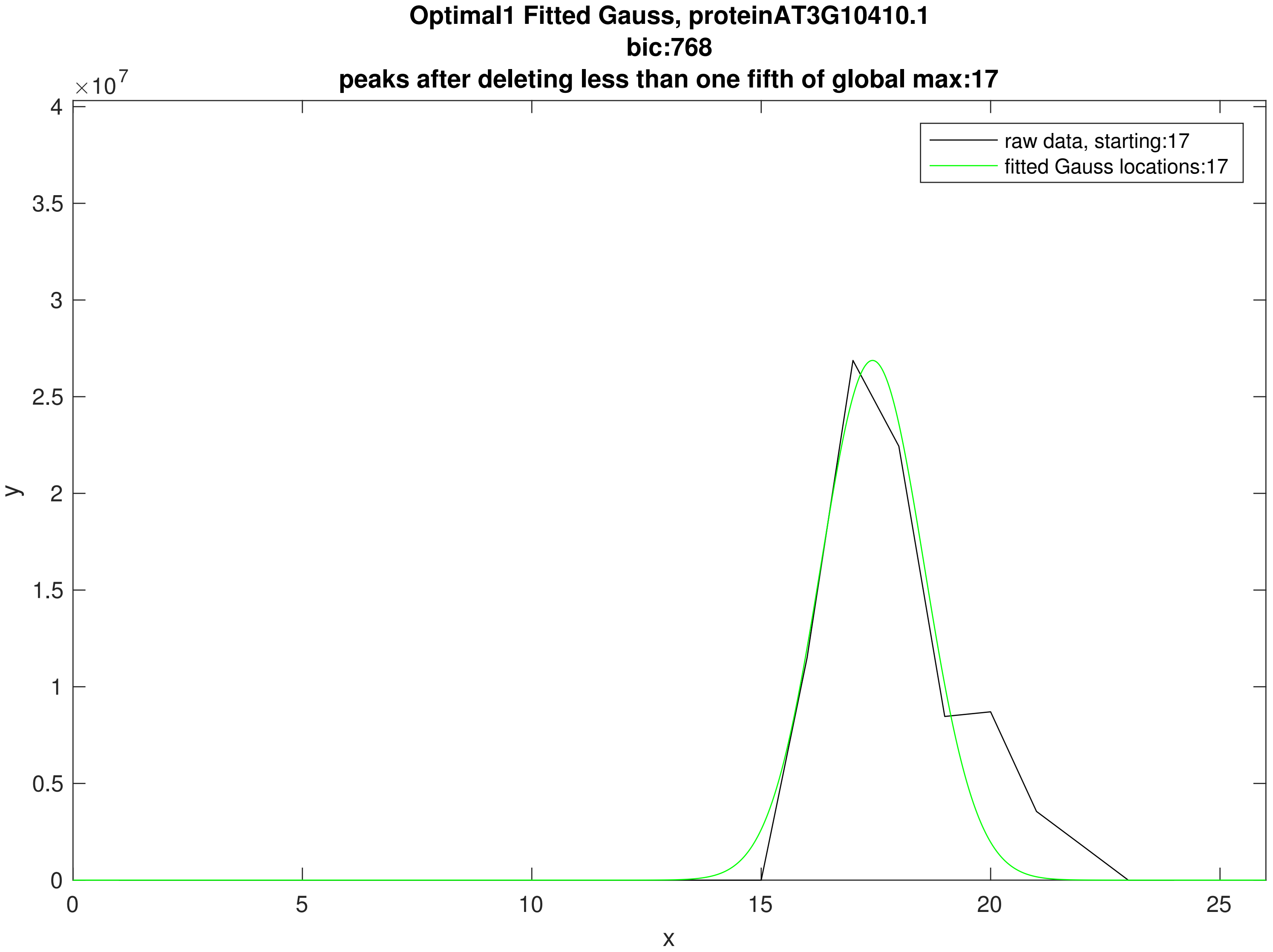

Optimal1 Fitted Gauss, proteinAT3G10410.1
bic:768
peaks after deleting less than one fifth of global max:17
×10^7
raw data, starting:17
fitted Gauss locations:17
y
x
0
0.5
1
1.5
2
2.5
3
3.5
4
0
5
10
15
20
25

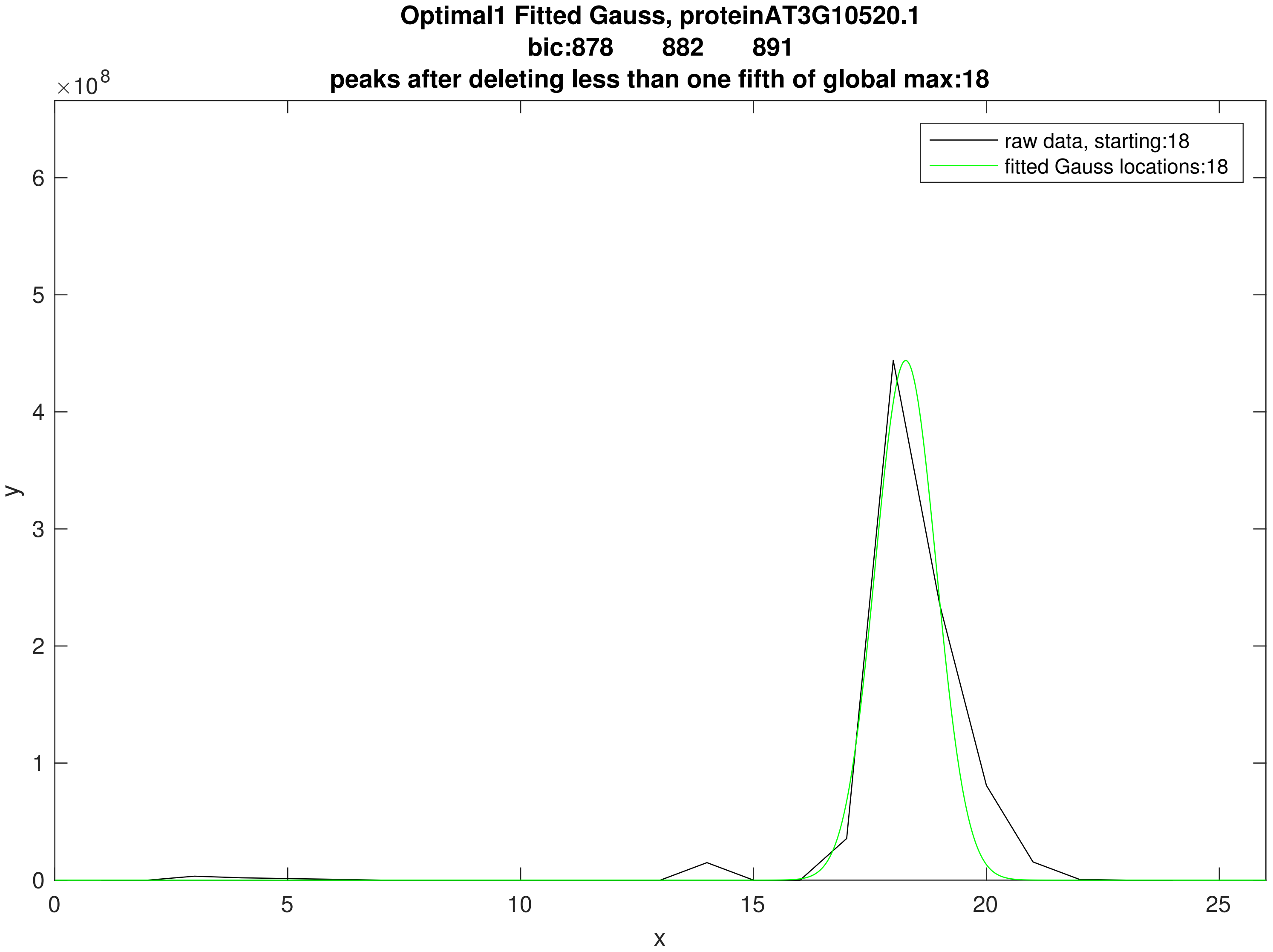

Optimal1 Fitted Gauss, proteinAT3G10520.1
bic:878 882 891
peaks after deleting less than one fifth of global max:18
×10^8
raw data, starting:18
fitted Gauss locations:18
6
5
4
3
2
1
0
y
0
5
10
15
20
25
x

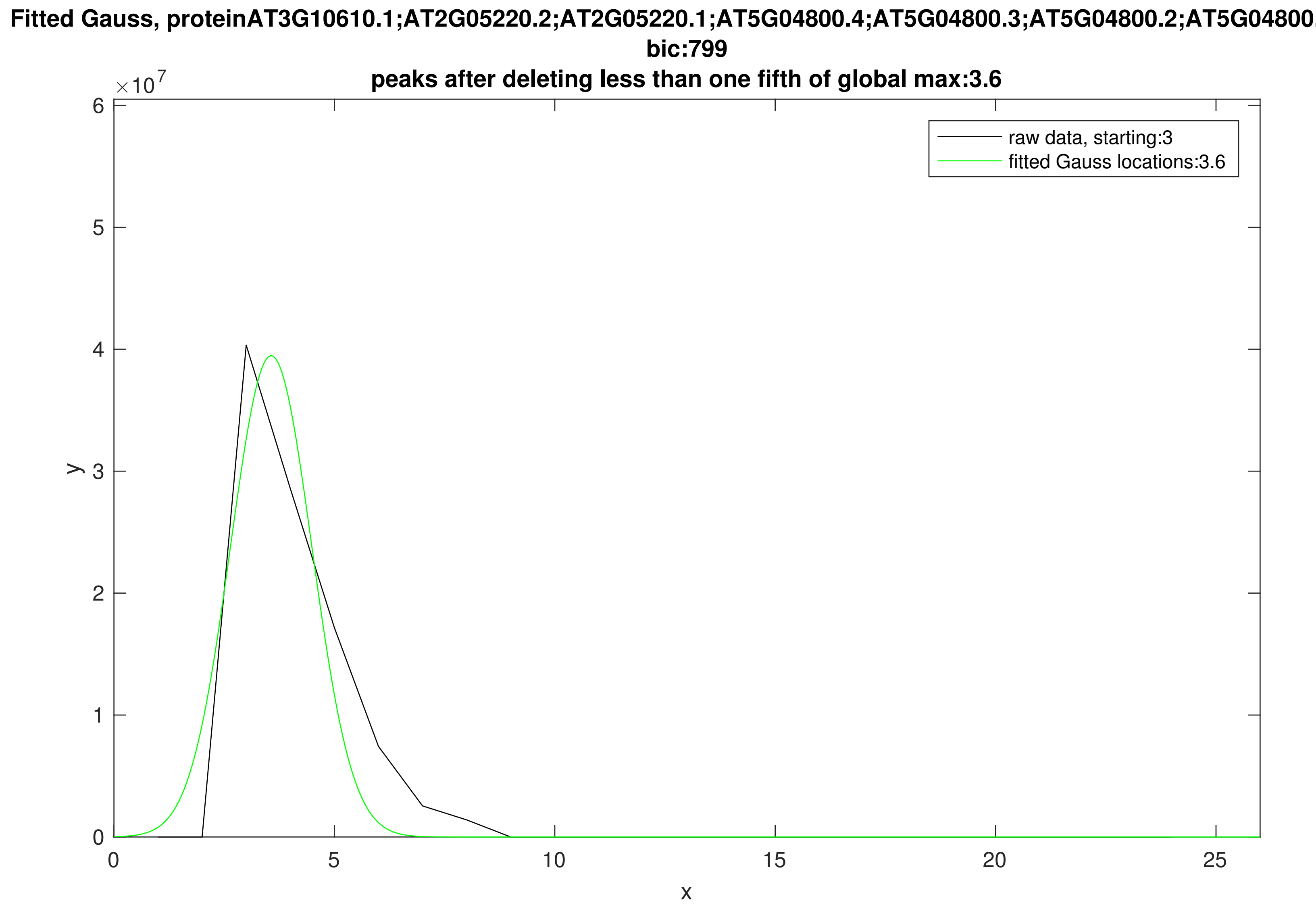

Fitted Gauss, proteinAT3G10610.1;AT2G05220.2;AT2G05220.1;AT5G04800.4;AT5G04800.3;AT5G04800.2;AT5G04800.
bic:799
peaks after deleting less than one fifth of global max:3.6
×10^7
raw data, starting:3
fitted Gauss locations:3.6
y
x
0
1
2
3
4
5
6
0
5
10
15
20
25

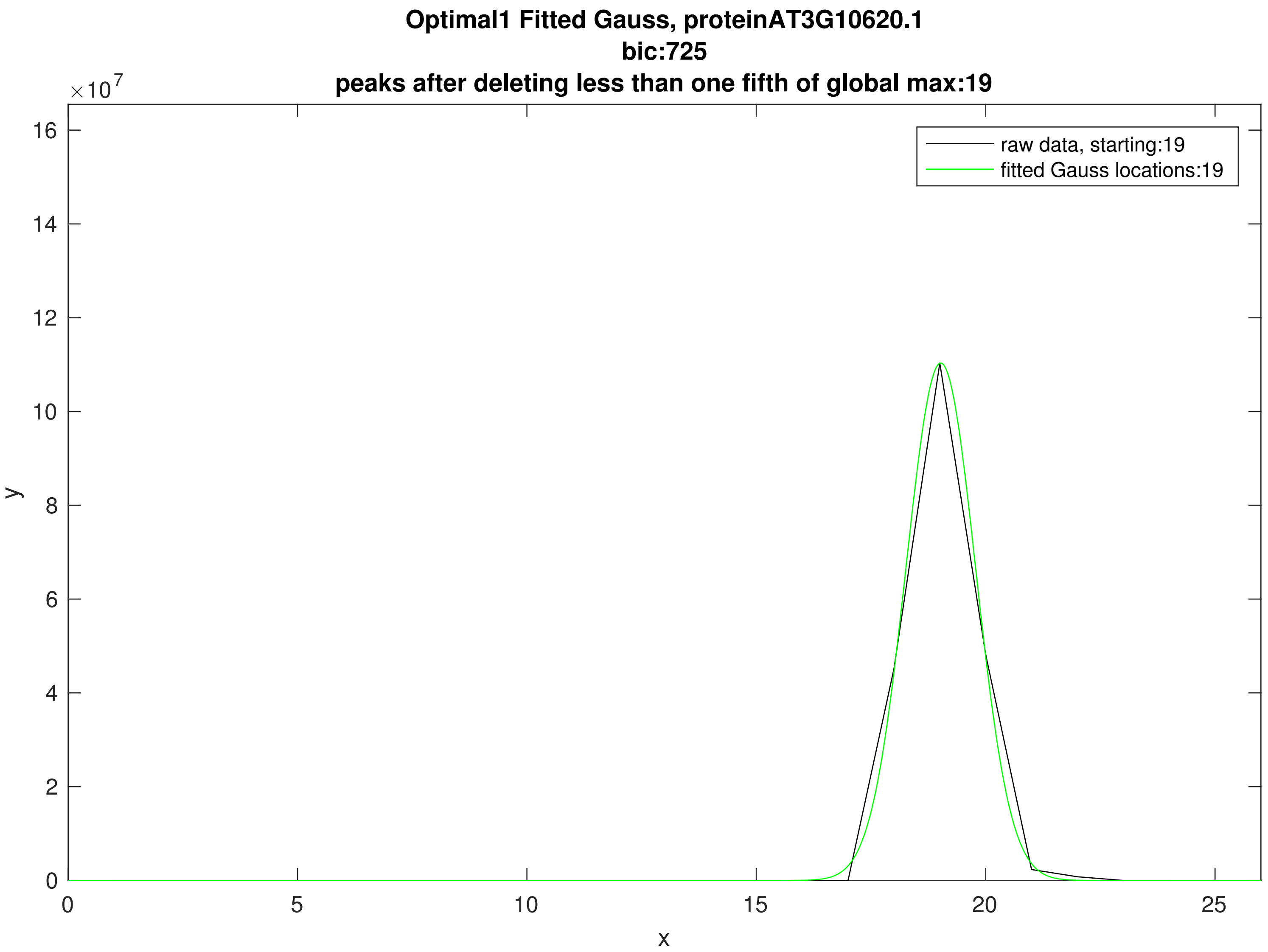
Optimal1 Fitted Gauss, proteinAT3G10620.1
bic:725
peaks after deleting less than one fifth of global max:19
×10^7
y
x
raw data, starting:19
fitted Gauss locations:19
0
2
4
6
8
10
12
14
16
0
5
10
15
20
25

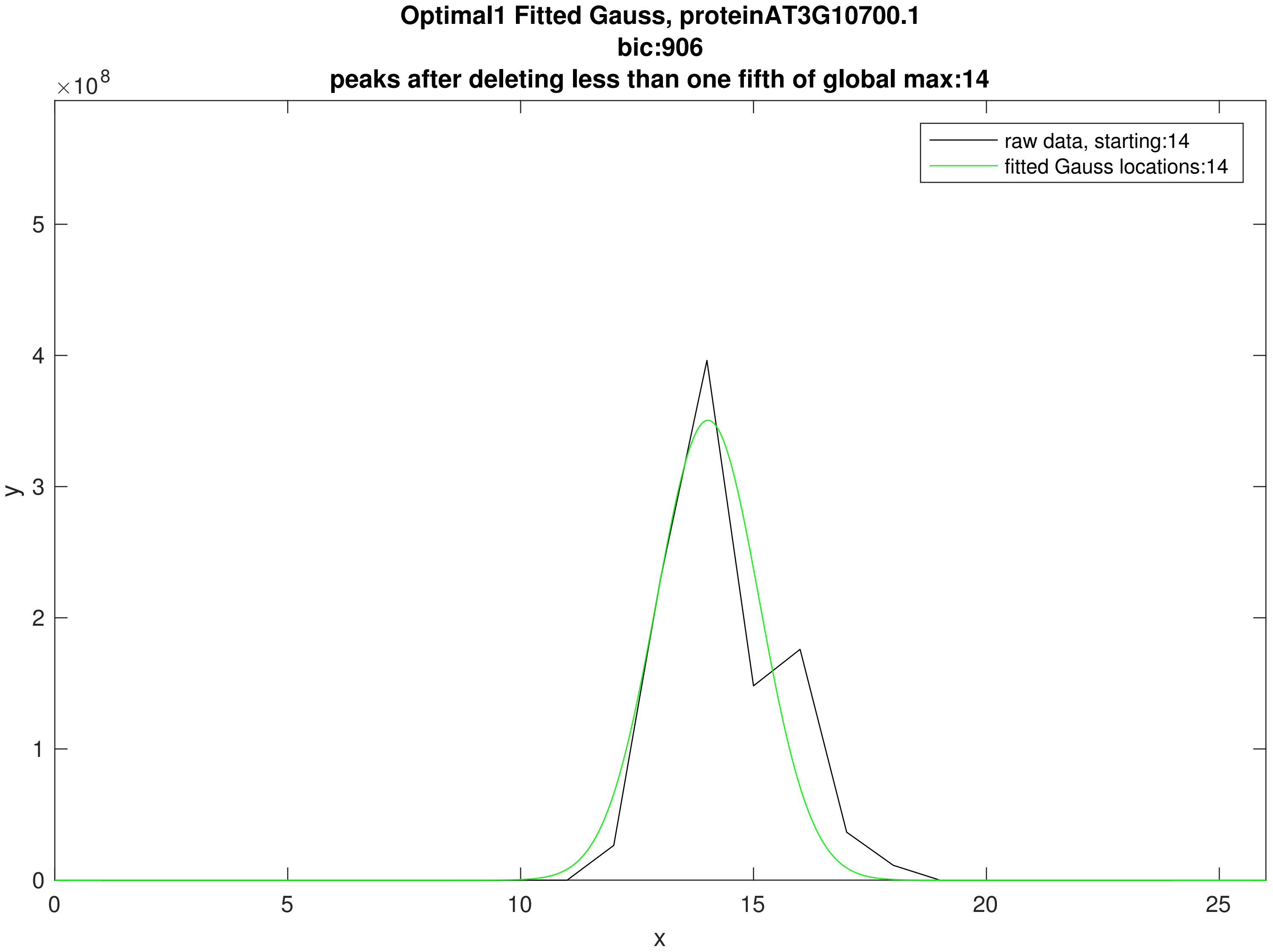
Optimal1 Fitted Gauss, proteinAT3G10700.1
bic:906
peaks after deleting less than one fifth of global max:14
×10^8
5
4
3
2
1
0
y
0
5
10
15
20
25
x
raw data, starting:14
fitted Gauss locations:14

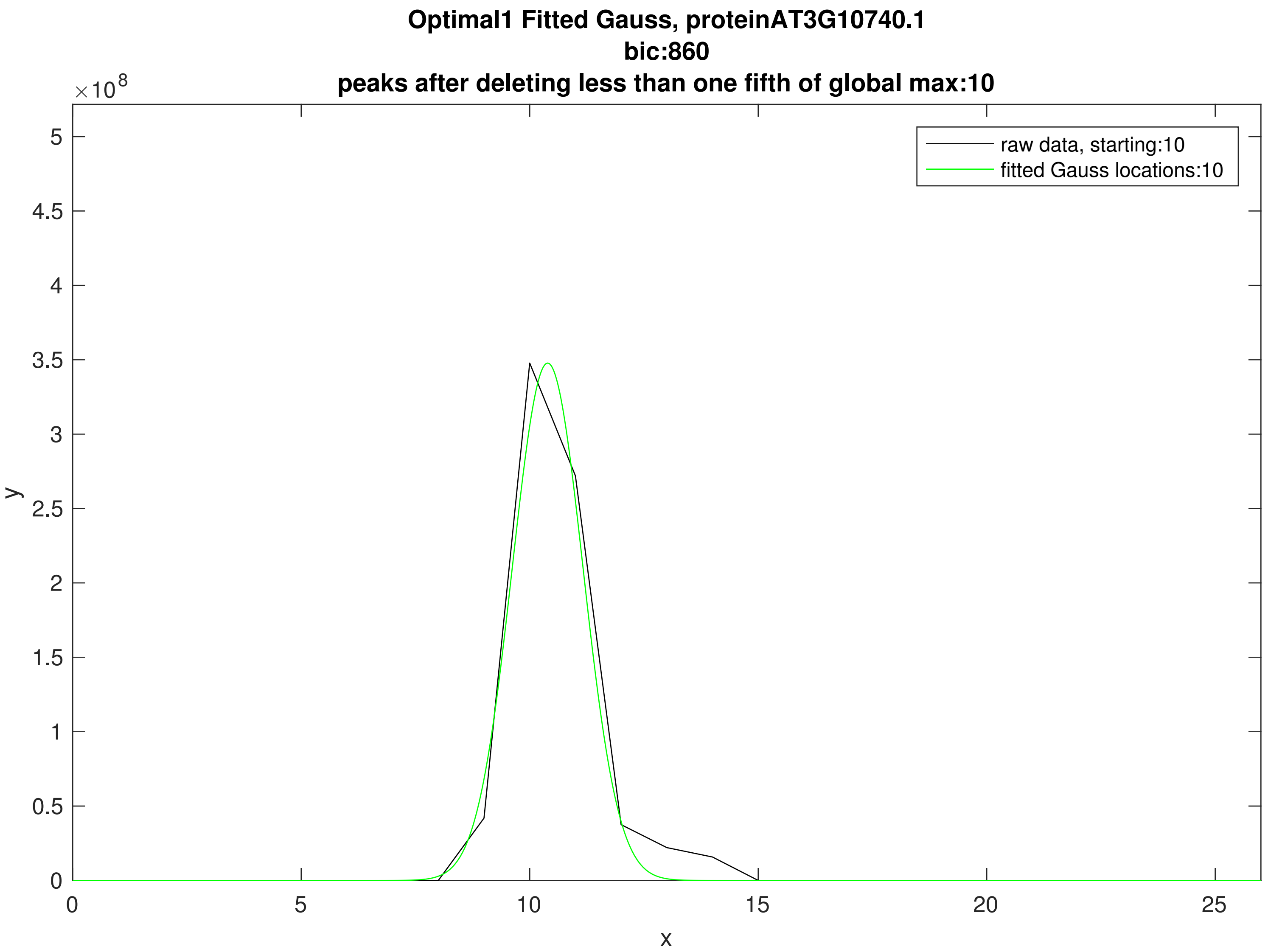

Optimal1 Fitted Gauss, proteinAT3G10740.1
bic:860
peaks after deleting less than one fifth of global max:10
×10^8
5
4.5
4
3.5
3
2.5
2
1.5
1
0.5
0
y
0
5
10
15
20
25
x
raw data, starting:10
fitted Gauss locations:10

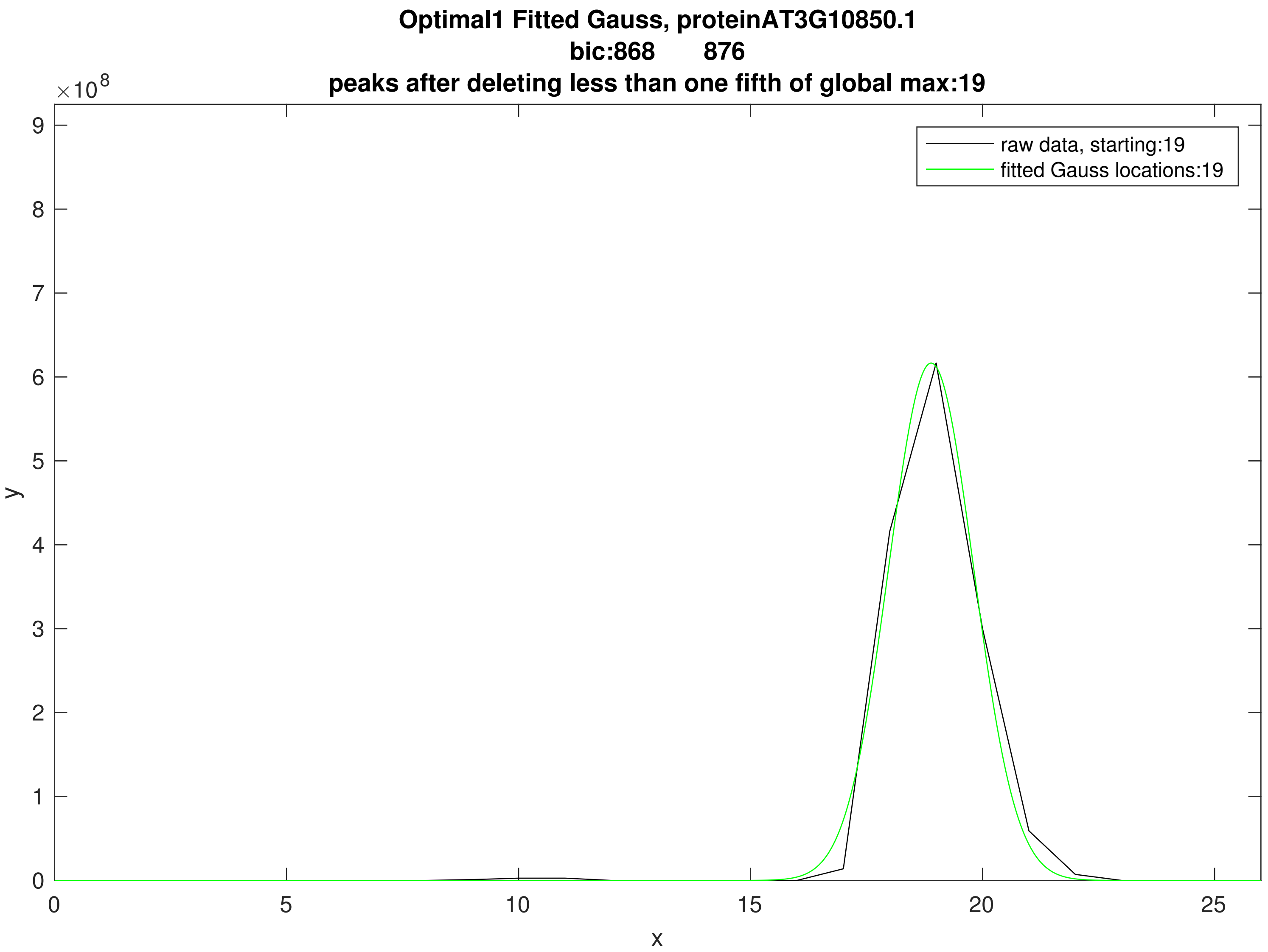

Optimal1 Fitted Gauss, proteinAT3G10850.1
bic:868 876
peaks after deleting less than one fifth of global max:19
×10^8
raw data, starting:19
fitted Gauss locations:19
y
x
0
1
2
3
4
5
6
7
8
9
0
5
10
15
20
25

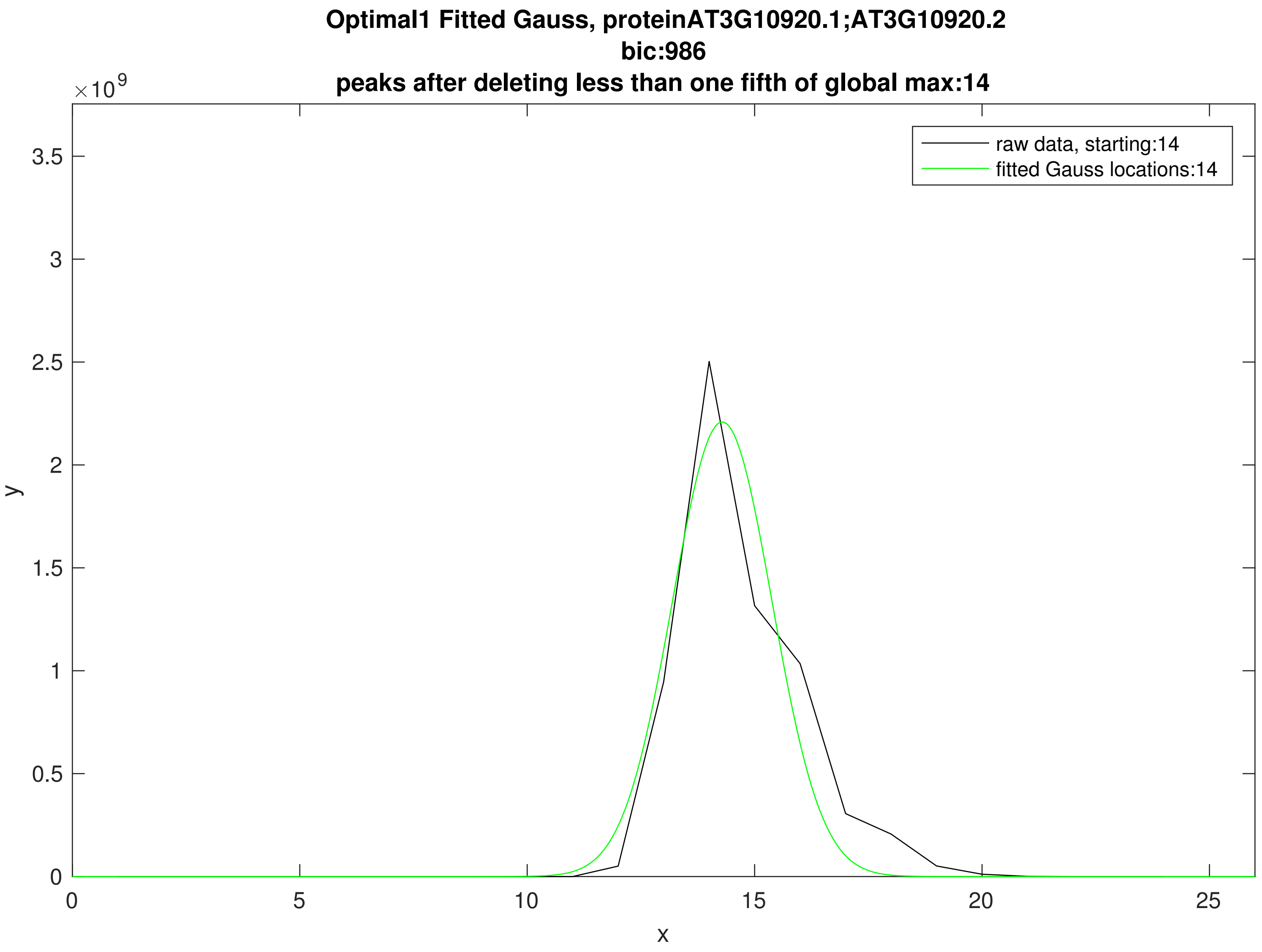

Optimal1 Fitted Gauss, proteinAT3G10920.1;AT3G10920.2
bic:986
peaks after deleting less than one fifth of global max:14
×10^9
3.5
3
2.5
2
1.5
1
0.5
0
y
0
5
10
15
20
25
x
raw data, starting:14
fitted Gauss locations:14

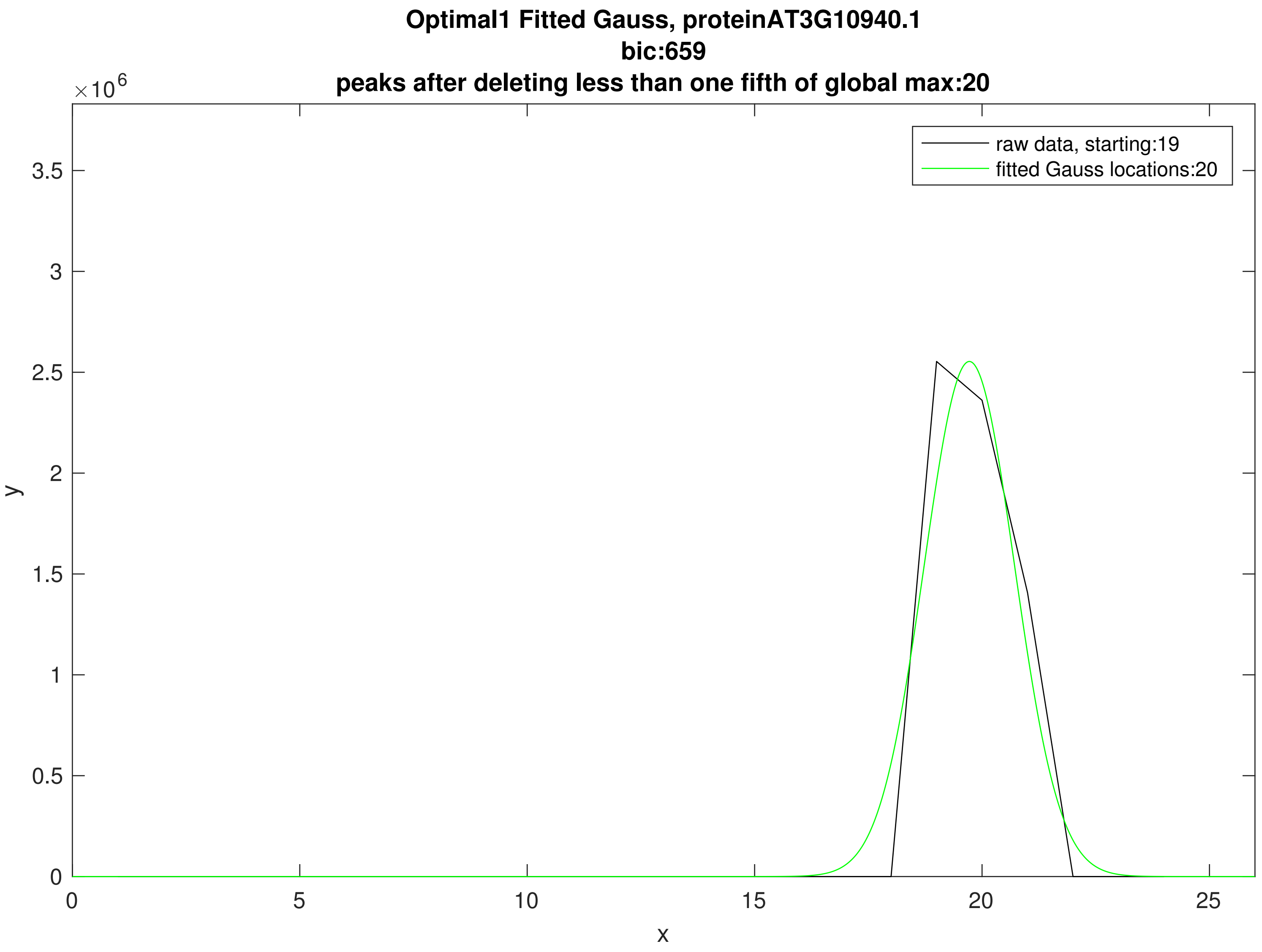
Optimal1 Fitted Gauss, proteinAT3G10940.1
bic:659
peaks after deleting less than one fifth of global max:20
×10^6
raw data, starting:19
fitted Gauss locations:20
3.5
3
2.5
2
1.5
1
0.5
0
y
0
5
10
15
20
25
x

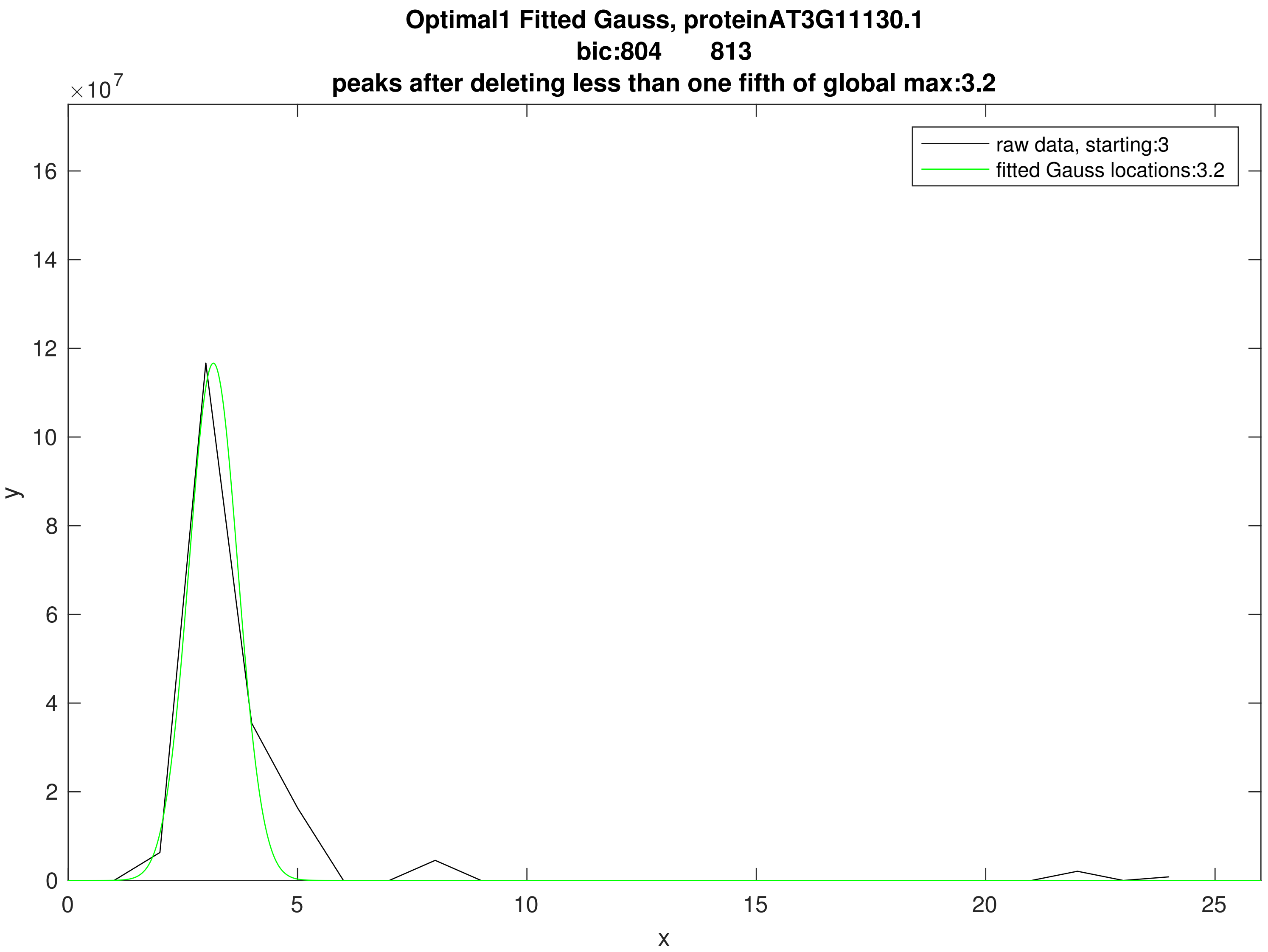

Optimal1 Fitted Gauss, proteinAT3G11130.1
bic:804 813
peaks after deleting less than one fifth of global max:3.2
×10^7
raw data, starting:3
fitted Gauss locations:3.2
16
14
12
10
8
6
4
2
0
y
0
5
10
15
20
25
x

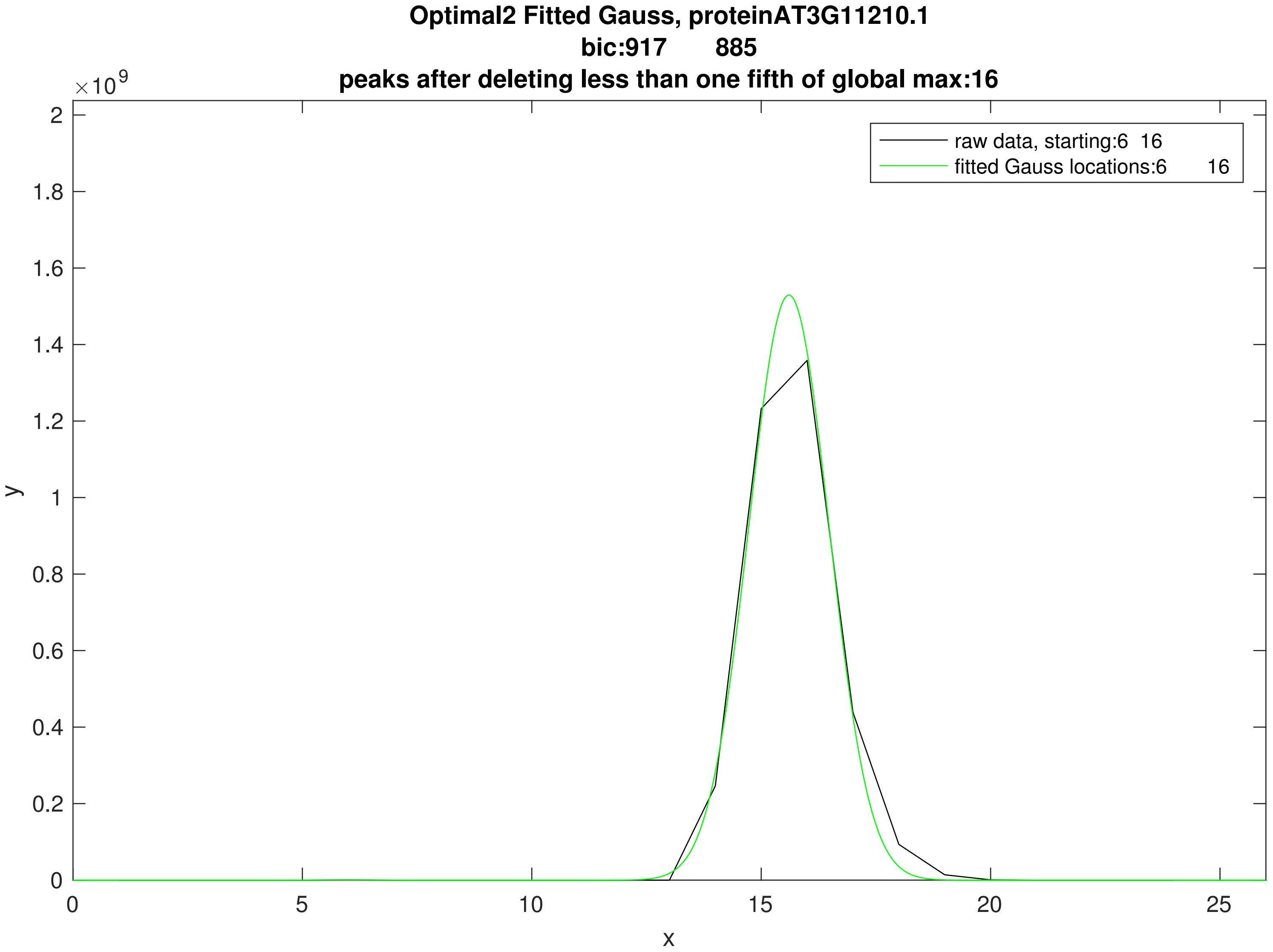
Optimal2 Fitted Gauss, proteinAT3G11210.1
bic:917 885
peaks after deleting less than one fifth of global max:16
raw data, starting:6 16
fitted Gauss locations:6 16
$\times 10^9$
2
1.8
1.6
1.4
1.2
1
0.8
0.6
0.4
0.2
0
y
0
5
10
15
20
25
x

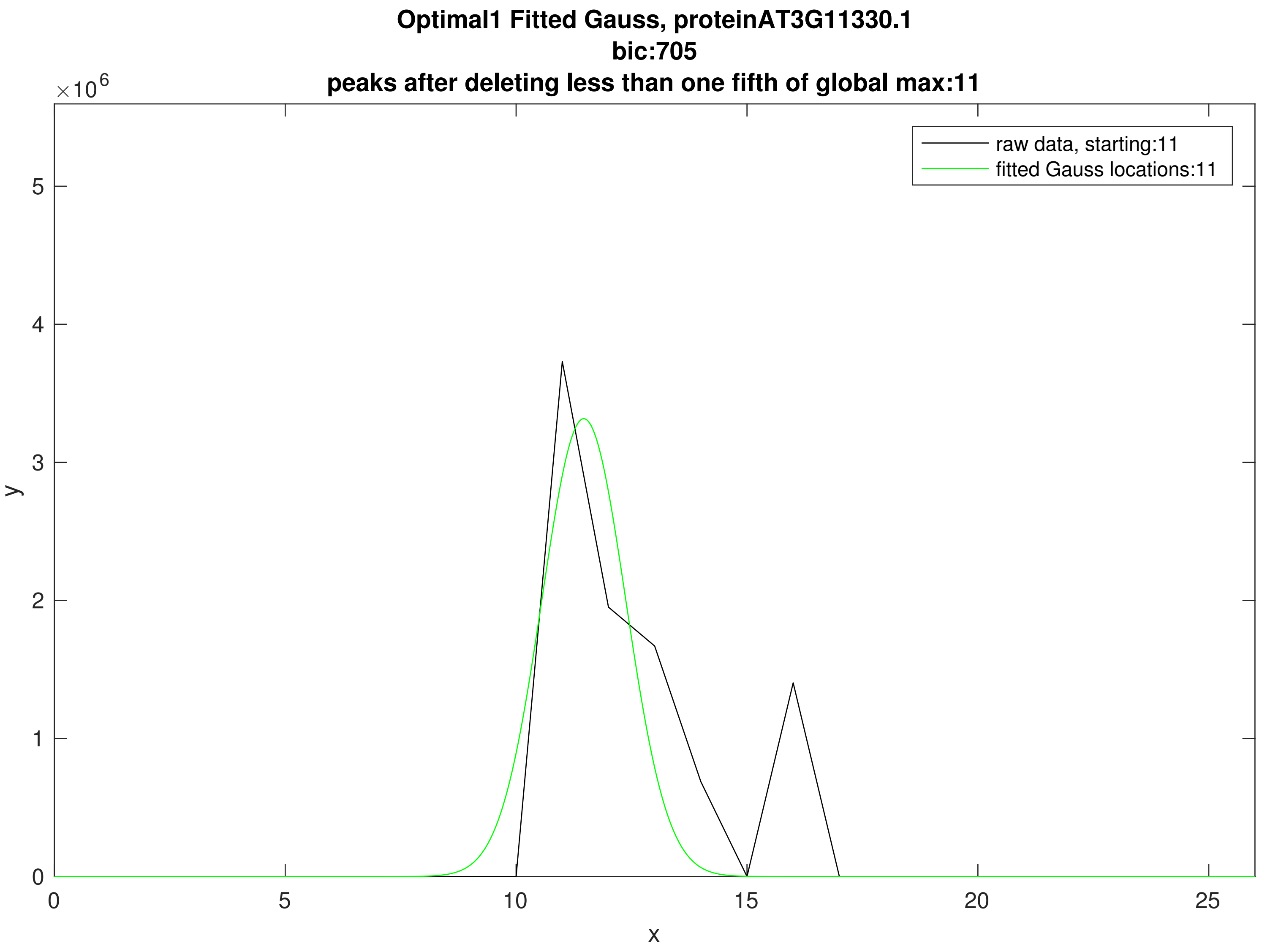

Optimal1 Fitted Gauss, proteinAT3G11330.1
bic:705
peaks after deleting less than one fifth of global max:11
×10^6
raw data, starting:11
fitted Gauss locations:11
5
4
3
2
1
0
y
0
5
10
15
20
25
x

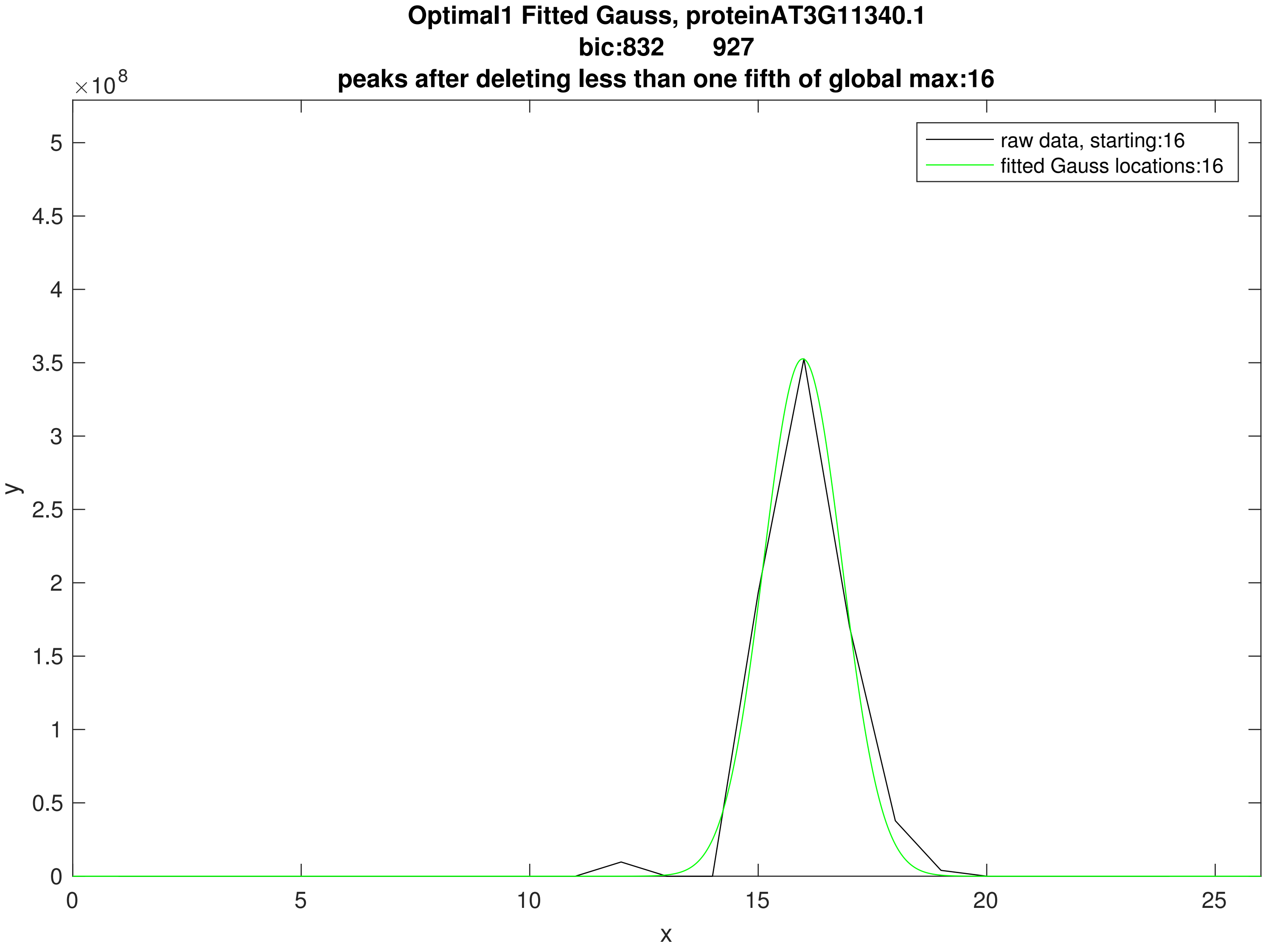
Optimal1 Fitted Gauss, proteinAT3G11340.1
bic:832 927
peaks after deleting less than one fifth of global max:16
×10^8
y
x
raw data, starting:16
fitted Gauss locations:16

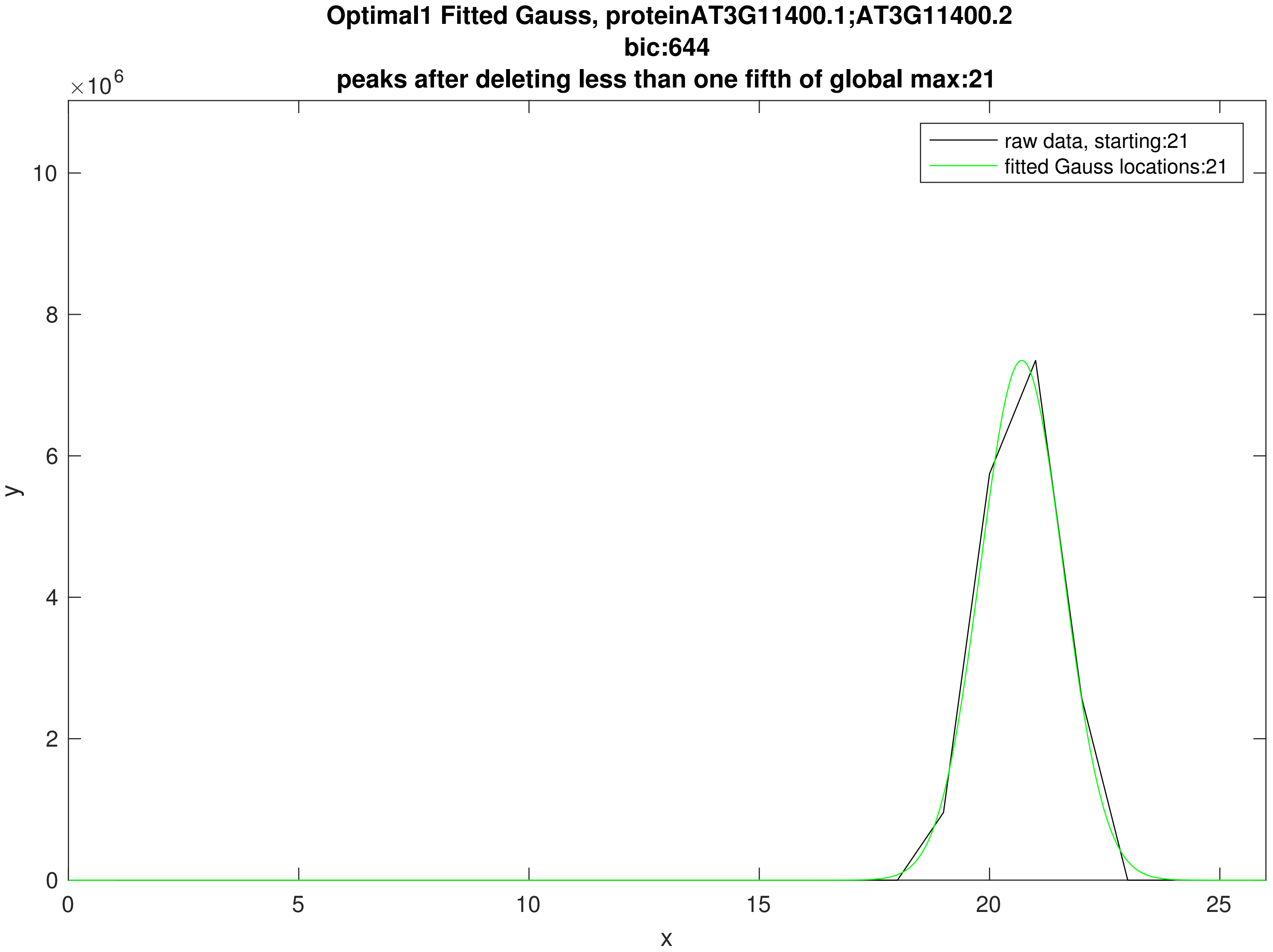

Optimal1 Fitted Gauss, proteinAT3G11400.1;AT3G11400.2
bic:644
peaks after deleting less than one fifth of global max:21
×10^6
raw data, starting:21
fitted Gauss locations:21
y
x
0
2
4
6
8
10
0
5
10
15
20
25

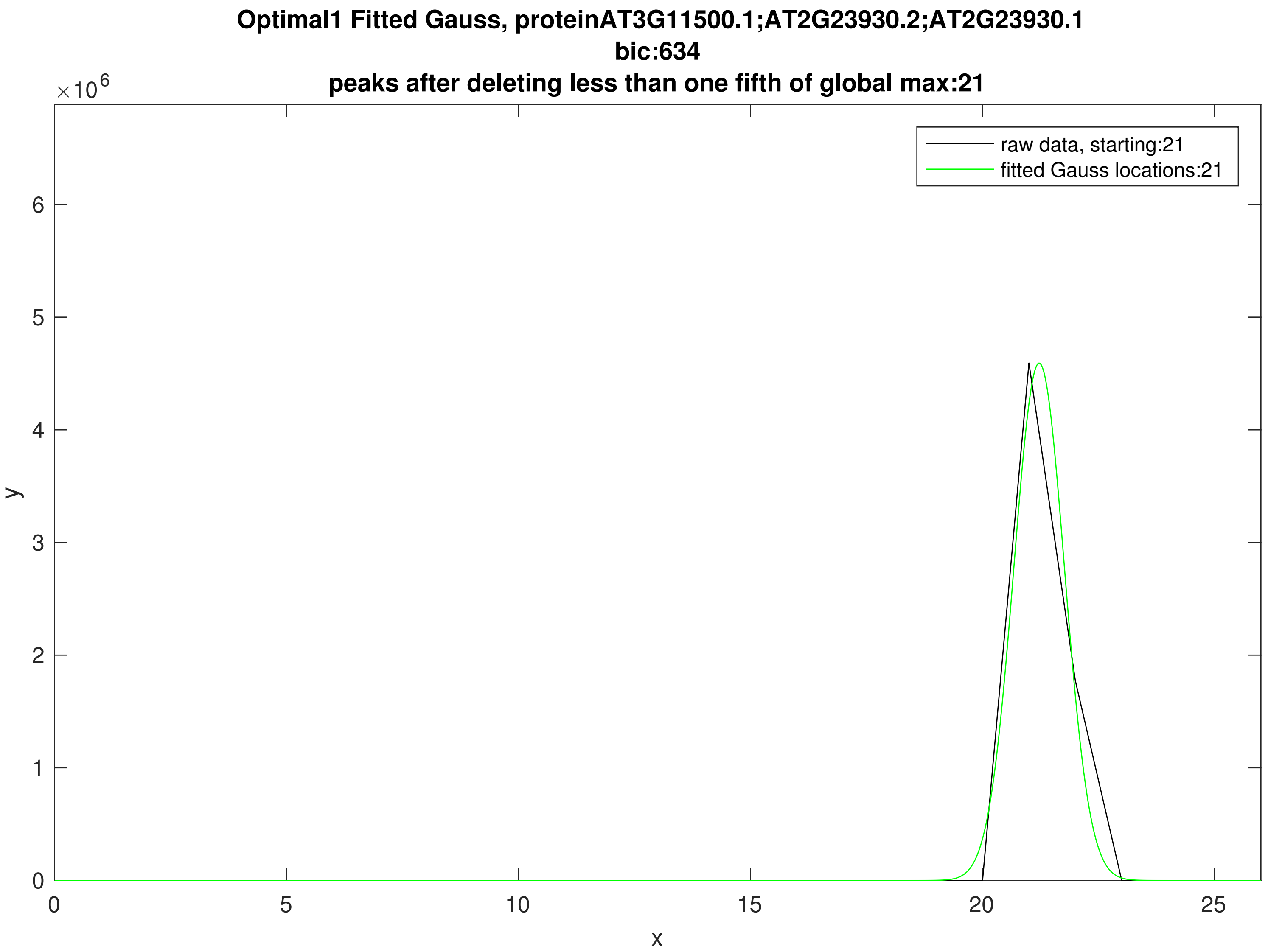
Optimal1 Fitted Gauss, proteinAT3G11500.1;AT2G23930.2;AT2G23930.1
bic:634
peaks after deleting less than one fifth of global max:21
×10^6
raw data, starting:21
fitted Gauss locations:21
y
x
0
1
2
3
4
5
6
0
5
10
15
20
25

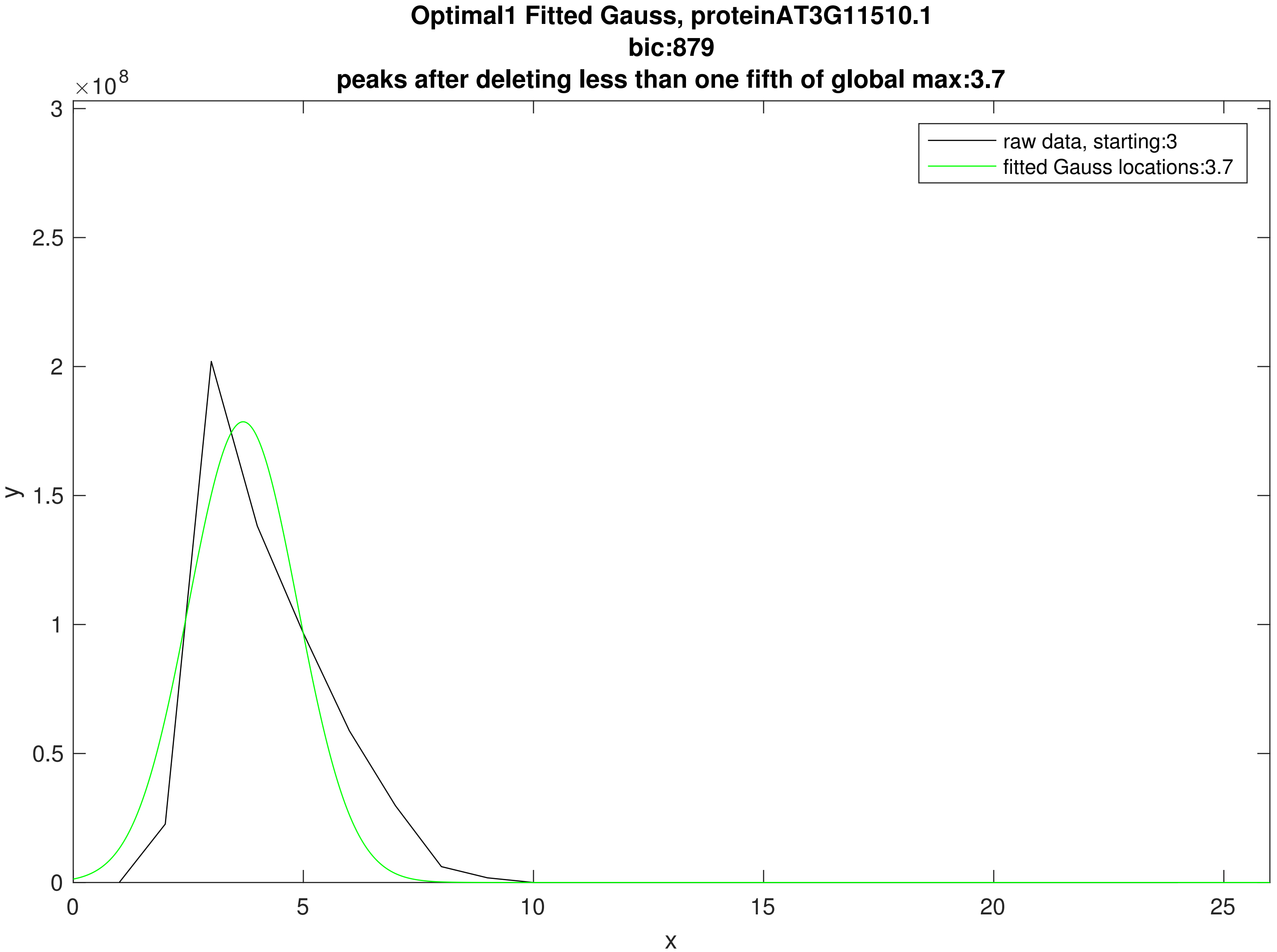
Optimal1 Fitted Gauss, proteinAT3G11510.1
bic:879
peaks after deleting less than one fifth of global max:3.7
×10^8
3
2.5
2
1.5
1
0.5
0
y
0
5
10
15
20
25
x
raw data, starting:3
fitted Gauss locations:3.7

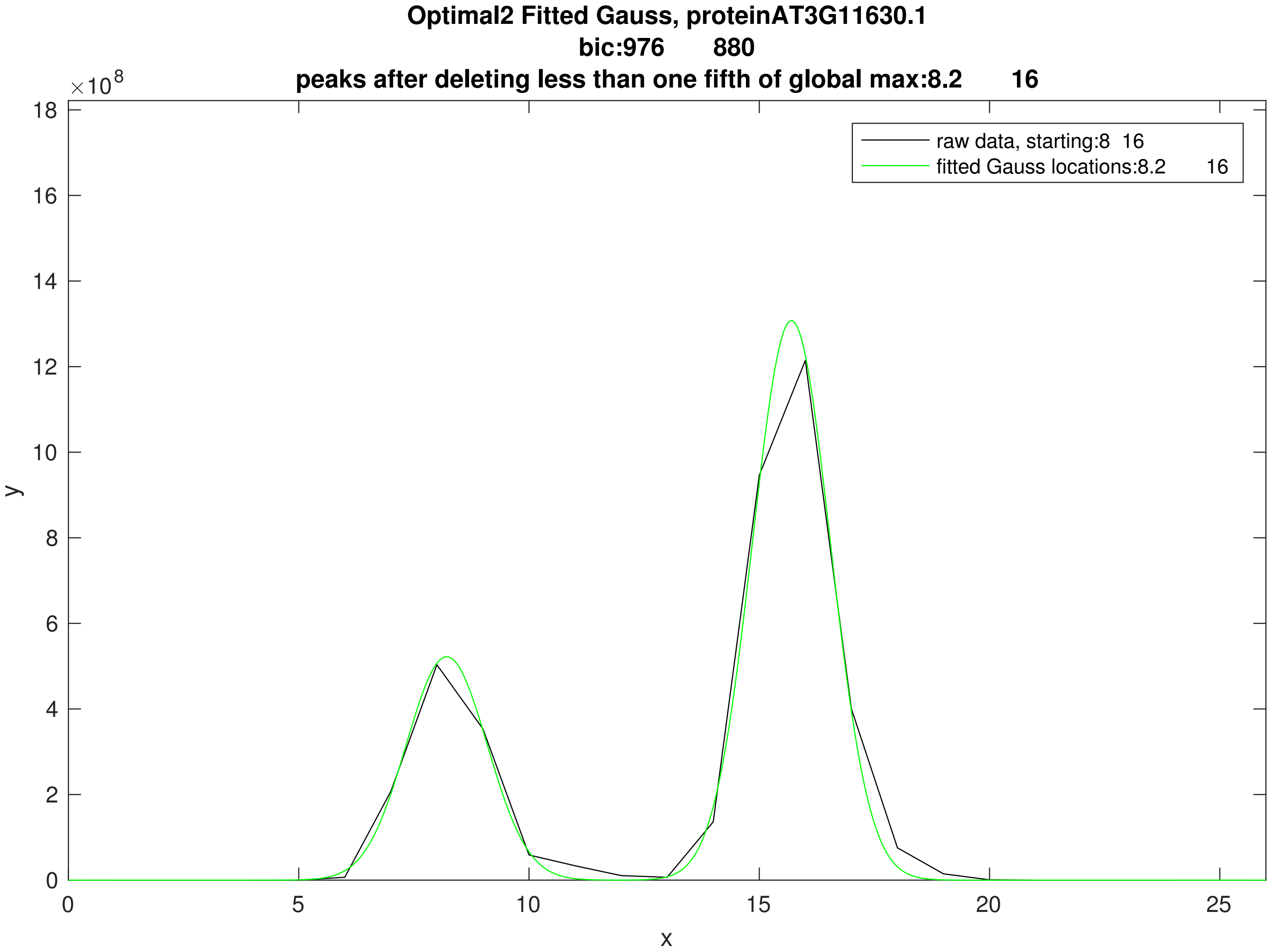
Optimal2 Fitted Gauss, proteinAT3G11630.1
bic:976 880
peaks after deleting less than one fifth of global max:8.2 16
×10^8
raw data, starting:8 16
fitted Gauss locations:8.2 16
y
x
0
2
4
6
8
10
12
14
16
18
5
10
15
20
25

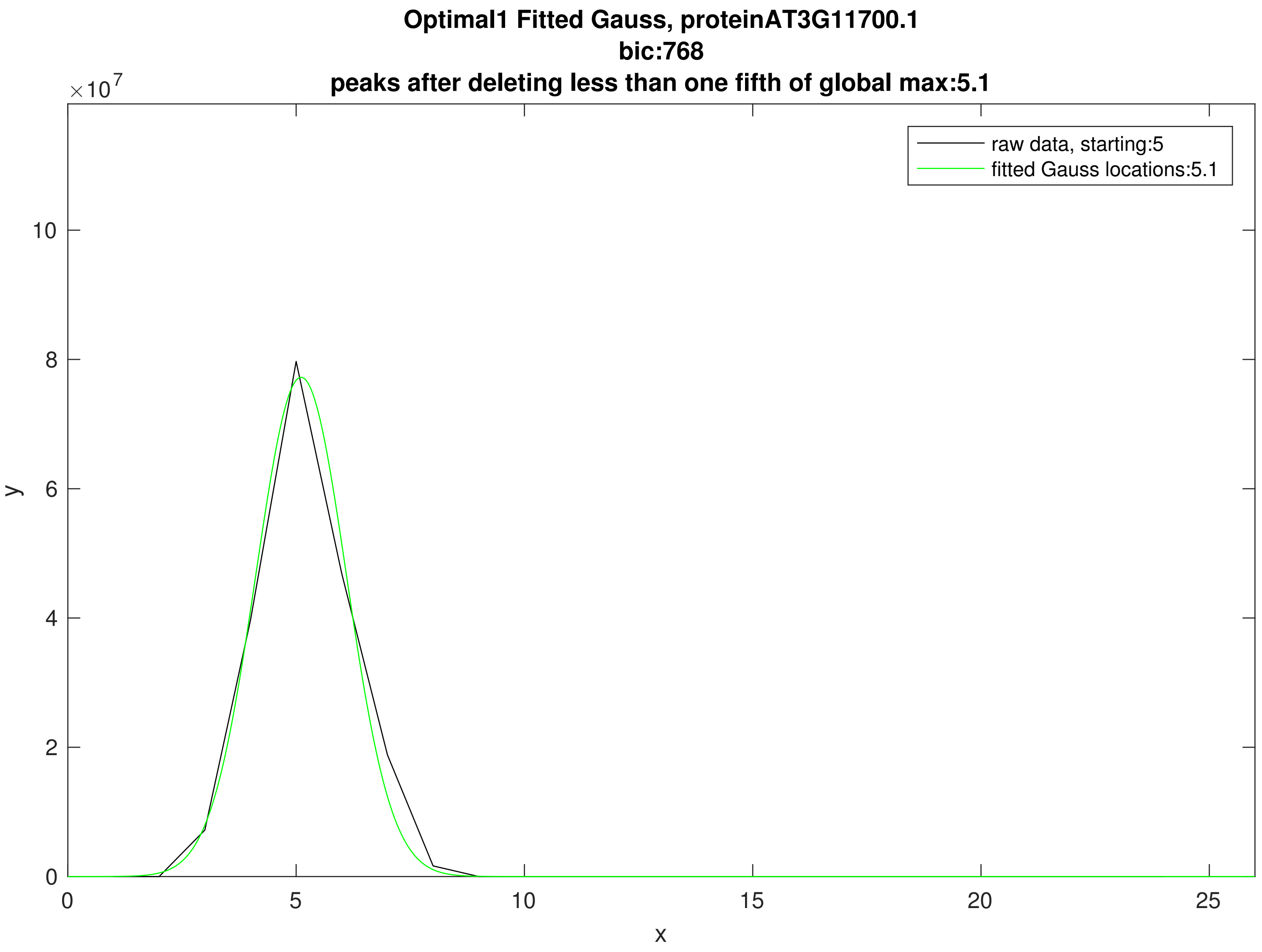

Optimal1 Fitted Gauss, proteinAT3G11700.1
bic:768
peaks after deleting less than one fifth of global max:5.1
×10^7
raw data, starting:5
fitted Gauss locations:5.1
y
x
0
2
4
6
8
10
5
15
20
25

**Optimal1 Fitted Gauss, proteinAT3G11770.1**
**bic:813**
**peaks after deleting less than one fifth of global max:15**

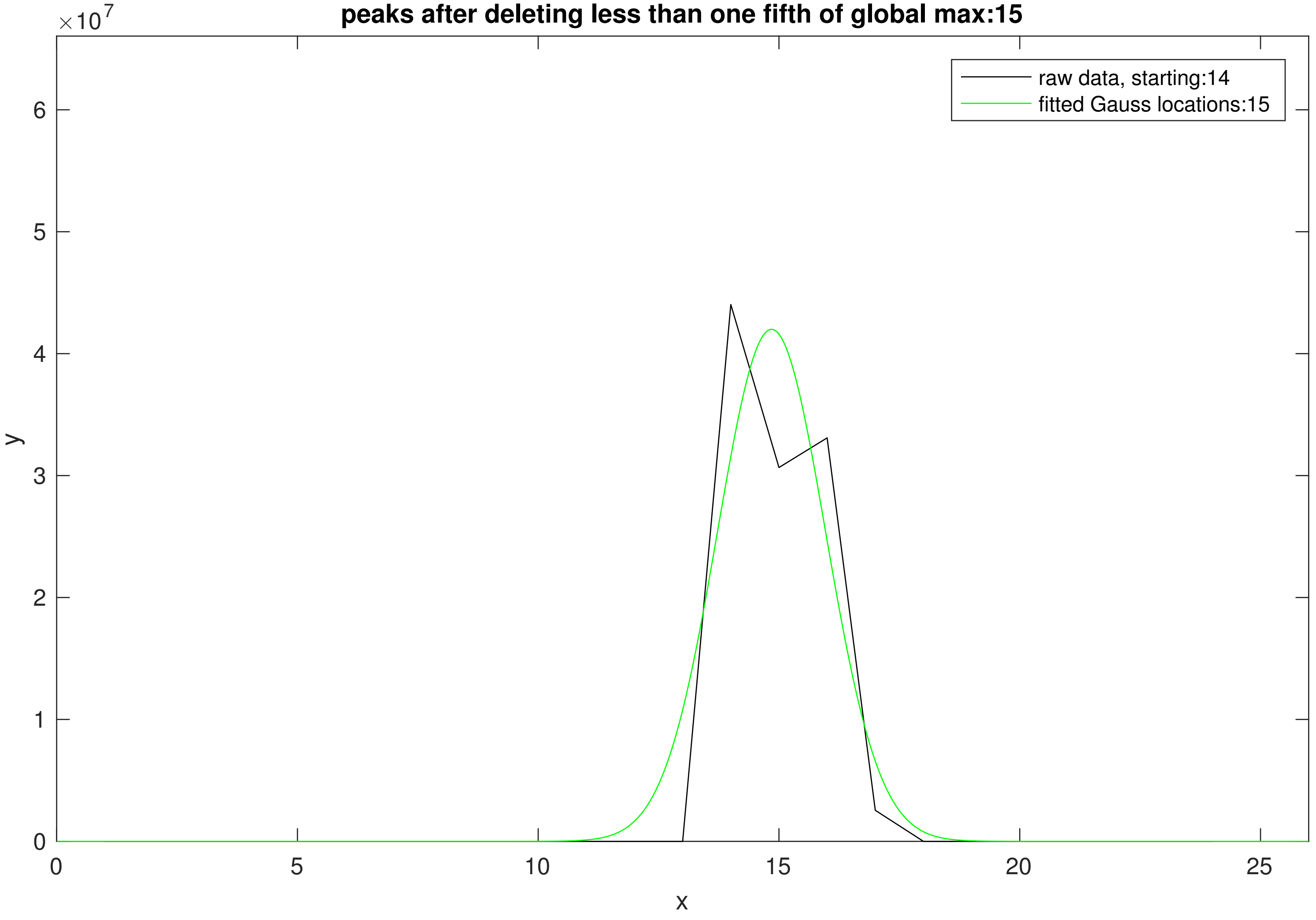

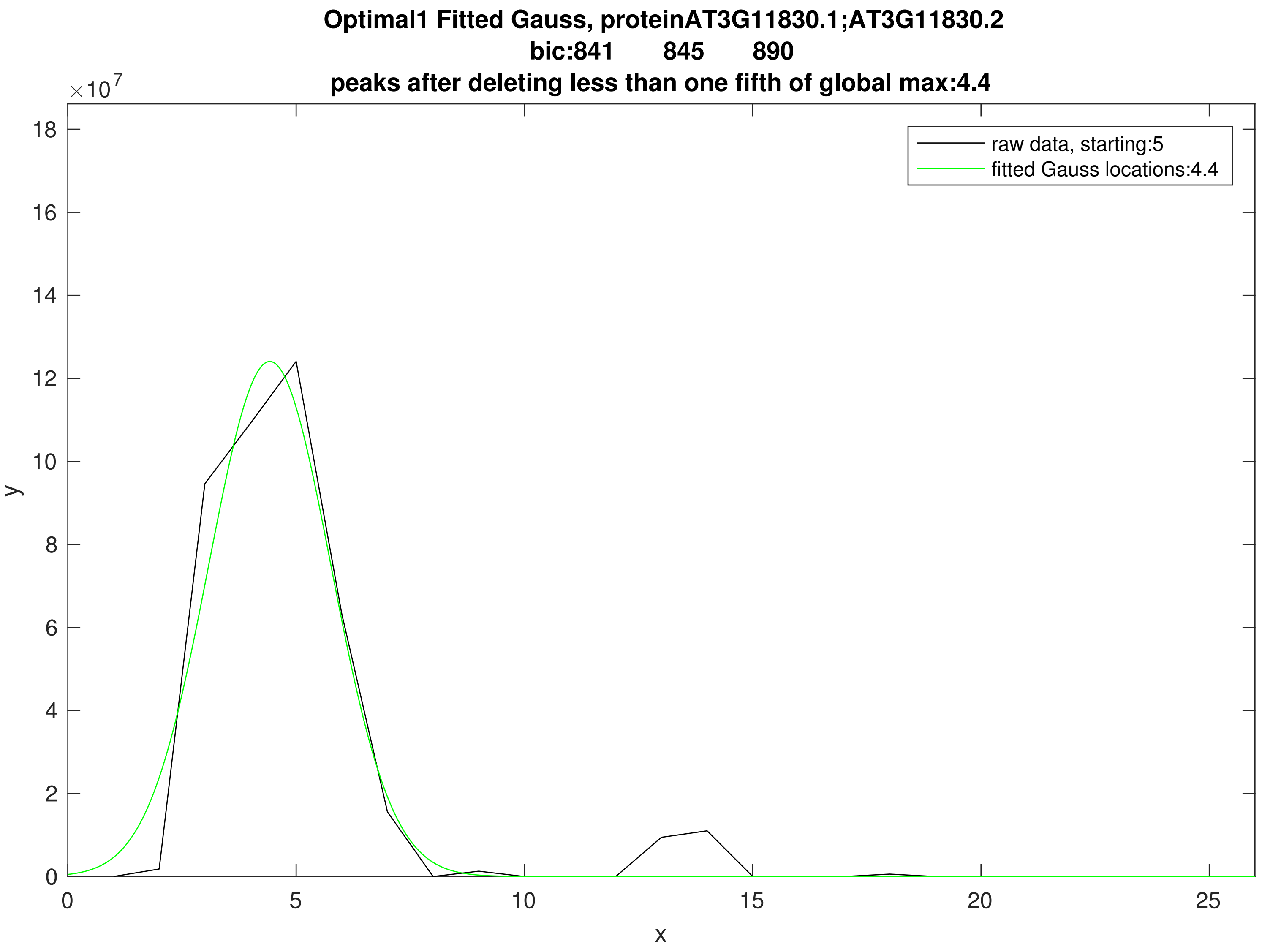
Optimal1 Fitted Gauss, proteinAT3G11830.1;AT3G11830.2
bic:841 845 890
peaks after deleting less than one fifth of global max:4.4
×10^7
raw data, starting:5
fitted Gauss locations:4.4
y
x
0
2
4
6
8
10
12
14
16
18
0
5
10
15
20
25

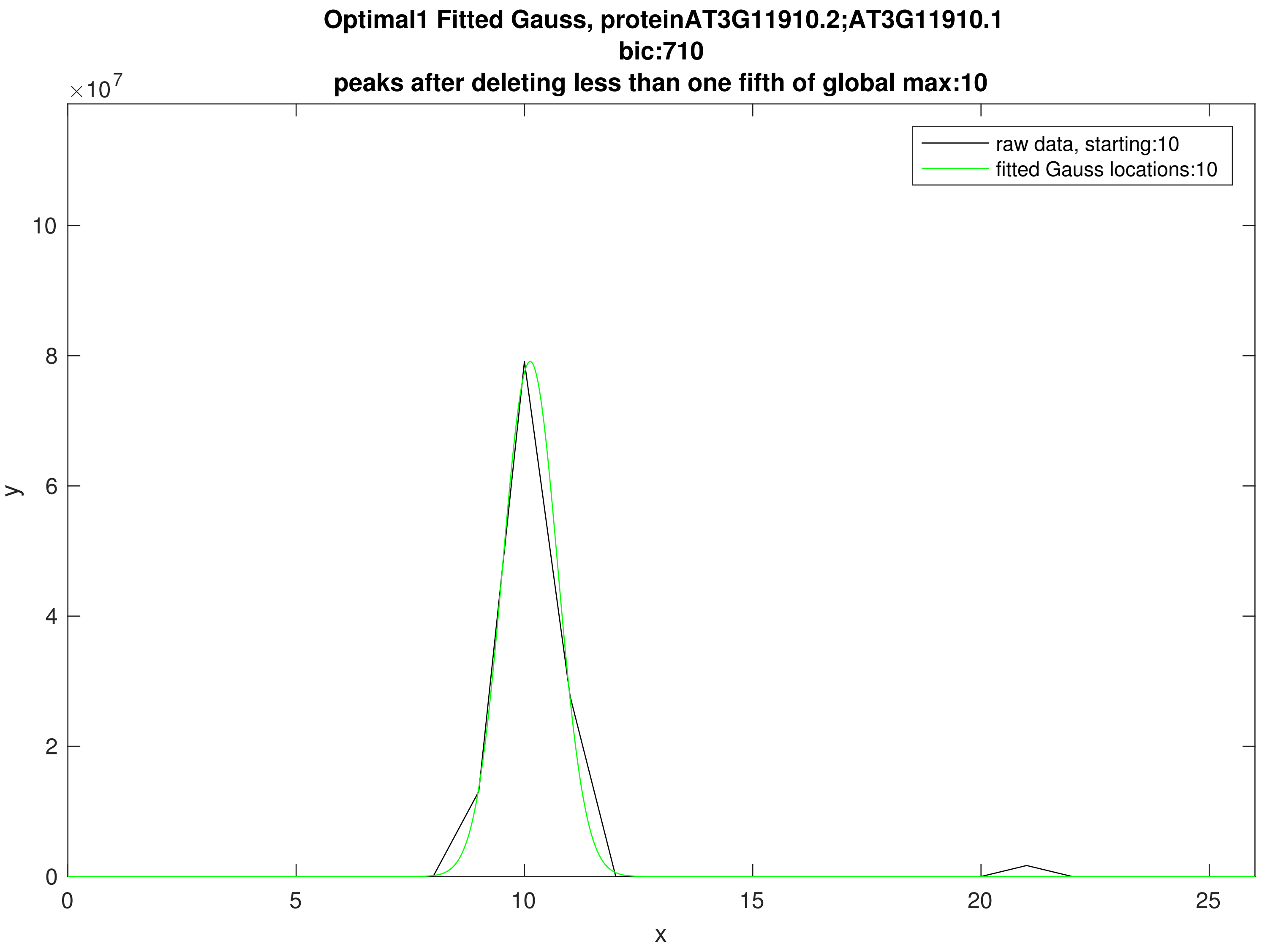
Optimal1 Fitted Gauss, proteinAT3G11910.2;AT3G11910.1
bic:710
peaks after deleting less than one fifth of global max:10
×10^7
raw data, starting:10
fitted Gauss locations:10
y
x
0
2
4
6
8
10
0
5
10
15
20
25

**Optimal1 Fitted Gauss, proteinAT3G11940.2;AT3G11940.1**
**bic:830**
**peaks after deleting less than one fifth of global max:3.7**

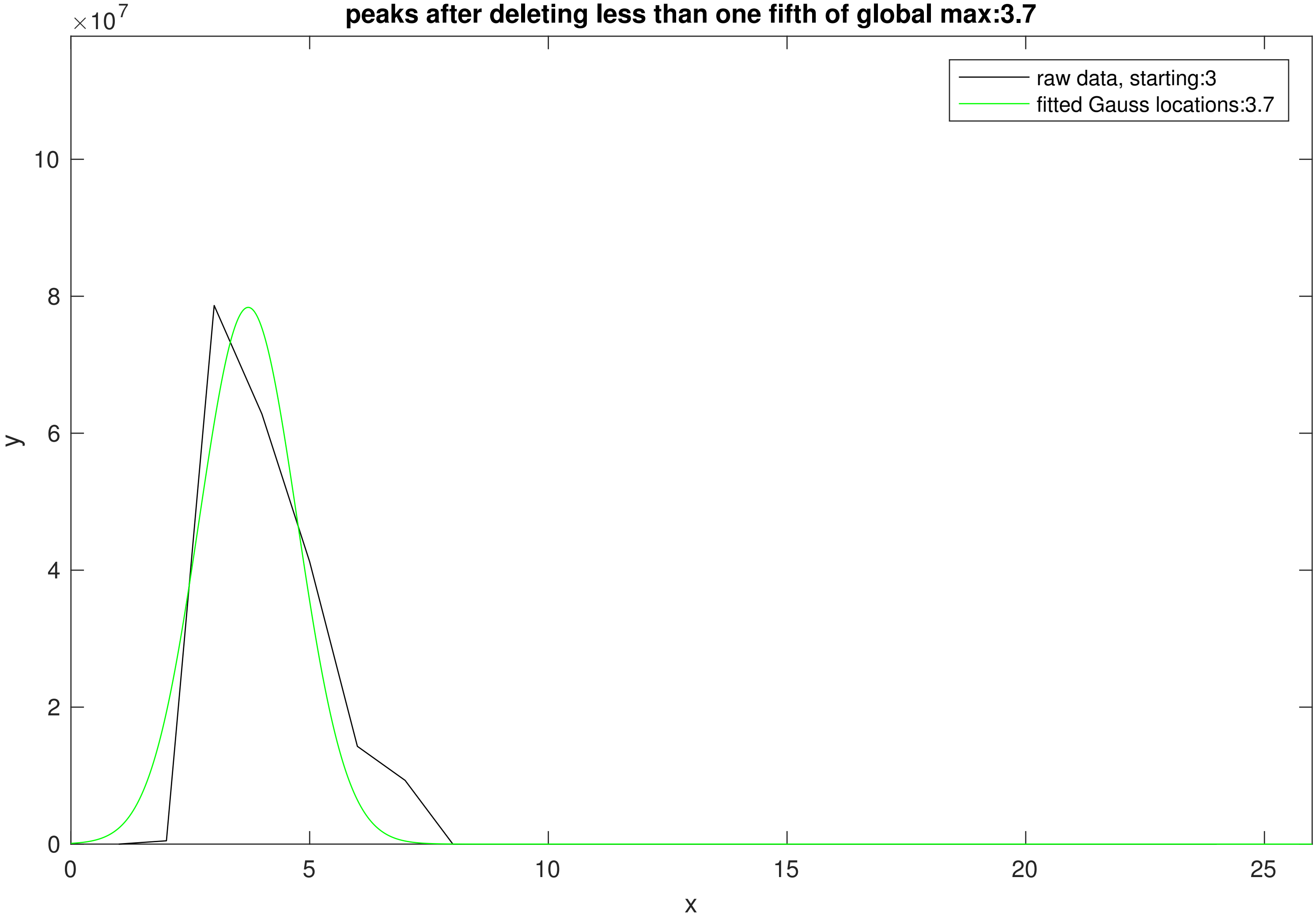

**Optimal2 Fitted Gauss, proteinAT3G12110.1**
**bic:822 813 818 823**
**peaks after deleting less than one fifth of global max:14**

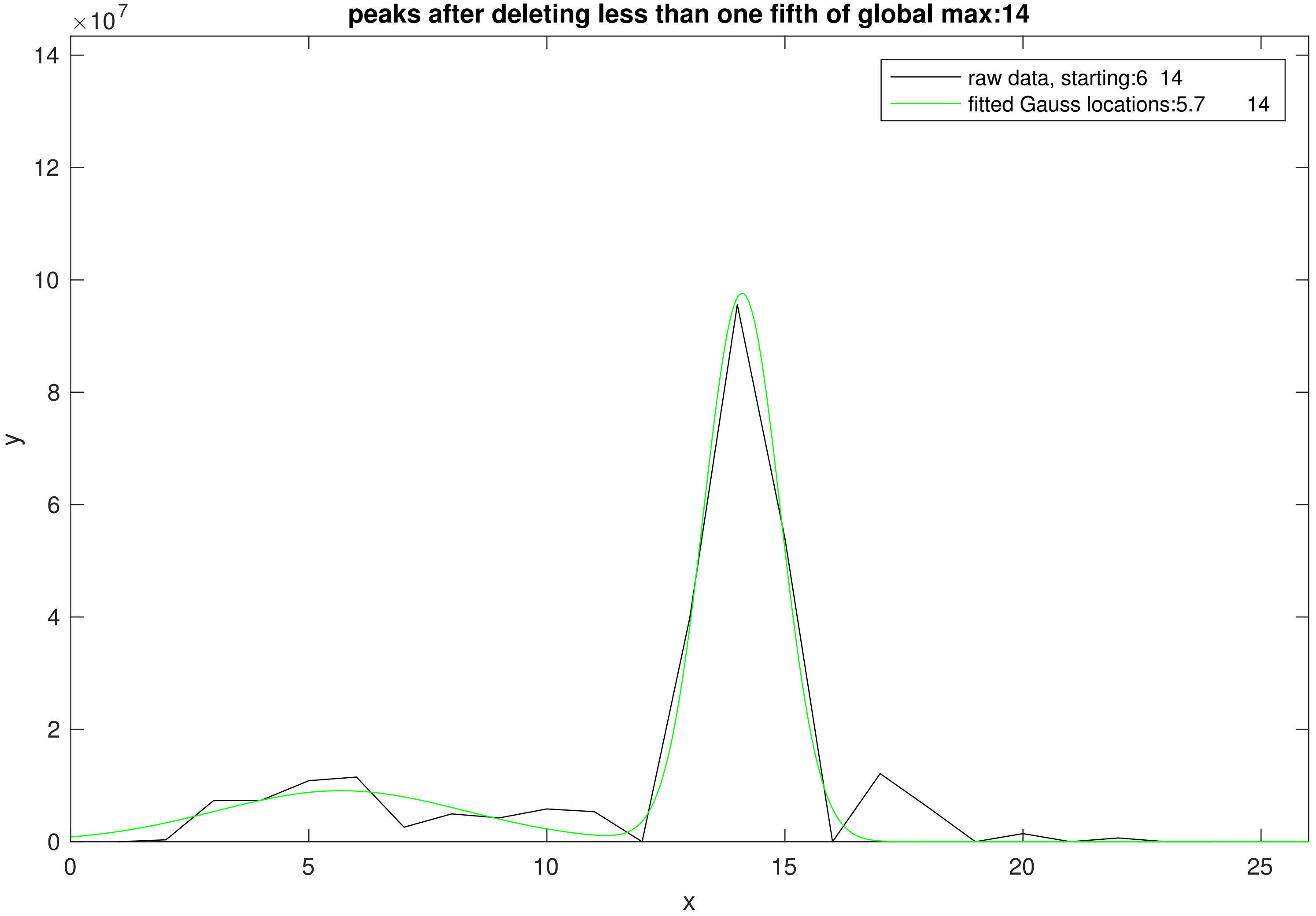

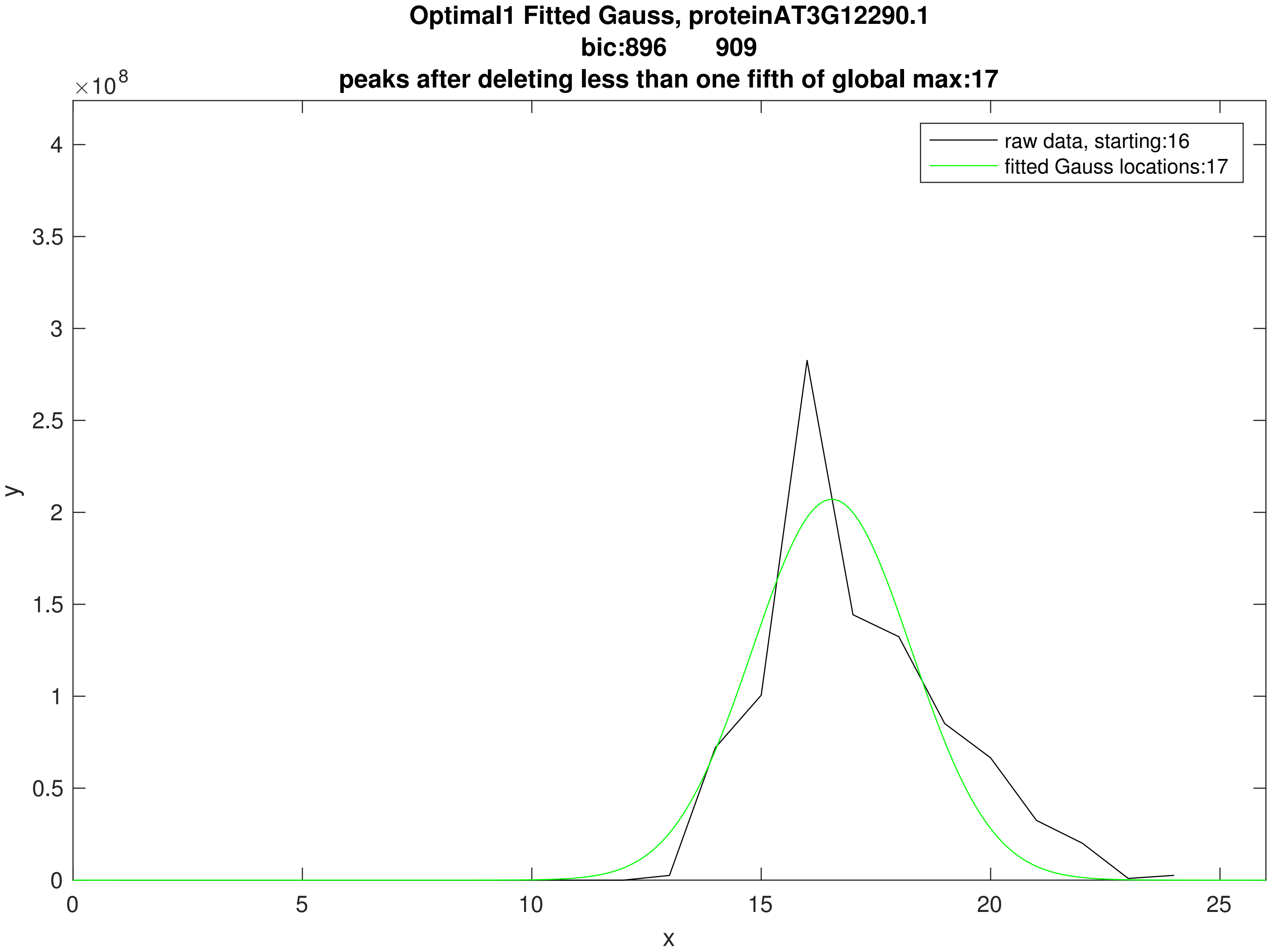
Optimal1 Fitted Gauss, proteinAT3G12290.1
bic:896 909
peaks after deleting less than one fifth of global max:17
×10^8
raw data, starting:16
fitted Gauss locations:17
y
x
0
0.5
1
1.5
2
2.5
3
3.5
4
0
5
10
15
20
25

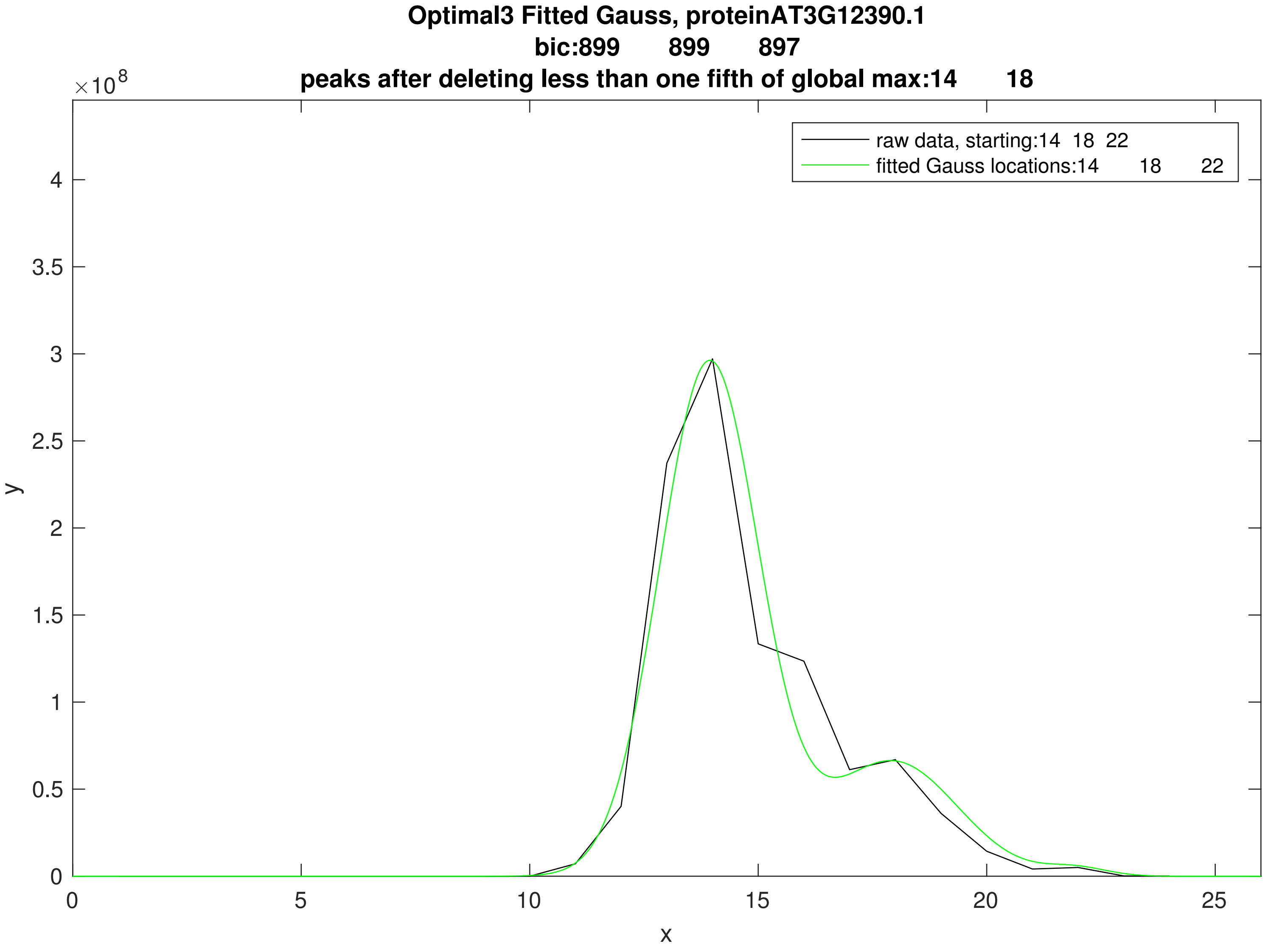
Optimal3 Fitted Gauss, proteinAT3G12390.1
bic:899 899 897
peaks after deleting less than one fifth of global max:14 18
raw data, starting:14 18 22
fitted Gauss locations:14 18 22
×10^8
y
x

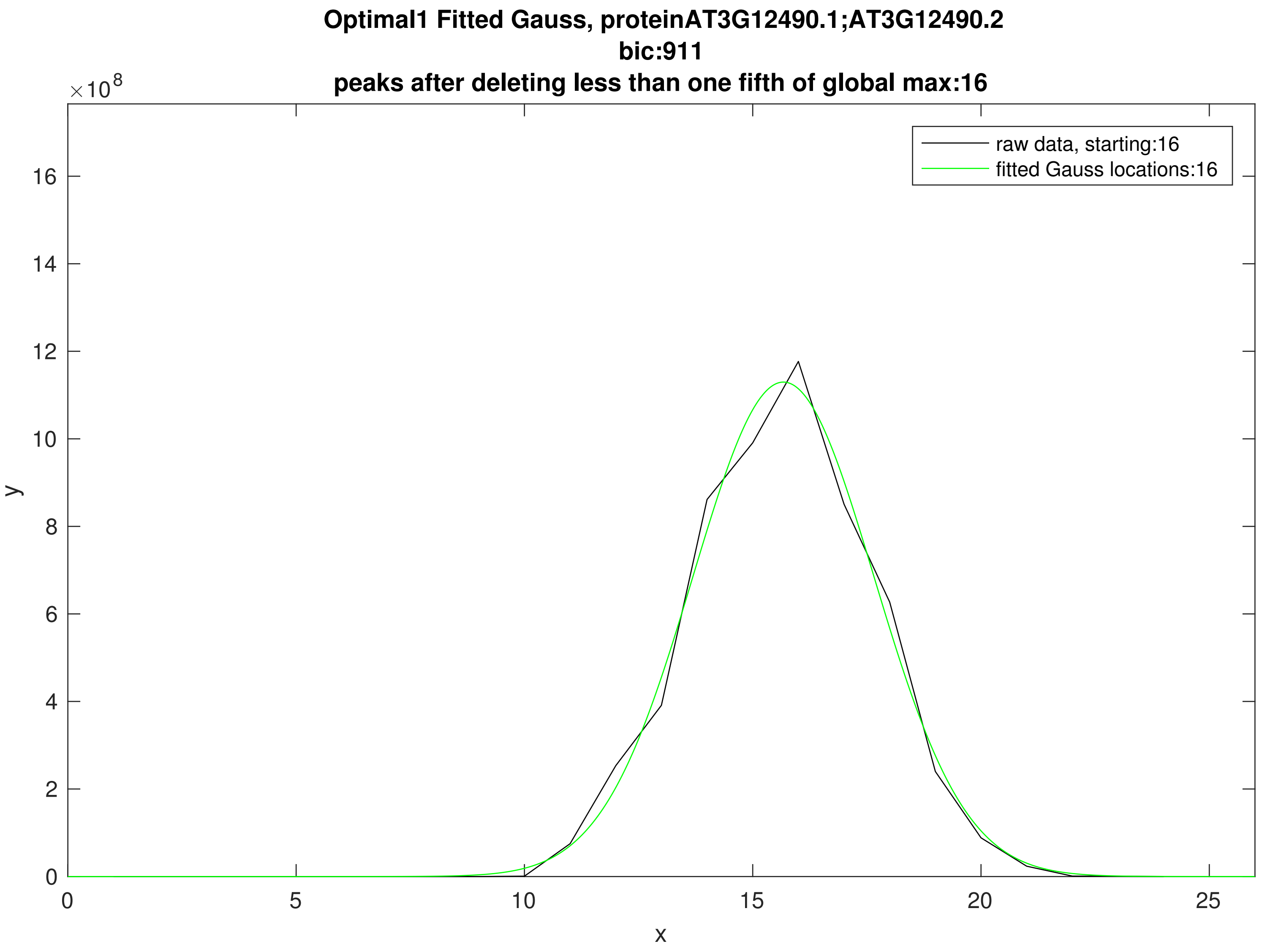

Optimal1 Fitted Gauss, proteinAT3G12490.1;AT3G12490.2
bic:911
peaks after deleting less than one fifth of global max:16
×10^8
raw data, starting:16
fitted Gauss locations:16
y
x
0
2
4
6
8
10
12
14
16
5
15
20
25

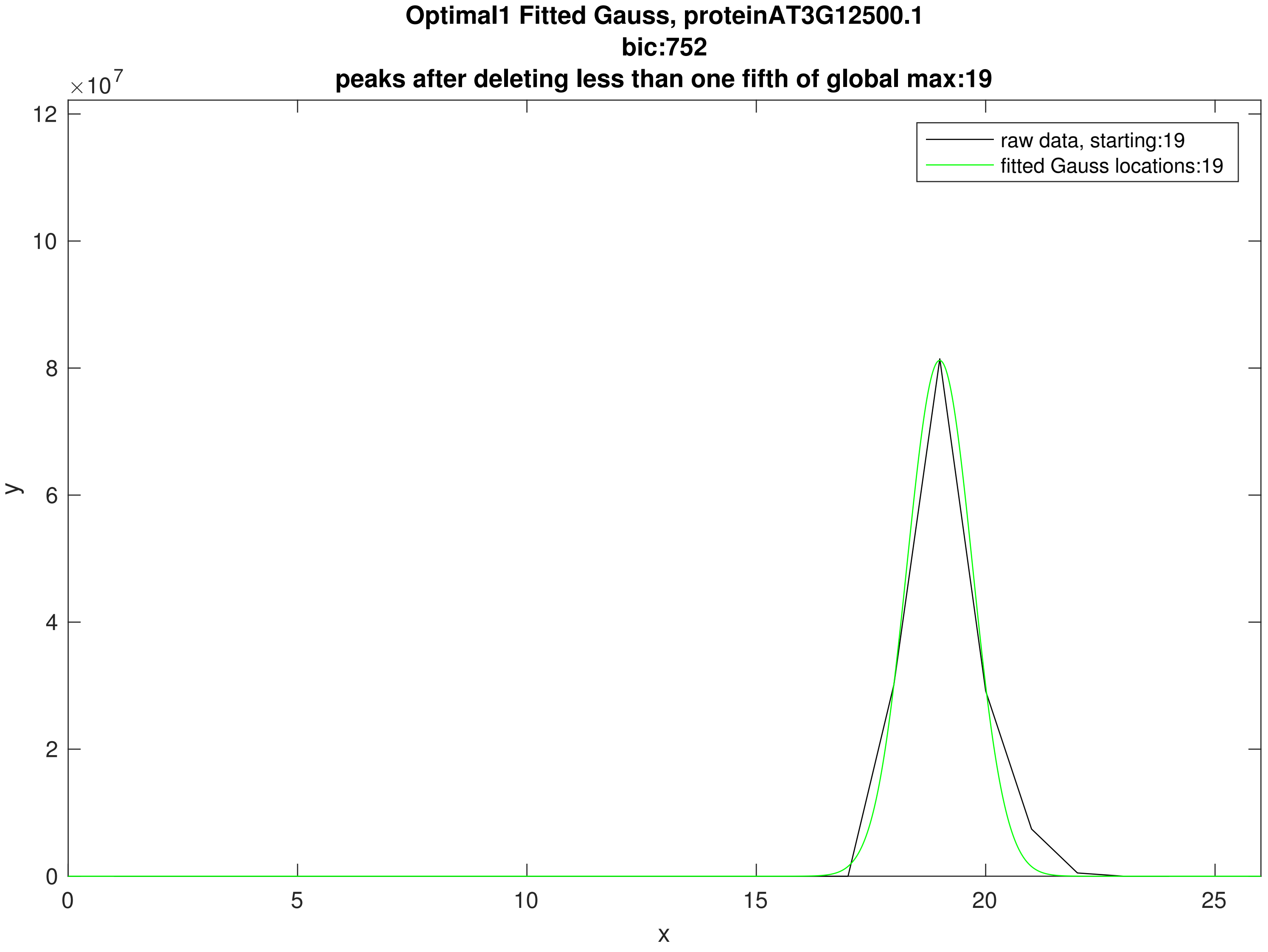

Optimal1 Fitted Gauss, proteinAT3G12500.1
bic:752
peaks after deleting less than one fifth of global max:19
×10^7
12
10
8
6
4
2
0
y
0
5
10
15
20
25
x
raw data, starting:19
fitted Gauss locations:19

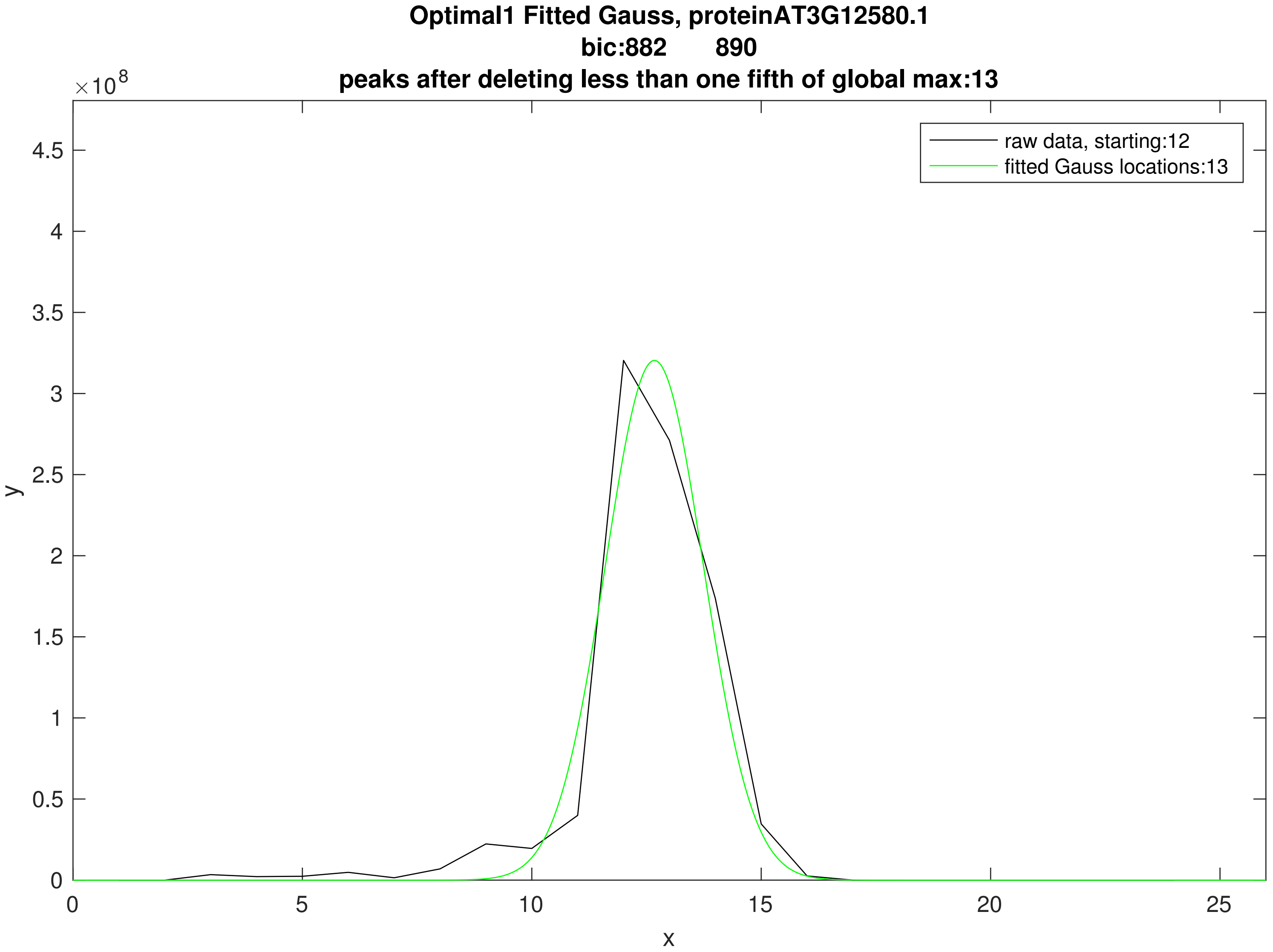
Optimal1 Fitted Gauss, proteinAT3G12580.1
bic:882 890
peaks after deleting less than one fifth of global max:13
×10^8
raw data, starting:12
fitted Gauss locations:13
y
x
0
0.5
1
1.5
2
2.5
3
3.5
4
4.5
0
5
10
15
20
25

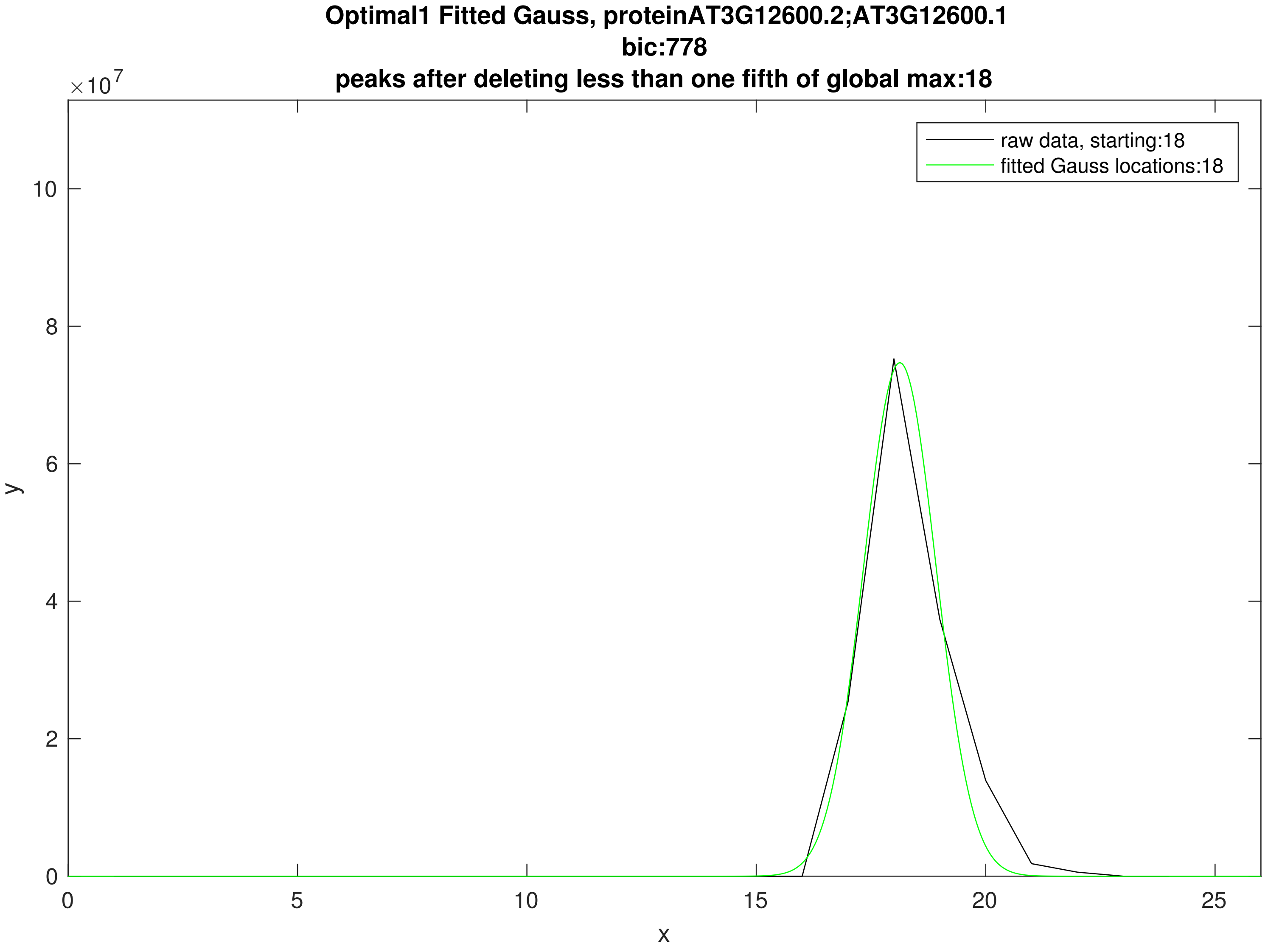
Optimal1 Fitted Gauss, proteinAT3G12600.2;AT3G12600.1
bic:778
peaks after deleting less than one fifth of global max:18
×10^7
raw data, starting:18
fitted Gauss locations:18
10
8
6
4
2
0
y
0
5
10
15
20
25
x

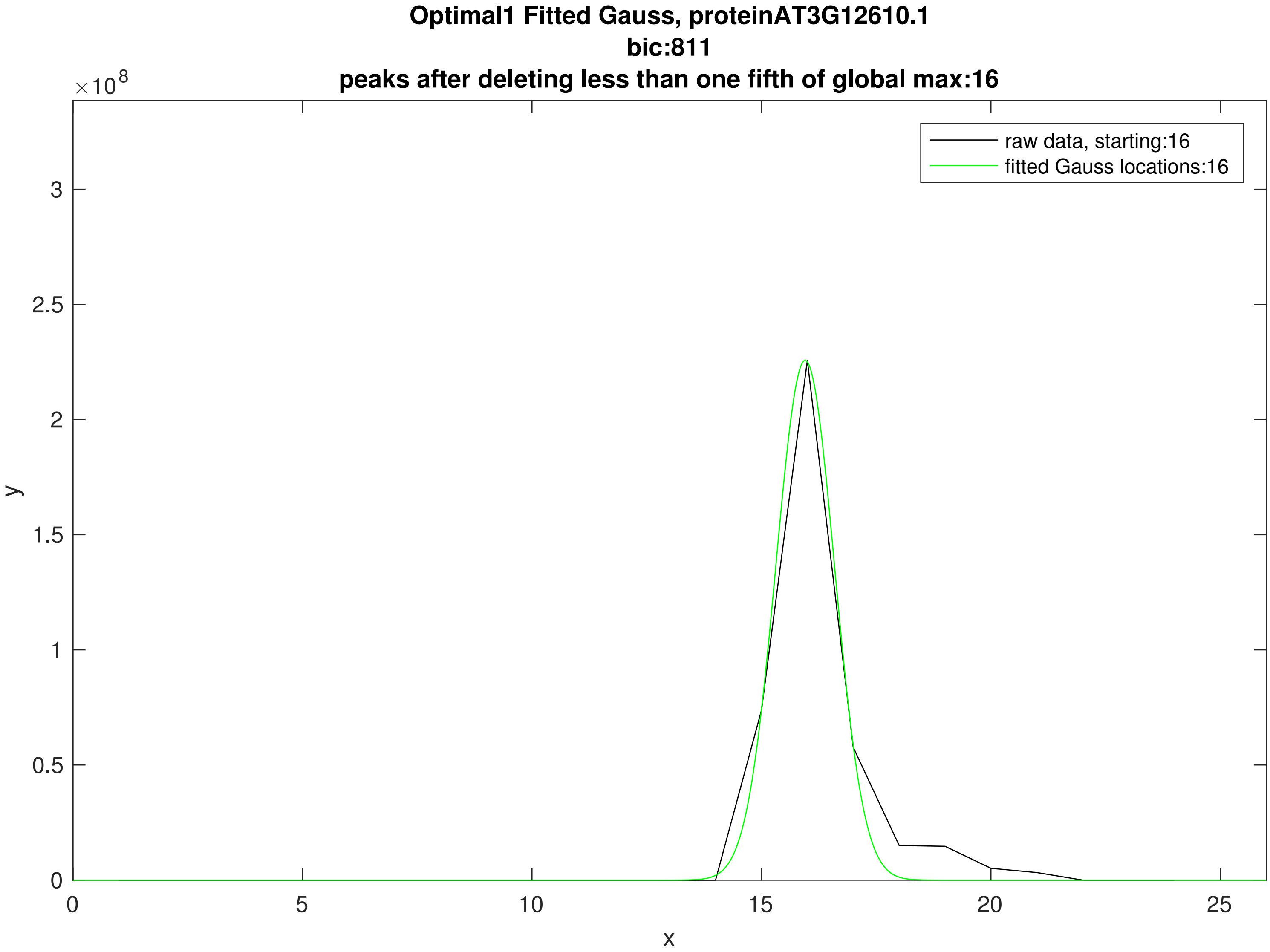

Optimal1 Fitted Gauss, proteinAT3G12610.1
bic:811
peaks after deleting less than one fifth of global max:16
×10^8
raw data, starting:16
fitted Gauss locations:16
3
2.5
2
1.5
1
0.5
0
y
0
5
10
15
20
25
x

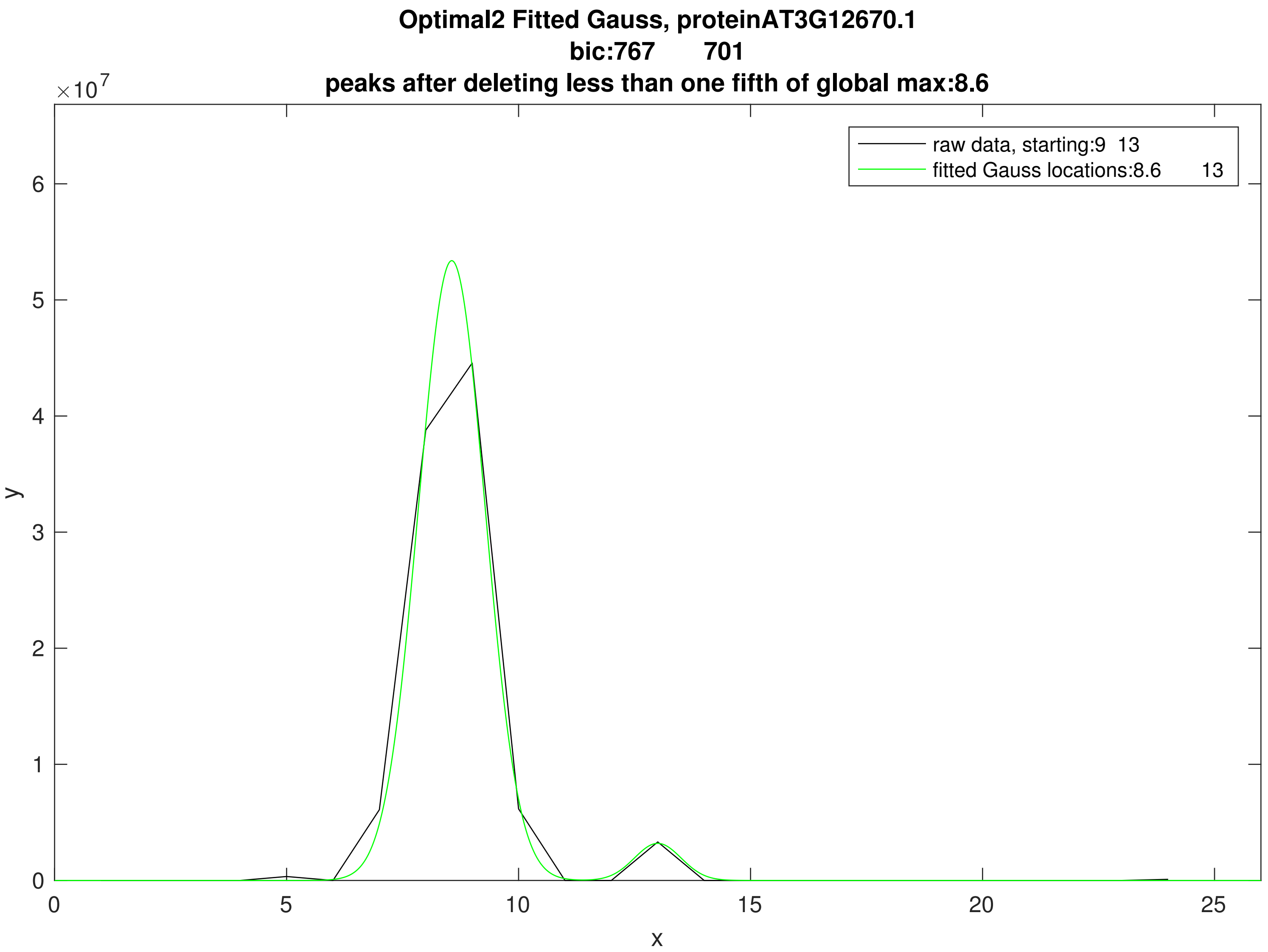

Optimal2 Fitted Gauss, proteinAT3G12670.1
bic:767 701
peaks after deleting less than one fifth of global max:8.6
×10^7
raw data, starting:9 13
fitted Gauss locations:8.6 13
y
x
0
1
2
3
4
5
6
0
5
10
15
20
25

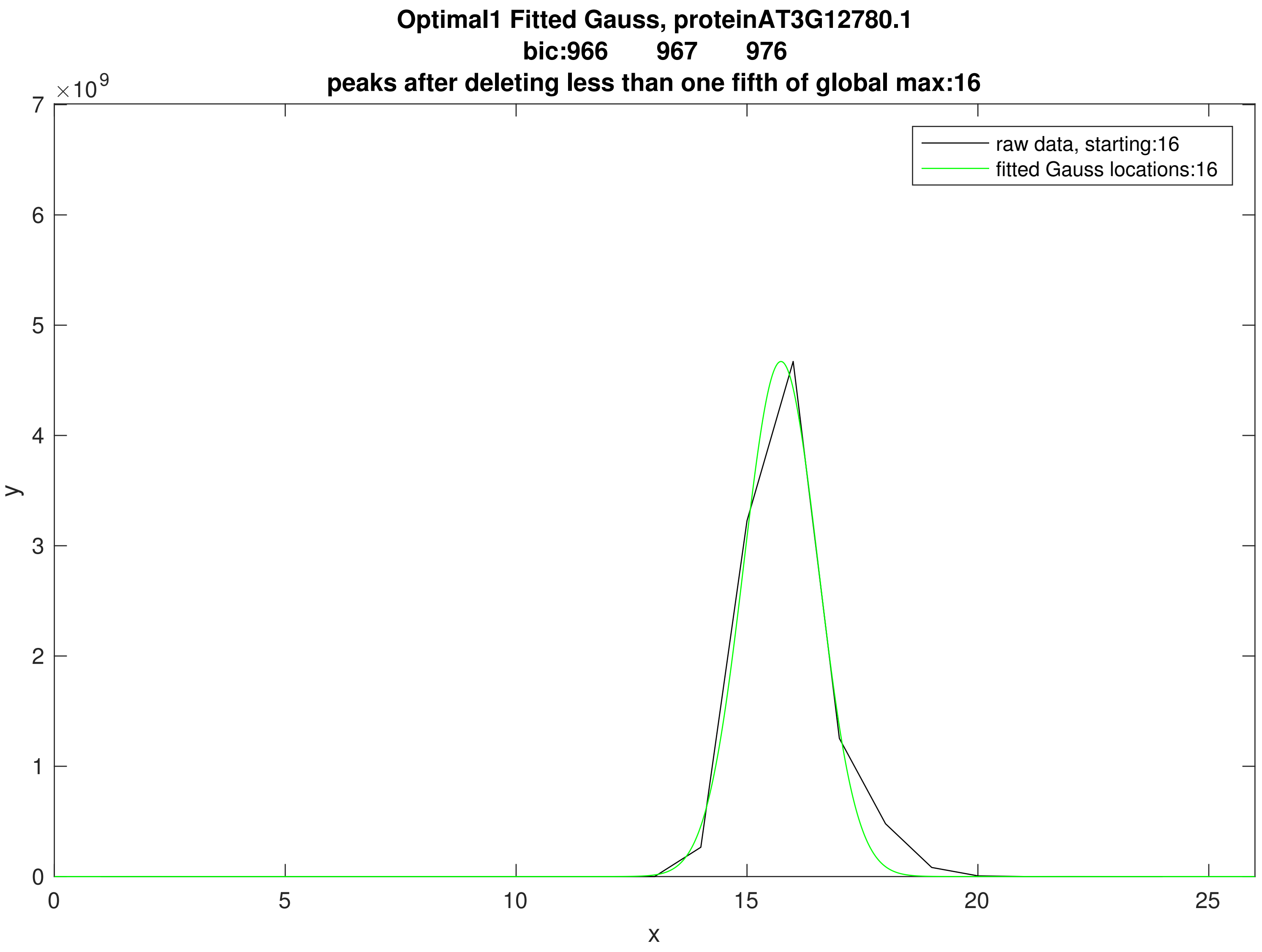

Optimal1 Fitted Gauss, proteinAT3G12780.1
bic:966 967 976
peaks after deleting less than one fifth of global max:16
×10^9
7
6
5
4
3
2
1
0
y
0
5
10
15
20
25
x
raw data, starting:16
fitted Gauss locations:16

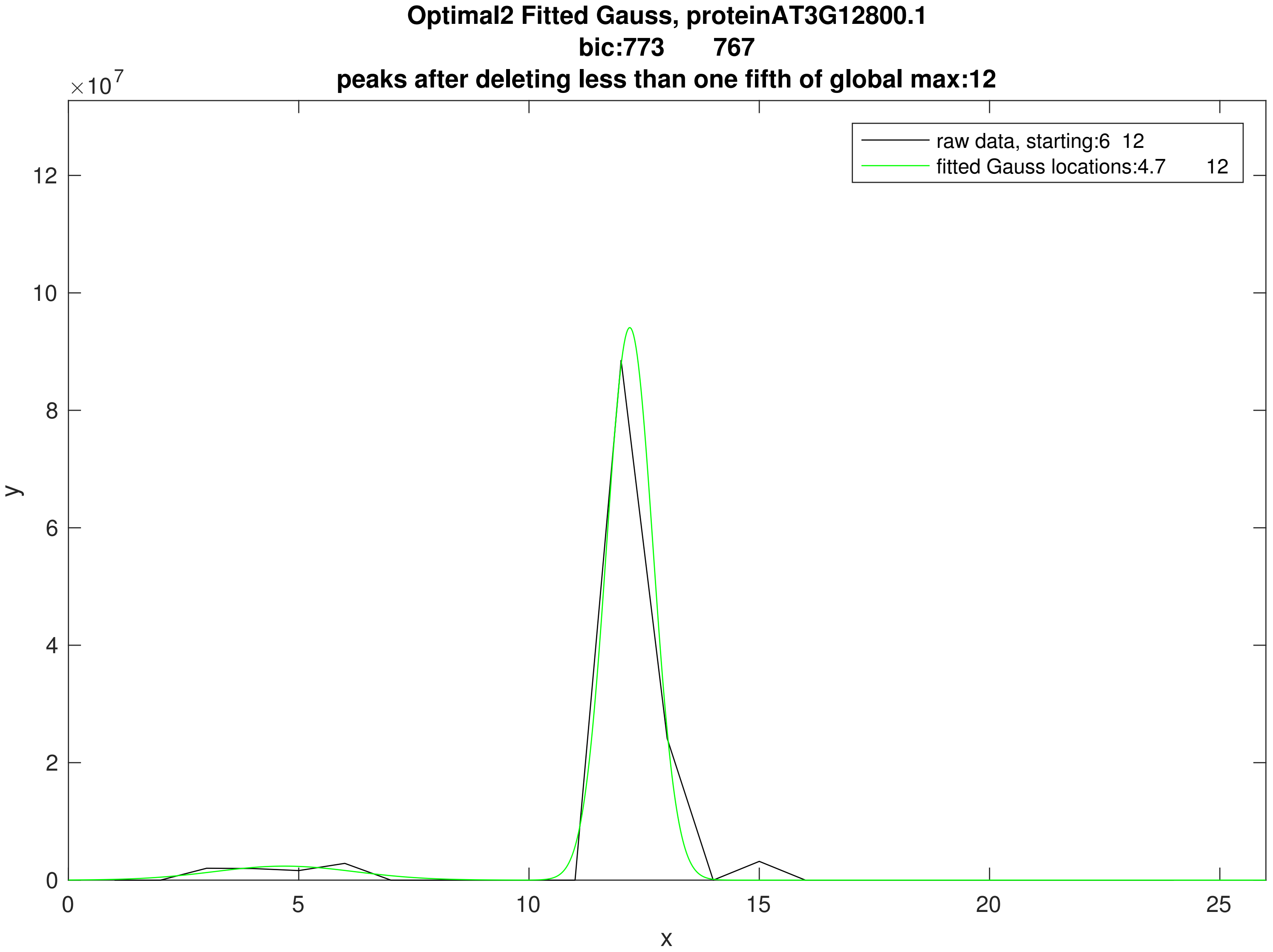

Optimal2 Fitted Gauss, proteinAT3G12800.1
bic:773 767
peaks after deleting less than one fifth of global max:12
×10^7
raw data, starting:6 12
fitted Gauss locations:4.7 12
y
x
0
2
4
6
8
10
12
5
15
20
25

**Optimal1 Fitted Gauss, proteinAT3G12915.1;AT3G12915.2**
**bic:812 820**
**peaks after deleting less than one fifth of global max:13**

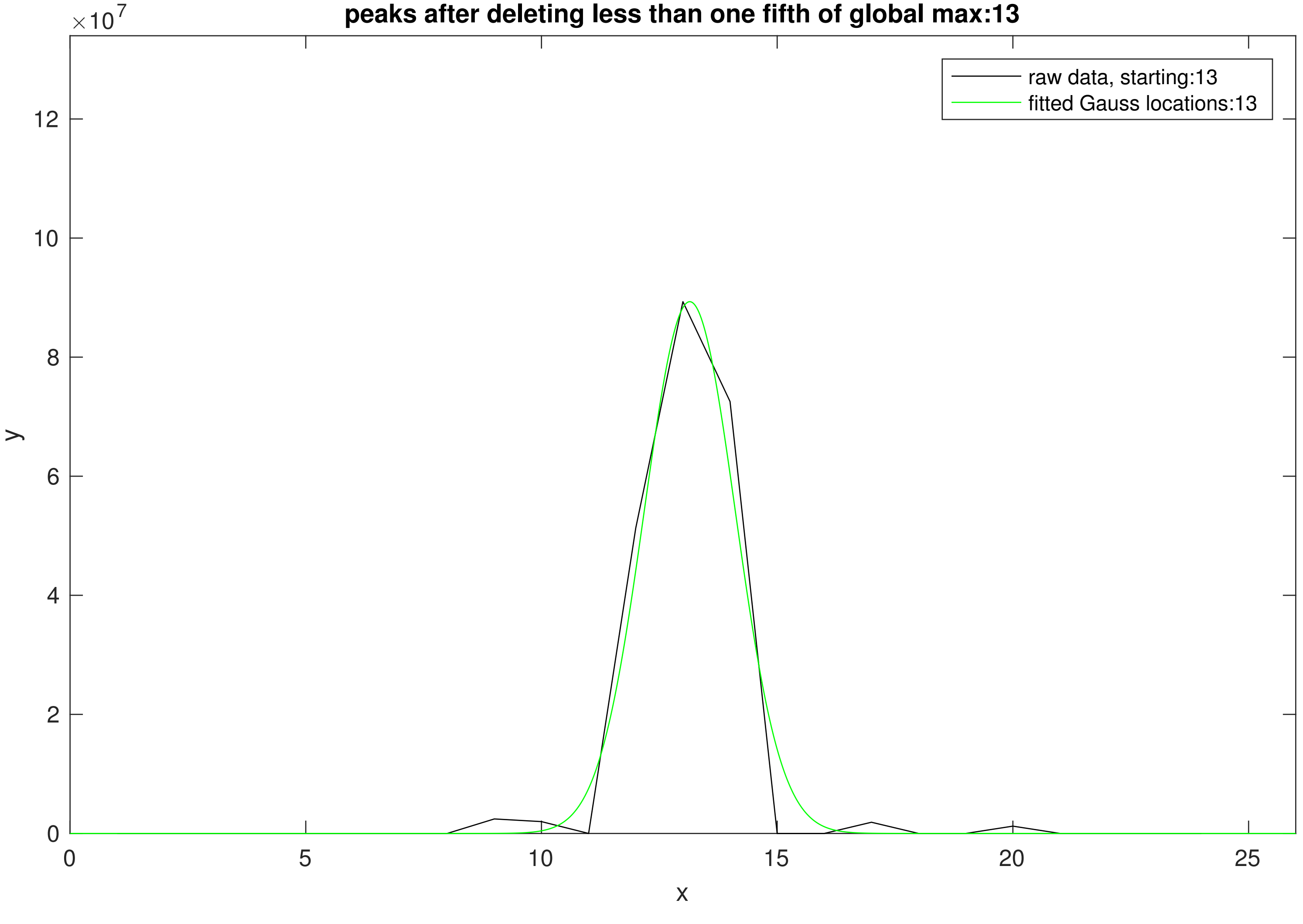

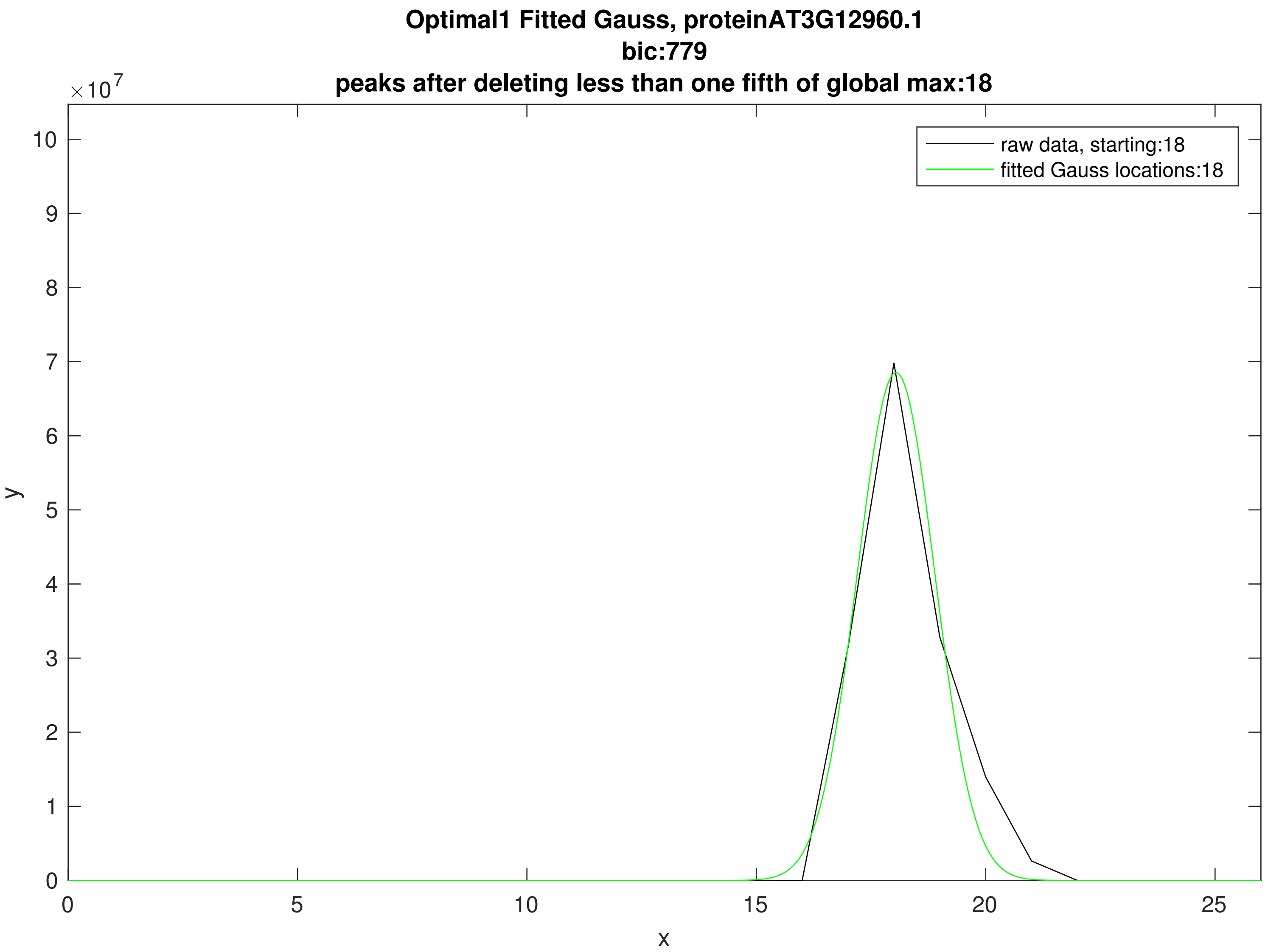

Optimal1 Fitted Gauss, proteinAT3G12960.1
bic:779
peaks after deleting less than one fifth of global max:18
×10^7
raw data, starting:18
fitted Gauss locations:18
y
x
0
1
2
3
4
5
6
7
8
9
10
0
5
10
15
20
25

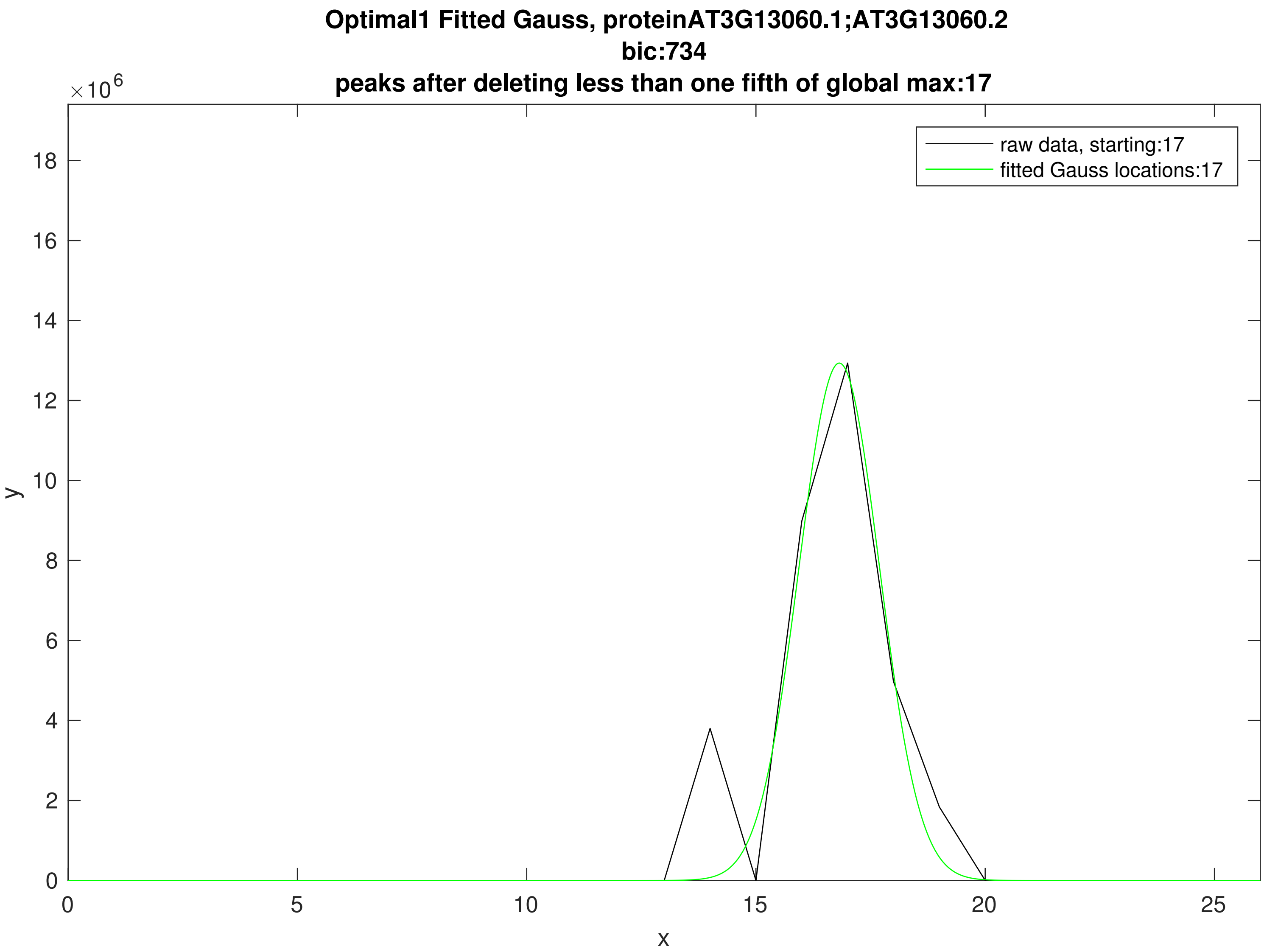

Optimal1 Fitted Gauss, proteinAT3G13060.1;AT3G13060.2
bic:734
peaks after deleting less than one fifth of global max:17
×10^6
raw data, starting:17
fitted Gauss locations:17
y
x
0
2
4
6
8
10
12
14
16
18
0
5
10
15
20
25

Optimal1 Fitted Gauss, proteinAT3G13110.1
bic:723
peaks after deleting less than one fifth of global max:10

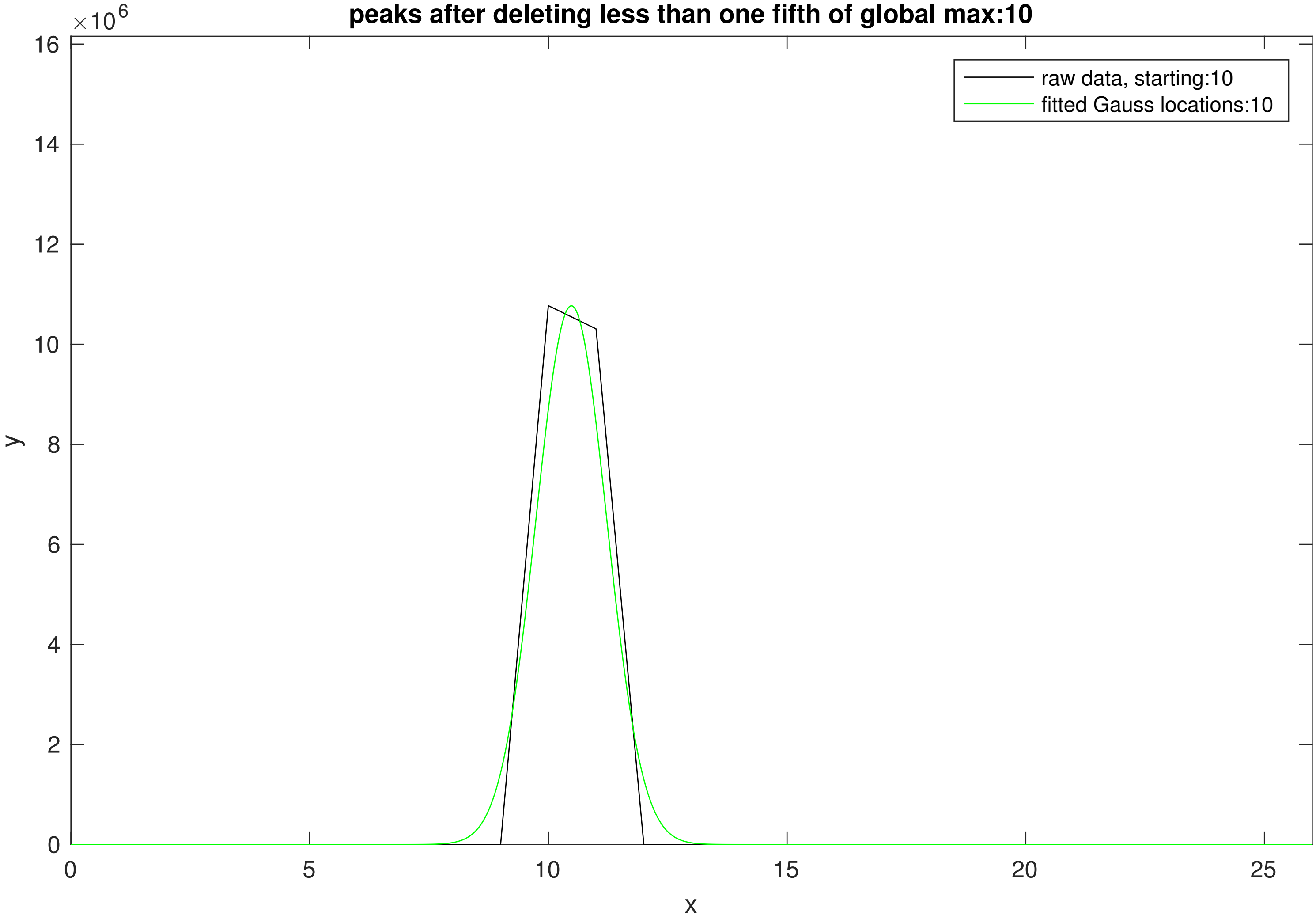

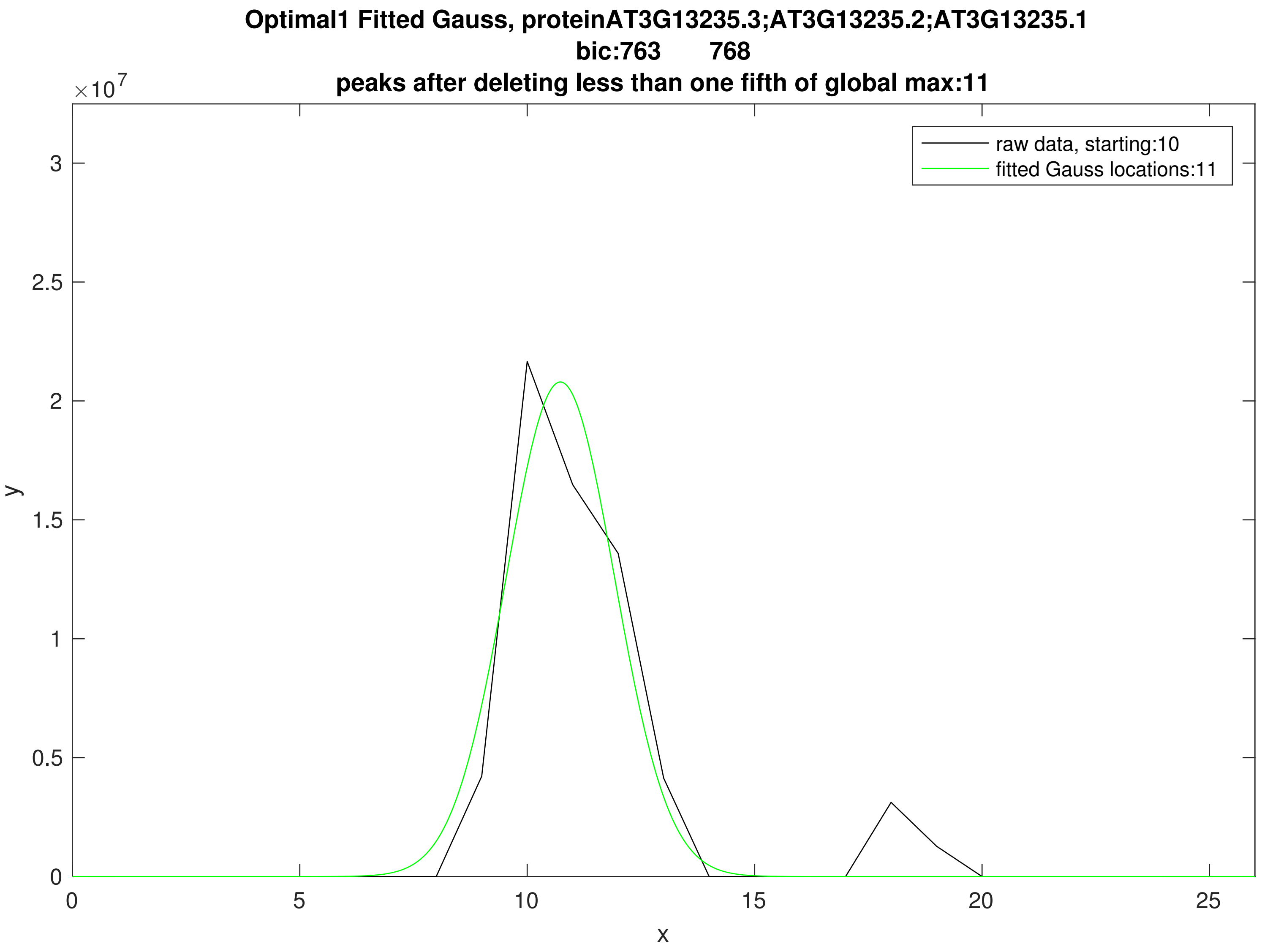
Optimal1 Fitted Gauss, proteinAT3G13235.3;AT3G13235.2;AT3G13235.1
bic:763 768
peaks after deleting less than one fifth of global max:11
×10^7
raw data, starting:10
fitted Gauss locations:11
y
x
0
0.5
1
1.5
2
2.5
3
0
5
10
15
20
25

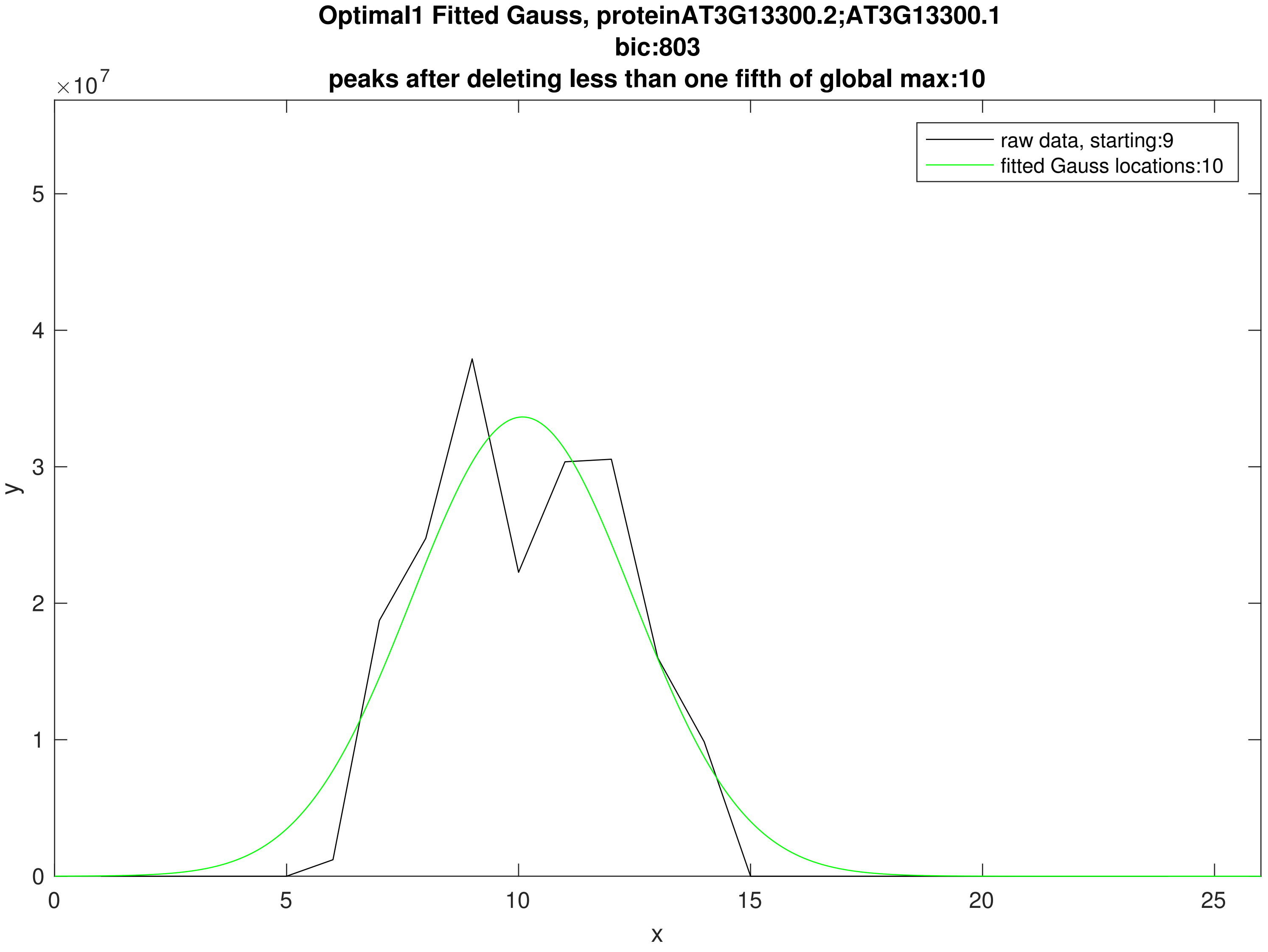

Optimal1 Fitted Gauss, proteinAT3G13300.2;AT3G13300.1
bic:803
peaks after deleting less than one fifth of global max:10
×10^7
raw data, starting:9
fitted Gauss locations:10
y
x
0
1
2
3
4
5
0
5
10
15
20
25

Optimal2 Fitted Gauss, proteinAT3G13330.1
bic:753 643
peaks after deleting less than one fifth of global max:4.5 9.6

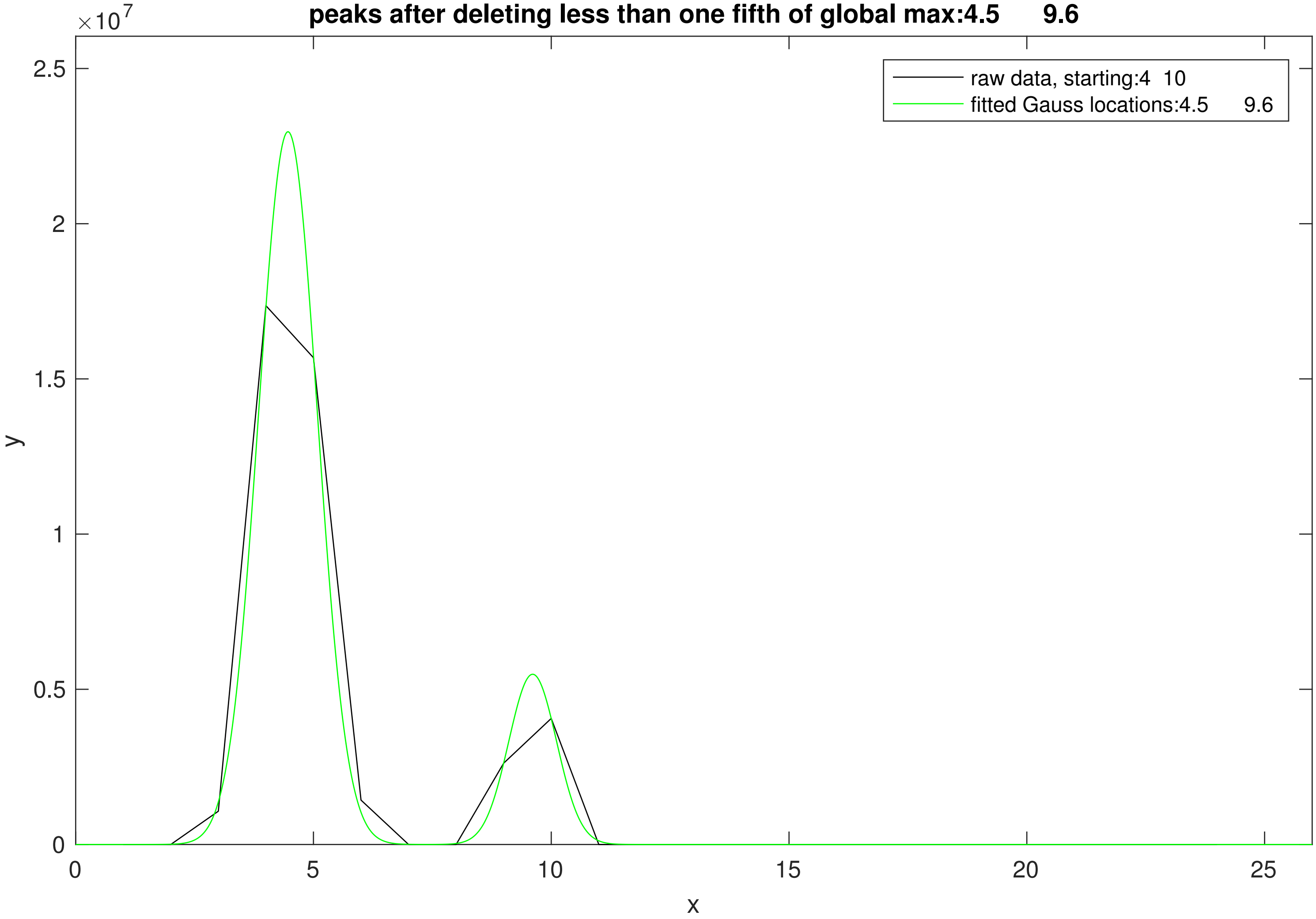

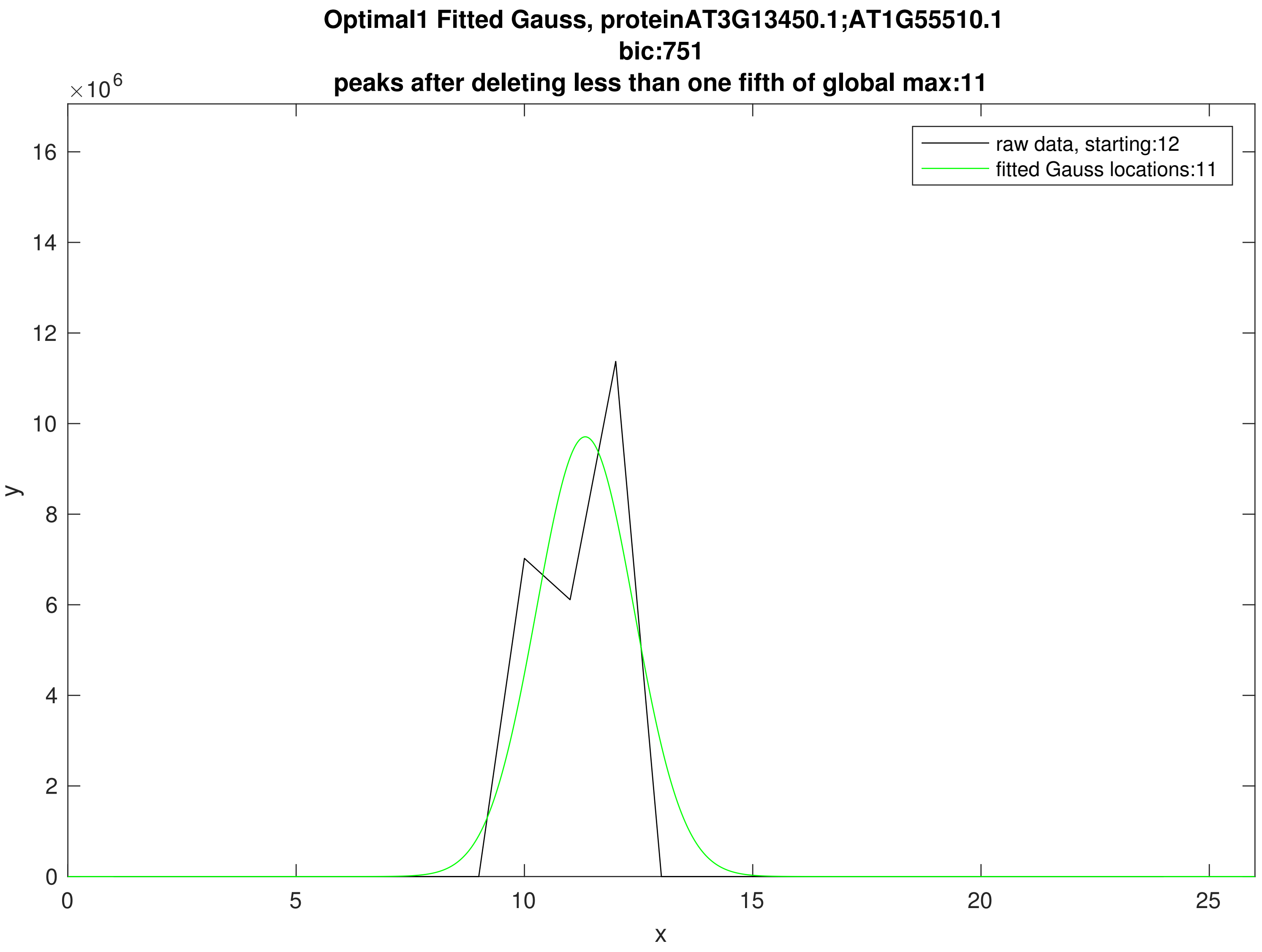
Optimal1 Fitted Gauss, proteinAT3G13450.1;AT1G55510.1
bic:751
peaks after deleting less than one fifth of global max:11
×10^6
raw data, starting:12
fitted Gauss locations:11
y
x
0
2
4
6
8
10
12
14
16
0
5
10
15
20
25

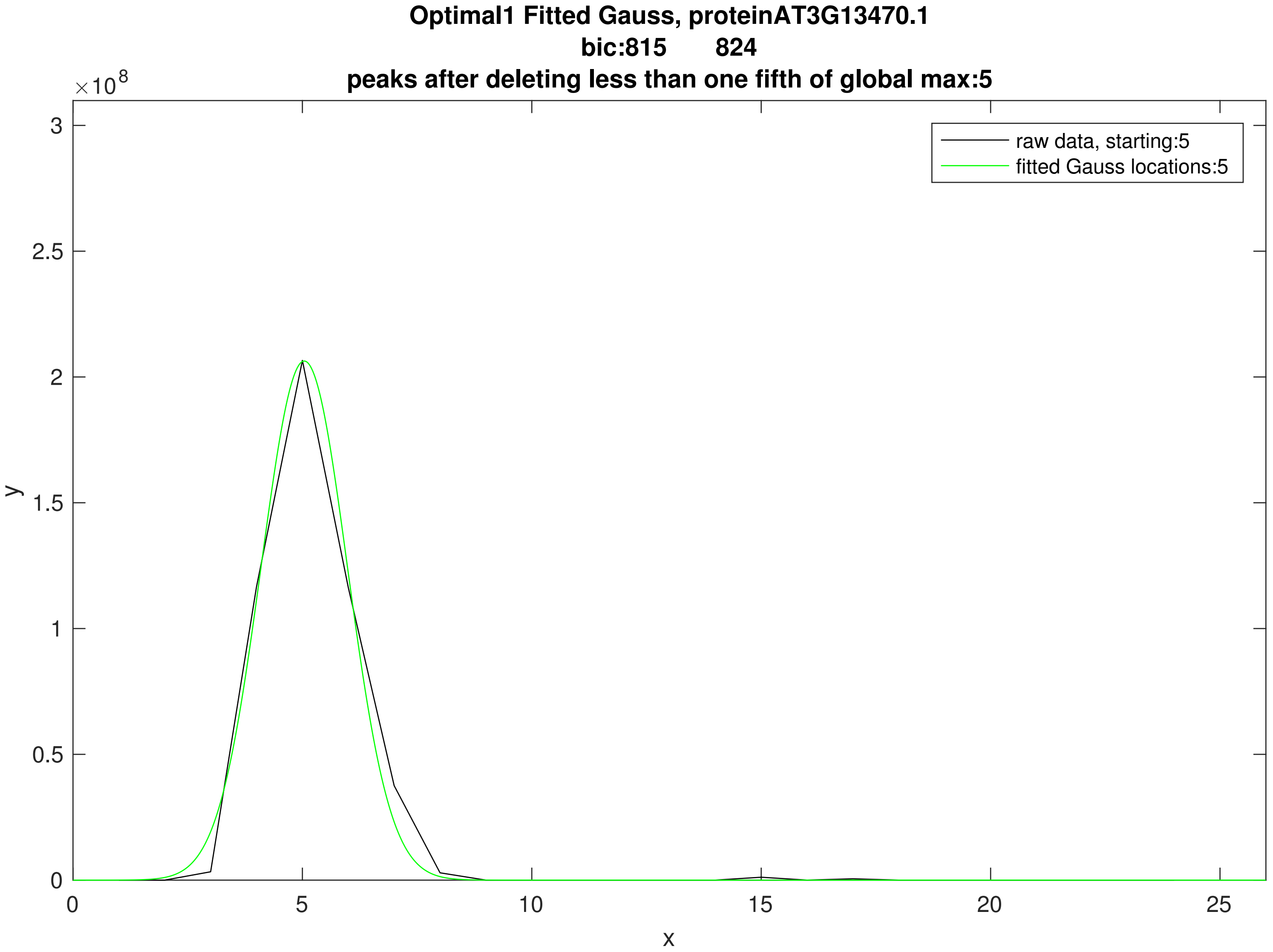
Optimal1 Fitted Gauss, proteinAT3G13470.1
bic:815 824
peaks after deleting less than one fifth of global max:5
×10^8
3
2.5
2
1.5
1
0.5
0
y
0
5
10
15
20
25
x
raw data, starting:5
fitted Gauss locations:5

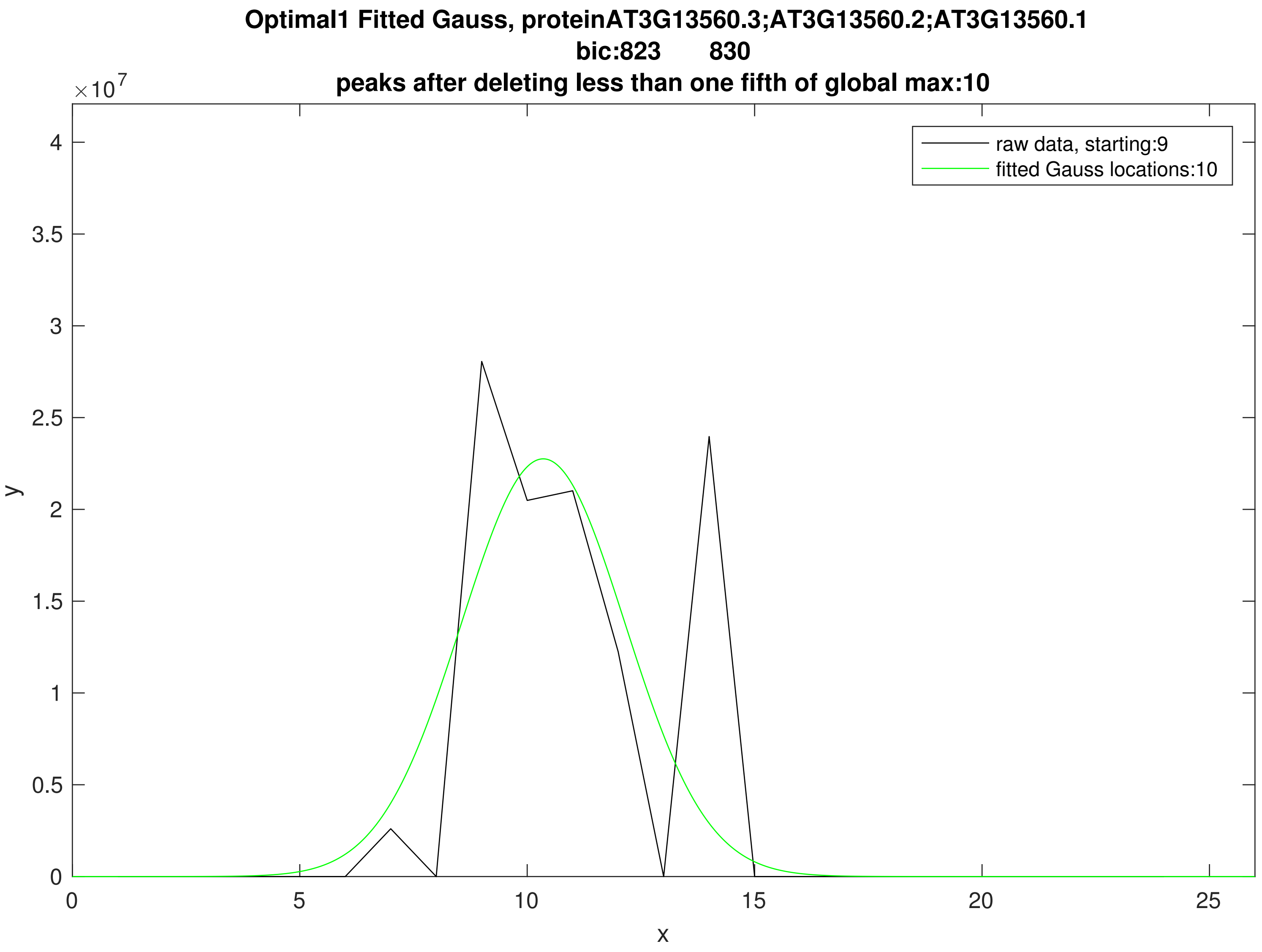

Optimal1 Fitted Gauss, proteinAT3G13560.3;AT3G13560.2;AT3G13560.1
bic:823 830
peaks after deleting less than one fifth of global max:10
×10^7
raw data, starting:9
fitted Gauss locations:10
y
x
0
0.5
1
1.5
2
2.5
3
3.5
4
0
5
10
15
20
25

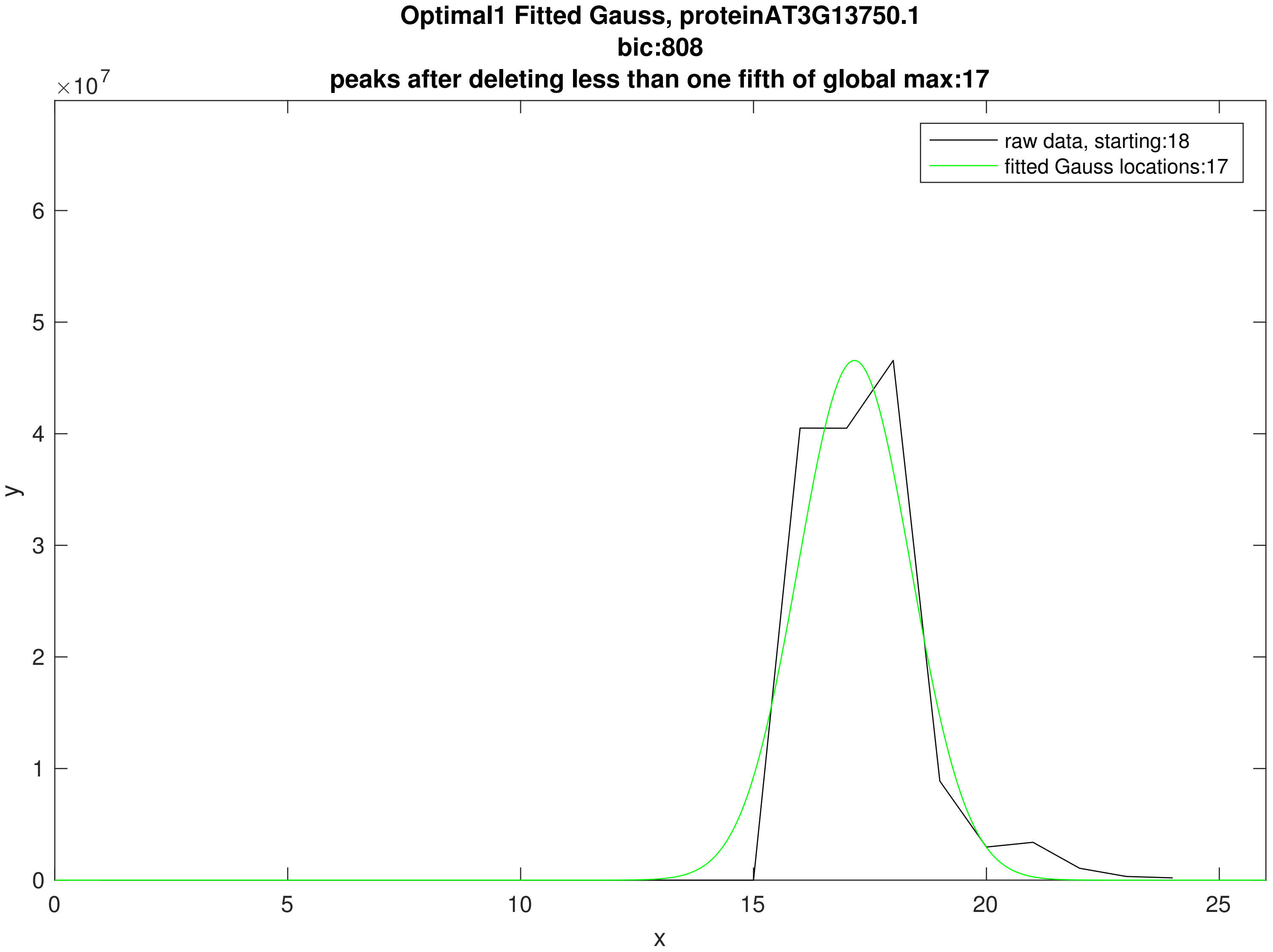

Optimal1 Fitted Gauss, proteinAT3G13750.1
bic:808
peaks after deleting less than one fifth of global max:17
×10^7
raw data, starting:18
fitted Gauss locations:17
y
x
0
1
2
3
4
5
6
0
5
10
15
20
25

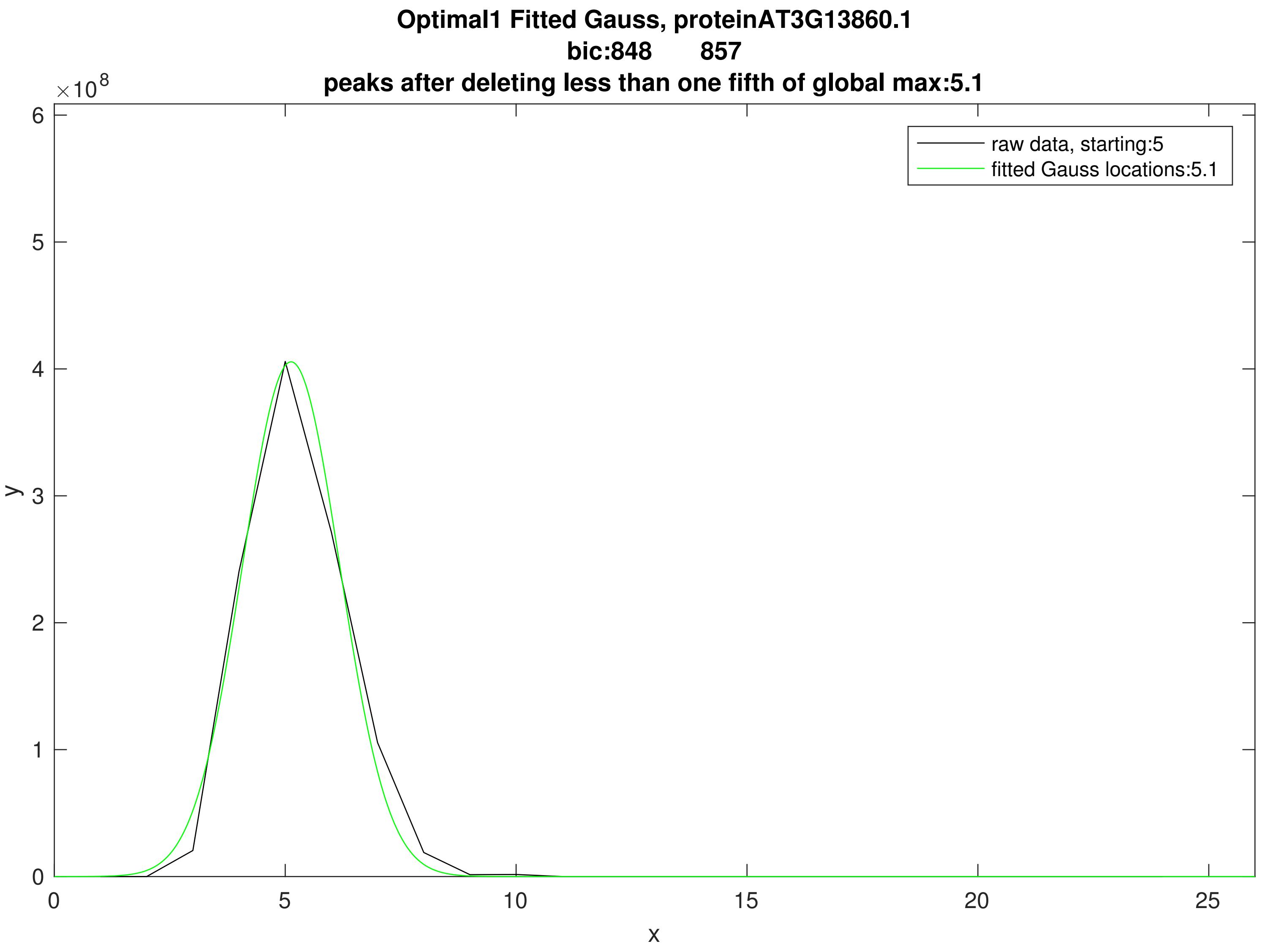

Optimal1 Fitted Gauss, proteinAT3G13860.1
bic:848 857
peaks after deleting less than one fifth of global max:5.1
×10^8
raw data, starting:5
fitted Gauss locations:5.1
y
x
0
1
2
3
4
5
6
0
5
10
15
20
25

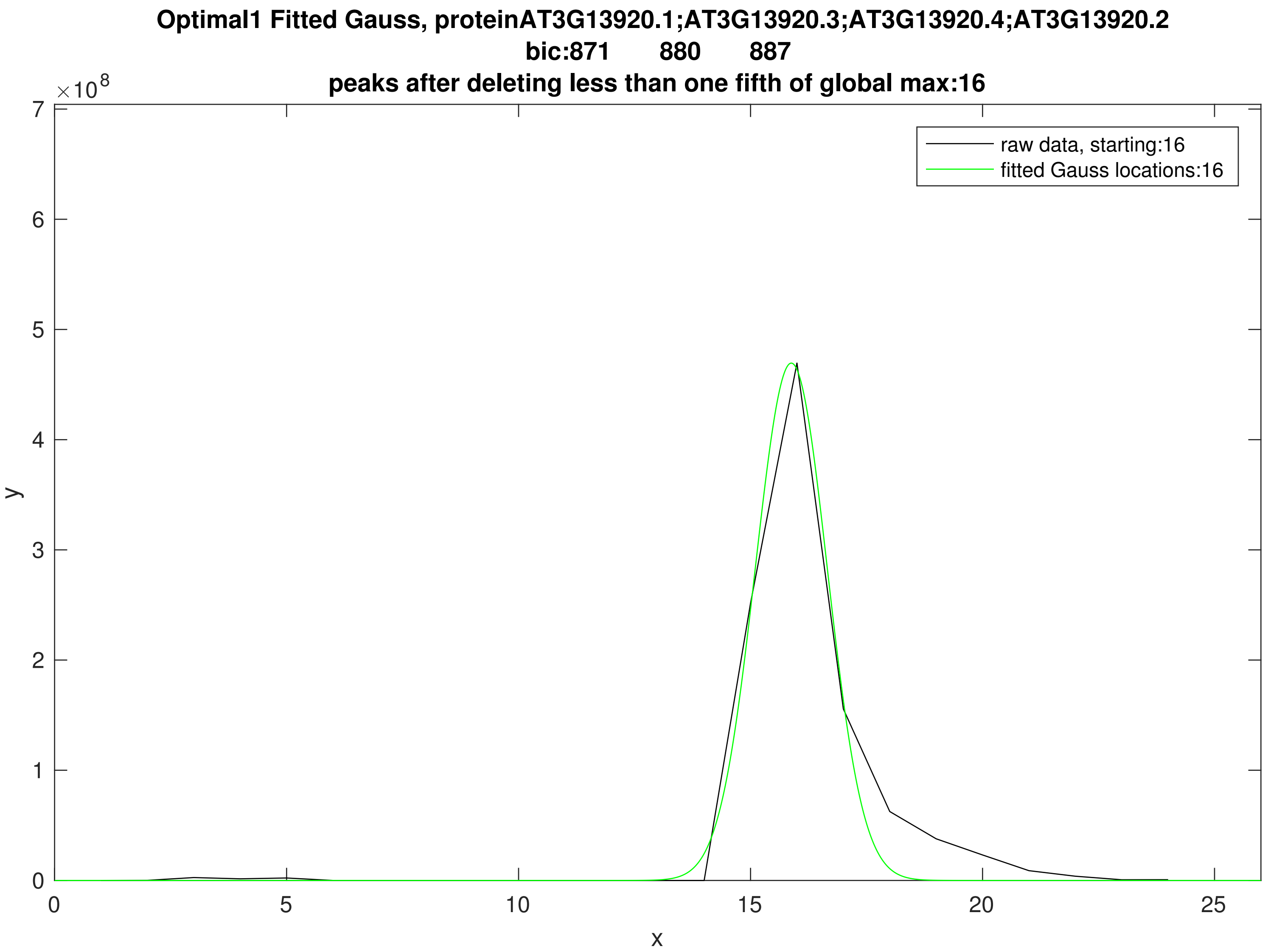

Optimal1 Fitted Gauss, proteinAT3G13920.1;AT3G13920.3;AT3G13920.4;AT3G13920.2
bic:871 880 887
peaks after deleting less than one fifth of global max:16
×10^8
y
x
raw data, starting:16
fitted Gauss locations:16

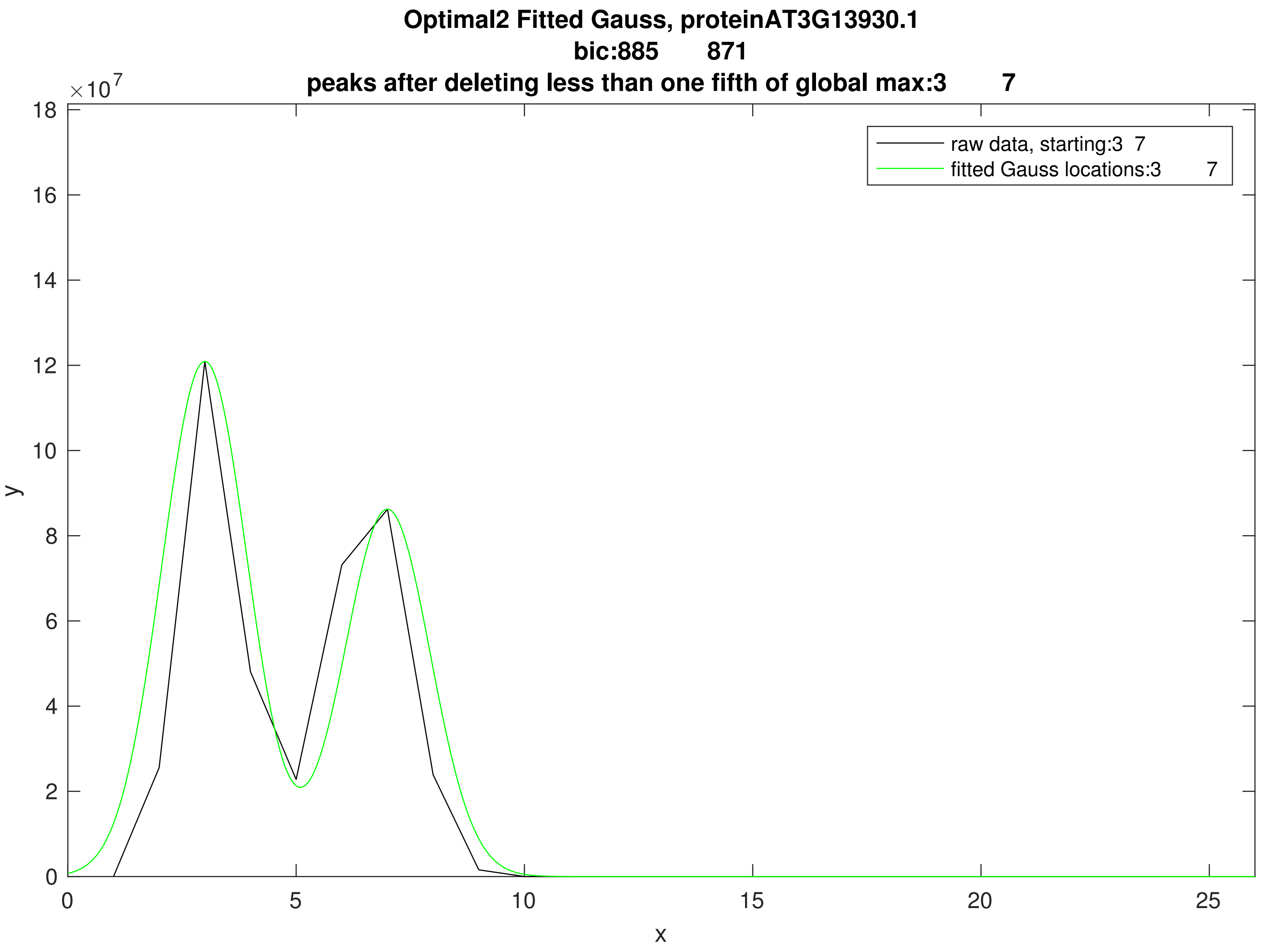

Optimal2 Fitted Gauss, proteinAT3G13930.1
bic:885 871
peaks after deleting less than one fifth of global max:3 7
×10^7
18
16
14
12
10
8
6
4
2
0
y
0
5
10
15
20
25
x
raw data, starting:3 7
fitted Gauss locations:3 7

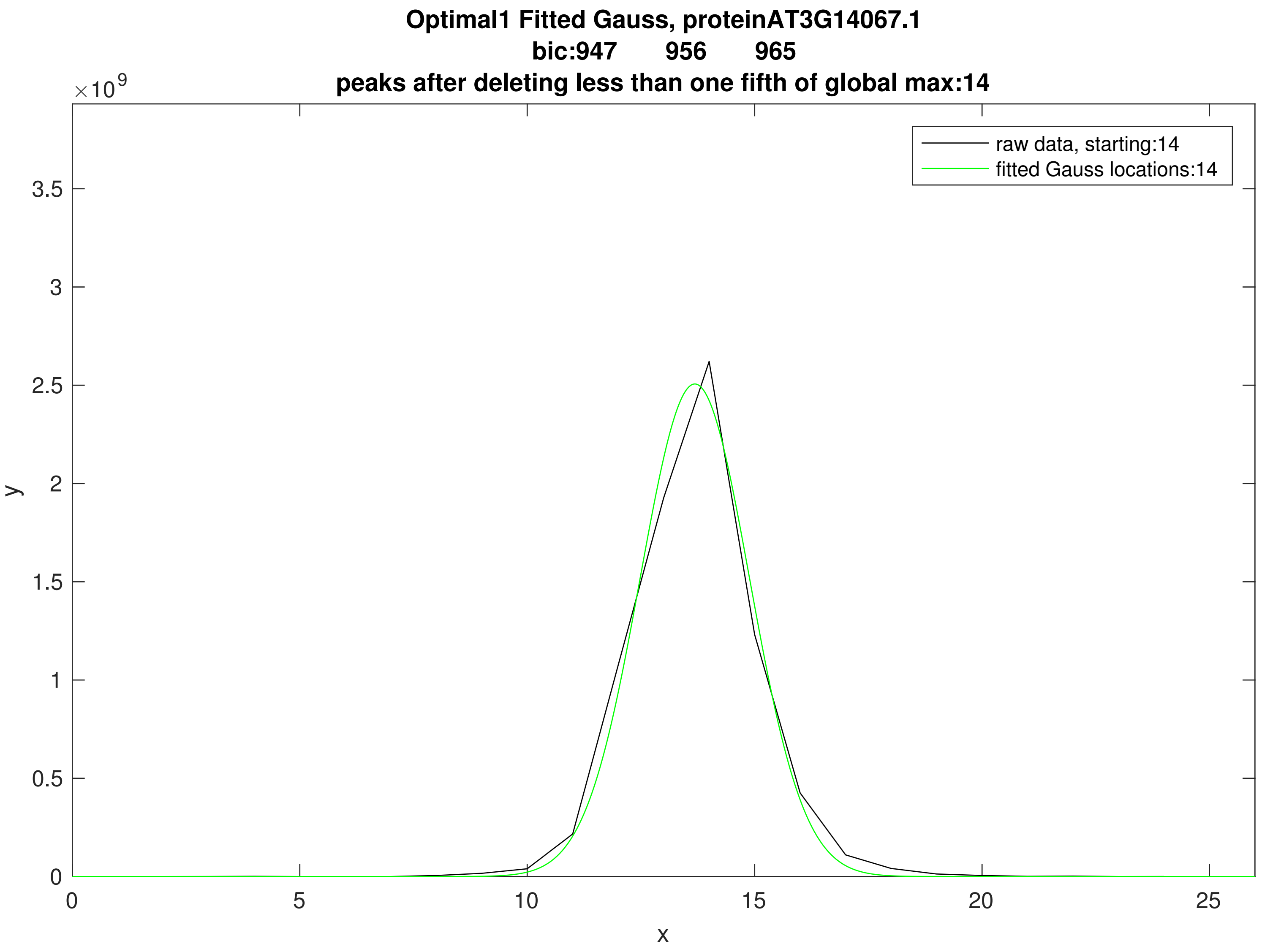

Optimal1 Fitted Gauss, proteinAT3G14067.1
bic:947 956 965
peaks after deleting less than one fifth of global max:14
×10^9
raw data, starting:14
fitted Gauss locations:14
3.5
3
2.5
2
1.5
1
0.5
0
y
0
5
10
15
20
25
x

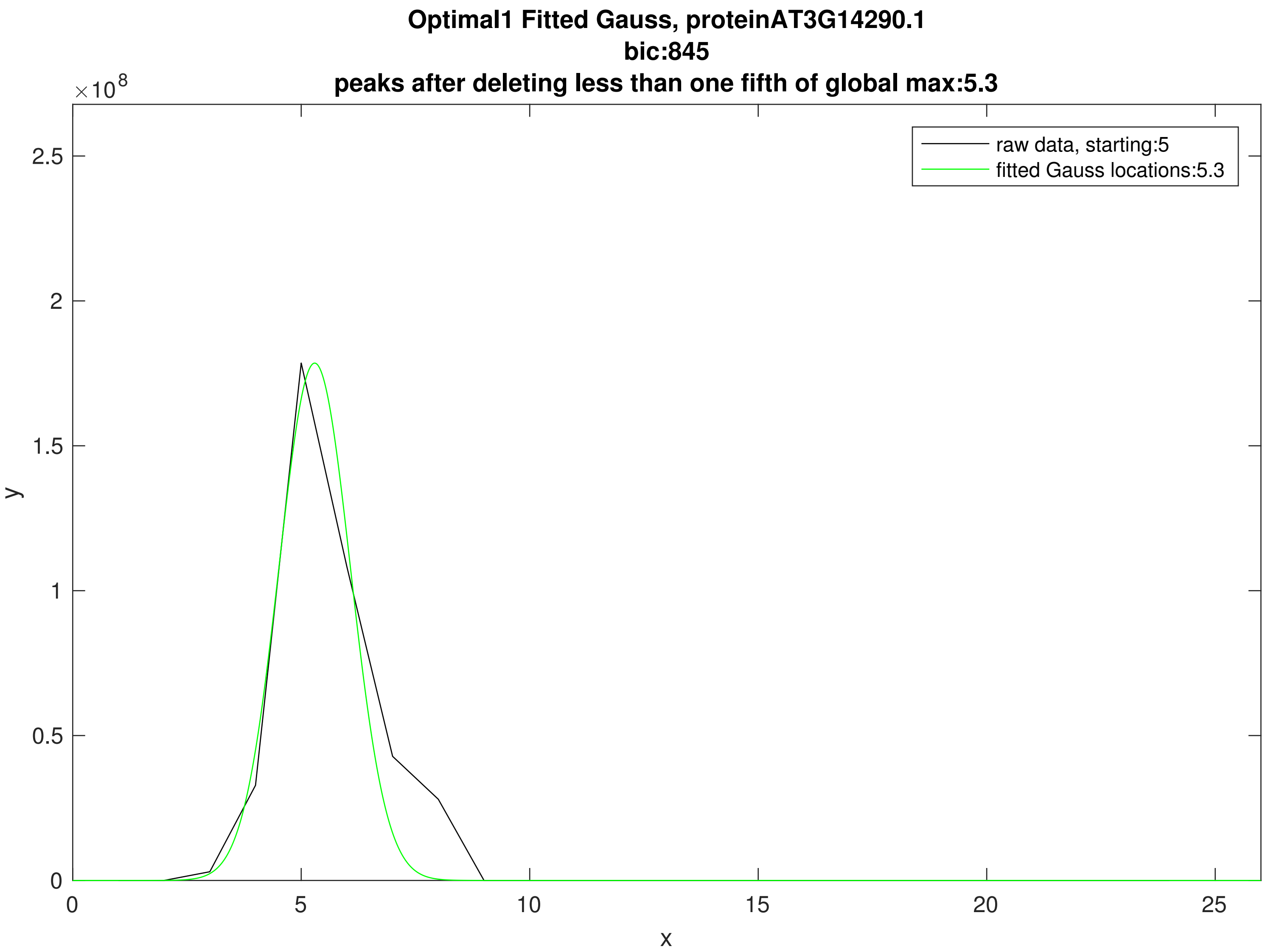

Optimal1 Fitted Gauss, proteinAT3G14290.1
bic:845
peaks after deleting less than one fifth of global max:5.3
×10^8
2.5
2
1.5
1
0.5
0
y
0
5
10
15
20
25
x
raw data, starting:5
fitted Gauss locations:5.3

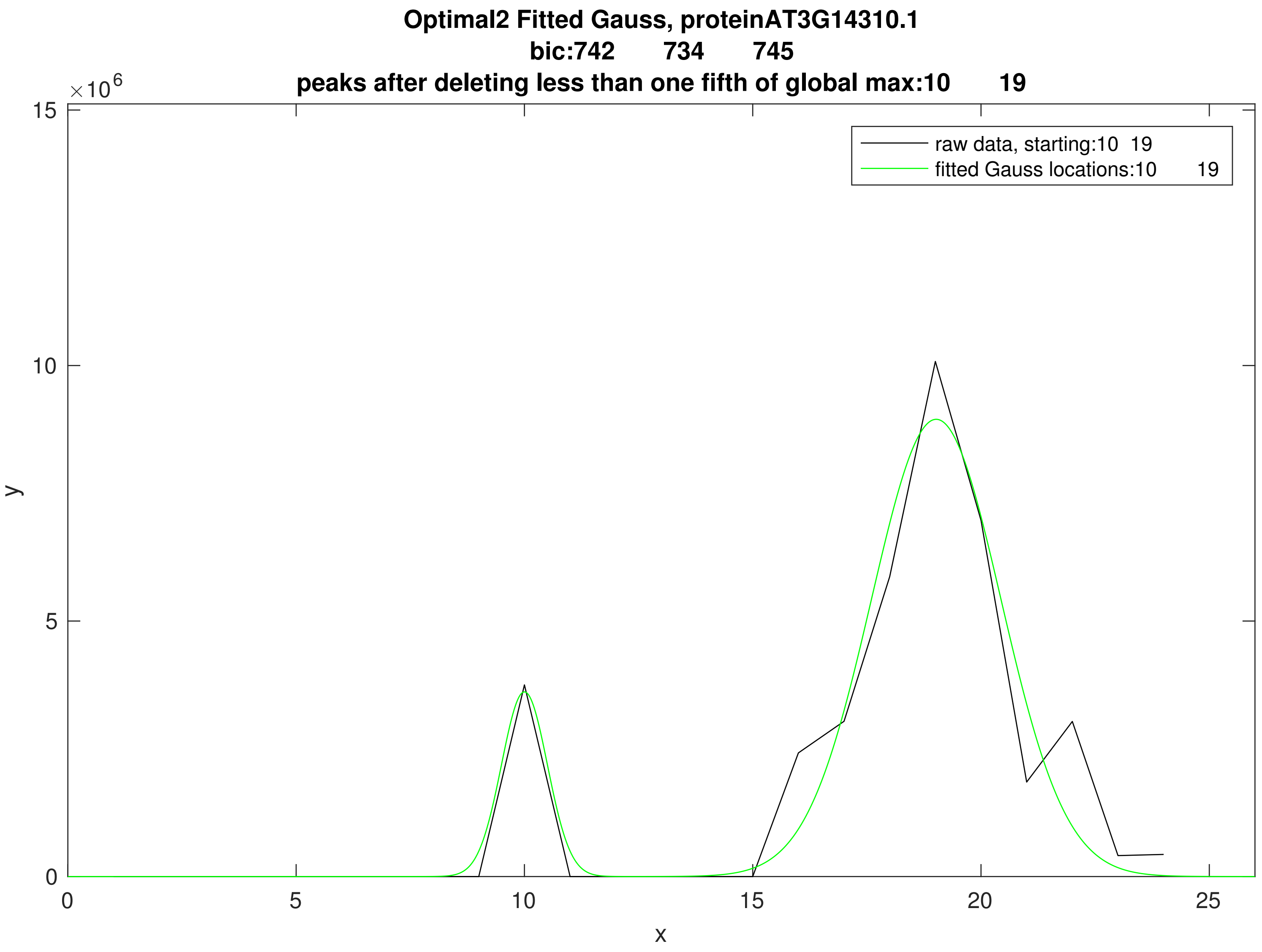
Optimal2 Fitted Gauss, proteinAT3G14310.1
bic:742 734 745
peaks after deleting less than one fifth of global max:10 19
×10^6
raw data, starting:10 19
fitted Gauss locations:10 19
15
10
5
0
y
0
5
10
15
20
25
x

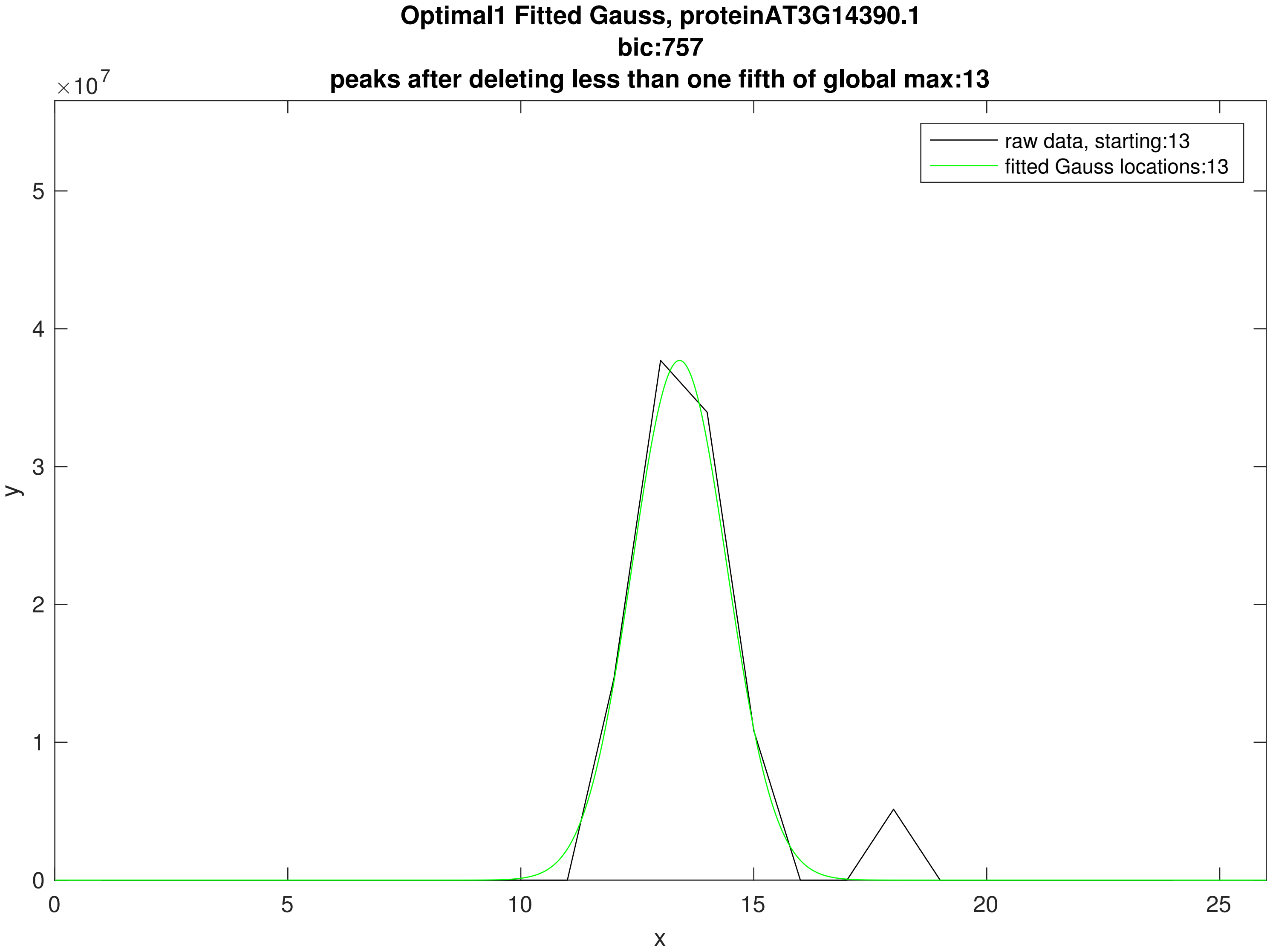

Optimal1 Fitted Gauss, proteinAT3G14390.1
bic:757
peaks after deleting less than one fifth of global max:13
×10^7
raw data, starting:13
fitted Gauss locations:13
y
x
0
1
2
3
4
5
0
5
10
15
20
25

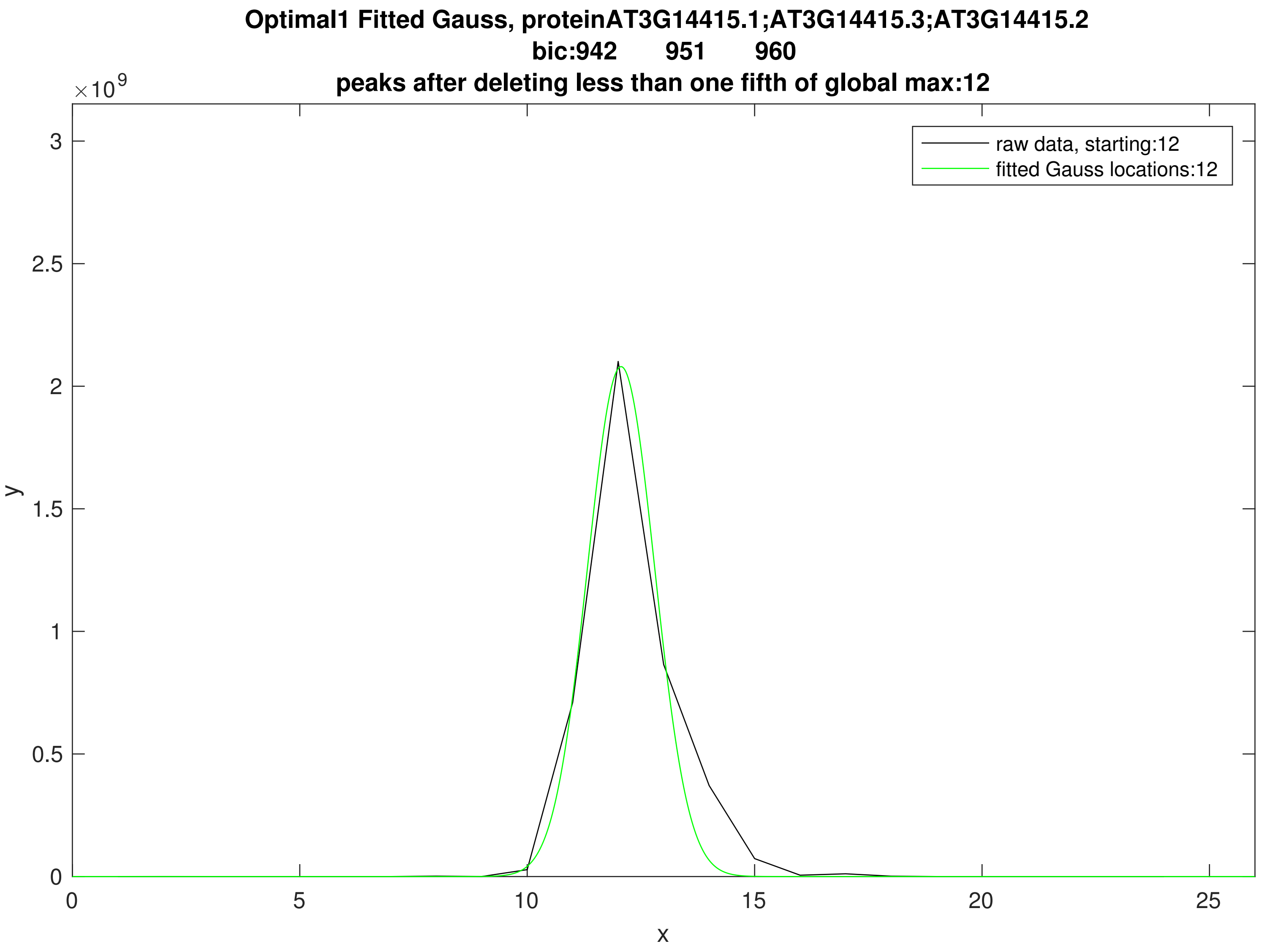
Optimal1 Fitted Gauss, proteinAT3G14415.1;AT3G14415.3;AT3G14415.2
bic:942 951 960
peaks after deleting less than one fifth of global max:12
×10^9
raw data, starting:12
fitted Gauss locations:12
y
x
0
0.5
1
1.5
2
2.5
3
5
10
15
20
25

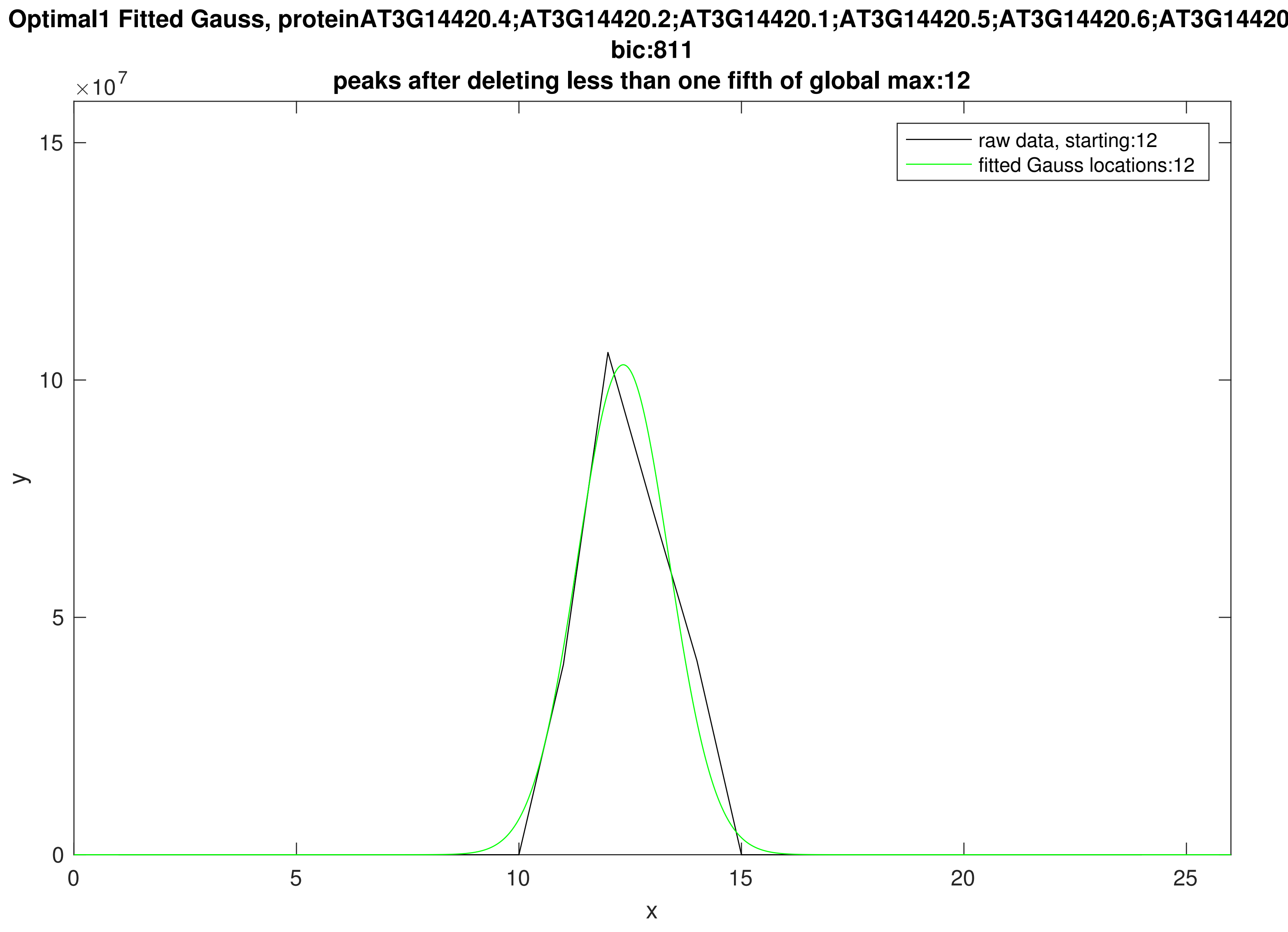
Optimal1 Fitted Gauss, proteinAT3G14420.4;AT3G14420.2;AT3G14420.1;AT3G14420.5;AT3G14420.6;AT3G14420
bic:811
peaks after deleting less than one fifth of global max:12
×10^7
raw data, starting:12
fitted Gauss locations:12
y
x
0
5
10
15
20
25

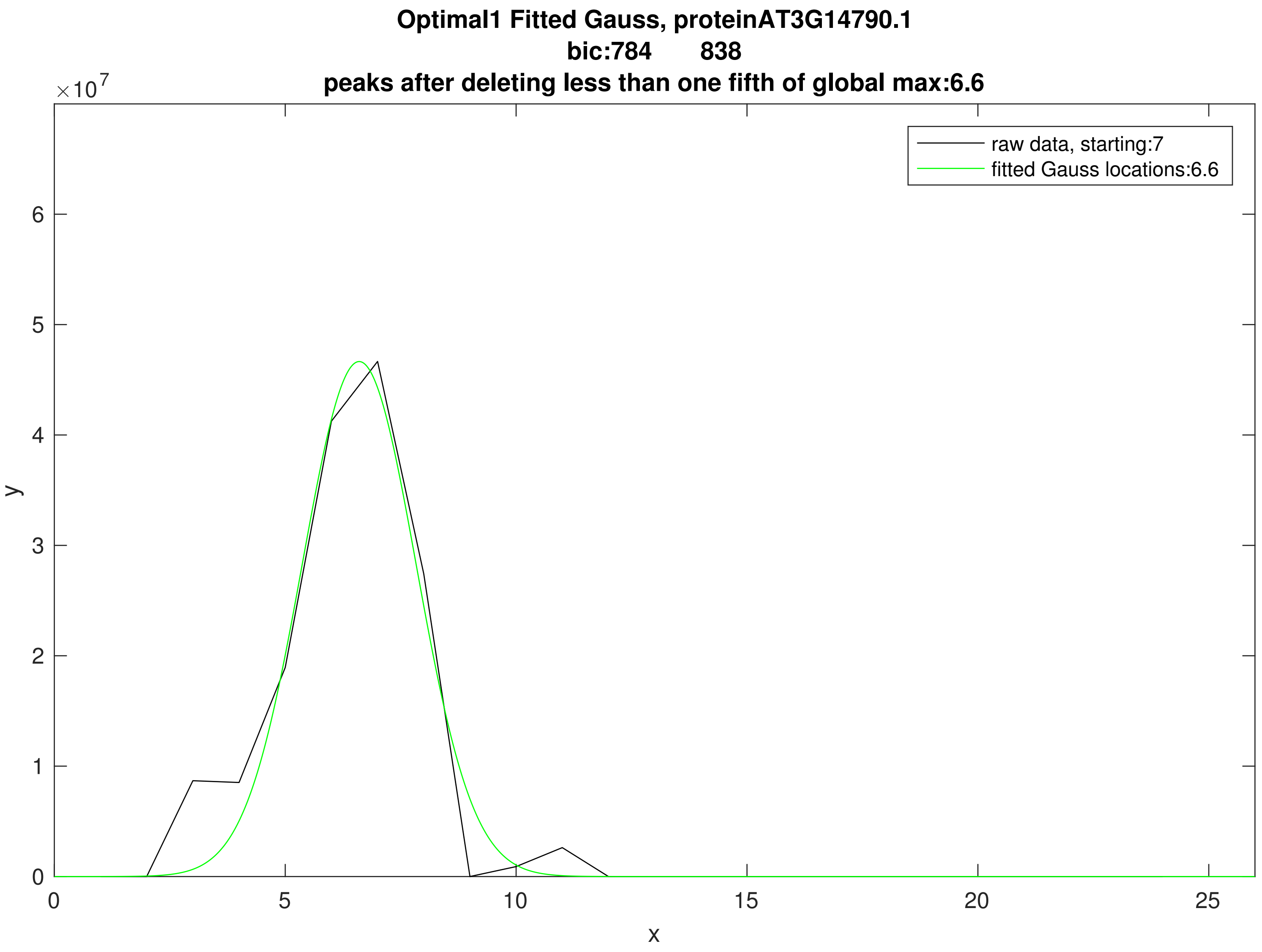

Optimal1 Fitted Gauss, proteinAT3G14790.1
bic:784 838
peaks after deleting less than one fifth of global max:6.6
×10^7
raw data, starting:7
fitted Gauss locations:6.6
y
x
0
1
2
3
4
5
6
0
5
10
15
20
25

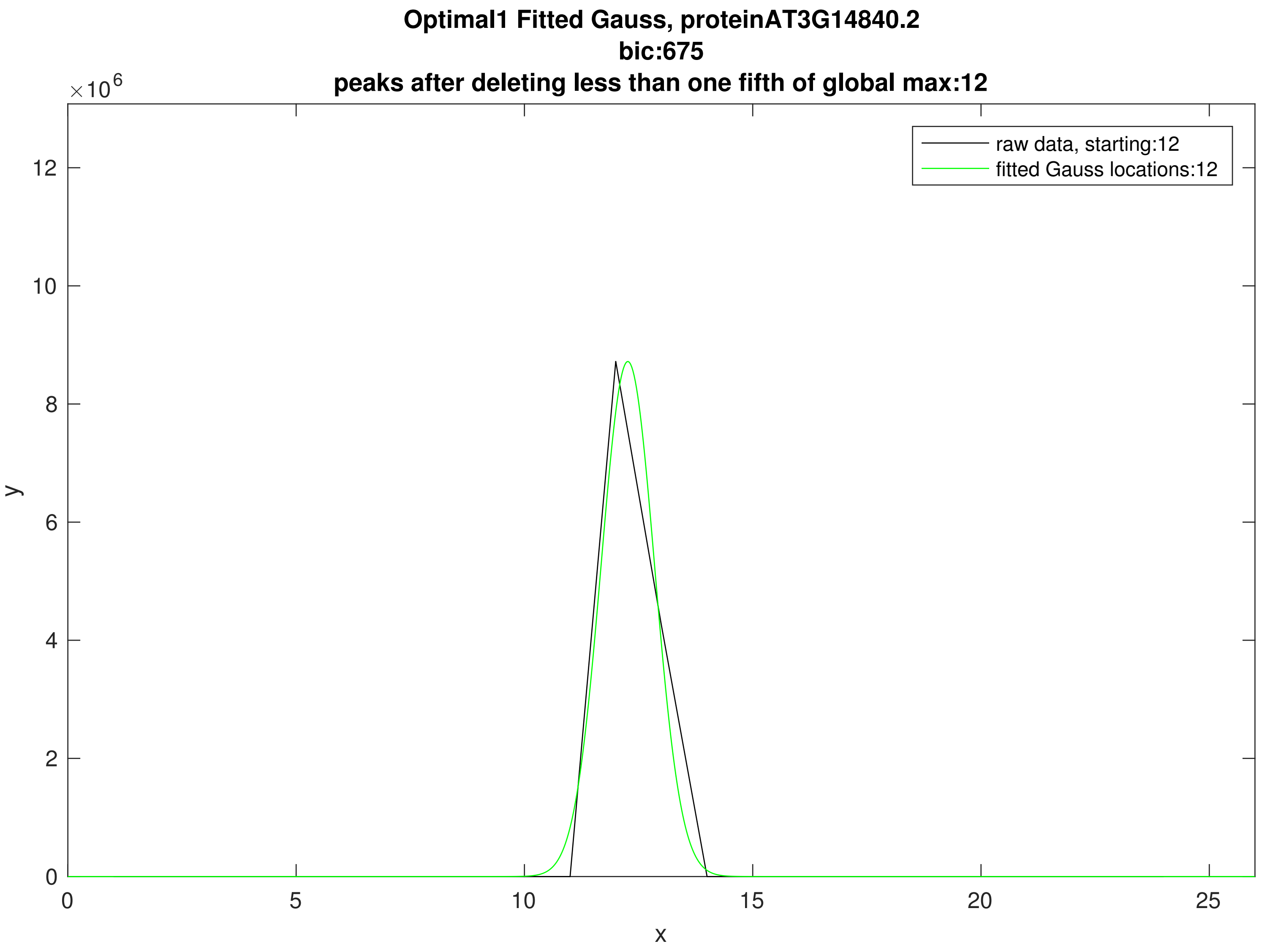
Optimal1 Fitted Gauss, proteinAT3G14840.2
bic:675
peaks after deleting less than one fifth of global max:12
×10^6
raw data, starting:12
fitted Gauss locations:12
y
x
0
2
4
6
8
10
12
0
5
10
15
20
25

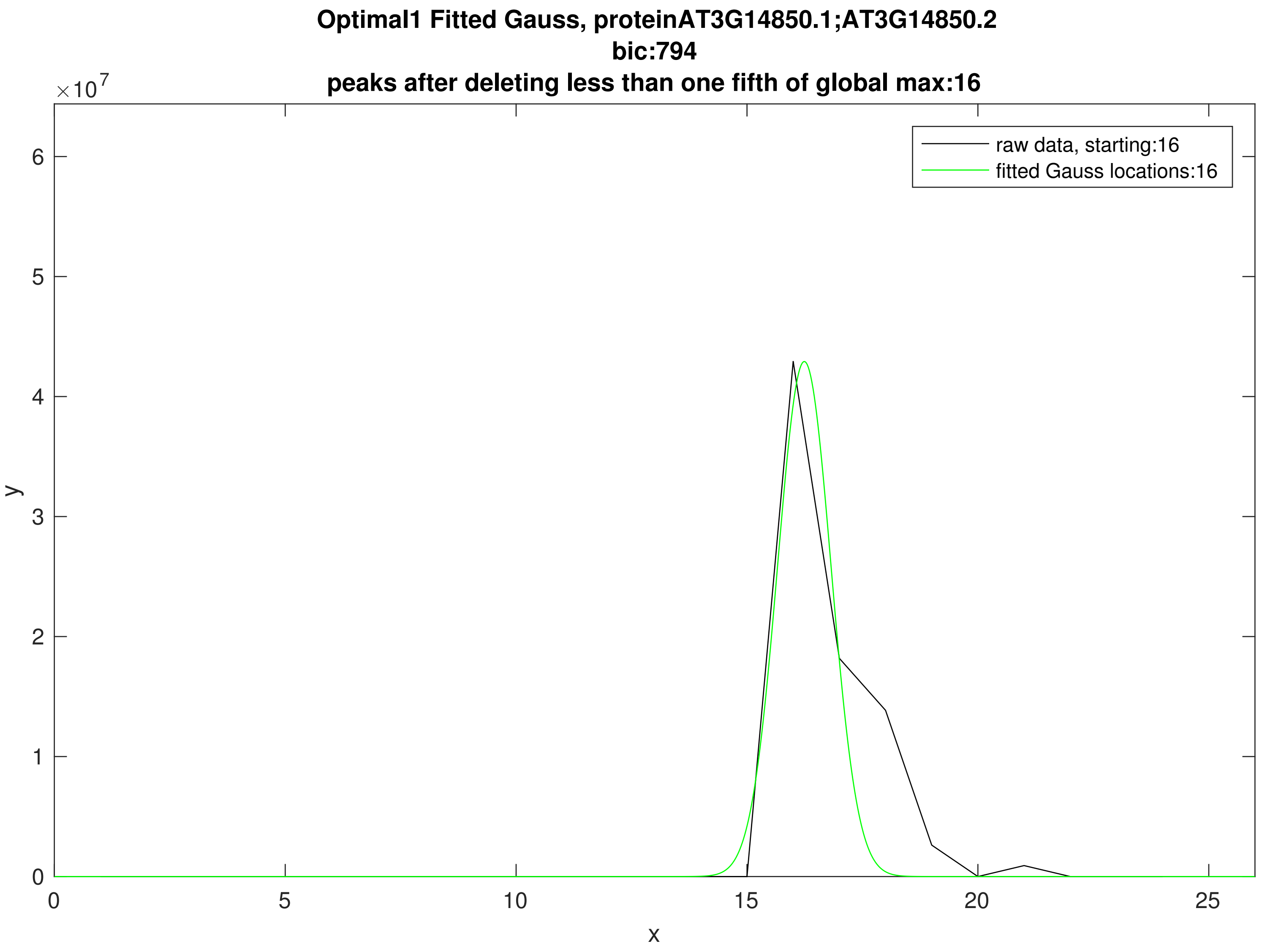
Optimal1 Fitted Gauss, proteinAT3G14850.1;AT3G14850.2
bic:794
peaks after deleting less than one fifth of global max:16
×10^7
raw data, starting:16
fitted Gauss locations:16
y
x
0
1
2
3
4
5
6
0
5
10
15
20
25

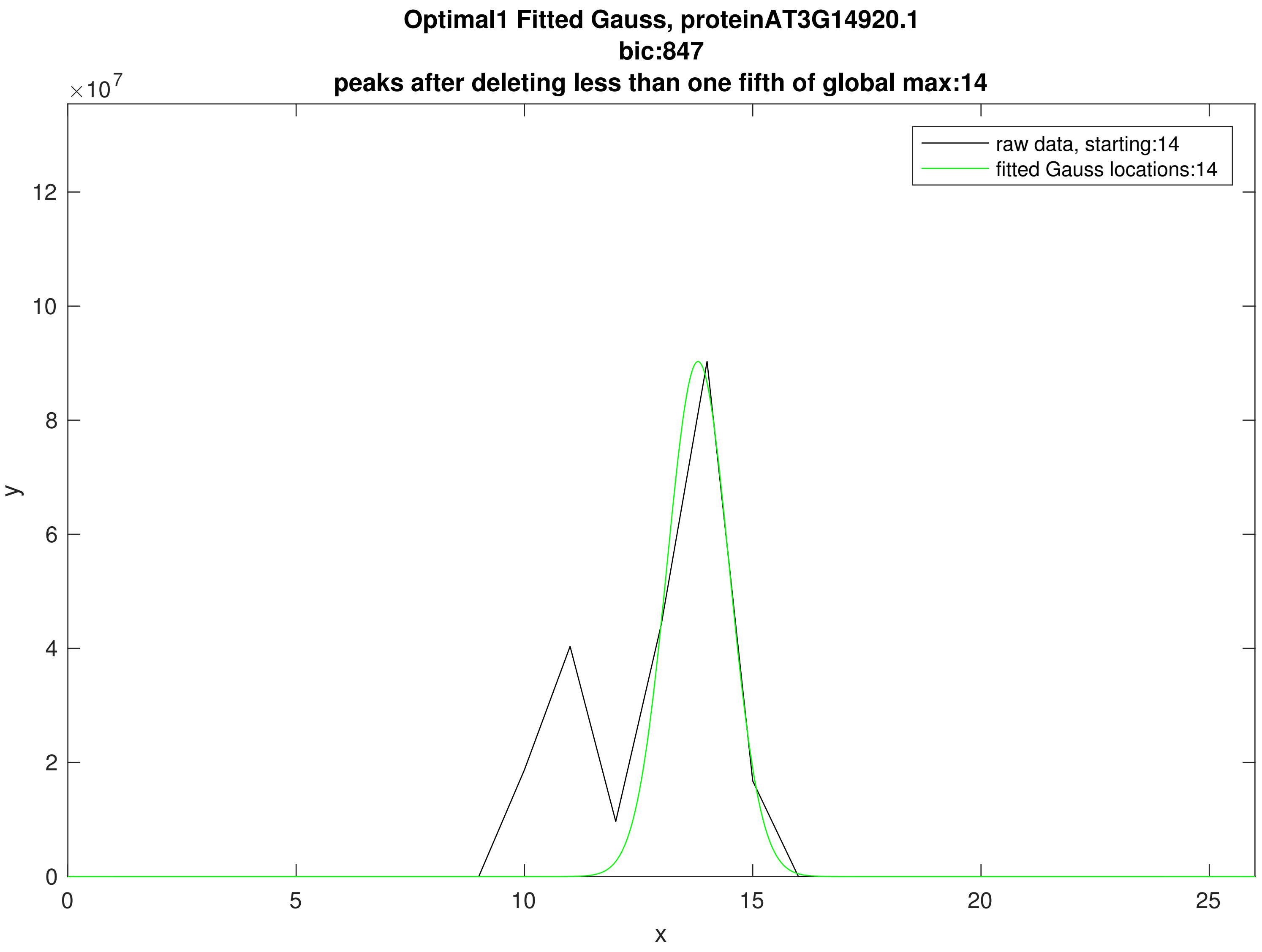
Optimal1 Fitted Gauss, proteinAT3G14920.1
bic:847
peaks after deleting less than one fifth of global max:14
×10^7
raw data, starting:14
fitted Gauss locations:14
12
10
8
6
4
2
0
y
0
5
10
15
20
25
x

**Optimal1 Fitted Gauss, proteinAT3G14940.1**
**bic:843 847 862**
**peaks after deleting less than one fifth of global max:7**

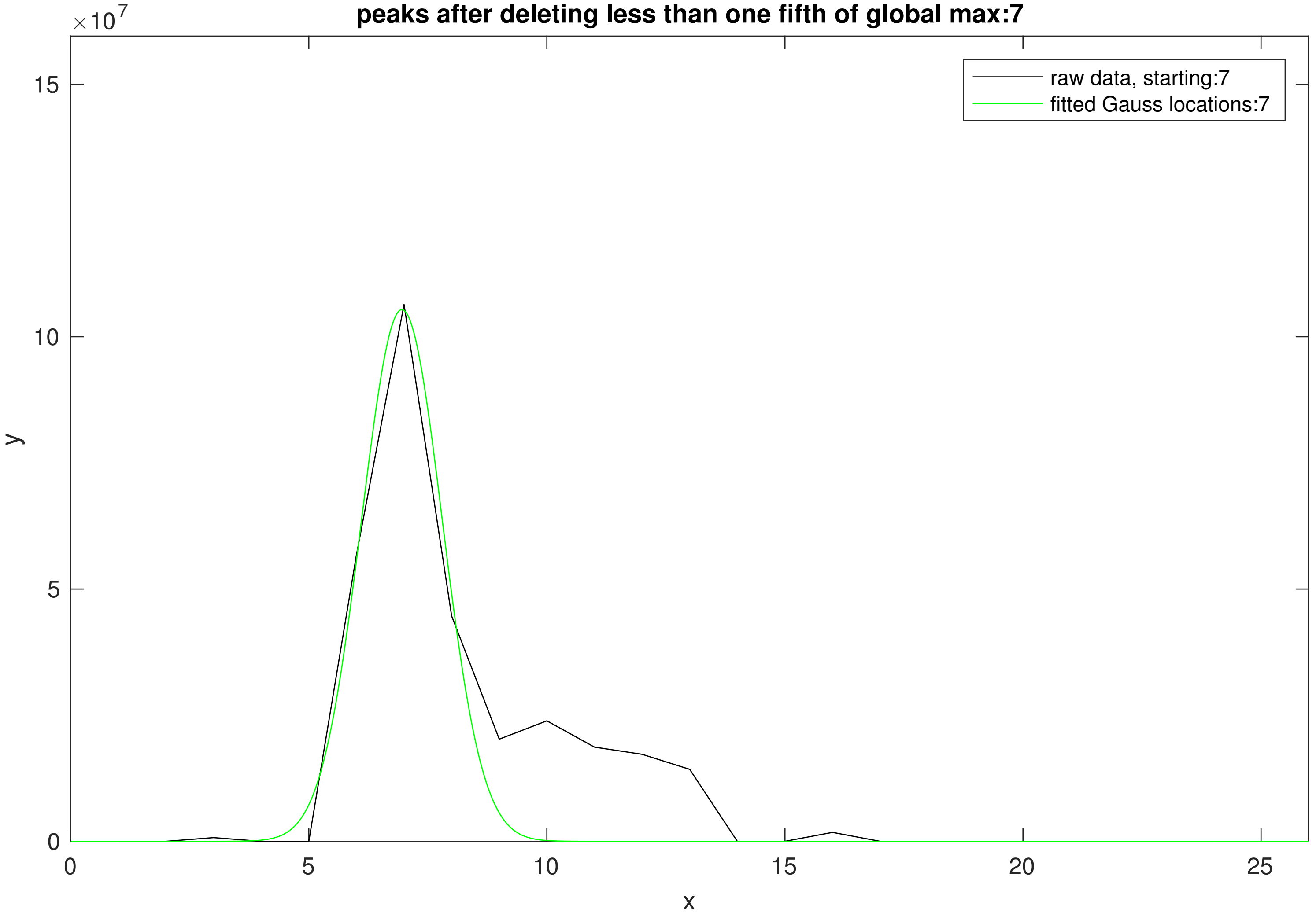

Optimal1 Fitted Gauss, proteinAT3G14990.1;AT3G14990.3;AT3G14990.2
bic:955
peaks after deleting less than one fifth of global max:17

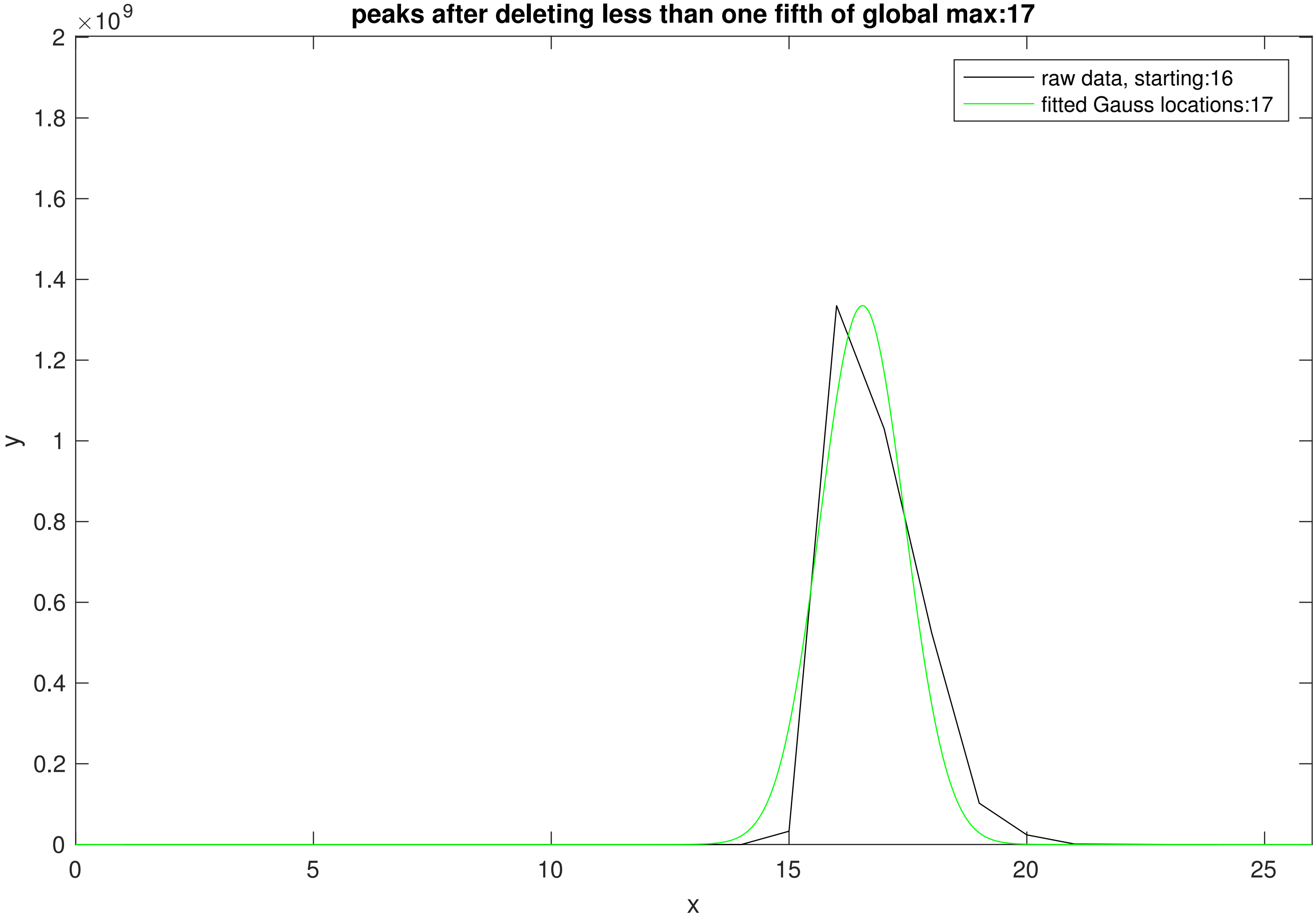

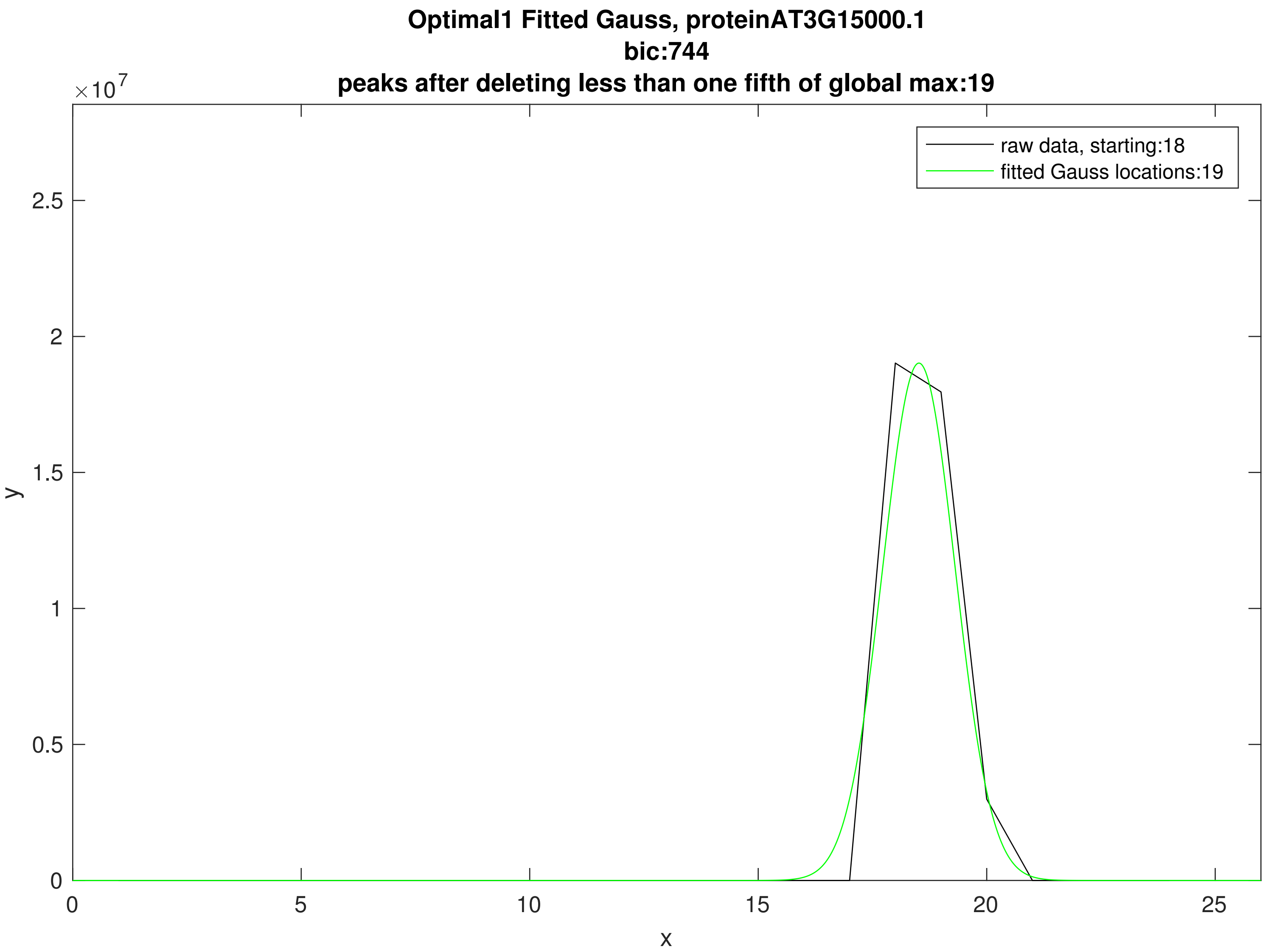

Optimal1 Fitted Gauss, proteinAT3G15000.1
bic:744
peaks after deleting less than one fifth of global max:19
×10^7
2.5
2
1.5
1
0.5
0
y
0
5
10
15
20
25
x
raw data, starting:18
fitted Gauss locations:19

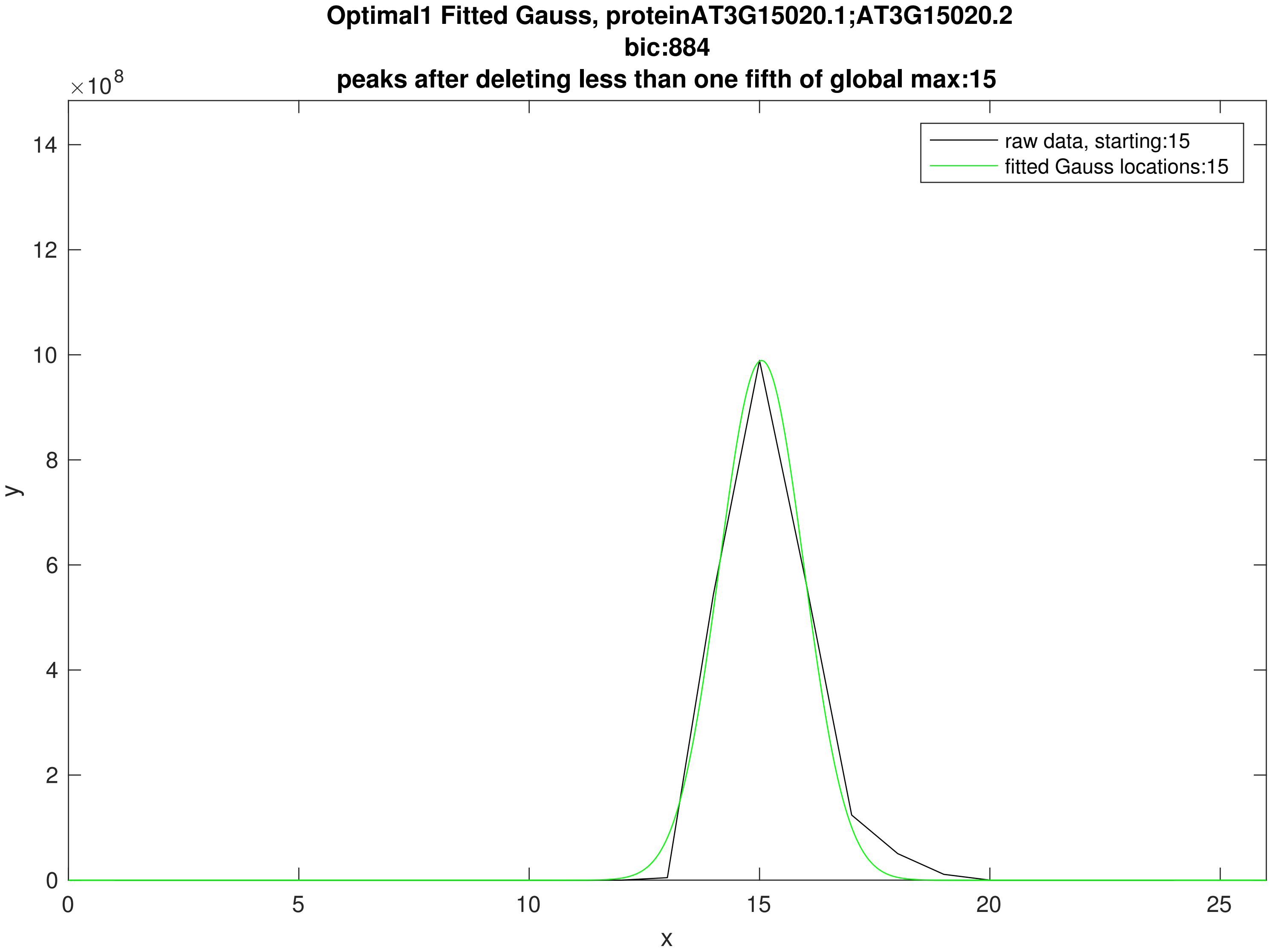

Optimal1 Fitted Gauss, proteinAT3G15020.1;AT3G15020.2
bic:884
peaks after deleting less than one fifth of global max:15
×10^8
14
12
10
8
6
4
2
0
y
0
5
10
15
20
25
x
raw data, starting:15
fitted Gauss locations:15

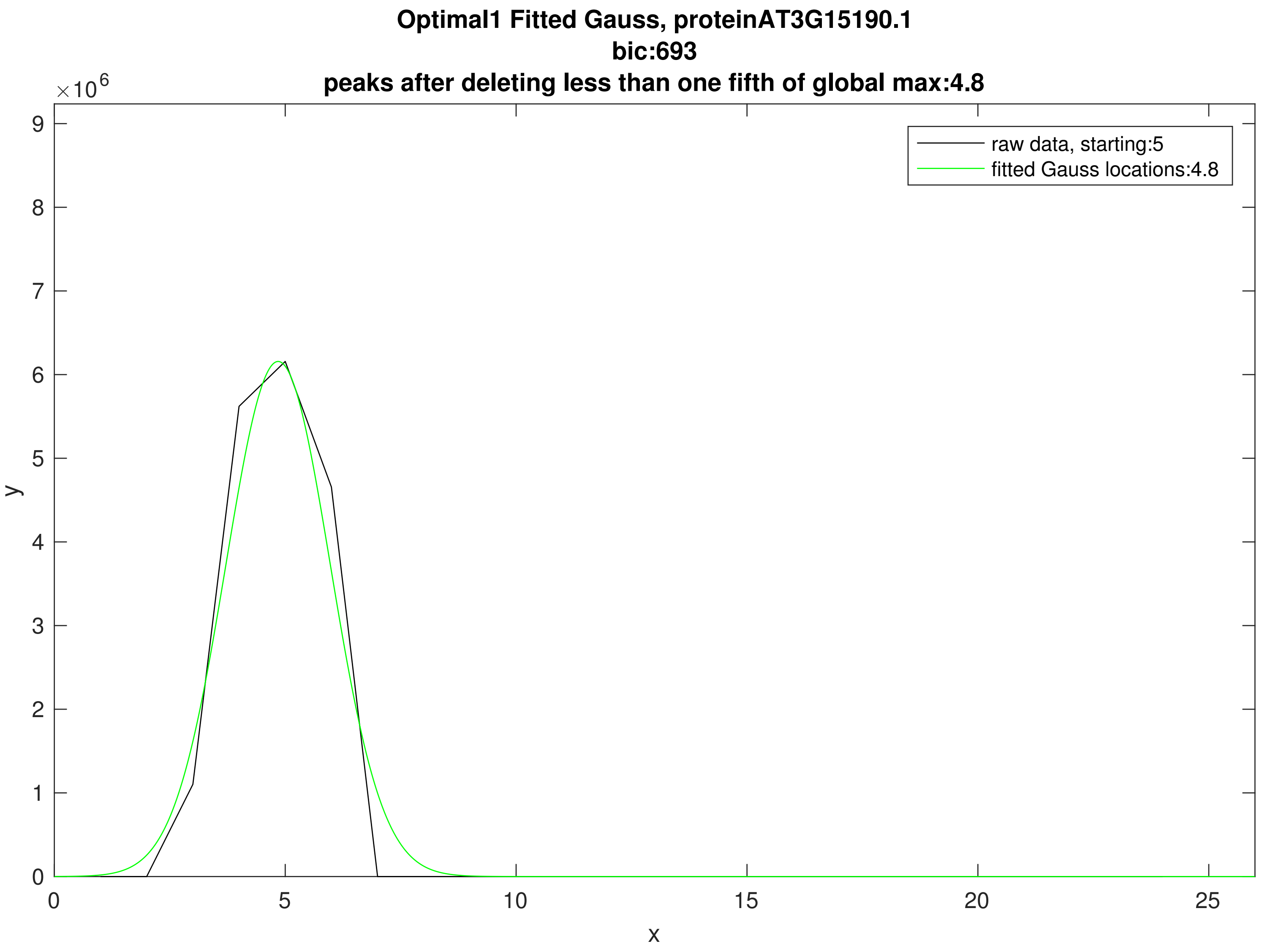
Optimal1 Fitted Gauss, proteinAT3G15190.1
bic:693
peaks after deleting less than one fifth of global max:4.8
×10^6
raw data, starting:5
fitted Gauss locations:4.8
y
x
0
1
2
3
4
5
6
7
8
9
0
5
10
15
20
25

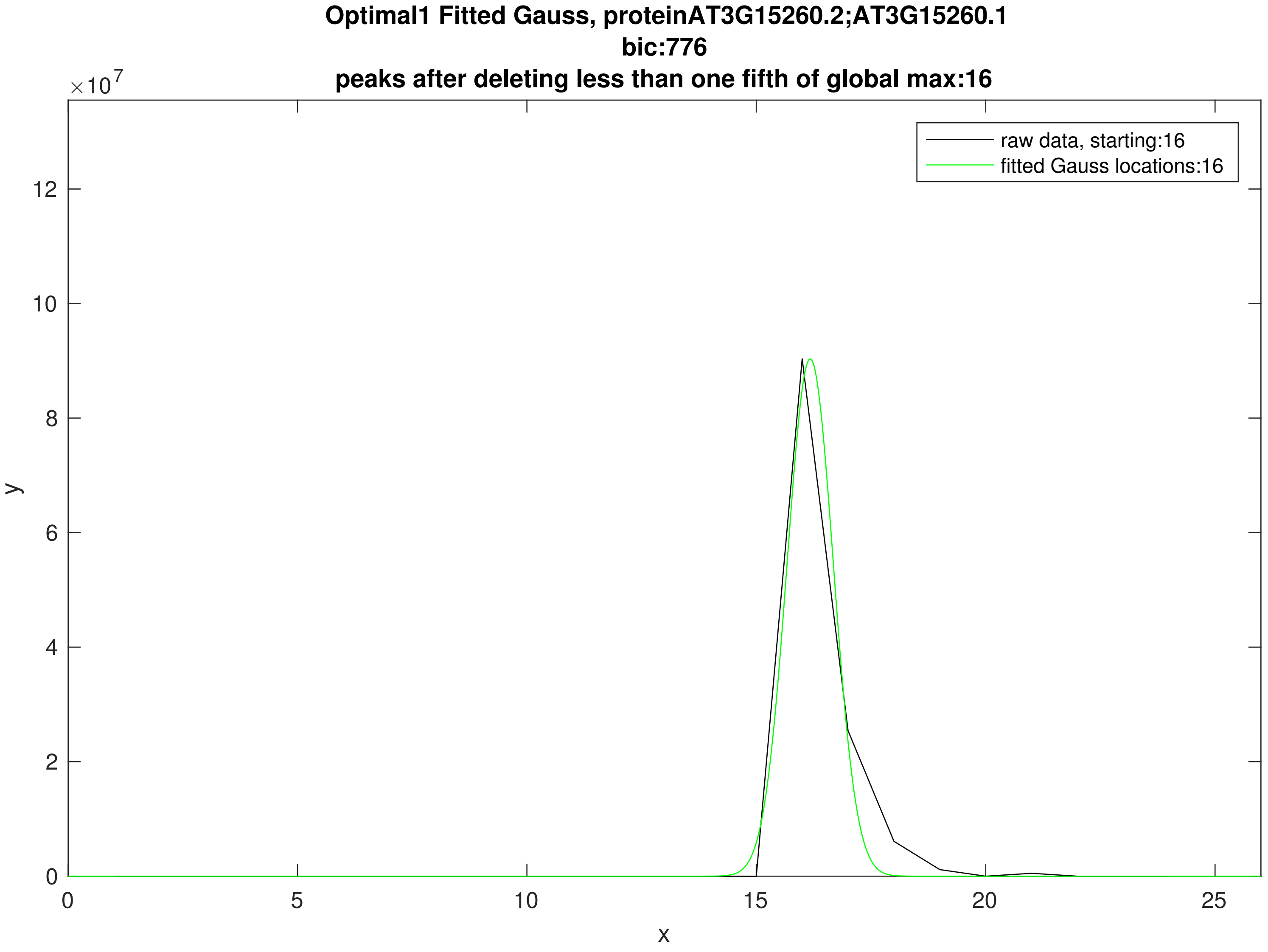
Optimal1 Fitted Gauss, proteinAT3G15260.2;AT3G15260.1
bic:776
peaks after deleting less than one fifth of global max:16
×10^7
raw data, starting:16
fitted Gauss locations:16
12
10
8
6
4
2
0
y
0
5
10
15
20
25
x

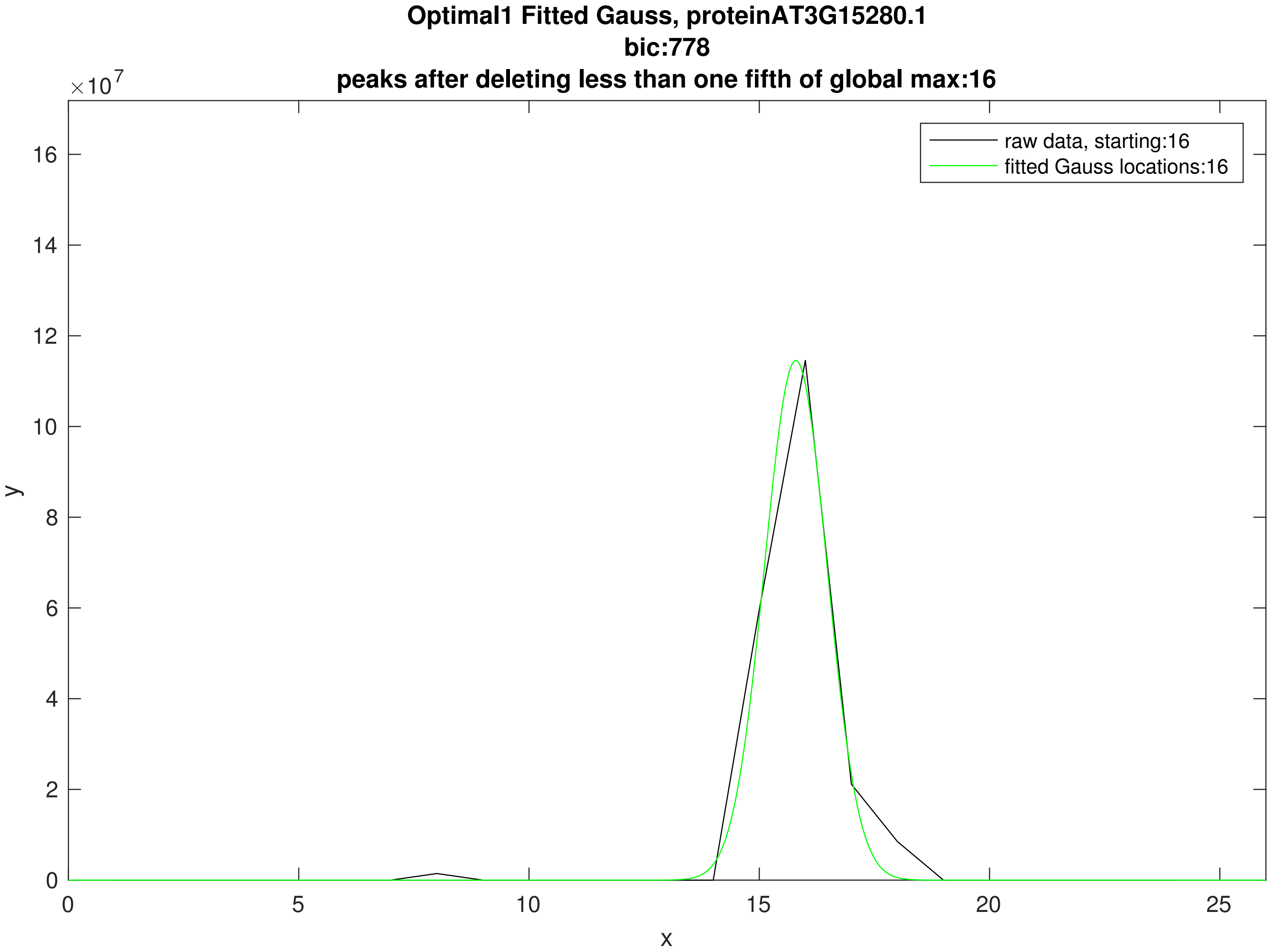

Optimal1 Fitted Gauss, proteinAT3G15280.1
bic:778
peaks after deleting less than one fifth of global max:16
×10^7
16
14
12
10
8
6
4
2
0
y
0
5
10
15
20
25
x
raw data, starting:16
fitted Gauss locations:16

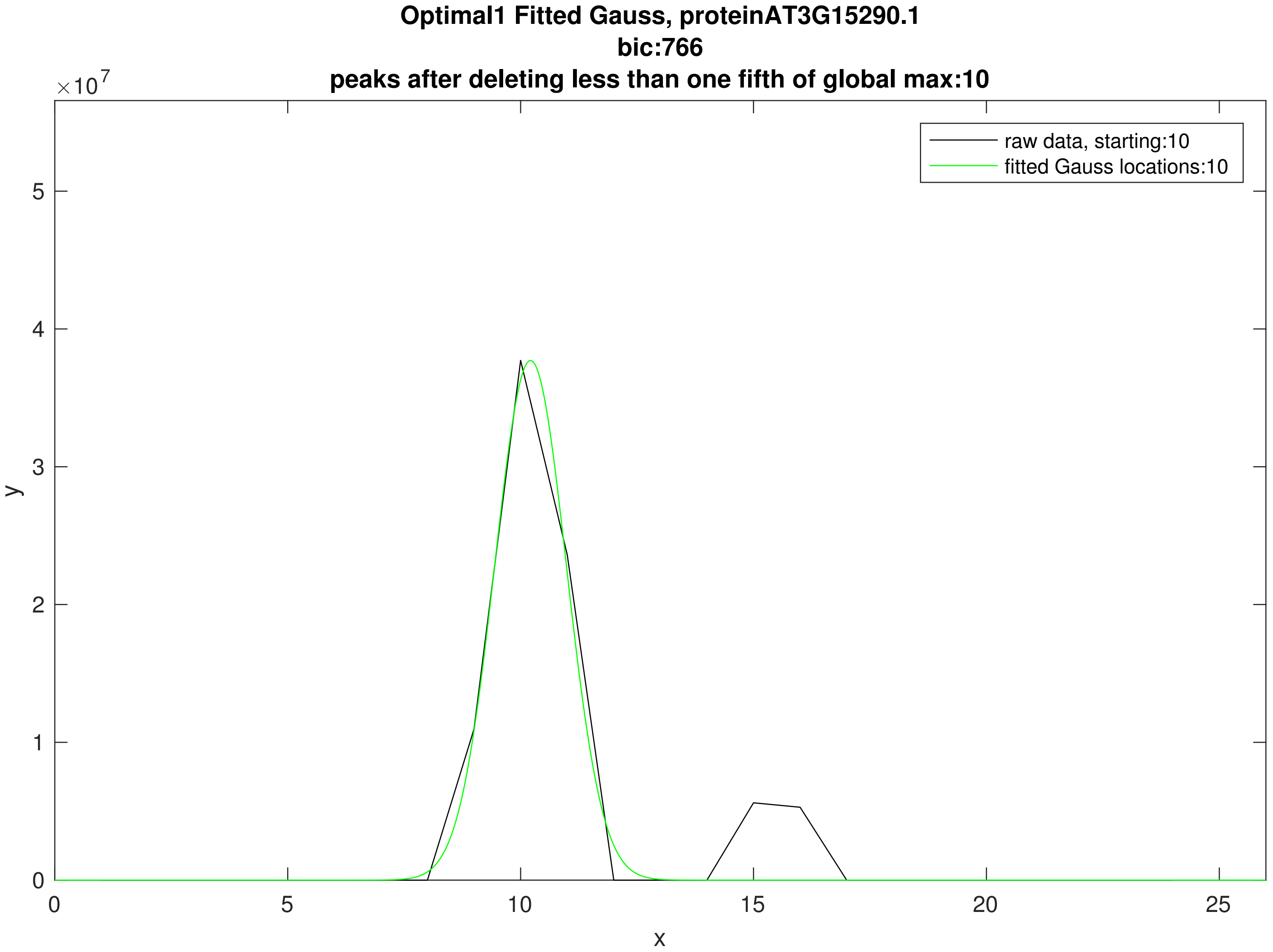
Optimal1 Fitted Gauss, proteinAT3G15290.1
bic:766
peaks after deleting less than one fifth of global max:10
×10^7
raw data, starting:10
fitted Gauss locations:10
y
x
0
1
2
3
4
5
0
5
10
15
20
25

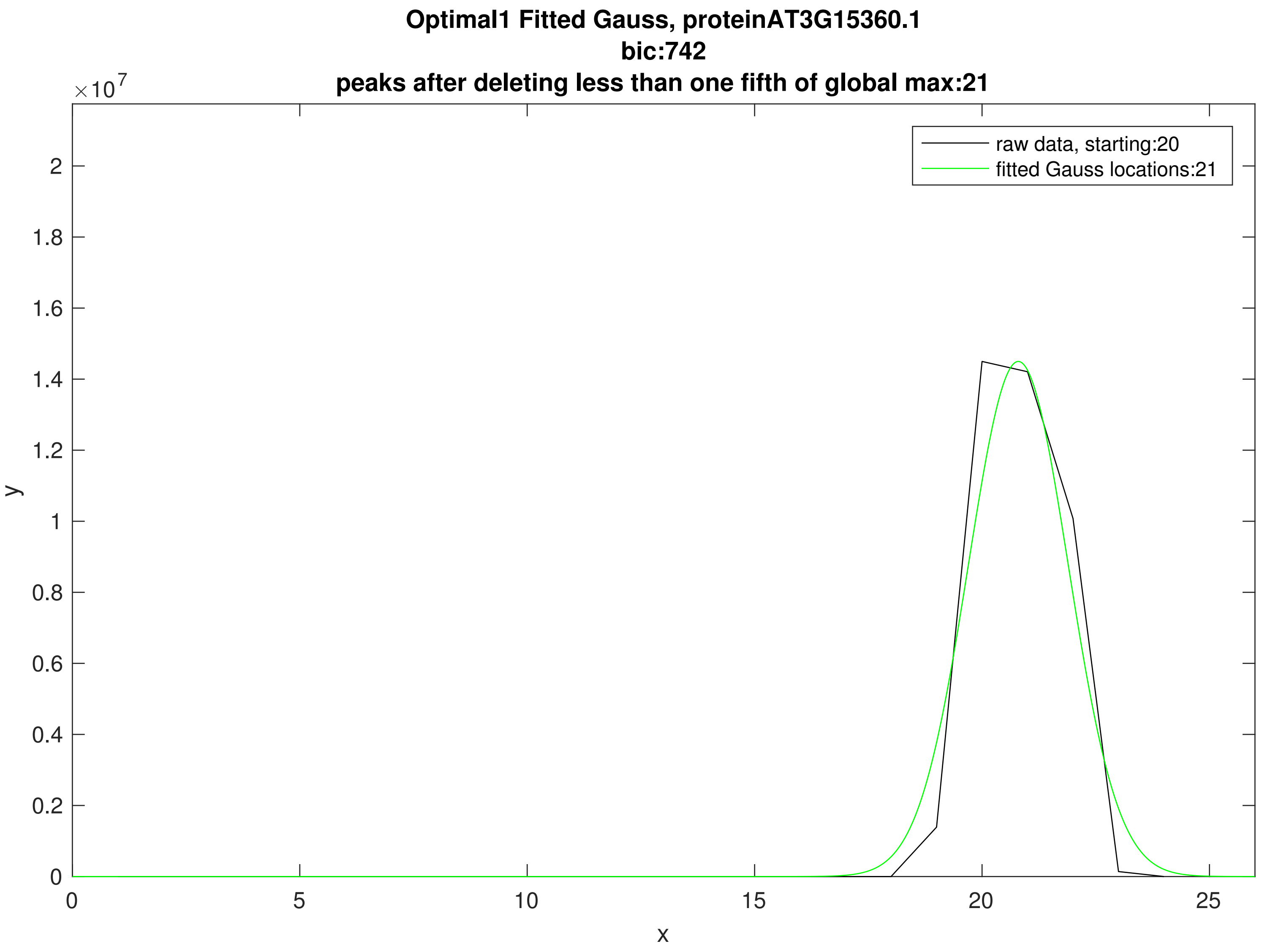

Optimal1 Fitted Gauss, proteinAT3G15360.1
bic:742
peaks after deleting less than one fifth of global max:21
×10^7
raw data, starting:20
fitted Gauss locations:21
2
1.8
1.6
1.4
1.2
1
0.8
0.6
0.4
0.2
0
y
0
5
10
15
20
25
x

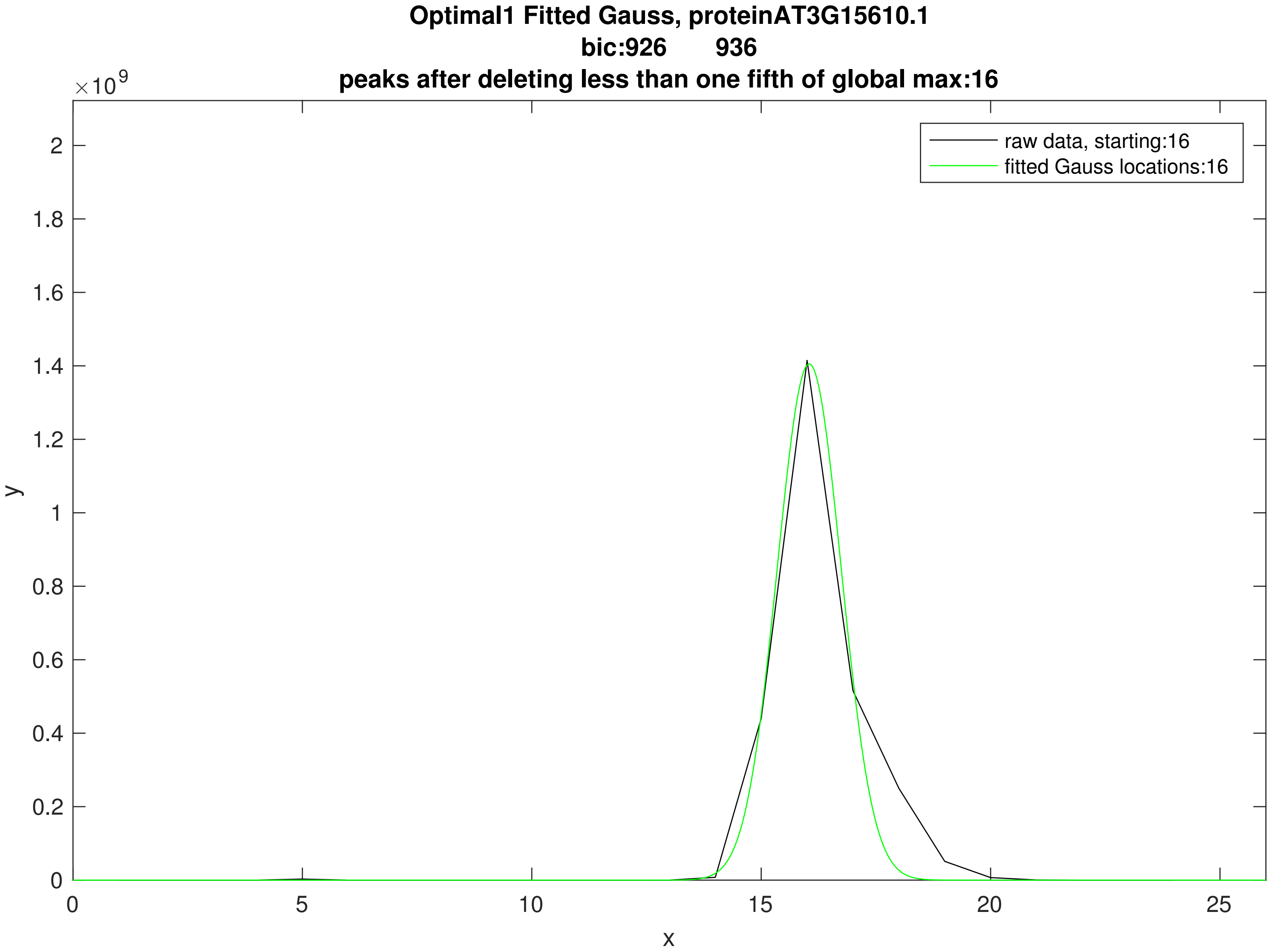

Optimal1 Fitted Gauss, proteinAT3G15610.1
bic:926 936
peaks after deleting less than one fifth of global max:16
×10^9
raw data, starting:16
fitted Gauss locations:16
y
x
0
0.2
0.4
0.6
0.8
1
1.2
1.4
1.6
1.8
2
5
10
15
20
25

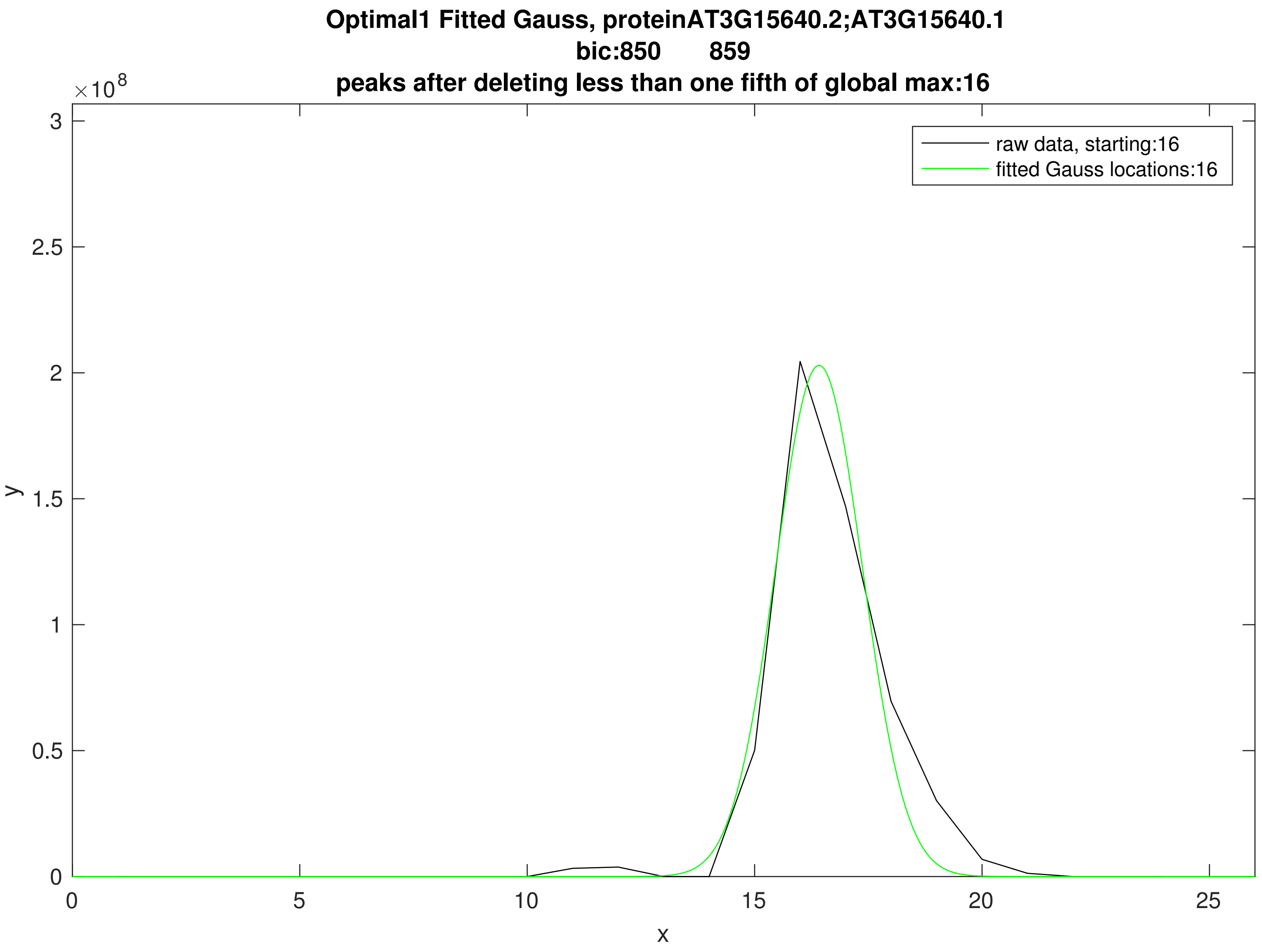
Optimal1 Fitted Gauss, proteinAT3G15640.2;AT3G15640.1
bic:850 859
peaks after deleting less than one fifth of global max:16
×10^8
3
2.5
2
1.5
1
0.5
0
y
0
5
10
15
20
25
x
raw data, starting:16
fitted Gauss locations:16

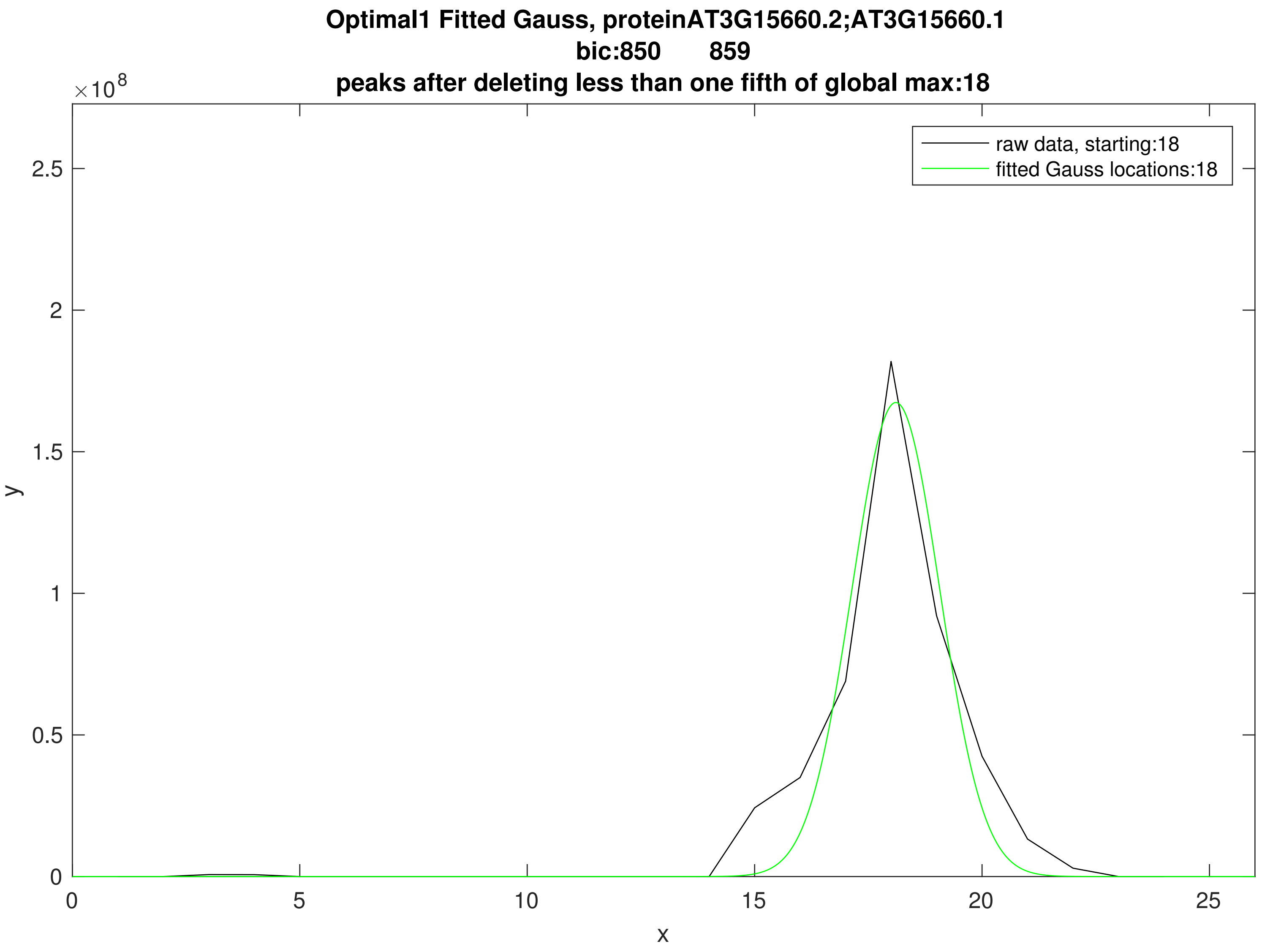
Optimal1 Fitted Gauss, proteinAT3G15660.2;AT3G15660.1
bic:850 859
peaks after deleting less than one fifth of global max:18
×10^8
2.5
2
1.5
1
0.5
0
y
0
5
10
15
20
25
x
raw data, starting:18
fitted Gauss locations:18

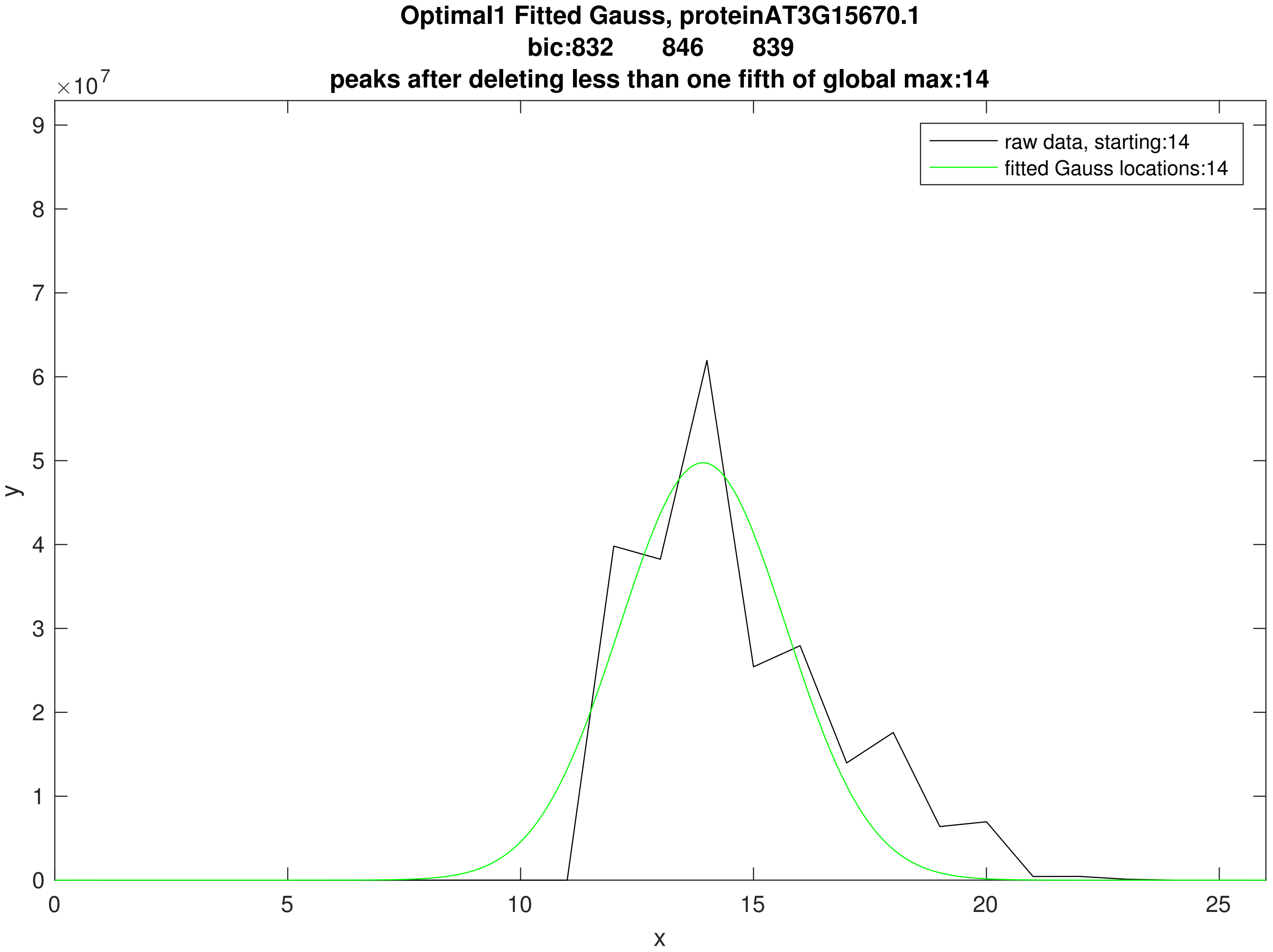
Optimal1 Fitted Gauss, proteinAT3G15670.1
bic:832 846 839
peaks after deleting less than one fifth of global max:14
×10^7
raw data, starting:14
fitted Gauss locations:14
y
x
0
1
2
3
4
5
6
7
8
9
0
5
10
15
20
25

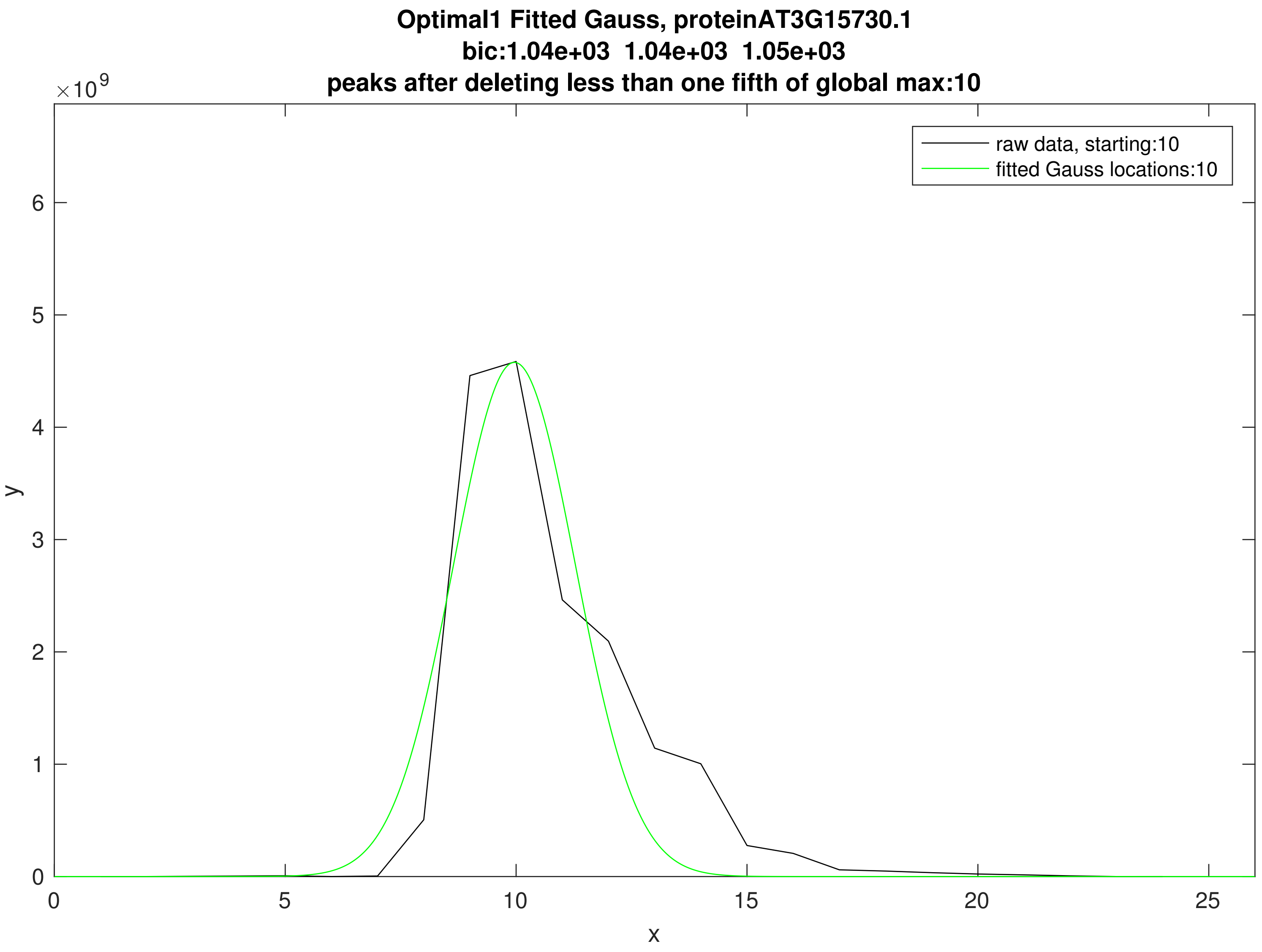
Optimal1 Fitted Gauss, proteinAT3G15730.1
bic:1.04e+03 1.04e+03 1.05e+03
peaks after deleting less than one fifth of global max:10
×10^9
raw data, starting:10
fitted Gauss locations:10
y
x
0
1
2
3
4
5
6
0
5
10
15
20
25

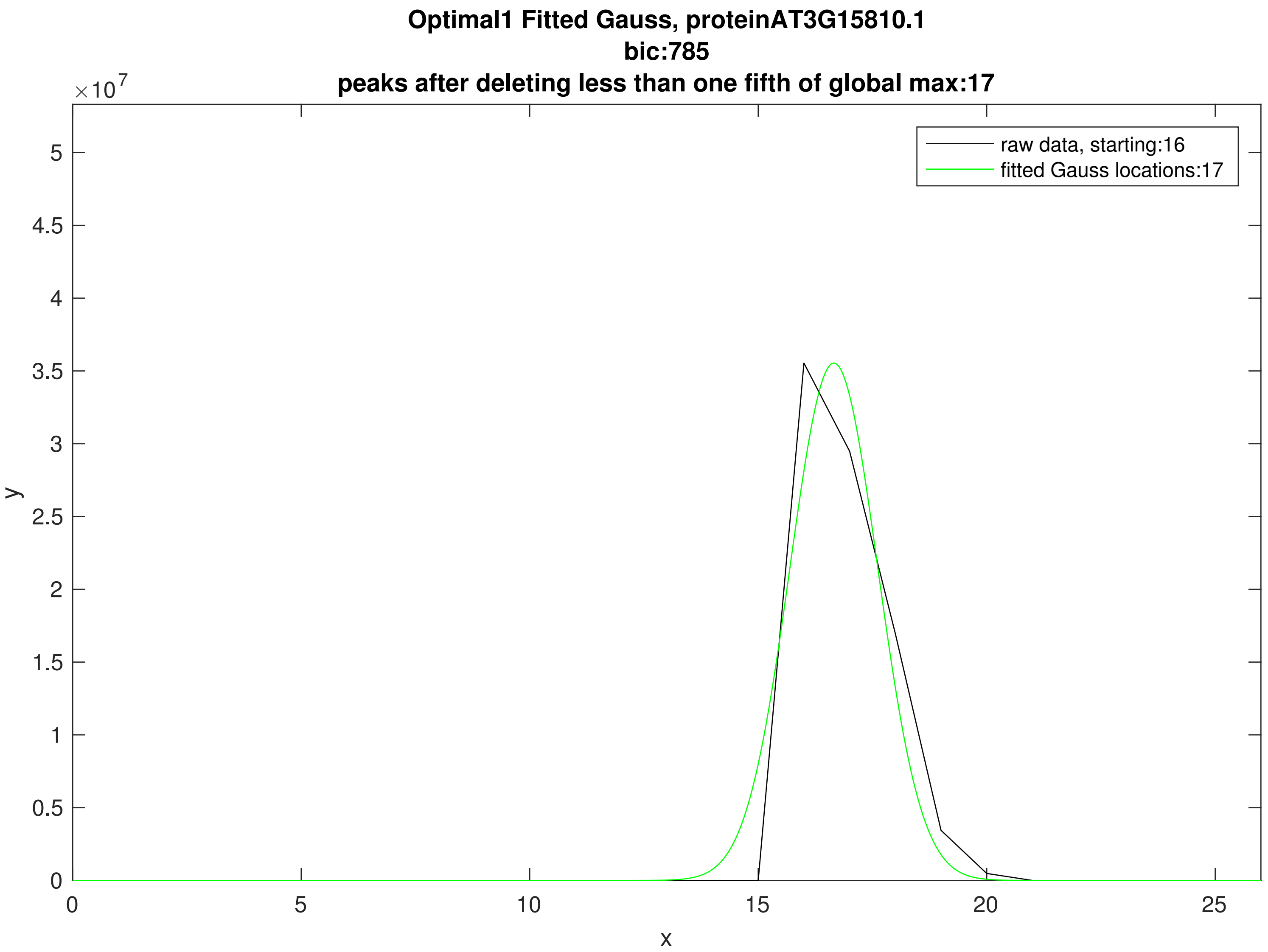
Optimal1 Fitted Gauss, proteinAT3G15810.1
bic:785
peaks after deleting less than one fifth of global max:17
×10^7
raw data, starting:16
fitted Gauss locations:17
y
x
0
0.5
1
1.5
2
2.5
3
3.5
4
4.5
5
0
5
10
15
20
25

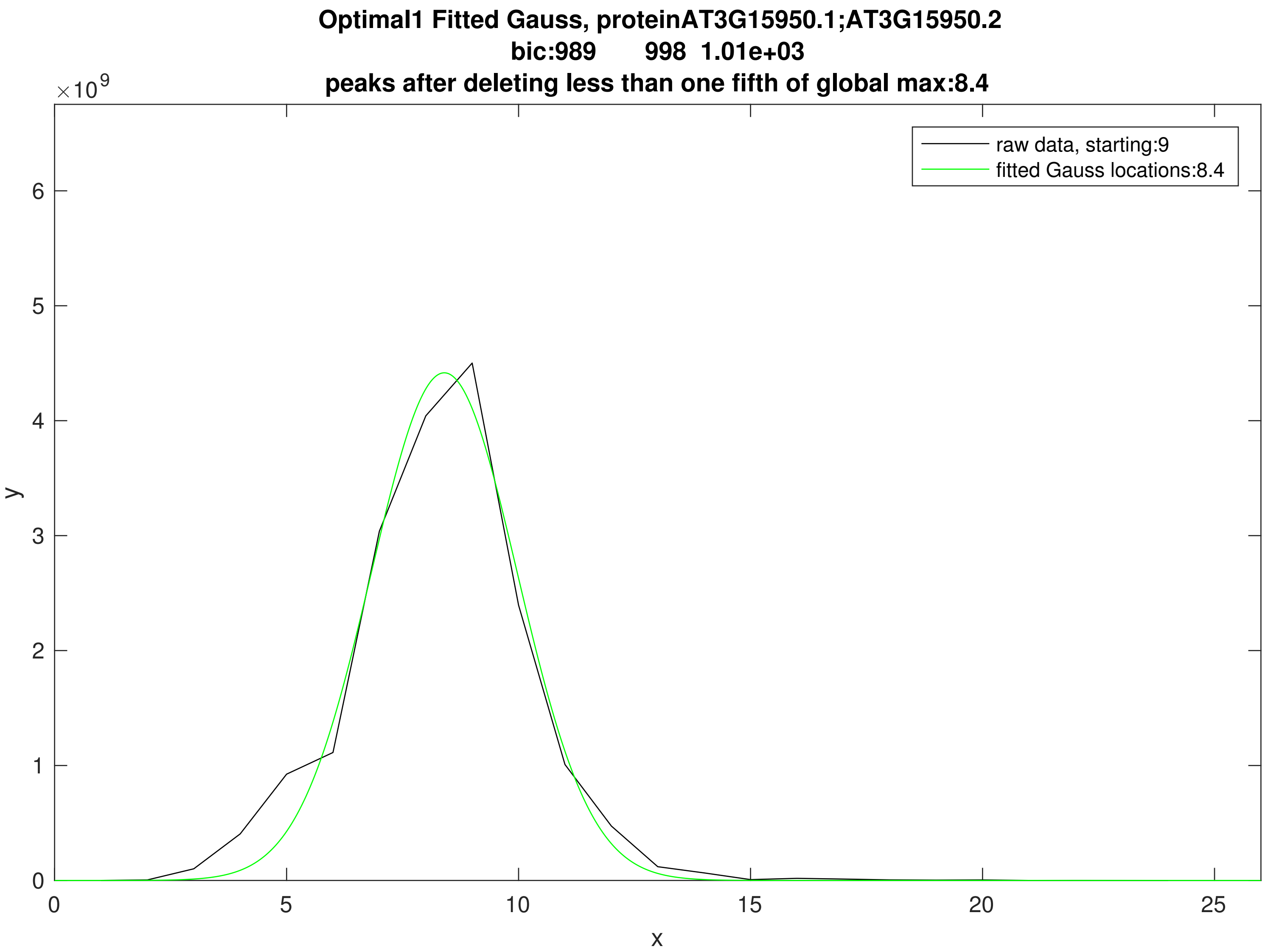
Optimal1 Fitted Gauss, proteinAT3G15950.1;AT3G15950.2
bic:989 998 1.01e+03
peaks after deleting less than one fifth of global max:8.4
×10^9
raw data, starting:9
fitted Gauss locations:8.4
y
x
0
1
2
3
4
5
6
0
5
10
15
20
25

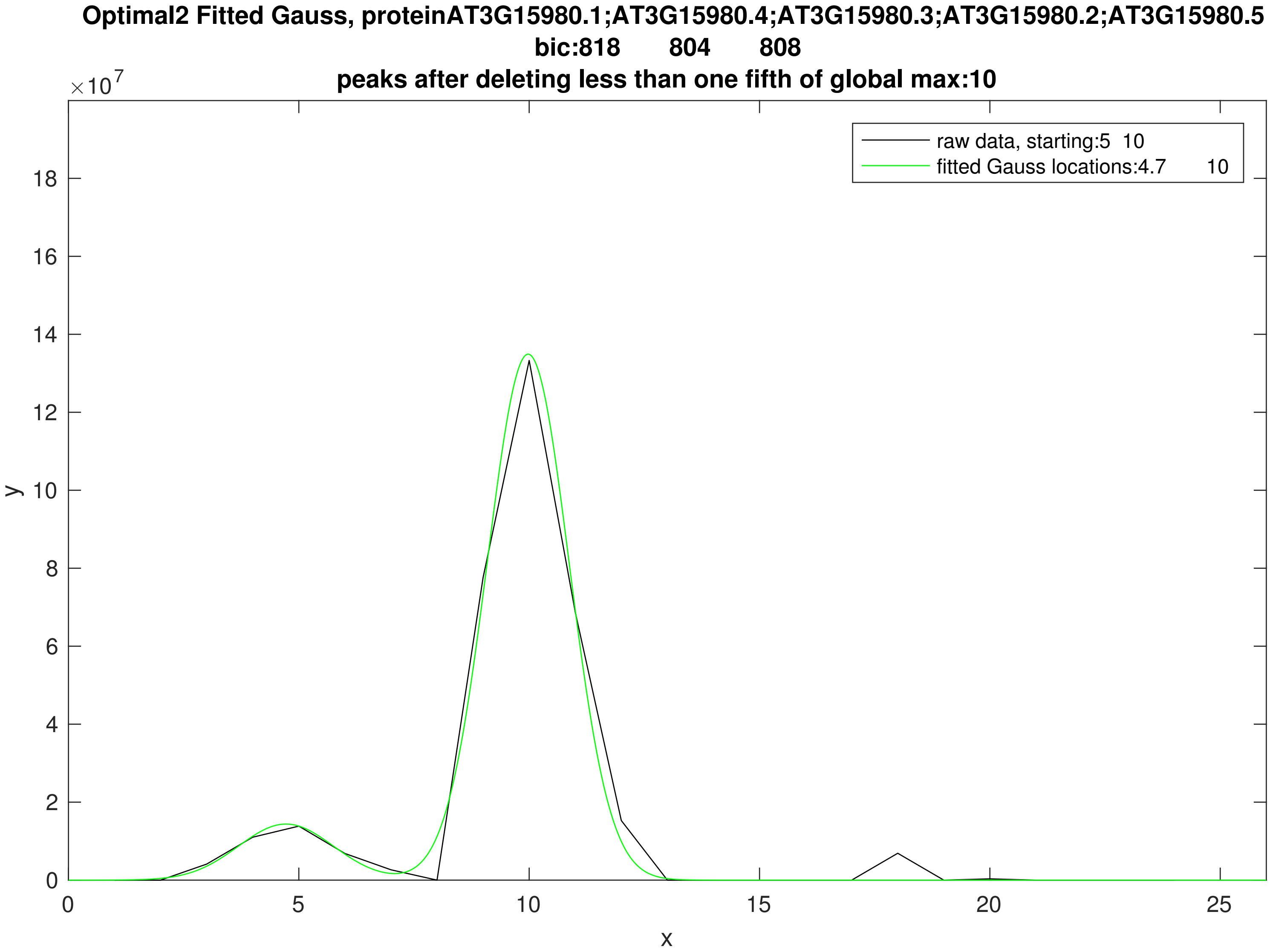
Optimal2 Fitted Gauss, proteinAT3G15980.1;AT3G15980.4;AT3G15980.3;AT3G15980.2;AT3G15980.5
bic:818 804 808
peaks after deleting less than one fifth of global max:10
×10^7
raw data, starting:5 10
fitted Gauss locations:4.7 10
y
x
0
2
4
6
8
10
12
14
16
18
0
5
10
15
20
25

**Optimal1 Fitted Gauss, proteinAT3G16050.1**
**bic:846**
**peaks after deleting less than one fifth of global max:6.5**

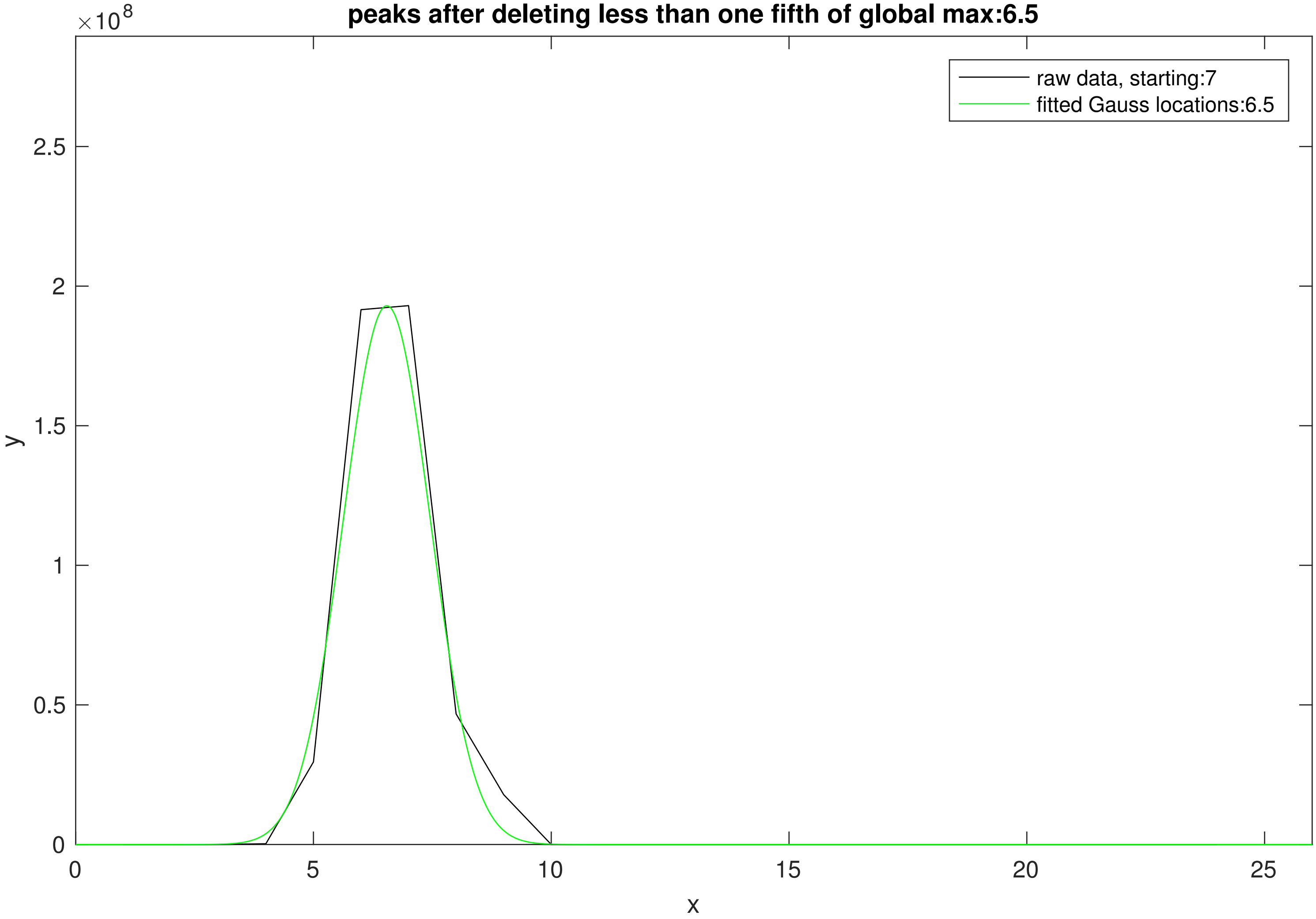

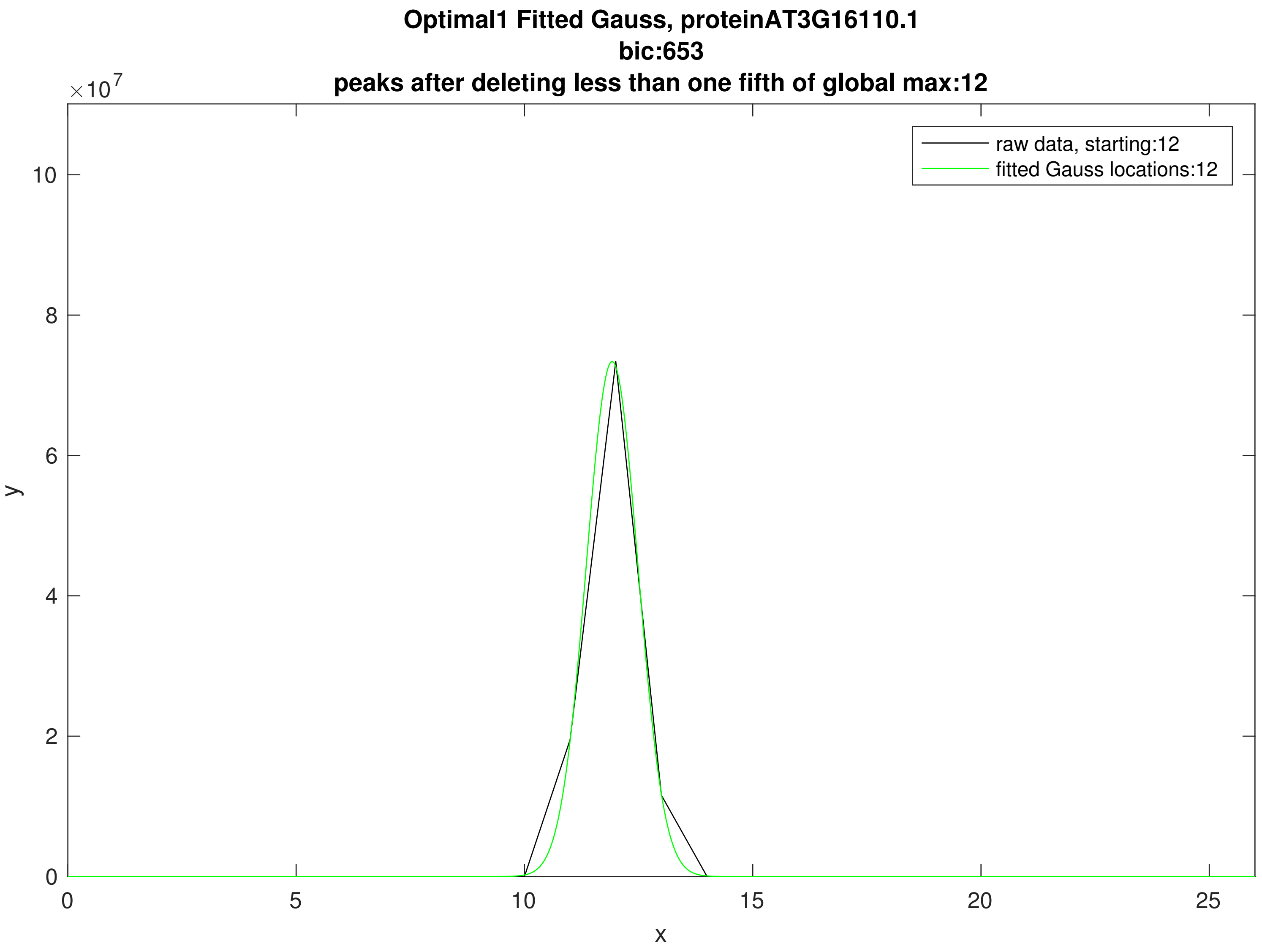
Optimal1 Fitted Gauss, proteinAT3G16110.1
bic:653
peaks after deleting less than one fifth of global max:12
×10^7
raw data, starting:12
fitted Gauss locations:12
y
x
0
2
4
6
8
10
0
5
10
15
20
25

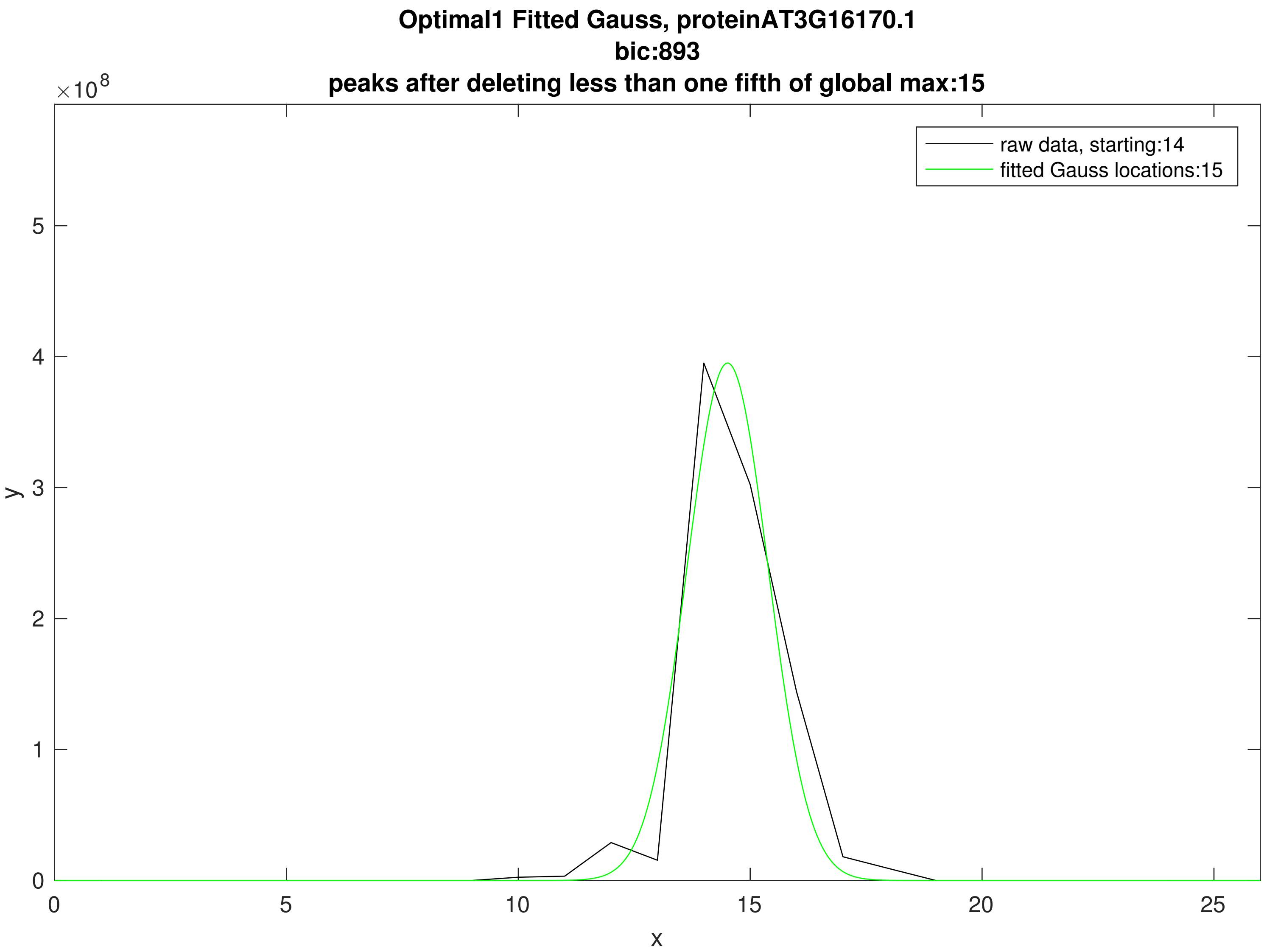

Optimal1 Fitted Gauss, proteinAT3G16170.1
bic:893
peaks after deleting less than one fifth of global max:15
×10^8
raw data, starting:14
fitted Gauss locations:15
5
4
3
2
1
0
y
0
5
10
15
20
25
x

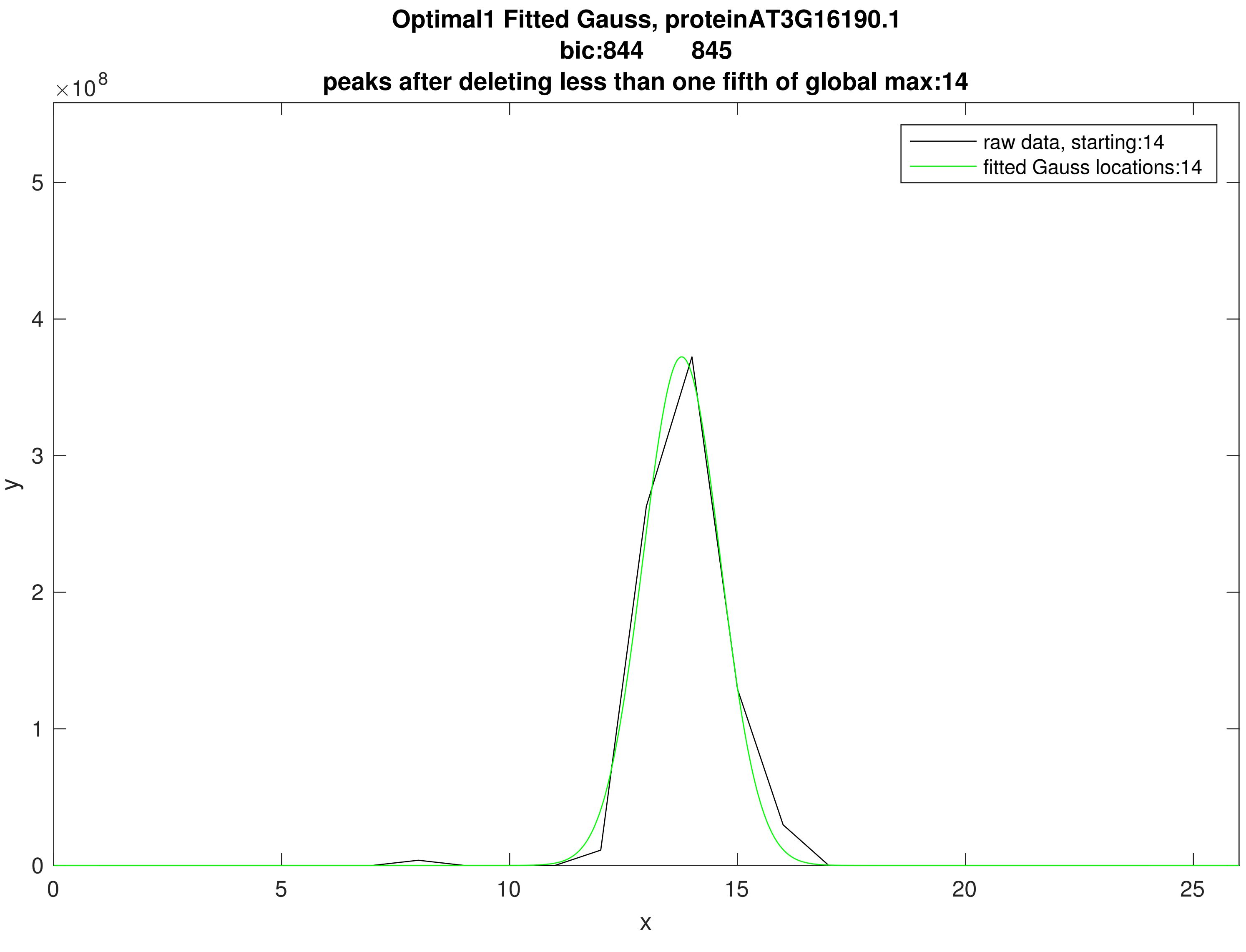
Optimal1 Fitted Gauss, proteinAT3G16190.1
bic:844 845
peaks after deleting less than one fifth of global max:14
×10^8
raw data, starting:14
fitted Gauss locations:14
y
x
0
1
2
3
4
5
0
5
10
15
20
25

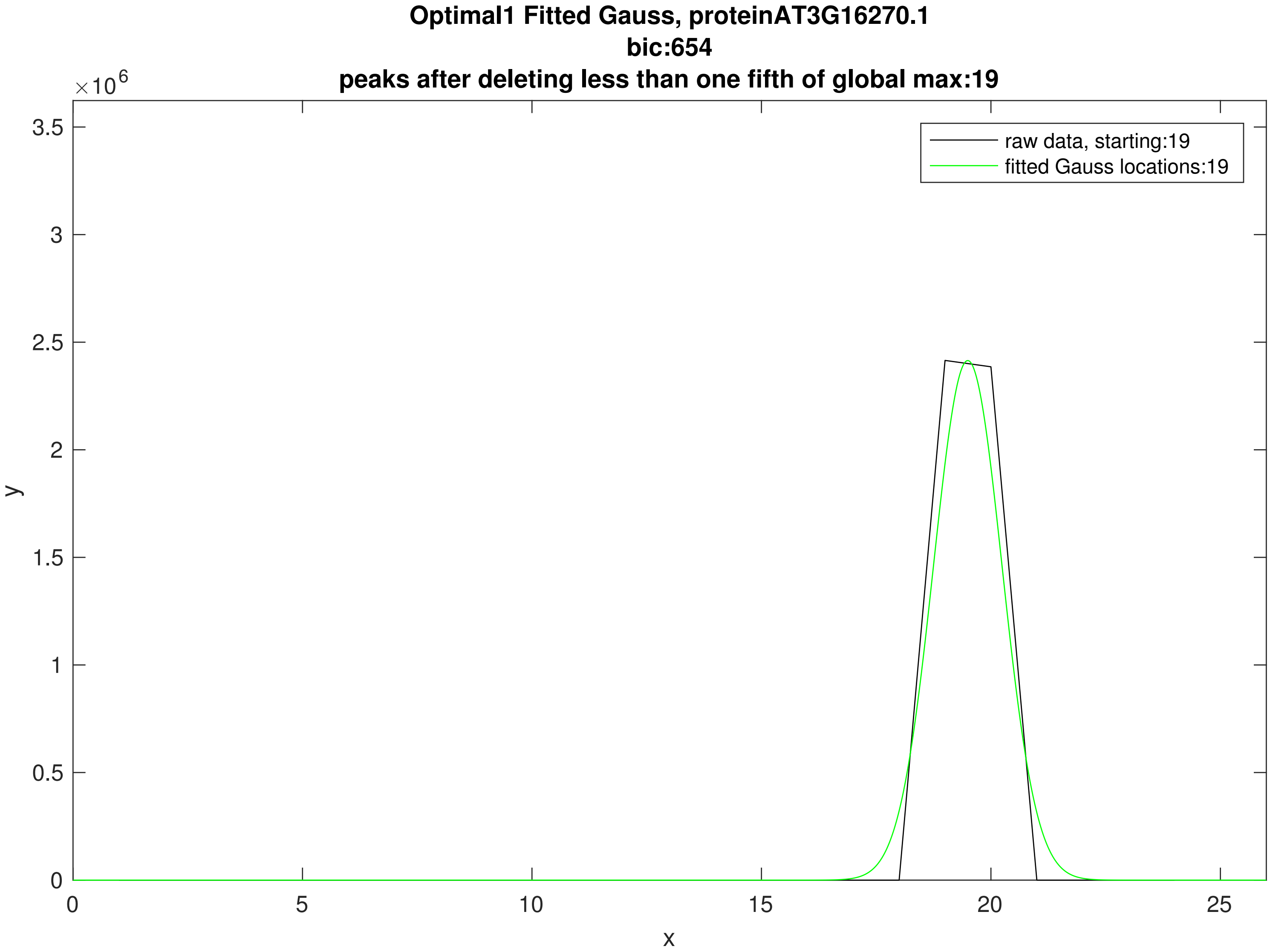

Optimal1 Fitted Gauss, proteinAT3G16270.1
bic:654
peaks after deleting less than one fifth of global max:19
×10^6
3.5
3
2.5
2
1.5
1
0.5
0
y
0
5
10
15
20
25
x
raw data, starting:19
fitted Gauss locations:19

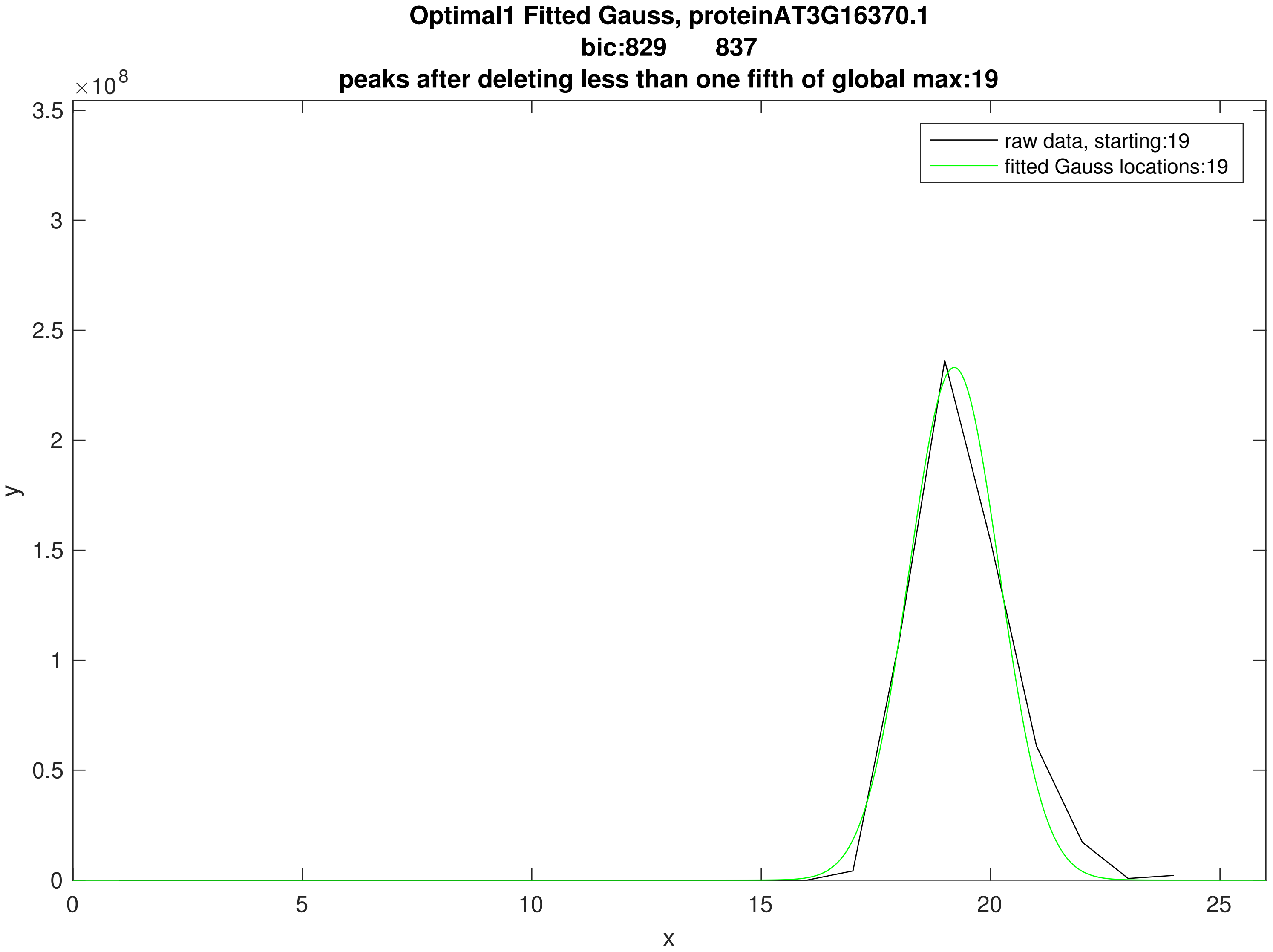
Optimal1 Fitted Gauss, proteinAT3G16370.1
bic:829 837
peaks after deleting less than one fifth of global max:19
×10^8
3.5
3
2.5
2
1.5
1
0.5
0
y
0
5
10
15
20
25
x
raw data, starting:19
fitted Gauss locations:19

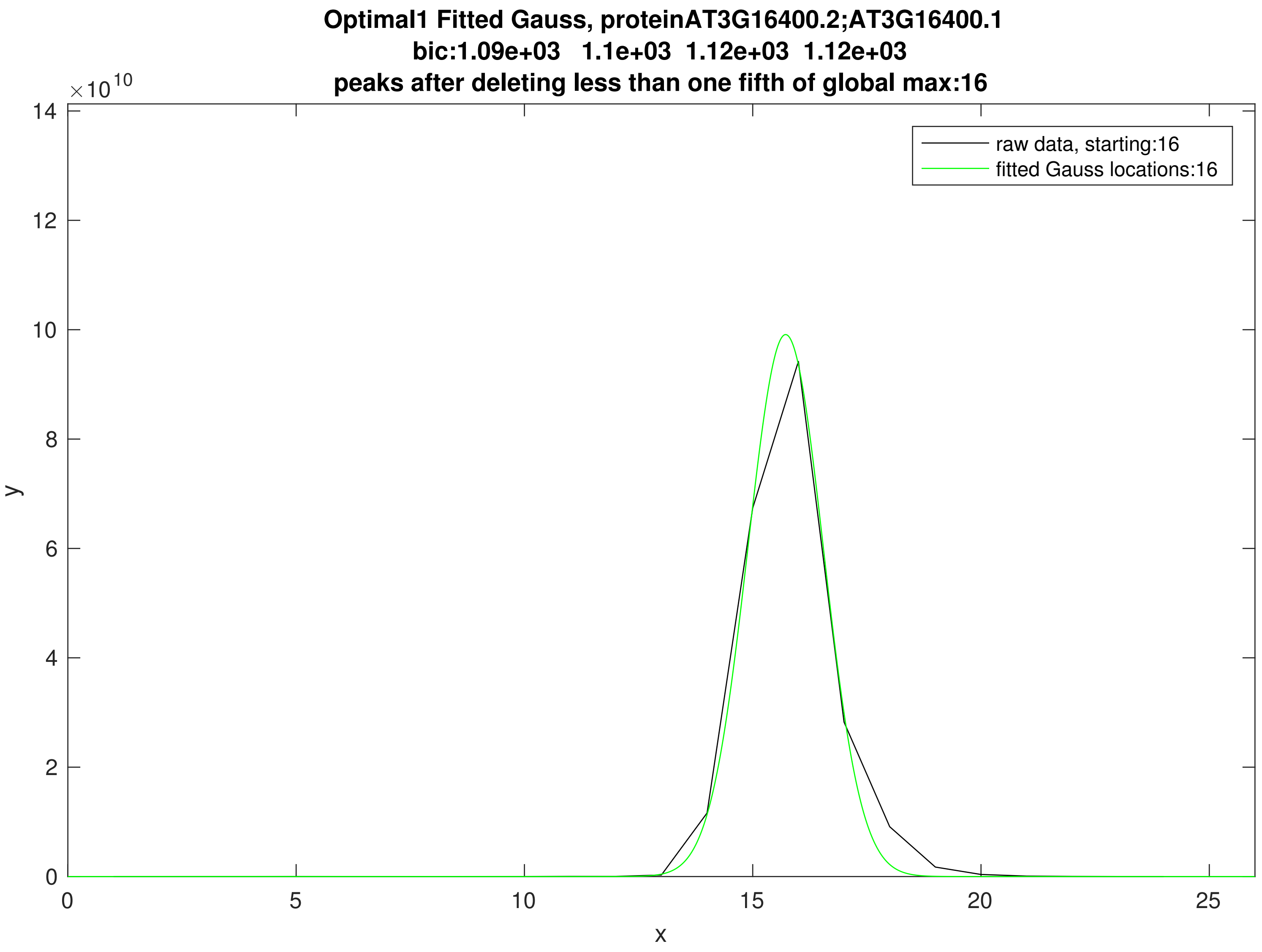
Optimal1 Fitted Gauss, proteinAT3G16400.2;AT3G16400.1
bic:1.09e+03 1.1e+03 1.12e+03 1.12e+03
peaks after deleting less than one fifth of global max:16
×10^10
raw data, starting:16
fitted Gauss locations:16
y
x
0
2
4
6
8
10
12
14
5
15
20
25

**Optimal1 Fitted Gauss, proteinAT3G16410.1**
**bic:952 958 967**
**peaks after deleting less than one fifth of global max:16**

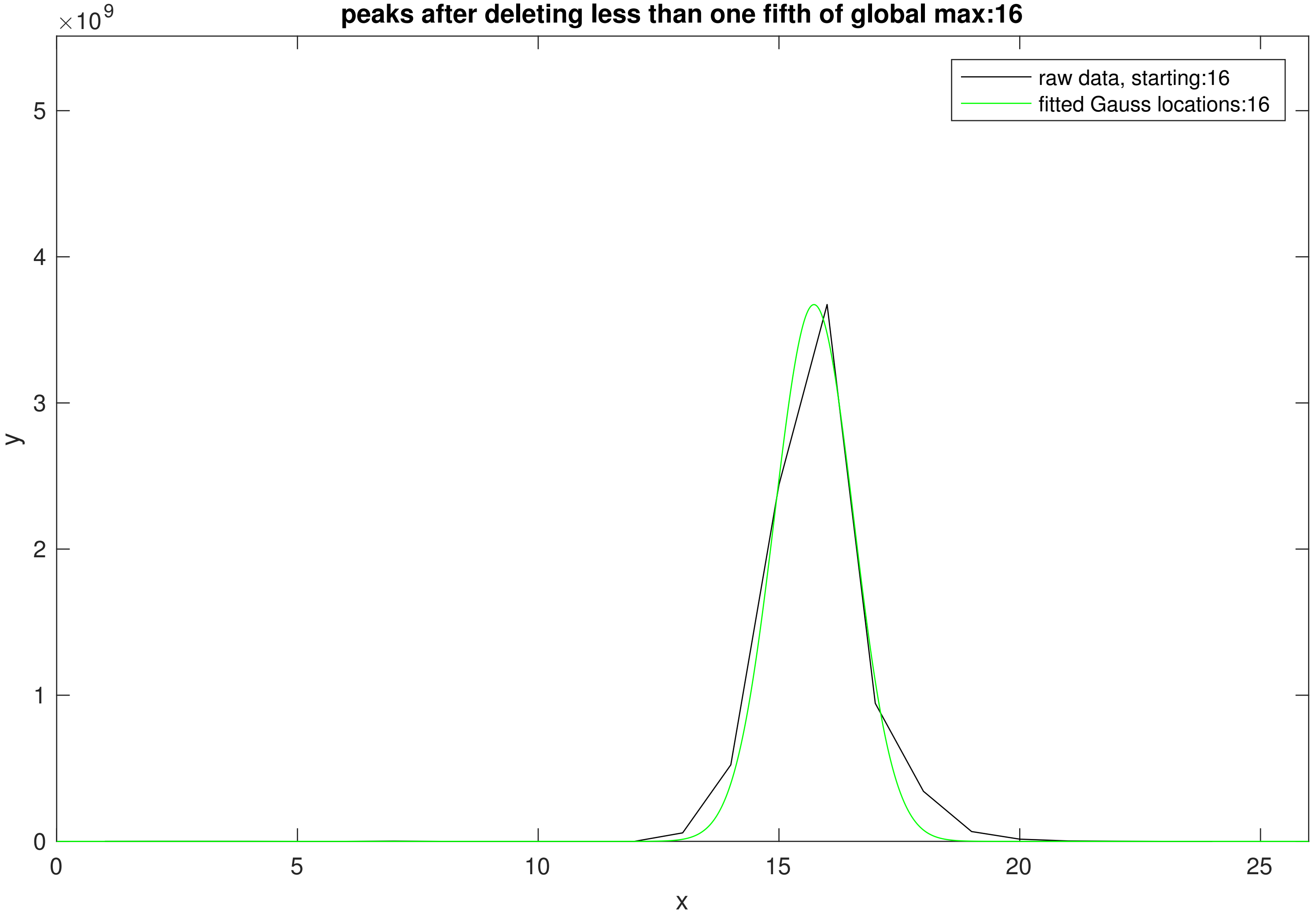

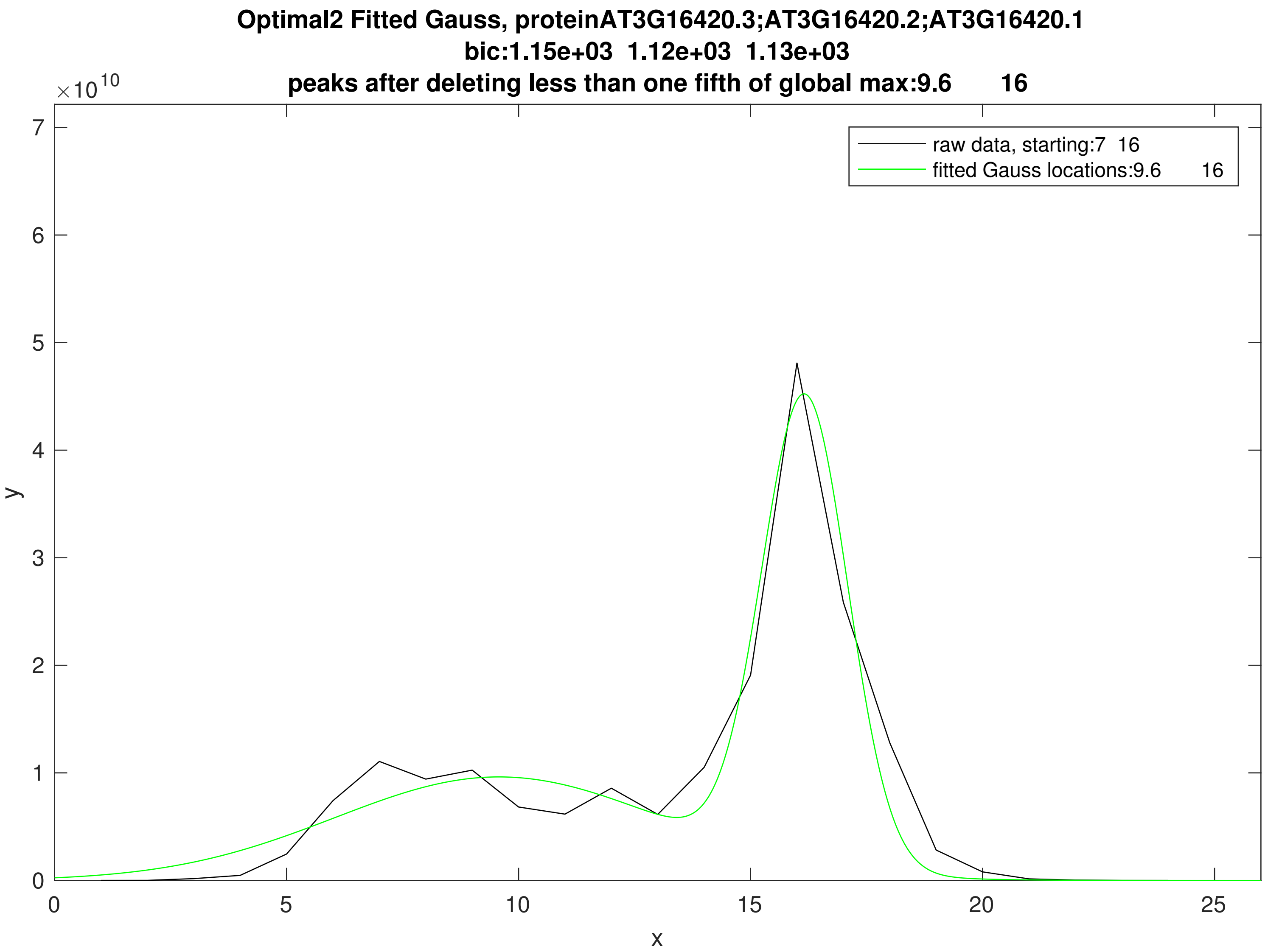

Optimal2 Fitted Gauss, proteinAT3G16420.3;AT3G16420.2;AT3G16420.1
bic:1.15e+03 1.12e+03 1.13e+03
peaks after deleting less than one fifth of global max:9.6 16
×10^10
raw data, starting:7 16
fitted Gauss locations:9.6 16
y
x
0
1
2
3
4
5
6
7
0
5
10
15
20
25

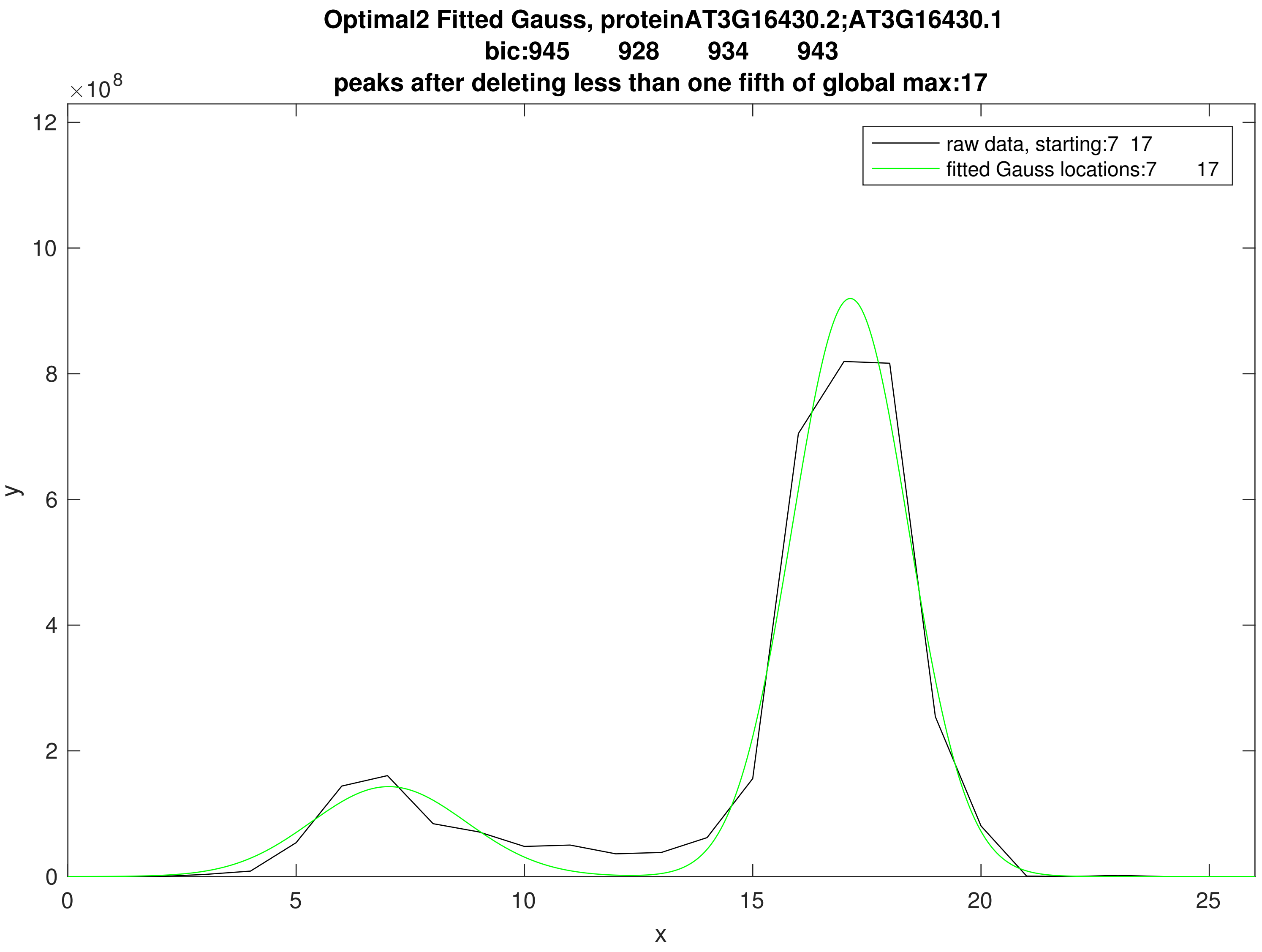
Optimal2 Fitted Gauss, proteinAT3G16430.2;AT3G16430.1
bic:945 928 934 943
peaks after deleting less than one fifth of global max:17
raw data, starting:7 17
fitted Gauss locations:7 17
×10^8
y
x
0
2
4
6
8
10
12
5
15
20
25

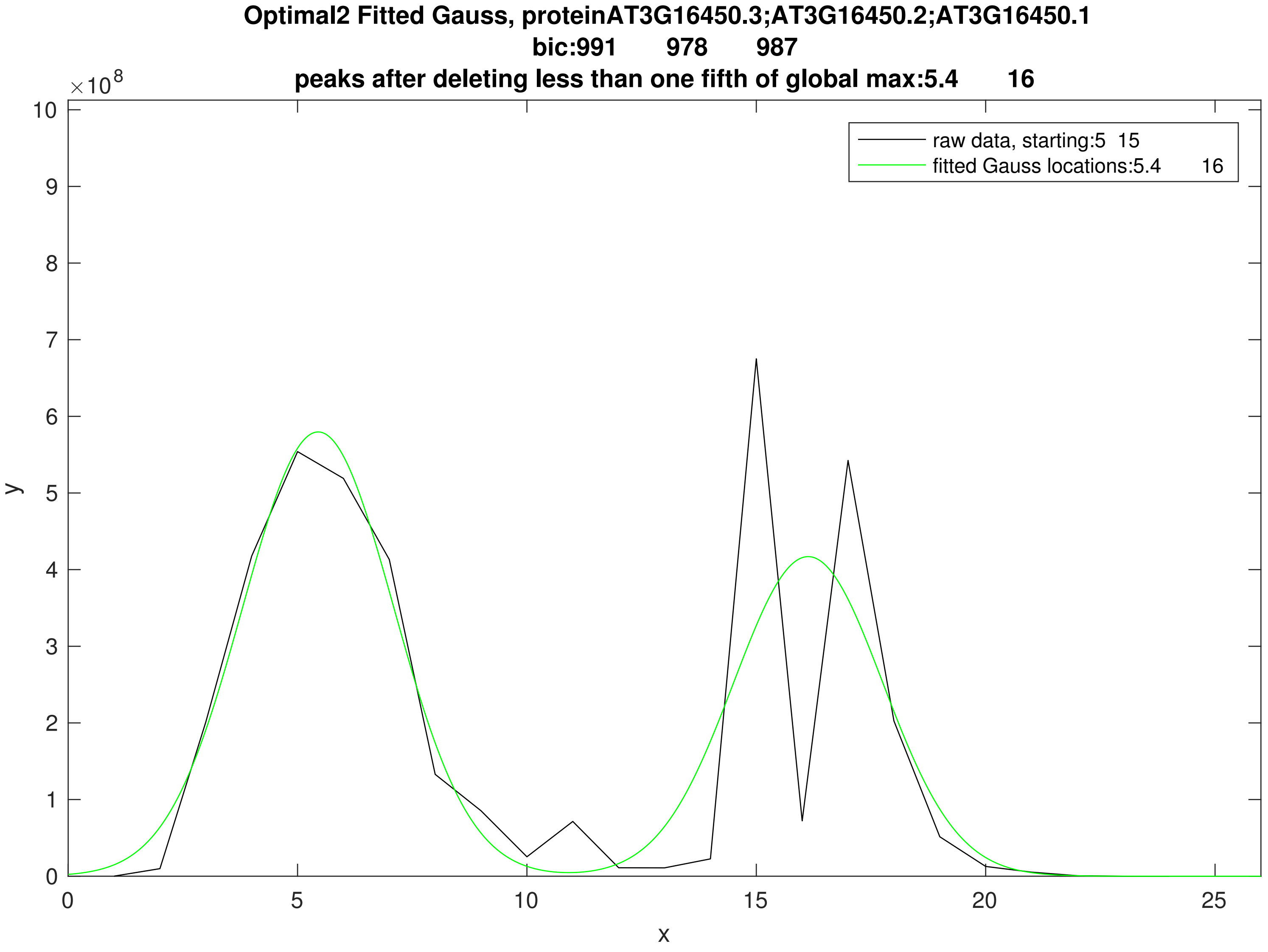

Optimal2 Fitted Gauss, proteinAT3G16450.3;AT3G16450.2;AT3G16450.1
bic:991 978 987
peaks after deleting less than one fifth of global max:5.4 16
×10^8
raw data, starting:5 15
fitted Gauss locations:5.4 16
y
x

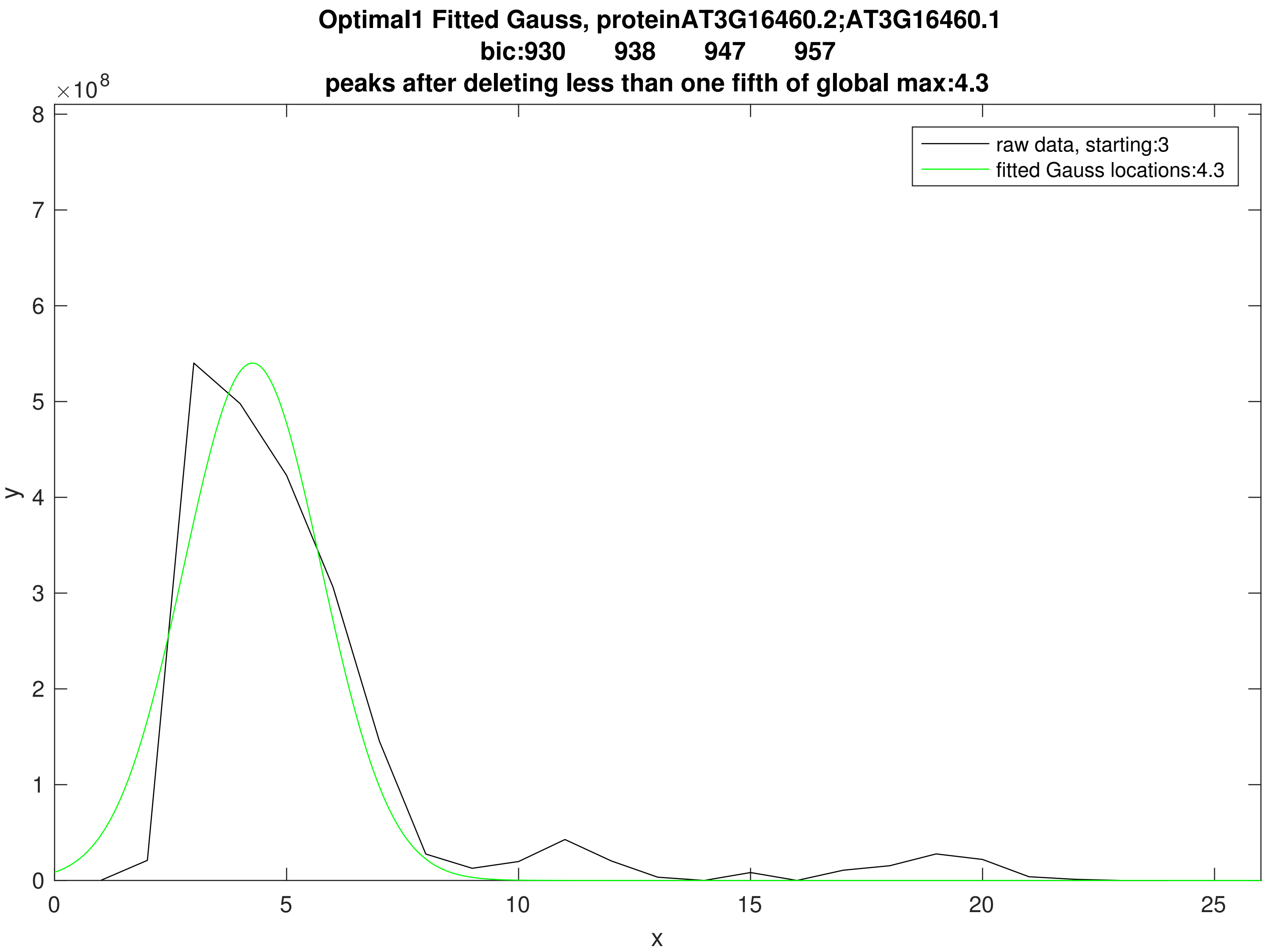

Optimal1 Fitted Gauss, proteinAT3G16460.2;AT3G16460.1
bic:930 938 947 957
peaks after deleting less than one fifth of global max:4.3
×10^8
raw data, starting:3
fitted Gauss locations:4.3
y
x
0
1
2
3
4
5
6
7
8
0
5
10
15
20
25

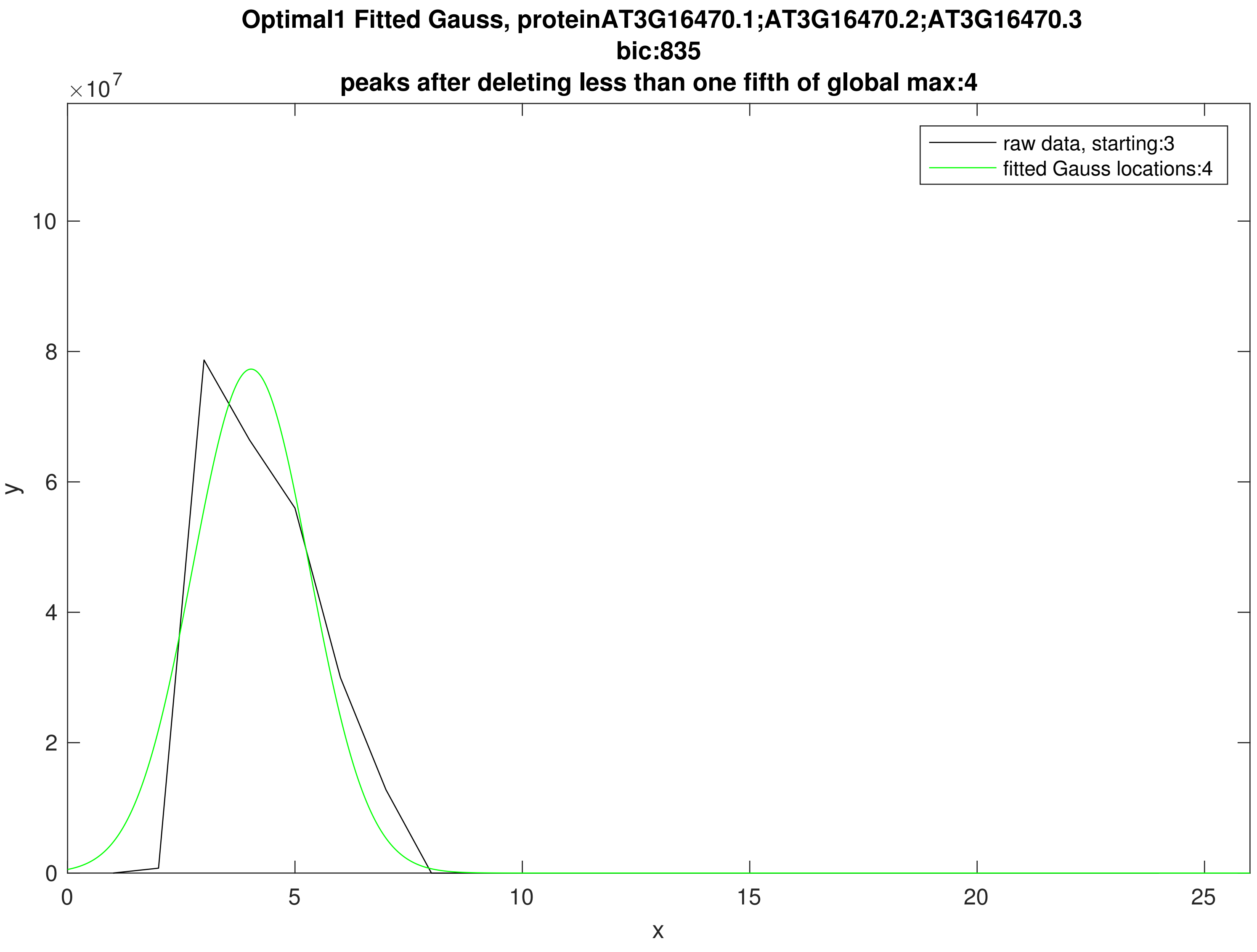
Optimal1 Fitted Gauss, proteinAT3G16470.1;AT3G16470.2;AT3G16470.3
bic:835
peaks after deleting less than one fifth of global max:4
×10^7
raw data, starting:3
fitted Gauss locations:4
10
8
6
4
2
0
y
0
5
10
15
20
25
x

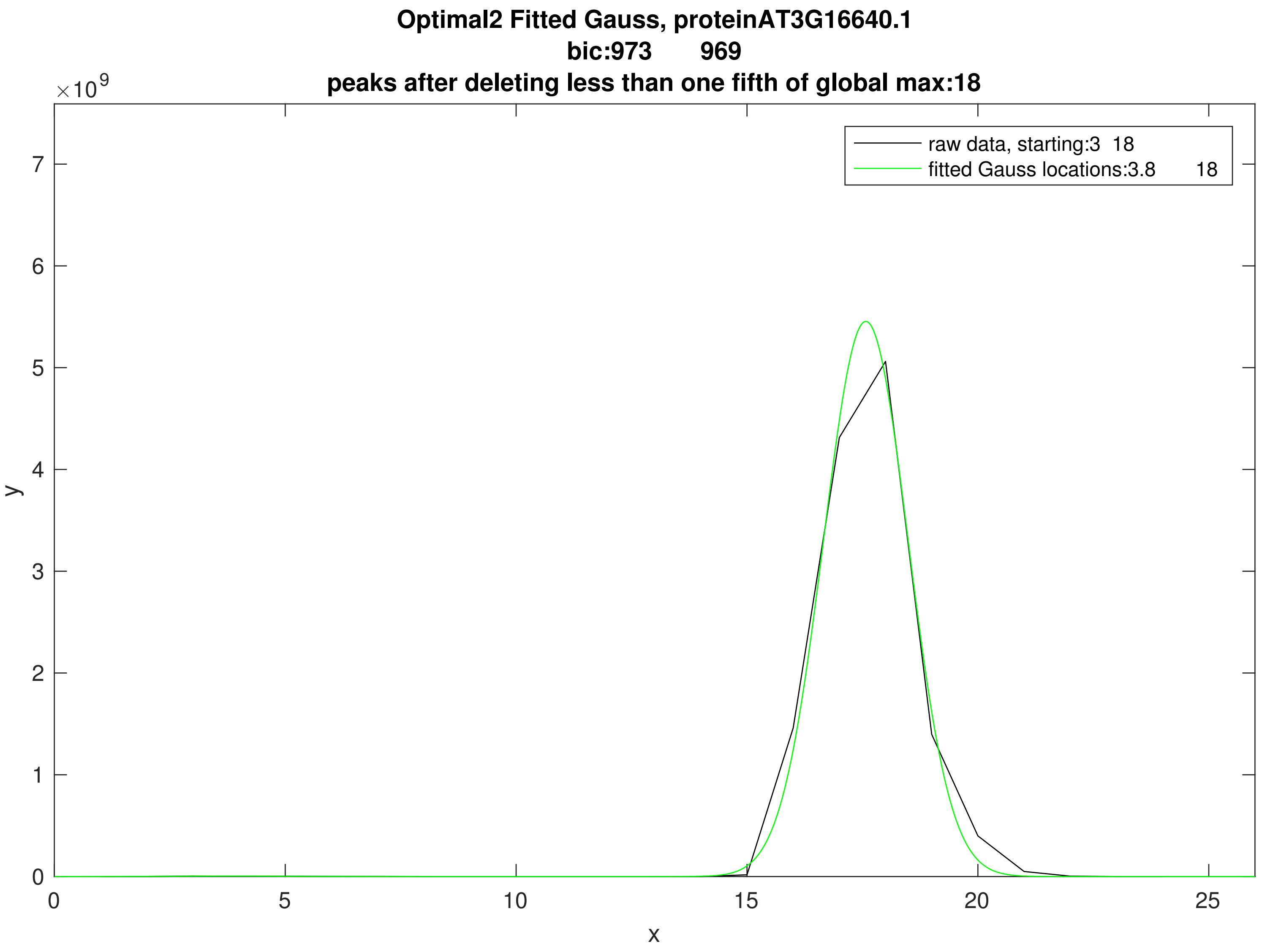

Optimal2 Fitted Gauss, proteinAT3G16640.1
bic:973 969
peaks after deleting less than one fifth of global max:18
×10^9
raw data, starting:3 18
fitted Gauss locations:3.8 18
y
x
0
1
2
3
4
5
6
7
0
5
10
15
20
25

**Optimal1 Fitted Gauss, proteinAT3G16780.1**
**bic:739**
**peaks after deleting less than one fifth of global max:9**

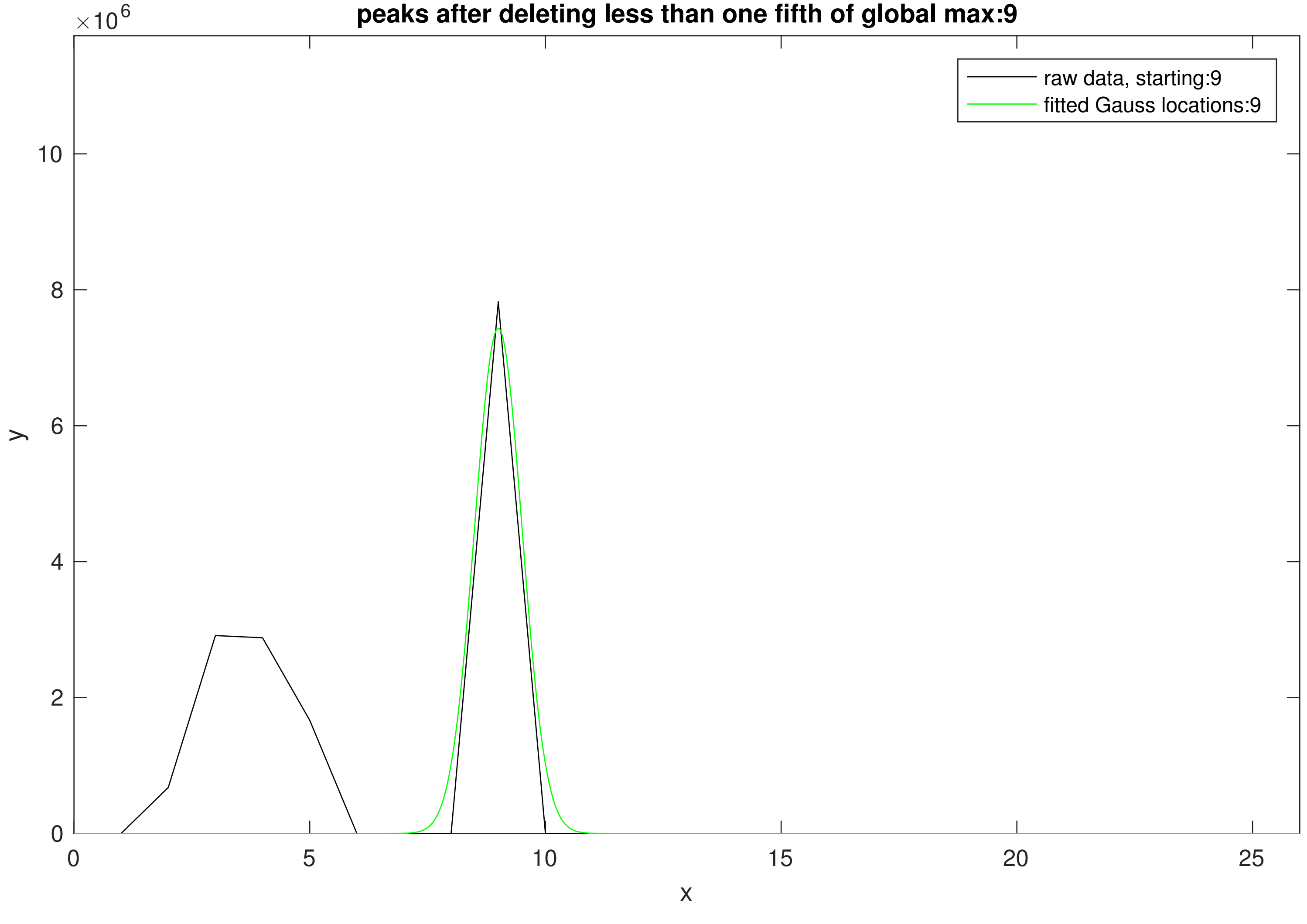

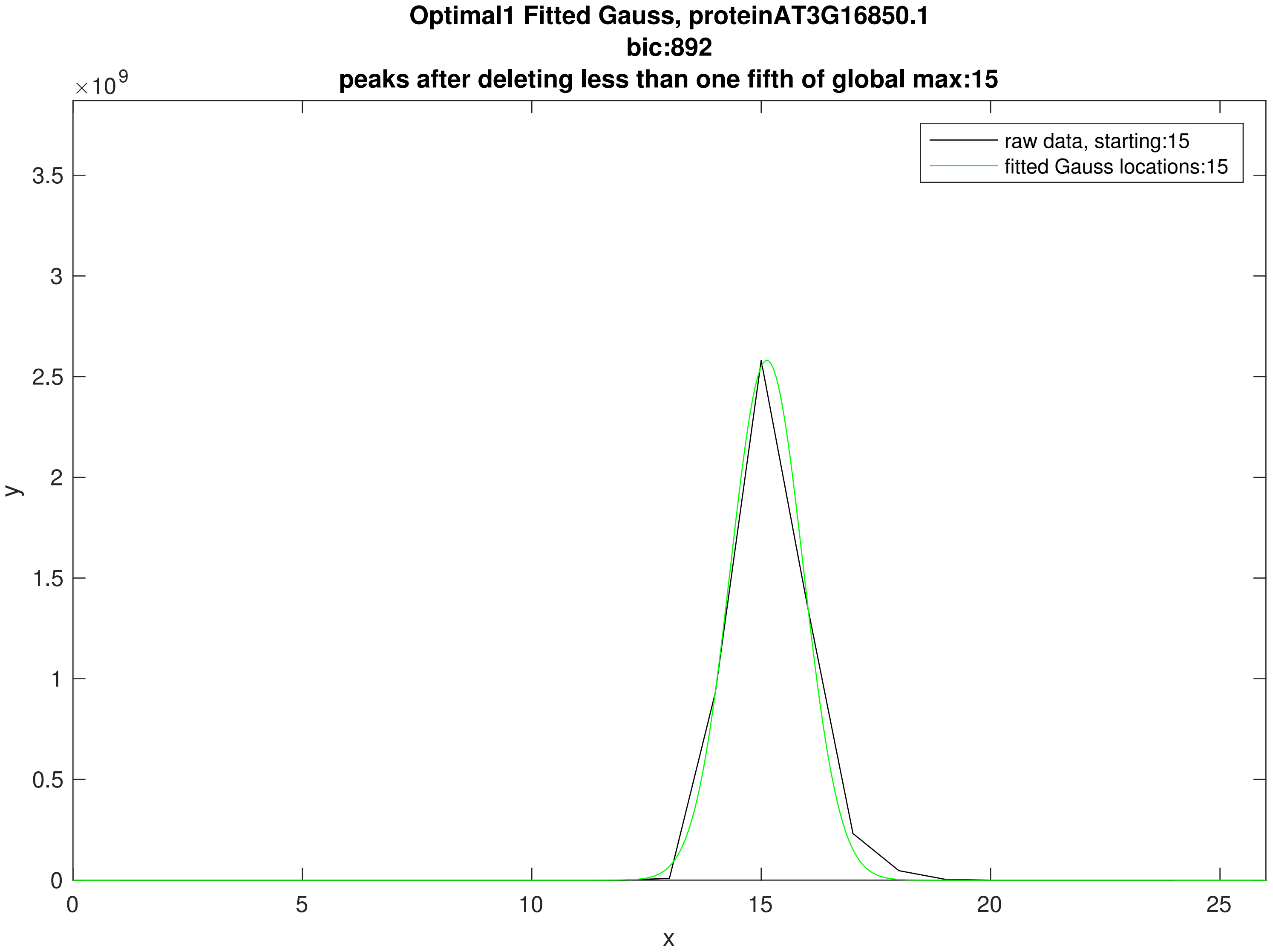
Optimal1 Fitted Gauss, proteinAT3G16850.1
bic:892
peaks after deleting less than one fifth of global max:15
×10^9
raw data, starting:15
fitted Gauss locations:15
3.5
3
2.5
2
1.5
1
0.5
0
y
0
5
10
15
20
25
x

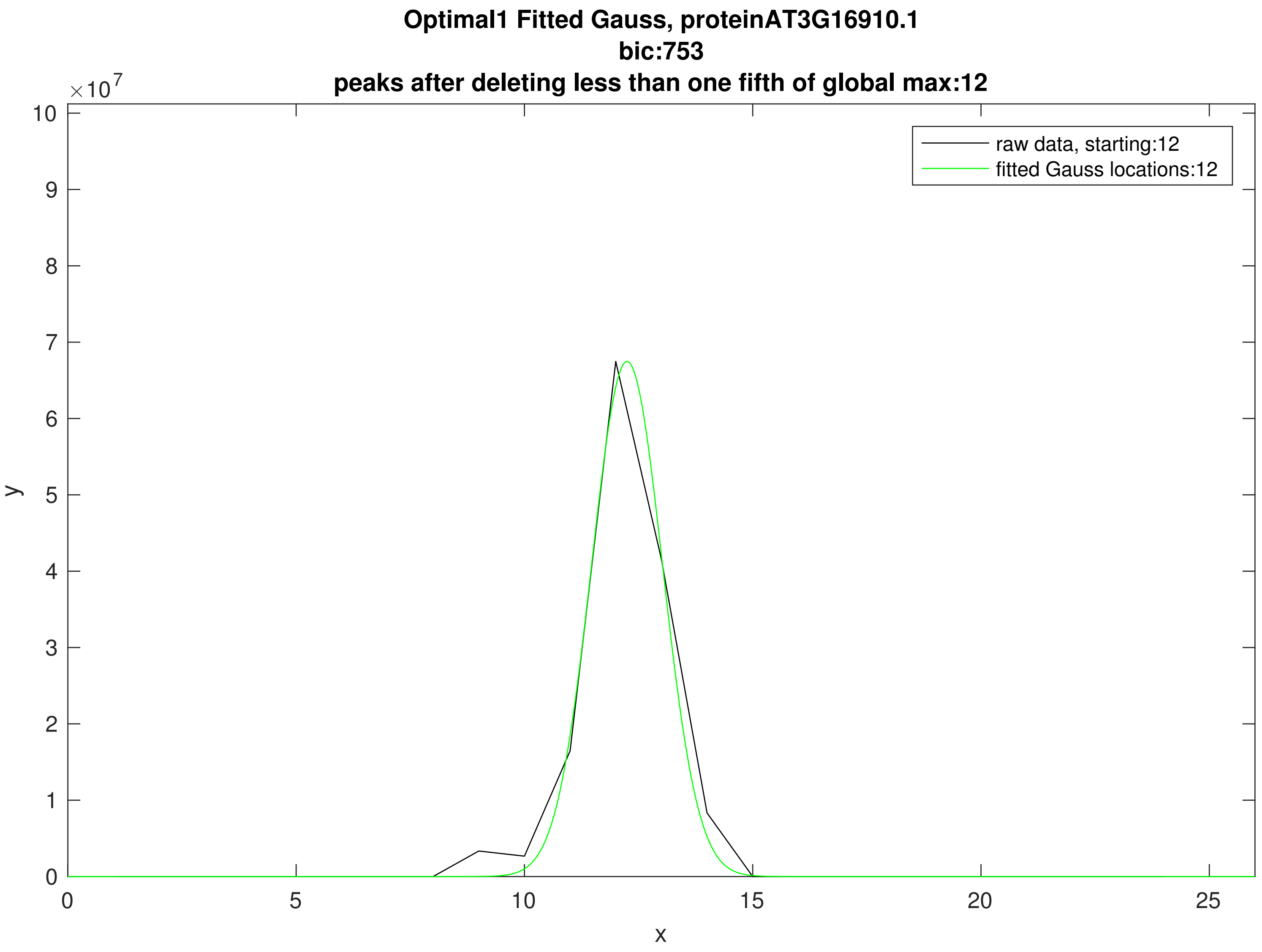
Optimal1 Fitted Gauss, proteinAT3G16910.1
bic:753
peaks after deleting less than one fifth of global max:12
×10^7
10
9
8
7
6
5
4
3
2
1
0
y
0
5
10
15
20
25
x
raw data, starting:12
fitted Gauss locations:12

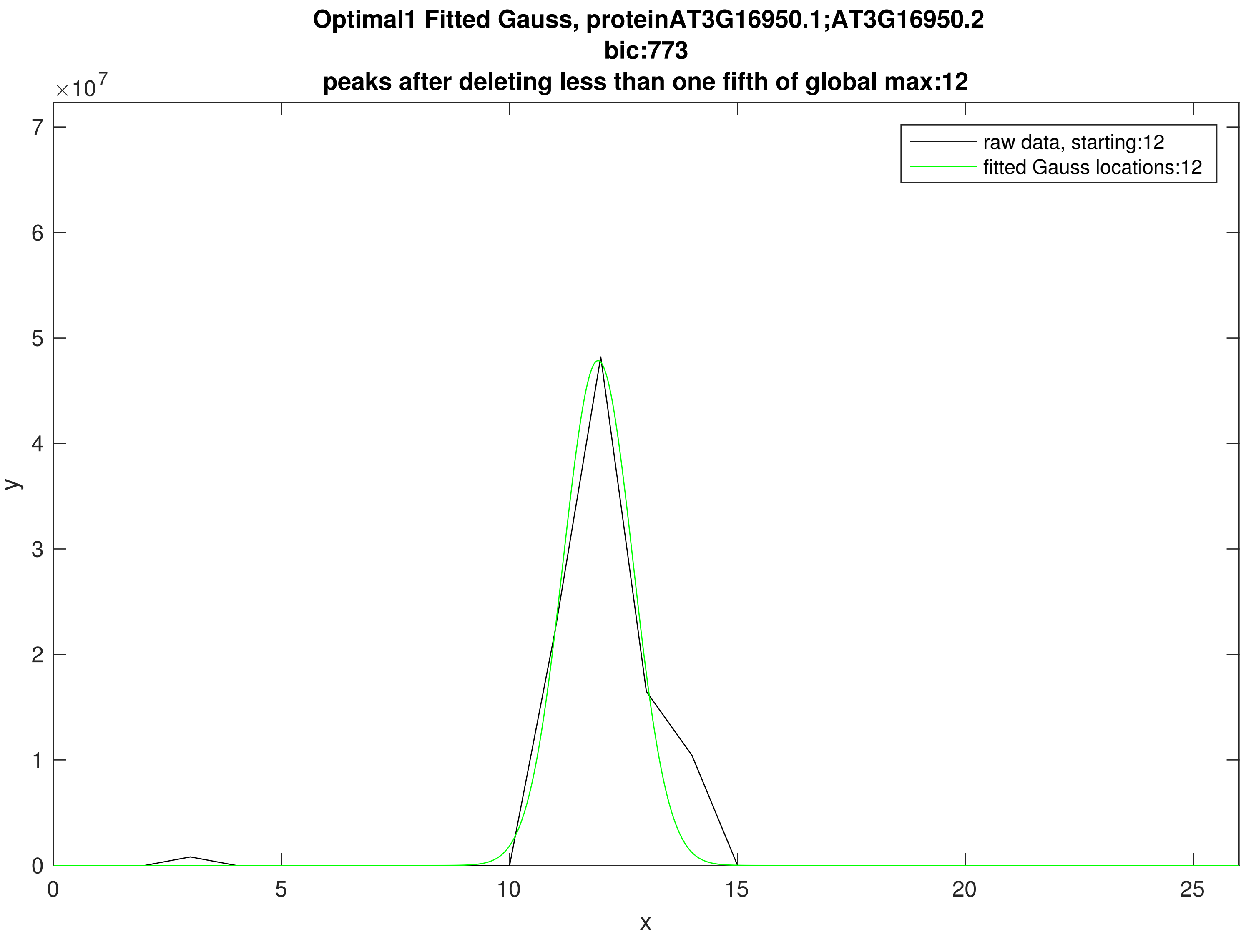
Optimal1 Fitted Gauss, proteinAT3G16950.1;AT3G16950.2
bic:773
peaks after deleting less than one fifth of global max:12
×10^7
raw data, starting:12
fitted Gauss locations:12
y
x
0
1
2
3
4
5
6
7
0
5
10
15
20
25

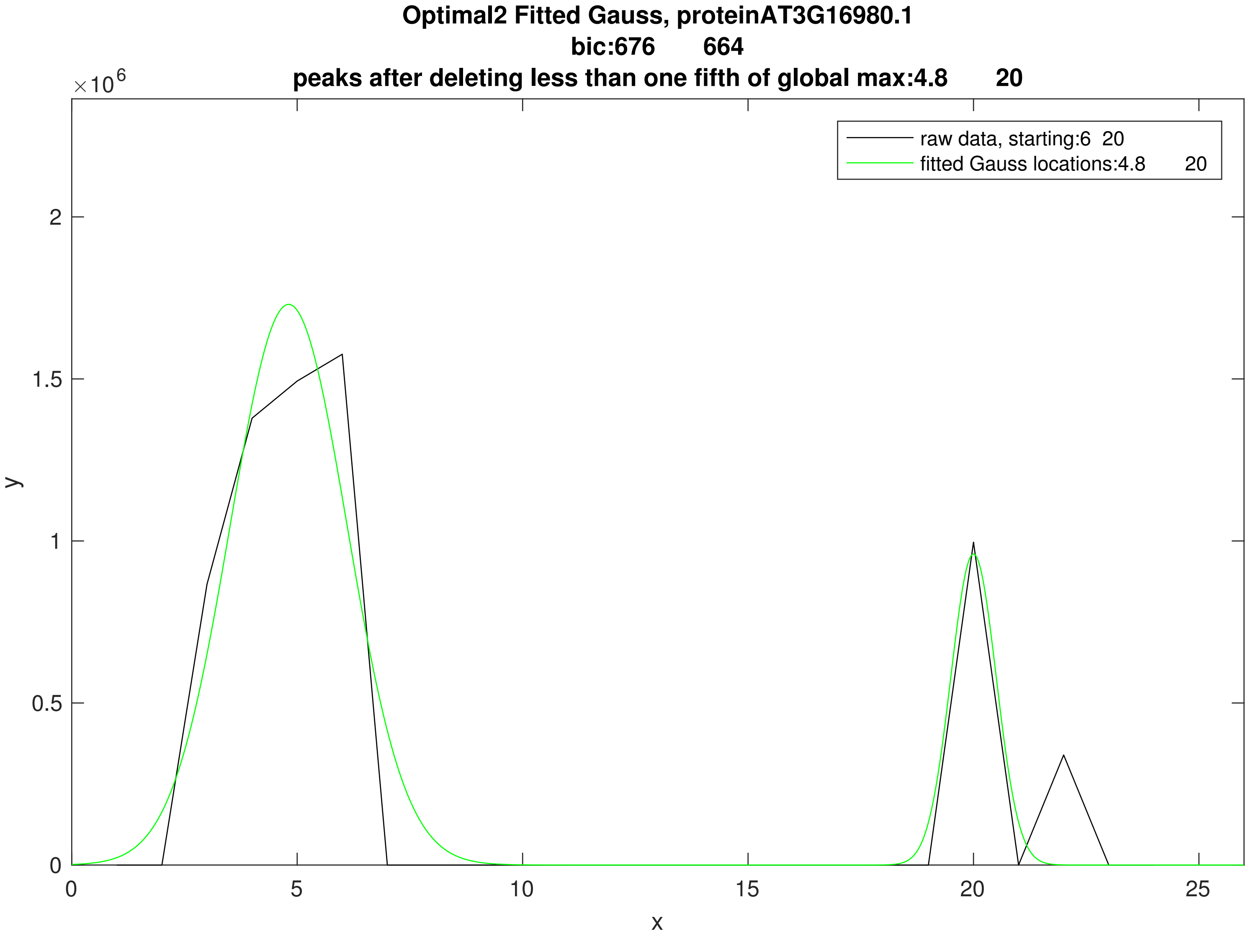

Optimal2 Fitted Gauss, proteinAT3G16980.1
bic:676 664
peaks after deleting less than one fifth of global max:4.8 20
×10^6
raw data, starting:6 20
fitted Gauss locations:4.8 20
2
1.5
1
0.5
0
y
0
5
10
15
20
25
x

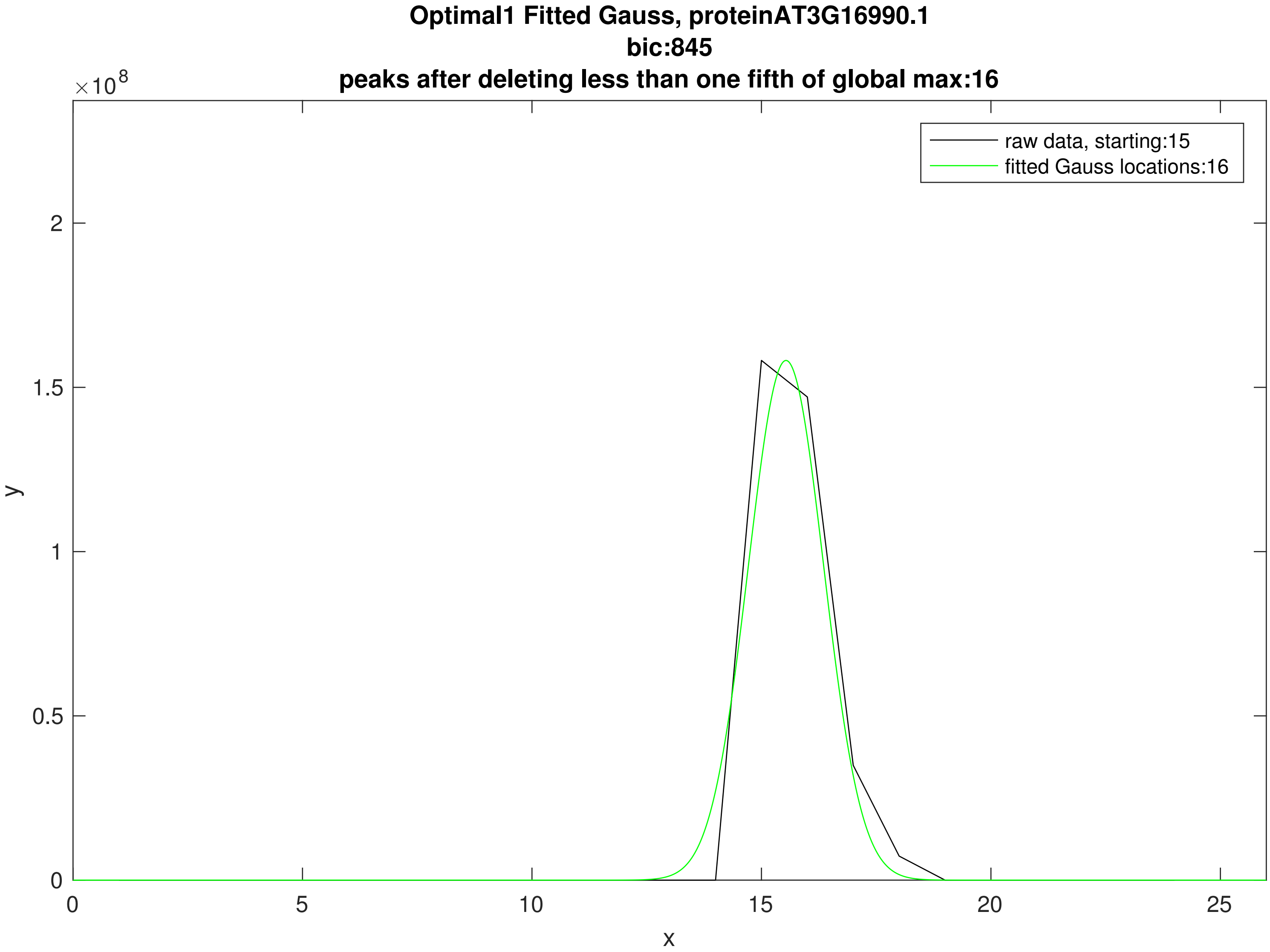
Optimal1 Fitted Gauss, proteinAT3G16990.1
bic:845
peaks after deleting less than one fifth of global max:16
×10^8
raw data, starting:15
fitted Gauss locations:16
y
x
0
0.5
1
1.5
2
0
5
10
15
20
25

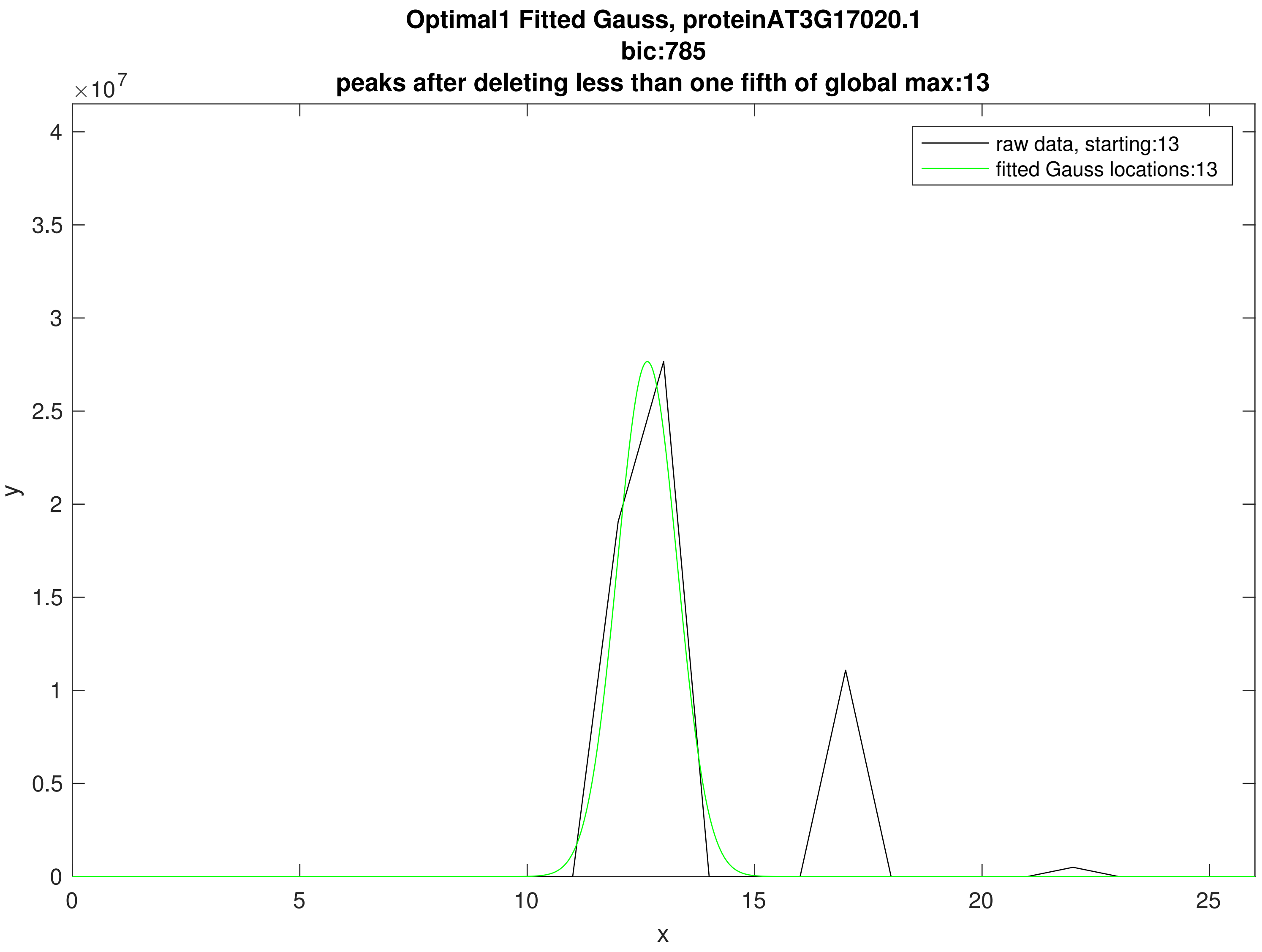

Optimal1 Fitted Gauss, proteinAT3G17020.1
bic:785
peaks after deleting less than one fifth of global max:13
×10^7
4
3.5
3
2.5
2
1.5
1
0.5
0
y
0
5
10
15
20
25
x
raw data, starting:13
fitted Gauss locations:13

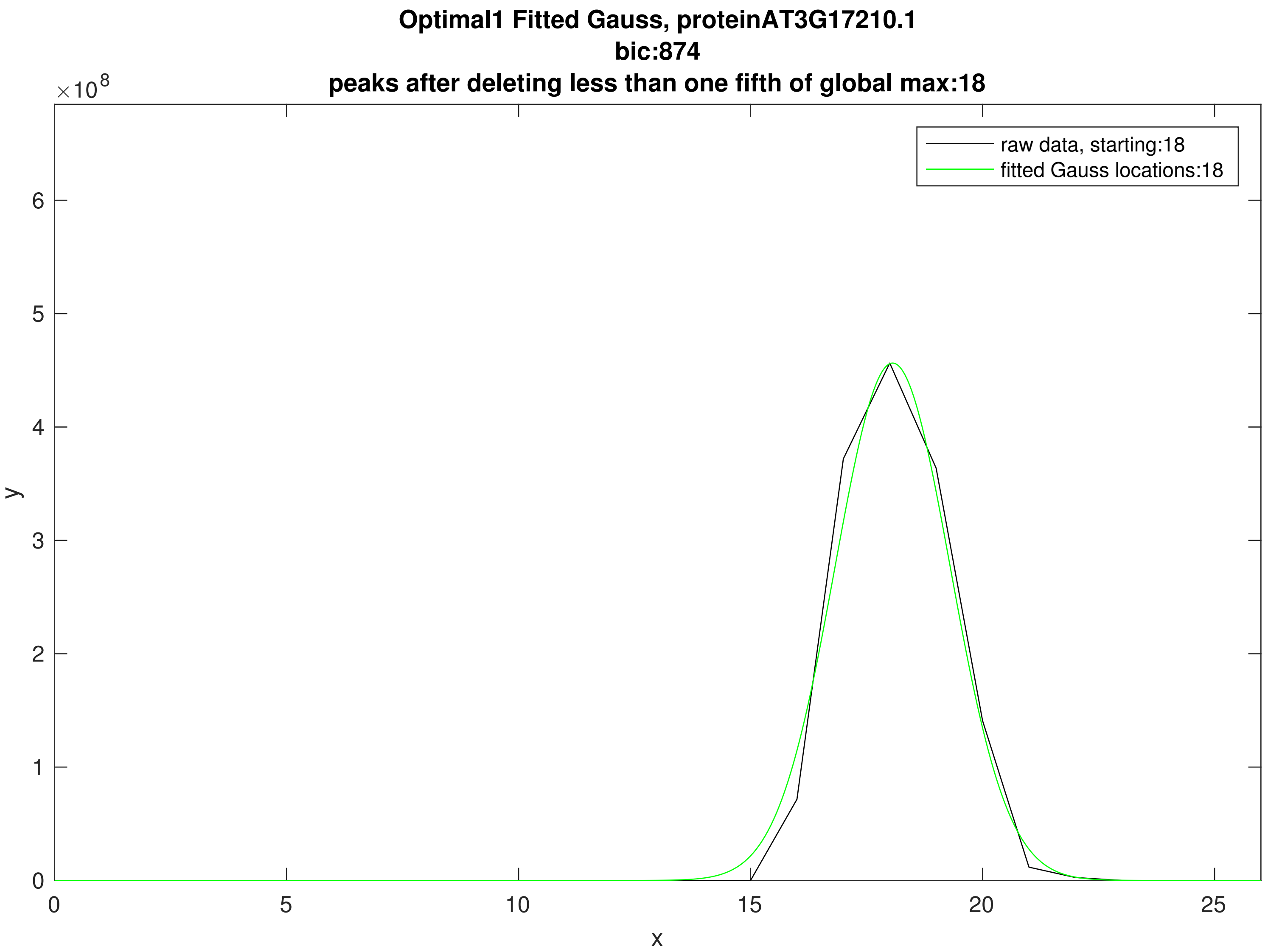
Optimal1 Fitted Gauss, proteinAT3G17210.1
bic:874
peaks after deleting less than one fifth of global max:18
×10^8
raw data, starting:18
fitted Gauss locations:18
y
x
0
1
2
3
4
5
6
0
5
10
15
20
25

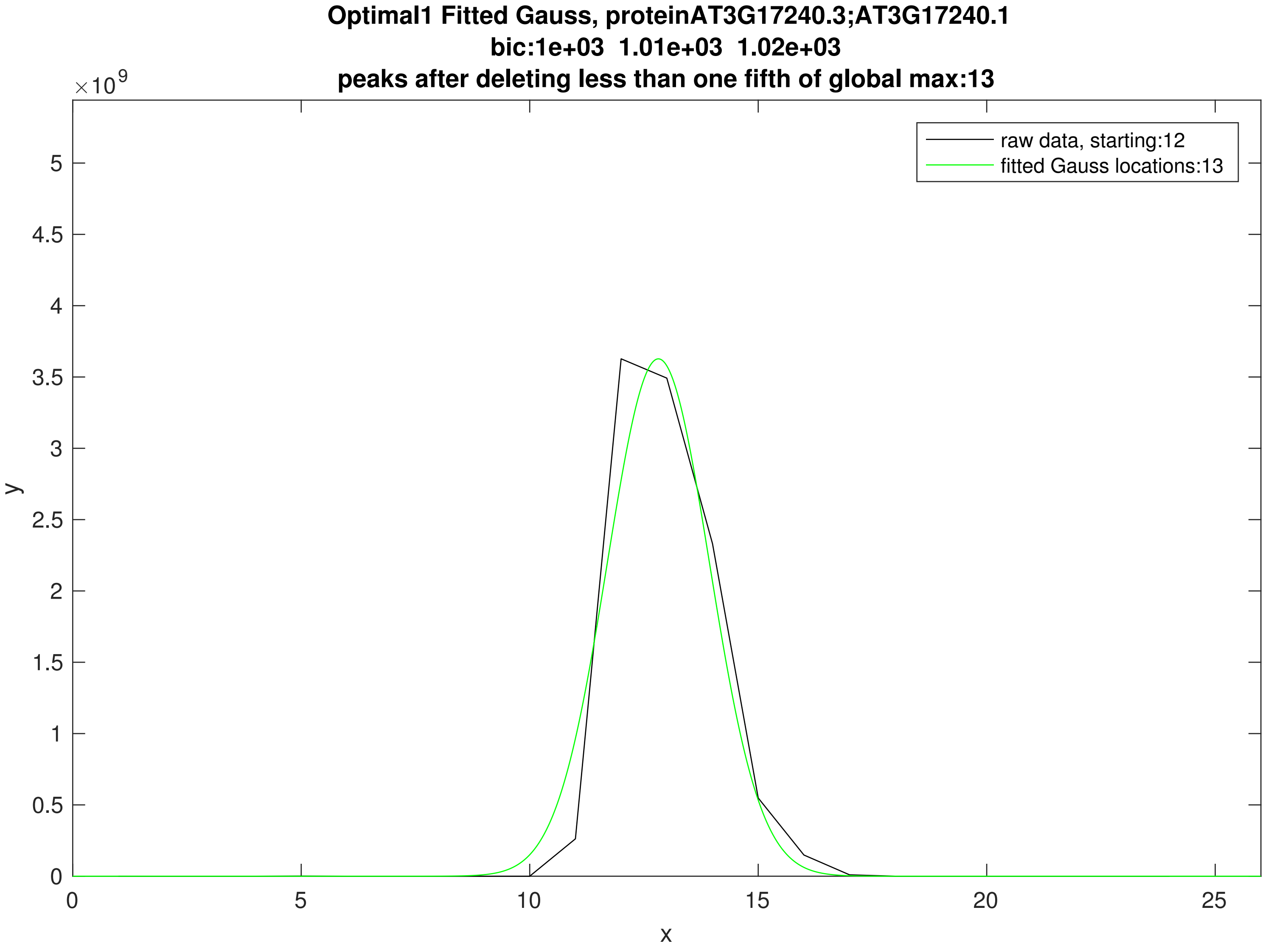
Optimal1 Fitted Gauss, proteinAT3G17240.3;AT3G17240.1
bic:1e+03 1.01e+03 1.02e+03
peaks after deleting less than one fifth of global max:13
×10^9
raw data, starting:12
fitted Gauss locations:13
y
x
0
0.5
1
1.5
2
2.5
3
3.5
4
4.5
5
0
5
10
15
20
25

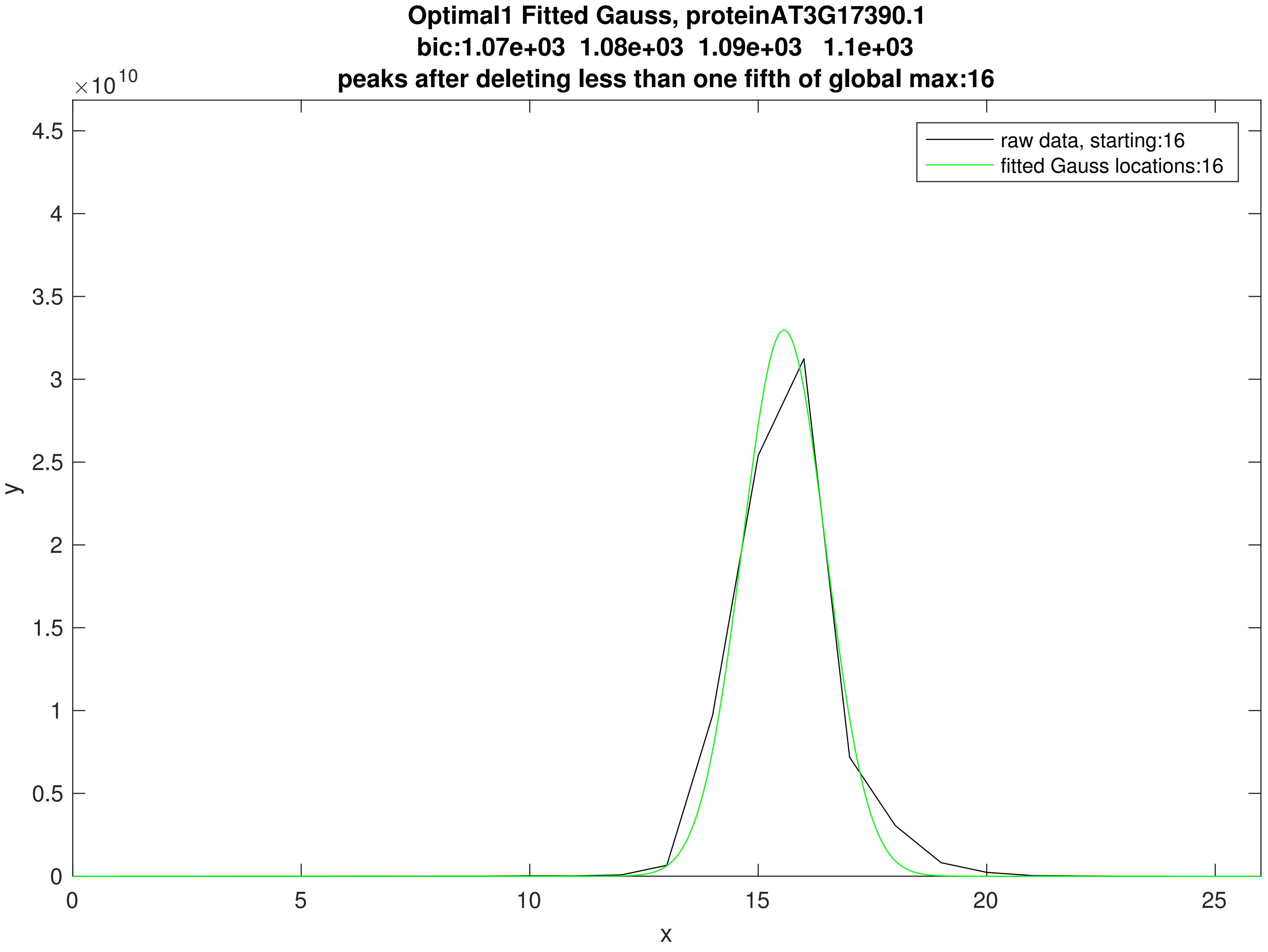
Optimal1 Fitted Gauss, proteinAT3G17390.1
bic:1.07e+03  1.08e+03  1.09e+03   1.1e+03
peaks after deleting less than one fifth of global max:16
×10^10
4.5
4
3.5
3
2.5
2
1.5
1
0.5
0
y
0
5
10
15
20
25
x
raw data, starting:16
fitted Gauss locations:16

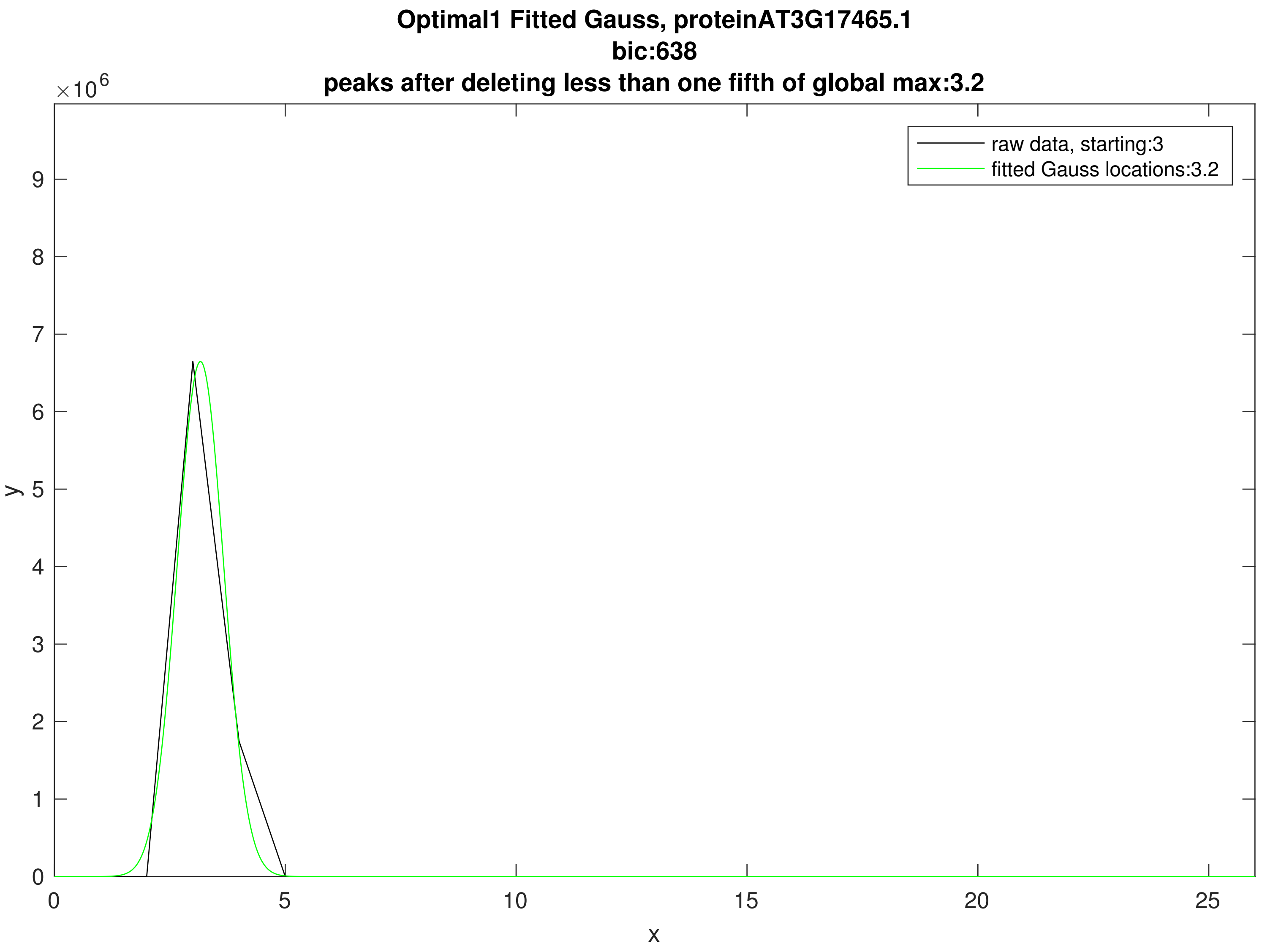
Optimal1 Fitted Gauss, proteinAT3G17465.1
bic:638
peaks after deleting less than one fifth of global max:3.2
×10^6
raw data, starting:3
fitted Gauss locations:3.2
y
x
0
1
2
3
4
5
6
7
8
9
0
5
10
15
20
25

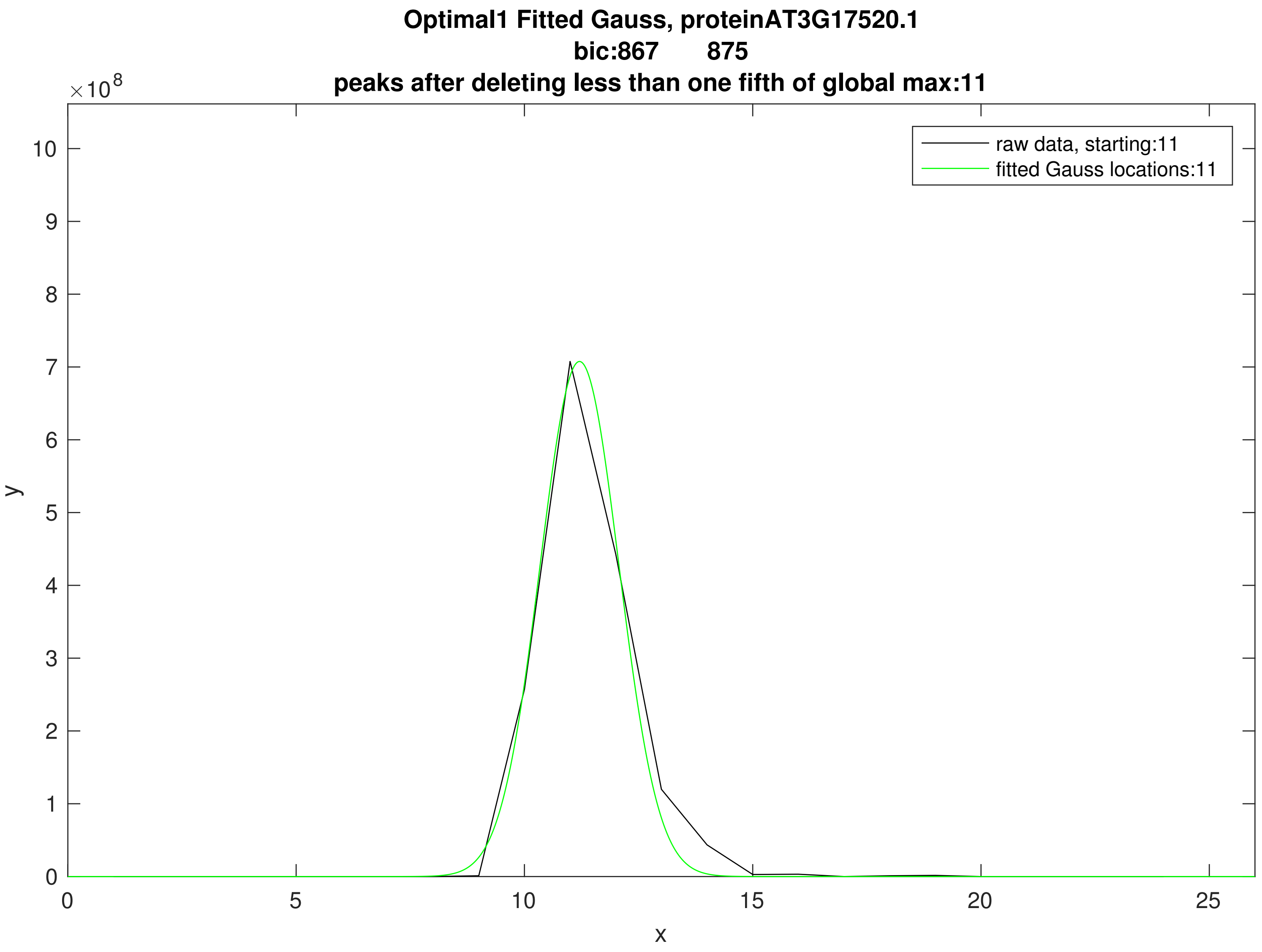
Optimal1 Fitted Gauss, proteinAT3G17520.1
bic:867 875
peaks after deleting less than one fifth of global max:11
×10^8
raw data, starting:11
fitted Gauss locations:11
y
x
0
1
2
3
4
5
6
7
8
9
10
0
5
10
15
20
25

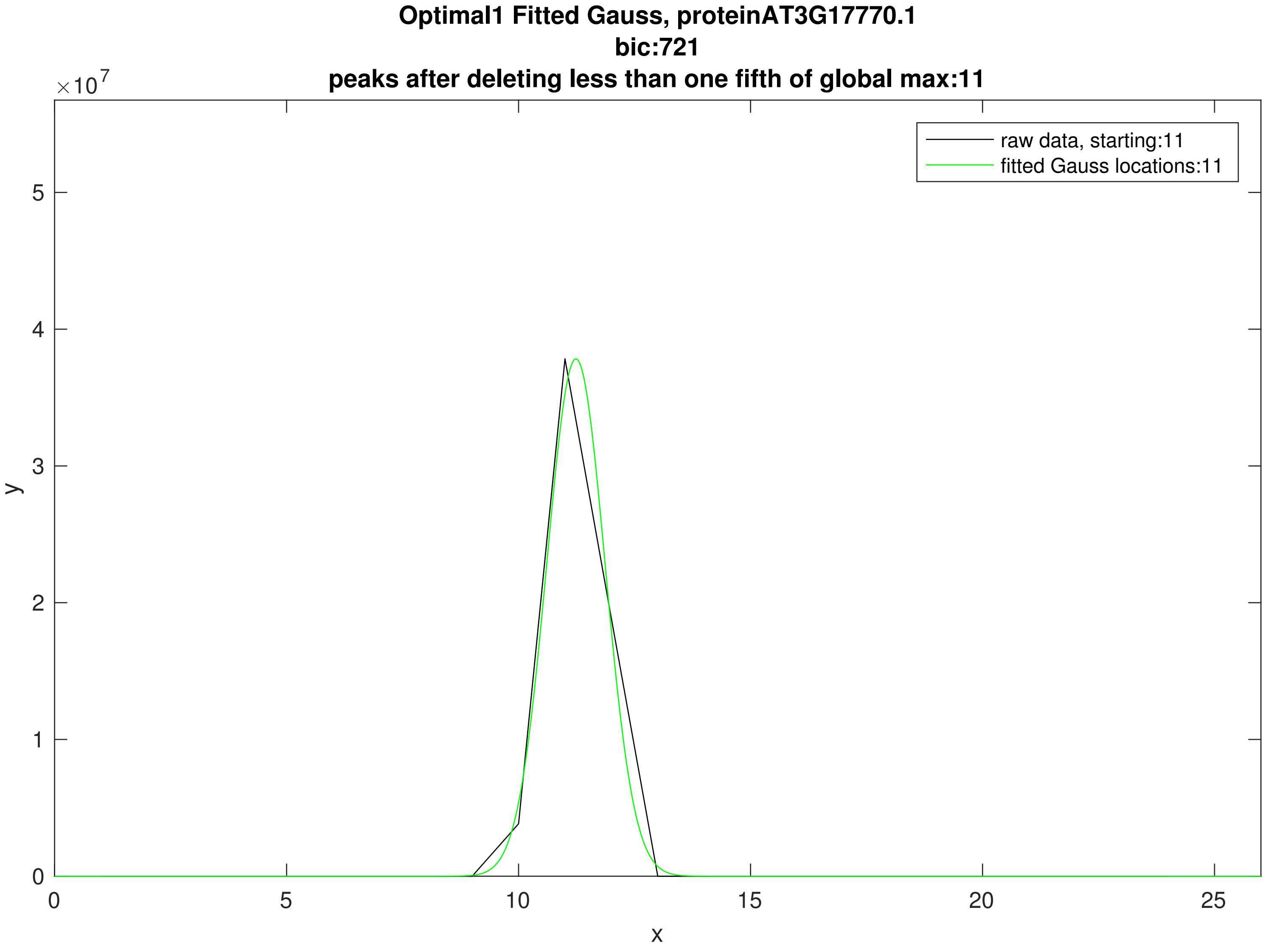

Optimal1 Fitted Gauss, proteinAT3G17770.1
bic:721
peaks after deleting less than one fifth of global max:11
×10^7
raw data, starting:11
fitted Gauss locations:11
y
x
0
1
2
3
4
5
0
5
10
15
20
25

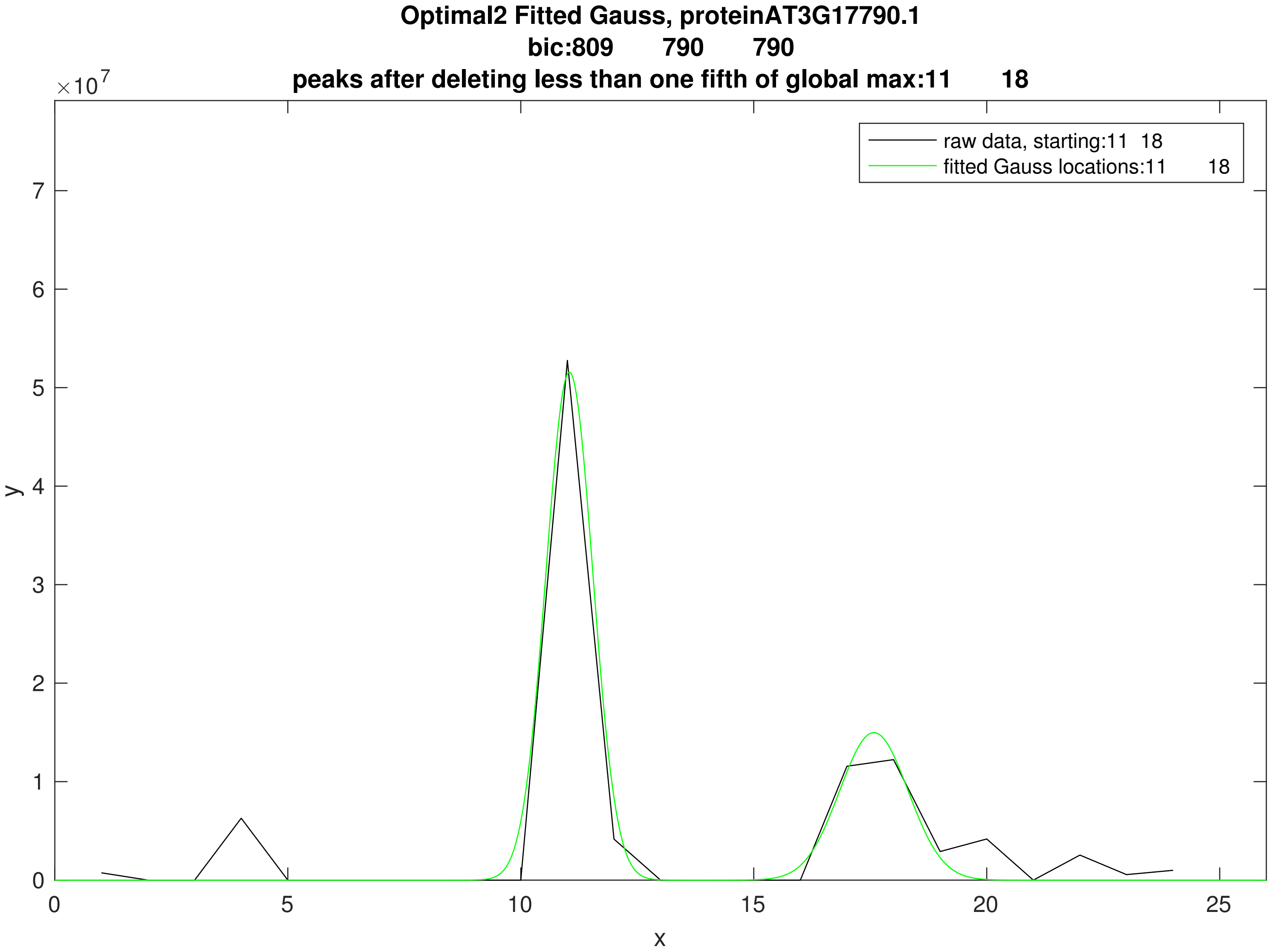
Optimal2 Fitted Gauss, proteinAT3G17790.1
bic:809 790 790
peaks after deleting less than one fifth of global max:11 18
×10^7
raw data, starting:11 18
fitted Gauss locations:11 18
y
x
0
1
2
3
4
5
6
7
0
5
10
15
20
25

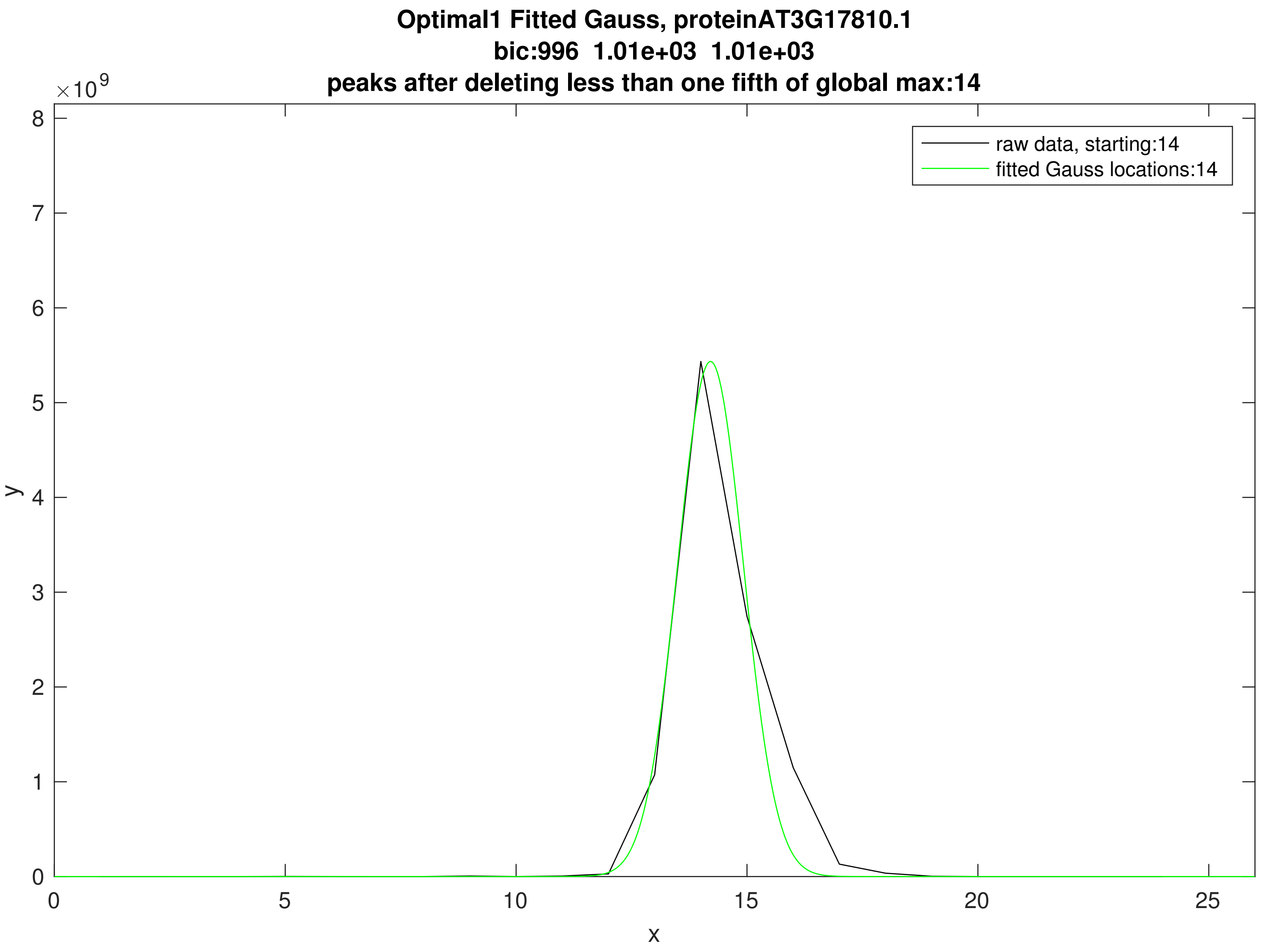
Optimal1 Fitted Gauss, proteinAT3G17810.1
bic:996  1.01e+03  1.01e+03
peaks after deleting less than one fifth of global max:14
×10^9
raw data, starting:14
fitted Gauss locations:14
y
x
0
1
2
3
4
5
6
7
8
0
5
10
15
20
25

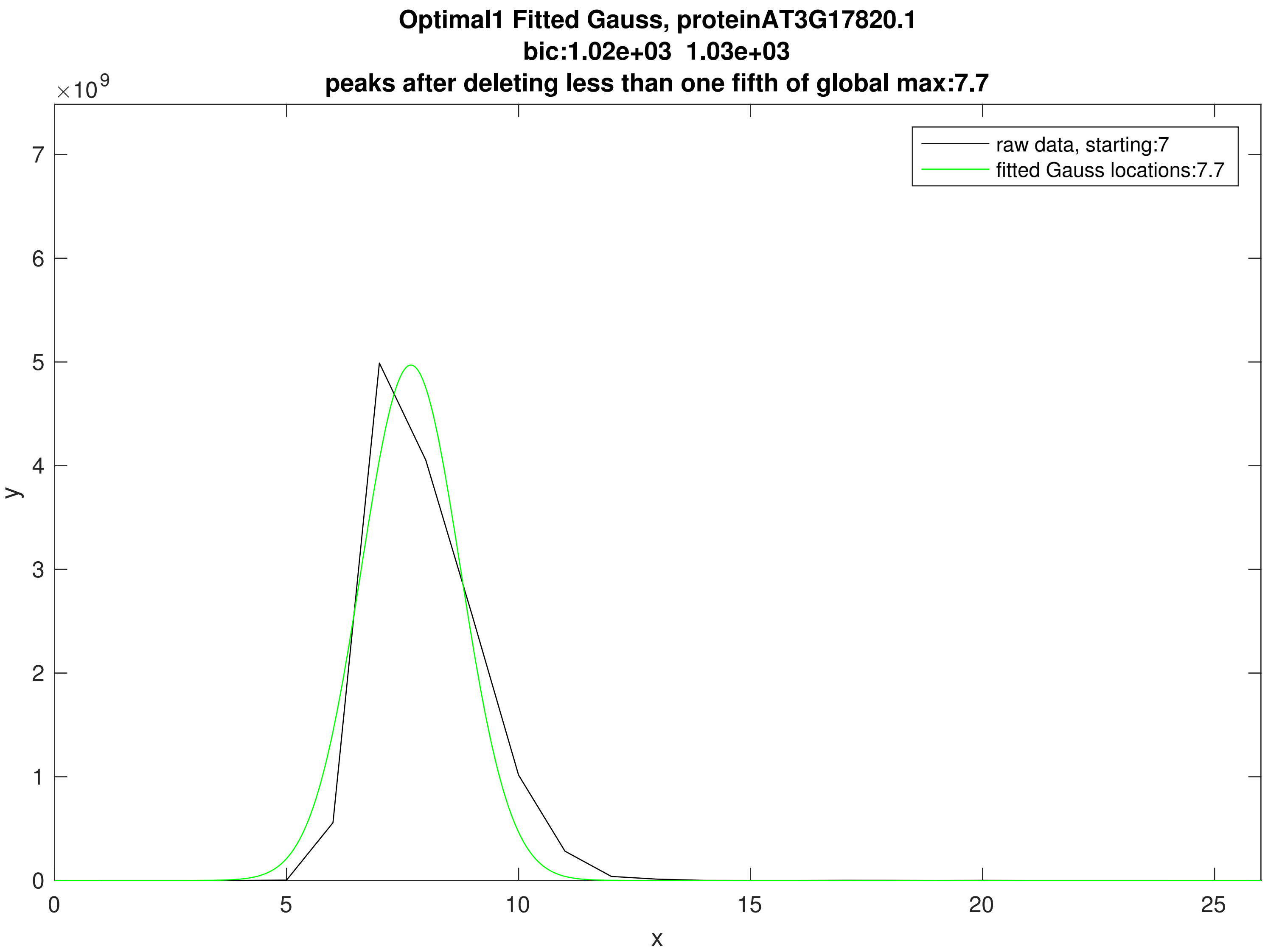
Optimal1 Fitted Gauss, proteinAT3G17820.1
bic:1.02e+03  1.03e+03
peaks after deleting less than one fifth of global max:7.7
×10^9
raw data, starting:7
fitted Gauss locations:7.7
y
x
0
1
2
3
4
5
6
7
0
5
10
15
20
25

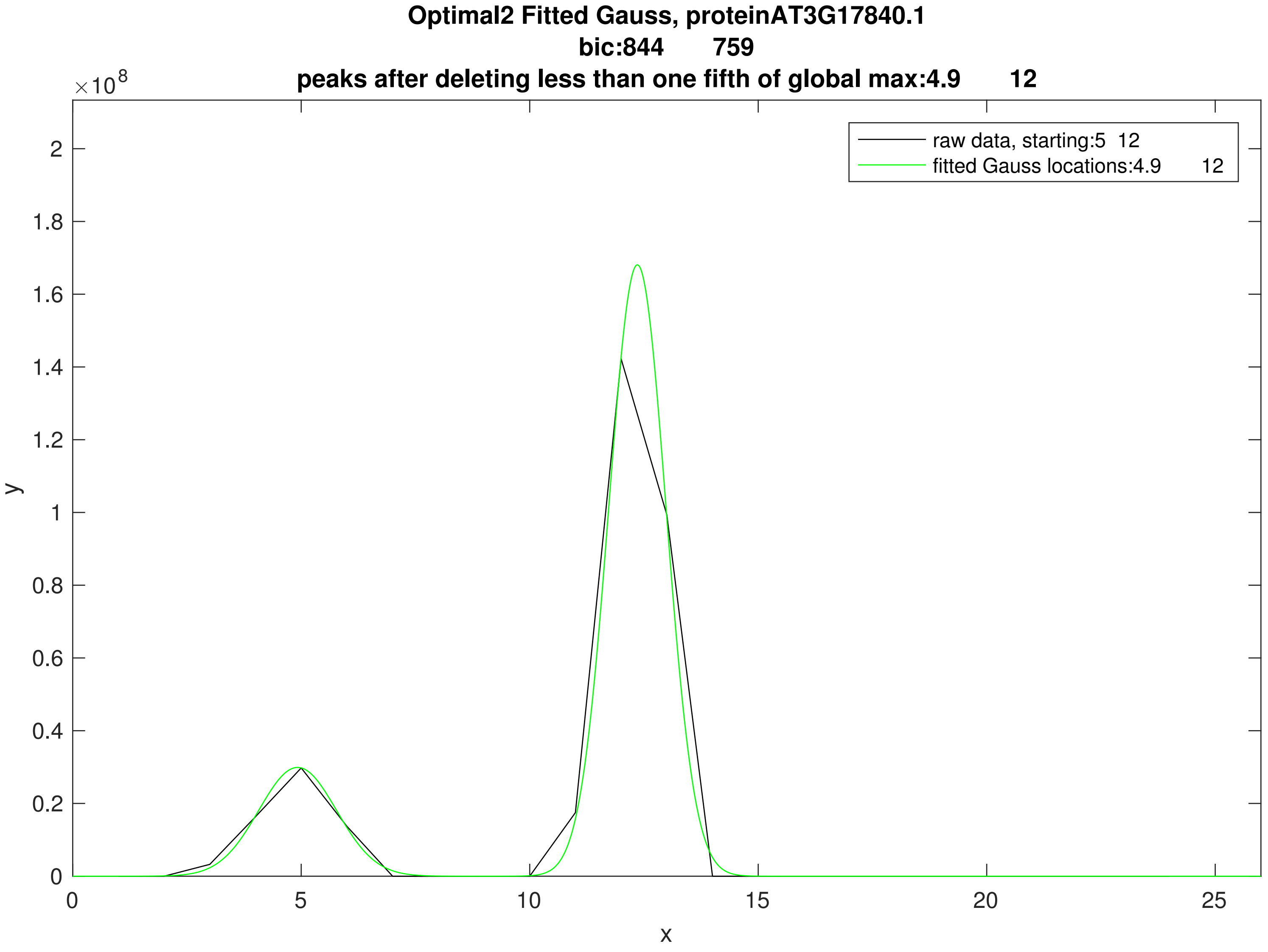
Optimal2 Fitted Gauss, proteinAT3G17840.1
bic:844 759
peaks after deleting less than one fifth of global max:4.9 12
raw data, starting:5 12
fitted Gauss locations:4.9 12
×10^8
y
x

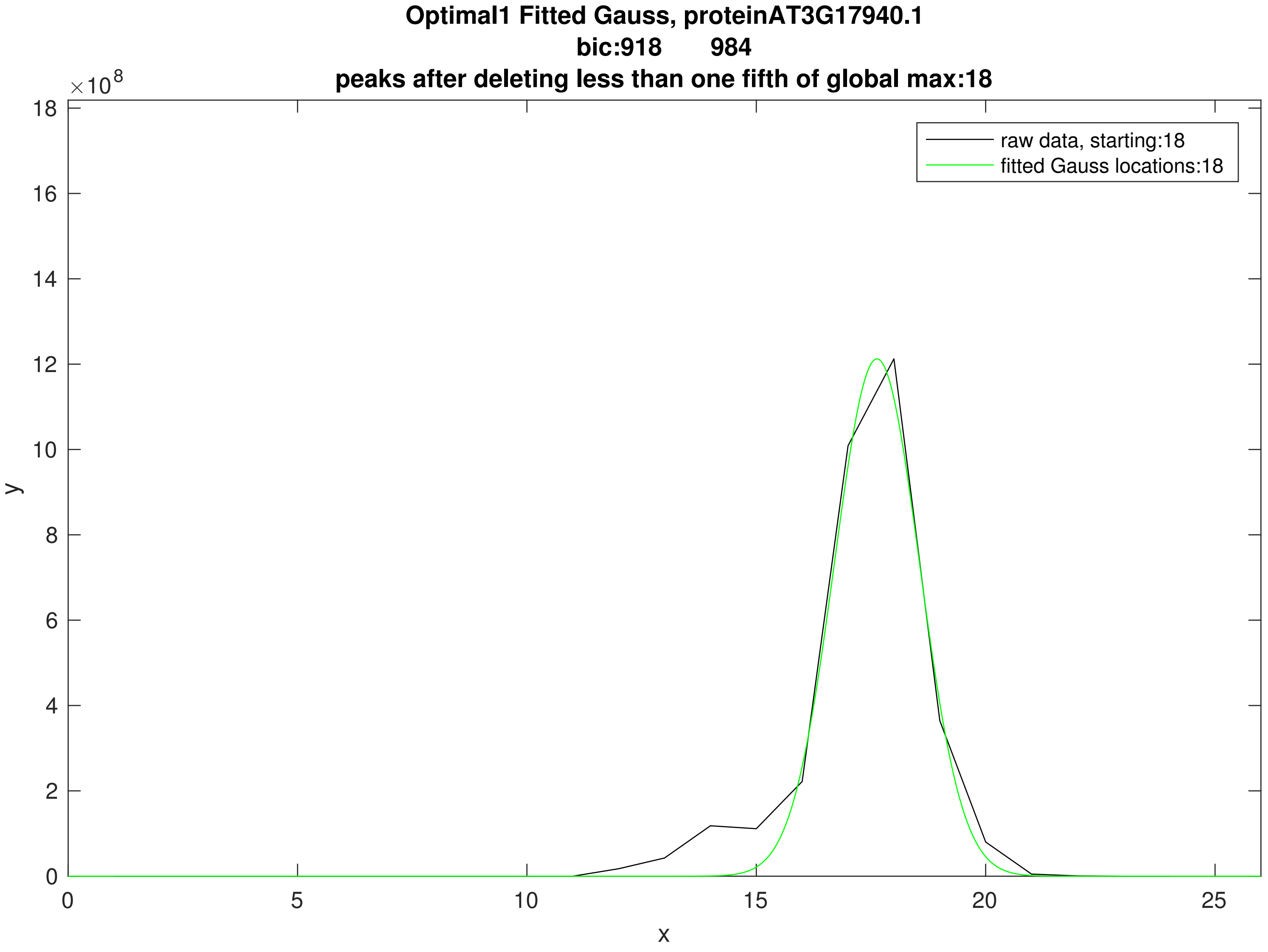

Optimal1 Fitted Gauss, proteinAT3G17940.1
bic:918 984
peaks after deleting less than one fifth of global max:18
×10^8
18
16
14
12
10
8
6
4
2
0
y
0
5
10
15
20
25
x
raw data, starting:18
fitted Gauss locations:18

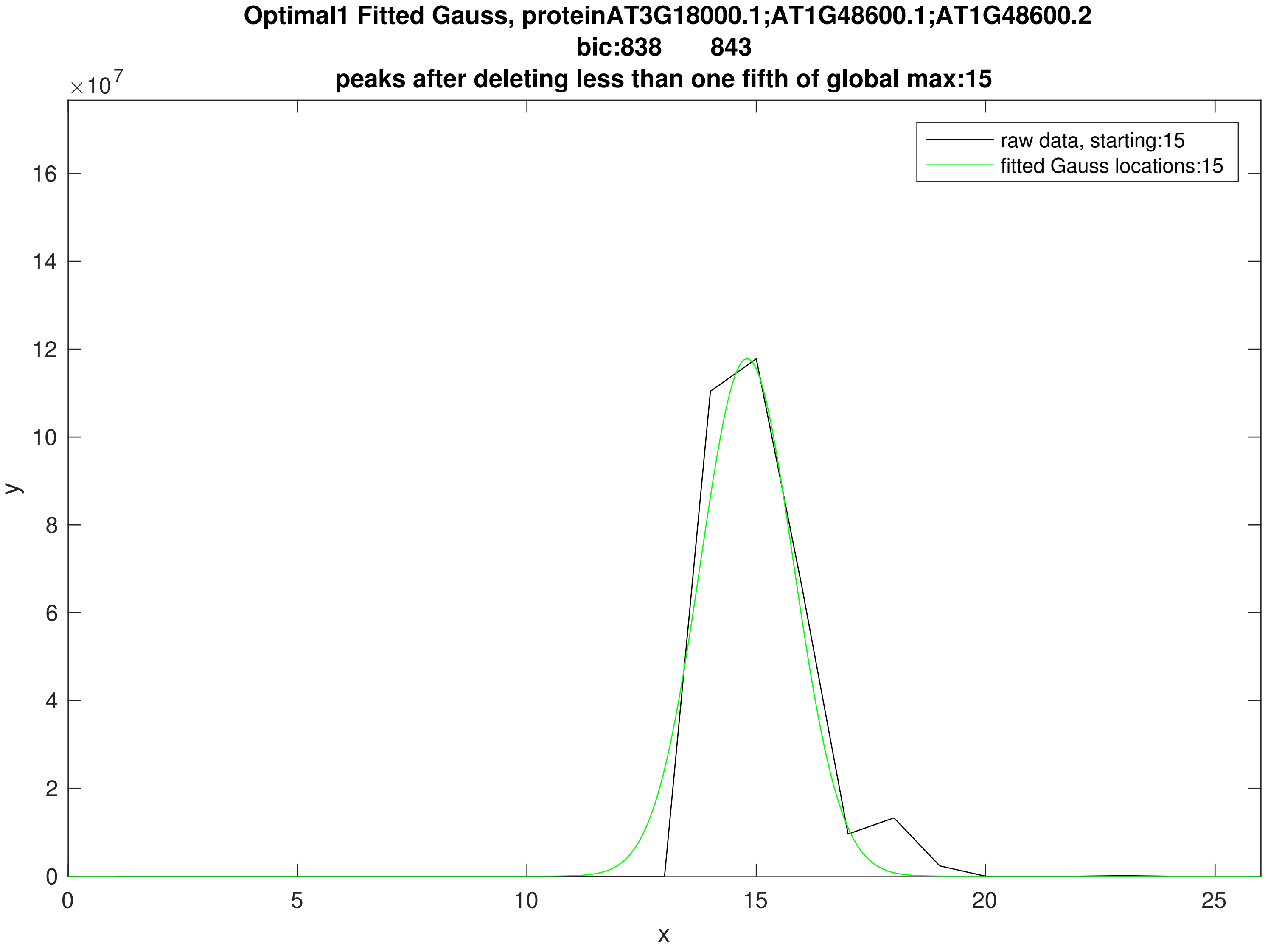
Optimal1 Fitted Gauss, proteinAT3G18000.1;AT1G48600.1;AT1G48600.2
bic:838 843
peaks after deleting less than one fifth of global max:15
×10^7
raw data, starting:15
fitted Gauss locations:15
y
x
0
2
4
6
8
10
12
14
16
0
5
10
15
20
25

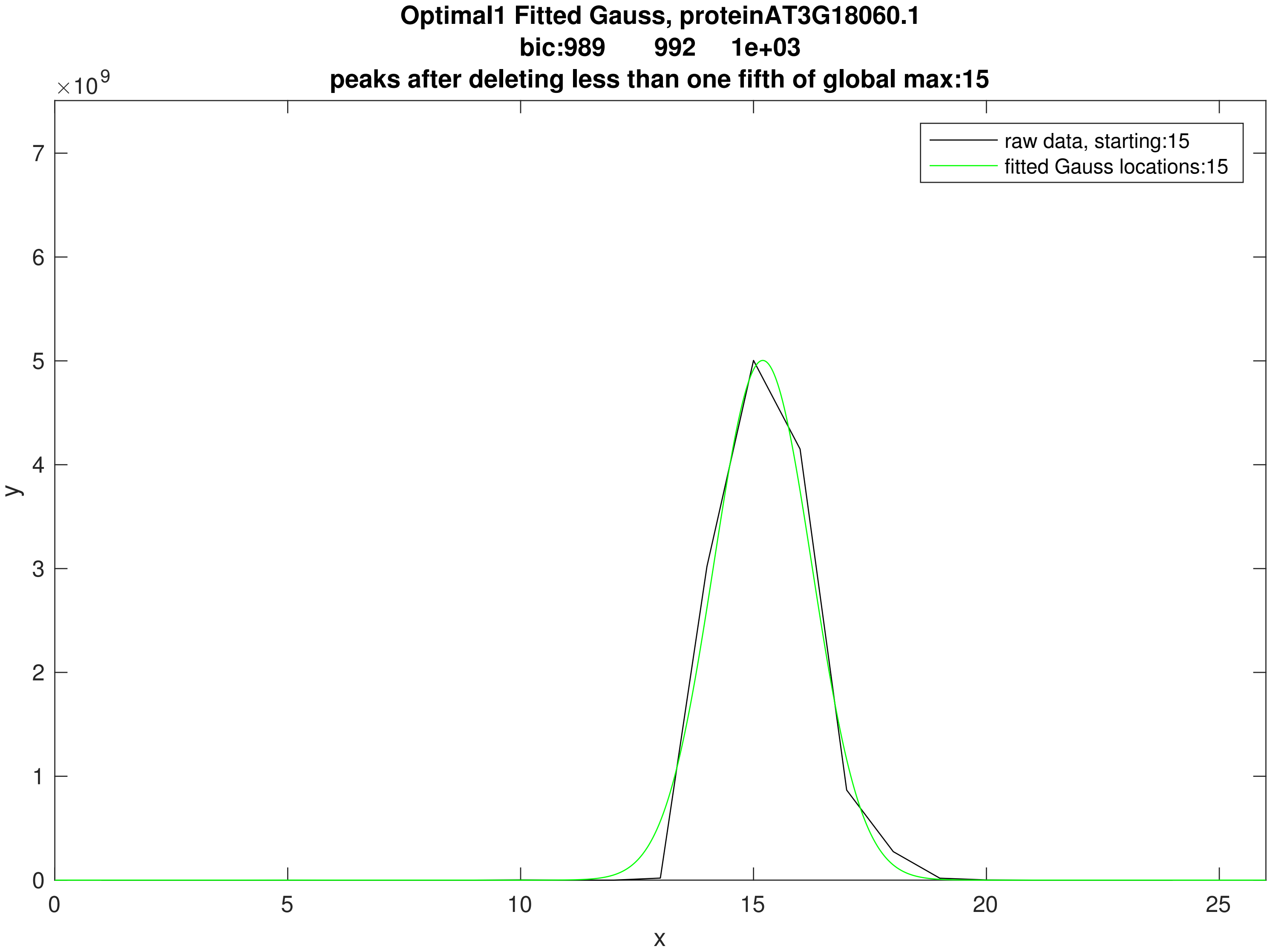
Optimal1 Fitted Gauss, proteinAT3G18060.1
bic:989 992 1e+03
peaks after deleting less than one fifth of global max:15
×10^9
raw data, starting:15
fitted Gauss locations:15
y
x
0
1
2
3
4
5
6
7
0
5
10
15
20
25

**Optimal1 Fitted Gauss, proteinAT3G18130.1**
**bic:832 837**
**peaks after deleting less than one fifth of global max:18**

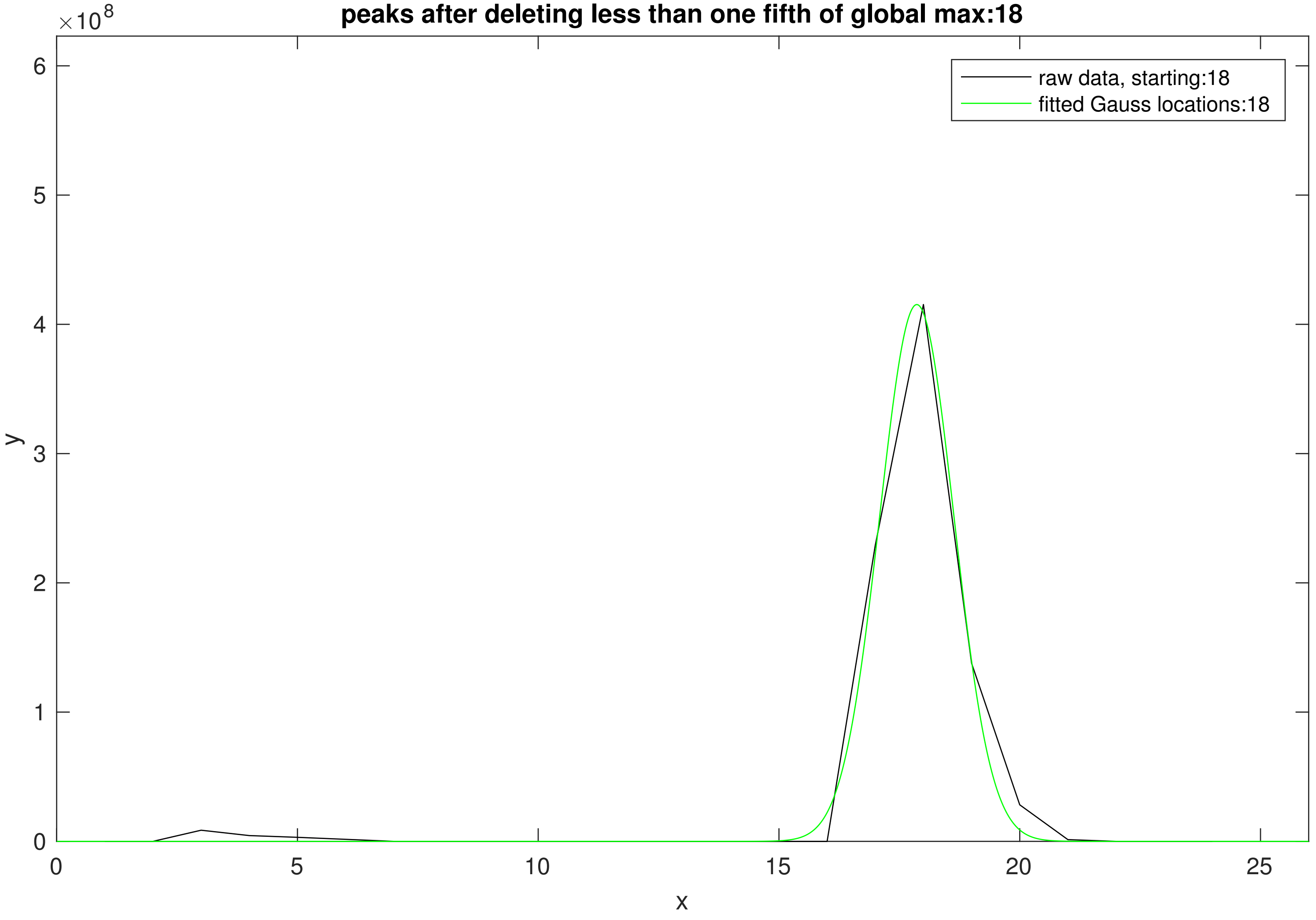

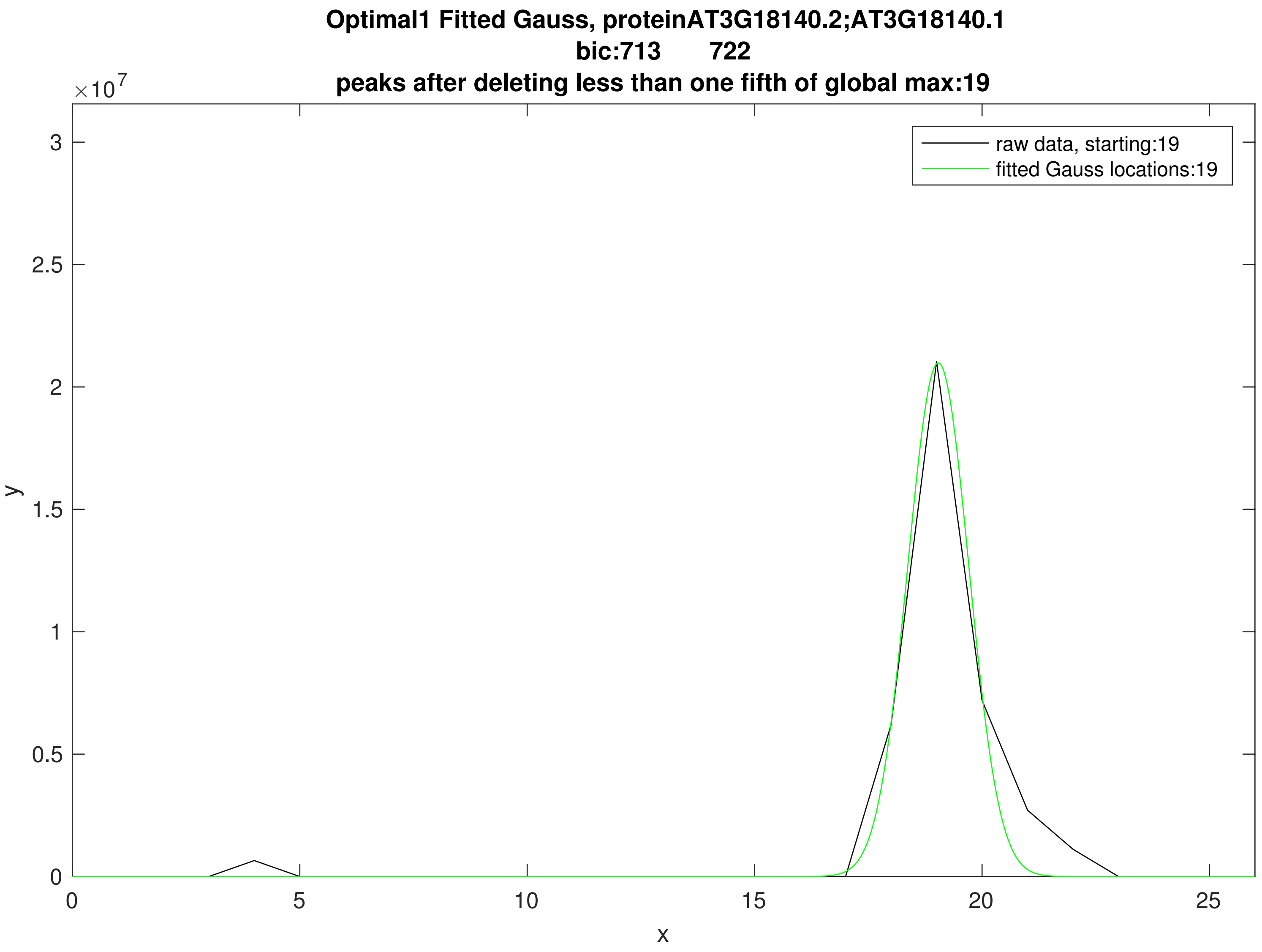

Optimal1 Fitted Gauss, proteinAT3G18140.2;AT3G18140.1
bic:713 722
peaks after deleting less than one fifth of global max:19
×10^7
3
2.5
2
1.5
1
0.5
0
y
0
5
10
15
20
25
x
raw data, starting:19
fitted Gauss locations:19

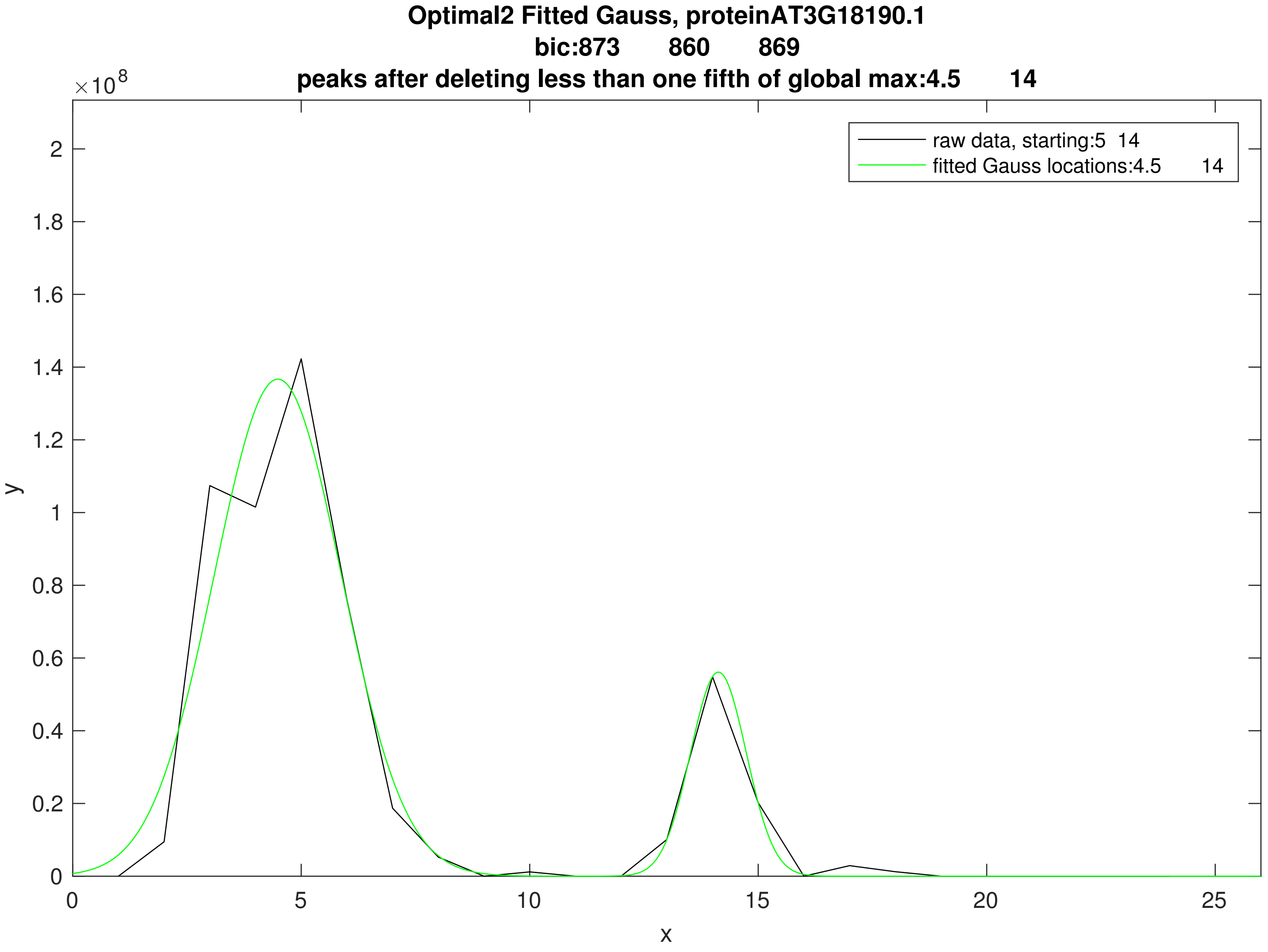

Optimal2 Fitted Gauss, proteinAT3G18190.1
bic:873 860 869
peaks after deleting less than one fifth of global max:4.5 14
×10^8
raw data, starting:5 14
fitted Gauss locations:4.5 14
y
x
0
5
10
15
20
25
0
0.2
0.4
0.6
0.8
1
1.2
1.4
1.6
1.8
2

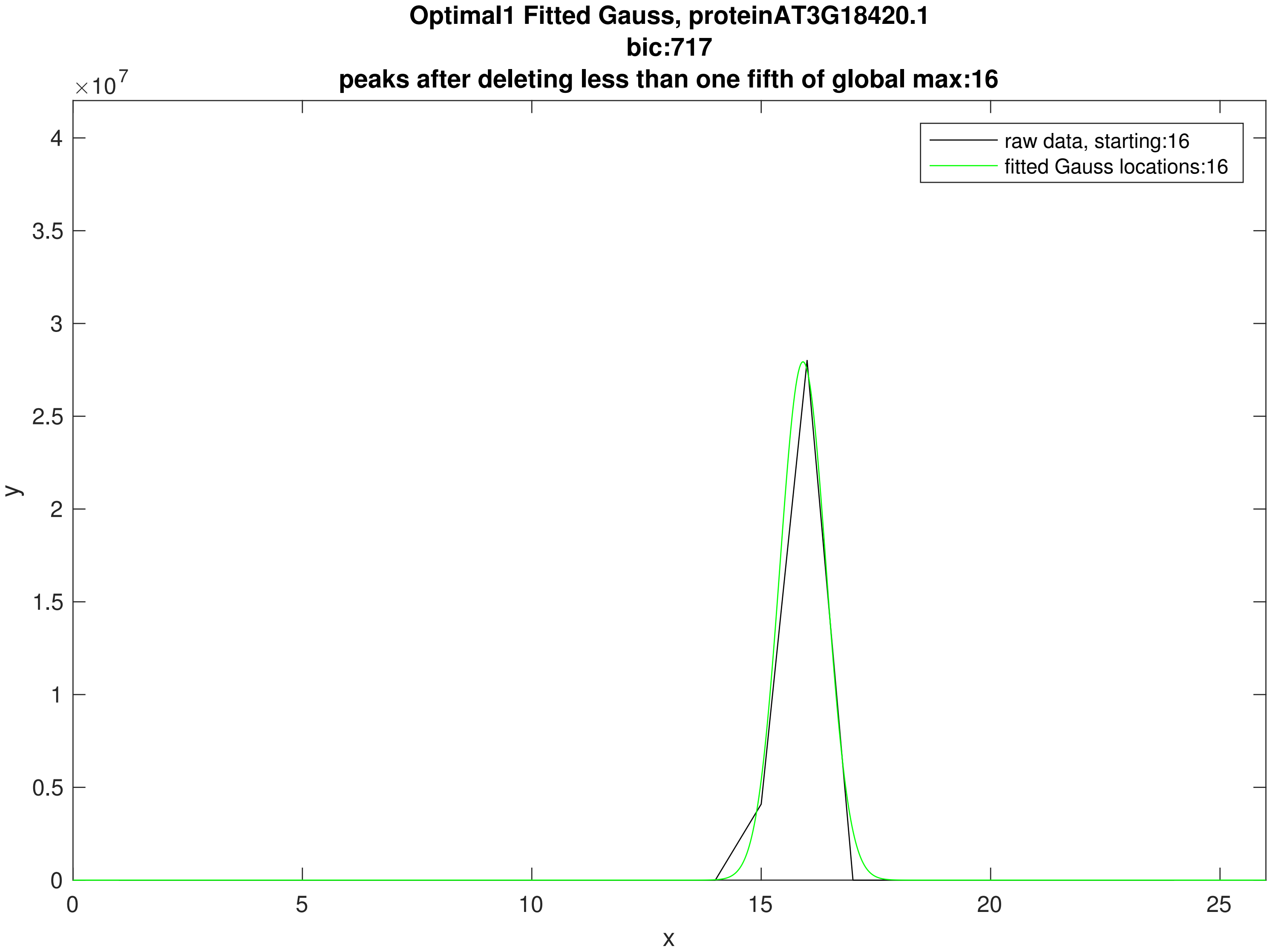

Optimal1 Fitted Gauss, proteinAT3G18420.1
bic:717
peaks after deleting less than one fifth of global max:16
×10^7
raw data, starting:16
fitted Gauss locations:16
y
x
0
0.5
1
1.5
2
2.5
3
3.5
4
0
5
10
15
20
25

**Optimal1 Fitted Gauss, proteinAT3G18490.1**
**bic:786 794**
**peaks after deleting less than one fifth of global max:17**

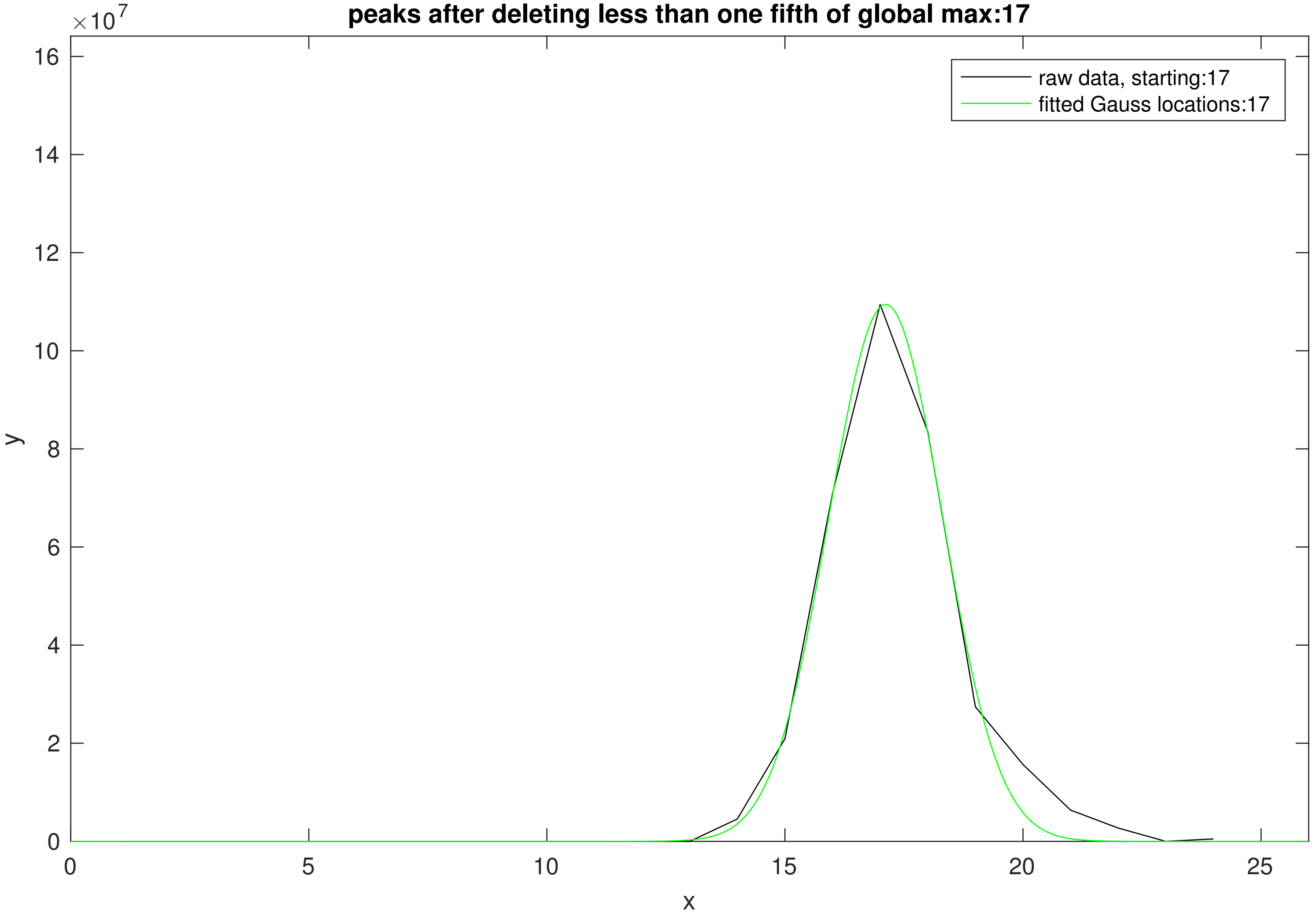

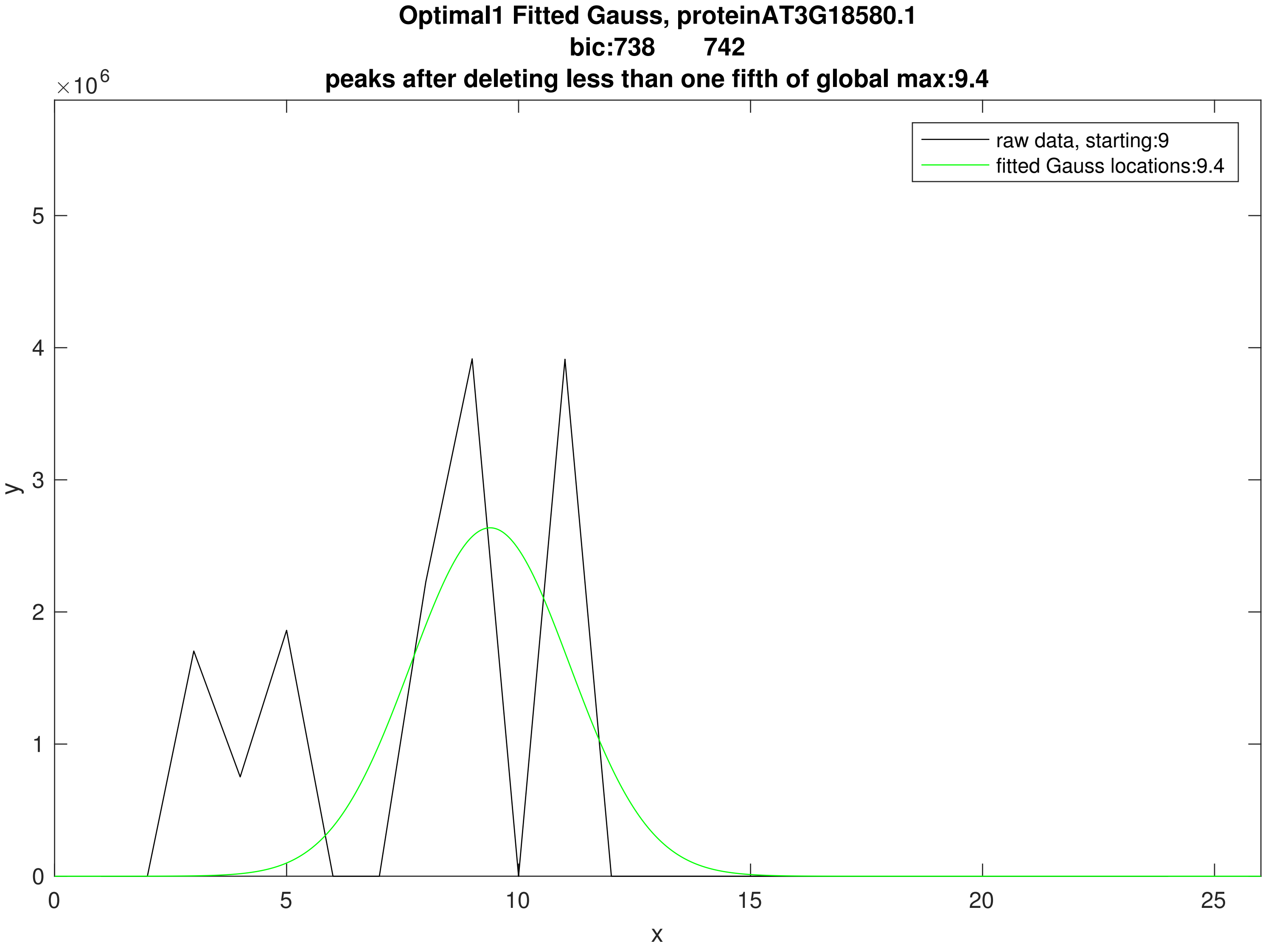

Optimal1 Fitted Gauss, proteinAT3G18580.1
bic:738 742
peaks after deleting less than one fifth of global max:9.4
×10^6
raw data, starting:9
fitted Gauss locations:9.4
y
x
0
1
2
3
4
5
0
5
10
15
20
25

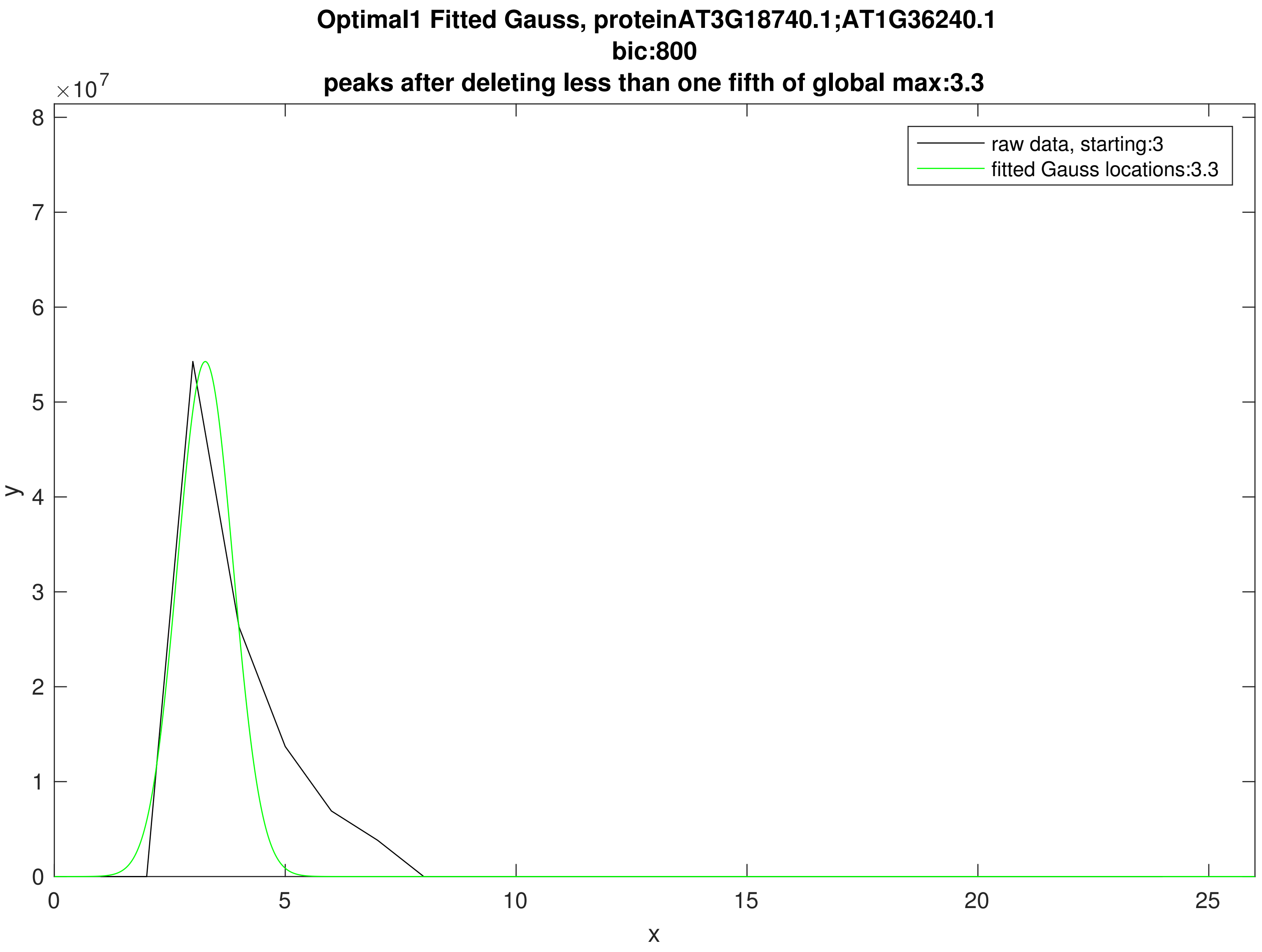

Optimal1 Fitted Gauss, proteinAT3G18740.1;AT1G36240.1
bic:800
peaks after deleting less than one fifth of global max:3.3
×10^7
raw data, starting:3
fitted Gauss locations:3.3
y
x
0
1
2
3
4
5
6
7
8
0
5
10
15
20
25

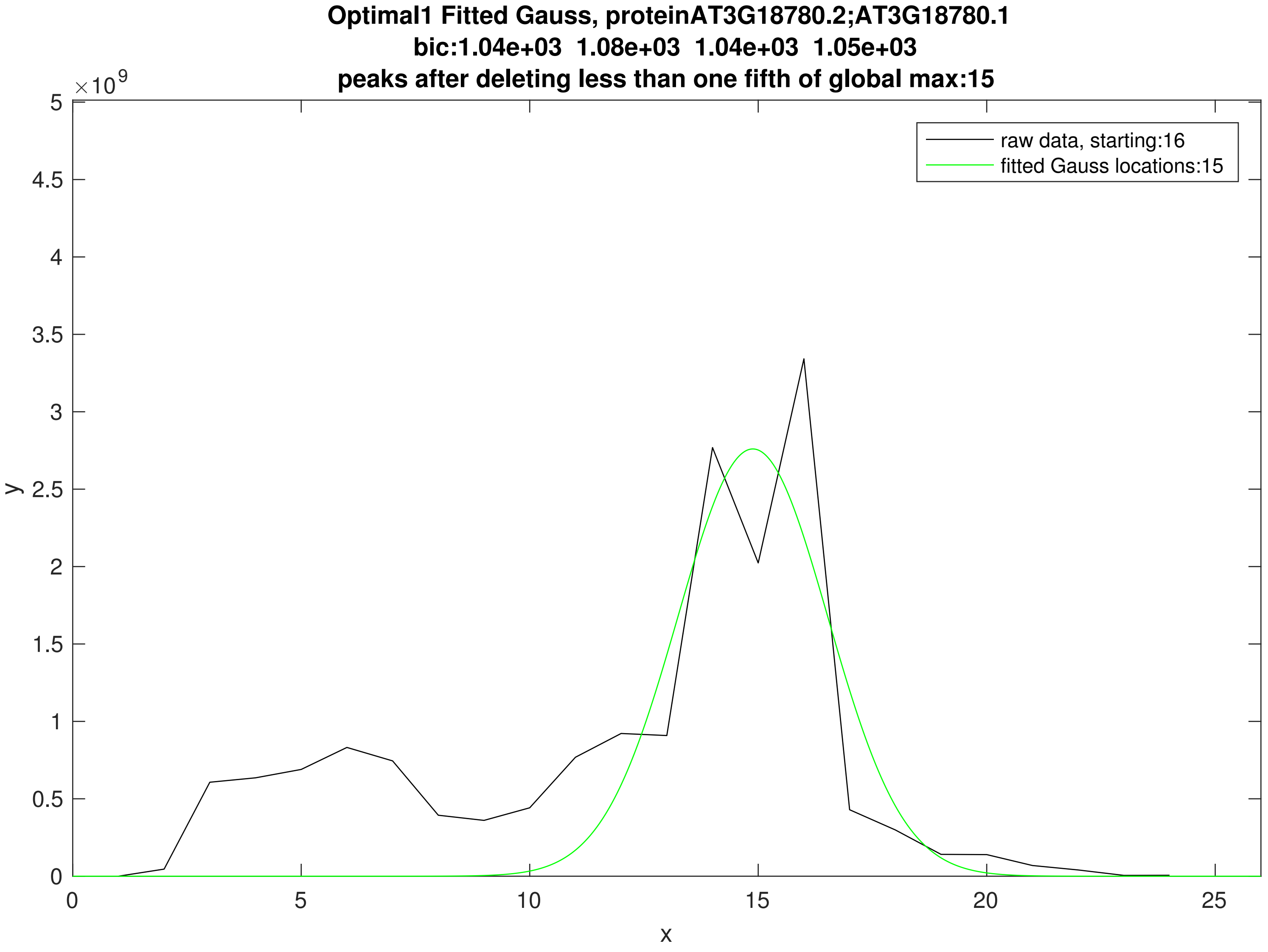
Optimal1 Fitted Gauss, proteinAT3G18780.2;AT3G18780.1
bic:1.04e+03 1.08e+03 1.04e+03 1.05e+03
peaks after deleting less than one fifth of global max:15
raw data, starting:16
fitted Gauss locations:15
×10^9
y
x

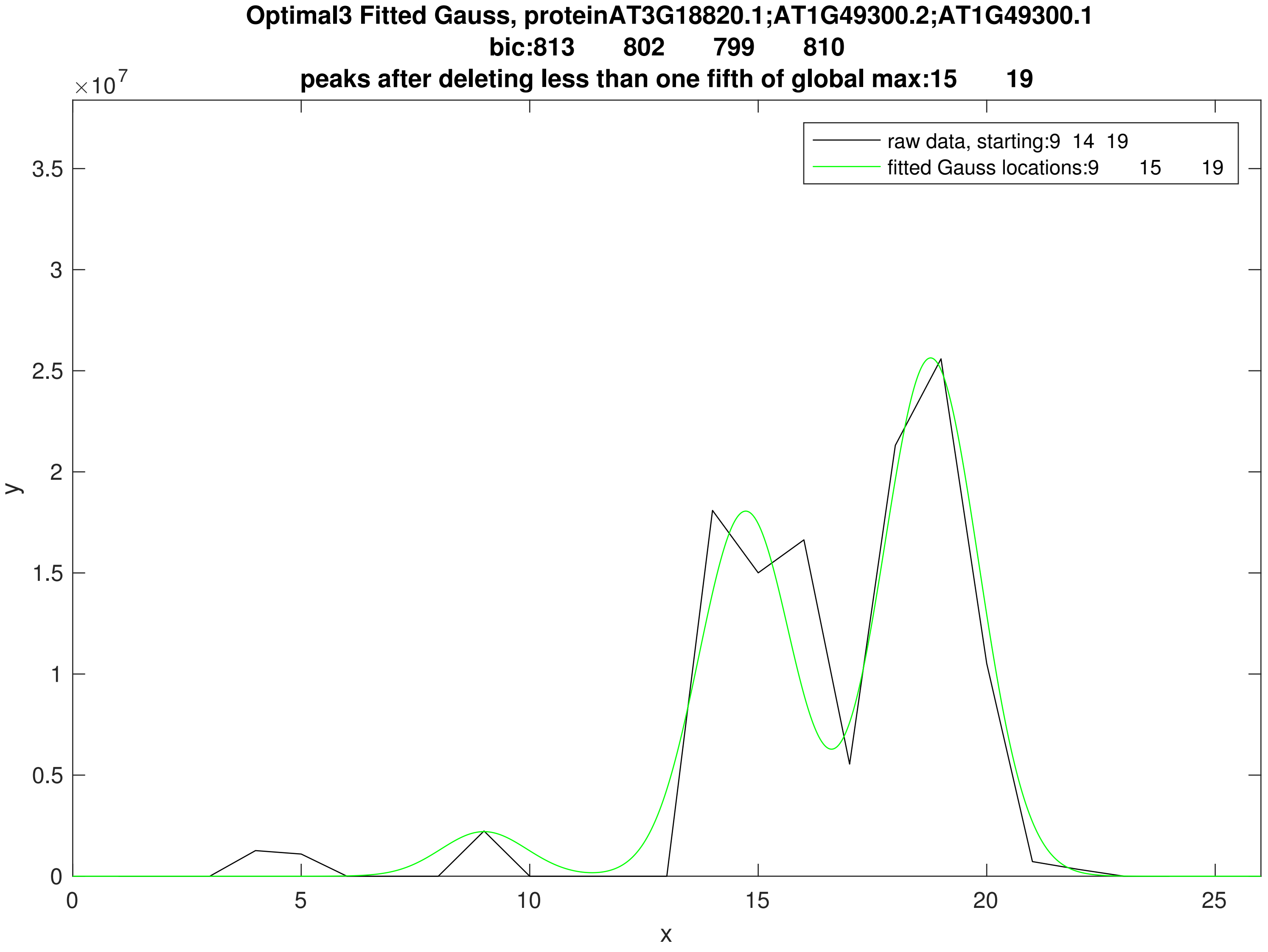

Optimal3 Fitted Gauss, proteinAT3G18820.1;AT1G49300.2;AT1G49300.1
bic:813 802 799 810
peaks after deleting less than one fifth of global max:15 19
×10^7
raw data, starting:9 14 19
fitted Gauss locations:9 15 19
3.5
3
2.5
2
1.5
1
0.5
0
y
0
5
10
15
20
25
x

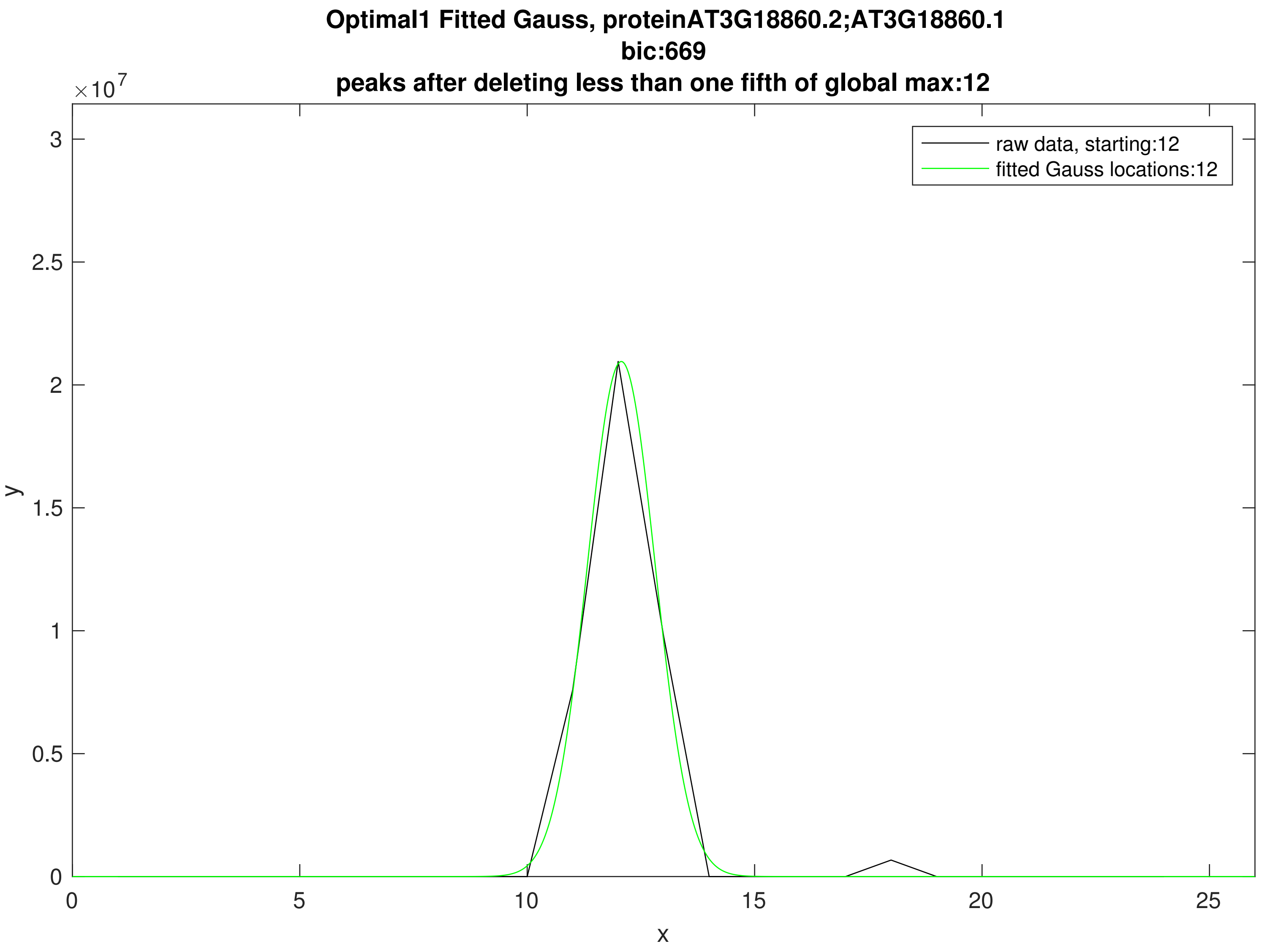

Optimal1 Fitted Gauss, proteinAT3G18860.2;AT3G18860.1
bic:669
peaks after deleting less than one fifth of global max:12
×10^7
raw data, starting:12
fitted Gauss locations:12
y
x
0
0.5
1
1.5
2
2.5
3
0
5
10
15
20
25

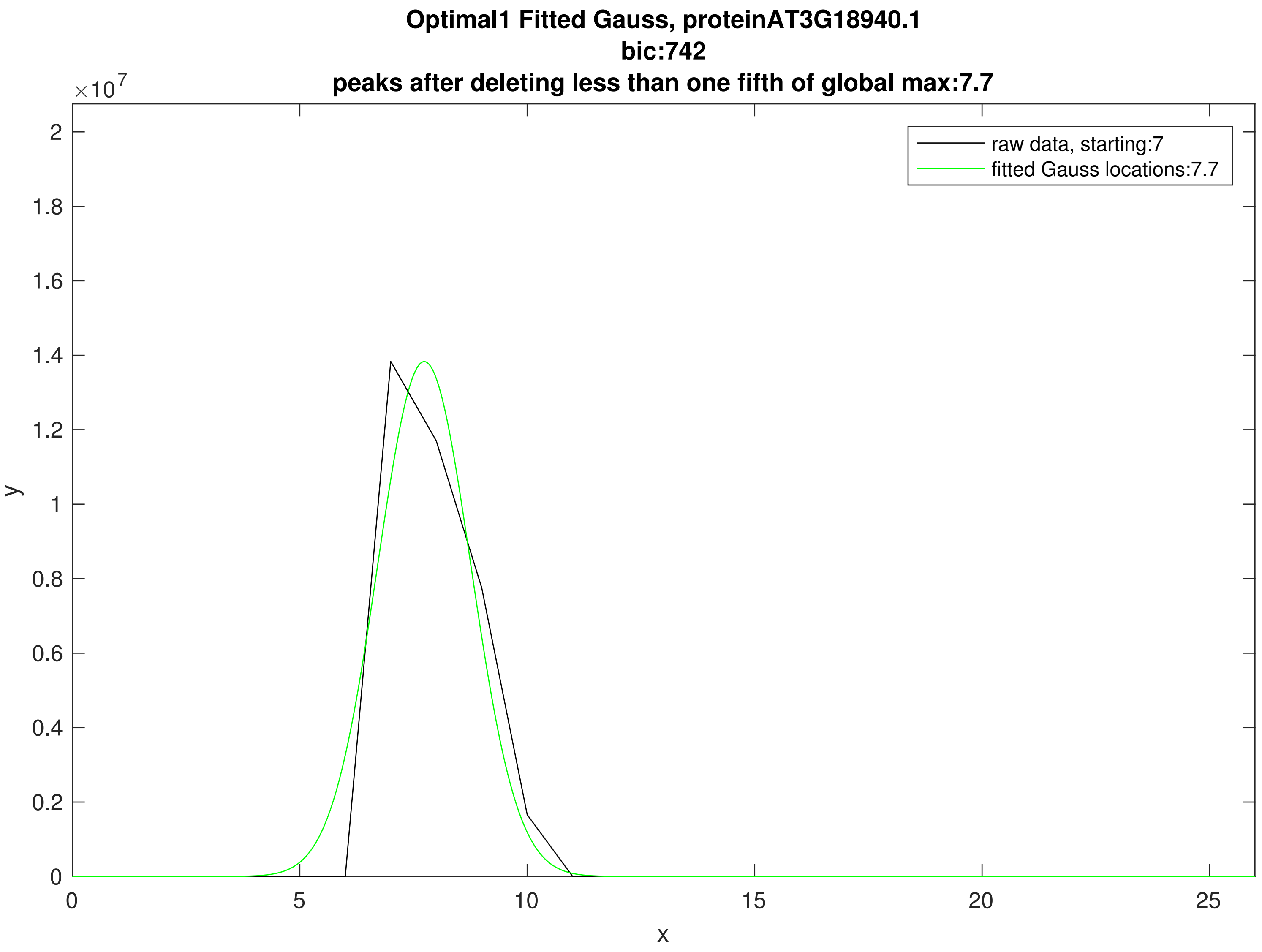

Optimal1 Fitted Gauss, proteinAT3G18940.1
bic:742
peaks after deleting less than one fifth of global max:7.7
×10^7
2
1.8
1.6
1.4
1.2
1
0.8
0.6
0.4
0.2
0
y
0
5
10
15
20
25
x
raw data, starting:7
fitted Gauss locations:7.7

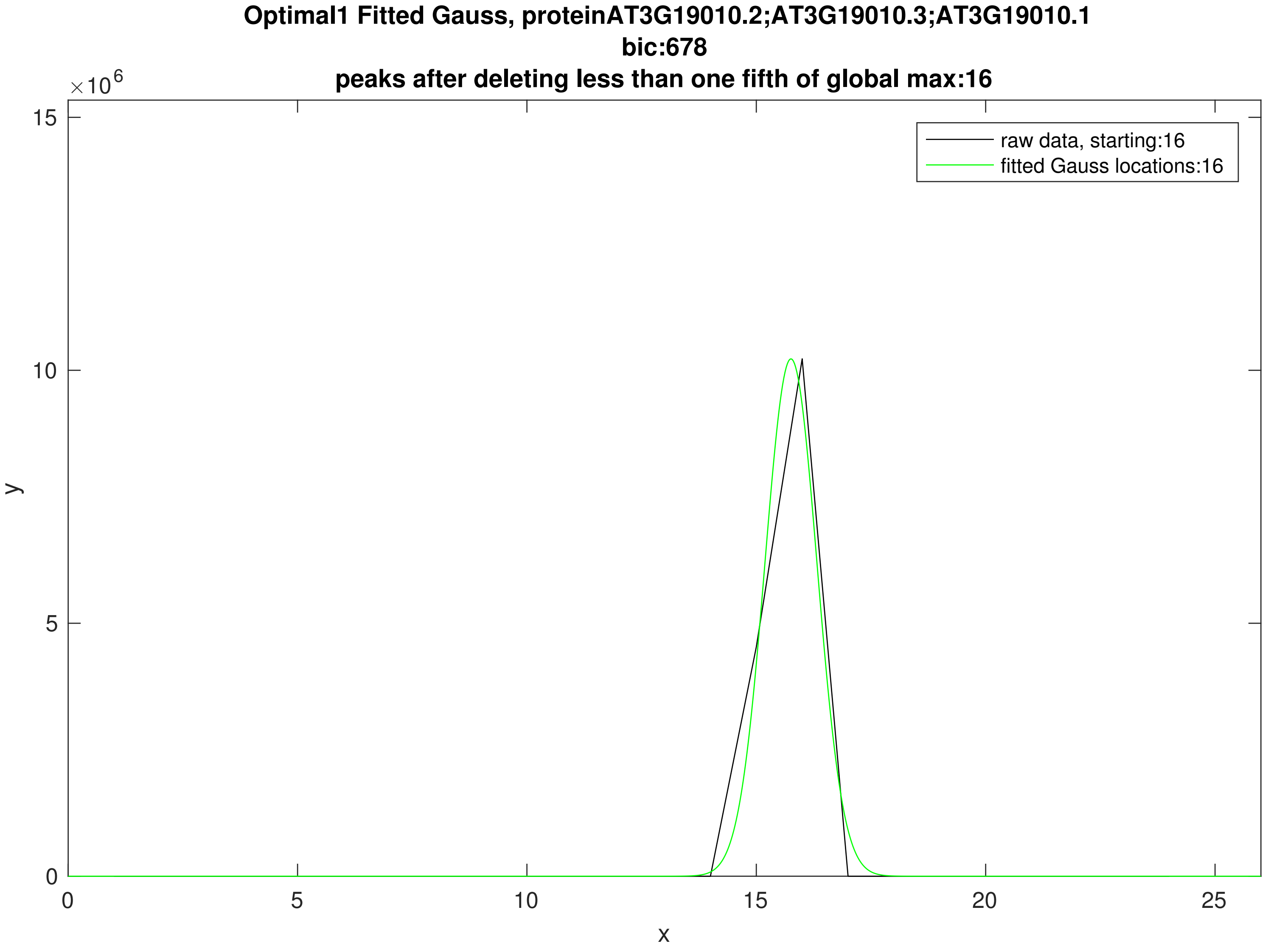

Optimal1 Fitted Gauss, proteinAT3G19010.2;AT3G19010.3;AT3G19010.1
bic:678
peaks after deleting less than one fifth of global max:16
×10^6
15
10
5
0
y
0
5
10
15
20
25
x
raw data, starting:16
fitted Gauss locations:16

**Optimal1 Fitted Gauss, proteinAT3G19170.1;AT3G19170.2**
**bic:952 960 969**
**peaks after deleting less than one fifth of global max:13**

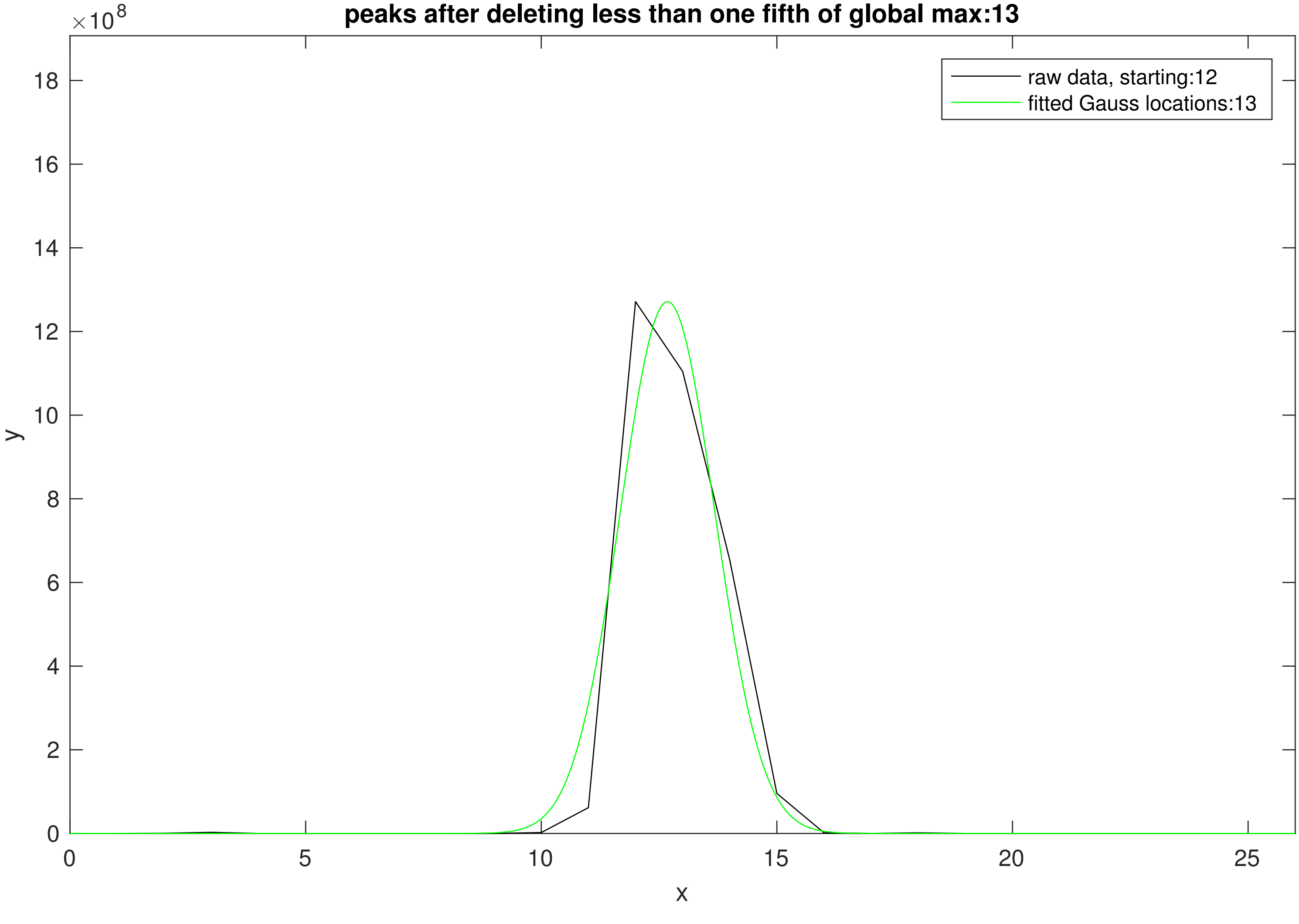

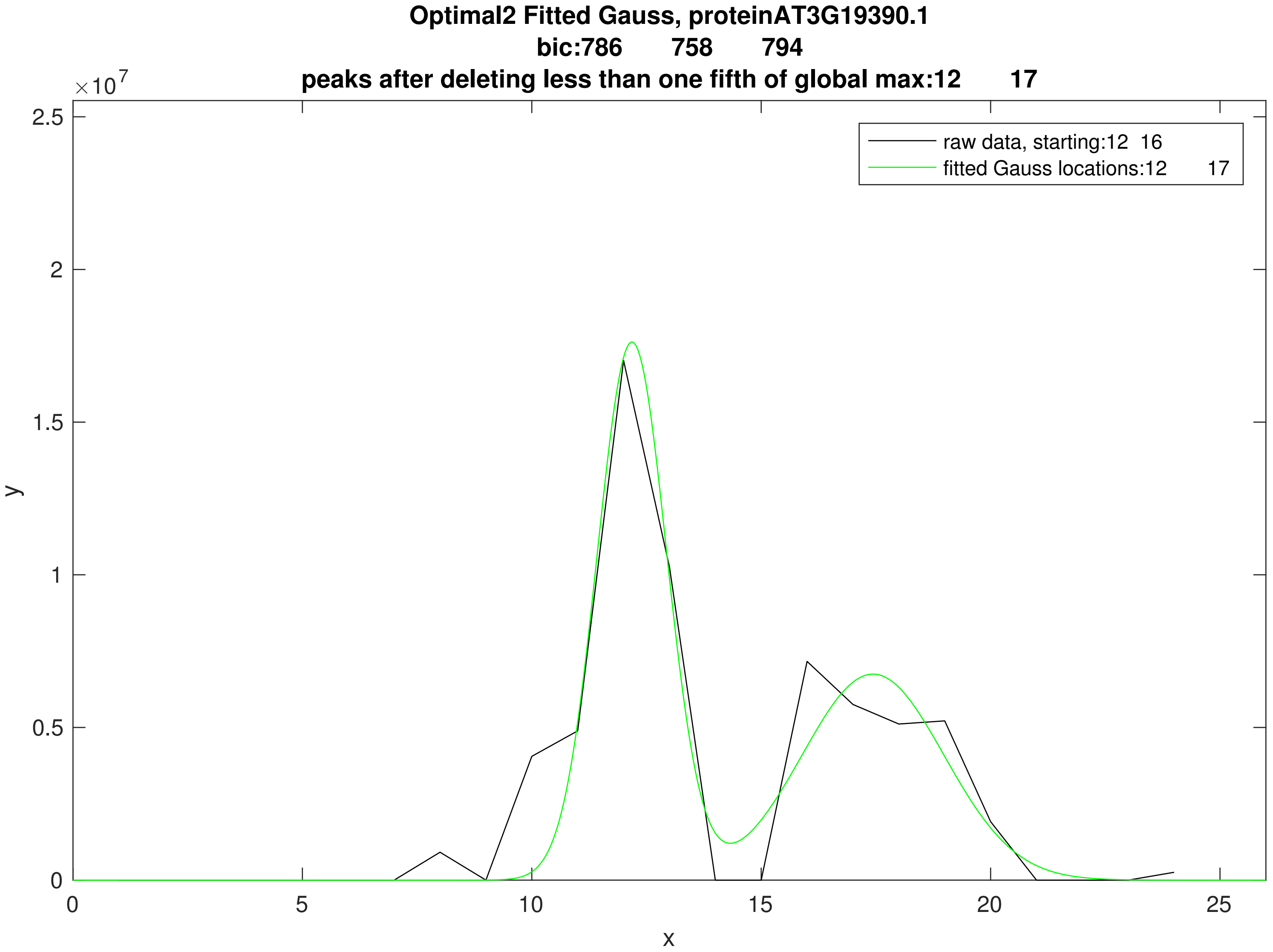
Optimal2 Fitted Gauss, proteinAT3G19390.1
bic:786 758 794
peaks after deleting less than one fifth of global max:12 17
×10^7
raw data, starting:12 16
fitted Gauss locations:12 17
y
x

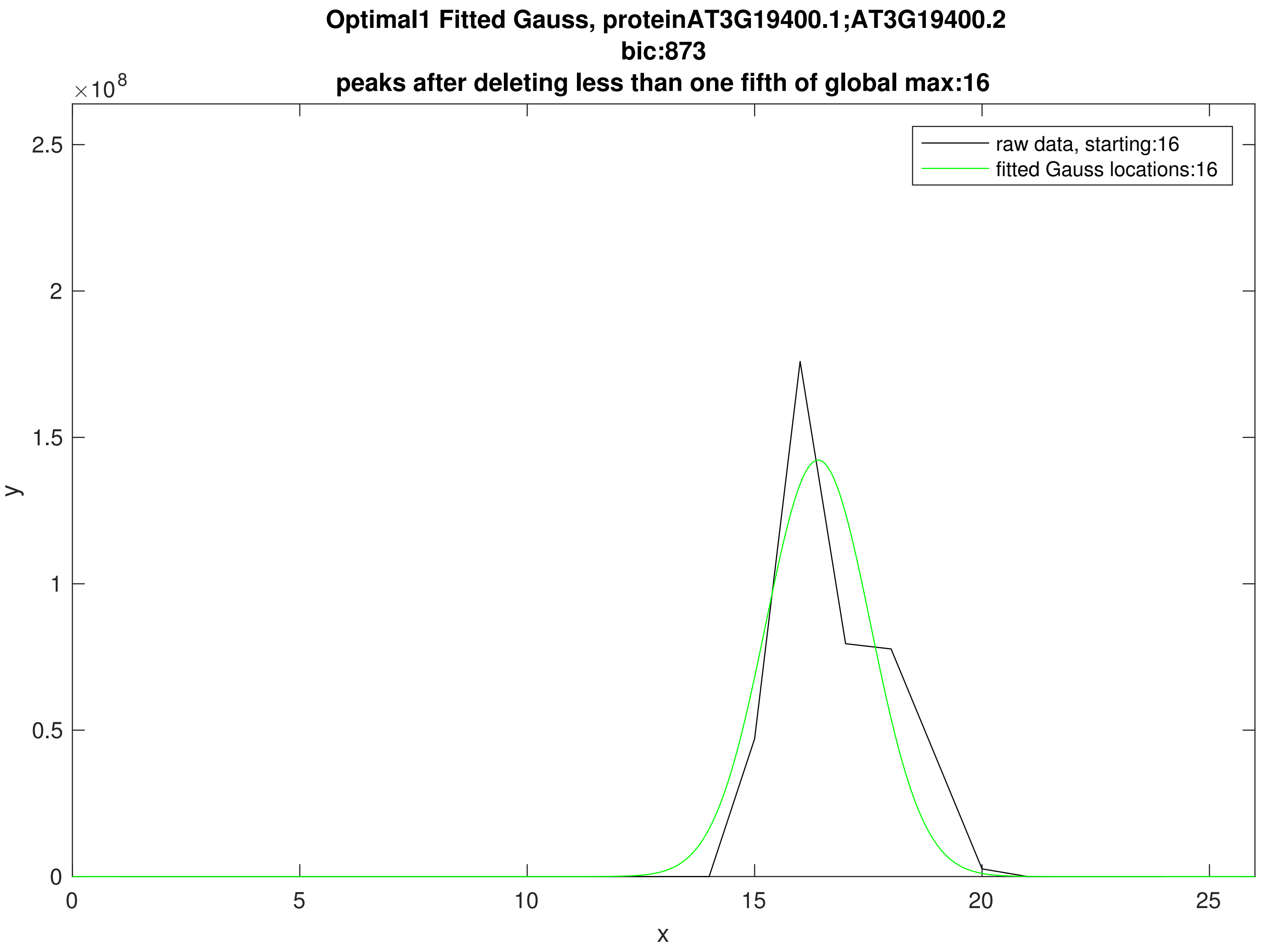

Optimal1 Fitted Gauss, proteinAT3G19400.1;AT3G19400.2
bic:873
peaks after deleting less than one fifth of global max:16
×10^8
2.5
2
1.5
1
0.5
0
y
0
5
10
15
20
25
x
raw data, starting:16
fitted Gauss locations:16

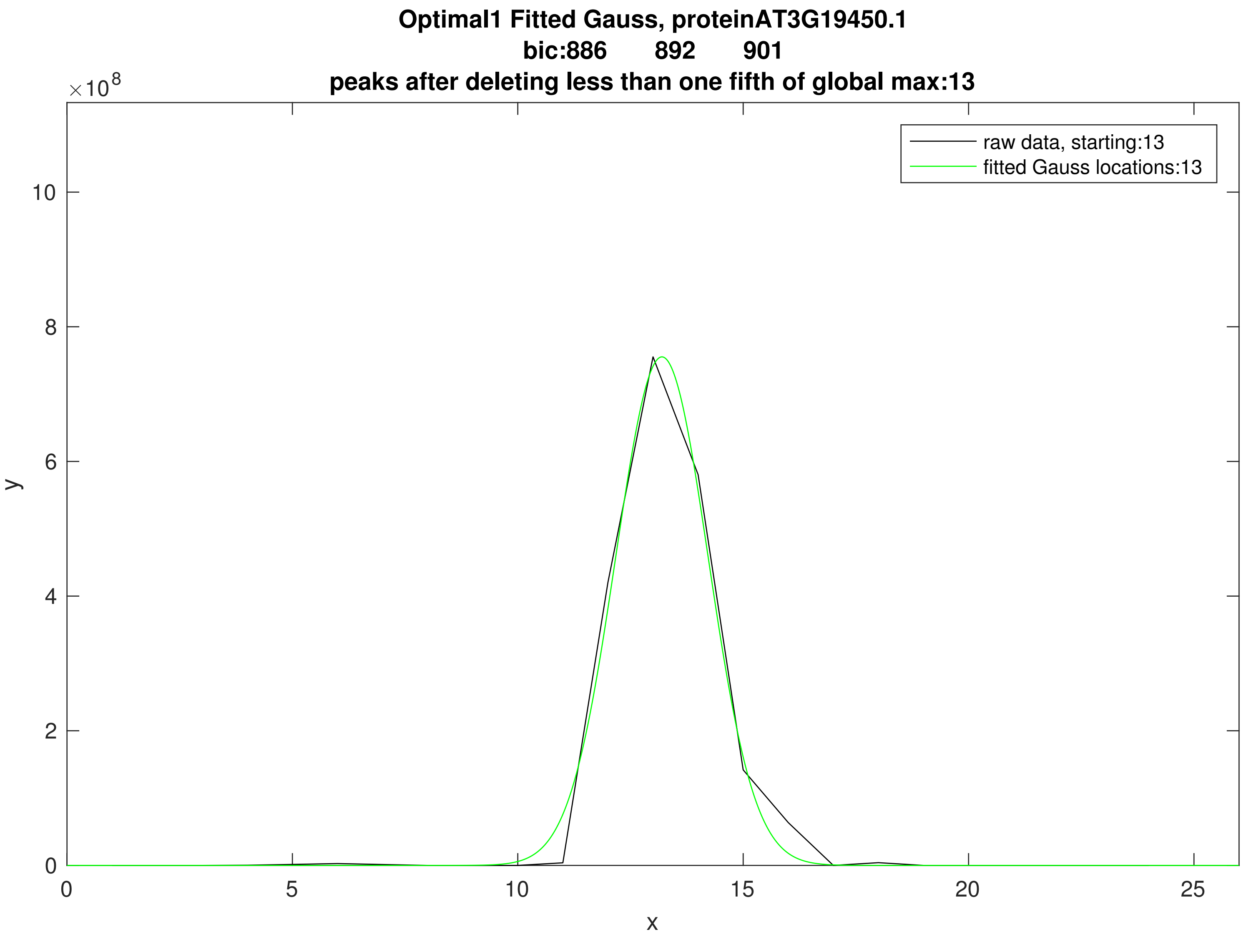

Optimal1 Fitted Gauss, proteinAT3G19450.1
bic:886 892 901
peaks after deleting less than one fifth of global max:13
×10^8
raw data, starting:13
fitted Gauss locations:13
y
x
0
2
4
6
8
10
0
5
10
15
20
25

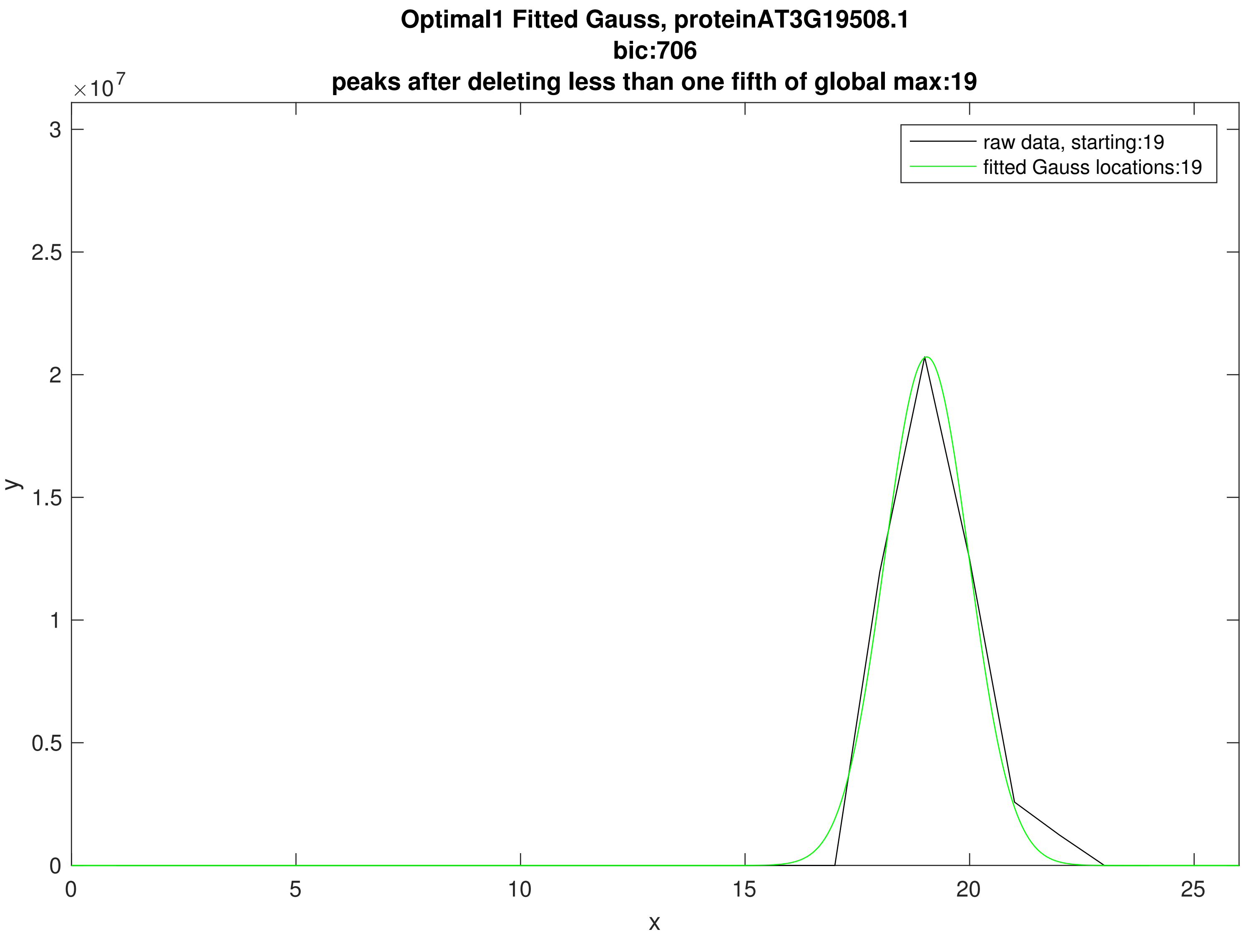

Optimal1 Fitted Gauss, proteinAT3G19508.1
bic:706
peaks after deleting less than one fifth of global max:19
×10^7
raw data, starting:19
fitted Gauss locations:19
y
x
0
0.5
1
1.5
2
2.5
3
0
5
10
15
20
25

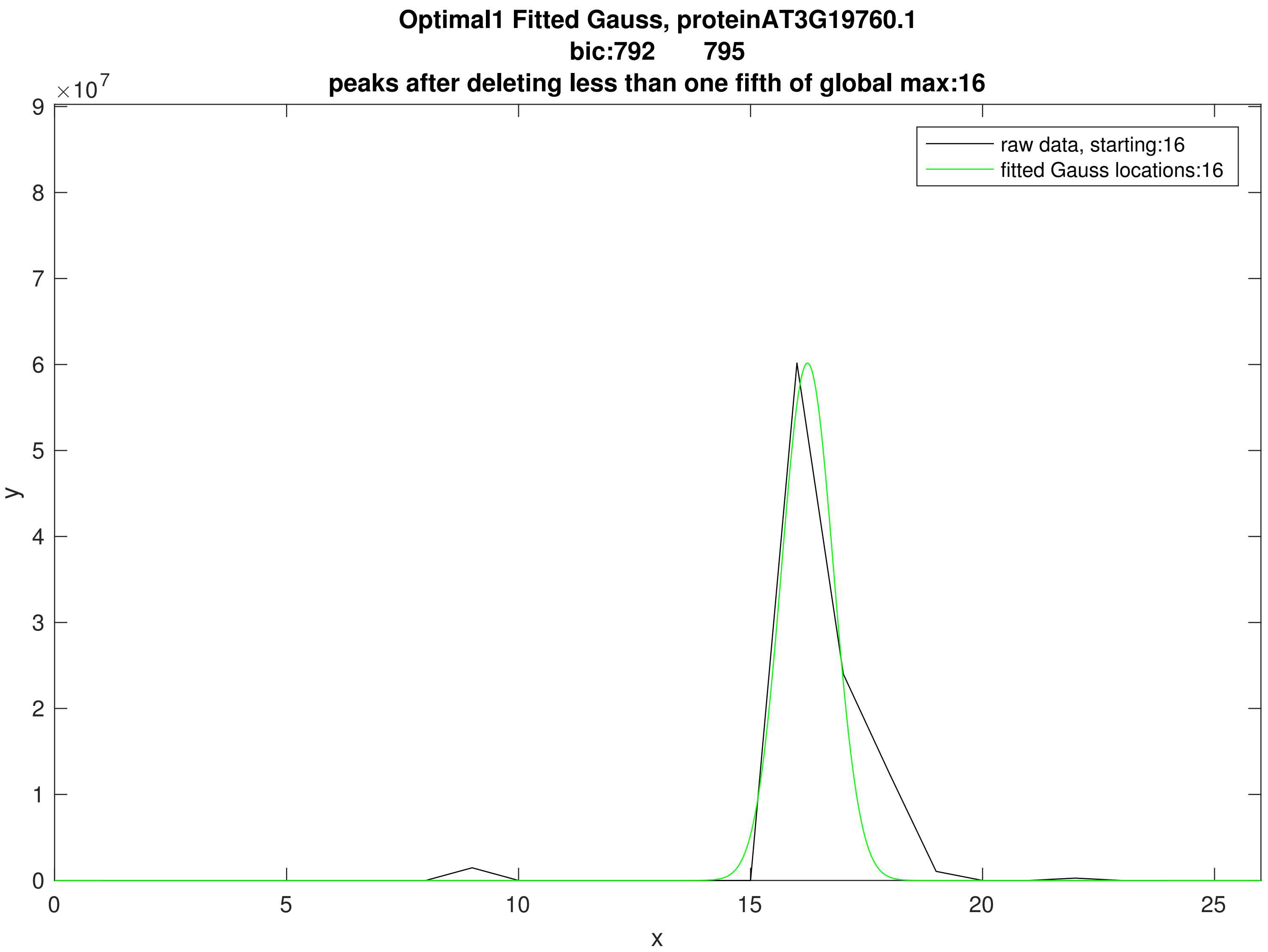
Optimal1 Fitted Gauss, proteinAT3G19760.1
bic:792 795
peaks after deleting less than one fifth of global max:16
×10^7
y
x
raw data, starting:16
fitted Gauss locations:16

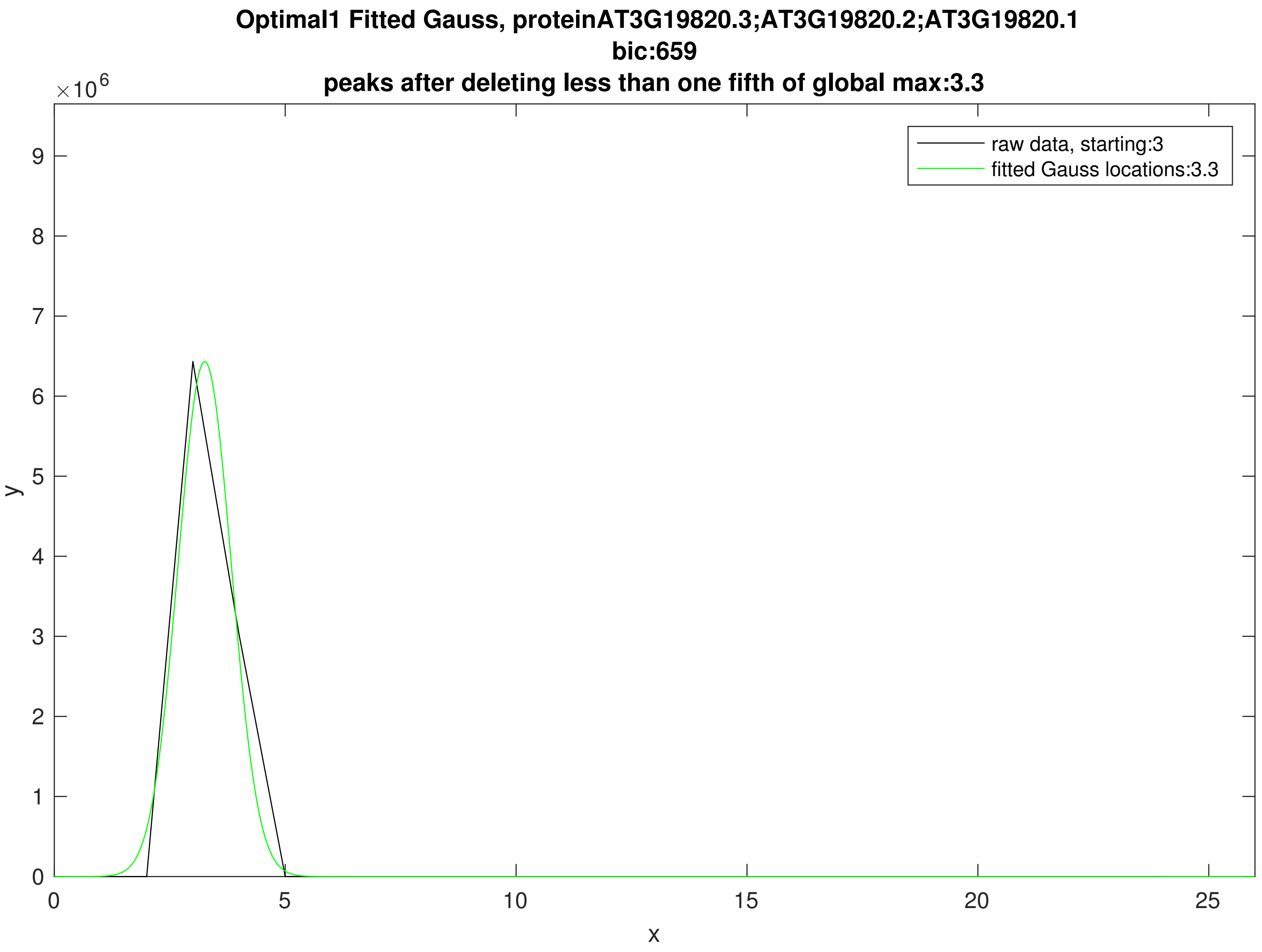
Optimal1 Fitted Gauss, proteinAT3G19820.3;AT3G19820.2;AT3G19820.1
bic:659
peaks after deleting less than one fifth of global max:3.3
×10^6
raw data, starting:3
fitted Gauss locations:3.3
y
x
0
1
2
3
4
5
6
7
8
9
0
5
10
15
20
25

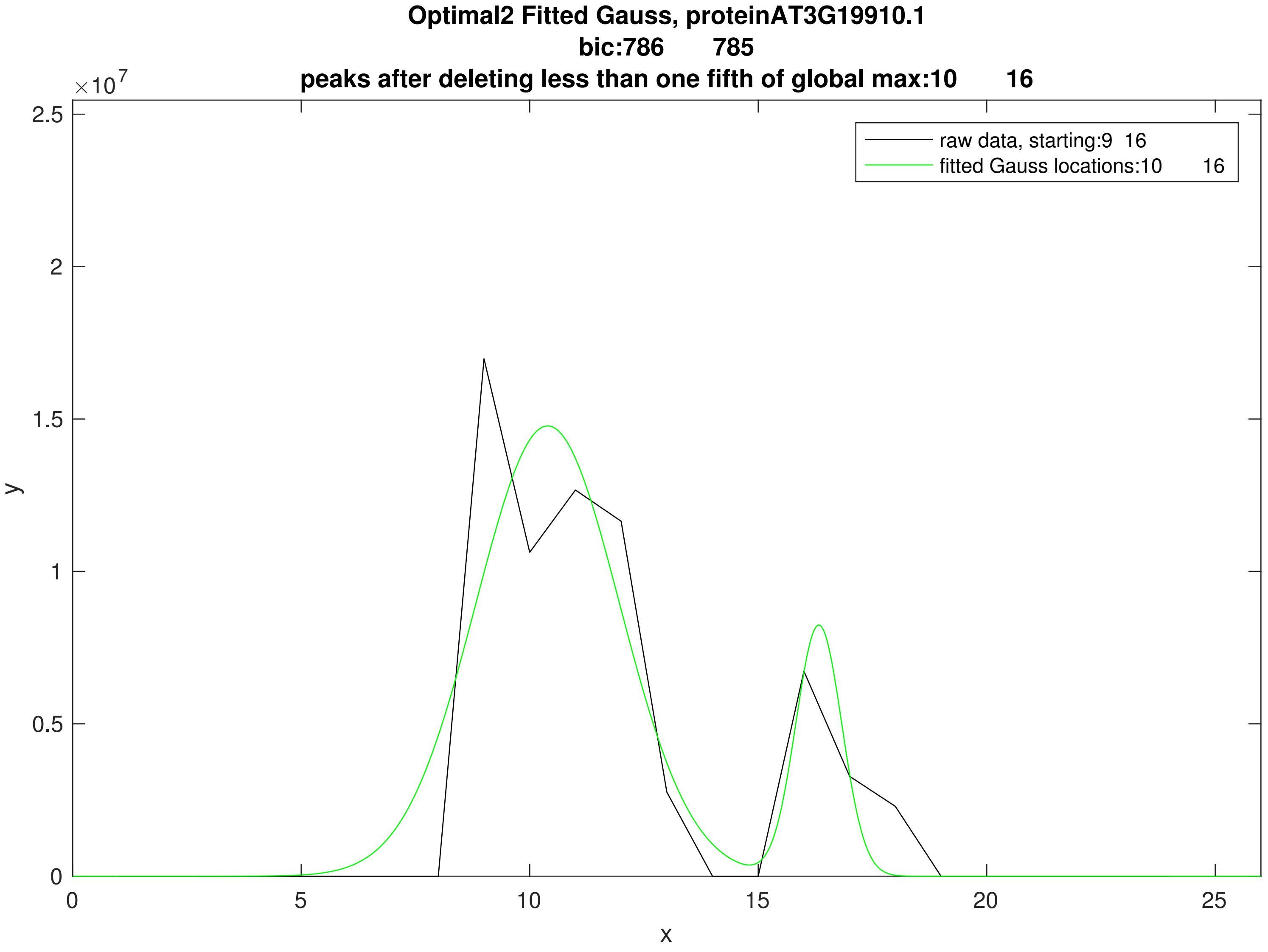

Optimal2 Fitted Gauss, proteinAT3G19910.1
bic:786 785
peaks after deleting less than one fifth of global max:10 16
×10^7
2.5
2
1.5
1
0.5
0
y
0
5
10
15
20
25
x
raw data, starting:9 16
fitted Gauss locations:10 16

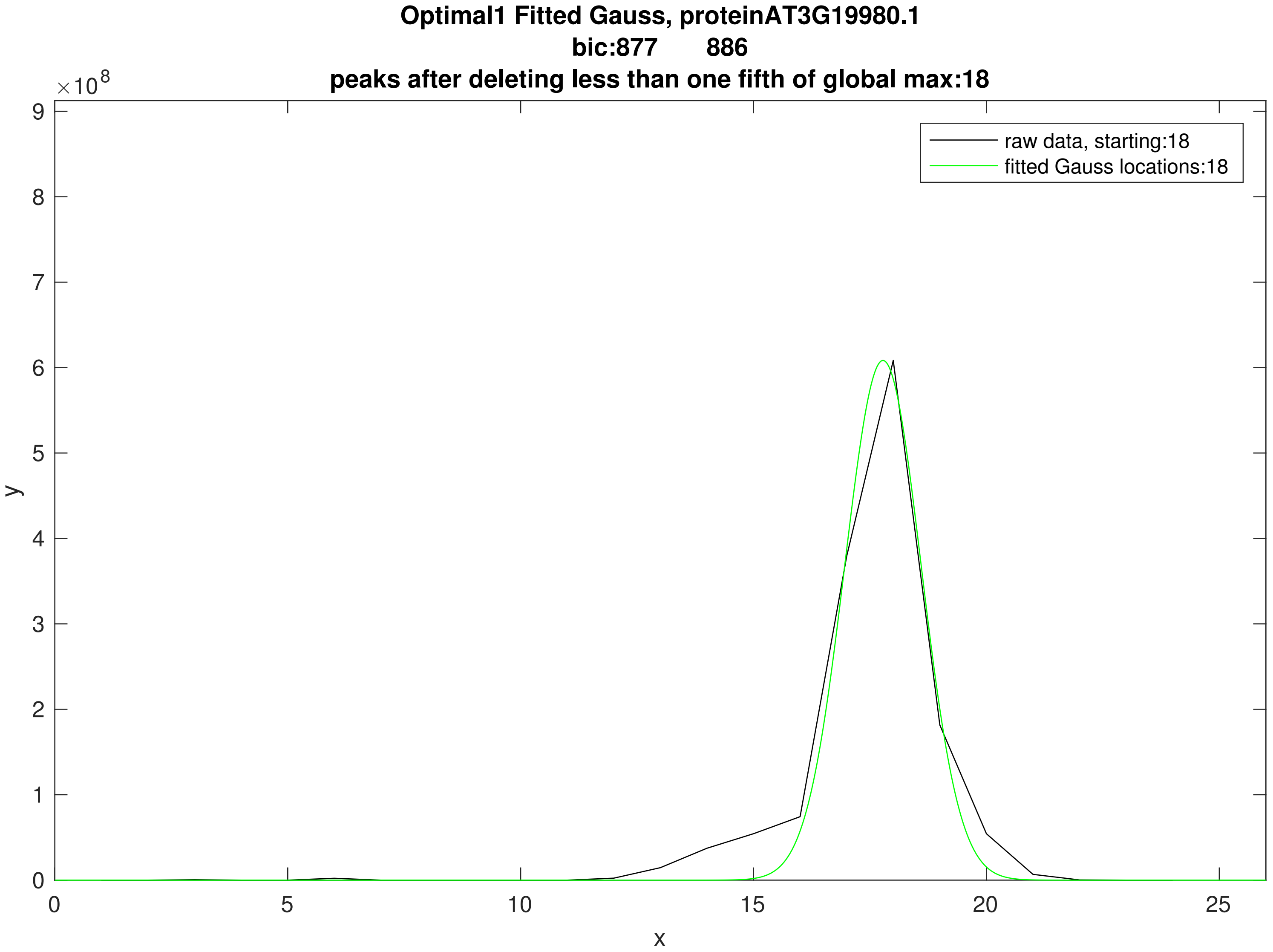

Optimal1 Fitted Gauss, proteinAT3G19980.1
bic:877 886
peaks after deleting less than one fifth of global max:18
×10^8
9
8
7
6
5
4
3
2
1
0
y
0
5
10
15
20
25
x
raw data, starting:18
fitted Gauss locations:18

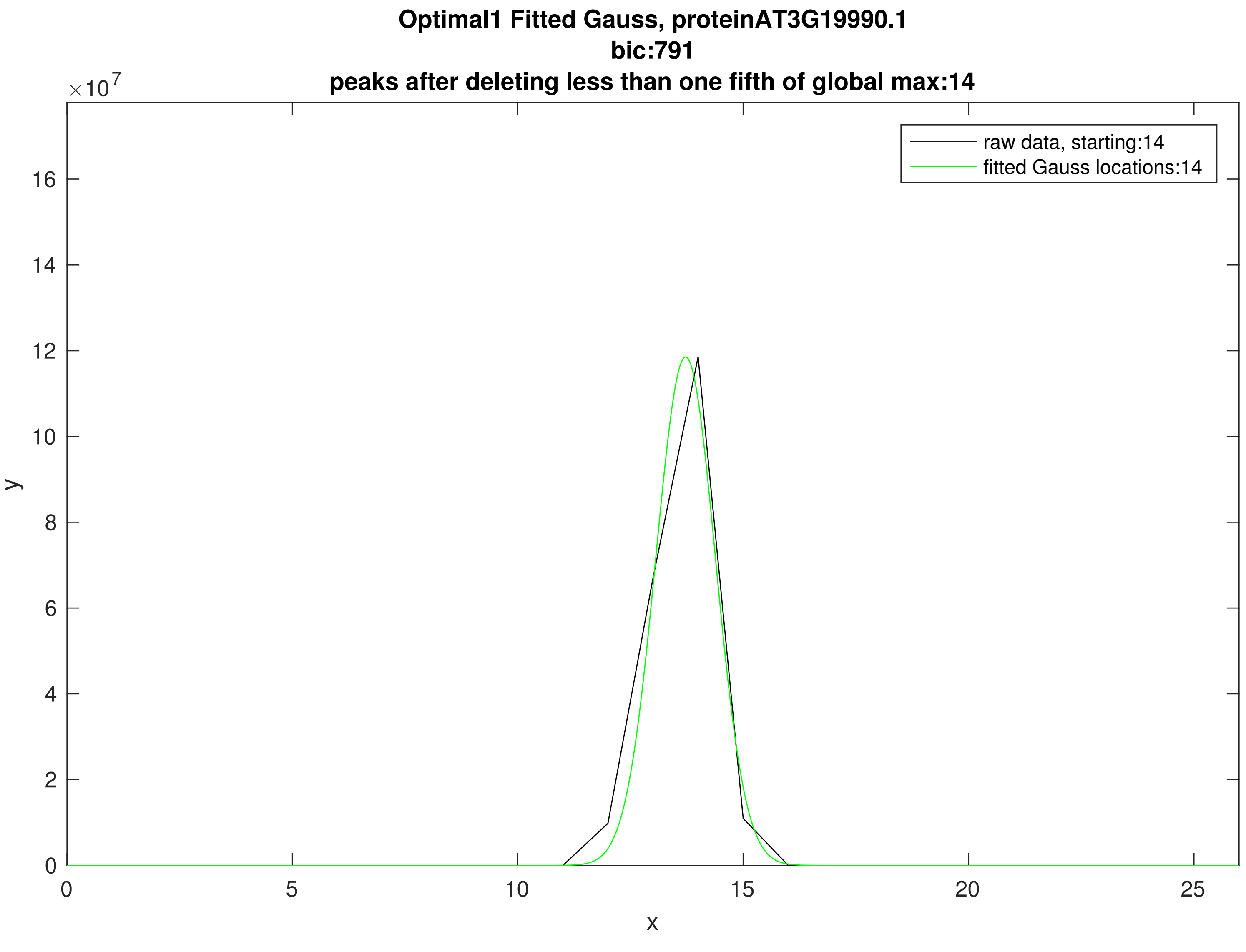
Optimal1 Fitted Gauss, proteinAT3G19990.1
bic:791
peaks after deleting less than one fifth of global max:14
×10^7
raw data, starting:14
fitted Gauss locations:14
y
x
0
2
4
6
8
10
12
14
16
0
5
10
15
20
25

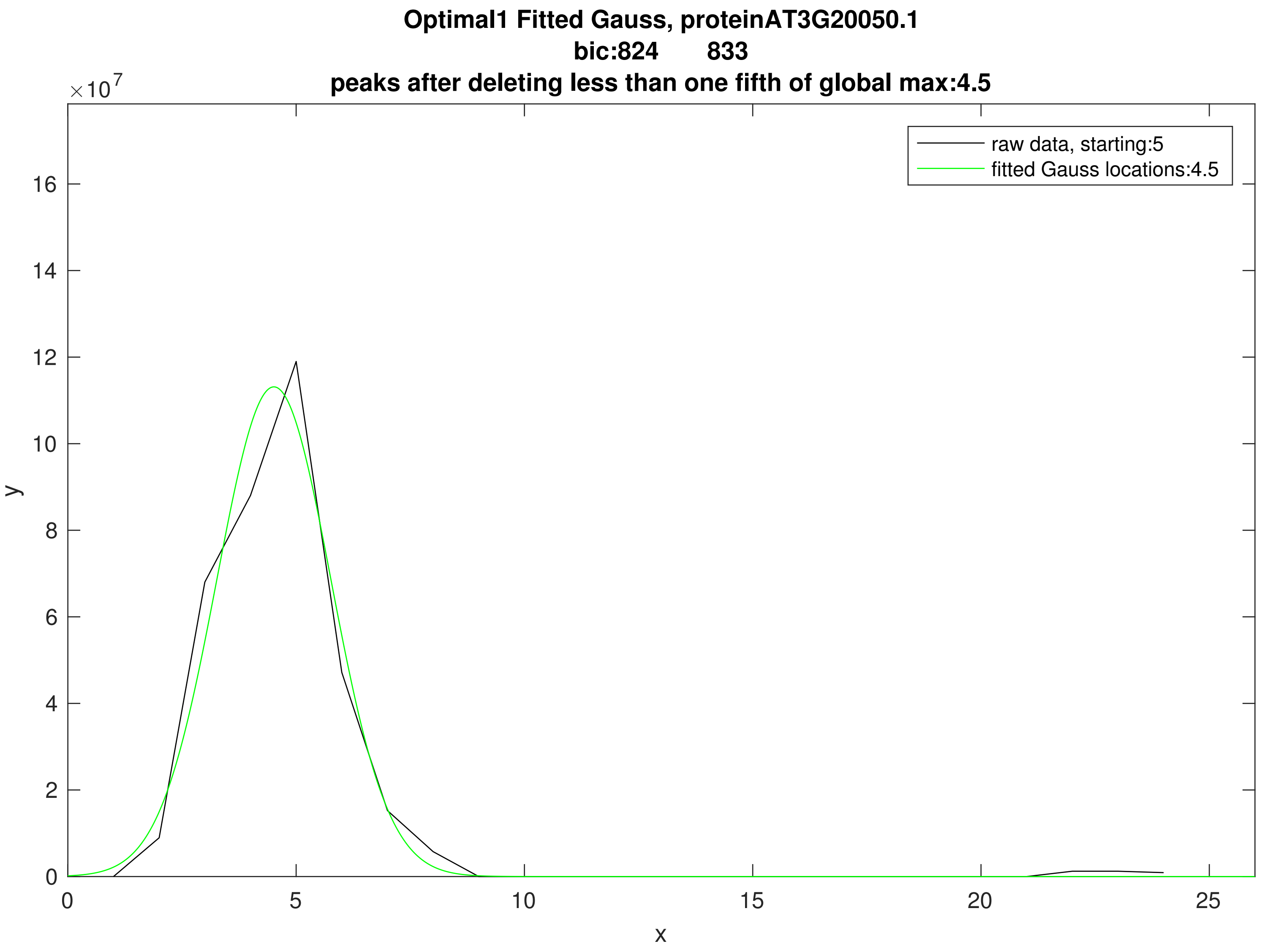
Optimal1 Fitted Gauss, proteinAT3G20050.1
bic:824 833
peaks after deleting less than one fifth of global max:4.5
×10^7
raw data, starting:5
fitted Gauss locations:4.5
y
x
0
2
4
6
8
10
12
14
16
0
5
10
15
20
25

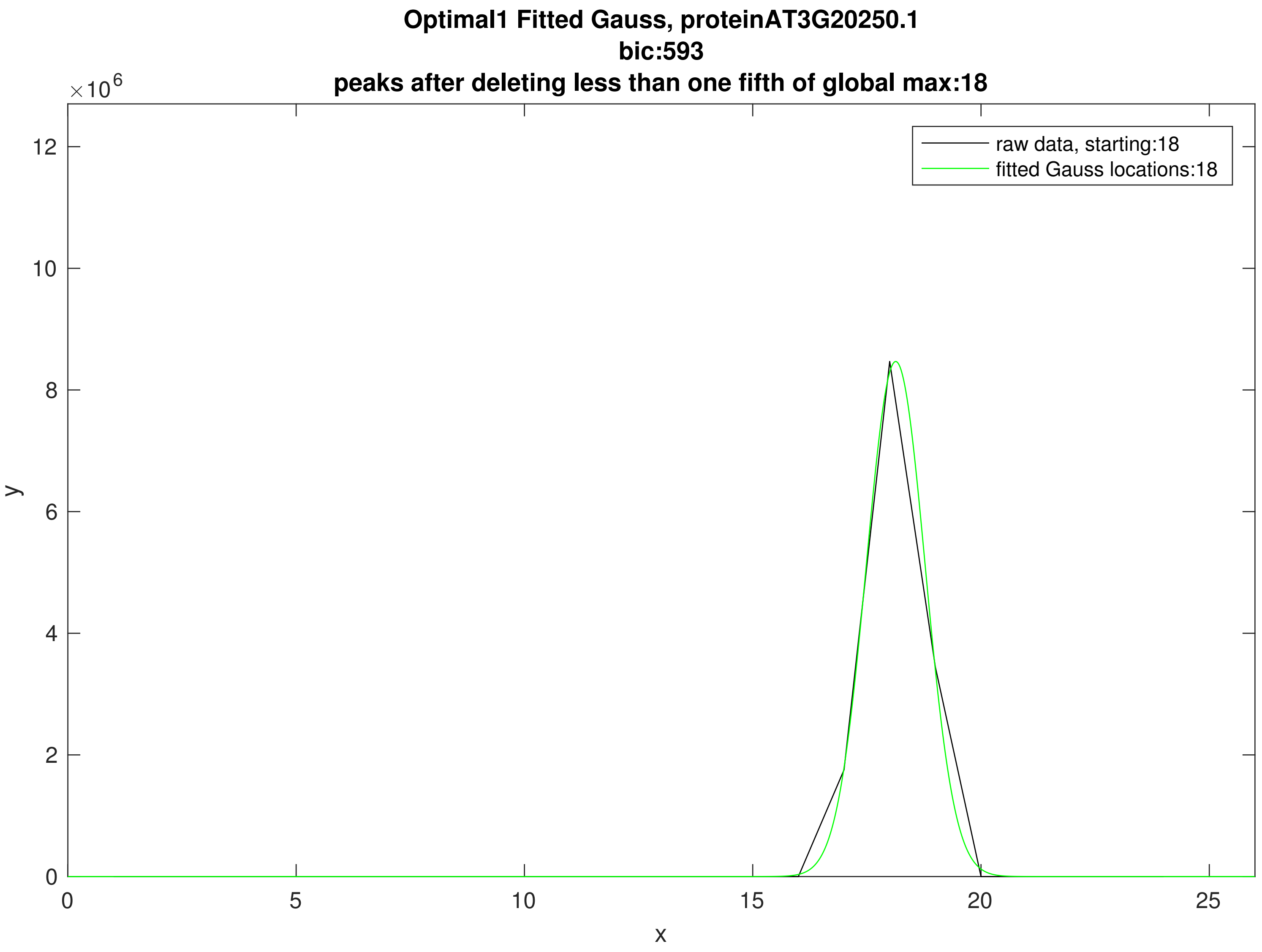

Optimal1 Fitted Gauss, proteinAT3G20250.1
bic:593
peaks after deleting less than one fifth of global max:18
×10^6
raw data, starting:18
fitted Gauss locations:18
y
x
0
2
4
6
8
10
12
0
5
10
15
20
25

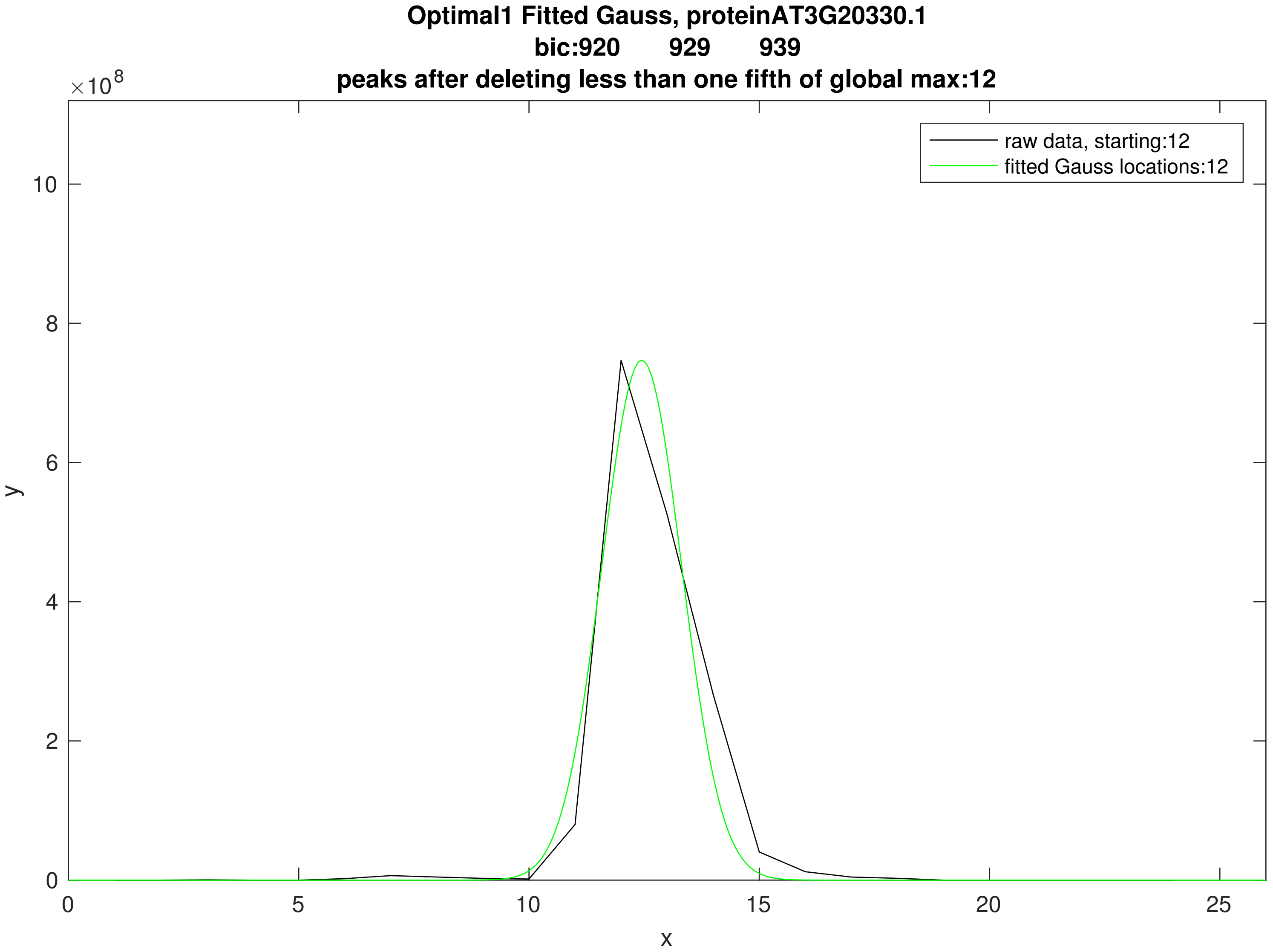
Optimal1 Fitted Gauss, proteinAT3G20330.1
bic:920 929 939
peaks after deleting less than one fifth of global max:12
×10^8
raw data, starting:12
fitted Gauss locations:12
y
x
0
2
4
6
8
10
0
5
10
15
20
25

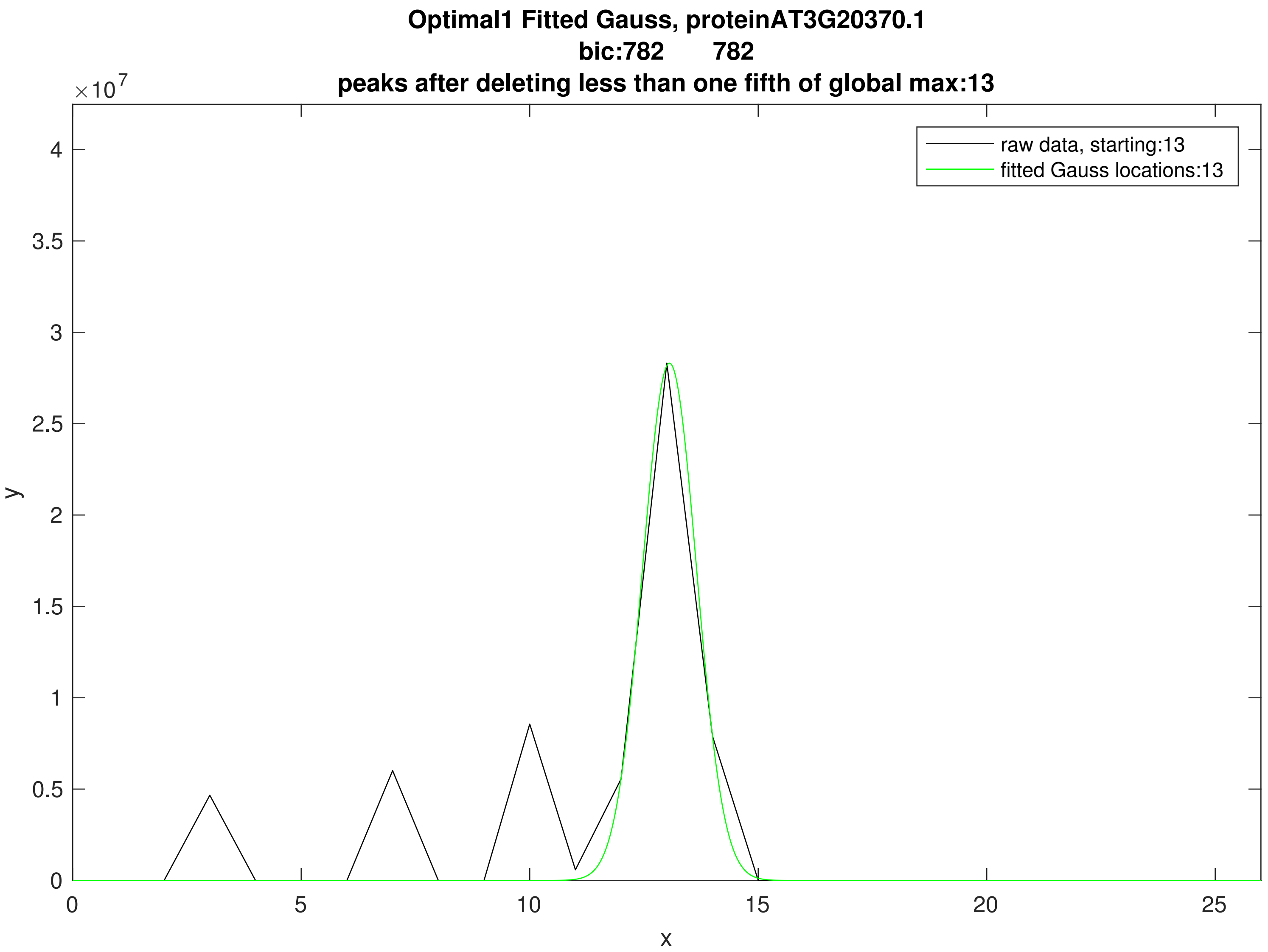
Optimal1 Fitted Gauss, proteinAT3G20370.1
bic:782 782
peaks after deleting less than one fifth of global max:13
×10^7
raw data, starting:13
fitted Gauss locations:13
y
x
0
0.5
1
1.5
2
2.5
3
3.5
4
5
10
15
20
25

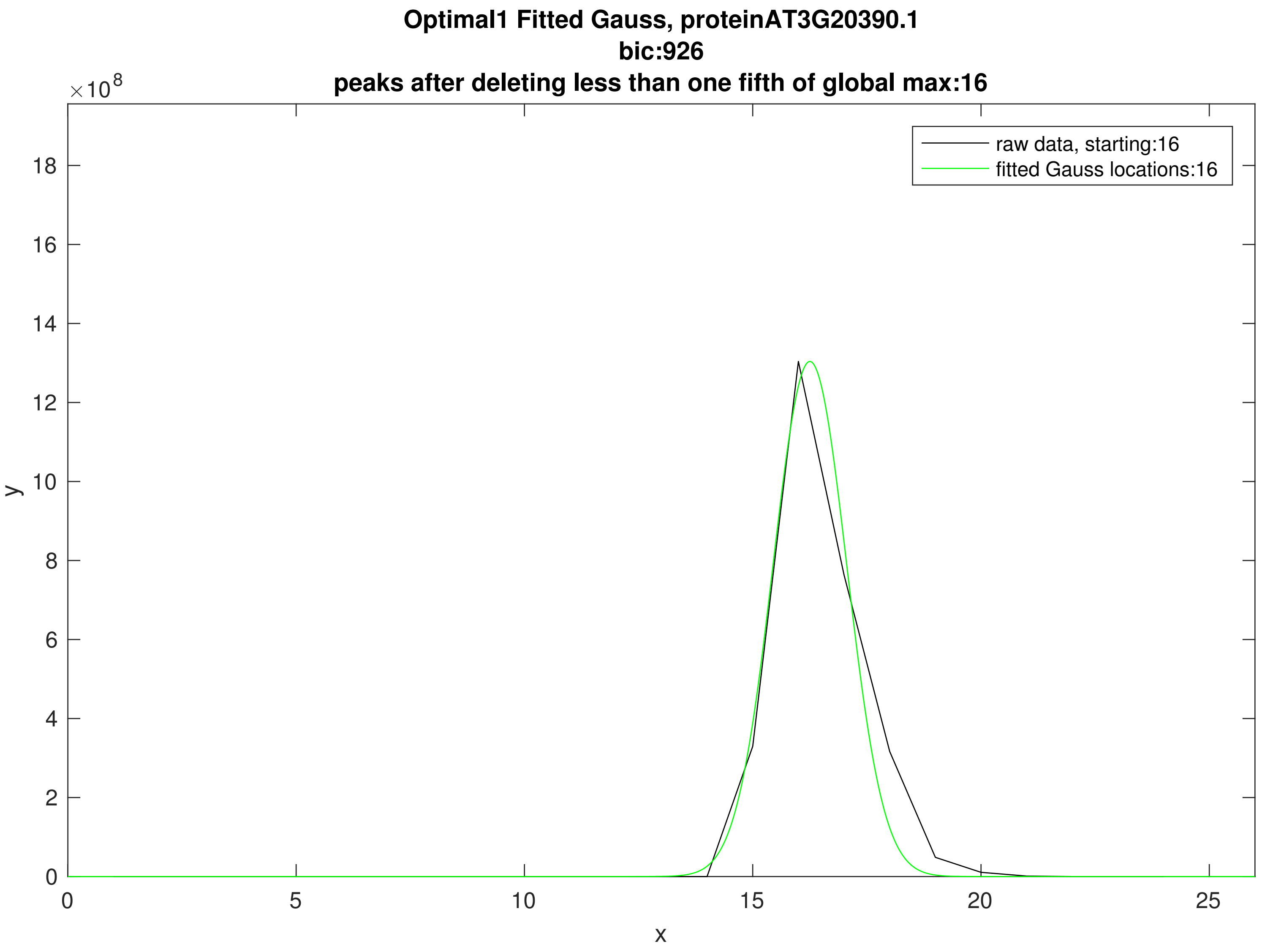
Optimal1 Fitted Gauss, proteinAT3G20390.1
bic:926
peaks after deleting less than one fifth of global max:16
×10^8
raw data, starting:16
fitted Gauss locations:16
y
x
0
2
4
6
8
10
12
14
16
18
0
5
10
15
20
25

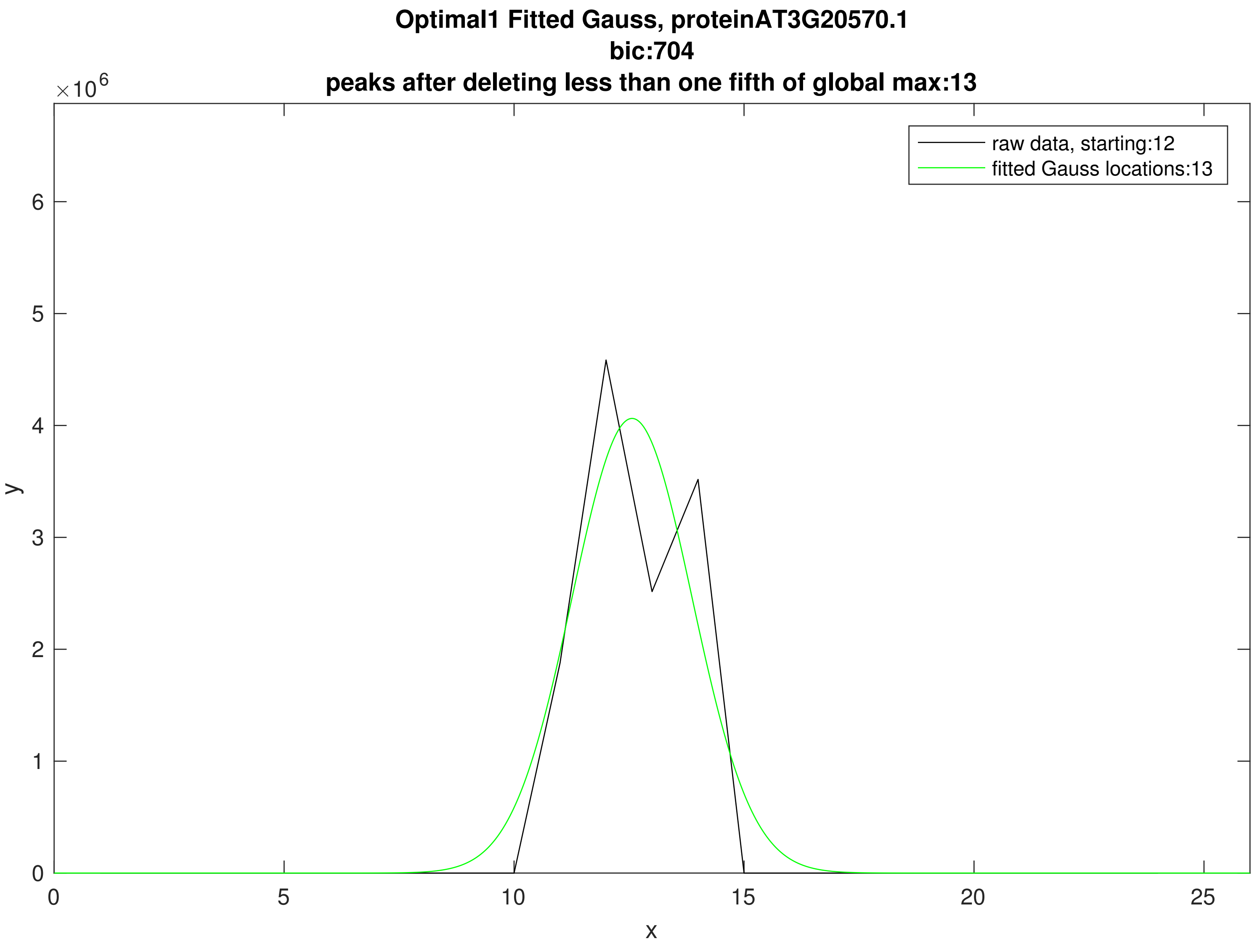

Optimal1 Fitted Gauss, proteinAT3G20570.1
bic:704
peaks after deleting less than one fifth of global max:13
×10^6
raw data, starting:12
fitted Gauss locations:13
y
x
0
1
2
3
4
5
6
0
5
10
15
20
25

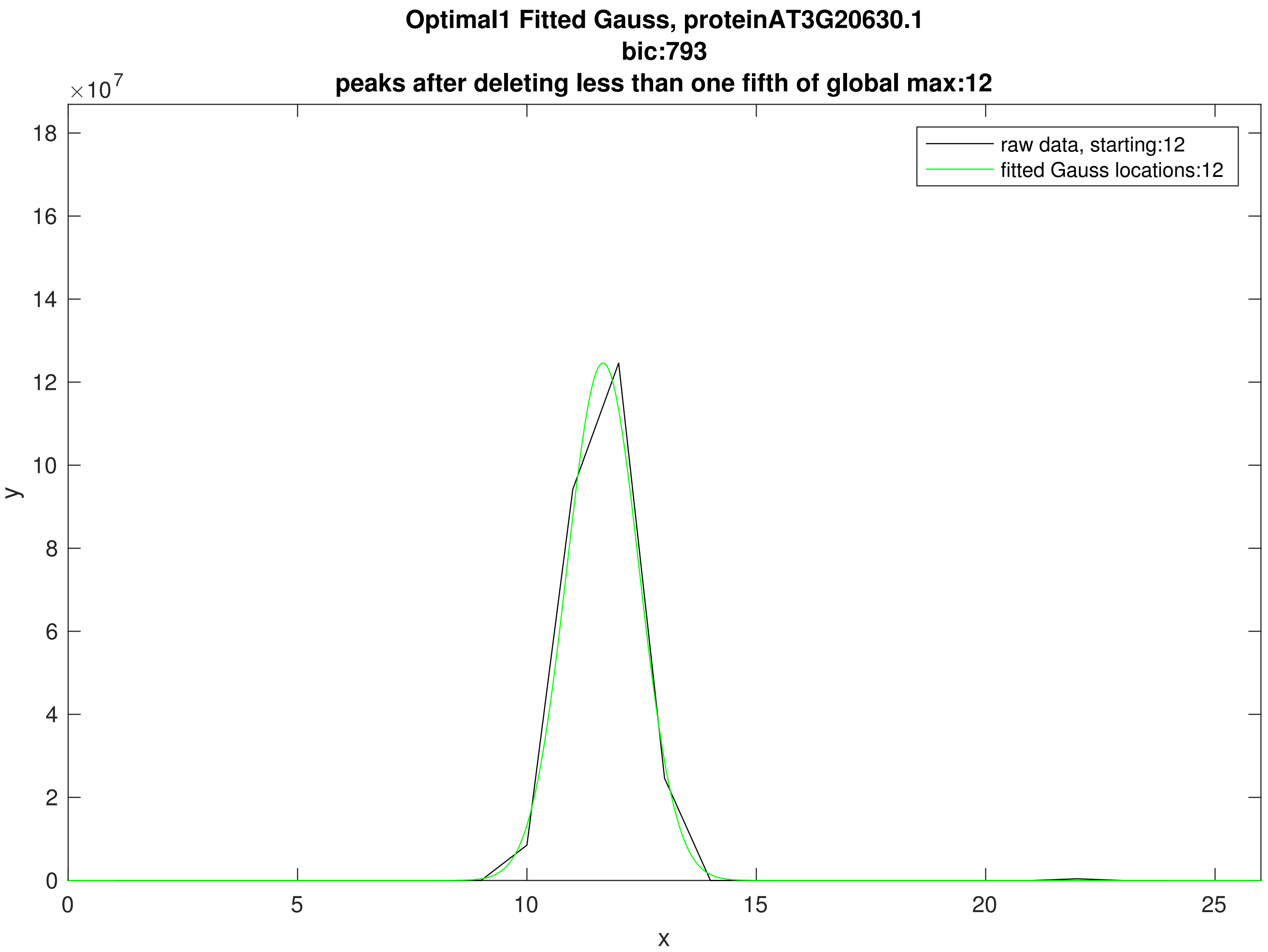
Optimal1 Fitted Gauss, proteinAT3G20630.1
bic:793
peaks after deleting less than one fifth of global max:12
×10^7
18
16
14
12
10
8
6
4
2
0
y
0
5
10
15
20
25
x
raw data, starting:12
fitted Gauss locations:12

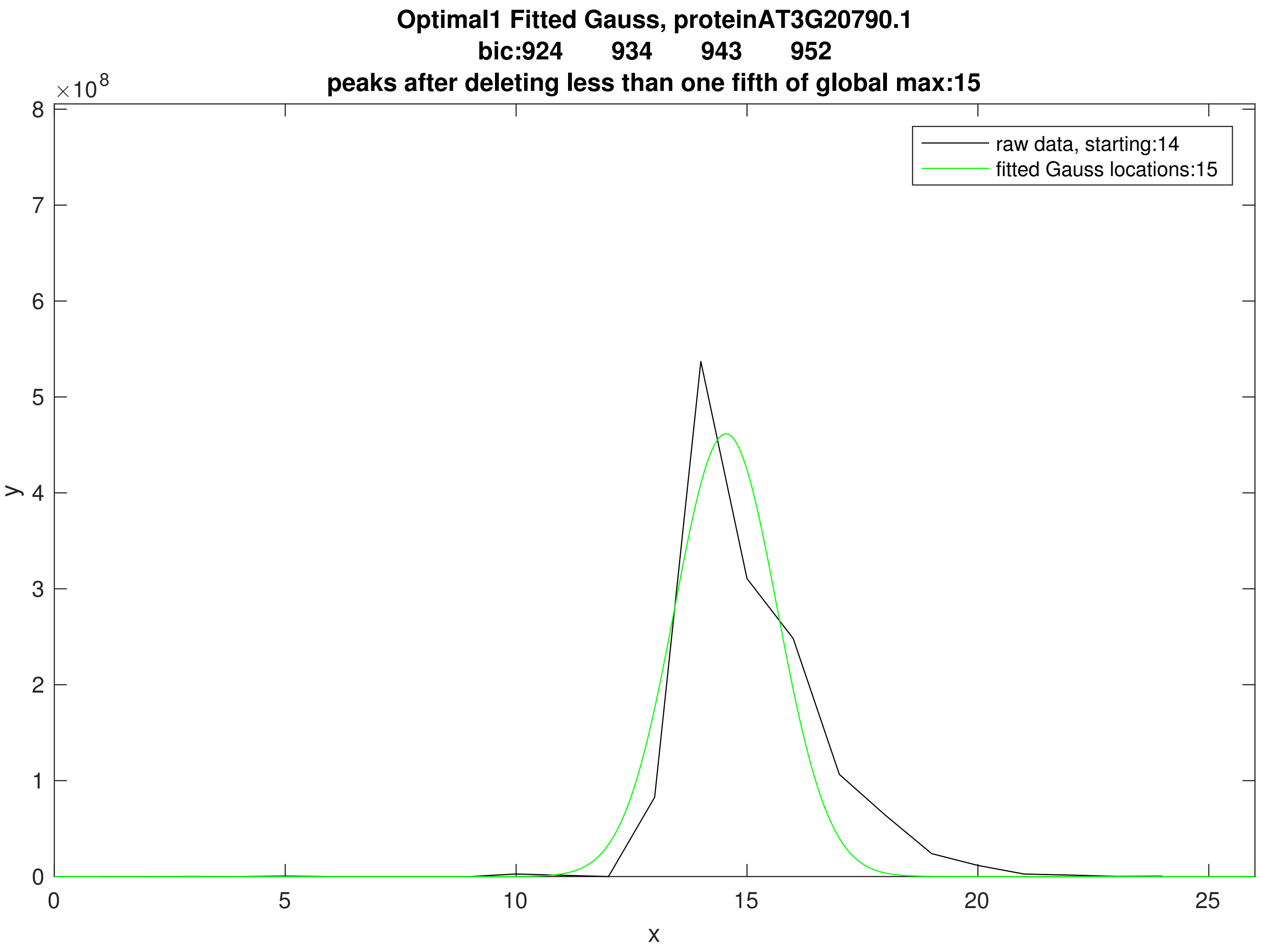

Optimal1 Fitted Gauss, proteinAT3G20790.1
bic:924 934 943 952
peaks after deleting less than one fifth of global max:15
×10^8
raw data, starting:14
fitted Gauss locations:15
y
x

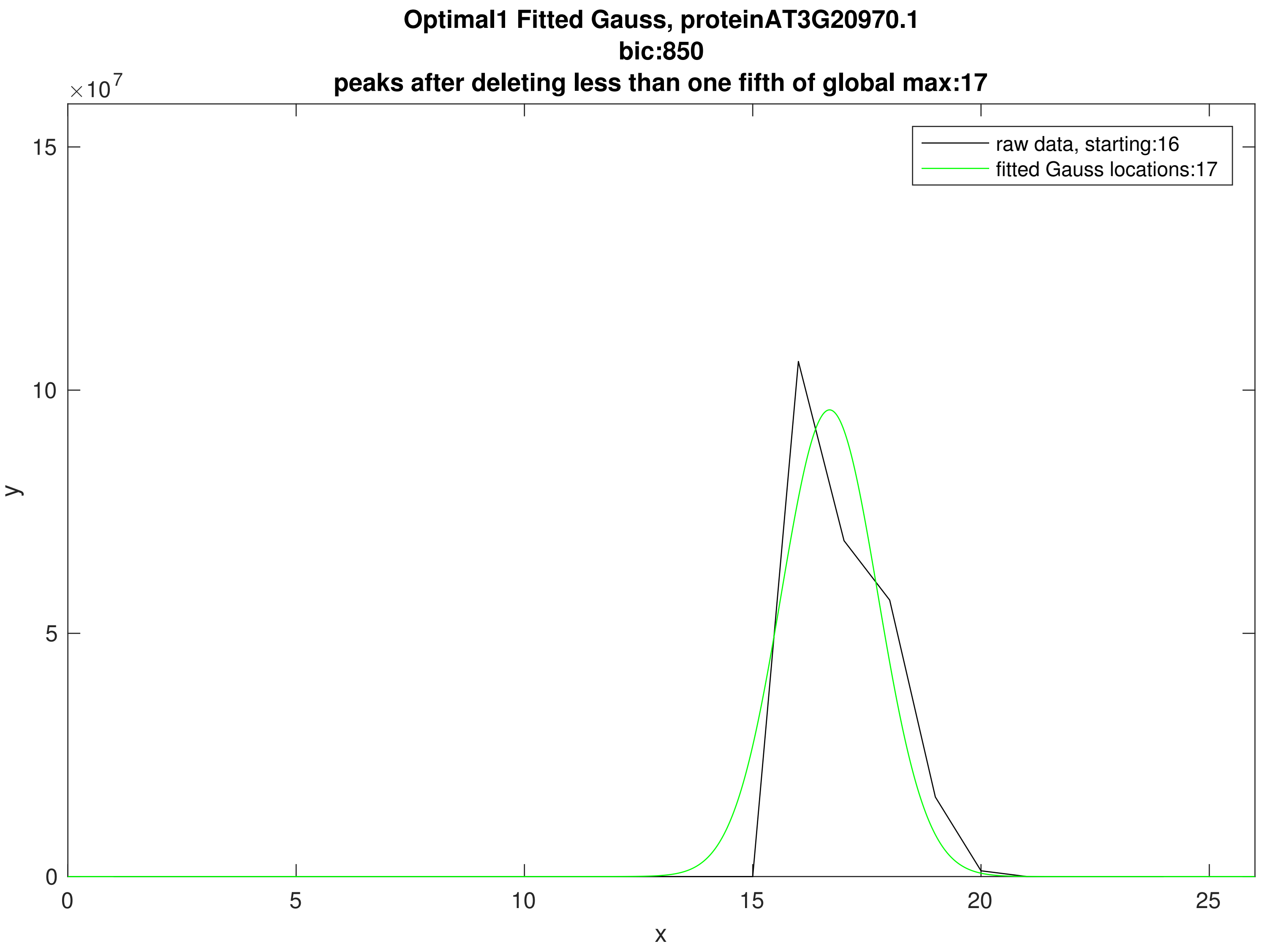

Optimal1 Fitted Gauss, proteinAT3G20970.1
bic:850
peaks after deleting less than one fifth of global max:17
×10^7
15
10
5
0
y
0
5
10
15
20
25
x
raw data, starting:16
fitted Gauss locations:17

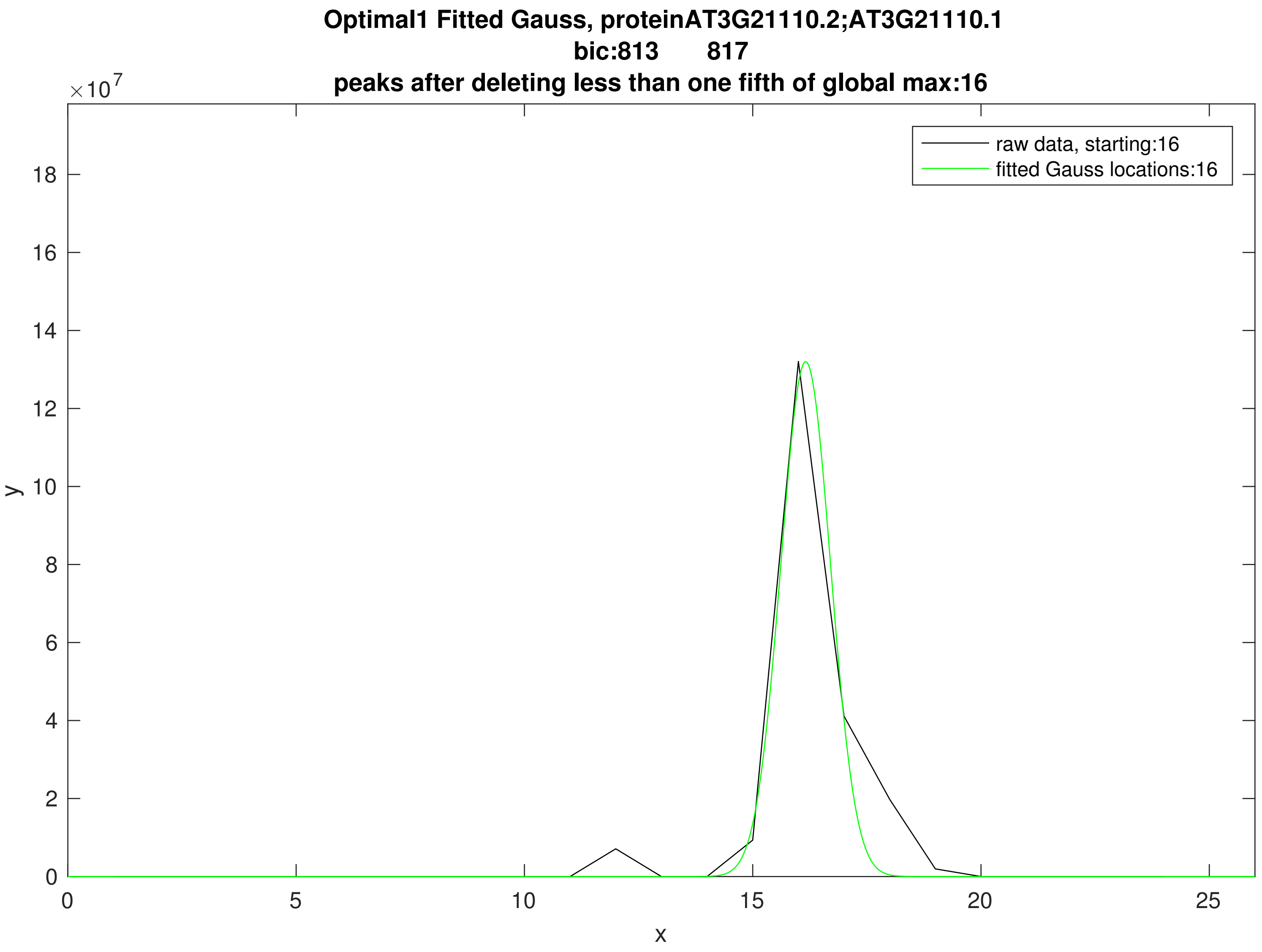

Optimal1 Fitted Gauss, proteinAT3G21110.2;AT3G21110.1
bic:813 817
peaks after deleting less than one fifth of global max:16
×10^7
18
16
14
12
10
8
6
4
2
0
y
0
5
10
15
20
25
x
raw data, starting:16
fitted Gauss locations:16

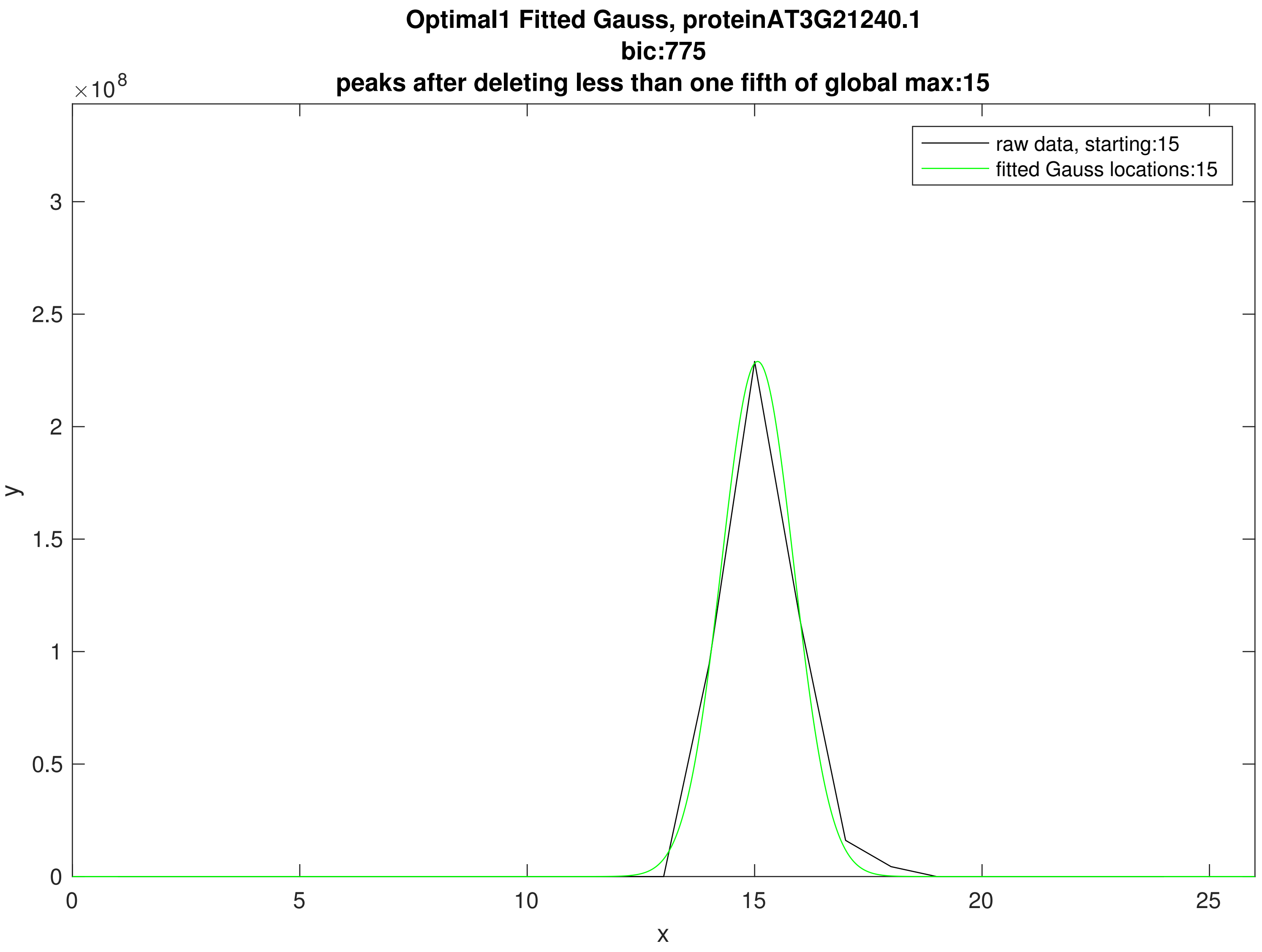
Optimal1 Fitted Gauss, proteinAT3G21240.1
bic:775
peaks after deleting less than one fifth of global max:15
×10^8
raw data, starting:15
fitted Gauss locations:15
y
x
0
0.5
1
1.5
2
2.5
3
0
5
10
15
20
25

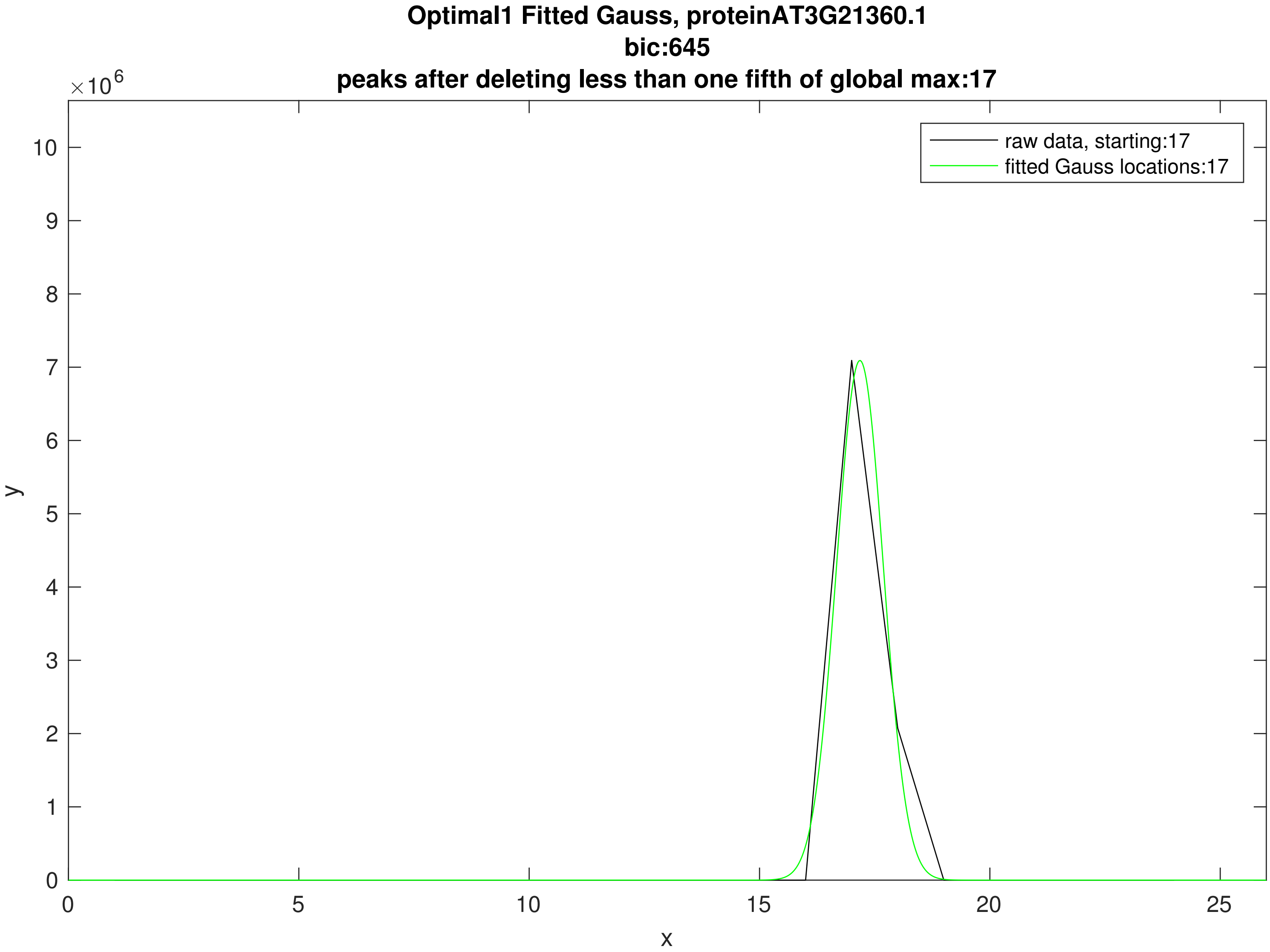
Optimal1 Fitted Gauss, proteinAT3G21360.1
bic:645
peaks after deleting less than one fifth of global max:17
×10^6
raw data, starting:17
fitted Gauss locations:17
y
x
0
1
2
3
4
5
6
7
8
9
10
0
5
10
15
20
25

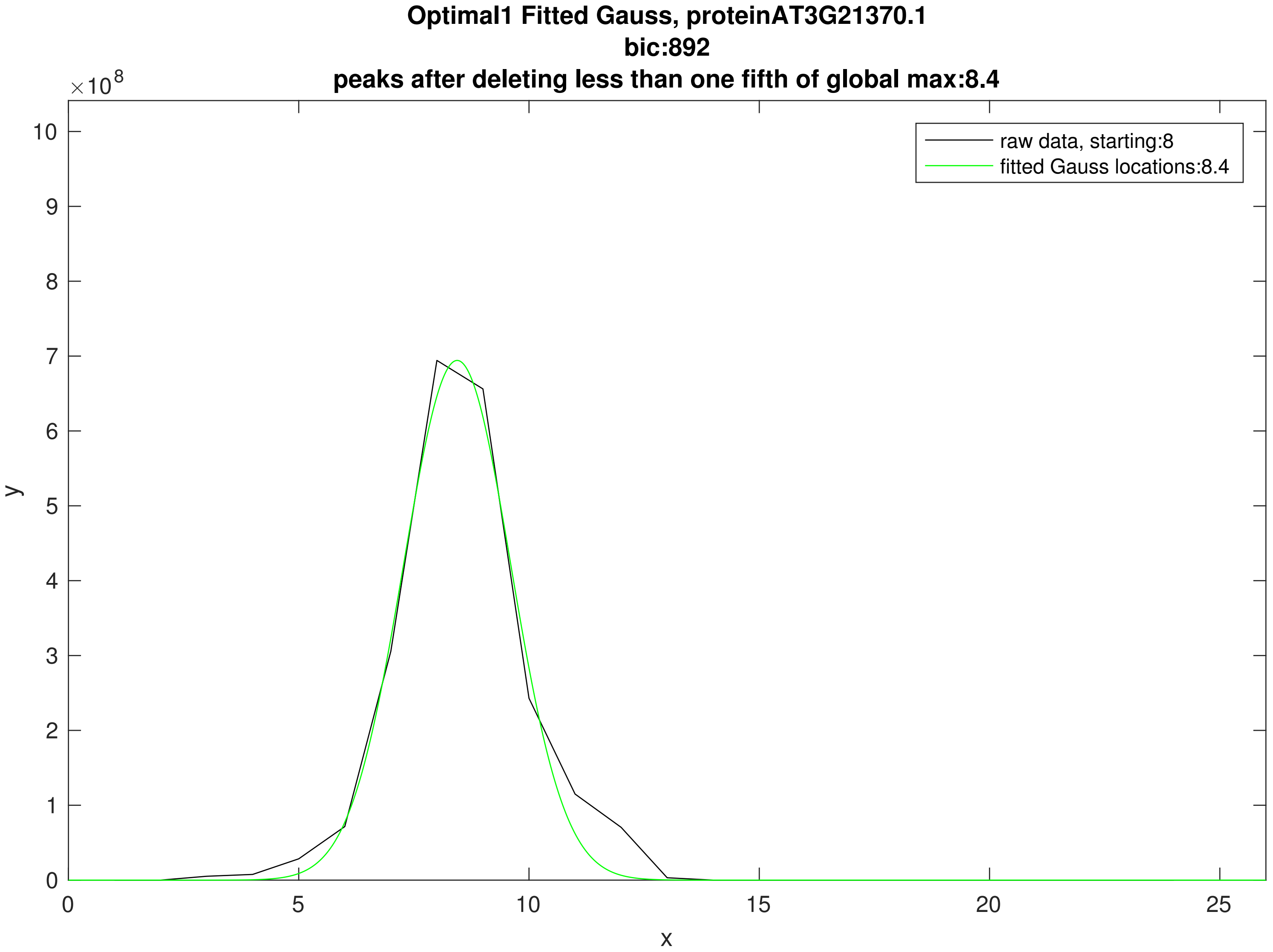
Optimal1 Fitted Gauss, proteinAT3G21370.1
bic:892
peaks after deleting less than one fifth of global max:8.4
×10^8
raw data, starting:8
fitted Gauss locations:8.4
y
x
0
1
2
3
4
5
6
7
8
9
10
0
5
10
15
20
25

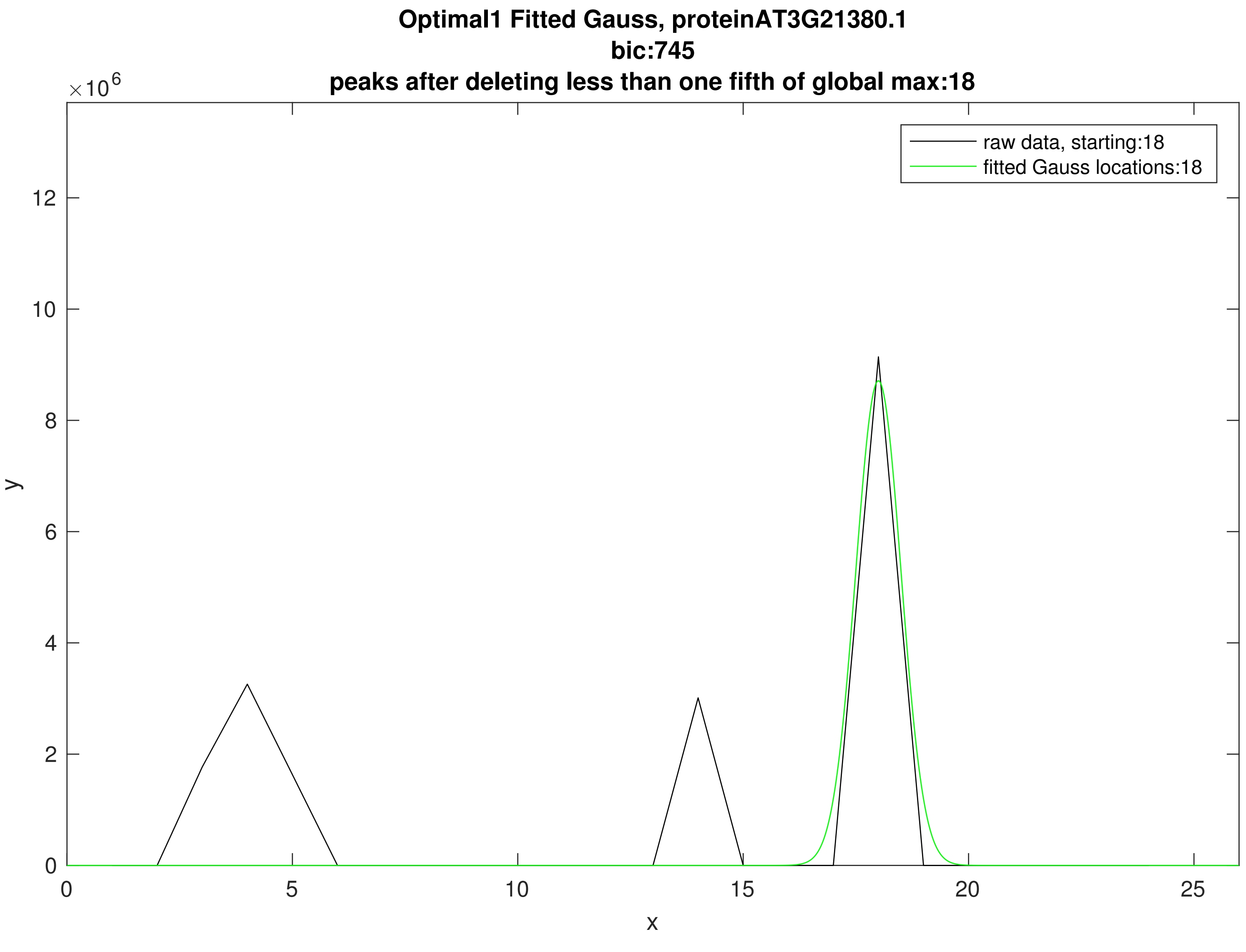
Optimal1 Fitted Gauss, proteinAT3G21380.1
bic:745
peaks after deleting less than one fifth of global max:18
×10^6
raw data, starting:18
fitted Gauss locations:18
y
x
0
2
4
6
8
10
12
0
5
10
15
20
25

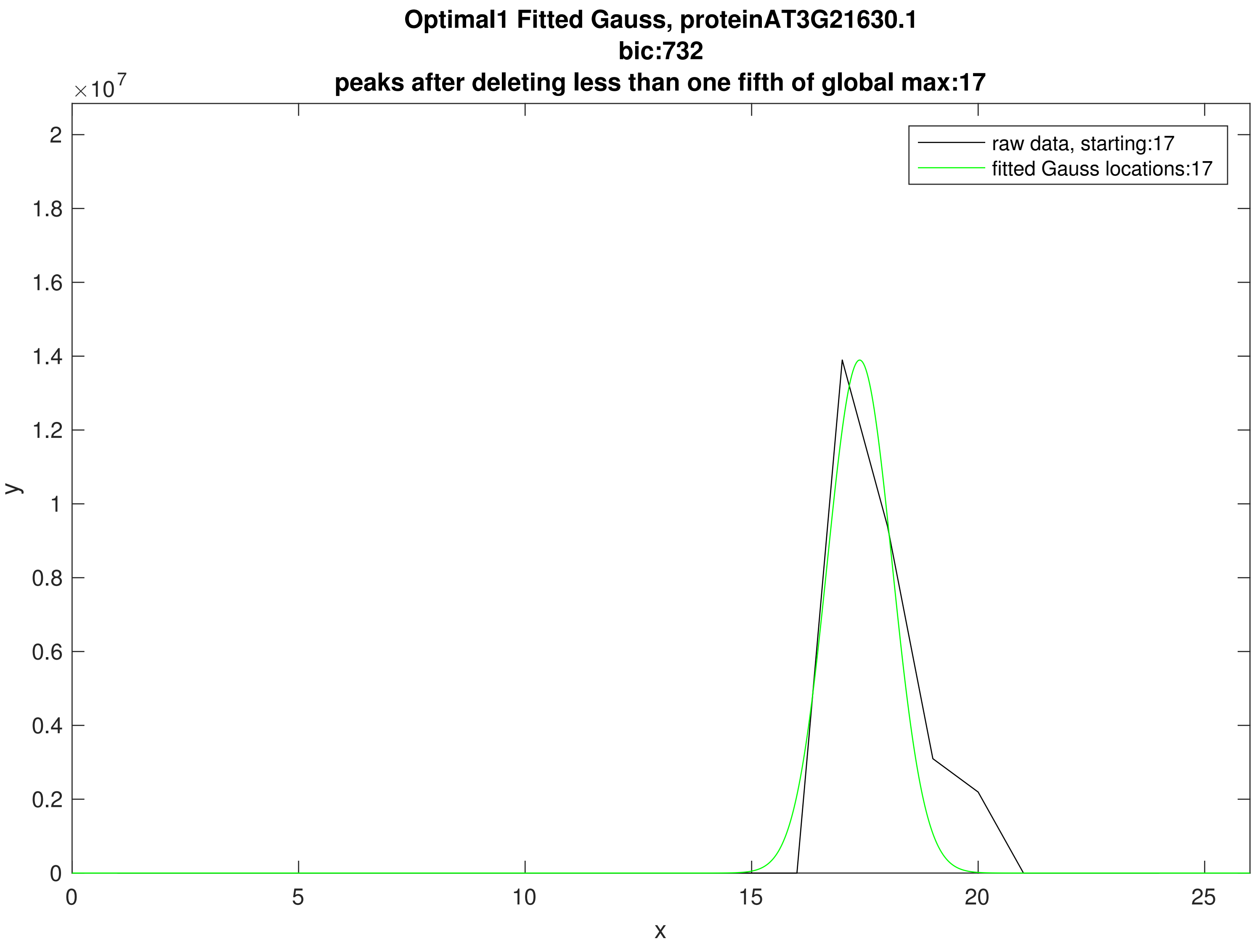
Optimal1 Fitted Gauss, proteinAT3G21630.1
bic:732
peaks after deleting less than one fifth of global max:17
×10^7
raw data, starting:17
fitted Gauss locations:17
y
x
0
0.2
0.4
0.6
0.8
1
1.2
1.4
1.6
1.8
2
0
5
10
15
20
25

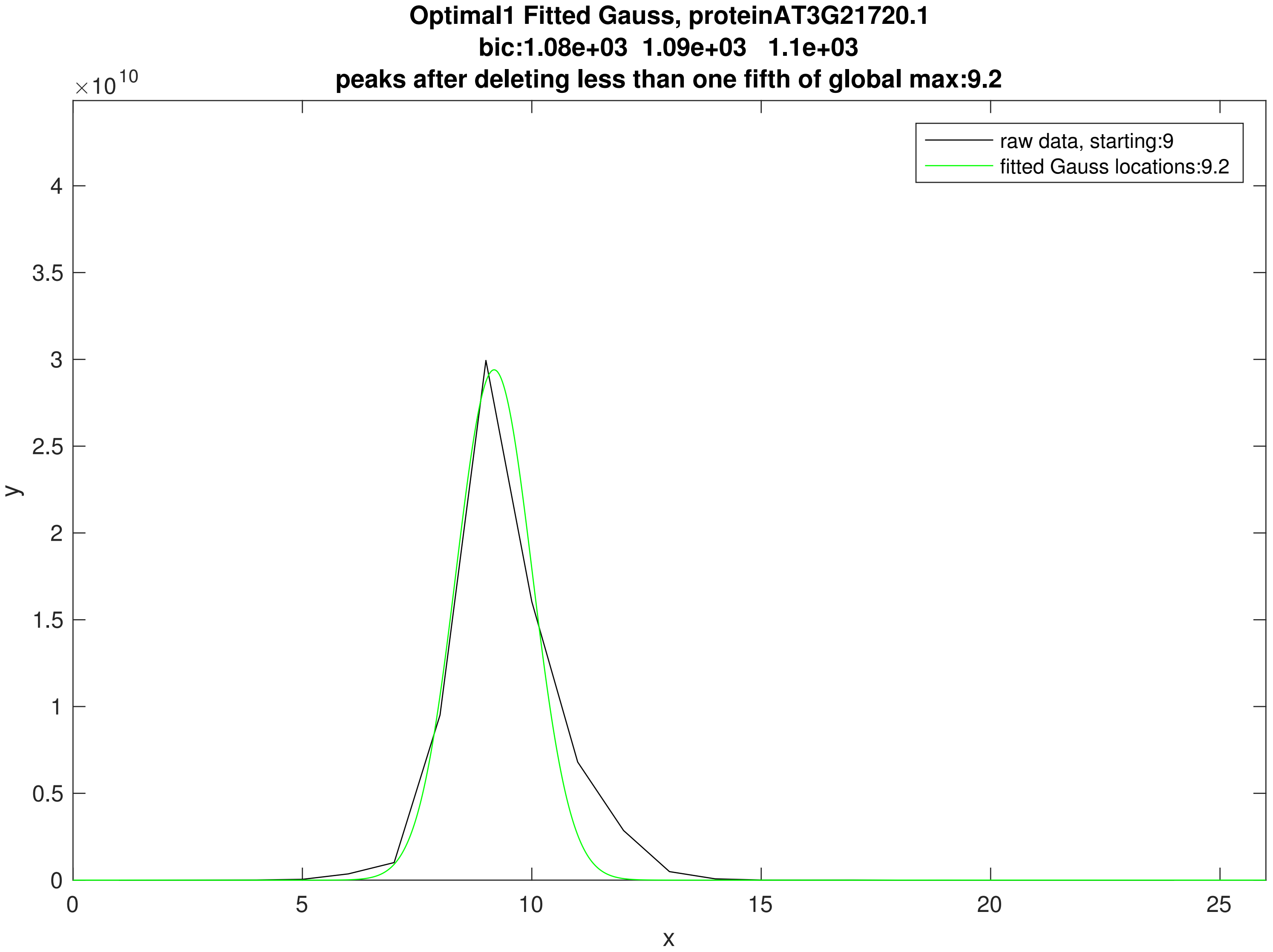
Optimal1 Fitted Gauss, proteinAT3G21720.1
bic:1.08e+03 1.09e+03 1.1e+03
peaks after deleting less than one fifth of global max:9.2
×10^10
raw data, starting:9
fitted Gauss locations:9.2
4
3.5
3
2.5
2
1.5
1
0.5
0
y
0
5
10
15
20
25
x

**Optimal1 Fitted Gauss, proteinAT3G22110.1**
**bic:914 923 932 942**
**peaks after deleting less than one fifth of global max:5.1**

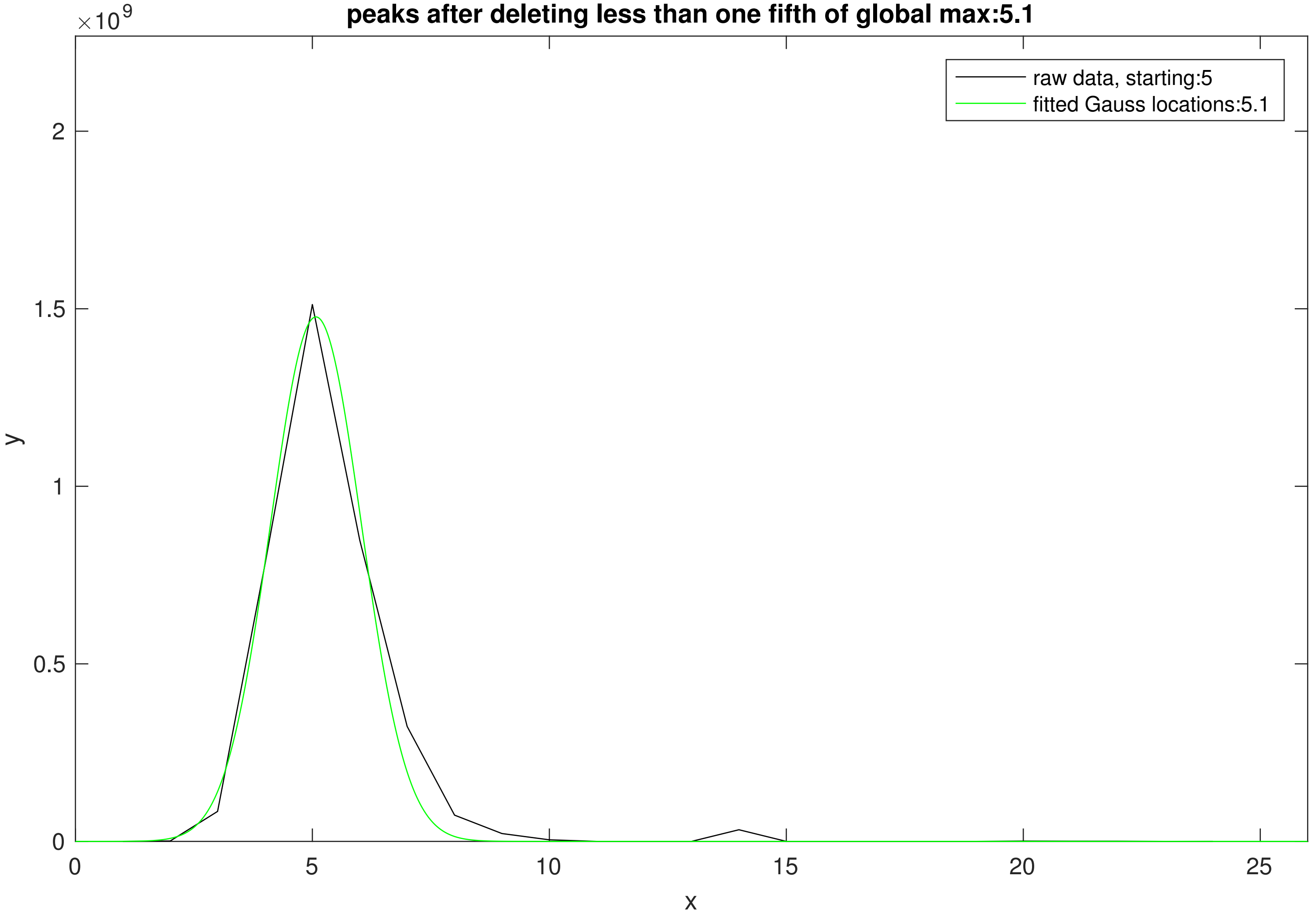

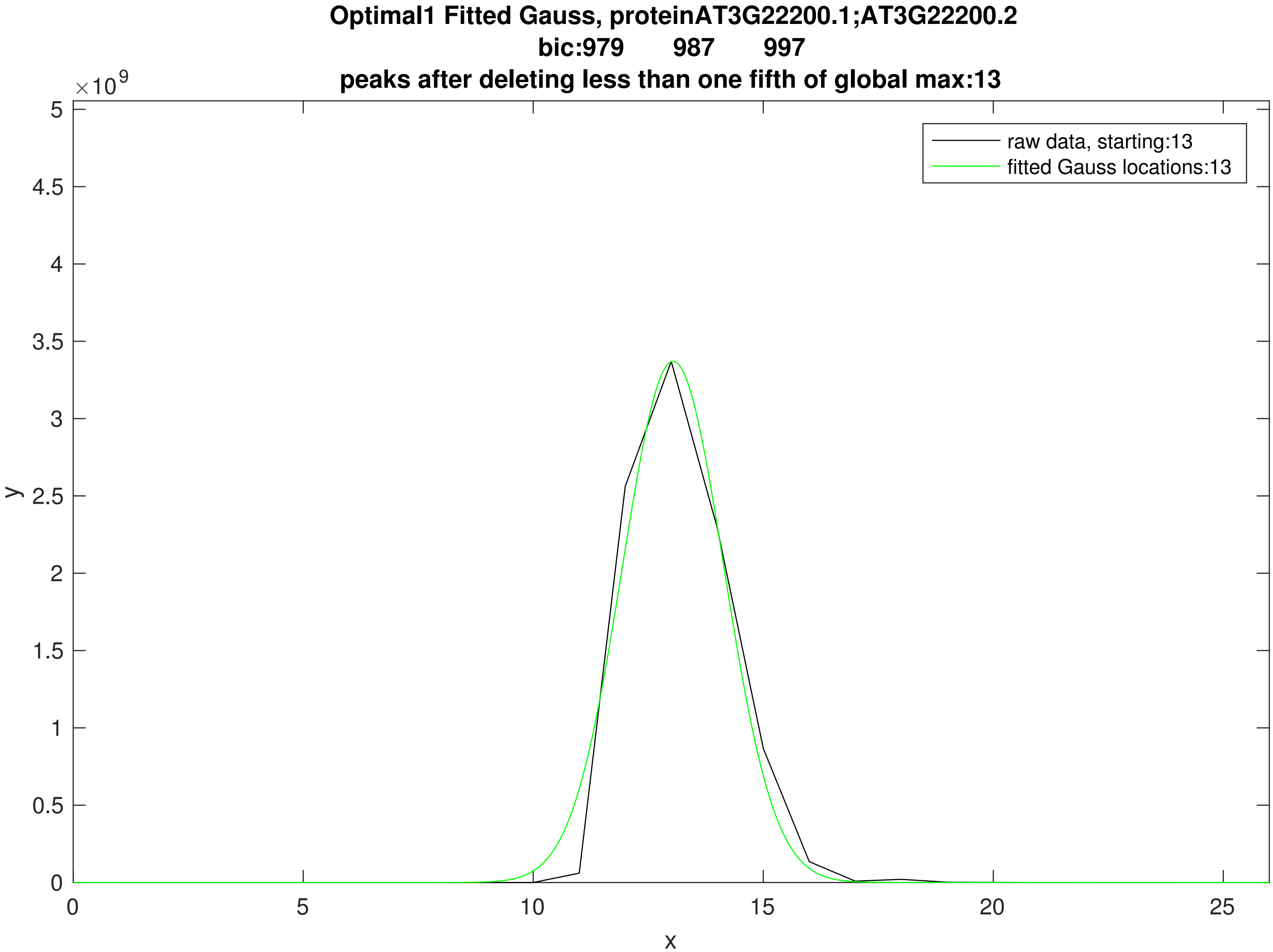
Optimal1 Fitted Gauss, proteinAT3G22200.1;AT3G22200.2
bic:979 987 997
peaks after deleting less than one fifth of global max:13
×10^9
5
4.5
4
3.5
3
2.5
2
1.5
1
0.5
0
y
0
5
10
15
20
25
x
raw data, starting:13
fitted Gauss locations:13

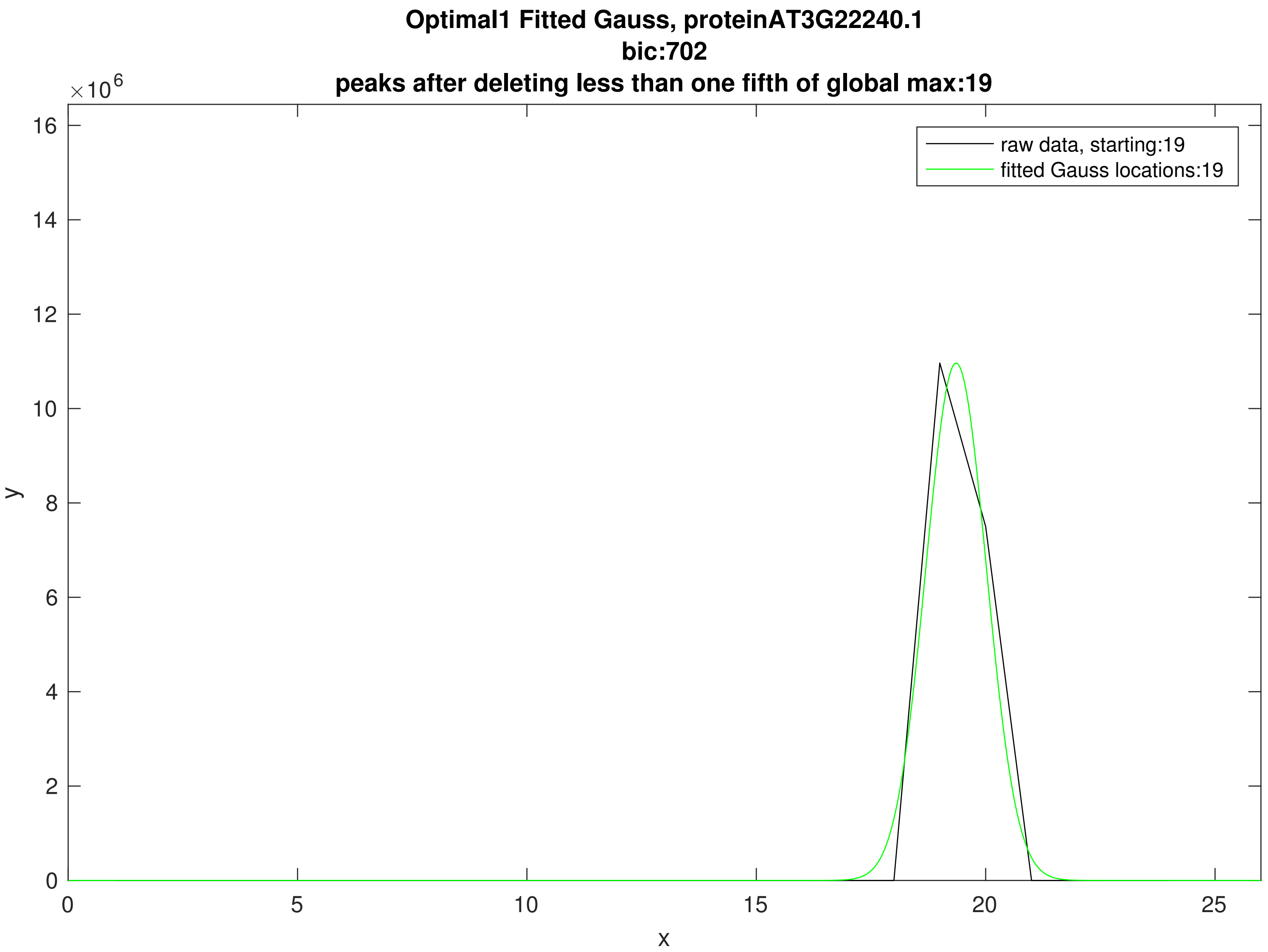
Optimal1 Fitted Gauss, proteinAT3G22240.1
bic:702
peaks after deleting less than one fifth of global max:19
×10^6
raw data, starting:19
fitted Gauss locations:19
y
x
0
2
4
6
8
10
12
14
16
0
5
10
15
20
25

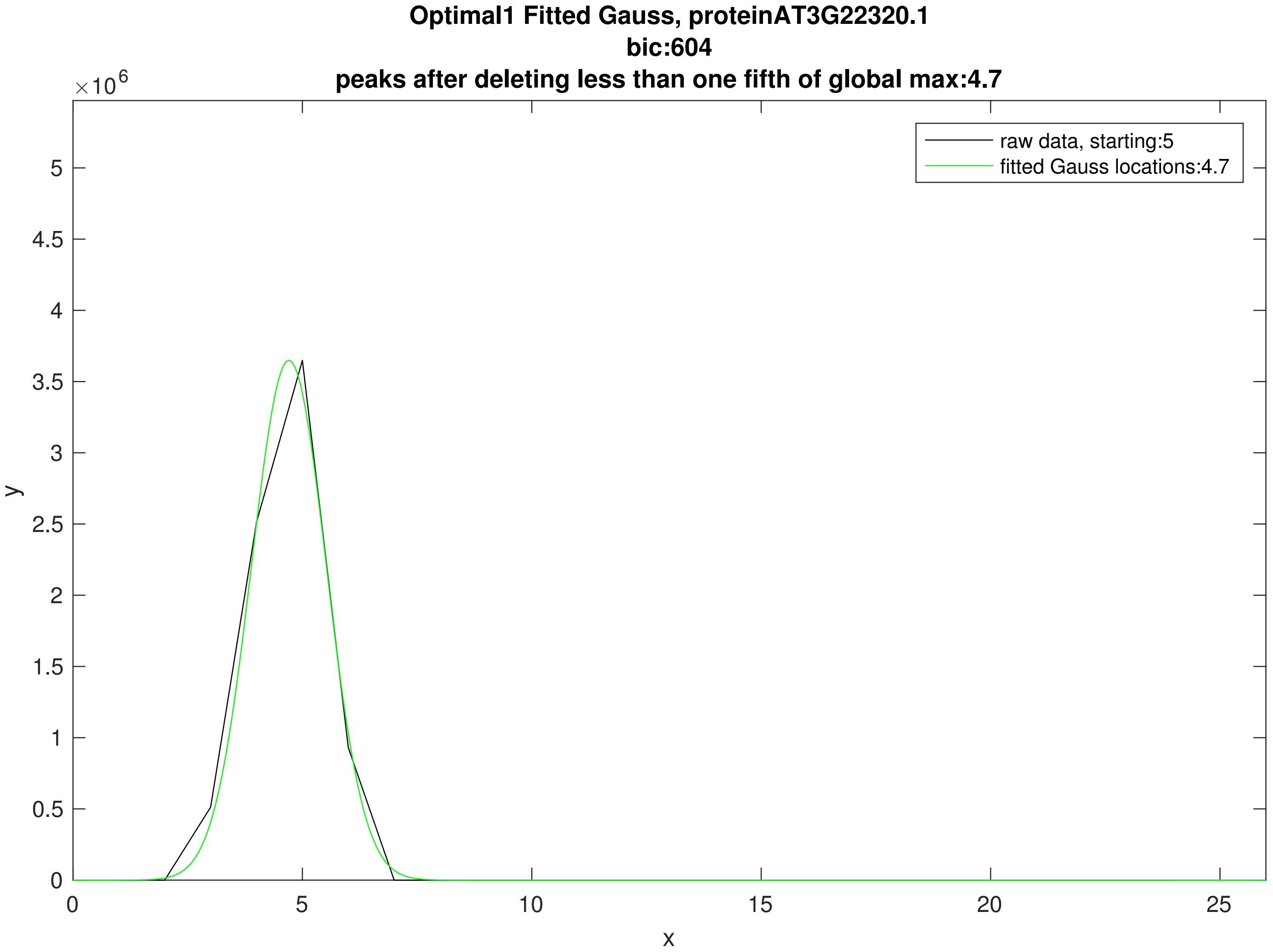
Optimal1 Fitted Gauss, proteinAT3G22320.1
bic:604
peaks after deleting less than one fifth of global max:4.7
×10^6
raw data, starting:5
fitted Gauss locations:4.7
y
x

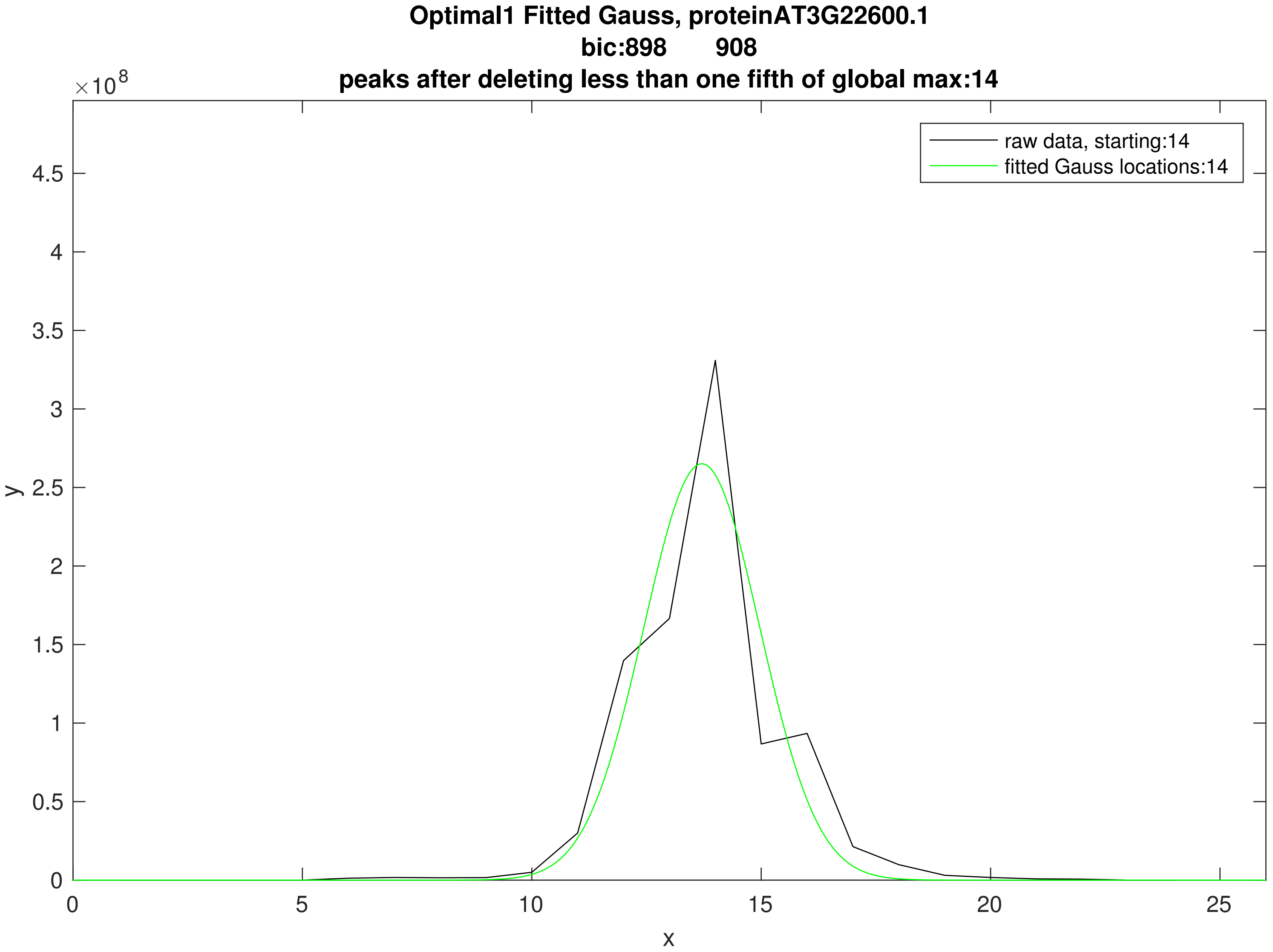
Optimal1 Fitted Gauss, proteinAT3G22600.1
bic:898 908
peaks after deleting less than one fifth of global max:14
×10^8
raw data, starting:14
fitted Gauss locations:14
y
x
0
0.5
1
1.5
2
2.5
3
3.5
4
4.5
0
5
10
15
20
25

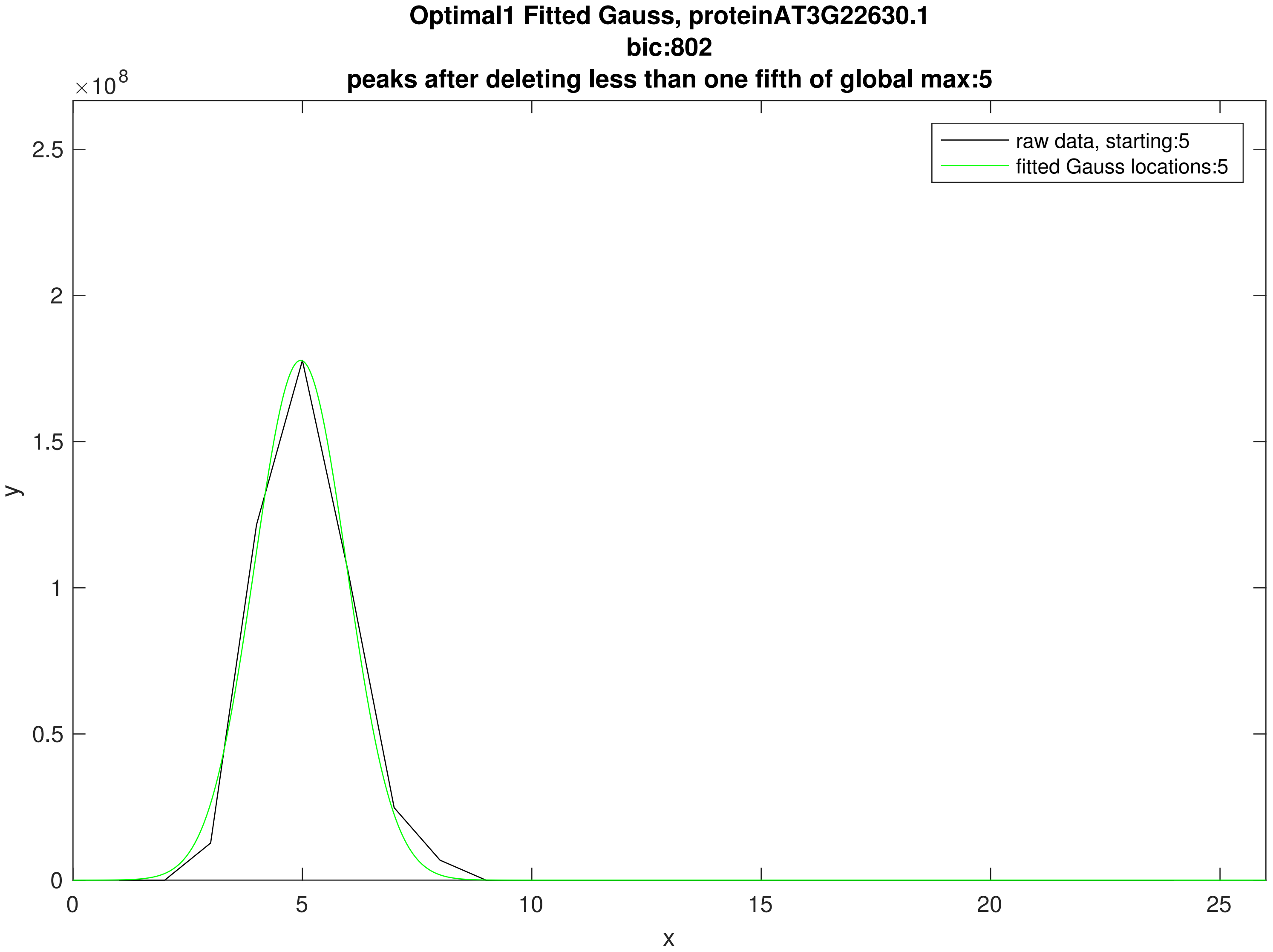
Optimal1 Fitted Gauss, proteinAT3G22630.1
bic:802
peaks after deleting less than one fifth of global max:5
×10^8
raw data, starting:5
fitted Gauss locations:5
y
x
0
0.5
1
1.5
2
2.5
5
10
15
20
25

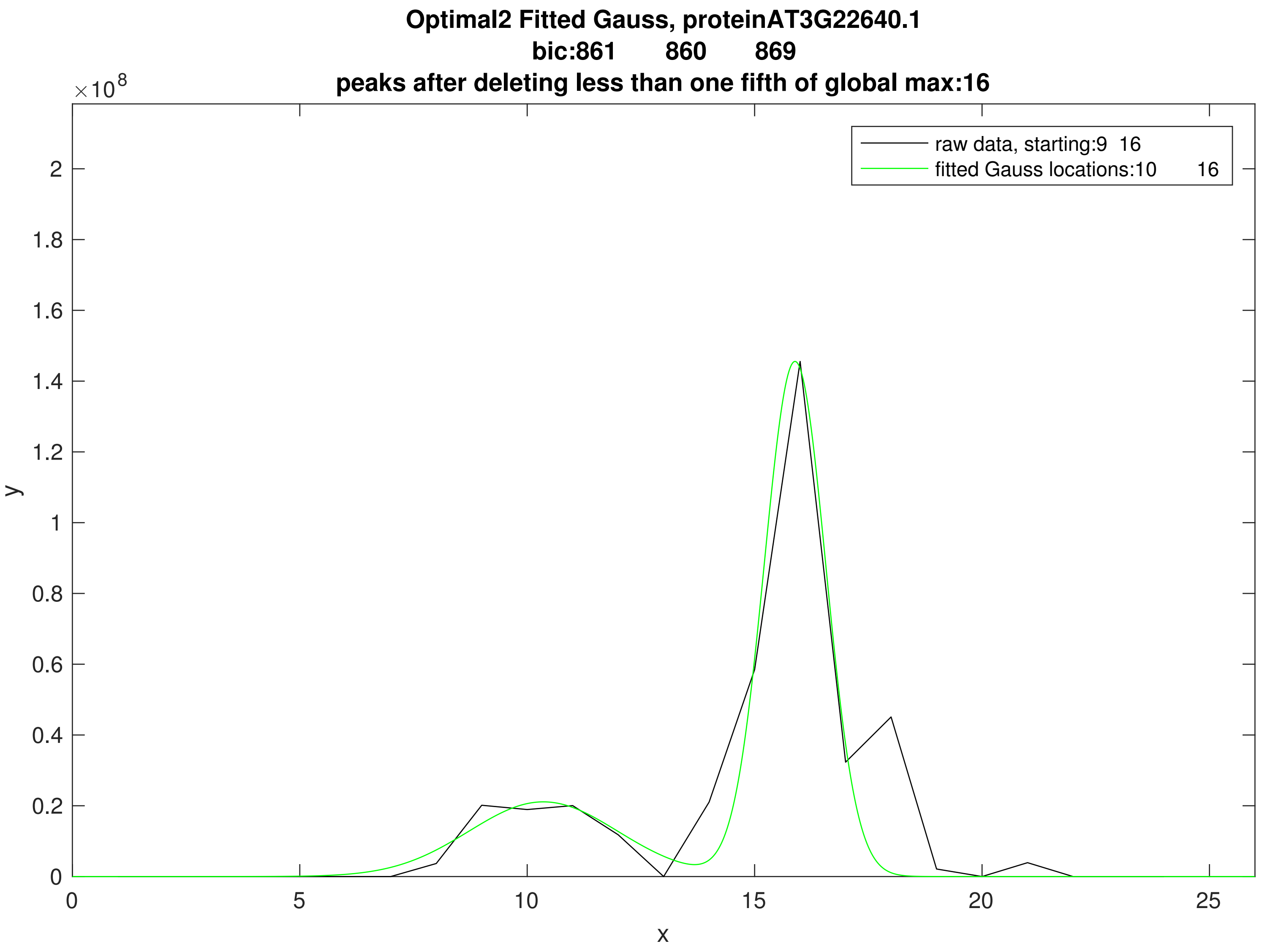
Optimal2 Fitted Gauss, proteinAT3G22640.1
bic:861 860 869
peaks after deleting less than one fifth of global max:16
×10^8
raw data, starting:9 16
fitted Gauss locations:10 16
y
x
0
0.2
0.4
0.6
0.8
1
1.2
1.4
1.6
1.8
2
0
5
10
15
20
25

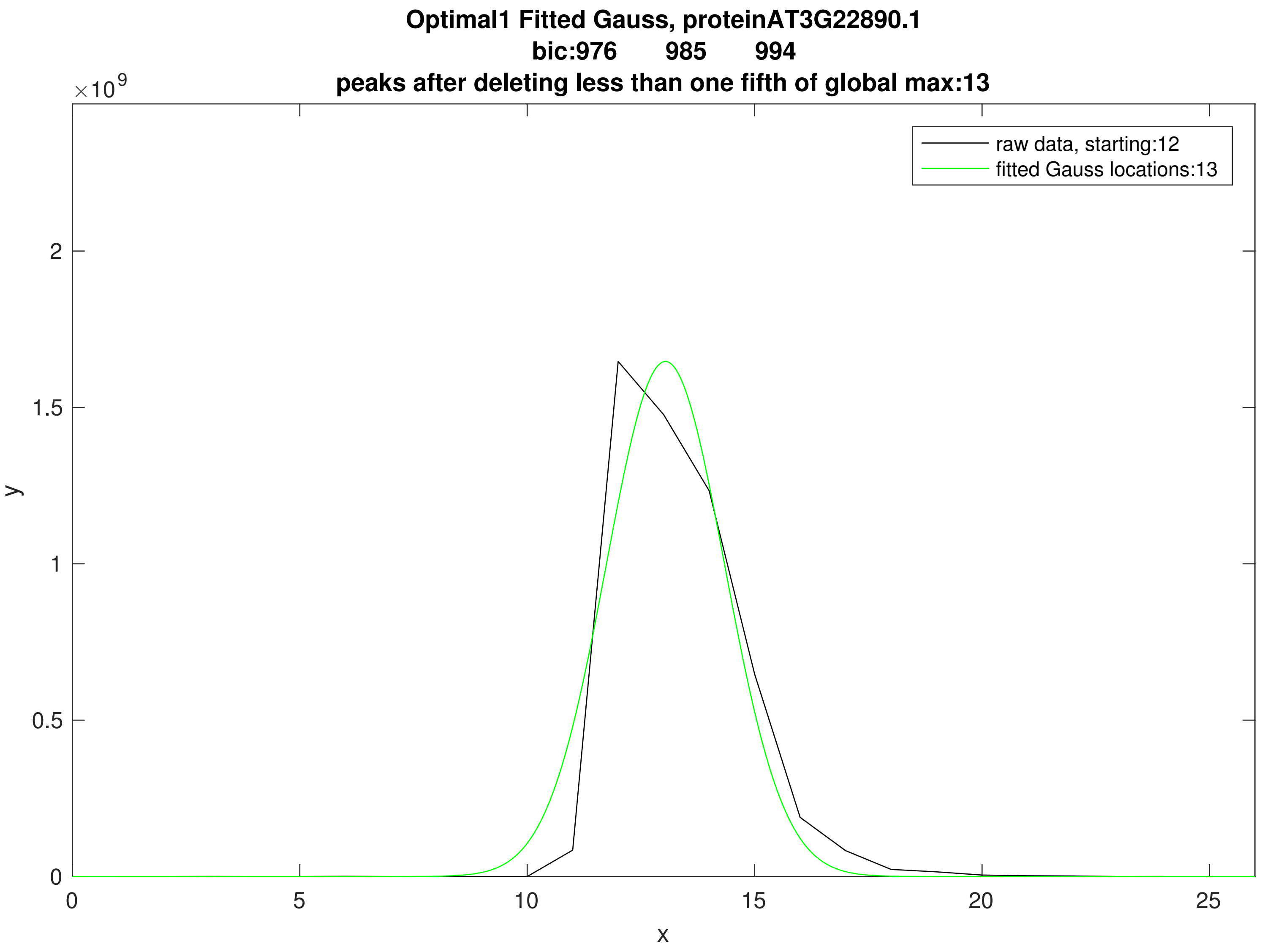
Optimal1 Fitted Gauss, proteinAT3G22890.1
bic:976 985 994
peaks after deleting less than one fifth of global max:13
×10^9
raw data, starting:12
fitted Gauss locations:13
y
x
0
0.5
1
1.5
2
0
5
10
15
20
25

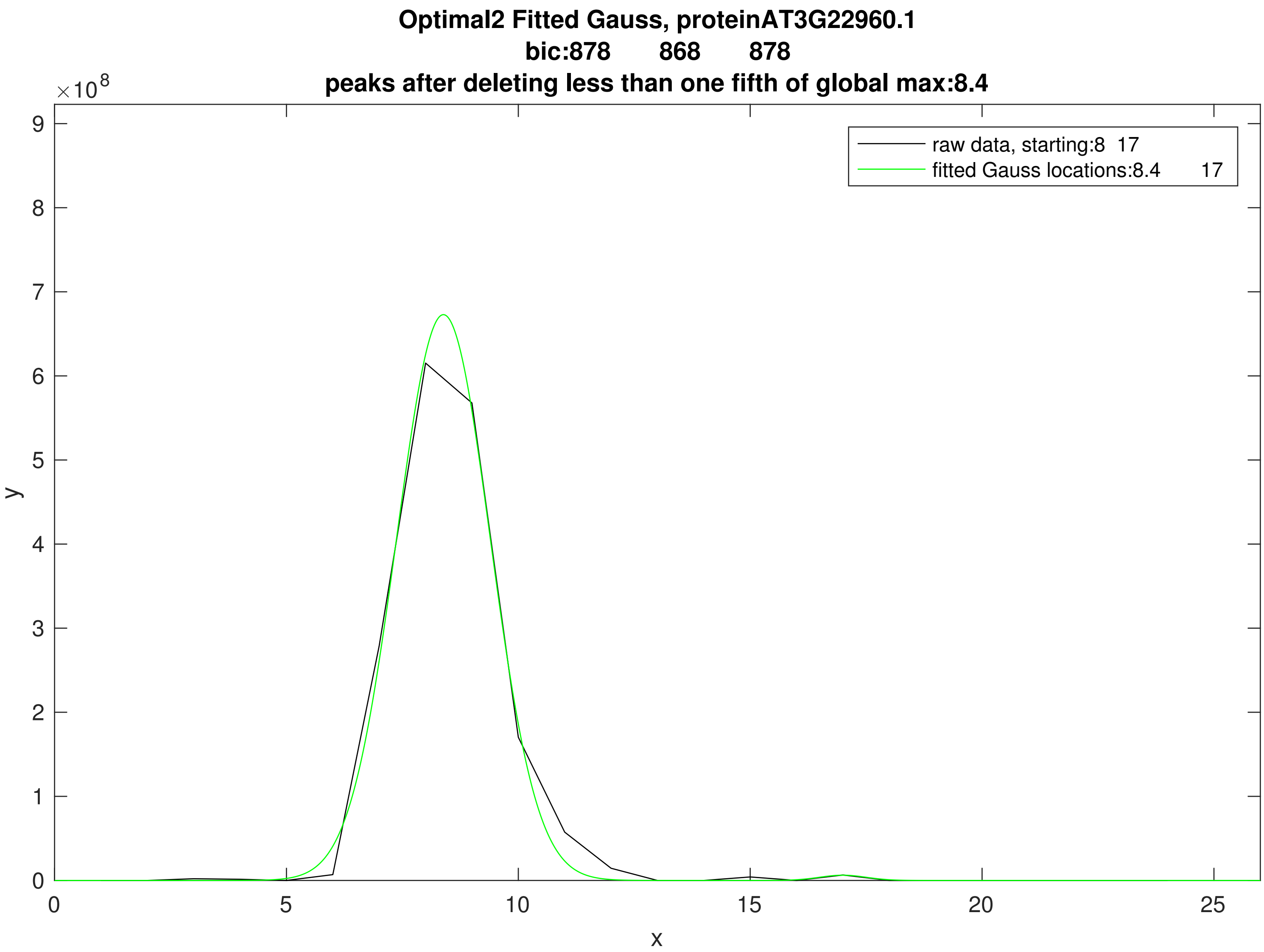
Optimal2 Fitted Gauss, proteinAT3G22960.1
bic:878 868 878
peaks after deleting less than one fifth of global max:8.4
×10^8
raw data, starting:8 17
fitted Gauss locations:8.4 17
y
x
0
1
2
3
4
5
6
7
8
9
0
5
10
15
20
25

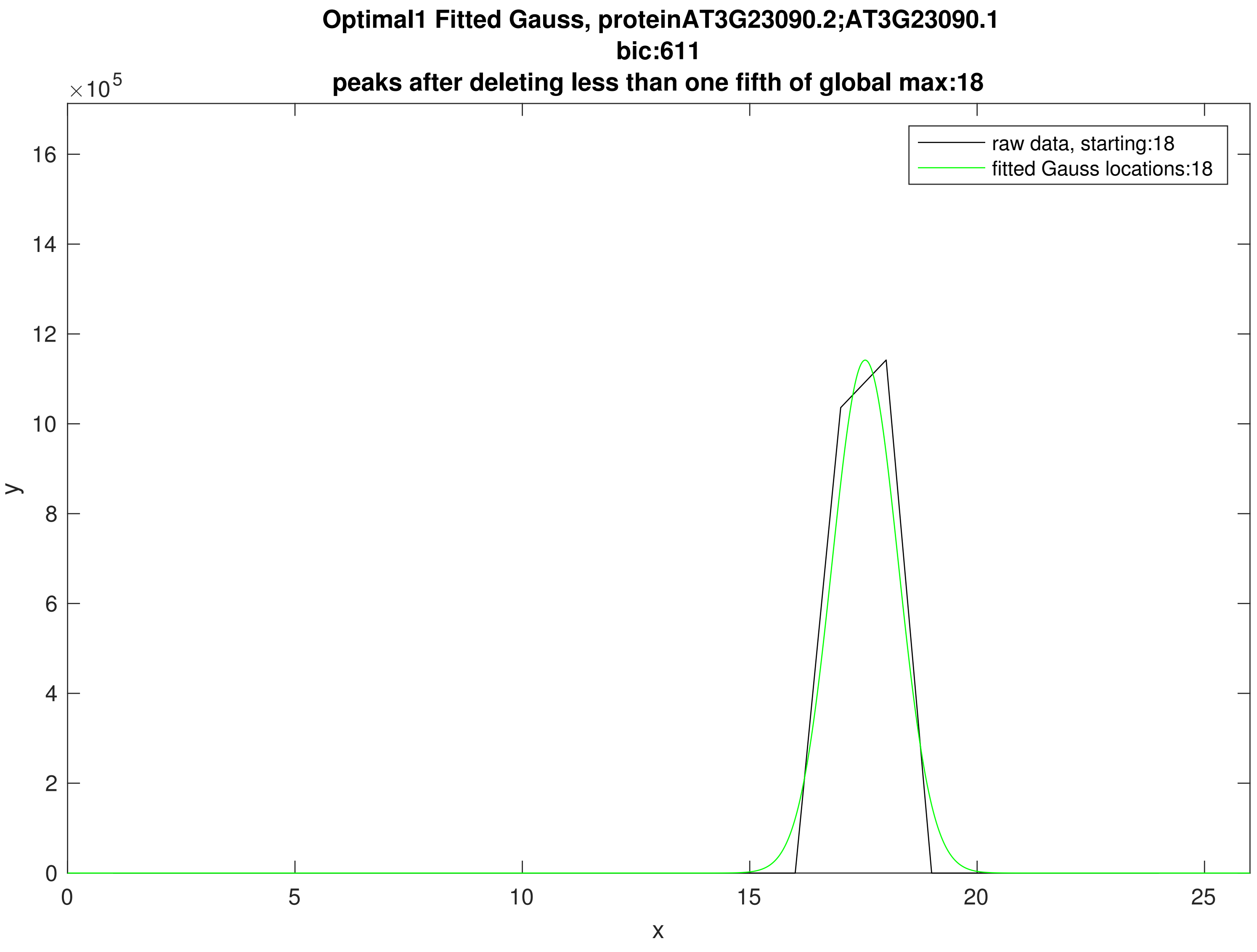

Optimal1 Fitted Gauss, proteinAT3G23090.2;AT3G23090.1
bic:611
peaks after deleting less than one fifth of global max:18
×10^5
raw data, starting:18
fitted Gauss locations:18
y
x
0
2
4
6
8
10
12
14
16
0
5
10
15
20
25

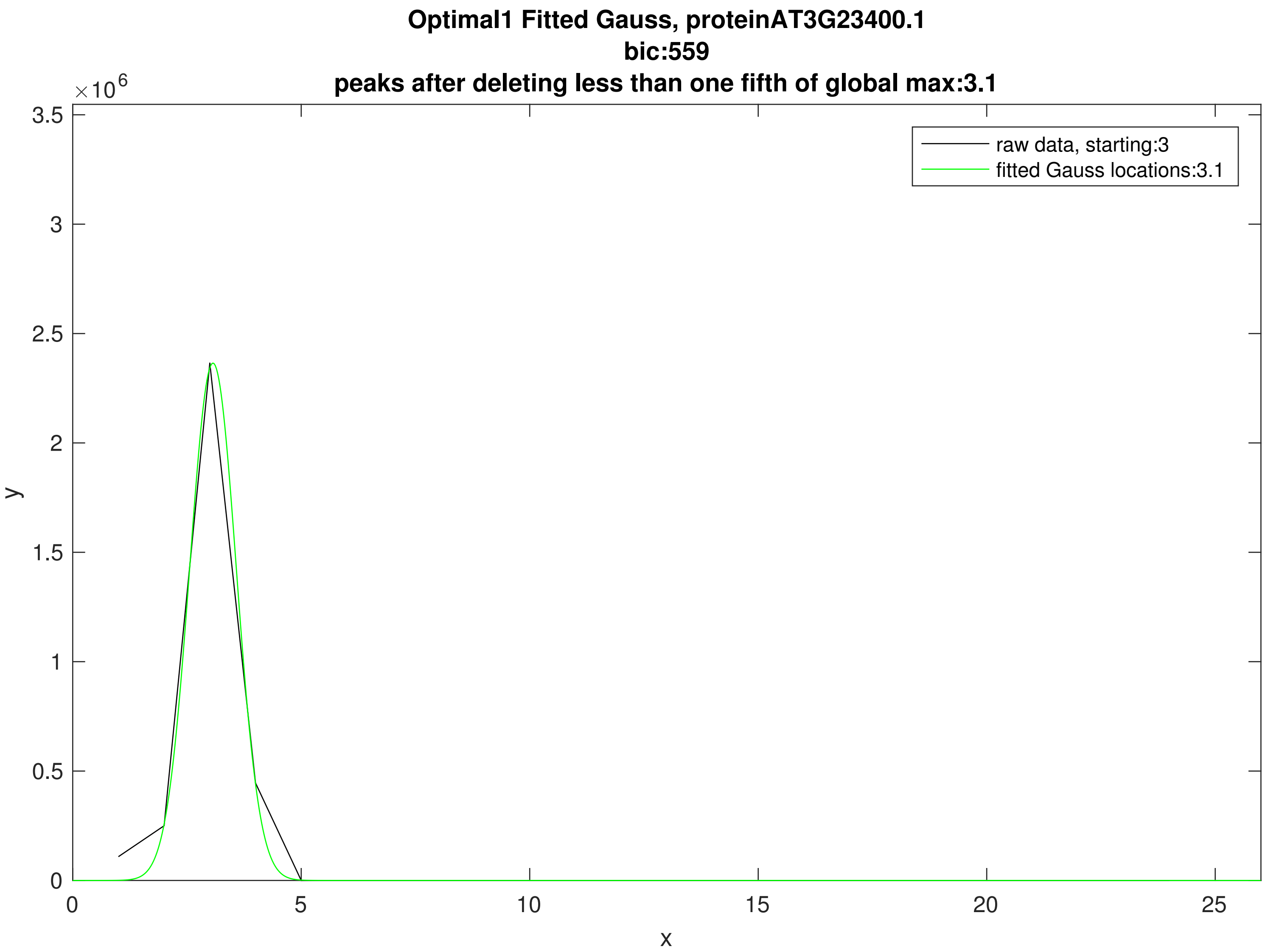
Optimal1 Fitted Gauss, proteinAT3G23400.1
bic:559
peaks after deleting less than one fifth of global max:3.1
×10^6
3.5
3
2.5
2
1.5
1
0.5
0
y
0
5
10
15
20
25
x
raw data, starting:3
fitted Gauss locations:3.1

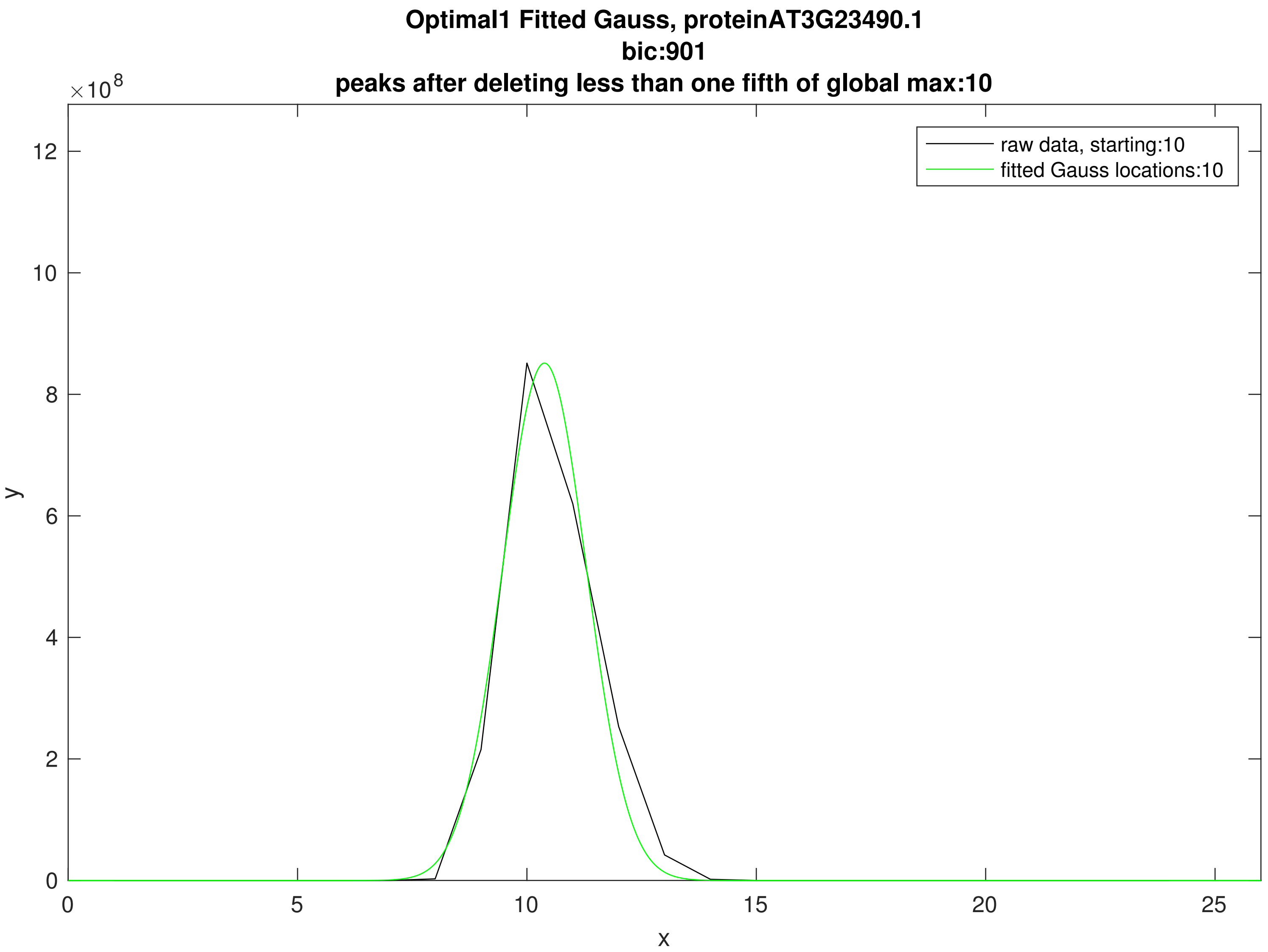

Optimal1 Fitted Gauss, proteinAT3G23490.1
bic:901
peaks after deleting less than one fifth of global max:10
×10^8
12
10
8
6
4
2
0
y
0
5
10
15
20
25
x
raw data, starting:10
fitted Gauss locations:10

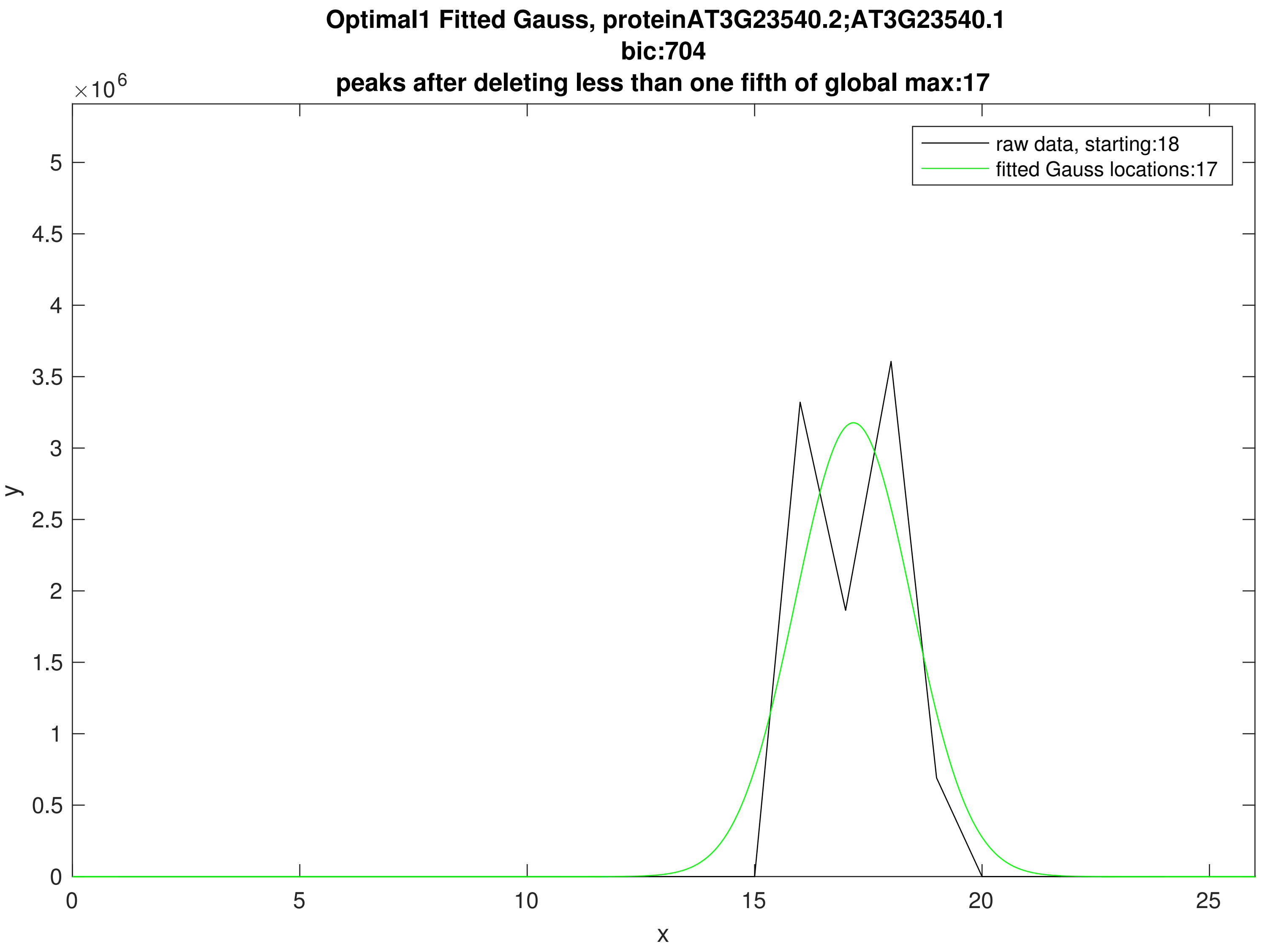

Optimal1 Fitted Gauss, proteinAT3G23540.2;AT3G23540.1
bic:704
peaks after deleting less than one fifth of global max:17
×10^6
raw data, starting:18
fitted Gauss locations:17
y
x

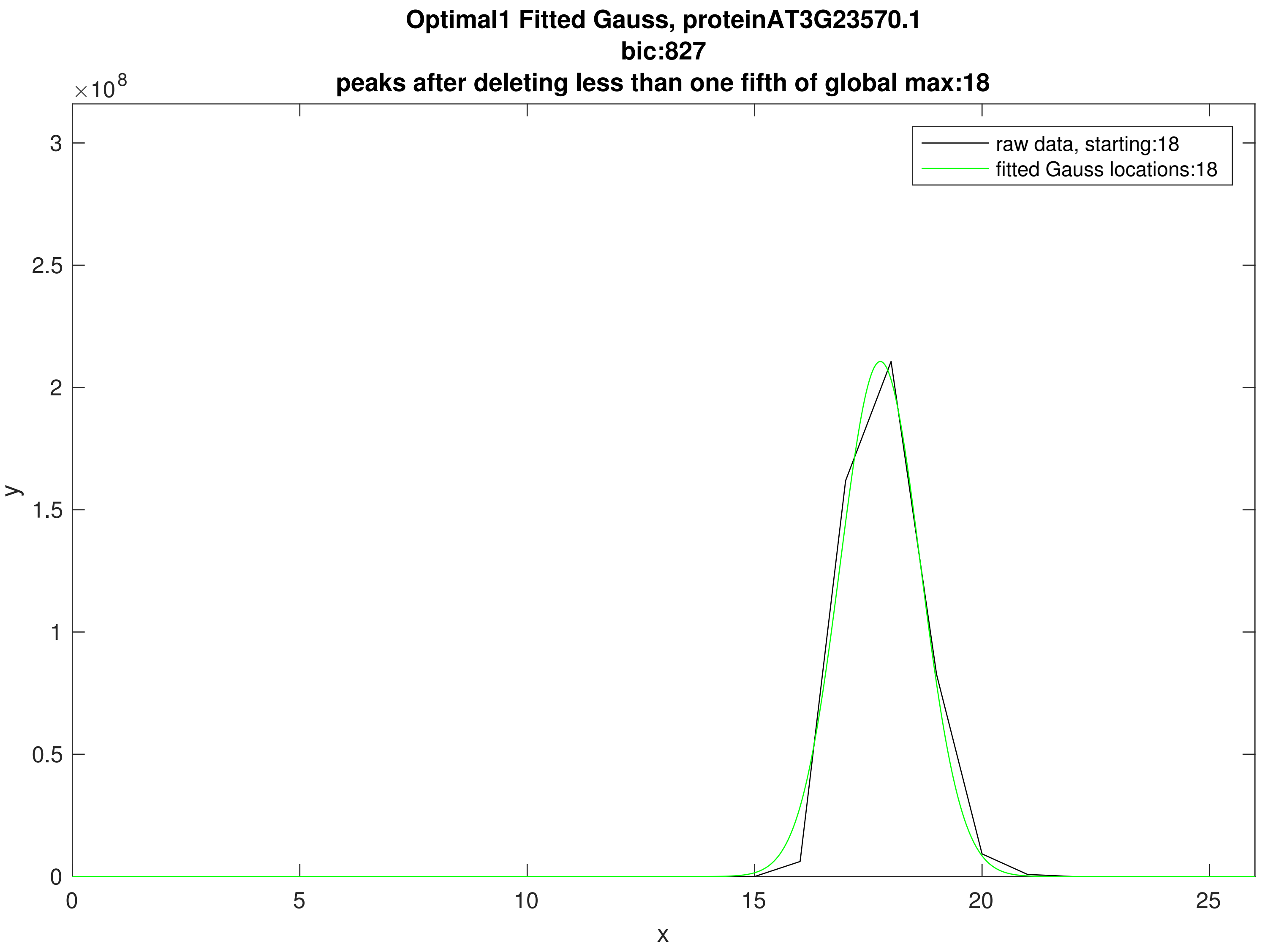
Optimal1 Fitted Gauss, proteinAT3G23570.1
bic:827
peaks after deleting less than one fifth of global max:18
×10^8
raw data, starting:18
fitted Gauss locations:18
y
x
0
0.5
1
1.5
2
2.5
3
0
5
10
15
20
25

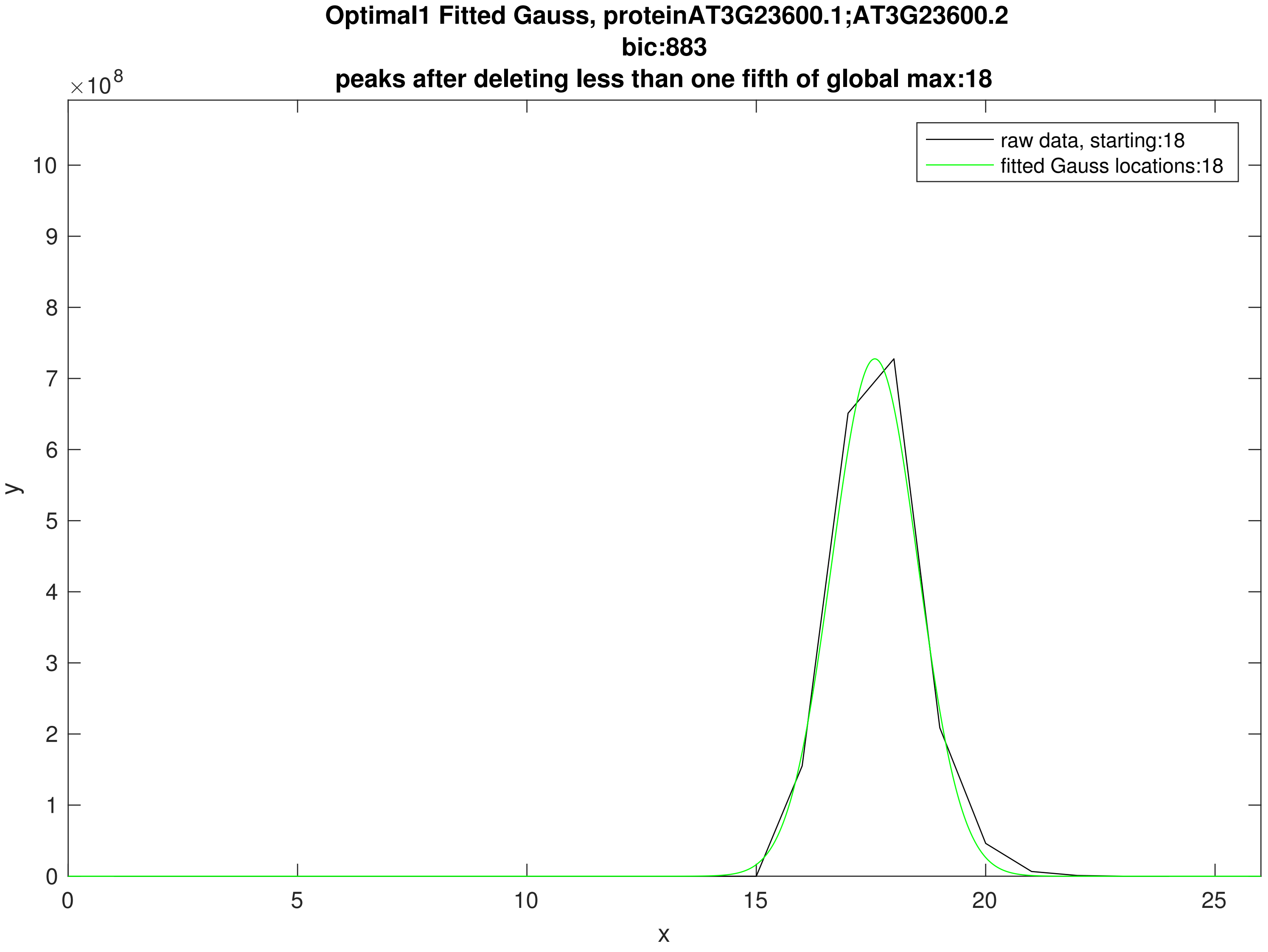

Optimal1 Fitted Gauss, proteinAT3G23600.1;AT3G23600.2
bic:883
peaks after deleting less than one fifth of global max:18
×10^8
raw data, starting:18
fitted Gauss locations:18
y
x
0
1
2
3
4
5
6
7
8
9
10
0
5
10
15
20
25

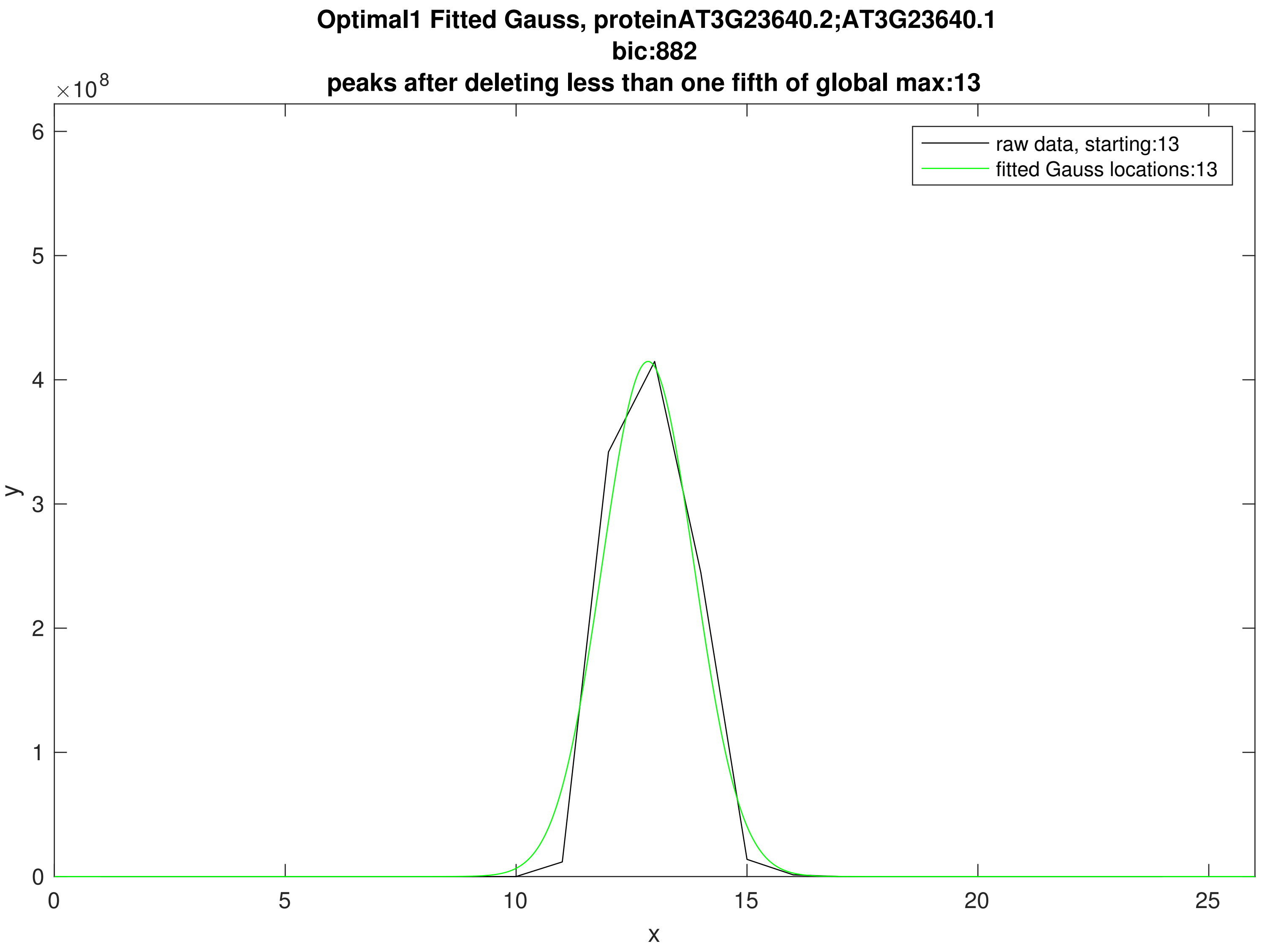

Optimal1 Fitted Gauss, proteinAT3G23640.2;AT3G23640.1
bic:882
peaks after deleting less than one fifth of global max:13
×10^8
y
x
raw data, starting:13
fitted Gauss locations:13

Optimal1 Fitted Gauss, proteinAT3G23660.1;AT3G23660.2
bic:647
peaks after deleting less than one fifth of global max:14

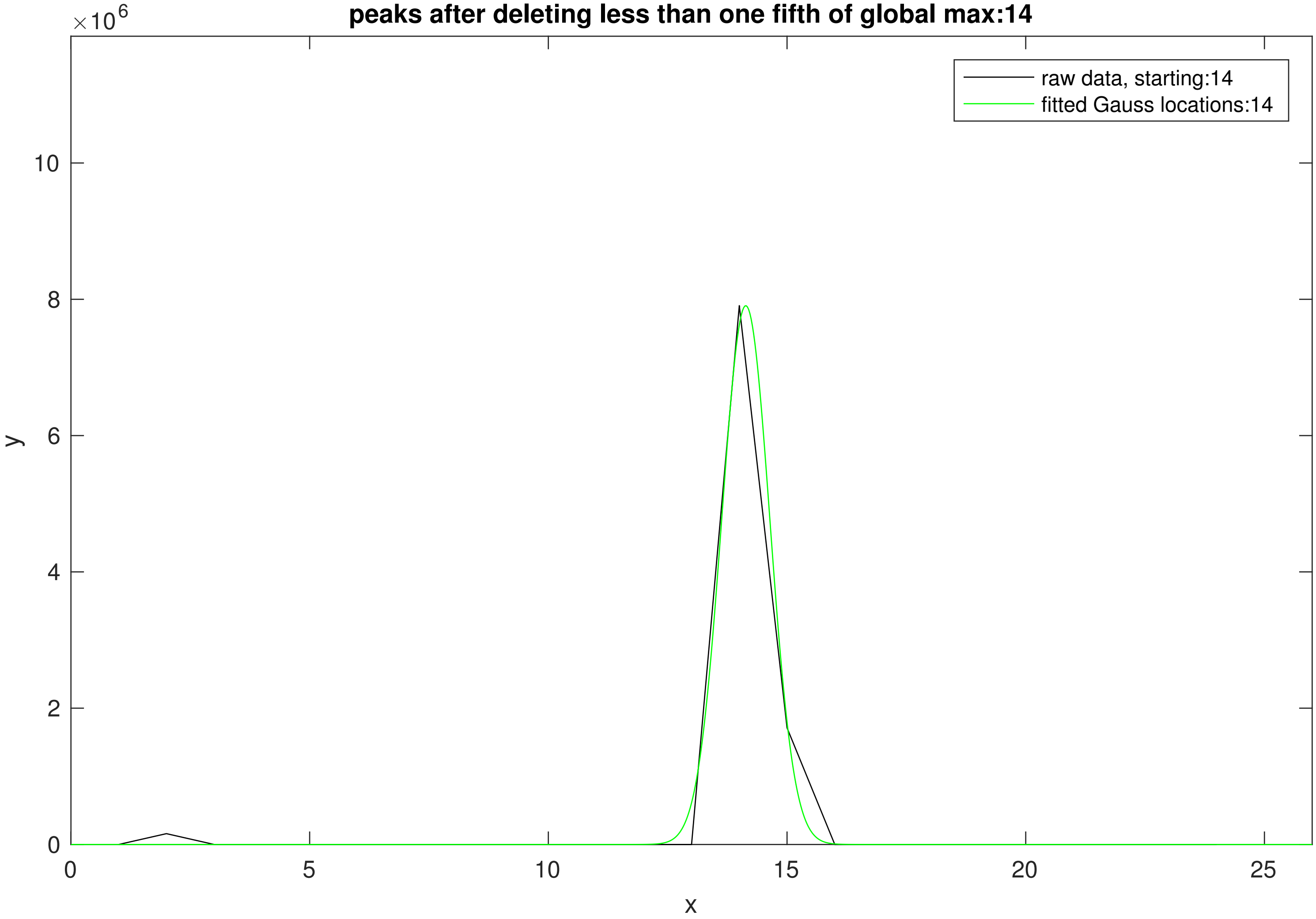

**Optimal1 Fitted Gauss, proteinAT3G23750.1**
**bic:966 973**
**peaks after deleting less than one fifth of global max:14**

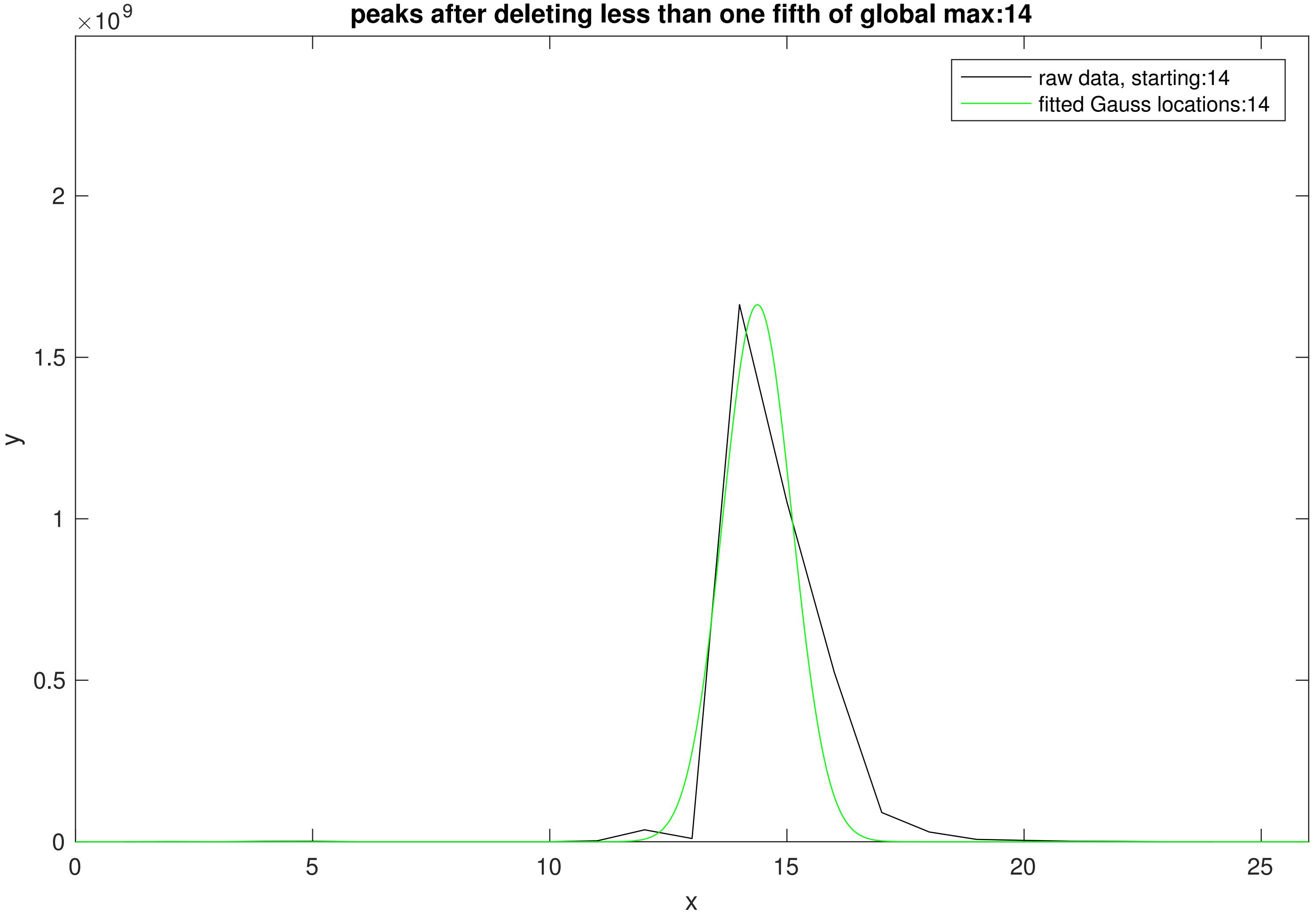

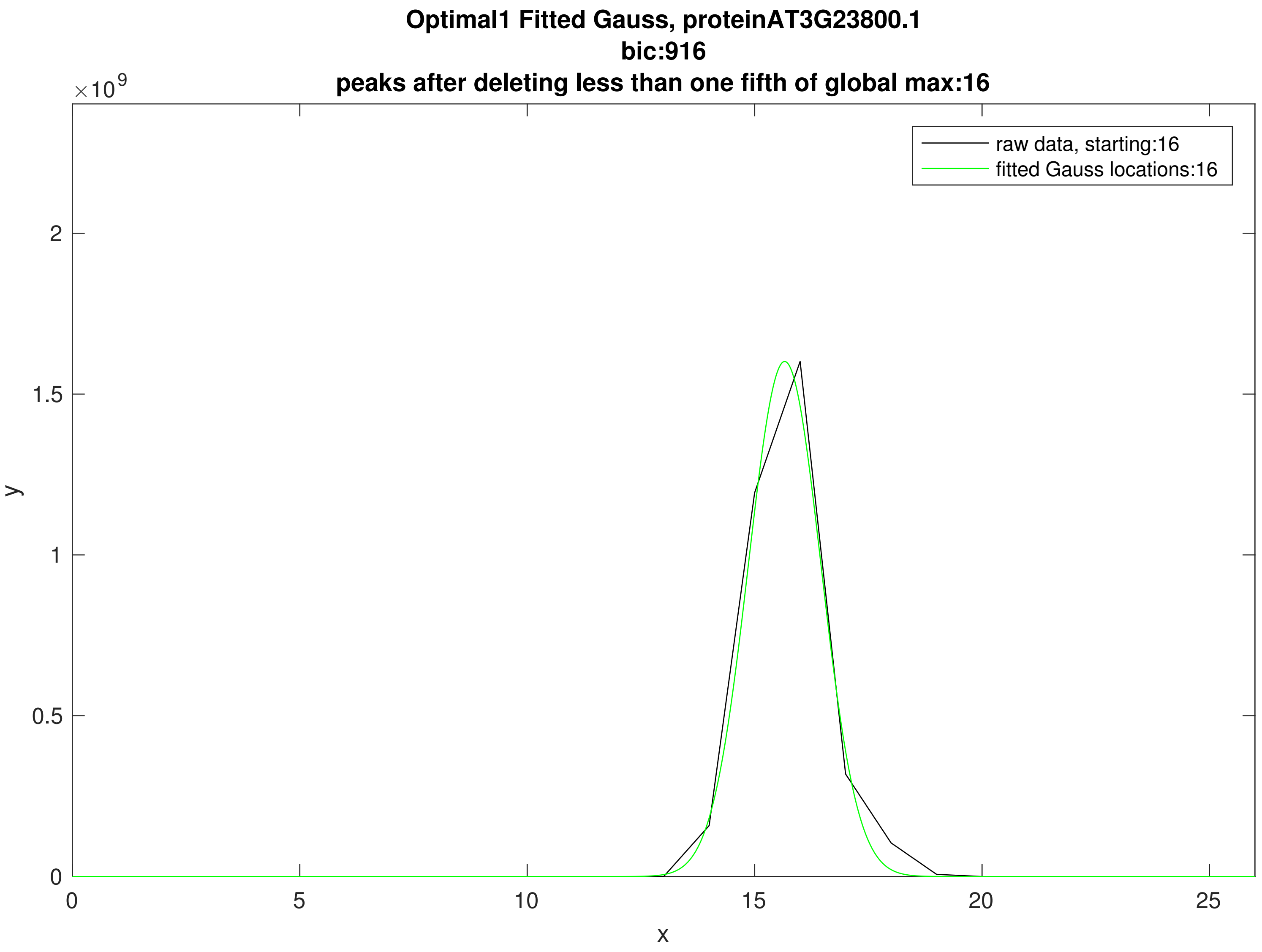
Optimal1 Fitted Gauss, proteinAT3G23800.1
bic:916
peaks after deleting less than one fifth of global max:16
×10^9
raw data, starting:16
fitted Gauss locations:16
y
x
0
0.5
1
1.5
2
0
5
10
15
20
25

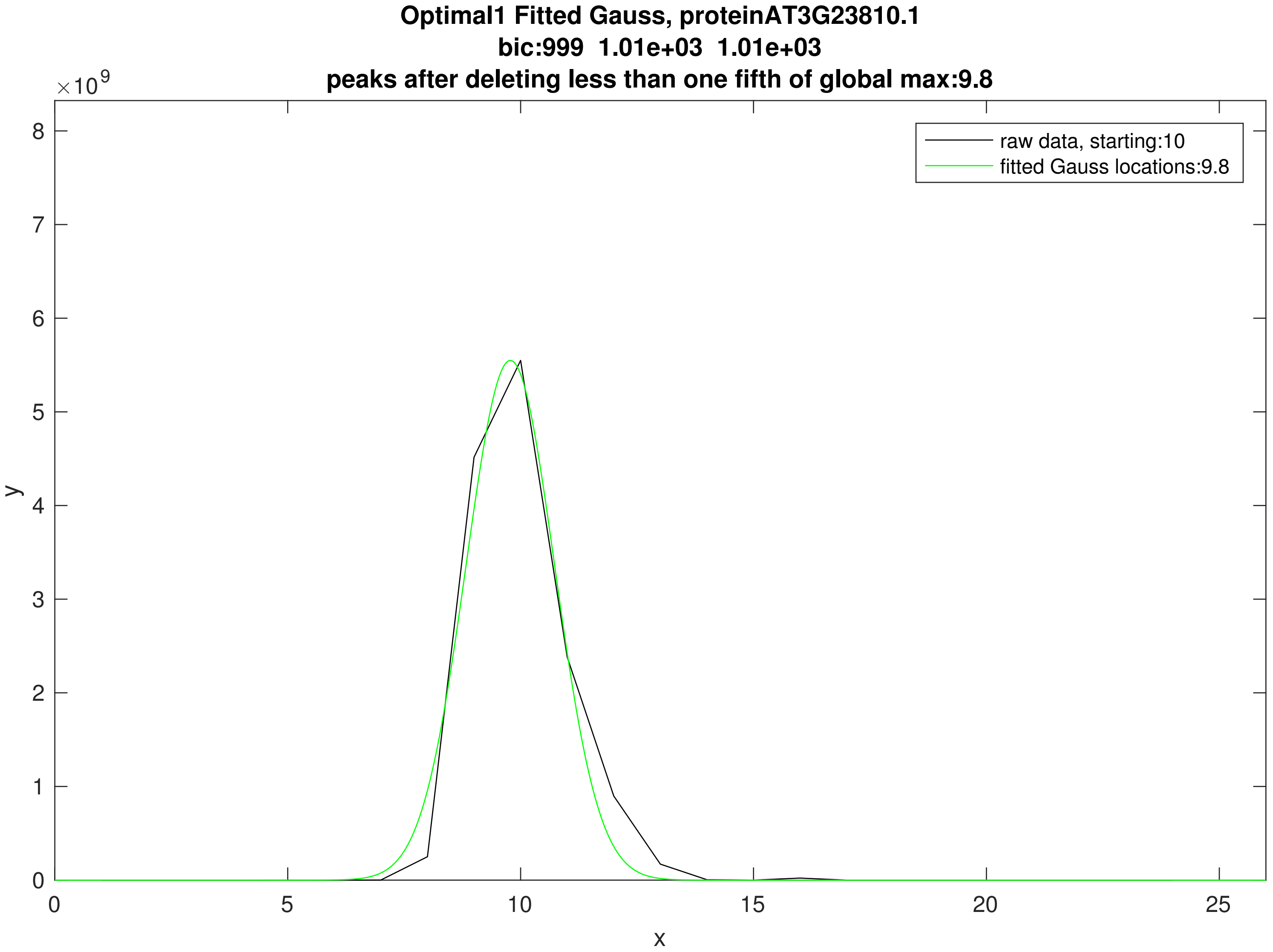
Optimal1 Fitted Gauss, proteinAT3G23810.1
bic:999 1.01e+03 1.01e+03
peaks after deleting less than one fifth of global max:9.8
×10^9
raw data, starting:10
fitted Gauss locations:9.8
y
x
0
1
2
3
4
5
6
7
8
0
5
10
15
20
25

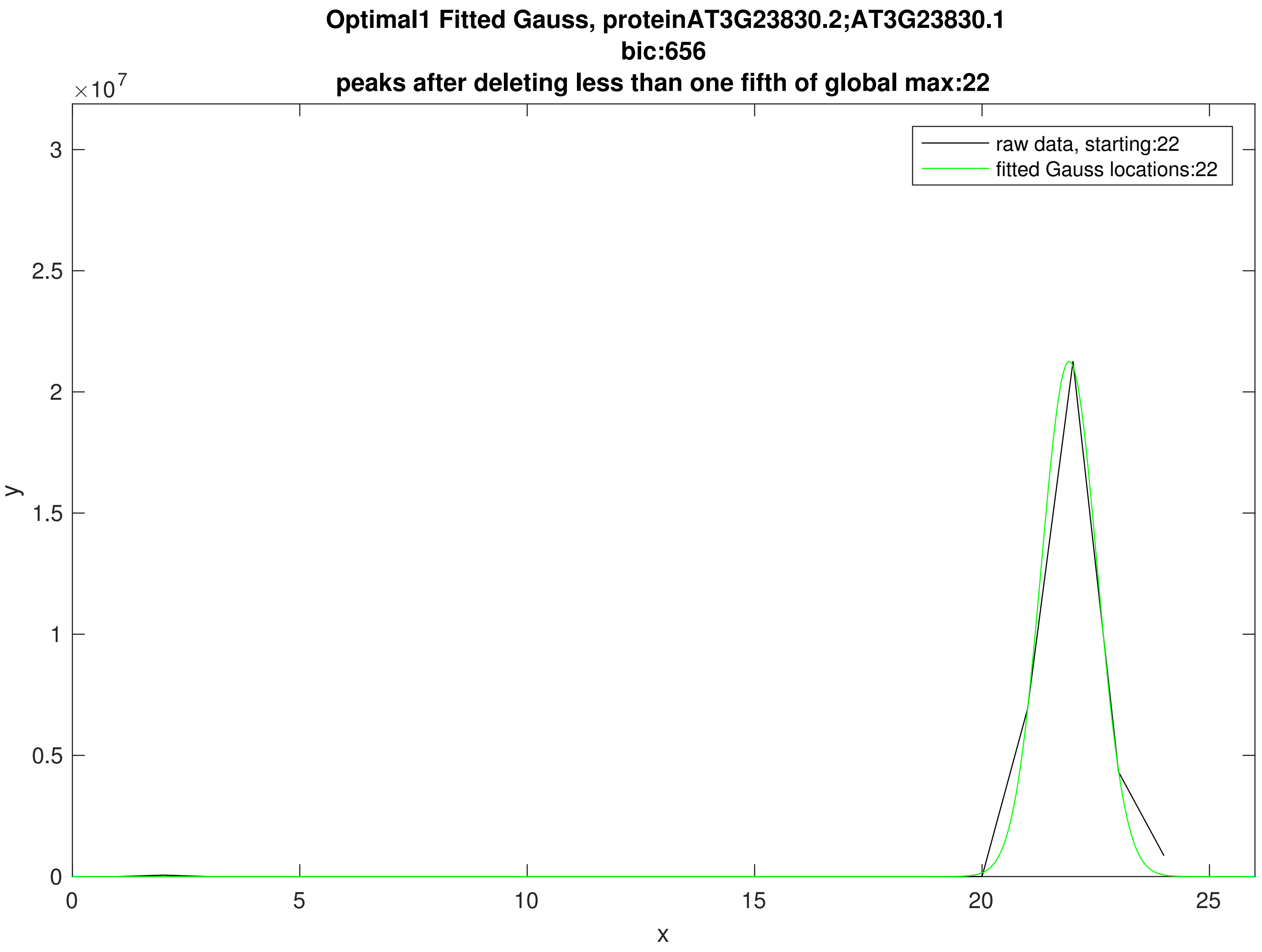
Optimal1 Fitted Gauss, proteinAT3G23830.2;AT3G23830.1
bic:656
peaks after deleting less than one fifth of global max:22
$\times10^7$
raw data, starting:22
fitted Gauss locations:22
y
x
0
0.5
1
1.5
2
2.5
3
0
5
10
15
20
25

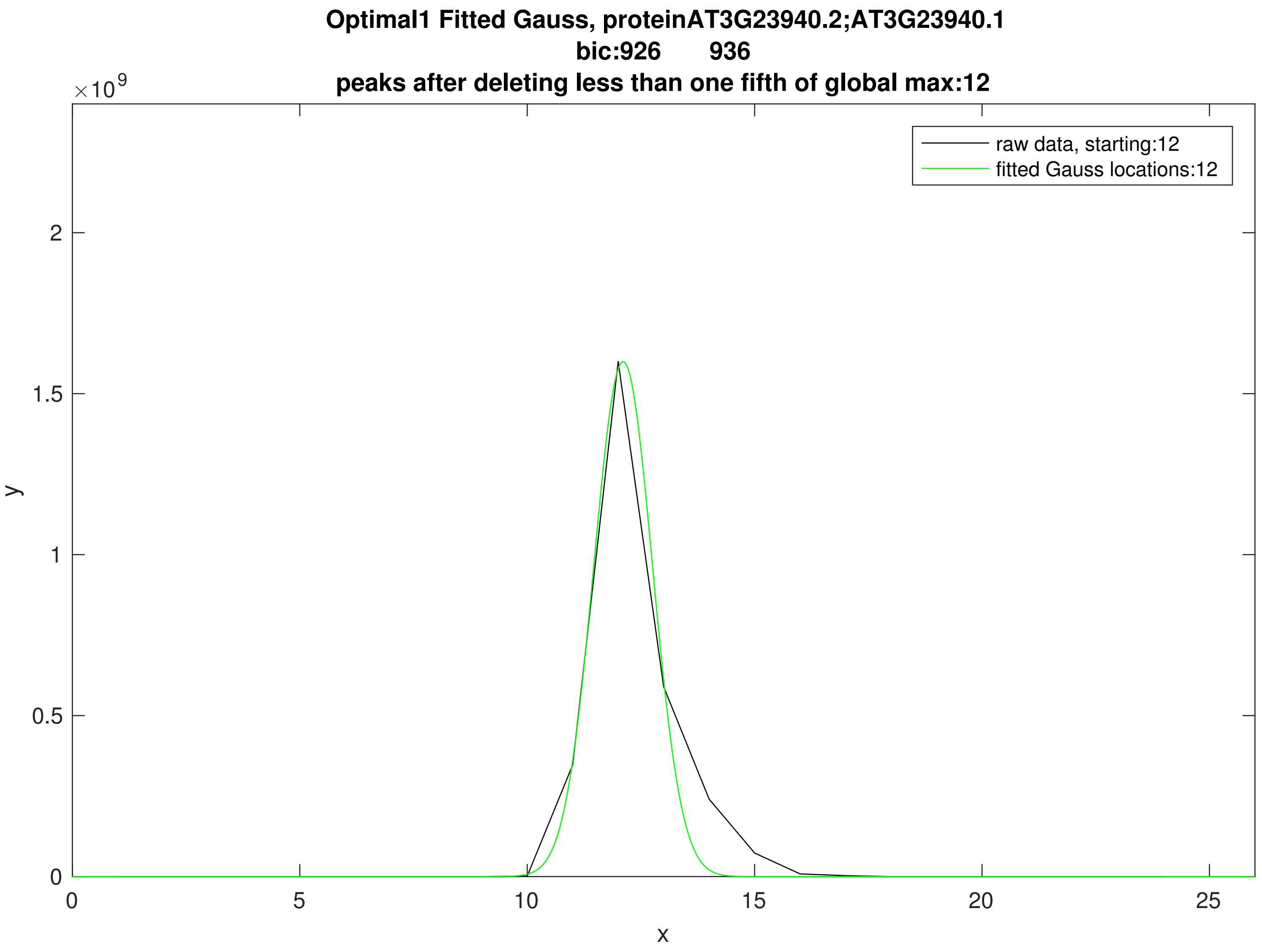
Optimal1 Fitted Gauss, proteinAT3G23940.2;AT3G23940.1
bic:926 936
peaks after deleting less than one fifth of global max:12
×10^9
raw data, starting:12
fitted Gauss locations:12
y
x
0
0.5
1
1.5
2
0
5
10
15
20
25

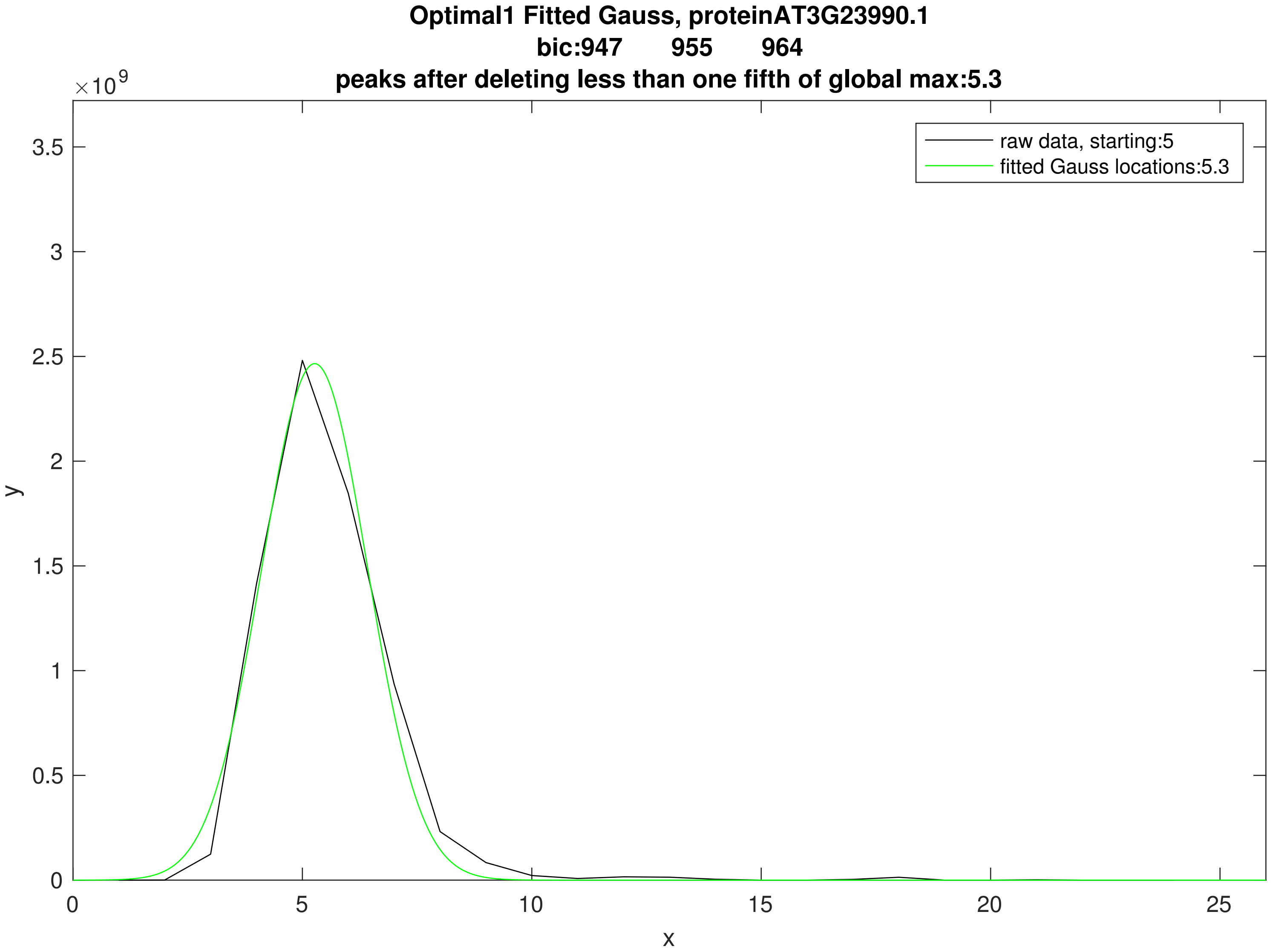
Optimal1 Fitted Gauss, proteinAT3G23990.1
bic:947 955 964
peaks after deleting less than one fifth of global max:5.3
×10^9
raw data, starting:5
fitted Gauss locations:5.3
y
x
0
0.5
1
1.5
2
2.5
3
3.5
5
10
15
20
25

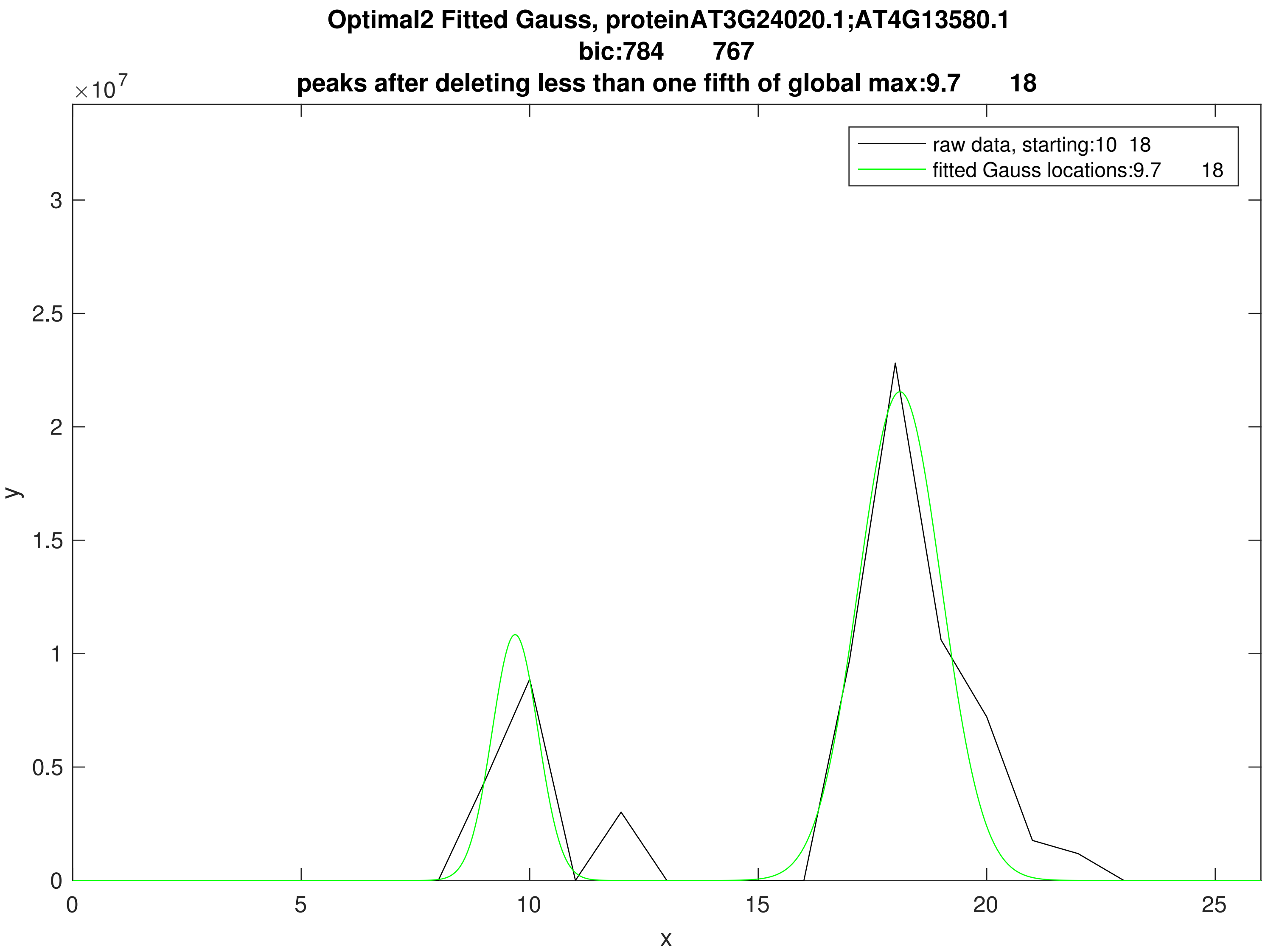
Optimal2 Fitted Gauss, proteinAT3G24020.1;AT4G13580.1
bic:784 767
peaks after deleting less than one fifth of global max:9.7 18
×10^7
raw data, starting:10 18
fitted Gauss locations:9.7 18
y
x
0
0.5
1
1.5
2
2.5
3
0
5
10
15
20
25

**Optimal1 Fitted Gauss, proteinAT3G24030.1**
**bic:694**
**peaks after deleting less than one fifth of global max:13**

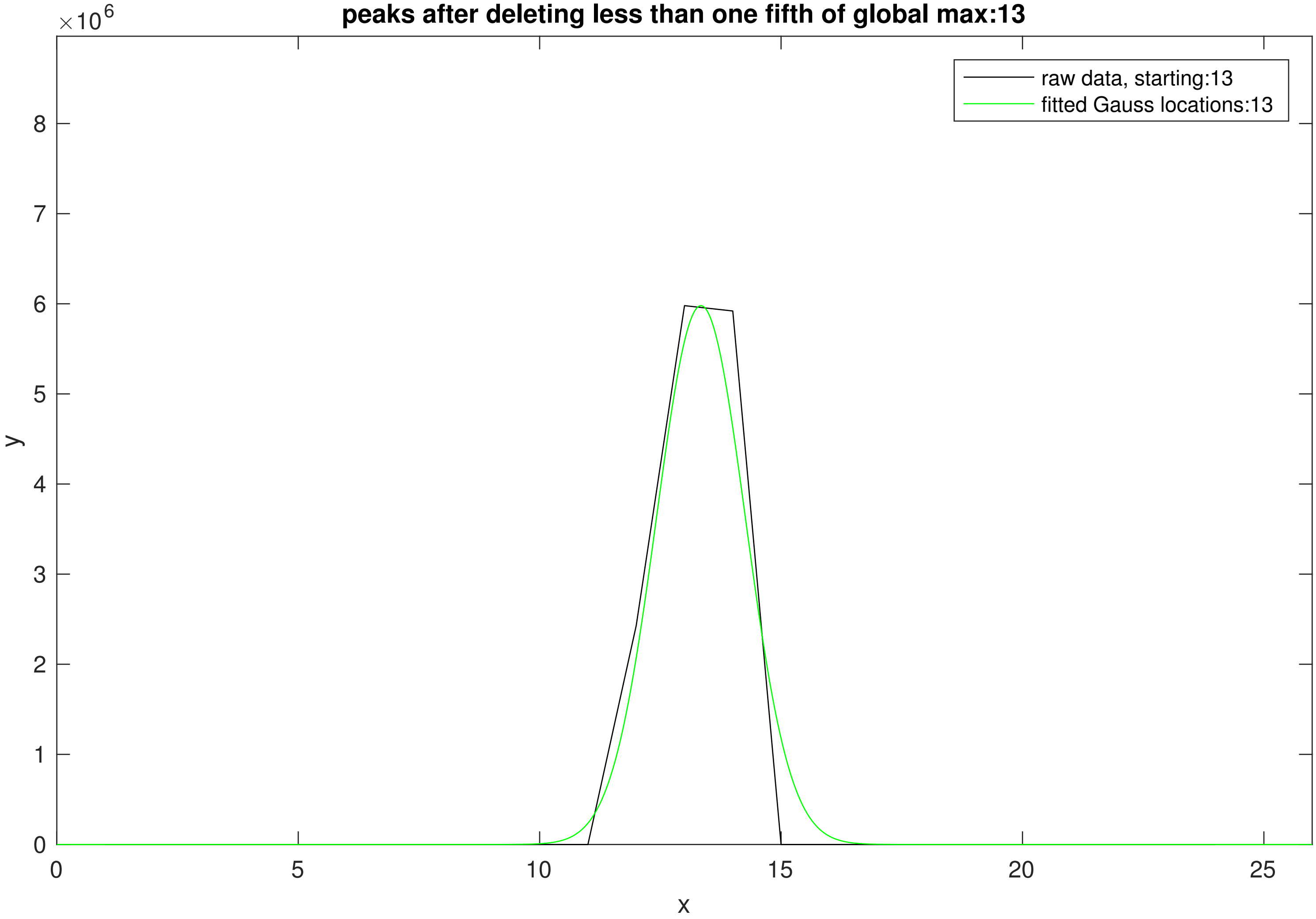

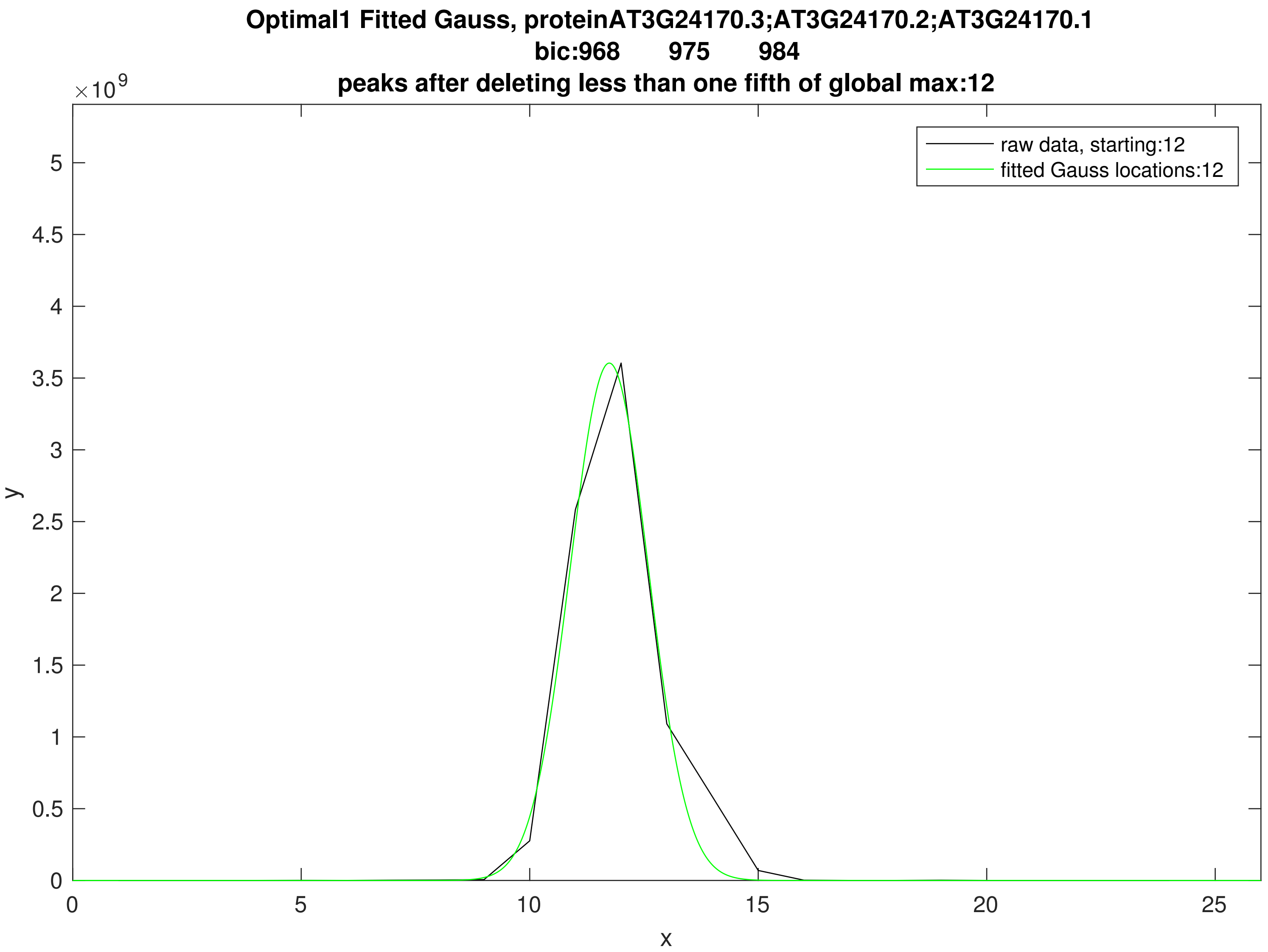

Optimal1 Fitted Gauss, proteinAT3G24170.3;AT3G24170.2;AT3G24170.1
bic:968 975 984
peaks after deleting less than one fifth of global max:12
×10^9
raw data, starting:12
fitted Gauss locations:12
5
4.5
4
3.5
3
2.5
2
1.5
1
0.5
0
y
0
5
10
15
20
25
x

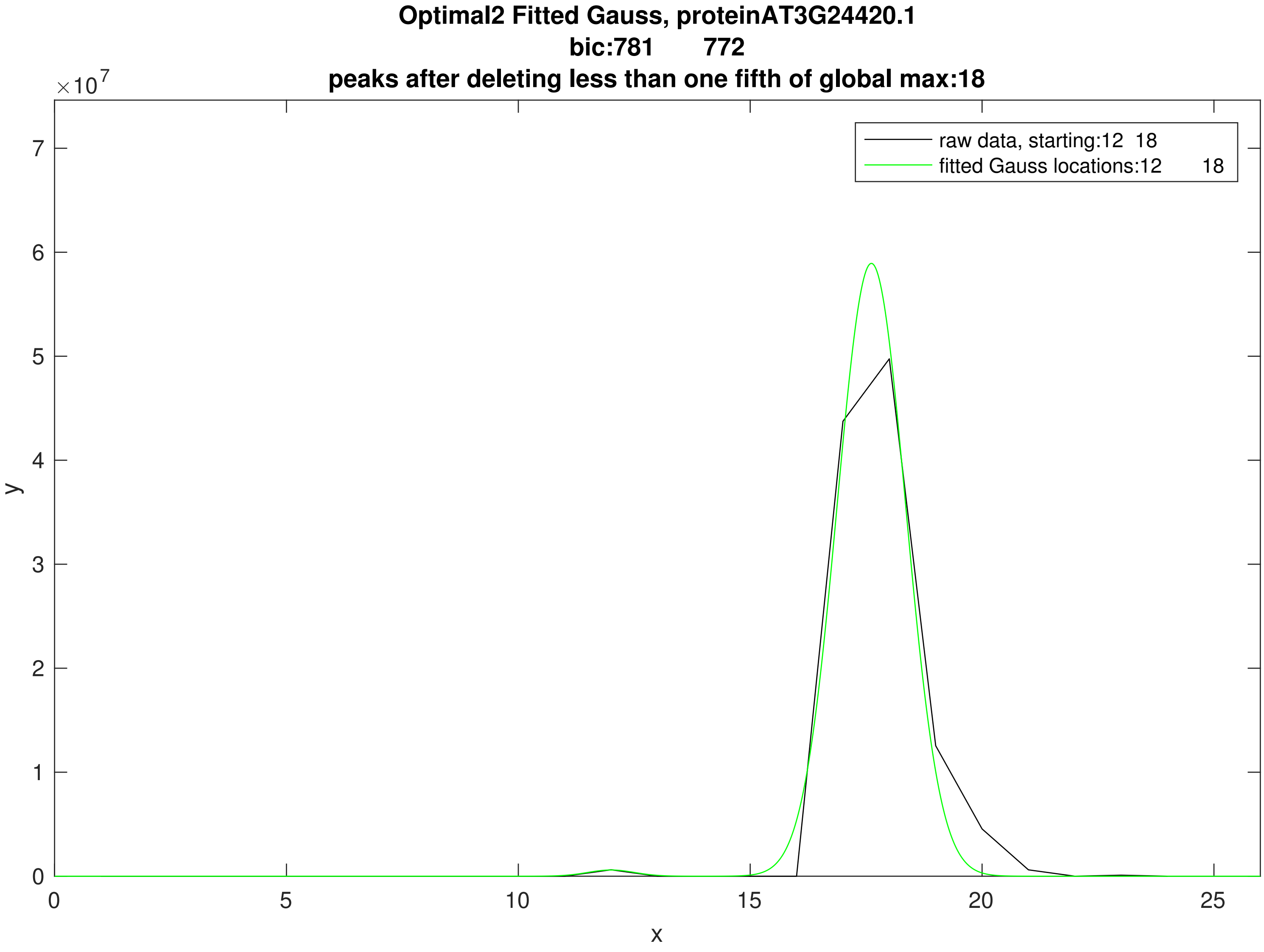

Optimal2 Fitted Gauss, proteinAT3G24420.1
bic:781 772
peaks after deleting less than one fifth of global max:18
×10^7
raw data, starting:12 18
fitted Gauss locations:12 18
y
x
0
1
2
3
4
5
6
7
0
5
10
15
20
25

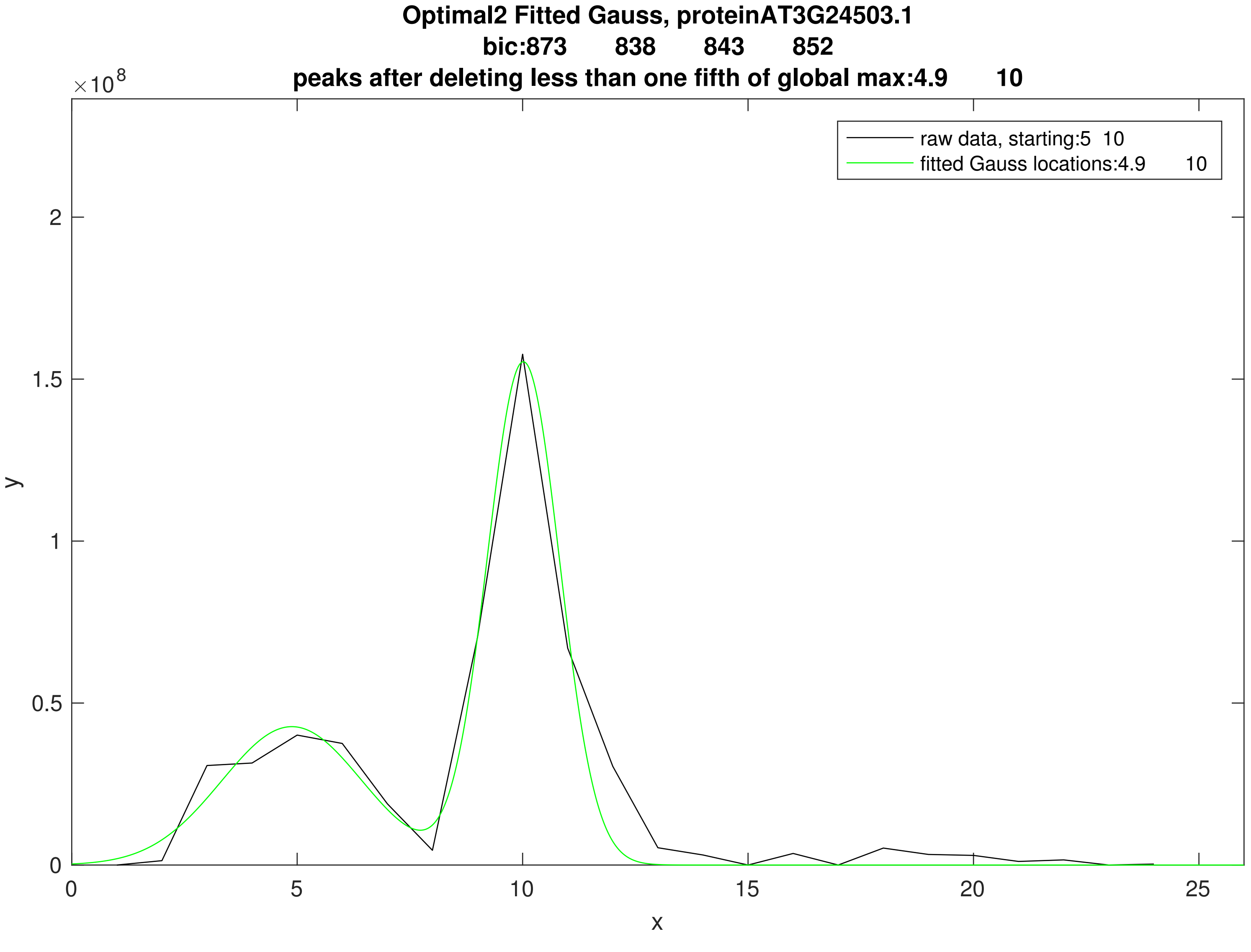

Optimal2 Fitted Gauss, proteinAT3G24503.1
bic:873 838 843 852
peaks after deleting less than one fifth of global max:4.9 10
×10^8
raw data, starting:5 10
fitted Gauss locations:4.9 10
y
x
0
0.5
1
1.5
2
5
10
15
20
25

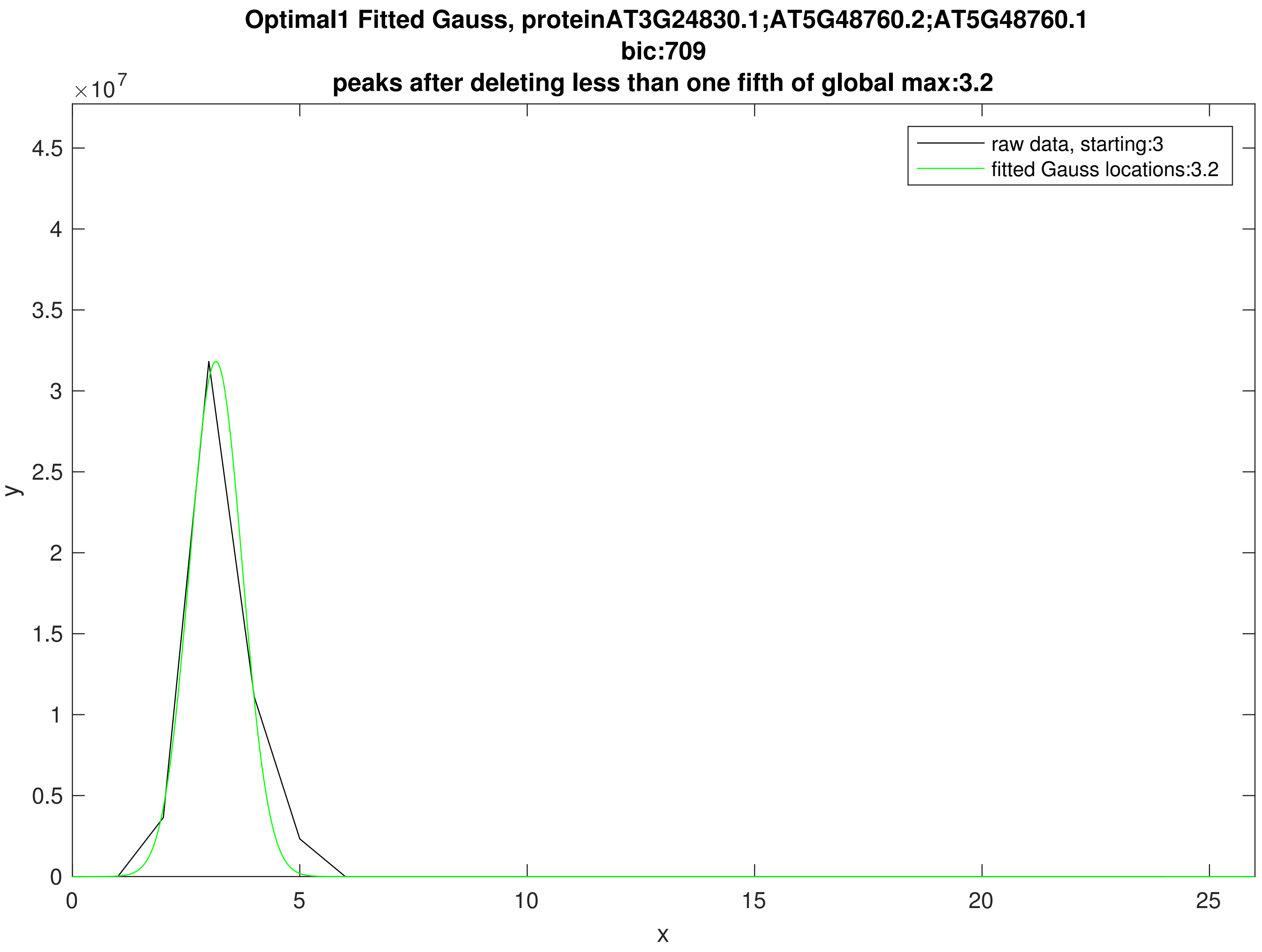

Optimal1 Fitted Gauss, proteinAT3G24830.1;AT5G48760.2;AT5G48760.1
bic:709
peaks after deleting less than one fifth of global max:3.2
×10^7
4.5
4
3.5
3
2.5
2
1.5
1
0.5
0
y
0
5
10
15
20
25
x
raw data, starting:3
fitted Gauss locations:3.2

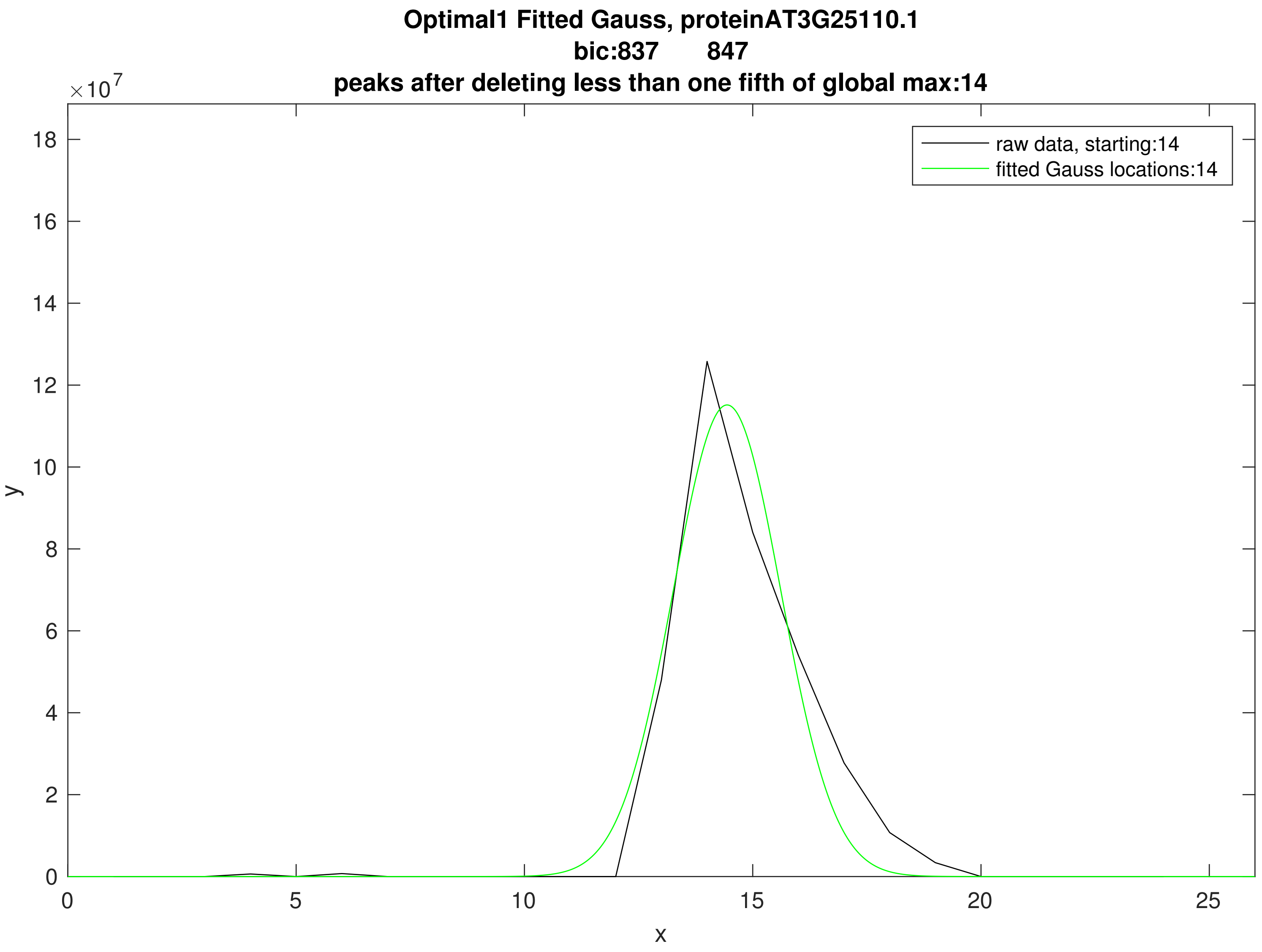

Optimal1 Fitted Gauss, proteinAT3G25110.1
bic:837 847
peaks after deleting less than one fifth of global max:14
×10^7
raw data, starting:14
fitted Gauss locations:14
y
x
0
2
4
6
8
10
12
14
16
18
0
5
10
15
20
25

**Optimal1 Fitted Gauss, proteinAT3G25220.1**
**bic:716**
**peaks after deleting less than one fifth of global max:21**

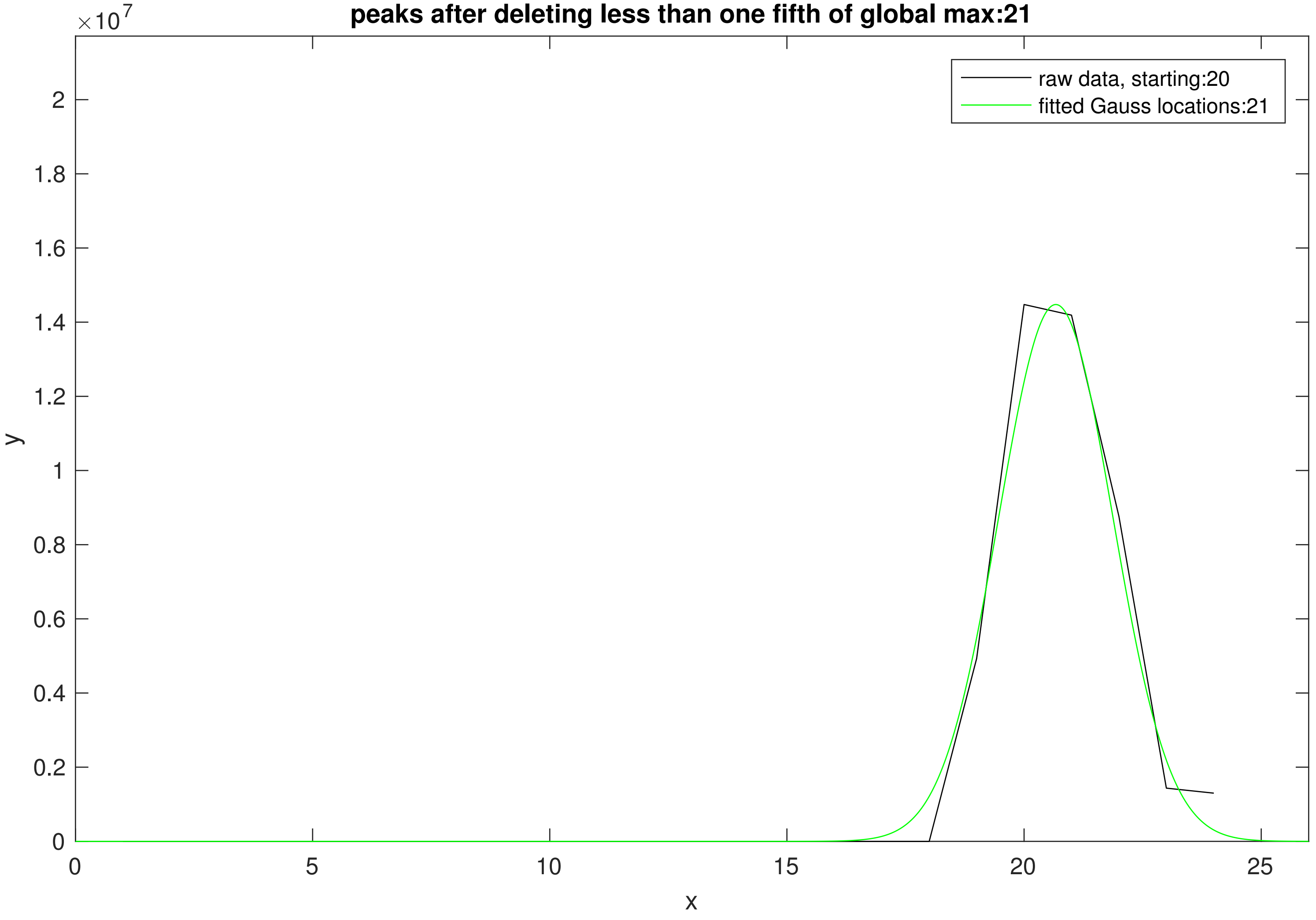

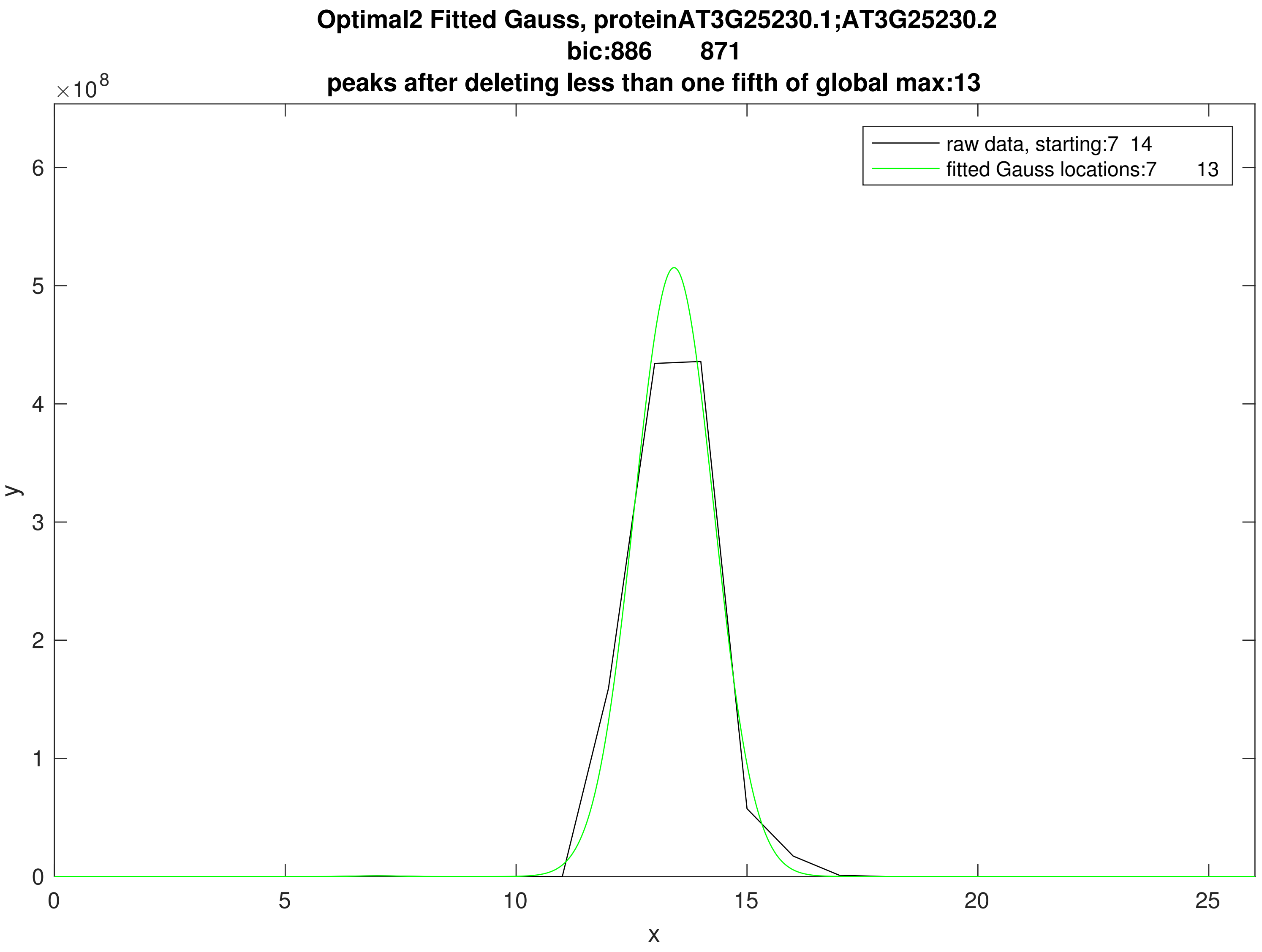

Optimal2 Fitted Gauss, proteinAT3G25230.1;AT3G25230.2
bic:886 871
peaks after deleting less than one fifth of global max:13
×10^8
raw data, starting:7 14
fitted Gauss locations:7 13
y
x
0
1
2
3
4
5
6
0
5
10
15
20
25

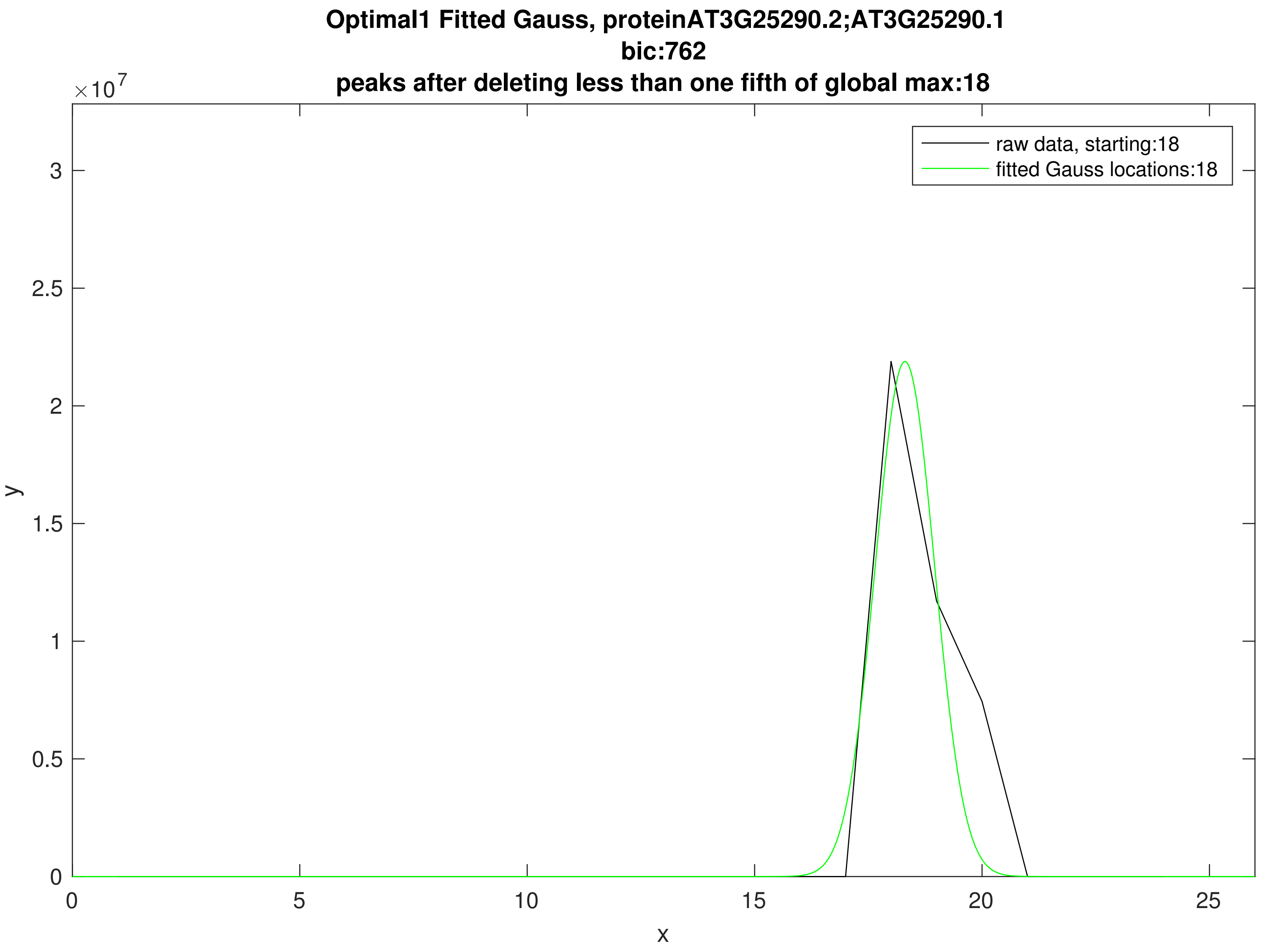

Optimal1 Fitted Gauss, proteinAT3G25290.2;AT3G25290.1
bic:762
peaks after deleting less than one fifth of global max:18
×10^7
3
2.5
2
1.5
1
0.5
0
y
0
5
10
15
20
25
x
raw data, starting:18
fitted Gauss locations:18

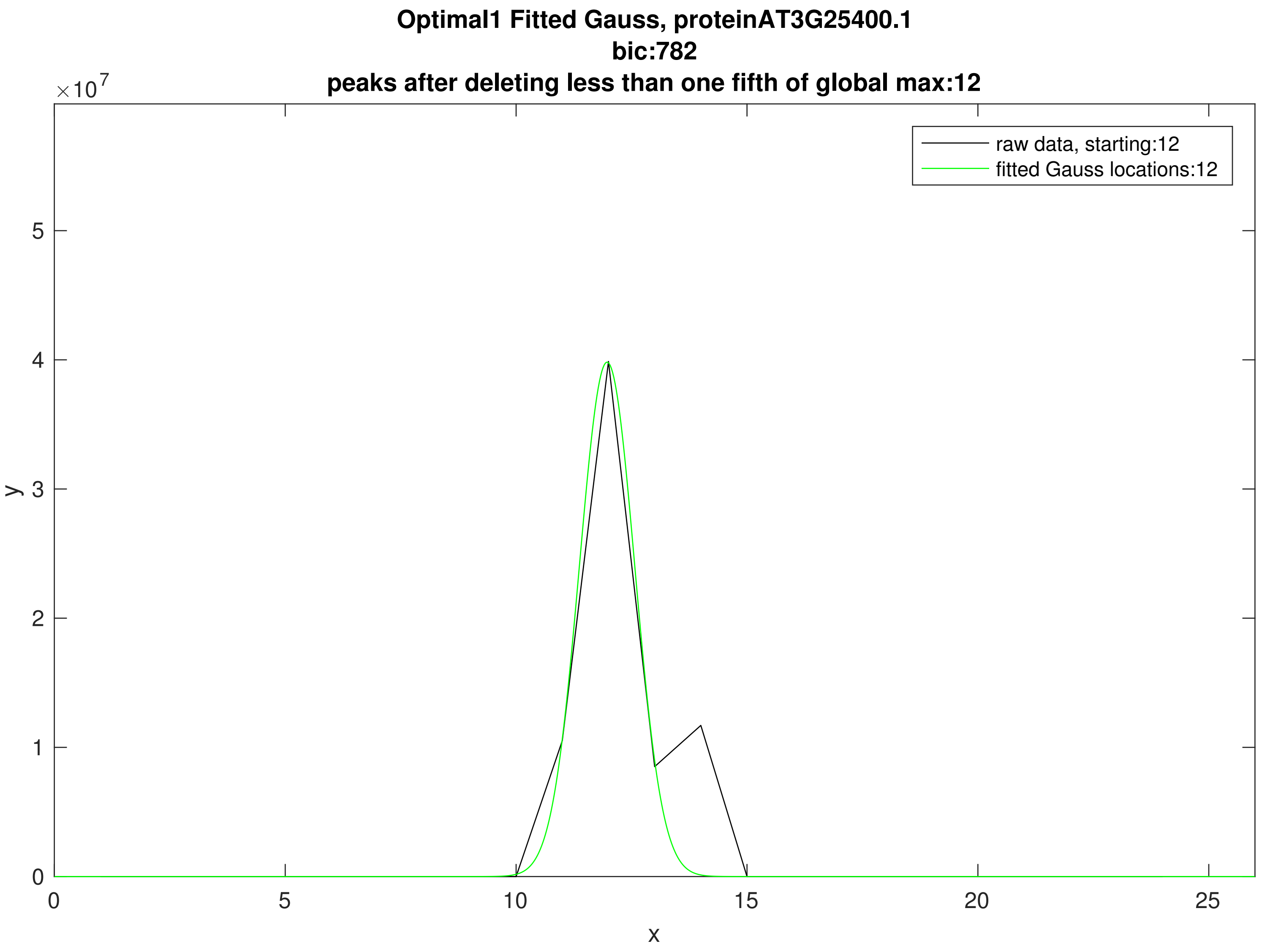

Optimal1 Fitted Gauss, proteinAT3G25400.1
bic:782
peaks after deleting less than one fifth of global max:12
×10^7
raw data, starting:12
fitted Gauss locations:12
y
x
0
1
2
3
4
5
0
5
10
15
20
25

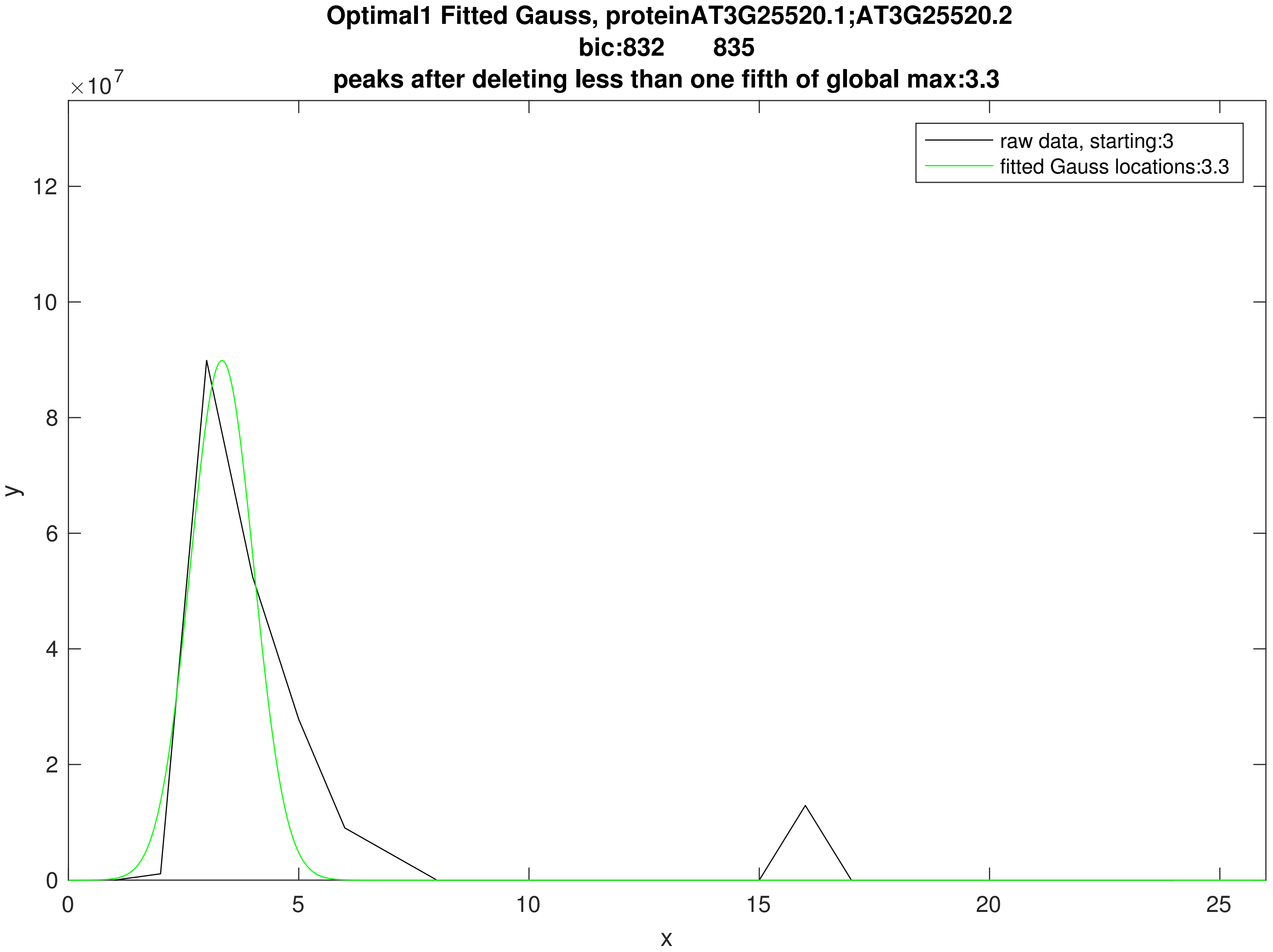
Optimal1 Fitted Gauss, proteinAT3G25520.1;AT3G25520.2
bic:832 835
peaks after deleting less than one fifth of global max:3.3
×10^7
raw data, starting:3
fitted Gauss locations:3.3
y
x
0
2
4
6
8
10
12
0
5
10
15
20
25

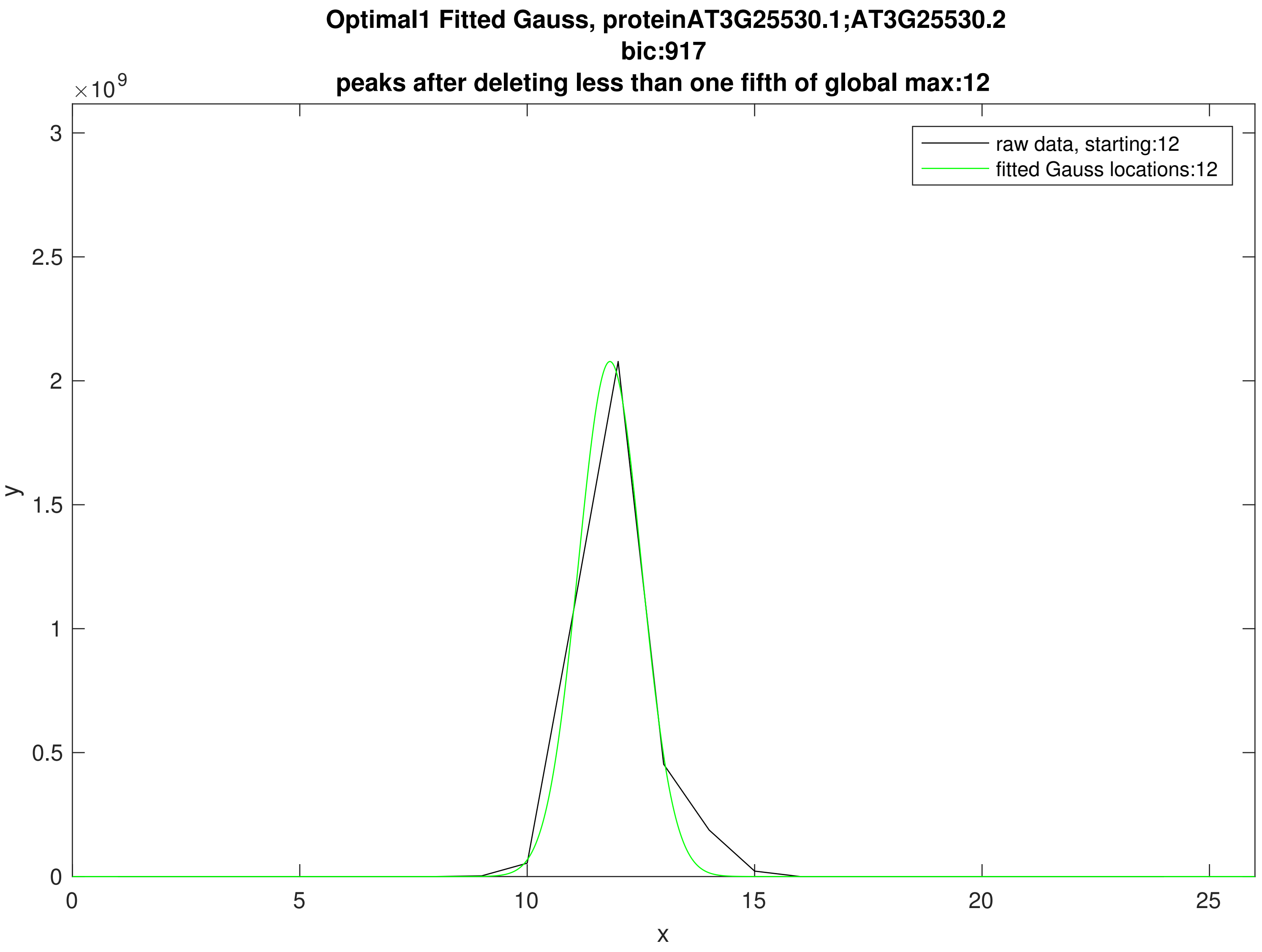
Optimal1 Fitted Gauss, proteinAT3G25530.1;AT3G25530.2
bic:917
peaks after deleting less than one fifth of global max:12
raw data, starting:12
fitted Gauss locations:12
x
y

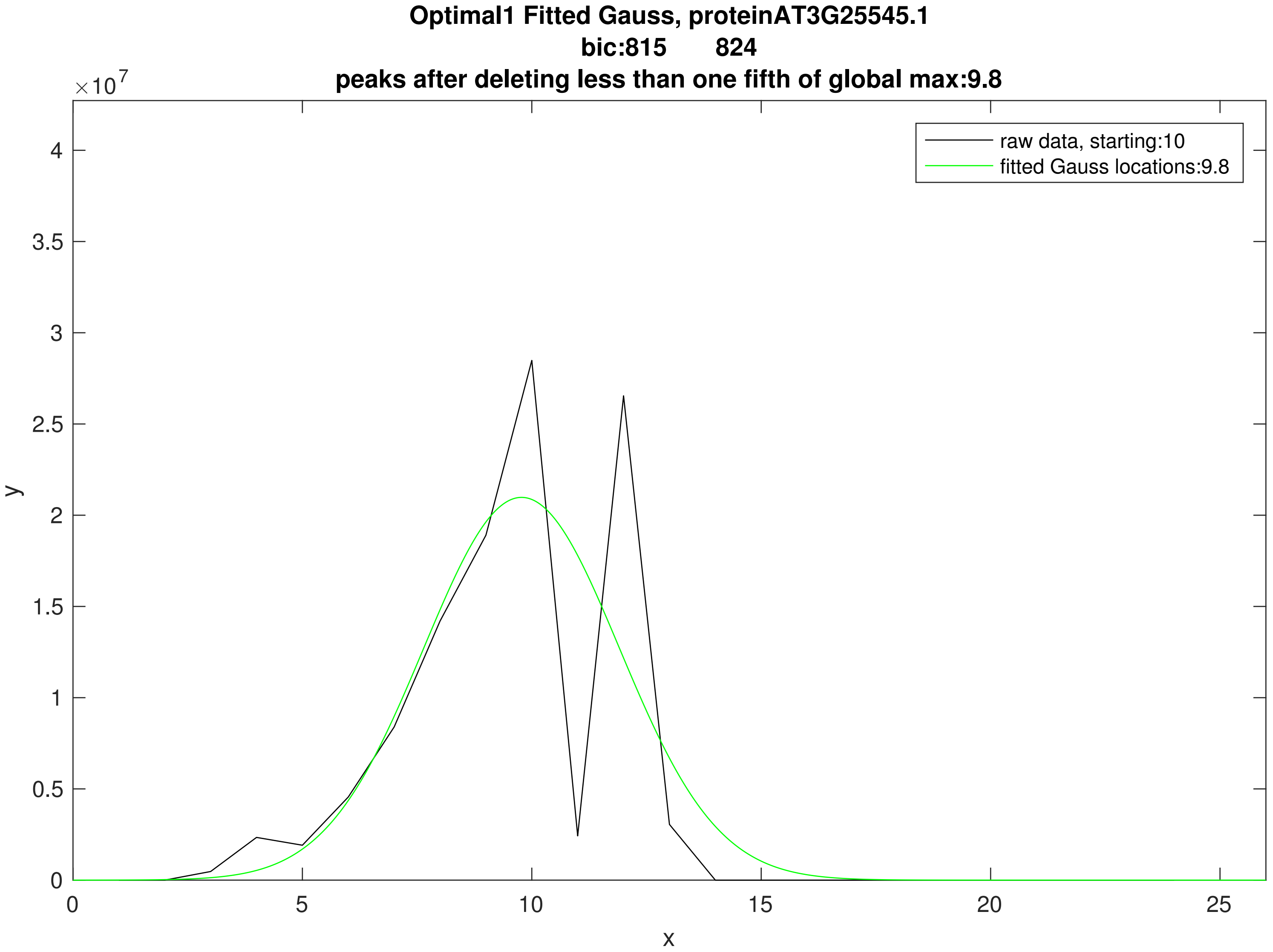
Optimal1 Fitted Gauss, proteinAT3G25545.1
bic:815 824
peaks after deleting less than one fifth of global max:9.8
×10^7
raw data, starting:10
fitted Gauss locations:9.8
4
3.5
3
2.5
2
1.5
1
0.5
0
y
0
5
10
15
20
25
x

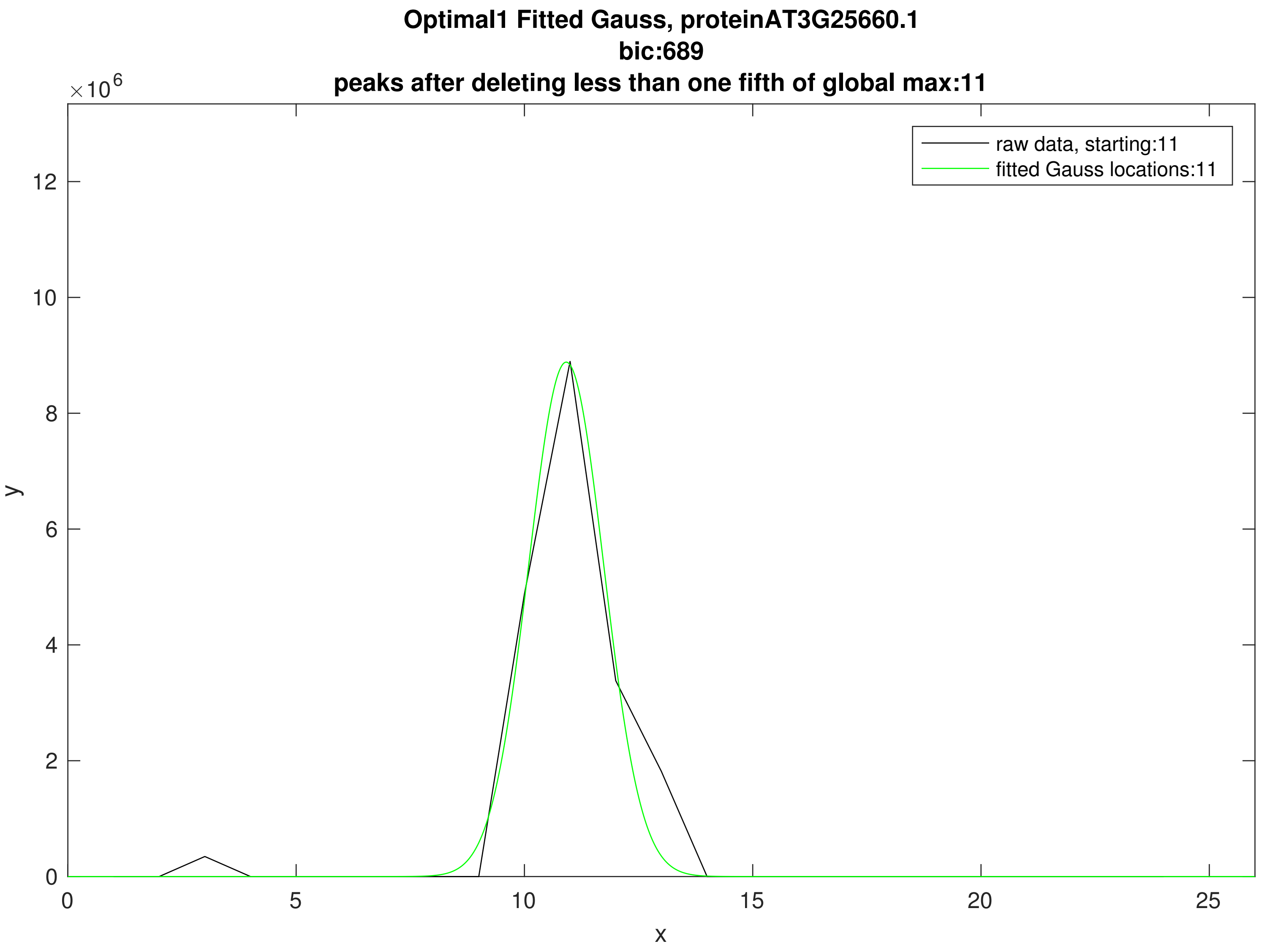

Optimal1 Fitted Gauss, proteinAT3G25660.1
bic:689
peaks after deleting less than one fifth of global max:11
×10^6
raw data, starting:11
fitted Gauss locations:11
12
10
8
6
4
2
0
y
0
5
10
15
20
25
x

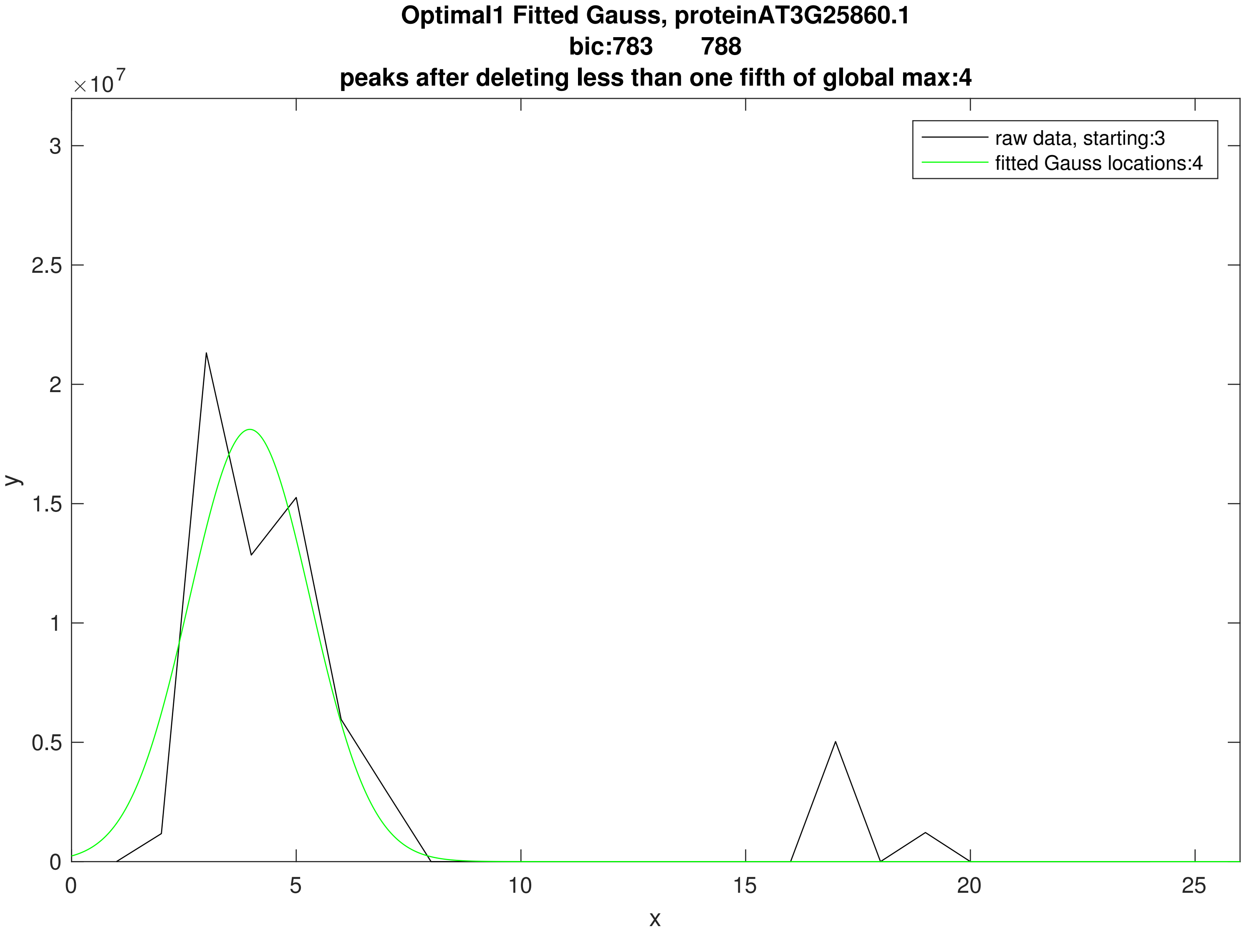

Optimal1 Fitted Gauss, proteinAT3G25860.1
bic:783 788
peaks after deleting less than one fifth of global max:4
×10^7
raw data, starting:3
fitted Gauss locations:4
y
x
0
0.5
1
1.5
2
2.5
3
5
10
15
20
25

#### Optimal1 Fitted Gauss, proteinAT3G25920.1
#### bic:774
#### peaks after deleting less than one fifth of global max:14

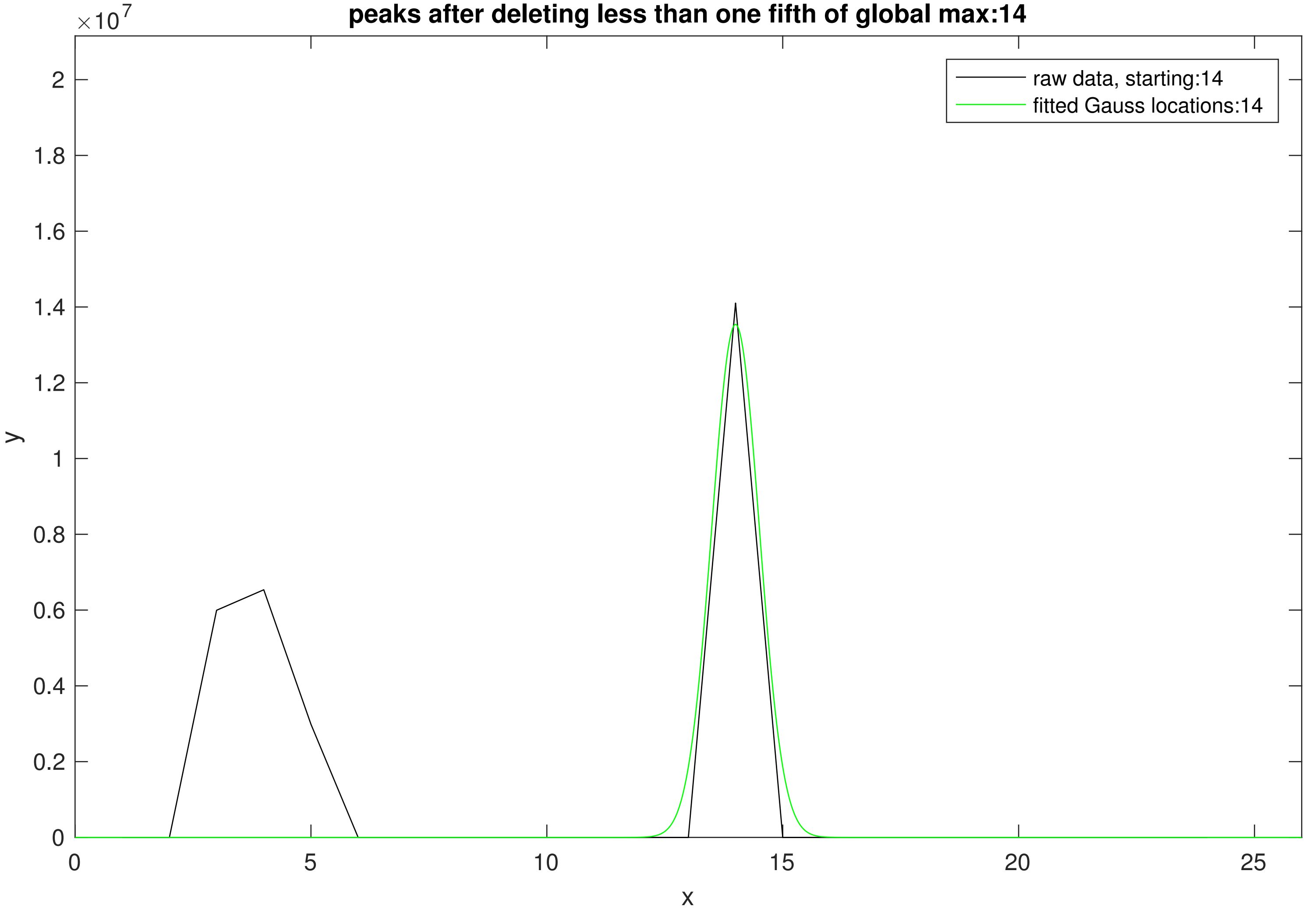

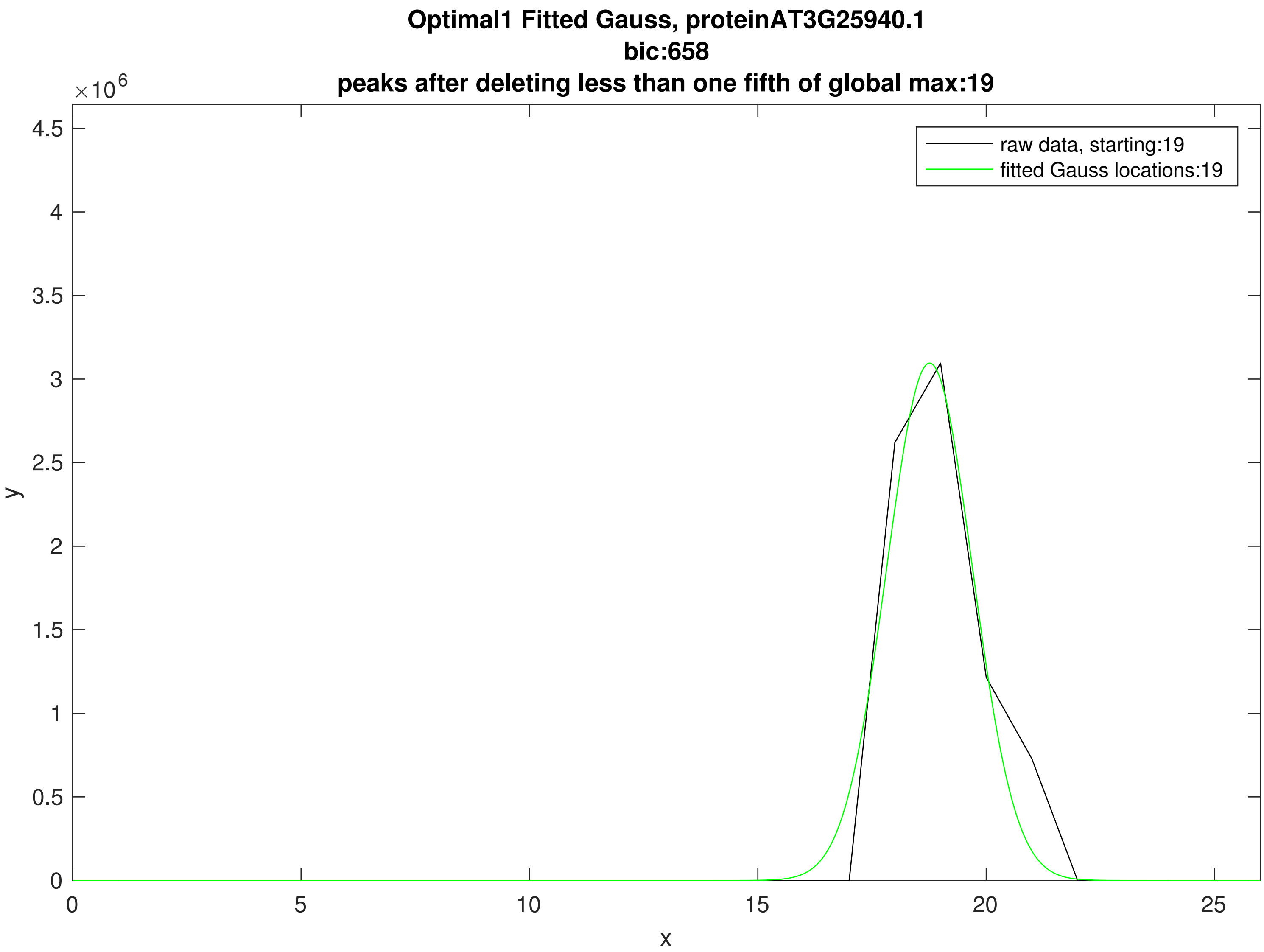
Optimal1 Fitted Gauss, proteinAT3G25940.1
bic:658
peaks after deleting less than one fifth of global max:19
×10^6
4.5
4
3.5
3
2.5
2
1.5
1
0.5
0
y
0
5
10
15
20
25
x
raw data, starting:19
fitted Gauss locations:19

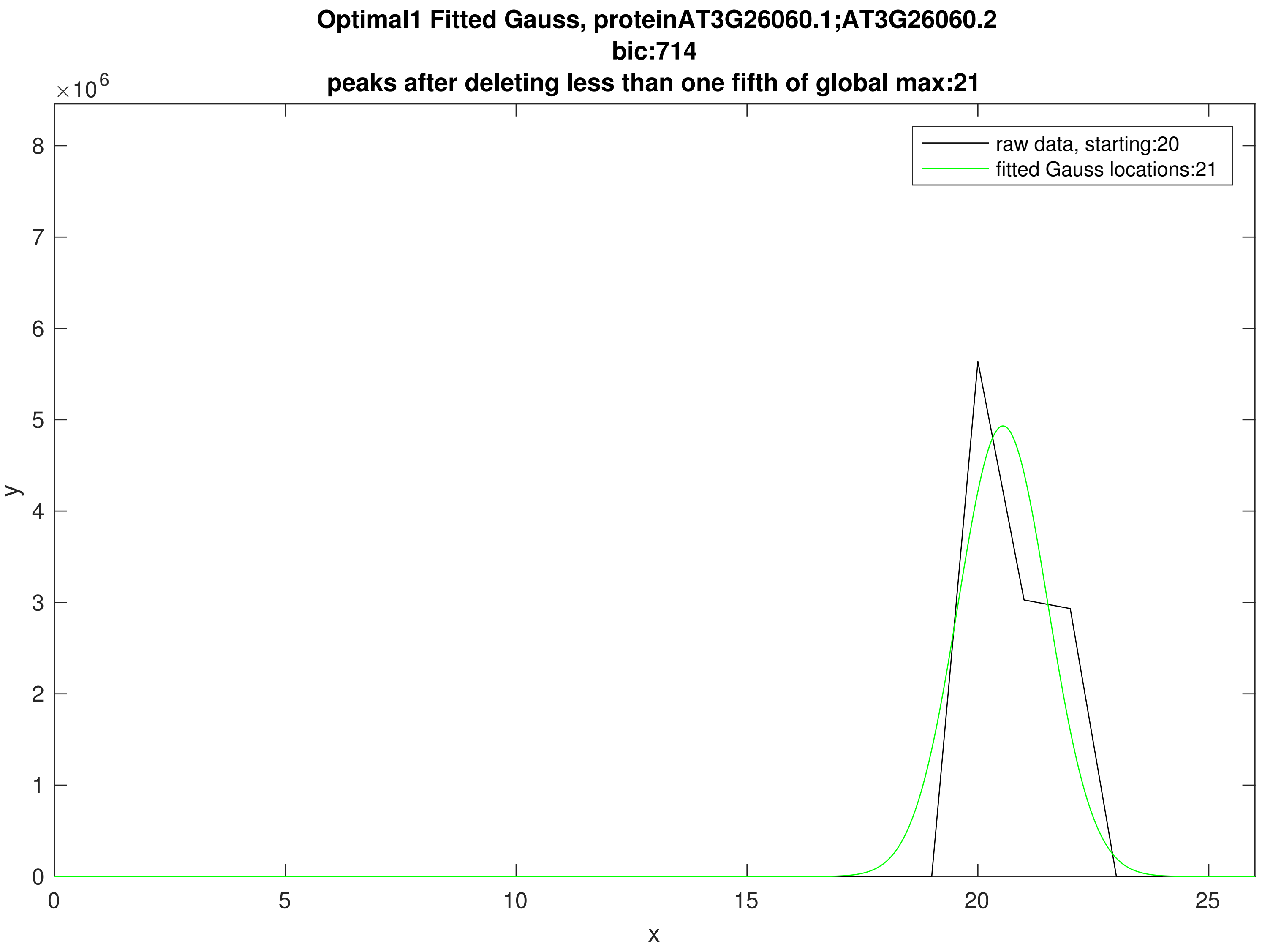
Optimal1 Fitted Gauss, proteinAT3G26060.1;AT3G26060.2
bic:714
peaks after deleting less than one fifth of global max:21
×10^6
raw data, starting:20
fitted Gauss locations:21
y
x
0
1
2
3
4
5
6
7
8
0
5
10
15
20
25

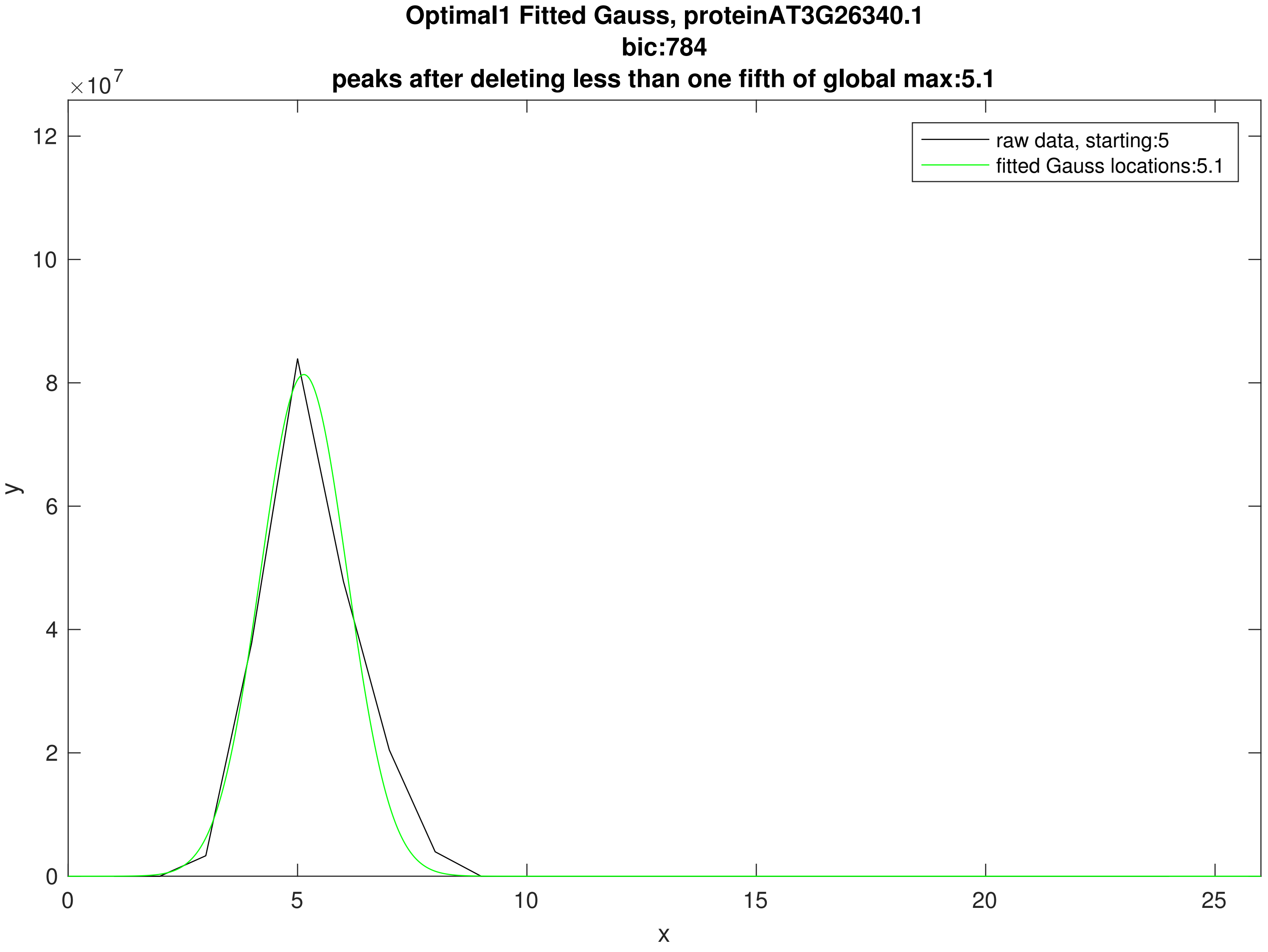

Optimal1 Fitted Gauss, proteinAT3G26340.1
bic:784
peaks after deleting less than one fifth of global max:5.1
×10^7
12
10
8
6
4
2
0
y
0
5
10
15
20
25
x
raw data, starting:5
fitted Gauss locations:5.1

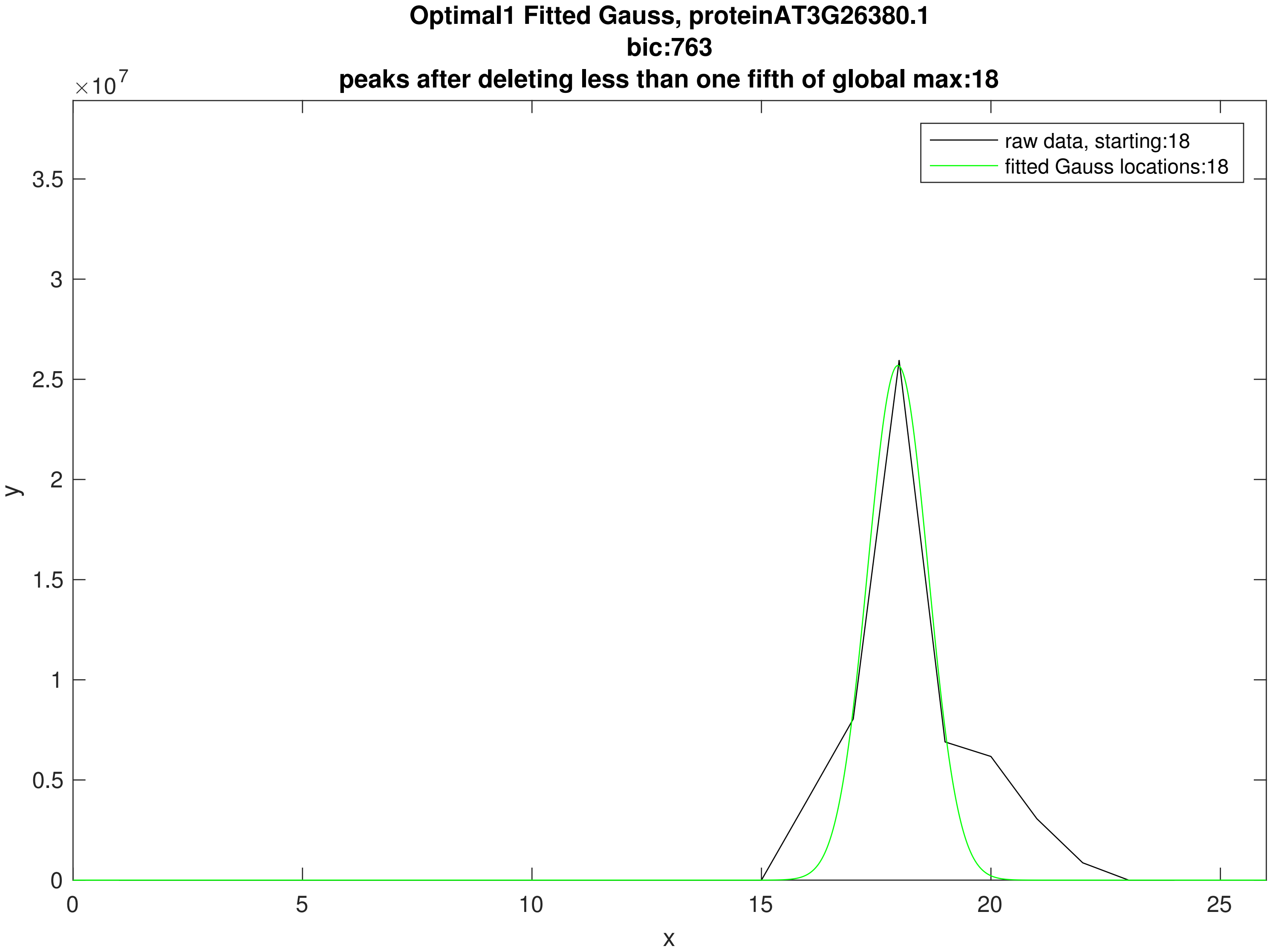
Optimal1 Fitted Gauss, proteinAT3G26380.1
bic:763
peaks after deleting less than one fifth of global max:18
×10^7
3.5
3
2.5
2
1.5
1
0.5
0
y
0
5
10
15
20
25
x
raw data, starting:18
fitted Gauss locations:18

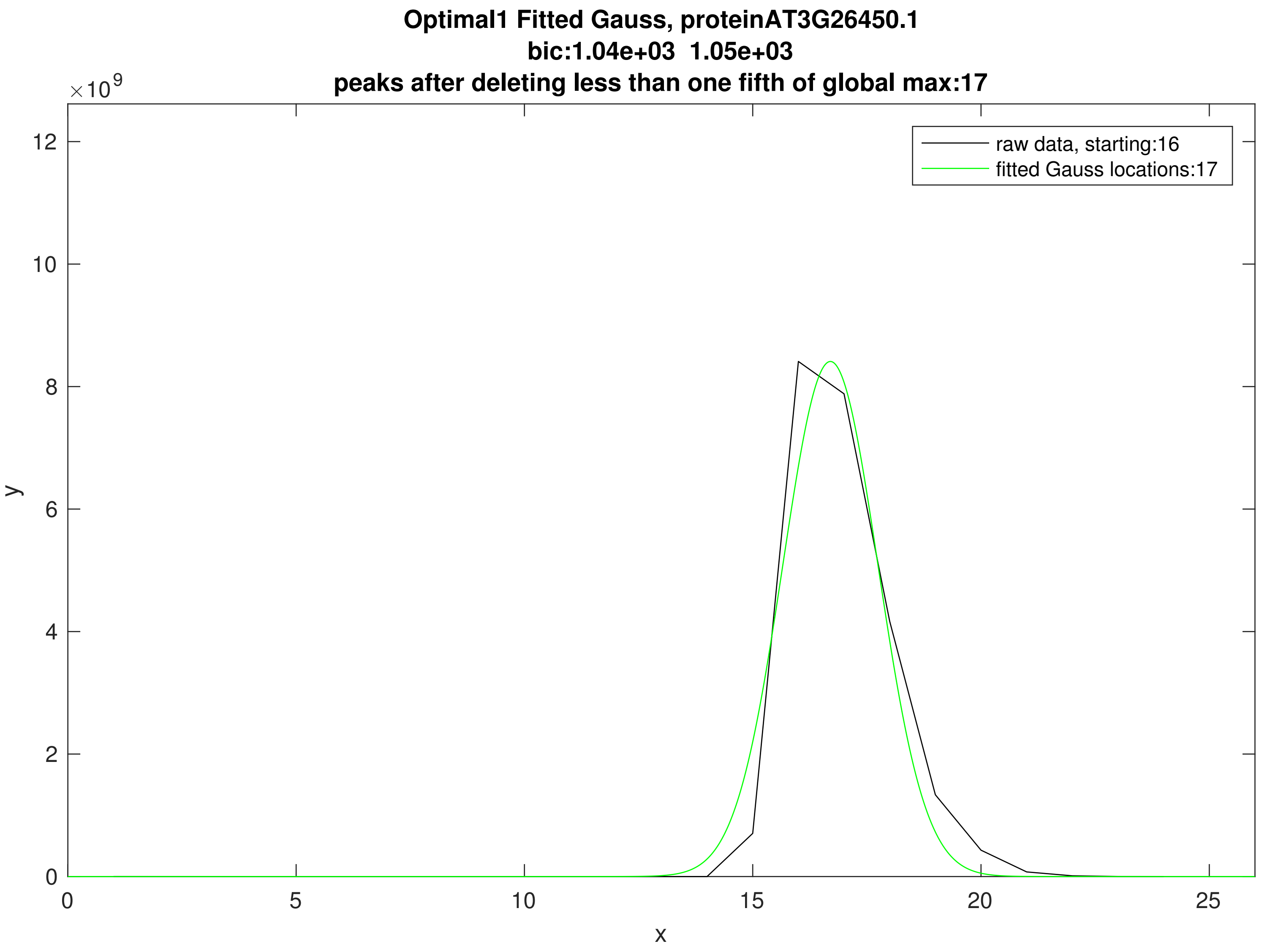
Optimal1 Fitted Gauss, proteinAT3G26450.1
bic:1.04e+03  1.05e+03
peaks after deleting less than one fifth of global max:17
×10^9
raw data, starting:16
fitted Gauss locations:17
y
x
0
2
4
6
8
10
12
0
5
10
15
20
25

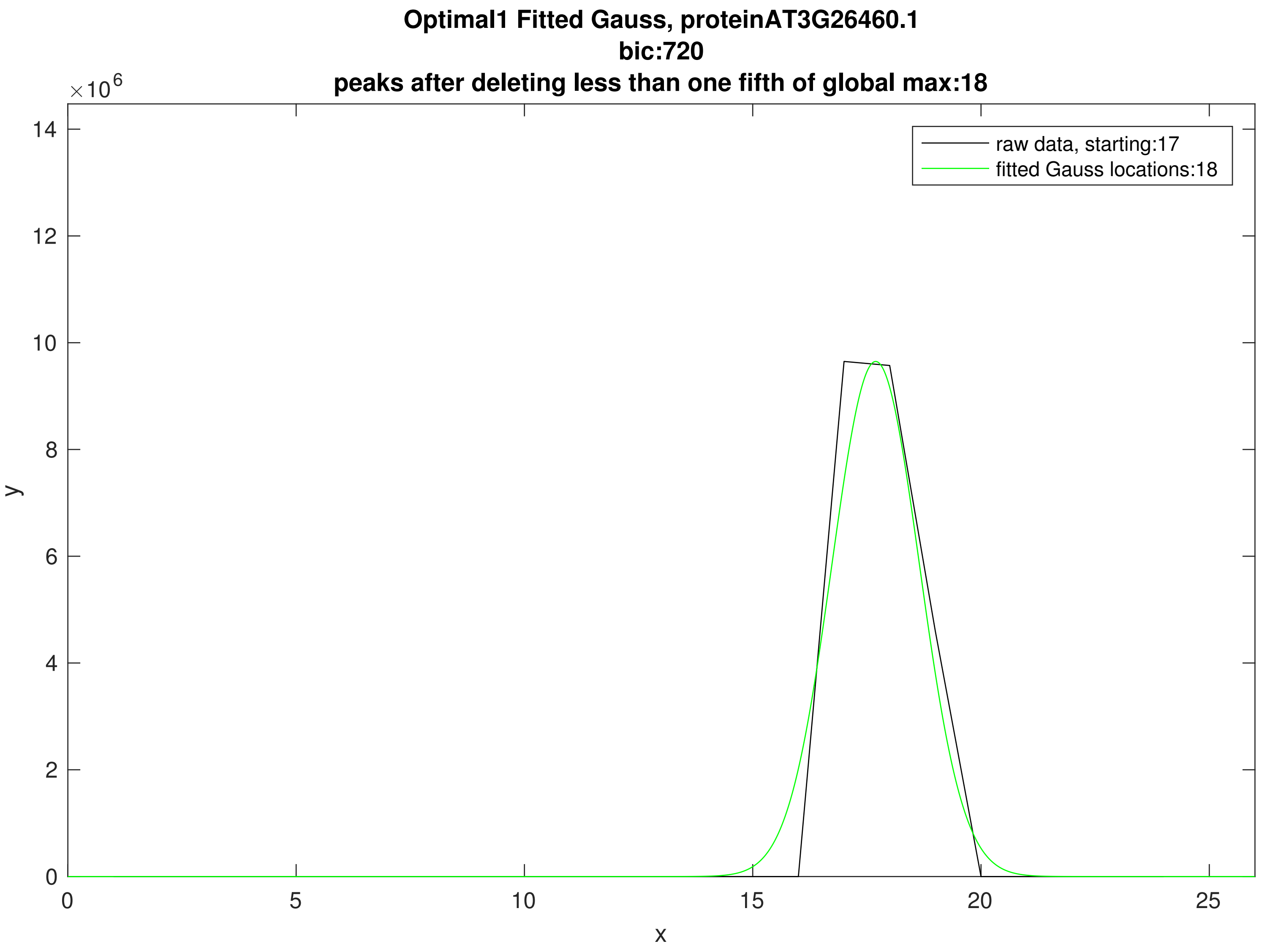
Optimal1 Fitted Gauss, proteinAT3G26460.1
bic:720
peaks after deleting less than one fifth of global max:18
×10^6
14
12
10
8
6
4
2
0
y
0
5
10
15
20
25
x
raw data, starting:17
fitted Gauss locations:18

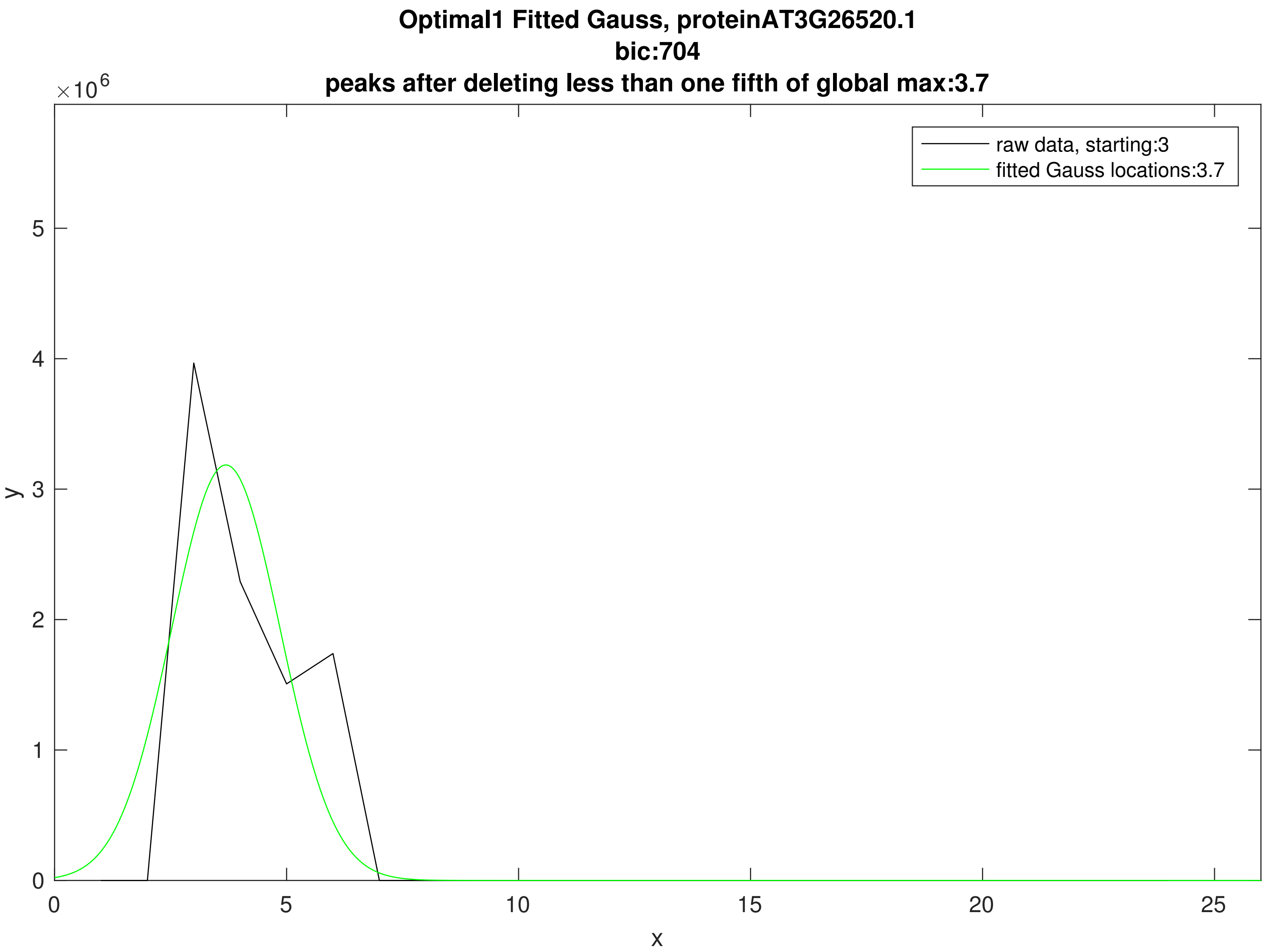
Optimal1 Fitted Gauss, proteinAT3G26520.1
bic:704
peaks after deleting less than one fifth of global max:3.7
×10^6
raw data, starting:3
fitted Gauss locations:3.7
y
x
0
1
2
3
4
5
0
5
10
15
20
25

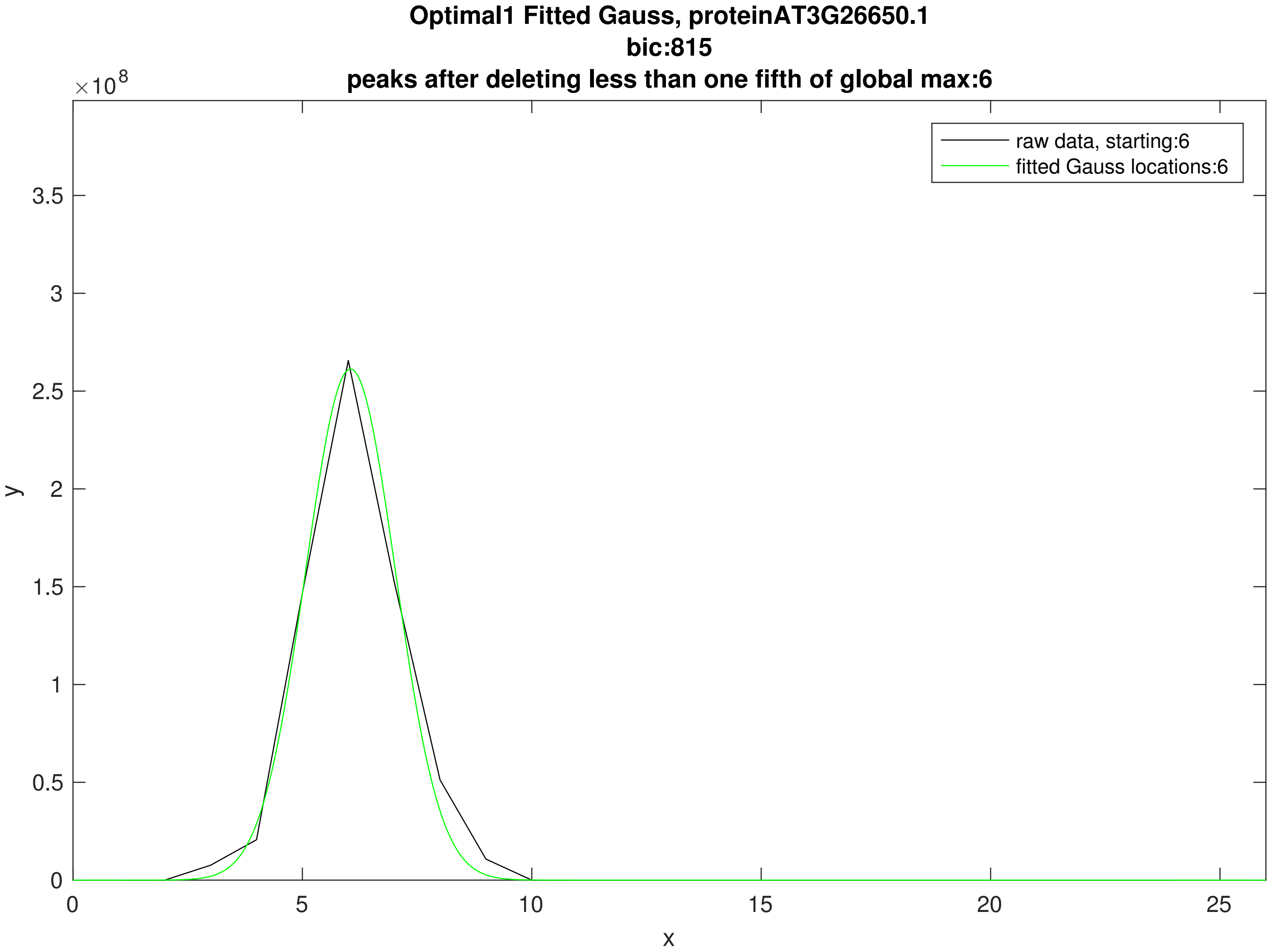

Optimal1 Fitted Gauss, proteinAT3G26650.1
bic:815
peaks after deleting less than one fifth of global max:6
×10^8
raw data, starting:6
fitted Gauss locations:6
3.5
3
2.5
2
1.5
1
0.5
0
y
0
5
10
15
20
25
x

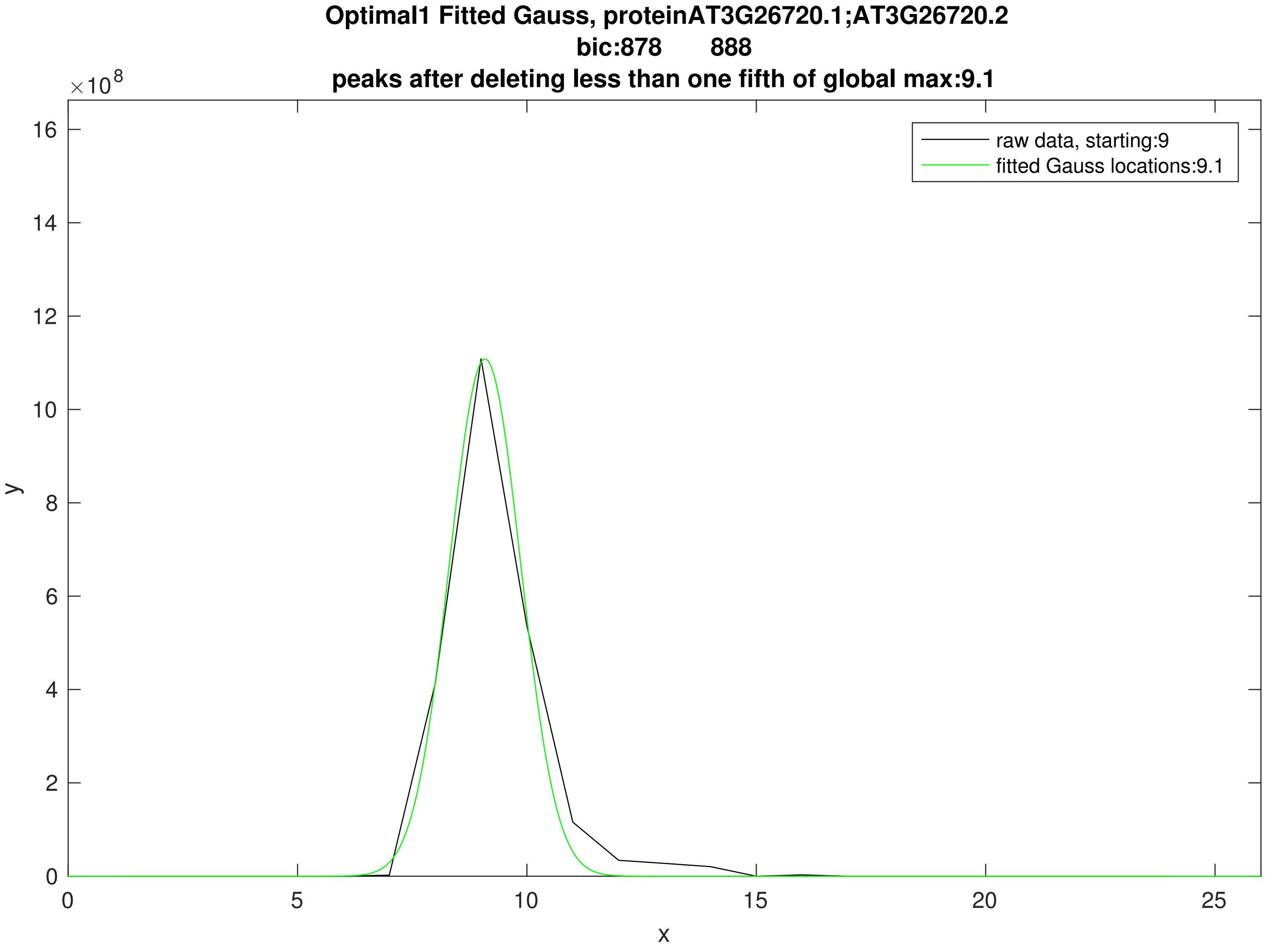
Optimal1 Fitted Gauss, proteinAT3G26720.1;AT3G26720.2
bic:878 888
peaks after deleting less than one fifth of global max:9.1
×10^8
16
14
12
10
8
6
4
2
0
y
0
5
10
15
20
25
x
raw data, starting:9
fitted Gauss locations:9.1

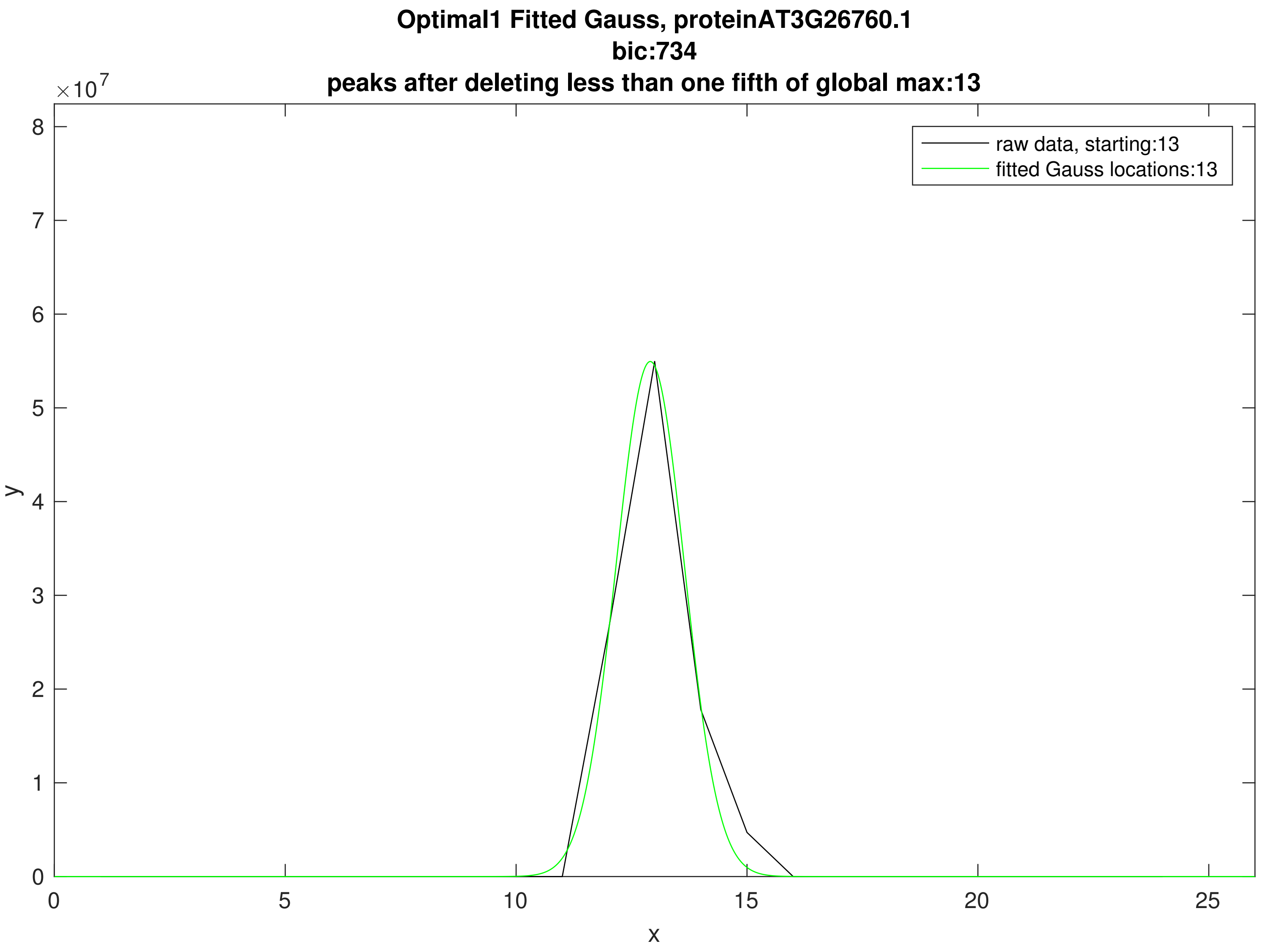
Optimal1 Fitted Gauss, proteinAT3G26760.1
bic:734
peaks after deleting less than one fifth of global max:13
×10^7
8
7
6
5
4
3
2
1
0
y
0
5
10
15
20
25
x
raw data, starting:13
fitted Gauss locations:13

Optimal1 Fitted Gauss, proteinAT3G26770.1
bic:837
peaks after deleting less than one fifth of global max:12

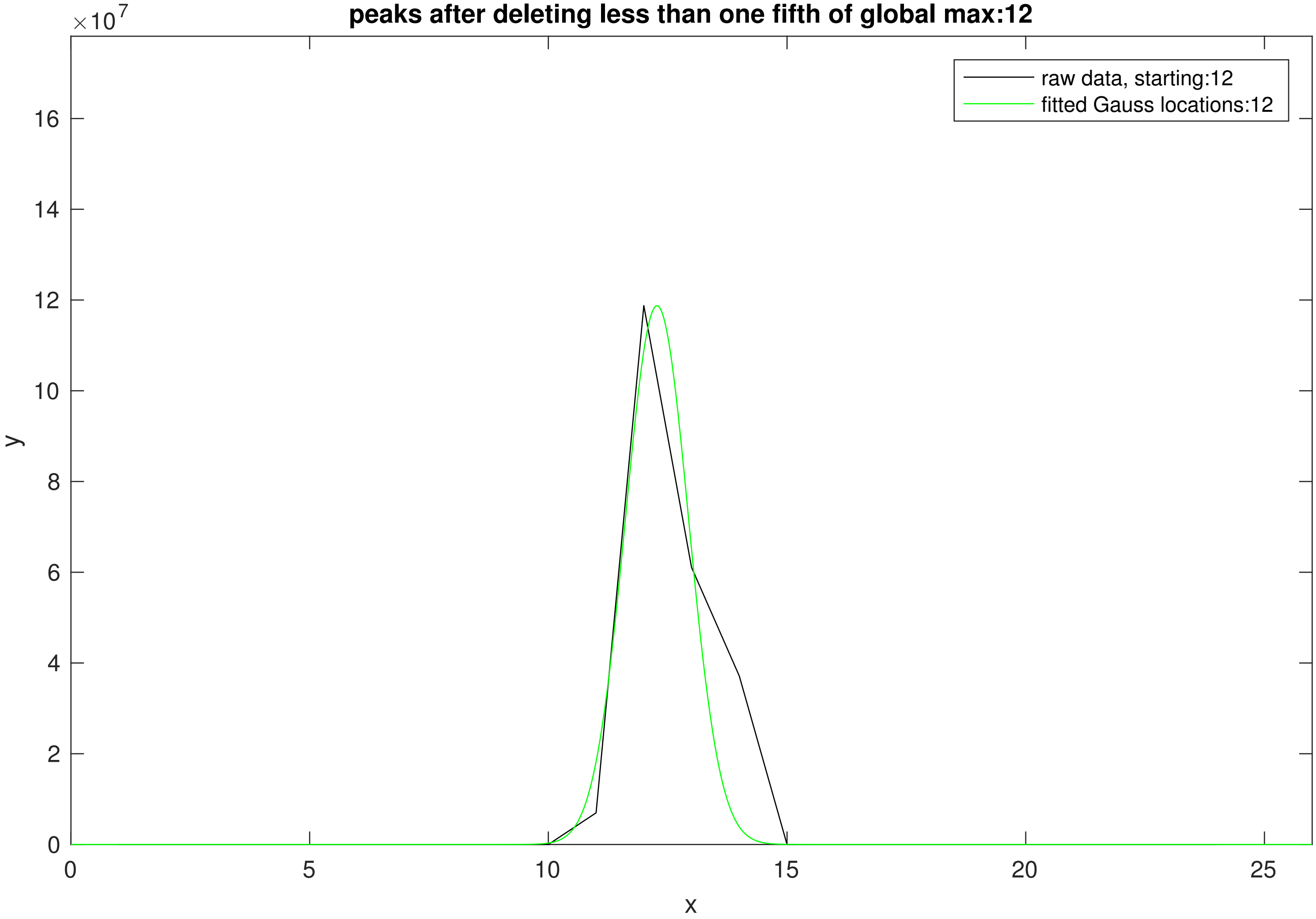

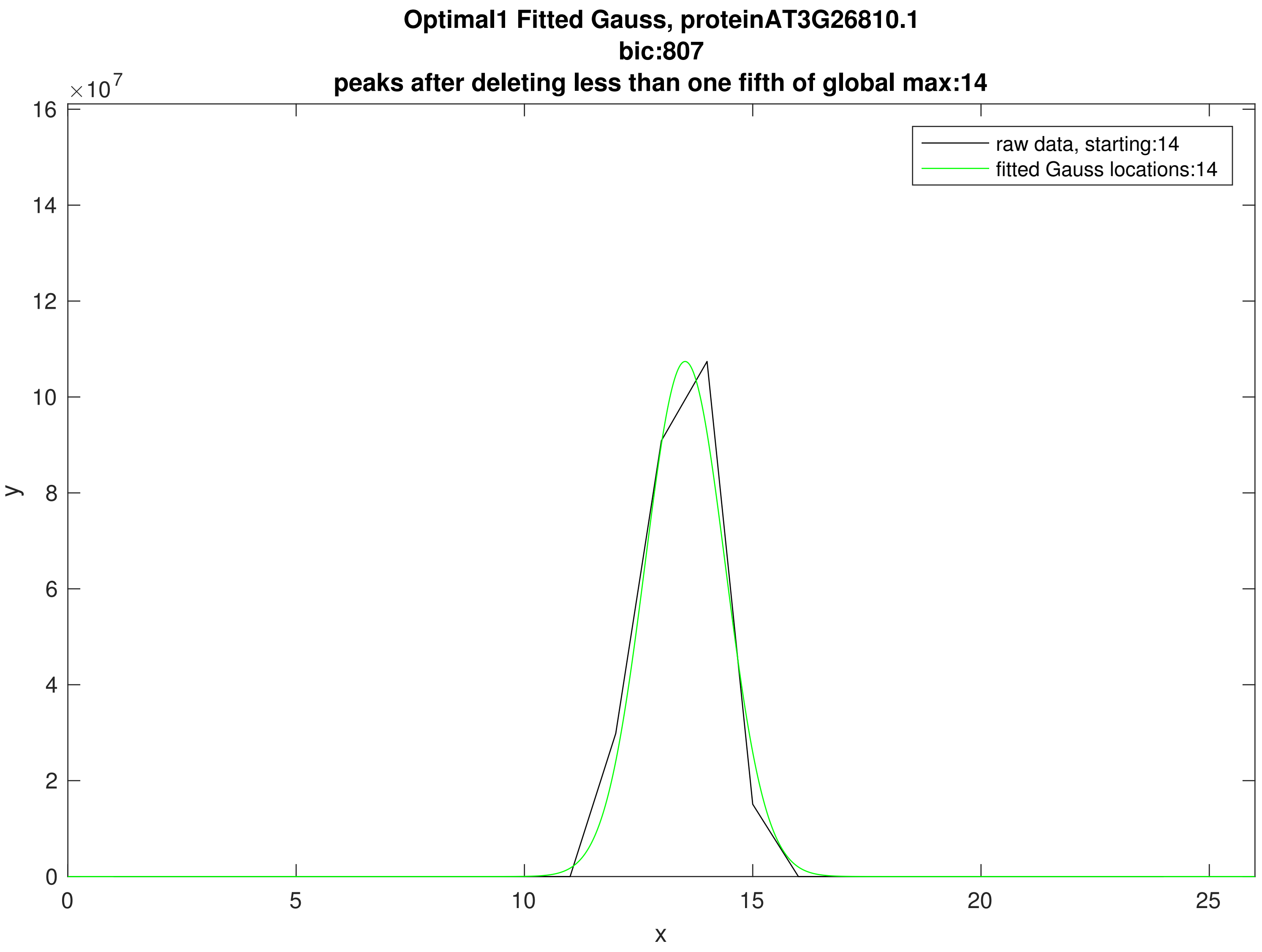
Optimal1 Fitted Gauss, proteinAT3G26810.1
bic:807
peaks after deleting less than one fifth of global max:14
×10^7
raw data, starting:14
fitted Gauss locations:14
y
x
0
2
4
6
8
10
12
14
16
0
5
10
15
20
25

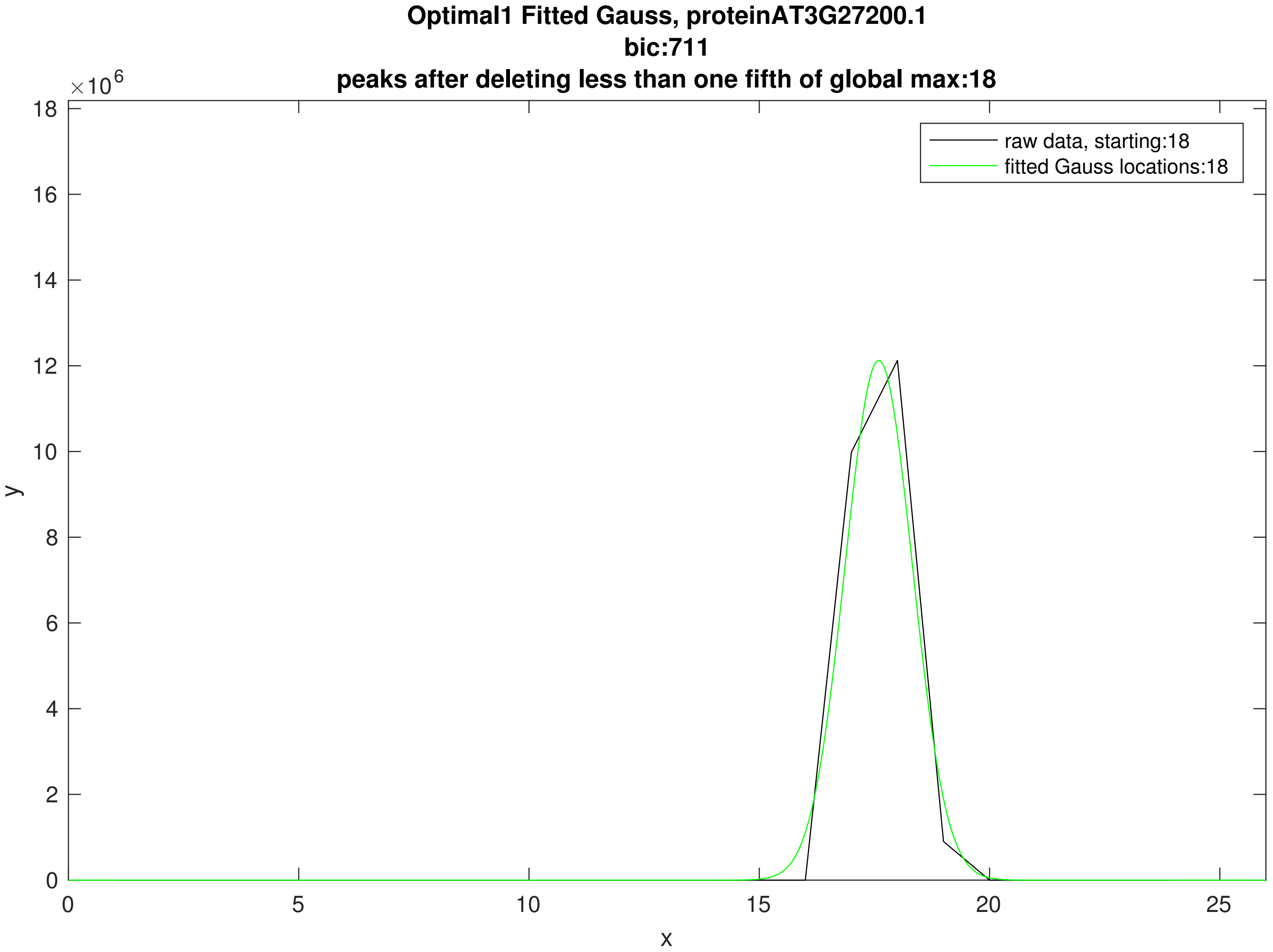
Optimal1 Fitted Gauss, proteinAT3G27200.1
bic:711
peaks after deleting less than one fifth of global max:18
raw data, starting:18
fitted Gauss locations:18
$\times 10^6$
y
x
0
2
4
6
8
10
12
14
16
18
0
5
10
15
20
25

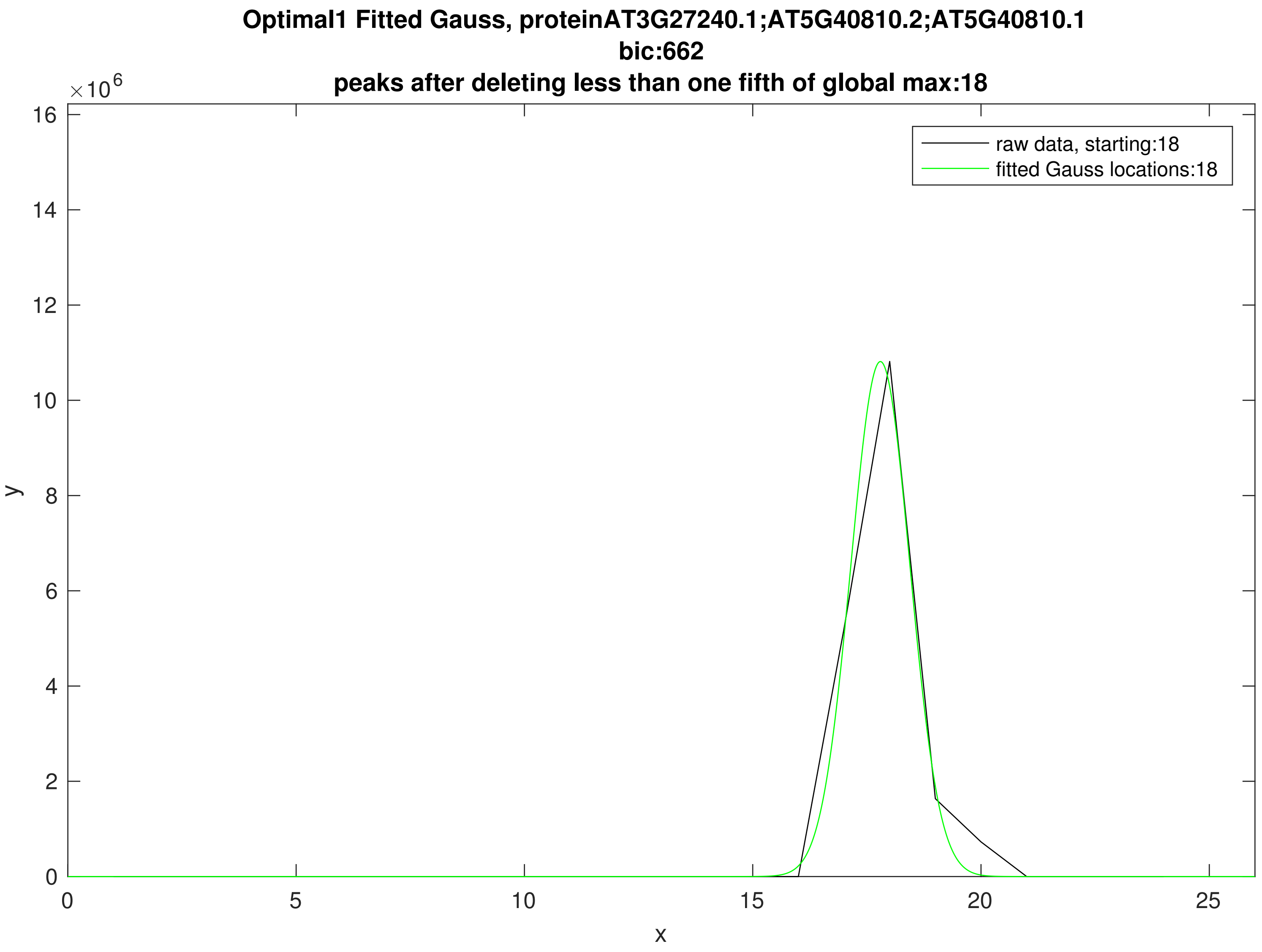
Optimal1 Fitted Gauss, proteinAT3G27240.1;AT5G40810.2;AT5G40810.1
bic:662
peaks after deleting less than one fifth of global max:18
$\times 10^6$
16
14
12
10
8
6
4
2
0
y
0
5
10
15
20
25
x
raw data, starting:18
fitted Gauss locations:18

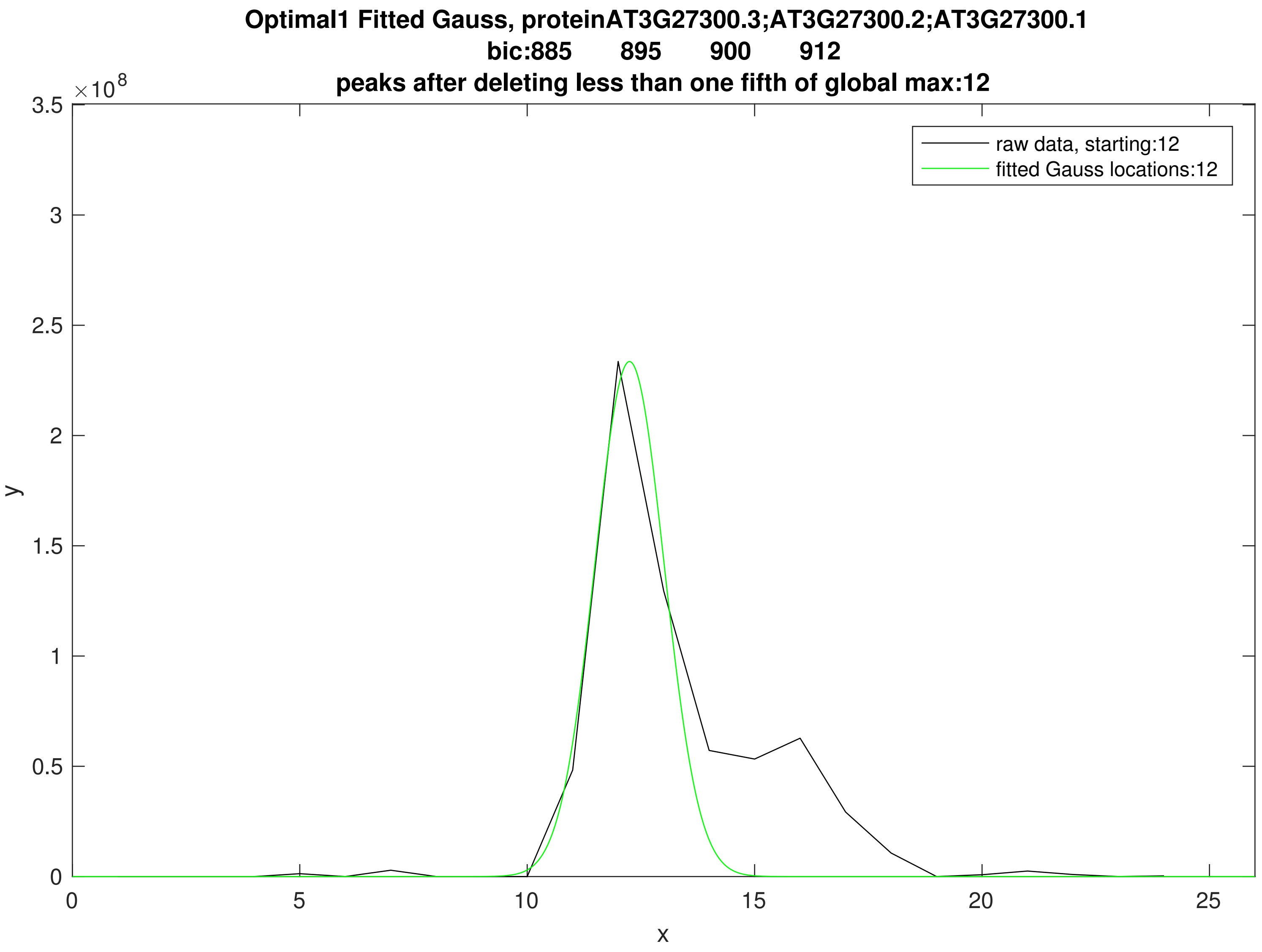
Optimal1 Fitted Gauss, proteinAT3G27300.3;AT3G27300.2;AT3G27300.1
bic:885 895 900 912
peaks after deleting less than one fifth of global max:12
×10^8
3.5
3
2.5
2
1.5
1
0.5
0
y
0
5
10
15
20
25
x
raw data, starting:12
fitted Gauss locations:12

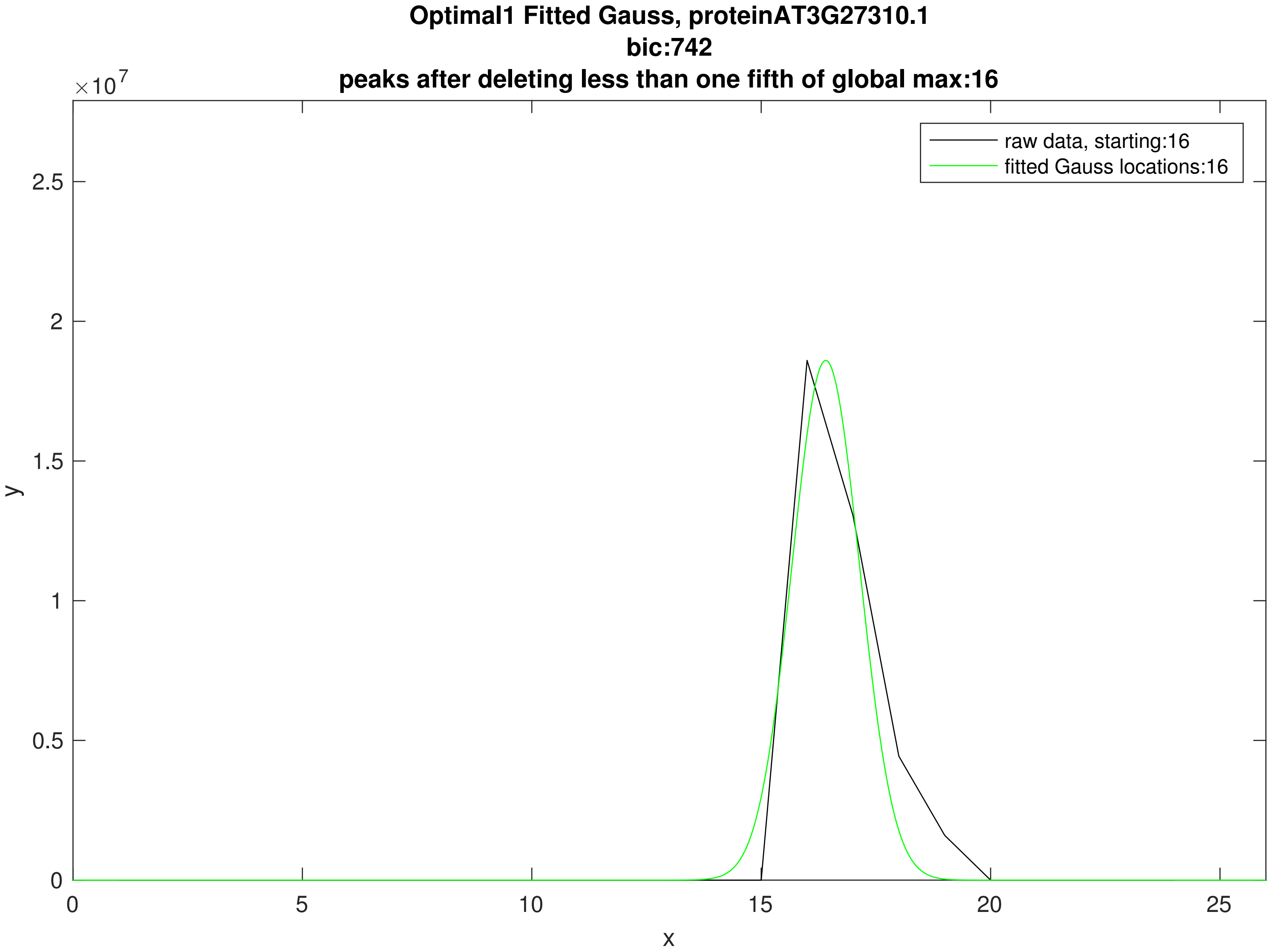
Optimal1 Fitted Gauss, proteinAT3G27310.1
bic:742
peaks after deleting less than one fifth of global max:16
×10^7
raw data, starting:16
fitted Gauss locations:16
2.5
2
1.5
1
0.5
0
y
0
5
10
15
20
25
x

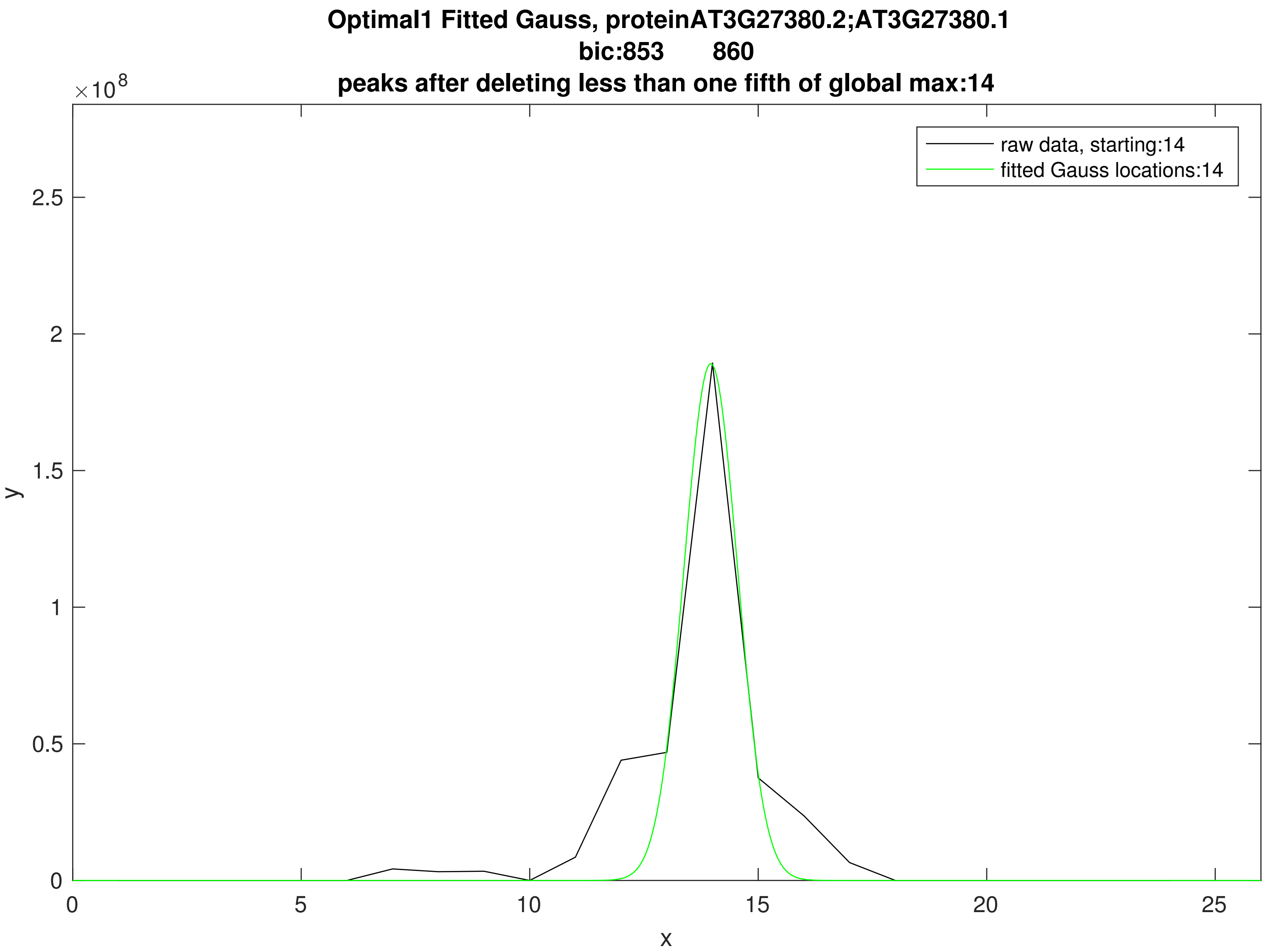

Optimal1 Fitted Gauss, proteinAT3G27380.2;AT3G27380.1
bic:853 860
peaks after deleting less than one fifth of global max:14
×10^8
raw data, starting:14
fitted Gauss locations:14
y
x

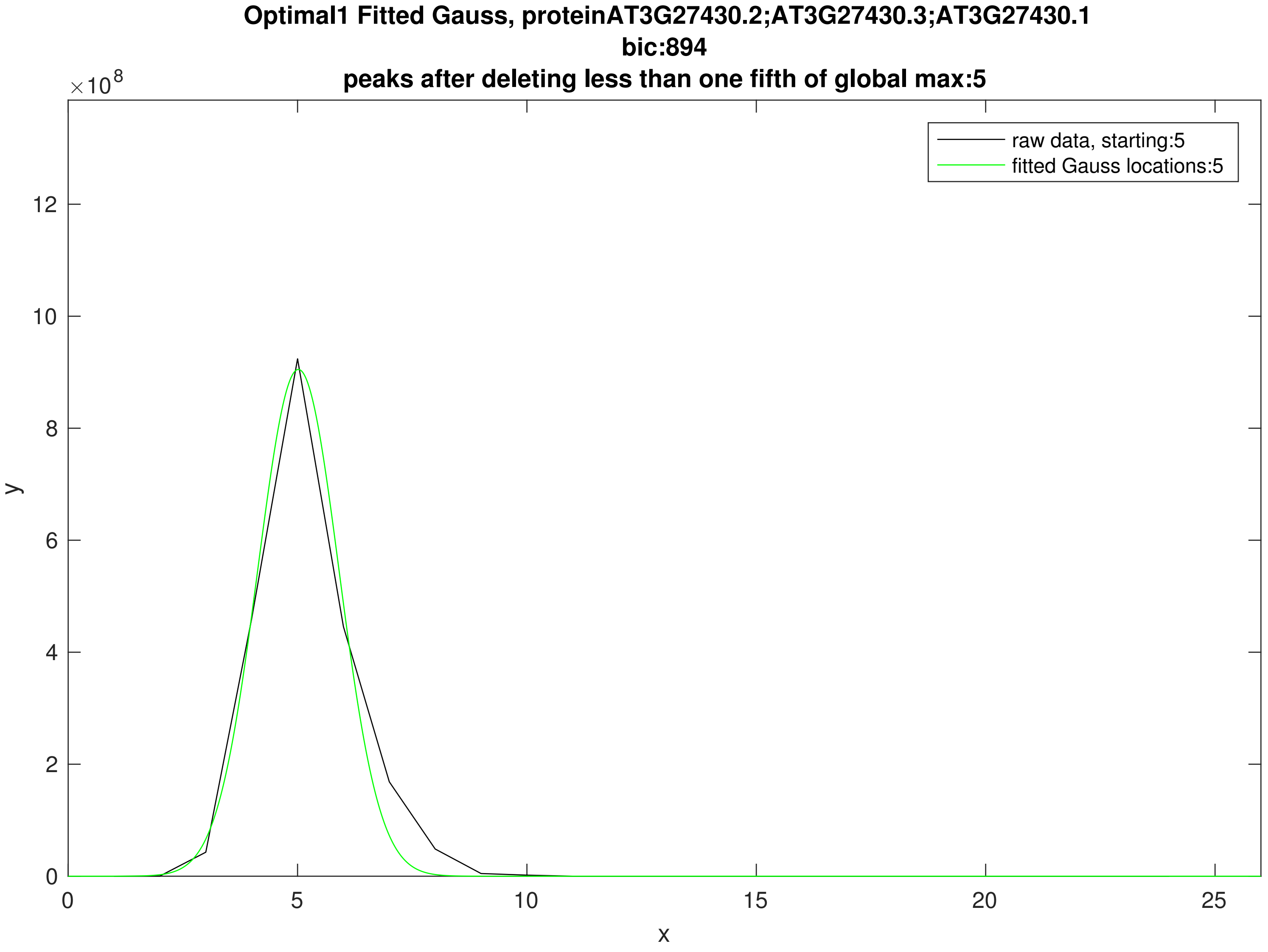

Optimal1 Fitted Gauss, proteinAT3G27430.2;AT3G27430.3;AT3G27430.1
bic:894
peaks after deleting less than one fifth of global max:5
×10^8
raw data, starting:5
fitted Gauss locations:5
y
x
0
2
4
6
8
10
12
5
15
20
25

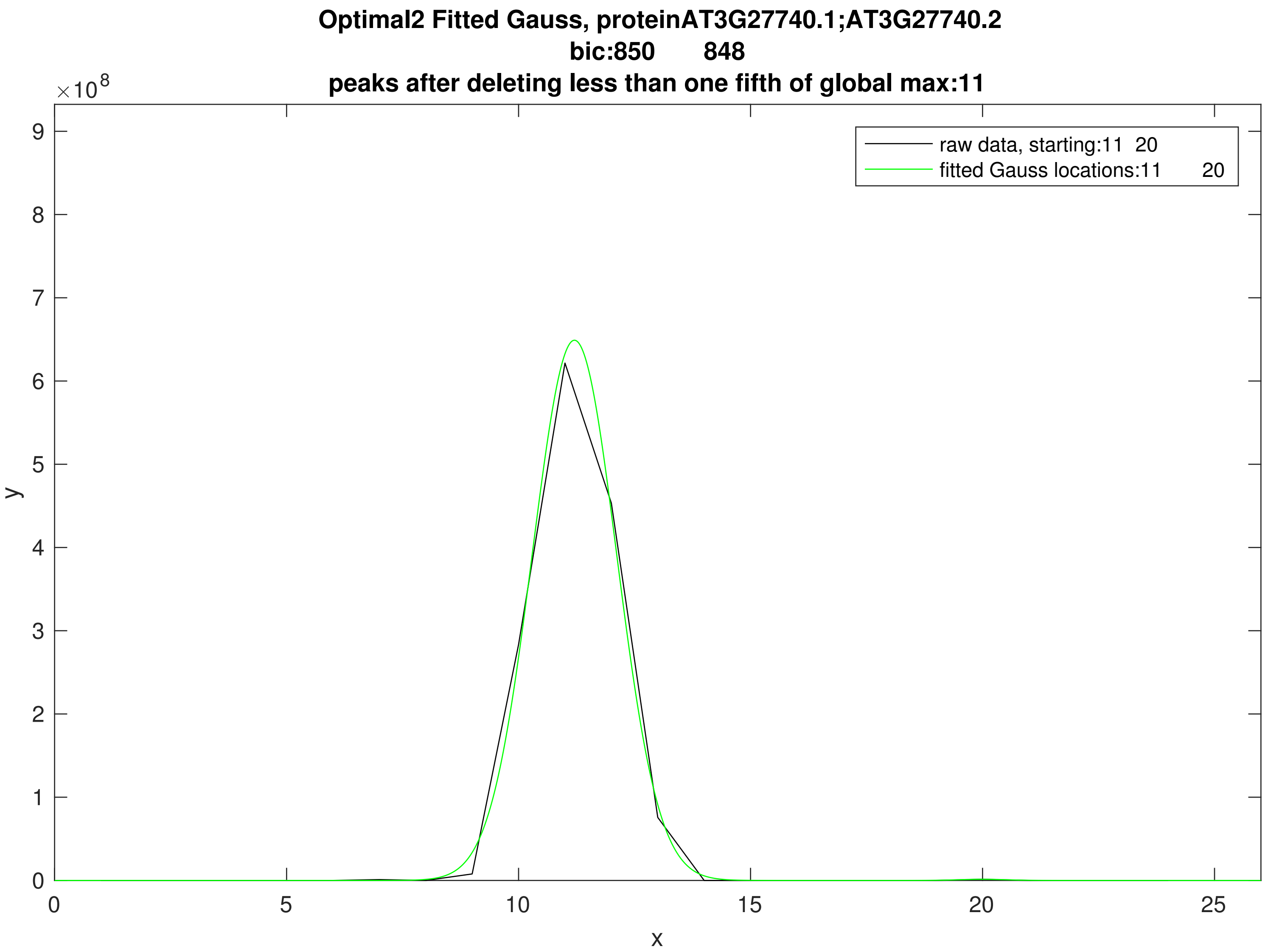
Optimal2 Fitted Gauss, proteinAT3G27740.1;AT3G27740.2
bic:850 848
peaks after deleting less than one fifth of global max:11
×10^8
raw data, starting:11 20
fitted Gauss locations:11 20
y
x
0
1
2
3
4
5
6
7
8
9
0
5
10
15
20
25

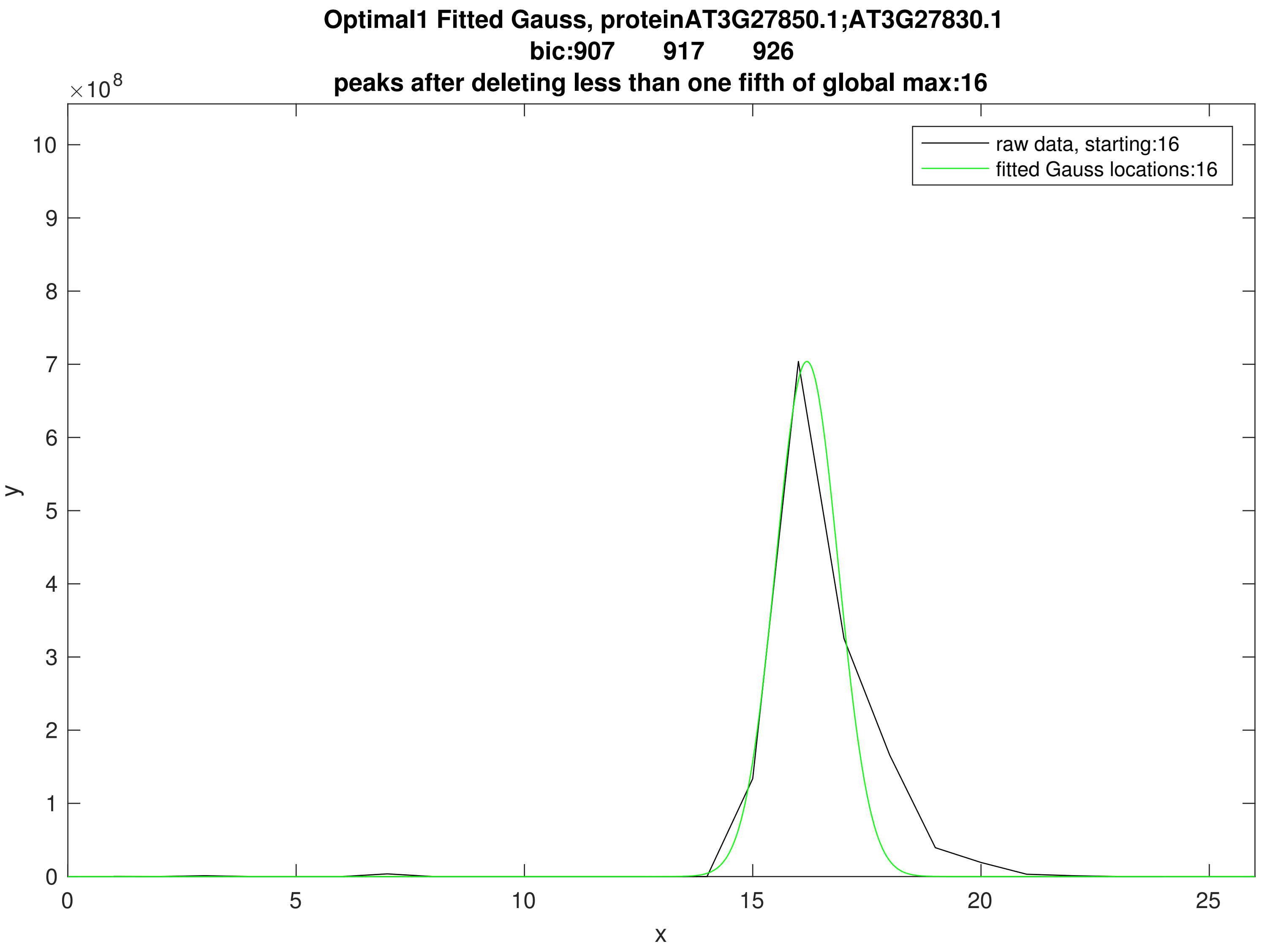

Optimal1 Fitted Gauss, proteinAT3G27850.1;AT3G27830.1
bic:907 917 926
peaks after deleting less than one fifth of global max:16
×10^8
raw data, starting:16
fitted Gauss locations:16
y
x
0
1
2
3
4
5
6
7
8
9
10
0
5
10
15
20
25

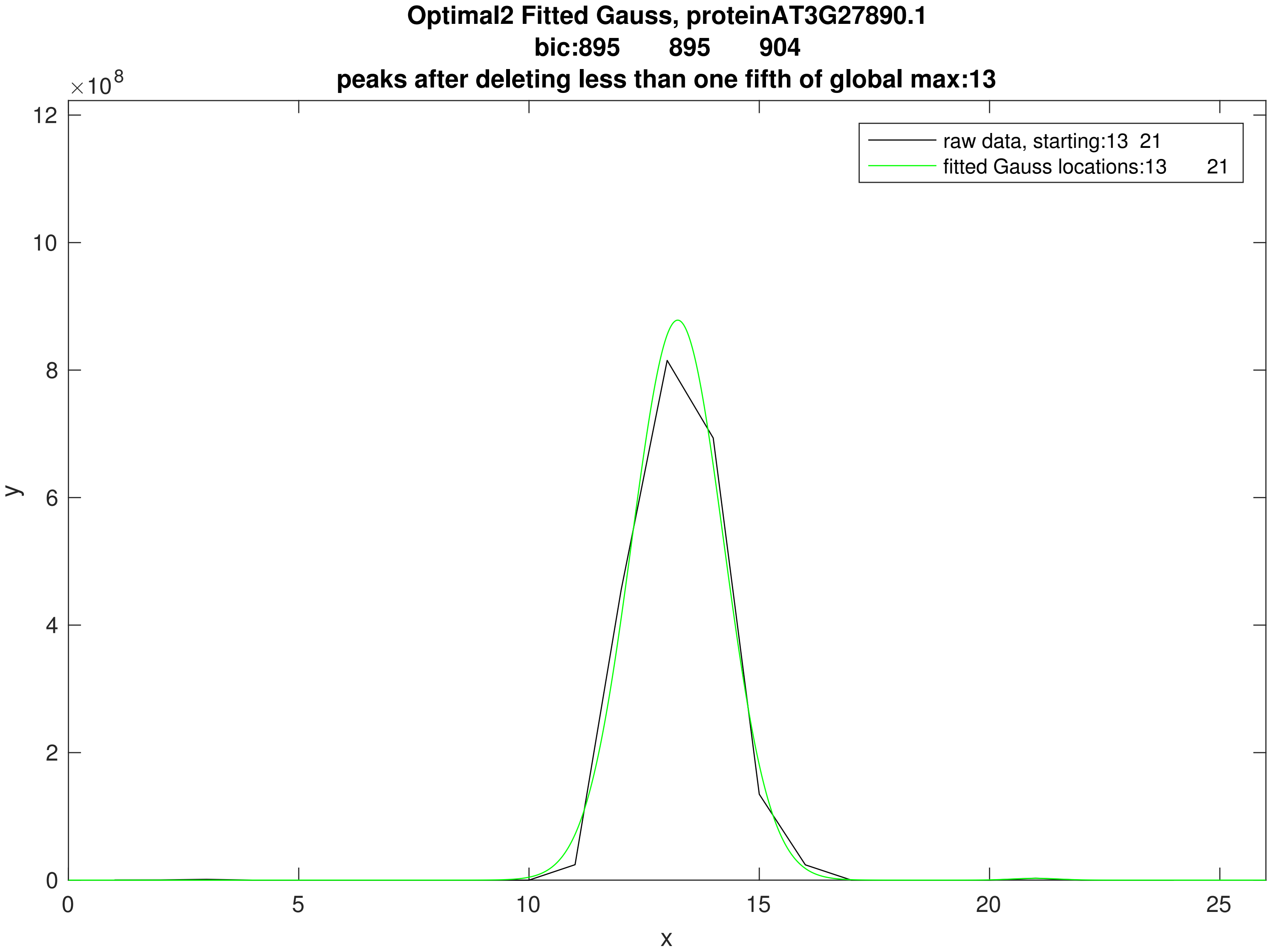
Optimal2 Fitted Gauss, proteinAT3G27890.1
bic:895 895 904
peaks after deleting less than one fifth of global max:13
×10^8
raw data, starting:13 21
fitted Gauss locations:13 21
y
x
0
2
4
6
8
10
12
0
5
10
15
20
25

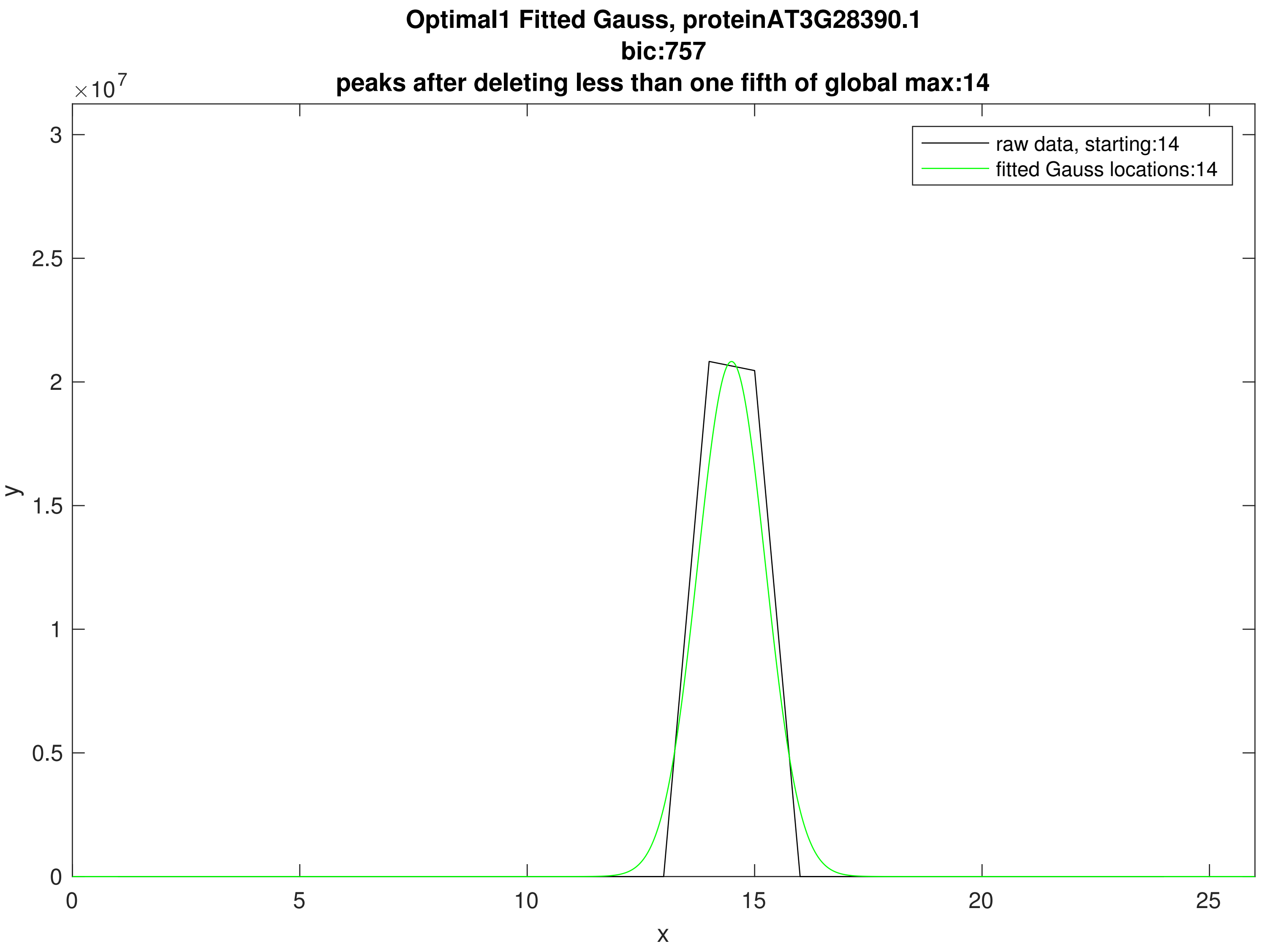

Optimal1 Fitted Gauss, proteinAT3G28390.1
bic:757
peaks after deleting less than one fifth of global max:14
×10^7
3
2.5
2
1.5
1
0.5
0
y
0
5
10
15
20
25
x
raw data, starting:14
fitted Gauss locations:14

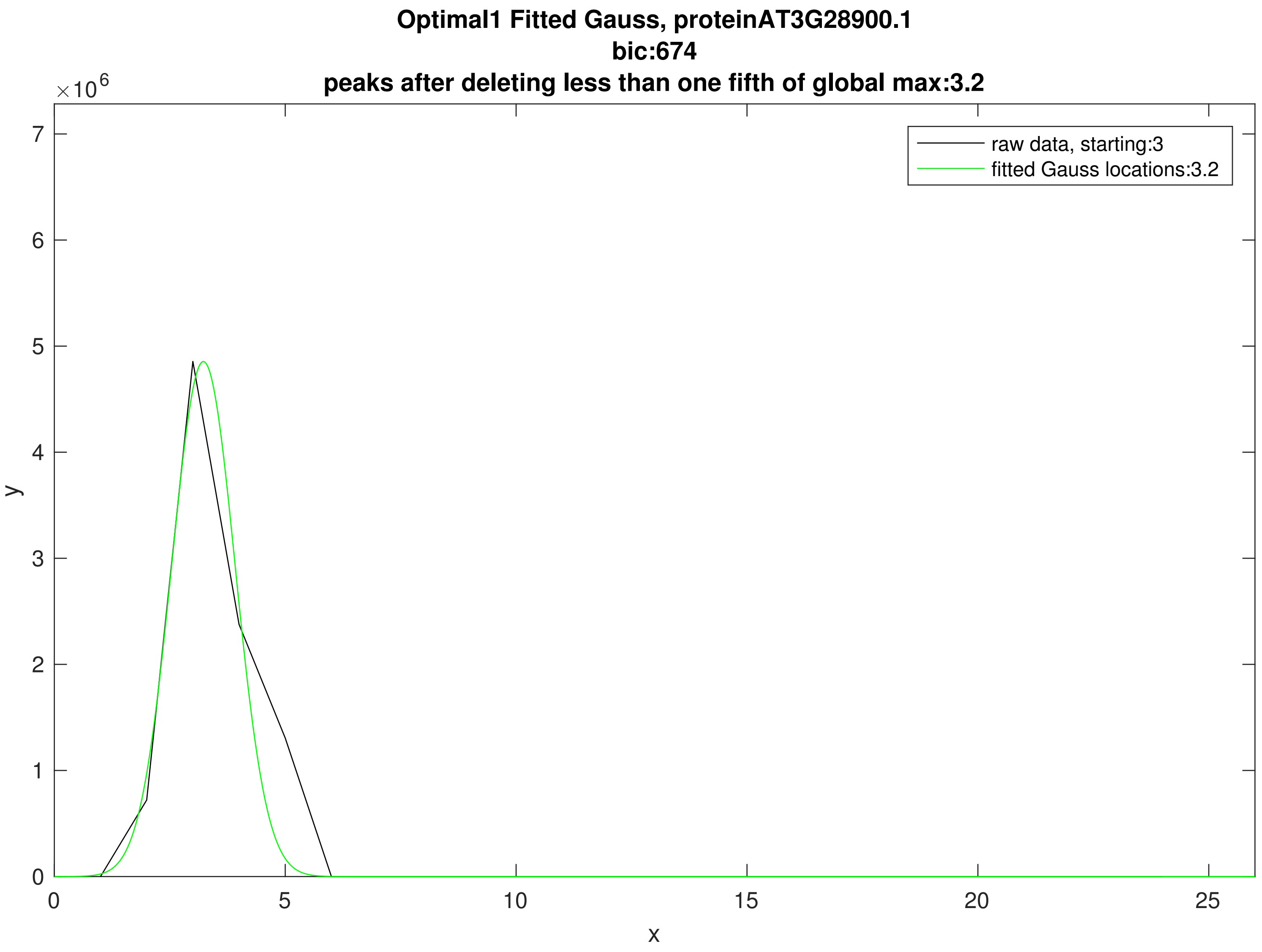

Optimal1 Fitted Gauss, proteinAT3G28900.1
bic:674
peaks after deleting less than one fifth of global max:3.2
×10^6
raw data, starting:3
fitted Gauss locations:3.2
y
x
0
1
2
3
4
5
6
7
0
5
10
15
20
25

Optimal1 Fitted Gauss, proteinAT3G28930.1
bic:755 760
peaks after deleting less than one fifth of global max:18

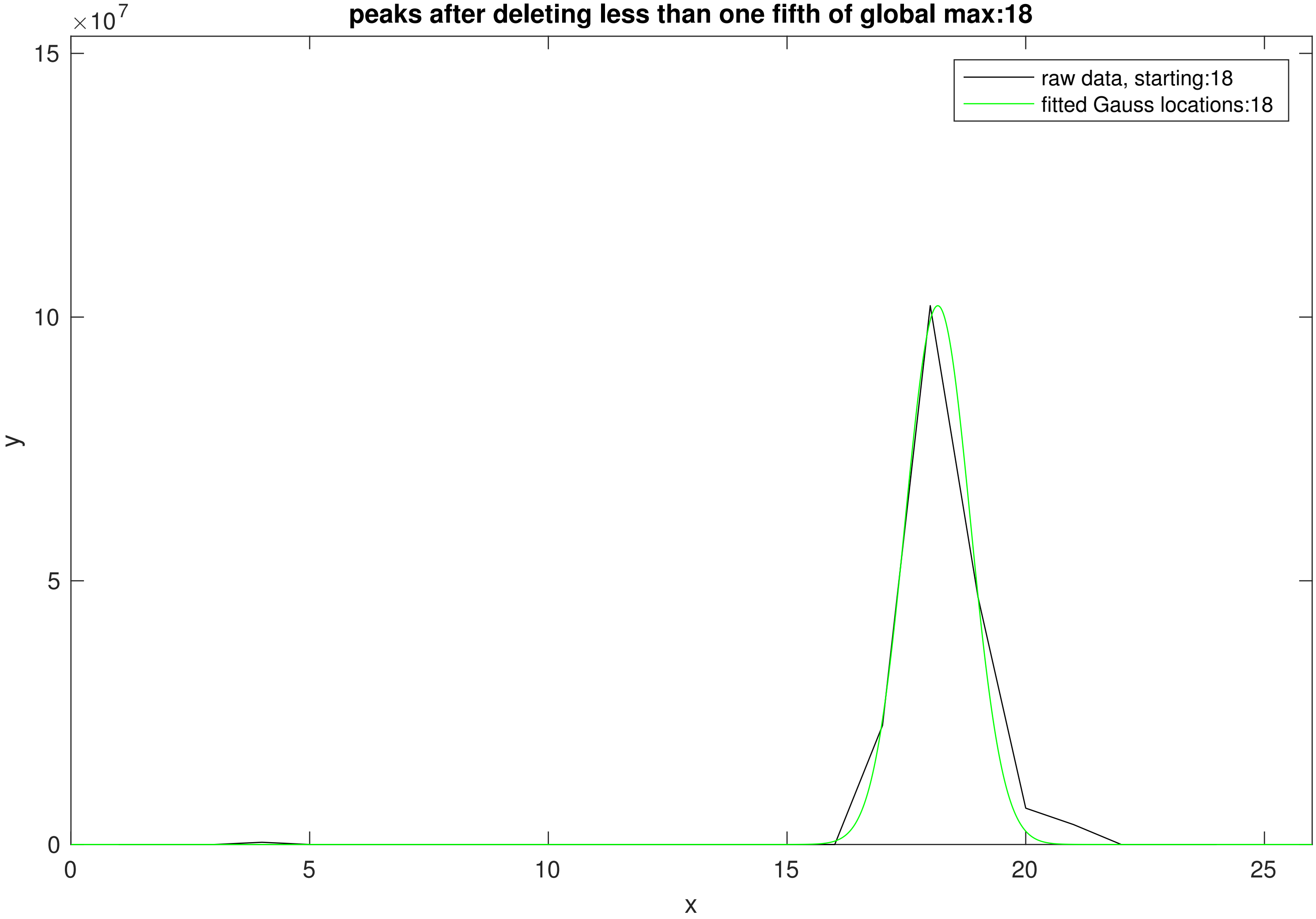

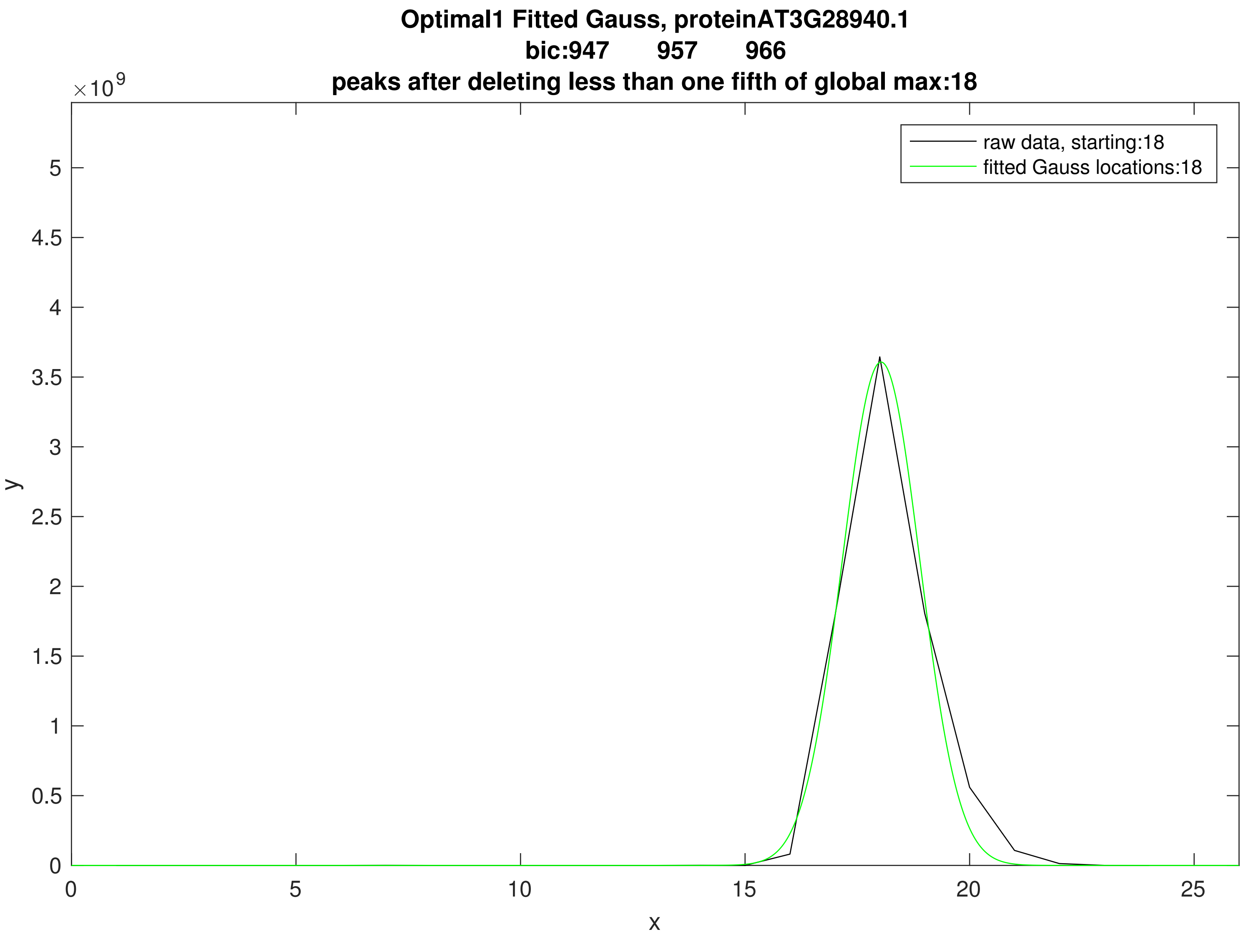
Optimal1 Fitted Gauss, proteinAT3G28940.1
bic:947 957 966
peaks after deleting less than one fifth of global max:18
×10^9
raw data, starting:18
fitted Gauss locations:18
y
x
0
0.5
1
1.5
2
2.5
3
3.5
4
4.5
5
0
5
10
15
20
25

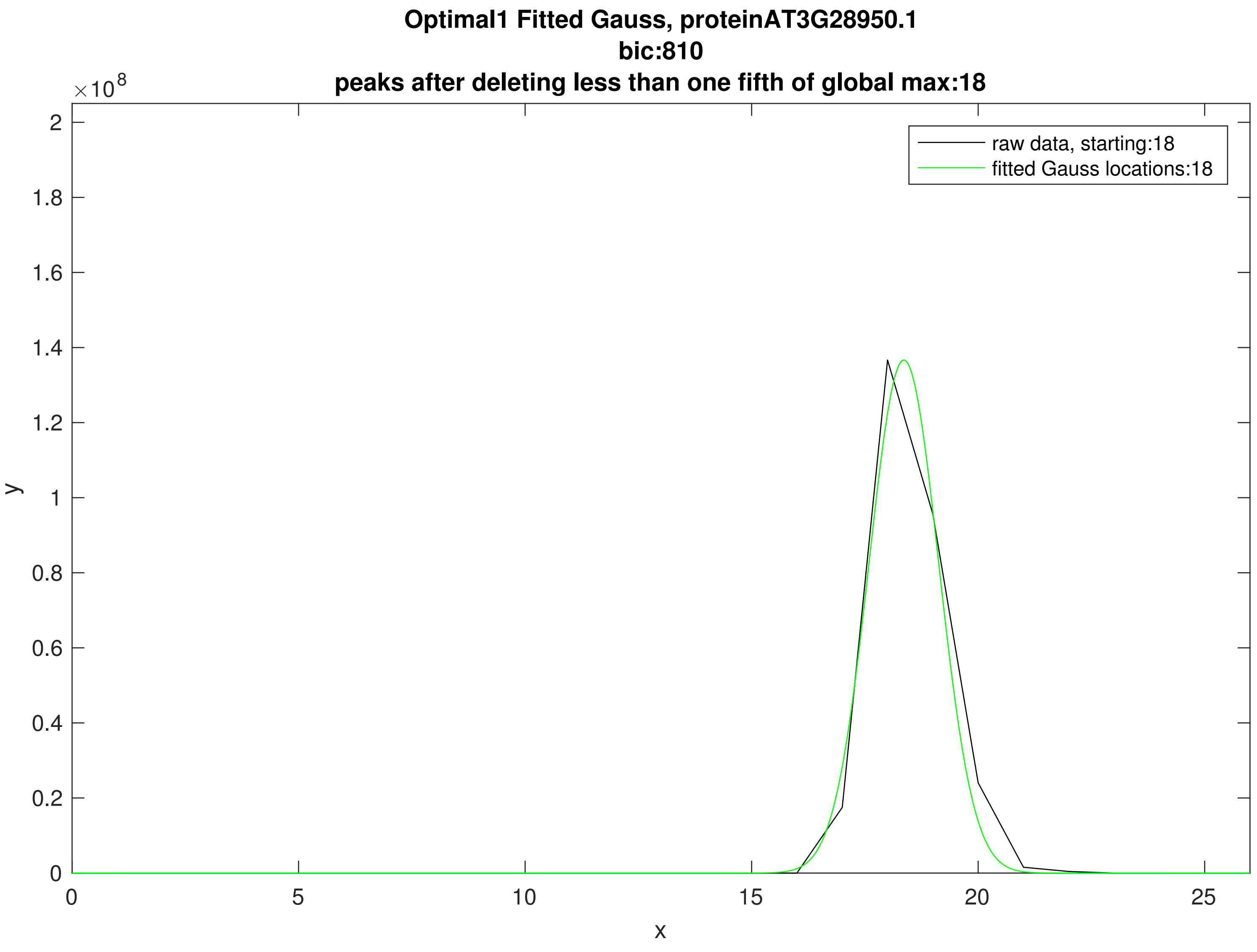
Optimal1 Fitted Gauss, proteinAT3G28950.1
bic:810
peaks after deleting less than one fifth of global max:18
×10^8
2
1.8
1.6
1.4
1.2
1
0.8
0.6
0.4
0.2
0
y
0
5
10
15
20
25
x
raw data, starting:18
fitted Gauss locations:18

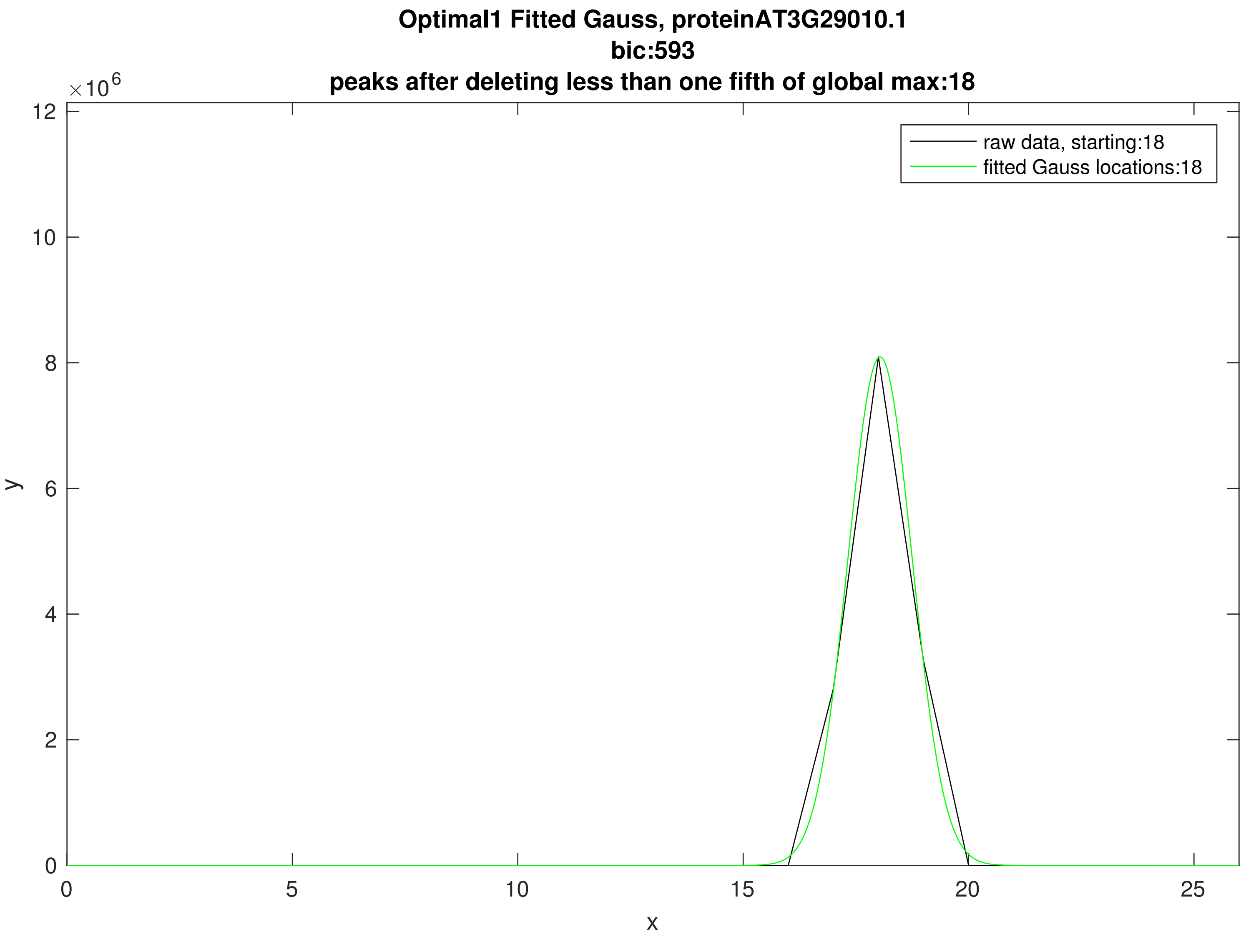
Optimal1 Fitted Gauss, proteinAT3G29010.1
bic:593
peaks after deleting less than one fifth of global max:18
×10^6
12
10
8
6
4
2
0
y
0
5
10
15
20
25
x
raw data, starting:18
fitted Gauss locations:18

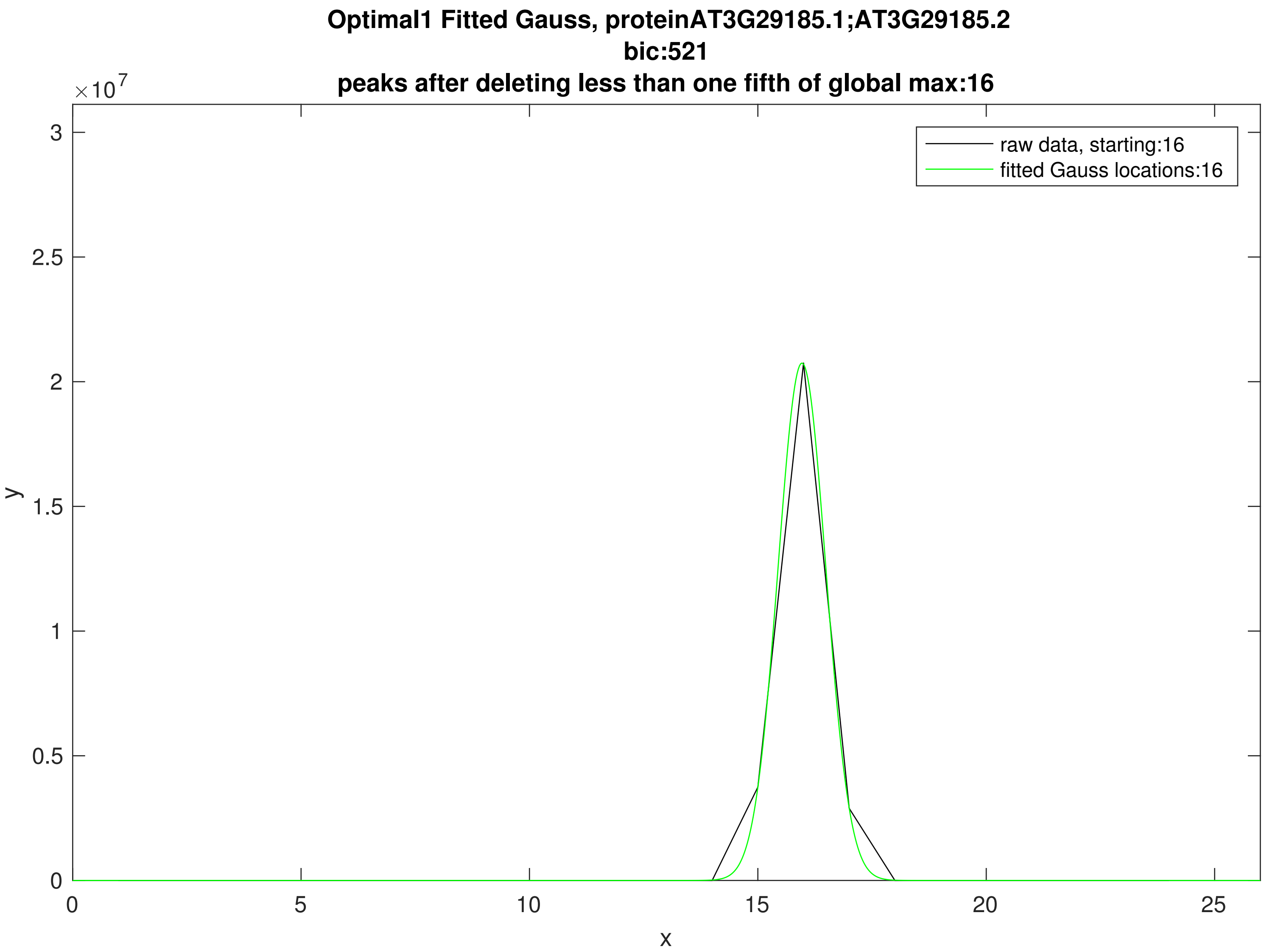
Optimal1 Fitted Gauss, proteinAT3G29185.1;AT3G29185.2
bic:521
peaks after deleting less than one fifth of global max:16
×10^7
raw data, starting:16
fitted Gauss locations:16
y
x
0
0.5
1
1.5
2
2.5
3
0
5
10
15
20
25

**Optimal1 Fitted Gauss, proteinAT3G29250.2;AT3G29250.1**
**bic:831 840 846**
**peaks after deleting less than one fifth of global max:14**

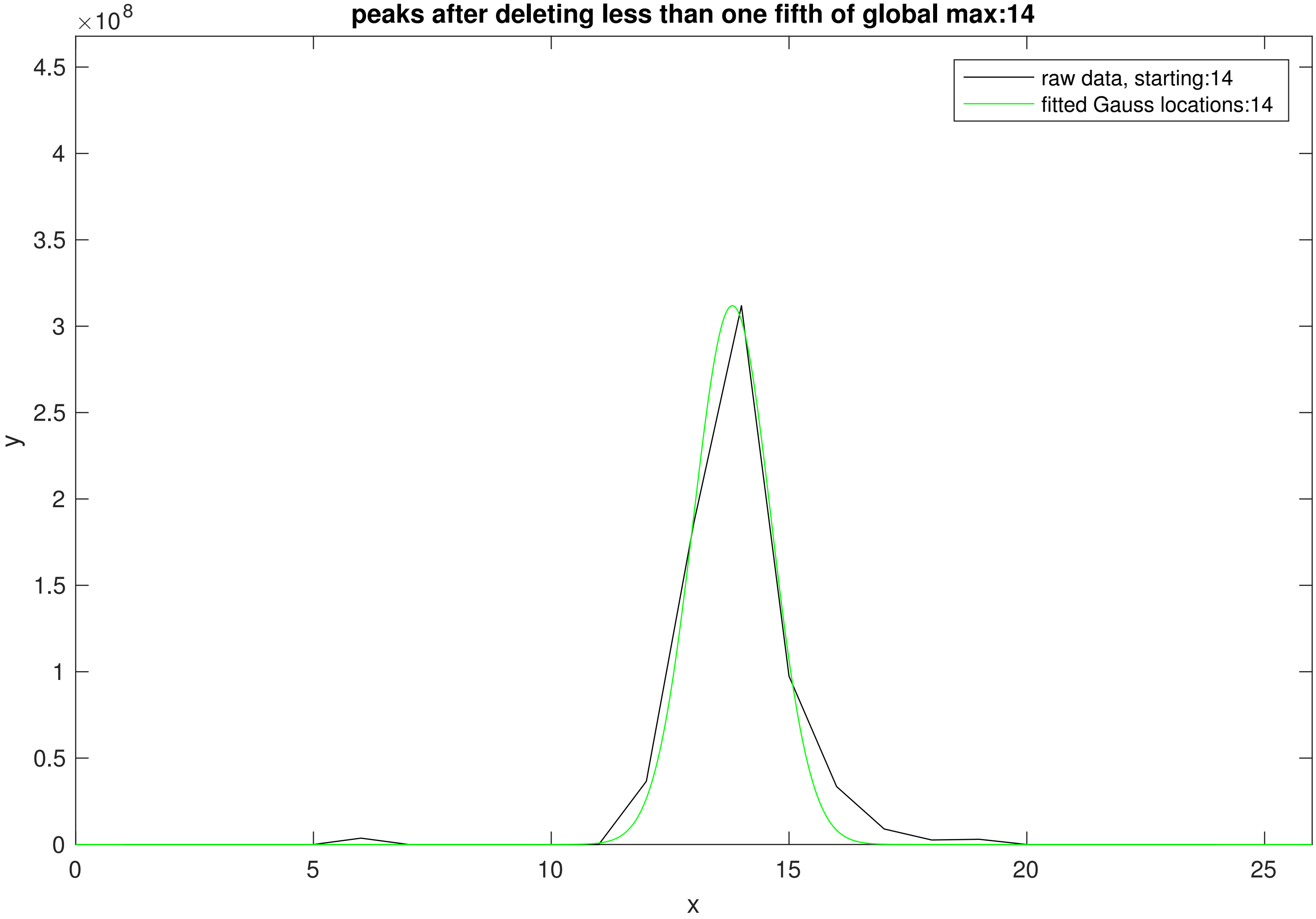

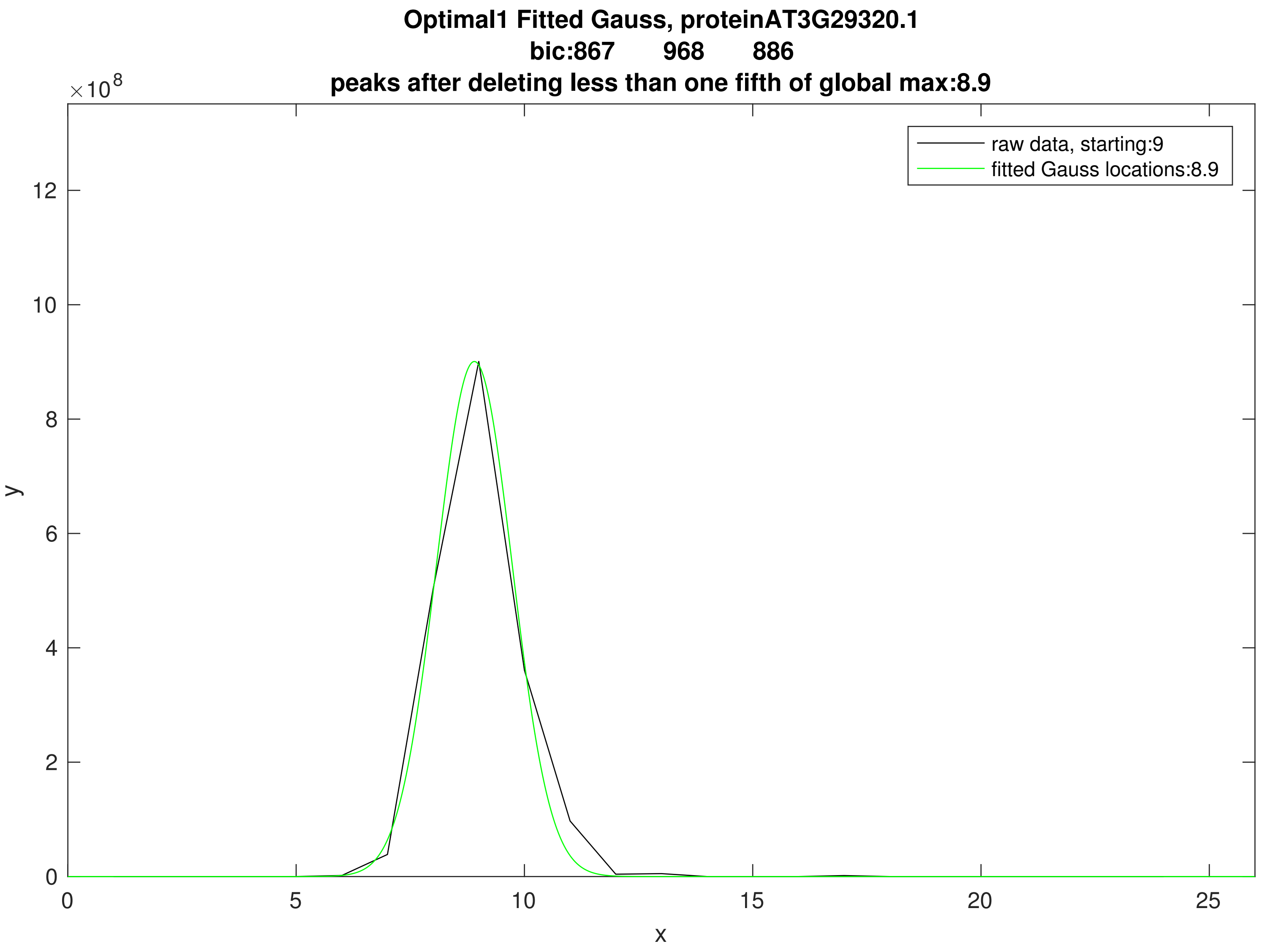
Optimal1 Fitted Gauss, proteinAT3G29320.1
bic:867 968 886
peaks after deleting less than one fifth of global max:8.9
×10^8
raw data, starting:9
fitted Gauss locations:8.9
y
x
0
2
4
6
8
10
12
0
5
10
15
20
25

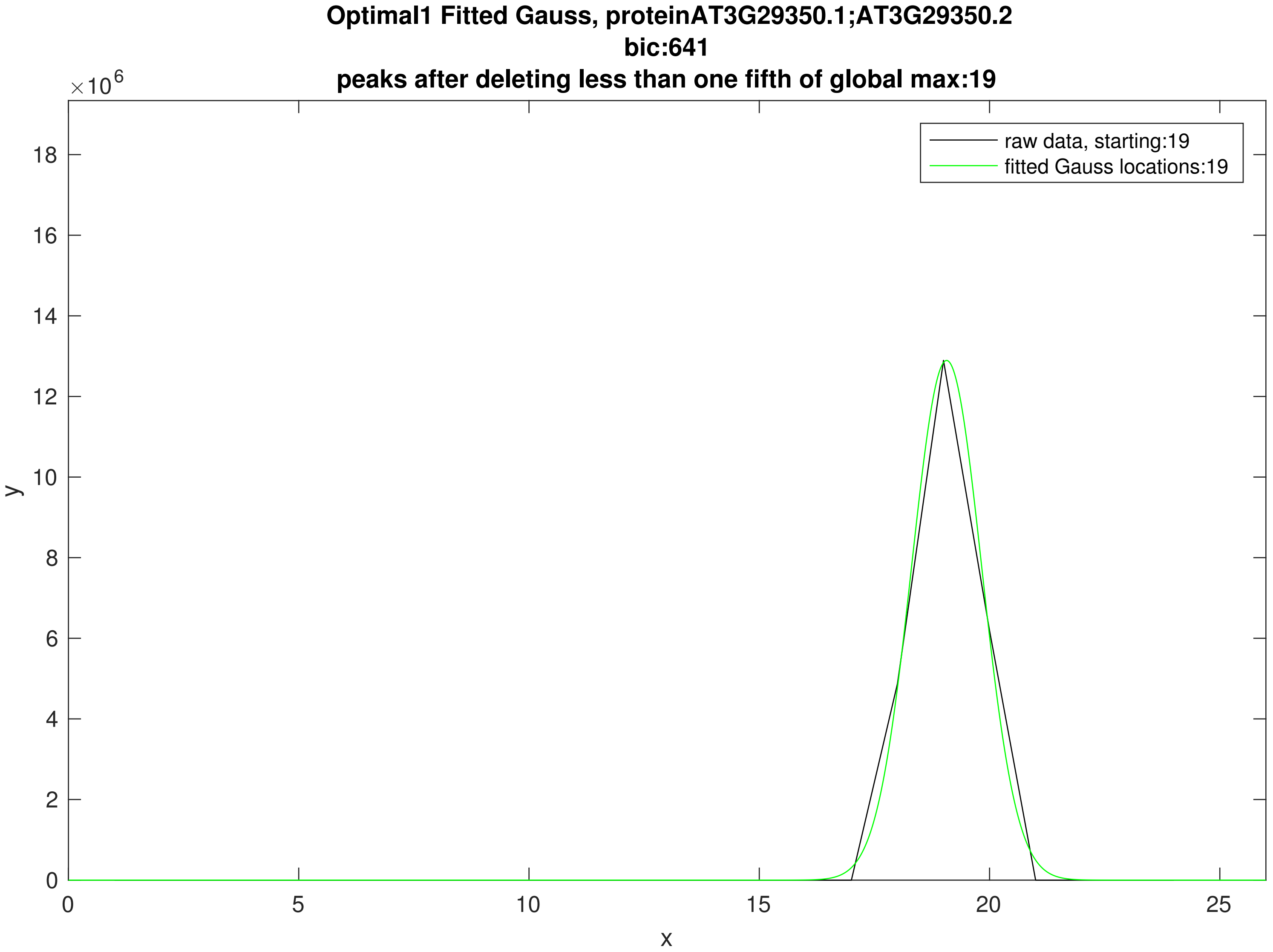
Optimal1 Fitted Gauss, proteinAT3G29350.1;AT3G29350.2
bic:641
peaks after deleting less than one fifth of global max:19
×10^6
raw data, starting:19
fitted Gauss locations:19
y
x
0
2
4
6
8
10
12
14
16
18
0
5
10
15
20
25

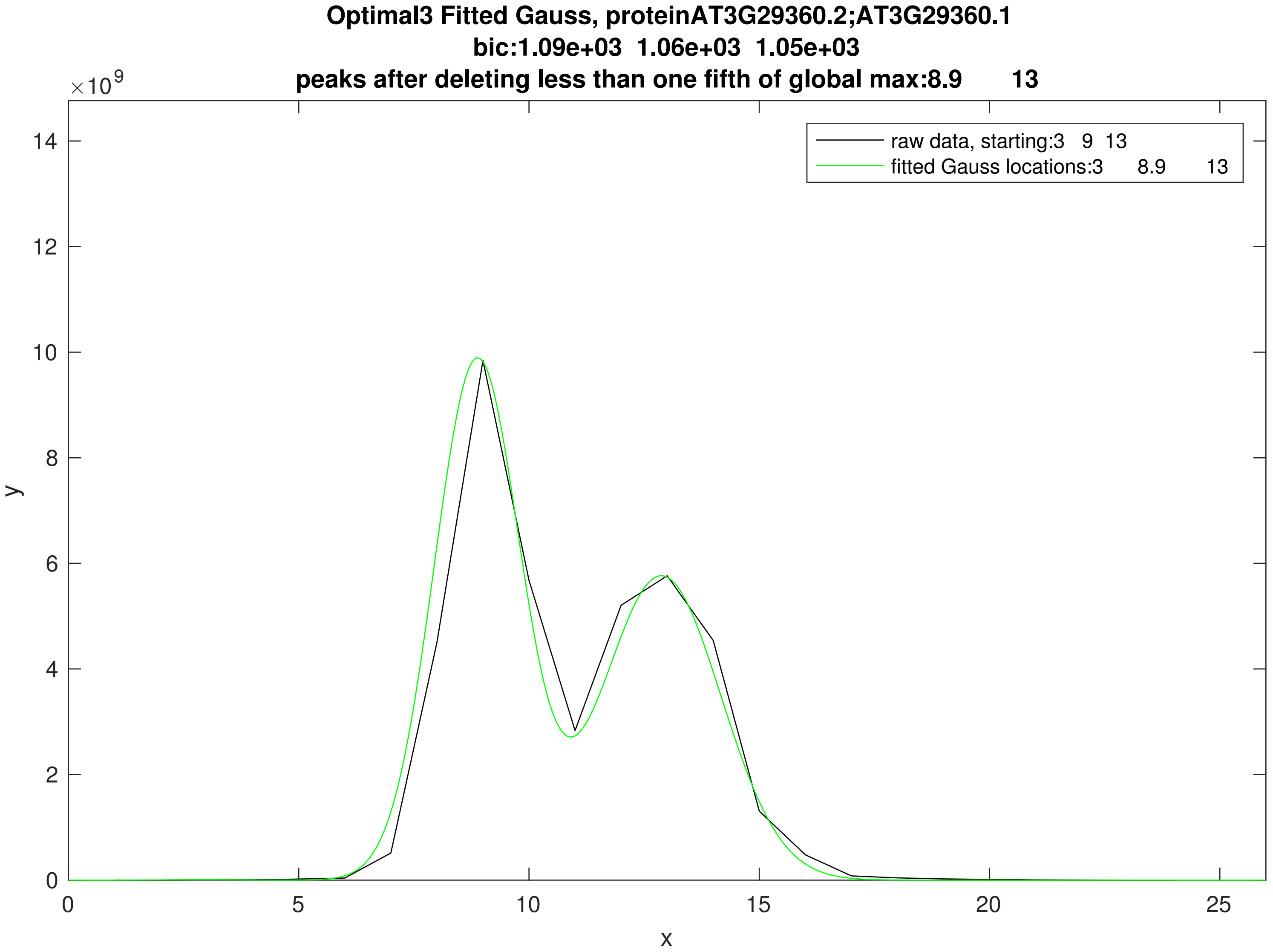
Optimal3 Fitted Gauss, proteinAT3G29360.2;AT3G29360.1
bic:1.09e+03 1.06e+03 1.05e+03
peaks after deleting less than one fifth of global max:8.9 13
raw data, starting:3 9 13
fitted Gauss locations:3 8.9 13
×10^9
y
x

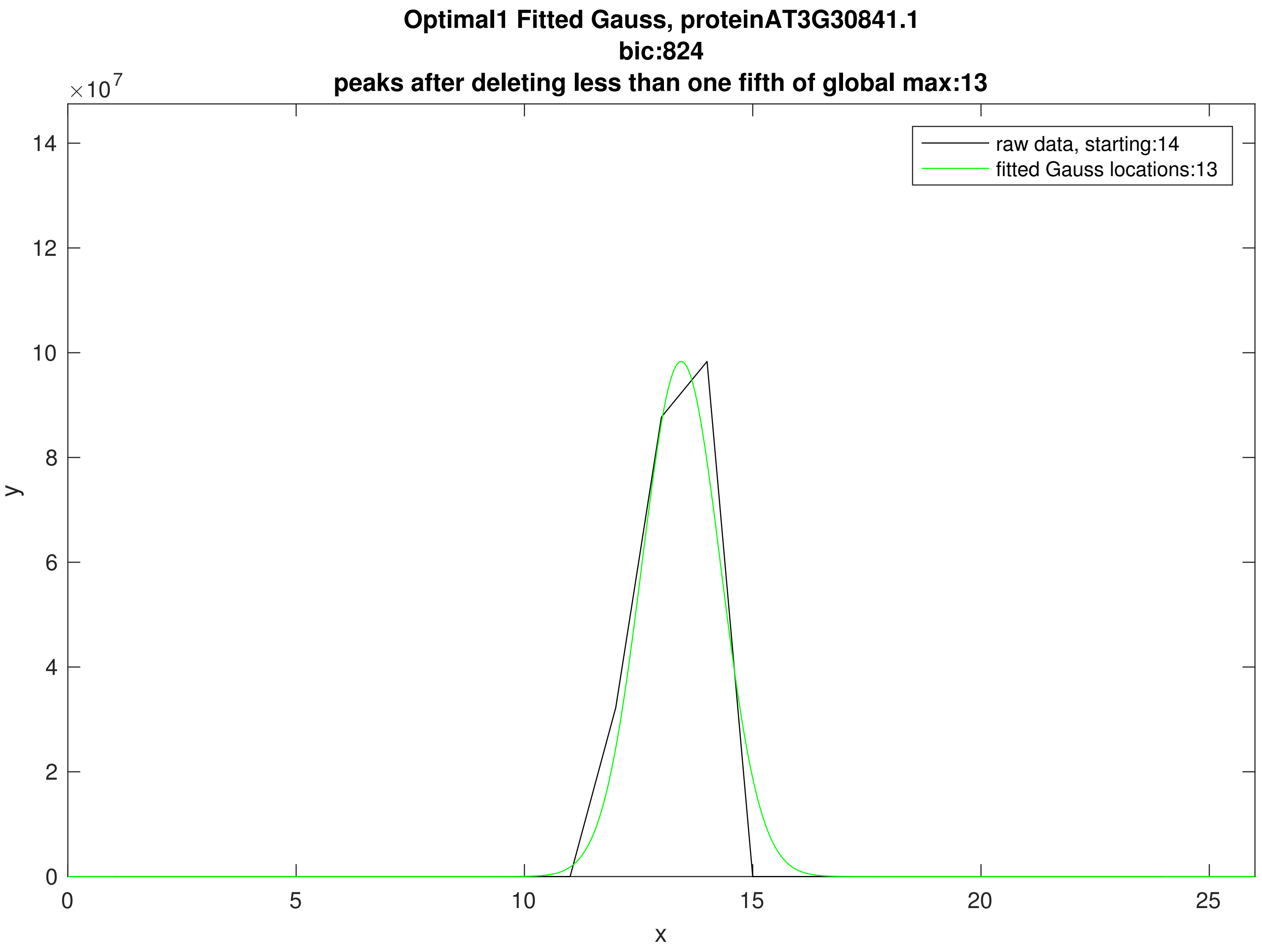
Optimal1 Fitted Gauss, proteinAT3G30841.1
bic:824
peaks after deleting less than one fifth of global max:13
×10^7
raw data, starting:14
fitted Gauss locations:13
y
x
0
2
4
6
8
10
12
14
0
5
10
15
20
25

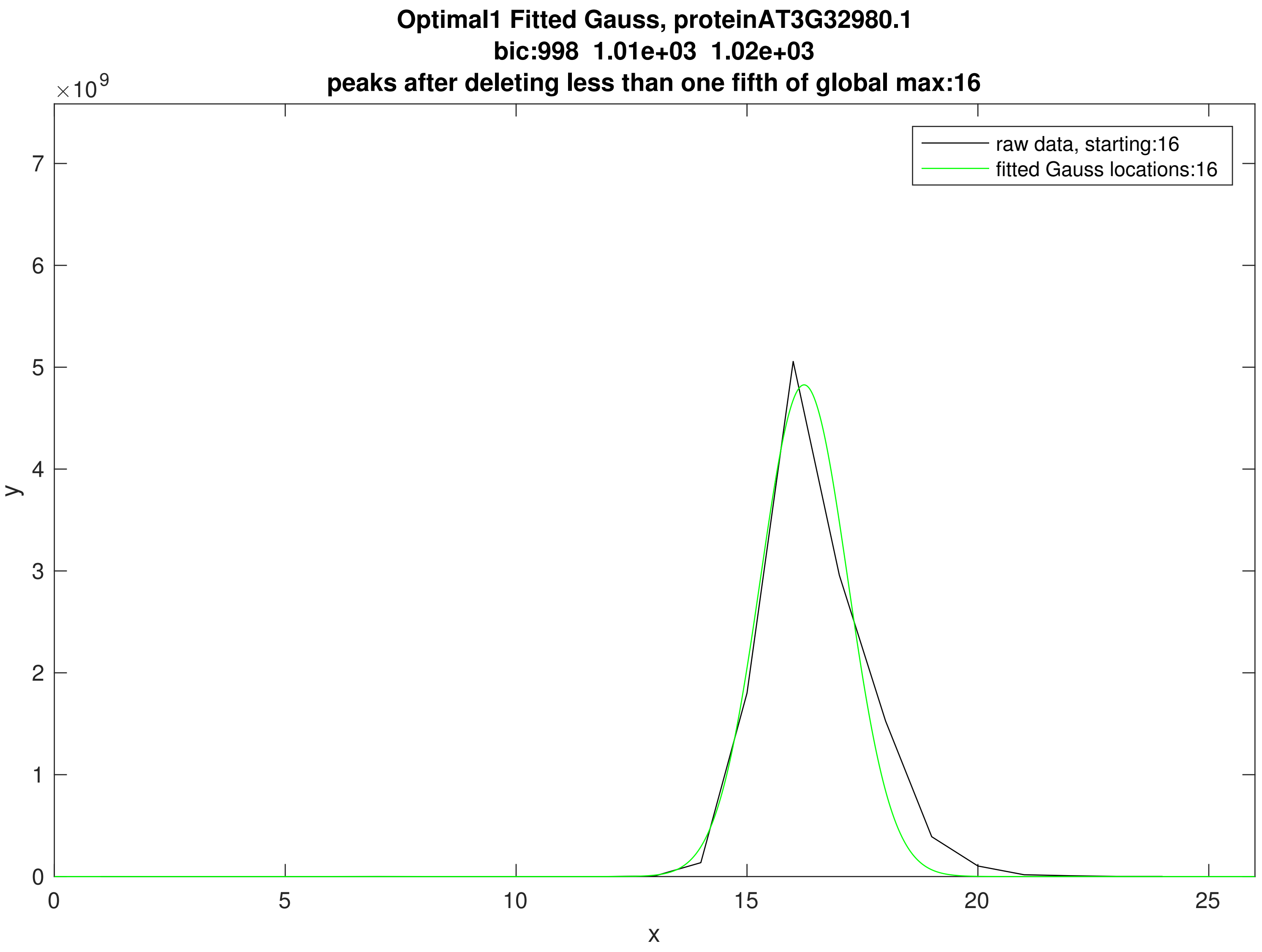

Optimal1 Fitted Gauss, proteinAT3G32980.1
bic:998 1.01e+03 1.02e+03
peaks after deleting less than one fifth of global max:16
×10^9
raw data, starting:16
fitted Gauss locations:16
y
x
0
1
2
3
4
5
6
7
0
5
10
15
20
25

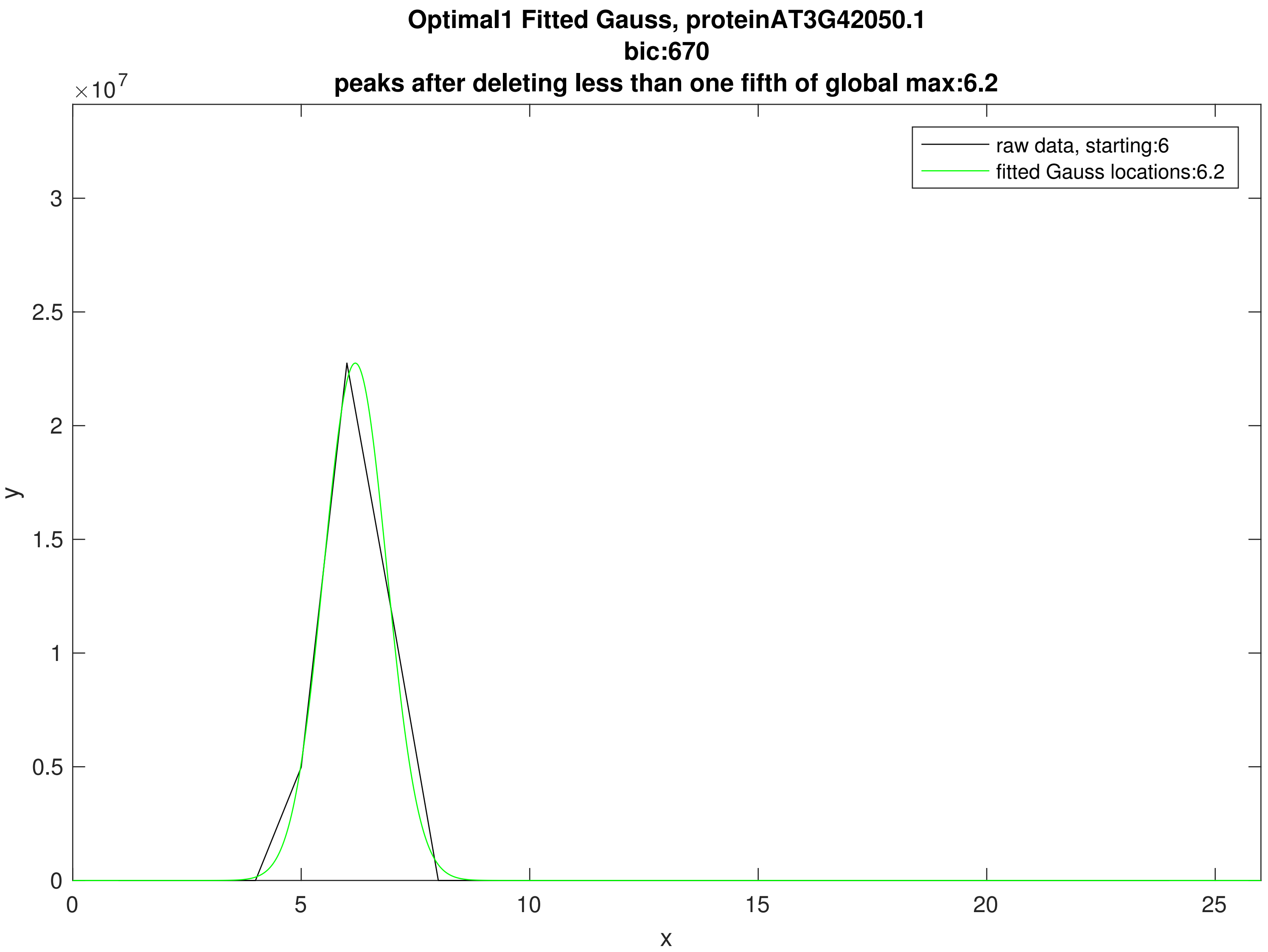
Optimal1 Fitted Gauss, proteinAT3G42050.1
bic:670
peaks after deleting less than one fifth of global max:6.2
×10^7
raw data, starting:6
fitted Gauss locations:6.2
y
x
0
0.5
1
1.5
2
2.5
3
0
5
10
15
20
25

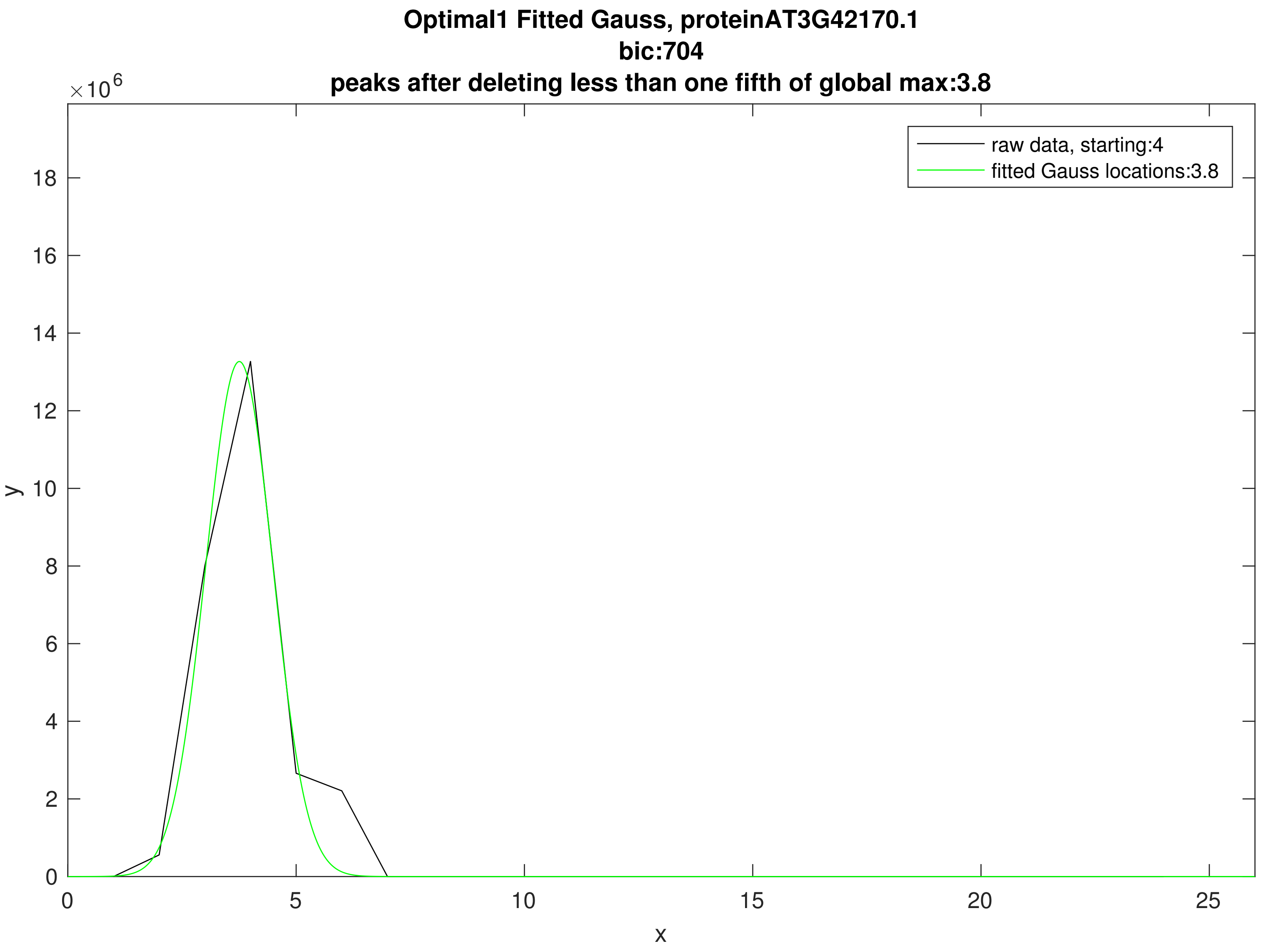
Optimal1 Fitted Gauss, proteinAT3G42170.1
bic:704
peaks after deleting less than one fifth of global max:3.8
×10^6
raw data, starting:4
fitted Gauss locations:3.8
y
x
0
2
4
6
8
10
12
14
16
18
0
5
10
15
20
25

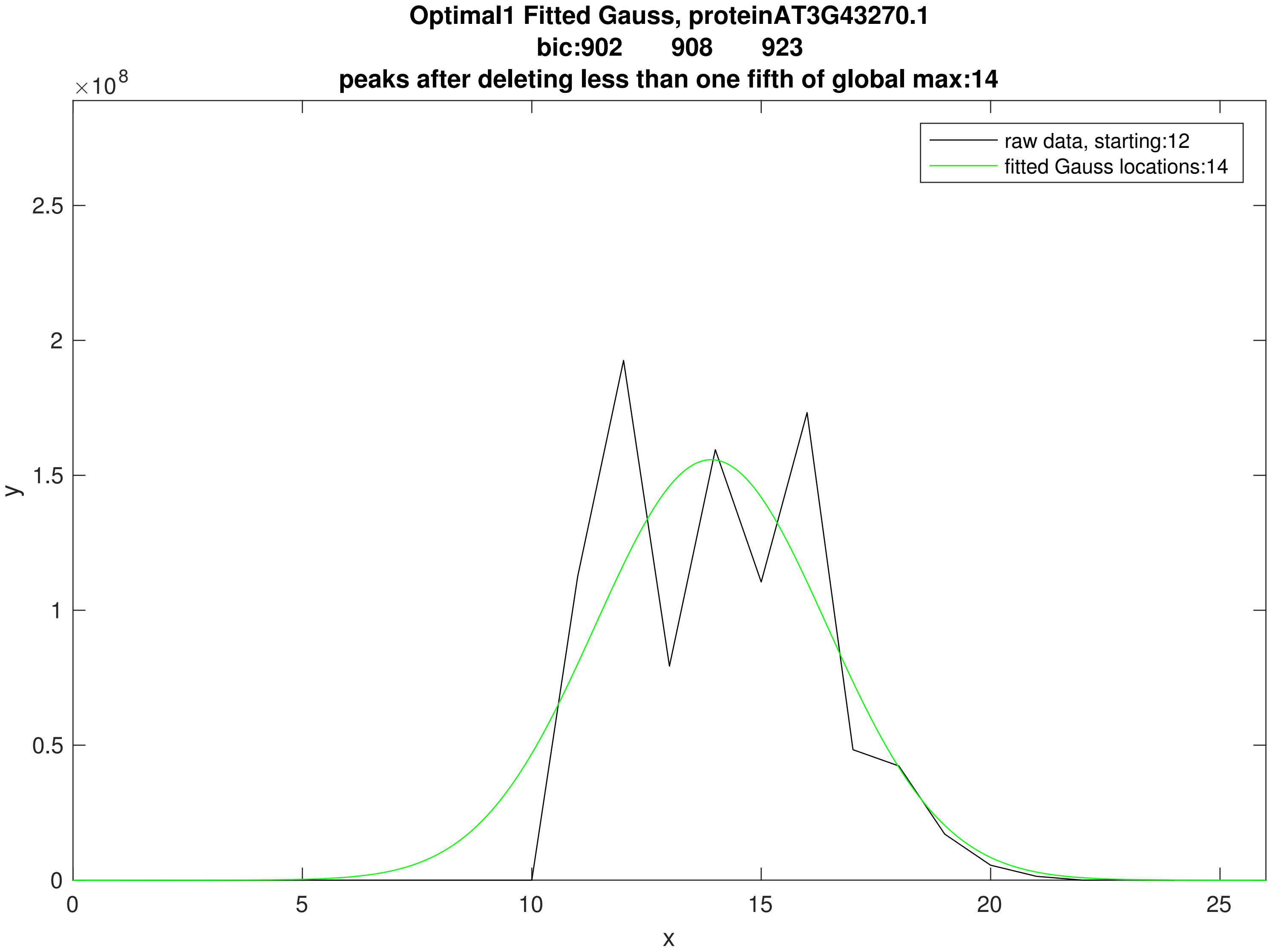
Optimal1 Fitted Gauss, proteinAT3G43270.1
bic:902 908 923
peaks after deleting less than one fifth of global max:14
×10^8
raw data, starting:12
fitted Gauss locations:14
y
x
0
0.5
1
1.5
2
2.5
0
5
10
15
20
25

**Optimal1 Fitted Gauss, proteinAT3G43600.1**
**bic:874**
**peaks after deleting less than one fifth of global max:8.5**

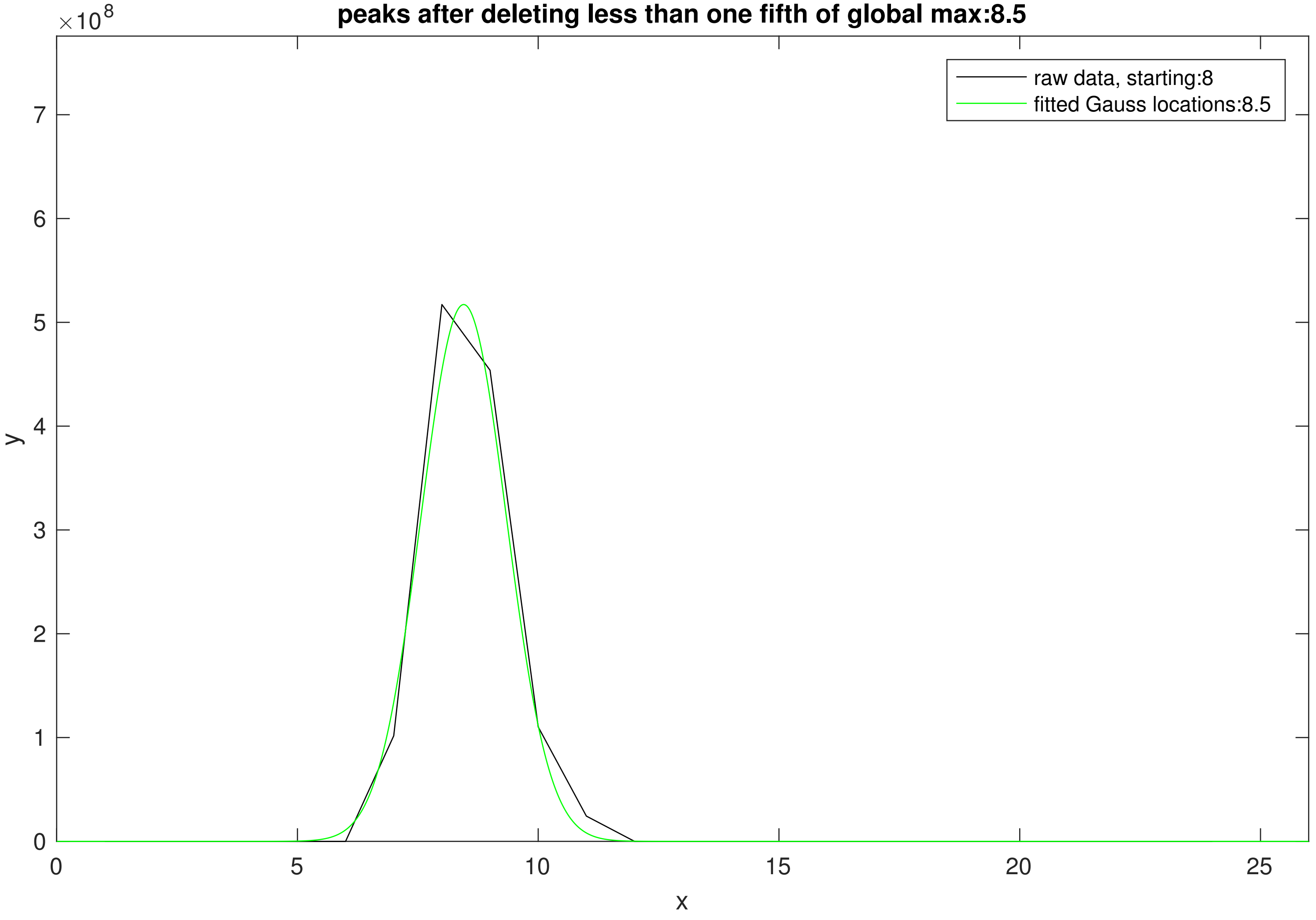

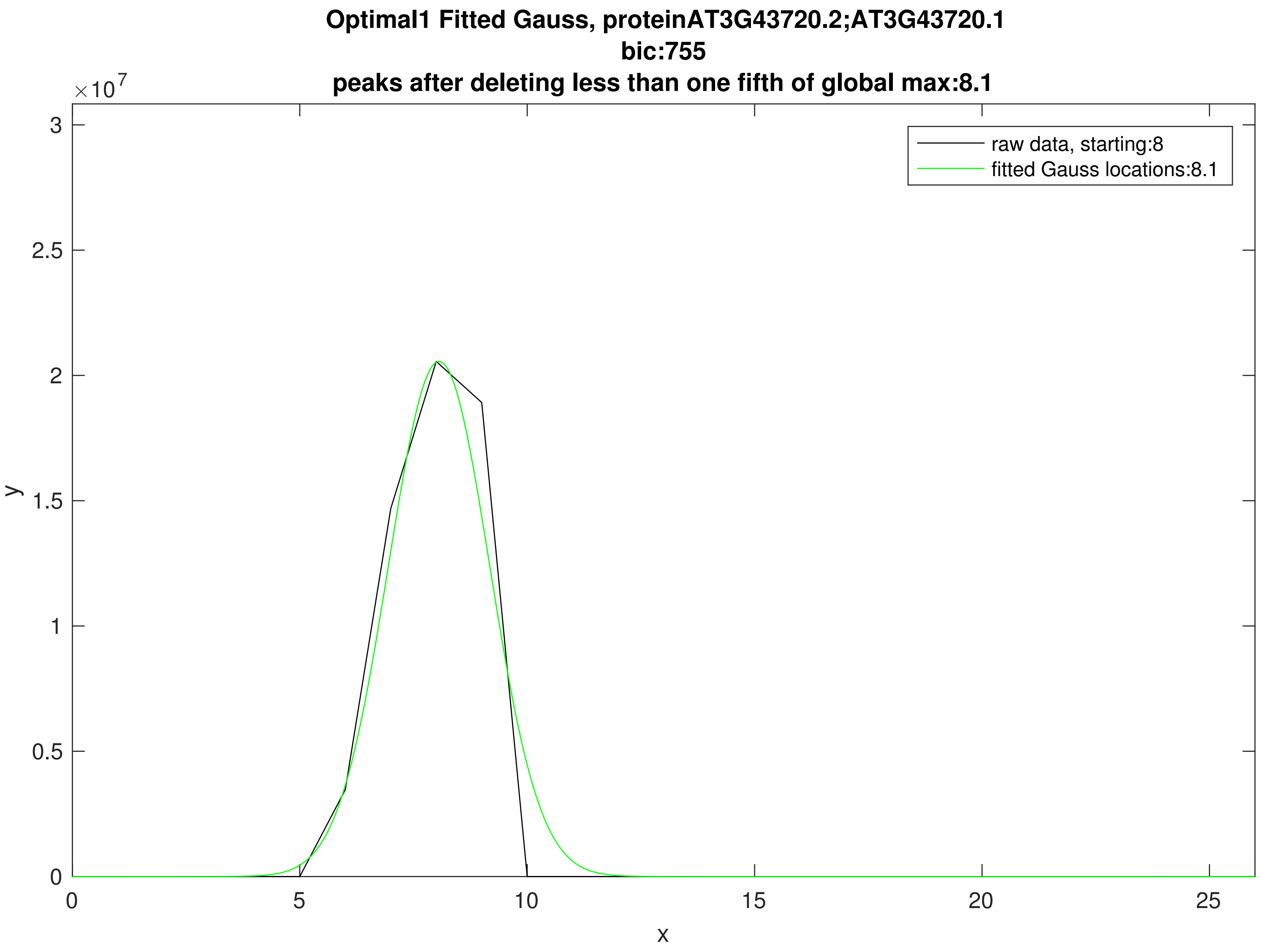
Optimal1 Fitted Gauss, proteinAT3G43720.2;AT3G43720.1
bic:755
peaks after deleting less than one fifth of global max:8.1
×10^7
3
2.5
2
1.5
1
0.5
0
y
0
5
10
15
20
25
x
raw data, starting:8
fitted Gauss locations:8.1

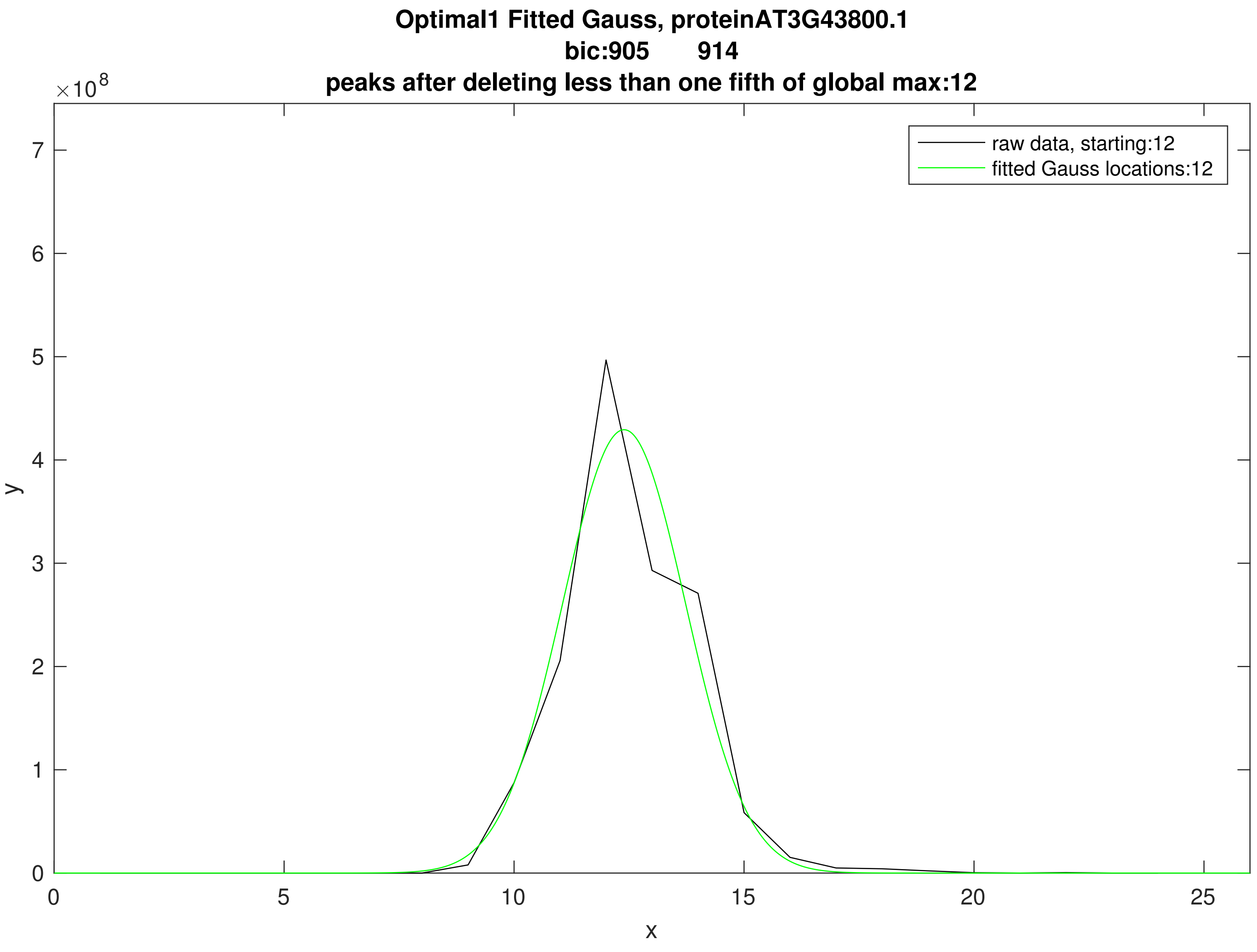
Optimal1 Fitted Gauss, proteinAT3G43800.1
bic:905 914
peaks after deleting less than one fifth of global max:12
×10^8
raw data, starting:12
fitted Gauss locations:12
y
x
0
1
2
3
4
5
6
7
0
5
10
15
20
25

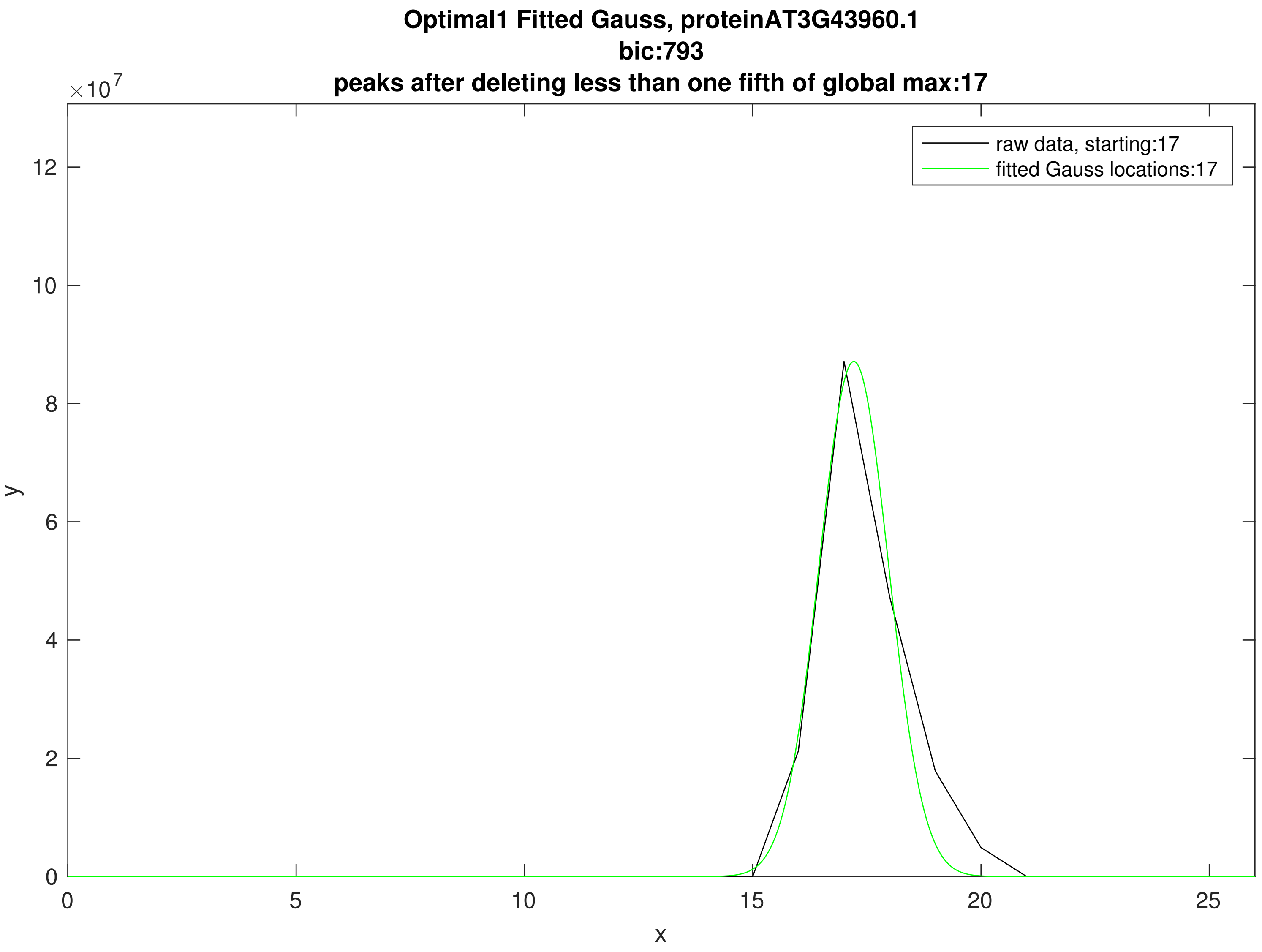
Optimal1 Fitted Gauss, proteinAT3G43960.1
bic:793
peaks after deleting less than one fifth of global max:17
×10^7
raw data, starting:17
fitted Gauss locations:17
y
x
0
2
4
6
8
10
12
0
5
10
15
20
25

**Optimal1 Fitted Gauss, proteinAT3G44100.1**
**bic:802 865**
**peaks after deleting less than one fifth of global max:21**

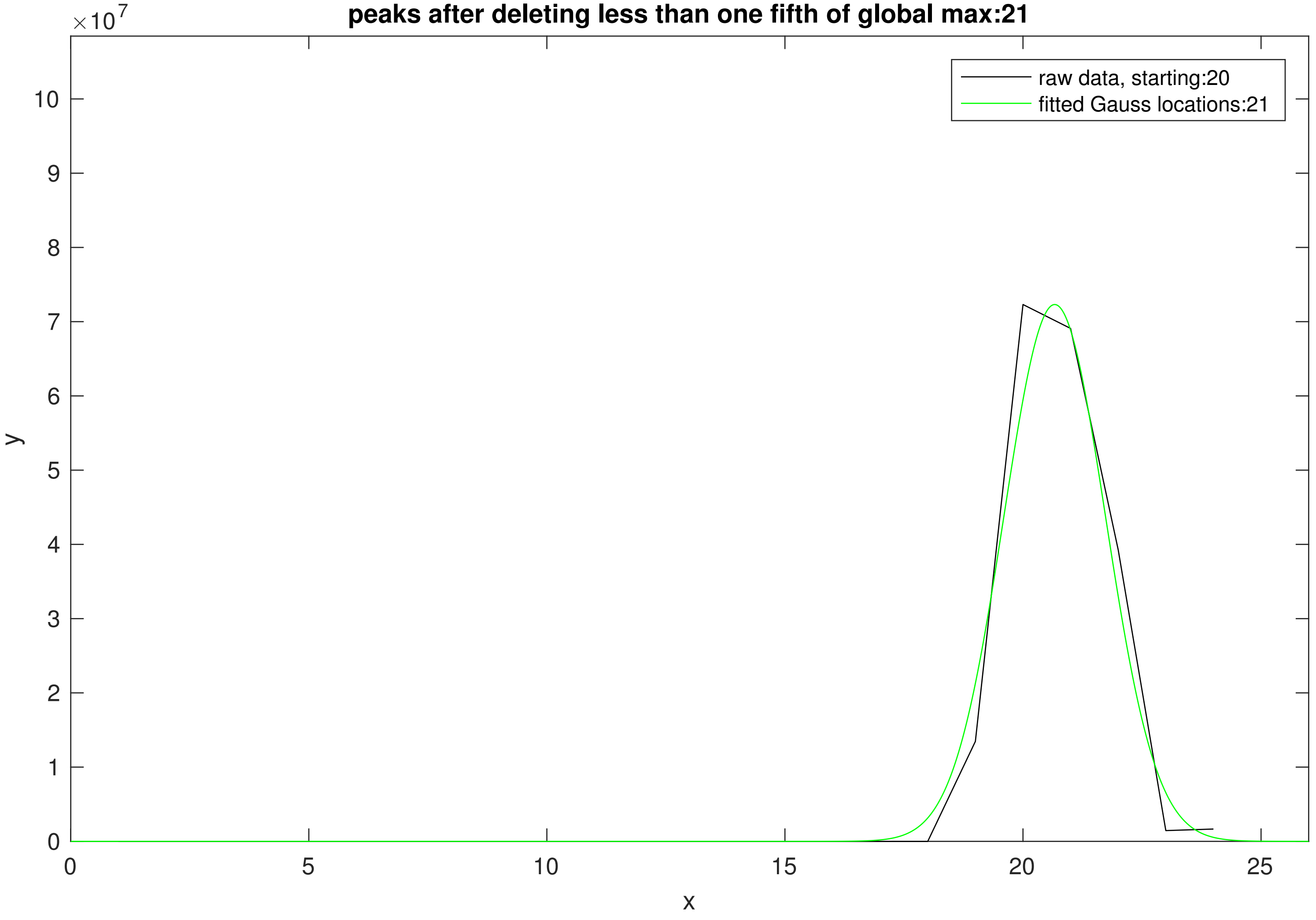

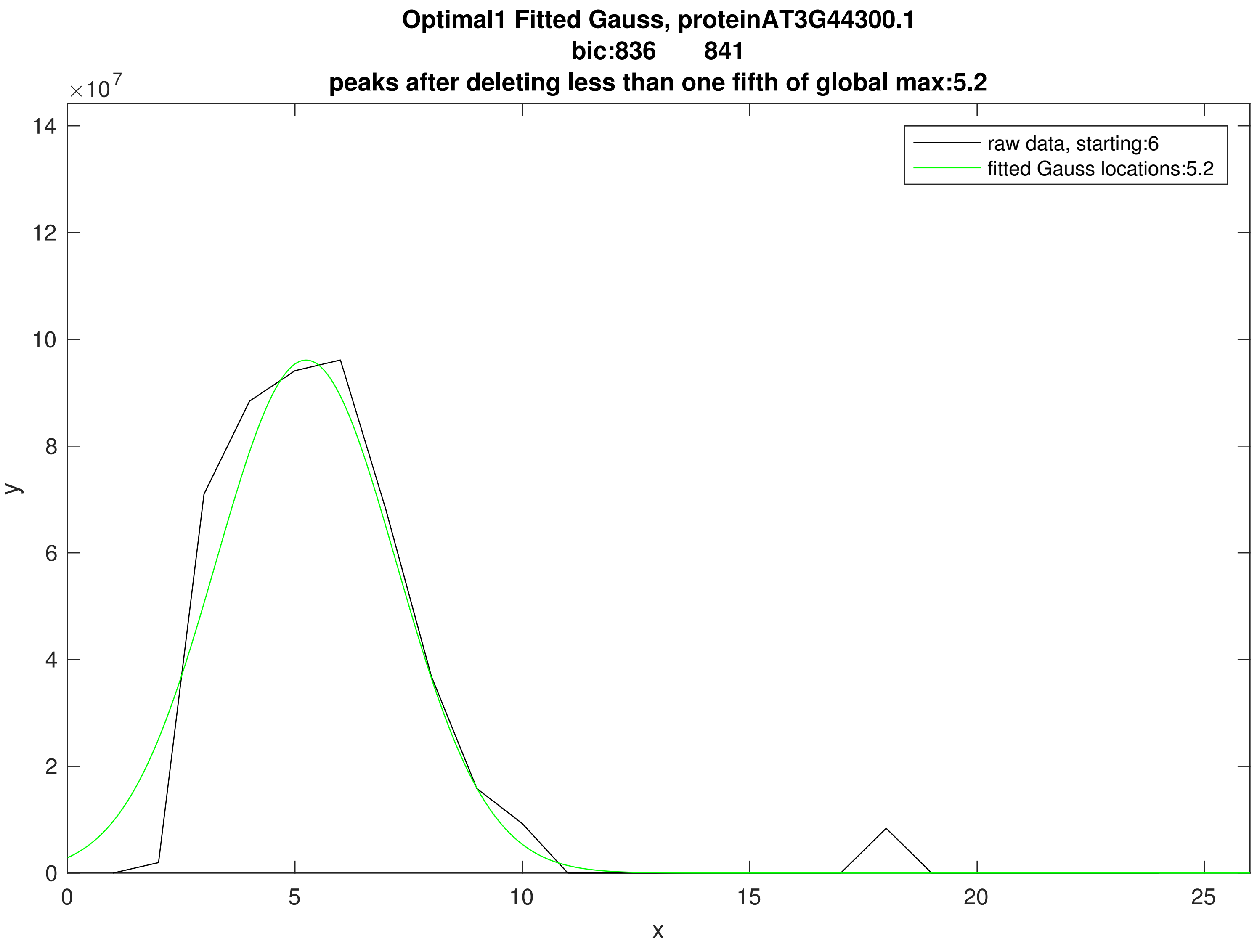

Optimal1 Fitted Gauss, proteinAT3G44300.1
bic:836 841
peaks after deleting less than one fifth of global max:5.2
×10^7
raw data, starting:6
fitted Gauss locations:5.2
y
x
0
2
4
6
8
10
12
14
0
5
10
15
20
25

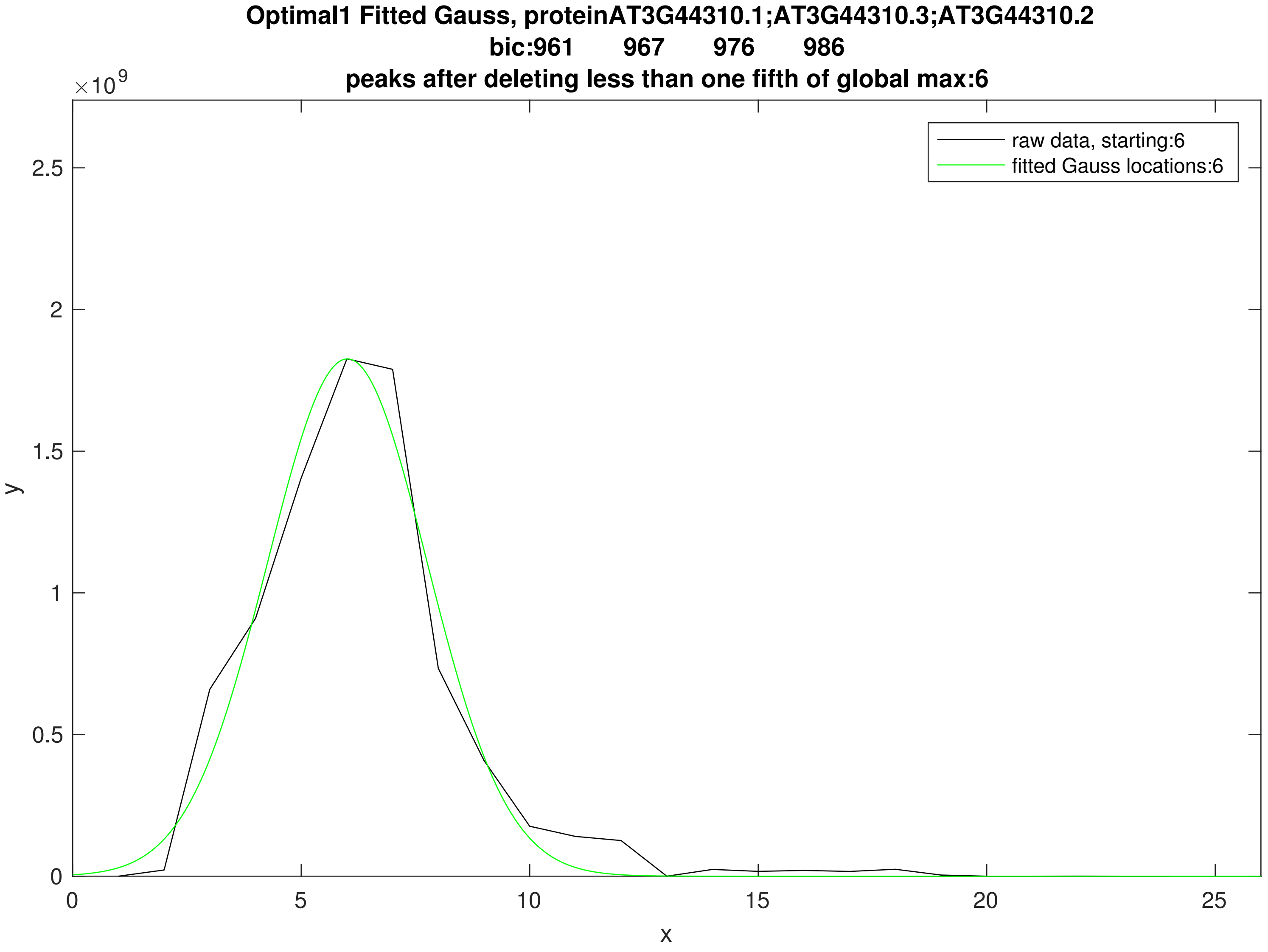

Optimal1 Fitted Gauss, proteinAT3G44310.1;AT3G44310.3;AT3G44310.2
bic:961 967 976 986
peaks after deleting less than one fifth of global max:6
×10^9
2.5
2
1.5
1
0.5
0
y
0
5
10
15
20
25
x
raw data, starting:6
fitted Gauss locations:6

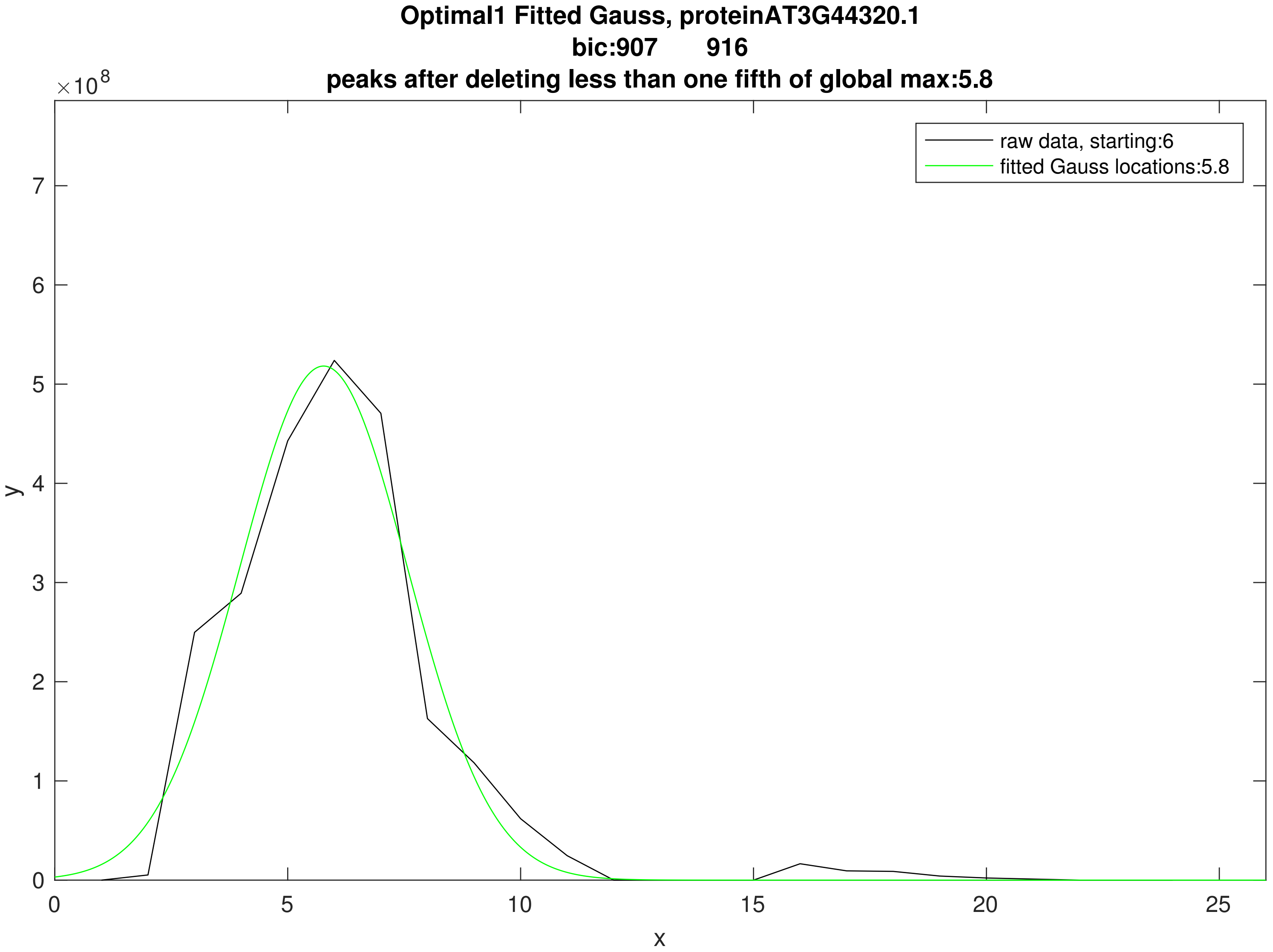

Optimal1 Fitted Gauss, proteinAT3G44320.1
bic:907 916
peaks after deleting less than one fifth of global max:5.8
×10^8
raw data, starting:6
fitted Gauss locations:5.8
y
x
0
1
2
3
4
5
6
7
0
5
10
15
20
25

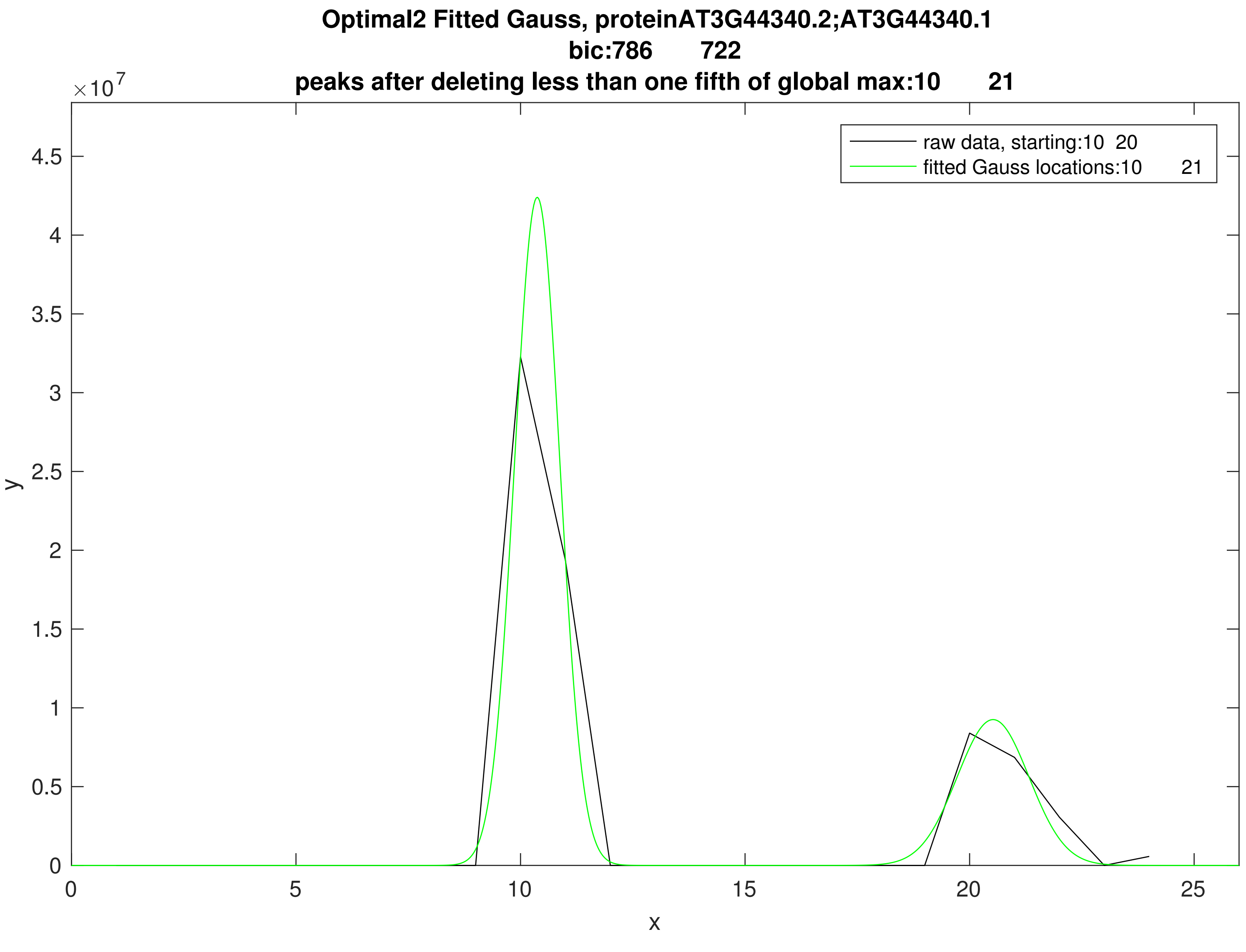

Optimal2 Fitted Gauss, proteinAT3G44340.2;AT3G44340.1
bic:786 722
peaks after deleting less than one fifth of global max:10 21
×10^7
raw data, starting:10 20
fitted Gauss locations:10 21
y
x
0
0.5
1
1.5
2
2.5
3
3.5
4
4.5
0
5
10
15
20
25

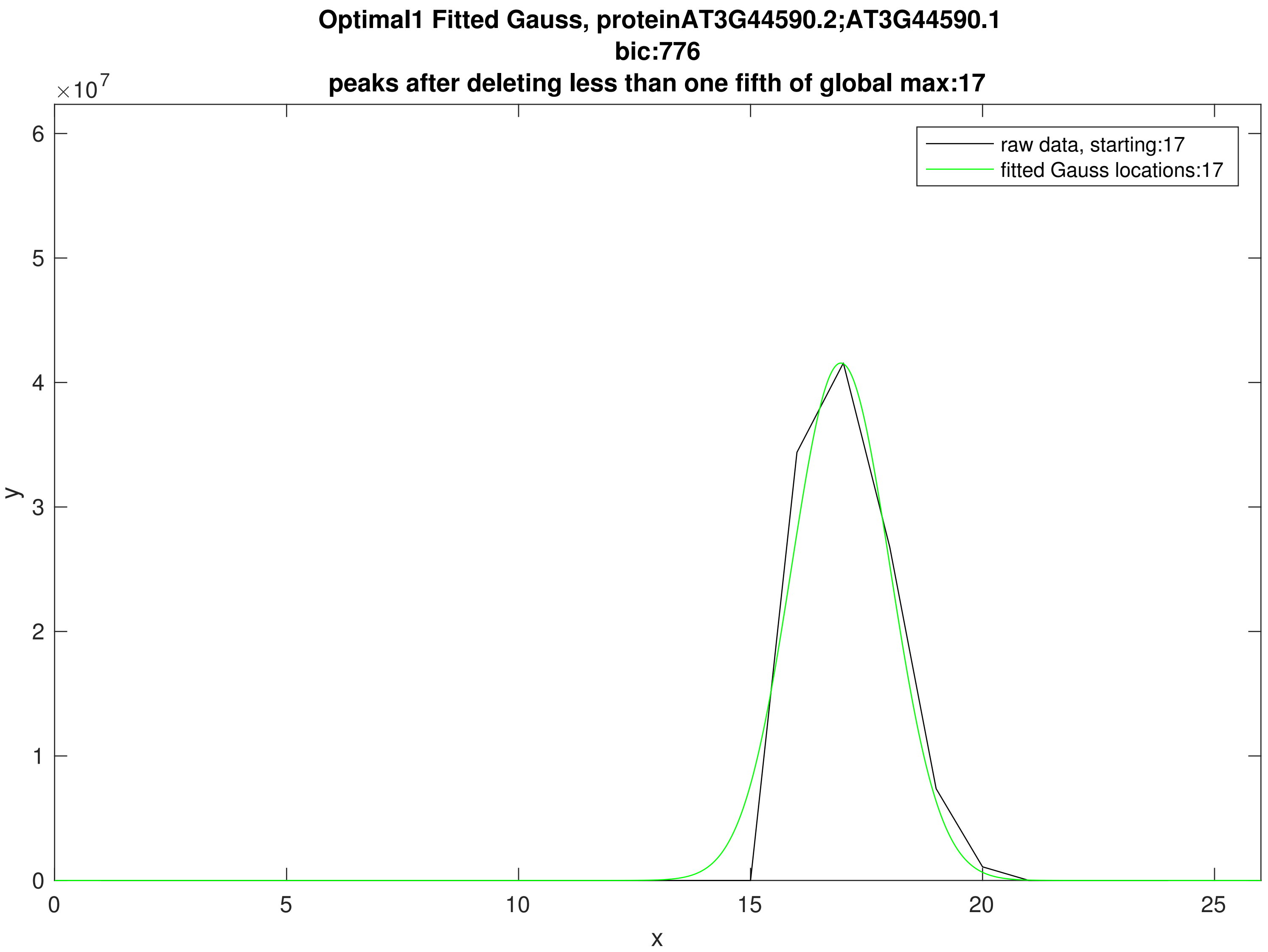
Optimal1 Fitted Gauss, proteinAT3G44590.2;AT3G44590.1
bic:776
peaks after deleting less than one fifth of global max:17
×10^7
raw data, starting:17
fitted Gauss locations:17
y
x
0
1
2
3
4
5
6
0
5
10
15
20
25

**Optimal1 Fitted Gauss, proteinAT3G44620.1;AT3G44620.2**
**bic:734**
**peaks after deleting less than one fifth of global max:20**

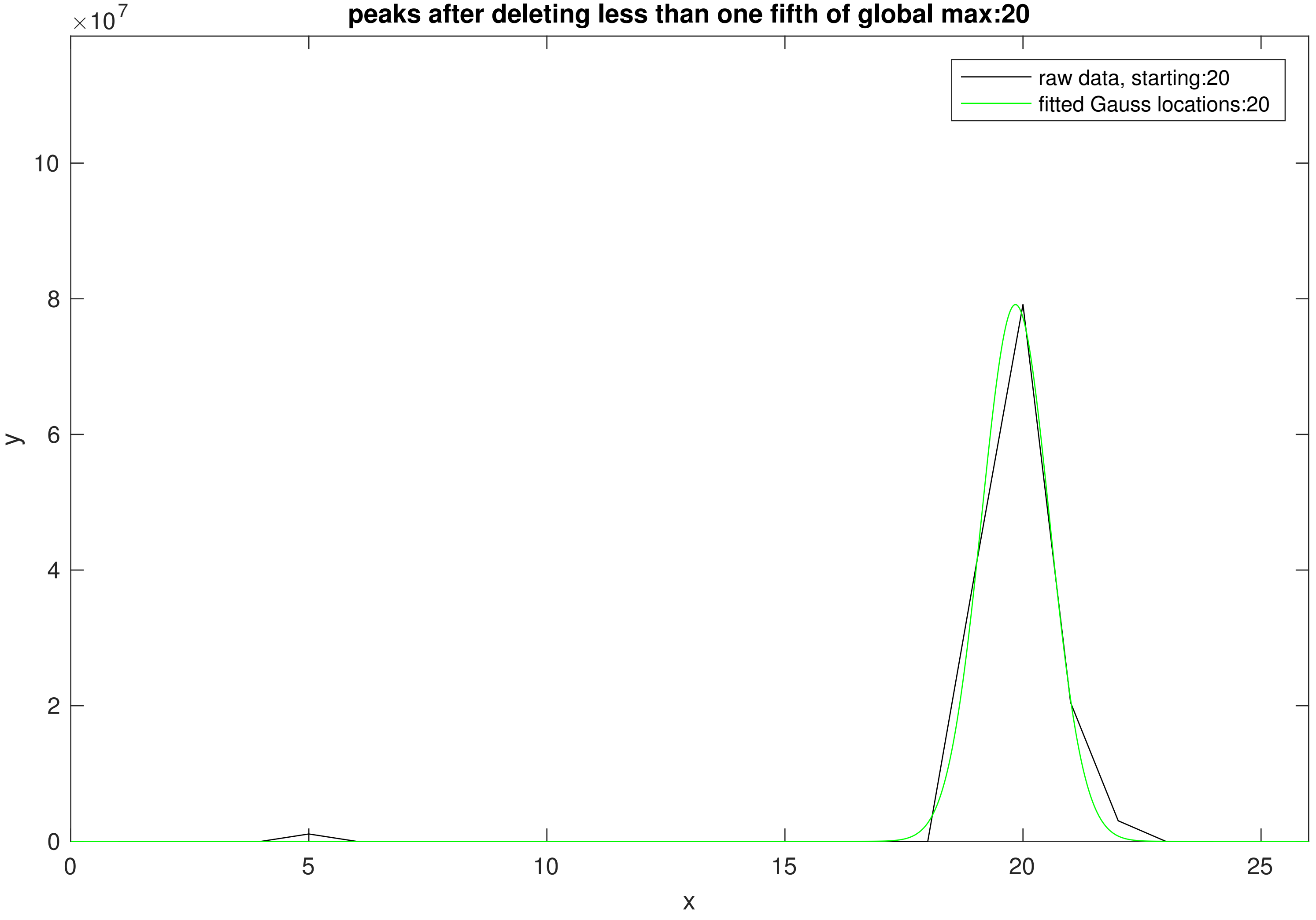

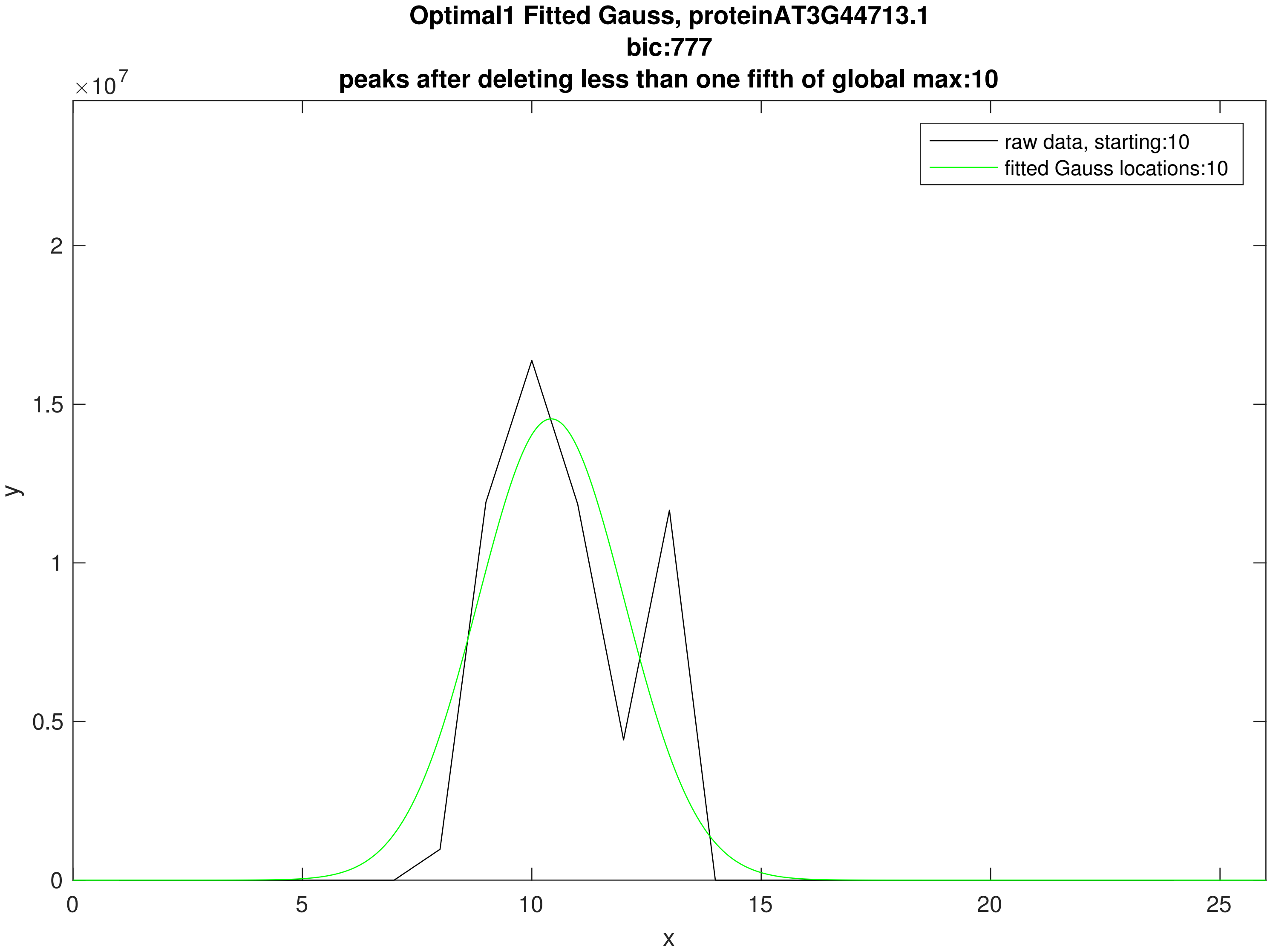
Optimal1 Fitted Gauss, proteinAT3G44713.1
bic:777
peaks after deleting less than one fifth of global max:10
×10^7
raw data, starting:10
fitted Gauss locations:10
y
x
0
0.5
1
1.5
2
0
5
10
15
20
25

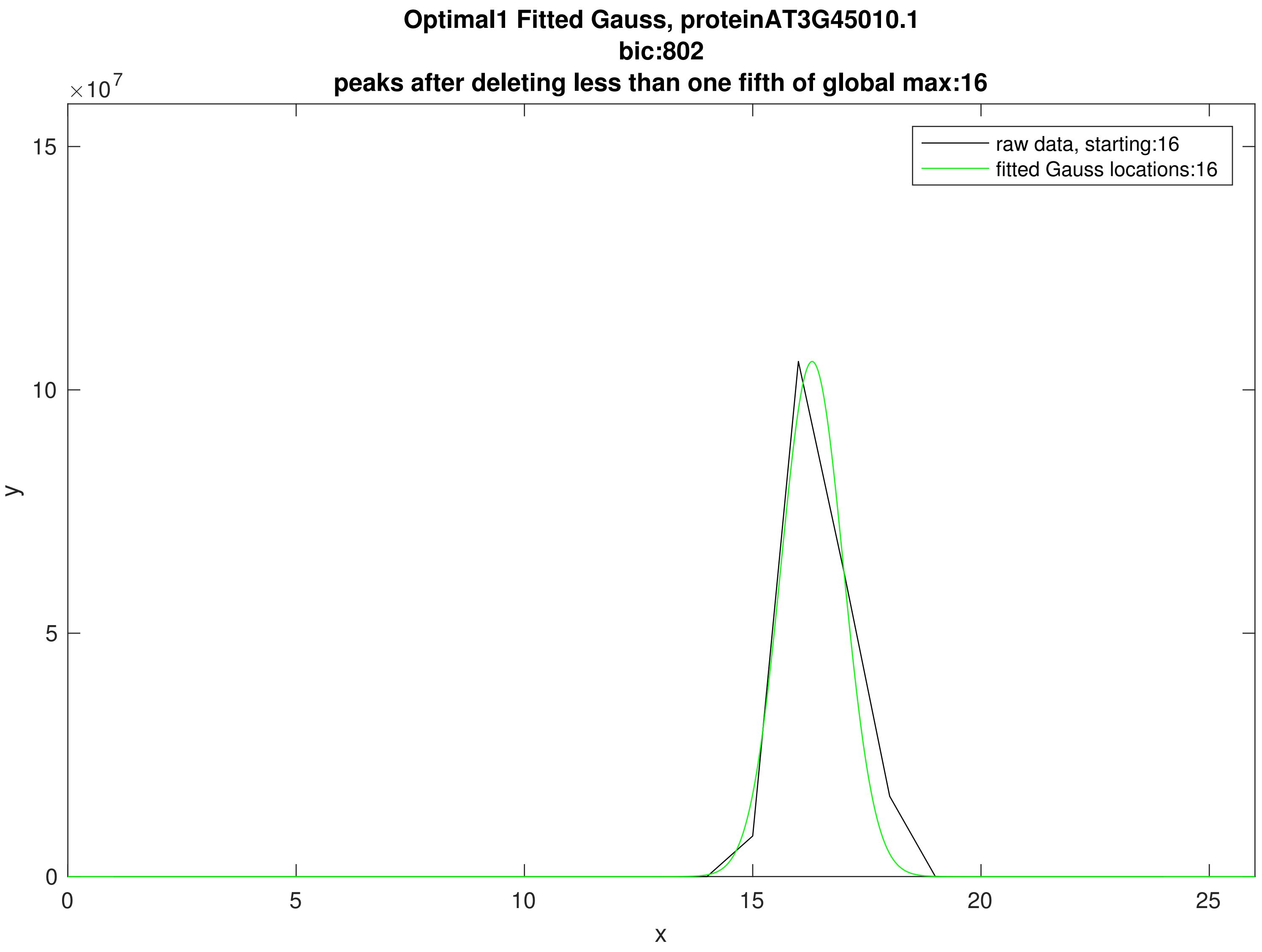
Optimal1 Fitted Gauss, proteinAT3G45010.1
bic:802
peaks after deleting less than one fifth of global max:16
×10^7
raw data, starting:16
fitted Gauss locations:16
y
x
0
5
10
15
20
25

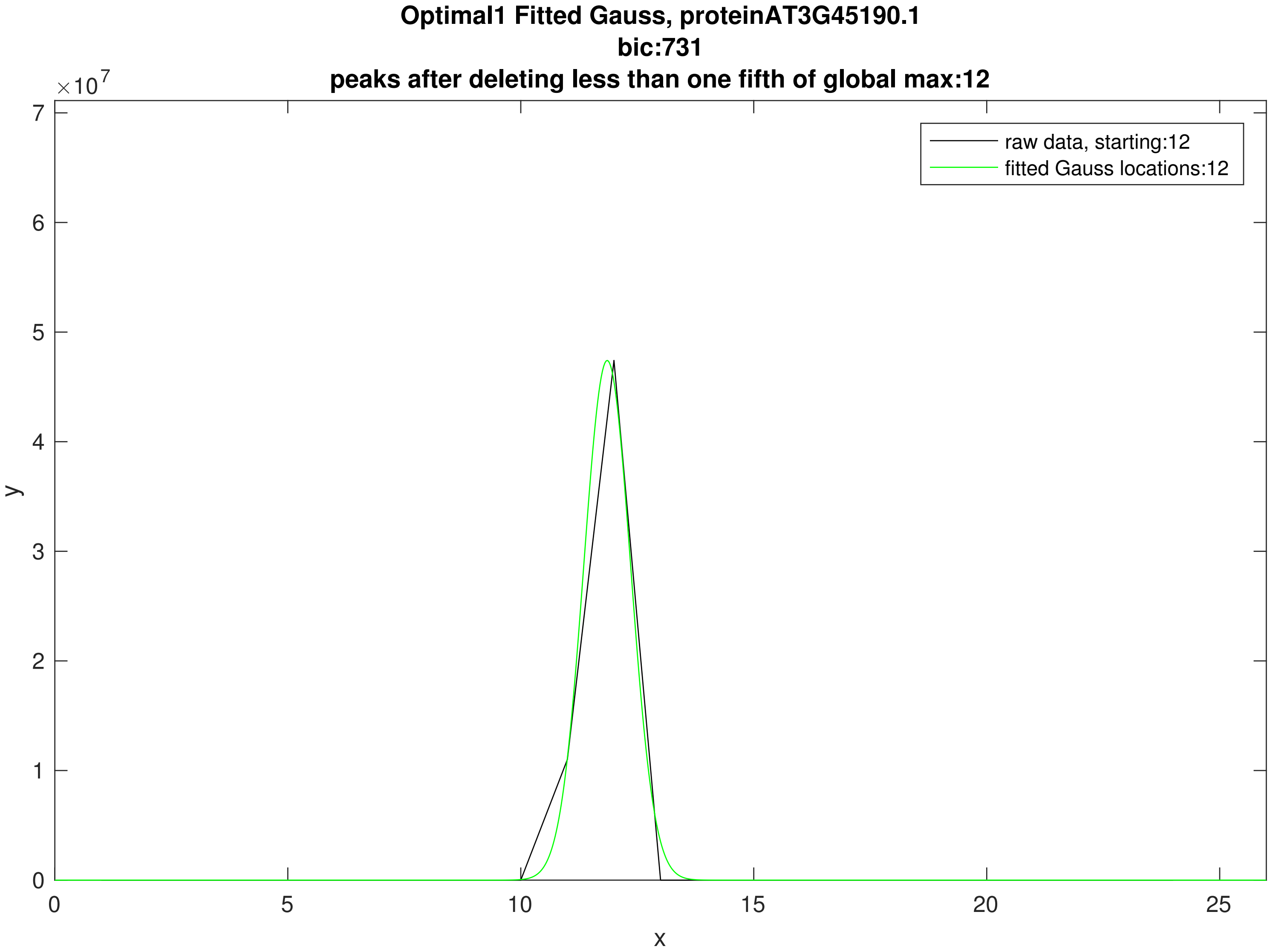

Optimal1 Fitted Gauss, proteinAT3G45190.1
bic:731
peaks after deleting less than one fifth of global max:12
×10^7
raw data, starting:12
fitted Gauss locations:12
y
x
0
1
2
3
4
5
6
7
0
5
10
15
20
25

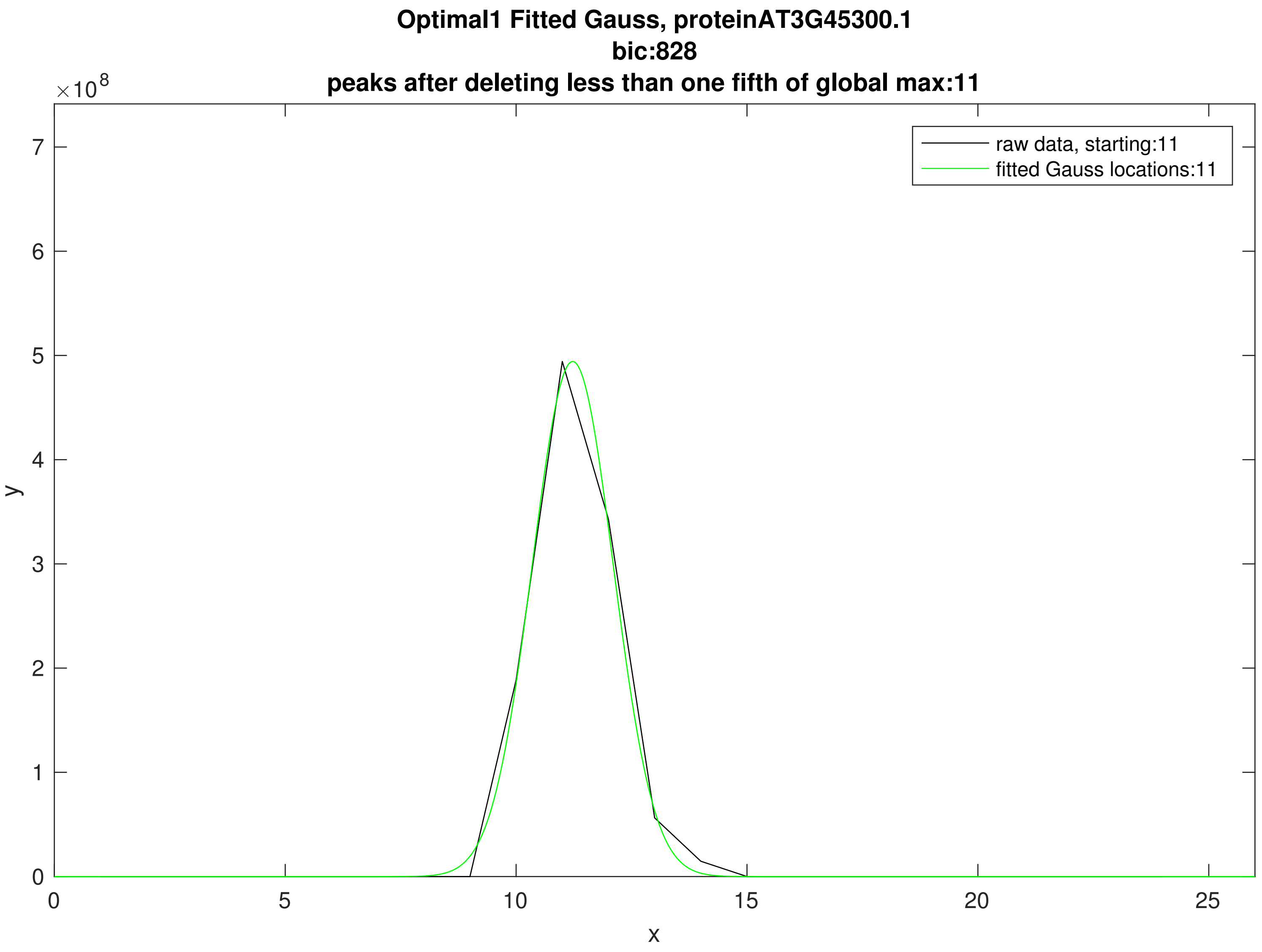

Optimal1 Fitted Gauss, proteinAT3G45300.1
bic:828
peaks after deleting less than one fifth of global max:11
×10^8
raw data, starting:11
fitted Gauss locations:11
y
x
0
1
2
3
4
5
6
7
0
5
10
15
20
25

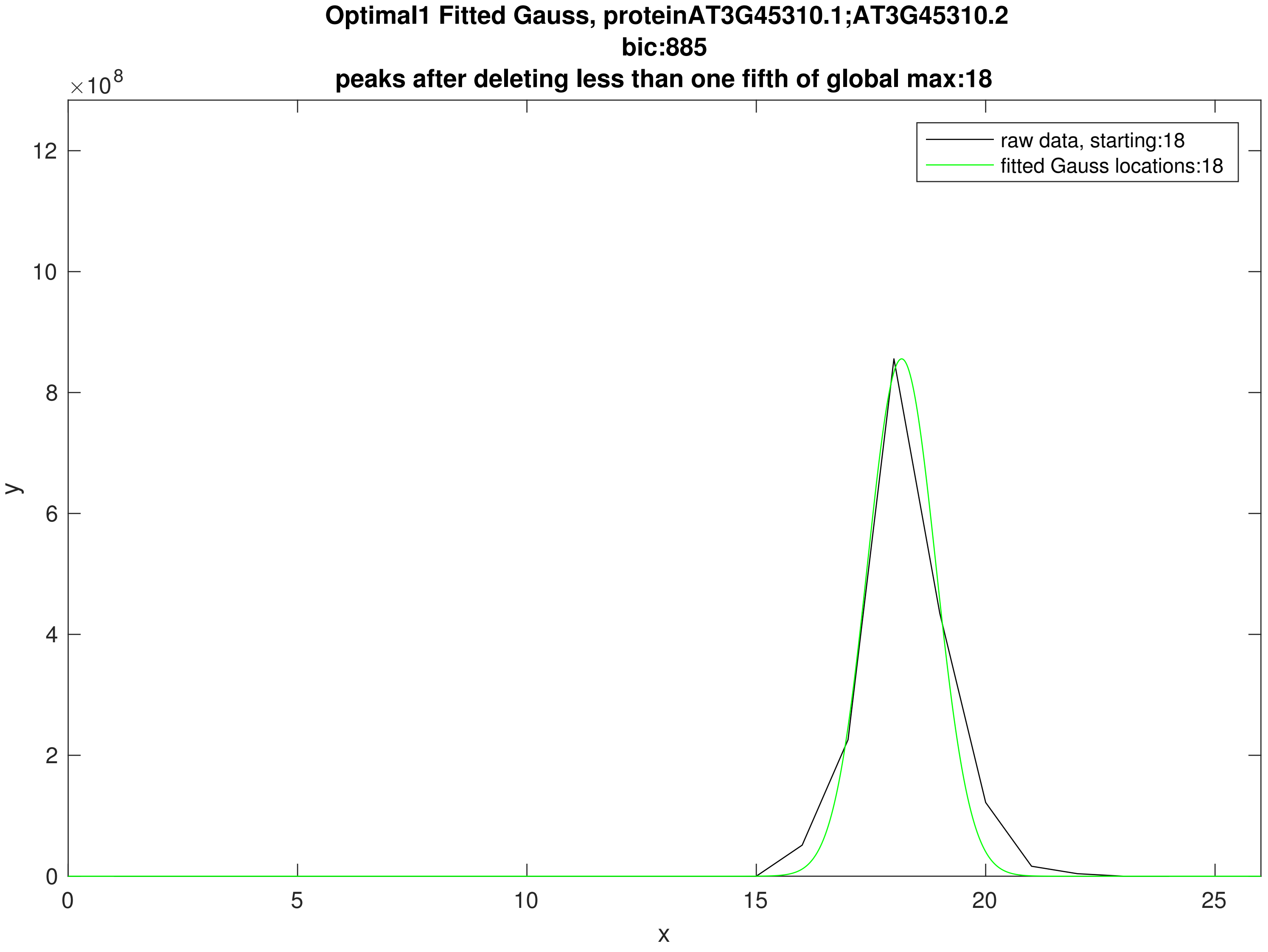

Optimal1 Fitted Gauss, proteinAT3G45310.1;AT3G45310.2
bic:885
peaks after deleting less than one fifth of global max:18
×10^8
raw data, starting:18
fitted Gauss locations:18
y
x
0
2
4
6
8
10
12
5
15
20
25

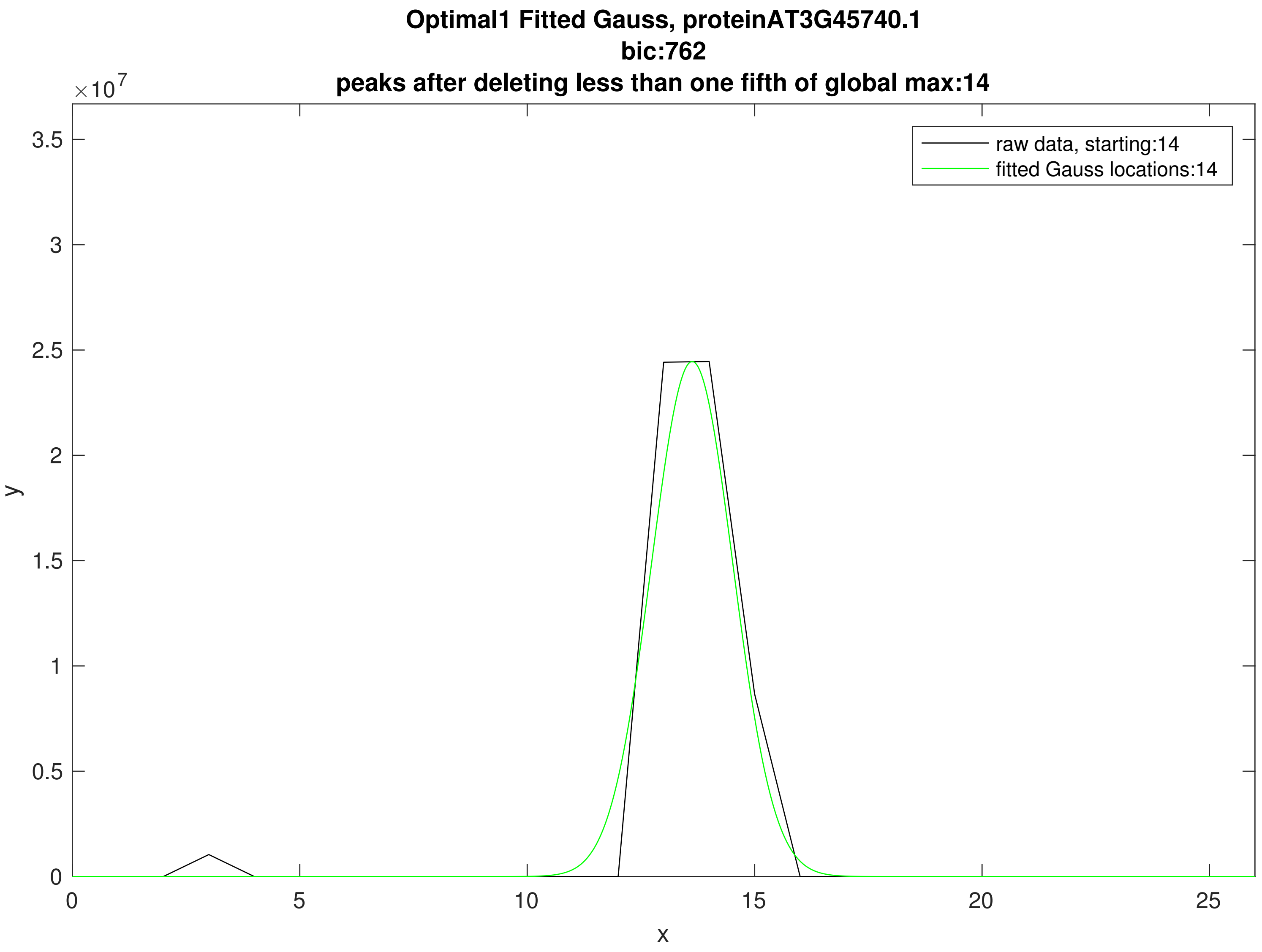

Optimal1 Fitted Gauss, proteinAT3G45740.1
bic:762
peaks after deleting less than one fifth of global max:14
$\times 10^7$
raw data, starting:14
fitted Gauss locations:14
3.5
3
2.5
2
1.5
1
0.5
0
y
0
5
10
15
20
25
x

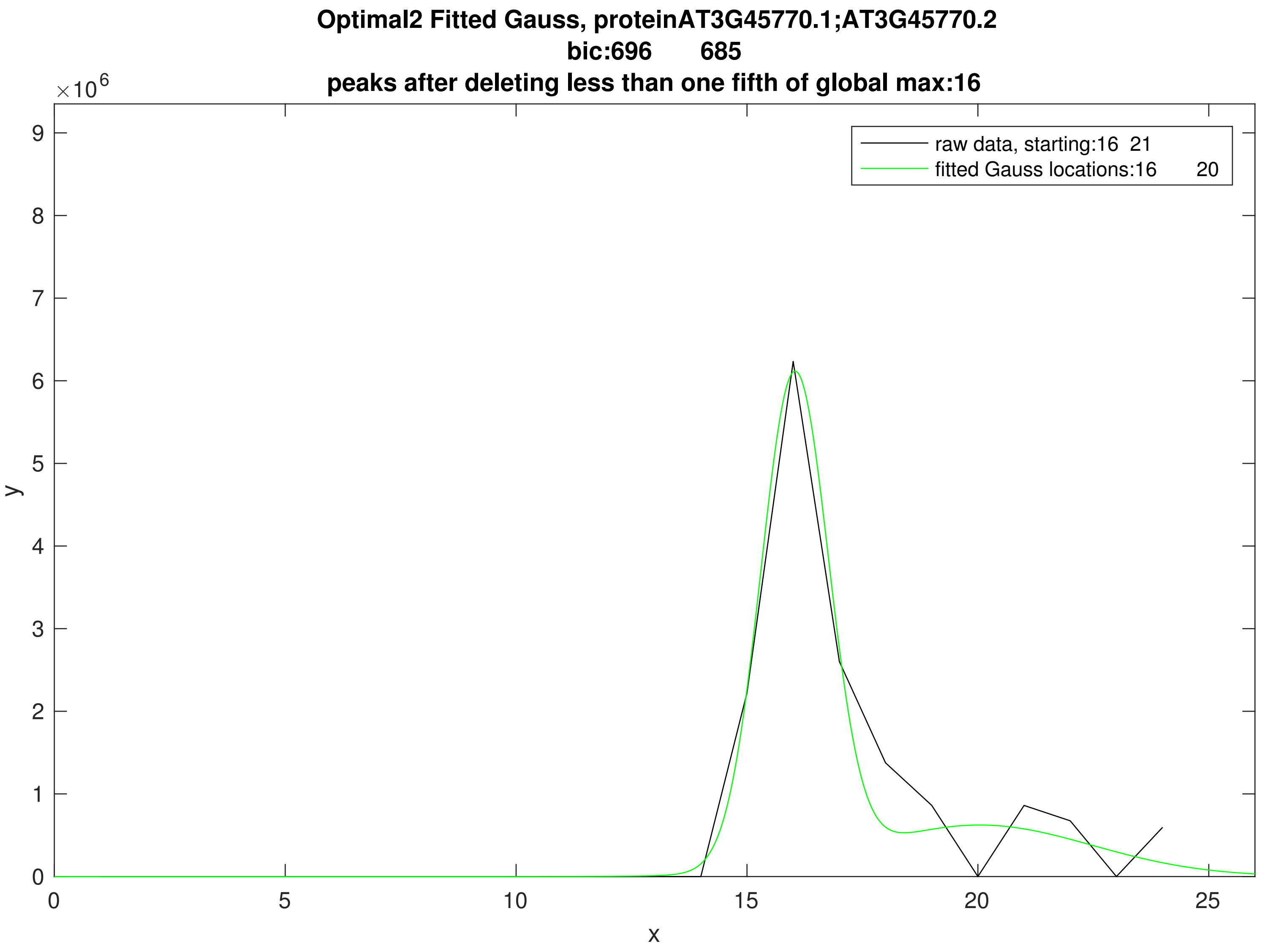
Optimal2 Fitted Gauss, proteinAT3G45770.1;AT3G45770.2
bic:696 685
peaks after deleting less than one fifth of global max:16
×10^6
raw data, starting:16 21
fitted Gauss locations:16 20
y
x
0
1
2
3
4
5
6
7
8
9
0
5
10
15
20
25

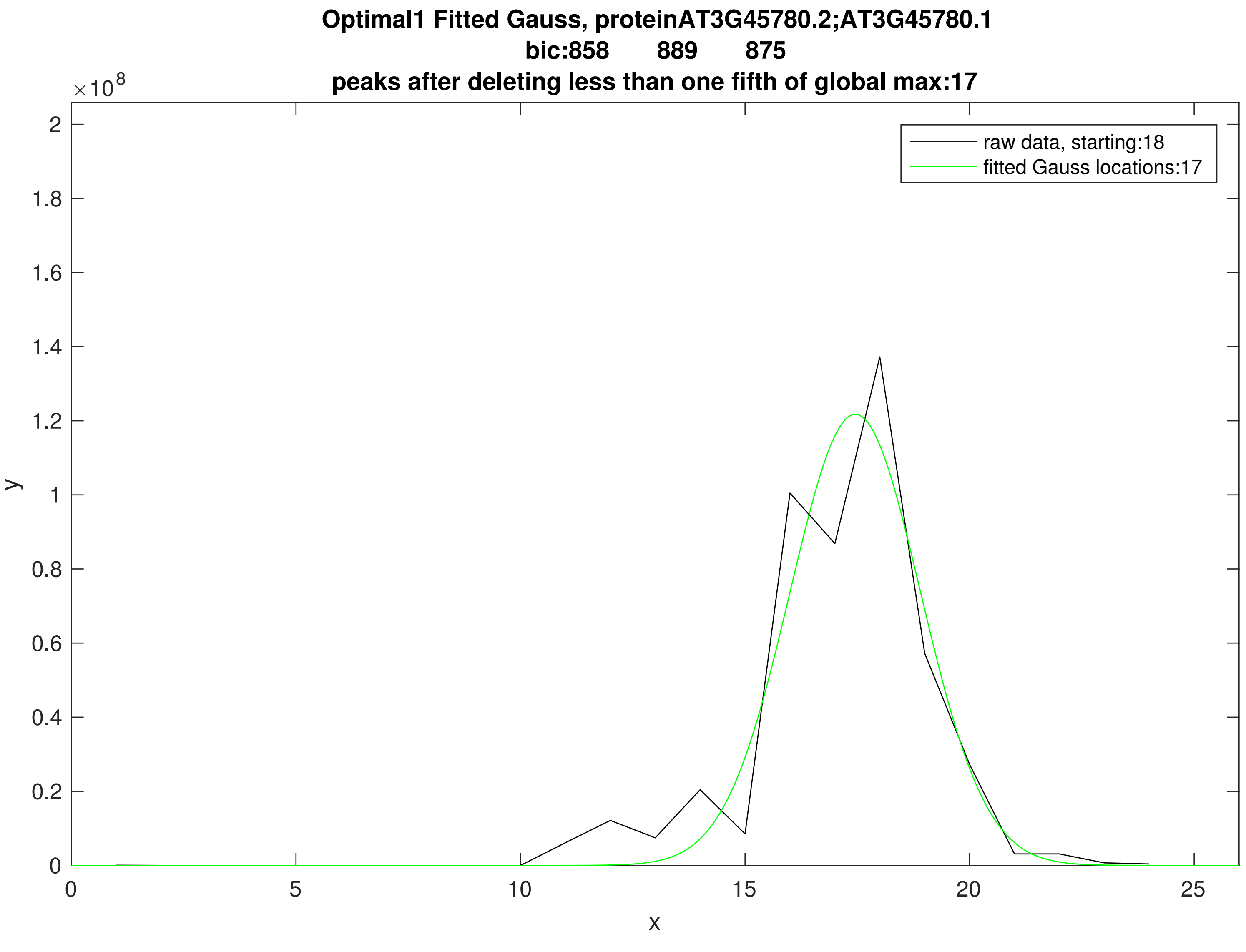
Optimal1 Fitted Gauss, proteinAT3G45780.2;AT3G45780.1
bic:858 889 875
peaks after deleting less than one fifth of global max:17
×10^8
raw data, starting:18
fitted Gauss locations:17
y
x
0
0.2
0.4
0.6
0.8
1
1.2
1.4
1.6
1.8
2
0
5
10
15
20
25

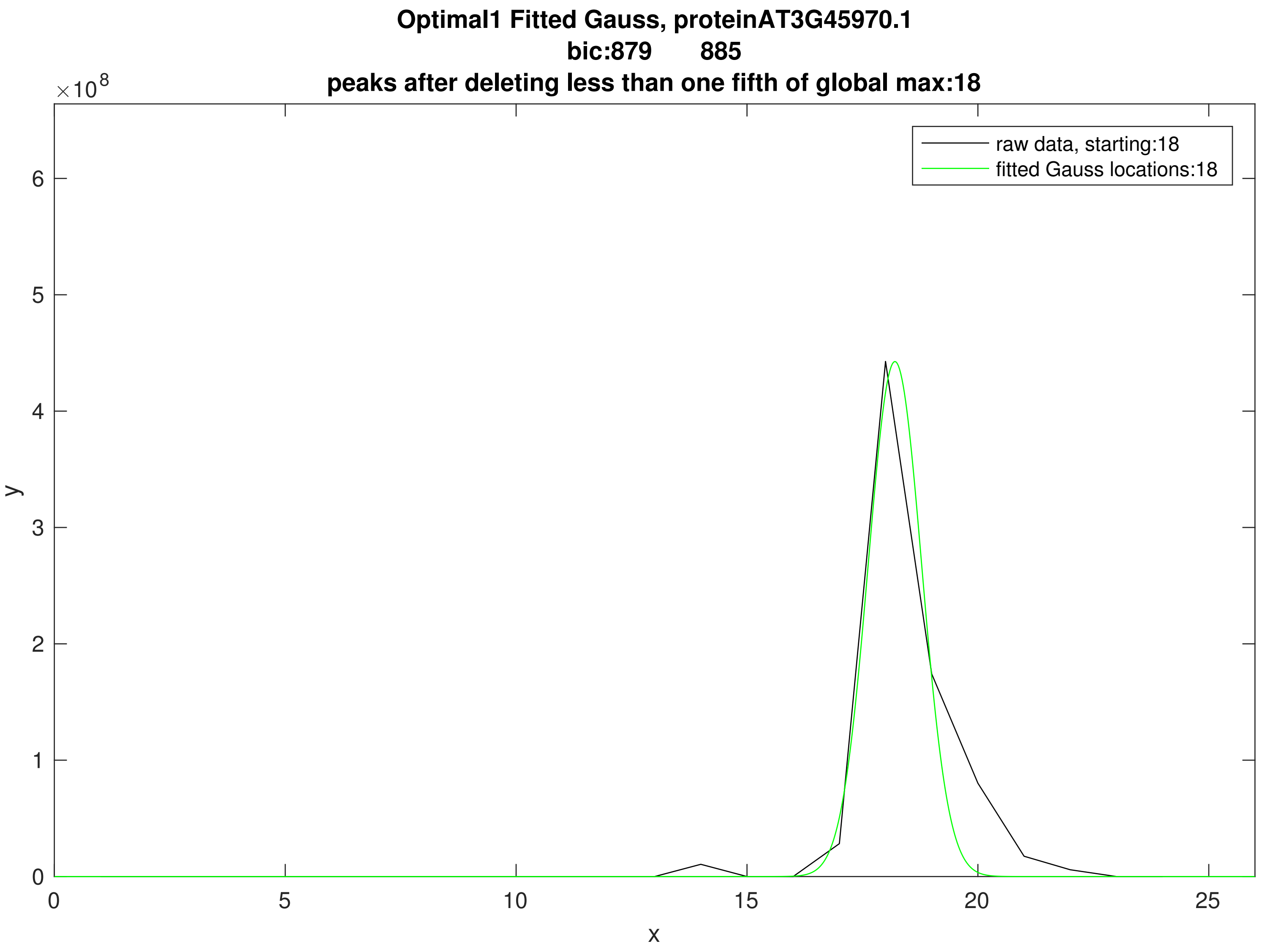

Optimal1 Fitted Gauss, proteinAT3G45970.1
bic:879 885
peaks after deleting less than one fifth of global max:18
×10^8
raw data, starting:18
fitted Gauss locations:18
y
x
0
1
2
3
4
5
6
0
5
10
15
20
25

**Optimal2 Fitted Gauss, proteinAT3G46000.1**
**bic:944 930 939**
**peaks after deleting less than one fifth of global max:19**

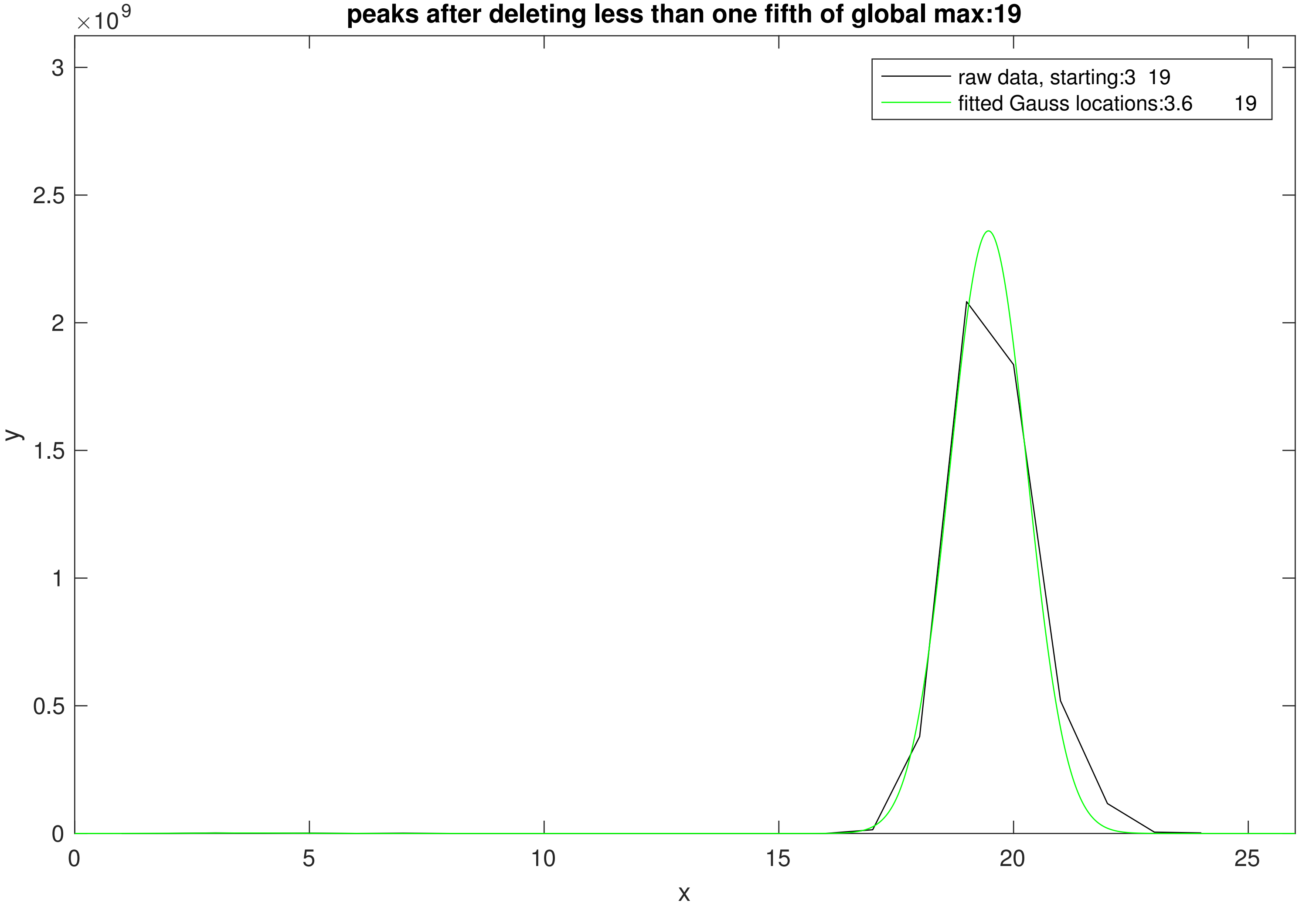

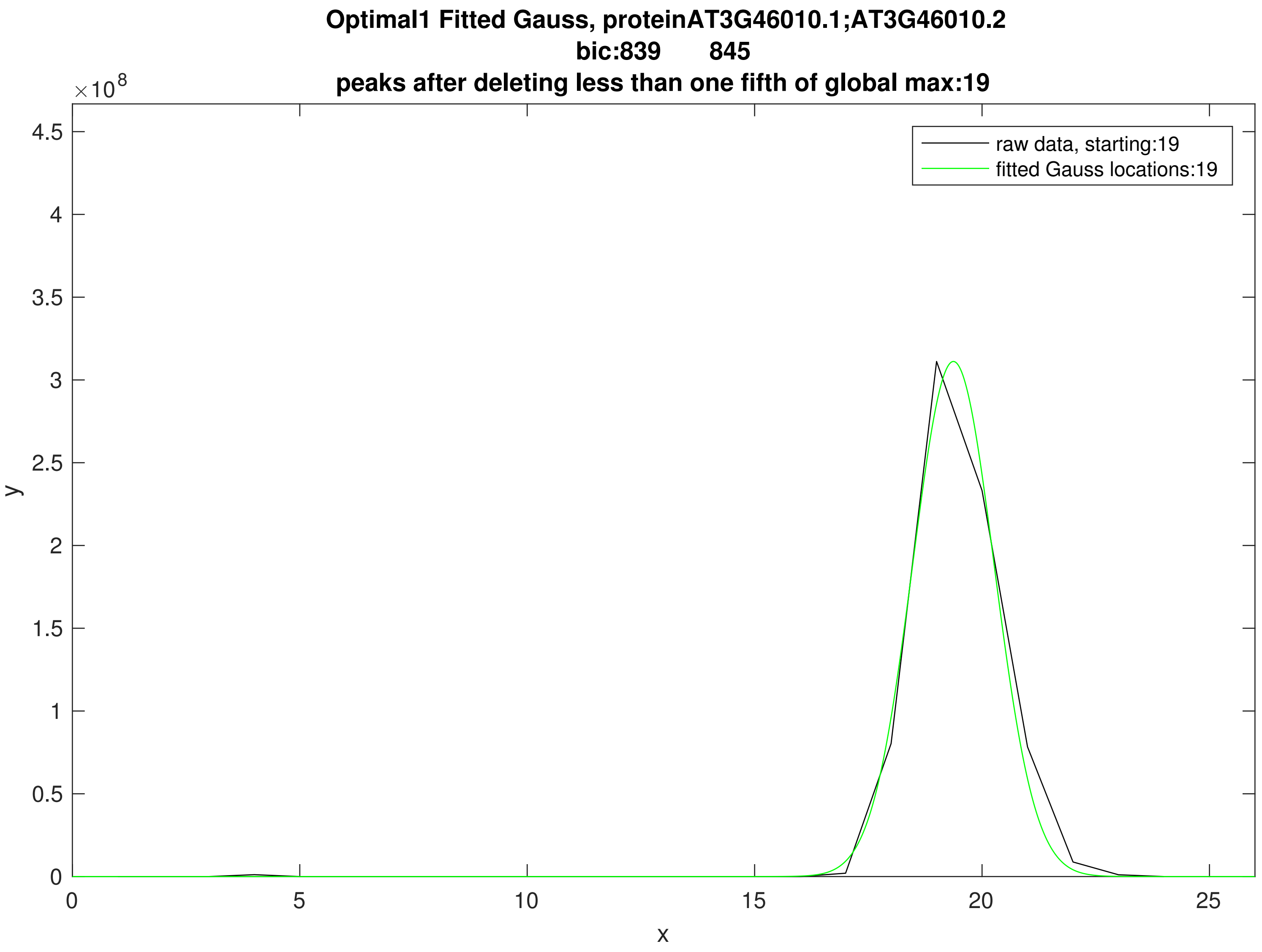
Optimal1 Fitted Gauss, proteinAT3G46010.1;AT3G46010.2
bic:839 845
peaks after deleting less than one fifth of global max:19
×10^8
4.5
4
3.5
3
2.5
2
1.5
1
0.5
0
y
0
5
10
15
20
25
x
raw data, starting:19
fitted Gauss locations:19

**Optimal1 Fitted Gauss, proteinAT3G46280.1**
**bic:739**
**peaks after deleting less than one fifth of global max:16**

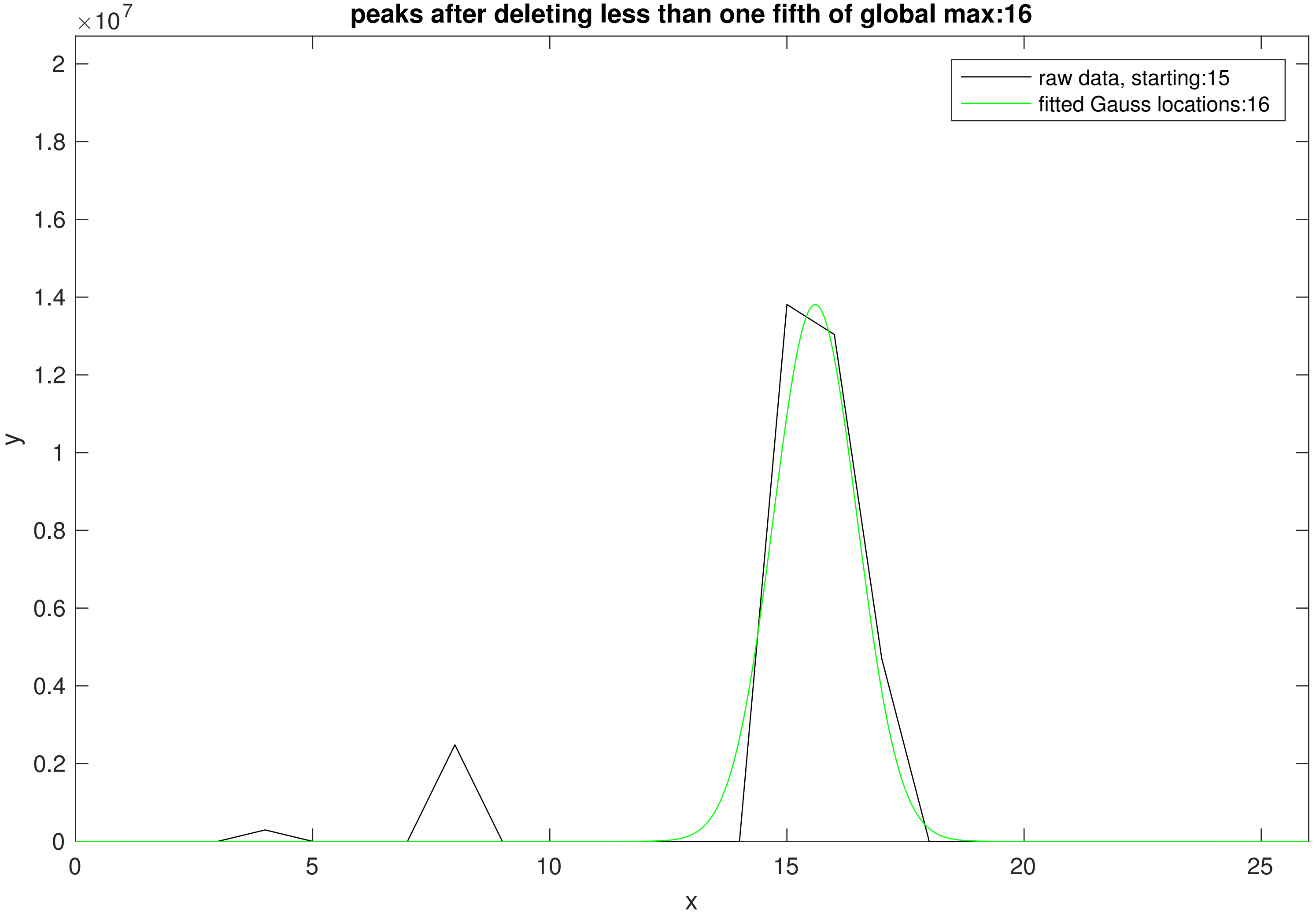

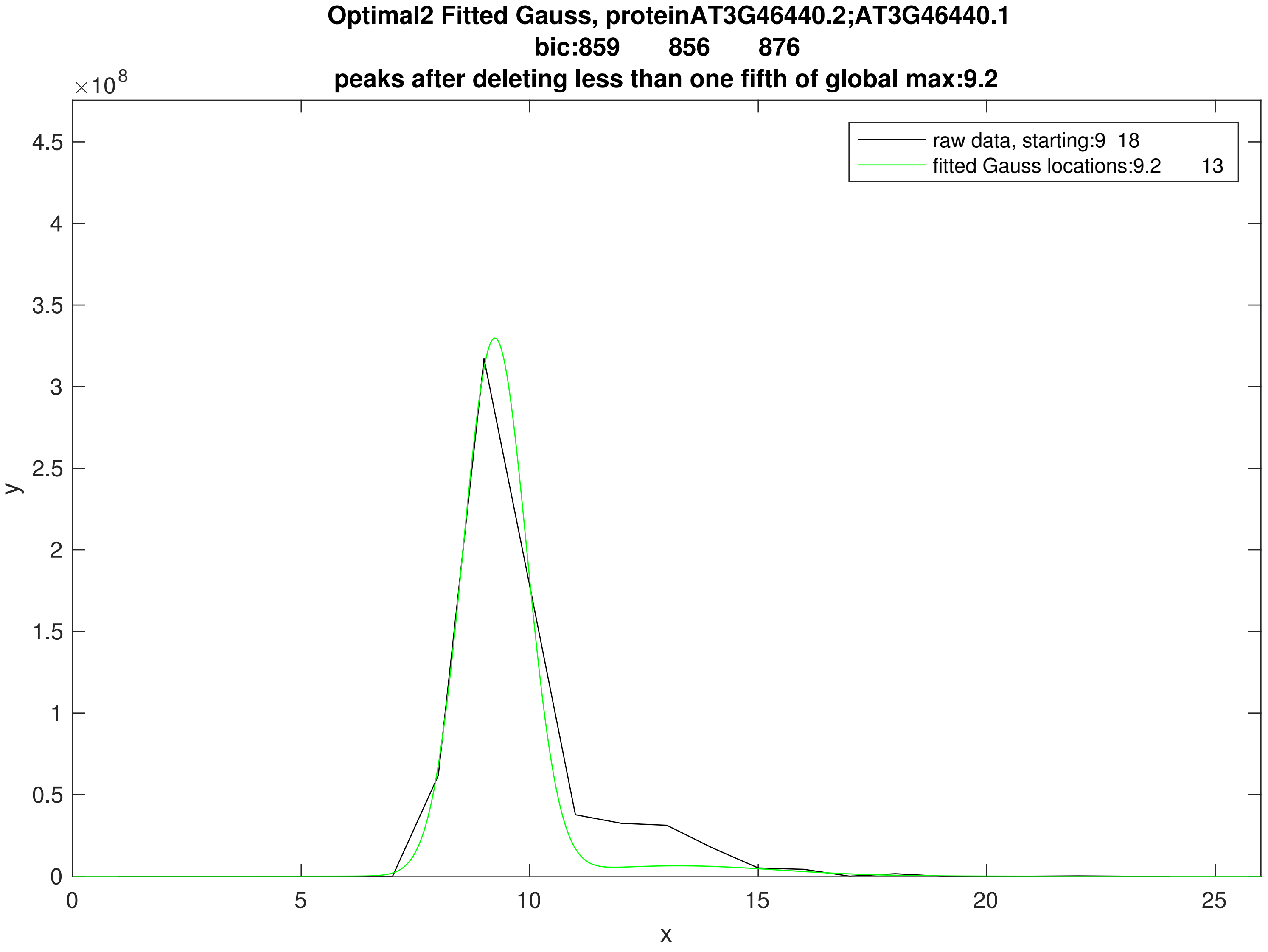
Optimal2 Fitted Gauss, proteinAT3G46440.2;AT3G46440.1
bic:859 856 876
peaks after deleting less than one fifth of global max:9.2
×10^8
raw data, starting:9 18
fitted Gauss locations:9.2 13
4.5
4
3.5
3
2.5
2
1.5
1
0.5
0
y
0
5
10
15
20
25
x

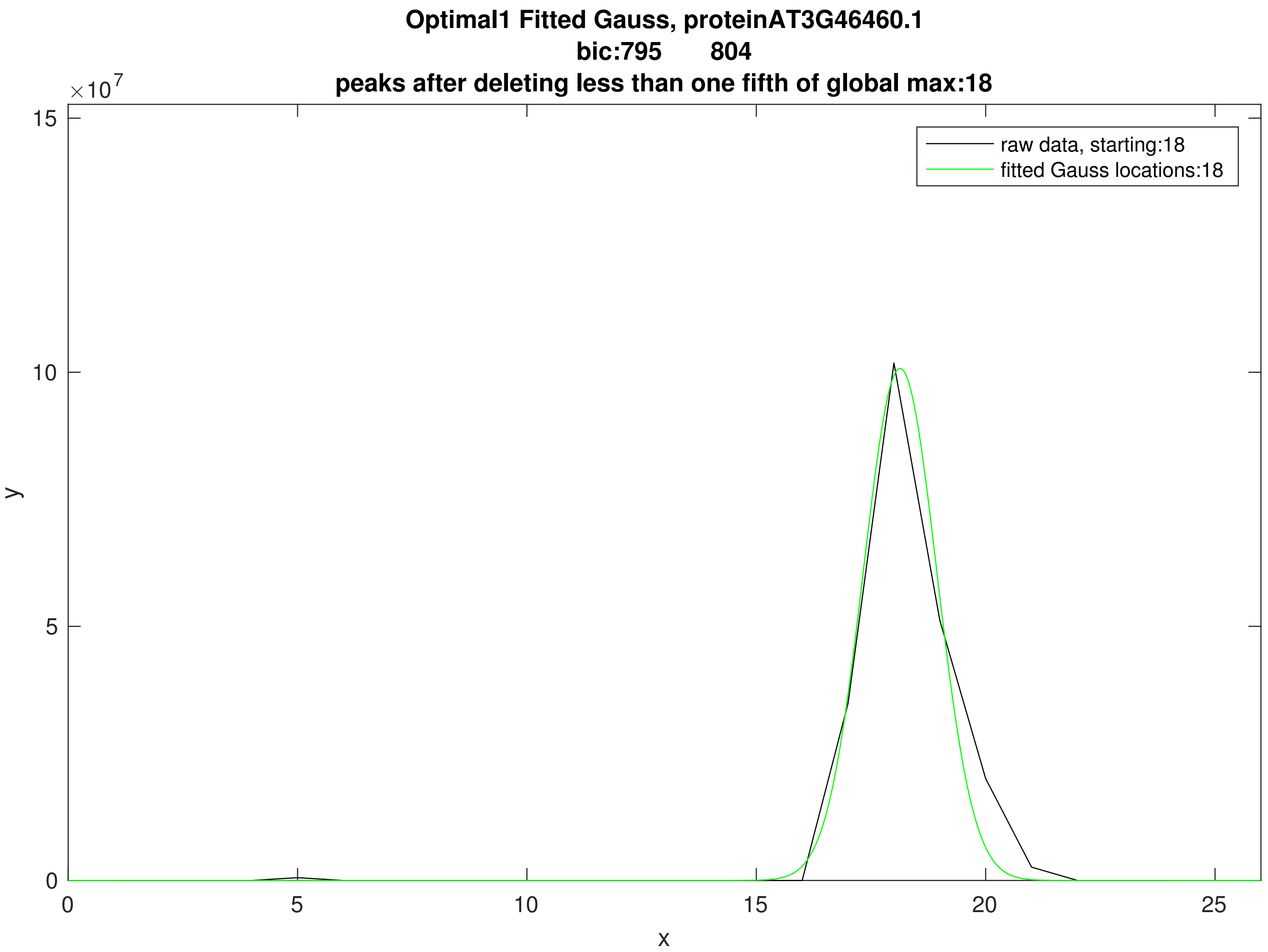

Optimal1 Fitted Gauss, proteinAT3G46460.1
bic:795 804
peaks after deleting less than one fifth of global max:18
×10^7
15
10
5
0
y
0
5
10
15
20
25
x
raw data, starting:18
fitted Gauss locations:18

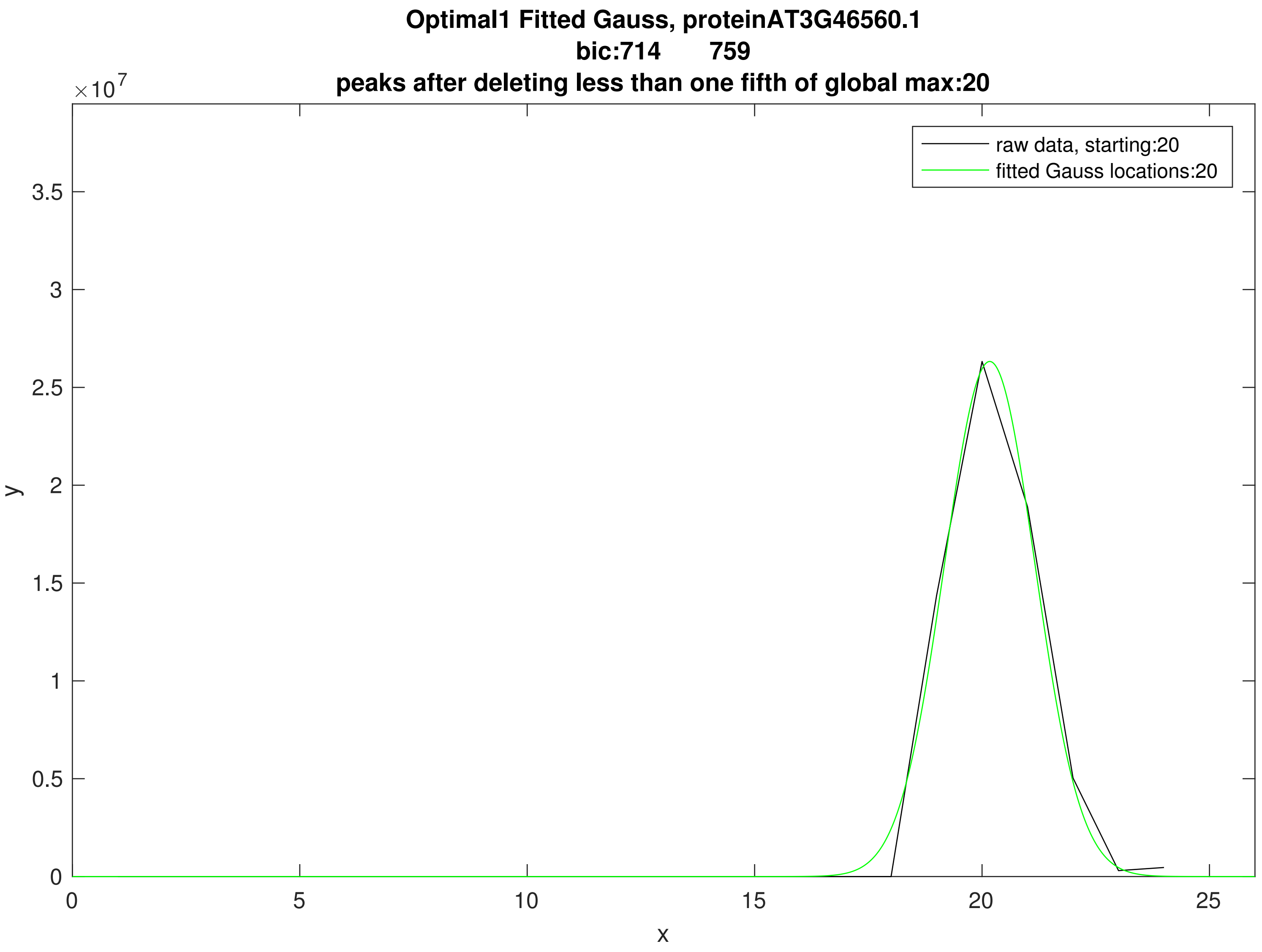
Optimal1 Fitted Gauss, proteinAT3G46560.1
bic:714 759
peaks after deleting less than one fifth of global max:20
×10^7
raw data, starting:20
fitted Gauss locations:20
3.5
3
2.5
2
1.5
1
0.5
0
y
0
5
10
15
20
25
x

**Optimal1 Fitted Gauss, proteinAT3G46940.1**
**bic:857**
**peaks after deleting less than one fifth of global max:14**

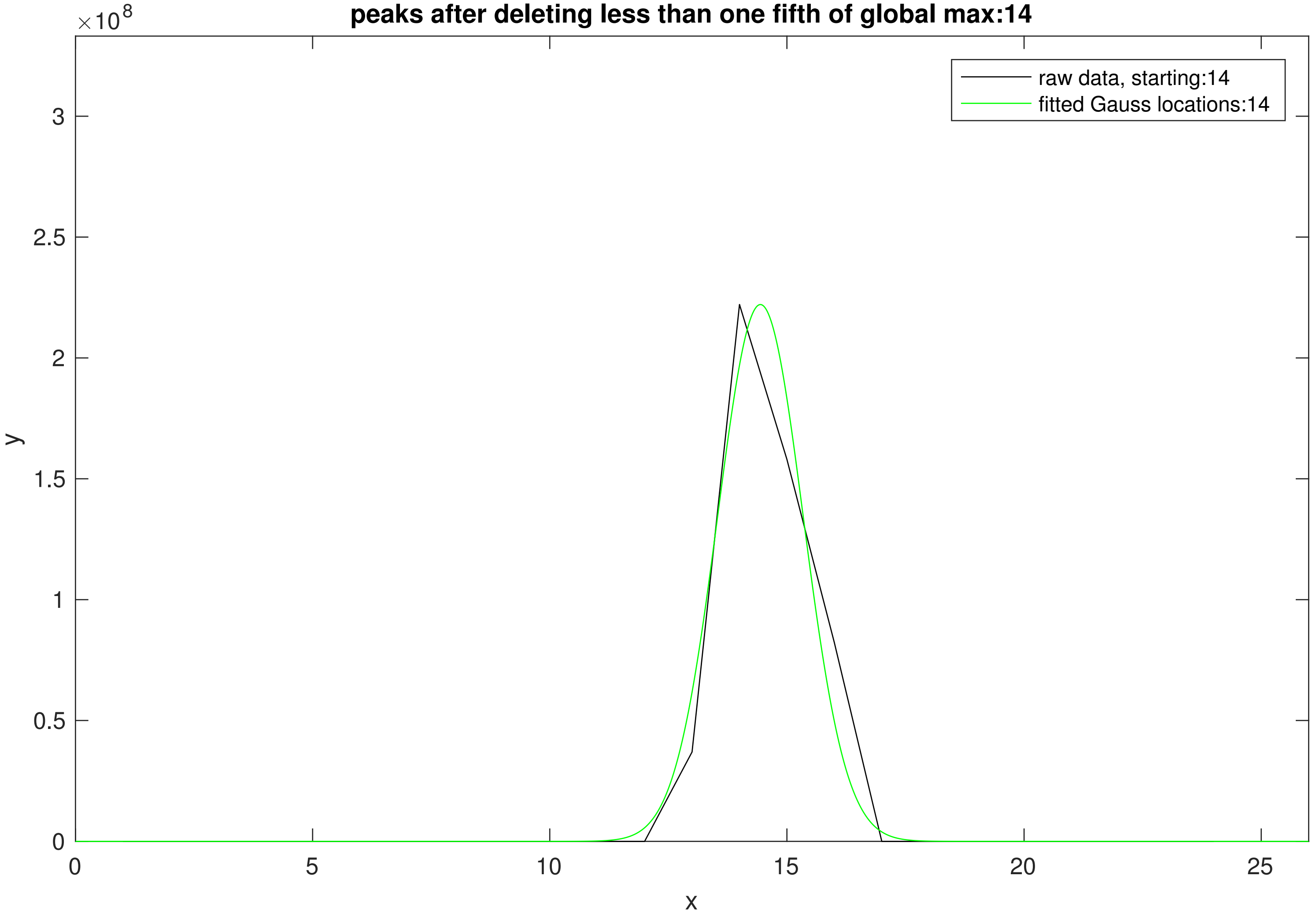

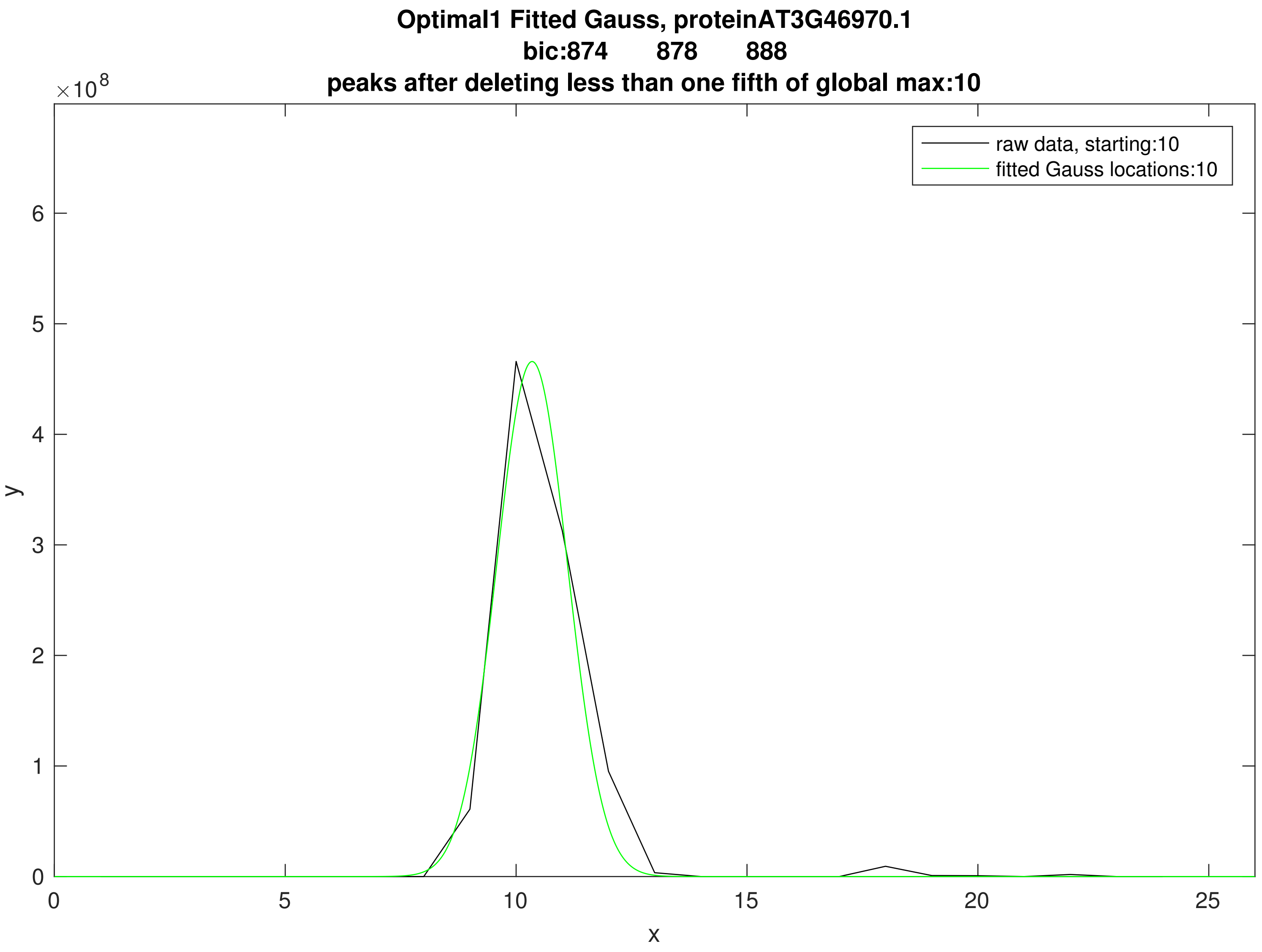

Optimal1 Fitted Gauss, proteinAT3G46970.1
bic:874 878 888
peaks after deleting less than one fifth of global max:10
×10^8
raw data, starting:10
fitted Gauss locations:10
y
x
0
1
2
3
4
5
6
0
5
10
15
20
25

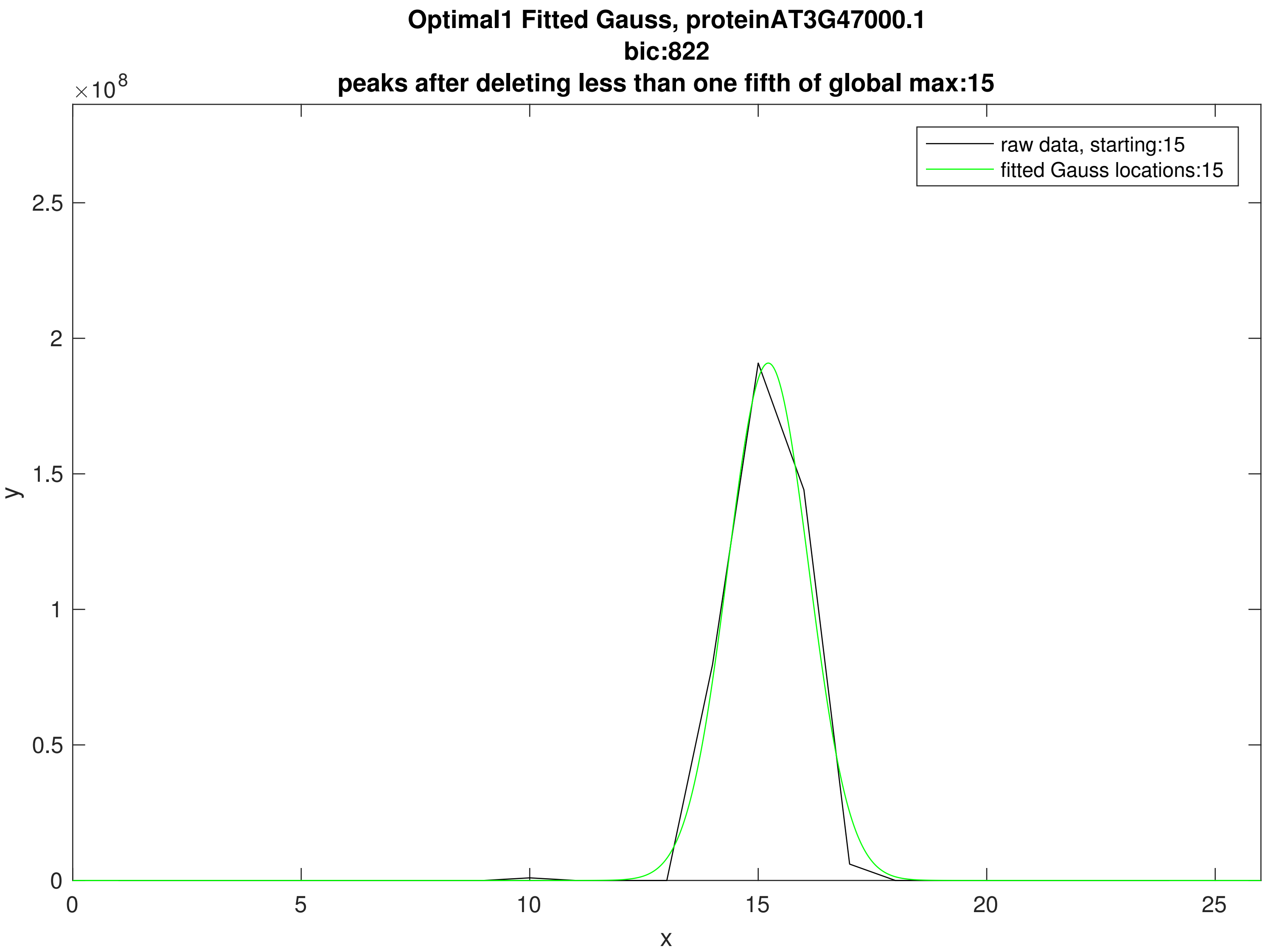
Optimal1 Fitted Gauss, proteinAT3G47000.1
bic:822
peaks after deleting less than one fifth of global max:15
×10^8
2.5
2
1.5
1
0.5
0
y
0
5
10
15
20
25
x
raw data, starting:15
fitted Gauss locations:15

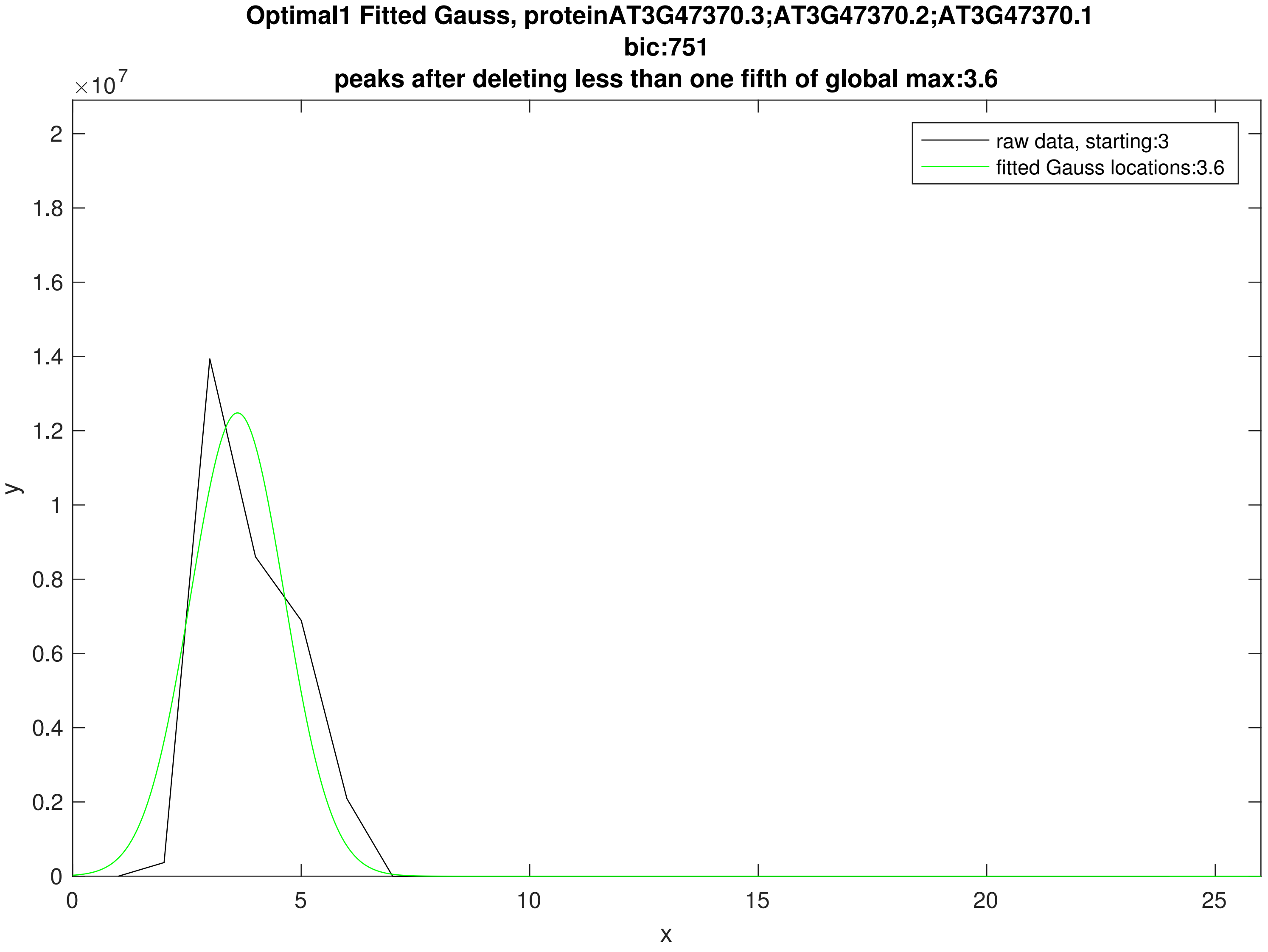
Optimal1 Fitted Gauss, proteinAT3G47370.3;AT3G47370.2;AT3G47370.1
bic:751
peaks after deleting less than one fifth of global max:3.6
×10^7
raw data, starting:3
fitted Gauss locations:3.6
y
x
0
0.2
0.4
0.6
0.8
1
1.2
1.4
1.6
1.8
2
5
10
15
20
25

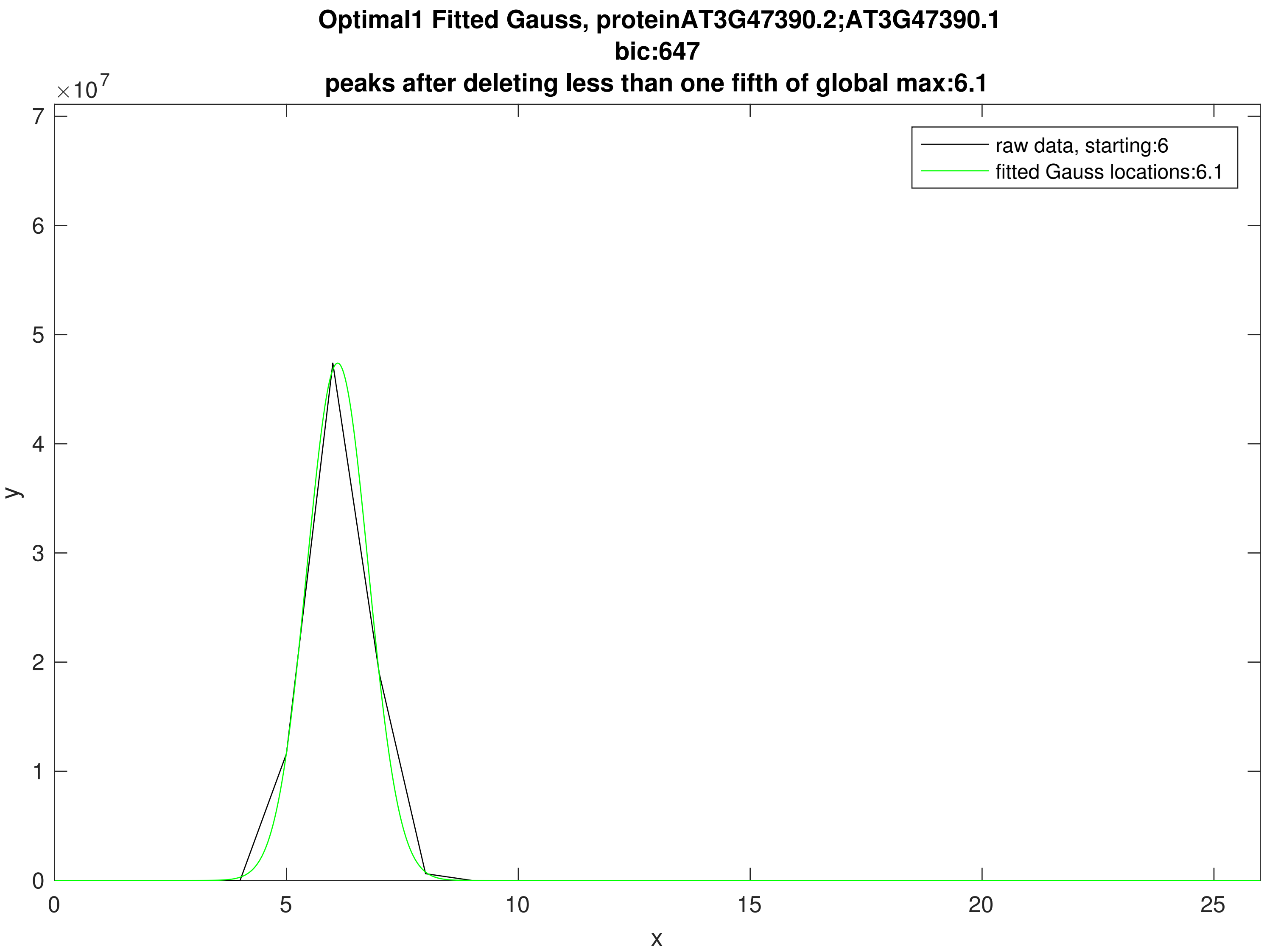
Optimal1 Fitted Gauss, proteinAT3G47390.2;AT3G47390.1
bic:647
peaks after deleting less than one fifth of global max:6.1
×10^7
raw data, starting:6
fitted Gauss locations:6.1
y
x
0
1
2
3
4
5
6
7
0
5
10
15
20
25

Optimal2 Fitted Gauss, proteinAT3G47460.1;AT5G62410.1
bic:859 842
peaks after deleting less than one fifth of global max:9.5 14

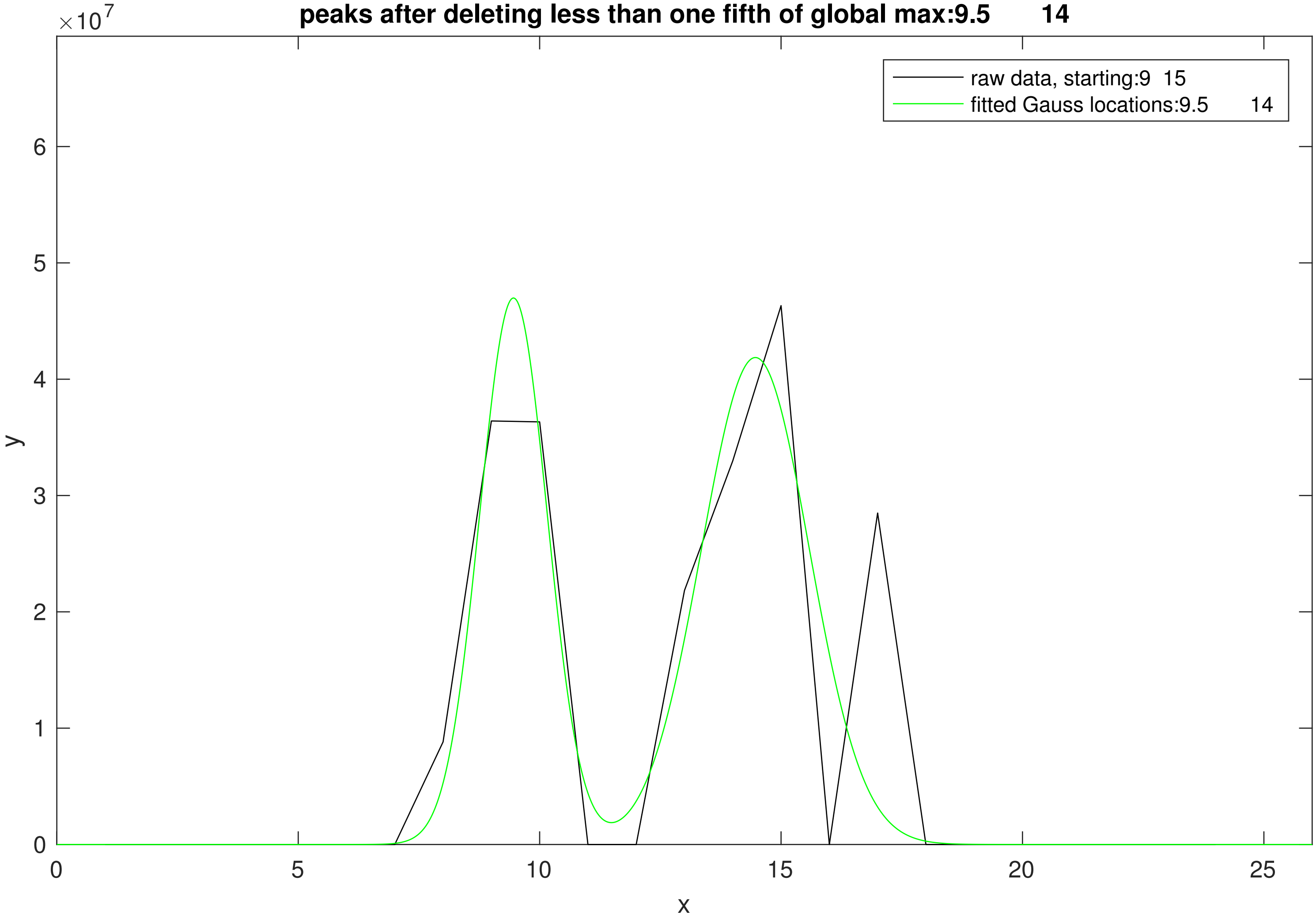

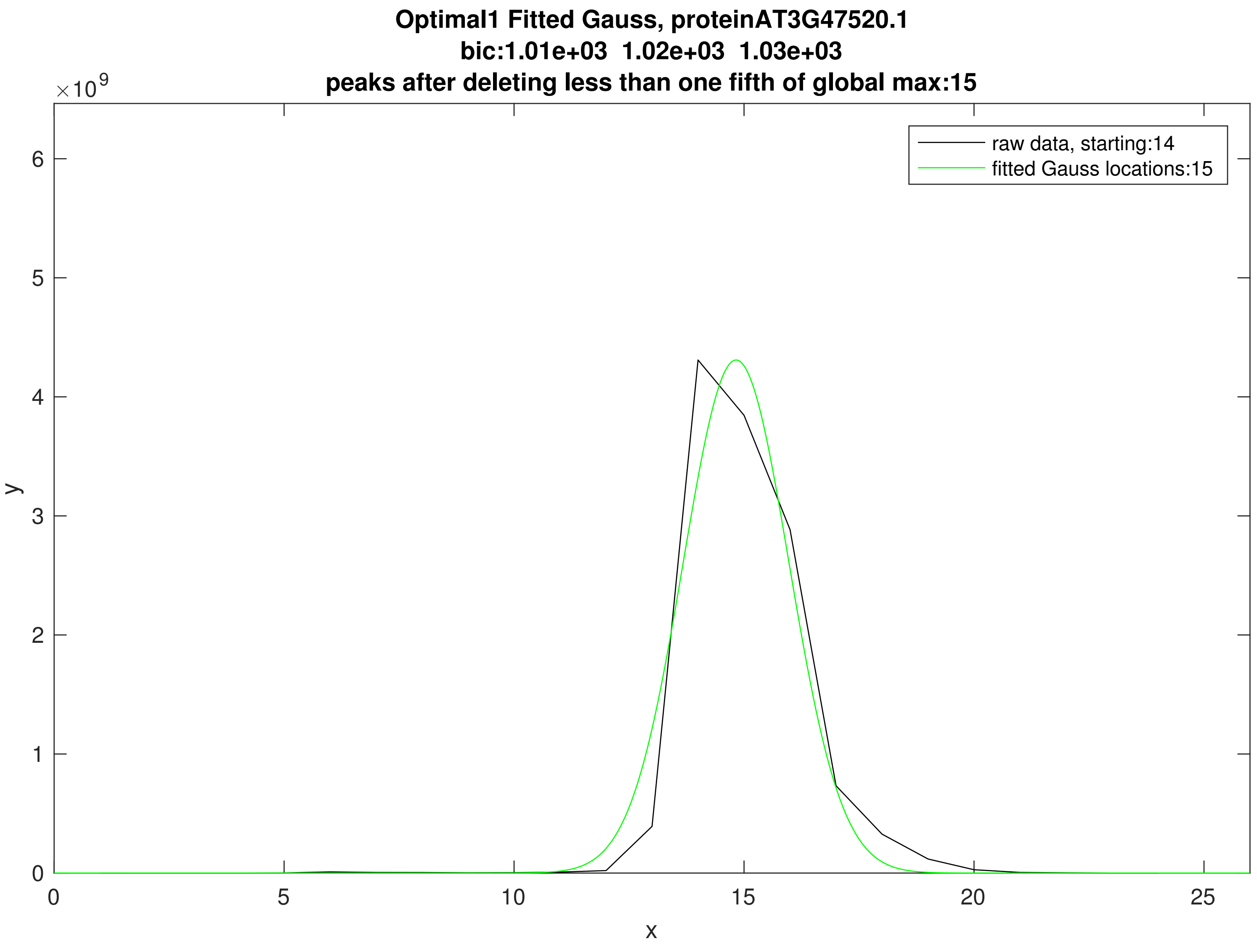
Optimal1 Fitted Gauss, proteinAT3G47520.1
bic:1.01e+03  1.02e+03  1.03e+03
peaks after deleting less than one fifth of global max:15
raw data, starting:14
fitted Gauss locations:15
×10^9
y
x

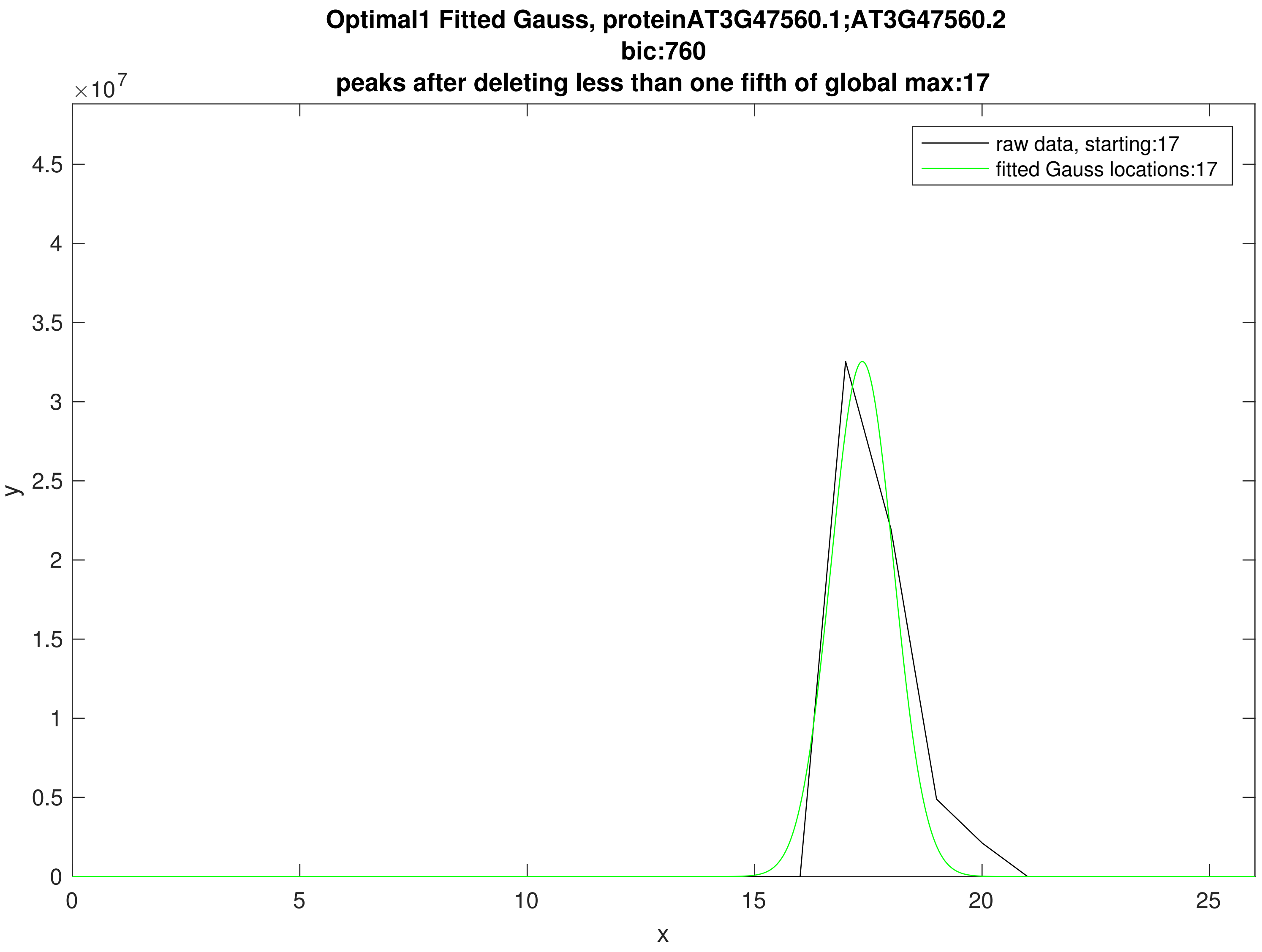
Optimal1 Fitted Gauss, proteinAT3G47560.1;AT3G47560.2
bic:760
peaks after deleting less than one fifth of global max:17
×10^7
4.5
4
3.5
3
2.5
2
1.5
1
0.5
0
y
0
5
10
15
20
25
x
raw data, starting:17
fitted Gauss locations:17

**Optimal1 Fitted Gauss, proteinAT3G47590.1**
**bic:845**
**peaks after deleting less than one fifth of global max:18**

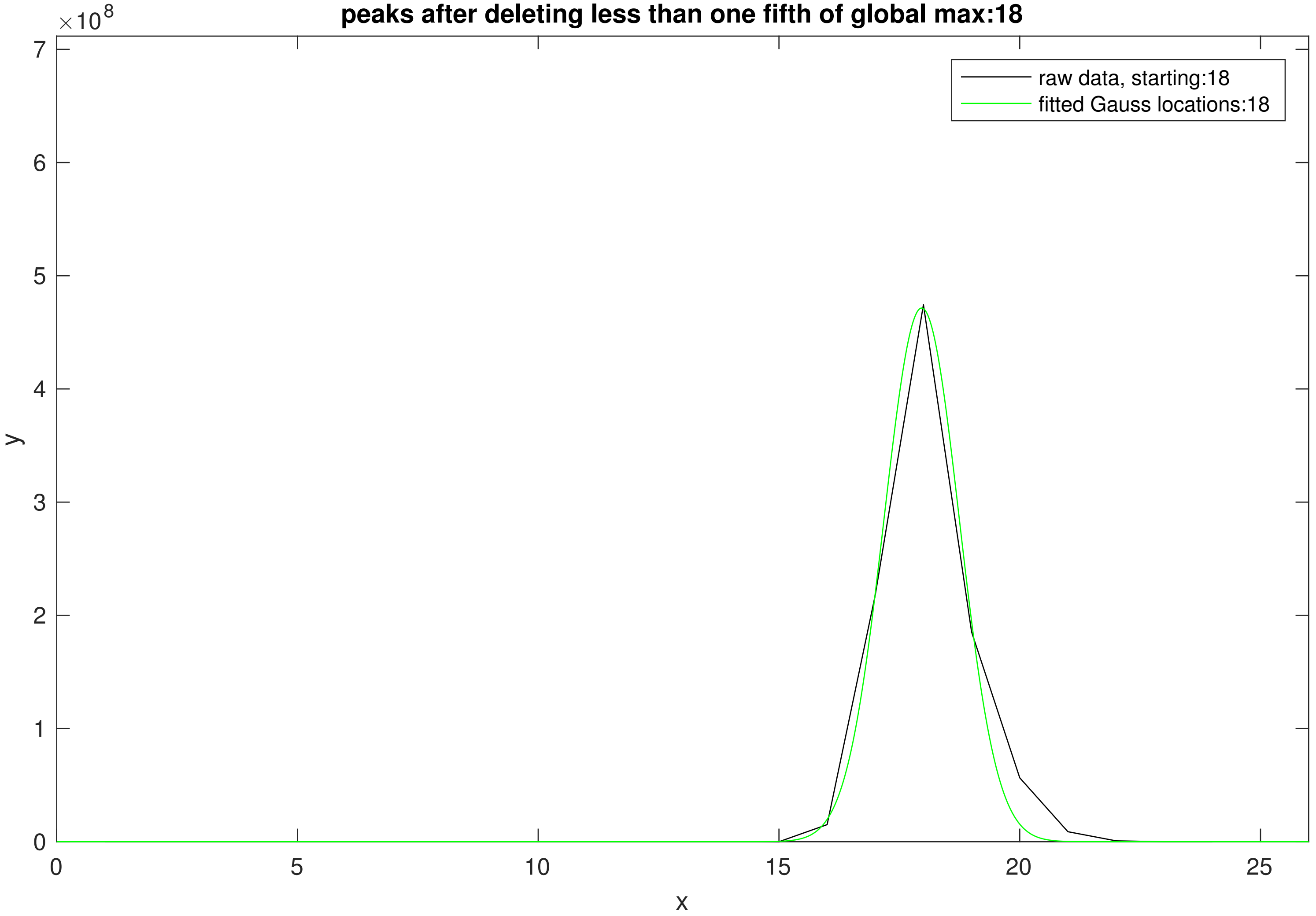

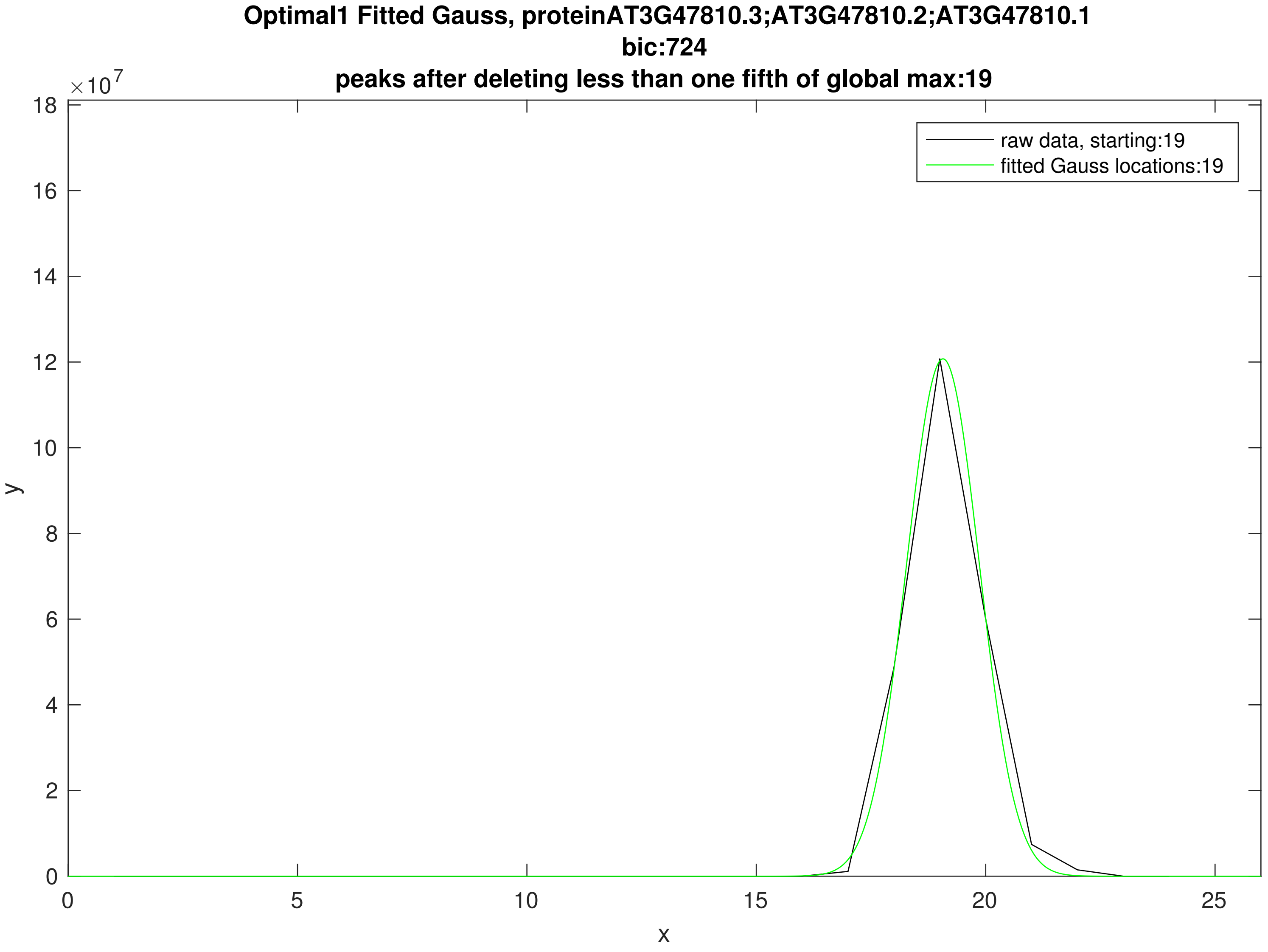
Optimal1 Fitted Gauss, proteinAT3G47810.3;AT3G47810.2;AT3G47810.1
bic:724
peaks after deleting less than one fifth of global max:19
$\times 10^7$
18
16
14
12
10
8
6
4
2
0
y
0
5
10
15
20
25
x
raw data, starting:19
fitted Gauss locations:19

Optimal1 Fitted Gauss, proteinAT3G48000.1
bic:954 963
peaks after deleting less than one fifth of global max:10

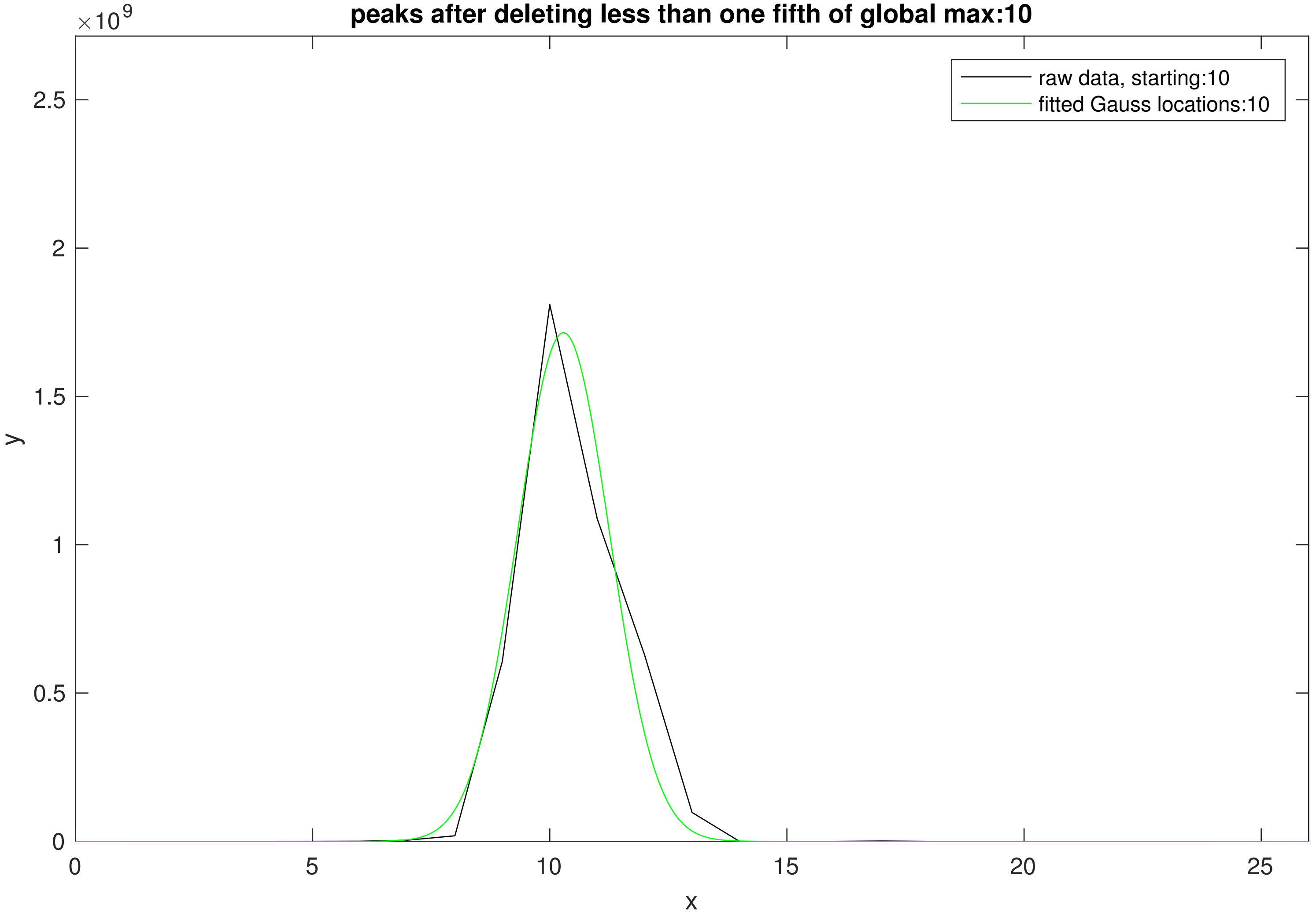

**Optimal1 Fitted Gauss, proteinAT3G48110.1**
**bic:851       853**
**peaks after deleting less than one fifth of global max:7.6**

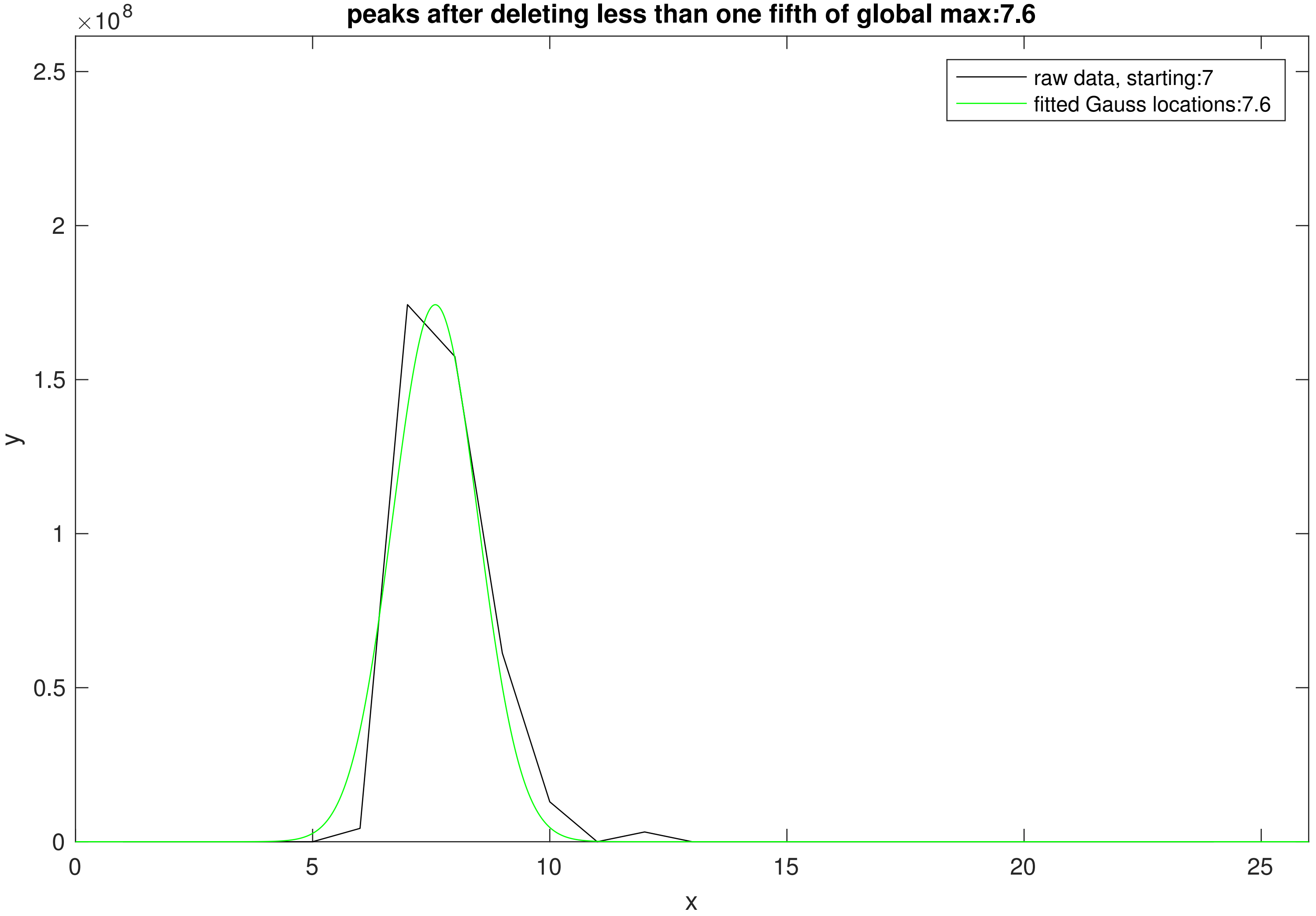

Optimal1 Fitted Gauss, proteinAT3G48170.1
bic:917
peaks after deleting less than one fifth of global max:13

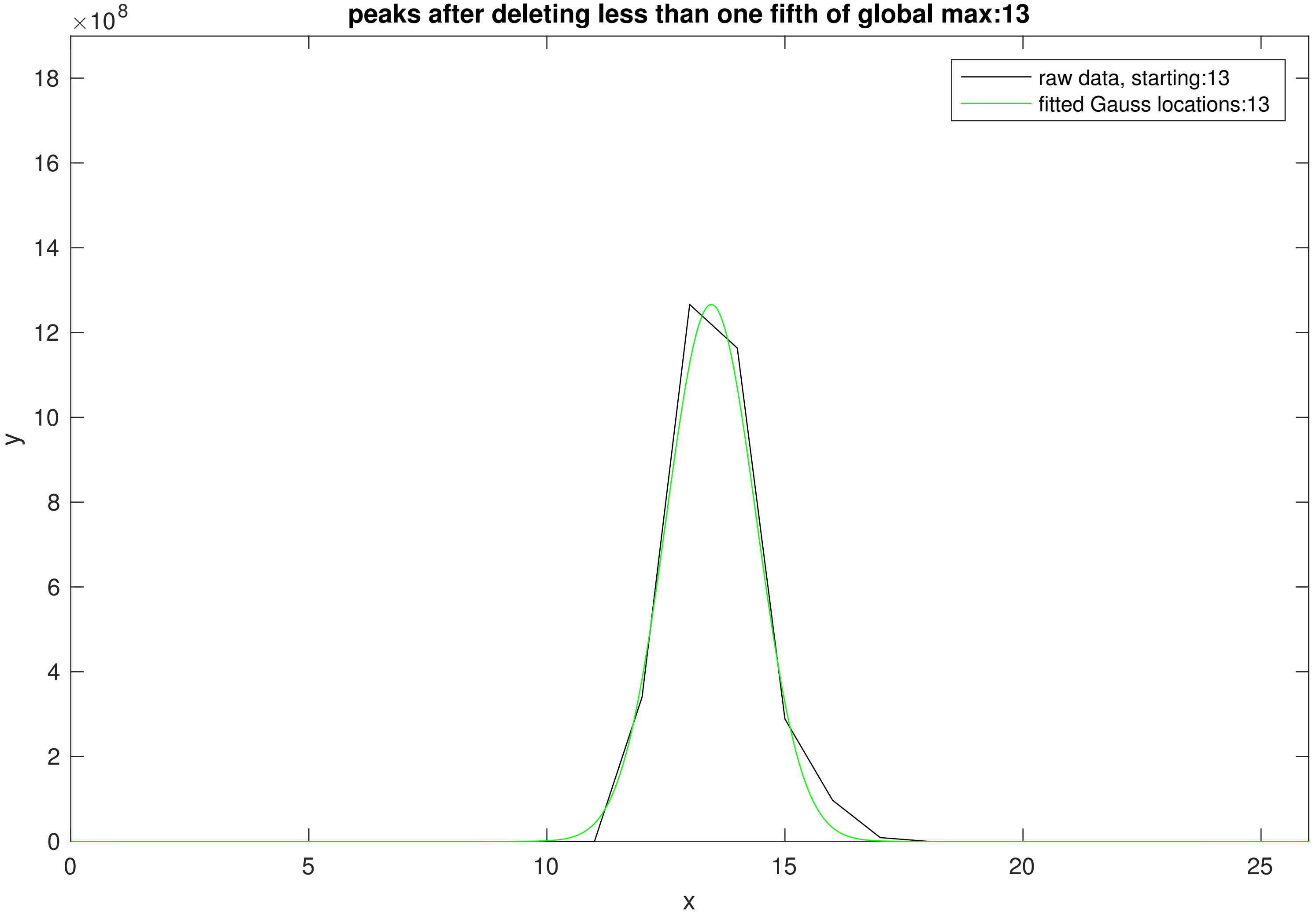

**Optimal1 Fitted Gauss, proteinAT3G48350.1**
**bic:776**
**peaks after deleting less than one fifth of global max:13**

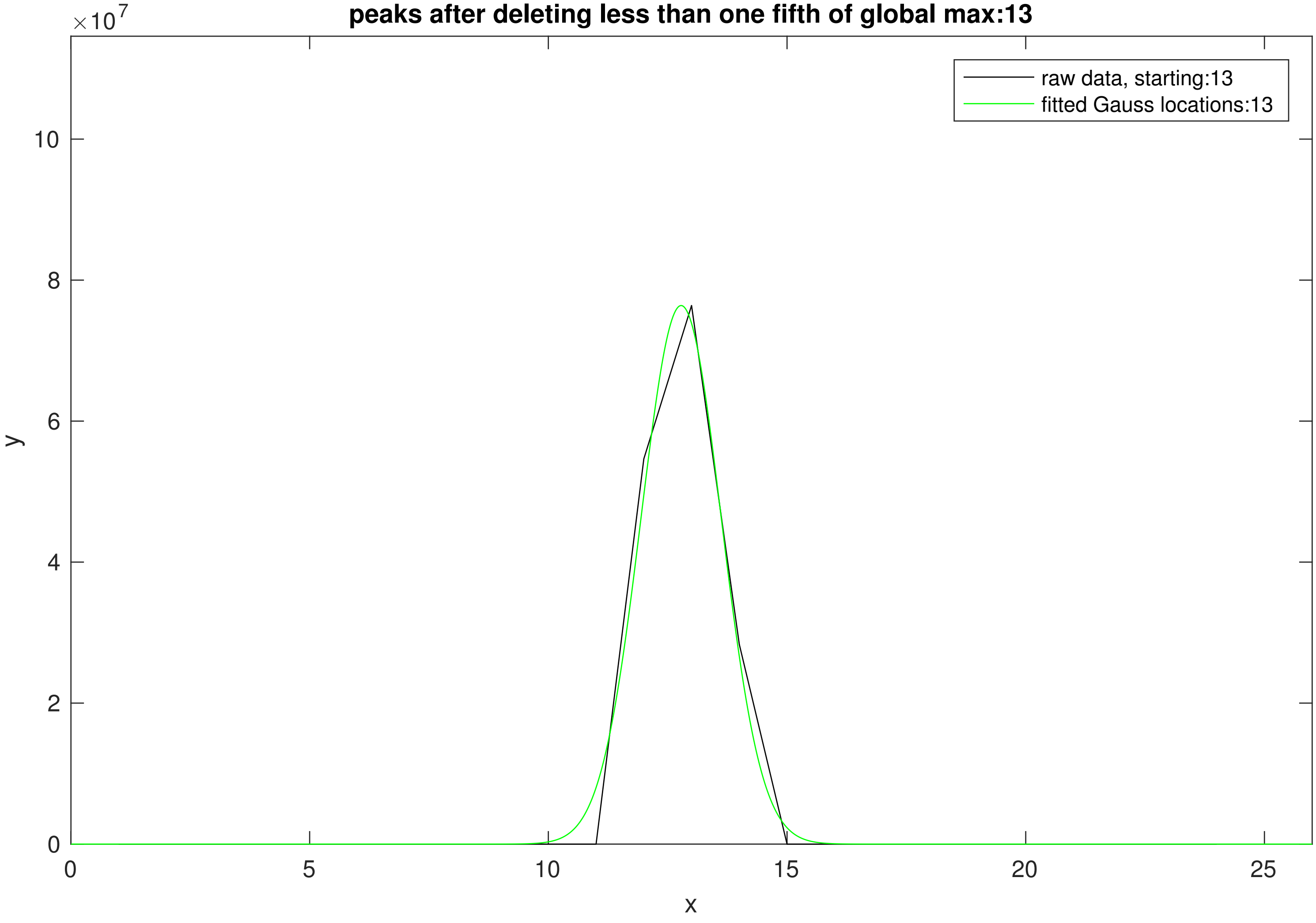

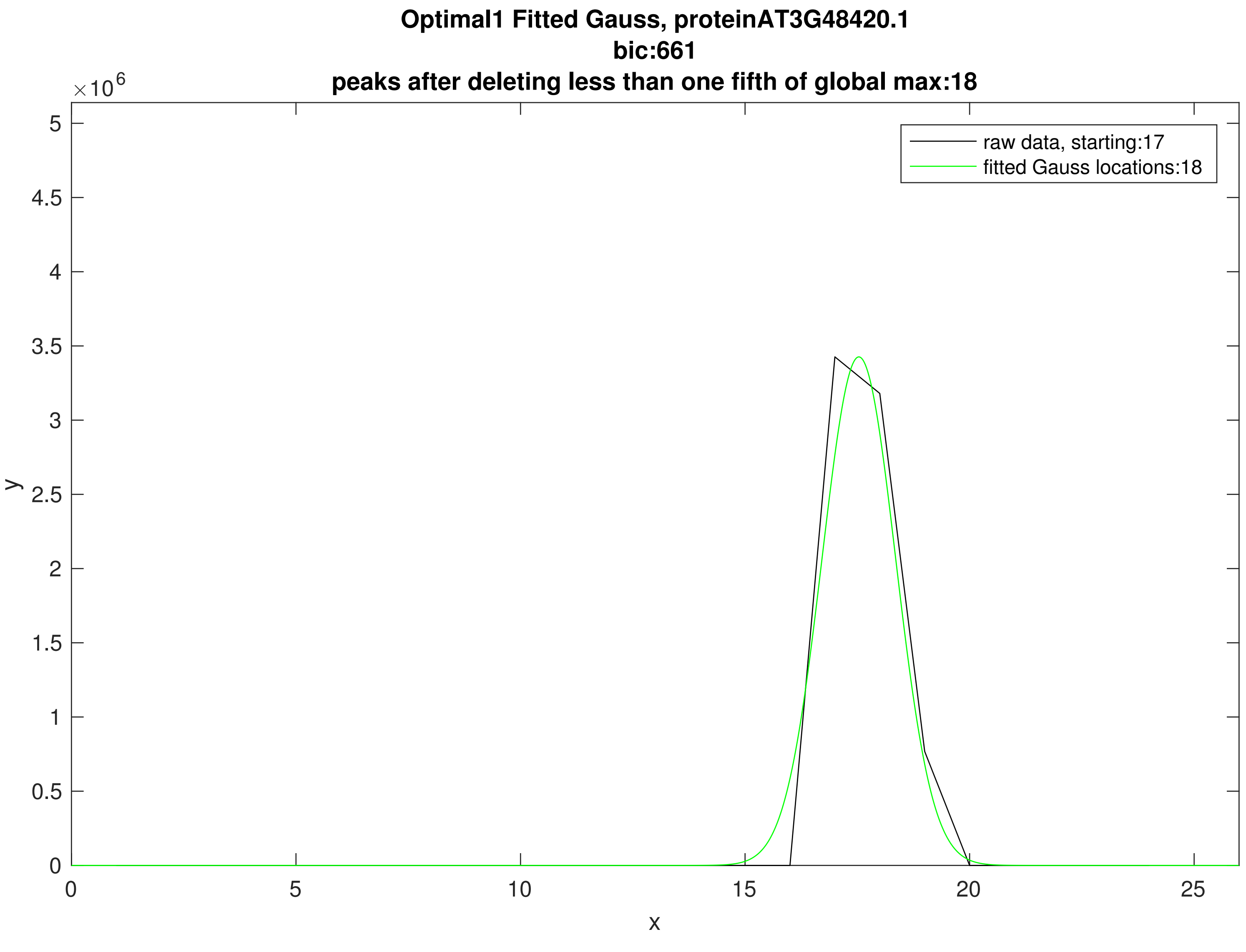

Optimal1 Fitted Gauss, proteinAT3G48420.1
bic:661
peaks after deleting less than one fifth of global max:18
×10^6
raw data, starting:17
fitted Gauss locations:18
y
x
0
0.5
1
1.5
2
2.5
3
3.5
4
4.5
5
0
5
10
15
20
25

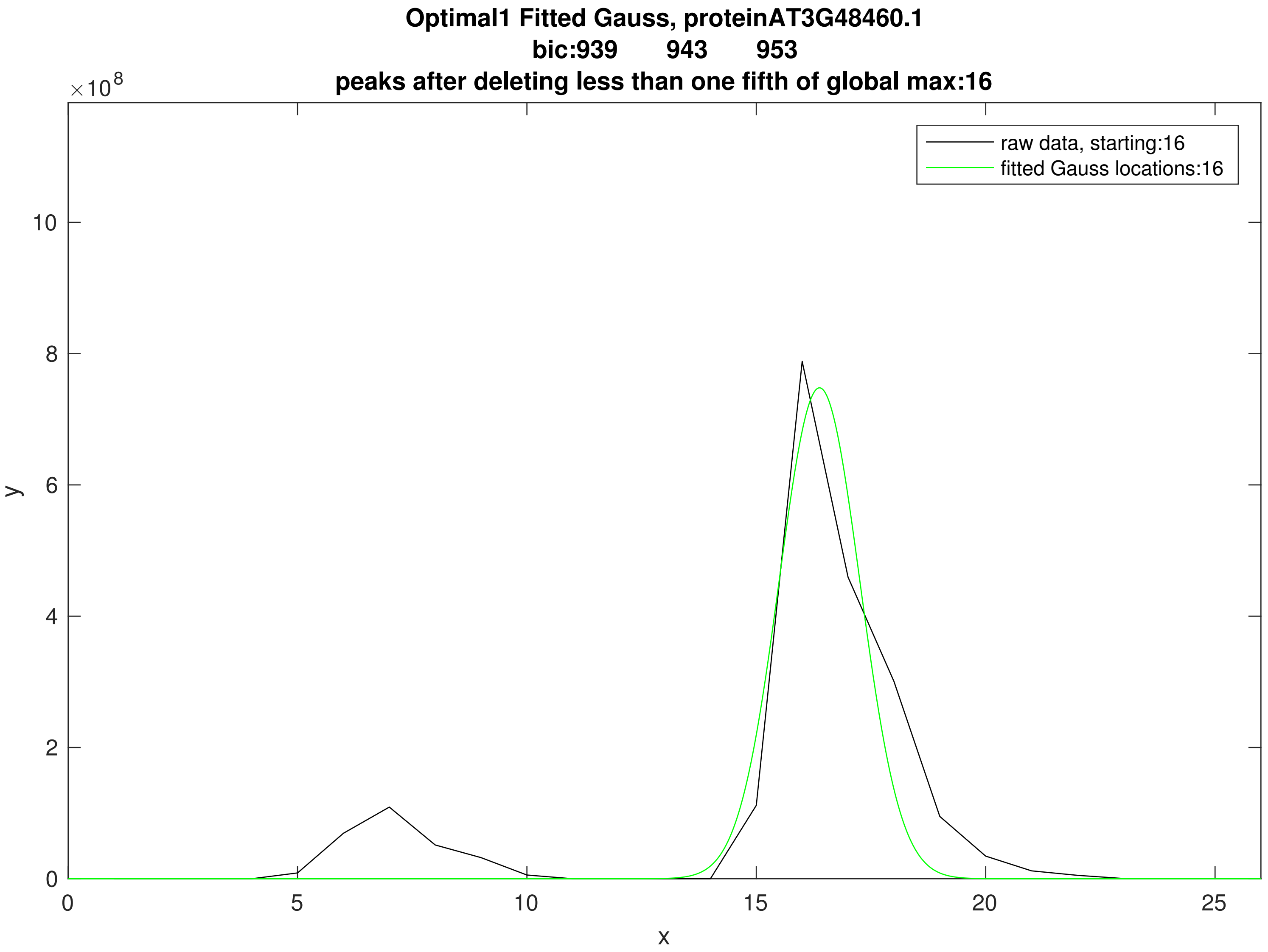
Optimal1 Fitted Gauss, proteinAT3G48460.1
bic:939 943 953
peaks after deleting less than one fifth of global max:16
raw data, starting:16
fitted Gauss locations:16
×10^8
y
x
0
2
4
6
8
10
0
5
10
15
20
25

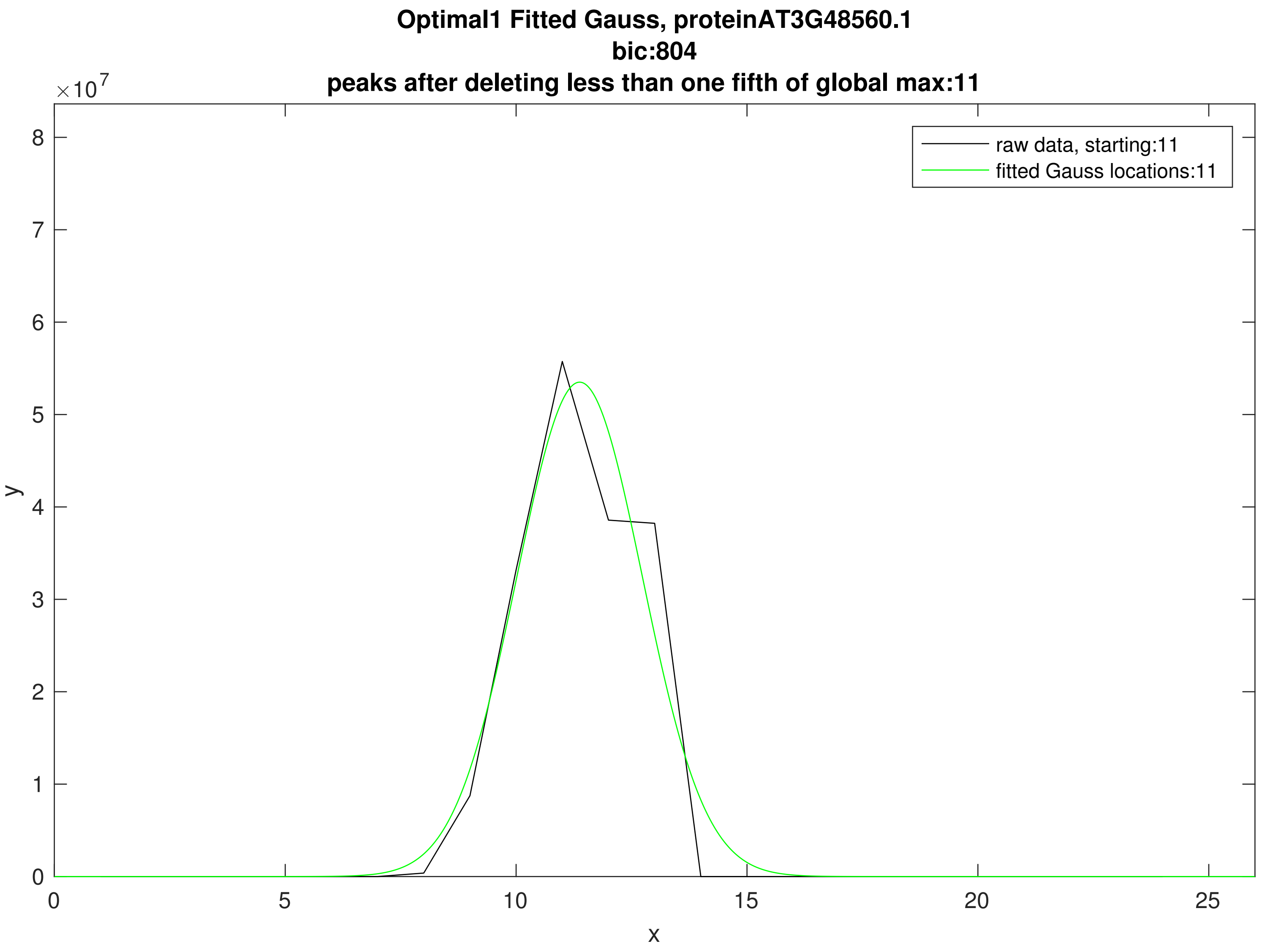

Optimal1 Fitted Gauss, proteinAT3G48560.1
bic:804
peaks after deleting less than one fifth of global max:11
×10^7
raw data, starting:11
fitted Gauss locations:11
y
x
0
1
2
3
4
5
6
7
8
0
5
10
15
20
25

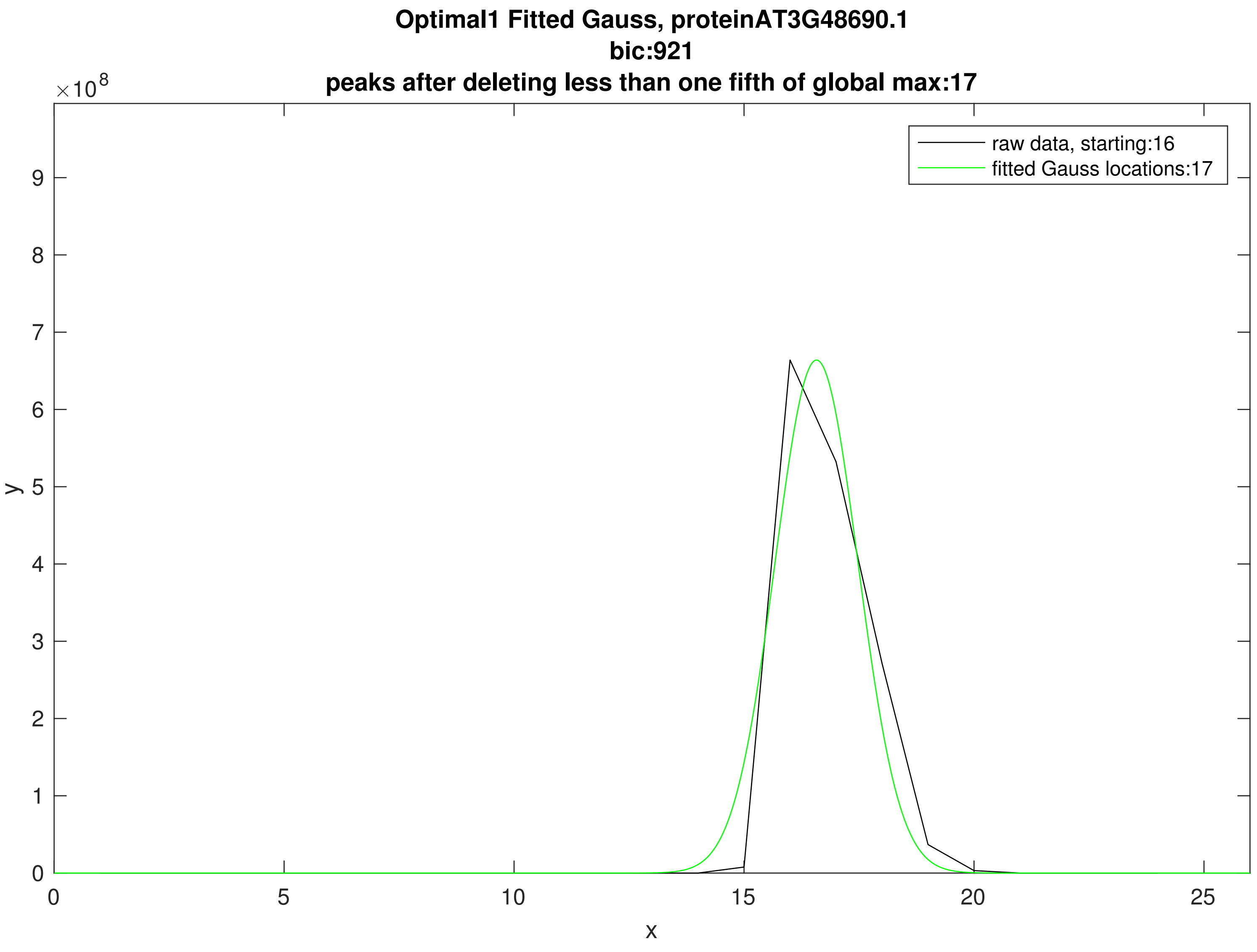
Optimal1 Fitted Gauss, proteinAT3G48690.1
bic:921
peaks after deleting less than one fifth of global max:17
×10^8
raw data, starting:16
fitted Gauss locations:17
y
x
0
1
2
3
4
5
6
7
8
9
0
5
10
15
20
25

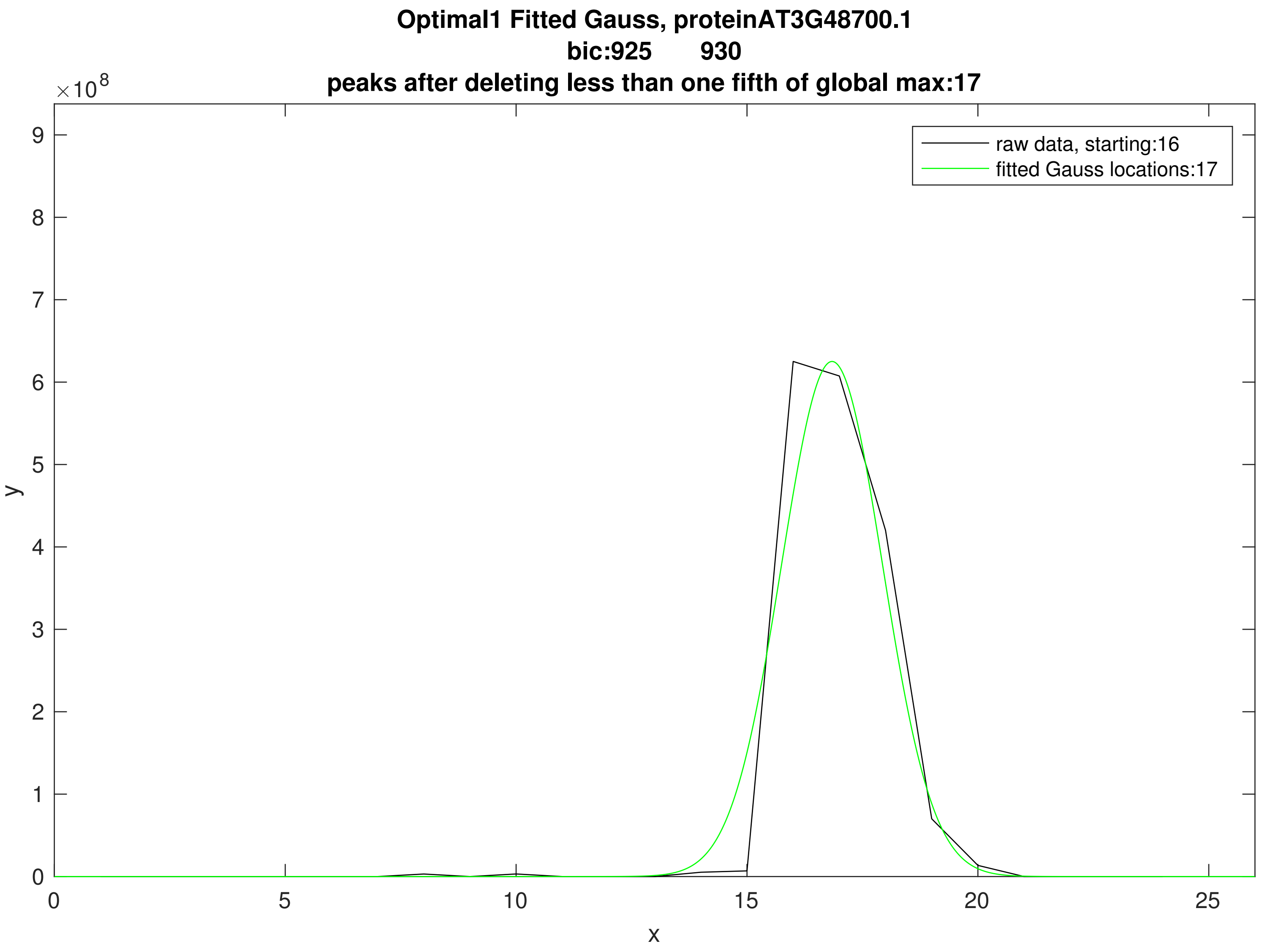

Optimal1 Fitted Gauss, proteinAT3G48700.1
bic:925 930
peaks after deleting less than one fifth of global max:17
×10^8
9
8
7
6
5
4
3
2
1
0
y
0
5
10
15
20
25
x
raw data, starting:16
fitted Gauss locations:17

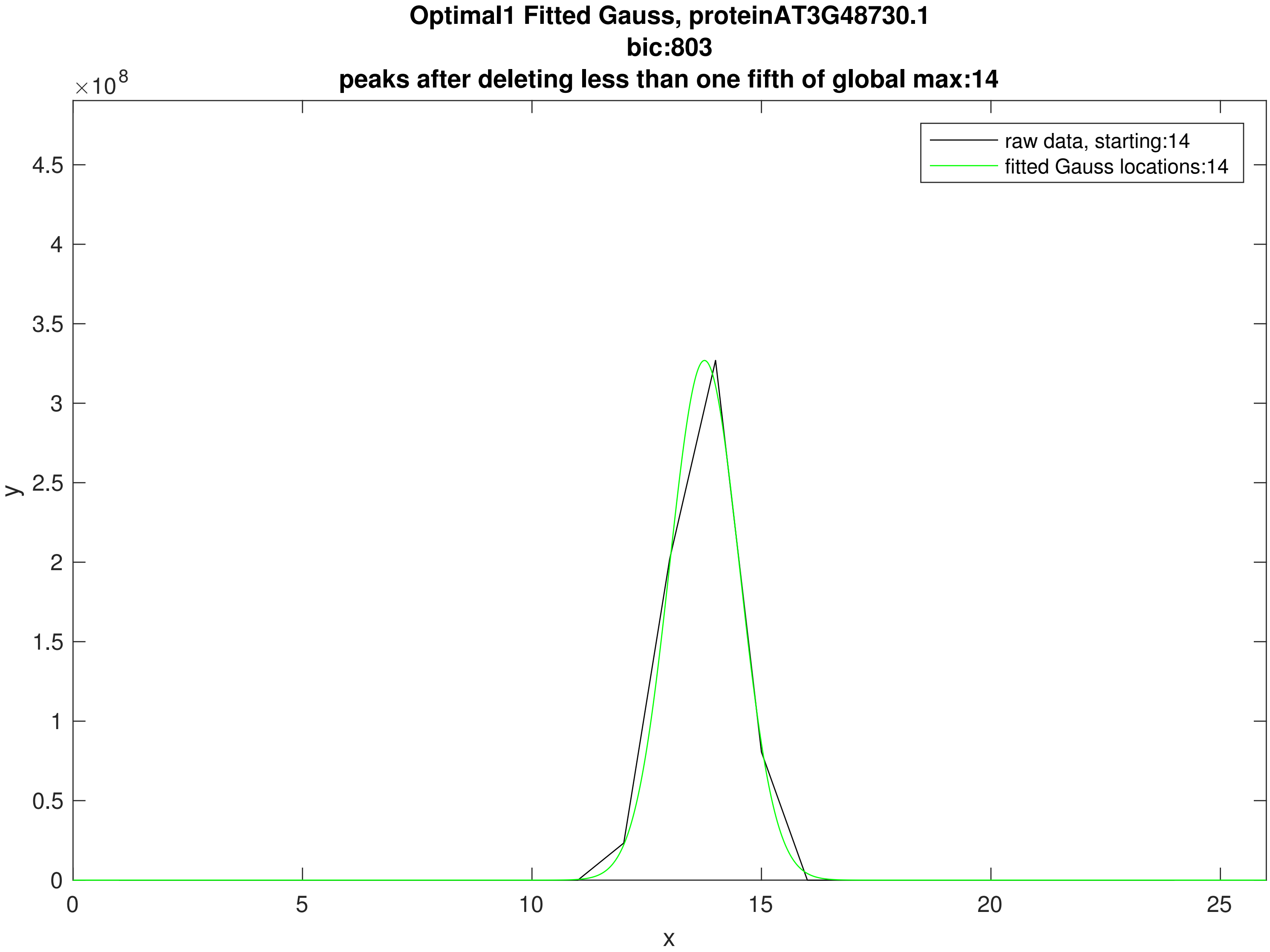

Optimal1 Fitted Gauss, proteinAT3G48730.1
bic:803
peaks after deleting less than one fifth of global max:14
×10^8
4.5
4
3.5
3
2.5
2
1.5
1
0.5
0
y
0
5
10
15
20
25
x
raw data, starting:14
fitted Gauss locations:14

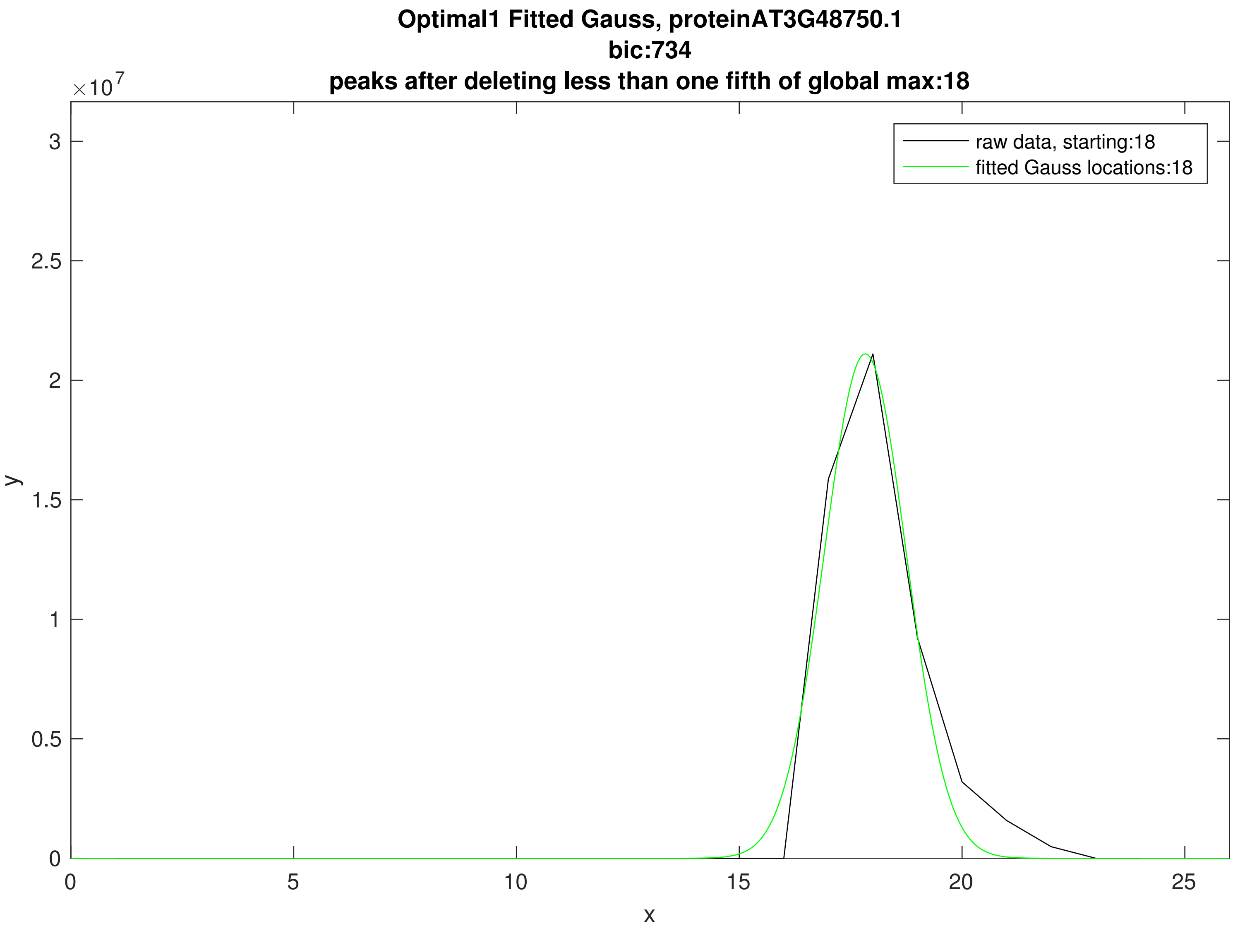

Optimal1 Fitted Gauss, proteinAT3G48750.1
bic:734
peaks after deleting less than one fifth of global max:18
×10^7
3
2.5
2
1.5
1
0.5
0
y
0
5
10
15
20
25
x
raw data, starting:18
fitted Gauss locations:18

**Optimal1 Fitted Gauss, proteinAT3G48930.1;AT4G30800.1**
**bic:850**
**peaks after deleting less than one fifth of global max:3.8**

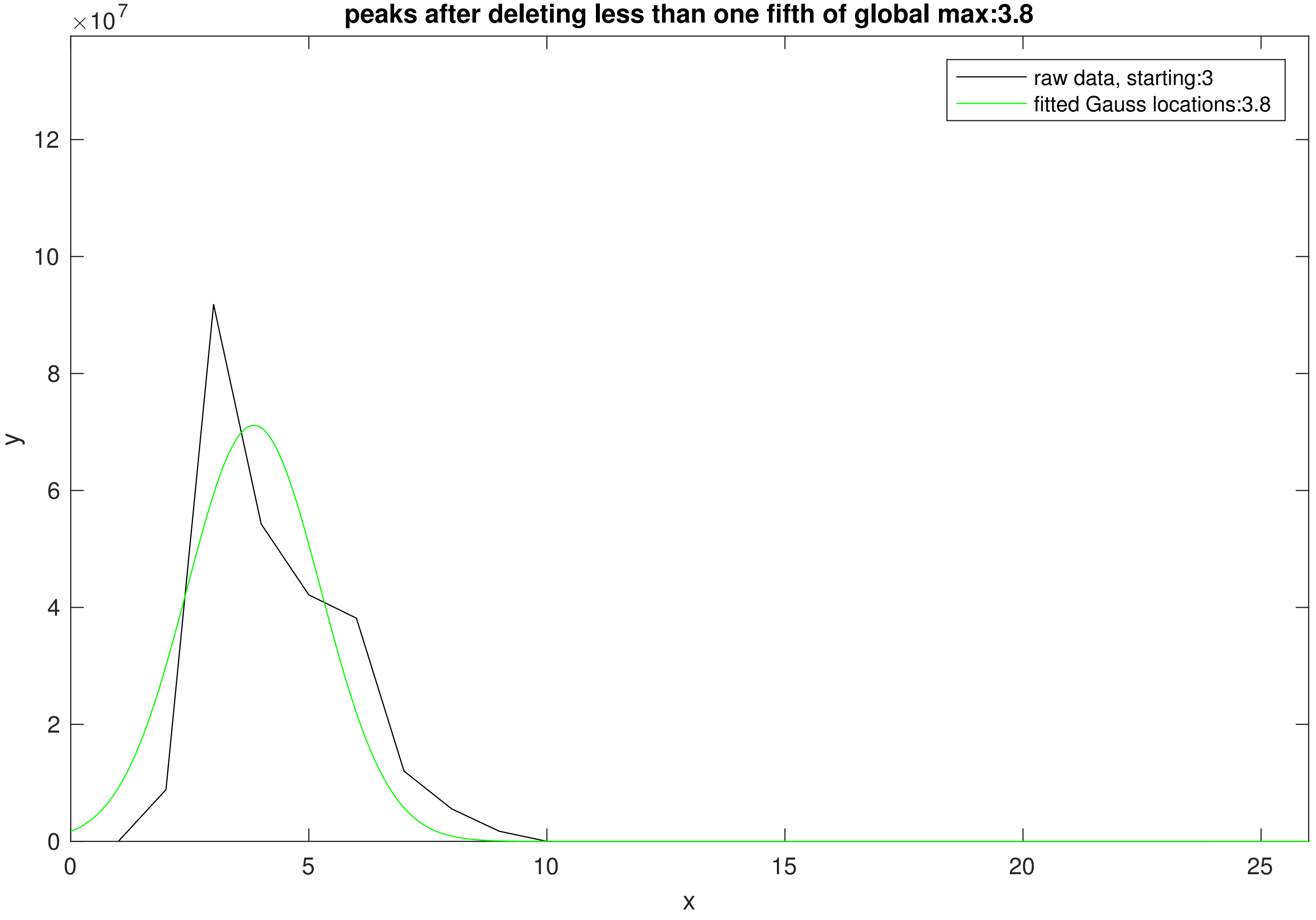

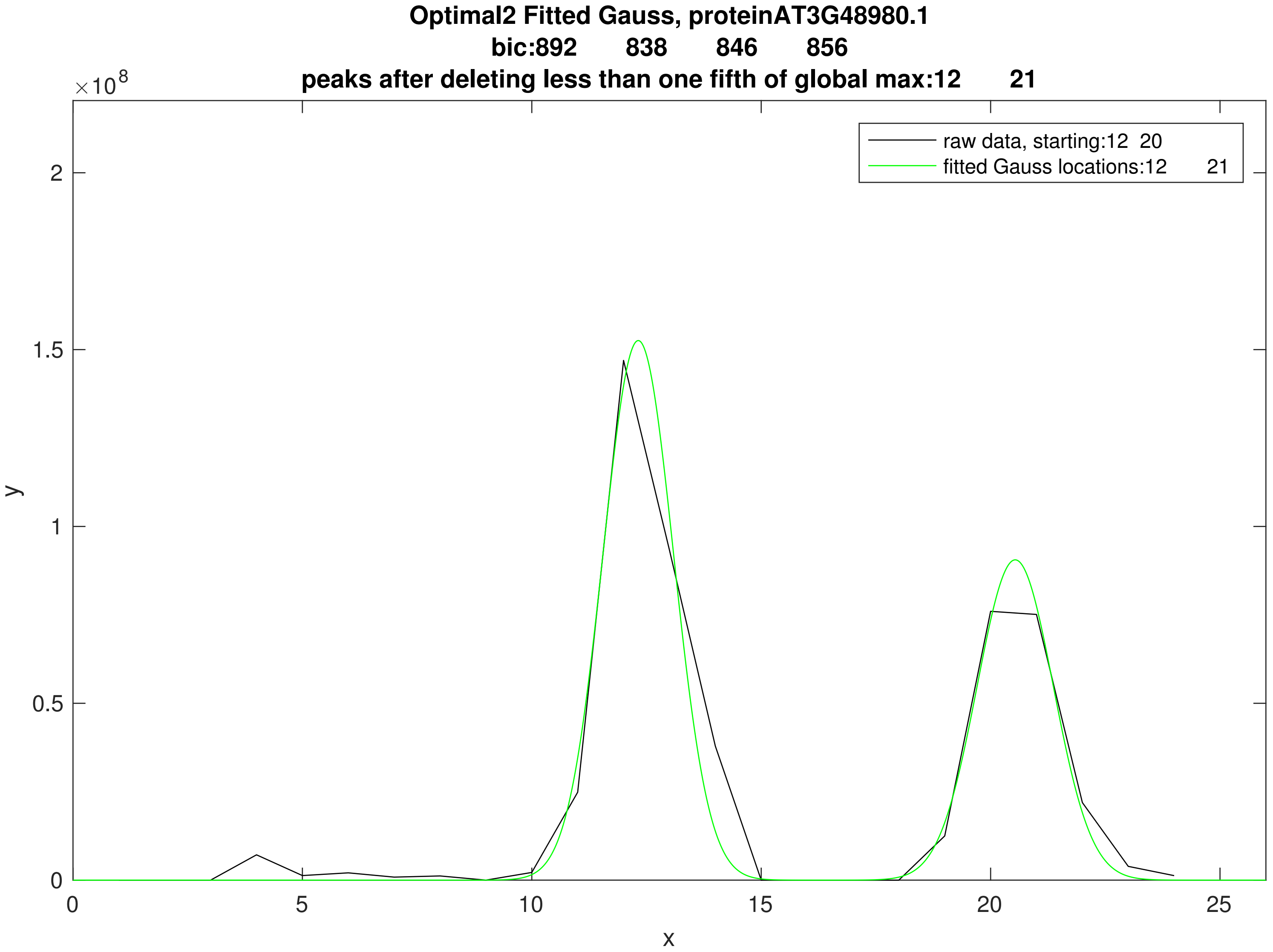

Optimal2 Fitted Gauss, proteinAT3G48980.1
bic:892 838 846 856
peaks after deleting less than one fifth of global max:12 21
×10^8
raw data, starting:12 20
fitted Gauss locations:12 21
y
x
0
0.5
1
1.5
2
5
10
15
20
25

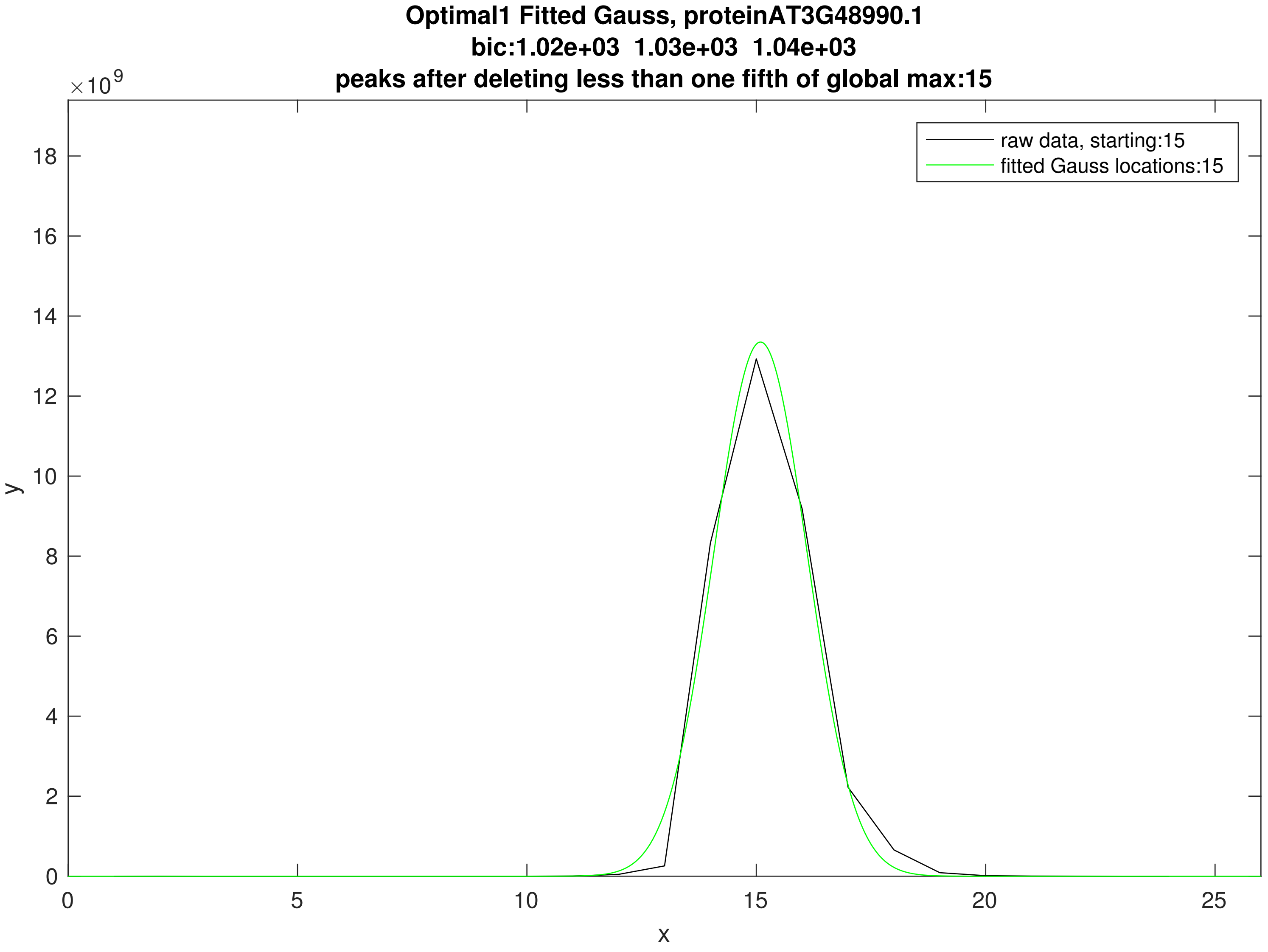

Optimal1 Fitted Gauss, proteinAT3G48990.1
bic:1.02e+03  1.03e+03  1.04e+03
peaks after deleting less than one fifth of global max:15
×10^9
y
x
raw data, starting:15
fitted Gauss locations:15
0
2
4
6
8
10
12
14
16
18
0
5
10
15
20
25

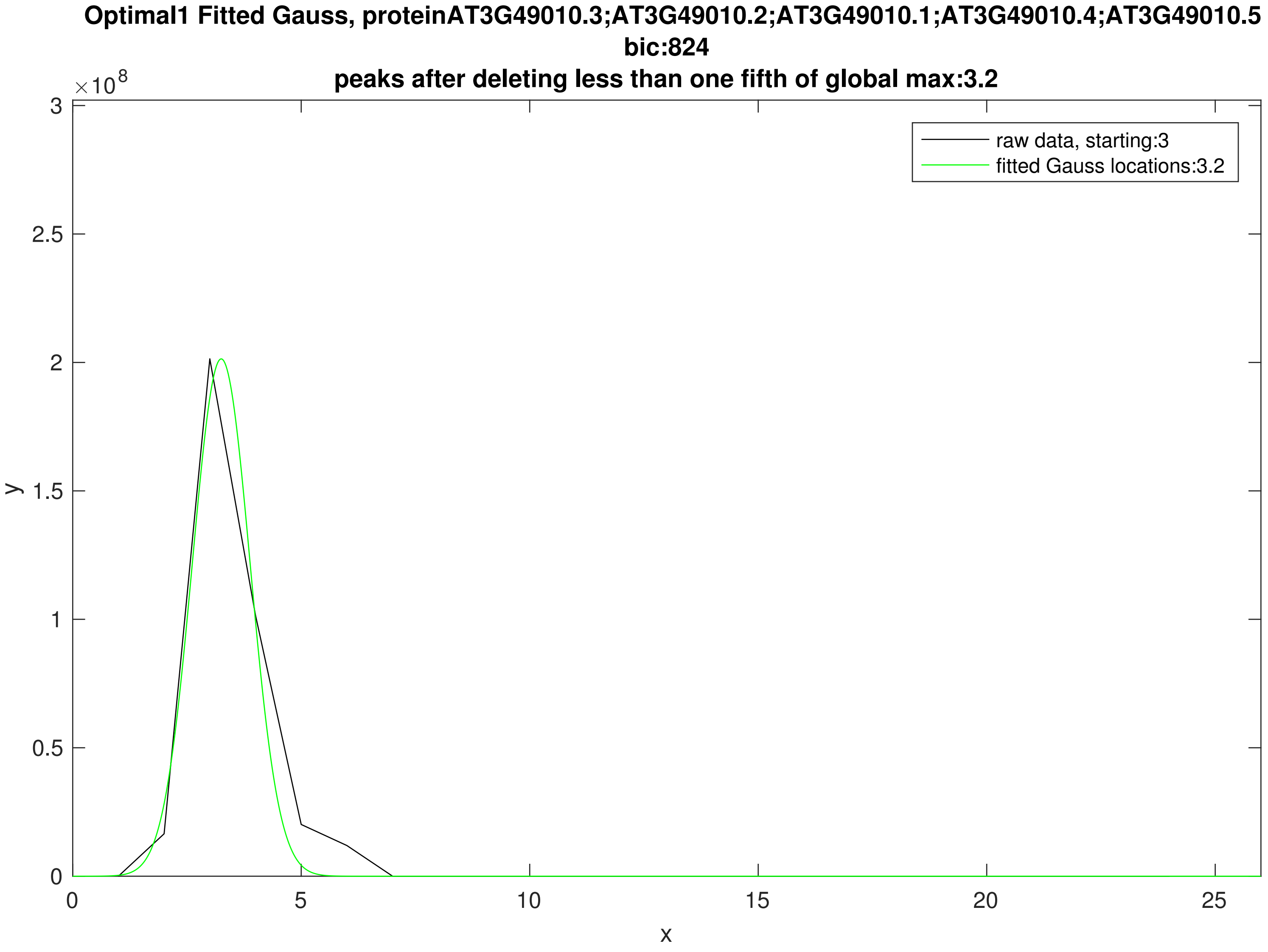
Optimal1 Fitted Gauss, proteinAT3G49010.3;AT3G49010.2;AT3G49010.1;AT3G49010.4;AT3G49010.5
bic:824
peaks after deleting less than one fifth of global max:3.2
×10^8
3
2.5
2
1.5
1
0.5
0
y
0
5
10
15
20
25
x
raw data, starting:3
fitted Gauss locations:3.2

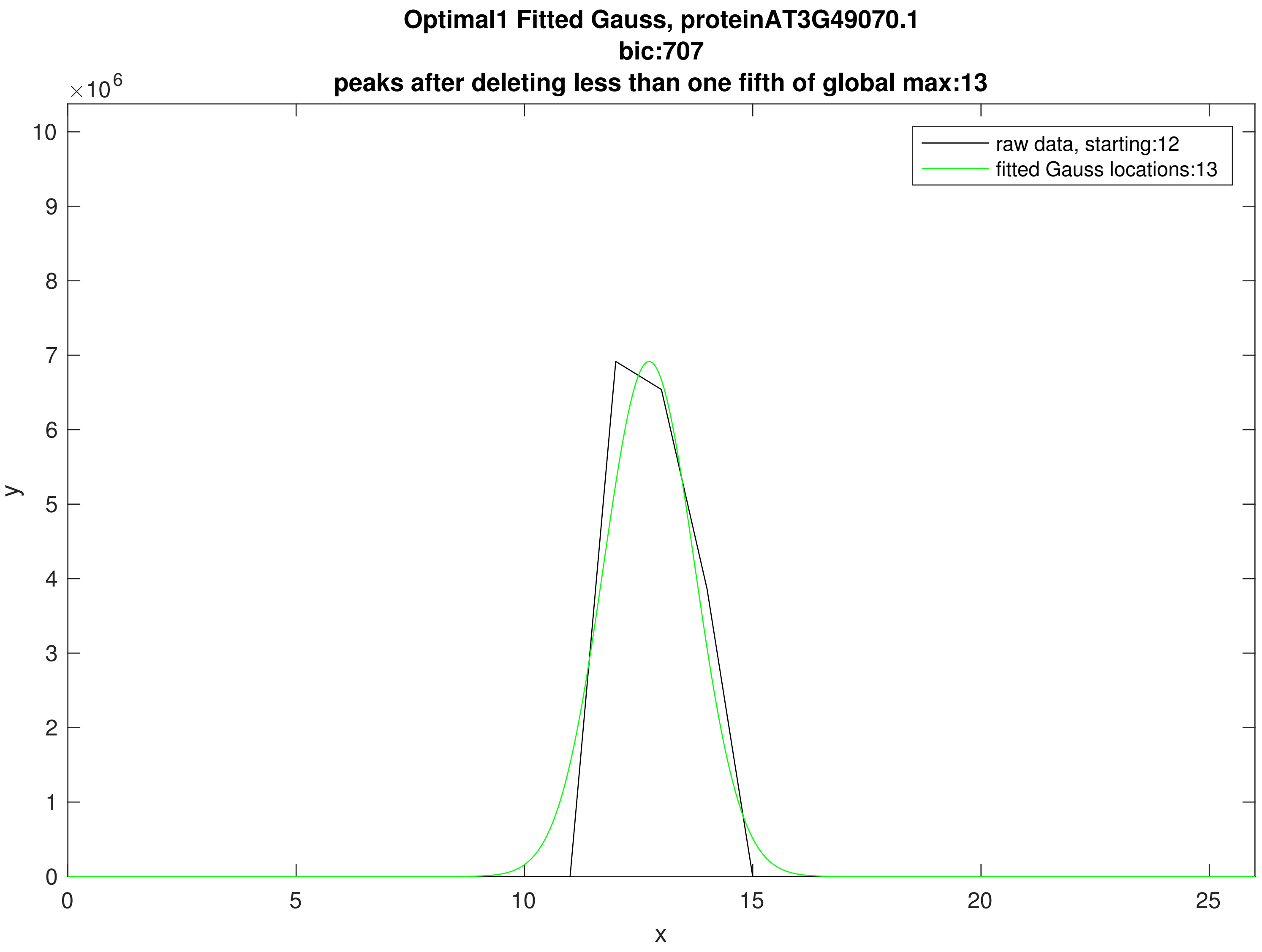
Optimal1 Fitted Gauss, proteinAT3G49070.1
bic:707
peaks after deleting less than one fifth of global max:13
×10^6
raw data, starting:12
fitted Gauss locations:13
y
x
0
1
2
3
4
5
6
7
8
9
10
0
5
10
15
20
25

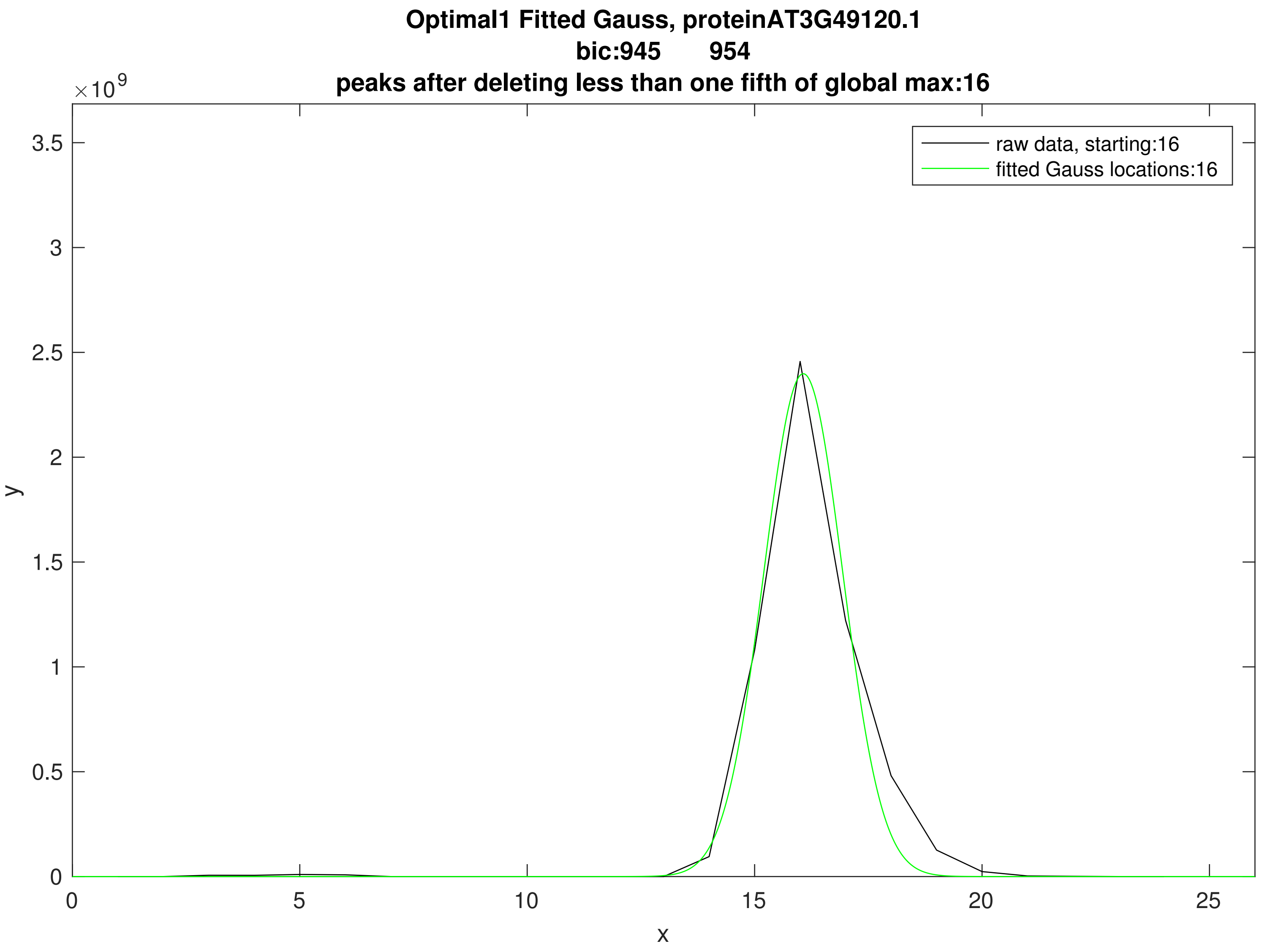

Optimal1 Fitted Gauss, proteinAT3G49120.1
bic:945 954
peaks after deleting less than one fifth of global max:16
×10^9
3.5
3
2.5
2
1.5
1
0.5
0
y
0
5
10
15
20
25
x
raw data, starting:16
fitted Gauss locations:16

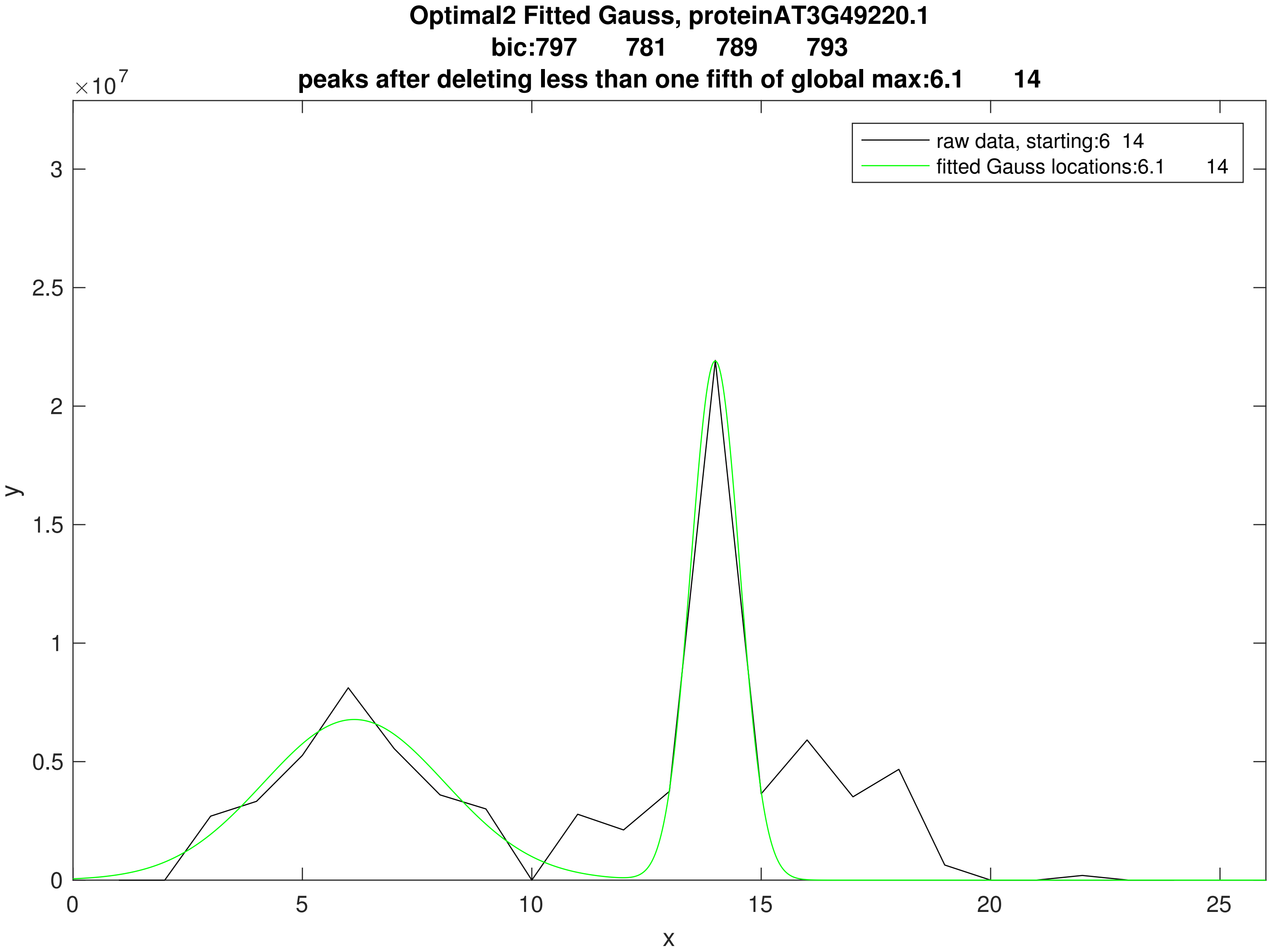

Optimal2 Fitted Gauss, proteinAT3G49220.1
bic:797 781 789 793
peaks after deleting less than one fifth of global max:6.1 14
×10^7
raw data, starting:6 14
fitted Gauss locations:6.1 14
y
x
0
0.5
1
1.5
2
2.5
3
5
10
15
20
25

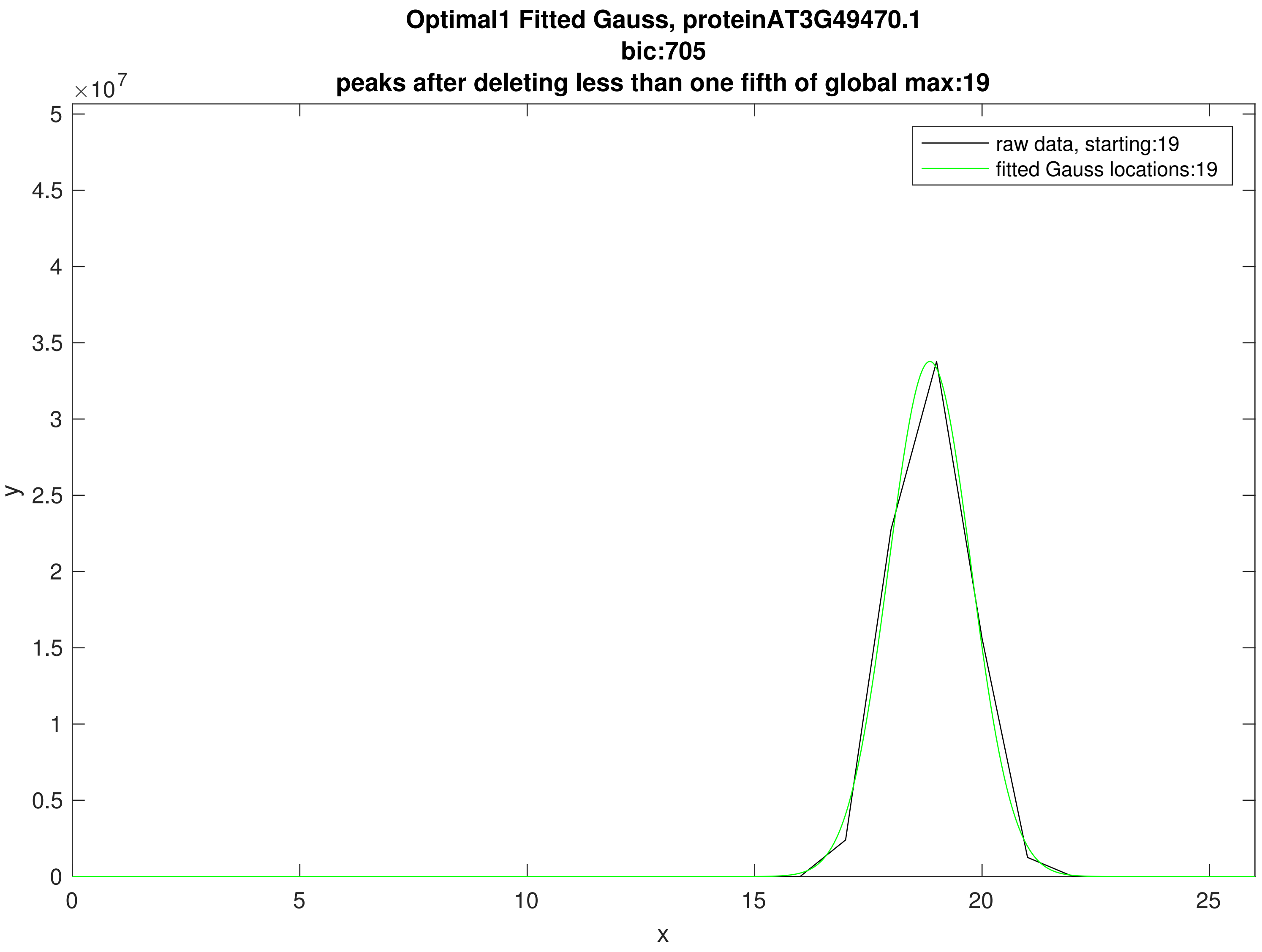

Optimal1 Fitted Gauss, proteinAT3G49470.1
bic:705
peaks after deleting less than one fifth of global max:19
×10^7
raw data, starting:19
fitted Gauss locations:19
y
x
0
0.5
1
1.5
2
2.5
3
3.5
4
4.5
5
0
5
10
15
20
25

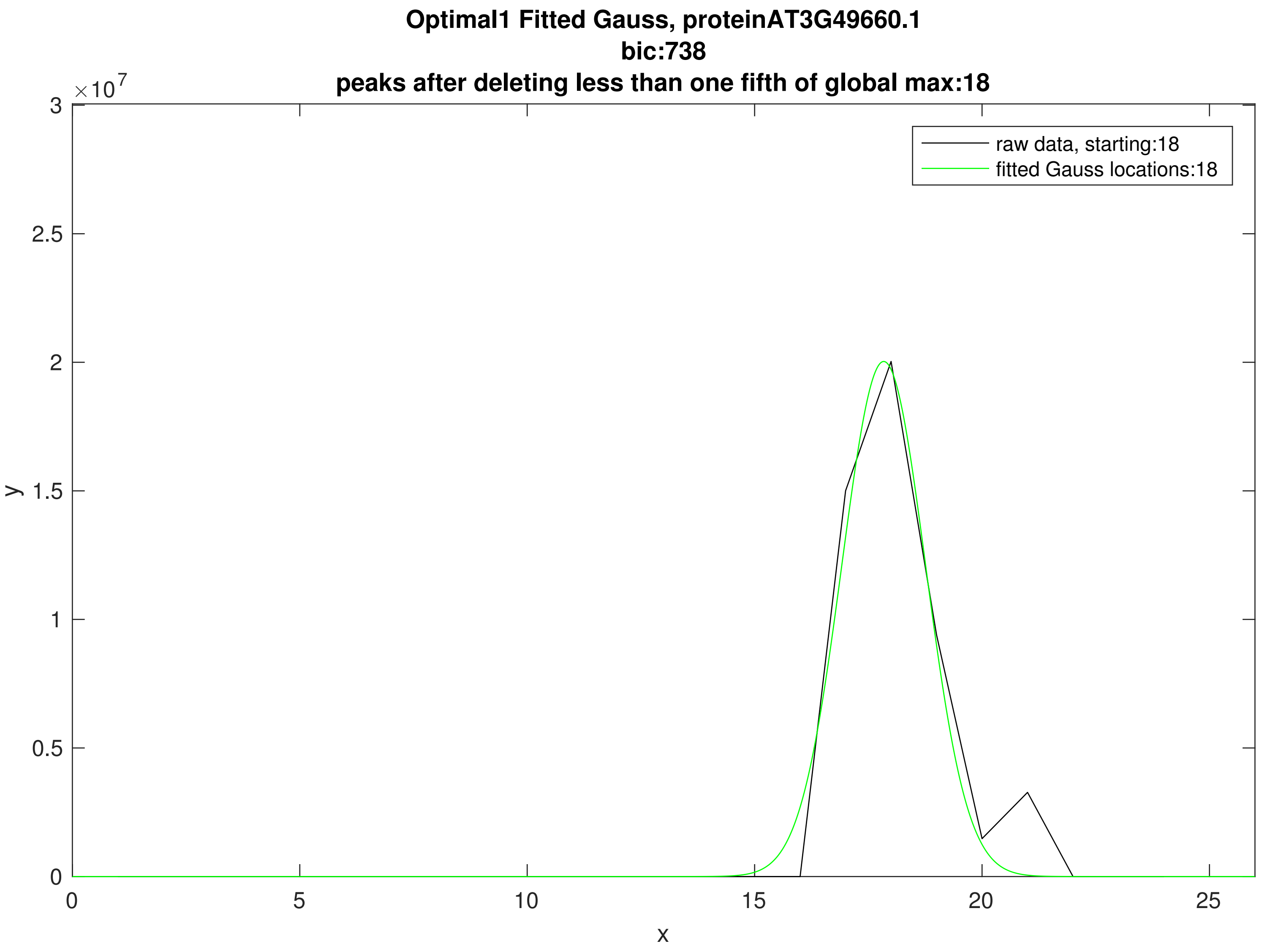
Optimal1 Fitted Gauss, proteinAT3G49660.1
bic:738
peaks after deleting less than one fifth of global max:18
×10^7
3
2.5
2
1.5
1
0.5
0
y
0
5
10
15
20
25
x
raw data, starting:18
fitted Gauss locations:18

**Optimal1 Fitted Gauss, proteinAT3G49680.2;AT3G49680.1**
**bic:871**
**peaks after deleting less than one fifth of global max:14**

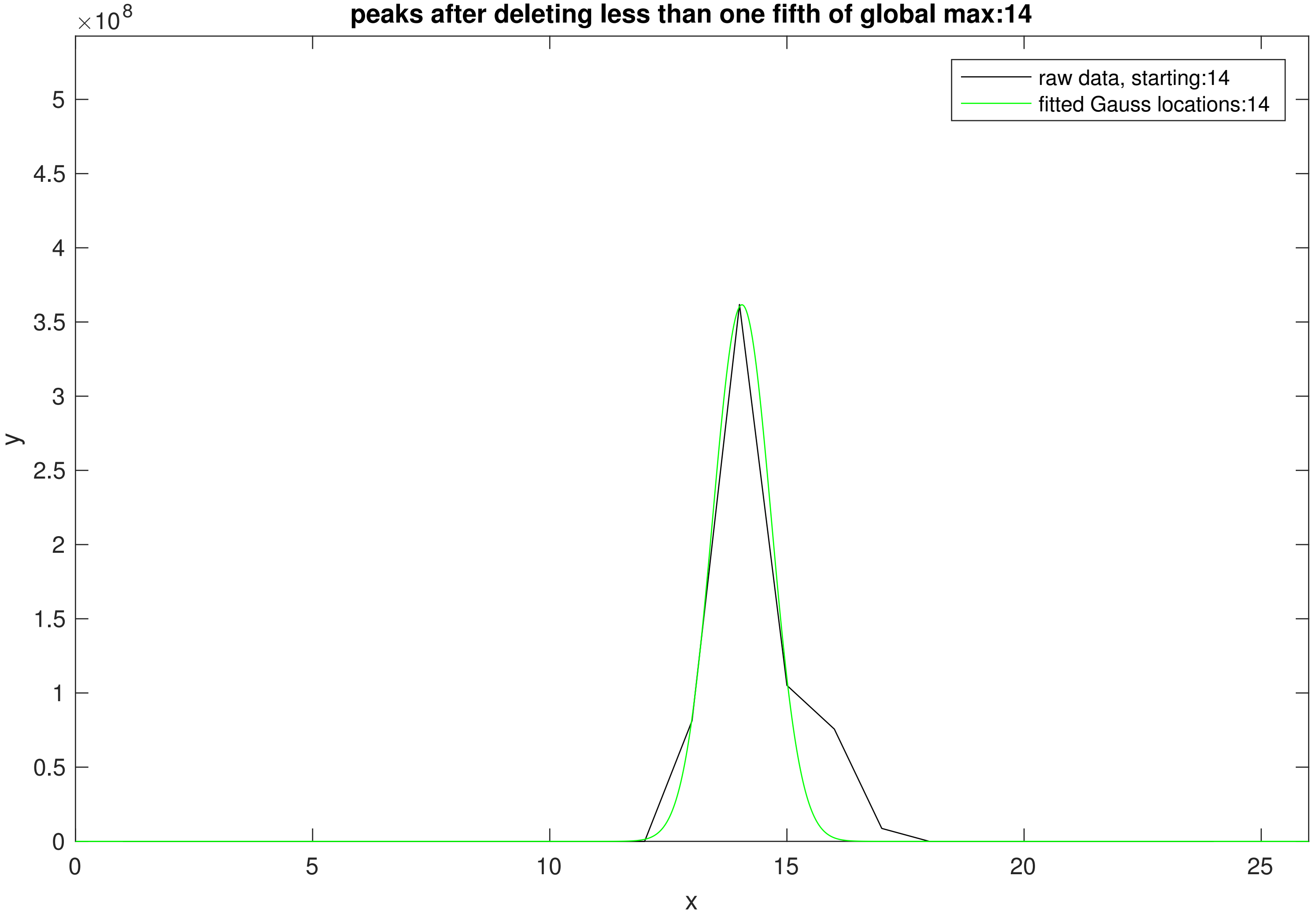

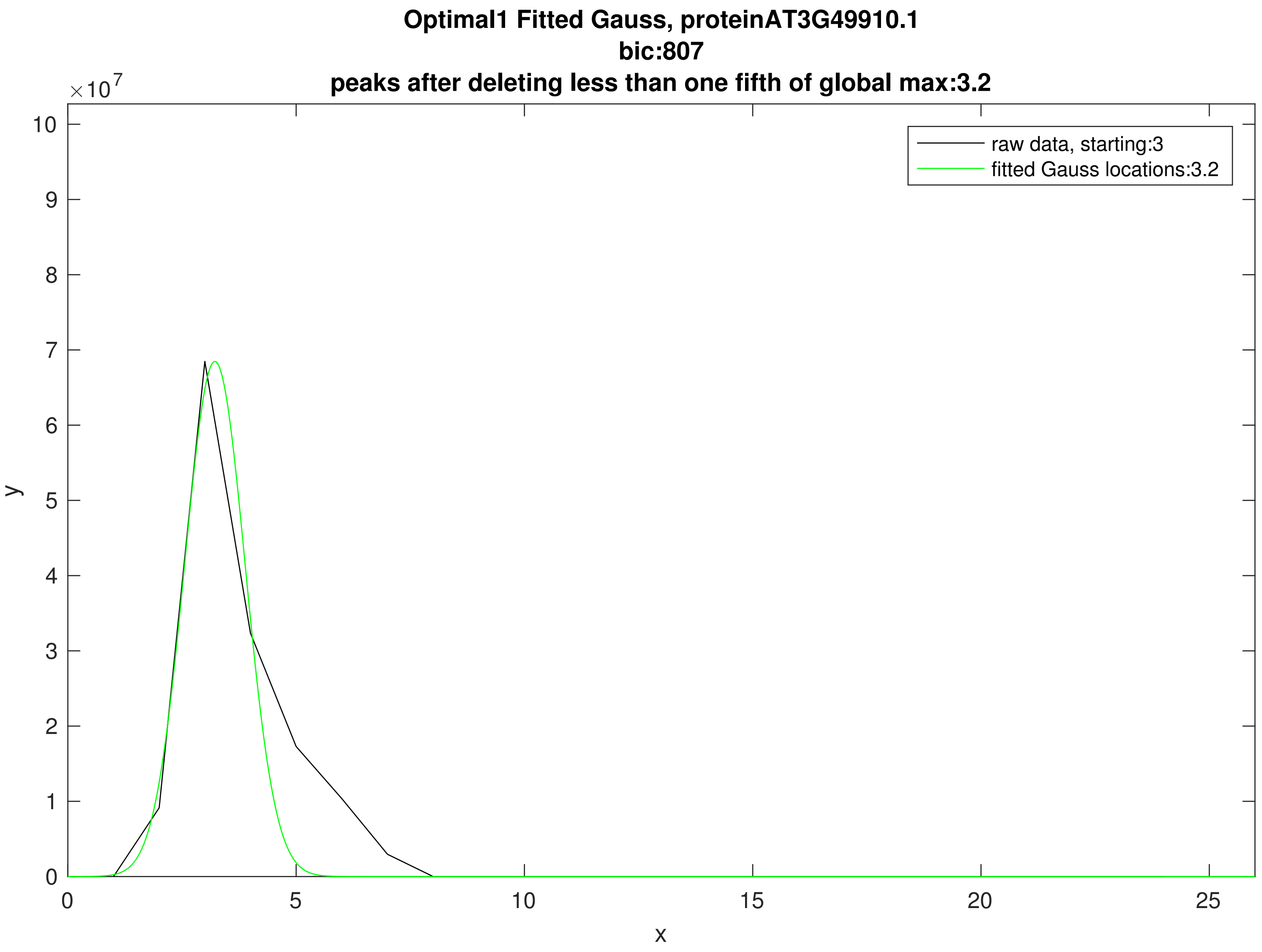

Optimal1 Fitted Gauss, proteinAT3G49910.1
bic:807
peaks after deleting less than one fifth of global max:3.2
×10^7
raw data, starting:3
fitted Gauss locations:3.2
y
x
0
1
2
3
4
5
6
7
8
9
10
0
5
10
15
20
25

Optimal1 Fitted Gauss, proteinAT3G50210.1;AT3G50210.3;AT3G50210.2
bic:787
peaks after deleting less than one fifth of global max:17

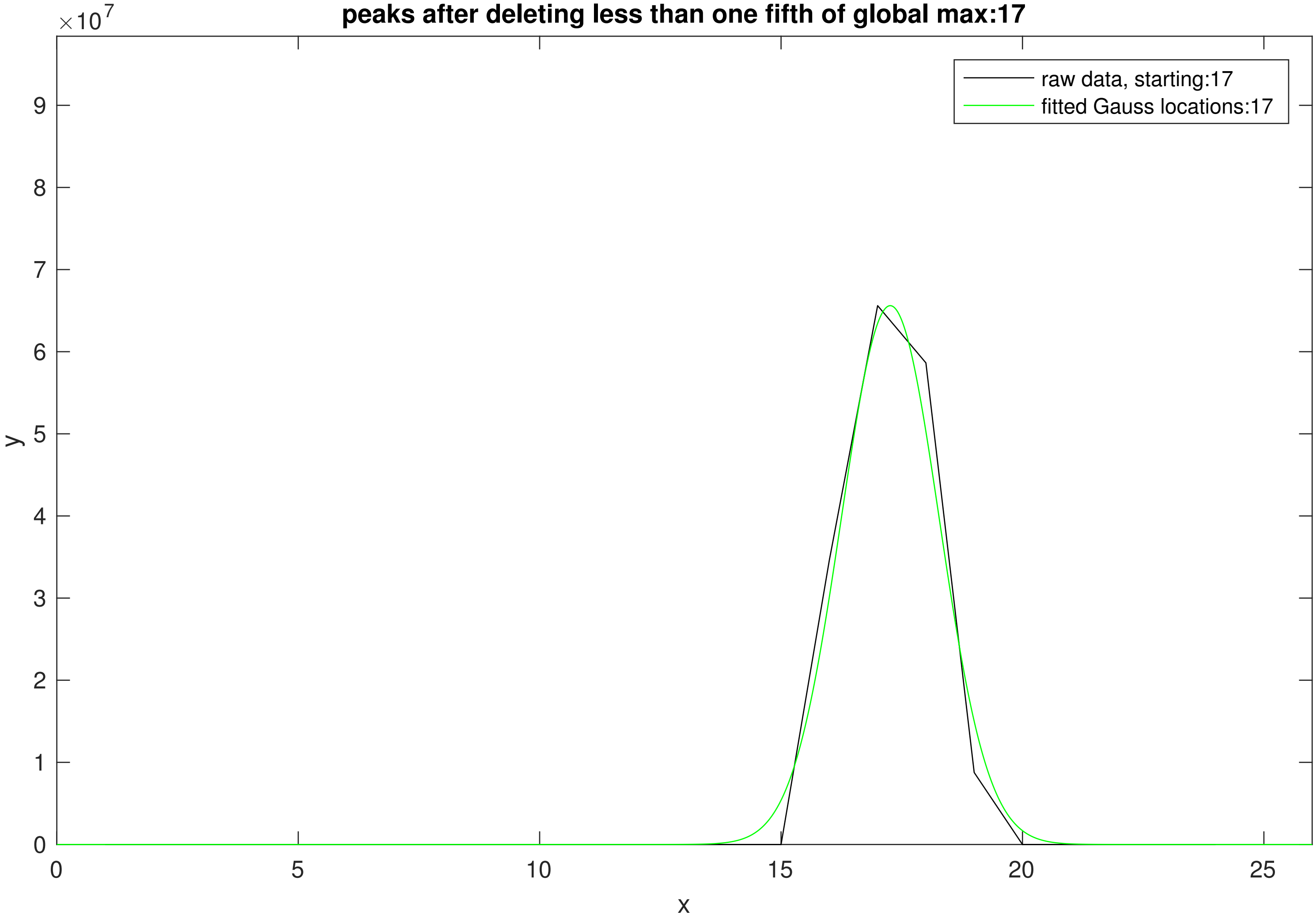

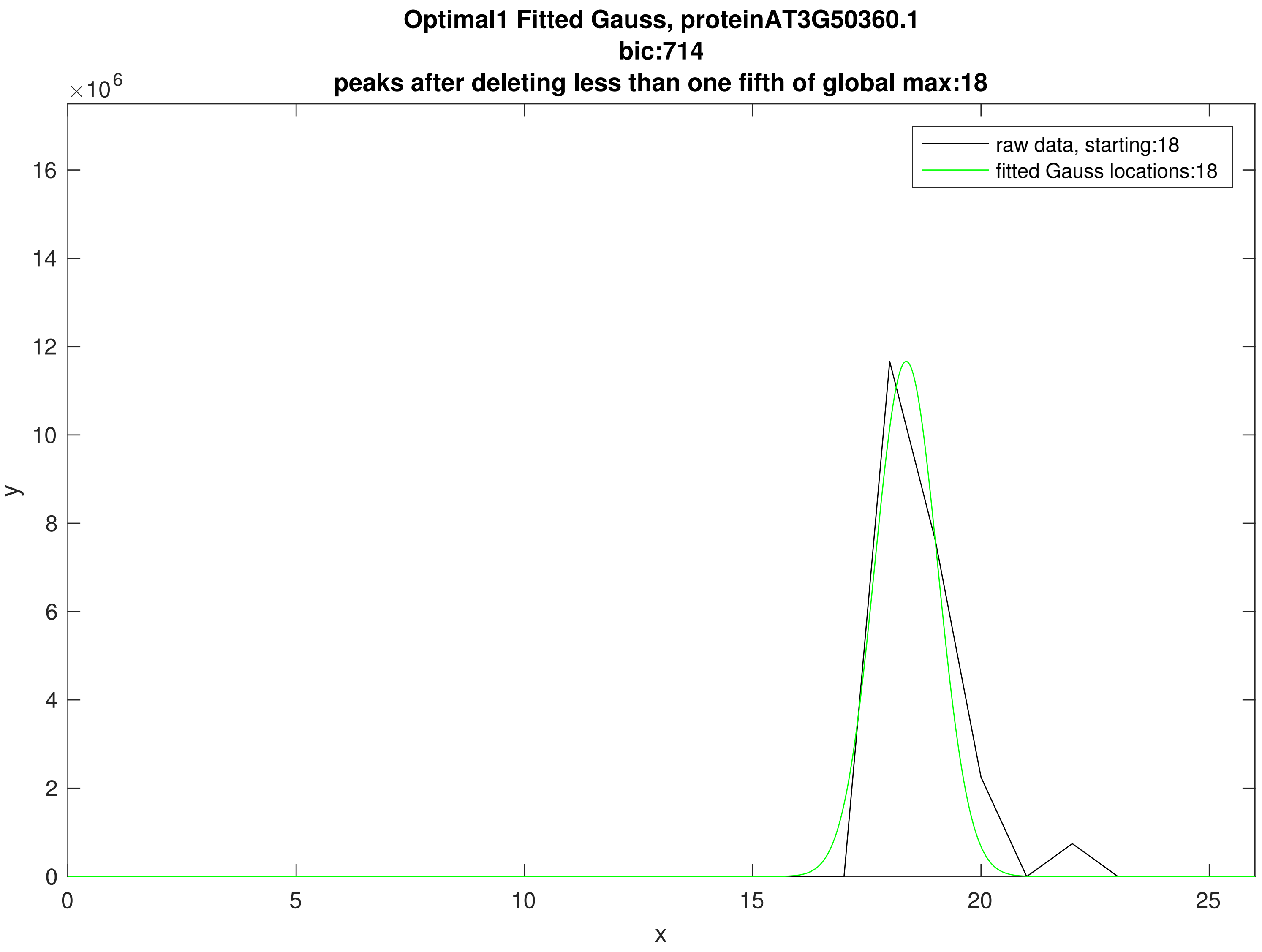
Optimal1 Fitted Gauss, proteinAT3G50360.1
bic:714
peaks after deleting less than one fifth of global max:18
×10^6
16
14
12
10
8
6
4
2
0
y
0
5
10
15
20
25
x
raw data, starting:18
fitted Gauss locations:18

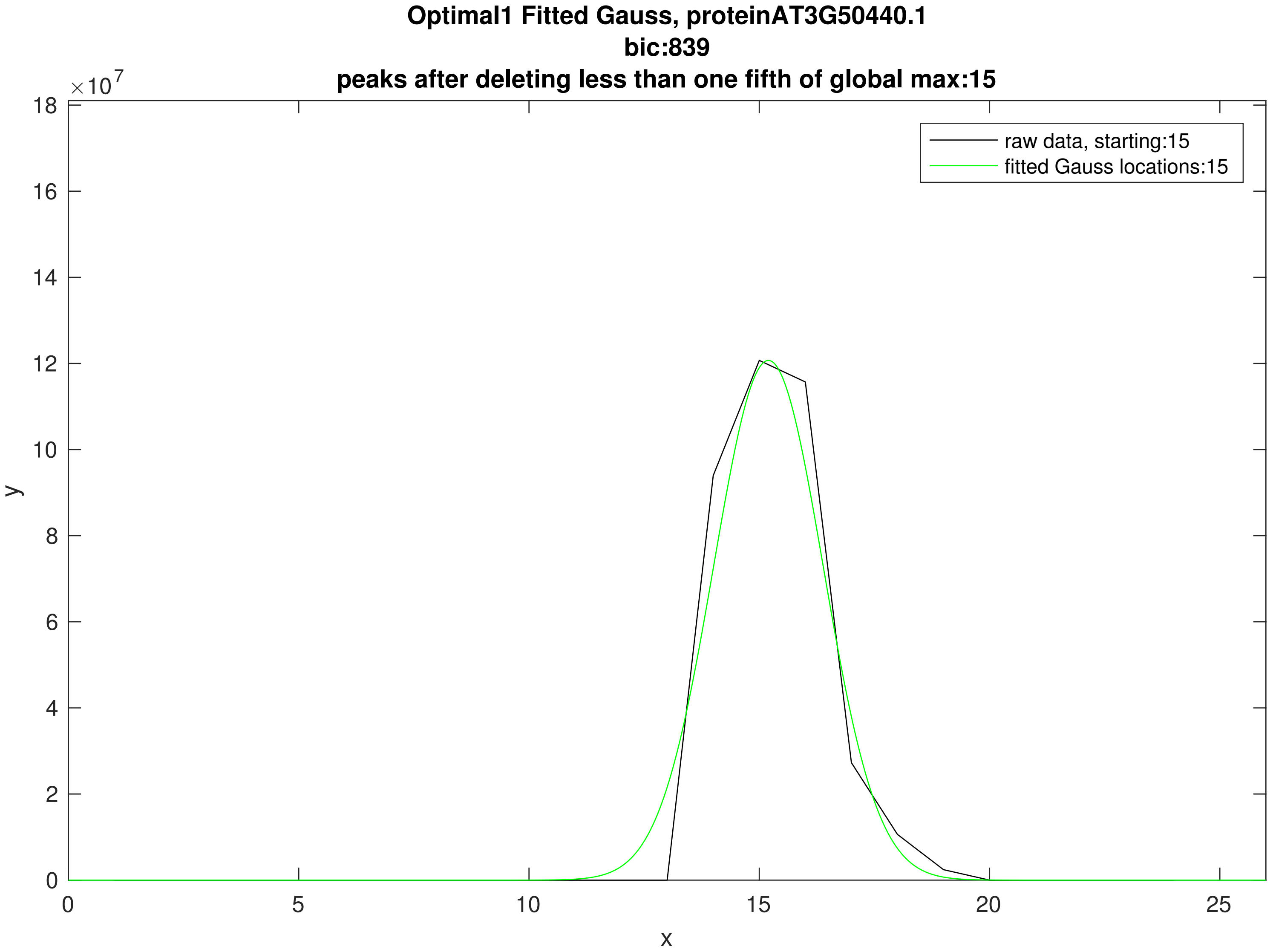

Optimal1 Fitted Gauss, proteinAT3G50440.1
bic:839
peaks after deleting less than one fifth of global max:15
×10^7
18
16
14
12
10
8
6
4
2
0
y
0
5
10
15
20
25
x
raw data, starting:15
fitted Gauss locations:15

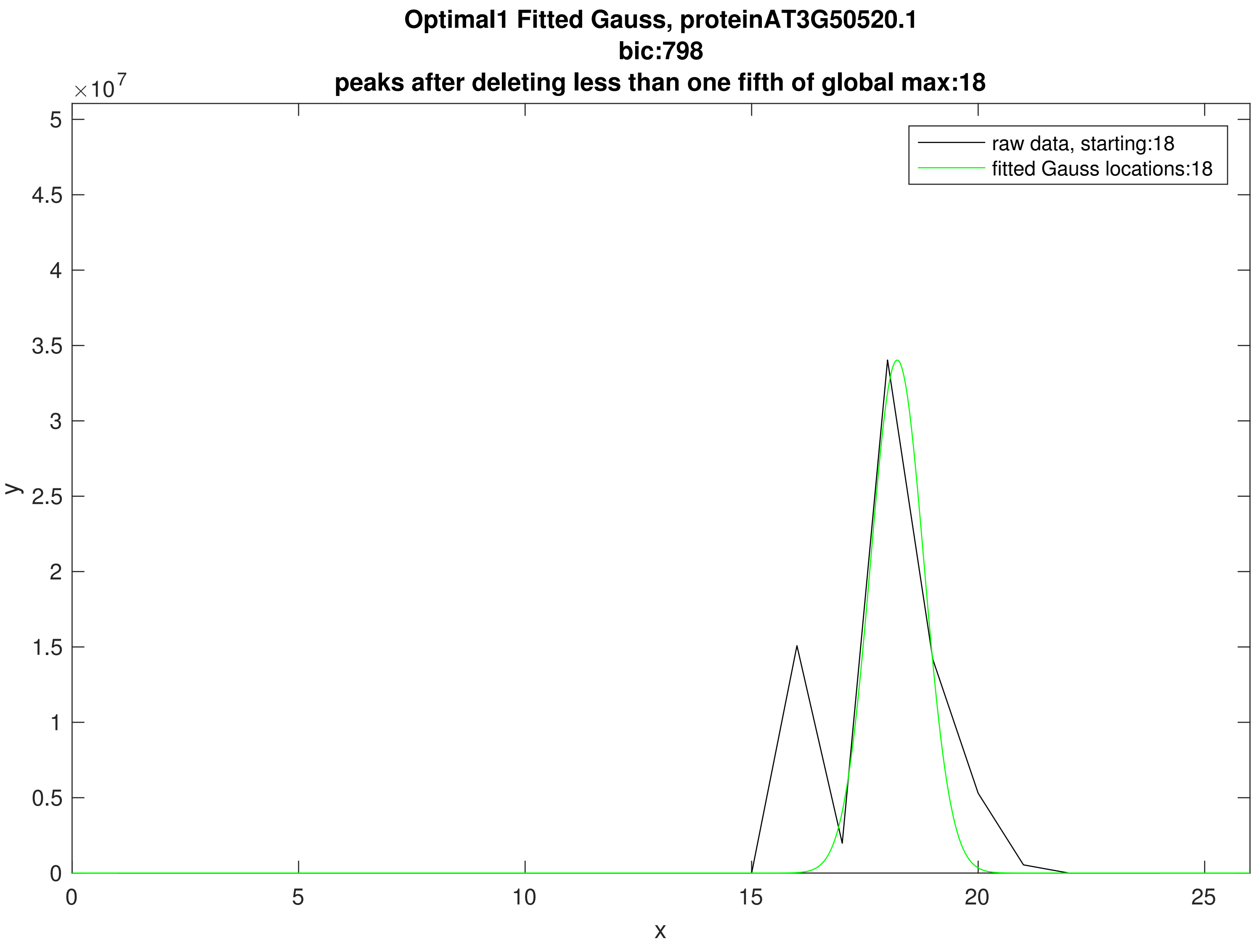

Optimal1 Fitted Gauss, proteinAT3G50520.1
bic:798
peaks after deleting less than one fifth of global max:18
×10^7
raw data, starting:18
fitted Gauss locations:18
y
x
0
0.5
1
1.5
2
2.5
3
3.5
4
4.5
5
0
5
10
15
20
25

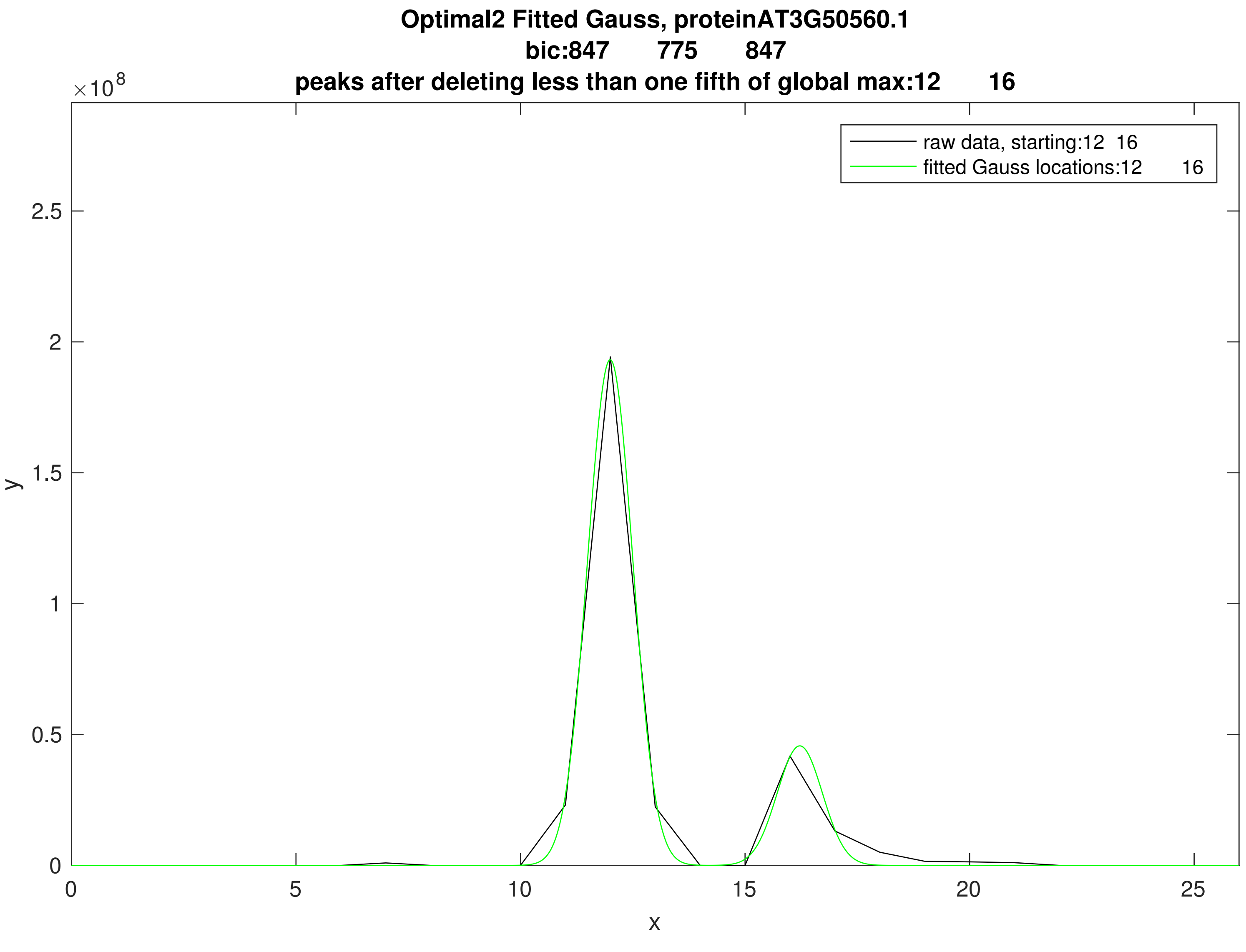

Optimal2 Fitted Gauss, proteinAT3G50560.1
bic:847 775 847
peaks after deleting less than one fifth of global max:12 16
×10^8
raw data, starting:12 16
fitted Gauss locations:12 16
2.5
2
1.5
1
0.5
0
y
0
5
10
15
20
25
x

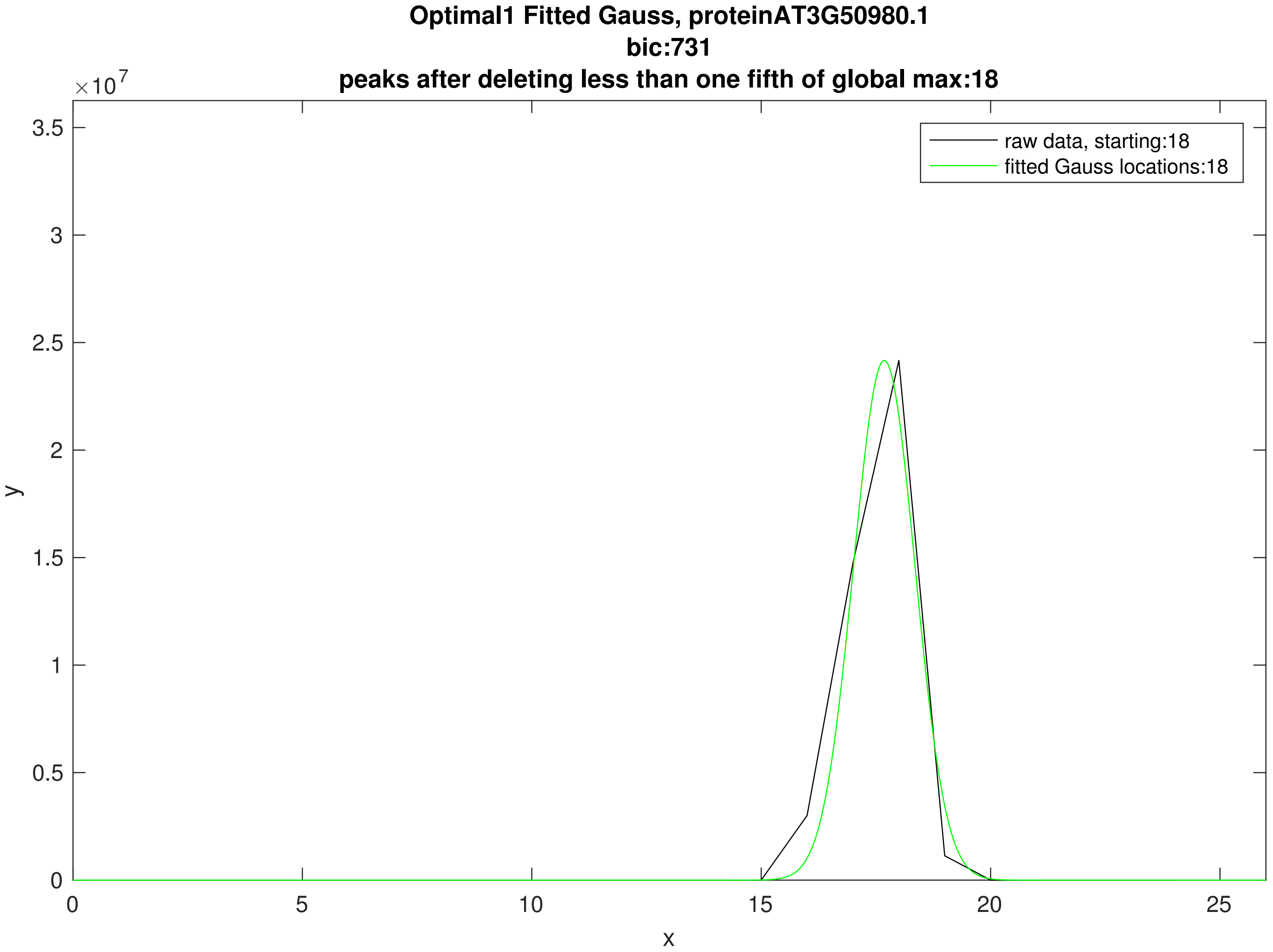
Optimal1 Fitted Gauss, proteinAT3G50980.1
bic:731
peaks after deleting less than one fifth of global max:18
×10^7
3.5
3
2.5
2
1.5
1
0.5
0
y
0
5
10
15
20
25
x
raw data, starting:18
fitted Gauss locations:18

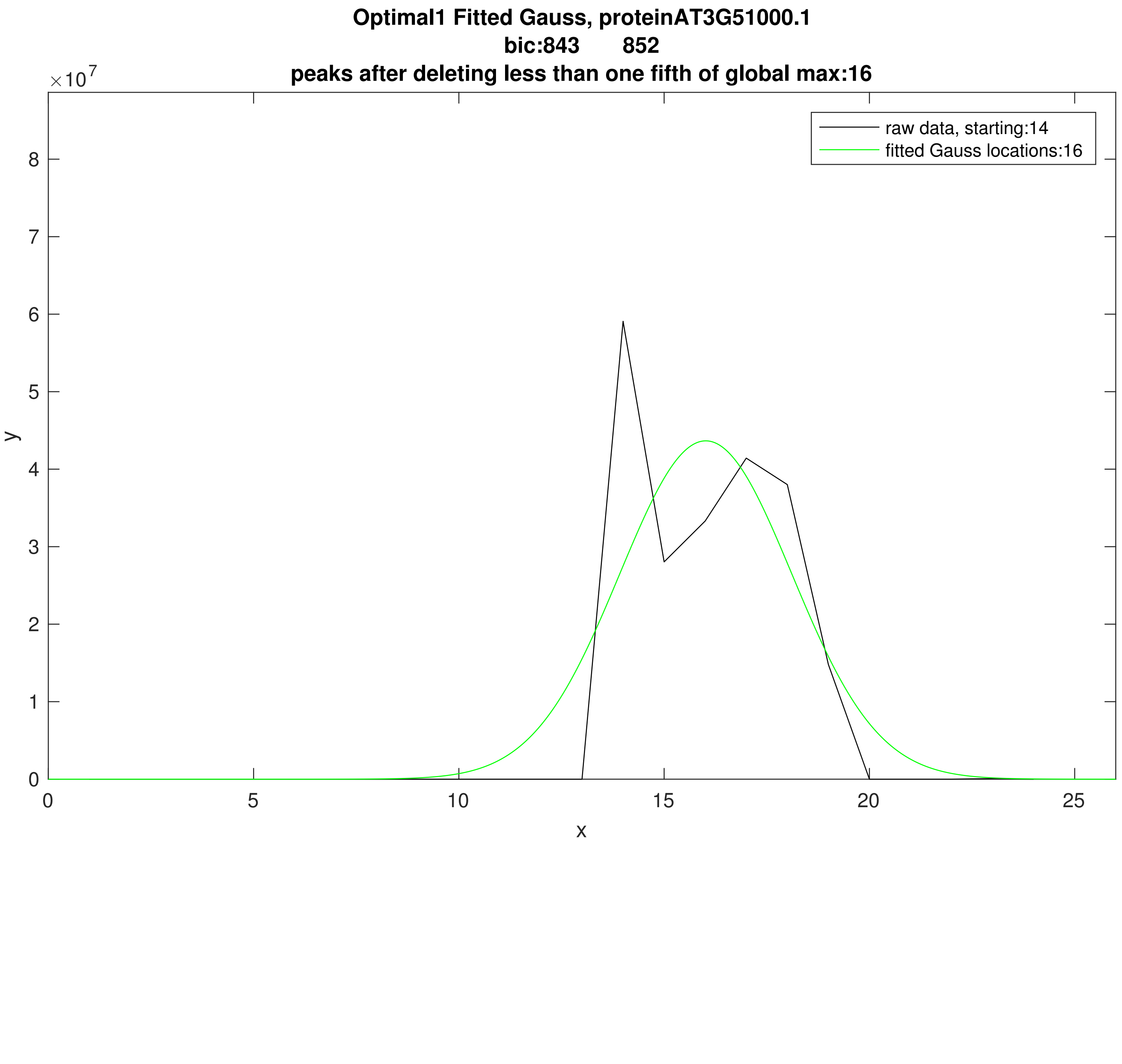
Optimal1 Fitted Gauss, proteinAT3G51000.1
bic:843 852
peaks after deleting less than one fifth of global max:16
×10^7
raw data, starting:14
fitted Gauss locations:16
y
x
0
1
2
3
4
5
6
7
8
0
5
10
15
20
25

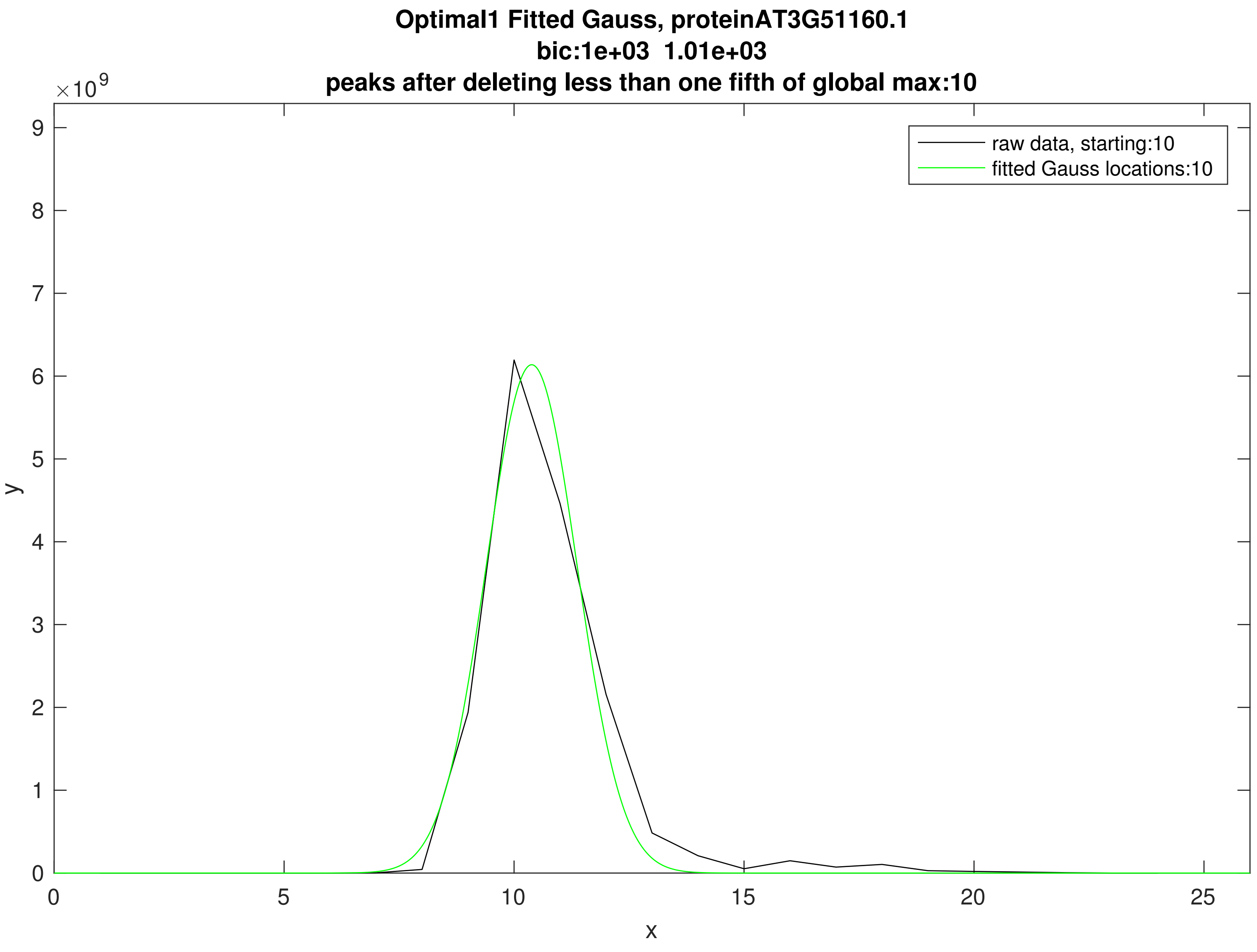
Optimal1 Fitted Gauss, proteinAT3G51160.1
bic:1e+03  1.01e+03
peaks after deleting less than one fifth of global max:10
×10^9
raw data, starting:10
fitted Gauss locations:10
y
x
0
1
2
3
4
5
6
7
8
9
0
5
10
15
20
25

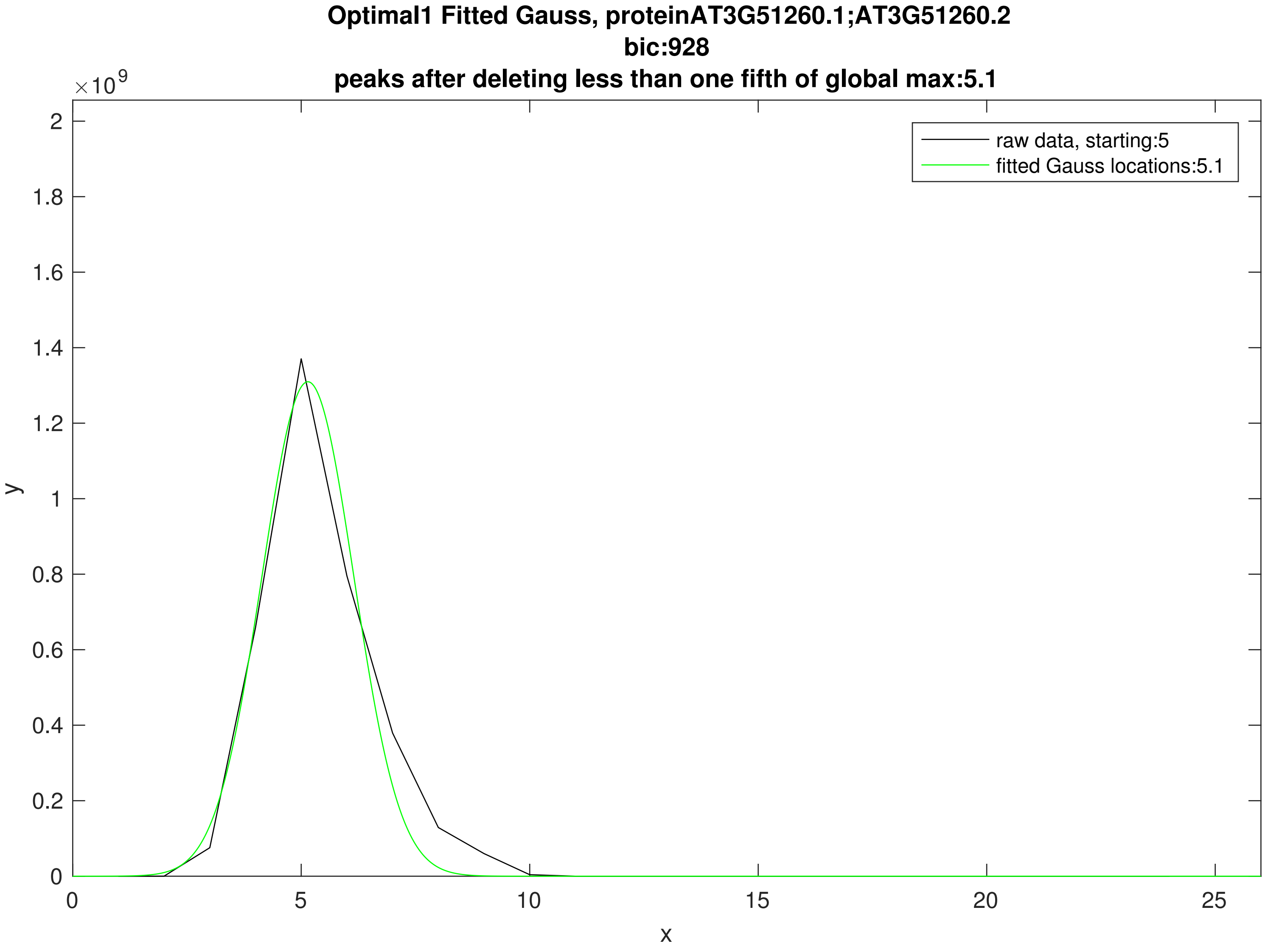
Optimal1 Fitted Gauss, proteinAT3G51260.1;AT3G51260.2
bic:928
peaks after deleting less than one fifth of global max:5.1
×10^9
2
1.8
1.6
1.4
1.2
1
0.8
0.6
0.4
0.2
0
y
0
5
10
15
20
25
x
raw data, starting:5
fitted Gauss locations:5.1

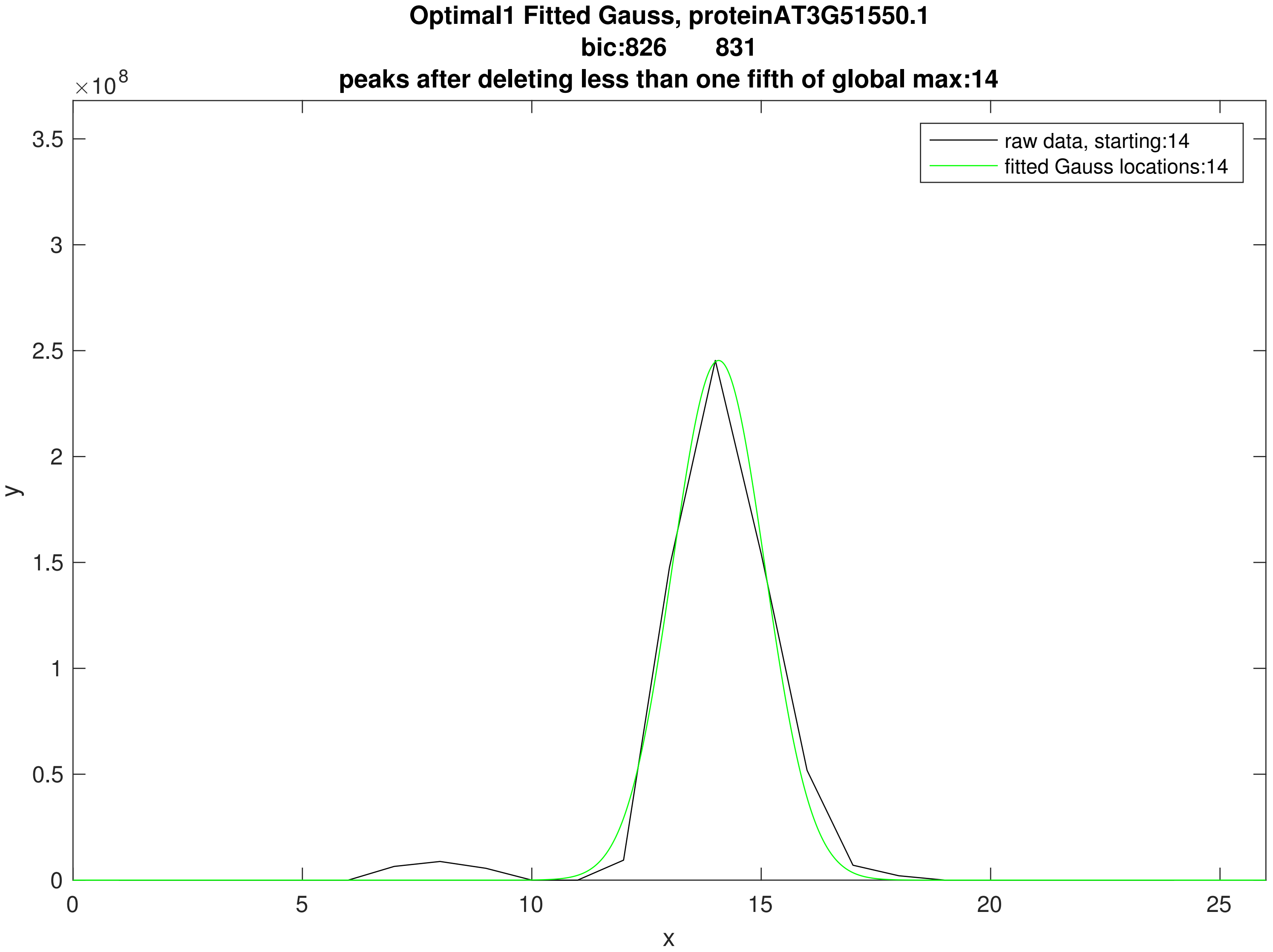

Optimal1 Fitted Gauss, proteinAT3G51550.1
bic:826 831
peaks after deleting less than one fifth of global max:14
×10^8
3.5
3
2.5
2
1.5
1
0.5
0
y
0
5
10
15
20
25
x
raw data, starting:14
fitted Gauss locations:14

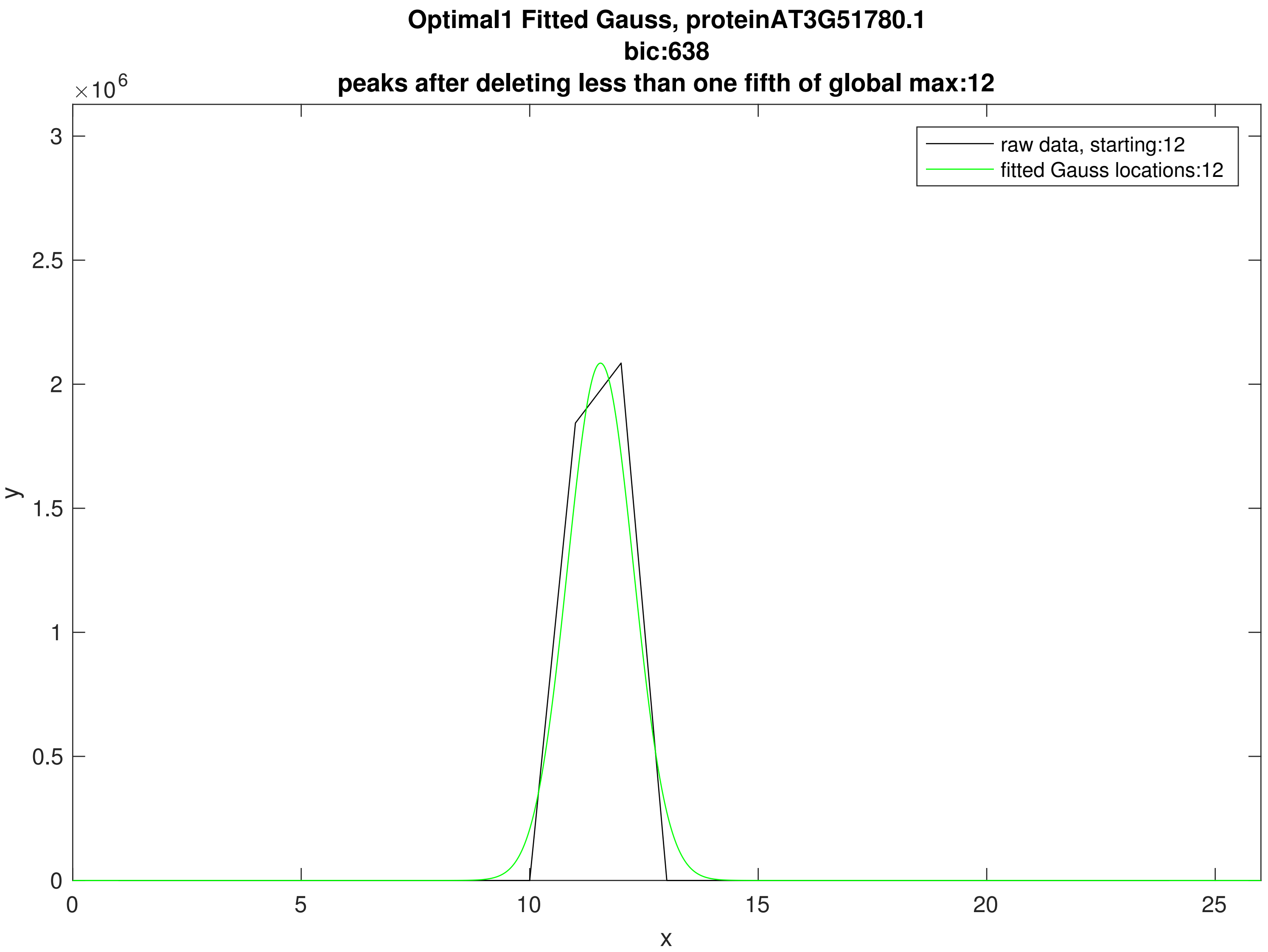
Optimal1 Fitted Gauss, proteinAT3G51780.1
bic:638
peaks after deleting less than one fifth of global max:12
×10^6
raw data, starting:12
fitted Gauss locations:12
y
x
0
0.5
1
1.5
2
2.5
3
0
5
10
15
20
25

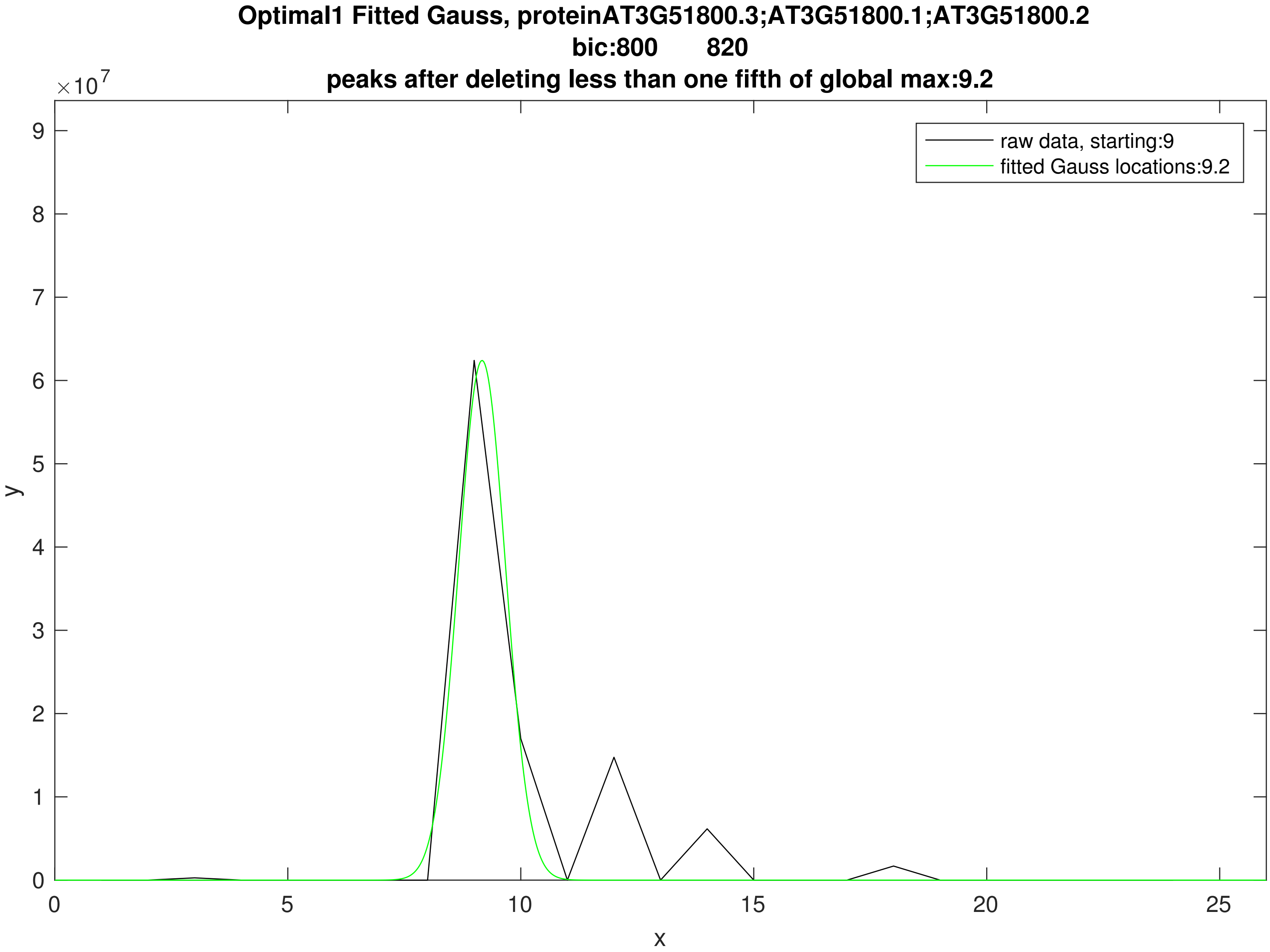

Optimal1 Fitted Gauss, proteinAT3G51800.3;AT3G51800.1;AT3G51800.2
bic:800 820
peaks after deleting less than one fifth of global max:9.2
×10^7
raw data, starting:9
fitted Gauss locations:9.2
y
x
0
1
2
3
4
5
6
7
8
9
0
5
10
15
20
25

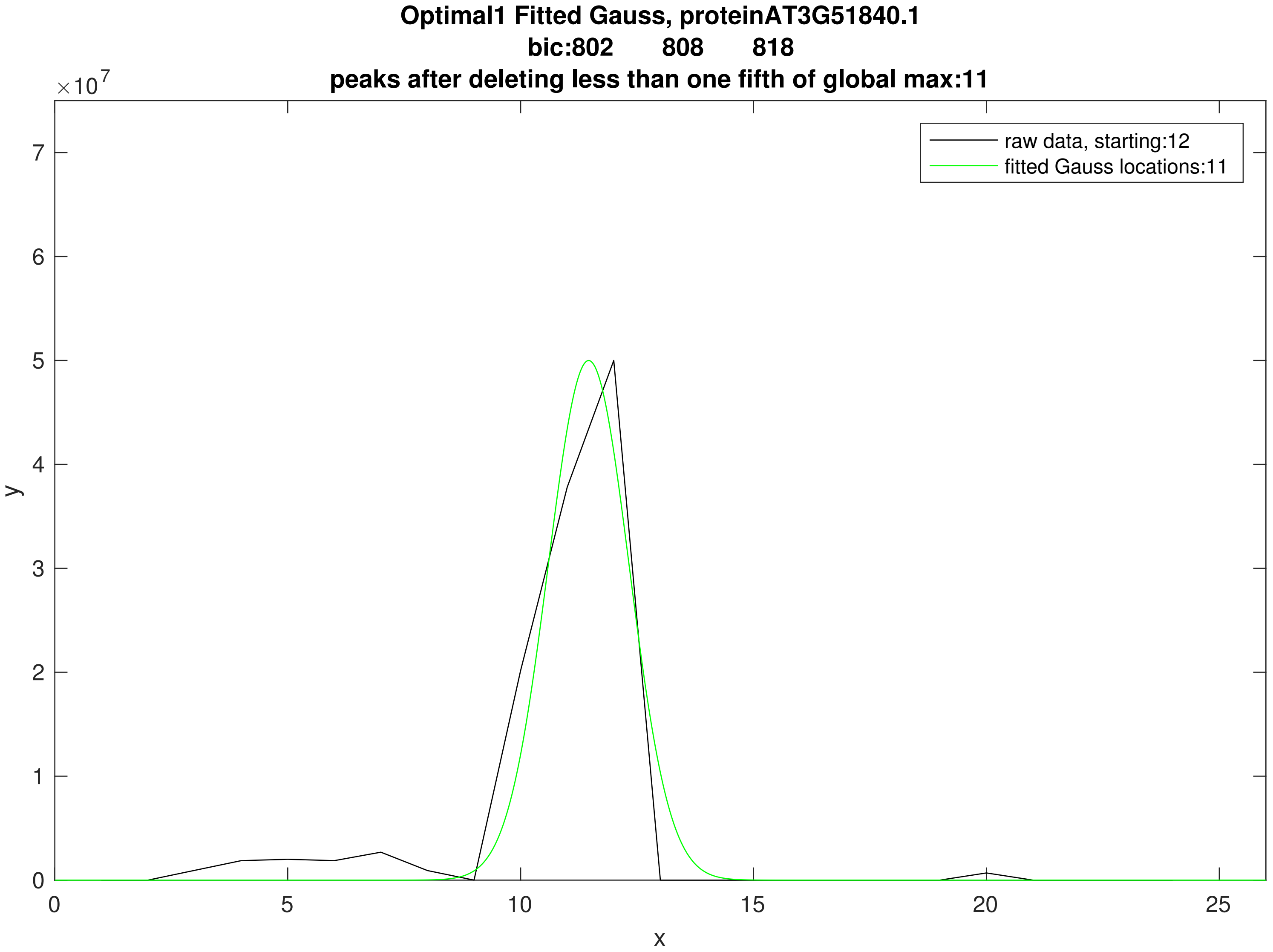

Optimal1 Fitted Gauss, proteinAT3G51840.1
bic:802 808 818
peaks after deleting less than one fifth of global max:11
×10^7
raw data, starting:12
fitted Gauss locations:11
y
x
0
1
2
3
4
5
6
7
10
15
20
25

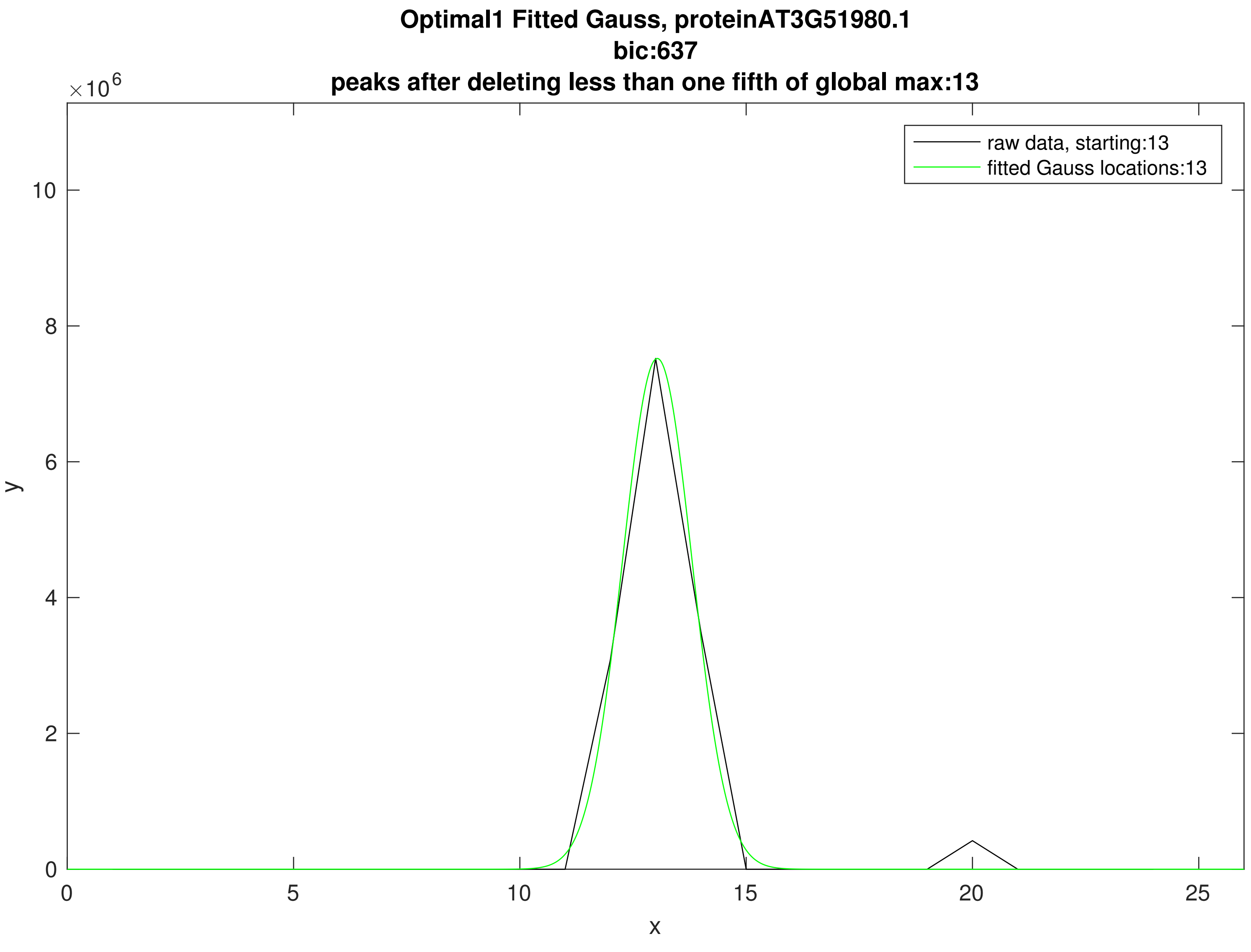

Optimal1 Fitted Gauss, proteinAT3G51980.1
bic:637
peaks after deleting less than one fifth of global max:13
×10^6
raw data, starting:13
fitted Gauss locations:13
y
x
0
2
4
6
8
10
0
5
10
15
20
25

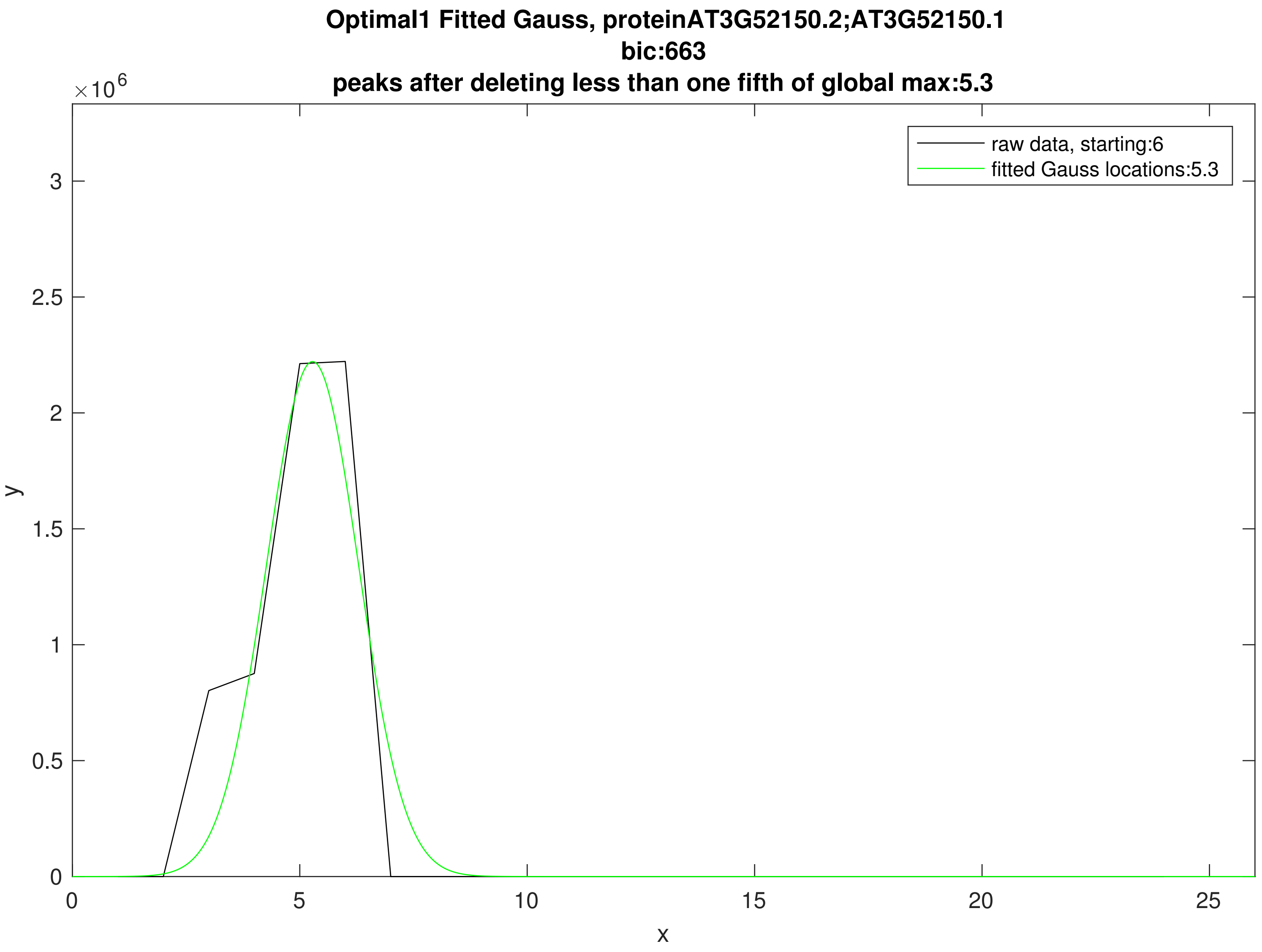
Optimal1 Fitted Gauss, proteinAT3G52150.2;AT3G52150.1
bic:663
peaks after deleting less than one fifth of global max:5.3
×10^6
raw data, starting:6
fitted Gauss locations:5.3
y
x
0
0.5
1
1.5
2
2.5
3
5
10
15
20
25

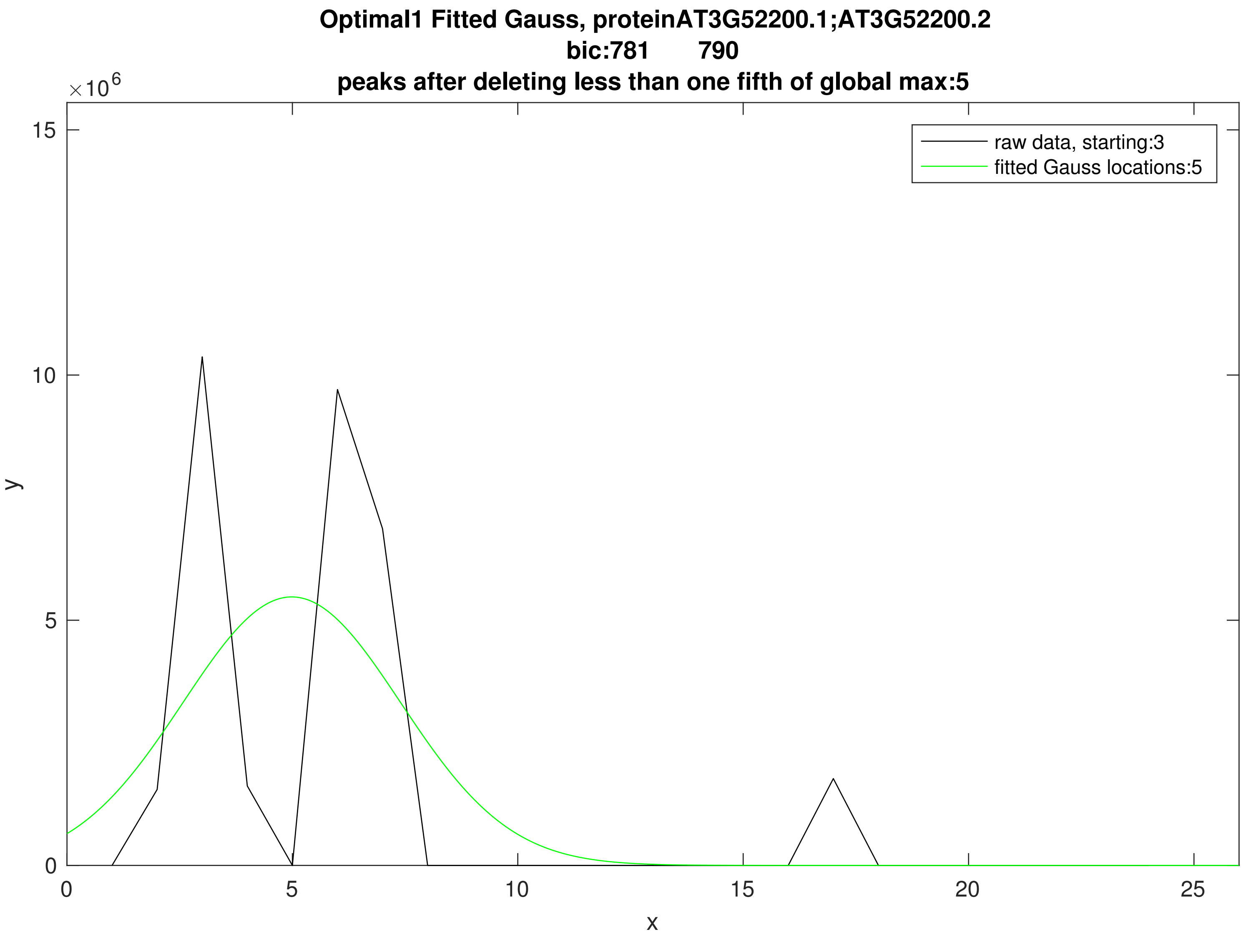

Optimal1 Fitted Gauss, proteinAT3G52200.1;AT3G52200.2
bic:781 790
peaks after deleting less than one fifth of global max:5
×10^6
15
10
5
0
y
0
5
10
15
20
25
x
raw data, starting:3
fitted Gauss locations:5

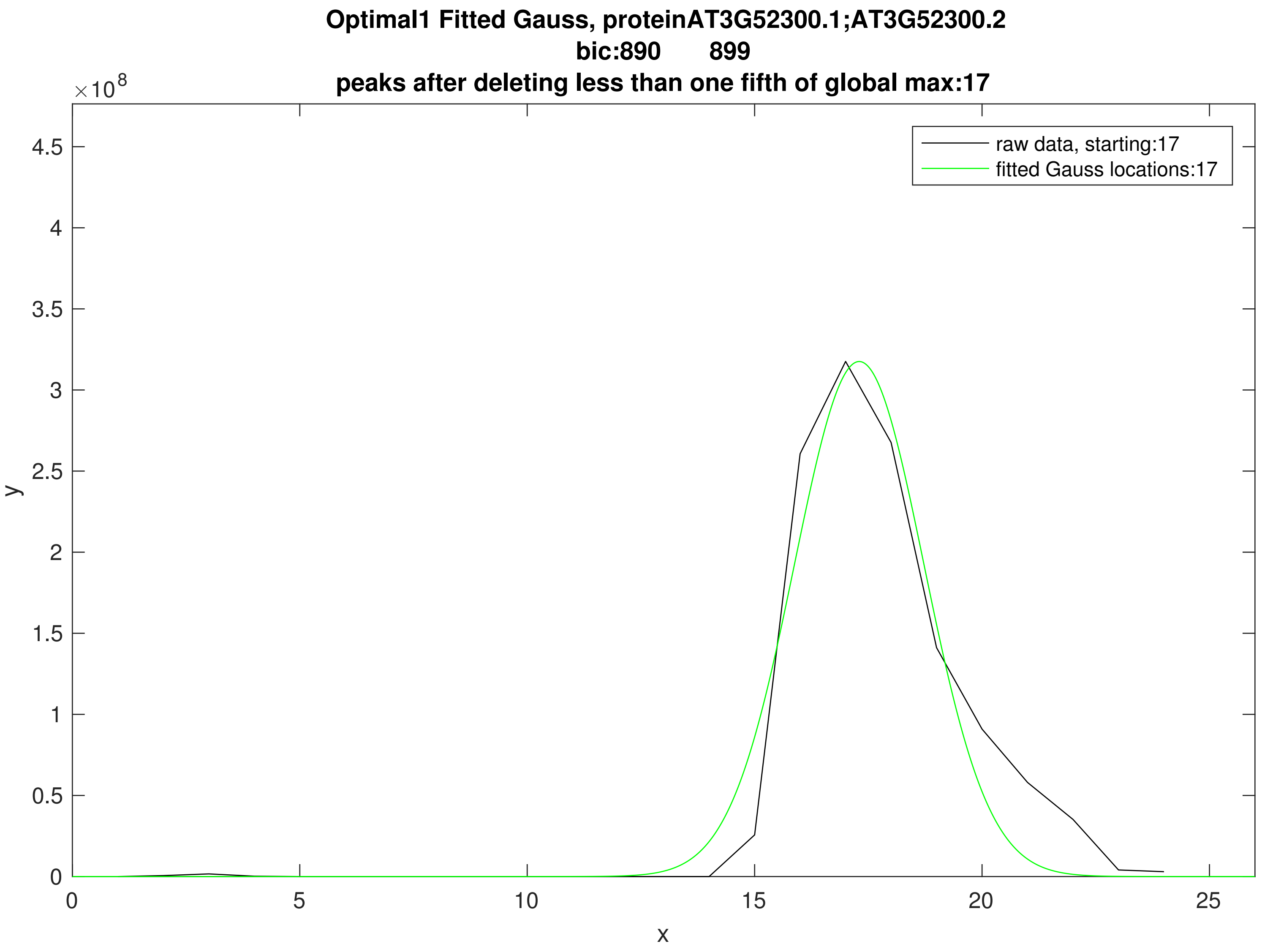

Optimal1 Fitted Gauss, proteinAT3G52300.1;AT3G52300.2
bic:890 899
peaks after deleting less than one fifth of global max:17
×10^8
raw data, starting:17
fitted Gauss locations:17
y
x
0
0.5
1
1.5
2
2.5
3
3.5
4
4.5
0
5
10
15
20
25

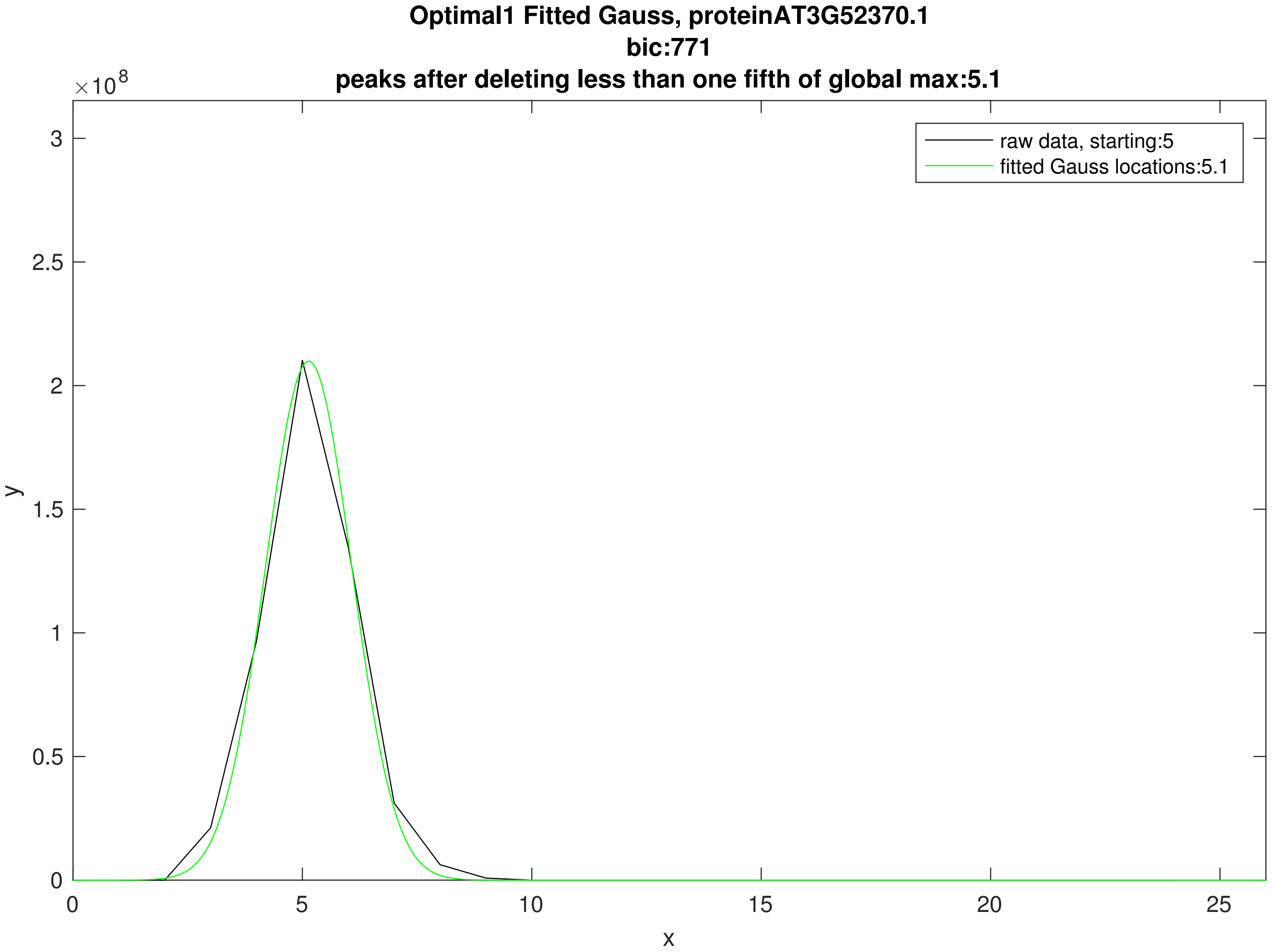

Optimal1 Fitted Gauss, proteinAT3G52370.1
bic:771
peaks after deleting less than one fifth of global max:5.1
×10^8
3
2.5
2
1.5
1
0.5
0
y
0
5
10
15
20
25
x
raw data, starting:5
fitted Gauss locations:5.1

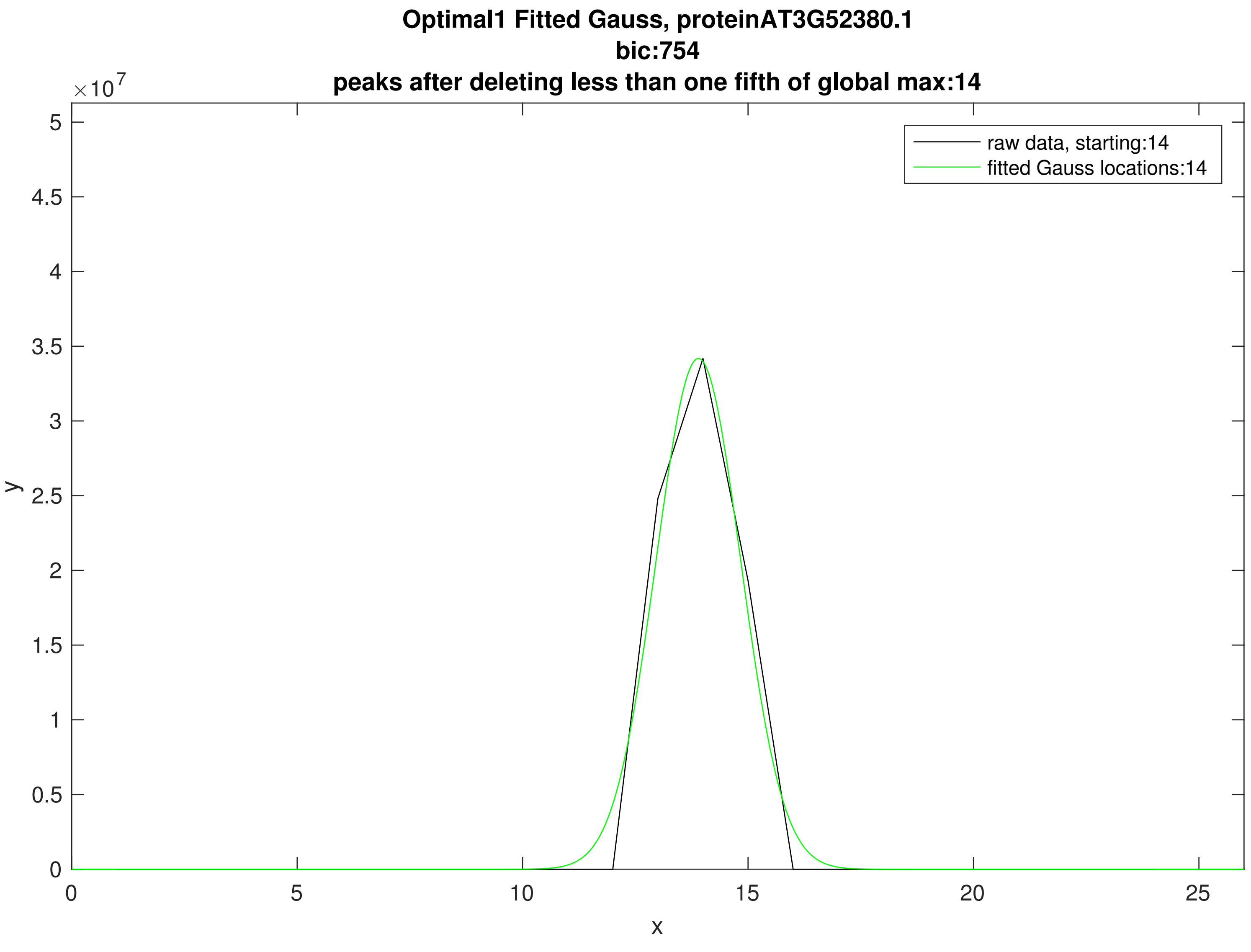

Optimal1 Fitted Gauss, proteinAT3G52380.1
bic:754
peaks after deleting less than one fifth of global max:14
×10^7
5
4.5
4
3.5
3
2.5
2
1.5
1
0.5
0
y
0
5
10
15
20
25
x
raw data, starting:14
fitted Gauss locations:14

Optimal1 Fitted Gauss, proteinAT3G52390.1;AT3G52390.2
bic:757
peaks after deleting less than one fifth of global max:18

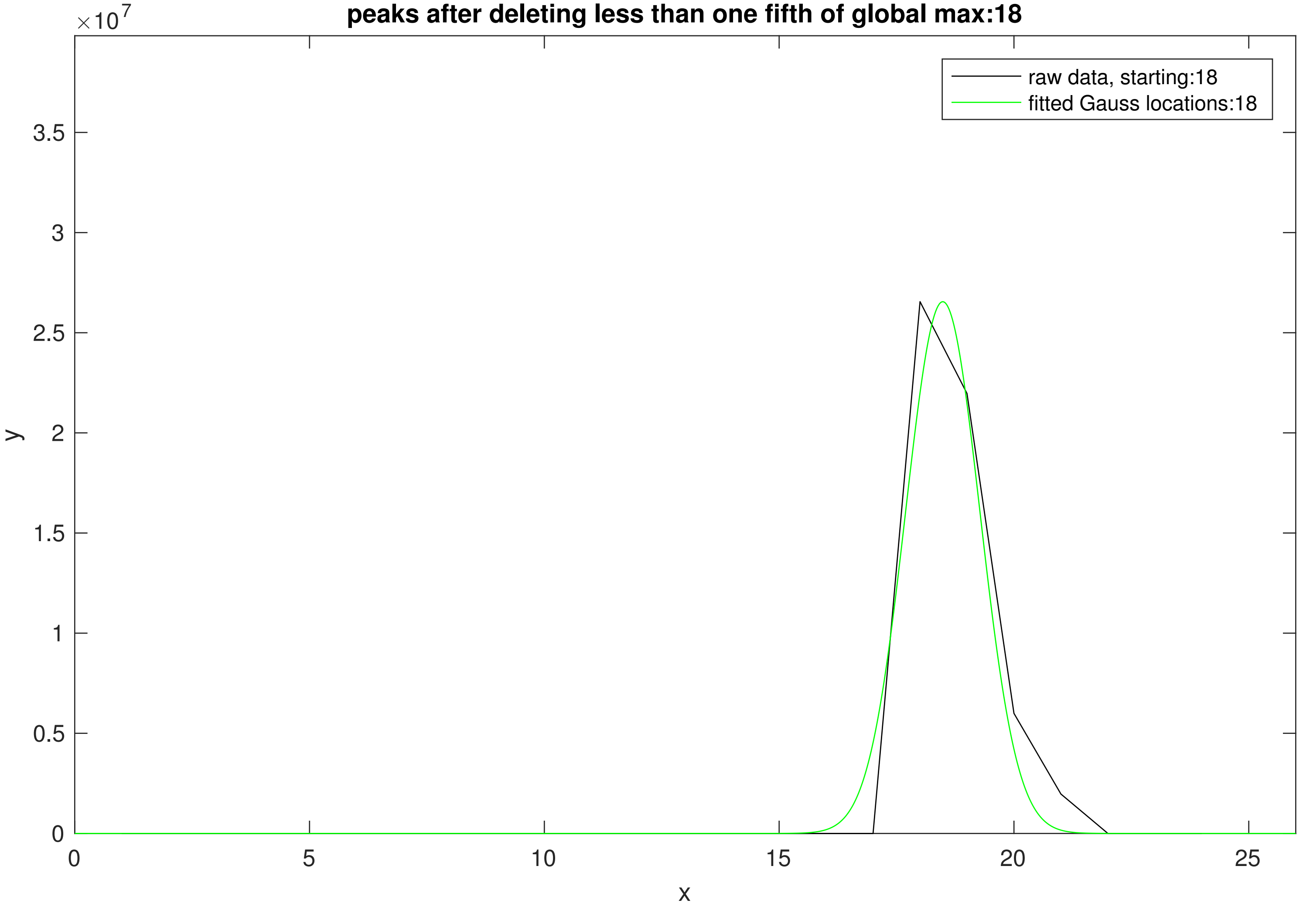

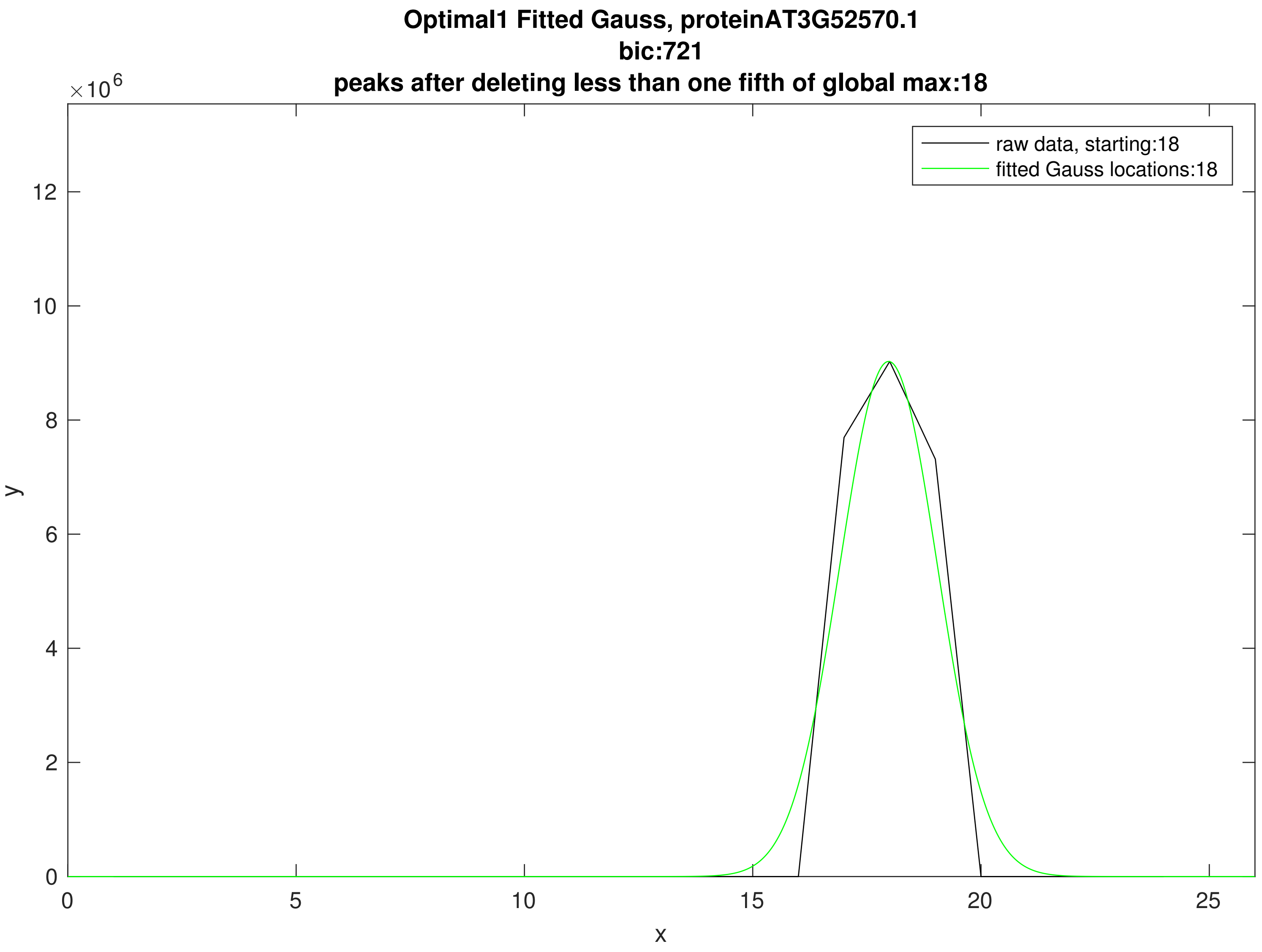
Optimal1 Fitted Gauss, proteinAT3G52570.1
bic:721
peaks after deleting less than one fifth of global max:18
×10^6
raw data, starting:18
fitted Gauss locations:18
y
x
0
2
4
6
8
10
12
0
5
10
15
20
25

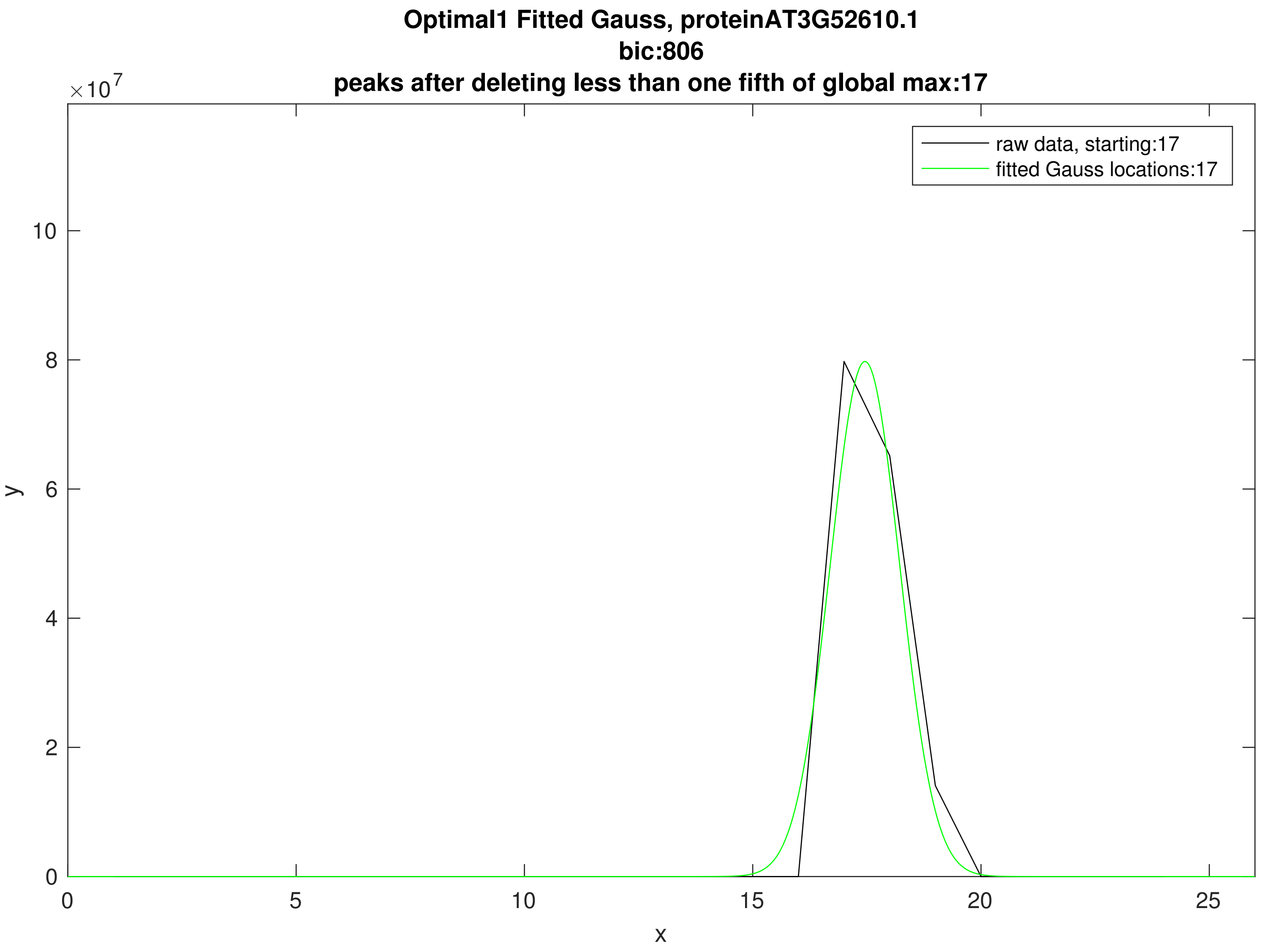

Optimal1 Fitted Gauss, proteinAT3G52610.1
bic:806
peaks after deleting less than one fifth of global max:17
×10^7
raw data, starting:17
fitted Gauss locations:17
10
8
6
4
2
0
y
0
5
10
15
20
25
x

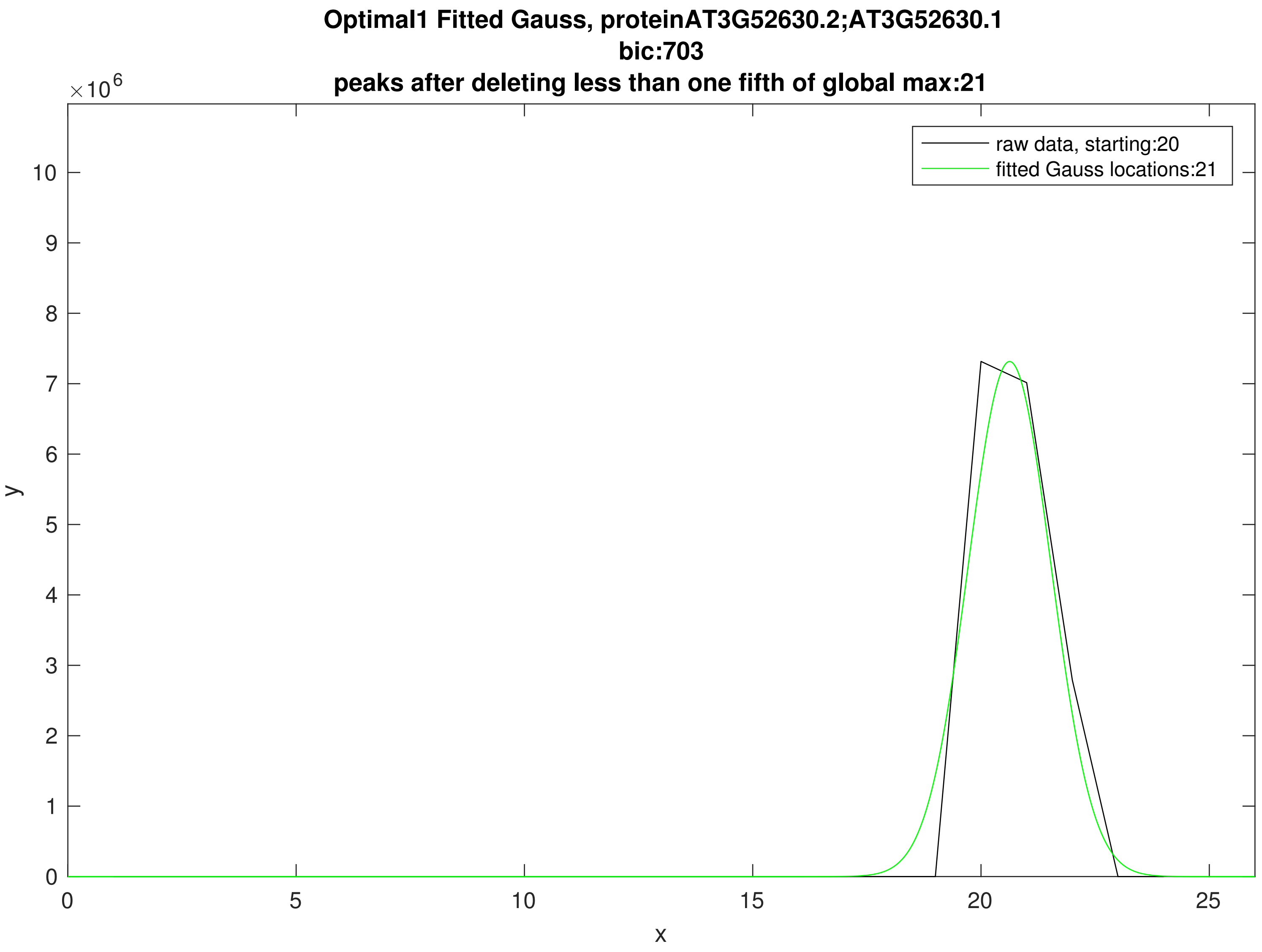

Optimal1 Fitted Gauss, proteinAT3G52630.2;AT3G52630.1
bic:703
peaks after deleting less than one fifth of global max:21
×10^6
raw data, starting:20
fitted Gauss locations:21
y
x
0
1
2
3
4
5
6
7
8
9
10
0
5
10
15
20
25

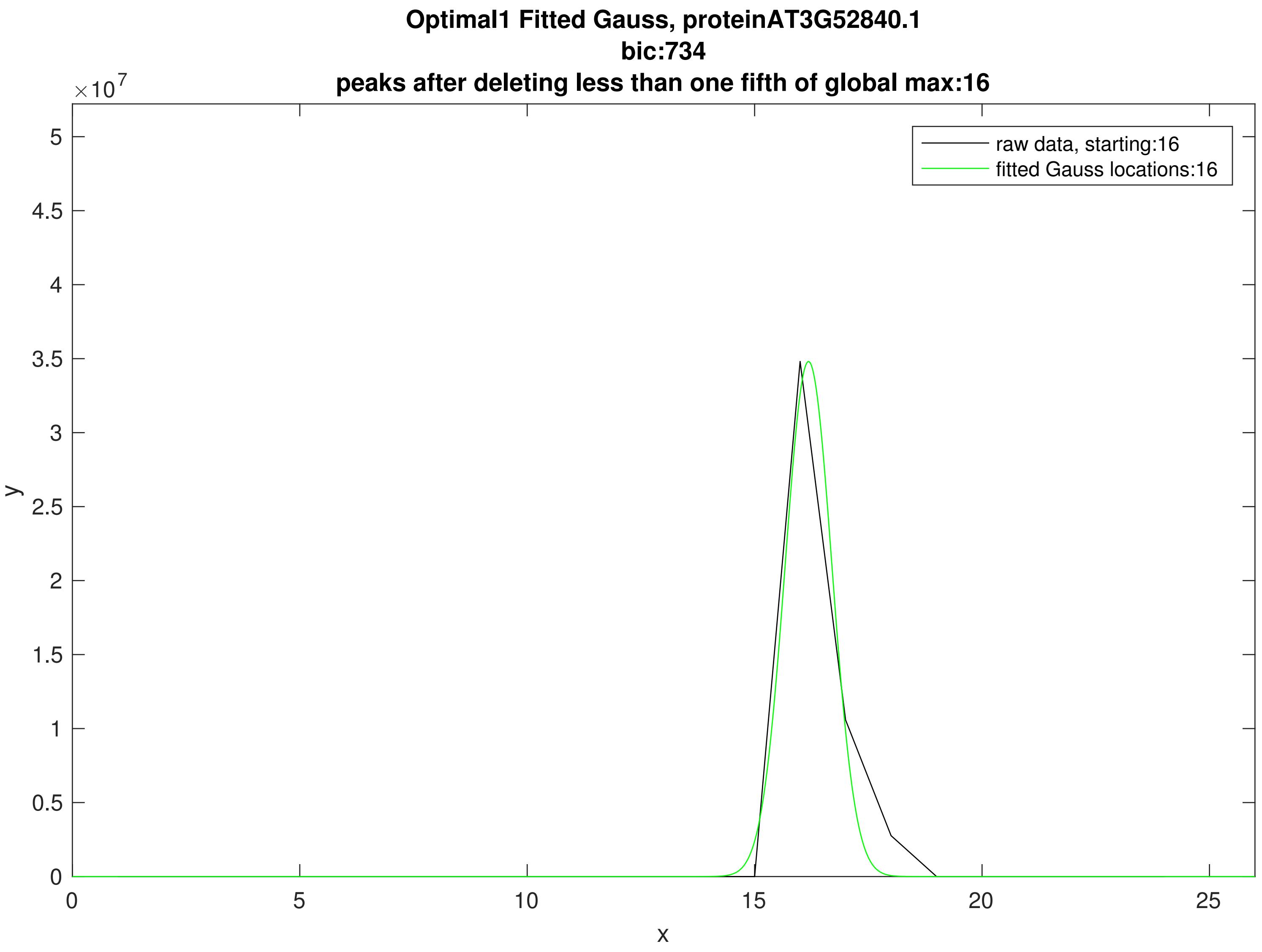
Optimal1 Fitted Gauss, proteinAT3G52840.1
bic:734
peaks after deleting less than one fifth of global max:16
×10^7
raw data, starting:16
fitted Gauss locations:16
y
x
0
0.5
1
1.5
2
2.5
3
3.5
4
4.5
5
0
5
10
15
20
25

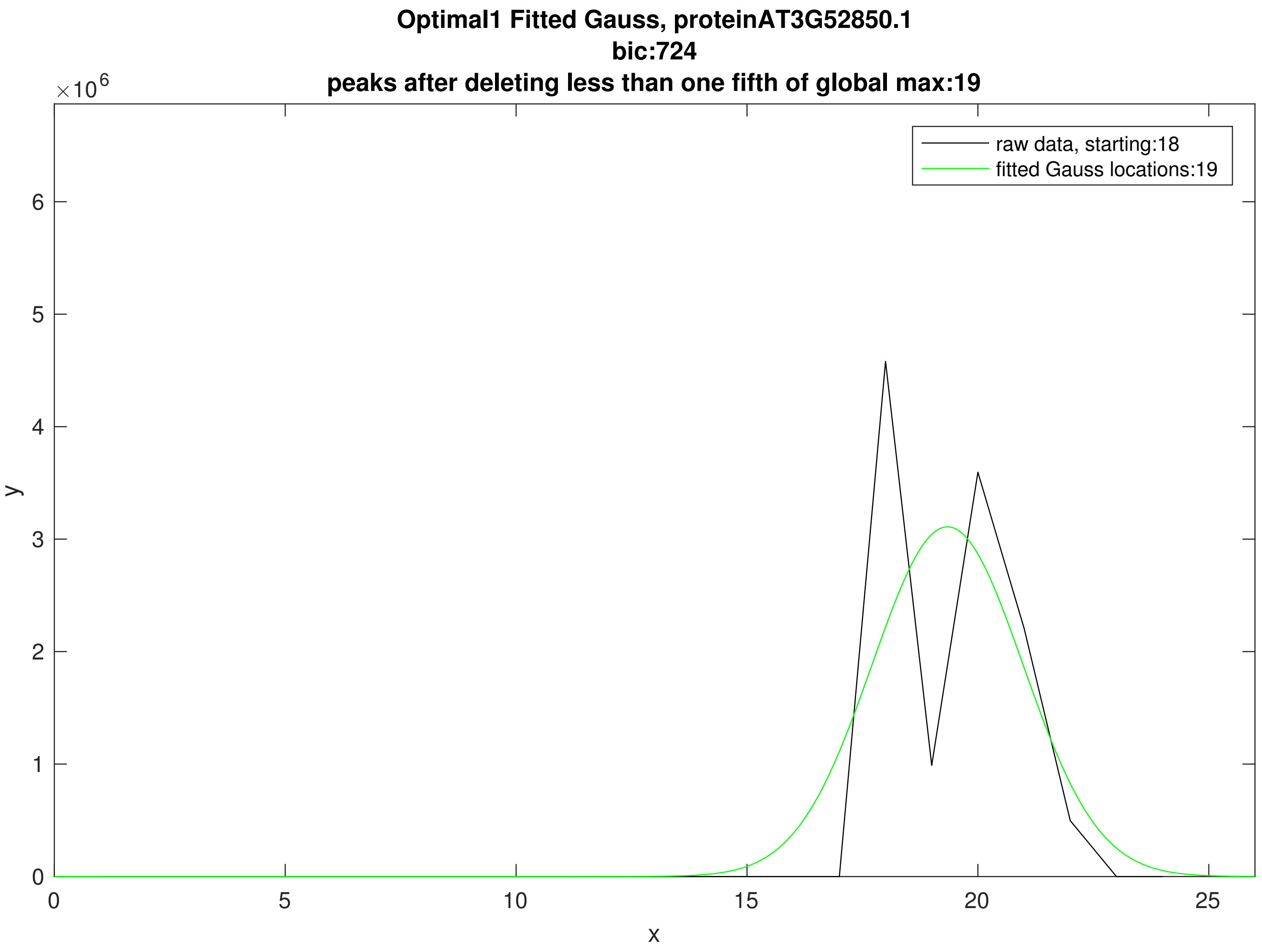

Optimal1 Fitted Gauss, proteinAT3G52850.1
bic:724
peaks after deleting less than one fifth of global max:19
×10^6
raw data, starting:18
fitted Gauss locations:19
y
x
0
1
2
3
4
5
6
0
5
10
15
20
25

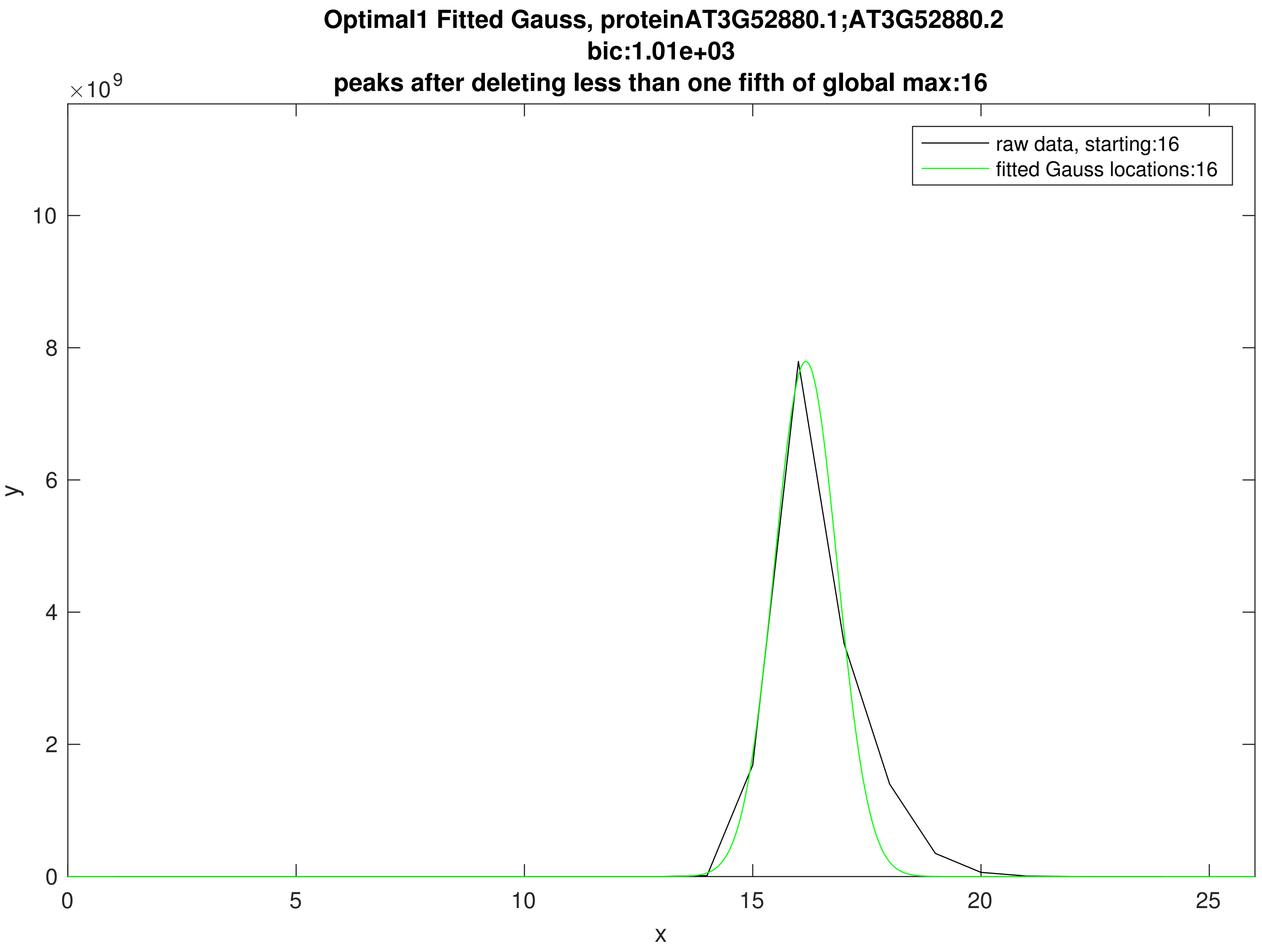
Optimal1 Fitted Gauss, proteinAT3G52880.1;AT3G52880.2
bic:1.01e+03
peaks after deleting less than one fifth of global max:16
×10^9
raw data, starting:16
fitted Gauss locations:16
y
x
0
2
4
6
8
10
0
5
10
15
20
25

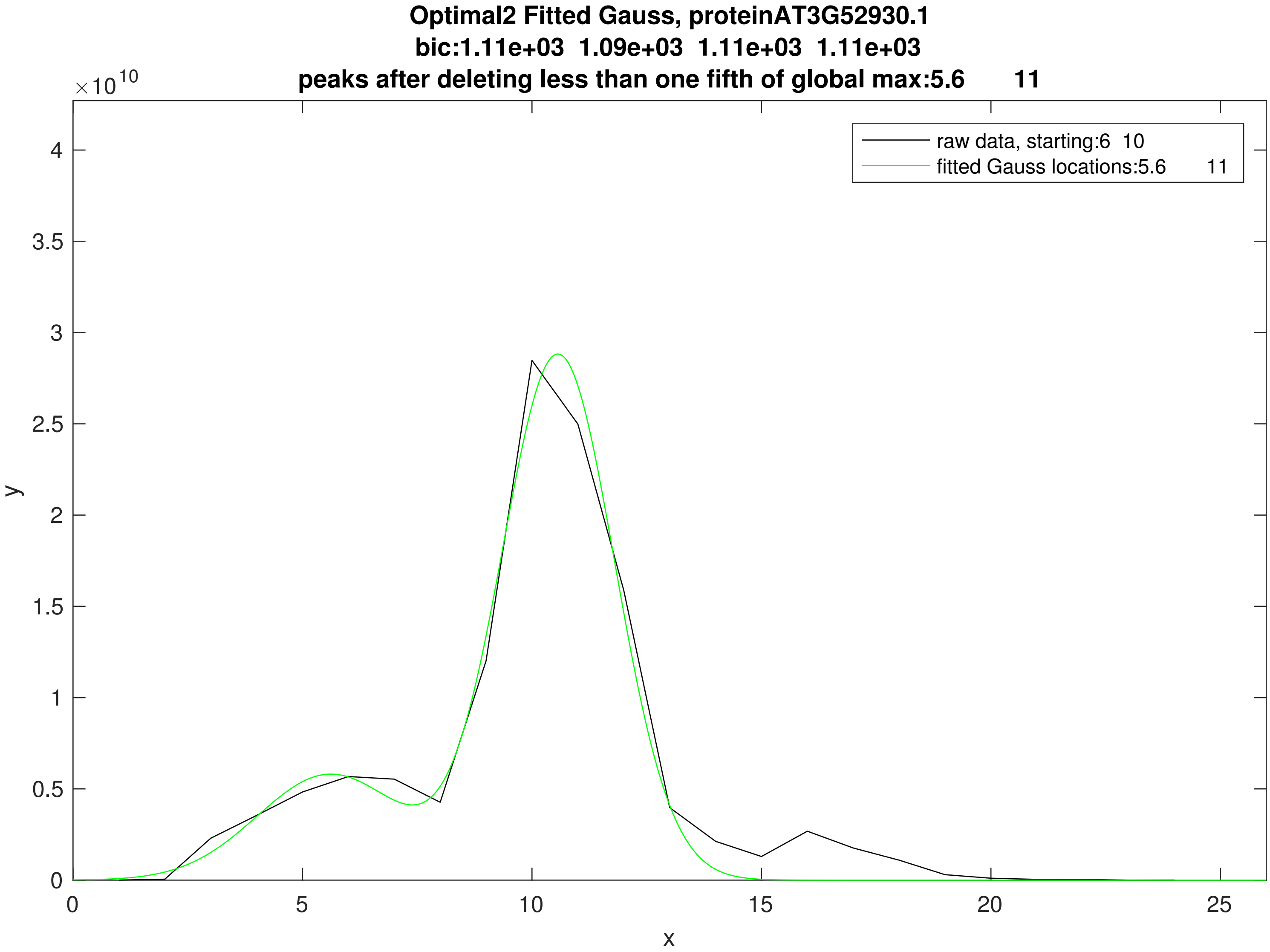
Optimal2 Fitted Gauss, proteinAT3G52930.1
bic:1.11e+03 1.09e+03 1.11e+03 1.11e+03
peaks after deleting less than one fifth of global max:5.6 11
×10^10
raw data, starting:6 10
fitted Gauss locations:5.6 11
y
x
0
0.5
1
1.5
2
2.5
3
3.5
4
5
10
15
20
25

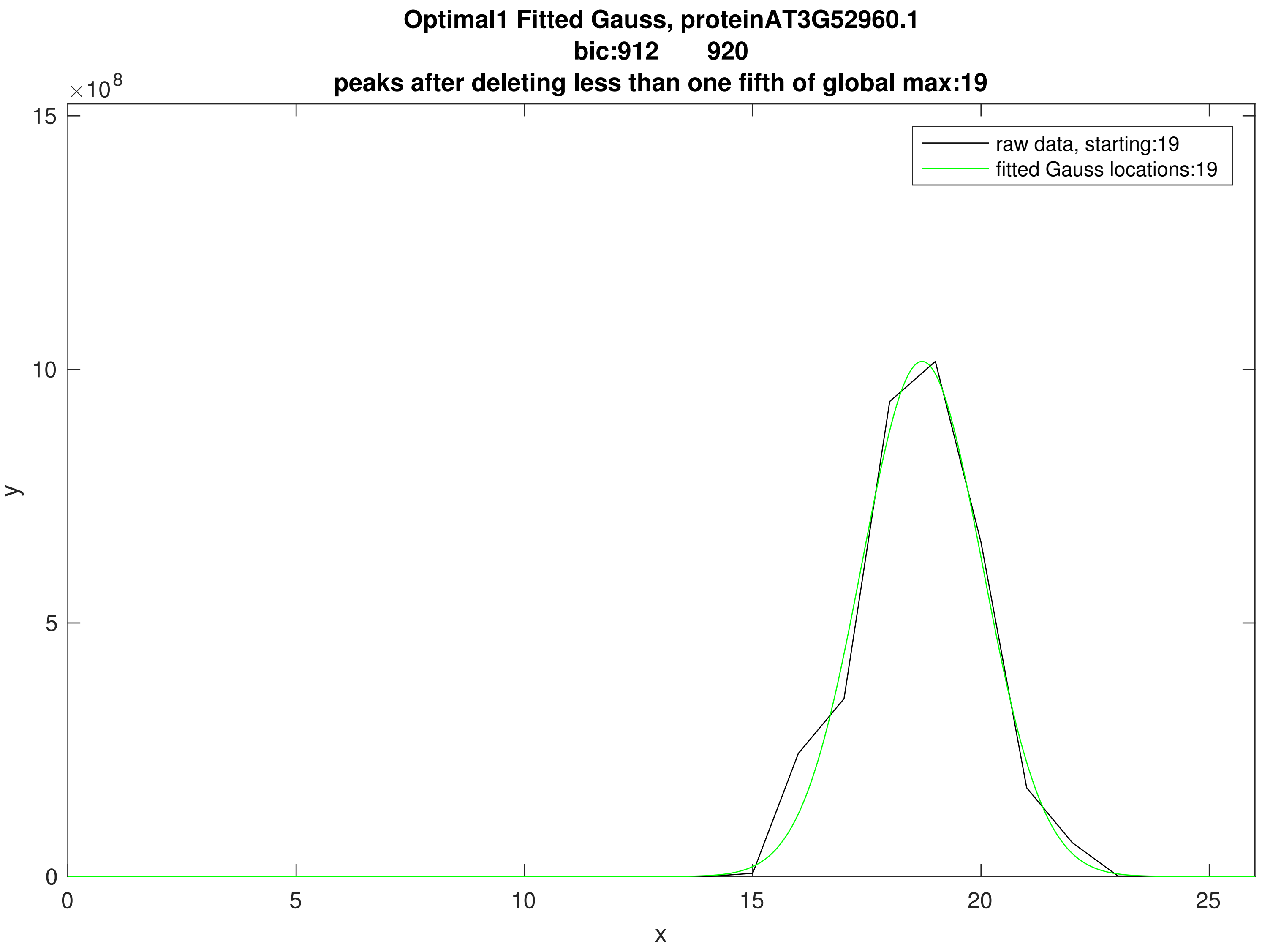
Optimal1 Fitted Gauss, proteinAT3G52960.1
bic:912 920
peaks after deleting less than one fifth of global max:19
×10^8
15
10
5
0
y
0
5
10
15
20
25
x
raw data, starting:19
fitted Gauss locations:19

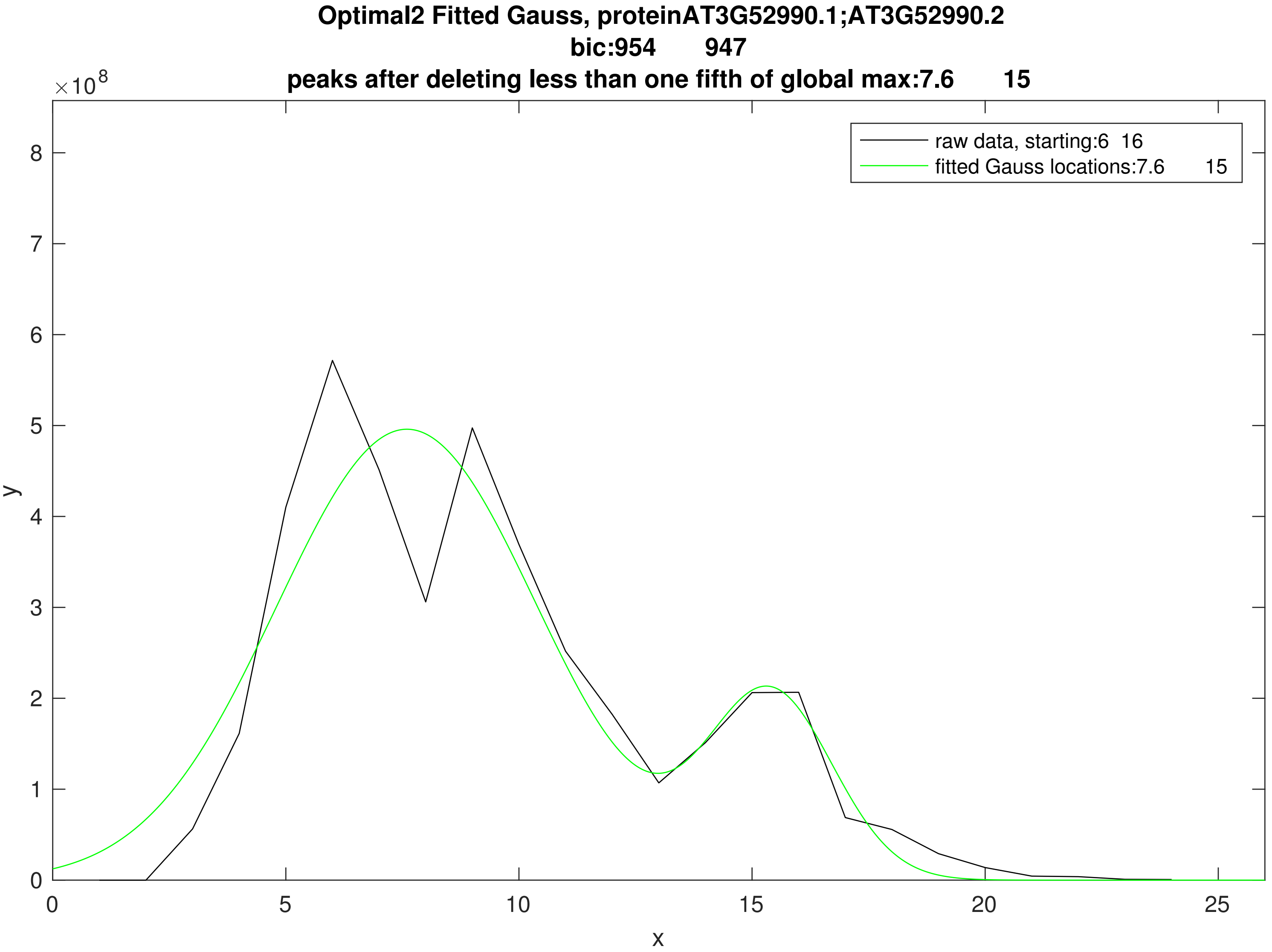

Optimal2 Fitted Gauss, proteinAT3G52990.1;AT3G52990.2
bic:954 947
peaks after deleting less than one fifth of global max:7.6 15
×10^8
raw data, starting:6 16
fitted Gauss locations:7.6 15
y
x
0
1
2
3
4
5
6
7
8
0
5
10
15
20
25

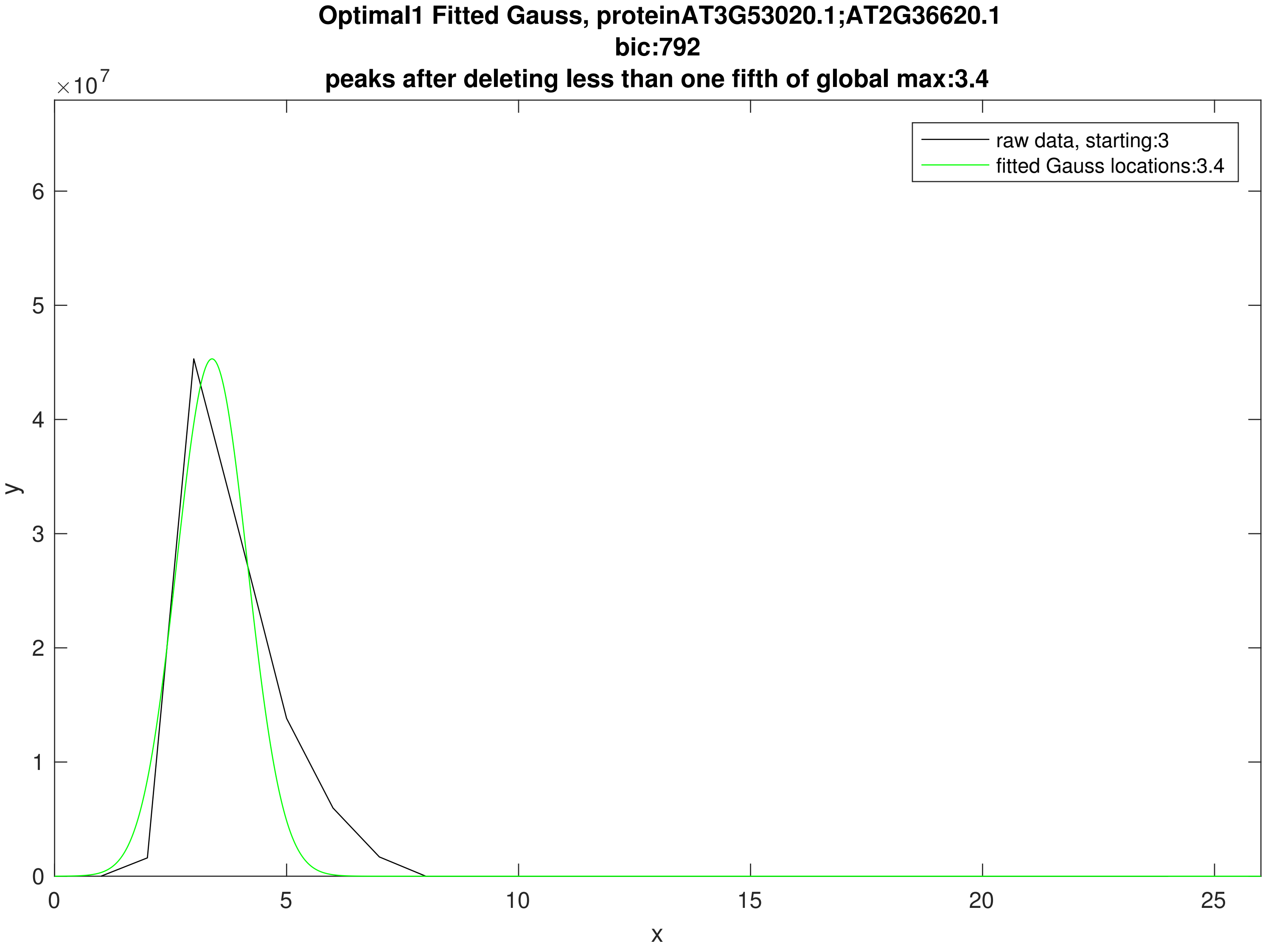
Optimal1 Fitted Gauss, proteinAT3G53020.1;AT2G36620.1
bic:792
peaks after deleting less than one fifth of global max:3.4
×10^7
raw data, starting:3
fitted Gauss locations:3.4
y
x
0
1
2
3
4
5
6
0
5
10
15
20
25

**Optimal1 Fitted Gauss, proteinAT3G53040.1**
**bic:929 935 944 953**
**peaks after deleting less than one fifth of global max:7.7**

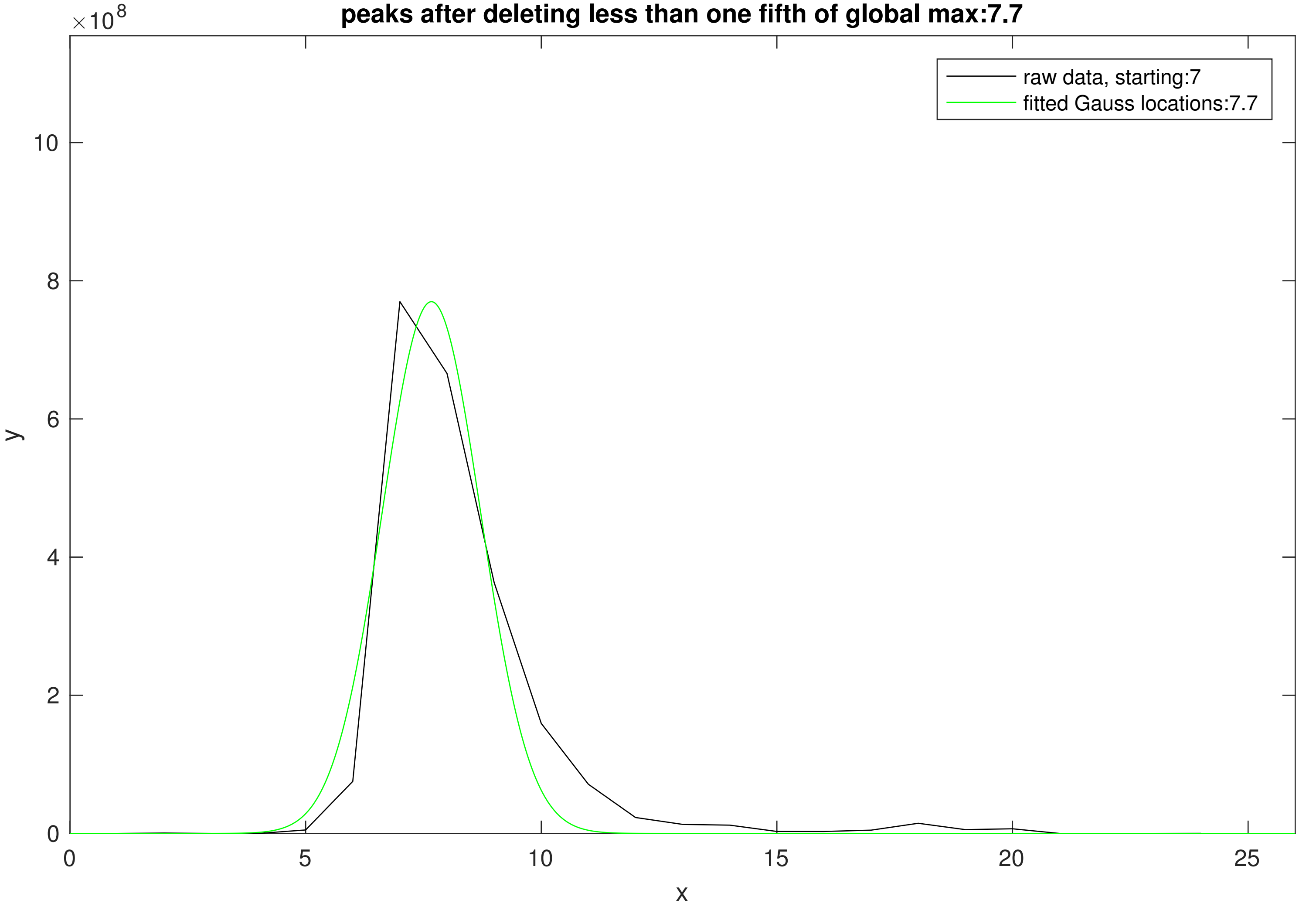

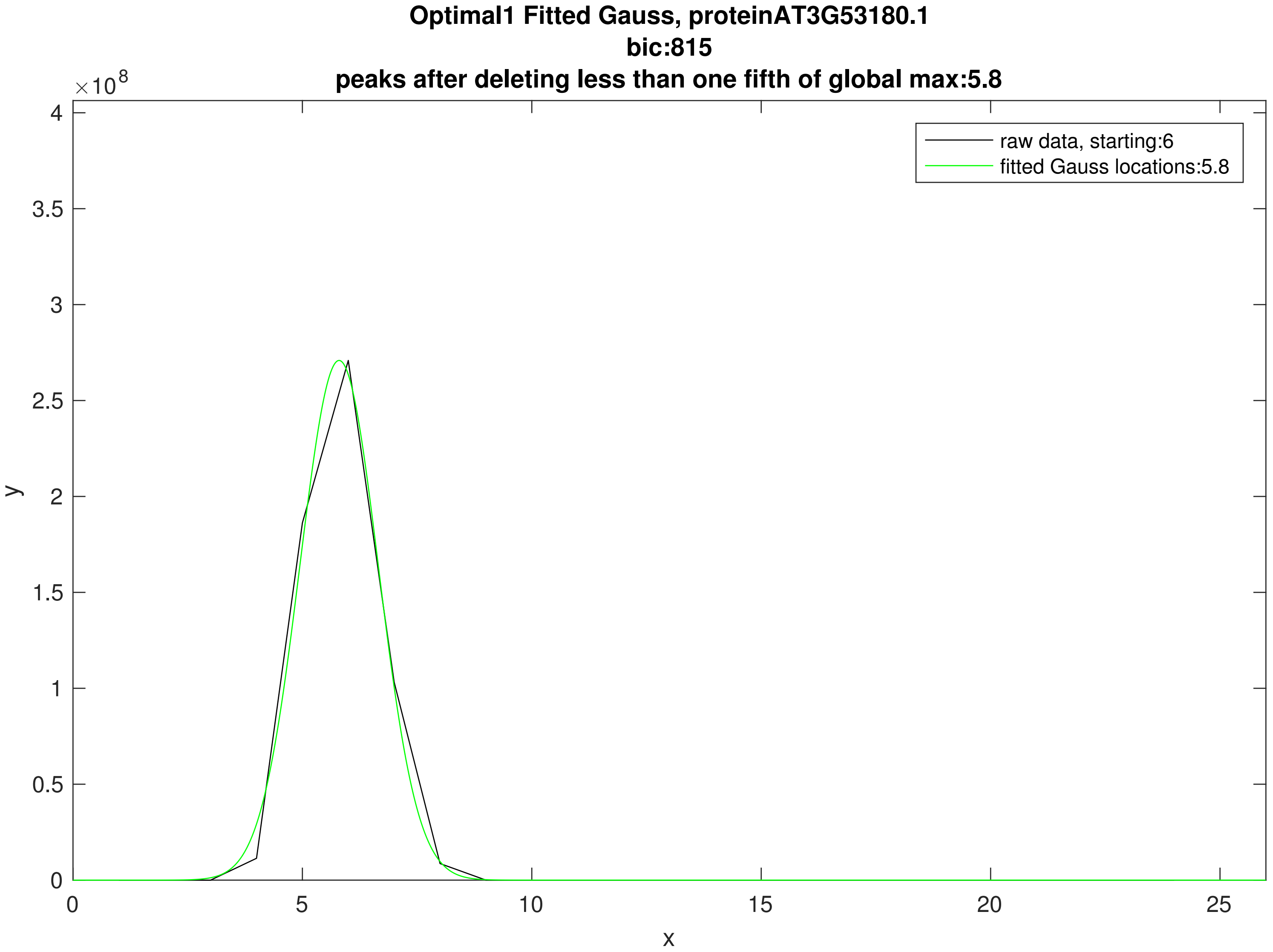

Optimal1 Fitted Gauss, proteinAT3G53180.1
bic:815
peaks after deleting less than one fifth of global max:5.8
×10^8
raw data, starting:6
fitted Gauss locations:5.8
y
x
0
0.5
1
1.5
2
2.5
3
3.5
4
0
5
10
15
20
25

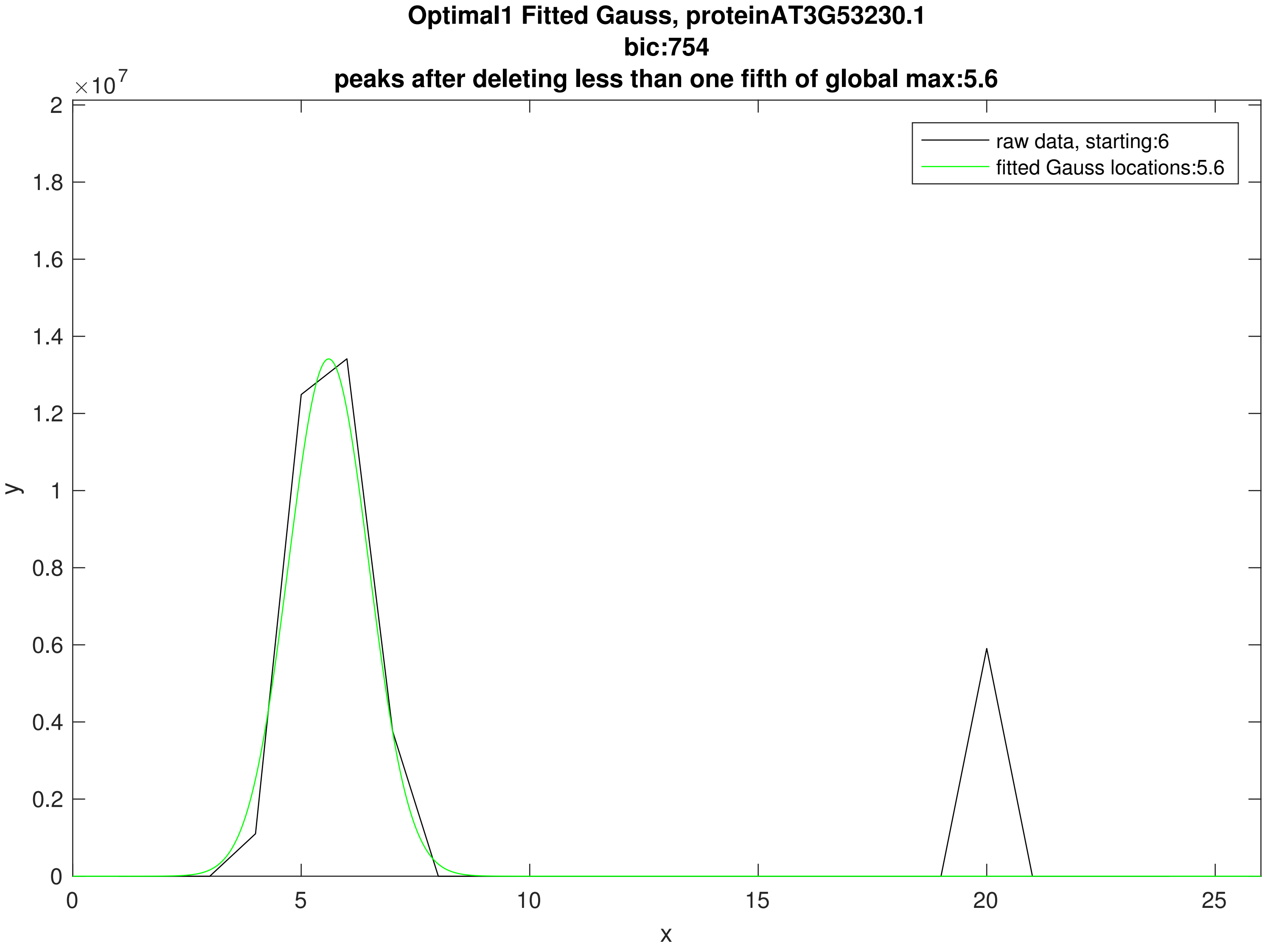

Optimal1 Fitted Gauss, proteinAT3G53230.1
bic:754
peaks after deleting less than one fifth of global max:5.6
×10^7
raw data, starting:6
fitted Gauss locations:5.6
y
x
0
0.2
0.4
0.6
0.8
1
1.2
1.4
1.6
1.8
2
0
5
10
15
20
25

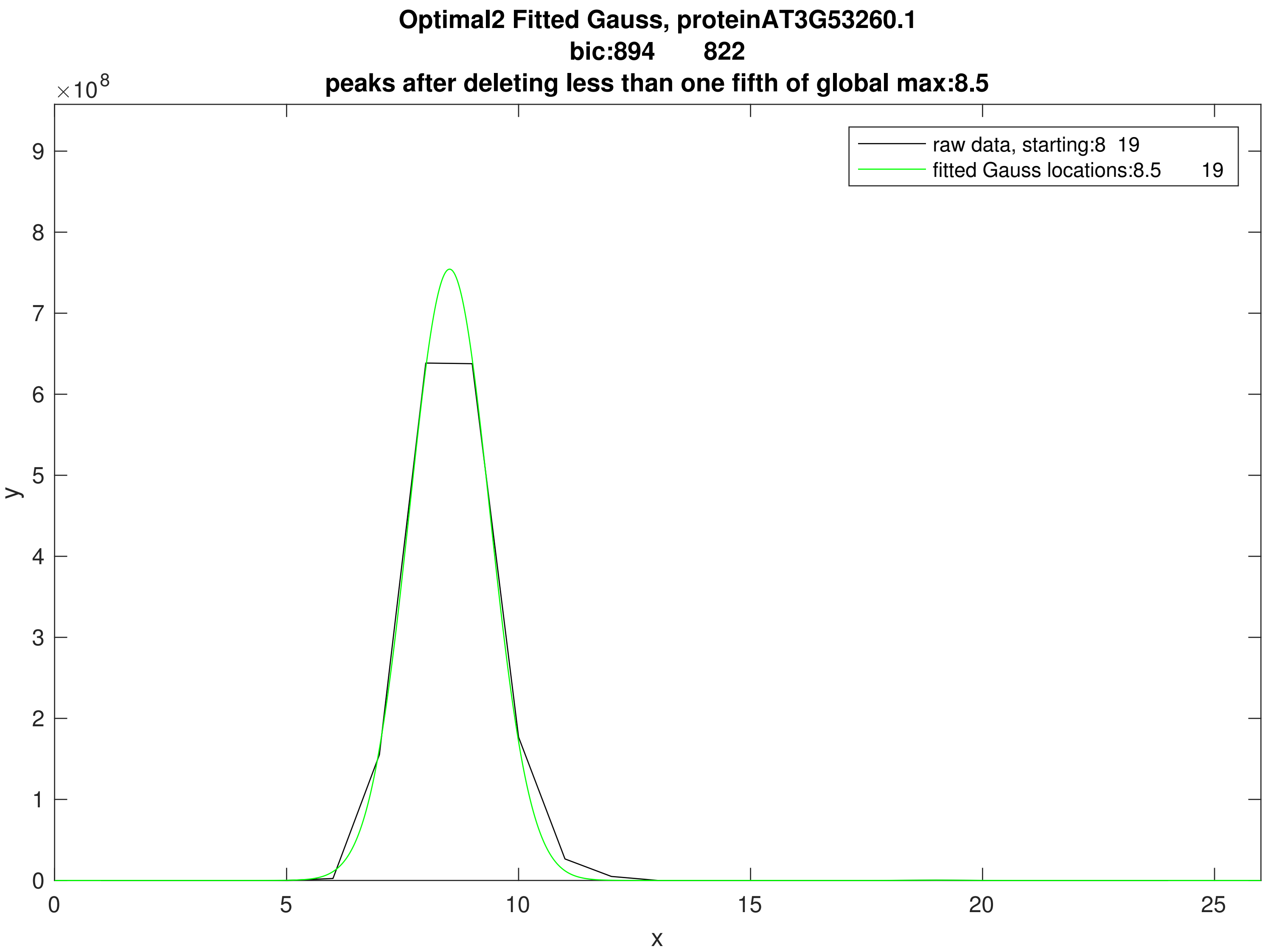

Optimal2 Fitted Gauss, proteinAT3G53260.1
bic:894 822
peaks after deleting less than one fifth of global max:8.5
×10^8
raw data, starting:8 19
fitted Gauss locations:8.5 19
y
x
0
1
2
3
4
5
6
7
8
9
0
5
10
15
20
25

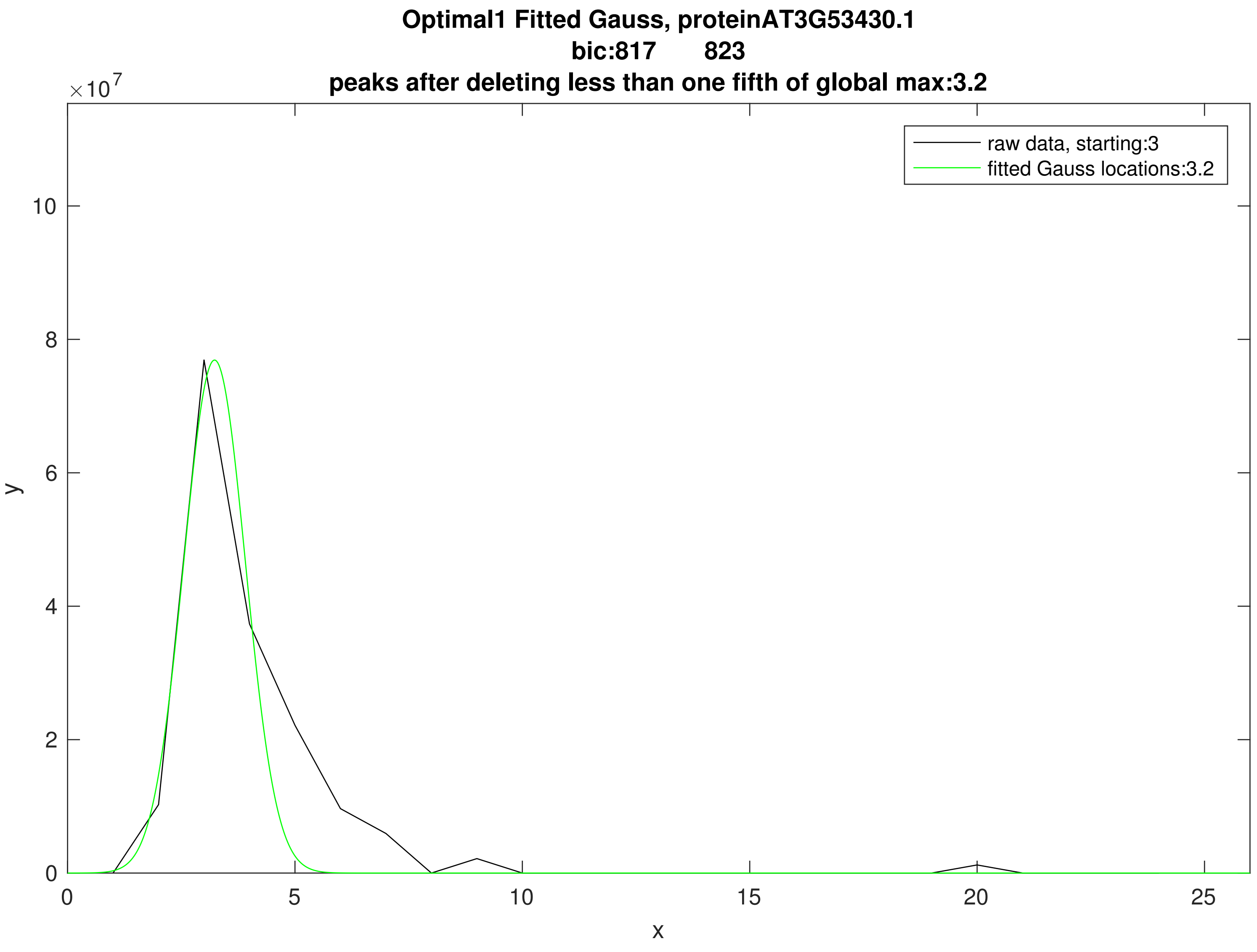
Optimal1 Fitted Gauss, proteinAT3G53430.1
bic:817 823
peaks after deleting less than one fifth of global max:3.2
×10^7
raw data, starting:3
fitted Gauss locations:3.2
y
x
0
2
4
6
8
10
0
5
10
15
20
25

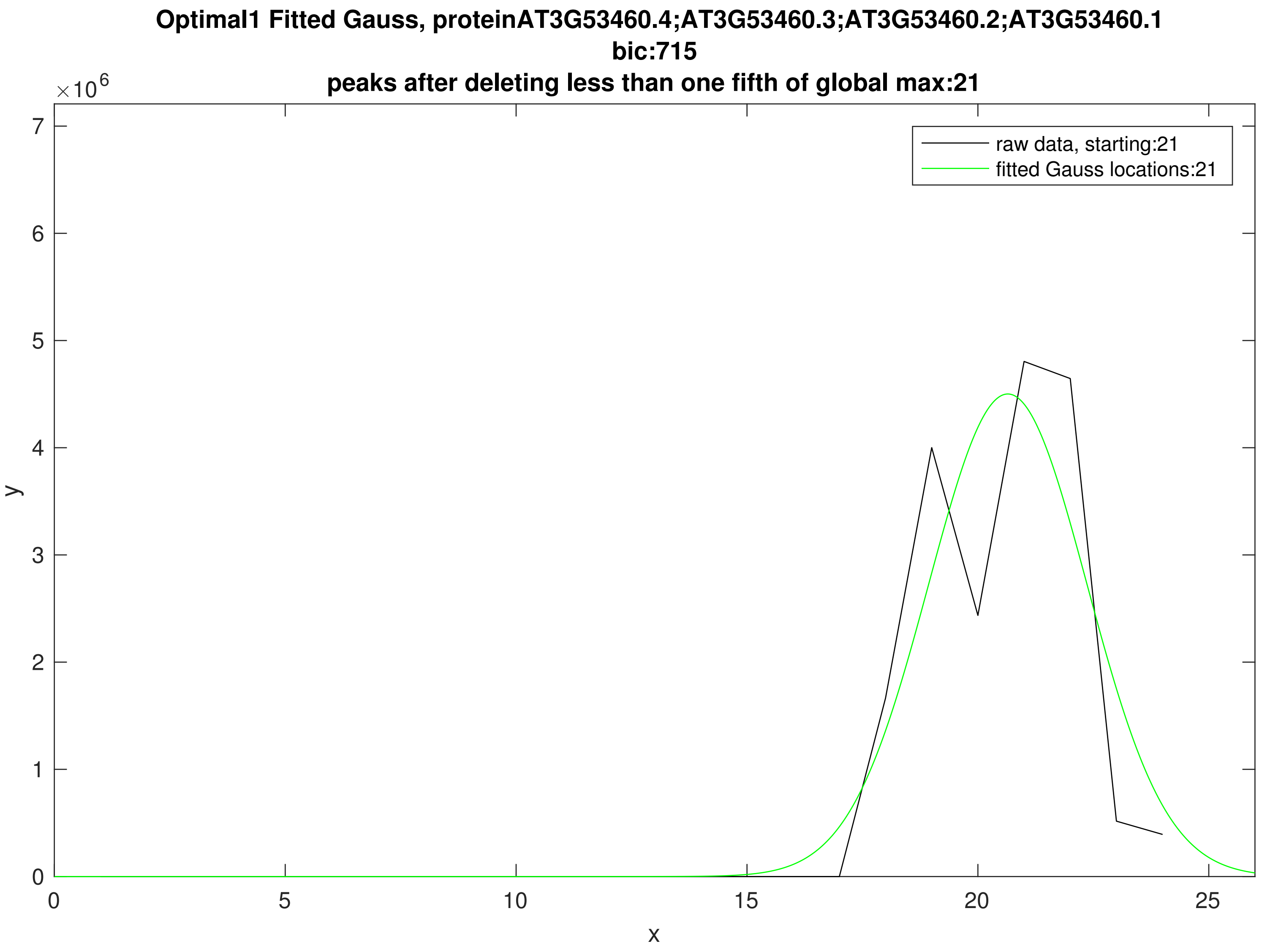
Optimal1 Fitted Gauss, proteinAT3G53460.4;AT3G53460.3;AT3G53460.2;AT3G53460.1
bic:715
peaks after deleting less than one fifth of global max:21
×10^6
raw data, starting:21
fitted Gauss locations:21
y
x
0
1
2
3
4
5
6
7
0
5
10
15
20
25

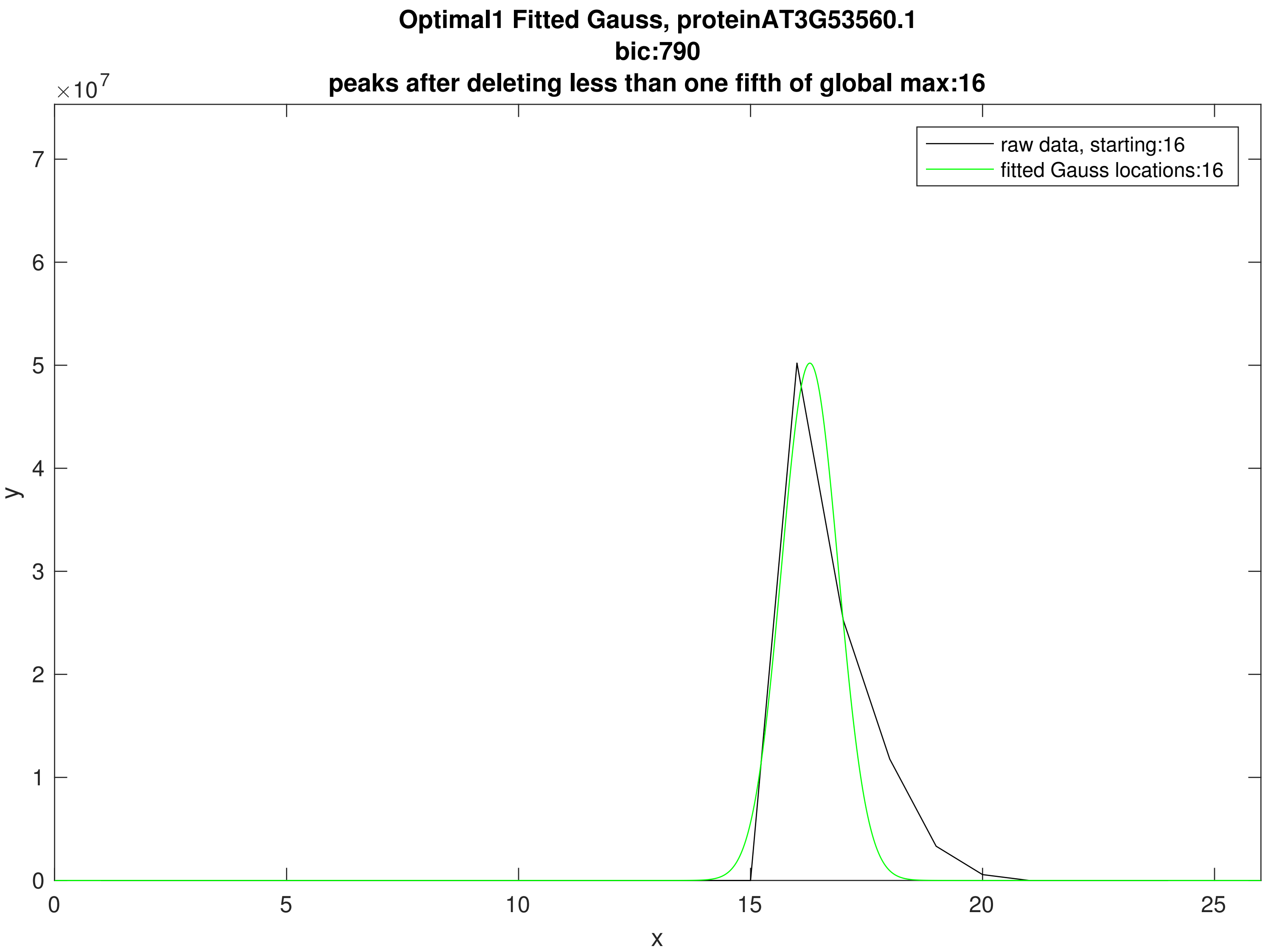

Optimal1 Fitted Gauss, proteinAT3G53560.1
bic:790
peaks after deleting less than one fifth of global max:16
×10^7
raw data, starting:16
fitted Gauss locations:16
y
x
0
1
2
3
4
5
6
7
0
5
10
15
20
25

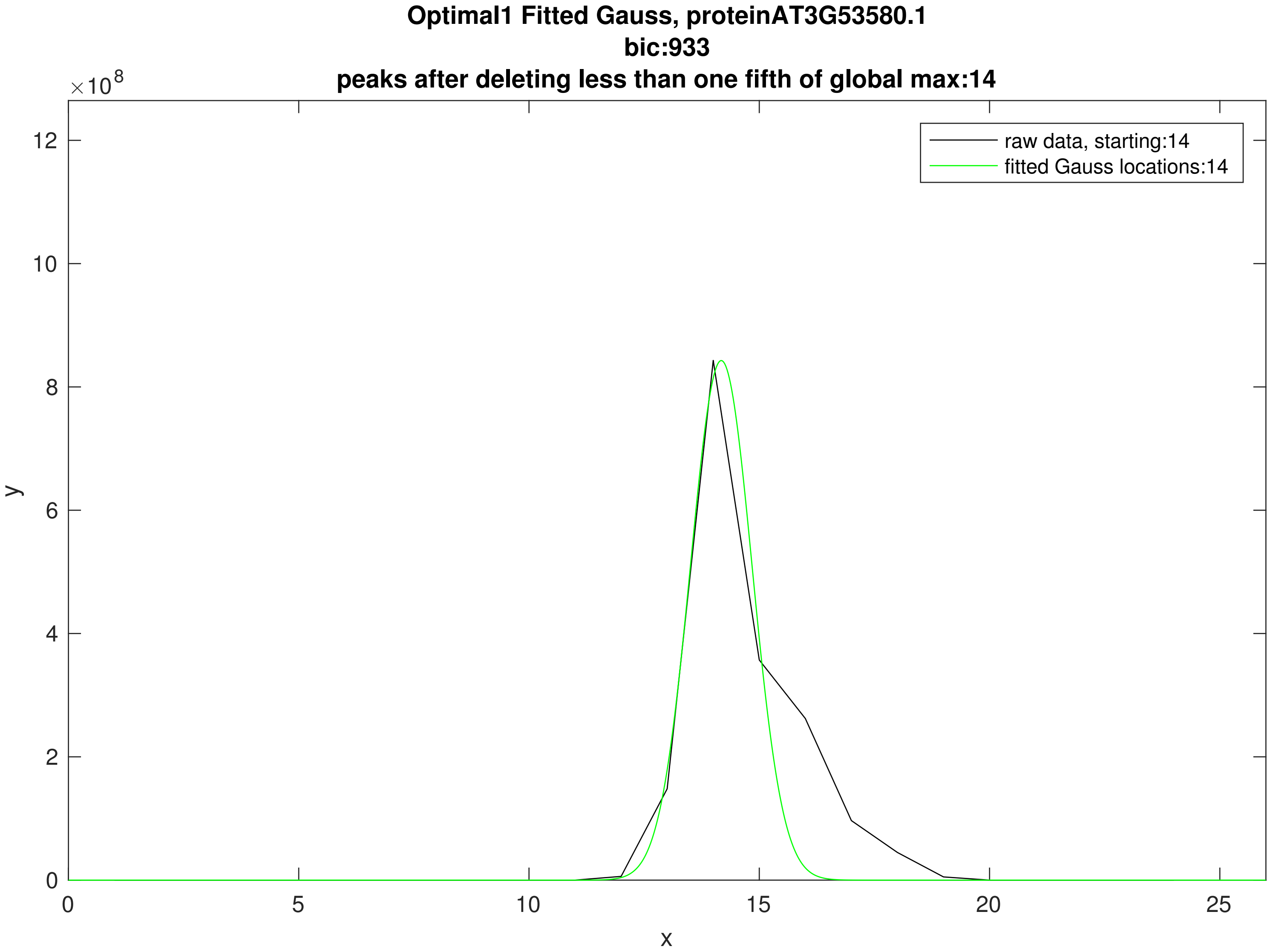

Optimal1 Fitted Gauss, proteinAT3G53580.1
bic:933
peaks after deleting less than one fifth of global max:14
×10^8
12
10
8
6
4
2
0
y
0
5
10
15
20
25
x
raw data, starting:14
fitted Gauss locations:14

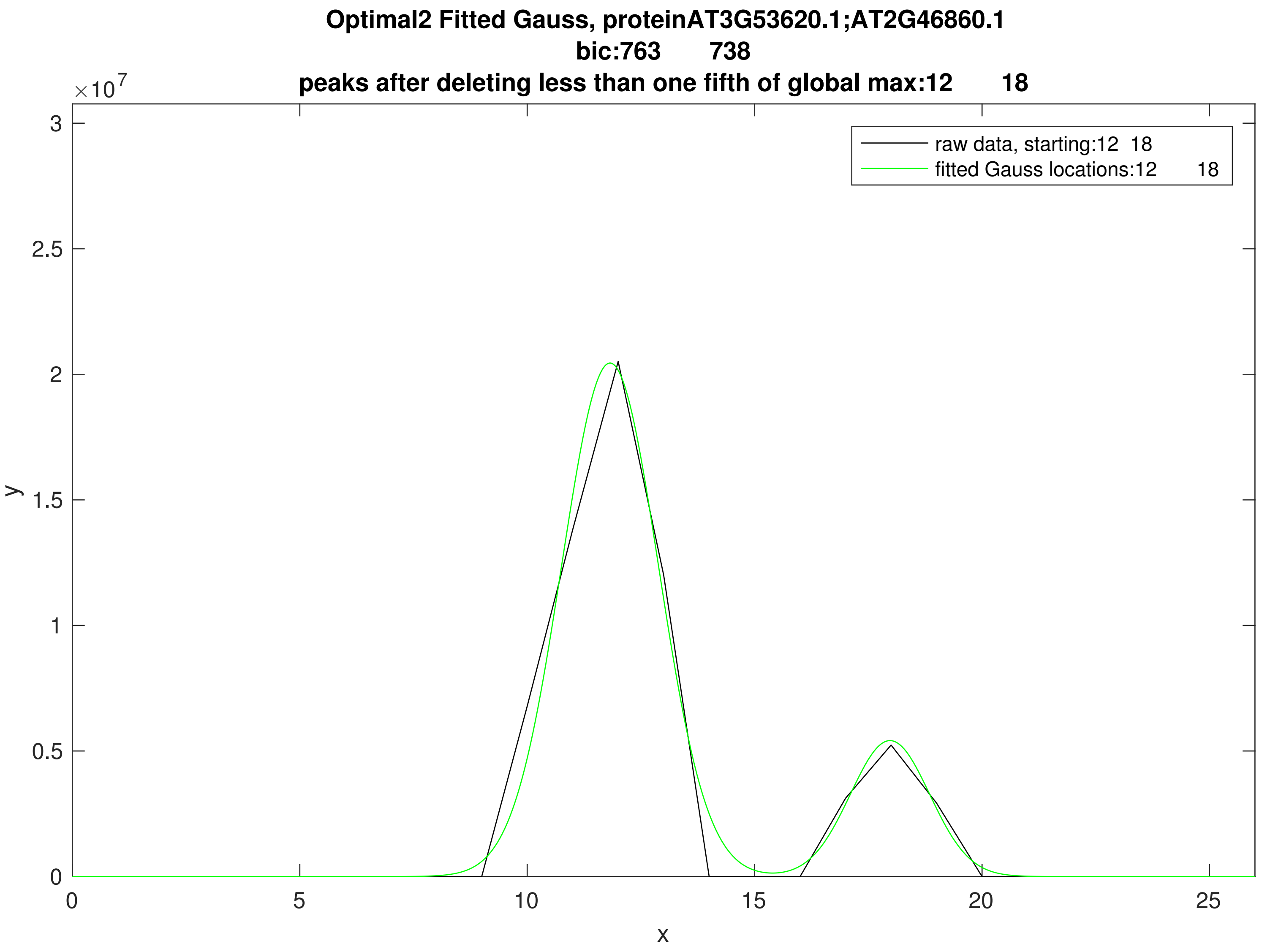

Optimal2 Fitted Gauss, proteinAT3G53620.1;AT2G46860.1
bic:763 738
peaks after deleting less than one fifth of global max:12 18
×10^7
raw data, starting:12 18
fitted Gauss locations:12 18
y
x
0
0.5
1
1.5
2
2.5
3
5
10
15
20
25

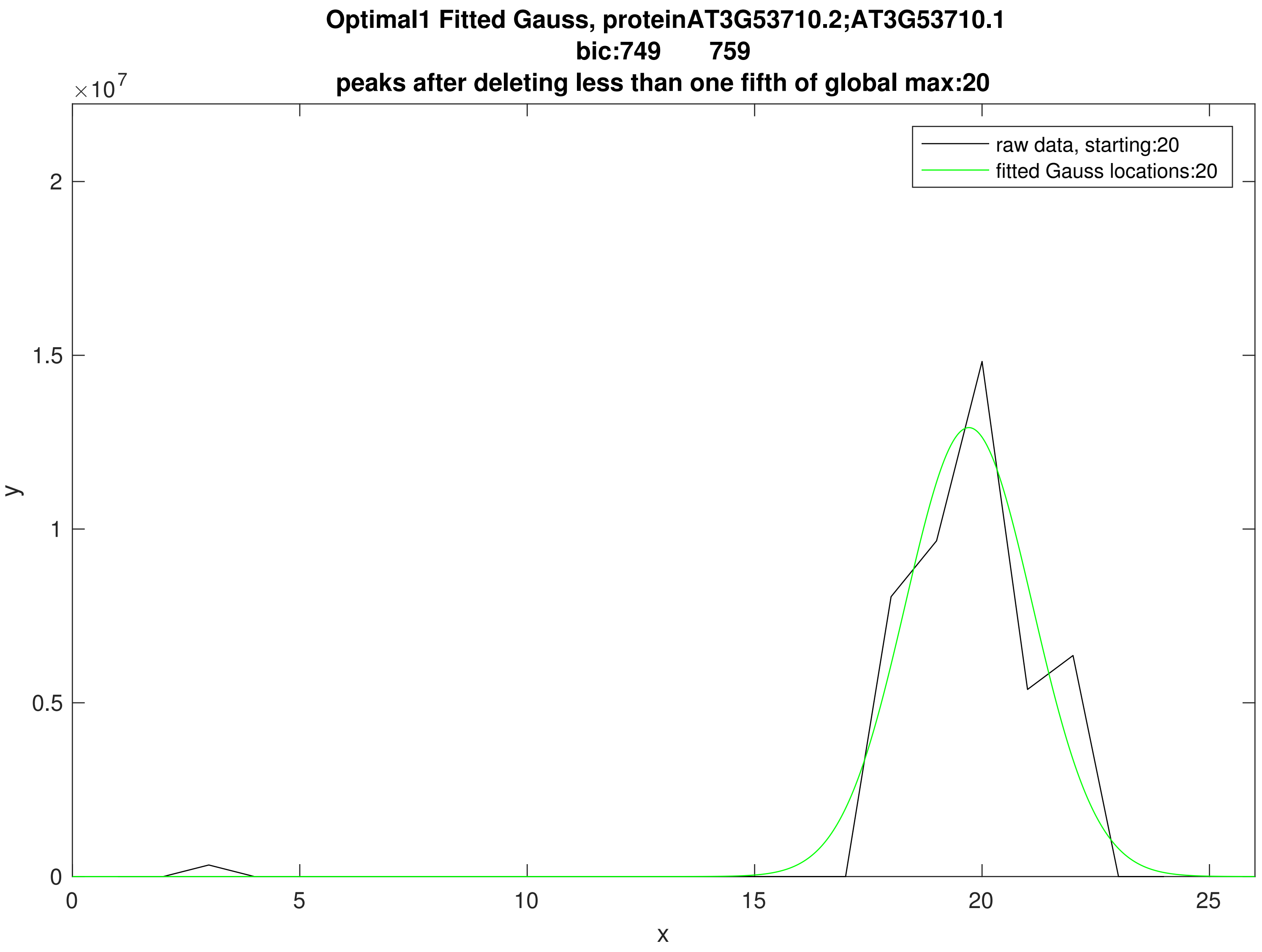

Optimal1 Fitted Gauss, proteinAT3G53710.2;AT3G53710.1
bic:749 759
peaks after deleting less than one fifth of global max:20
×10^7
raw data, starting:20
fitted Gauss locations:20
y
x
0
0.5
1
1.5
2
5
10
15
20
25

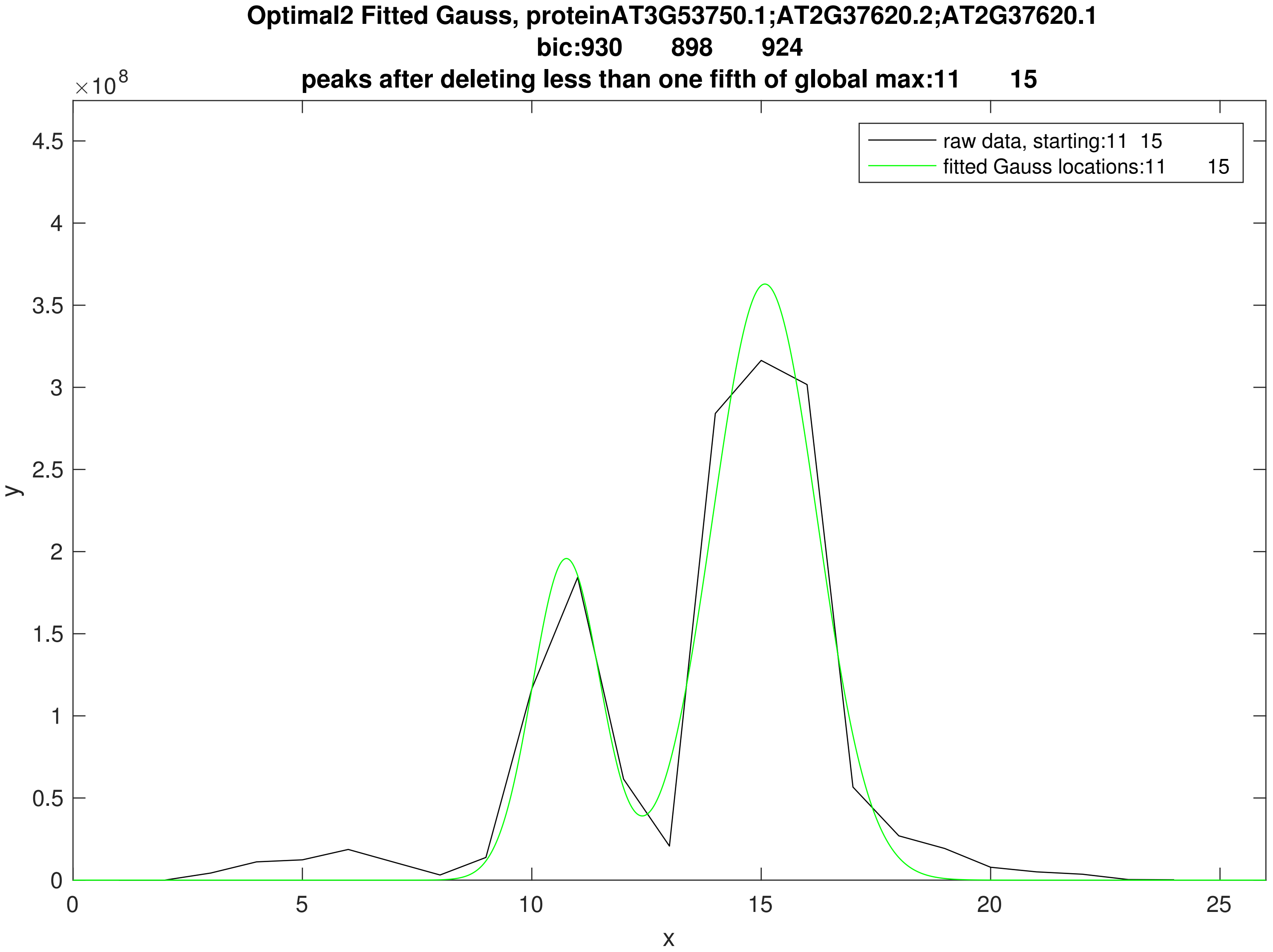
Optimal2 Fitted Gauss, proteinAT3G53750.1;AT2G37620.2;AT2G37620.1
bic:930 898 924
peaks after deleting less than one fifth of global max:11 15
×10^8
raw data, starting:11 15
fitted Gauss locations:11 15
y
x
0
0.5
1
1.5
2
2.5
3
3.5
4
4.5
5
10
15
20
25

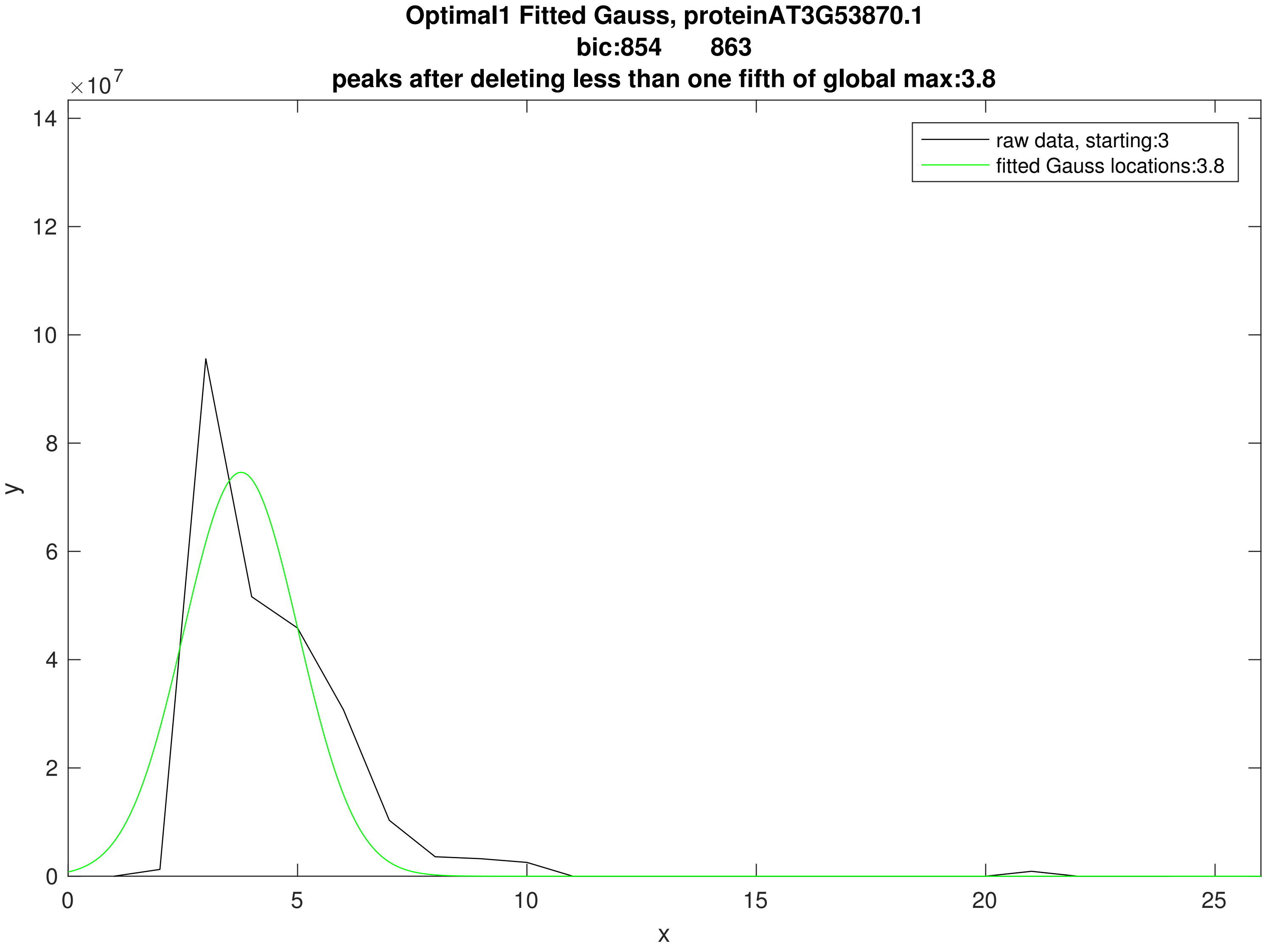
Optimal1 Fitted Gauss, proteinAT3G53870.1
bic:854 863
peaks after deleting less than one fifth of global max:3.8
×10^7
raw data, starting:3
fitted Gauss locations:3.8
y
x
0
2
4
6
8
10
12
14
0
5
10
15
20
25

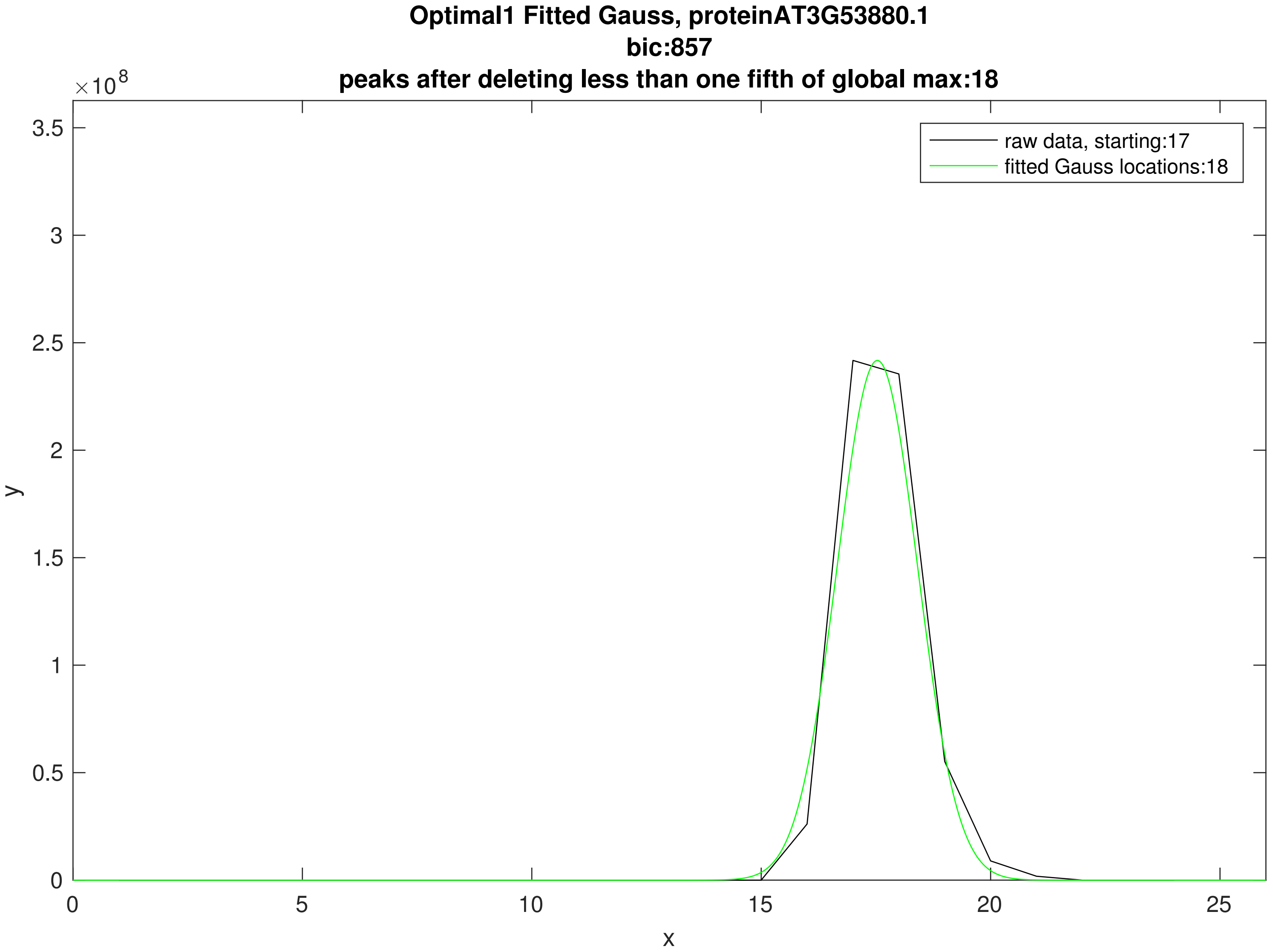
Optimal1 Fitted Gauss, proteinAT3G53880.1
bic:857
peaks after deleting less than one fifth of global max:18
×10^8
3.5
3
2.5
2
1.5
1
0.5
0
y
0
5
10
15
20
25
x
raw data, starting:17
fitted Gauss locations:18

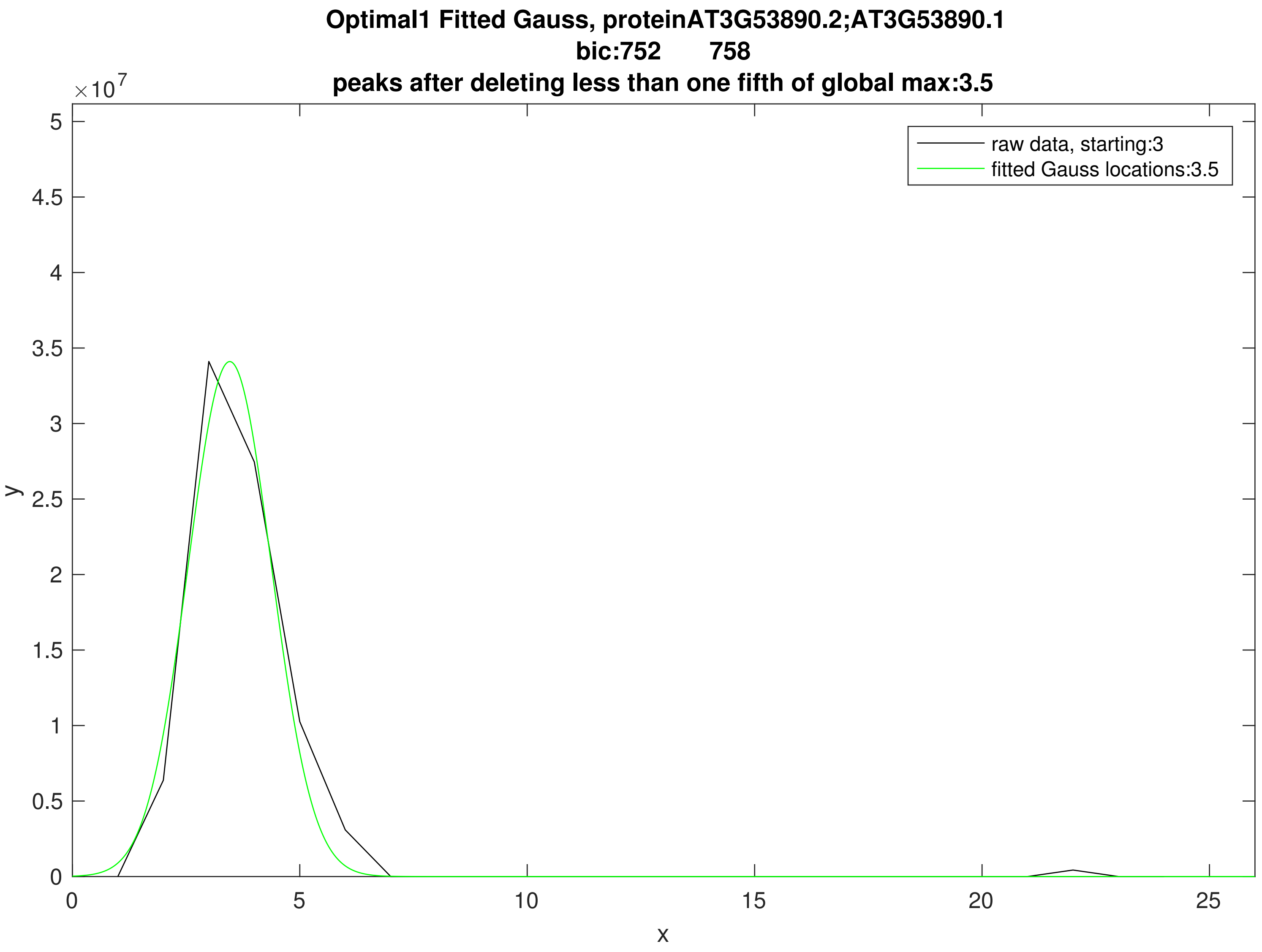
Optimal1 Fitted Gauss, proteinAT3G53890.2;AT3G53890.1
bic:752 758
peaks after deleting less than one fifth of global max:3.5
×10^7
raw data, starting:3
fitted Gauss locations:3.5
y
x
0
0.5
1
1.5
2
2.5
3
3.5
4
4.5
5
0
5
10
15
20
25

**Optimal1 Fitted Gauss, proteinAT3G53900.1;AT3G53900.2**
**bic:894 904 931**
**peaks after deleting less than one fifth of global max:14**

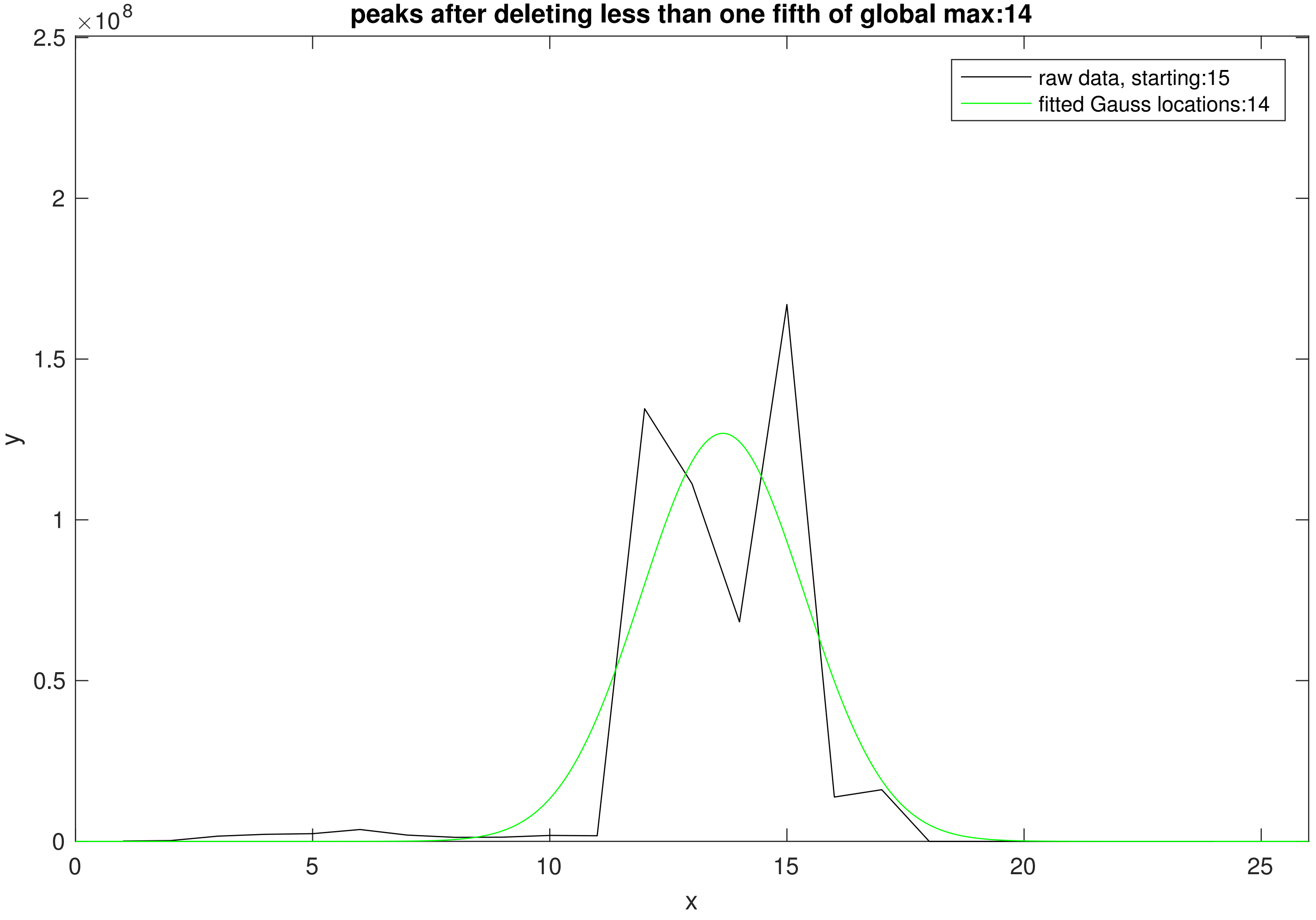

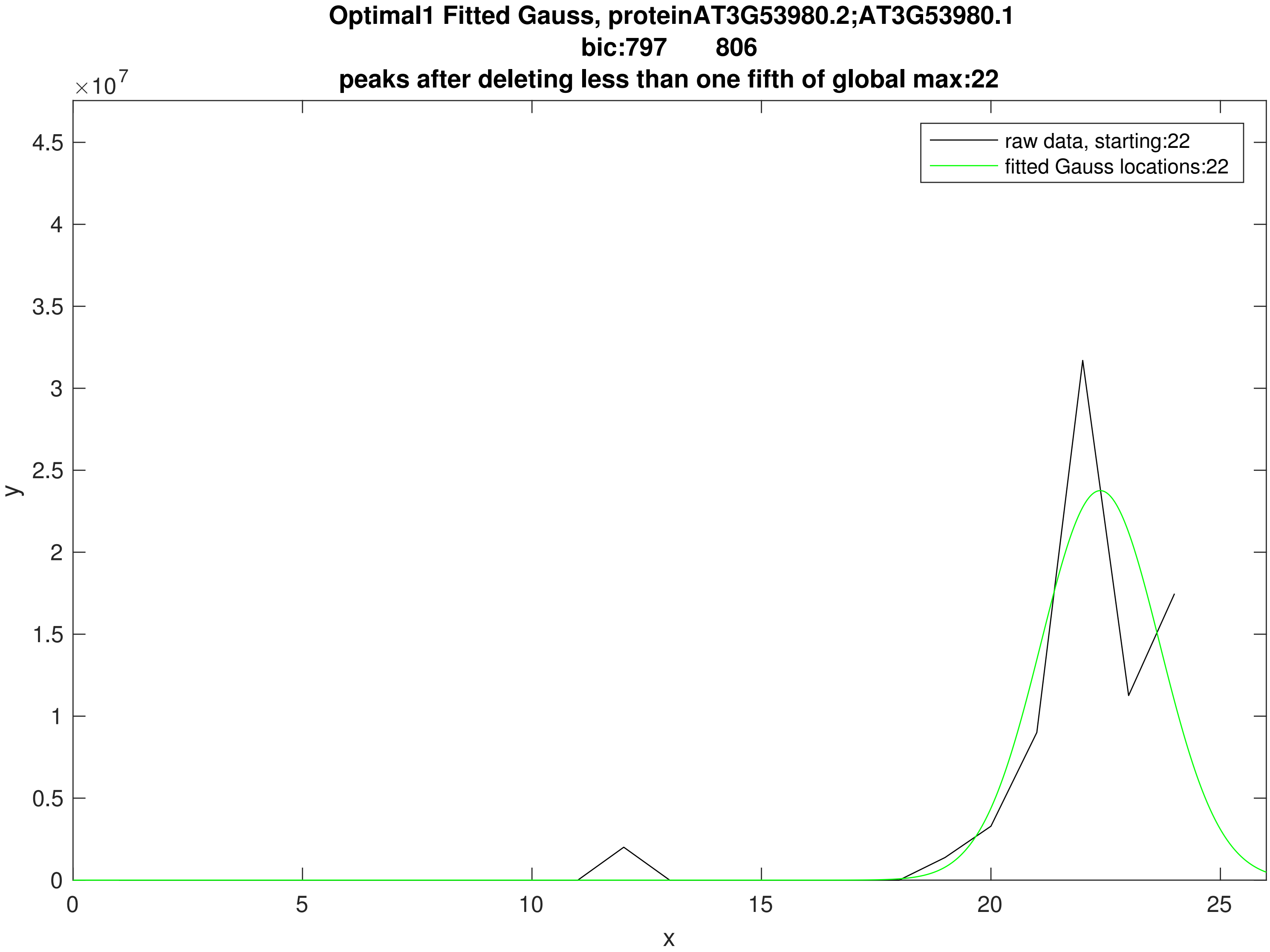
Optimal1 Fitted Gauss, proteinAT3G53980.2;AT3G53980.1
bic:797 806
peaks after deleting less than one fifth of global max:22
raw data, starting:22
fitted Gauss locations:22
y
x
×10^7

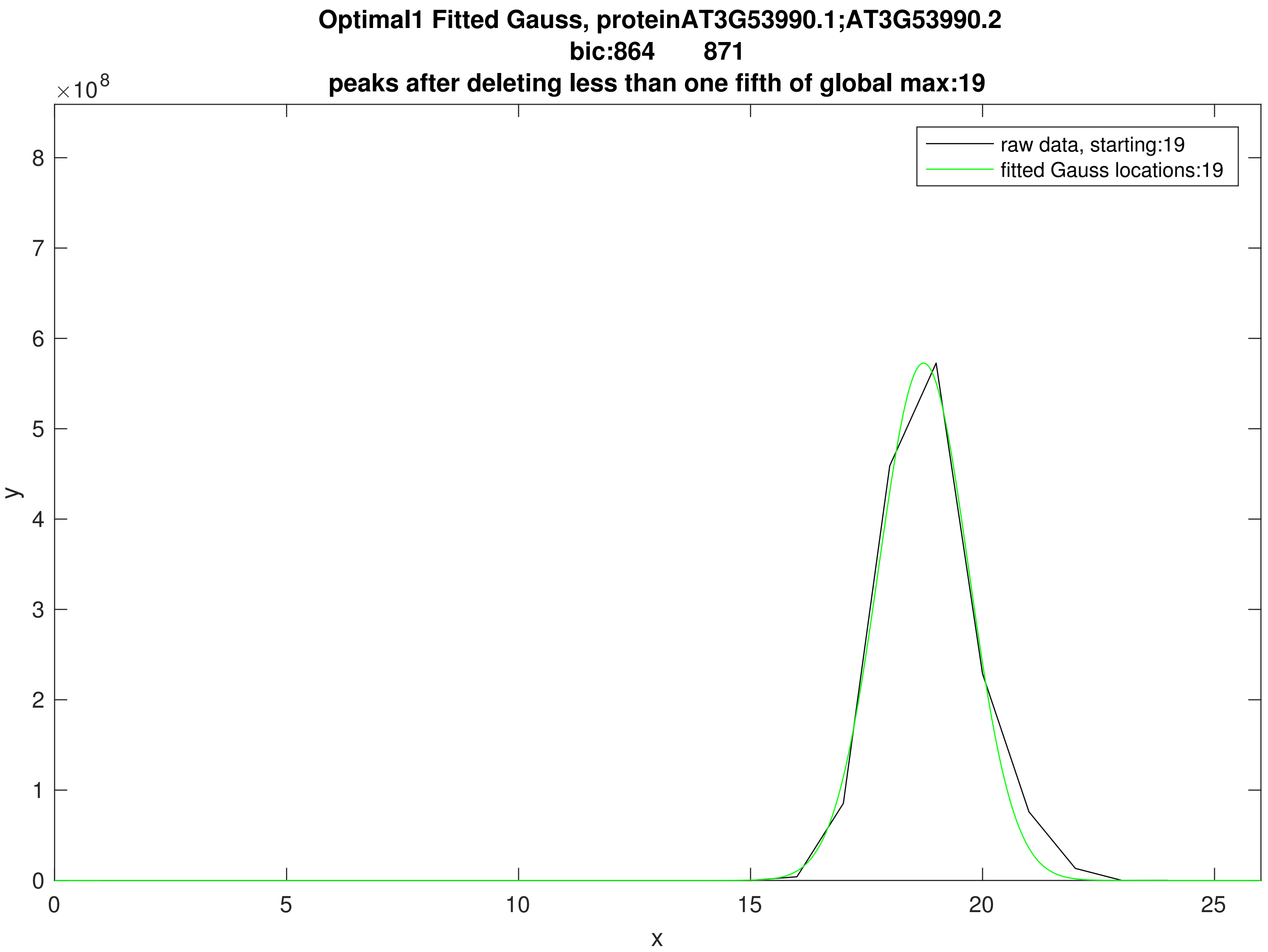

Optimal1 Fitted Gauss, proteinAT3G53990.1;AT3G53990.2
bic:864 871
peaks after deleting less than one fifth of global max:19
×10^8
raw data, starting:19
fitted Gauss locations:19
y
x
0
1
2
3
4
5
6
7
8
0
5
10
15
20
25

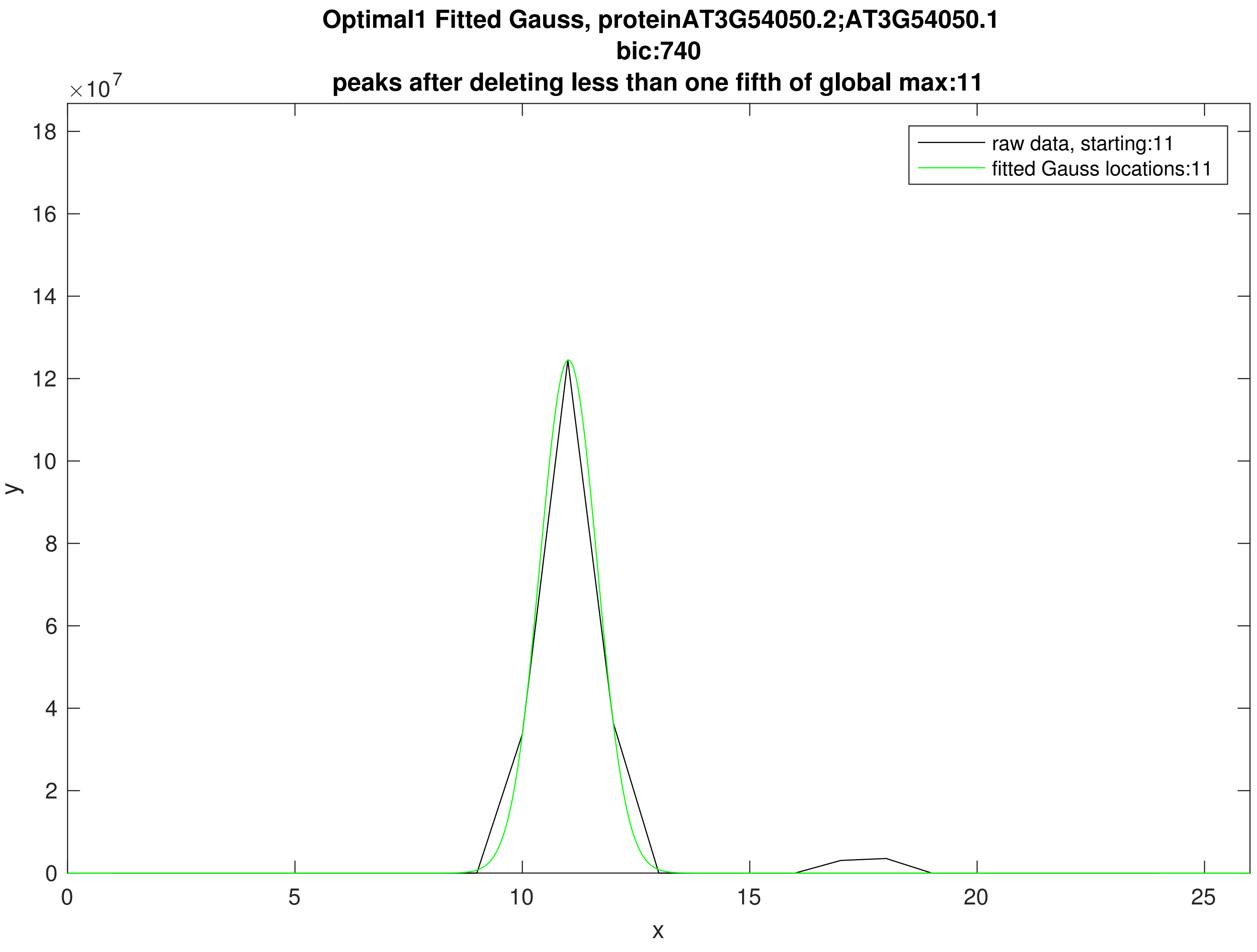
Optimal1 Fitted Gauss, proteinAT3G54050.2;AT3G54050.1
bic:740
peaks after deleting less than one fifth of global max:11
×10^7
raw data, starting:11
fitted Gauss locations:11
y
x
0
2
4
6
8
10
12
14
16
18
0
5
10
15
20
25

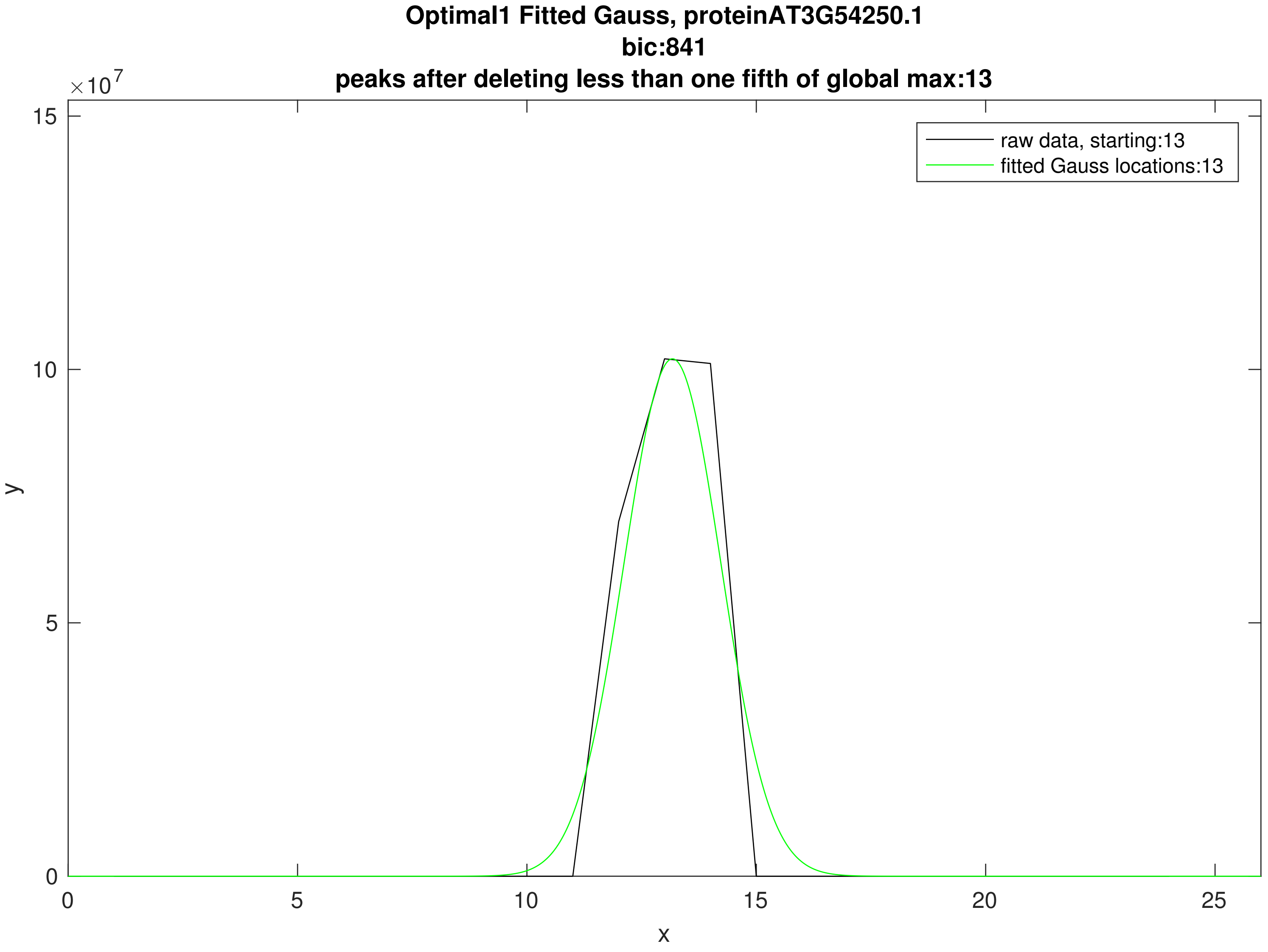

Optimal1 Fitted Gauss, proteinAT3G54250.1
bic:841
peaks after deleting less than one fifth of global max:13
×10^7
15
10
5
0
y
0
5
10
15
20
25
x
raw data, starting:13
fitted Gauss locations:13

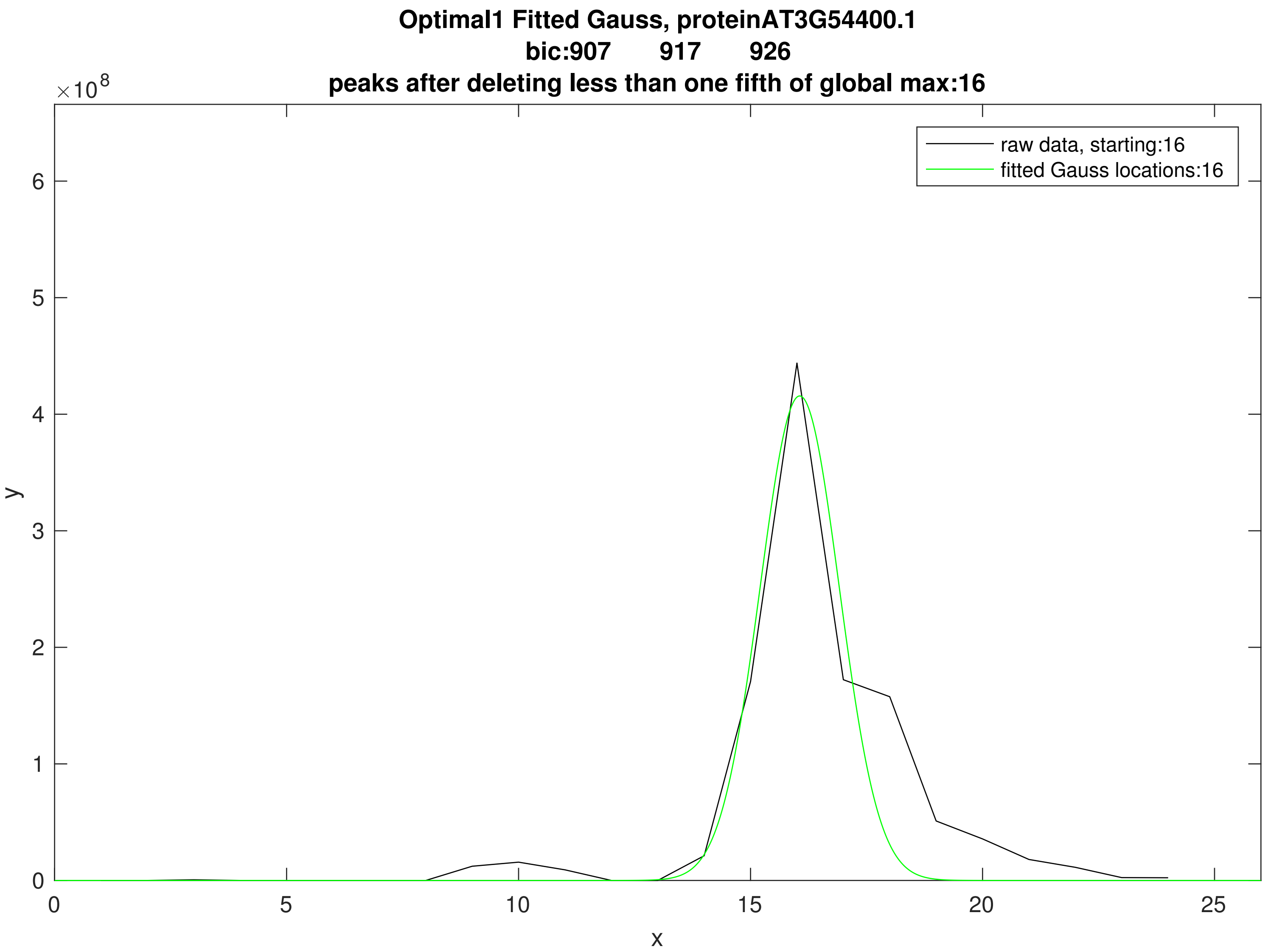
Optimal1 Fitted Gauss, proteinAT3G54400.1
bic:907 917 926
peaks after deleting less than one fifth of global max:16
×10^8
raw data, starting:16
fitted Gauss locations:16
y
x
0
1
2
3
4
5
6
0
5
10
15
20
25

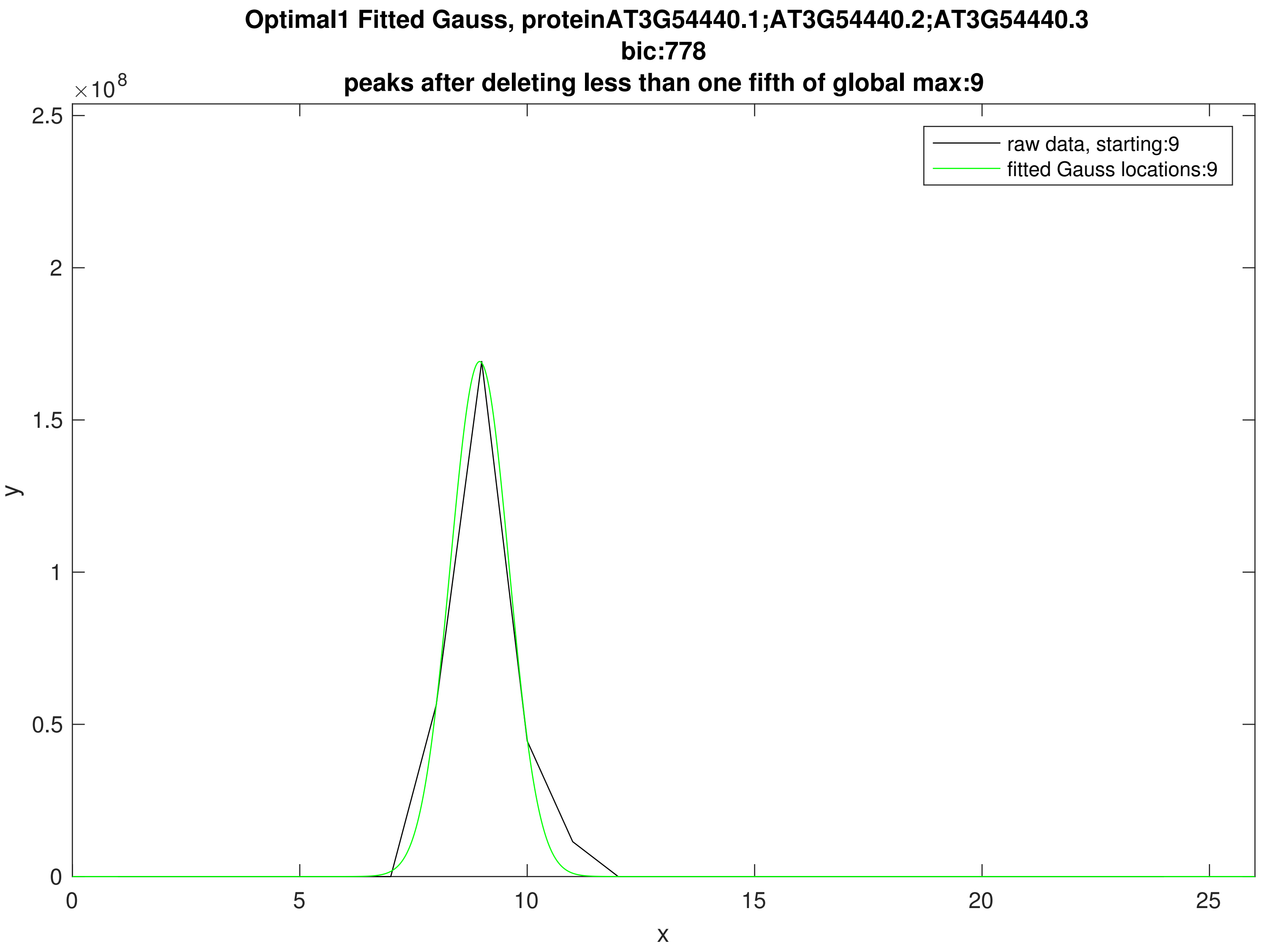

Optimal1 Fitted Gauss, proteinAT3G54440.1;AT3G54440.2;AT3G54440.3
bic:778
peaks after deleting less than one fifth of global max:9
×10^8
2.5
2
1.5
1
0.5
0
y
0
5
10
15
20
25
x
raw data, starting:9
fitted Gauss locations:9

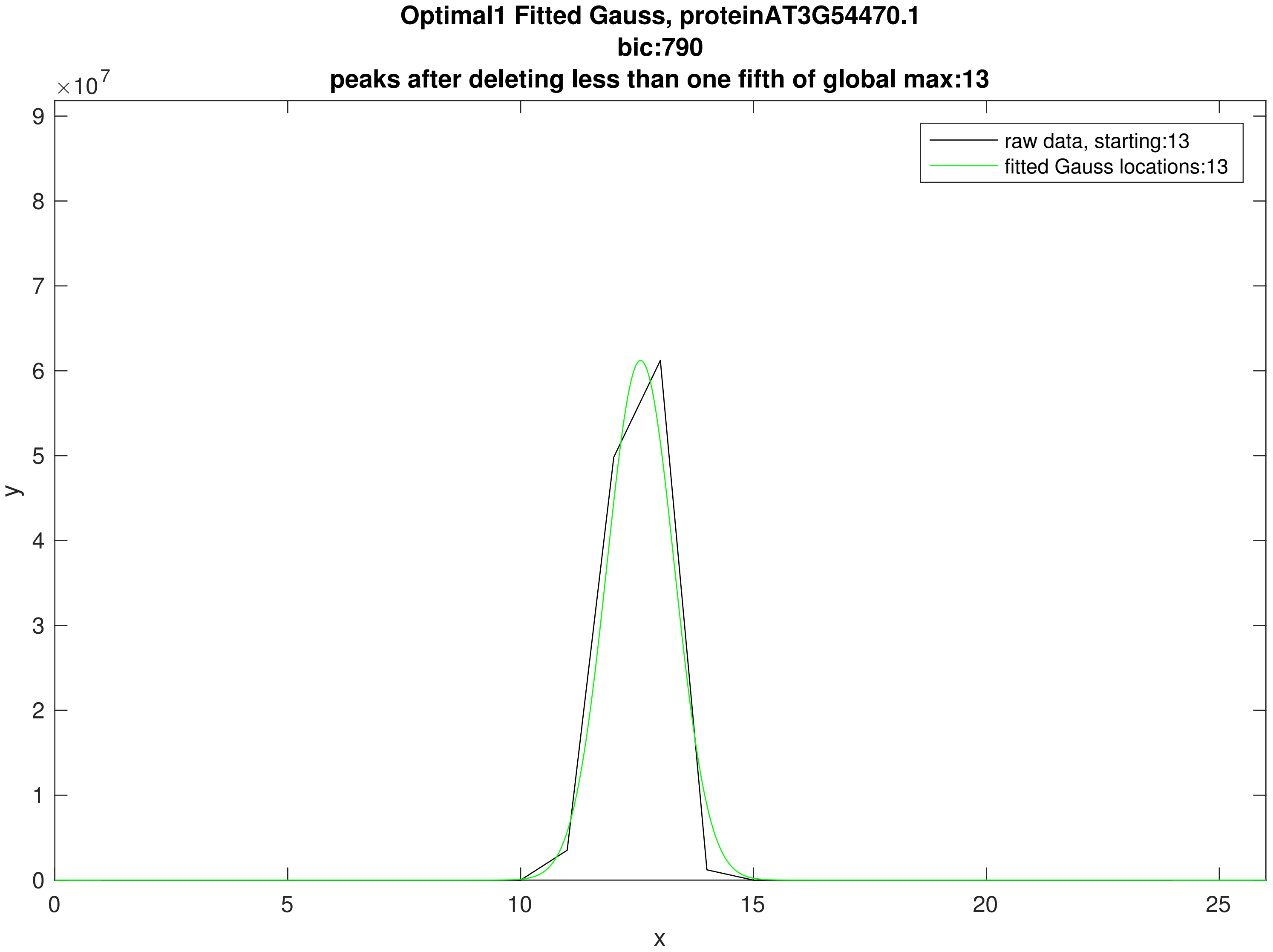
Optimal1 Fitted Gauss, proteinAT3G54470.1
bic:790
peaks after deleting less than one fifth of global max:13
×10^7
raw data, starting:13
fitted Gauss locations:13
y
x
0
1
2
3
4
5
6
7
8
9
0
5
10
15
20
25

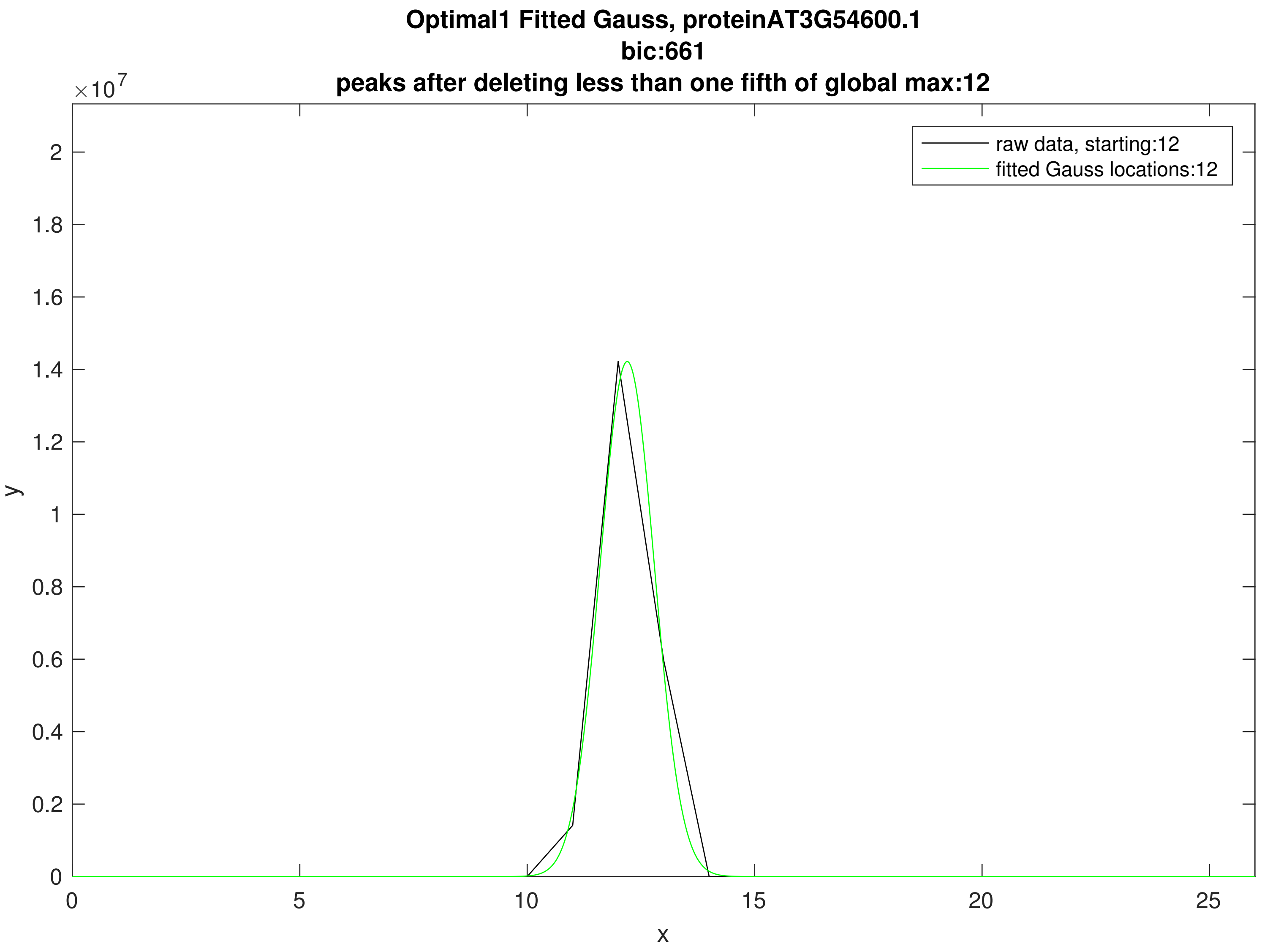

Optimal1 Fitted Gauss, proteinAT3G54600.1
bic:661
peaks after deleting less than one fifth of global max:12
×10^7
raw data, starting:12
fitted Gauss locations:12
y
x
0
0.2
0.4
0.6
0.8
1
1.2
1.4
1.6
1.8
2
0
5
10
15
20
25

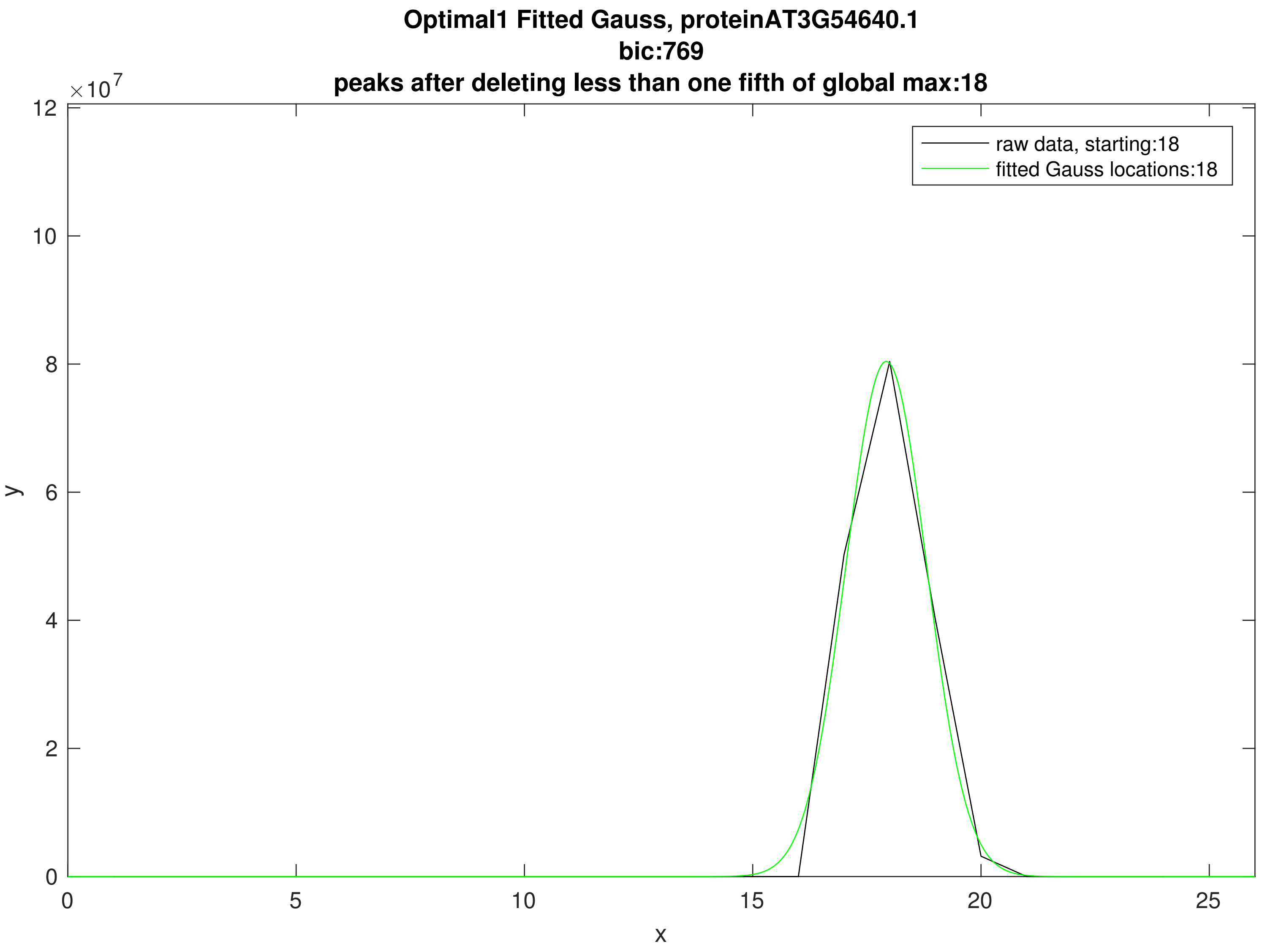
Optimal1 Fitted Gauss, proteinAT3G54640.1
bic:769
peaks after deleting less than one fifth of global max:18
×10^7
raw data, starting:18
fitted Gauss locations:18
y
x
0
2
4
6
8
10
12
0
5
10
15
20
25

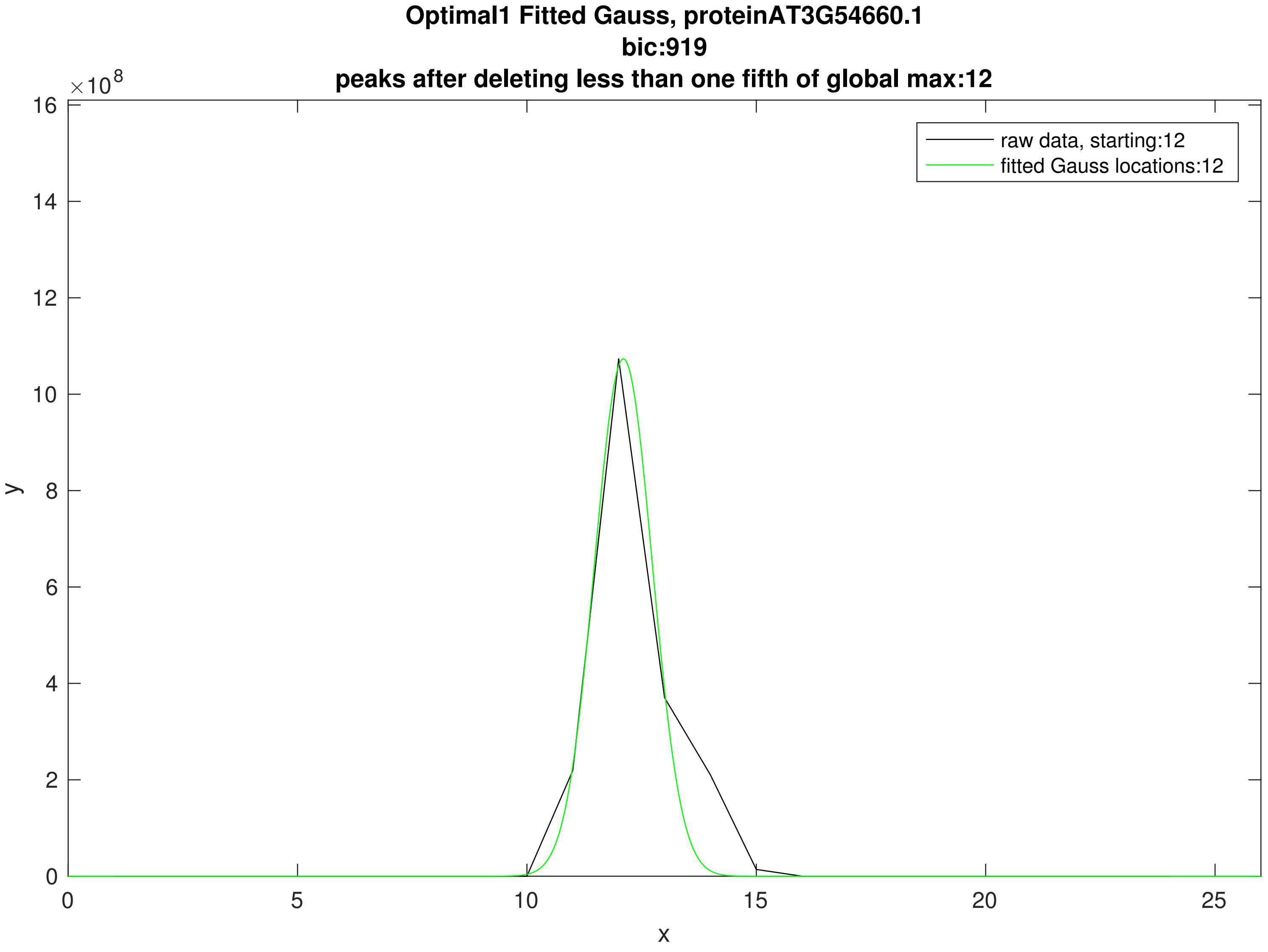
Optimal1 Fitted Gauss, proteinAT3G54660.1
bic:919
peaks after deleting less than one fifth of global max:12
×10^8
16
14
12
10
8
6
4
2
0
y
0
5
10
15
20
25
x
raw data, starting:12
fitted Gauss locations:12

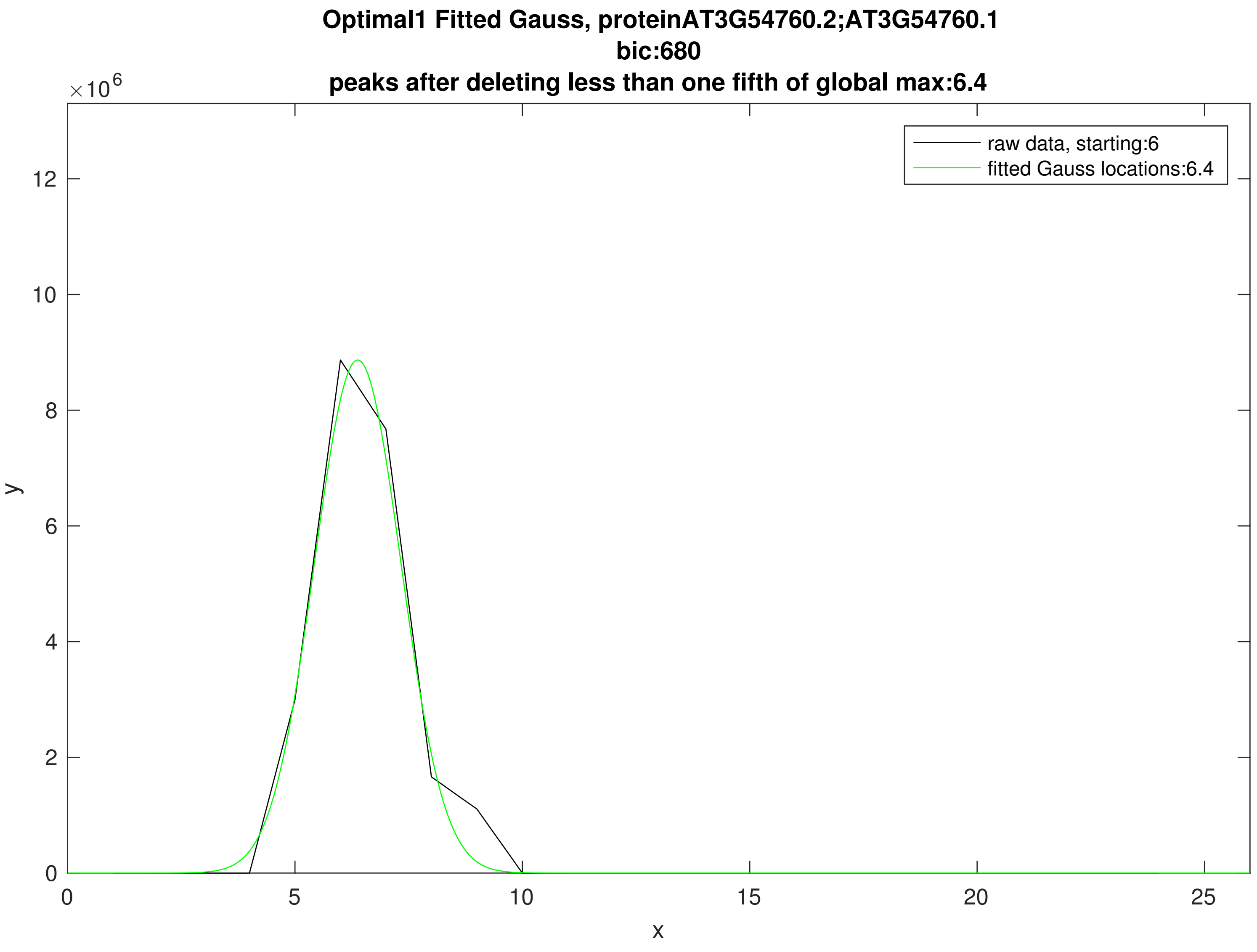
Optimal1 Fitted Gauss, proteinAT3G54760.2;AT3G54760.1
bic:680
peaks after deleting less than one fifth of global max:6.4
×10^6
raw data, starting:6
fitted Gauss locations:6.4
y
x
0
2
4
6
8
10
12
0
5
10
15
20
25

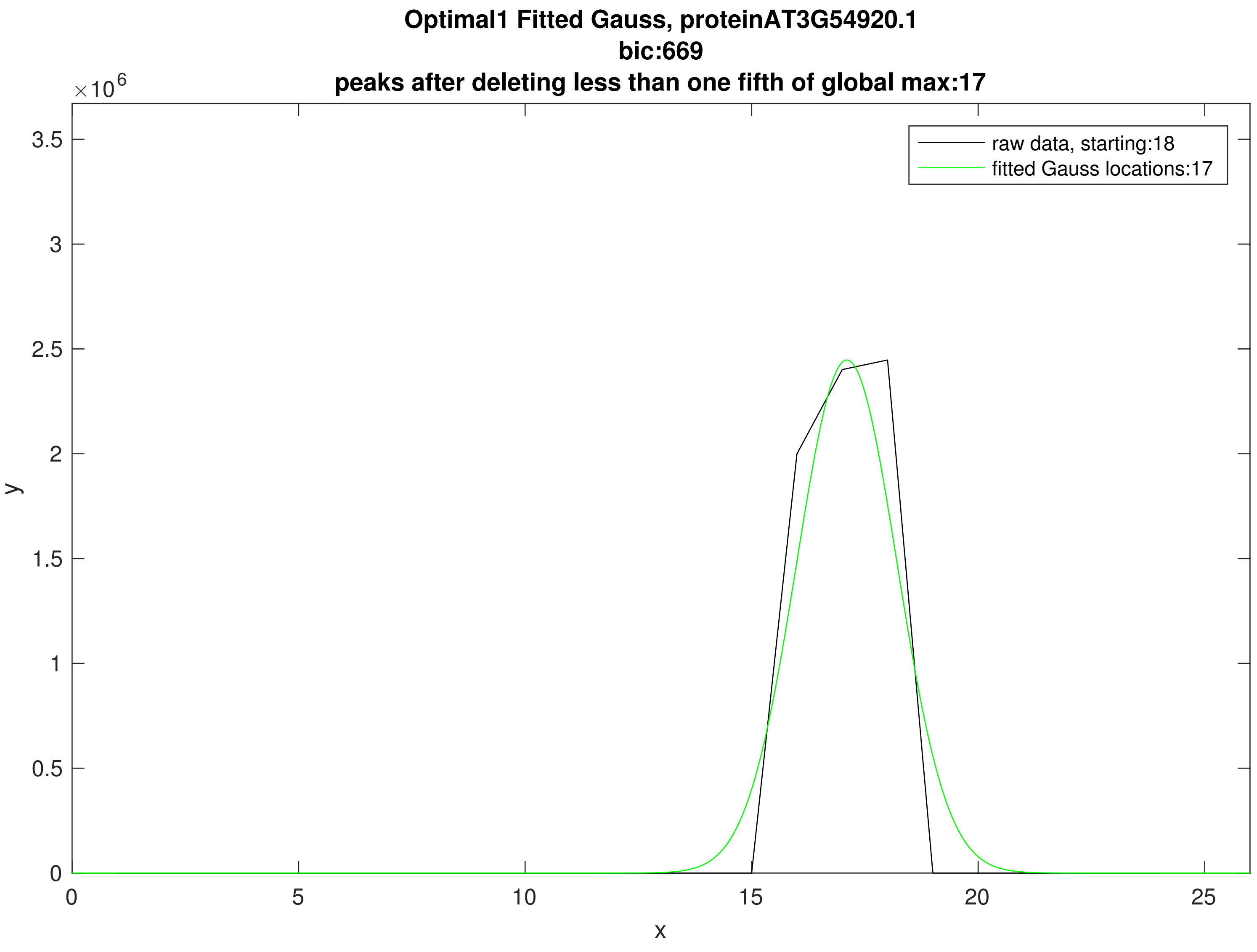
Optimal1 Fitted Gauss, proteinAT3G54920.1
bic:669
peaks after deleting less than one fifth of global max:17
×10^6
raw data, starting:18
fitted Gauss locations:17
y
x
0
0.5
1
1.5
2
2.5
3
3.5
0
5
10
15
20
25

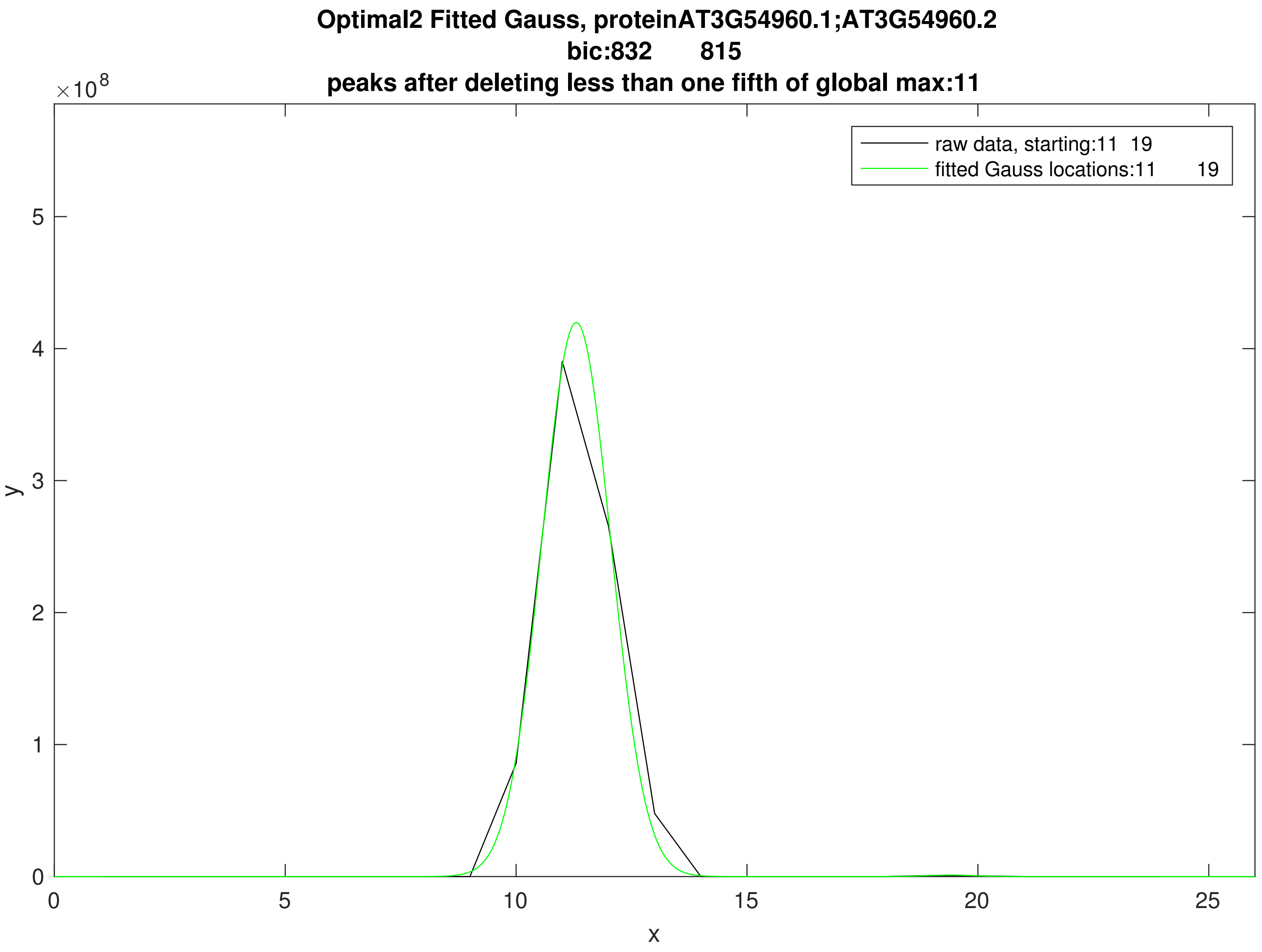
Optimal2 Fitted Gauss, proteinAT3G54960.1;AT3G54960.2
bic:832 815
peaks after deleting less than one fifth of global max:11
×10^8
raw data, starting:11 19
fitted Gauss locations:11 19
y
x
0
1
2
3
4
5
10
15
20
25

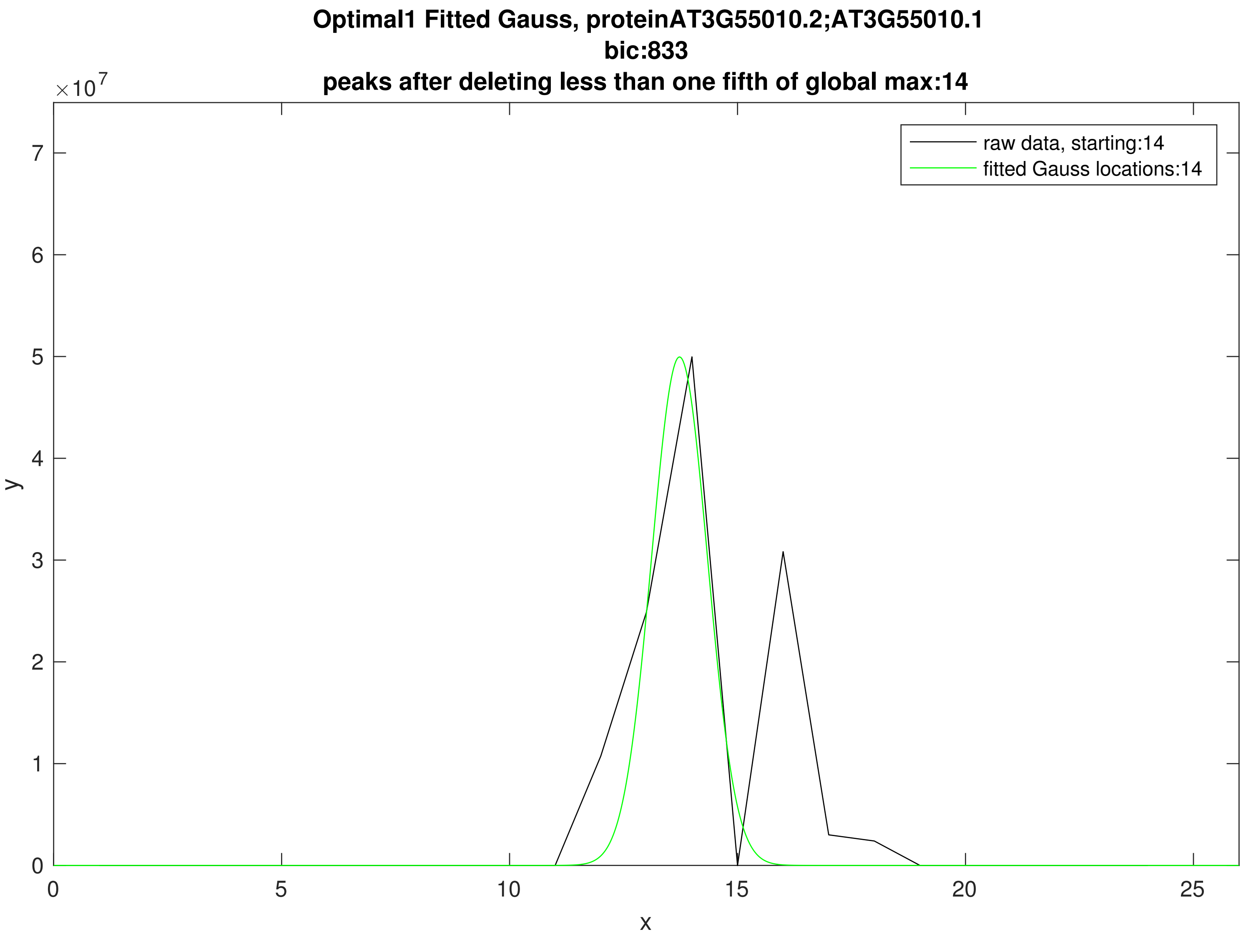
Optimal1 Fitted Gauss, proteinAT3G55010.2;AT3G55010.1
bic:833
peaks after deleting less than one fifth of global max:14
×10^7
raw data, starting:14
fitted Gauss locations:14
y
x
0
1
2
3
4
5
6
7
0
5
10
15
20
25

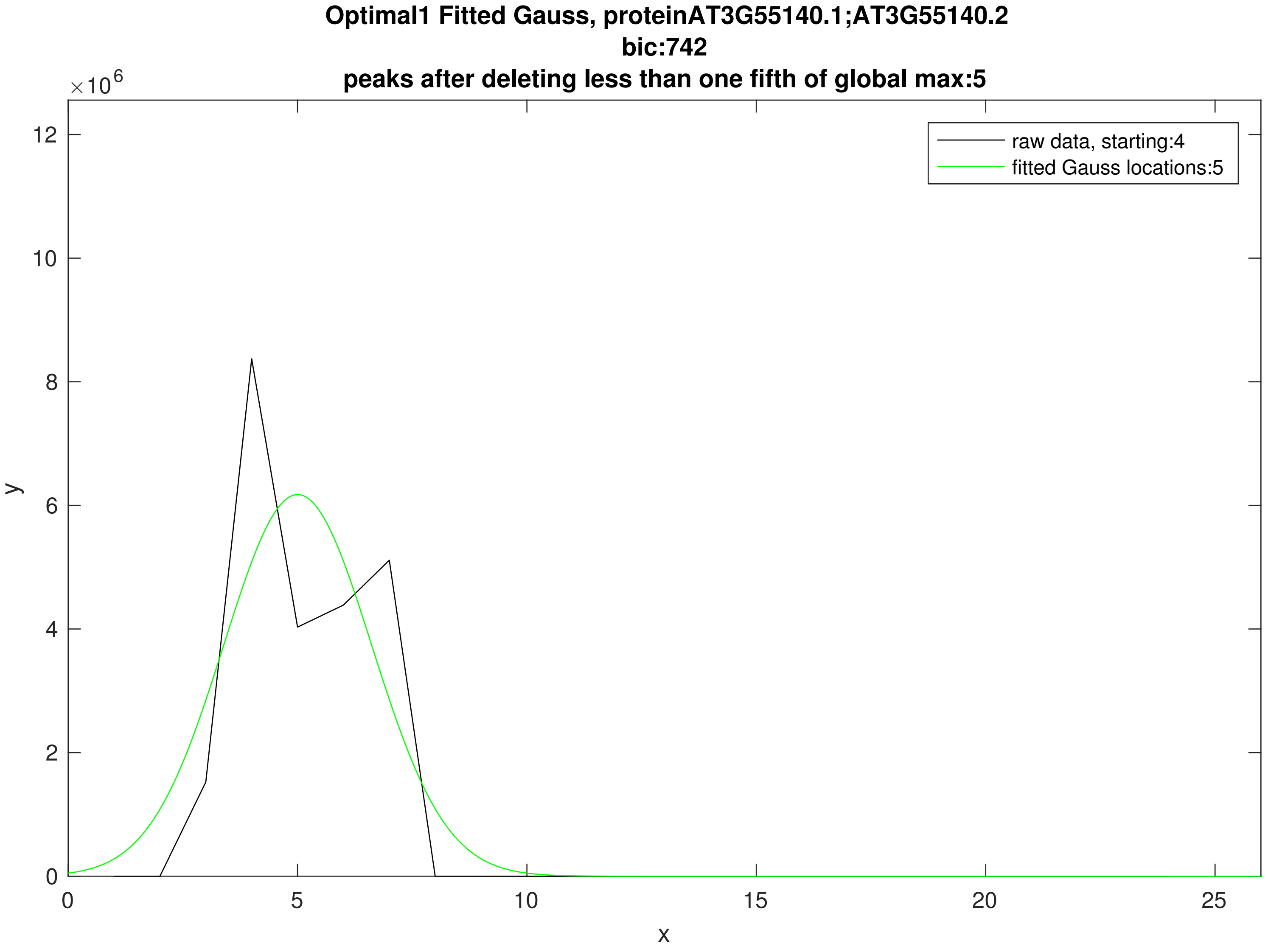
Optimal1 Fitted Gauss, proteinAT3G55140.1;AT3G55140.2
bic:742
peaks after deleting less than one fifth of global max:5
×10^6
raw data, starting:4
fitted Gauss locations:5
y
x
0
2
4
6
8
10
12
0
5
10
15
20
25

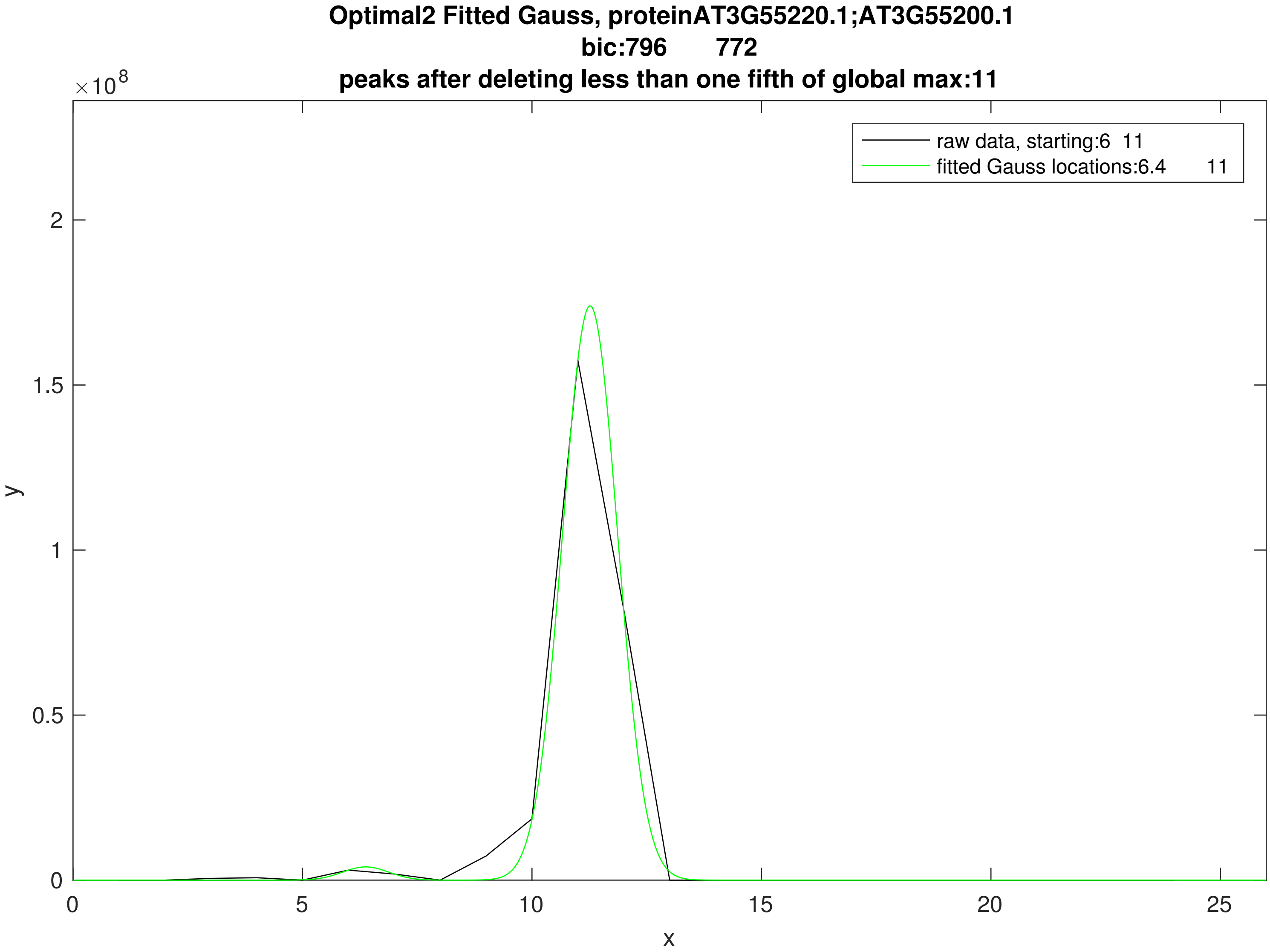
Optimal2 Fitted Gauss, proteinAT3G55220.1;AT3G55200.1
bic:796 772
peaks after deleting less than one fifth of global max:11
×10^8
raw data, starting:6 11
fitted Gauss locations:6.4 11
y
x
0
0.5
1
1.5
2
5
10
15
20
25

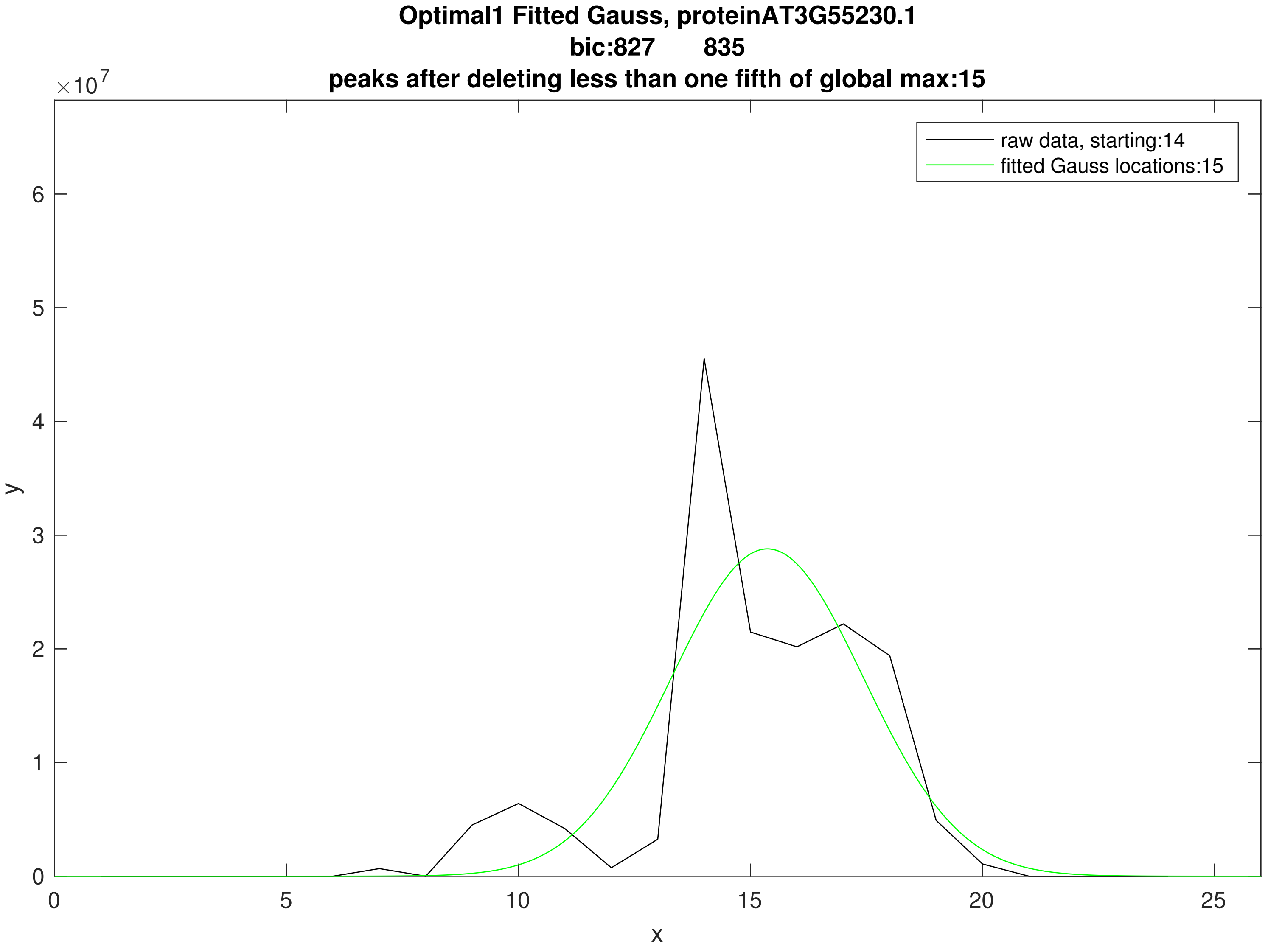

Optimal1 Fitted Gauss, proteinAT3G55230.1
bic:827 835
peaks after deleting less than one fifth of global max:15
×10^7
raw data, starting:14
fitted Gauss locations:15
y
x
0
1
2
3
4
5
6
0
5
10
15
20
25

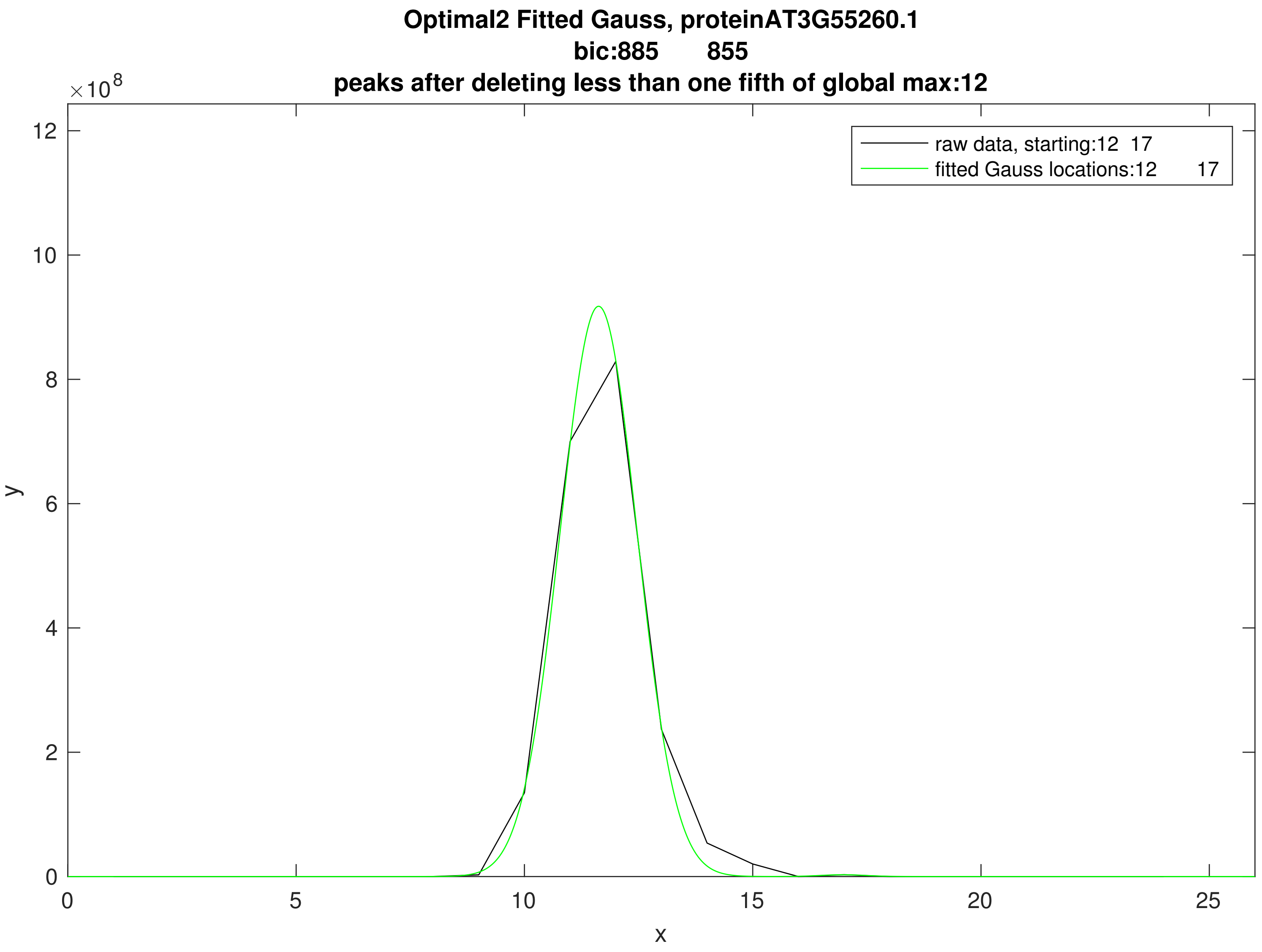
Optimal2 Fitted Gauss, proteinAT3G55260.1
bic:885 855
peaks after deleting less than one fifth of global max:12
×10^8
raw data, starting:12 17
fitted Gauss locations:12 17
y
x
0
2
4
6
8
10
12
5
15
20
25

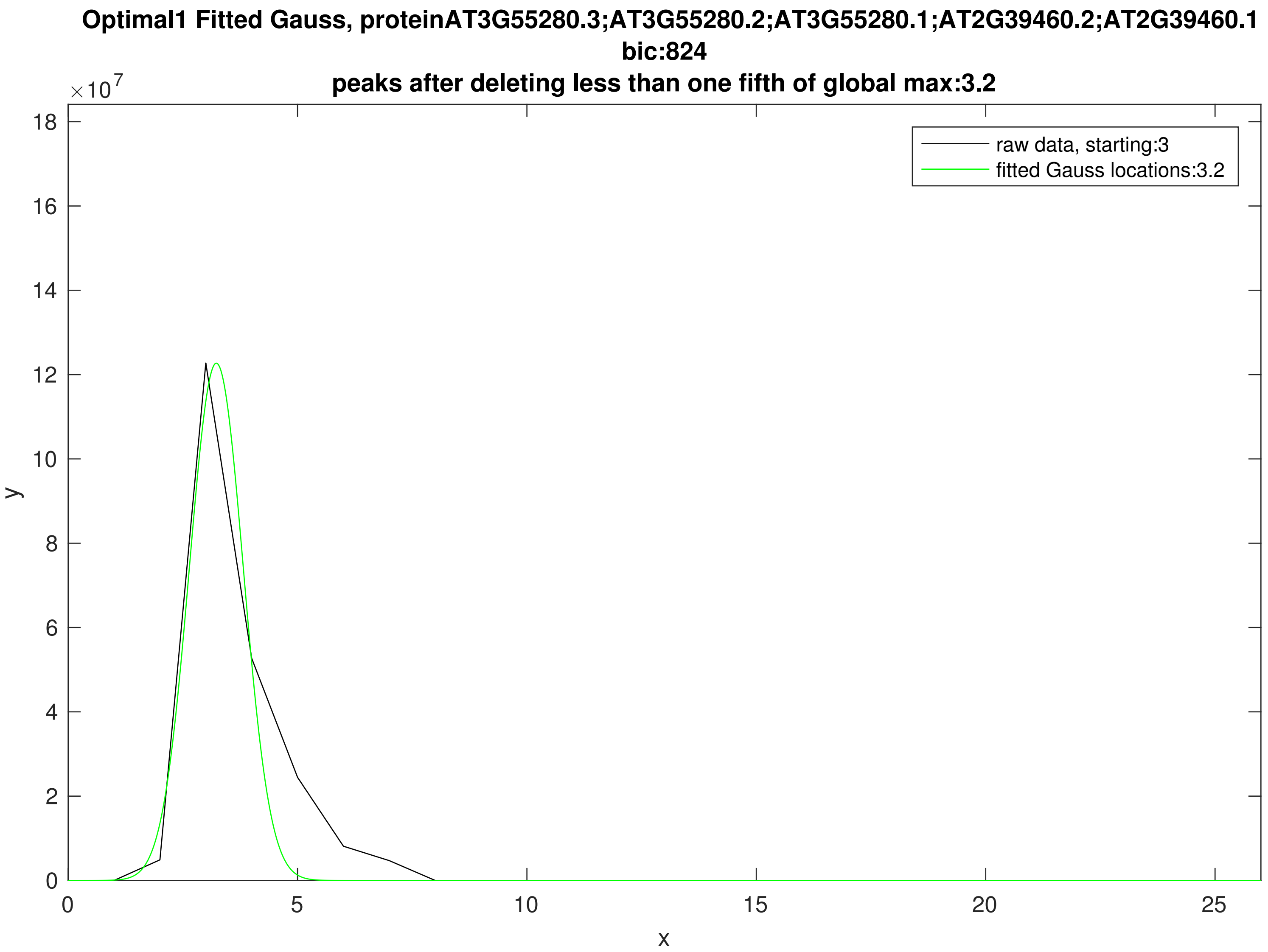
Optimal1 Fitted Gauss, proteinAT3G55280.3;AT3G55280.2;AT3G55280.1;AT2G39460.2;AT2G39460.1
bic:824
peaks after deleting less than one fifth of global max:3.2
×10^7
raw data, starting:3
fitted Gauss locations:3.2
y
x
0
2
4
6
8
10
12
14
16
18
0
5
10
15
20
25

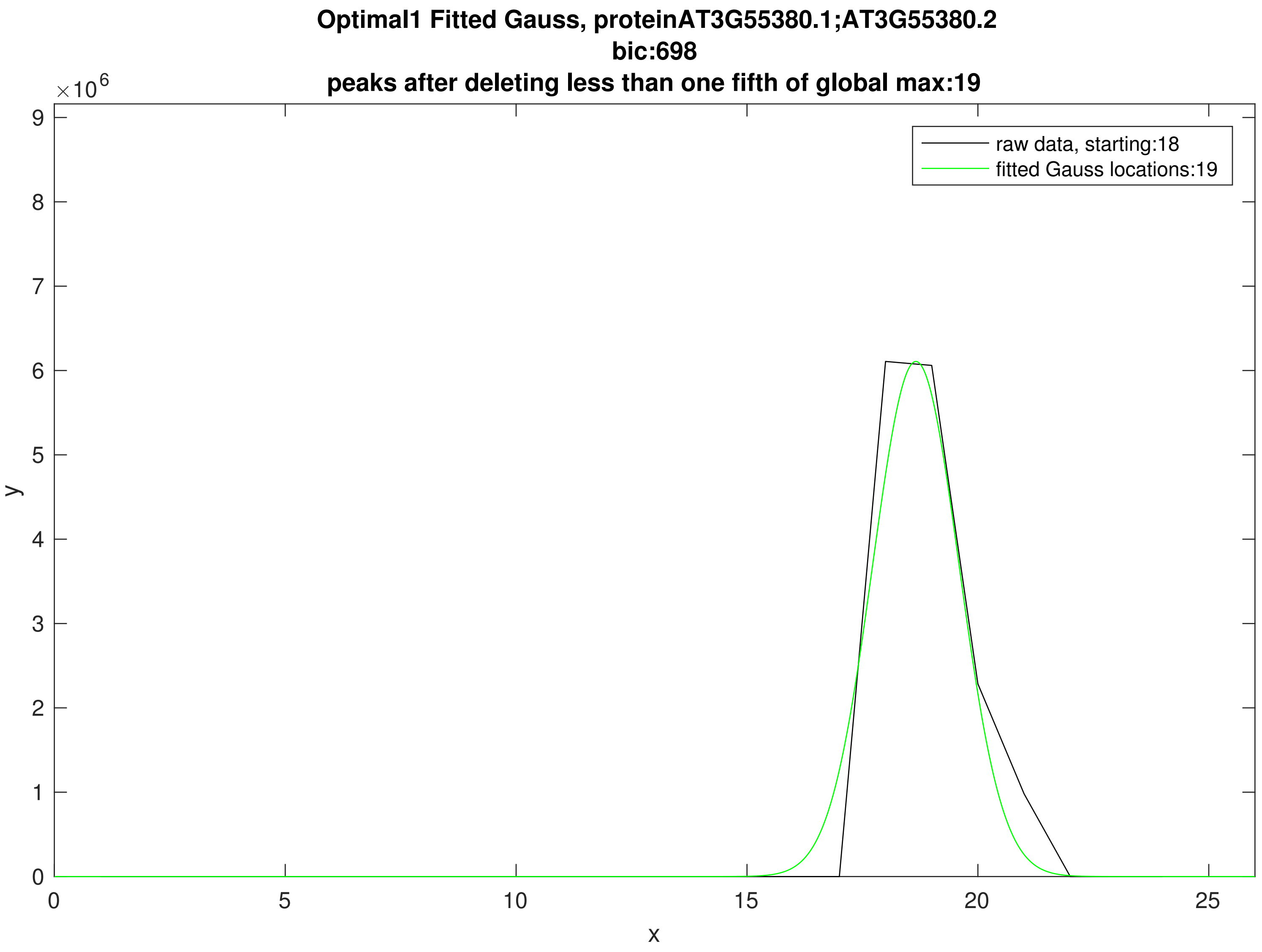

Optimal1 Fitted Gauss, proteinAT3G55380.1;AT3G55380.2
bic:698
peaks after deleting less than one fifth of global max:19
×10^6
y
x
0
1
2
3
4
5
6
7
8
9
0
5
10
15
20
25
raw data, starting:18
fitted Gauss locations:19

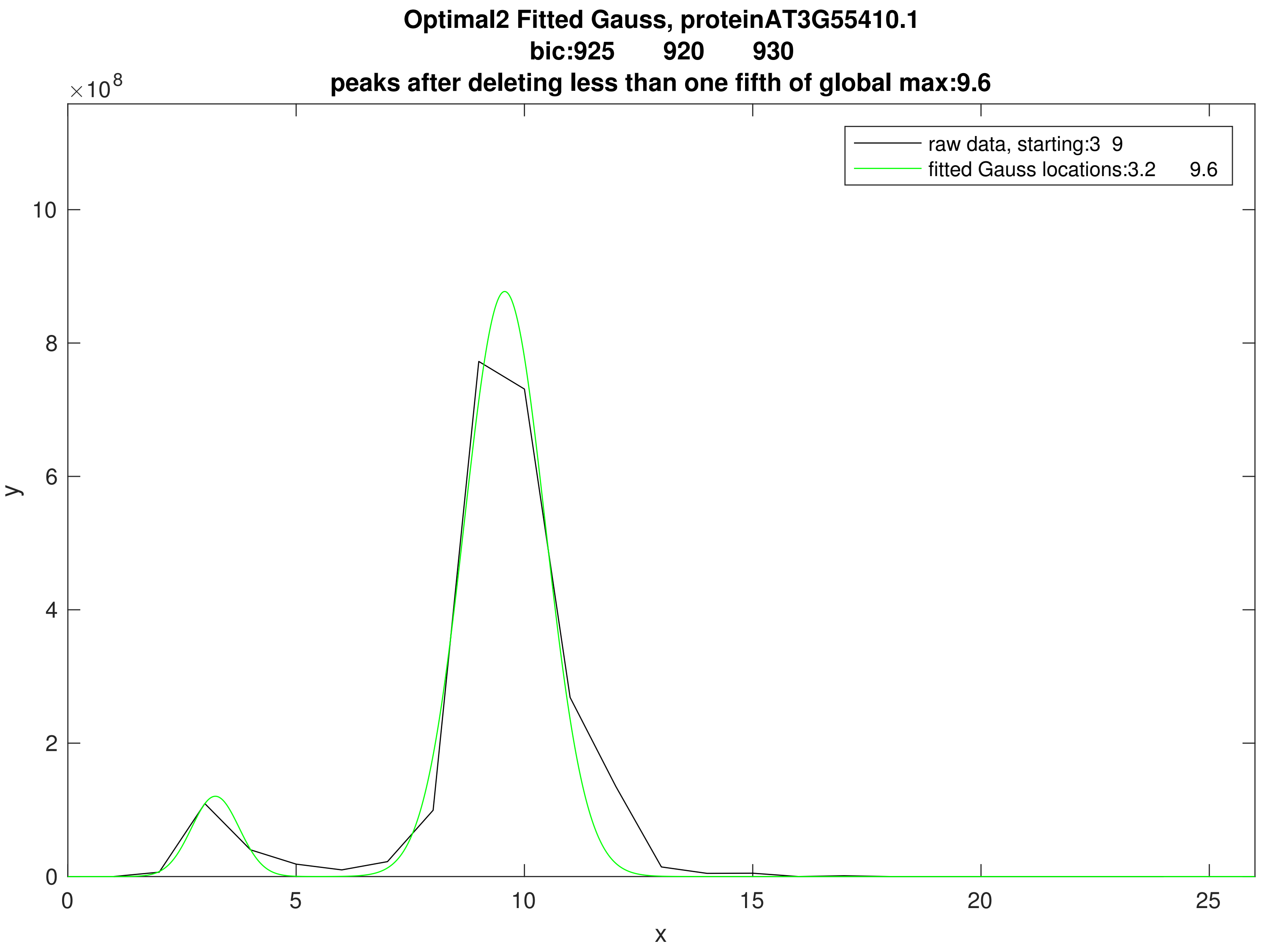
Optimal2 Fitted Gauss, proteinAT3G55410.1
bic:925 920 930
peaks after deleting less than one fifth of global max:9.6
×10^8
raw data, starting:3 9
fitted Gauss locations:3.2 9.6
y
x
0
2
4
6
8
10
0
5
10
15
20
25

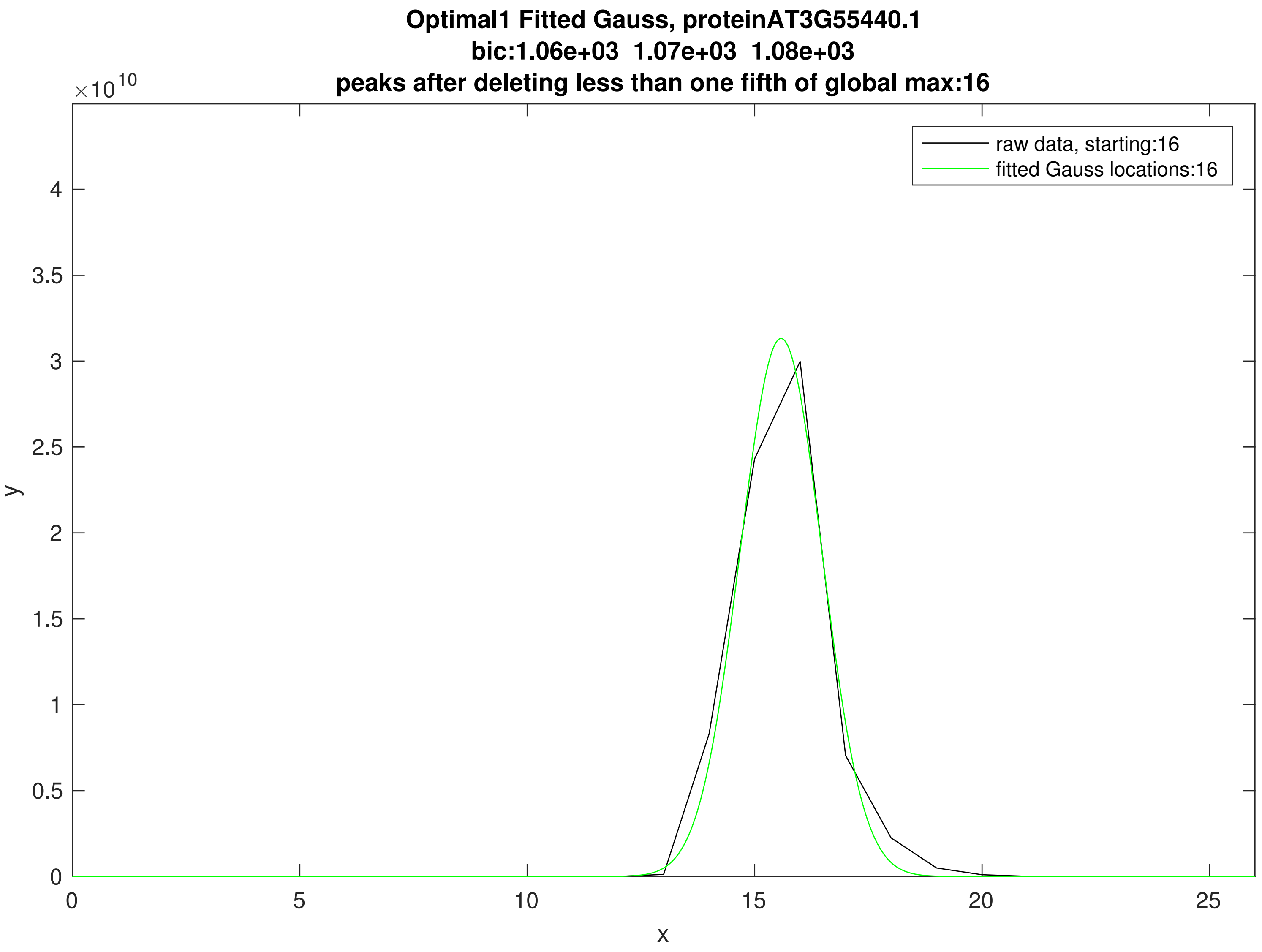
Optimal1 Fitted Gauss, proteinAT3G55440.1
bic:1.06e+03  1.07e+03  1.08e+03
peaks after deleting less than one fifth of global max:16
×10^10
raw data, starting:16
fitted Gauss locations:16
4
3.5
3
2.5
2
1.5
1
0.5
0
y
0
5
10
15
20
25
x

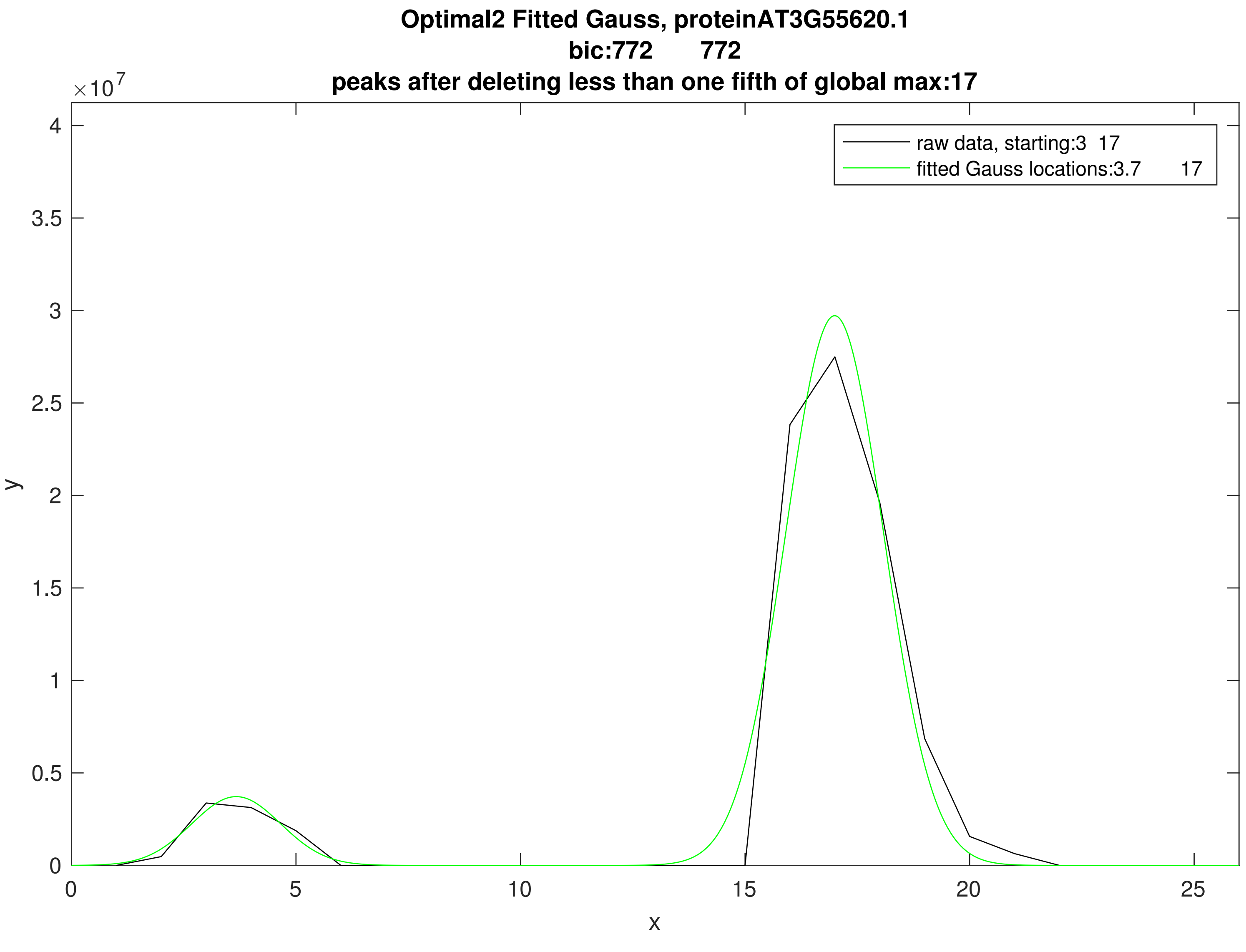
Optimal2 Fitted Gauss, proteinAT3G55620.1
bic:772 772
peaks after deleting less than one fifth of global max:17
×10^7
raw data, starting:3 17
fitted Gauss locations:3.7 17
y
x
0
0.5
1
1.5
2
2.5
3
3.5
4
5
10
15
20
25

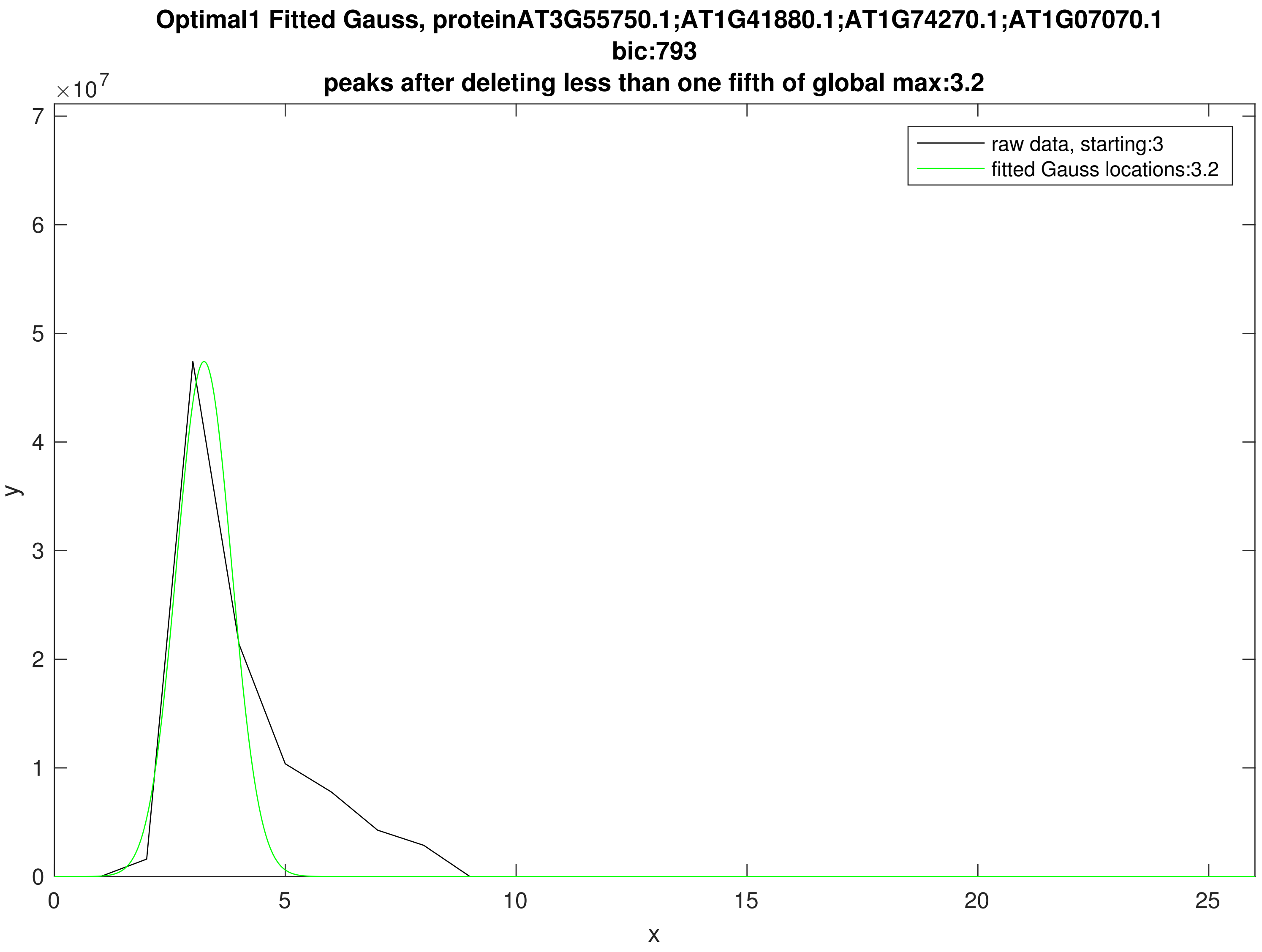

Optimal1 Fitted Gauss, proteinAT3G55750.1;AT1G41880.1;AT1G74270.1;AT1G07070.1
bic:793
peaks after deleting less than one fifth of global max:3.2
×10^7
7
6
5
4
3
2
1
0
y
0
5
10
15
20
25
x
raw data, starting:3
fitted Gauss locations:3.2

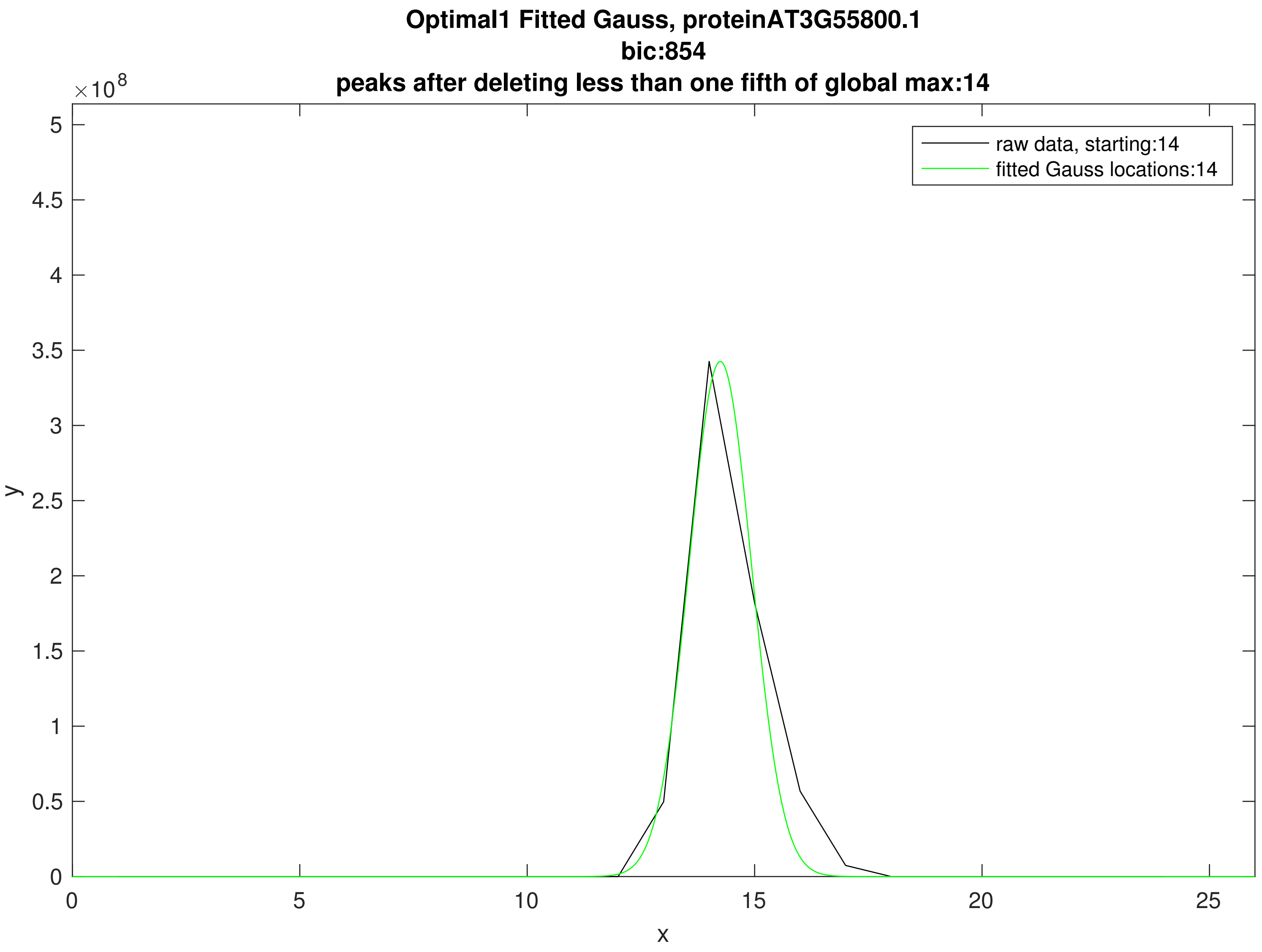
Optimal1 Fitted Gauss, proteinAT3G55800.1
bic:854
peaks after deleting less than one fifth of global max:14
×10^8
5
4.5
4
3.5
3
2.5
2
1.5
1
0.5
0
y
0
5
10
15
20
25
x
raw data, starting:14
fitted Gauss locations:14

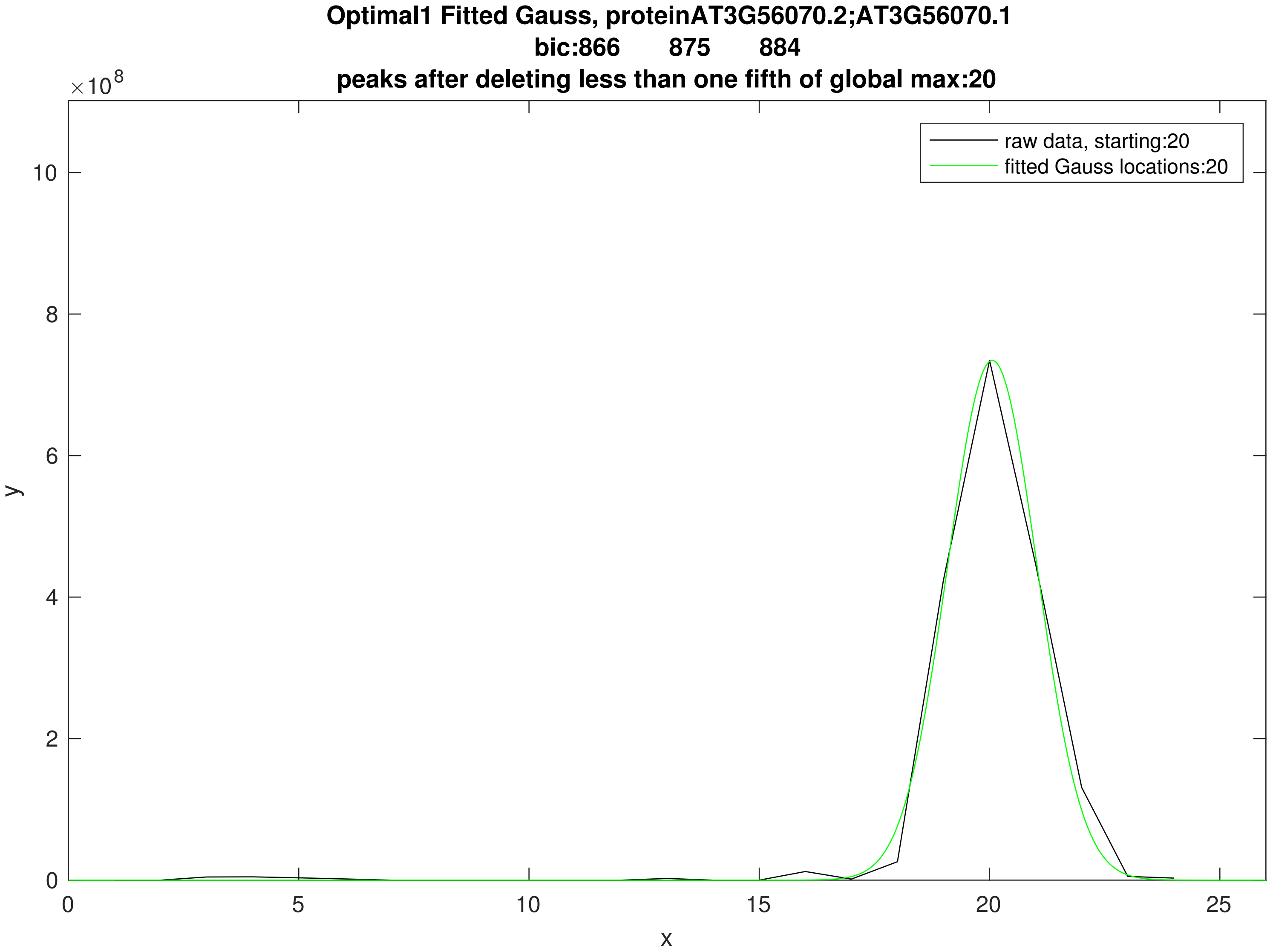
Optimal1 Fitted Gauss, proteinAT3G56070.2;AT3G56070.1
bic:866 875 884
peaks after deleting less than one fifth of global max:20
×10^8
raw data, starting:20
fitted Gauss locations:20
y
x
0
2
4
6
8
10
0
5
10
15
20
25

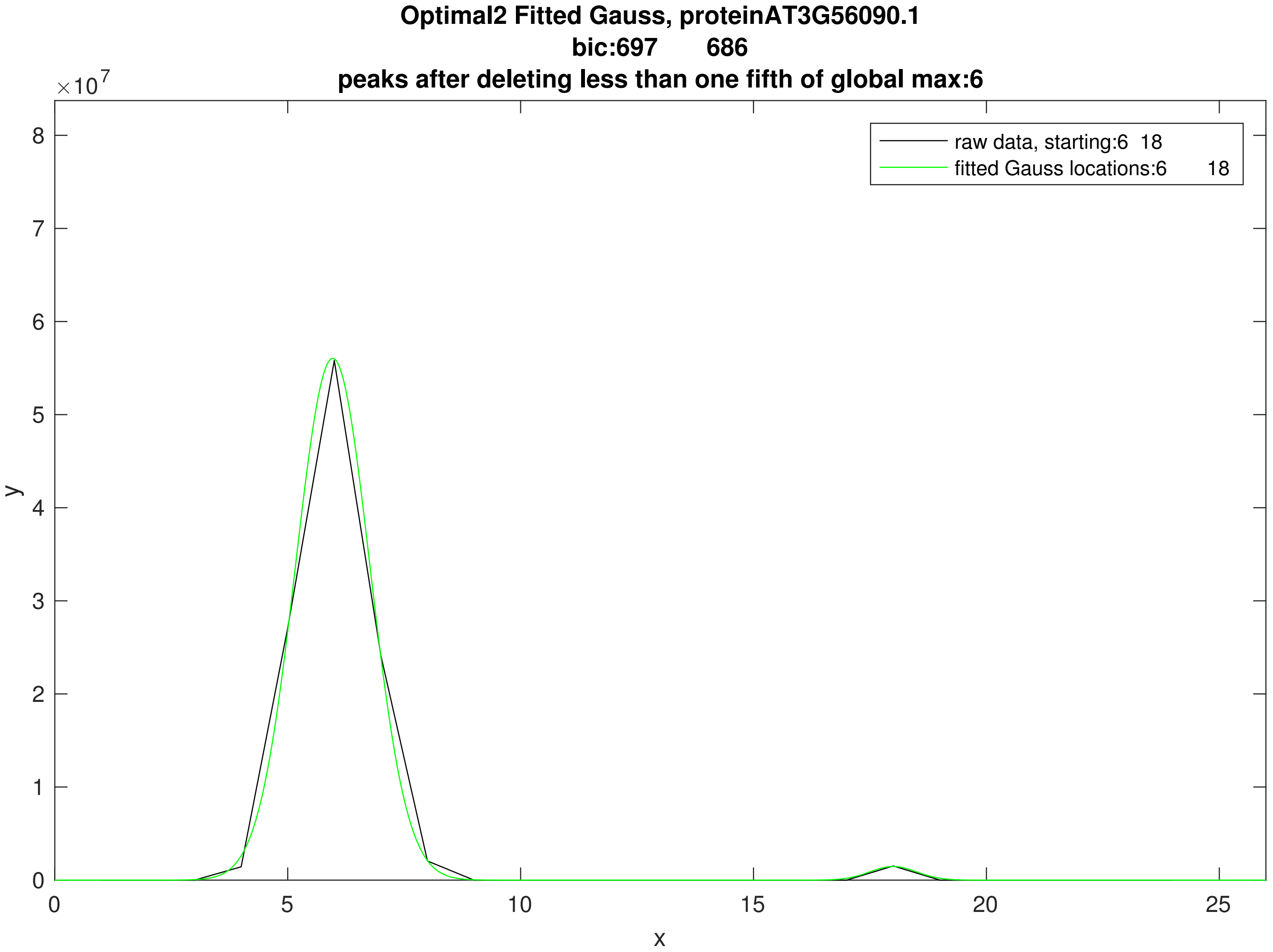
Optimal2 Fitted Gauss, proteinAT3G56090.1
bic:697 686
peaks after deleting less than one fifth of global max:6
×10^7
raw data, starting:6 18
fitted Gauss locations:6 18
0
1
2
3
4
5
6
7
8
y
0
5
10
15
20
25
x

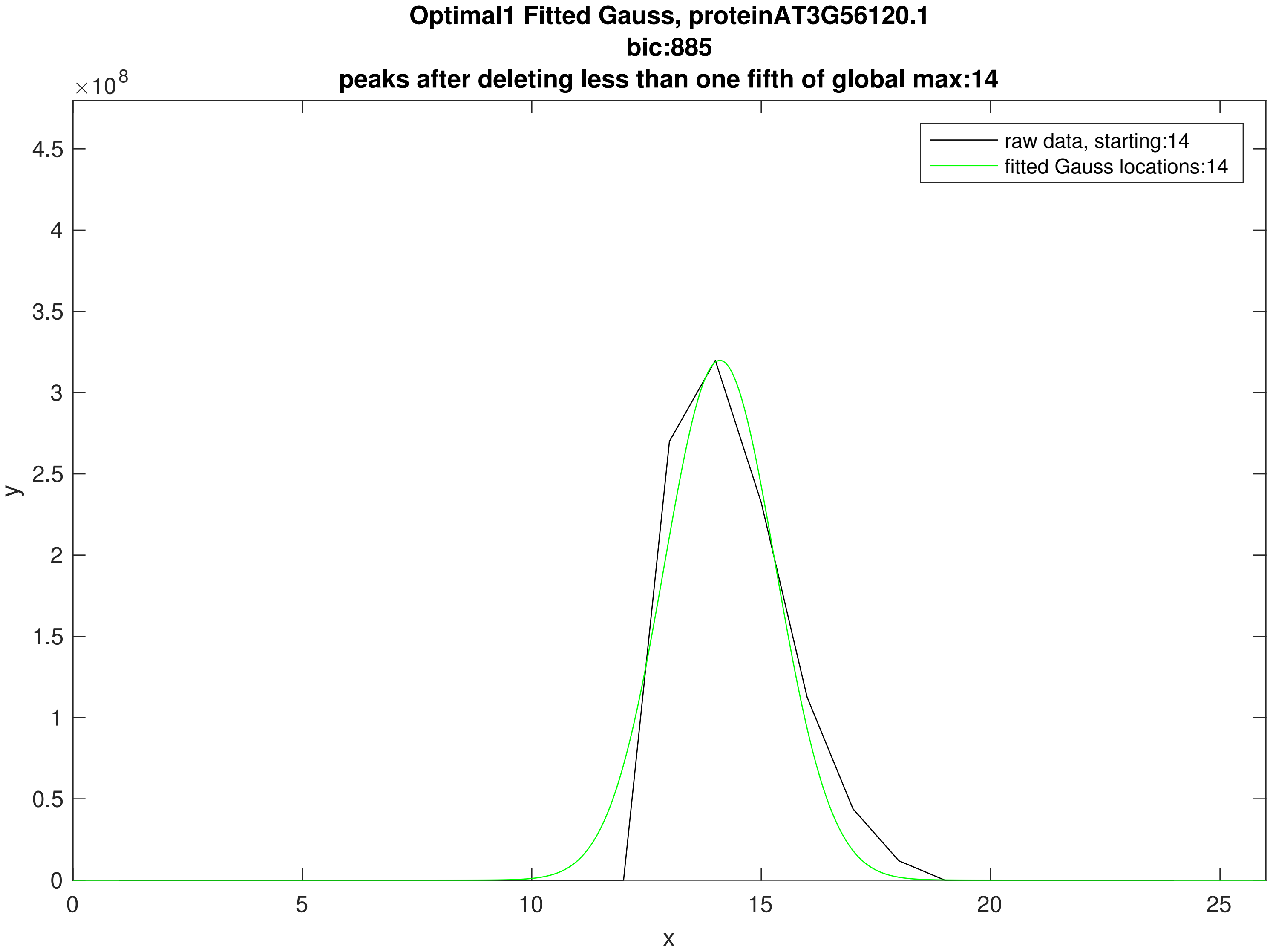

Optimal1 Fitted Gauss, proteinAT3G56120.1
bic:885
peaks after deleting less than one fifth of global max:14
×10^8
4.5
4
3.5
3
2.5
2
1.5
1
0.5
0
y
0
5
10
15
20
25
x
raw data, starting:14
fitted Gauss locations:14

**Optimal2 Fitted Gauss, proteinAT3G56130.4;AT3G56130.1;AT3G56130.3;AT3G56130.2**
**bic:748 734**
**peaks after deleting less than one fifth of global max:14 18**

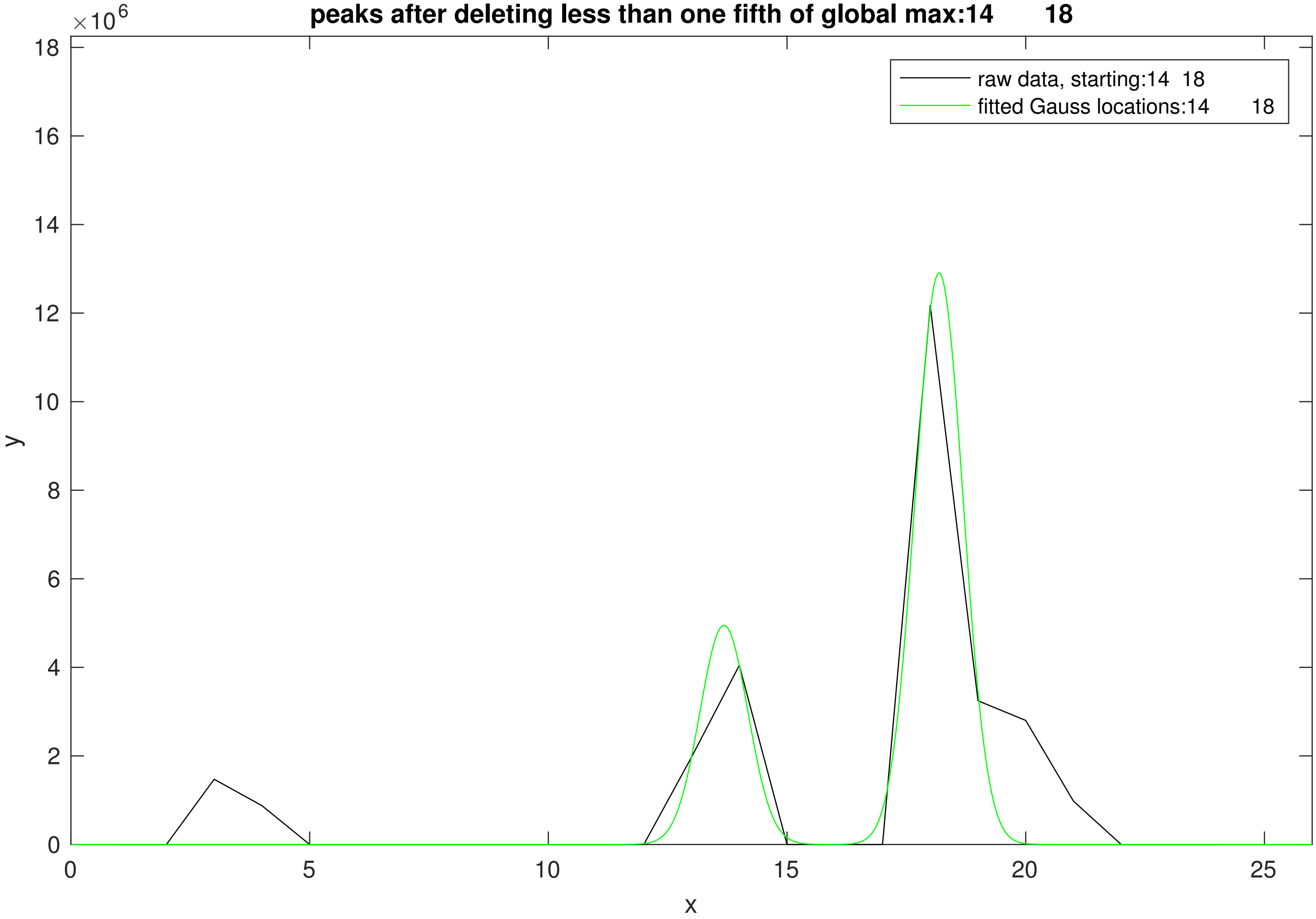

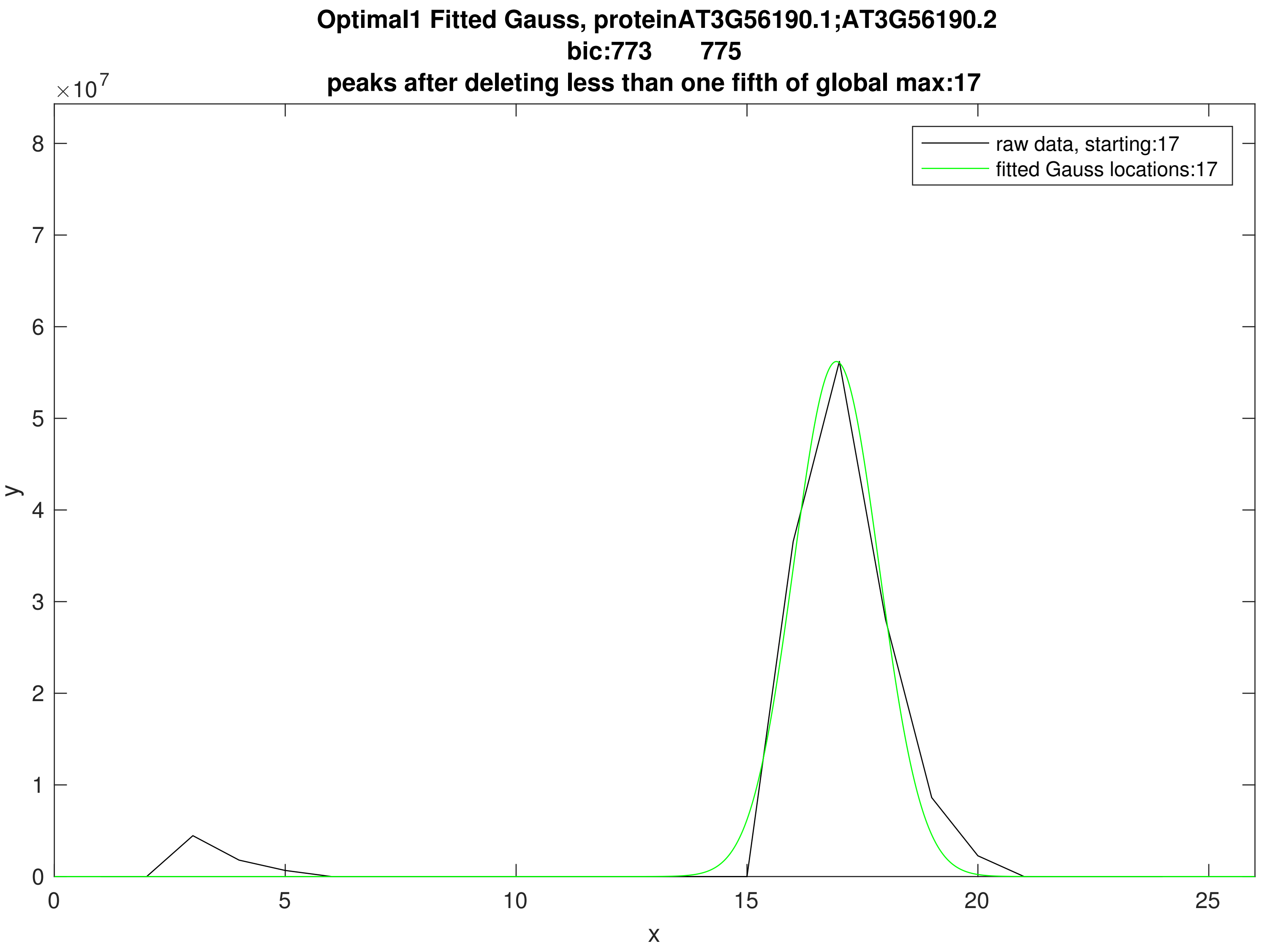
Optimal1 Fitted Gauss, proteinAT3G56190.1;AT3G56190.2
bic:773 775
peaks after deleting less than one fifth of global max:17
×10^7
raw data, starting:17
fitted Gauss locations:17
y
x
0
1
2
3
4
5
6
7
8
0
5
10
15
20
25

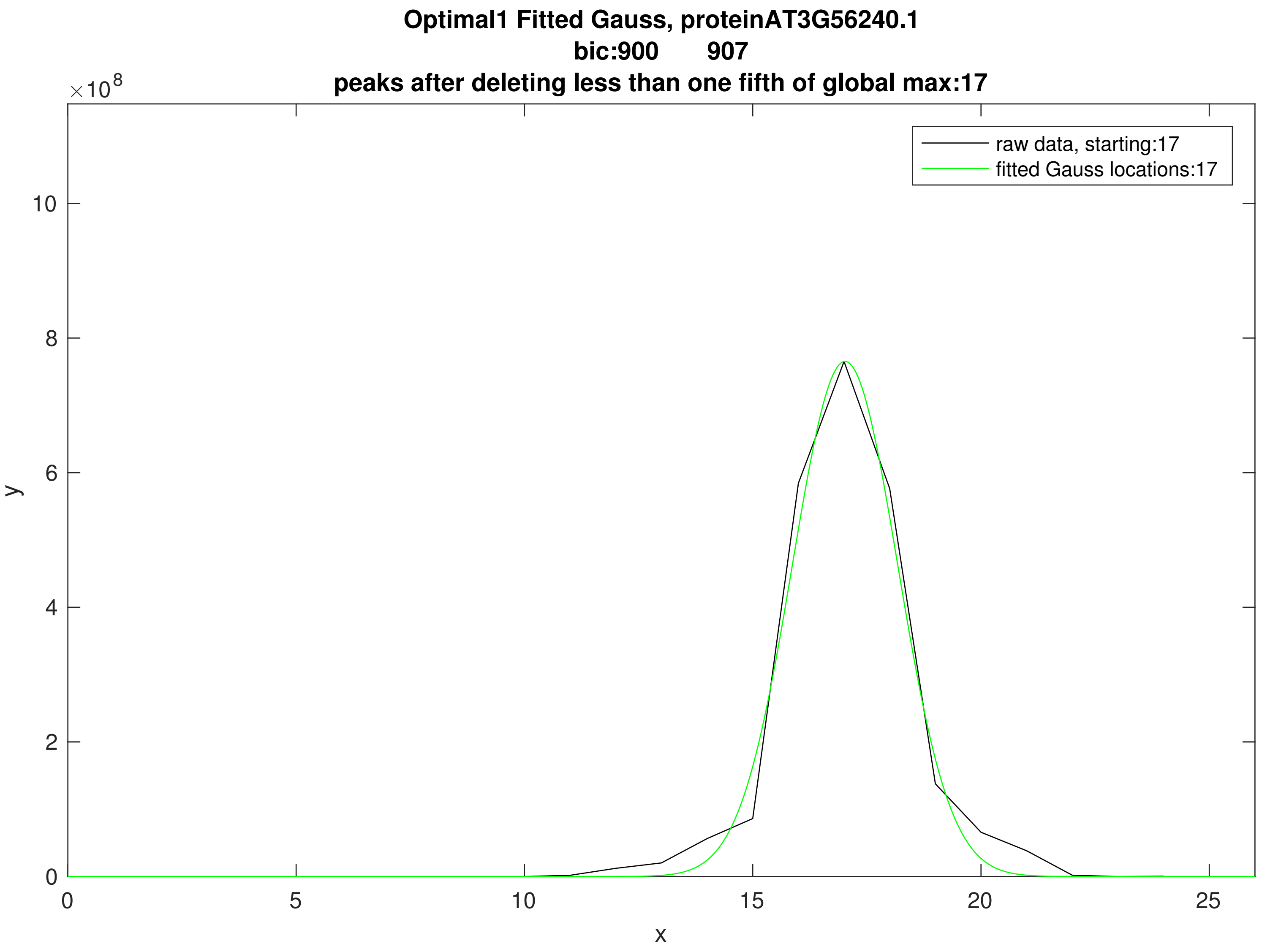
Optimal1 Fitted Gauss, proteinAT3G56240.1
bic:900 907
peaks after deleting less than one fifth of global max:17
×10^8
raw data, starting:17
fitted Gauss locations:17
y
x
0
2
4
6
8
10
5
15
20
25

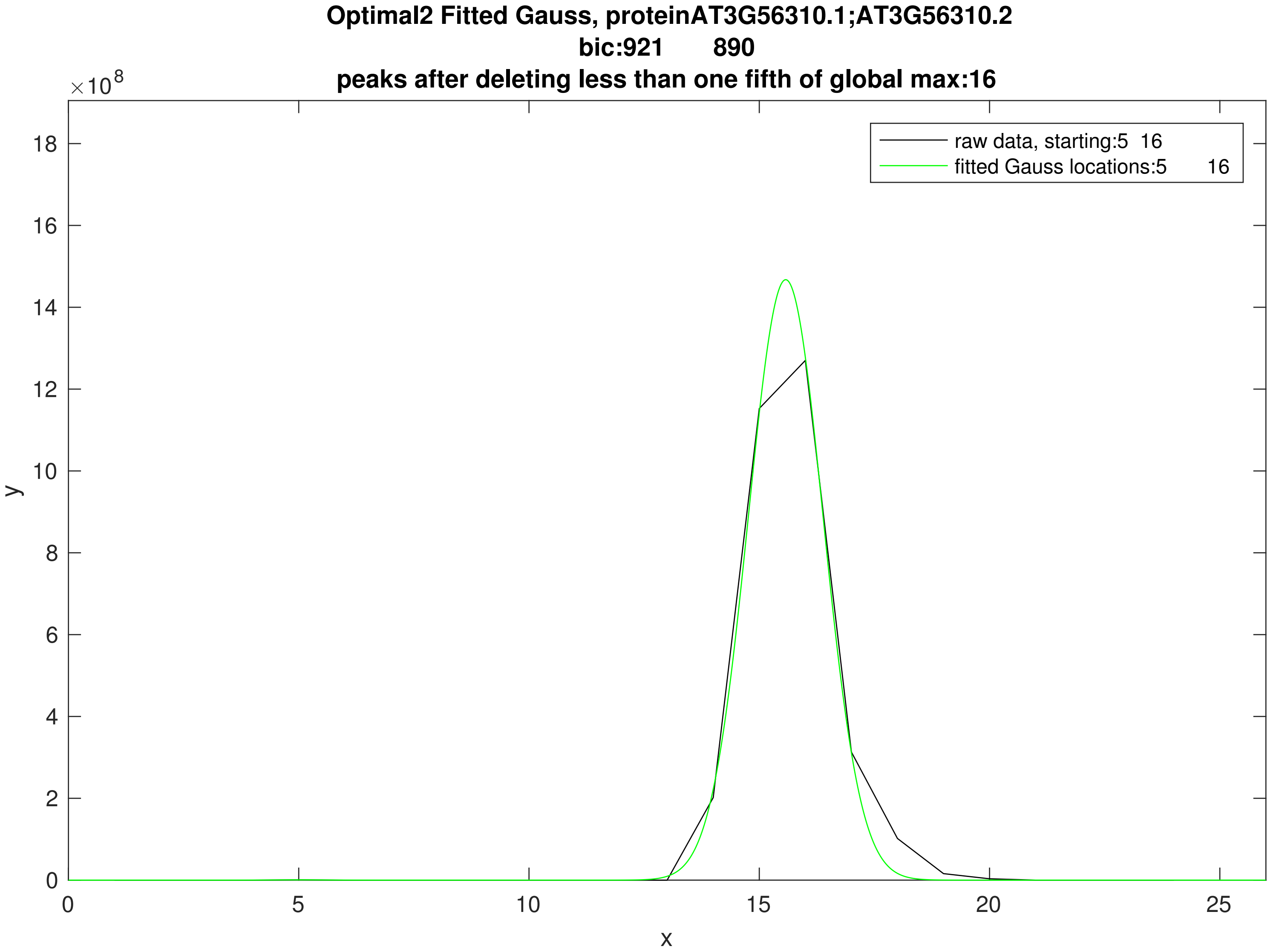

Optimal2 Fitted Gauss, proteinAT3G56310.1;AT3G56310.2
bic:921 890
peaks after deleting less than one fifth of global max:16
×10^8
raw data, starting:5 16
fitted Gauss locations:5 16
y
x
0
2
4
6
8
10
12
14
16
18
0
5
10
15
20
25

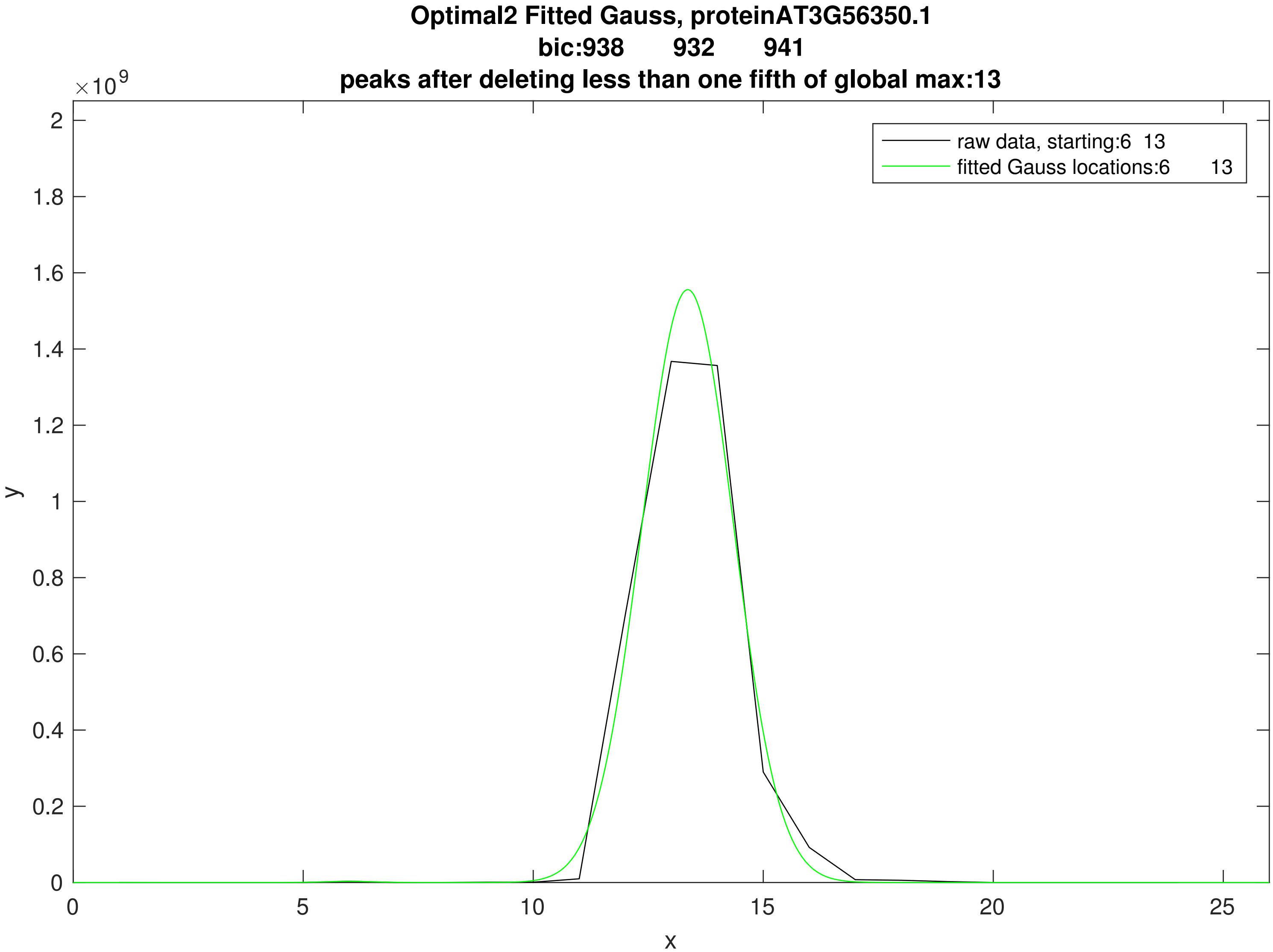
Optimal2 Fitted Gauss, proteinAT3G56350.1
bic:938 932 941
peaks after deleting less than one fifth of global max:13
×10^9
raw data, starting:6 13
fitted Gauss locations:6 13
y
x
0
0.2
0.4
0.6
0.8
1
1.2
1.4
1.6
1.8
2
5
10
15
20
25

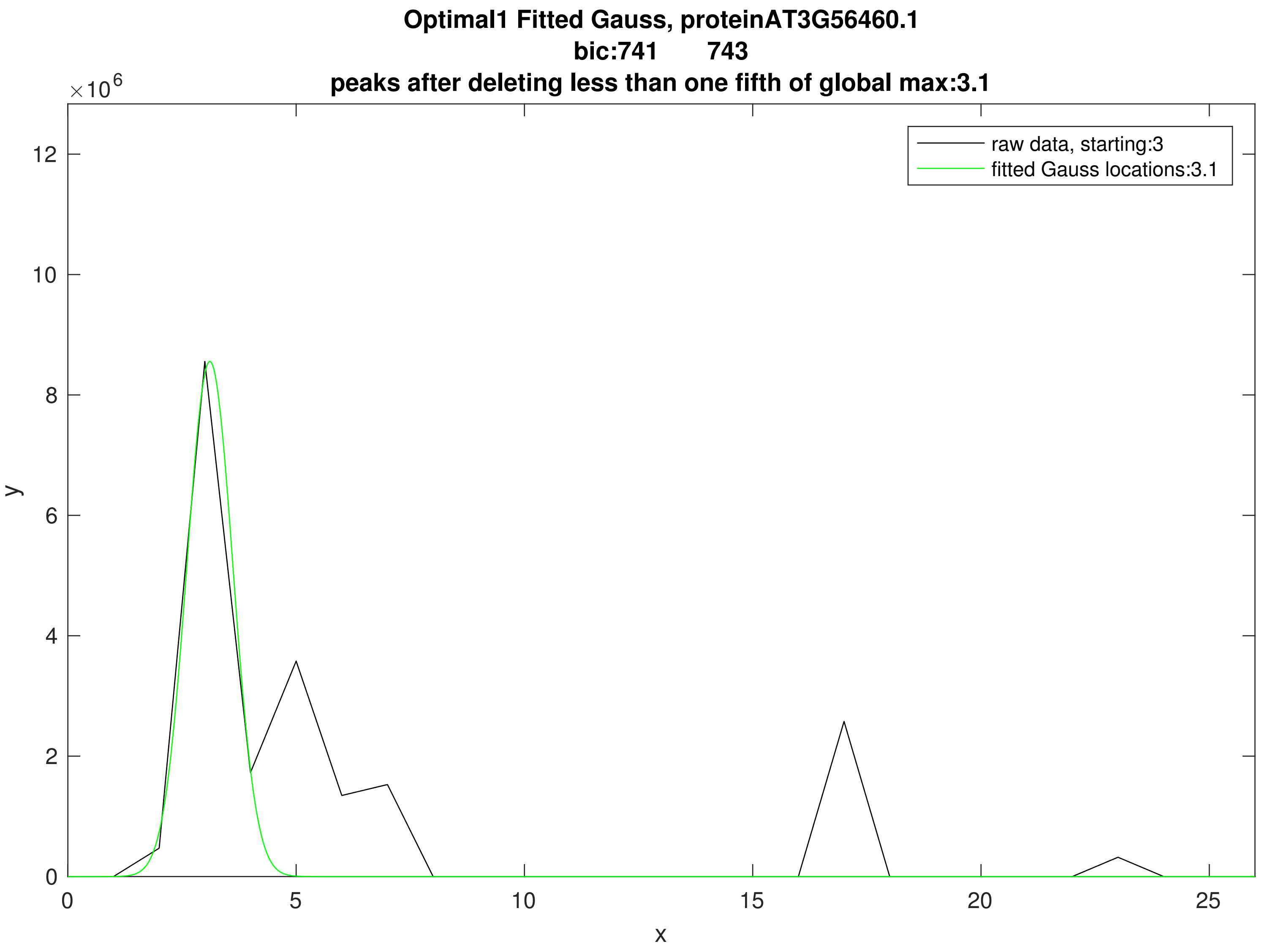
Optimal1 Fitted Gauss, proteinAT3G56460.1
bic:741 743
peaks after deleting less than one fifth of global max:3.1
×10^6
raw data, starting:3
fitted Gauss locations:3.1
y
x
0
2
4
6
8
10
12
5
15
20
25

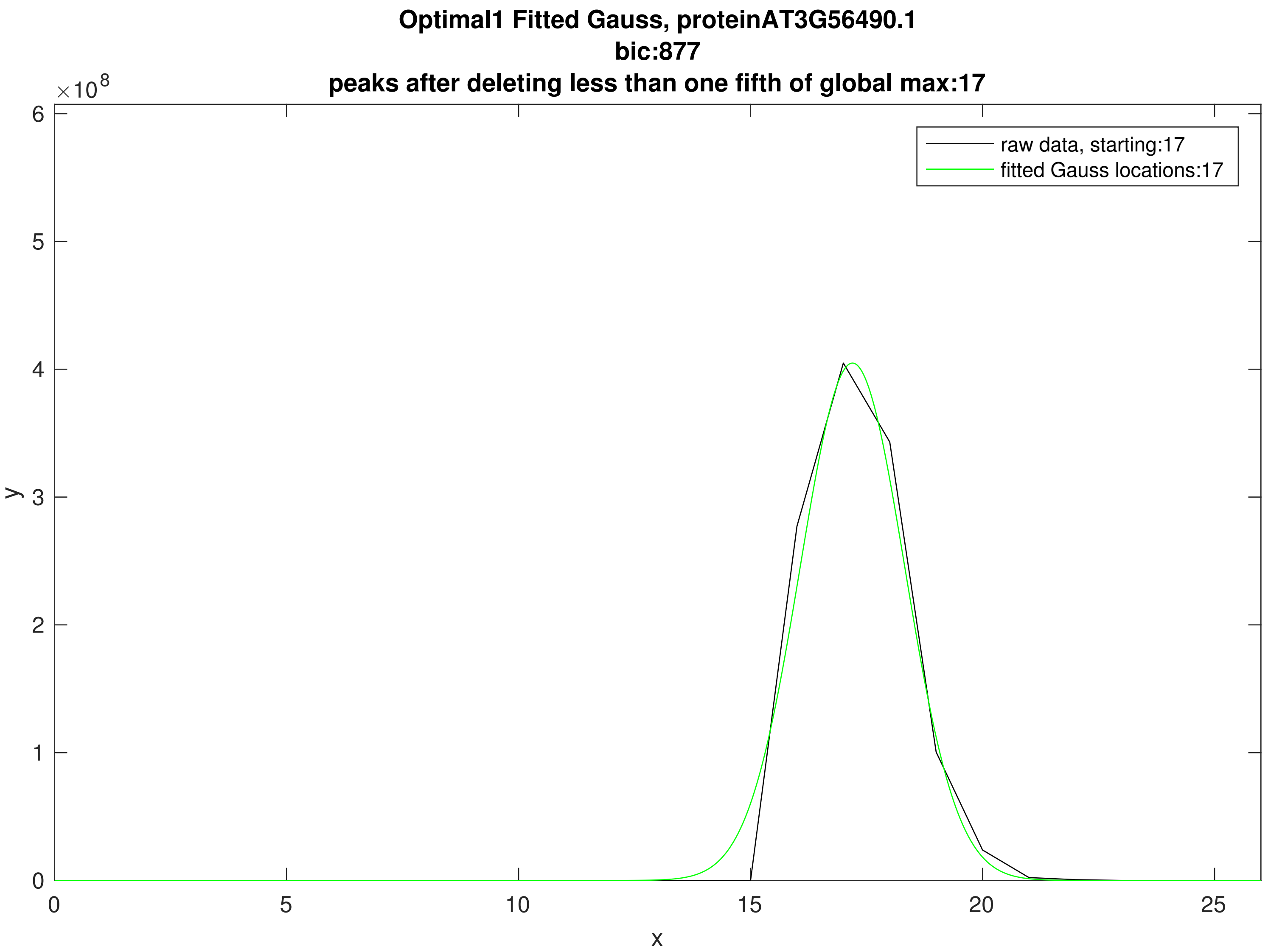
Optimal1 Fitted Gauss, proteinAT3G56490.1
bic:877
peaks after deleting less than one fifth of global max:17
×10^8
6
5
4
3
2
1
0
y
0
5
10
15
20
25
x
raw data, starting:17
fitted Gauss locations:17

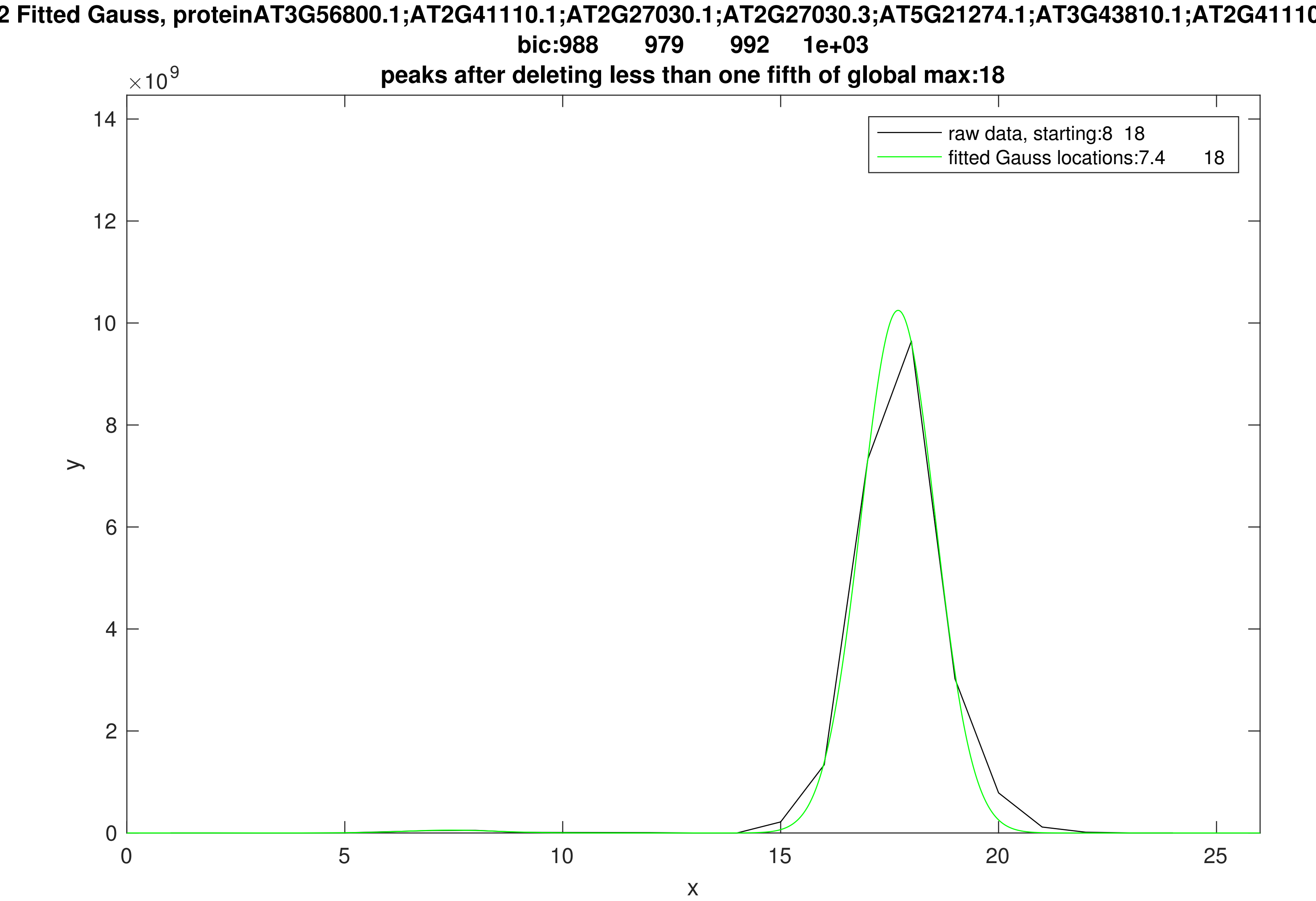

2 Fitted Gauss, proteinAT3G56800.1;AT2G41110.1;AT2G27030.1;AT2G27030.3;AT5G21274.1;AT3G43810.1;AT2G4111
bic:988 979 992 1e+03
peaks after deleting less than one fifth of global max:18
×10^9
14
12
10
8
6
4
2
0
y
0
5
10
15
20
25
x
raw data, starting:8 18
fitted Gauss locations:7.4 18

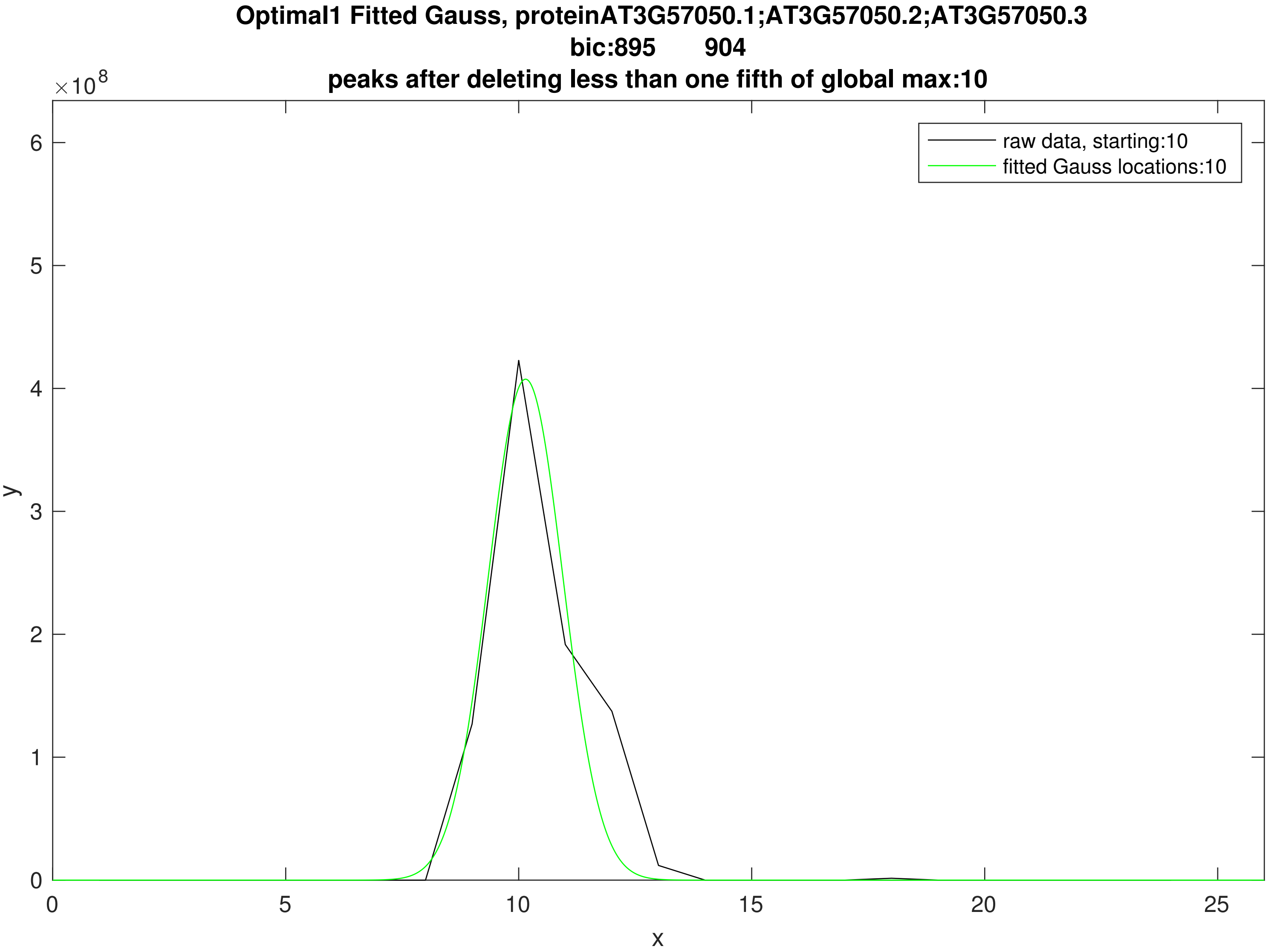
Optimal1 Fitted Gauss, proteinAT3G57050.1;AT3G57050.2;AT3G57050.3
bic:895 904
peaks after deleting less than one fifth of global max:10
×10^8
raw data, starting:10
fitted Gauss locations:10
y
x
0
1
2
3
4
5
6
0
5
10
15
20
25

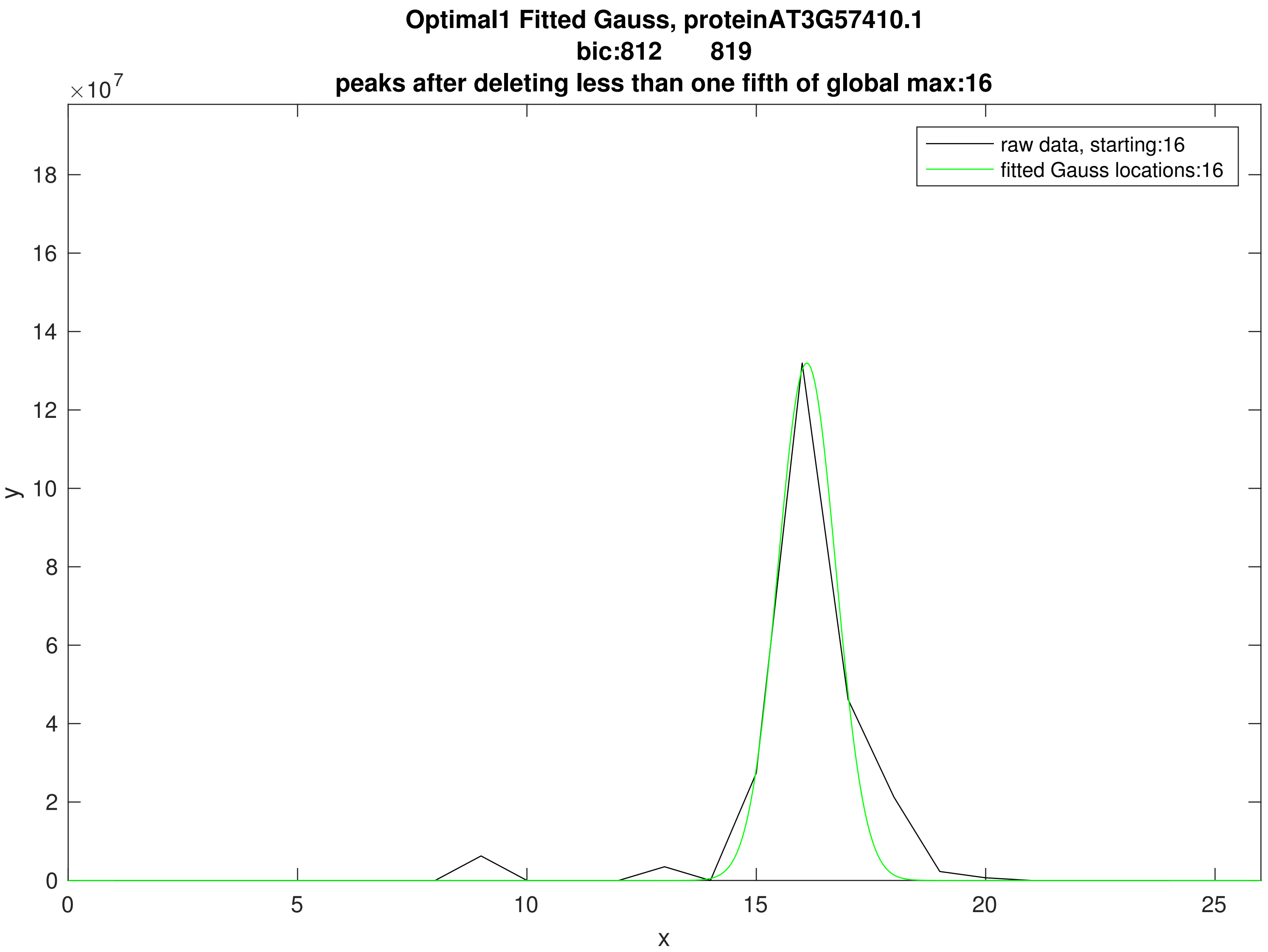

Optimal1 Fitted Gauss, proteinAT3G57410.1
bic:812 819
peaks after deleting less than one fifth of global max:16
×10^7
raw data, starting:16
fitted Gauss locations:16
y
x
0
2
4
6
8
10
12
14
16
18
0
5
10
15
20
25

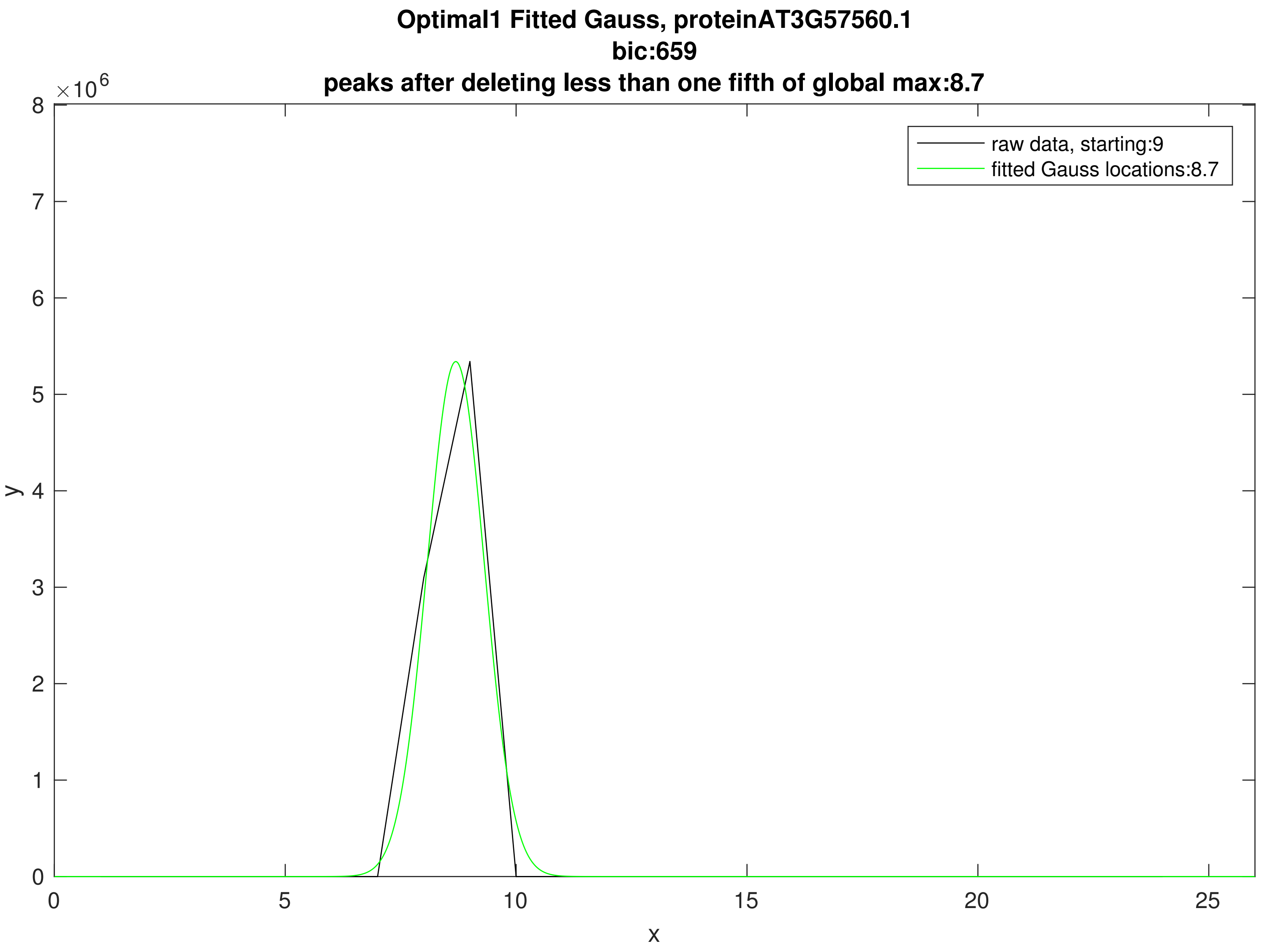
Optimal1 Fitted Gauss, proteinAT3G57560.1
bic:659
peaks after deleting less than one fifth of global max:8.7
×10^6
raw data, starting:9
fitted Gauss locations:8.7
y
x
0
1
2
3
4
5
6
7
8
0
5
10
15
20
25

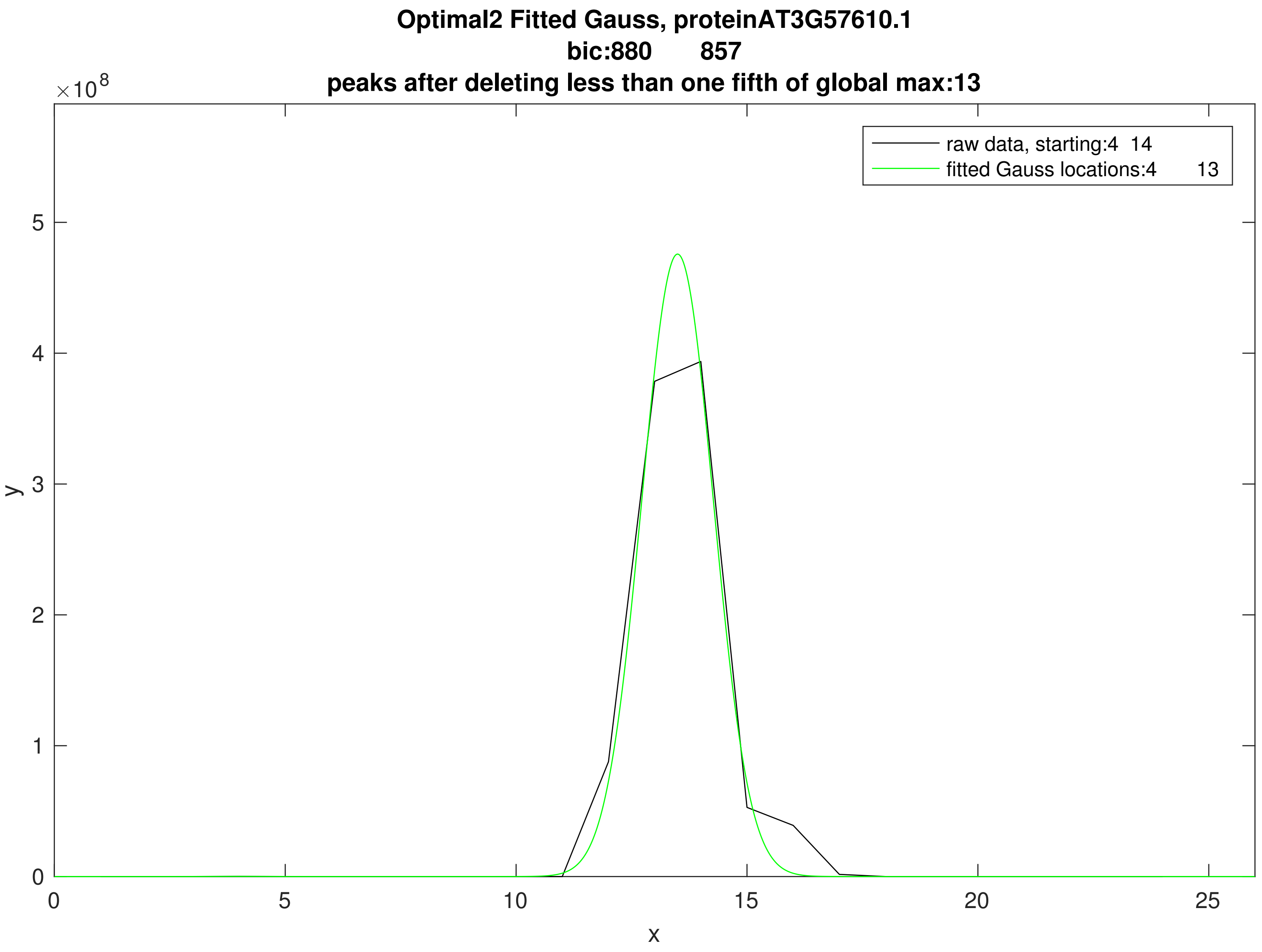

Optimal2 Fitted Gauss, proteinAT3G57610.1
bic:880 857
peaks after deleting less than one fifth of global max:13
×10^8
raw data, starting:4 14
fitted Gauss locations:4 13
y
x
0
1
2
3
4
5
0
5
10
15
20
25

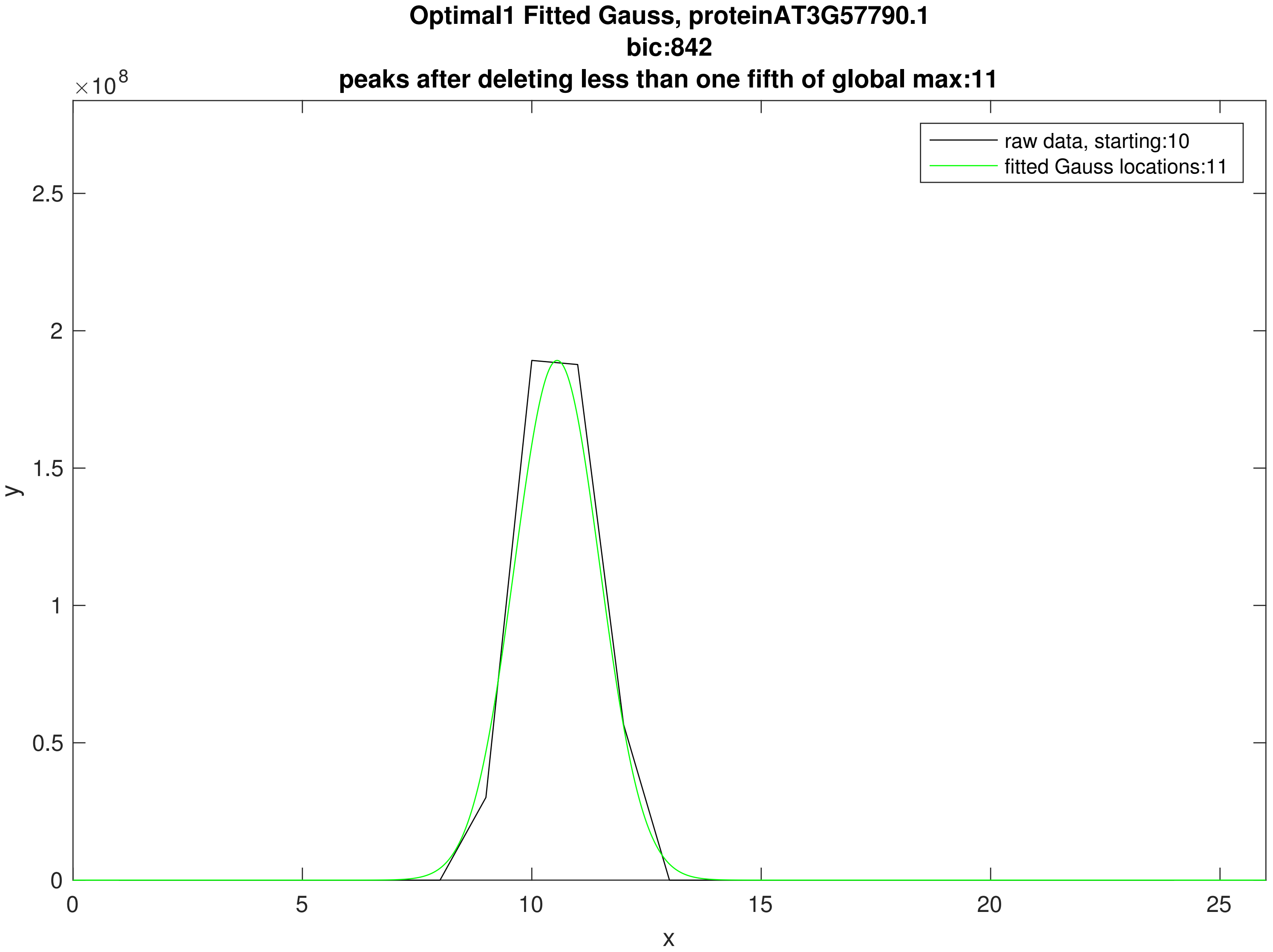
Optimal1 Fitted Gauss, proteinAT3G57790.1
bic:842
peaks after deleting less than one fifth of global max:11
×10^8
raw data, starting:10
fitted Gauss locations:11
2.5
2
1.5
1
0.5
0
y
0
5
10
15
20
25
x

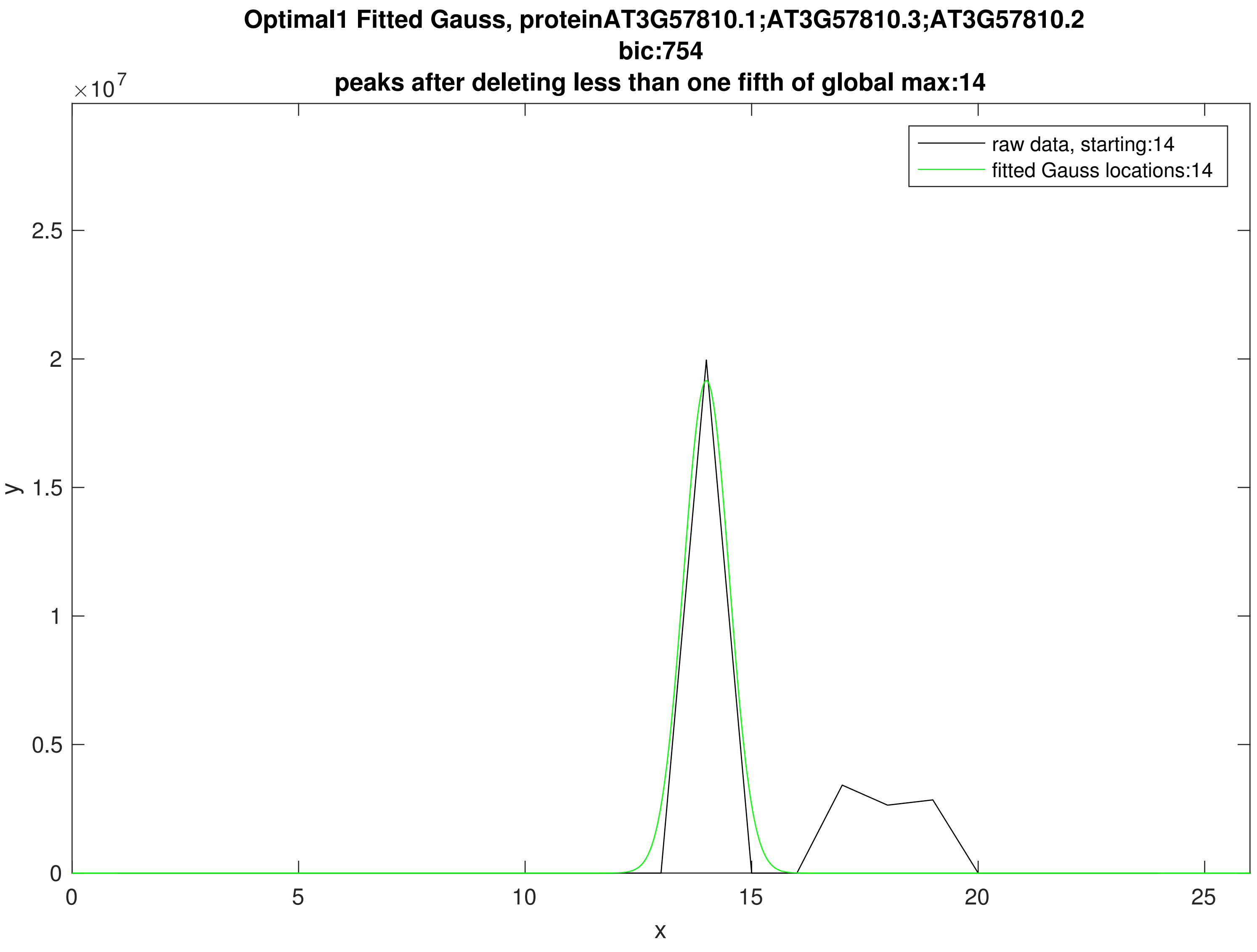

Optimal1 Fitted Gauss, proteinAT3G57810.1;AT3G57810.3;AT3G57810.2
bic:754
peaks after deleting less than one fifth of global max:14
×10^7
raw data, starting:14
fitted Gauss locations:14
2.5
2
1.5
1
0.5
0
y
0
5
10
15
20
25
x

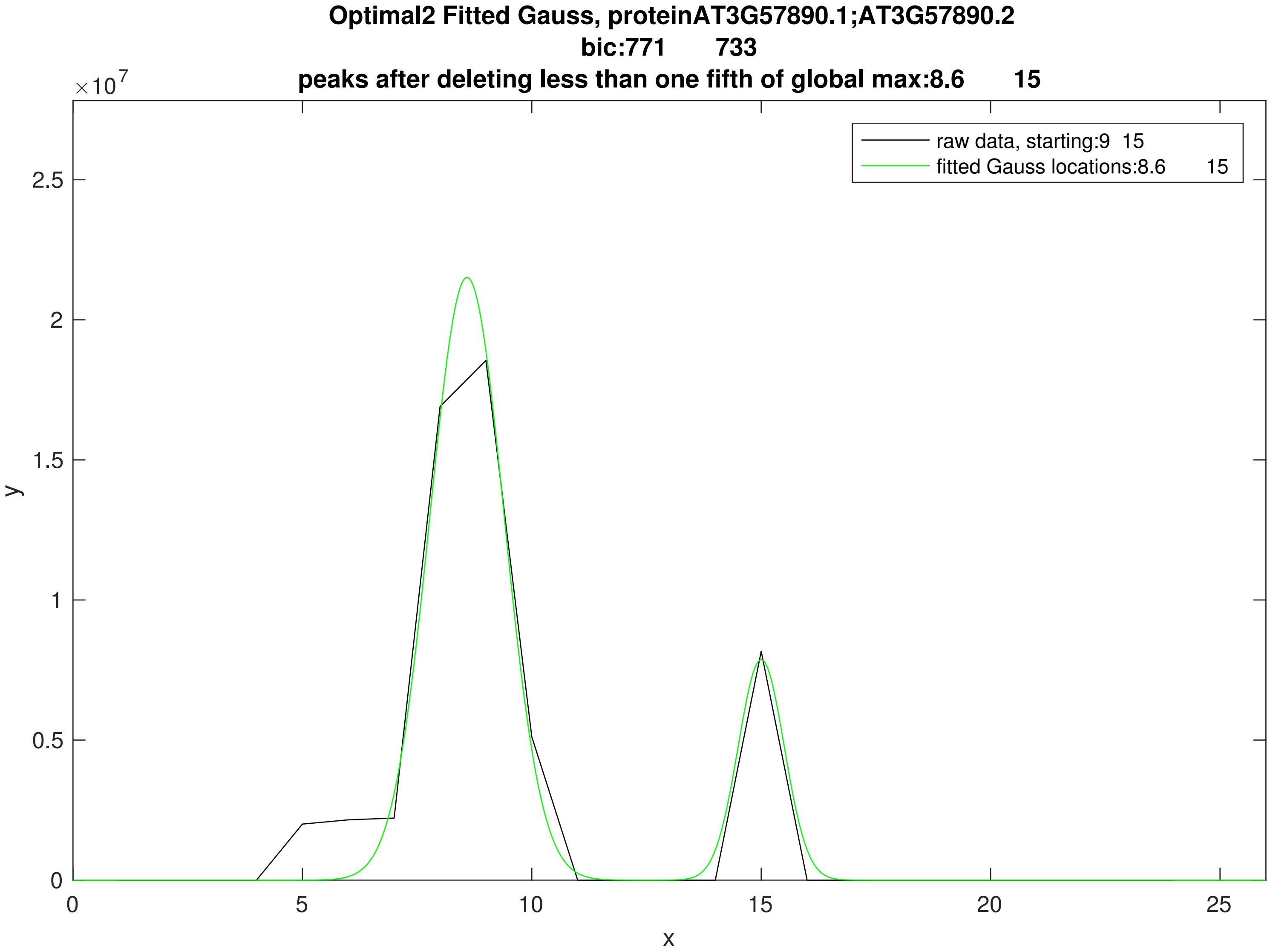
Optimal2 Fitted Gauss, proteinAT3G57890.1;AT3G57890.2
bic:771 733
peaks after deleting less than one fifth of global max:8.6 15
×10^7
raw data, starting:9 15
fitted Gauss locations:8.6 15
y
x
0
0.5
1
1.5
2
2.5
5
10
15
20
25

Optimal1 Fitted Gauss, proteinAT3G58140.1
bic:871
peaks after deleting less than one fifth of global max:16

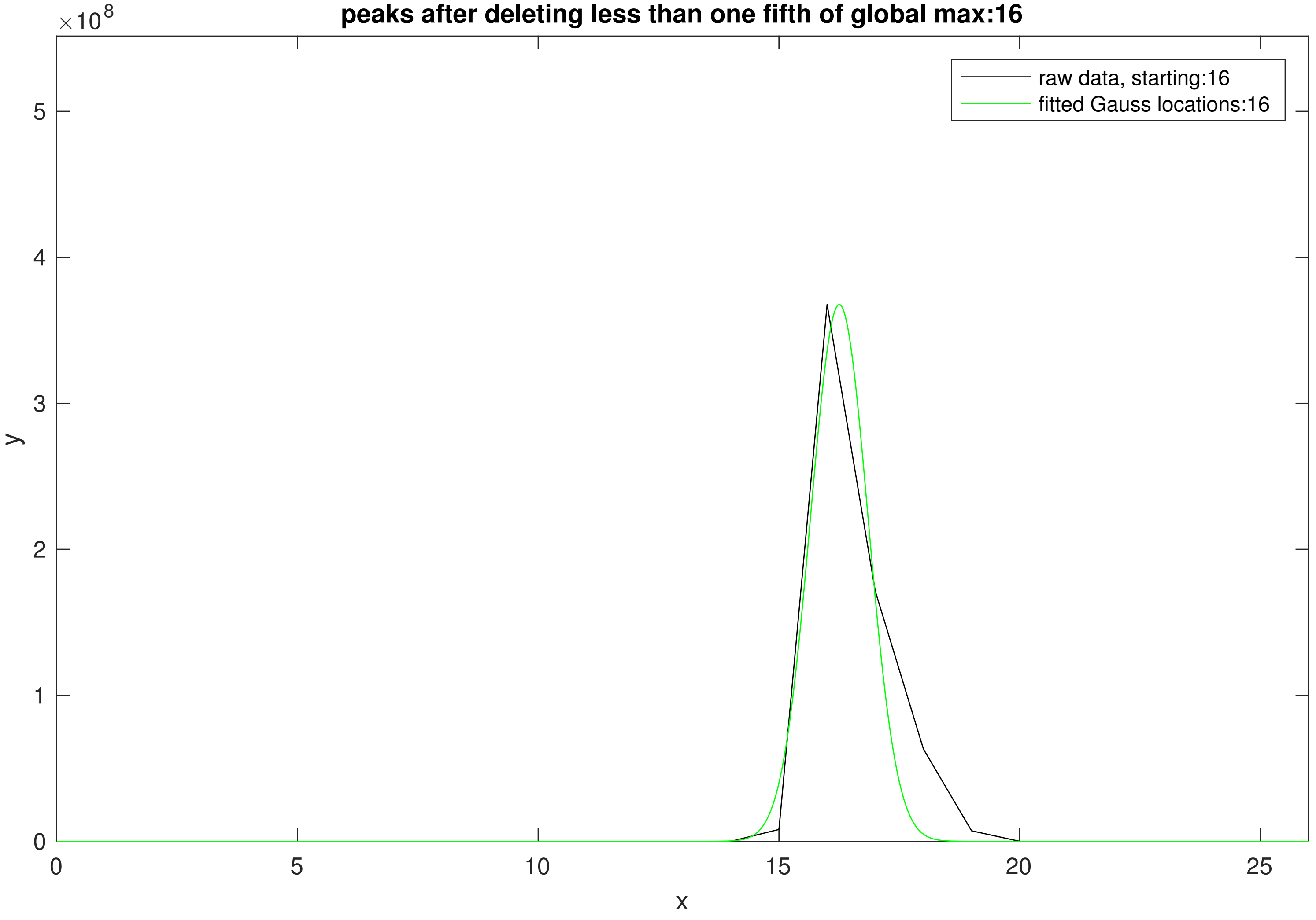

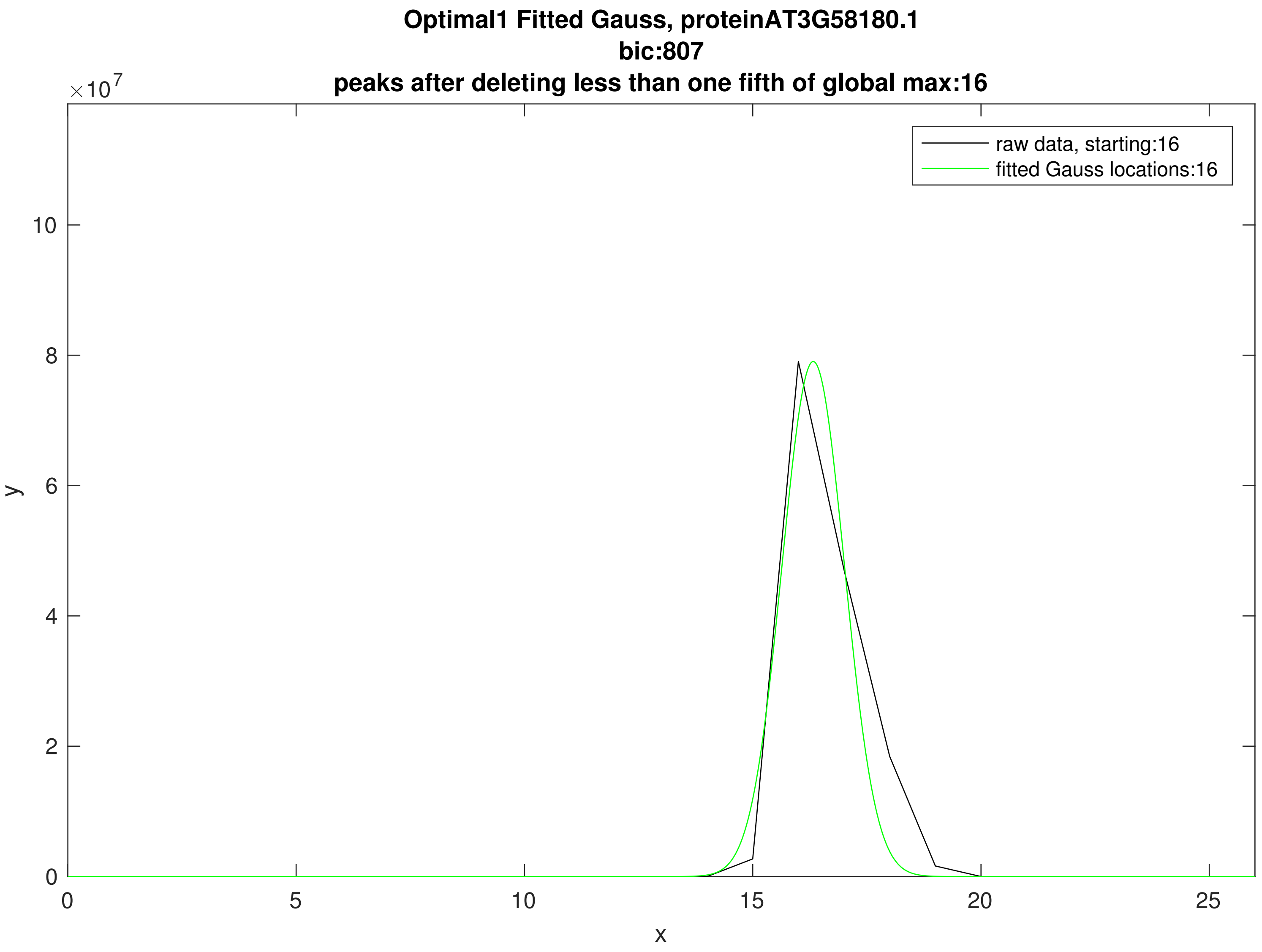
Optimal1 Fitted Gauss, proteinAT3G58180.1
bic:807
peaks after deleting less than one fifth of global max:16
×10^7
raw data, starting:16
fitted Gauss locations:16
y
x
0
2
4
6
8
10
0
5
10
15
20
25

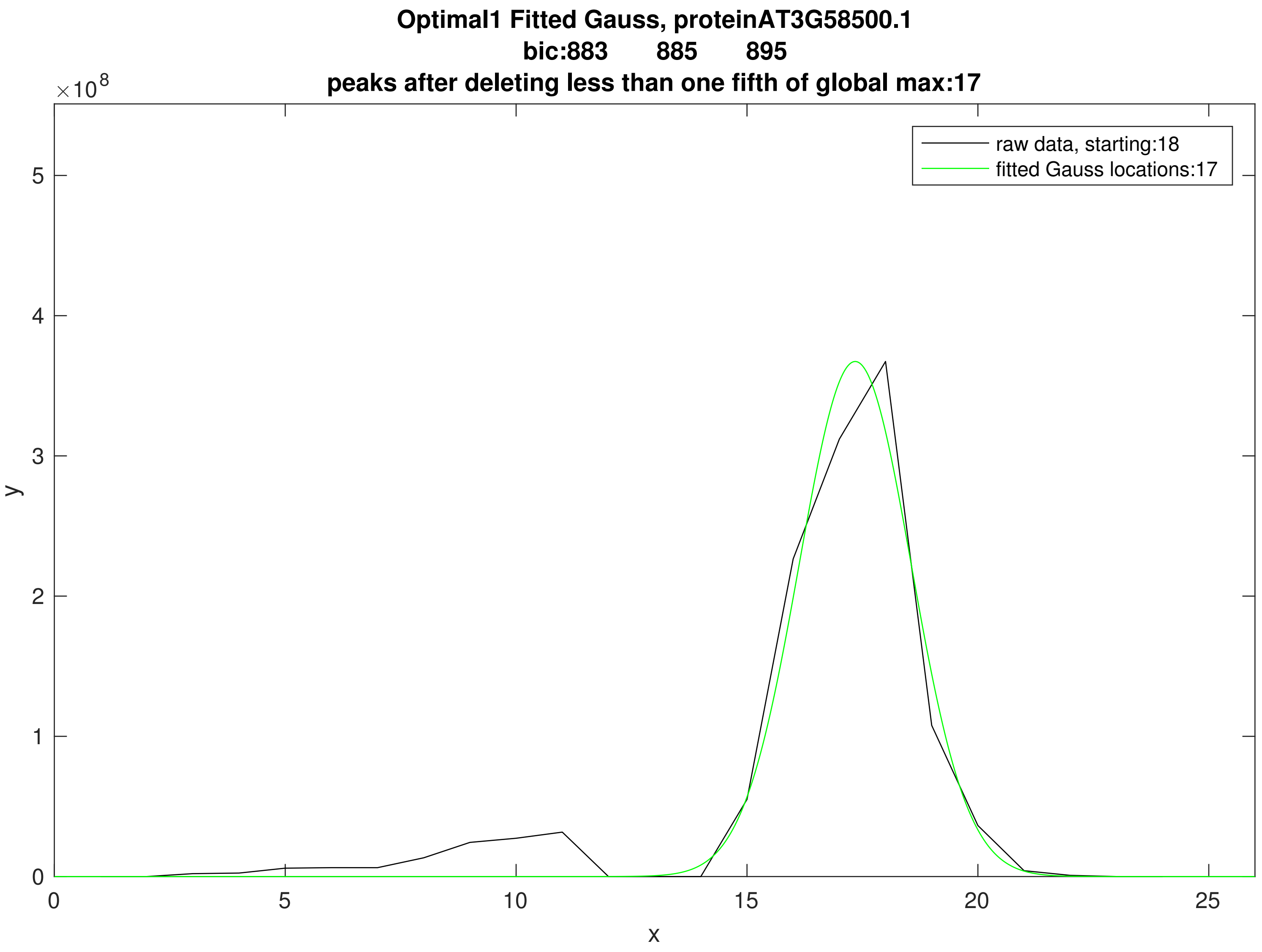
Optimal1 Fitted Gauss, proteinAT3G58500.1
bic:883 885 895
peaks after deleting less than one fifth of global max:17
×10^8
raw data, starting:18
fitted Gauss locations:17
y
x
0
1
2
3
4
5
0
5
10
15
20
25

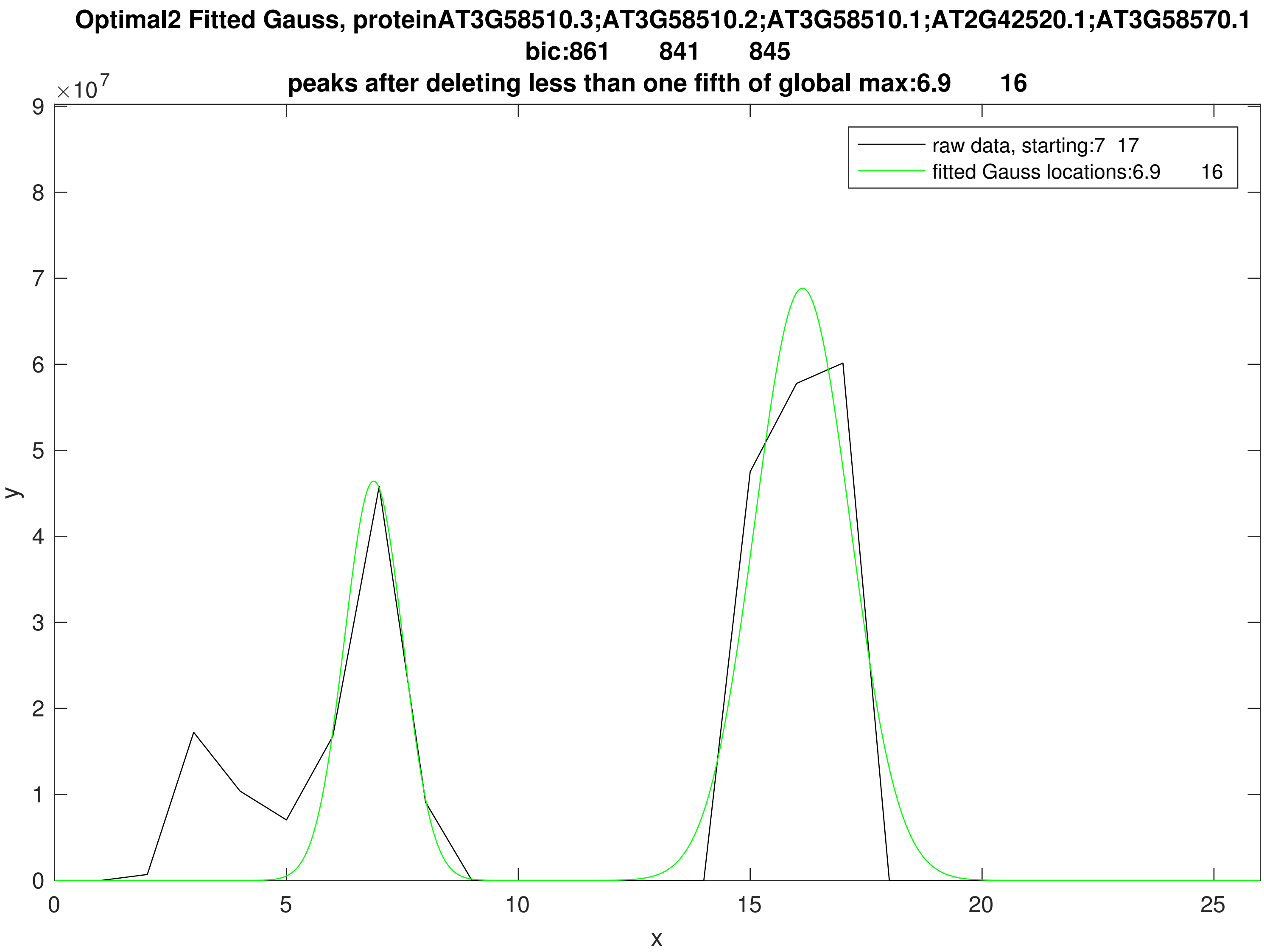
Optimal2 Fitted Gauss, proteinAT3G58510.3;AT3G58510.2;AT3G58510.1;AT2G42520.1;AT3G58570.1
bic:861 841 845
peaks after deleting less than one fifth of global max:6.9 16
raw data, starting:7 17
fitted Gauss locations:6.9 16
$\times 10^7$
y
x

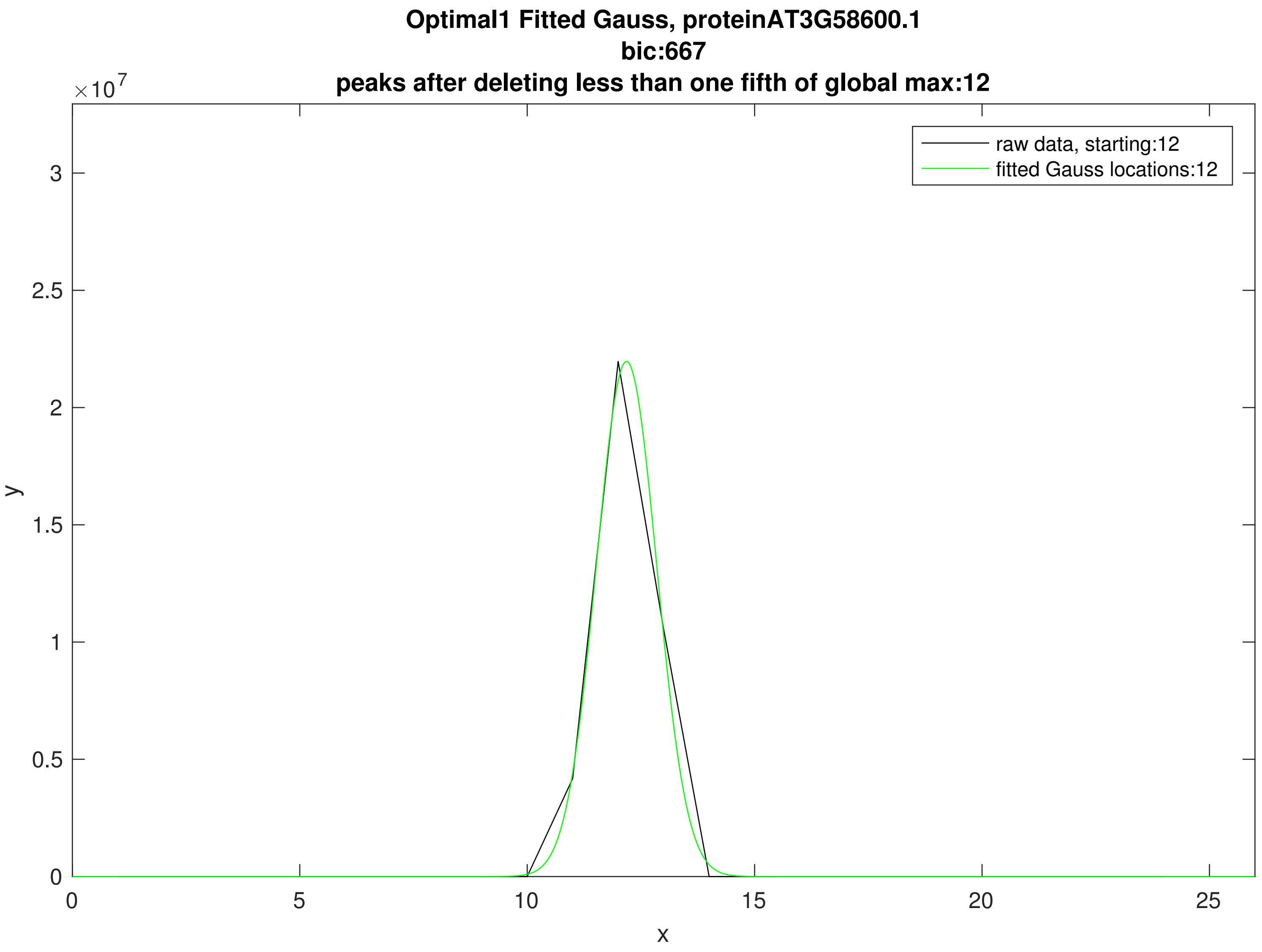
Optimal1 Fitted Gauss, proteinAT3G58600.1
bic:667
peaks after deleting less than one fifth of global max:12
×10^7
3
2.5
2
1.5
1
0.5
0
y
0
5
10
15
20
25
x
raw data, starting:12
fitted Gauss locations:12

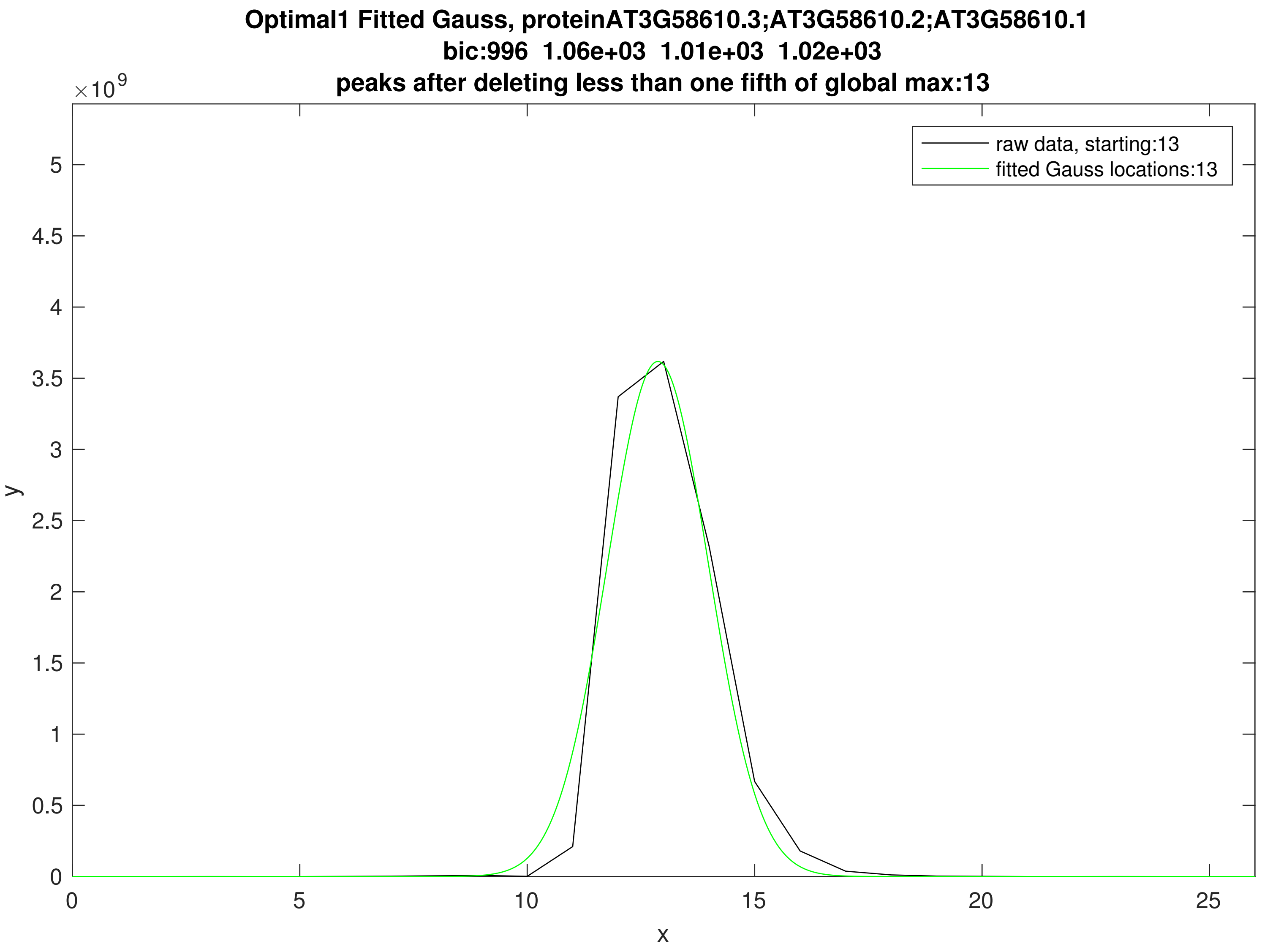

Optimal1 Fitted Gauss, proteinAT3G58610.3;AT3G58610.2;AT3G58610.1
bic:996 1.06e+03 1.01e+03 1.02e+03
peaks after deleting less than one fifth of global max:13
×10^9
raw data, starting:13
fitted Gauss locations:13
y
x
0
0.5
1
1.5
2
2.5
3
3.5
4
4.5
5
0
5
10
15
20
25

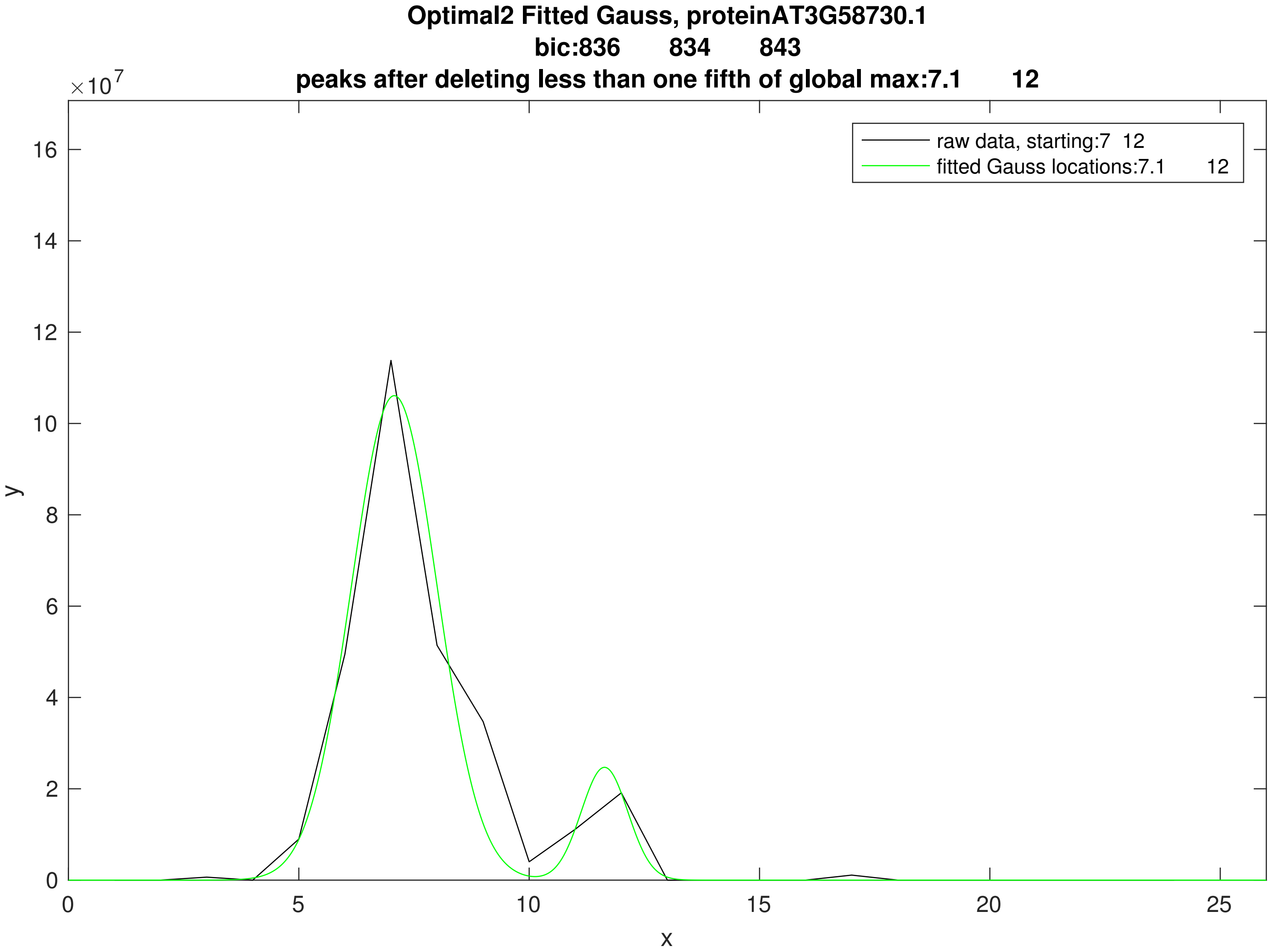

Optimal2 Fitted Gauss, proteinAT3G58730.1
bic:836 834 843
peaks after deleting less than one fifth of global max:7.1 12
×10^7
raw data, starting:7 12
fitted Gauss locations:7.1 12
y
x
0
2
4
6
8
10
12
14
16
5
10
15
20
25

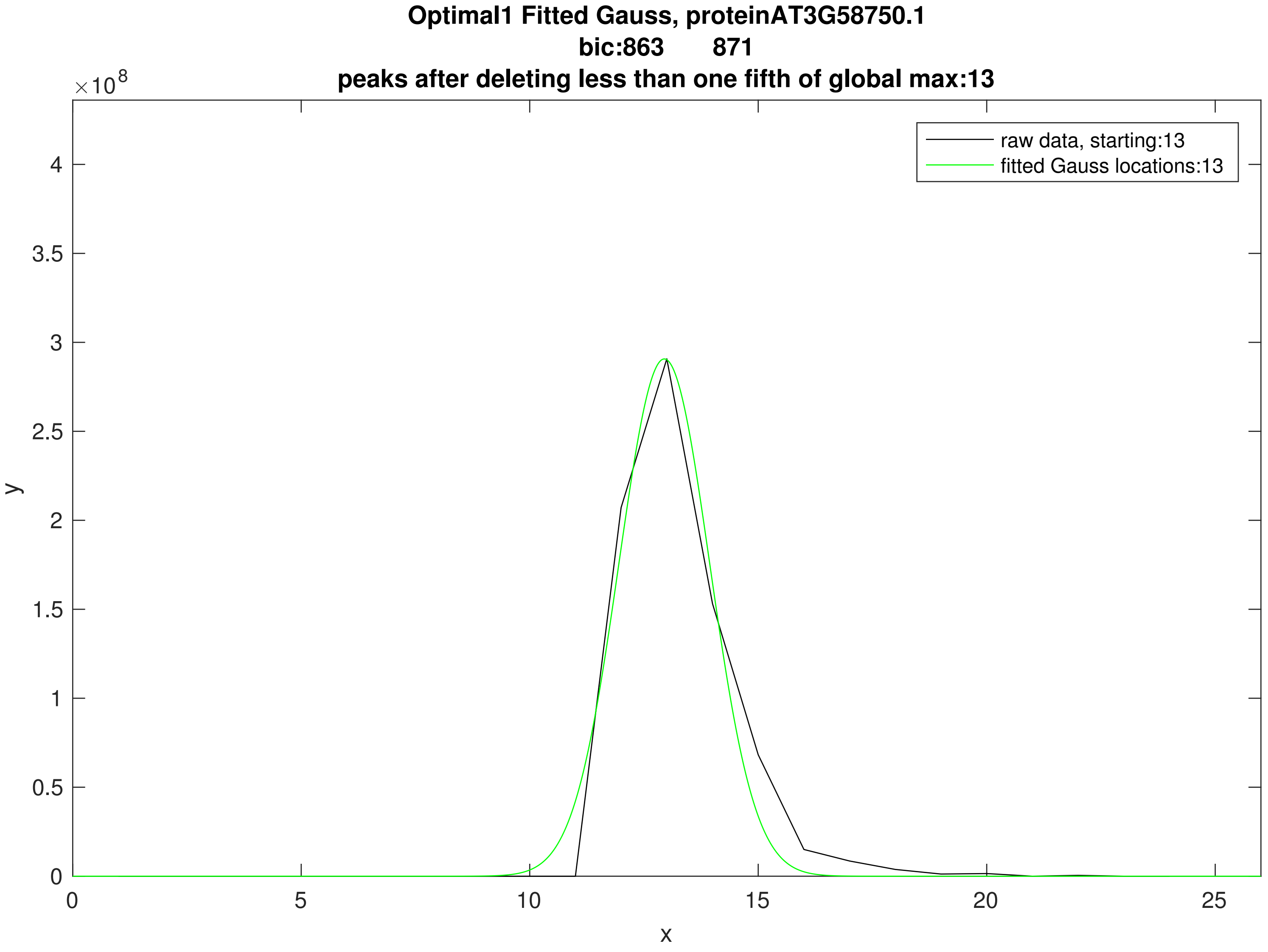

Optimal1 Fitted Gauss, proteinAT3G58750.1
bic:863 871
peaks after deleting less than one fifth of global max:13
×10^8
raw data, starting:13
fitted Gauss locations:13
4
3.5
3
2.5
2
1.5
1
0.5
0
y
0
5
10
15
20
25
x

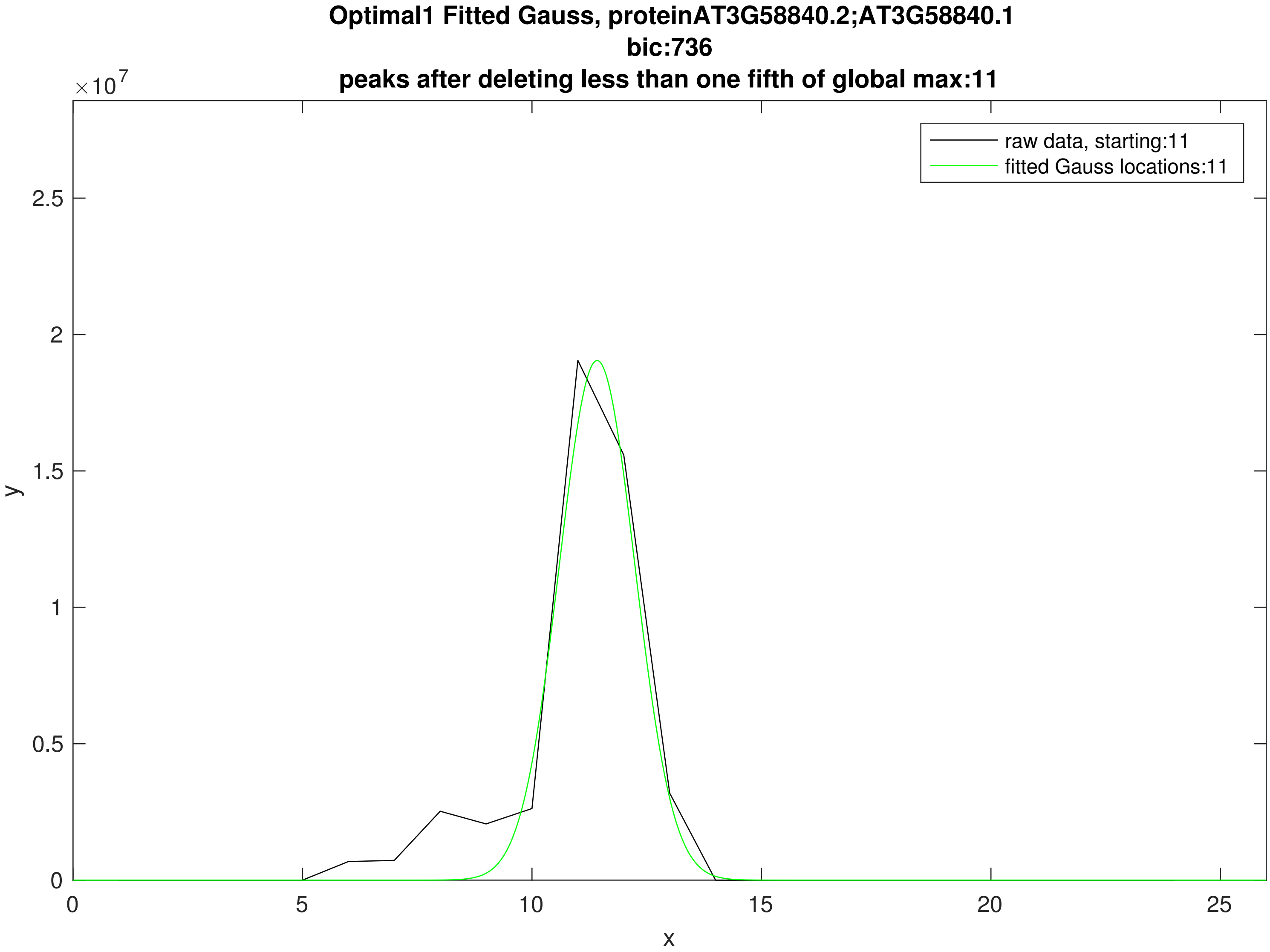

Optimal1 Fitted Gauss, proteinAT3G58840.2;AT3G58840.1
bic:736
peaks after deleting less than one fifth of global max:11
×10^7
raw data, starting:11
fitted Gauss locations:11
2.5
2
1.5
1
0.5
0
y
0
5
10
15
20
25
x

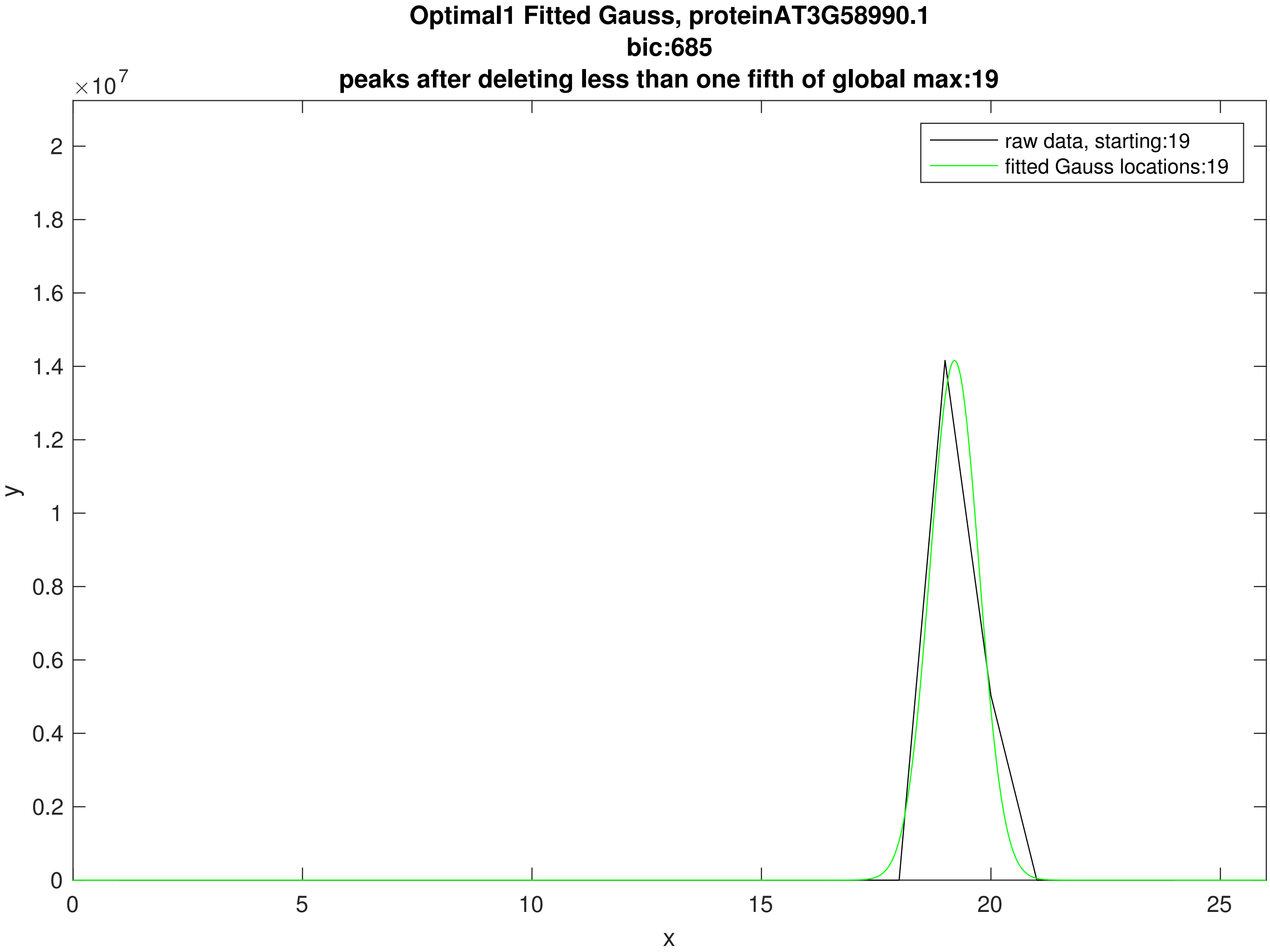

Optimal1 Fitted Gauss, proteinAT3G58990.1
bic:685
peaks after deleting less than one fifth of global max:19
×10^7
raw data, starting:19
fitted Gauss locations:19
y
x
0
0.2
0.4
0.6
0.8
1
1.2
1.4
1.6
1.8
2
0
5
10
15
20
25

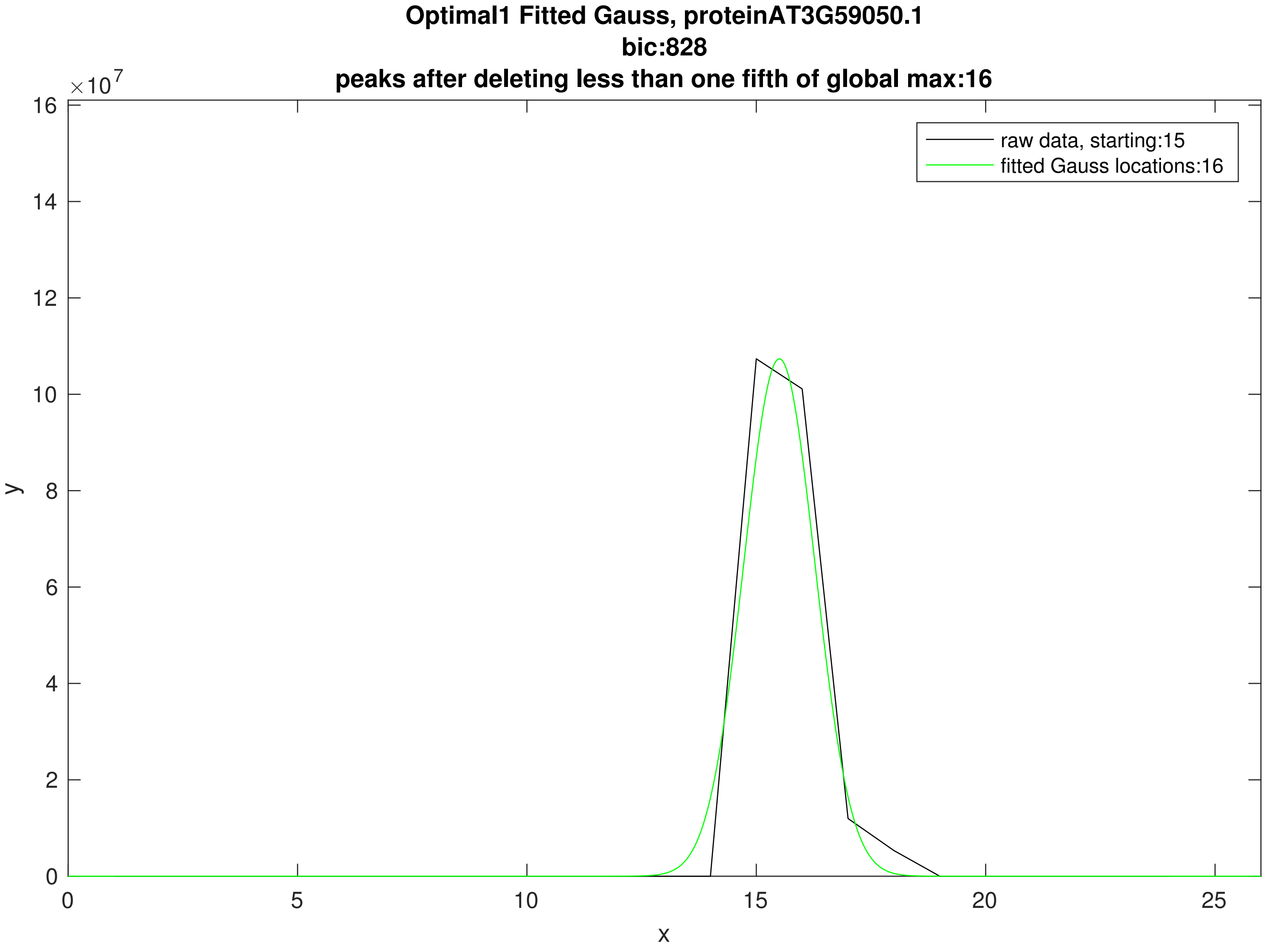
Optimal1 Fitted Gauss, proteinAT3G59050.1
bic:828
peaks after deleting less than one fifth of global max:16
×10^7
16
14
12
10
8
6
4
2
0
y
0
5
10
15
20
25
x
raw data, starting:15
fitted Gauss locations:16

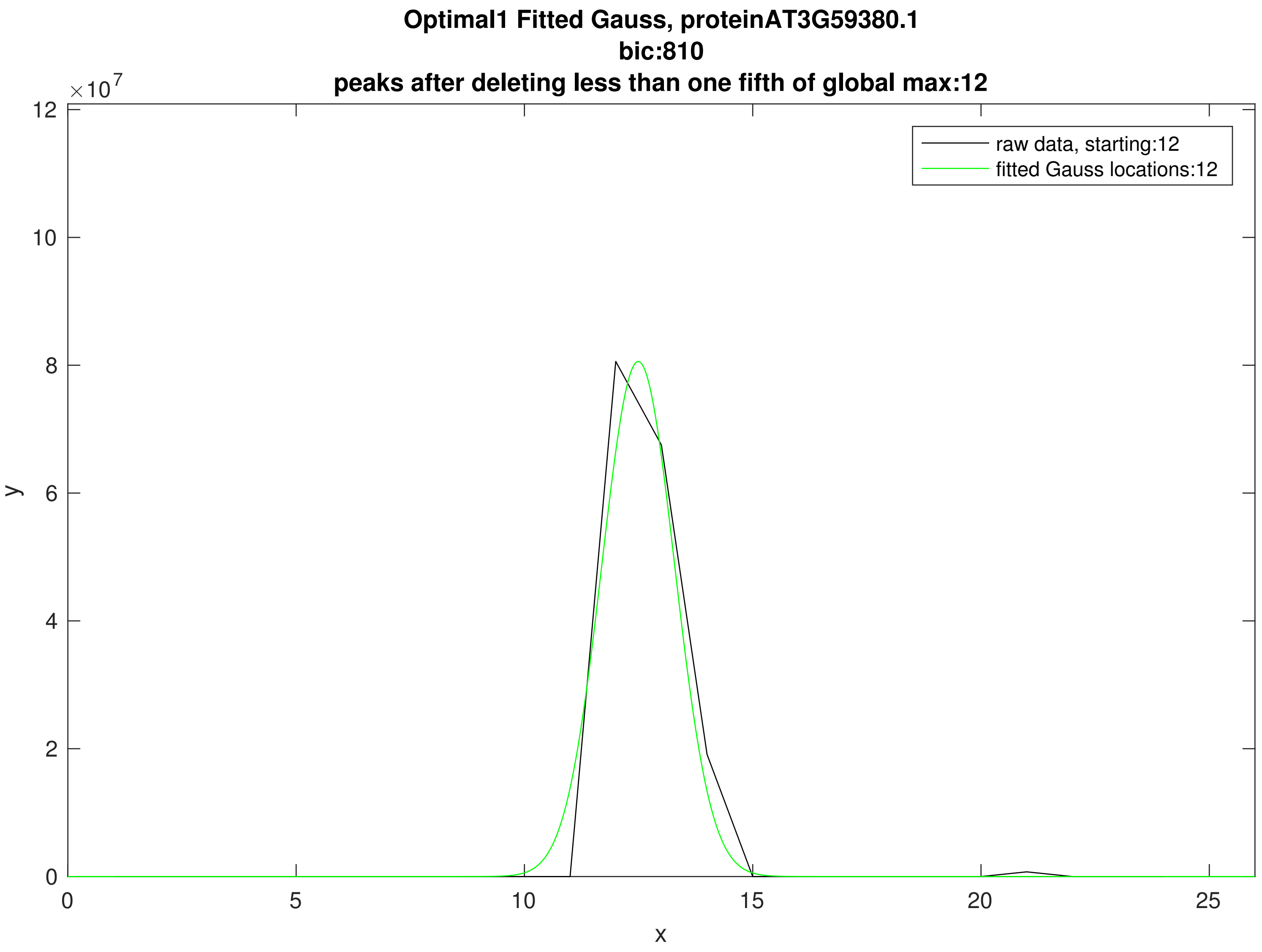

Optimal1 Fitted Gauss, proteinAT3G59380.1
bic:810
peaks after deleting less than one fifth of global max:12
×10^7
raw data, starting:12
fitted Gauss locations:12
y
x
0
2
4
6
8
10
12
0
5
10
15
20
25

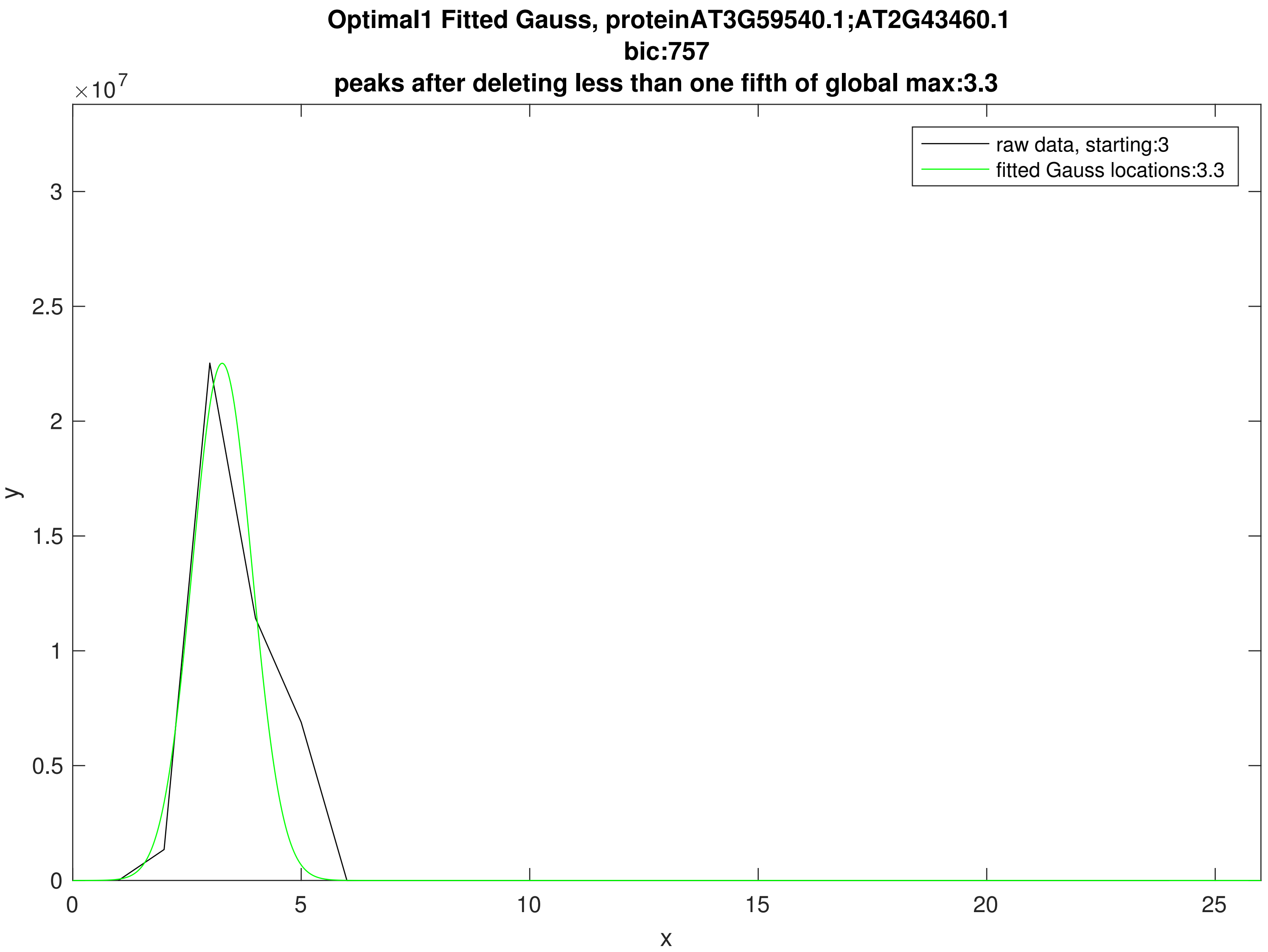

Optimal1 Fitted Gauss, proteinAT3G59540.1;AT2G43460.1
bic:757
peaks after deleting less than one fifth of global max:3.3
×10^7
raw data, starting:3
fitted Gauss locations:3.3
3
2.5
2
1.5
1
0.5
0
y
0
5
10
15
20
25
x

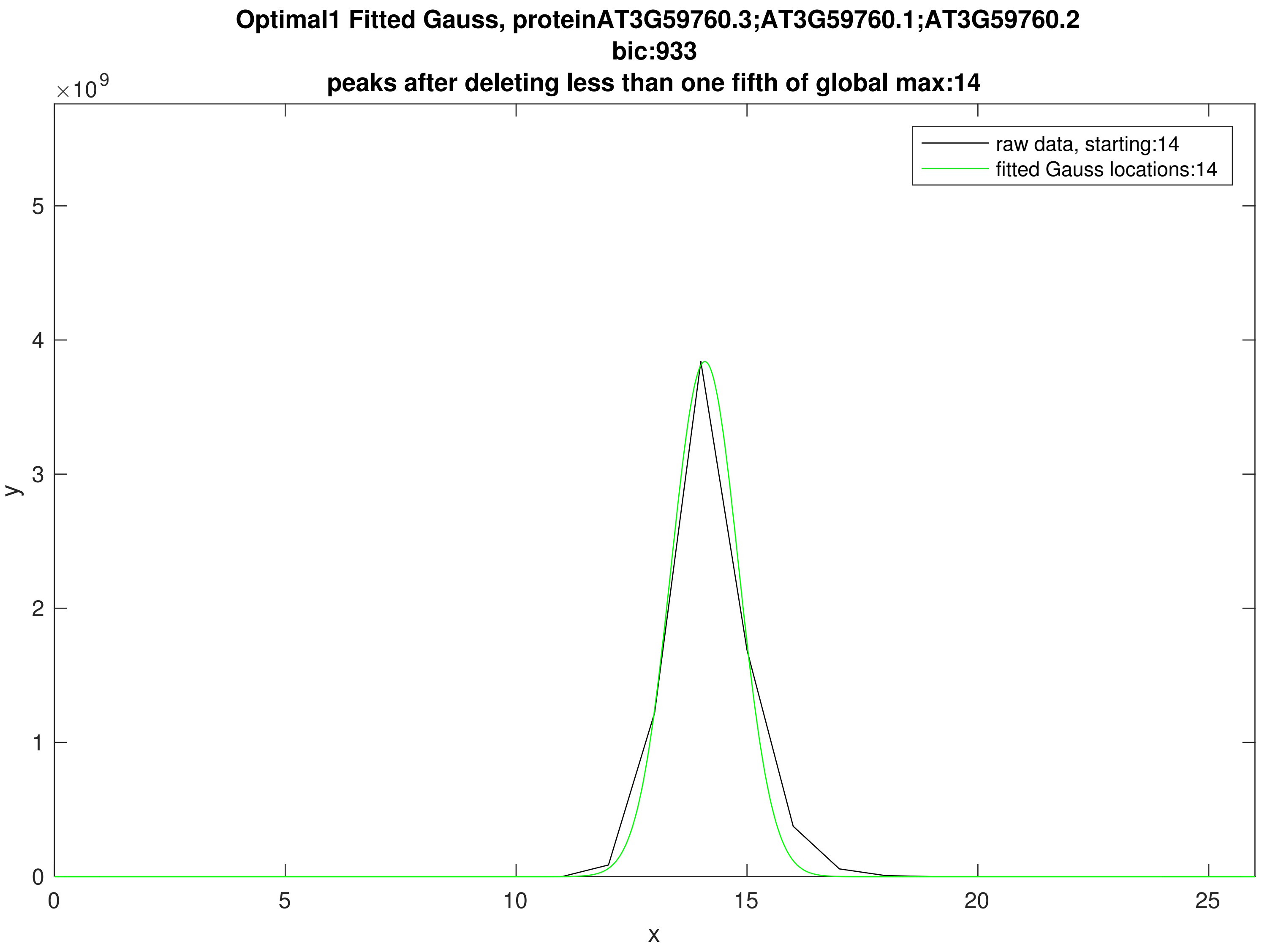

Optimal1 Fitted Gauss, proteinAT3G59760.3;AT3G59760.1;AT3G59760.2
bic:933
peaks after deleting less than one fifth of global max:14
×10^9
raw data, starting:14
fitted Gauss locations:14
y
x
0
1
2
3
4
5
0
5
10
15
20
25

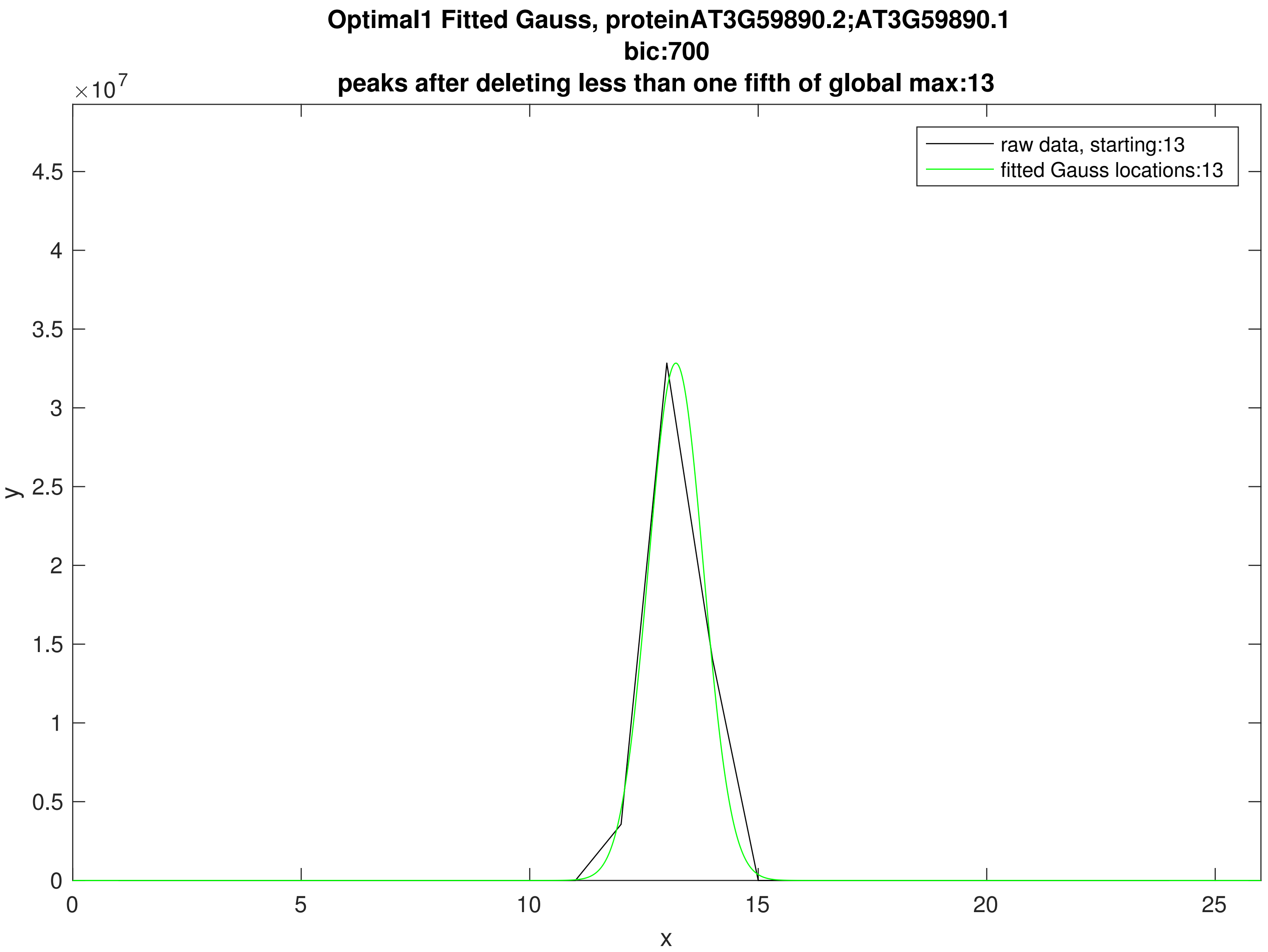

Optimal1 Fitted Gauss, proteinAT3G59890.2;AT3G59890.1
bic:700
peaks after deleting less than one fifth of global max:13
×10^7
raw data, starting:13
fitted Gauss locations:13
y
x
0
0.5
1
1.5
2
2.5
3
3.5
4
4.5
5
10
15
20
25

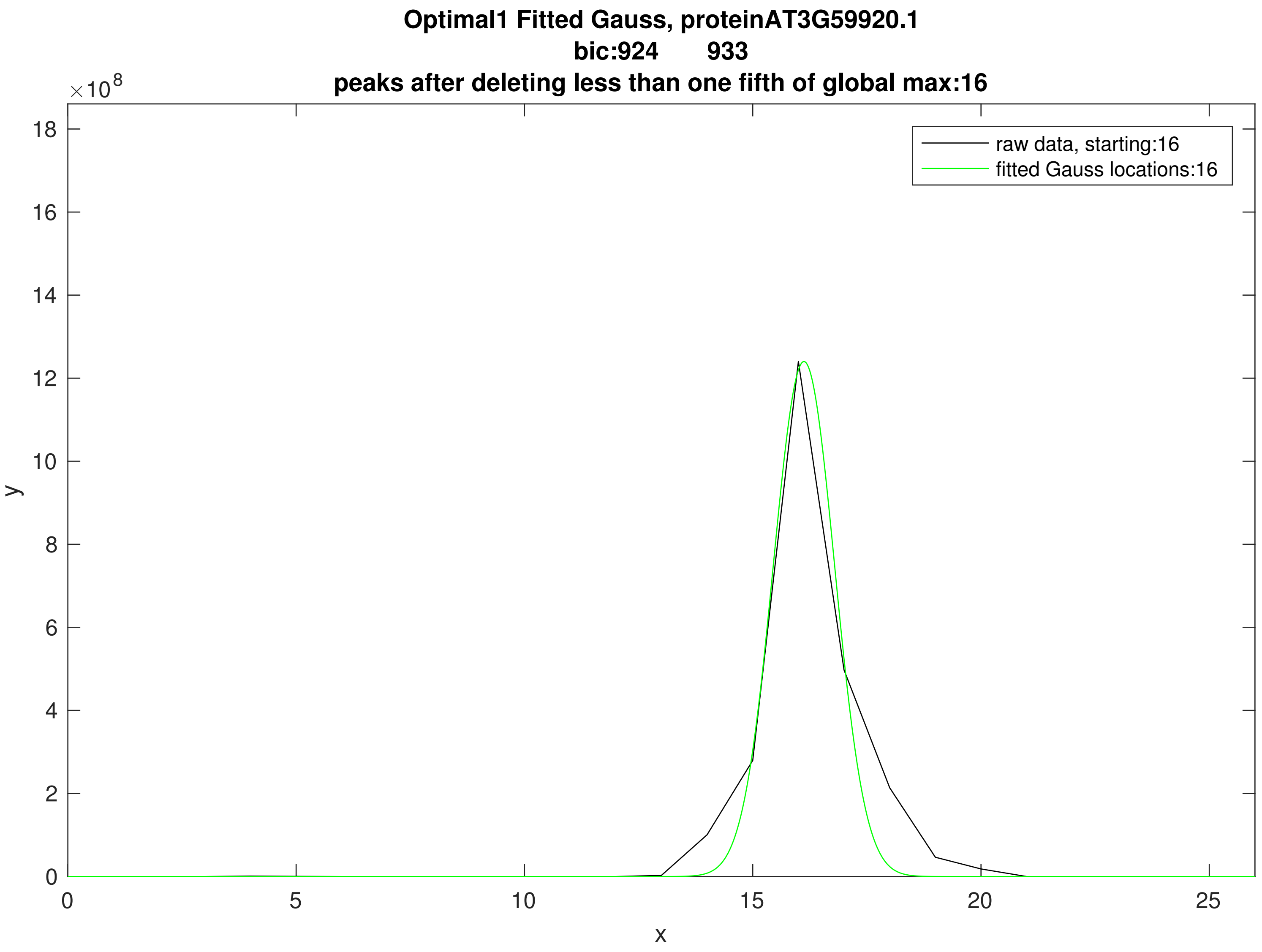

Optimal1 Fitted Gauss, proteinAT3G59920.1
bic:924 933
peaks after deleting less than one fifth of global max:16
×10^8
18
16
14
12
10
8
6
4
2
0
y
0
5
10
15
20
25
x
raw data, starting:16
fitted Gauss locations:16

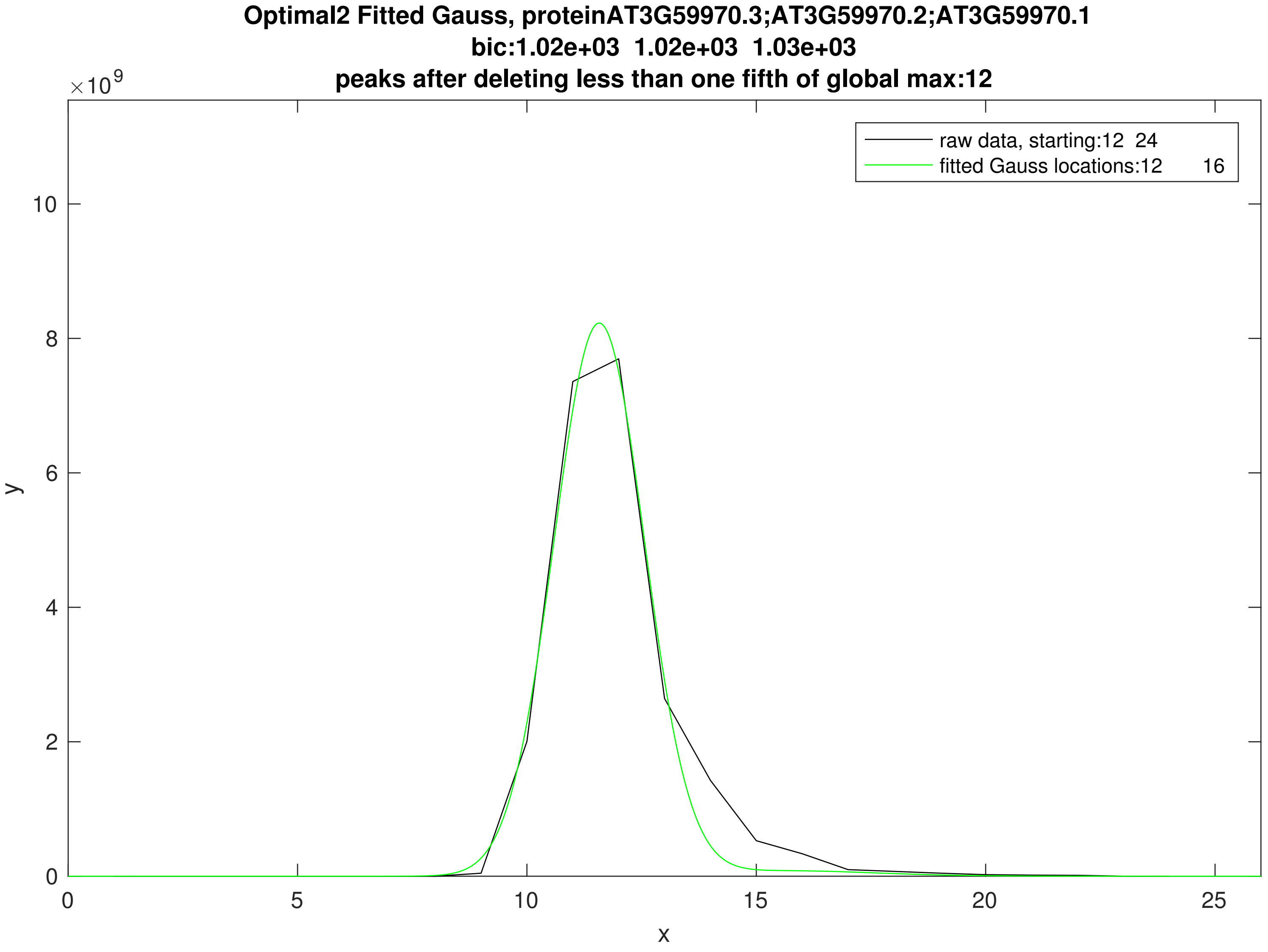
Optimal2 Fitted Gauss, proteinAT3G59970.3;AT3G59970.2;AT3G59970.1
bic:1.02e+03 1.02e+03 1.03e+03
peaks after deleting less than one fifth of global max:12
×10^9
raw data, starting:12 24
fitted Gauss locations:12 16
10
8
6
4
2
0
y
0
5
10
15
20
25
x

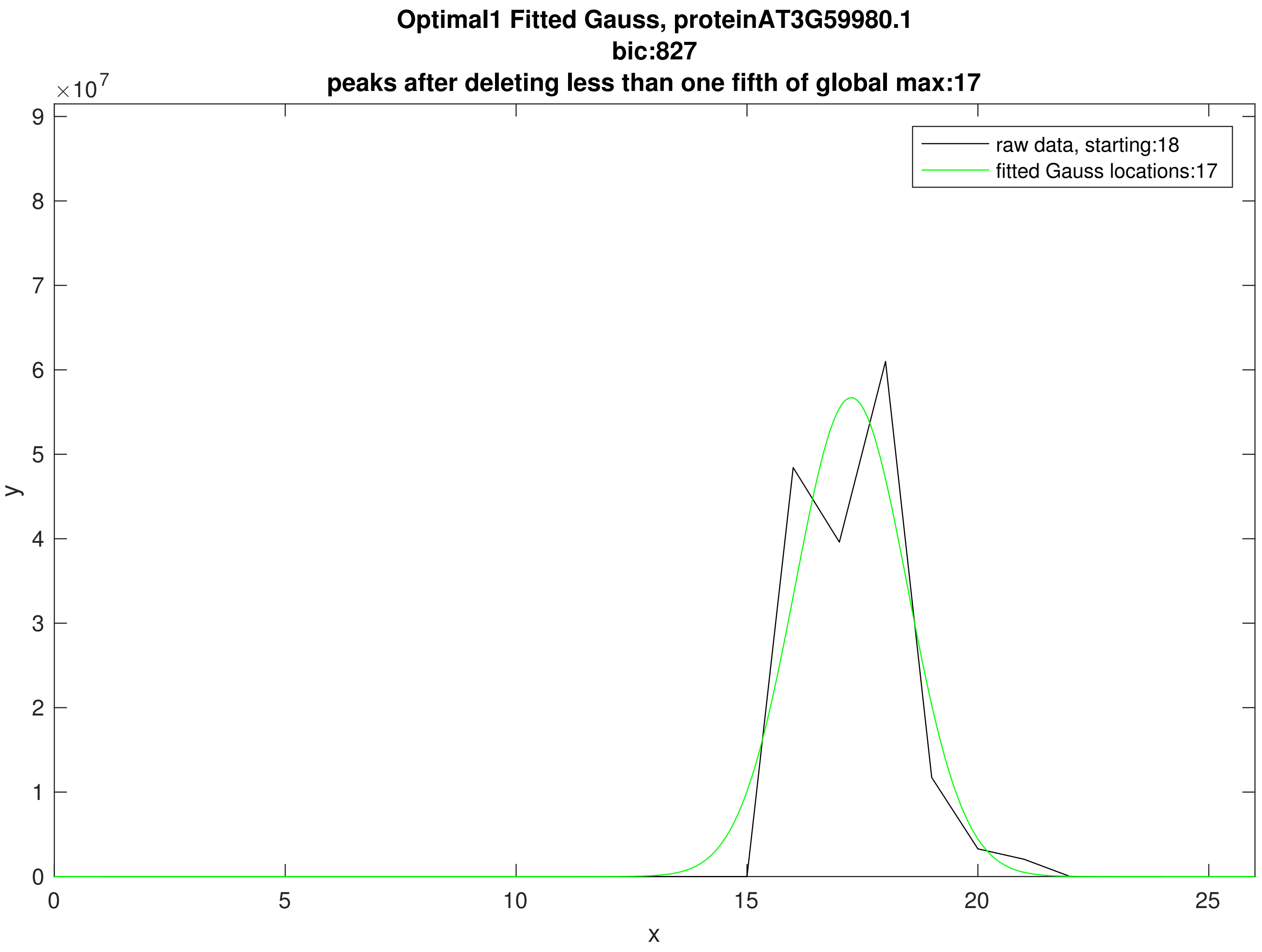

Optimal1 Fitted Gauss, proteinAT3G59980.1
bic:827
peaks after deleting less than one fifth of global max:17
×10^7
raw data, starting:18
fitted Gauss locations:17
y
x
0
1
2
3
4
5
6
7
8
9
0
5
10
15
20
25

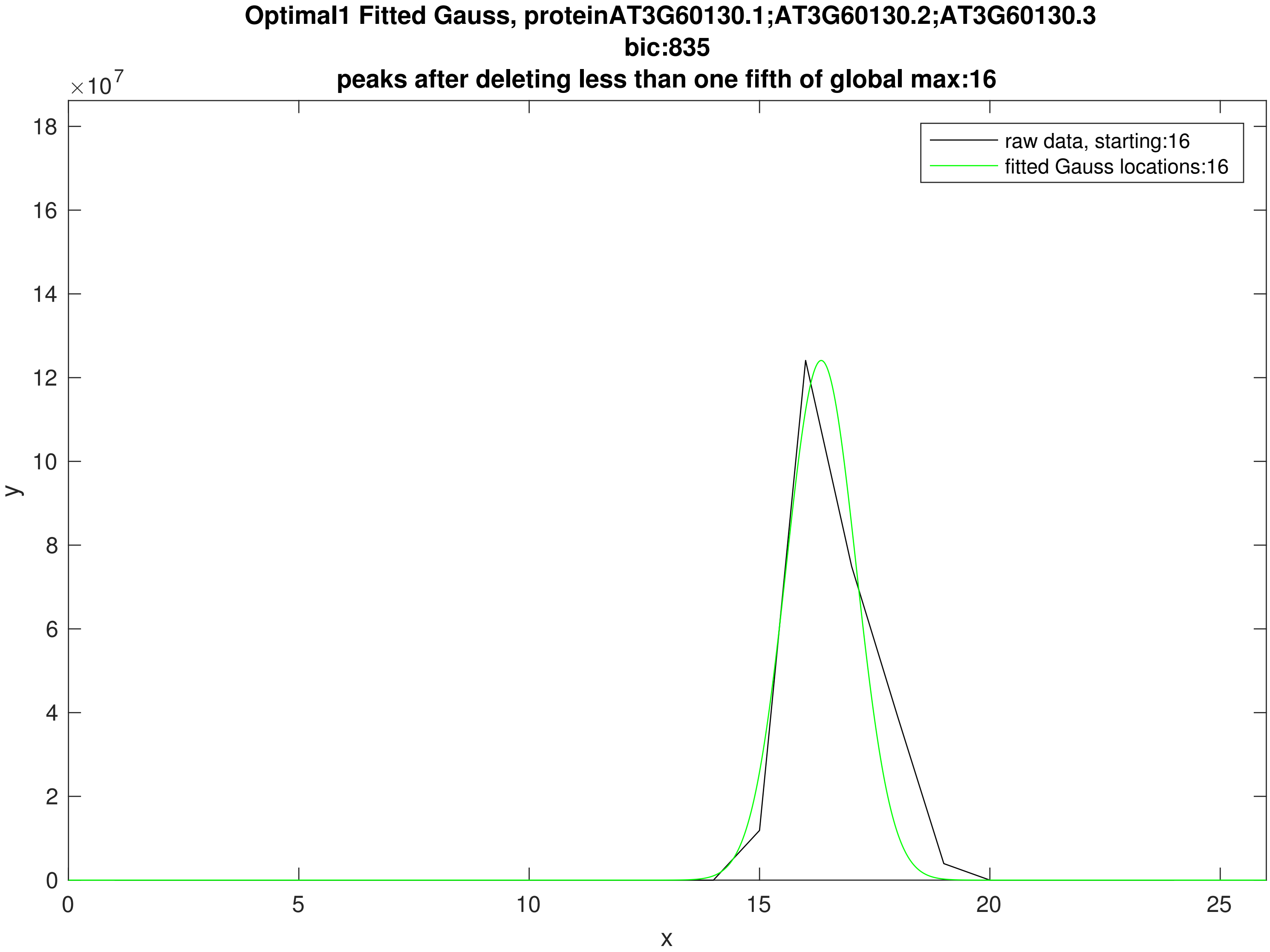
Optimal1 Fitted Gauss, proteinAT3G60130.1;AT3G60130.2;AT3G60130.3
bic:835
peaks after deleting less than one fifth of global max:16
×10^7
y
x
raw data, starting:16
fitted Gauss locations:16
0
2
4
6
8
10
12
14
16
18
0
5
10
15
20
25

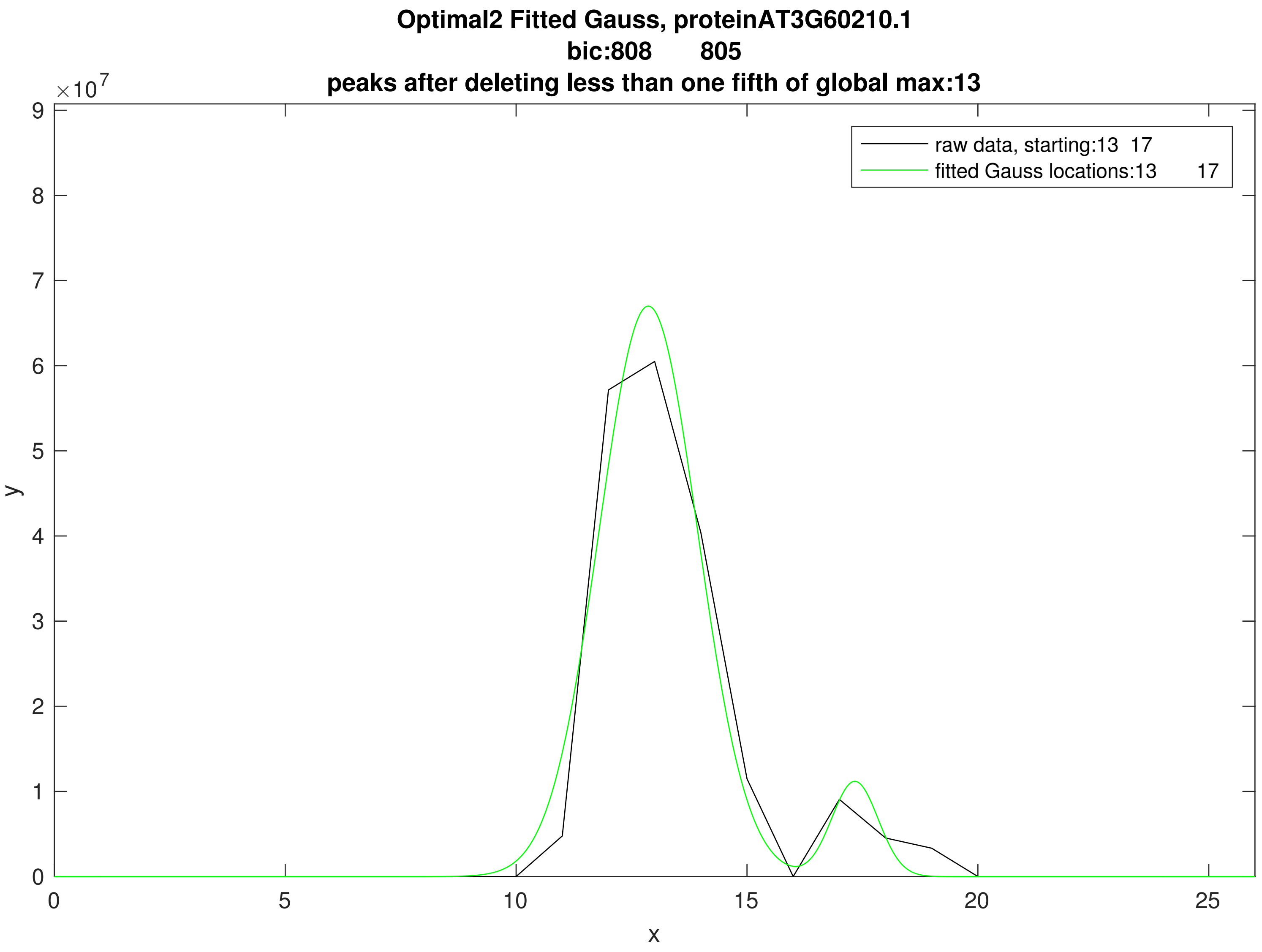
Optimal2 Fitted Gauss, proteinAT3G60210.1
bic:808 805
peaks after deleting less than one fifth of global max:13
×10^7
9
8
7
6
5
4
3
2
1
0
y
0
5
10
15
20
25
x
raw data, starting:13 17
fitted Gauss locations:13 17

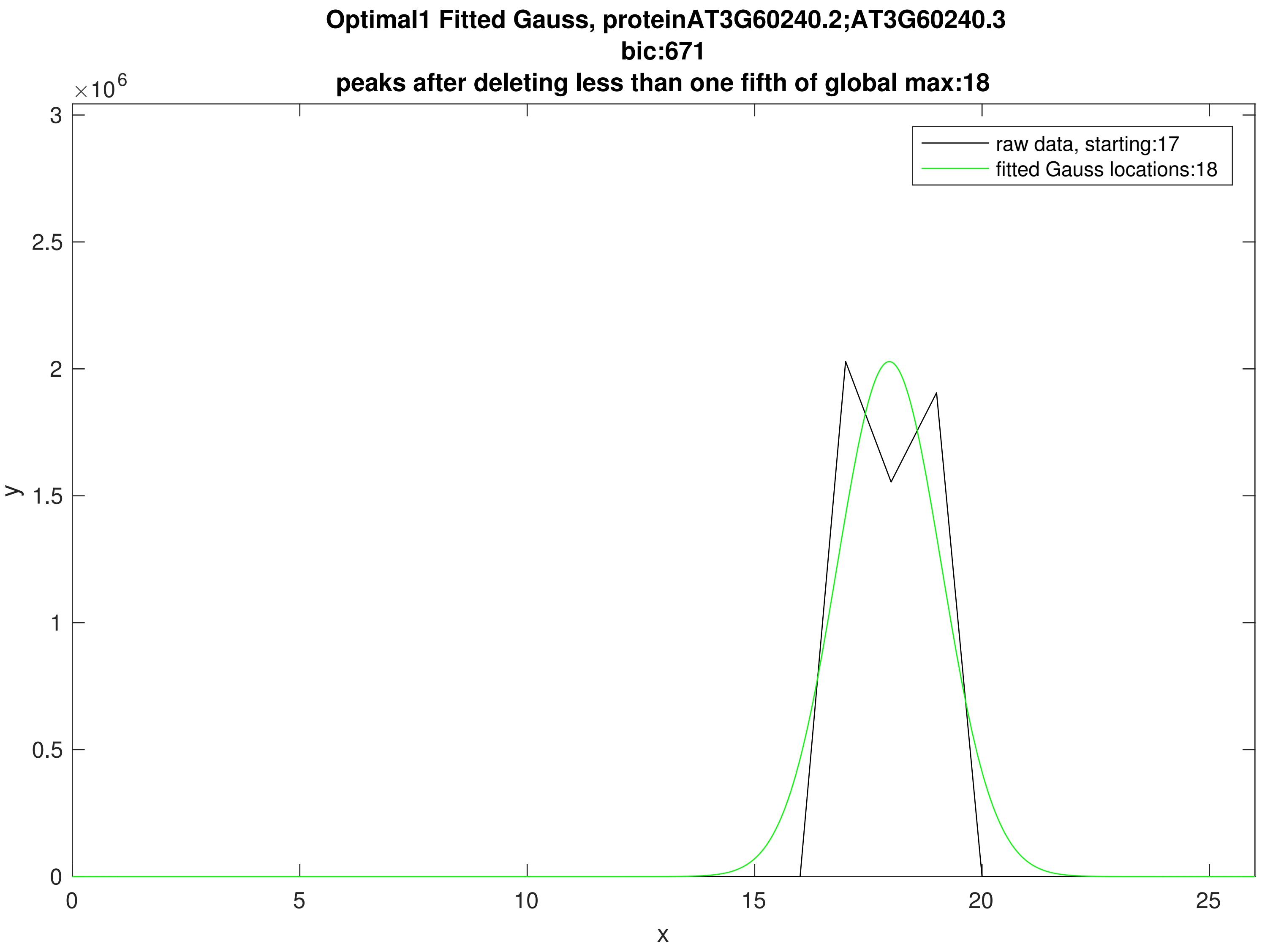
Optimal1 Fitted Gauss, proteinAT3G60240.2;AT3G60240.3
bic:671
peaks after deleting less than one fifth of global max:18
×10^6
raw data, starting:17
fitted Gauss locations:18
y
x
0
0.5
1
1.5
2
2.5
3
0
5
10
15
20
25

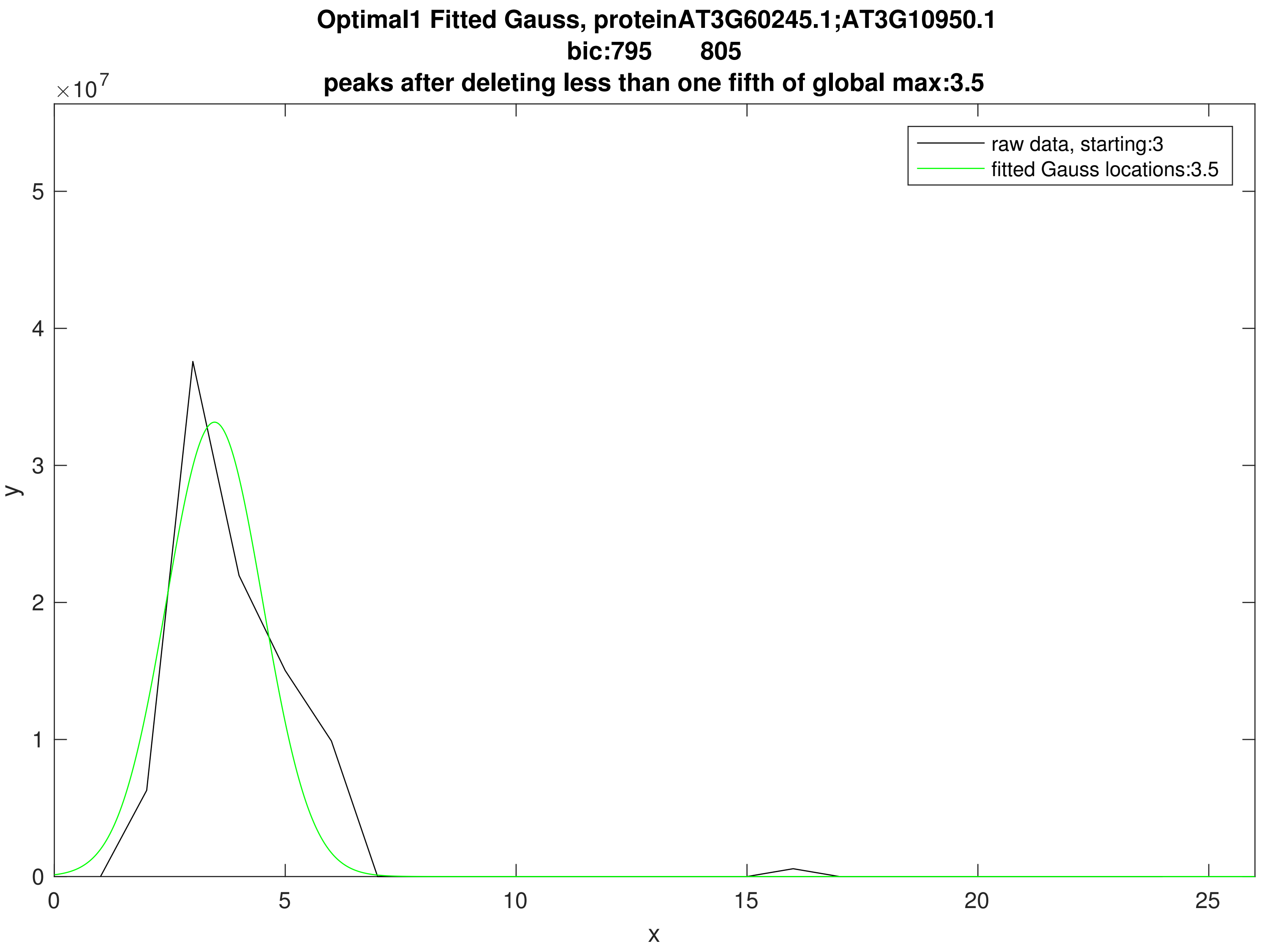
Optimal1 Fitted Gauss, proteinAT3G60245.1;AT3G10950.1
bic:795 805
peaks after deleting less than one fifth of global max:3.5
×10^7
raw data, starting:3
fitted Gauss locations:3.5
y
x
0
1
2
3
4
5
0
5
10
15
20
25

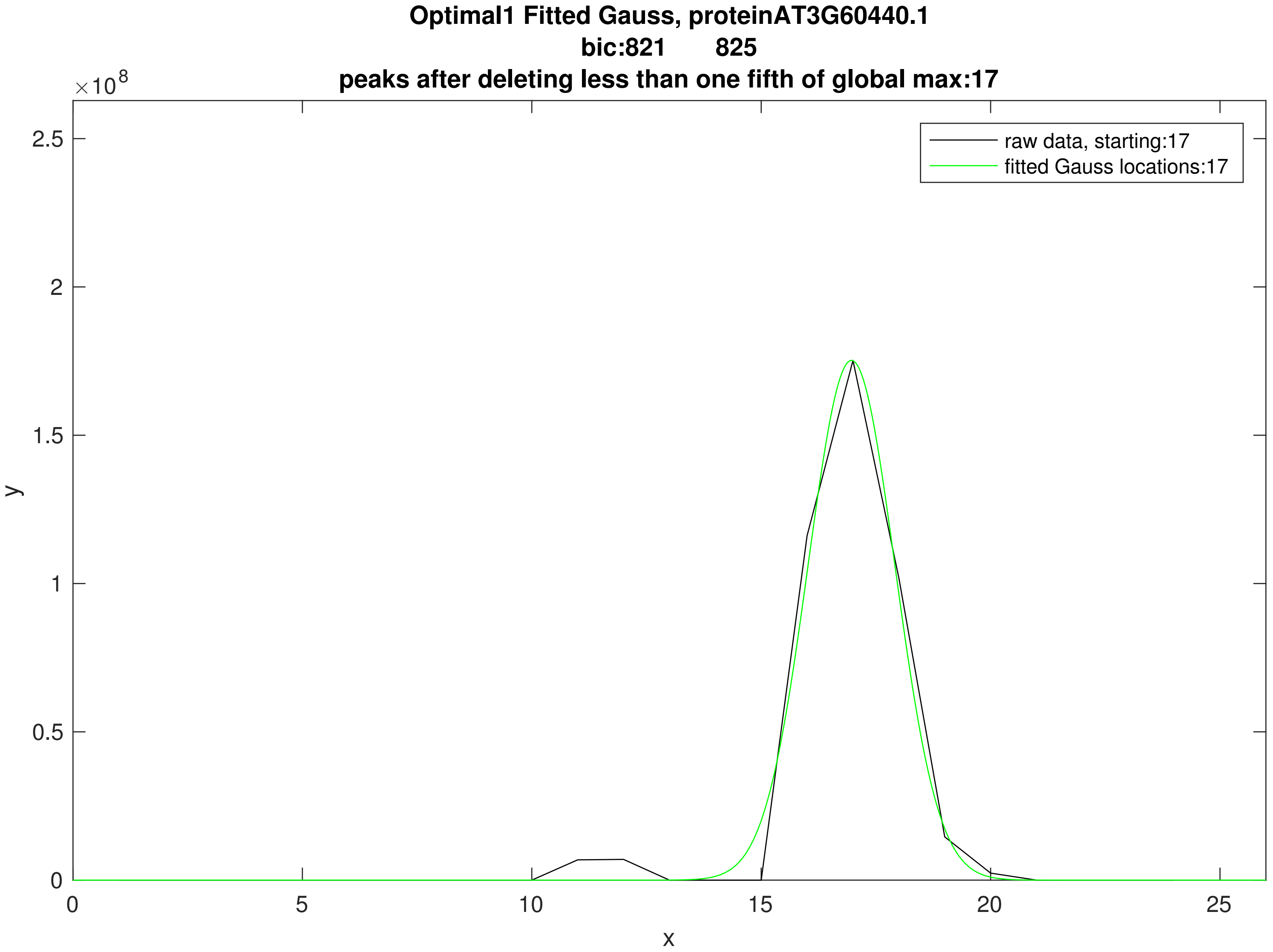
Optimal1 Fitted Gauss, proteinAT3G60440.1
bic:821 825
peaks after deleting less than one fifth of global max:17
raw data, starting:17
fitted Gauss locations:17
×10^8
2.5
2
1.5
1
0.5
0
y
0
5
10
15
20
25
x

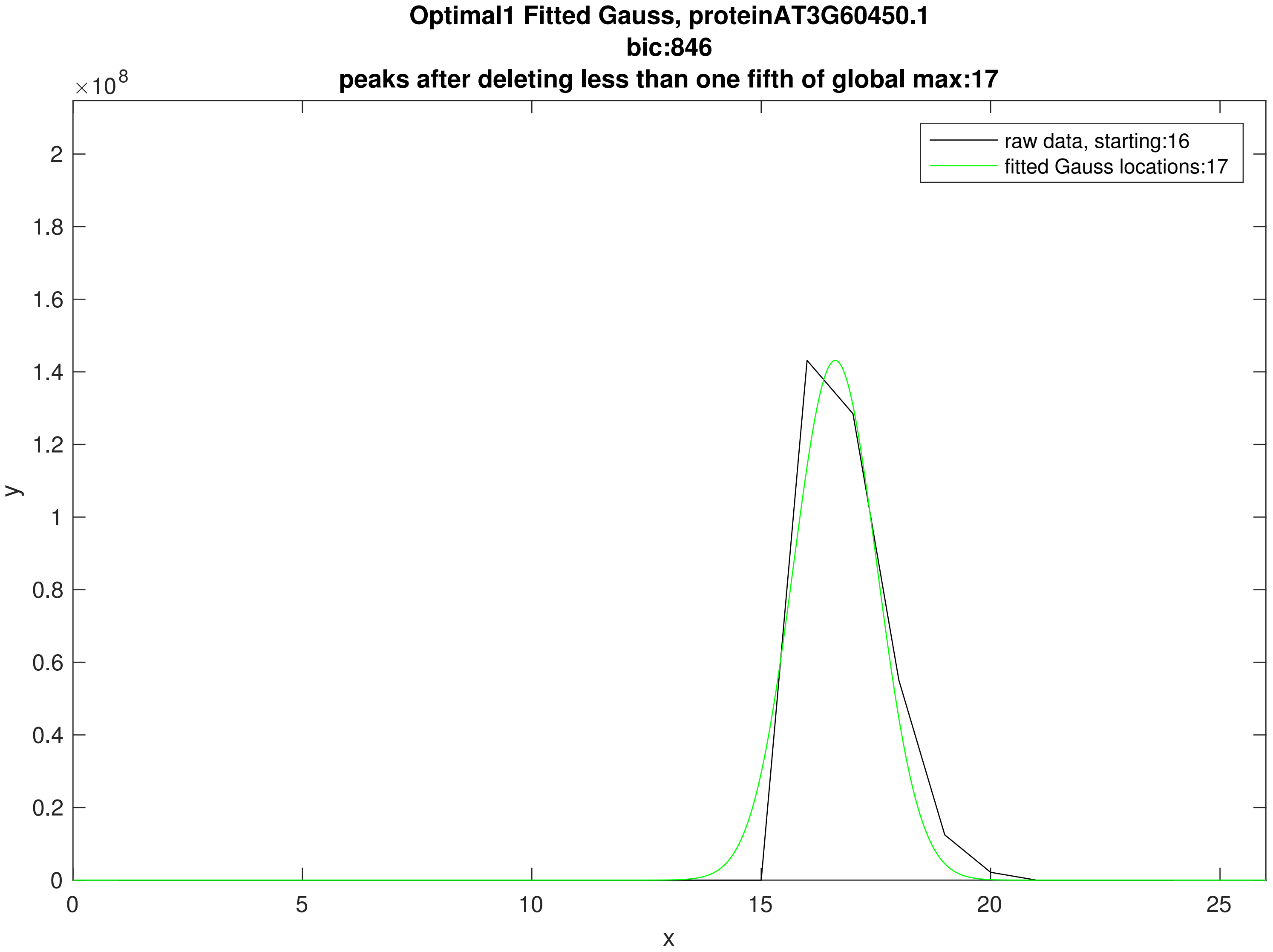
Optimal1 Fitted Gauss, proteinAT3G60450.1
bic:846
peaks after deleting less than one fifth of global max:17
×10^8
raw data, starting:16
fitted Gauss locations:17
y
x
0
5
10
15
20
25
0
0.2
0.4
0.6
0.8
1
1.2
1.4
1.6
1.8
2

**Optimal1 Fitted Gauss, proteinAT3G60750.1;AT3G60750.2**
**bic:1.01e+03  1.02e+03  1.03e+03**
**peaks after deleting less than one fifth of global max:12**

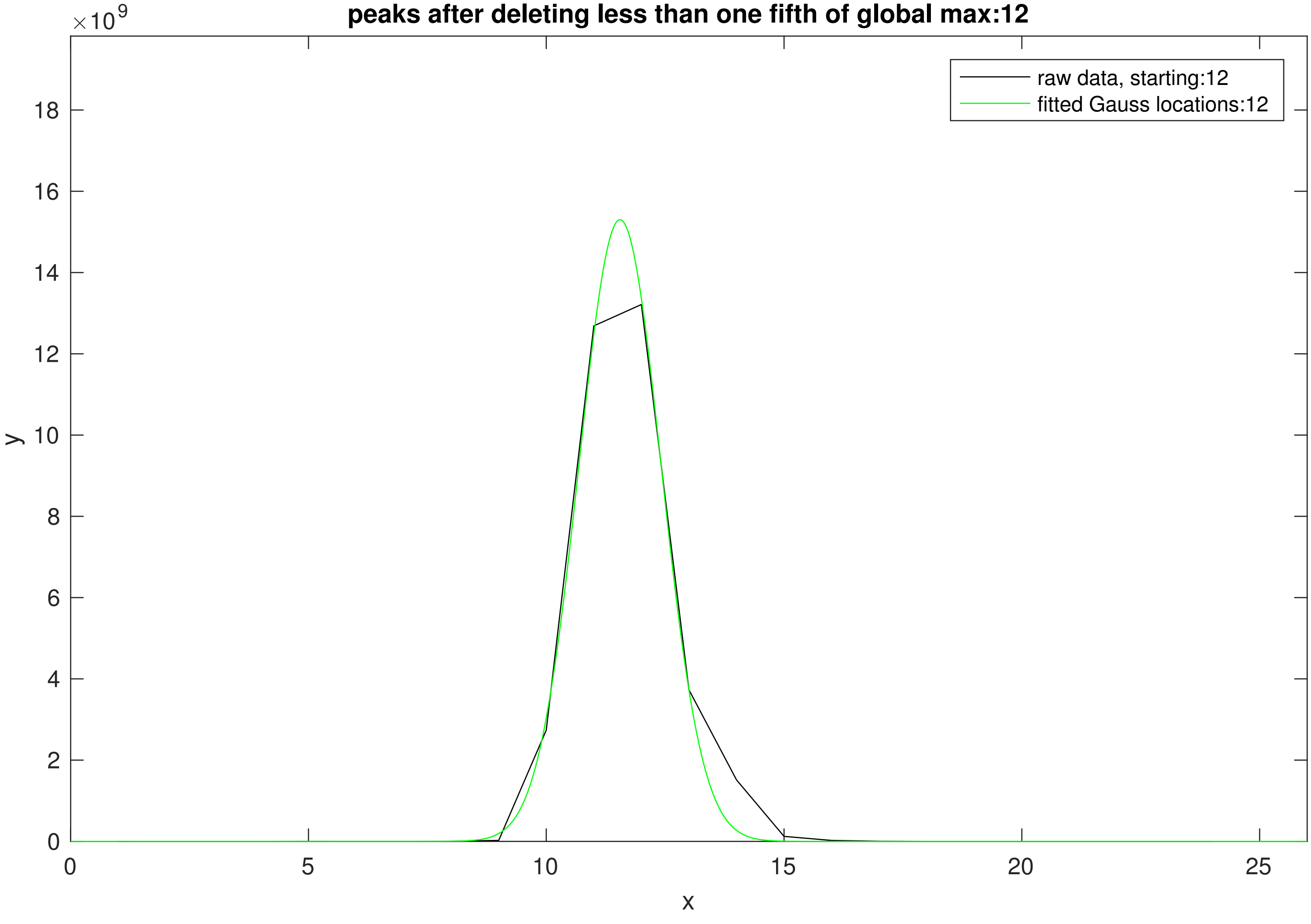

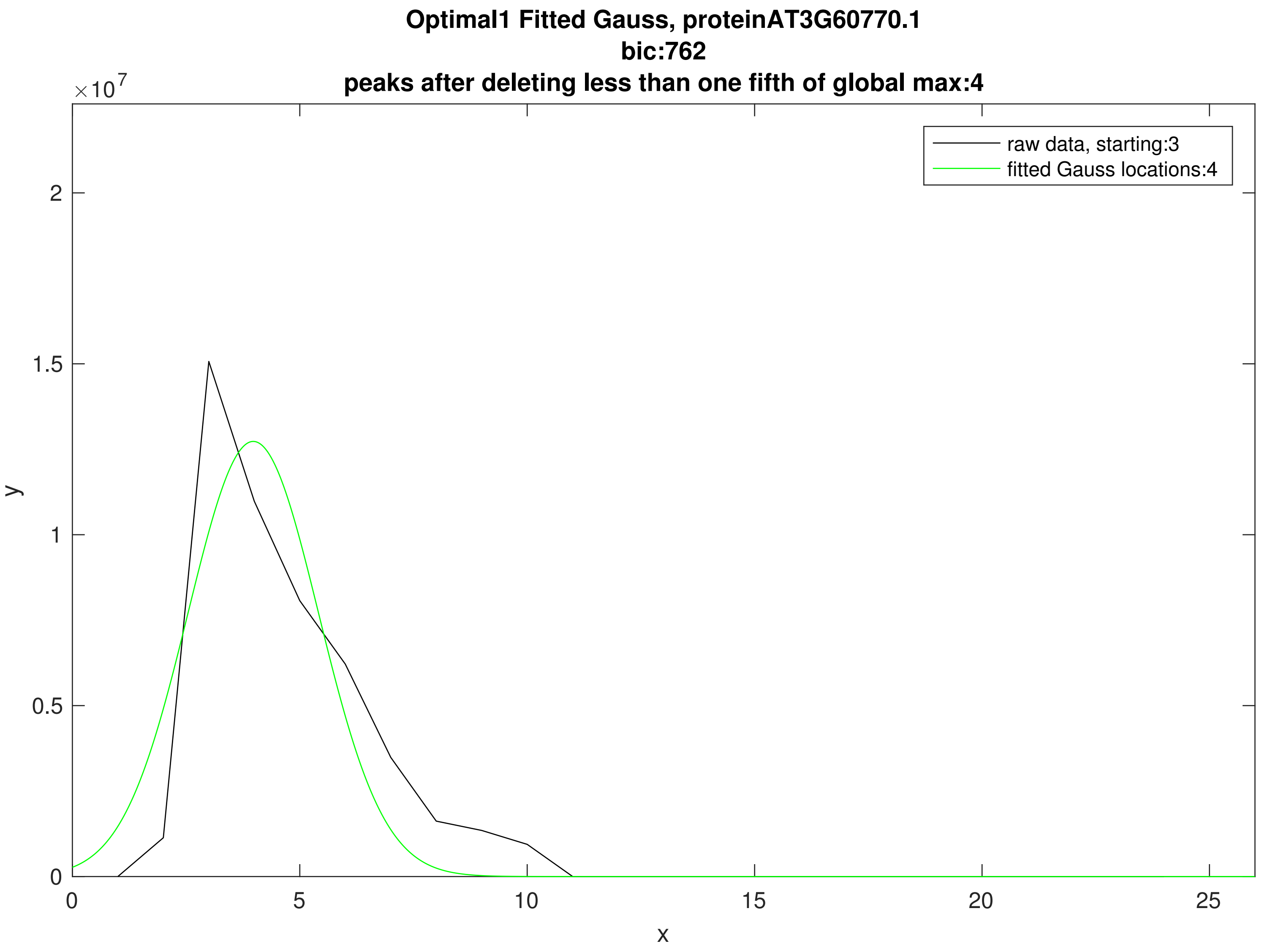

Optimal1 Fitted Gauss, proteinAT3G60770.1
bic:762
peaks after deleting less than one fifth of global max:4
×10^7
raw data, starting:3
fitted Gauss locations:4
0
0.5
1
1.5
2
y
0
5
10
15
20
25
x

Optimal1 Fitted Gauss, proteinAT3G60820.1;AT3G60820.3;AT3G60820.2
bic:921 931 940
peaks after deleting less than one fifth of global max:5

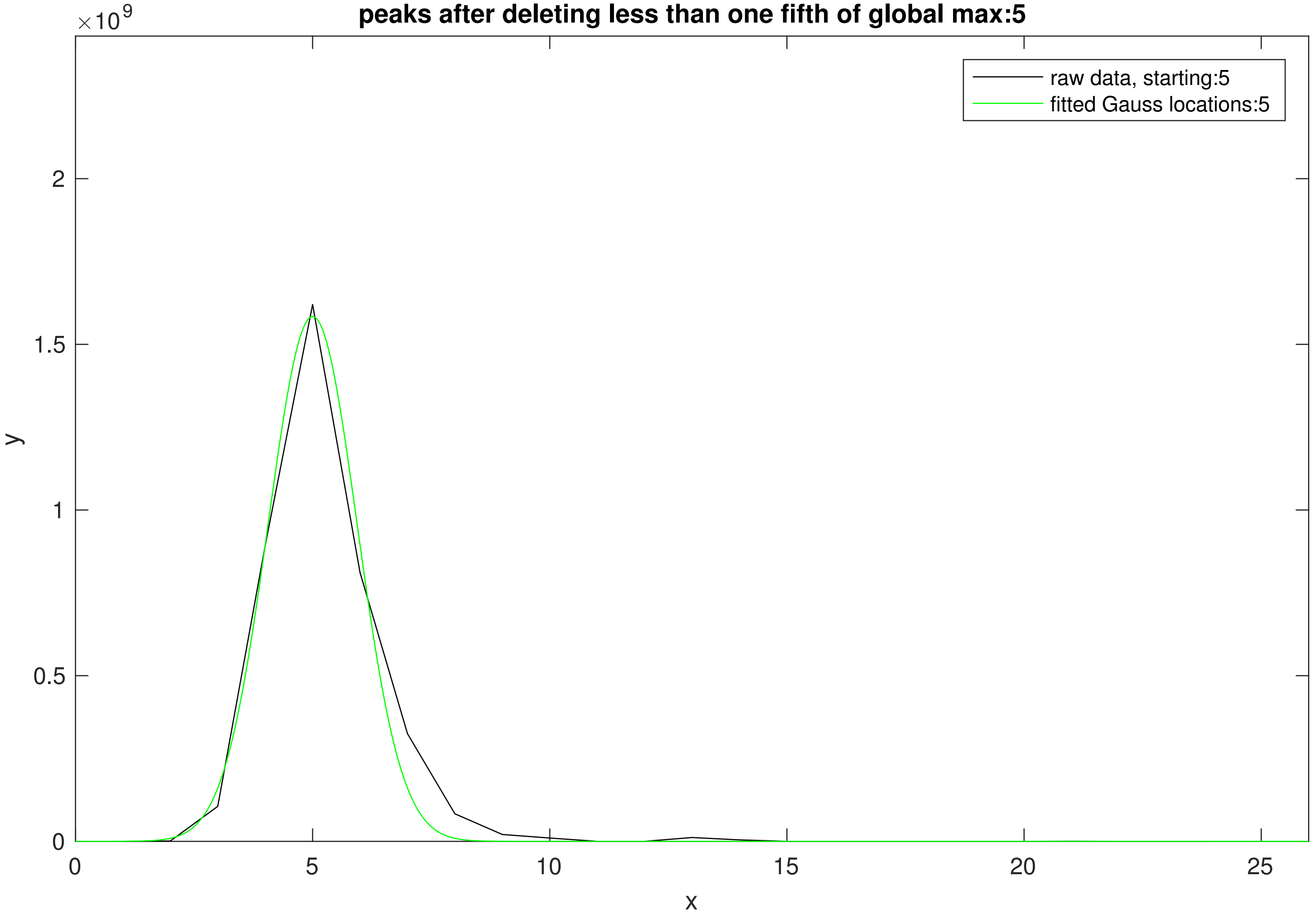

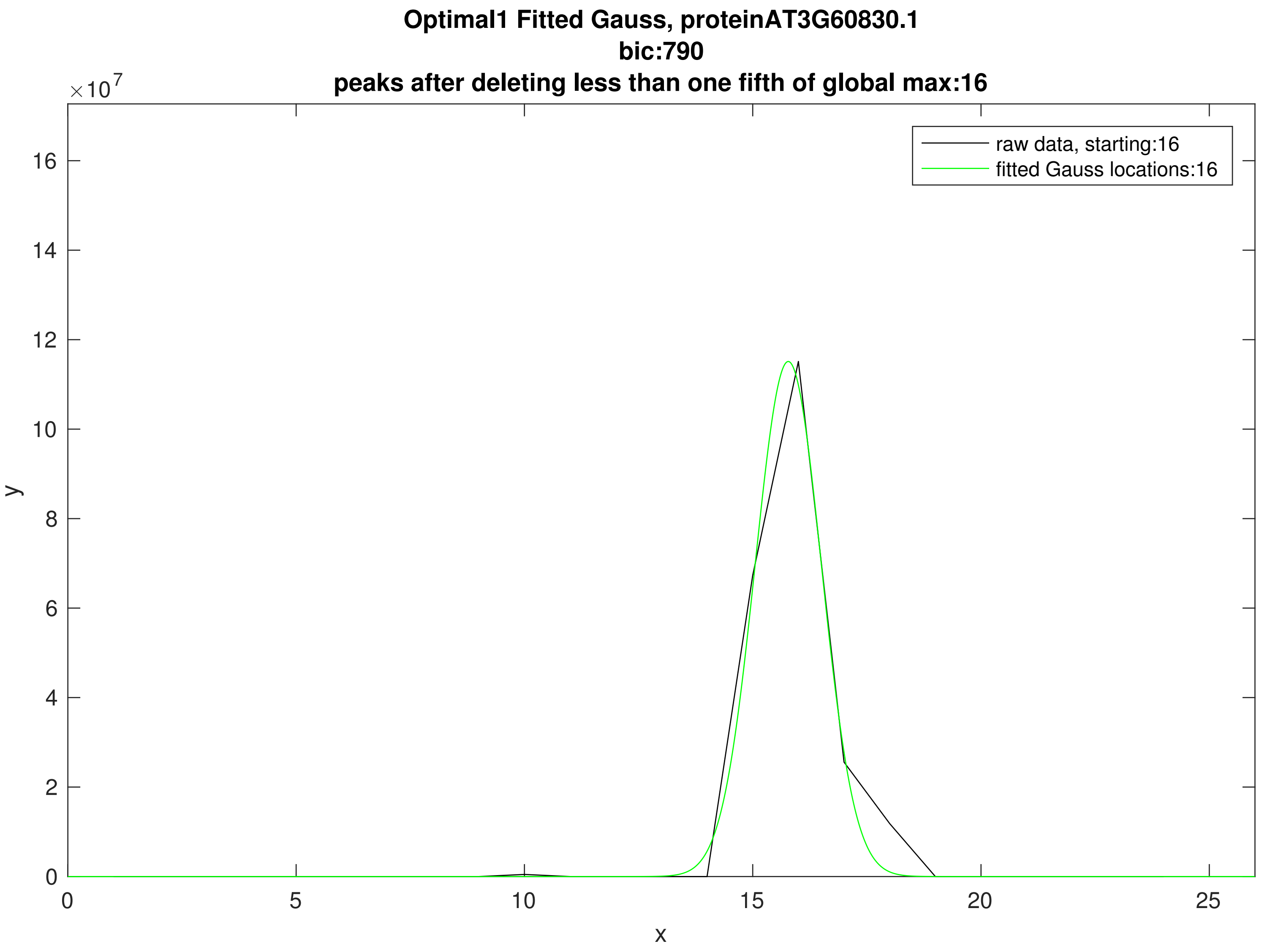
Optimal1 Fitted Gauss, proteinAT3G60830.1
bic:790
peaks after deleting less than one fifth of global max:16
×10^7
raw data, starting:16
fitted Gauss locations:16
y
x
0
2
4
6
8
10
12
14
16
0
5
10
15
20
25

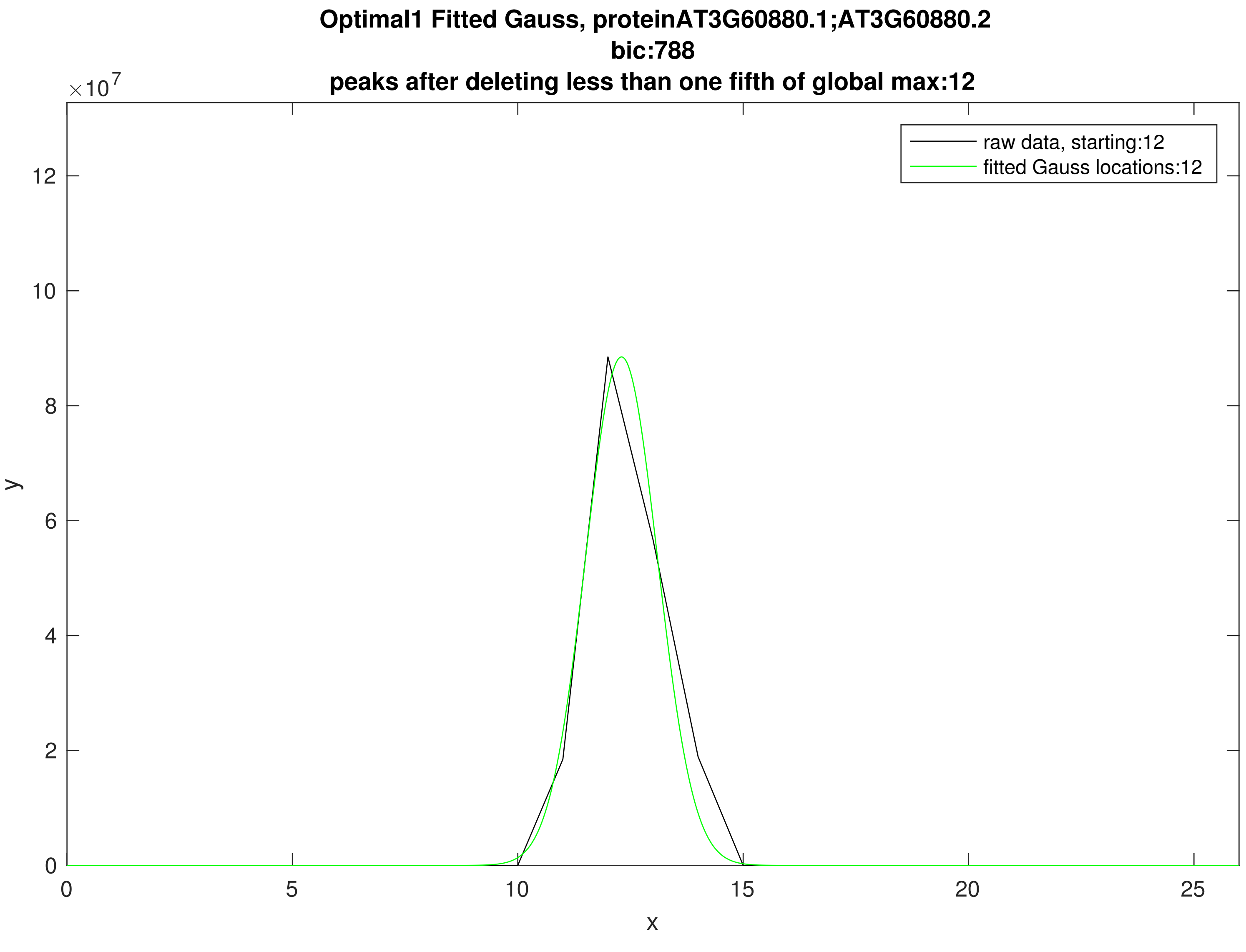
Optimal1 Fitted Gauss, proteinAT3G60880.1;AT3G60880.2
bic:788
peaks after deleting less than one fifth of global max:12
×10^7
12
10
8
6
4
2
0
y
0
5
10
15
20
25
x
raw data, starting:12
fitted Gauss locations:12

**Optimal1 Fitted Gauss, proteinAT3G61220.1;AT3G61220.2**
**bic:879 887 897**
**peaks after deleting less than one fifth of global max:13**

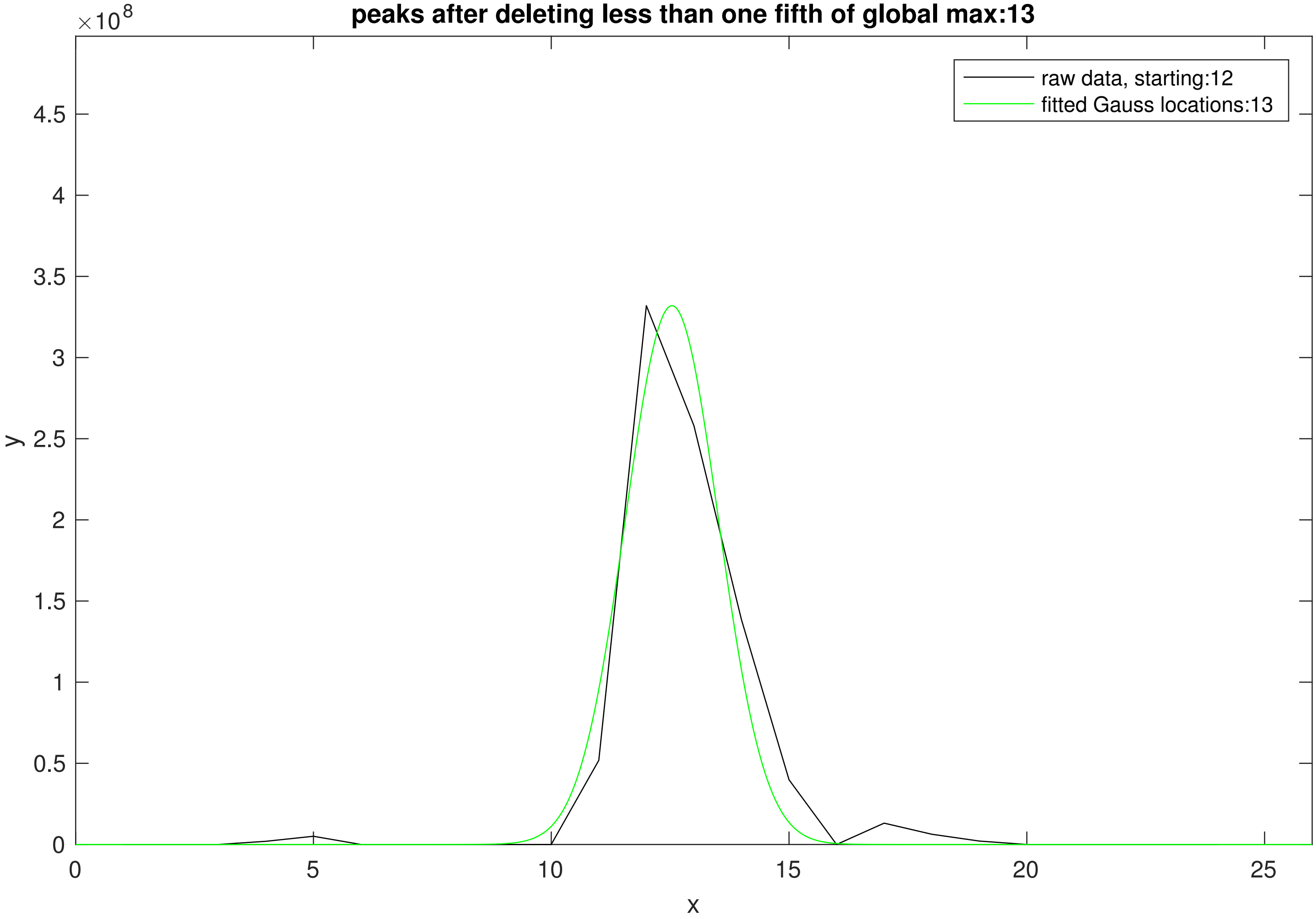

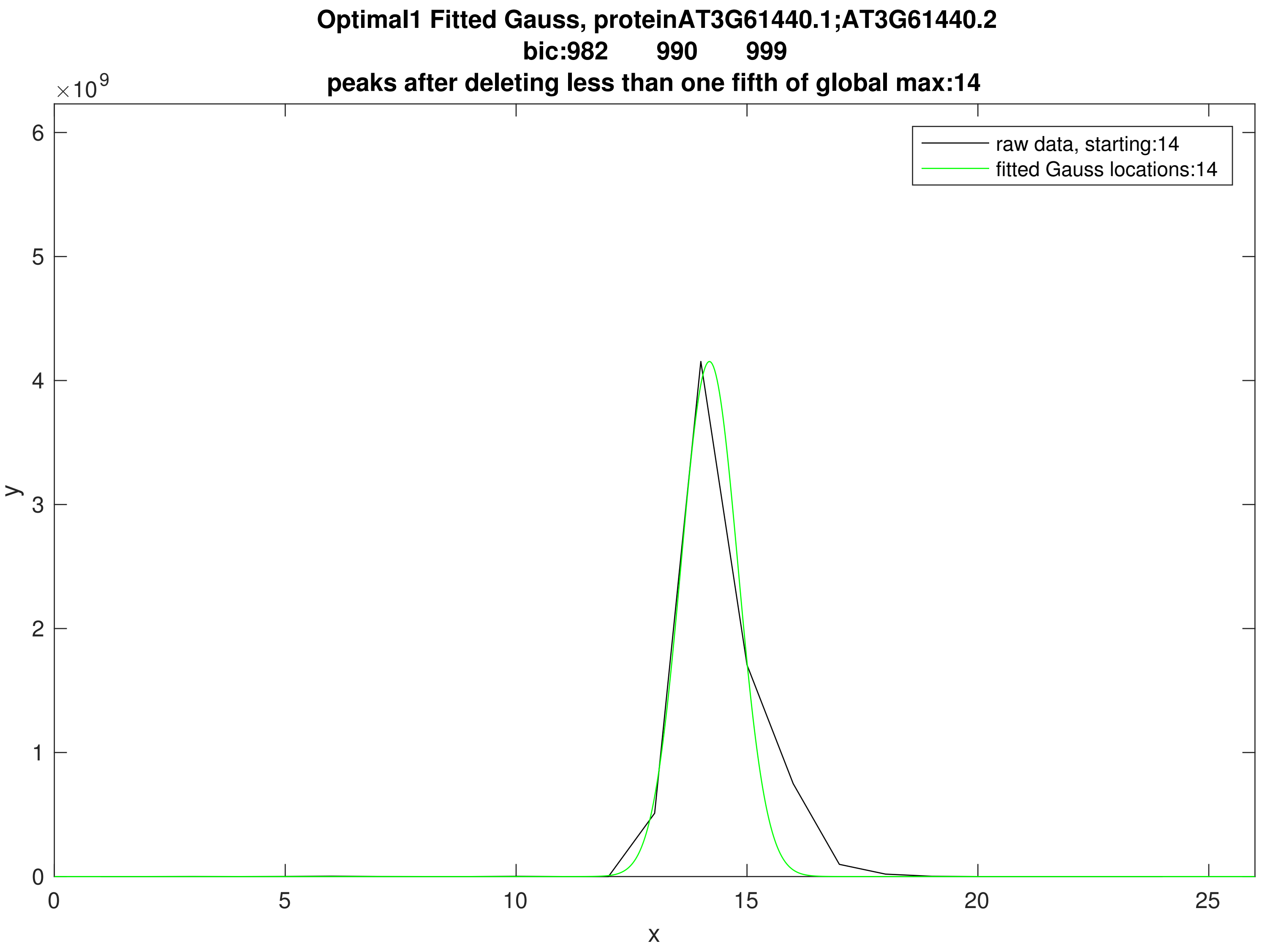

Optimal1 Fitted Gauss, proteinAT3G61440.1;AT3G61440.2
bic:982 990 999
peaks after deleting less than one fifth of global max:14
×10^9
raw data, starting:14
fitted Gauss locations:14
y
x
0
1
2
3
4
5
6
0
5
10
15
20
25

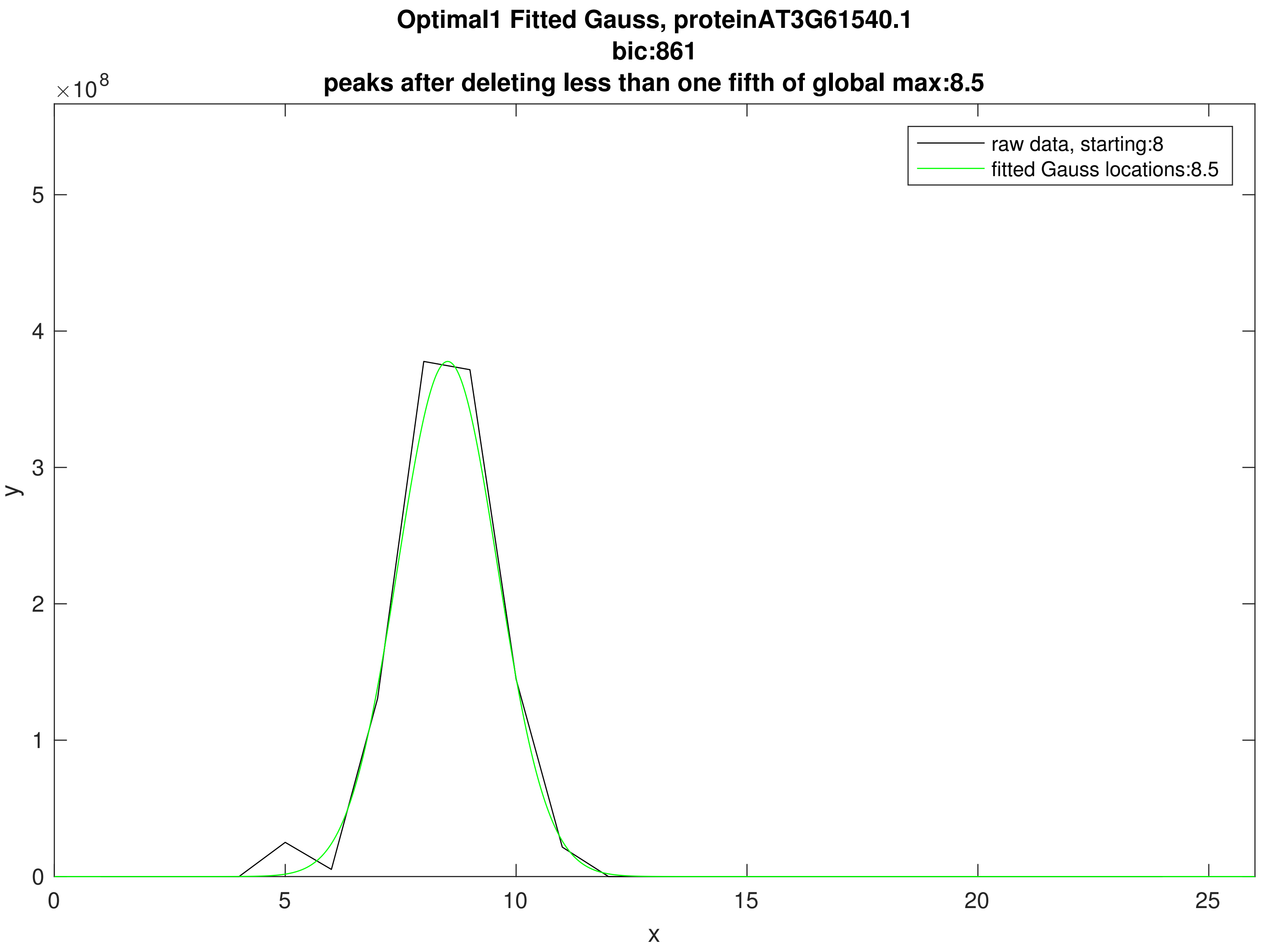

Optimal1 Fitted Gauss, proteinAT3G61540.1
bic:861
peaks after deleting less than one fifth of global max:8.5
×10^8
raw data, starting:8
fitted Gauss locations:8.5
y
x
0
1
2
3
4
5
0
5
10
15
20
25

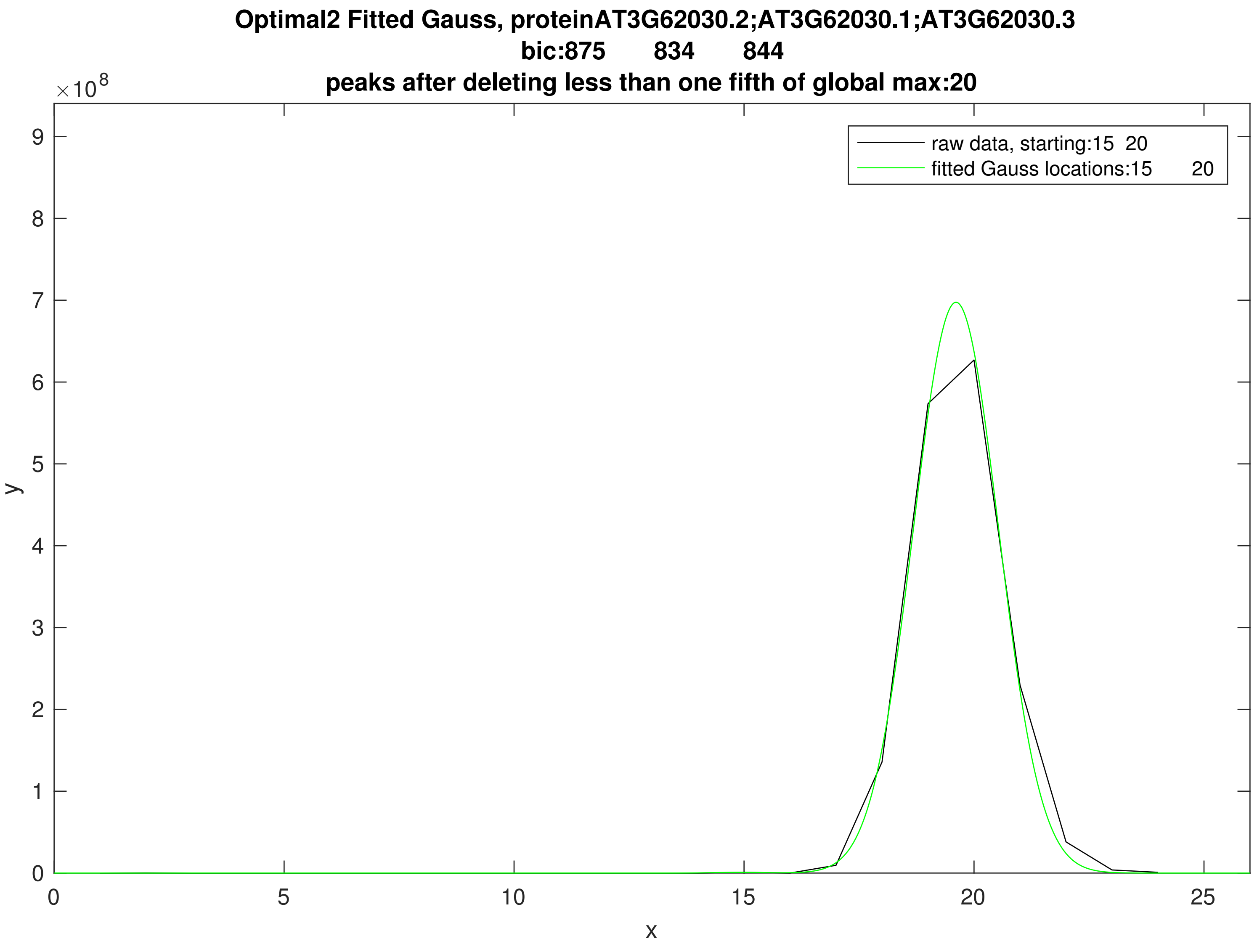
Optimal2 Fitted Gauss, proteinAT3G62030.2;AT3G62030.1;AT3G62030.3
bic:875 834 844
peaks after deleting less than one fifth of global max:20
×10^8
raw data, starting:15 20
fitted Gauss locations:15 20
y
x
0
1
2
3
4
5
6
7
8
9
0
5
10
15
20
25

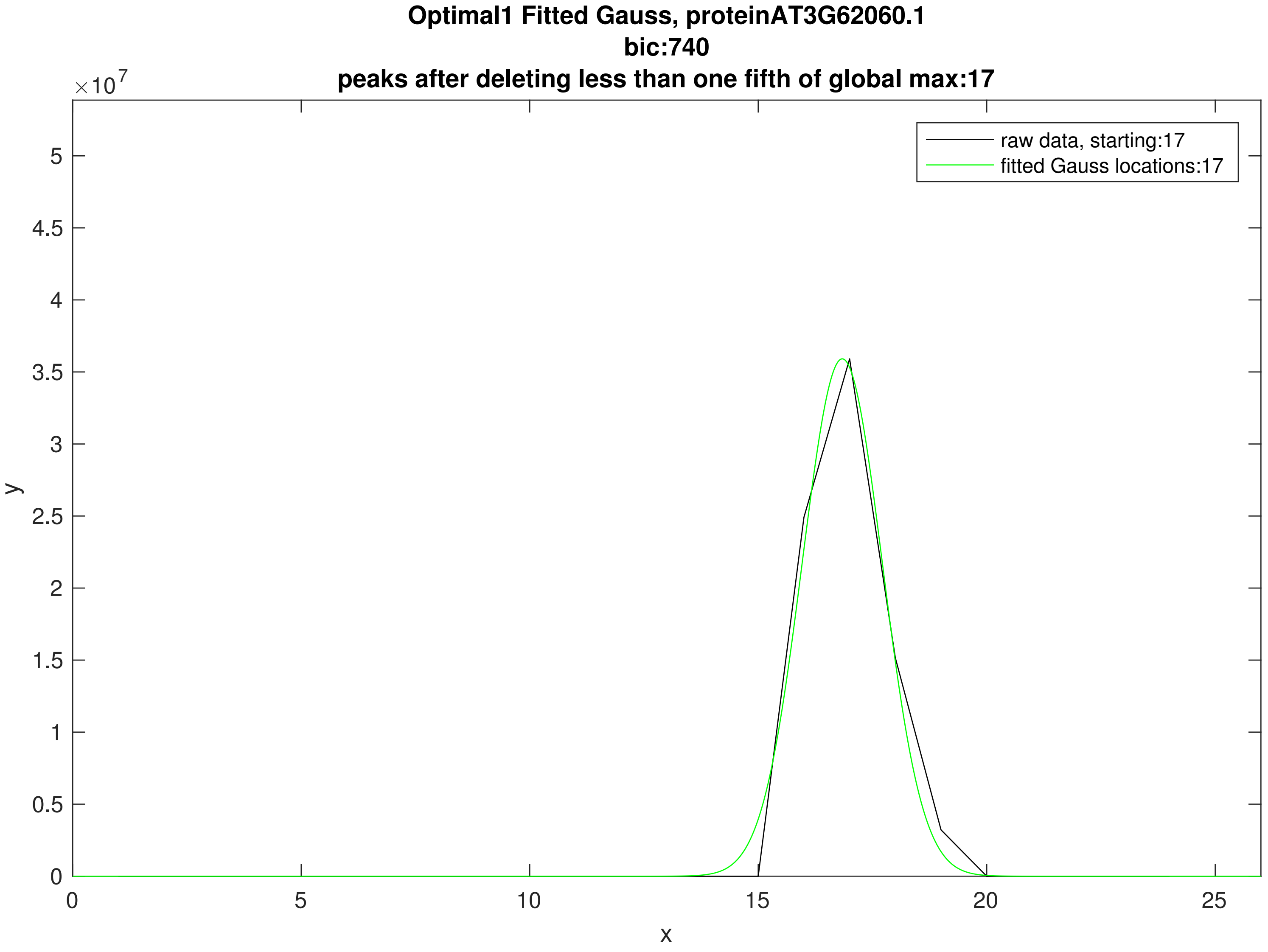
Optimal1 Fitted Gauss, proteinAT3G62060.1
bic:740
peaks after deleting less than one fifth of global max:17
×10^7
raw data, starting:17
fitted Gauss locations:17
y
x
0
0.5
1
1.5
2
2.5
3
3.5
4
4.5
5
0
5
10
15
20
25

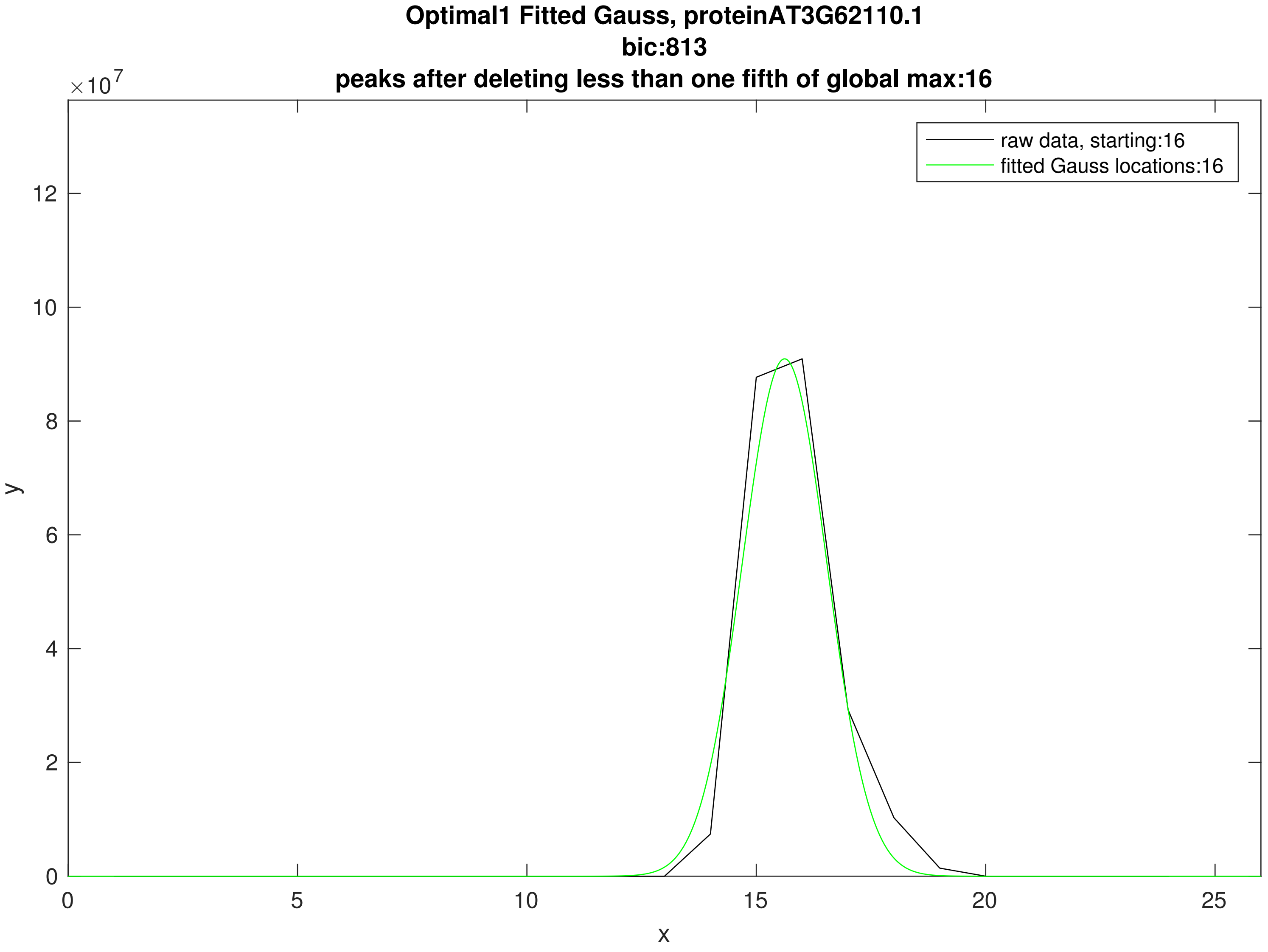

Optimal1 Fitted Gauss, proteinAT3G62110.1
bic:813
peaks after deleting less than one fifth of global max:16
×10^7
raw data, starting:16
fitted Gauss locations:16
12
10
8
6
4
2
0
y
0
5
10
15
20
25
x

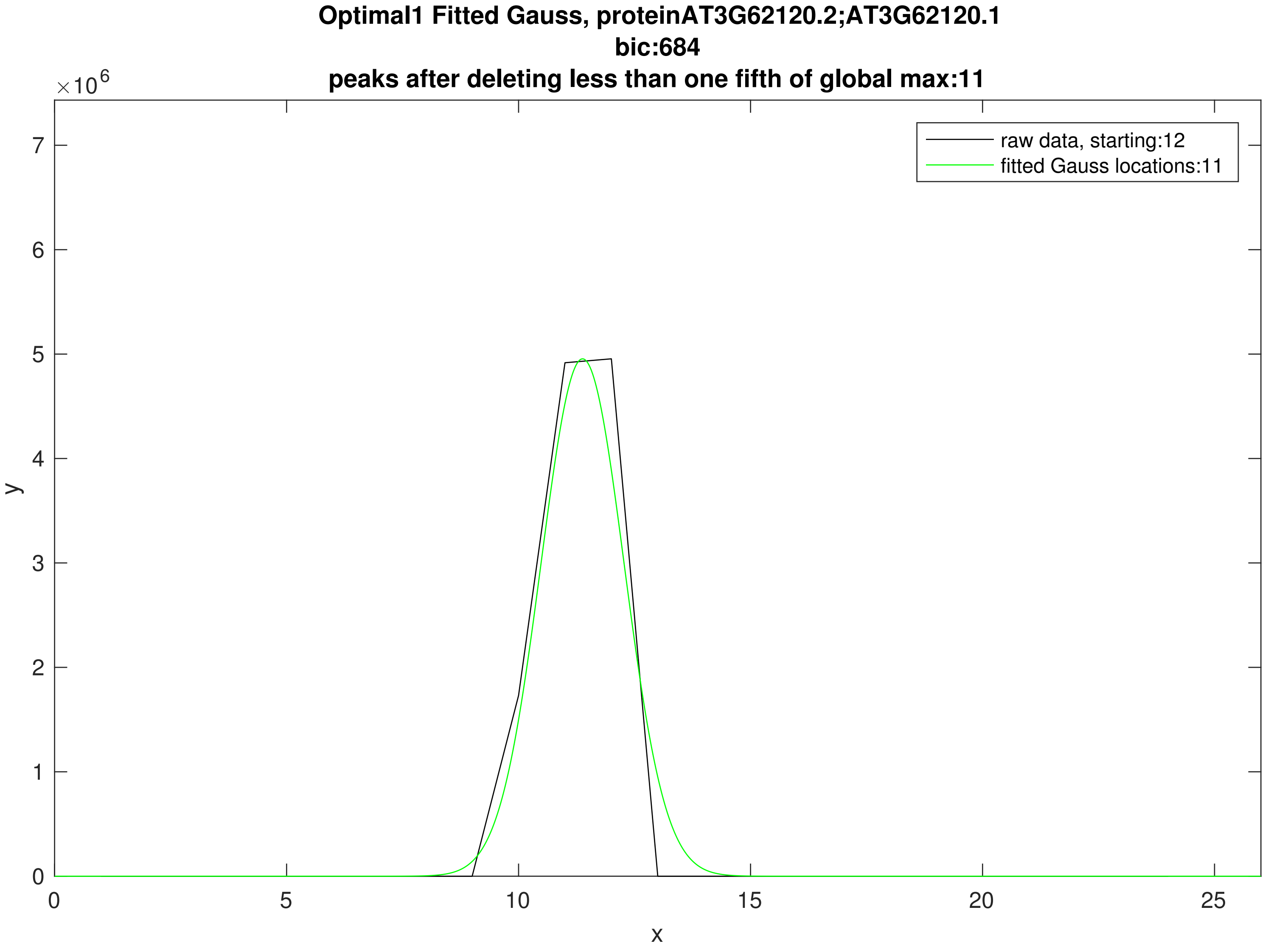

Optimal1 Fitted Gauss, proteinAT3G62120.2;AT3G62120.1
bic:684
peaks after deleting less than one fifth of global max:11
×10^6
raw data, starting:12
fitted Gauss locations:11
y
x
0
1
2
3
4
5
6
7
0
5
10
15
20
25

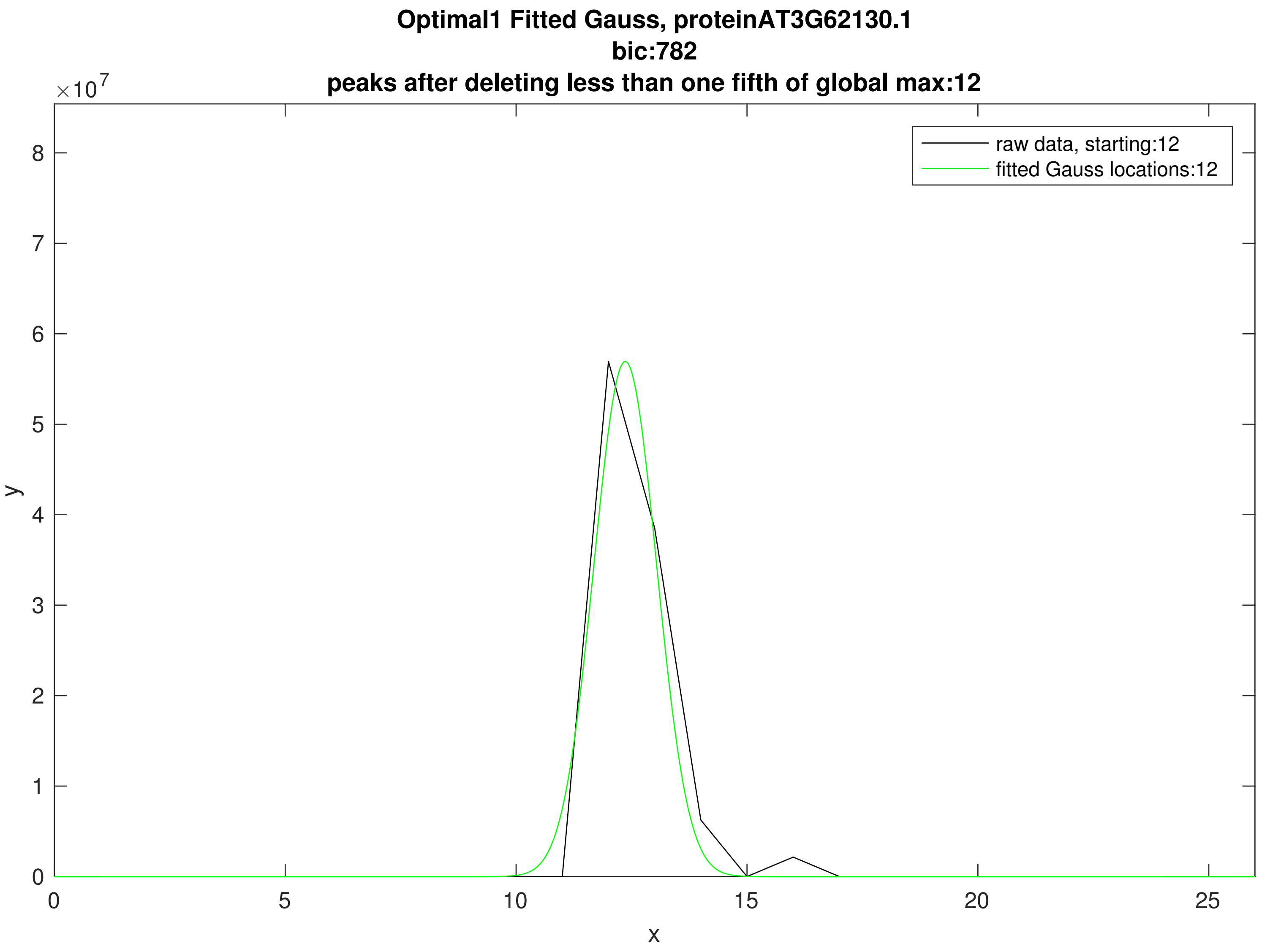

Optimal1 Fitted Gauss, proteinAT3G62130.1
bic:782
peaks after deleting less than one fifth of global max:12
×10^7
raw data, starting:12
fitted Gauss locations:12
8
7
6
5
4
3
2
1
0
y
0
5
10
15
20
25
x

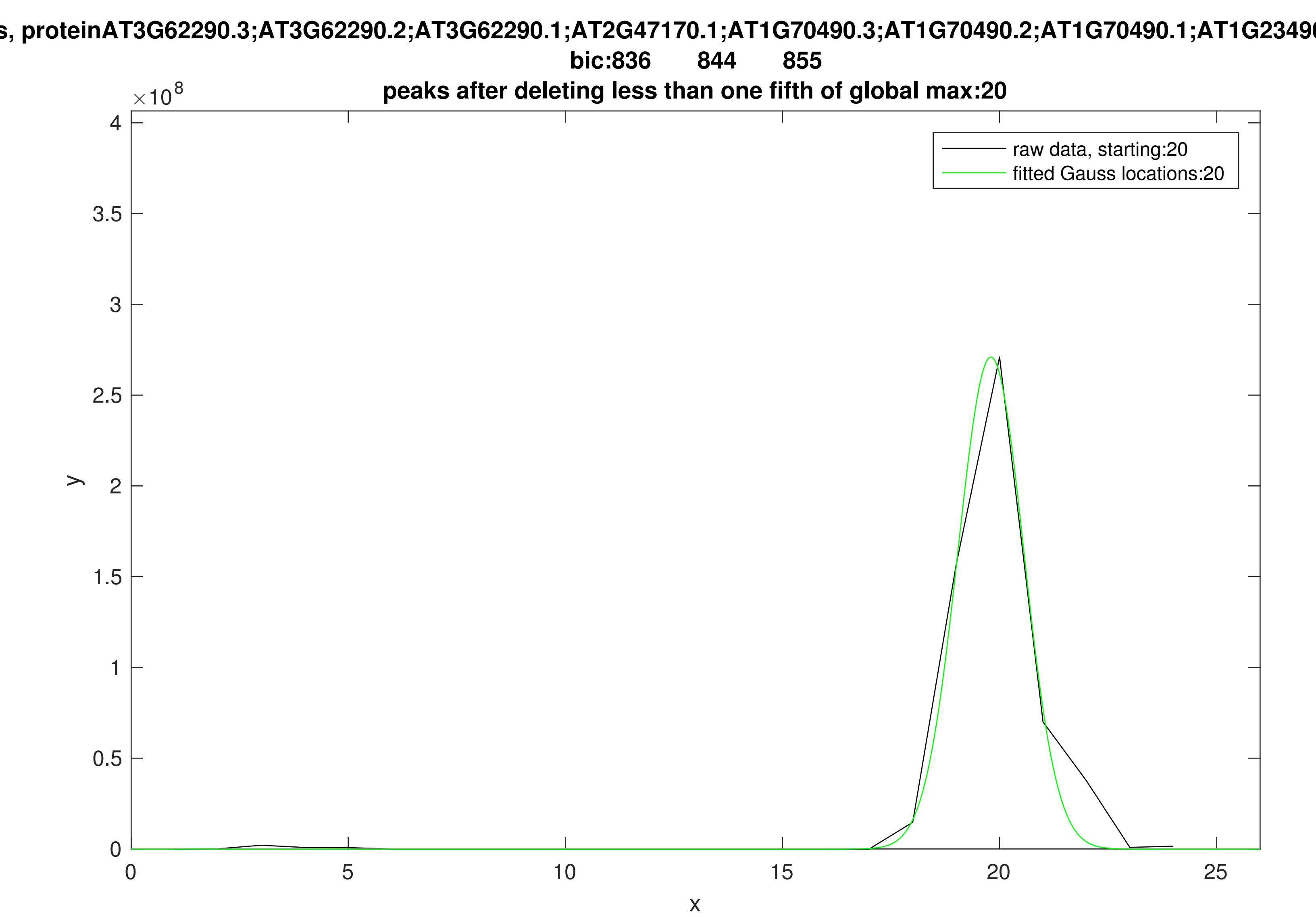
s, proteinAT3G62290.3;AT3G62290.2;AT3G62290.1;AT2G47170.1;AT1G70490.3;AT1G70490.2;AT1G70490.1;AT1G2349
bic:836 844 855
peaks after deleting less than one fifth of global max:20
×10^8
4
3.5
3
2.5
2
1.5
1
0.5
0
y
0
5
10
15
20
25
x
raw data, starting:20
fitted Gauss locations:20

Optimal1 Fitted Gauss, proteinAT3G62360.1
bic:642
peaks after deleting less than one fifth of global max:19

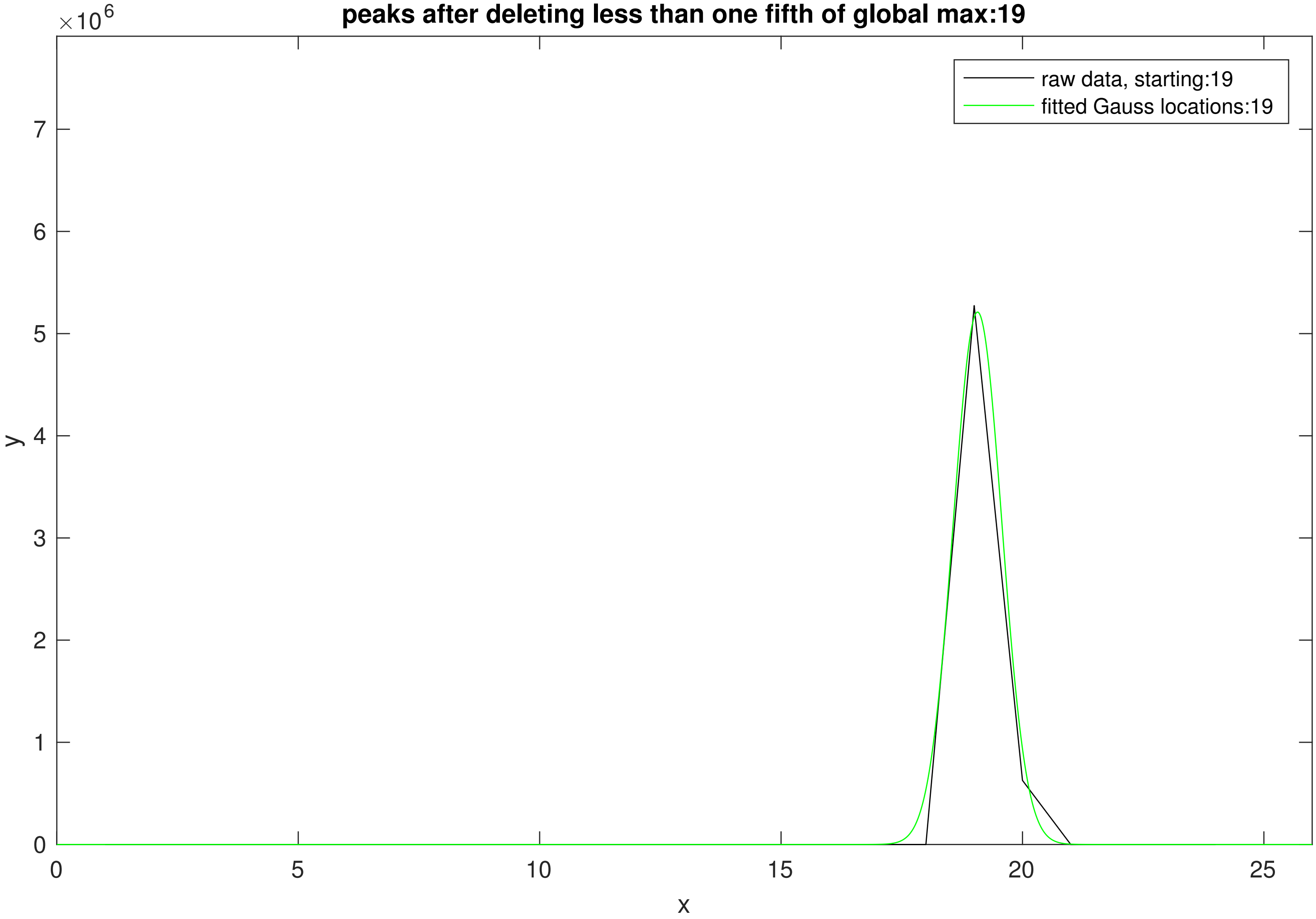

**Optimal1 Fitted Gauss, proteinAT3G62530.1**
**bic:809**
**peaks after deleting less than one fifth of global max:18**

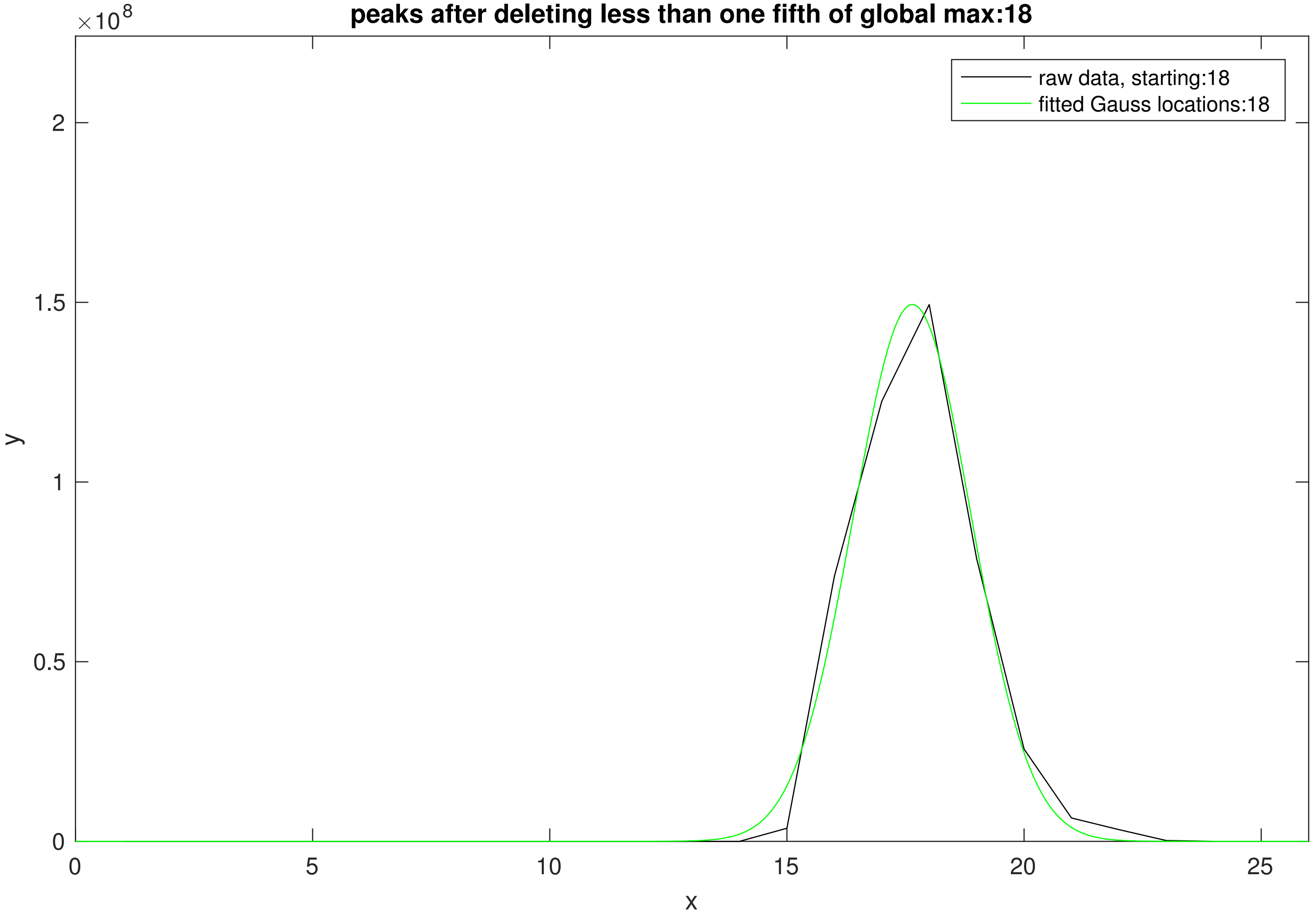

**Optimal1 Fitted Gauss, proteinAT3G62810.1**
**bic:837**
**peaks after deleting less than one fifth of global max:17**

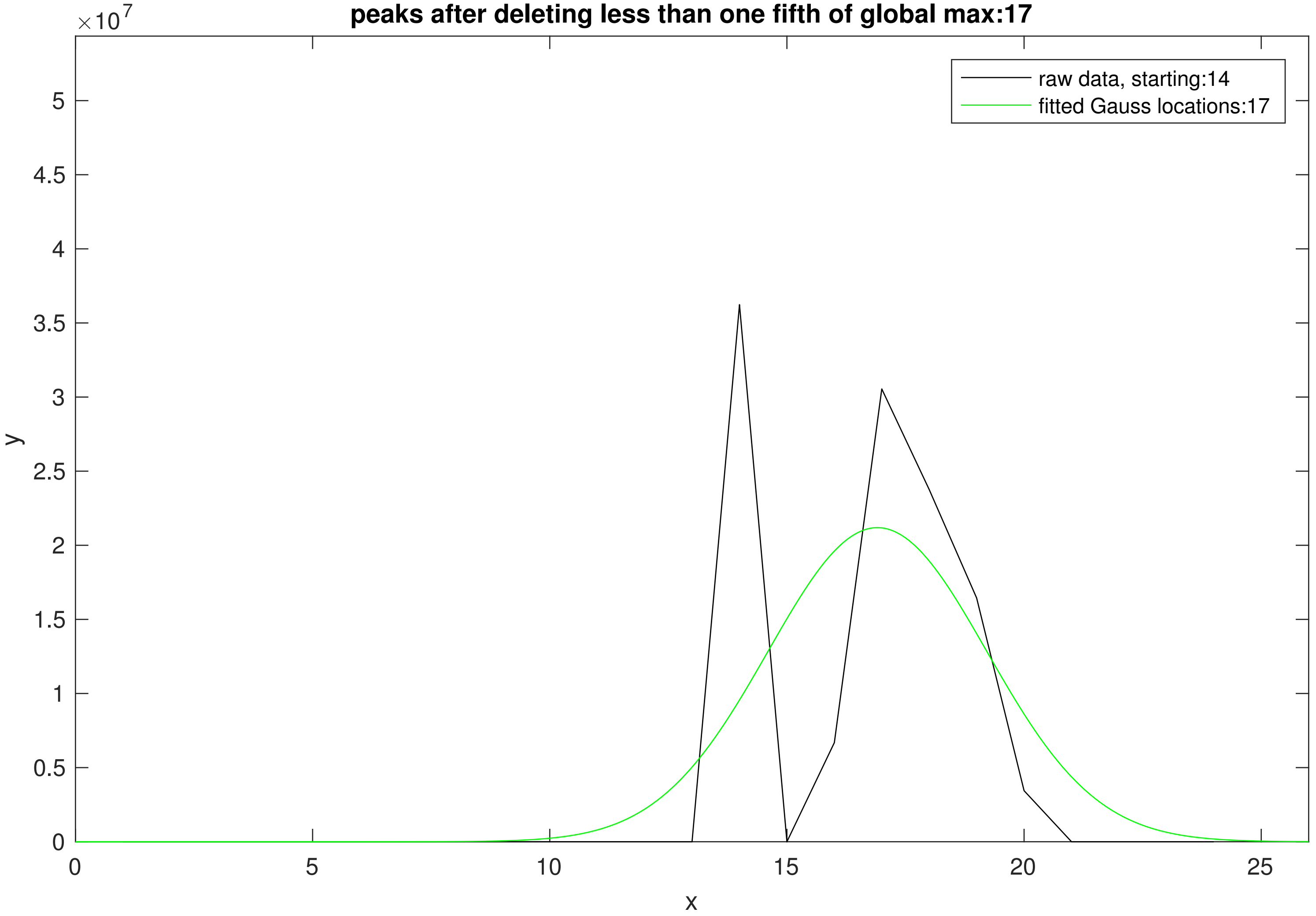

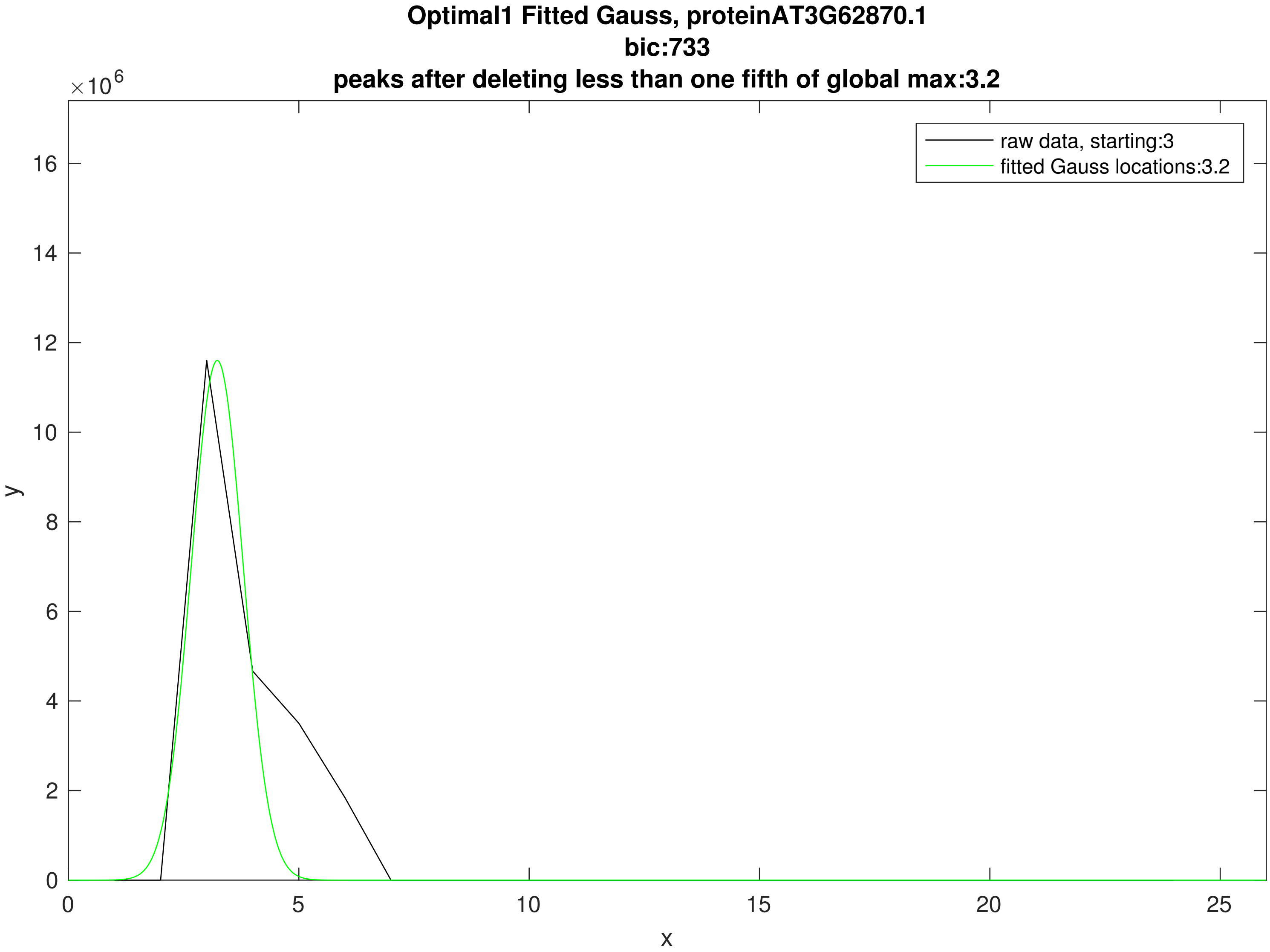

Optimal1 Fitted Gauss, proteinAT3G62870.1
bic:733
peaks after deleting less than one fifth of global max:3.2
×10^6
raw data, starting:3
fitted Gauss locations:3.2
y
x
0
2
4
6
8
10
12
14
16
0
5
10
15
20
25

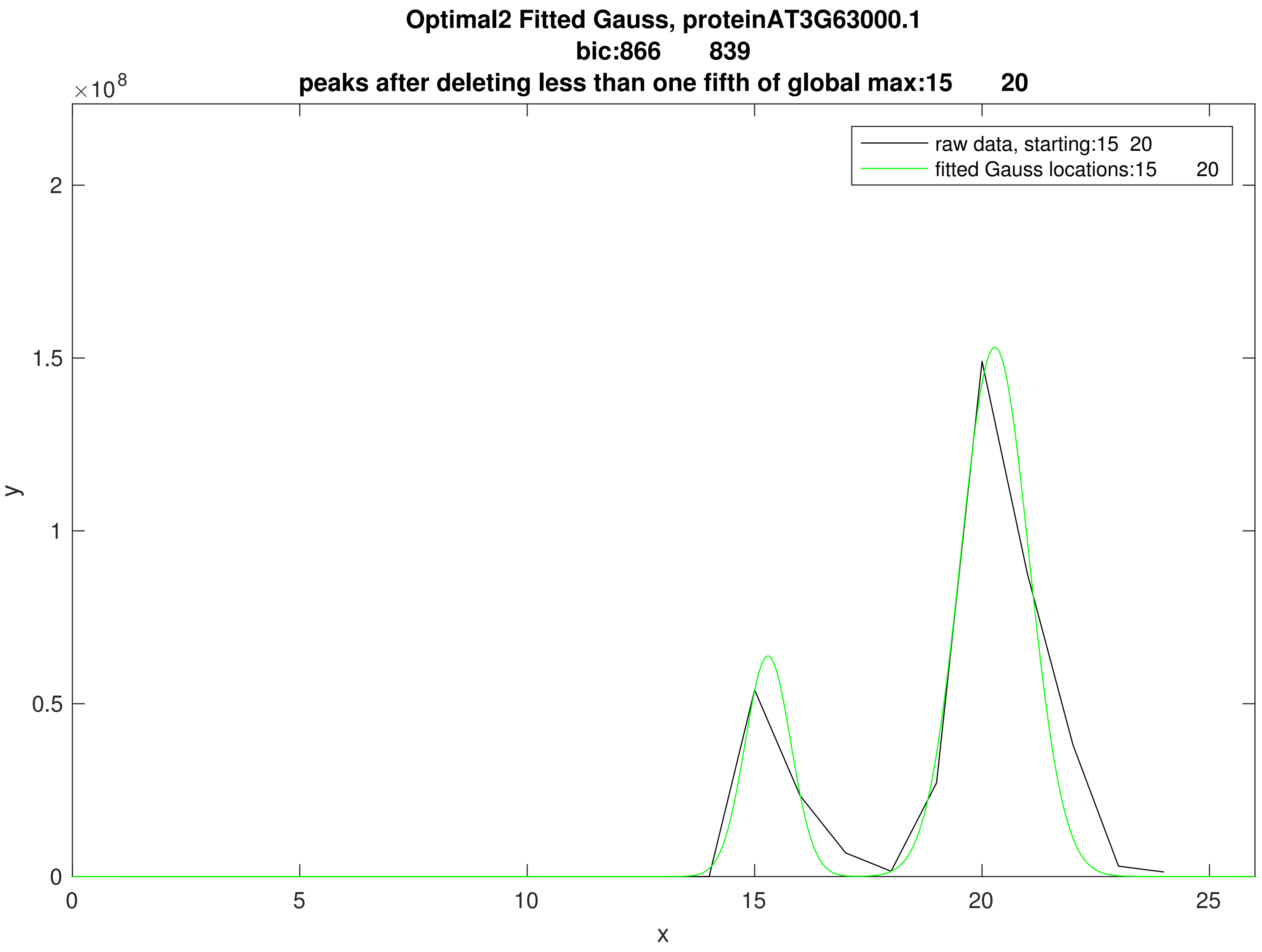
Optimal2 Fitted Gauss, proteinAT3G63000.1
bic:866 839
peaks after deleting less than one fifth of global max:15 20
×10^8
raw data, starting:15 20
fitted Gauss locations:15 20
2
1.5
1
0.5
0
y
0
5
10
15
20
25
x

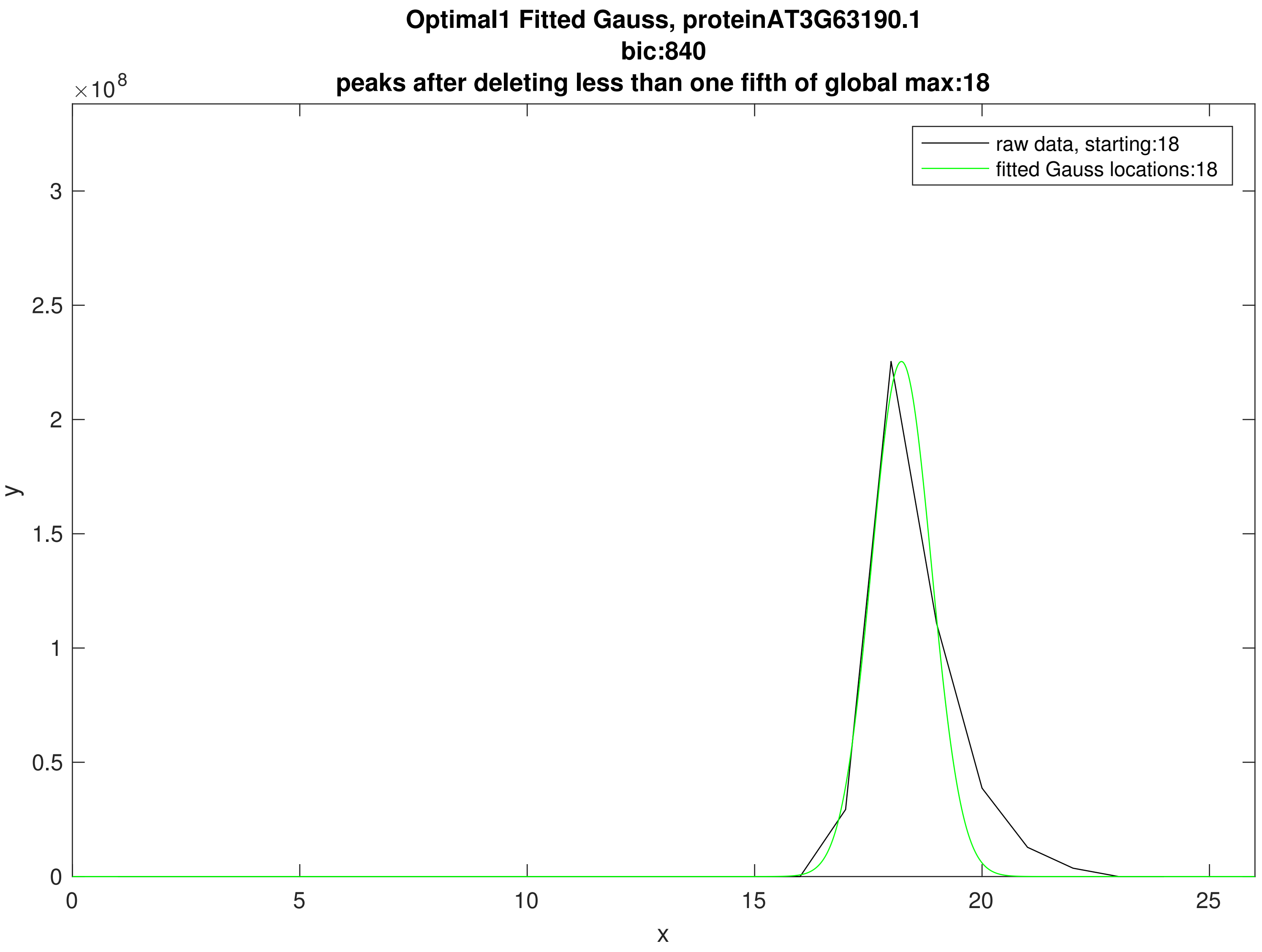
Optimal1 Fitted Gauss, proteinAT3G63190.1
bic:840
peaks after deleting less than one fifth of global max:18
×10^8
raw data, starting:18
fitted Gauss locations:18
y
x
0
0.5
1
1.5
2
2.5
3
0
5
10
15
20
25

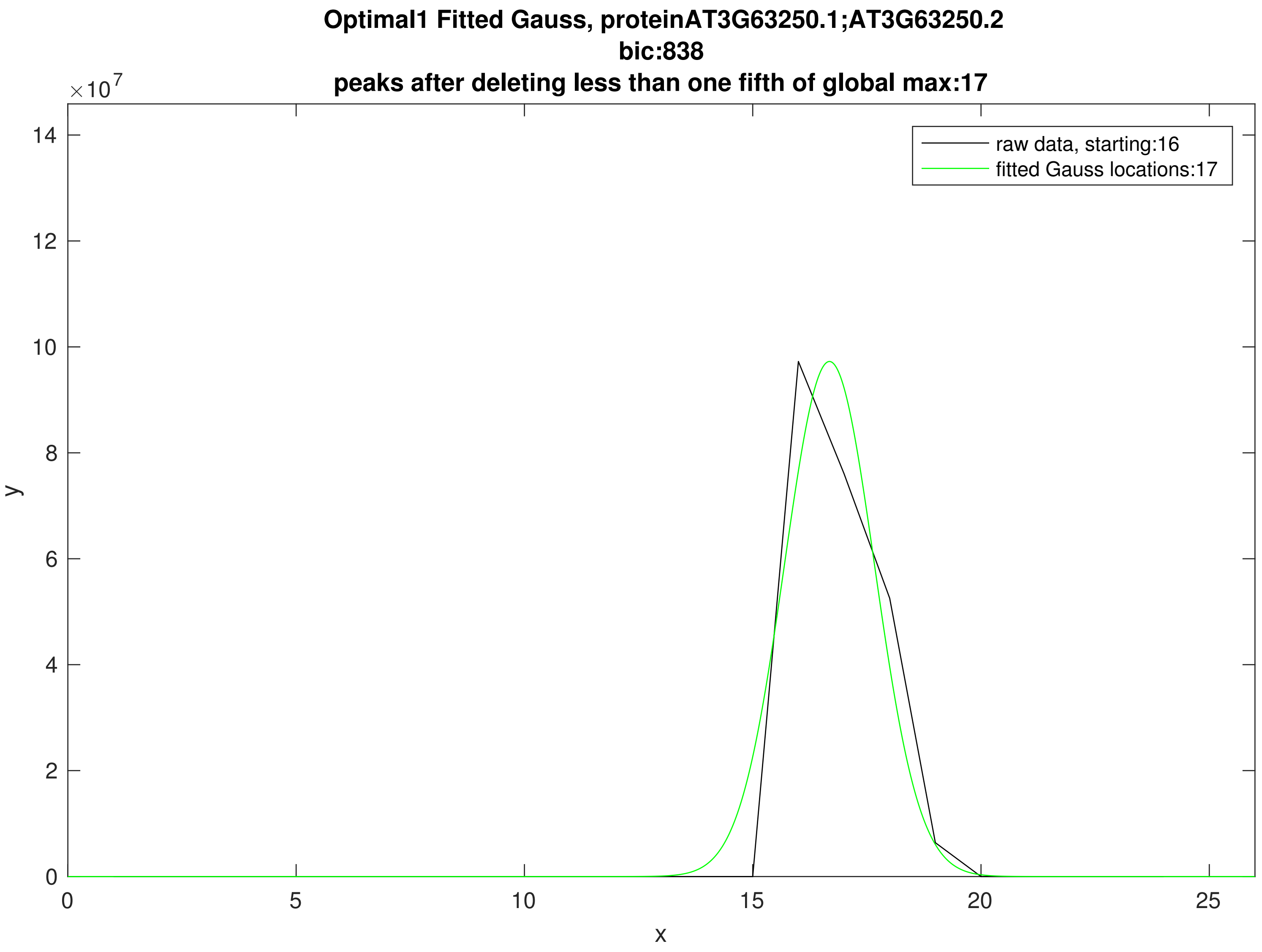

Optimal1 Fitted Gauss, proteinAT3G63250.1;AT3G63250.2
bic:838
peaks after deleting less than one fifth of global max:17
×10^7
14
12
10
8
6
4
2
0
y
0
5
10
15
20
25
x
raw data, starting:16
fitted Gauss locations:17

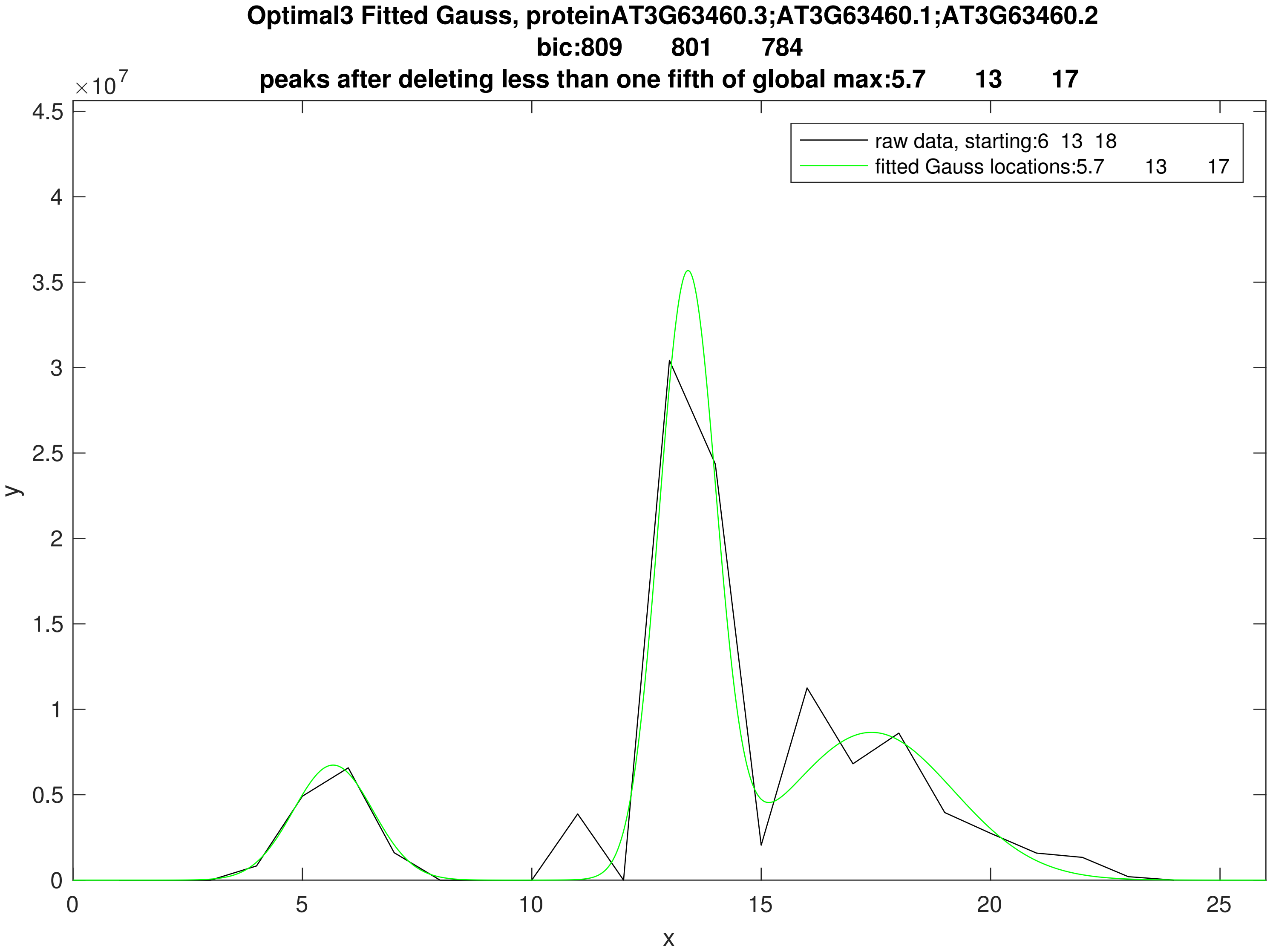

Optimal3 Fitted Gauss, proteinAT3G63460.3;AT3G63460.1;AT3G63460.2
bic:809 801 784
peaks after deleting less than one fifth of global max:5.7 13 17
×10^7
raw data, starting:6 13 18
fitted Gauss locations:5.7 13 17
y
x
0
0.5
1
1.5
2
2.5
3
3.5
4
4.5
5
10
15
20
25

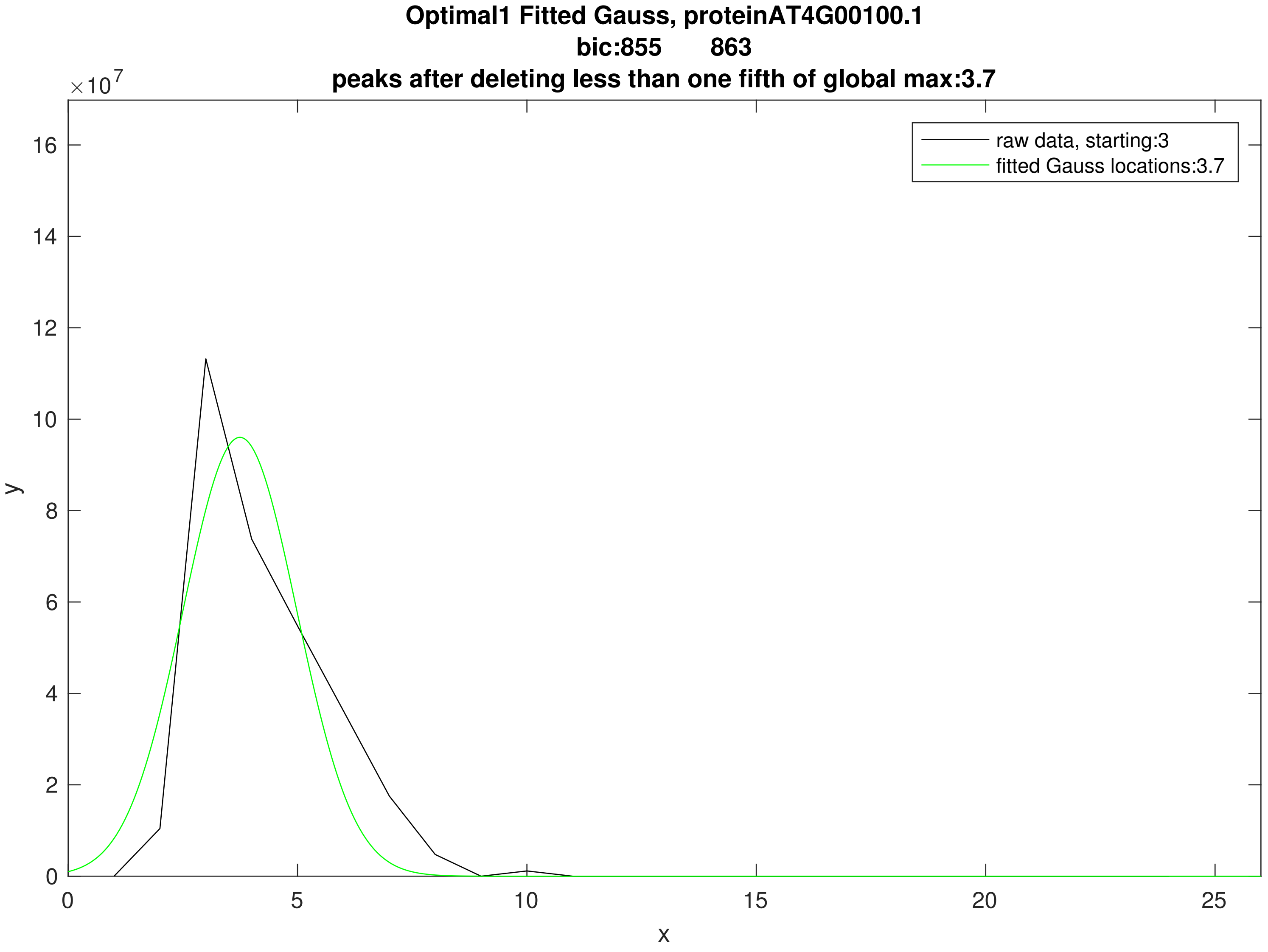

Optimal1 Fitted Gauss, proteinAT4G00100.1
bic:855 863
peaks after deleting less than one fifth of global max:3.7
×10^7
raw data, starting:3
fitted Gauss locations:3.7
y
x
0
2
4
6
8
10
12
14
16
0
5
10
15
20
25

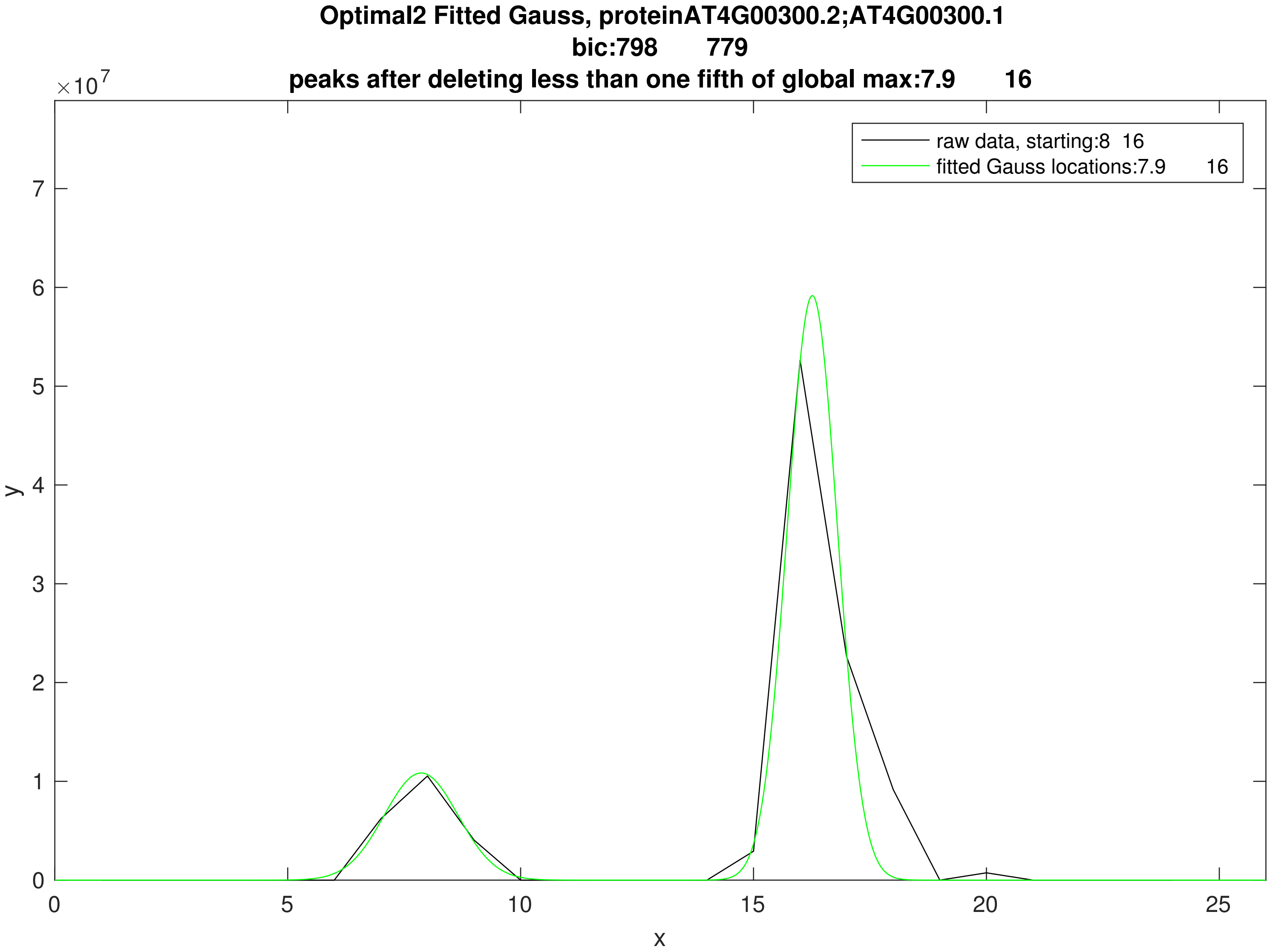

Optimal2 Fitted Gauss, proteinAT4G00300.2;AT4G00300.1
bic:798 779
peaks after deleting less than one fifth of global max:7.9 16
×10^7
raw data, starting:8 16
fitted Gauss locations:7.9 16
y
x
0
1
2
3
4
5
6
7
0
5
10
15
20
25

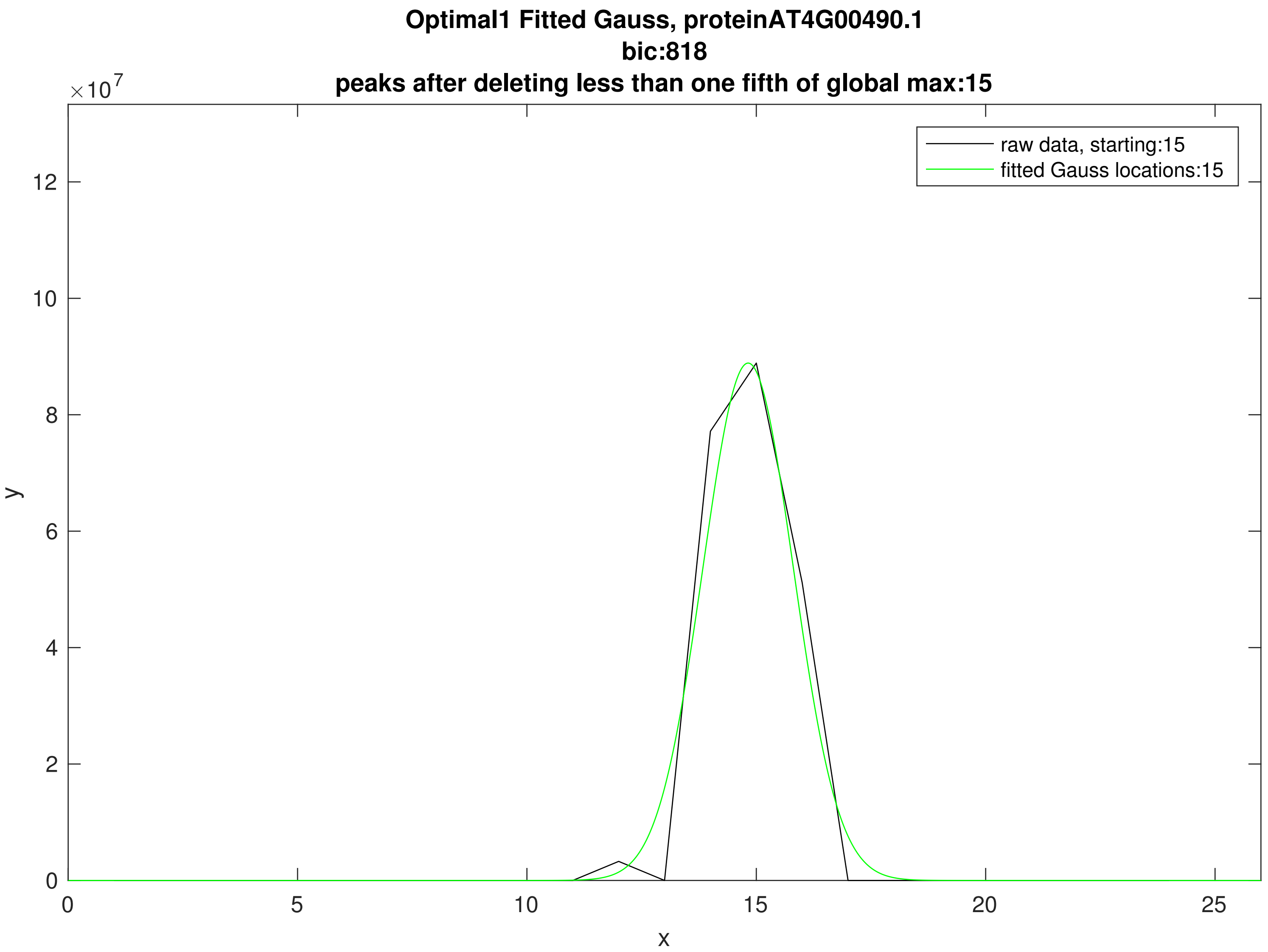
Optimal1 Fitted Gauss, proteinAT4G00490.1
bic:818
peaks after deleting less than one fifth of global max:15
×10^7
raw data, starting:15
fitted Gauss locations:15
y
x
0
2
4
6
8
10
12
0
5
10
15
20
25

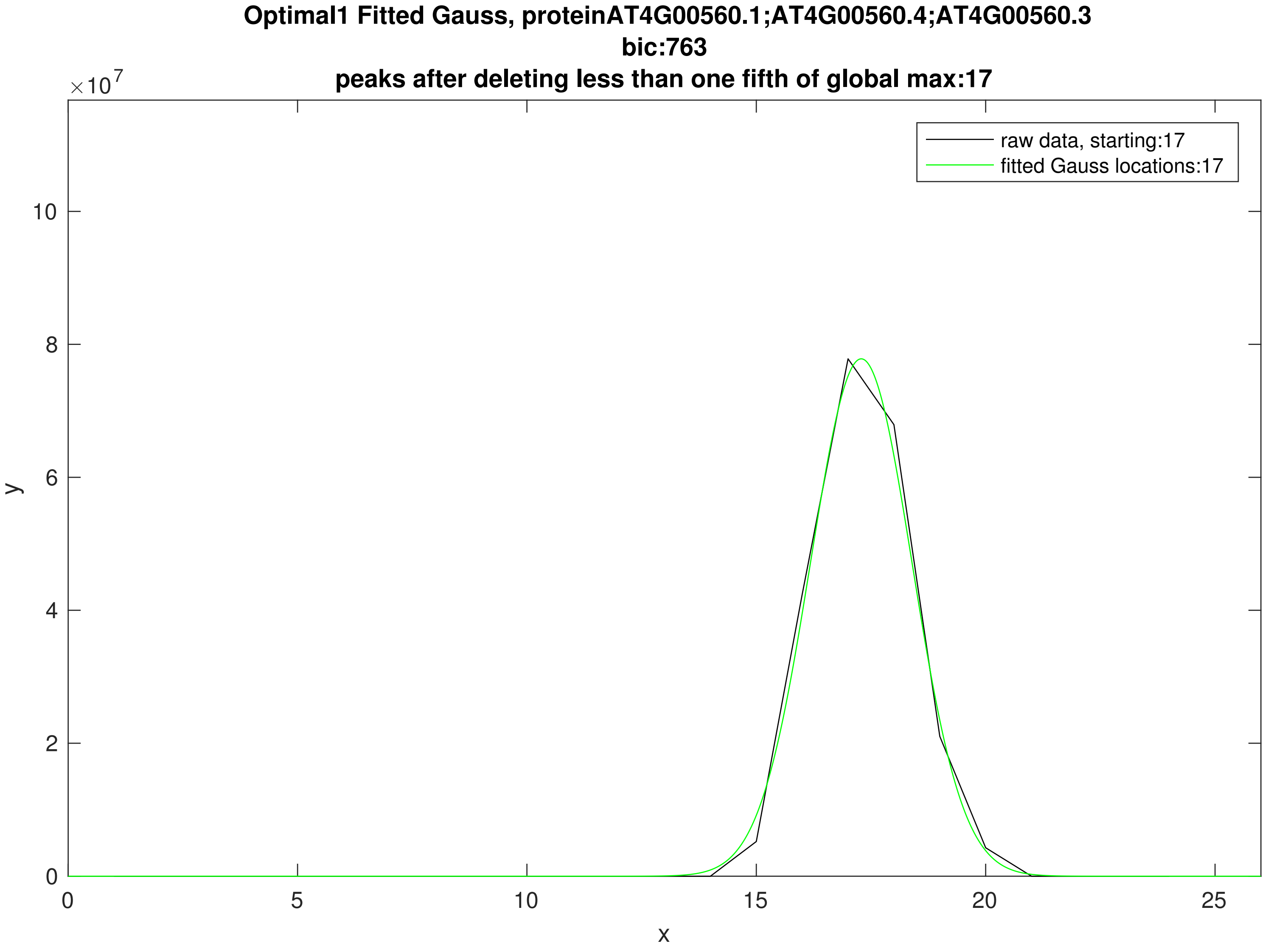
Optimal1 Fitted Gauss, proteinAT4G00560.1;AT4G00560.4;AT4G00560.3
bic:763
peaks after deleting less than one fifth of global max:17
×10^7
raw data, starting:17
fitted Gauss locations:17
y
x
0
2
4
6
8
10
0
5
10
15
20
25

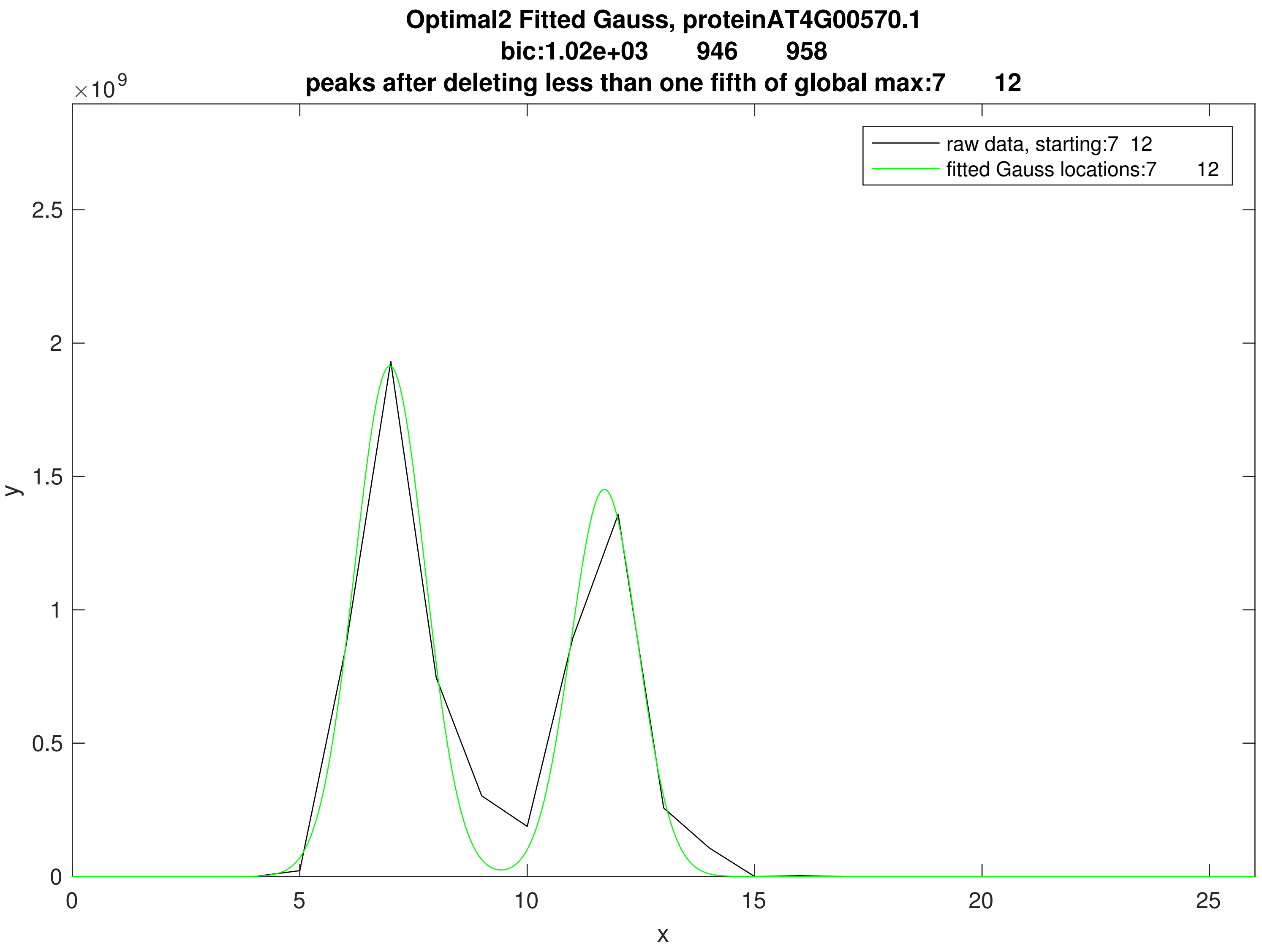

Optimal2 Fitted Gauss, proteinAT4G00570.1
bic:1.02e+03 946 958
peaks after deleting less than one fifth of global max:7 12
×10^9
raw data, starting:7 12
fitted Gauss locations:7 12
y
x

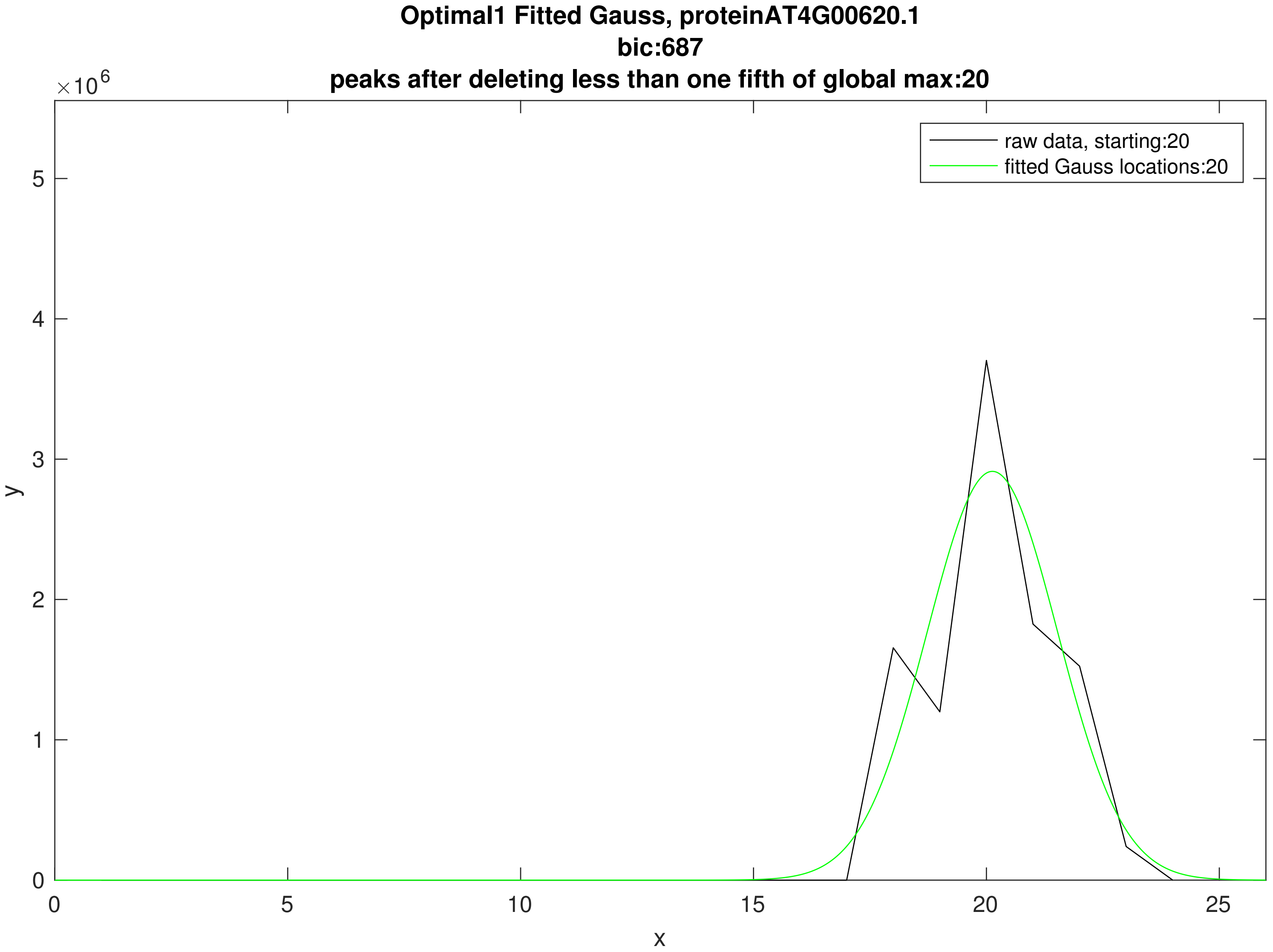
Optimal1 Fitted Gauss, proteinAT4G00620.1
bic:687
peaks after deleting less than one fifth of global max:20
×10^6
raw data, starting:20
fitted Gauss locations:20
y
x
0
1
2
3
4
5
0
5
10
15
20
25

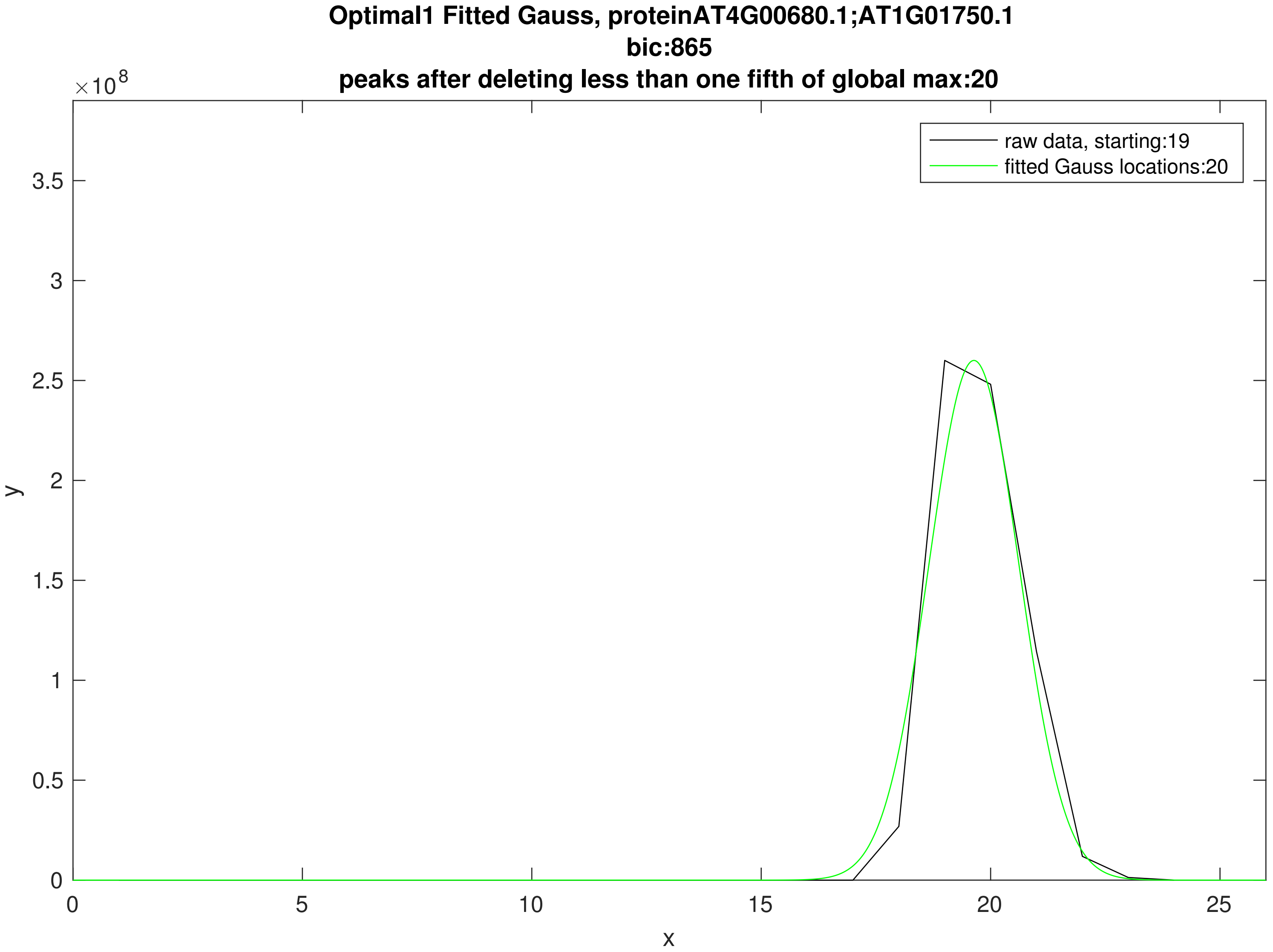
Optimal1 Fitted Gauss, proteinAT4G00680.1;AT1G01750.1
bic:865
peaks after deleting less than one fifth of global max:20
×10^8
3.5
3
2.5
2
1.5
1
0.5
0
y
0
5
10
15
20
25
x
raw data, starting:19
fitted Gauss locations:20

Optimal1 Fitted Gauss, proteinAT4G00810.2;AT4G00810.1
bic:655
peaks after deleting less than one fifth of global max:18

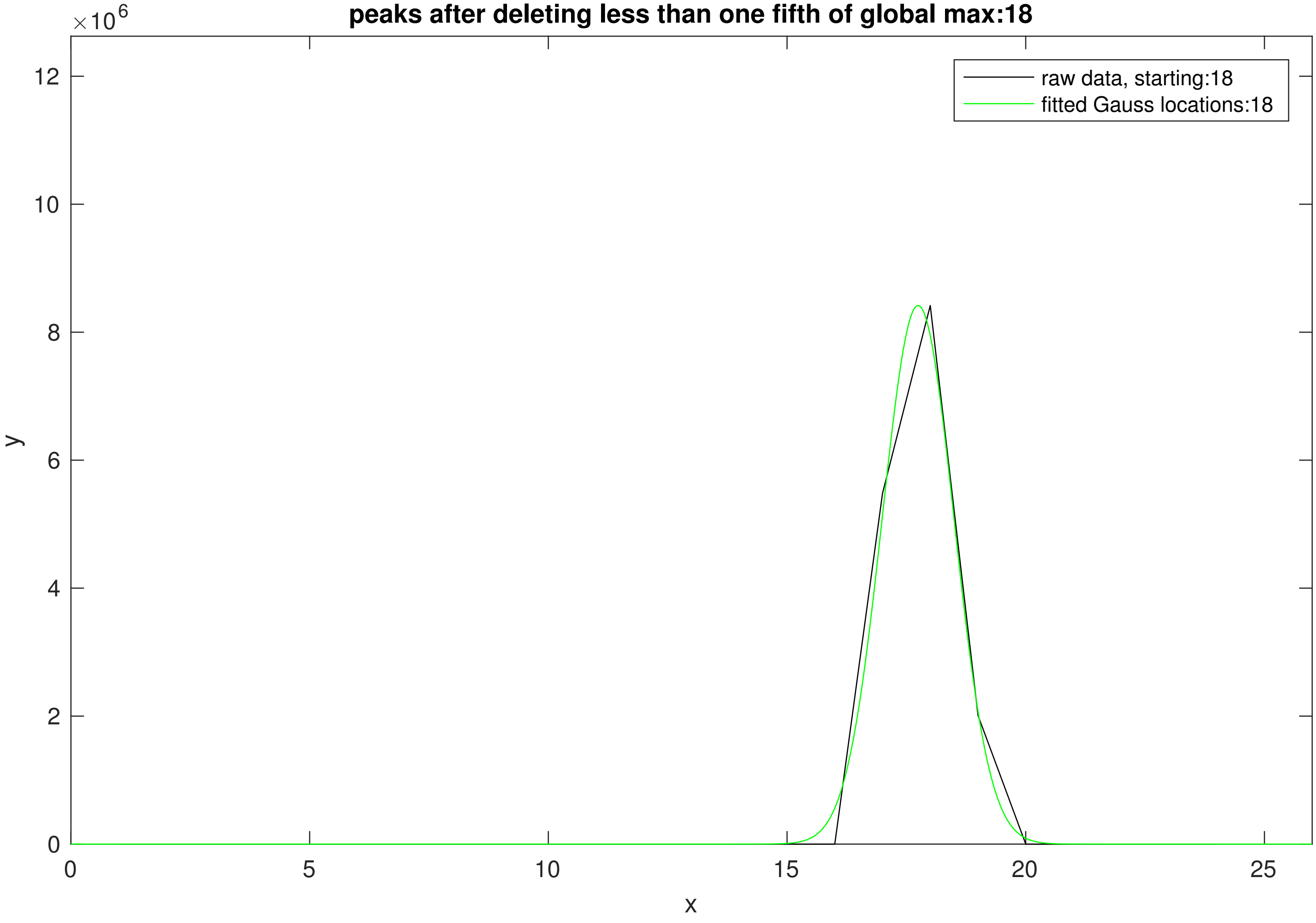

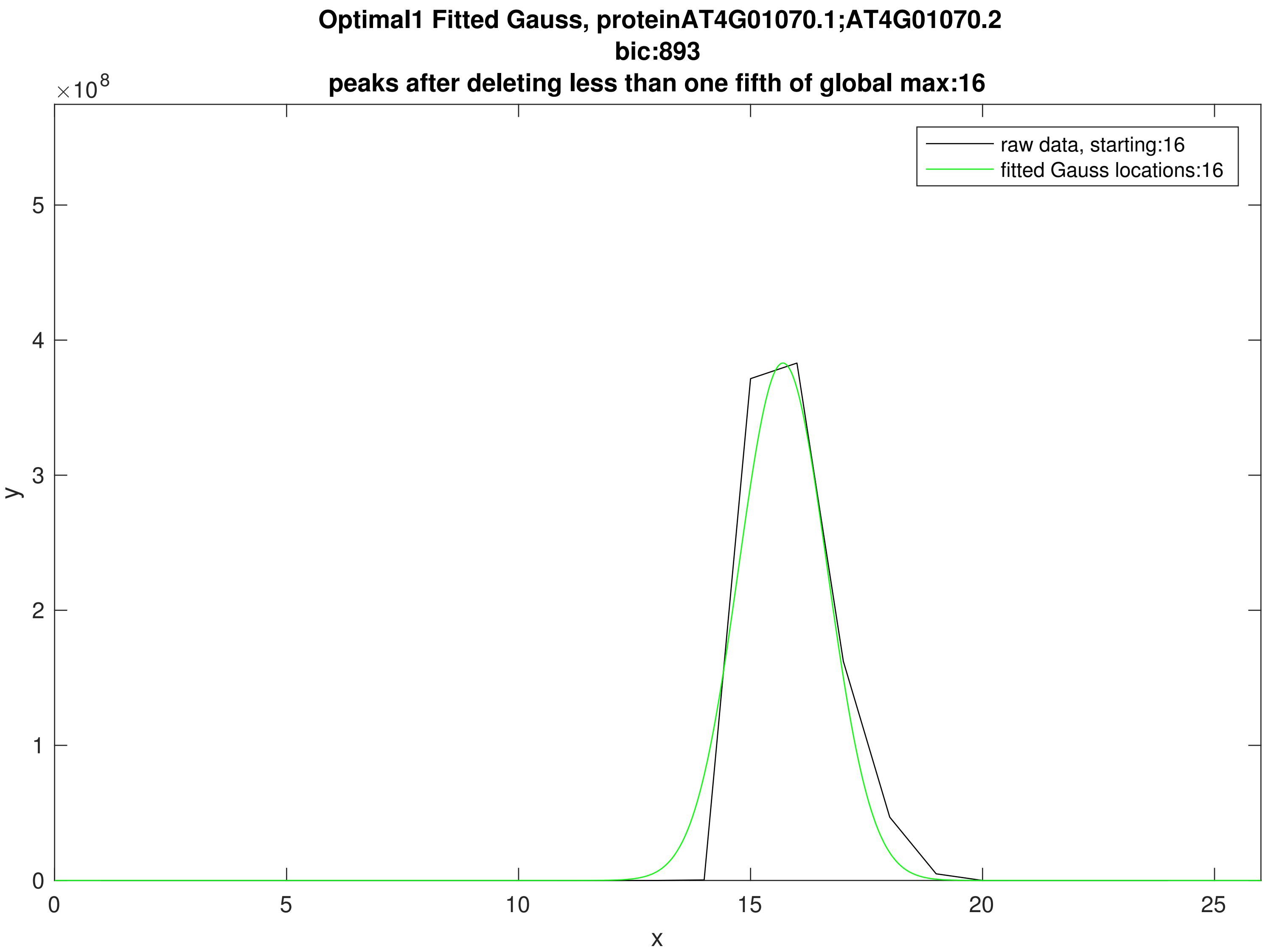
Optimal1 Fitted Gauss, proteinAT4G01070.1;AT4G01070.2
bic:893
peaks after deleting less than one fifth of global max:16
×10^8
raw data, starting:16
fitted Gauss locations:16
y
x
0
1
2
3
4
5
0
5
10
15
20
25

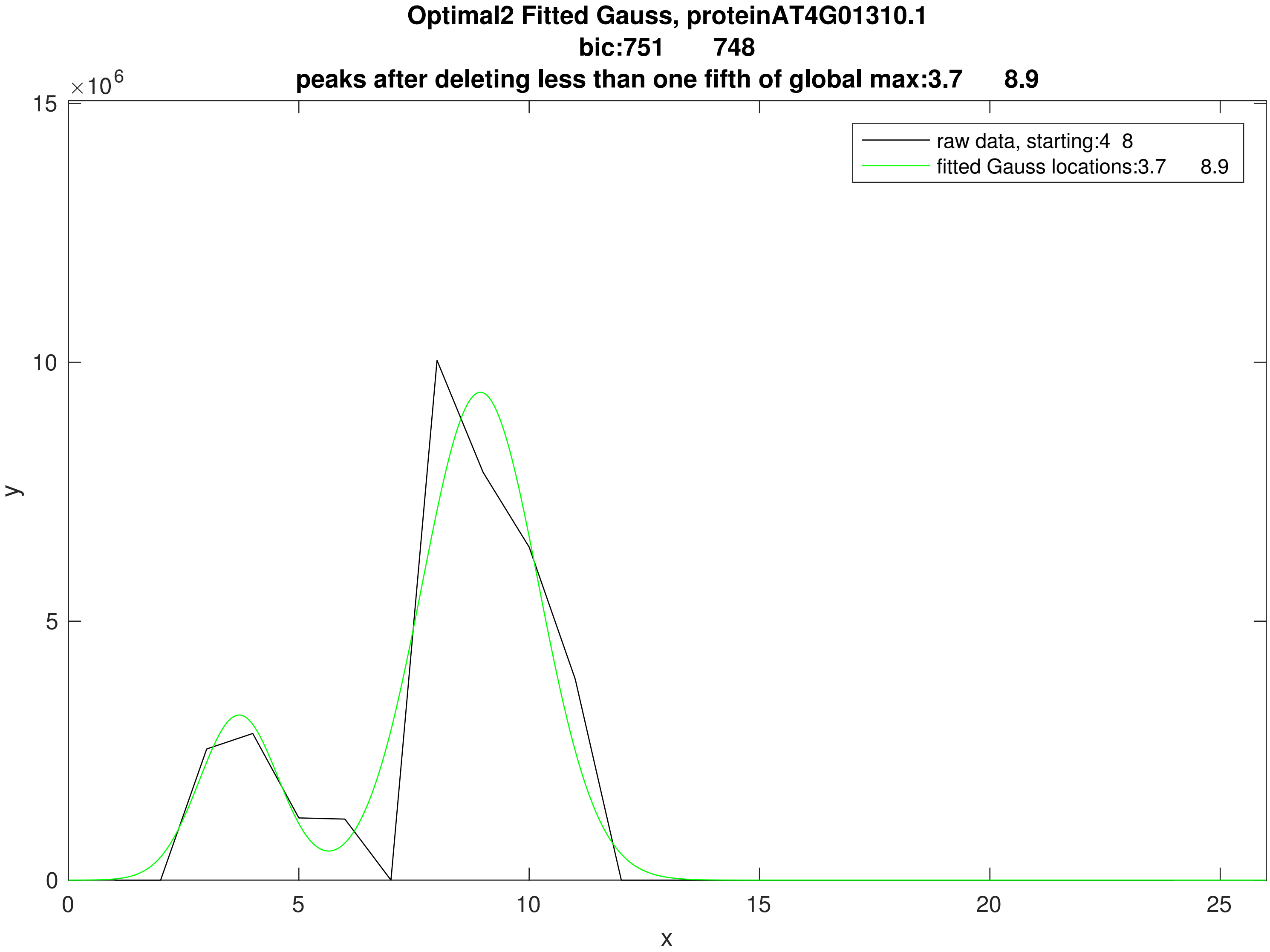

Optimal2 Fitted Gauss, proteinAT4G01310.1
bic:751 748
peaks after deleting less than one fifth of global max:3.7 8.9
×10^6
15
10
5
0
y
0
5
10
15
20
25
x
raw data, starting:4 8
fitted Gauss locations:3.7 8.9

#### Optimal1 Fitted Gauss, proteinAT4G01370.1
#### bic:727
#### peaks after deleting less than one fifth of global max:18

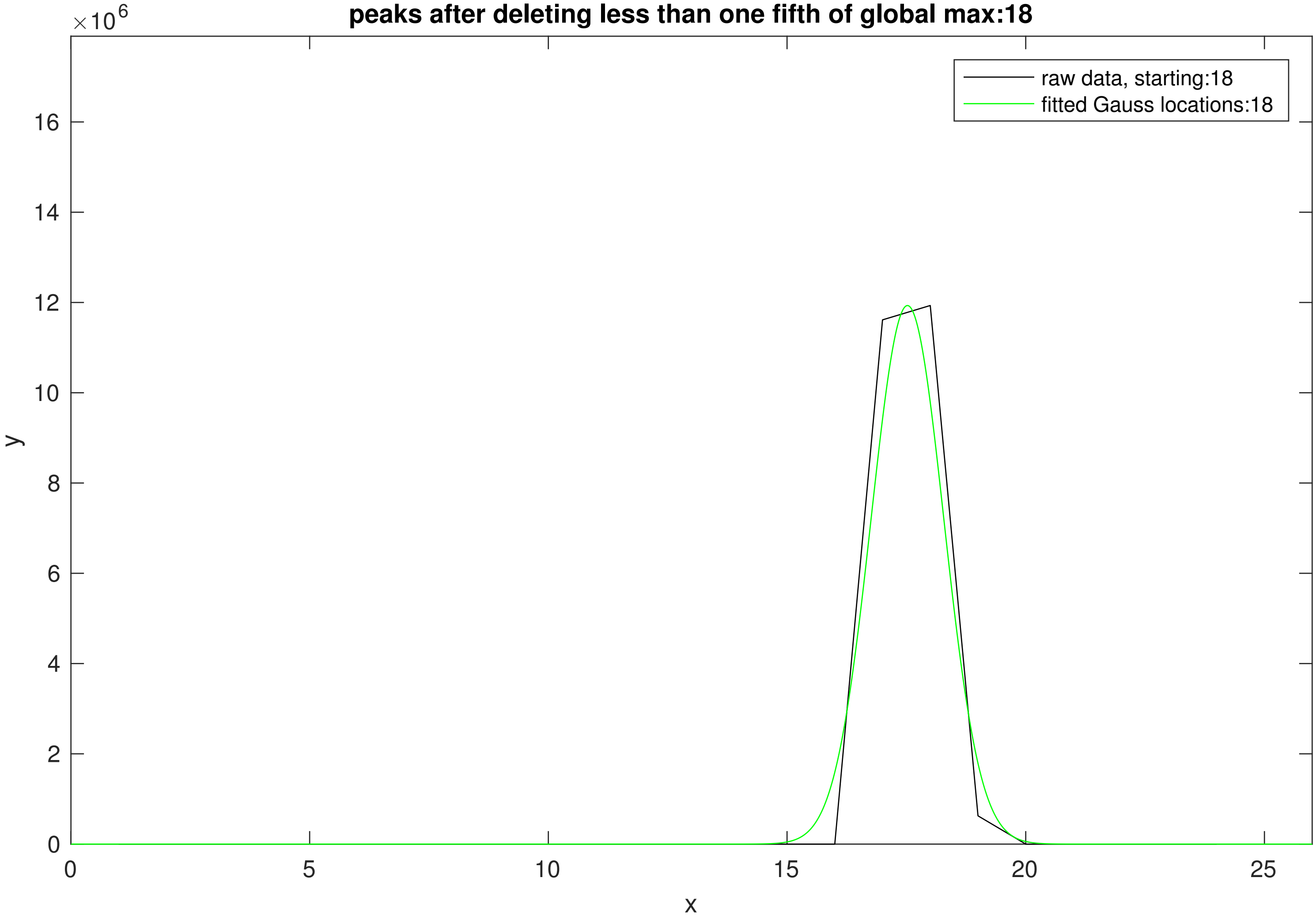

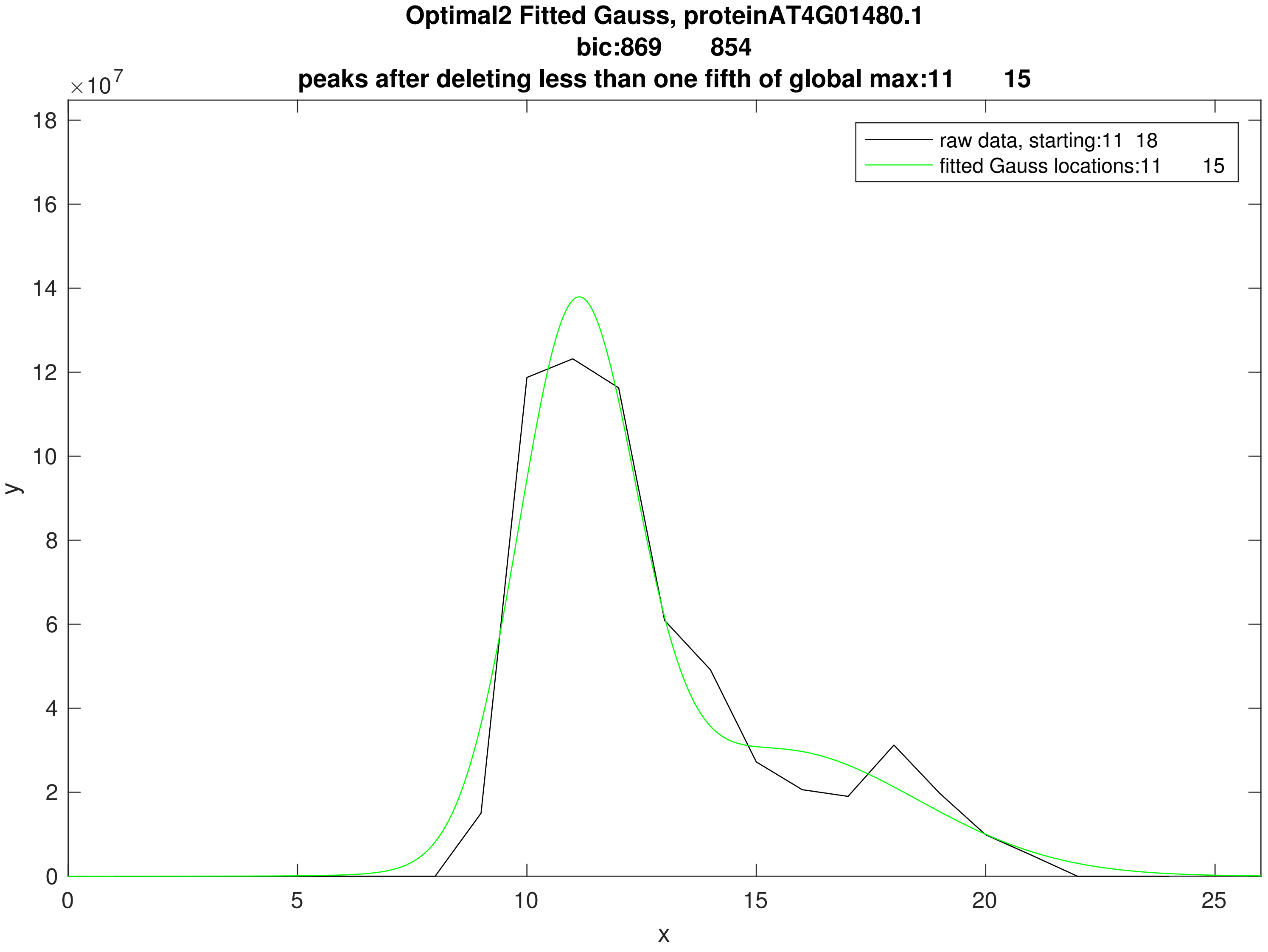

Optimal2 Fitted Gauss, proteinAT4G01480.1
bic:869 854
peaks after deleting less than one fifth of global max:11 15
×10^7
raw data, starting:11 18
fitted Gauss locations:11 15
y
x
0
2
4
6
8
10
12
14
16
18
0
5
10
15
20
25

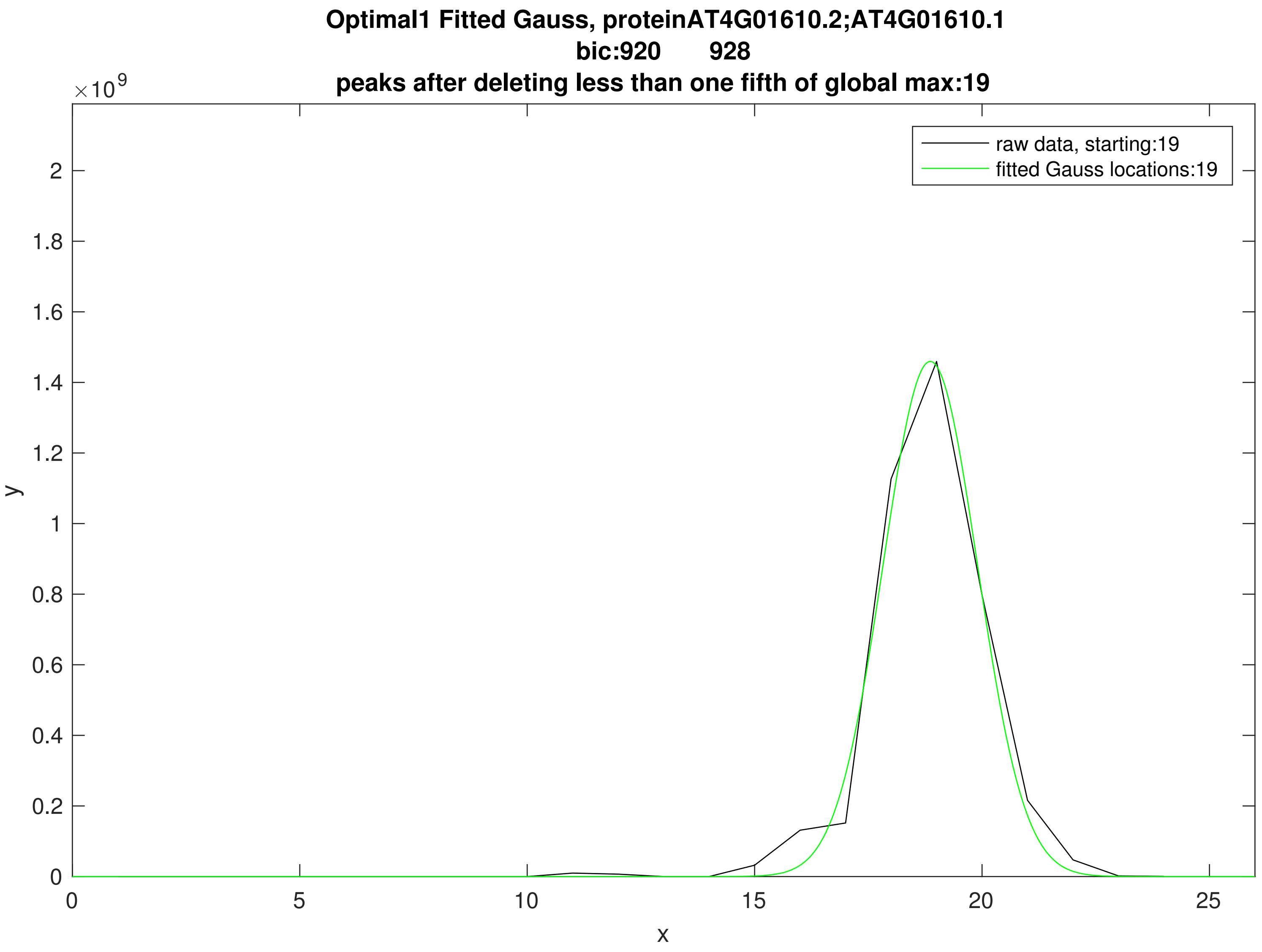
Optimal1 Fitted Gauss, proteinAT4G01610.2;AT4G01610.1
bic:920 928
peaks after deleting less than one fifth of global max:19
×10^9
raw data, starting:19
fitted Gauss locations:19
y
x
0
0.2
0.4
0.6
0.8
1
1.2
1.4
1.6
1.8
2
0
5
10
15
20
25

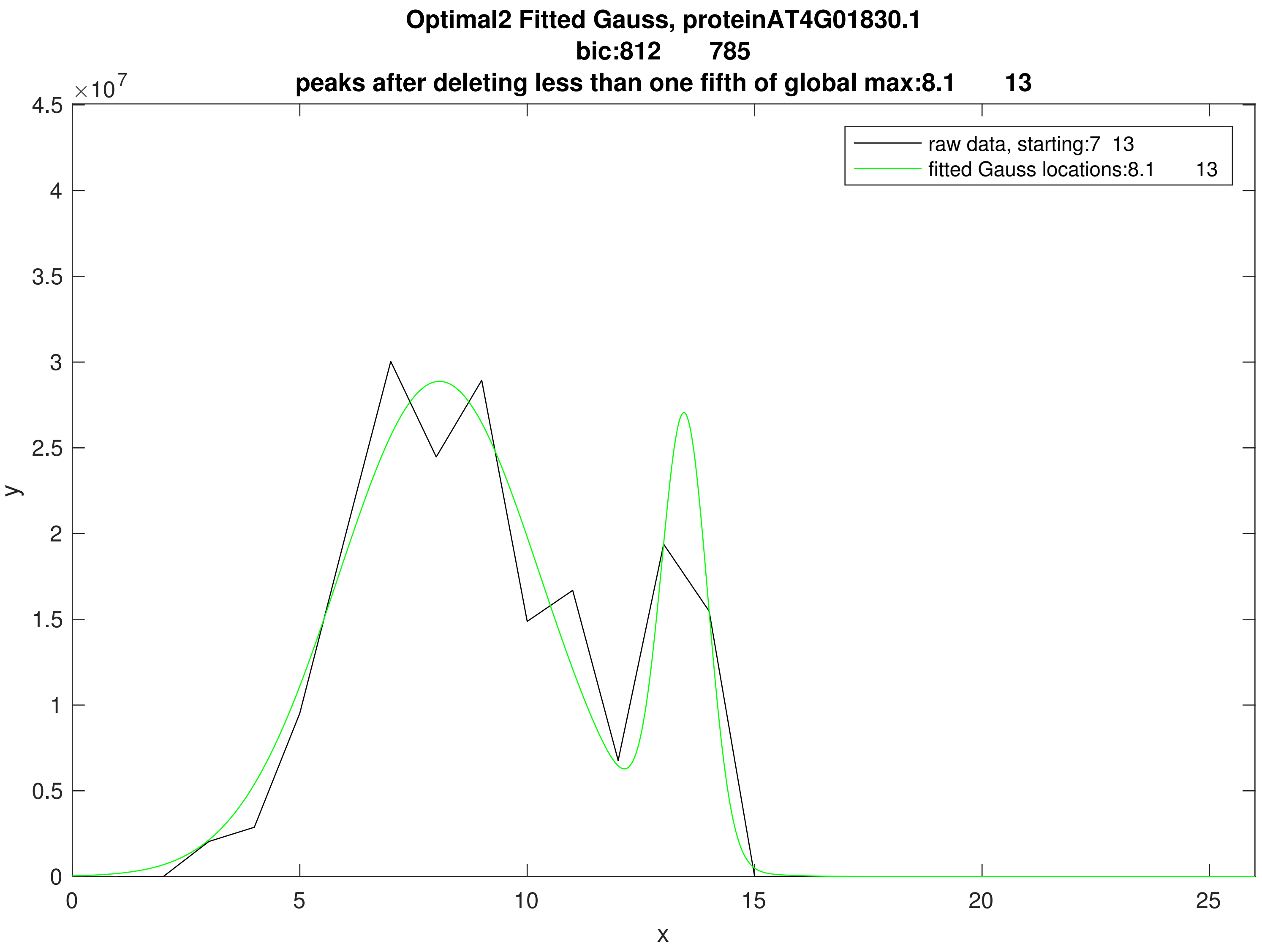
Optimal2 Fitted Gauss, proteinAT4G01830.1
bic:812 785
peaks after deleting less than one fifth of global max:8.1 13
×10^7
4.5
4
3.5
3
2.5
2
1.5
1
0.5
0
y
0
5
10
15
20
25
x
raw data, starting:7 13
fitted Gauss locations:8.1 13

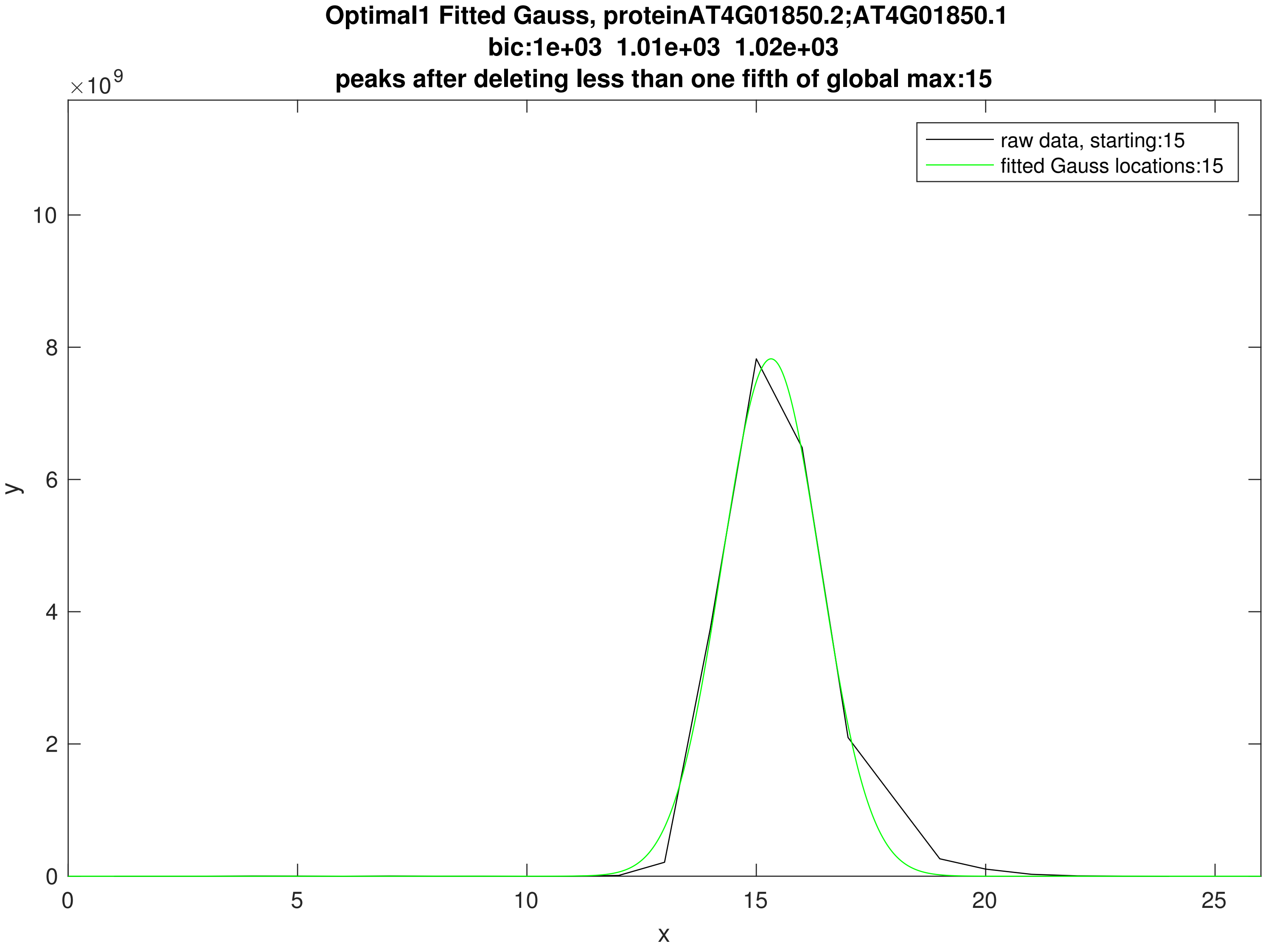

Optimal1 Fitted Gauss, proteinAT4G01850.2;AT4G01850.1
bic:1e+03 1.01e+03 1.02e+03
peaks after deleting less than one fifth of global max:15
×10^9
raw data, starting:15
fitted Gauss locations:15
0
2
4
6
8
10
y
0
5
10
15
20
25
x

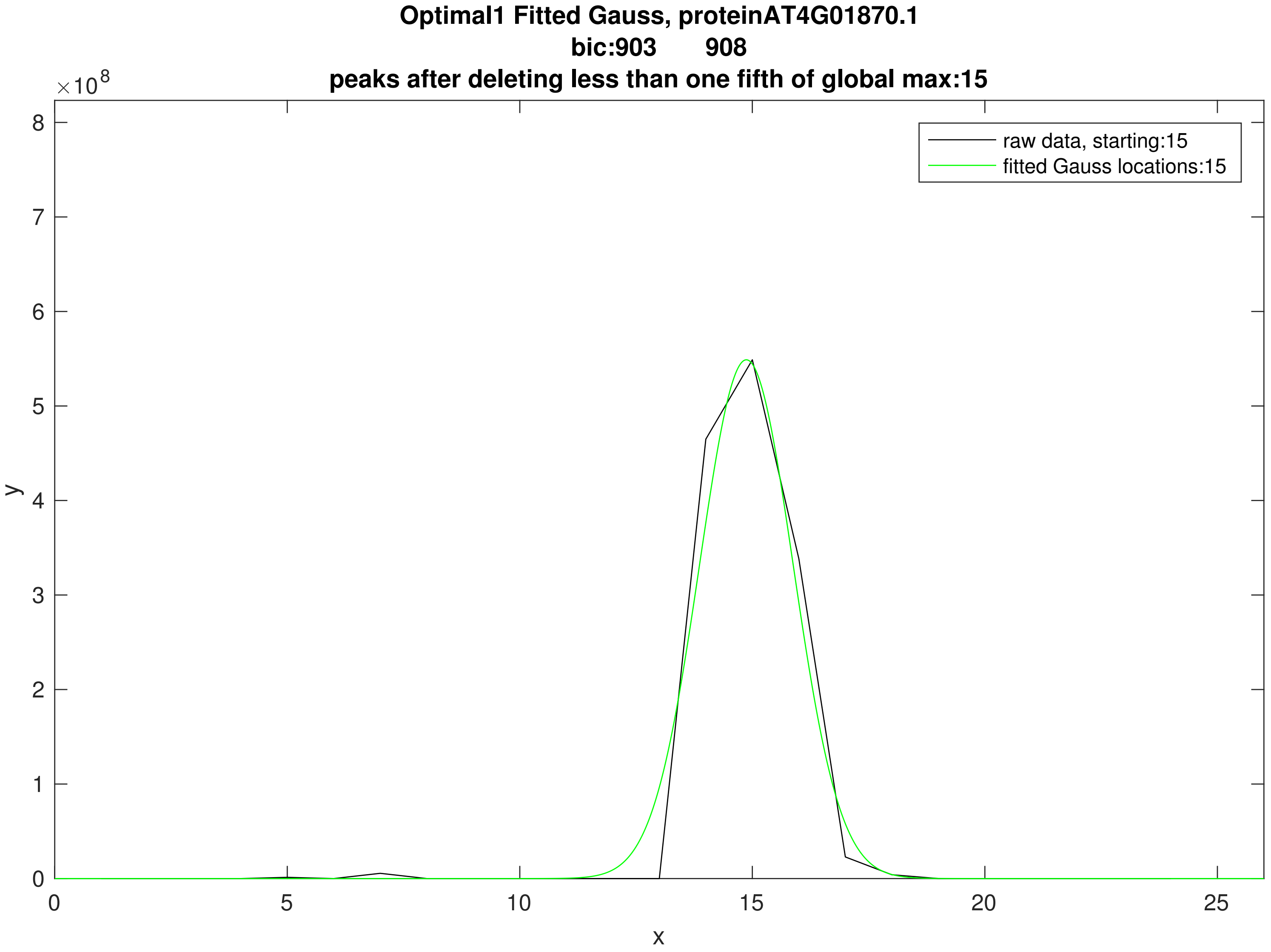
Optimal1 Fitted Gauss, proteinAT4G01870.1
bic:903 908
peaks after deleting less than one fifth of global max:15
×10^8
y
x
raw data, starting:15
fitted Gauss locations:15

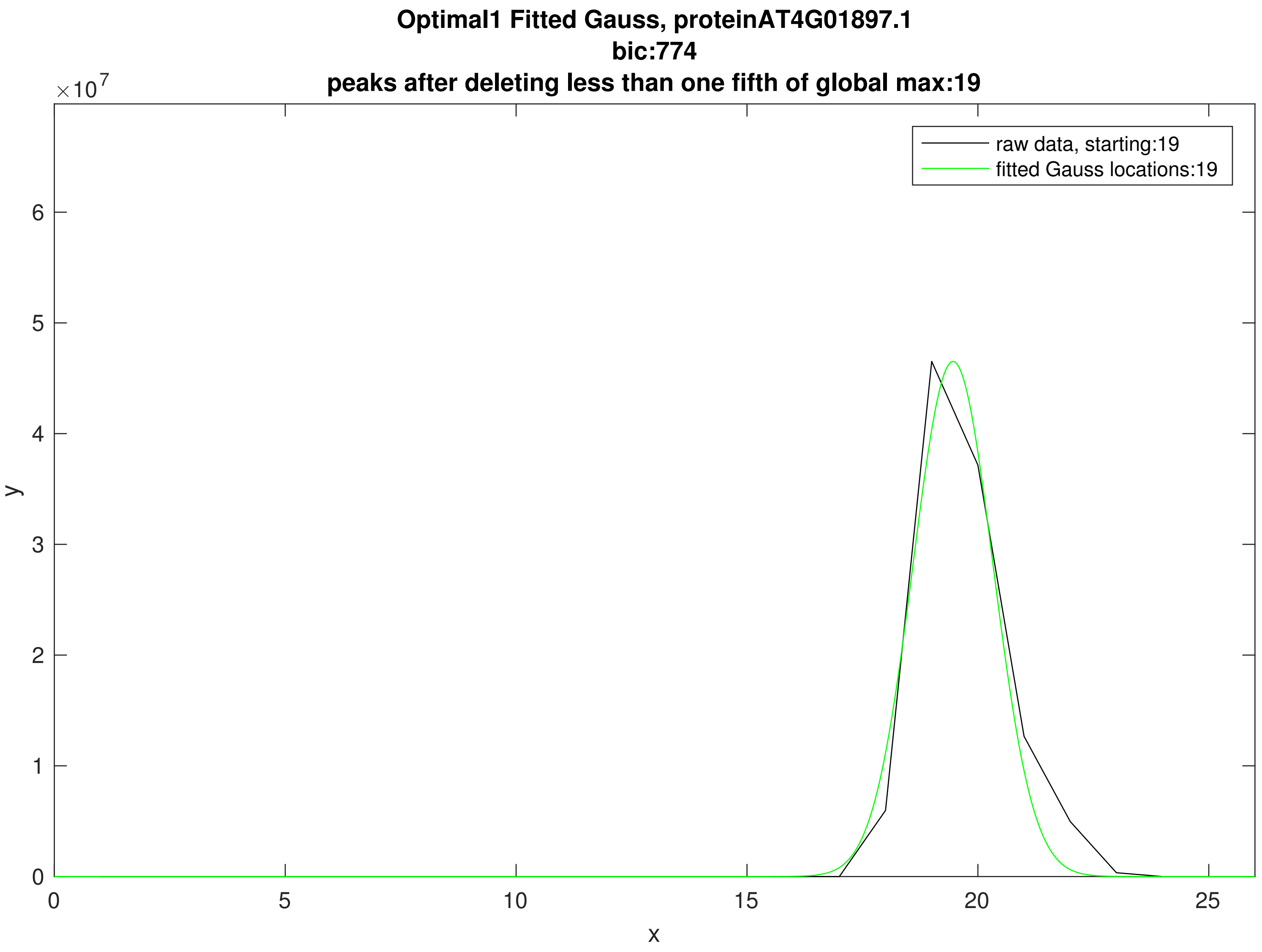

Optimal1 Fitted Gauss, proteinAT4G01897.1
bic:774
peaks after deleting less than one fifth of global max:19
×10^7
raw data, starting:19
fitted Gauss locations:19
y
x
0
1
2
3
4
5
6
0
5
10
15
20
25

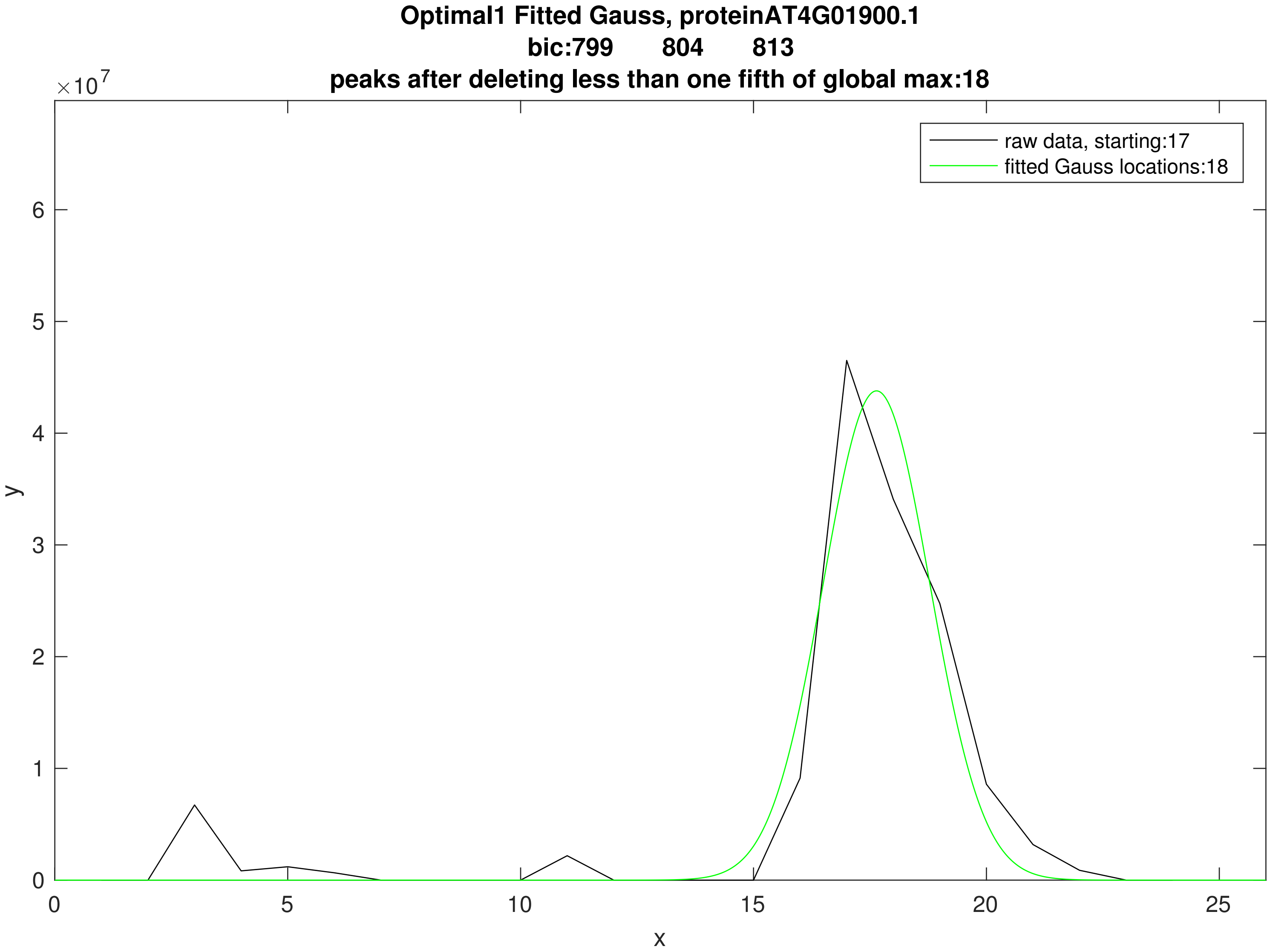
Optimal1 Fitted Gauss, proteinAT4G01900.1
bic:799 804 813
peaks after deleting less than one fifth of global max:18
×10^7
raw data, starting:17
fitted Gauss locations:18
y
x
0
1
2
3
4
5
6
0
5
10
15
20
25

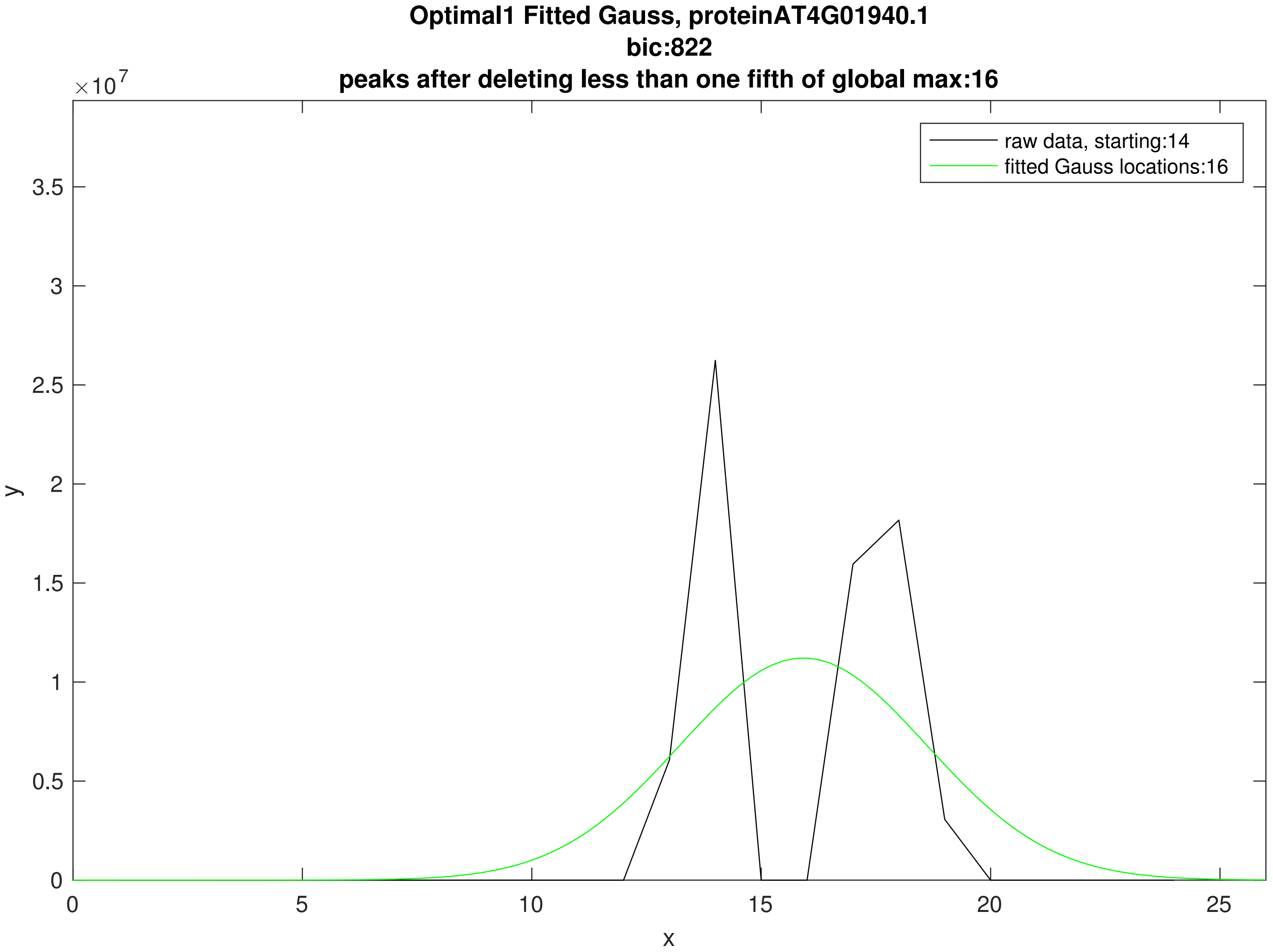
Optimal1 Fitted Gauss, proteinAT4G01940.1
bic:822
peaks after deleting less than one fifth of global max:16
×10^7
raw data, starting:14
fitted Gauss locations:16
y
x
0
0.5
1
1.5
2
2.5
3
3.5
0
5
10
15
20
25

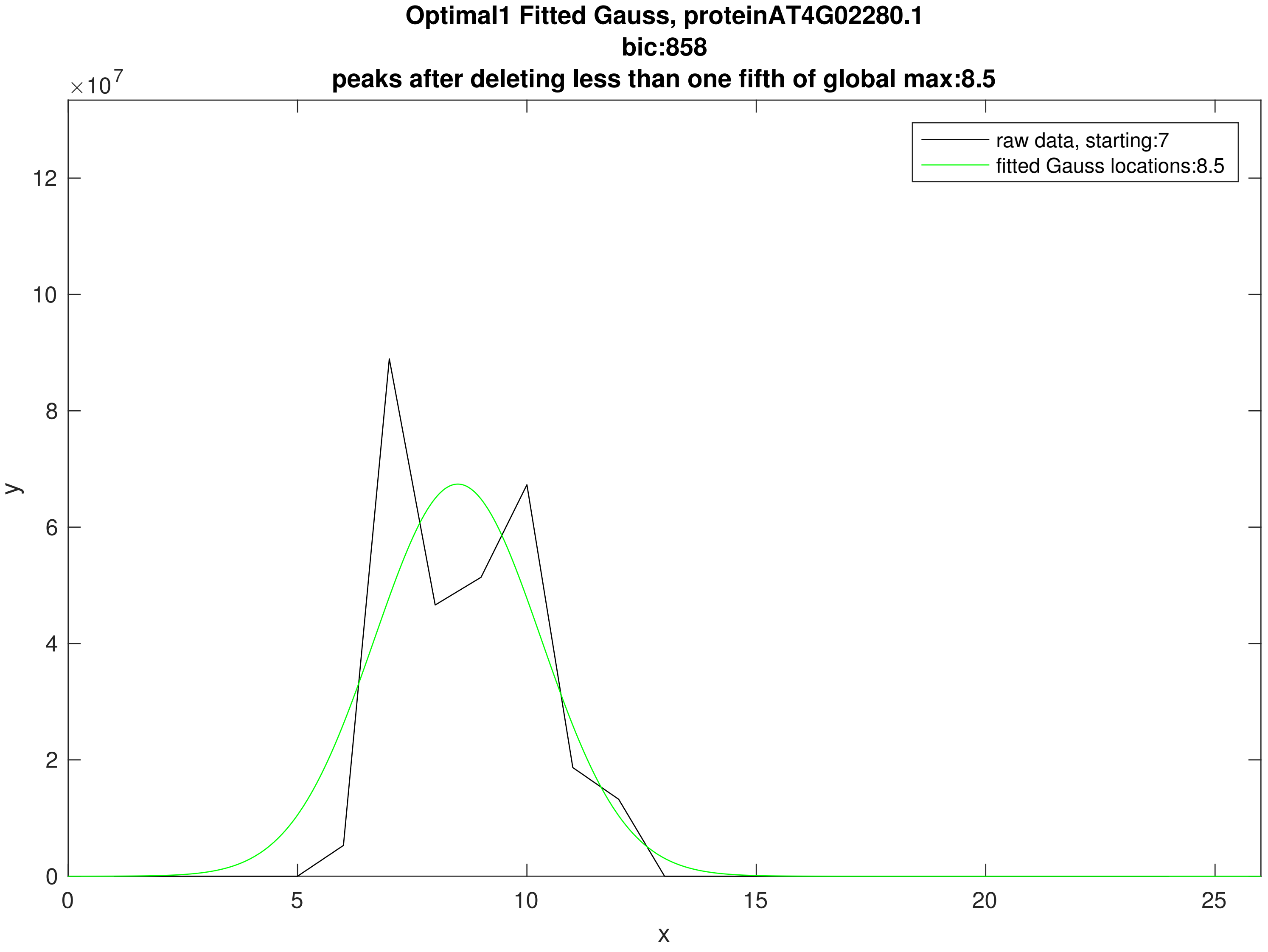
Optimal1 Fitted Gauss, proteinAT4G02280.1
bic:858
peaks after deleting less than one fifth of global max:8.5
×10^7
raw data, starting:7
fitted Gauss locations:8.5
y
x
0
2
4
6
8
10
12
5
15
20
25

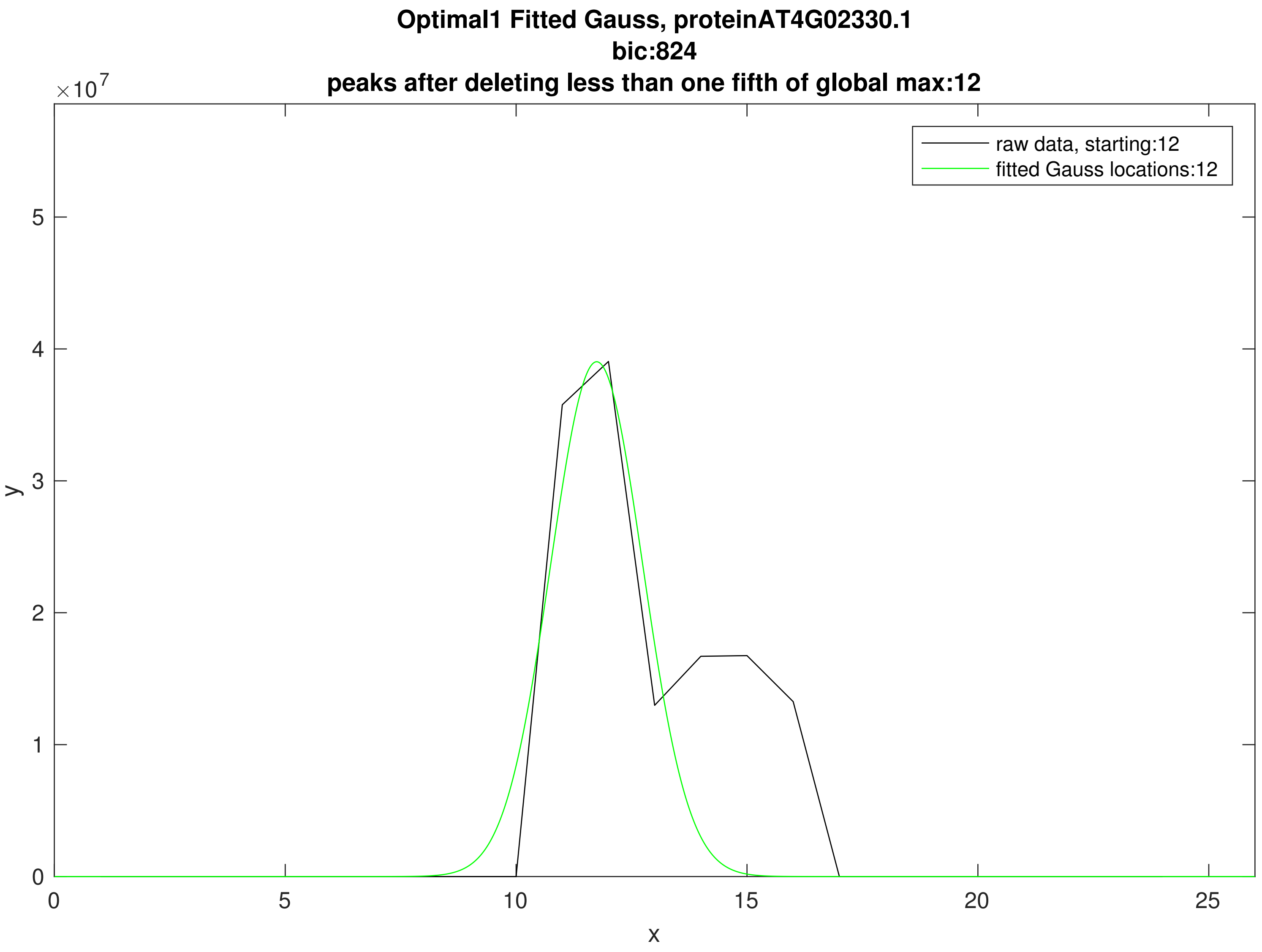
Optimal1 Fitted Gauss, proteinAT4G02330.1
bic:824
peaks after deleting less than one fifth of global max:12
×10^7
raw data, starting:12
fitted Gauss locations:12
y
x
0
1
2
3
4
5
0
5
10
15
20
25

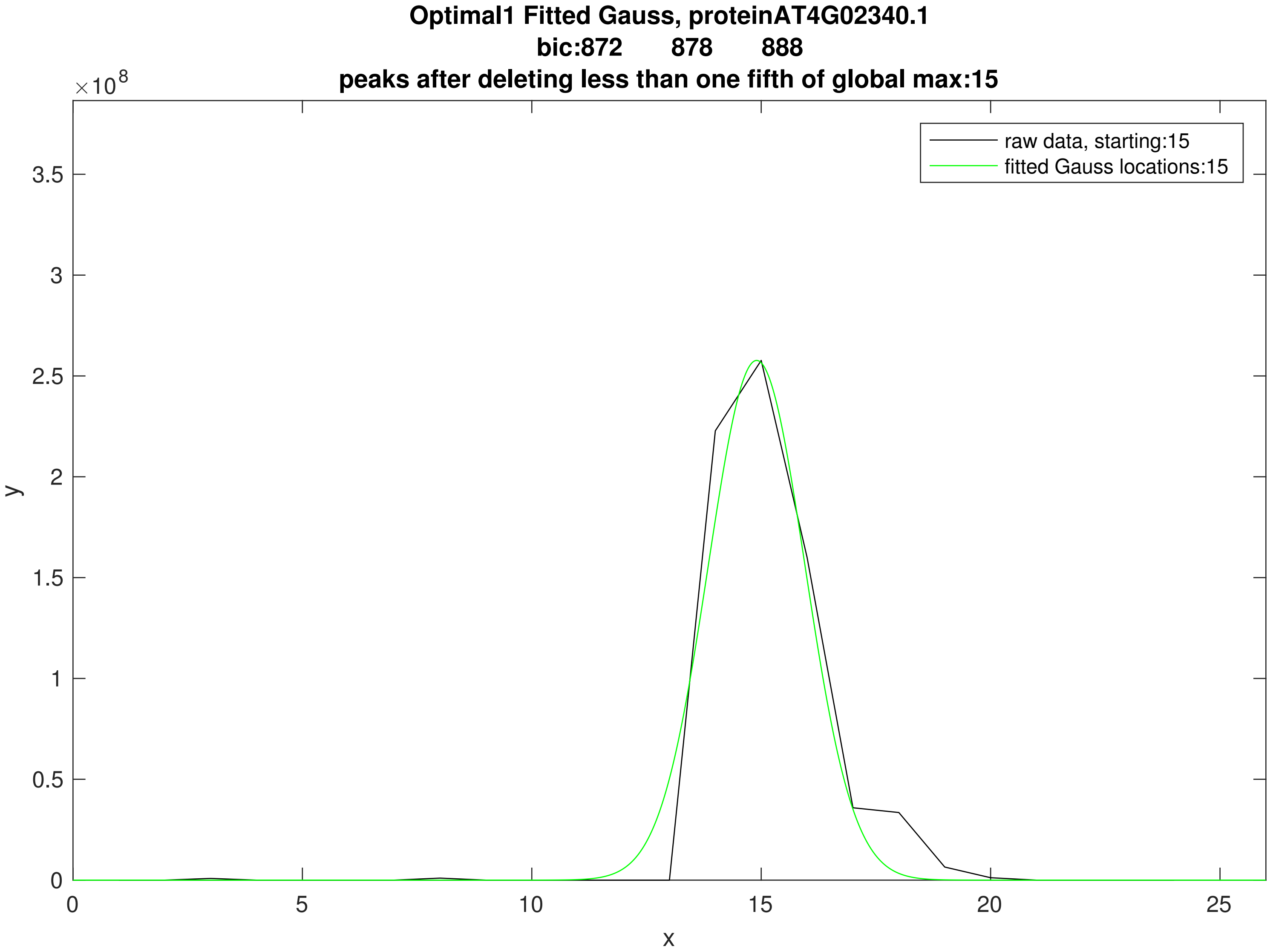
Optimal1 Fitted Gauss, proteinAT4G02340.1
bic:872 878 888
peaks after deleting less than one fifth of global max:15
×10^8
raw data, starting:15
fitted Gauss locations:15
y
x
0
0.5
1
1.5
2
2.5
3
3.5
0
5
10
15
20
25

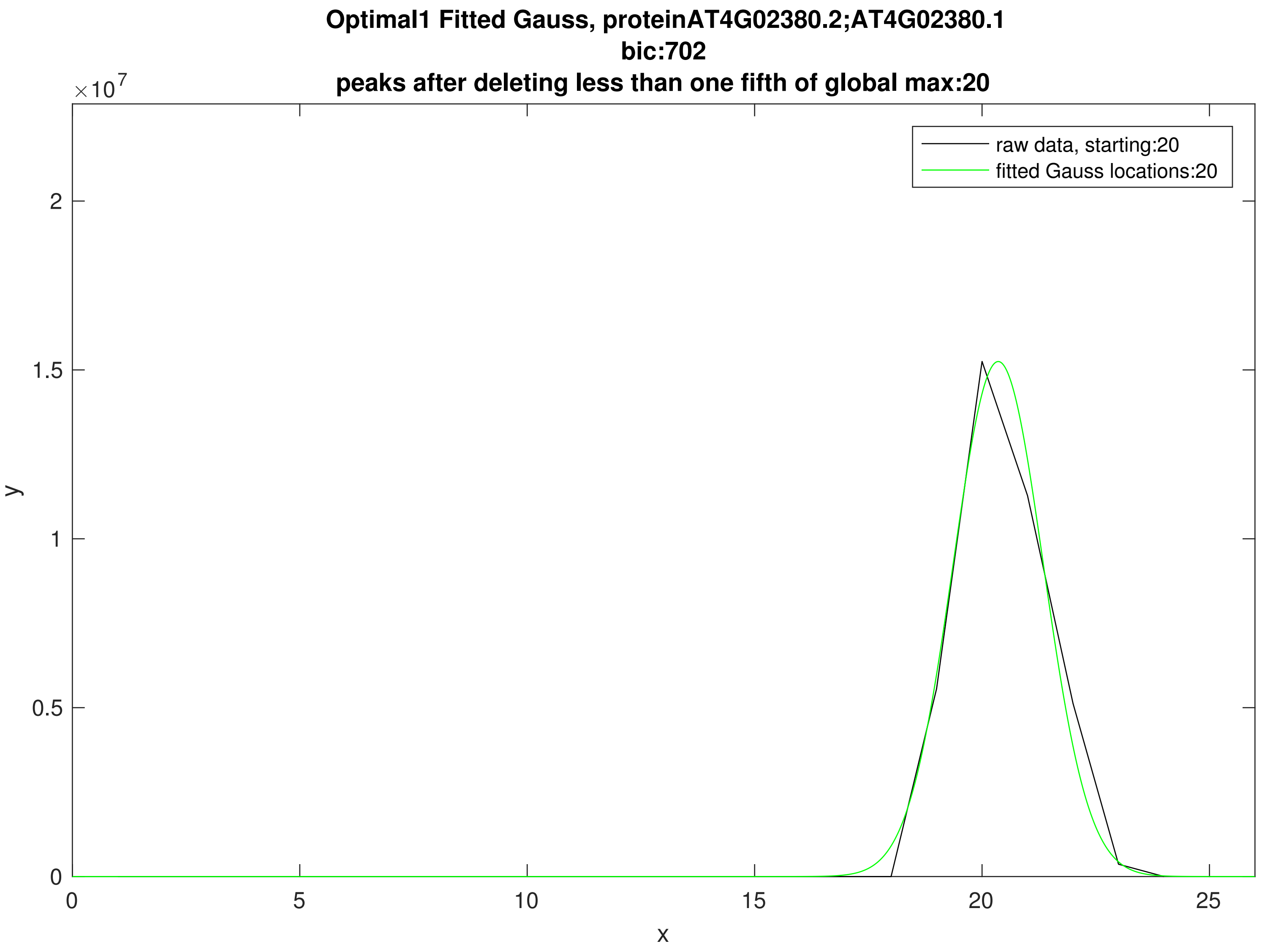
Optimal1 Fitted Gauss, proteinAT4G02380.2;AT4G02380.1
bic:702
peaks after deleting less than one fifth of global max:20
×10^7
raw data, starting:20
fitted Gauss locations:20
y
x
0
0.5
1
1.5
2
5
10
15
20
25

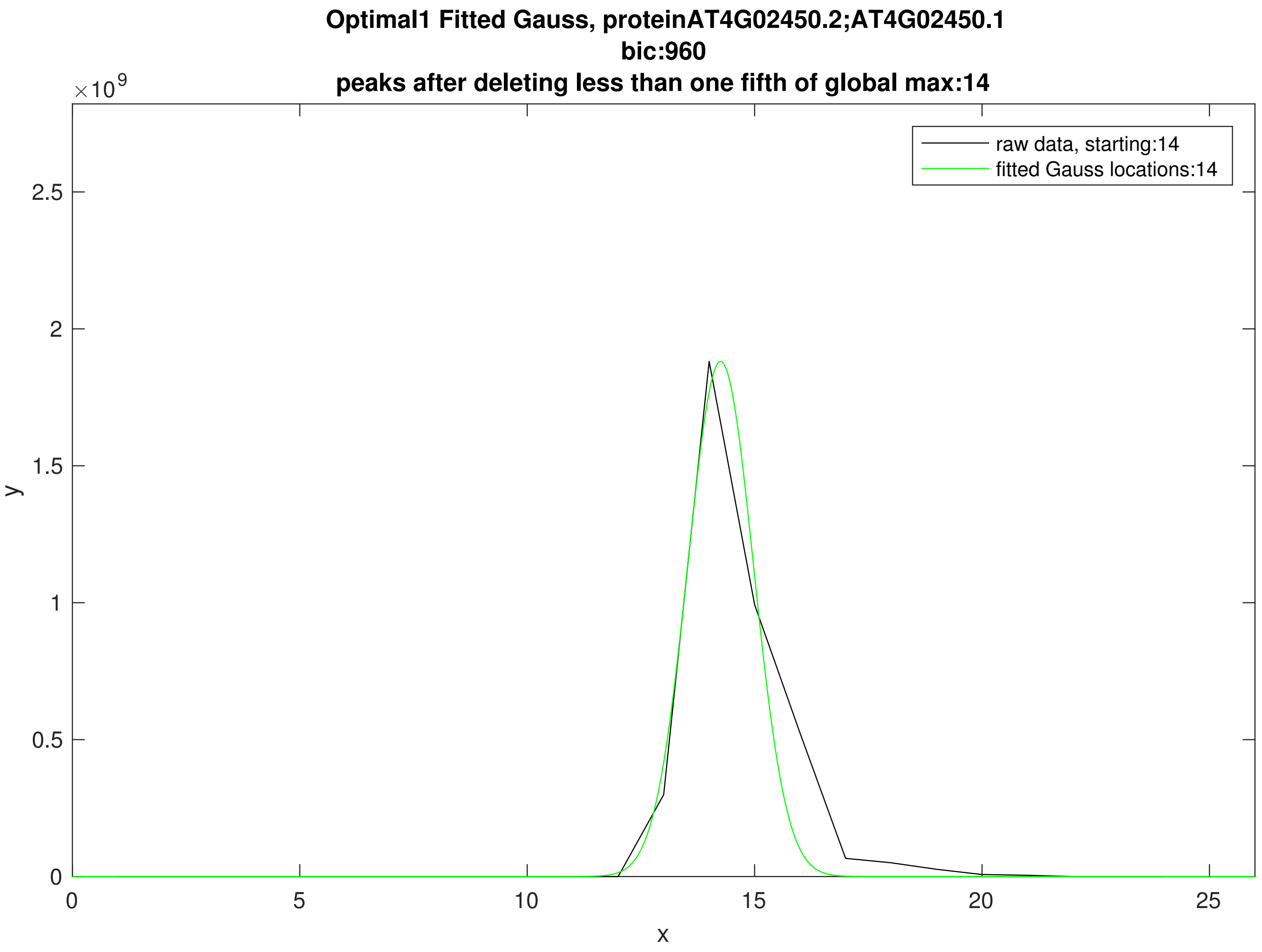
Optimal1 Fitted Gauss, proteinAT4G02450.2;AT4G02450.1
bic:960
peaks after deleting less than one fifth of global max:14
×10^9
raw data, starting:14
fitted Gauss locations:14
y
x
0
0.5
1
1.5
2
2.5
5
10
15
20
25

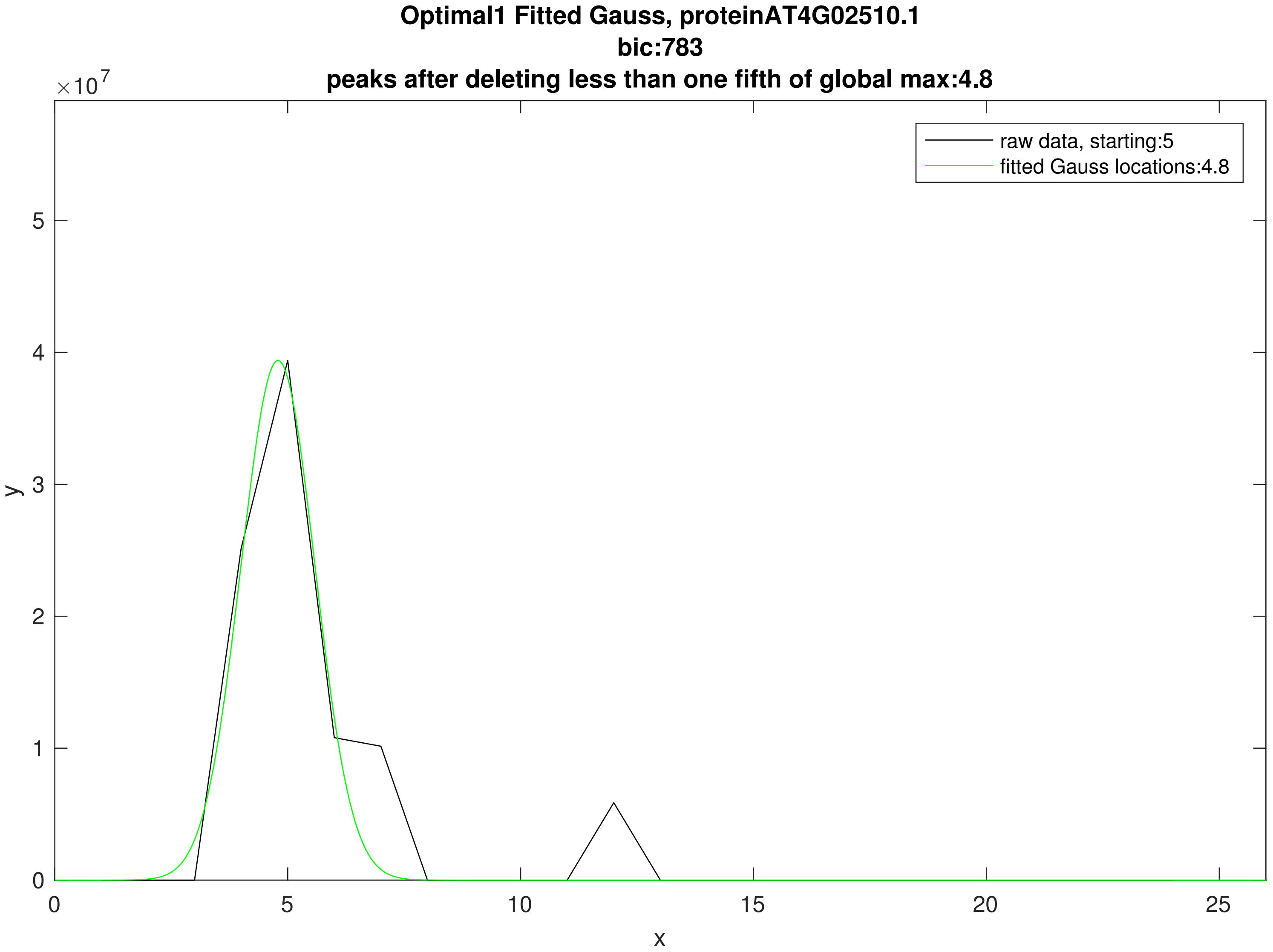
Optimal1 Fitted Gauss, proteinAT4G02510.1
bic:783
peaks after deleting less than one fifth of global max:4.8
×10^7
raw data, starting:5
fitted Gauss locations:4.8
y
x
0
1
2
3
4
5
10
15
20
25

**Optimal1 Fitted Gauss, proteinAT4G02520.1**
**bic:1.01e+03  1.02e+03  1.03e+03  1.04e+03**
**peaks after deleting less than one fifth of global max:17**

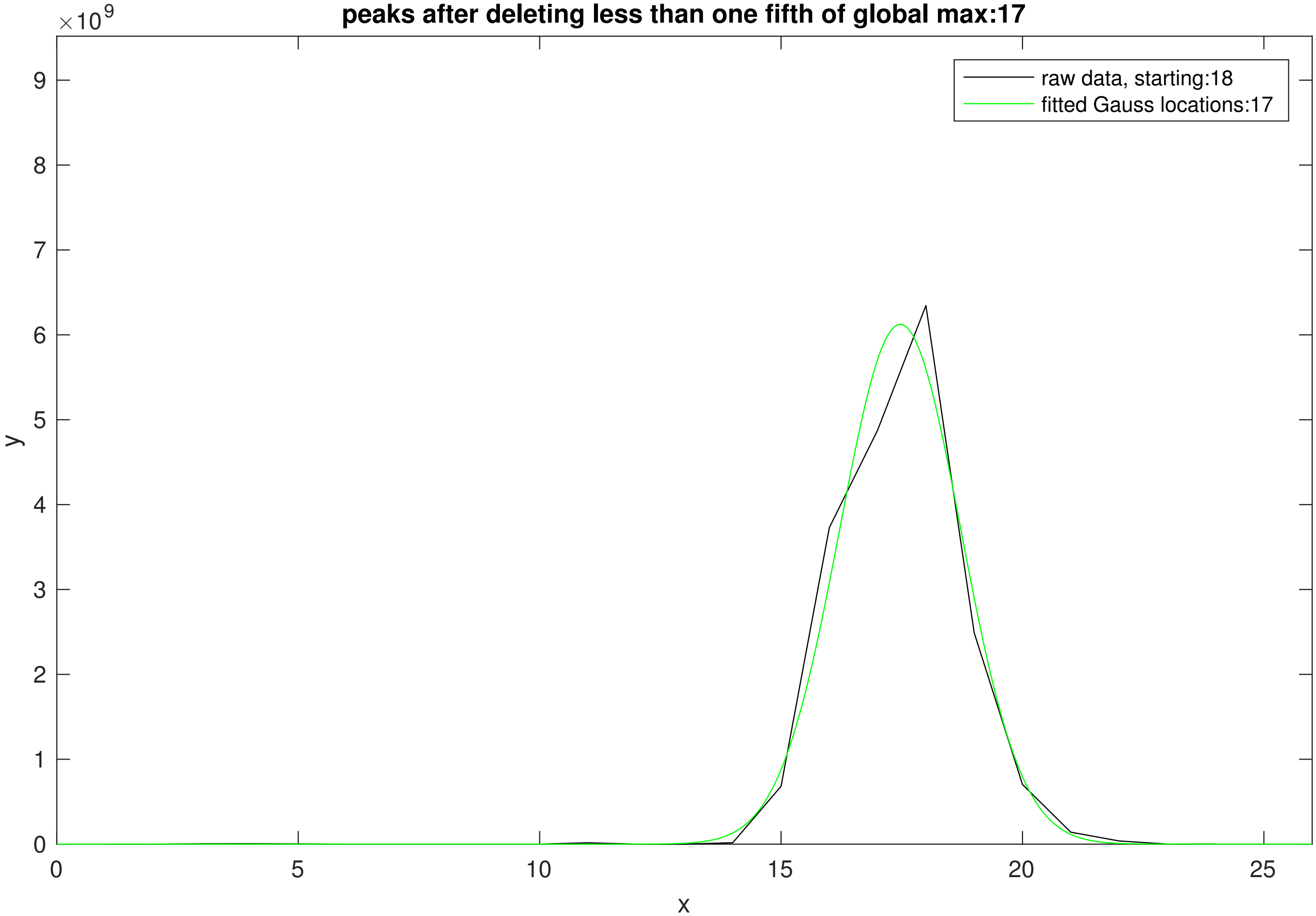

**Optimal1 Fitted Gauss, proteinAT4G02530.1**
**bic:726 735**
**peaks after deleting less than one fifth of global max:18**

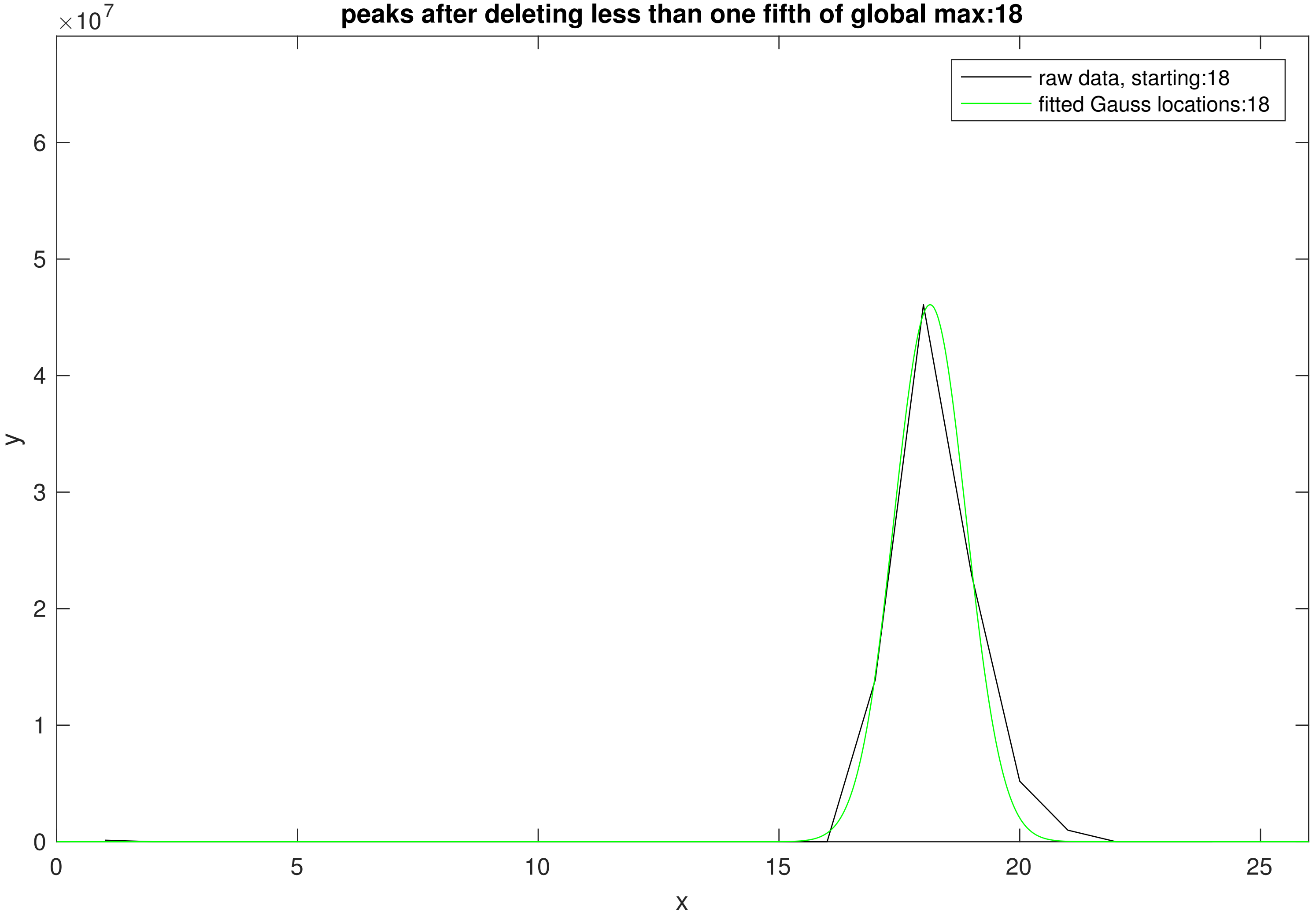

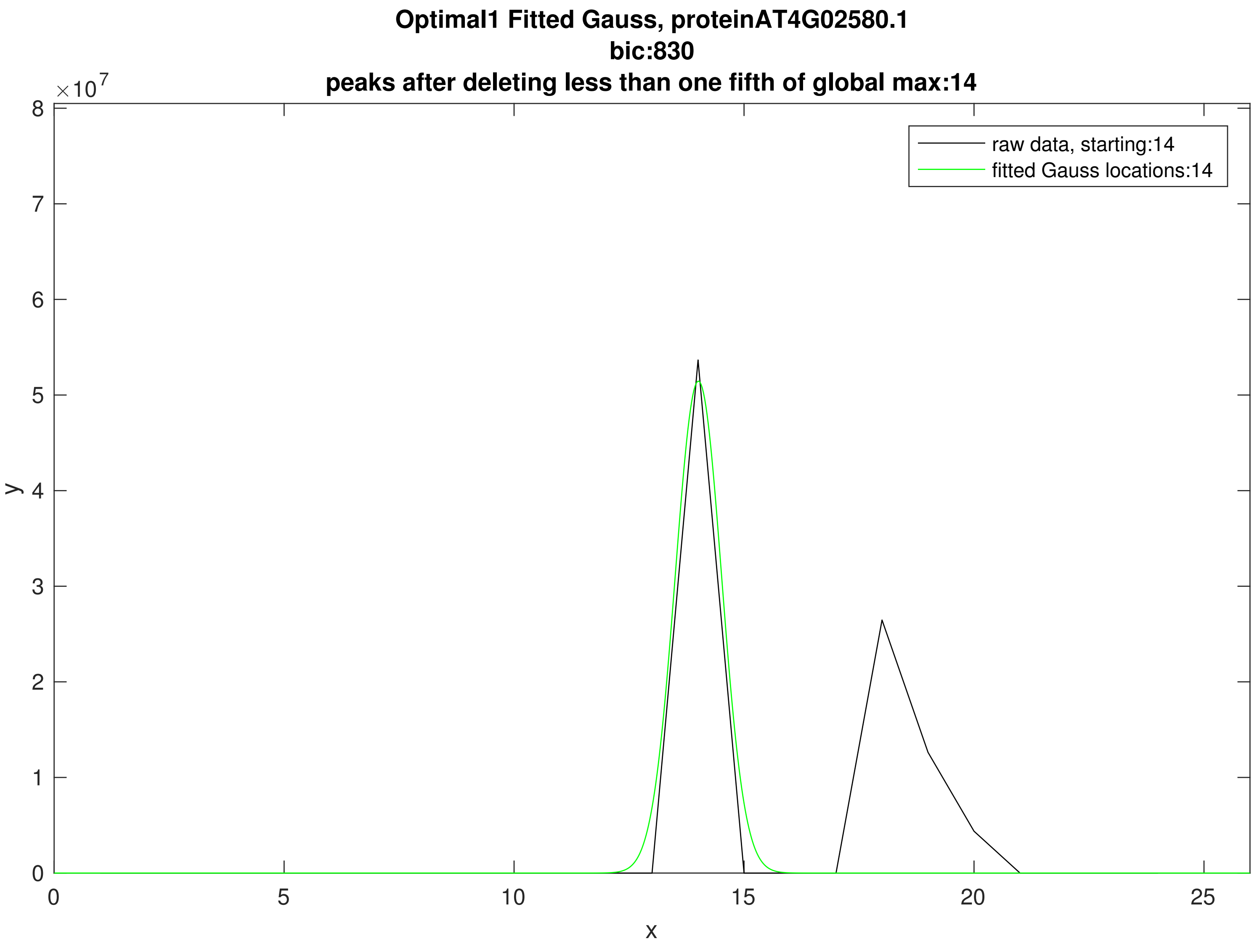

Optimal1 Fitted Gauss, proteinAT4G02580.1
bic:830
peaks after deleting less than one fifth of global max:14
×10^7
raw data, starting:14
fitted Gauss locations:14
y
x
0
1
2
3
4
5
6
7
8
0
5
10
15
20
25

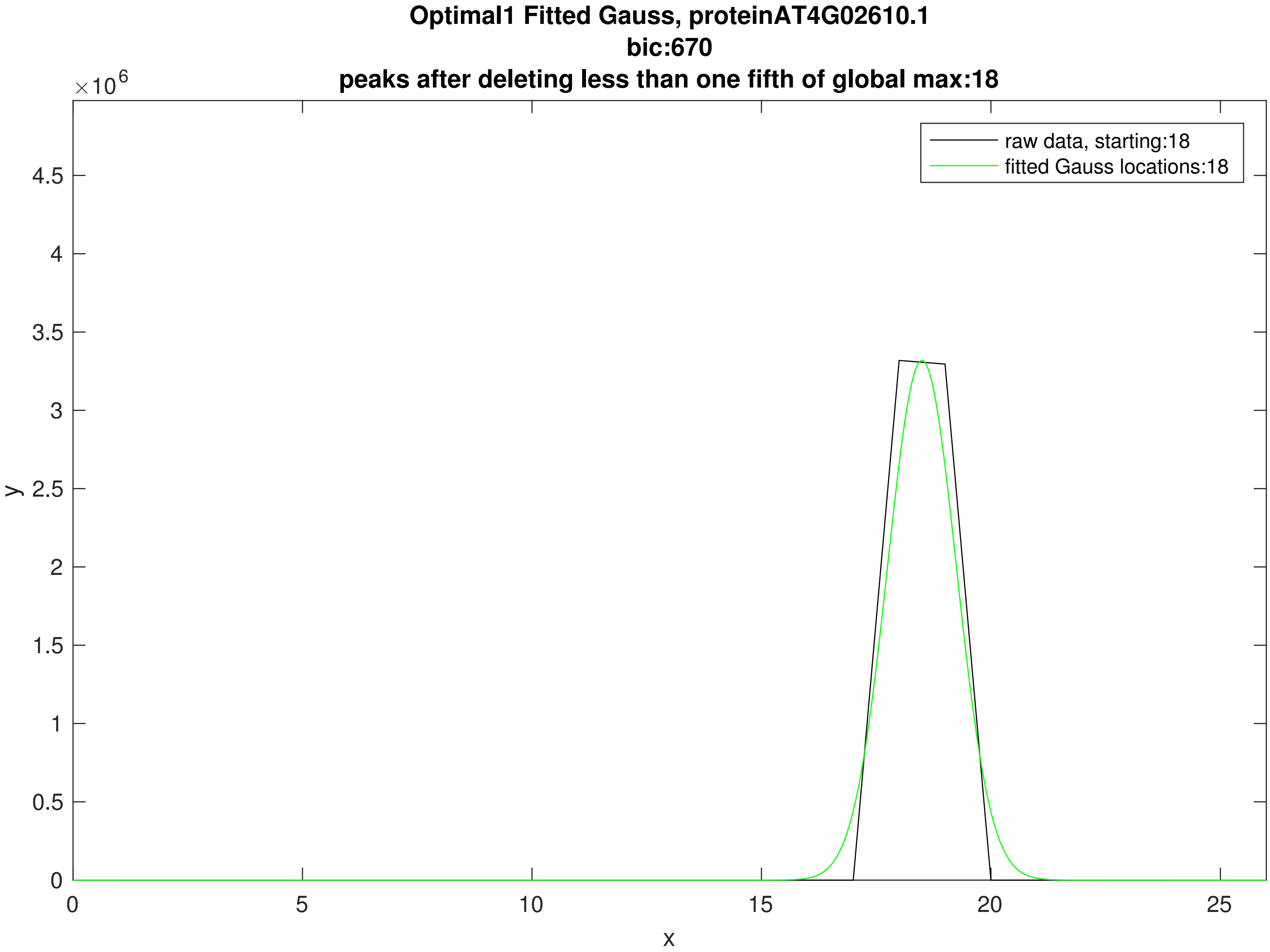
Optimal1 Fitted Gauss, proteinAT4G02610.1
bic:670
peaks after deleting less than one fifth of global max:18
$\times 10^6$
4.5
4
3.5
3
2.5
2
1.5
1
0.5
0
y
0
5
10
15
20
25
x
raw data, starting:18
fitted Gauss locations:18

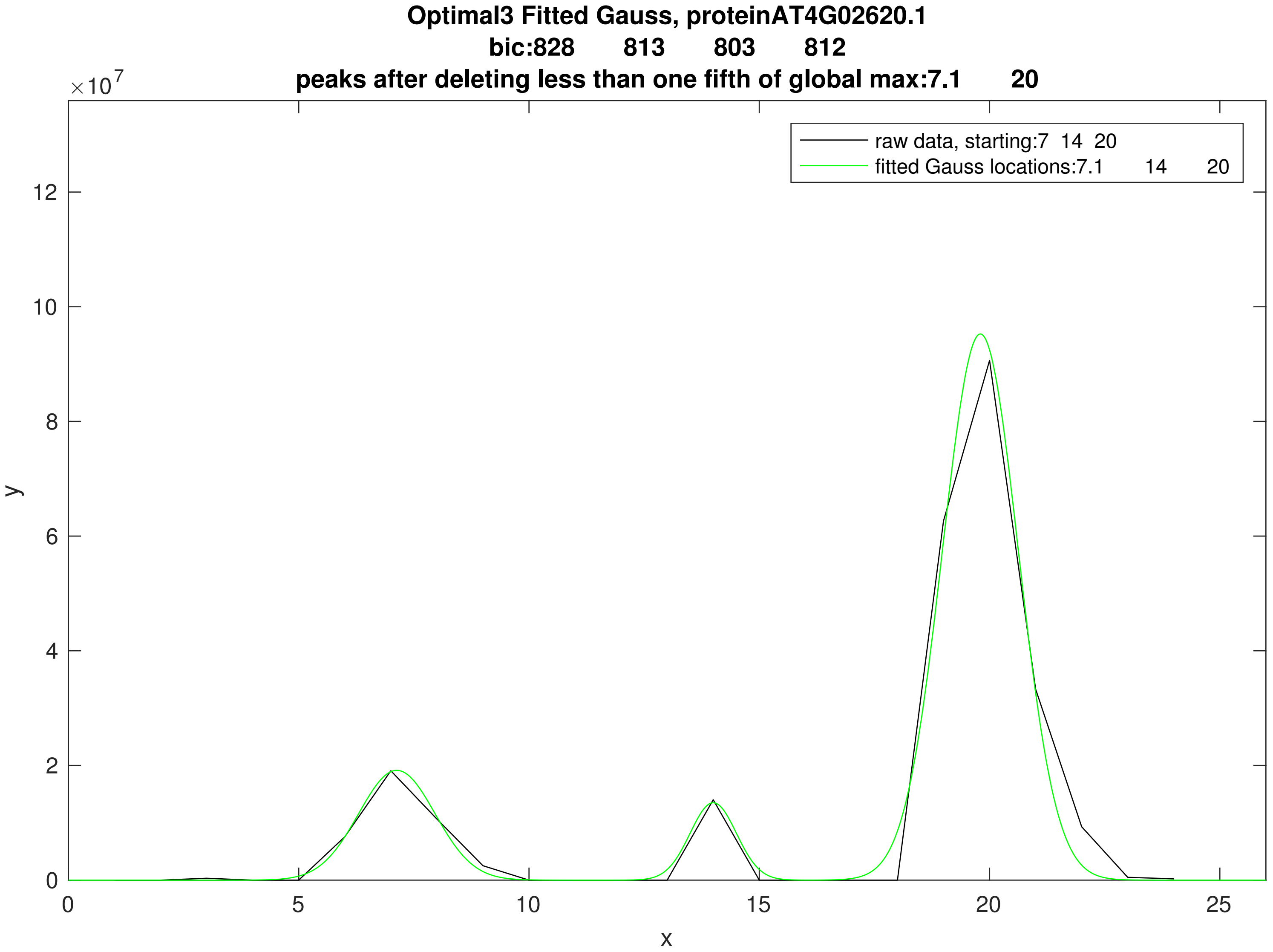

Optimal3 Fitted Gauss, proteinAT4G02620.1
bic:828 813 803 812
peaks after deleting less than one fifth of global max:7.1 20
×10^7
raw data, starting:7 14 20
fitted Gauss locations:7.1 14 20
y
x
0
2
4
6
8
10
12
5
15
20
25

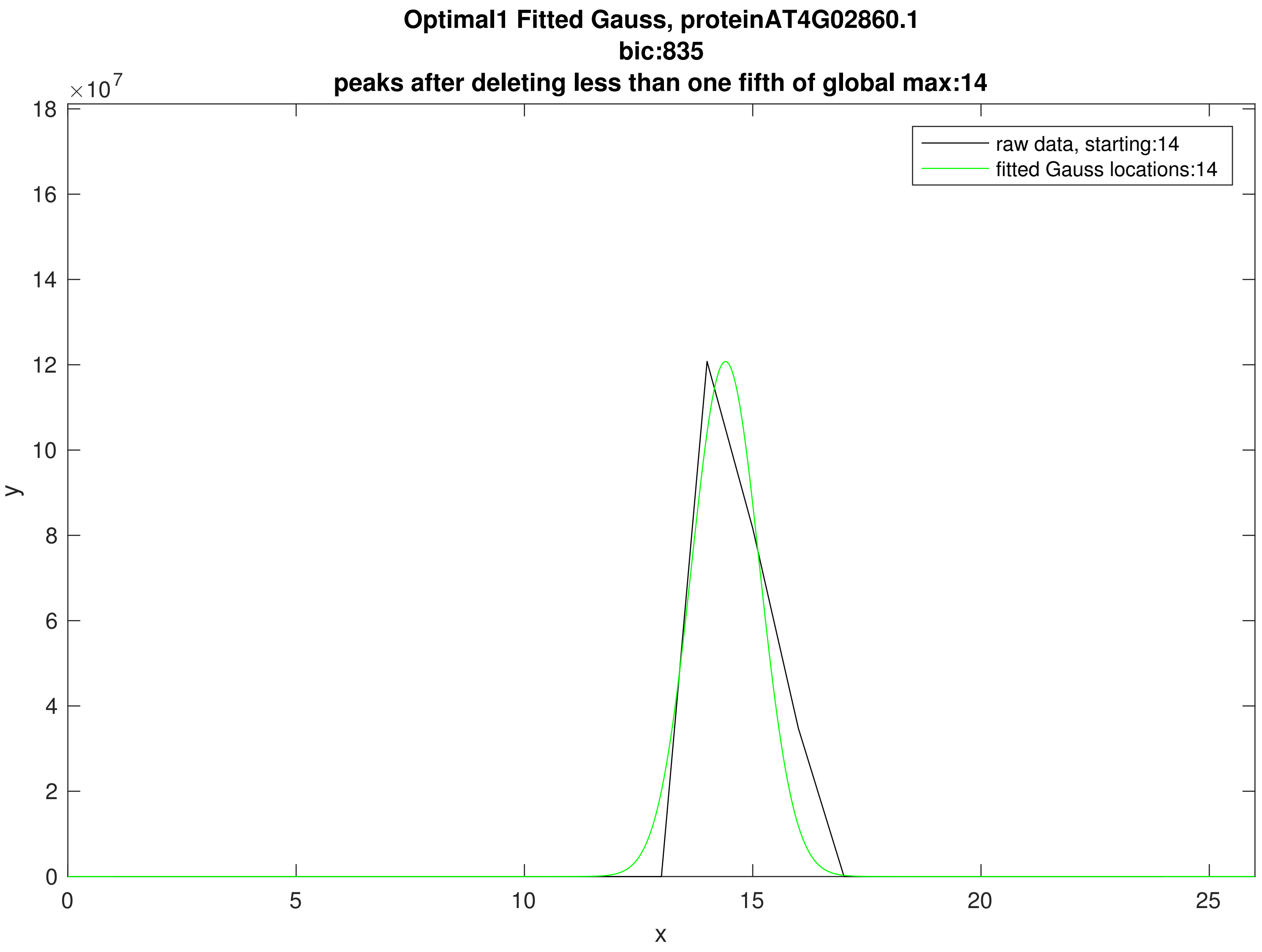
Optimal1 Fitted Gauss, proteinAT4G02860.1
bic:835
peaks after deleting less than one fifth of global max:14
×10^7
18
16
14
12
10
8
6
4
2
0
y
0
5
10
15
20
25
x
raw data, starting:14
fitted Gauss locations:14

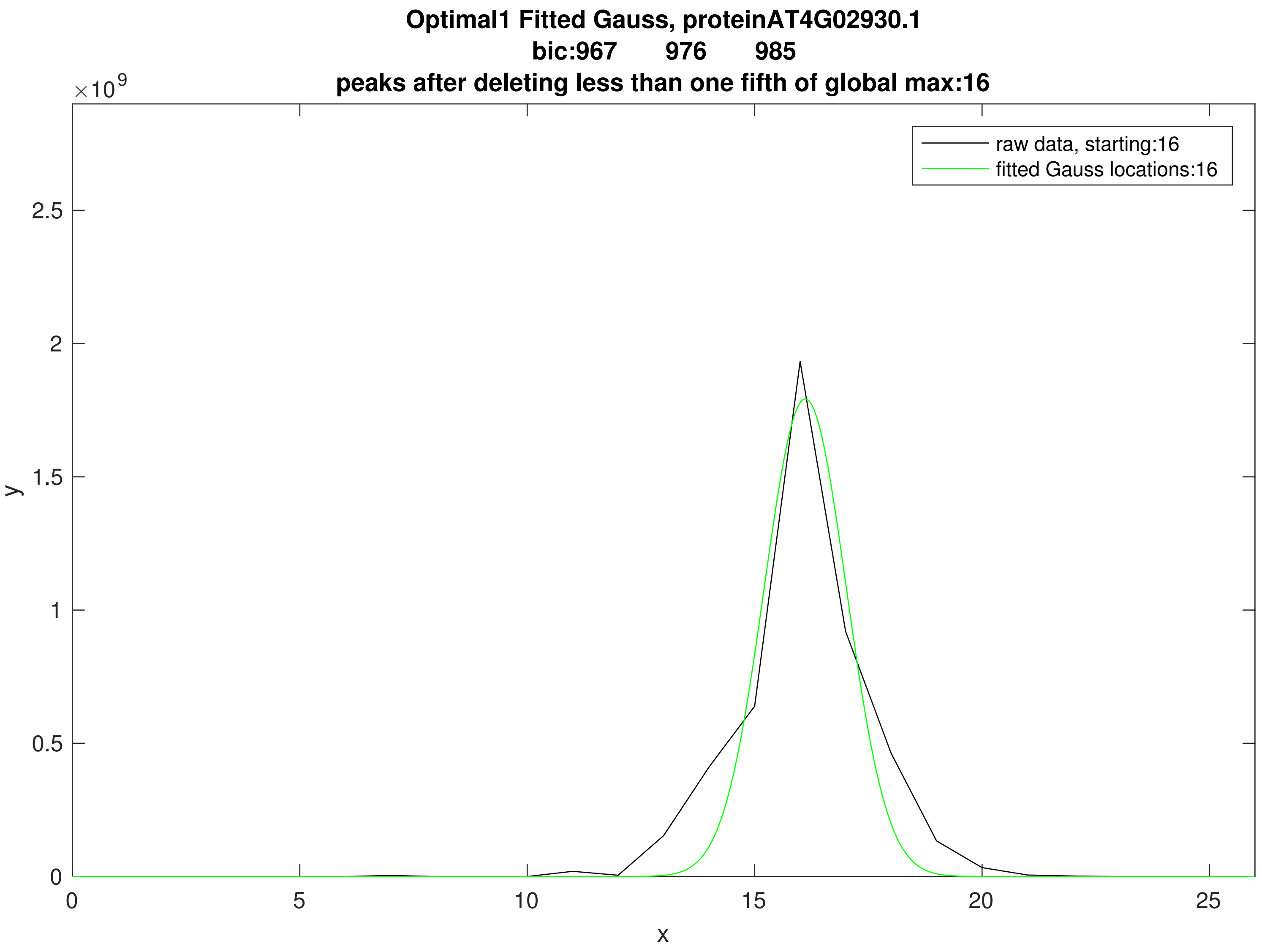
Optimal1 Fitted Gauss, proteinAT4G02930.1
bic:967 976 985
peaks after deleting less than one fifth of global max:16
×10^9
raw data, starting:16
fitted Gauss locations:16
2.5
2
1.5
1
0.5
0
y
0
5
10
15
20
25
x

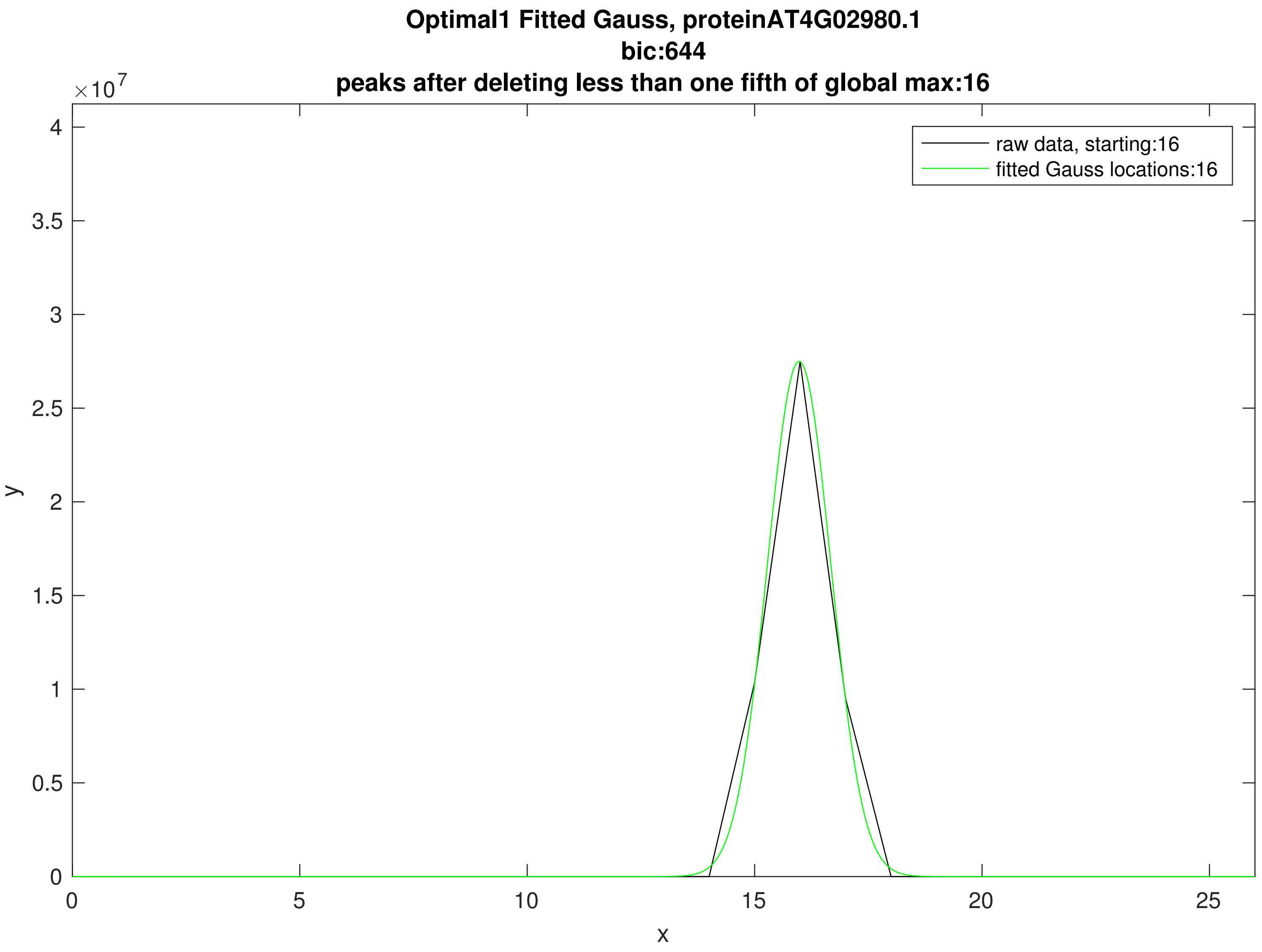
Optimal1 Fitted Gauss, proteinAT4G02980.1
bic:644
peaks after deleting less than one fifth of global max:16
×10^7
raw data, starting:16
fitted Gauss locations:16
y
x
0
0.5
1
1.5
2
2.5
3
3.5
4
5
10
15
20
25

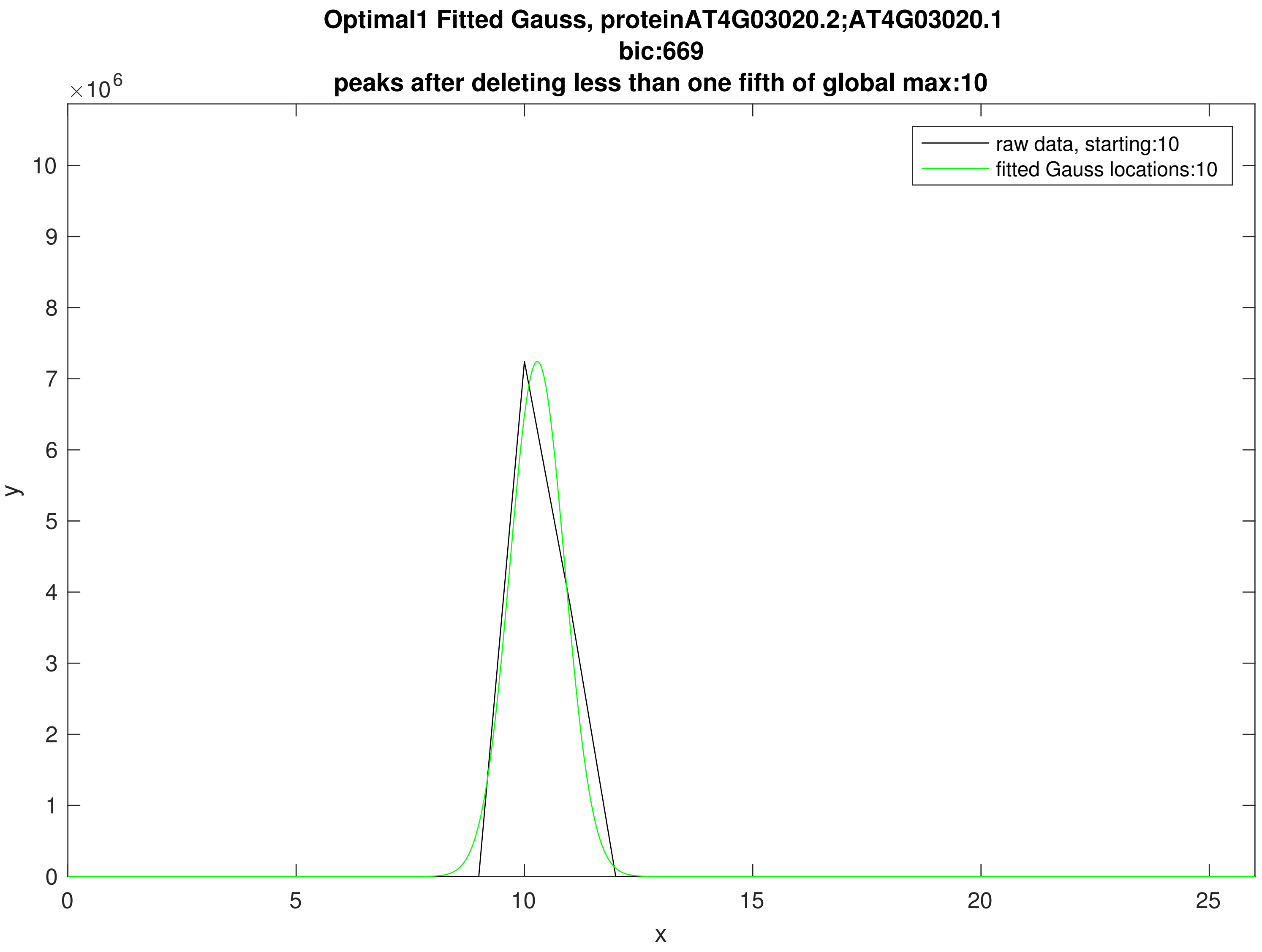

Optimal1 Fitted Gauss, proteinAT4G03020.2;AT4G03020.1
bic:669
peaks after deleting less than one fifth of global max:10
×10^6
raw data, starting:10
fitted Gauss locations:10
y
x
0
1
2
3
4
5
6
7
8
9
10
0
5
10
15
20
25

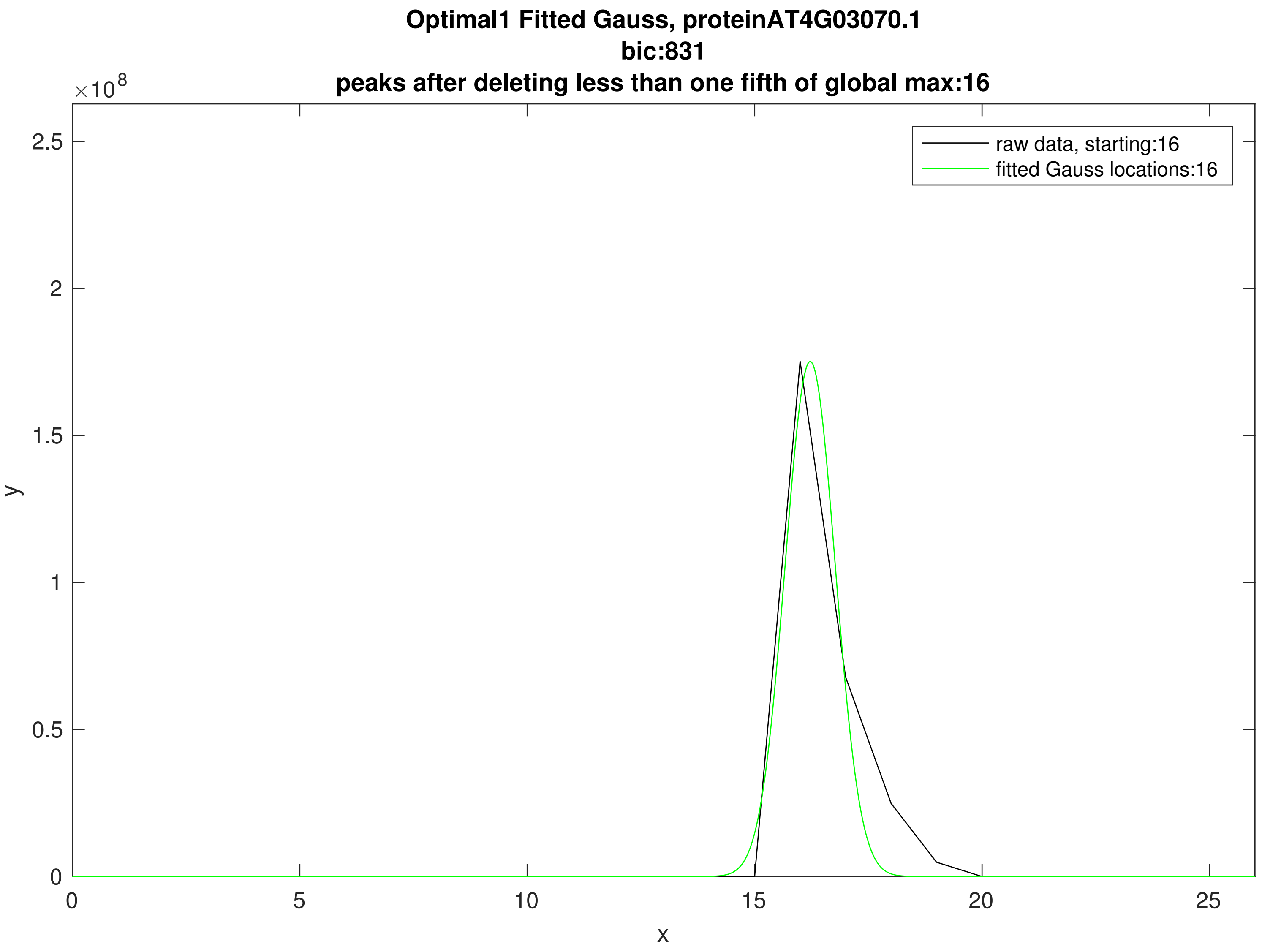

Optimal1 Fitted Gauss, proteinAT4G03070.1
bic:831
peaks after deleting less than one fifth of global max:16
×10^8
2.5
2
1.5
1
0.5
0
y
0
5
10
15
20
25
x
raw data, starting:16
fitted Gauss locations:16

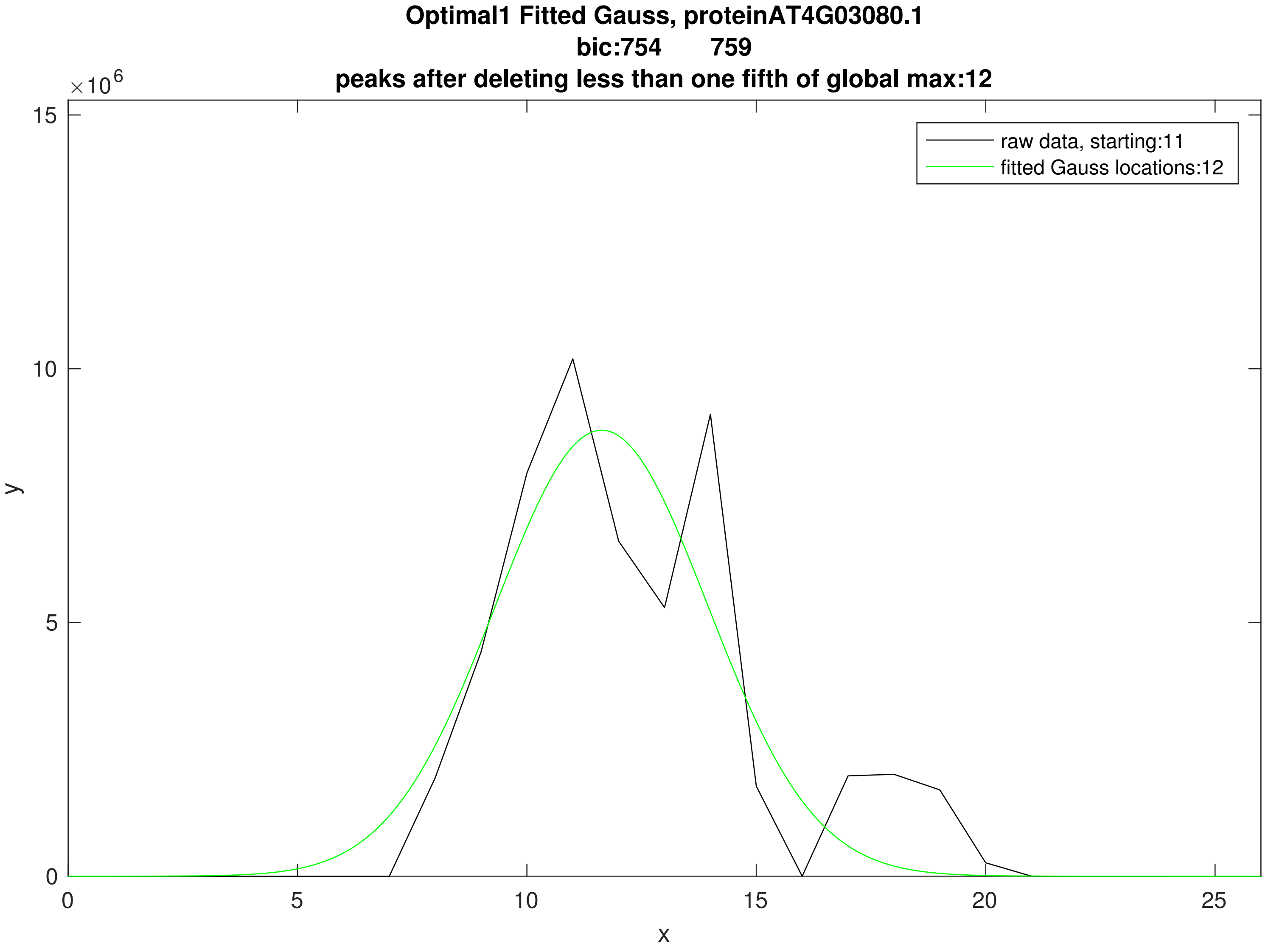

Optimal1 Fitted Gauss, proteinAT4G03080.1
bic:754 759
peaks after deleting less than one fifth of global max:12
×10^6
15
10
5
0
y
0
5
10
15
20
25
x
raw data, starting:11
fitted Gauss locations:12

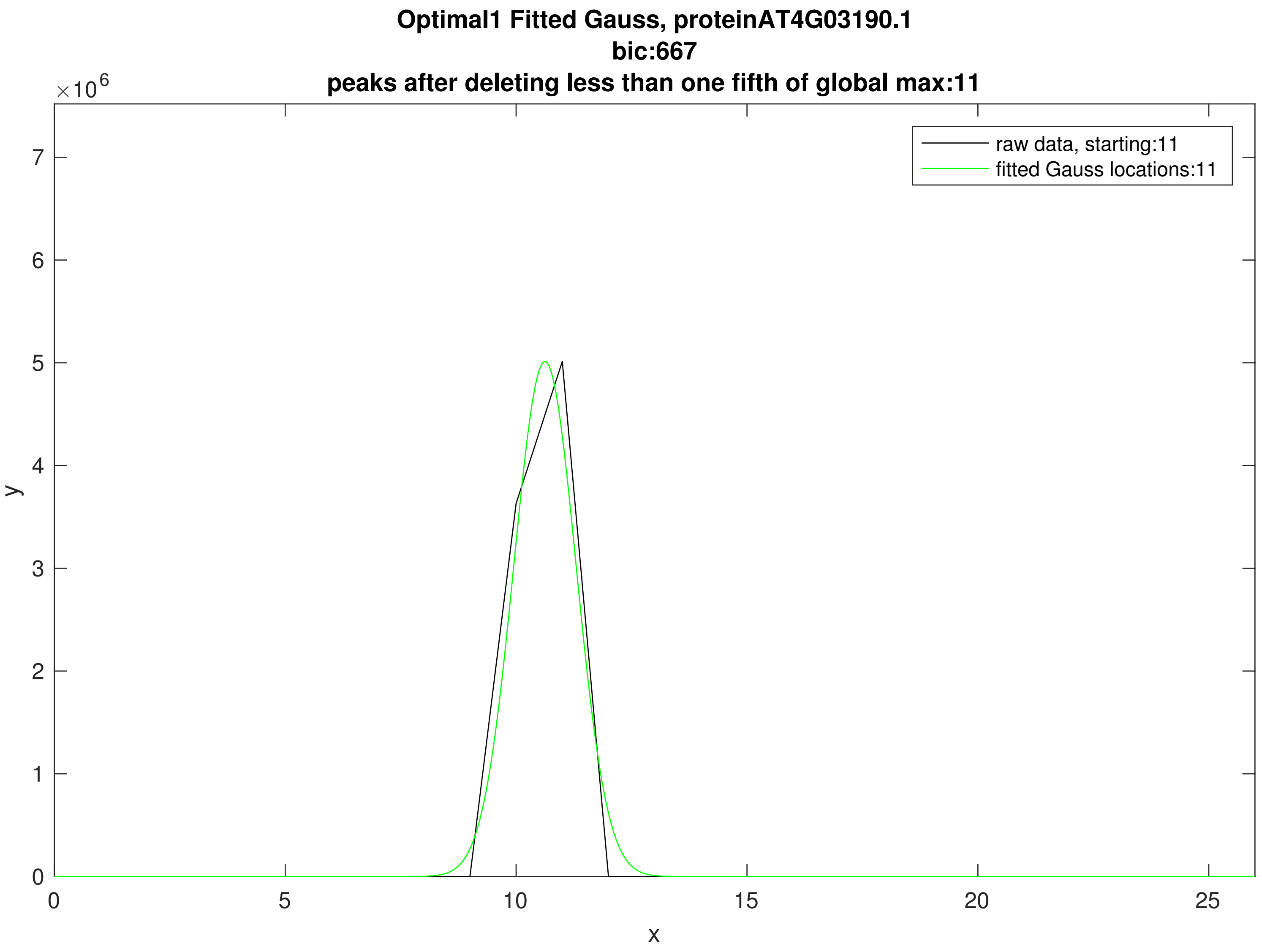

Optimal1 Fitted Gauss, proteinAT4G03190.1
bic:667
peaks after deleting less than one fifth of global max:11
×10^6
raw data, starting:11
fitted Gauss locations:11
y
x
0
1
2
3
4
5
6
7
0
5
10
15
20
25

**Optimal1 Fitted Gauss, proteinAT4G03200.2;AT4G03200.1**
**bic:697**
**peaks after deleting less than one fifth of global max:14**

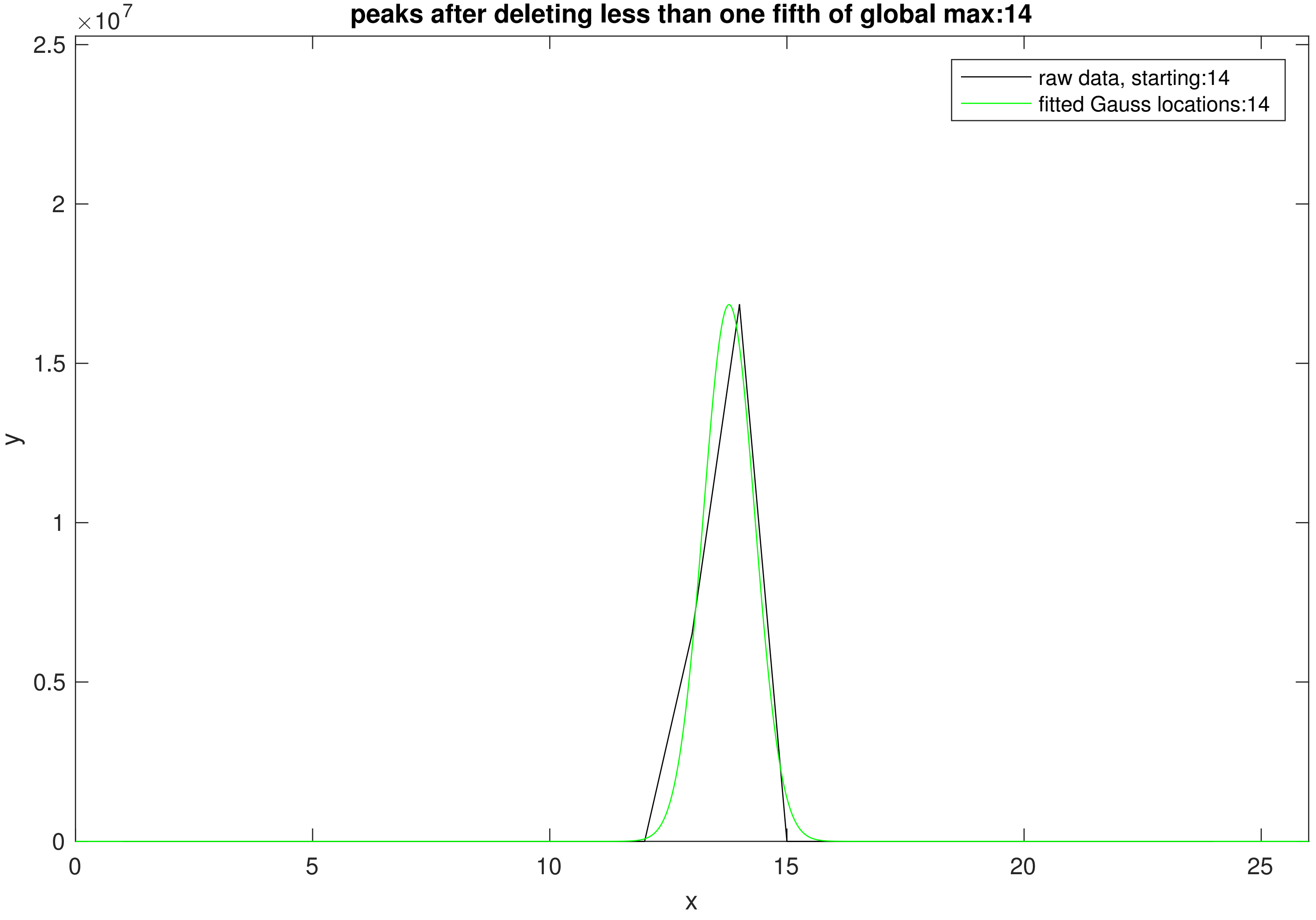

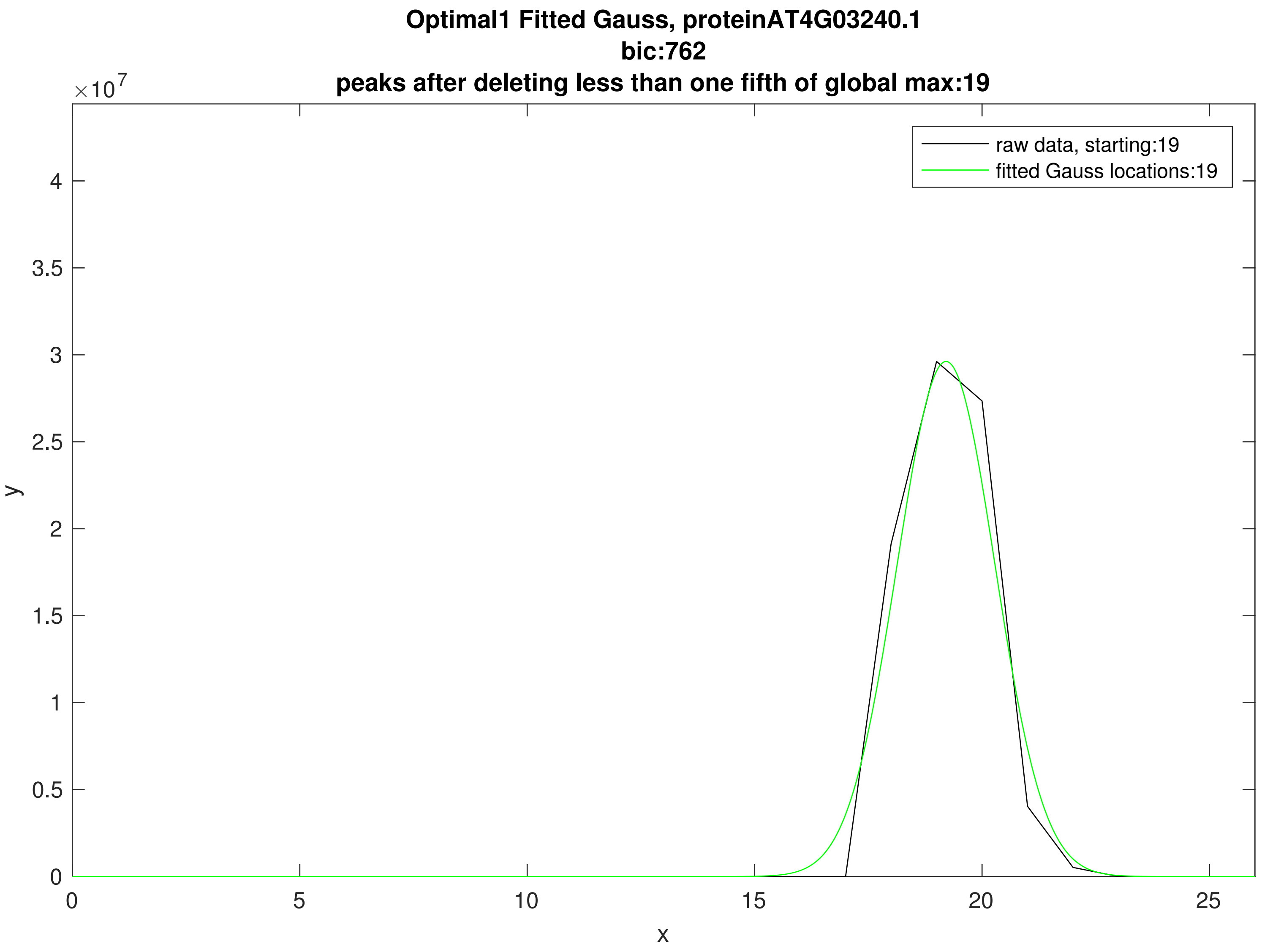
Optimal1 Fitted Gauss, proteinAT4G03240.1
bic:762
peaks after deleting less than one fifth of global max:19
×10^7
raw data, starting:19
fitted Gauss locations:19
0
0.5
1
1.5
2
2.5
3
3.5
4
0
5
10
15
20
25
x
y

Optimal1 Fitted Gauss, proteinAT4G03520.1;AT4G03520.2
bic:788
peaks after deleting less than one fifth of global max:21

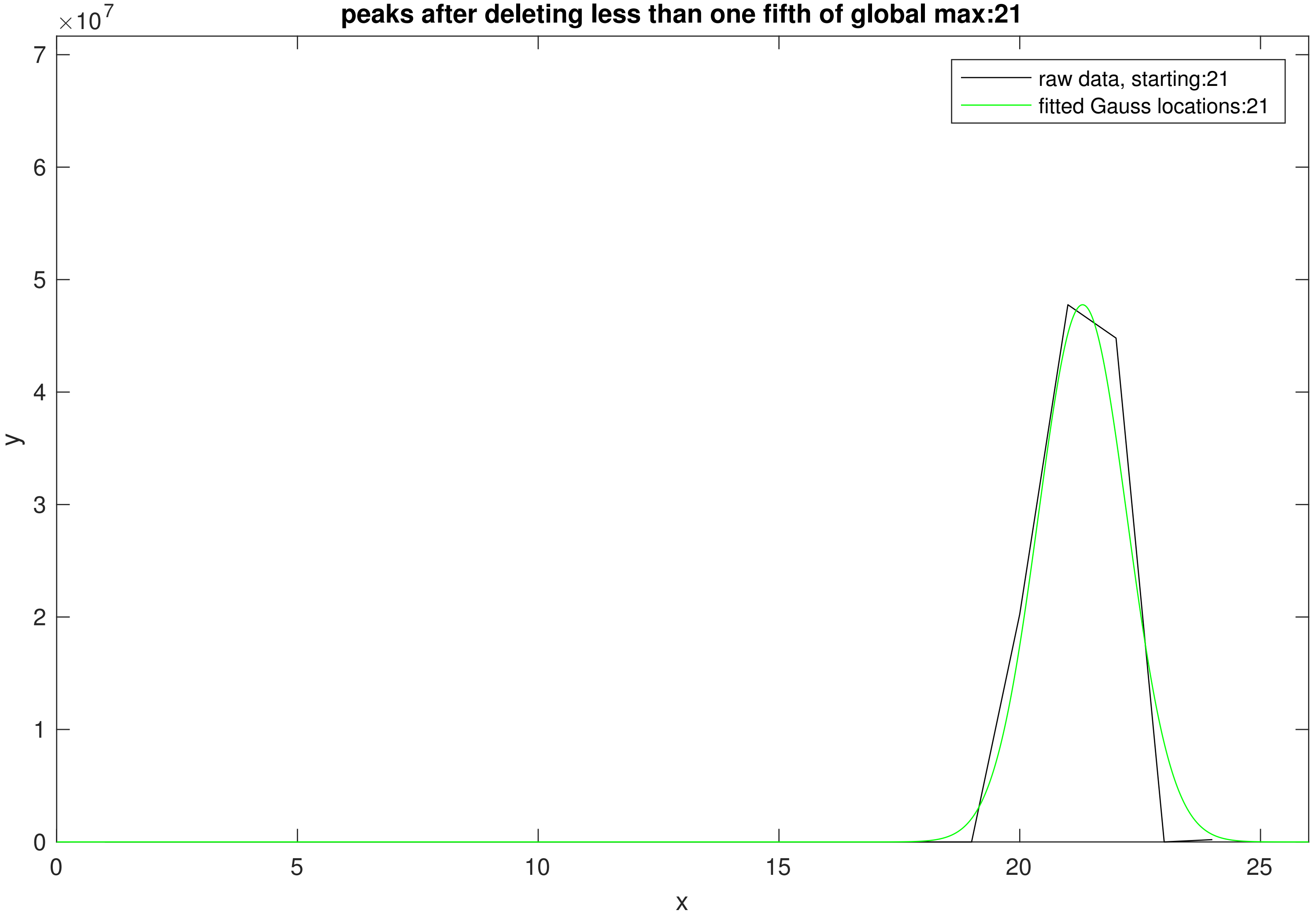

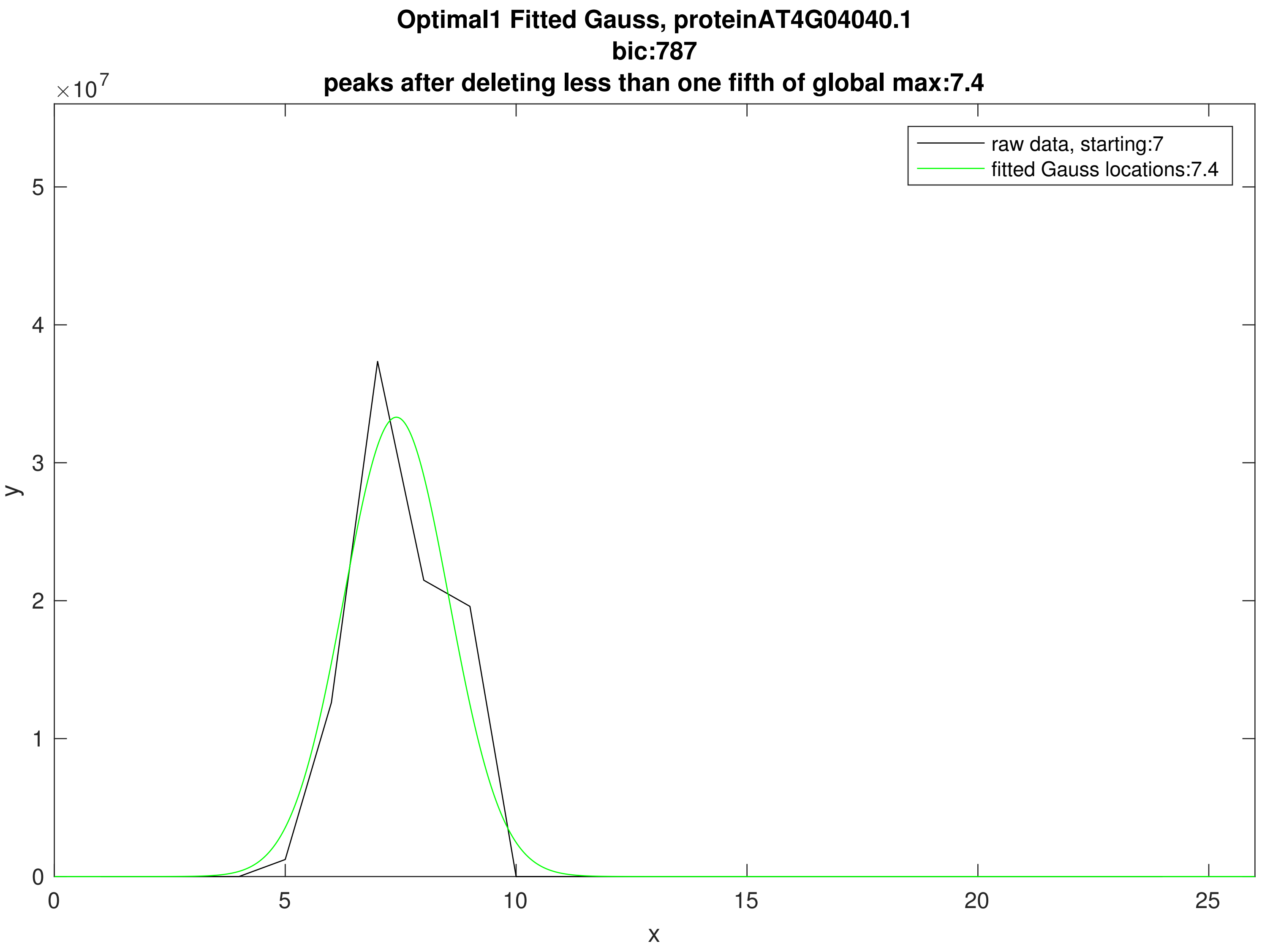
Optimal1 Fitted Gauss, proteinAT4G04040.1
bic:787
peaks after deleting less than one fifth of global max:7.4
×10^7
raw data, starting:7
fitted Gauss locations:7.4
y
x
0
1
2
3
4
5
0
5
10
15
20
25

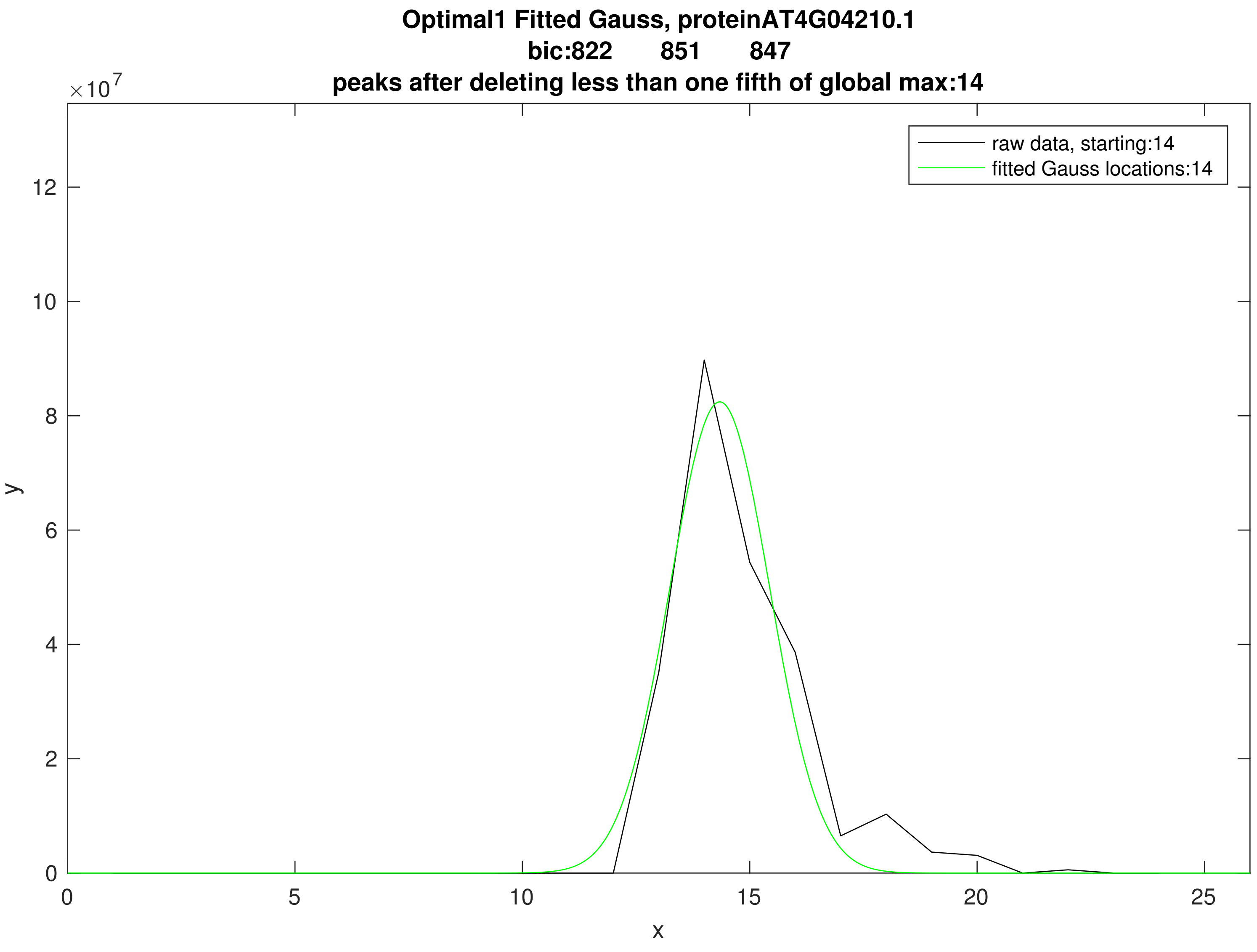
Optimal1 Fitted Gauss, proteinAT4G04210.1
bic:822 851 847
peaks after deleting less than one fifth of global max:14
×10^7
raw data, starting:14
fitted Gauss locations:14
y
x
0
2
4
6
8
10
12
0
5
10
15
20
25

**Optimal1 Fitted Gauss, proteinAT4G04640.1**
**bic:820 827 837**
**peaks after deleting less than one fifth of global max:7.3**

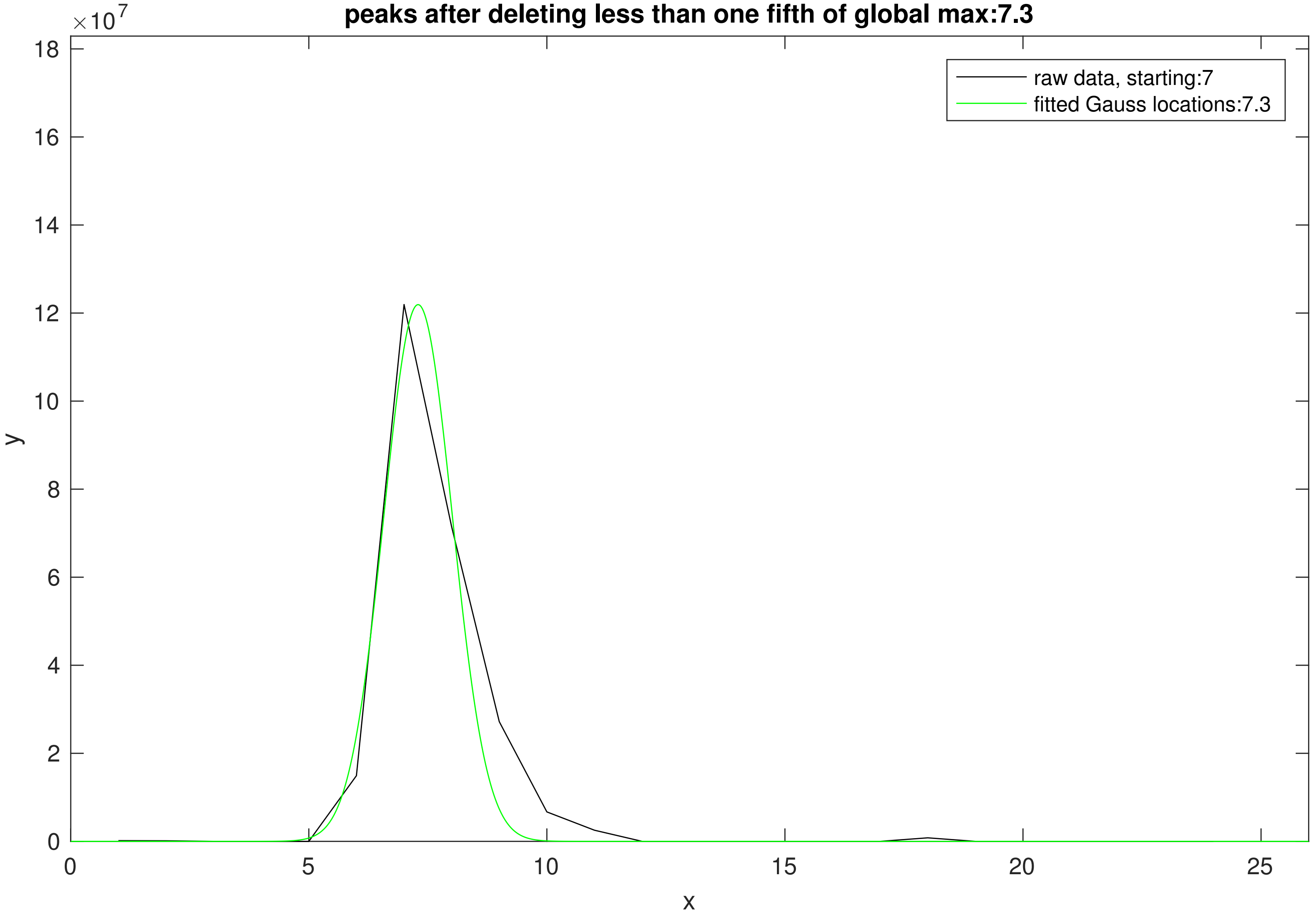

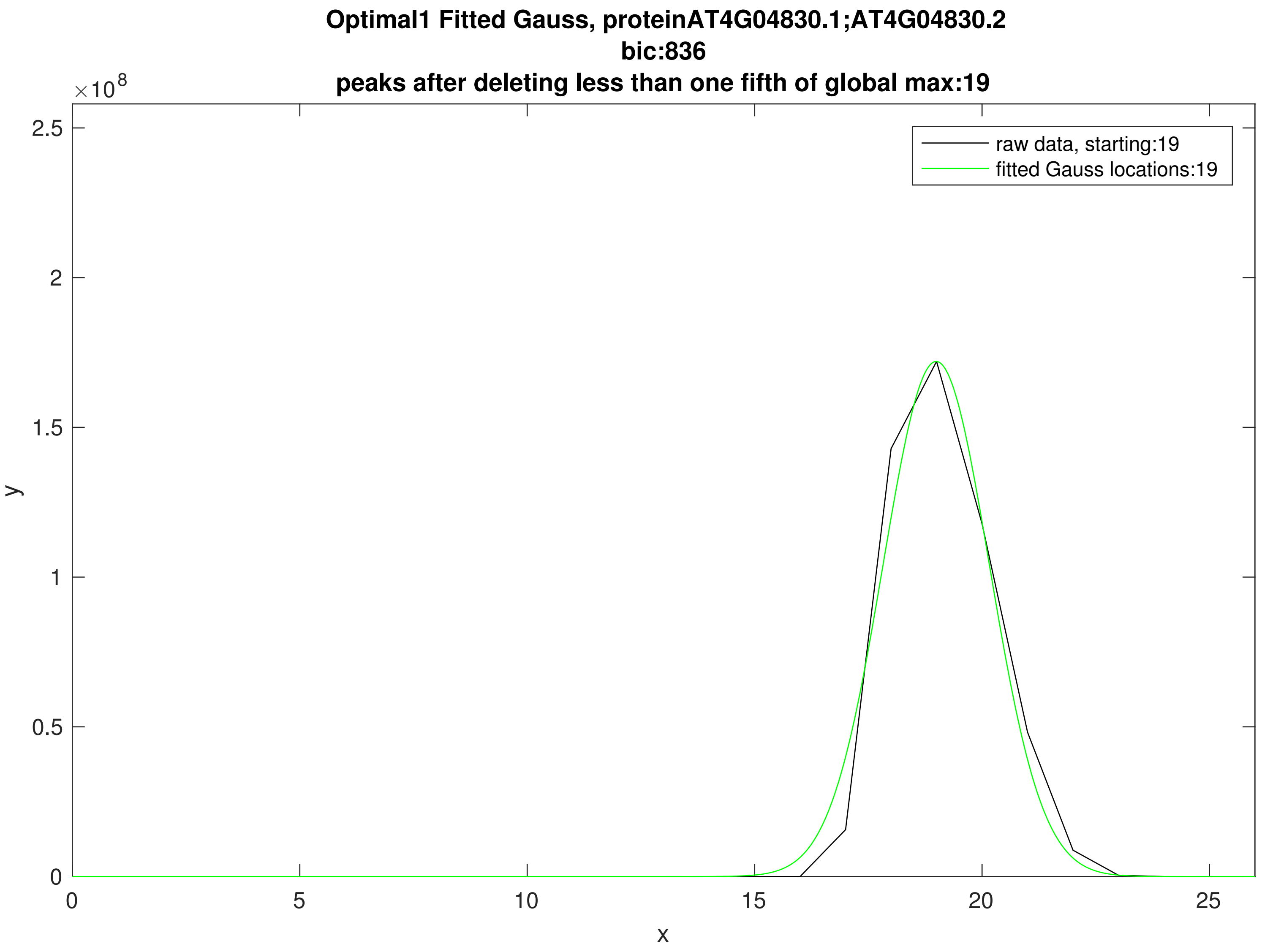

Optimal1 Fitted Gauss, proteinAT4G04830.1;AT4G04830.2
bic:836
peaks after deleting less than one fifth of global max:19
×10^8
2.5
2
1.5
1
0.5
0
y
0
5
10
15
20
25
x
raw data, starting:19
fitted Gauss locations:19

**Optimal1 Fitted Gauss, proteinAT4G04880.1**
**bic:679**
**peaks after deleting less than one fifth of global max:17**

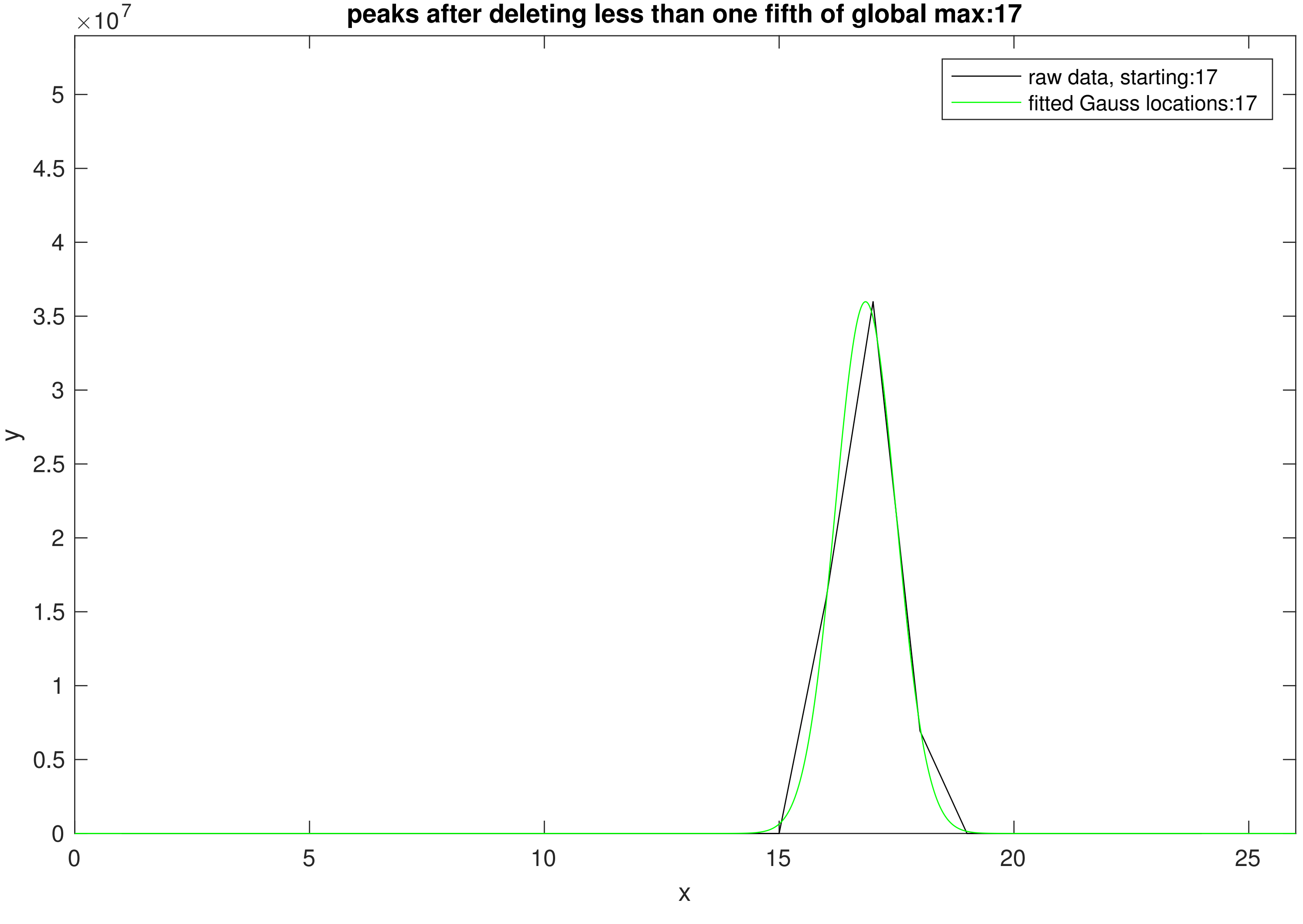

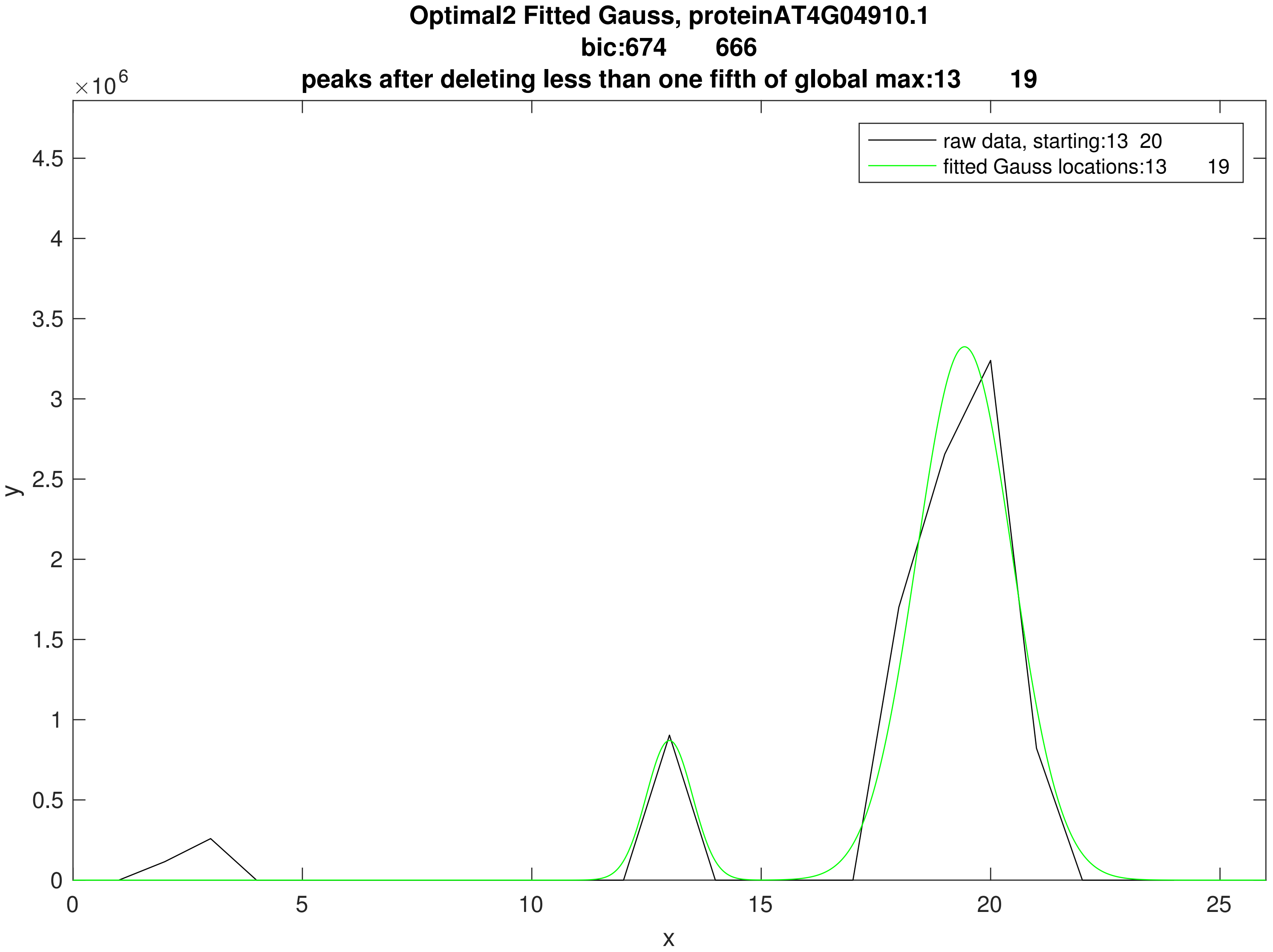
Optimal2 Fitted Gauss, proteinAT4G04910.1
bic:674 666
peaks after deleting less than one fifth of global max:13 19
raw data, starting:13 20
fitted Gauss locations:13 19
×10^6
y
x
0
0.5
1
1.5
2
2.5
3
3.5
4
4.5
5
10
15
20
25

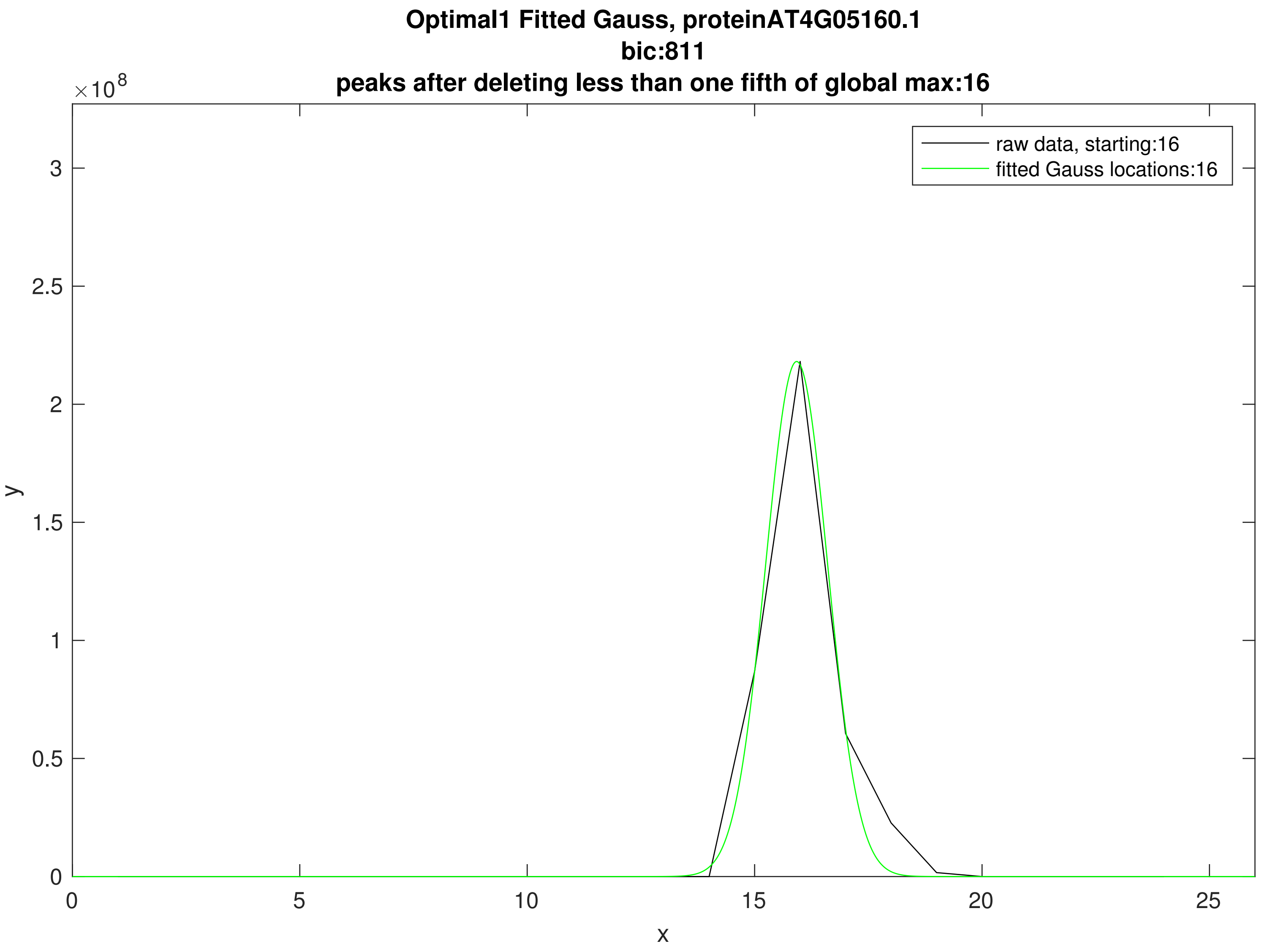
Optimal1 Fitted Gauss, proteinAT4G05160.1
bic:811
peaks after deleting less than one fifth of global max:16
×10^8
raw data, starting:16
fitted Gauss locations:16
y
x
0
0.5
1
1.5
2
2.5
3
0
5
10
15
20
25

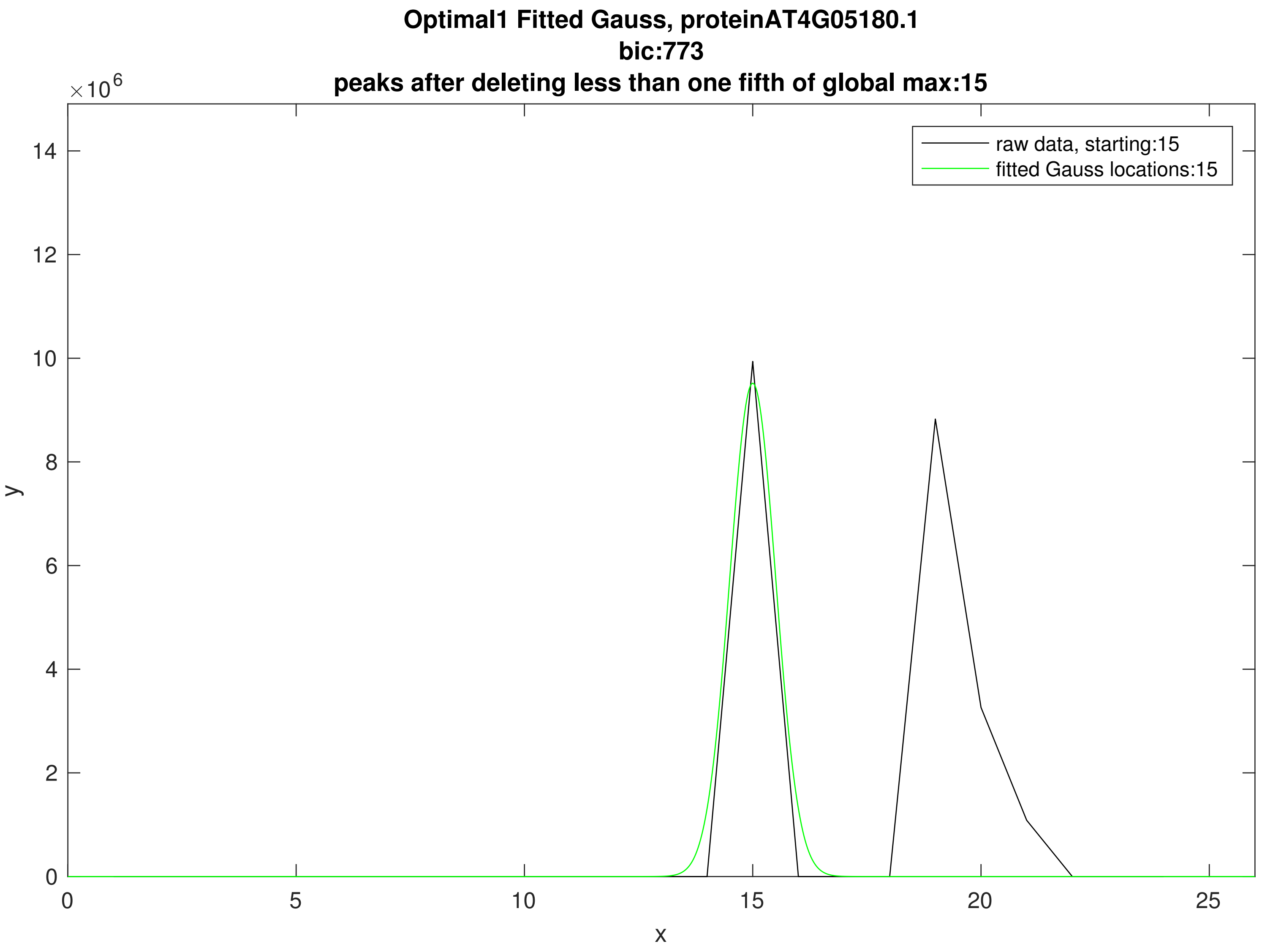

Optimal1 Fitted Gauss, proteinAT4G05180.1
bic:773
peaks after deleting less than one fifth of global max:15
×10^6
raw data, starting:15
fitted Gauss locations:15
14
12
10
8
6
4
2
0
y
0
5
10
15
20
25
x

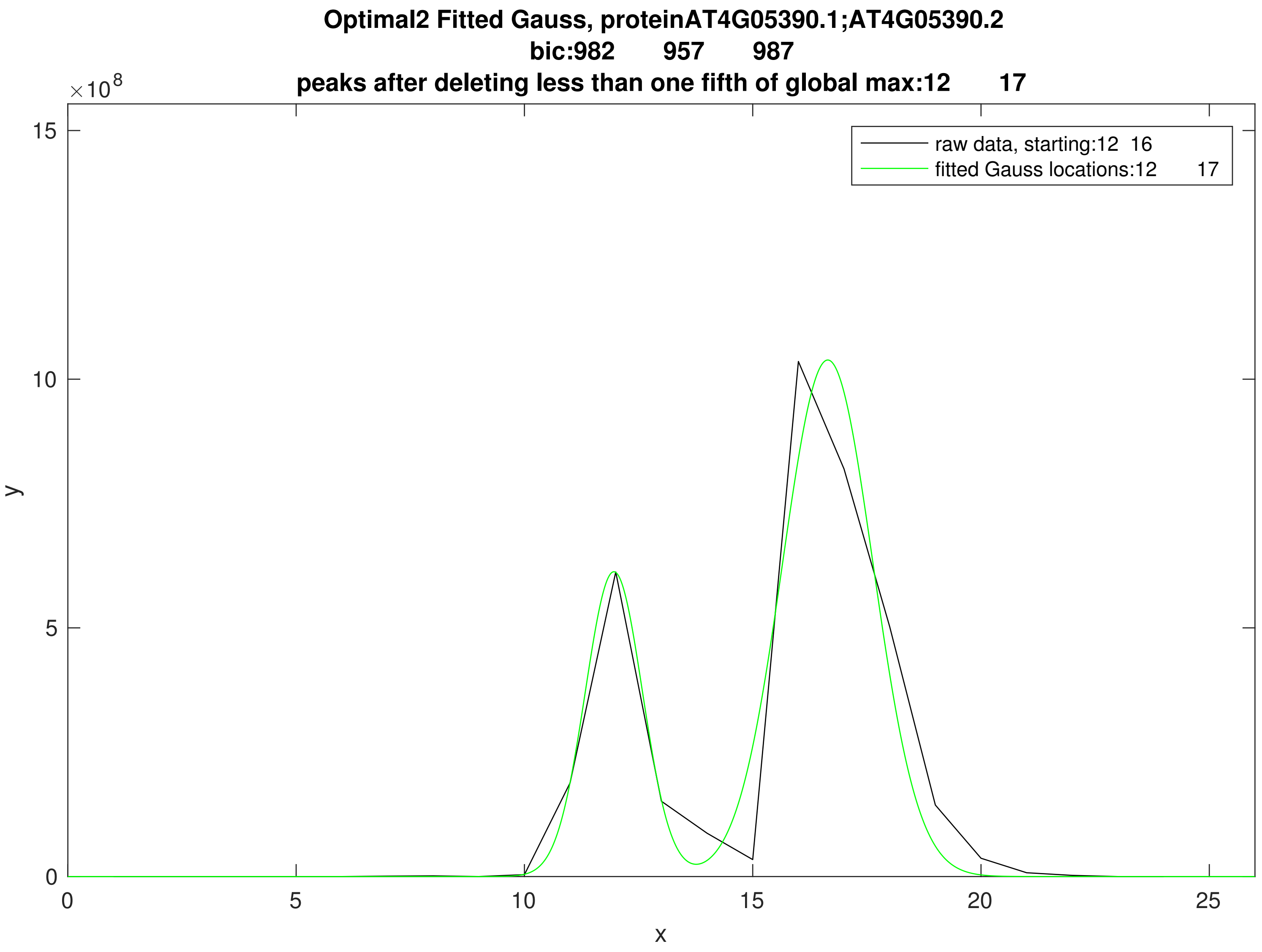

Optimal2 Fitted Gauss, proteinAT4G05390.1;AT4G05390.2
bic:982 957 987
peaks after deleting less than one fifth of global max:12 17
×10^8
15
10
5
0
y
0
5
10
15
20
25
x
raw data, starting:12 16
fitted Gauss locations:12 17

**Optimal1 Fitted Gauss, proteinAT4G05400.2;AT4G05400.1**
**bic:665**
**peaks after deleting less than one fifth of global max:3.1**

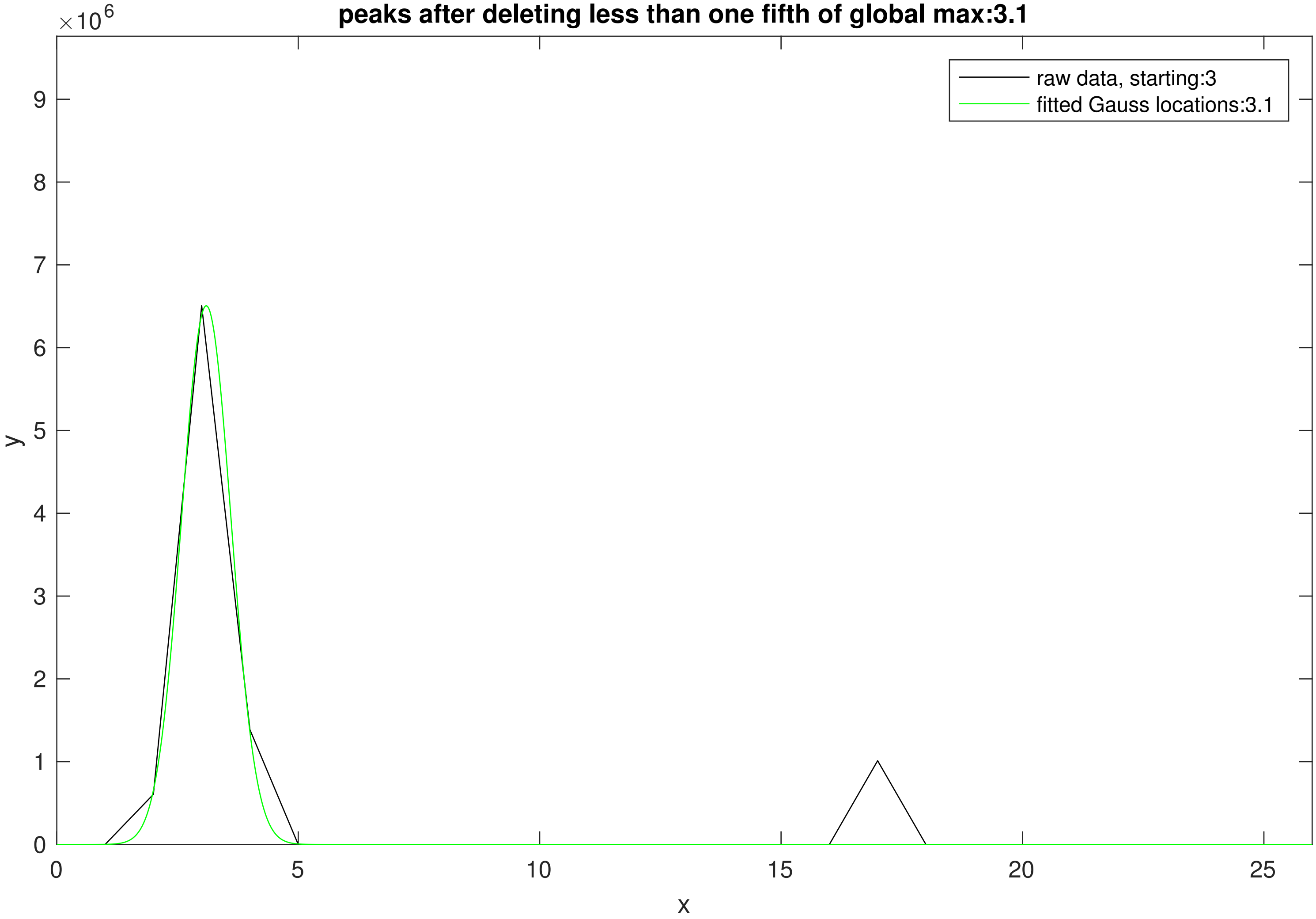

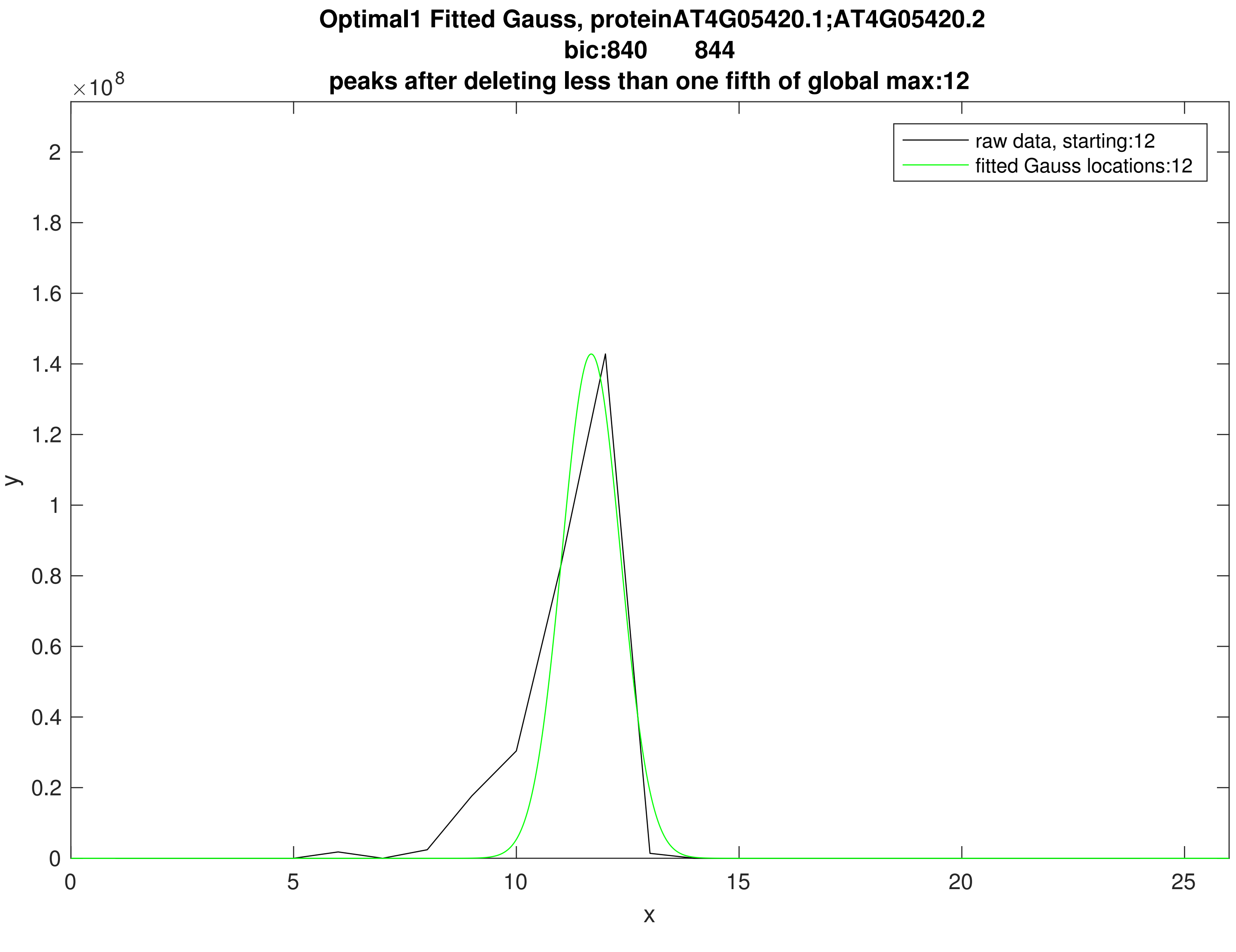
Optimal1 Fitted Gauss, proteinAT4G05420.1;AT4G05420.2
bic:840 844
peaks after deleting less than one fifth of global max:12
×10^8
raw data, starting:12
fitted Gauss locations:12
y
x

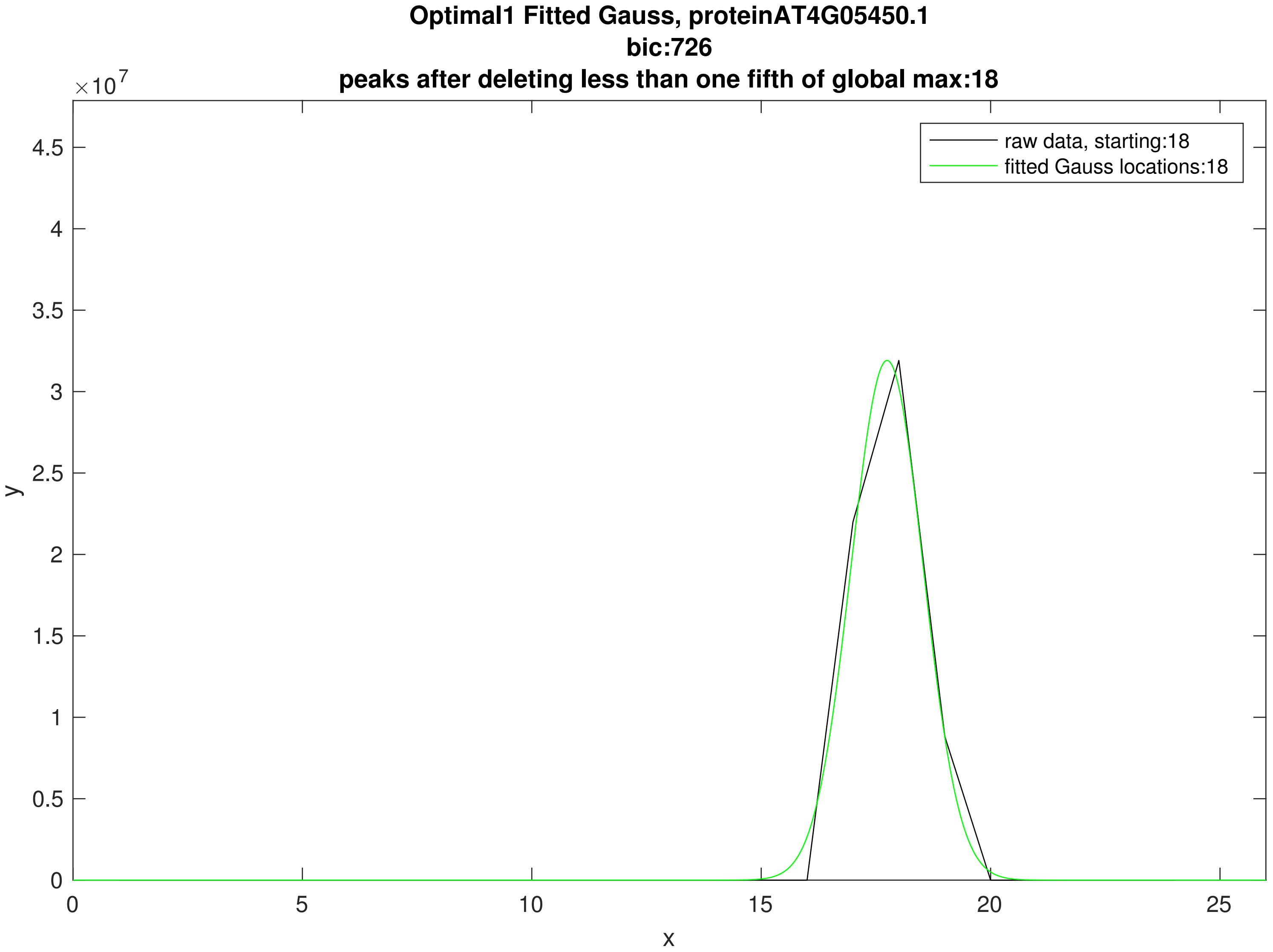

Optimal1 Fitted Gauss, proteinAT4G05450.1
bic:726
peaks after deleting less than one fifth of global max:18
×10^7
4.5
4
3.5
3
2.5
2
1.5
1
0.5
0
y
0
5
10
15
20
25
x
raw data, starting:18
fitted Gauss locations:18

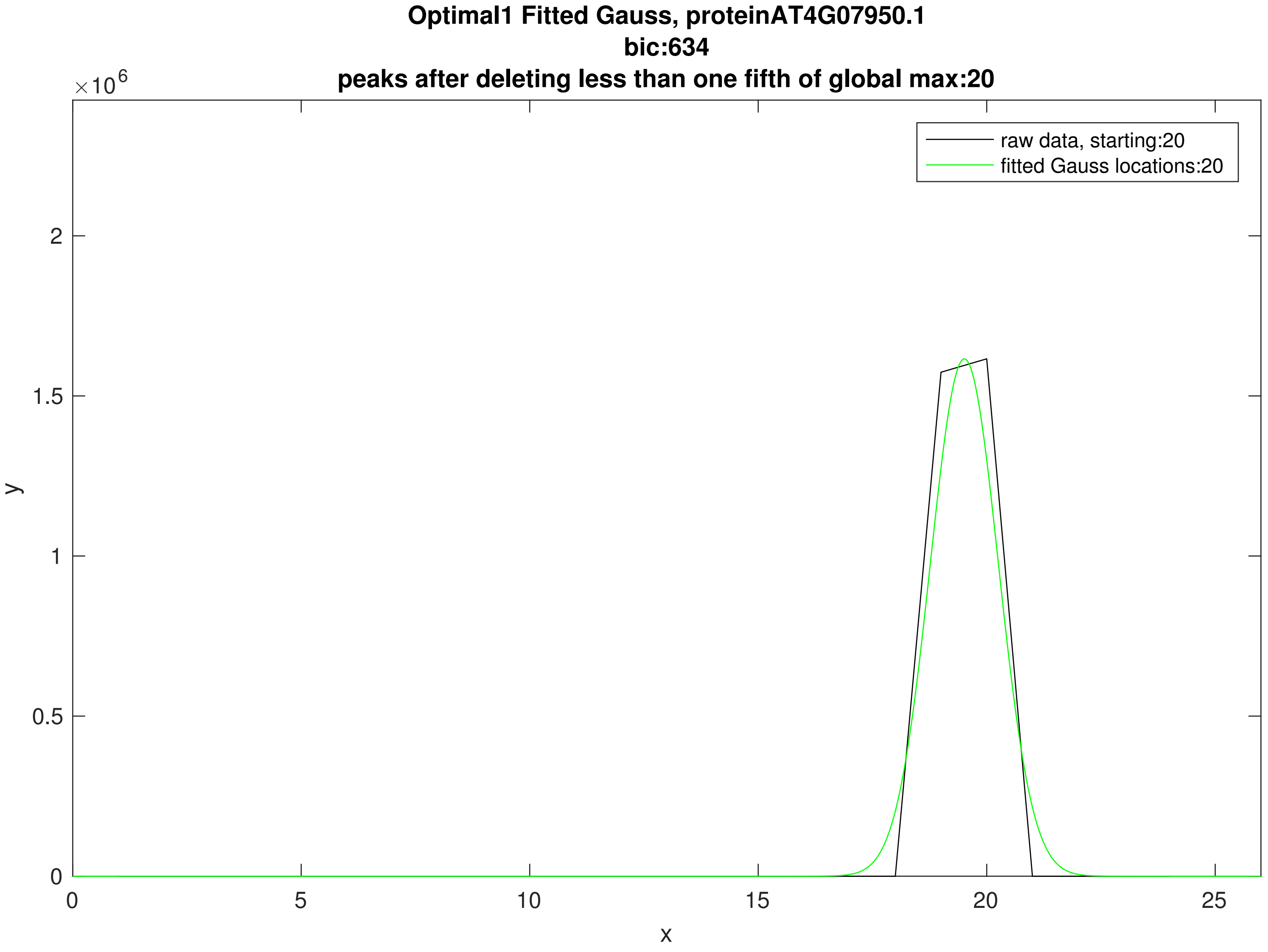
Optimal1 Fitted Gauss, proteinAT4G07950.1
bic:634
peaks after deleting less than one fifth of global max:20
×10^6
raw data, starting:20
fitted Gauss locations:20
y
x
0
0.5
1
1.5
2
5
10
15
20
25

**Optimal1 Fitted Gauss, proteinAT4G08390.3;AT4G08390.2;AT4G08390.1;AT4G08390.4**
**bic:940**
**peaks after deleting less than one fifth of global max:18**

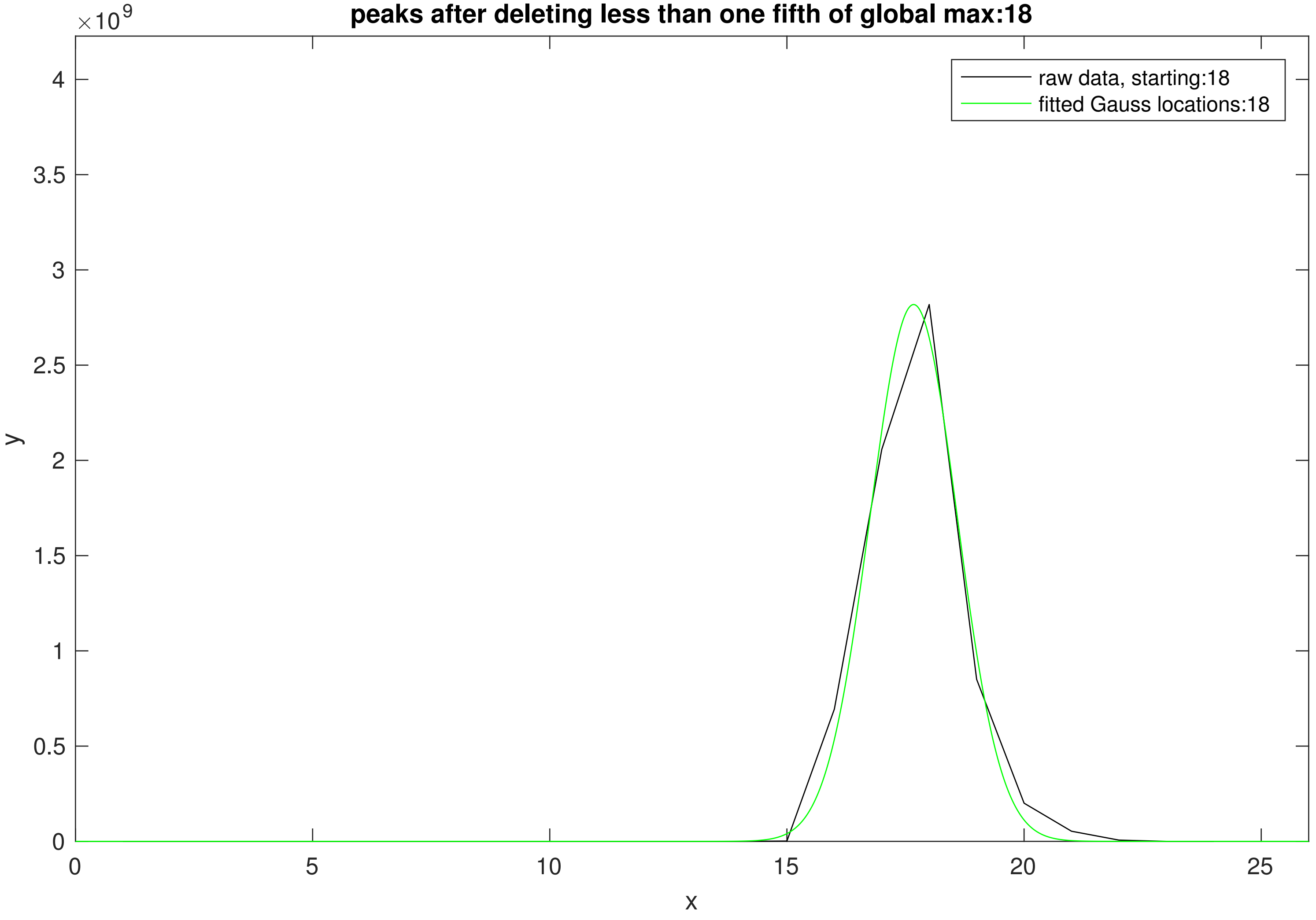

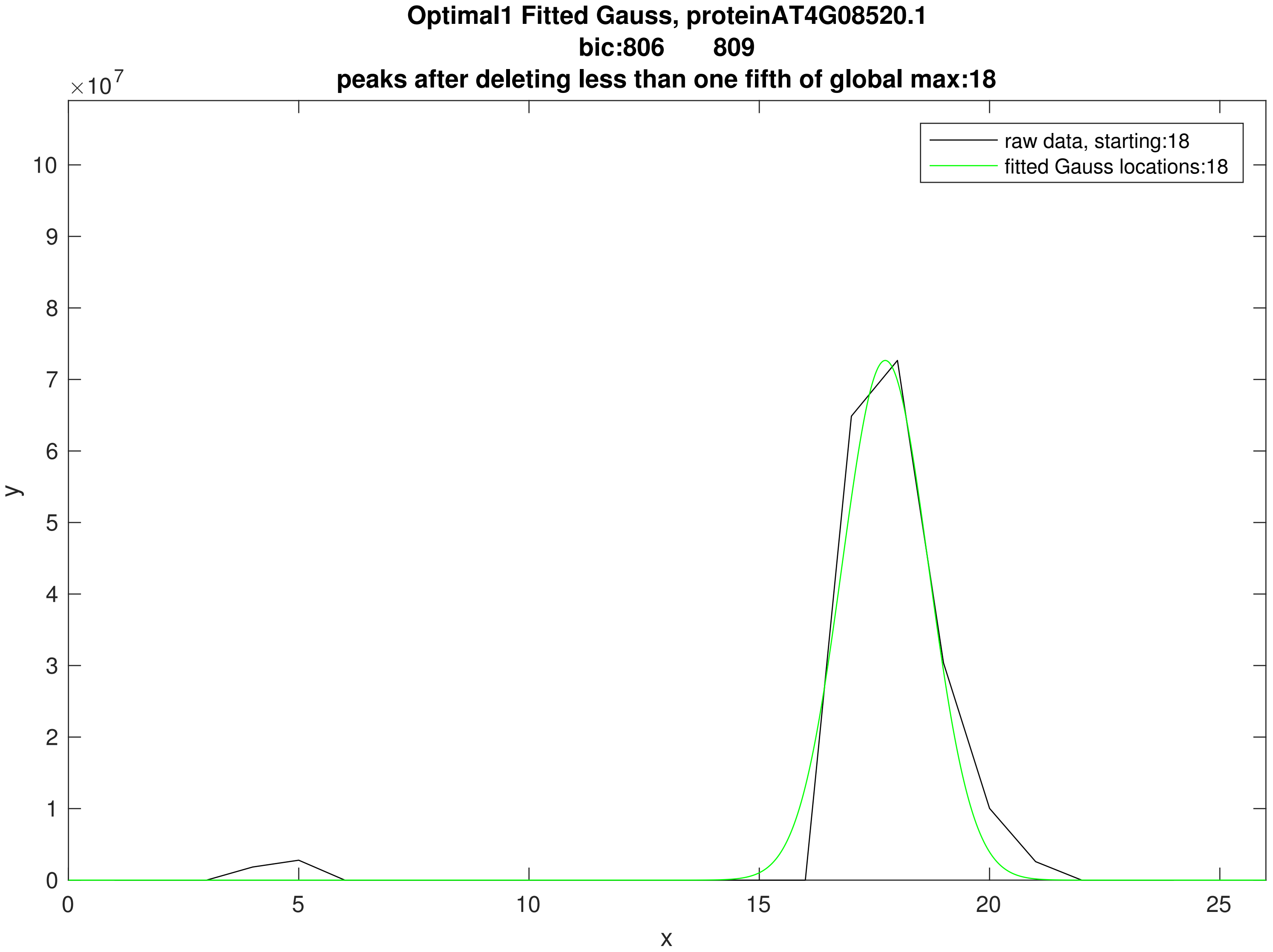

Optimal1 Fitted Gauss, proteinAT4G08520.1
bic:806 809
peaks after deleting less than one fifth of global max:18
×10^7
raw data, starting:18
fitted Gauss locations:18
y
x
0
1
2
3
4
5
6
7
8
9
10
0
5
10
15
20
25

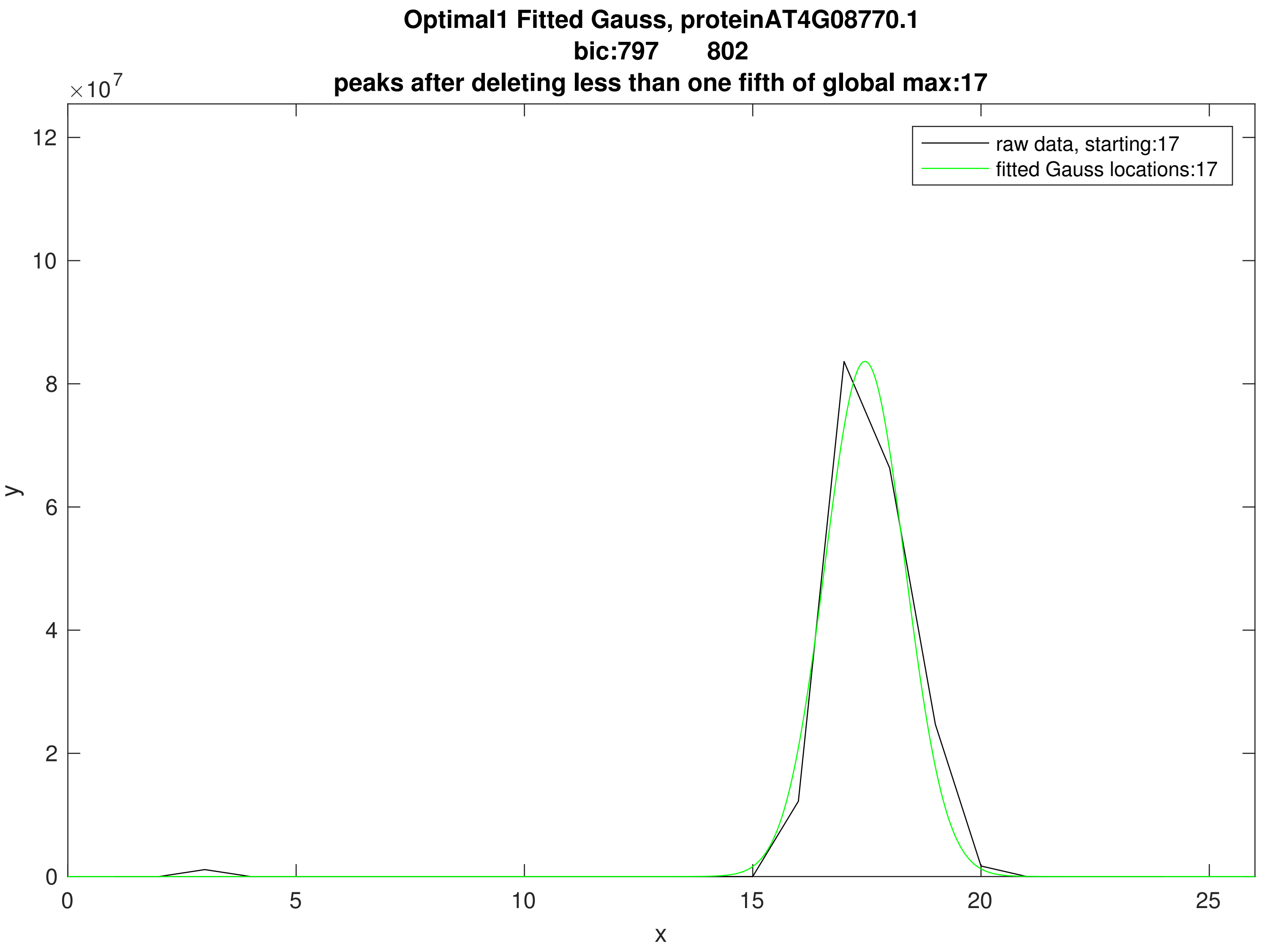
Optimal1 Fitted Gauss, proteinAT4G08770.1
bic:797 802
peaks after deleting less than one fifth of global max:17
×10^7
raw data, starting:17
fitted Gauss locations:17
y
x
0
2
4
6
8
10
12
5
15
20
25

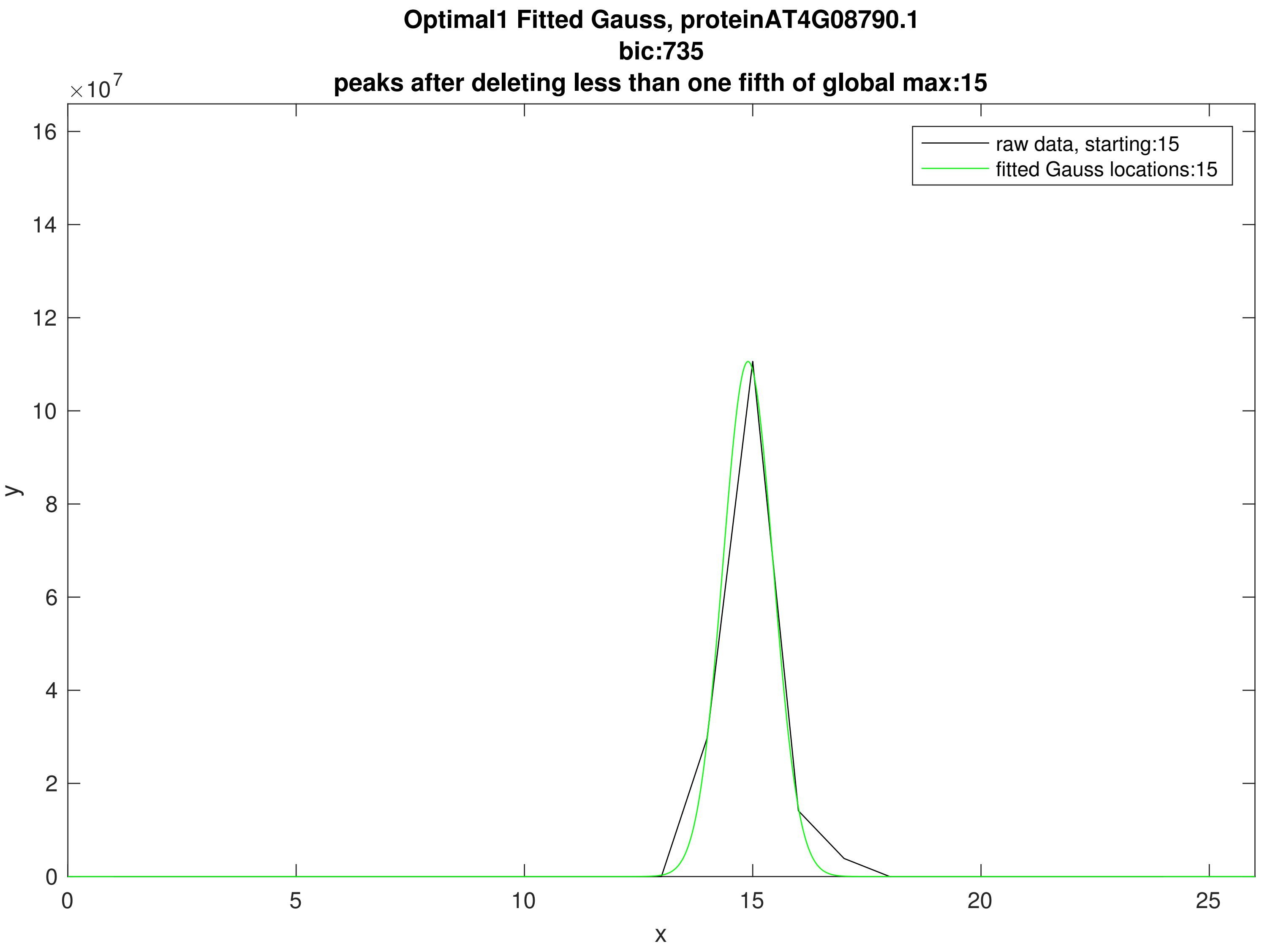
Optimal1 Fitted Gauss, proteinAT4G08790.1
bic:735
peaks after deleting less than one fifth of global max:15
×10^7
raw data, starting:15
fitted Gauss locations:15
y
x
0
2
4
6
8
10
12
14
16
0
5
10
15
20
25

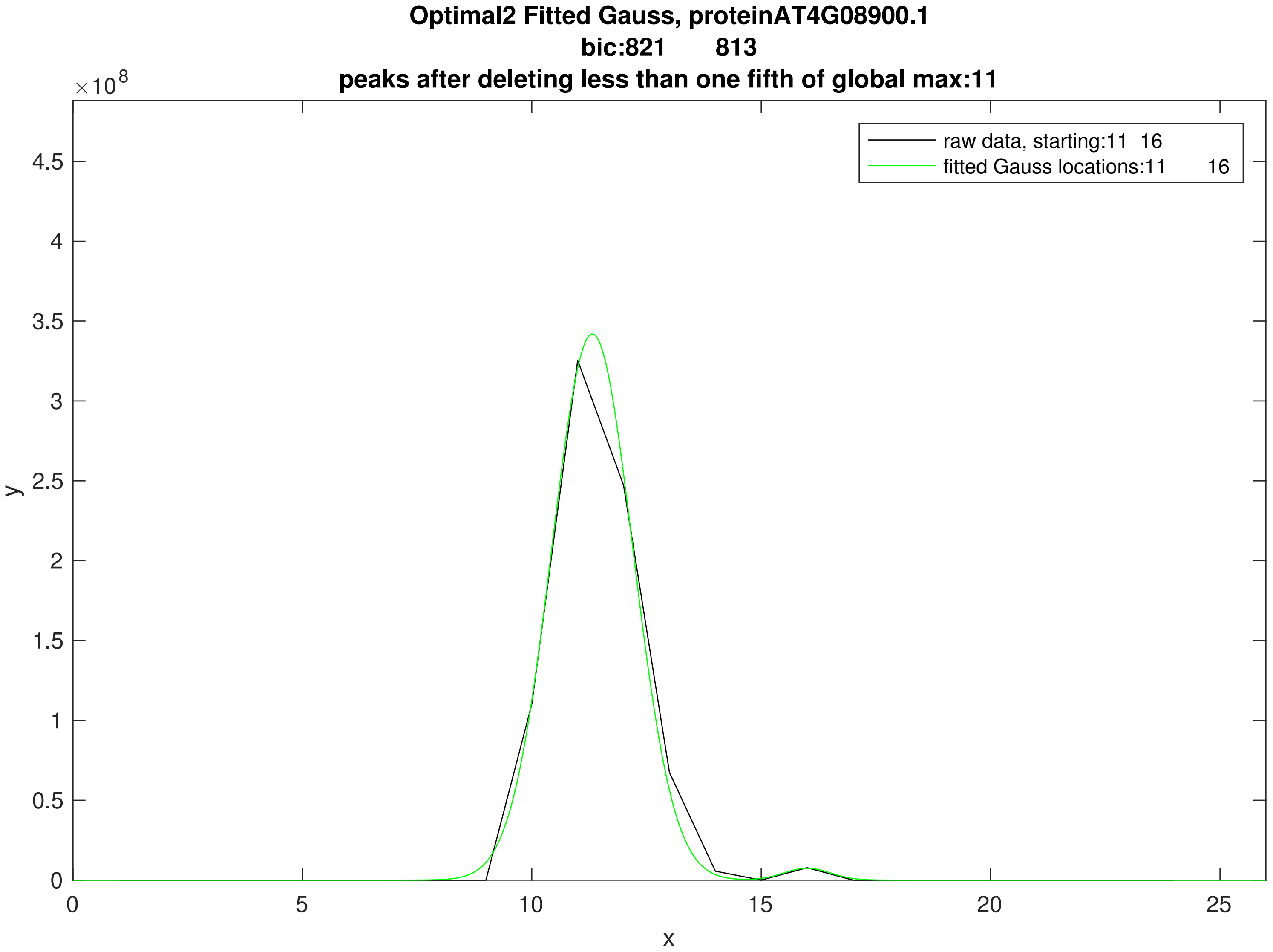
Optimal2 Fitted Gauss, proteinAT4G08900.1
bic:821 813
peaks after deleting less than one fifth of global max:11
×10^8
raw data, starting:11 16
fitted Gauss locations:11 16
y
x
0
0.5
1
1.5
2
2.5
3
3.5
4
4.5
0
5
10
15
20
25

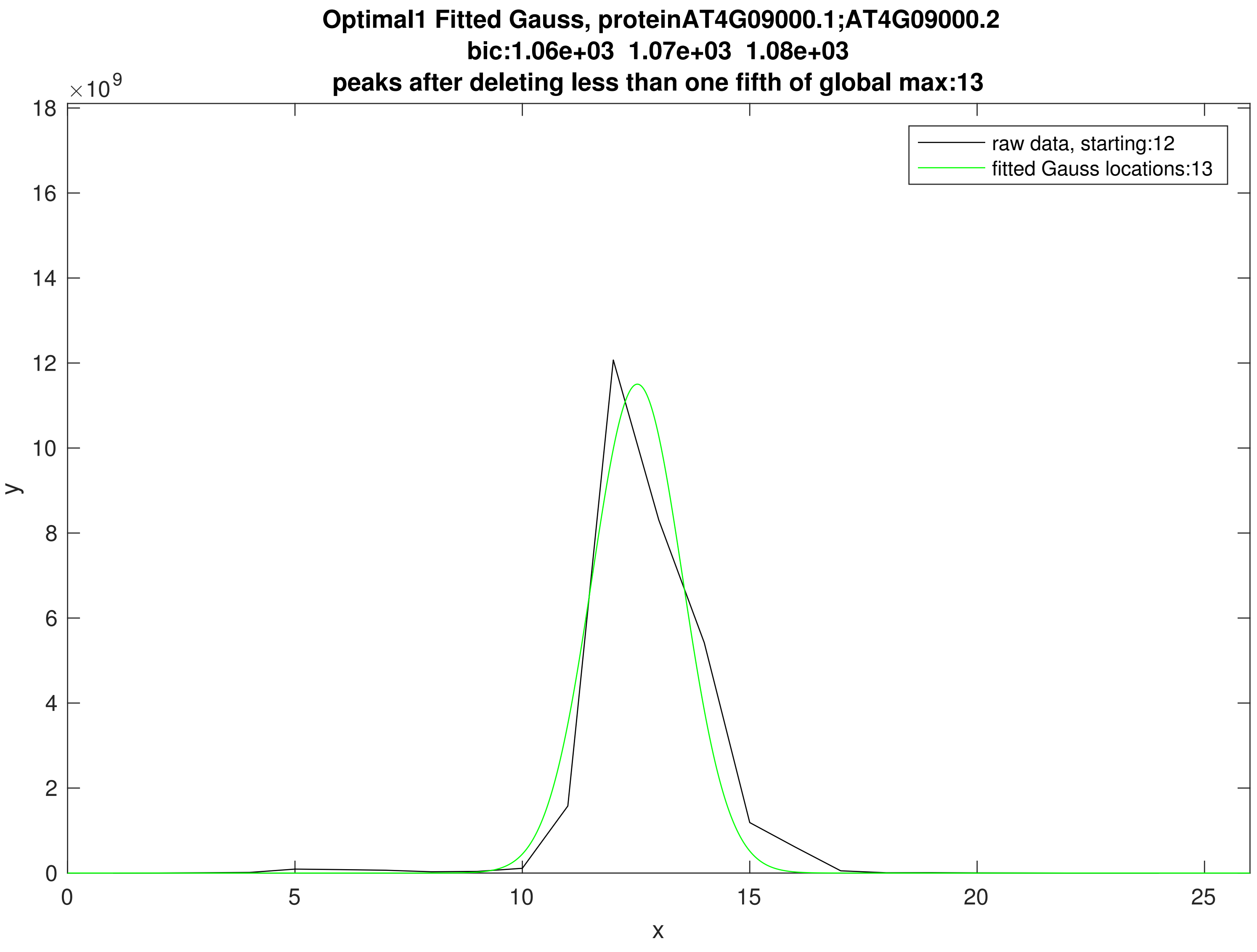

Optimal1 Fitted Gauss, proteinAT4G09000.1;AT4G09000.2
bic:1.06e+03  1.07e+03  1.08e+03
peaks after deleting less than one fifth of global max:13
×10^9
raw data, starting:12
fitted Gauss locations:13
y
x
0
2
4
6
8
10
12
14
16
18
0
5
10
15
20
25

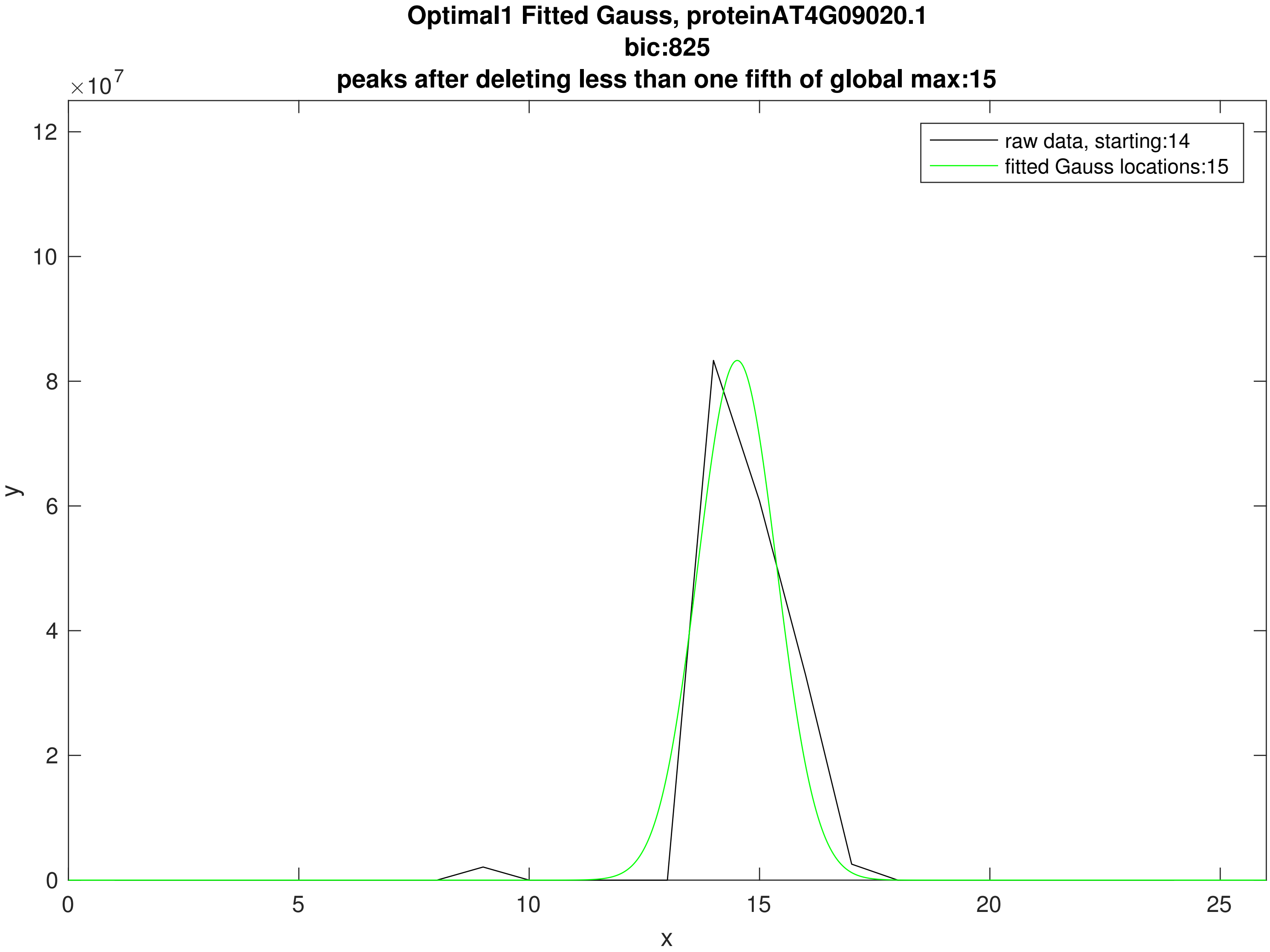
Optimal1 Fitted Gauss, proteinAT4G09020.1
bic:825
peaks after deleting less than one fifth of global max:15
×10^7
12
10
8
6
4
2
0
y
0
5
10
15
20
25
x
raw data, starting:14
fitted Gauss locations:15

**Optimal1 Fitted Gauss, proteinAT4G09160.1**
**bic:677**
**peaks after deleting less than one fifth of global max:9.2**

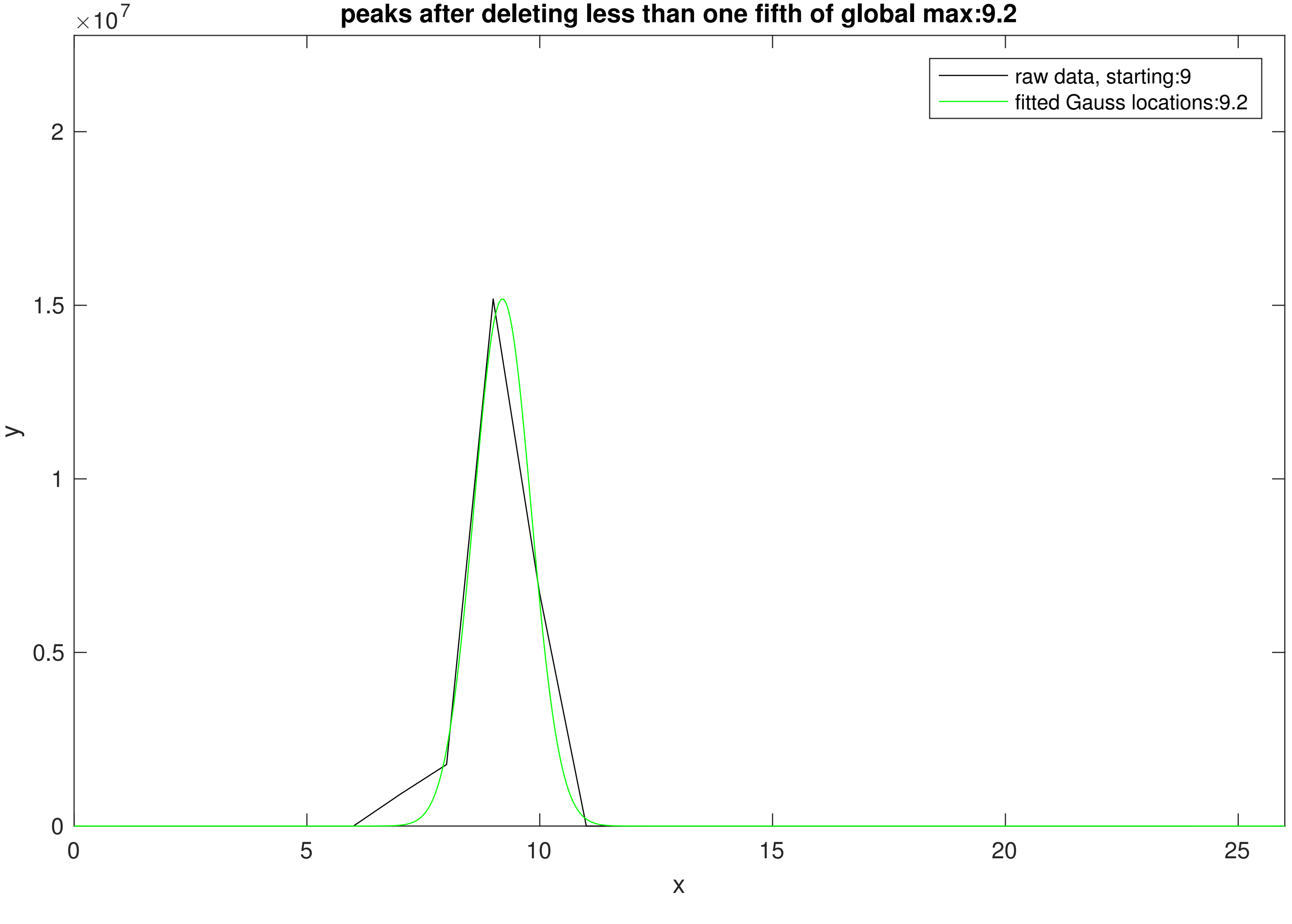

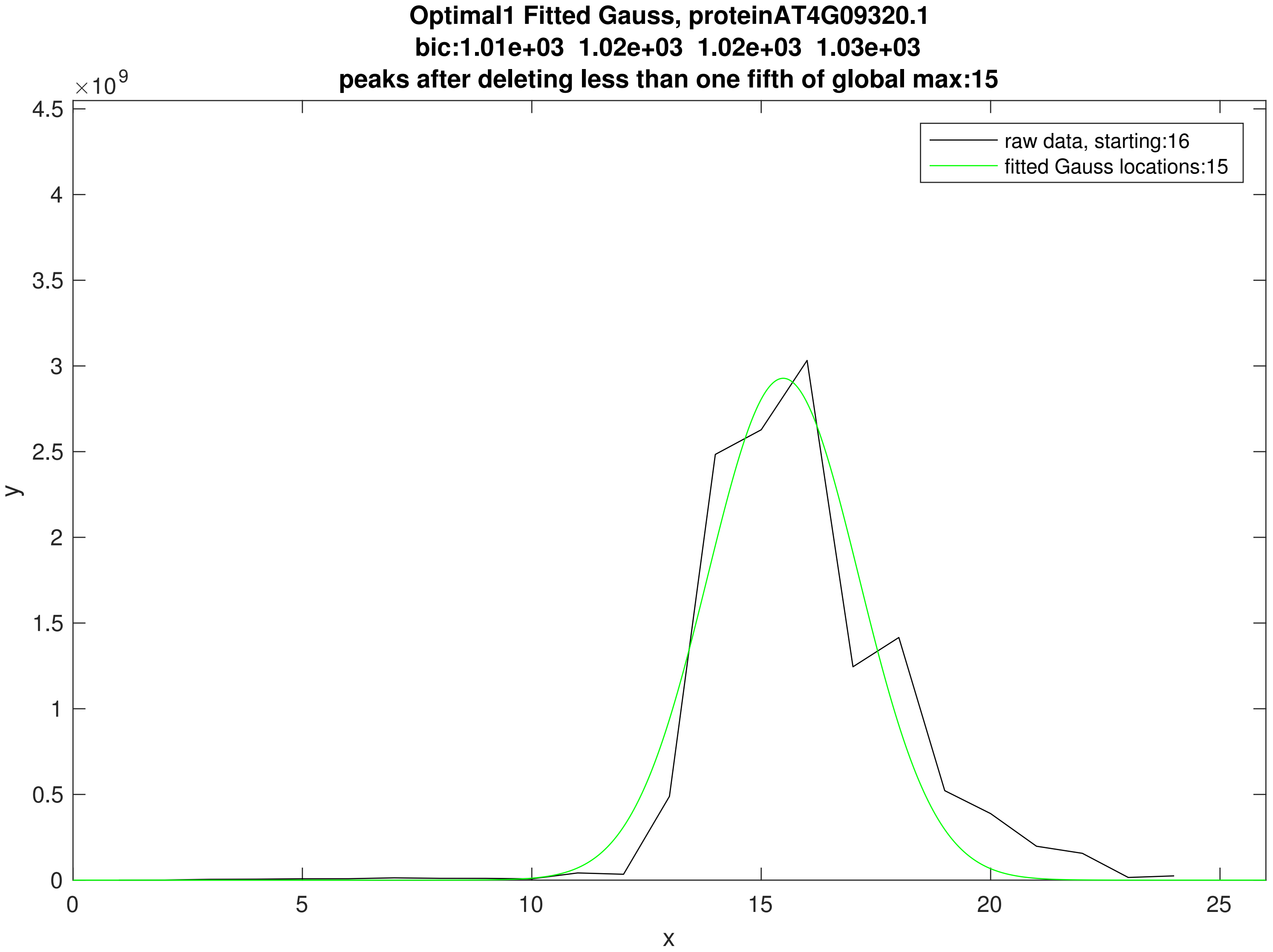

Optimal1 Fitted Gauss, proteinAT4G09320.1
bic:1.01e+03 1.02e+03 1.02e+03 1.03e+03
peaks after deleting less than one fifth of global max:15
×10^9
4.5
4
3.5
3
2.5
2
1.5
1
0.5
0
y
0
5
10
15
20
25
x
raw data, starting:16
fitted Gauss locations:15

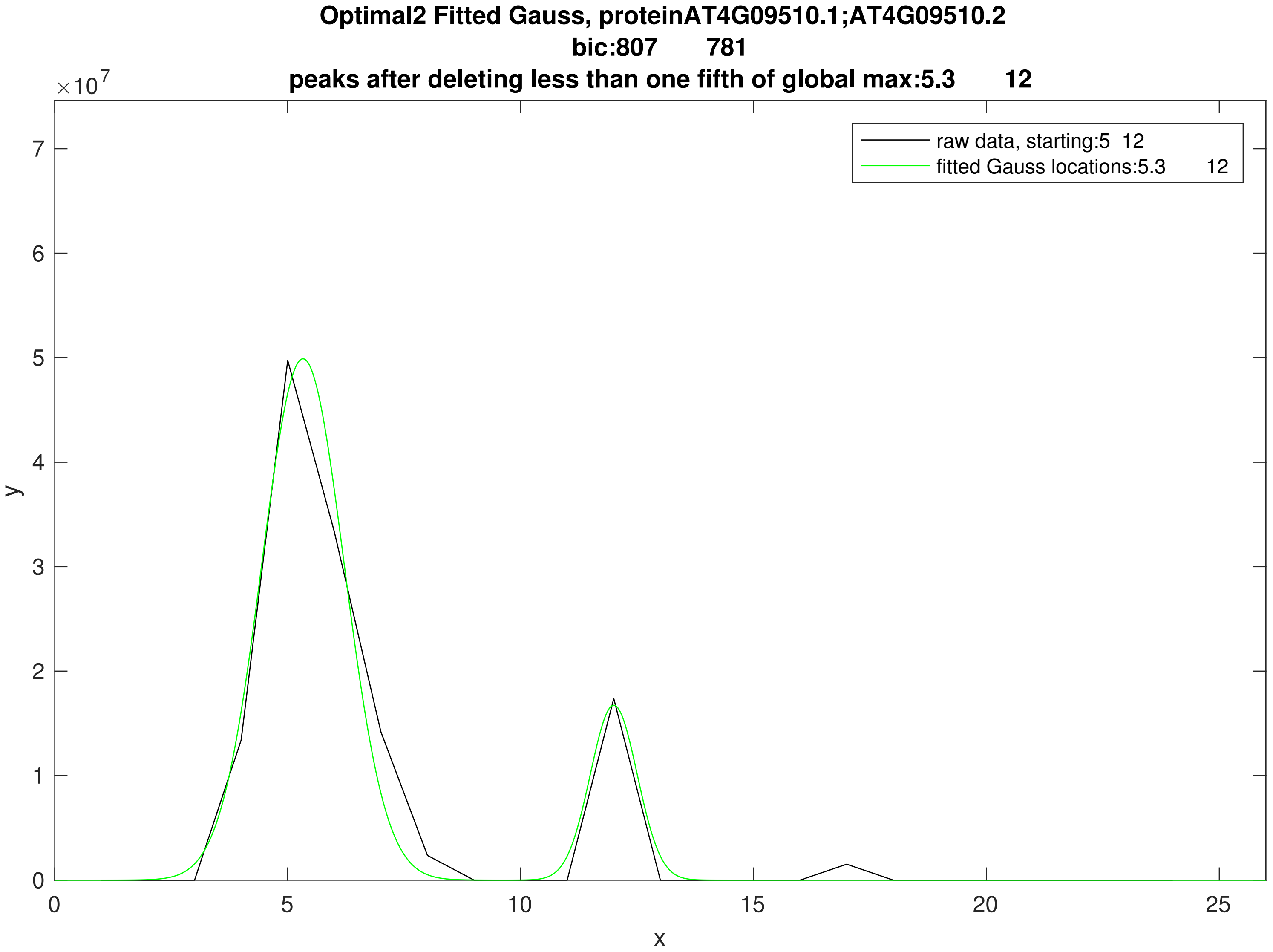

Optimal2 Fitted Gauss, proteinAT4G09510.1;AT4G09510.2
bic:807 781
peaks after deleting less than one fifth of global max:5.3 12
×10^7
raw data, starting:5 12
fitted Gauss locations:5.3 12
y
x
0
1
2
3
4
5
6
7
10
15
20
25

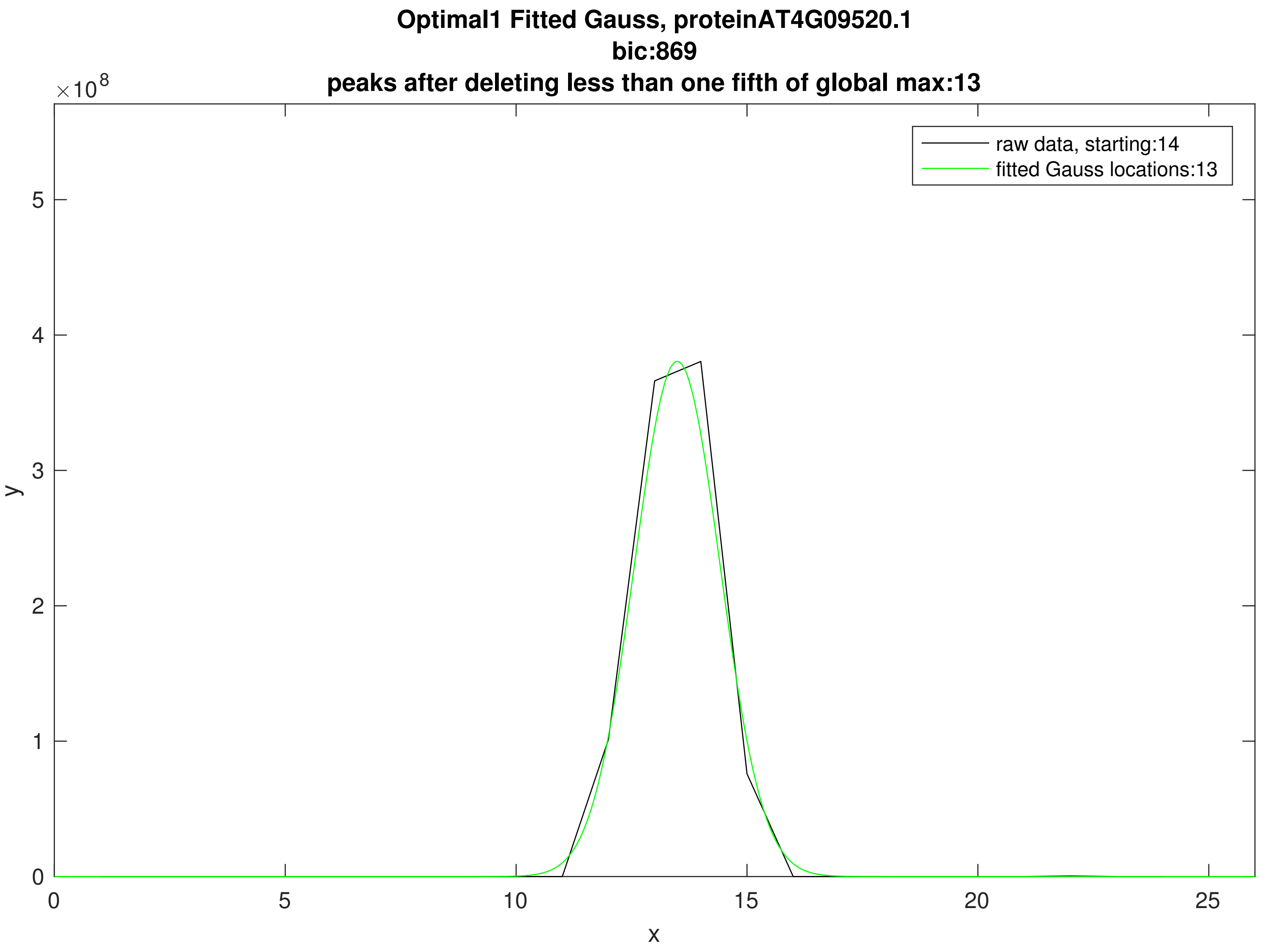

Optimal1 Fitted Gauss, proteinAT4G09520.1
bic:869
peaks after deleting less than one fifth of global max:13
×10^8
raw data, starting:14
fitted Gauss locations:13
y
x
0
1
2
3
4
5
0
5
10
15
20
25

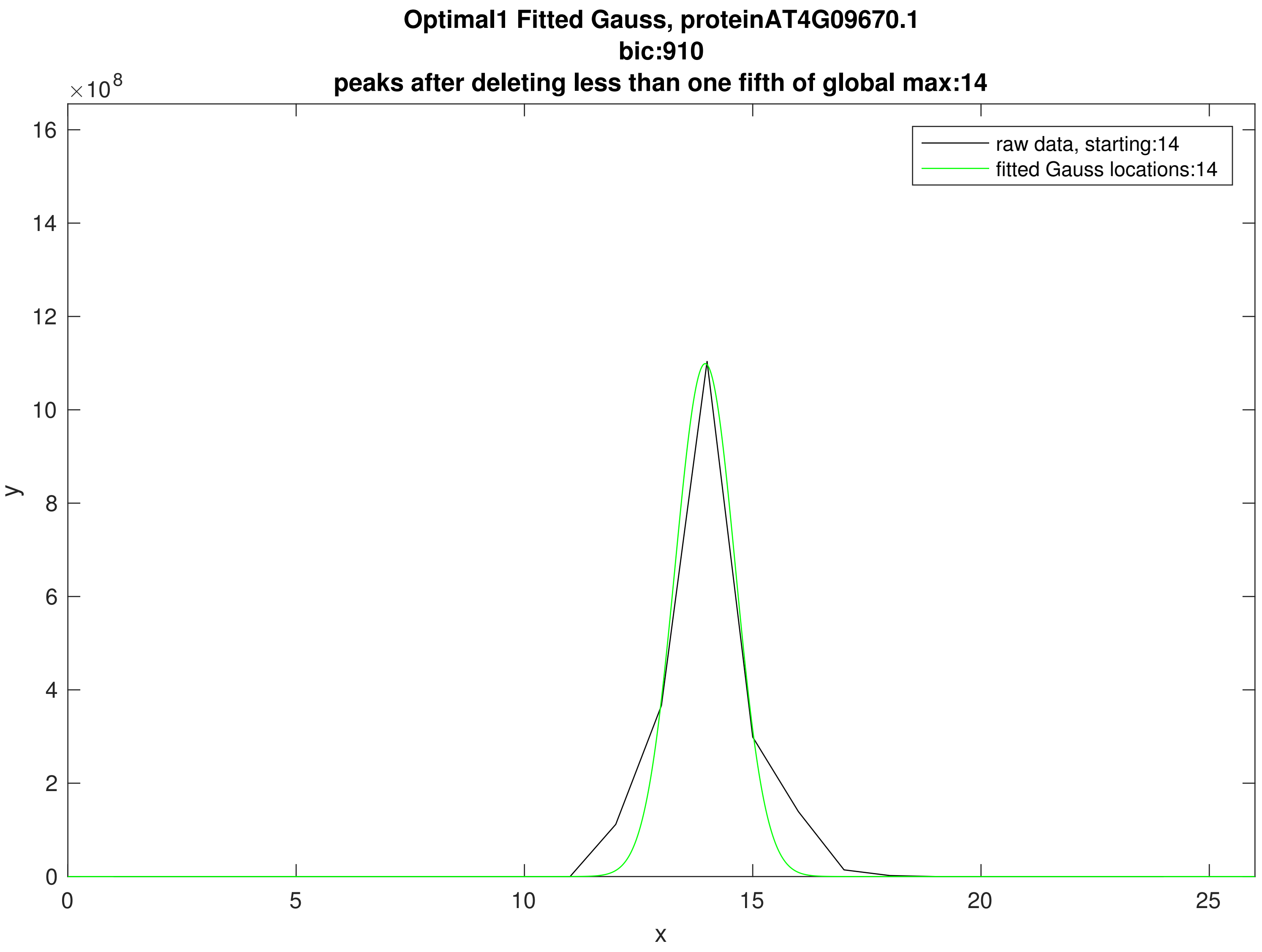

Optimal1 Fitted Gauss, proteinAT4G09670.1
bic:910
peaks after deleting less than one fifth of global max:14
×10^8
raw data, starting:14
fitted Gauss locations:14
y
x
0
2
4
6
8
10
12
14
16
0
5
10
15
20
25

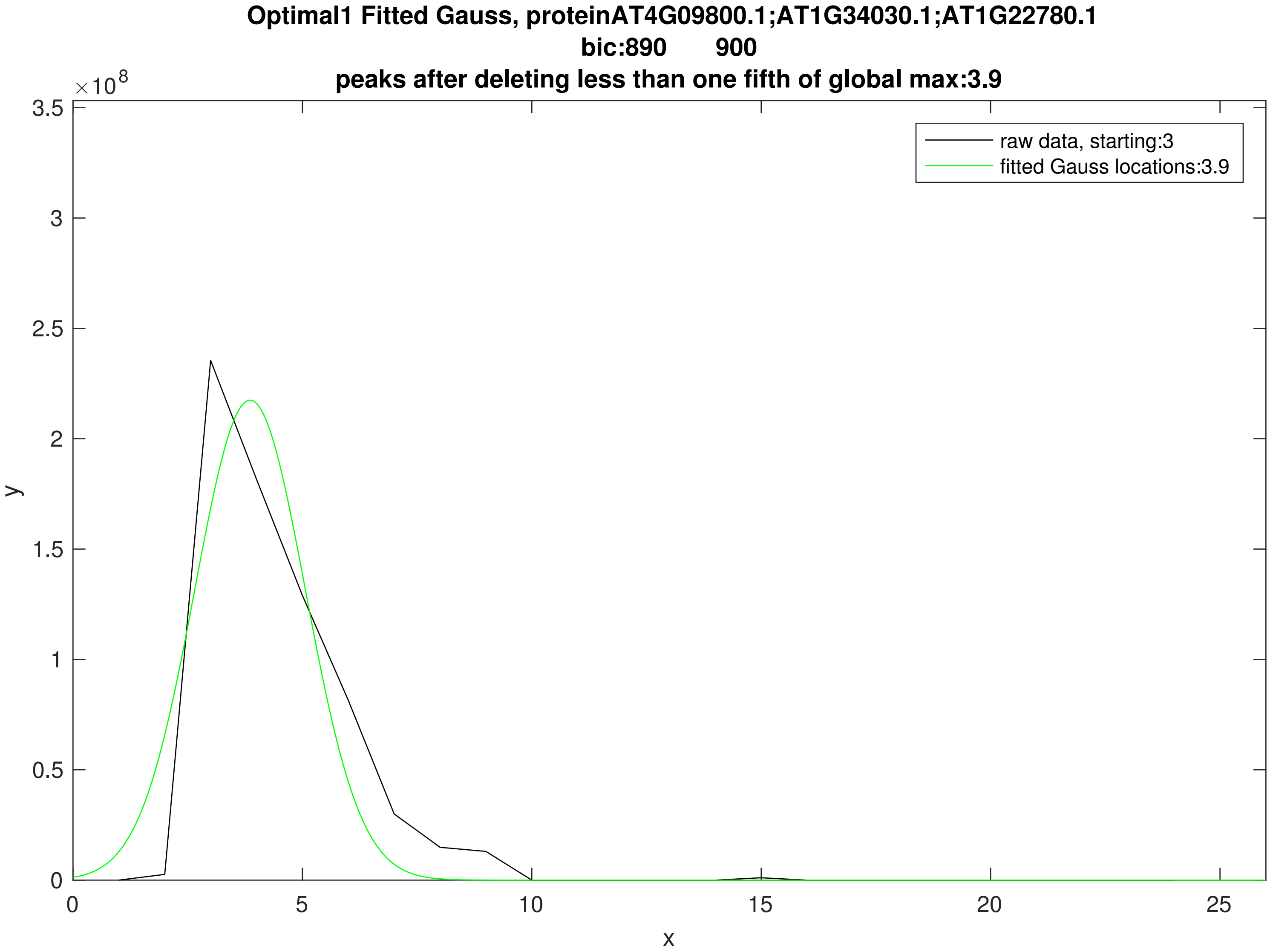

Optimal1 Fitted Gauss, proteinAT4G09800.1;AT1G34030.1;AT1G22780.1
bic:890 900
peaks after deleting less than one fifth of global max:3.9
×10^8
3.5
3
2.5
2
1.5
1
0.5
0
y
0
5
10
15
20
25
x
raw data, starting:3
fitted Gauss locations:3.9

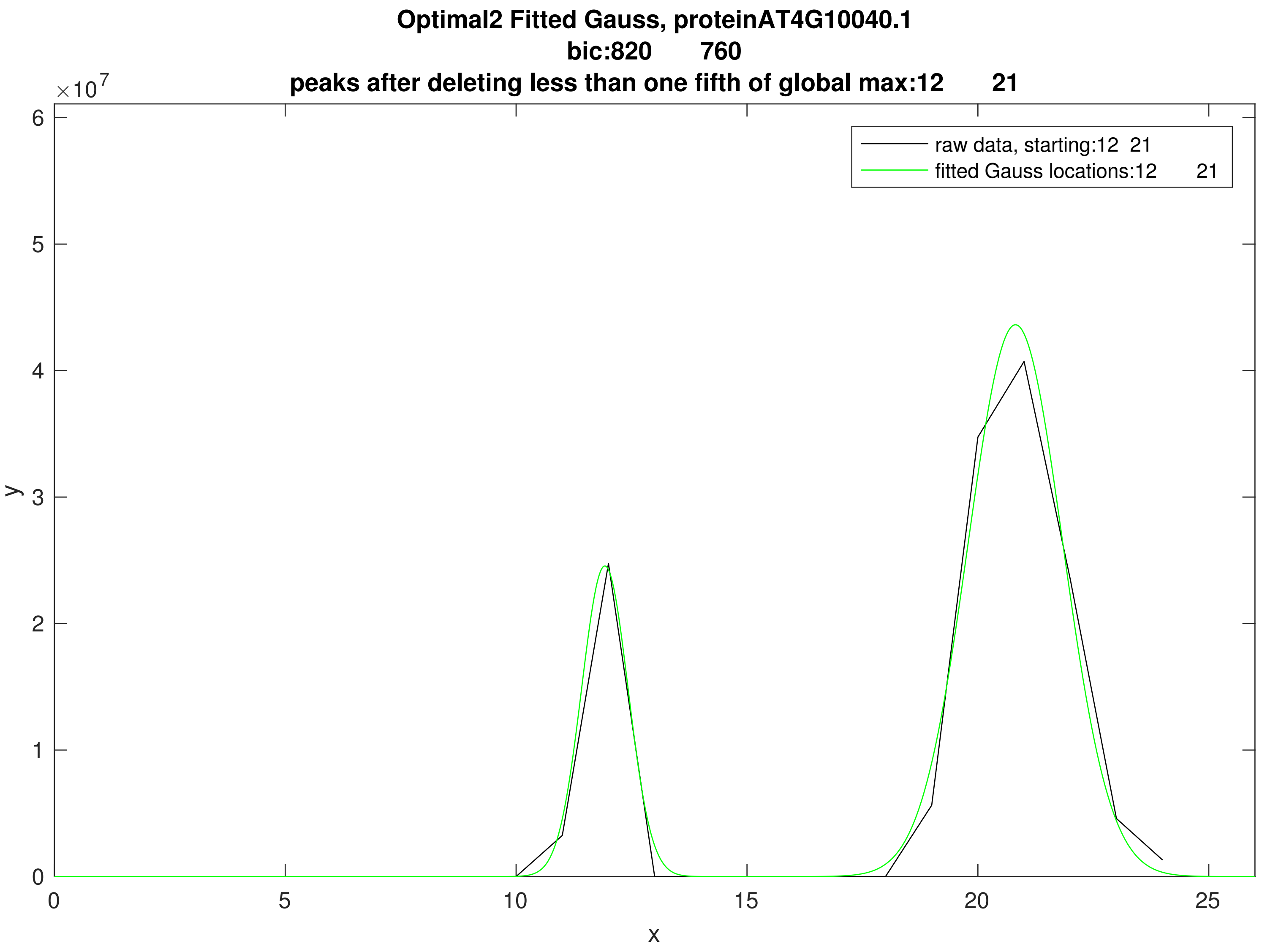

Optimal2 Fitted Gauss, proteinAT4G10040.1
bic:820 760
peaks after deleting less than one fifth of global max:12 21
×10^7
y
x
raw data, starting:12 21
fitted Gauss locations:12 21
0
1
2
3
4
5
6
0
5
10
15
20
25

Optimal1 Fitted Gauss, proteinAT4G10100.3;AT4G10100.2;AT4G10100.1
bic:743
peaks after deleting less than one fifth of global max:19

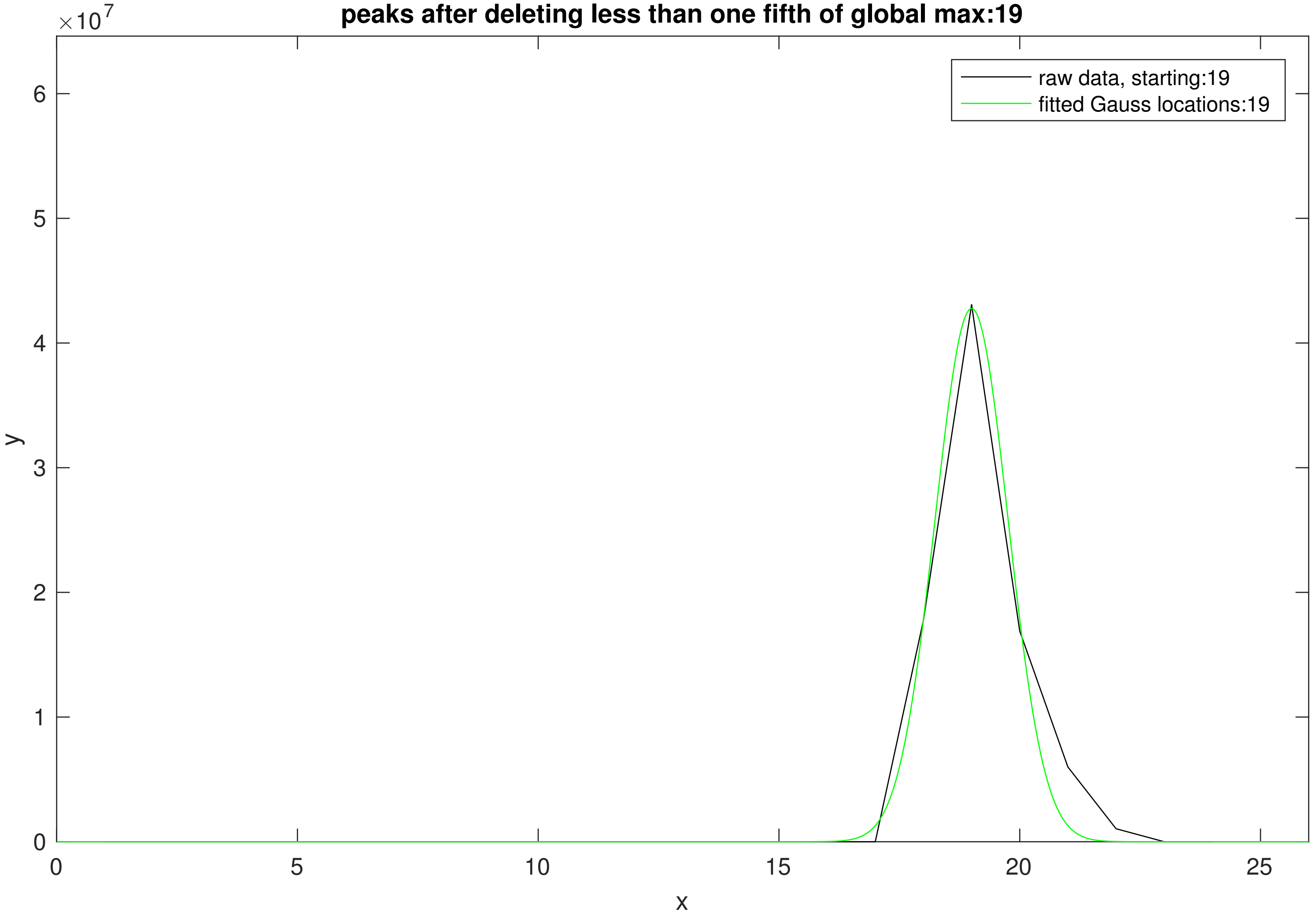

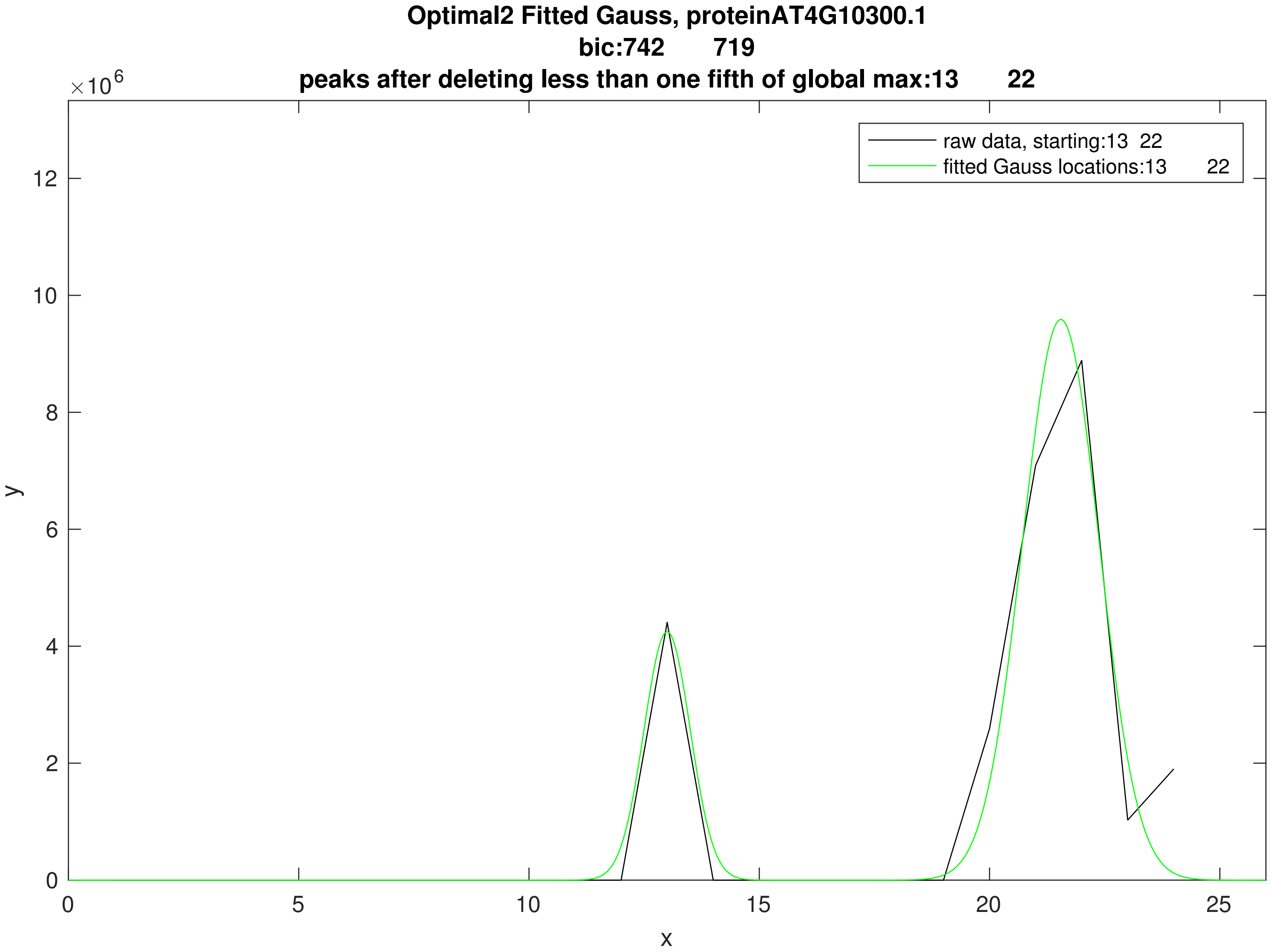
Optimal2 Fitted Gauss, proteinAT4G10300.1
bic:742 719
peaks after deleting less than one fifth of global max:13 22
×10^6
raw data, starting:13 22
fitted Gauss locations:13 22
y
x
0
2
4
6
8
10
12
0
5
10
15
20
25

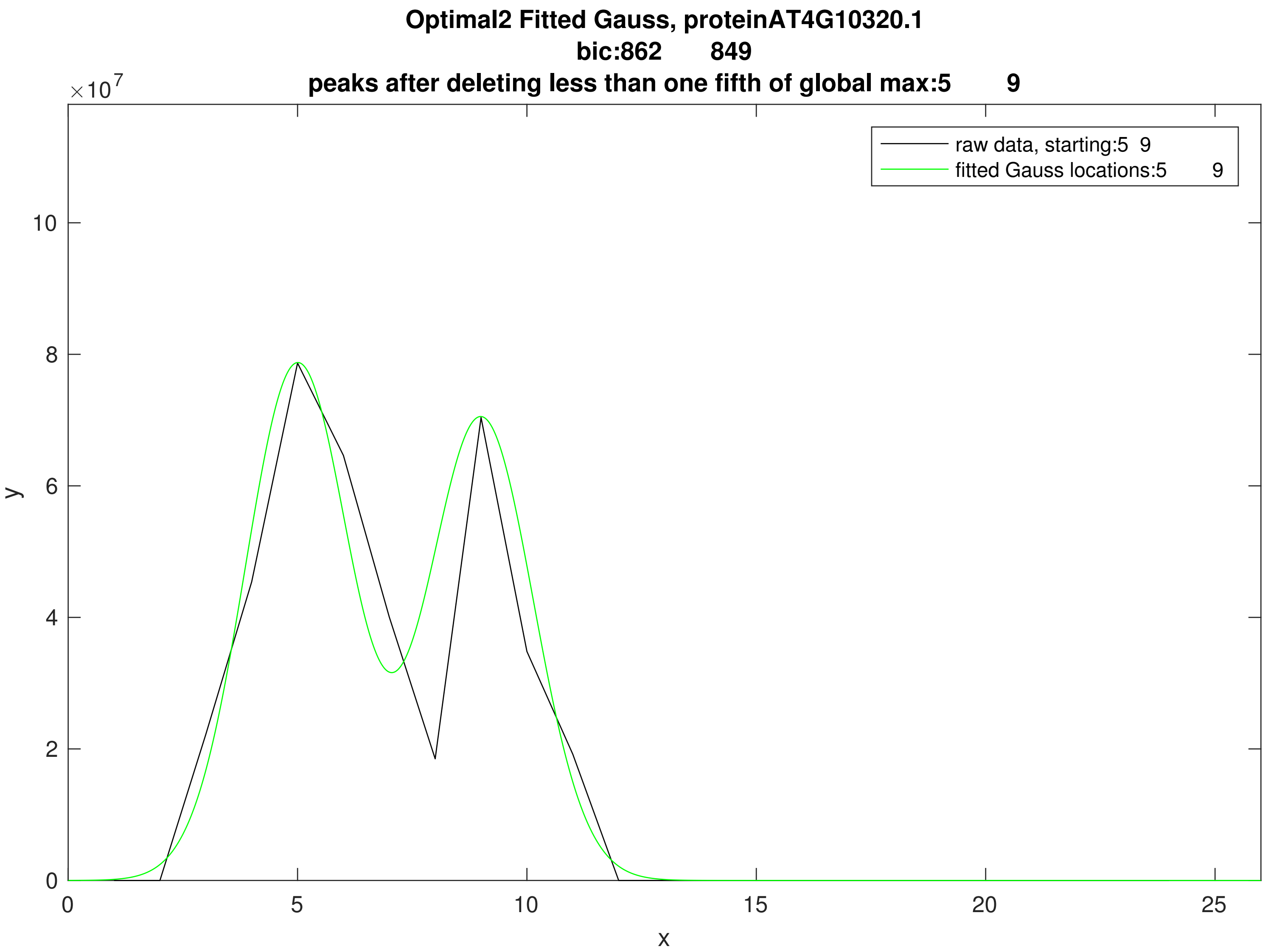
Optimal2 Fitted Gauss, proteinAT4G10320.1
bic:862 849
peaks after deleting less than one fifth of global max:5 9
×10^7
raw data, starting:5 9
fitted Gauss locations:5 9
y
x
0
2
4
6
8
10
0
5
10
15
20
25

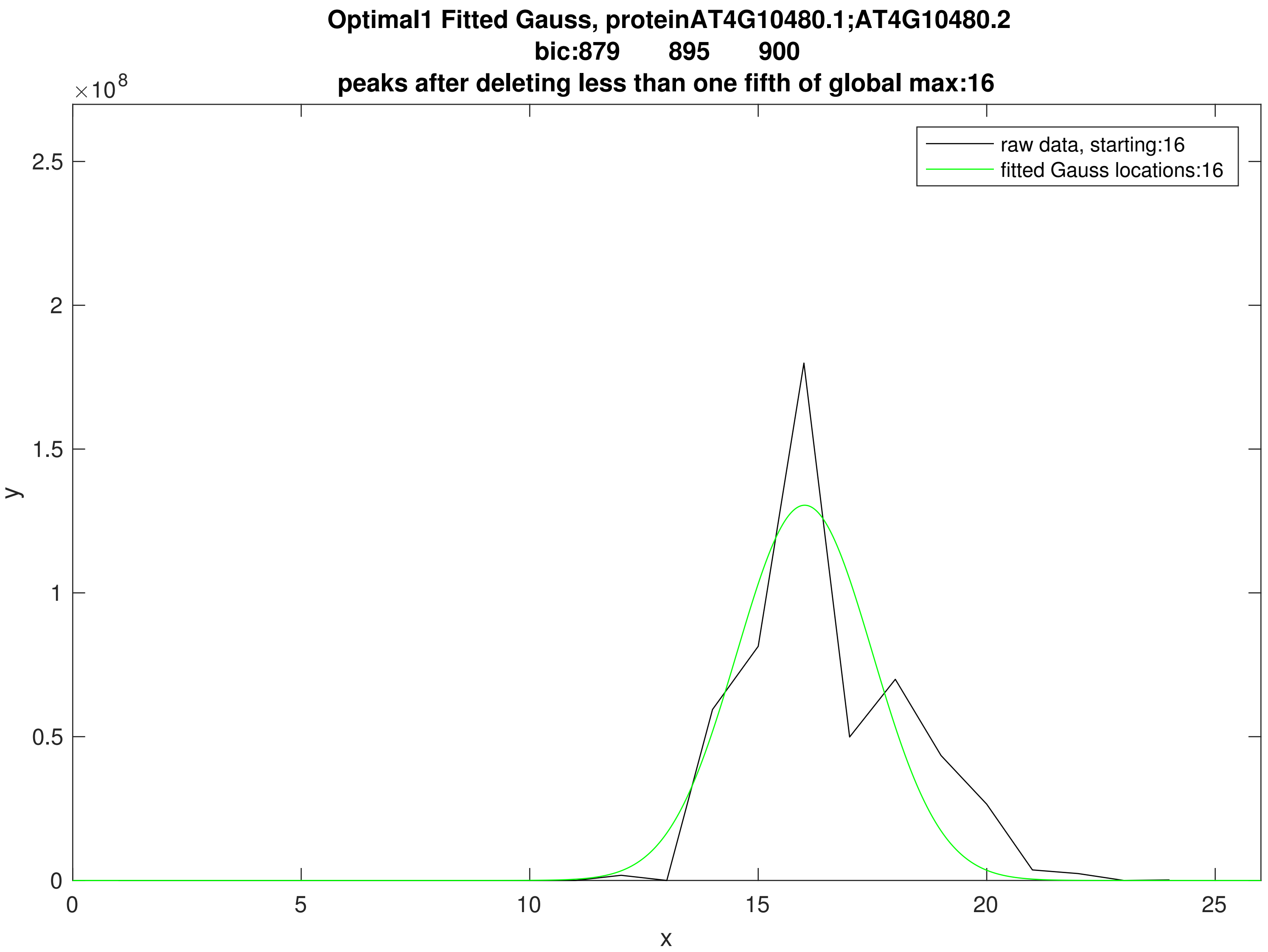

Optimal1 Fitted Gauss, proteinAT4G10480.1;AT4G10480.2
bic:879 895 900
peaks after deleting less than one fifth of global max:16
×10^8
raw data, starting:16
fitted Gauss locations:16
y
x
0
0.5
1
1.5
2
2.5
0
5
10
15
20
25

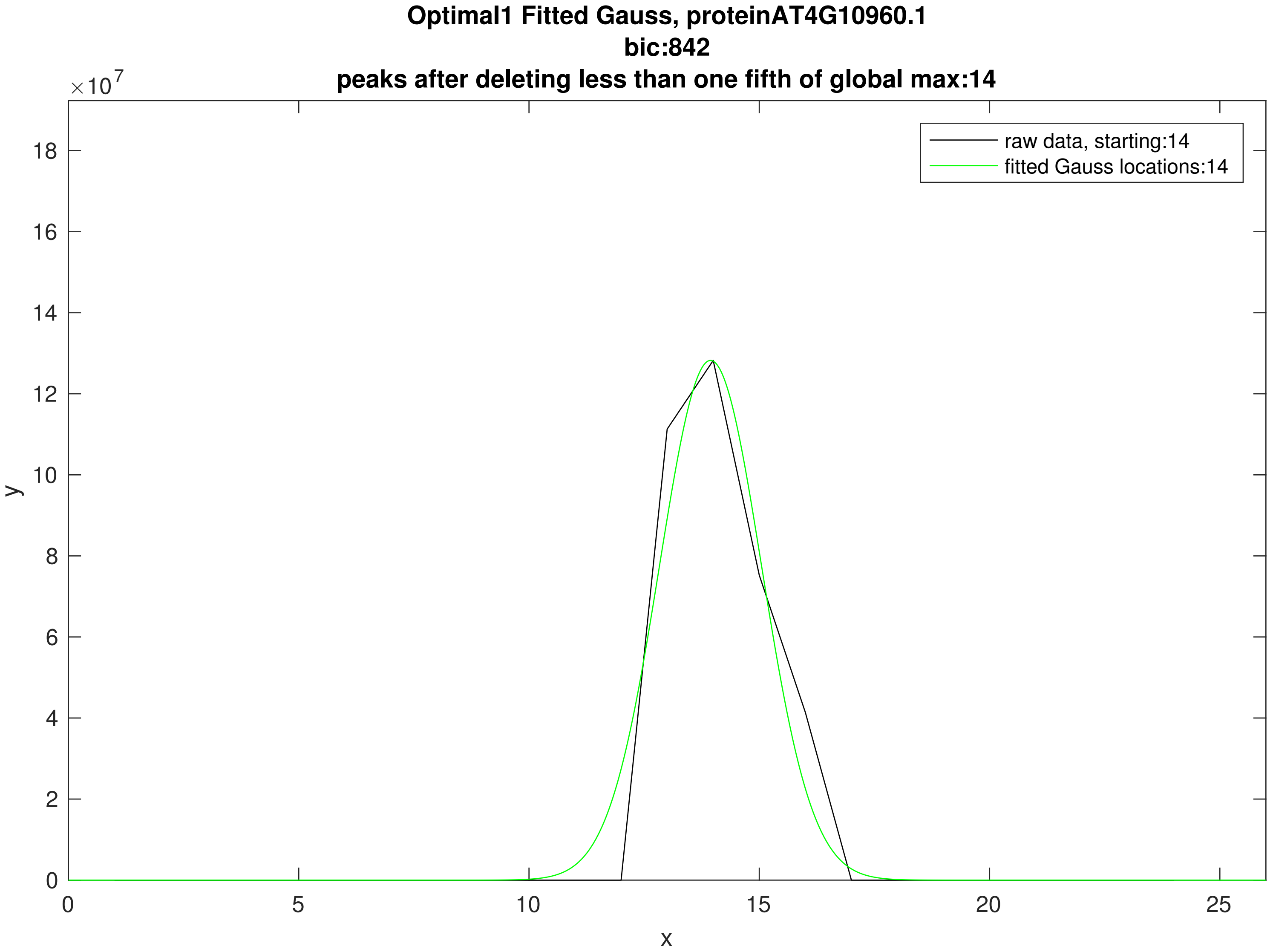
Optimal1 Fitted Gauss, proteinAT4G10960.1
bic:842
peaks after deleting less than one fifth of global max:14
×10^7
raw data, starting:14
fitted Gauss locations:14
y
x
0
2
4
6
8
10
12
14
16
18
0
5
10
15
20
25

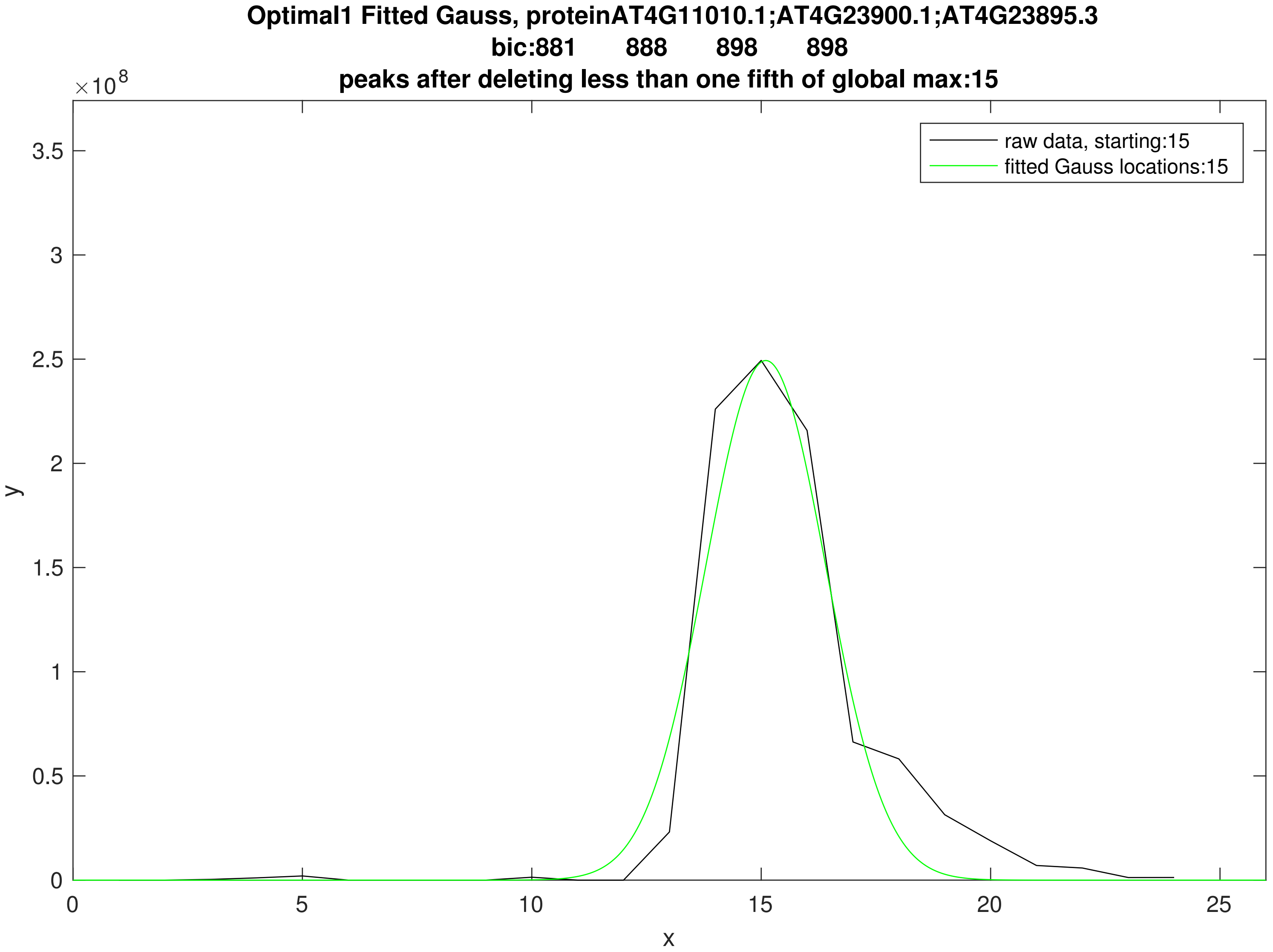

Optimal1 Fitted Gauss, proteinAT4G11010.1;AT4G23900.1;AT4G23895.3
bic:881 888 898 898
peaks after deleting less than one fifth of global max:15
×10^8
3.5
3
2.5
2
1.5
1
0.5
0
y
0
5
10
15
20
25
x
raw data, starting:15
fitted Gauss locations:15

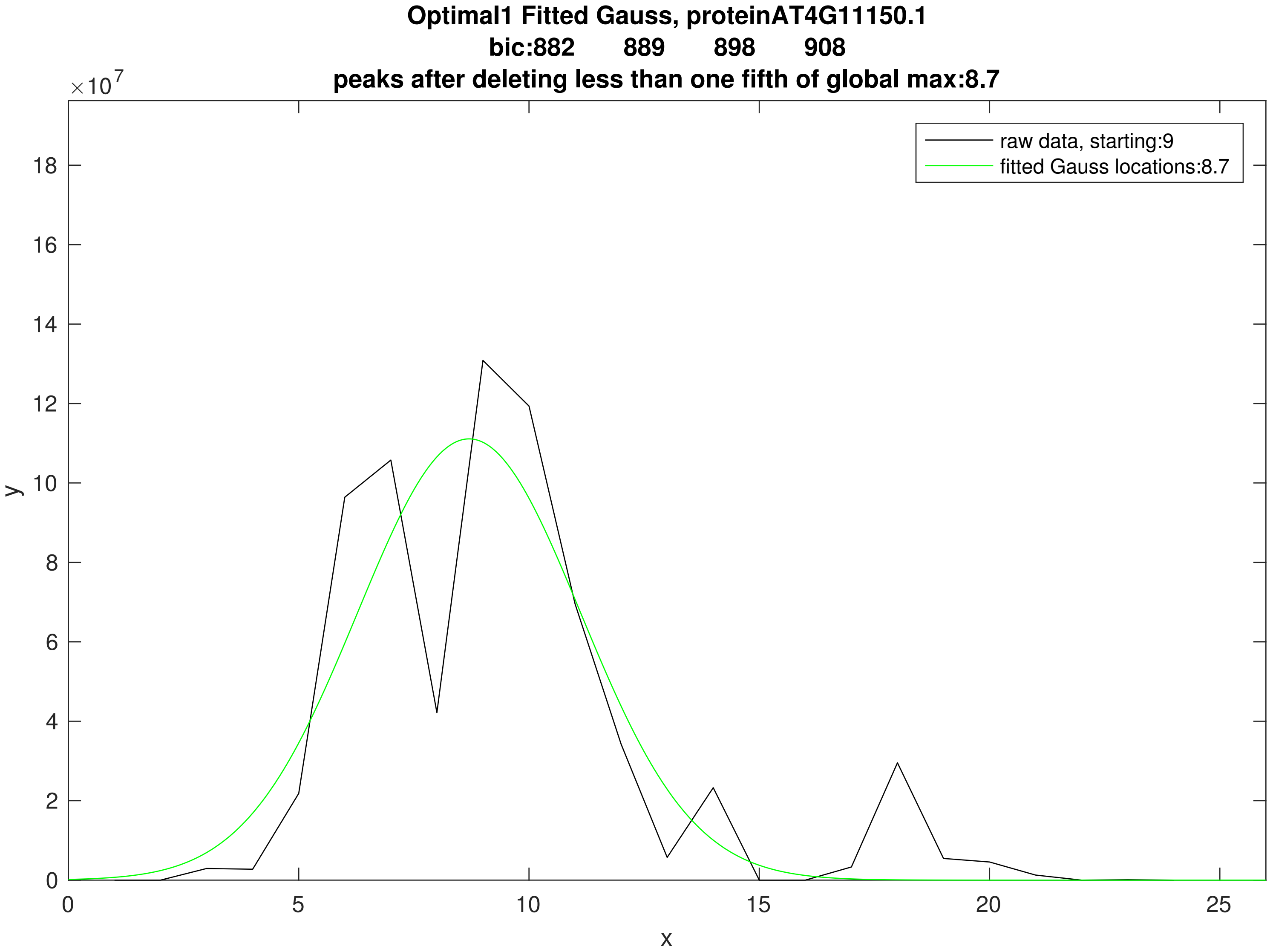
Optimal1 Fitted Gauss, proteinAT4G11150.1
bic:882 889 898 908
peaks after deleting less than one fifth of global max:8.7
×10^7
raw data, starting:9
fitted Gauss locations:8.7
y
x
0
2
4
6
8
10
12
14
16
18
0
5
10
15
20
25

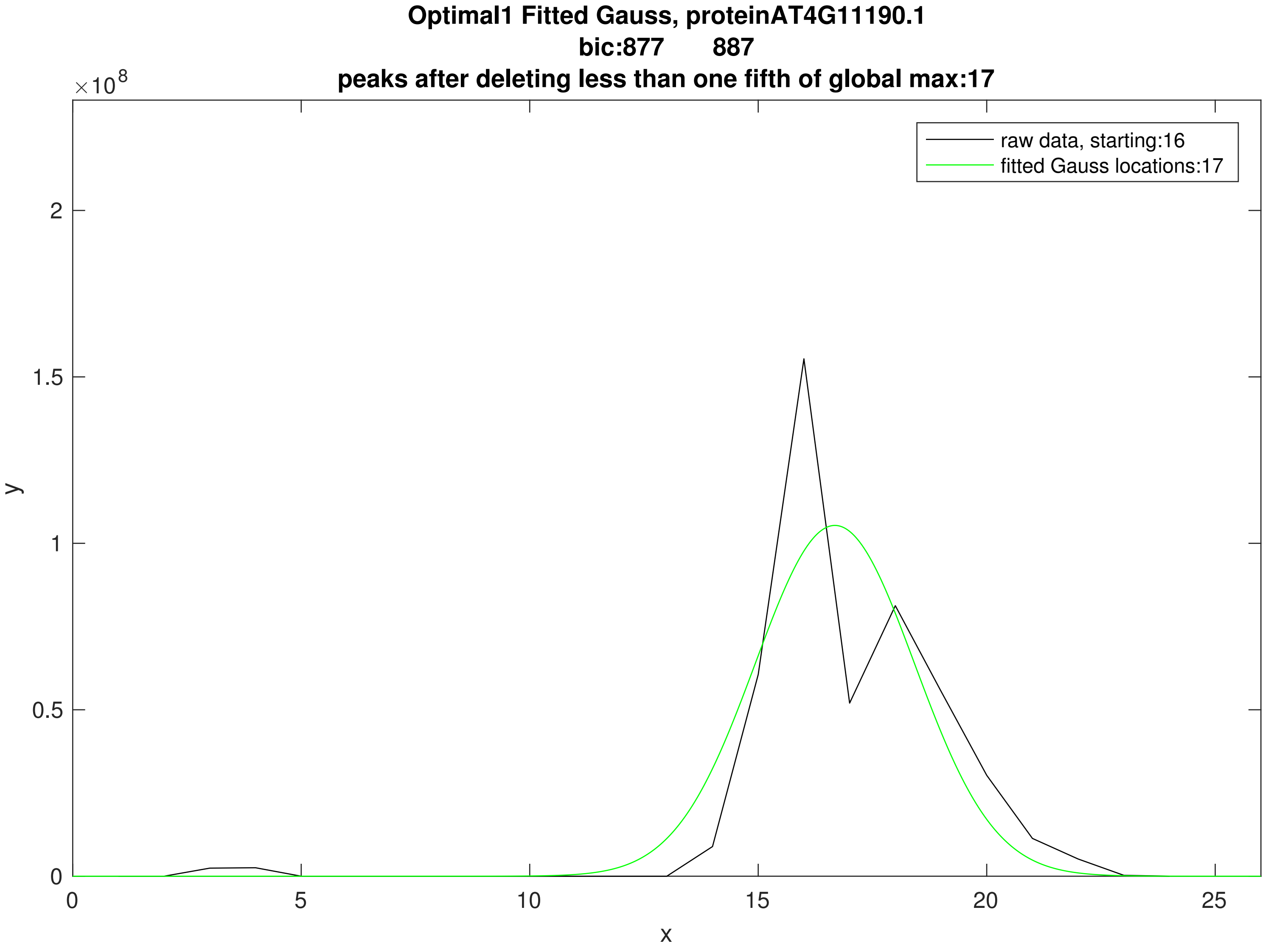
Optimal1 Fitted Gauss, proteinAT4G11190.1
bic:877 887
peaks after deleting less than one fifth of global max:17
×10^8
raw data, starting:16
fitted Gauss locations:17
y
x
0
0.5
1
1.5
2
0
5
10
15
20
25

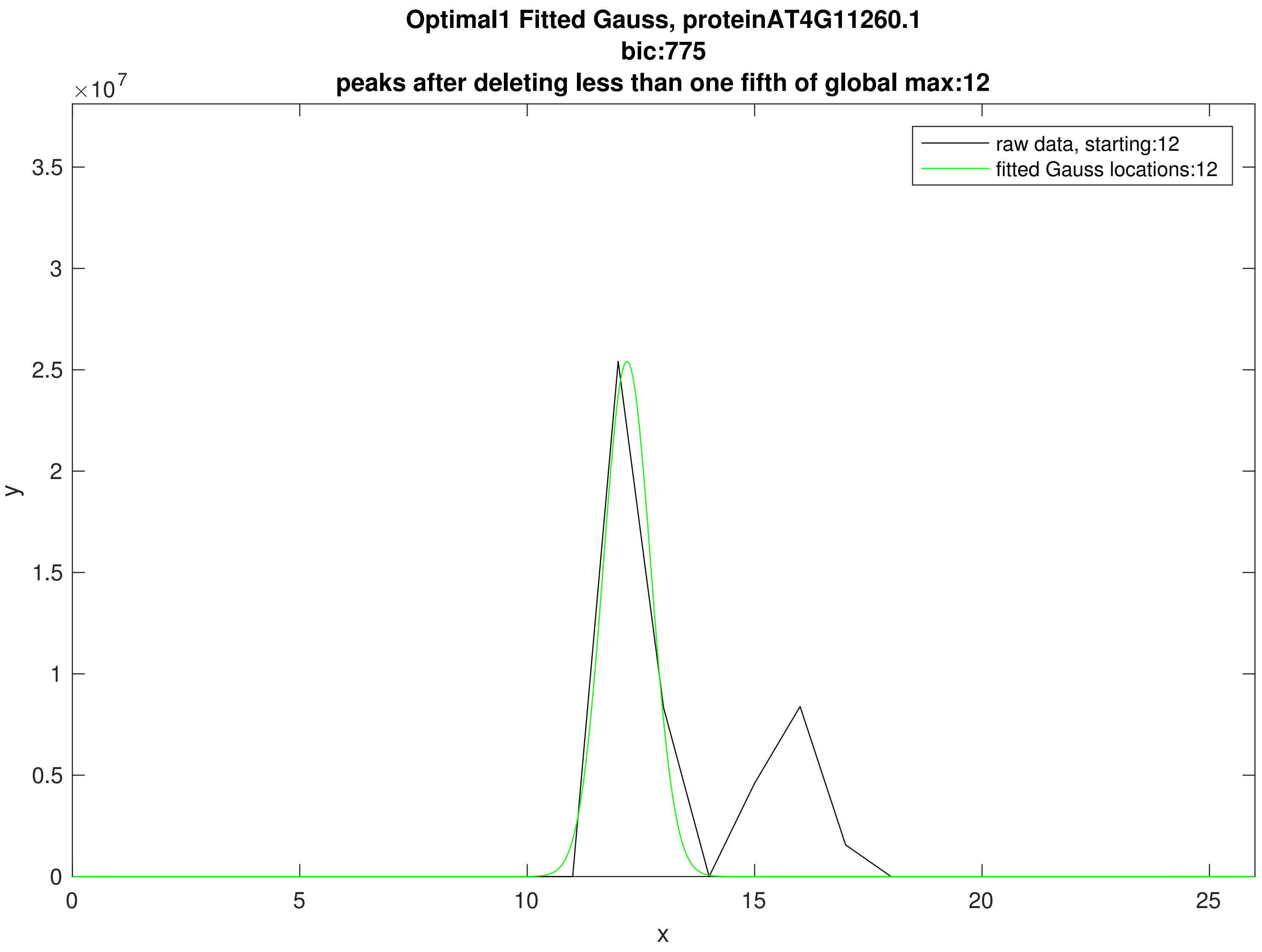
Optimal1 Fitted Gauss, proteinAT4G11260.1
bic:775
peaks after deleting less than one fifth of global max:12
×10^7
raw data, starting:12
fitted Gauss locations:12
y
x
0
0.5
1
1.5
2
2.5
3
3.5
5
10
15
20
25

Optimal1 Fitted Gauss, proteinAT4G11290.1
bic:943 1.04e+03 952
peaks after deleting less than one fifth of global max:17

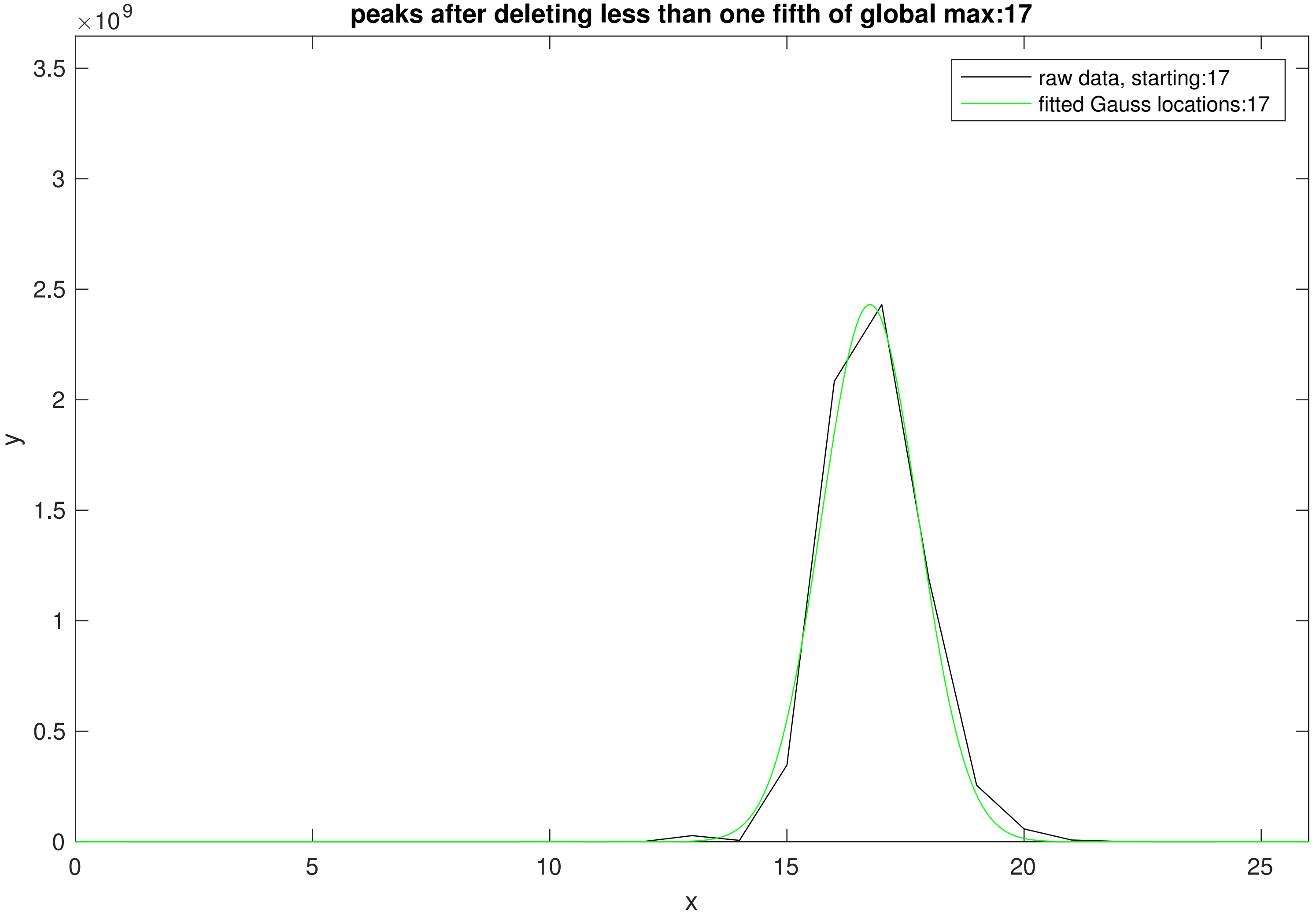

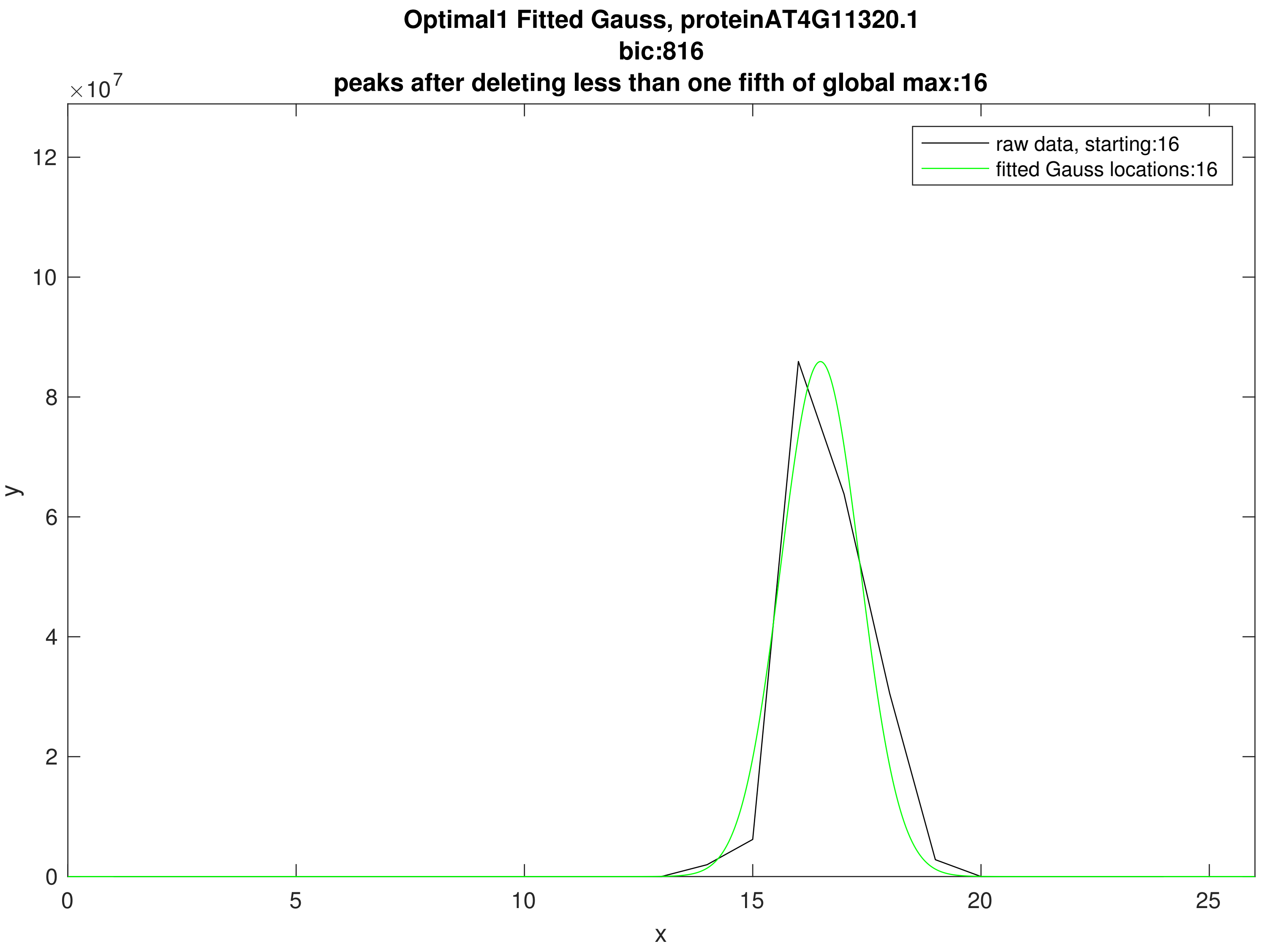
Optimal1 Fitted Gauss, proteinAT4G11320.1
bic:816
peaks after deleting less than one fifth of global max:16
×10^7
raw data, starting:16
fitted Gauss locations:16
y
x
0
2
4
6
8
10
12
0
5
10
15
20
25

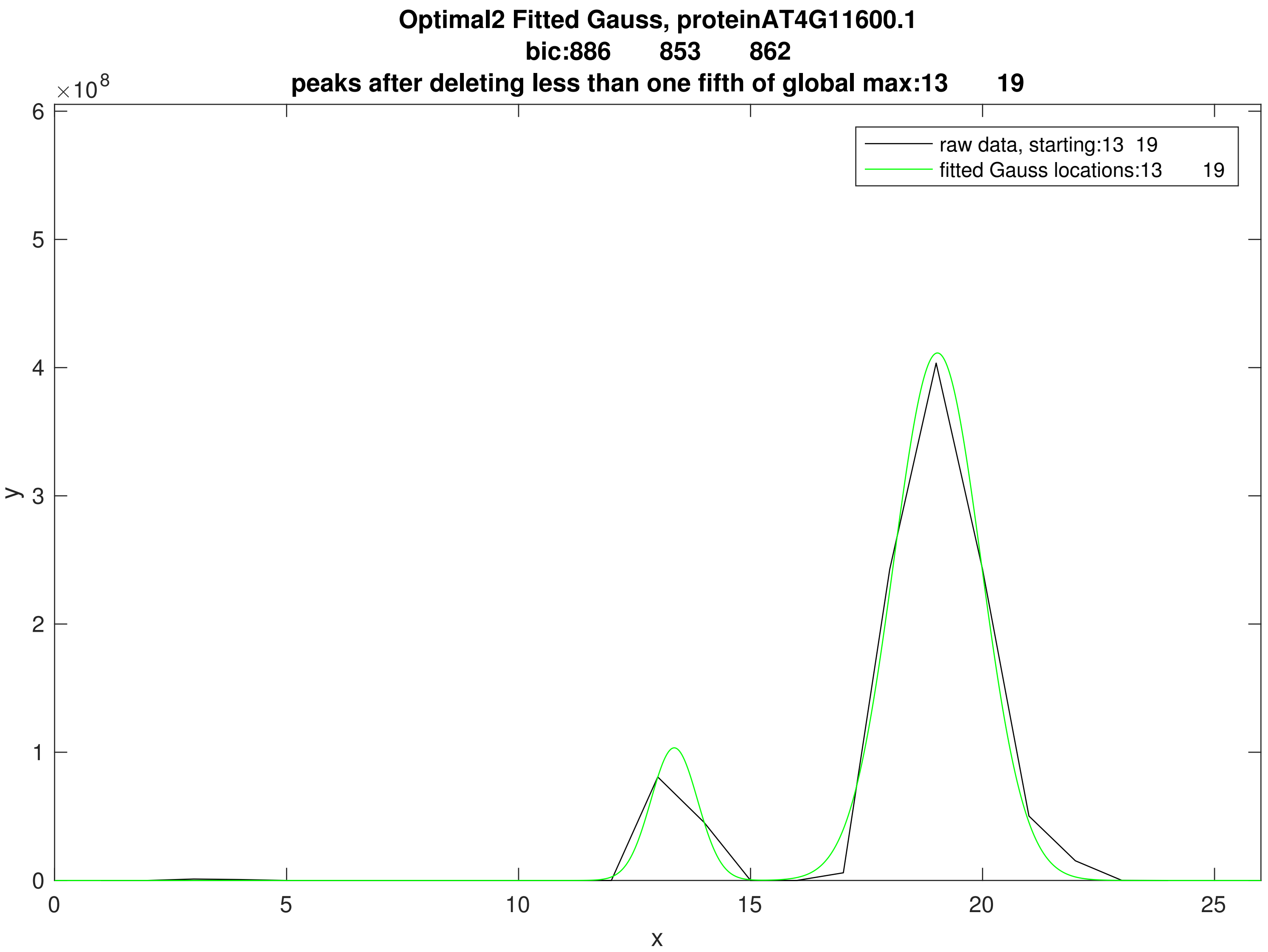
Optimal2 Fitted Gauss, proteinAT4G11600.1
bic:886 853 862
peaks after deleting less than one fifth of global max:13 19
×10^8
6
5
4
3
2
1
0
y
0
5
10
15
20
25
x
raw data, starting:13 19
fitted Gauss locations:13 19

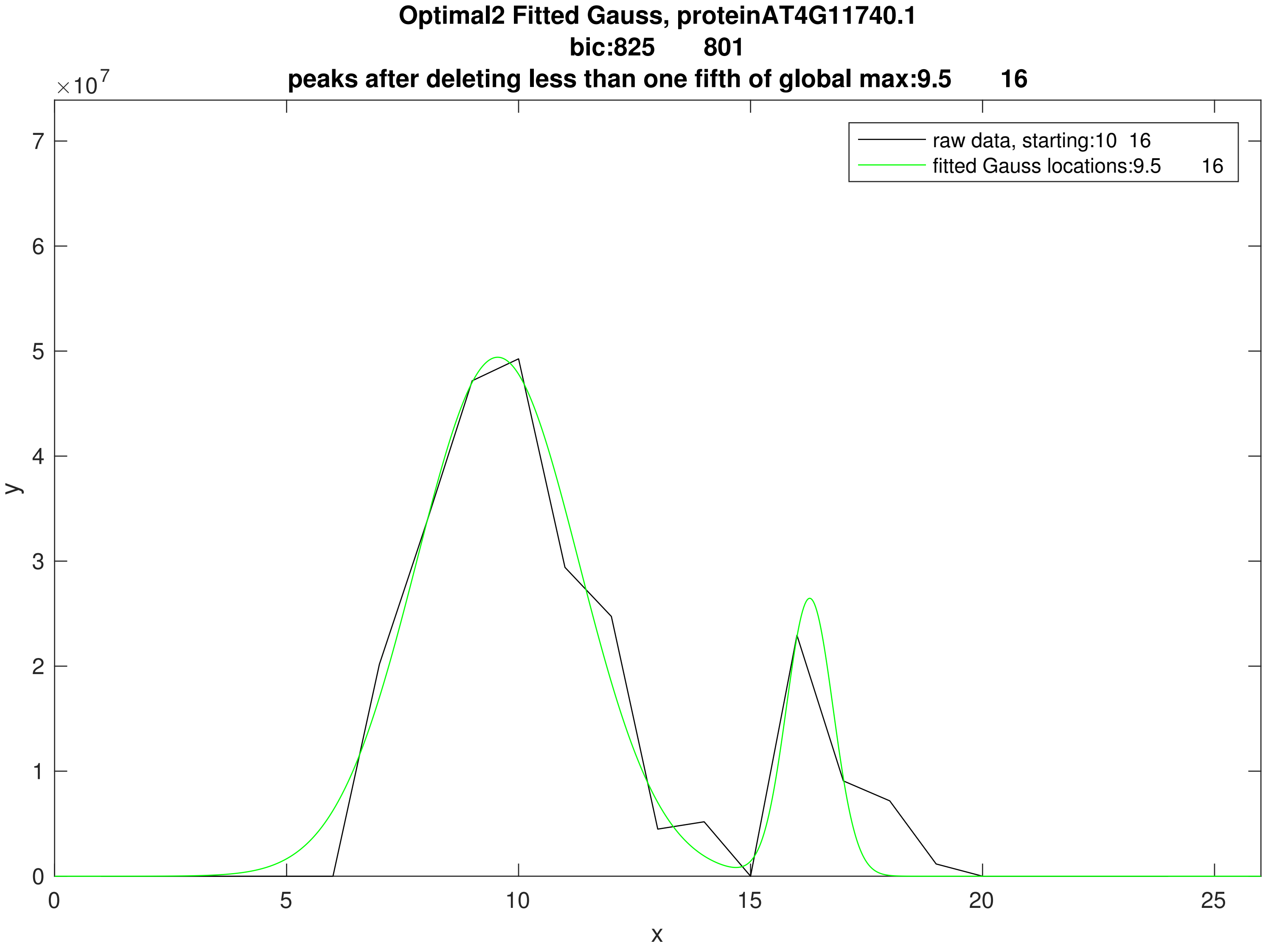

Optimal2 Fitted Gauss, proteinAT4G11740.1
bic:825 801
peaks after deleting less than one fifth of global max:9.5 16
×10^7
raw data, starting:10 16
fitted Gauss locations:9.5 16
y
x
0
1
2
3
4
5
6
7
0
5
10
15
20
25

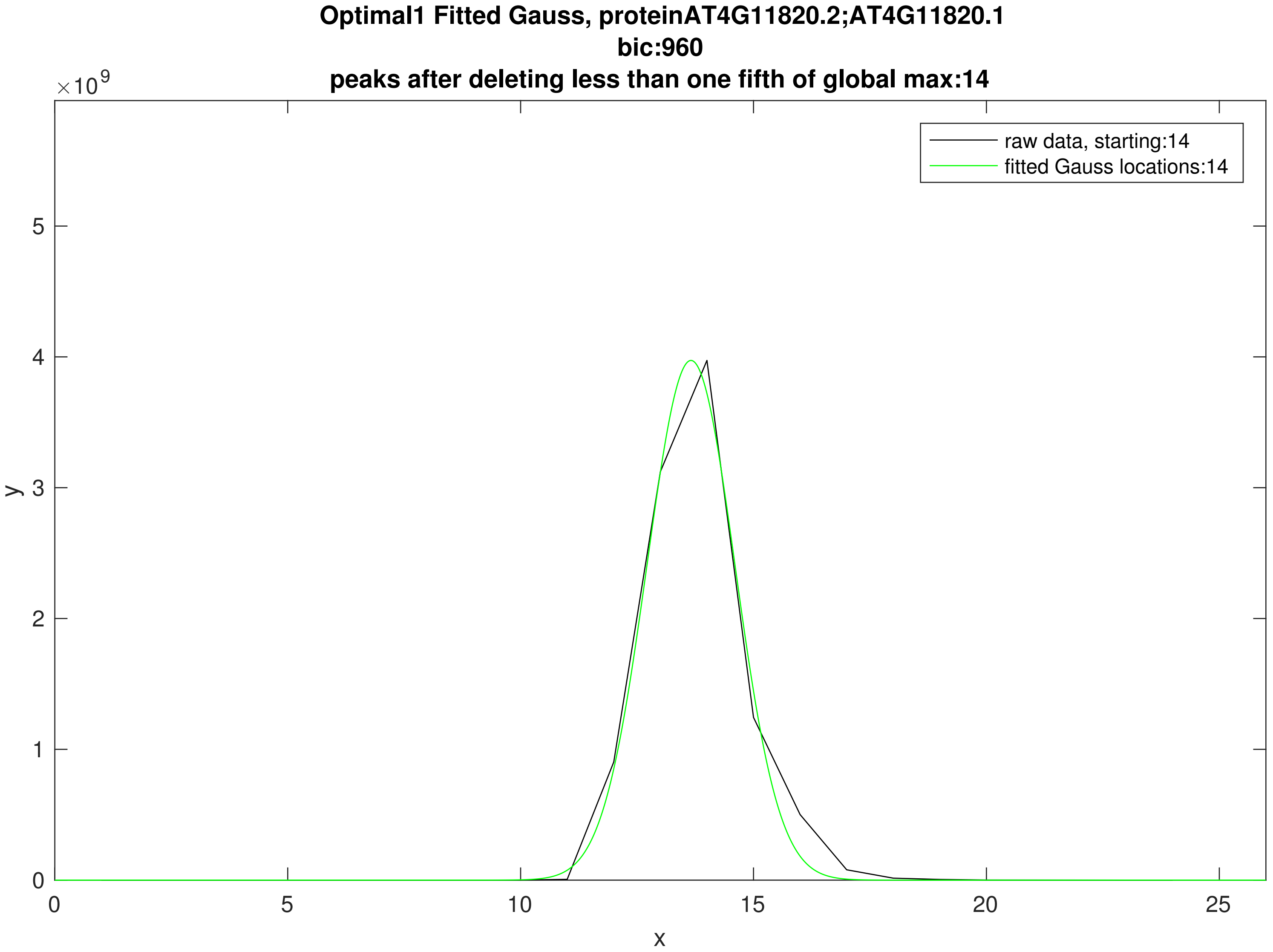

Optimal1 Fitted Gauss, proteinAT4G11820.2;AT4G11820.1
bic:960
peaks after deleting less than one fifth of global max:14
×10^9
raw data, starting:14
fitted Gauss locations:14
y
x
0
1
2
3
4
5
0
5
10
15
20
25

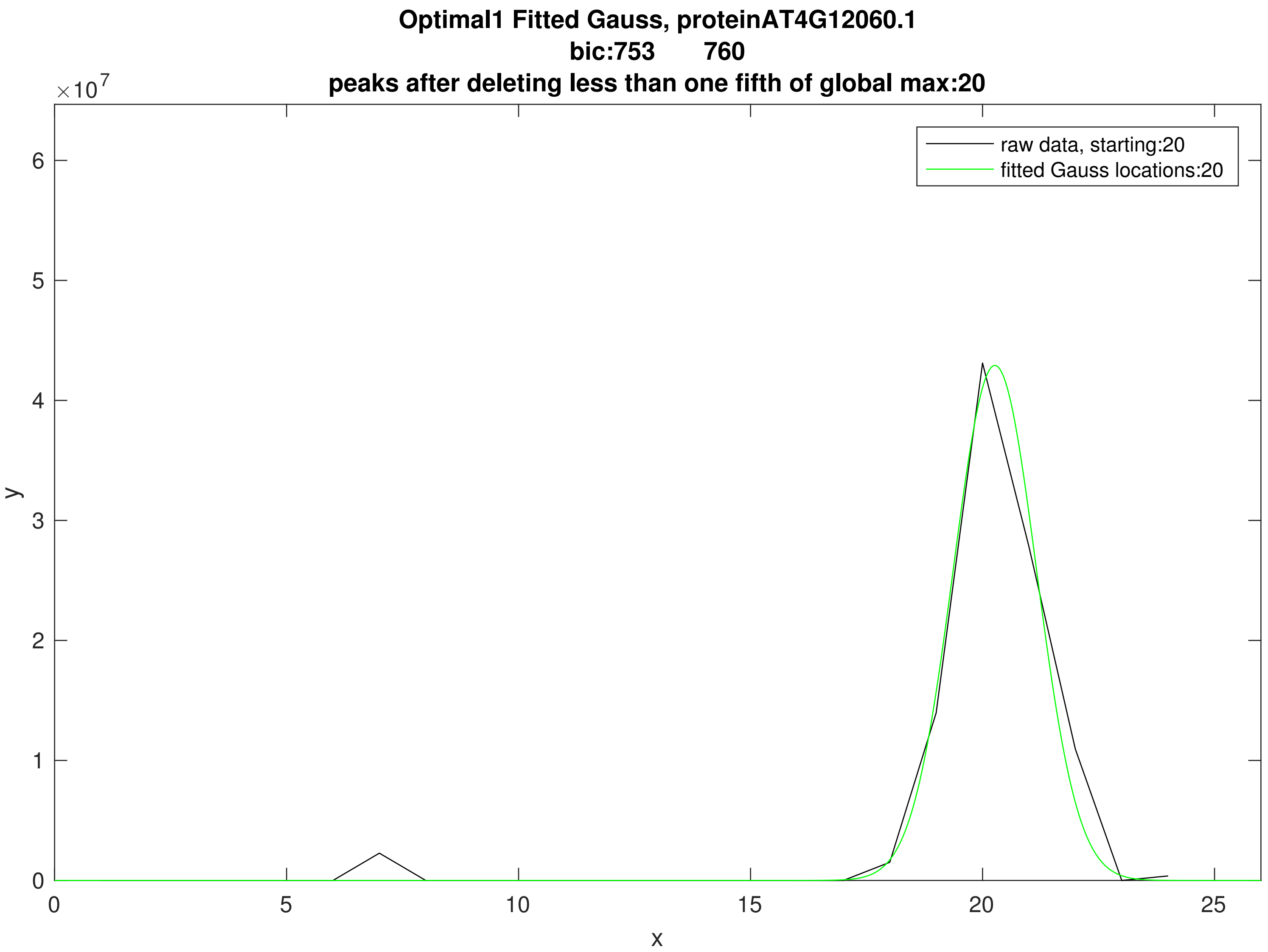
Optimal1 Fitted Gauss, proteinAT4G12060.1
bic:753 760
peaks after deleting less than one fifth of global max:20
×10^7
raw data, starting:20
fitted Gauss locations:20
y
x
0
1
2
3
4
5
6
0
5
10
15
20
25

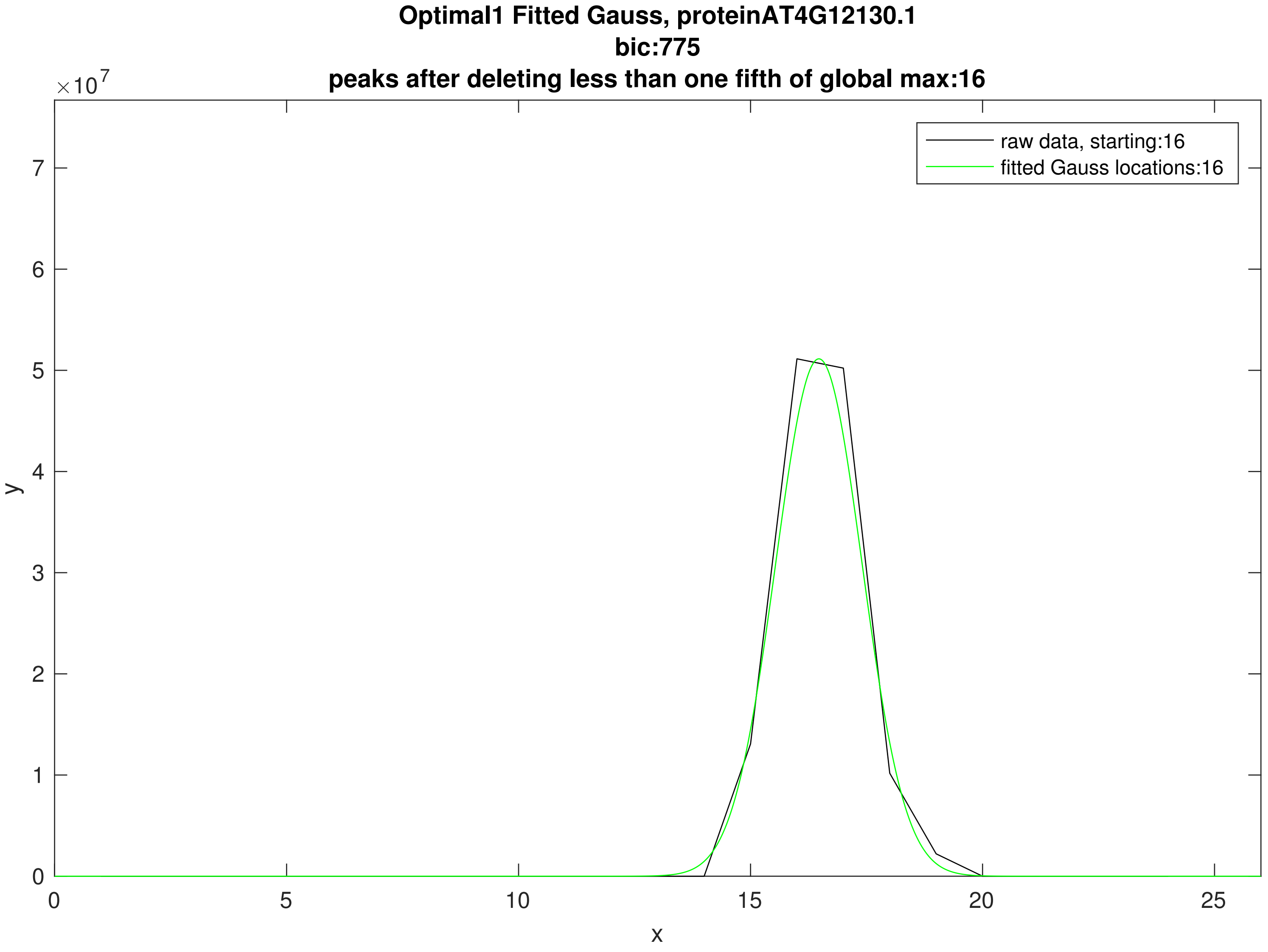

Optimal1 Fitted Gauss, proteinAT4G12130.1
bic:775
peaks after deleting less than one fifth of global max:16
×10^7
raw data, starting:16
fitted Gauss locations:16
y
x
0
1
2
3
4
5
6
7
0
5
10
15
20
25

**Optimal1 Fitted Gauss, proteinAT4G12290.1**
**bic:763**
**peaks after deleting less than one fifth of global max:11**

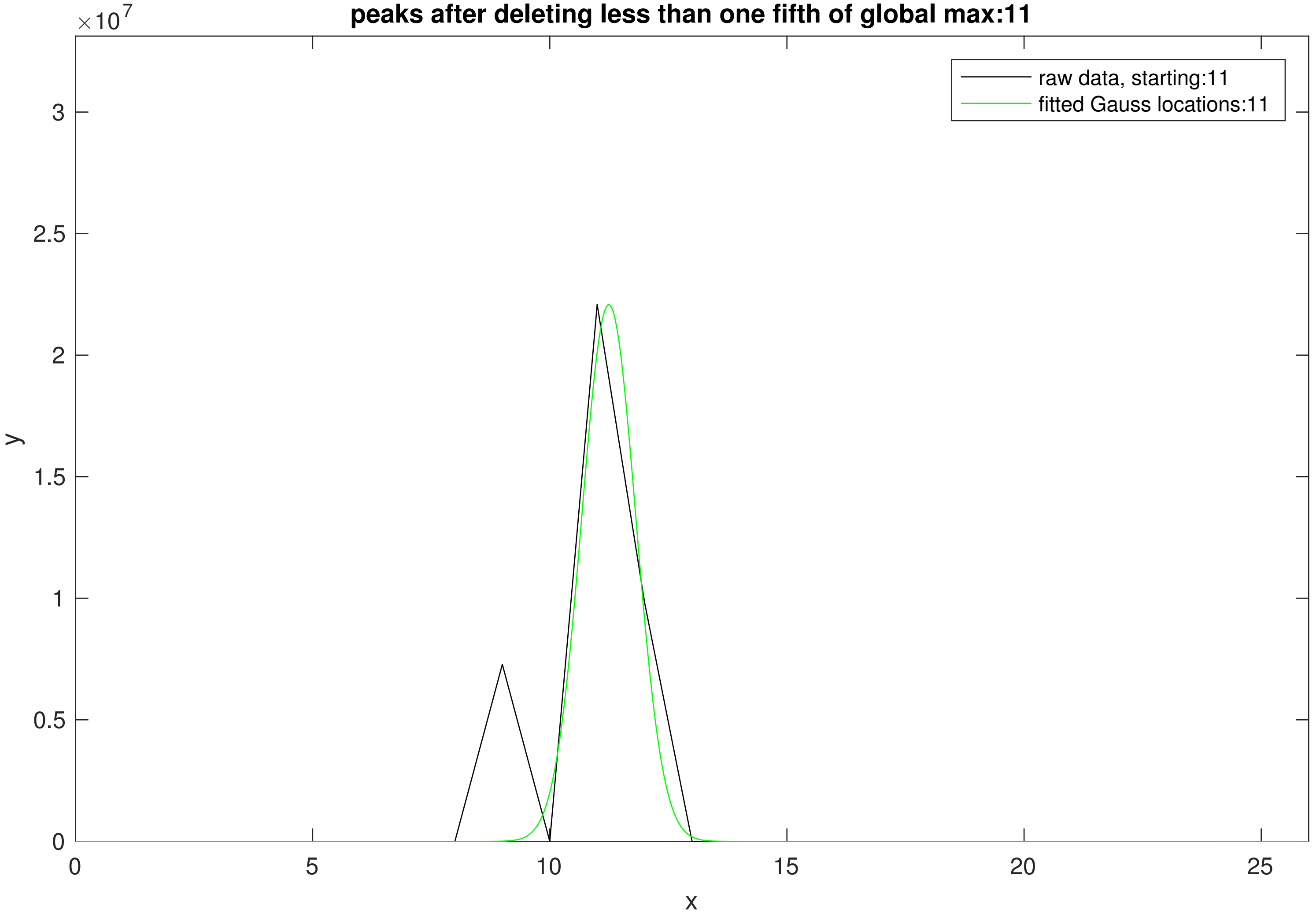

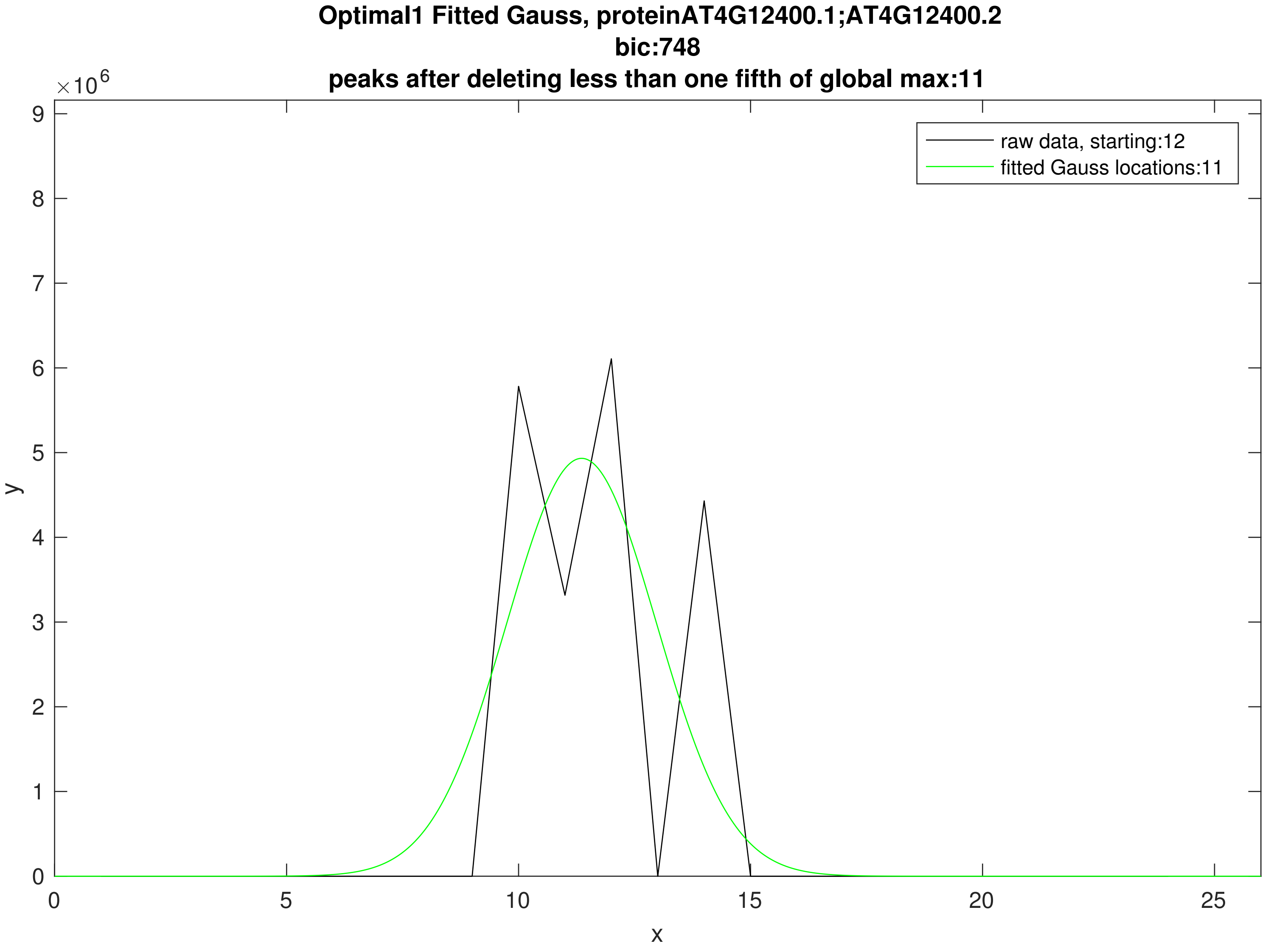
Optimal1 Fitted Gauss, proteinAT4G12400.1;AT4G12400.2
bic:748
peaks after deleting less than one fifth of global max:11
×10^6
9
8
7
6
5
4
3
2
1
0
y
0
5
10
15
20
25
x
raw data, starting:12
fitted Gauss locations:11

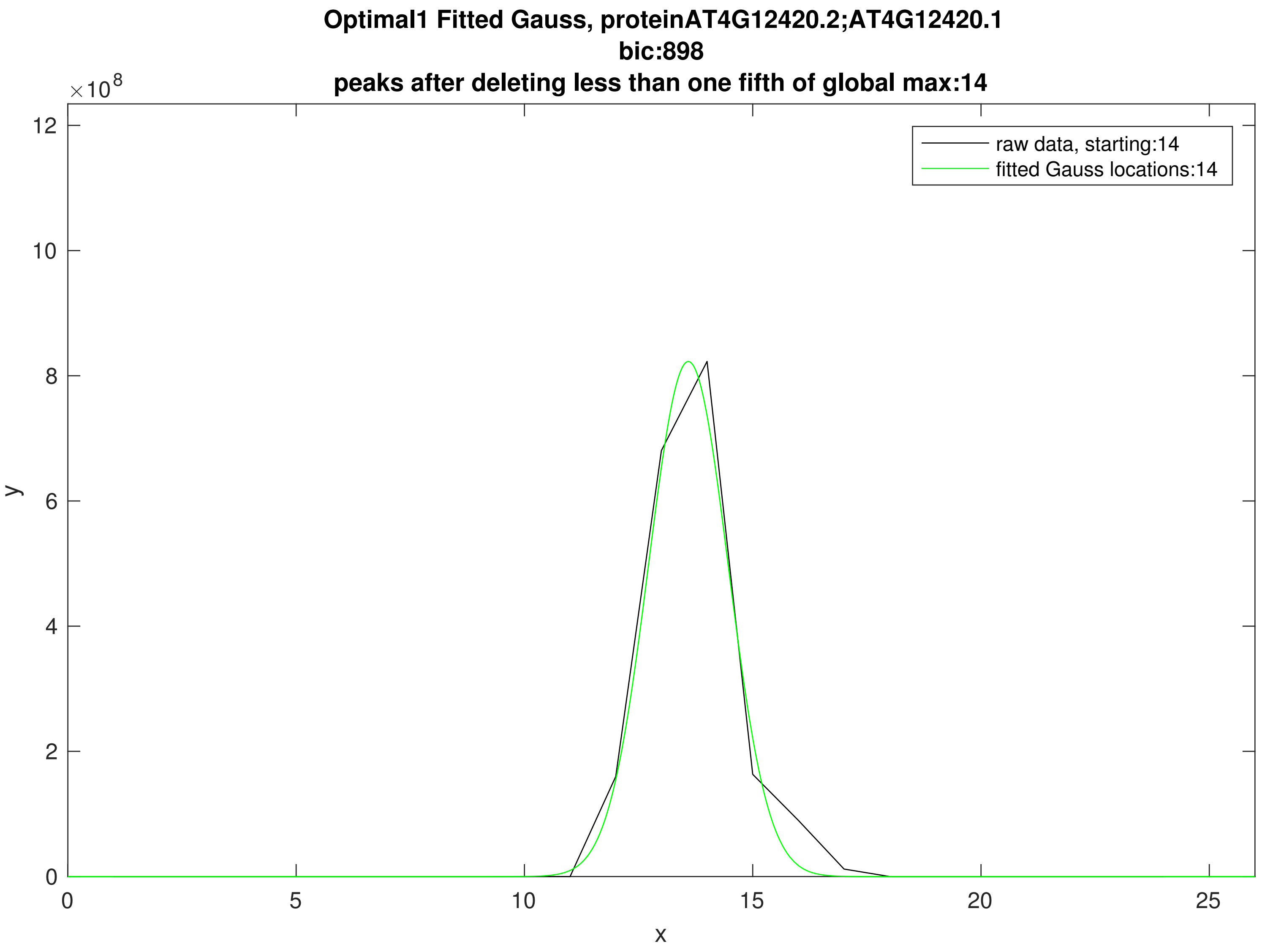
Optimal1 Fitted Gauss, proteinAT4G12420.2;AT4G12420.1
bic:898
peaks after deleting less than one fifth of global max:14
×10^8
12
10
8
6
4
2
0
y
0
5
10
15
20
25
x
raw data, starting:14
fitted Gauss locations:14

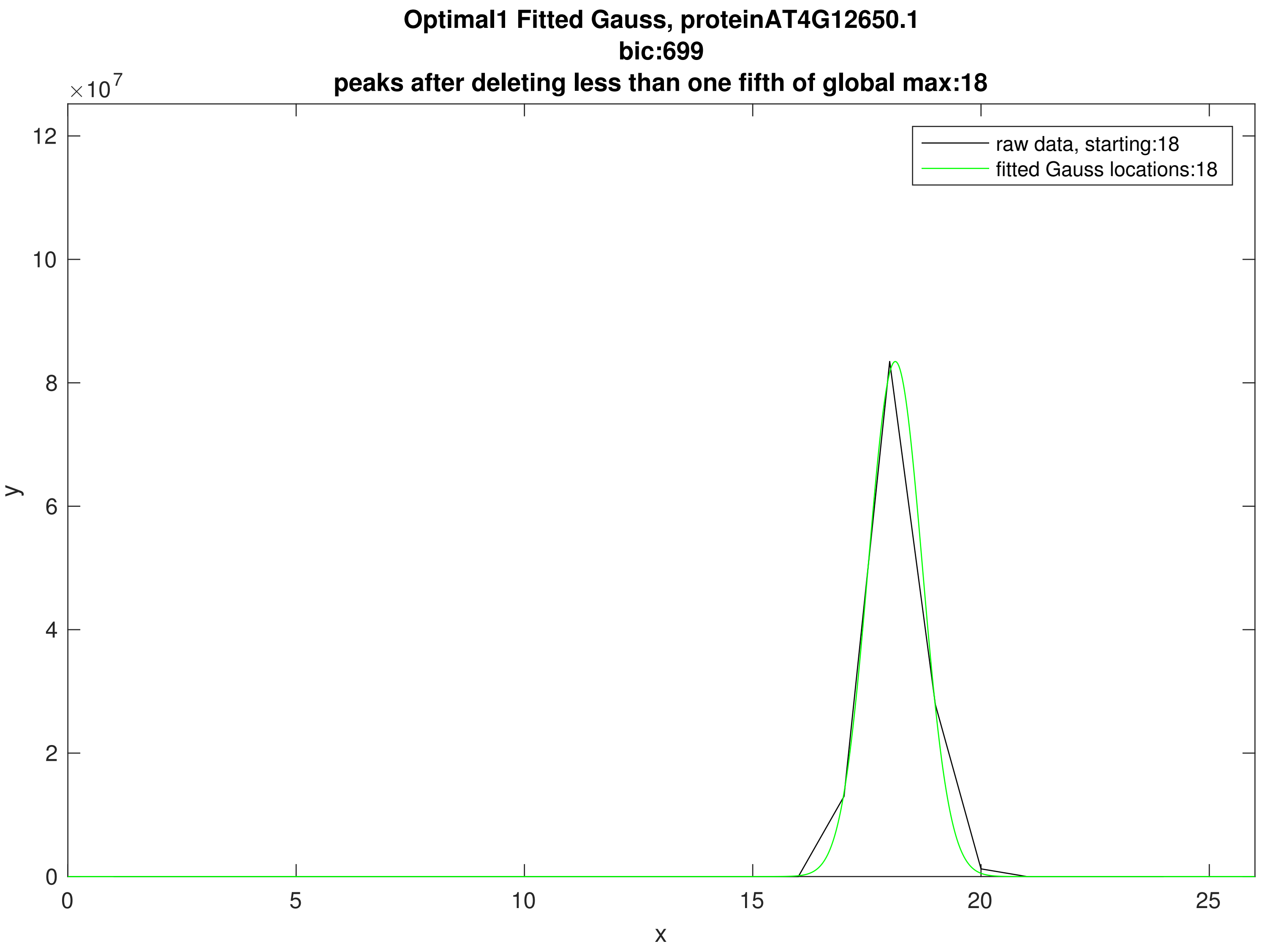
Optimal1 Fitted Gauss, proteinAT4G12650.1
bic:699
peaks after deleting less than one fifth of global max:18
raw data, starting:18
fitted Gauss locations:18
y
x
×10^7

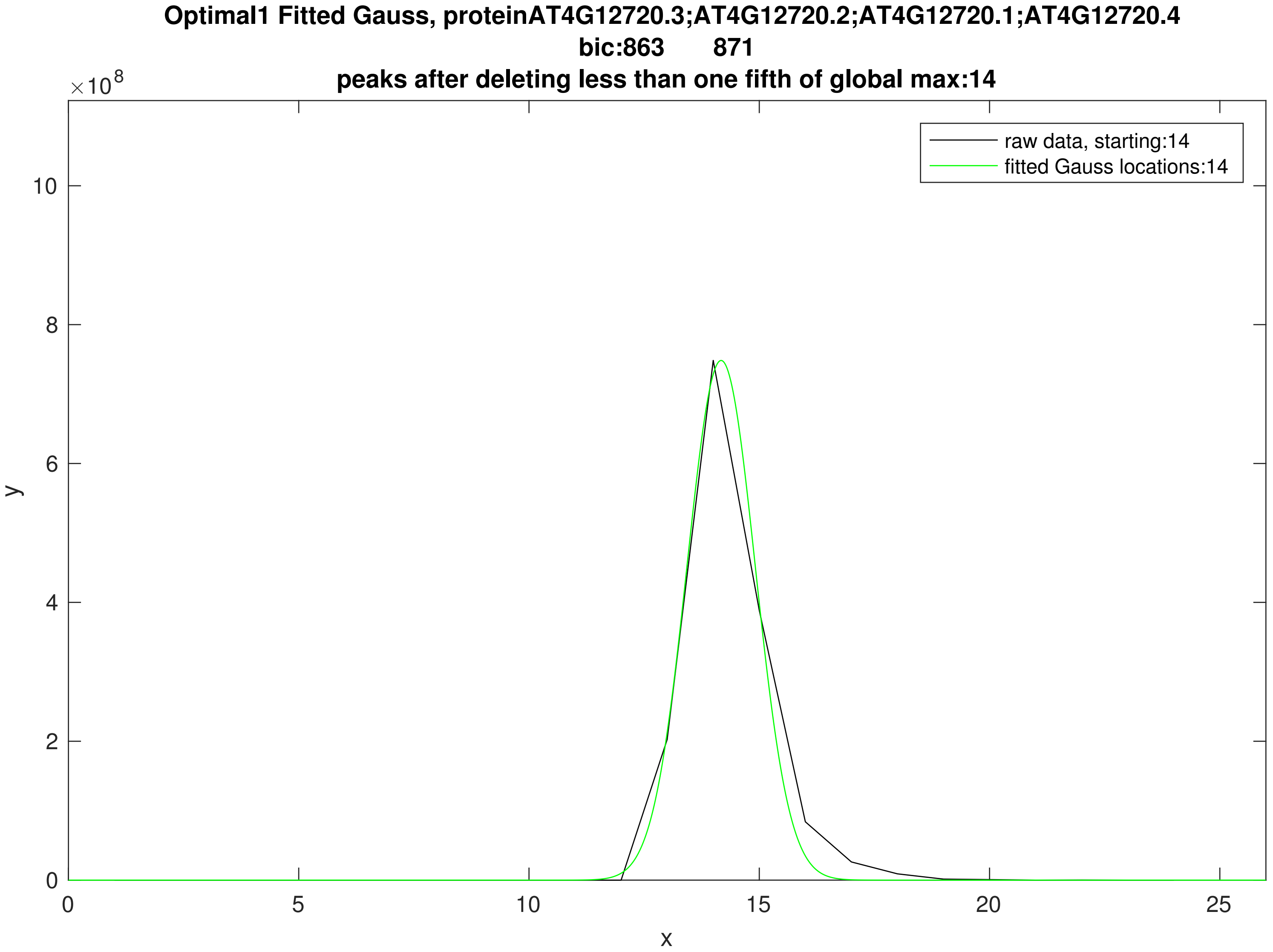
Optimal1 Fitted Gauss, proteinAT4G12720.3;AT4G12720.2;AT4G12720.1;AT4G12720.4
bic:863 871
peaks after deleting less than one fifth of global max:14
×10^8
raw data, starting:14
fitted Gauss locations:14
y
x
0
2
4
6
8
10
0
5
10
15
20
25

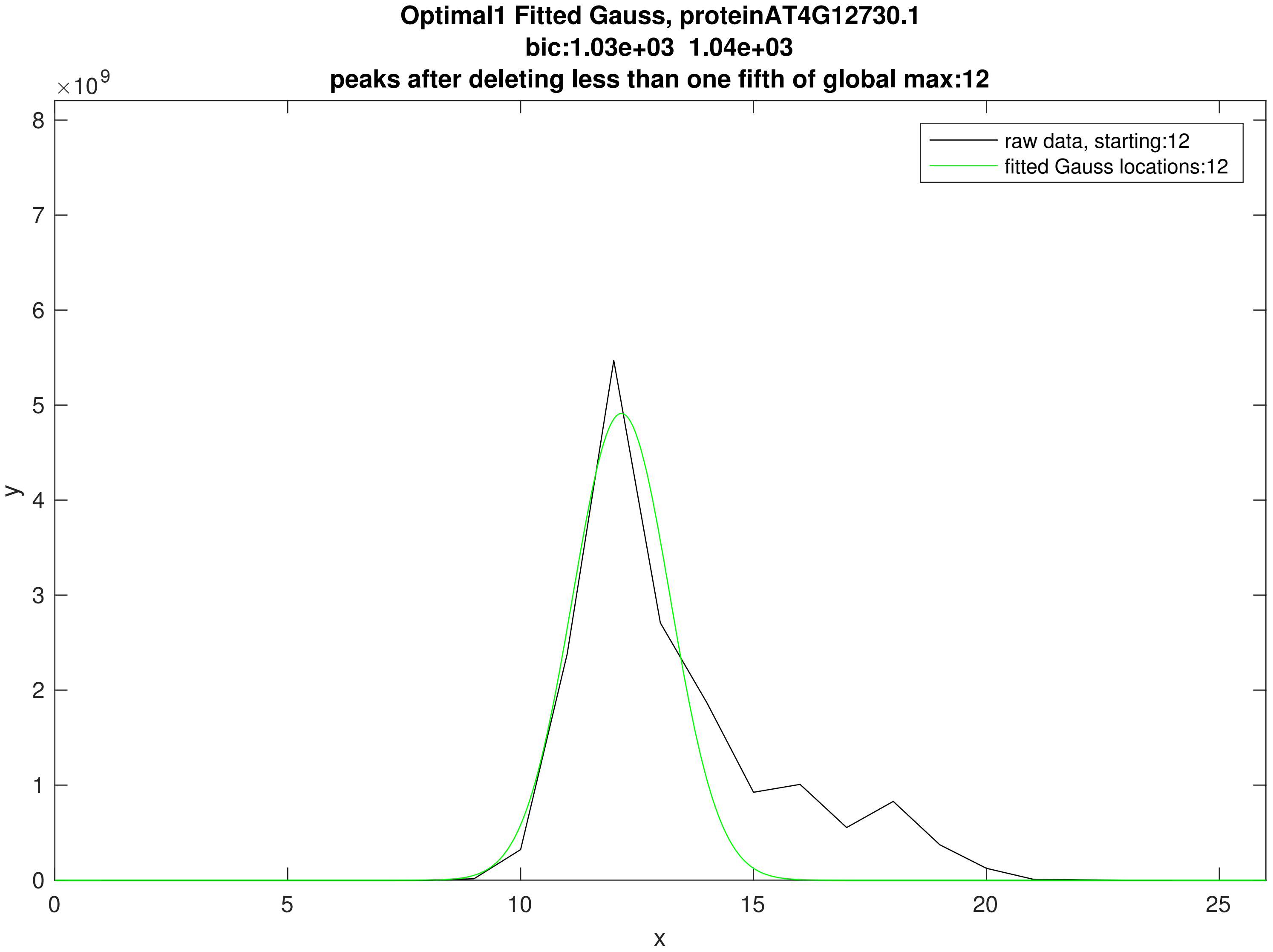
Optimal1 Fitted Gauss, proteinAT4G12730.1
bic:1.03e+03 1.04e+03
peaks after deleting less than one fifth of global max:12
×10^9
raw data, starting:12
fitted Gauss locations:12
y
x

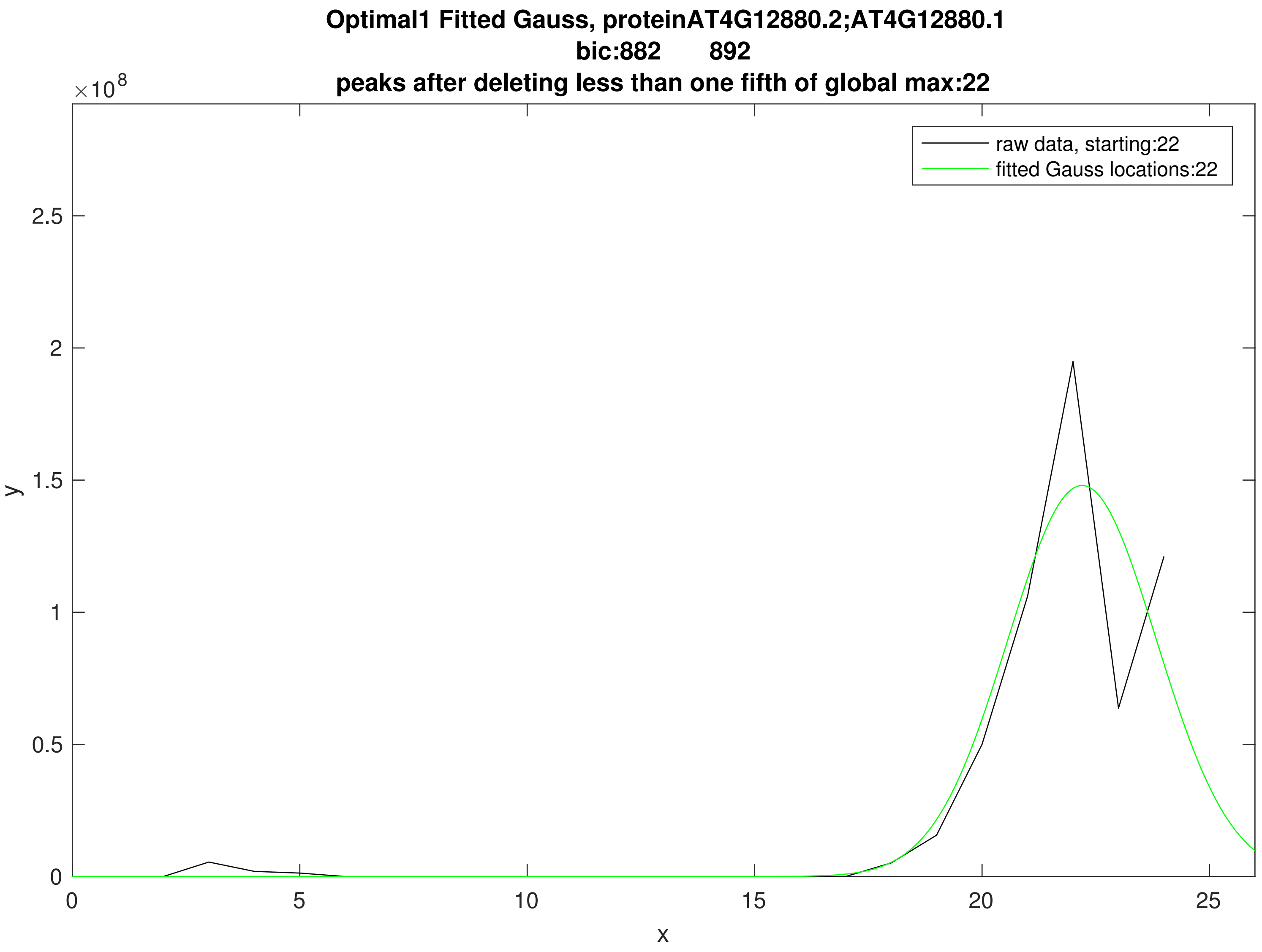

Optimal1 Fitted Gauss, proteinAT4G12880.2;AT4G12880.1
bic:882 892
peaks after deleting less than one fifth of global max:22
raw data, starting:22
fitted Gauss locations:22
×10^8
2.5
2
1.5
1
0.5
0
y
0
5
10
15
20
25
x

#### Optimal1 Fitted Gauss, proteinAT4G12910.1
#### bic:730
#### peaks after deleting less than one fifth of global max:13

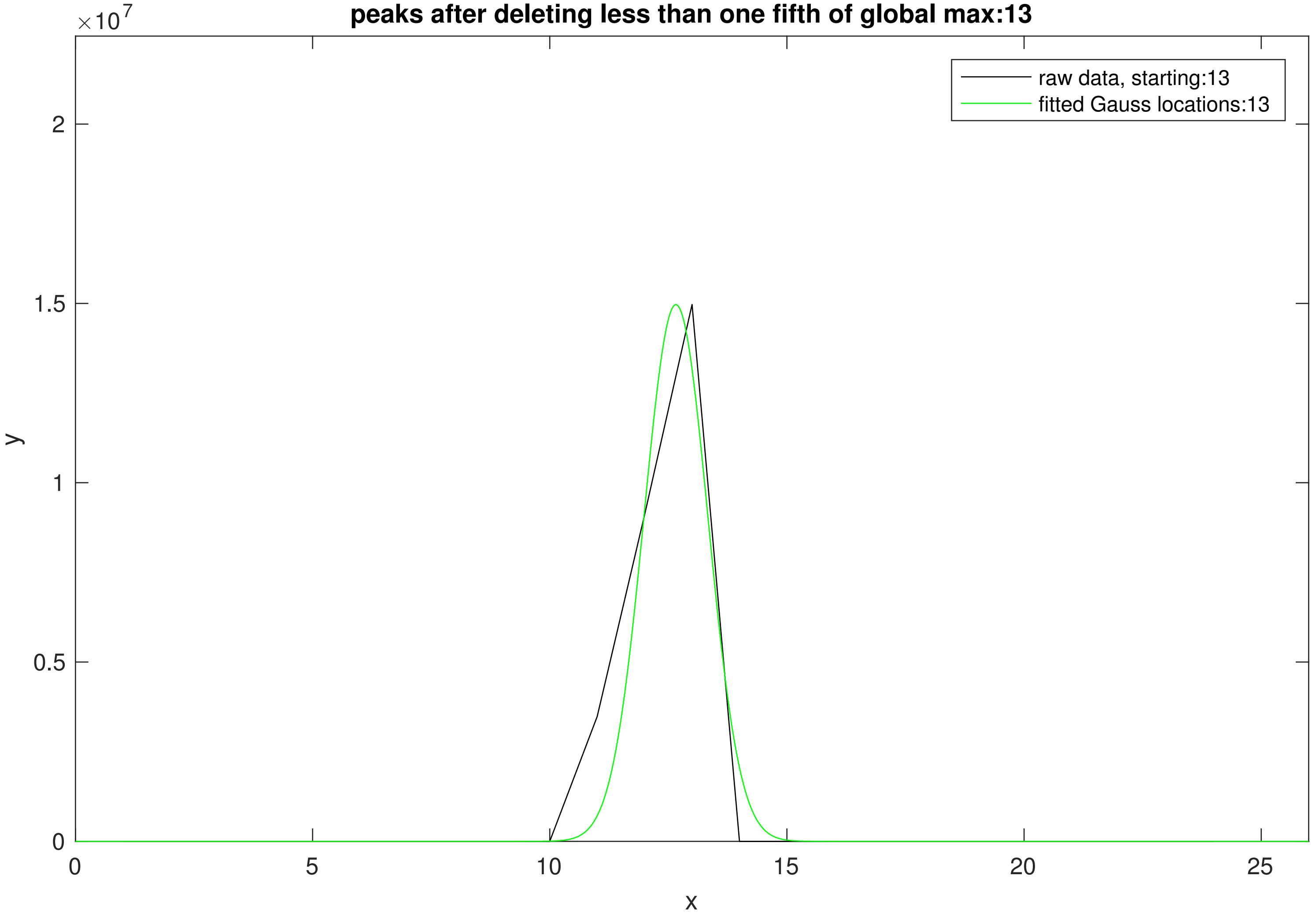

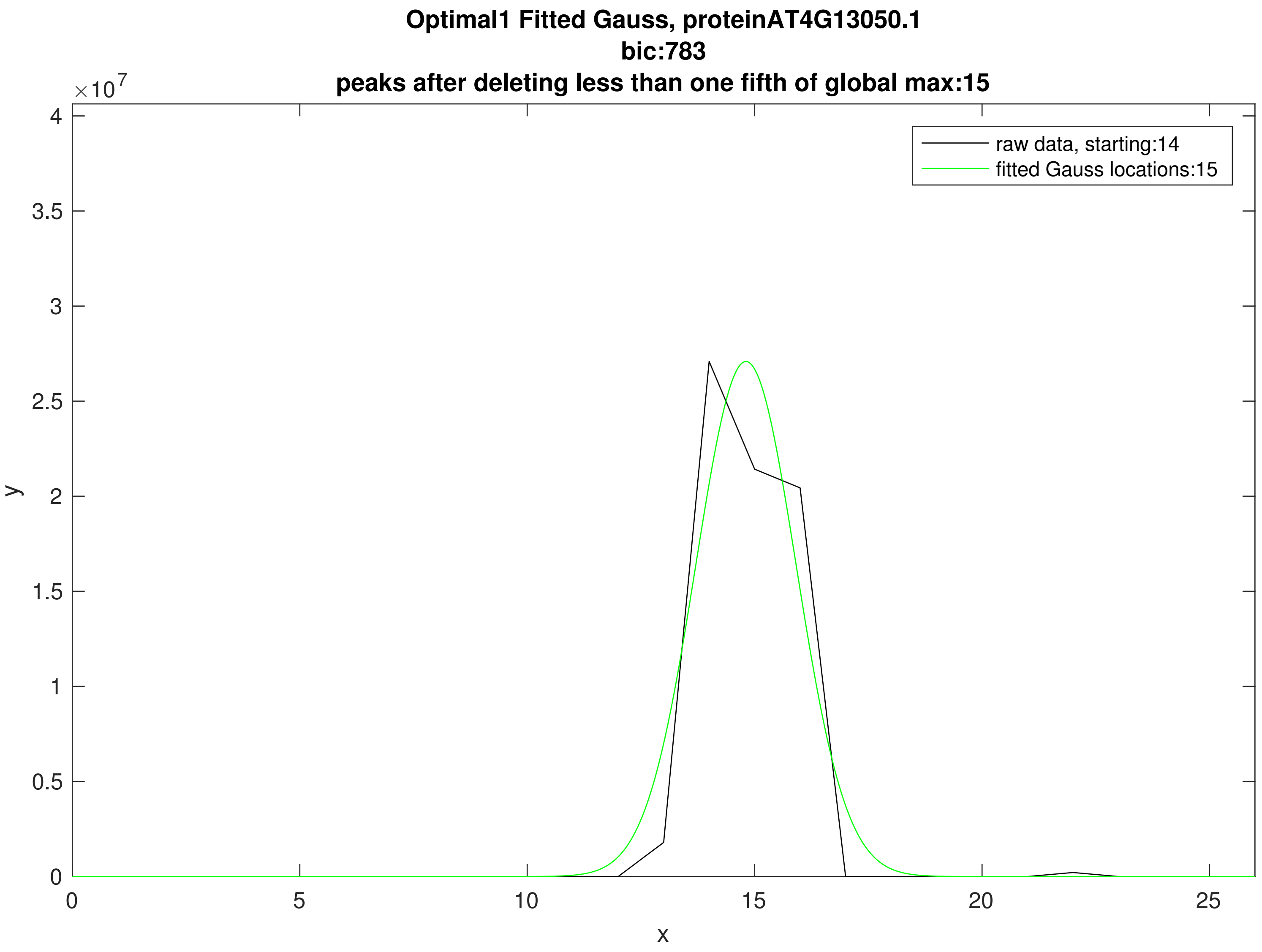

Optimal1 Fitted Gauss, proteinAT4G13050.1
bic:783
peaks after deleting less than one fifth of global max:15
×10^7
4
3.5
3
2.5
2
1.5
1
0.5
0
y
0
5
10
15
20
25
x
raw data, starting:14
fitted Gauss locations:15

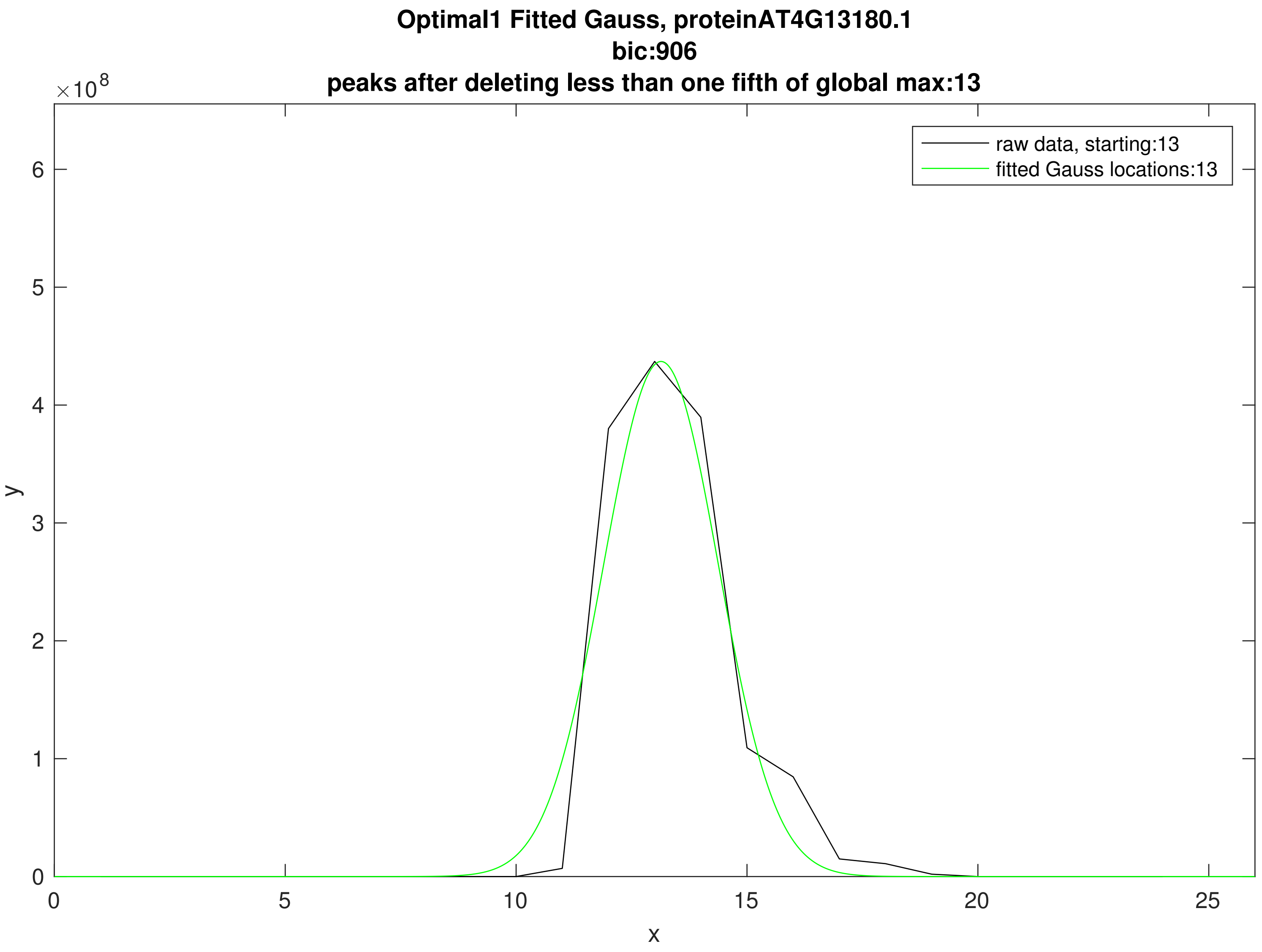
Optimal1 Fitted Gauss, proteinAT4G13180.1
bic:906
peaks after deleting less than one fifth of global max:13
×10^8
raw data, starting:13
fitted Gauss locations:13
y
x
0
1
2
3
4
5
6
0
5
10
15
20
25

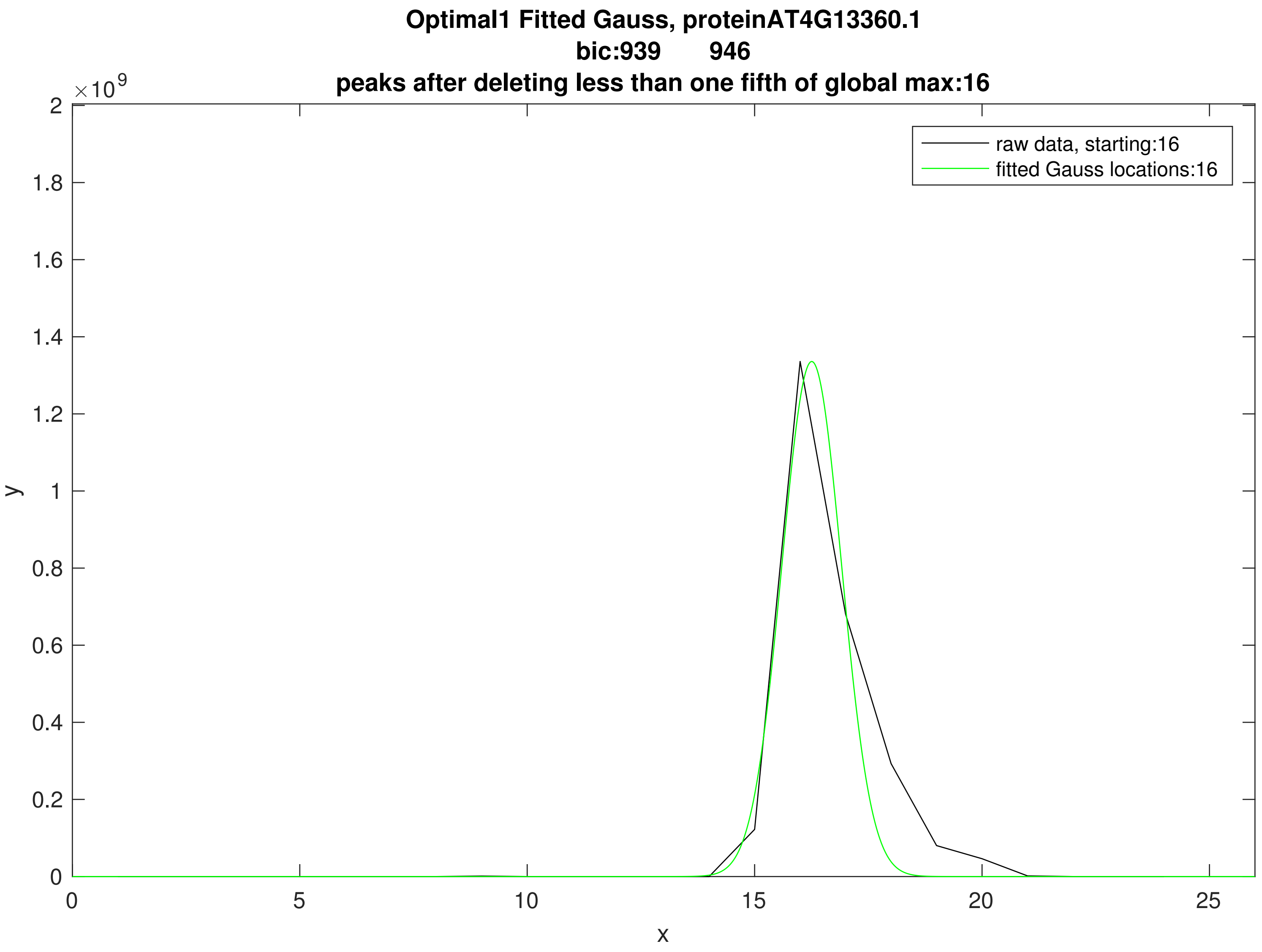

Optimal1 Fitted Gauss, proteinAT4G13360.1
bic:939 946
peaks after deleting less than one fifth of global max:16
×10^9
raw data, starting:16
fitted Gauss locations:16
y
x
0
0.2
0.4
0.6
0.8
1
1.2
1.4
1.6
1.8
2
5
10
15
20
25

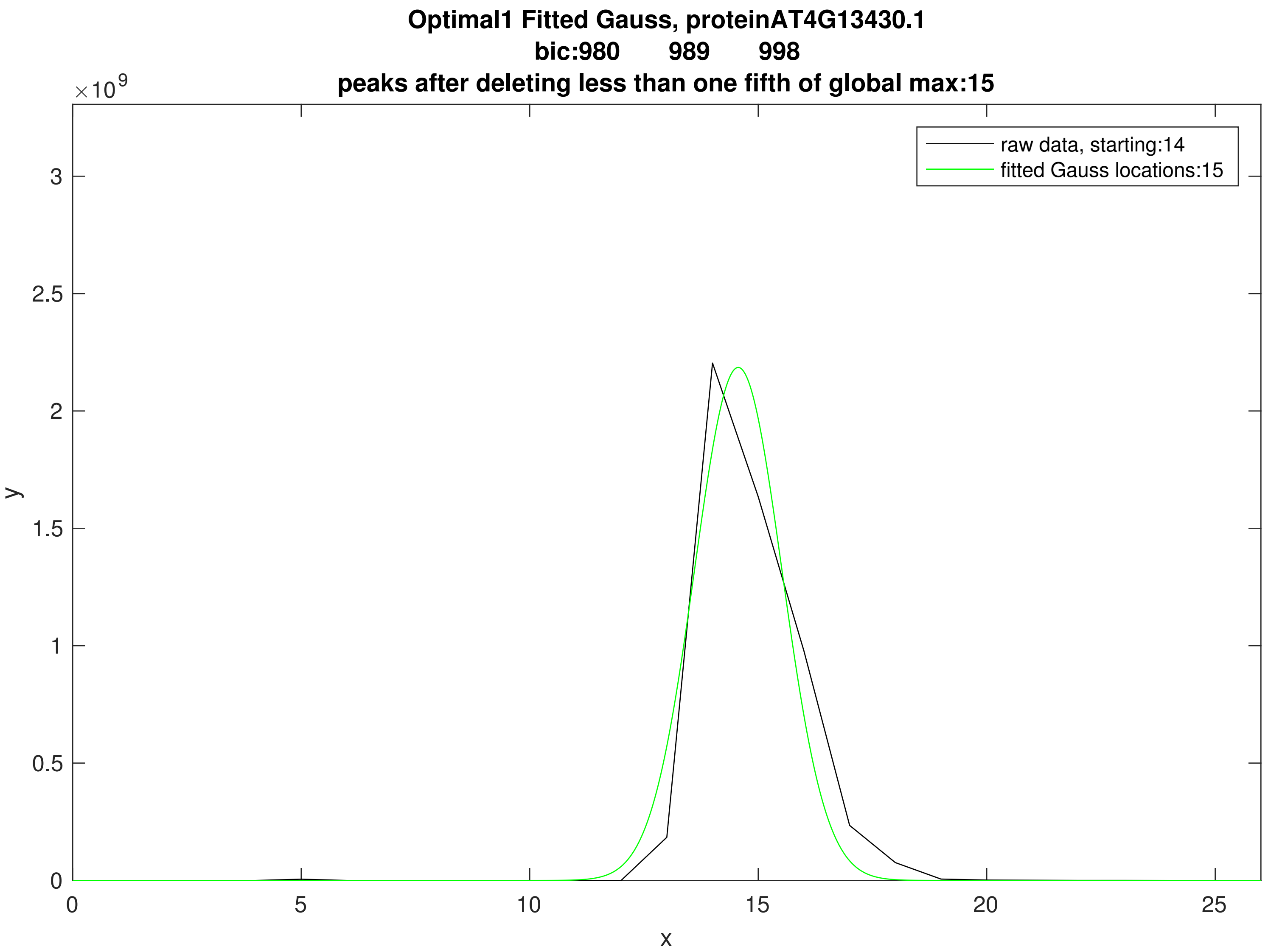

Optimal1 Fitted Gauss, proteinAT4G13430.1
bic:980 989 998
peaks after deleting less than one fifth of global max:15
×10^9
raw data, starting:14
fitted Gauss locations:15
3
2.5
2
1.5
1
0.5
0
y
0
5
10
15
20
25
x

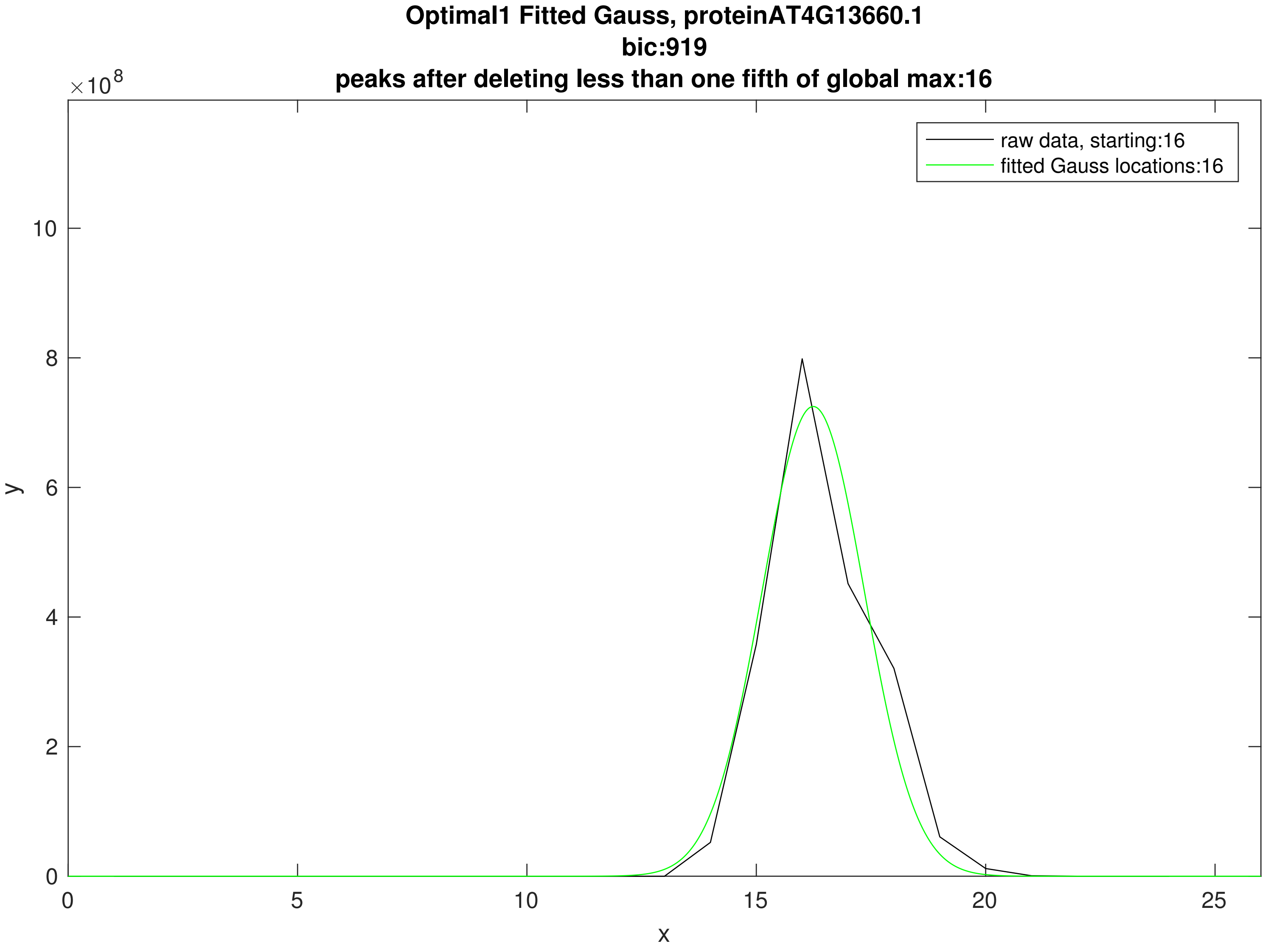
Optimal1 Fitted Gauss, proteinAT4G13660.1
bic:919
peaks after deleting less than one fifth of global max:16
×10^8
raw data, starting:16
fitted Gauss locations:16
10
8
6
4
2
0
y
0
5
10
15
20
25
x

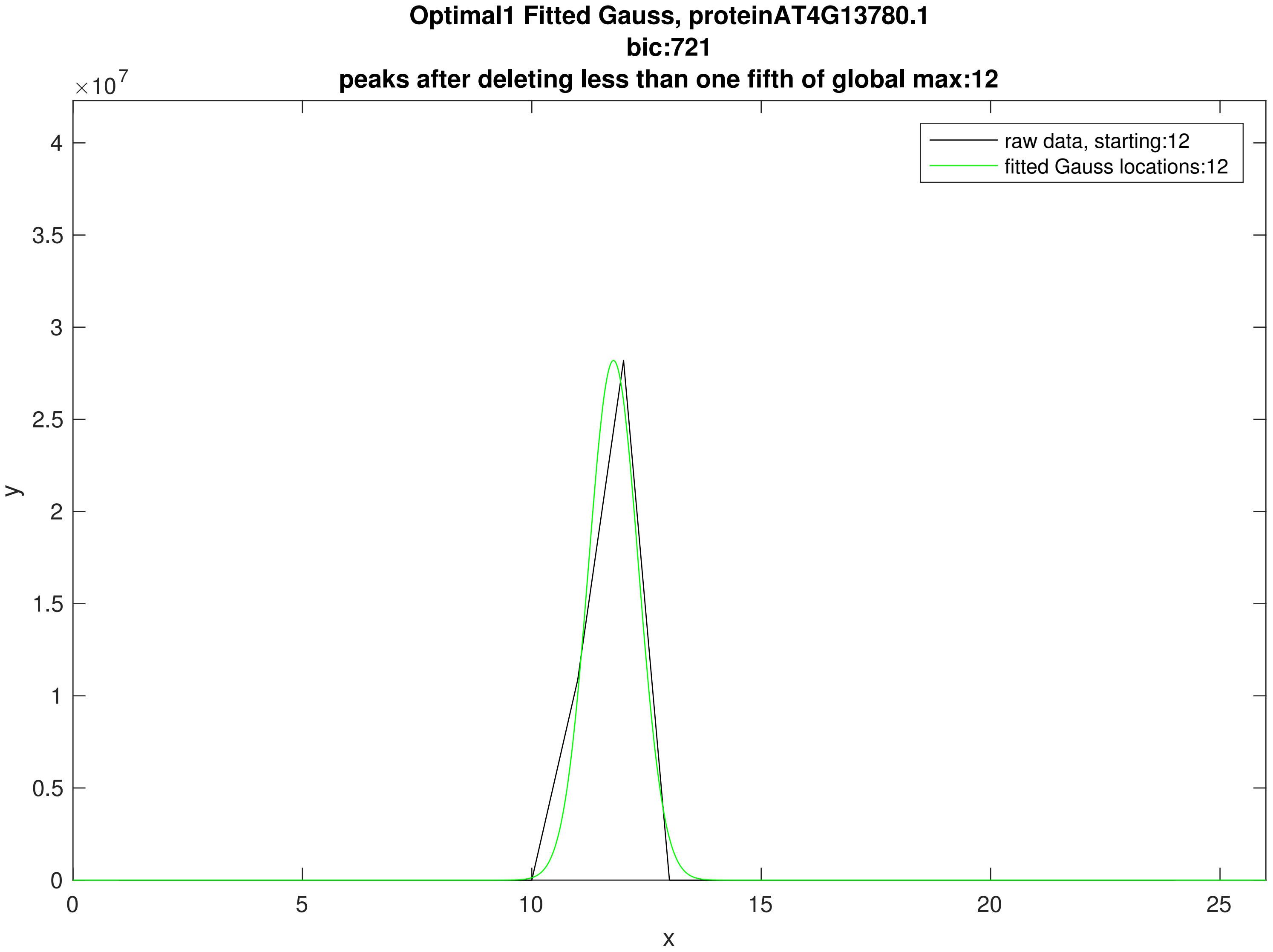
Optimal1 Fitted Gauss, proteinAT4G13780.1
bic:721
peaks after deleting less than one fifth of global max:12
×10^7
raw data, starting:12
fitted Gauss locations:12
y
x
0
0.5
1
1.5
2
2.5
3
3.5
4
0
5
10
15
20
25

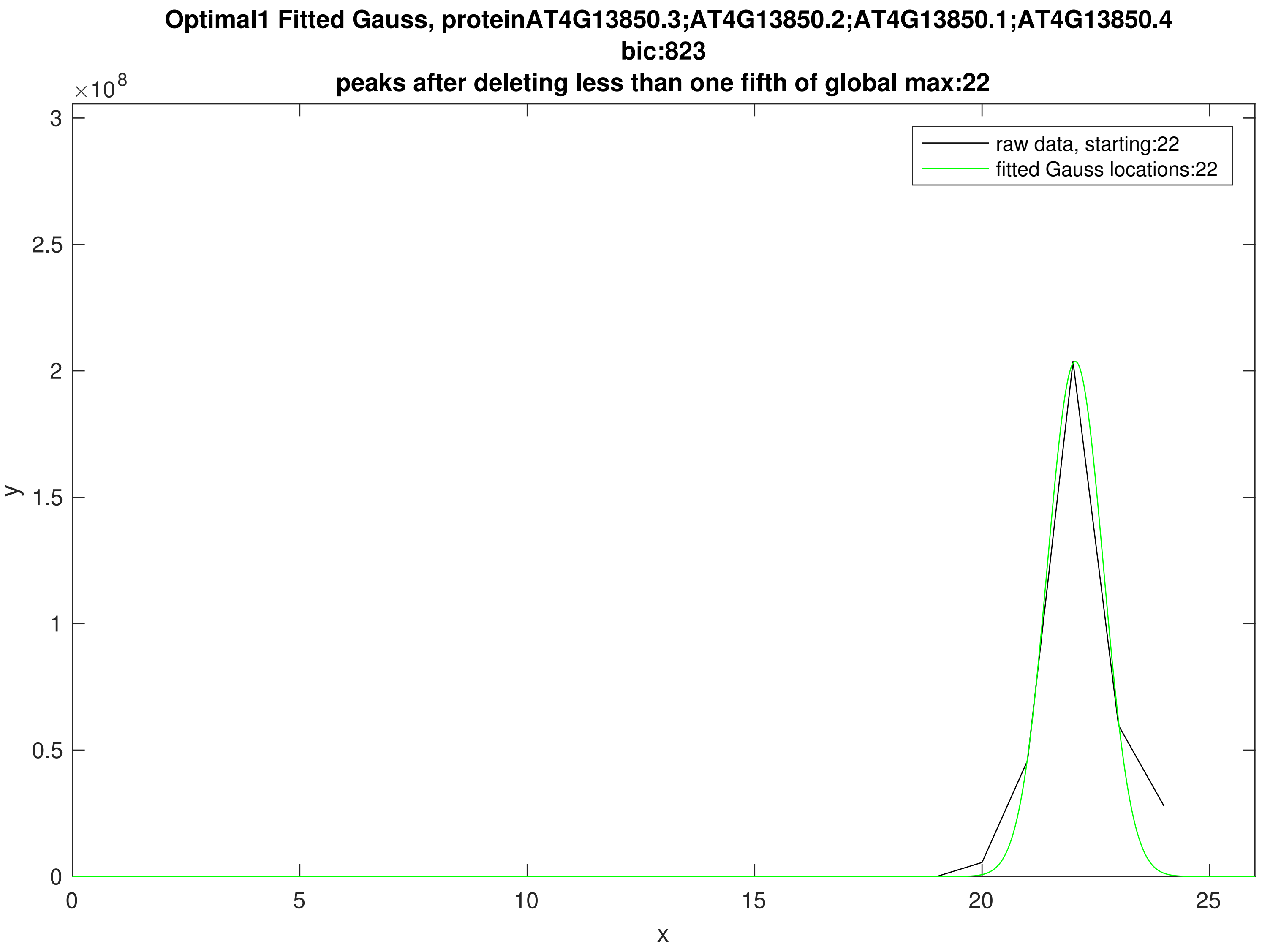

Optimal1 Fitted Gauss, proteinAT4G13850.3;AT4G13850.2;AT4G13850.1;AT4G13850.4
bic:823
peaks after deleting less than one fifth of global max:22
×10^8
3
2.5
2
1.5
1
0.5
0
y
0
5
10
15
20
25
x
raw data, starting:22
fitted Gauss locations:22

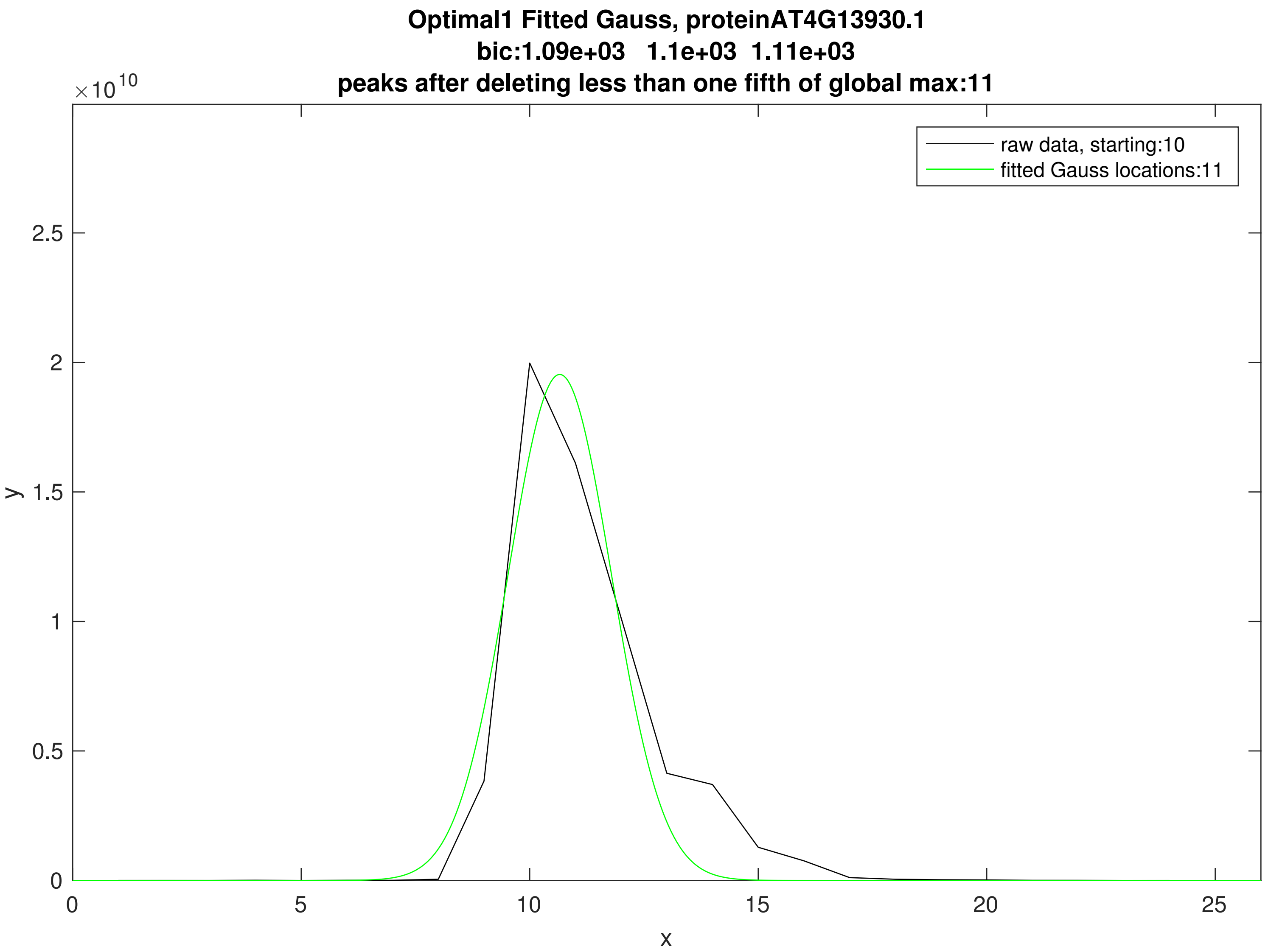
Optimal1 Fitted Gauss, proteinAT4G13930.1
bic:1.09e+03 1.1e+03 1.11e+03
peaks after deleting less than one fifth of global max:11
×10^10
raw data, starting:10
fitted Gauss locations:11
0
0.5
1
1.5
2
2.5
y
0
5
10
15
20
25
x

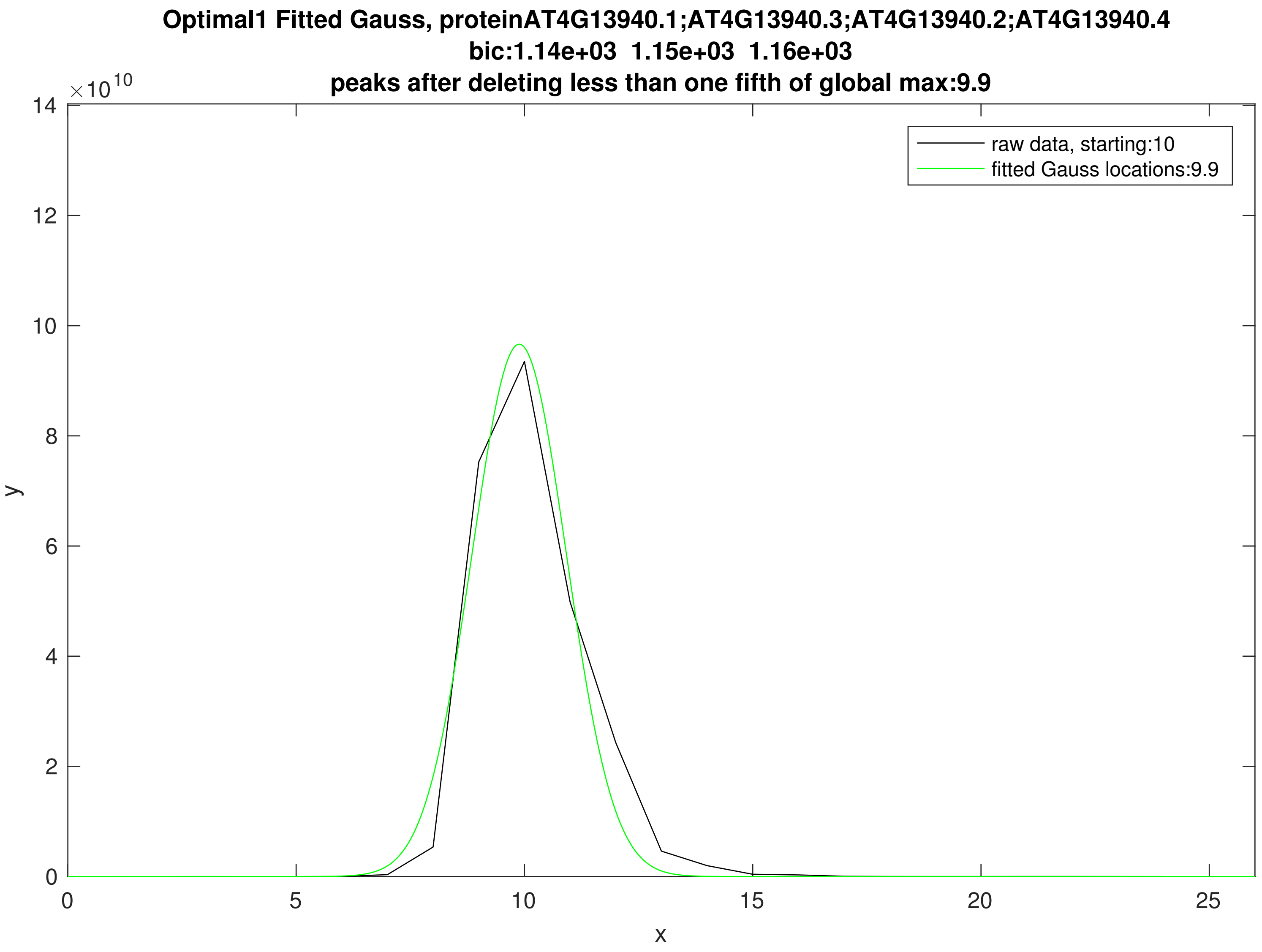
Optimal1 Fitted Gauss, proteinAT4G13940.1;AT4G13940.3;AT4G13940.2;AT4G13940.4
bic:1.14e+03 1.15e+03 1.16e+03
peaks after deleting less than one fifth of global max:9.9
×10^10
raw data, starting:10
fitted Gauss locations:9.9
y
x
0
2
4
6
8
10
12
14
5
15
20
25

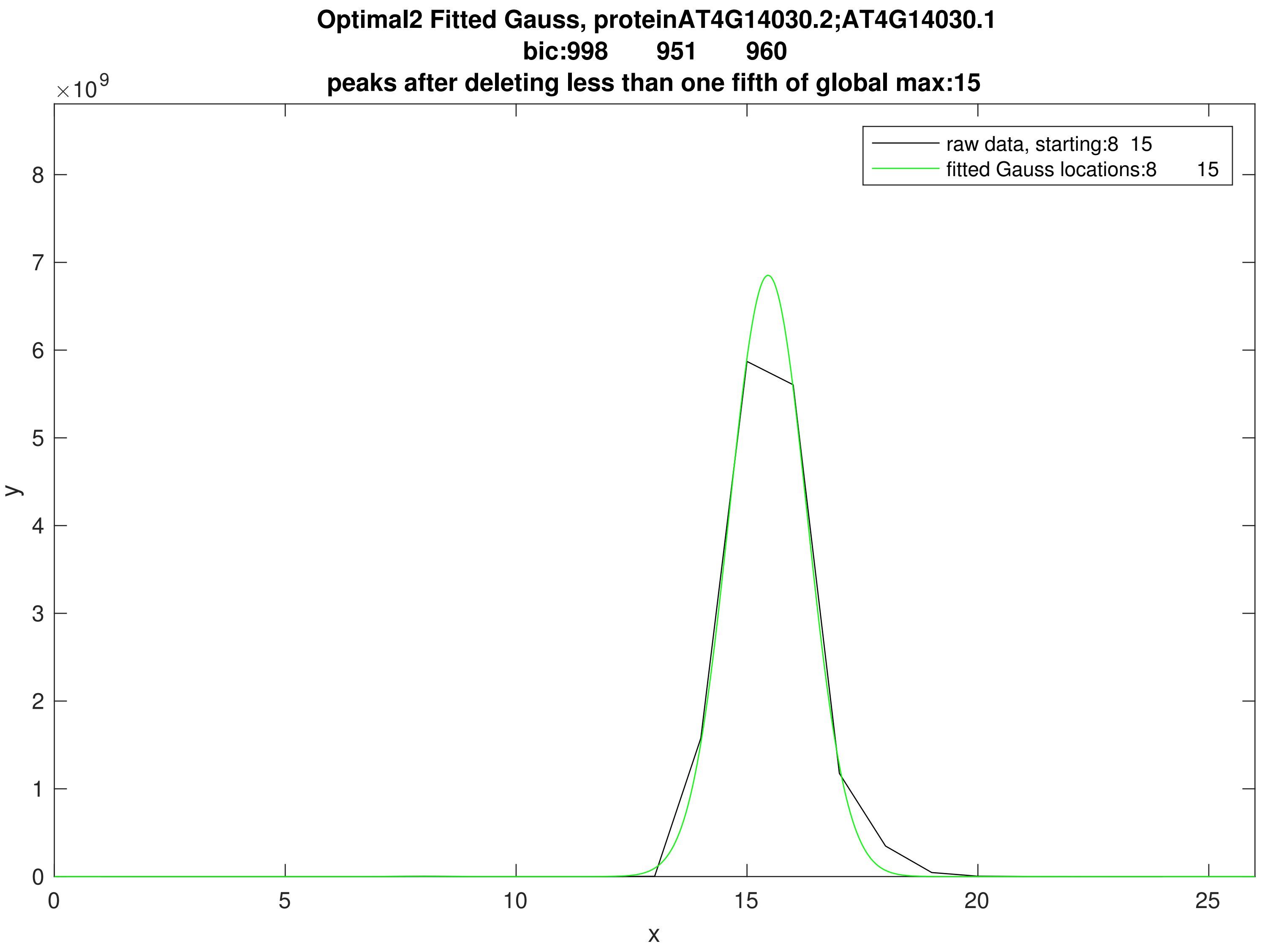
Optimal2 Fitted Gauss, proteinAT4G14030.2;AT4G14030.1
bic:998 951 960
peaks after deleting less than one fifth of global max:15
×10^9
raw data, starting:8 15
fitted Gauss locations:8 15
y
x
0
1
2
3
4
5
6
7
8
0
5
10
15
20
25

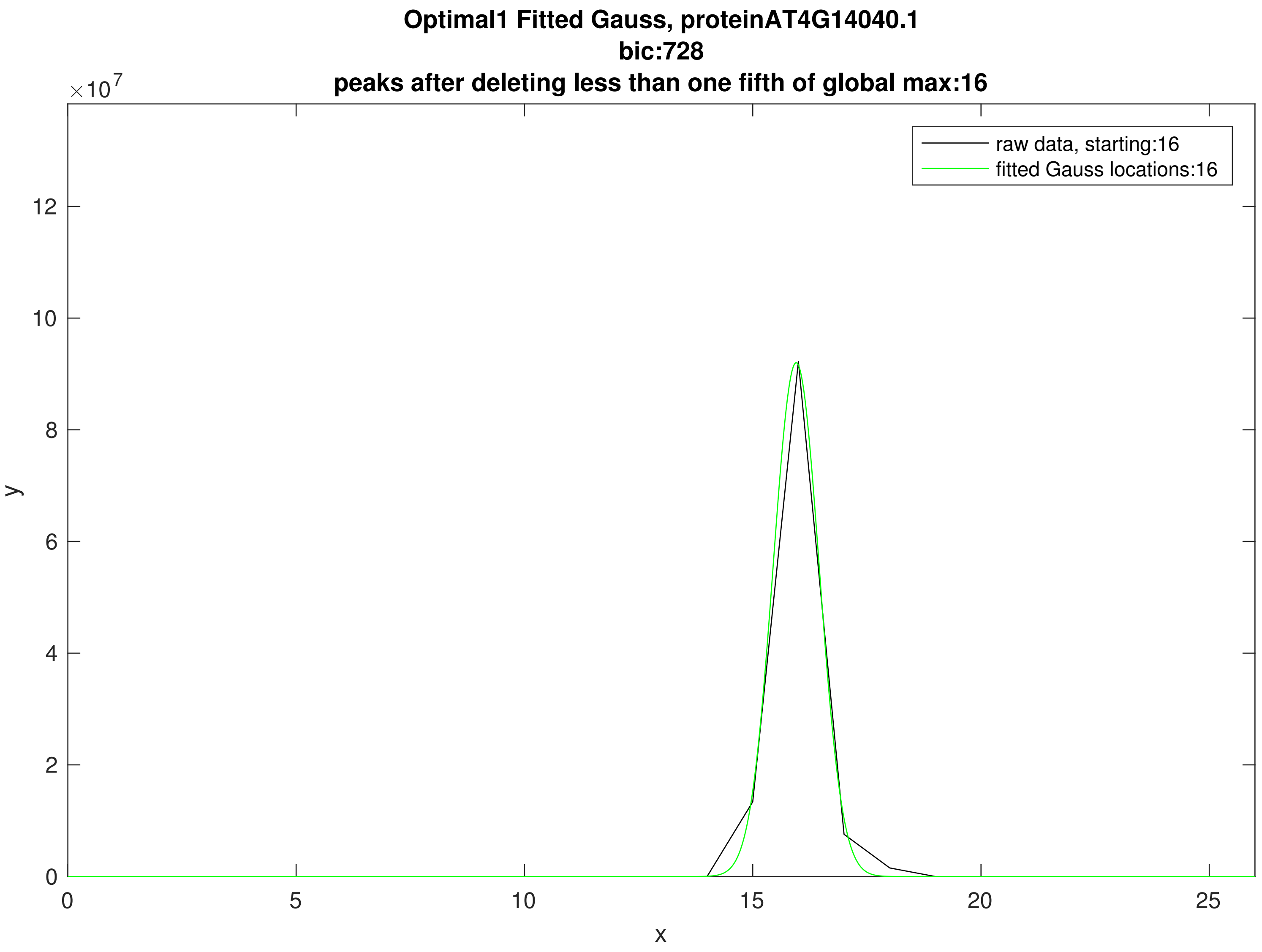
Optimal1 Fitted Gauss, proteinAT4G14040.1
bic:728
peaks after deleting less than one fifth of global max:16
×10^7
raw data, starting:16
fitted Gauss locations:16
y
x
0
2
4
6
8
10
12
0
5
10
15
20
25

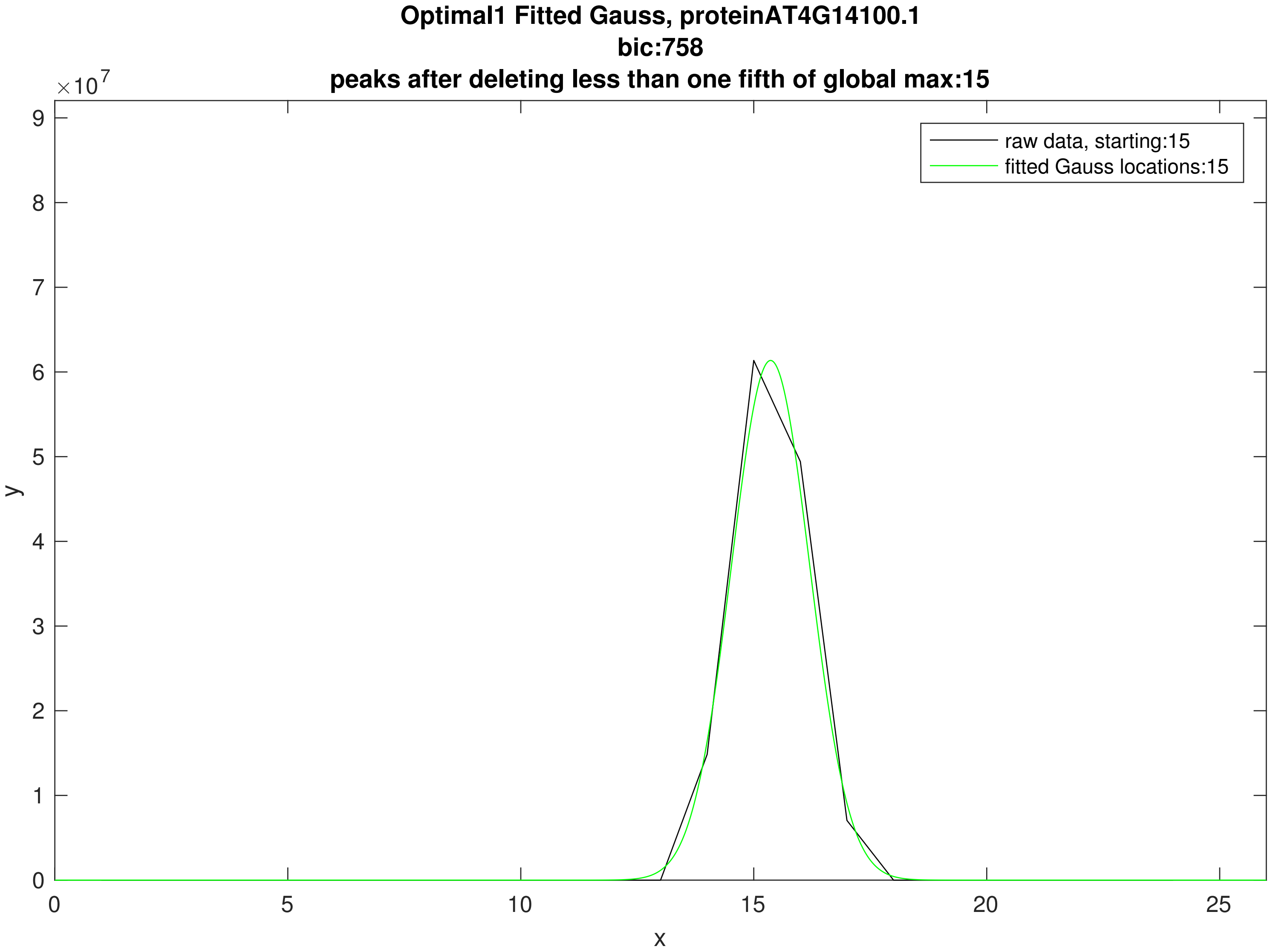
Optimal1 Fitted Gauss, proteinAT4G14100.1
bic:758
peaks after deleting less than one fifth of global max:15
×10^7
raw data, starting:15
fitted Gauss locations:15
y
x

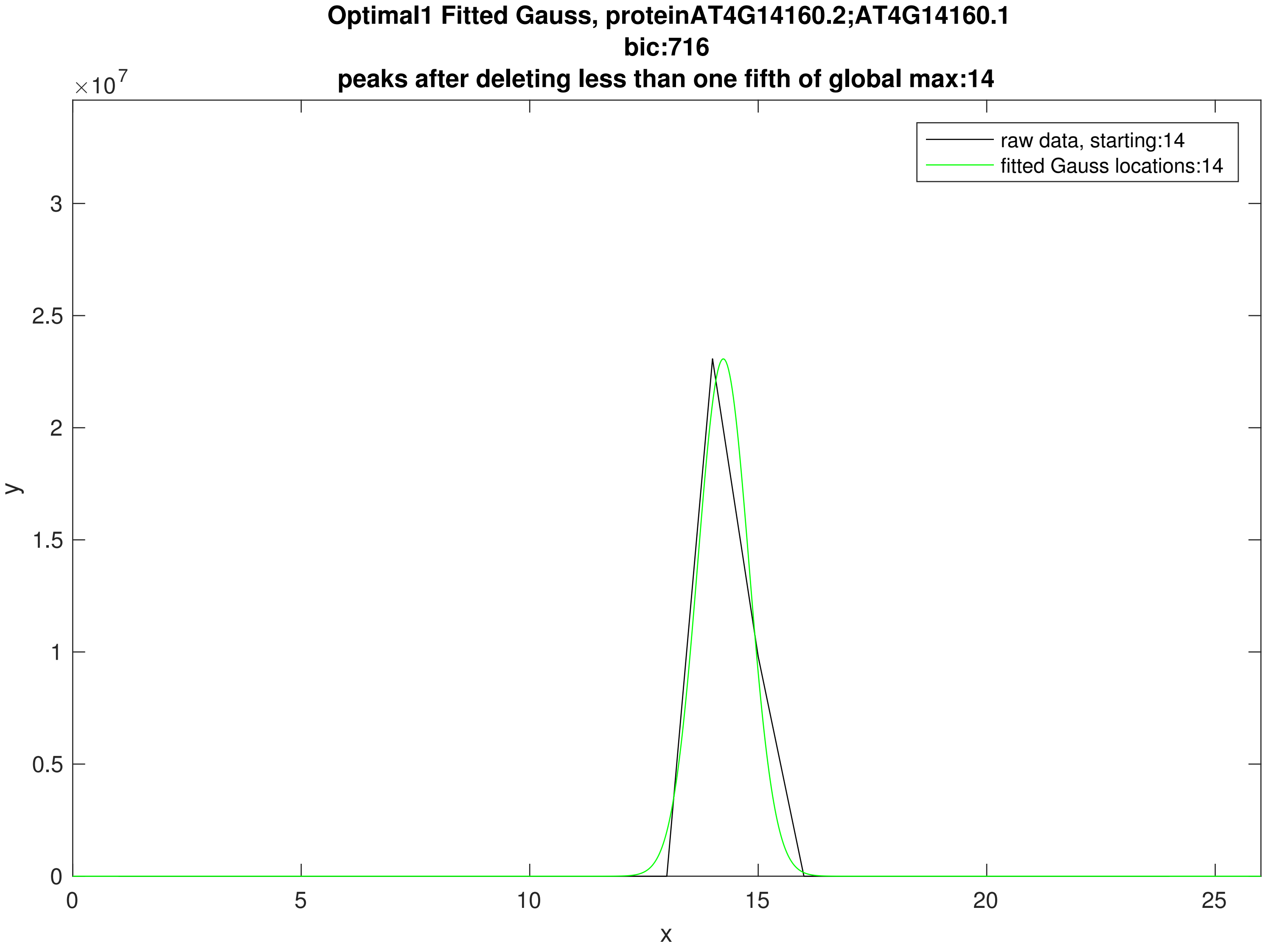
Optimal1 Fitted Gauss, proteinAT4G14160.2;AT4G14160.1
bic:716
peaks after deleting less than one fifth of global max:14
×10^7
raw data, starting:14
fitted Gauss locations:14
y
x
0
0.5
1
1.5
2
2.5
3
0
5
10
15
20
25

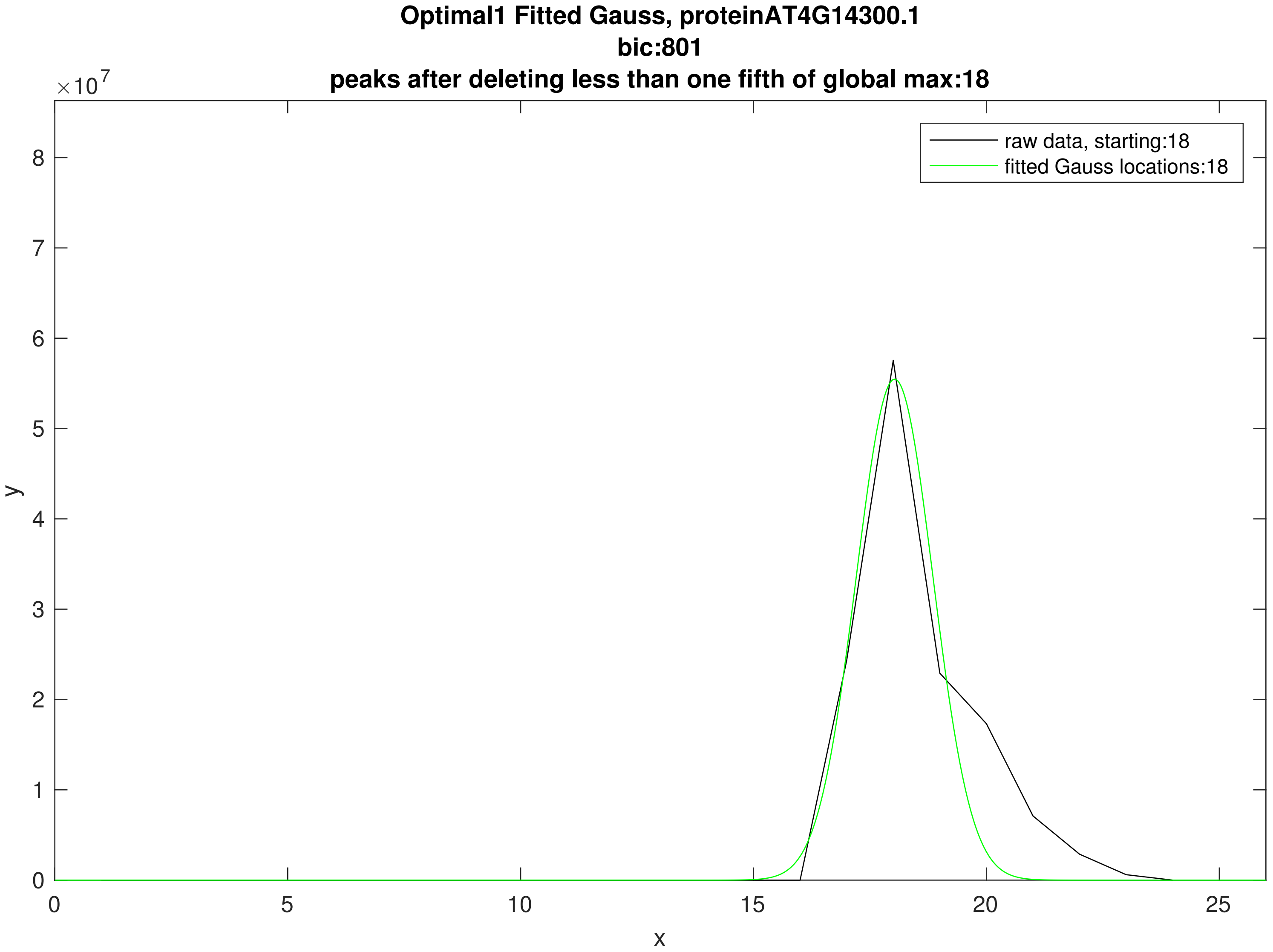

Optimal1 Fitted Gauss, proteinAT4G14300.1
bic:801
peaks after deleting less than one fifth of global max:18
×10^7
raw data, starting:18
fitted Gauss locations:18
y
x
0
1
2
3
4
5
6
7
8
0
5
10
15
20
25

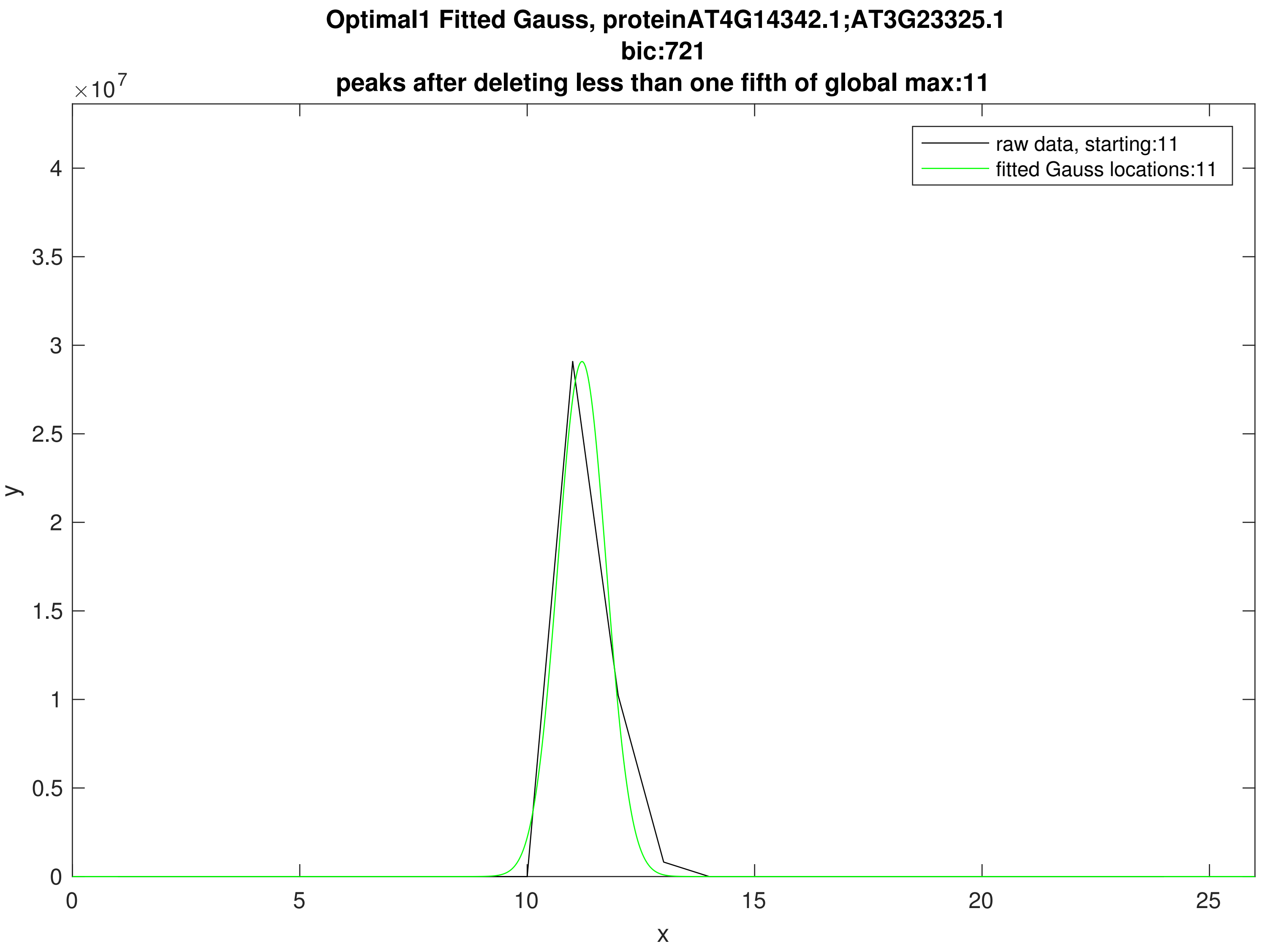
Optimal1 Fitted Gauss, proteinAT4G14342.1;AT3G23325.1
bic:721
peaks after deleting less than one fifth of global max:11
×10^7
raw data, starting:11
fitted Gauss locations:11
y
x
0
0.5
1
1.5
2
2.5
3
3.5
4
0
5
10
15
20
25

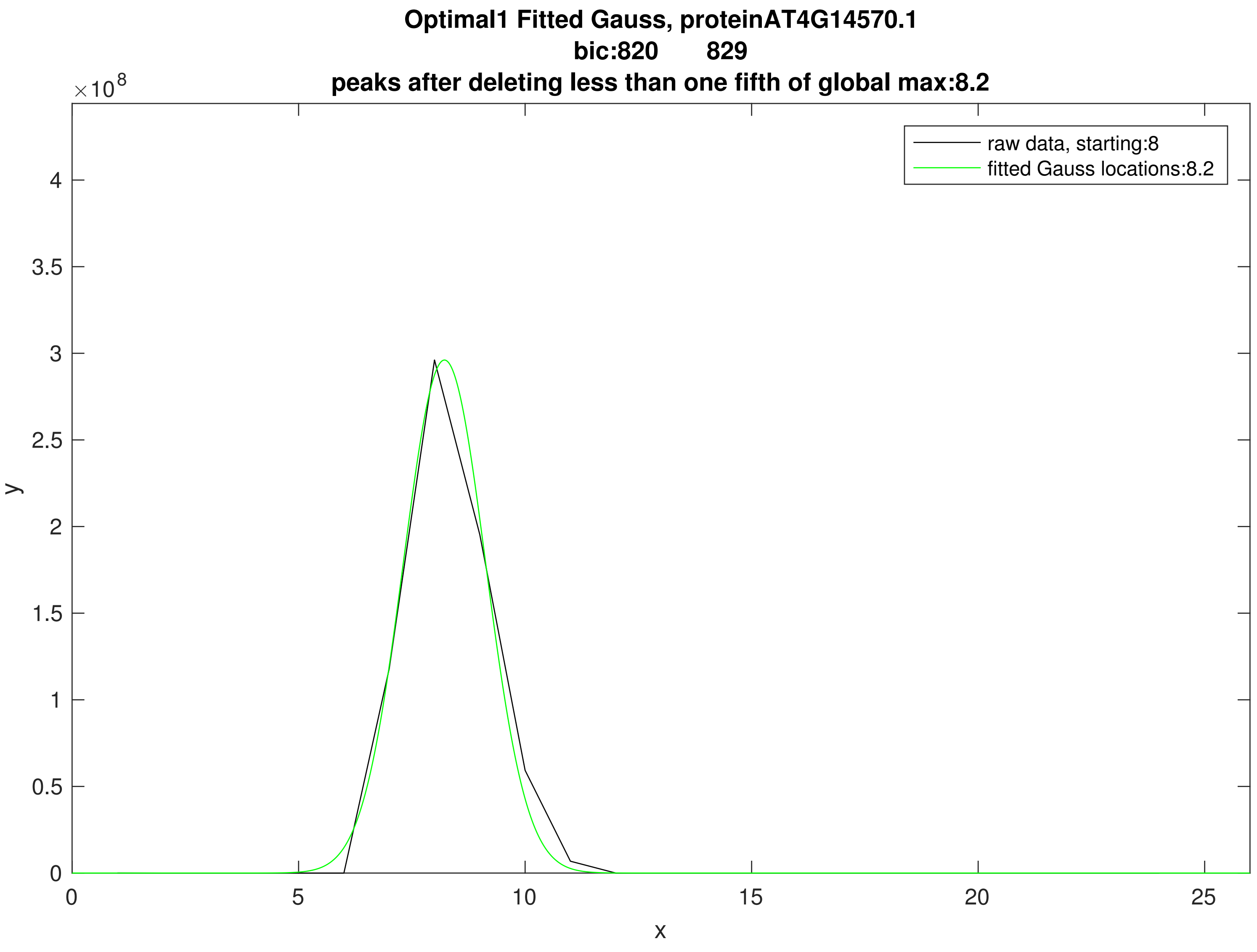
Optimal1 Fitted Gauss, proteinAT4G14570.1
bic:820 829
peaks after deleting less than one fifth of global max:8.2
×10^8
raw data, starting:8
fitted Gauss locations:8.2
y
x
0
0.5
1
1.5
2
2.5
3
3.5
4
0
5
10
15
20
25

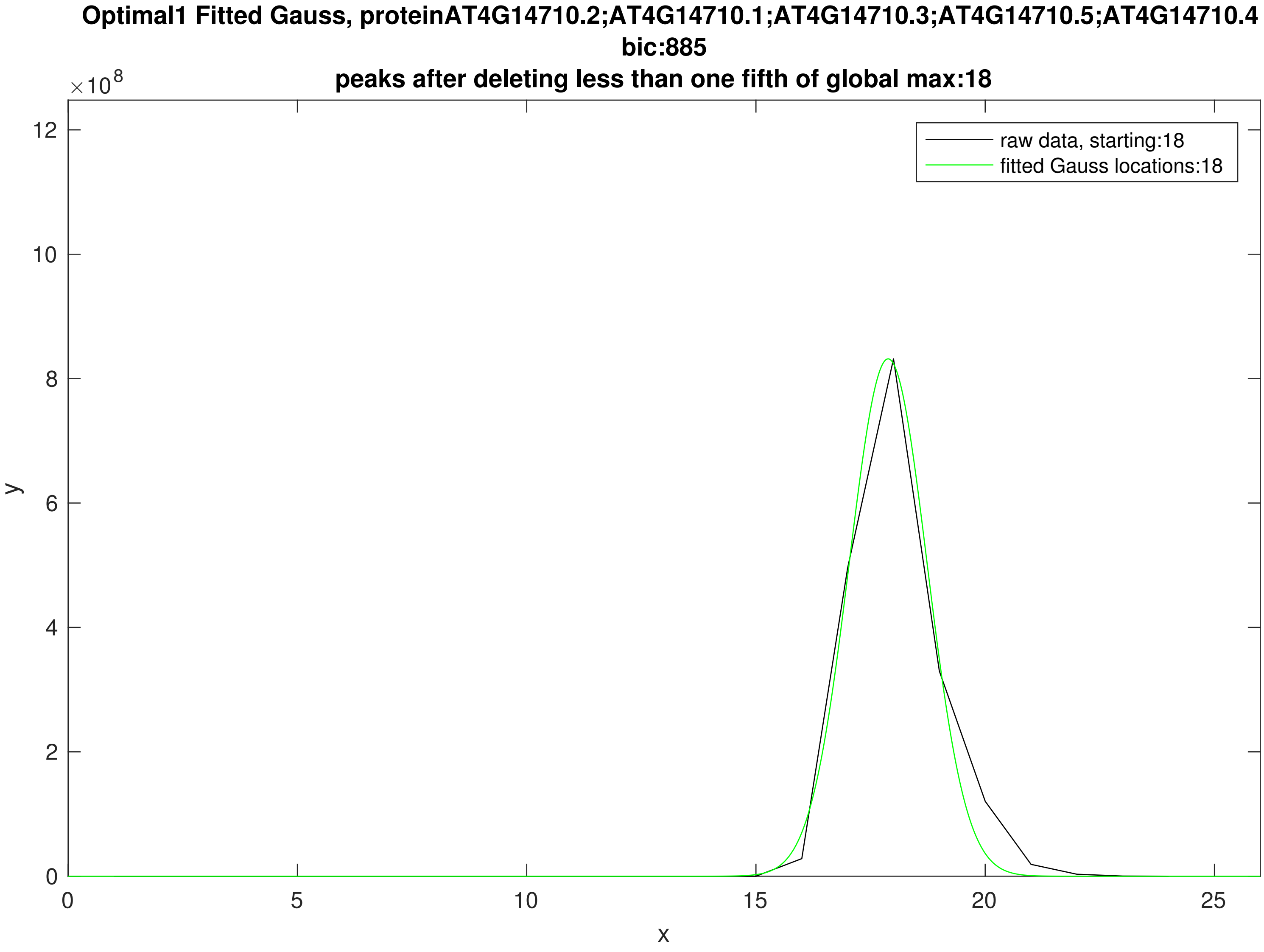
Optimal1 Fitted Gauss, proteinAT4G14710.2;AT4G14710.1;AT4G14710.3;AT4G14710.5;AT4G14710.4
bic:885
peaks after deleting less than one fifth of global max:18
×10^8
12
10
8
6
4
2
0
y
0
5
10
15
20
25
x
raw data, starting:18
fitted Gauss locations:18

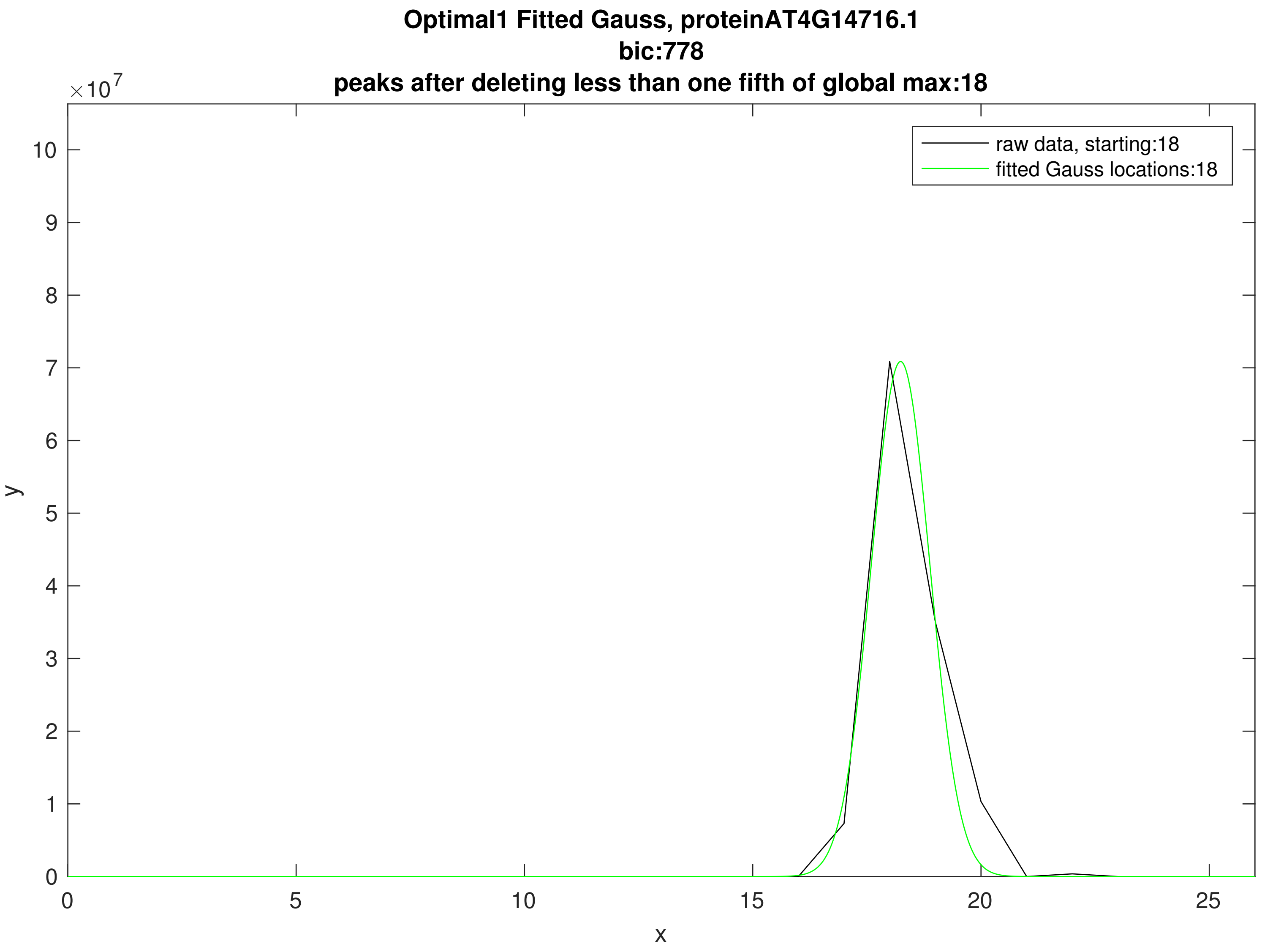
Optimal1 Fitted Gauss, proteinAT4G14716.1
bic:778
peaks after deleting less than one fifth of global max:18
×10^7
raw data, starting:18
fitted Gauss locations:18
y
x
0
1
2
3
4
5
6
7
8
9
10
0
5
10
15
20
25

Optimal1 Fitted Gauss, proteinAT4G14760.1
bic:779 784
peaks after deleting less than one fifth of global max:18

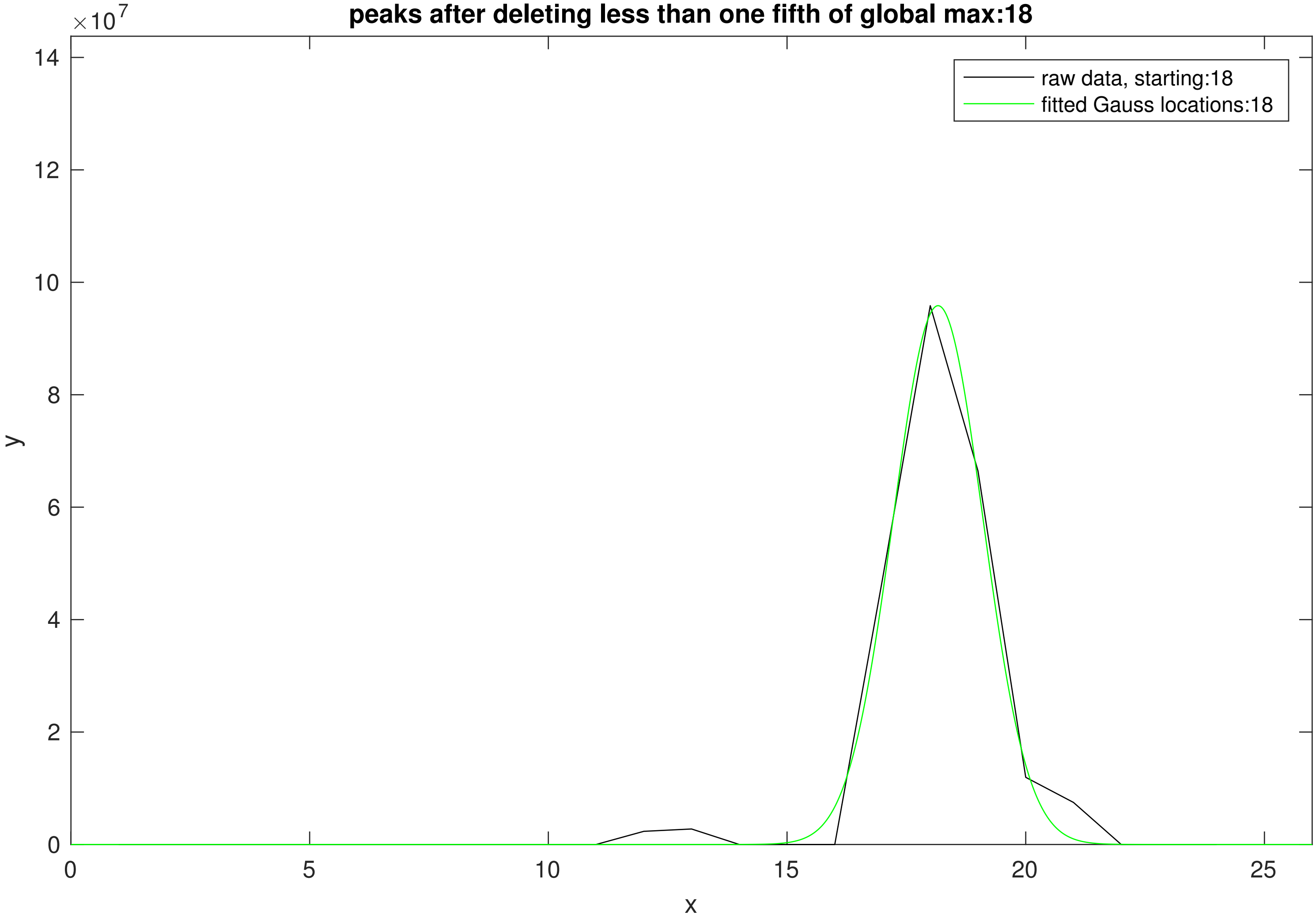

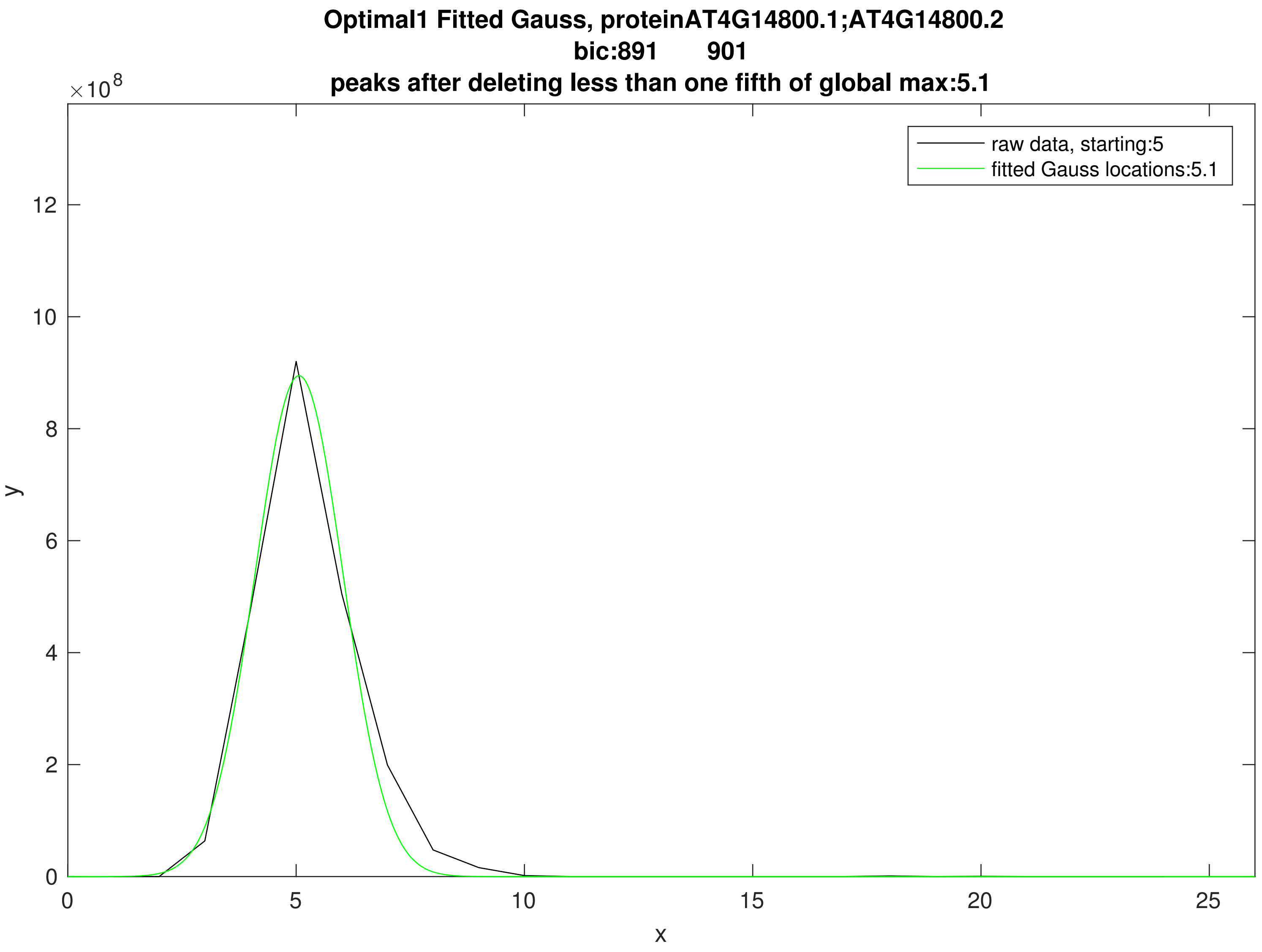

Optimal1 Fitted Gauss, proteinAT4G14800.1;AT4G14800.2
bic:891 901
peaks after deleting less than one fifth of global max:5.1
×10^8
raw data, starting:5
fitted Gauss locations:5.1
y
x
0
2
4
6
8
10
12
5
15
20
25

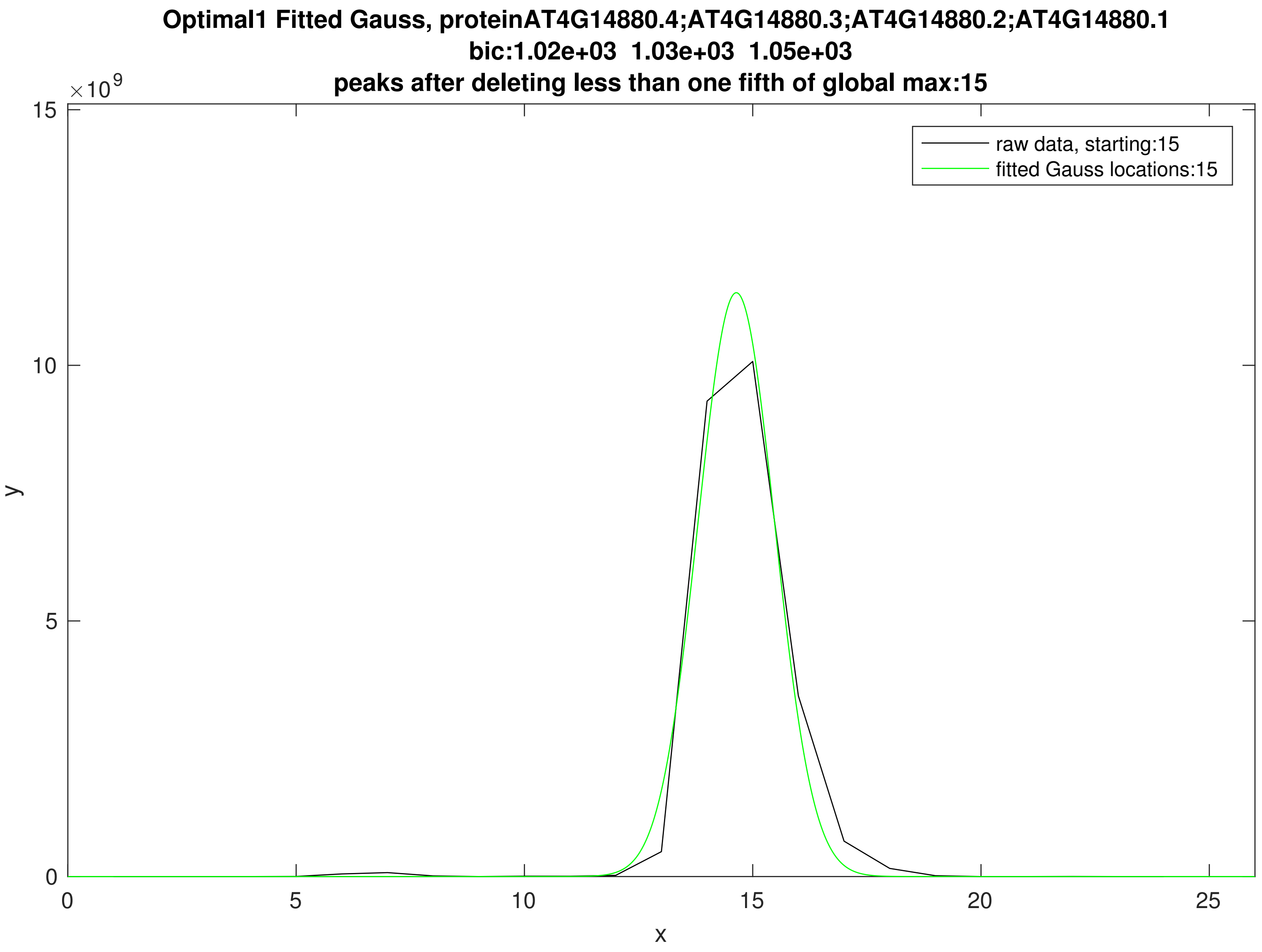

Optimal1 Fitted Gauss, proteinAT4G14880.4;AT4G14880.3;AT4G14880.2;AT4G14880.1
bic:1.02e+03 1.03e+03 1.05e+03
peaks after deleting less than one fifth of global max:15
×10^9
15
10
5
0
y
0
5
10
15
20
25
x
raw data, starting:15
fitted Gauss locations:15

#### Optimal1 Fitted Gauss, proteinAT4G14890.1
#### bic:728
#### peaks after deleting less than one fifth of global max:19

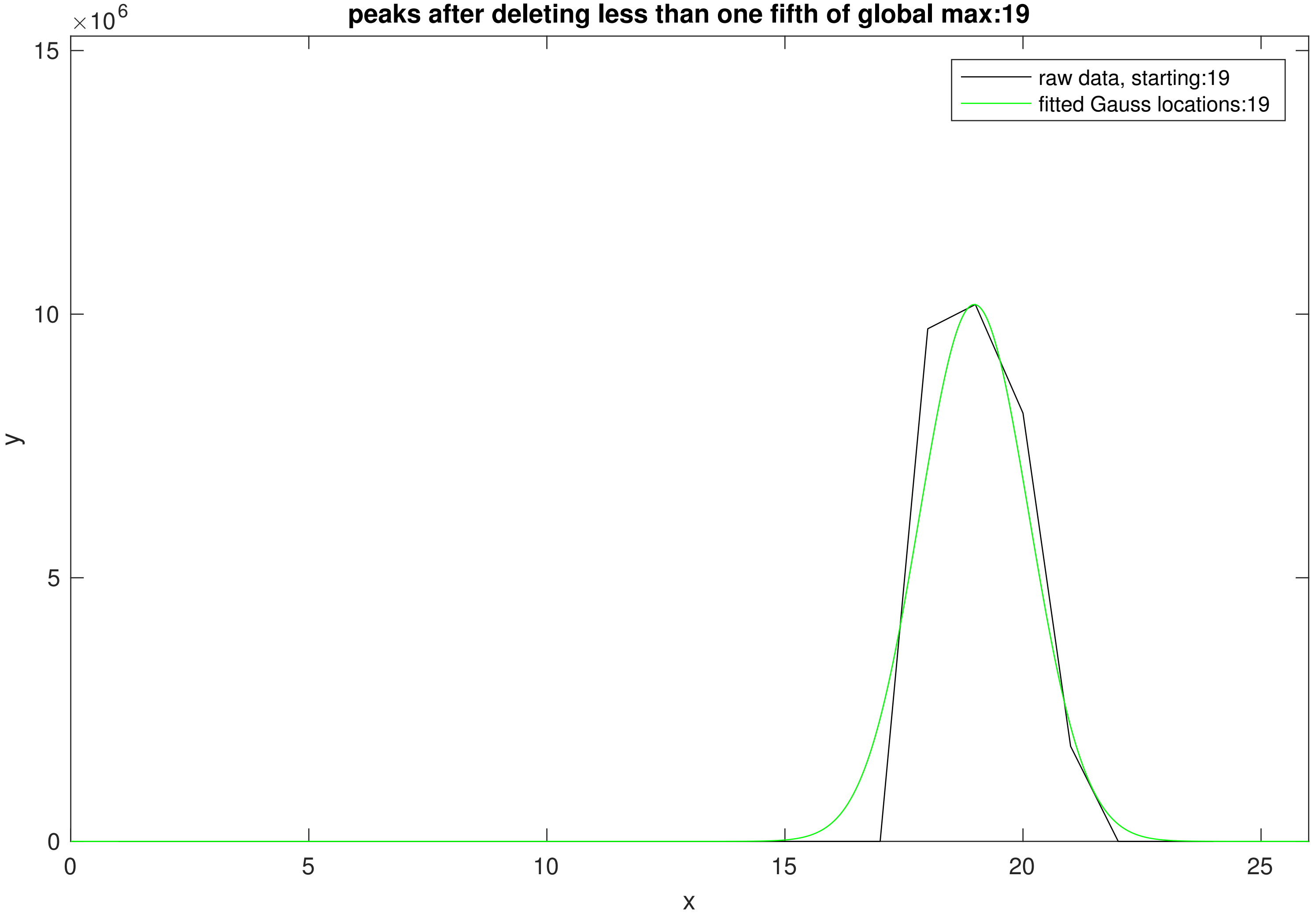

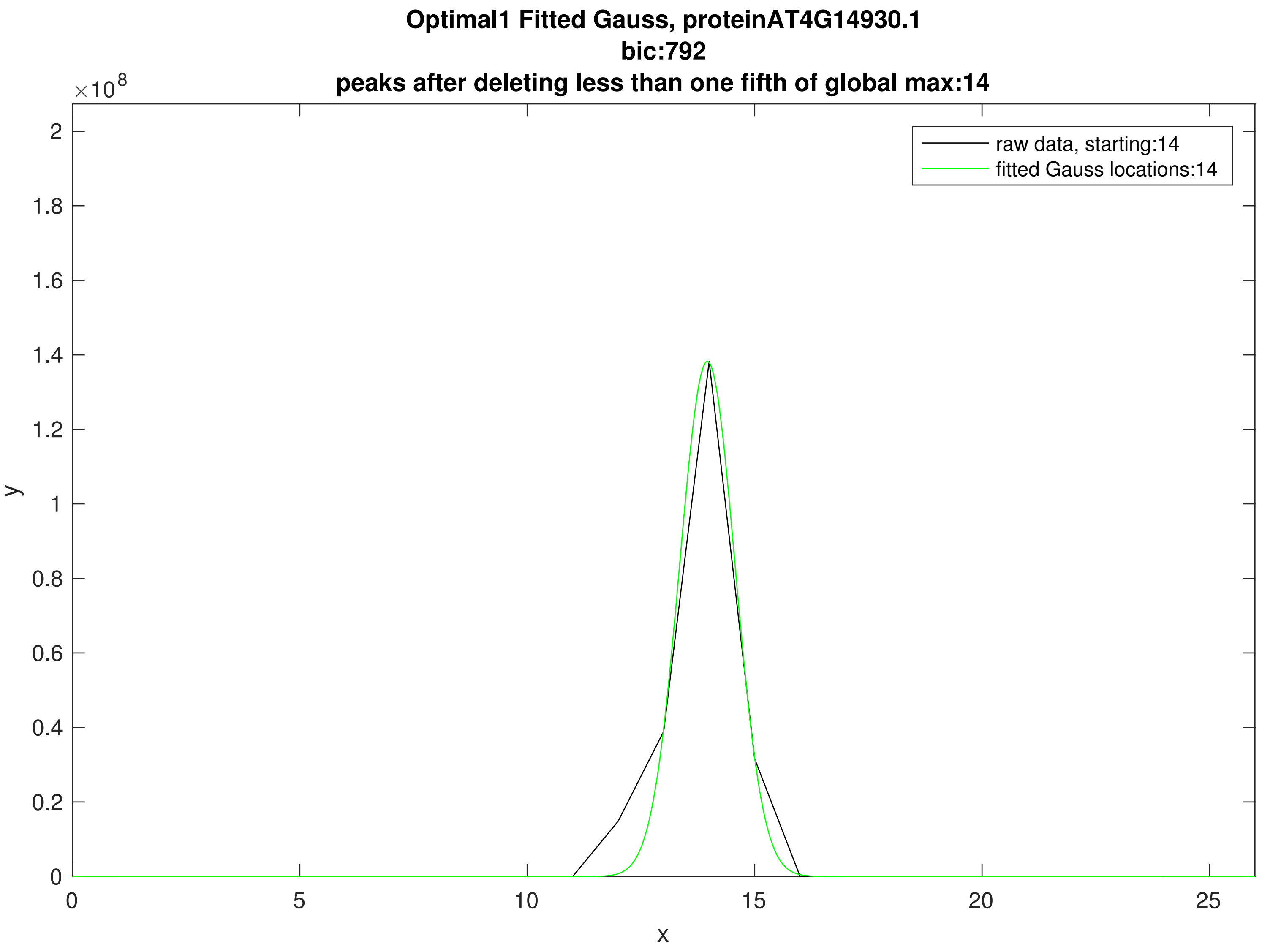

Optimal1 Fitted Gauss, proteinAT4G14930.1
bic:792
peaks after deleting less than one fifth of global max:14
×10^8
raw data, starting:14
fitted Gauss locations:14
y
x
0
0.2
0.4
0.6
0.8
1
1.2
1.4
1.6
1.8
2
0
5
10
15
20
25

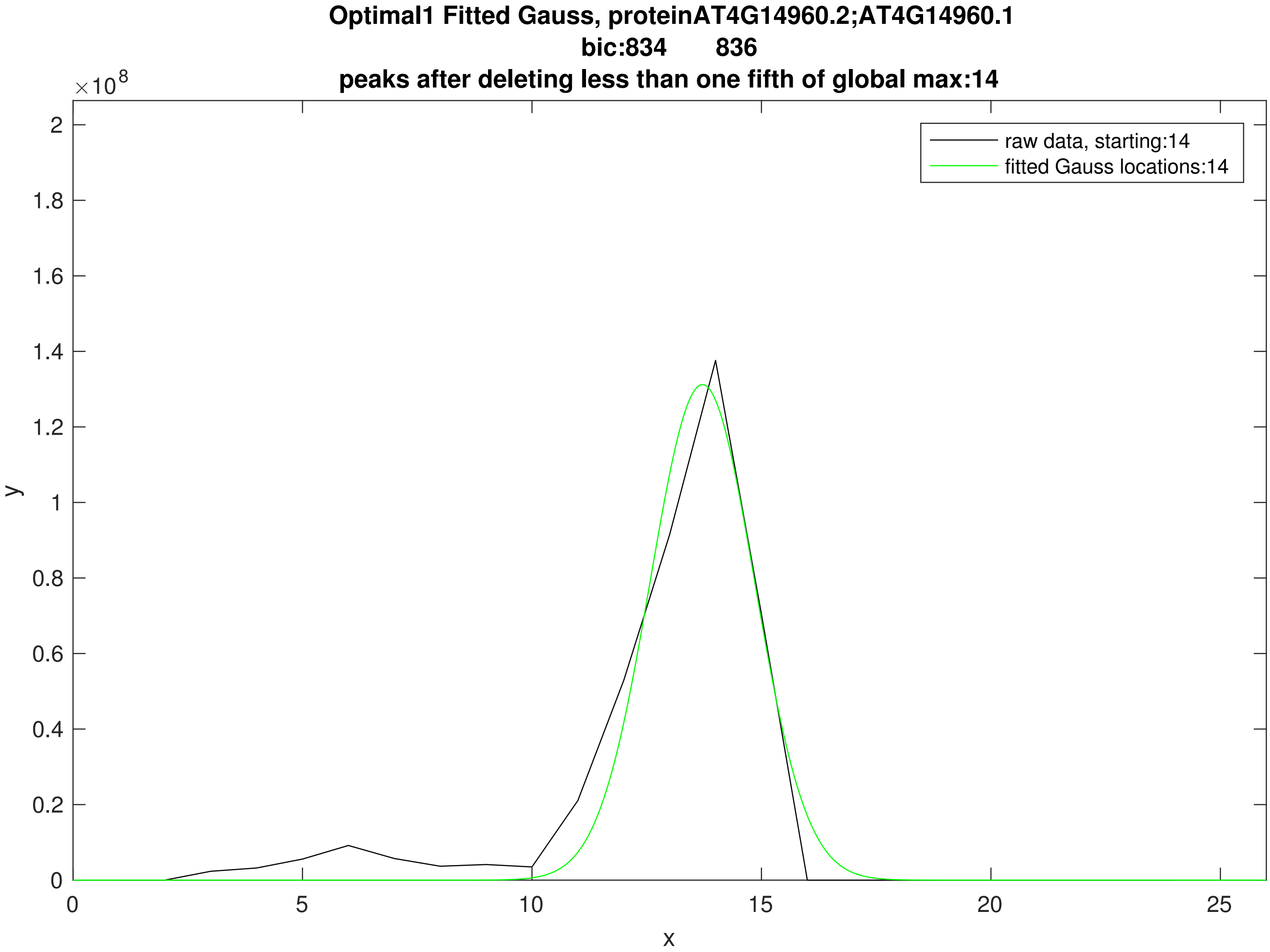
Optimal1 Fitted Gauss, proteinAT4G14960.2;AT4G14960.1
bic:834 836
peaks after deleting less than one fifth of global max:14
×10^8
2
1.8
1.6
1.4
1.2
1
0.8
0.6
0.4
0.2
0
y
0
5
10
15
20
25
x
raw data, starting:14
fitted Gauss locations:14

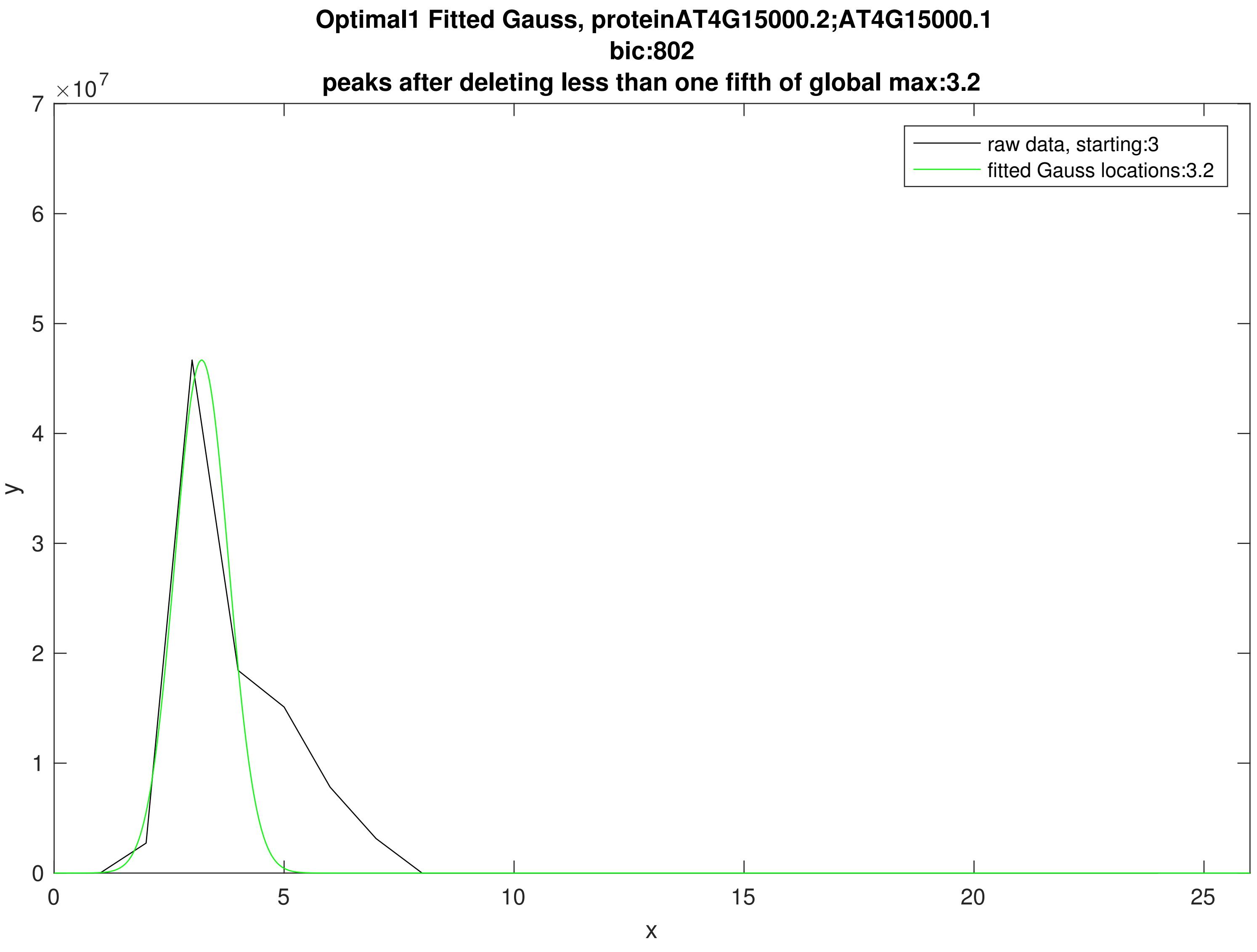

Optimal1 Fitted Gauss, proteinAT4G15000.2;AT4G15000.1
bic:802
peaks after deleting less than one fifth of global max:3.2
×10^7
7
6
5
4
3
2
1
0
y
0
5
10
15
20
25
x
raw data, starting:3
fitted Gauss locations:3.2

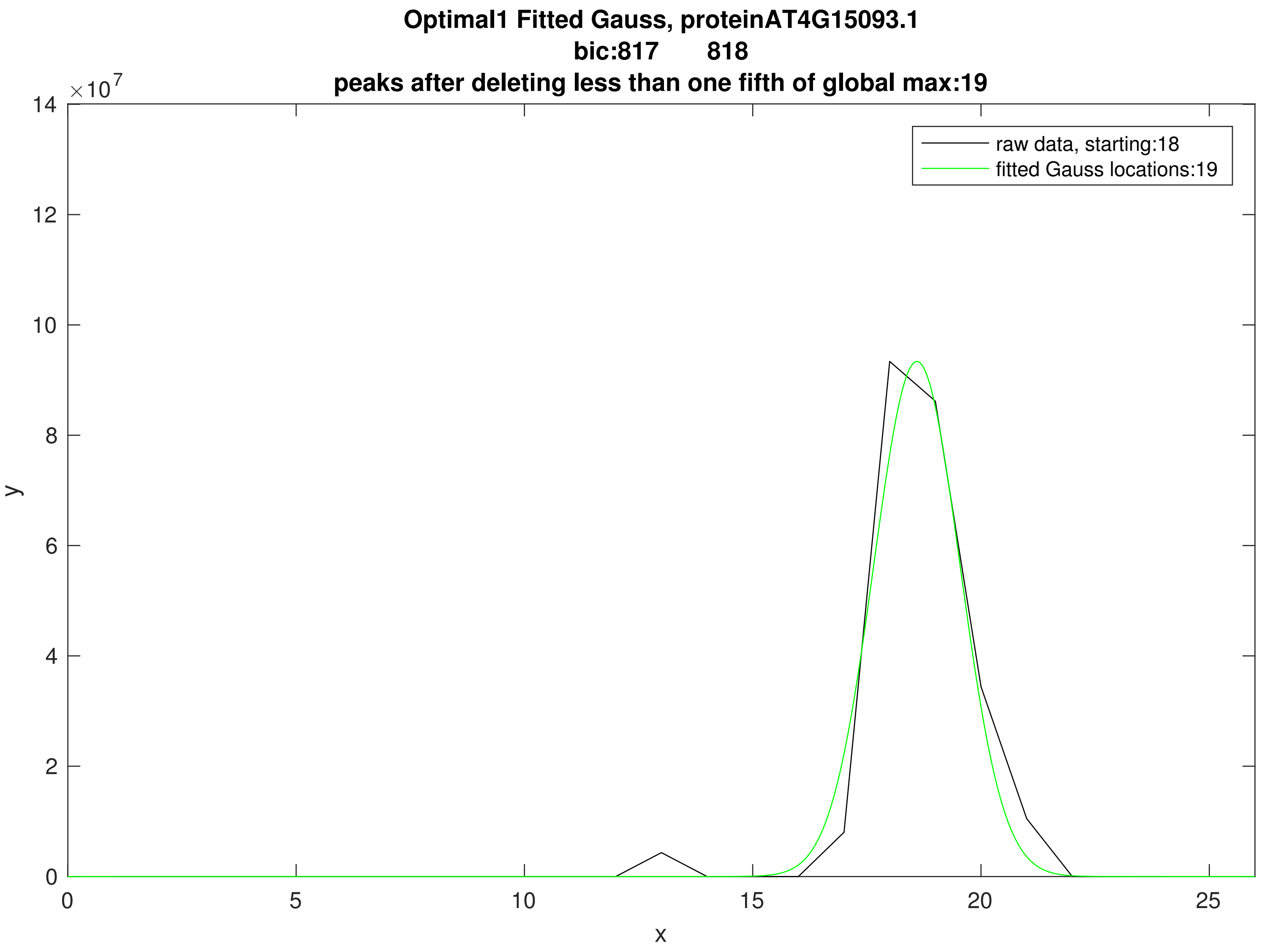

Optimal1 Fitted Gauss, proteinAT4G15093.1
bic:817 818
peaks after deleting less than one fifth of global max:19
×10^7
14
12
10
8
6
4
2
0
y
0
5
10
15
20
25
x
raw data, starting:18
fitted Gauss locations:19

Optimal1 Fitted Gauss, proteinAT4G15210.1;AT4G15210.2
bic:716
peaks after deleting less than one fifth of global max:16

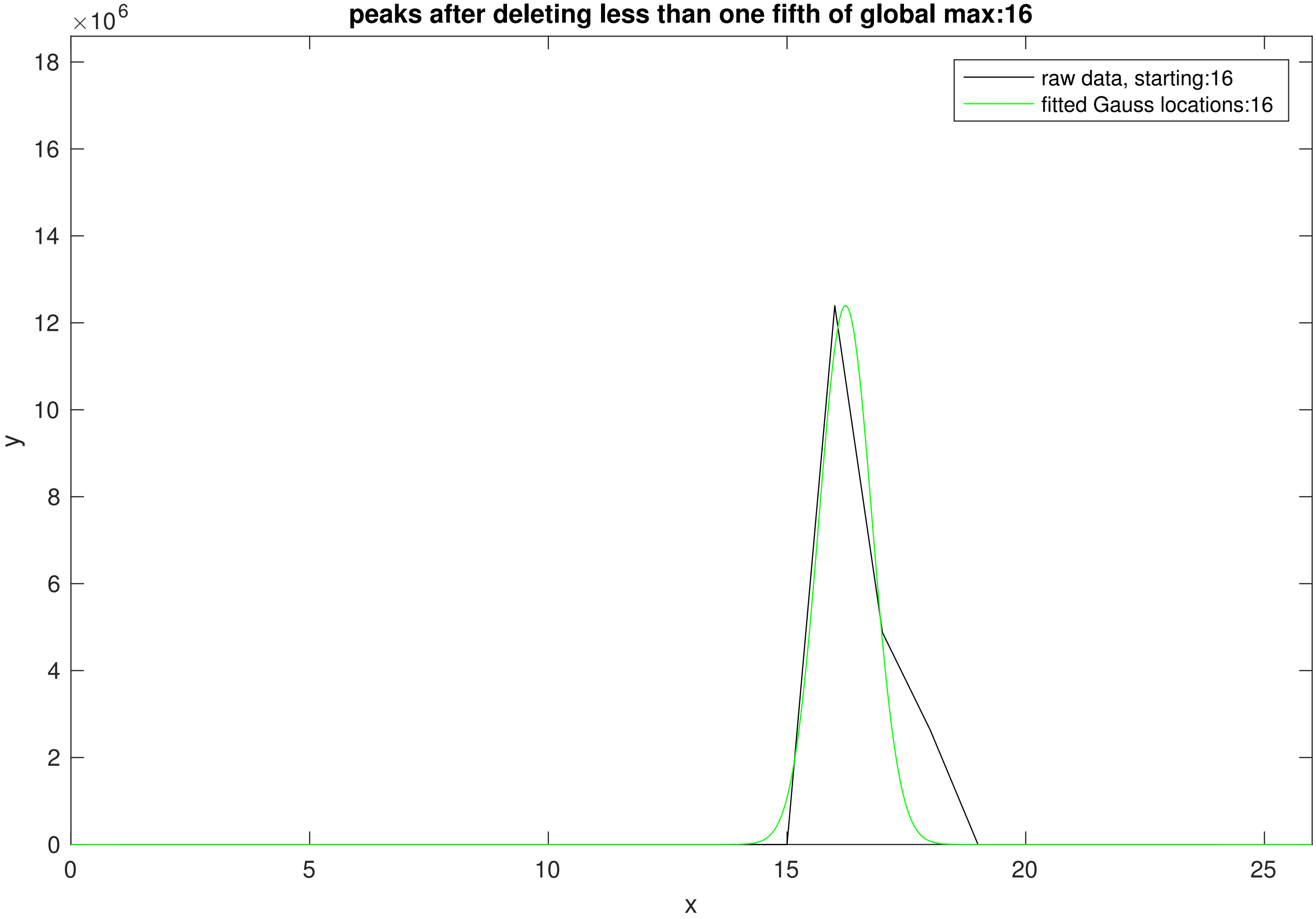

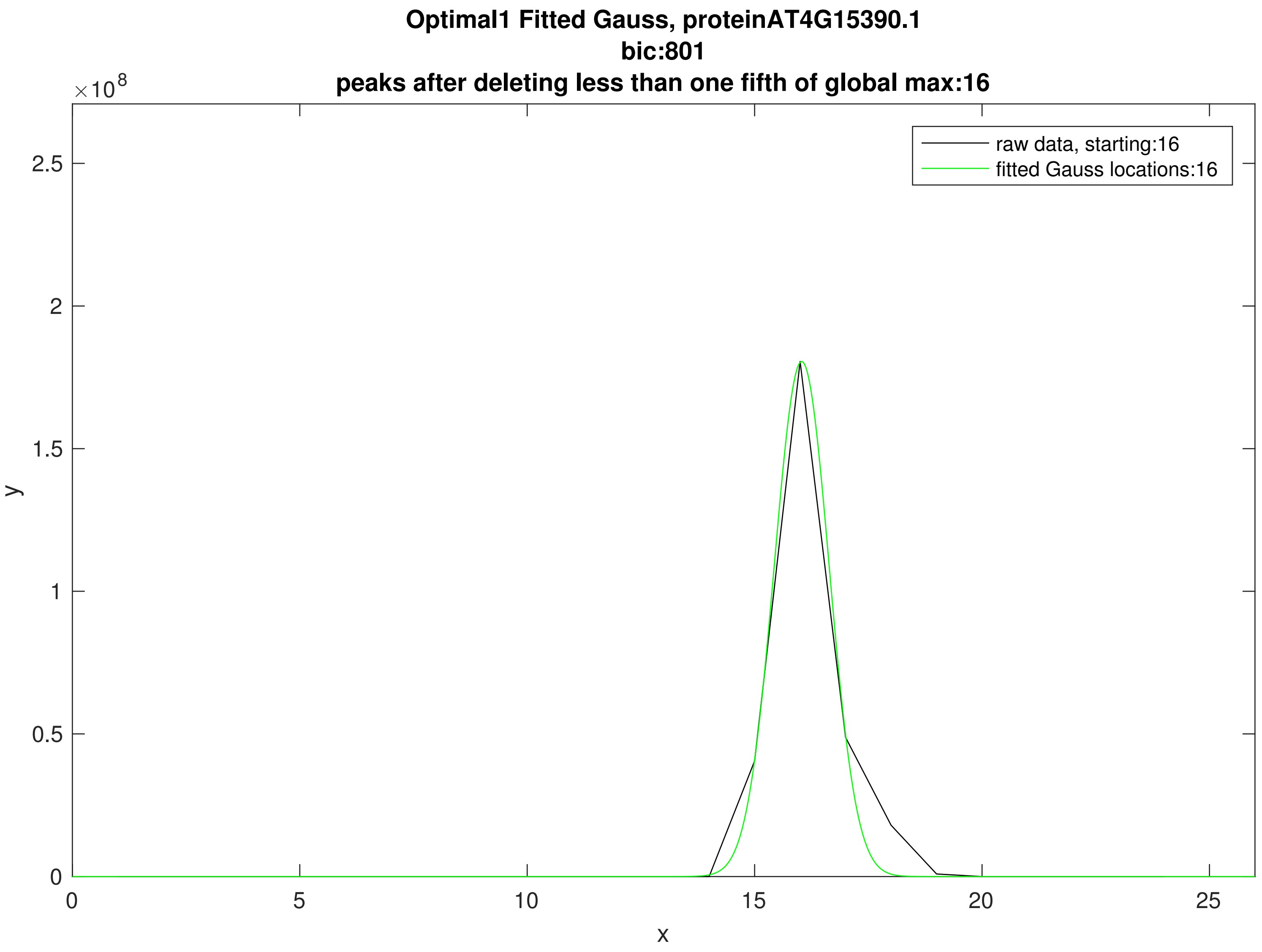
Optimal1 Fitted Gauss, proteinAT4G15390.1
bic:801
peaks after deleting less than one fifth of global max:16
×10^8
raw data, starting:16
fitted Gauss locations:16
y
x
0
0.5
1
1.5
2
2.5
5
10
15
20
25

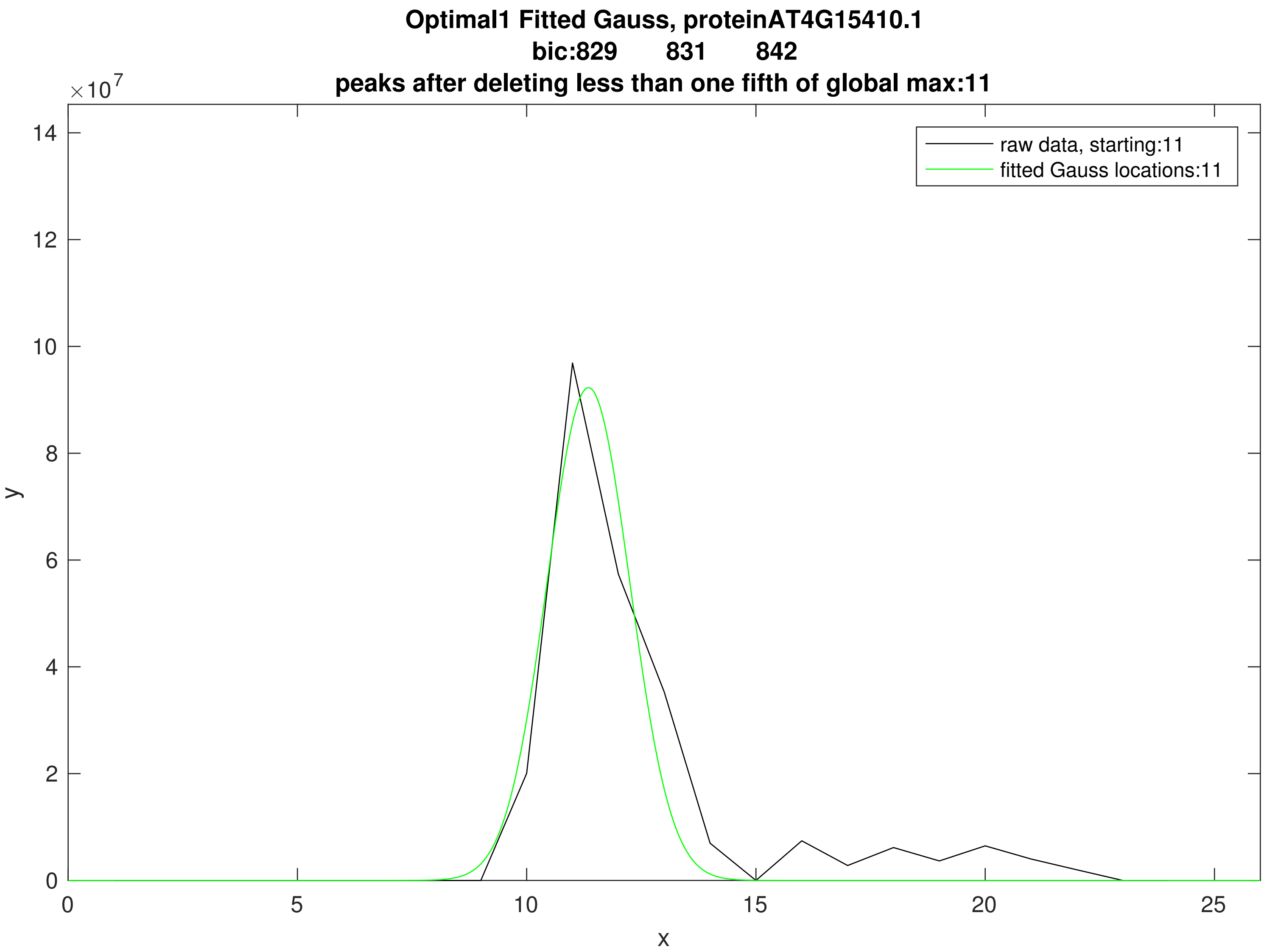
Optimal1 Fitted Gauss, proteinAT4G15410.1
bic:829 831 842
peaks after deleting less than one fifth of global max:11
×10^7
raw data, starting:11
fitted Gauss locations:11
y
x
0
2
4
6
8
10
12
14
0
5
10
15
20
25

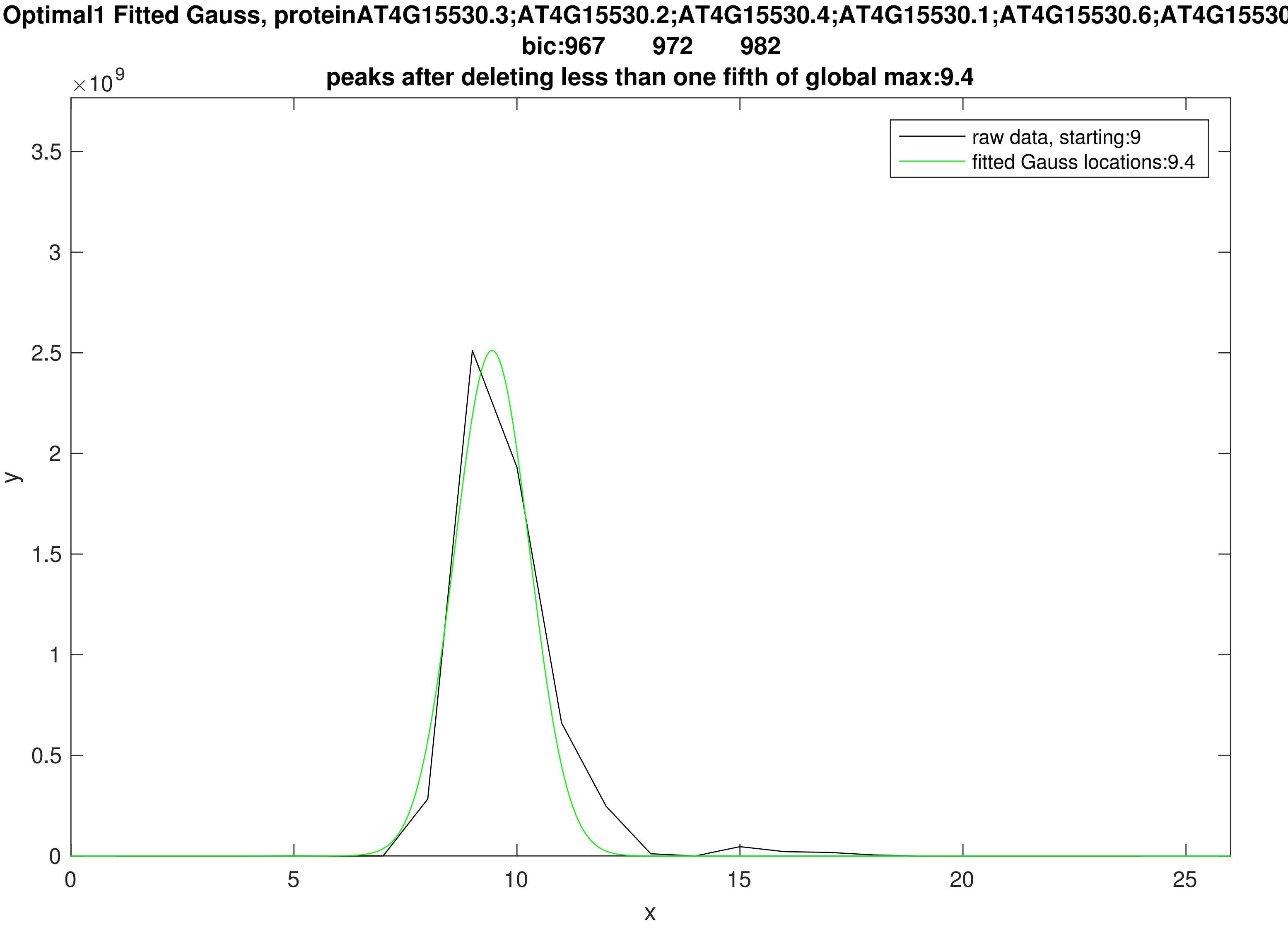
Optimal1 Fitted Gauss, proteinAT4G15530.3;AT4G15530.2;AT4G15530.4;AT4G15530.1;AT4G15530.6;AT4G15530
bic:967 972 982
peaks after deleting less than one fifth of global max:9.4
×10^9
3.5
3
2.5
2
1.5
1
0.5
0
y
0
5
10
15
20
25
x
raw data, starting:9
fitted Gauss locations:9.4

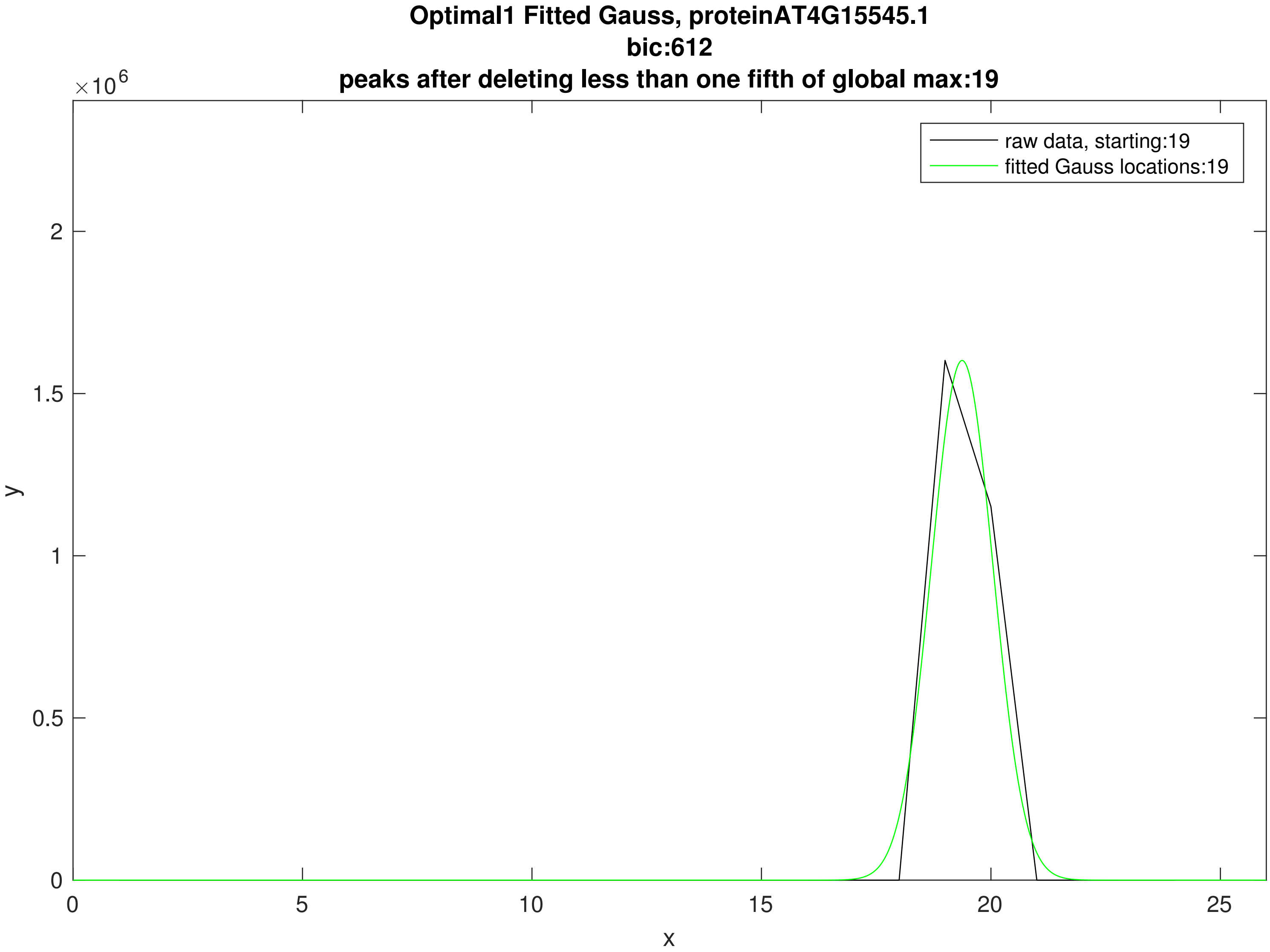
Optimal1 Fitted Gauss, proteinAT4G15545.1
bic:612
peaks after deleting less than one fifth of global max:19
×10^6
raw data, starting:19
fitted Gauss locations:19
y
x
0
0.5
1
1.5
2
0
5
10
15
20
25

**Optimal1 Fitted Gauss, proteinAT4G15550.1**
**bic:754**
**peaks after deleting less than one fifth of global max:16**

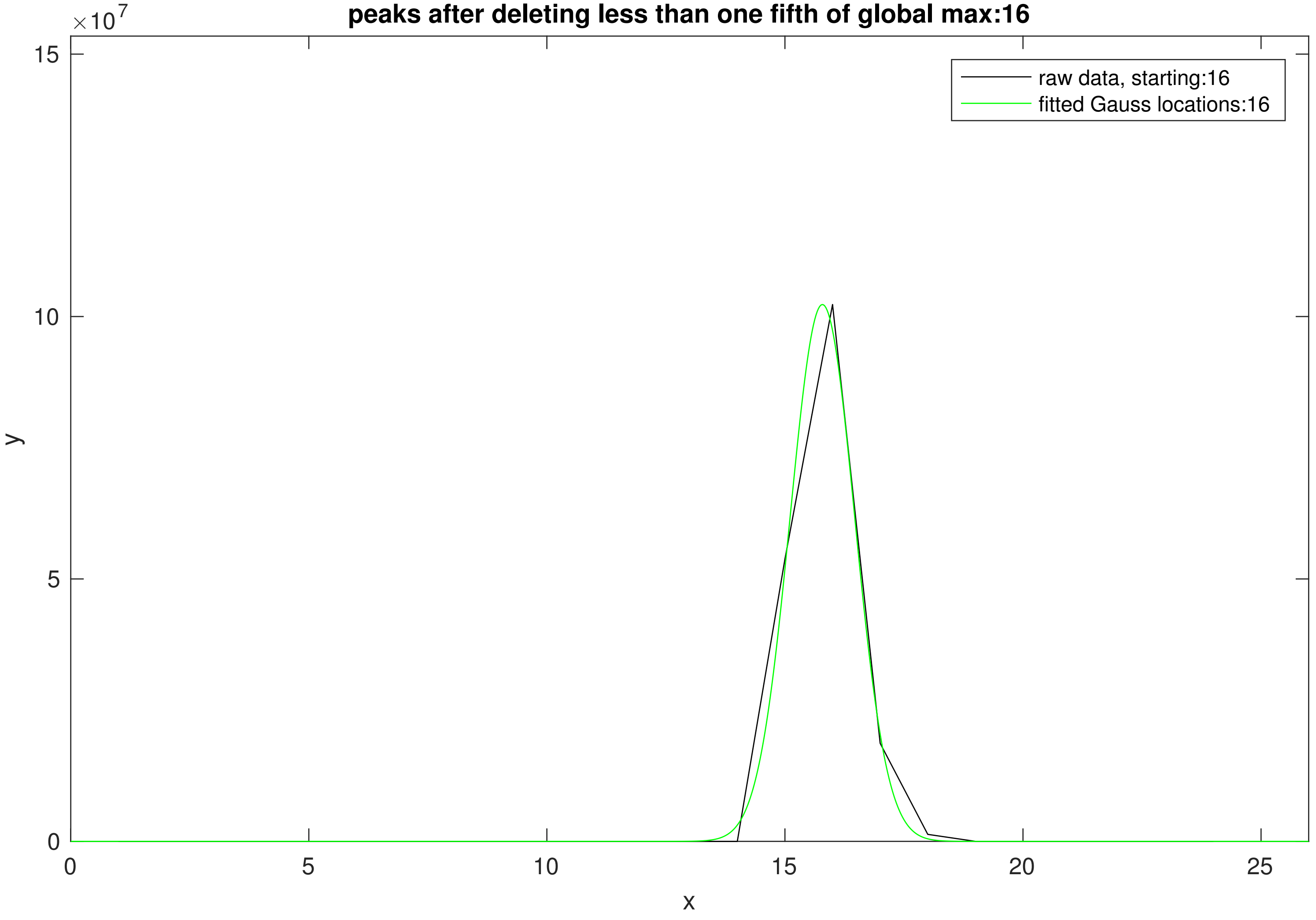

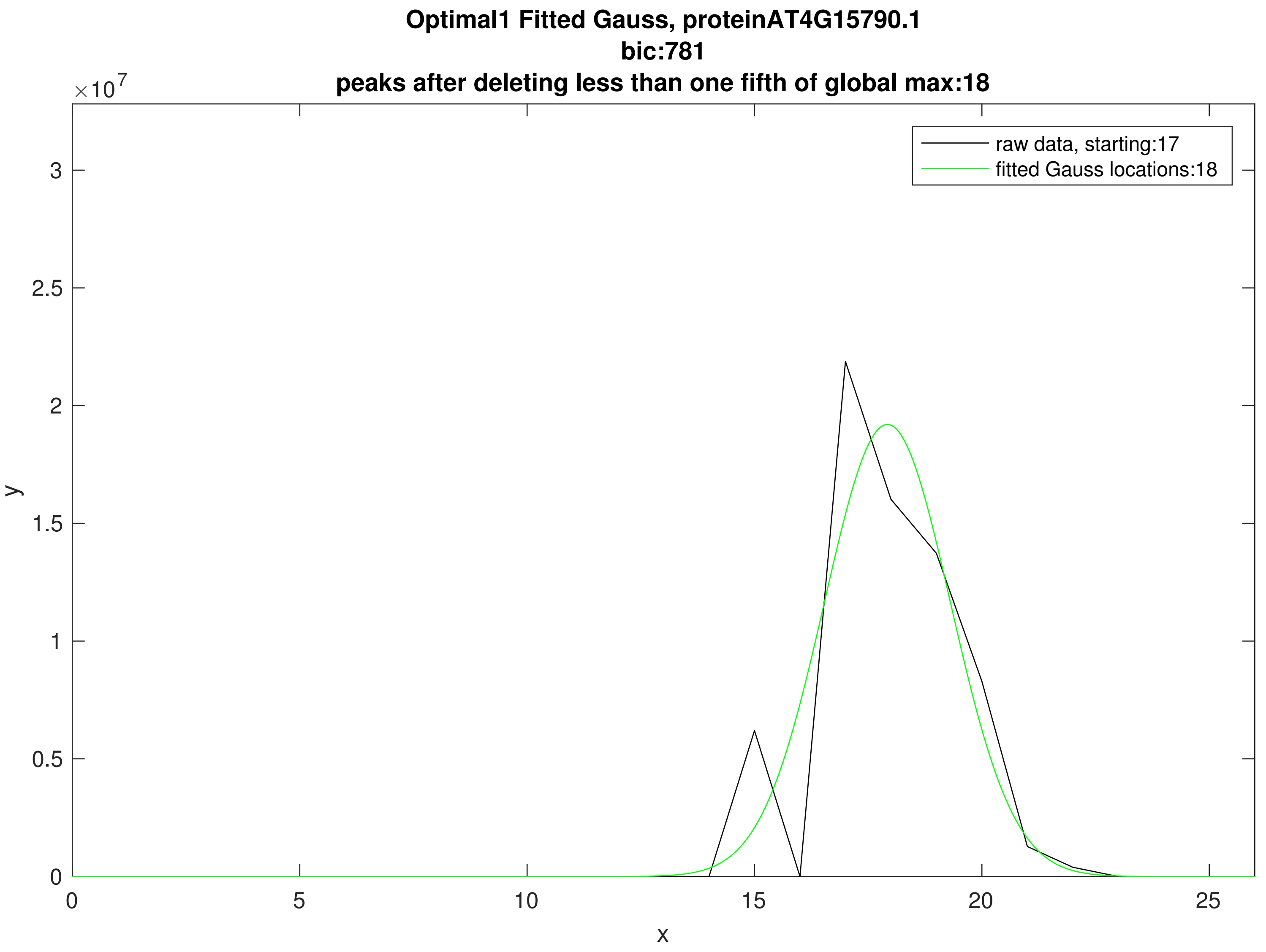

Optimal1 Fitted Gauss, proteinAT4G15790.1
bic:781
peaks after deleting less than one fifth of global max:18
×10^7
raw data, starting:17
fitted Gauss locations:18
y
x
0
0.5
1
1.5
2
2.5
3
0
5
10
15
20
25

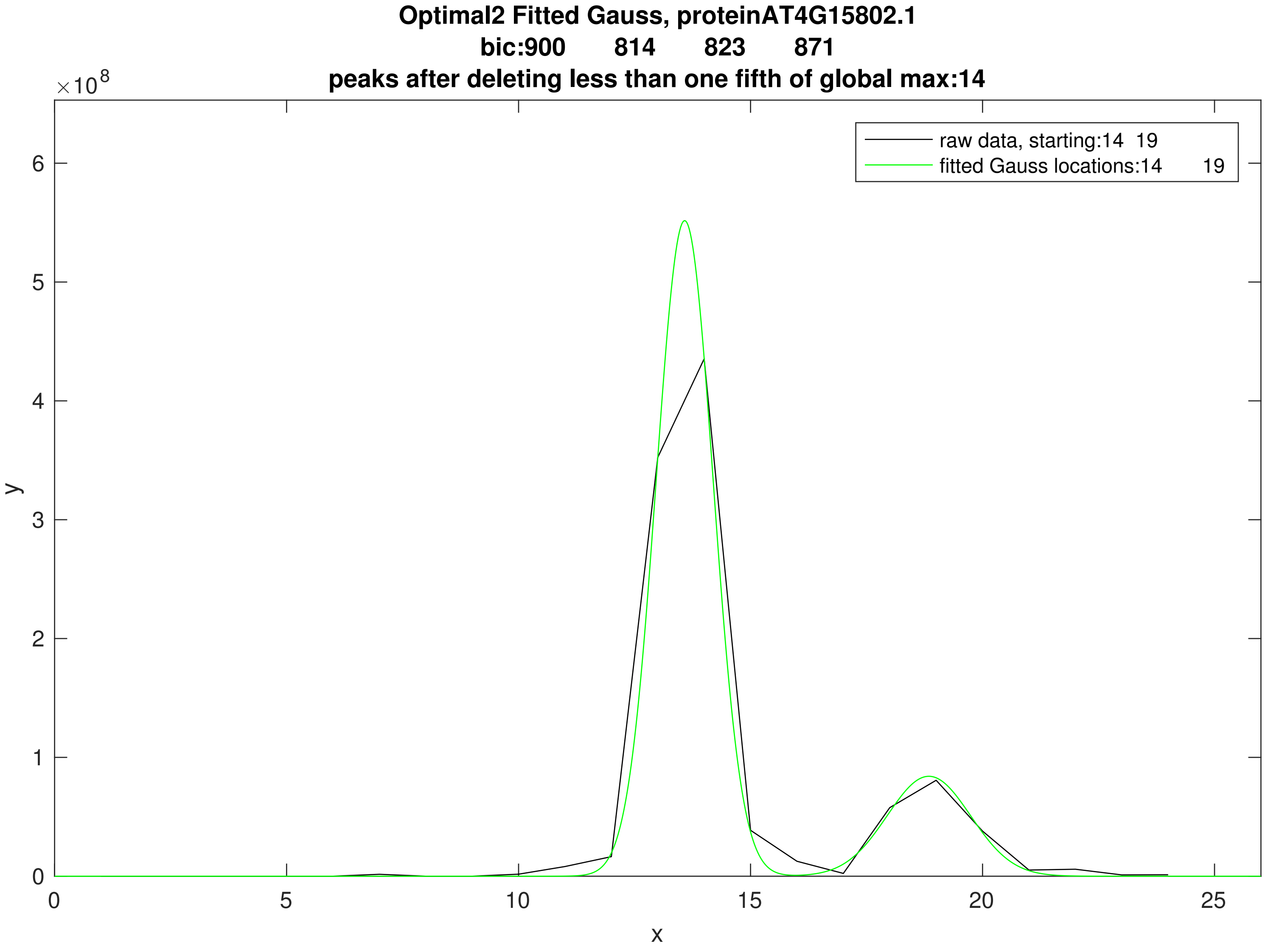

Optimal2 Fitted Gauss, proteinAT4G15802.1
bic:900 814 823 871
peaks after deleting less than one fifth of global max:14
×10^8
raw data, starting:14 19
fitted Gauss locations:14 19
y
x
0
1
2
3
4
5
6
0
5
10
15
20
25

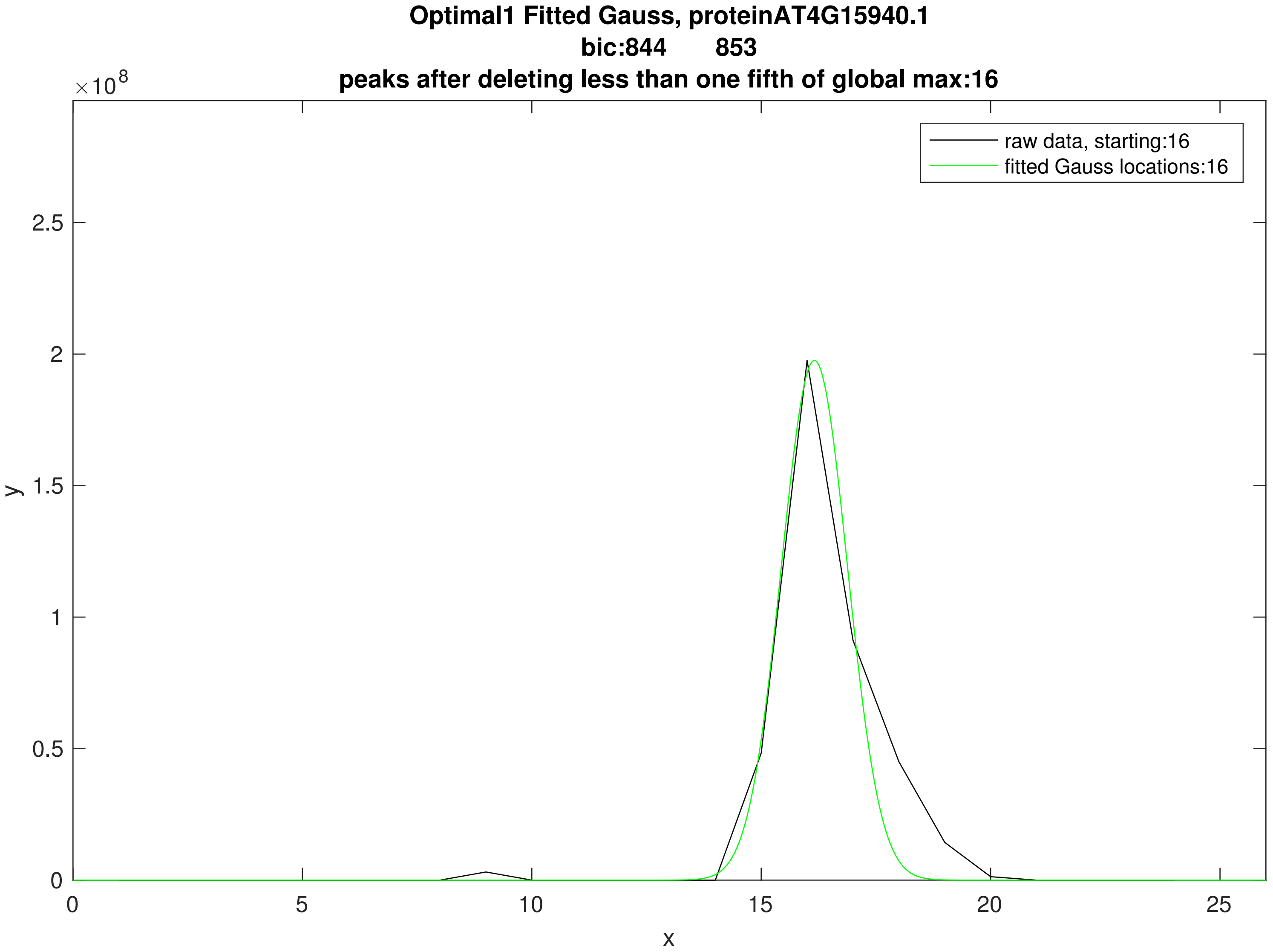

Optimal1 Fitted Gauss, proteinAT4G15940.1
bic:844 853
peaks after deleting less than one fifth of global max:16
×10^8
raw data, starting:16
fitted Gauss locations:16
2.5
2
1.5
1
0.5
0
y
0
5
10
15
20
25
x

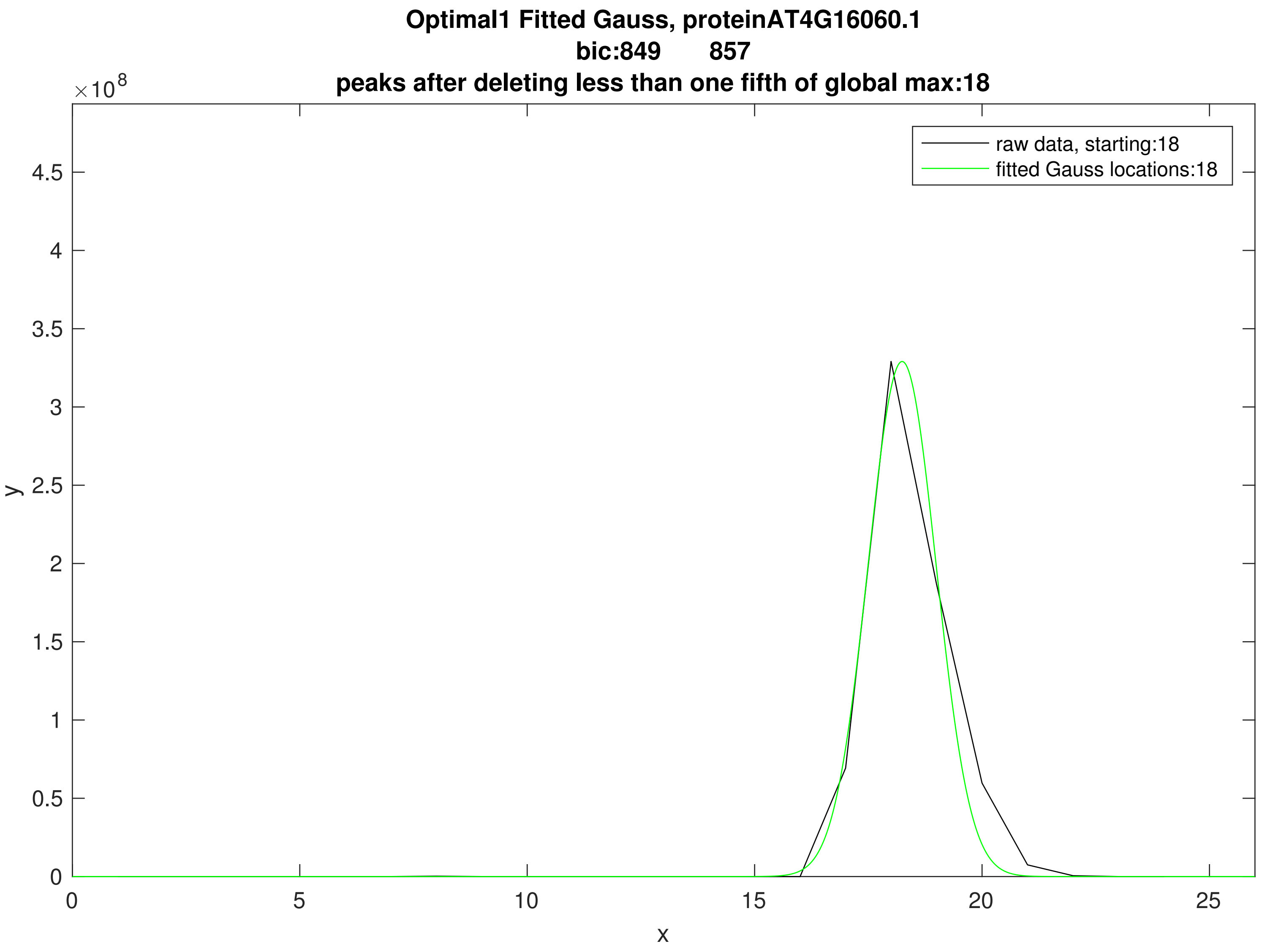

Optimal1 Fitted Gauss, proteinAT4G16060.1
bic:849 857
peaks after deleting less than one fifth of global max:18
×10^8
4.5
4
3.5
3
2.5
2
1.5
1
0.5
0
y
0
5
10
15
20
25
x
raw data, starting:18
fitted Gauss locations:18

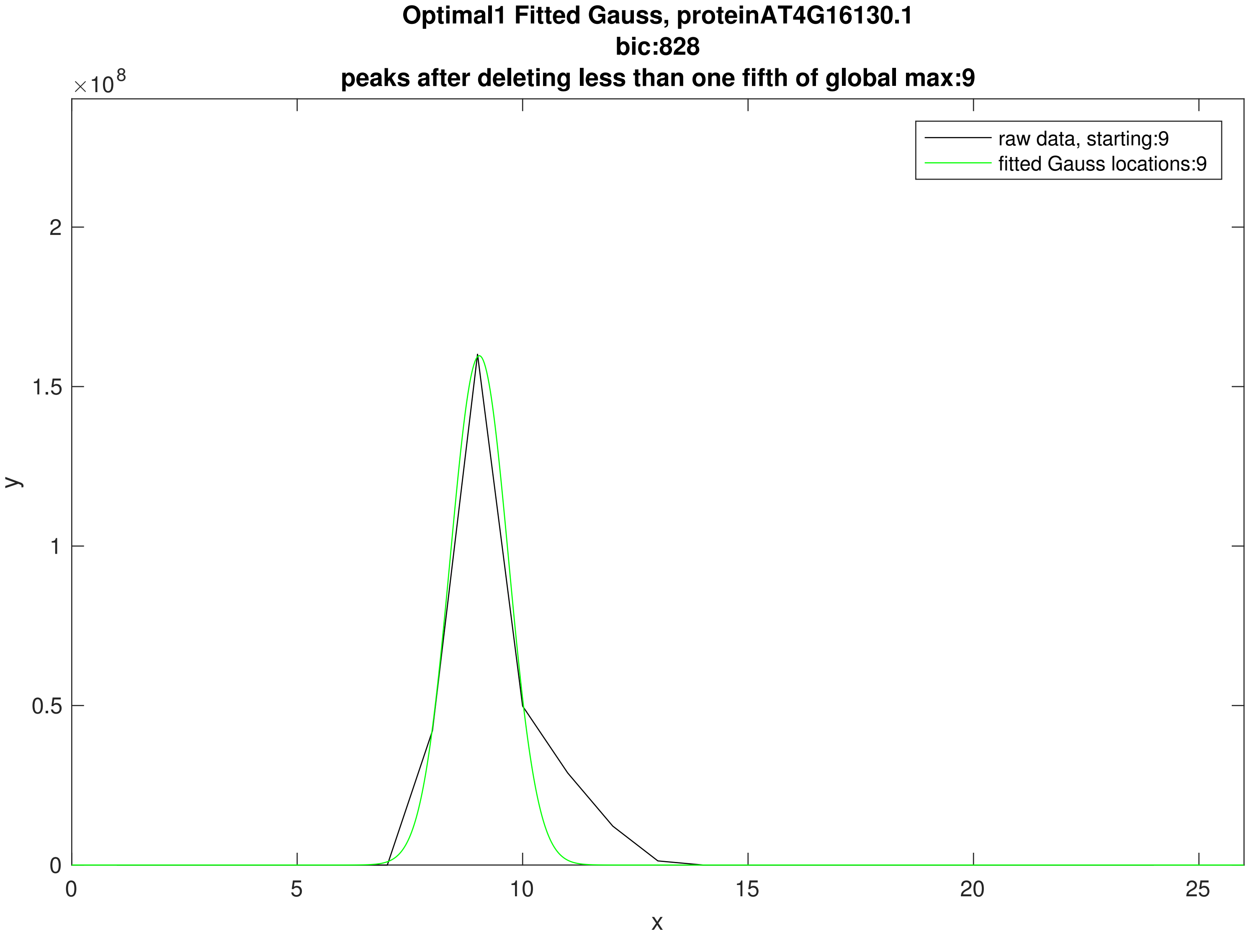

Optimal1 Fitted Gauss, proteinAT4G16130.1
bic:828
peaks after deleting less than one fifth of global max:9
×10^8
raw data, starting:9
fitted Gauss locations:9
y
x
0
0.5
1
1.5
2
0
5
10
15
20
25

**Optimal1 Fitted Gauss, proteinAT4G16155.1**
**bic:831 840**
**peaks after deleting less than one fifth of global max:12**

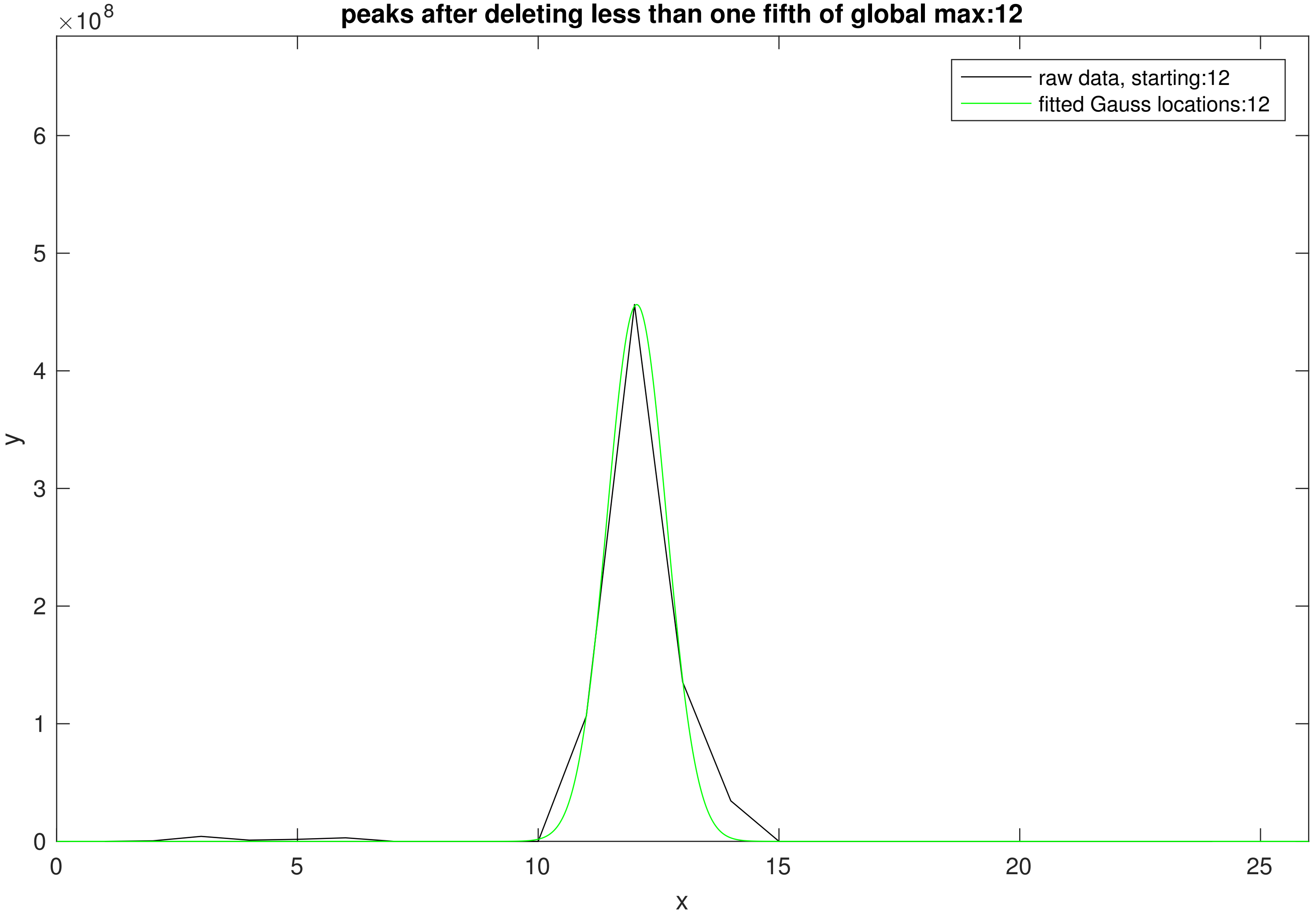

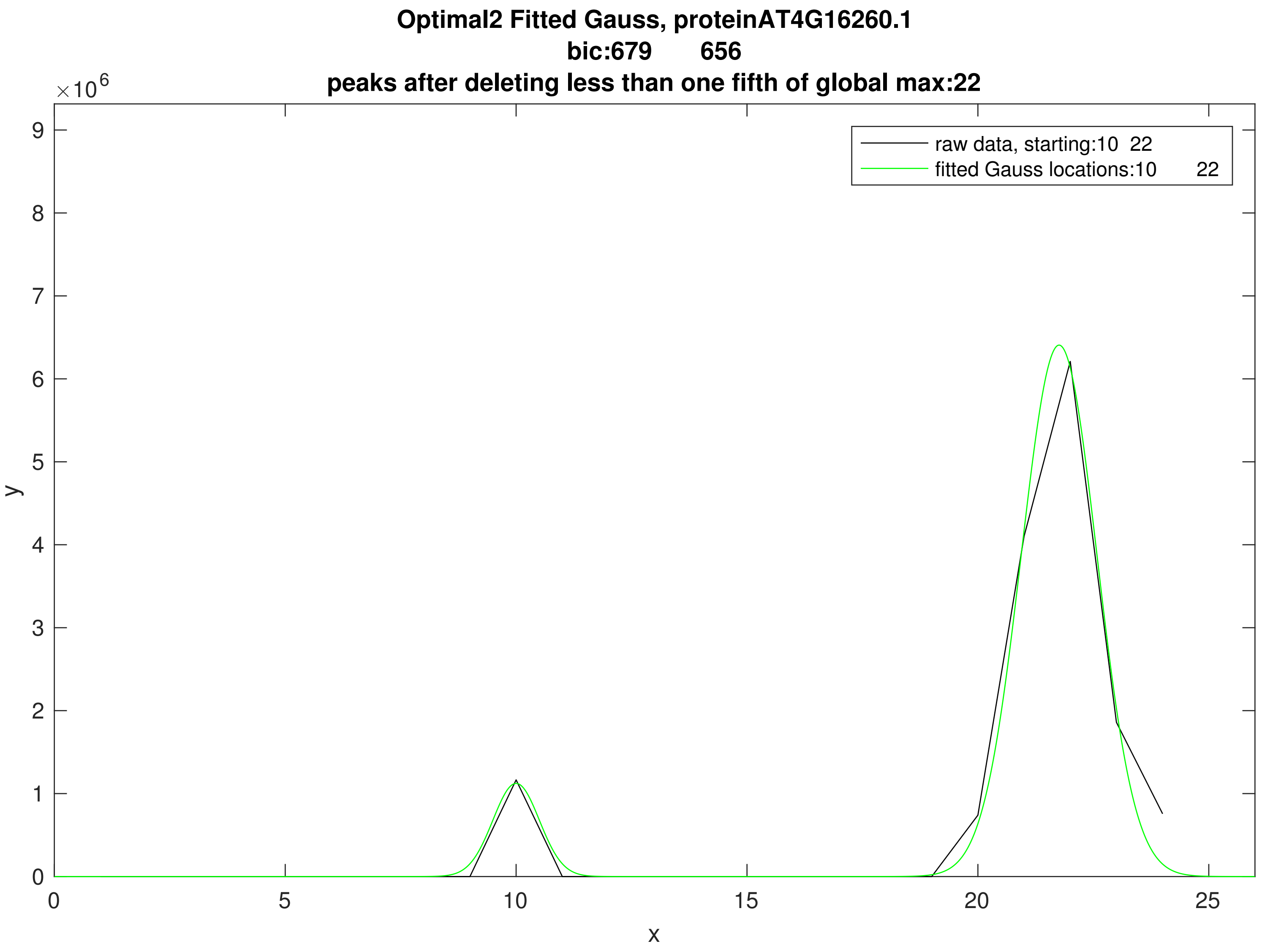

Optimal2 Fitted Gauss, proteinAT4G16260.1
bic:679 656
peaks after deleting less than one fifth of global max:22
×10^6
9
8
7
6
5
4
3
2
1
0
y
0
5
10
15
20
25
x
raw data, starting:10 22
fitted Gauss locations:10 22

Optimal2 Fitted Gauss, proteinAT4G16500.1
bic:840 824
peaks after deleting less than one fifth of global max:16 21

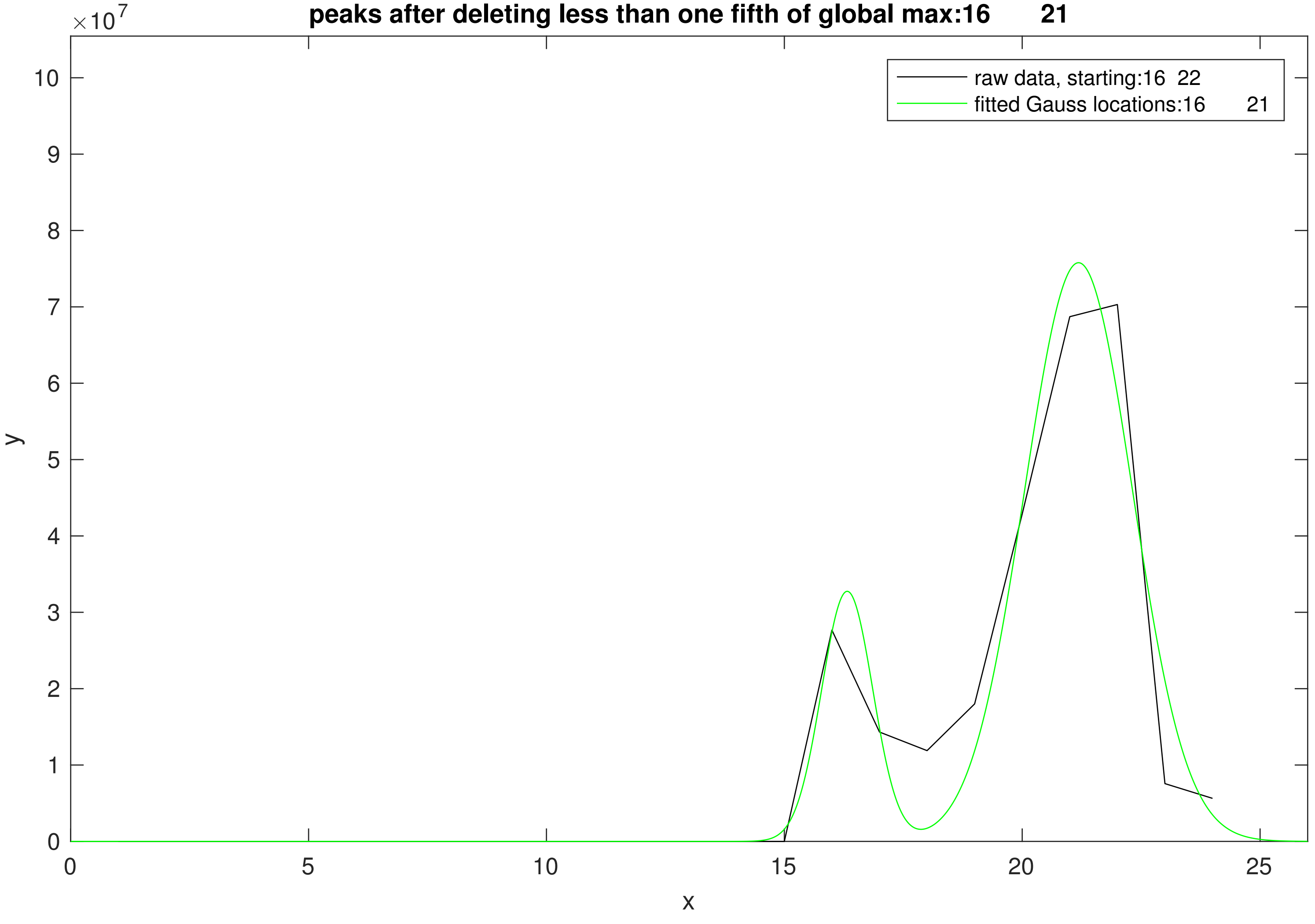

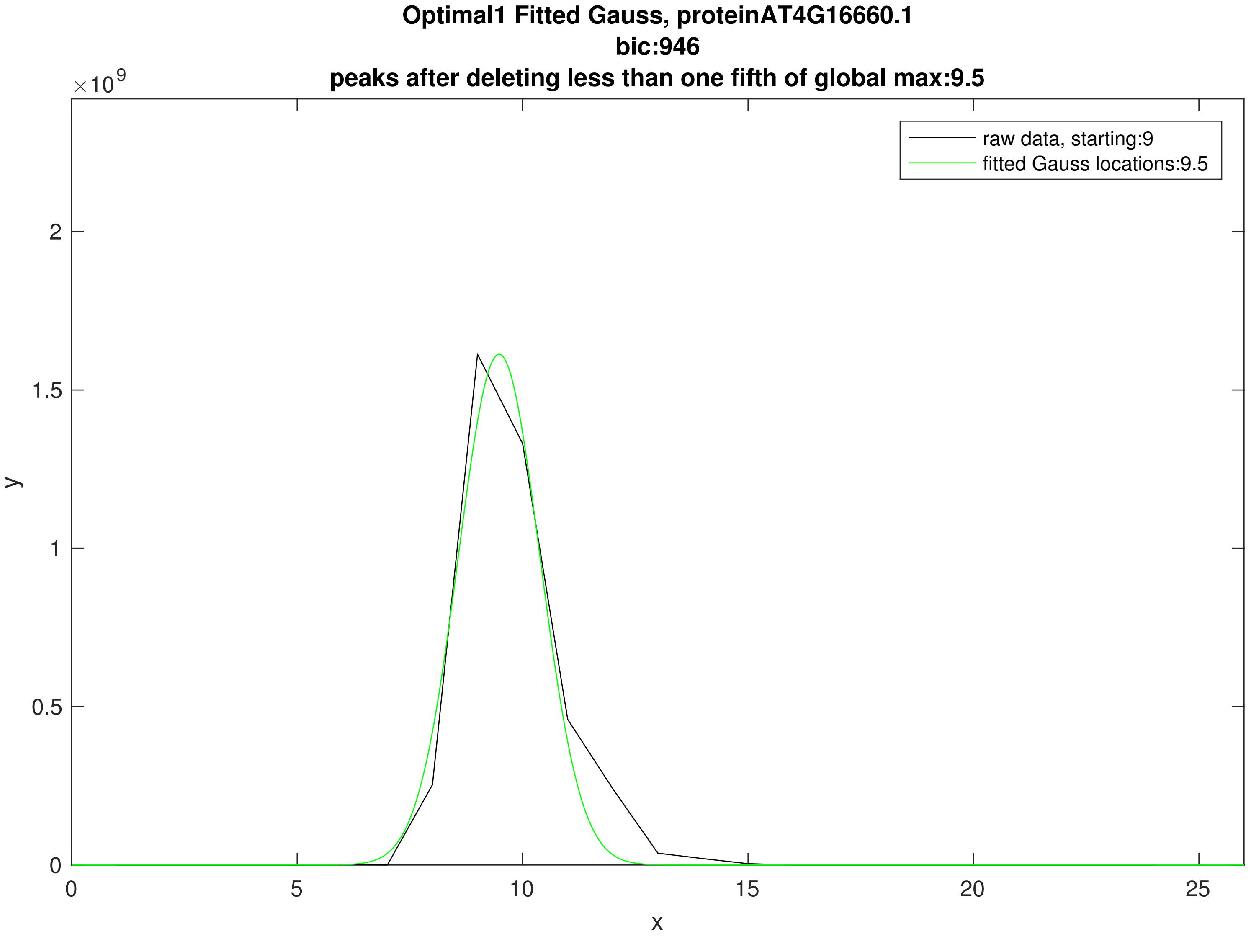
Optimal1 Fitted Gauss, proteinAT4G16660.1
bic:946
peaks after deleting less than one fifth of global max:9.5
raw data, starting:9
fitted Gauss locations:9.5
y
x

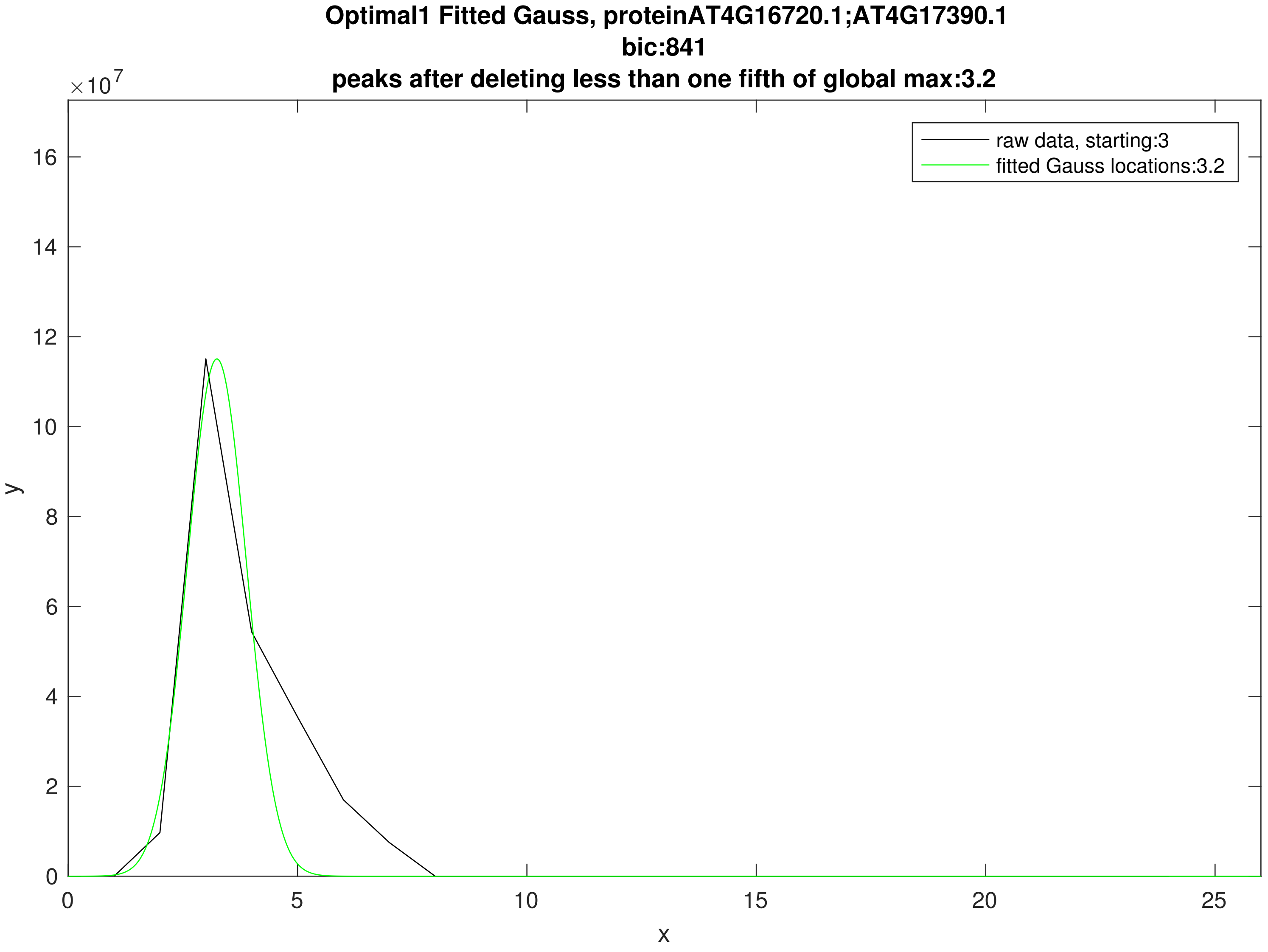
Optimal1 Fitted Gauss, proteinAT4G16720.1;AT4G17390.1
bic:841
peaks after deleting less than one fifth of global max:3.2
×10^7
raw data, starting:3
fitted Gauss locations:3.2
y
x
0
2
4
6
8
10
12
14
16
0
5
10
15
20
25

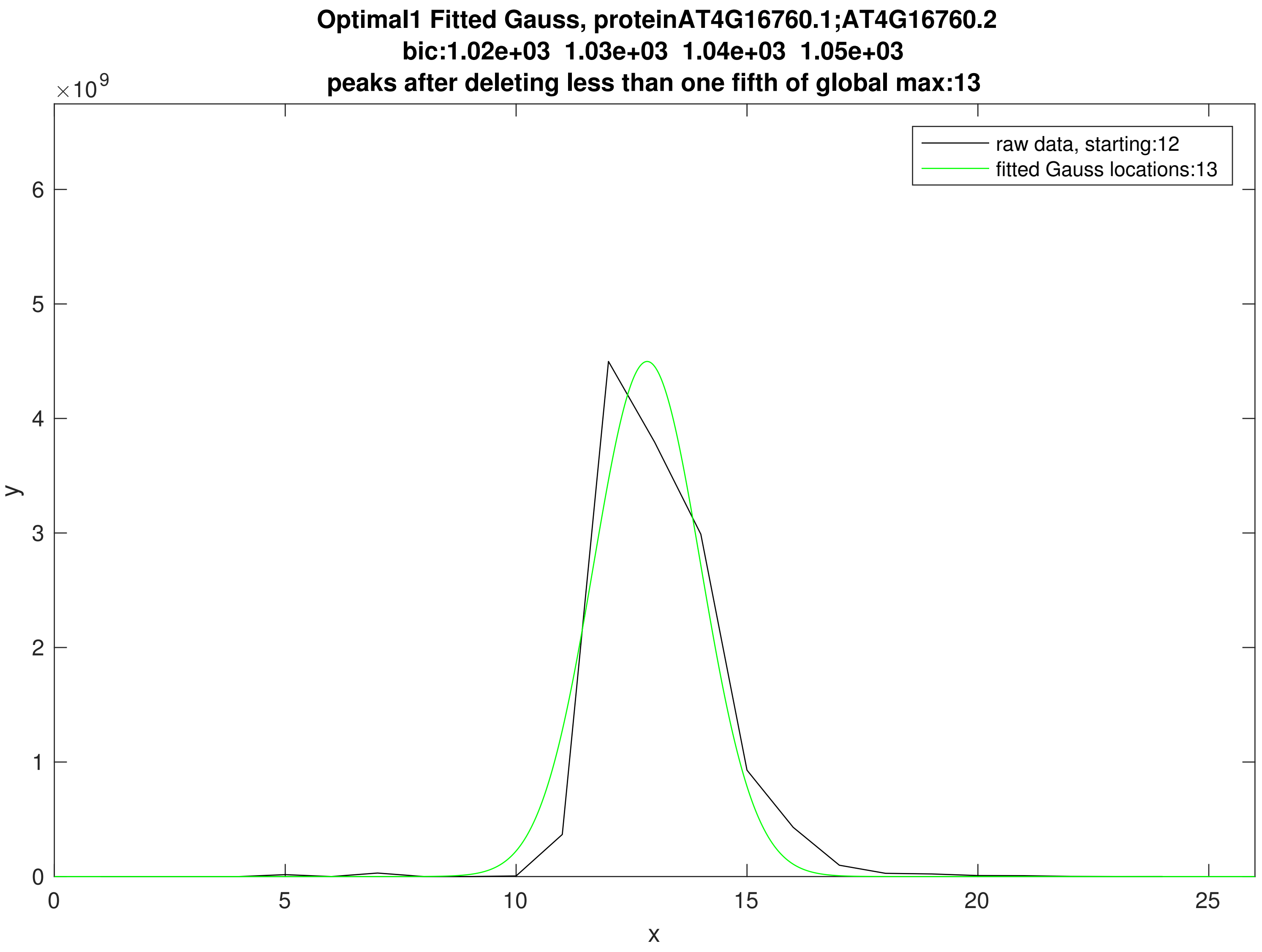

Optimal1 Fitted Gauss, proteinAT4G16760.1;AT4G16760.2
bic:1.02e+03 1.03e+03 1.04e+03 1.05e+03
peaks after deleting less than one fifth of global max:13
×10^9
raw data, starting:12
fitted Gauss locations:13
y
x
0
1
2
3
4
5
6
0
5
10
15
20
25

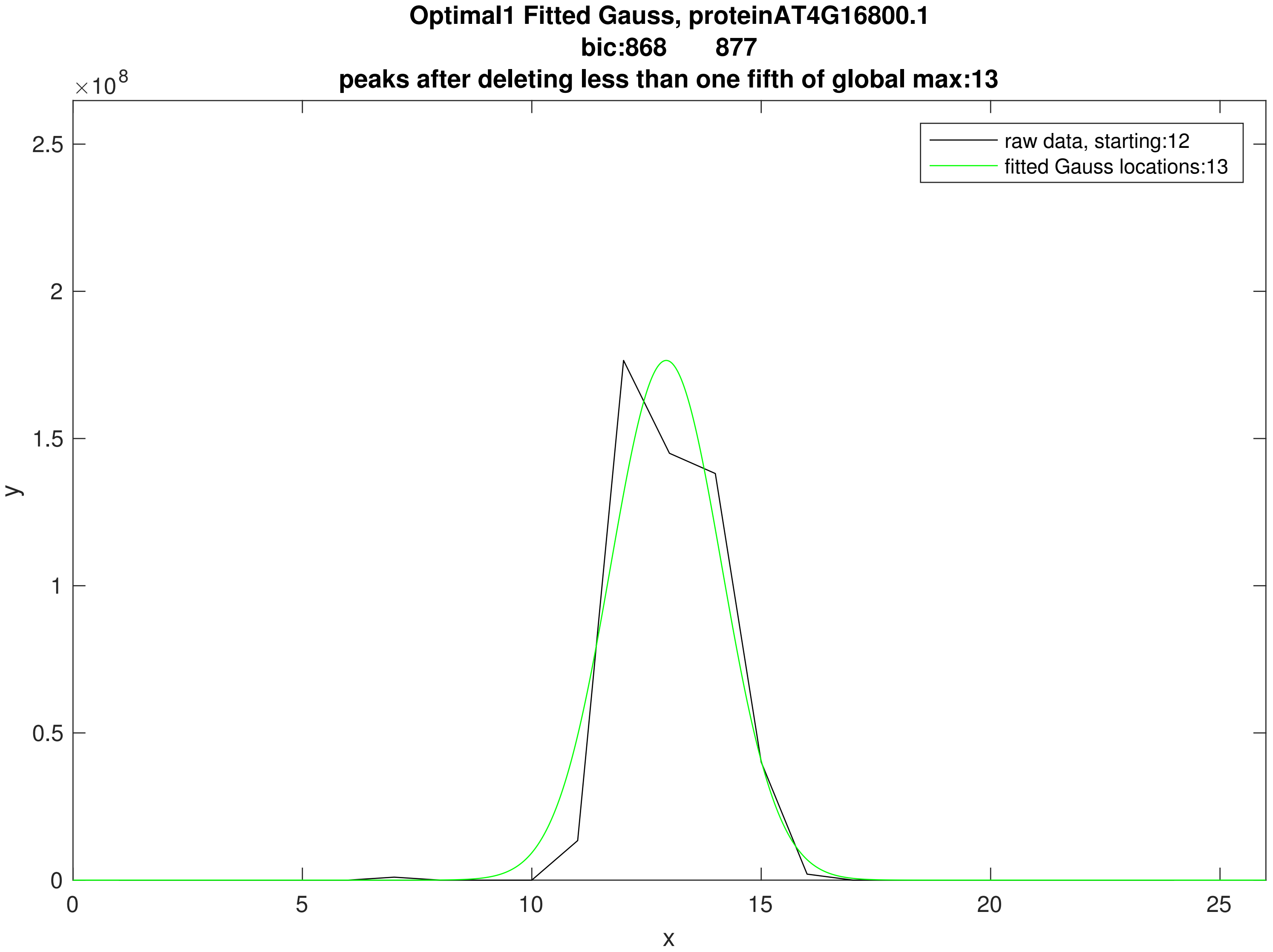
Optimal1 Fitted Gauss, proteinAT4G16800.1
bic:868 877
peaks after deleting less than one fifth of global max:13
×10^8
2.5
2
1.5
1
0.5
0
y
0
5
10
15
20
25
x
raw data, starting:12
fitted Gauss locations:13

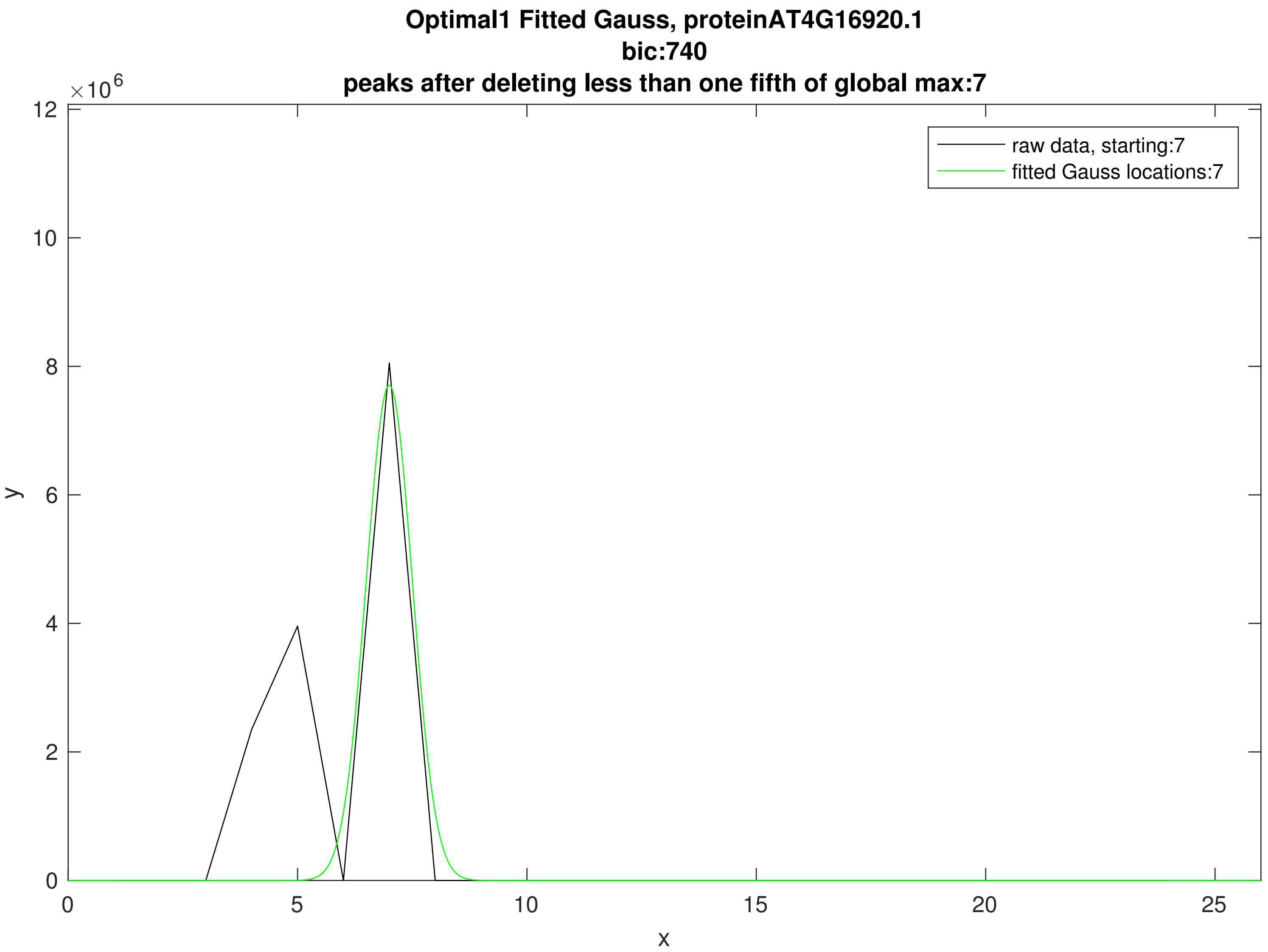
Optimal1 Fitted Gauss, proteinAT4G16920.1
bic:740
peaks after deleting less than one fifth of global max:7
×10^6
raw data, starting:7
fitted Gauss locations:7
y
x
0
2
4
6
8
10
12
0
5
10
15
20
25

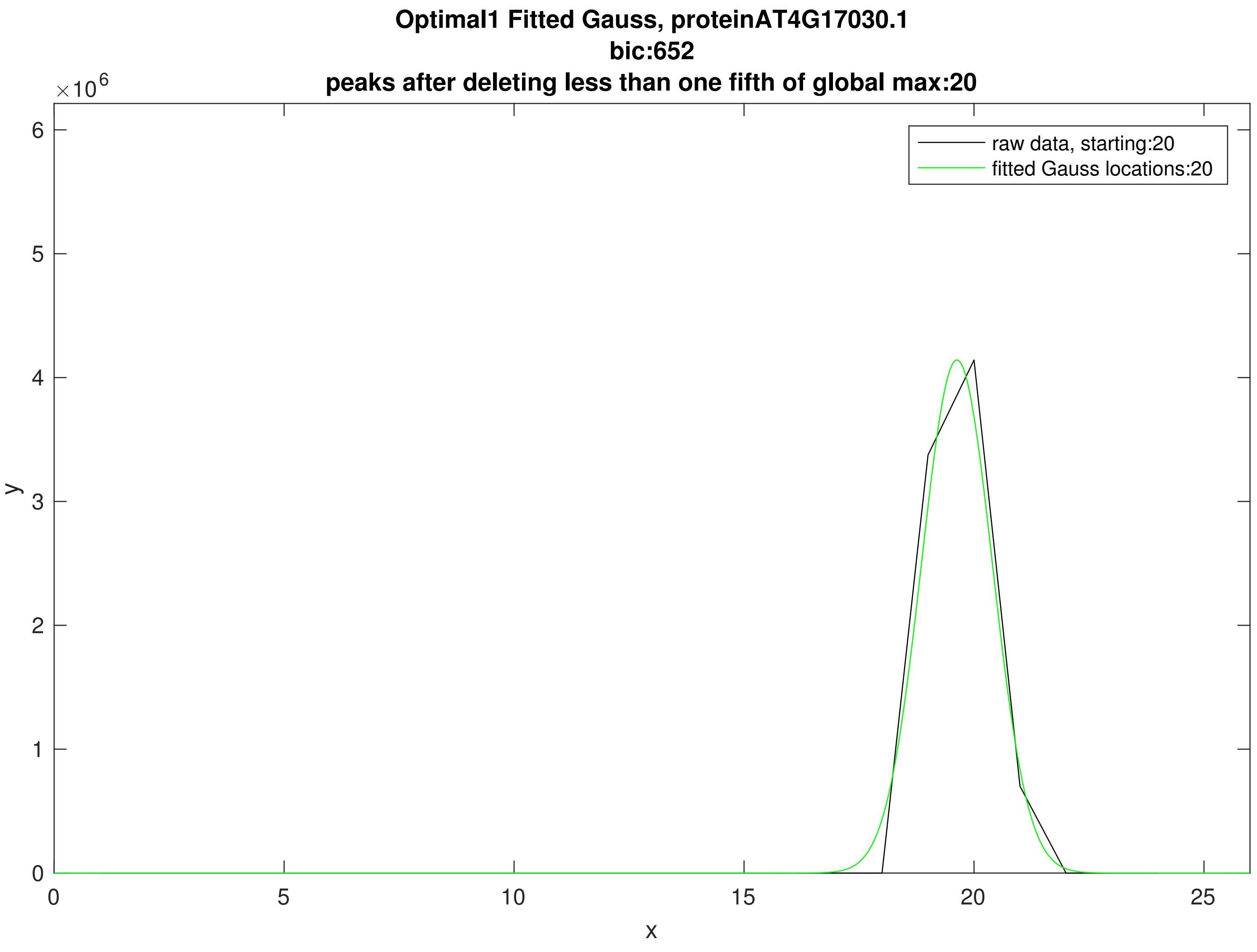
Optimal1 Fitted Gauss, proteinAT4G17030.1
bic:652
peaks after deleting less than one fifth of global max:20
×10^6
raw data, starting:20
fitted Gauss locations:20
y
x
0
1
2
3
4
5
6
0
5
10
15
20
25

**Optimal1 Fitted Gauss, proteinAT4G17040.1**
**bic:817**
**peaks after deleting less than one fifth of global max:9.5**

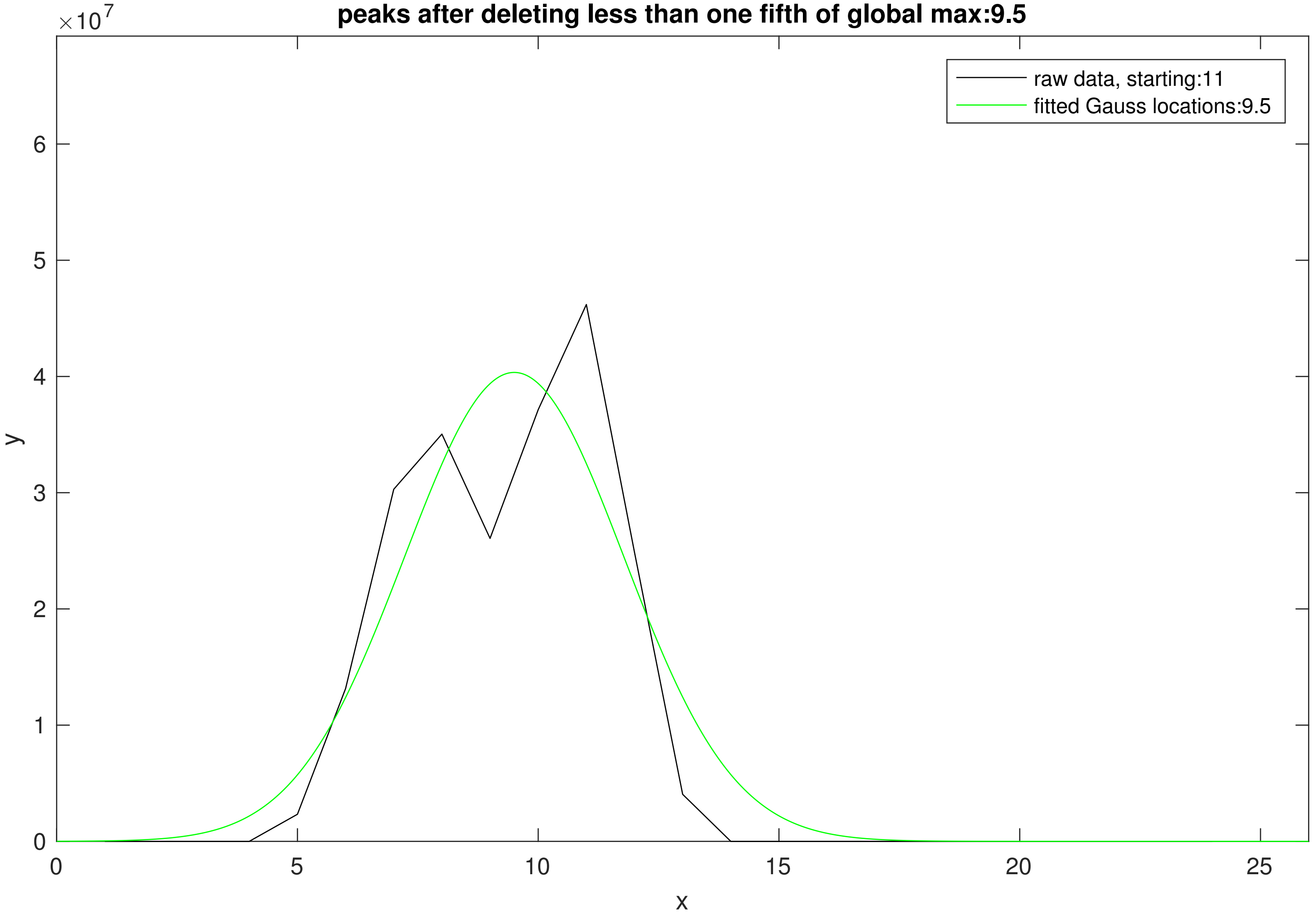

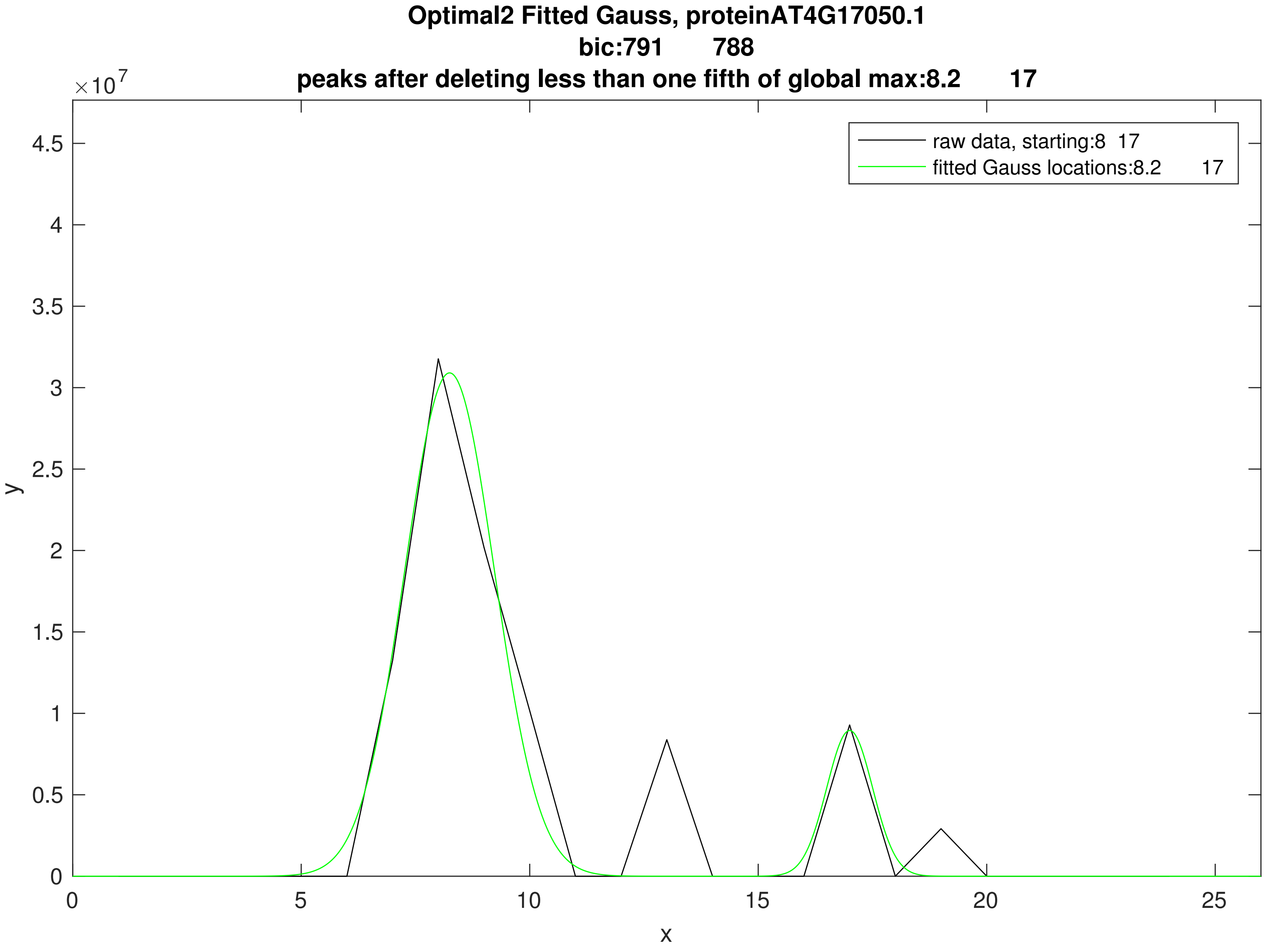

Optimal2 Fitted Gauss, proteinAT4G17050.1
bic:791 788
peaks after deleting less than one fifth of global max:8.2 17
×10^7
4.5
4
3.5
3
2.5
2
1.5
1
0.5
0
y
0
5
10
15
20
25
x
raw data, starting:8 17
fitted Gauss locations:8.2 17

#### Optimal1 Fitted Gauss, proteinAT4G17100.1;AT4G17100.2
#### bic:940 949 959
#### peaks after deleting less than one fifth of global max:14

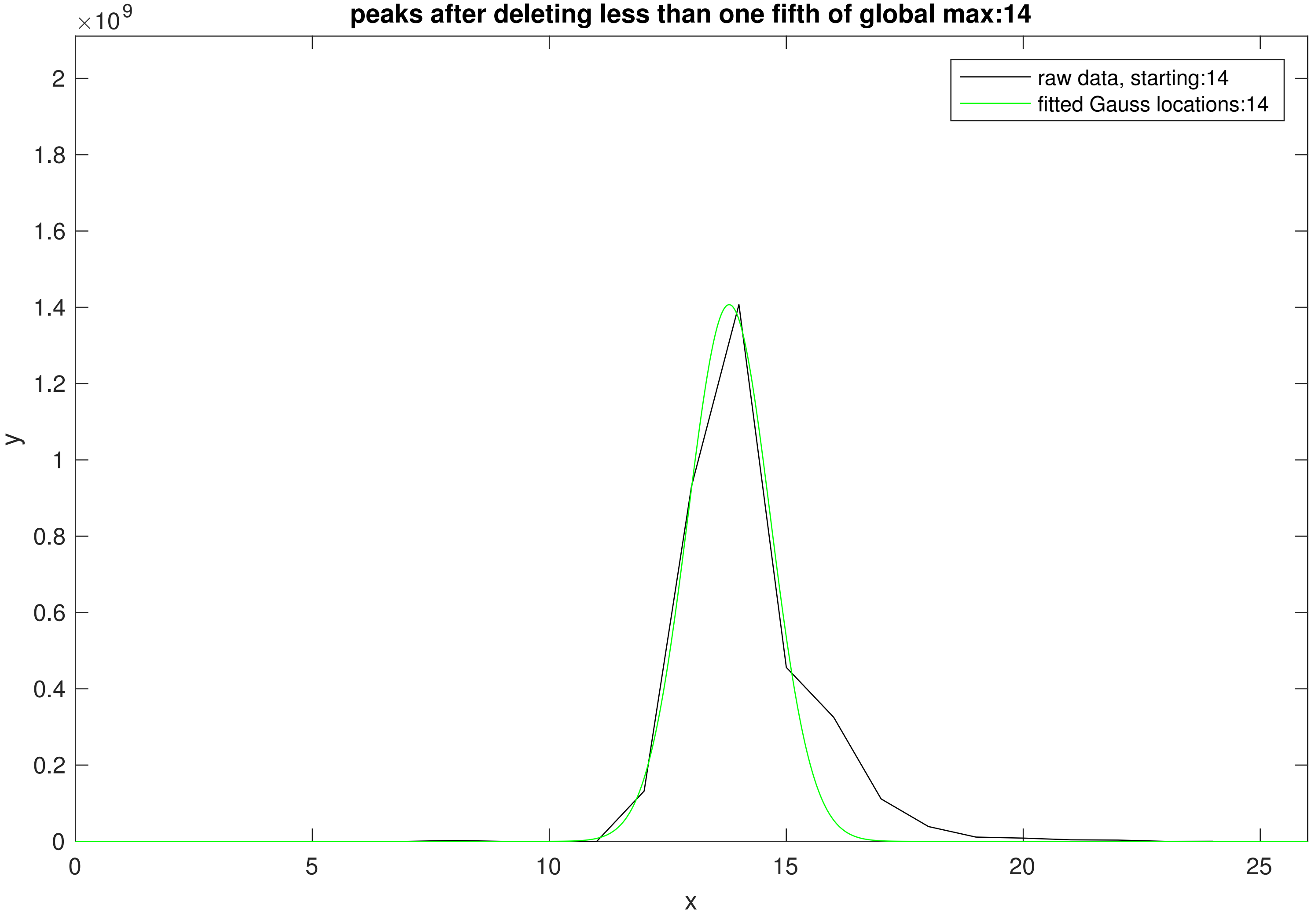

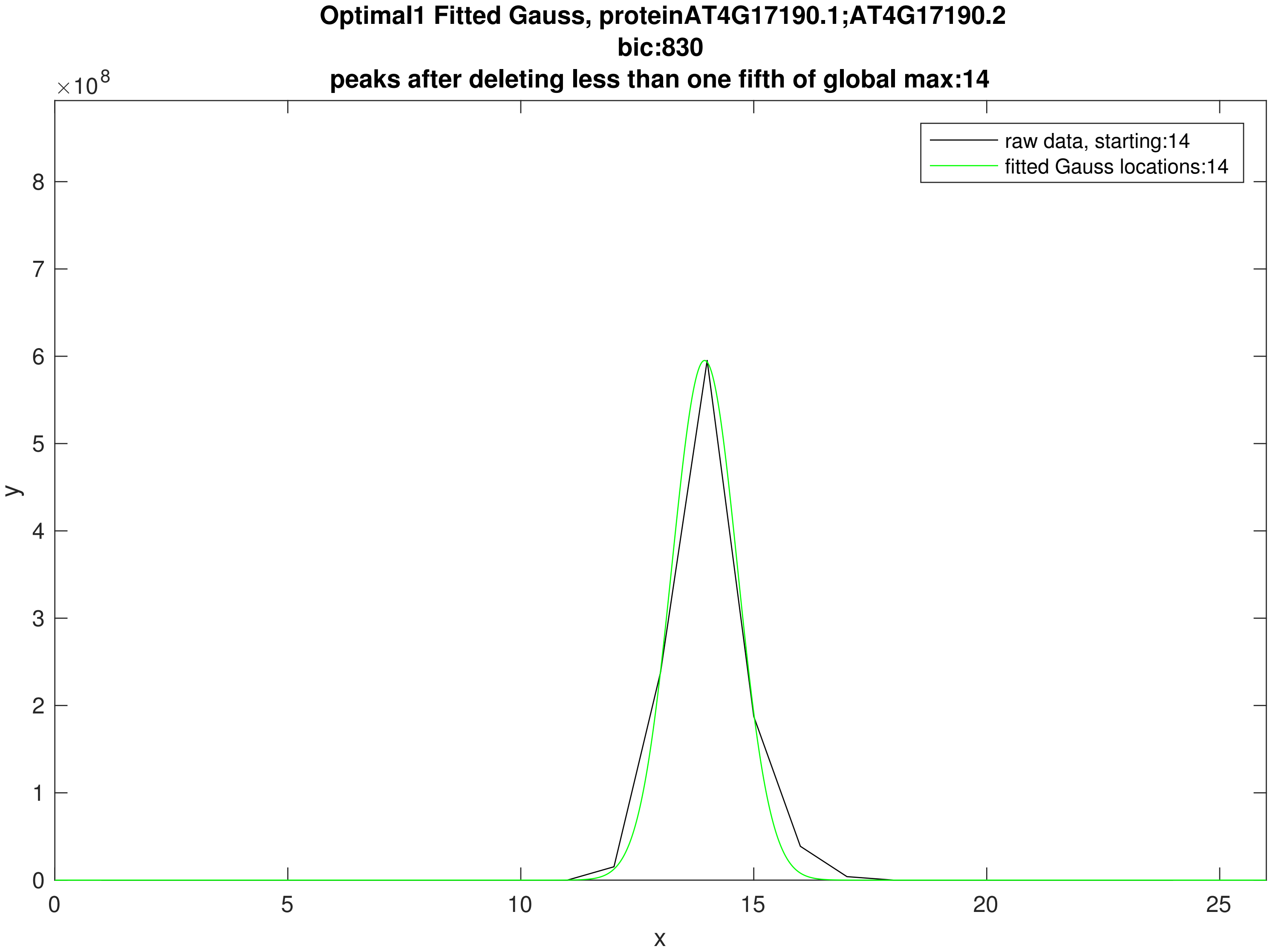

Optimal1 Fitted Gauss, proteinAT4G17190.1;AT4G17190.2
bic:830
peaks after deleting less than one fifth of global max:14
×10^8
raw data, starting:14
fitted Gauss locations:14
y
x
0
1
2
3
4
5
6
7
8
0
5
10
15
20
25

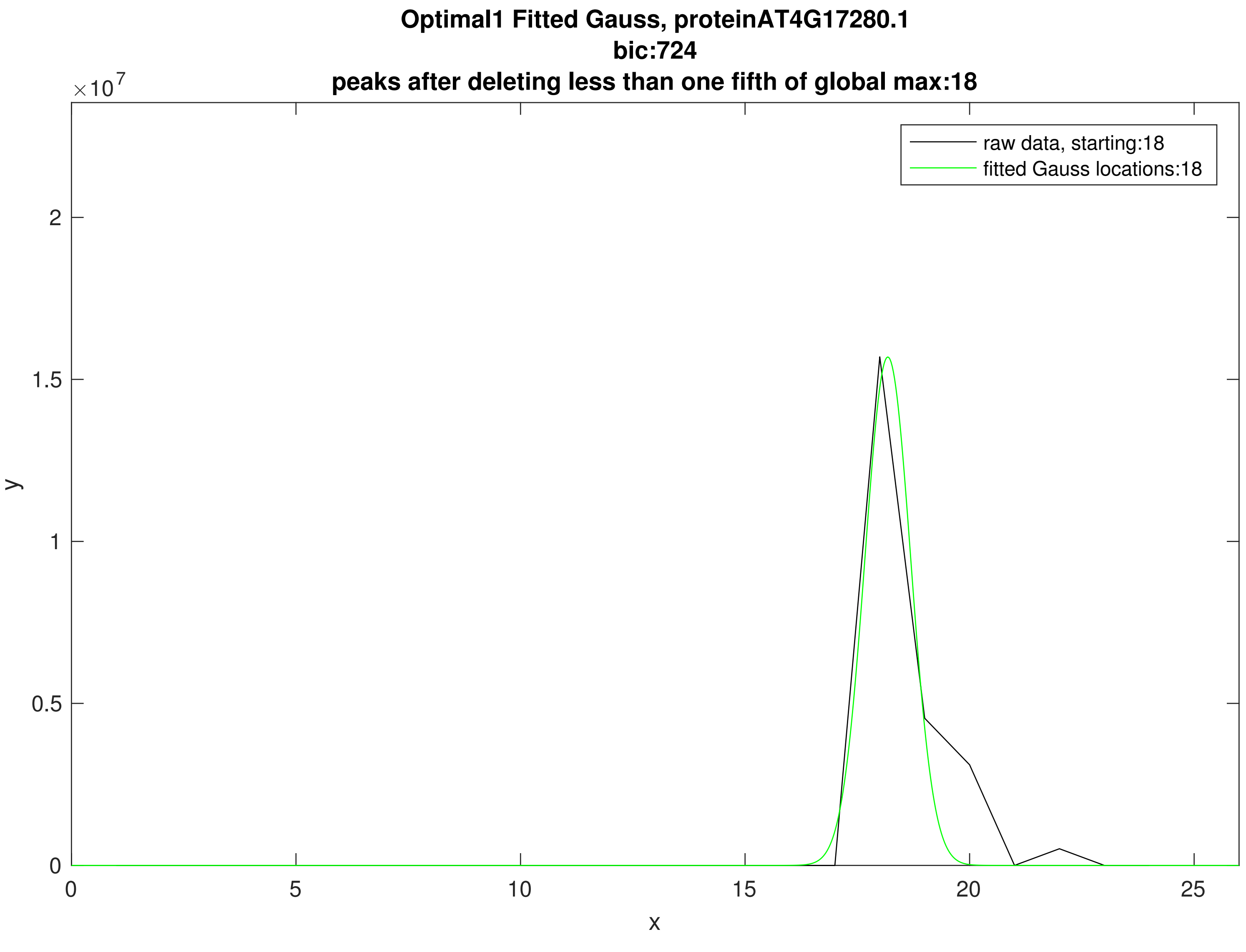
Optimal1 Fitted Gauss, proteinAT4G17280.1
bic:724
peaks after deleting less than one fifth of global max:18
×10^7
raw data, starting:18
fitted Gauss locations:18
y
x
0
0.5
1
1.5
2
0
5
10
15
20
25

Optimal1 Fitted Gauss, proteinAT4G17300.1
bic:824
peaks after deleting less than one fifth of global max:13

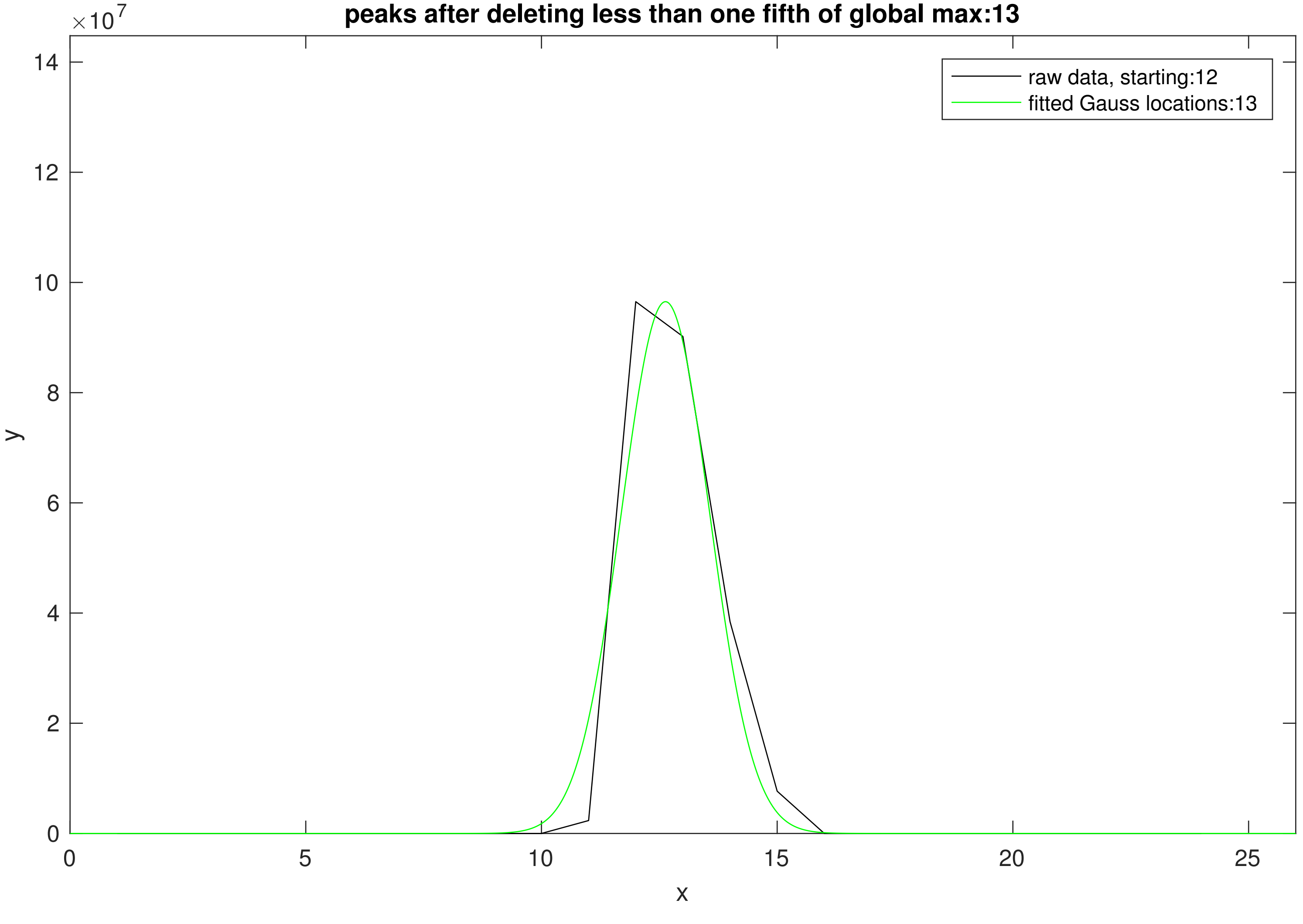

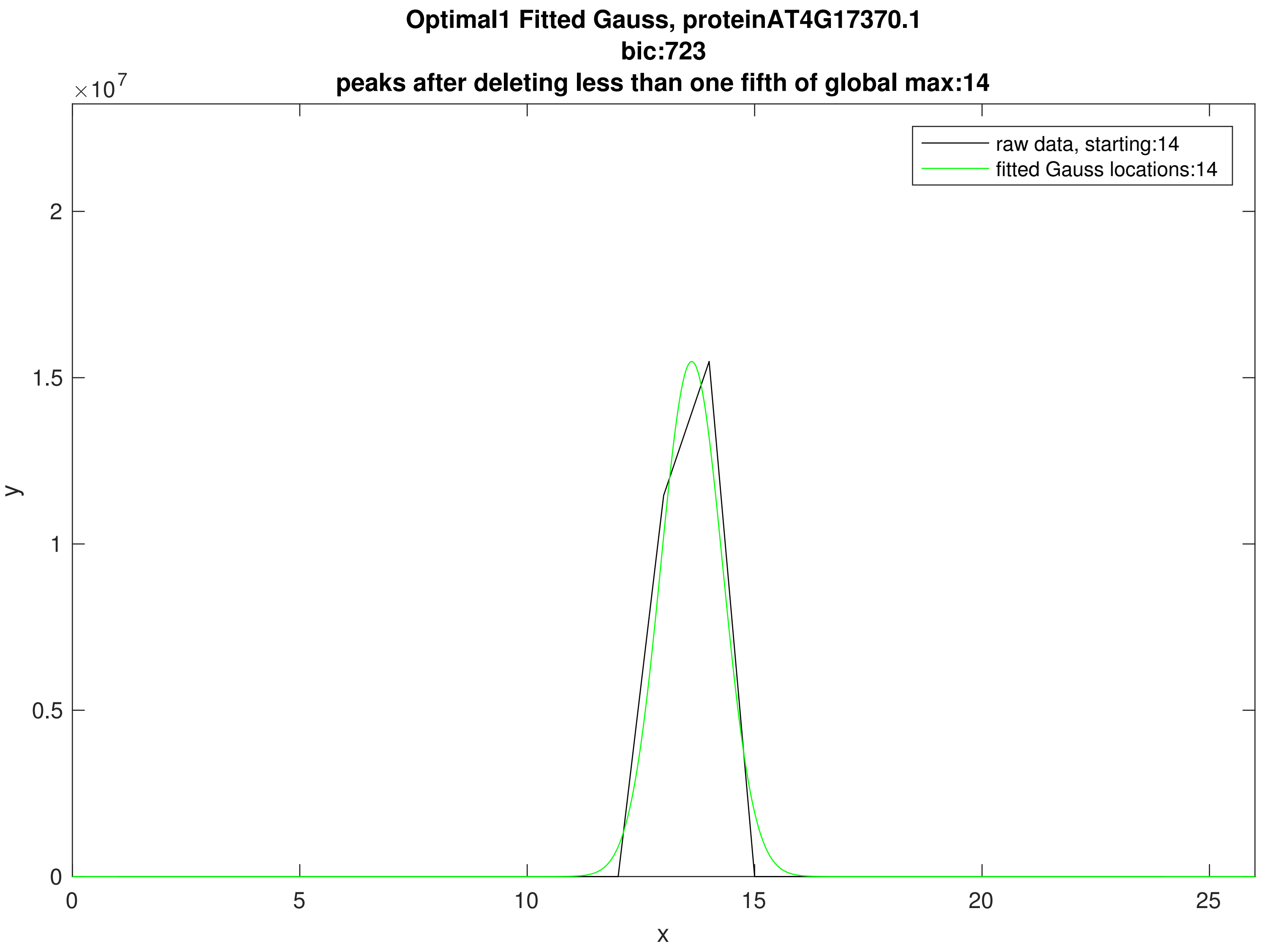
Optimal1 Fitted Gauss, proteinAT4G17370.1
bic:723
peaks after deleting less than one fifth of global max:14
×10^7
raw data, starting:14
fitted Gauss locations:14
y
x
0
0.5
1
1.5
2
0
5
10
15
20
25

**Optimal1 Fitted Gauss, proteinAT4G17470.3;AT4G17470.2;AT4G17470.1**
**bic:811**
**peaks after deleting less than one fifth of global max:18**

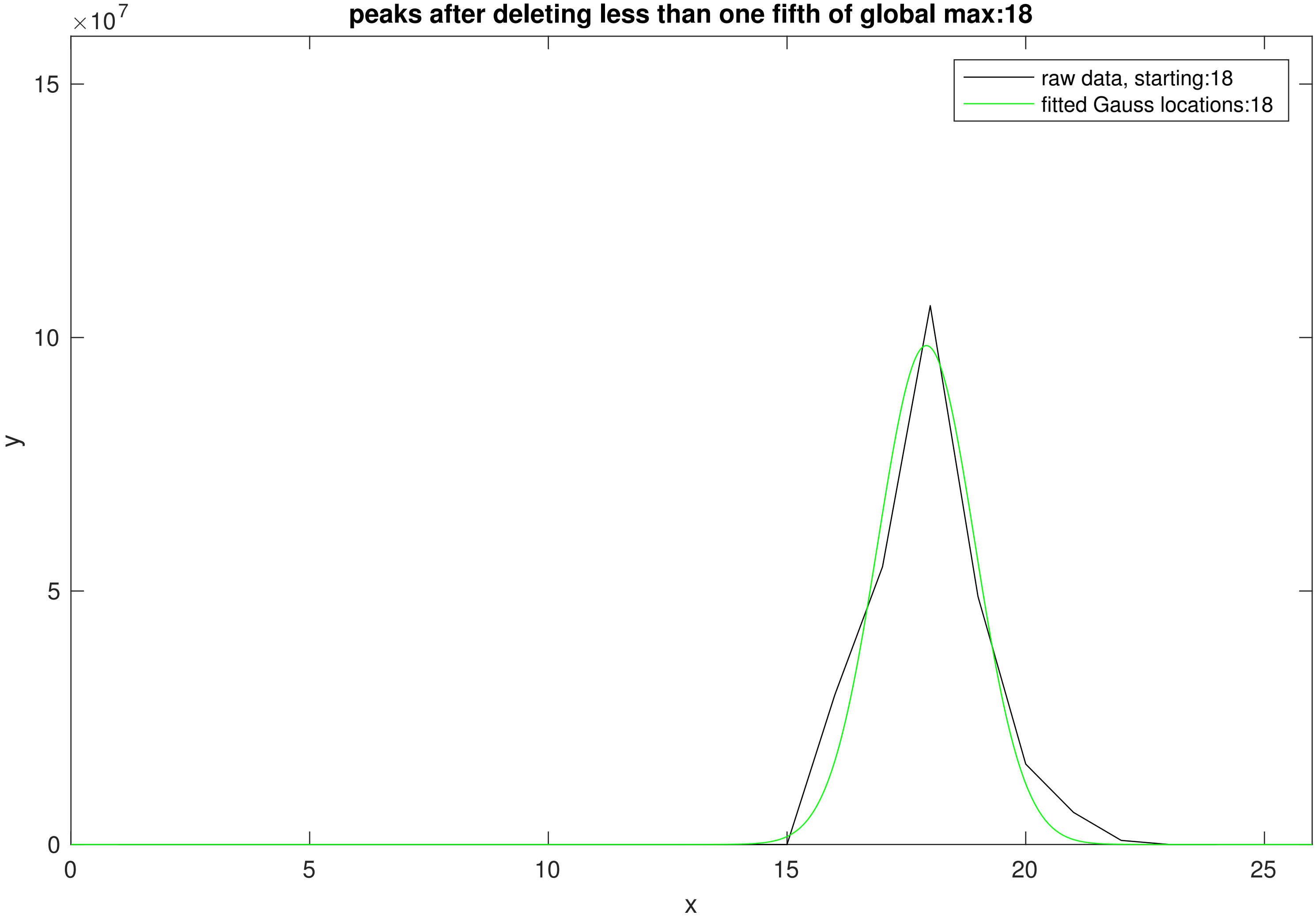

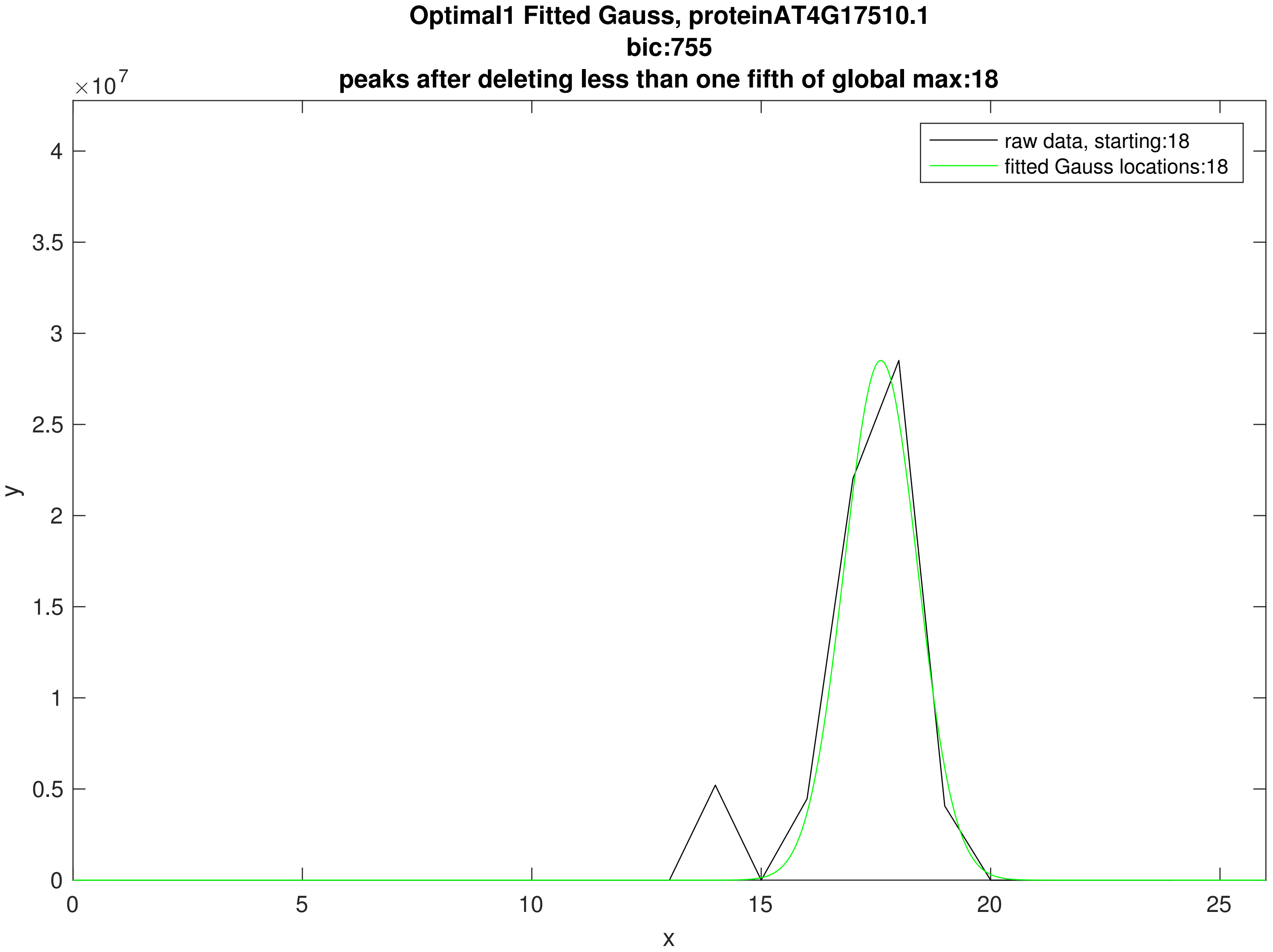
Optimal1 Fitted Gauss, proteinAT4G17510.1
bic:755
peaks after deleting less than one fifth of global max:18
×10^7
raw data, starting:18
fitted Gauss locations:18
y
x
0
0.5
1
1.5
2
2.5
3
3.5
4
5
10
15
20
25

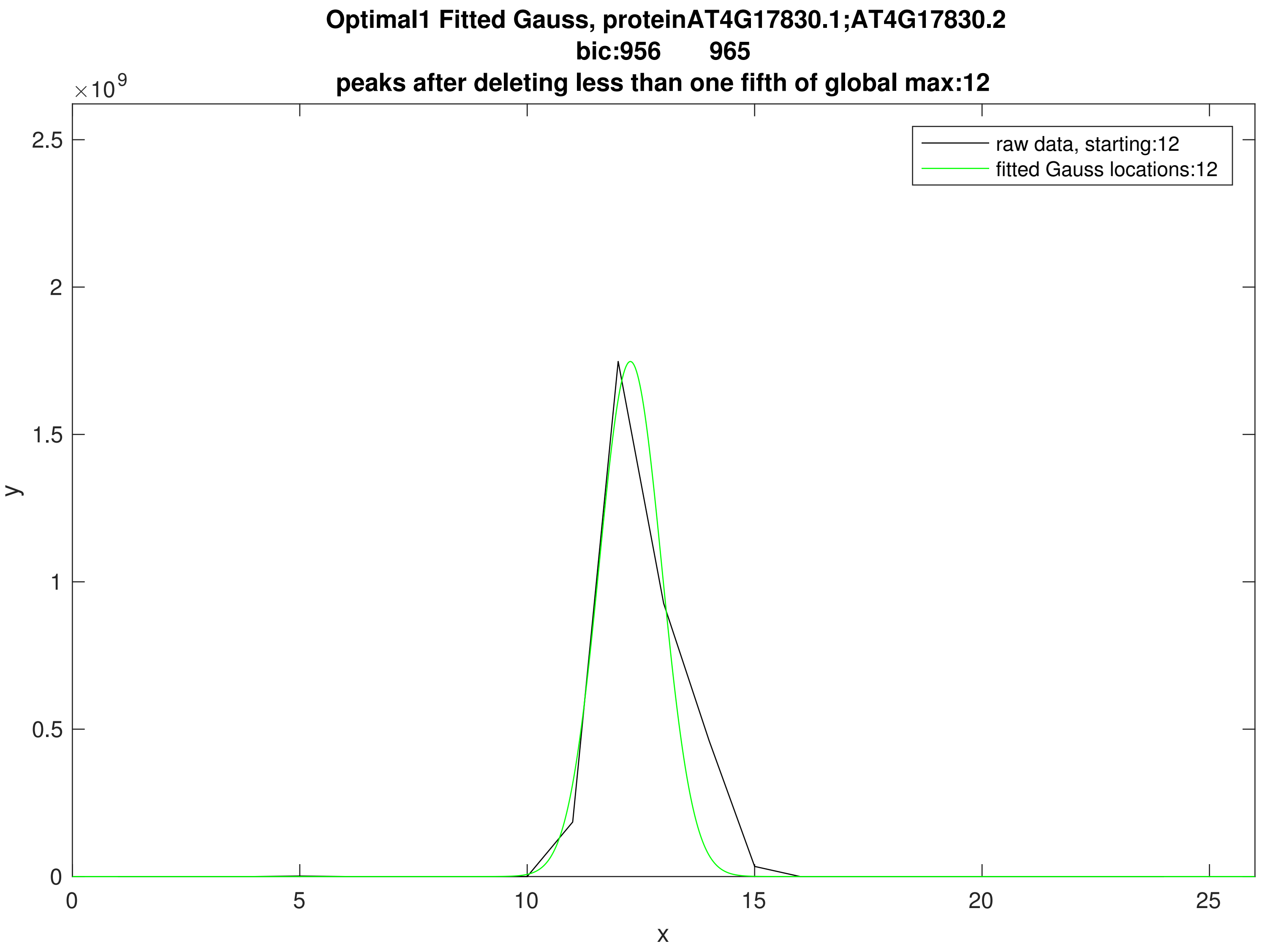
Optimal1 Fitted Gauss, proteinAT4G17830.1;AT4G17830.2
bic:956 965
peaks after deleting less than one fifth of global max:12
×10^9
2.5
2
1.5
1
0.5
0
y
0
5
10
15
20
25
x
raw data, starting:12
fitted Gauss locations:12

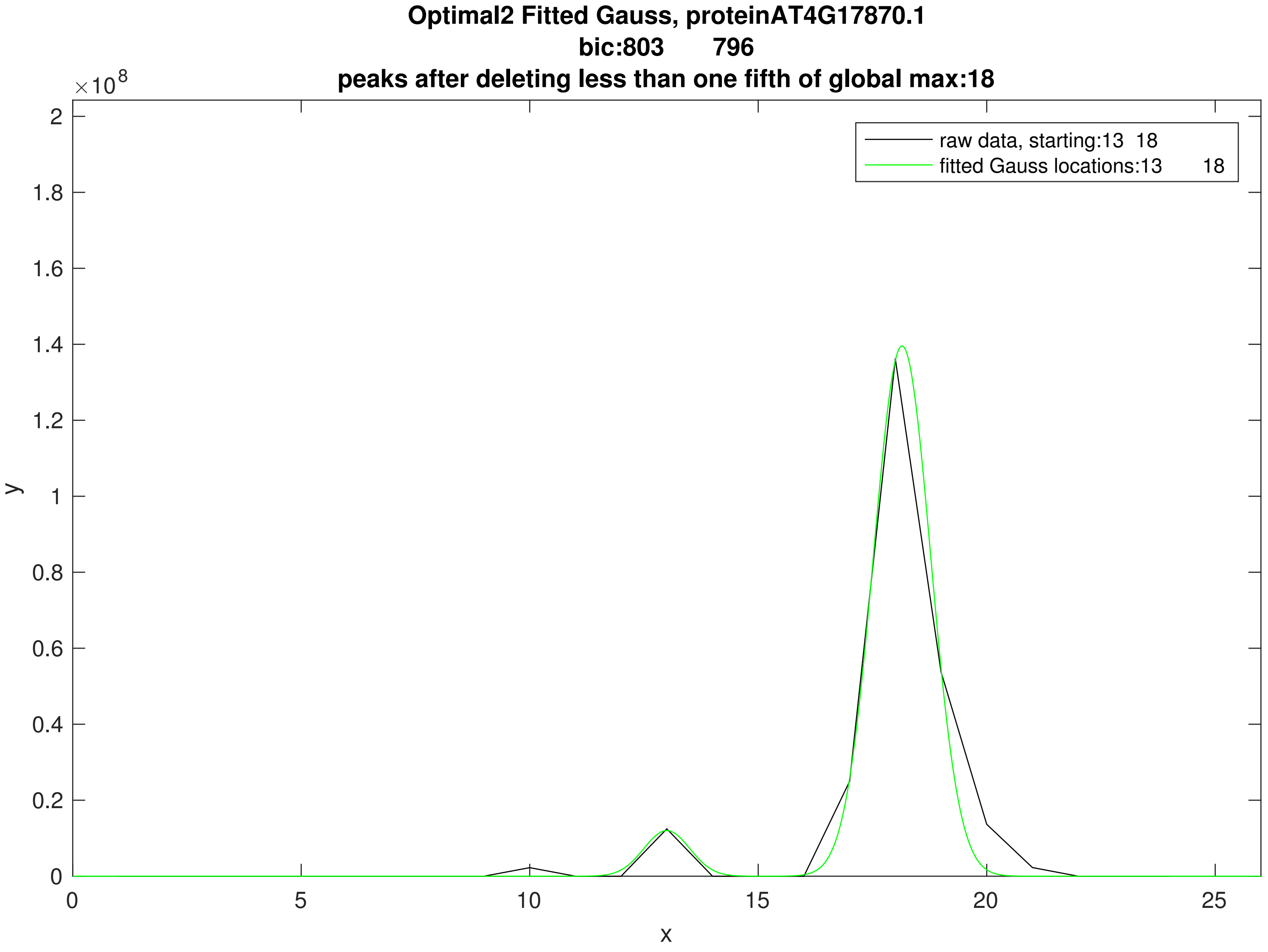
Optimal2 Fitted Gauss, proteinAT4G17870.1
bic:803 796
peaks after deleting less than one fifth of global max:18
×10^8
raw data, starting:13 18
fitted Gauss locations:13 18
y
x
0
0.2
0.4
0.6
0.8
1
1.2
1.4
1.6
1.8
2
0
5
10
15
20
25

**Optimal1 Fitted Gauss, proteinAT4G17890.2;AT4G17890.1**
**bic:749**
**peaks after deleting less than one fifth of global max:19**

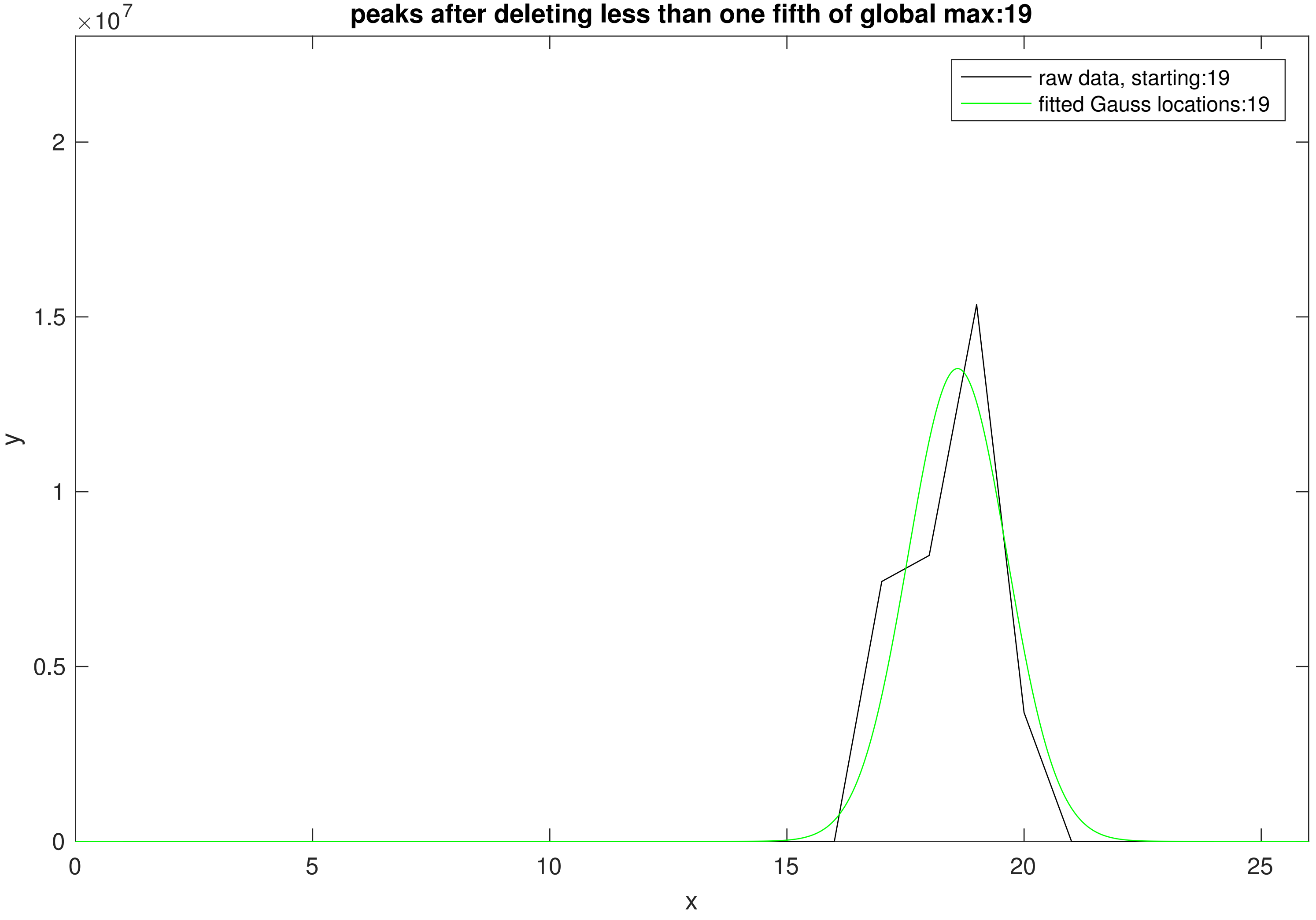

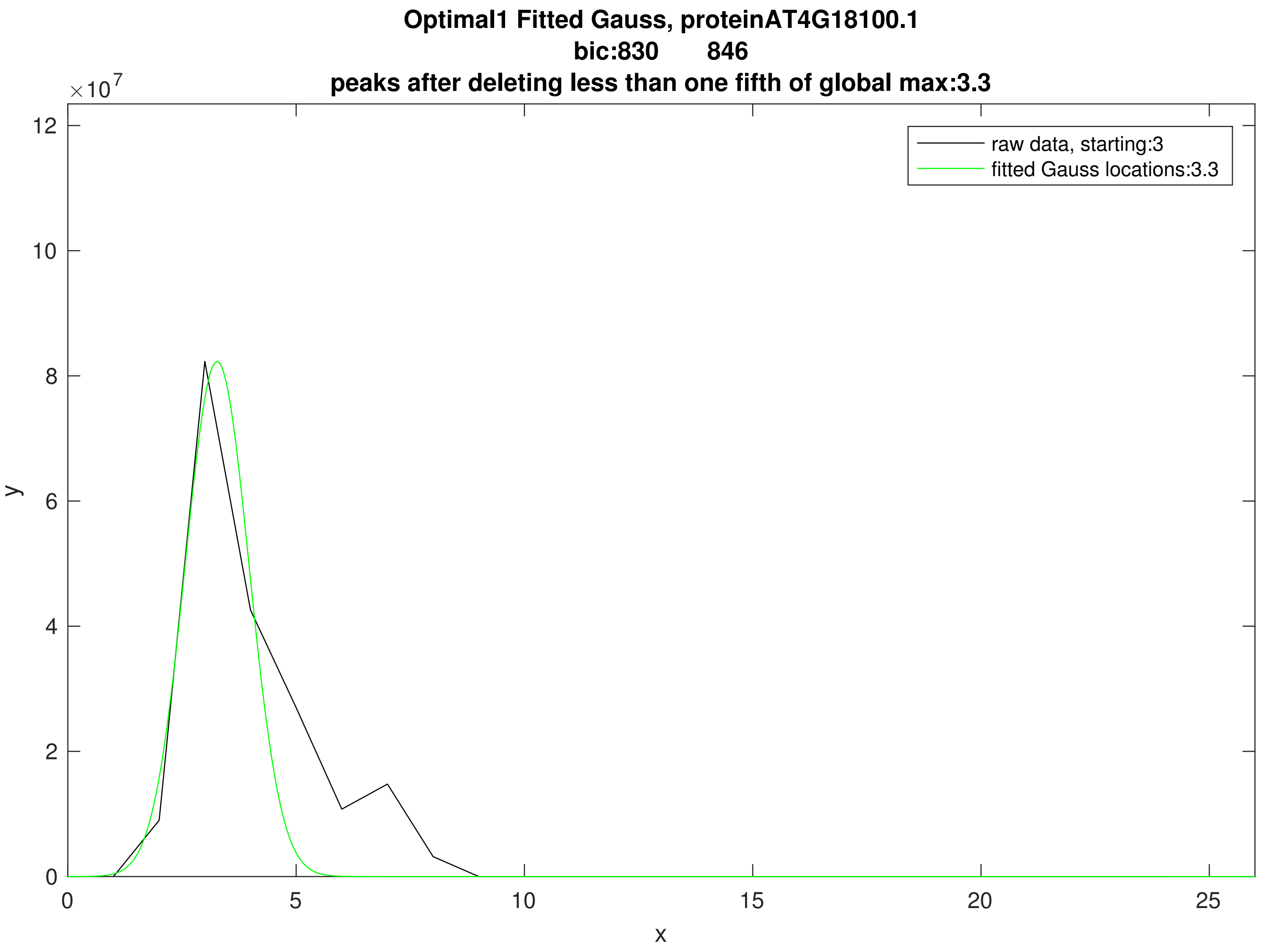

Optimal1 Fitted Gauss, proteinAT4G18100.1
bic:830 846
peaks after deleting less than one fifth of global max:3.3
×10^7
12
10
8
6
4
2
0
y
0
5
10
15
20
25
x
raw data, starting:3
fitted Gauss locations:3.3

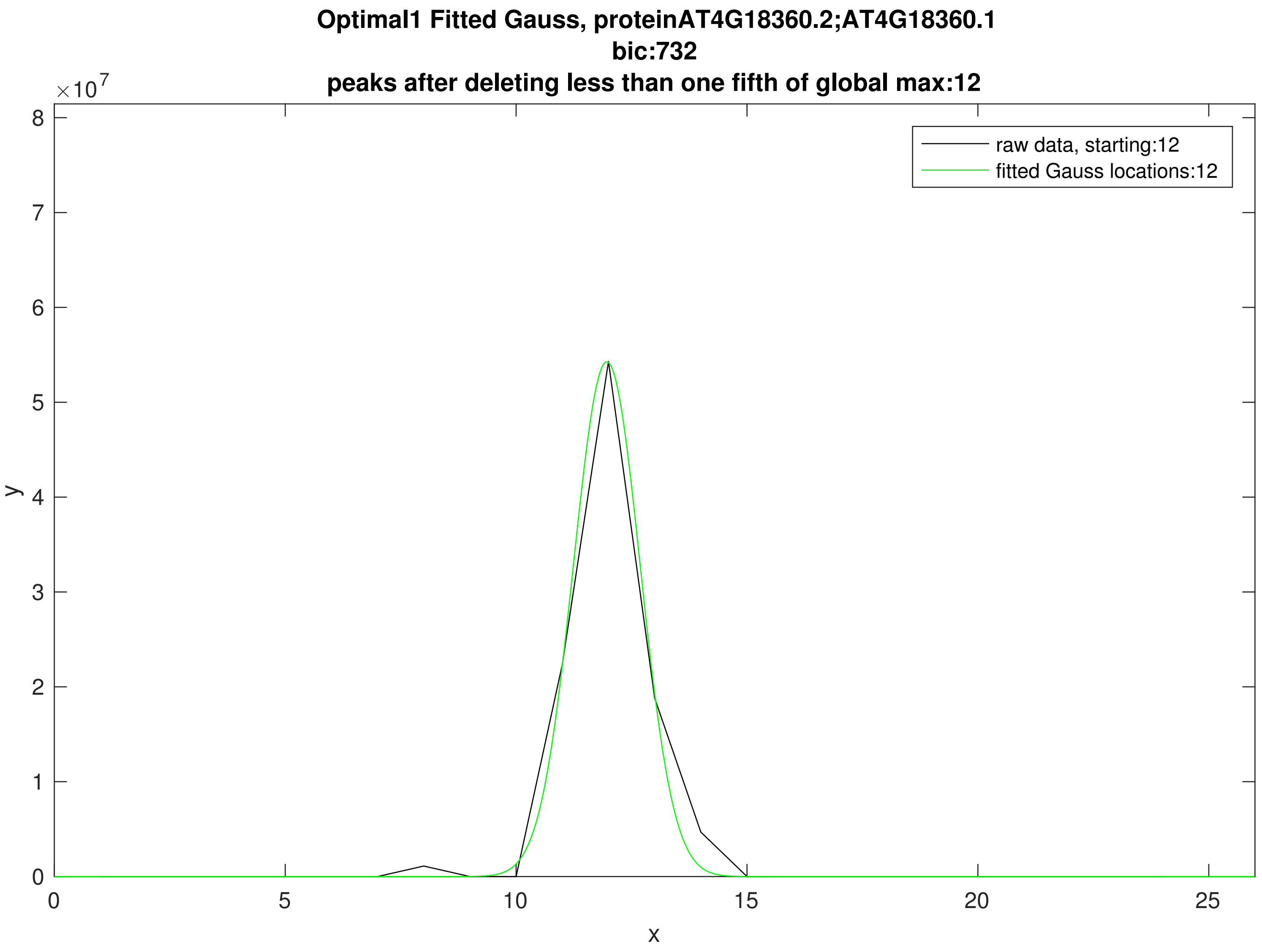
Optimal1 Fitted Gauss, proteinAT4G18360.2;AT4G18360.1
bic:732
peaks after deleting less than one fifth of global max:12
×10^7
raw data, starting:12
fitted Gauss locations:12
y
x
0
1
2
3
4
5
6
7
8
0
5
10
15
20
25

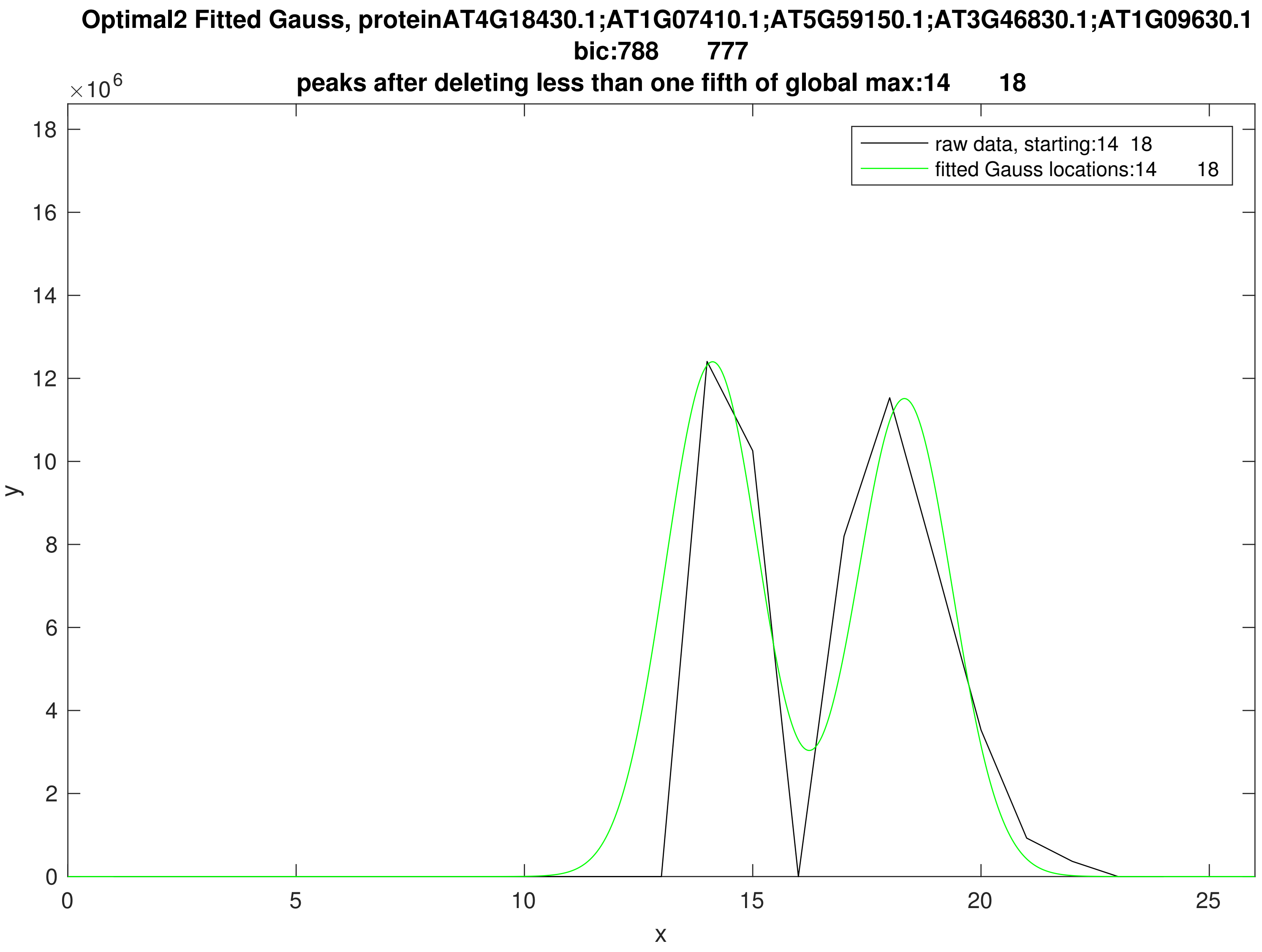
Optimal2 Fitted Gauss, proteinAT4G18430.1;AT1G07410.1;AT5G59150.1;AT3G46830.1;AT1G09630.1
bic:788 777
peaks after deleting less than one fifth of global max:14 18
×10^6
raw data, starting:14 18
fitted Gauss locations:14 18
y
x

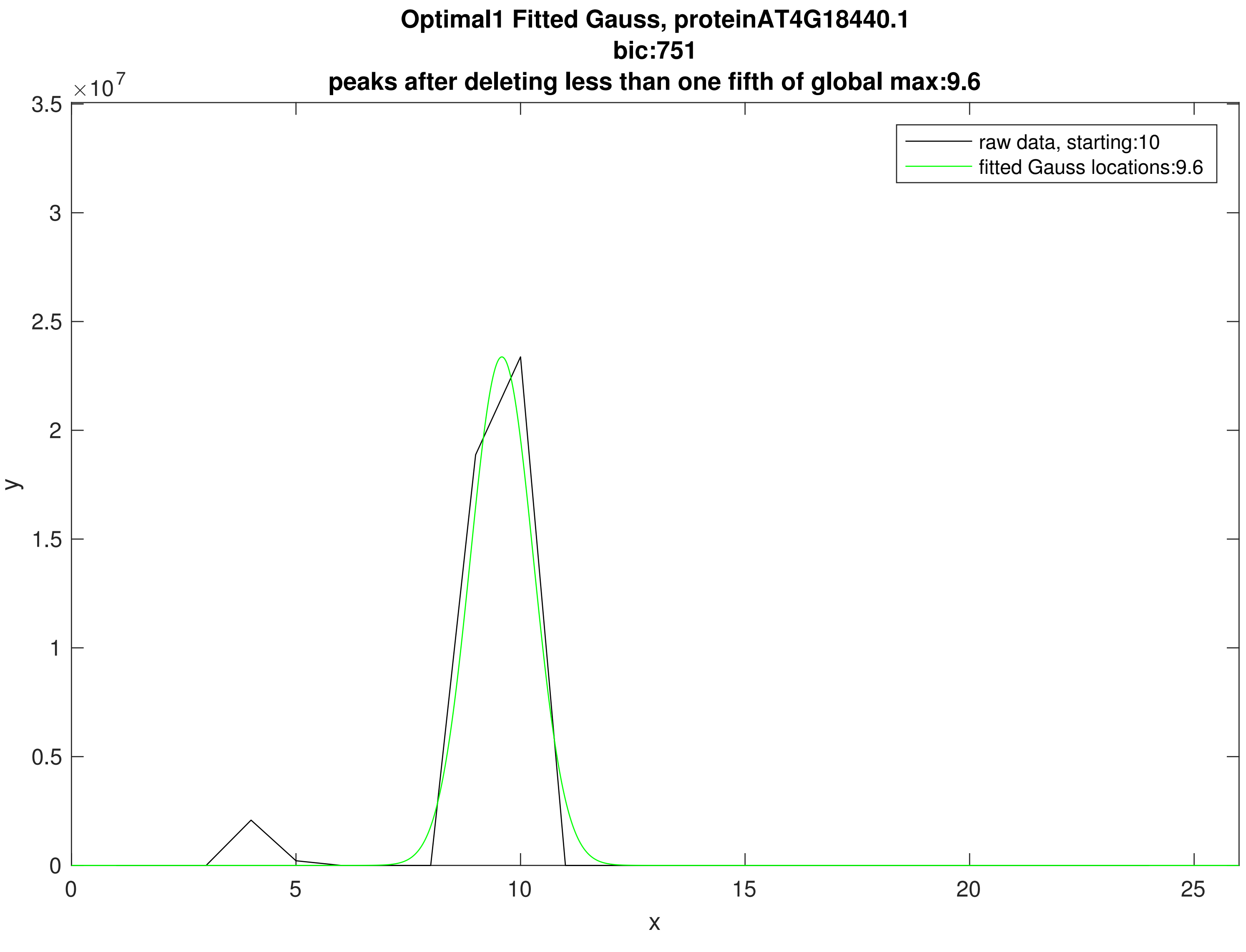
Optimal1 Fitted Gauss, proteinAT4G18440.1
bic:751
peaks after deleting less than one fifth of global max:9.6
raw data, starting:10
fitted Gauss locations:9.6
×10^7
y
x

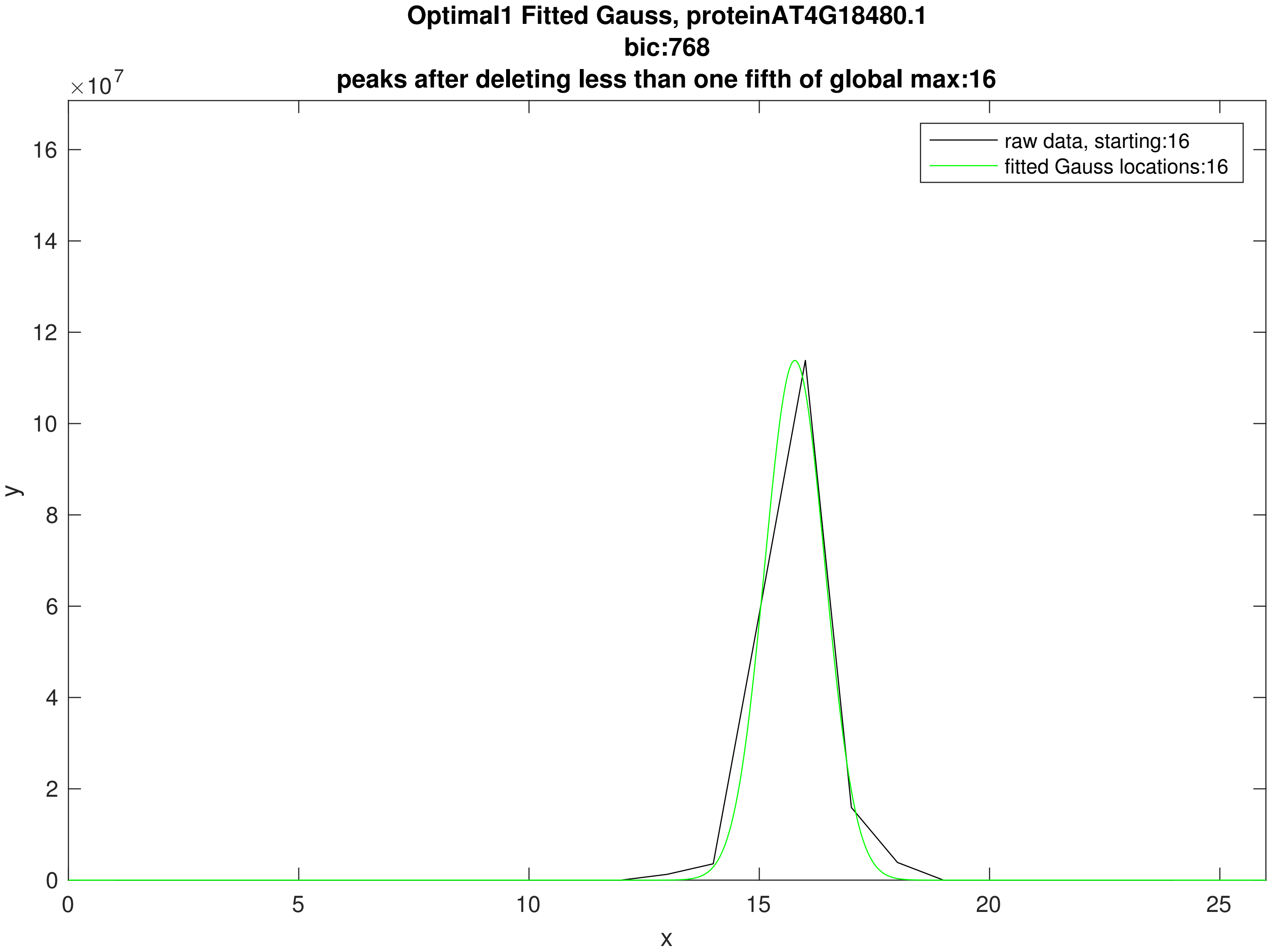
Optimal1 Fitted Gauss, proteinAT4G18480.1
bic:768
peaks after deleting less than one fifth of global max:16
×10^7
raw data, starting:16
fitted Gauss locations:16
y
x
0
2
4
6
8
10
12
14
16
0
5
10
15
20
25

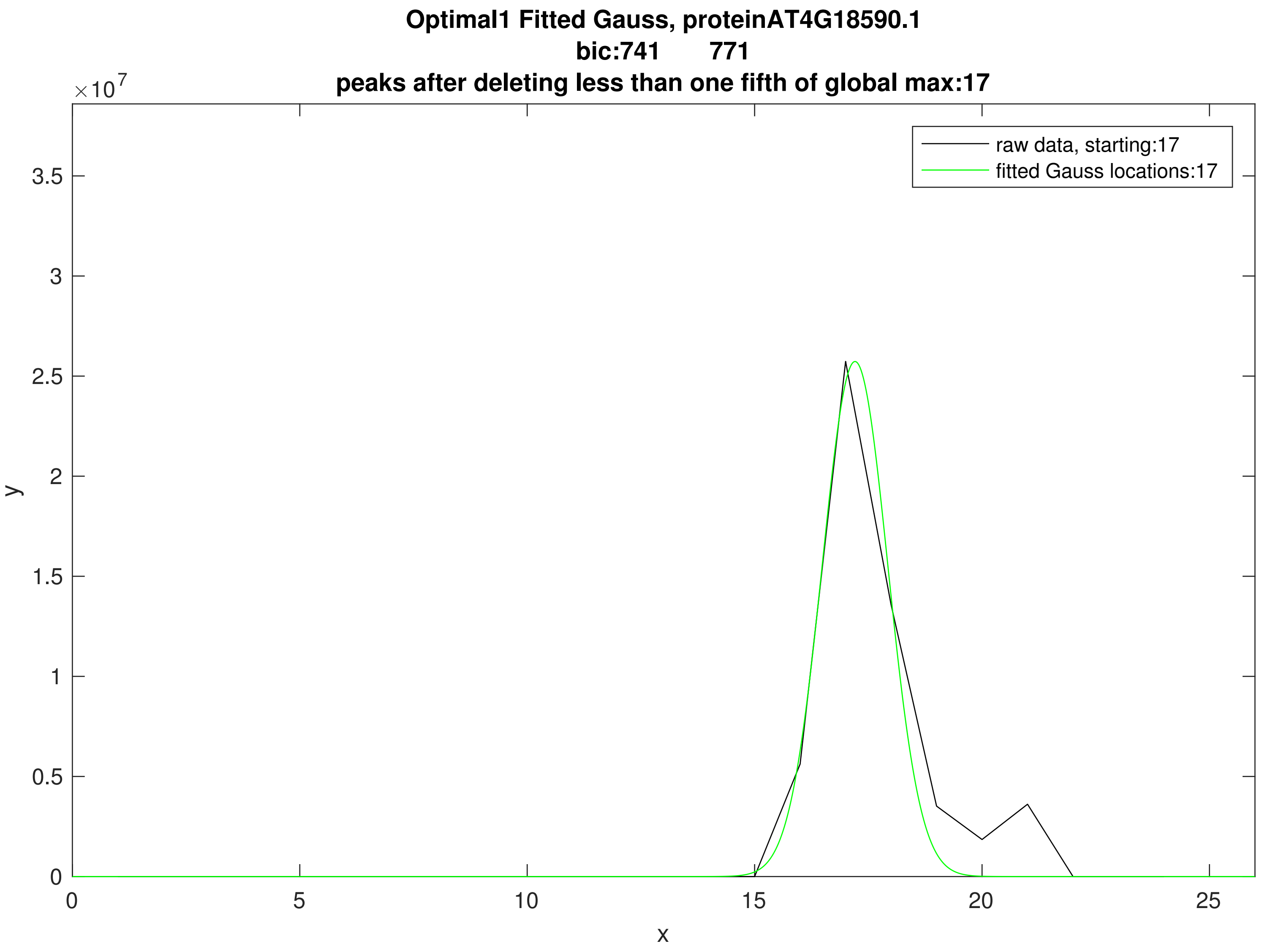

Optimal1 Fitted Gauss, proteinAT4G18590.1
bic:741 771
peaks after deleting less than one fifth of global max:17
×10^7
3.5
3
2.5
2
1.5
1
0.5
0
y
0
5
10
15
20
25
x
raw data, starting:17
fitted Gauss locations:17

Optimal2 Fitted Gauss, proteinAT4G18870.1
bic:898 870
peaks after deleting less than one fifth of global max:13 18

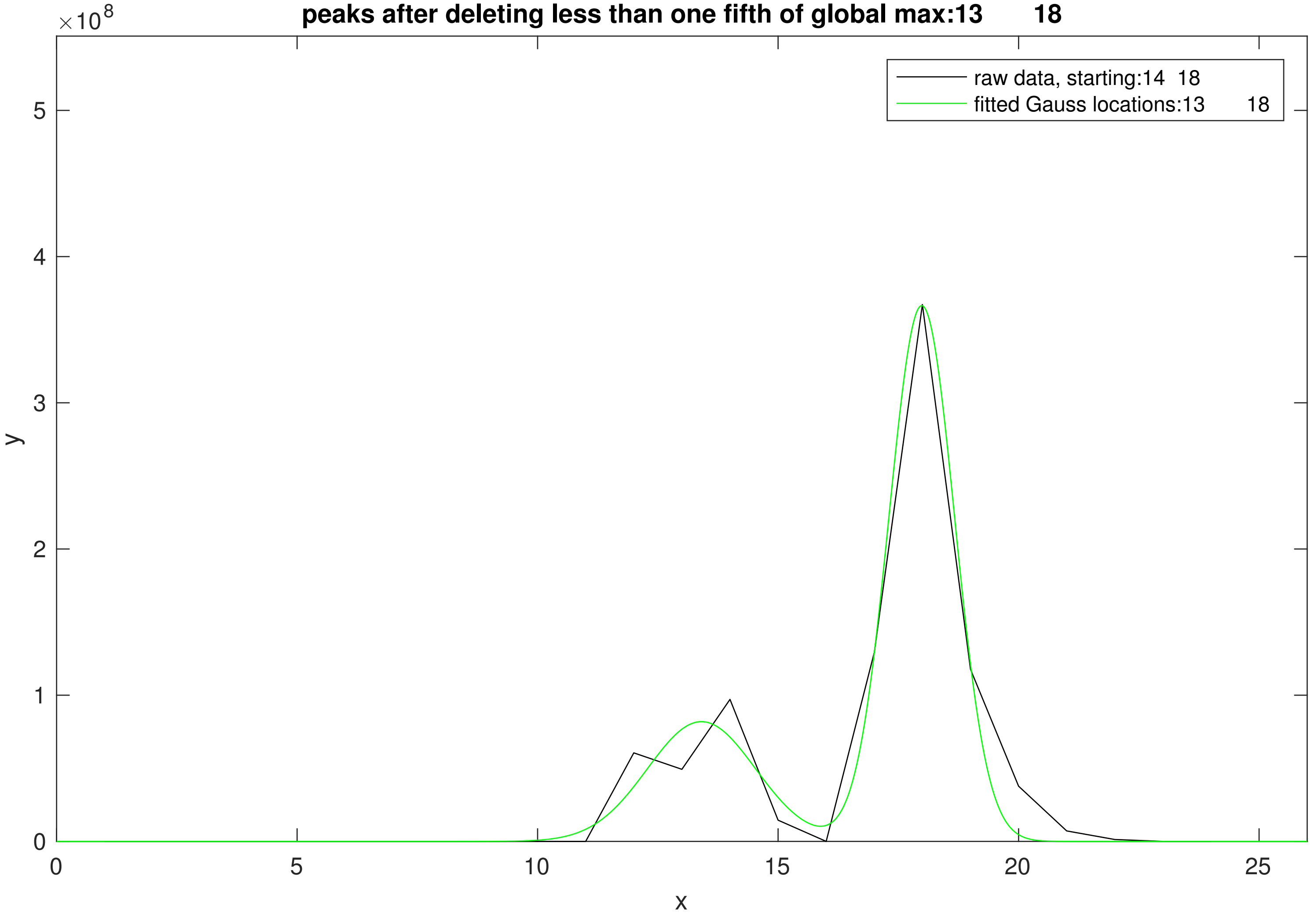

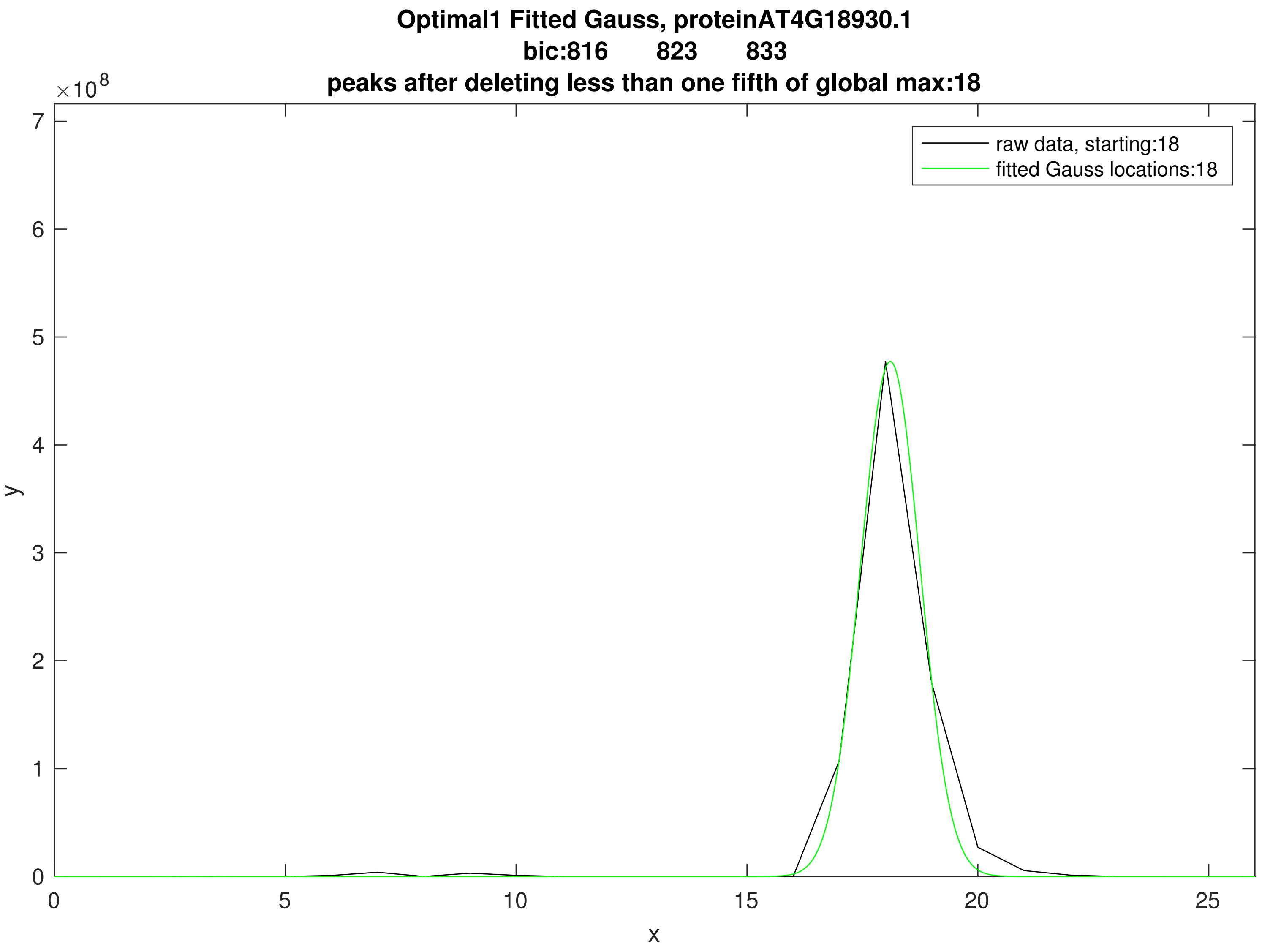
Optimal1 Fitted Gauss, proteinAT4G18930.1
bic:816 823 833
peaks after deleting less than one fifth of global max:18
×10^8
raw data, starting:18
fitted Gauss locations:18
y
x
0
1
2
3
4
5
6
7
0
5
10
15
20
25

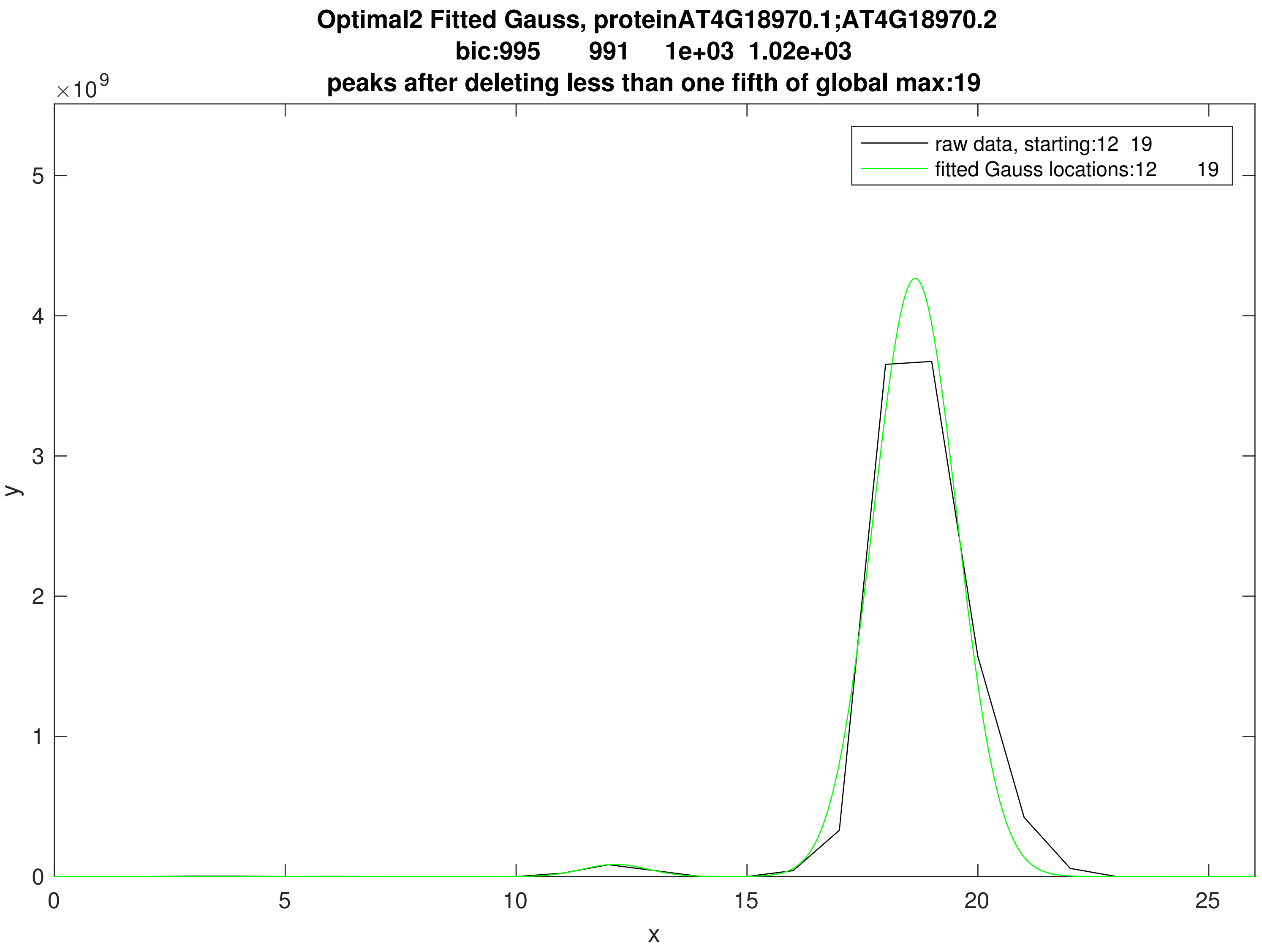

Optimal2 Fitted Gauss, proteinAT4G18970.1;AT4G18970.2
bic:995 991 1e+03 1.02e+03
peaks after deleting less than one fifth of global max:19
×10^9
raw data, starting:12 19
fitted Gauss locations:12 19
y
x
0
1
2
3
4
5
0
5
10
15
20
25

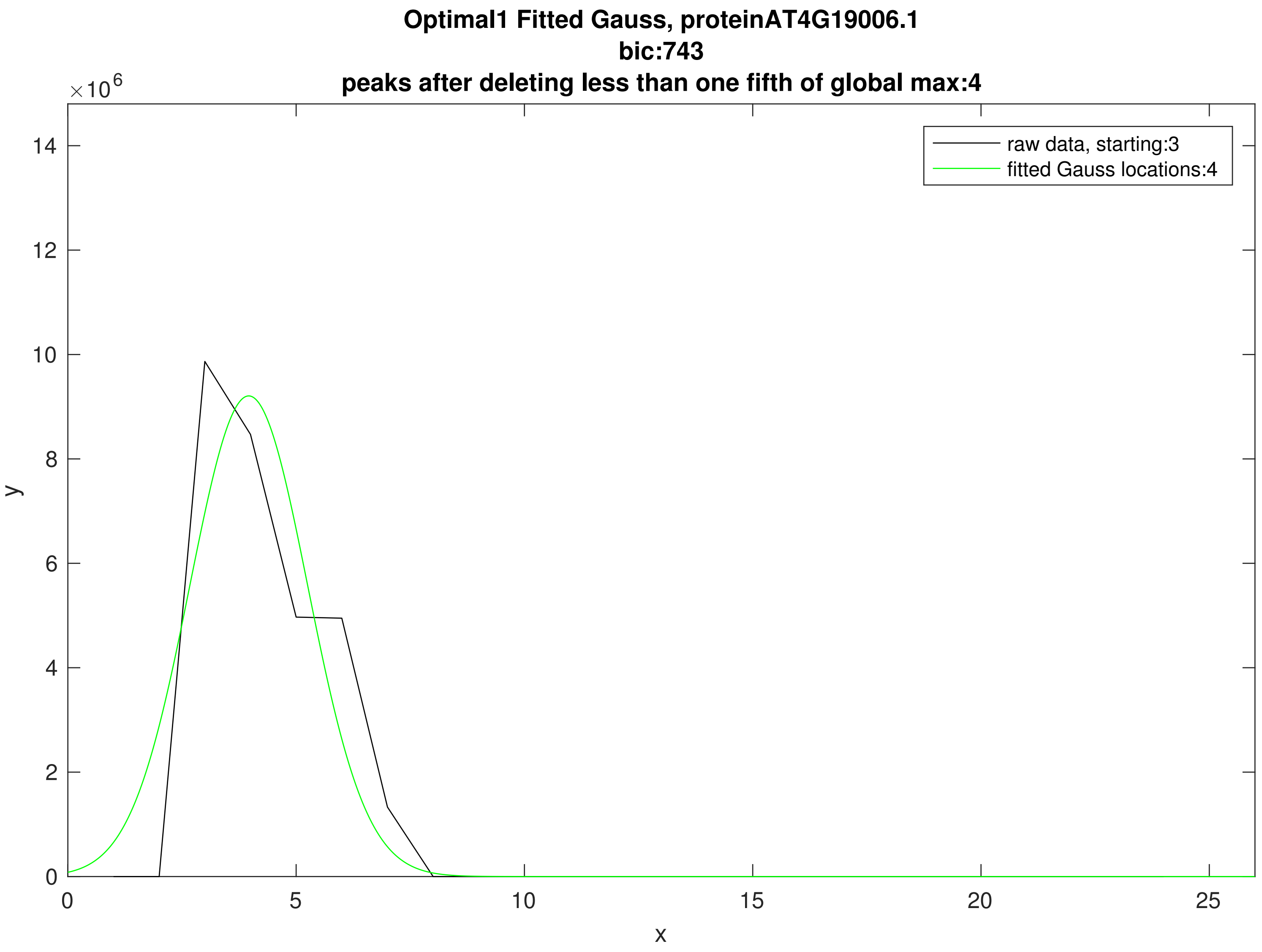
Optimal1 Fitted Gauss, proteinAT4G19006.1
bic:743
peaks after deleting less than one fifth of global max:4
×10^6
raw data, starting:3
fitted Gauss locations:4
y
x
0
2
4
6
8
10
12
14
0
5
10
15
20
25

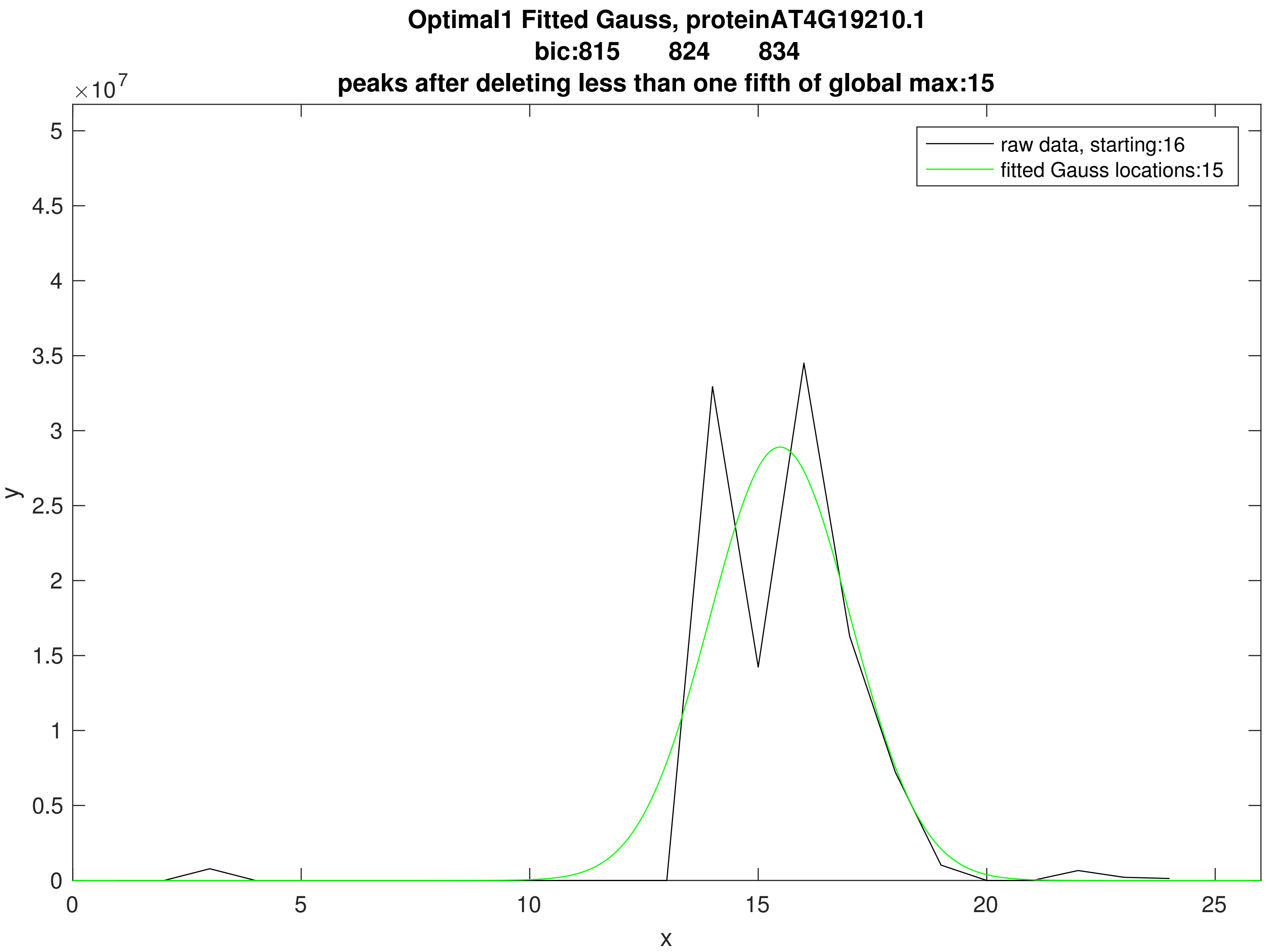
Optimal1 Fitted Gauss, proteinAT4G19210.1
bic:815 824 834
peaks after deleting less than one fifth of global max:15
×10^7
raw data, starting:16
fitted Gauss locations:15
y
x
0
0.5
1
1.5
2
2.5
3
3.5
4
4.5
5
0
5
10
15
20
25

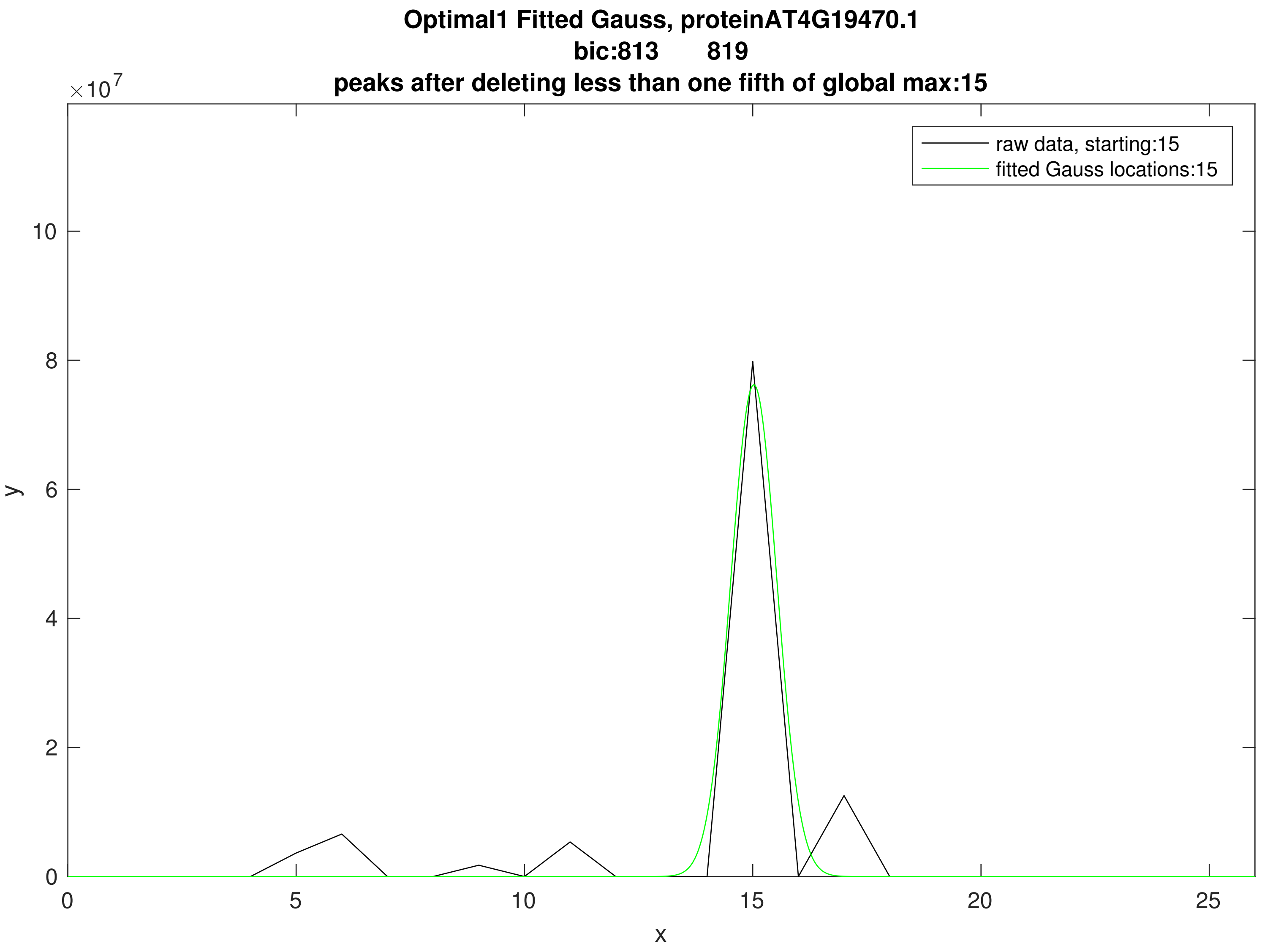
Optimal1 Fitted Gauss, proteinAT4G19470.1
bic:813 819
peaks after deleting less than one fifth of global max:15
×10^7
raw data, starting:15
fitted Gauss locations:15
y
x
0
2
4
6
8
10
0
5
10
15
20
25

Optimal1 Fitted Gauss, proteinAT4G19710.2;AT4G19710.1
bic:768 776
peaks after deleting less than one fifth of global max:11

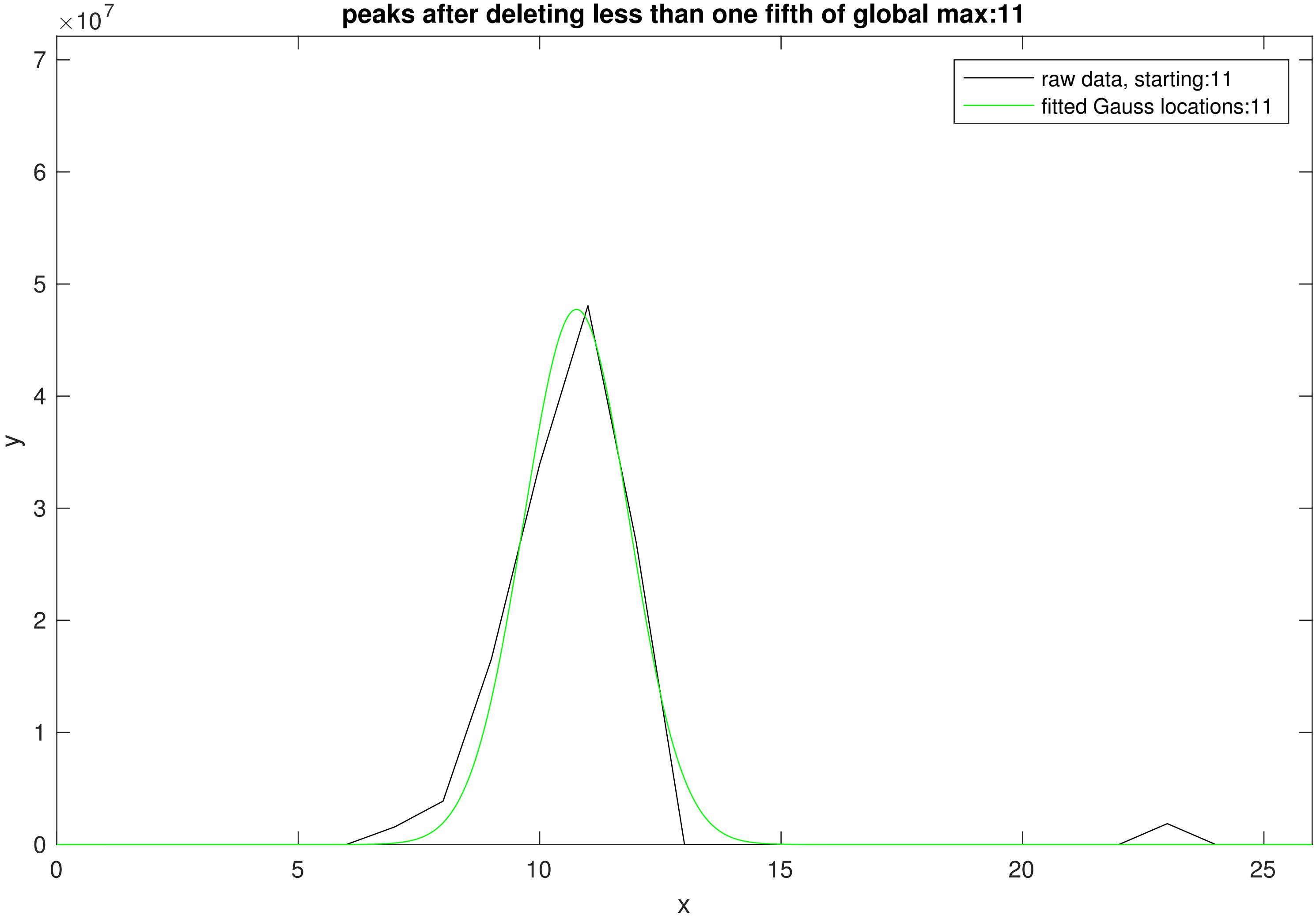

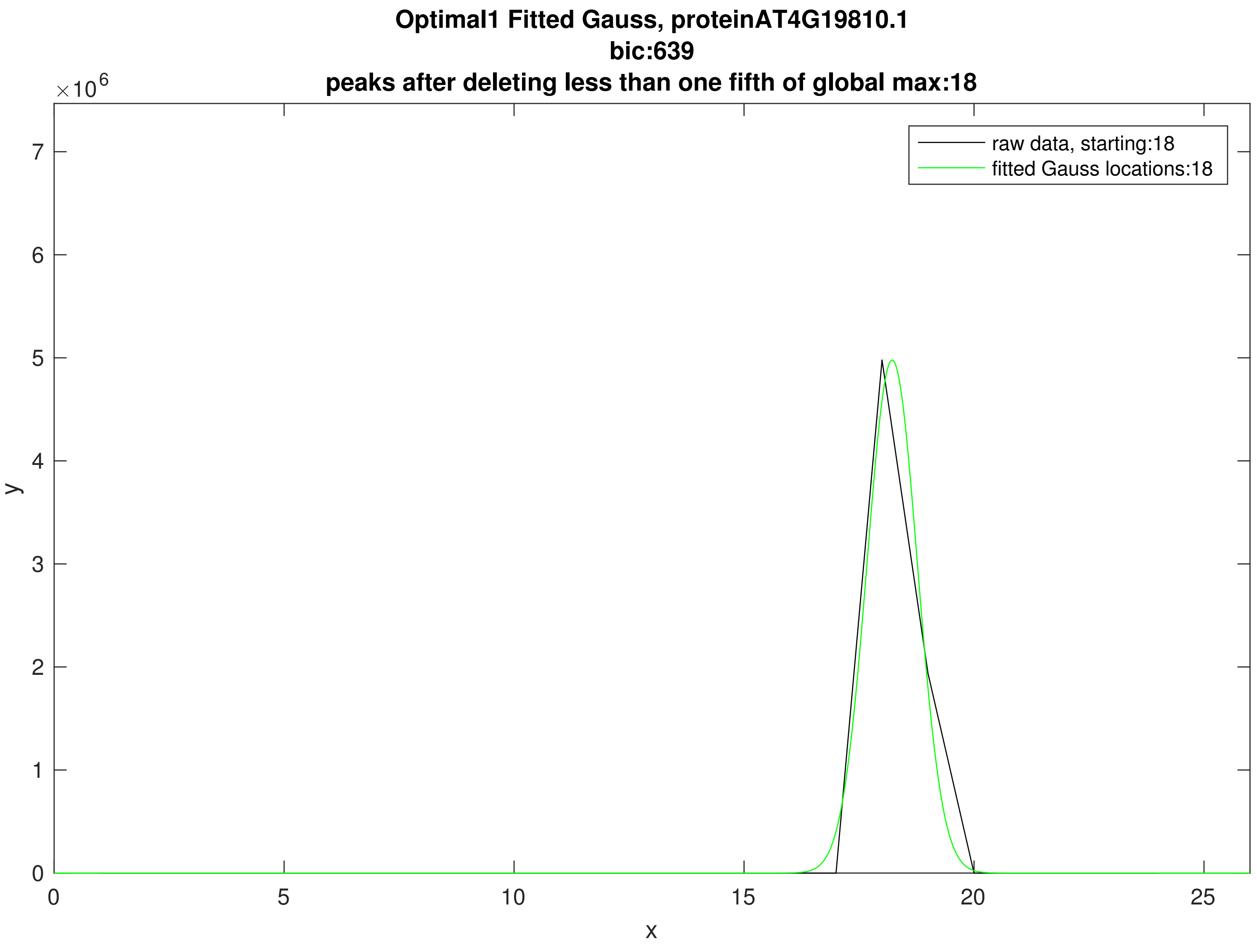
Optimal1 Fitted Gauss, proteinAT4G19810.1
bic:639
peaks after deleting less than one fifth of global max:18
×10^6
raw data, starting:18
fitted Gauss locations:18
y
x
0
1
2
3
4
5
6
7
0
5
10
15
20
25

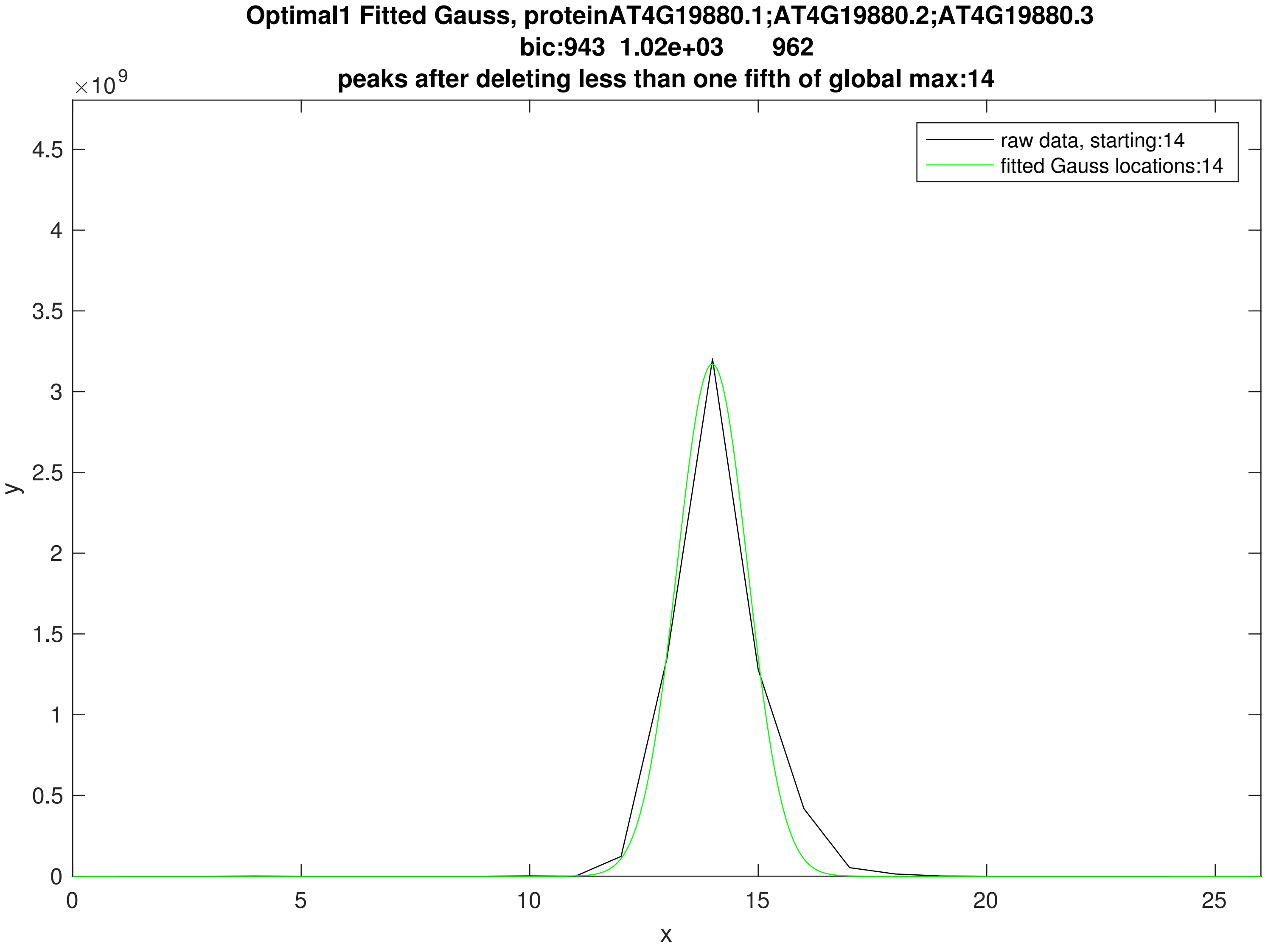
Optimal1 Fitted Gauss, proteinAT4G19880.1;AT4G19880.2;AT4G19880.3
bic:943 1.02e+03 962
peaks after deleting less than one fifth of global max:14
×10^9
4.5
4
3.5
3
2.5
2
1.5
1
0.5
0
y
0
5
10
15
20
25
x
raw data, starting:14
fitted Gauss locations:14

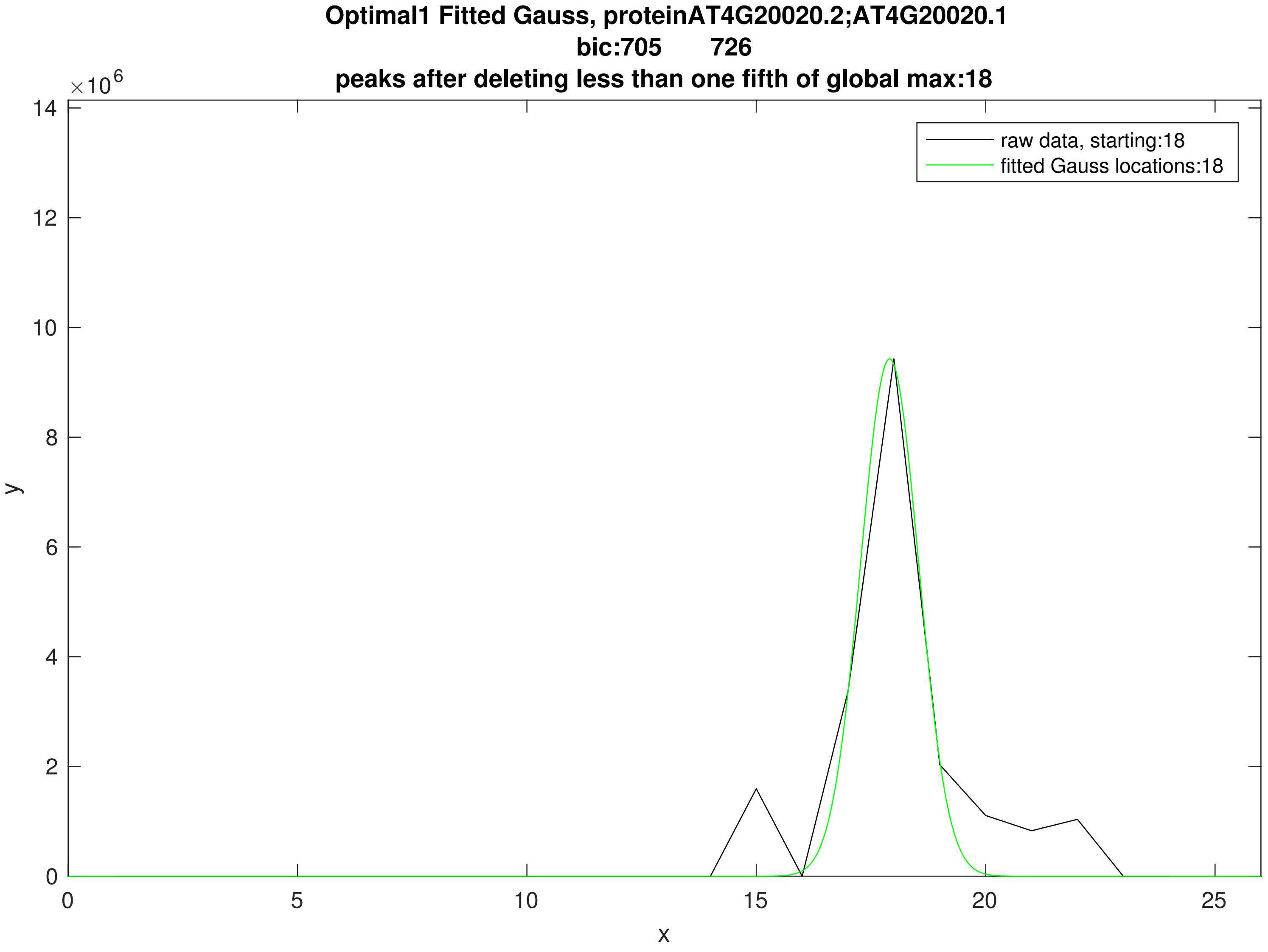
Optimal1 Fitted Gauss, proteinAT4G20020.2;AT4G20020.1
bic:705 726
peaks after deleting less than one fifth of global max:18
×10^6
14
12
10
8
6
4
2
0
y
0
5
10
15
20
25
x
raw data, starting:18
fitted Gauss locations:18

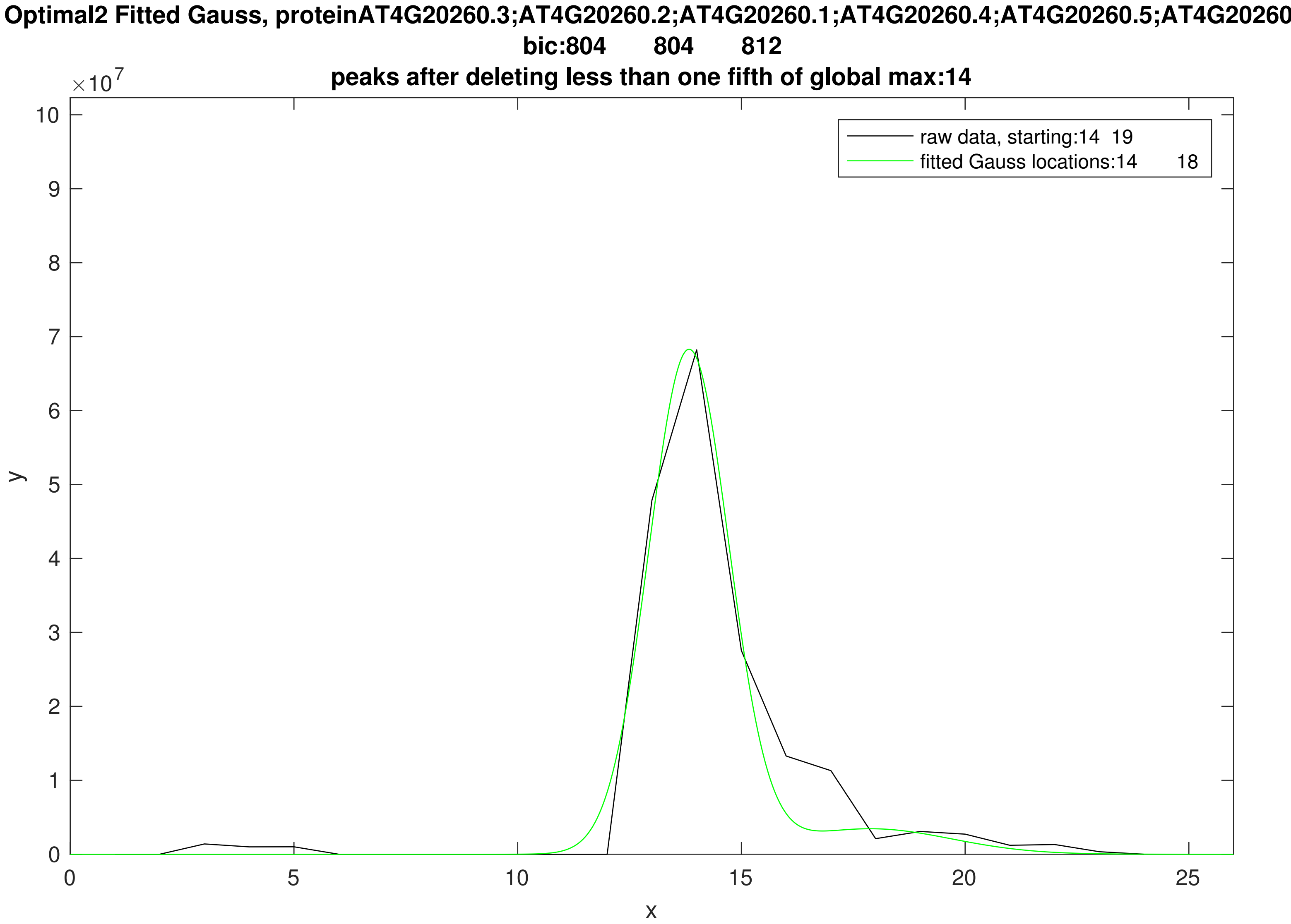
Optimal2 Fitted Gauss, proteinAT4G20260.3;AT4G20260.2;AT4G20260.1;AT4G20260.4;AT4G20260.5;AT4G20260
bic:804 804 812
peaks after deleting less than one fifth of global max:14
×10^7
raw data, starting:14 19
fitted Gauss locations:14 18
y
x
0
1
2
3
4
5
6
7
8
9
10
0
5
10
15
20
25

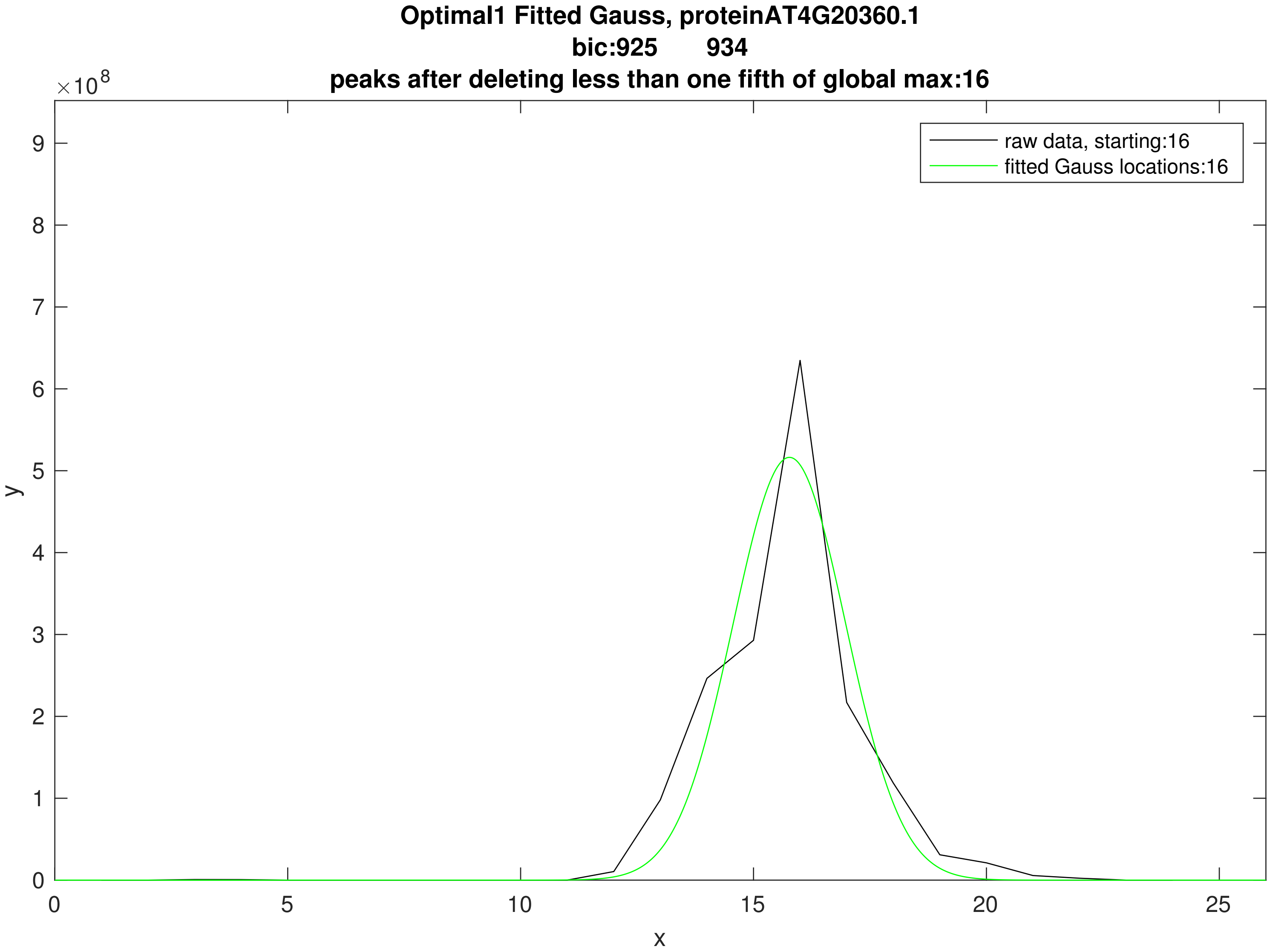
Optimal1 Fitted Gauss, proteinAT4G20360.1
bic:925 934
peaks after deleting less than one fifth of global max:16
×10^8
raw data, starting:16
fitted Gauss locations:16
y
x
0
1
2
3
4
5
6
7
8
9
0
5
10
15
20
25

#### Optimal1 Fitted Gauss, proteinAT4G20780.1
#### bic:724
#### peaks after deleting less than one fifth of global max:18

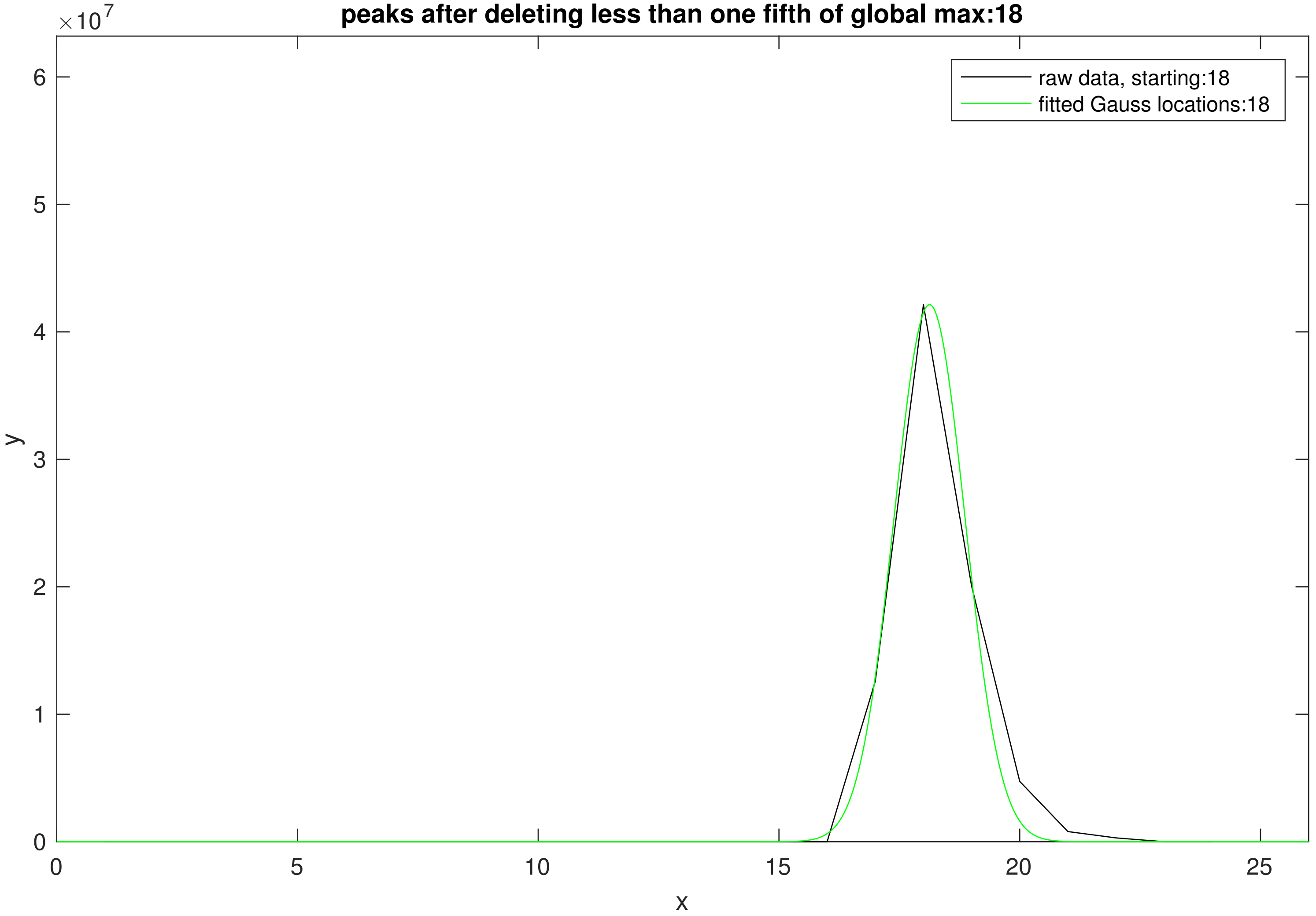

**Optimal1 Fitted Gauss, proteinAT4G20840.1**
**bic:806**
**peaks after deleting less than one fifth of global max:15**

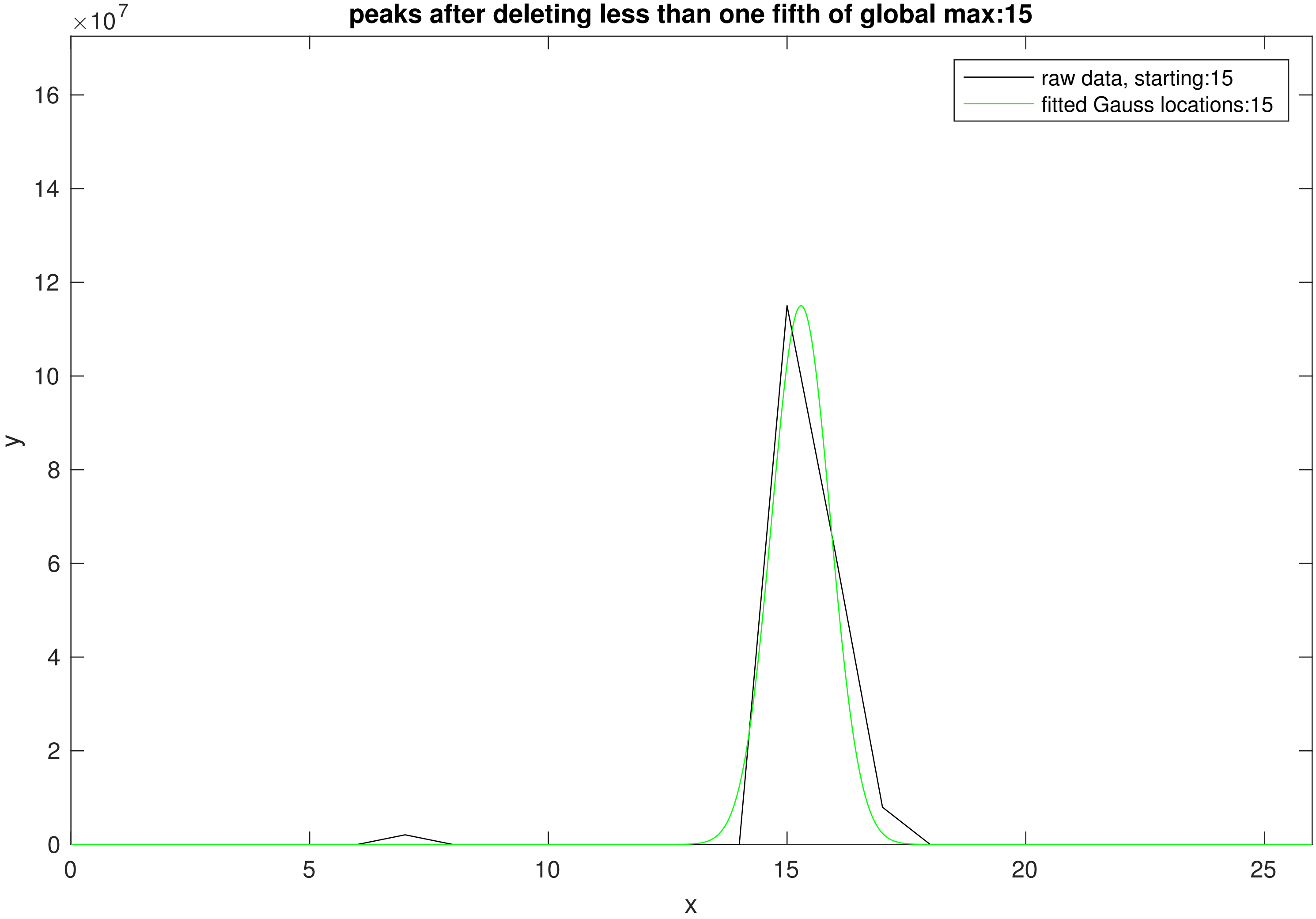

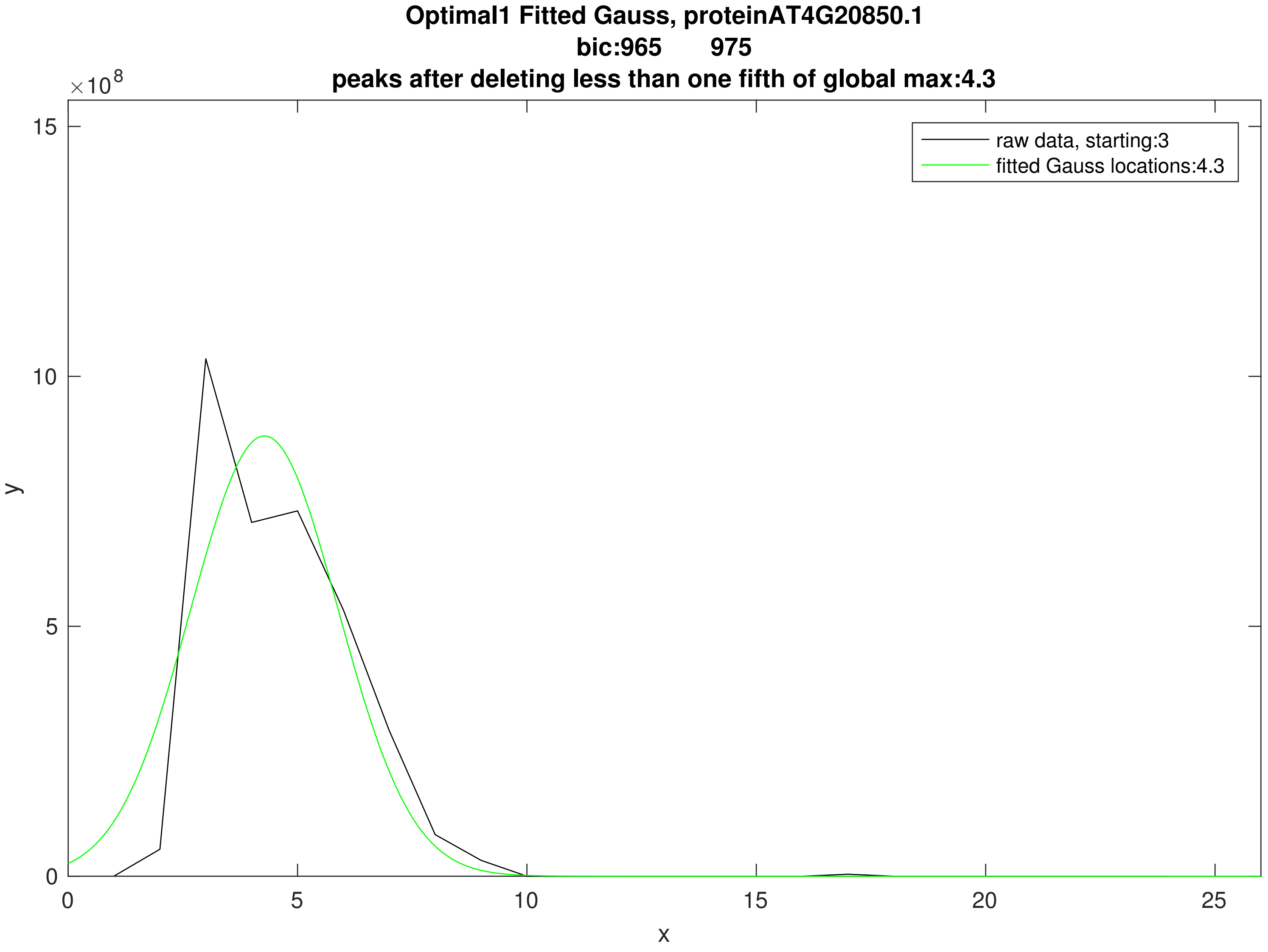
Optimal1 Fitted Gauss, proteinAT4G20850.1
bic:965 975
peaks after deleting less than one fifth of global max:4.3
×10^8
15
10
5
0
y
0
5
10
15
20
25
x
raw data, starting:3
fitted Gauss locations:4.3

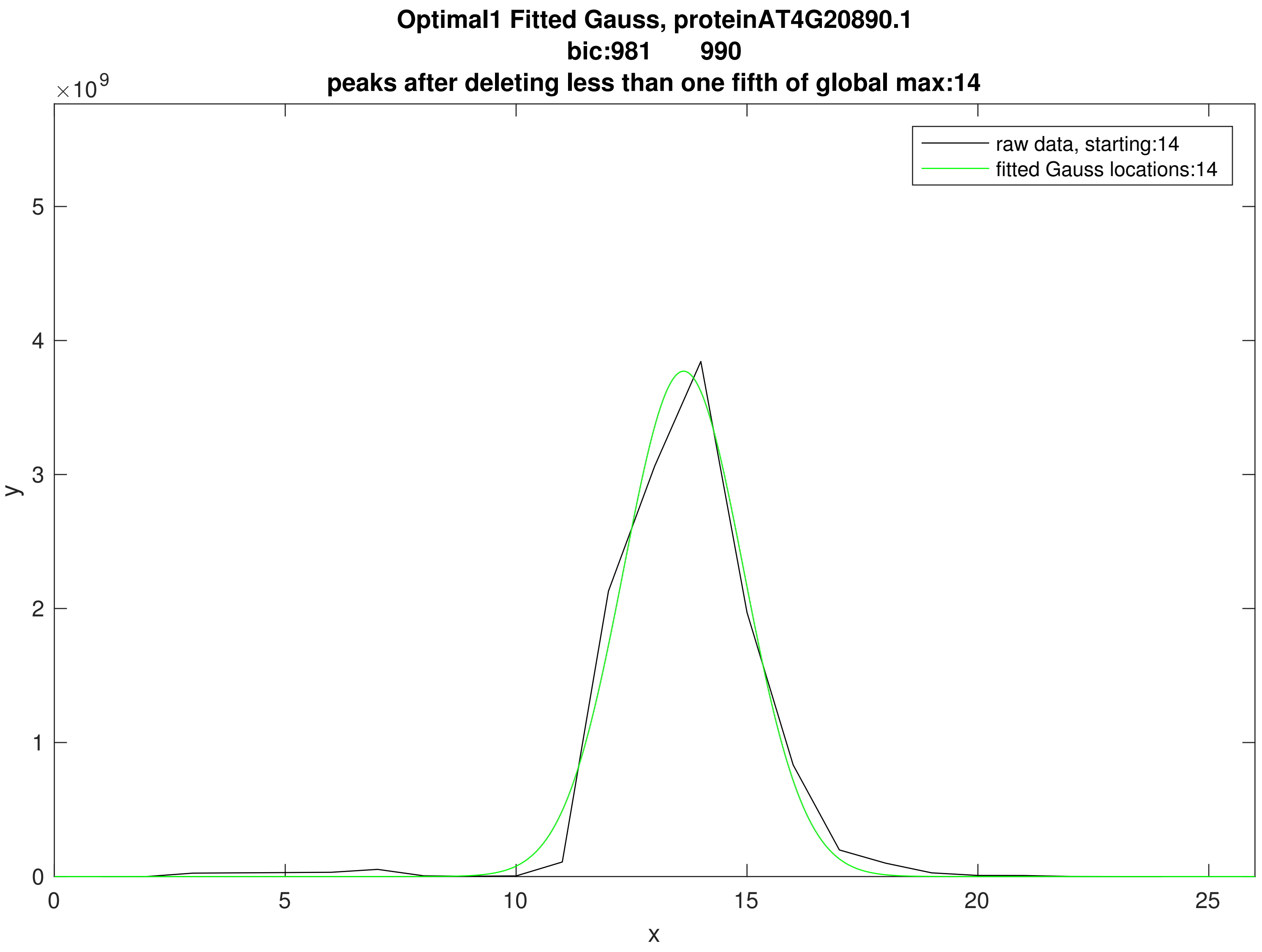
Optimal1 Fitted Gauss, proteinAT4G20890.1
bic:981 990
peaks after deleting less than one fifth of global max:14
×10^9
raw data, starting:14
fitted Gauss locations:14
y
x
0
1
2
3
4
5
10
15
20
25

#### Optimal1 Fitted Gauss, proteinAT4G21020.1
#### bic:899 904
#### peaks after deleting less than one fifth of global max:12

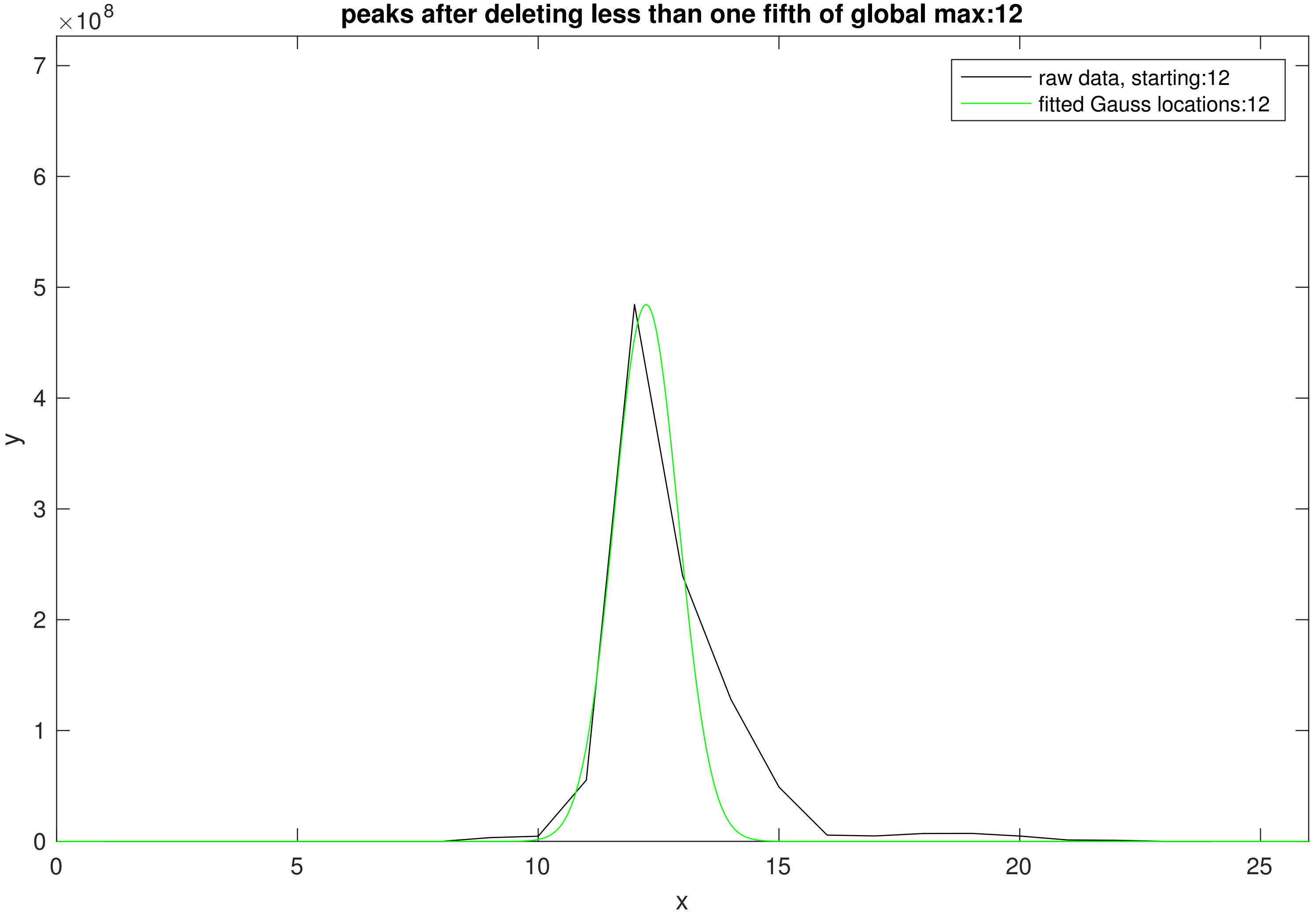

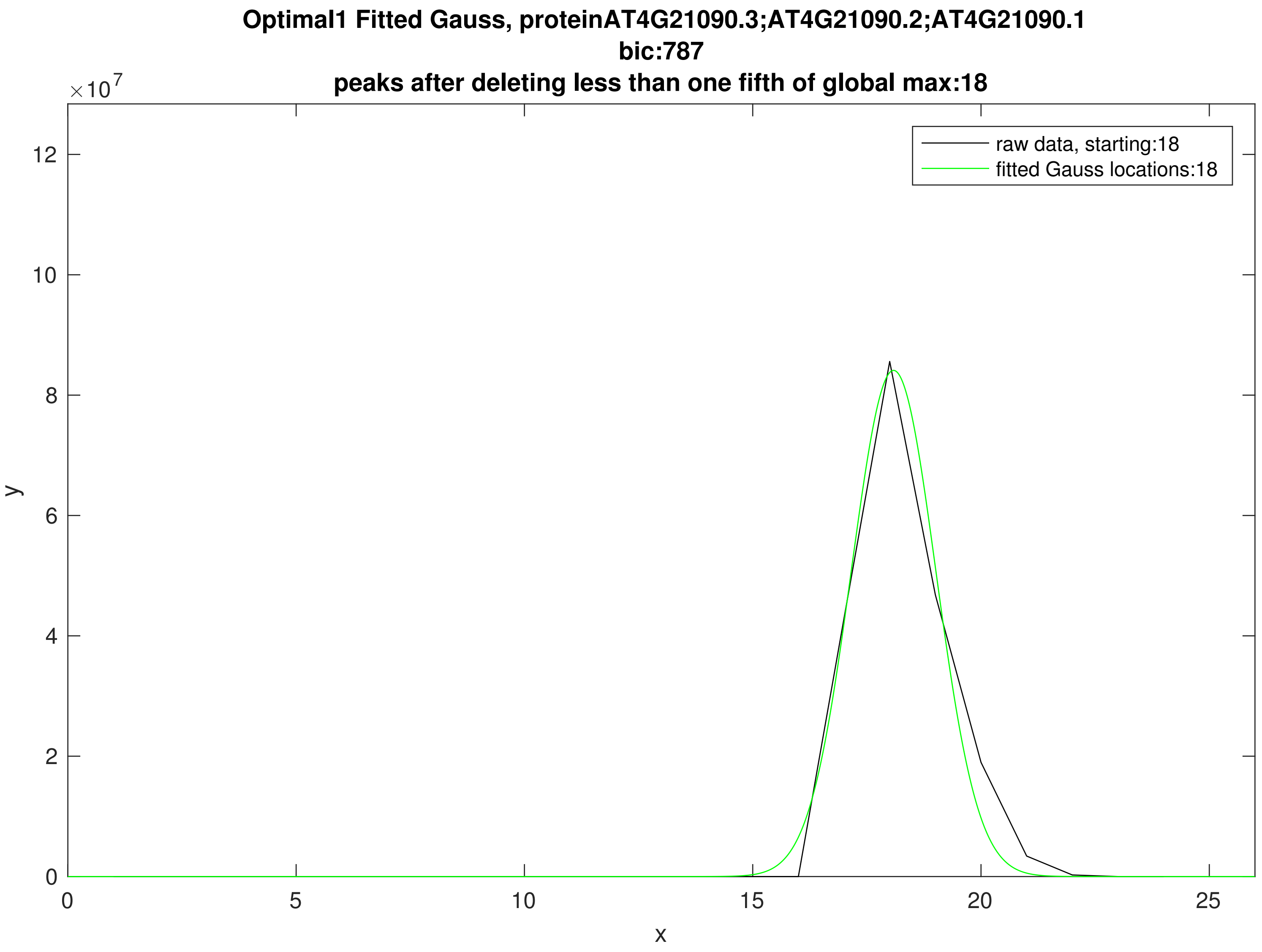
Optimal1 Fitted Gauss, proteinAT4G21090.3;AT4G21090.2;AT4G21090.1
bic:787
peaks after deleting less than one fifth of global max:18
×10^7
12
10
8
6
4
2
0
y
0
5
10
15
20
25
x
raw data, starting:18
fitted Gauss locations:18

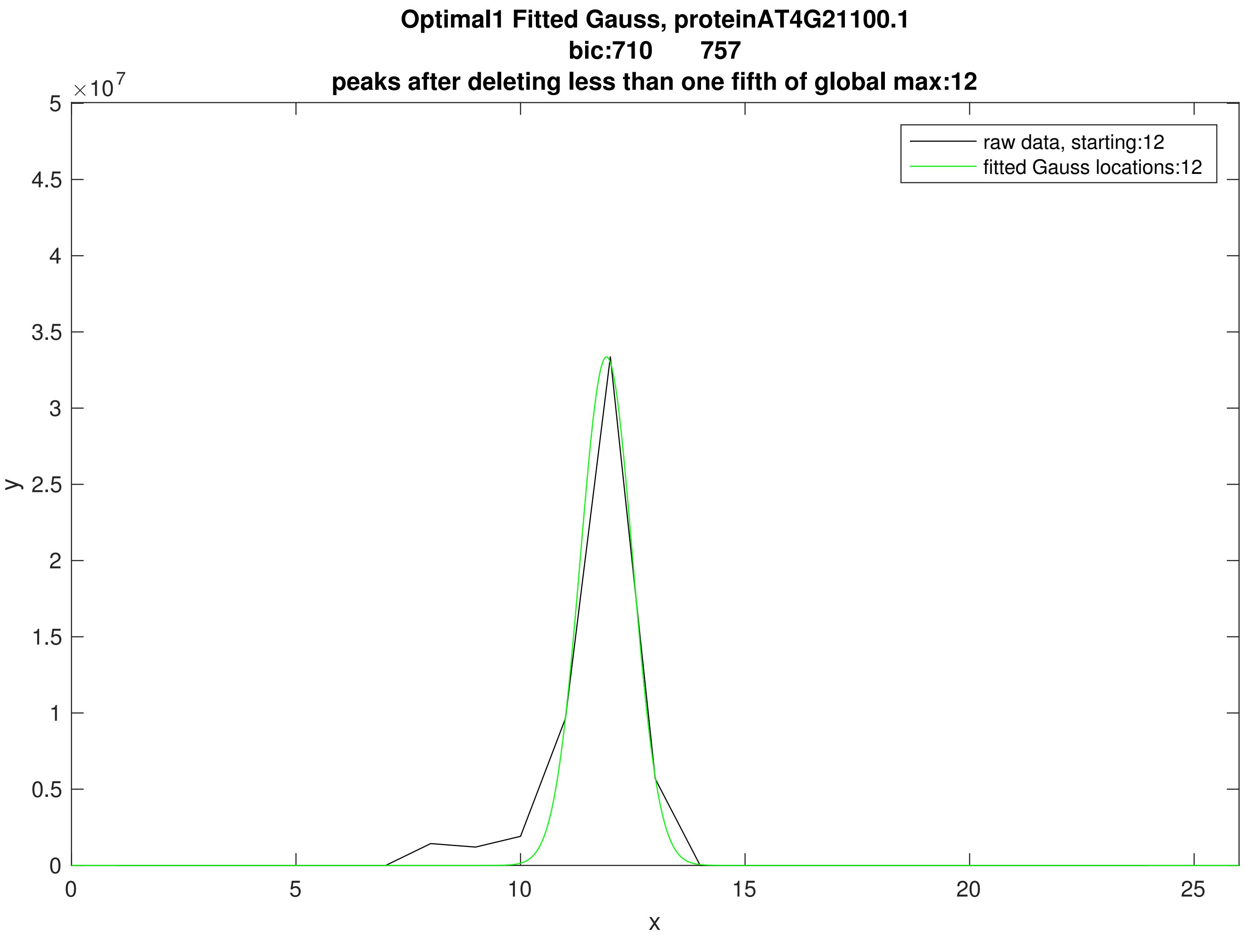
Optimal1 Fitted Gauss, proteinAT4G21100.1
bic:710 757
peaks after deleting less than one fifth of global max:12
×10^7
y
x
raw data, starting:12
fitted Gauss locations:12

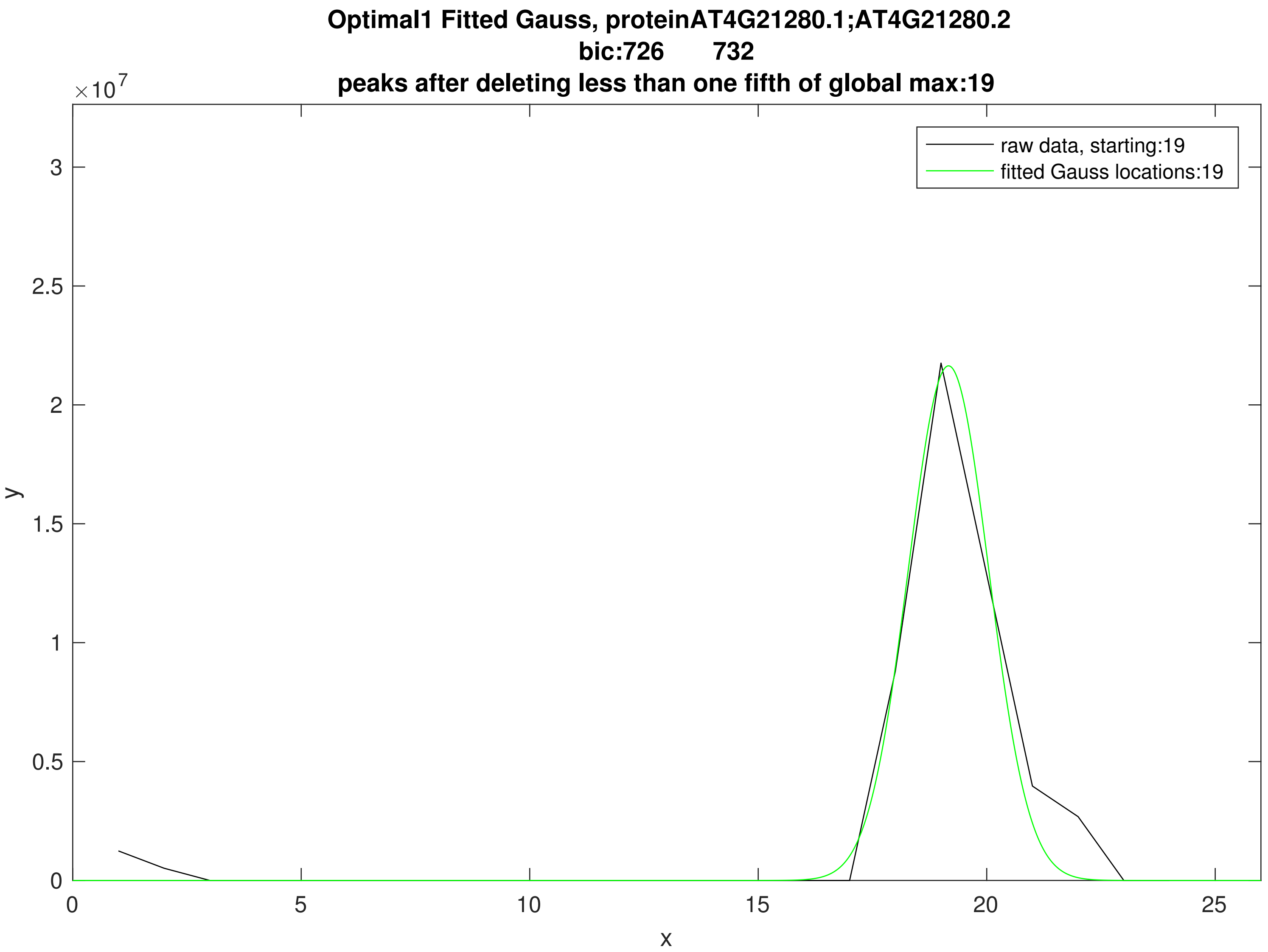

Optimal1 Fitted Gauss, proteinAT4G21280.1;AT4G21280.2
bic:726 732
peaks after deleting less than one fifth of global max:19
×10^7
3
2.5
2
1.5
1
0.5
0
y
0
5
10
15
20
25
x
raw data, starting:19
fitted Gauss locations:19

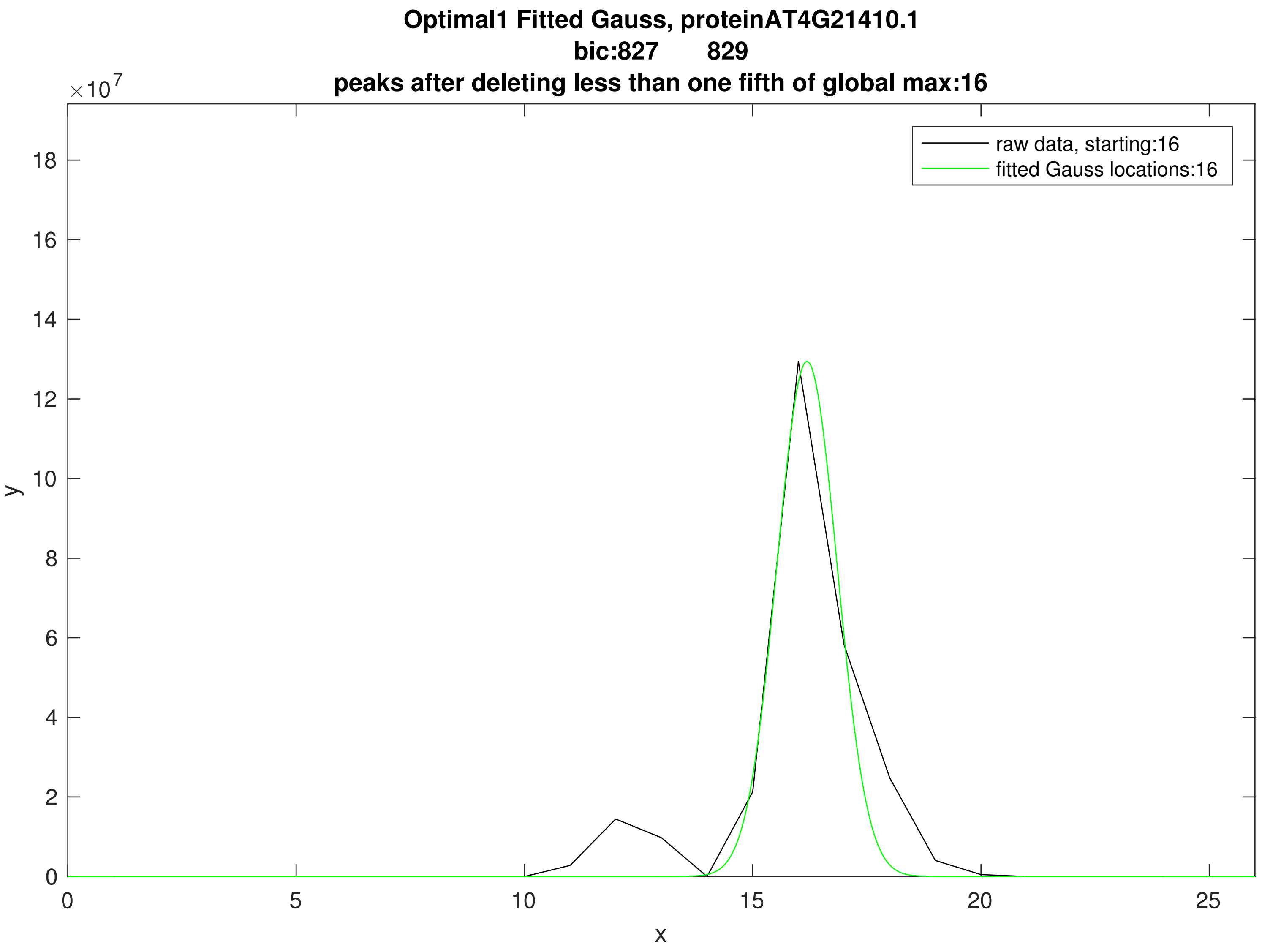

Optimal1 Fitted Gauss, proteinAT4G21410.1
bic:827 829
peaks after deleting less than one fifth of global max:16
×10^7
raw data, starting:16
fitted Gauss locations:16
y
x
0
2
4
6
8
10
12
14
16
18
0
5
10
15
20
25

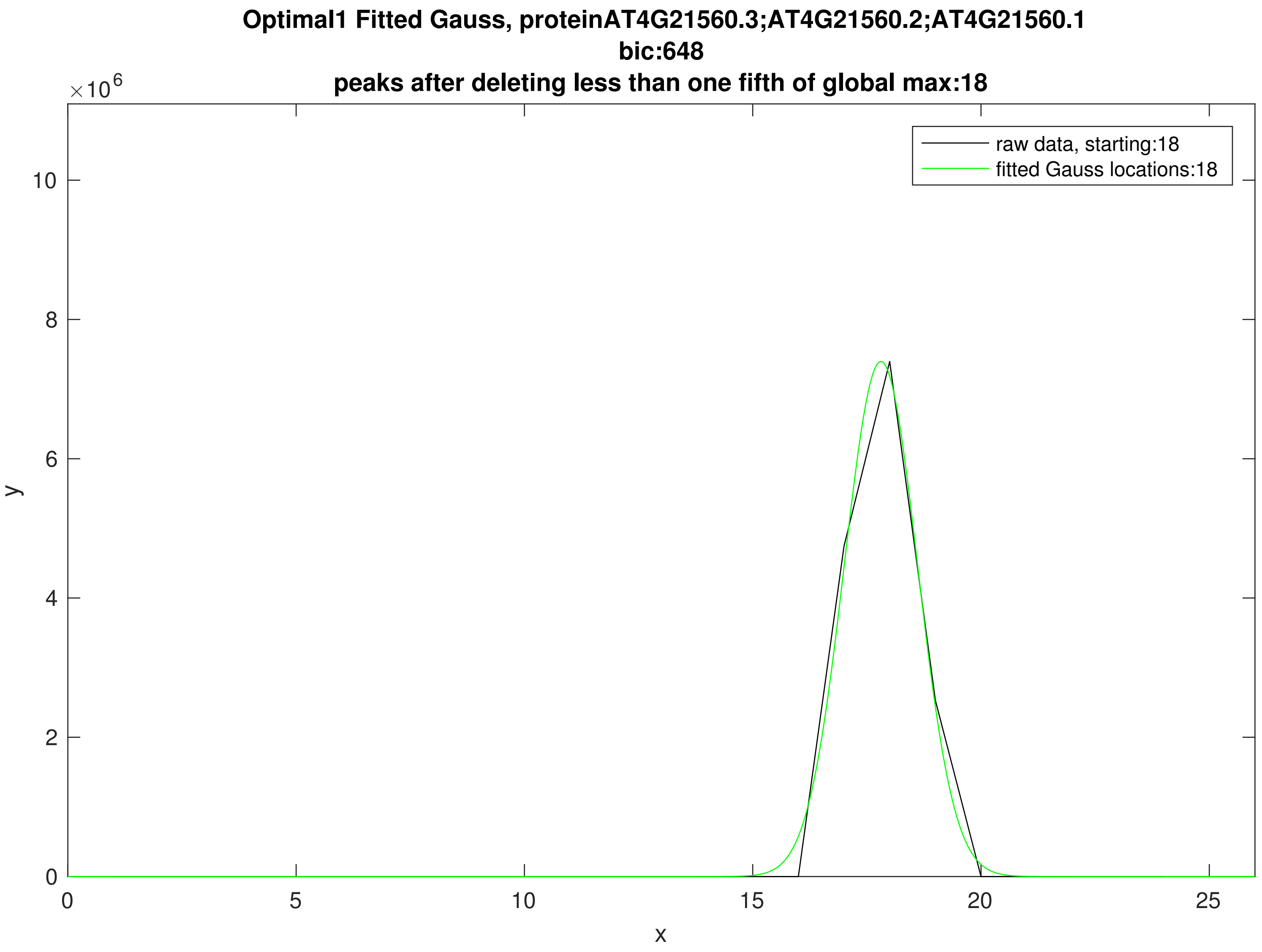

Optimal1 Fitted Gauss, proteinAT4G21560.3;AT4G21560.2;AT4G21560.1
bic:648
peaks after deleting less than one fifth of global max:18
×10^6
raw data, starting:18
fitted Gauss locations:18
10
8
6
4
2
0
y
0
5
10
15
20
25
x

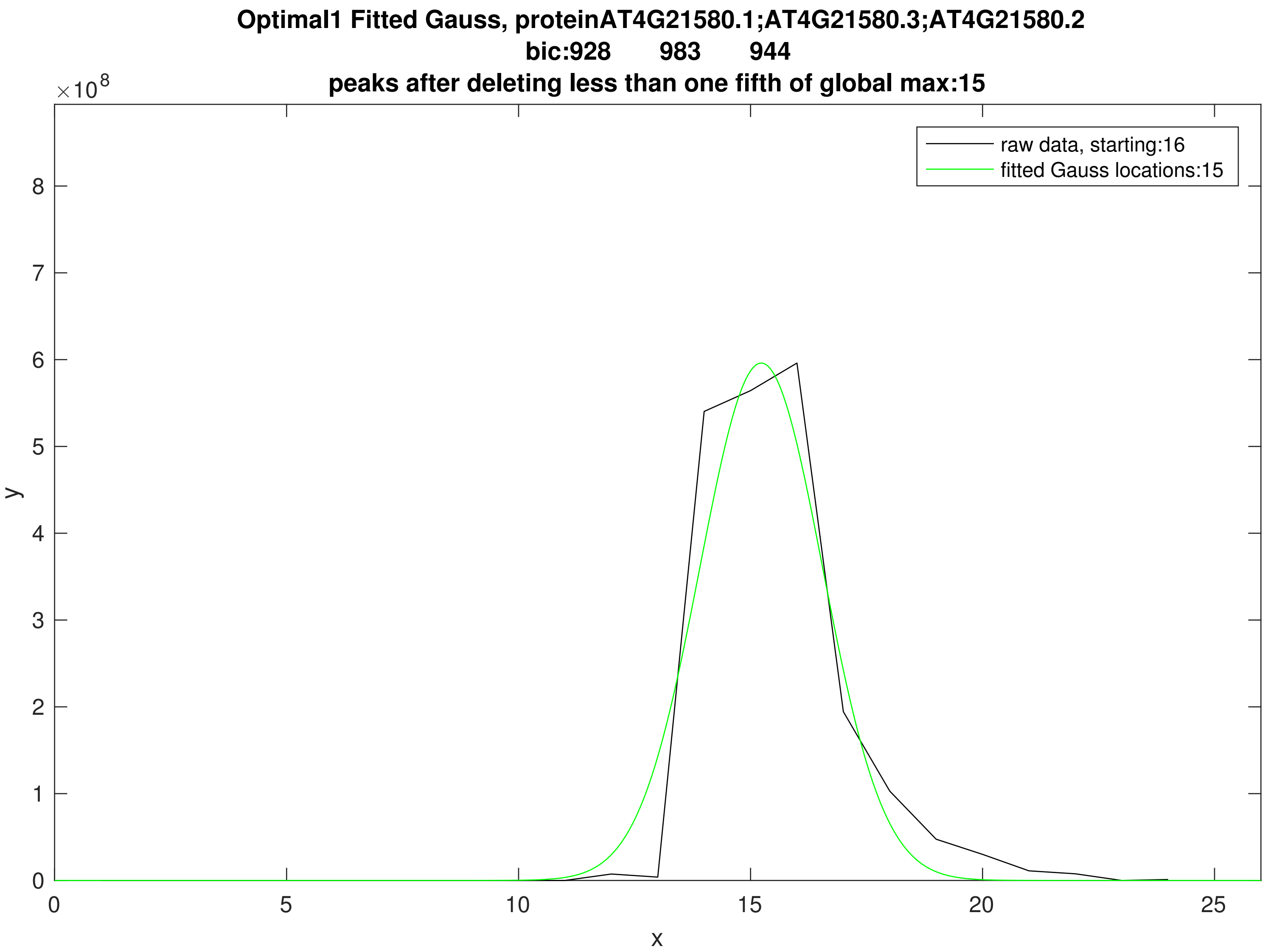
Optimal1 Fitted Gauss, proteinAT4G21580.1;AT4G21580.3;AT4G21580.2
bic:928 983 944
peaks after deleting less than one fifth of global max:15
×10^8
raw data, starting:16
fitted Gauss locations:15
y
x

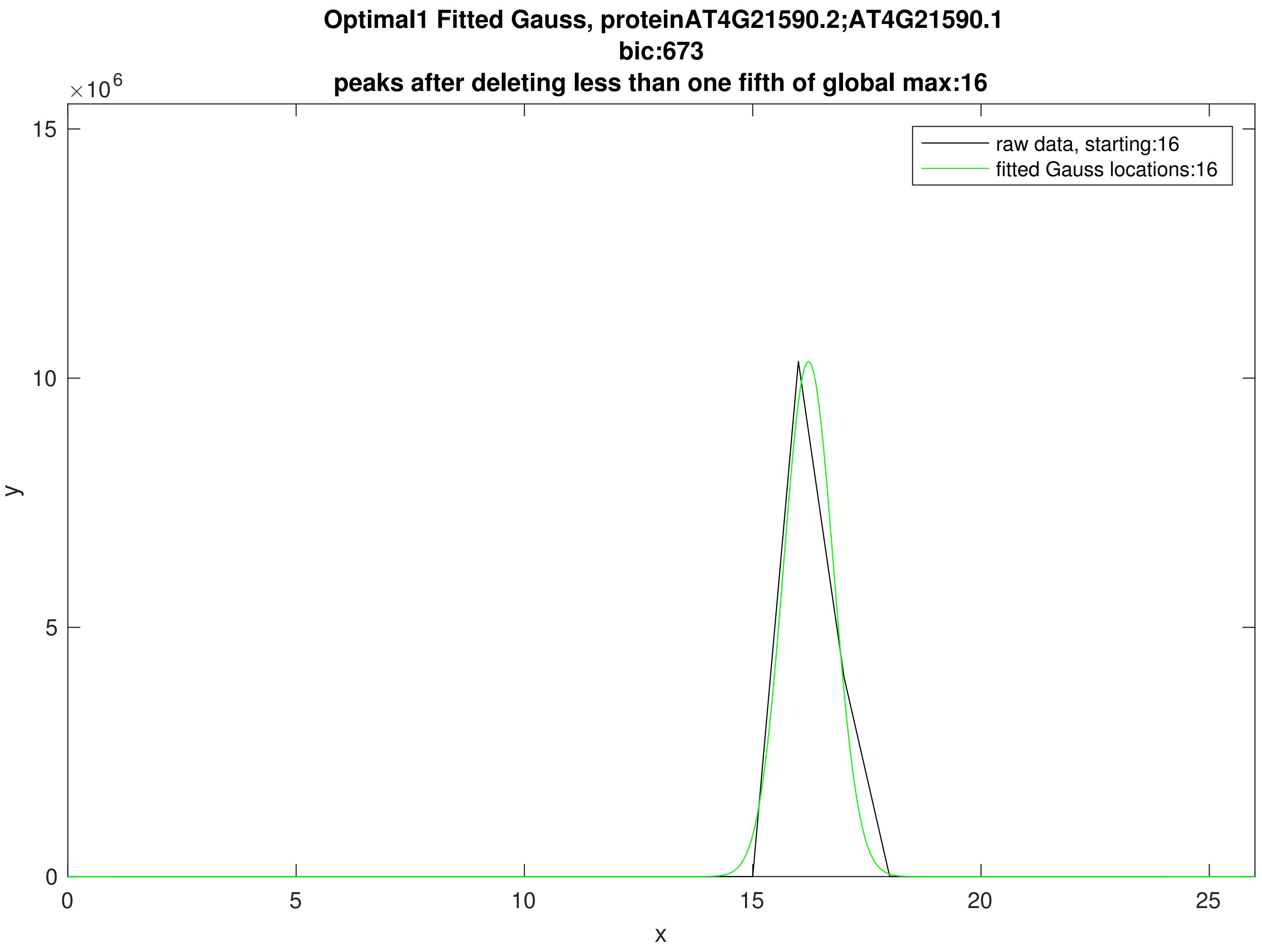

Optimal1 Fitted Gauss, proteinAT4G21590.2;AT4G21590.1
bic:673
peaks after deleting less than one fifth of global max:16
×10^6
15
10
5
0
y
0
5
10
15
20
25
x
raw data, starting:16
fitted Gauss locations:16

**Optimal1 Fitted Gauss, proteinAT4G21600.1**
**bic:879**
**peaks after deleting less than one fifth of global max:17**

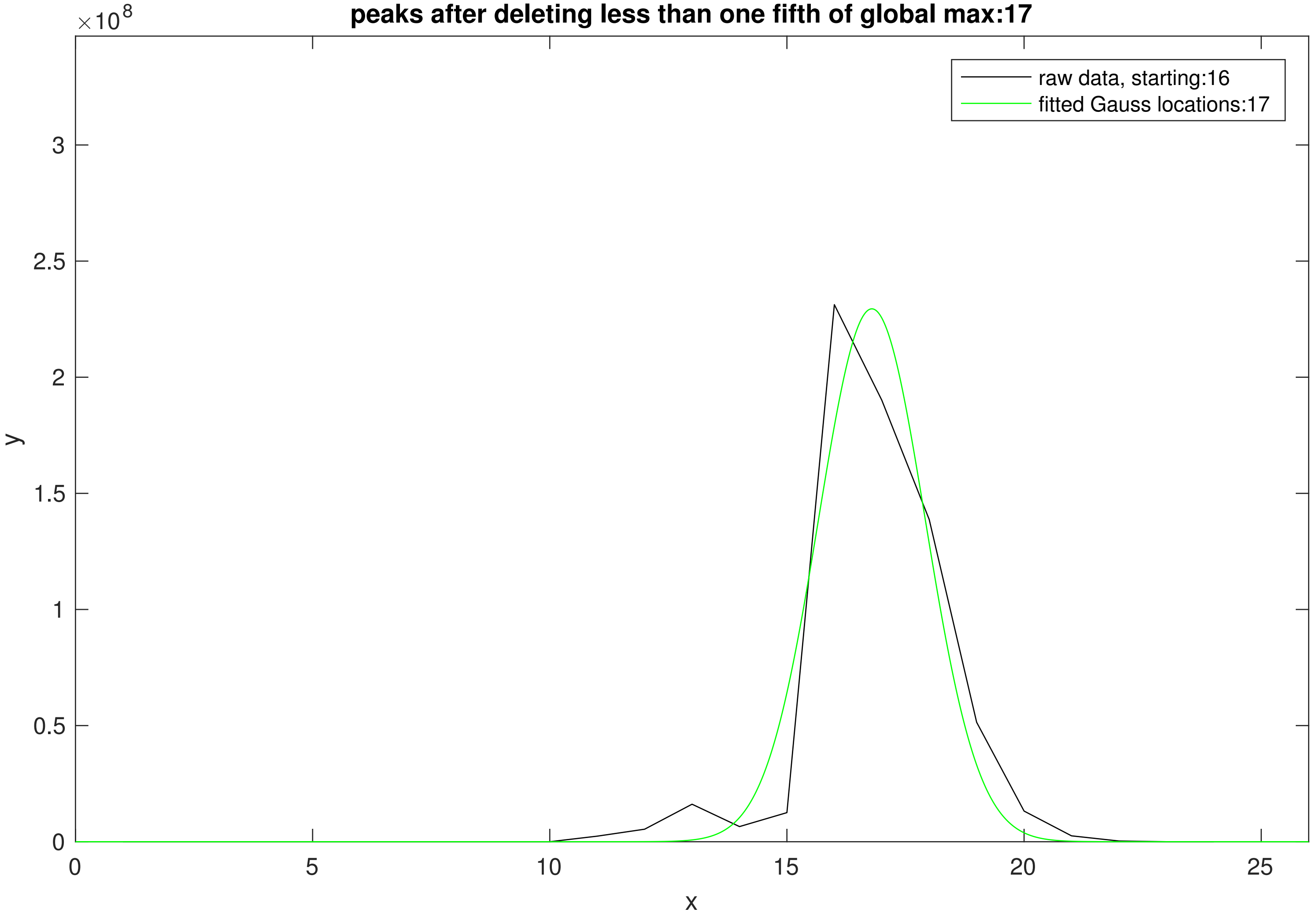

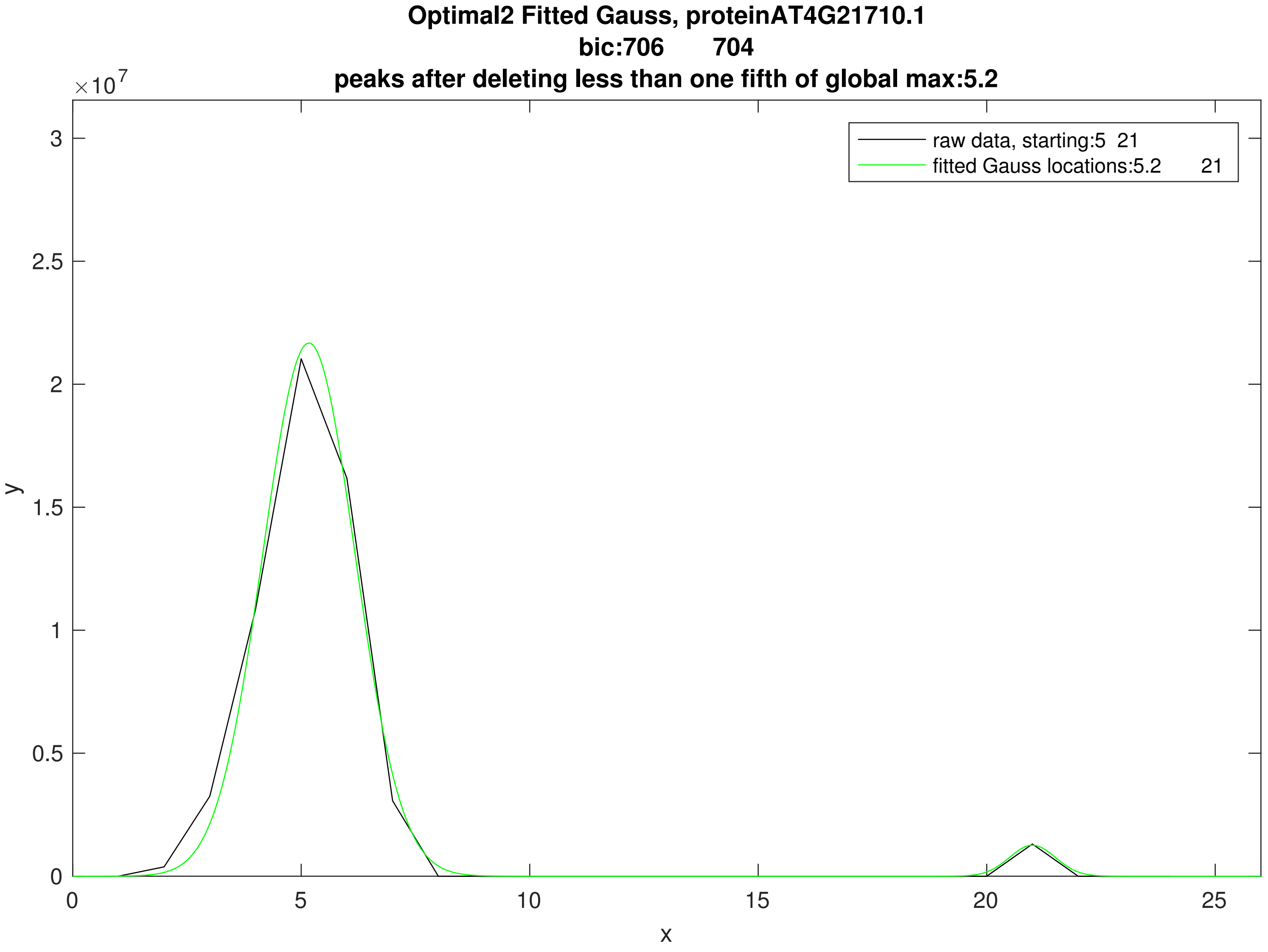

Optimal2 Fitted Gauss, proteinAT4G21710.1
bic:706 704
peaks after deleting less than one fifth of global max:5.2
×10^7
raw data, starting:5 21
fitted Gauss locations:5.2 21
y
x
0
0.5
1
1.5
2
2.5
3
0
5
10
15
20
25

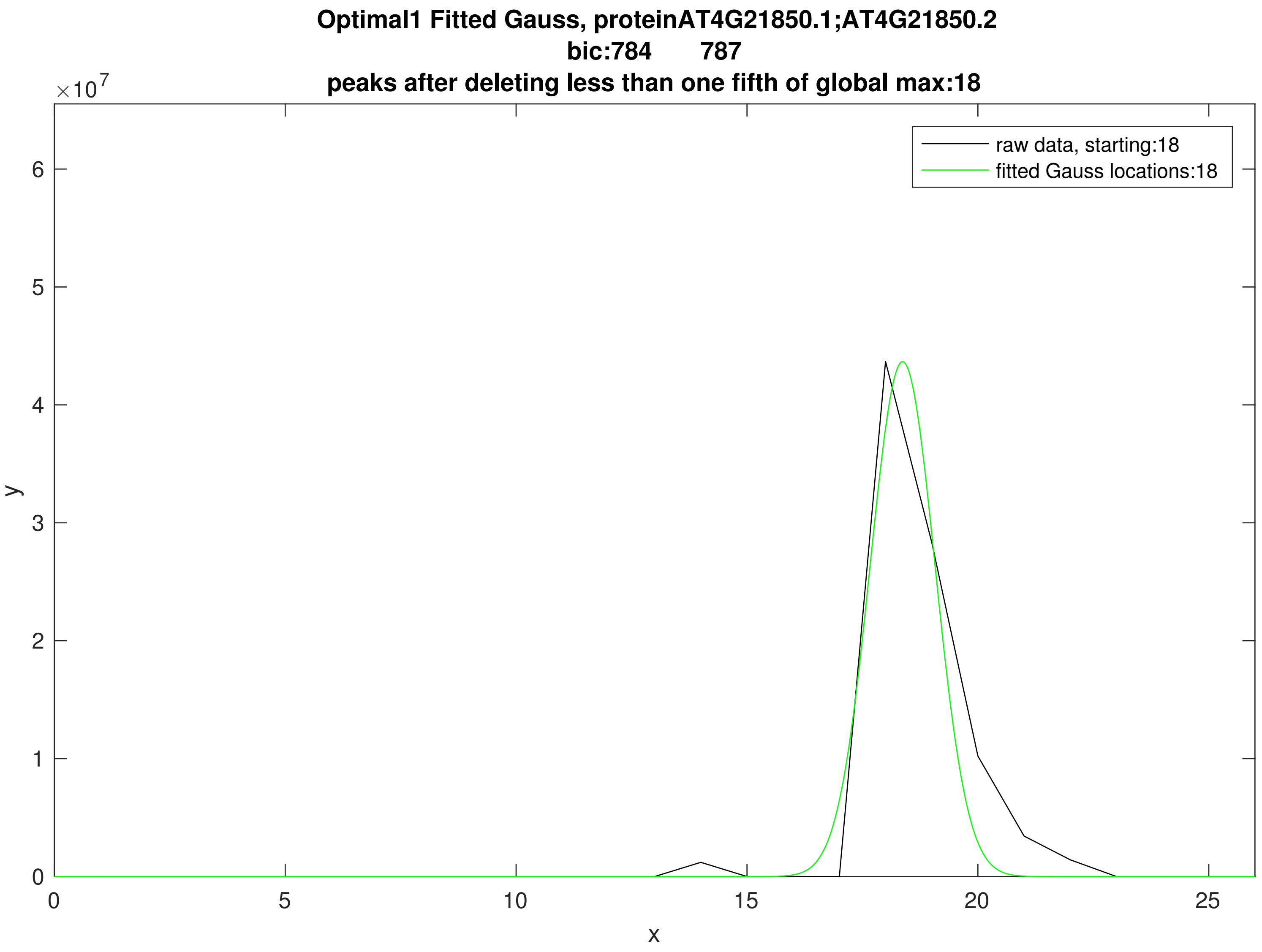

Optimal1 Fitted Gauss, proteinAT4G21850.1;AT4G21850.2
bic:784 787
peaks after deleting less than one fifth of global max:18
×10^7
raw data, starting:18
fitted Gauss locations:18
y
x
0
1
2
3
4
5
6
0
5
10
15
20
25

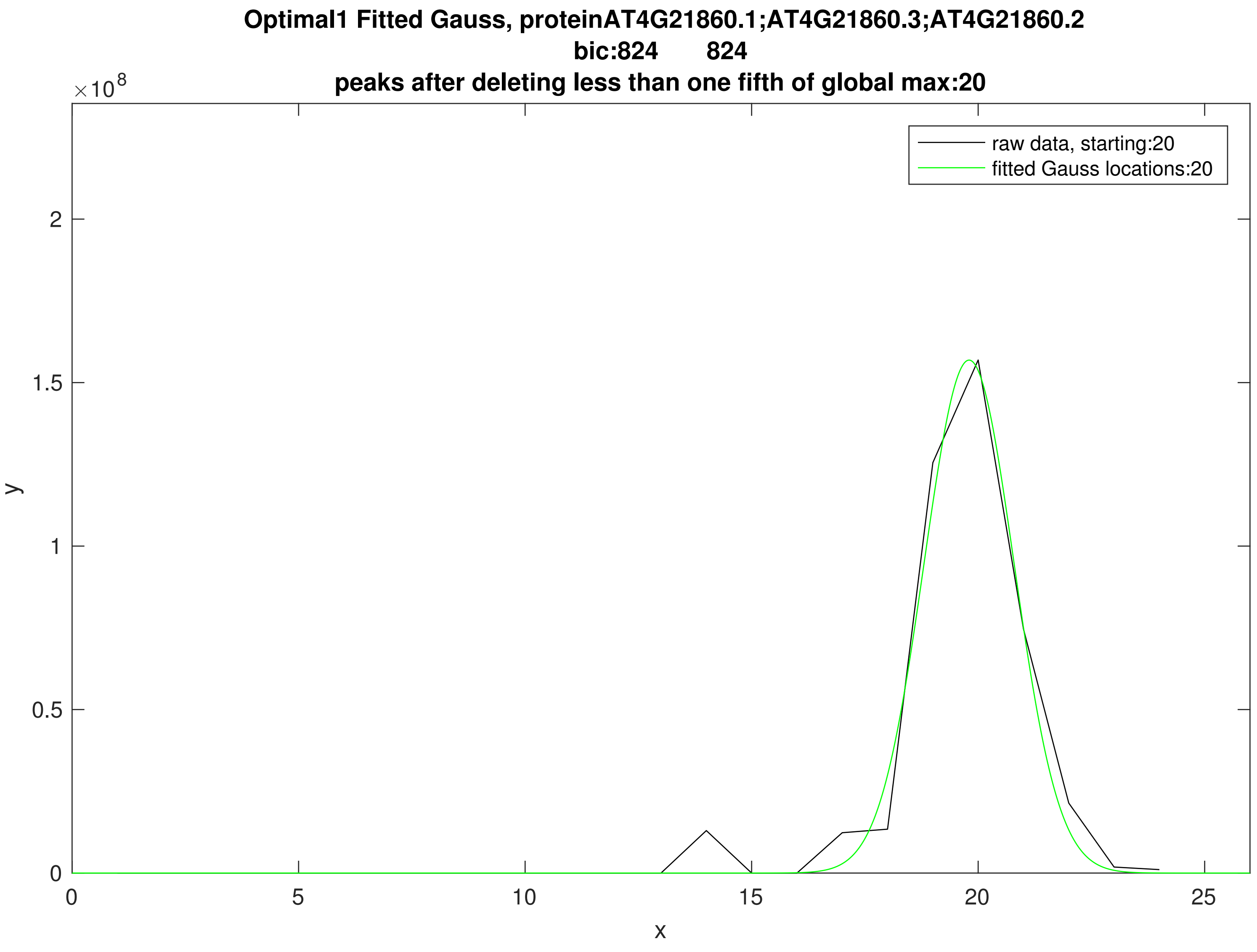
Optimal1 Fitted Gauss, proteinAT4G21860.1;AT4G21860.3;AT4G21860.2
bic:824 824
peaks after deleting less than one fifth of global max:20
×10^8
raw data, starting:20
fitted Gauss locations:20
y
x
0
0.5
1
1.5
2
5
10
15
20
25

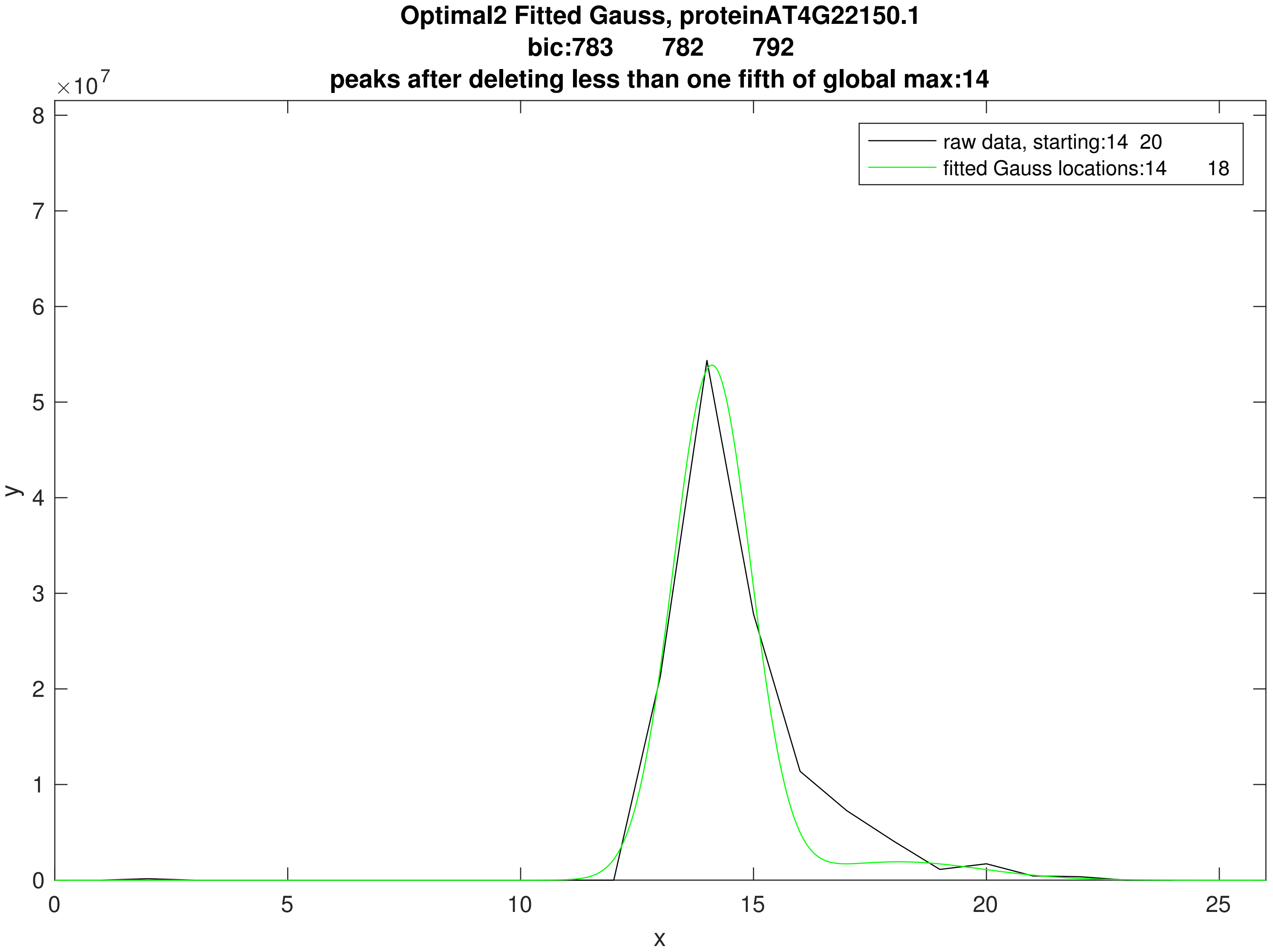
Optimal2 Fitted Gauss, proteinAT4G22150.1
bic:783 782 792
peaks after deleting less than one fifth of global max:14
×10^7
raw data, starting:14 20
fitted Gauss locations:14 18
y
x
0
1
2
3
4
5
6
7
8
0
5
10
15
20
25

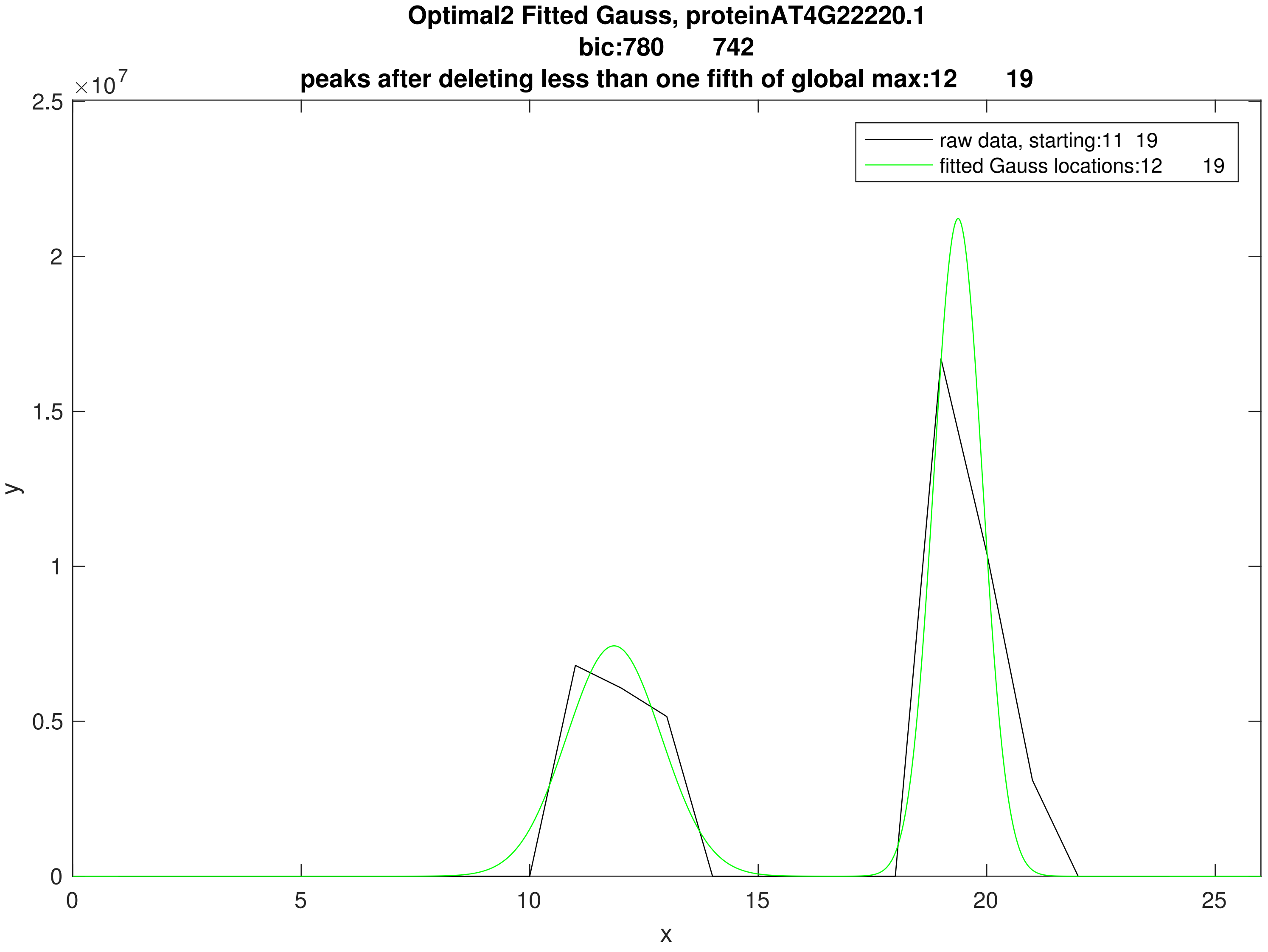

Optimal2 Fitted Gauss, proteinAT4G22220.1
bic:780 742
peaks after deleting less than one fifth of global max:12 19
×10^7
raw data, starting:11 19
fitted Gauss locations:12 19
y
x
0
0.5
1
1.5
2
2.5
0
5
10
15
20
25

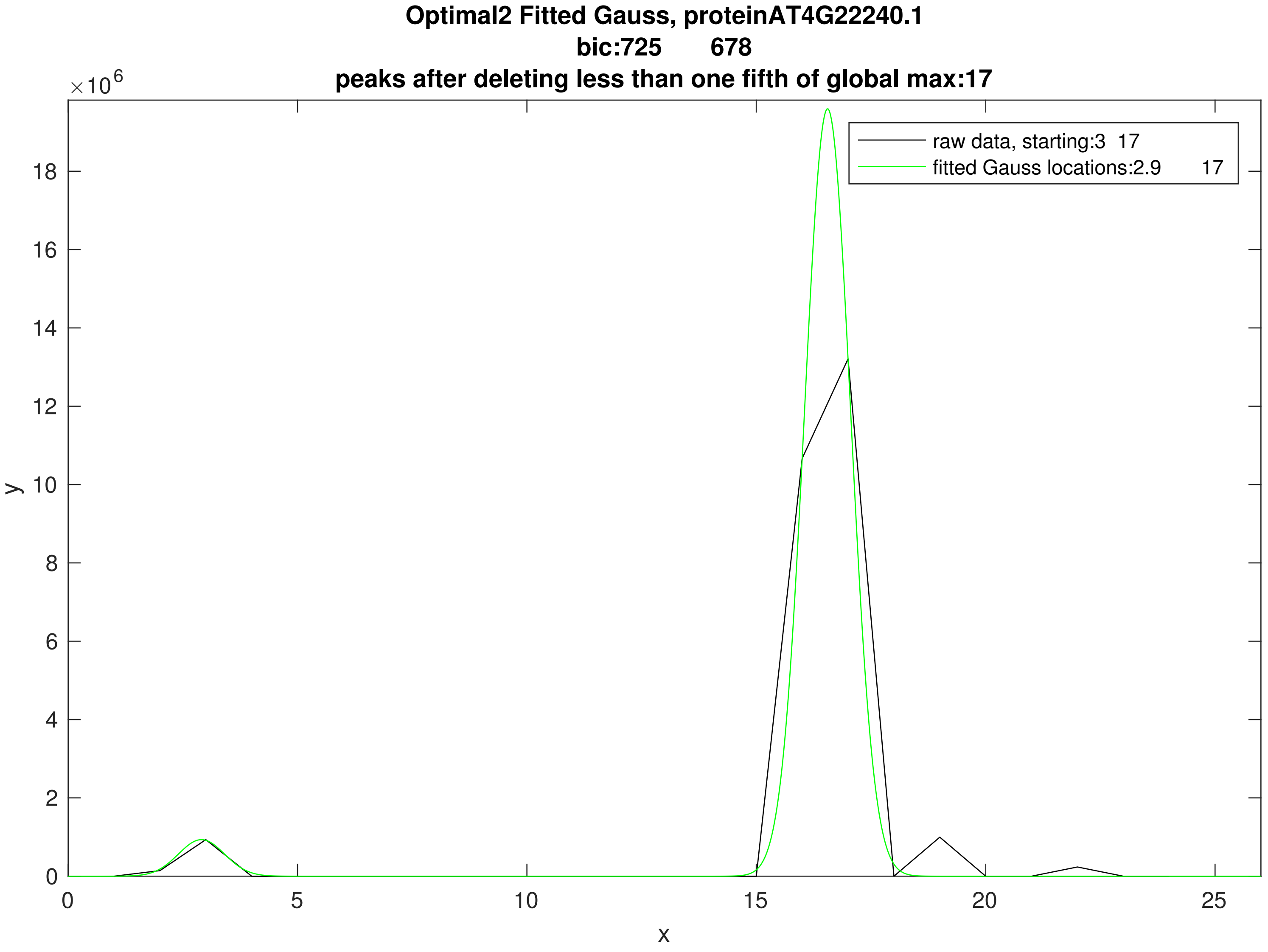

Optimal2 Fitted Gauss, proteinAT4G22240.1
bic:725 678
peaks after deleting less than one fifth of global max:17
×10^6
raw data, starting:3 17
fitted Gauss locations:2.9 17
y
x
0
2
4
6
8
10
12
14
16
18
5
15
20
25

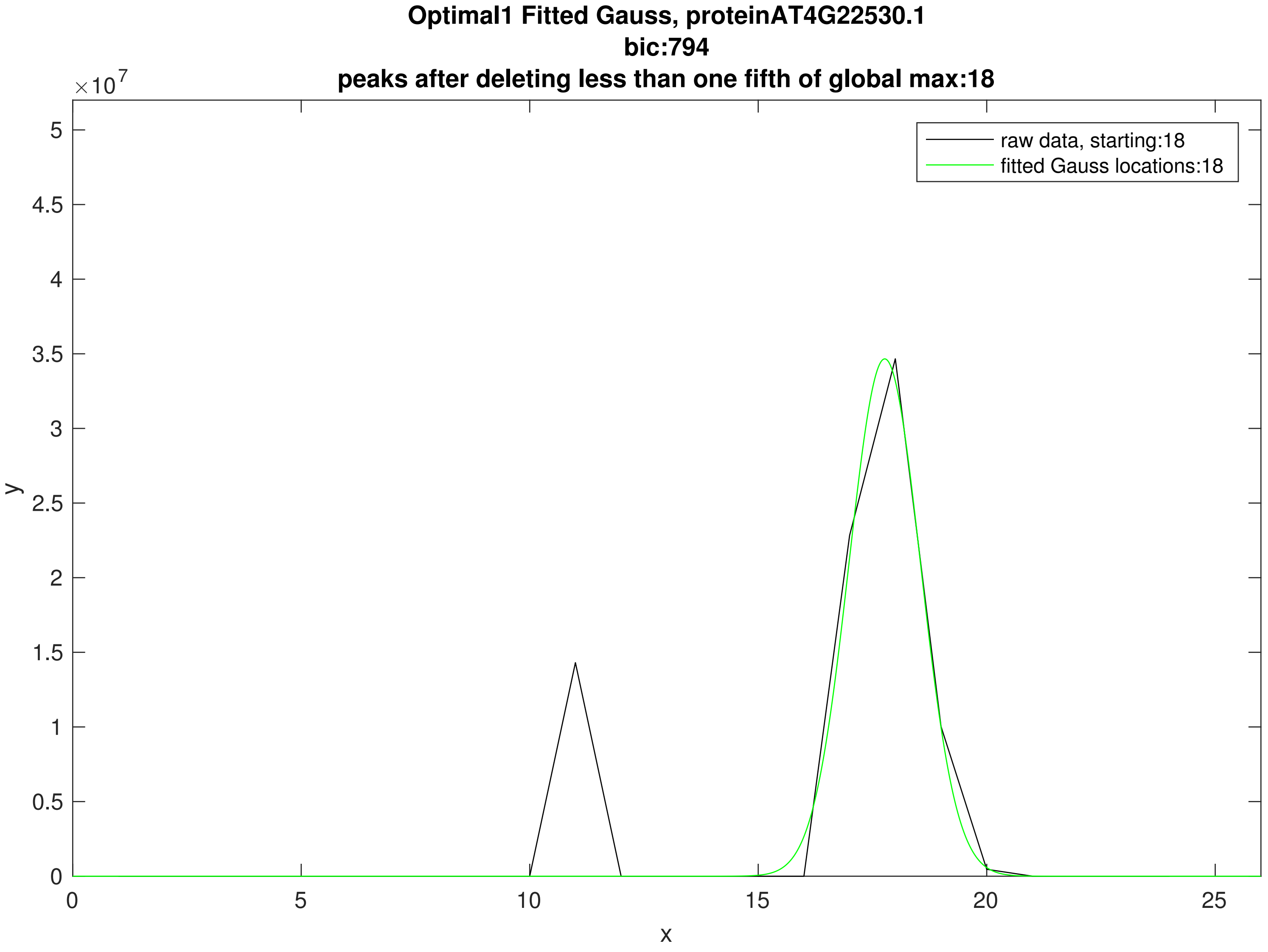
Optimal1 Fitted Gauss, proteinAT4G22530.1
bic:794
peaks after deleting less than one fifth of global max:18
×10^7
raw data, starting:18
fitted Gauss locations:18
y
x
0
0.5
1
1.5
2
2.5
3
3.5
4
4.5
5
0
5
10
15
20
25

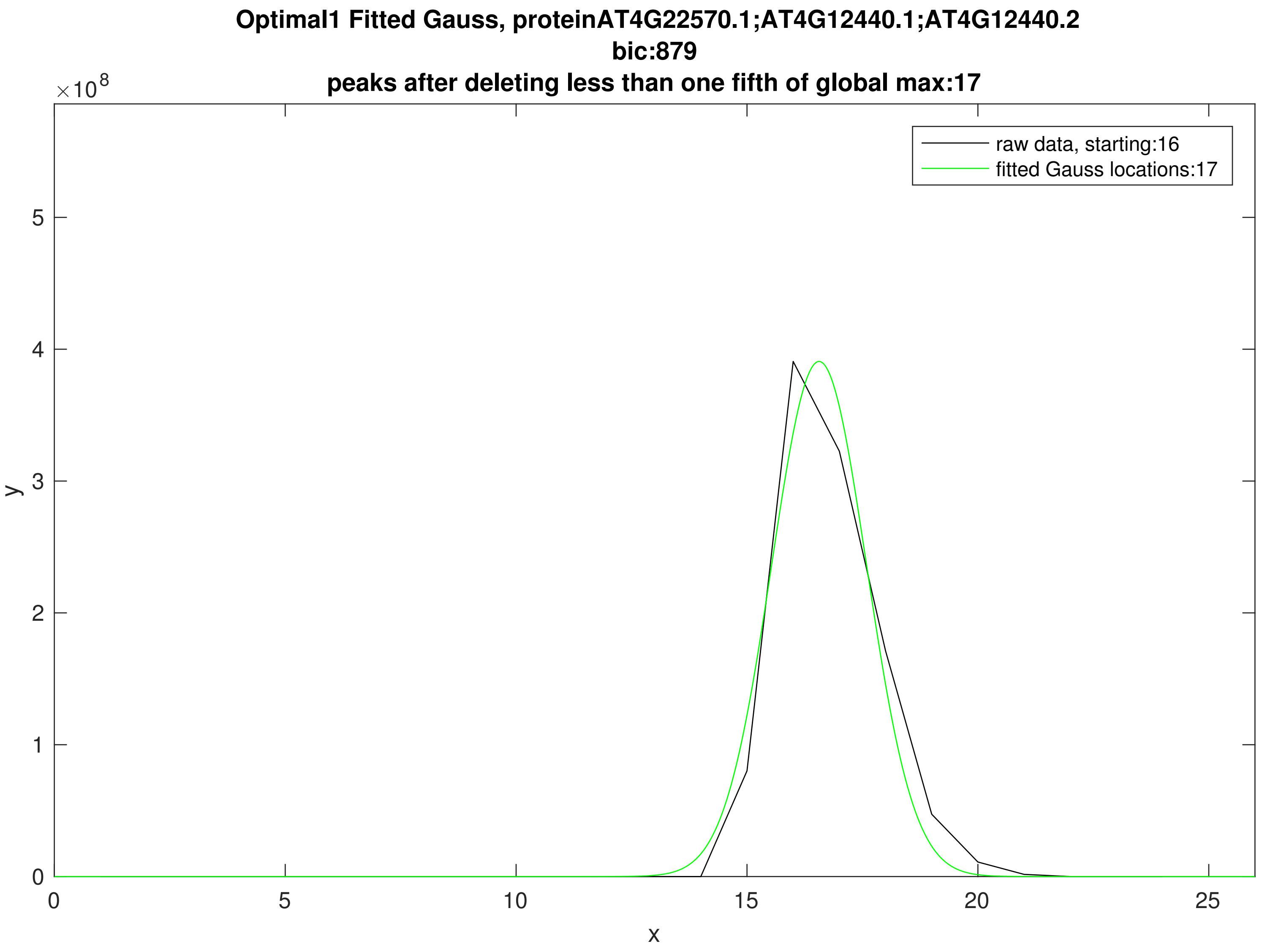
Optimal1 Fitted Gauss, proteinAT4G22570.1;AT4G12440.1;AT4G12440.2
bic:879
peaks after deleting less than one fifth of global max:17
×10^8
raw data, starting:16
fitted Gauss locations:17
0
1
2
3
4
5
y
0
5
10
15
20
25
x

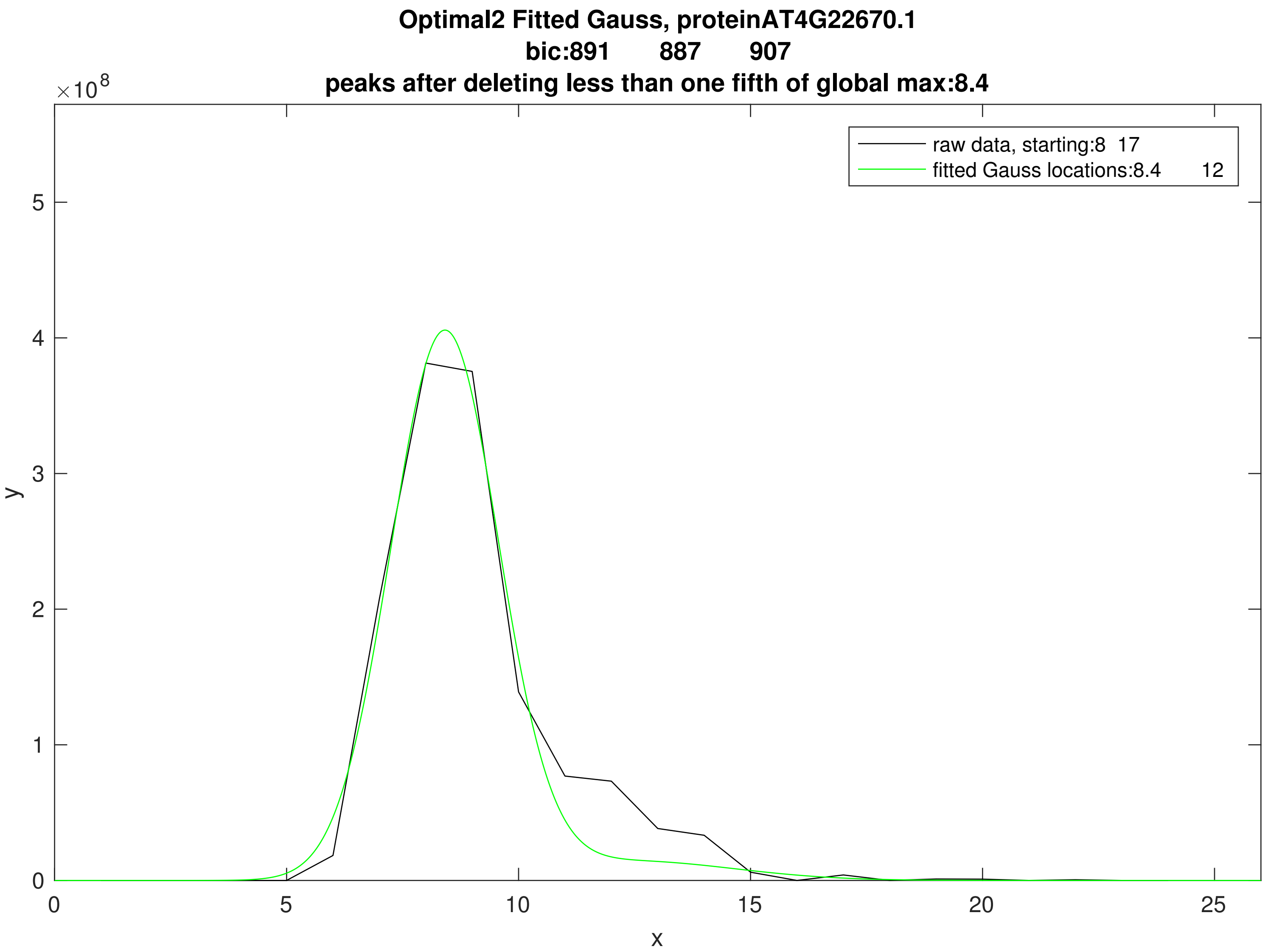
Optimal2 Fitted Gauss, proteinAT4G22670.1
bic:891 887 907
peaks after deleting less than one fifth of global max:8.4
×10^8
raw data, starting:8 17
fitted Gauss locations:8.4 12
y
x
0
1
2
3
4
5
0
5
10
15
20
25

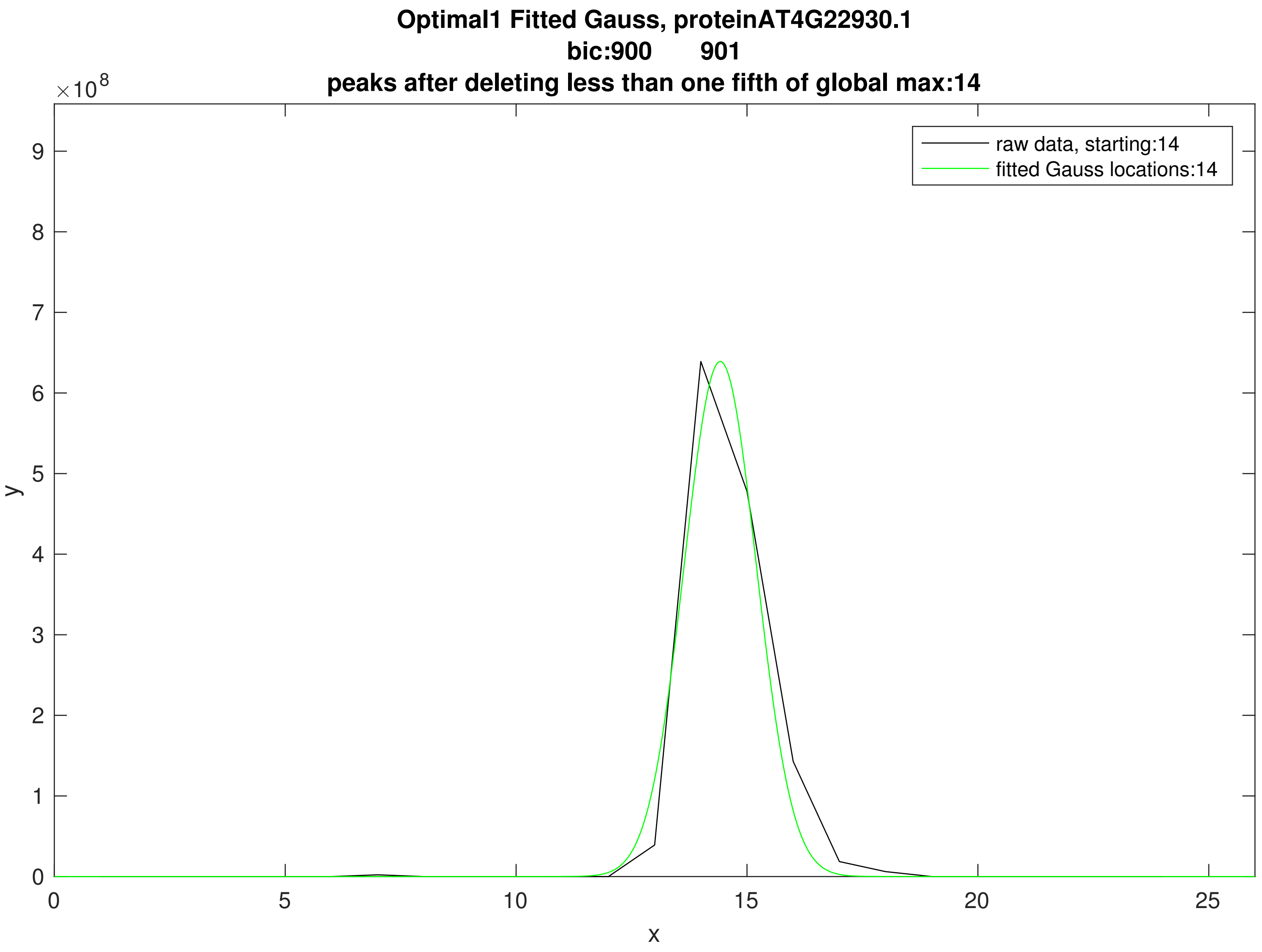
Optimal1 Fitted Gauss, proteinAT4G22930.1
bic:900 901
peaks after deleting less than one fifth of global max:14
×10^8
raw data, starting:14
fitted Gauss locations:14
y
x
0
1
2
3
4
5
6
7
8
9
0
5
10
15
20
25

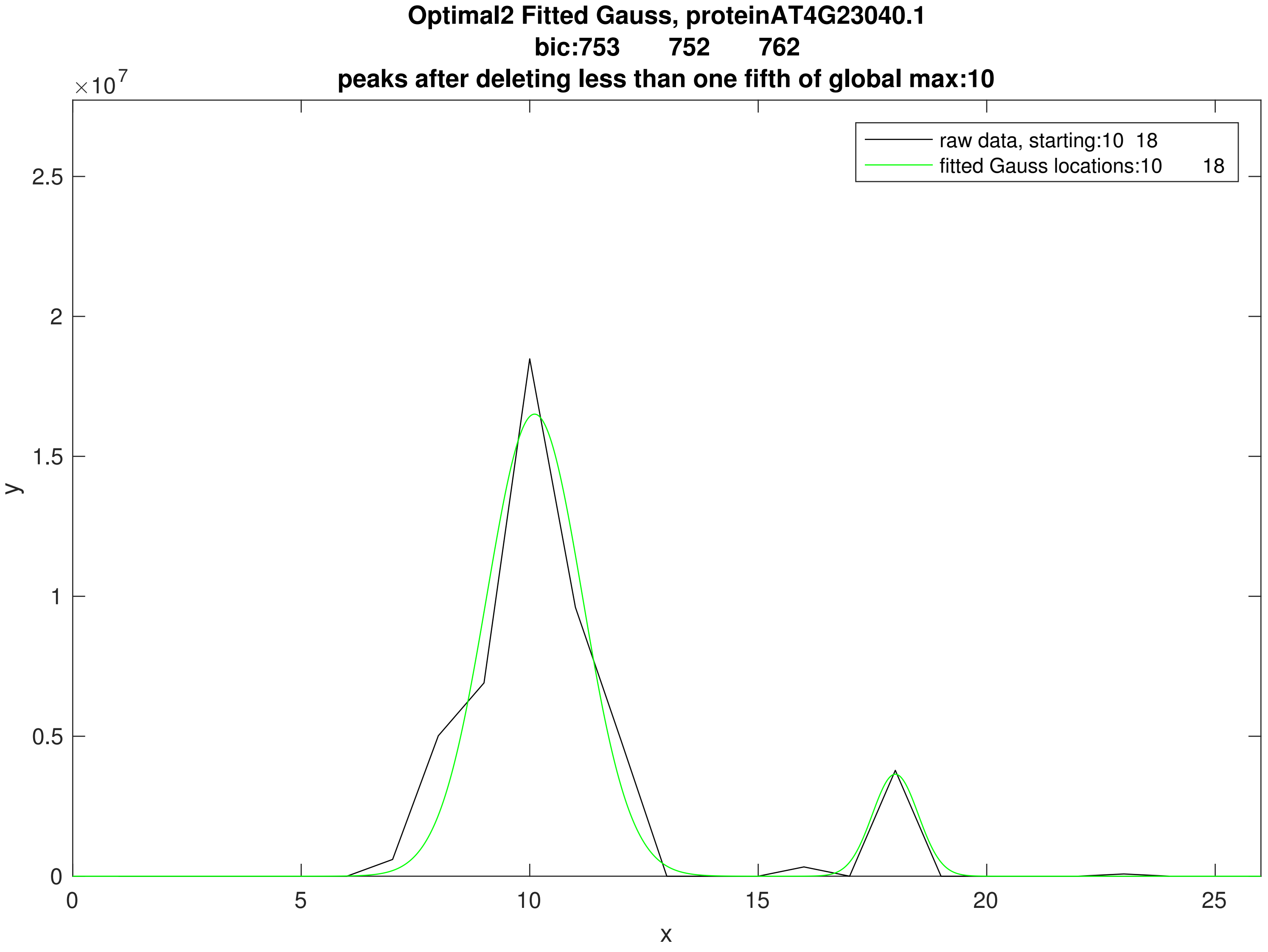

Optimal2 Fitted Gauss, proteinAT4G23040.1
bic:753 752 762
peaks after deleting less than one fifth of global max:10
×10^7
raw data, starting:10 18
fitted Gauss locations:10 18
y
x
0
0.5
1
1.5
2
2.5
5
10
15
20
25

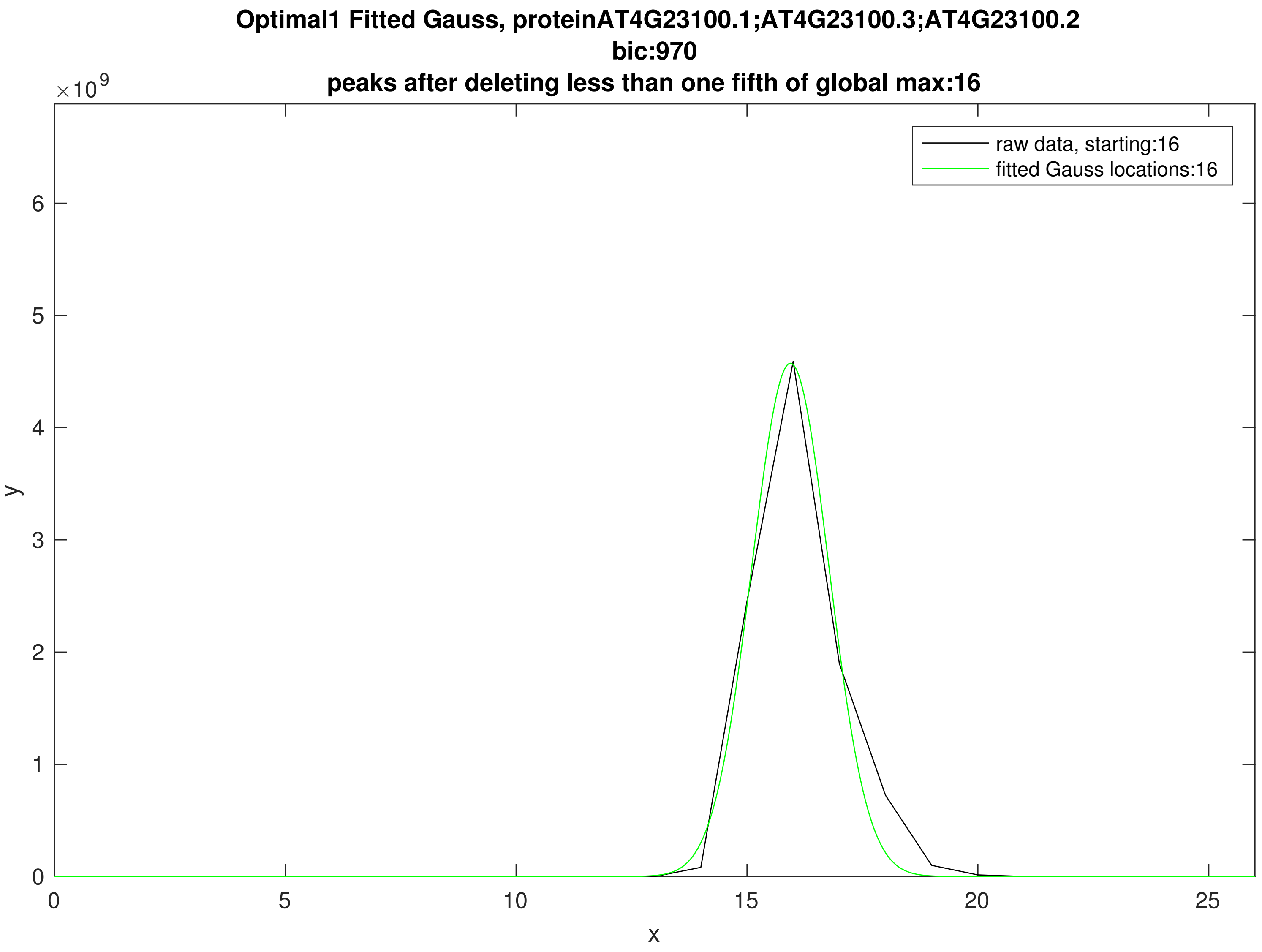
Optimal1 Fitted Gauss, proteinAT4G23100.1;AT4G23100.3;AT4G23100.2
bic:970
peaks after deleting less than one fifth of global max:16
×10^9
raw data, starting:16
fitted Gauss locations:16
y
x
0
1
2
3
4
5
6
0
5
10
15
20
25

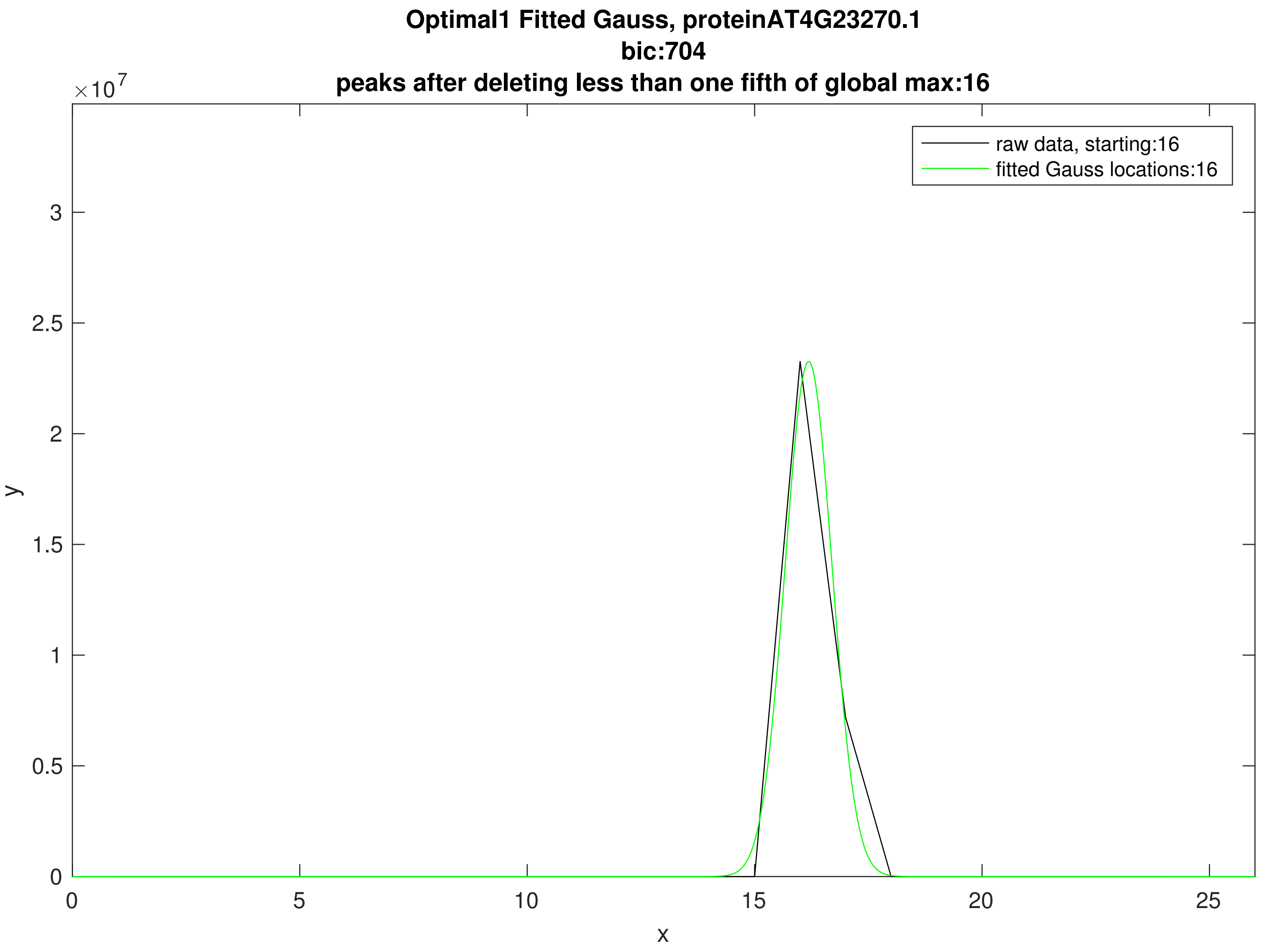
Optimal1 Fitted Gauss, proteinAT4G23270.1
bic:704
peaks after deleting less than one fifth of global max:16
×10^7
raw data, starting:16
fitted Gauss locations:16
3
2.5
2
1.5
1
0.5
0
y
0
5
10
15
20
25
x

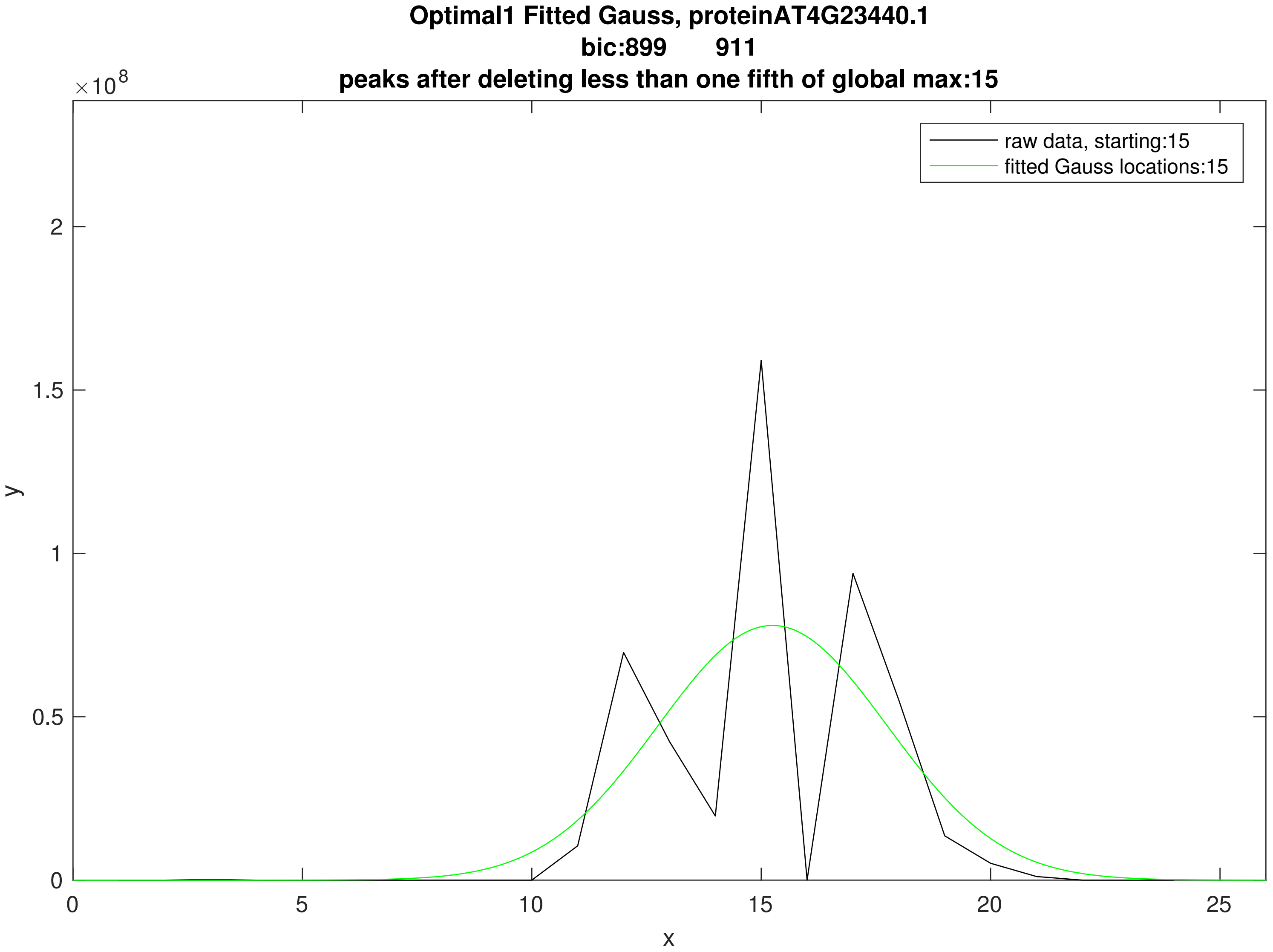
Optimal1 Fitted Gauss, proteinAT4G23440.1
bic:899 911
peaks after deleting less than one fifth of global max:15
×10^8
raw data, starting:15
fitted Gauss locations:15
y
x
0
0.5
1
1.5
2
0
5
10
15
20
25

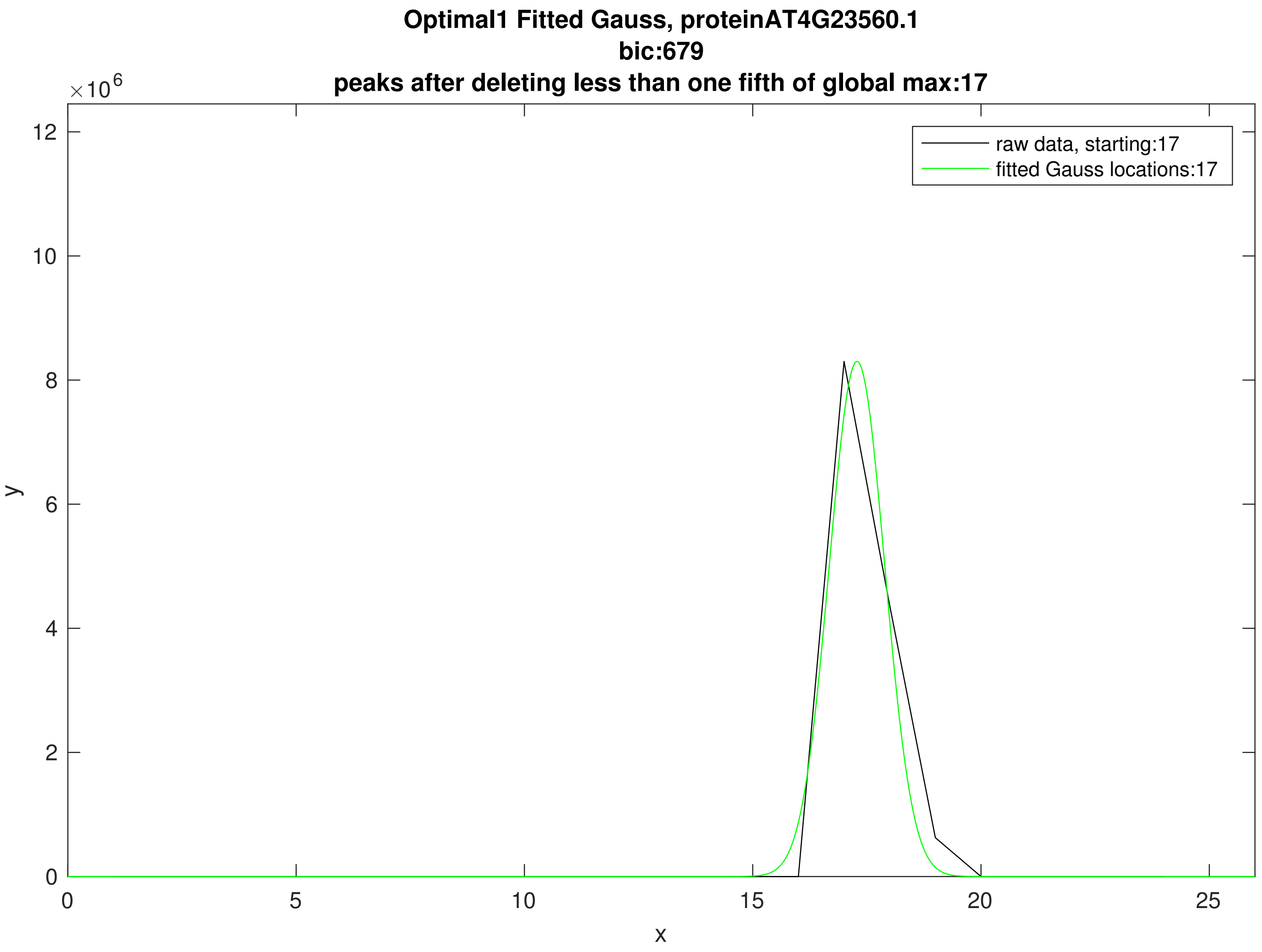

Optimal1 Fitted Gauss, proteinAT4G23560.1
bic:679
peaks after deleting less than one fifth of global max:17
×10^6
raw data, starting:17
fitted Gauss locations:17
y
x
0
2
4
6
8
10
12
0
5
10
15
20
25

#### Optimal1 Fitted Gauss, proteinAT4G23630.1
#### bic:626
#### peaks after deleting less than one fifth of global max:19

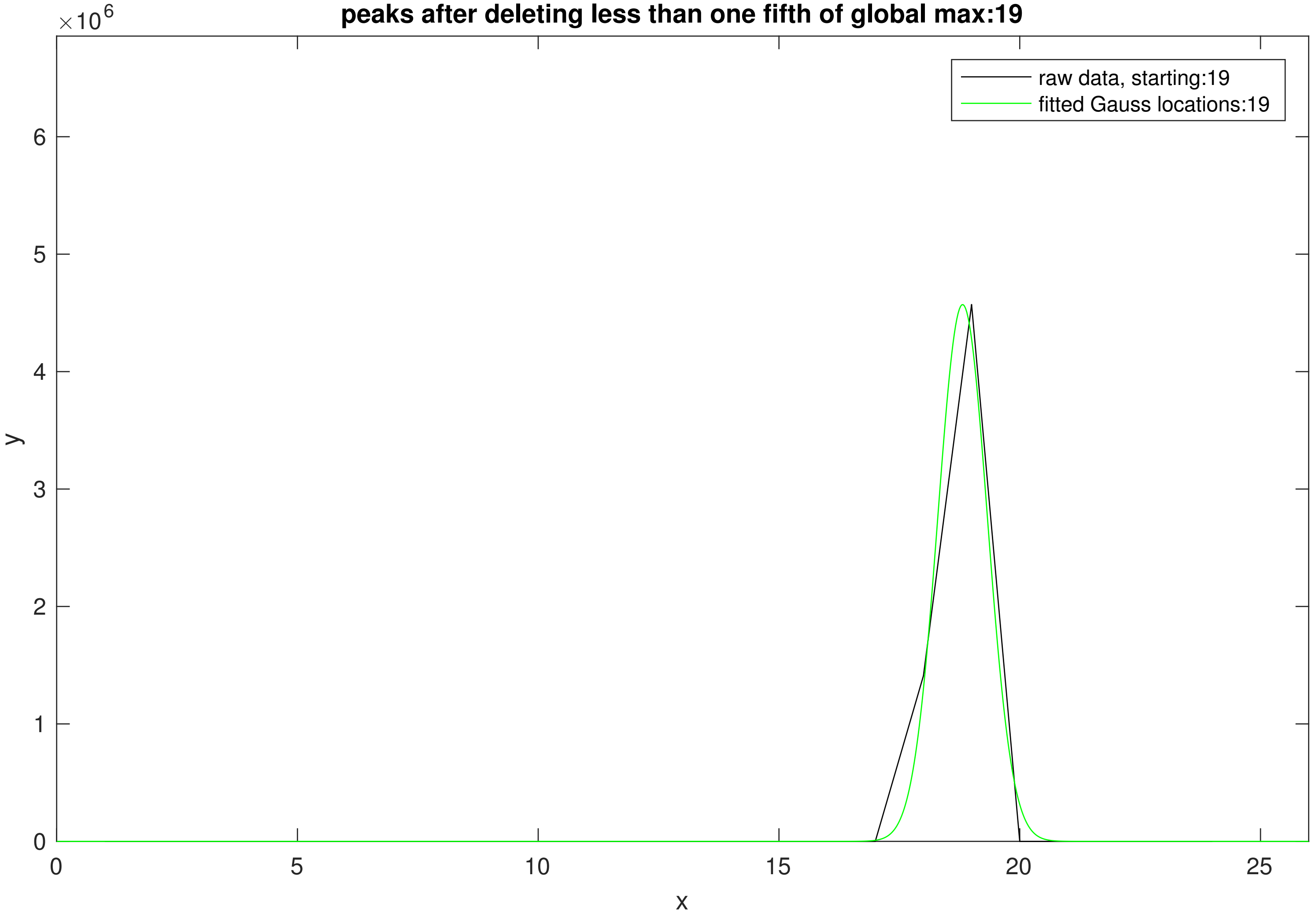

**Optimal1 Fitted Gauss, proteinAT4G23650.1**
**bic:727 736**
**peaks after deleting less than one fifth of global max:16**

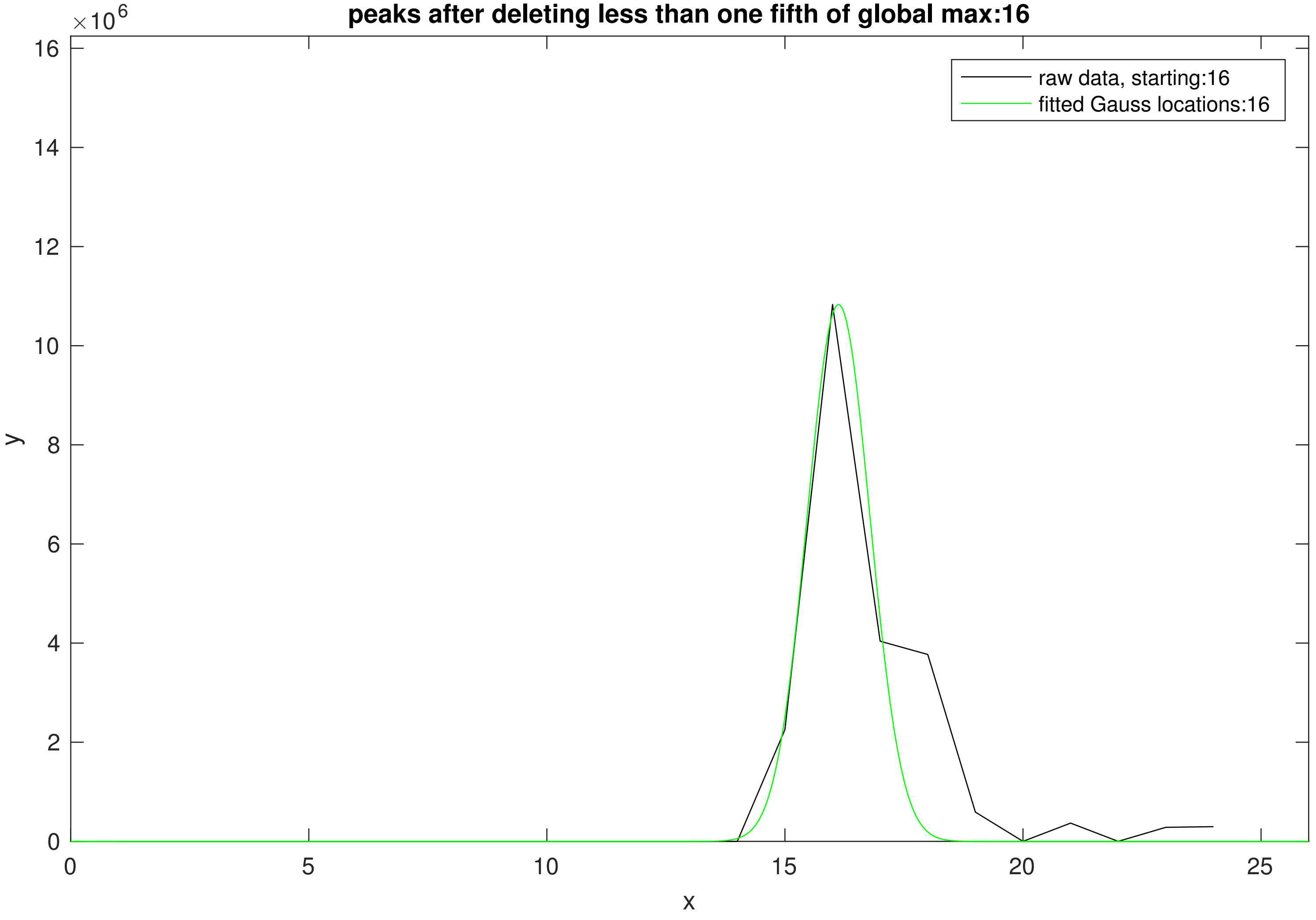

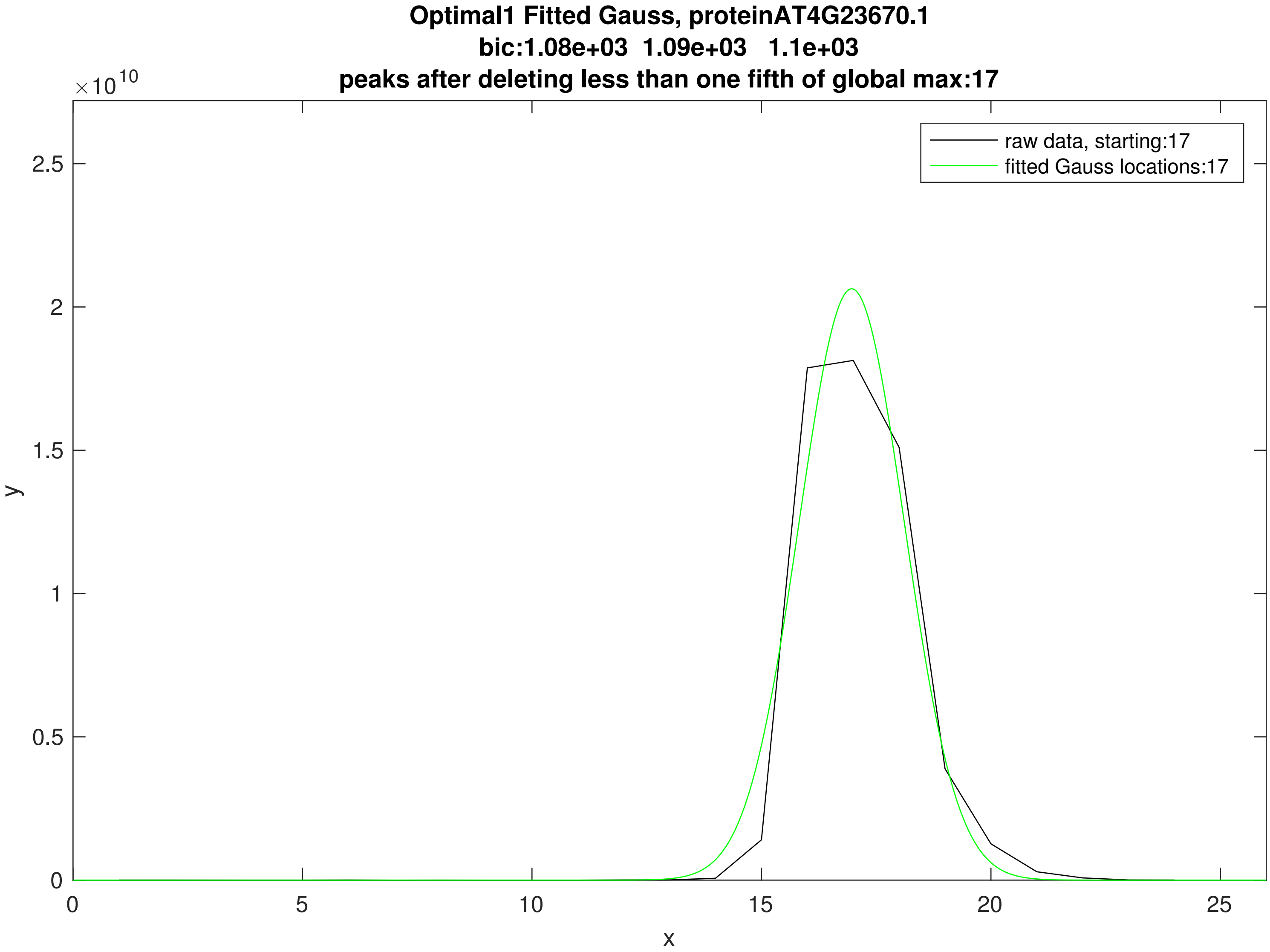
Optimal1 Fitted Gauss, proteinAT4G23670.1
bic:1.08e+03 1.09e+03 1.1e+03
peaks after deleting less than one fifth of global max:17
×10^10
raw data, starting:17
fitted Gauss locations:17
y
x
0
0.5
1
1.5
2
2.5
5
10
15
20
25

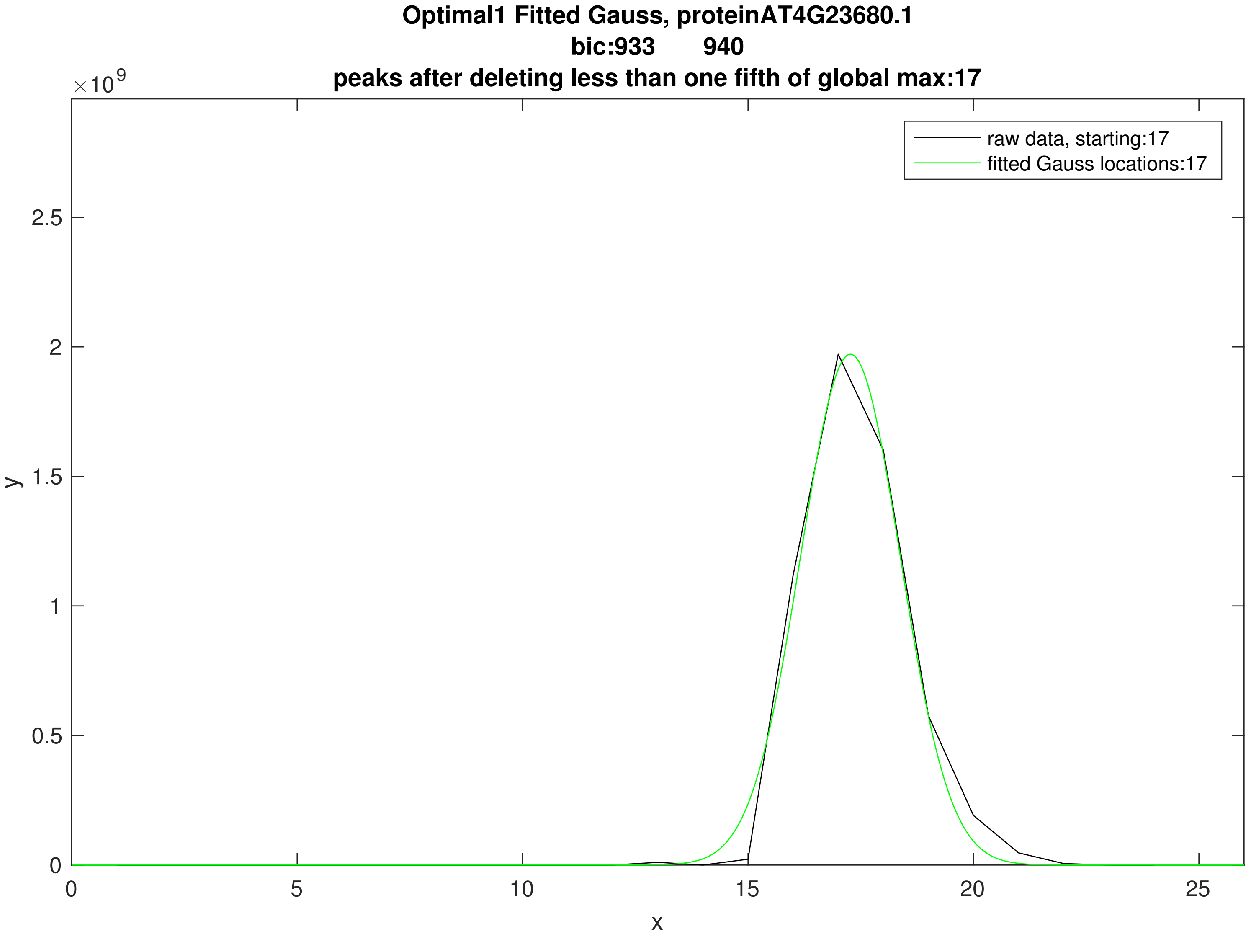
Optimal1 Fitted Gauss, proteinAT4G23680.1
bic:933 940
peaks after deleting less than one fifth of global max:17
×10^9
raw data, starting:17
fitted Gauss locations:17
y
x
0
0.5
1
1.5
2
2.5
0
5
10
15
20
25

Optimal1 Fitted Gauss, proteinAT4G23690.1;AT1G64160.1
bic:735
peaks after deleting less than one fifth of global max:15

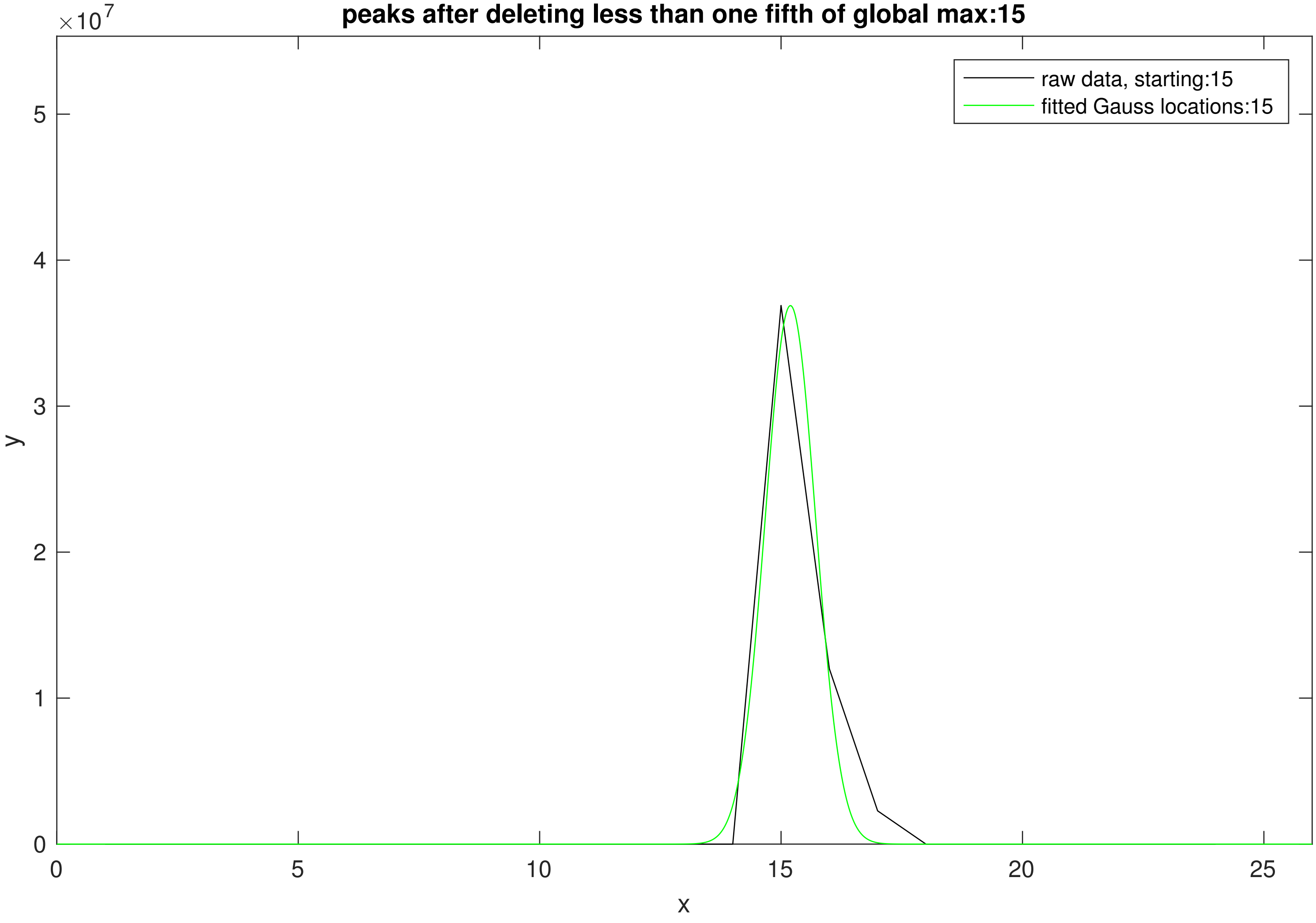

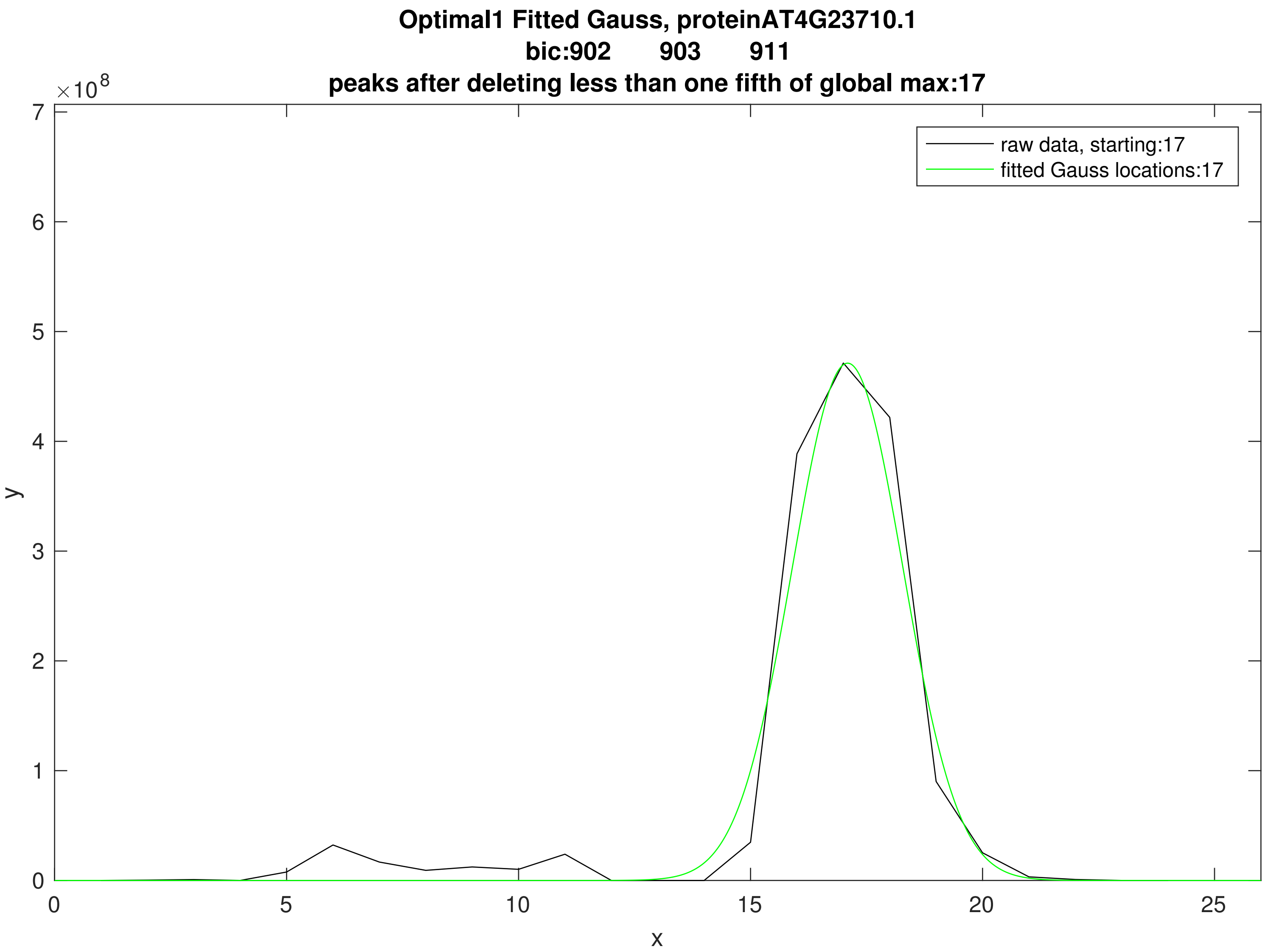

Optimal1 Fitted Gauss, proteinAT4G23710.1
bic:902 903 911
peaks after deleting less than one fifth of global max:17
×10^8
raw data, starting:17
fitted Gauss locations:17
y
x
0
1
2
3
4
5
6
7
0
5
10
15
20
25

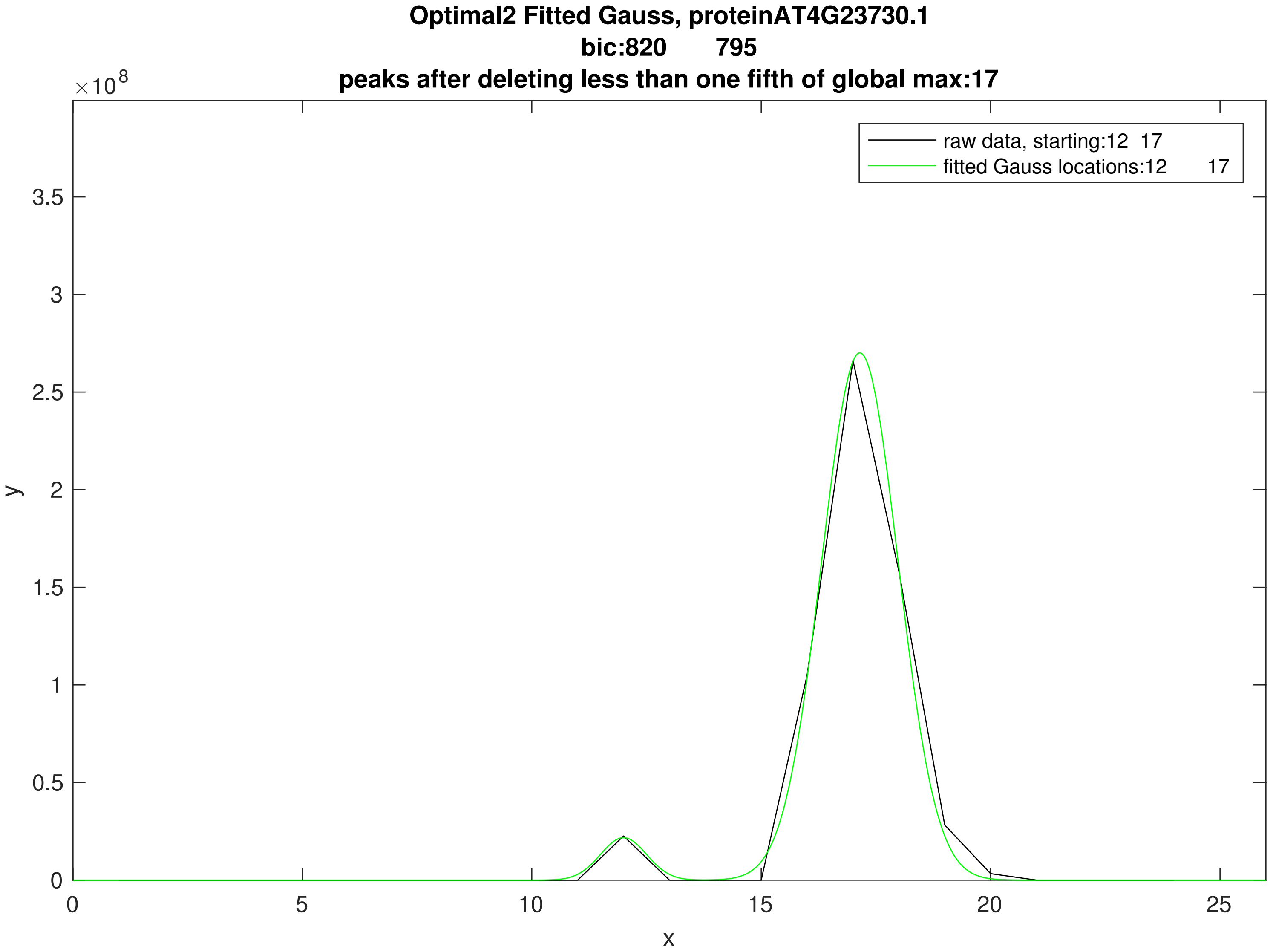

Optimal2 Fitted Gauss, proteinAT4G23730.1
bic:820 795
peaks after deleting less than one fifth of global max:17
×10^8
raw data, starting:12 17
fitted Gauss locations:12 17
3.5
3
2.5
2
1.5
1
0.5
0
y
0
5
10
15
20
25
x

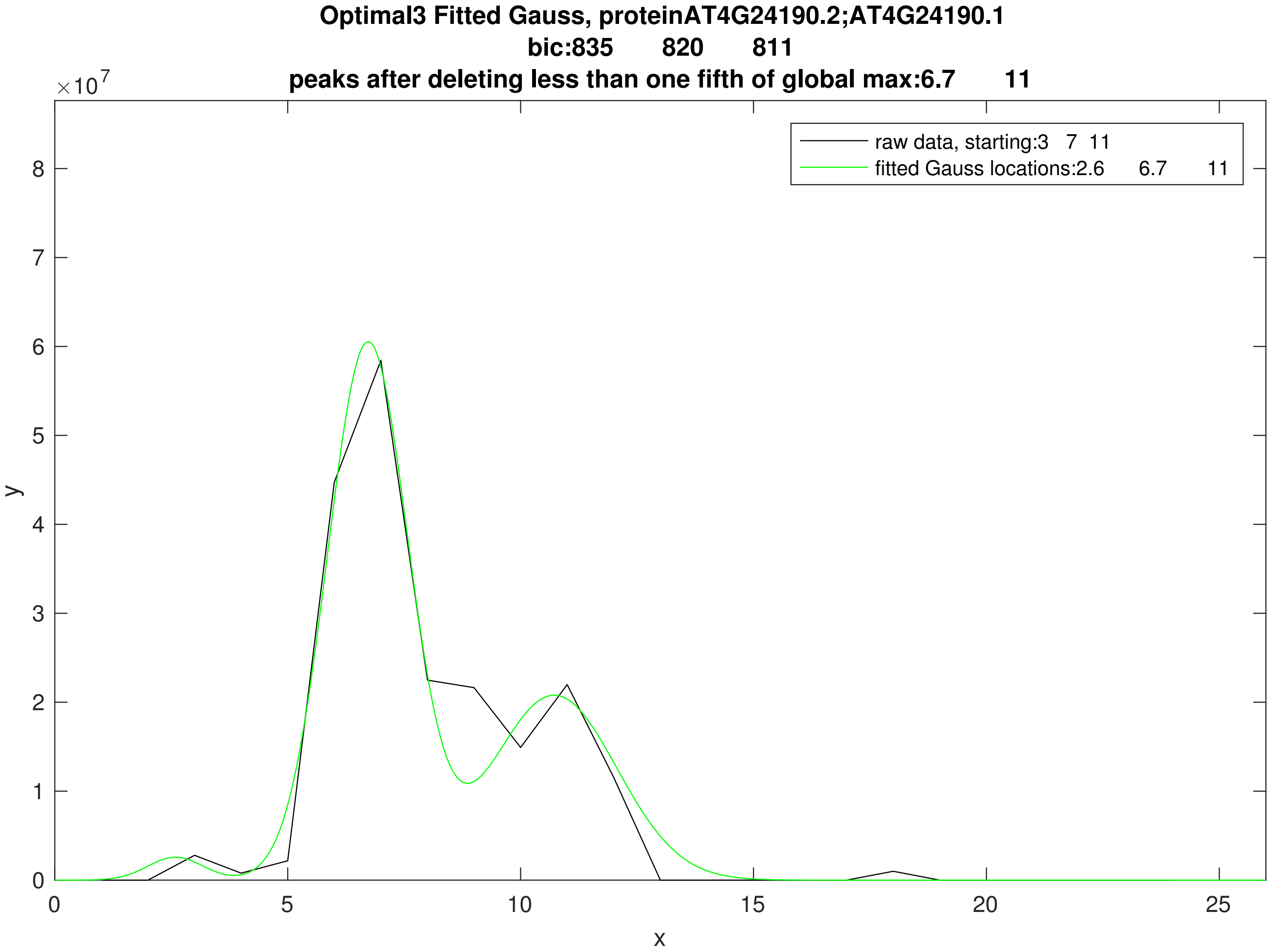

Optimal3 Fitted Gauss, proteinAT4G24190.2;AT4G24190.1
bic:835 820 811
peaks after deleting less than one fifth of global max:6.7 11
×10^7
raw data, starting:3 7 11
fitted Gauss locations:2.6 6.7 11
y
x
0
1
2
3
4
5
6
7
8
0
5
10
15
20
25

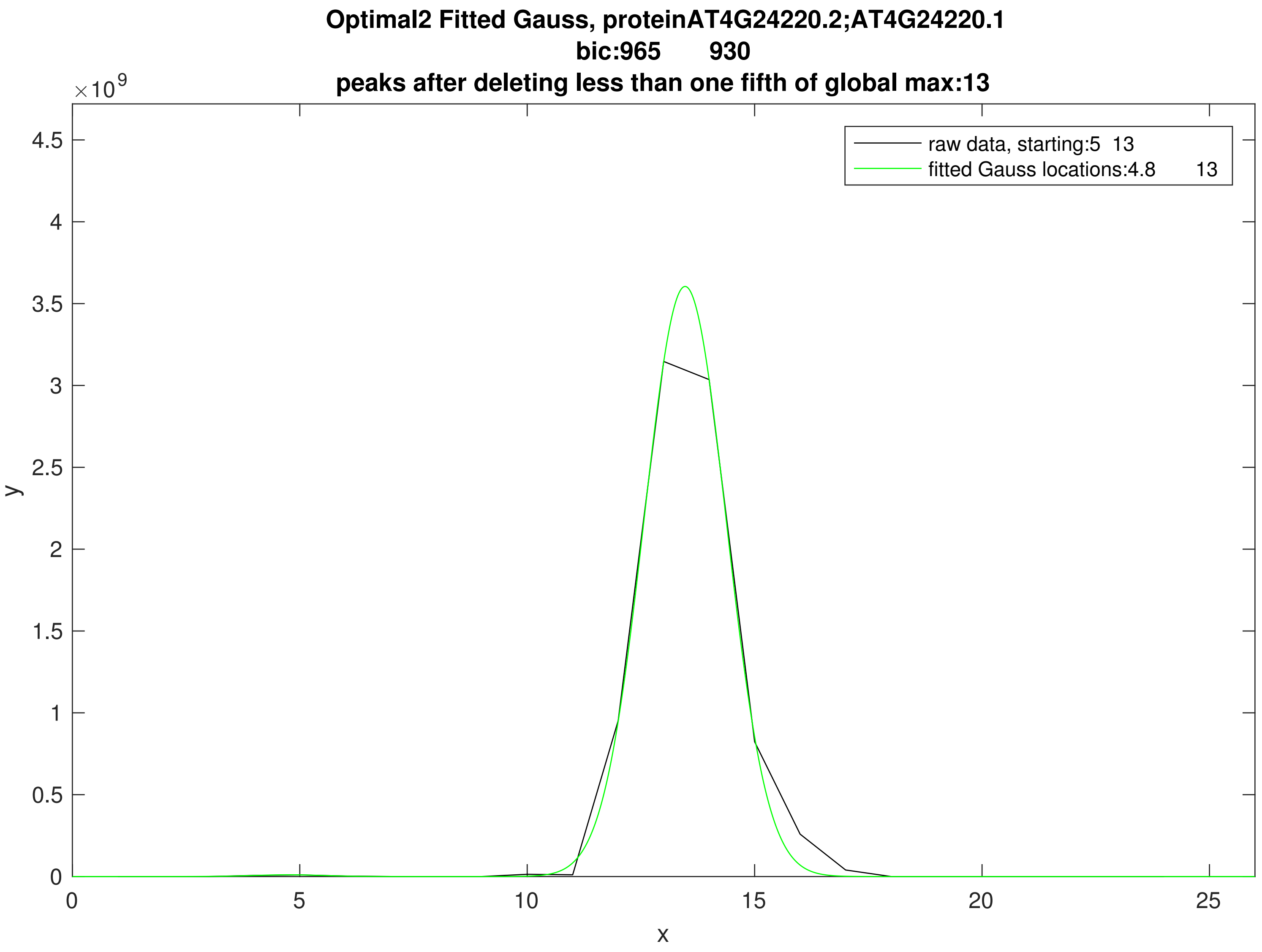
Optimal2 Fitted Gauss, proteinAT4G24220.2;AT4G24220.1
bic:965 930
peaks after deleting less than one fifth of global max:13
×10^9
raw data, starting:5 13
fitted Gauss locations:4.8 13
y
x
0
0.5
1
1.5
2
2.5
3
3.5
4
4.5
0
5
10
15
20
25

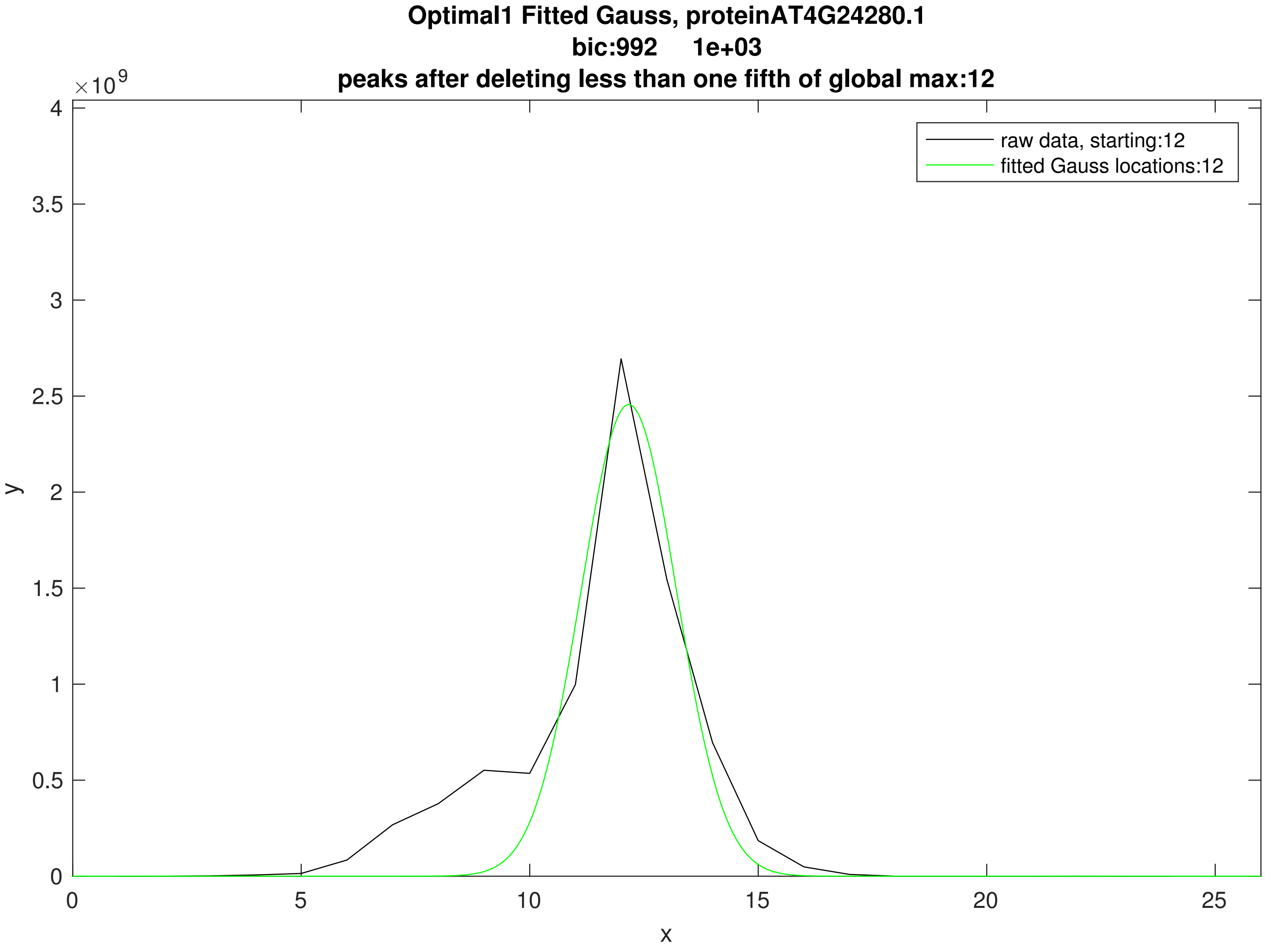

Optimal1 Fitted Gauss, proteinAT4G24280.1
bic:992 1e+03
peaks after deleting less than one fifth of global max:12
×10^9
raw data, starting:12
fitted Gauss locations:12
y
x
0
0.5
1
1.5
2
2.5
3
3.5
4
5
10
15
20
25

**Optimal1 Fitted Gauss, proteinAT4G24440.2;AT4G24440.1**
**bic:777**
**peaks after deleting less than one fifth of global max:11**

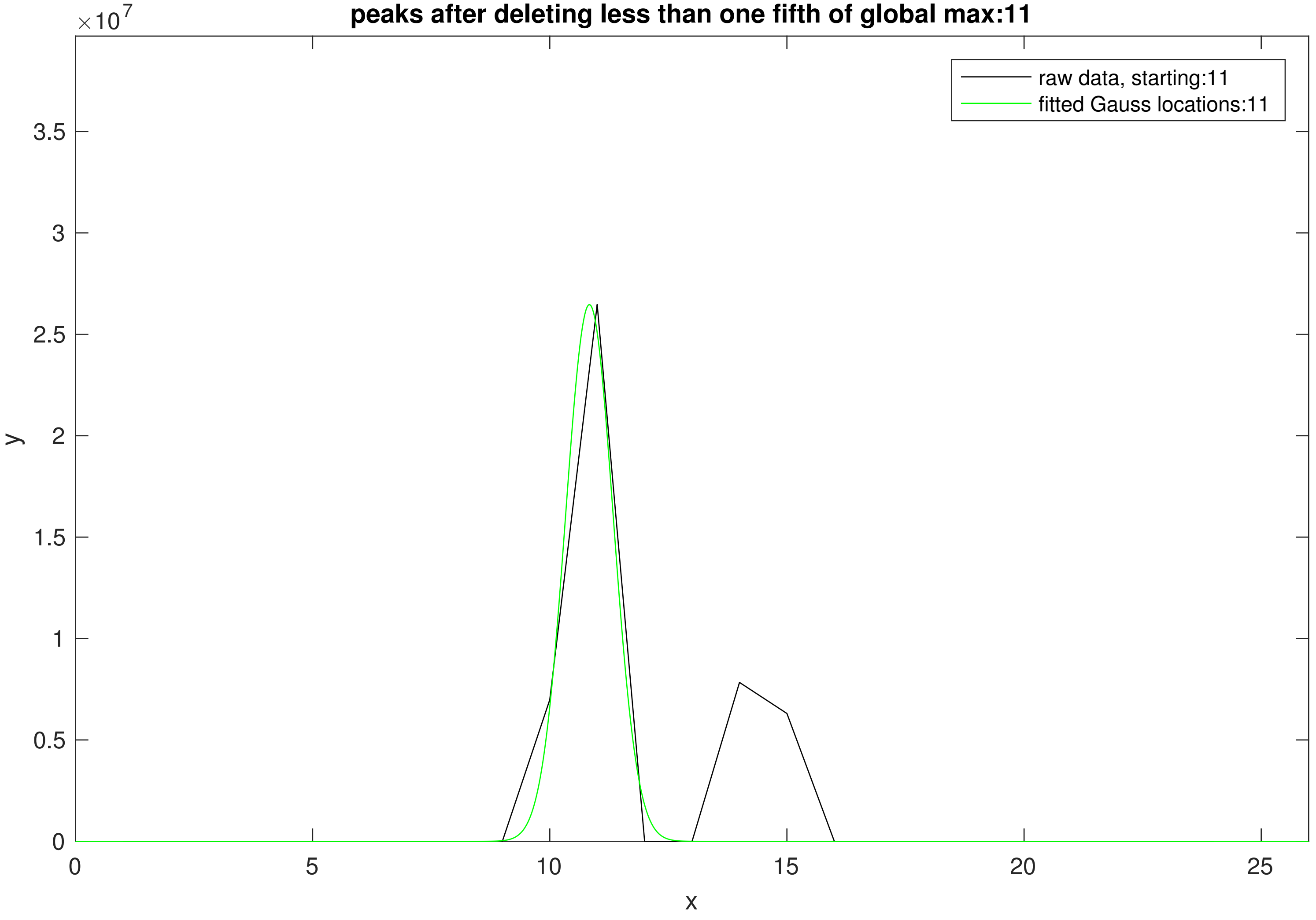

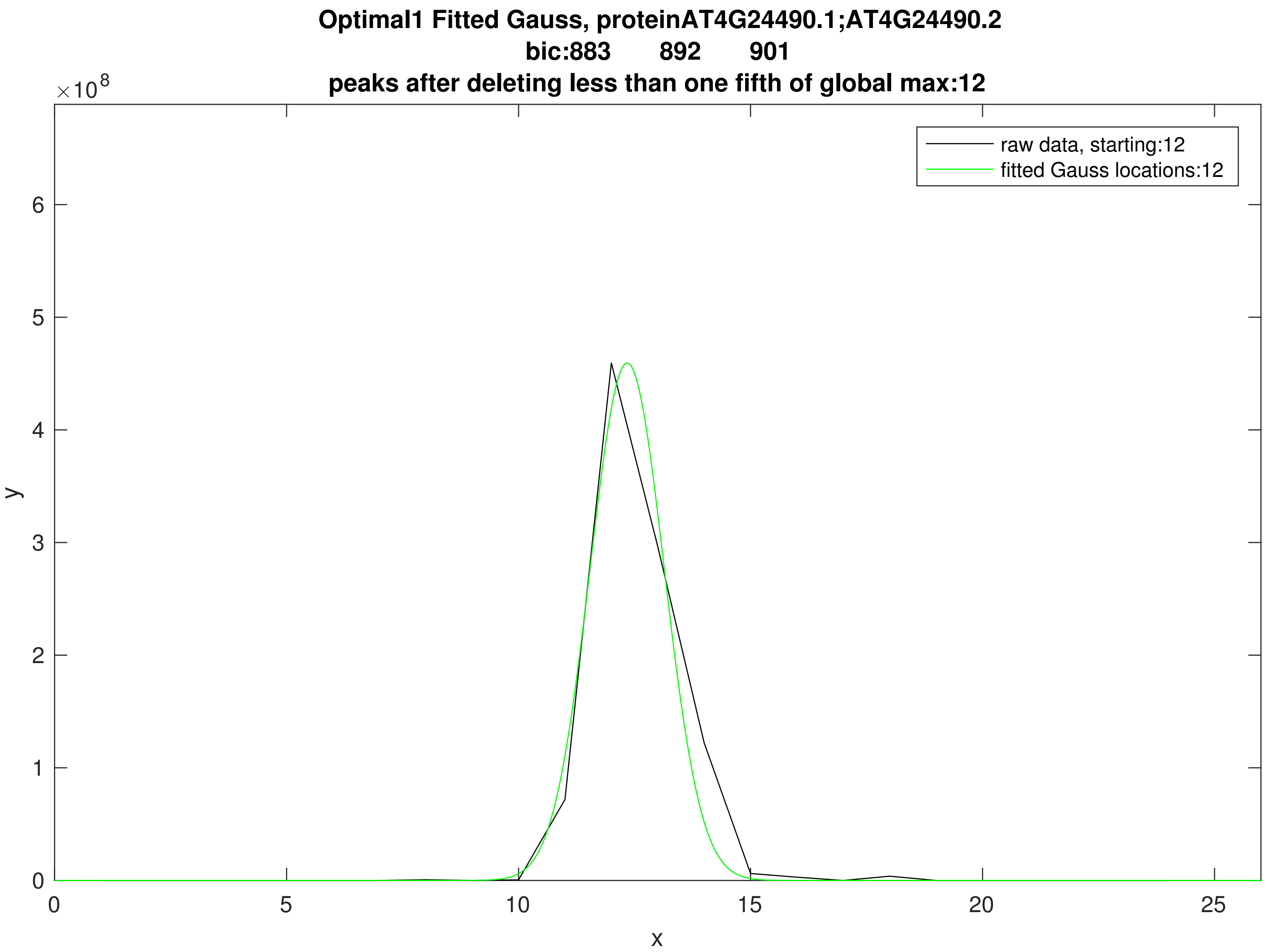
Optimal1 Fitted Gauss, proteinAT4G24490.1;AT4G24490.2
bic:883 892 901
peaks after deleting less than one fifth of global max:12
×10^8
raw data, starting:12
fitted Gauss locations:12
y
x
0
1
2
3
4
5
6
0
5
10
15
20
25

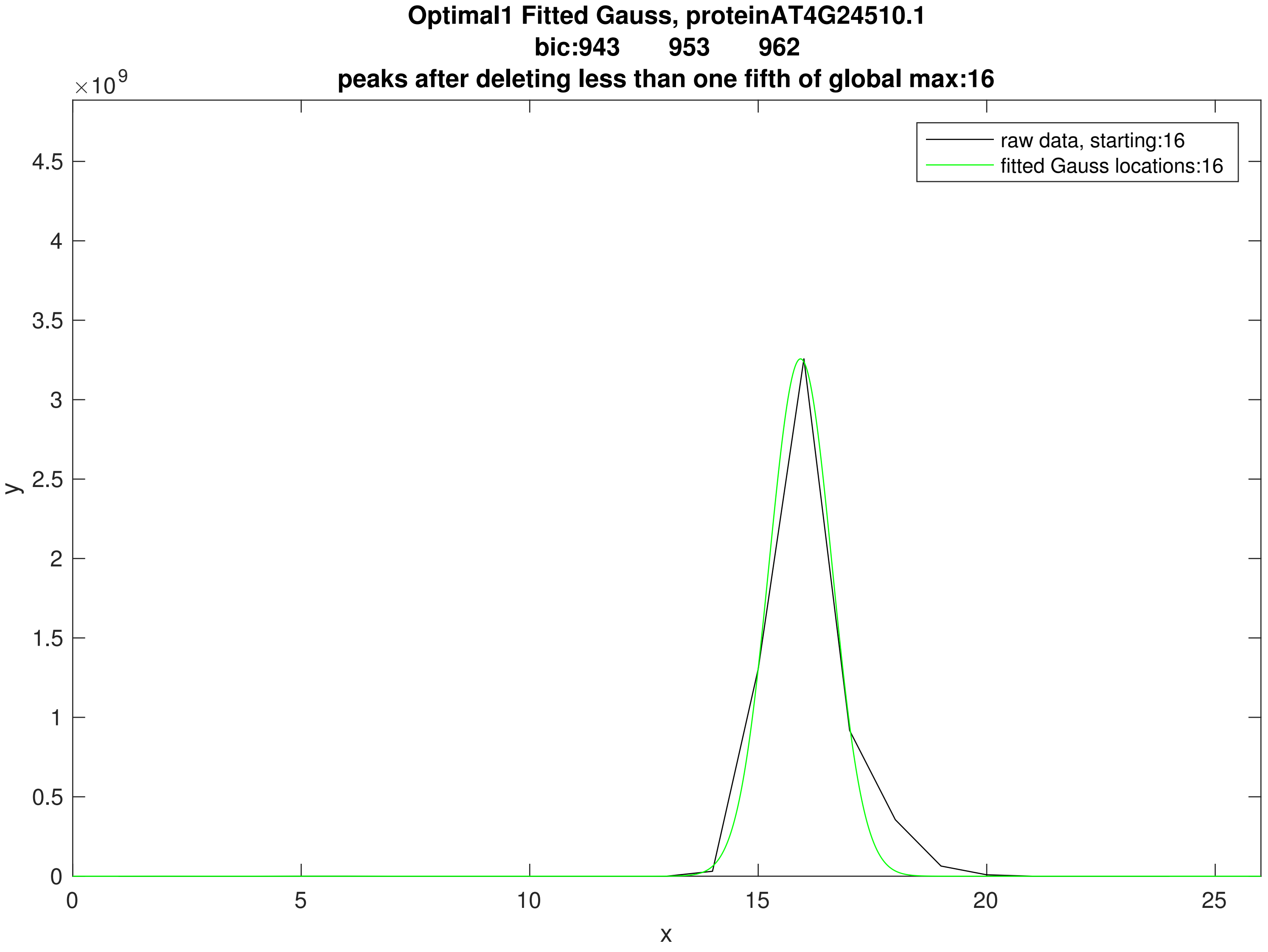
Optimal1 Fitted Gauss, proteinAT4G24510.1
bic:943 953 962
peaks after deleting less than one fifth of global max:16
×10^9
raw data, starting:16
fitted Gauss locations:16
4.5
4
3.5
3
2.5
2
1.5
1
0.5
0
y
0
5
10
15
20
25
x

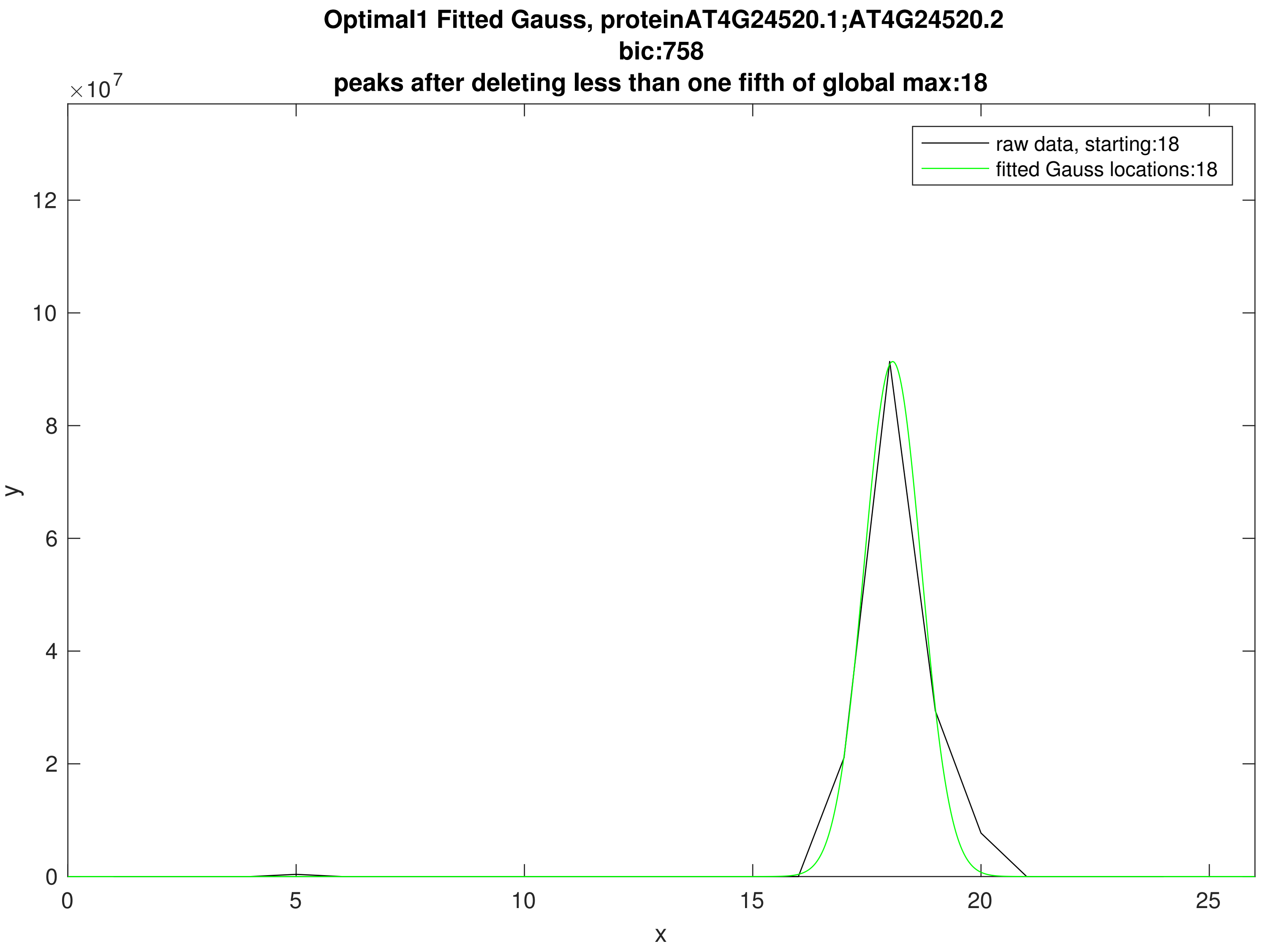
Optimal1 Fitted Gauss, proteinAT4G24520.1;AT4G24520.2
bic:758
peaks after deleting less than one fifth of global max:18
×10^7
raw data, starting:18
fitted Gauss locations:18
y
x
0
2
4
6
8
10
12
0
5
10
15
20
25

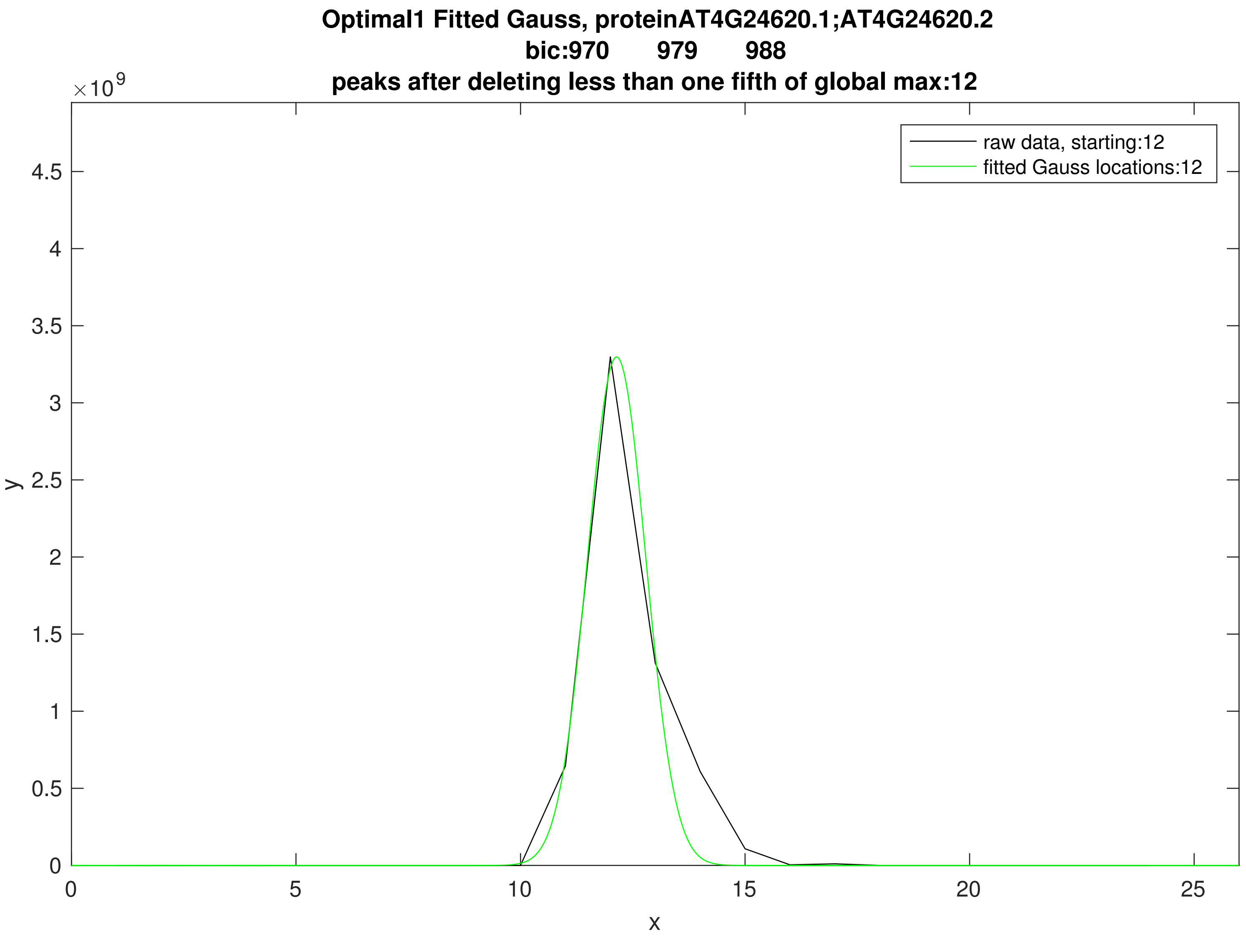
Optimal1 Fitted Gauss, proteinAT4G24620.1;AT4G24620.2
bic:970 979 988
peaks after deleting less than one fifth of global max:12
×10^9
4.5
4
3.5
3
2.5
2
1.5
1
0.5
0
y
0
5
10
15
20
25
x
raw data, starting:12
fitted Gauss locations:12

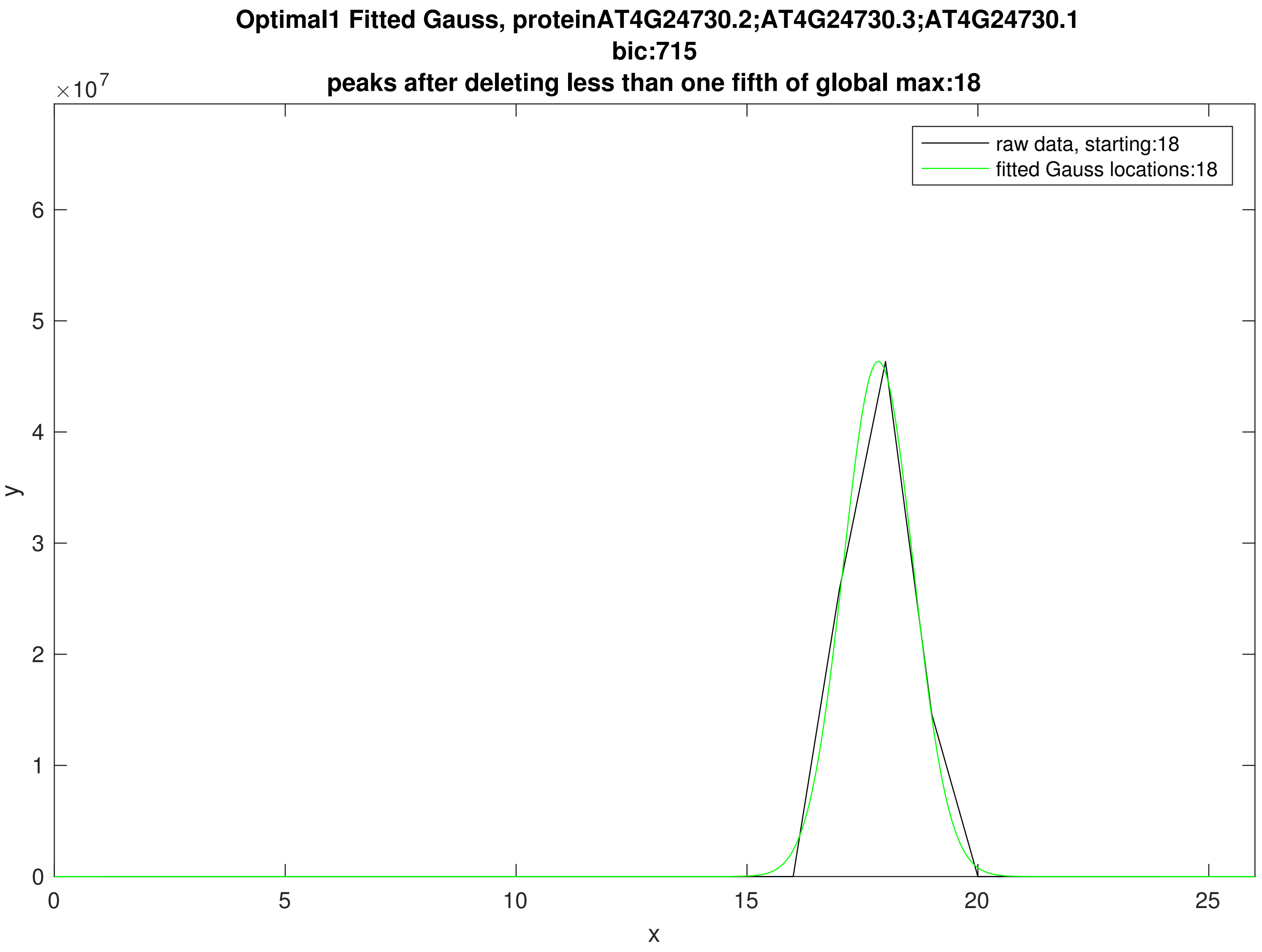

Optimal1 Fitted Gauss, proteinAT4G24730.2;AT4G24730.3;AT4G24730.1
bic:715
peaks after deleting less than one fifth of global max:18
×10^7
raw data, starting:18
fitted Gauss locations:18
y
x
0
1
2
3
4
5
6
0
5
10
15
20
25

**Optimal1 Fitted Gauss, proteinAT4G24770.1**
**bic:932 960 974 984**
**peaks after deleting less than one fifth of global max:15**

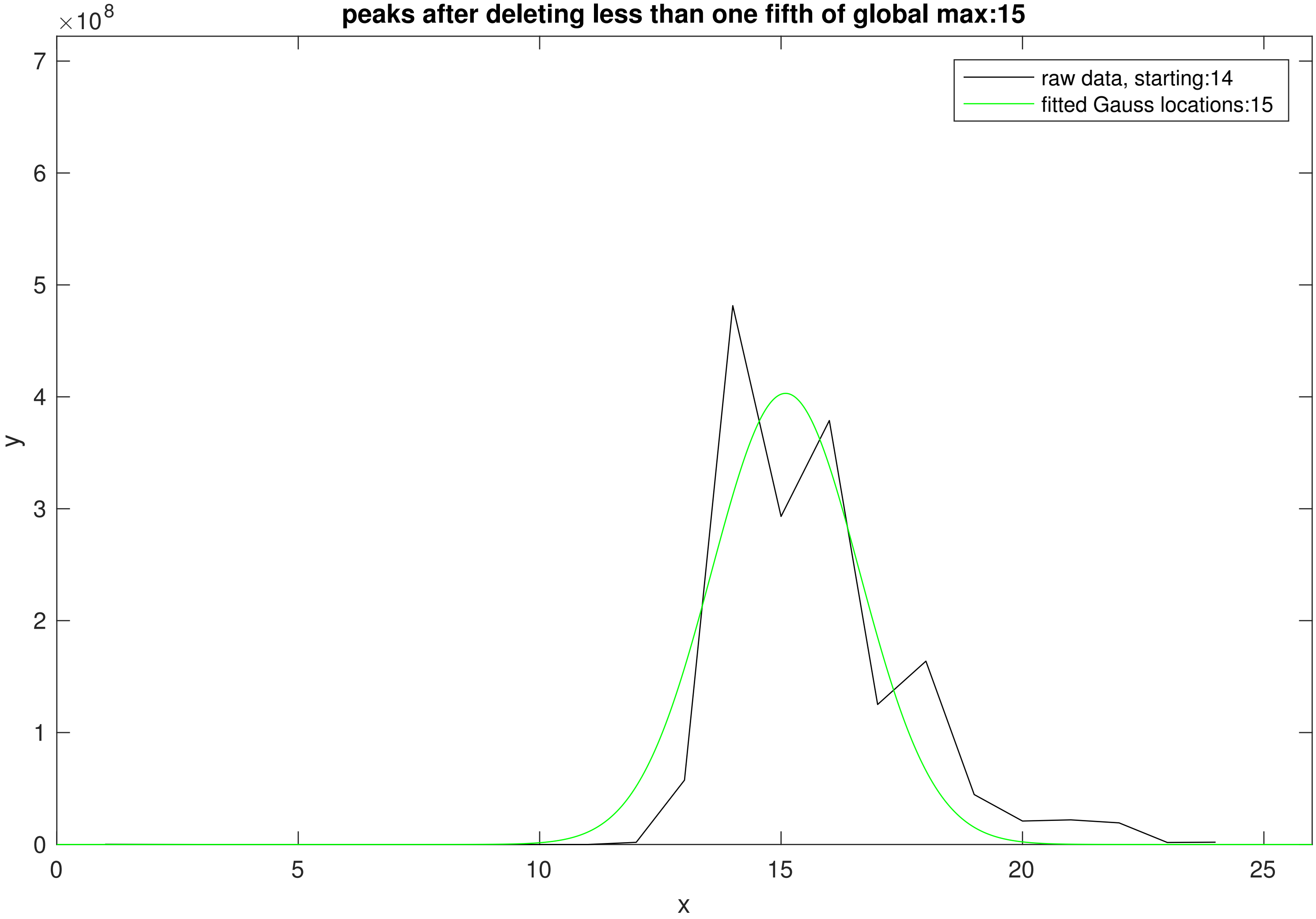

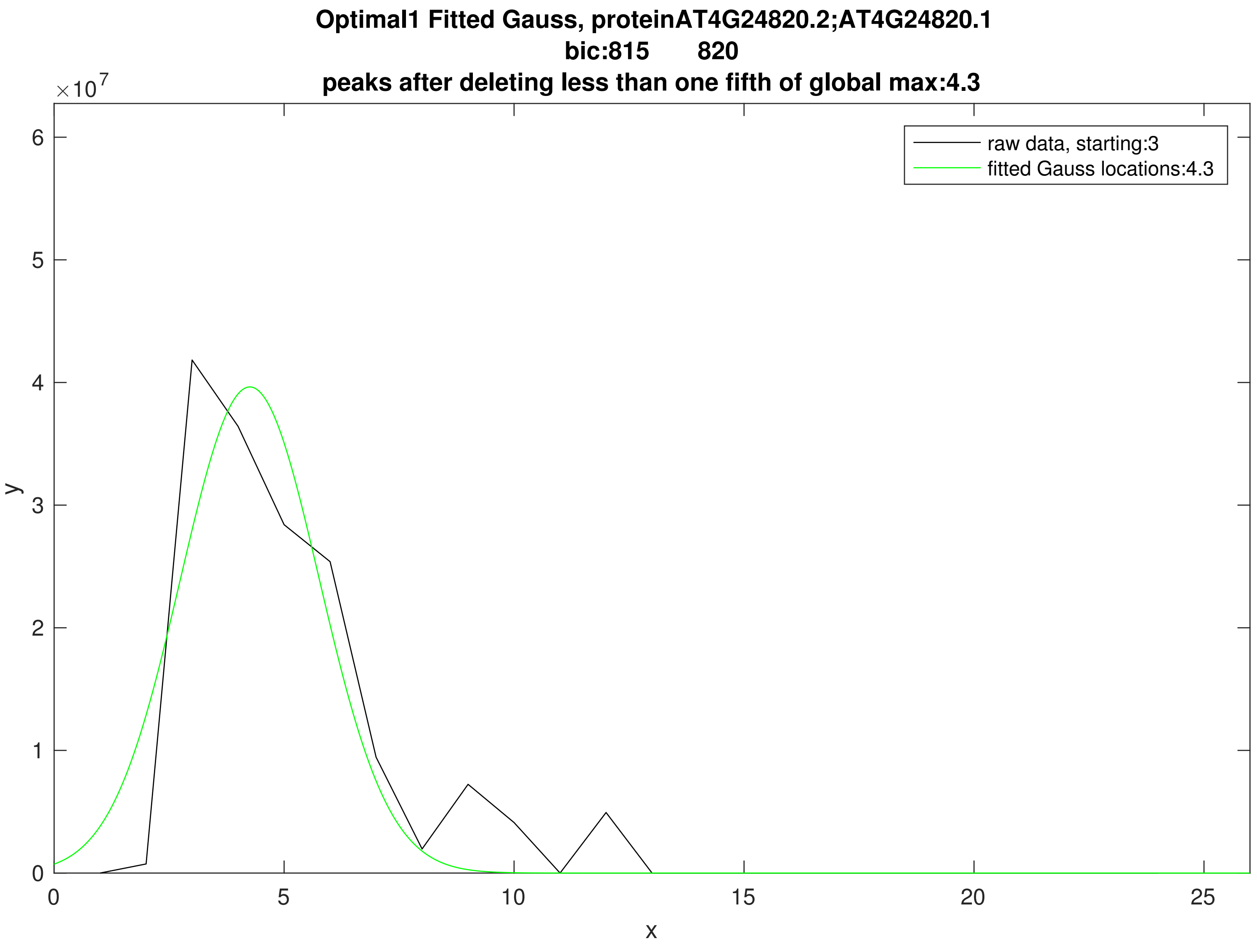
Optimal1 Fitted Gauss, proteinAT4G24820.2;AT4G24820.1
bic:815 820
peaks after deleting less than one fifth of global max:4.3
×10^7
raw data, starting:3
fitted Gauss locations:4.3
y
x
0
1
2
3
4
5
6
0
5
10
15
20
25

#### Optimal1 Fitted Gauss, proteinAT4G24830.1;AT4G24830.2
#### bic:793 801
#### peaks after deleting less than one fifth of global max:9.9

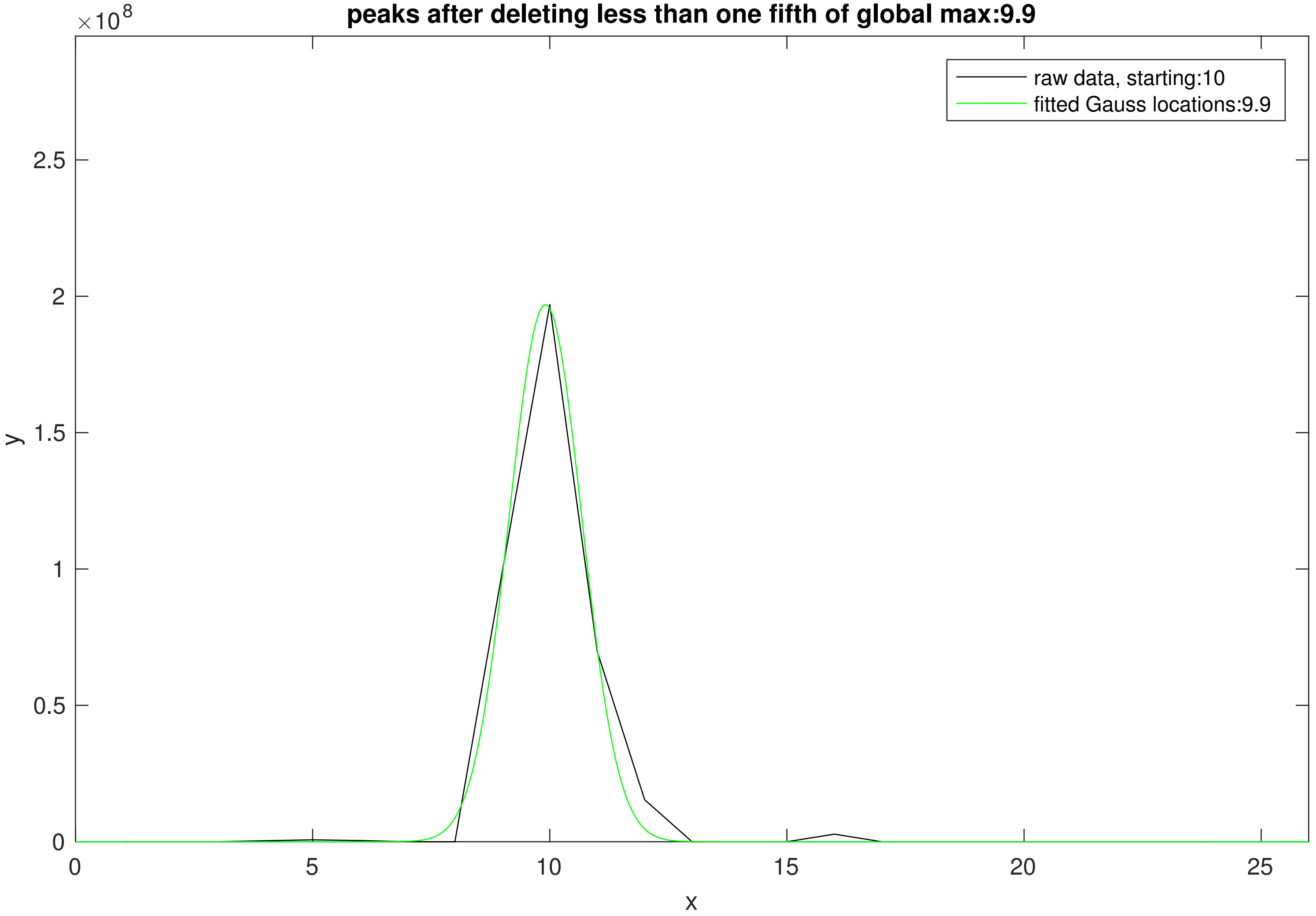

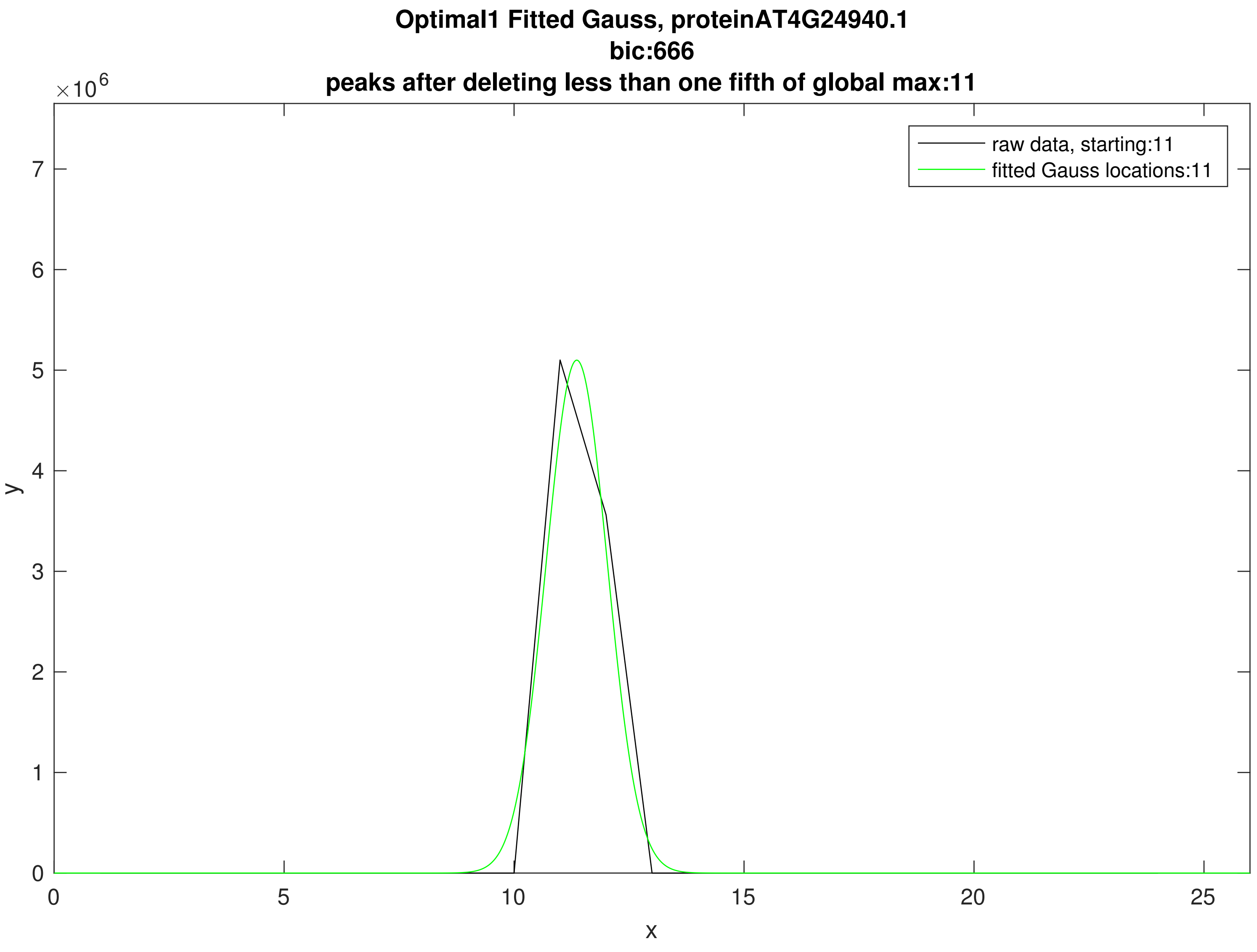
Optimal1 Fitted Gauss, proteinAT4G24940.1
bic:666
peaks after deleting less than one fifth of global max:11
×10^6
raw data, starting:11
fitted Gauss locations:11
y
x
0
1
2
3
4
5
6
7
0
5
10
15
20
25

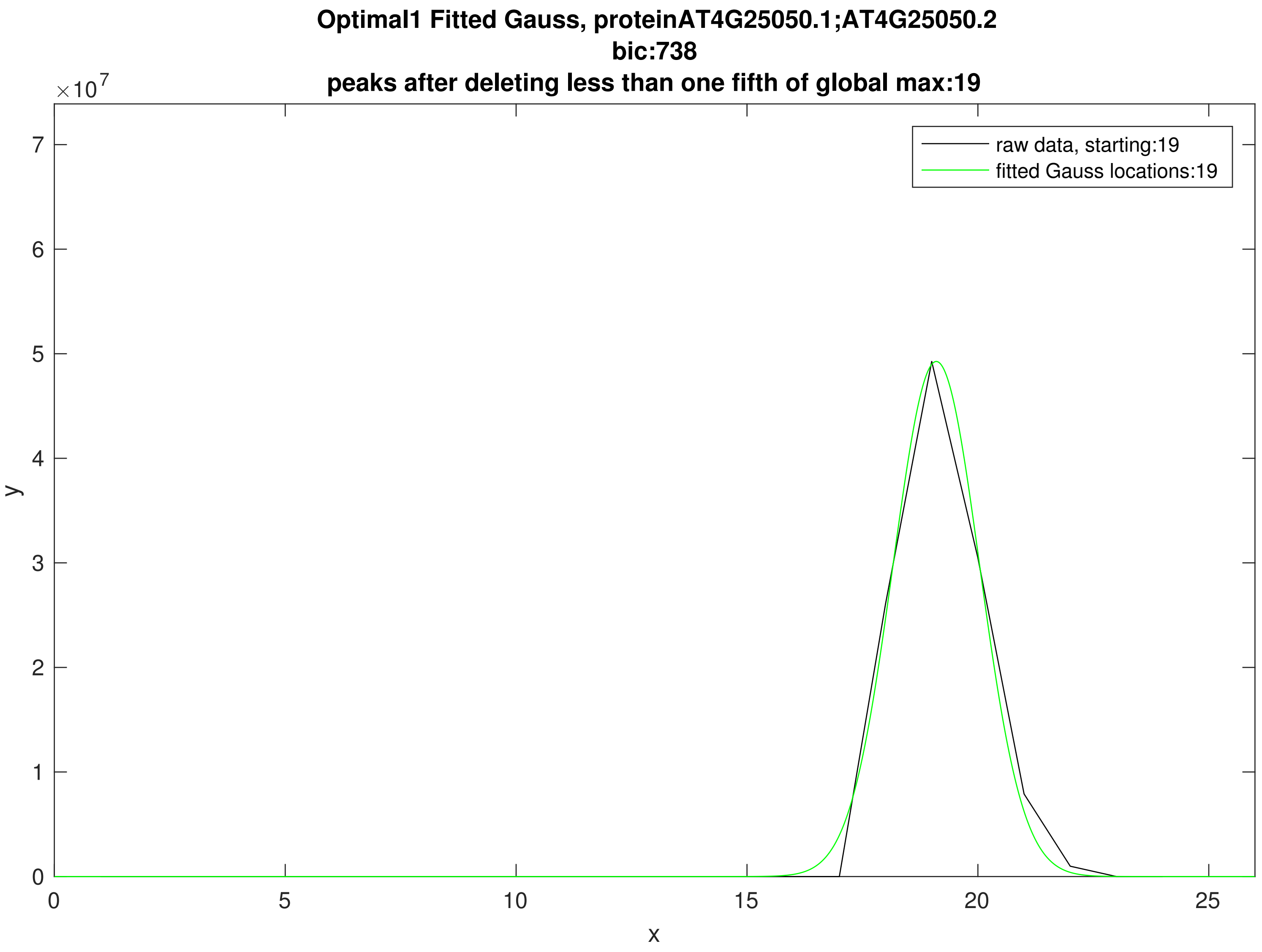
Optimal1 Fitted Gauss, proteinAT4G25050.1;AT4G25050.2
bic:738
peaks after deleting less than one fifth of global max:19
×10^7
raw data, starting:19
fitted Gauss locations:19
y
x
0
1
2
3
4
5
6
7
0
5
10
15
20
25

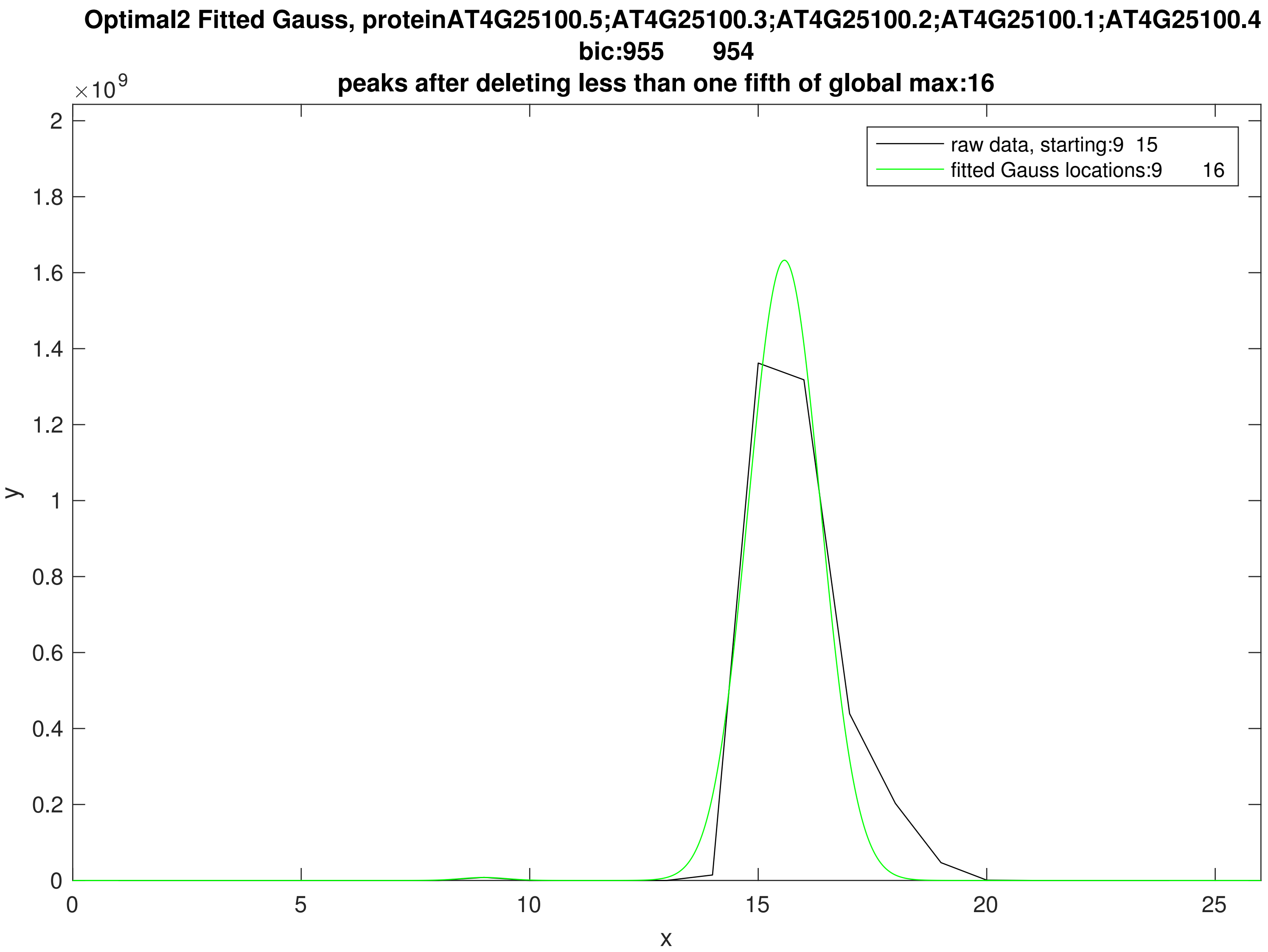
Optimal2 Fitted Gauss, proteinAT4G25100.5;AT4G25100.3;AT4G25100.2;AT4G25100.1;AT4G25100.4
bic:955 954
peaks after deleting less than one fifth of global max:16
×10^9
raw data, starting:9 15
fitted Gauss locations:9 16
y
x
0
0.2
0.4
0.6
0.8
1
1.2
1.4
1.6
1.8
2
0
5
10
15
20
25

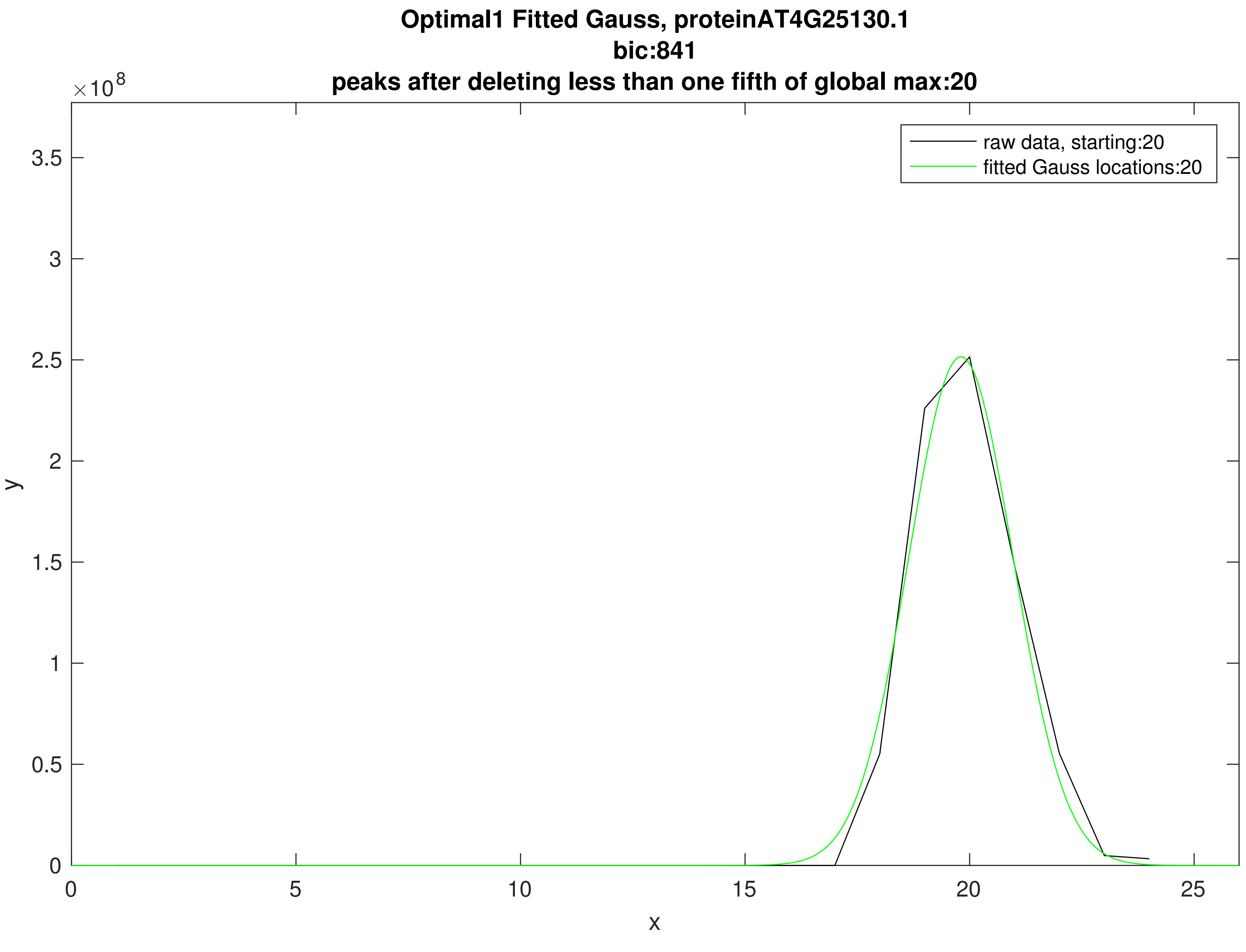

Optimal1 Fitted Gauss, proteinAT4G25130.1
bic:841
peaks after deleting less than one fifth of global max:20
×10^8
3.5
3
2.5
2
1.5
1
0.5
0
y
0
5
10
15
20
25
x
raw data, starting:20
fitted Gauss locations:20

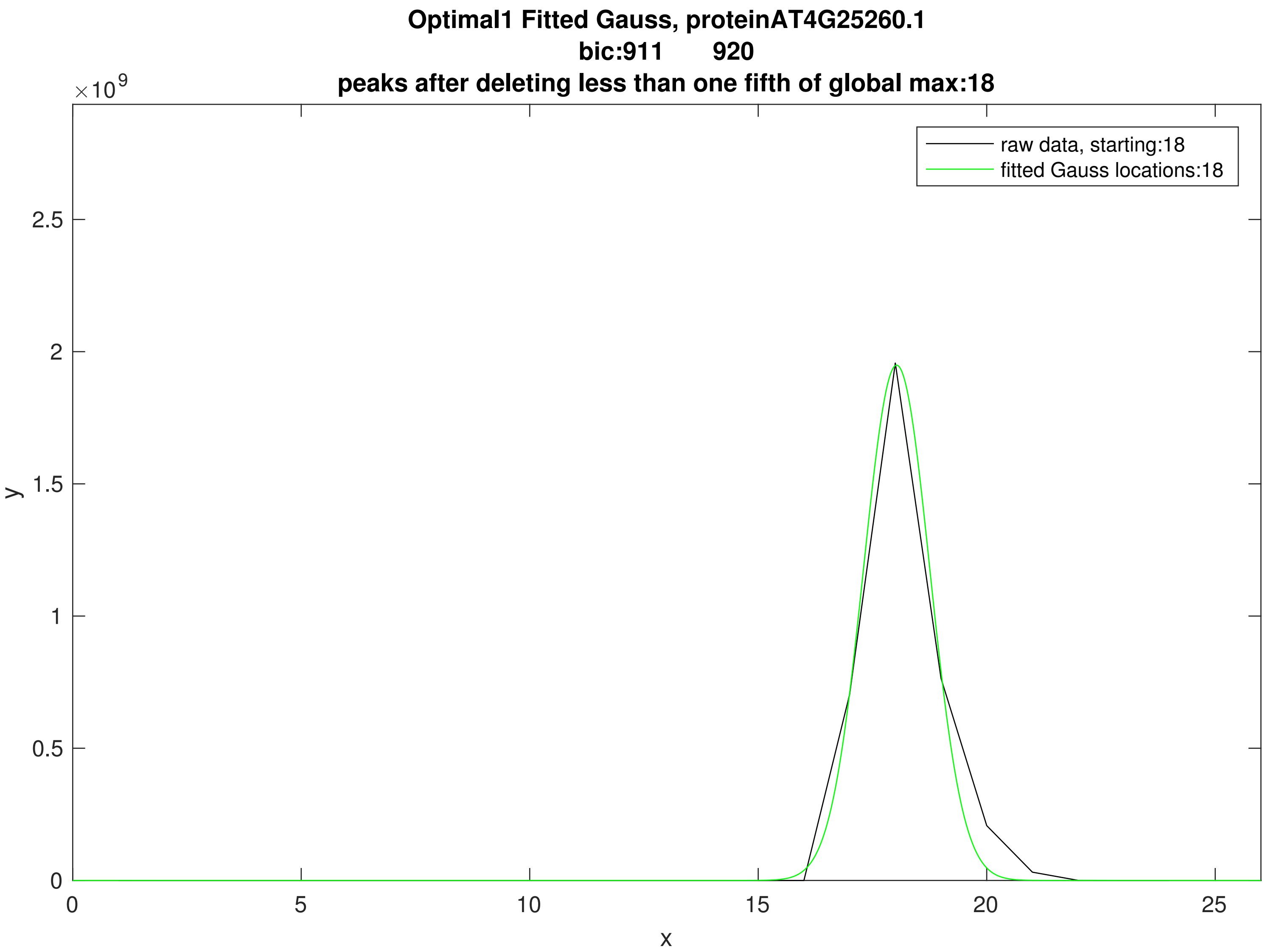
Optimal1 Fitted Gauss, proteinAT4G25260.1
bic:911 920
peaks after deleting less than one fifth of global max:18
×10^9
raw data, starting:18
fitted Gauss locations:18
y
x
0
0.5
1
1.5
2
2.5
0
5
10
15
20
25

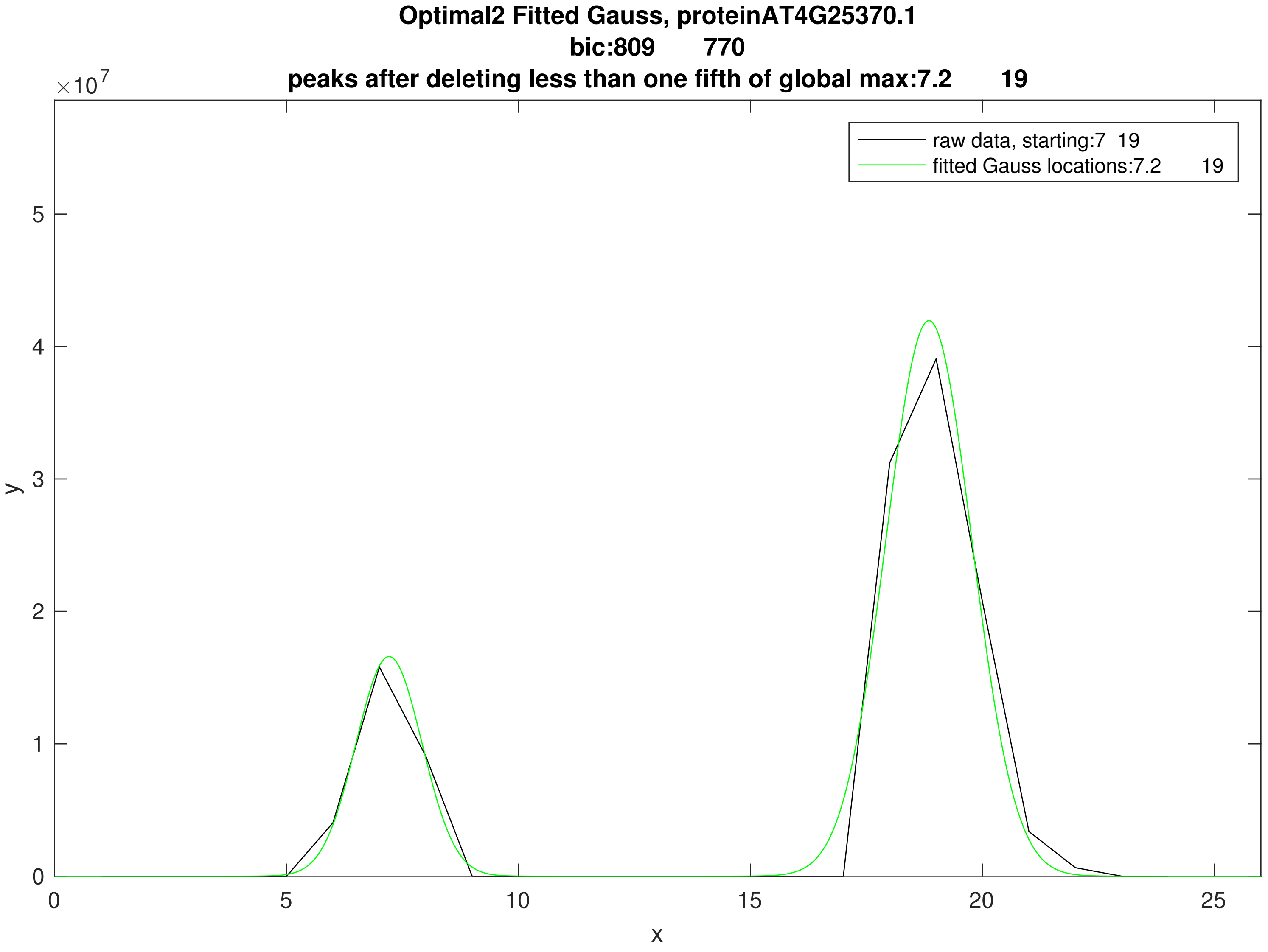

Optimal2 Fitted Gauss, proteinAT4G25370.1
bic:809 770
peaks after deleting less than one fifth of global max:7.2 19
×10^7
raw data, starting:7 19
fitted Gauss locations:7.2 19
y
x
0
1
2
3
4
5
0
5
10
15
20
25

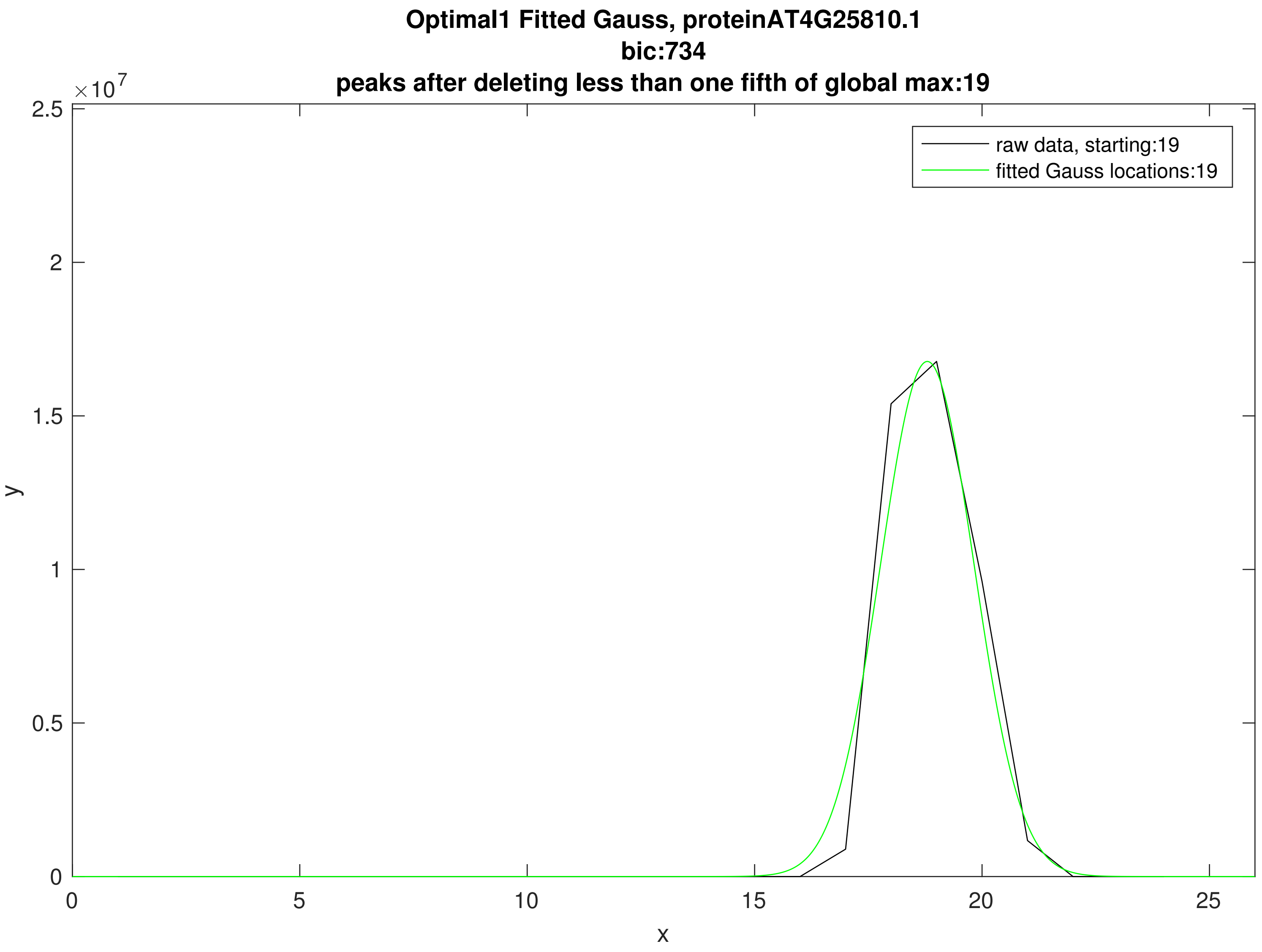

Optimal1 Fitted Gauss, proteinAT4G25810.1
bic:734
peaks after deleting less than one fifth of global max:19
$\times 10^7$
2.5
2
1.5
1
0.5
0
y
0
5
10
15
20
25
x
raw data, starting:19
fitted Gauss locations:19

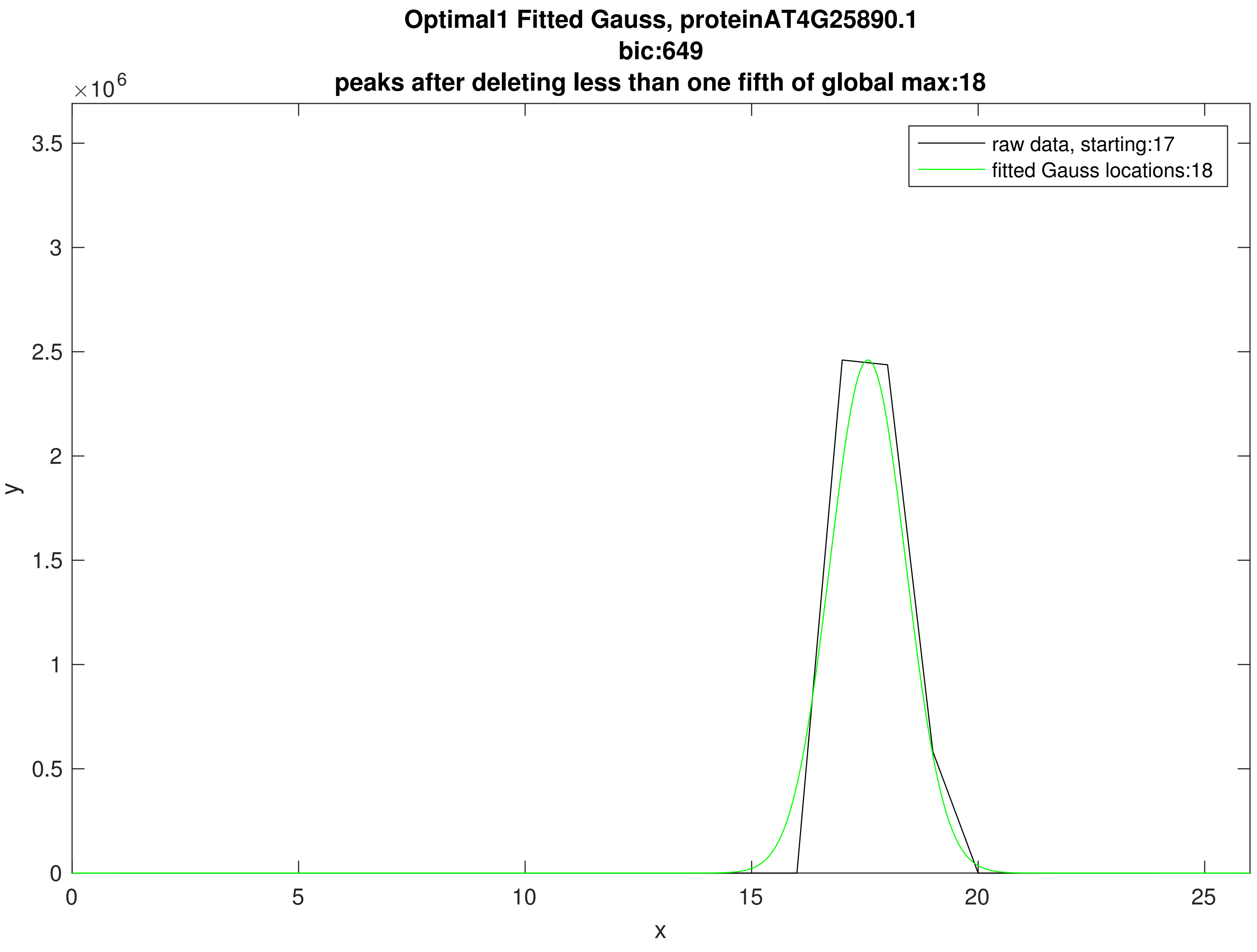
Optimal1 Fitted Gauss, proteinAT4G25890.1
bic:649
peaks after deleting less than one fifth of global max:18
×10^6
raw data, starting:17
fitted Gauss locations:18
3.5
3
2.5
2
1.5
1
0.5
0
y
0
5
10
15
20
25
x

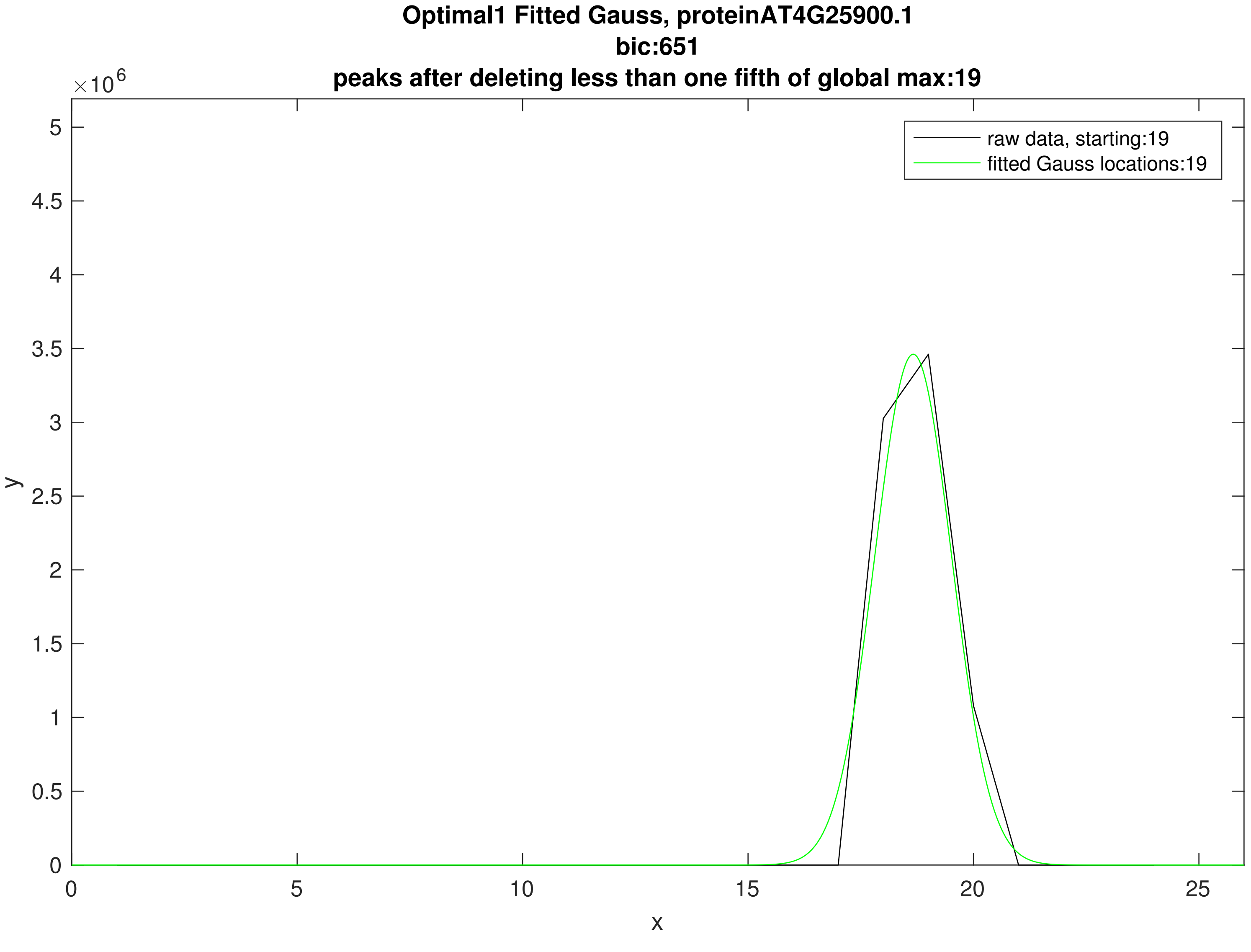
Optimal1 Fitted Gauss, proteinAT4G25900.1
bic:651
peaks after deleting less than one fifth of global max:19
×10^6
raw data, starting:19
fitted Gauss locations:19
y
x
0
0.5
1
1.5
2
2.5
3
3.5
4
4.5
5
0
5
10
15
20
25

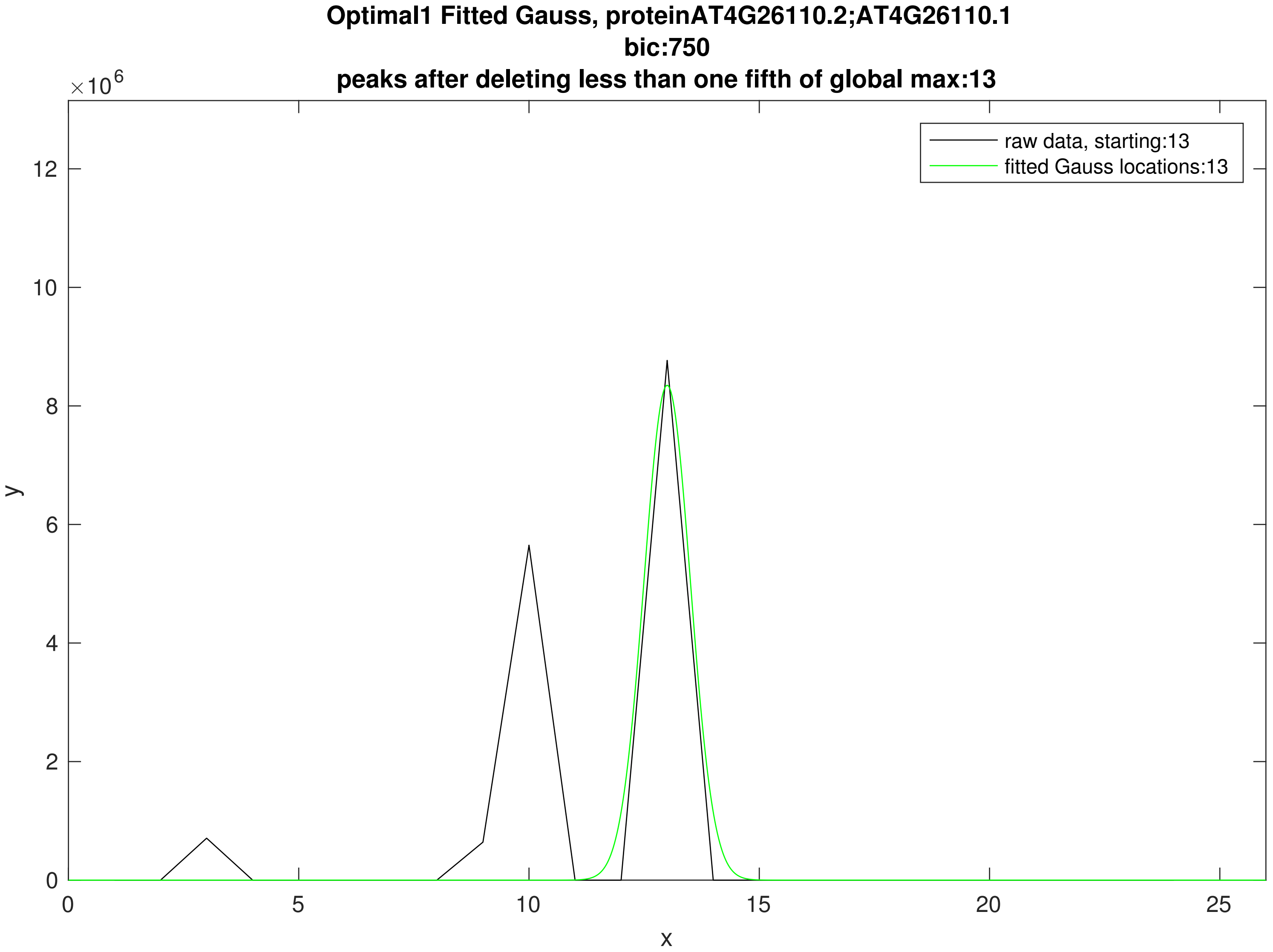
Optimal1 Fitted Gauss, proteinAT4G26110.2;AT4G26110.1
bic:750
peaks after deleting less than one fifth of global max:13
×10^6
raw data, starting:13
fitted Gauss locations:13
y
x
0
2
4
6
8
10
12
0
5
10
15
20
25

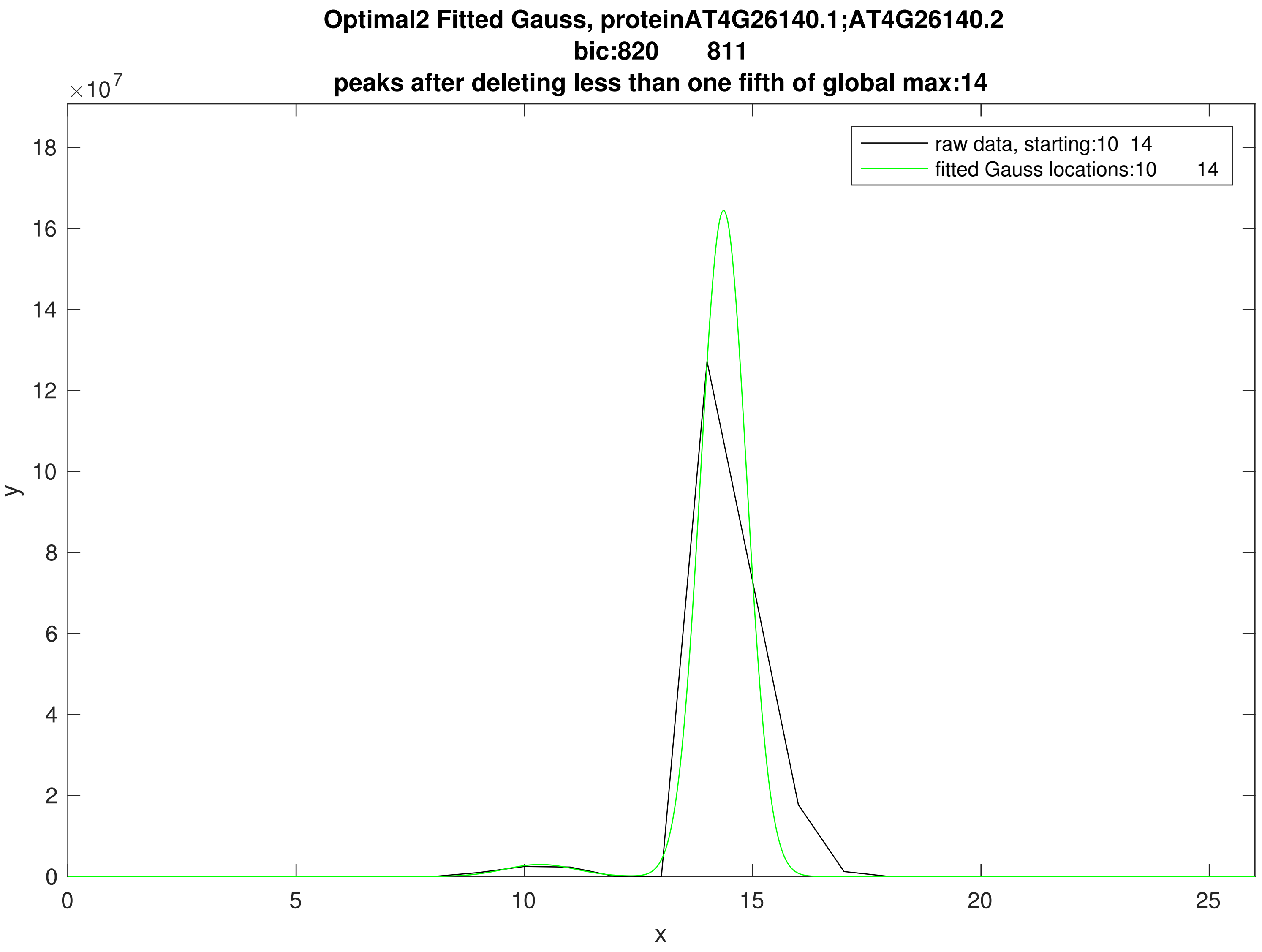
Optimal2 Fitted Gauss, proteinAT4G26140.1;AT4G26140.2
bic:820 811
peaks after deleting less than one fifth of global max:14
×10^7
raw data, starting:10 14
fitted Gauss locations:10 14
18
16
14
12
10
8
6
4
2
0
y
0
5
10
15
20
25
x

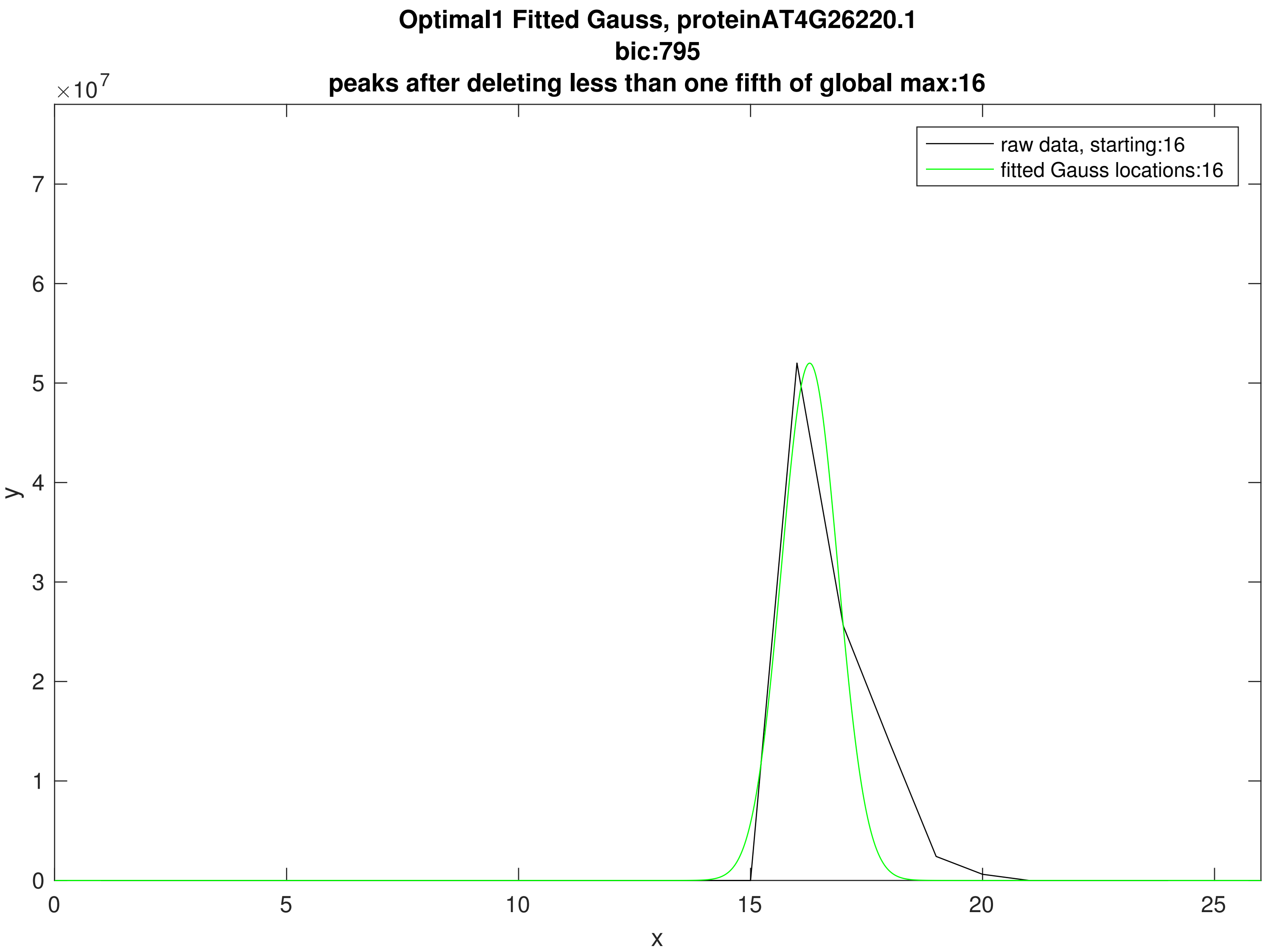

Optimal1 Fitted Gauss, proteinAT4G26220.1
bic:795
peaks after deleting less than one fifth of global max:16
×10^7
raw data, starting:16
fitted Gauss locations:16
y
x
0
1
2
3
4
5
6
7
0
5
10
15
20
25

Optimal1 Fitted Gauss, proteinAT4G26300.1
bic:870 872
peaks after deleting less than one fifth of global max:14

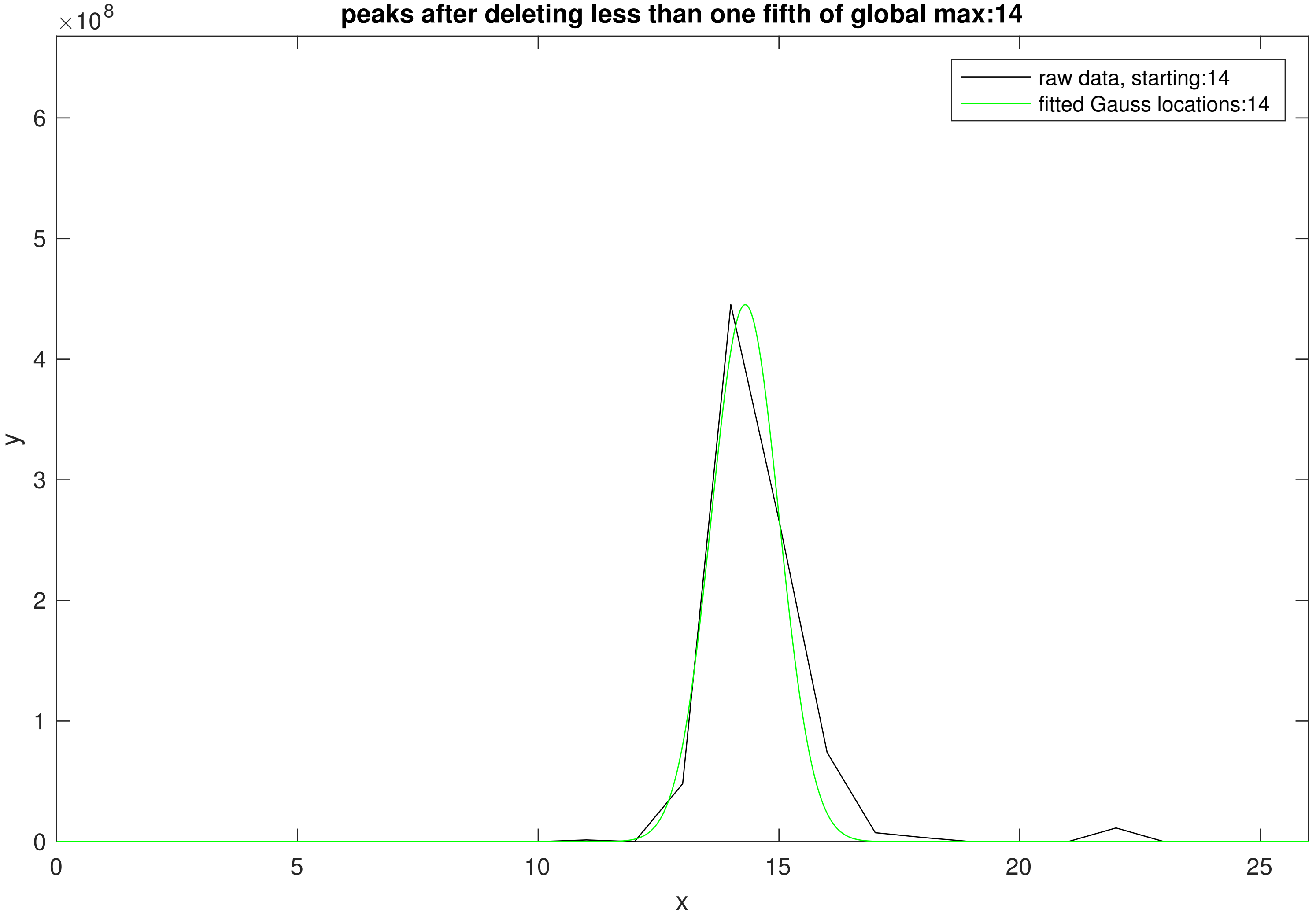

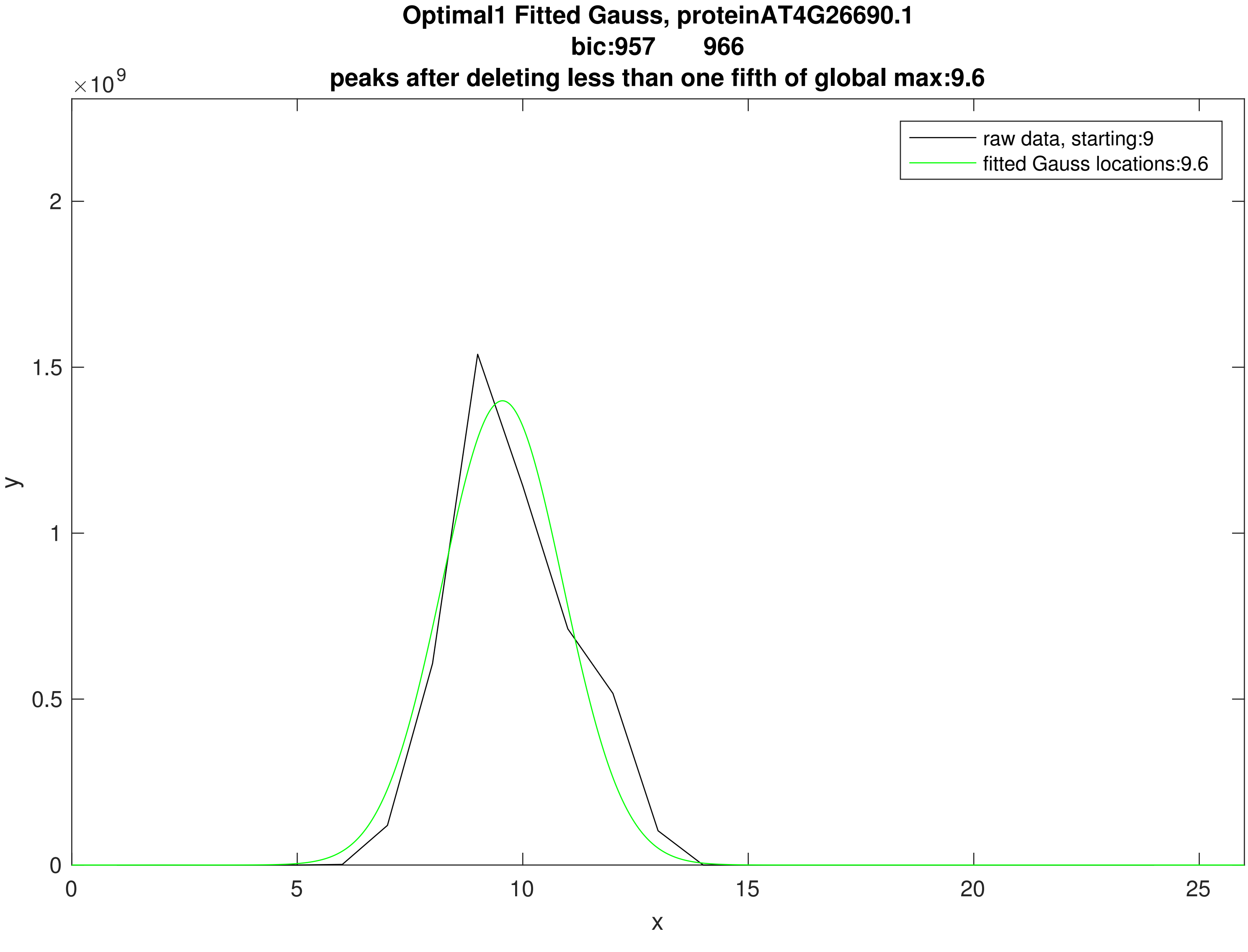
Optimal1 Fitted Gauss, proteinAT4G26690.1
bic:957 966
peaks after deleting less than one fifth of global max:9.6
×10^9
raw data, starting:9
fitted Gauss locations:9.6
y
x
0
0.5
1
1.5
2
5
10
15
20
25

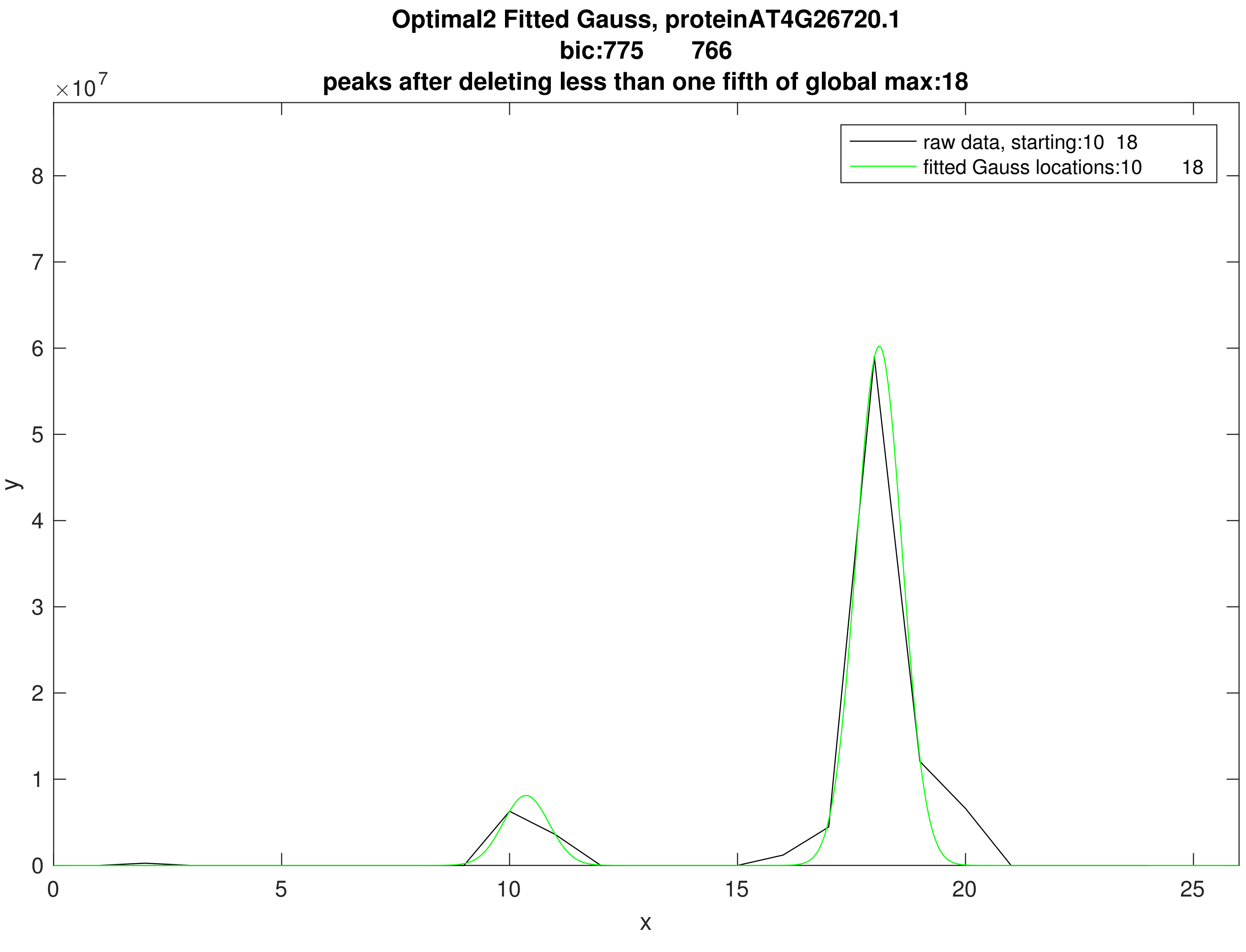
Optimal2 Fitted Gauss, proteinAT4G26720.1
bic:775 766
peaks after deleting less than one fifth of global max:18
×10^7
raw data, starting:10 18
fitted Gauss locations:10 18
y
x
0
1
2
3
4
5
6
7
8
0
5
10
15
20
25

**Optimal1 Fitted Gauss, proteinAT4G26780.1**
**bic:783 792**
**peaks after deleting less than one fifth of global max:8.9**

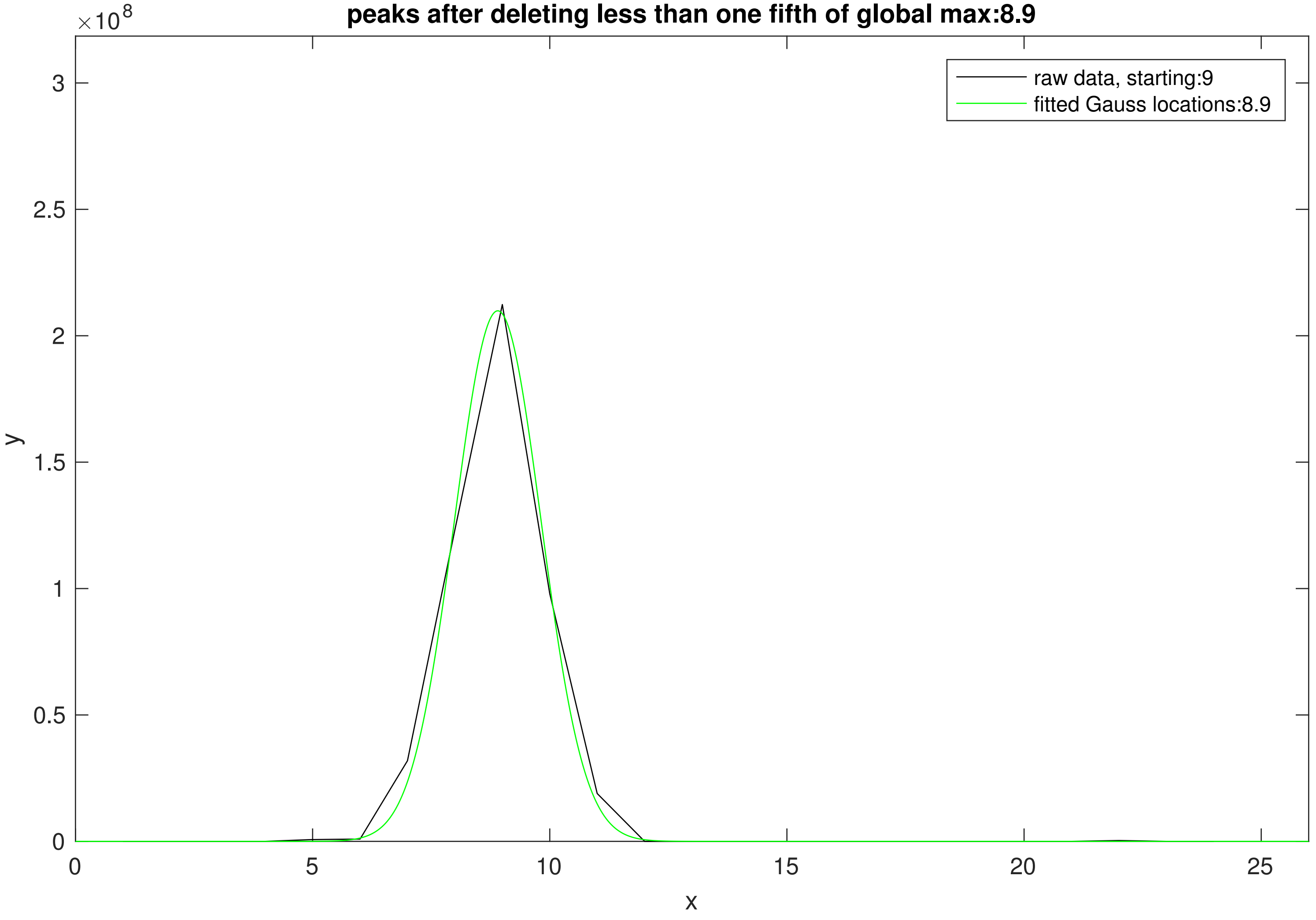

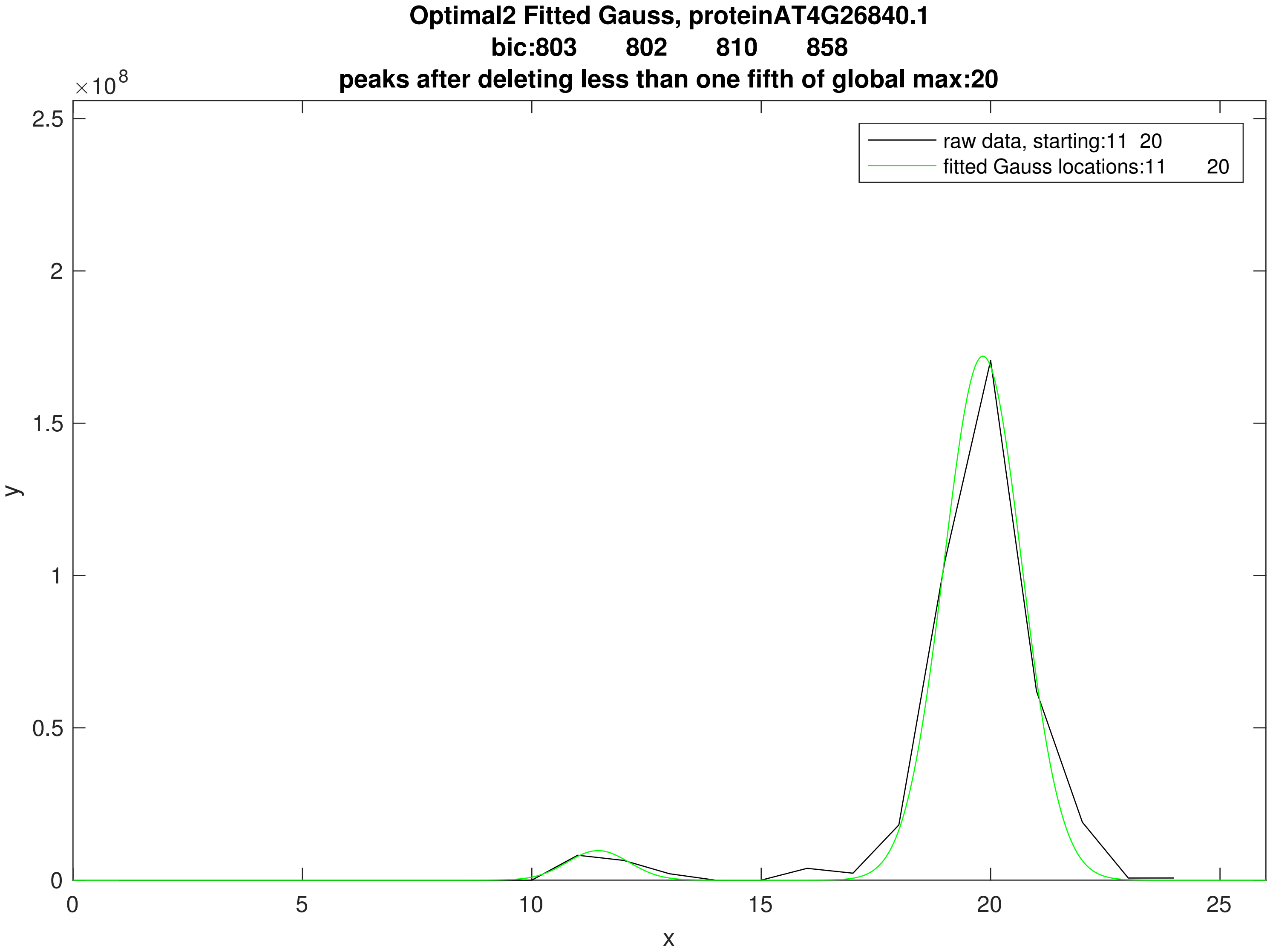

Optimal2 Fitted Gauss, proteinAT4G26840.1
bic:803 802 810 858
peaks after deleting less than one fifth of global max:20
×10^8
2.5
2
1.5
1
0.5
0
y
0
5
10
15
20
25
x
raw data, starting:11 20
fitted Gauss locations:11 20

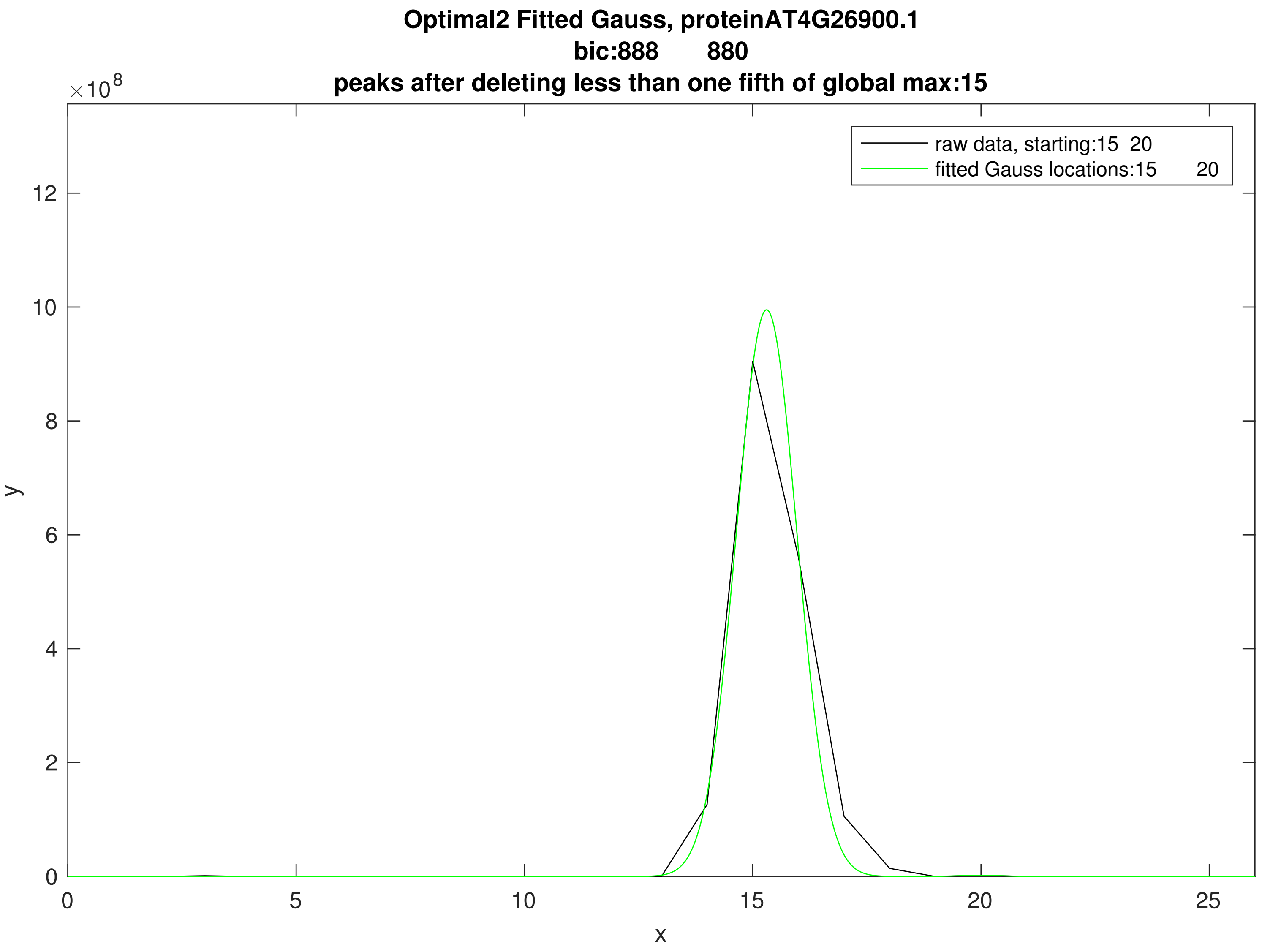
Optimal2 Fitted Gauss, proteinAT4G26900.1
bic:888 880
peaks after deleting less than one fifth of global max:15
×10^8
raw data, starting:15 20
fitted Gauss locations:15 20
0
2
4
6
8
10
12
y
0
5
10
15
20
25
x

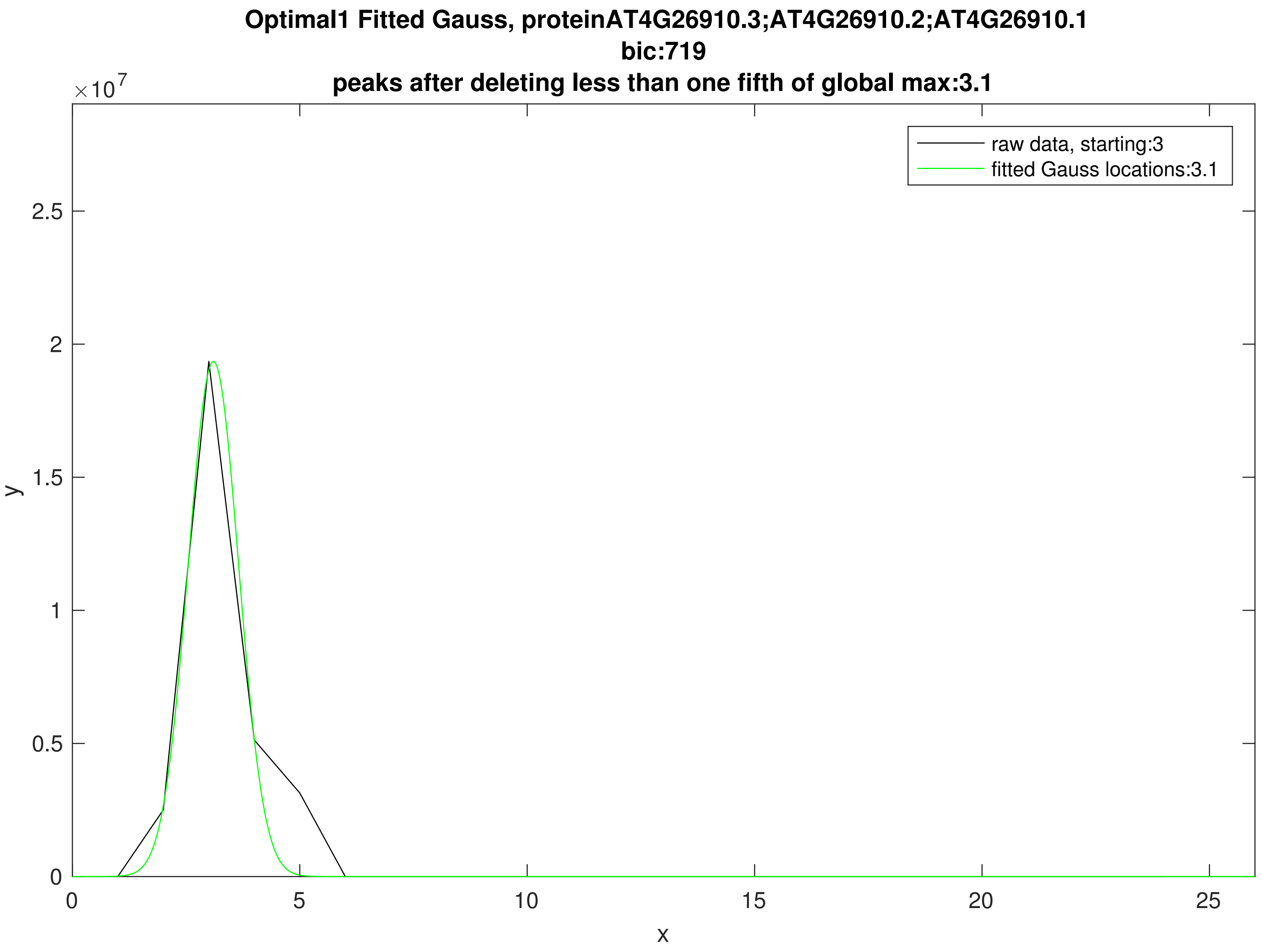
Optimal1 Fitted Gauss, proteinAT4G26910.3;AT4G26910.2;AT4G26910.1
bic:719
peaks after deleting less than one fifth of global max:3.1
×10^7
raw data, starting:3
fitted Gauss locations:3.1
y
x
0
0.5
1
1.5
2
2.5
0
5
10
15
20
25

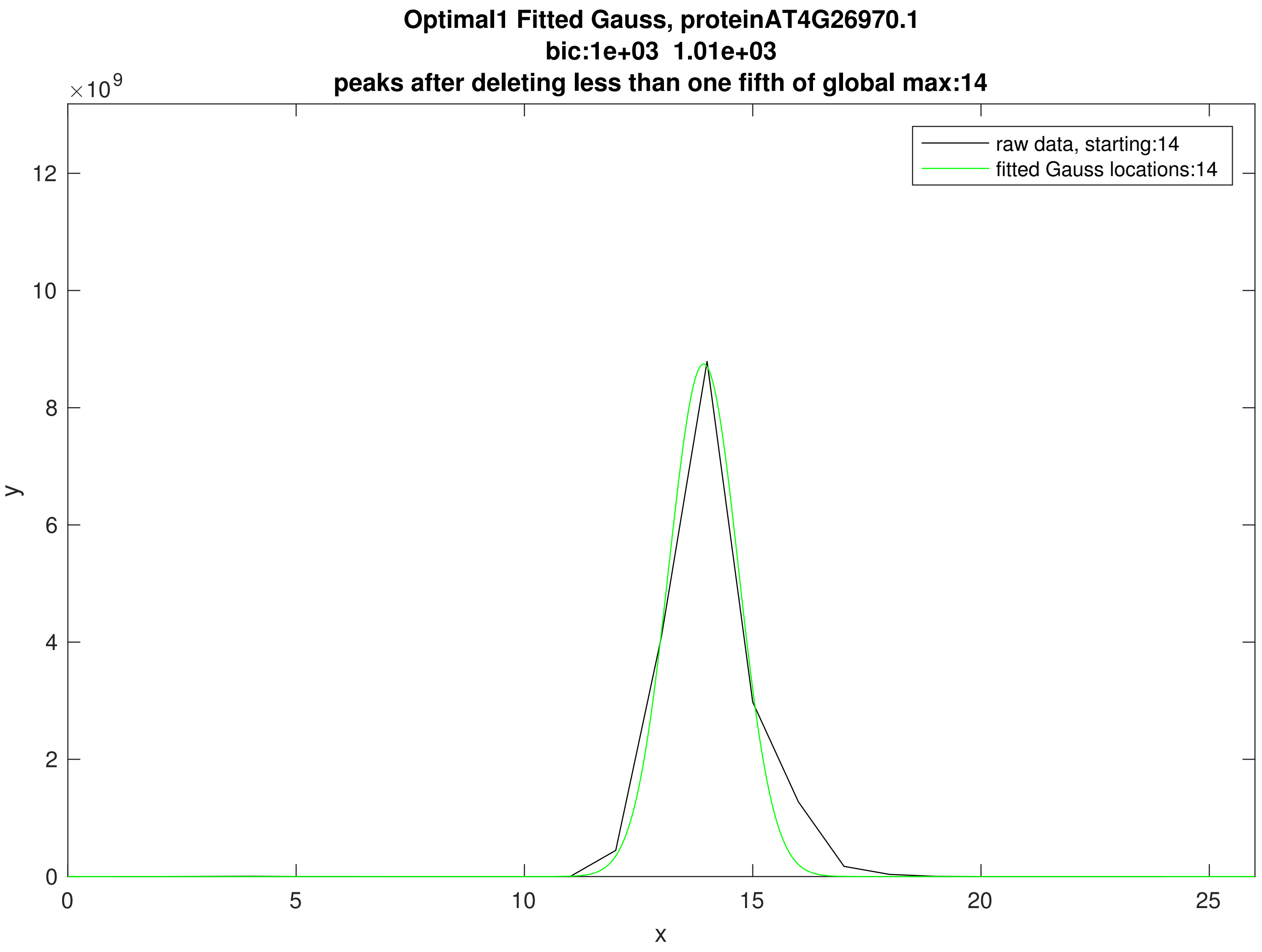

Optimal1 Fitted Gauss, proteinAT4G26970.1
bic:1e+03  1.01e+03
peaks after deleting less than one fifth of global max:14
×10^9
12
10
8
6
4
2
0
y
0
5
10
15
20
25
x
raw data, starting:14
fitted Gauss locations:14

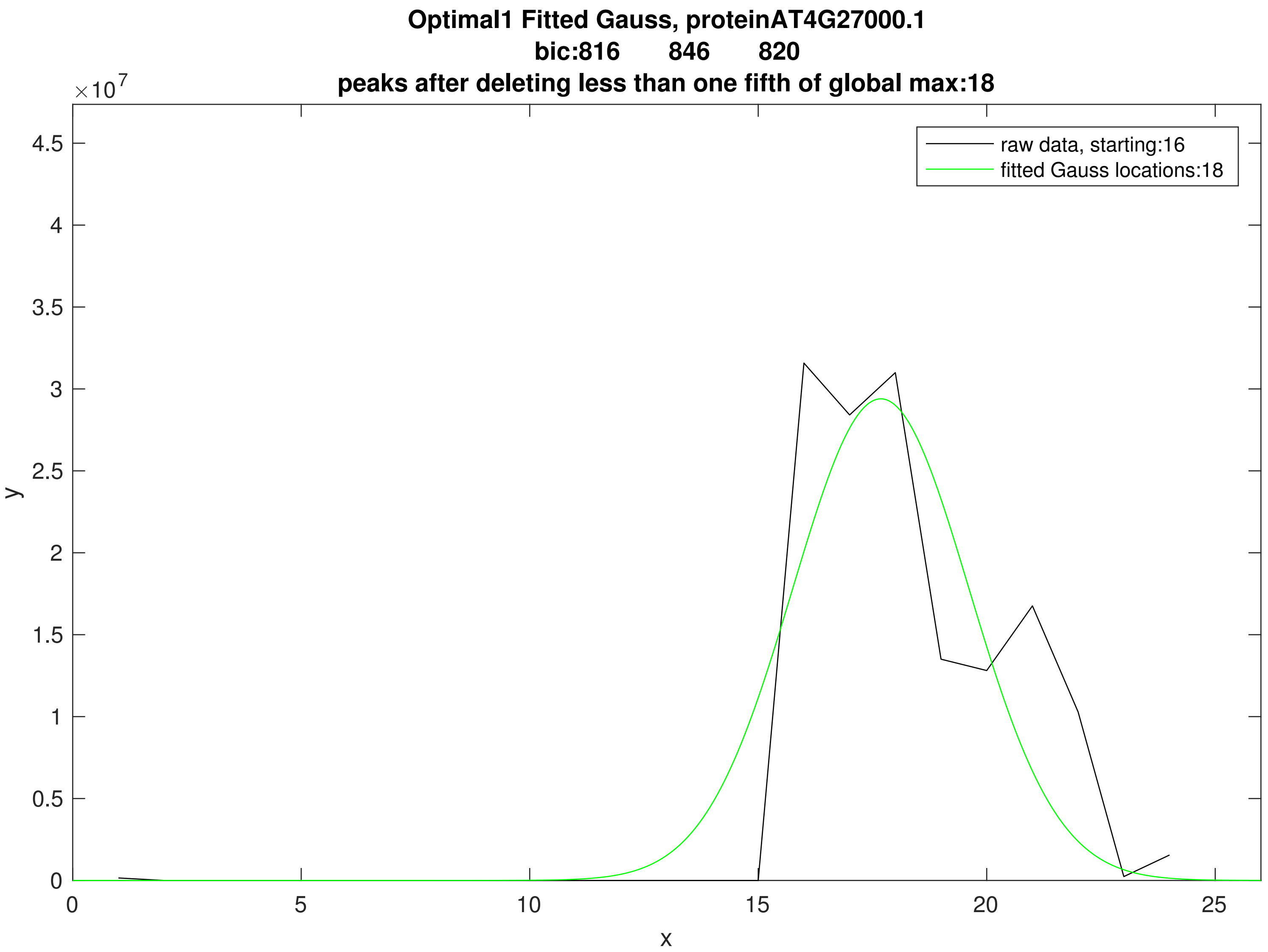

Optimal1 Fitted Gauss, proteinAT4G27000.1
bic:816 846 820
peaks after deleting less than one fifth of global max:18
×10^7
raw data, starting:16
fitted Gauss locations:18
y
x

**Optimal1 Fitted Gauss, proteinAT4G27070.1**
**bic:738**
**peaks after deleting less than one fifth of global max:12**

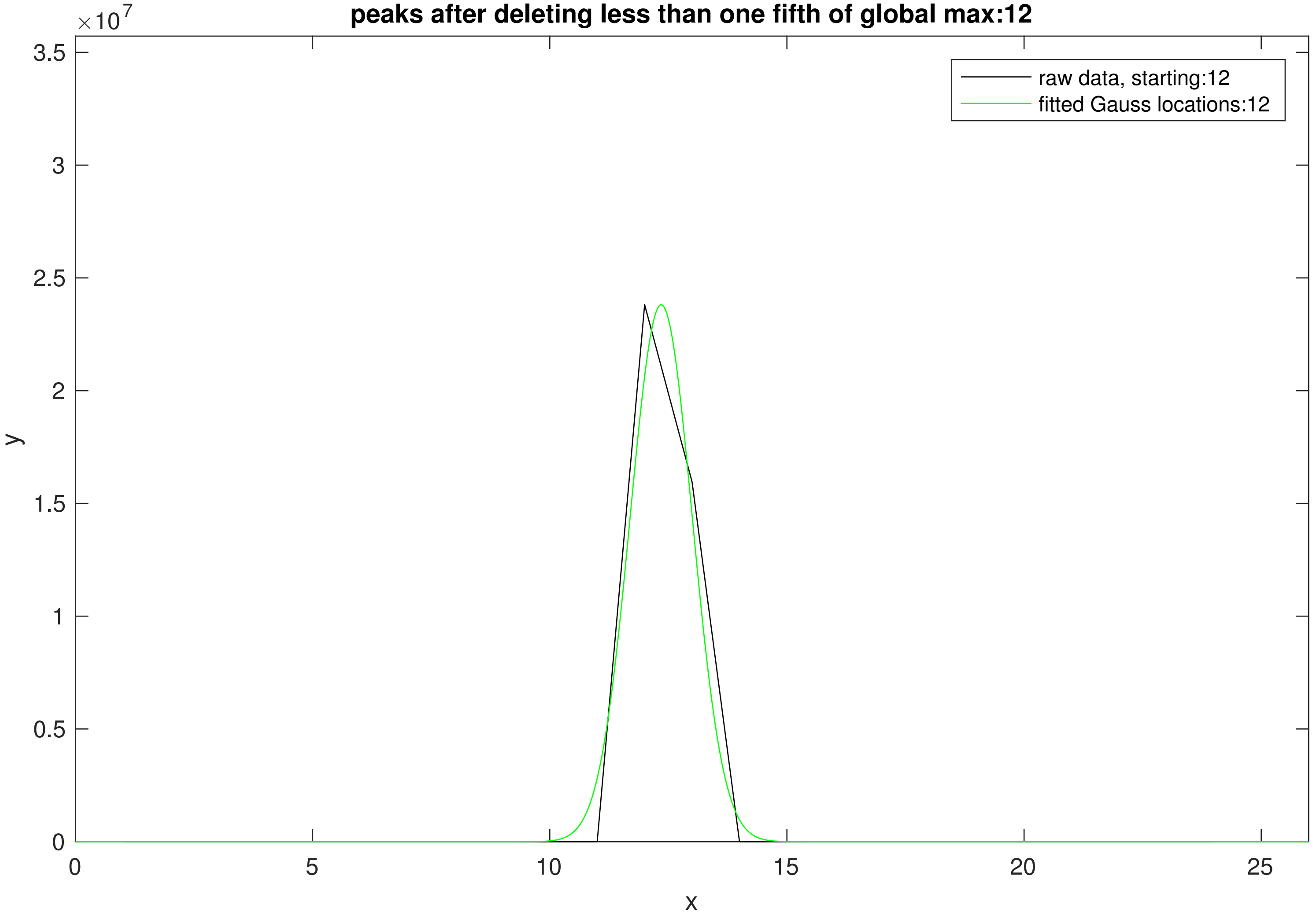

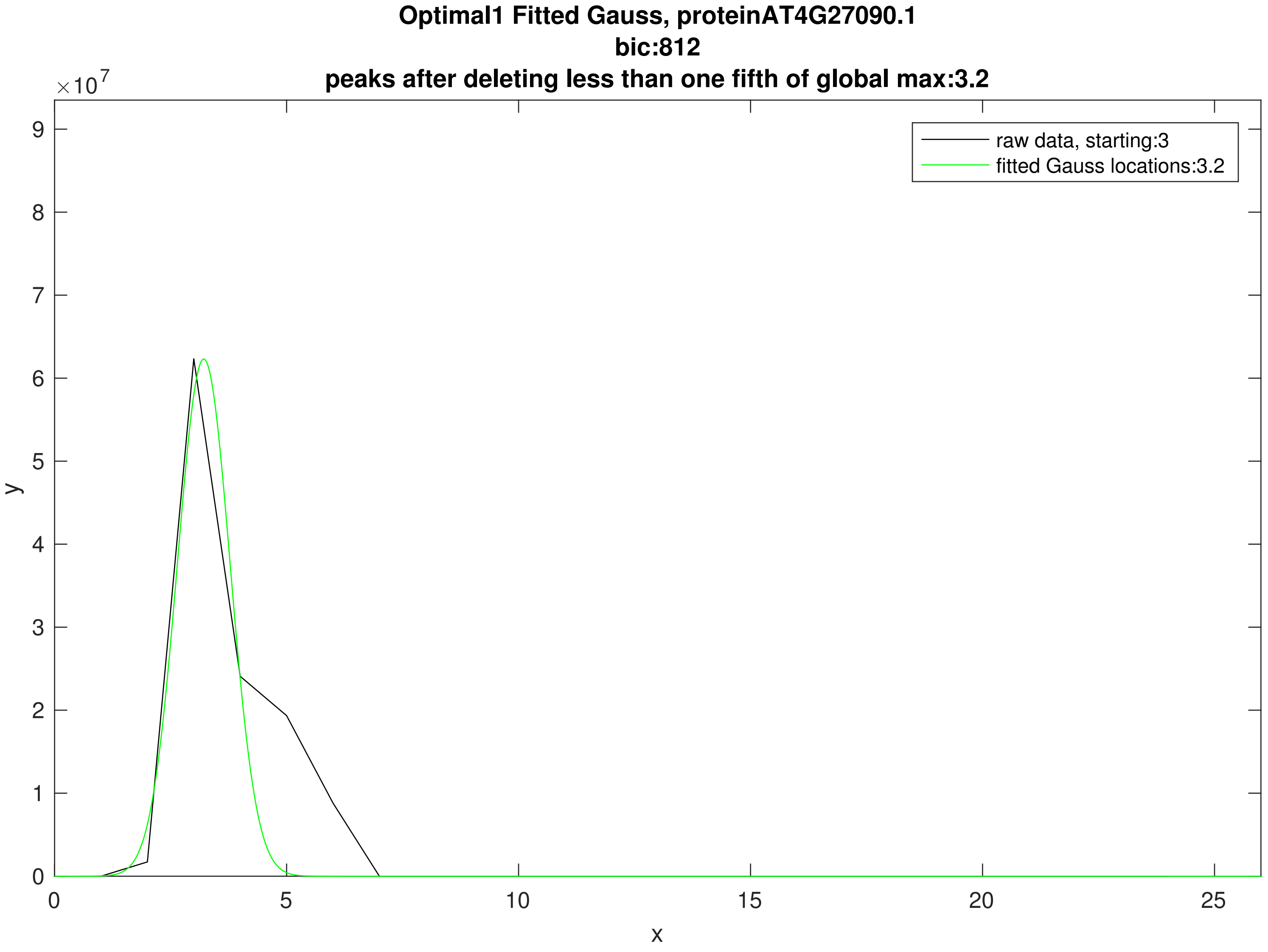
Optimal1 Fitted Gauss, proteinAT4G27090.1
bic:812
peaks after deleting less than one fifth of global max:3.2
×10^7
raw data, starting:3
fitted Gauss locations:3.2
y
x
0
1
2
3
4
5
6
7
8
9
0
5
10
15
20
25

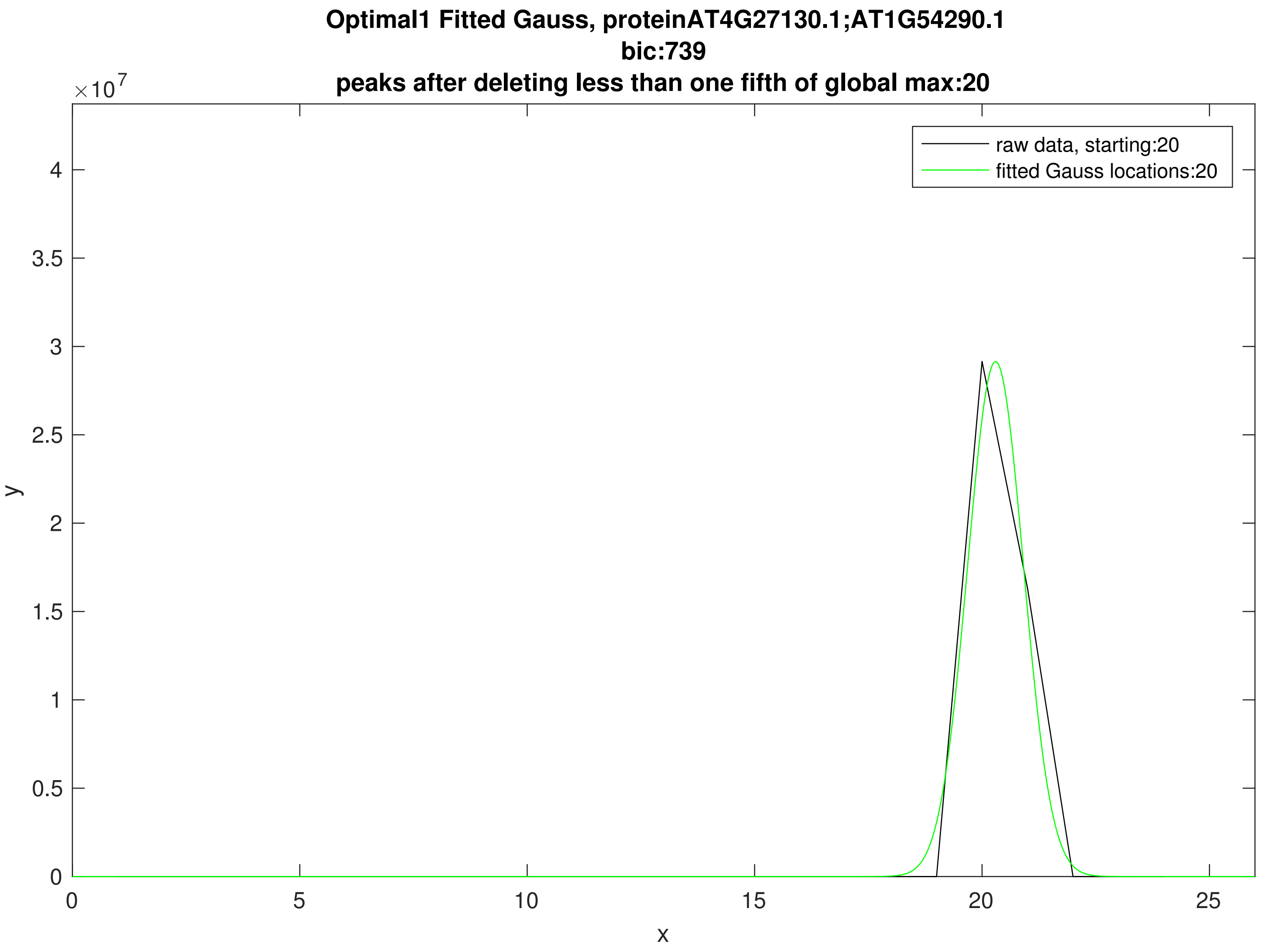
Optimal1 Fitted Gauss, proteinAT4G27130.1;AT1G54290.1
bic:739
peaks after deleting less than one fifth of global max:20
×10^7
raw data, starting:20
fitted Gauss locations:20
y
x
0
0.5
1
1.5
2
2.5
3
3.5
4
5
10
15
20
25

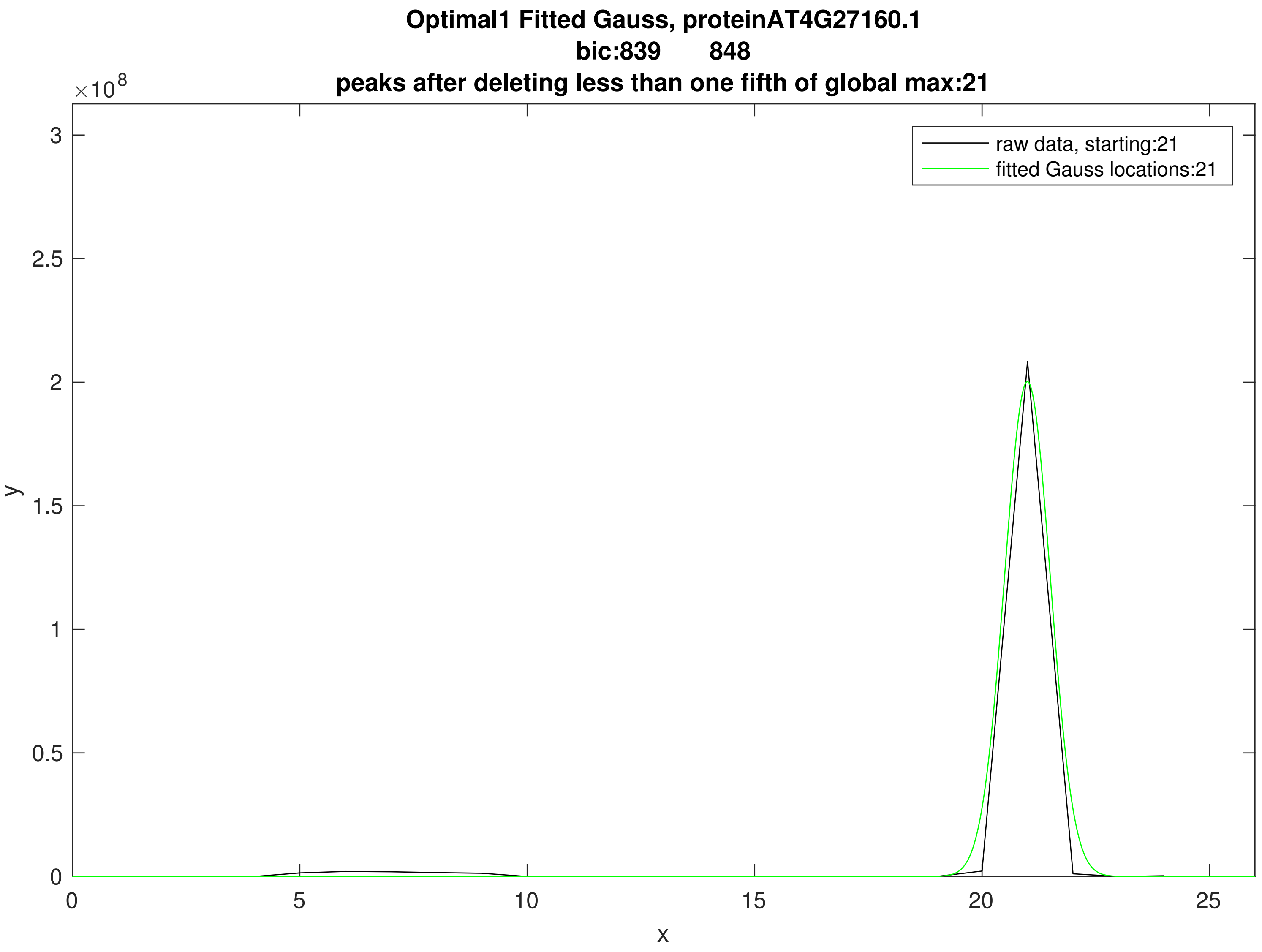
Optimal1 Fitted Gauss, proteinAT4G27160.1
bic:839 848
peaks after deleting less than one fifth of global max:21
×10^8
3
2.5
2
1.5
1
0.5
0
y
0
5
10
15
20
25
x
raw data, starting:21
fitted Gauss locations:21

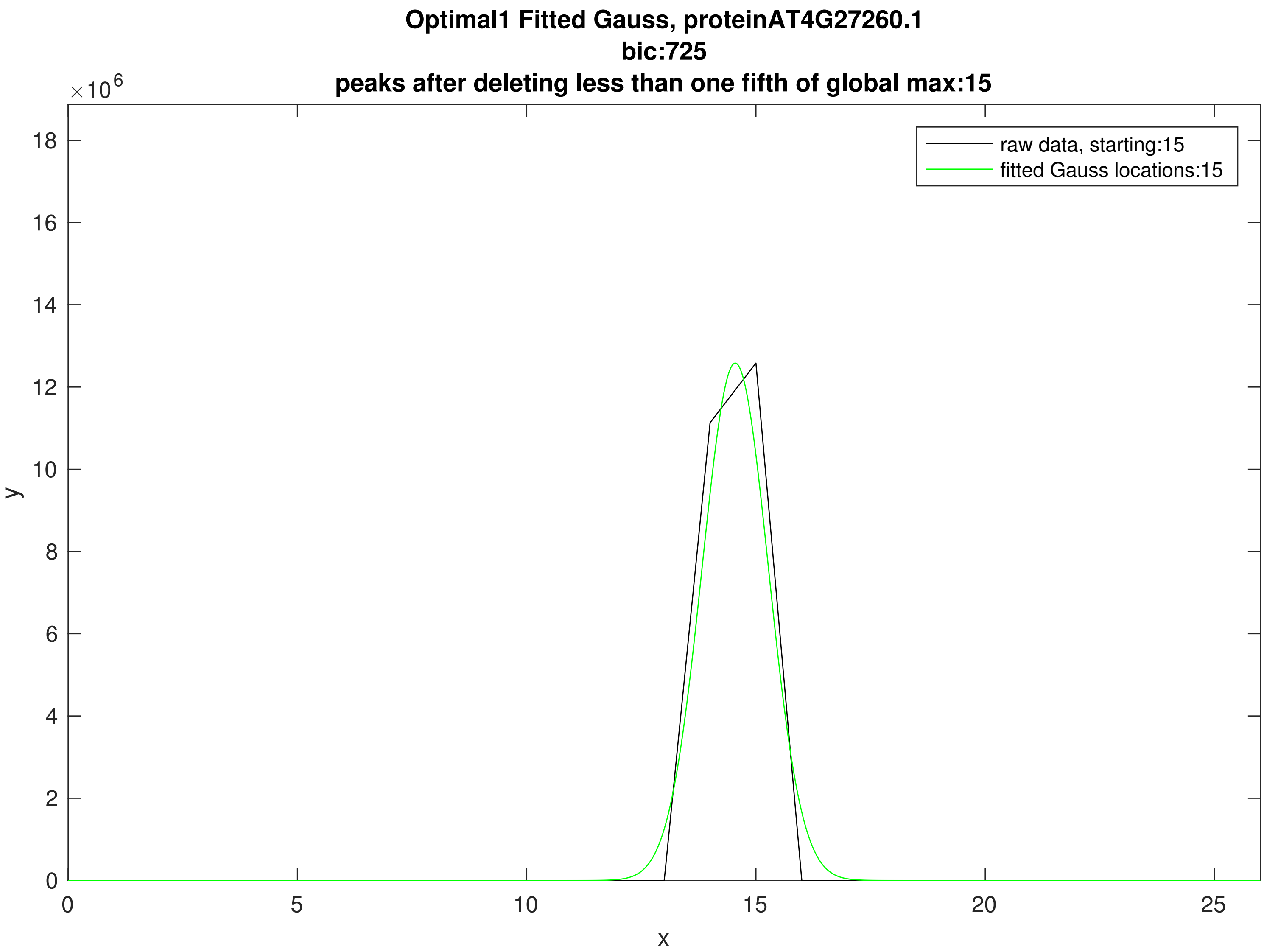

Optimal1 Fitted Gauss, proteinAT4G27260.1
bic:725
peaks after deleting less than one fifth of global max:15
×10^6
raw data, starting:15
fitted Gauss locations:15
y
x
0
2
4
6
8
10
12
14
16
18
0
5
10
15
20
25

**Optimal1 Fitted Gauss, proteinAT4G27320.1**
**bic:796**
**peaks after deleting less than one fifth of global max:15**

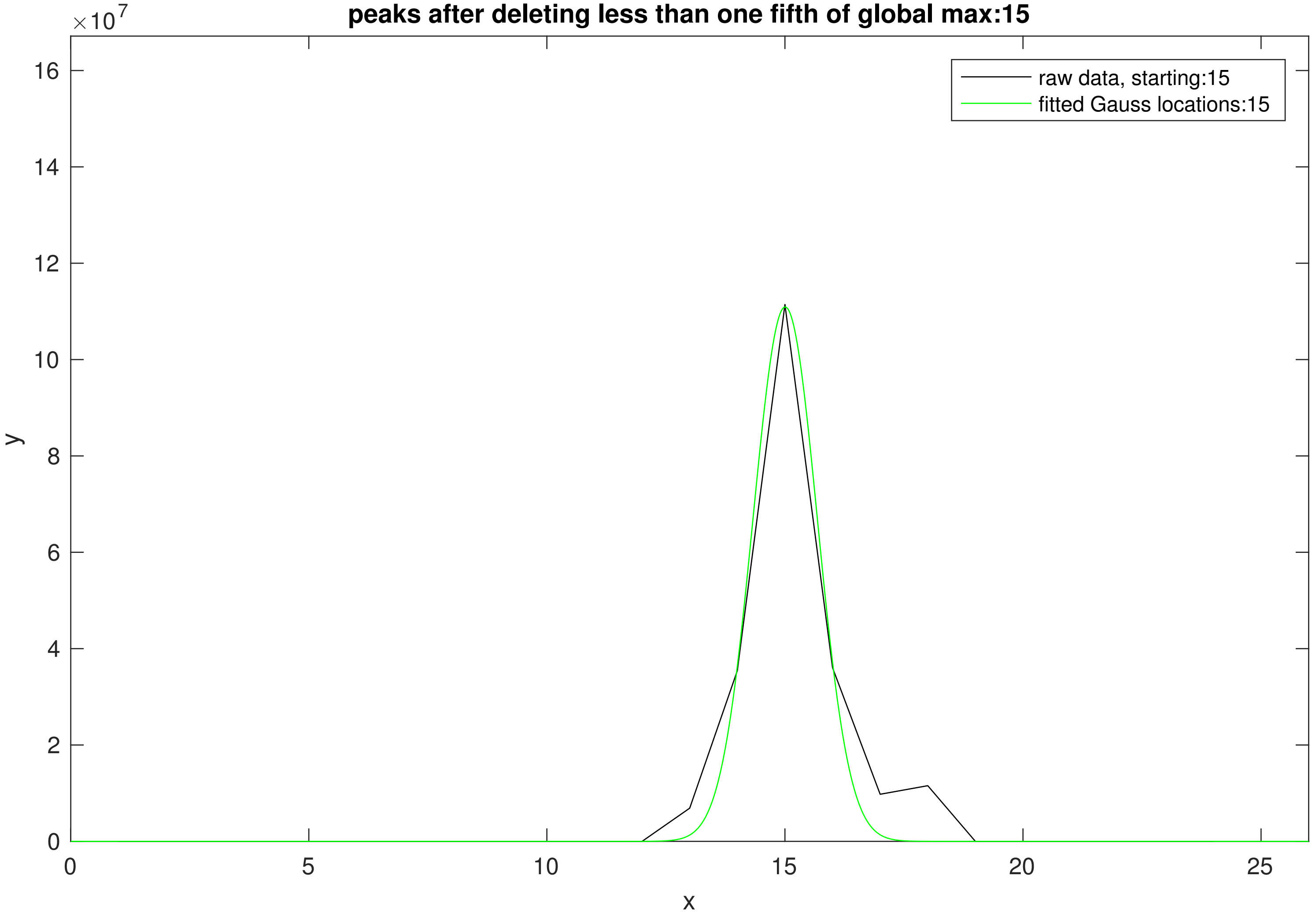

Optimal1 Fitted Gauss, proteinAT4G27520.1
bic:738
peaks after deleting less than one fifth of global max:6.1

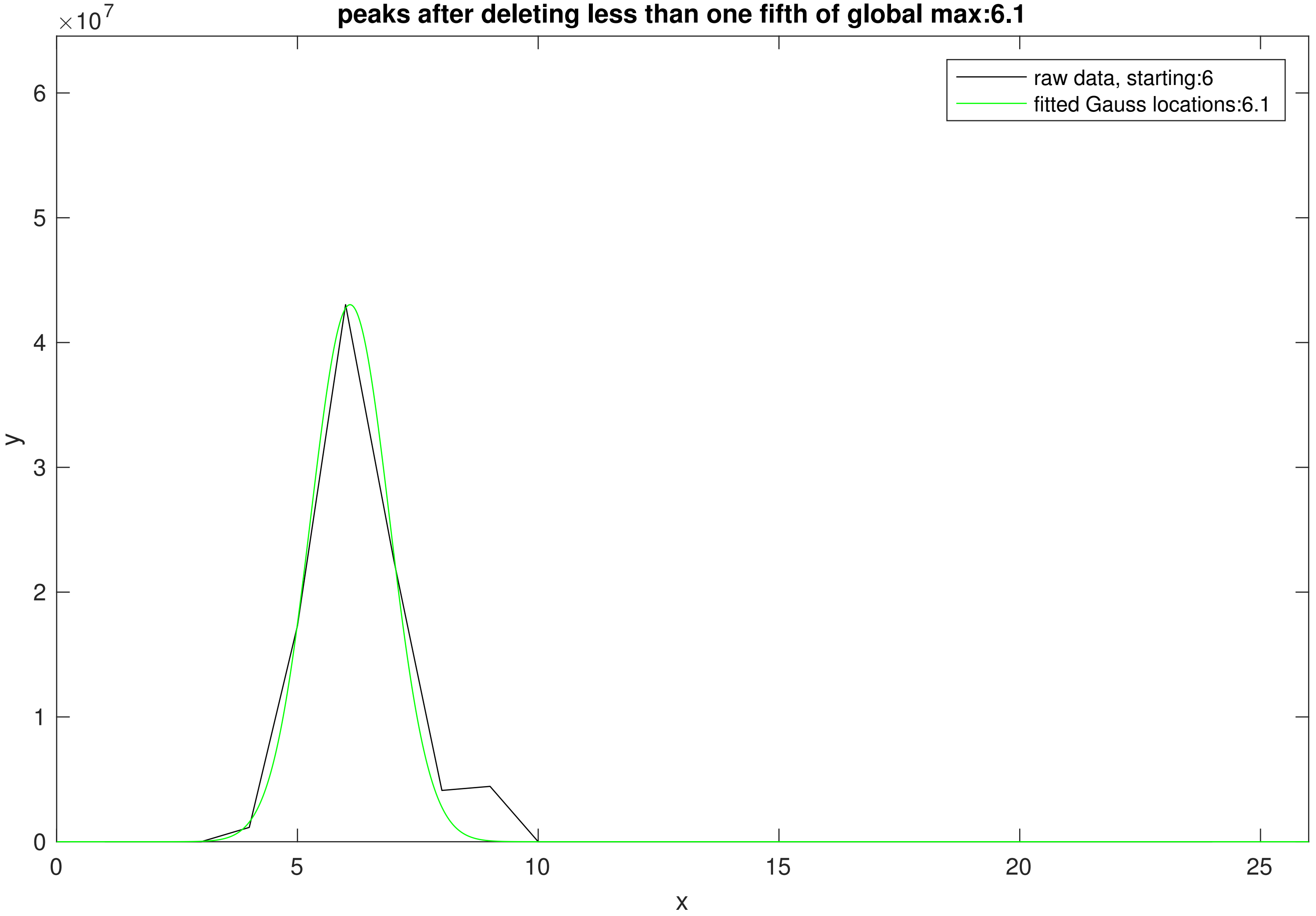

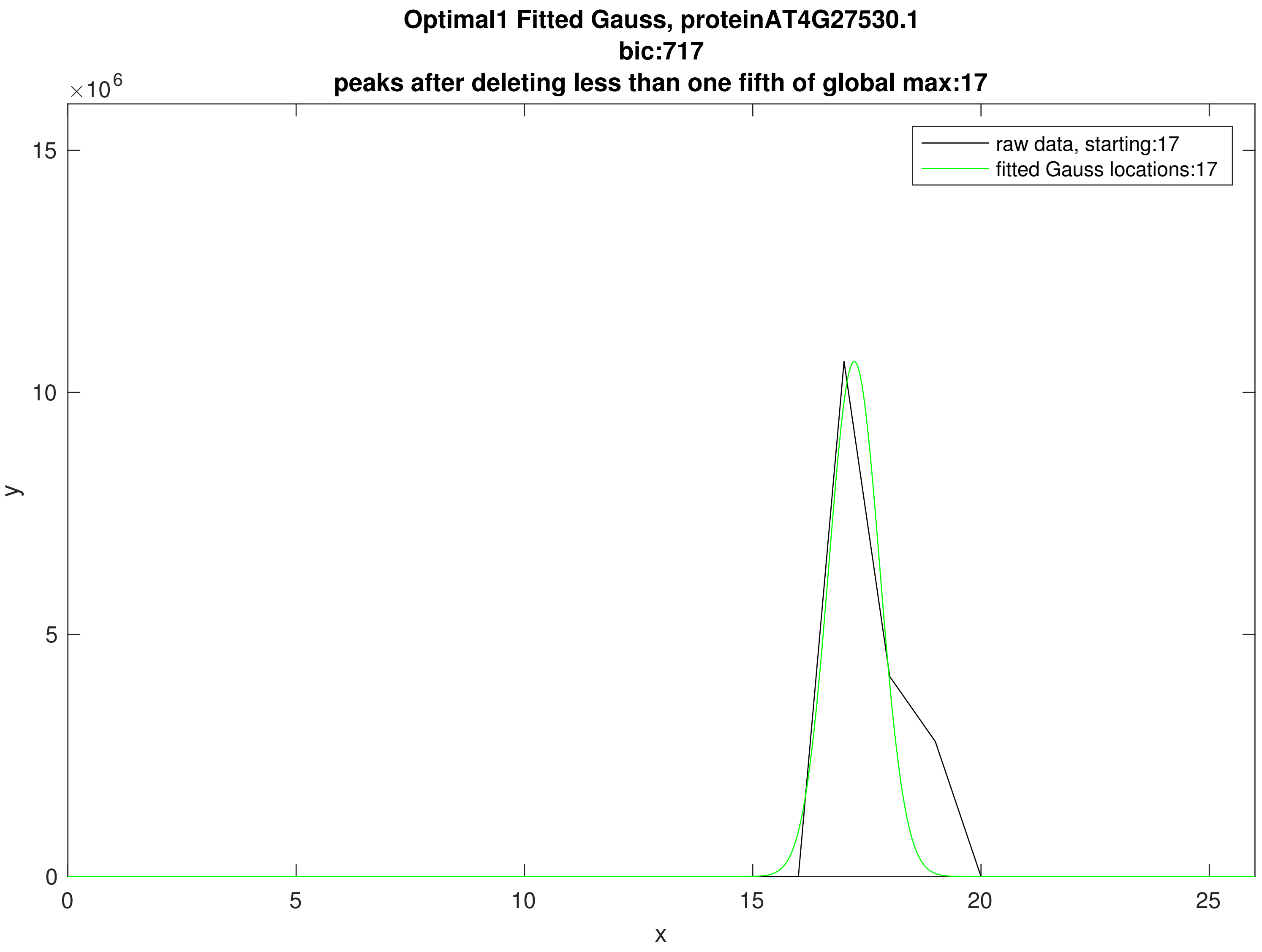
Optimal1 Fitted Gauss, proteinAT4G27530.1
bic:717
peaks after deleting less than one fifth of global max:17
×10^6
raw data, starting:17
fitted Gauss locations:17
y
x
0
5
10
15
20
25

#### Optimal1 Fitted Gauss, proteinAT4G27690.2;AT4G27690.1
#### bic:807 814
#### peaks after deleting less than one fifth of global max:18

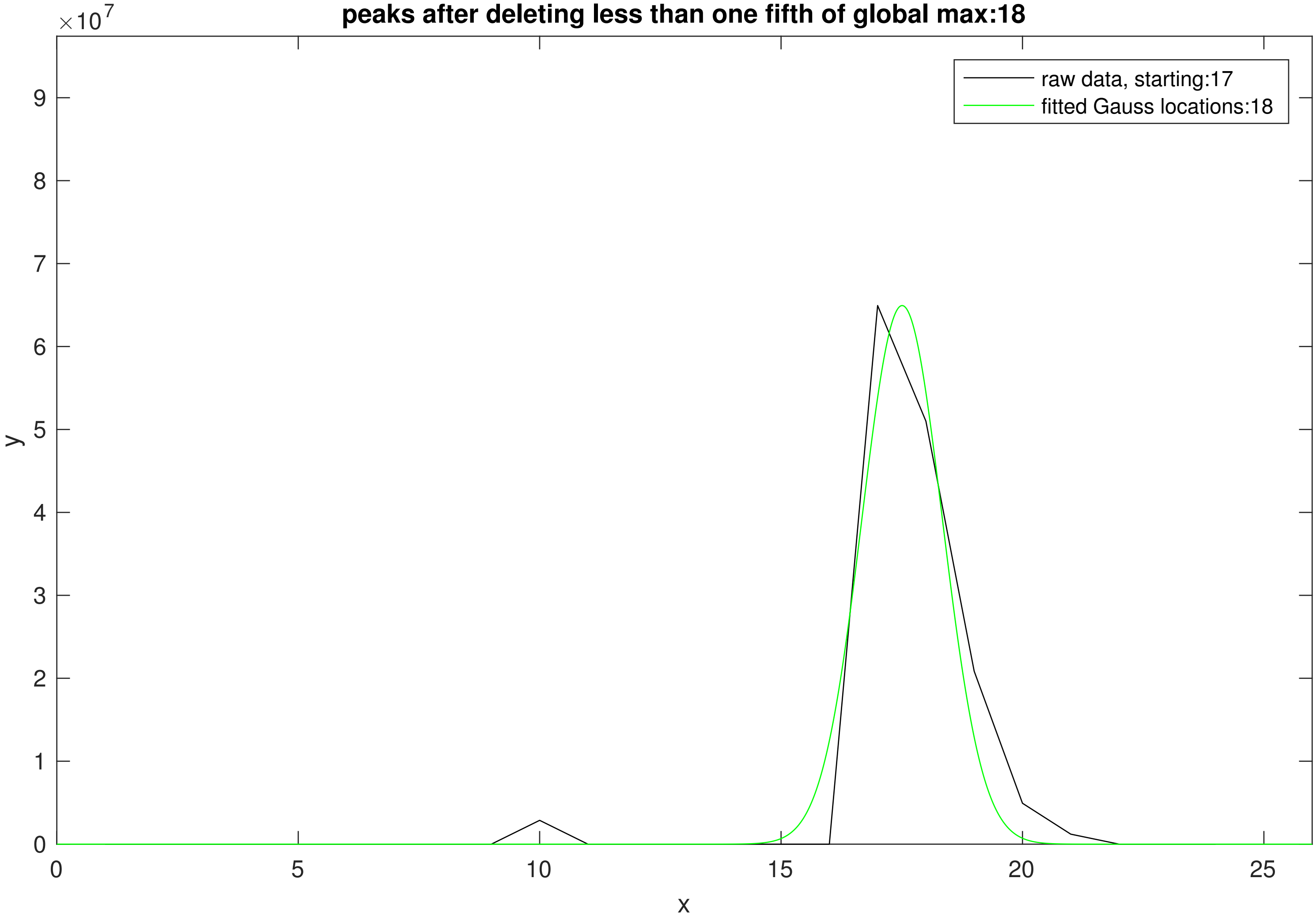

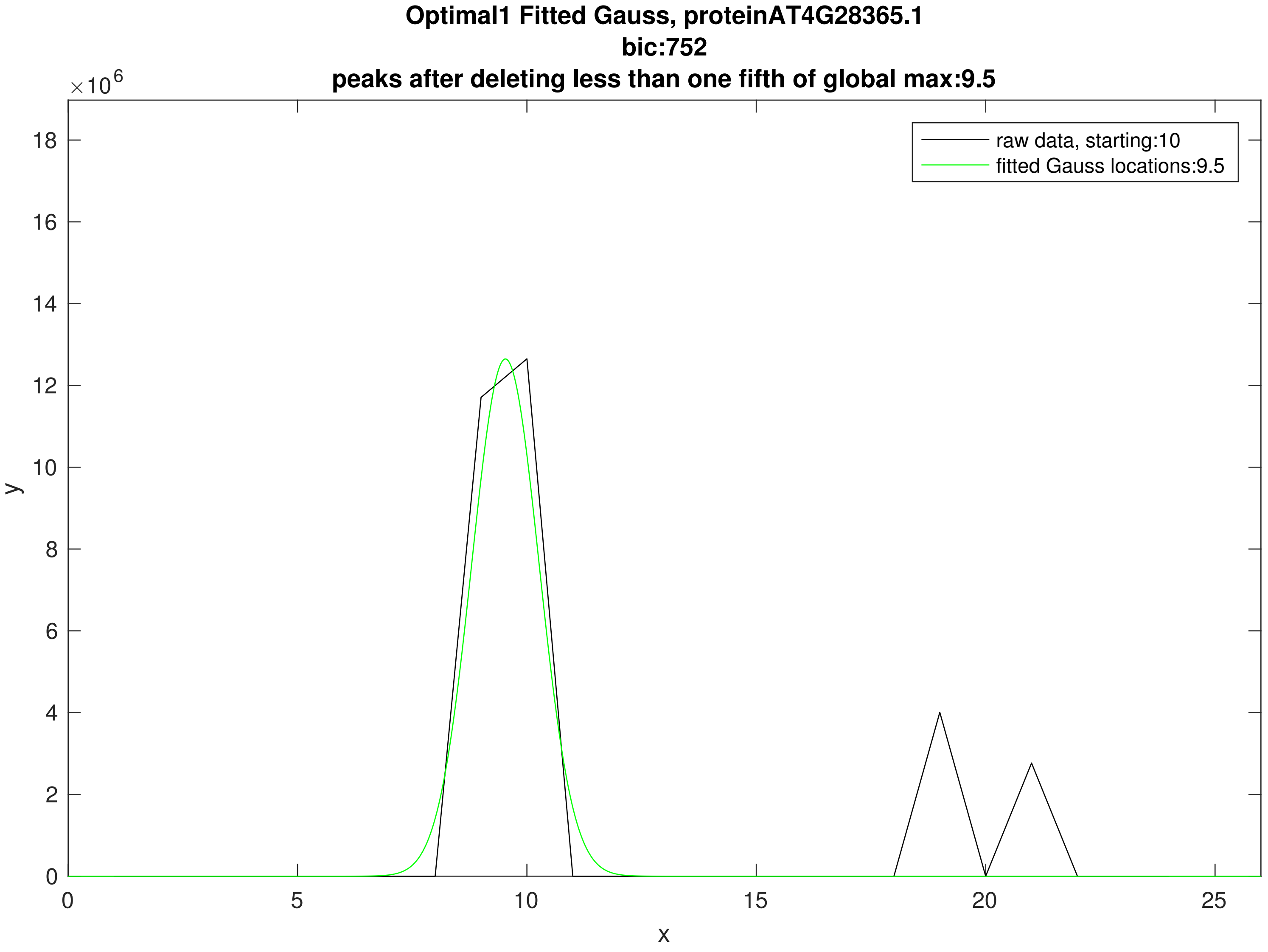

Optimal1 Fitted Gauss, proteinAT4G28365.1
bic:752
peaks after deleting less than one fifth of global max:9.5
×10^6
raw data, starting:10
fitted Gauss locations:9.5
y
x
0
2
4
6
8
10
12
14
16
18
0
5
10
15
20
25

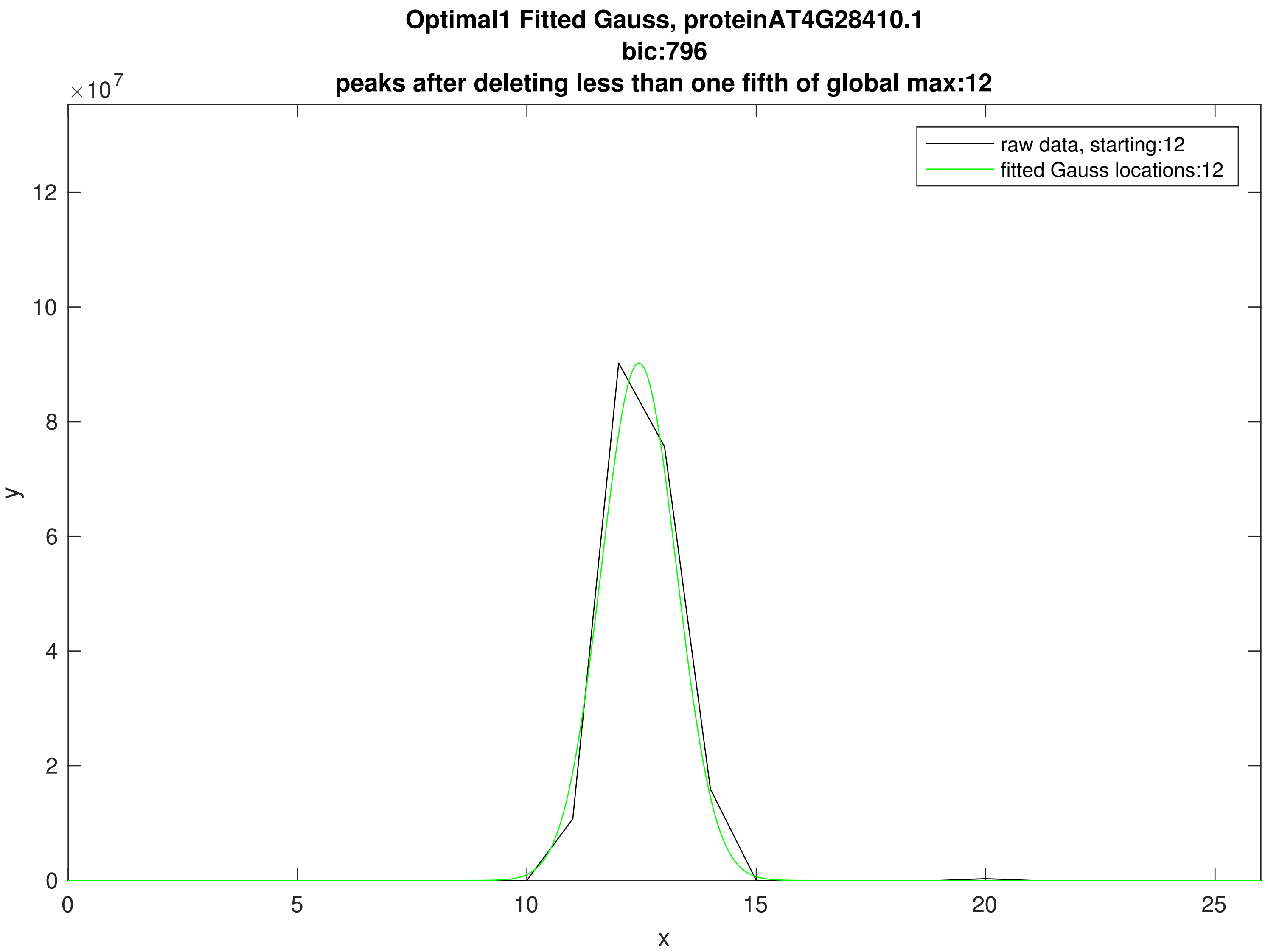
Optimal1 Fitted Gauss, proteinAT4G28410.1
bic:796
peaks after deleting less than one fifth of global max:12
×10^7
raw data, starting:12
fitted Gauss locations:12
y
x
0
2
4
6
8
10
12
0
5
10
15
20
25

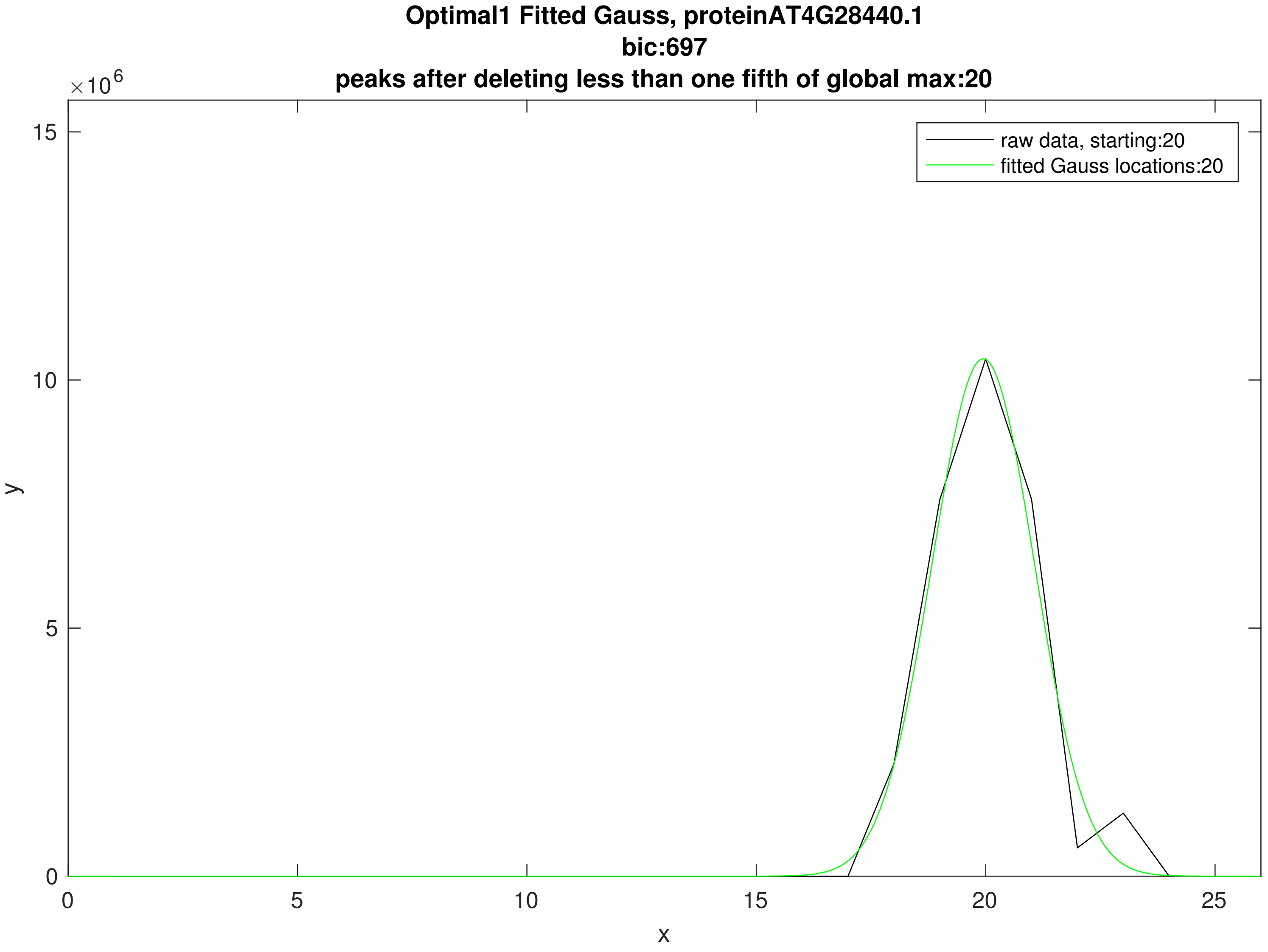
Optimal1 Fitted Gauss, proteinAT4G28440.1
bic:697
peaks after deleting less than one fifth of global max:20
×10^6
15
10
5
0
y
0
5
10
15
20
25
x
raw data, starting:20
fitted Gauss locations:20

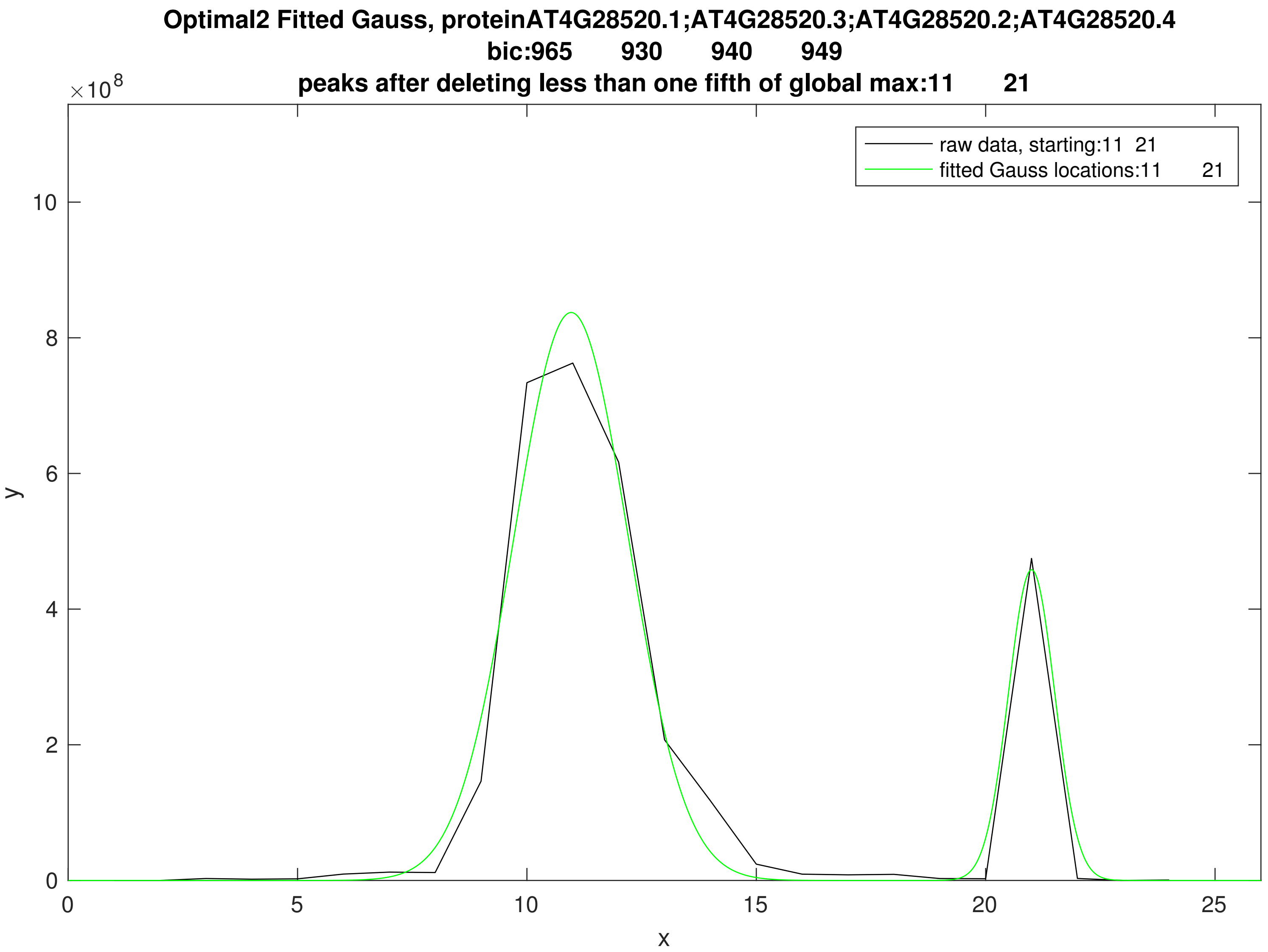

Optimal2 Fitted Gauss, proteinAT4G28520.1;AT4G28520.3;AT4G28520.2;AT4G28520.4
bic:965 930 940 949
peaks after deleting less than one fifth of global max:11 21
×10^8
raw data, starting:11 21
fitted Gauss locations:11 21
y
x
0
2
4
6
8
10
5
15
20
25

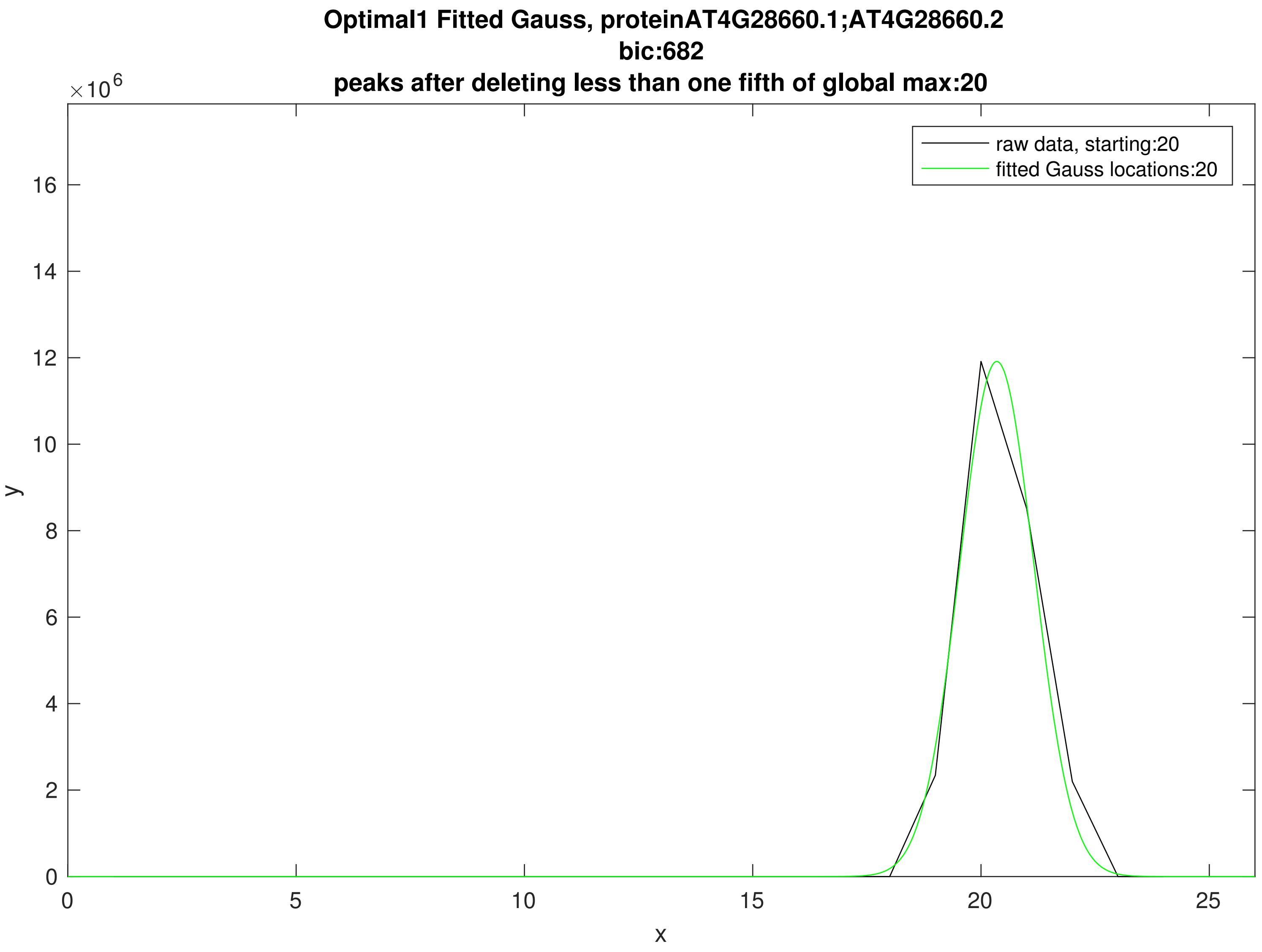
Optimal1 Fitted Gauss, proteinAT4G28660.1;AT4G28660.2
bic:682
peaks after deleting less than one fifth of global max:20
raw data, starting:20
fitted Gauss locations:20
$\times 10^6$
y
x
0
2
4
6
8
10
12
14
16
5
15
20
25

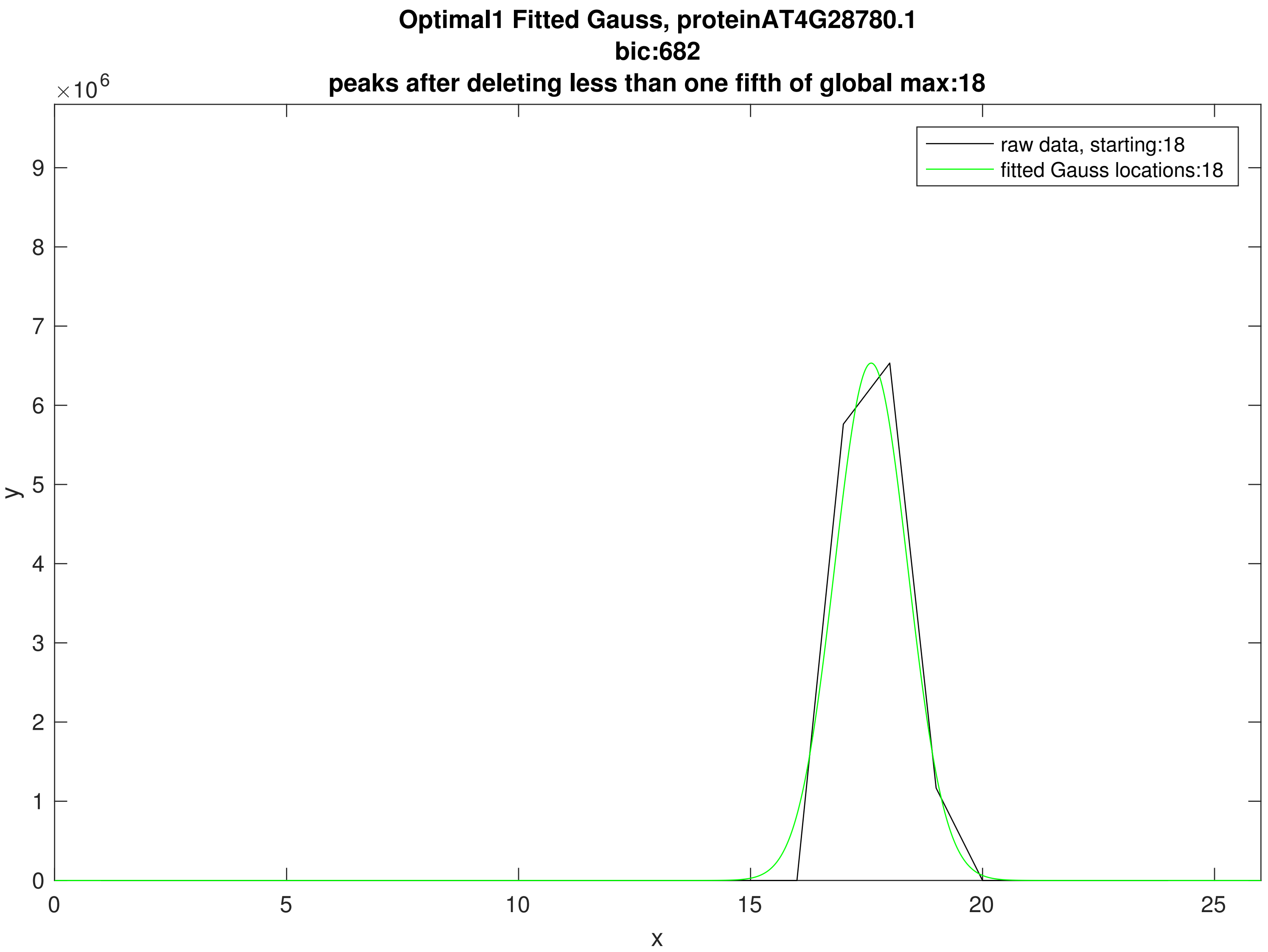

Optimal1 Fitted Gauss, proteinAT4G28780.1
bic:682
peaks after deleting less than one fifth of global max:18
×10^6
raw data, starting:18
fitted Gauss locations:18
y
x
0
1
2
3
4
5
6
7
8
9
0
5
10
15
20
25

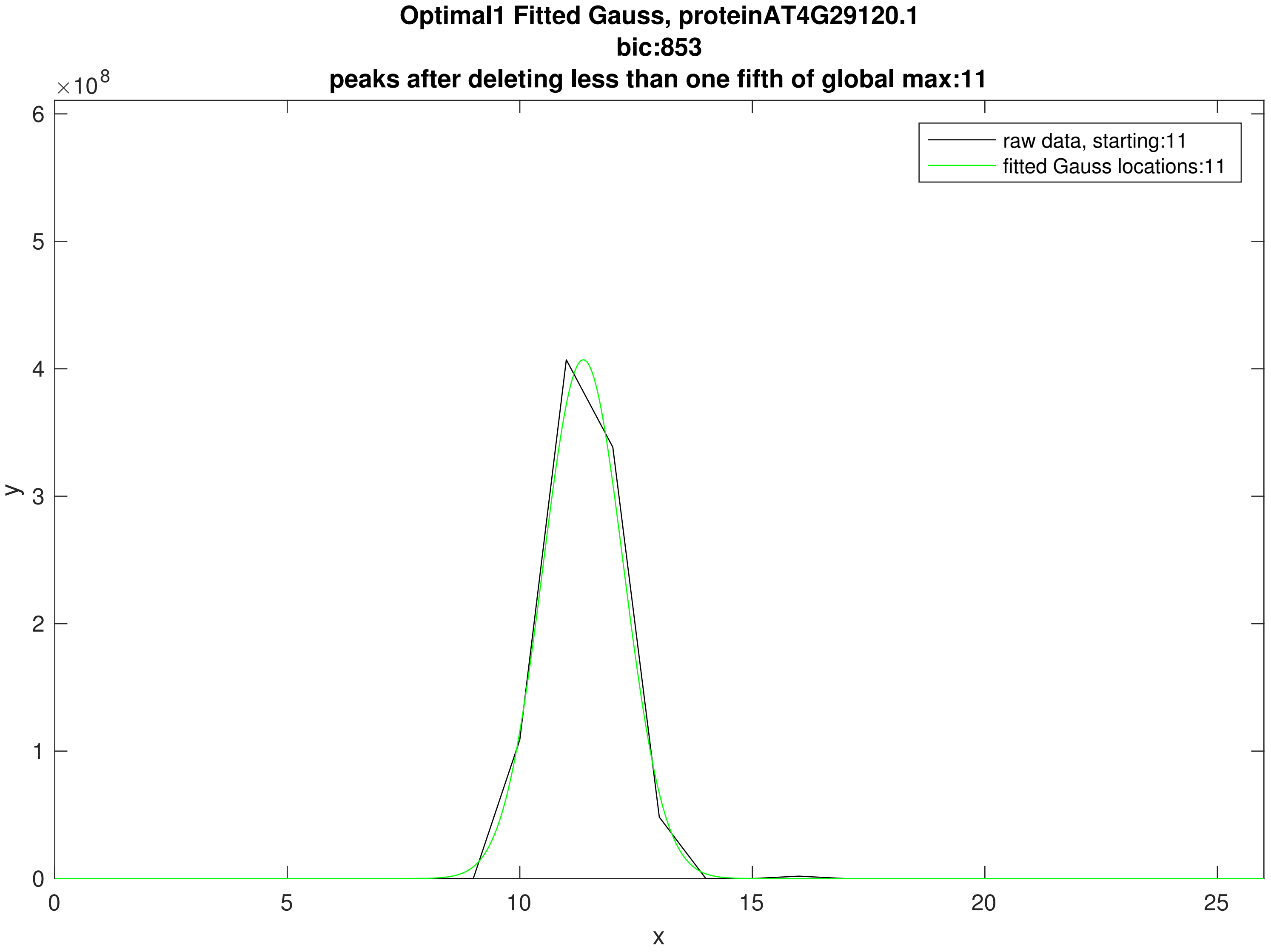
Optimal1 Fitted Gauss, proteinAT4G29120.1
bic:853
peaks after deleting less than one fifth of global max:11
×10^8
raw data, starting:11
fitted Gauss locations:11
y
x
0
1
2
3
4
5
6
0
5
10
15
20
25

Optimal1 Fitted Gauss, proteinAT4G29160.2;AT4G29160.3;AT4G29160.1
bic:806 814
peaks after deleting less than one fifth of global max:16

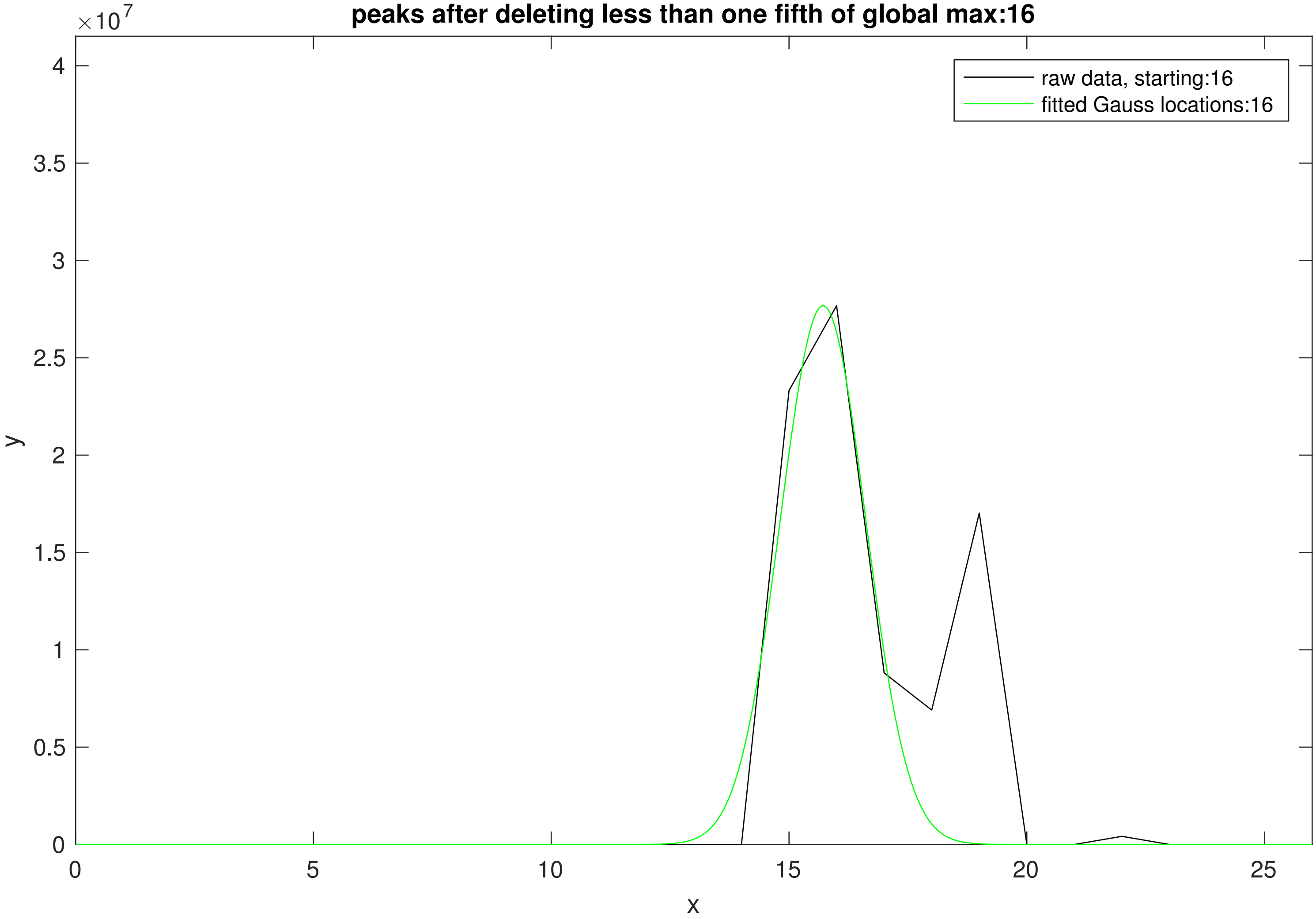

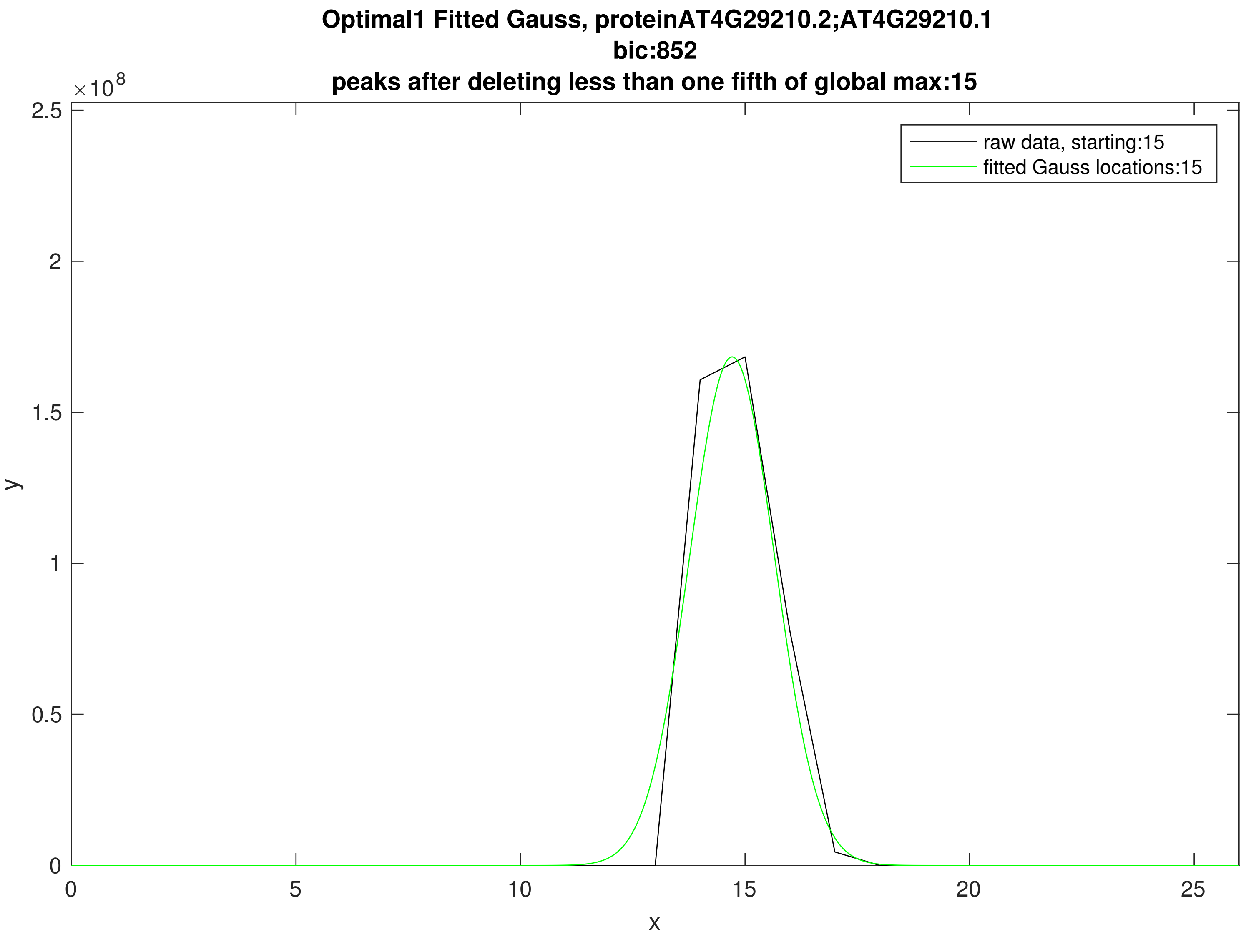
Optimal1 Fitted Gauss, proteinAT4G29210.2;AT4G29210.1
bic:852
peaks after deleting less than one fifth of global max:15
×10^8
2.5
2
1.5
1
0.5
0
y
0
5
10
15
20
25
x
raw data, starting:15
fitted Gauss locations:15

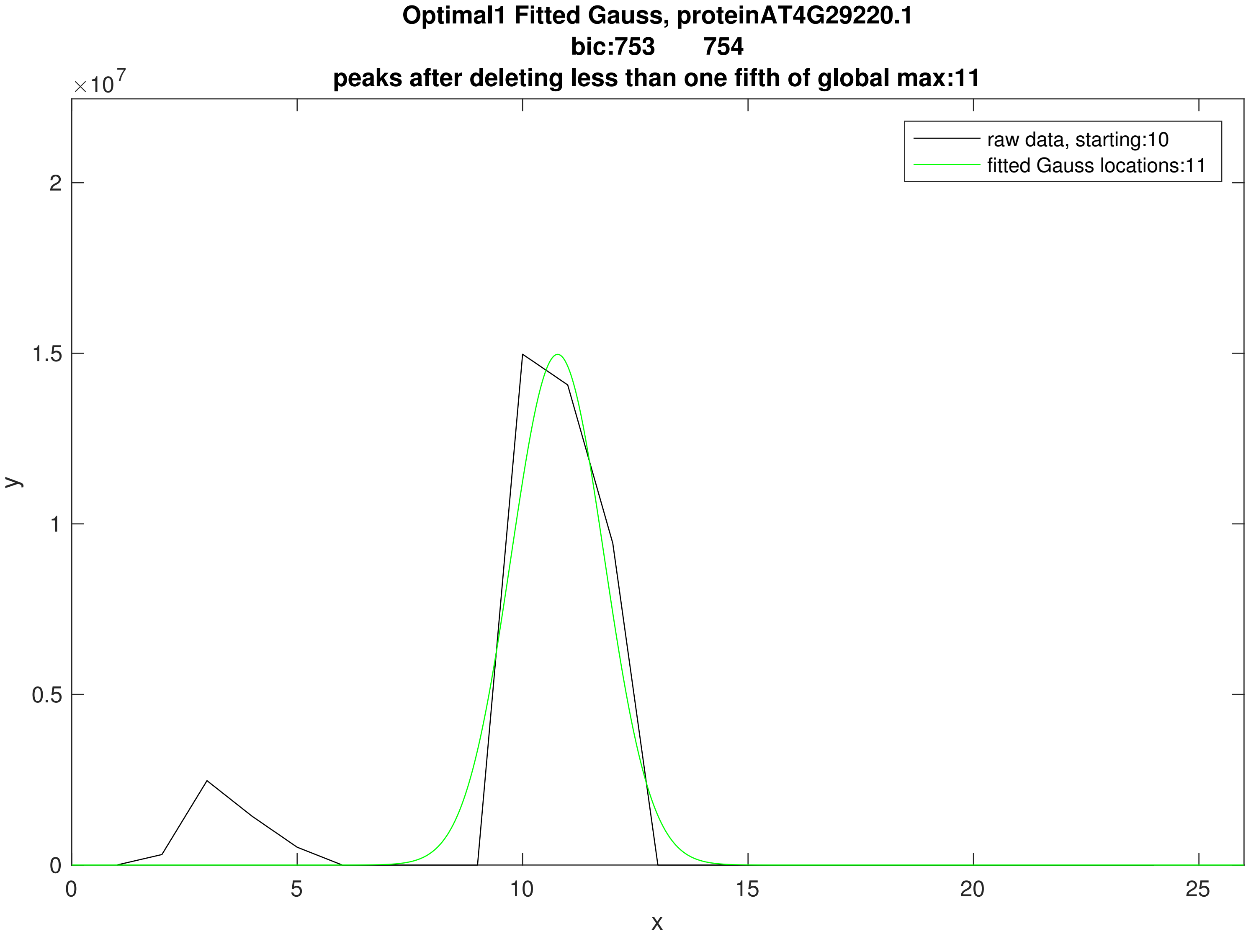

Optimal1 Fitted Gauss, proteinAT4G29220.1
bic:753 754
peaks after deleting less than one fifth of global max:11
×10^7
raw data, starting:10
fitted Gauss locations:11
y
x
0
0.5
1
1.5
2
5
10
15
20
25

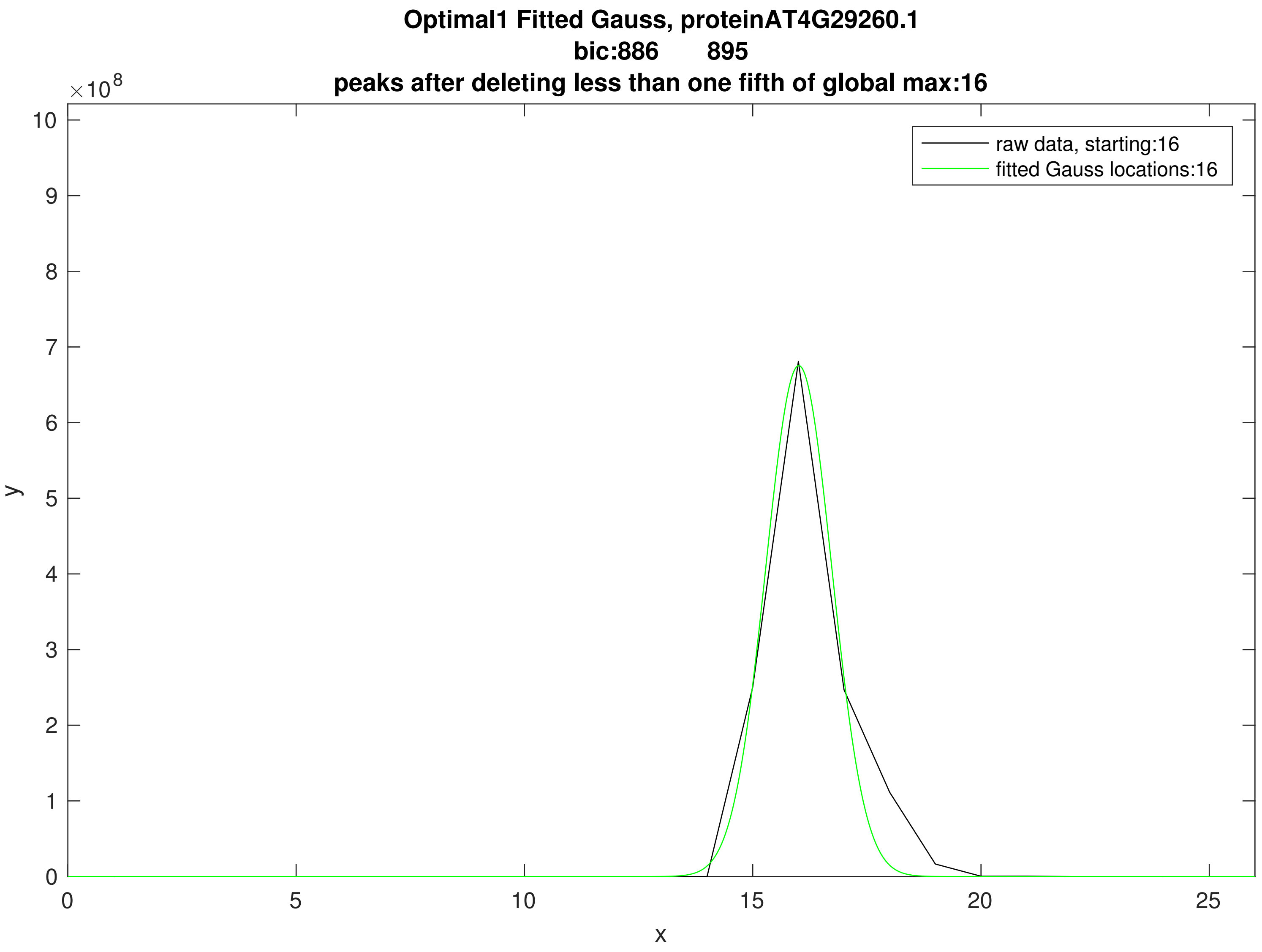
Optimal1 Fitted Gauss, proteinAT4G29260.1
bic:886 895
peaks after deleting less than one fifth of global max:16
×10^8
10
9
8
7
6
5
4
3
2
1
0
y
0
5
10
15
20
25
x
raw data, starting:16
fitted Gauss locations:16

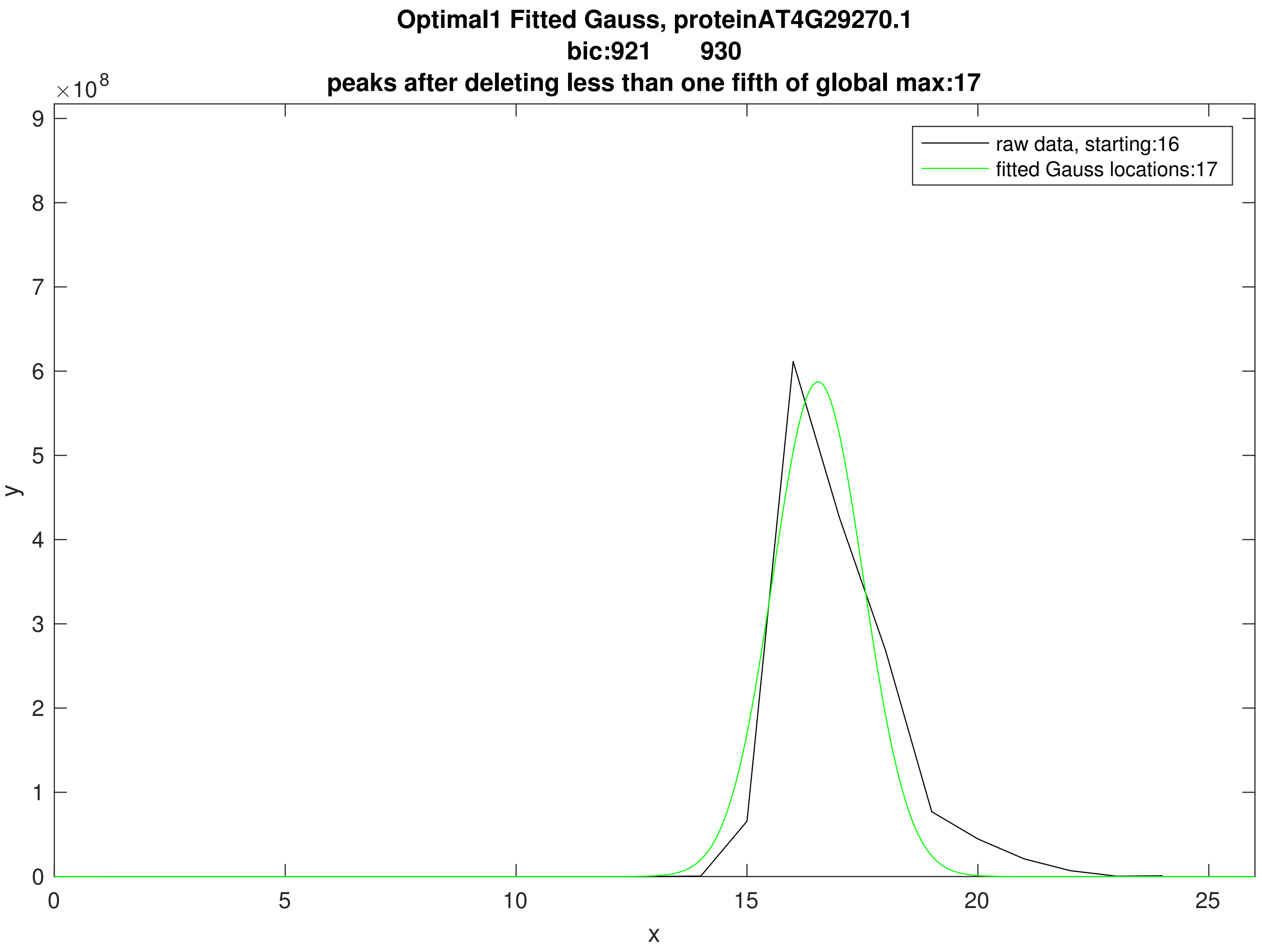
Optimal1 Fitted Gauss, proteinAT4G29270.1
bic:921 930
peaks after deleting less than one fifth of global max:17
×10^8
9
8
7
6
5
4
3
2
1
0
y
0
5
10
15
20
25
x
raw data, starting:16
fitted Gauss locations:17

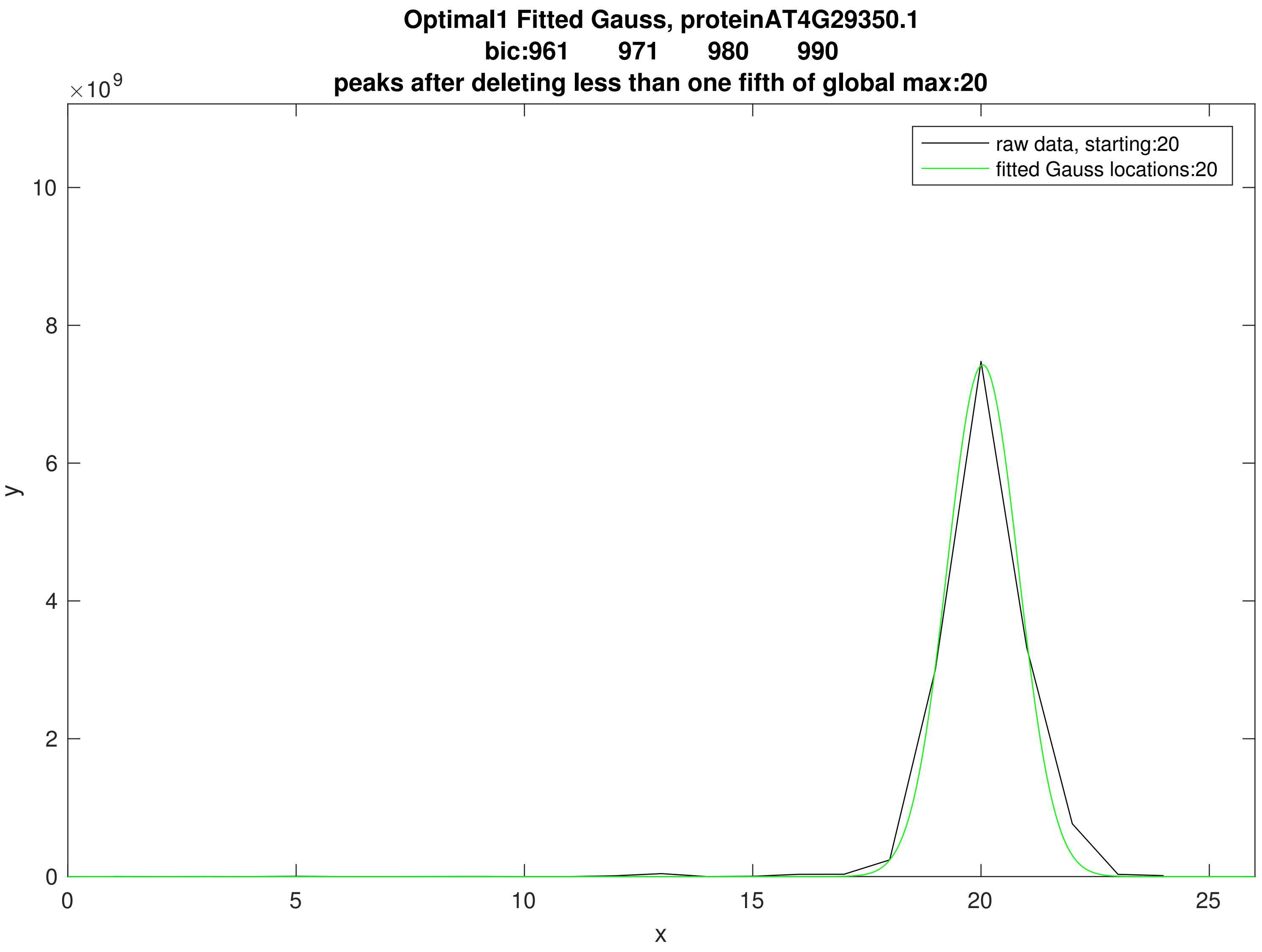

Optimal1 Fitted Gauss, proteinAT4G29350.1
bic:961 971 980 990
peaks after deleting less than one fifth of global max:20
×10^9
raw data, starting:20
fitted Gauss locations:20
10
8
6
4
2
0
y
0
5
10
15
20
25
x

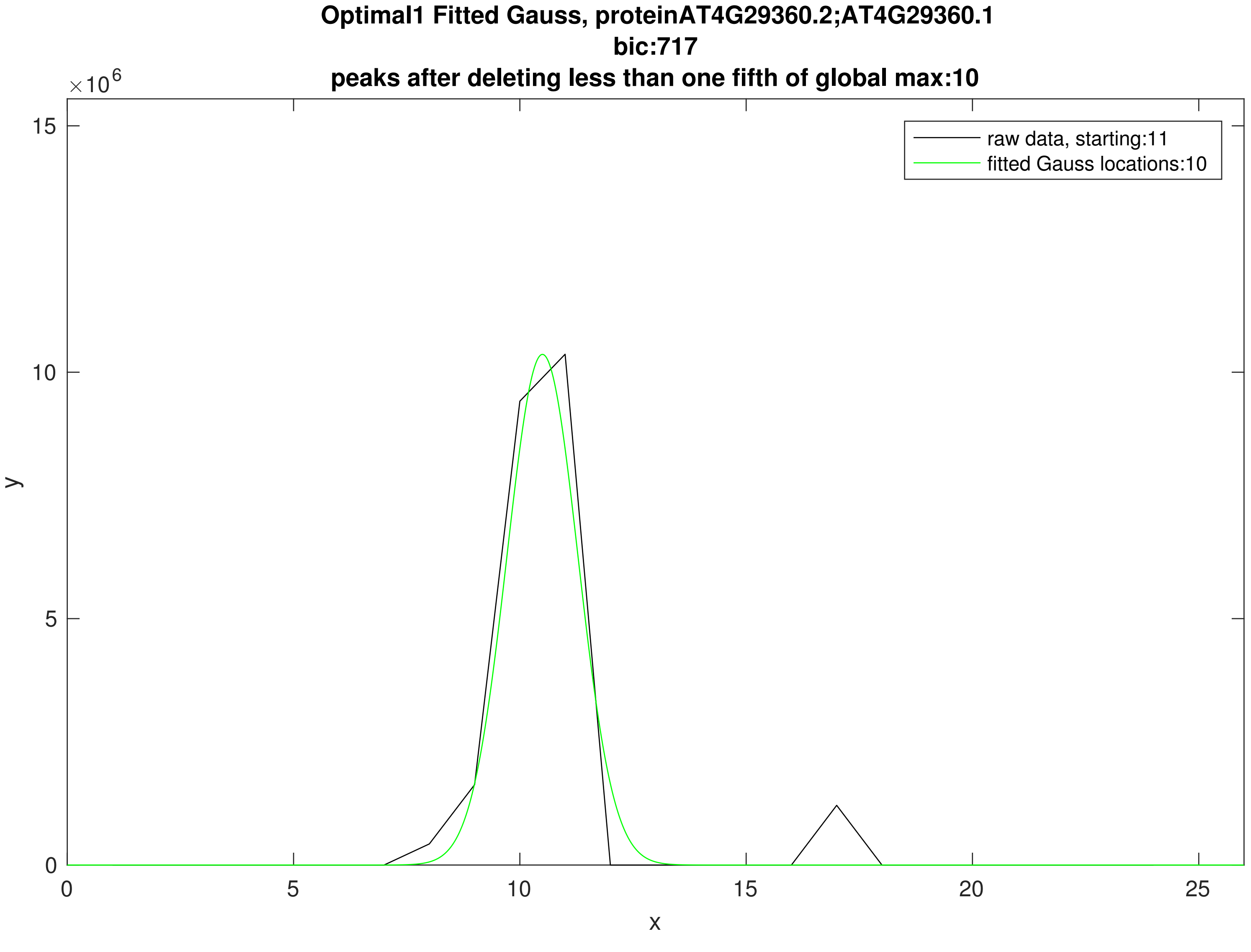
Optimal1 Fitted Gauss, proteinAT4G29360.2;AT4G29360.1
bic:717
peaks after deleting less than one fifth of global max:10
×10^6
15
10
5
0
y
0
5
10
15
20
25
x
raw data, starting:11
fitted Gauss locations:10

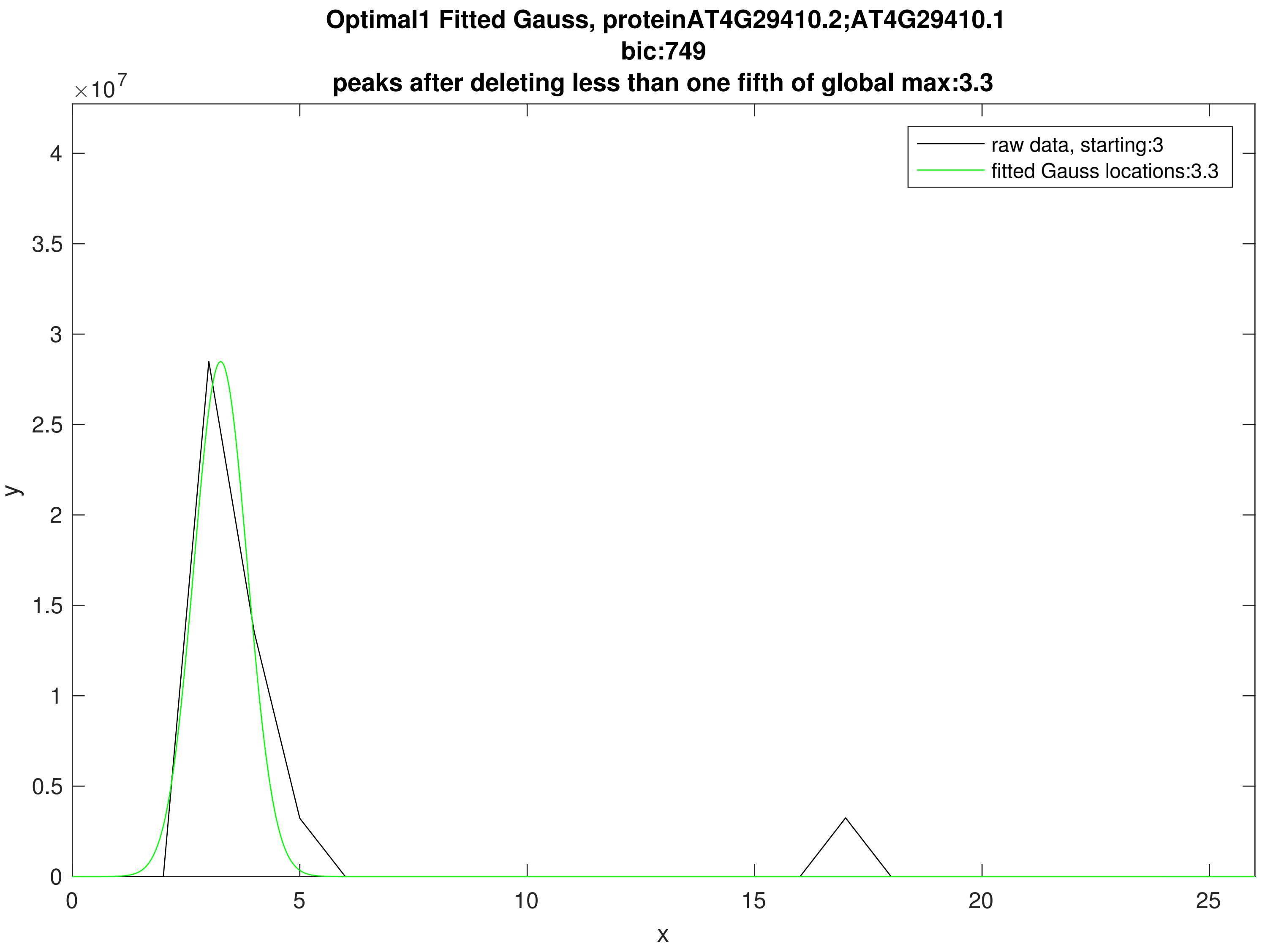
Optimal1 Fitted Gauss, proteinAT4G29410.2;AT4G29410.1
bic:749
peaks after deleting less than one fifth of global max:3.3
×10^7
raw data, starting:3
fitted Gauss locations:3.3
y
x
0
0.5
1
1.5
2
2.5
3
3.5
4
5
10
15
20
25

Optimal1 Fitted Gauss, proteinAT4G29490.1
bic:778
peaks after deleting less than one fifth of global max:13

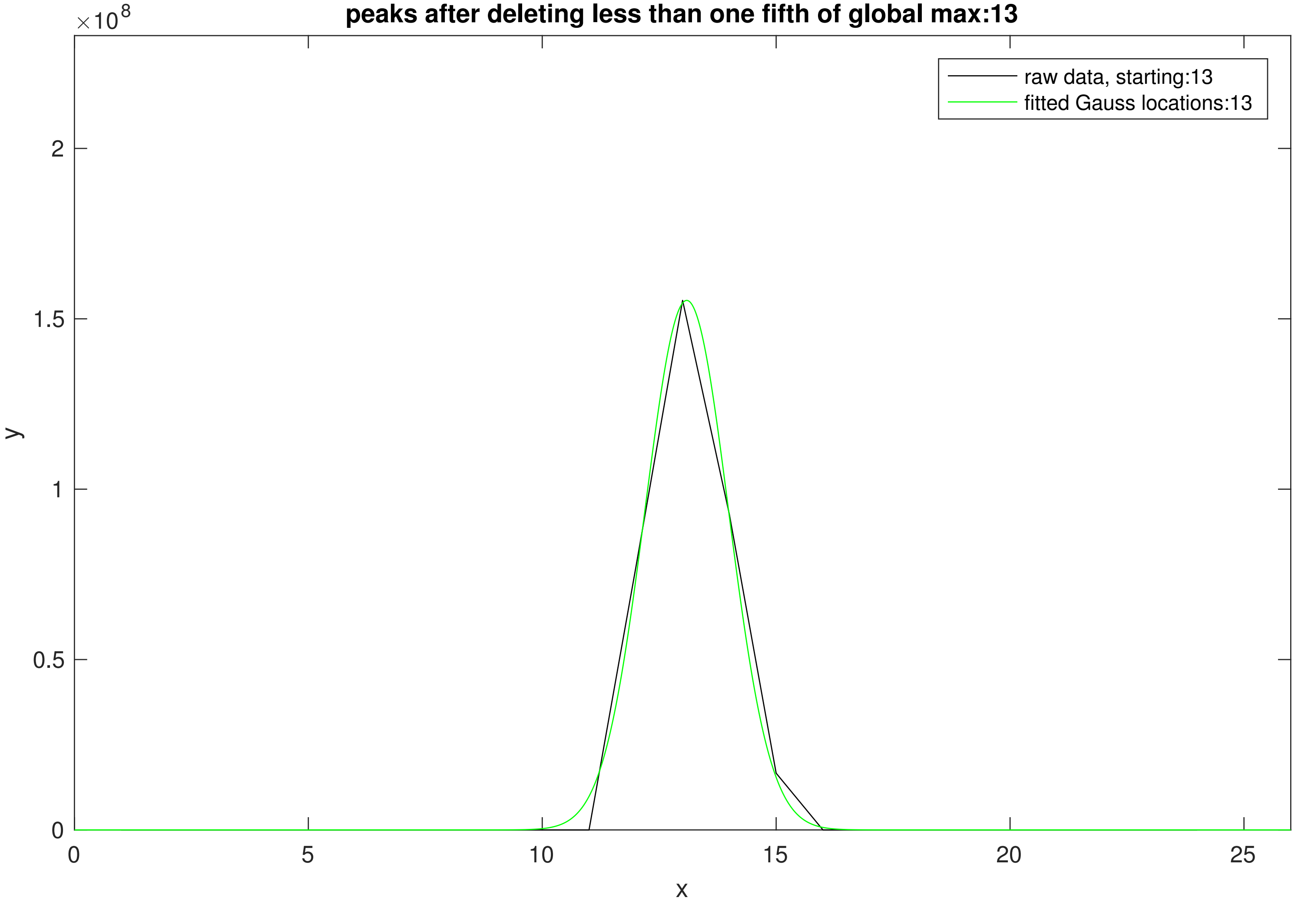

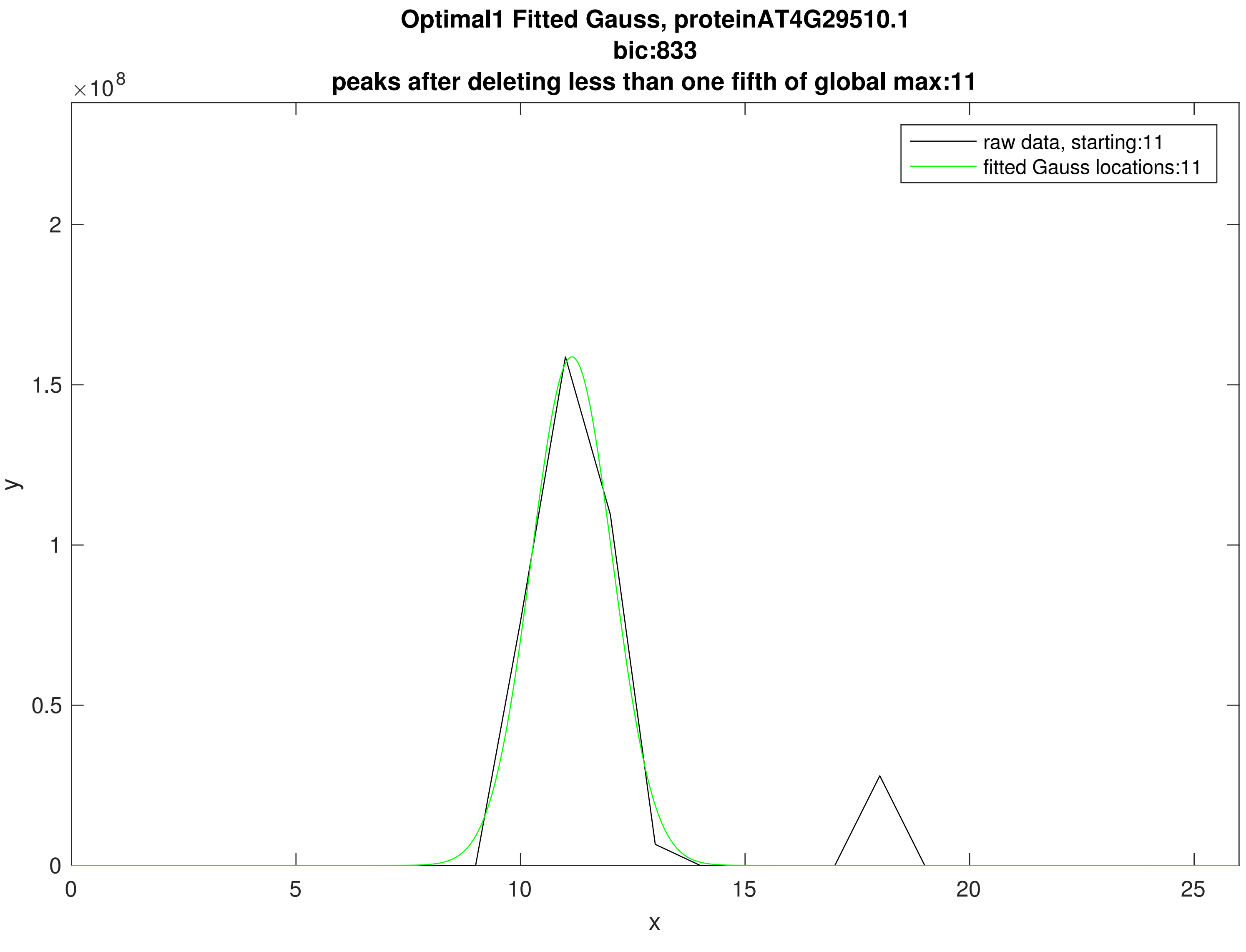
Optimal1 Fitted Gauss, proteinAT4G29510.1
bic:833
peaks after deleting less than one fifth of global max:11
×10^8
raw data, starting:11
fitted Gauss locations:11
y
x
0
0.5
1
1.5
2
0
5
10
15
20
25

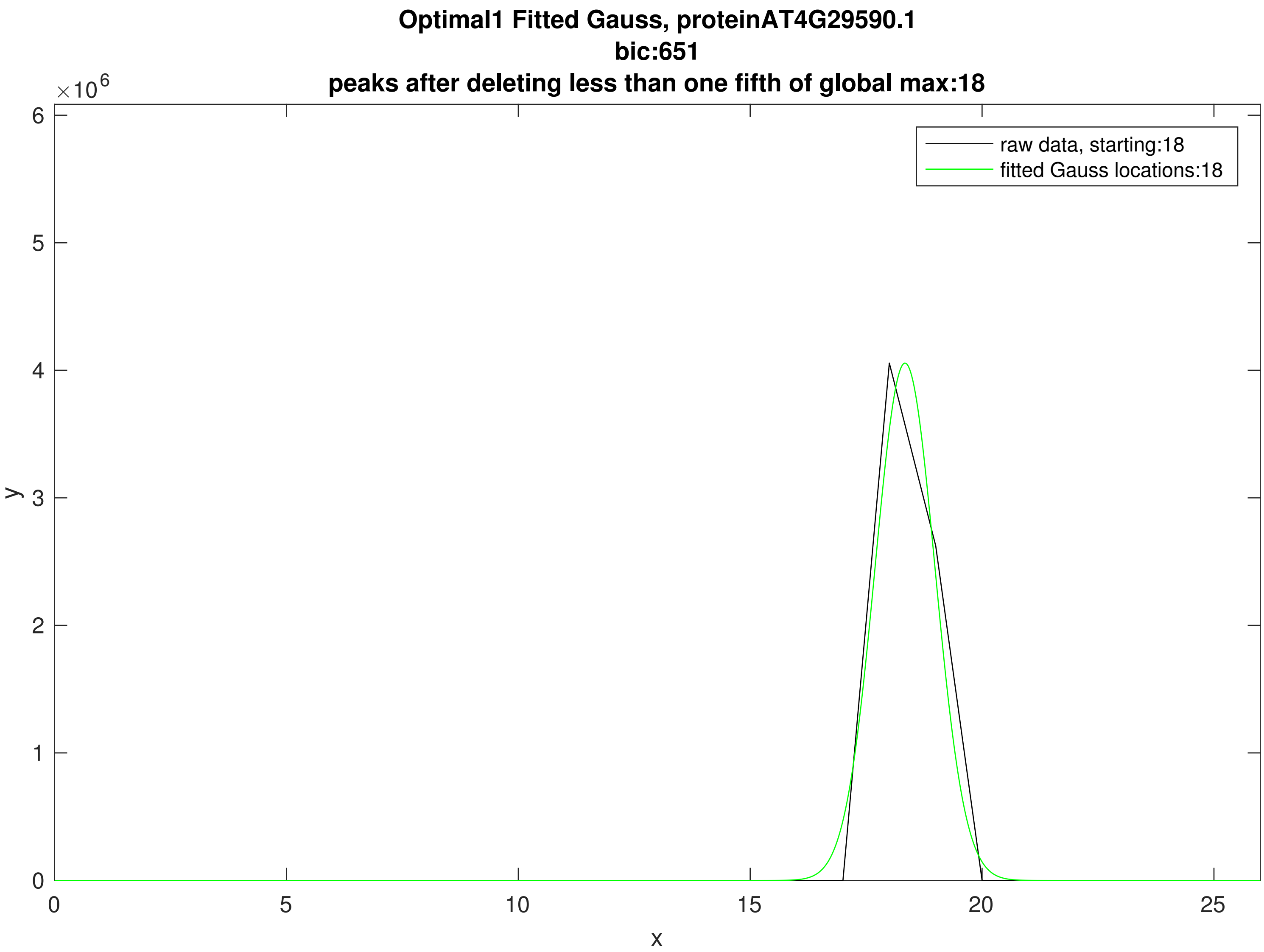
Optimal1 Fitted Gauss, proteinAT4G29590.1
bic:651
peaks after deleting less than one fifth of global max:18
×10^6
6
5
4
3
2
1
0
y
0
5
10
15
20
25
x
raw data, starting:18
fitted Gauss locations:18

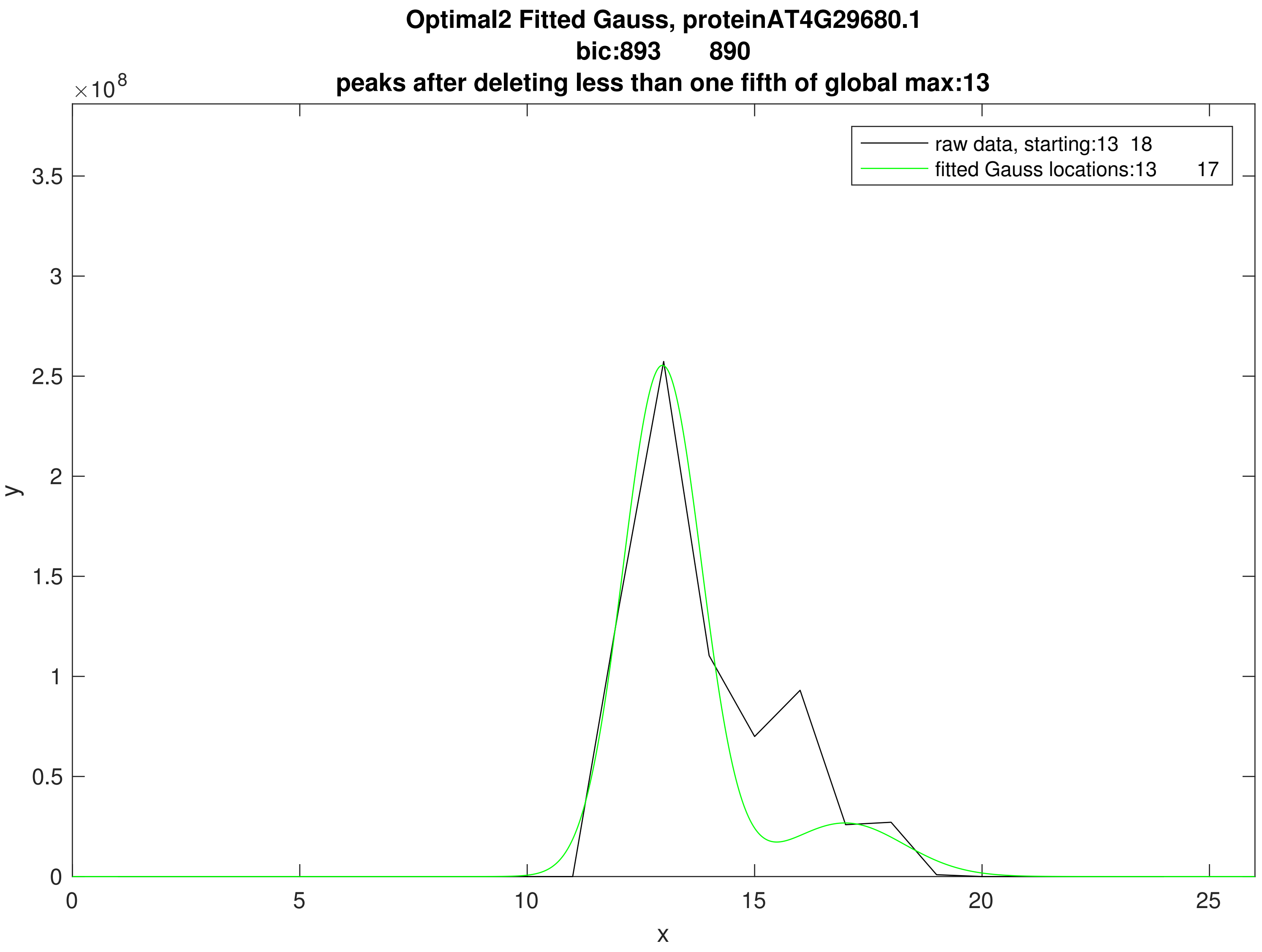

Optimal2 Fitted Gauss, proteinAT4G29680.1
bic:893 890
peaks after deleting less than one fifth of global max:13
×10^8
3.5
3
2.5
2
1.5
1
0.5
0
y
0
5
10
15
20
25
x
raw data, starting:13 18
fitted Gauss locations:13 17

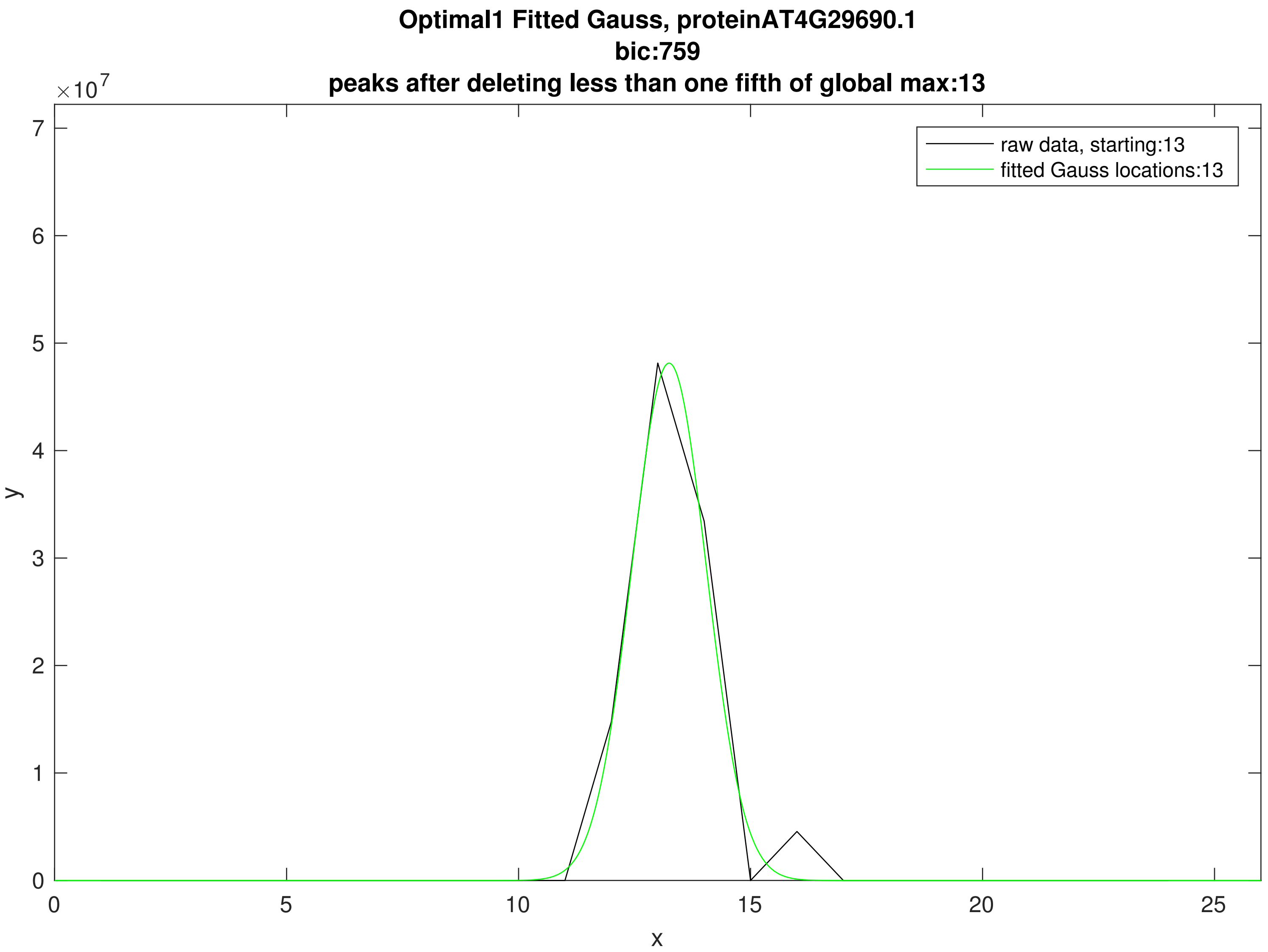

Optimal1 Fitted Gauss, proteinAT4G29690.1
bic:759
peaks after deleting less than one fifth of global max:13
×10^7
raw data, starting:13
fitted Gauss locations:13
y
x
0
1
2
3
4
5
6
7
0
5
10
15
20
25

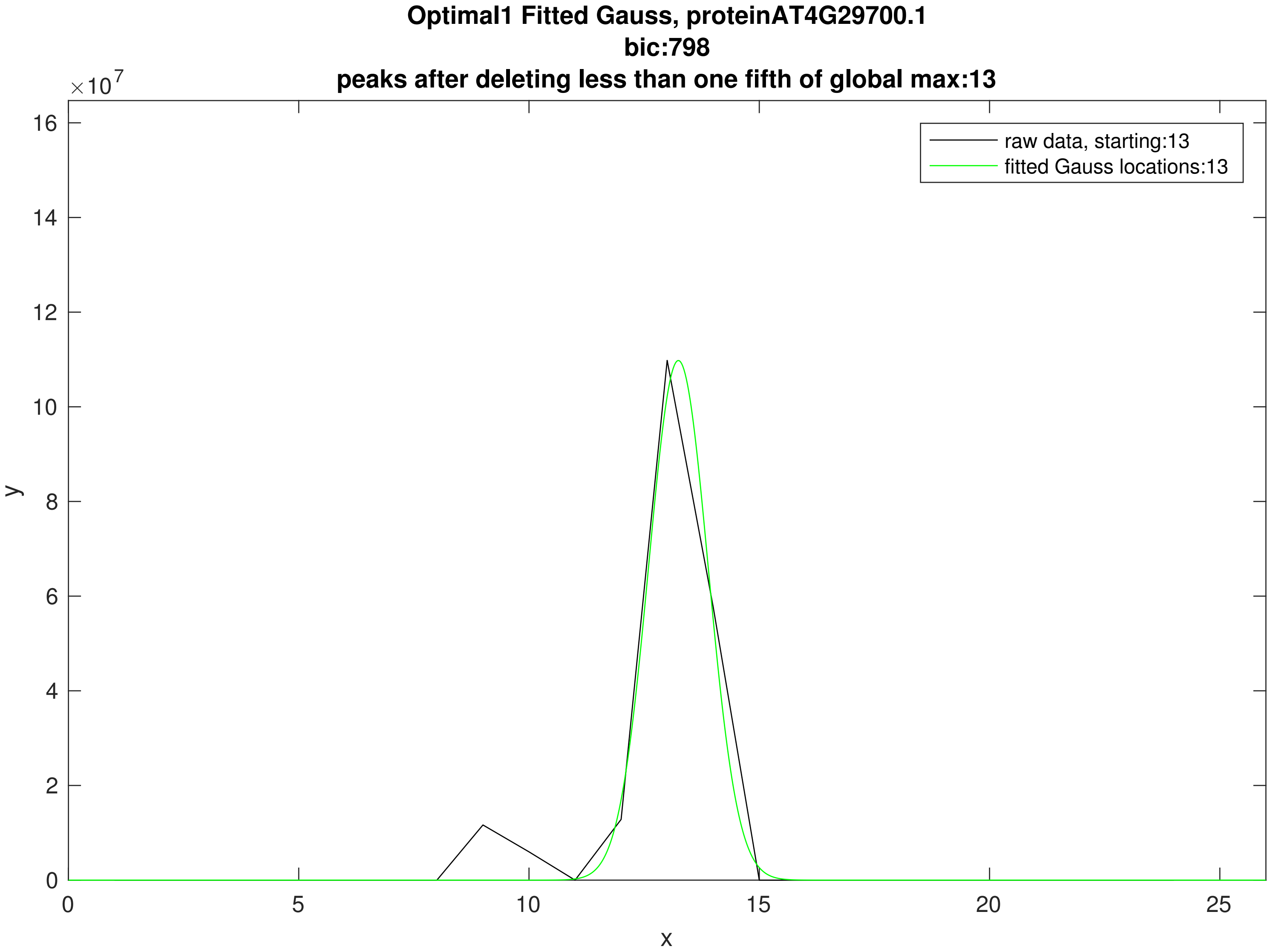

Optimal1 Fitted Gauss, proteinAT4G29700.1
bic:798
peaks after deleting less than one fifth of global max:13
×10^7
raw data, starting:13
fitted Gauss locations:13
y
x
0
2
4
6
8
10
12
14
16
0
5
10
15
20
25

**Optimal1 Fitted Gauss, proteinAT4G29830.1**
**bic:810**
**peaks after deleting less than one fifth of global max:18**

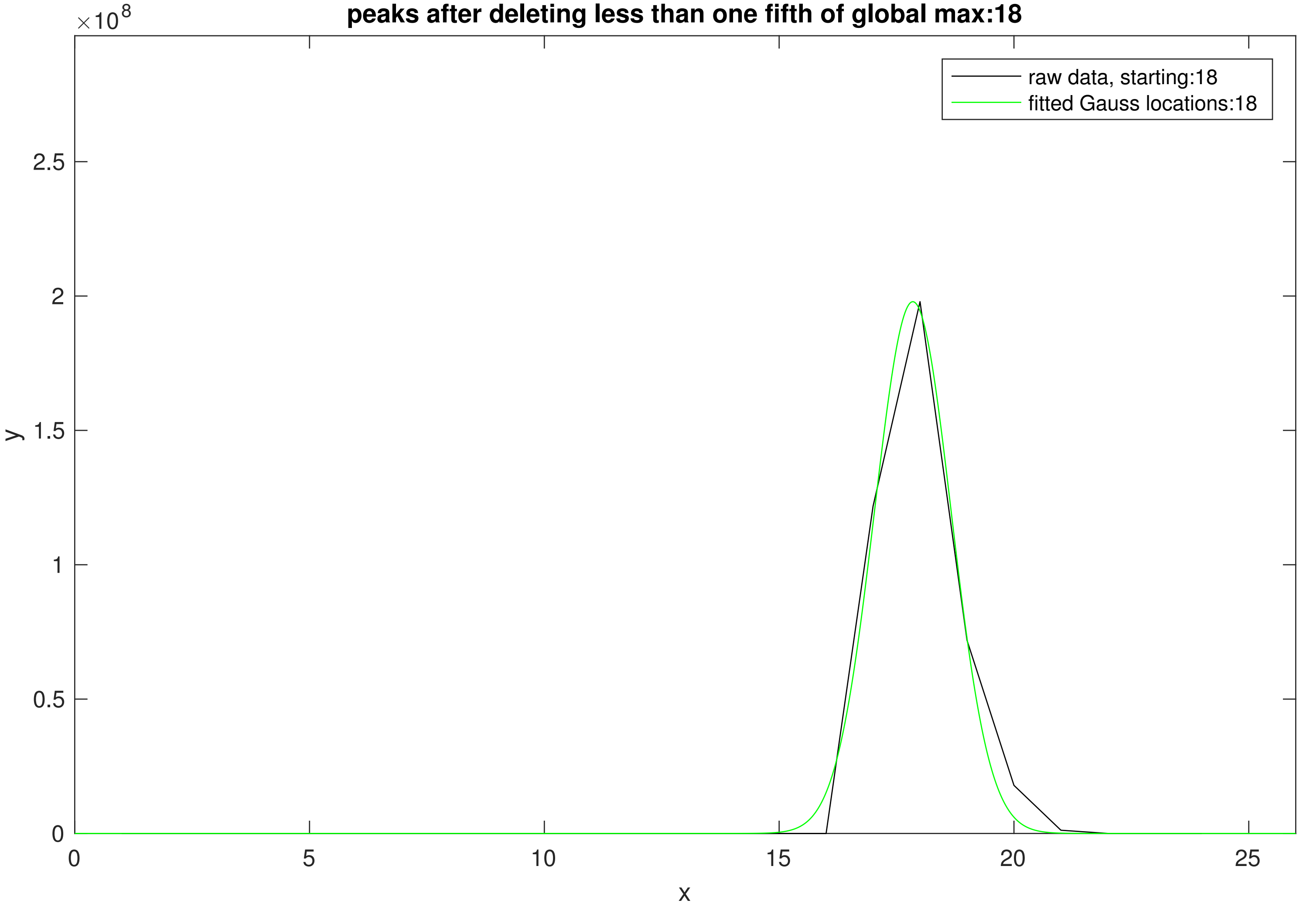

**Optimal1 Fitted Gauss, proteinAT4G29840.1**
**bic:924 933**
**peaks after deleting less than one fifth of global max:13**

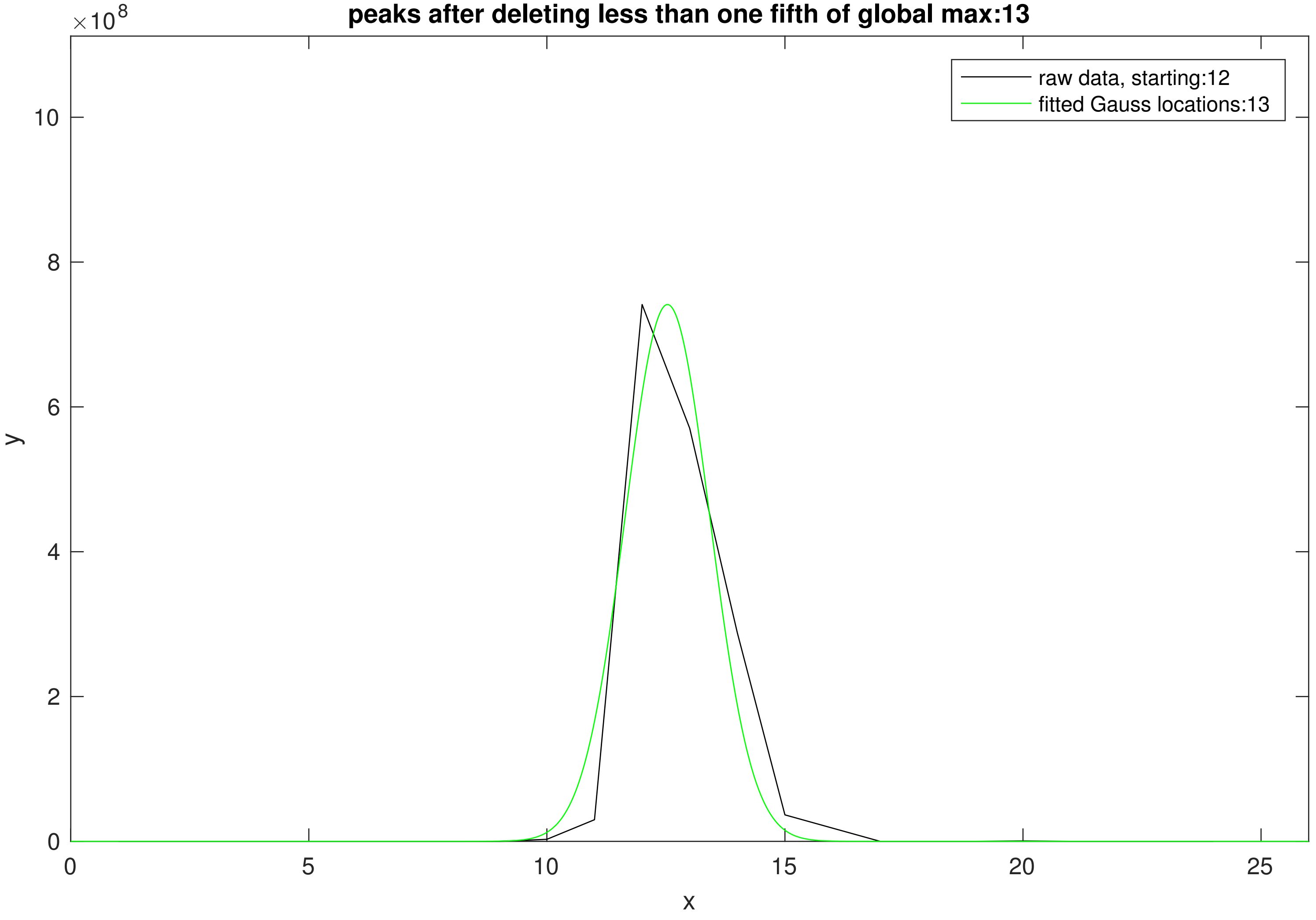

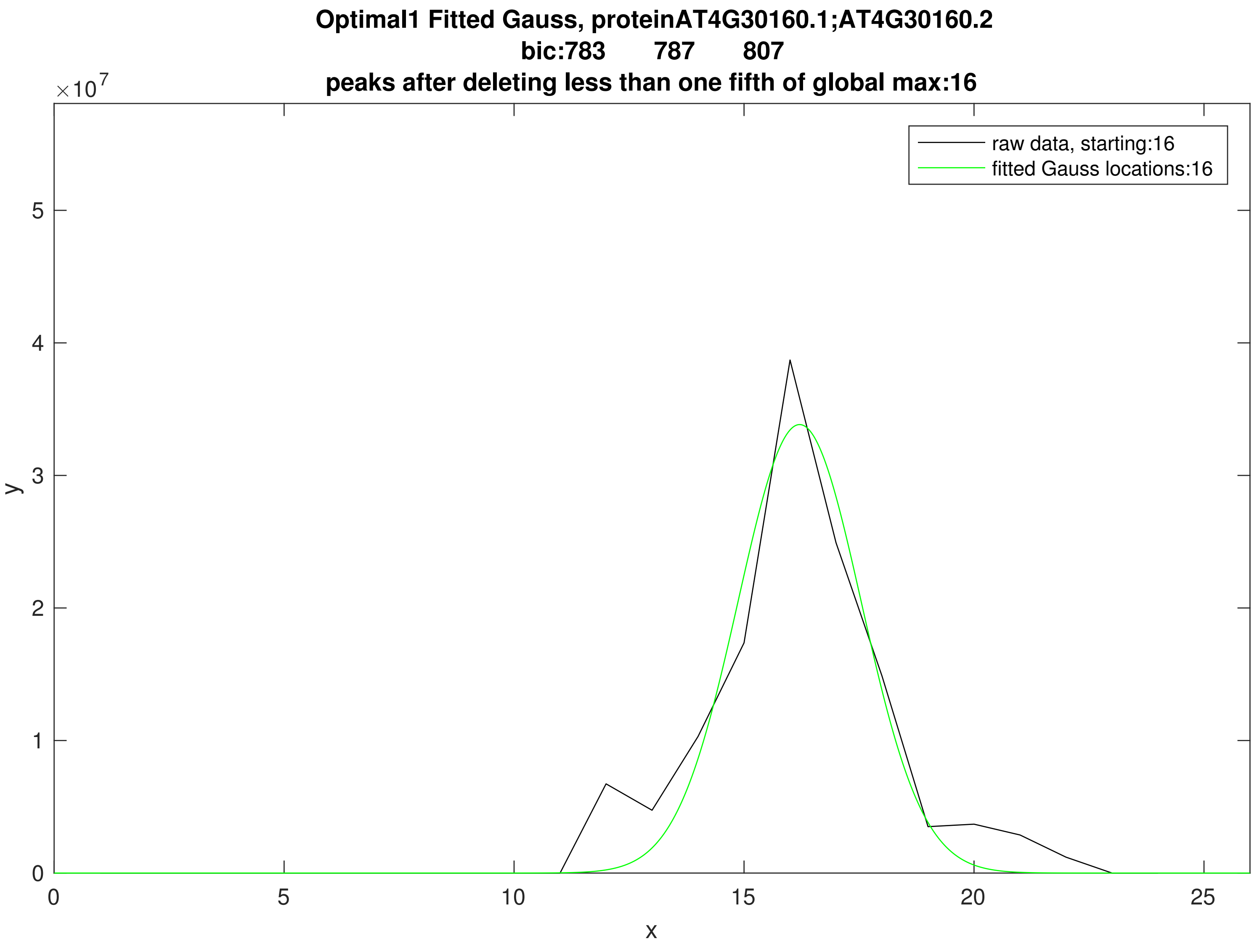

Optimal1 Fitted Gauss, proteinAT4G30160.1;AT4G30160.2
bic:783 787 807
peaks after deleting less than one fifth of global max:16
×10^7
raw data, starting:16
fitted Gauss locations:16
y
x
0
1
2
3
4
5
0
5
10
15
20
25

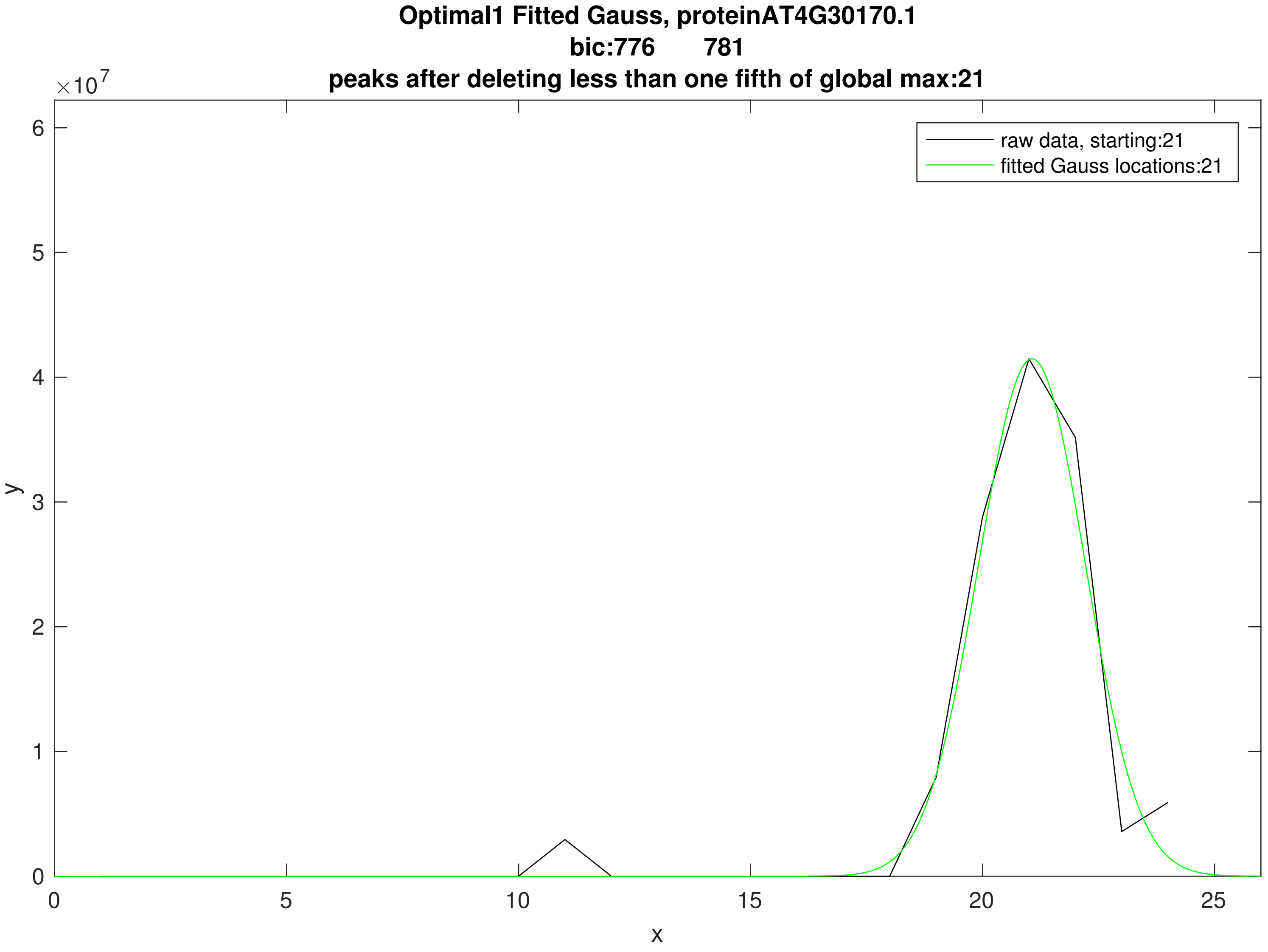
Optimal1 Fitted Gauss, proteinAT4G30170.1
bic:776 781
peaks after deleting less than one fifth of global max:21
×10^7
raw data, starting:21
fitted Gauss locations:21
y
x
0
1
2
3
4
5
6
0
5
10
15
20
25

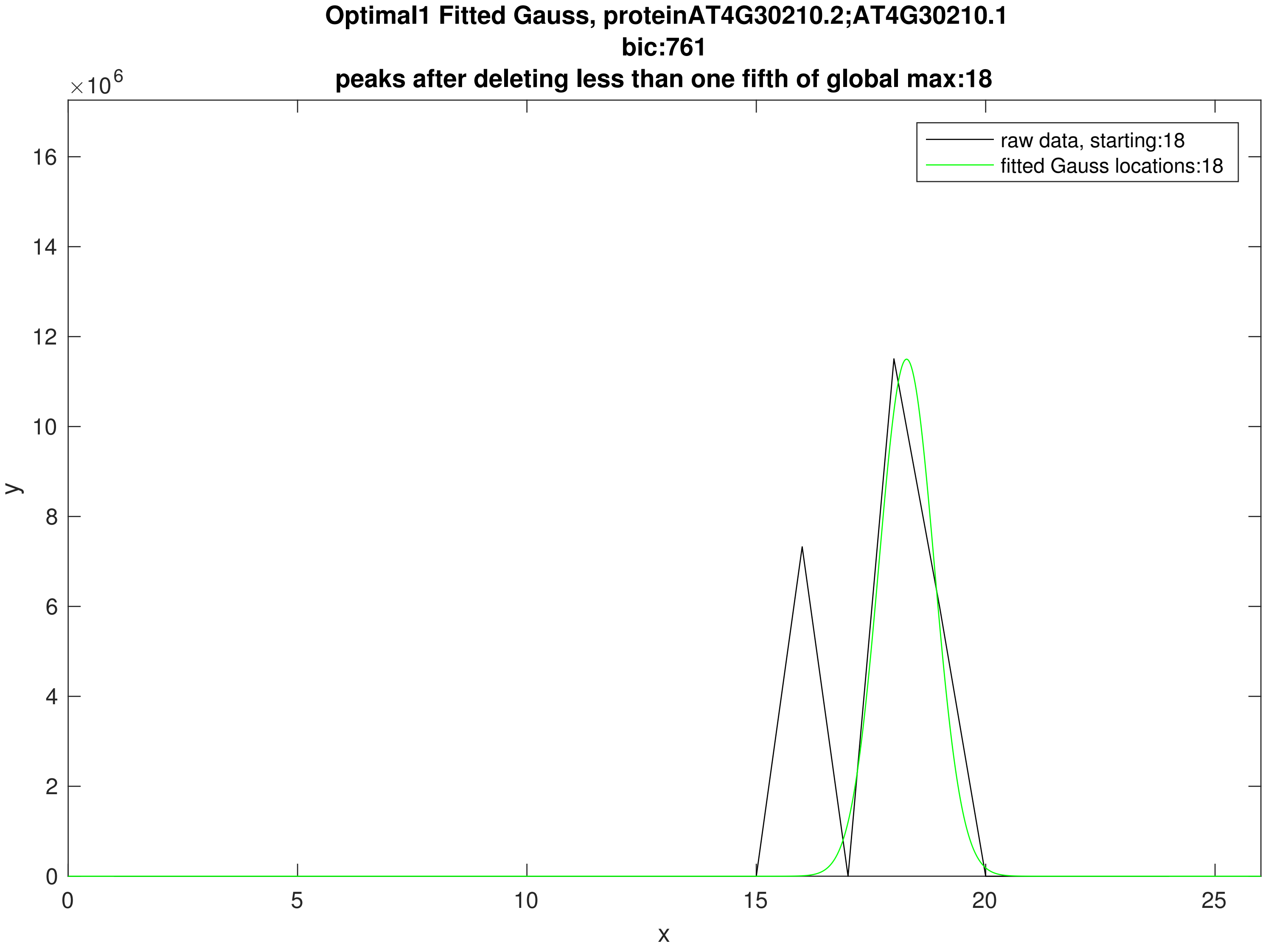

Optimal1 Fitted Gauss, proteinAT4G30210.2;AT4G30210.1
bic:761
peaks after deleting less than one fifth of global max:18
×10^6
raw data, starting:18
fitted Gauss locations:18
y
x
0
2
4
6
8
10
12
14
16
0
5
10
15
20
25

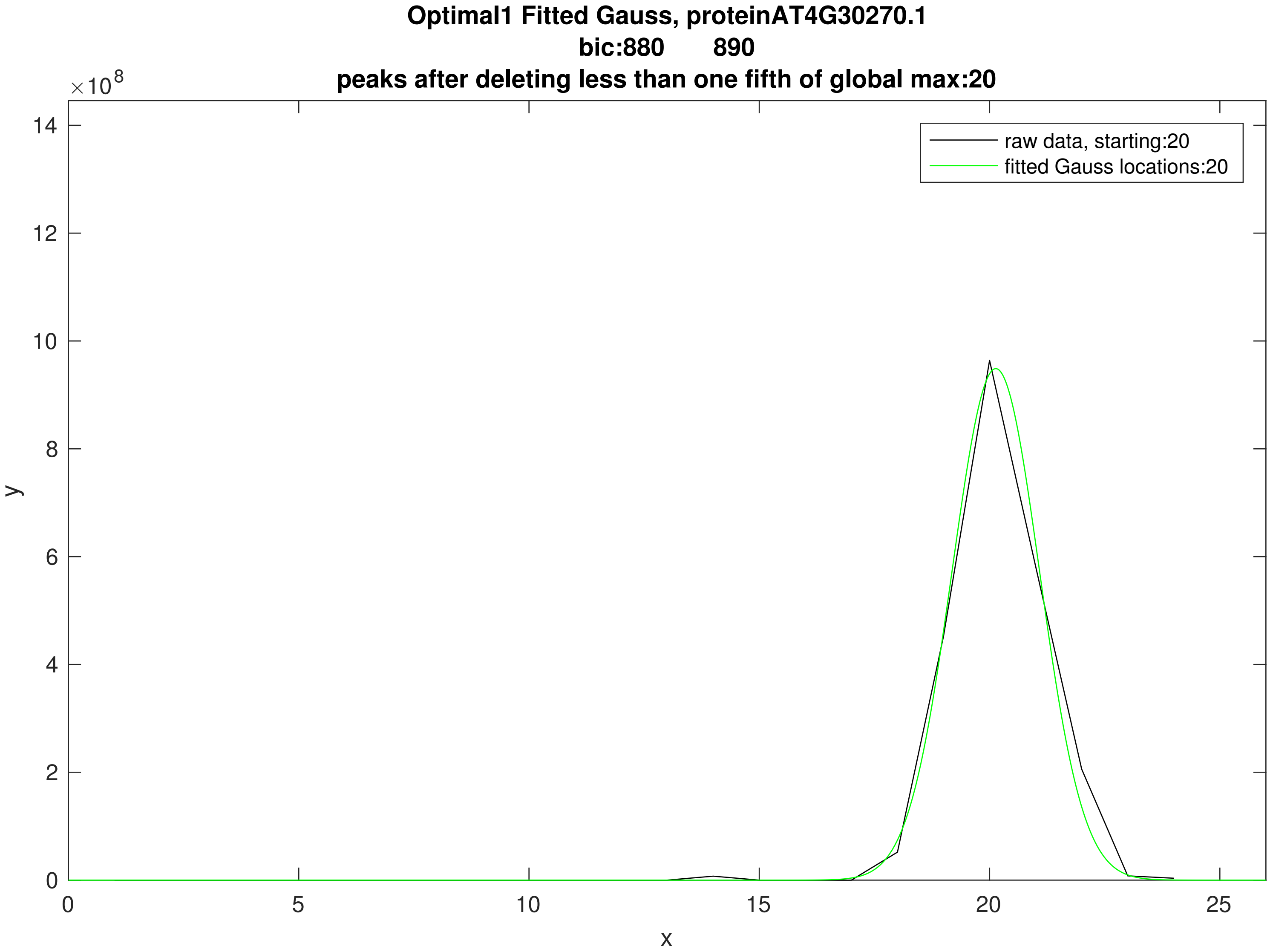
Optimal1 Fitted Gauss, proteinAT4G30270.1
bic:880 890
peaks after deleting less than one fifth of global max:20
×10^8
raw data, starting:20
fitted Gauss locations:20
14
12
10
8
6
4
2
0
y
0
5
10
15
20
25
x

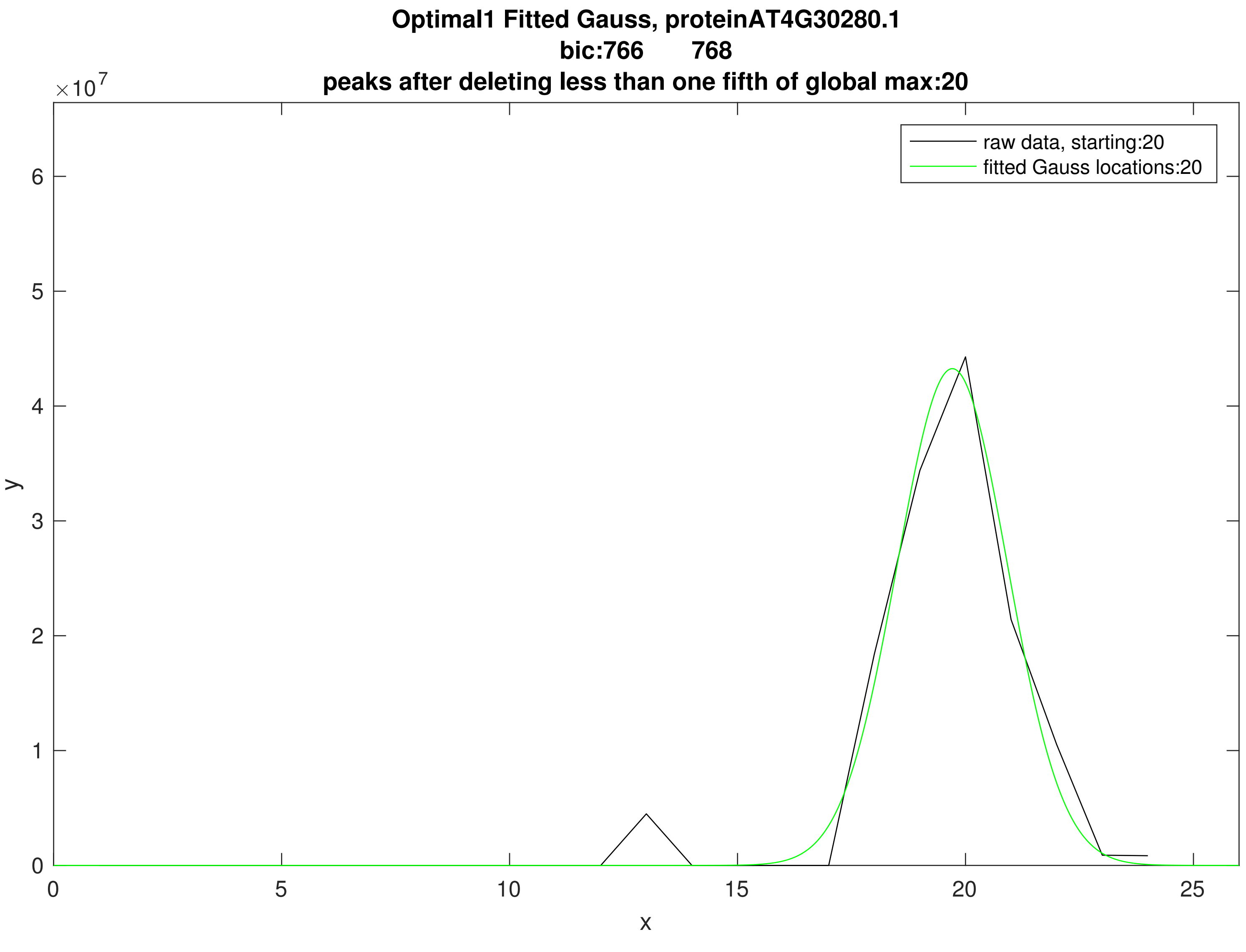
Optimal1 Fitted Gauss, proteinAT4G30280.1
bic:766 768
peaks after deleting less than one fifth of global max:20
×10^7
raw data, starting:20
fitted Gauss locations:20
y
x
0
1
2
3
4
5
6
0
5
10
15
20
25

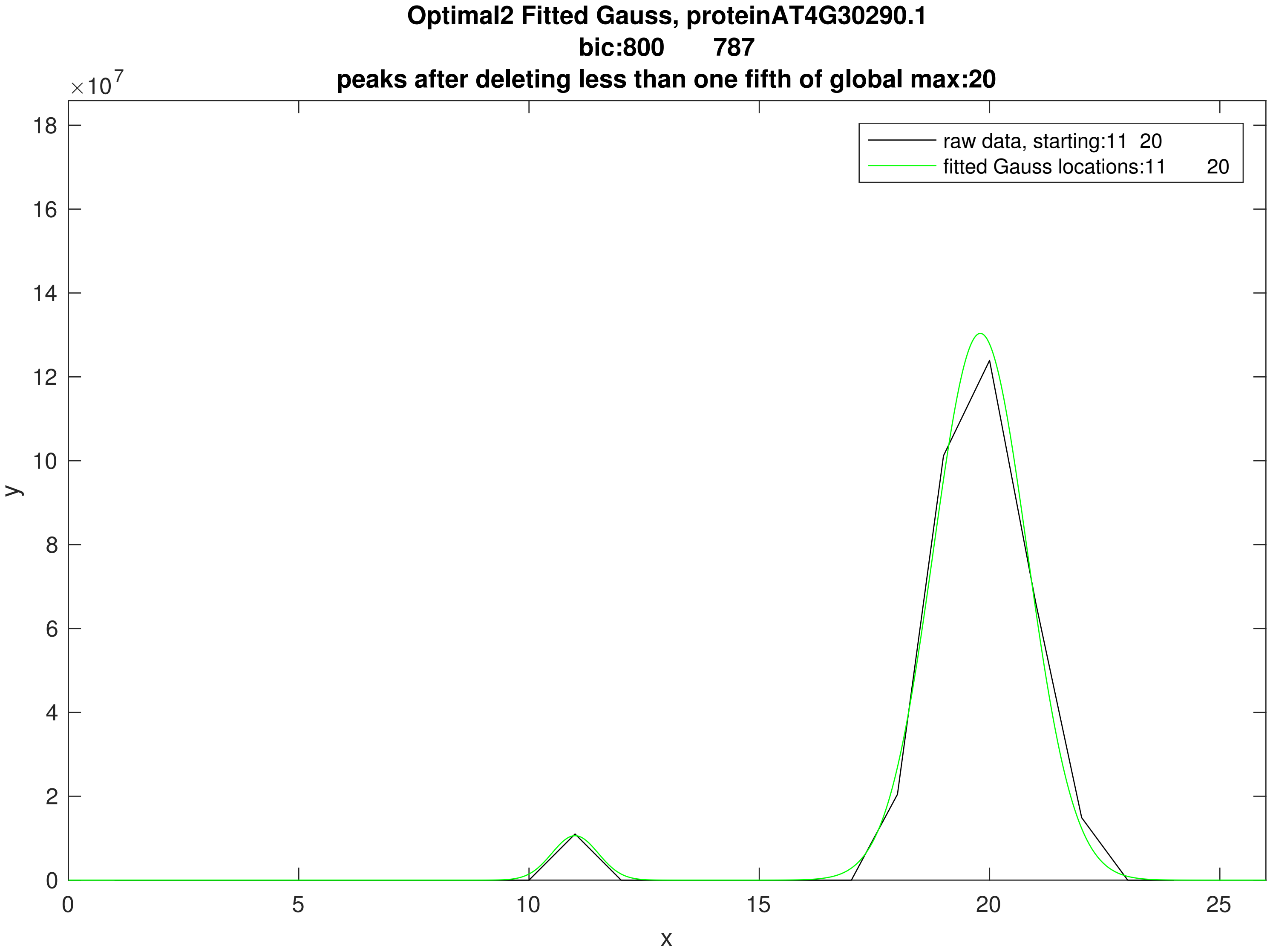

Optimal2 Fitted Gauss, proteinAT4G30290.1
bic:800 787
peaks after deleting less than one fifth of global max:20
×10^7
raw data, starting:11 20
fitted Gauss locations:11 20
y
x
0
2
4
6
8
10
12
14
16
18
5
15
20
25

Optimal1 Fitted Gauss, proteinAT4G30310.2;AT4G30310.3;AT4G30310.1
bic:856
peaks after deleting less than one fifth of global max:12

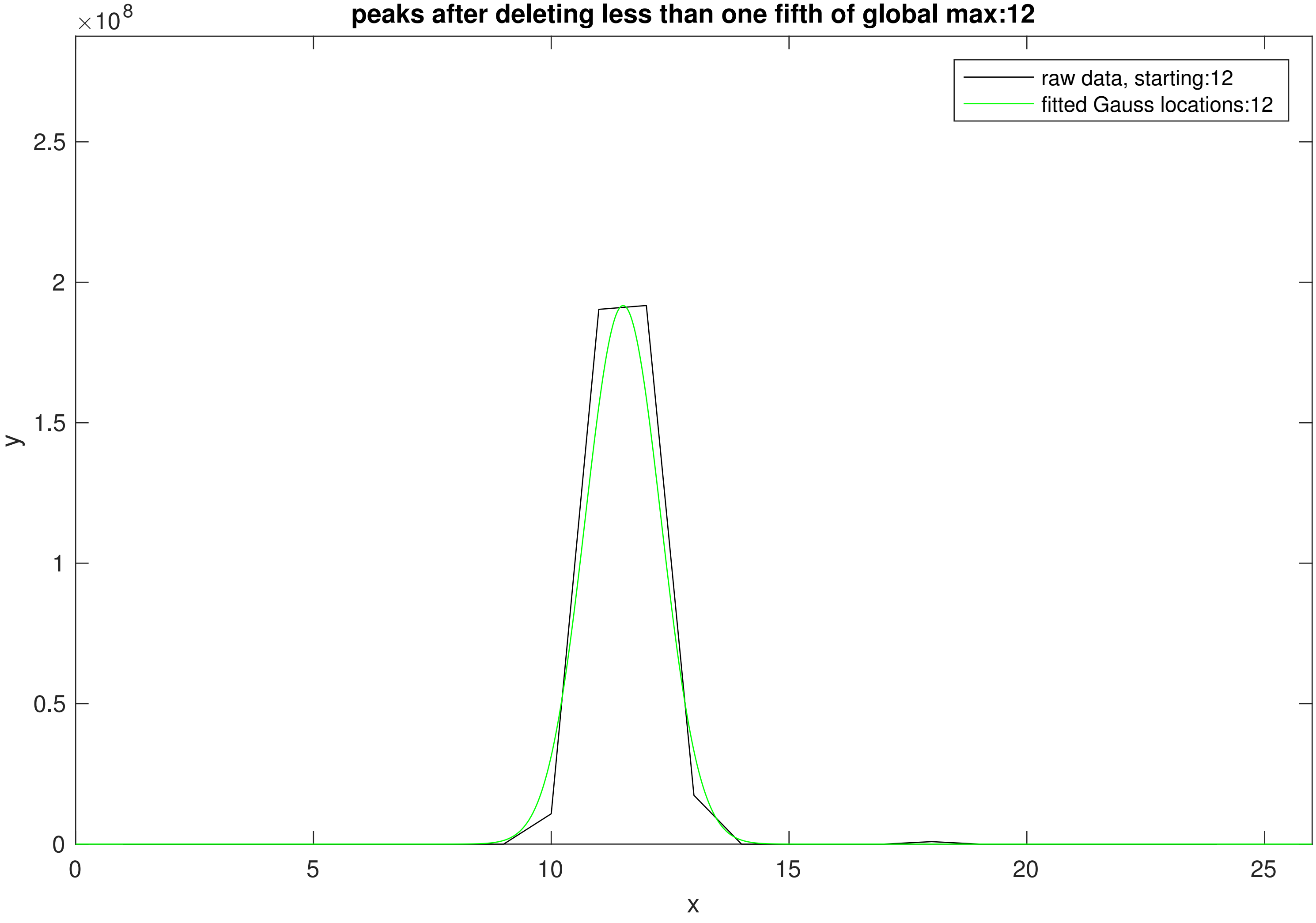

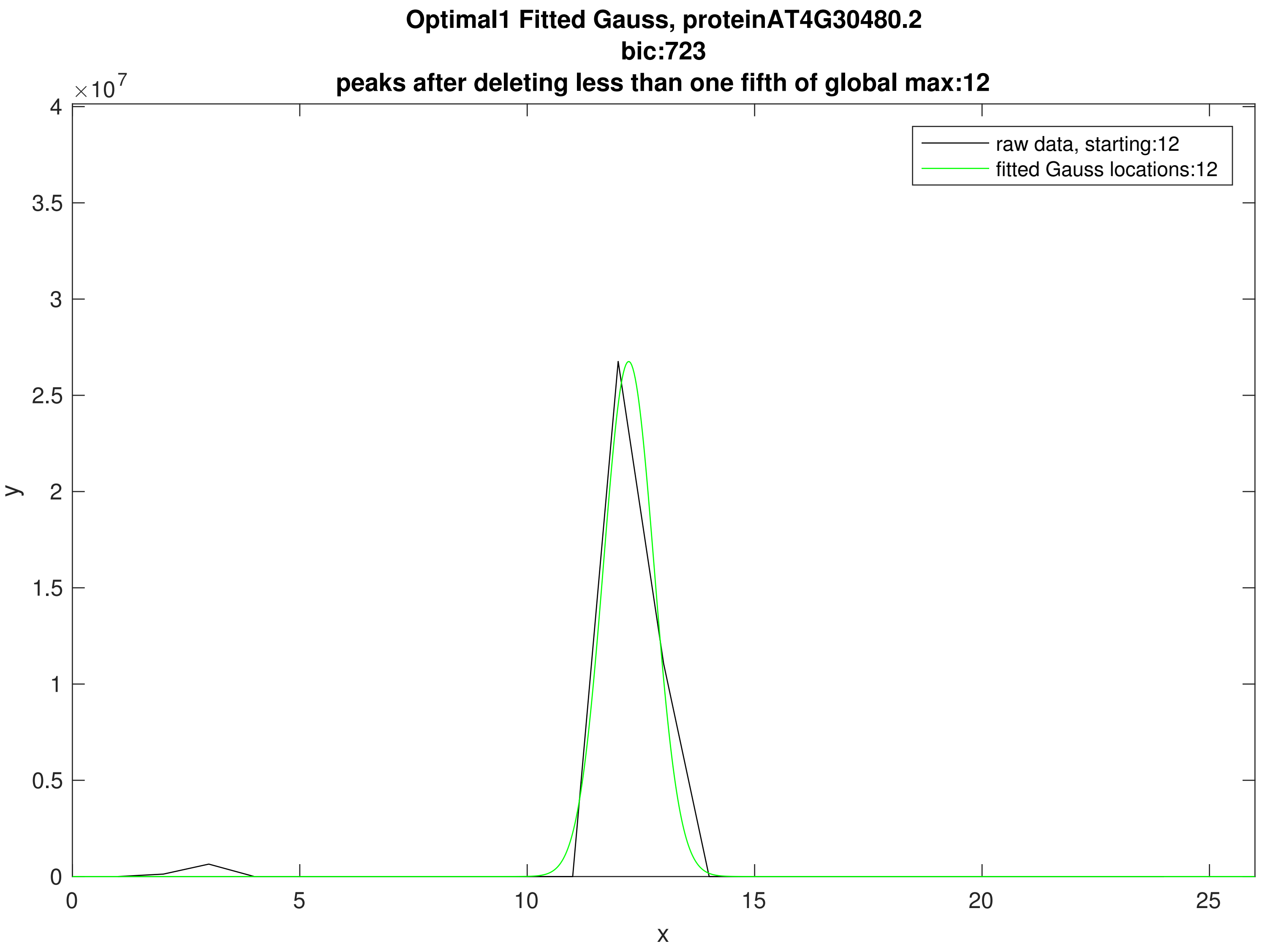
Optimal1 Fitted Gauss, proteinAT4G30480.2
bic:723
peaks after deleting less than one fifth of global max:12
×10^7
raw data, starting:12
fitted Gauss locations:12
y
x
0
0.5
1
1.5
2
2.5
3
3.5
4
5
10
15
20
25

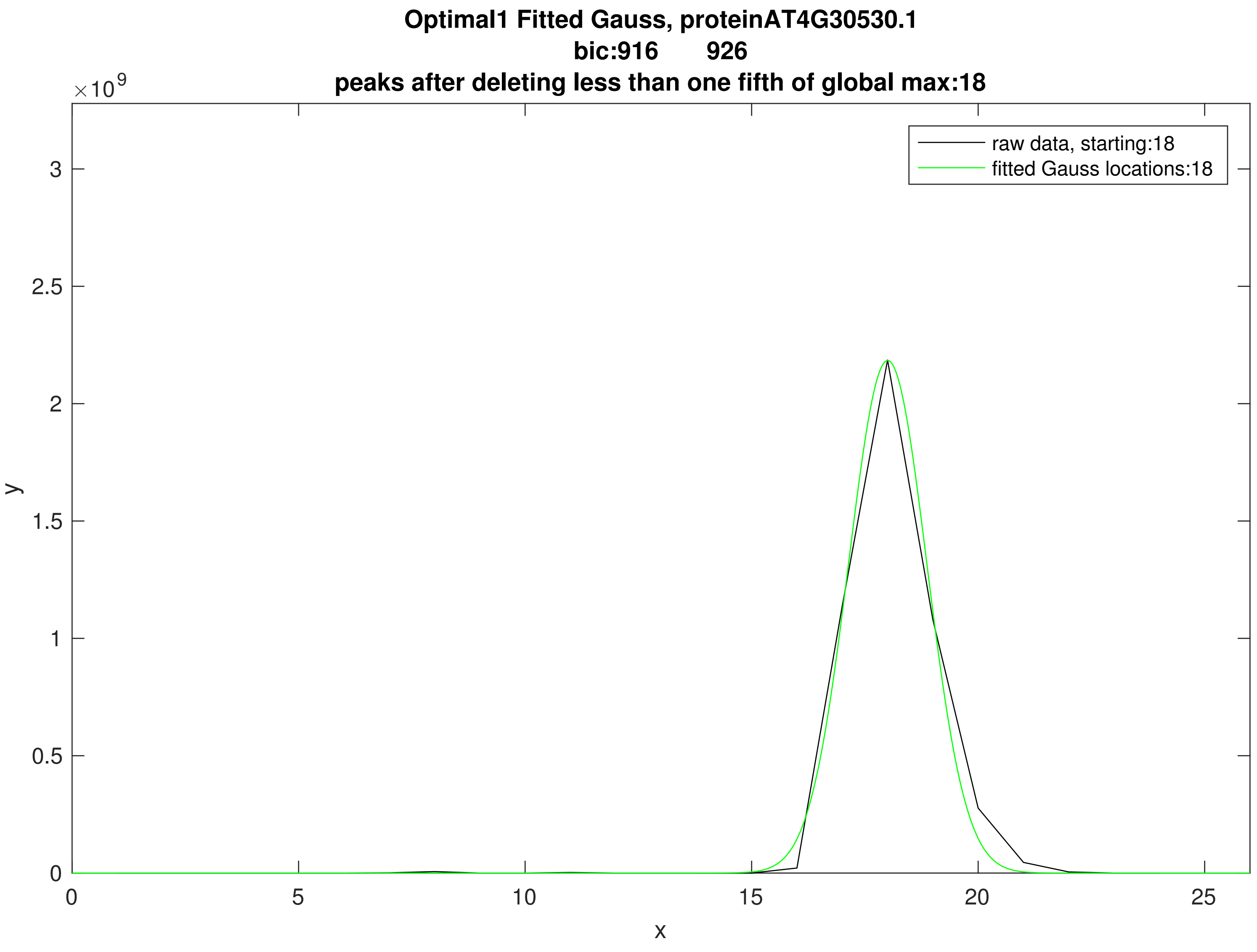

Optimal1 Fitted Gauss, proteinAT4G30530.1
bic:916 926
peaks after deleting less than one fifth of global max:18
×10^9
raw data, starting:18
fitted Gauss locations:18
3
2.5
2
1.5
1
0.5
0
y
0
5
10
15
20
25
x

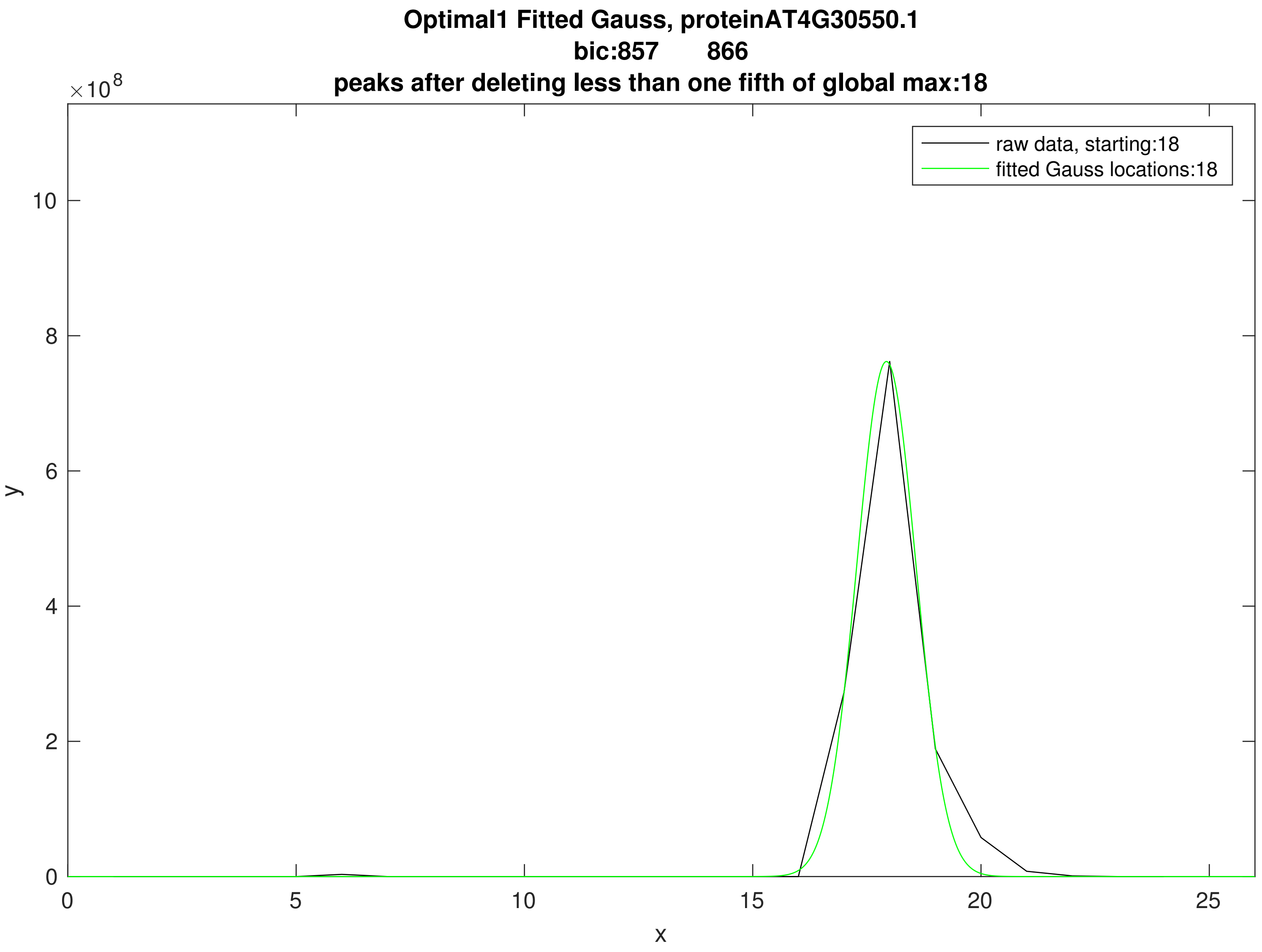

Optimal1 Fitted Gauss, proteinAT4G30550.1
bic:857 866
peaks after deleting less than one fifth of global max:18
×10^8
raw data, starting:18
fitted Gauss locations:18
10
8
6
4
2
0
y
0
5
10
15
20
25
x

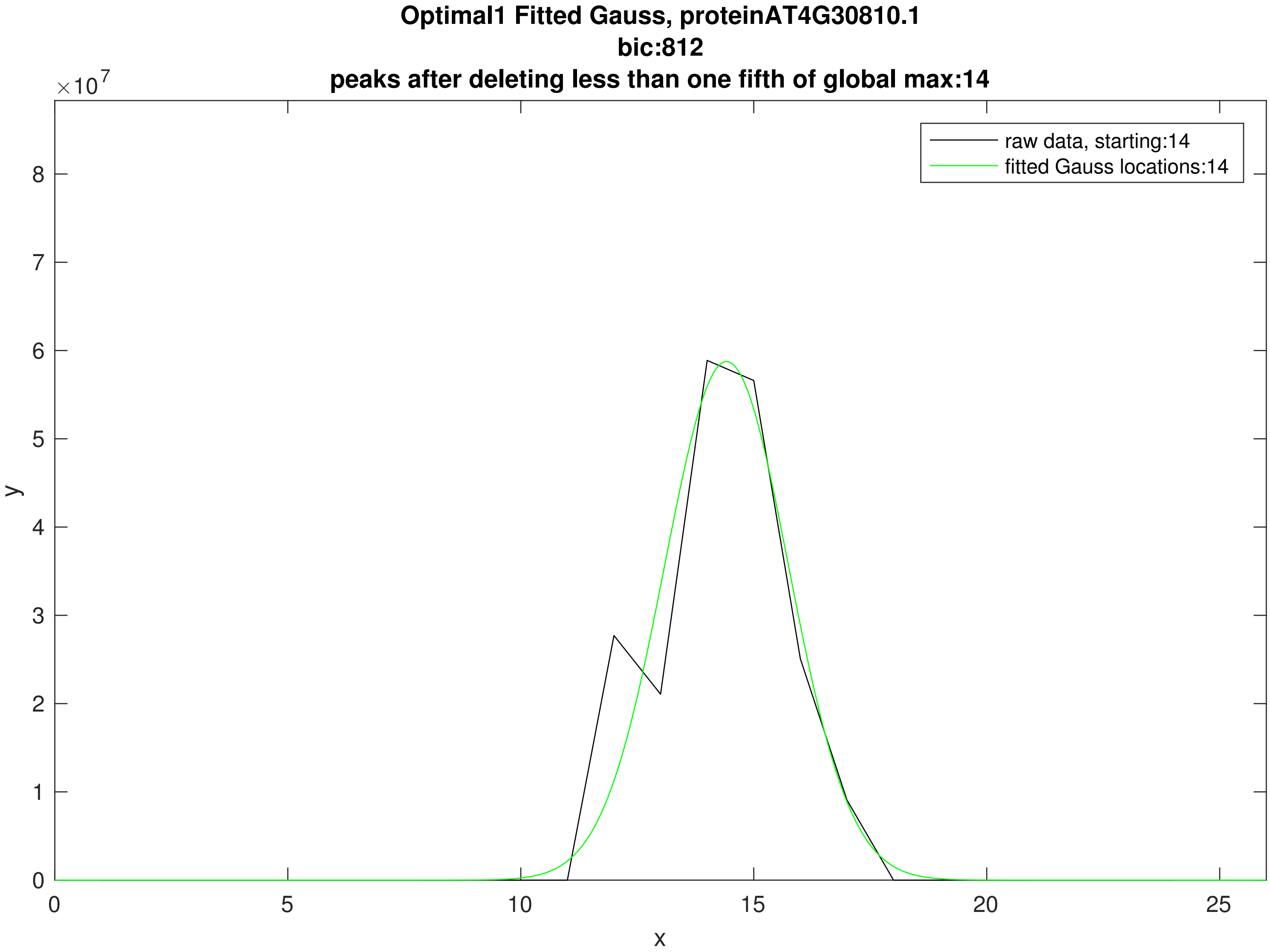
Optimal1 Fitted Gauss, proteinAT4G30810.1
bic:812
peaks after deleting less than one fifth of global max:14
×10^7
raw data, starting:14
fitted Gauss locations:14
y
x
0
1
2
3
4
5
6
7
8
0
5
10
15
20
25

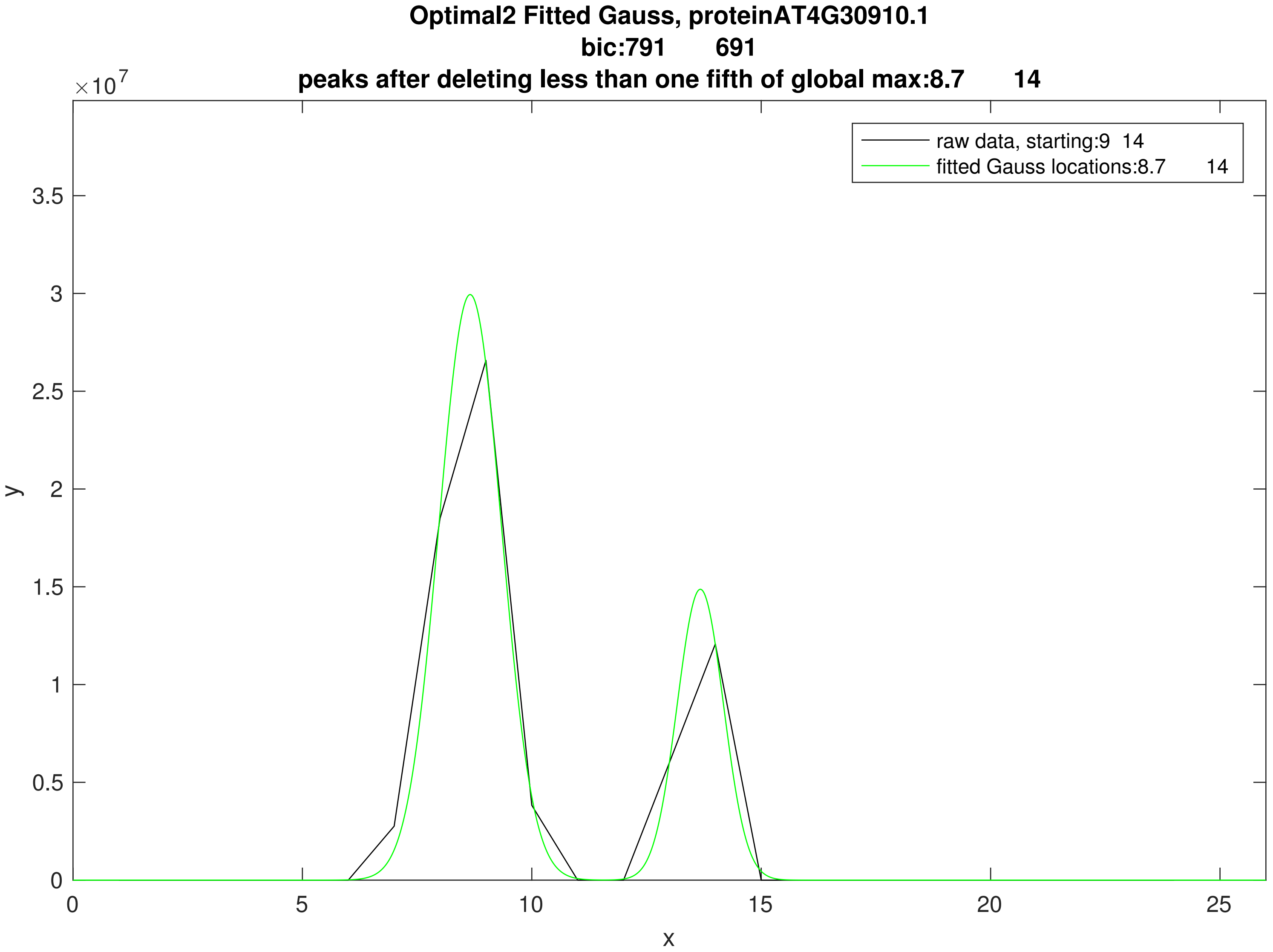

Optimal2 Fitted Gauss, proteinAT4G30910.1
bic:791 691
peaks after deleting less than one fifth of global max:8.7 14
×10^7
raw data, starting:9 14
fitted Gauss locations:8.7 14
y
x
0
0.5
1
1.5
2
2.5
3
3.5
0
5
10
15
20
25

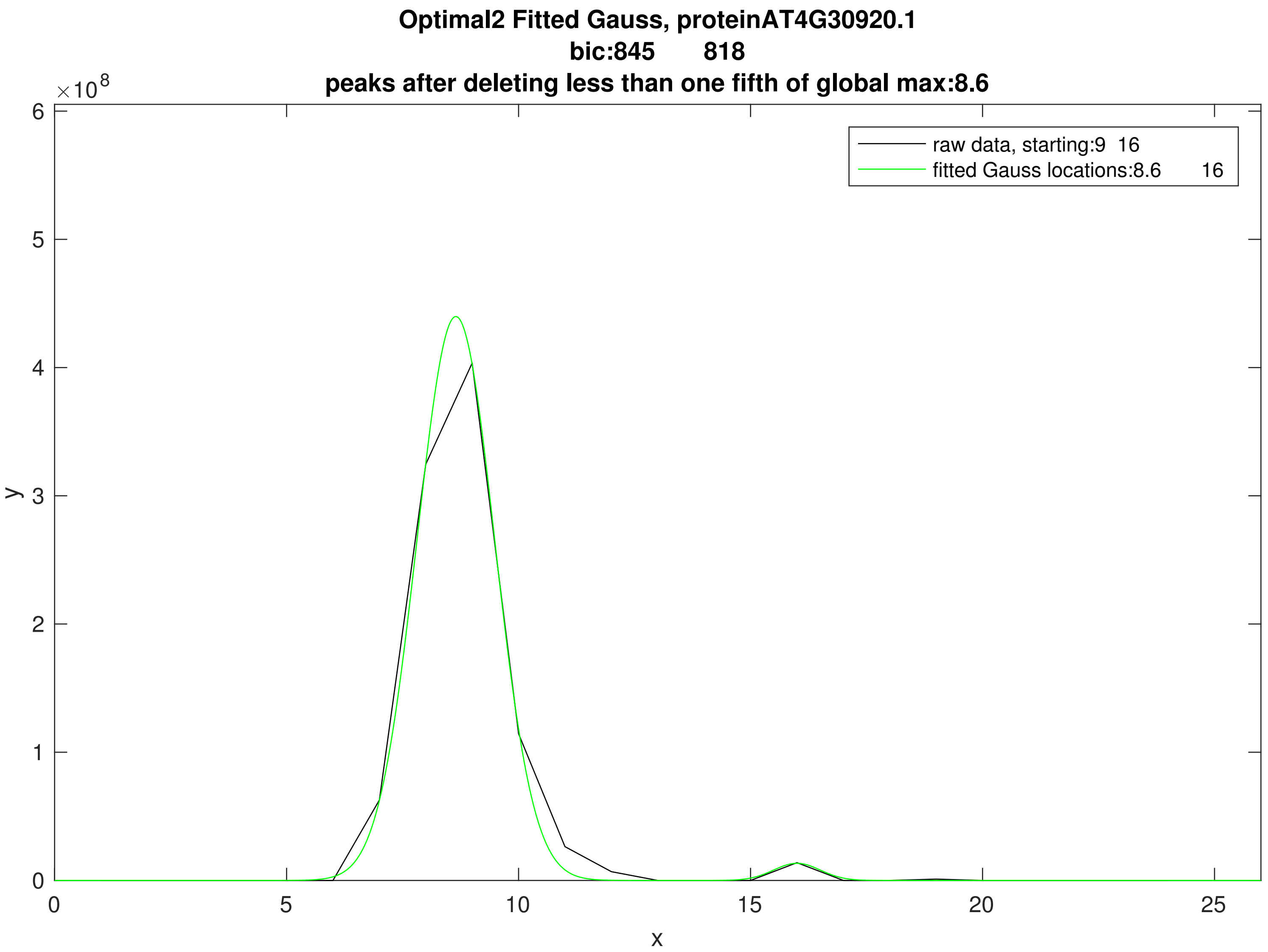

Optimal2 Fitted Gauss, proteinAT4G30920.1
bic:845 818
peaks after deleting less than one fifth of global max:8.6
×10^8
raw data, starting:9 16
fitted Gauss locations:8.6 16
y
x
0
1
2
3
4
5
6
0
5
10
15
20
25

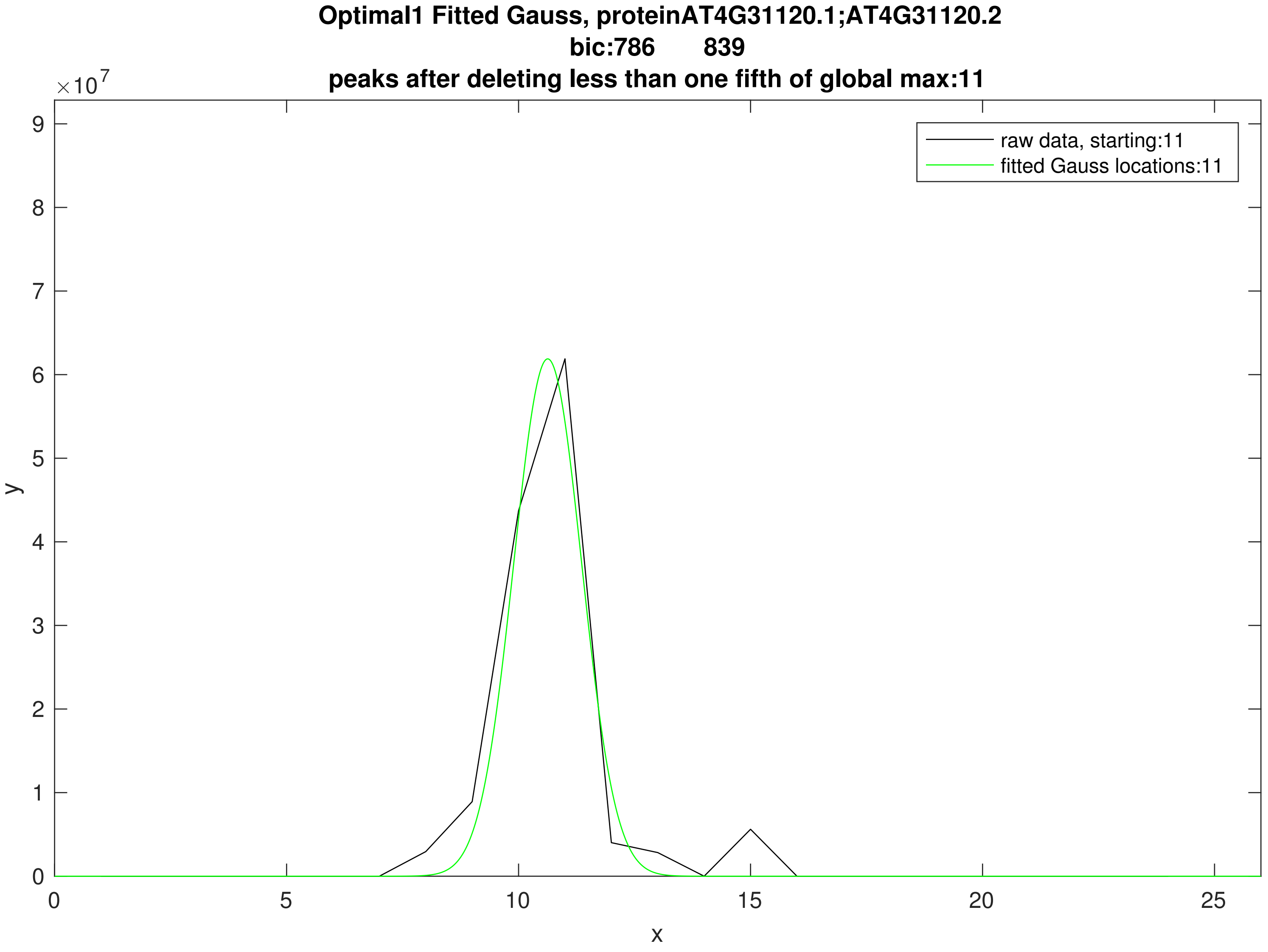

Optimal1 Fitted Gauss, proteinAT4G31120.1;AT4G31120.2
bic:786 839
peaks after deleting less than one fifth of global max:11
×10^7
raw data, starting:11
fitted Gauss locations:11
y
x
0
1
2
3
4
5
6
7
8
9
0
5
10
15
20
25

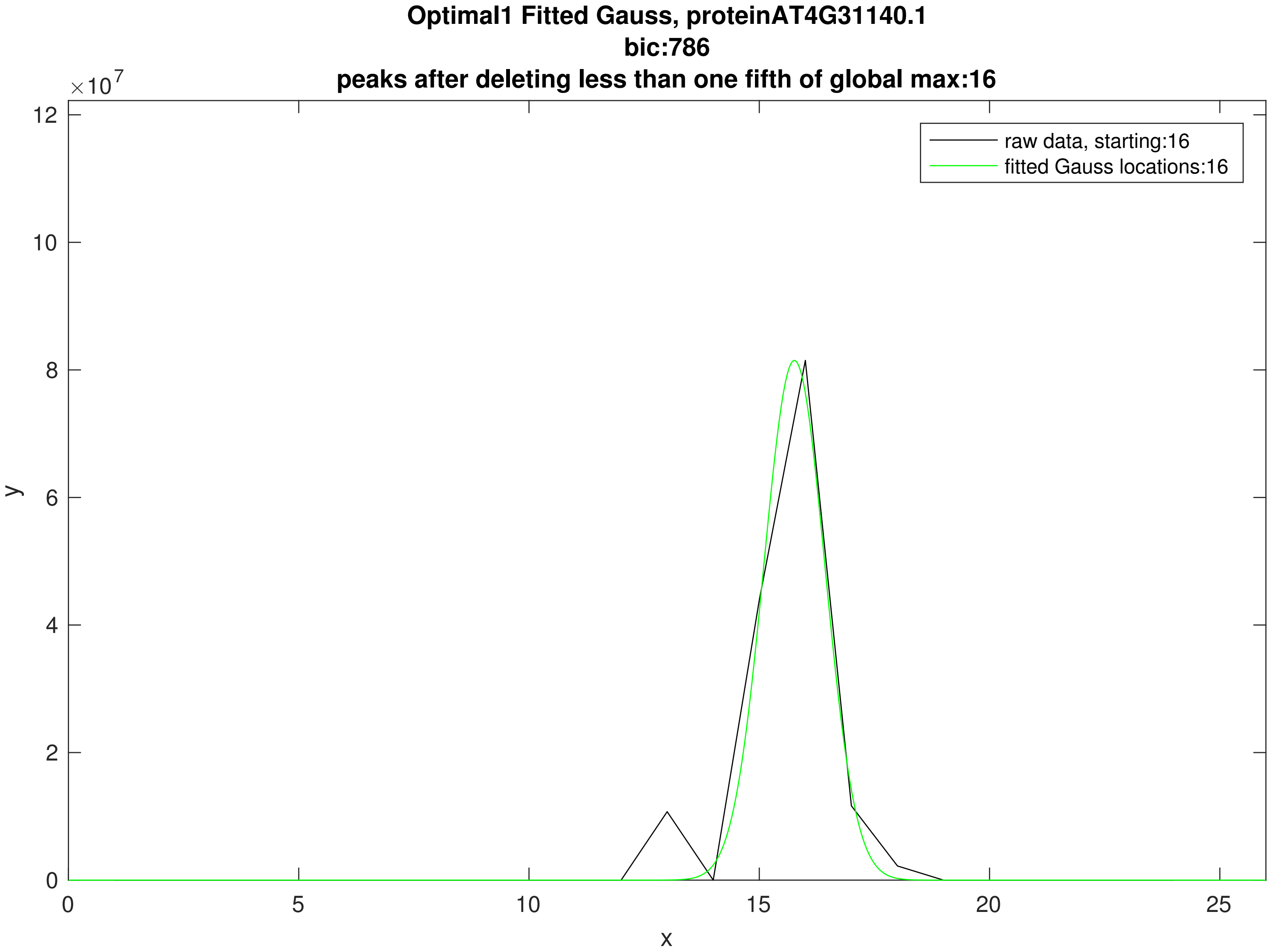

Optimal1 Fitted Gauss, proteinAT4G31140.1
bic:786
peaks after deleting less than one fifth of global max:16
×10^7
raw data, starting:16
fitted Gauss locations:16
y
x
0
2
4
6
8
10
12
0
5
10
15
20
25

**Optimal1 Fitted Gauss, proteinAT4G31300.1;AT4G31300.3;AT4G31300.2**
**bic:912 922**
**peaks after deleting less than one fifth of global max:5.2**

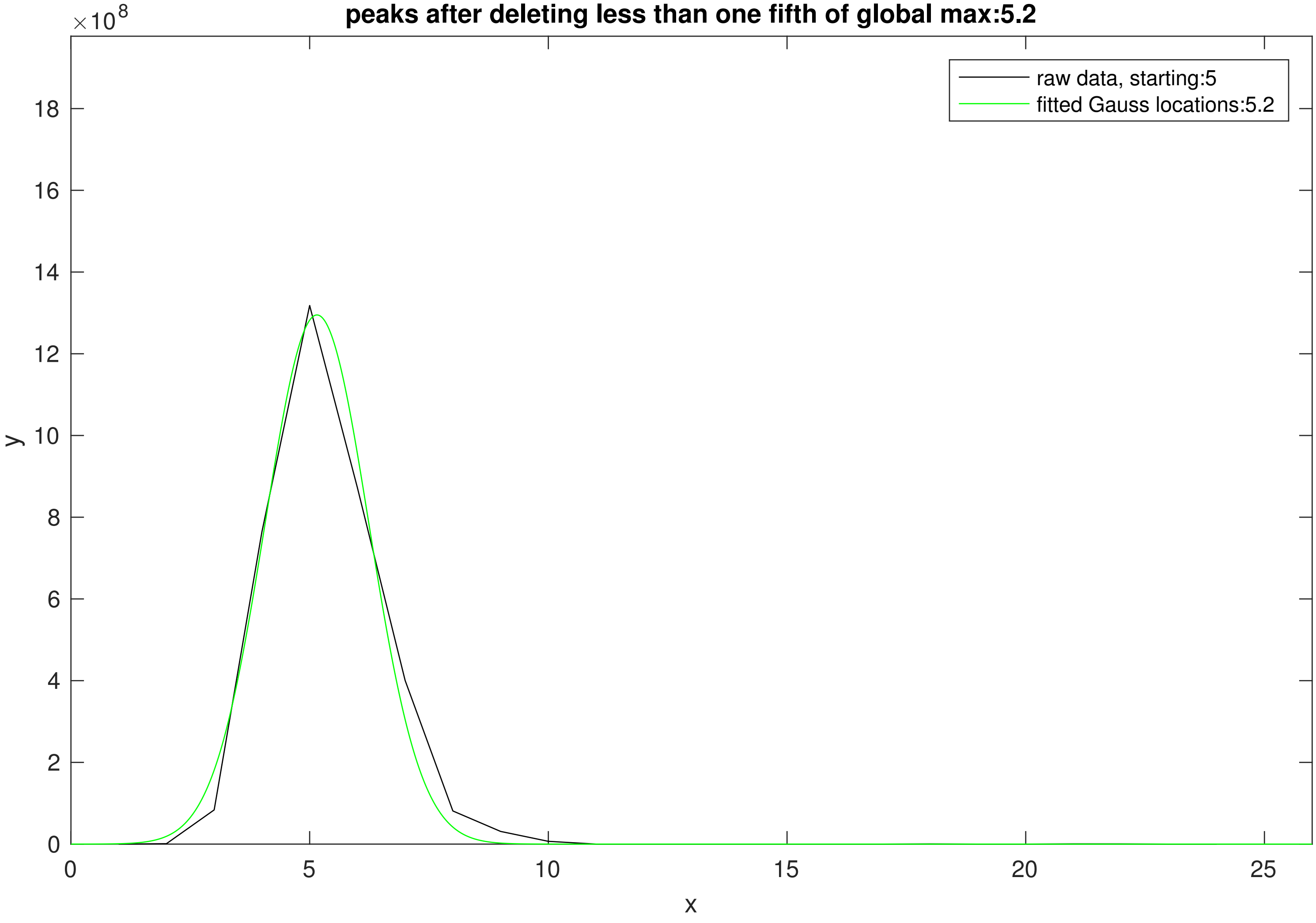

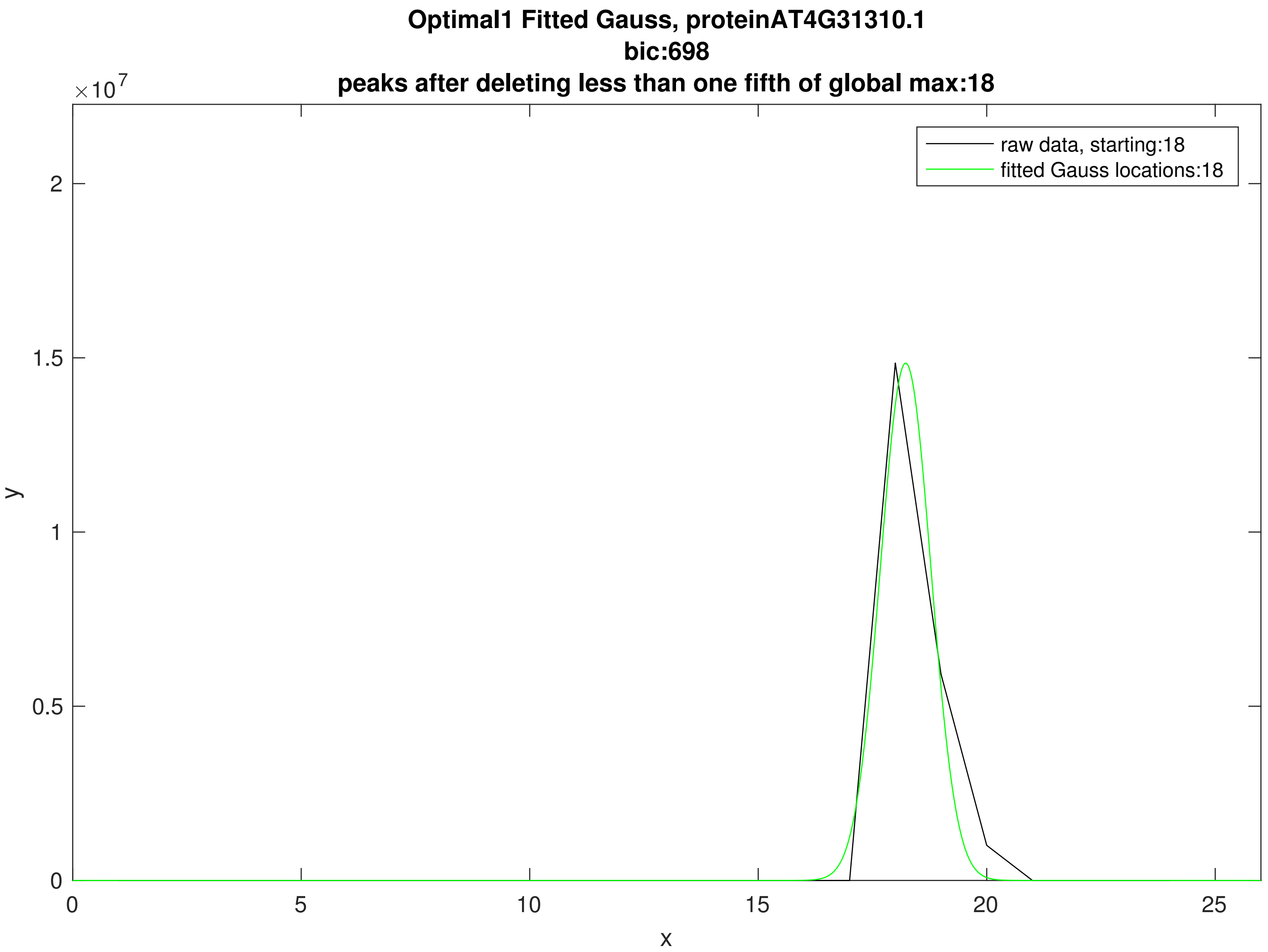
Optimal1 Fitted Gauss, proteinAT4G31310.1
bic:698
peaks after deleting less than one fifth of global max:18
×10^7
raw data, starting:18
fitted Gauss locations:18
y
x
0
0.5
1
1.5
2
5
10
15
20
25

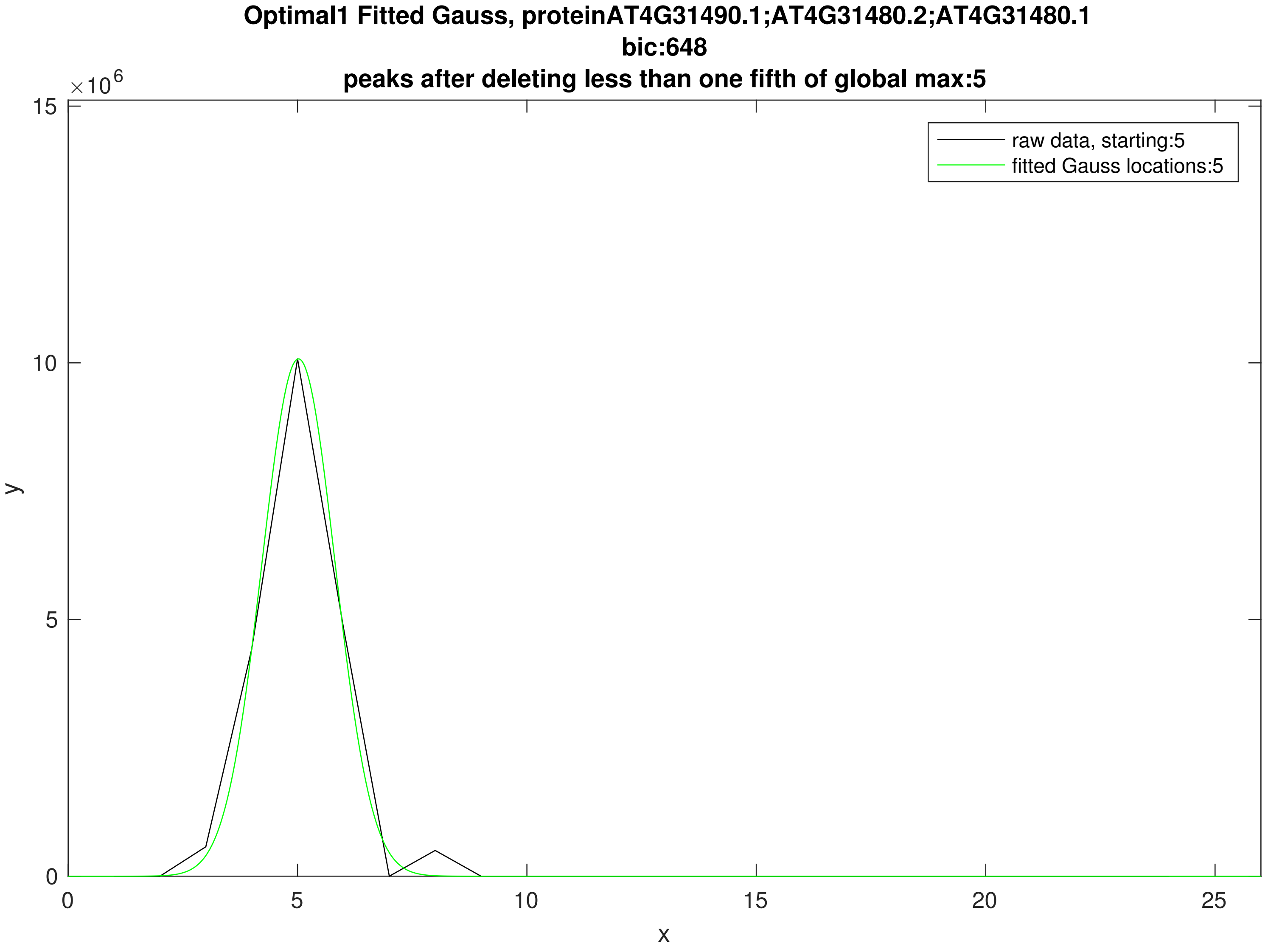
Optimal1 Fitted Gauss, proteinAT4G31490.1;AT4G31480.2;AT4G31480.1
bic:648
peaks after deleting less than one fifth of global max:5
×10^6
15
10
5
0
y
0
5
10
15
20
25
x
raw data, starting:5
fitted Gauss locations:5

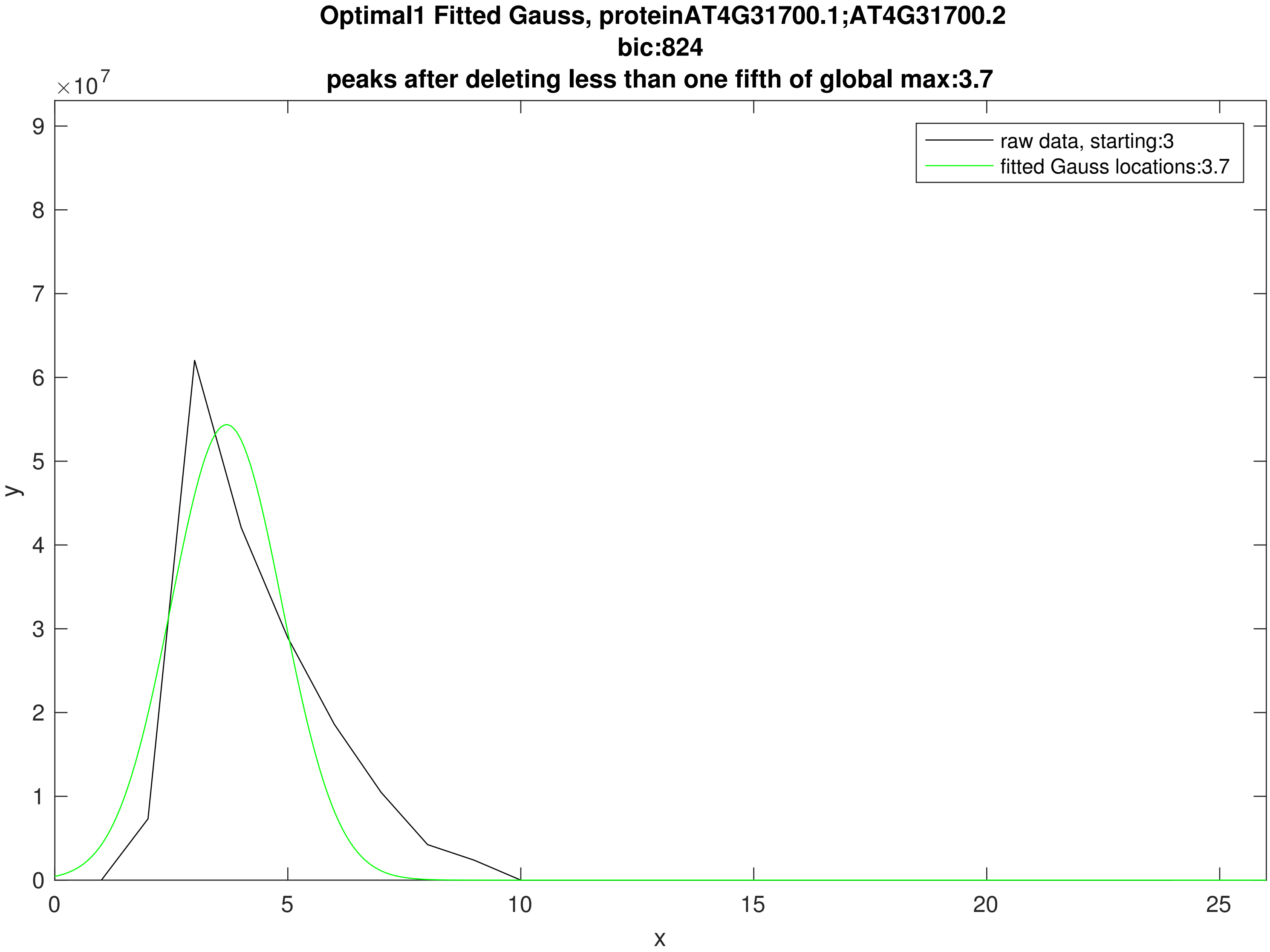
Optimal1 Fitted Gauss, proteinAT4G31700.1;AT4G31700.2
bic:824
peaks after deleting less than one fifth of global max:3.7
×10^7
raw data, starting:3
fitted Gauss locations:3.7
y
x
0
1
2
3
4
5
6
7
8
9
0
5
10
15
20
25

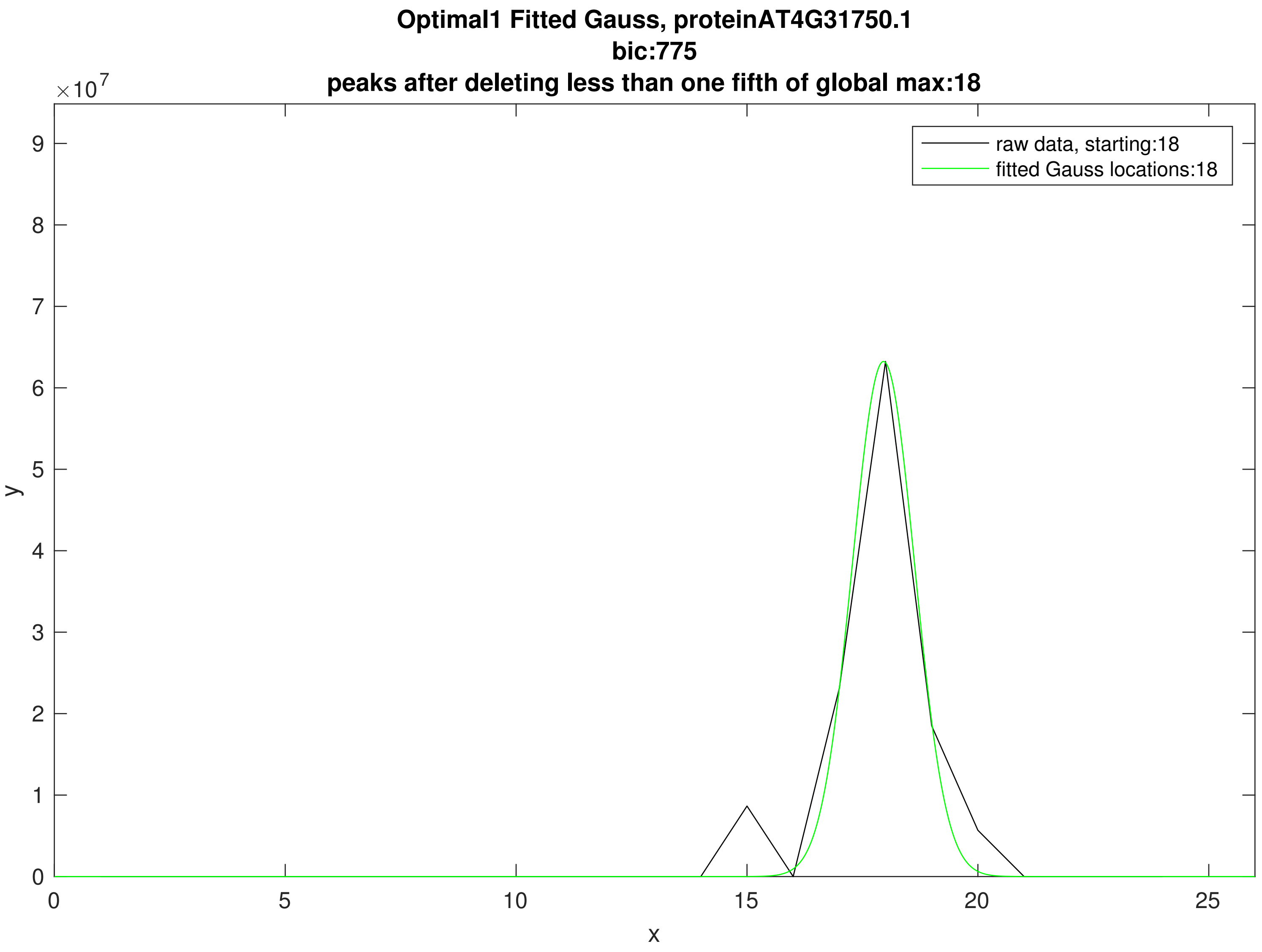
Optimal1 Fitted Gauss, proteinAT4G31750.1
bic:775
peaks after deleting less than one fifth of global max:18
×10^7
raw data, starting:18
fitted Gauss locations:18
y
x
0
1
2
3
4
5
6
7
8
9
0
5
10
15
20
25

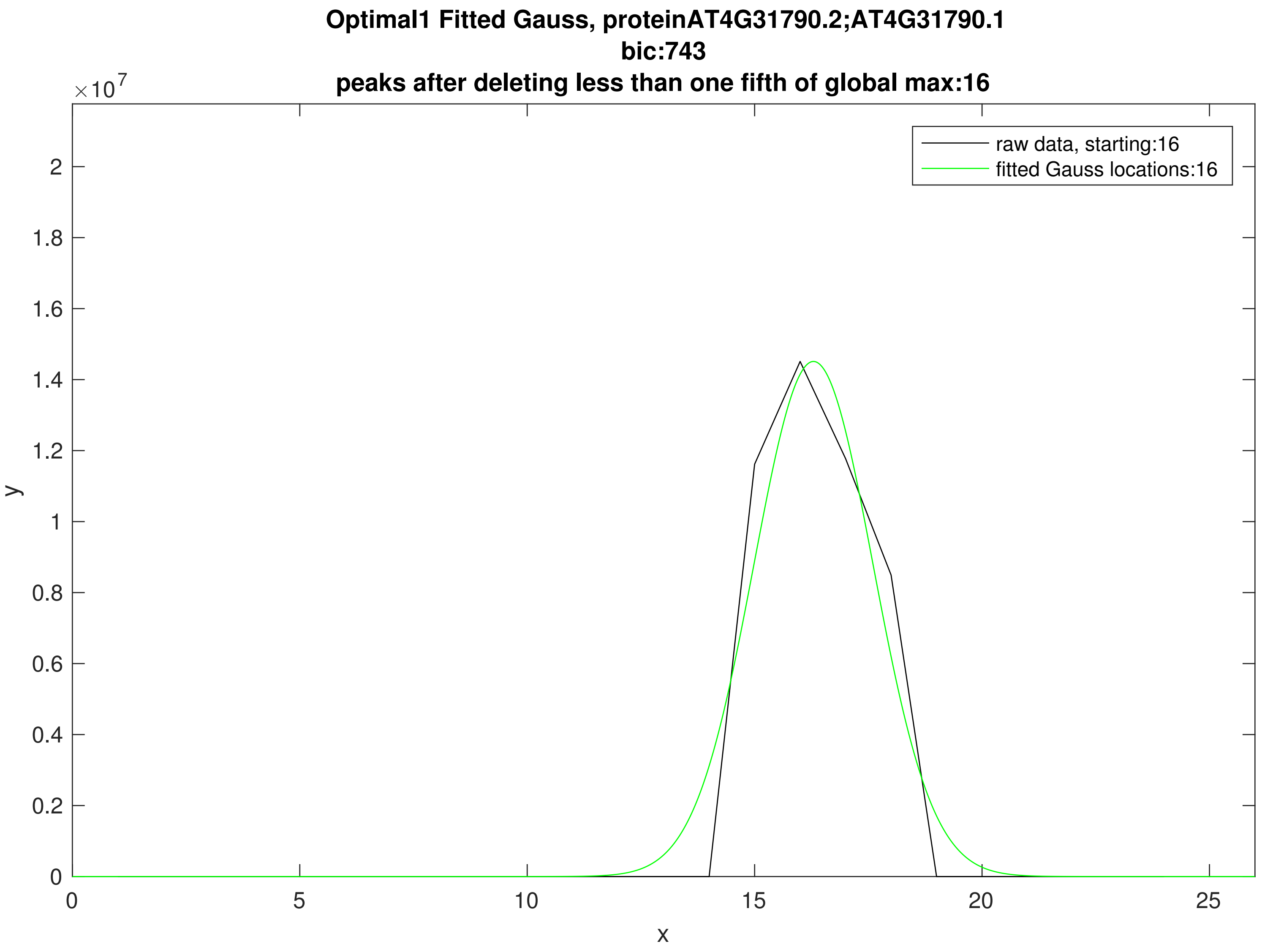
Optimal1 Fitted Gauss, proteinAT4G31790.2;AT4G31790.1
bic:743
peaks after deleting less than one fifth of global max:16
×10^7
raw data, starting:16
fitted Gauss locations:16
2
1.8
1.6
1.4
1.2
1
0.8
0.6
0.4
0.2
0
y
0
5
10
15
20
25
x

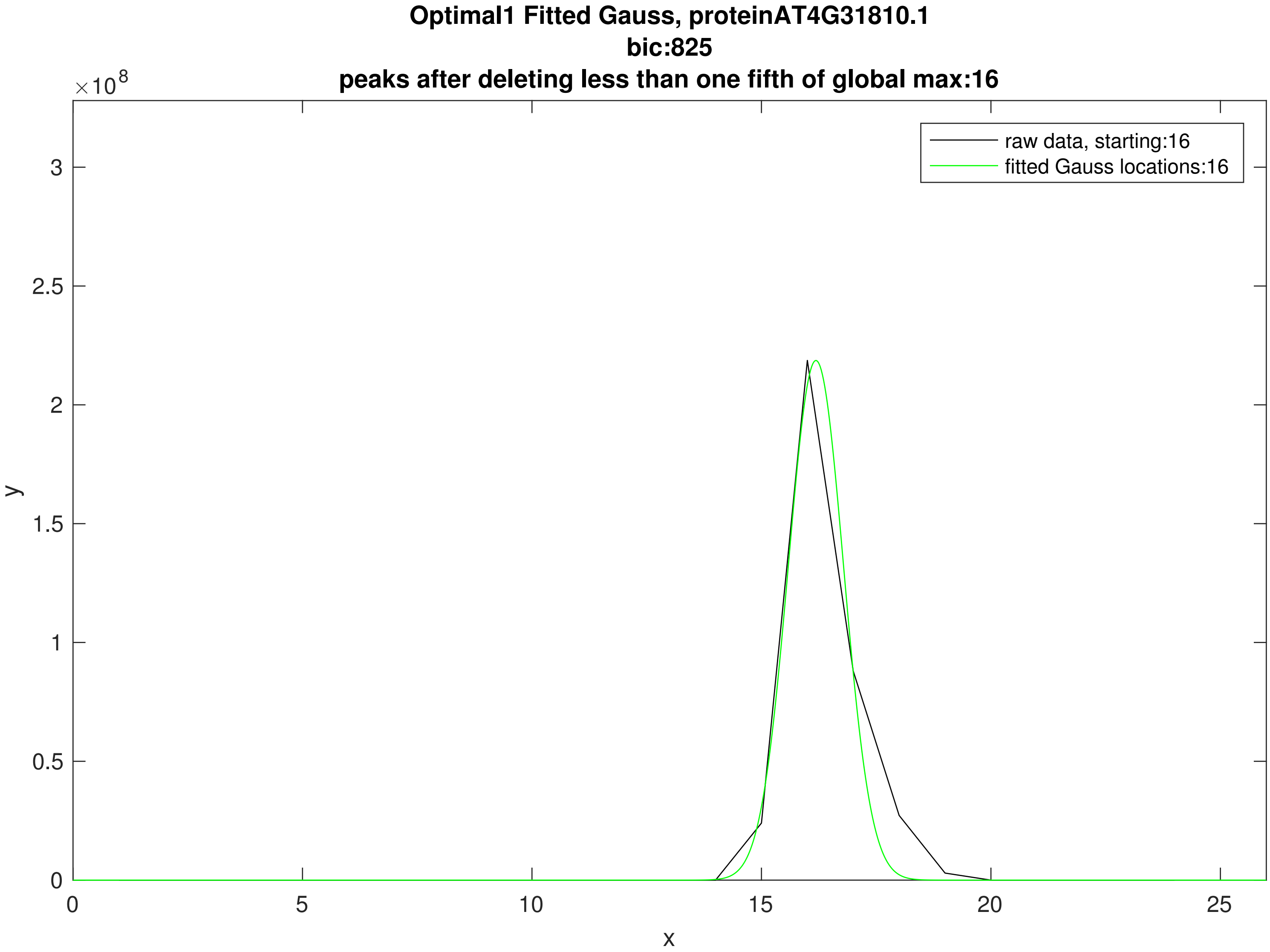
Optimal1 Fitted Gauss, proteinAT4G31810.1
bic:825
peaks after deleting less than one fifth of global max:16
×10^8
raw data, starting:16
fitted Gauss locations:16
3
2.5
2
1.5
1
0.5
0
y
0
5
10
15
20
25
x

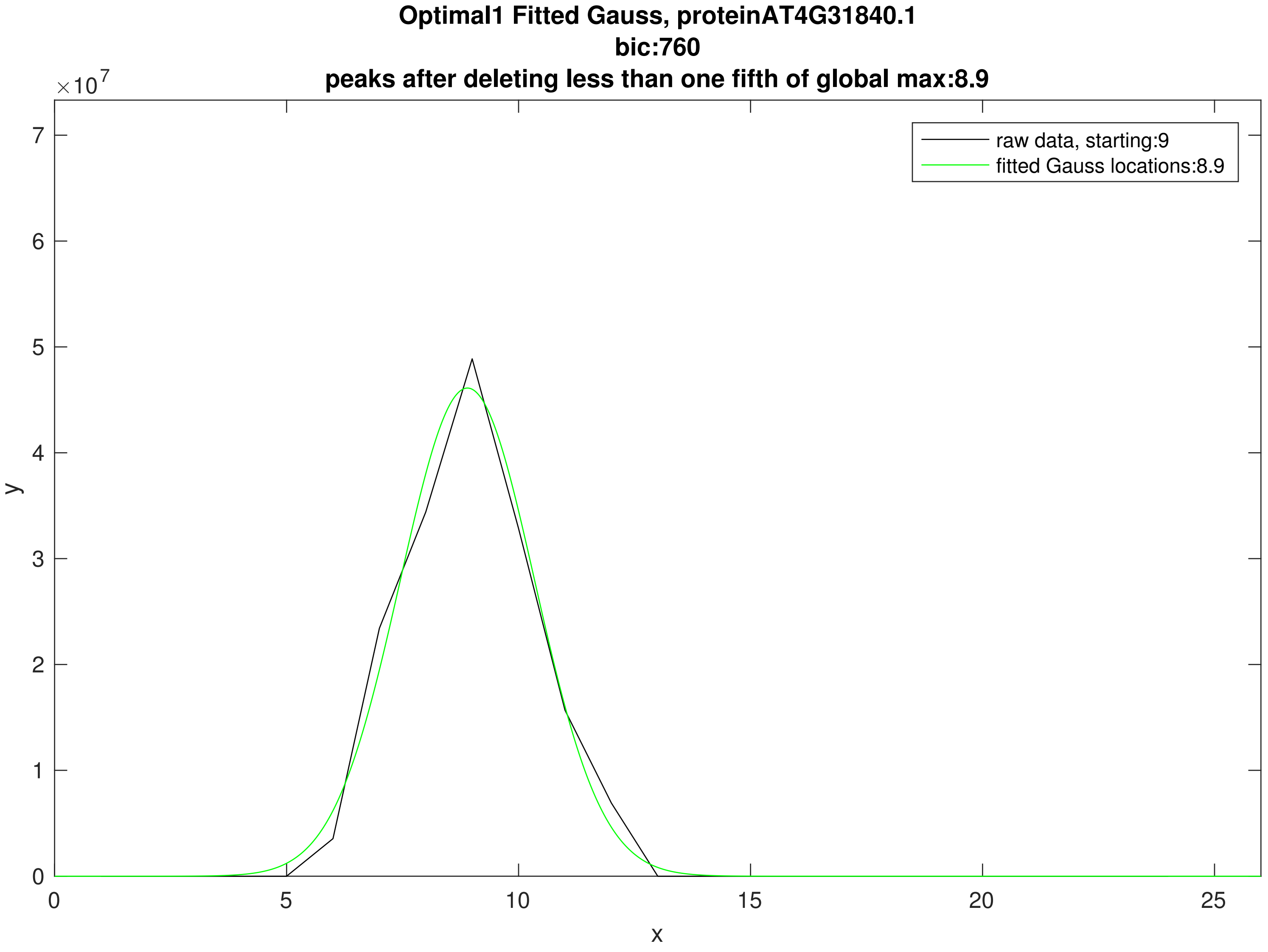

Optimal1 Fitted Gauss, proteinAT4G31840.1
bic:760
peaks after deleting less than one fifth of global max:8.9
×10^7
raw data, starting:9
fitted Gauss locations:8.9
y
x
0
1
2
3
4
5
6
7
0
5
10
15
20
25

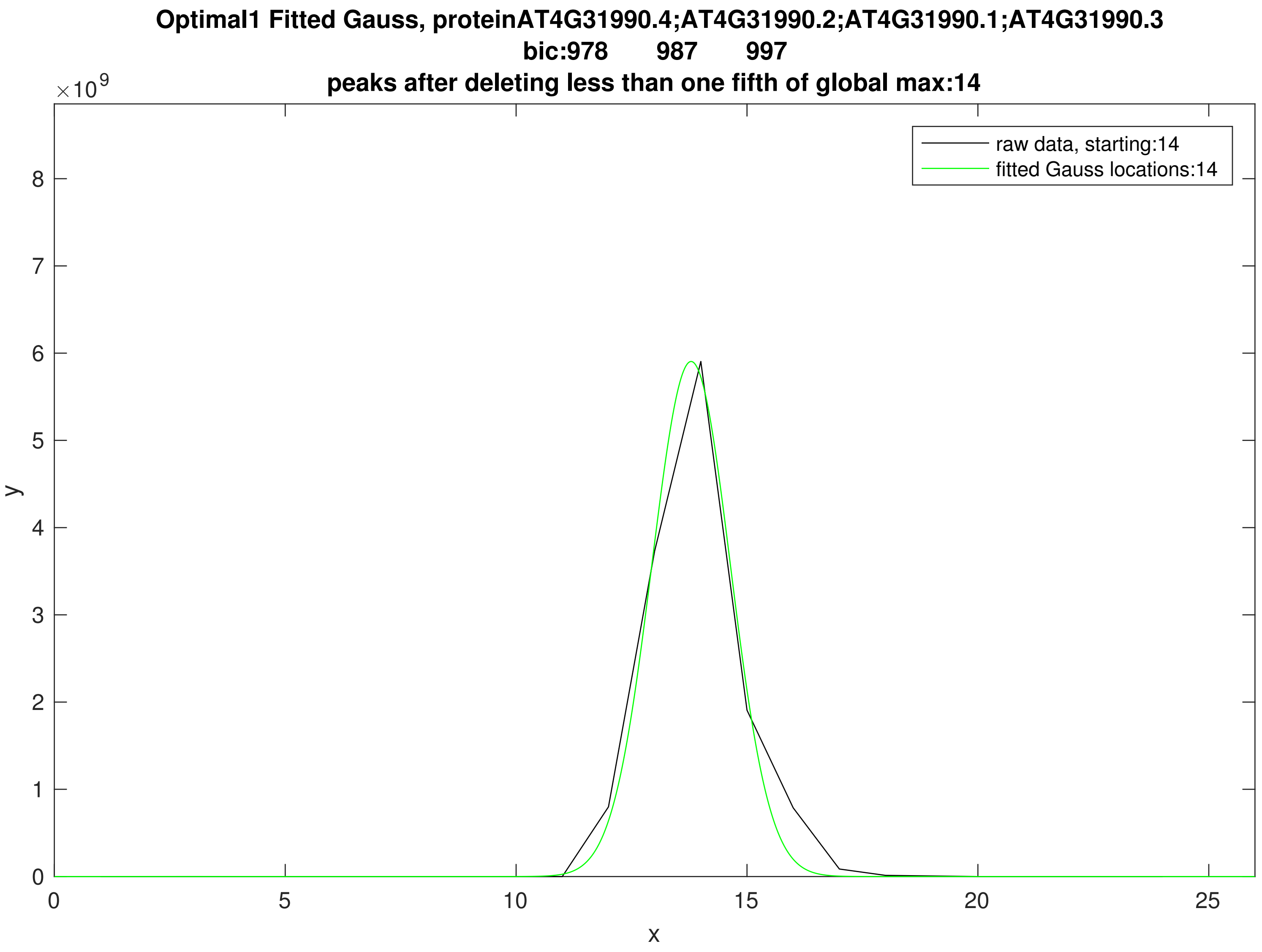

Optimal1 Fitted Gauss, proteinAT4G31990.4;AT4G31990.2;AT4G31990.1;AT4G31990.3
bic:978 987 997
peaks after deleting less than one fifth of global max:14
×10^9
raw data, starting:14
fitted Gauss locations:14
y
x
0
1
2
3
4
5
6
7
8
0
5
10
15
20
25

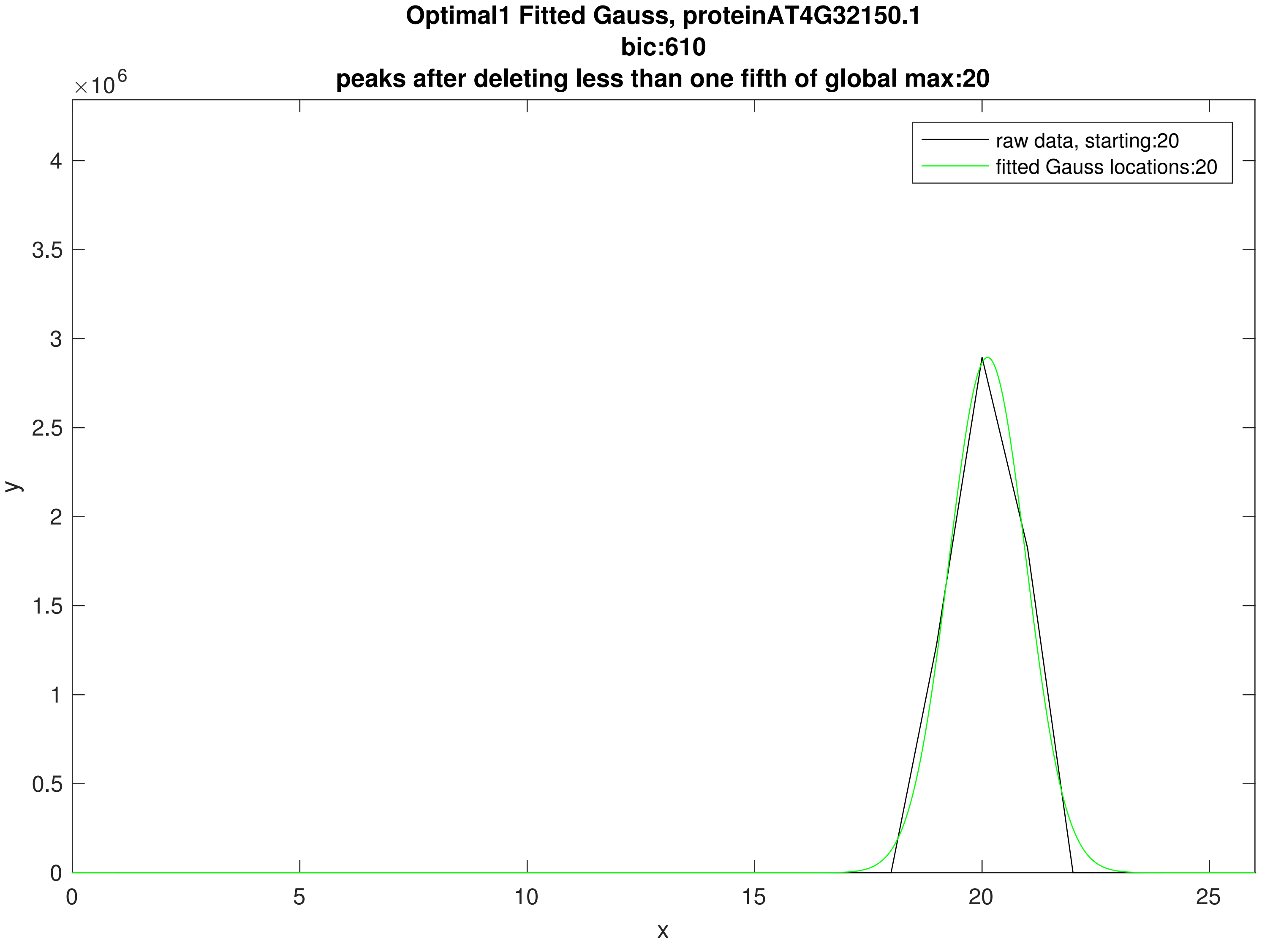
Optimal1 Fitted Gauss, proteinAT4G32150.1
bic:610
peaks after deleting less than one fifth of global max:20
×10^6
4
3.5
3
2.5
2
1.5
1
0.5
0
y
0
5
10
15
20
25
x
raw data, starting:20
fitted Gauss locations:20

Optimal1 Fitted Gauss, proteinAT4G32272.1;AT4G32272.2
bic:953  1.02e+03  1.02e+03
peaks after deleting less than one fifth of global max:13

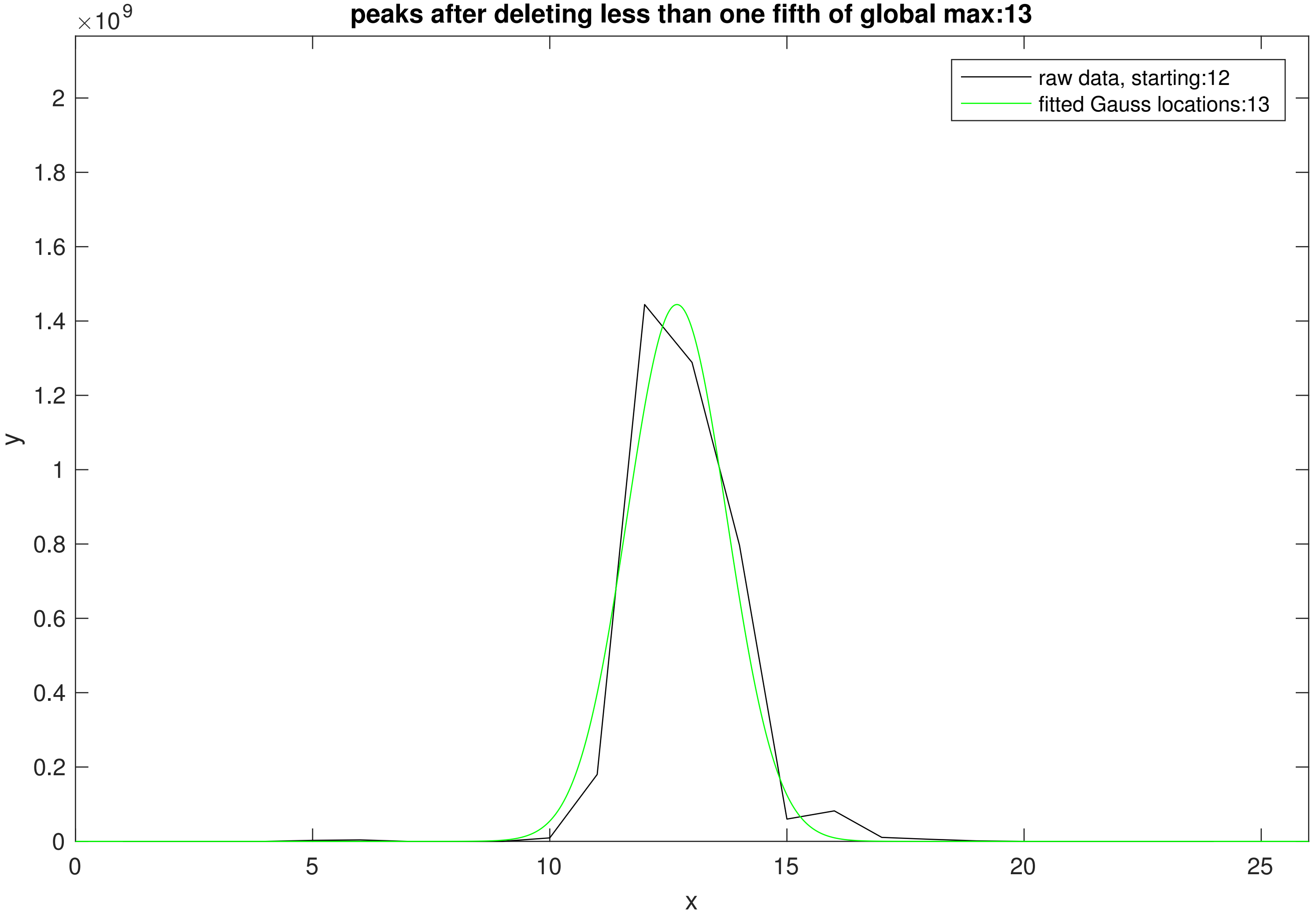

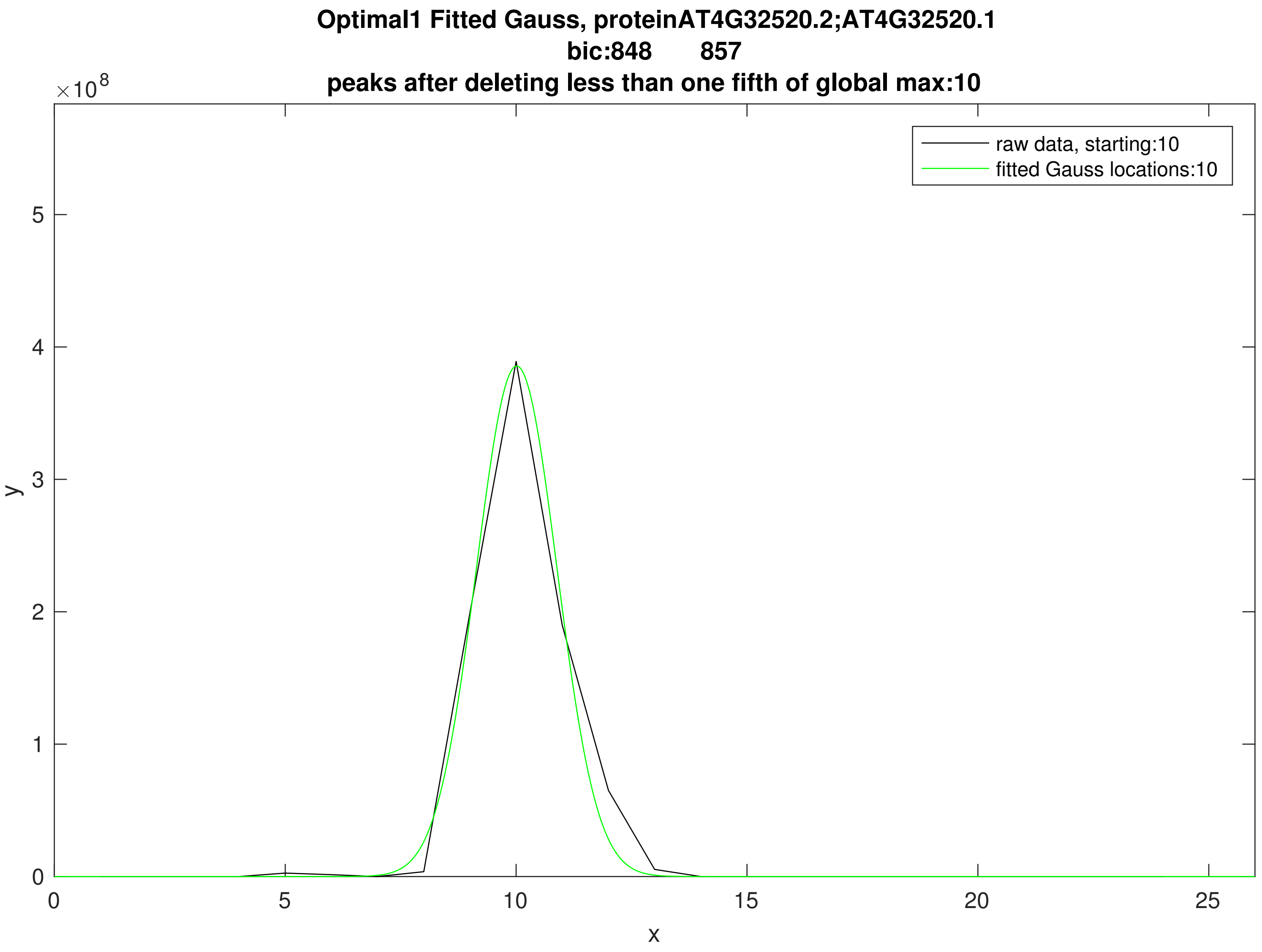
Optimal1 Fitted Gauss, proteinAT4G32520.2;AT4G32520.1
bic:848 857
peaks after deleting less than one fifth of global max:10
×10^8
raw data, starting:10
fitted Gauss locations:10
y
x
0
1
2
3
4
5
0
5
10
15
20
25

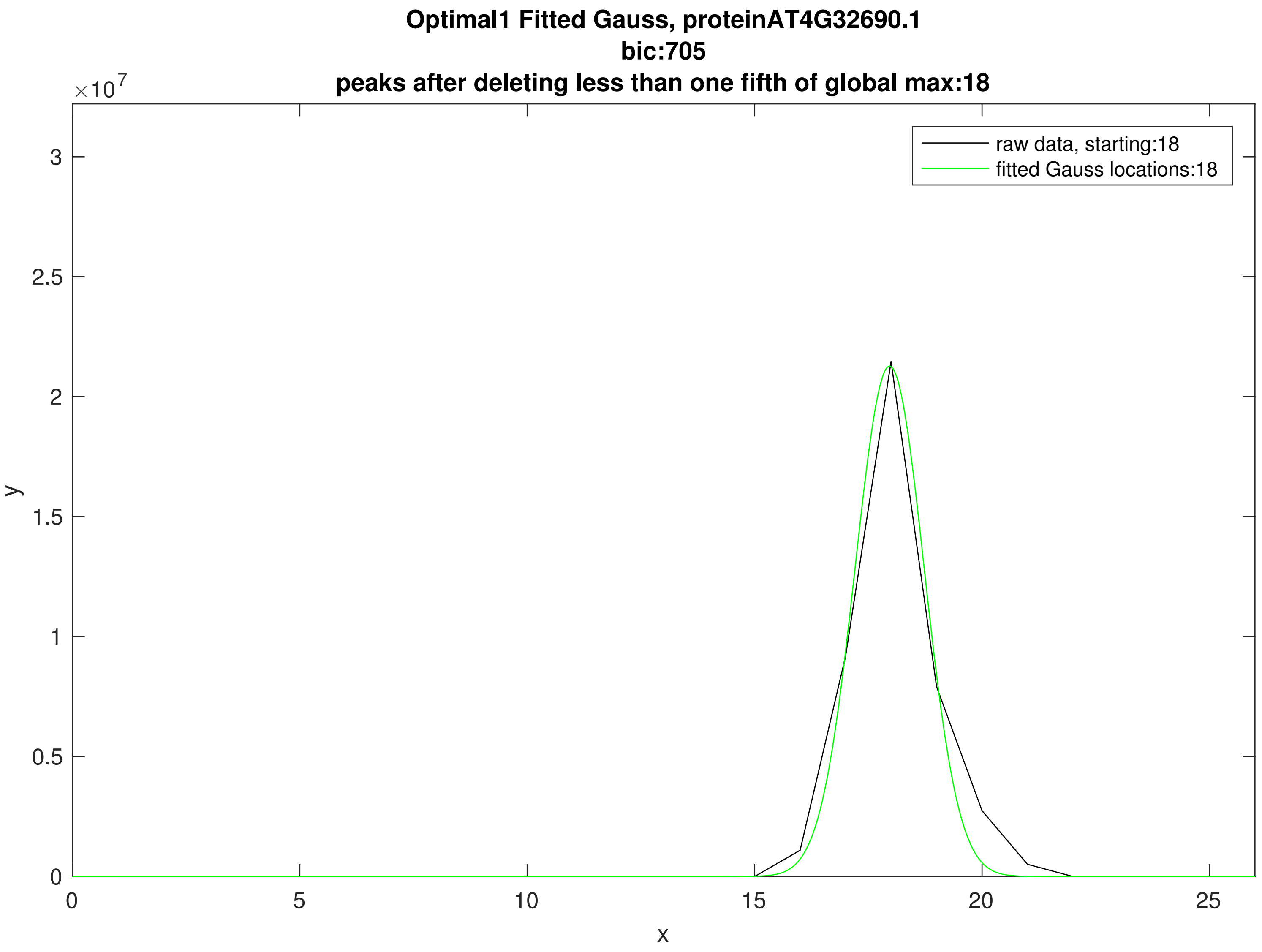
Optimal1 Fitted Gauss, proteinAT4G32690.1
bic:705
peaks after deleting less than one fifth of global max:18
×10^7
raw data, starting:18
fitted Gauss locations:18
3
2.5
2
1.5
1
0.5
0
y
0
5
10
15
20
25
x

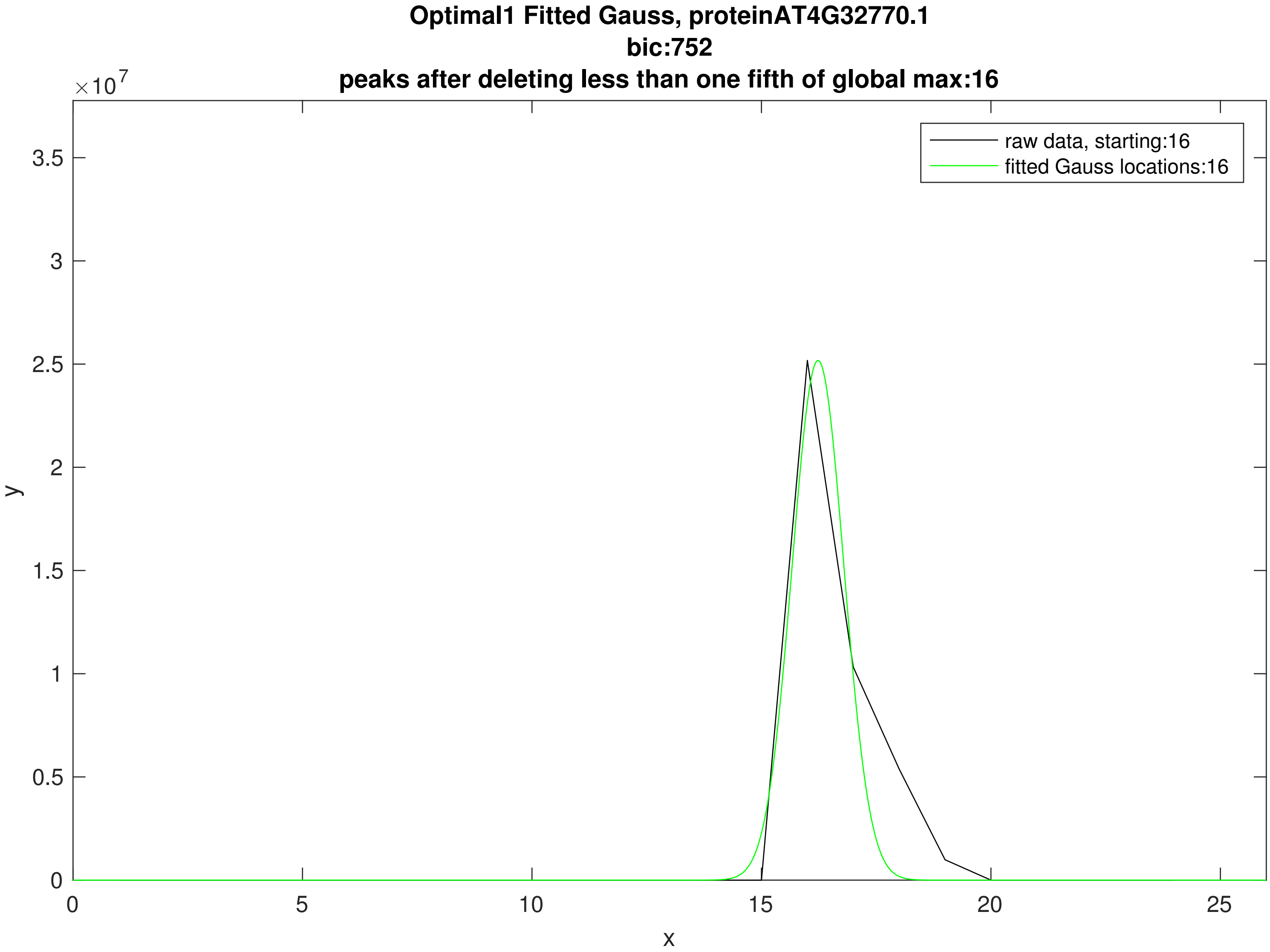

Optimal1 Fitted Gauss, proteinAT4G32770.1
bic:752
peaks after deleting less than one fifth of global max:16
×10^7
3.5
3
2.5
2
1.5
1
0.5
0
y
0
5
10
15
20
25
x
raw data, starting:16
fitted Gauss locations:16

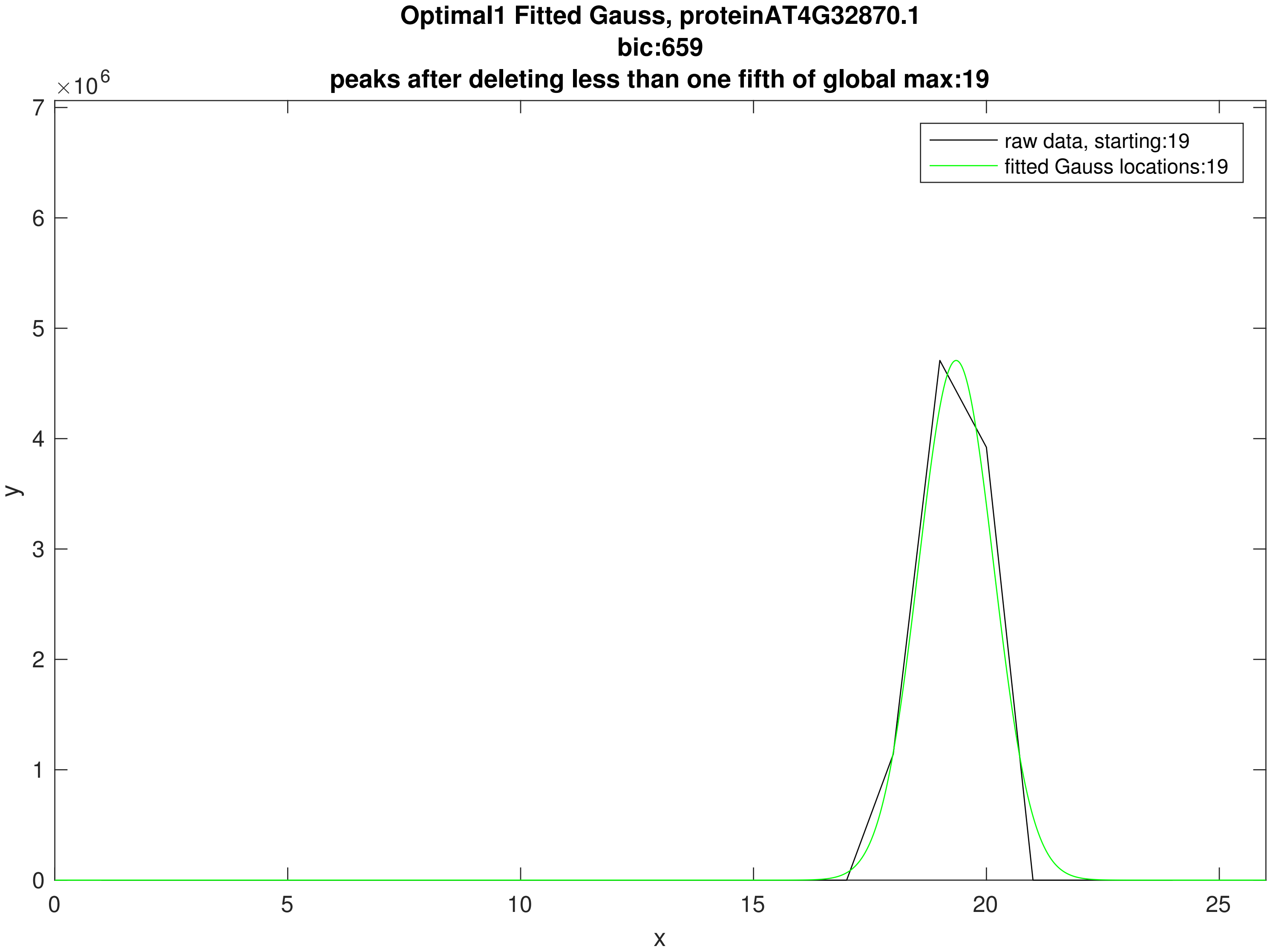
Optimal1 Fitted Gauss, proteinAT4G32870.1
bic:659
peaks after deleting less than one fifth of global max:19
×10^6
raw data, starting:19
fitted Gauss locations:19
y
x
0
1
2
3
4
5
6
7
0
5
10
15
20
25

**Optimal1 Fitted Gauss, proteinAT4G32915.1**
**bic:735**
**peaks after deleting less than one fifth of global max:18**

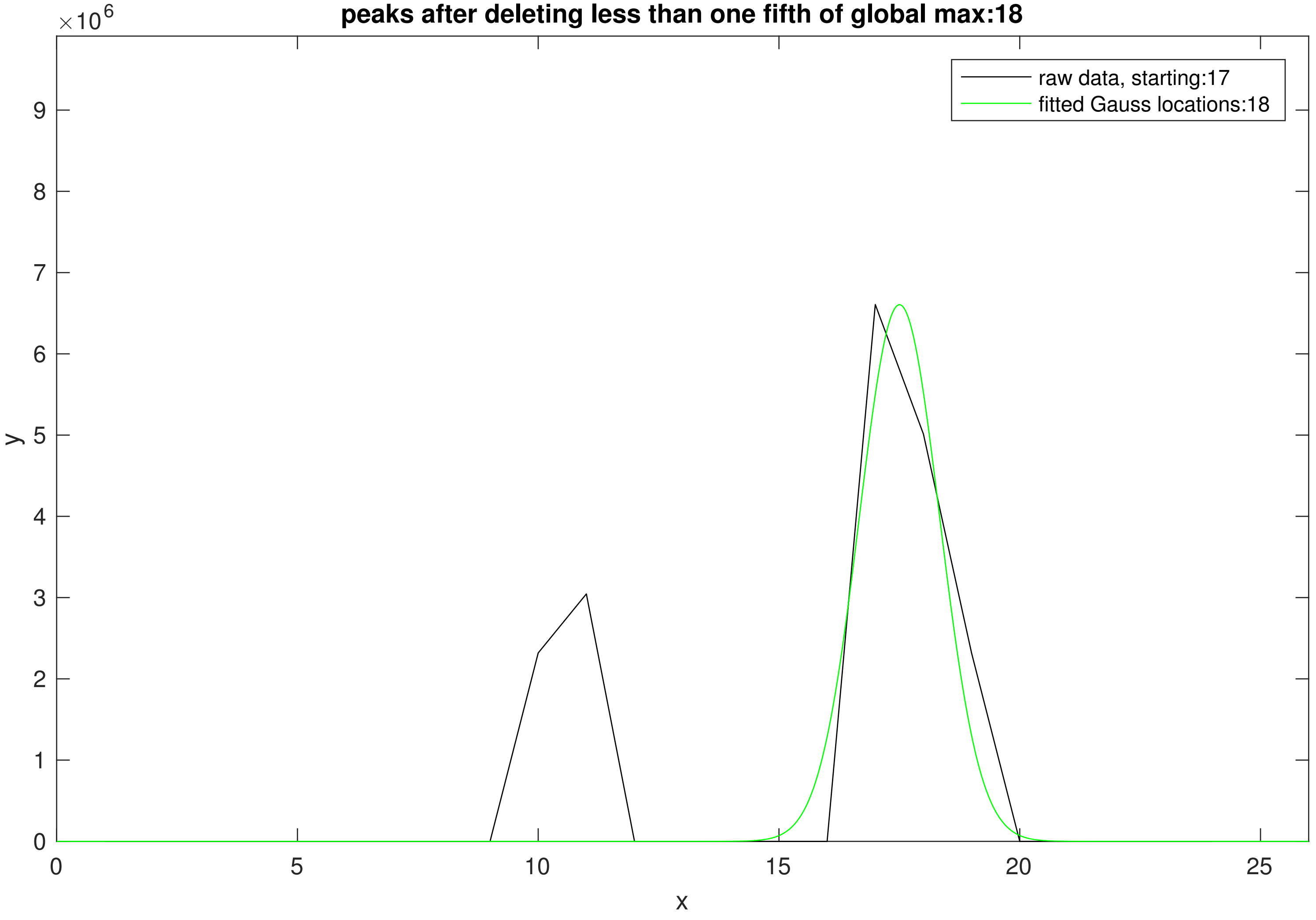

**Optimal1 Fitted Gauss, proteinAT4G32930.1;AT4G32930.2**
**bic:868 950**
**peaks after deleting less than one fifth of global max:19**

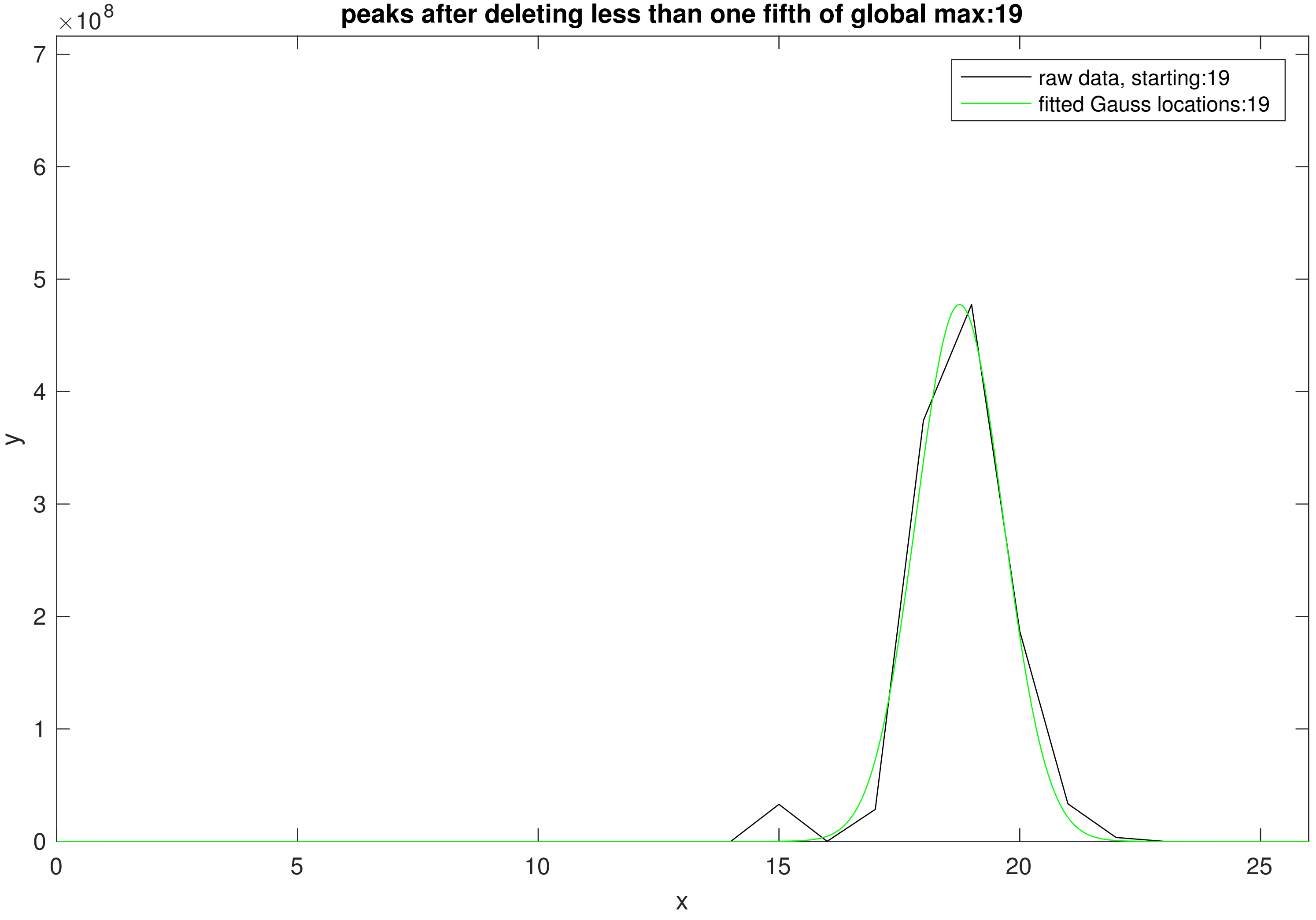

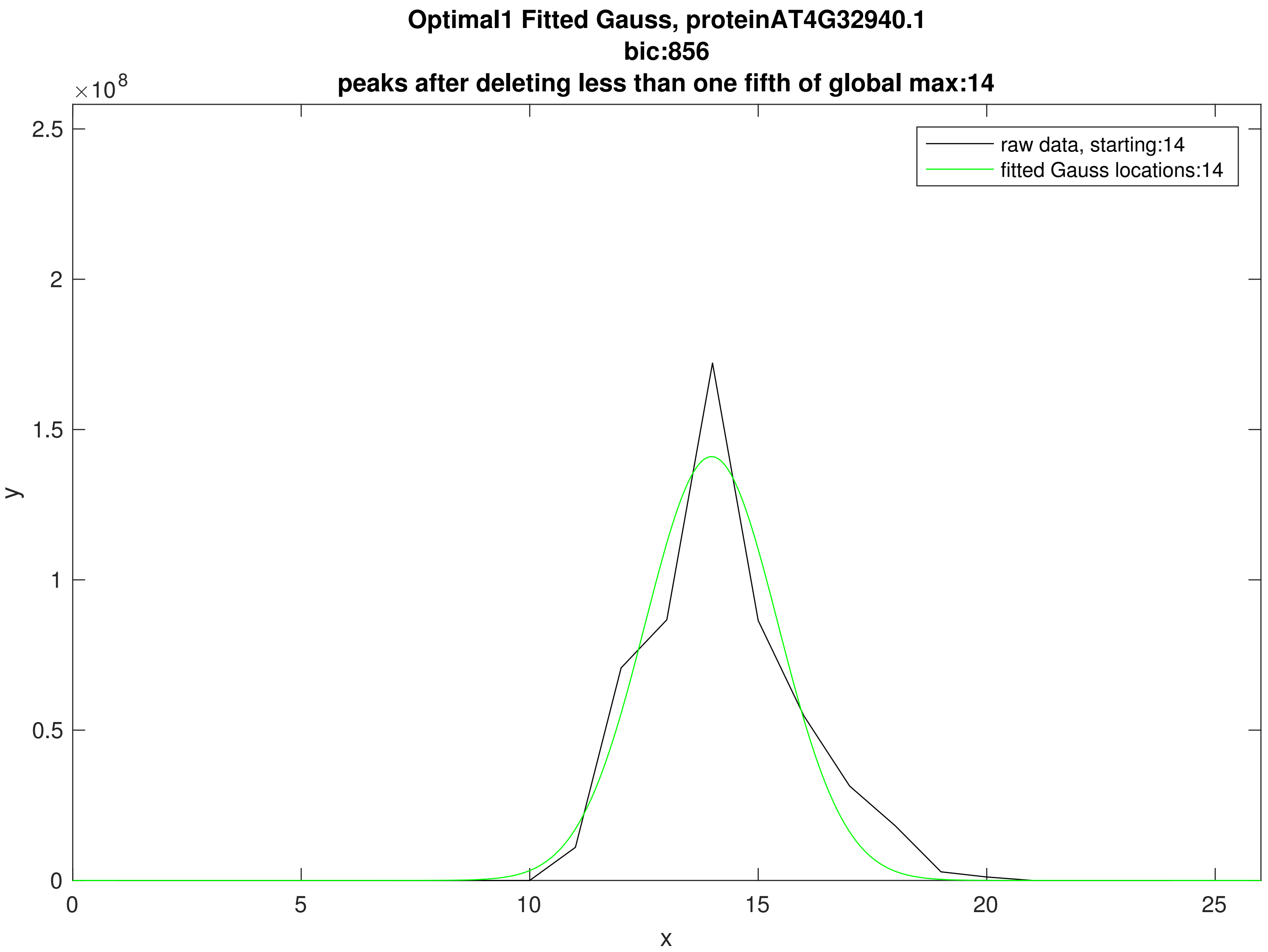
Optimal1 Fitted Gauss, proteinAT4G32940.1
bic:856
peaks after deleting less than one fifth of global max:14
×10^8
2.5
2
1.5
1
0.5
0
y
0
5
10
15
20
25
x
raw data, starting:14
fitted Gauss locations:14

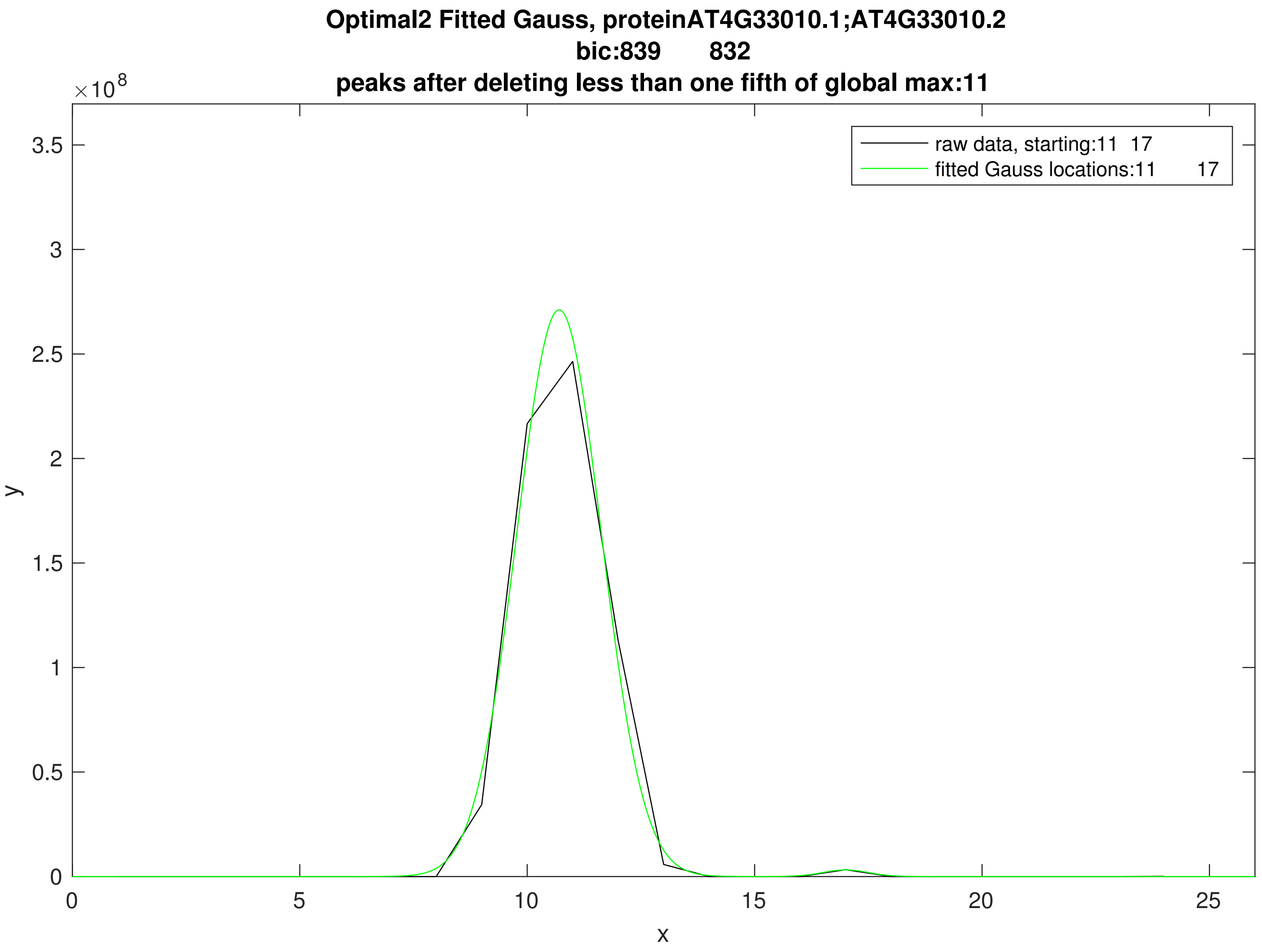
Optimal2 Fitted Gauss, proteinAT4G33010.1;AT4G33010.2
bic:839 832
peaks after deleting less than one fifth of global max:11
×10^8
raw data, starting:11 17
fitted Gauss locations:11 17
y
x
0
0.5
1
1.5
2
2.5
3
3.5
0
5
10
15
20
25

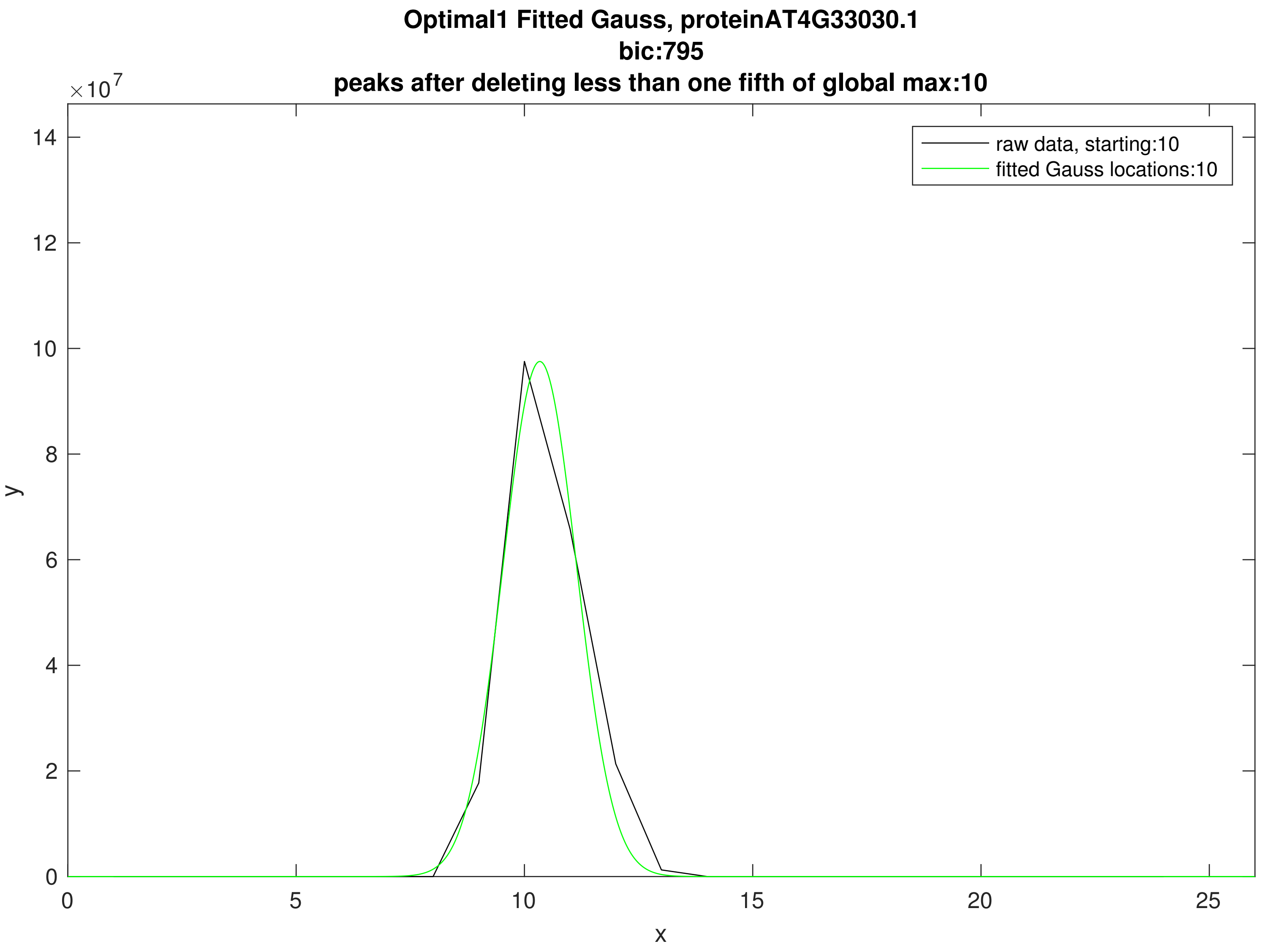
Optimal1 Fitted Gauss, proteinAT4G33030.1
bic:795
peaks after deleting less than one fifth of global max:10
×10^7
14
12
10
8
6
4
2
0
y
0
5
10
15
20
25
x
raw data, starting:10
fitted Gauss locations:10

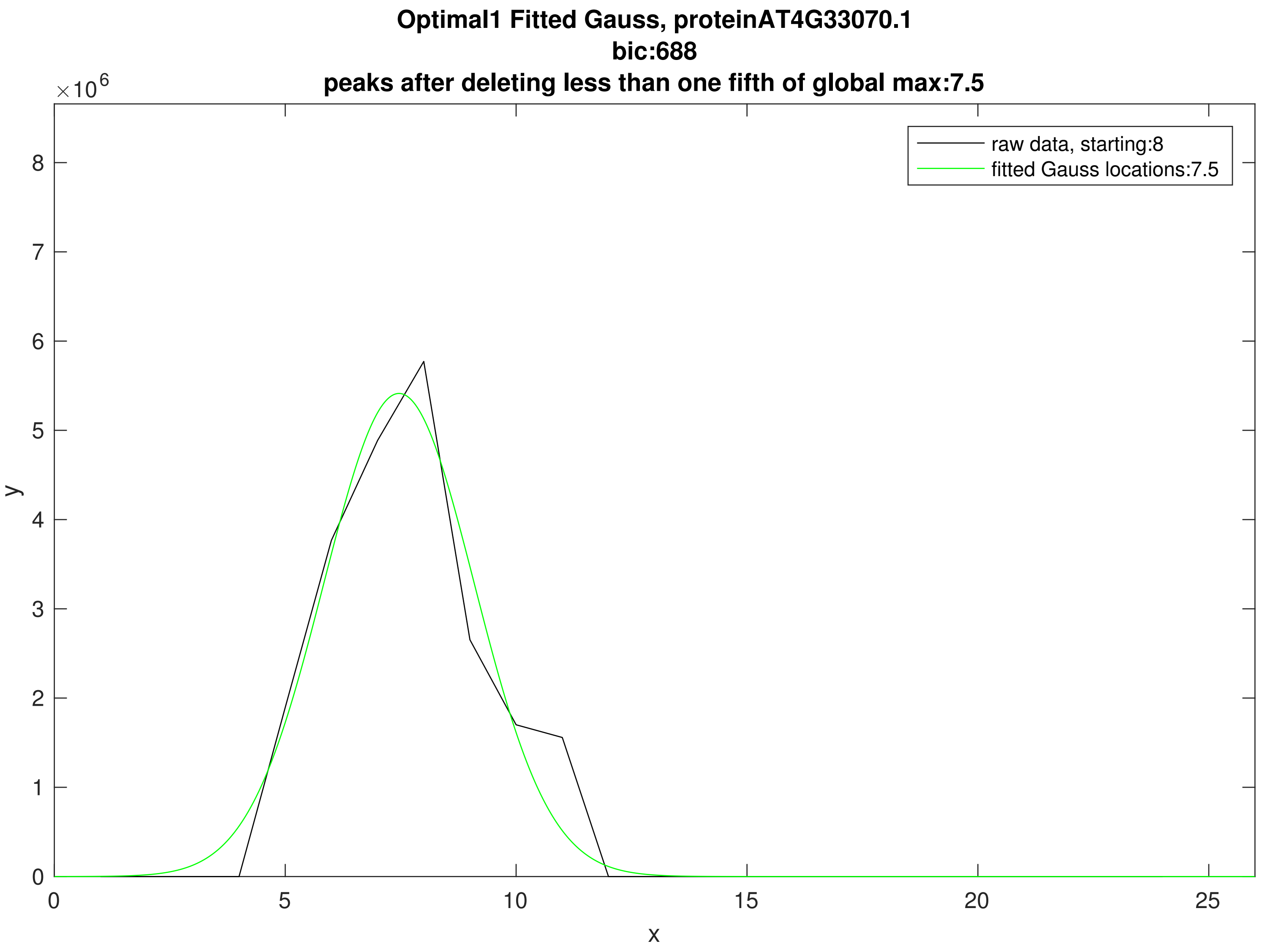

Optimal1 Fitted Gauss, proteinAT4G33070.1
bic:688
peaks after deleting less than one fifth of global max:7.5
×10^6
raw data, starting:8
fitted Gauss locations:7.5
y
x
0
1
2
3
4
5
6
7
8
0
5
10
15
20
25

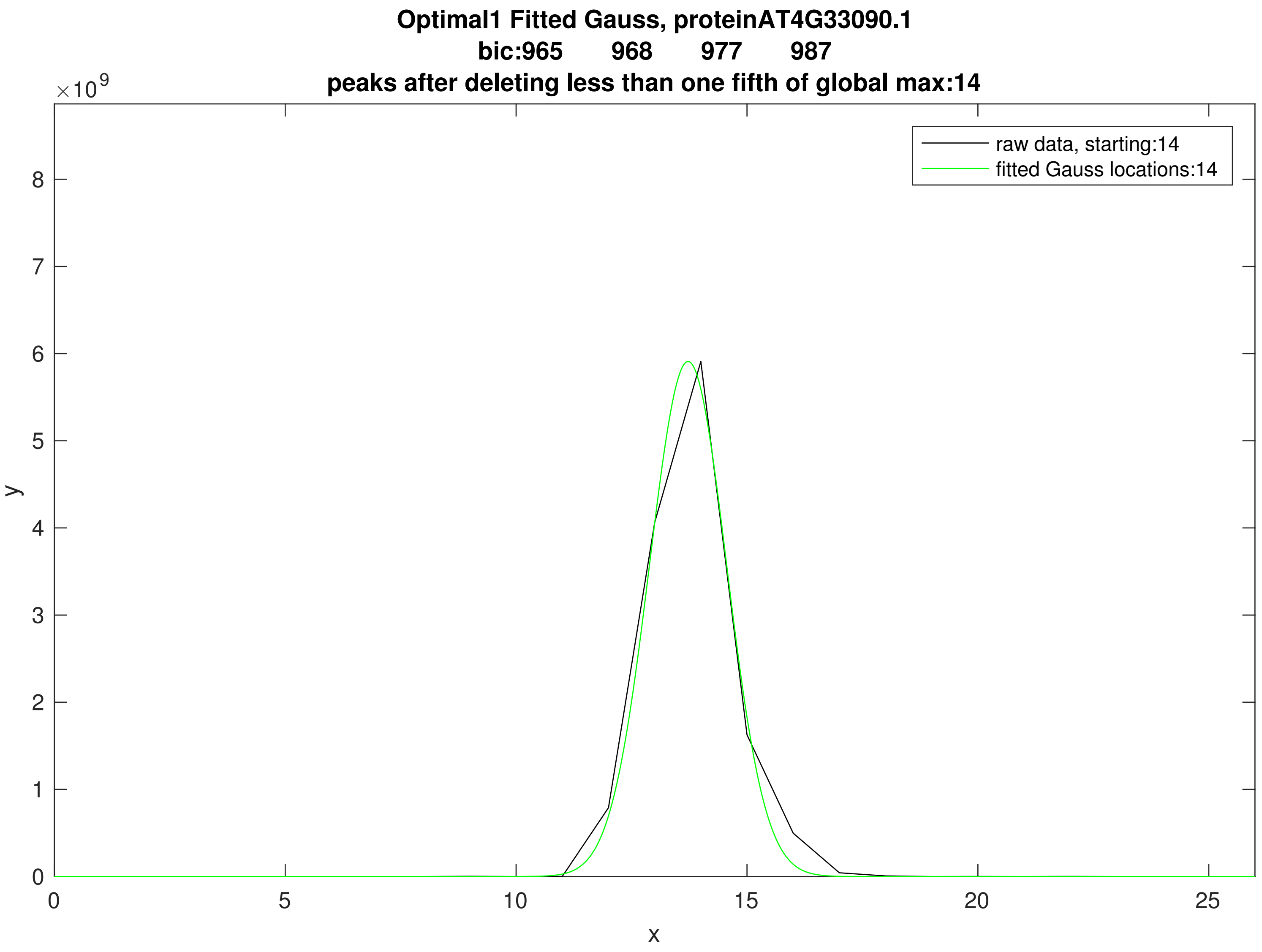
Optimal1 Fitted Gauss, proteinAT4G33090.1
bic:965 968 977 987
peaks after deleting less than one fifth of global max:14
×10^9
raw data, starting:14
fitted Gauss locations:14
y
x
0
1
2
3
4
5
6
7
8
0
5
10
15
20
25

Optimal2 Fitted Gauss, proteinAT4G33150.1;AT4G33150.2;AT4G33150.3
bic:845 749
peaks after deleting less than one fifth of global max:16

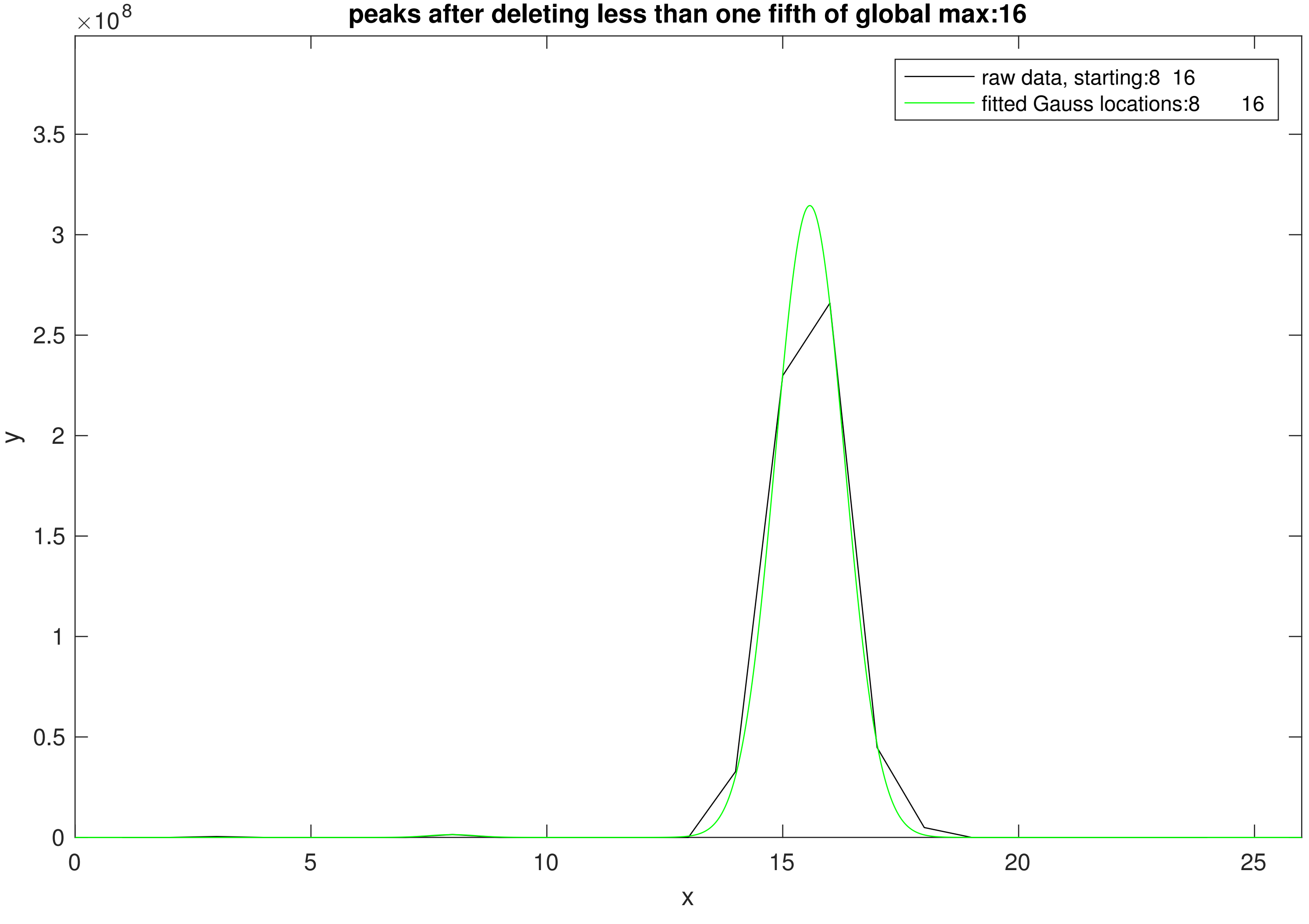

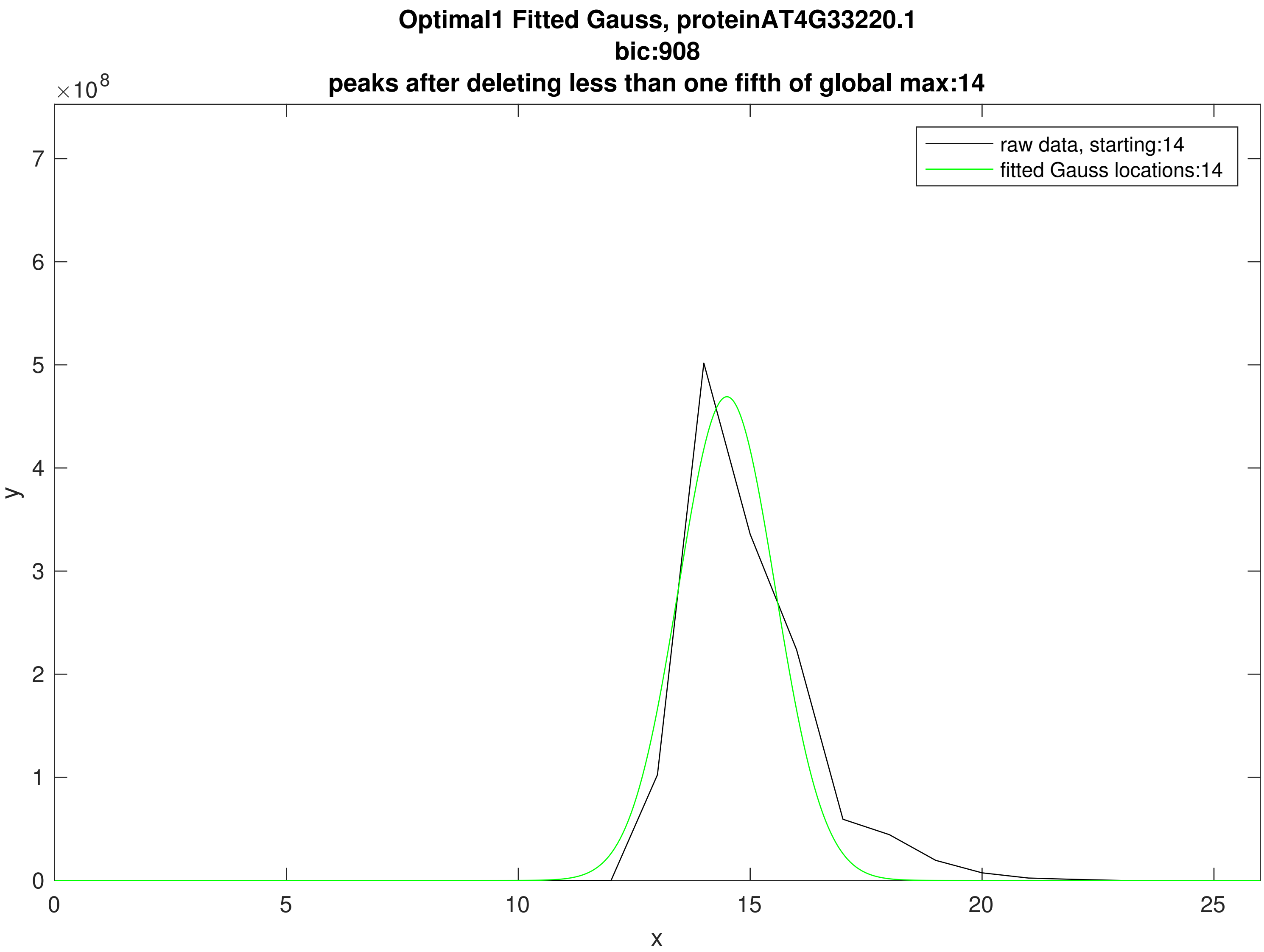
Optimal1 Fitted Gauss, proteinAT4G33220.1
bic:908
peaks after deleting less than one fifth of global max:14
×10^8
raw data, starting:14
fitted Gauss locations:14
y
x
0
1
2
3
4
5
6
7
0
5
10
15
20
25

**Optimal1 Fitted Gauss, proteinAT4G33250.1**
**bic:757**
**peaks after deleting less than one fifth of global max:17**

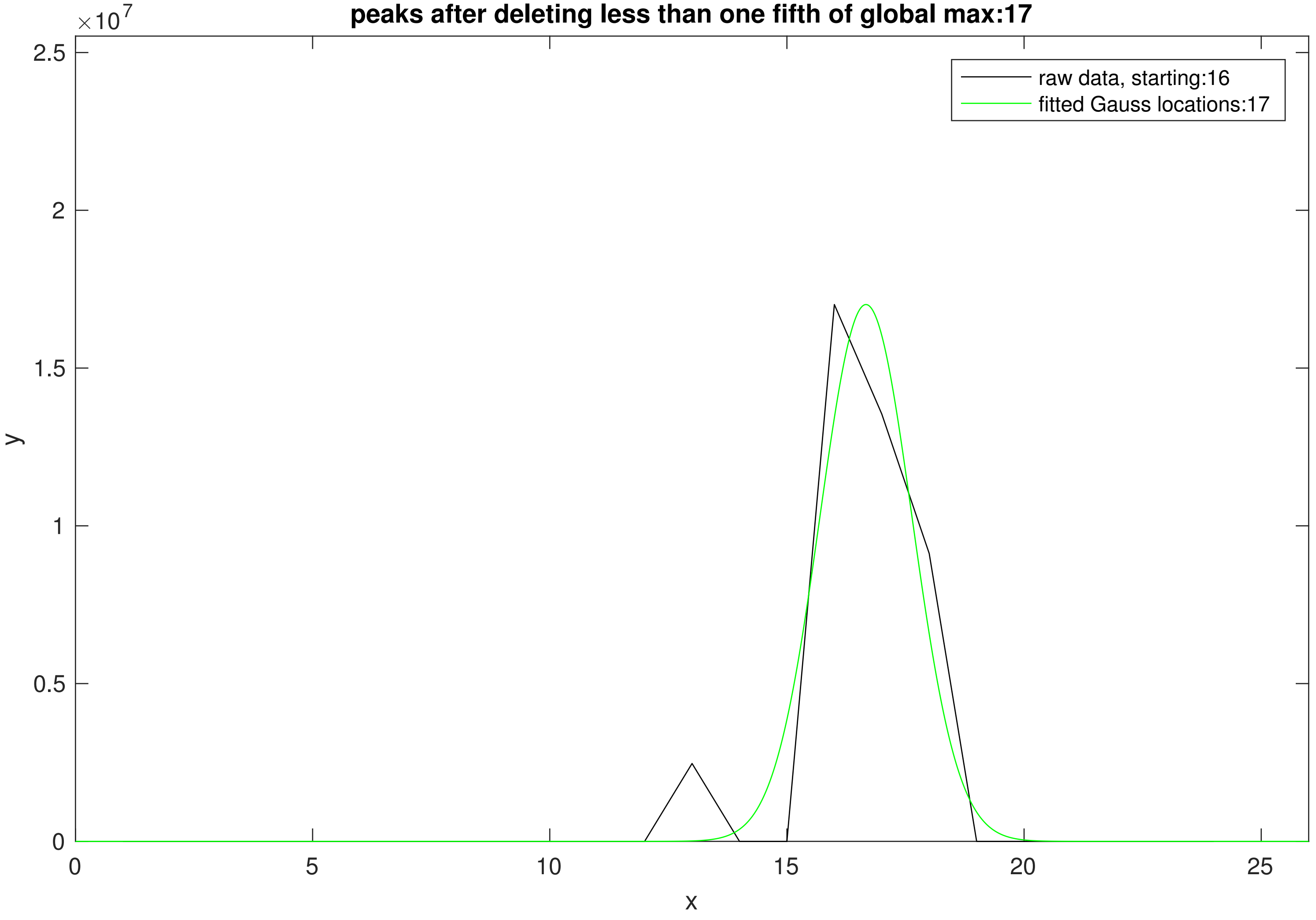

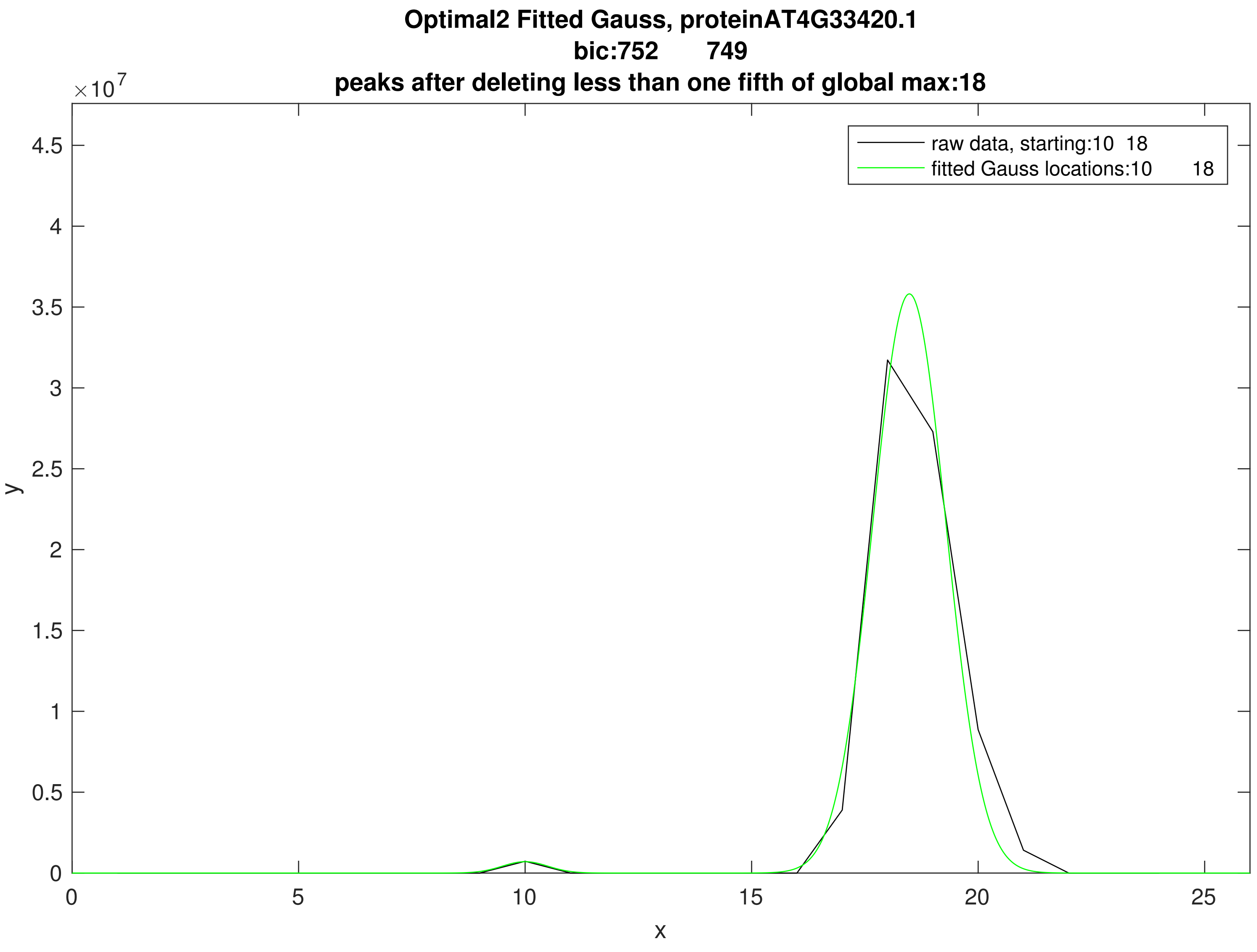
Optimal2 Fitted Gauss, proteinAT4G33420.1
bic:752 749
peaks after deleting less than one fifth of global max:18
×10^7
4.5
4
3.5
3
2.5
2
1.5
1
0.5
0
y
0
5
10
15
20
25
x
raw data, starting:10 18
fitted Gauss locations:10 18

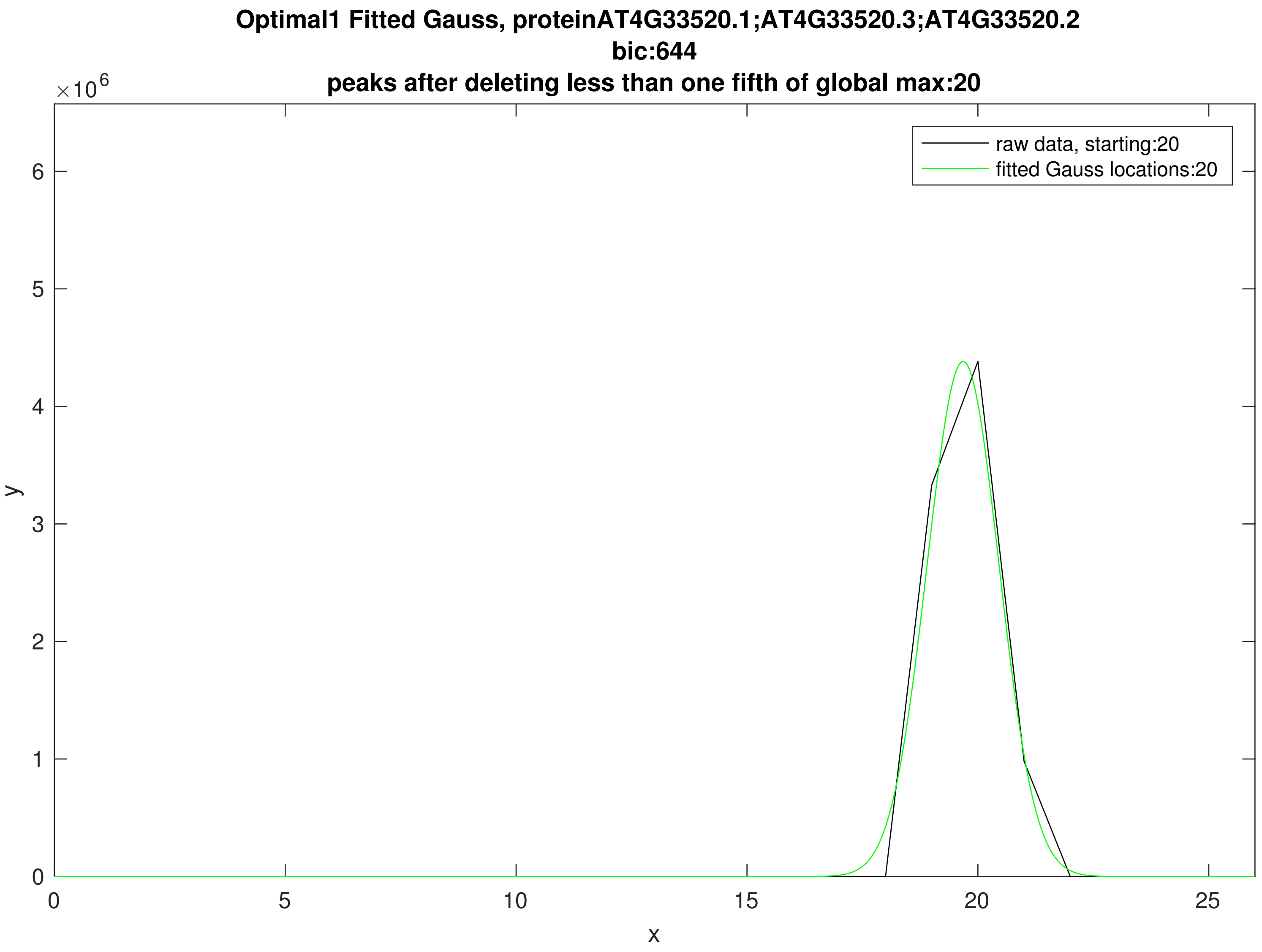
Optimal1 Fitted Gauss, proteinAT4G33520.1;AT4G33520.3;AT4G33520.2
bic:644
peaks after deleting less than one fifth of global max:20
×10^6
raw data, starting:20
fitted Gauss locations:20
y
x
0
1
2
3
4
5
6
0
5
10
15
20
25

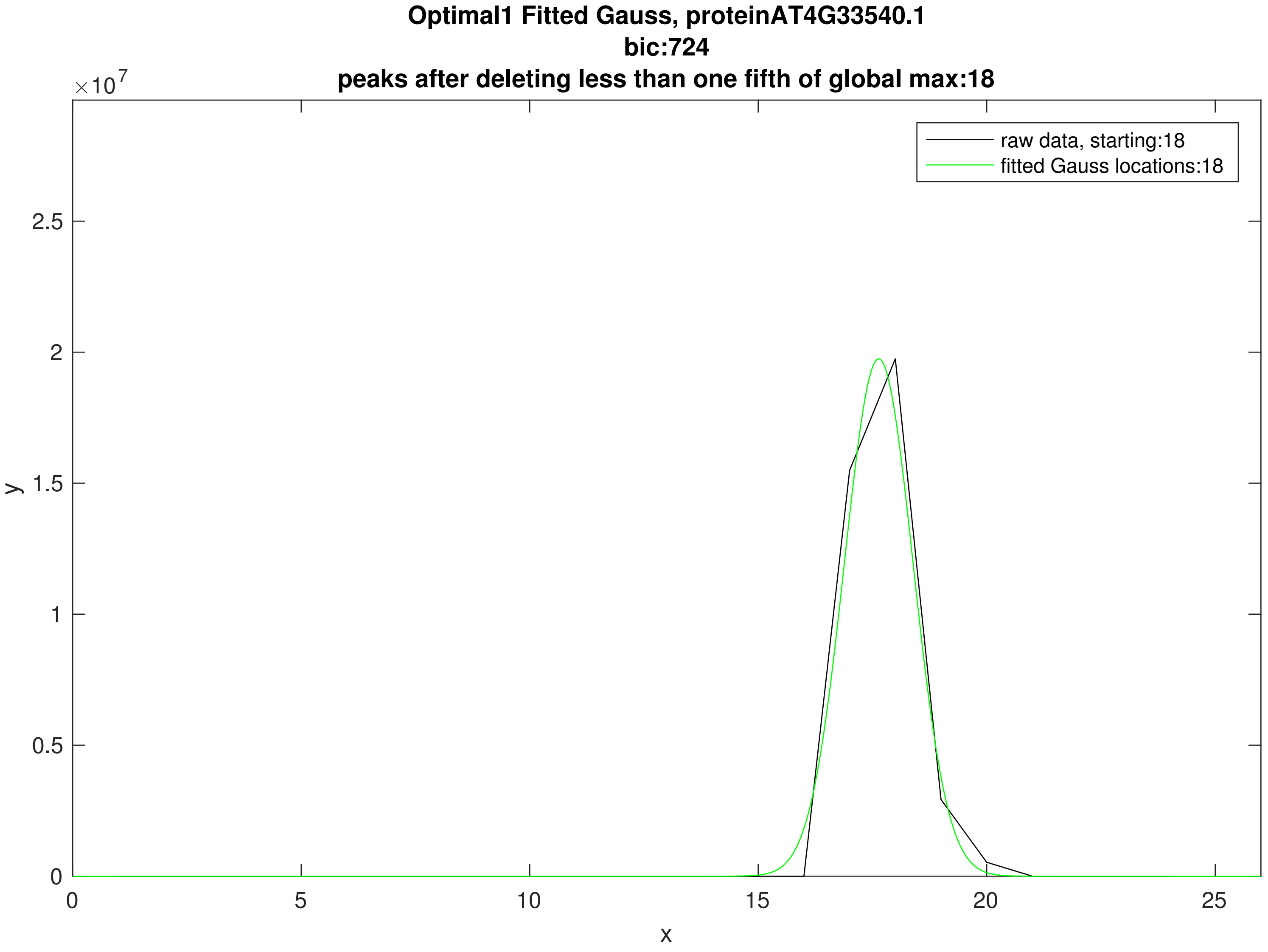

Optimal1 Fitted Gauss, proteinAT4G33540.1
bic:724
peaks after deleting less than one fifth of global max:18
×10^7
raw data, starting:18
fitted Gauss locations:18
2.5
2
1.5
1
0.5
0
y
0
5
10
15
20
25
x

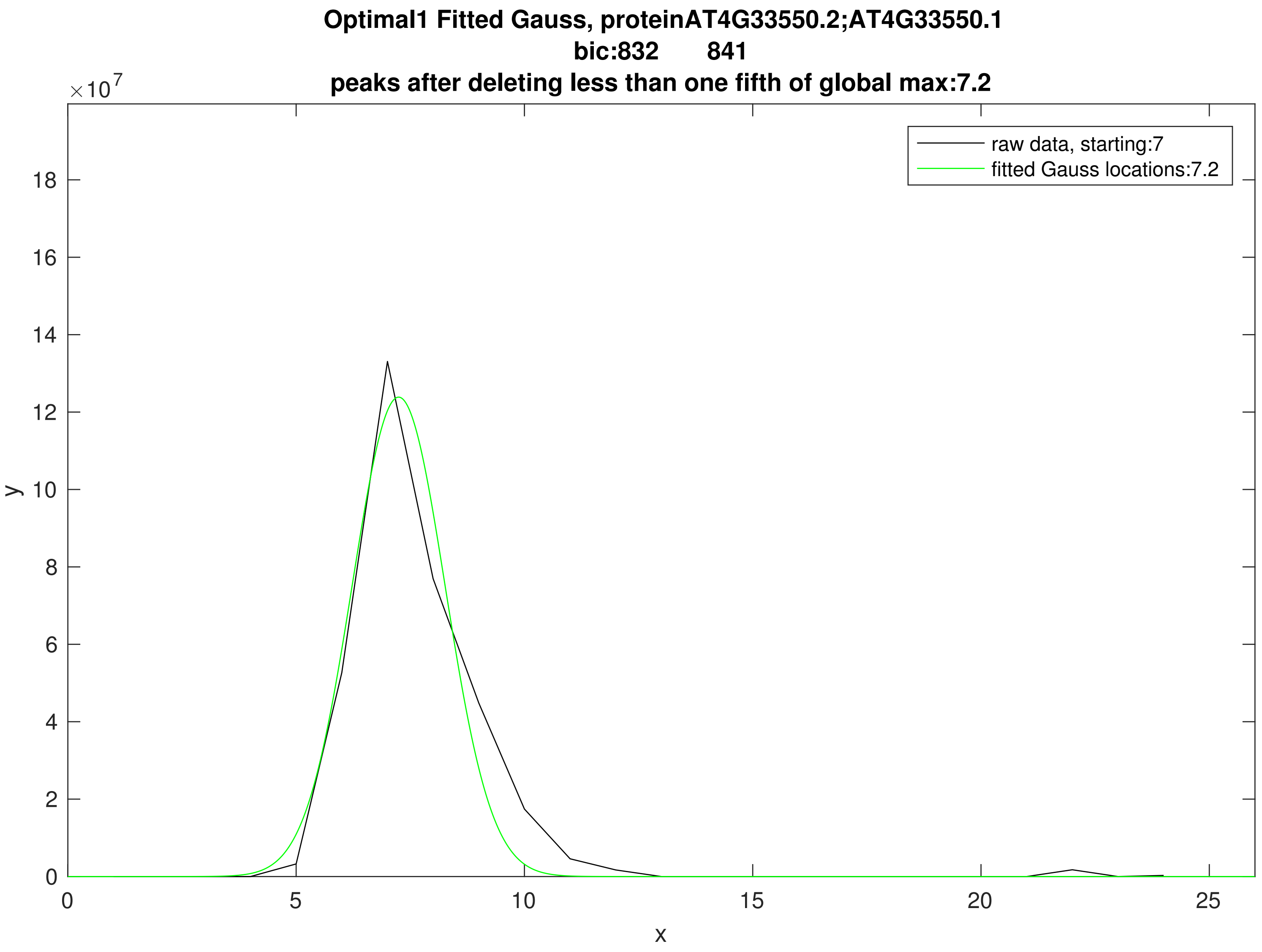
Optimal1 Fitted Gauss, proteinAT4G33550.2;AT4G33550.1
bic:832 841
peaks after deleting less than one fifth of global max:7.2
×10^7
raw data, starting:7
fitted Gauss locations:7.2
y
x
0
2
4
6
8
10
12
14
16
18
0
5
10
15
20
25

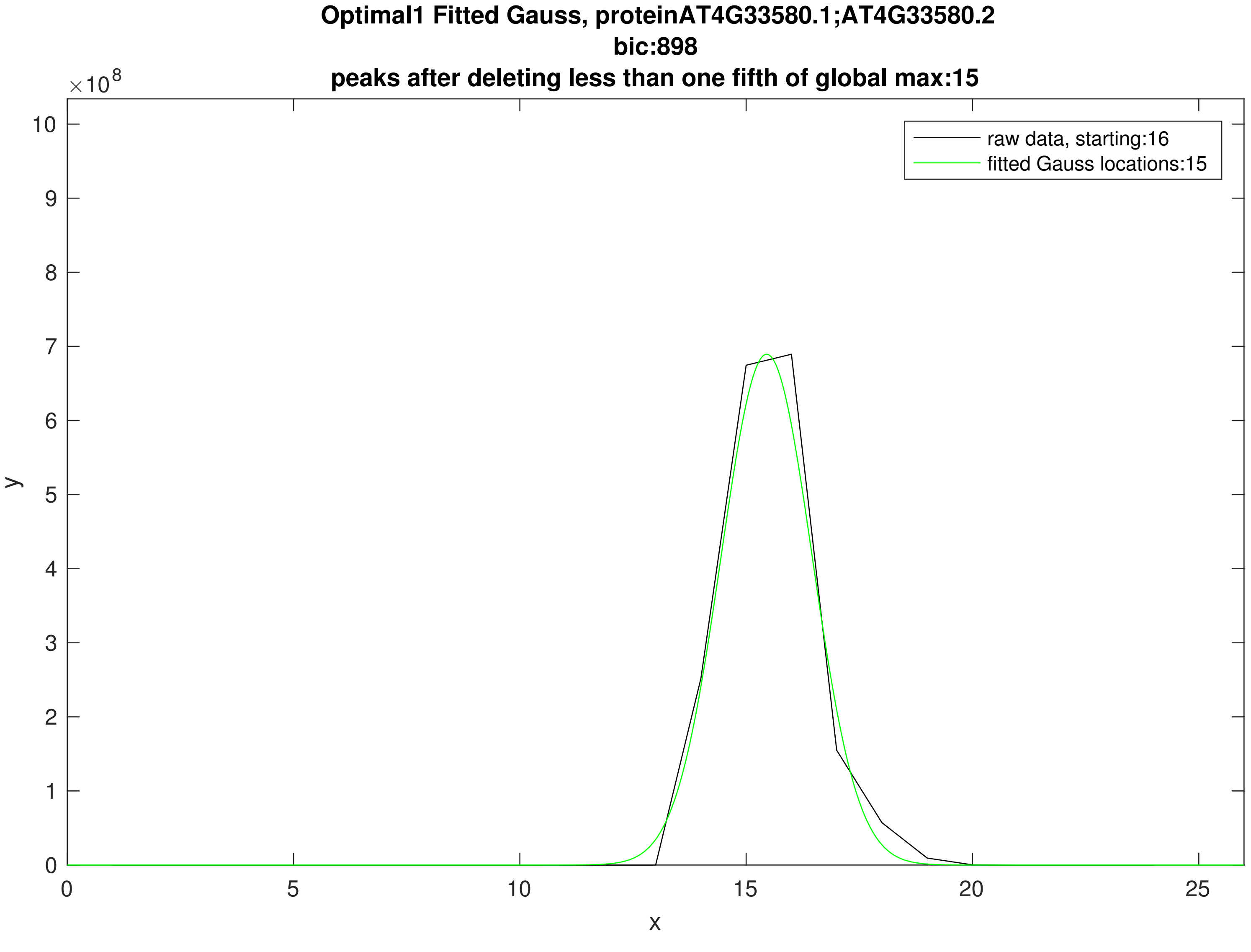
Optimal1 Fitted Gauss, proteinAT4G33580.1;AT4G33580.2
bic:898
peaks after deleting less than one fifth of global max:15
×10^8
raw data, starting:16
fitted Gauss locations:15
y
x
0
1
2
3
4
5
6
7
8
9
10
0
5
10
15
20
25

Optimal1 Fitted Gauss, proteinAT4G33640.1
bic:821
peaks after deleting less than one fifth of global max:20

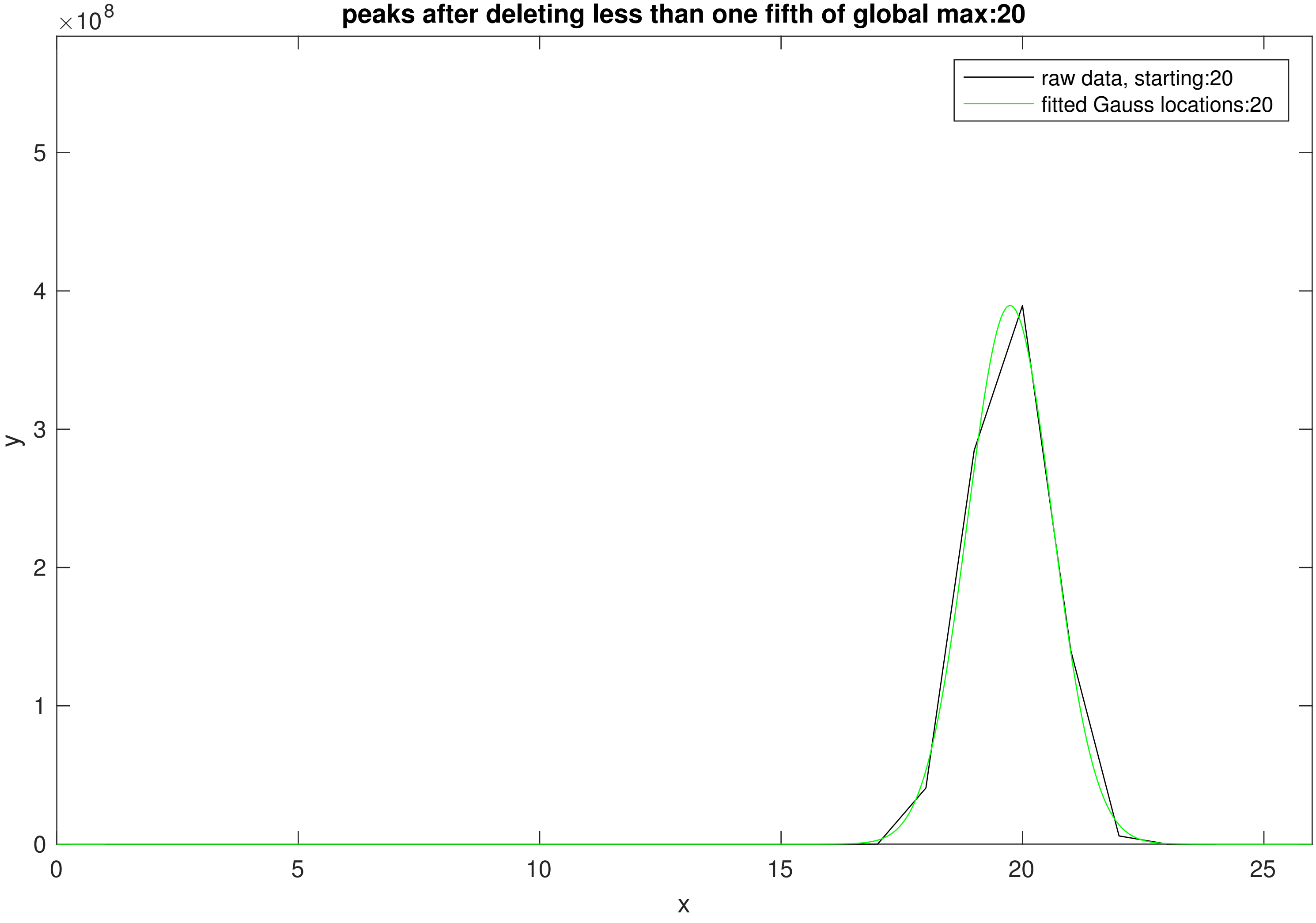

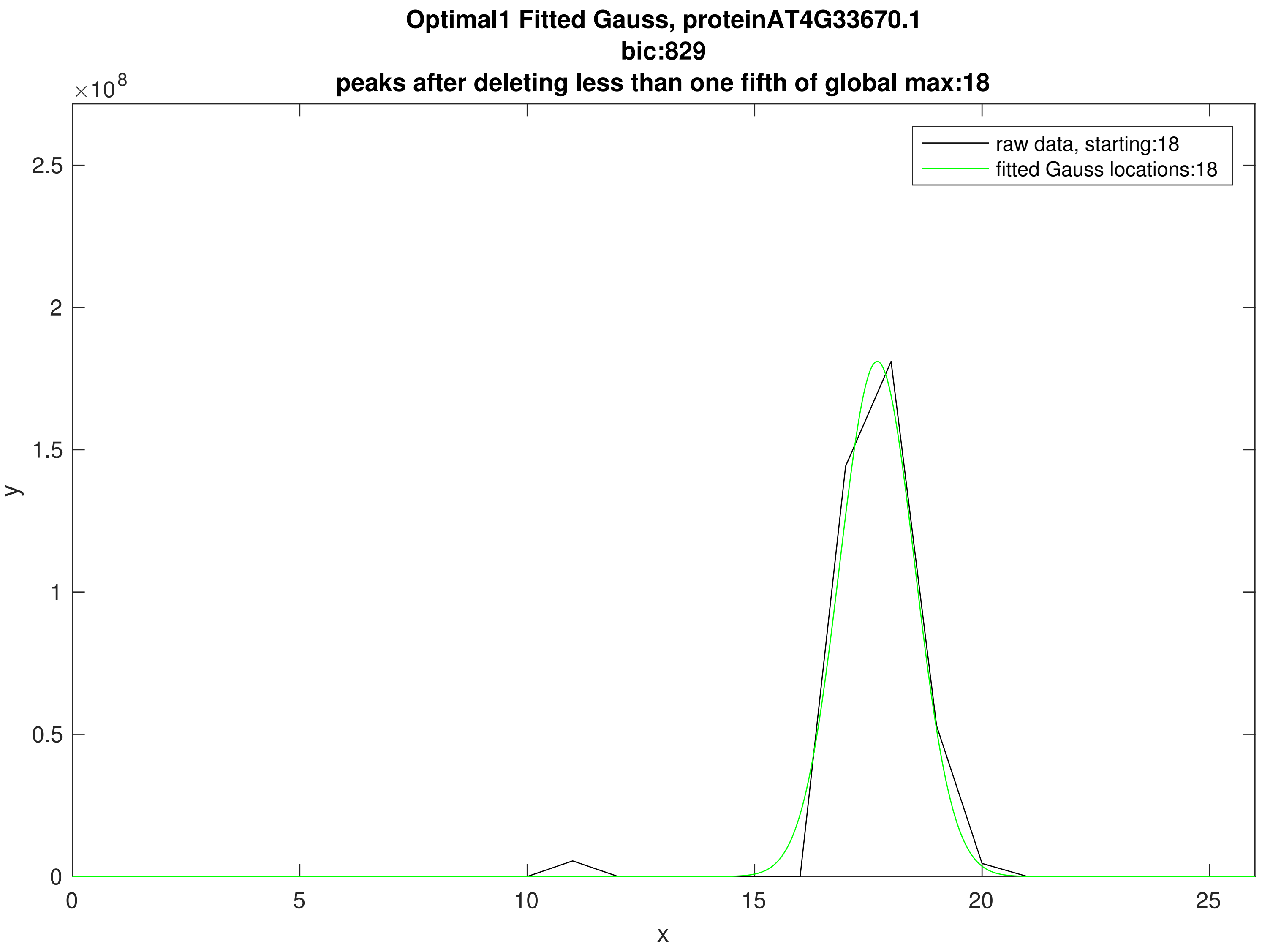
Optimal1 Fitted Gauss, proteinAT4G33670.1
bic:829
peaks after deleting less than one fifth of global max:18
×10^8
raw data, starting:18
fitted Gauss locations:18
2.5
2
1.5
1
0.5
0
y
0
5
10
15
20
25
x

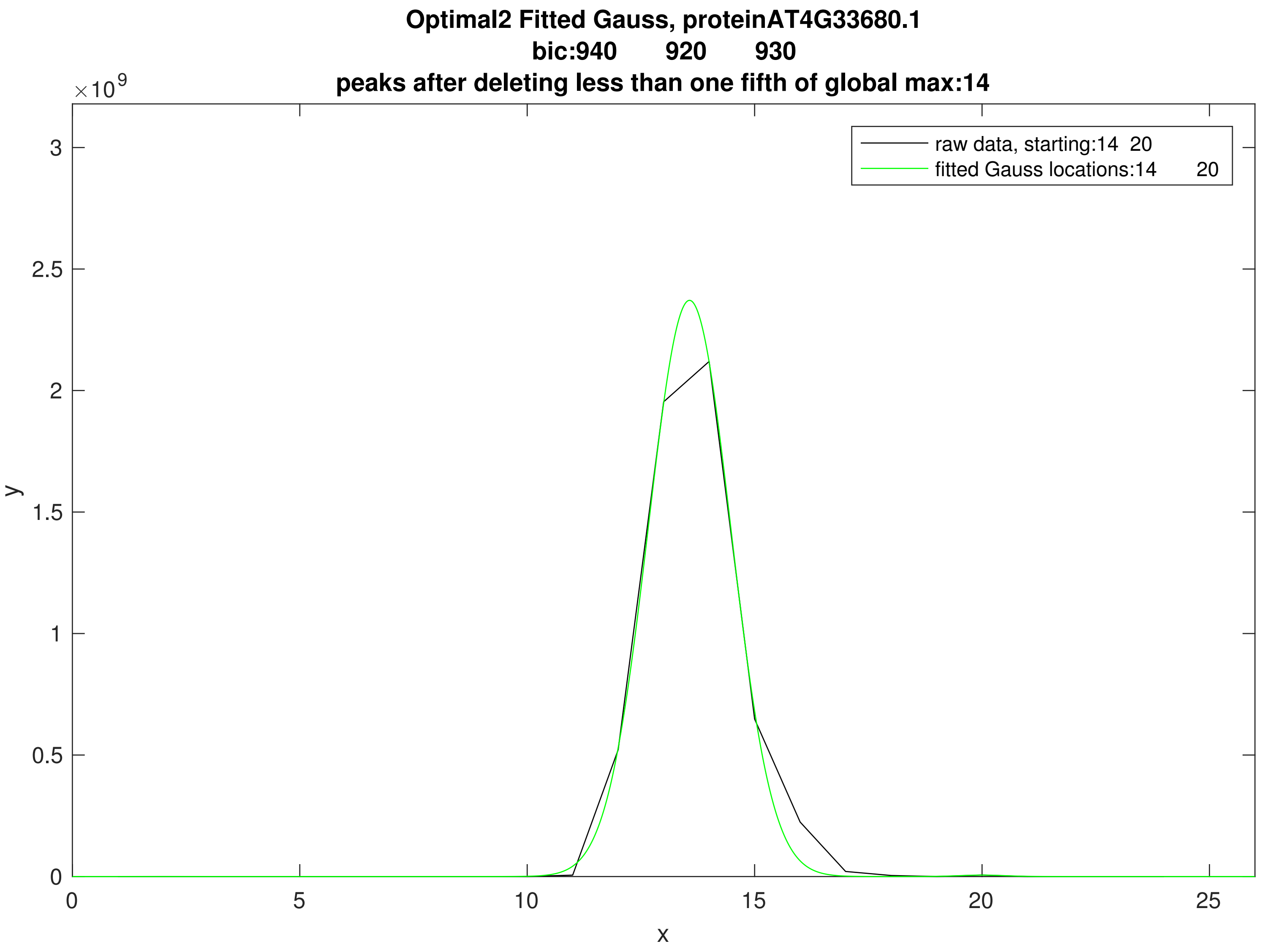
Optimal2 Fitted Gauss, proteinAT4G33680.1
bic:940 920 930
peaks after deleting less than one fifth of global max:14
×10^9
raw data, starting:14 20
fitted Gauss locations:14 20
y
x
0
0.5
1
1.5
2
2.5
3
0
5
10
15
20
25

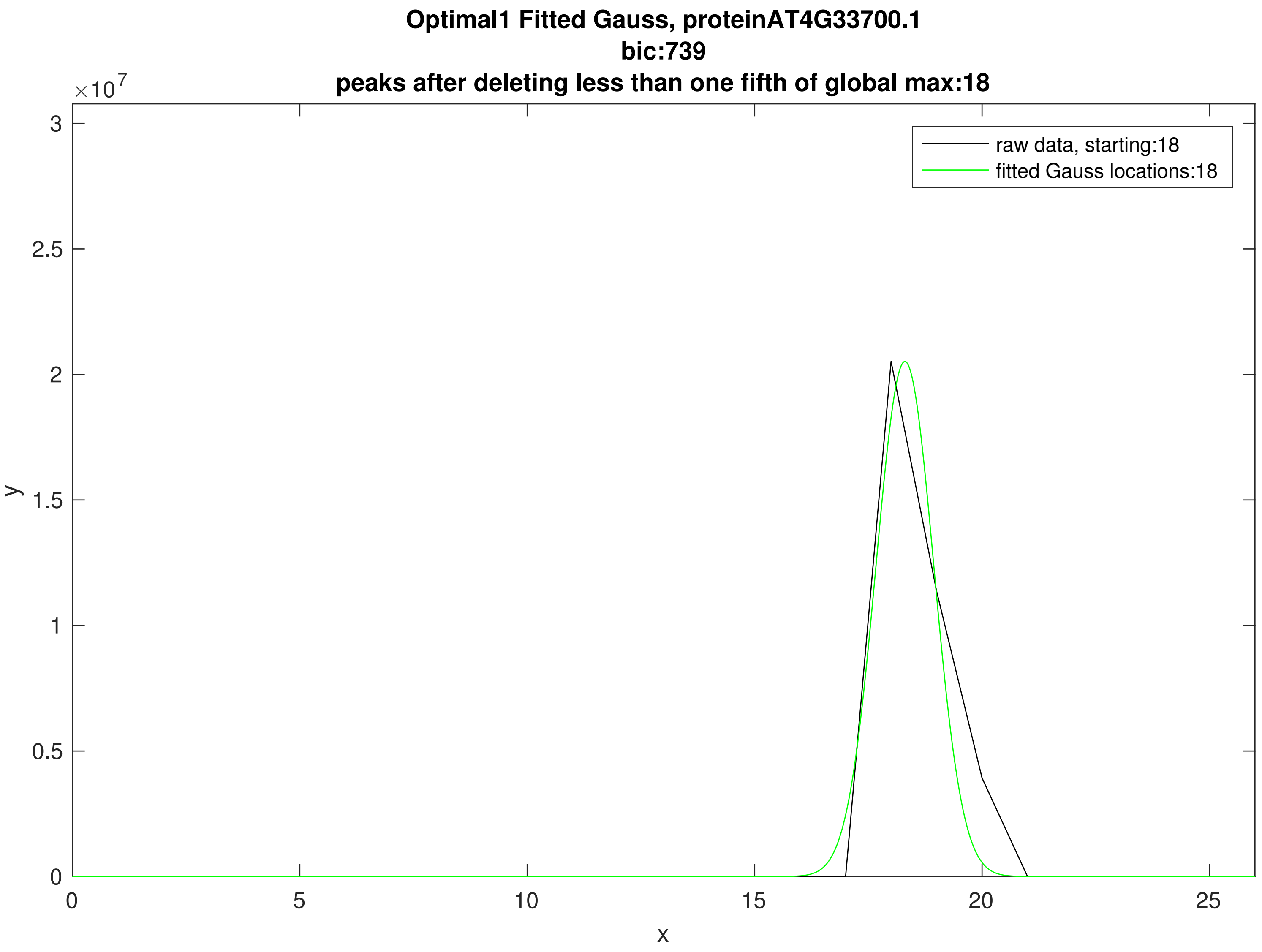
Optimal1 Fitted Gauss, proteinAT4G33700.1
bic:739
peaks after deleting less than one fifth of global max:18
×10^7
raw data, starting:18
fitted Gauss locations:18
y
x
0
0.5
1
1.5
2
2.5
3
5
10
15
20
25

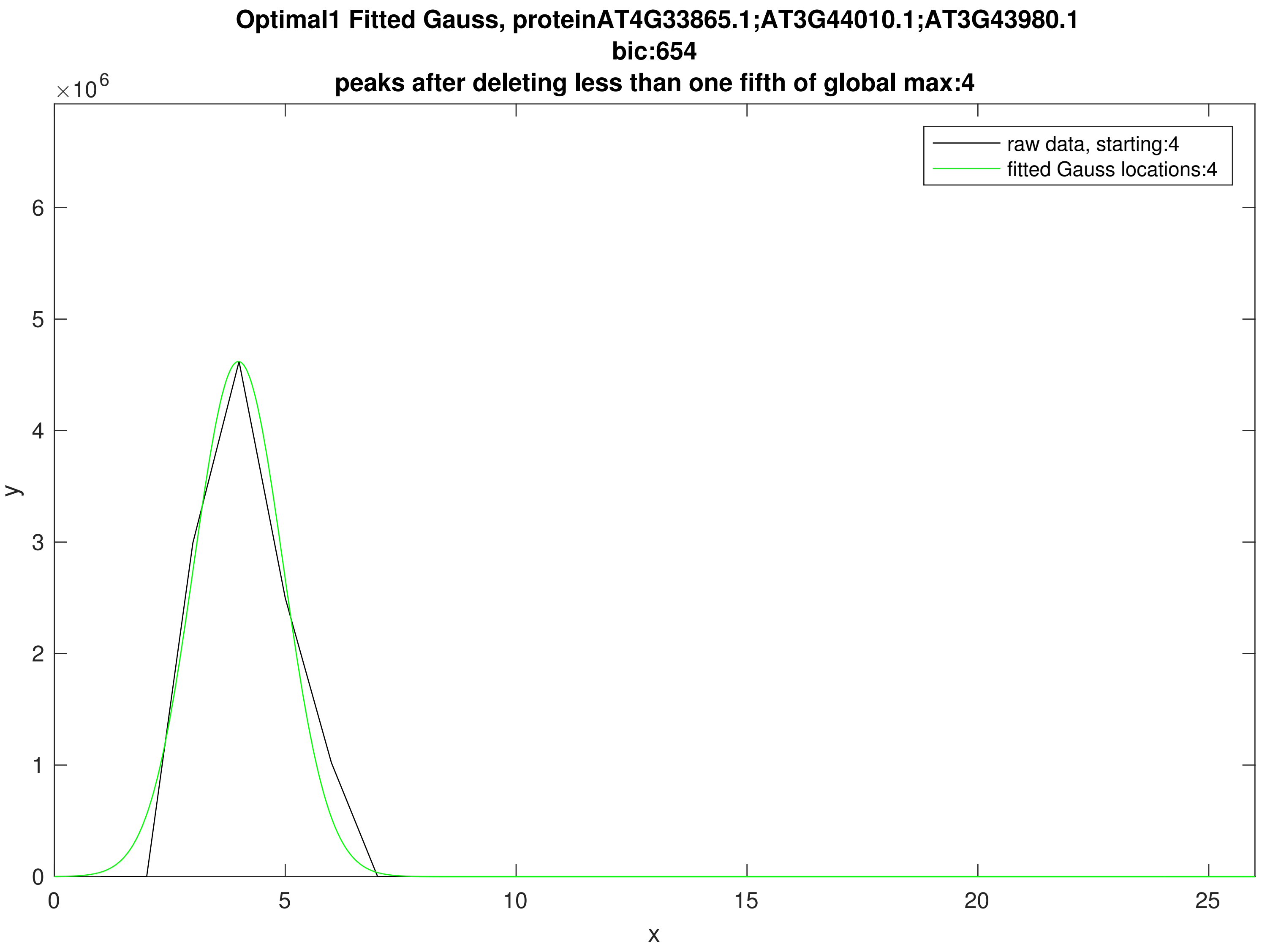

Optimal1 Fitted Gauss, proteinAT4G33865.1;AT3G44010.1;AT3G43980.1
bic:654
peaks after deleting less than one fifth of global max:4
×10^6
raw data, starting:4
fitted Gauss locations:4
y
x
0
1
2
3
4
5
6
0
5
10
15
20
25

**Optimal1 Fitted Gauss, proteinAT4G34030.1**
**bic:705**
**peaks after deleting less than one fifth of global max:5.1**

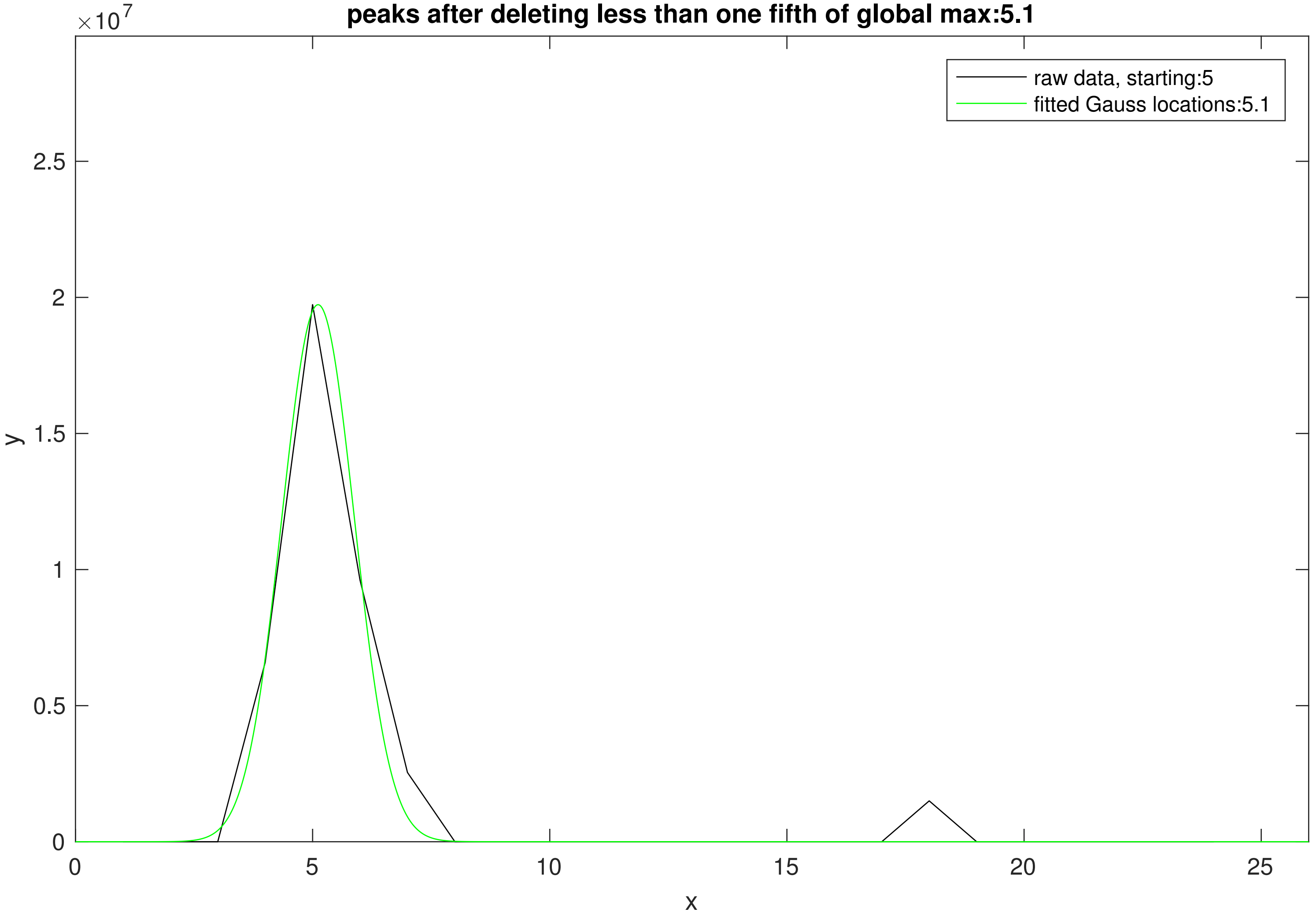

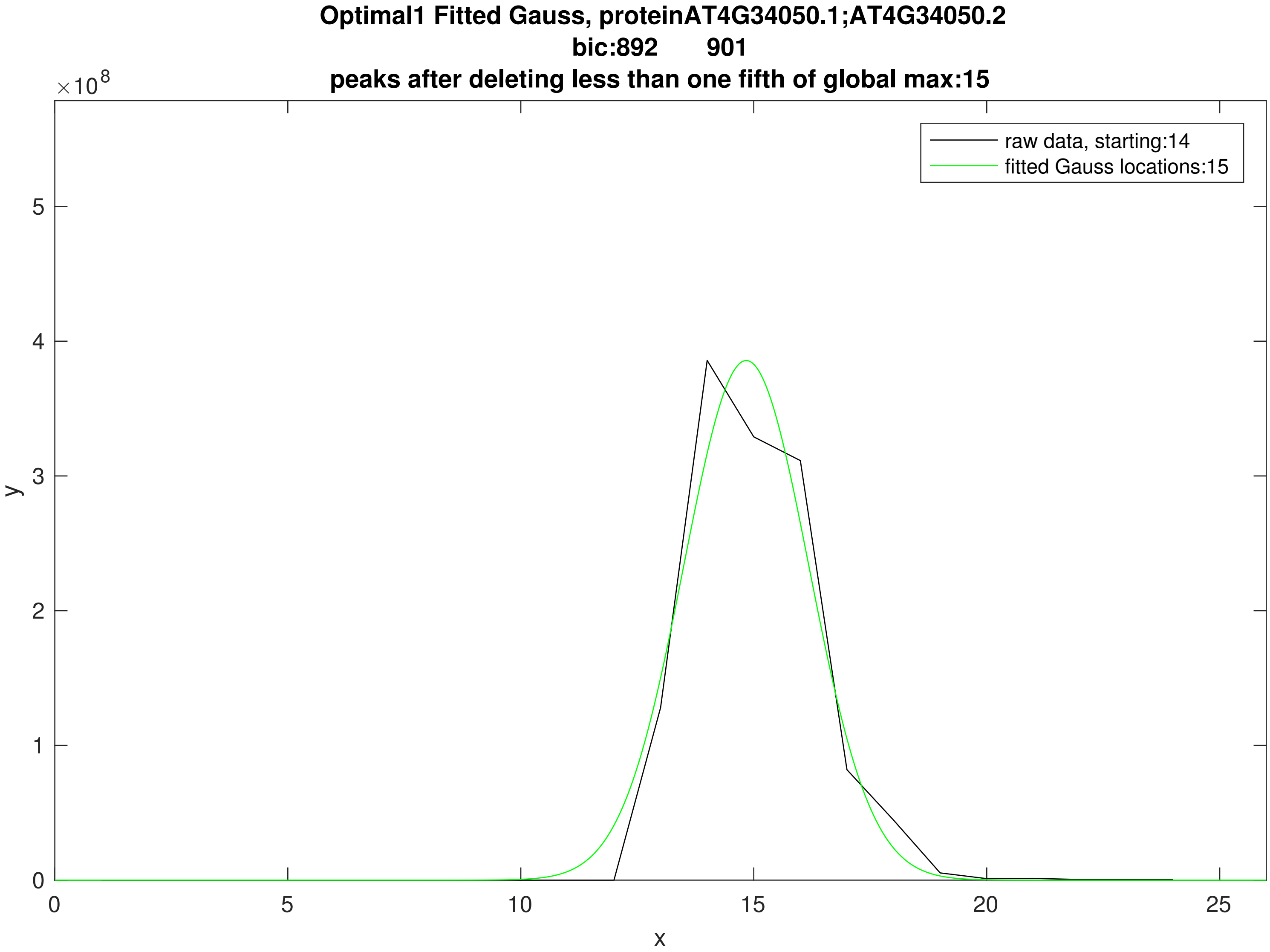
Optimal1 Fitted Gauss, proteinAT4G34050.1;AT4G34050.2
bic:892 901
peaks after deleting less than one fifth of global max:15
×10^8
raw data, starting:14
fitted Gauss locations:15
y
x
0
1
2
3
4
5
0
5
10
15
20
25

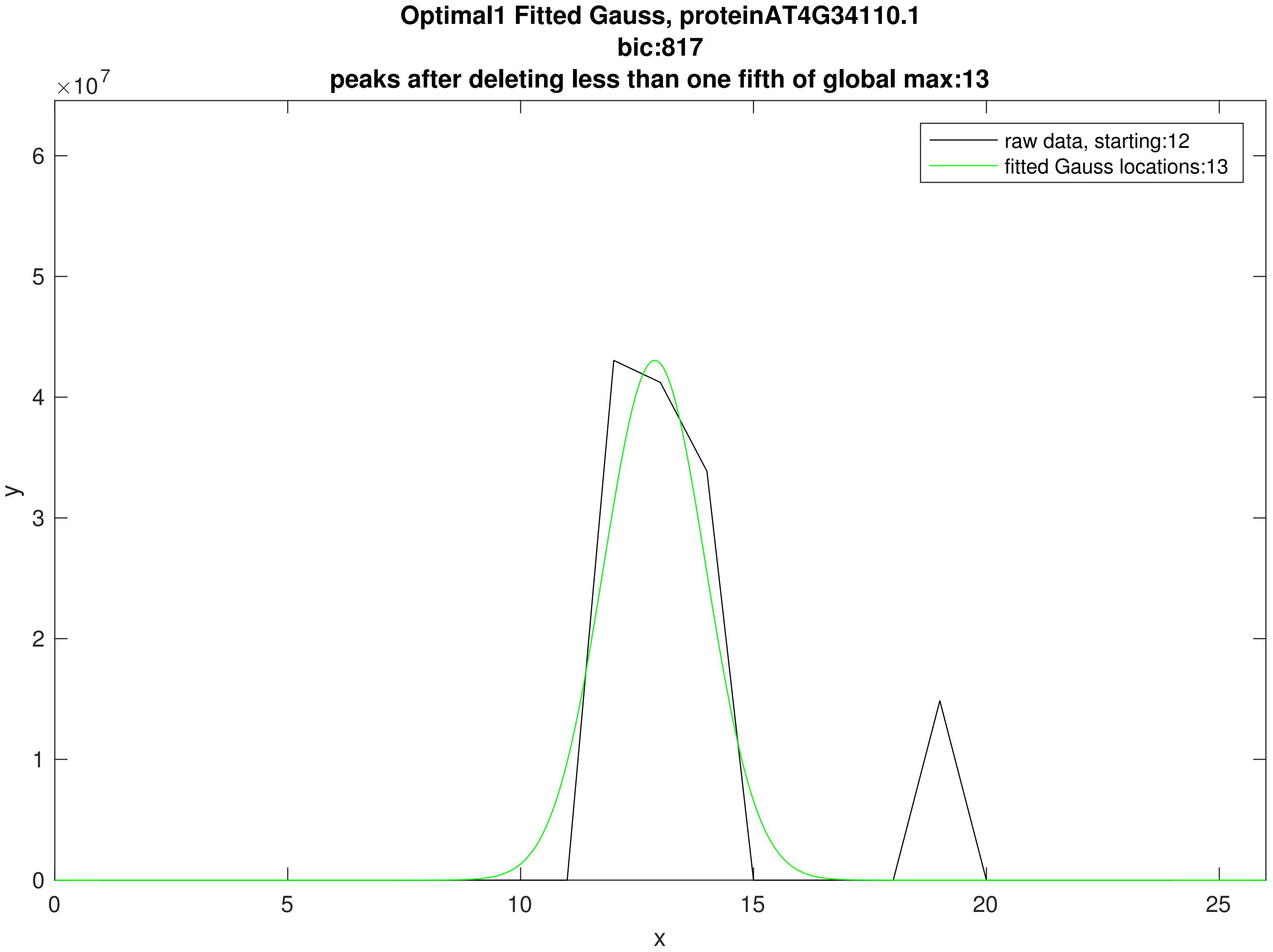
Optimal1 Fitted Gauss, proteinAT4G34110.1
bic:817
peaks after deleting less than one fifth of global max:13
×10^7
raw data, starting:12
fitted Gauss locations:13
y
x
0
1
2
3
4
5
6
0
5
10
15
20
25

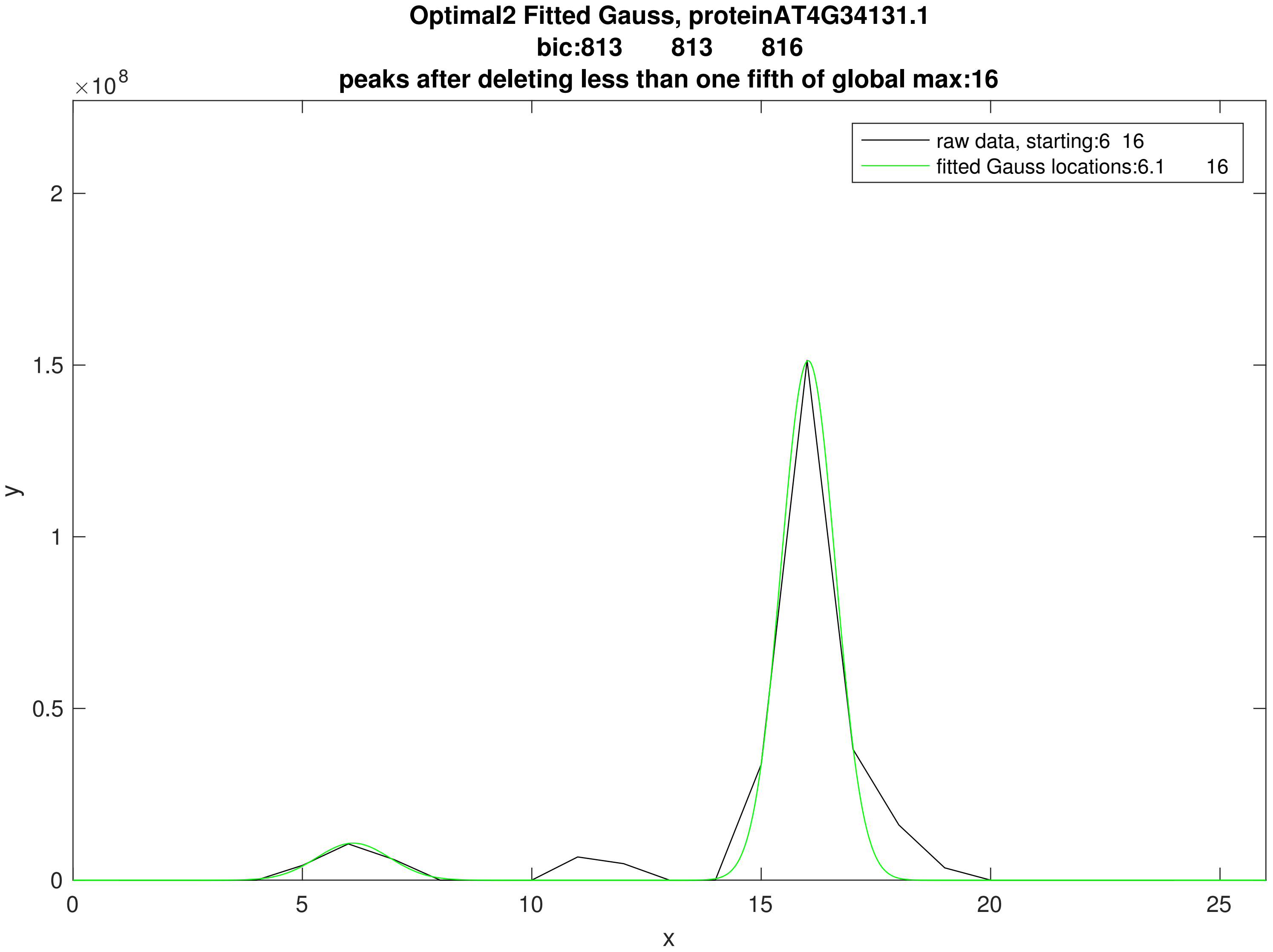

Optimal2 Fitted Gauss, proteinAT4G34131.1
bic:813 813 816
peaks after deleting less than one fifth of global max:16
×10^8
raw data, starting:6 16
fitted Gauss locations:6.1 16
y
x
0
0.5
1
1.5
2
0
5
10
15
20
25

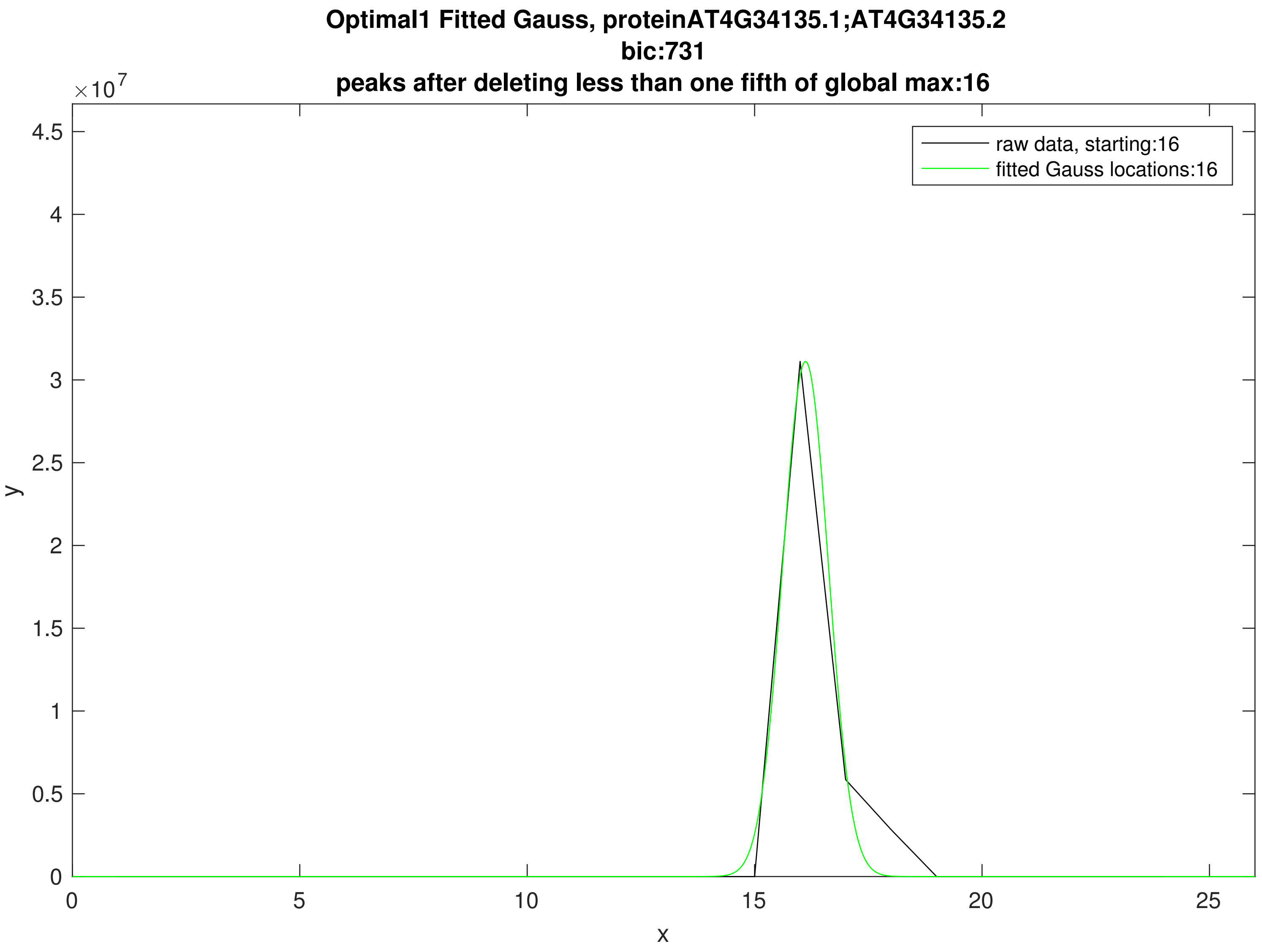
Optimal1 Fitted Gauss, proteinAT4G34135.1;AT4G34135.2
bic:731
peaks after deleting less than one fifth of global max:16
×10^7
4.5
4
3.5
3
2.5
2
1.5
1
0.5
0
y
0
5
10
15
20
25
x
raw data, starting:16
fitted Gauss locations:16

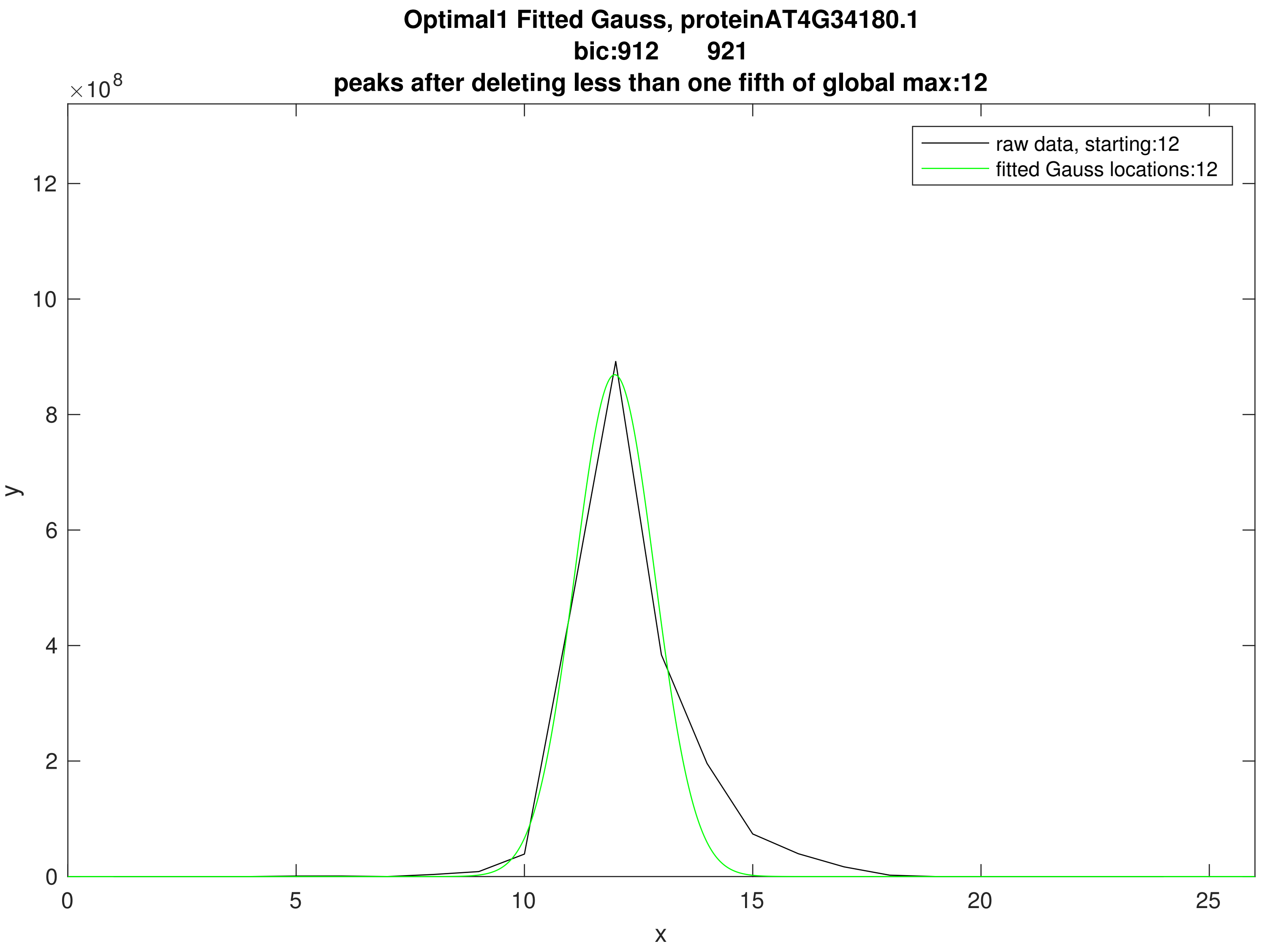

Optimal1 Fitted Gauss, proteinAT4G34180.1
bic:912 921
peaks after deleting less than one fifth of global max:12
×10^8
raw data, starting:12
fitted Gauss locations:12
12
10
8
6
4
2
0
y
0
5
10
15
20
25
x

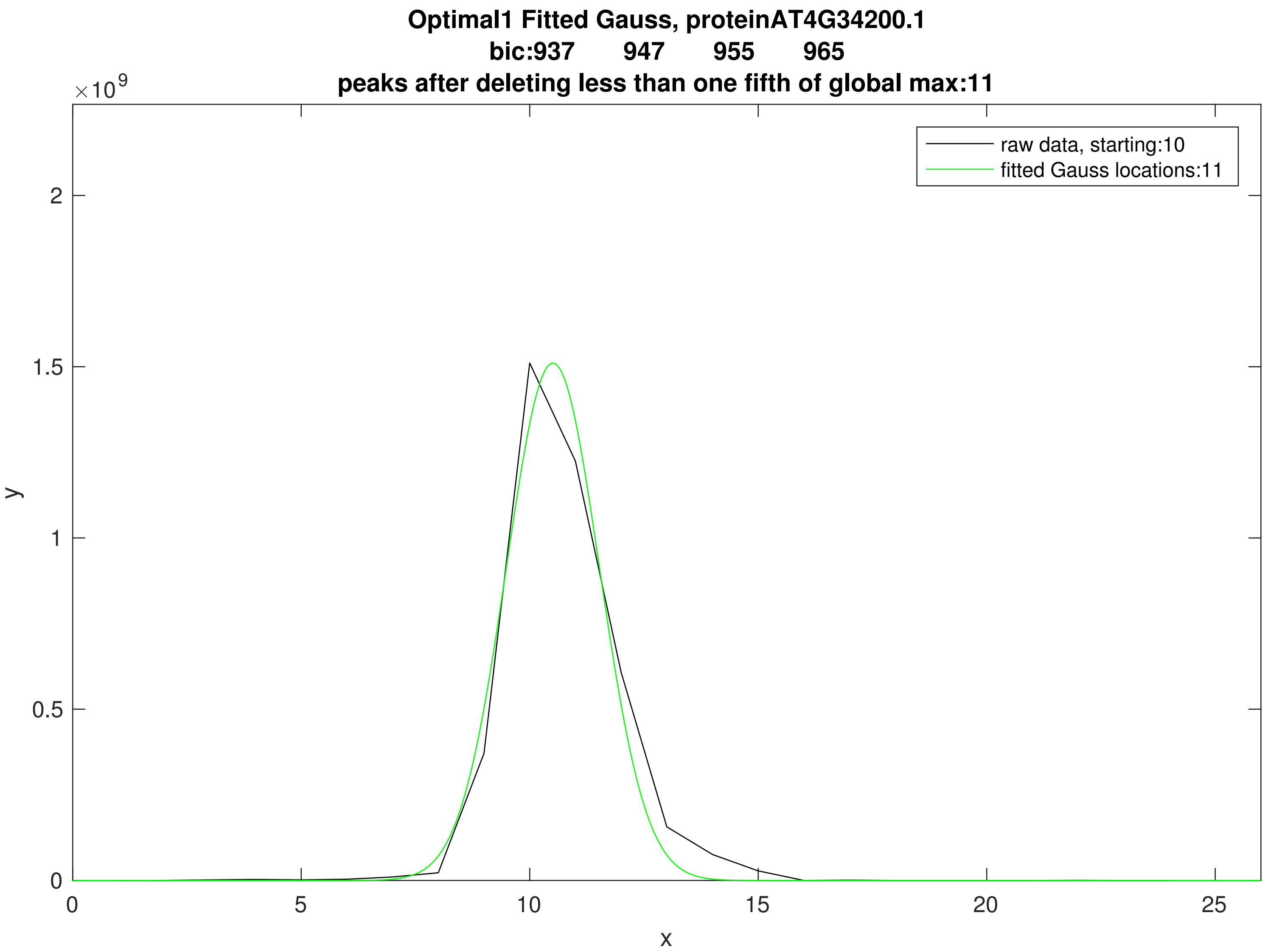
Optimal1 Fitted Gauss, proteinAT4G34200.1
bic:937 947 955 965
peaks after deleting less than one fifth of global max:11
×10^9
raw data, starting:10
fitted Gauss locations:11
y
x
0
0.5
1
1.5
2
0
5
10
15
20
25

#### Optimal1 Fitted Gauss, proteinAT4G34215.2;AT4G34215.1
#### bic:771
#### peaks after deleting less than one fifth of global max:18

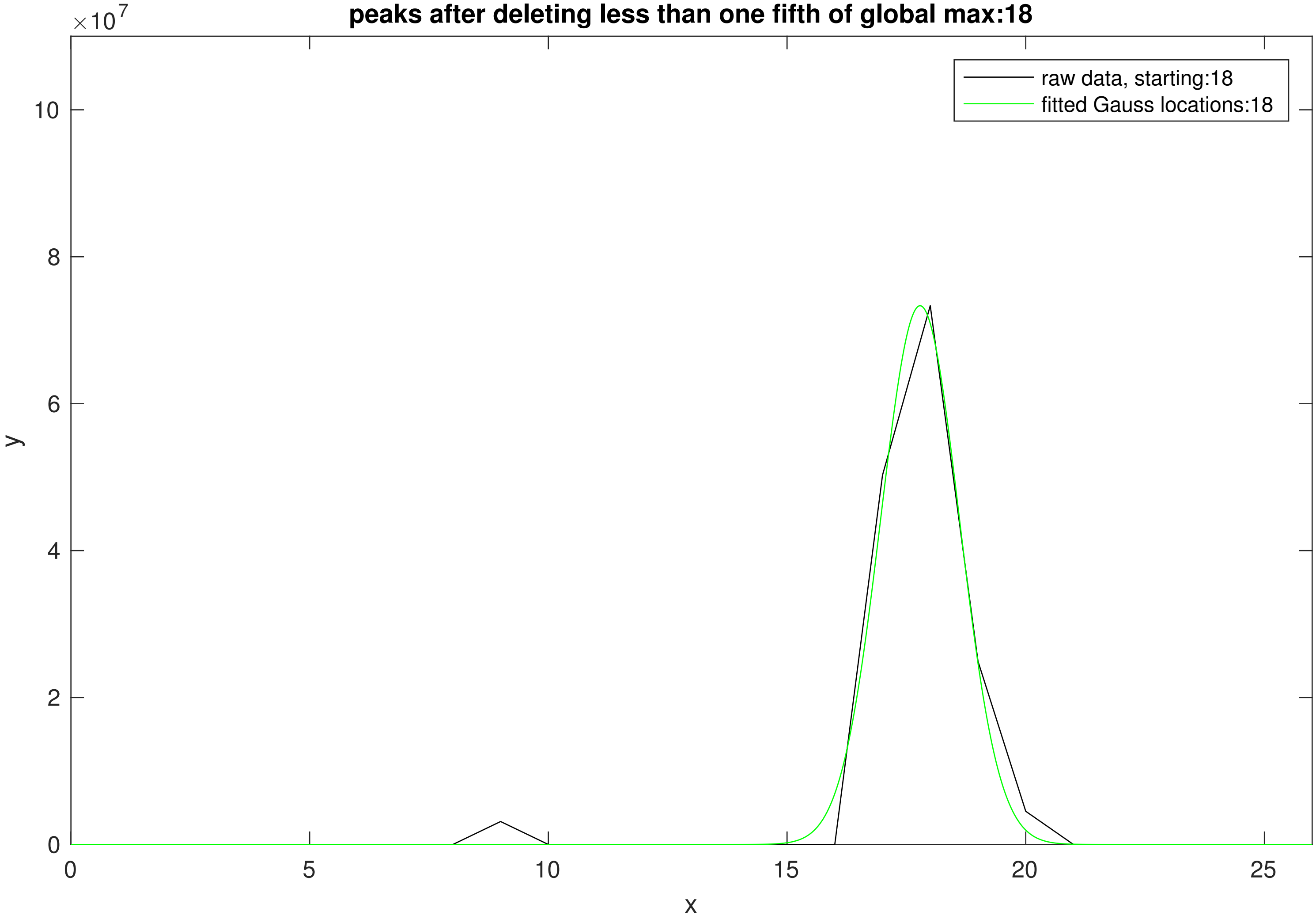

#### Optimal1 Fitted Gauss, proteinAT4G34230.1;AT4G34230.2
#### bic:972 982
#### peaks after deleting less than one fifth of global max:14

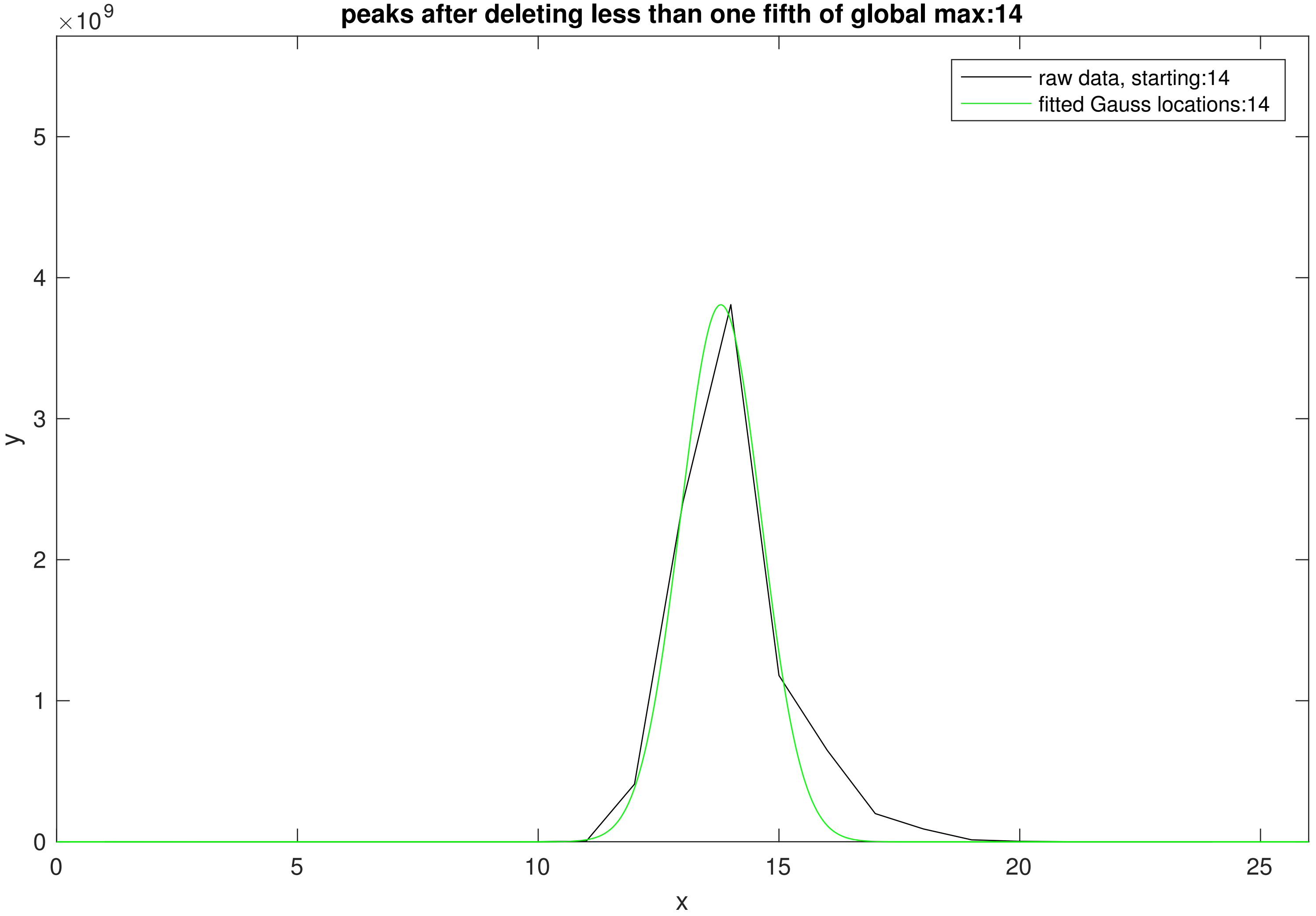

#### Optimal1 Fitted Gauss, proteinAT4G34260.1
#### bic:906
#### peaks after deleting less than one fifth of global max:14

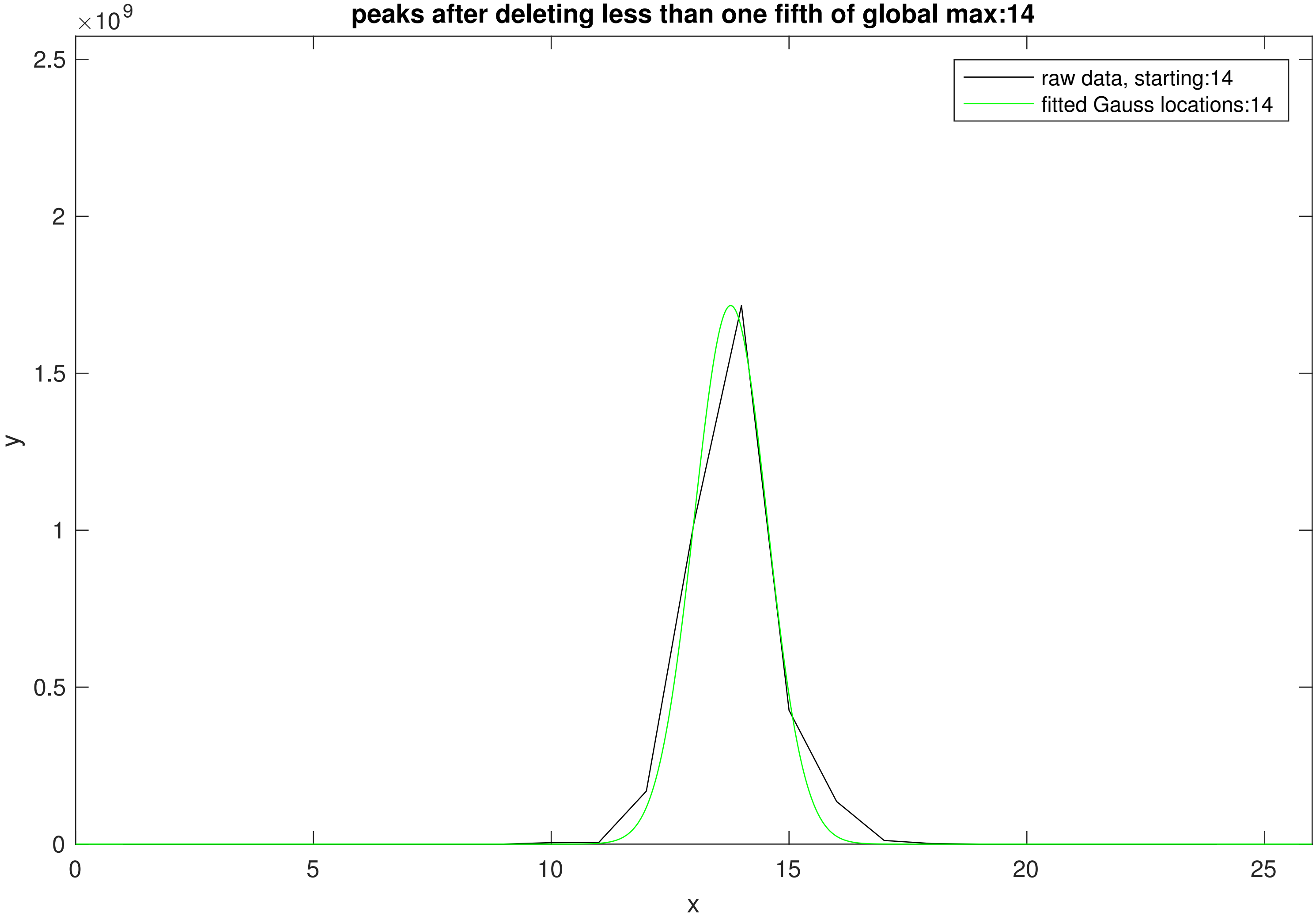

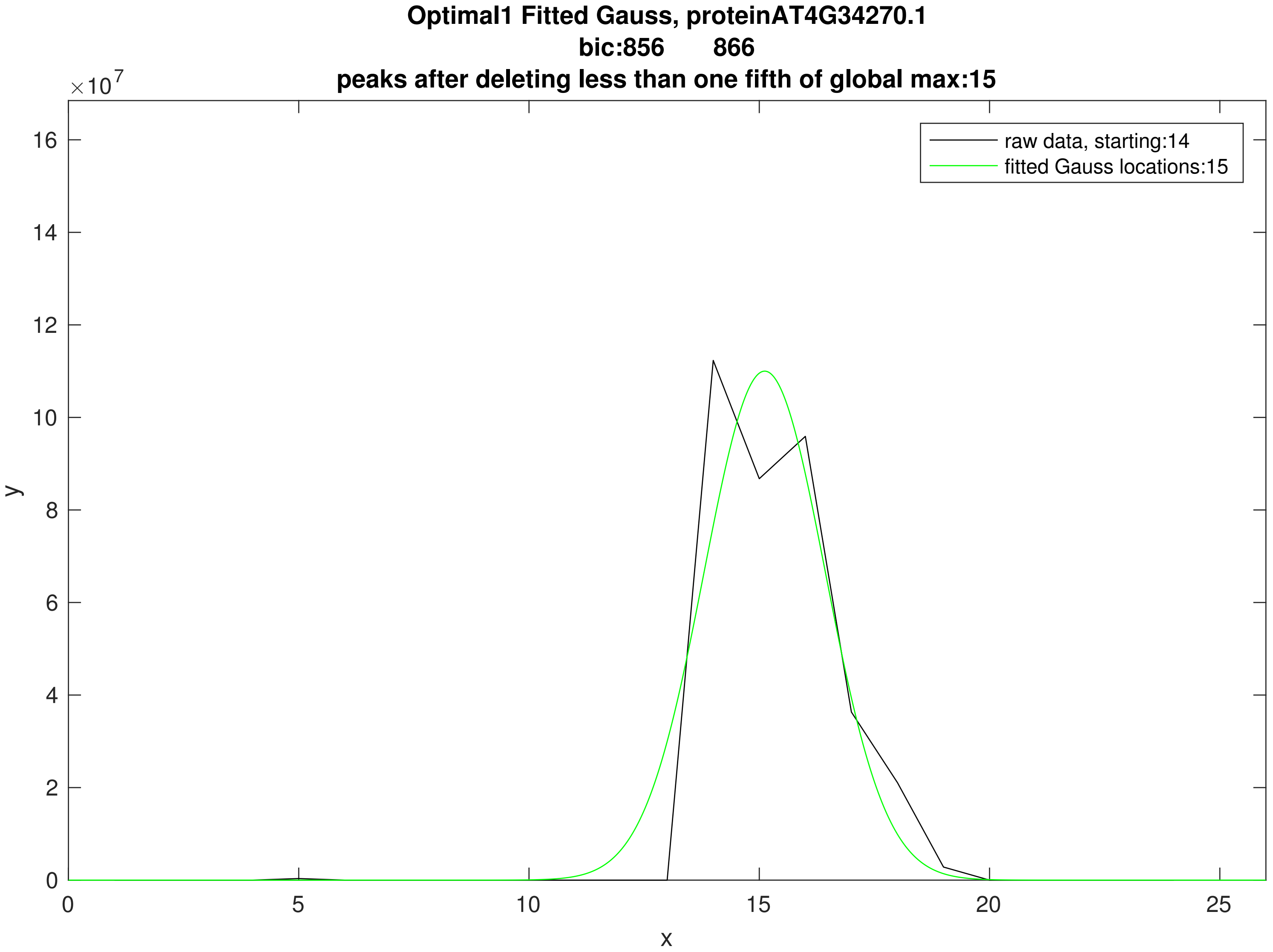
Optimal1 Fitted Gauss, proteinAT4G34270.1
bic:856 866
peaks after deleting less than one fifth of global max:15
×10^7
16
14
12
10
8
6
4
2
0
y
0
5
10
15
20
25
x
raw data, starting:14
fitted Gauss locations:15

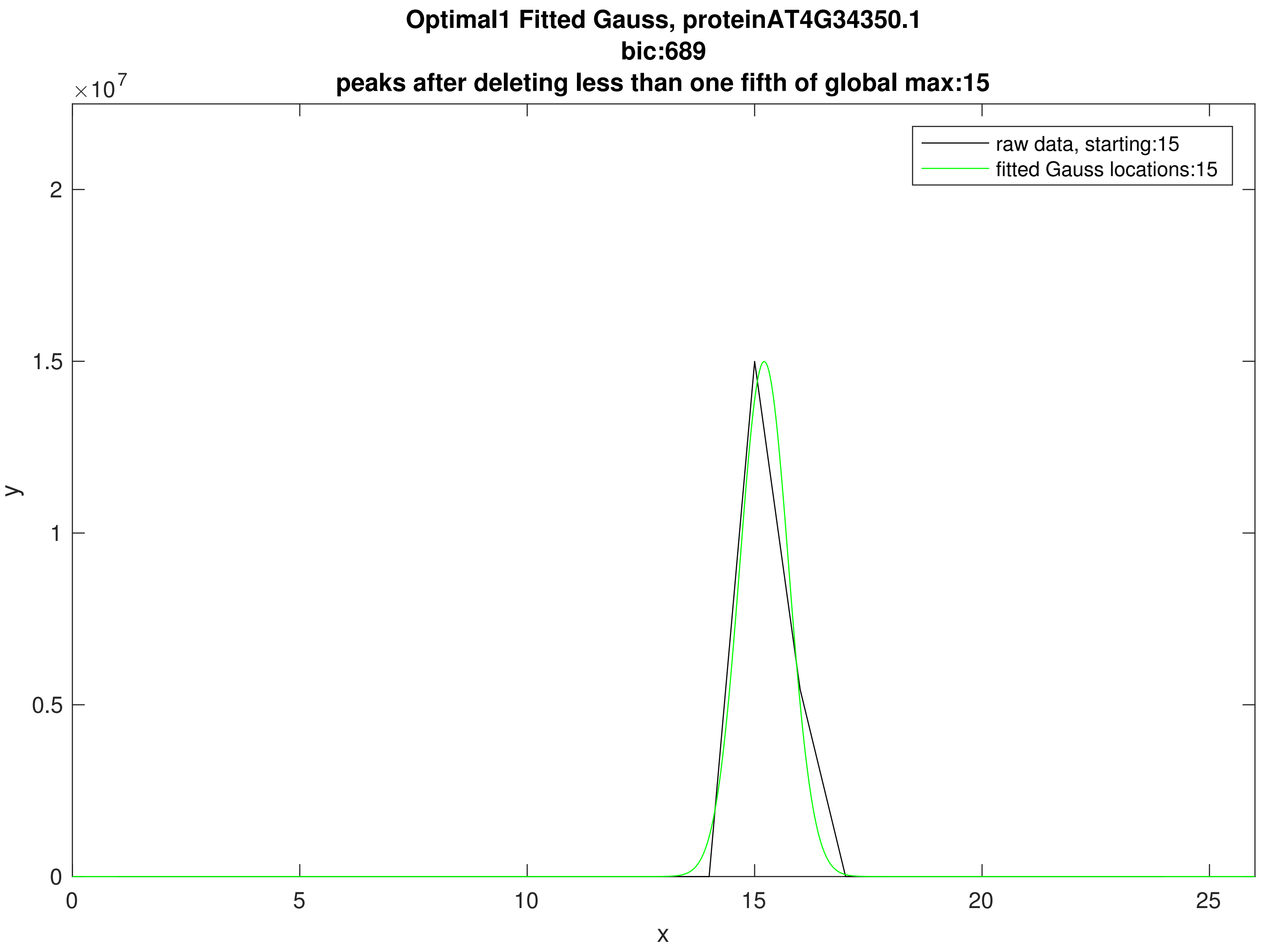
Optimal1 Fitted Gauss, proteinAT4G34350.1
bic:689
peaks after deleting less than one fifth of global max:15
×10^7
raw data, starting:15
fitted Gauss locations:15
y
x
0
0.5
1
1.5
2
0
5
10
15
20
25

#### Optimal1 Fitted Gauss, proteinAT4G34450.1
#### bic:800 806 804
#### peaks after deleting less than one fifth of global max:7.4

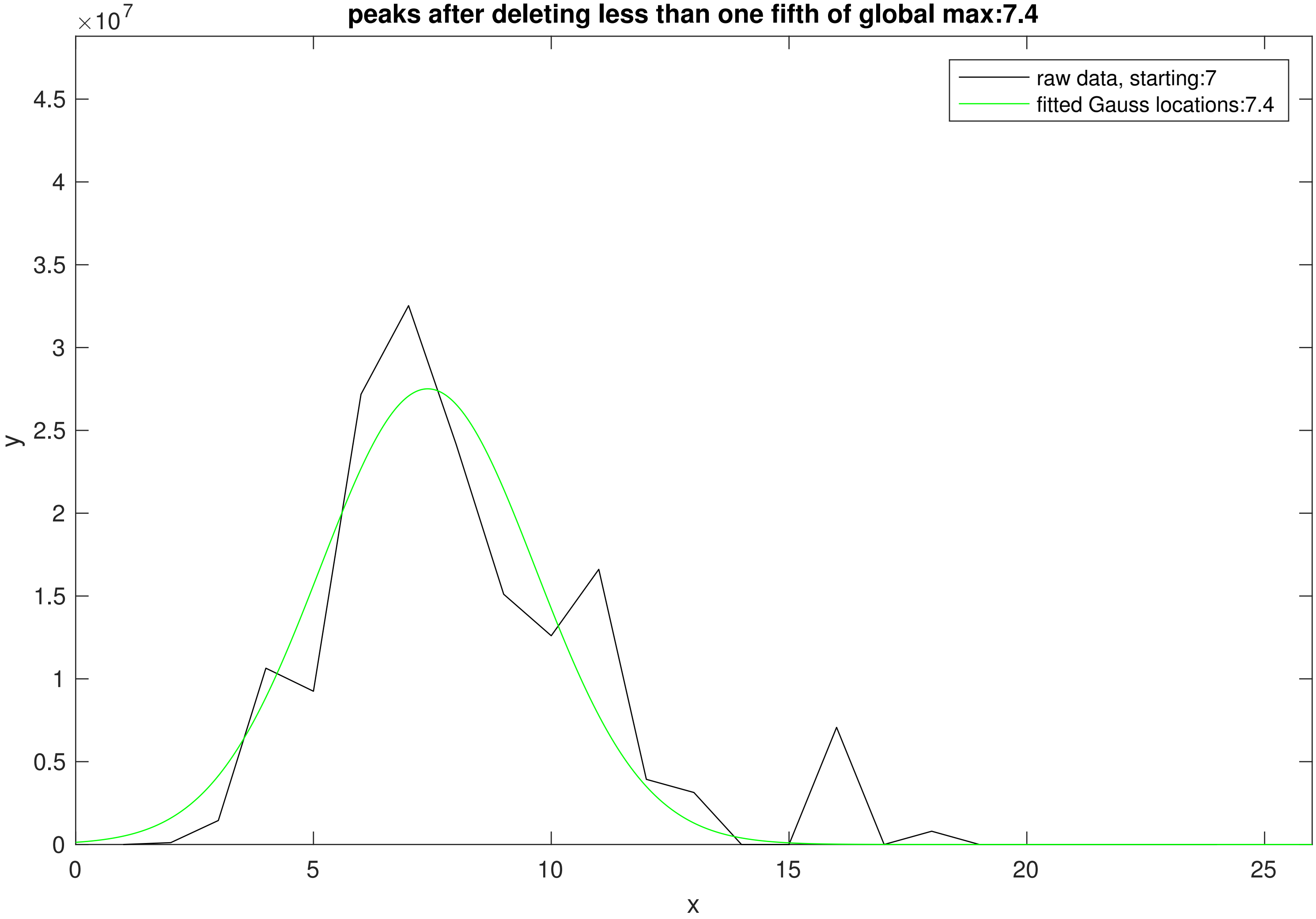

**Optimal1 Fitted Gauss, proteinAT4G34480.1**
**bic:797**
**peaks after deleting less than one fifth of global max:16**

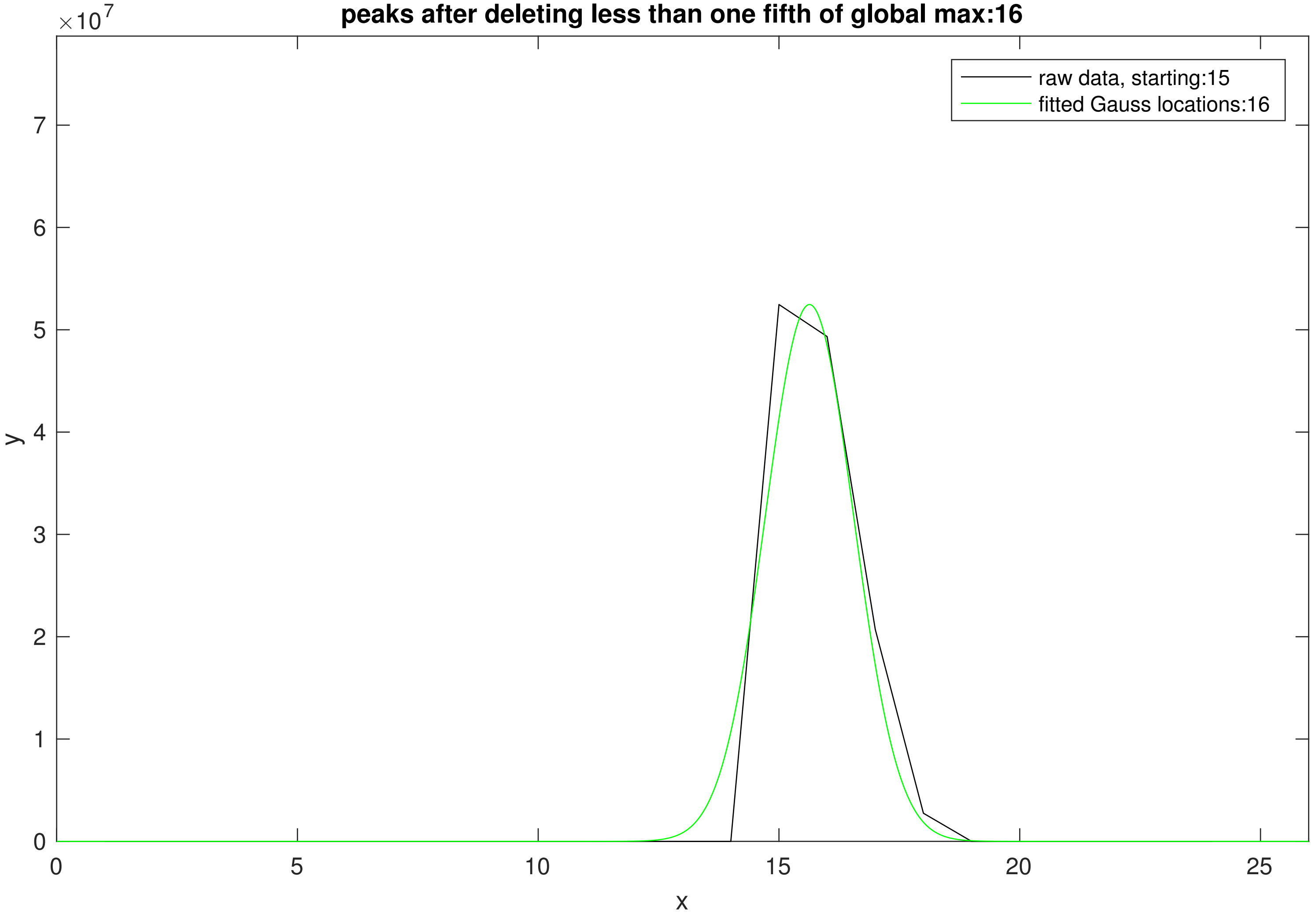

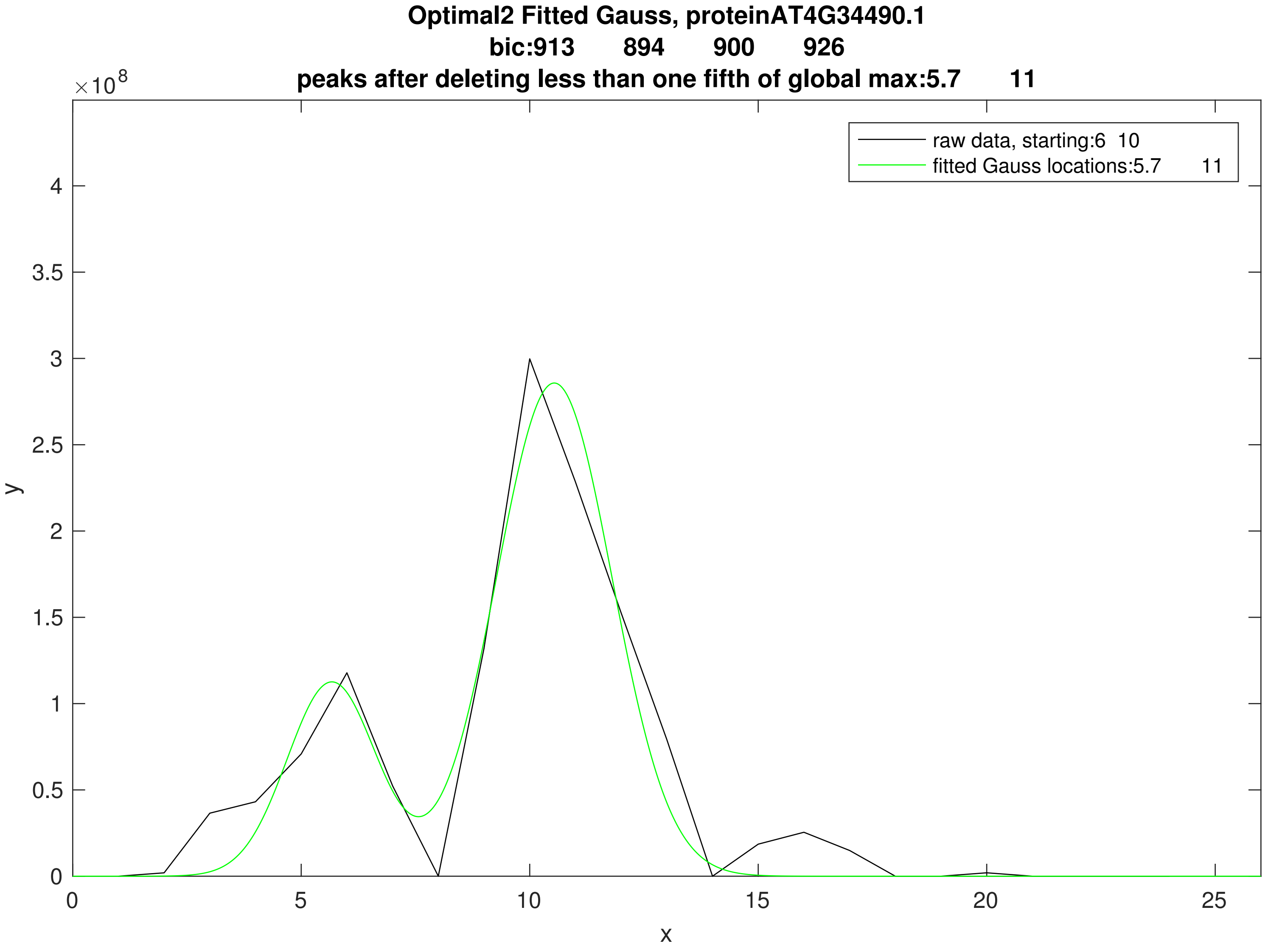
Optimal2 Fitted Gauss, proteinAT4G34490.1
bic:913 894 900 926
peaks after deleting less than one fifth of global max:5.7 11
×10^8
raw data, starting:6 10
fitted Gauss locations:5.7 11
y
x
0
0.5
1
1.5
2
2.5
3
3.5
4
0
5
10
15
20
25

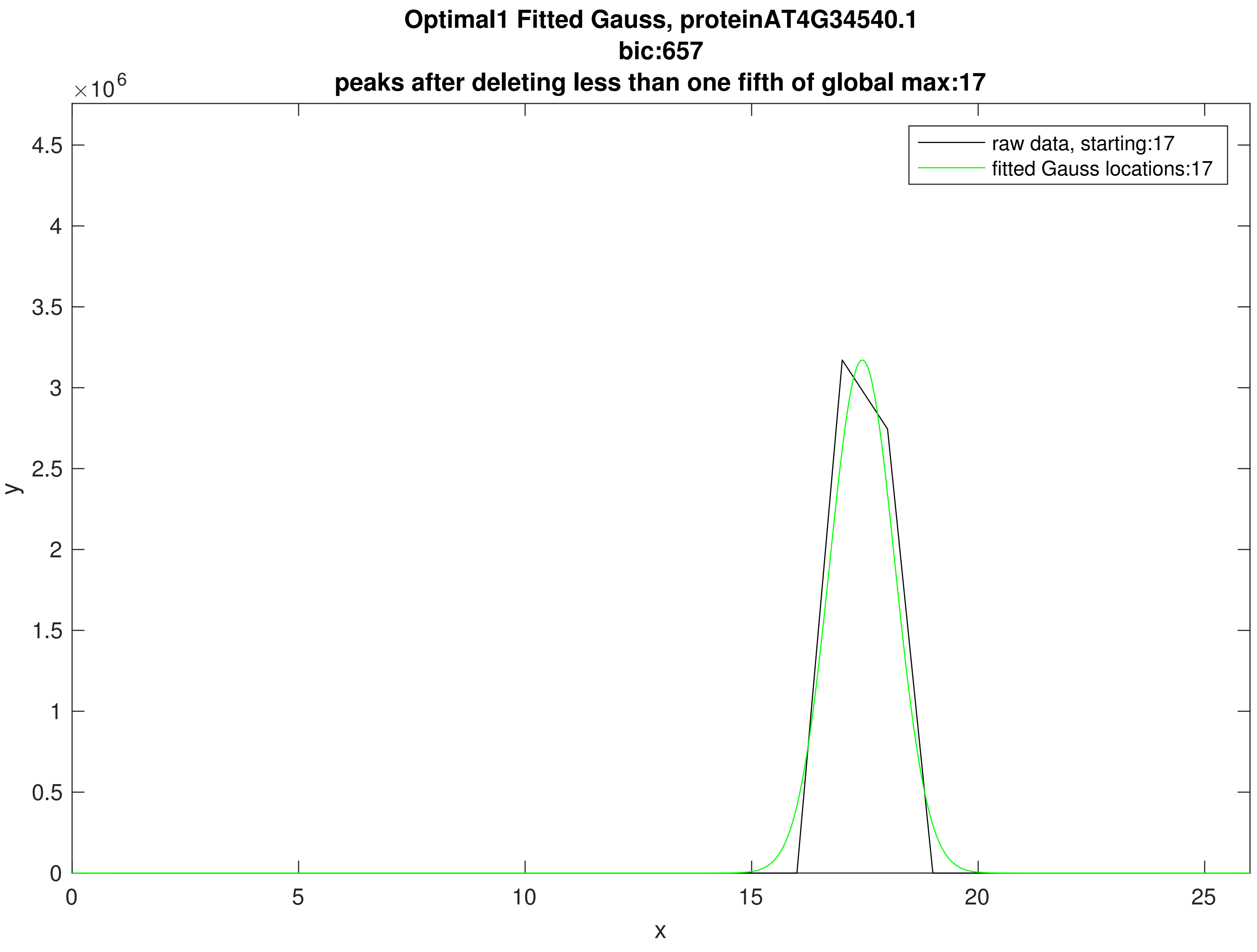
Optimal1 Fitted Gauss, proteinAT4G34540.1
bic:657
peaks after deleting less than one fifth of global max:17
×10^6
4.5
4
3.5
3
2.5
2
1.5
1
0.5
0
y
0
5
10
15
20
25
x
raw data, starting:17
fitted Gauss locations:17

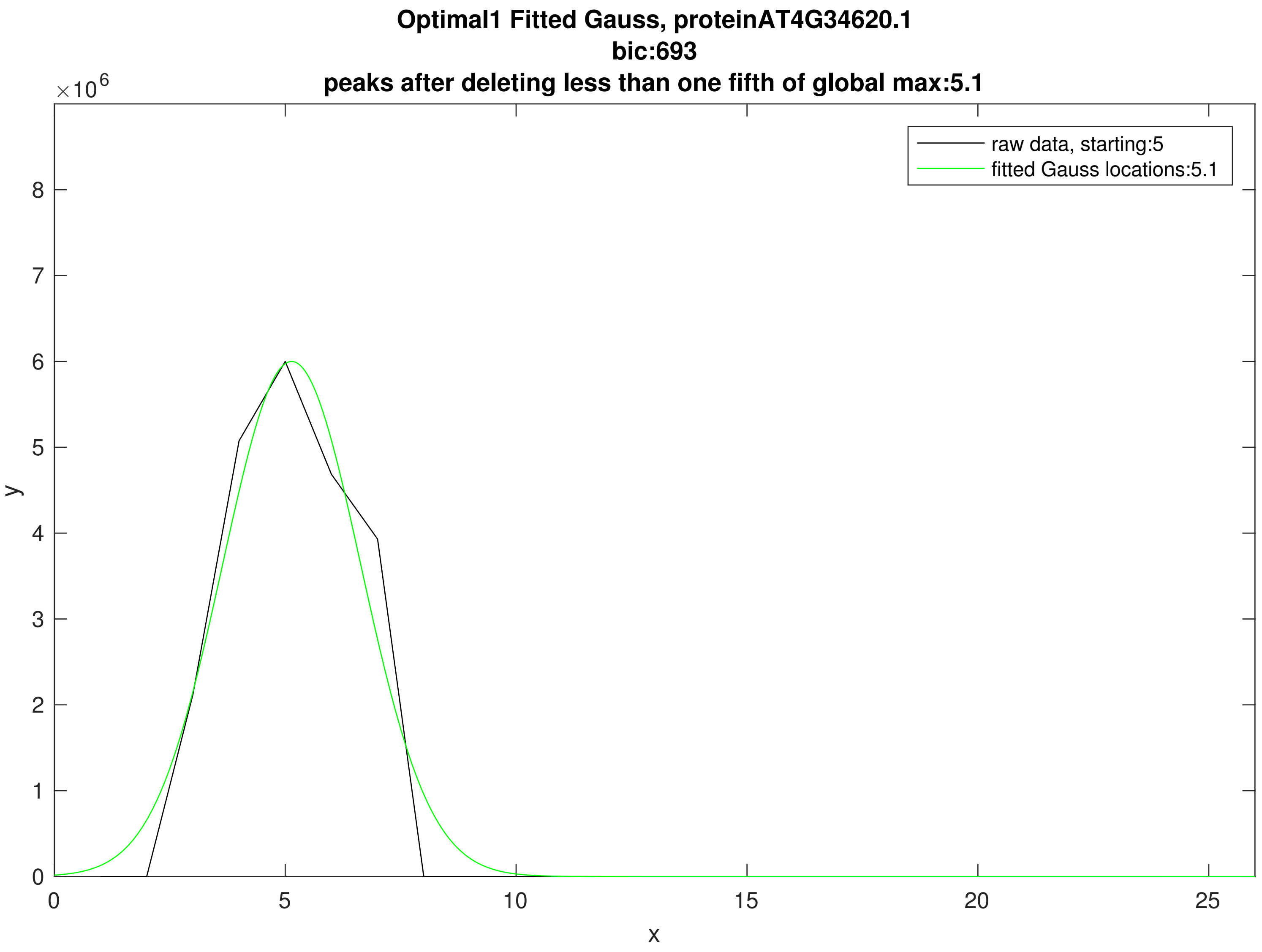
Optimal1 Fitted Gauss, proteinAT4G34620.1
bic:693
peaks after deleting less than one fifth of global max:5.1
×10^6
raw data, starting:5
fitted Gauss locations:5.1
y
x
0
1
2
3
4
5
6
7
8
0
5
10
15
20
25

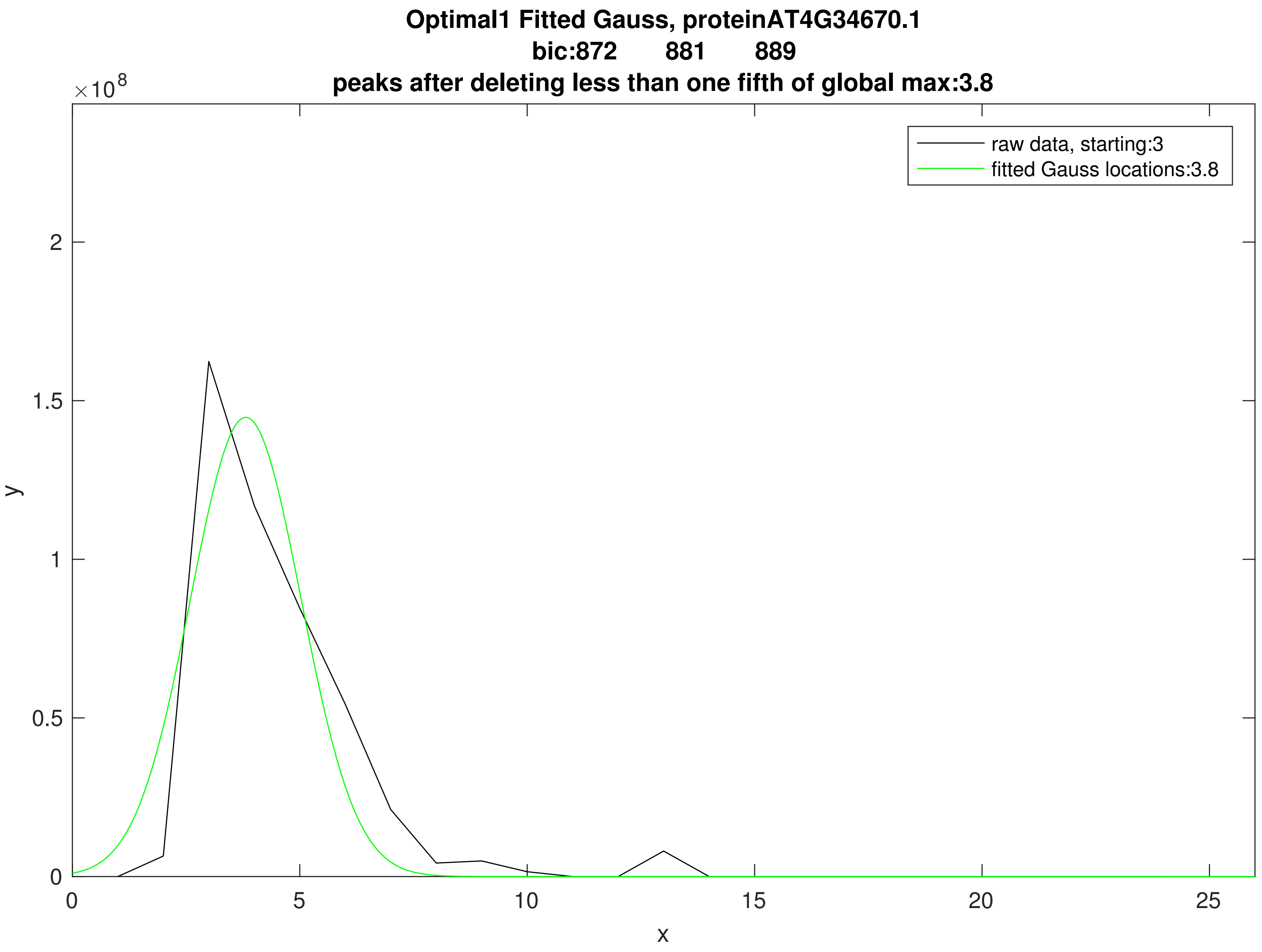
Optimal1 Fitted Gauss, proteinAT4G34670.1
bic:872 881 889
peaks after deleting less than one fifth of global max:3.8
×10^8
raw data, starting:3
fitted Gauss locations:3.8
y
x
0
0.5
1
1.5
2
0
5
10
15
20
25

Optimal1 Fitted Gauss, proteinAT4G34740.1
bic:776
peaks after deleting less than one fifth of global max:16

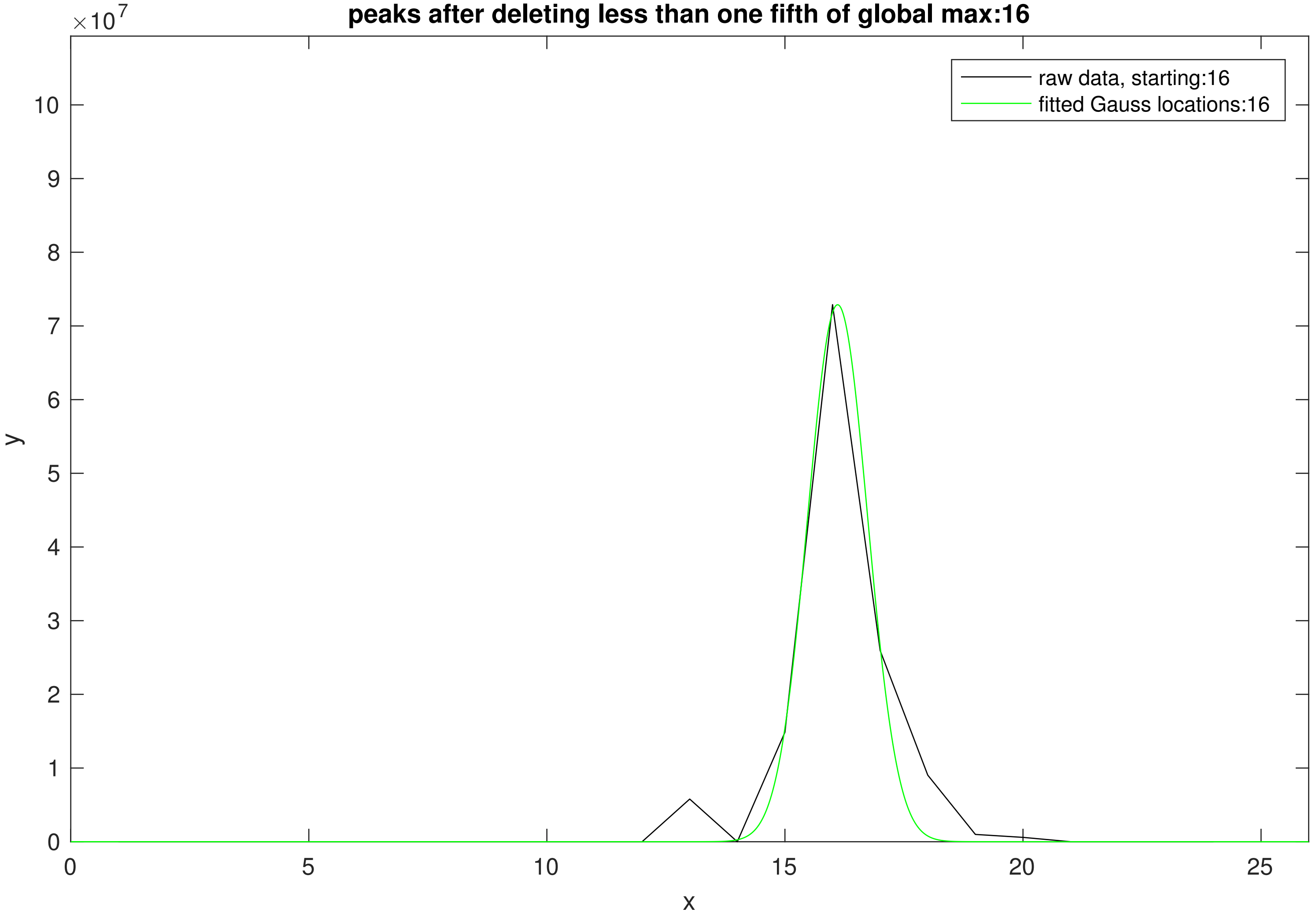

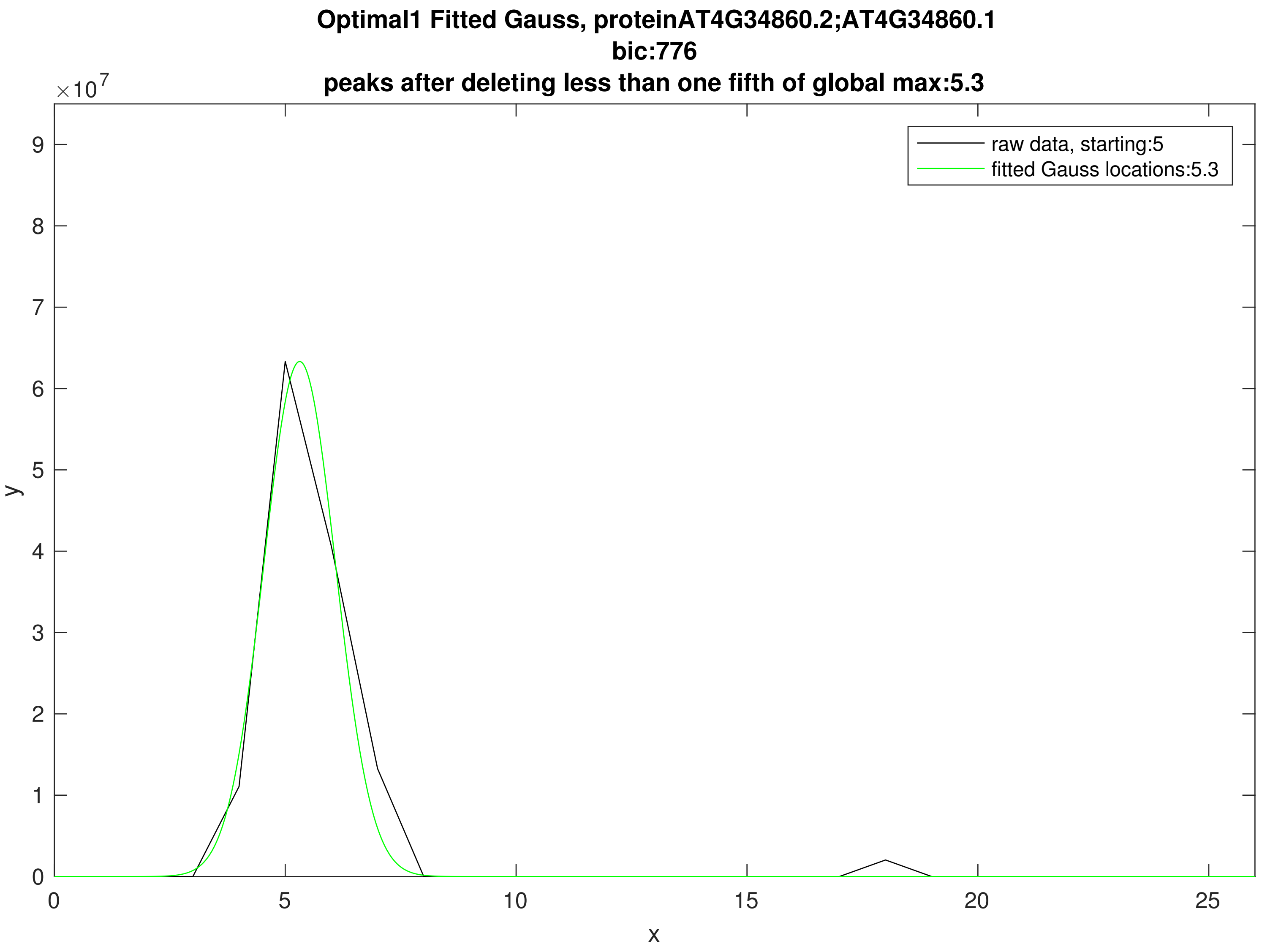

Optimal1 Fitted Gauss, proteinAT4G34860.2;AT4G34860.1
bic:776
peaks after deleting less than one fifth of global max:5.3
×10^7
raw data, starting:5
fitted Gauss locations:5.3
y
x
0
1
2
3
4
5
6
7
8
9
0
5
10
15
20
25

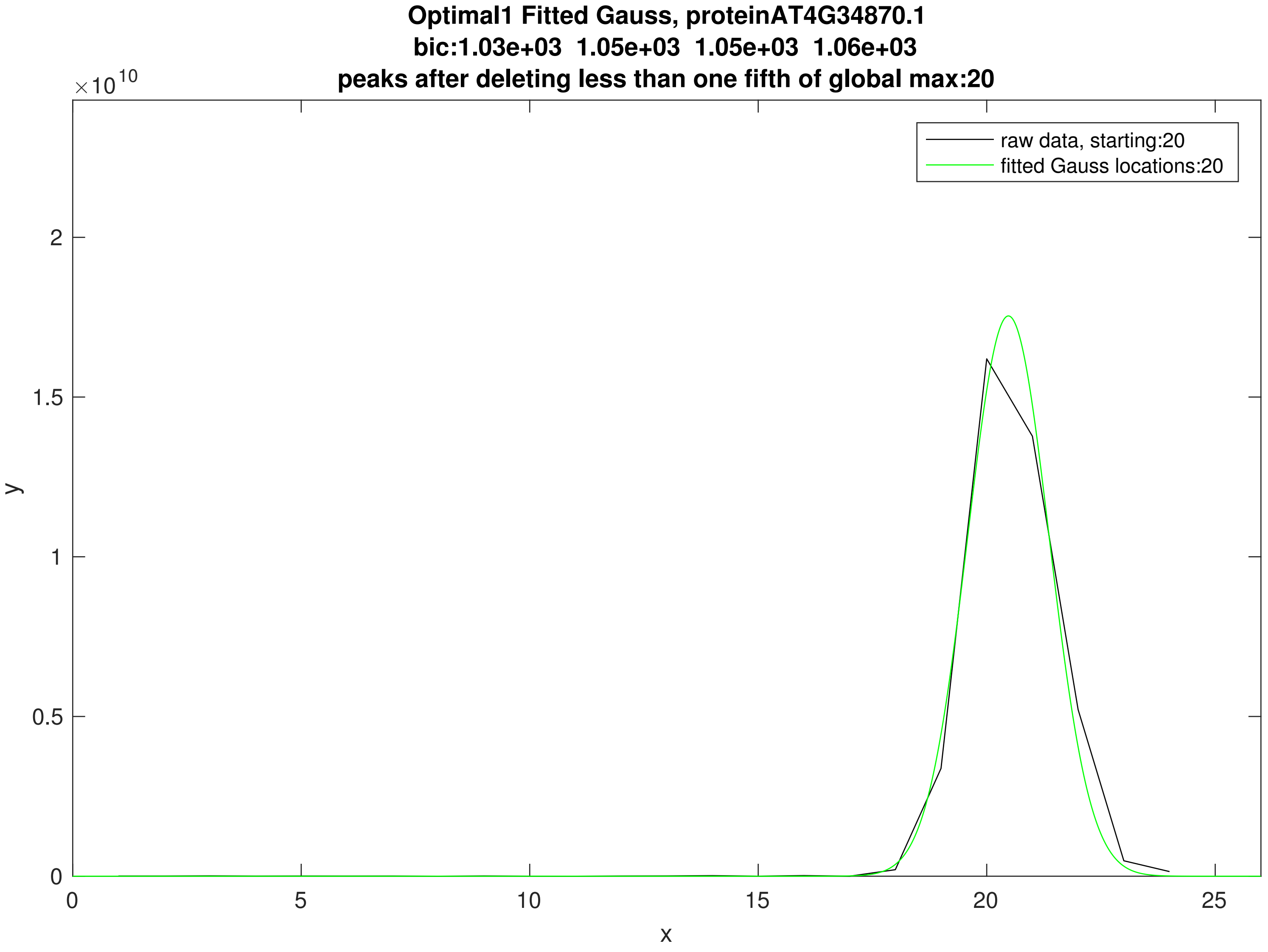
Optimal1 Fitted Gauss, proteinAT4G34870.1
bic:1.03e+03  1.05e+03  1.05e+03  1.06e+03
peaks after deleting less than one fifth of global max:20
×10^10
raw data, starting:20
fitted Gauss locations:20
y
x
0
0.5
1
1.5
2
5
10
15
20
25

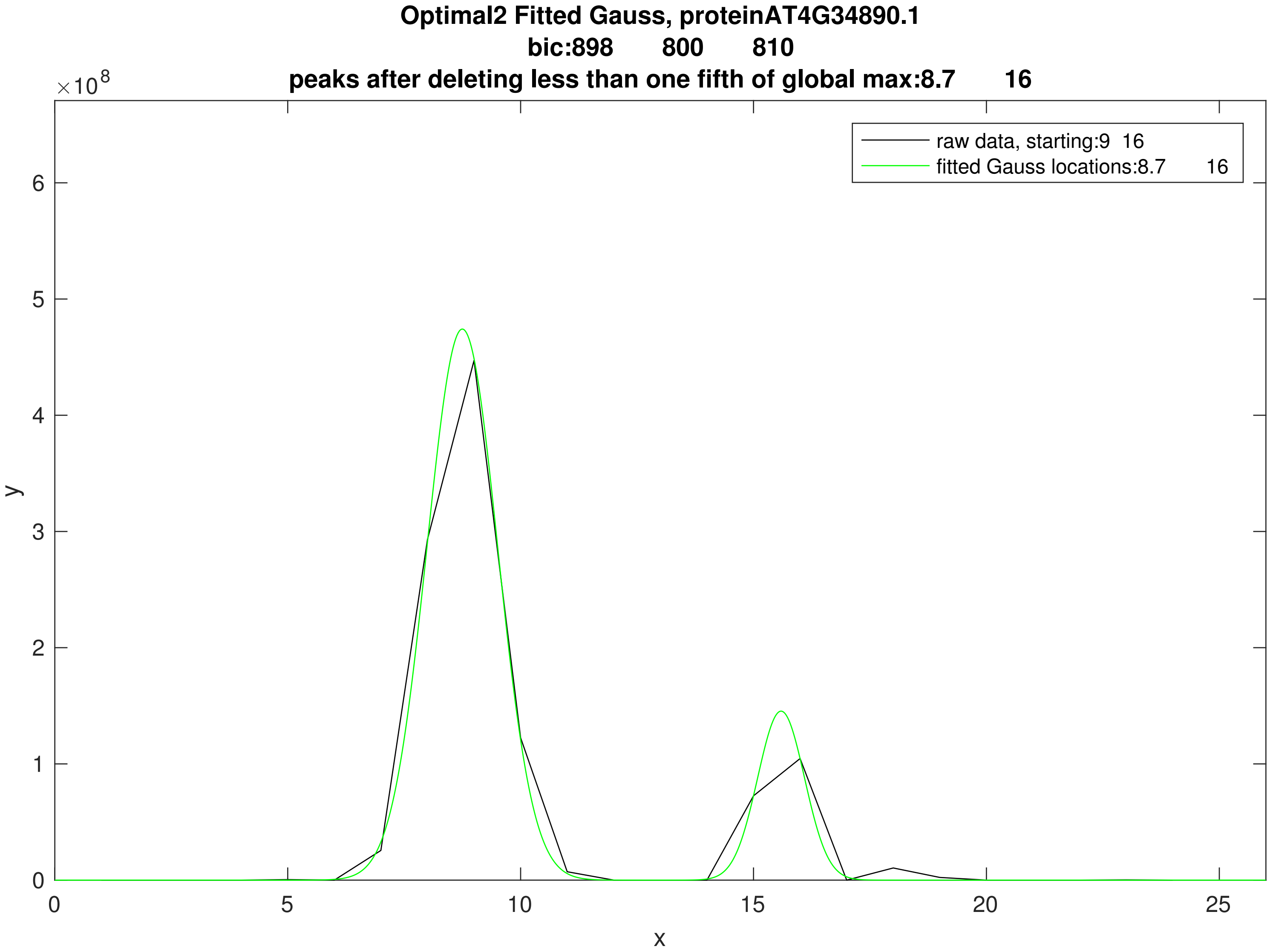

Optimal2 Fitted Gauss, proteinAT4G34890.1
bic:898 800 810
peaks after deleting less than one fifth of global max:8.7 16
×10^8
raw data, starting:9 16
fitted Gauss locations:8.7 16
y
x
0
1
2
3
4
5
6
0
5
10
15
20
25

Optimal1 Fitted Gauss, proteinAT4G34920.1
bic:779
peaks after deleting less than one fifth of global max:18

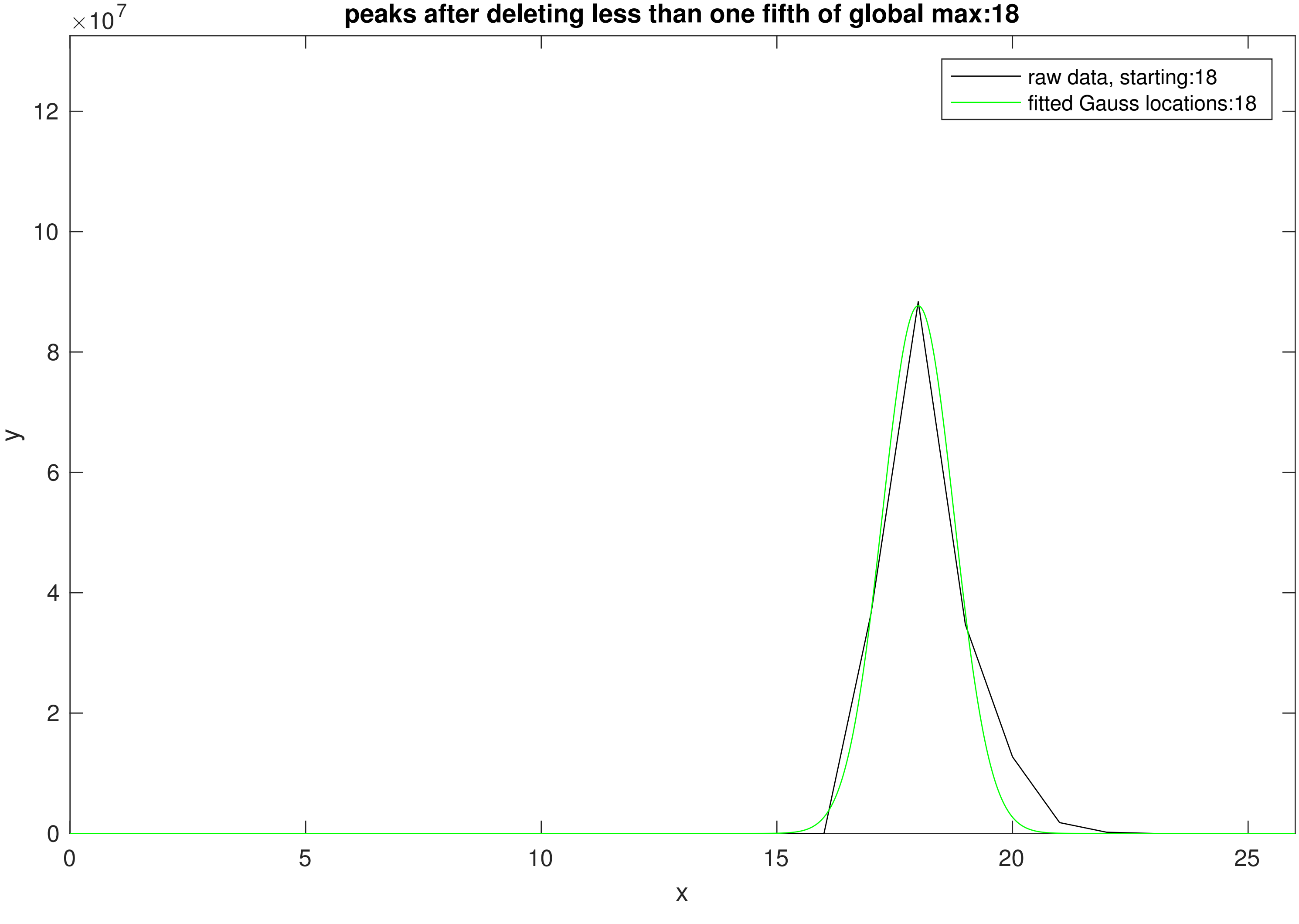

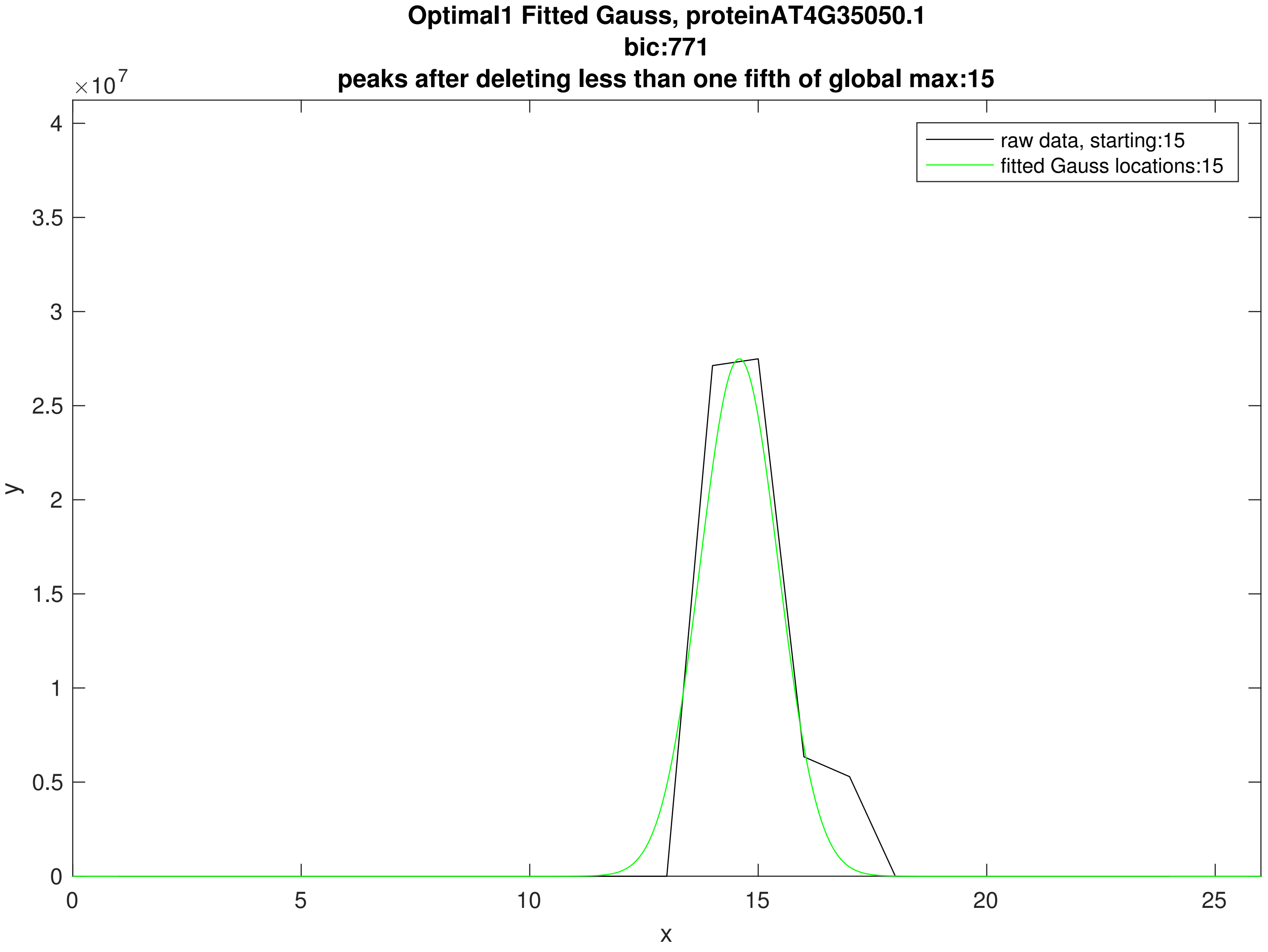

Optimal1 Fitted Gauss, proteinAT4G35050.1
bic:771
peaks after deleting less than one fifth of global max:15
×10^7
raw data, starting:15
fitted Gauss locations:15
y
x

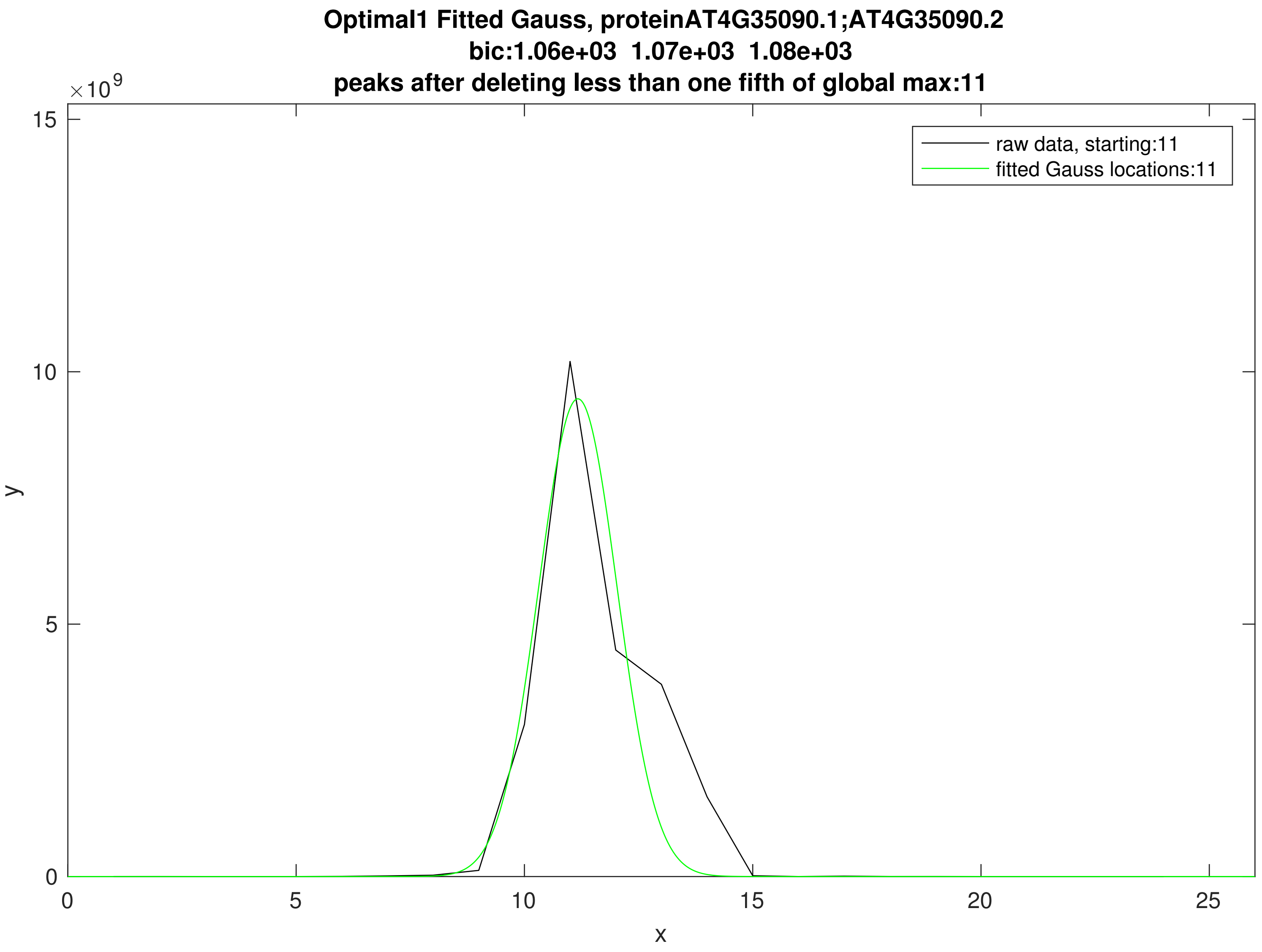
Optimal1 Fitted Gauss, proteinAT4G35090.1;AT4G35090.2
bic:1.06e+03 1.07e+03 1.08e+03
peaks after deleting less than one fifth of global max:11
×10^9
15
10
5
0
y
0
5
10
15
20
25
x
raw data, starting:11
fitted Gauss locations:11

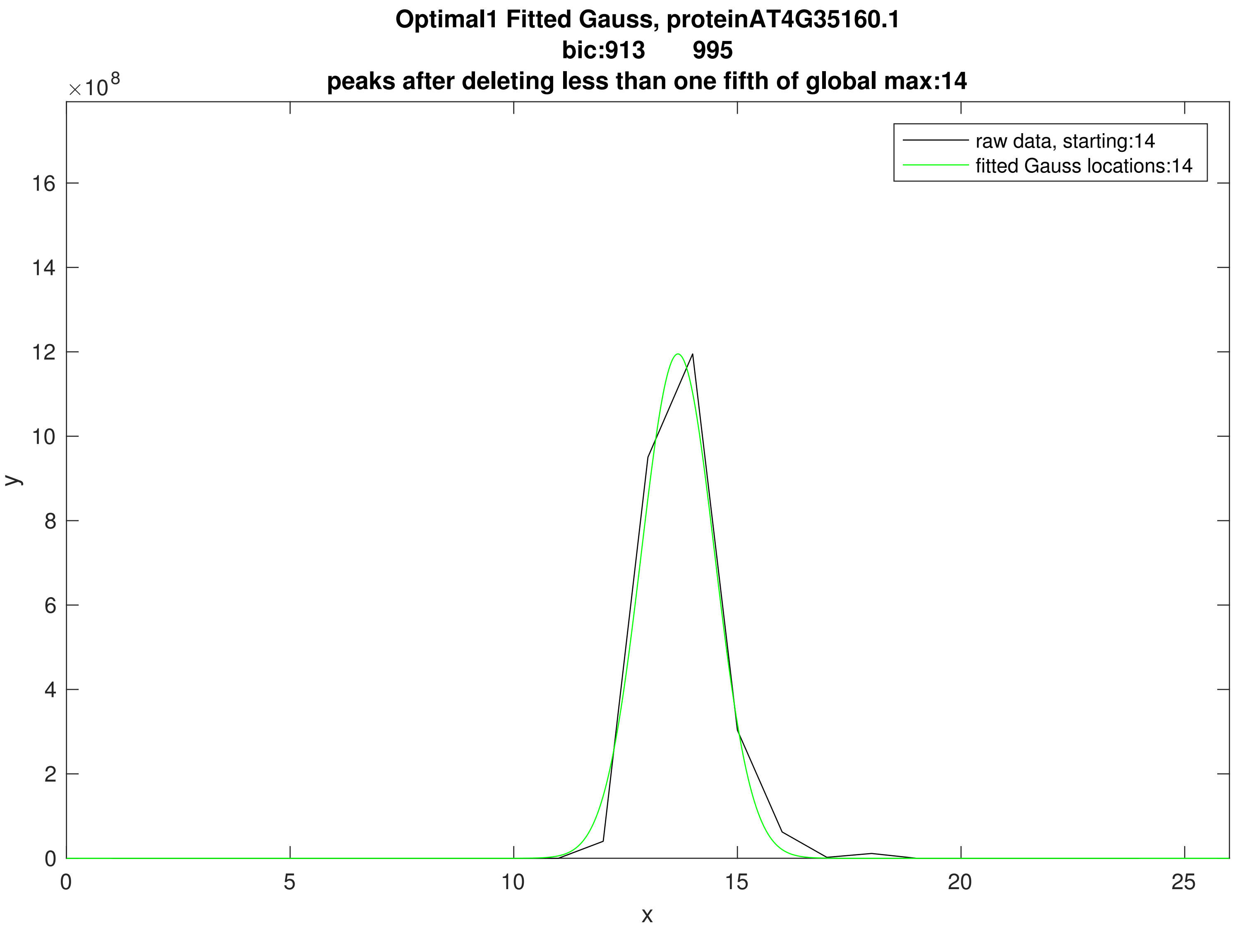
Optimal1 Fitted Gauss, proteinAT4G35160.1
bic:913 995
peaks after deleting less than one fifth of global max:14
×10^8
raw data, starting:14
fitted Gauss locations:14
16
14
12
10
8
6
4
2
0
y
0
5
10
15
20
25
x

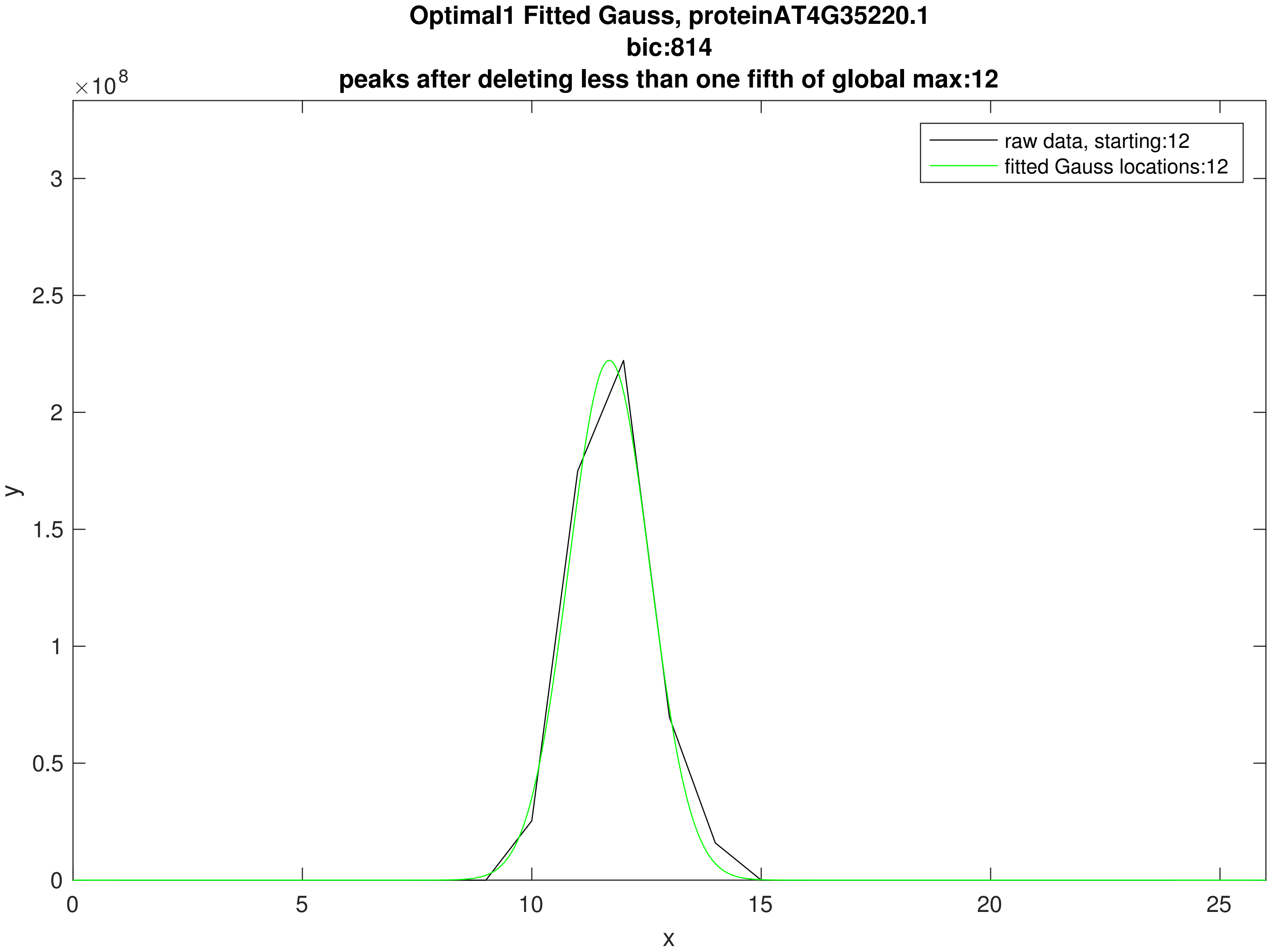
Optimal1 Fitted Gauss, proteinAT4G35220.1
bic:814
peaks after deleting less than one fifth of global max:12
×10^8
raw data, starting:12
fitted Gauss locations:12
y
x
0
0.5
1
1.5
2
2.5
3
0
5
10
15
20
25

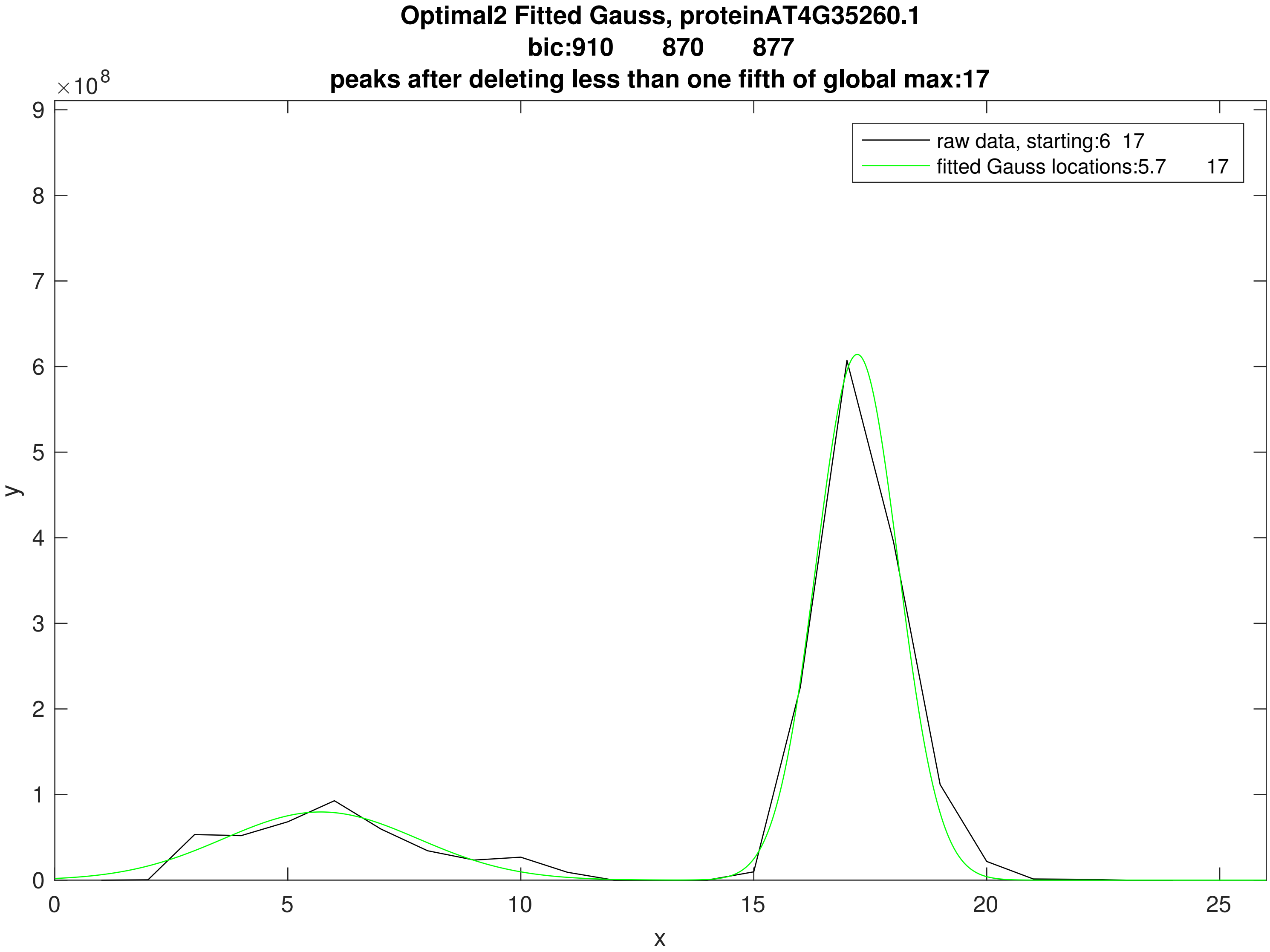
Optimal2 Fitted Gauss, proteinAT4G35260.1
bic:910 870 877
peaks after deleting less than one fifth of global max:17
×10^8
raw data, starting:6 17
fitted Gauss locations:5.7 17
y
x
0
1
2
3
4
5
6
7
8
9
0
5
10
15
20
25

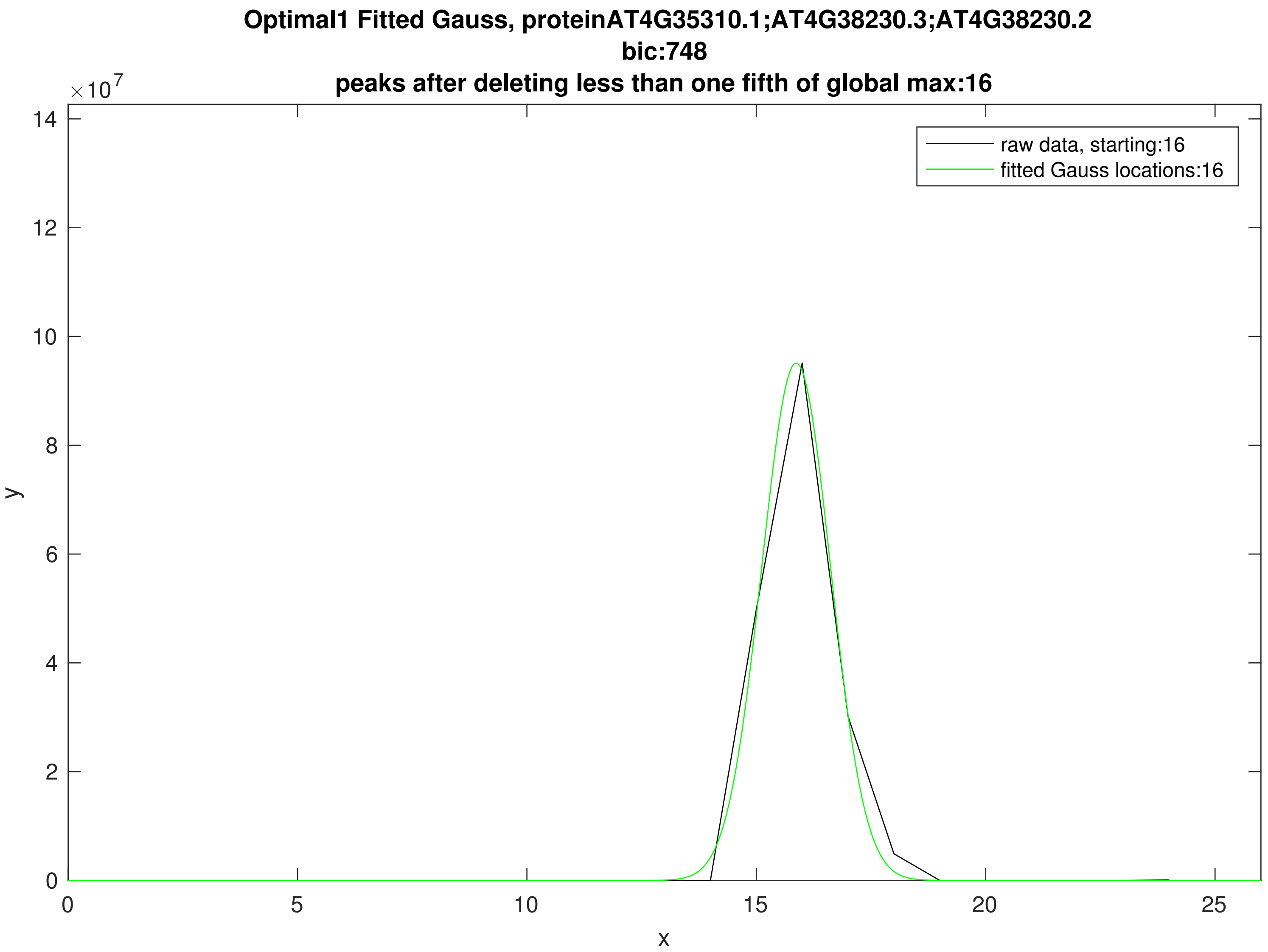

Optimal1 Fitted Gauss, proteinAT4G35310.1;AT4G38230.3;AT4G38230.2
bic:748
peaks after deleting less than one fifth of global max:16
×10^7
14
12
10
8
6
4
2
0
y
0
5
10
15
20
25
x
raw data, starting:16
fitted Gauss locations:16

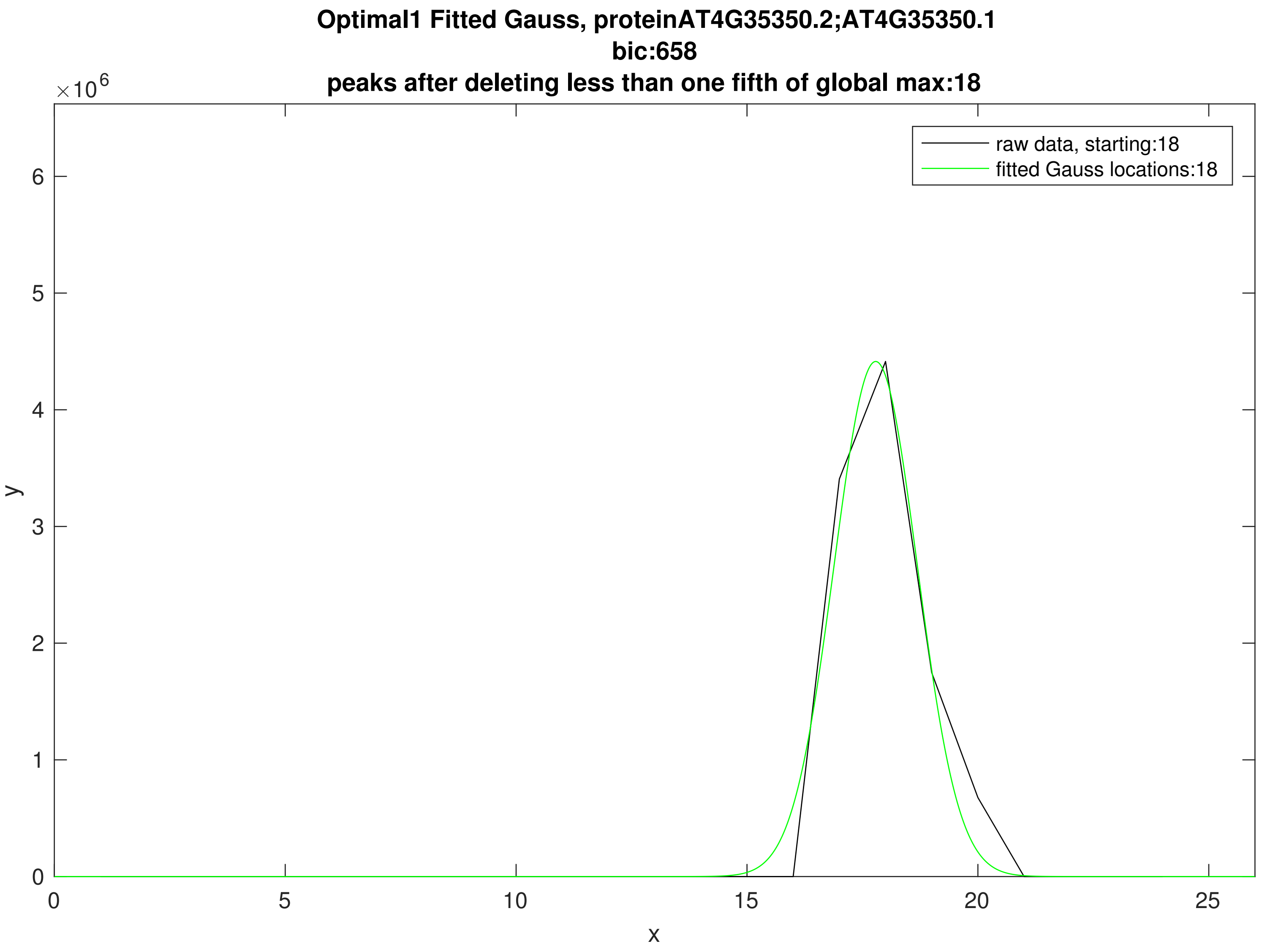
Optimal1 Fitted Gauss, proteinAT4G35350.2;AT4G35350.1
bic:658
peaks after deleting less than one fifth of global max:18
×10^6
y
x
raw data, starting:18
fitted Gauss locations:18
0
1
2
3
4
5
6
0
5
10
15
20
25

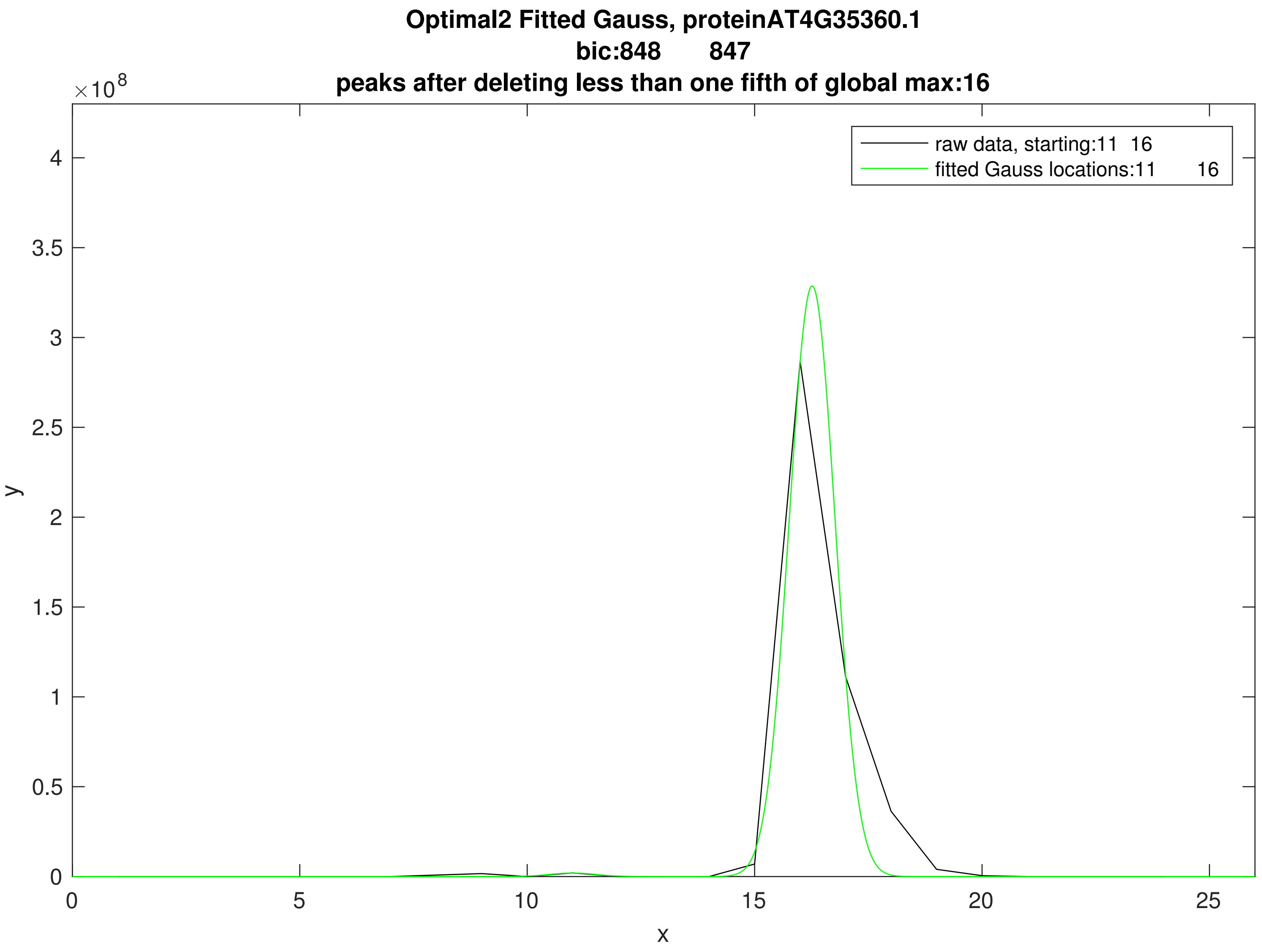

Optimal2 Fitted Gauss, proteinAT4G35360.1
bic:848 847
peaks after deleting less than one fifth of global max:16
×10^8
raw data, starting:11 16
fitted Gauss locations:11 16
y
x
0
0.5
1
1.5
2
2.5
3
3.5
4
5
10
15
20
25

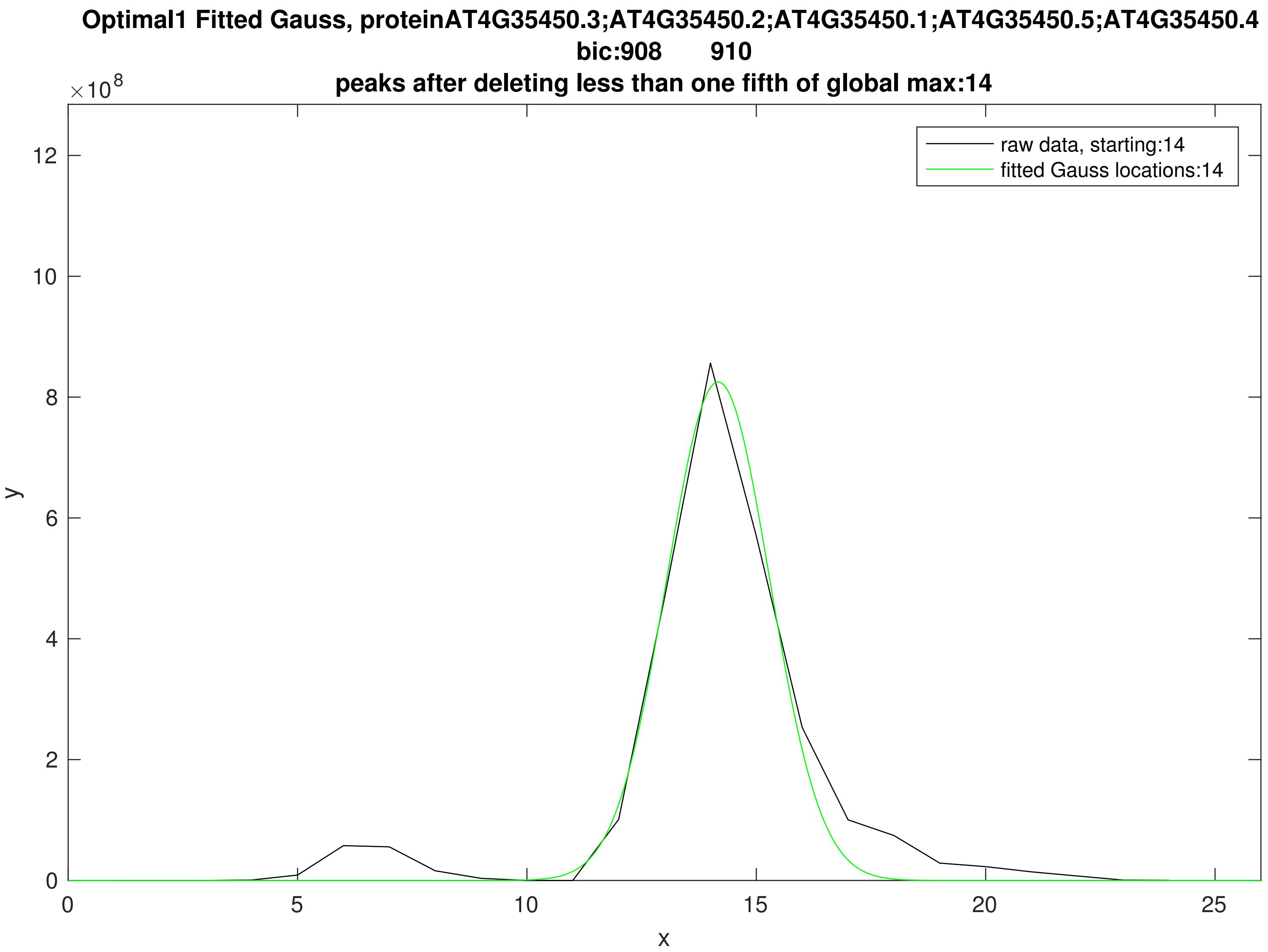
Optimal1 Fitted Gauss, proteinAT4G35450.3;AT4G35450.2;AT4G35450.1;AT4G35450.5;AT4G35450.4
bic:908 910
peaks after deleting less than one fifth of global max:14
×10^8
raw data, starting:14
fitted Gauss locations:14
y
x
0
2
4
6
8
10
12
0
5
10
15
20
25

**Optimal1 Fitted Gauss, proteinAT4G35460.1**
**bic:872**
**peaks after deleting less than one fifth of global max:14**

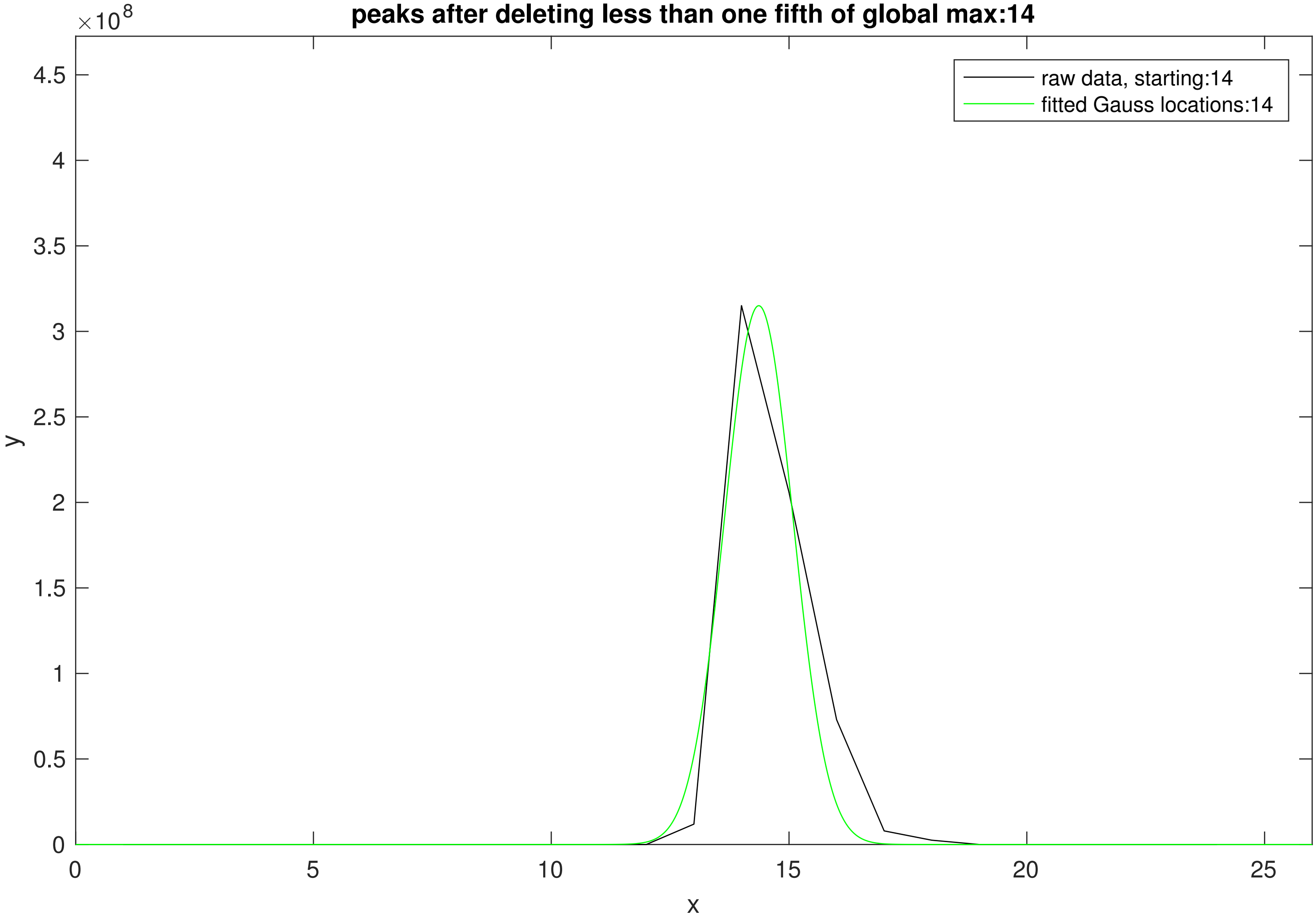

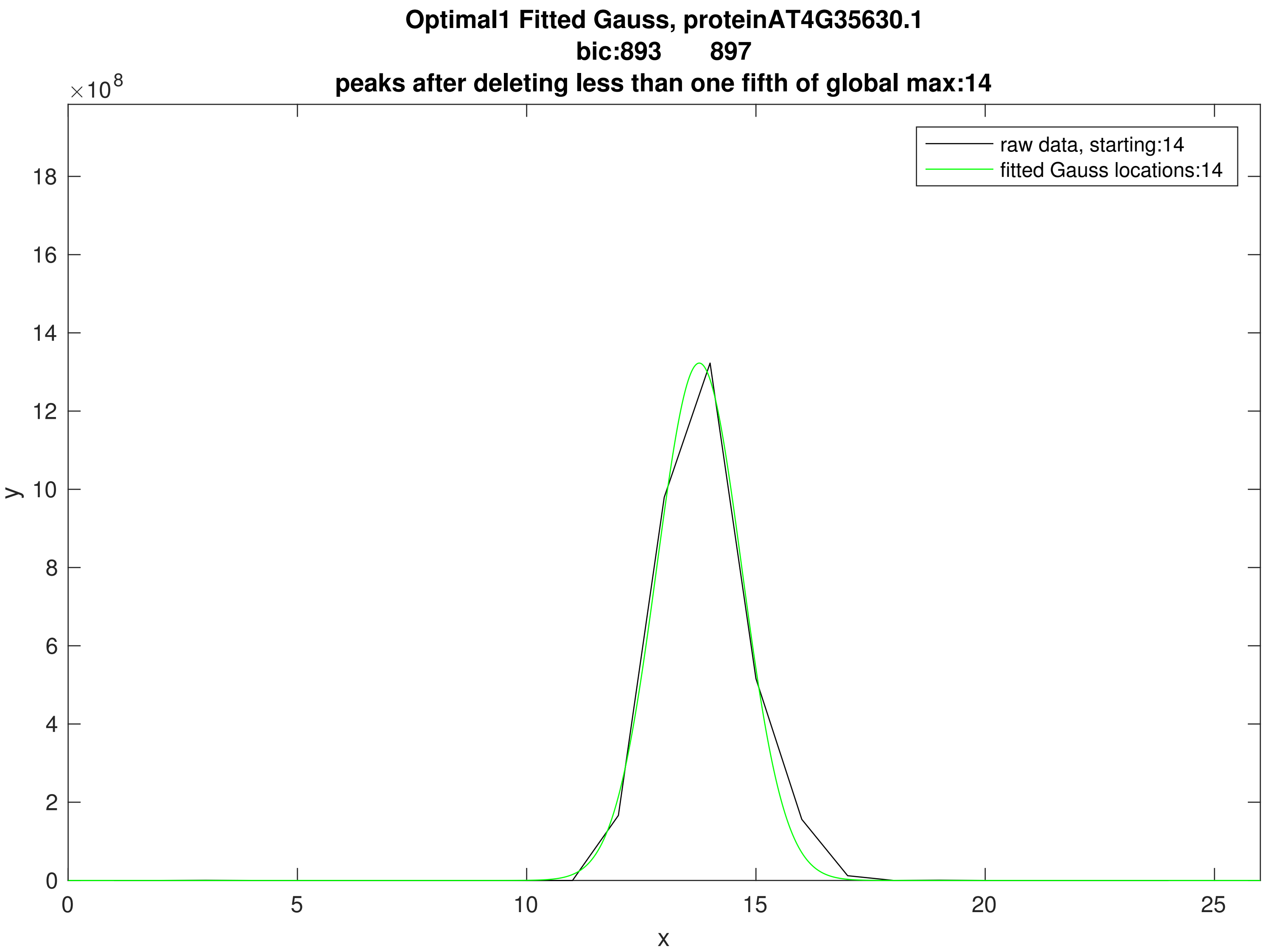

Optimal1 Fitted Gauss, proteinAT4G35630.1
bic:893 897
peaks after deleting less than one fifth of global max:14
×10^8
raw data, starting:14
fitted Gauss locations:14
y
x
0
2
4
6
8
10
12
14
16
18
5
15
20
25

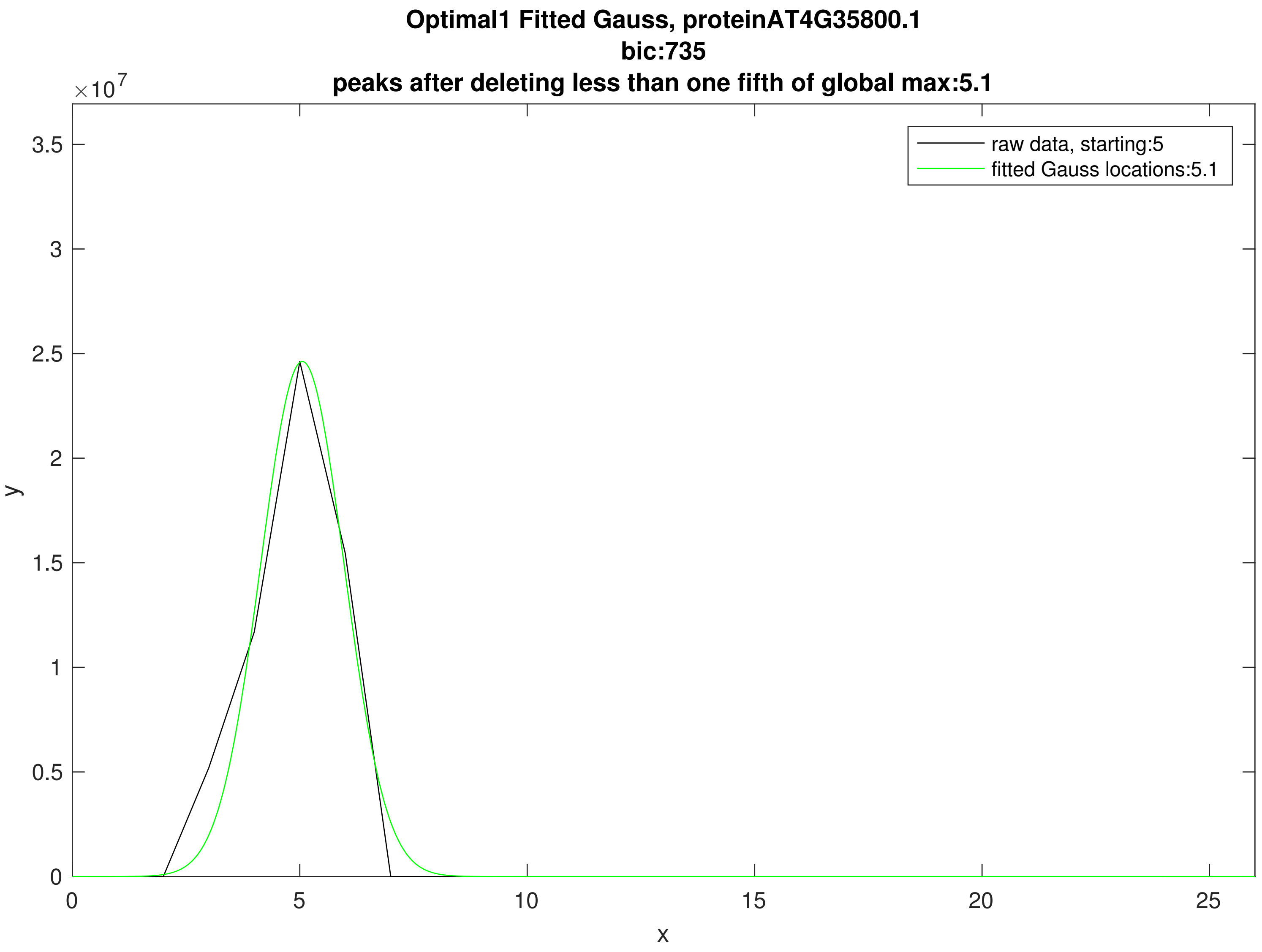
Optimal1 Fitted Gauss, proteinAT4G35800.1
bic:735
peaks after deleting less than one fifth of global max:5.1
×10^7
3.5
3
2.5
2
1.5
1
0.5
0
y
0
5
10
15
20
25
x
raw data, starting:5
fitted Gauss locations:5.1

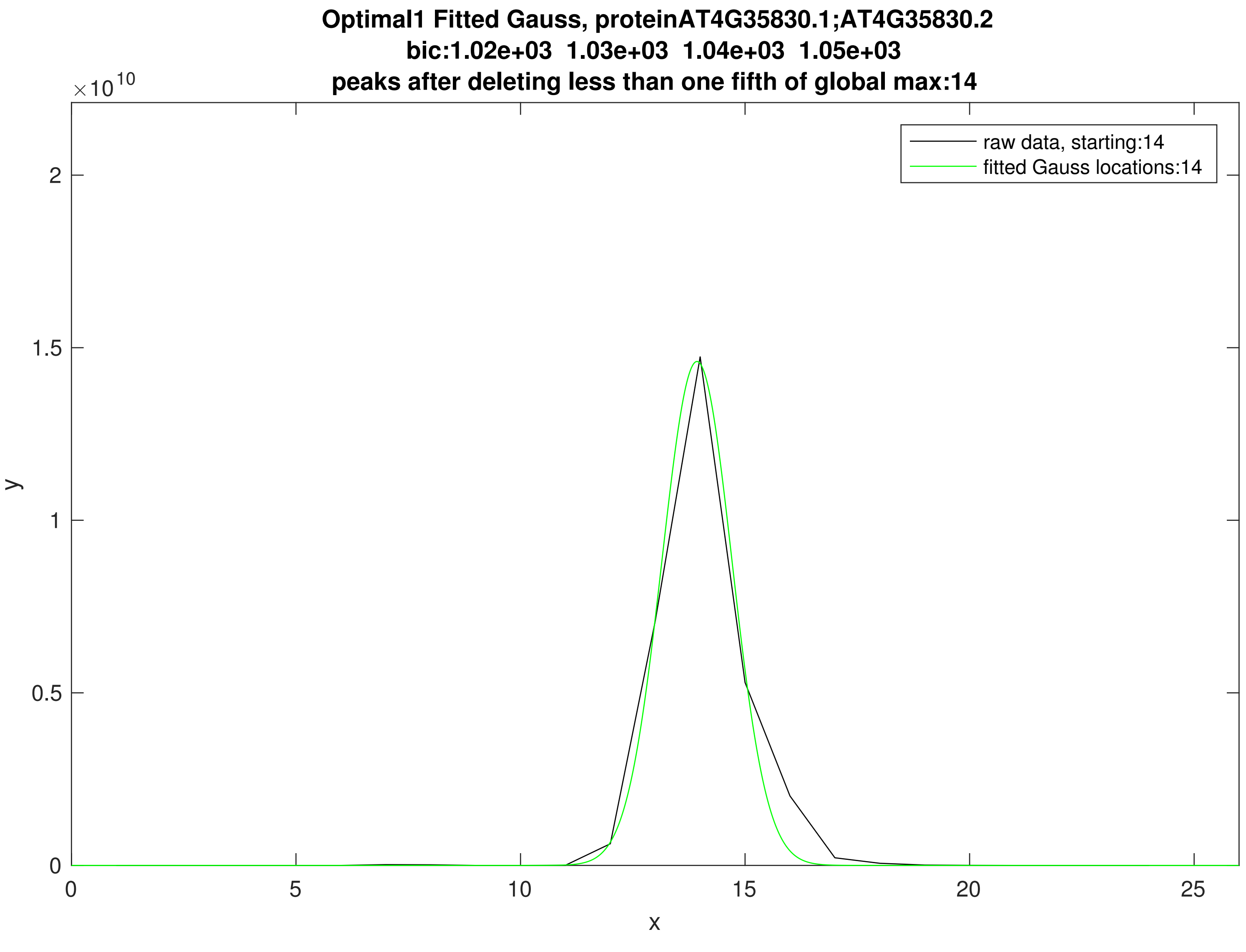
Optimal1 Fitted Gauss, proteinAT4G35830.1;AT4G35830.2
bic:1.02e+03 1.03e+03 1.04e+03 1.05e+03
peaks after deleting less than one fifth of global max:14
×10^10
raw data, starting:14
fitted Gauss locations:14
2
1.5
1
0.5
0
y
0
5
10
15
20
25
x

Optimal2 Fitted Gauss, proteinAT4G36010.2;AT4G36010.1
bic:725 724
peaks after deleting less than one fifth of global max:18

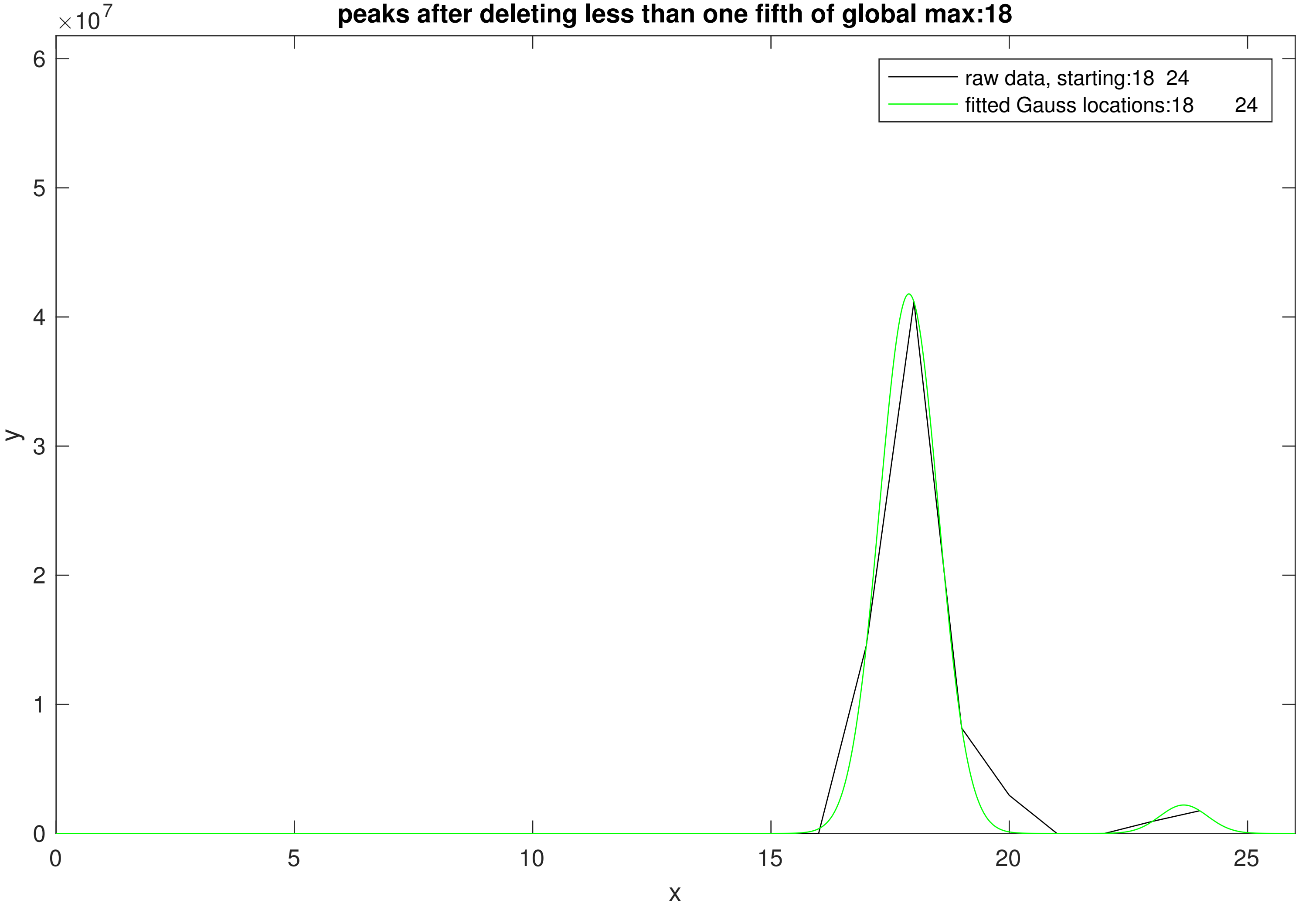

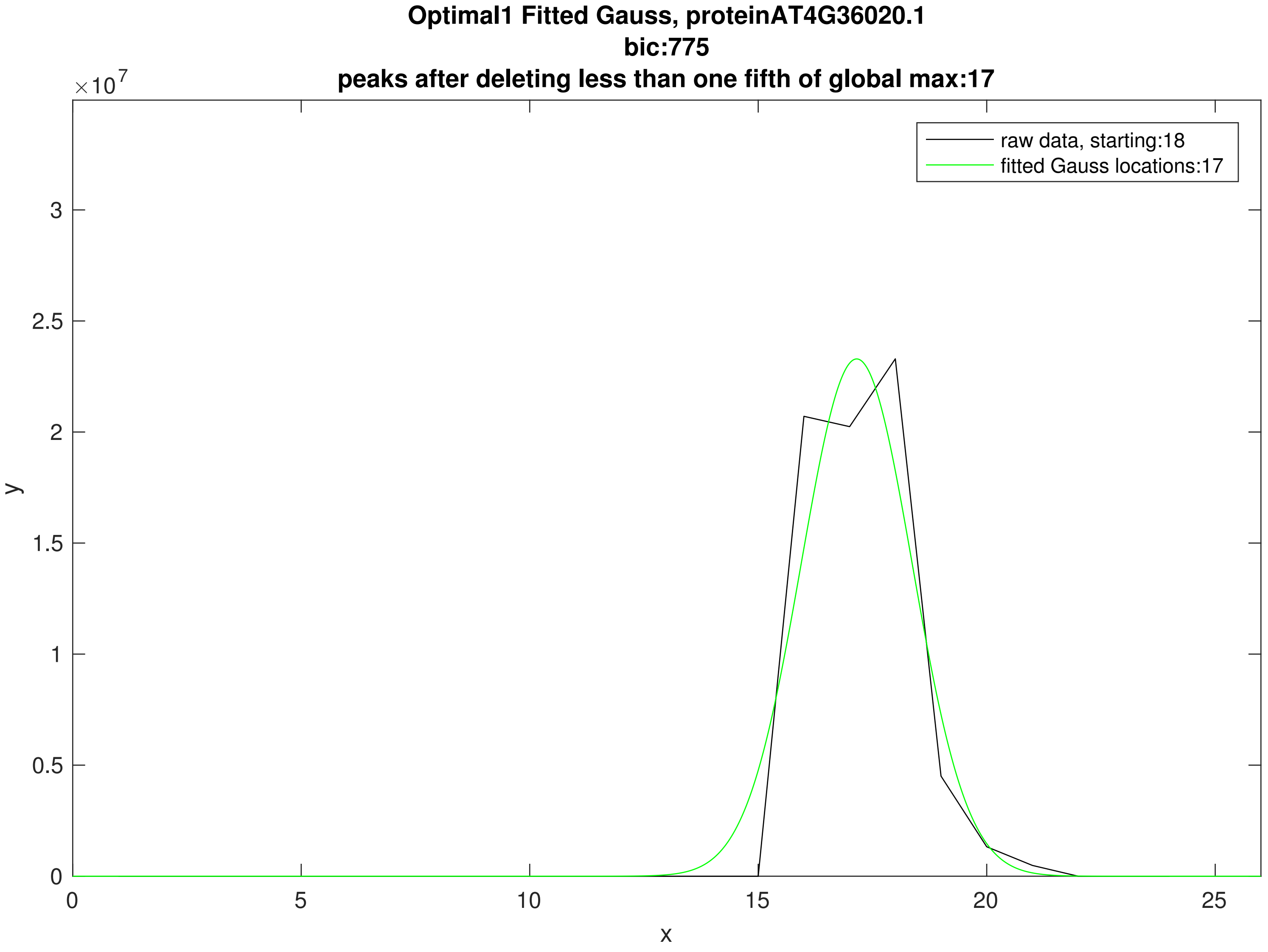
Optimal1 Fitted Gauss, proteinAT4G36020.1
bic:775
peaks after deleting less than one fifth of global max:17
×10^7
raw data, starting:18
fitted Gauss locations:17
3
2.5
2
1.5
1
0.5
0
5
10
15
20
25
x
y

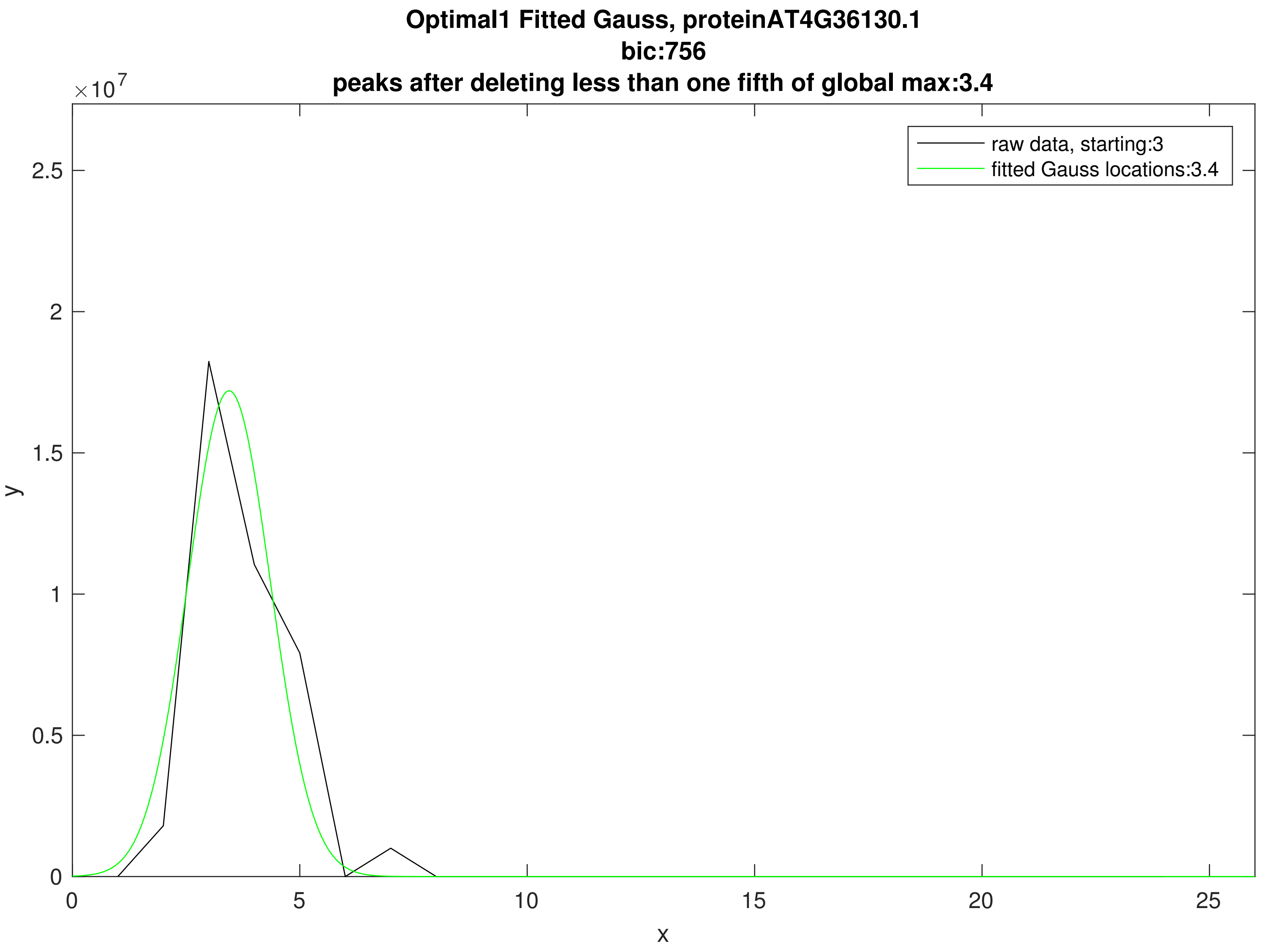
Optimal1 Fitted Gauss, proteinAT4G36130.1
bic:756
peaks after deleting less than one fifth of global max:3.4
×10^7
raw data, starting:3
fitted Gauss locations:3.4
y
x
0
0.5
1
1.5
2
2.5
5
10
15
20
25

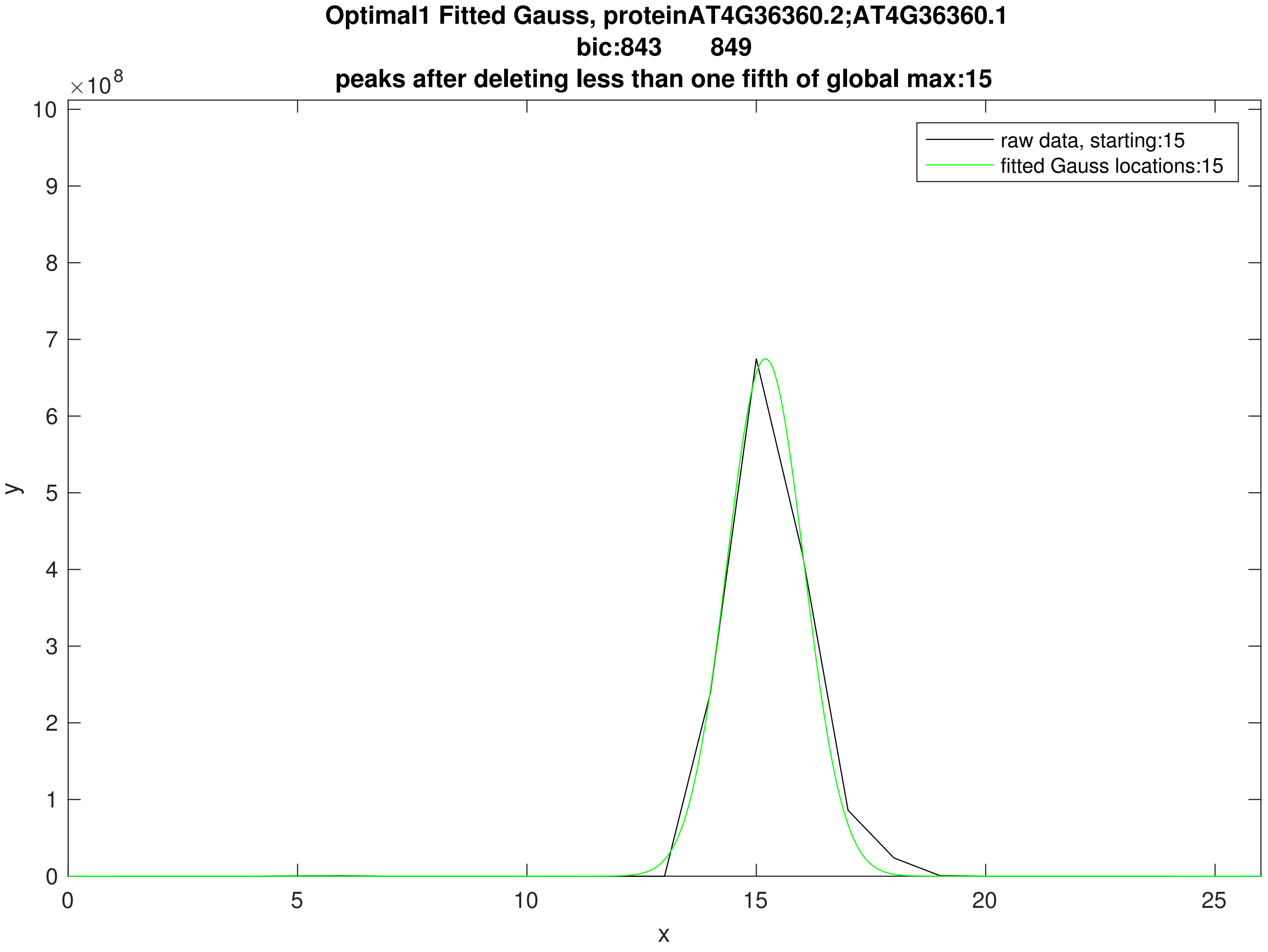

Optimal1 Fitted Gauss, proteinAT4G36360.2;AT4G36360.1
bic:843 849
peaks after deleting less than one fifth of global max:15
×10^8
raw data, starting:15
fitted Gauss locations:15
y
x
0
1
2
3
4
5
6
7
8
9
10
0
5
10
15
20
25

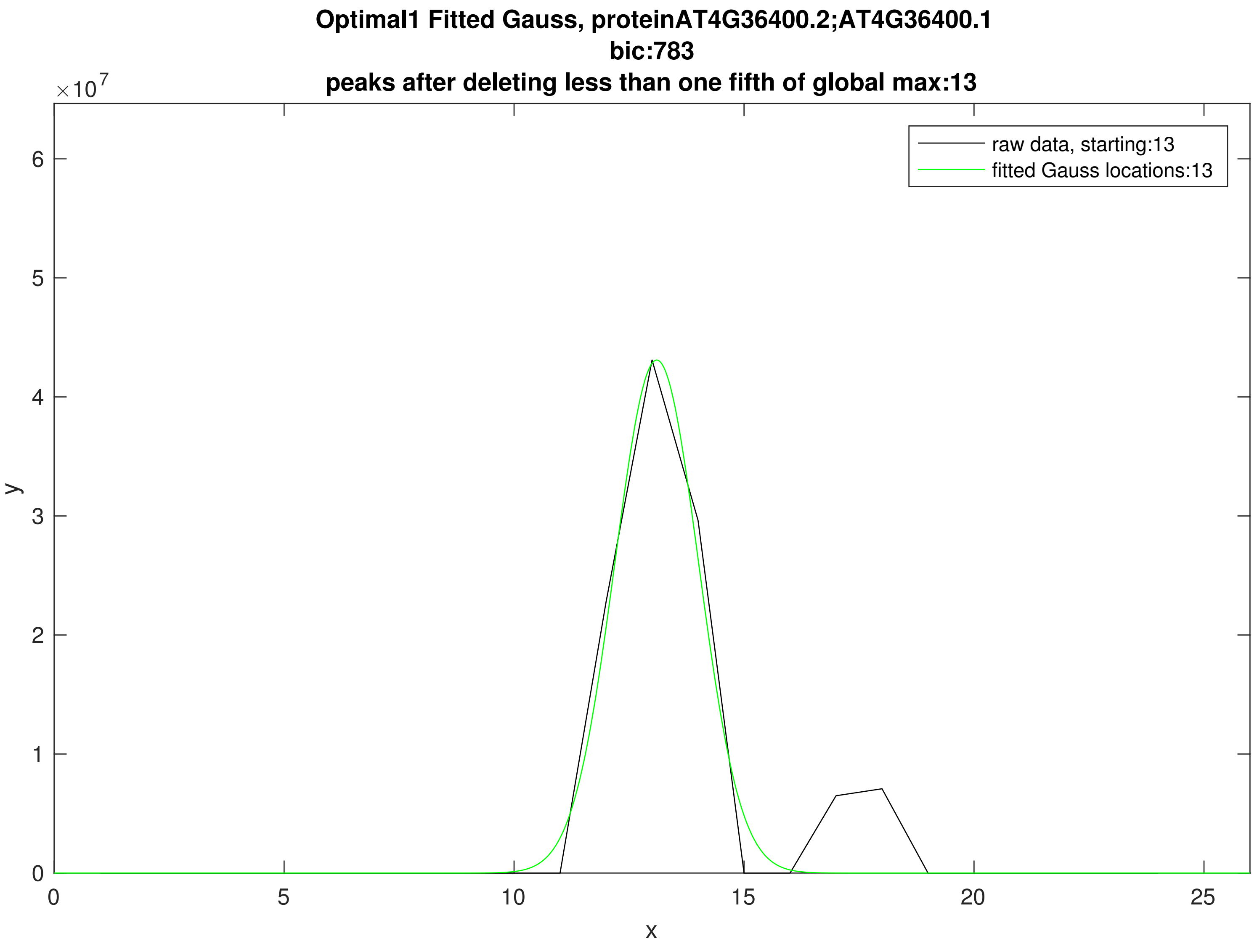
Optimal1 Fitted Gauss, proteinAT4G36400.2;AT4G36400.1
bic:783
peaks after deleting less than one fifth of global max:13
×10^7
raw data, starting:13
fitted Gauss locations:13
y
x
0
1
2
3
4
5
6
0
5
10
15
20
25

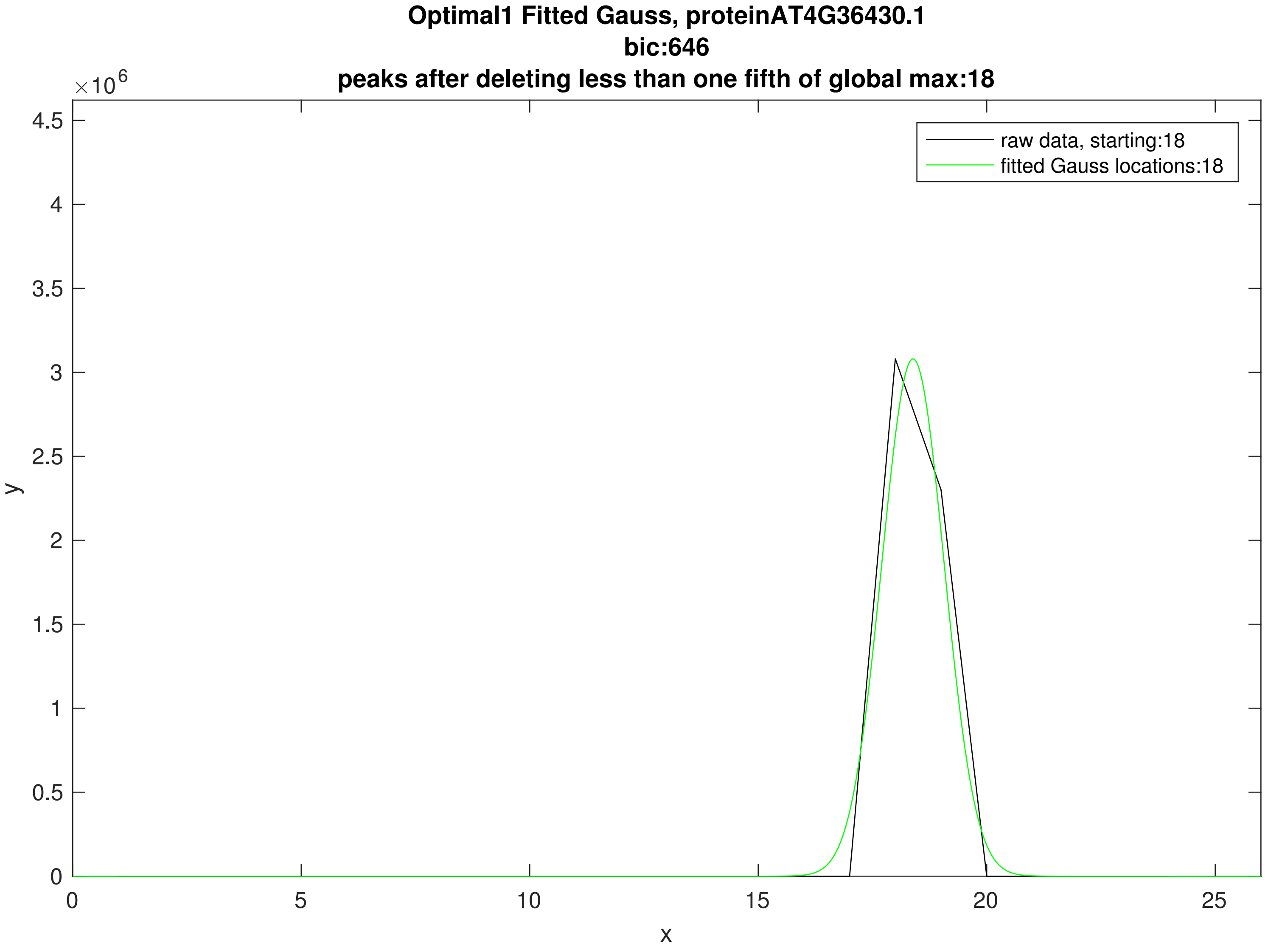

Optimal1 Fitted Gauss, proteinAT4G36430.1
bic:646
peaks after deleting less than one fifth of global max:18
×10^6
4.5
4
3.5
3
2.5
2
1.5
1
0.5
0
y
0
5
10
15
20
25
x
raw data, starting:18
fitted Gauss locations:18

**Optimal3 Fitted Gauss, proteinAT4G36700.1**
**bic:870 830 829**
**peaks after deleting less than one fifth of global max:5.9**

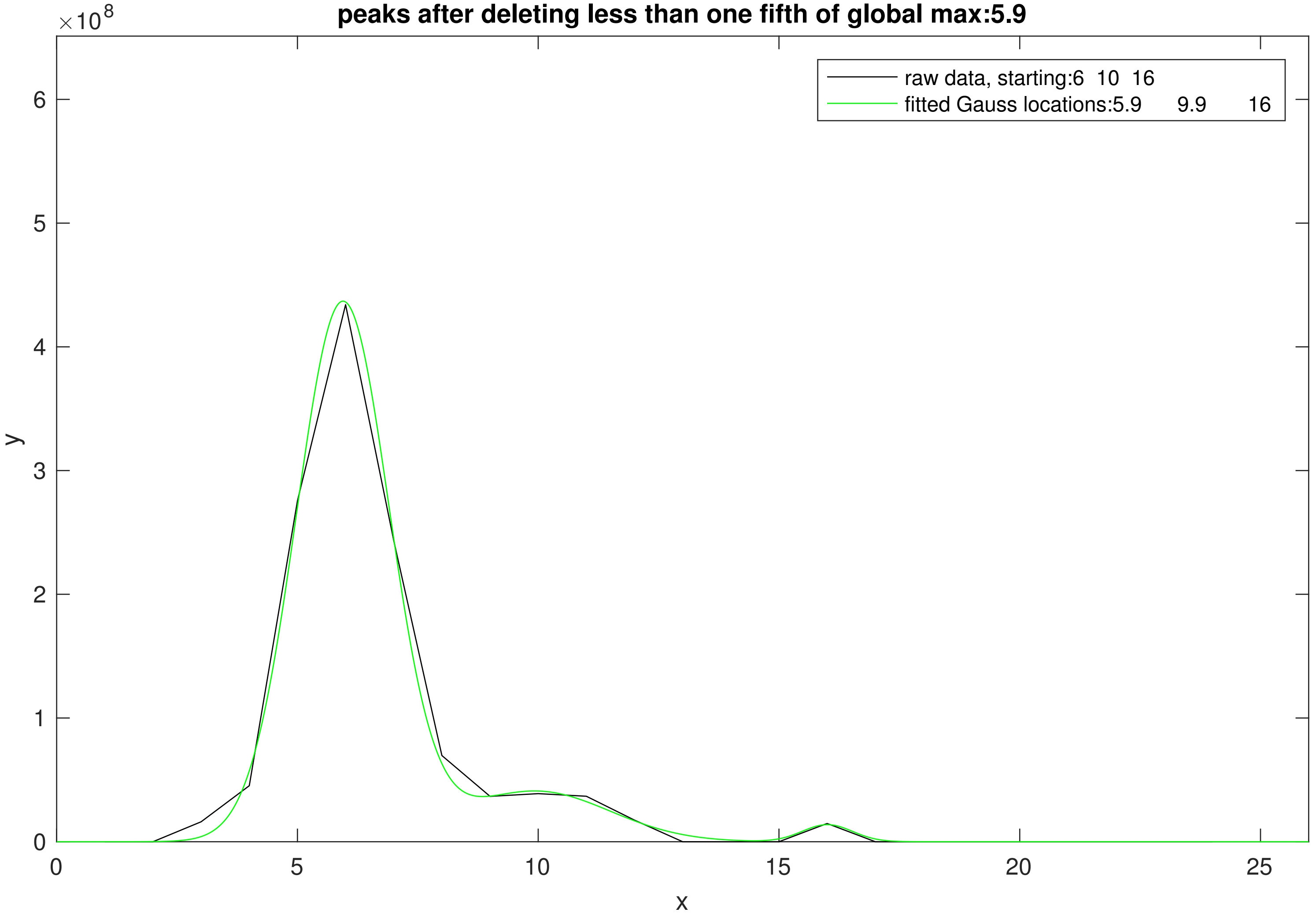

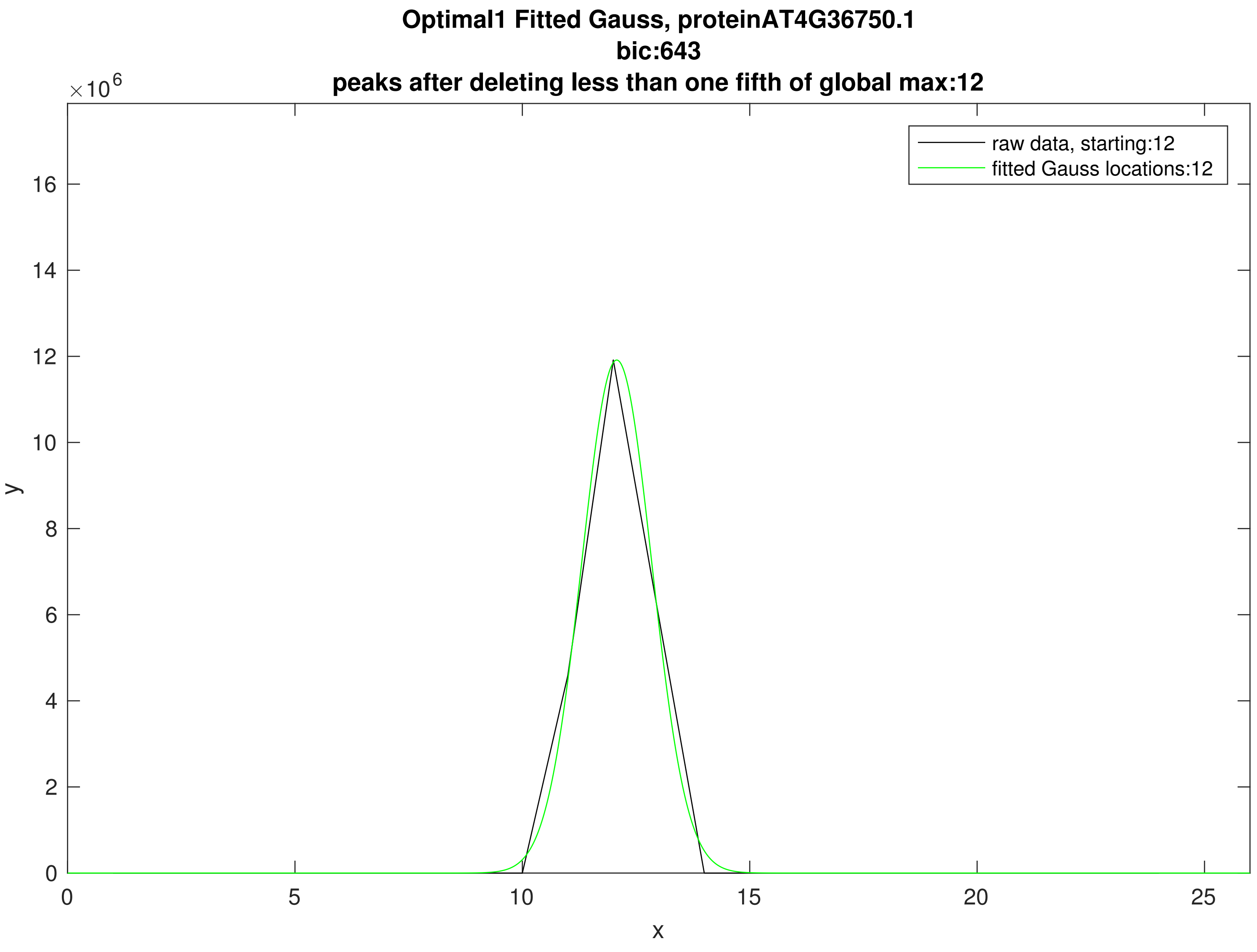

Optimal1 Fitted Gauss, proteinAT4G36750.1
bic:643
peaks after deleting less than one fifth of global max:12
×10^6
raw data, starting:12
fitted Gauss locations:12
16
14
12
10
8
6
4
2
0
y
0
5
10
15
20
25
x

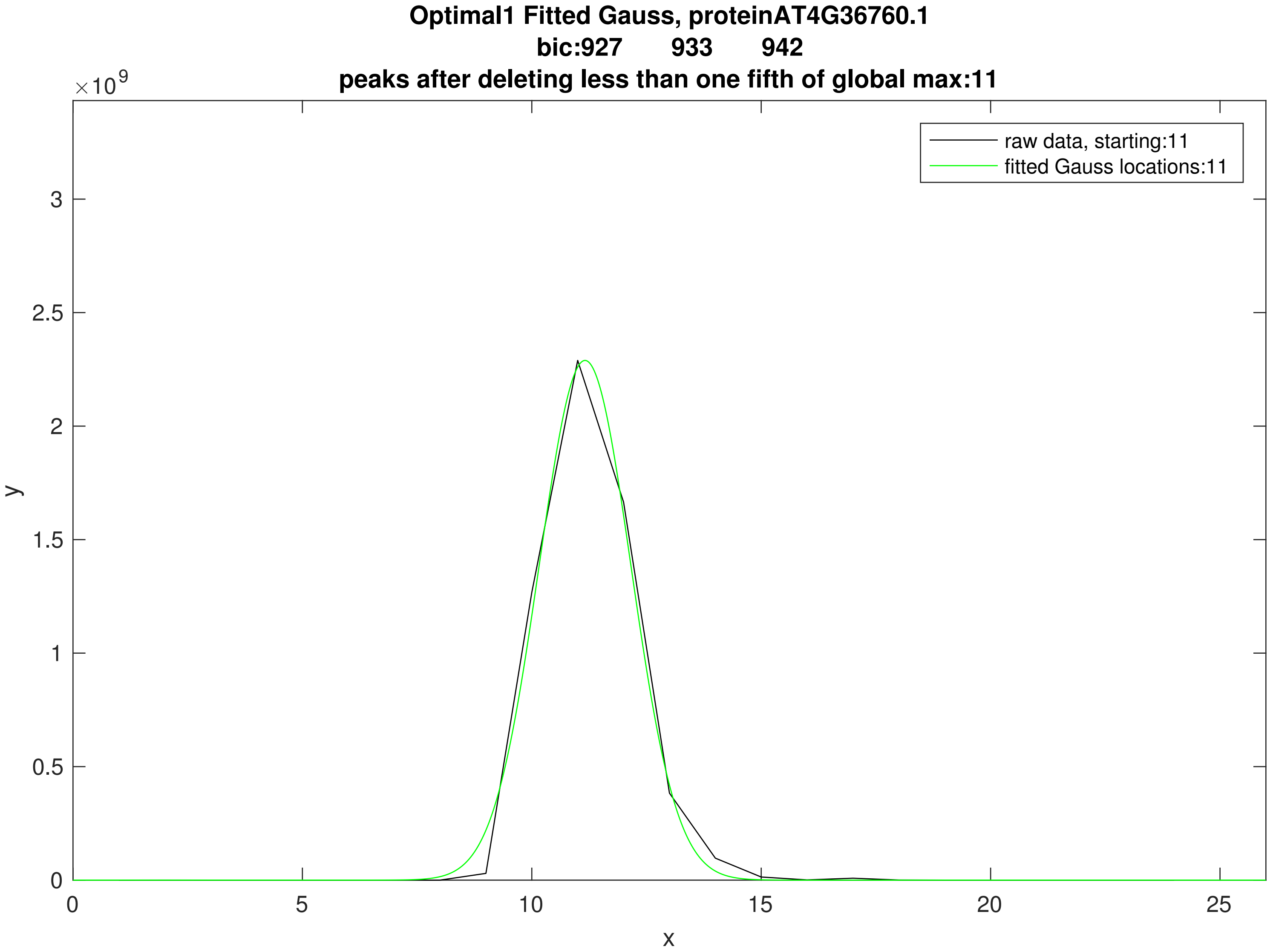

Optimal1 Fitted Gauss, proteinAT4G36760.1
bic:927 933 942
peaks after deleting less than one fifth of global max:11
×10^9
raw data, starting:11
fitted Gauss locations:11
y
x
0
0.5
1
1.5
2
2.5
3
0
5
10
15
20
25

**Optimal1 Fitted Gauss, proteinAT4G36880.1**
**bic:749**
**peaks after deleting less than one fifth of global max:19**

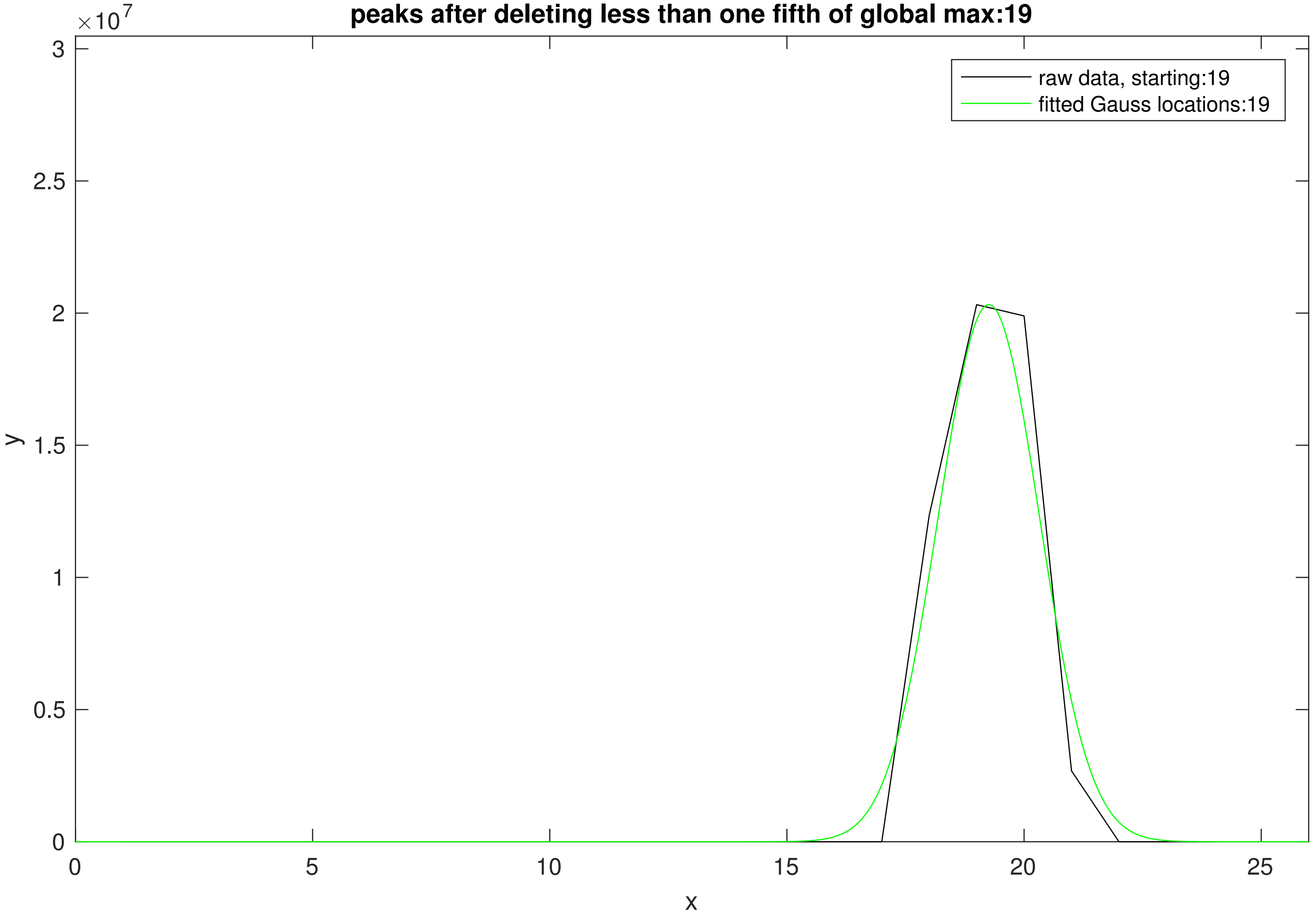

**Optimal1 Fitted Gauss, proteinAT4G36910.1**
**bic:791 791**
**peaks after deleting less than one fifth of global max:19**

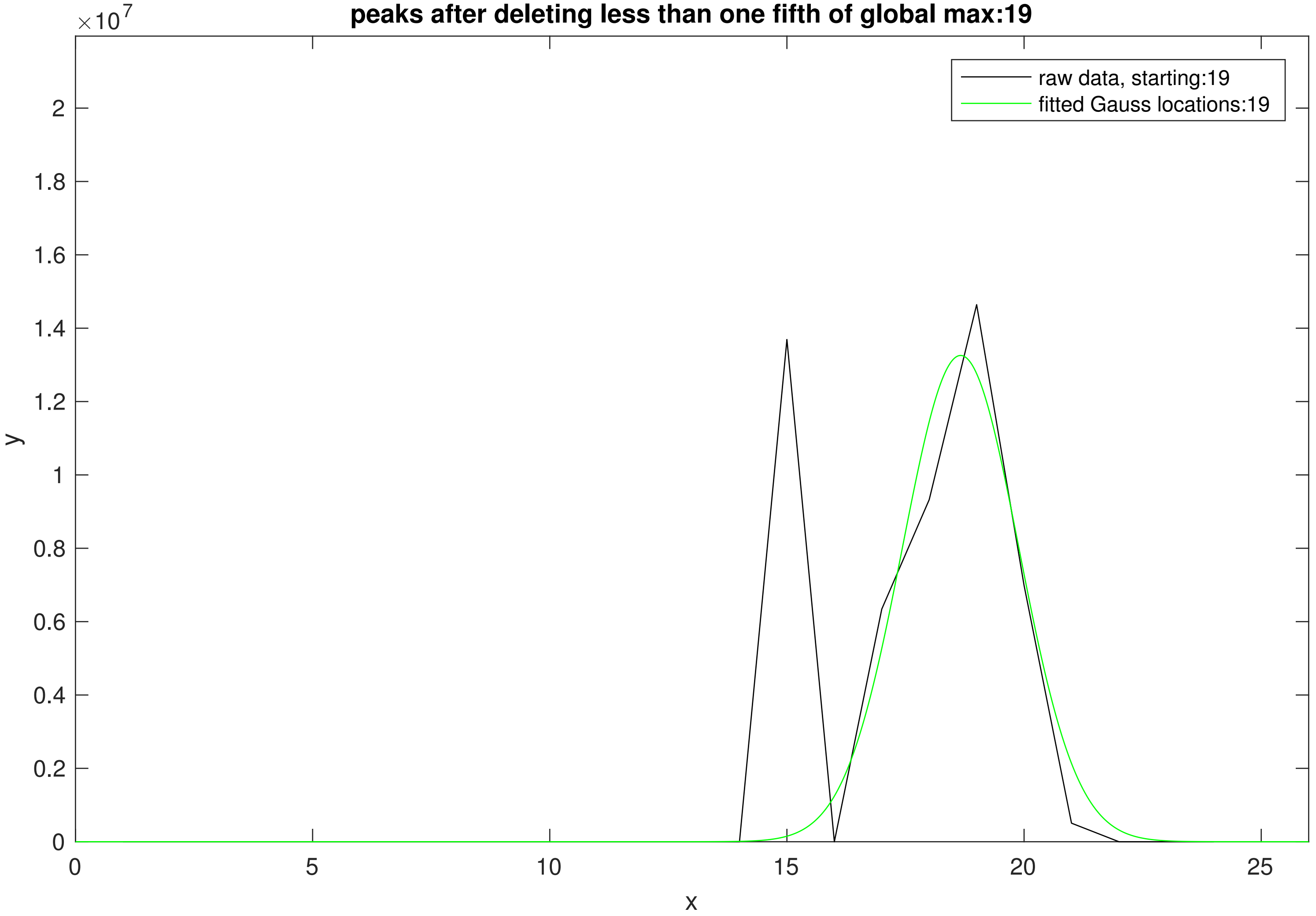

**Optimal1 Fitted Gauss, proteinAT4G36940.1**
**bic:810**
**peaks after deleting less than one fifth of global max:12**

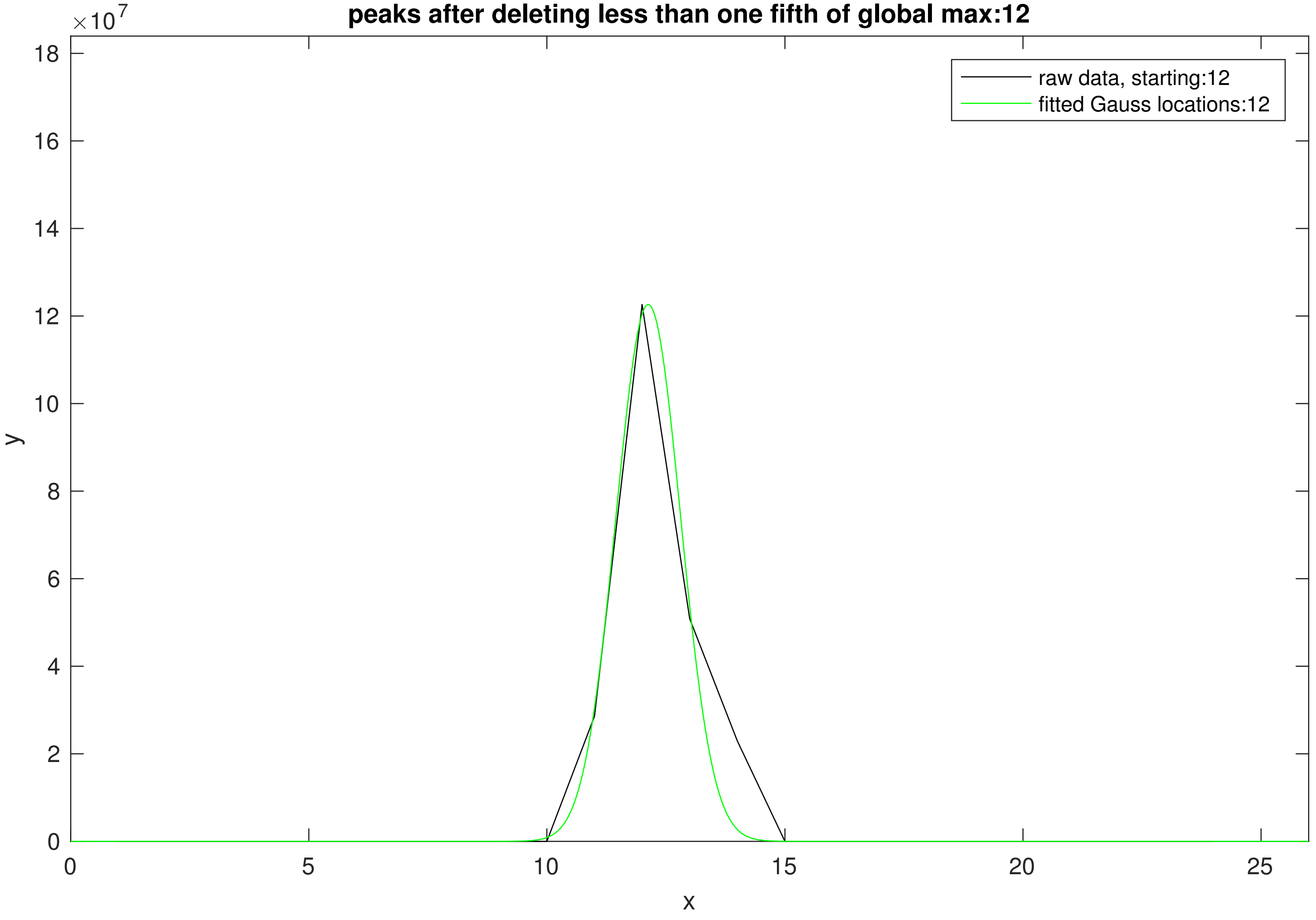

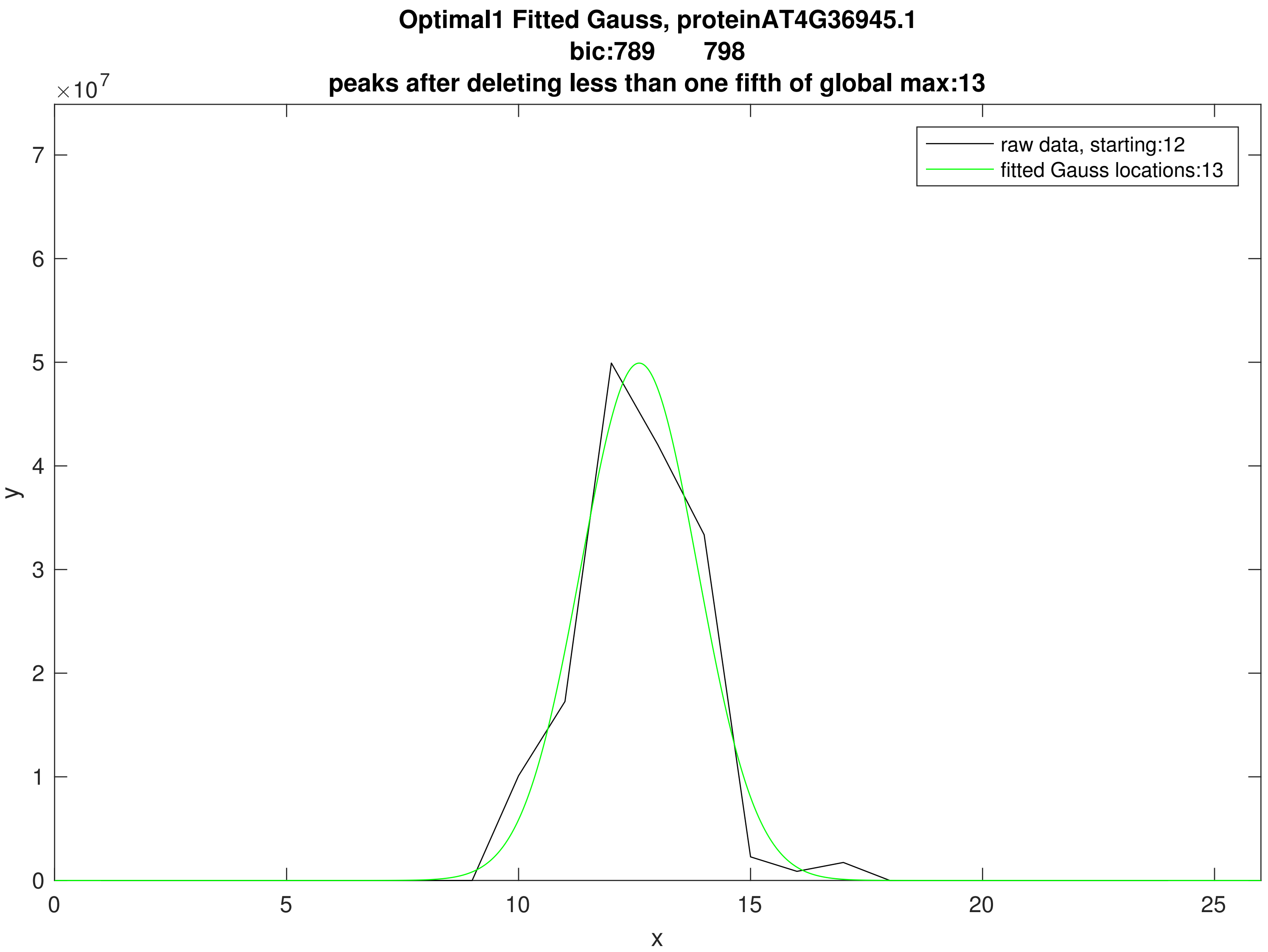
Optimal1 Fitted Gauss, proteinAT4G36945.1
bic:789 798
peaks after deleting less than one fifth of global max:13
×10^7
raw data, starting:12
fitted Gauss locations:13
y
x
0
1
2
3
4
5
6
7
0
5
10
15
20
25

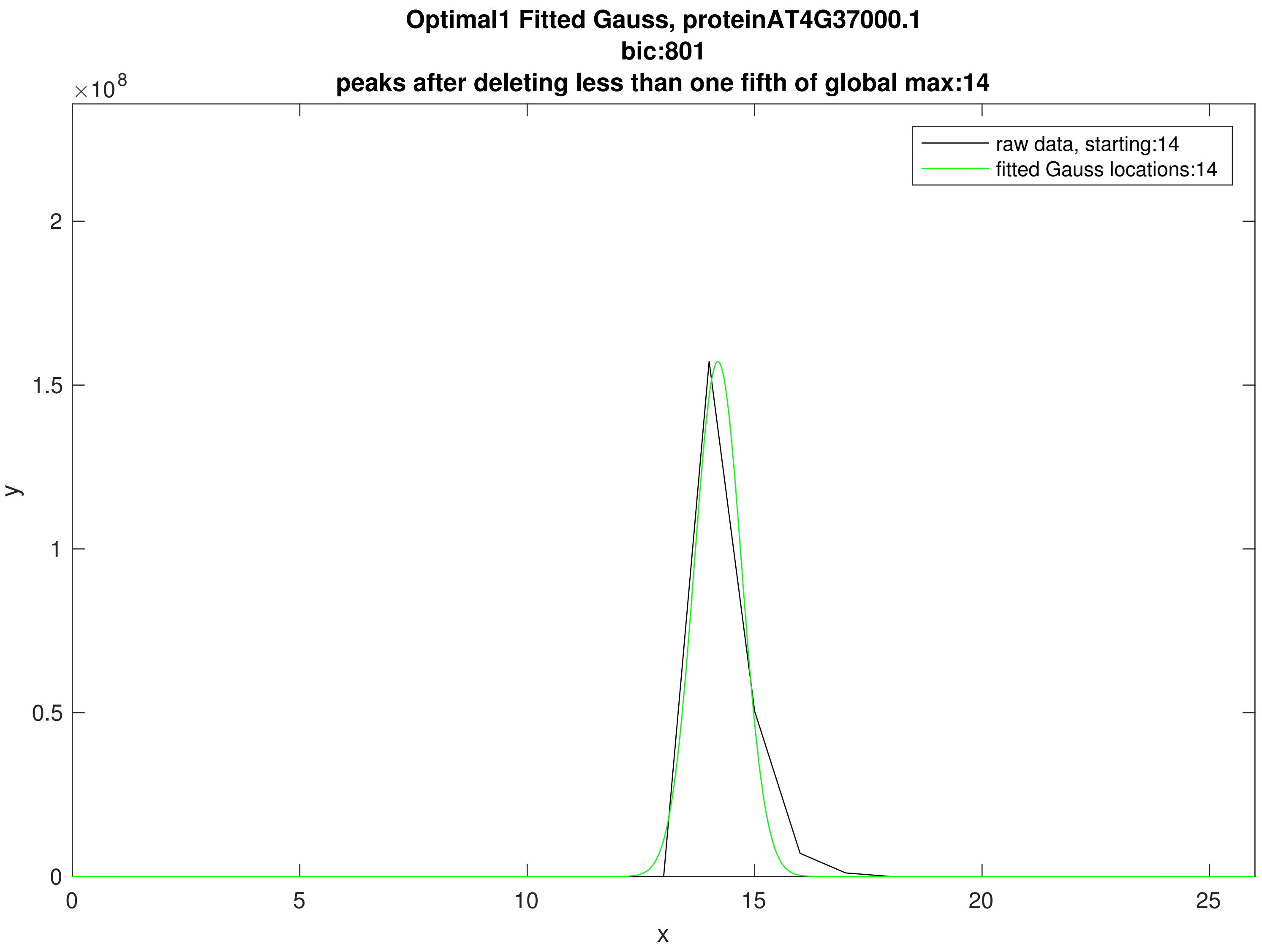
Optimal1 Fitted Gauss, proteinAT4G37000.1
bic:801
peaks after deleting less than one fifth of global max:14
×10^8
raw data, starting:14
fitted Gauss locations:14
2
1.5
1
0.5
0
y
0
5
10
15
20
25
x

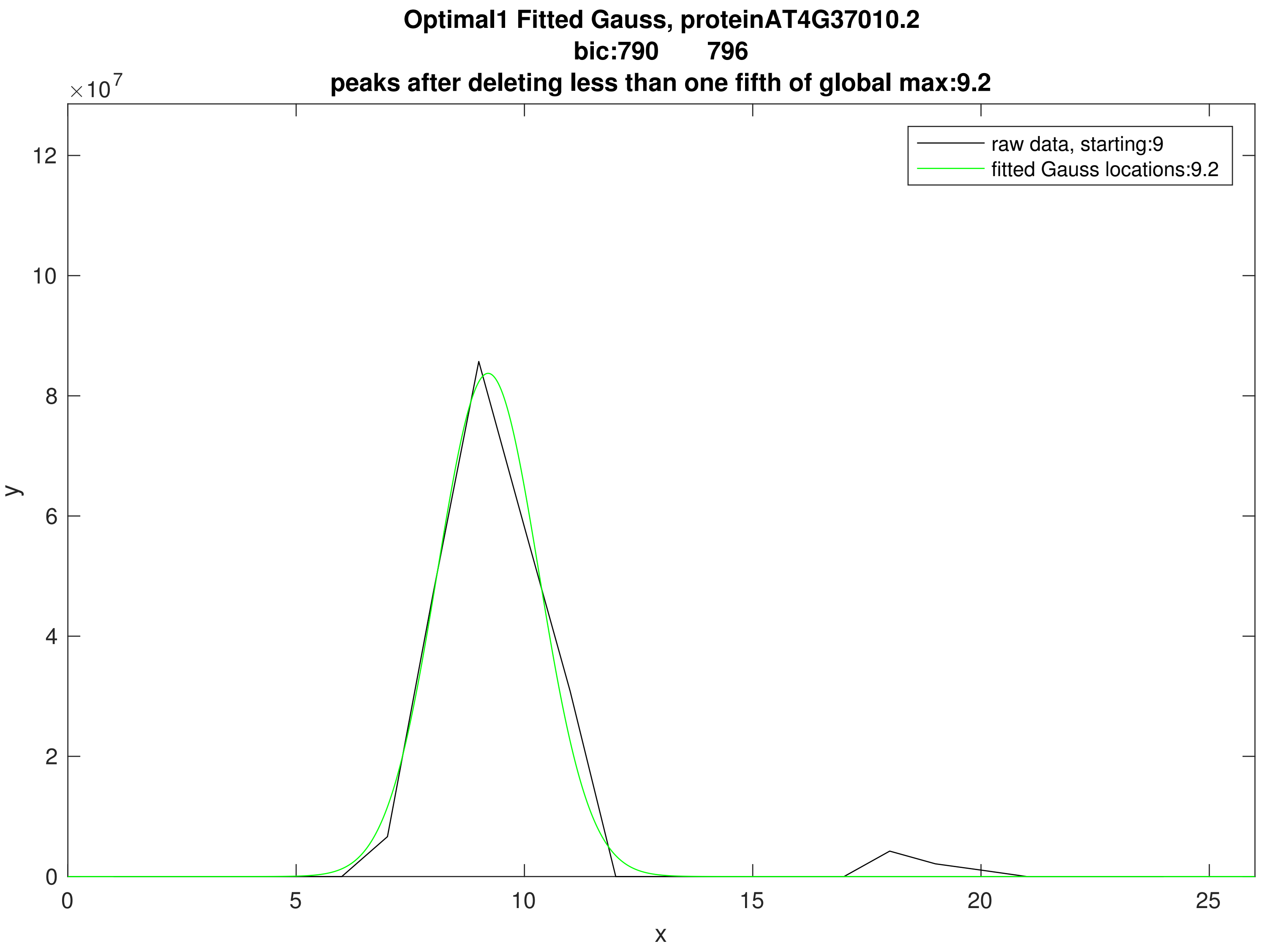
Optimal1 Fitted Gauss, proteinAT4G37010.2
bic:790 796
peaks after deleting less than one fifth of global max:9.2
×10^7
12
10
8
6
4
2
0
y
0
5
10
15
20
25
x
raw data, starting:9
fitted Gauss locations:9.2

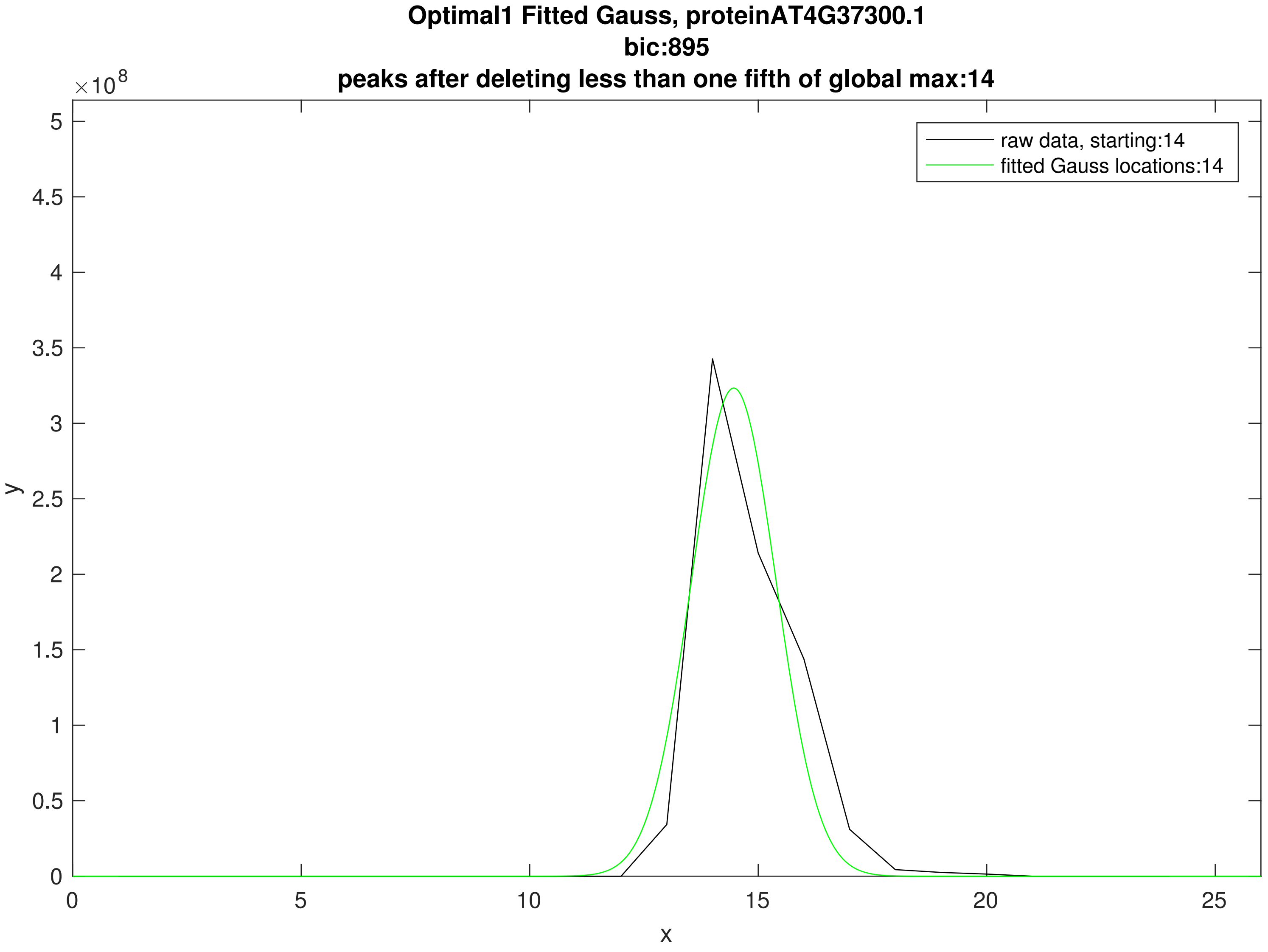
Optimal1 Fitted Gauss, proteinAT4G37300.1
bic:895
peaks after deleting less than one fifth of global max:14
×10^8
5
4.5
4
3.5
3
2.5
2
1.5
1
0.5
0
y
0
5
10
15
20
25
x
raw data, starting:14
fitted Gauss locations:14

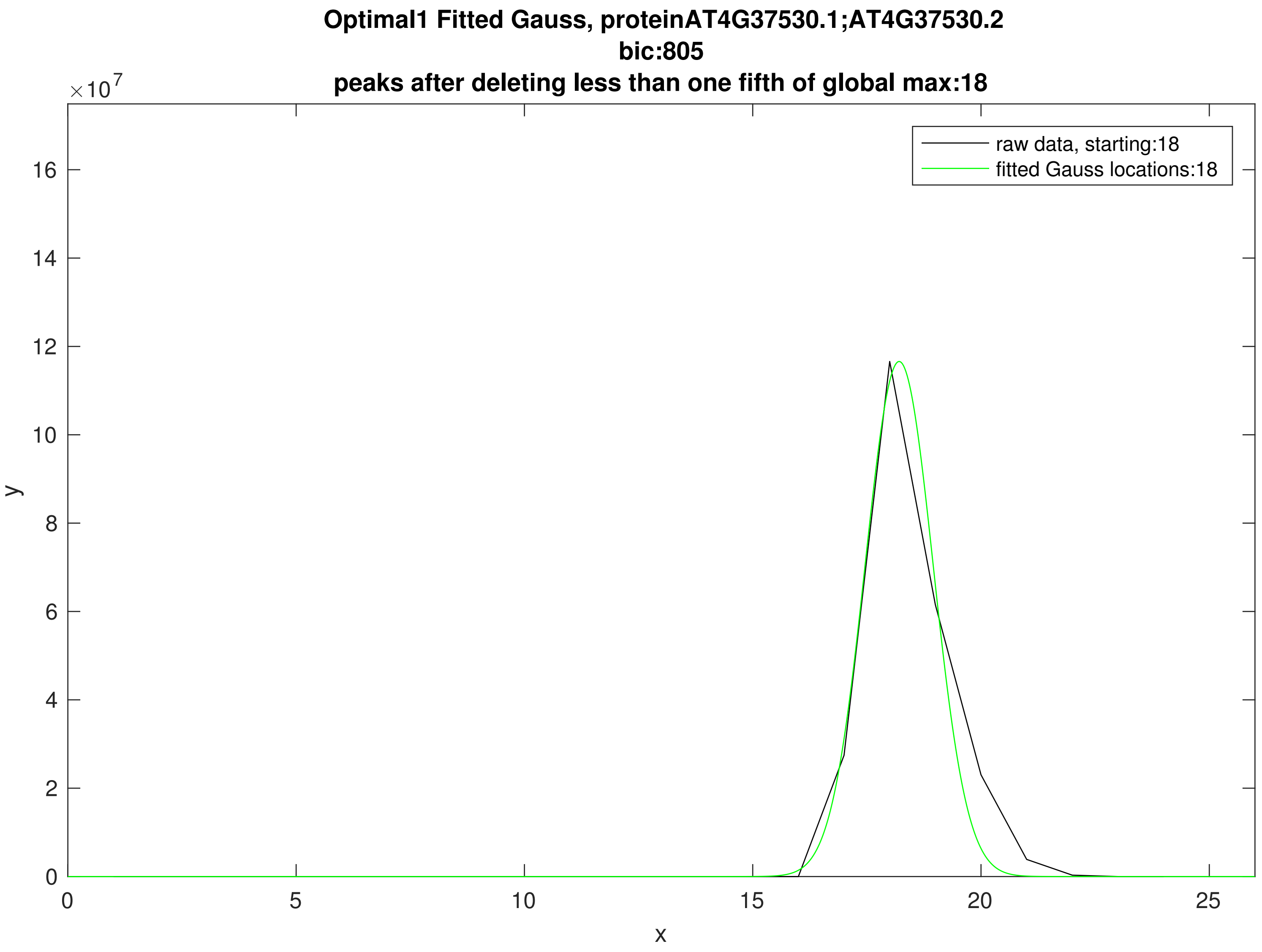

Optimal1 Fitted Gauss, proteinAT4G37530.1;AT4G37530.2
bic:805
peaks after deleting less than one fifth of global max:18
$\times 10^7$
raw data, starting:18
fitted Gauss locations:18
y
x
0
2
4
6
8
10
12
14
16
0
5
10
15
20
25

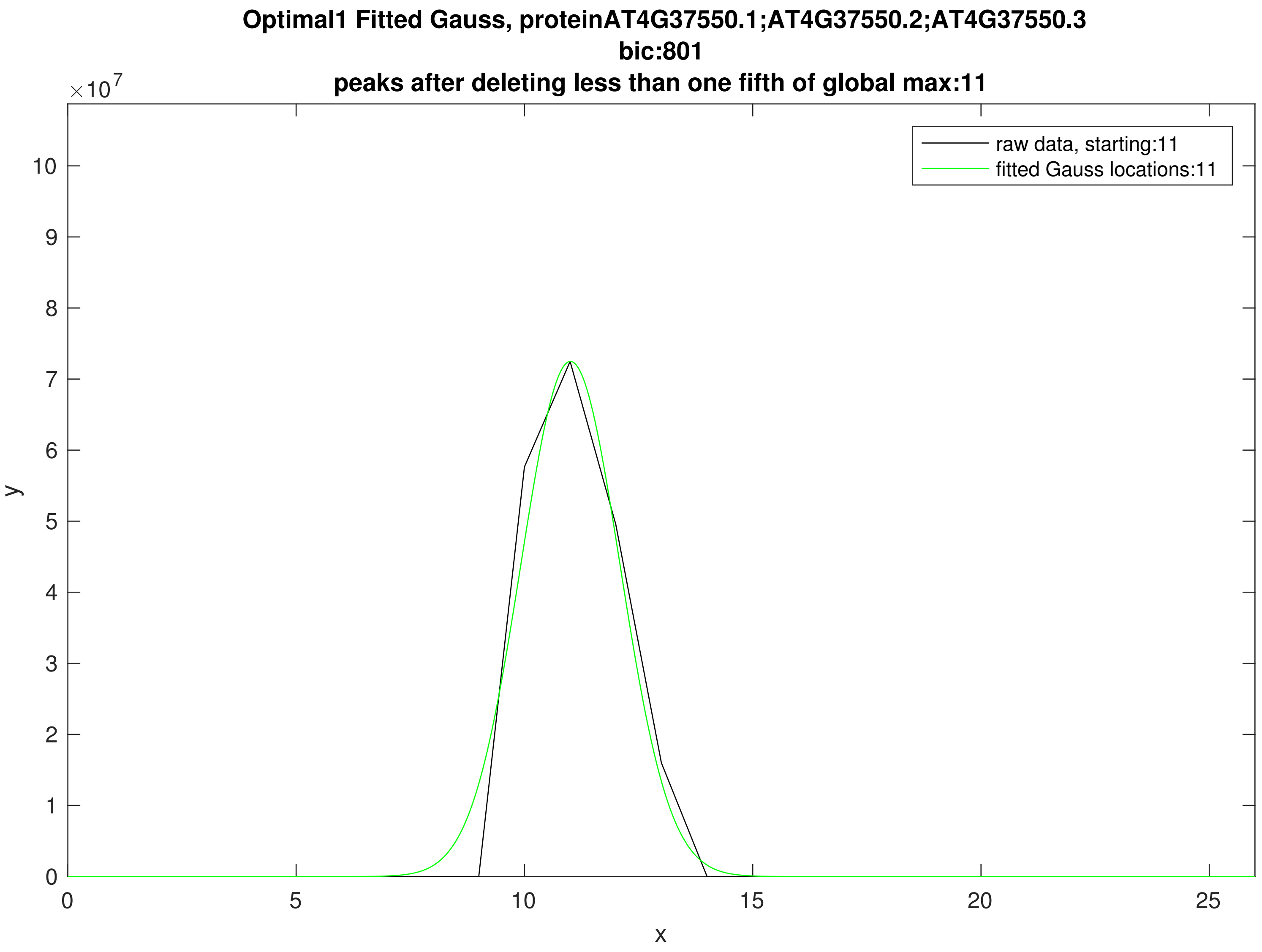
Optimal1 Fitted Gauss, proteinAT4G37550.1;AT4G37550.2;AT4G37550.3
bic:801
peaks after deleting less than one fifth of global max:11
×10^7
raw data, starting:11
fitted Gauss locations:11
y
x
0
1
2
3
4
5
6
7
8
9
10
0
5
10
15
20
25

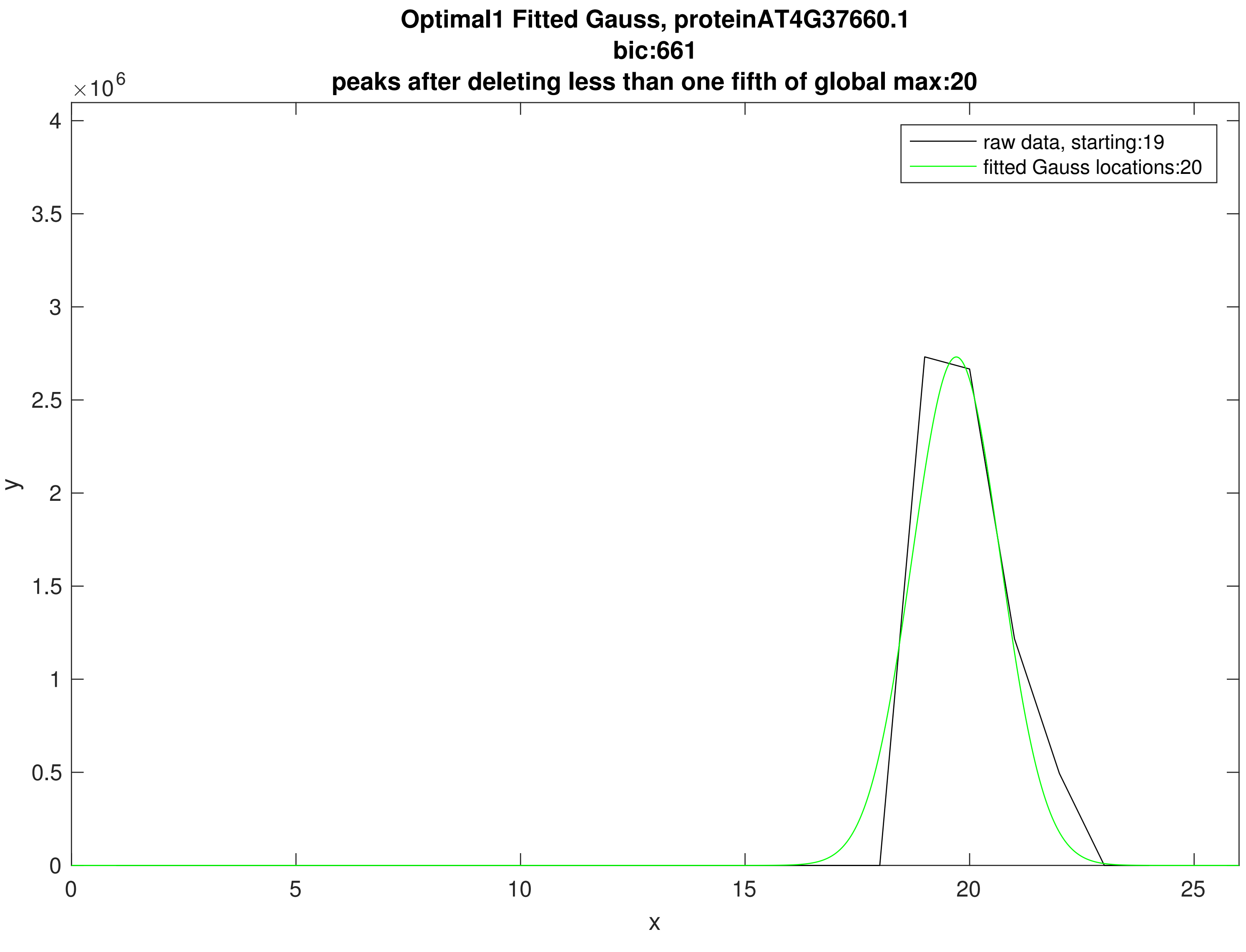
Optimal1 Fitted Gauss, proteinAT4G37660.1
bic:661
peaks after deleting less than one fifth of global max:20
×10^6
raw data, starting:19
fitted Gauss locations:20
y
x
0
0.5
1
1.5
2
2.5
3
3.5
4
0
5
10
15
20
25

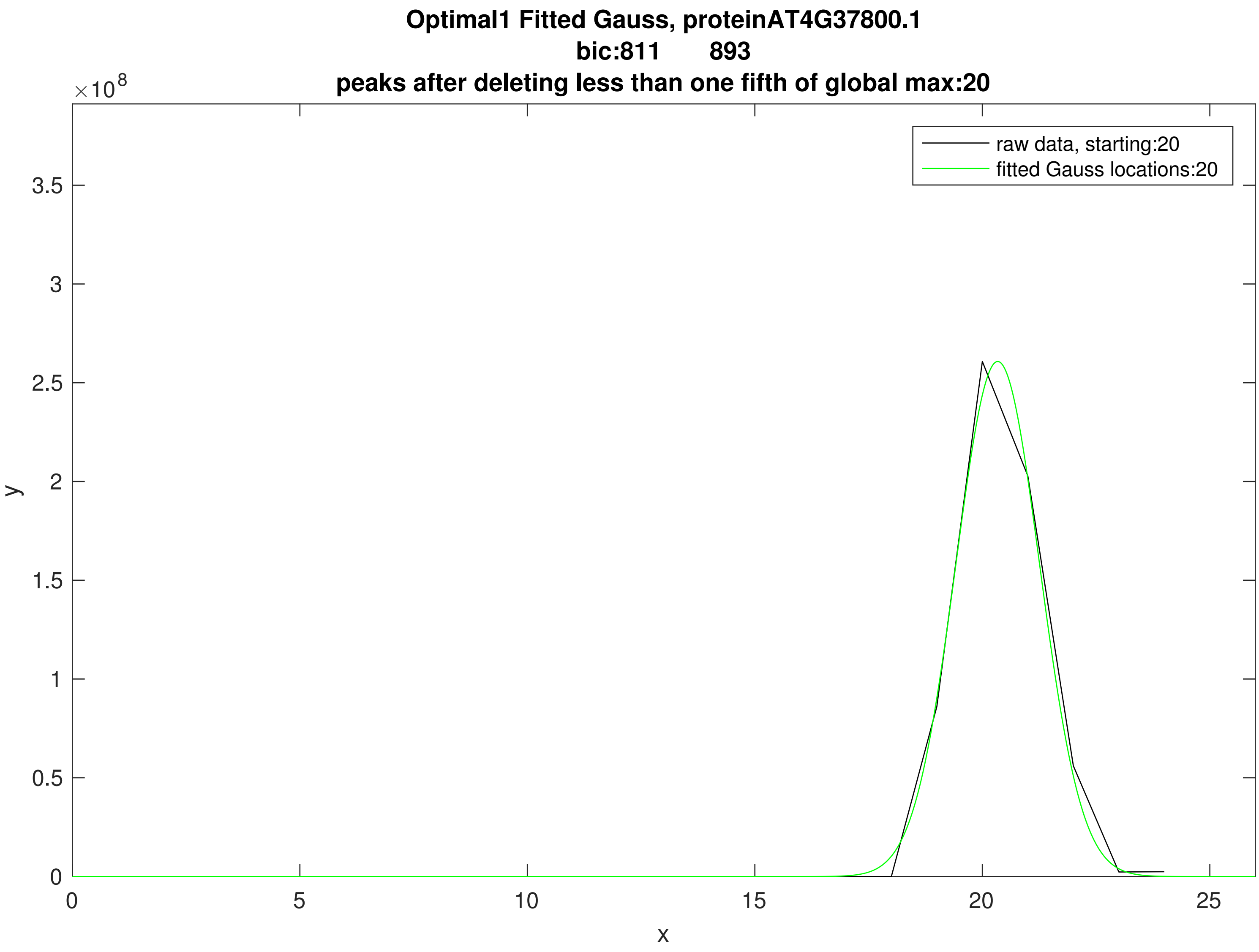
Optimal1 Fitted Gauss, proteinAT4G37800.1
bic:811 893
peaks after deleting less than one fifth of global max:20
×10^8
raw data, starting:20
fitted Gauss locations:20
y
x
0
0.5
1
1.5
2
2.5
3
3.5
0
5
10
15
20
25

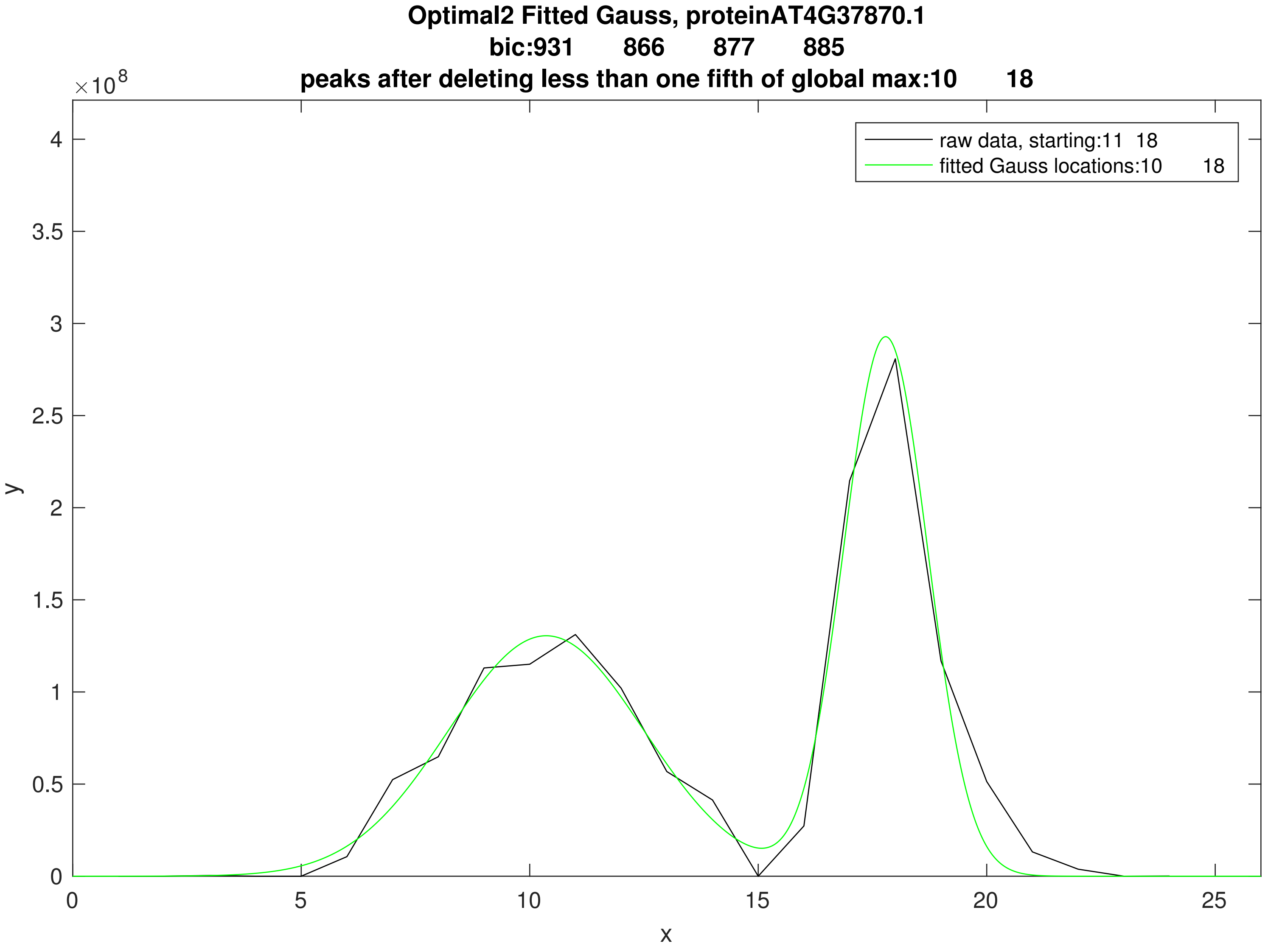
Optimal2 Fitted Gauss, proteinAT4G37870.1
bic:931 866 877 885
peaks after deleting less than one fifth of global max:10 18
×10^8
raw data, starting:11 18
fitted Gauss locations:10 18
y
x
0
0.5
1
1.5
2
2.5
3
3.5
4
0
5
10
15
20
25

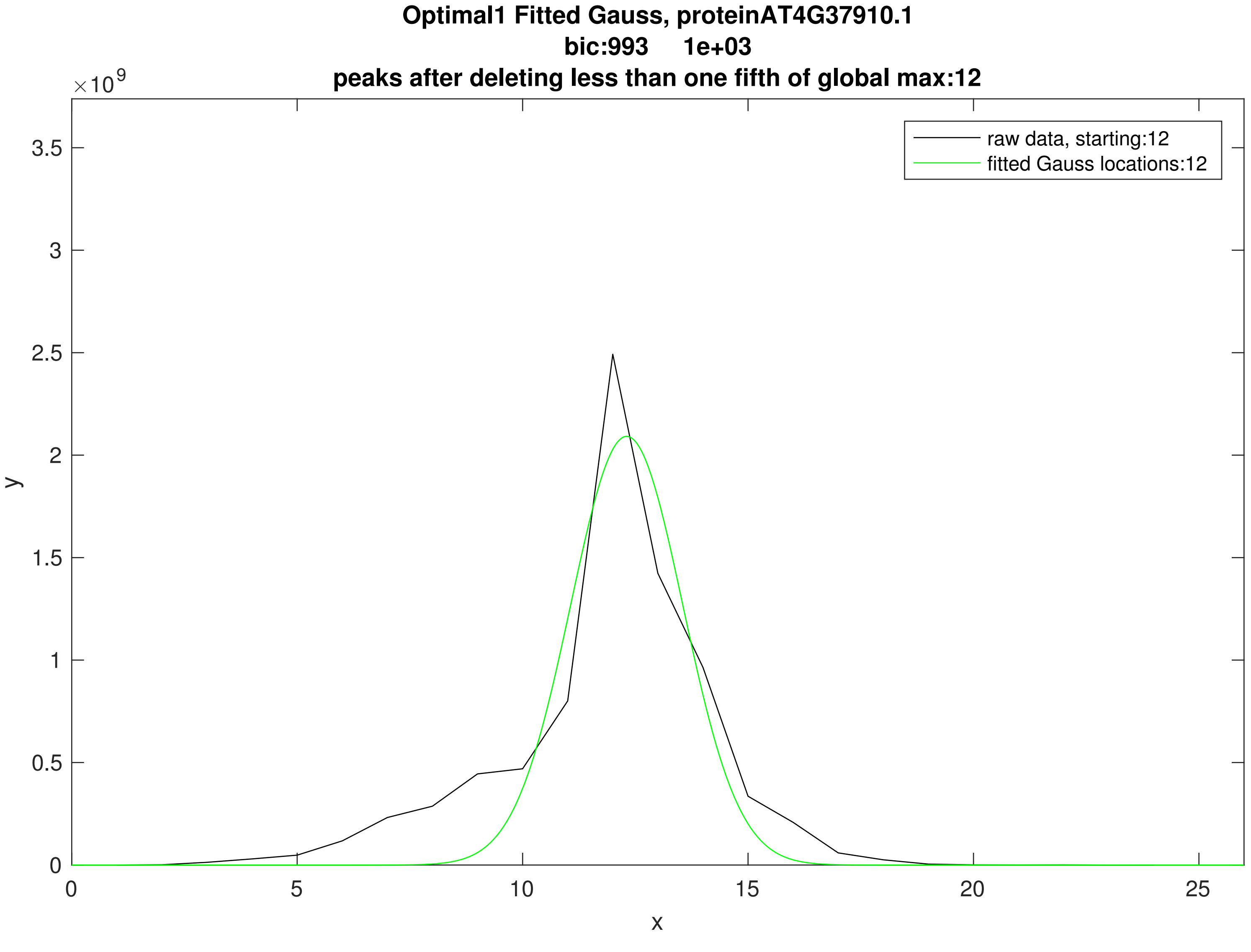
Optimal1 Fitted Gauss, proteinAT4G37910.1
bic:993 1e+03
peaks after deleting less than one fifth of global max:12
×10^9
raw data, starting:12
fitted Gauss locations:12
y
x
0
0.5
1
1.5
2
2.5
3
3.5
0
5
10
15
20
25

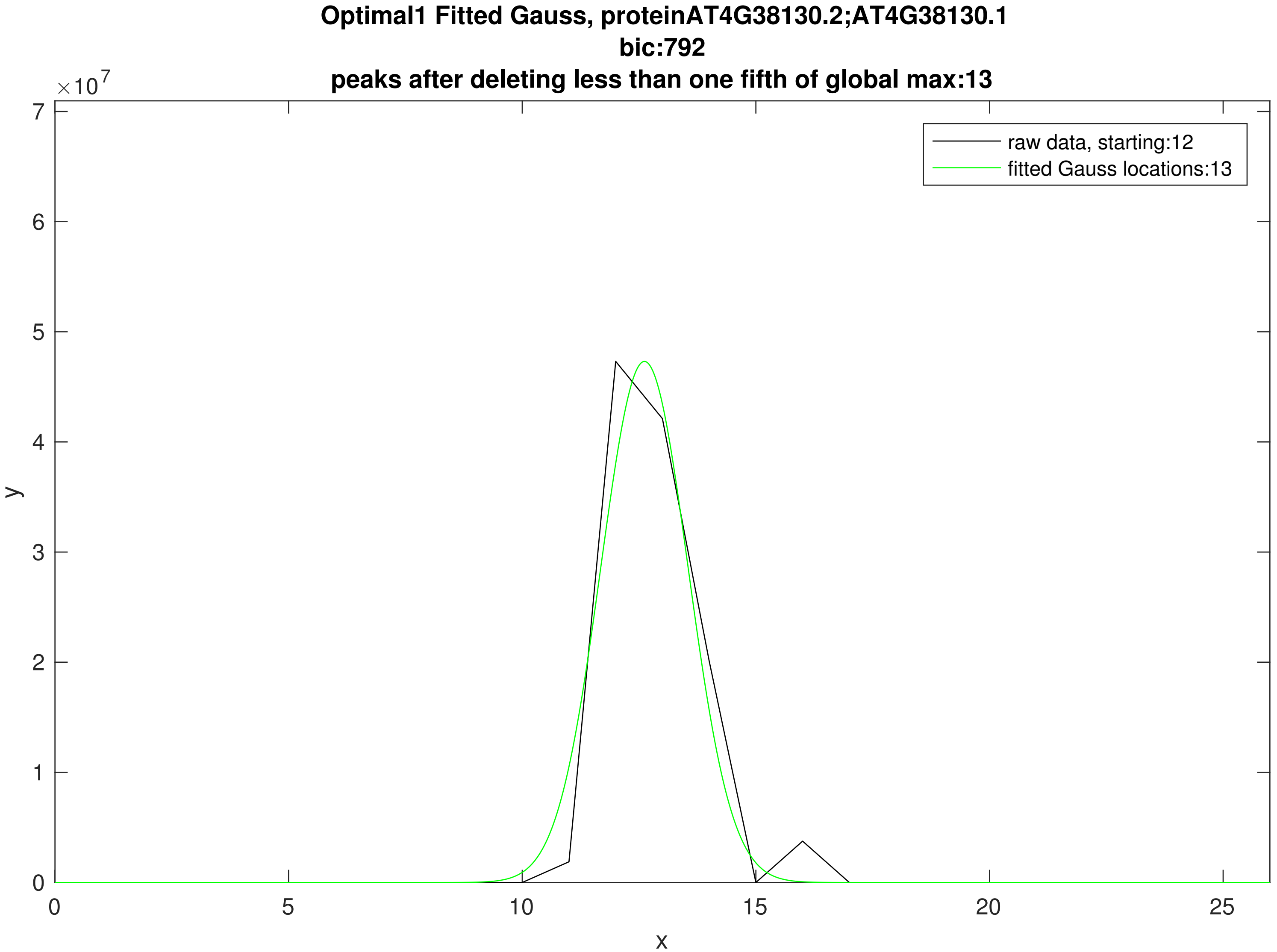

Optimal1 Fitted Gauss, proteinAT4G38130.2;AT4G38130.1
bic:792
peaks after deleting less than one fifth of global max:13
×10^7
7
6
5
4
3
2
1
0
y
0
5
10
15
20
25
x
raw data, starting:12
fitted Gauss locations:13

Optimal1 Fitted Gauss, proteinAT4G38220.1;AT4G38220.2
bic:946 949
peaks after deleting less than one fifth of global max:9.7

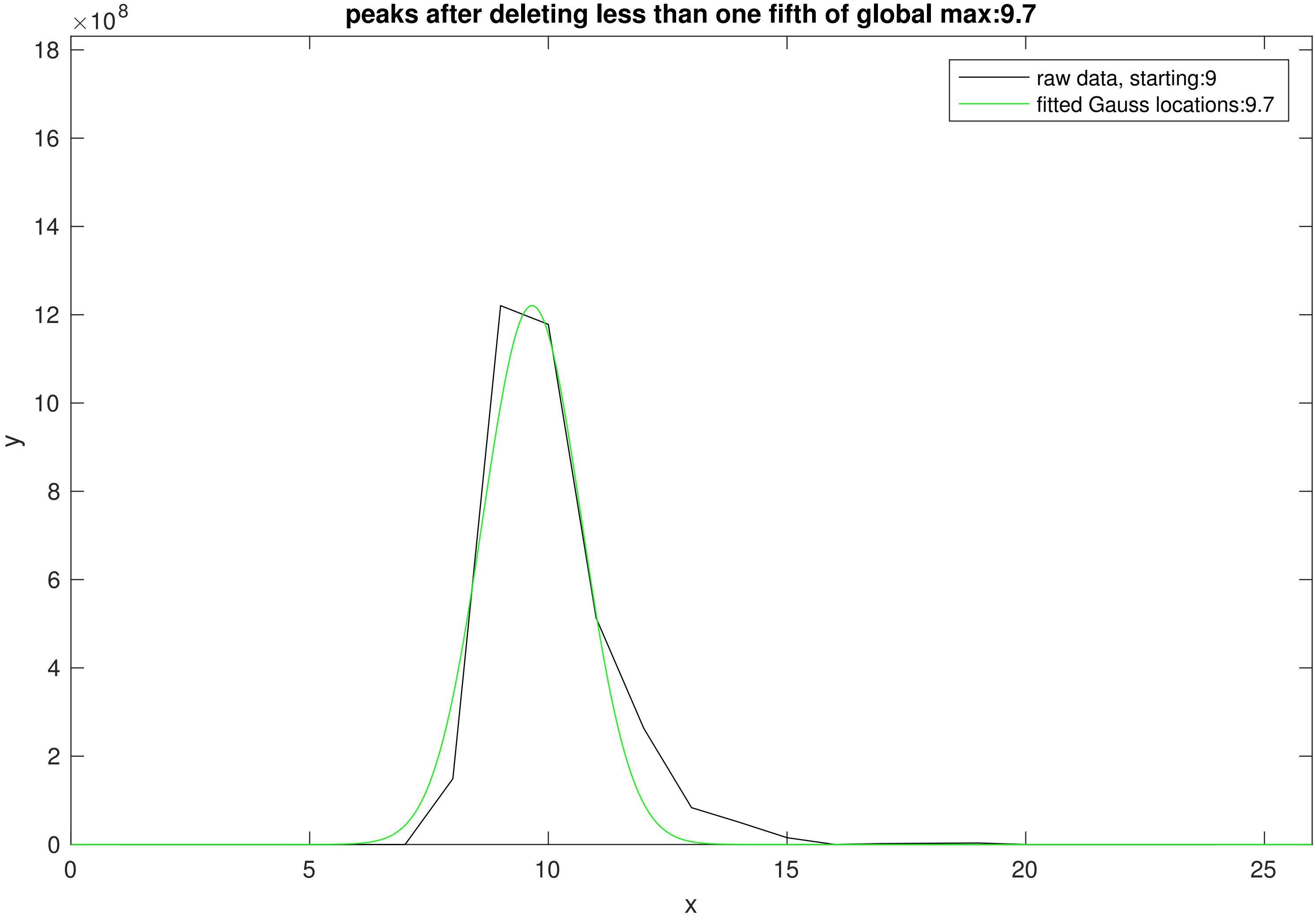

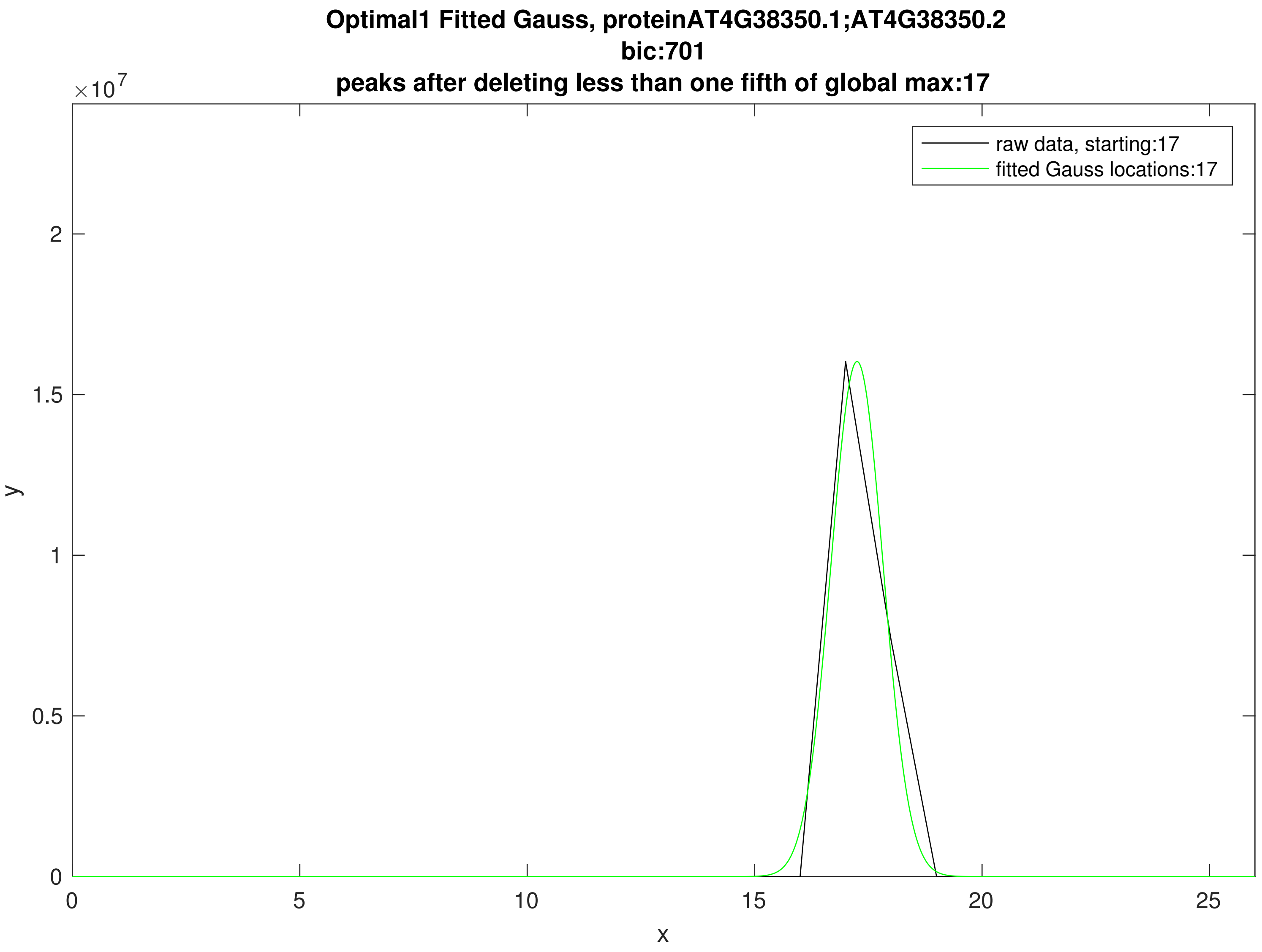

Optimal1 Fitted Gauss, proteinAT4G38350.1;AT4G38350.2
bic:701
peaks after deleting less than one fifth of global max:17
×10^7
raw data, starting:17
fitted Gauss locations:17
2
1.5
1
0.5
0
y
0
5
10
15
20
25
x

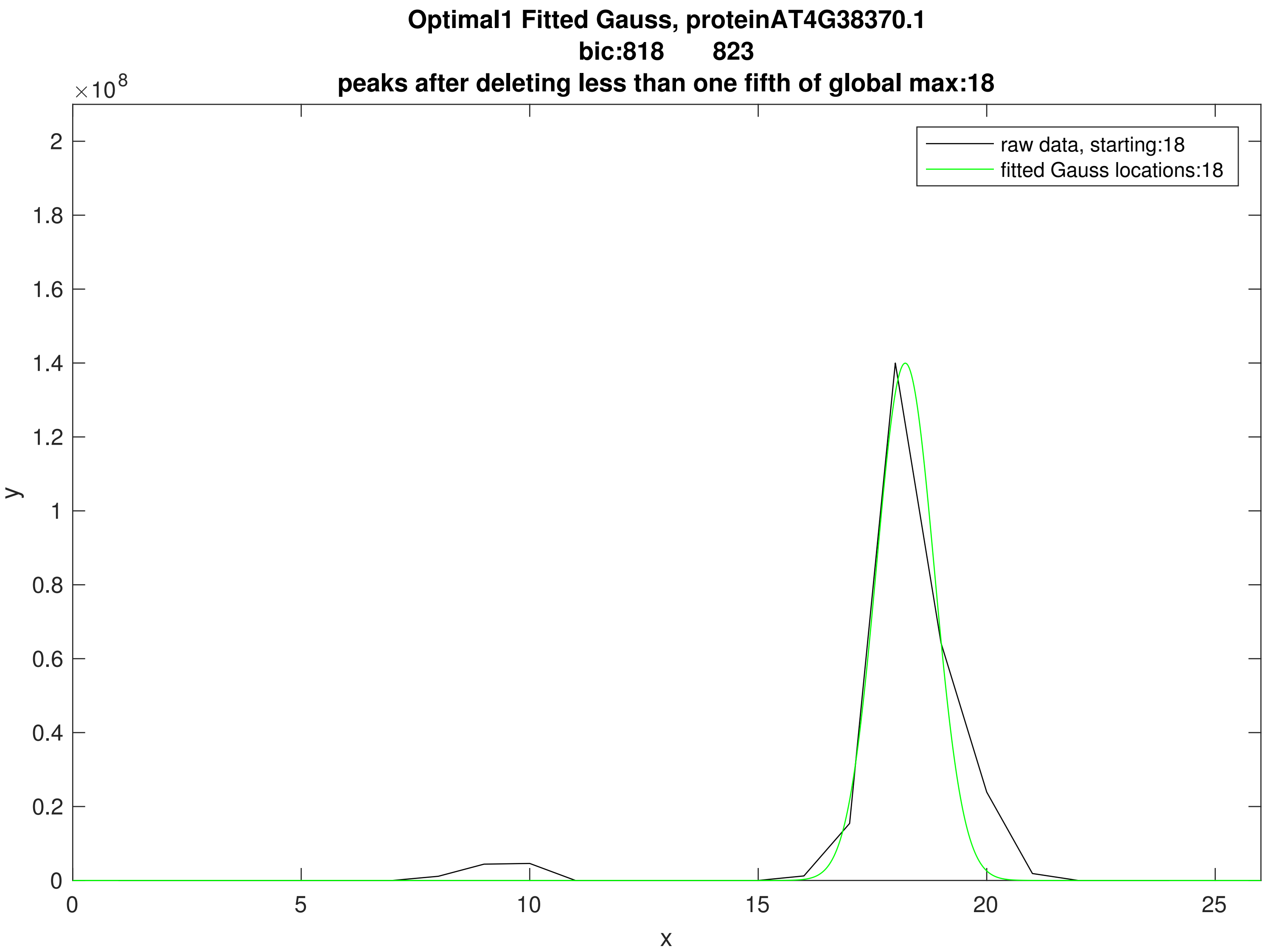
Optimal1 Fitted Gauss, proteinAT4G38370.1
bic:818 823
peaks after deleting less than one fifth of global max:18
×10^8
raw data, starting:18
fitted Gauss locations:18
y
x
0
0.2
0.4
0.6
0.8
1
1.2
1.4
1.6
1.8
2
0
5
10
15
20
25

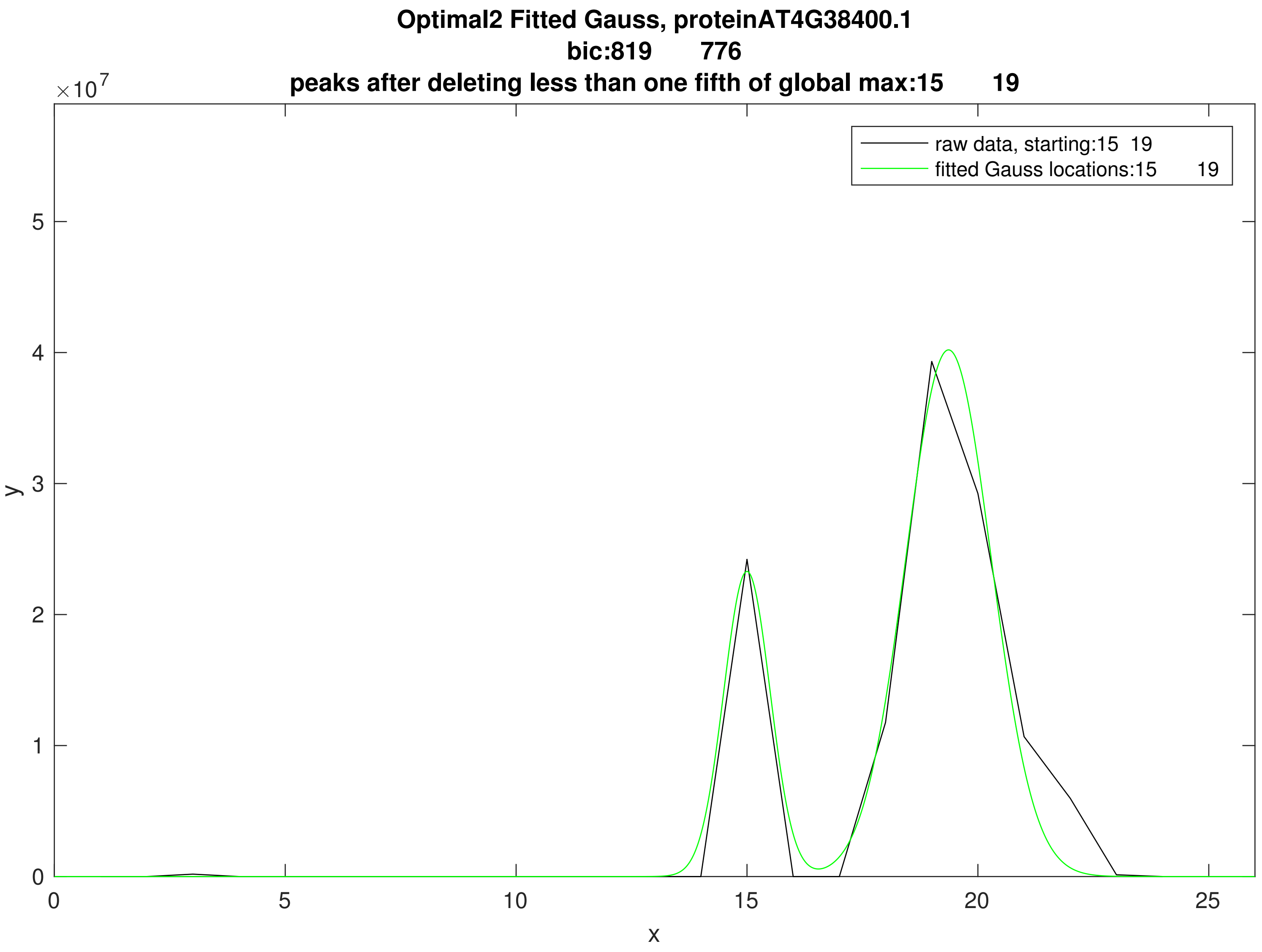
Optimal2 Fitted Gauss, proteinAT4G38400.1
bic:819 776
peaks after deleting less than one fifth of global max:15 19
×10^7
raw data, starting:15 19
fitted Gauss locations:15 19
5
4
3
2
1
0
y
0
5
10
15
20
25
x

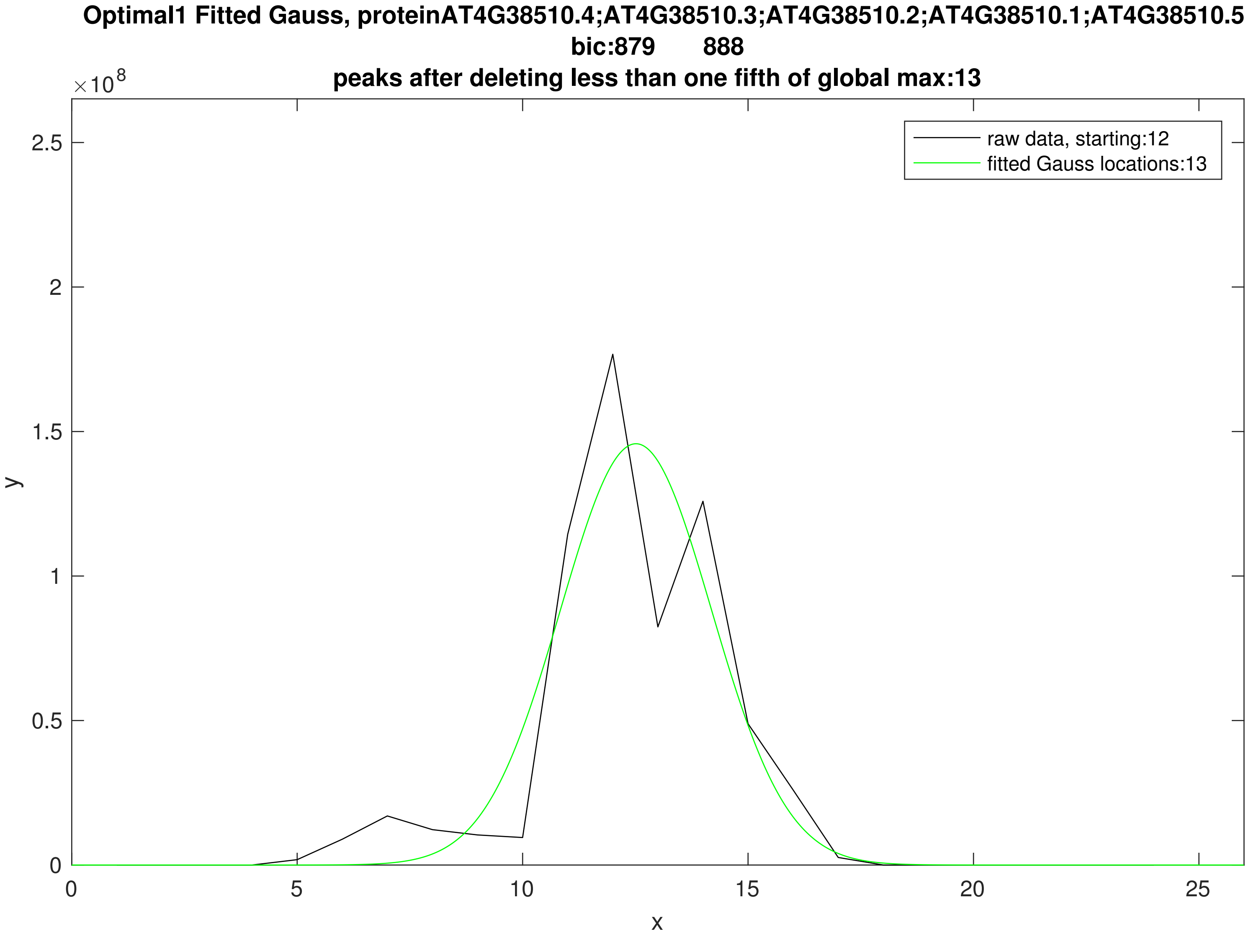

Optimal1 Fitted Gauss, proteinAT4G38510.4;AT4G38510.3;AT4G38510.2;AT4G38510.1;AT4G38510.5
bic:879 888
peaks after deleting less than one fifth of global max:13
×10^8
2.5
2
1.5
1
0.5
0
y
0
5
10
15
20
25
x
raw data, starting:12
fitted Gauss locations:13

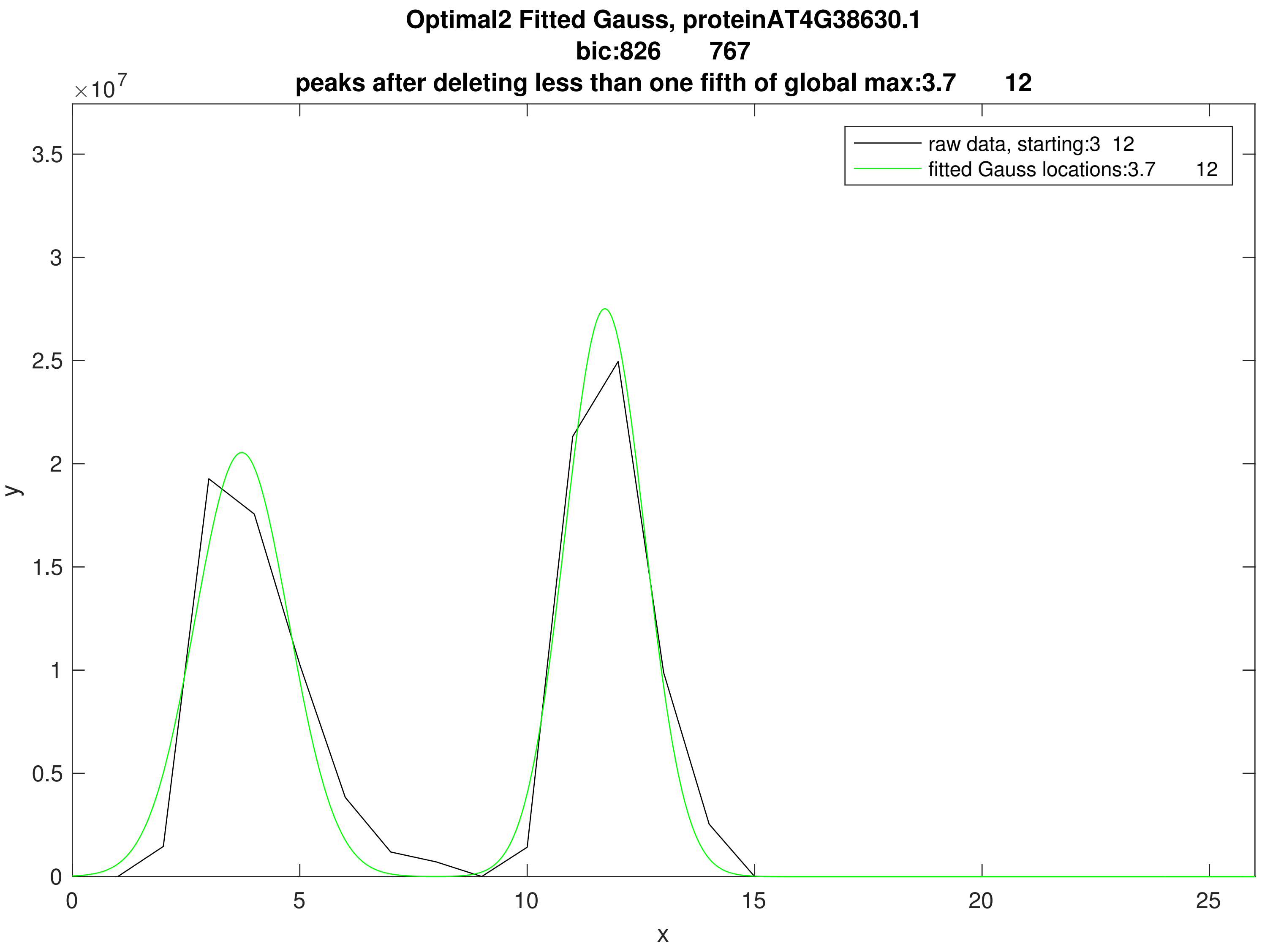

Optimal2 Fitted Gauss, proteinAT4G38630.1
bic:826 767
peaks after deleting less than one fifth of global max:3.7 12
×10^7
3.5
3
2.5
2
1.5
1
0.5
0
y
0
5
10
15
20
25
x
raw data, starting:3 12
fitted Gauss locations:3.7 12

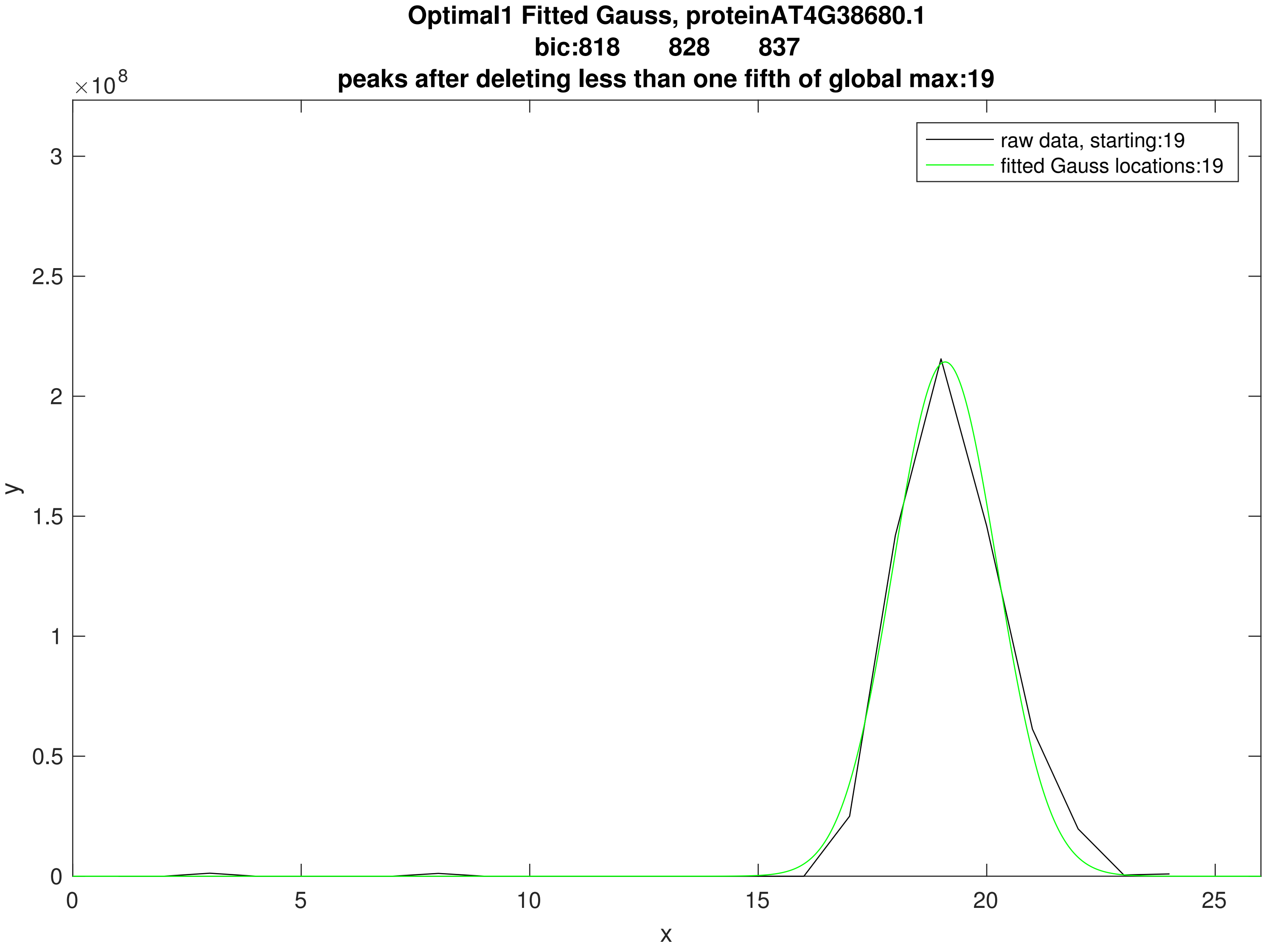
Optimal1 Fitted Gauss, proteinAT4G38680.1
bic:818 828 837
peaks after deleting less than one fifth of global max:19
×10^8
raw data, starting:19
fitted Gauss locations:19
3
2.5
2
1.5
1
0.5
0
y
0
5
10
15
20
25
x

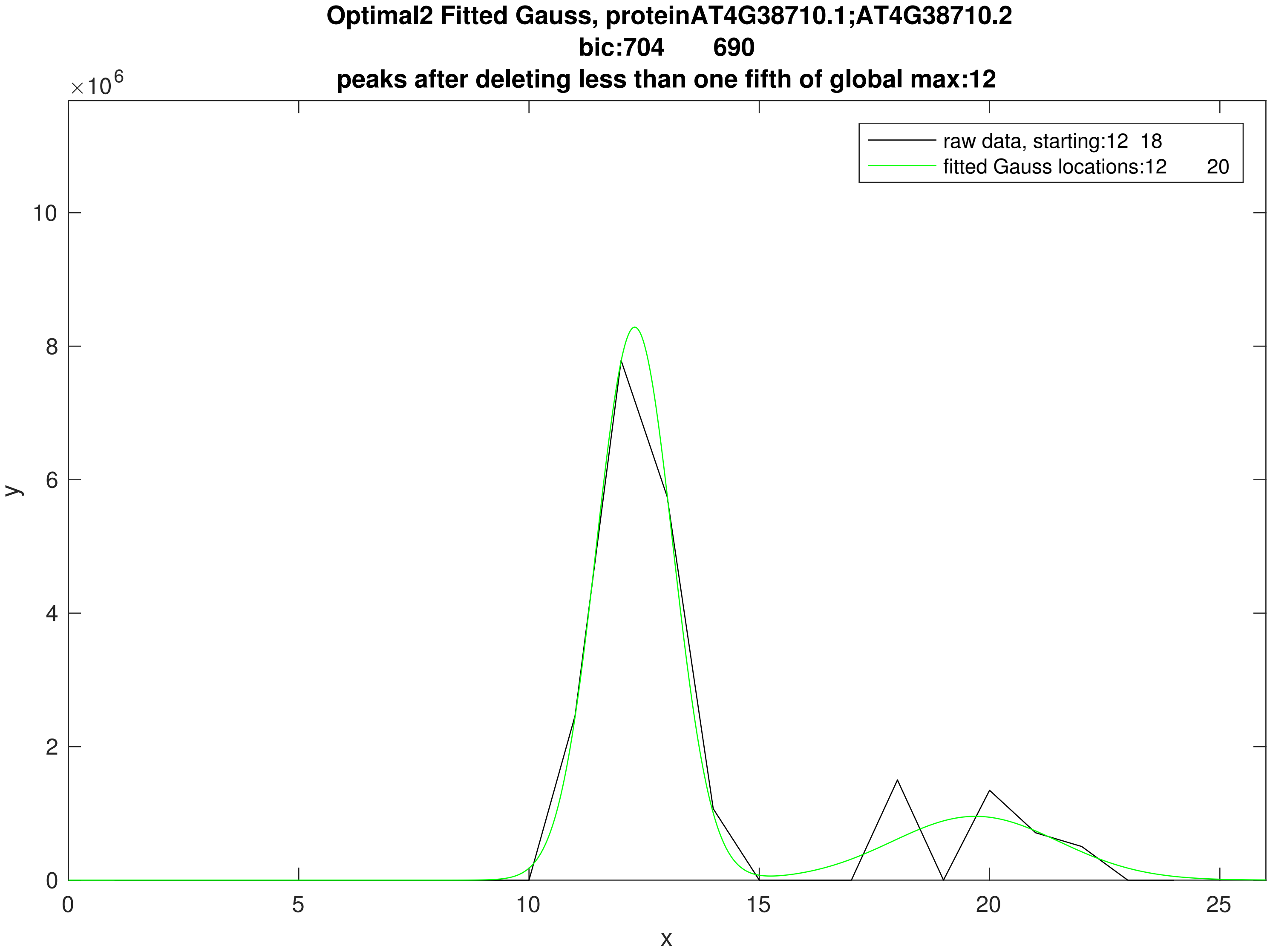

Optimal2 Fitted Gauss, proteinAT4G38710.1;AT4G38710.2
bic:704 690
peaks after deleting less than one fifth of global max:12
×10^6
raw data, starting:12 18
fitted Gauss locations:12 20
y
x
0
2
4
6
8
10
0
5
10
15
20
25

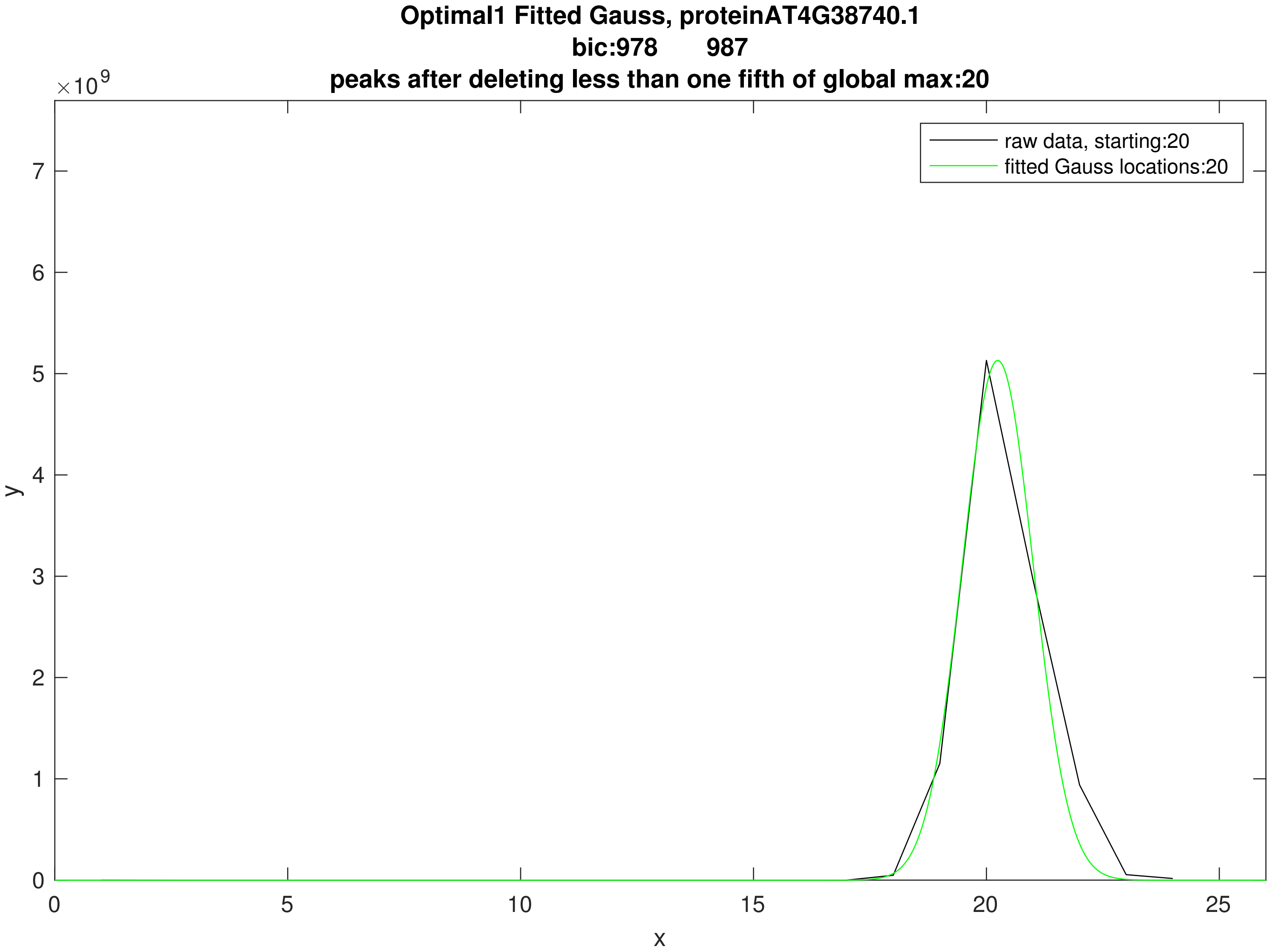
Optimal1 Fitted Gauss, proteinAT4G38740.1
bic:978 987
peaks after deleting less than one fifth of global max:20
×10^9
raw data, starting:20
fitted Gauss locations:20
y
x
0
1
2
3
4
5
6
7
0
5
10
15
20
25

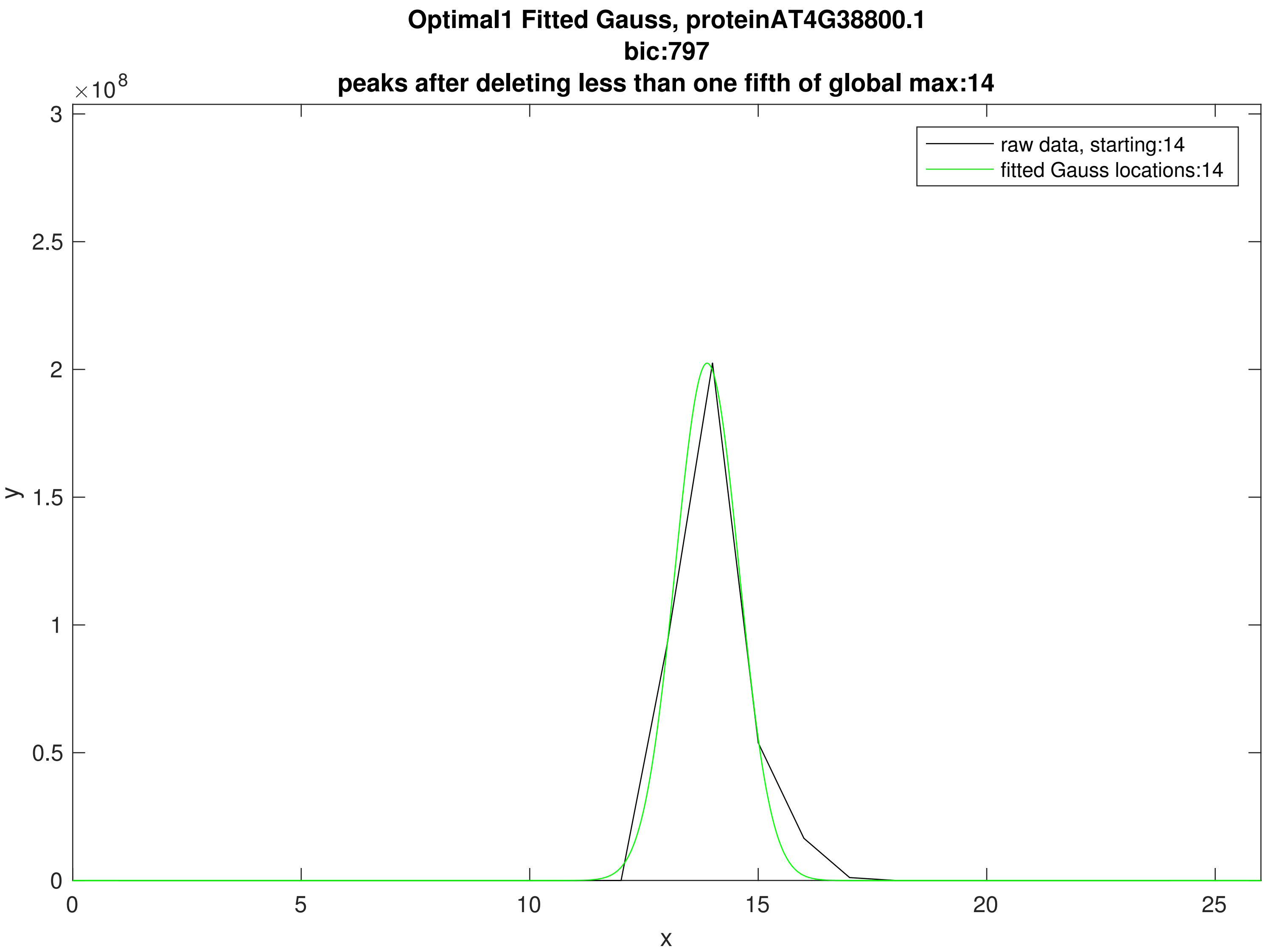
Optimal1 Fitted Gauss, proteinAT4G38800.1
bic:797
peaks after deleting less than one fifth of global max:14
×10^8
3
2.5
2
1.5
1
0.5
0
y
0
5
10
15
20
25
x
raw data, starting:14
fitted Gauss locations:14

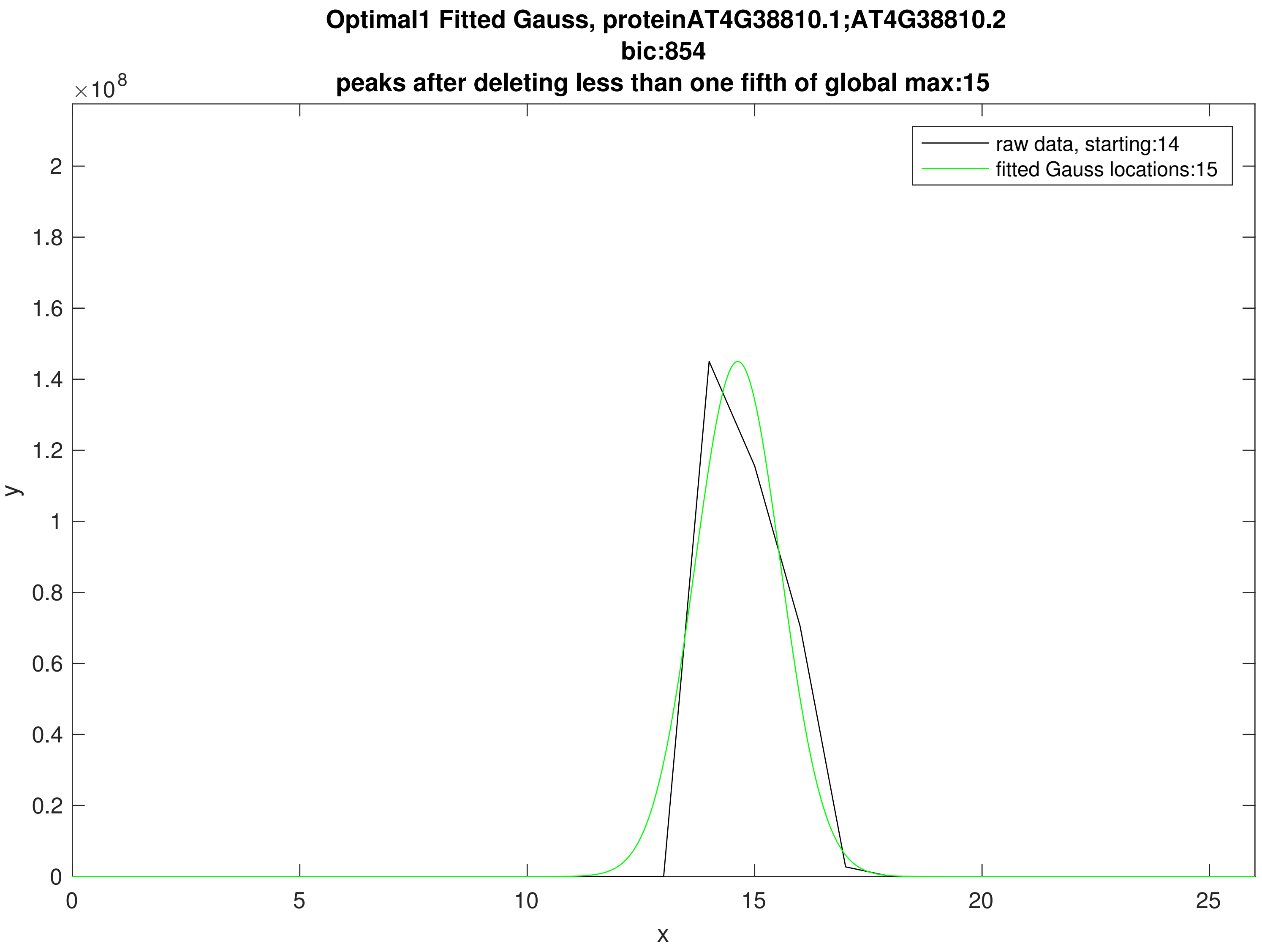
Optimal1 Fitted Gauss, proteinAT4G38810.1;AT4G38810.2
bic:854
peaks after deleting less than one fifth of global max:15
×10^8
raw data, starting:14
fitted Gauss locations:15
y
x
0
0.2
0.4
0.6
0.8
1
1.2
1.4
1.6
1.8
2
0
5
10
15
20
25

**Optimal2 Fitted Gauss, proteinAT4G38970.1;AT4G38970.2**
**bic:854 838**
**peaks after deleting less than one fifth of global max:10**

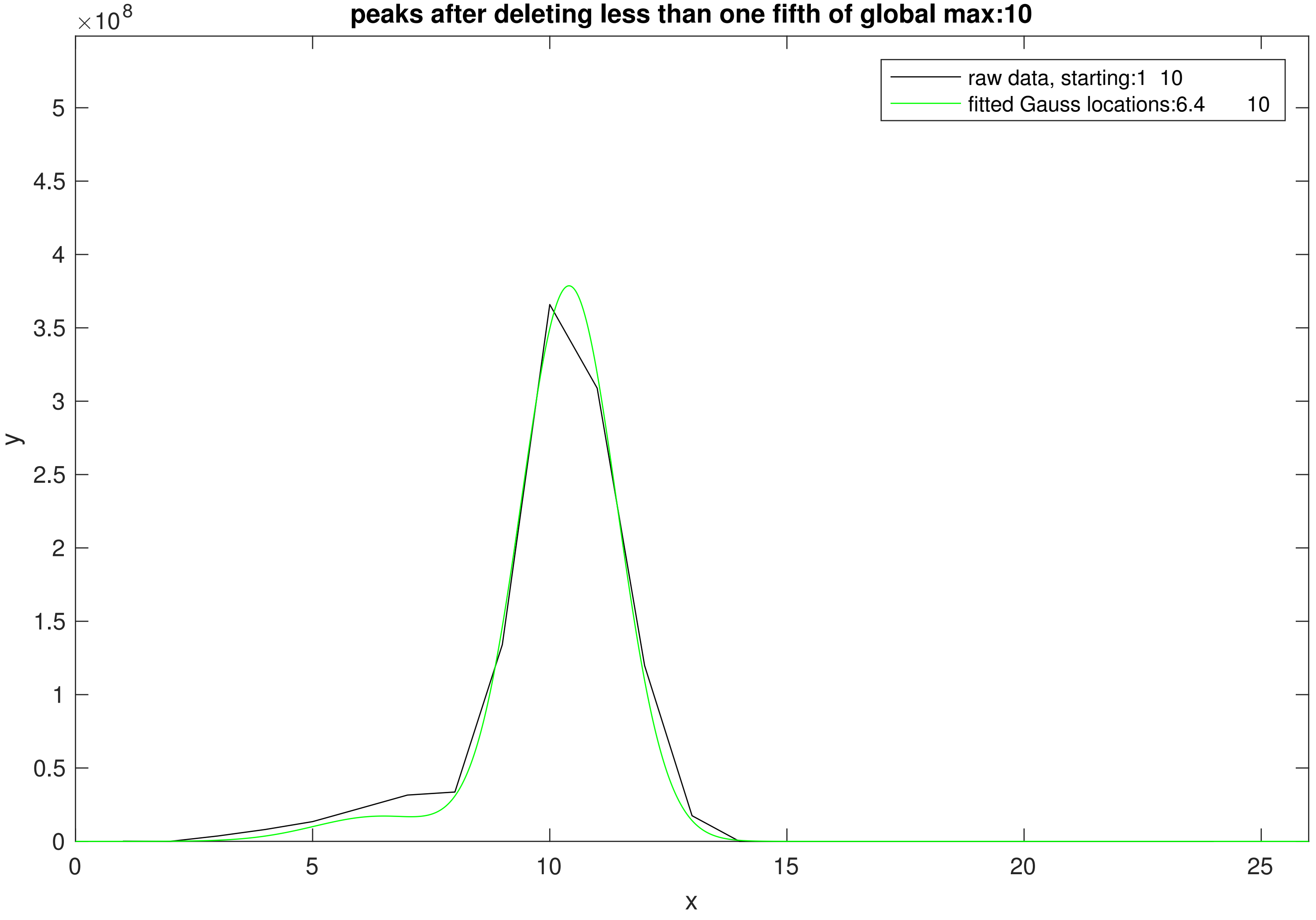

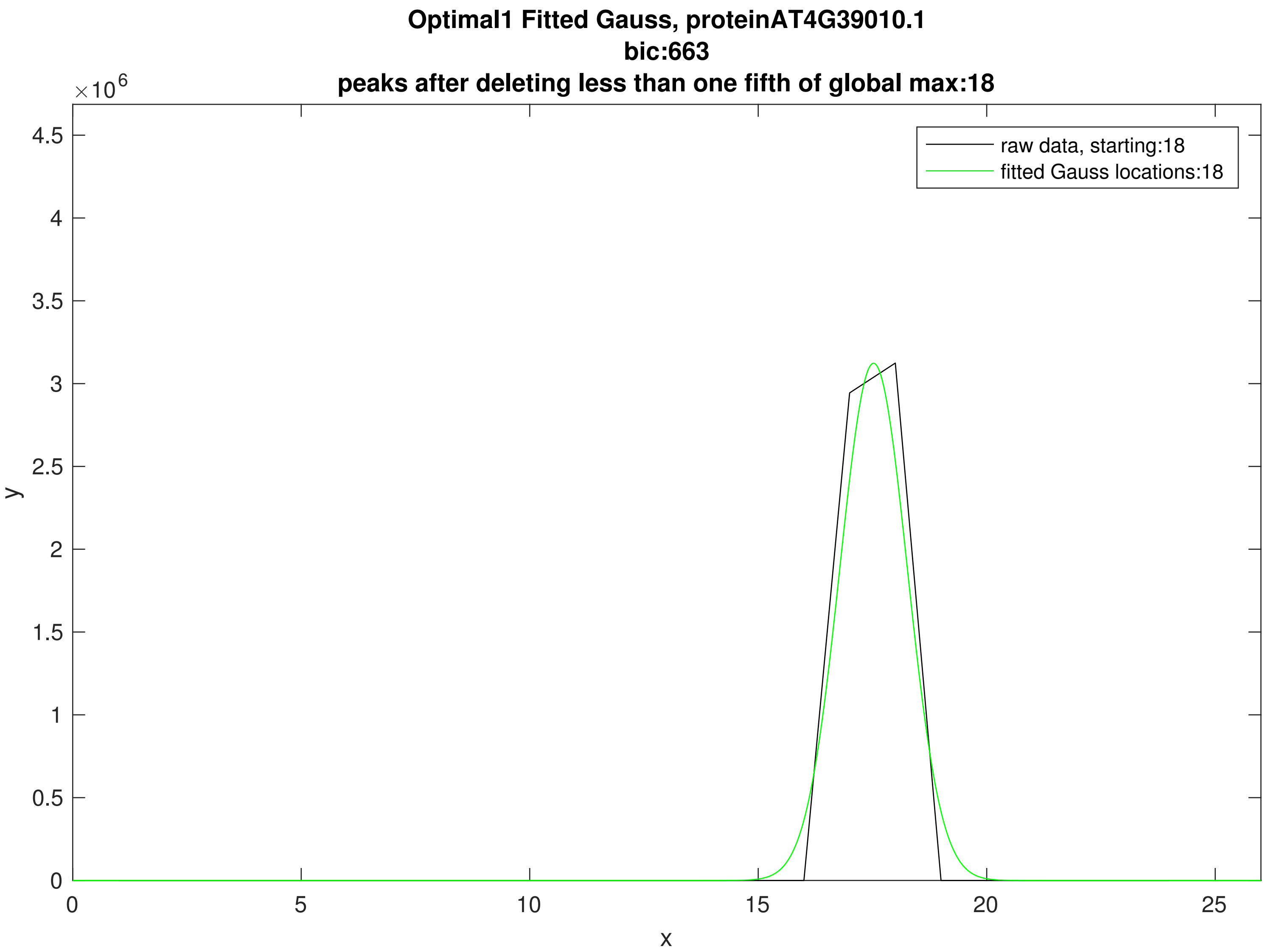

Optimal1 Fitted Gauss, proteinAT4G39010.1
bic:663
peaks after deleting less than one fifth of global max:18
×10^6
4.5
4
3.5
3
2.5
2
1.5
1
0.5
0
y
0
5
10
15
20
25
x
raw data, starting:18
fitted Gauss locations:18

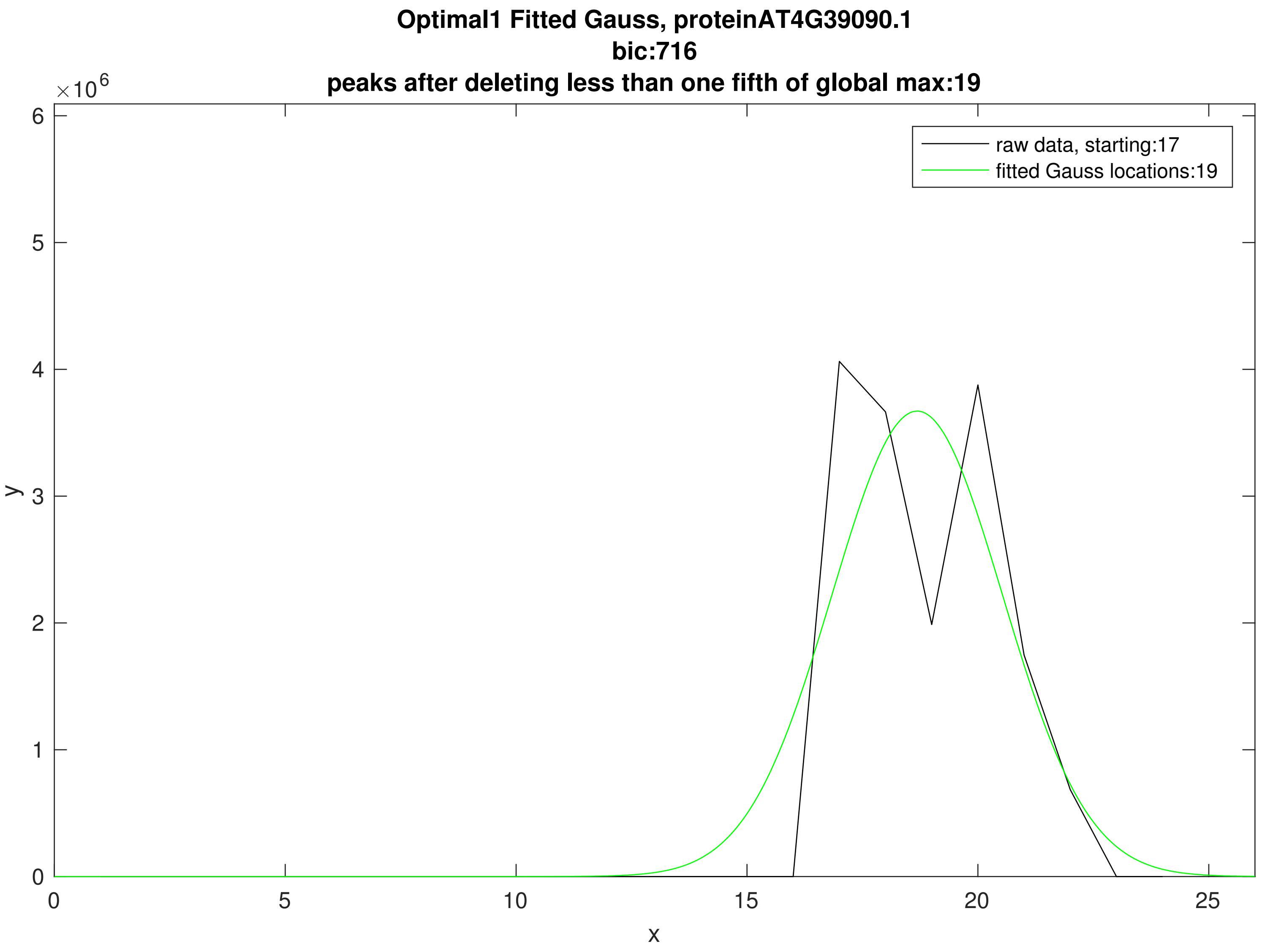
Optimal1 Fitted Gauss, proteinAT4G39090.1
bic:716
peaks after deleting less than one fifth of global max:19
×10^6
raw data, starting:17
fitted Gauss locations:19
y
x
0
1
2
3
4
5
6
0
5
10
15
20
25

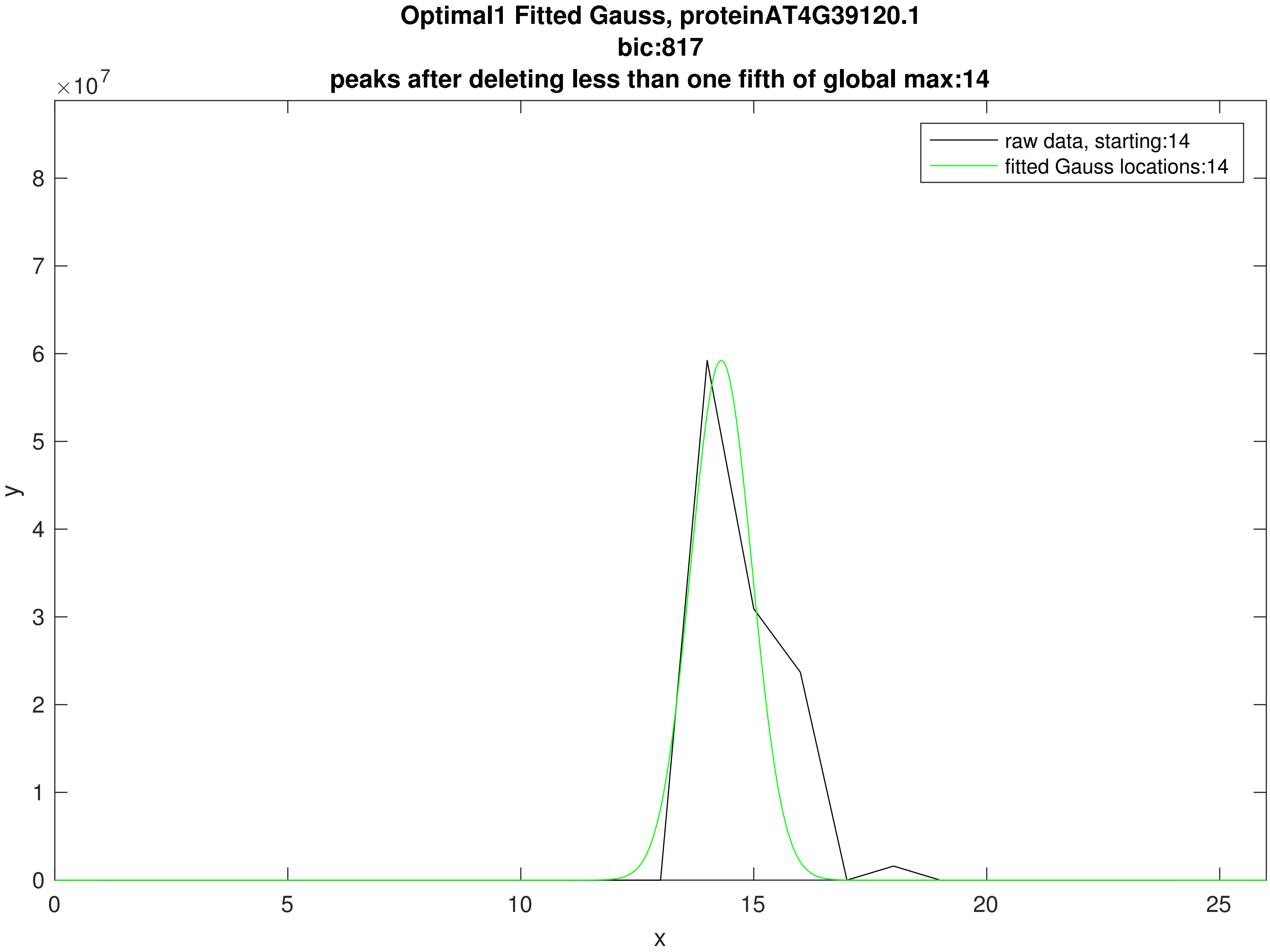

Optimal1 Fitted Gauss, proteinAT4G39120.1
bic:817
peaks after deleting less than one fifth of global max:14
×10^7
raw data, starting:14
fitted Gauss locations:14
y
x
0
1
2
3
4
5
6
7
8
0
5
10
15
20
25

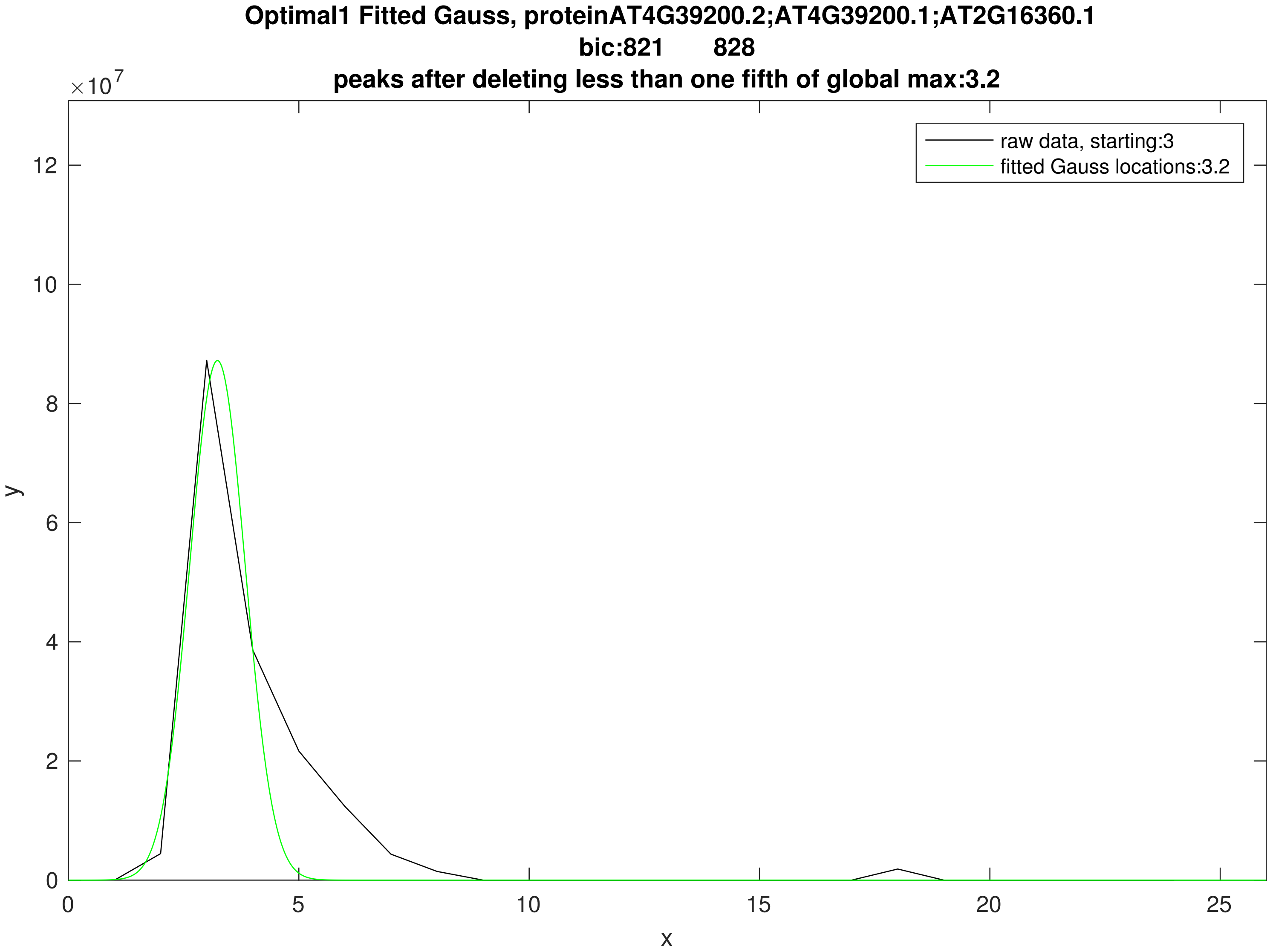
Optimal1 Fitted Gauss, proteinAT4G39200.2;AT4G39200.1;AT2G16360.1
bic:821 828
peaks after deleting less than one fifth of global max:3.2
×10^7
raw data, starting:3
fitted Gauss locations:3.2
y
x
0
2
4
6
8
10
12
0
5
10
15
20
25

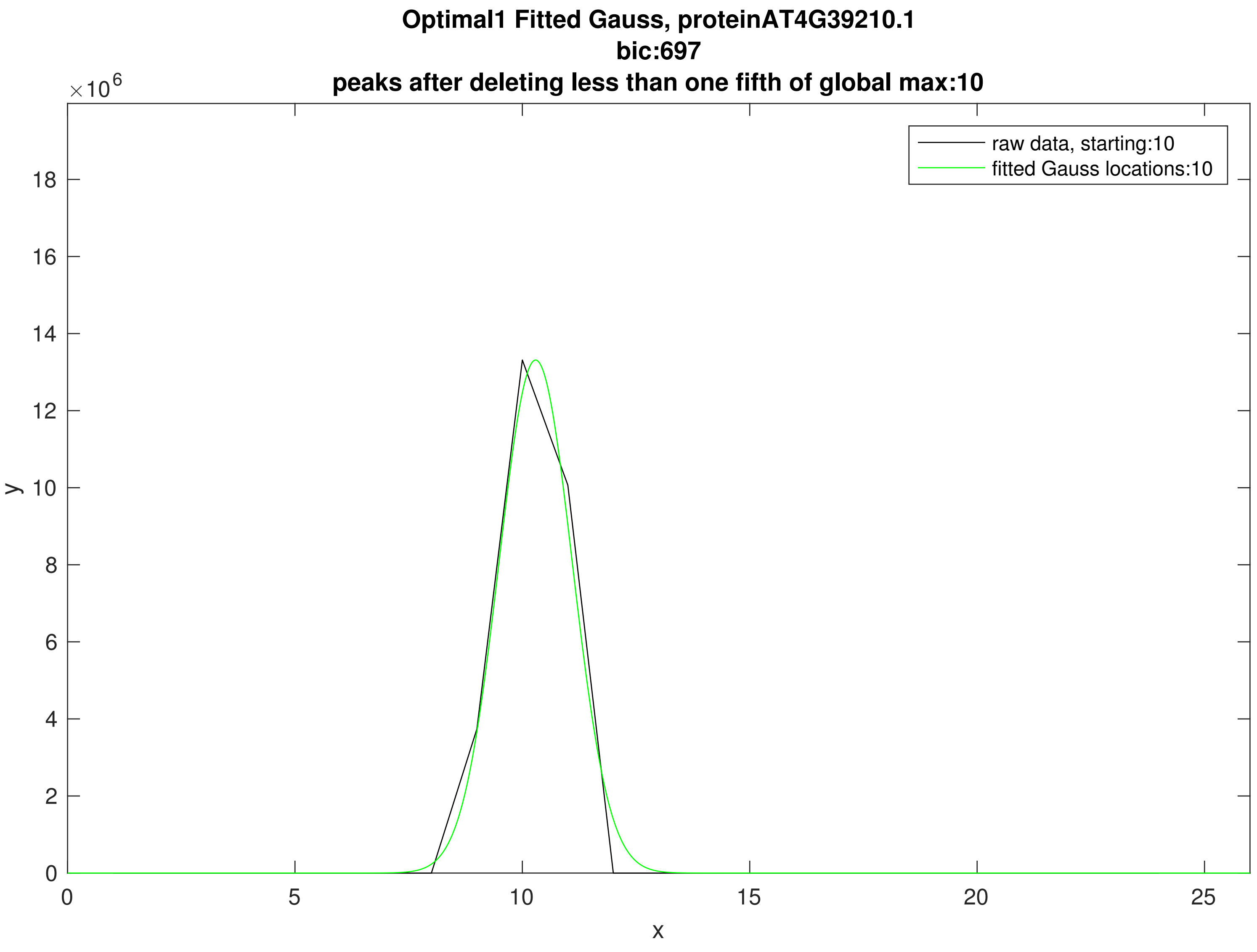
Optimal1 Fitted Gauss, proteinAT4G39210.1
bic:697
peaks after deleting less than one fifth of global max:10
×10^6
raw data, starting:10
fitted Gauss locations:10
y
x
0
2
4
6
8
10
12
14
16
18
0
5
10
15
20
25

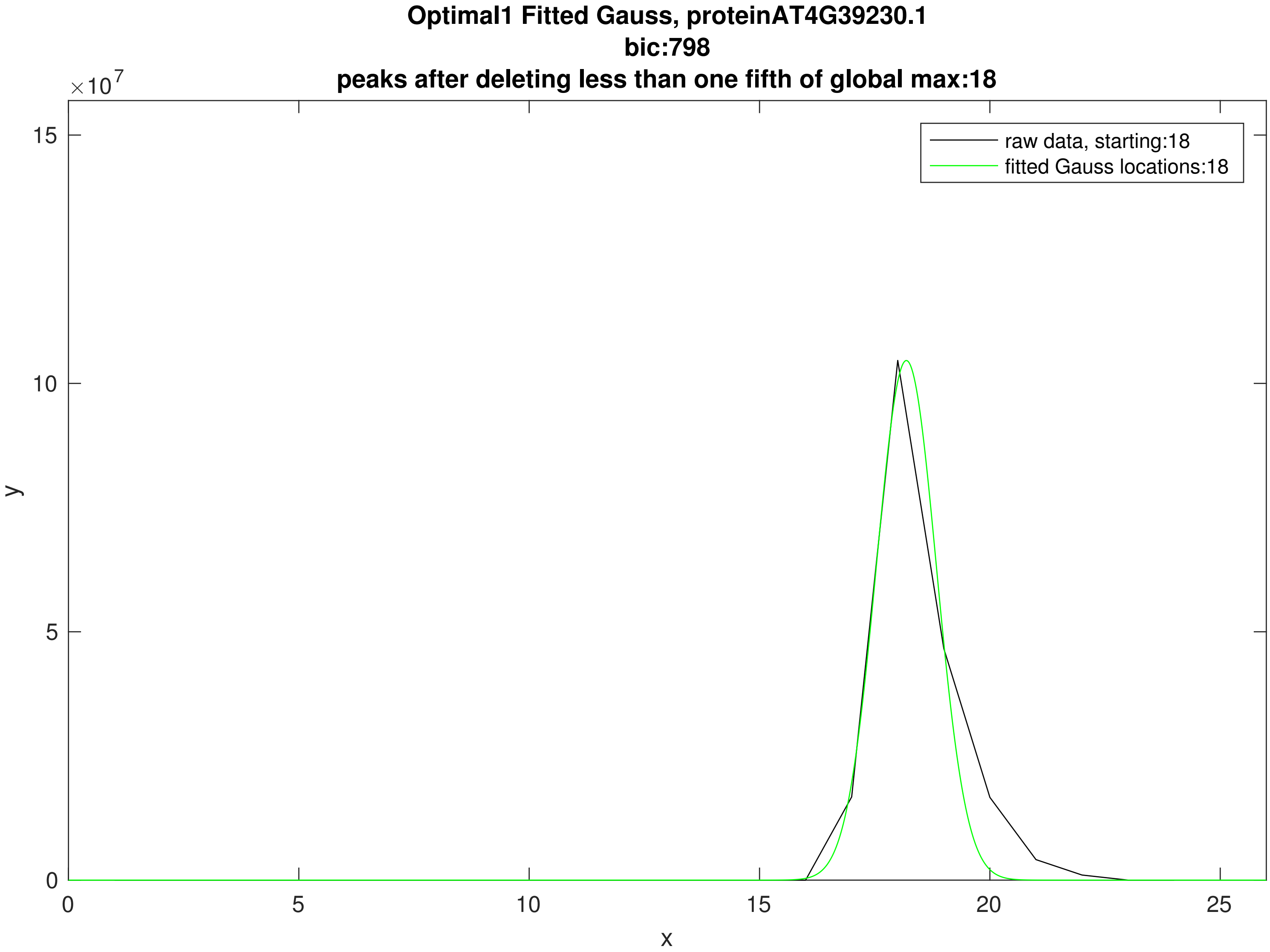

Optimal1 Fitted Gauss, proteinAT4G39230.1
bic:798
peaks after deleting less than one fifth of global max:18
×10^7
15
10
5
0
y
0
5
10
15
20
25
x
raw data, starting:18
fitted Gauss locations:18

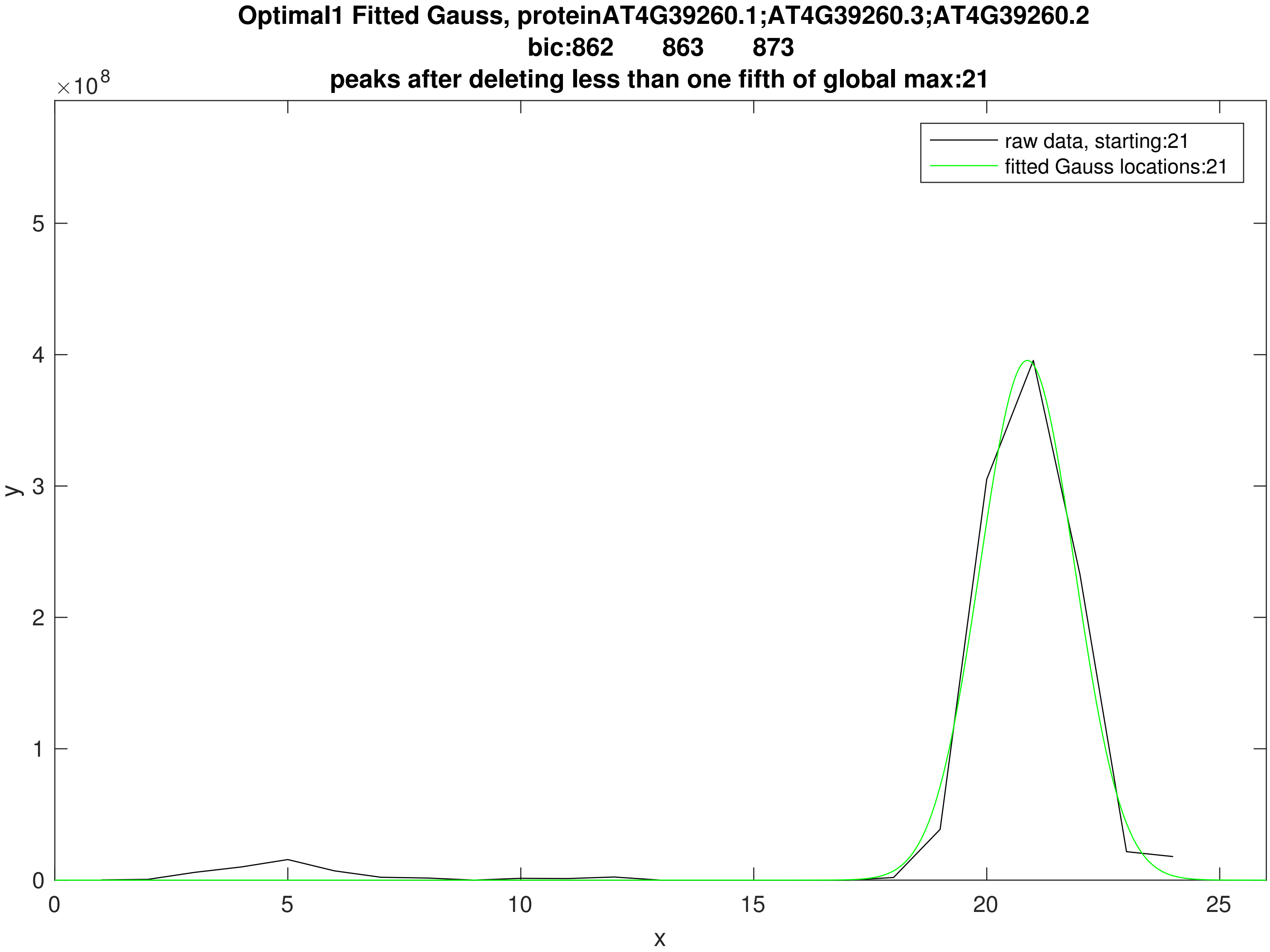

Optimal1 Fitted Gauss, proteinAT4G39260.1;AT4G39260.3;AT4G39260.2
bic:862 863 873
peaks after deleting less than one fifth of global max:21
×10^8
raw data, starting:21
fitted Gauss locations:21
y
x
0
1
2
3
4
5
10
15
20
25

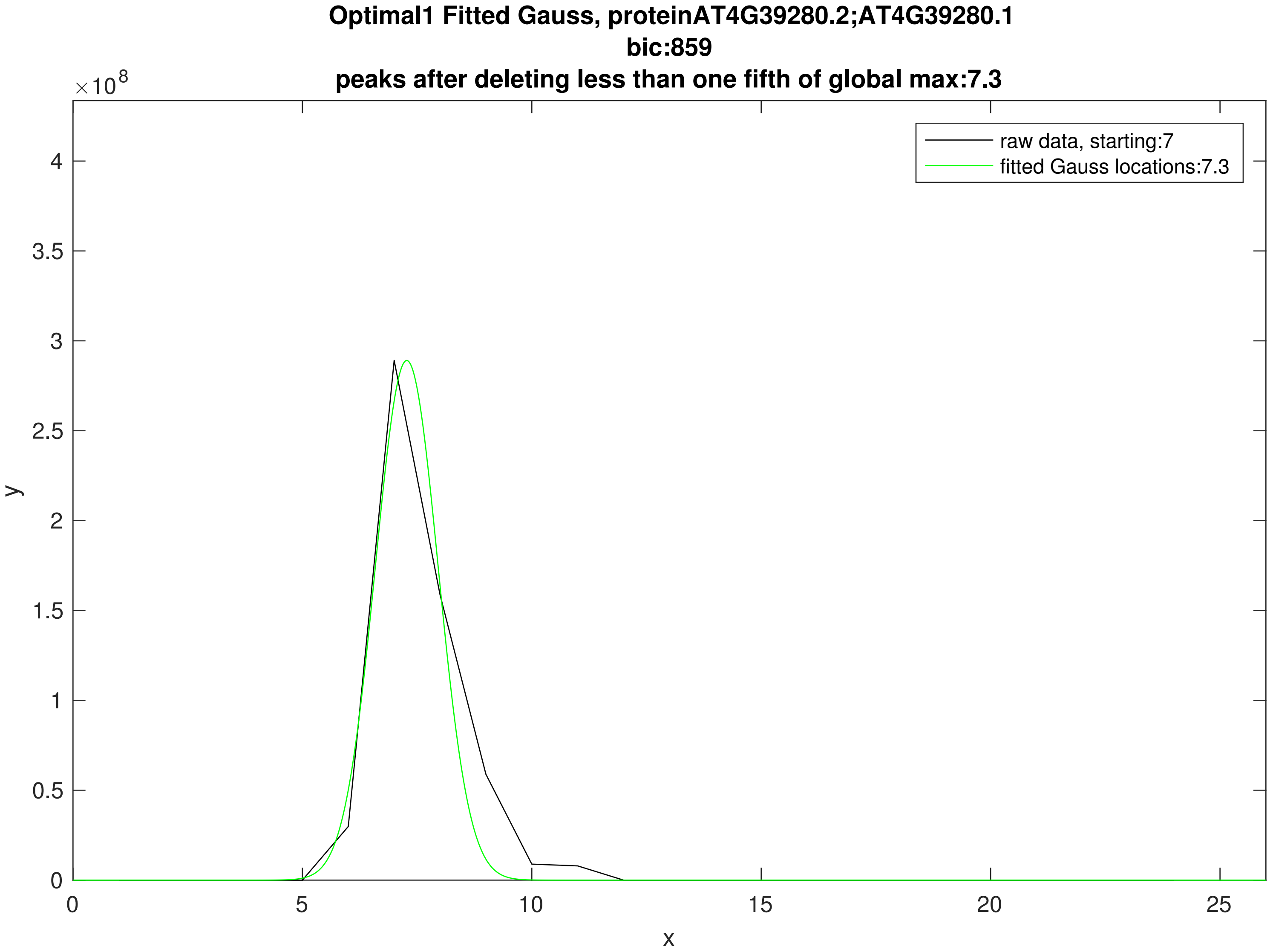

Optimal1 Fitted Gauss, proteinAT4G39280.2;AT4G39280.1
bic:859
peaks after deleting less than one fifth of global max:7.3
×10^8
raw data, starting:7
fitted Gauss locations:7.3
y
x
0
0.5
1
1.5
2
2.5
3
3.5
4
0
5
10
15
20
25

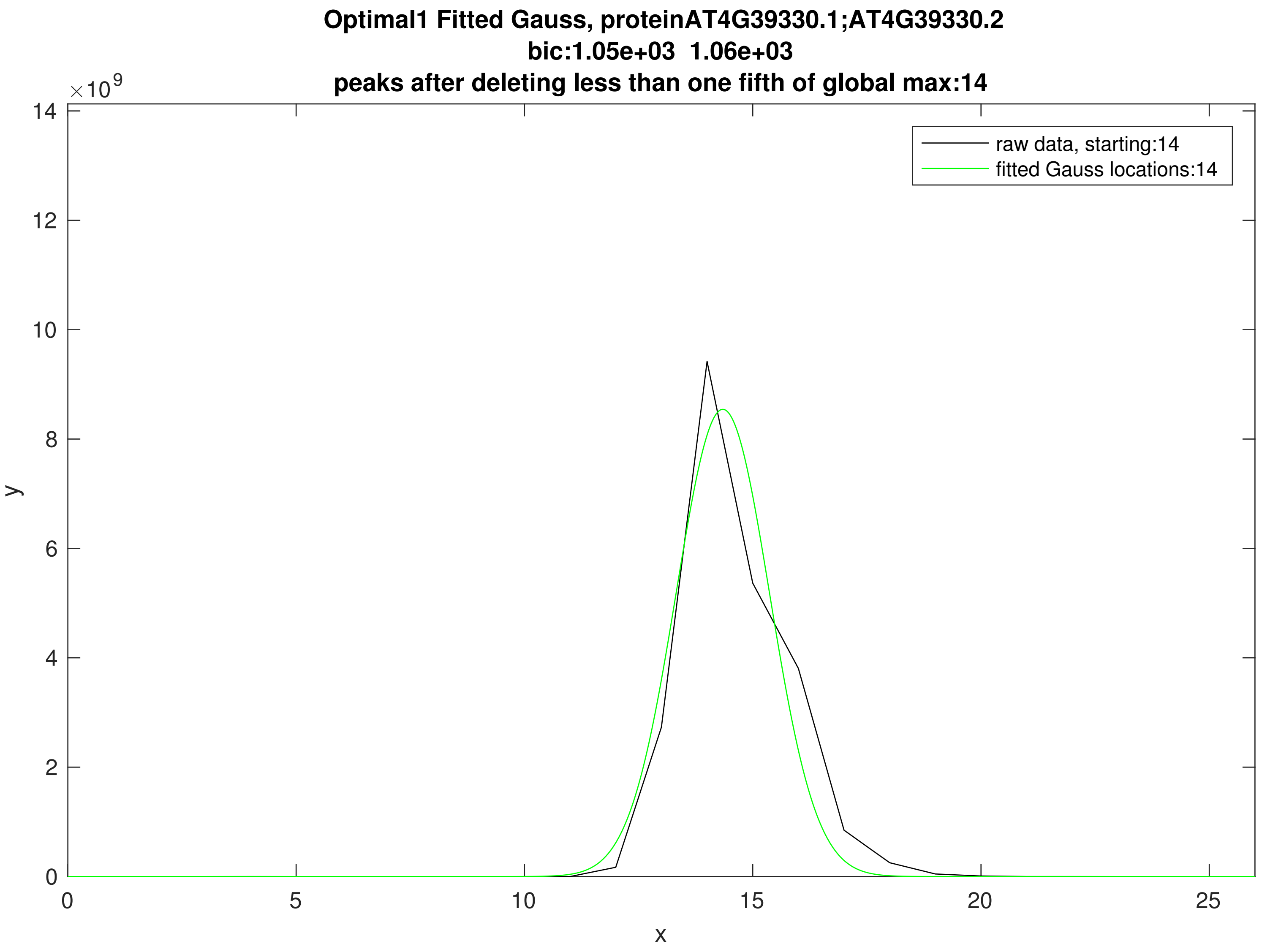
Optimal1 Fitted Gauss, proteinAT4G39330.1;AT4G39330.2
bic:1.05e+03  1.06e+03
peaks after deleting less than one fifth of global max:14
×10^9
raw data, starting:14
fitted Gauss locations:14
y
x
0
2
4
6
8
10
12
14
0
5
10
15
20
25

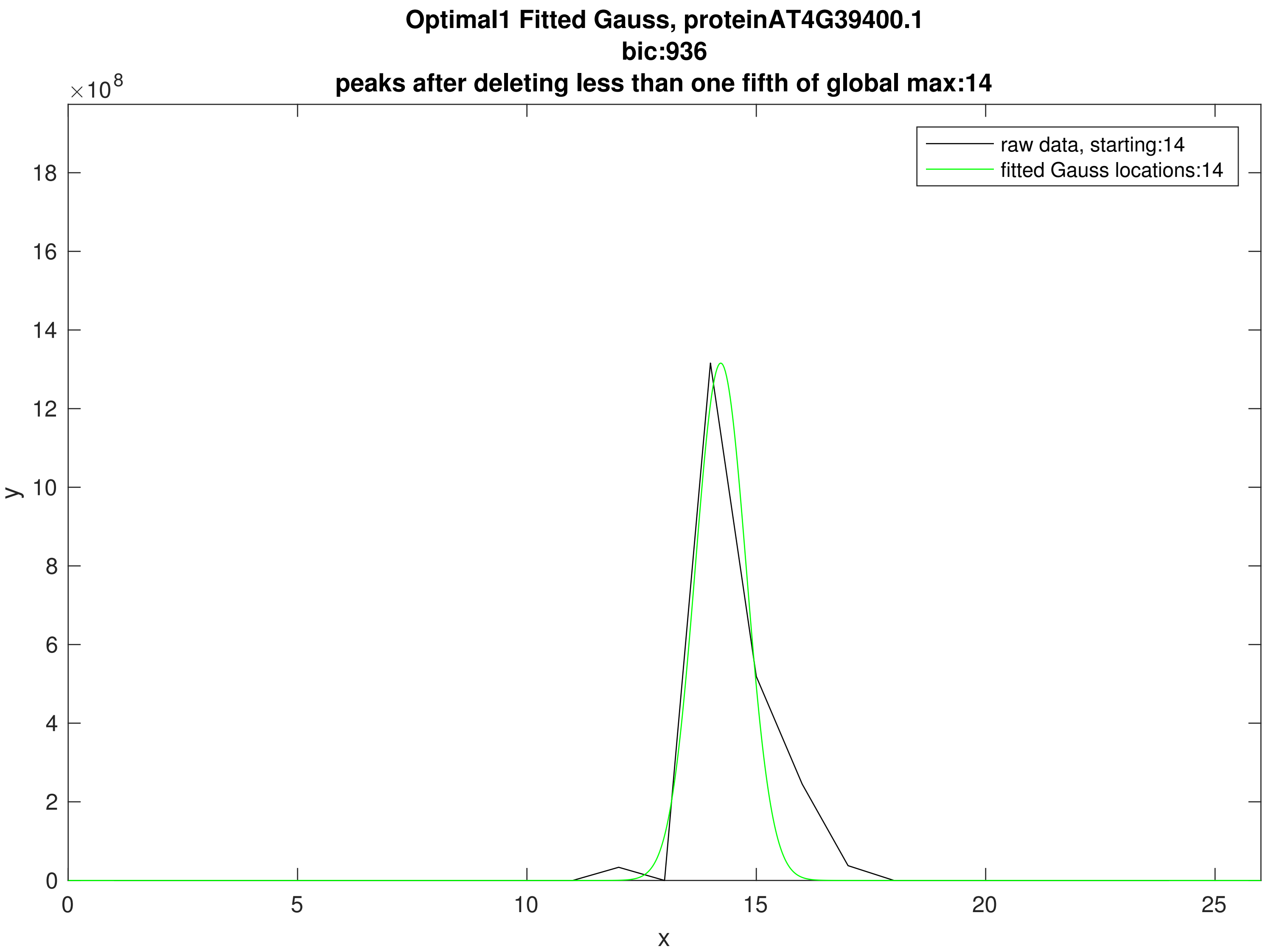
Optimal1 Fitted Gauss, proteinAT4G39400.1
bic:936
peaks after deleting less than one fifth of global max:14
×10^8
raw data, starting:14
fitted Gauss locations:14
y
x
0
2
4
6
8
10
12
14
16
18
0
5
10
15
20
25

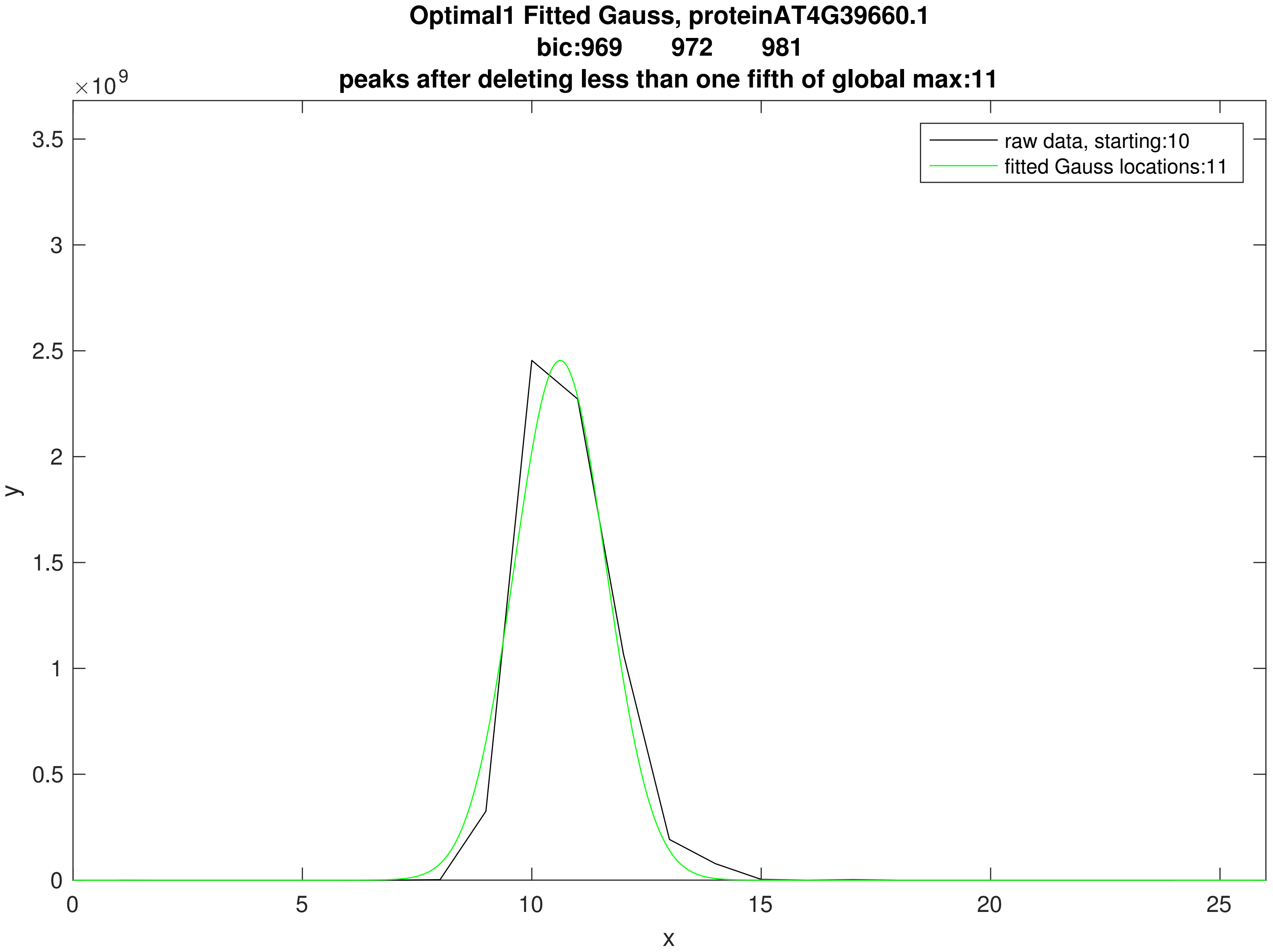
Optimal1 Fitted Gauss, proteinAT4G39660.1
bic:969 972 981
peaks after deleting less than one fifth of global max:11
×10^9
3.5
3
2.5
2
1.5
1
0.5
0
y
0
5
10
15
20
25
x
raw data, starting:10
fitted Gauss locations:11

Optimal1 Fitted Gauss, proteinAT4G39680.2;AT4G39680.1
bic:768
peaks after deleting less than one fifth of global max:7.9

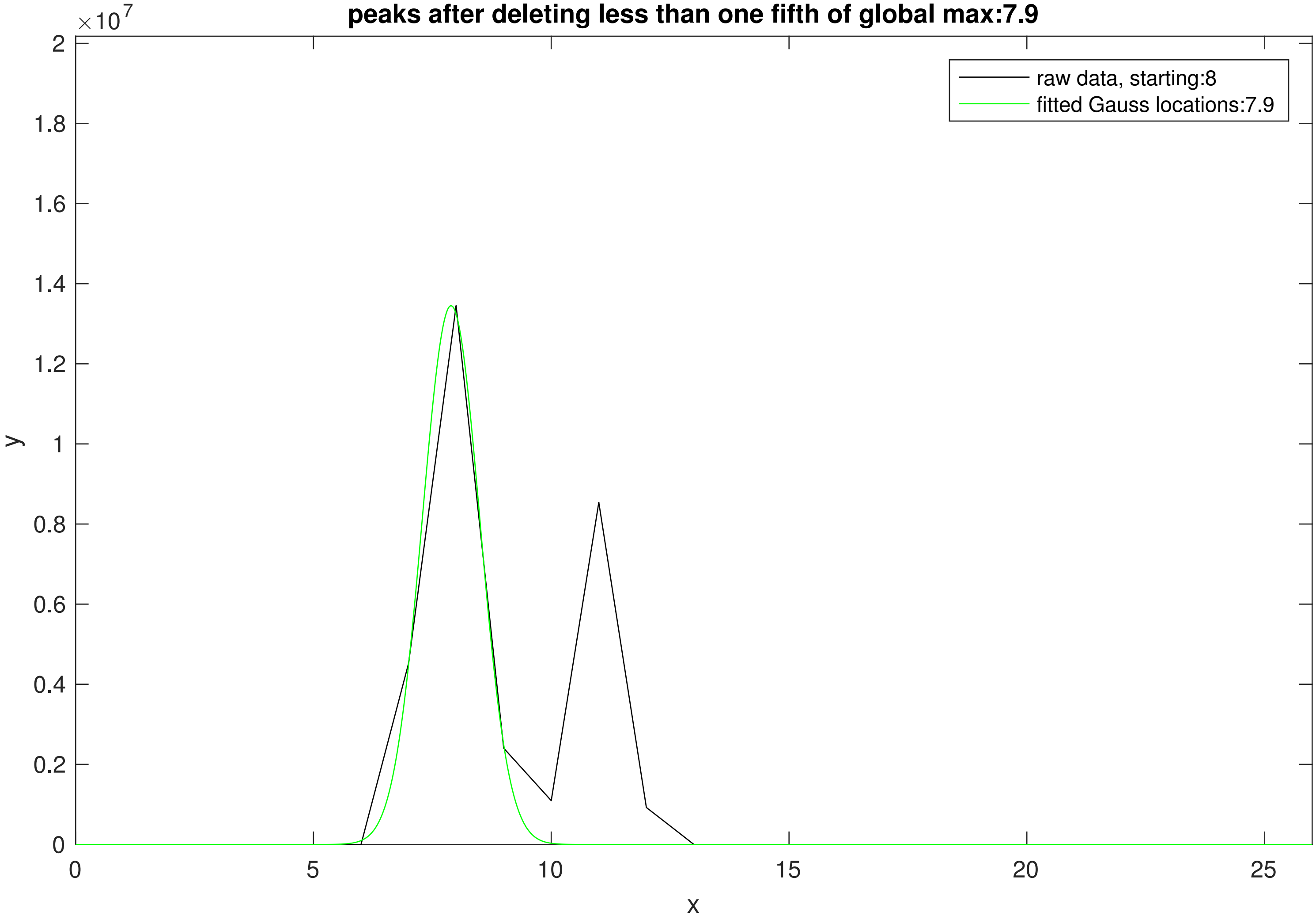

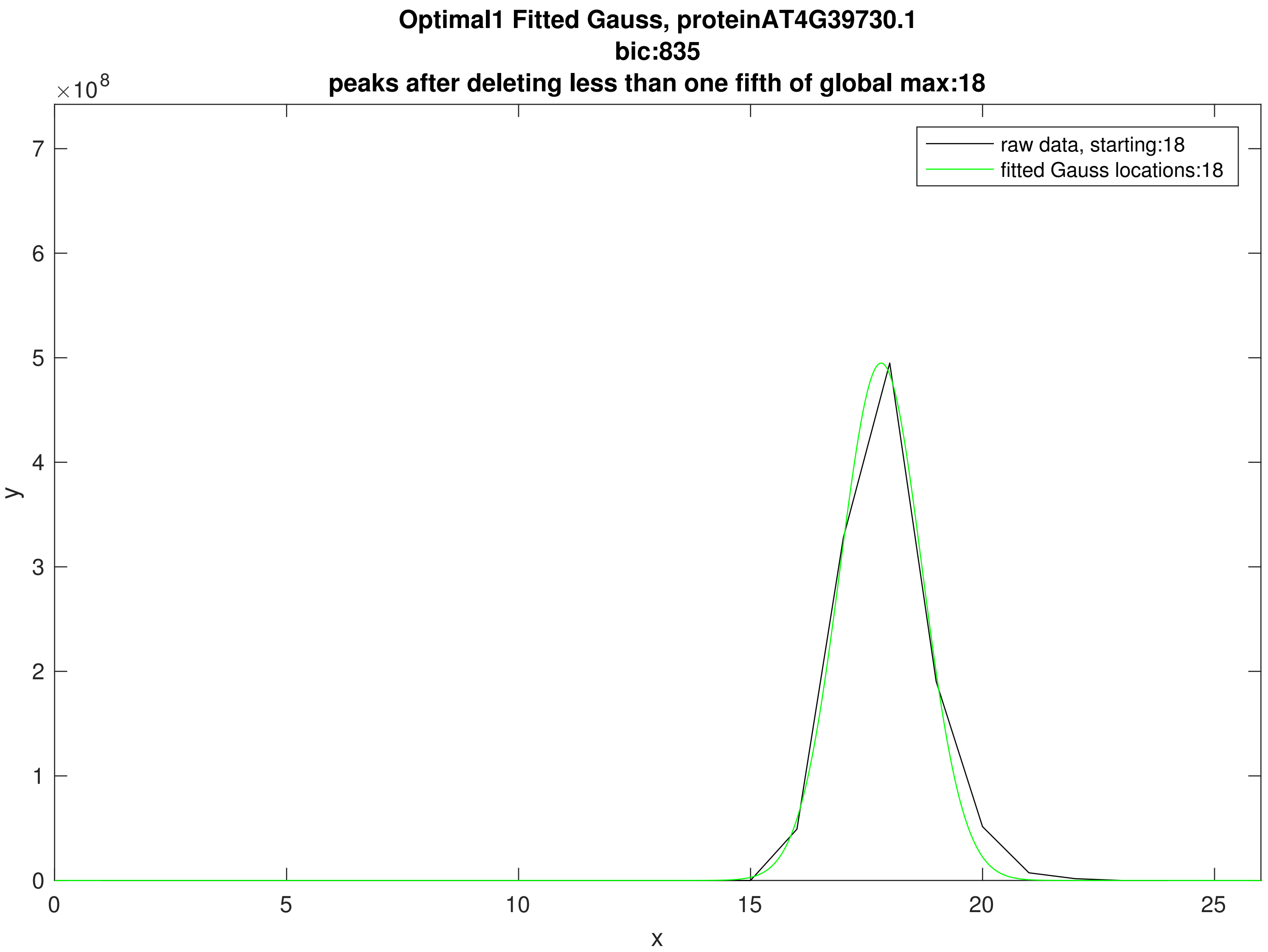
Optimal1 Fitted Gauss, proteinAT4G39730.1
bic:835
peaks after deleting less than one fifth of global max:18
×10^8
y
x
raw data, starting:18
fitted Gauss locations:18

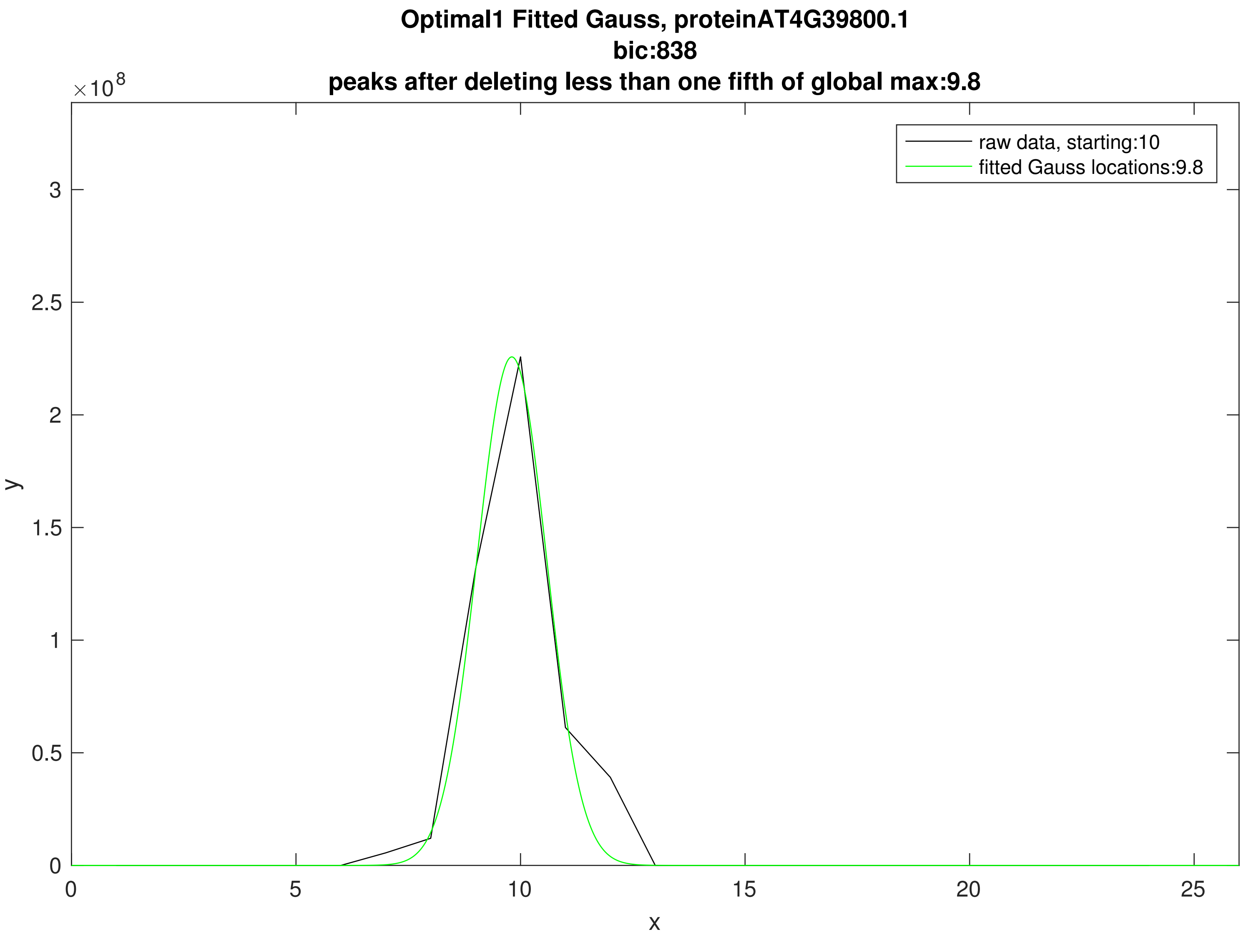
Optimal1 Fitted Gauss, proteinAT4G39800.1
bic:838
peaks after deleting less than one fifth of global max:9.8
×10^8
raw data, starting:10
fitted Gauss locations:9.8
y
x
0
0.5
1
1.5
2
2.5
3
5
10
15
20
25

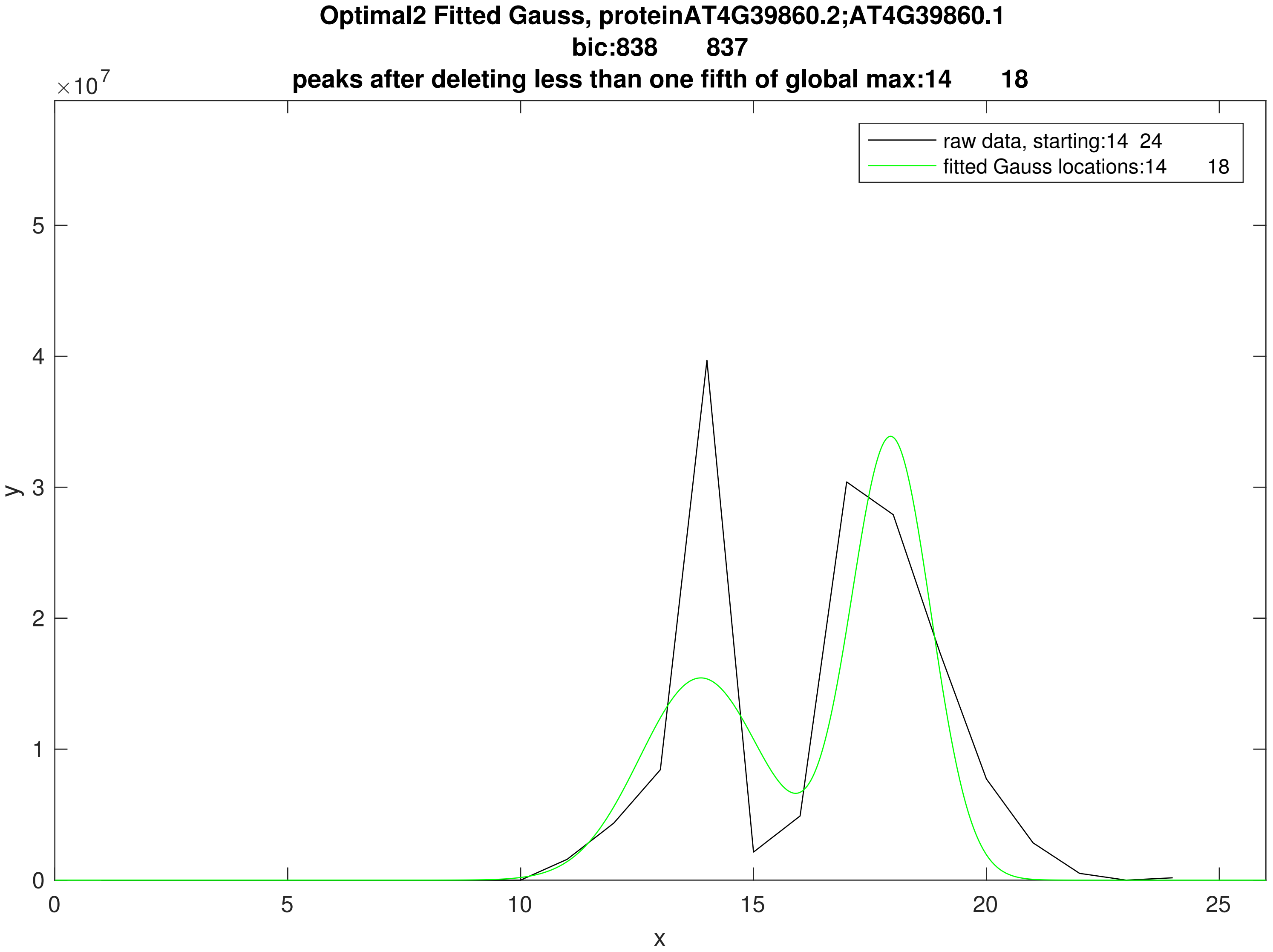
Optimal2 Fitted Gauss, proteinAT4G39860.2;AT4G39860.1
bic:838 837
peaks after deleting less than one fifth of global max:14 18
×10^7
raw data, starting:14 24
fitted Gauss locations:14 18
y
x
0
1
2
3
4
5
0
5
10
15
20
25

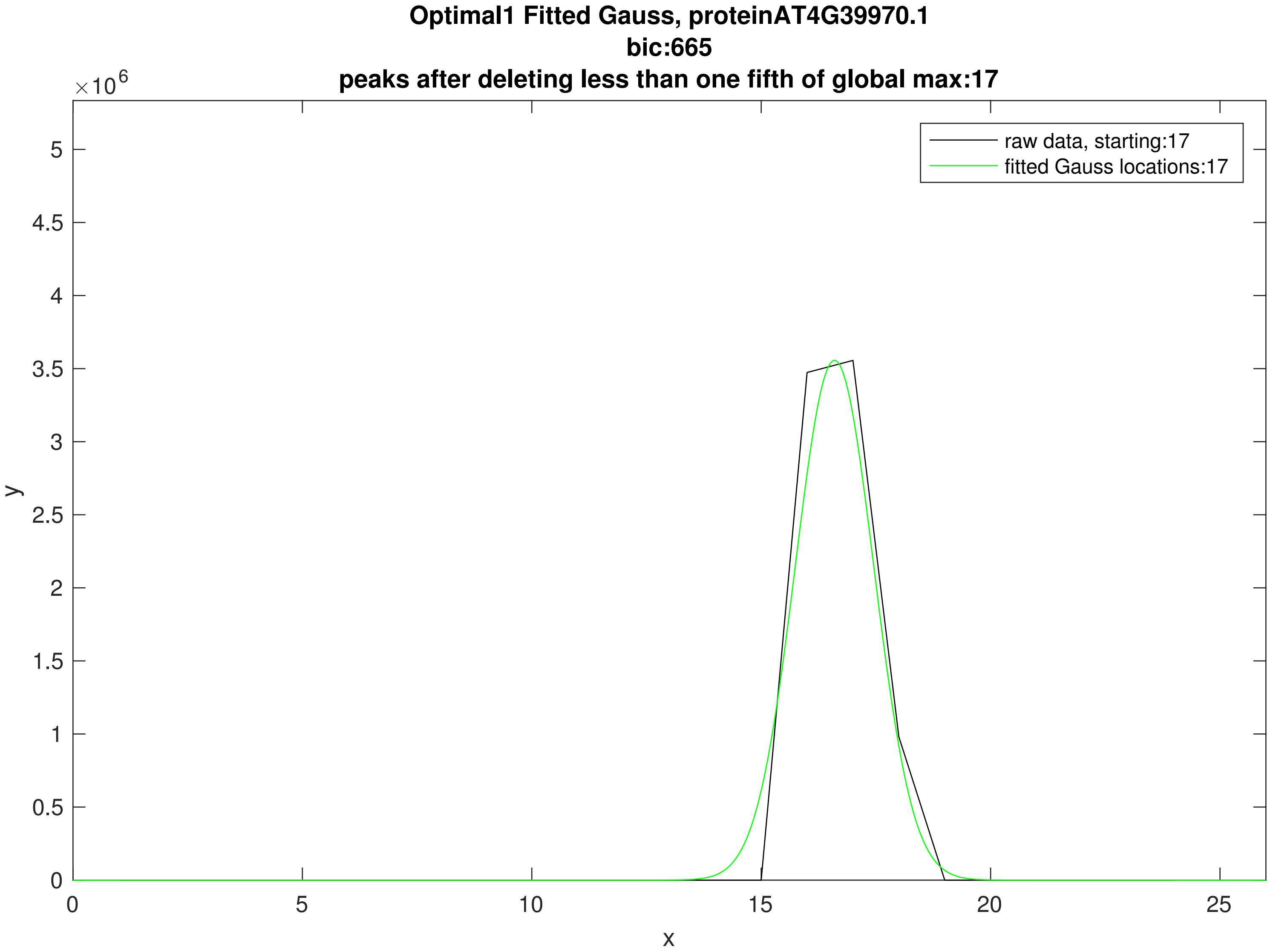
Optimal1 Fitted Gauss, proteinAT4G39970.1
bic:665
peaks after deleting less than one fifth of global max:17
×10^6
raw data, starting:17
fitted Gauss locations:17
y
x
0
0.5
1
1.5
2
2.5
3
3.5
4
4.5
5
0
5
10
15
20
25

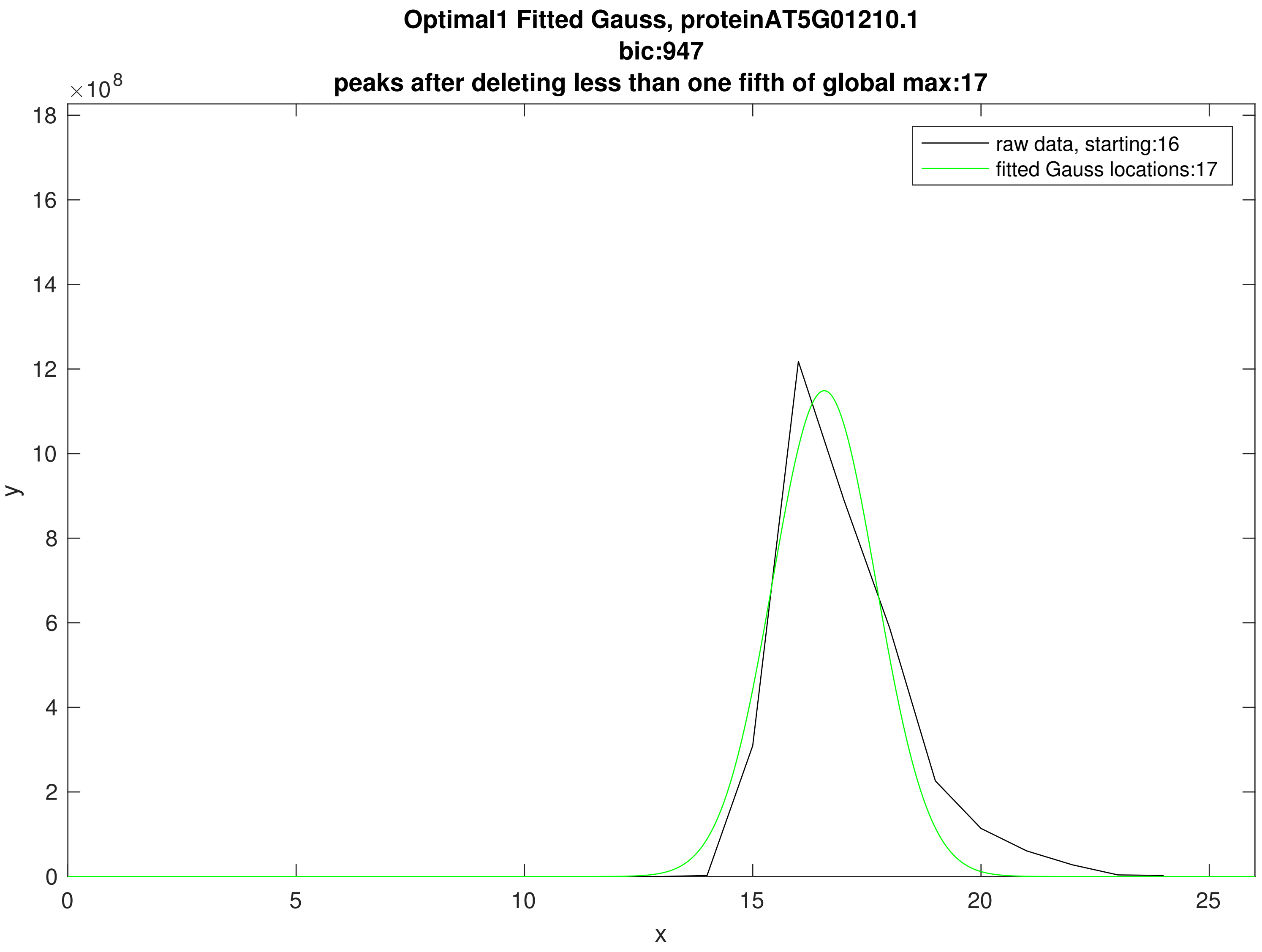
Optimal1 Fitted Gauss, proteinAT5G01210.1
bic:947
peaks after deleting less than one fifth of global max:17
×10^8
raw data, starting:16
fitted Gauss locations:17
y
x
0
2
4
6
8
10
12
14
16
18
0
5
10
15
20
25

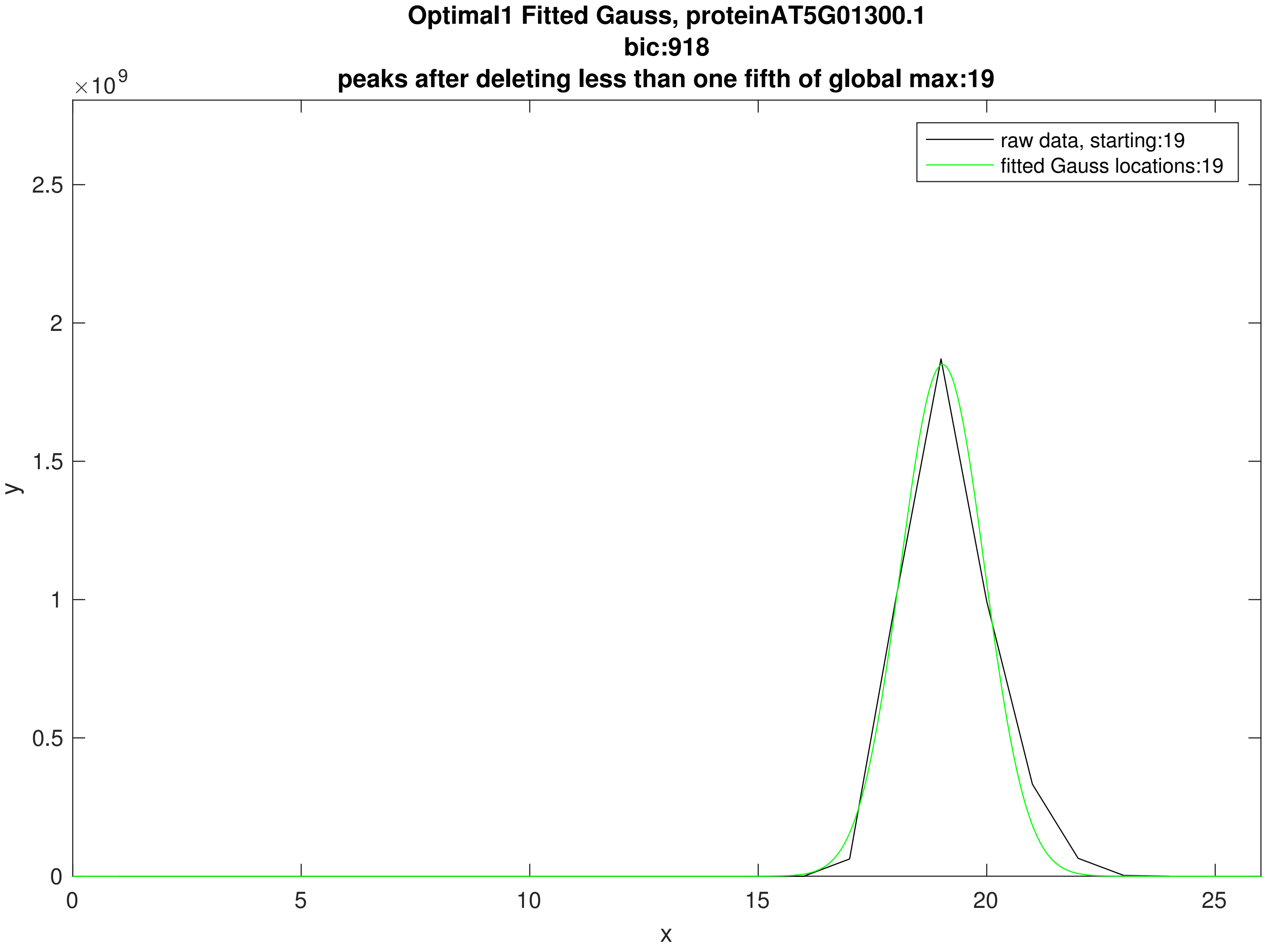
Optimal1 Fitted Gauss, proteinAT5G01300.1
bic:918
peaks after deleting less than one fifth of global max:19
×10^9
raw data, starting:19
fitted Gauss locations:19
2.5
2
1.5
1
0.5
0
y
0
5
10
15
20
25
x

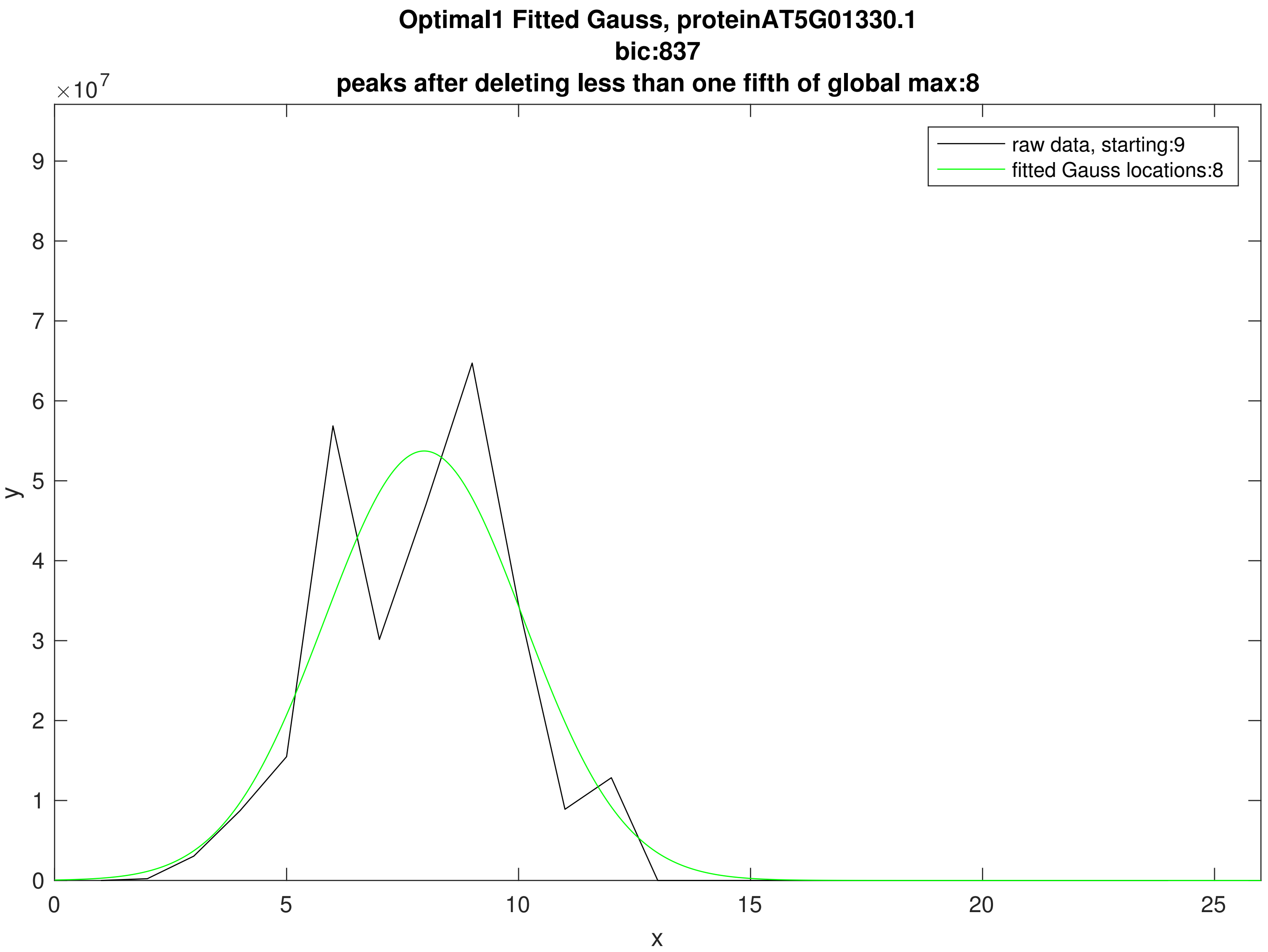
Optimal1 Fitted Gauss, proteinAT5G01330.1
bic:837
peaks after deleting less than one fifth of global max:8
×10^7
raw data, starting:9
fitted Gauss locations:8
y
x
0
1
2
3
4
5
6
7
8
9
0
5
10
15
20
25

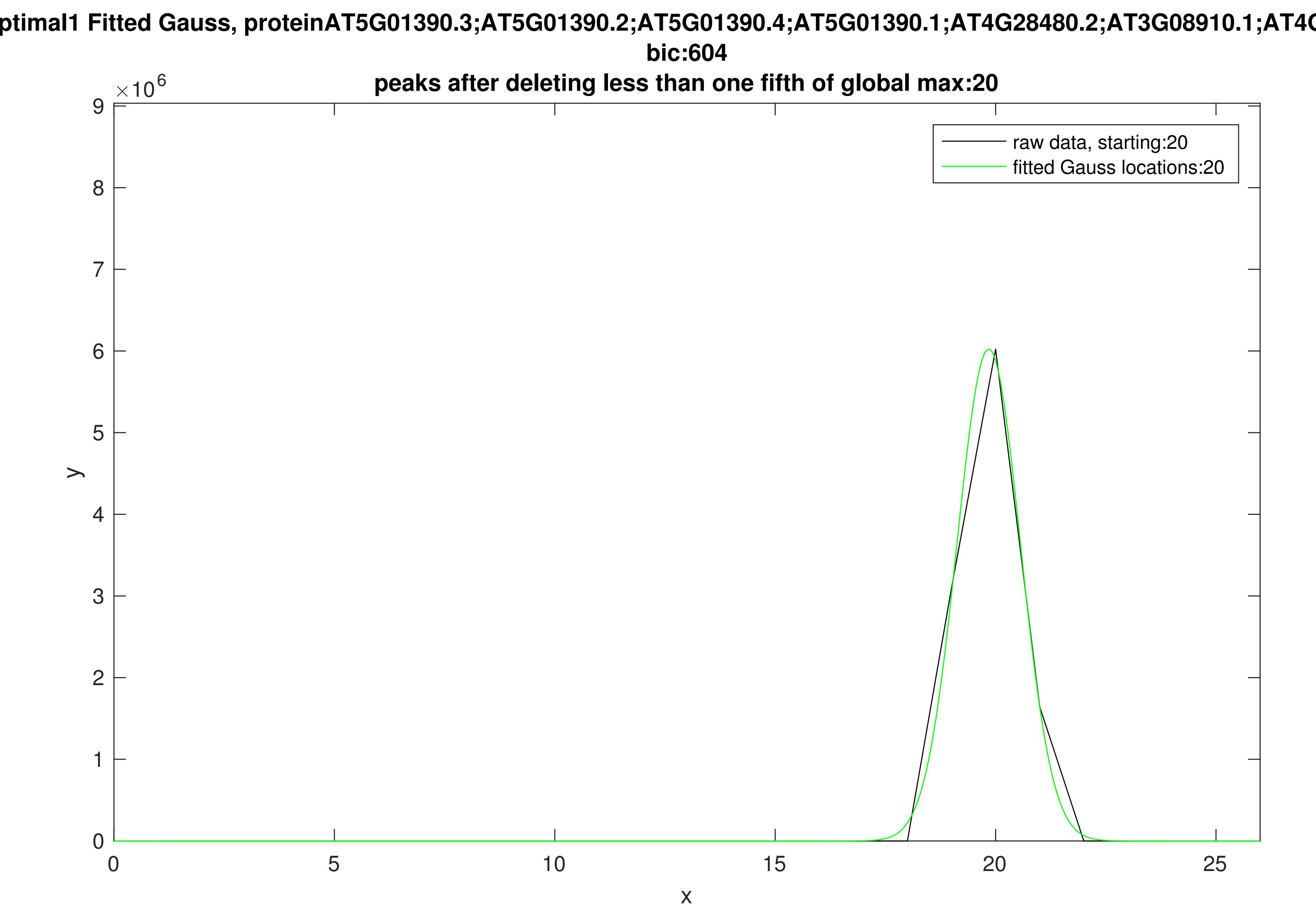
ptimal1 Fitted Gauss, proteinAT5G01390.3;AT5G01390.2;AT5G01390.4;AT5G01390.1;AT4G28480.2;AT3G08910.1;AT4C
bic:604
peaks after deleting less than one fifth of global max:20
×10^6
raw data, starting:20
fitted Gauss locations:20
y
x
0
1
2
3
4
5
6
7
8
9
0
5
10
15
20
25

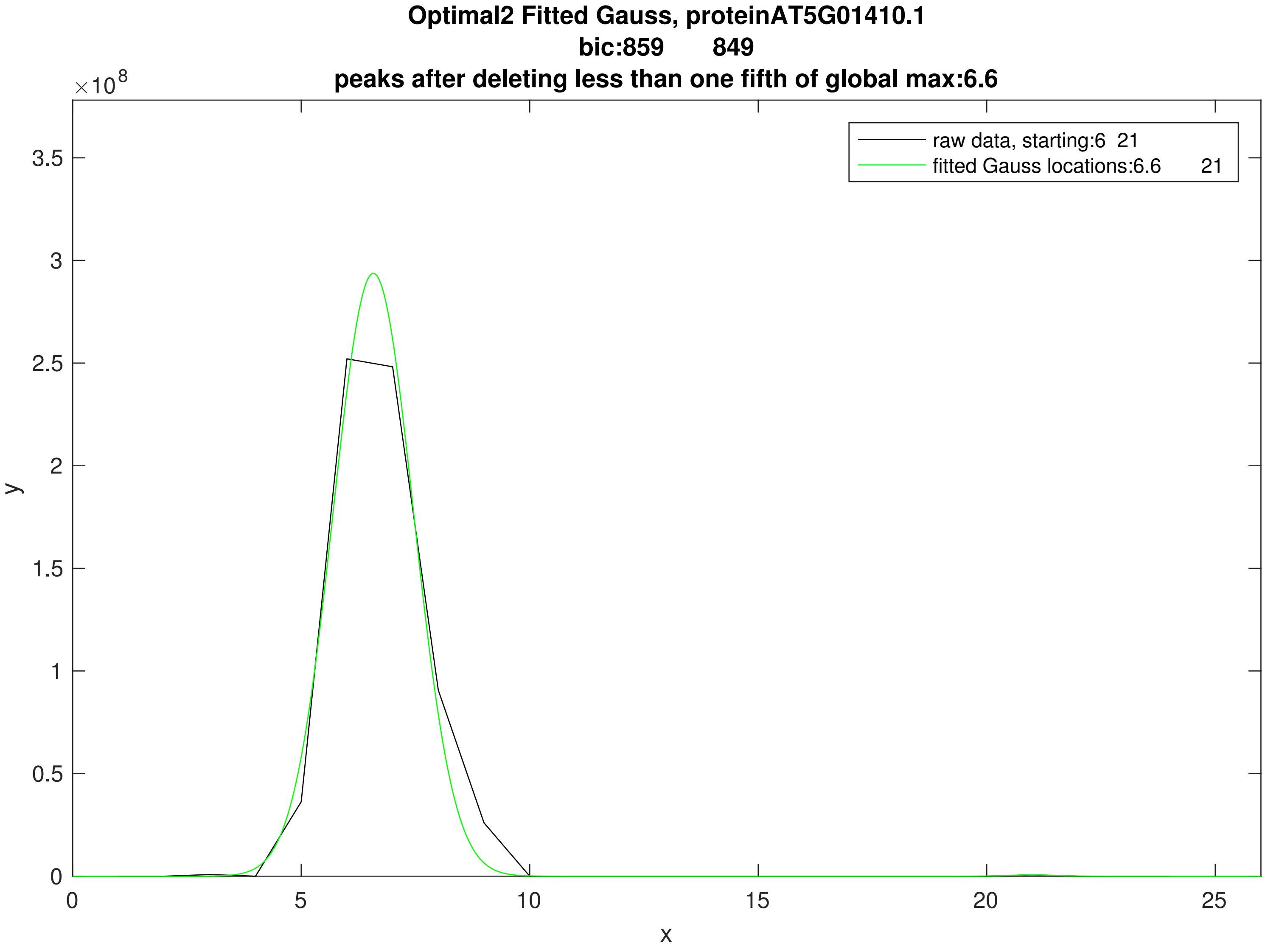

Optimal2 Fitted Gauss, proteinAT5G01410.1
bic:859 849
peaks after deleting less than one fifth of global max:6.6
×10^8
raw data, starting:6 21
fitted Gauss locations:6.6 21
3.5
3
2.5
2
1.5
1
0.5
0
y
0
5
10
15
20
25
x

Optimal1 Fitted Gauss, proteinAT5G01530.1
bic:654
peaks after deleting less than one fifth of global max:1

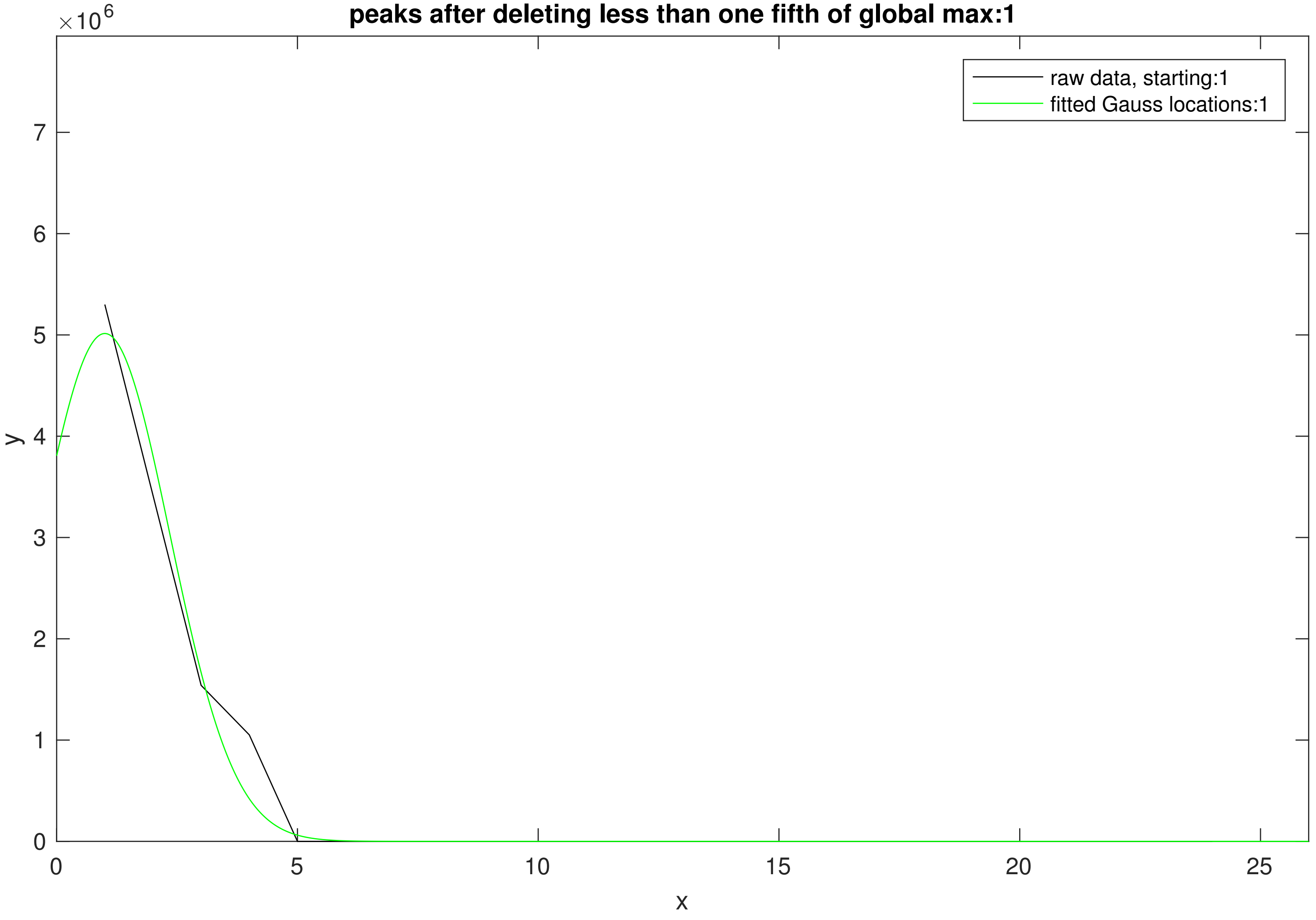

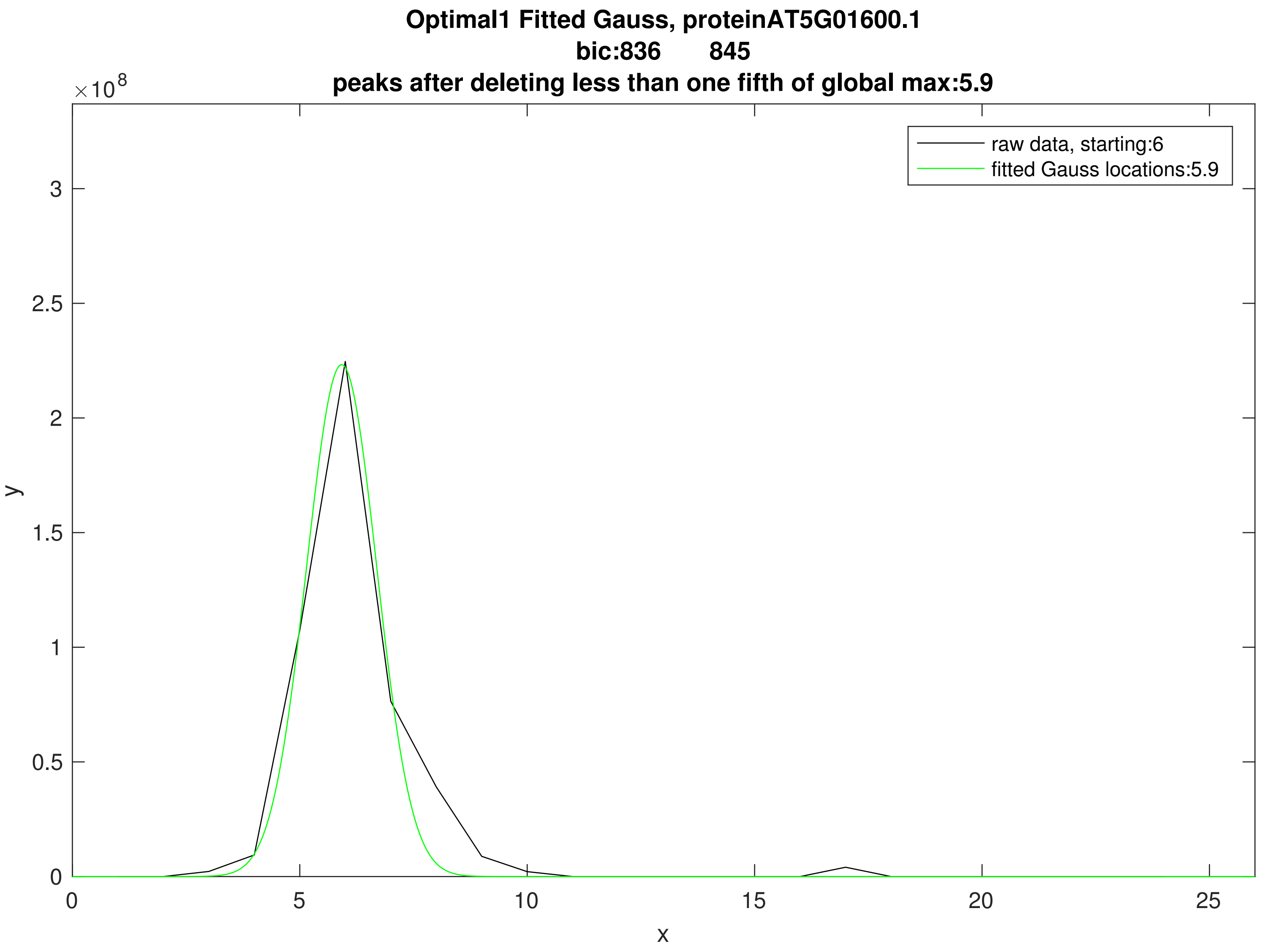
Optimal1 Fitted Gauss, proteinAT5G01600.1
bic:836 845
peaks after deleting less than one fifth of global max:5.9
×10^8
raw data, starting:6
fitted Gauss locations:5.9
y
x
0
0.5
1
1.5
2
2.5
3
0
5
10
15
20
25

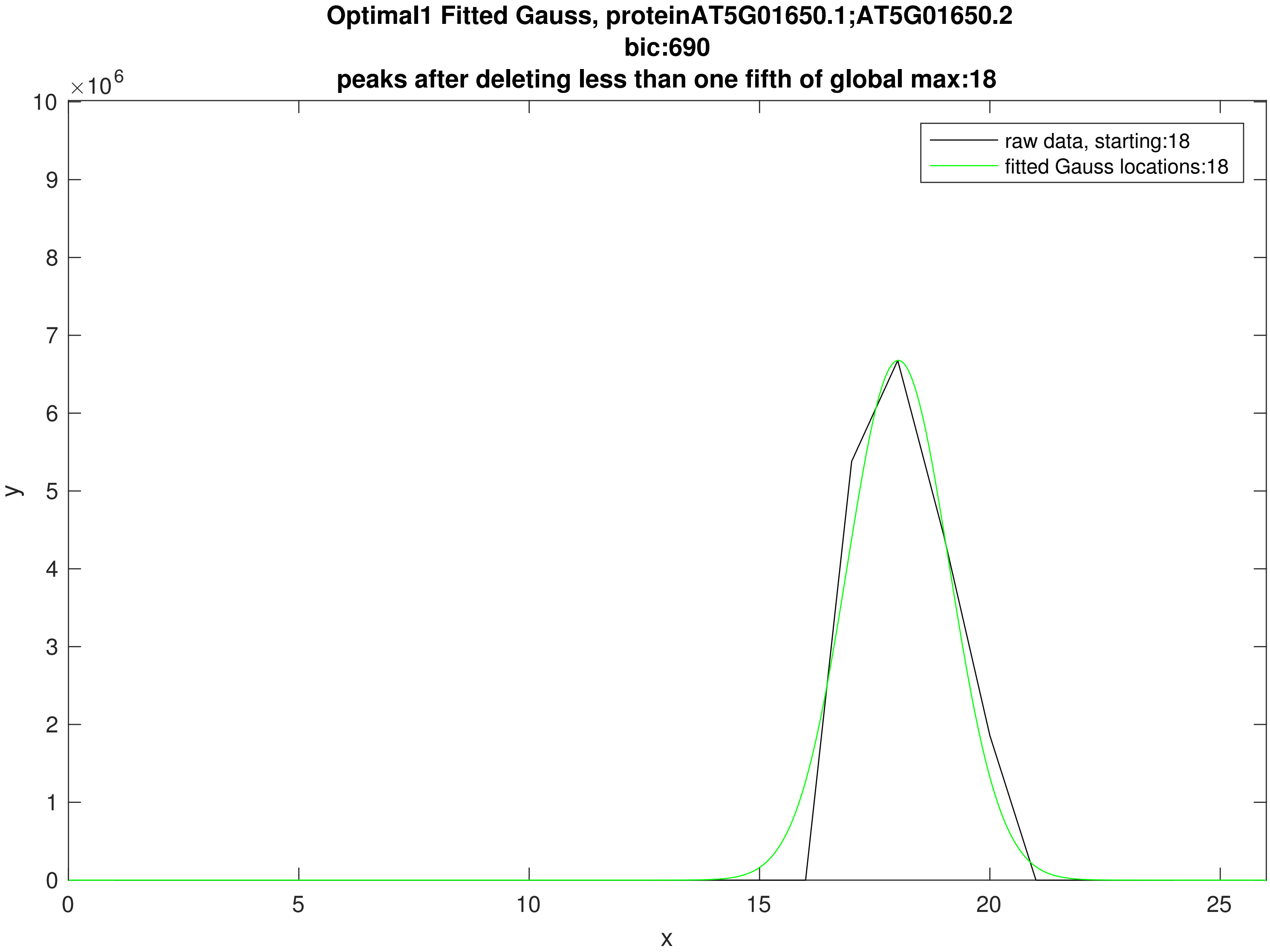
Optimal1 Fitted Gauss, proteinAT5G01650.1;AT5G01650.2
bic:690
peaks after deleting less than one fifth of global max:18
raw data, starting:18
fitted Gauss locations:18
y
x

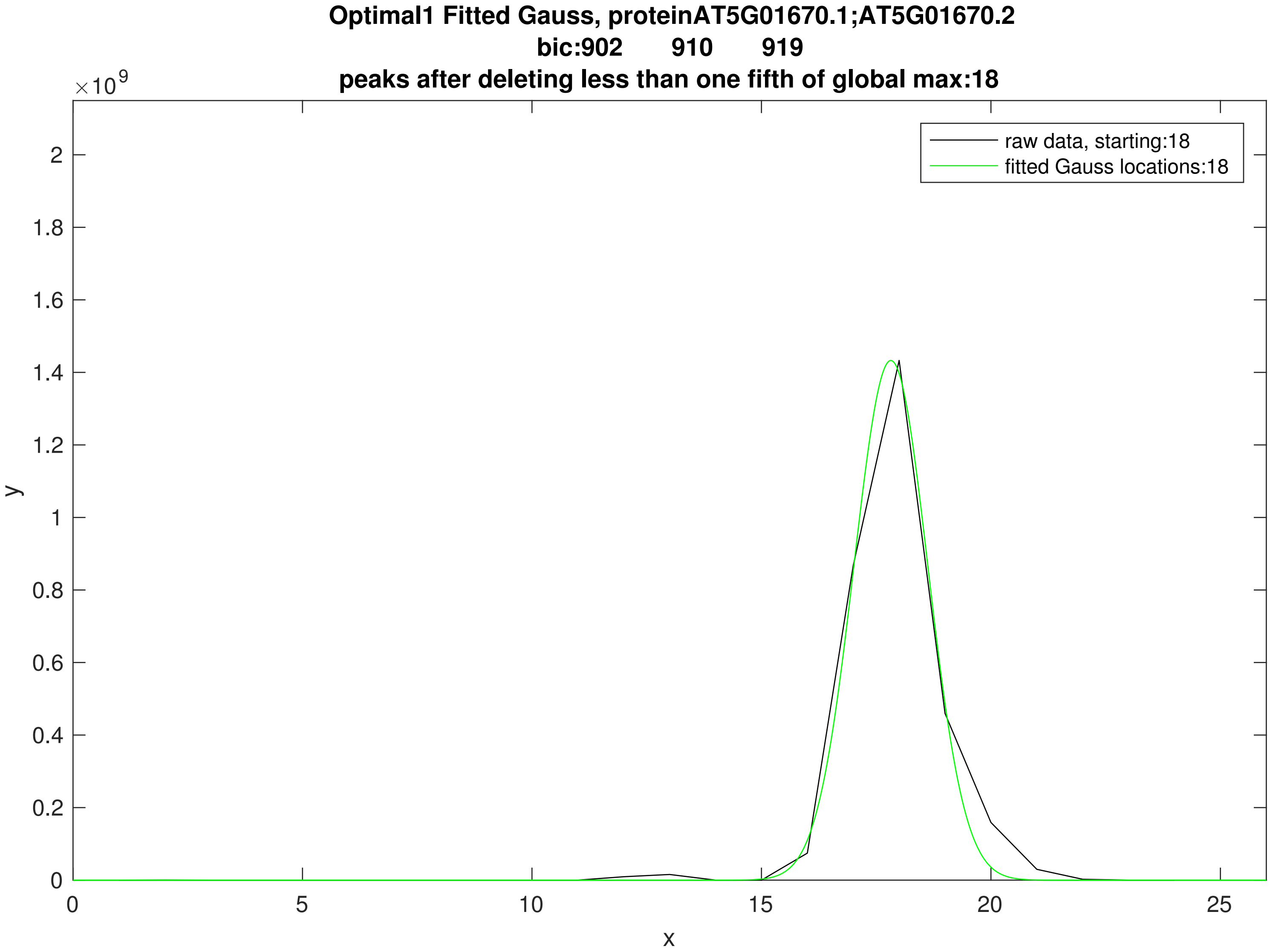
Optimal1 Fitted Gauss, proteinAT5G01670.1;AT5G01670.2
bic:902 910 919
peaks after deleting less than one fifth of global max:18
×10^9
raw data, starting:18
fitted Gauss locations:18
y
x
0
0.2
0.4
0.6
0.8
1
1.2
1.4
1.6
1.8
2
0
5
10
15
20
25

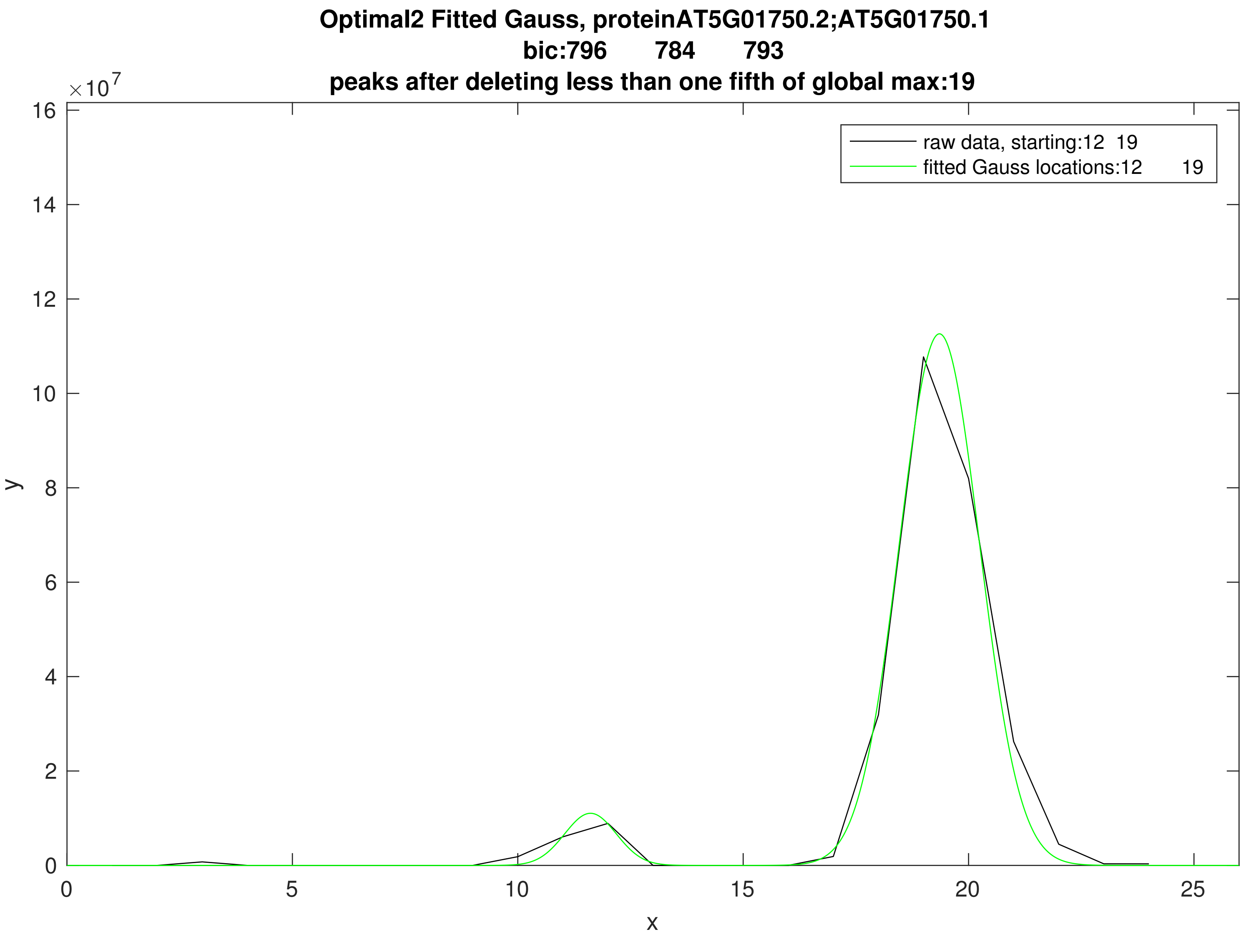
Optimal2 Fitted Gauss, proteinAT5G01750.2;AT5G01750.1
bic:796 784 793
peaks after deleting less than one fifth of global max:19
×10^7
raw data, starting:12 19
fitted Gauss locations:12 19
y
x
0
2
4
6
8
10
12
14
16
5
15
20
25

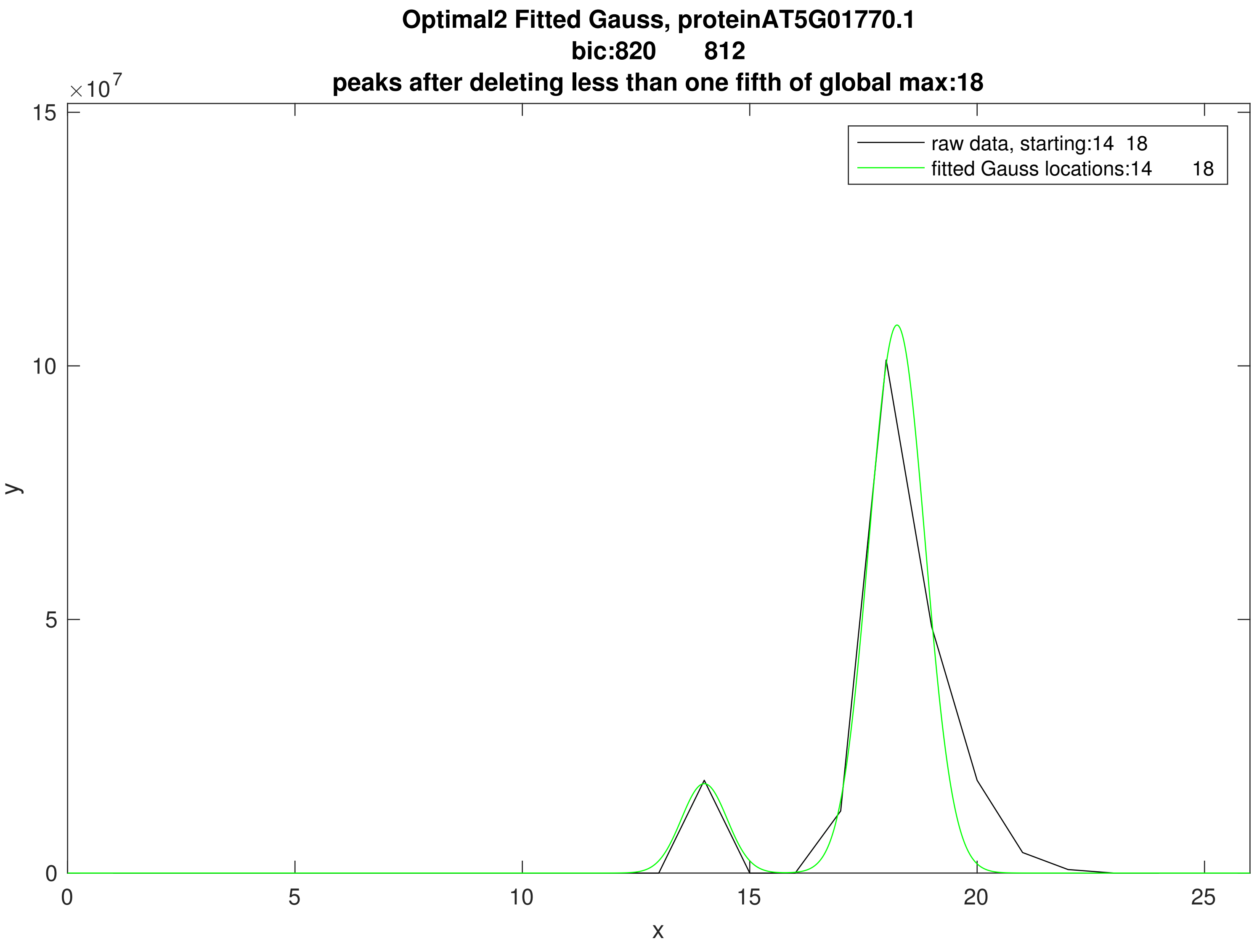
Optimal2 Fitted Gauss, proteinAT5G01770.1
bic:820 812
peaks after deleting less than one fifth of global max:18
×10^7
15
10
5
0
y
0
5
10
15
20
25
x
raw data, starting:14 18
fitted Gauss locations:14 18

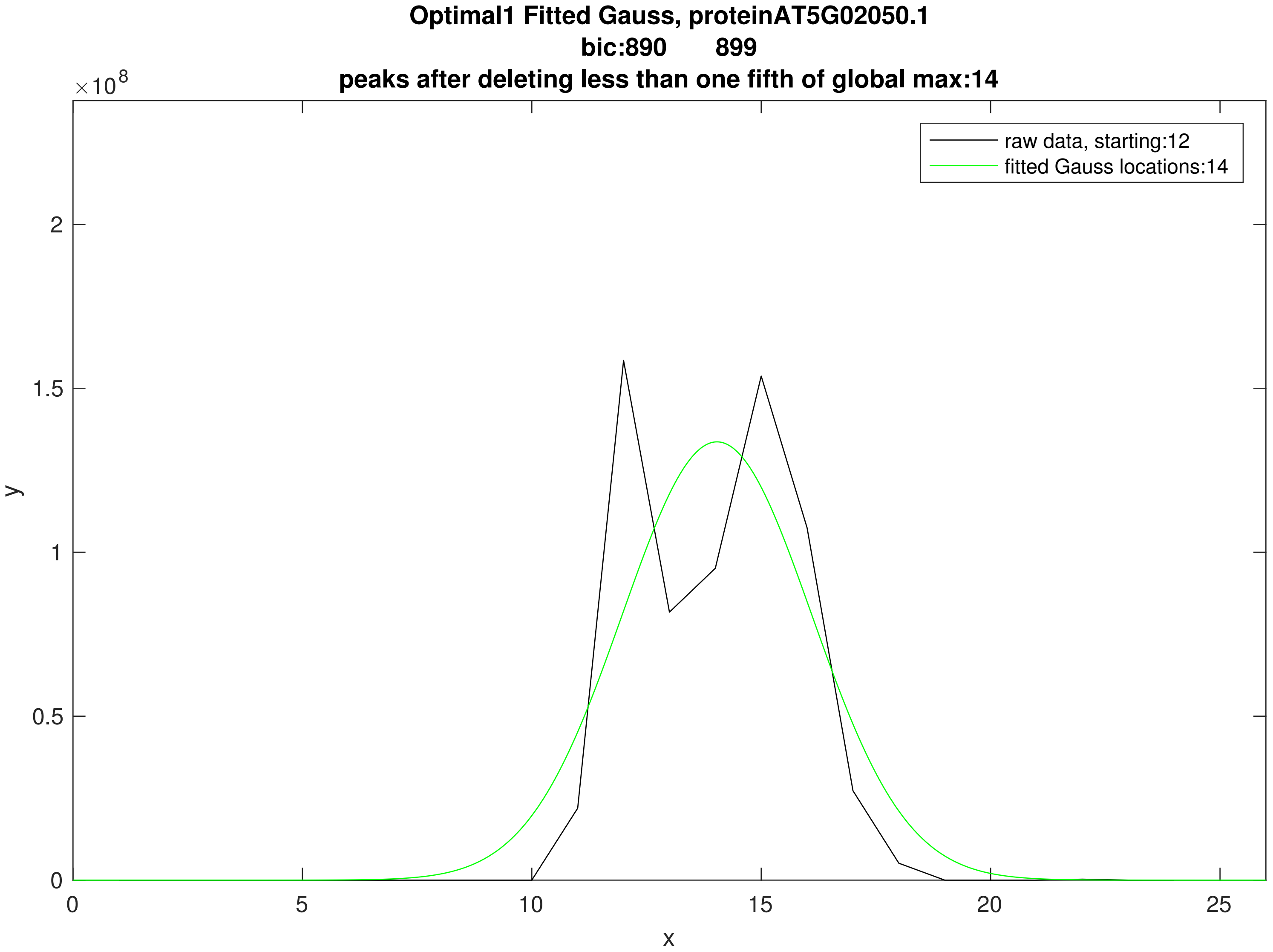

Optimal1 Fitted Gauss, proteinAT5G02050.1
bic:890 899
peaks after deleting less than one fifth of global max:14
×10^8
raw data, starting:12
fitted Gauss locations:14
y
x
0
0.5
1
1.5
2
0
5
10
15
20
25

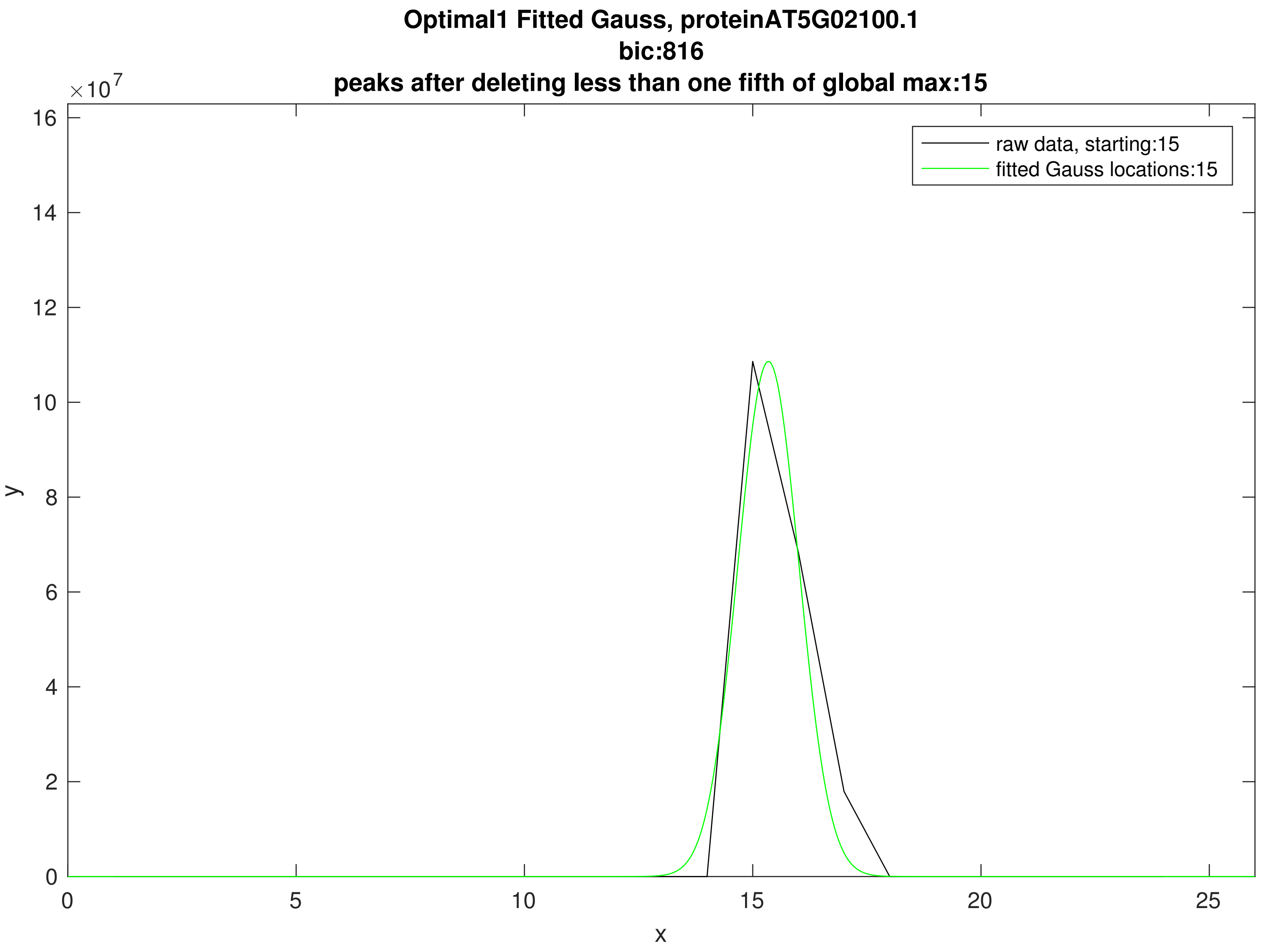
Optimal1 Fitted Gauss, proteinAT5G02100.1
bic:816
peaks after deleting less than one fifth of global max:15
×10^7
raw data, starting:15
fitted Gauss locations:15
y
x
0
2
4
6
8
10
12
14
16
0
5
10
15
20
25

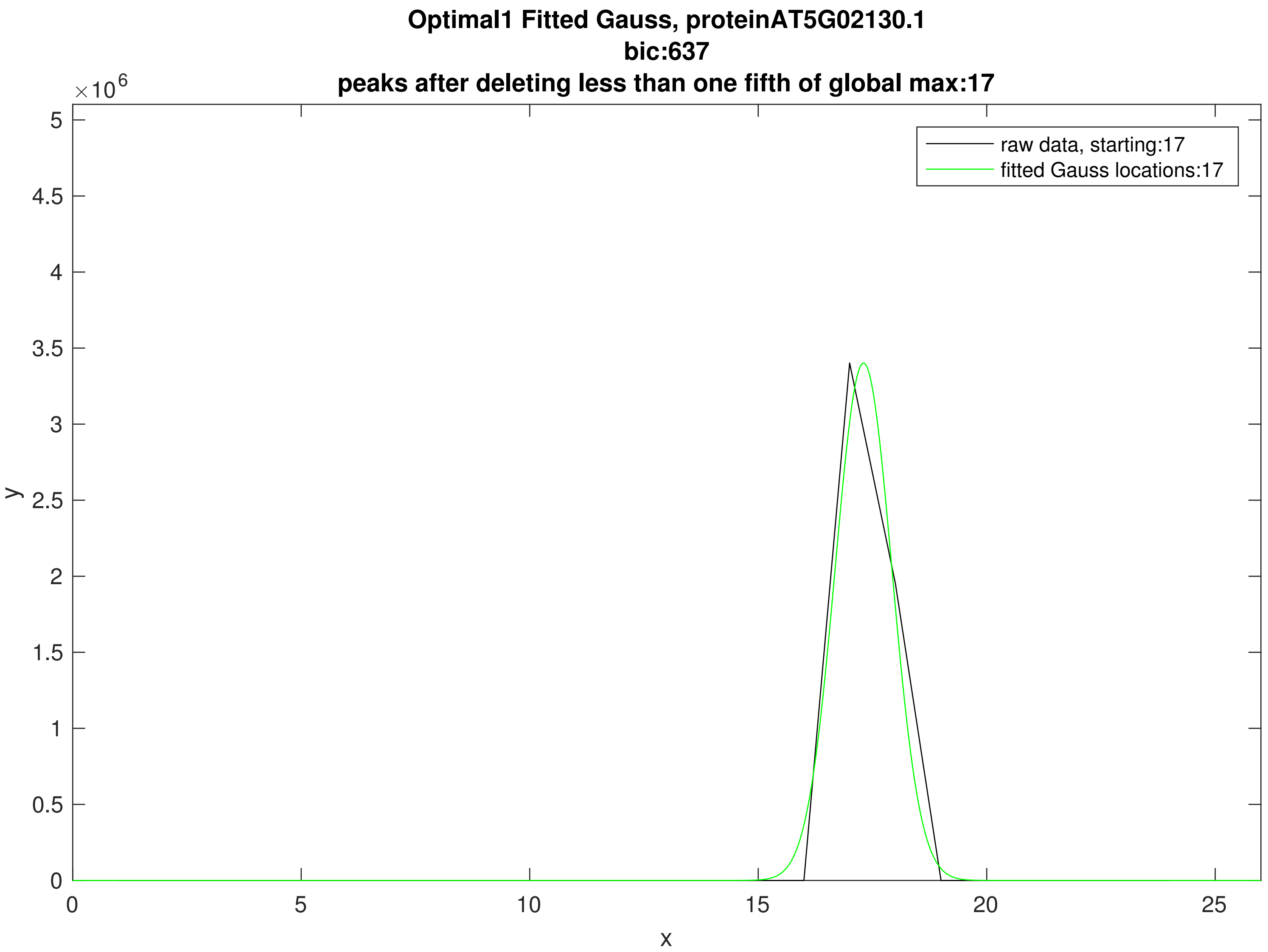

Optimal1 Fitted Gauss, proteinAT5G02130.1
bic:637
peaks after deleting less than one fifth of global max:17
×10^6
raw data, starting:17
fitted Gauss locations:17
y
x
0
0.5
1
1.5
2
2.5
3
3.5
4
4.5
5
0
5
10
15
20
25

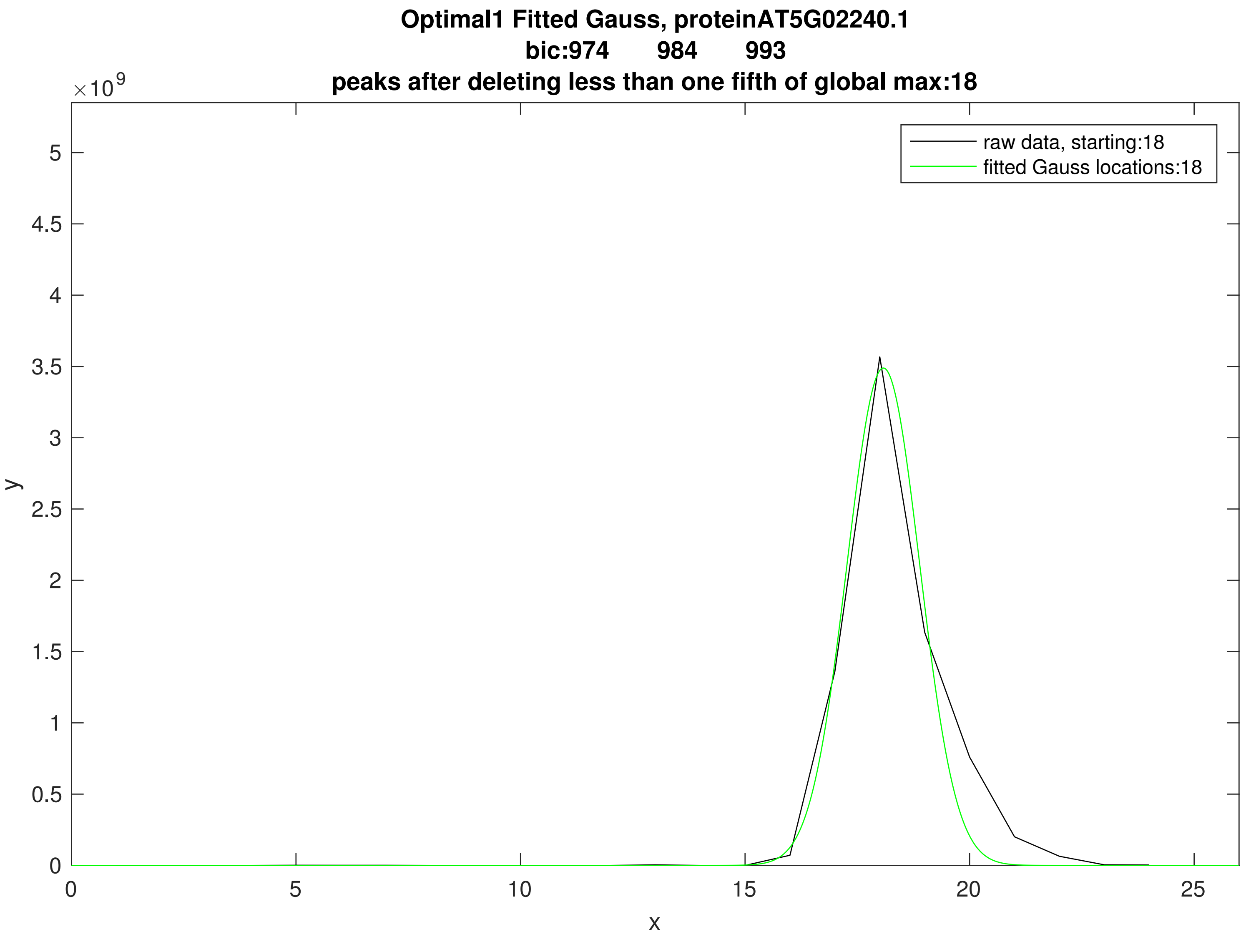
Optimal1 Fitted Gauss, proteinAT5G02240.1
bic:974 984 993
peaks after deleting less than one fifth of global max:18
×10^9
raw data, starting:18
fitted Gauss locations:18
y
x
0
0.5
1
1.5
2
2.5
3
3.5
4
4.5
5
0
5
10
15
20
25

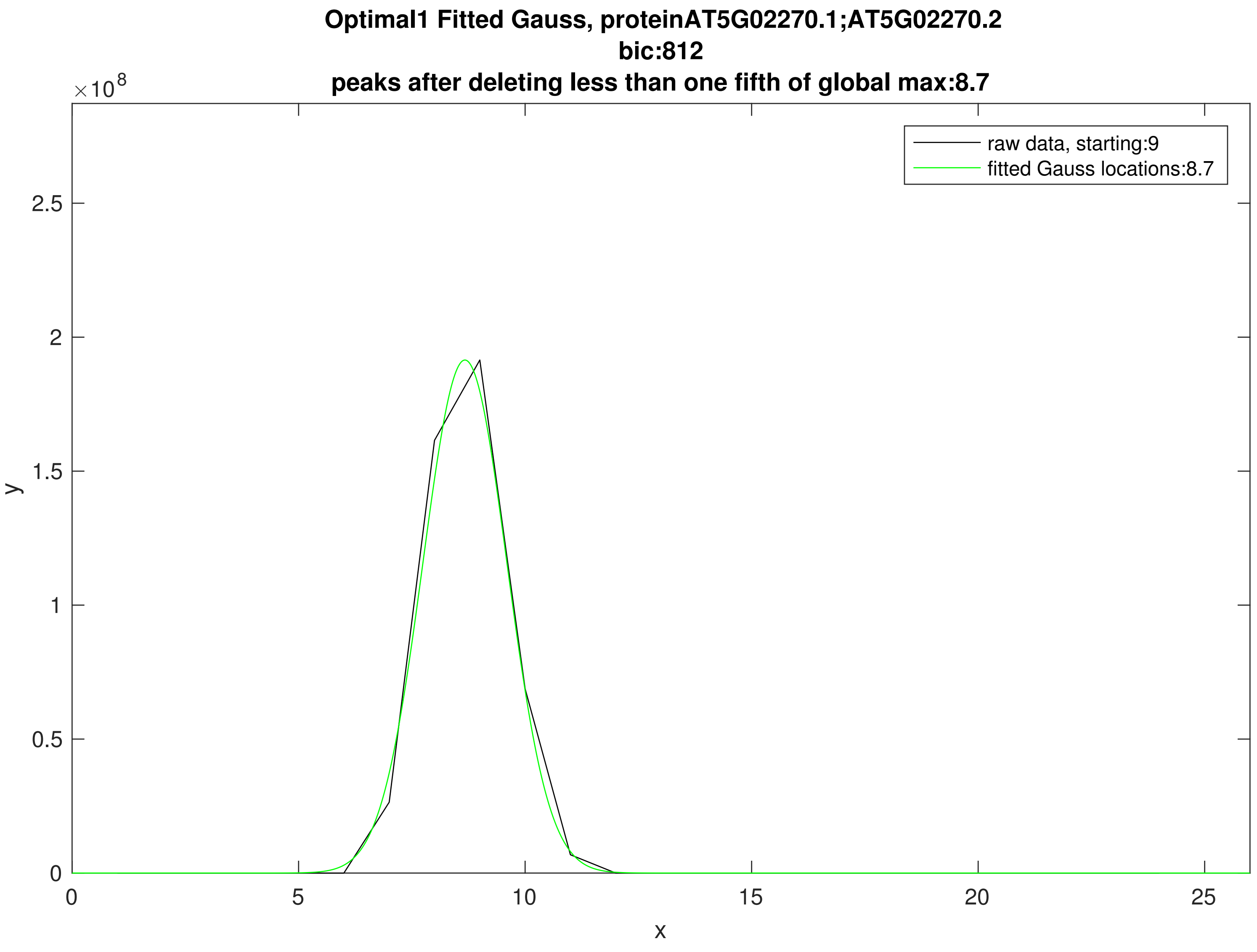
Optimal1 Fitted Gauss, proteinAT5G02270.1;AT5G02270.2
bic:812
peaks after deleting less than one fifth of global max:8.7
×10^8
raw data, starting:9
fitted Gauss locations:8.7
y
x
0
0.5
1
1.5
2
2.5
5
10
15
20
25

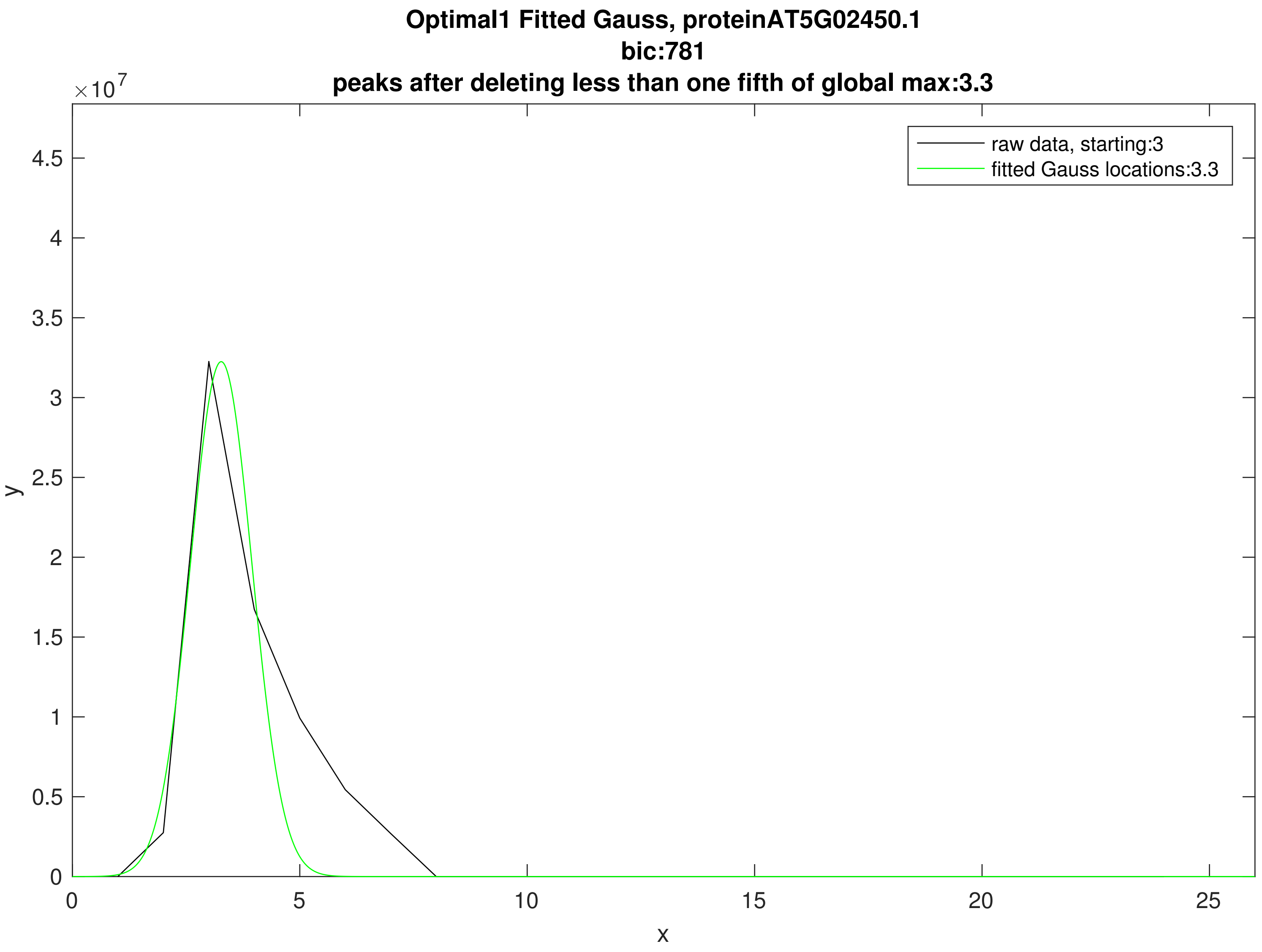
Optimal1 Fitted Gauss, proteinAT5G02450.1
bic:781
peaks after deleting less than one fifth of global max:3.3
×10^7
raw data, starting:3
fitted Gauss locations:3.3
y
x
0
0.5
1
1.5
2
2.5
3
3.5
4
4.5
5
10
15
20
25

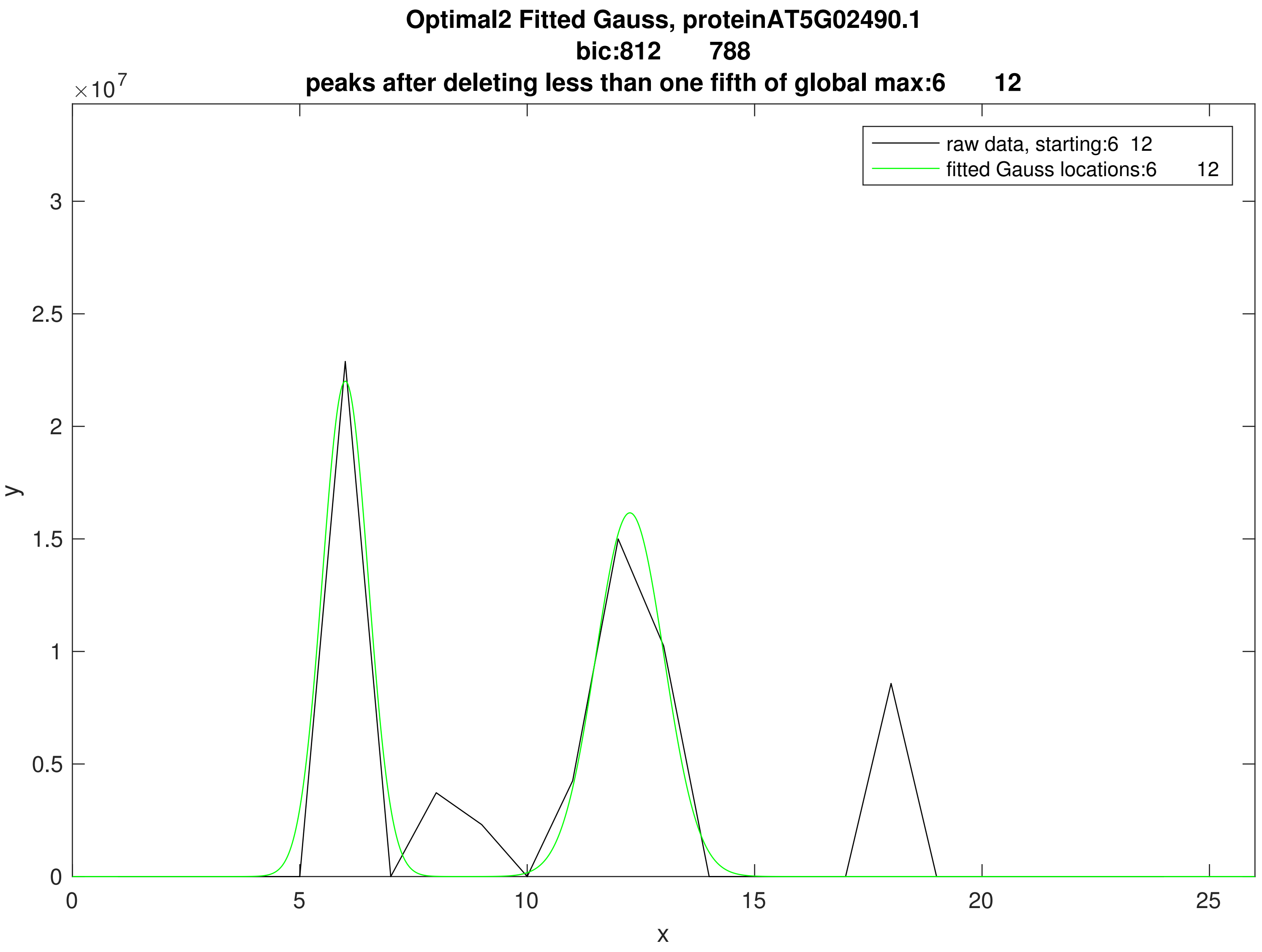
Optimal2 Fitted Gauss, proteinAT5G02490.1
bic:812 788
peaks after deleting less than one fifth of global max:6 12
raw data, starting:6 12
fitted Gauss locations:6 12
×10^7
y
x
0
0.5
1
1.5
2
2.5
3
5
10
15
20
25

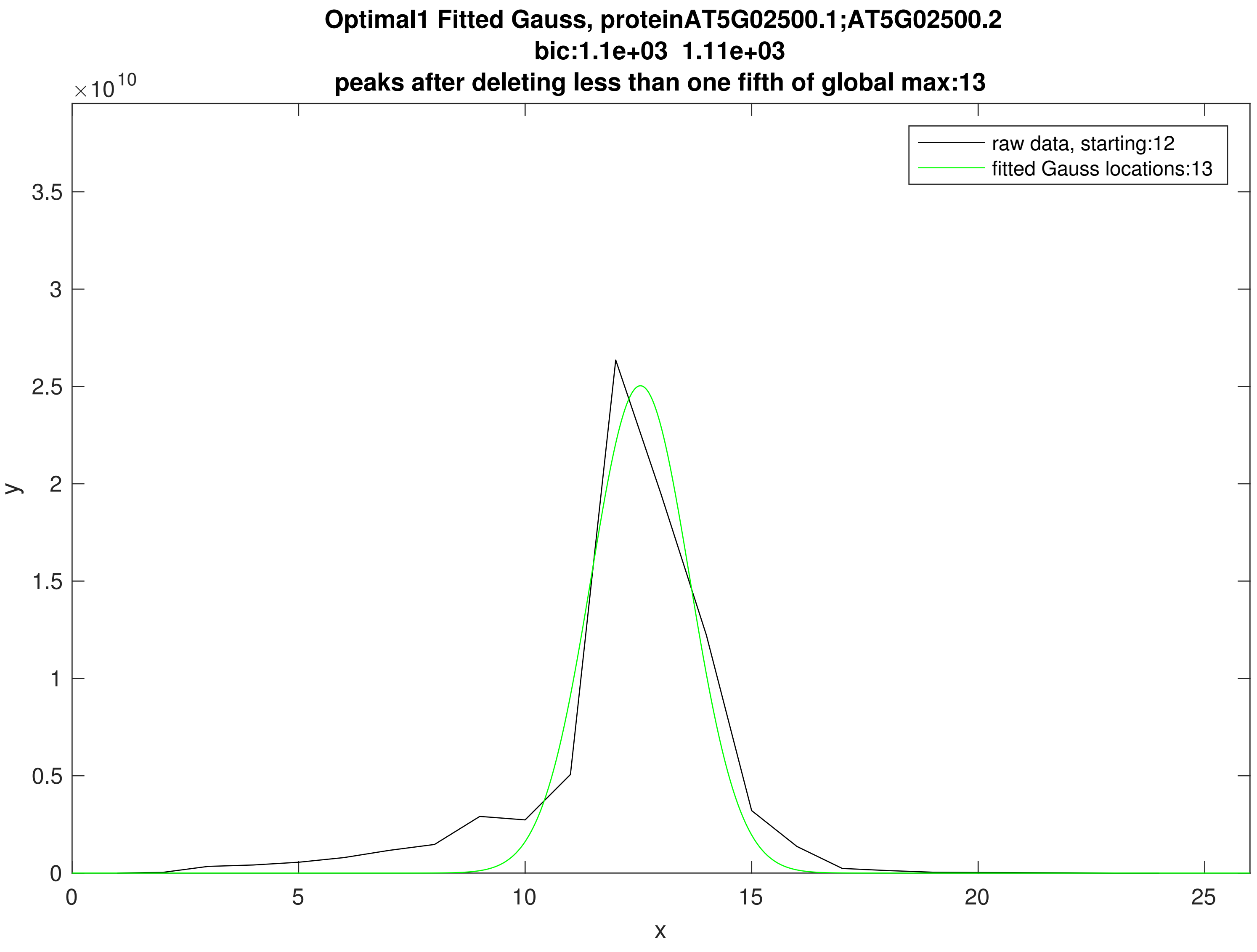
Optimal1 Fitted Gauss, proteinAT5G02500.1;AT5G02500.2
bic:1.1e+03  1.11e+03
peaks after deleting less than one fifth of global max:13
×10^10
raw data, starting:12
fitted Gauss locations:13
y
x
0
0.5
1
1.5
2
2.5
3
3.5
5
10
15
20
25

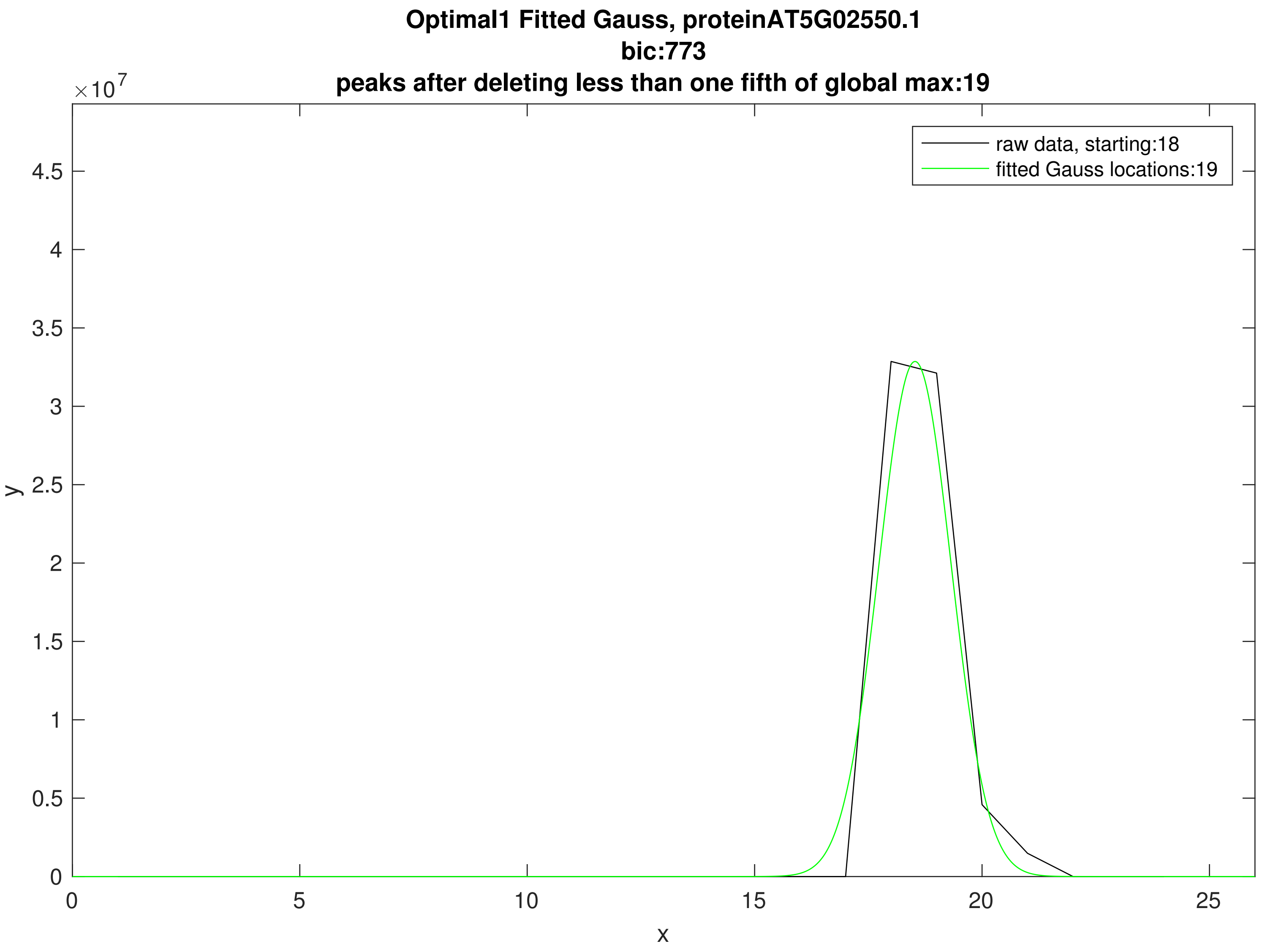
Optimal1 Fitted Gauss, proteinAT5G02550.1
bic:773
peaks after deleting less than one fifth of global max:19
×10^7
4.5
4
3.5
3
2.5
2
1.5
1
0.5
0
y
0
5
10
15
20
25
x
raw data, starting:18
fitted Gauss locations:19

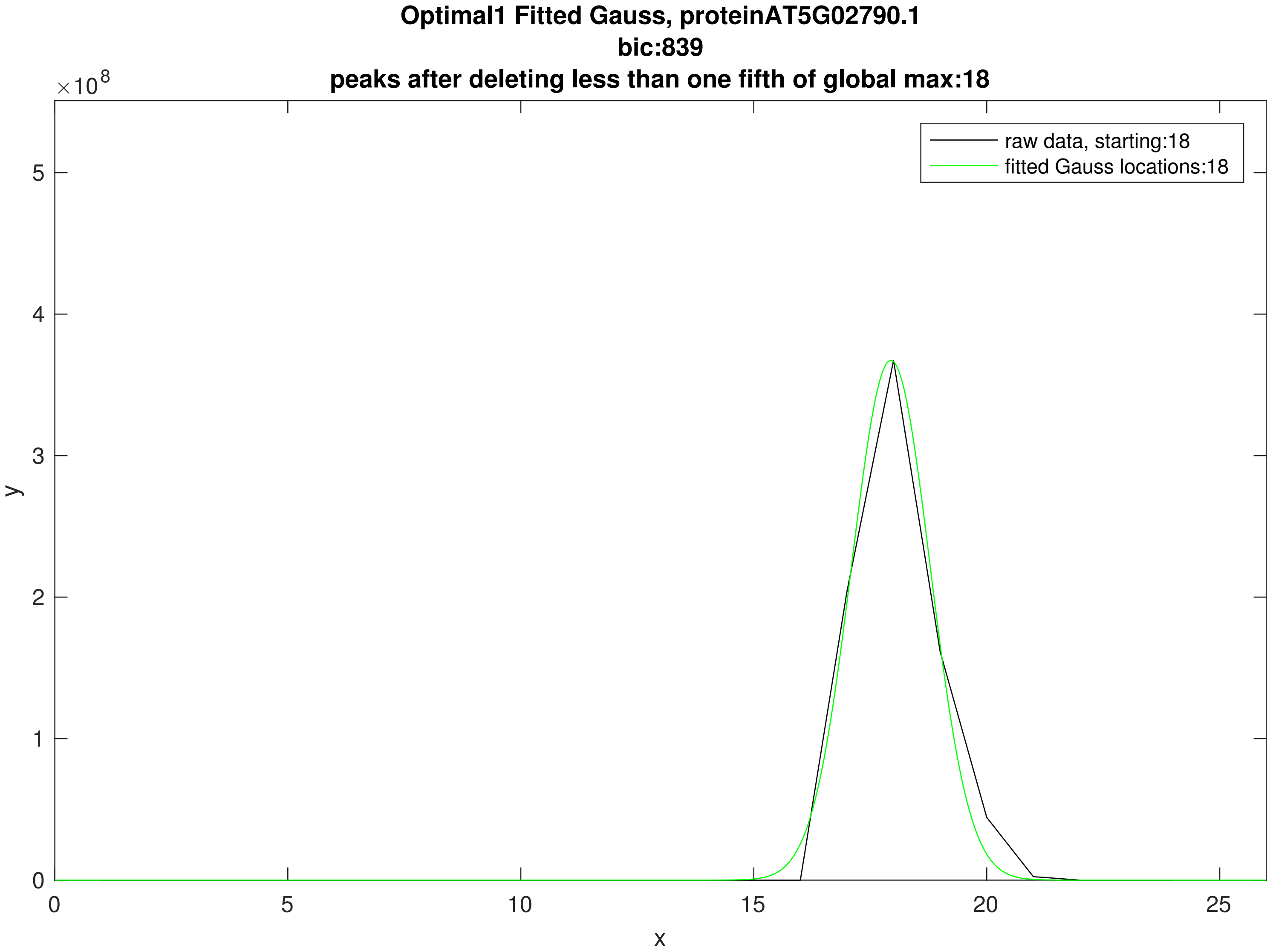

Optimal1 Fitted Gauss, proteinAT5G02790.1
bic:839
peaks after deleting less than one fifth of global max:18
×10^8
raw data, starting:18
fitted Gauss locations:18
y
x
0
1
2
3
4
5
0
5
10
15
20
25

Optimal1 Fitted Gauss, proteinAT5G02870.1;AT5G02870.2
bic:802
peaks after deleting less than one fifth of global max:3.2

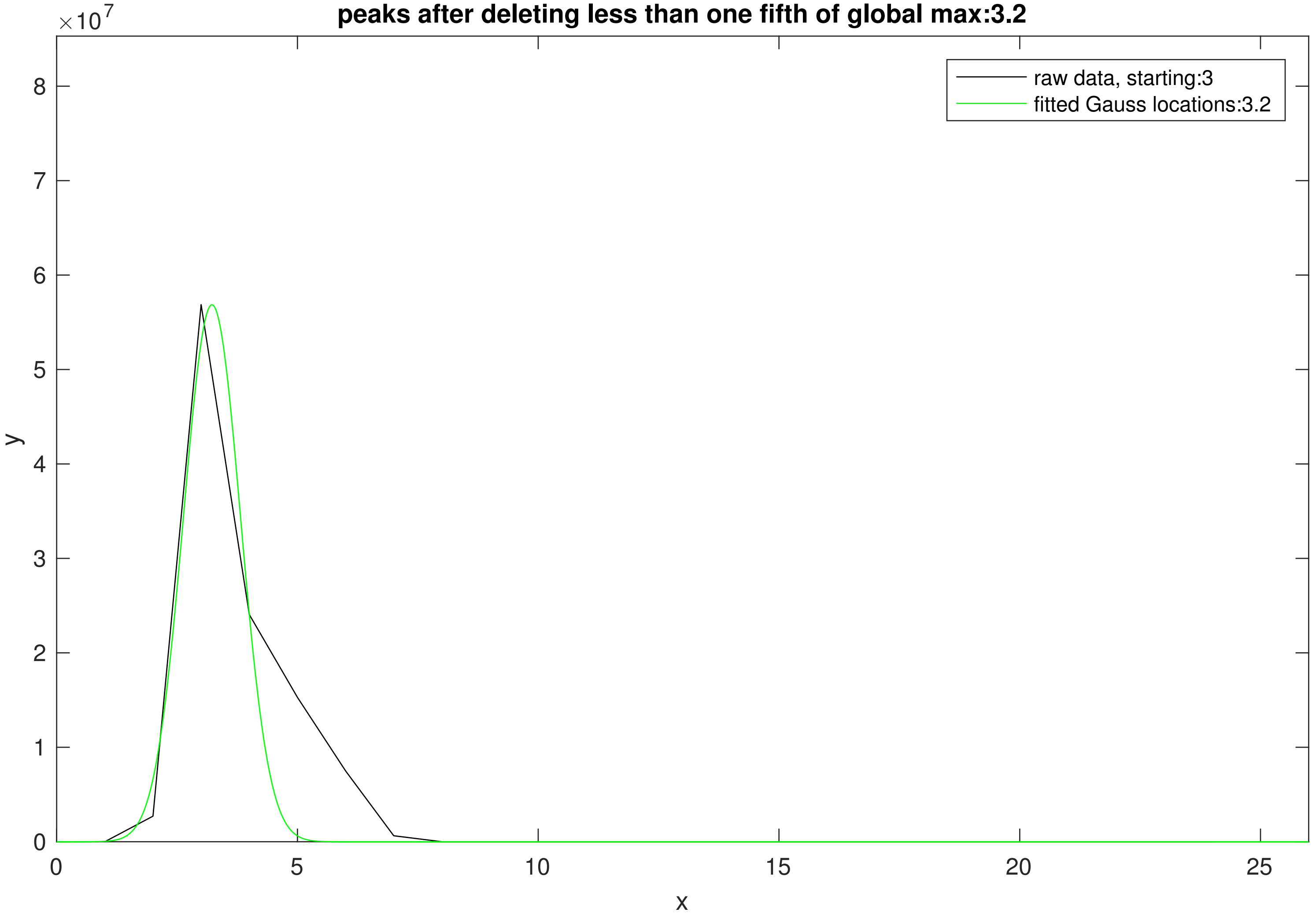

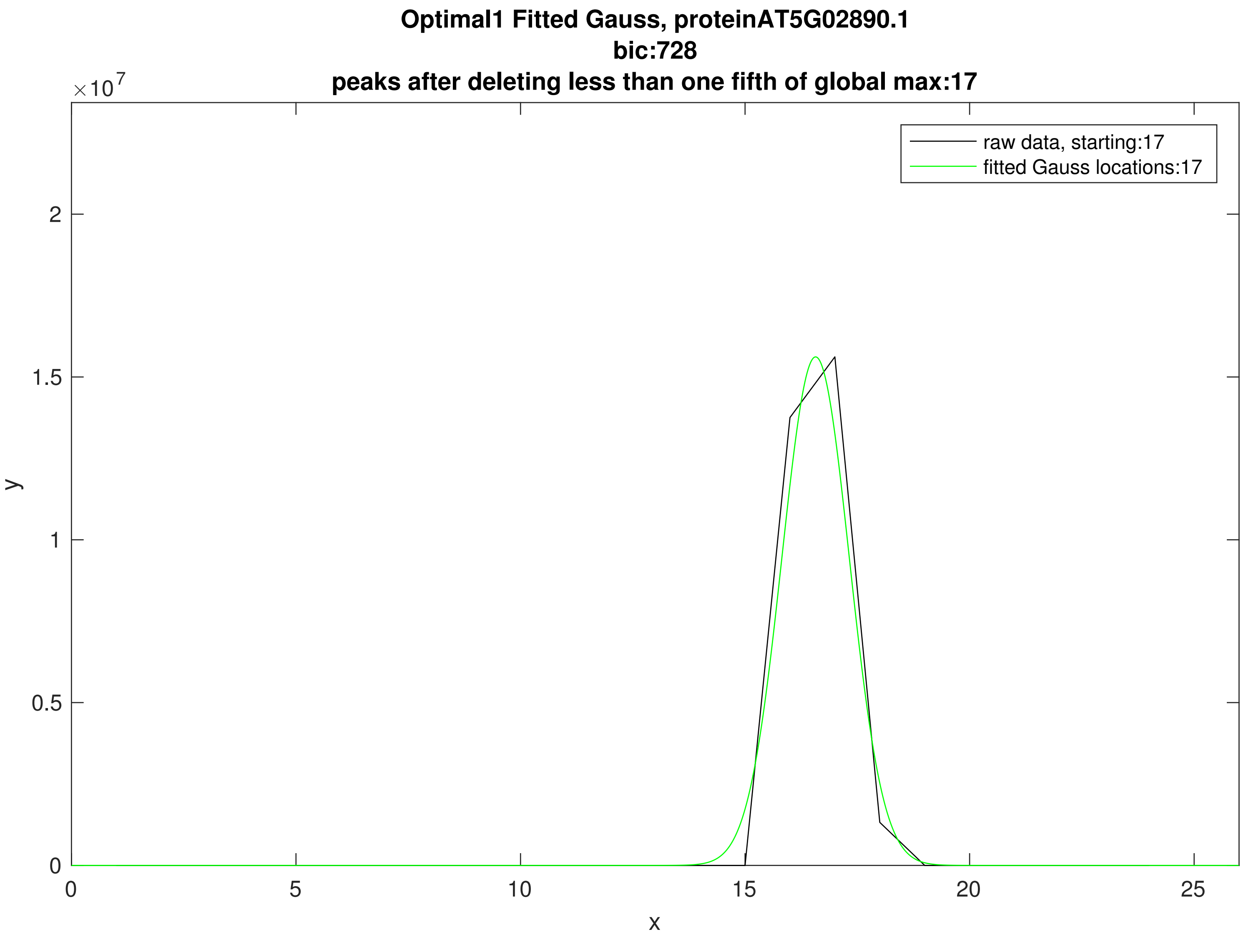

Optimal1 Fitted Gauss, proteinAT5G02890.1
bic:728
peaks after deleting less than one fifth of global max:17
×10^7
raw data, starting:17
fitted Gauss locations:17
2
1.5
1
0.5
0
y
0
5
10
15
20
25
x

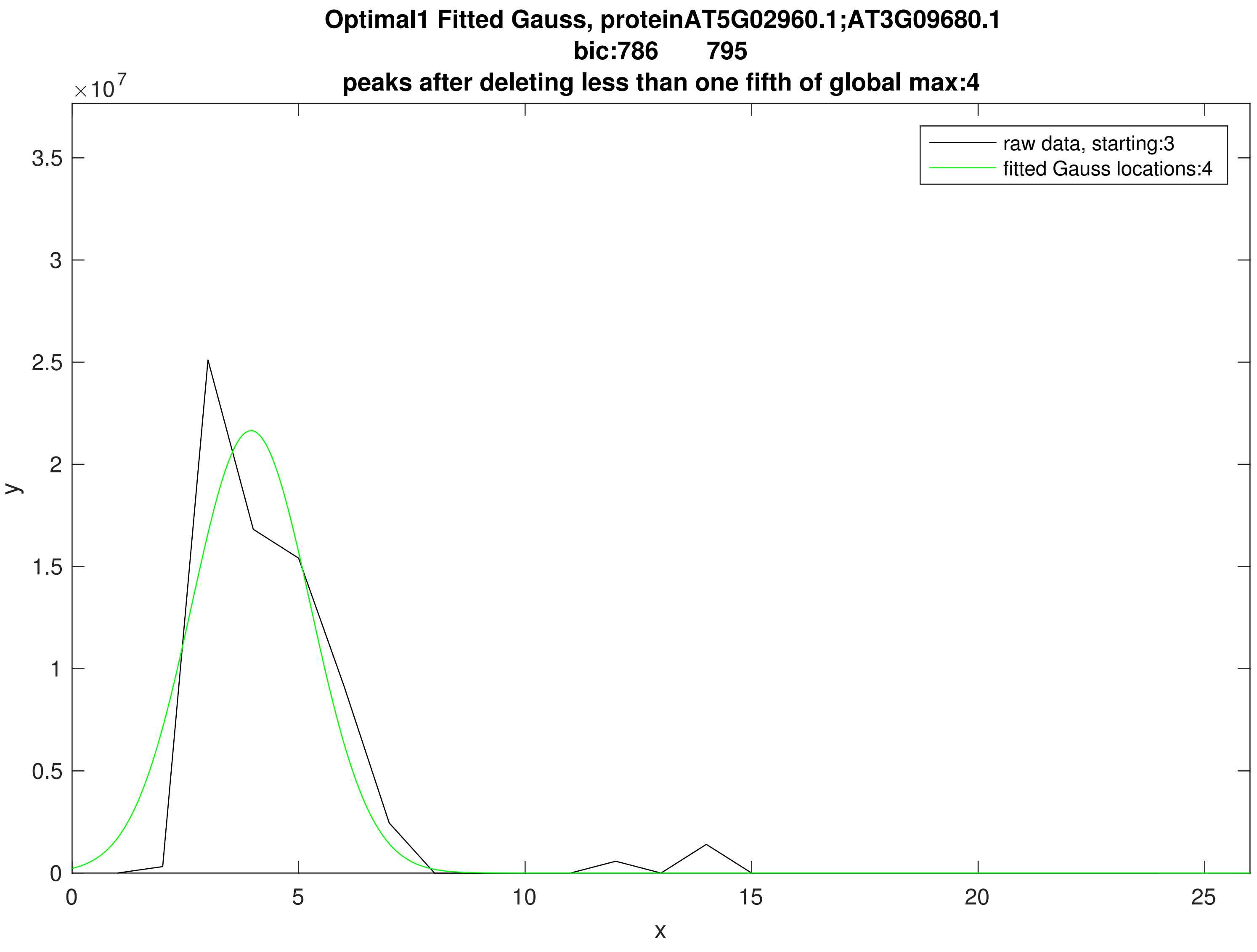

Optimal1 Fitted Gauss, proteinAT5G02960.1;AT3G09680.1
bic:786 795
peaks after deleting less than one fifth of global max:4
×10^7
3.5
3
2.5
2
1.5
1
0.5
0
y
0
5
10
15
20
25
x
raw data, starting:3
fitted Gauss locations:4

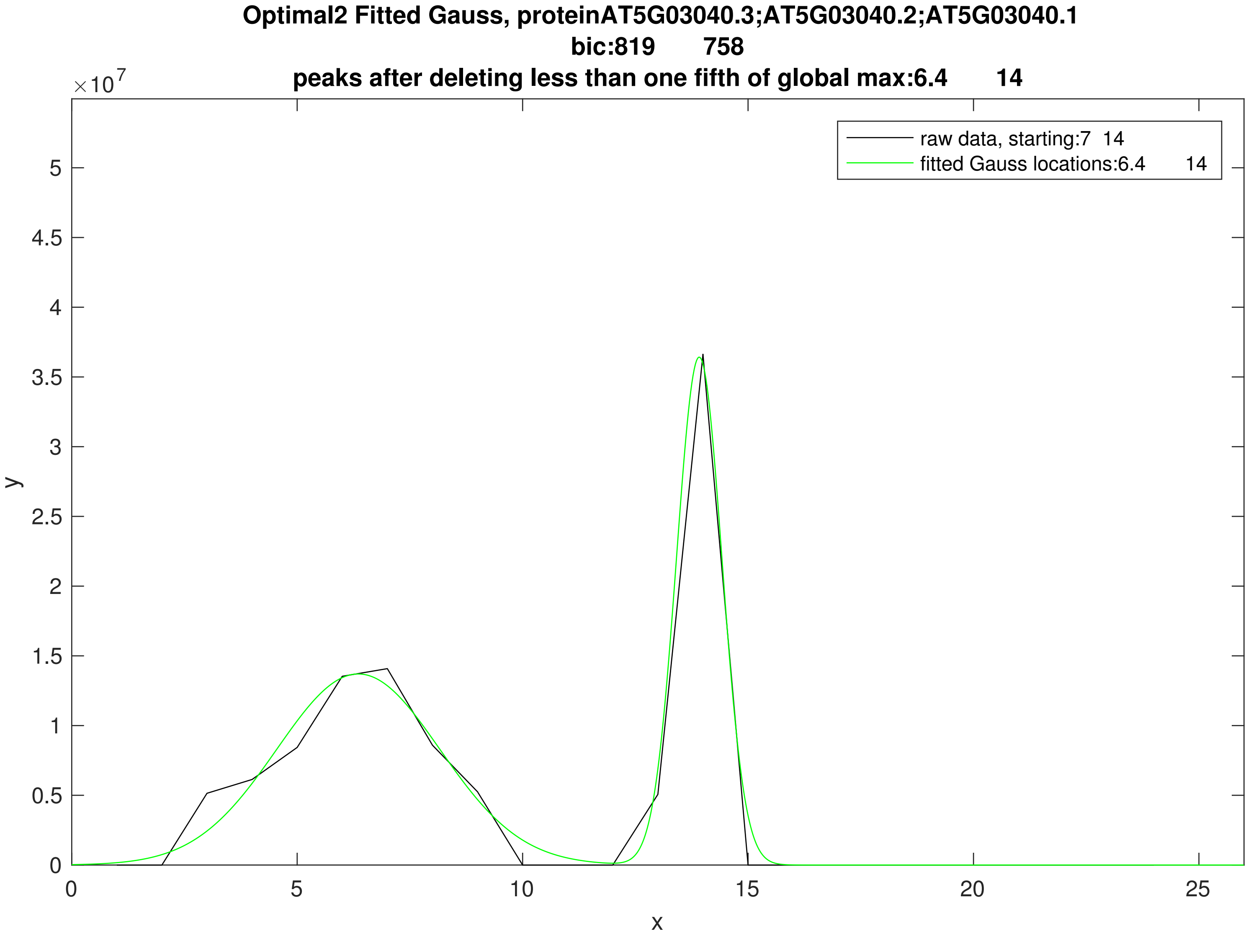
Optimal2 Fitted Gauss, proteinAT5G03040.3;AT5G03040.2;AT5G03040.1
bic:819 758
peaks after deleting less than one fifth of global max:6.4 14
×10^7
raw data, starting:7 14
fitted Gauss locations:6.4 14
y
x
0
0.5
1
1.5
2
2.5
3
3.5
4
4.5
5
0
5
10
15
20
25

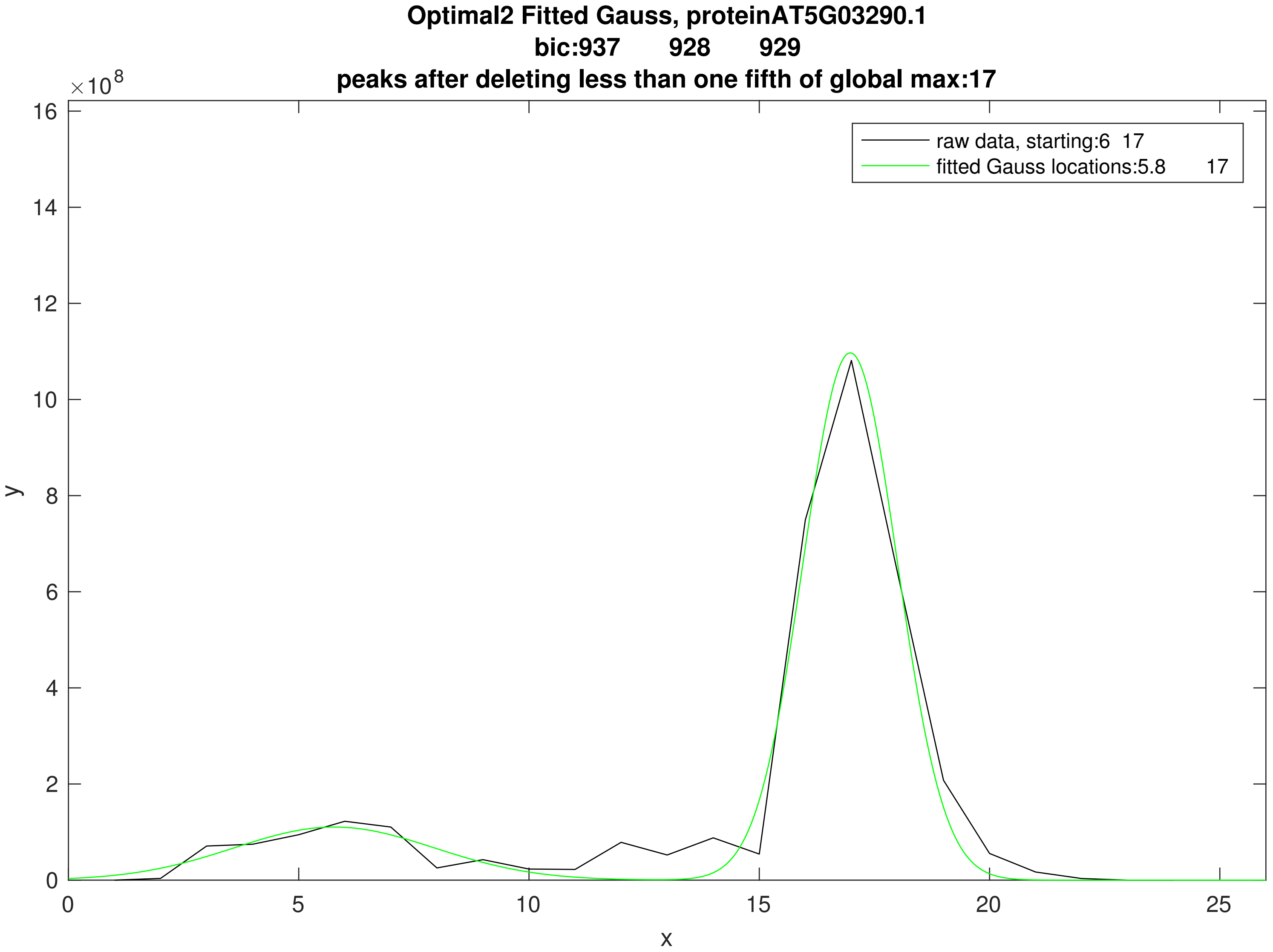

Optimal2 Fitted Gauss, proteinAT5G03290.1
bic:937 928 929
peaks after deleting less than one fifth of global max:17
×10^8
raw data, starting:6 17
fitted Gauss locations:5.8 17
y
x
0
2
4
6
8
10
12
14
16
0
5
10
15
20
25

**Optimal1 Fitted Gauss, proteinAT5G03300.1**
**bic:980 989 999**
**peaks after deleting less than one fifth of global max:14**

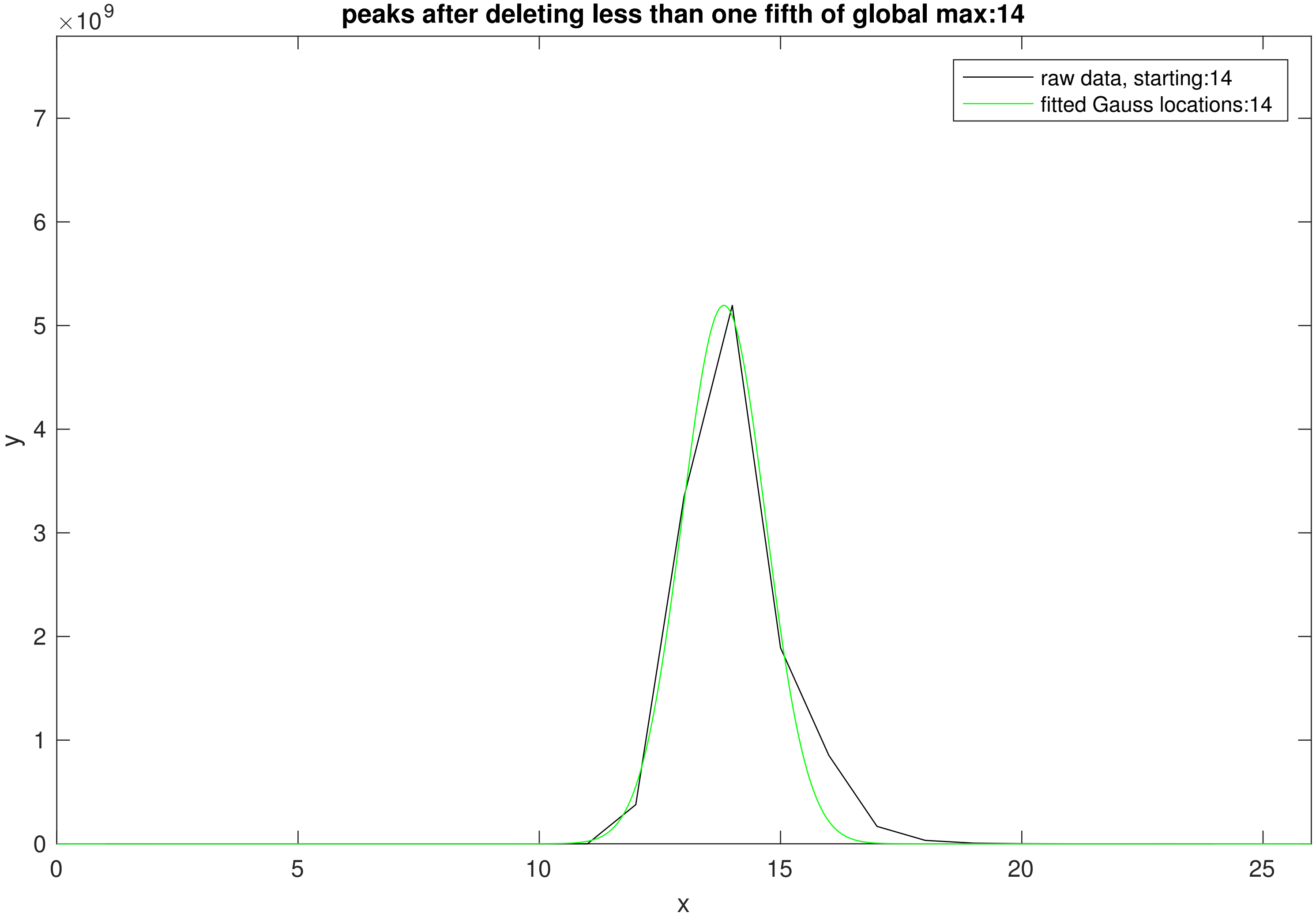

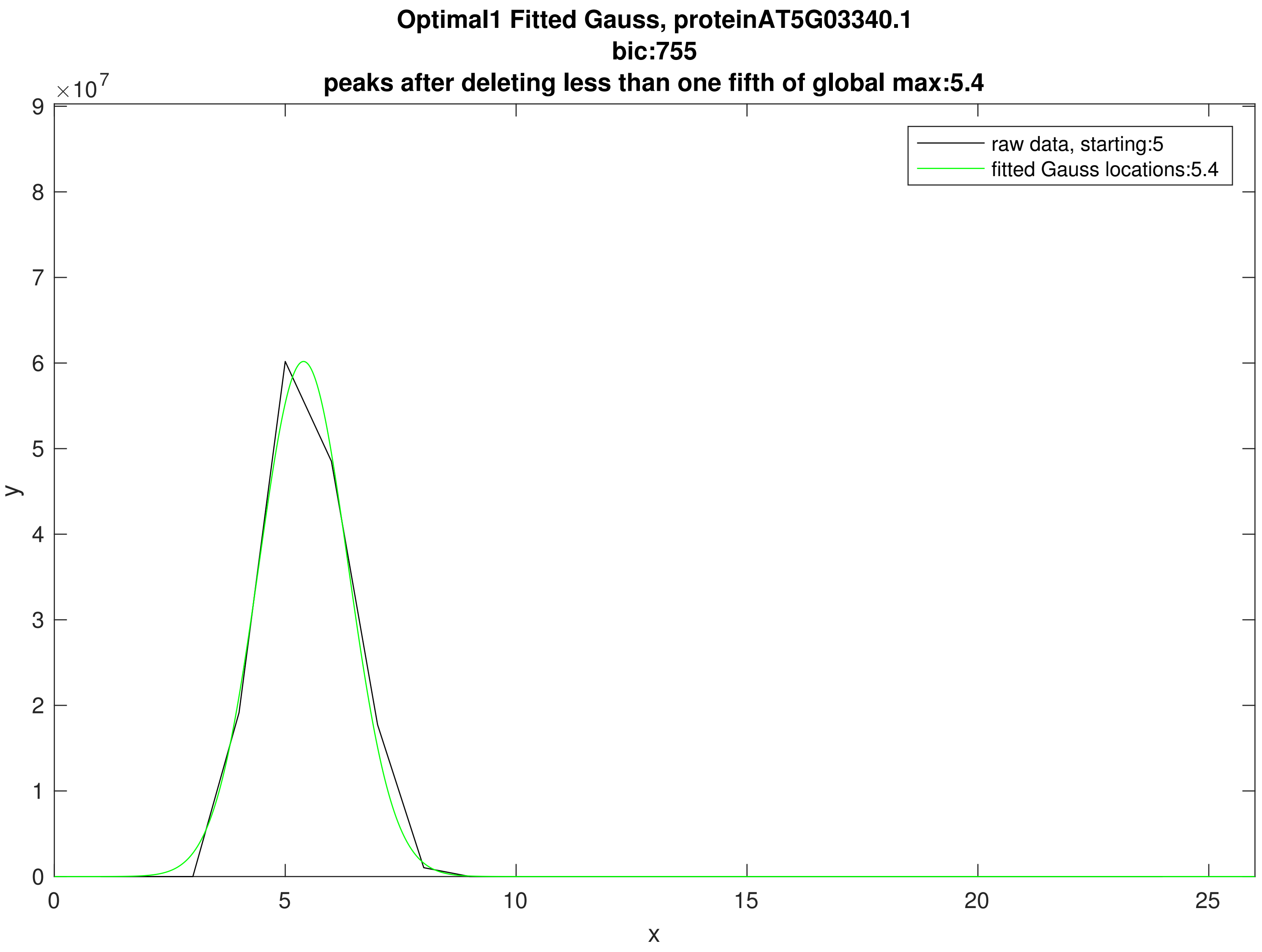

Optimal1 Fitted Gauss, proteinAT5G03340.1
bic:755
peaks after deleting less than one fifth of global max:5.4
×10^7
9
8
7
6
5
4
3
2
1
0
y
0
5
10
15
20
25
x
raw data, starting:5
fitted Gauss locations:5.4

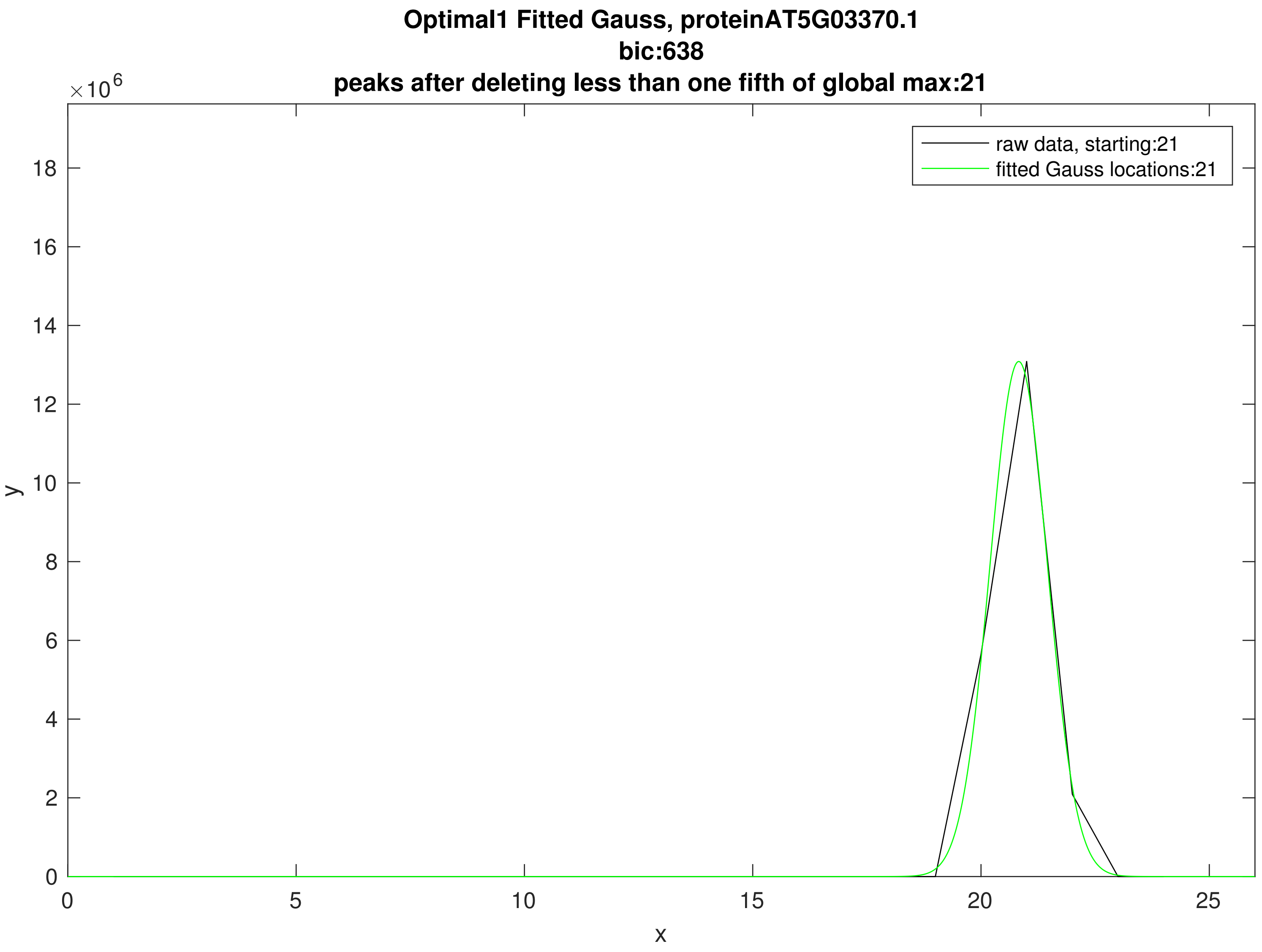
Optimal1 Fitted Gauss, proteinAT5G03370.1
bic:638
peaks after deleting less than one fifth of global max:21
$\times 10^6$
raw data, starting:21
fitted Gauss locations:21
y
x
0
2
4
6
8
10
12
14
16
18
0
5
10
15
20
25

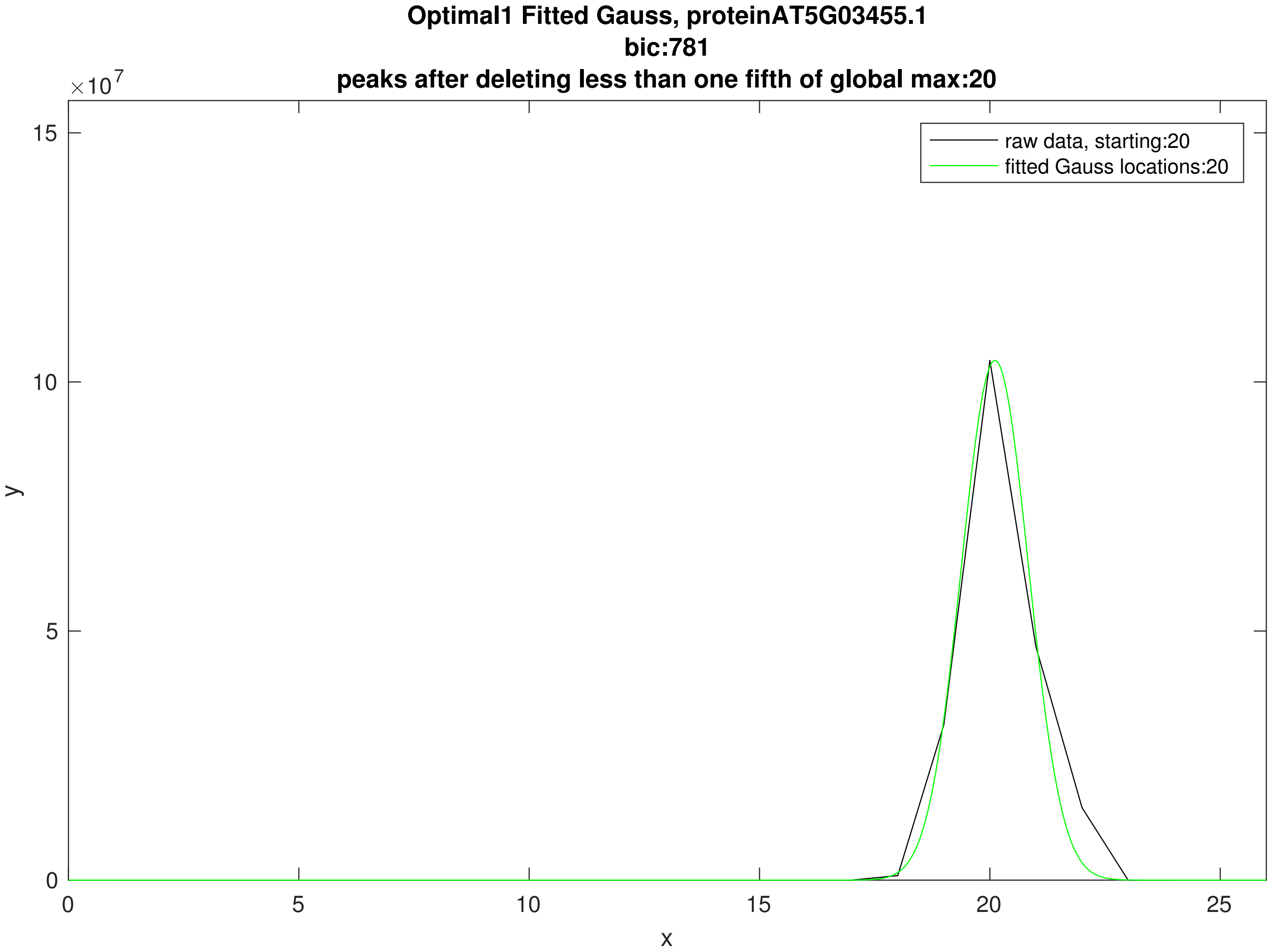

Optimal1 Fitted Gauss, proteinAT5G03455.1
bic:781
peaks after deleting less than one fifth of global max:20
×10^7
15
10
5
0
y
0
5
10
15
20
25
x
raw data, starting:20
fitted Gauss locations:20

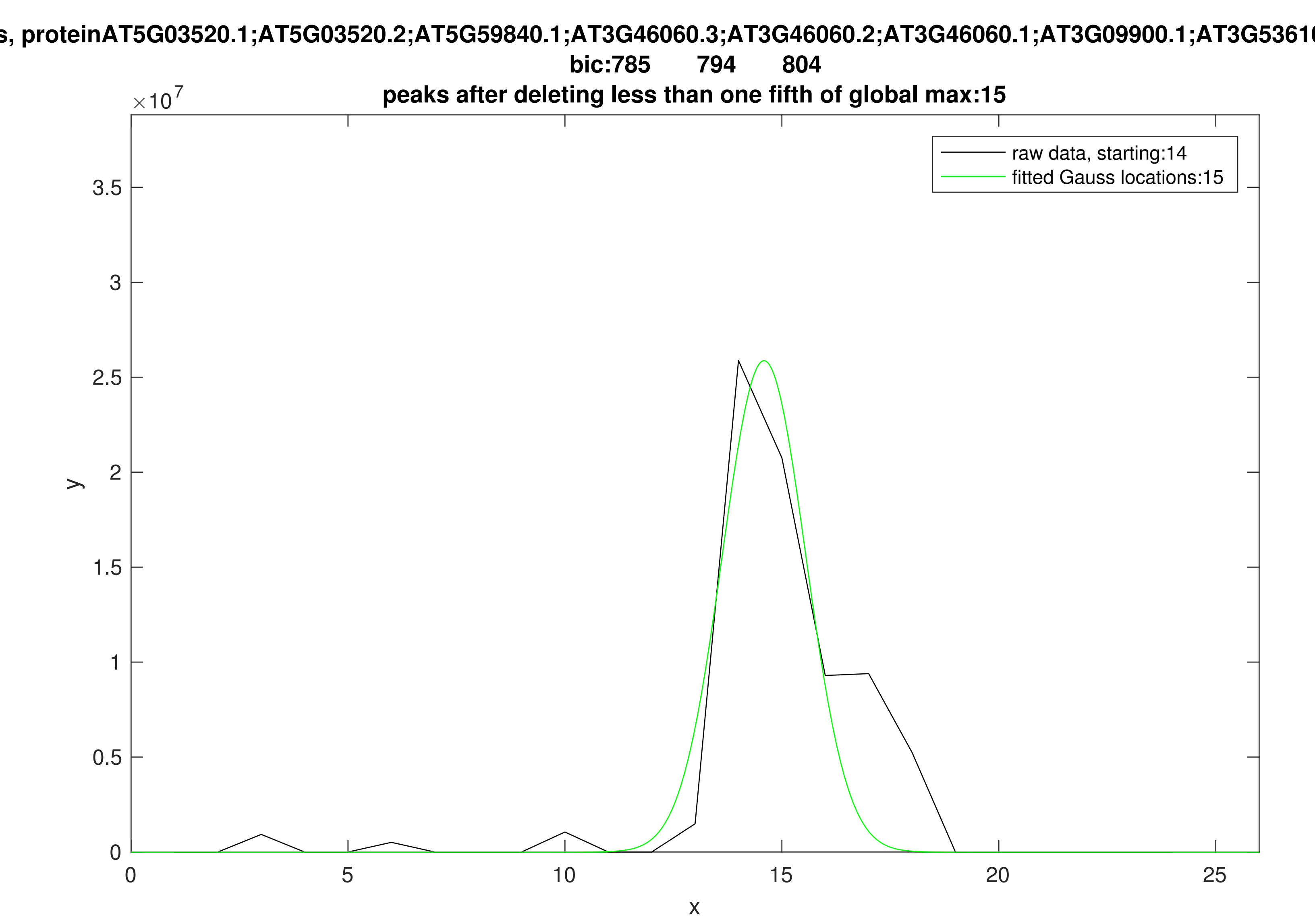
s, proteinAT5G03520.1;AT5G03520.2;AT5G59840.1;AT3G46060.3;AT3G46060.2;AT3G46060.1;AT3G09900.1;AT3G5361
bic:785 794 804
peaks after deleting less than one fifth of global max:15
×10^7
raw data, starting:14
fitted Gauss locations:15
y
x
0
0.5
1
1.5
2
2.5
3
3.5
5
10
15
20
25

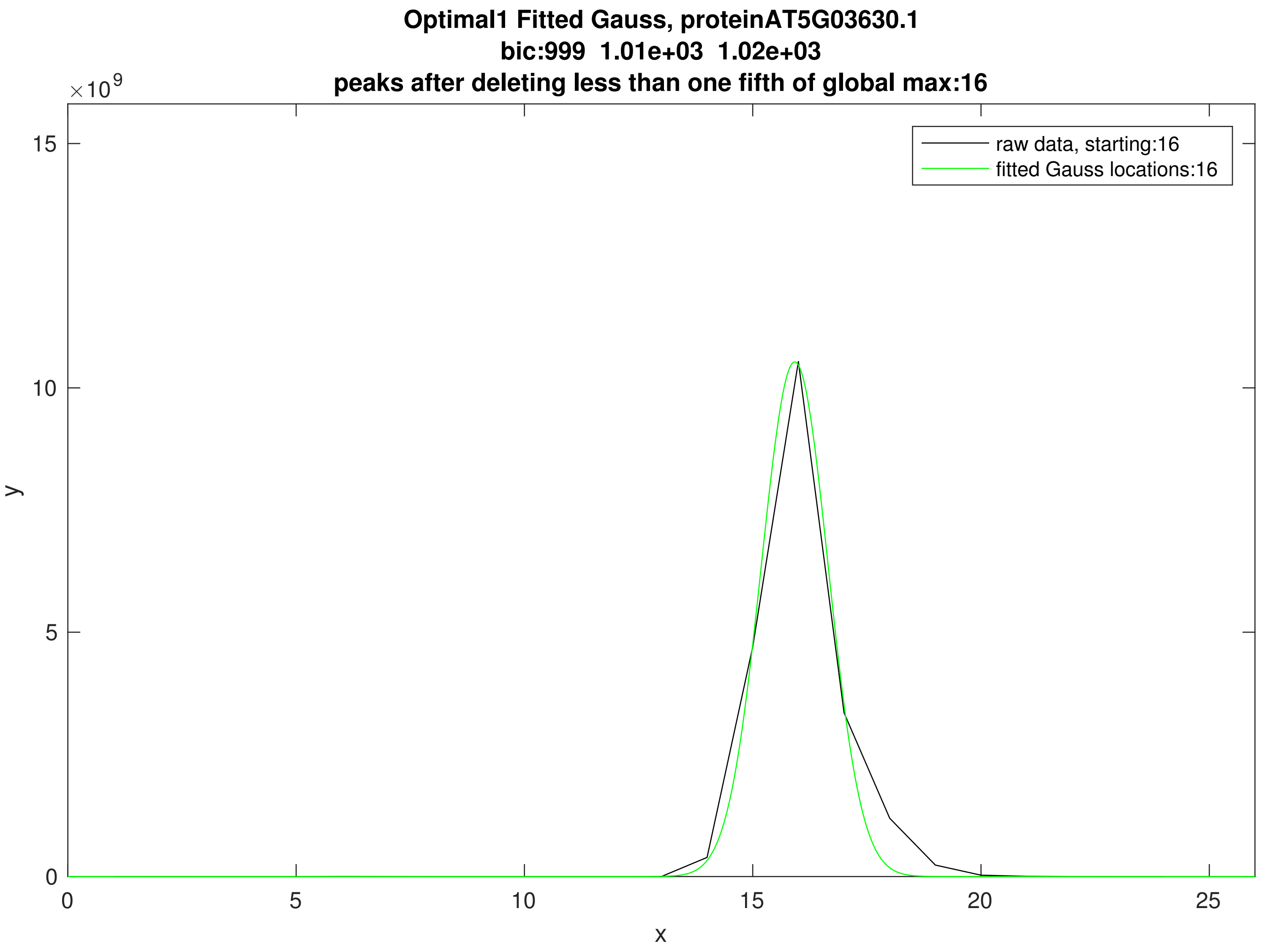
Optimal1 Fitted Gauss, proteinAT5G03630.1
bic:999 1.01e+03 1.02e+03
peaks after deleting less than one fifth of global max:16
×10^9
15
10
5
0
y
0
5
10
15
20
25
x
raw data, starting:16
fitted Gauss locations:16

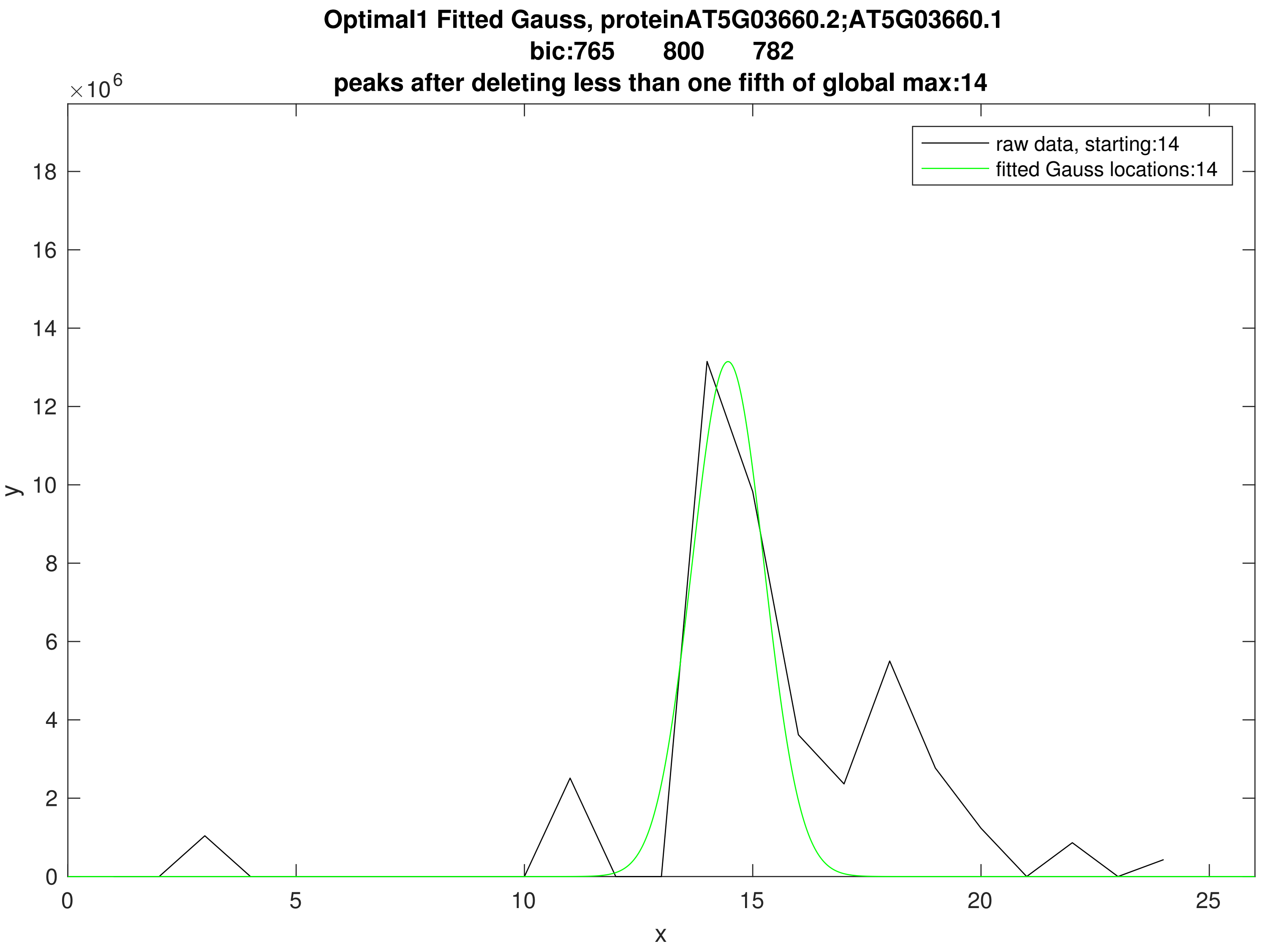

Optimal1 Fitted Gauss, proteinAT5G03660.2;AT5G03660.1
bic:765 800 782
peaks after deleting less than one fifth of global max:14
×10^6
raw data, starting:14
fitted Gauss locations:14
y
x
0
2
4
6
8
10
12
14
16
18
0
5
10
15
20
25

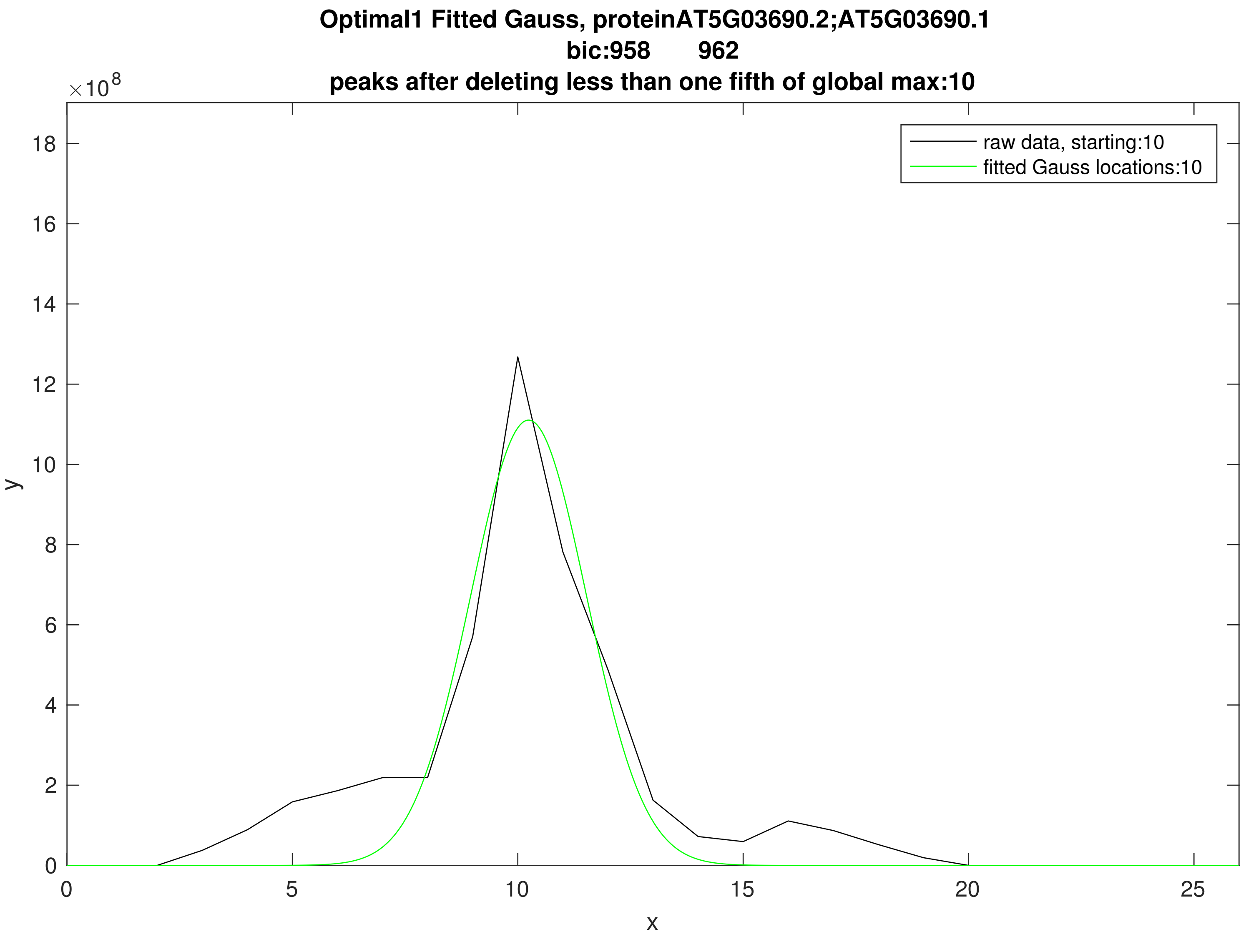

Optimal1 Fitted Gauss, proteinAT5G03690.2;AT5G03690.1
bic:958 962
peaks after deleting less than one fifth of global max:10
×10^8
raw data, starting:10
fitted Gauss locations:10
y
x
0
2
4
6
8
10
12
14
16
18
0
5
10
15
20
25

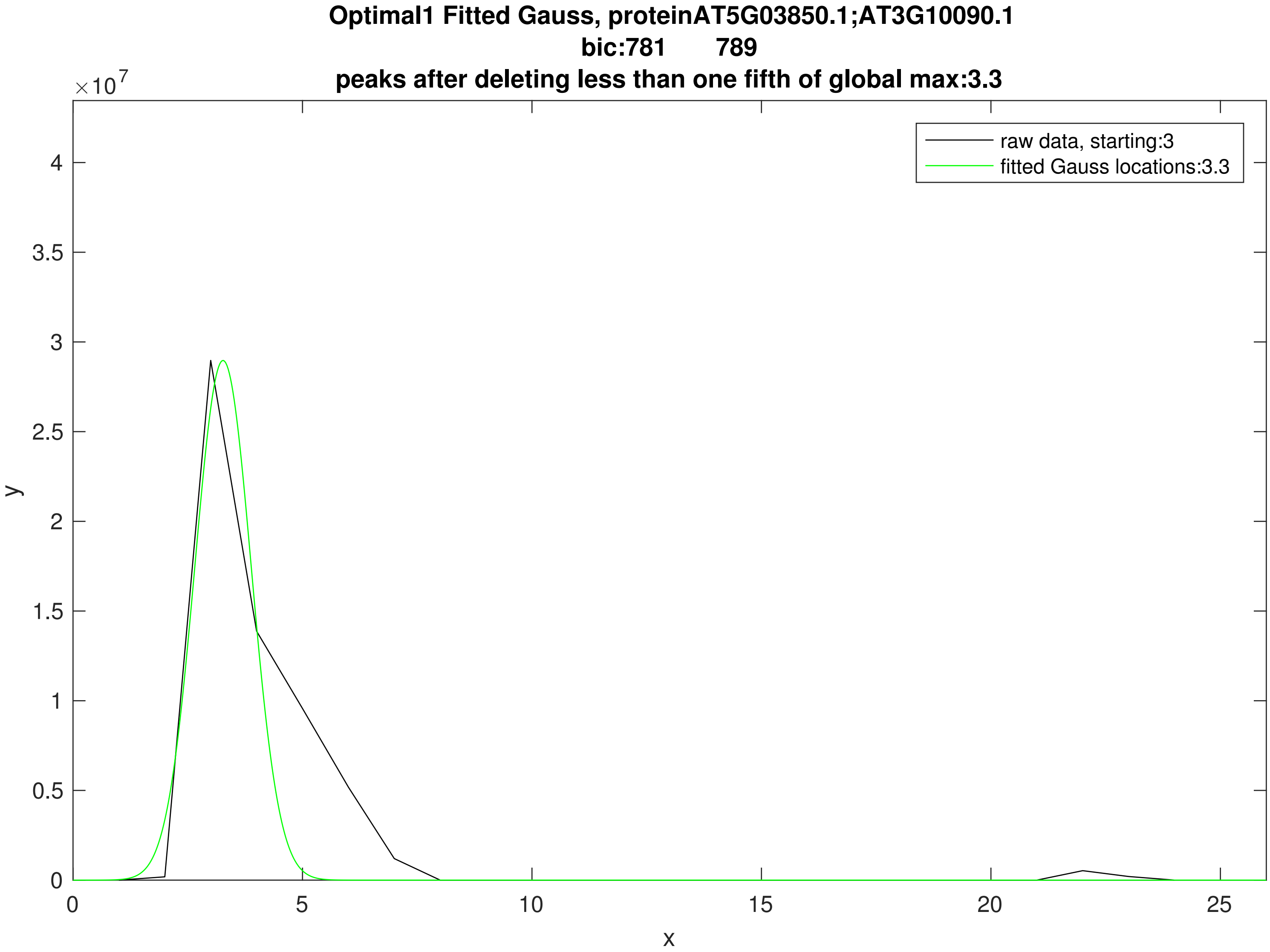
Optimal1 Fitted Gauss, proteinAT5G03850.1;AT3G10090.1
bic:781 789
peaks after deleting less than one fifth of global max:3.3
×10^7
raw data, starting:3
fitted Gauss locations:3.3
y
x
0
0.5
1
1.5
2
2.5
3
3.5
4
0
5
10
15
20
25

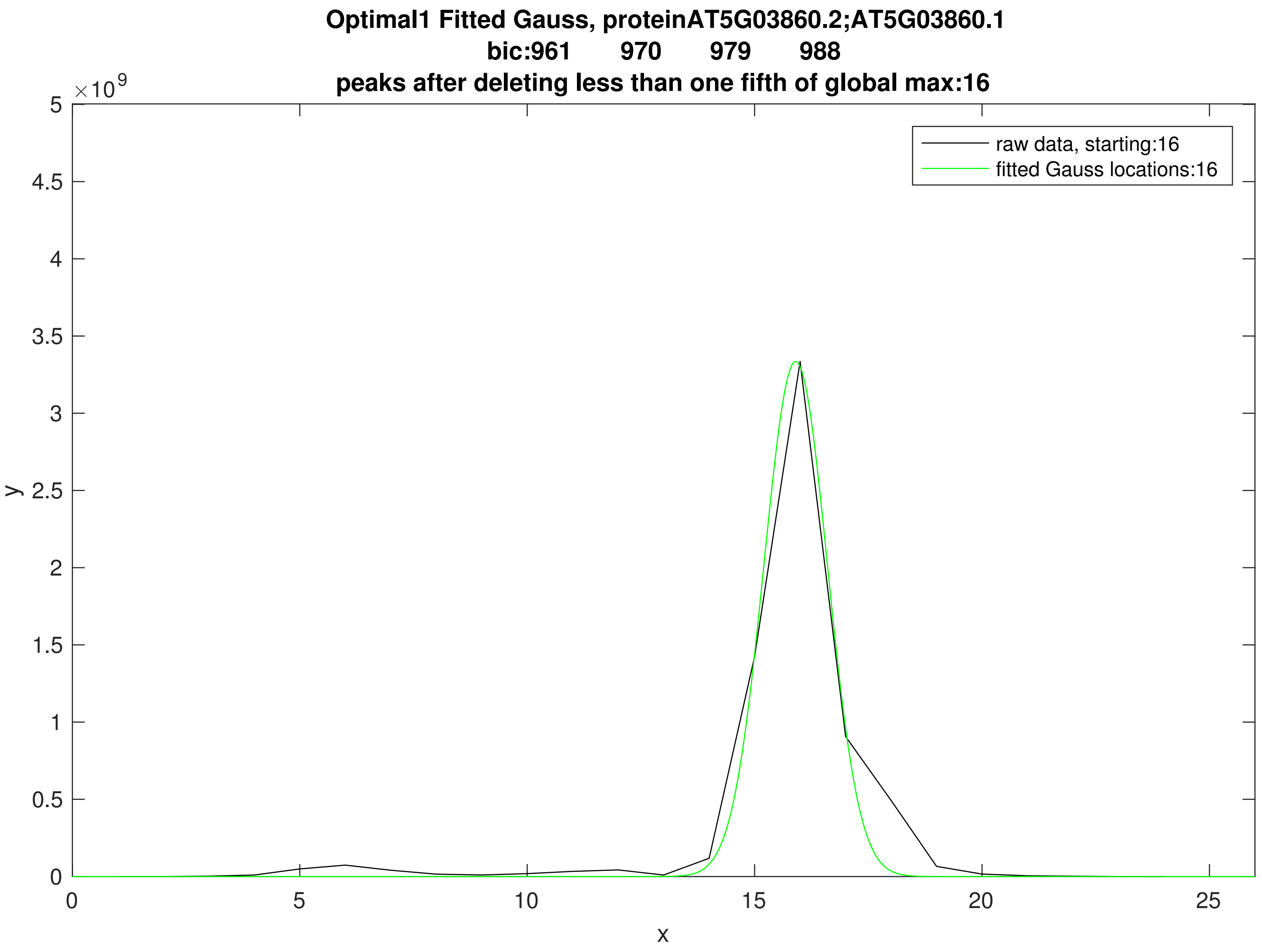

Optimal1 Fitted Gauss, proteinAT5G03860.2;AT5G03860.1
bic:961 970 979 988
peaks after deleting less than one fifth of global max:16
×10^9
5
4.5
4
3.5
3
2.5
2
1.5
1
0.5
0
y
0
5
10
15
20
25
x
raw data, starting:16
fitted Gauss locations:16

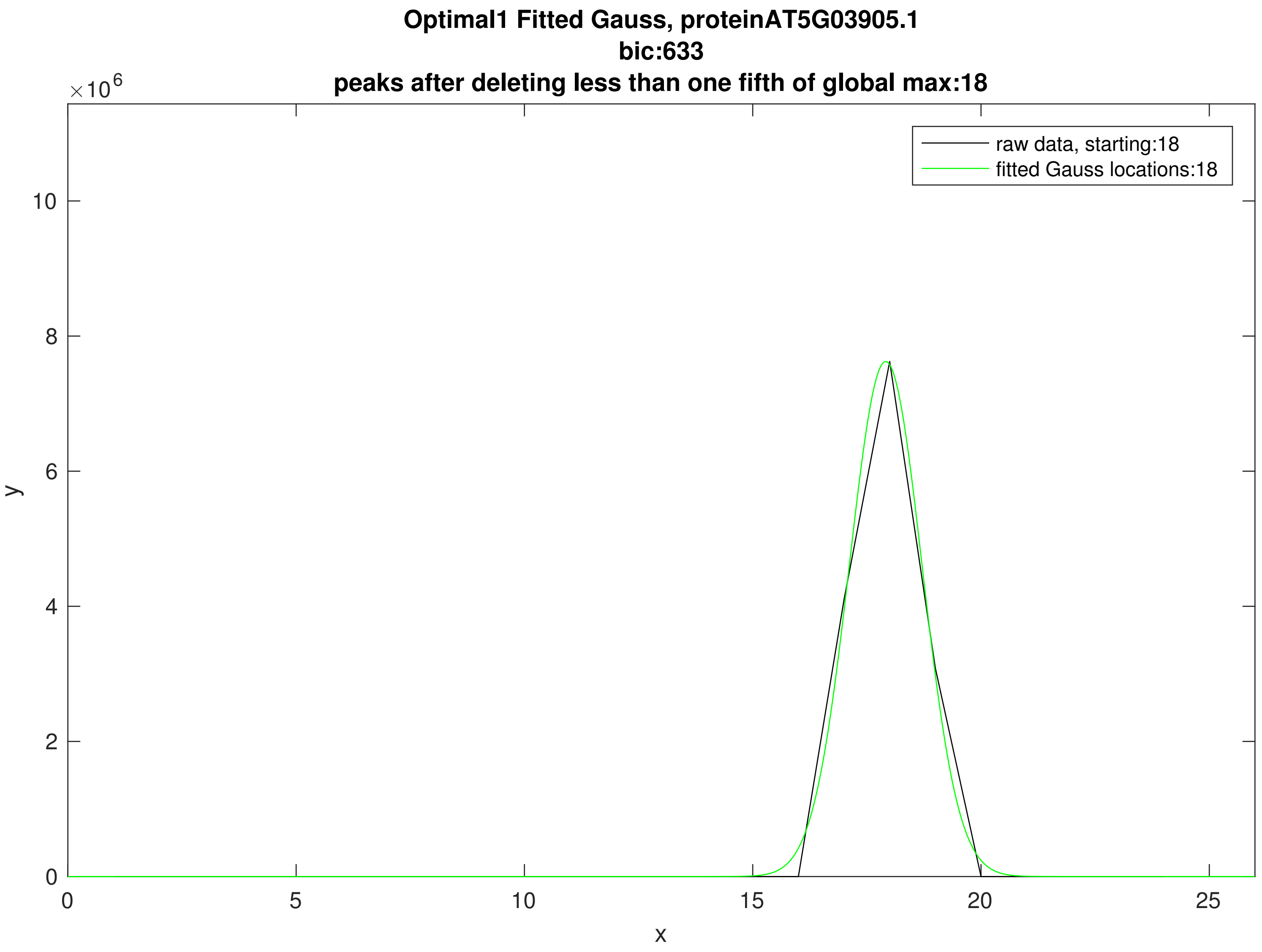

Optimal1 Fitted Gauss, proteinAT5G03905.1
bic:633
peaks after deleting less than one fifth of global max:18
×10^6
raw data, starting:18
fitted Gauss locations:18
10
8
6
4
2
0
y
0
5
10
15
20
25
x

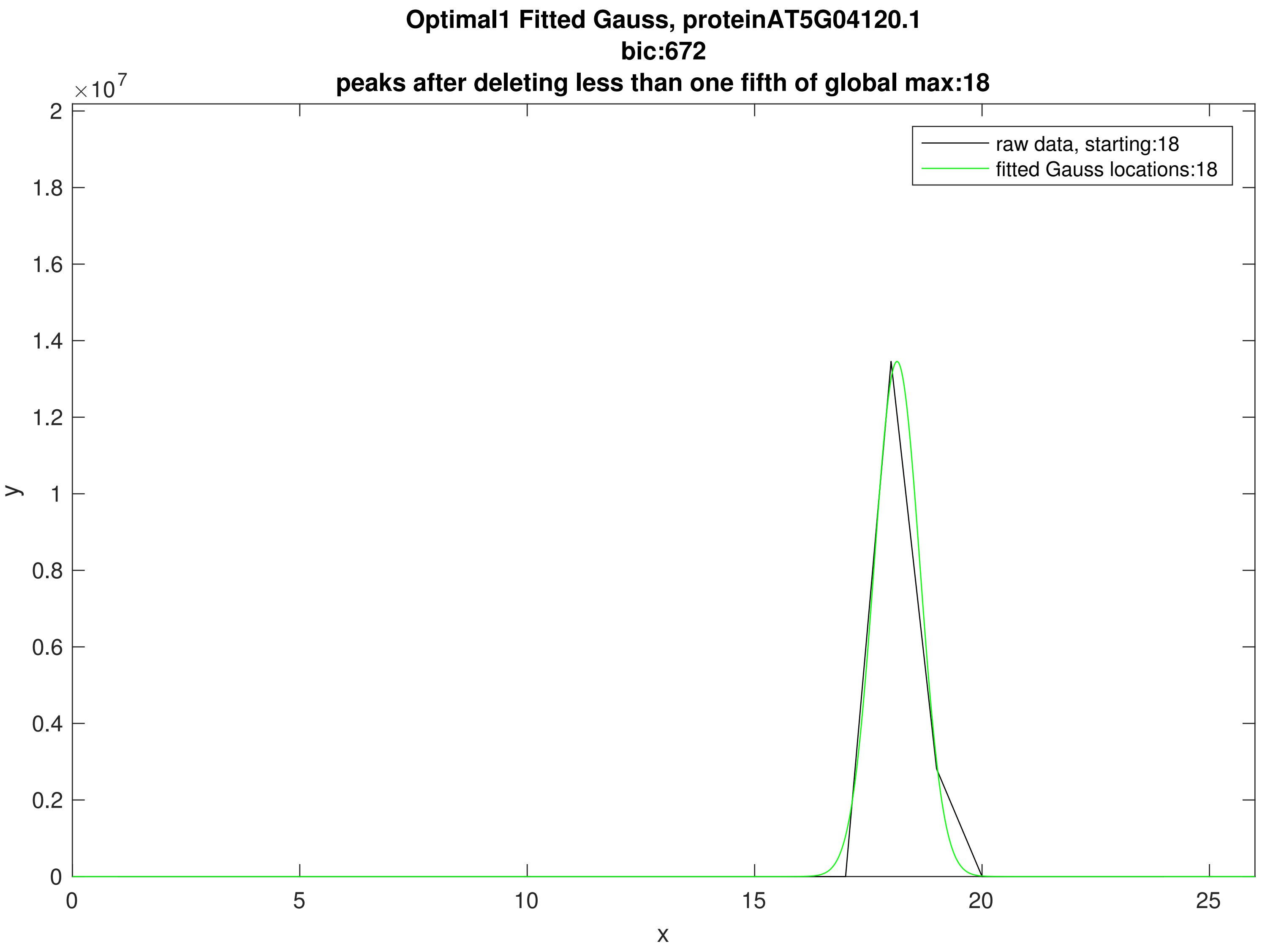
Optimal1 Fitted Gauss, proteinAT5G04120.1
bic:672
peaks after deleting less than one fifth of global max:18
×10^7
2
1.8
1.6
1.4
1.2
1
0.8
0.6
0.4
0.2
0
y
0
5
10
15
20
25
x
raw data, starting:18
fitted Gauss locations:18

Optimal1 Fitted Gauss, proteinAT5G04140.1;AT5G04140.2
bic:678
peaks after deleting less than one fifth of global max:5.1

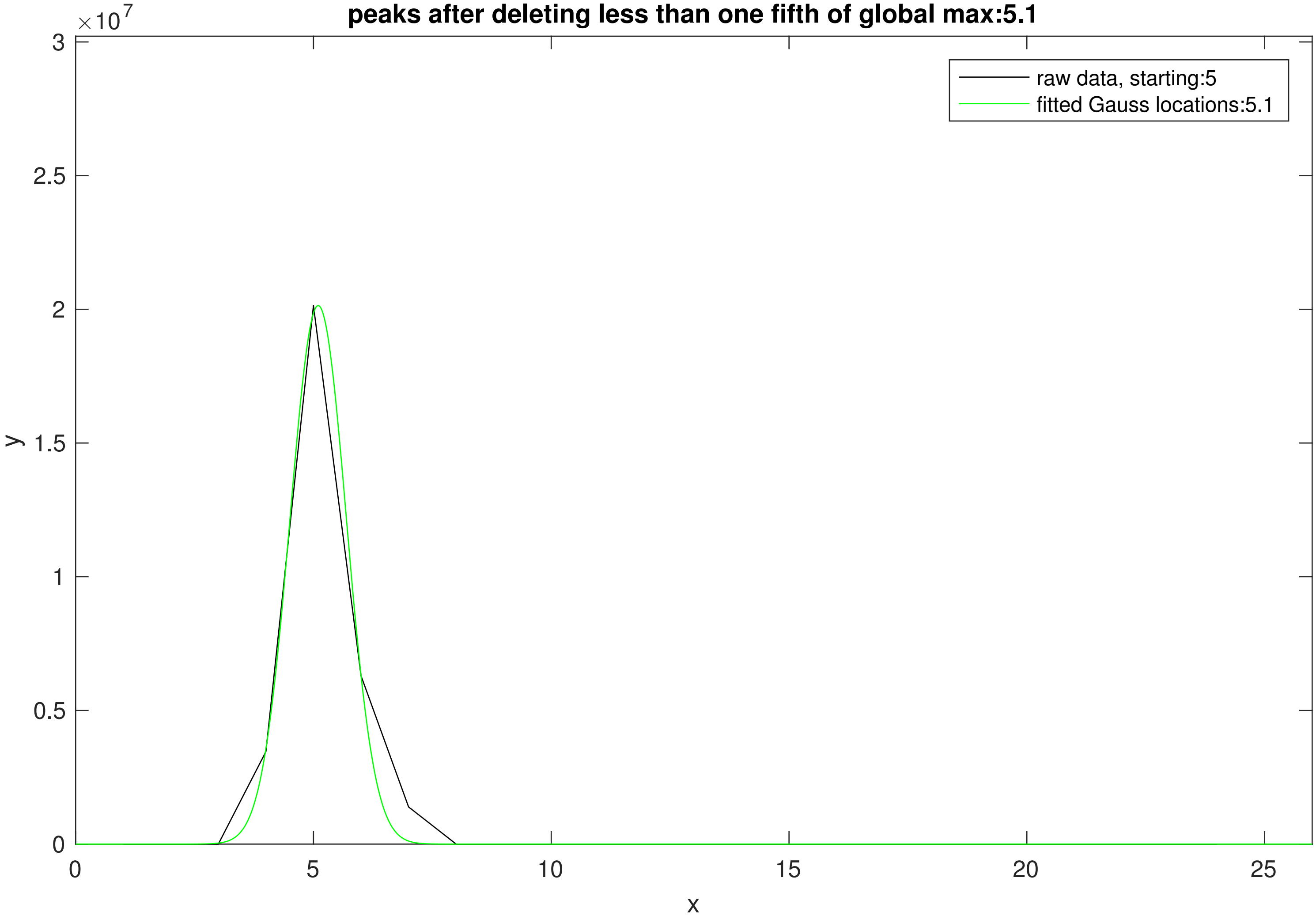

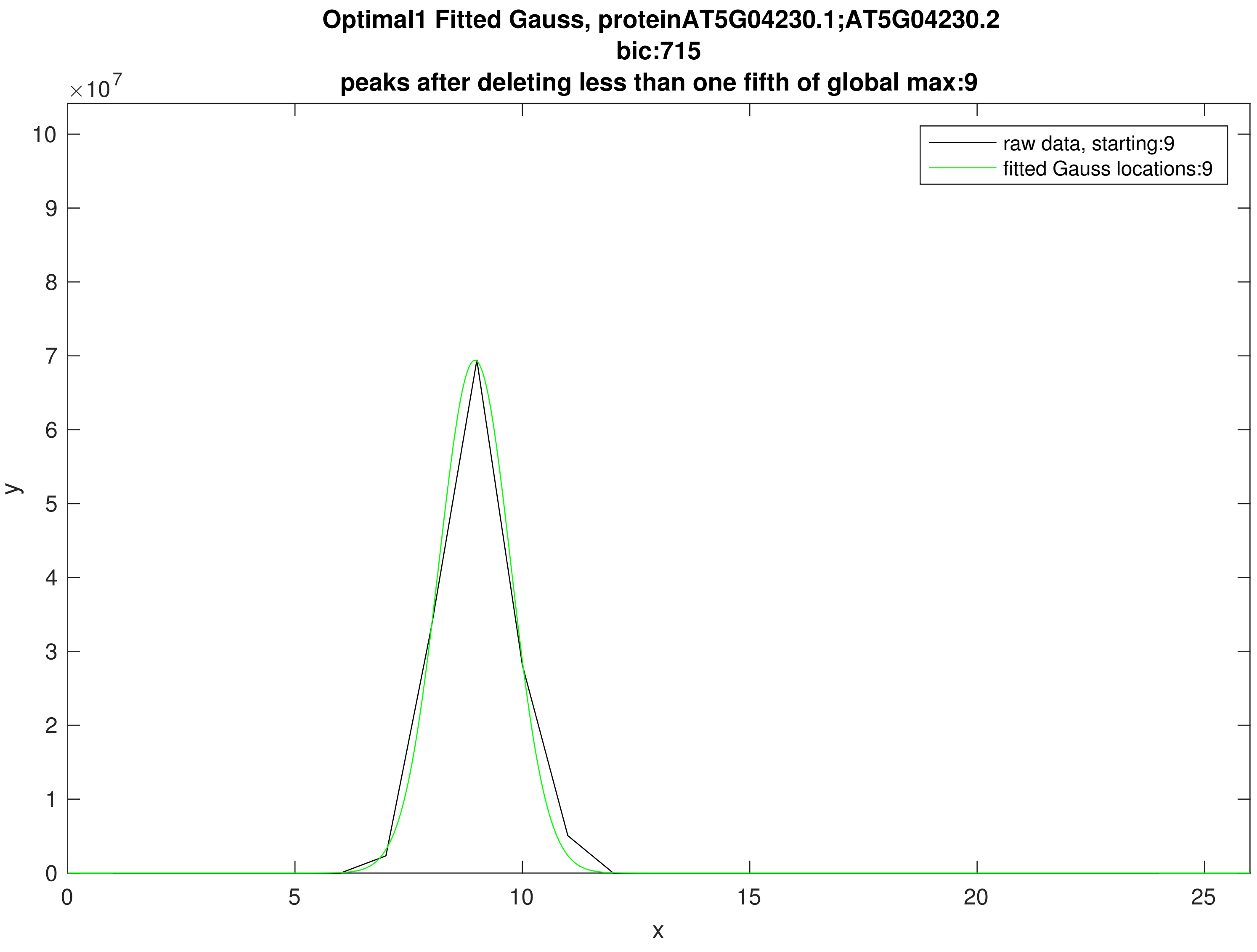
Optimal1 Fitted Gauss, proteinAT5G04230.1;AT5G04230.2
bic:715
peaks after deleting less than one fifth of global max:9
×10^7
raw data, starting:9
fitted Gauss locations:9
y
x
0
1
2
3
4
5
6
7
8
9
10
0
5
10
15
20
25

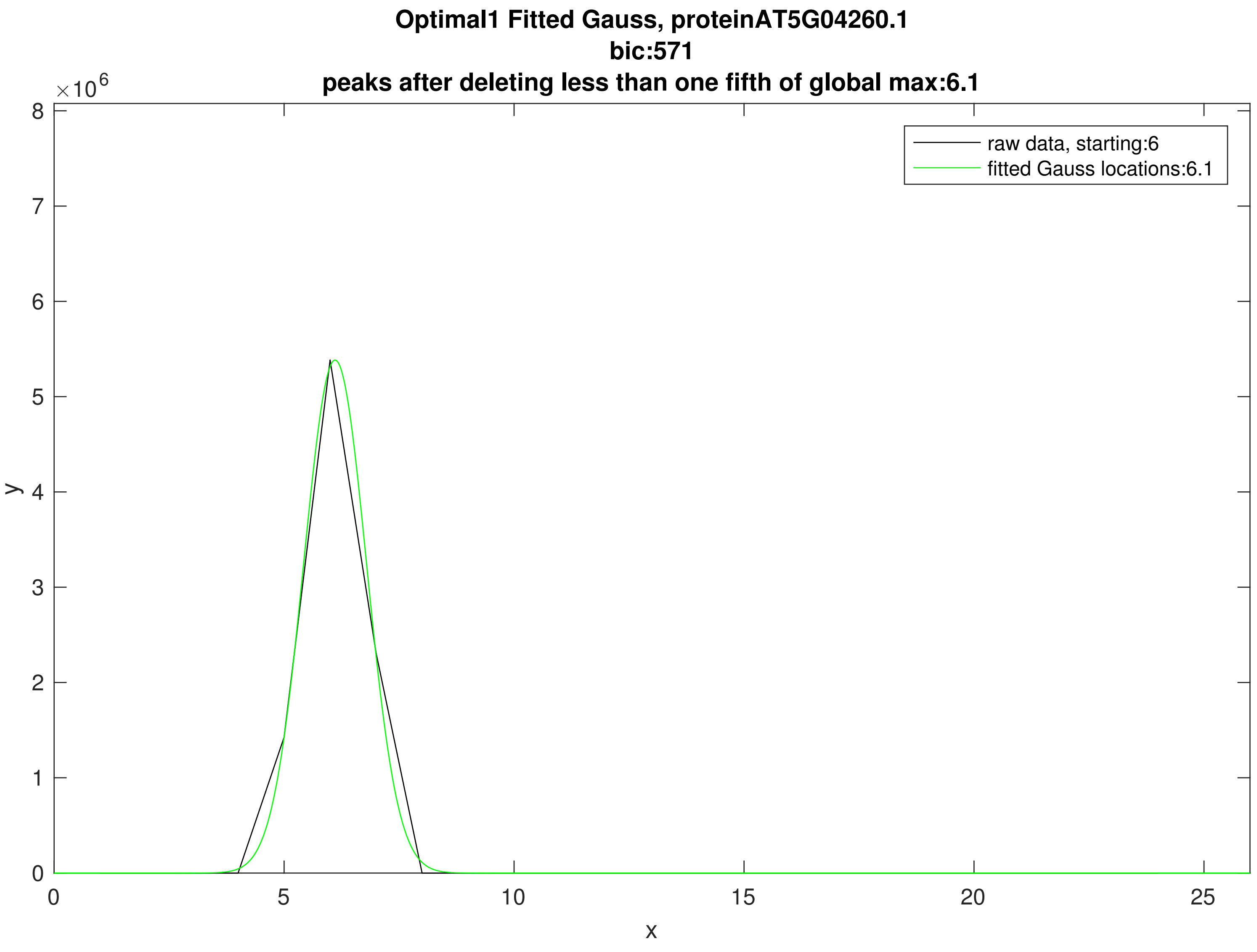
Optimal1 Fitted Gauss, proteinAT5G04260.1
bic:571
peaks after deleting less than one fifth of global max:6.1
×10^6
raw data, starting:6
fitted Gauss locations:6.1
y
x
0
1
2
3
4
5
6
7
8
0
5
10
15
20
25

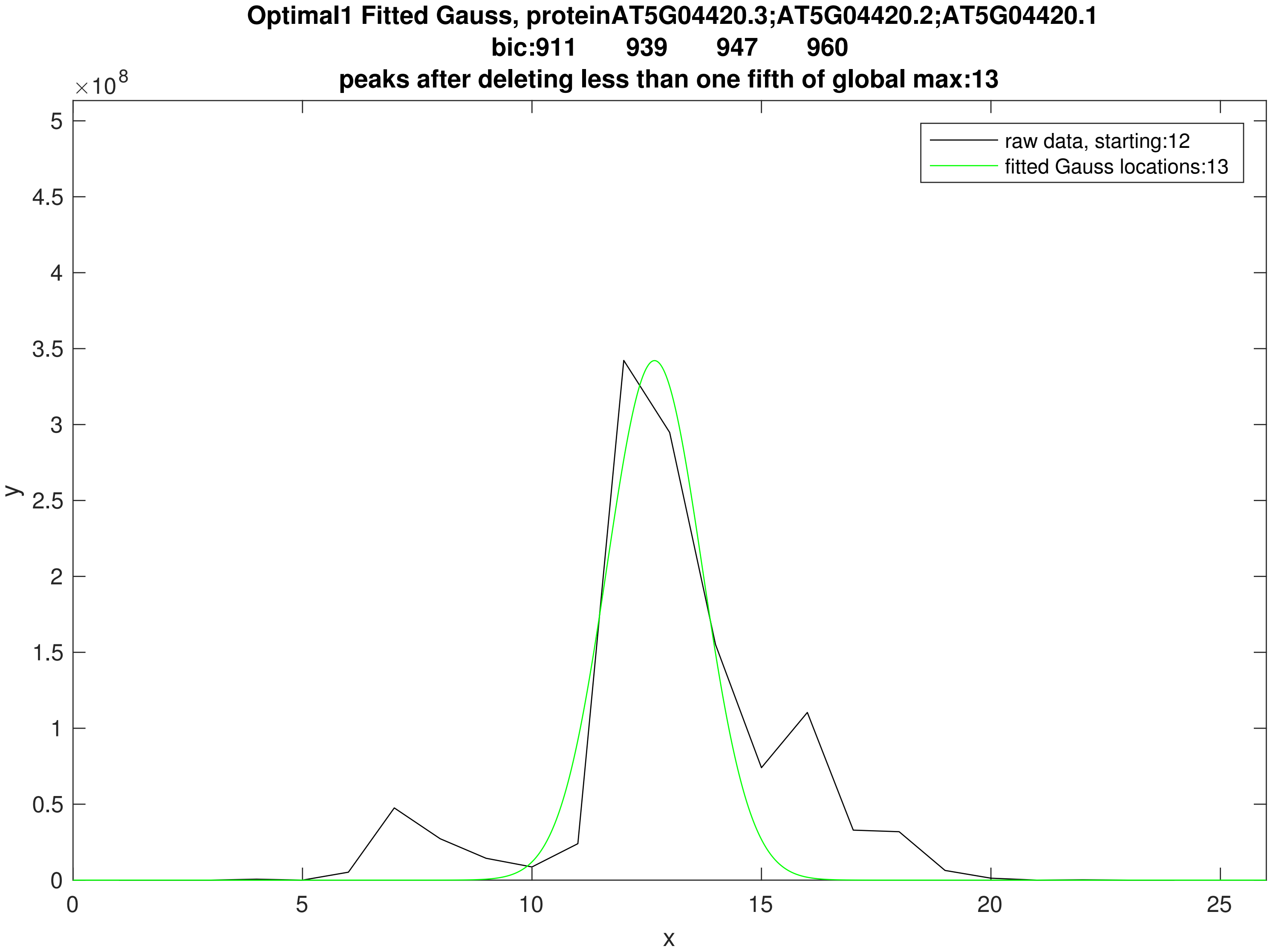
Optimal1 Fitted Gauss, proteinAT5G04420.3;AT5G04420.2;AT5G04420.1
bic:911 939 947 960
peaks after deleting less than one fifth of global max:13
×10^8
raw data, starting:12
fitted Gauss locations:13
y
x
0
0.5
1
1.5
2
2.5
3
3.5
4
4.5
5
0
5
10
15
20
25

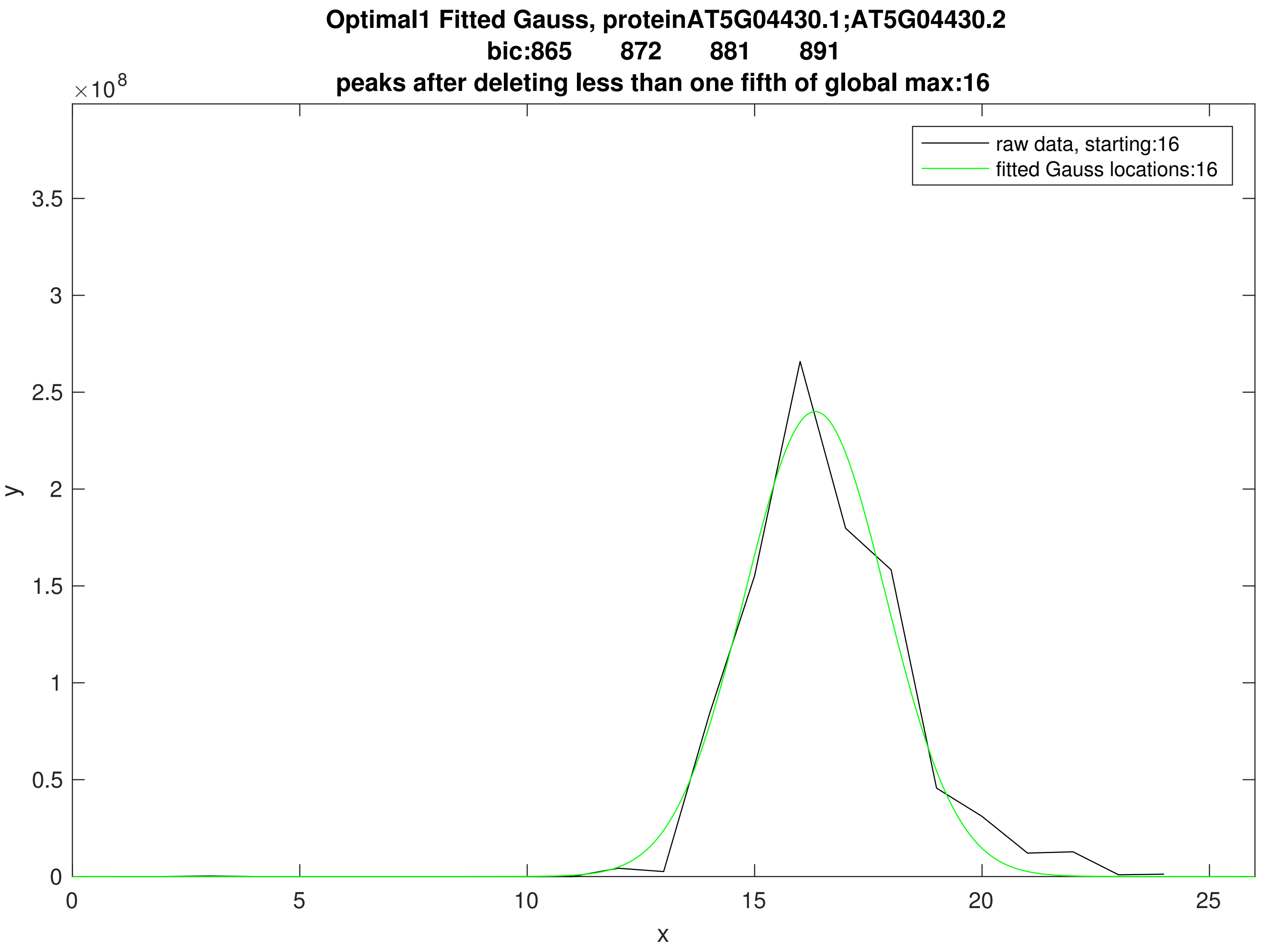
Optimal1 Fitted Gauss, proteinAT5G04430.1;AT5G04430.2
bic:865 872 881 891
peaks after deleting less than one fifth of global max:16
×10^8
raw data, starting:16
fitted Gauss locations:16
y
x
0
0.5
1
1.5
2
2.5
3
3.5
5
10
15
20
25

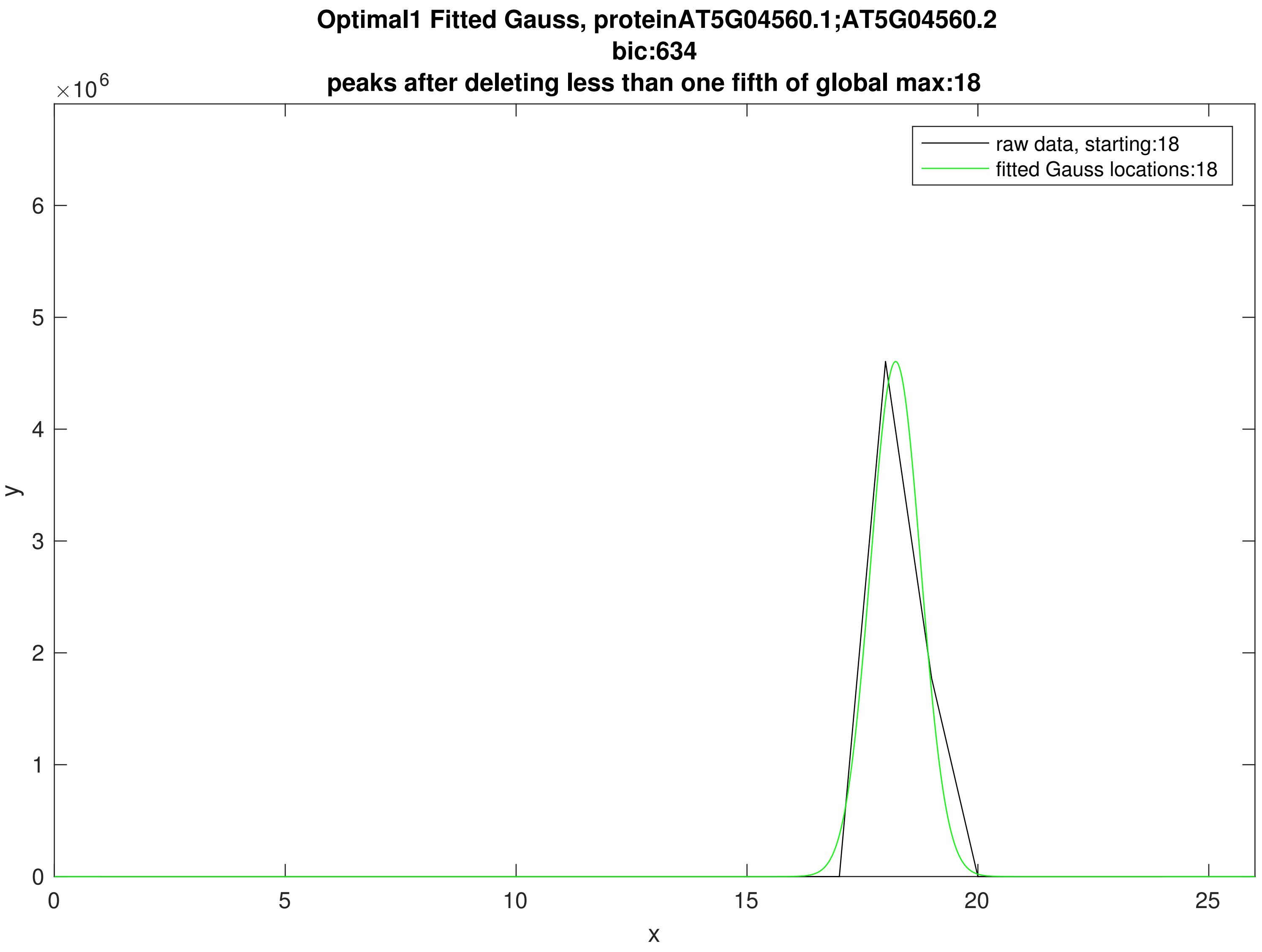
Optimal1 Fitted Gauss, proteinAT5G04560.1;AT5G04560.2
bic:634
peaks after deleting less than one fifth of global max:18
×10^6
raw data, starting:18
fitted Gauss locations:18
y
x
0
1
2
3
4
5
6
0
5
10
15
20
25

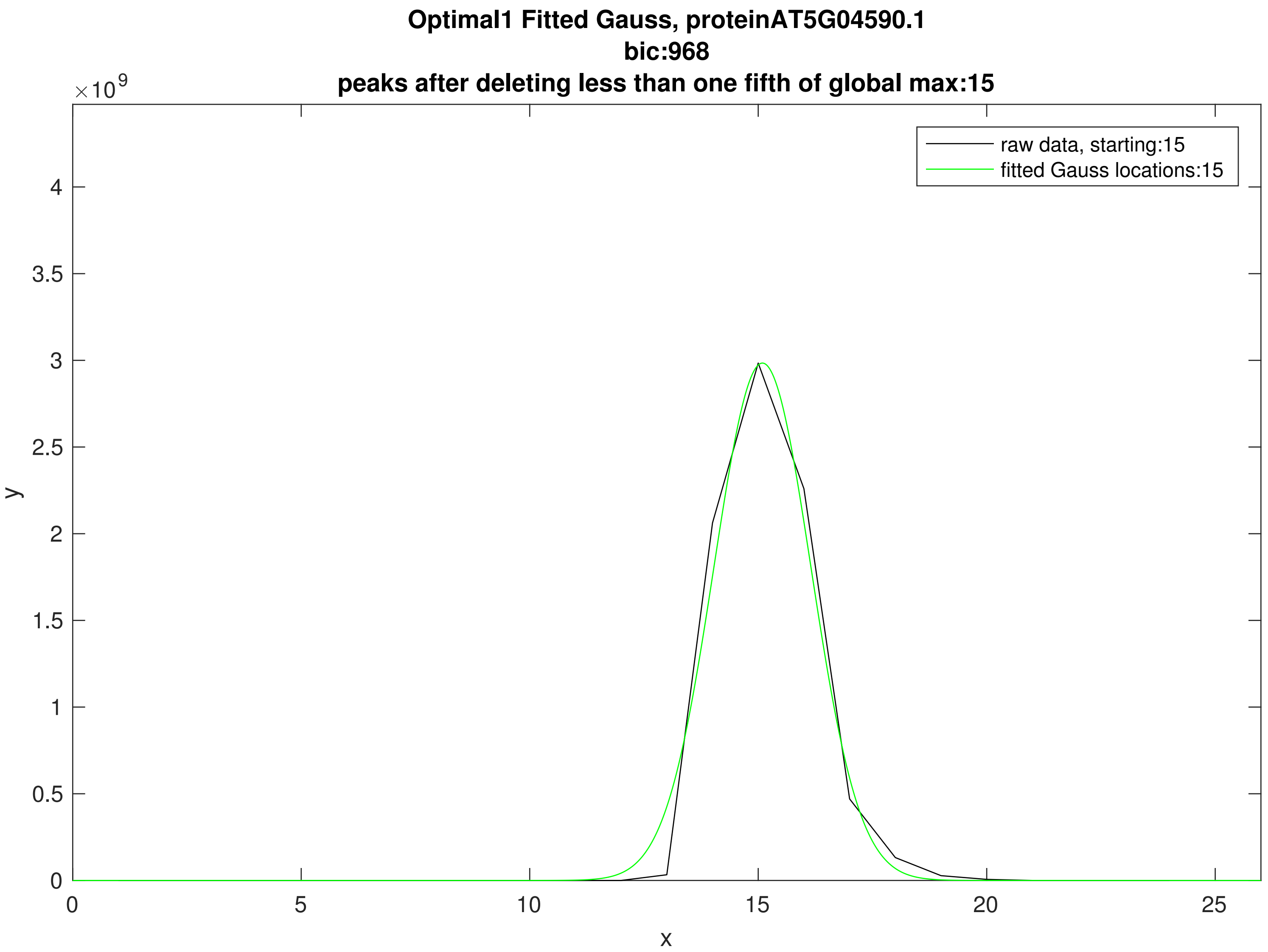

Optimal1 Fitted Gauss, proteinAT5G04590.1
bic:968
peaks after deleting less than one fifth of global max:15
×10^9
raw data, starting:15
fitted Gauss locations:15
4
3.5
3
2.5
2
1.5
1
0.5
0
y
0
5
10
15
20
25
x

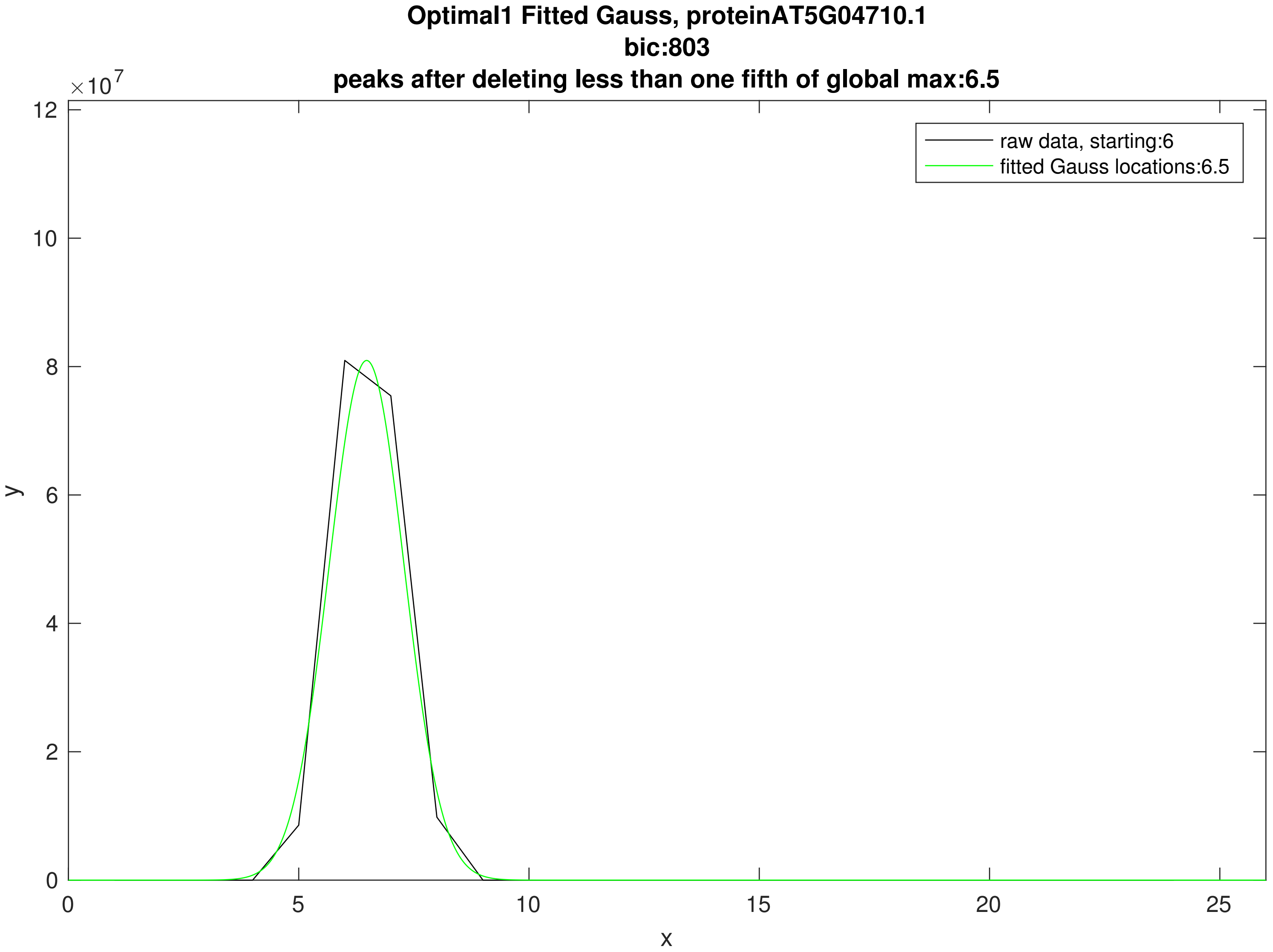
Optimal1 Fitted Gauss, proteinAT5G04710.1
bic:803
peaks after deleting less than one fifth of global max:6.5
×10^7
12
10
8
6
4
2
0
y
0
5
10
15
20
25
x
raw data, starting:6
fitted Gauss locations:6.5

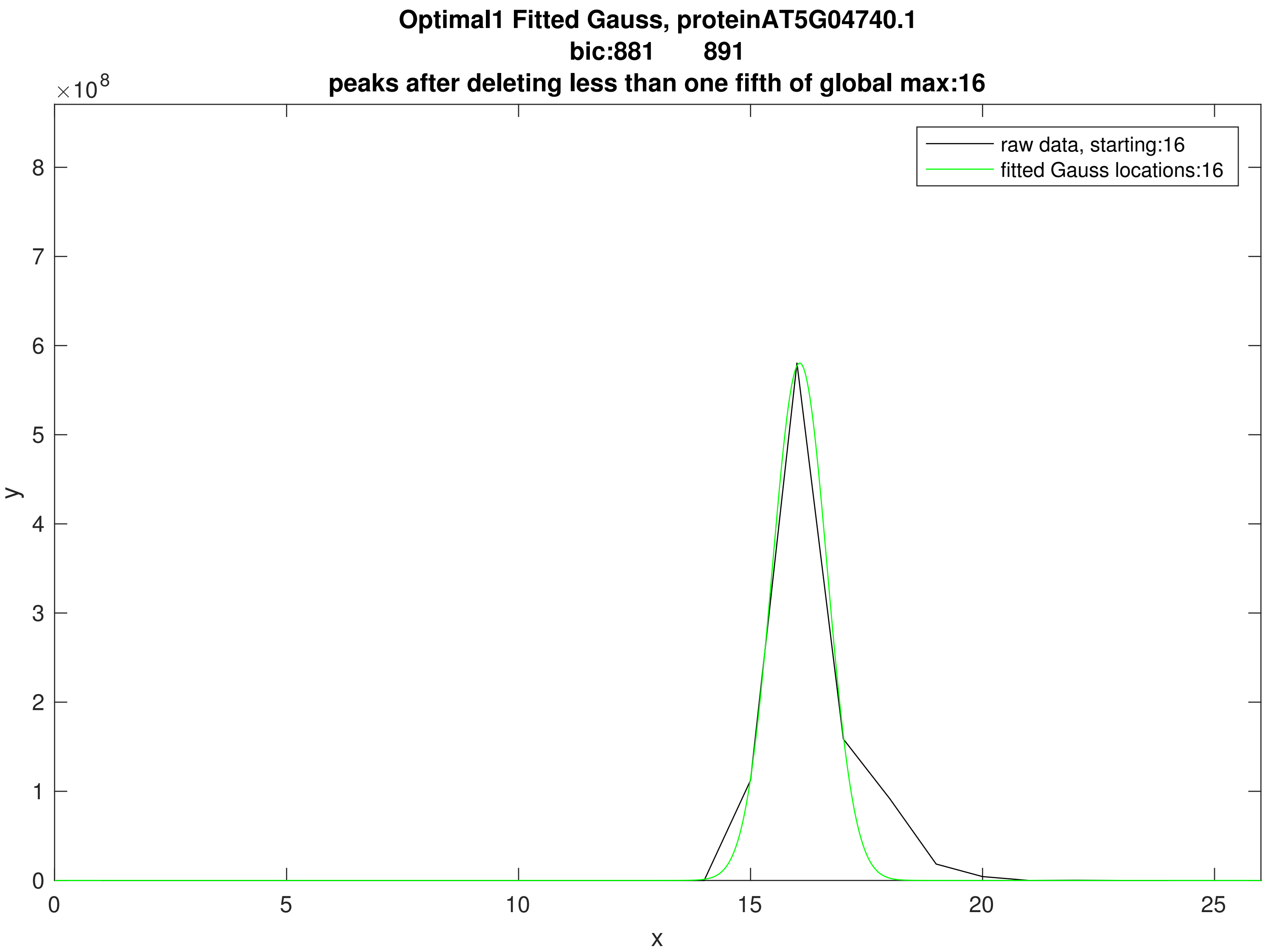
Optimal1 Fitted Gauss, proteinAT5G04740.1
bic:881 891
peaks after deleting less than one fifth of global max:16
×10^8
raw data, starting:16
fitted Gauss locations:16
y
x
0
1
2
3
4
5
6
7
8
0
5
10
15
20
25

**Optimal1 Fitted Gauss, proteinAT5G04750.1;AT5G04750.2**
**bic:786**
**peaks after deleting less than one fifth of global max:20**

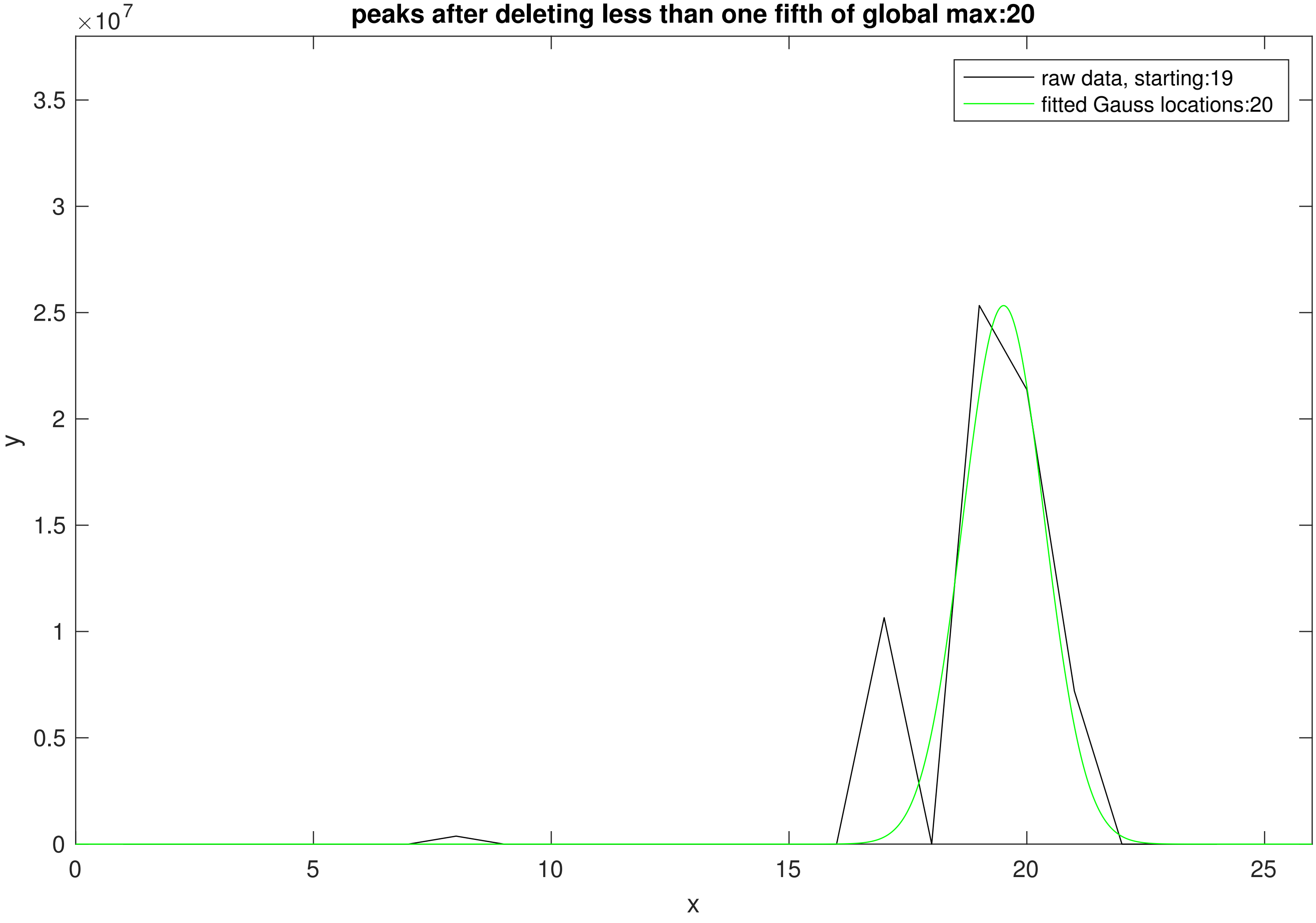

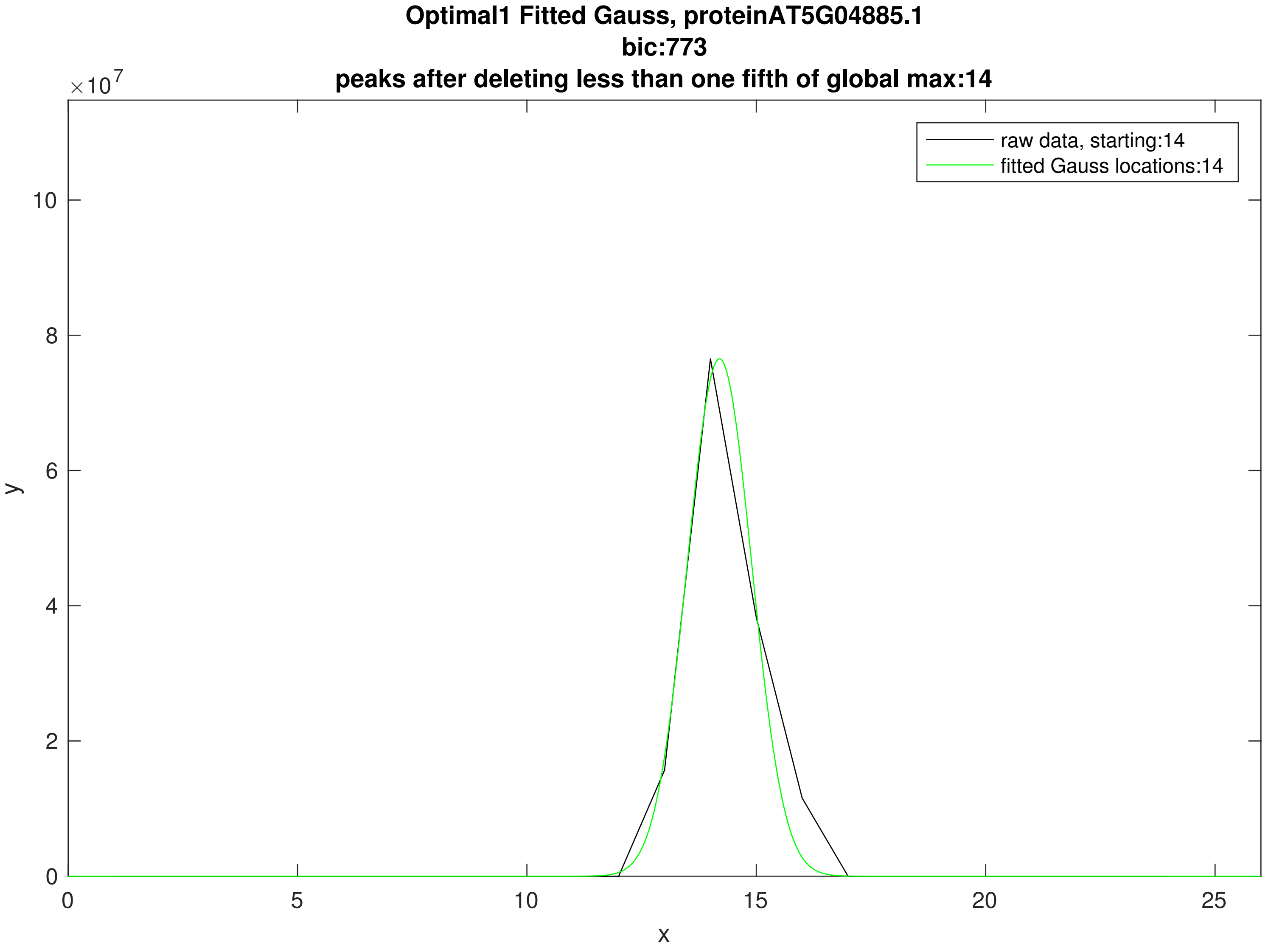
Optimal1 Fitted Gauss, proteinAT5G04885.1
bic:773
peaks after deleting less than one fifth of global max:14
×10^7
raw data, starting:14
fitted Gauss locations:14
y
x
0
2
4
6
8
10
5
15
20
25

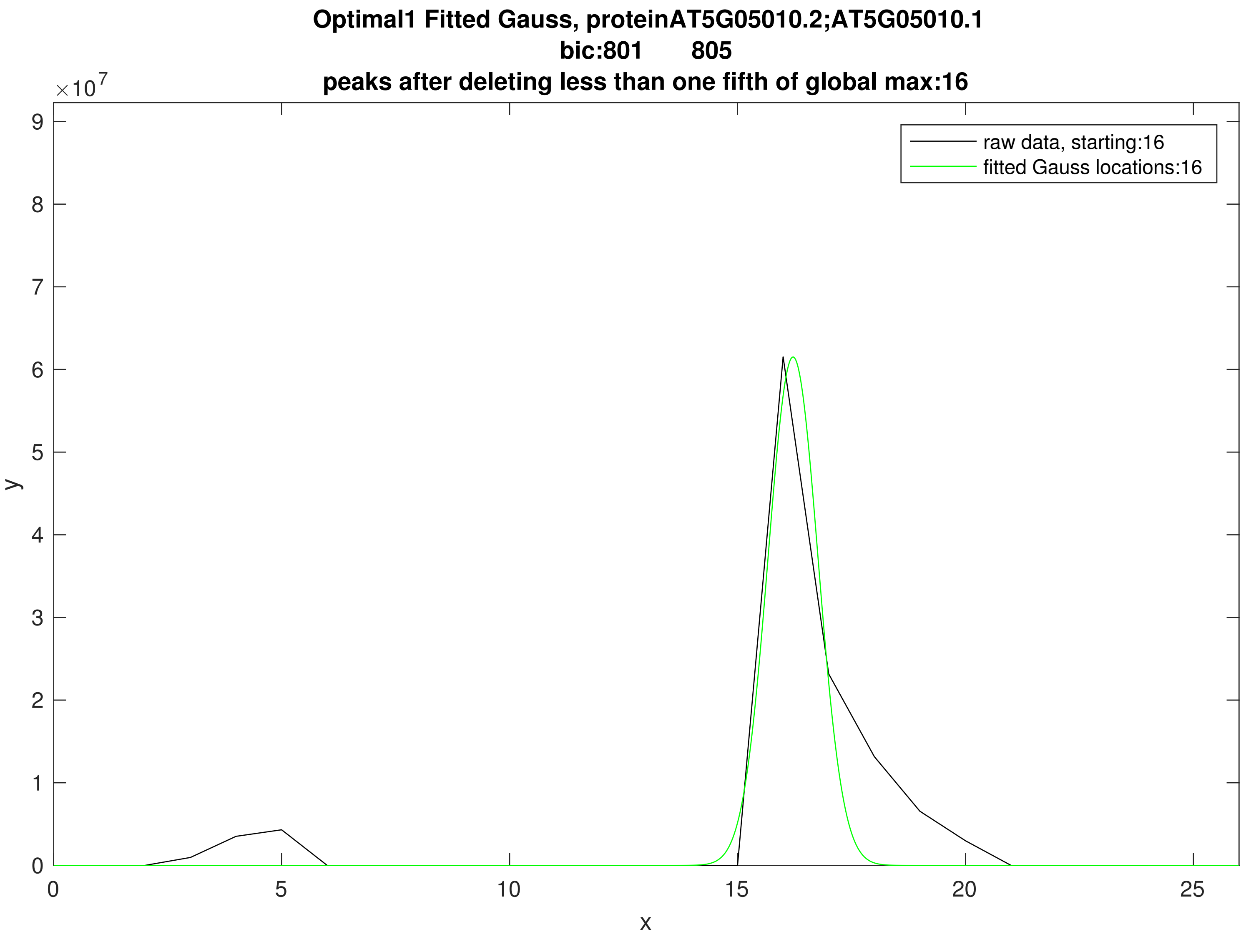
Optimal1 Fitted Gauss, proteinAT5G05010.2;AT5G05010.1
bic:801 805
peaks after deleting less than one fifth of global max:16
×10^7
raw data, starting:16
fitted Gauss locations:16
y
x
0
1
2
3
4
5
6
7
8
9
0
5
10
15
20
25

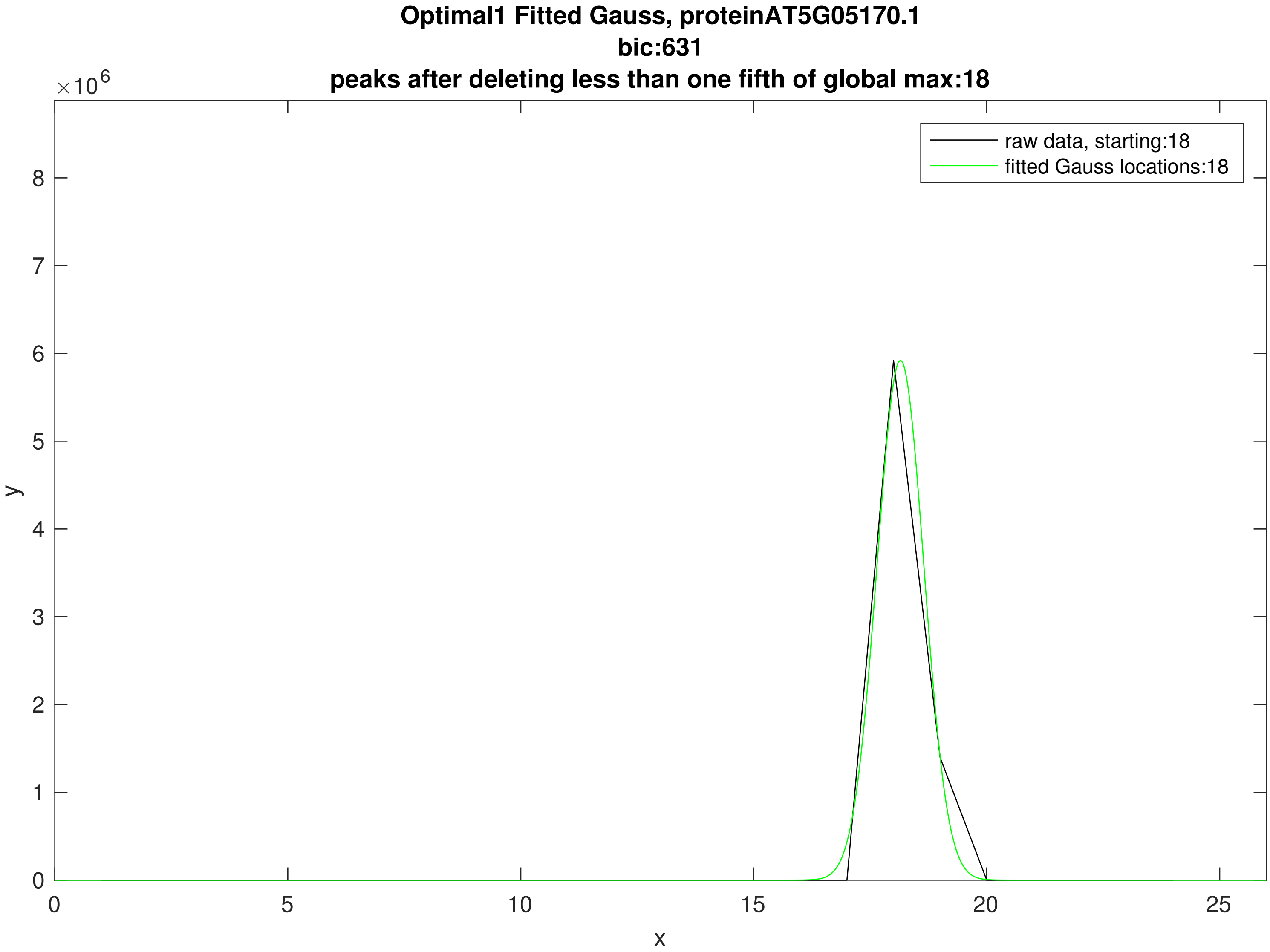
Optimal1 Fitted Gauss, proteinAT5G05170.1
bic:631
peaks after deleting less than one fifth of global max:18
×10^6
raw data, starting:18
fitted Gauss locations:18
y
x

#### Optimal1 Fitted Gauss, proteinAT5G05340.1
#### bic:703
#### peaks after deleting less than one fifth of global max:18

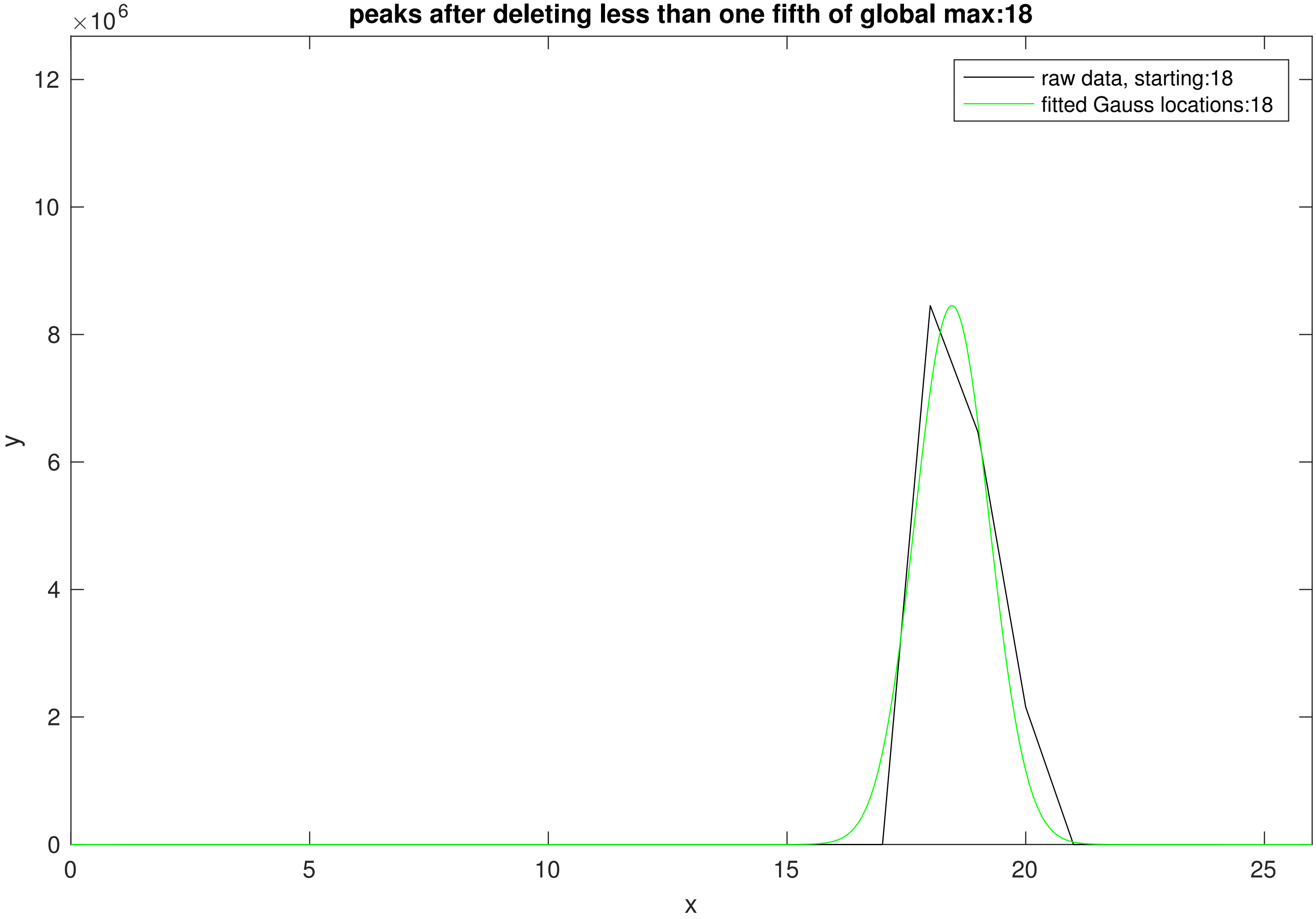

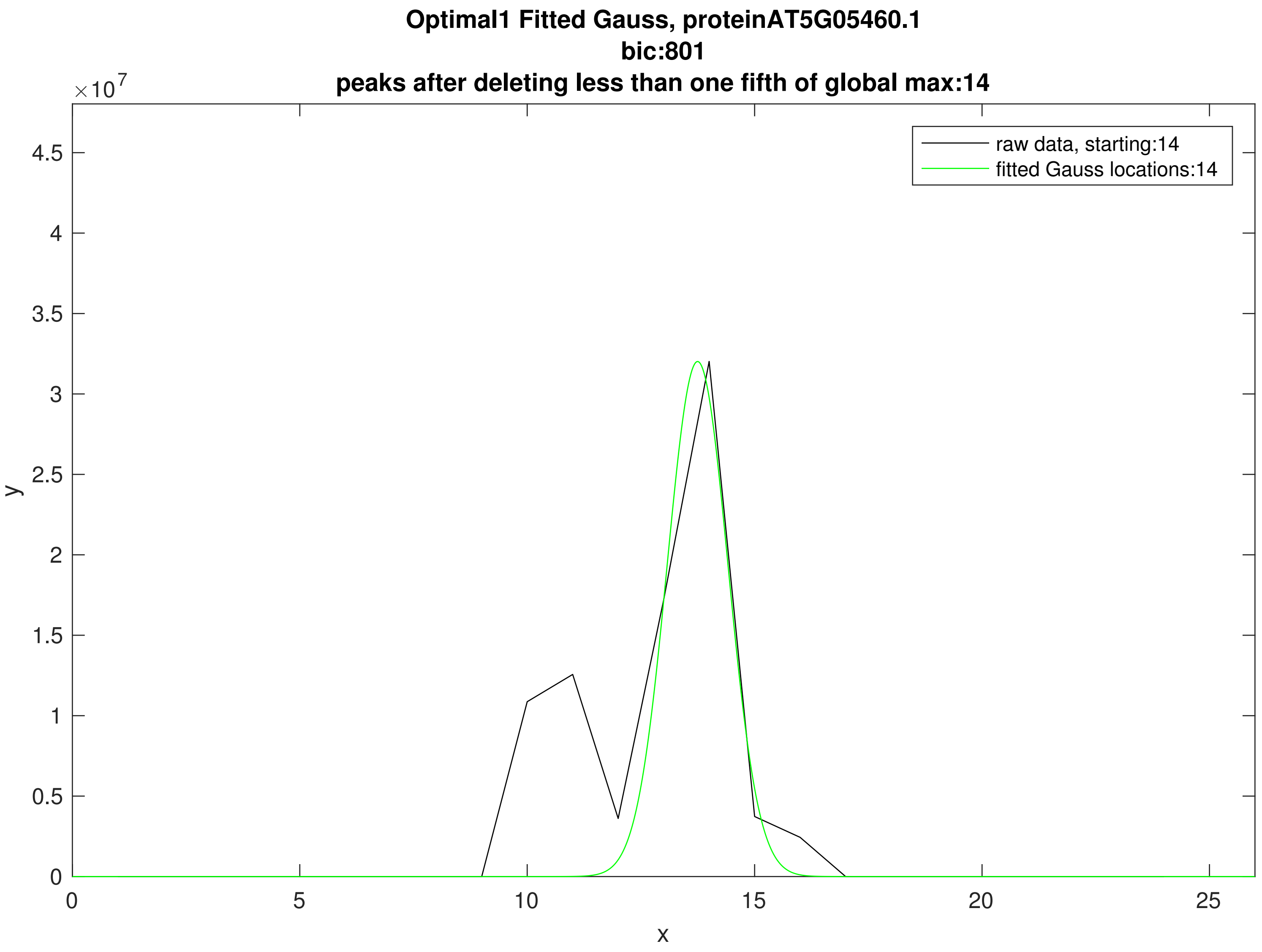
Optimal1 Fitted Gauss, proteinAT5G05460.1
bic:801
peaks after deleting less than one fifth of global max:14
×10^7
4.5
4
3.5
3
2.5
2
1.5
1
0.5
0
y
0
5
10
15
20
25
x
raw data, starting:14
fitted Gauss locations:14

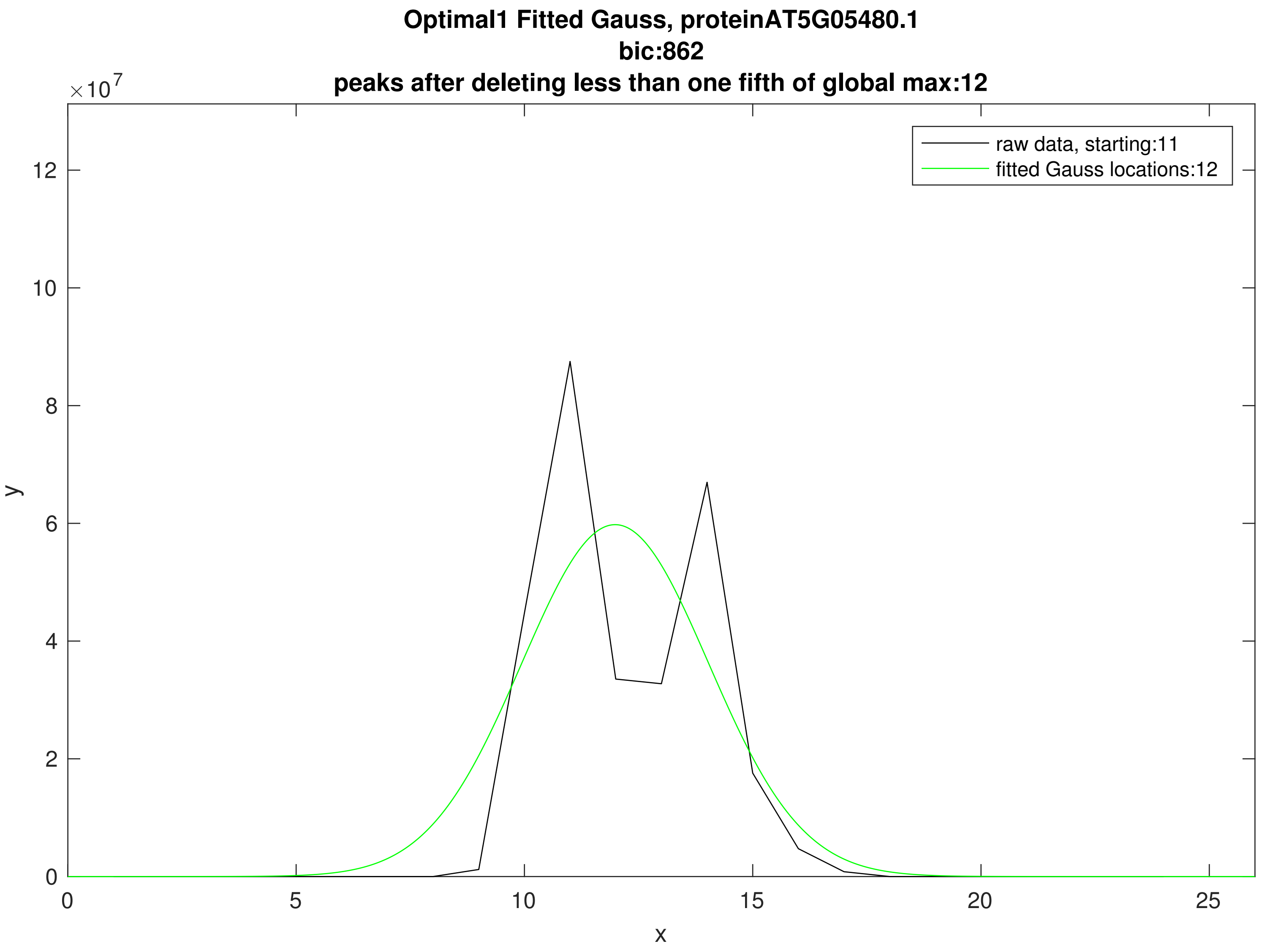
Optimal1 Fitted Gauss, proteinAT5G05480.1
bic:862
peaks after deleting less than one fifth of global max:12
×10^7
raw data, starting:11
fitted Gauss locations:12
12
10
8
6
4
2
0
y
0
5
10
15
20
25
x

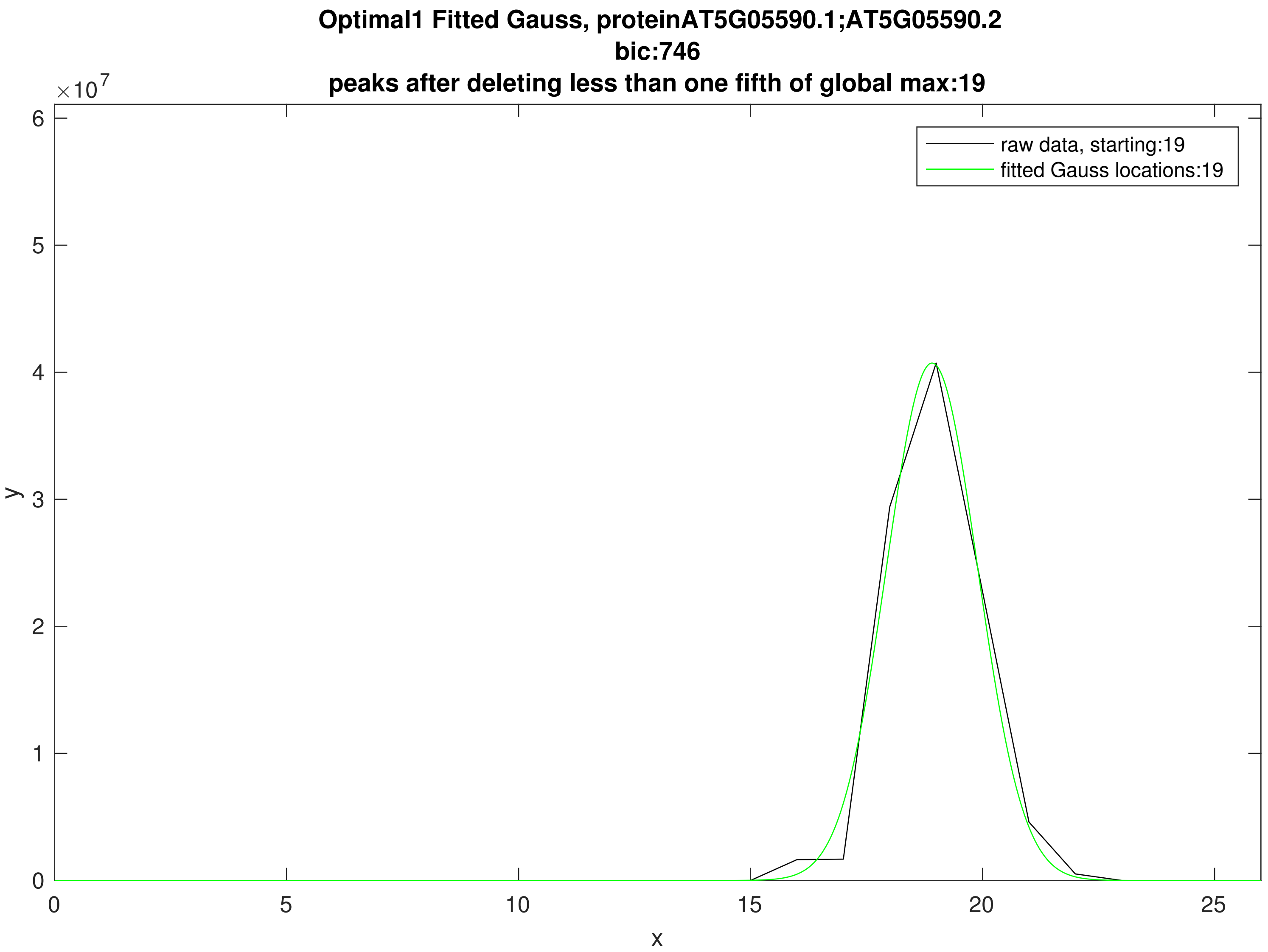
Optimal1 Fitted Gauss, proteinAT5G05590.1;AT5G05590.2
bic:746
peaks after deleting less than one fifth of global max:19
raw data, starting:19
fitted Gauss locations:19
y
x

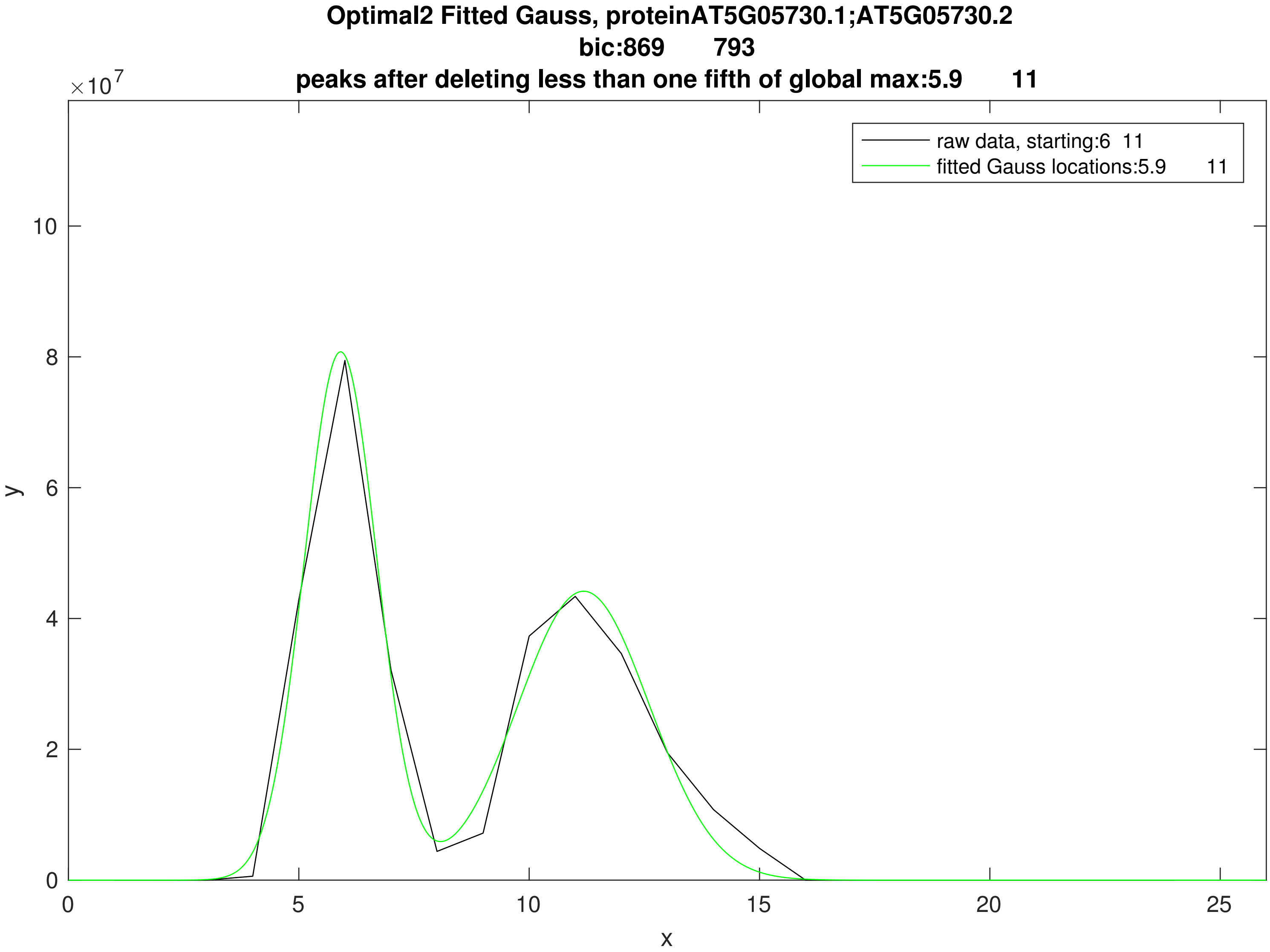
Optimal2 Fitted Gauss, proteinAT5G05730.1;AT5G05730.2
bic:869 793
peaks after deleting less than one fifth of global max:5.9 11
×10^7
raw data, starting:6 11
fitted Gauss locations:5.9 11
10
8
6
4
2
0
y
0
5
10
15
20
25
x

#### Optimal1 Fitted Gauss, proteinAT5G05780.1;AT5G05780.2;AT3G11270.1;AT3G11270.2
#### bic:756
#### peaks after deleting less than one fifth of global max:4.2

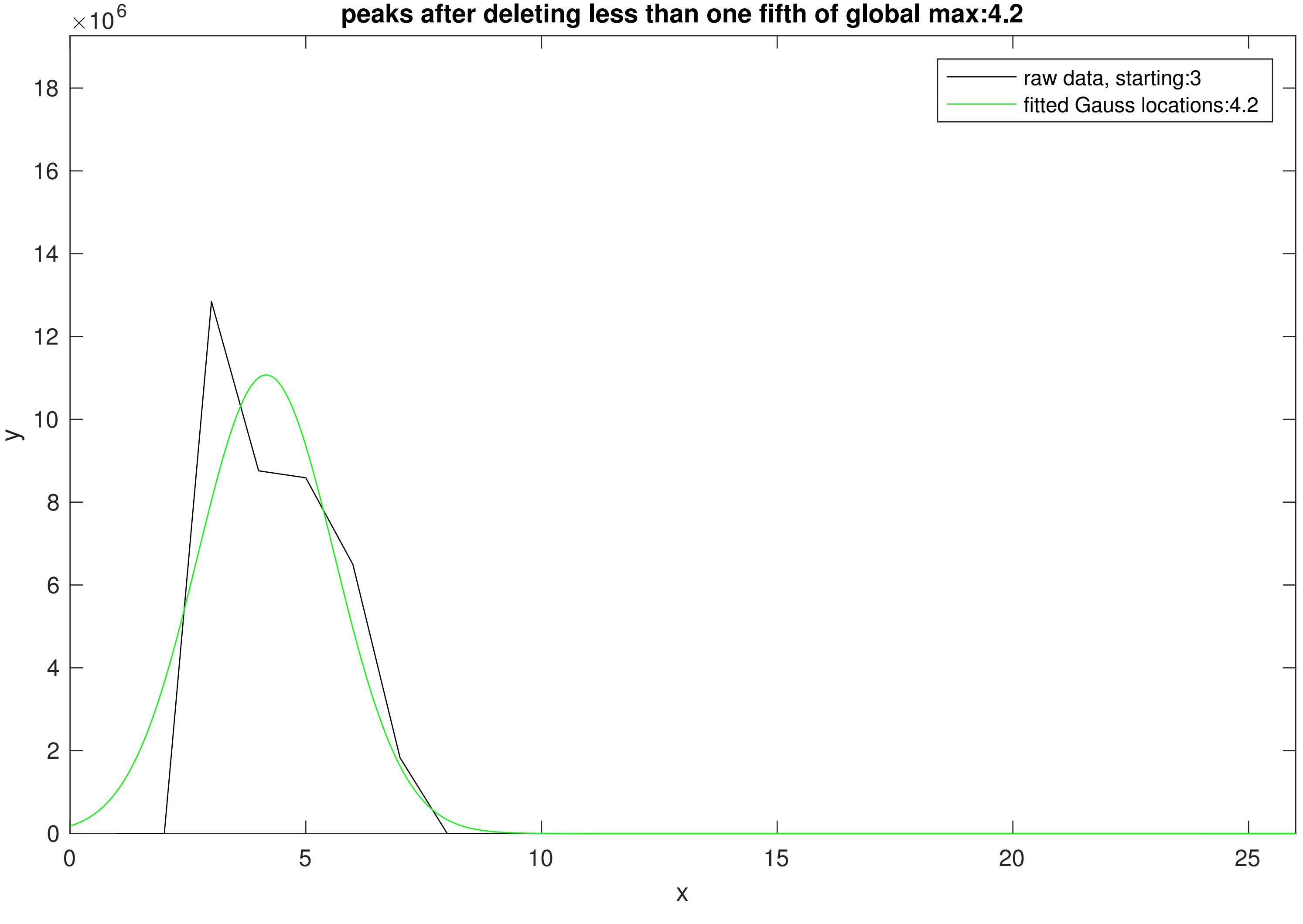

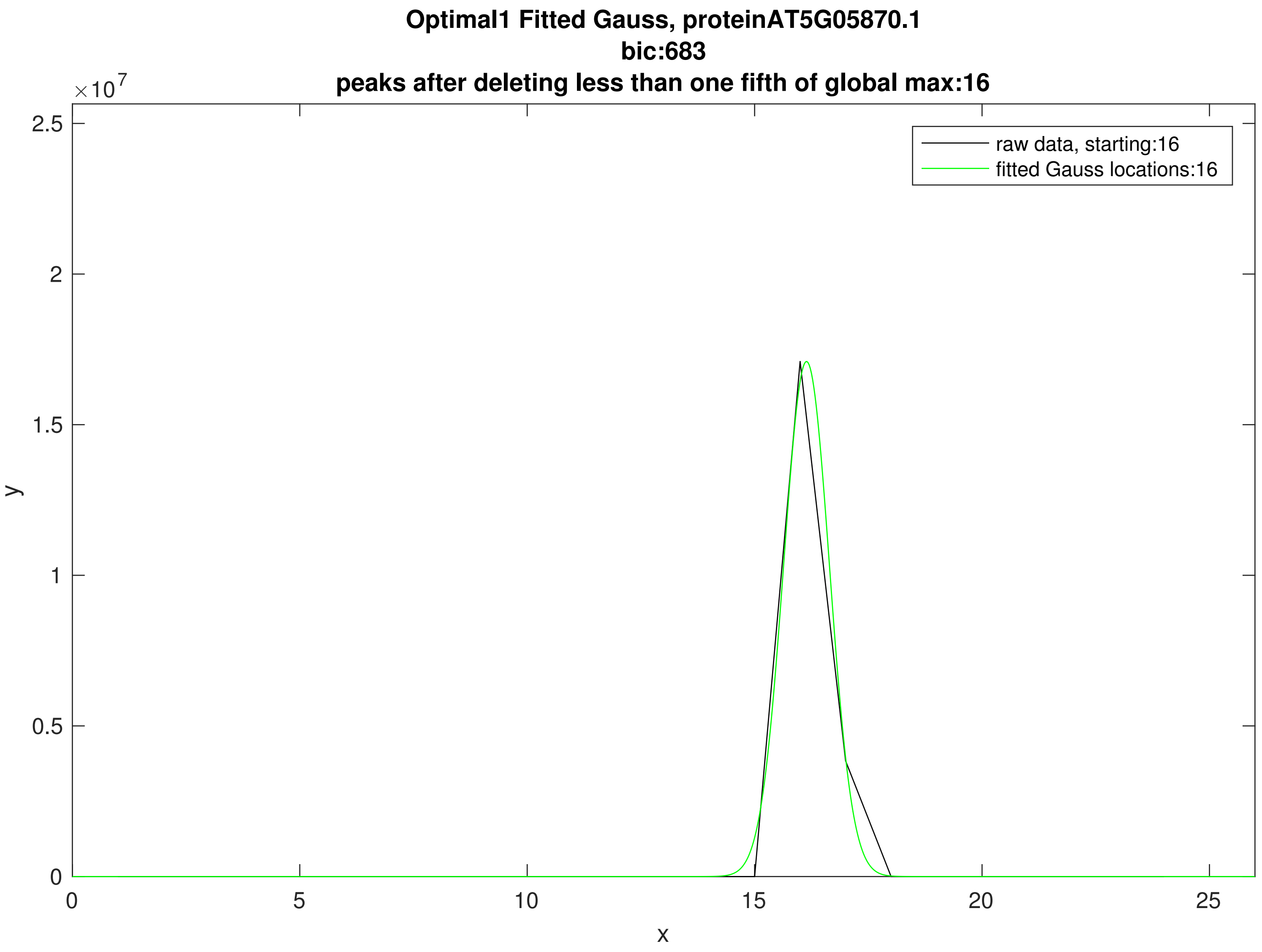

Optimal1 Fitted Gauss, proteinAT5G05870.1
bic:683
peaks after deleting less than one fifth of global max:16
×10^7
2.5
2
1.5
1
0.5
0
y
0
5
10
15
20
25
x
raw data, starting:16
fitted Gauss locations:16

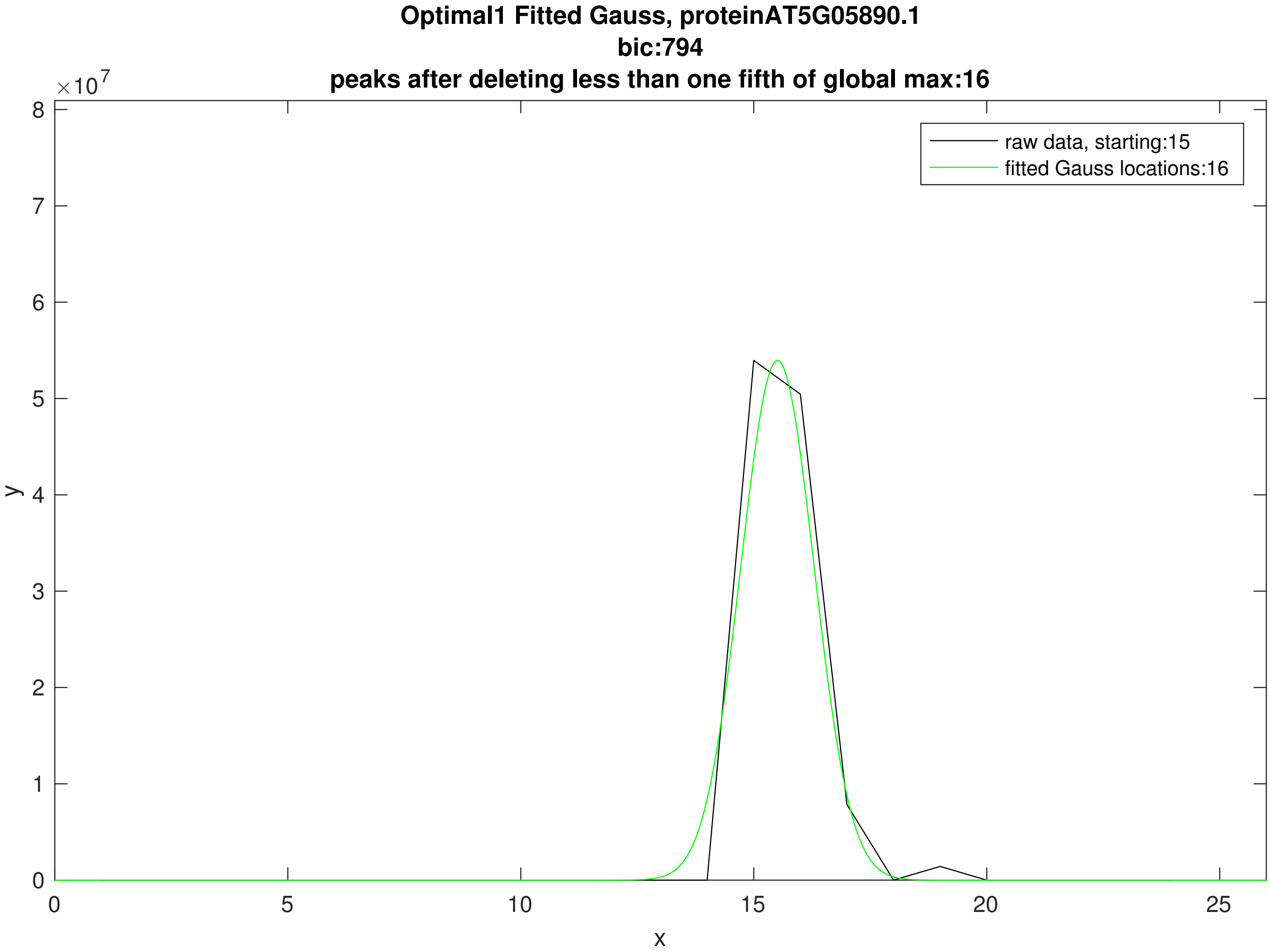
Optimal1 Fitted Gauss, proteinAT5G05890.1
bic:794
peaks after deleting less than one fifth of global max:16
×10^7
8
7
6
5
4
3
2
1
0
y
0
5
10
15
20
25
x
raw data, starting:15
fitted Gauss locations:16

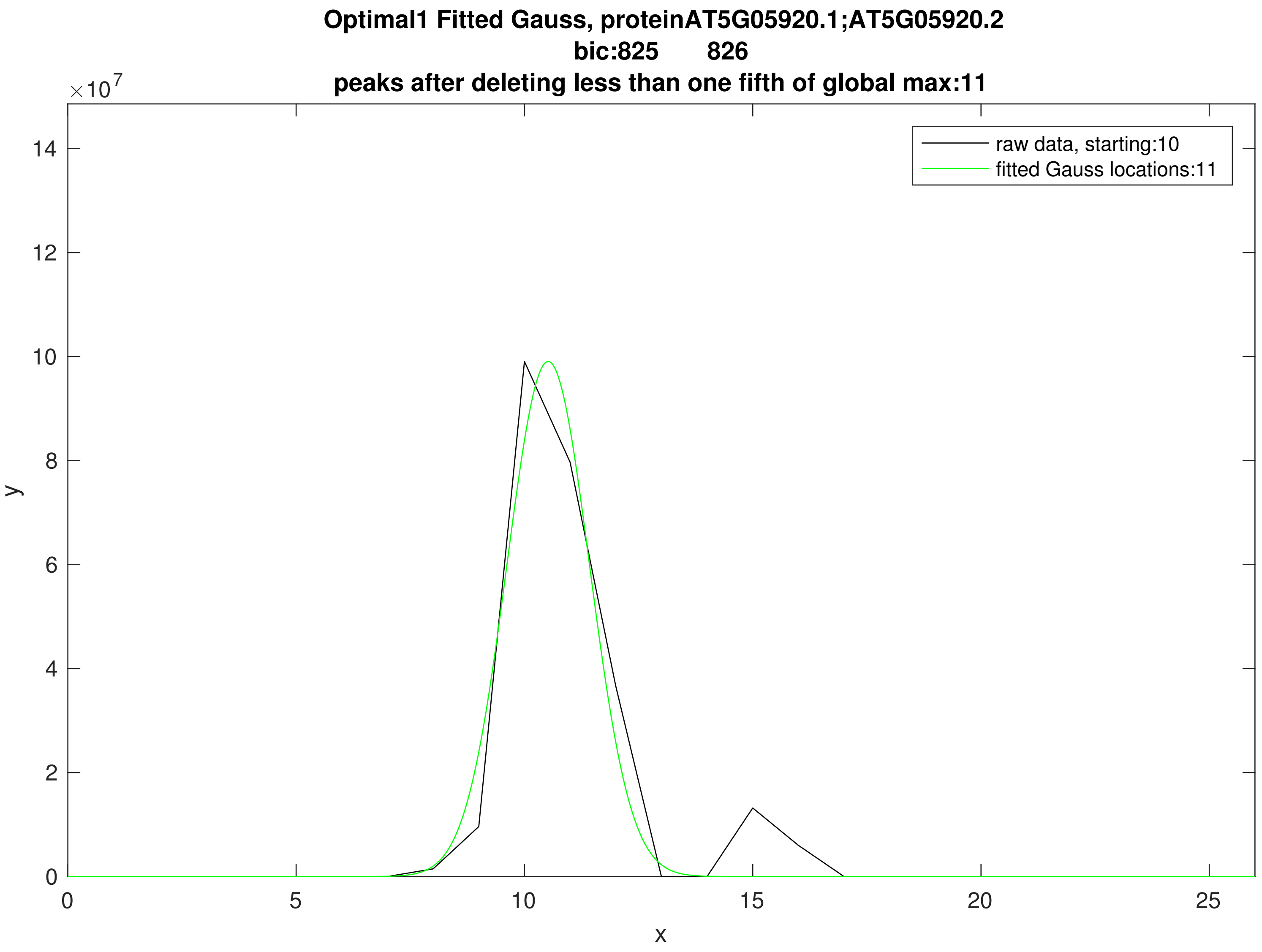
Optimal1 Fitted Gauss, proteinAT5G05920.1;AT5G05920.2
bic:825 826
peaks after deleting less than one fifth of global max:11
×10^7
raw data, starting:10
fitted Gauss locations:11
y
x
0
2
4
6
8
10
12
14
0
5
10
15
20
25

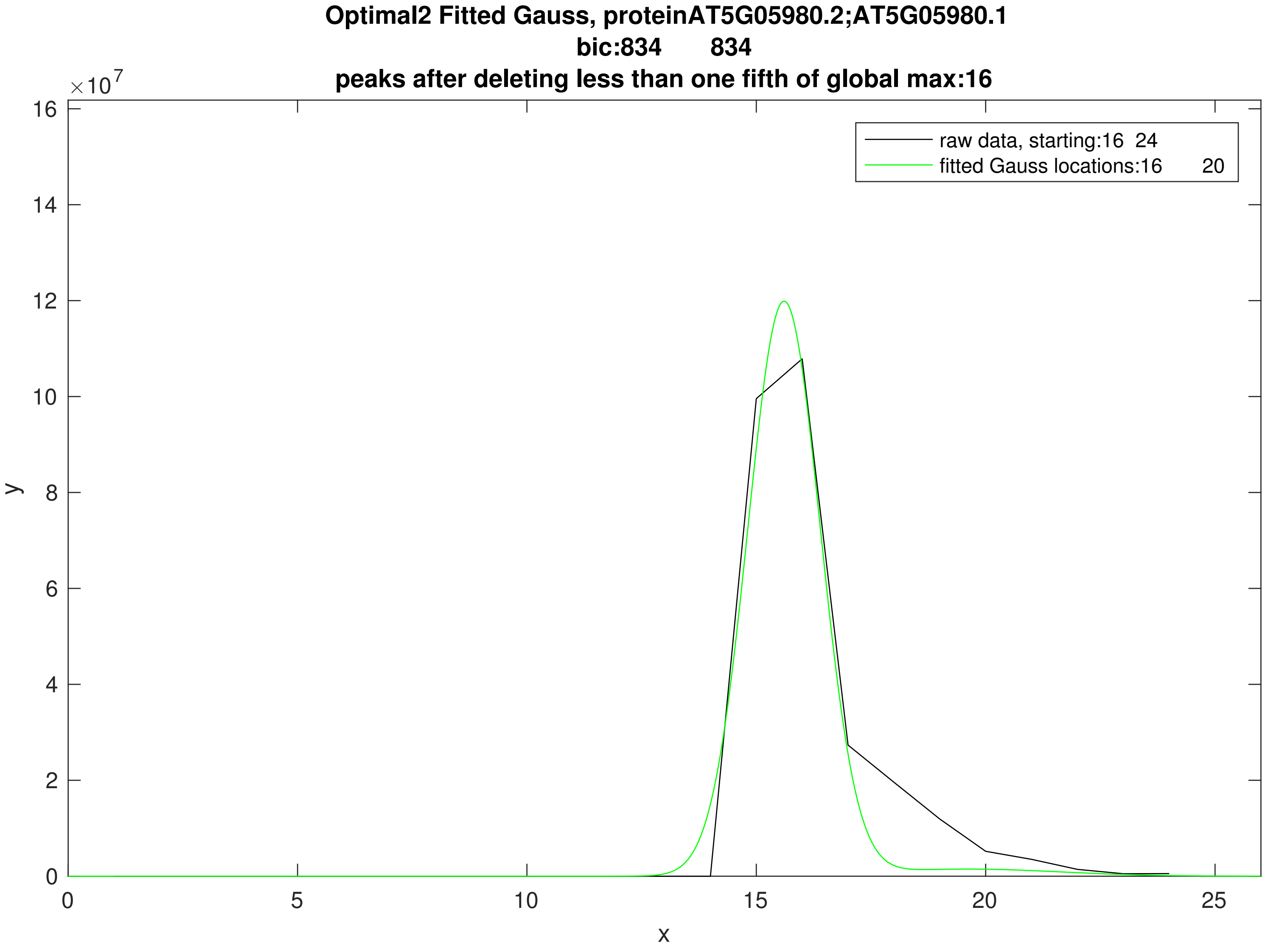

Optimal2 Fitted Gauss, proteinAT5G05980.2;AT5G05980.1
bic:834 834
peaks after deleting less than one fifth of global max:16
×10^7
16
14
12
10
8
6
4
2
0
y
raw data, starting:16 24
fitted Gauss locations:16 20
0
5
10
15
20
25
x

**Optimal1 Fitted Gauss, proteinAT5G06060.1**
**bic:858       944**
**peaks after deleting less than one fifth of global max:14**

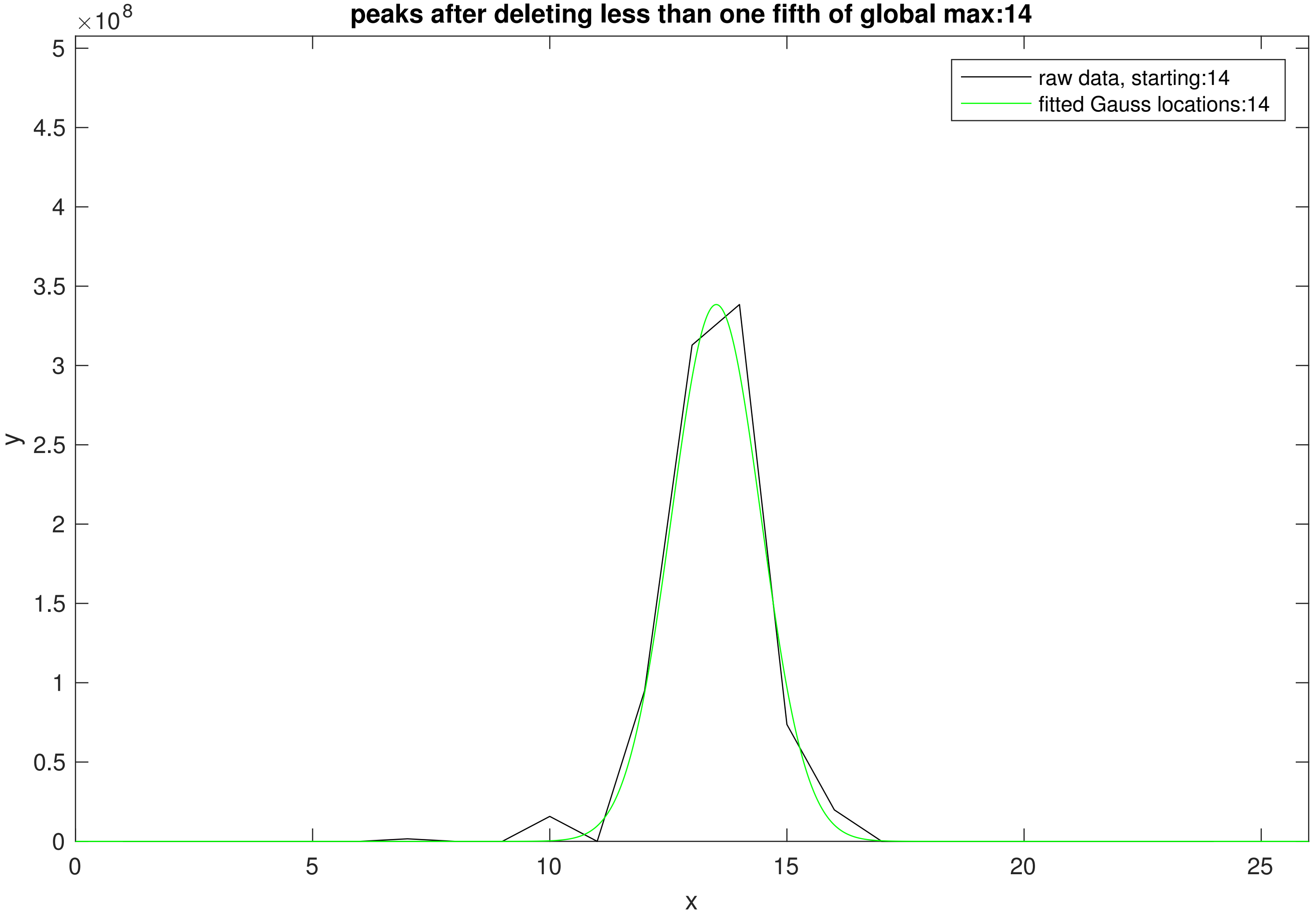

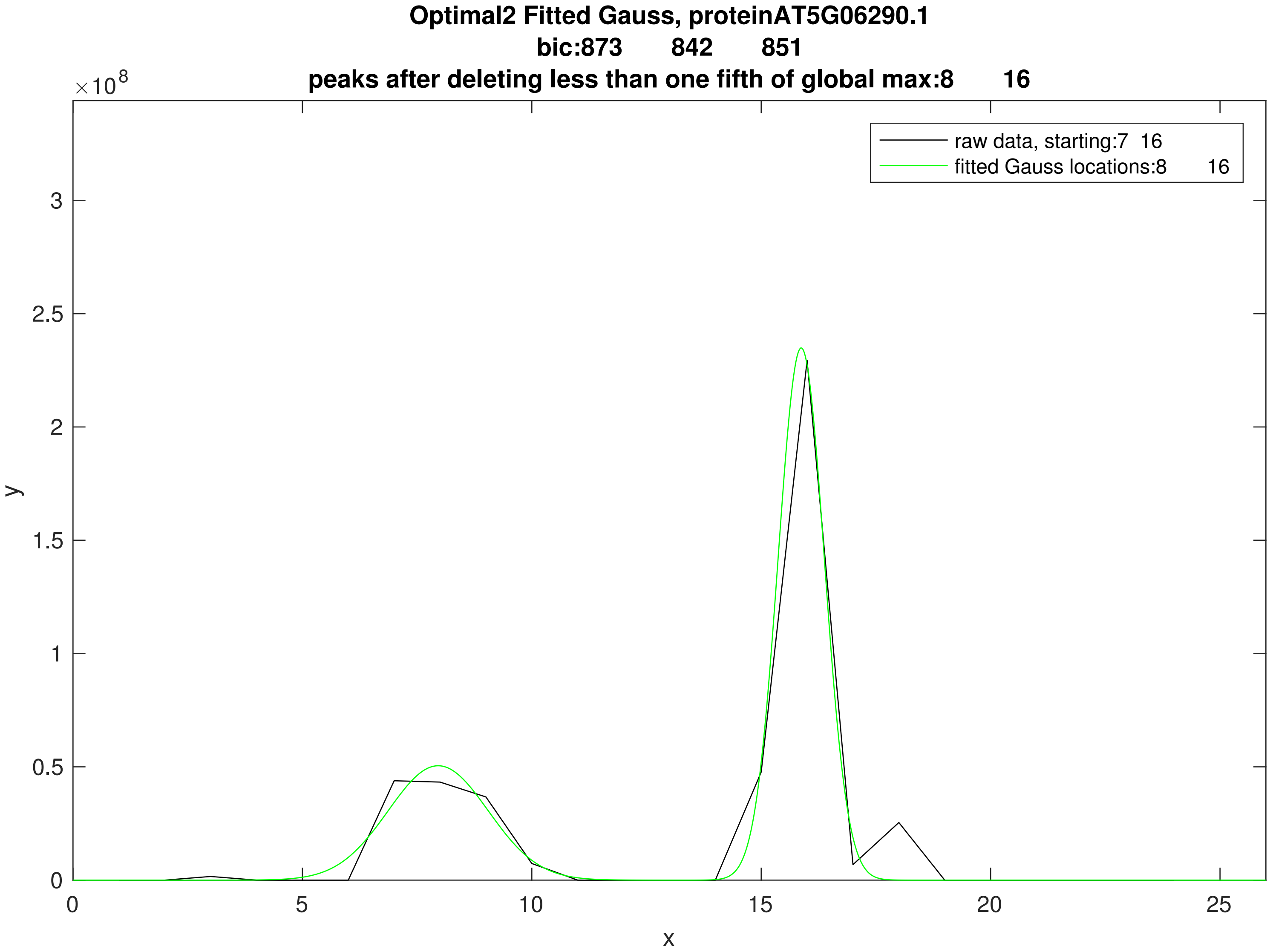

Optimal2 Fitted Gauss, proteinAT5G06290.1
bic:873 842 851
peaks after deleting less than one fifth of global max:8 16
×10^8
raw data, starting:7 16
fitted Gauss locations:8 16
3
2.5
2
1.5
1
0.5
0
y
0
5
10
15
20
25
x

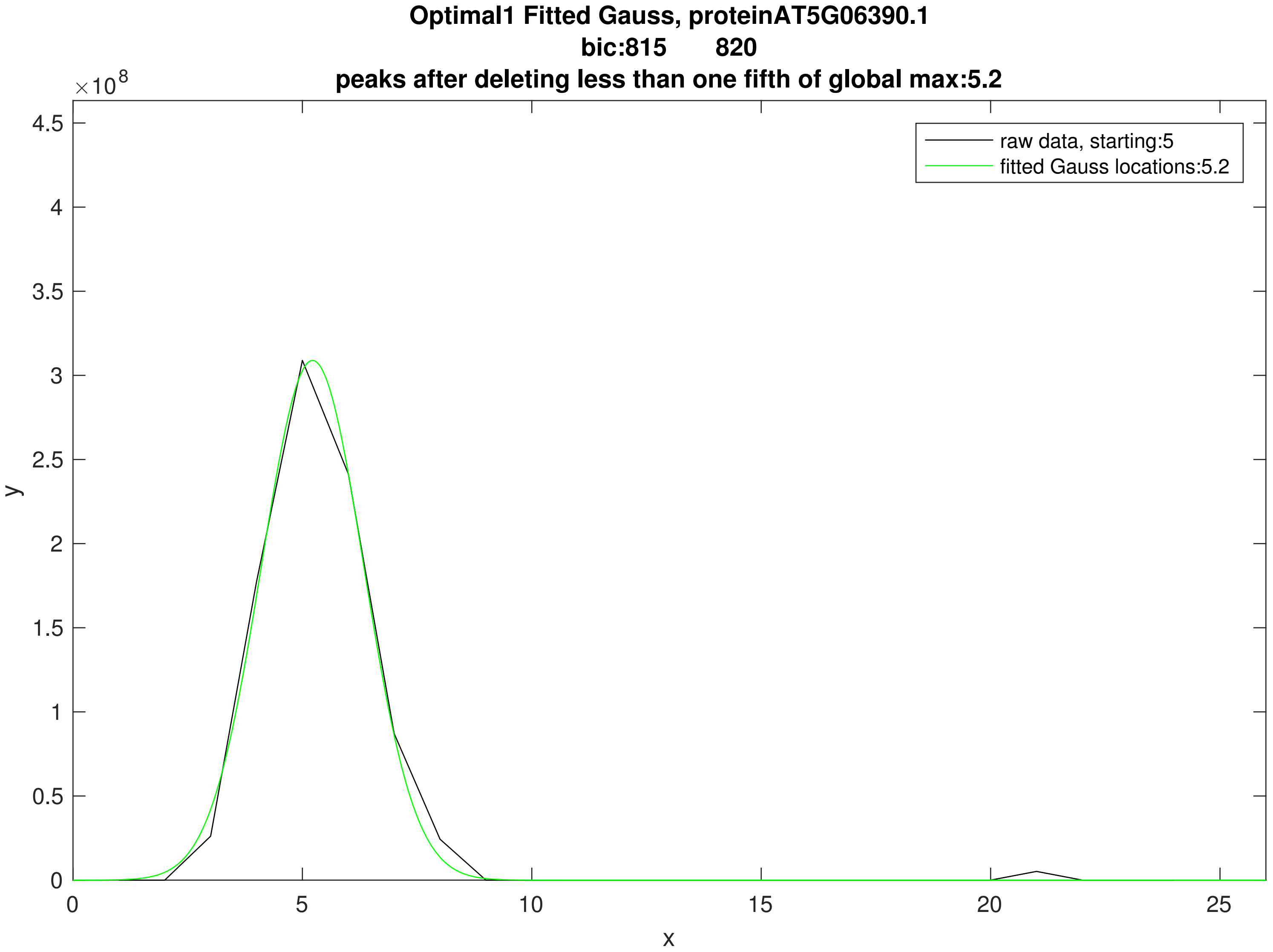

Optimal1 Fitted Gauss, proteinAT5G06390.1
bic:815 820
peaks after deleting less than one fifth of global max:5.2
×10^8
4.5
4
3.5
3
2.5
2
1.5
1
0.5
0
y
0
5
10
15
20
25
x
raw data, starting:5
fitted Gauss locations:5.2

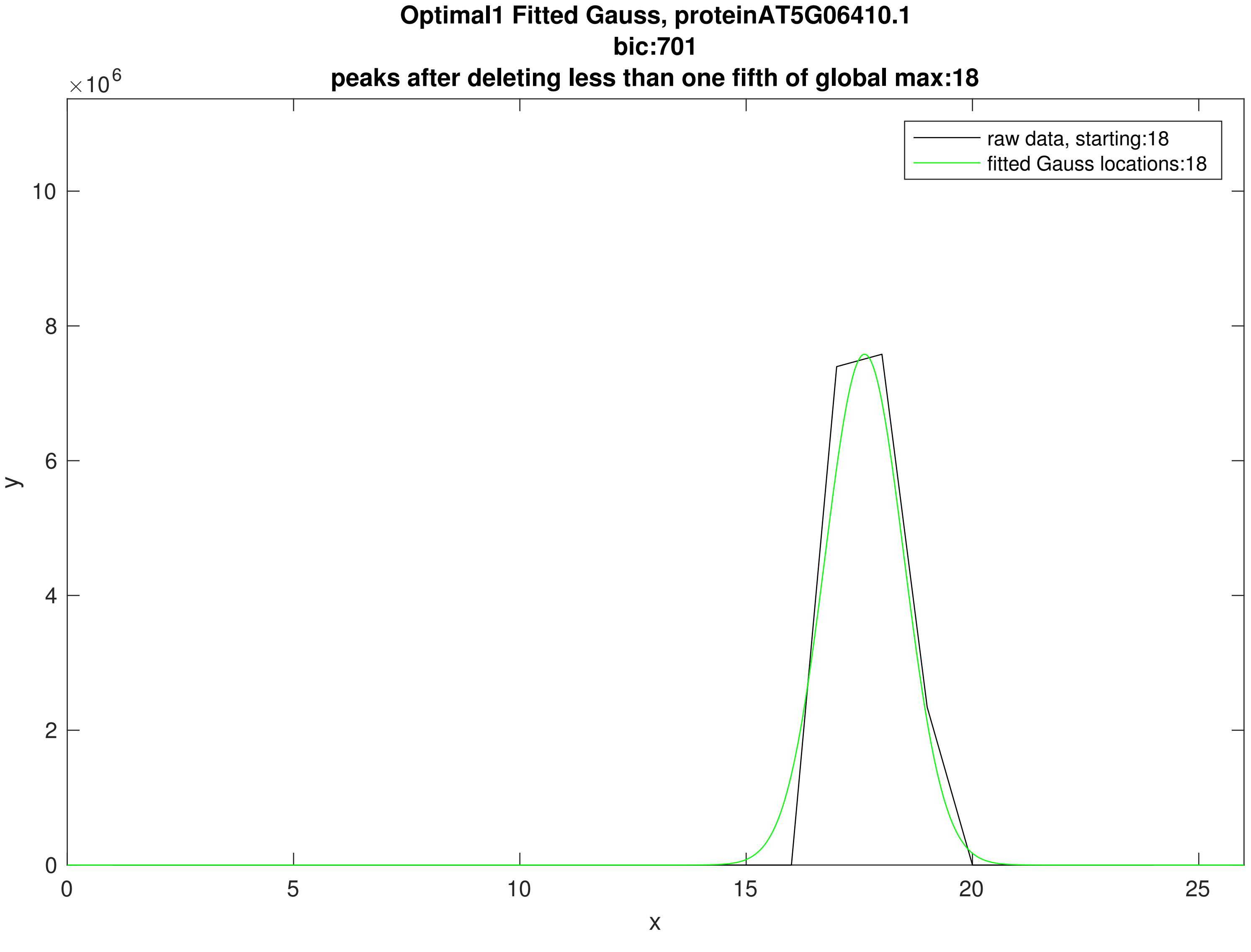
Optimal1 Fitted Gauss, proteinAT5G06410.1
bic:701
peaks after deleting less than one fifth of global max:18
×10^6
raw data, starting:18
fitted Gauss locations:18
10
8
6
4
2
0
y
0
5
10
15
20
25
x

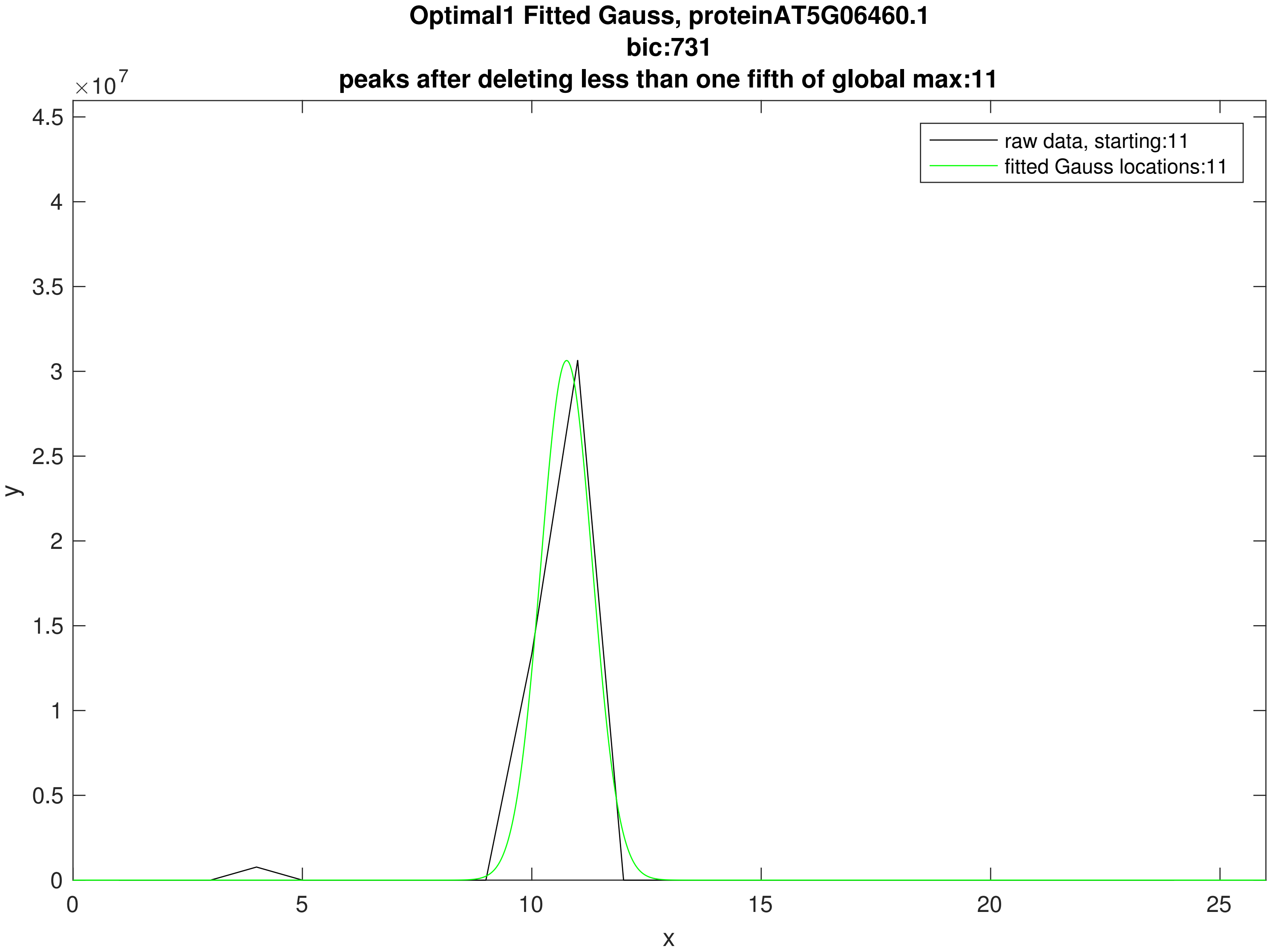
Optimal1 Fitted Gauss, proteinAT5G06460.1
bic:731
peaks after deleting less than one fifth of global max:11
×10^7
4.5
4
3.5
3
2.5
2
1.5
1
0.5
0
y
0
5
10
15
20
25
x
raw data, starting:11
fitted Gauss locations:11

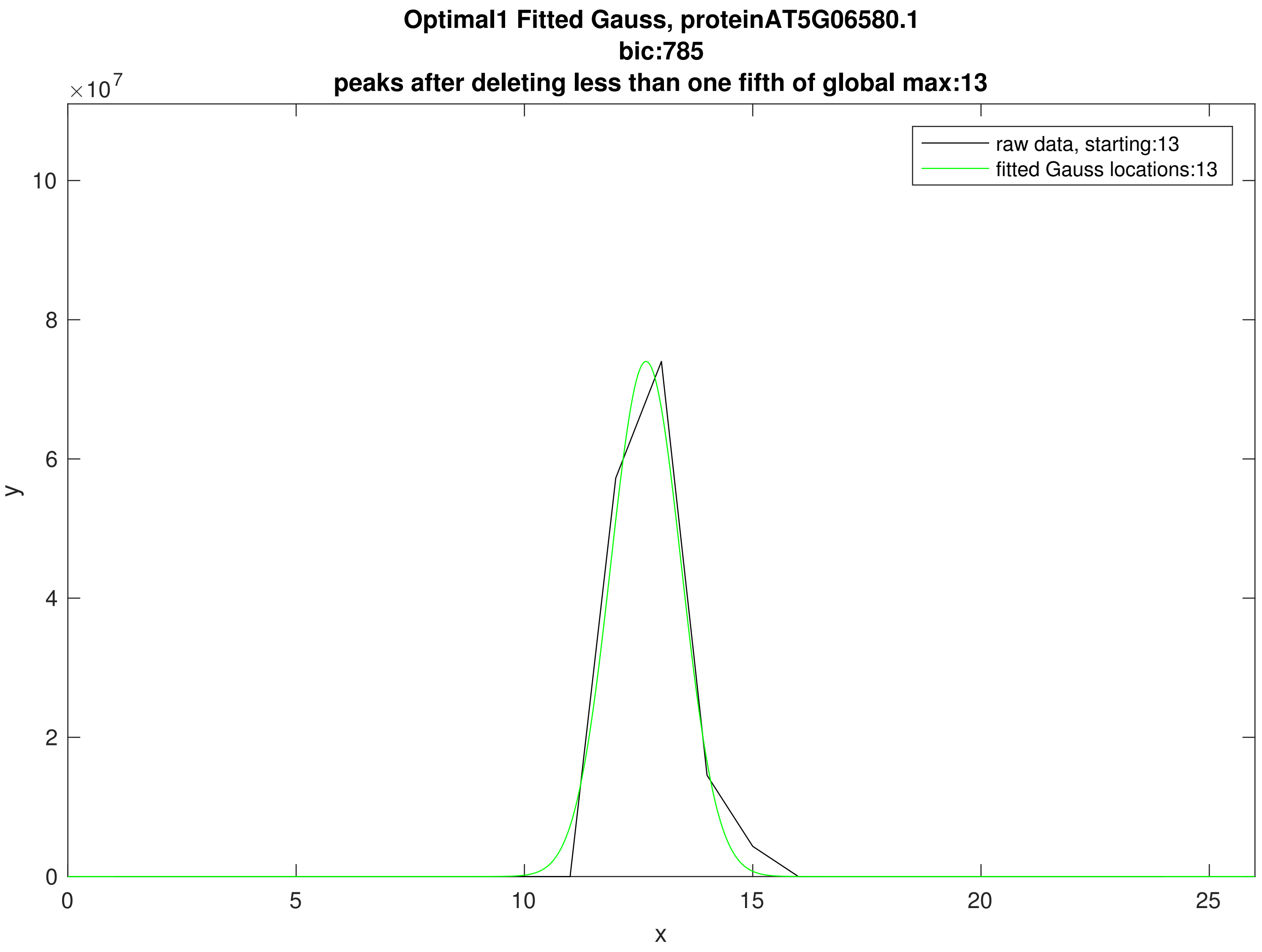
Optimal1 Fitted Gauss, proteinAT5G06580.1
bic:785
peaks after deleting less than one fifth of global max:13
×10^7
raw data, starting:13
fitted Gauss locations:13
10
8
6
4
2
0
y
0
5
10
15
20
25
x

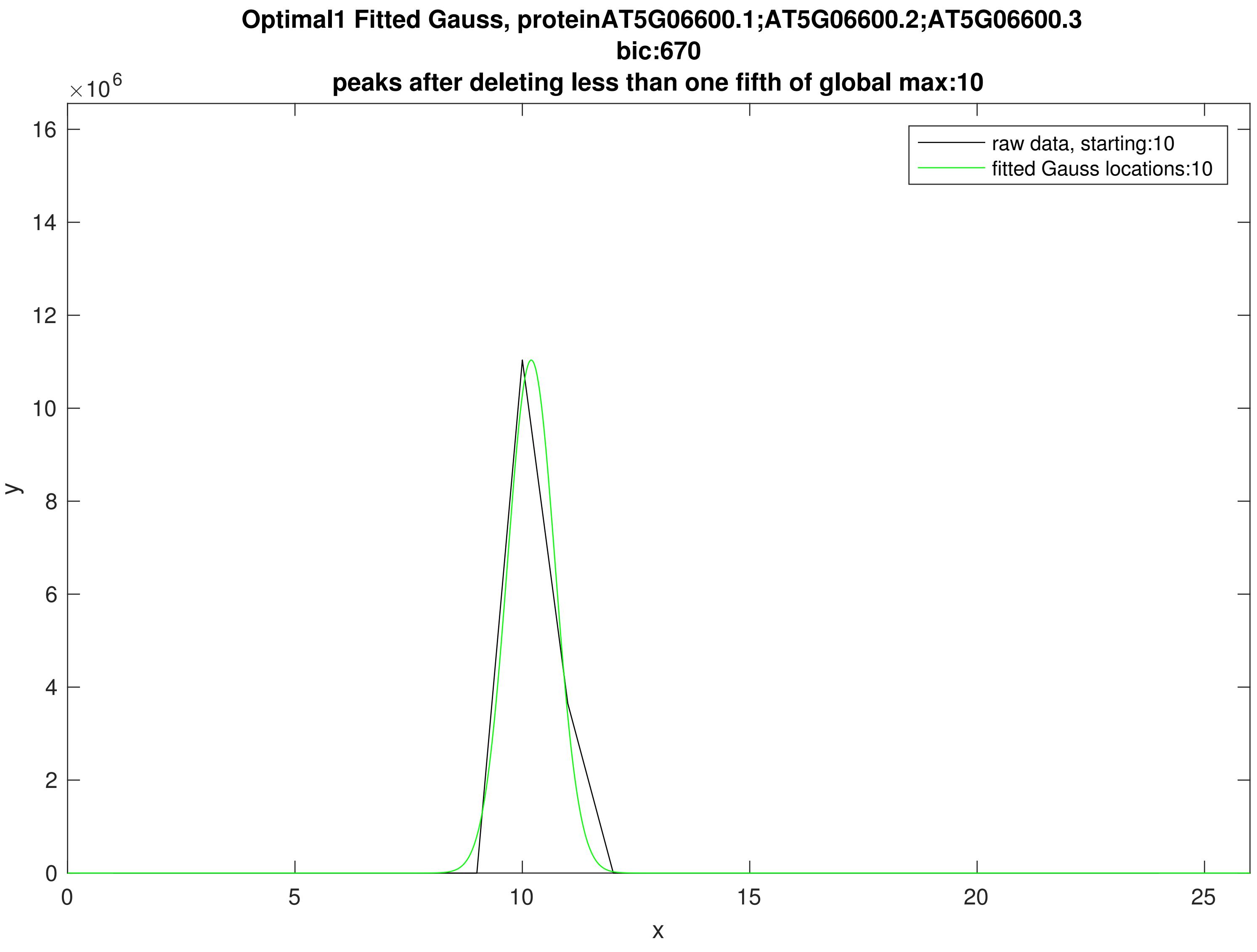
Optimal1 Fitted Gauss, proteinAT5G06600.1;AT5G06600.2;AT5G06600.3
bic:670
peaks after deleting less than one fifth of global max:10
×10^6
raw data, starting:10
fitted Gauss locations:10
y
x
0
2
4
6
8
10
12
14
16
0
5
10
15
20
25

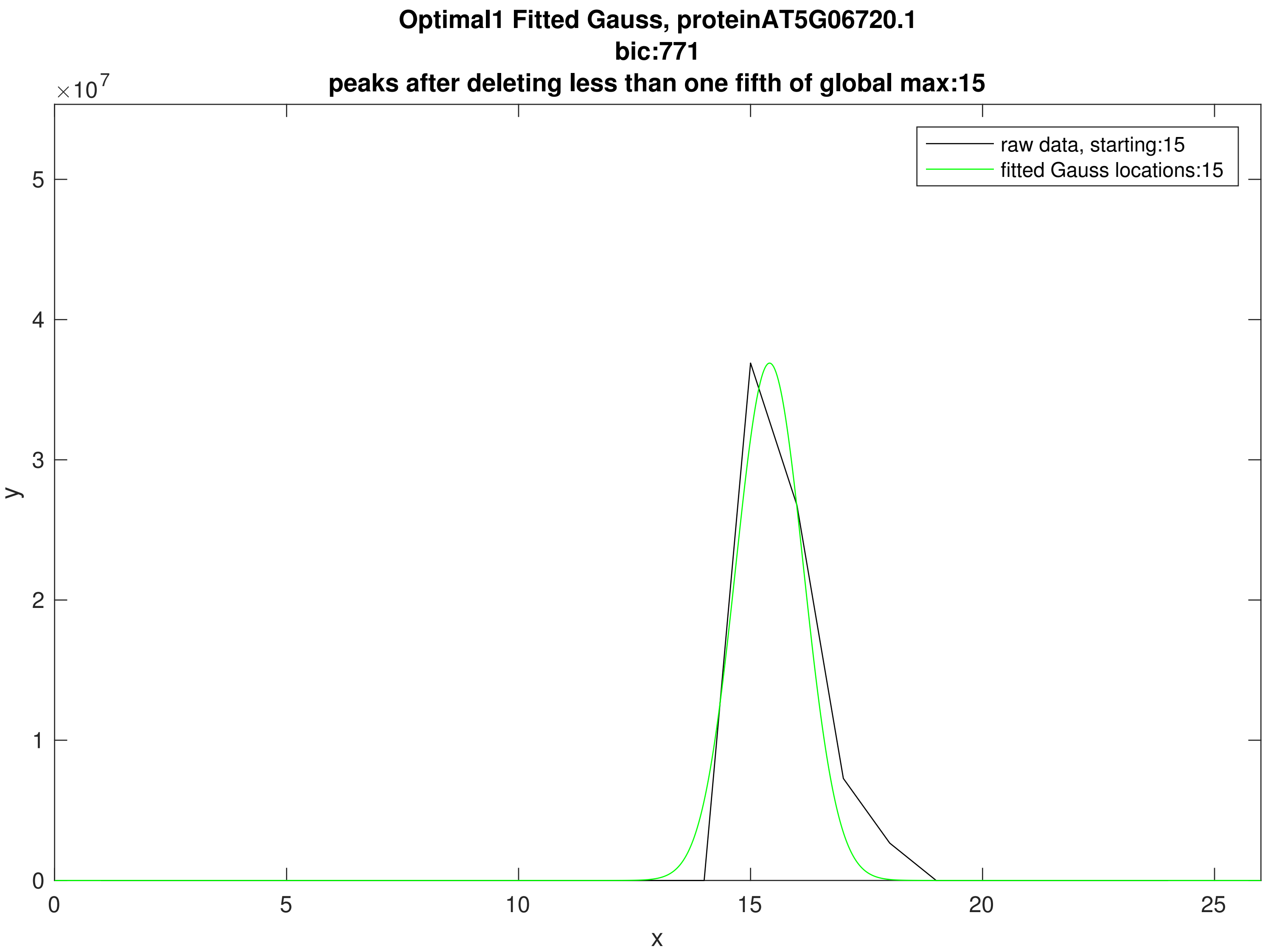

Optimal1 Fitted Gauss, proteinAT5G06720.1
bic:771
peaks after deleting less than one fifth of global max:15
×10^7
raw data, starting:15
fitted Gauss locations:15
y
x
0
1
2
3
4
5
0
5
10
15
20
25

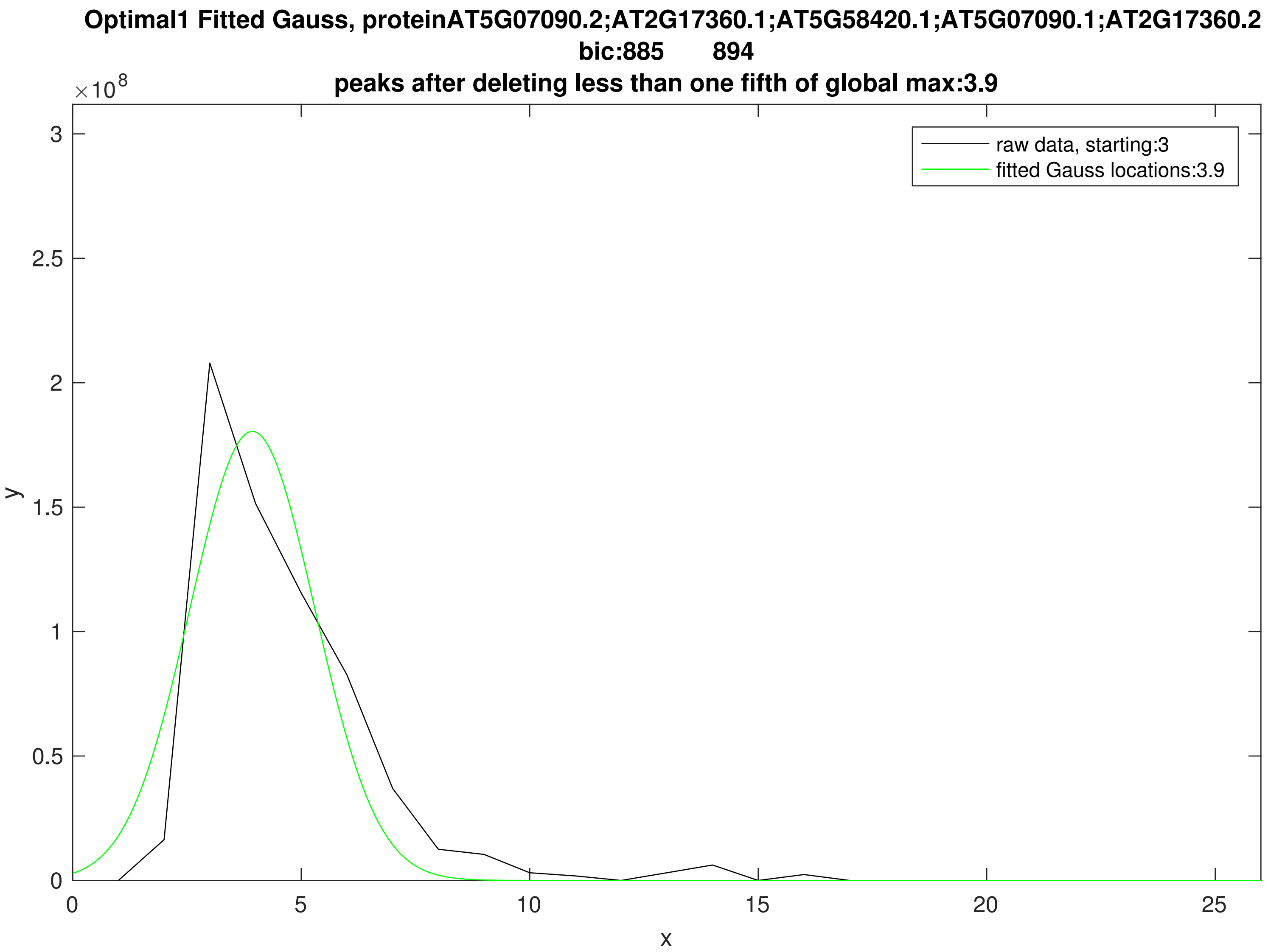

Optimal1 Fitted Gauss, proteinAT5G07090.2;AT2G17360.1;AT5G58420.1;AT5G07090.1;AT2G17360.2
bic:885 894
peaks after deleting less than one fifth of global max:3.9
×10^8
3
2.5
2
1.5
1
0.5
0
y
0
5
10
15
20
25
x
raw data, starting:3
fitted Gauss locations:3.9

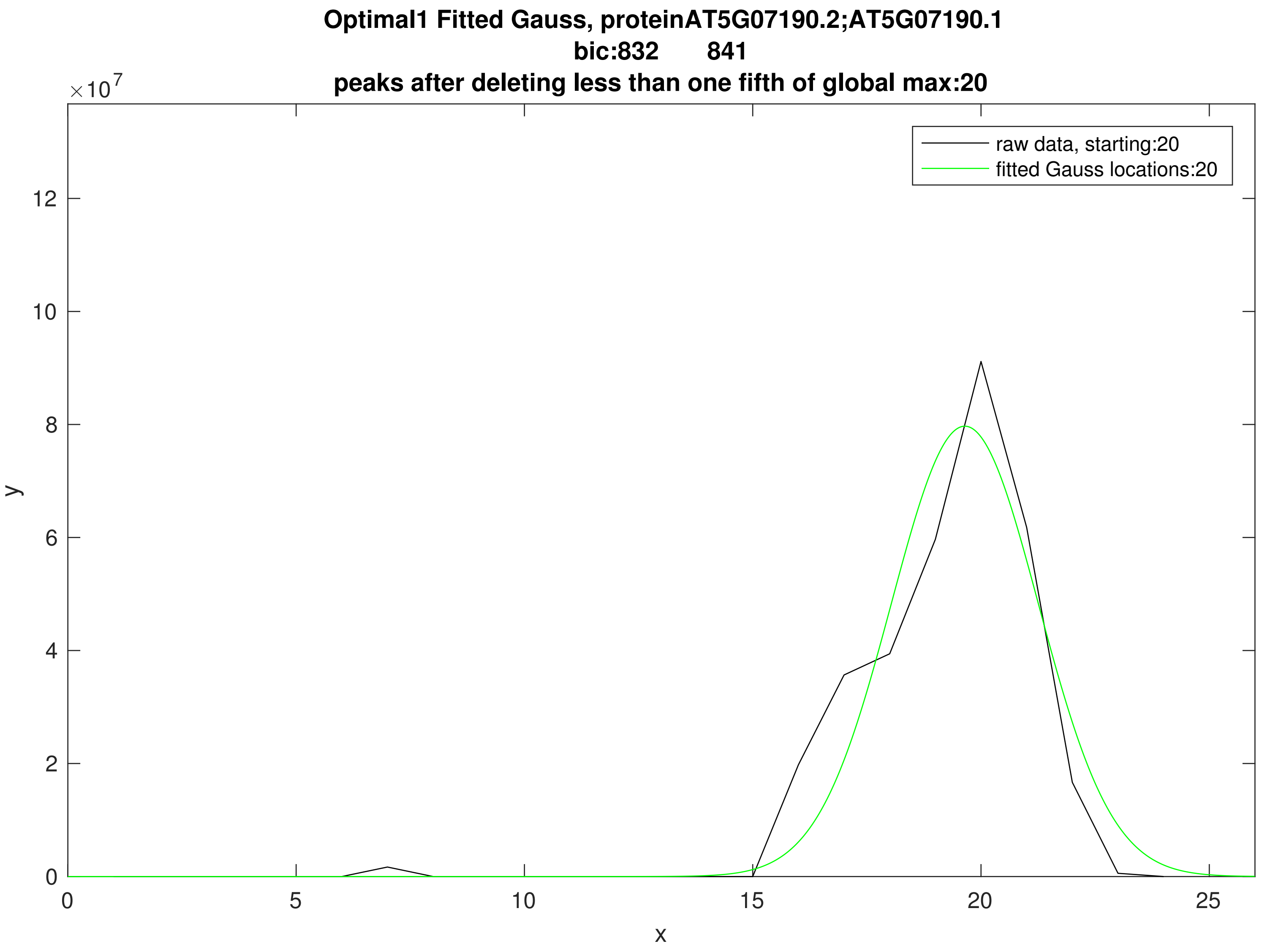

Optimal1 Fitted Gauss, proteinAT5G07190.2;AT5G07190.1
bic:832 841
peaks after deleting less than one fifth of global max:20
×10^7
raw data, starting:20
fitted Gauss locations:20
0
2
4
6
8
10
12
y
0
5
10
15
20
25
x

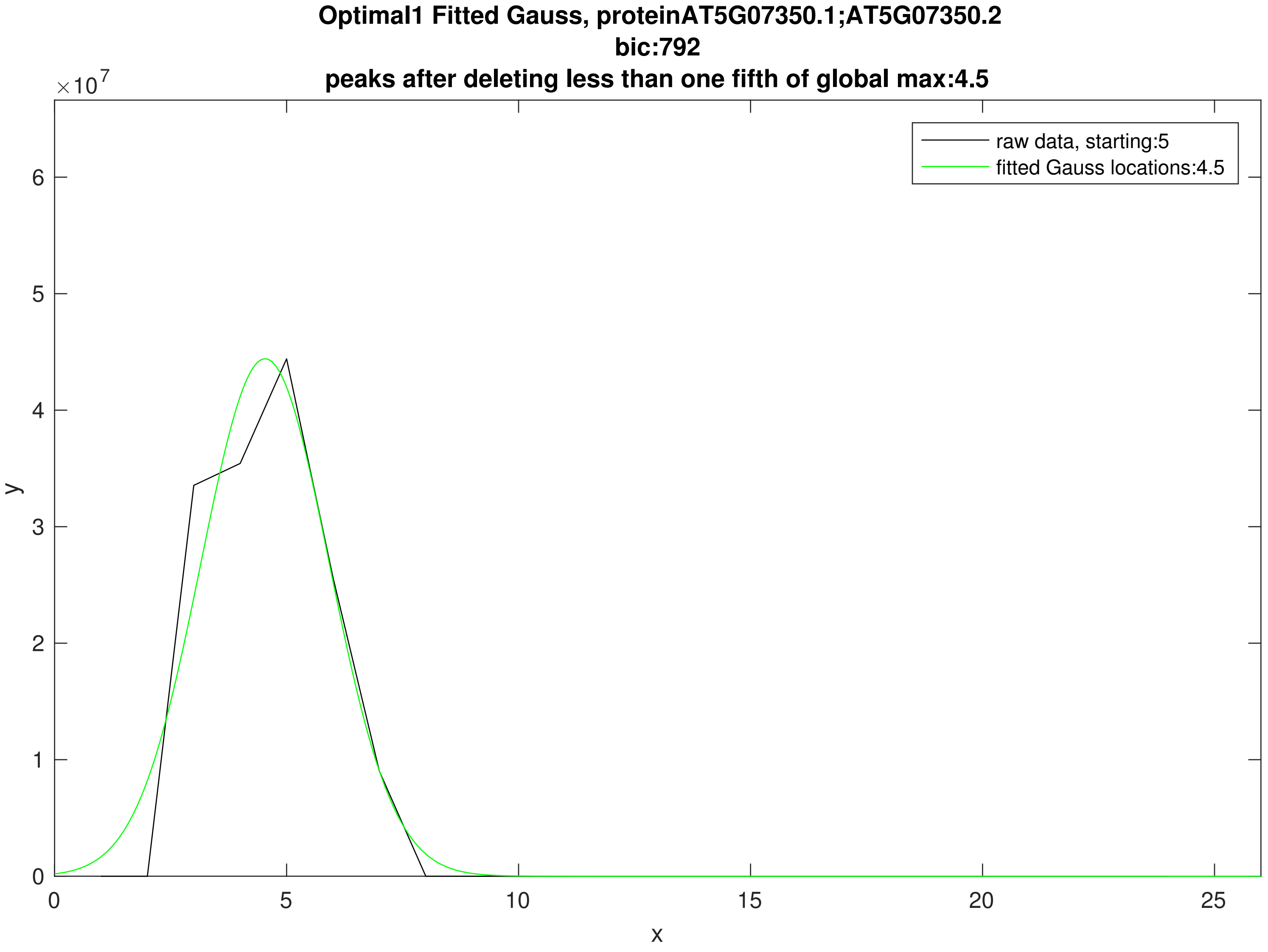
Optimal1 Fitted Gauss, proteinAT5G07350.1;AT5G07350.2
bic:792
peaks after deleting less than one fifth of global max:4.5
×10^7
raw data, starting:5
fitted Gauss locations:4.5
y
x
0
1
2
3
4
5
6
10
15
20
25

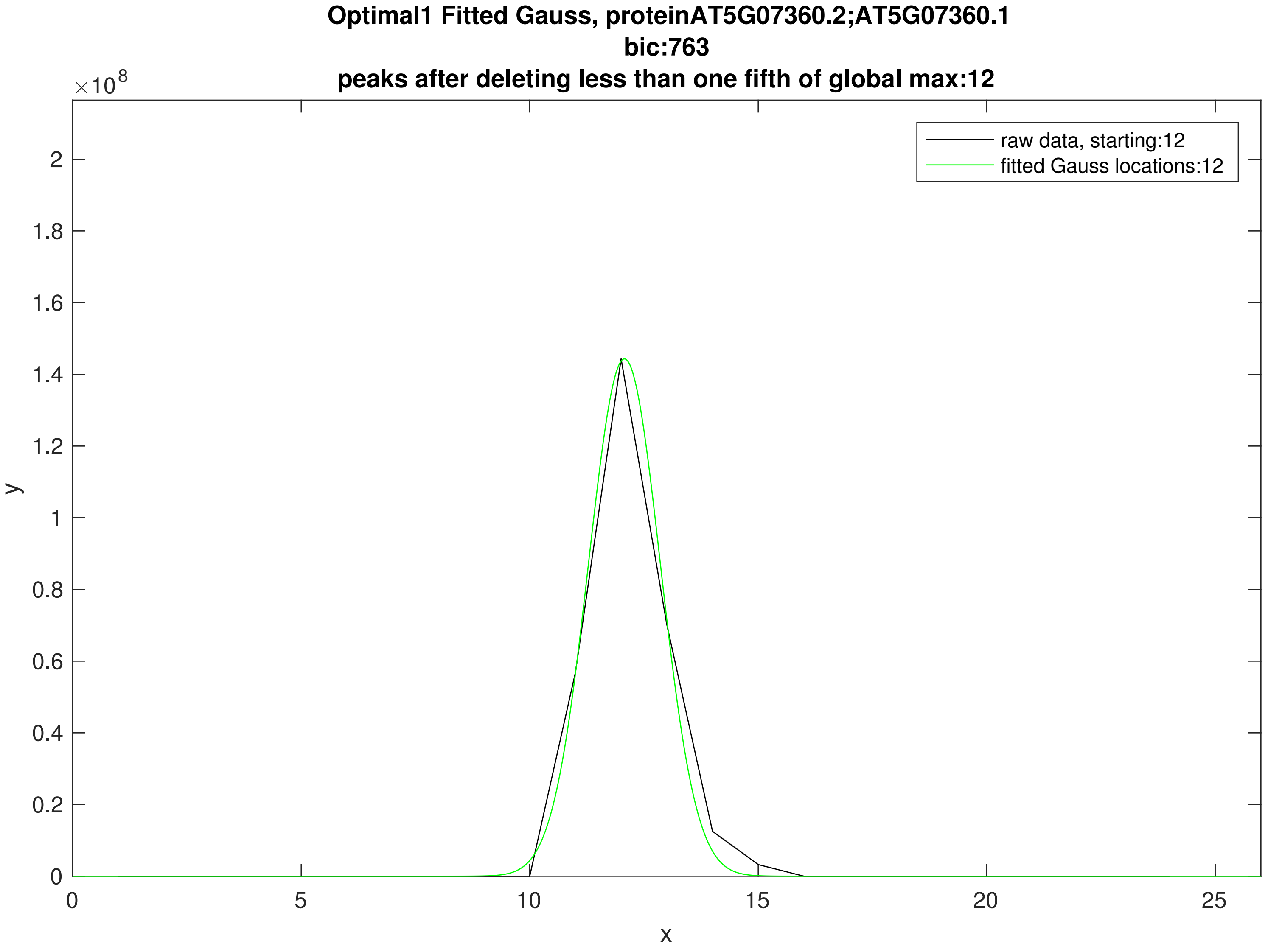
Optimal1 Fitted Gauss, proteinAT5G07360.2;AT5G07360.1
bic:763
peaks after deleting less than one fifth of global max:12
×10^8
raw data, starting:12
fitted Gauss locations:12
y
x
0
0.2
0.4
0.6
0.8
1
1.2
1.4
1.6
1.8
2
5
10
15
20
25

**Optimal1 Fitted Gauss, proteinAT5G07440.1;AT5G07440.2;AT5G07440.3**
**bic:978 988 998 1.01e+03**
**peaks after deleting less than one fifth of global max:9.1**

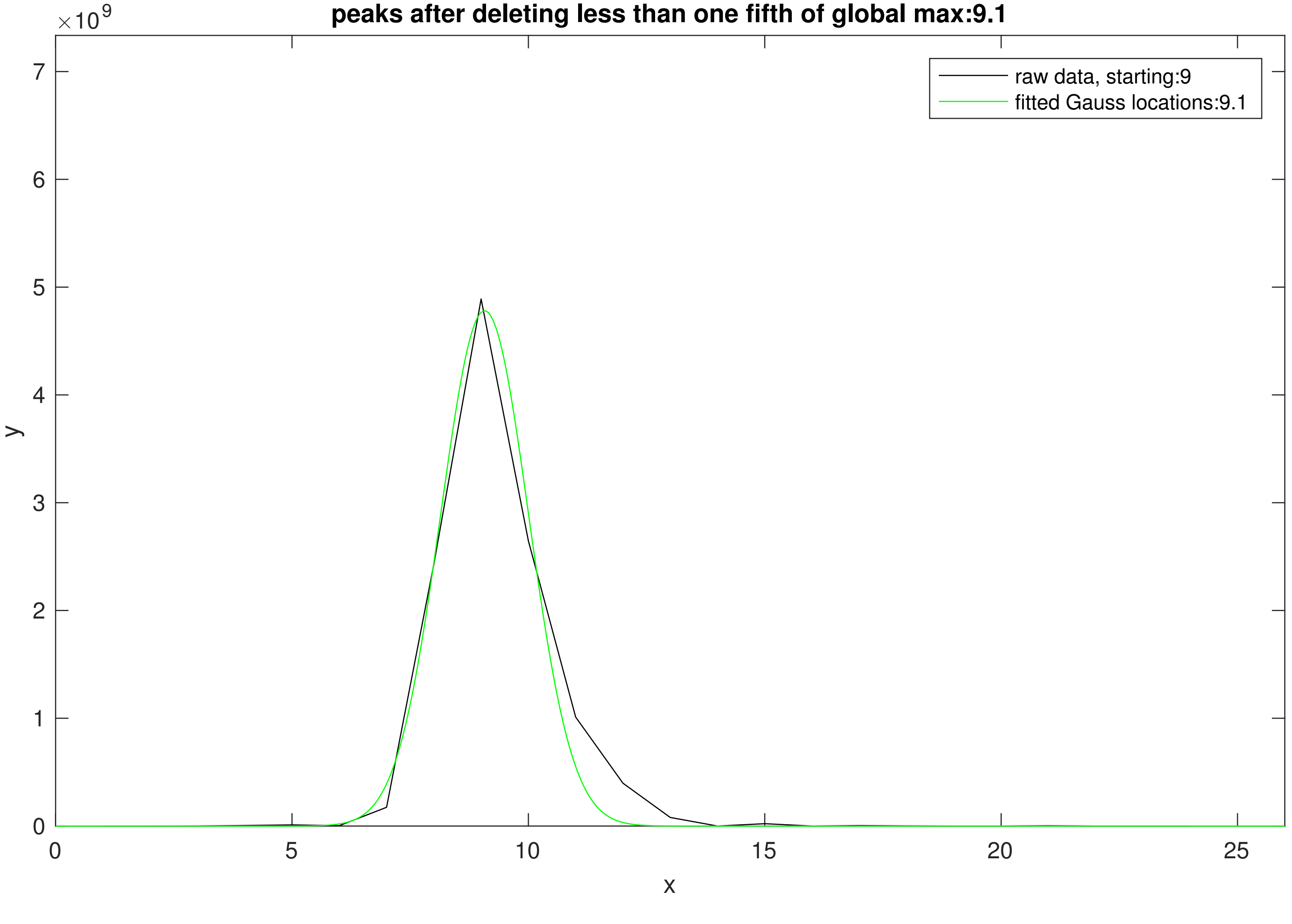

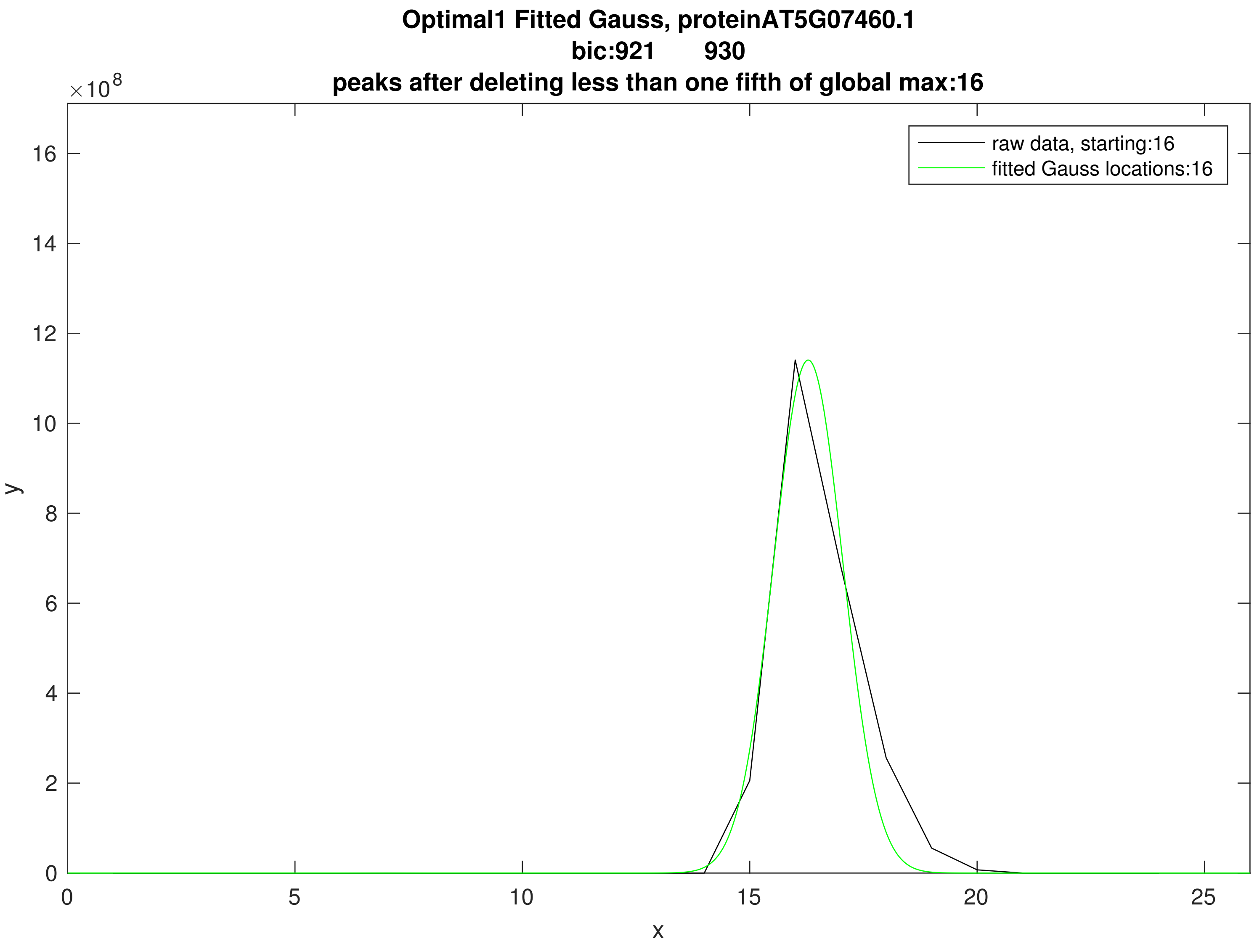
Optimal1 Fitted Gauss, proteinAT5G07460.1
bic:921 930
peaks after deleting less than one fifth of global max:16
×10^8
raw data, starting:16
fitted Gauss locations:16
y
x
0
2
4
6
8
10
12
14
16
0
5
10
15
20
25

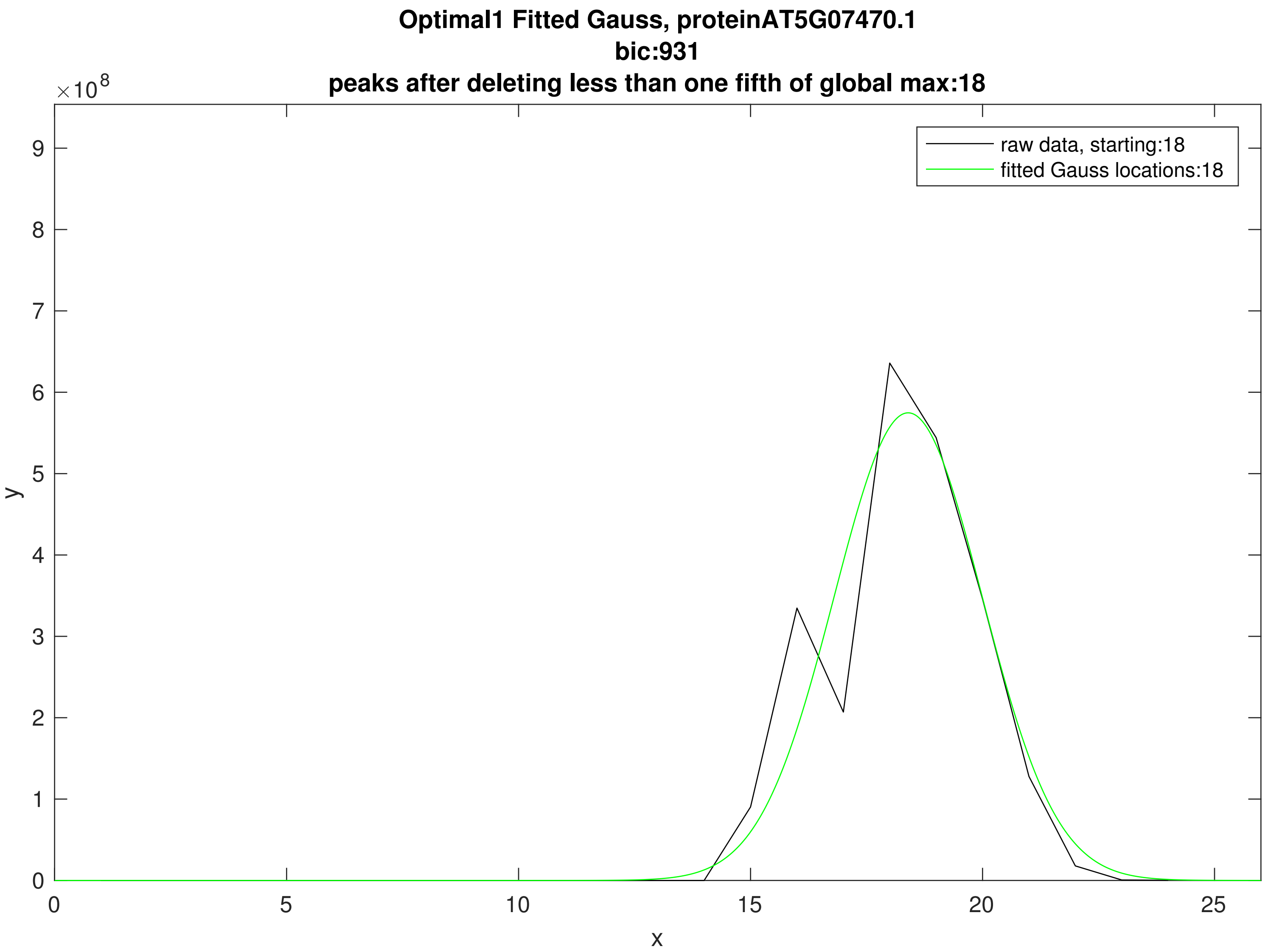
Optimal1 Fitted Gauss, proteinAT5G07470.1
bic:931
peaks after deleting less than one fifth of global max:18
×10^8
raw data, starting:18
fitted Gauss locations:18
y
x
0
1
2
3
4
5
6
7
8
9
0
5
10
15
20
25

**Optimal1 Fitted Gauss, proteinAT5G07830.1**
**bic:677**
**peaks after deleting less than one fifth of global max:19**

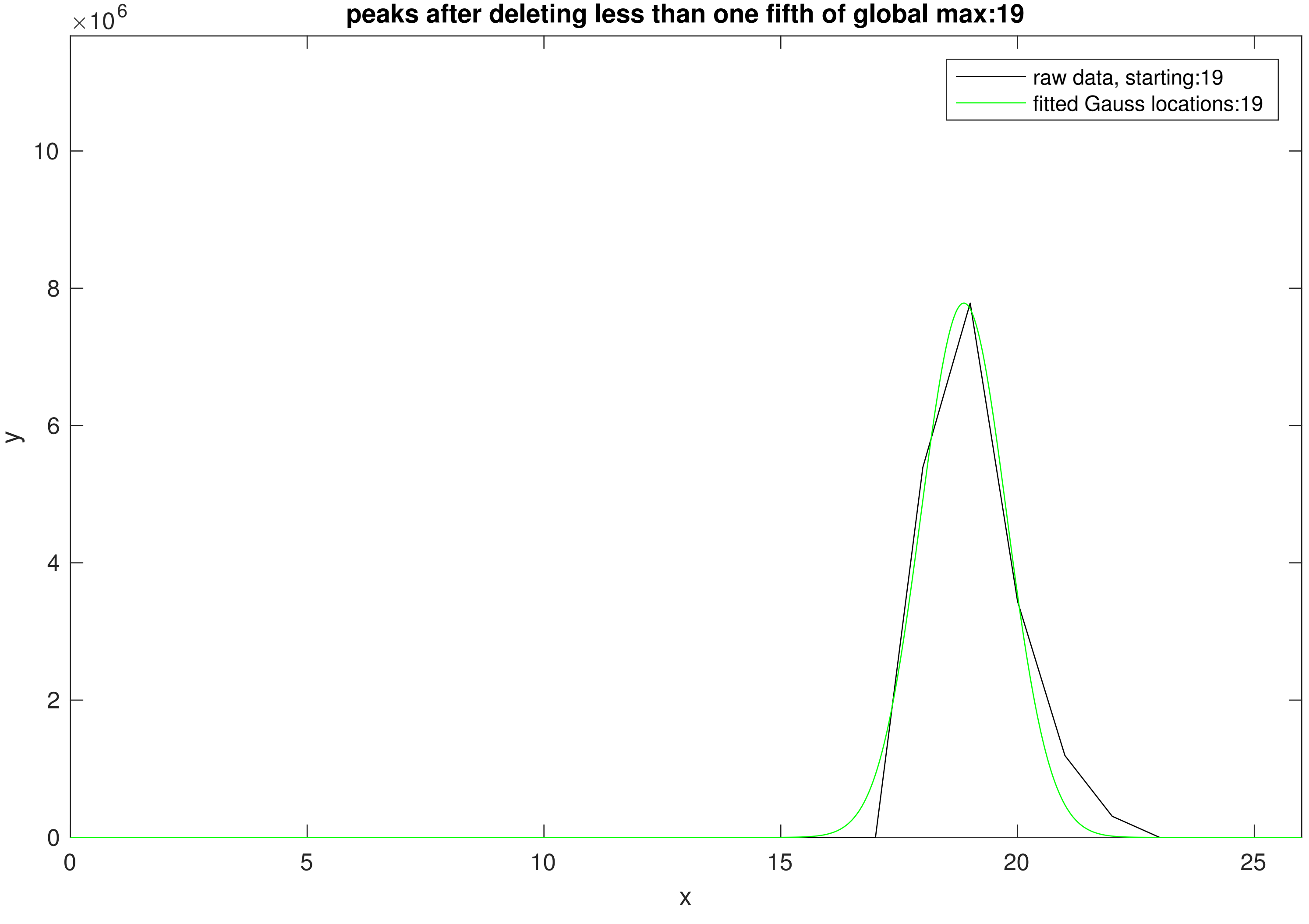

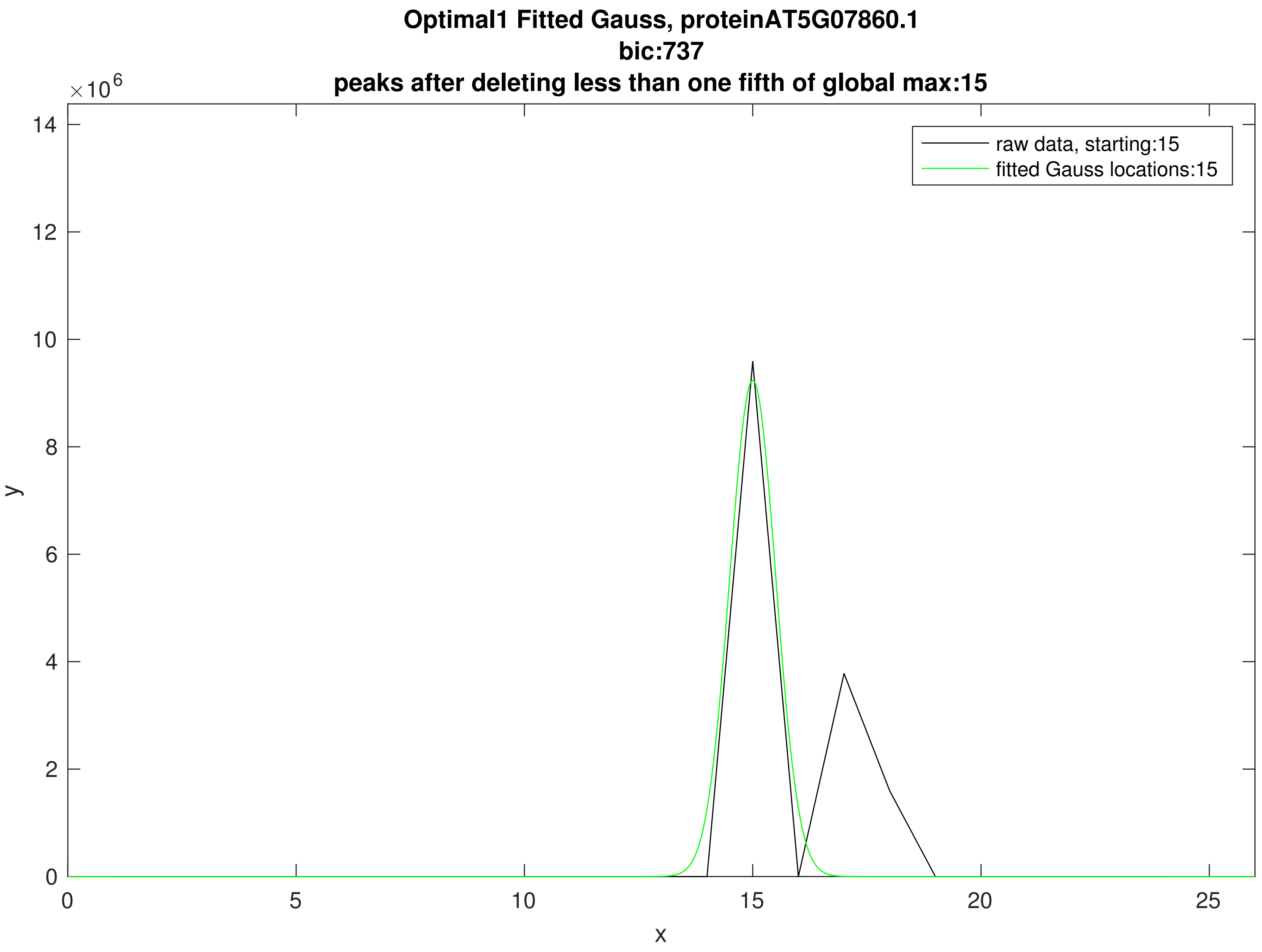
Optimal1 Fitted Gauss, proteinAT5G07860.1
bic:737
peaks after deleting less than one fifth of global max:15
×10^6
raw data, starting:15
fitted Gauss locations:15
y
x
0
2
4
6
8
10
12
14
0
5
10
15
20
25

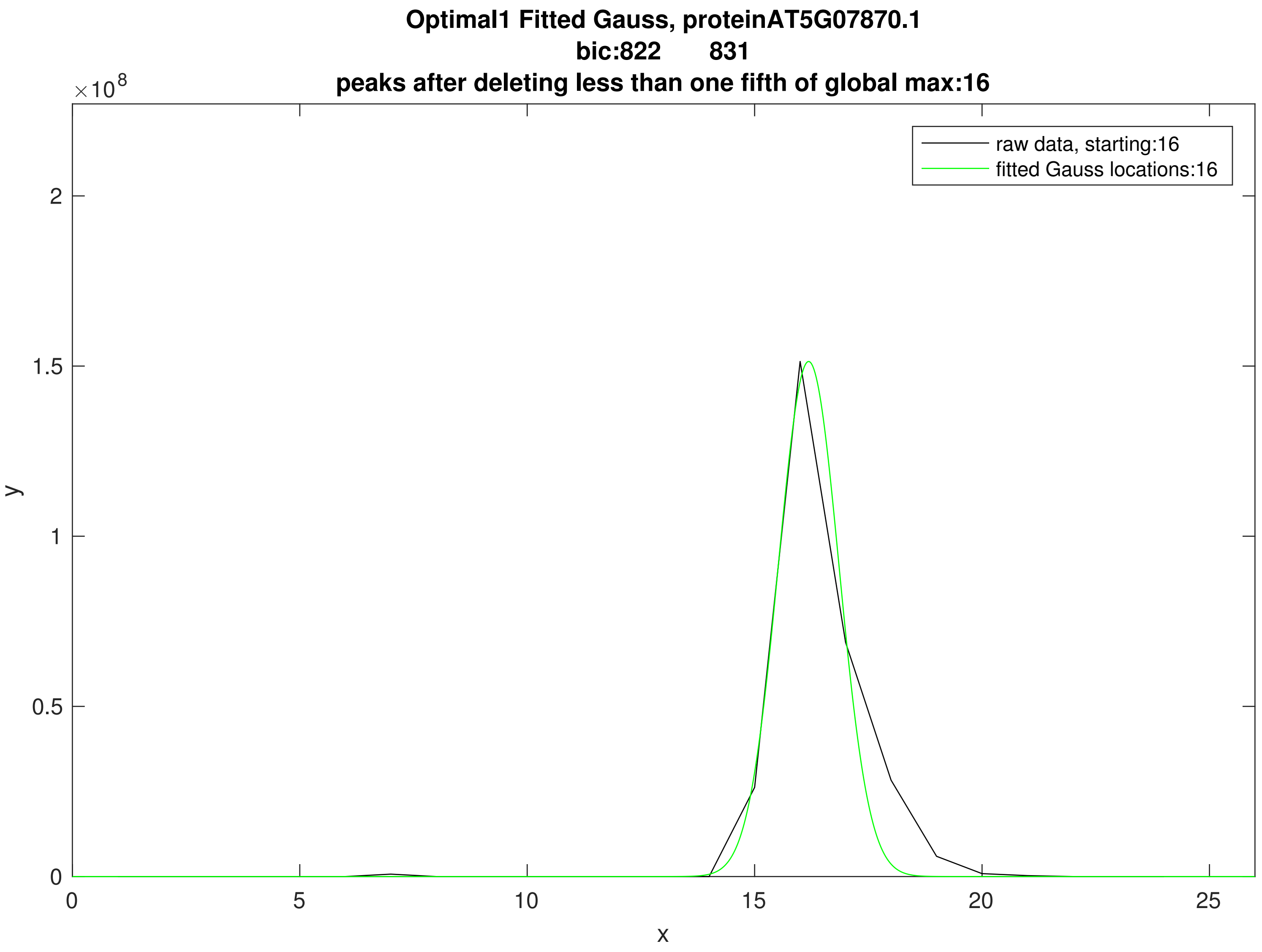
Optimal1 Fitted Gauss, proteinAT5G07870.1
bic:822 831
peaks after deleting less than one fifth of global max:16
×10^8
raw data, starting:16
fitted Gauss locations:16
y
x
0
0.5
1
1.5
2
0
5
10
15
20
25

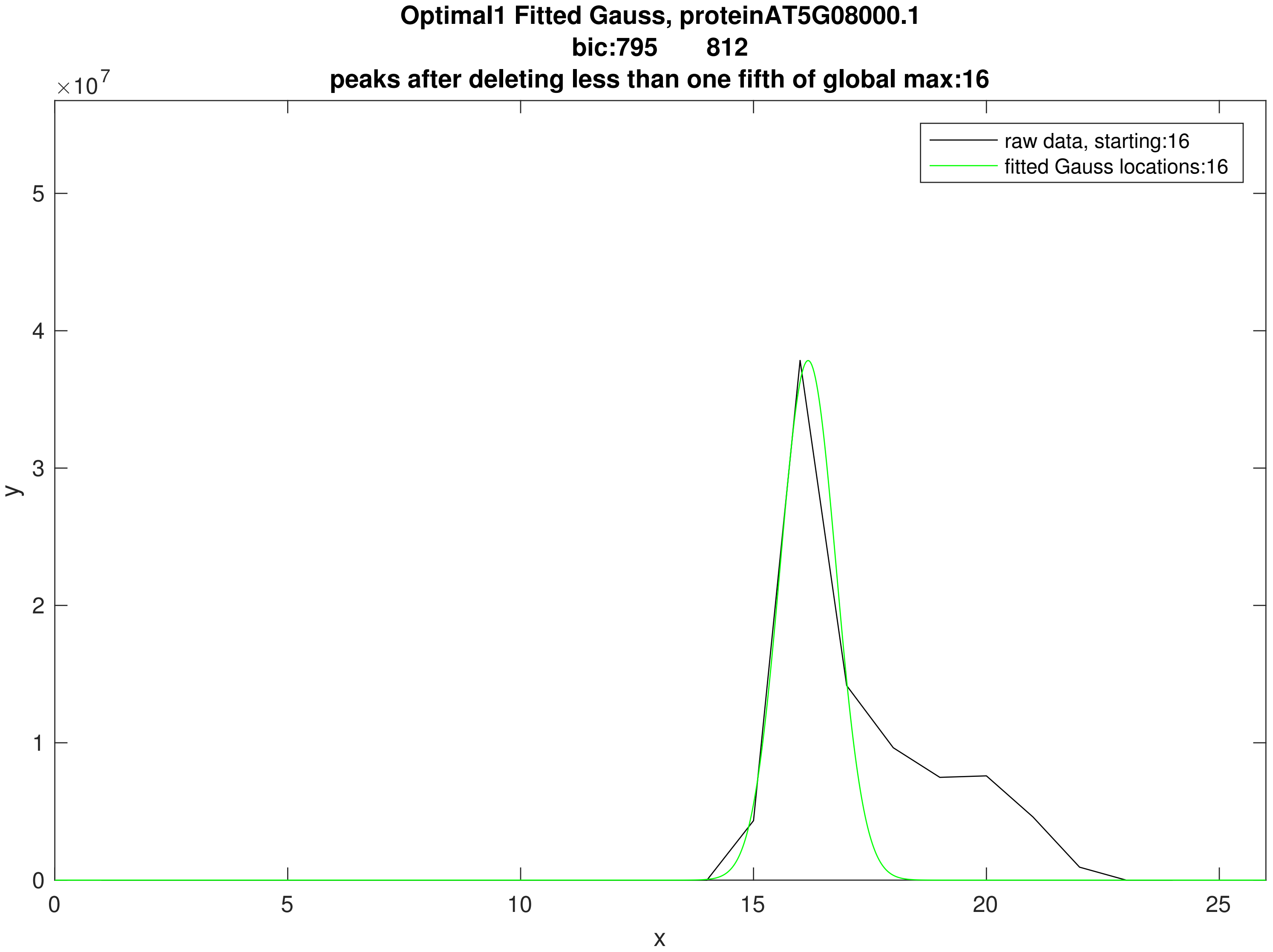
Optimal1 Fitted Gauss, proteinAT5G08000.1
bic:795 812
peaks after deleting less than one fifth of global max:16
×10^7
5
4
3
2
1
0
y
0
5
10
15
20
25
x
raw data, starting:16
fitted Gauss locations:16

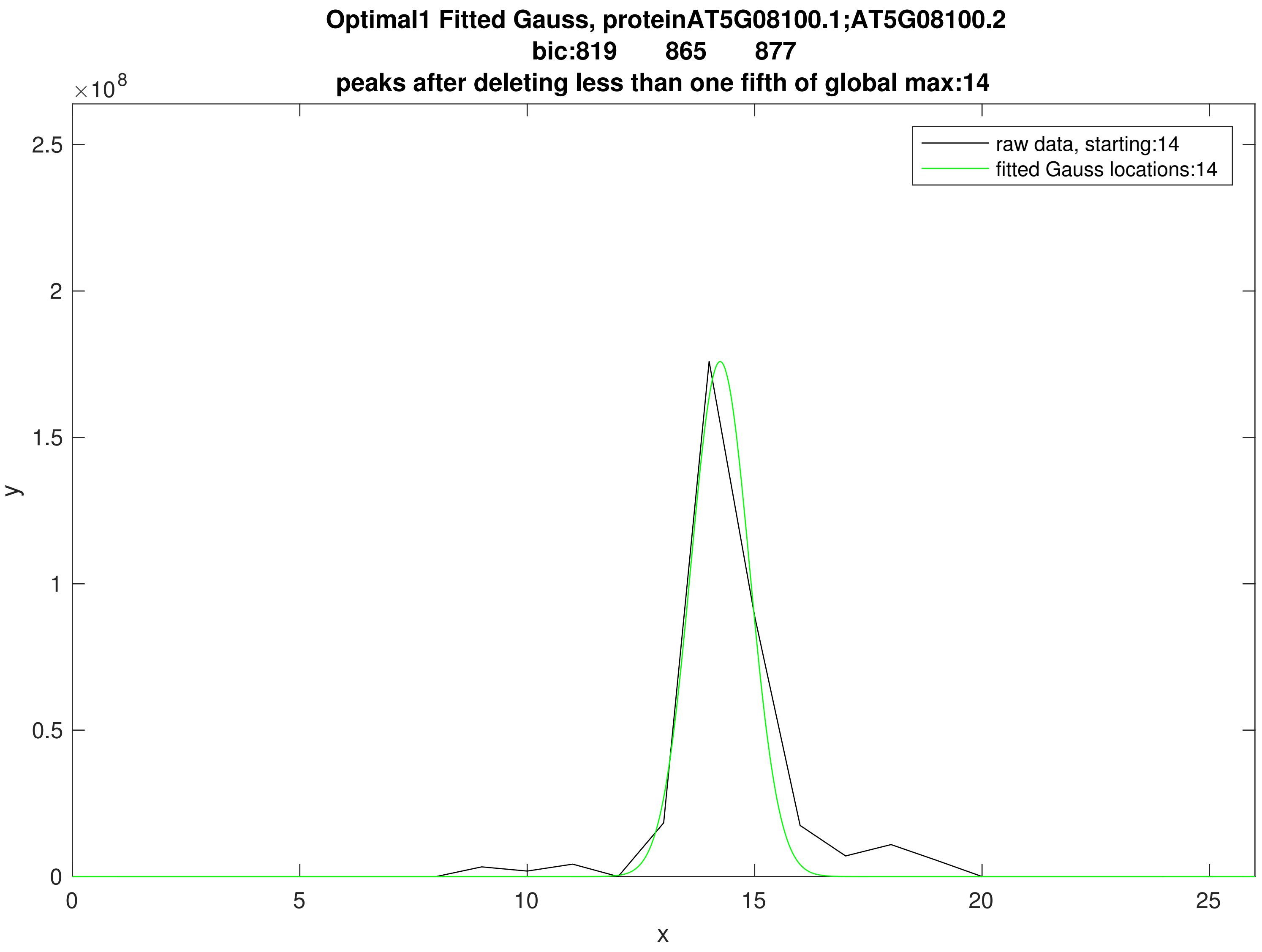

Optimal1 Fitted Gauss, proteinAT5G08100.1;AT5G08100.2
bic:819 865 877
peaks after deleting less than one fifth of global max:14
×10^8
2.5
2
1.5
1
0.5
0
y
0
5
10
15
20
25
x
raw data, starting:14
fitted Gauss locations:14

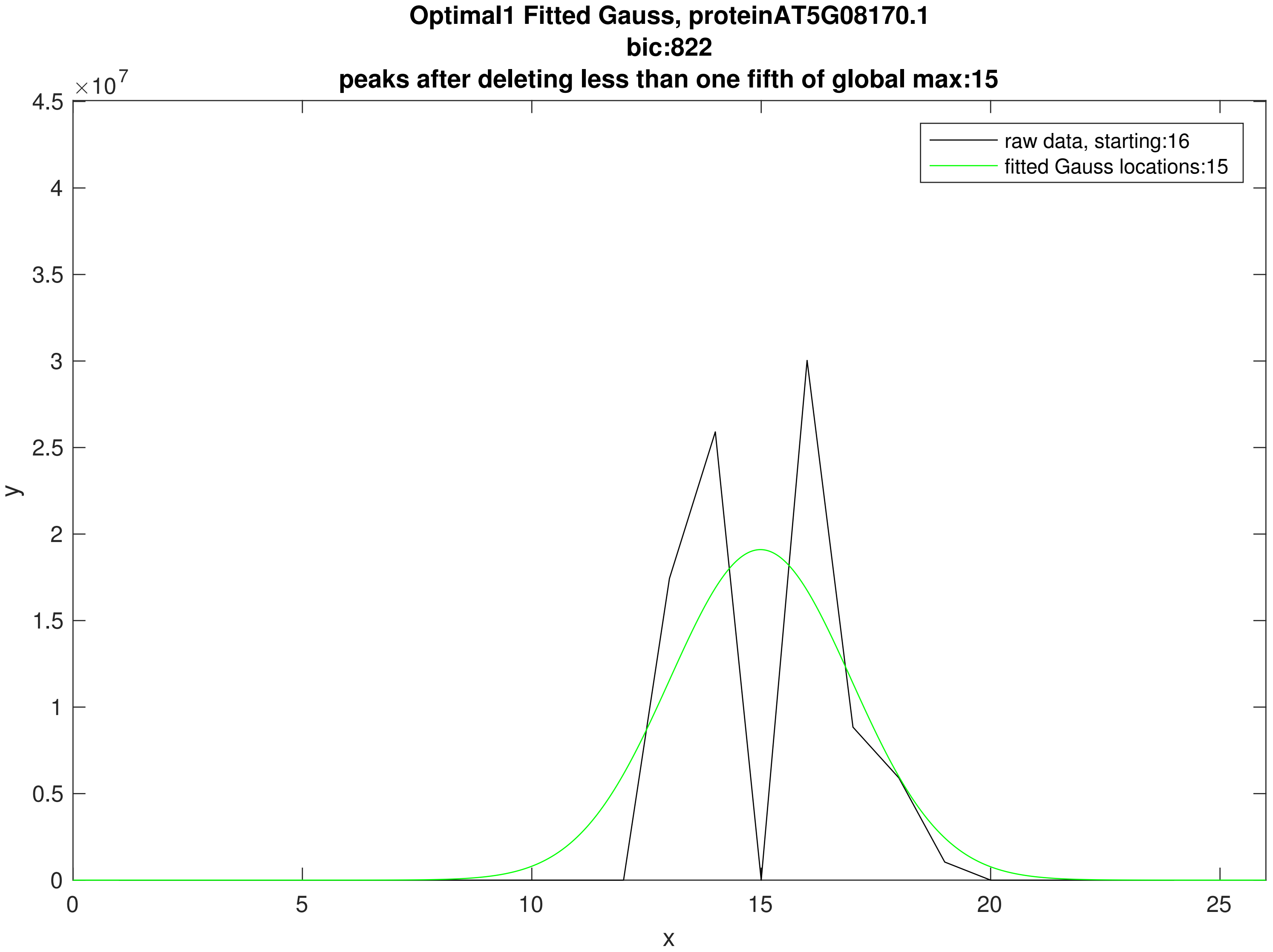
Optimal1 Fitted Gauss, proteinAT5G08170.1
bic:822
peaks after deleting less than one fifth of global max:15
×10^7
4.5
4
3.5
3
2.5
2
1.5
1
0.5
0
y
0
5
10
15
20
25
x
raw data, starting:16
fitted Gauss locations:15

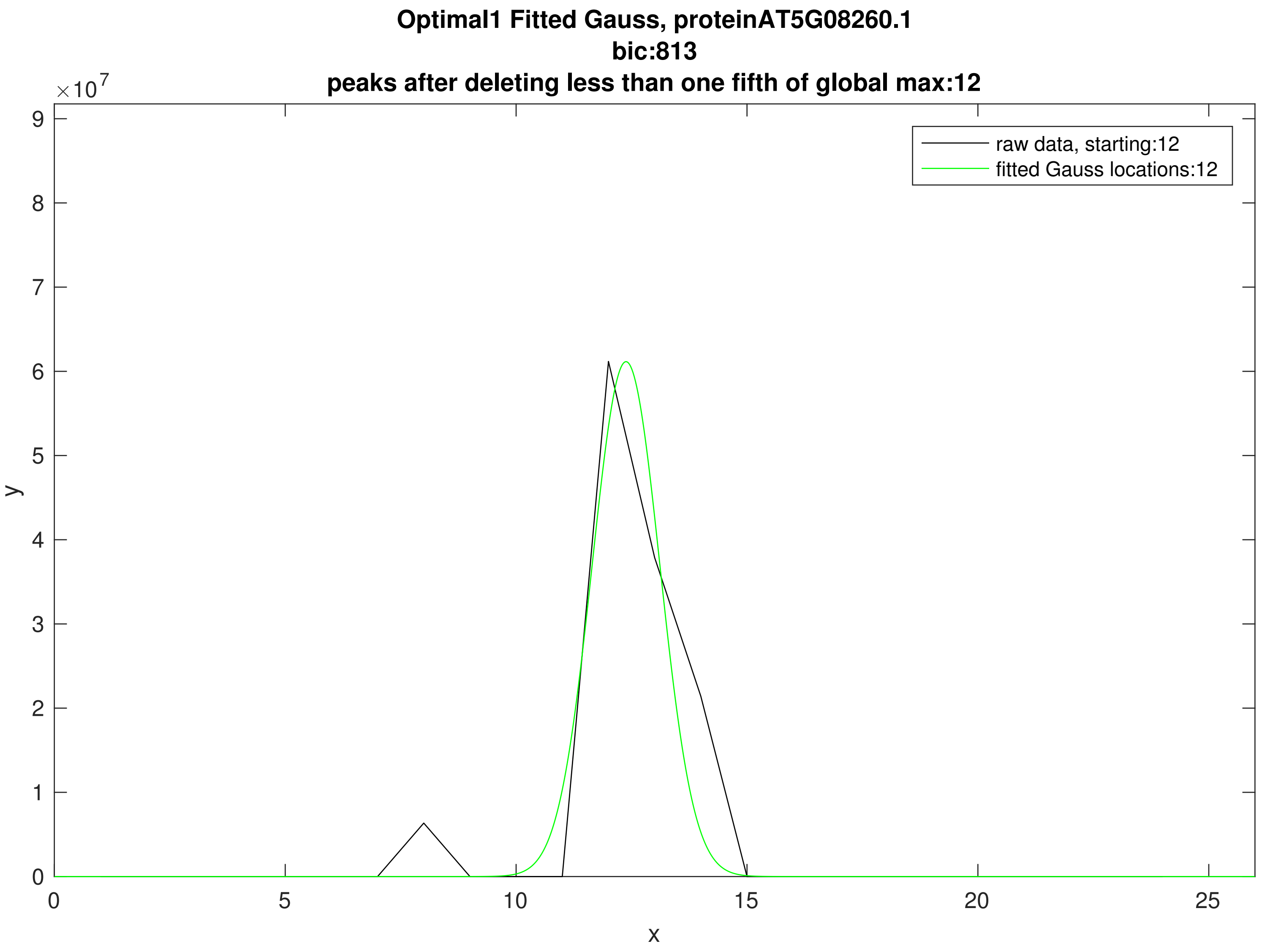
Optimal1 Fitted Gauss, proteinAT5G08260.1
bic:813
peaks after deleting less than one fifth of global max:12
×10^7
raw data, starting:12
fitted Gauss locations:12
y
x
0
1
2
3
4
5
6
7
8
9
0
5
10
15
20
25

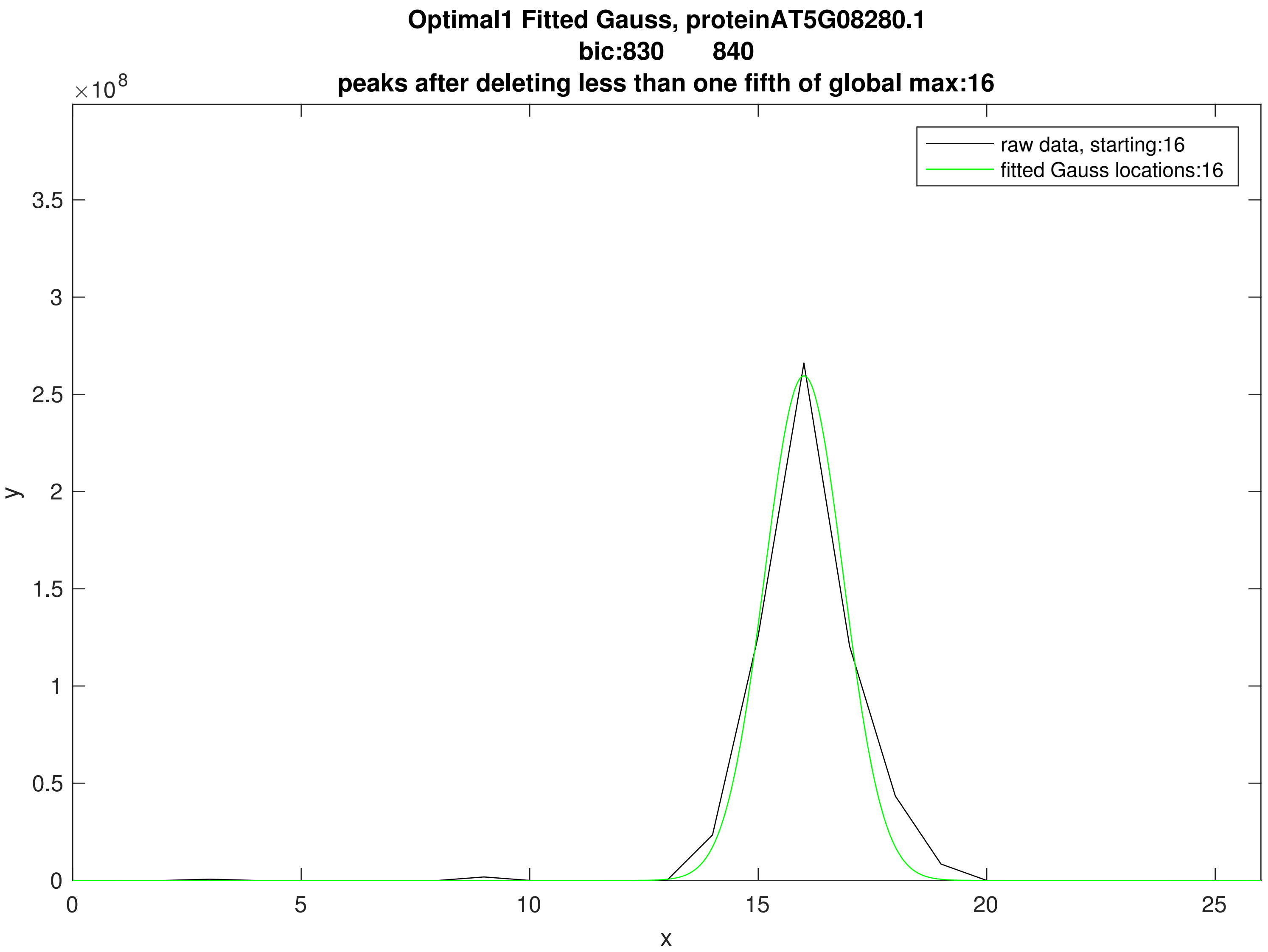
Optimal1 Fitted Gauss, proteinAT5G08280.1
bic:830 840
peaks after deleting less than one fifth of global max:16
×10^8
raw data, starting:16
fitted Gauss locations:16
y
x
0
0.5
1
1.5
2
2.5
3
3.5
0
5
10
15
20
25

**Optimal1 Fitted Gauss, proteinAT5G08300.1**
**bic:996**
**peaks after deleting less than one fifth of global max:14**

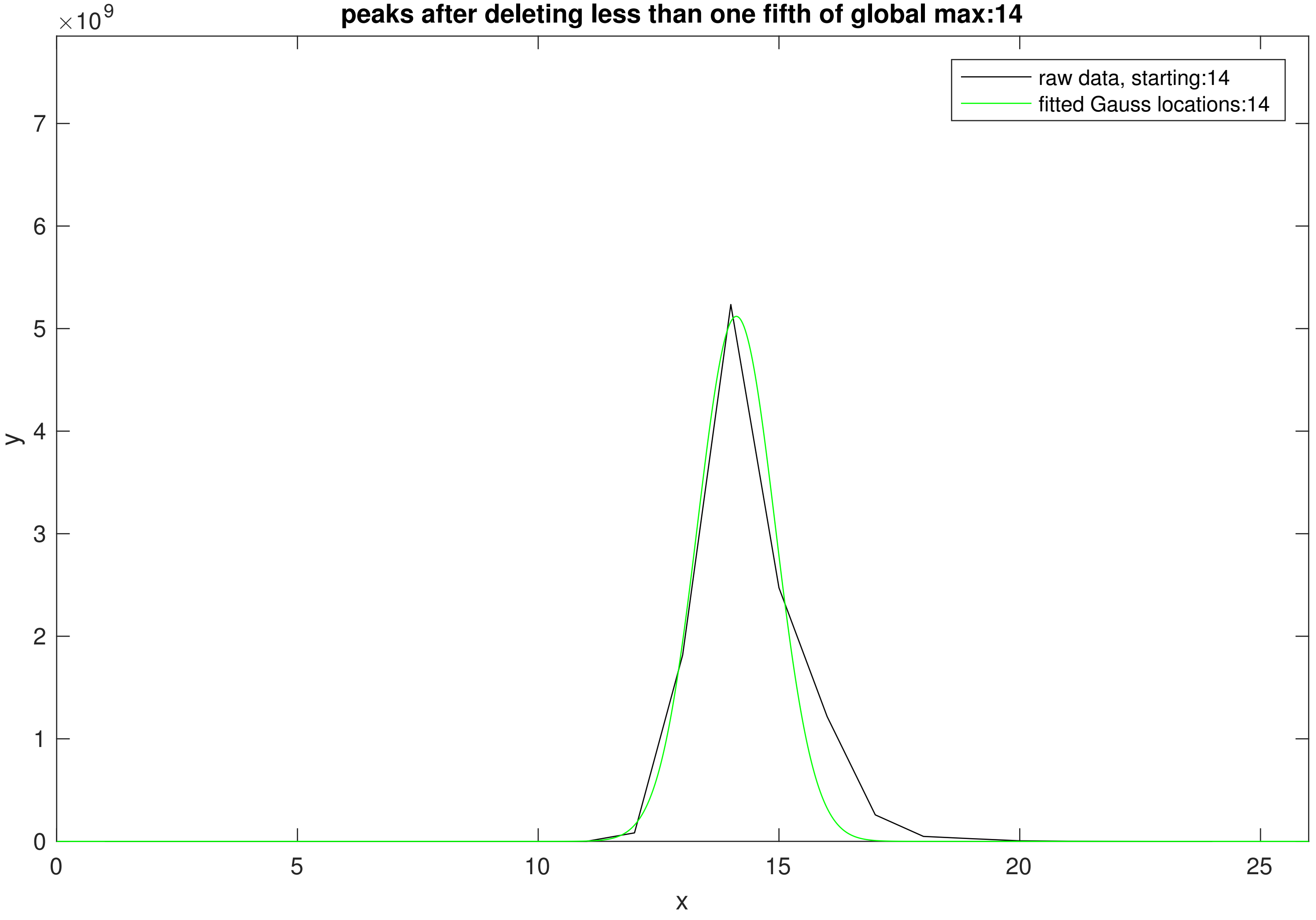

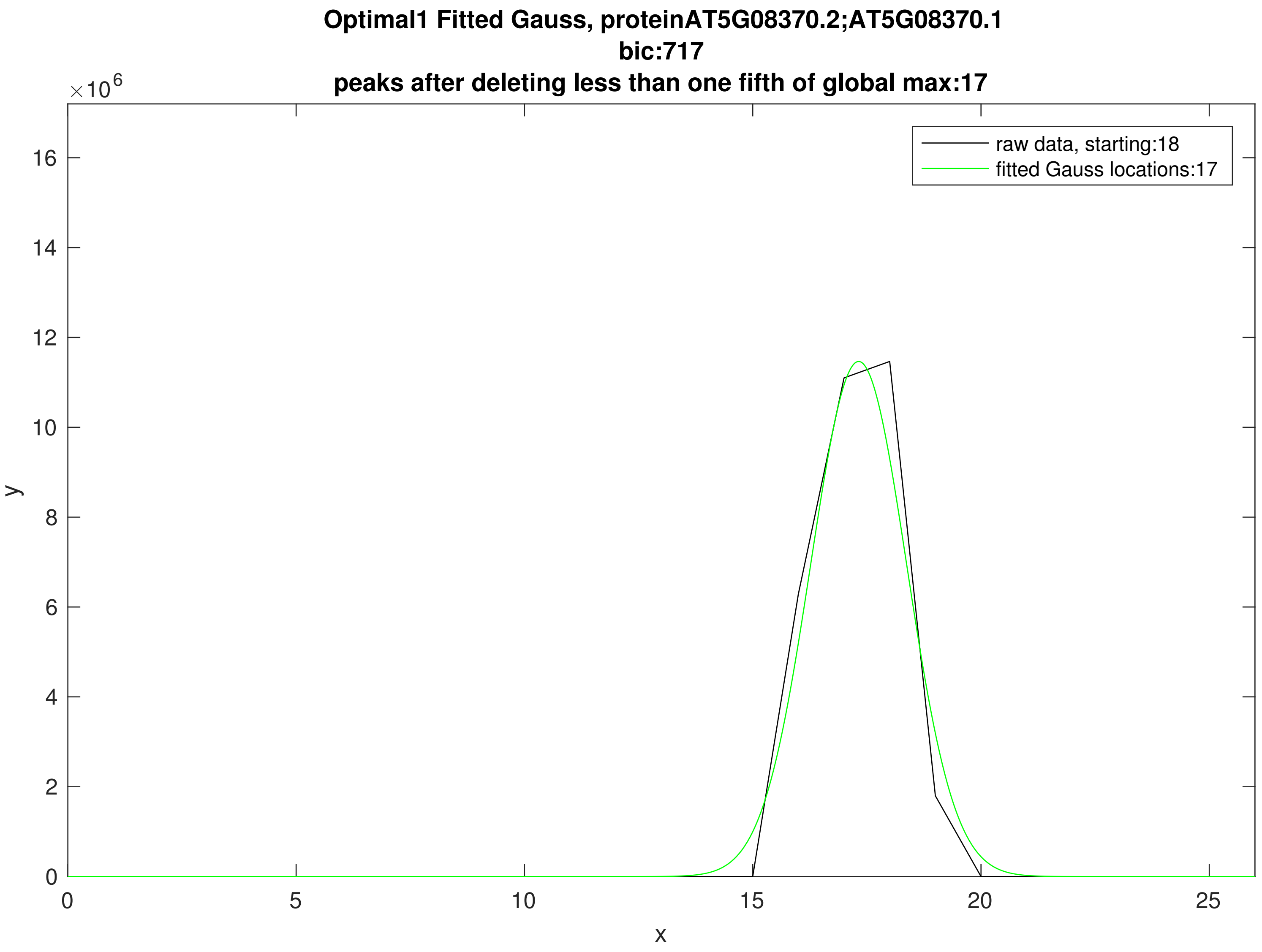

Optimal1 Fitted Gauss, proteinAT5G08370.2;AT5G08370.1
bic:717
peaks after deleting less than one fifth of global max:17
×10^6
raw data, starting:18
fitted Gauss locations:17
y
x
0
2
4
6
8
10
12
14
16
0
5
10
15
20
25

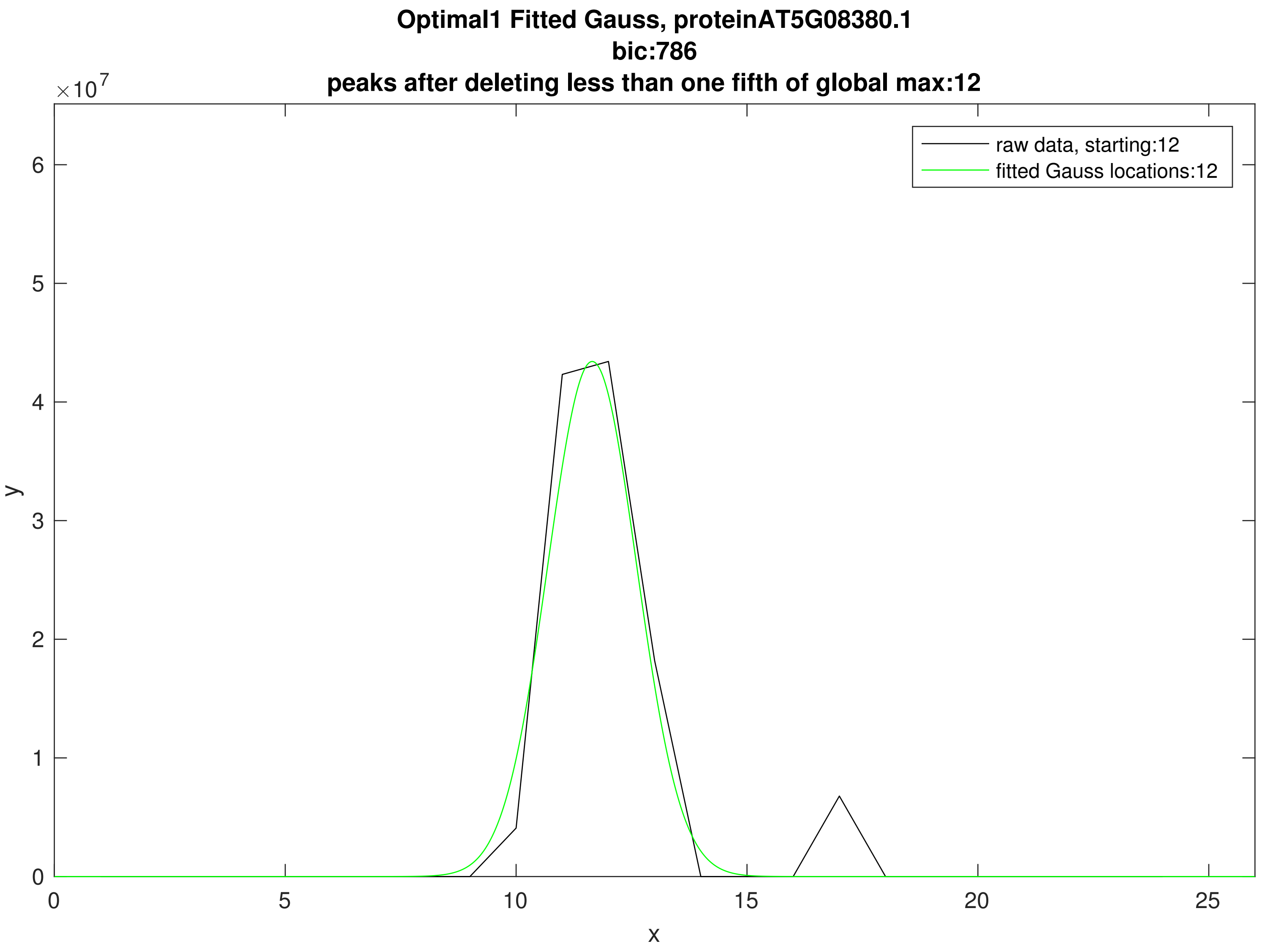
Optimal1 Fitted Gauss, proteinAT5G08380.1
bic:786
peaks after deleting less than one fifth of global max:12
×10^7
raw data, starting:12
fitted Gauss locations:12
y
x
0
1
2
3
4
5
6
0
5
10
15
20
25

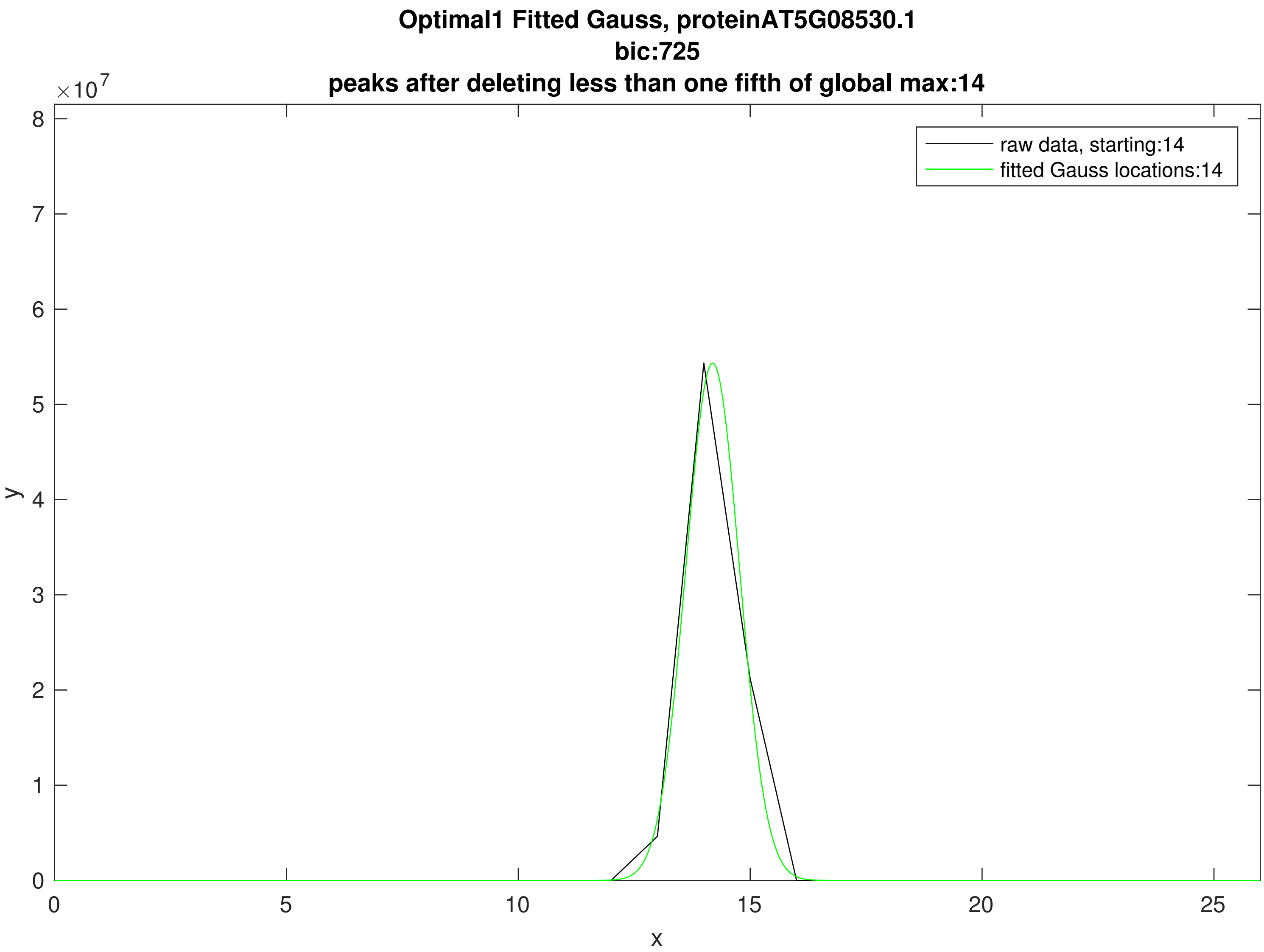

Optimal1 Fitted Gauss, proteinAT5G08530.1
bic:725
peaks after deleting less than one fifth of global max:14
×10^7
8
7
6
5
4
3
2
1
0
y
0
5
10
15
20
25
x
raw data, starting:14
fitted Gauss locations:14

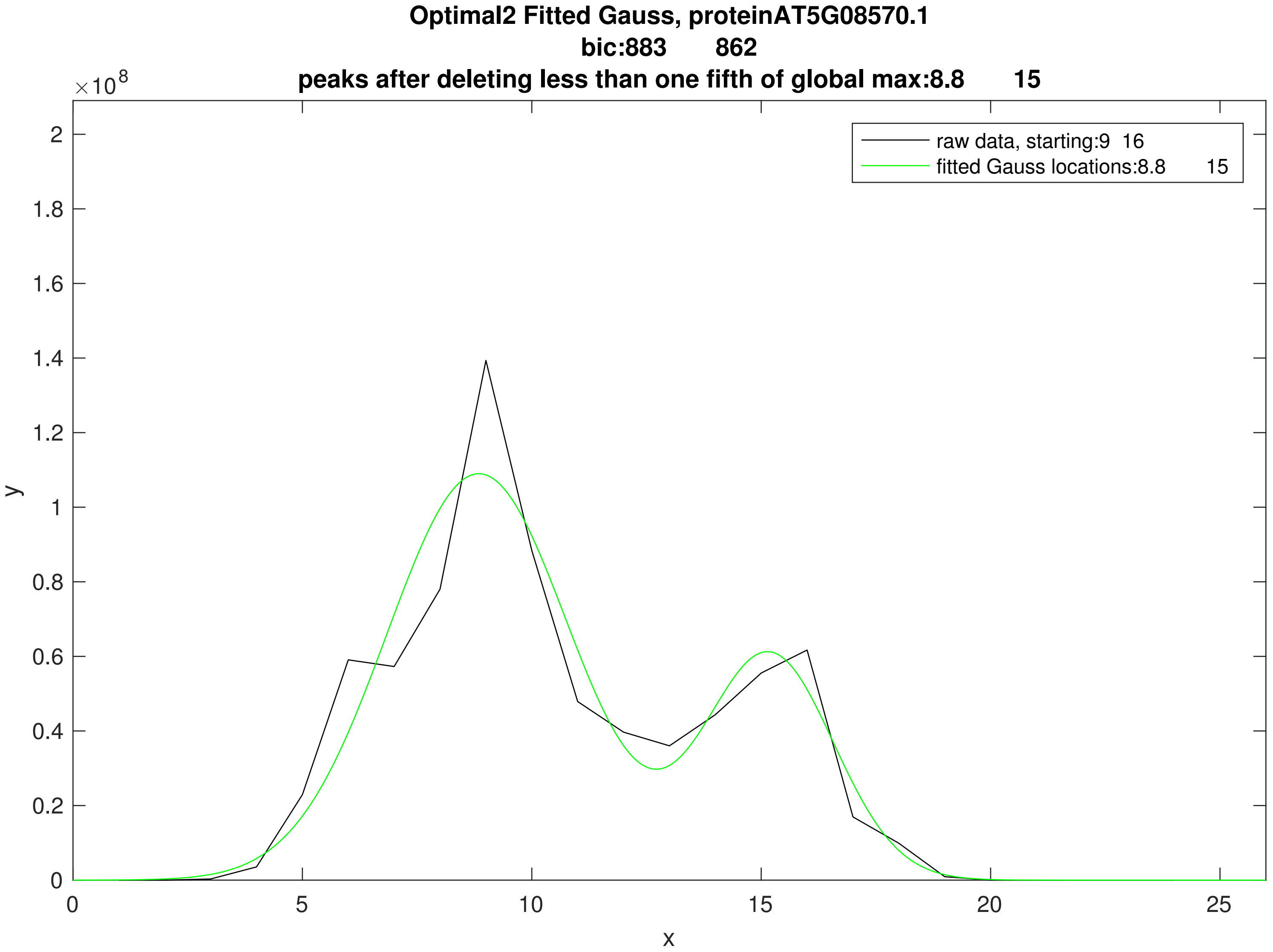

Optimal2 Fitted Gauss, proteinAT5G08570.1
bic:883 862
peaks after deleting less than one fifth of global max:8.8 15
×10^8
raw data, starting:9 16
fitted Gauss locations:8.8 15
y
x
0
0.2
0.4
0.6
0.8
1
1.2
1.4
1.6
1.8
2
0
5
10
15
20
25

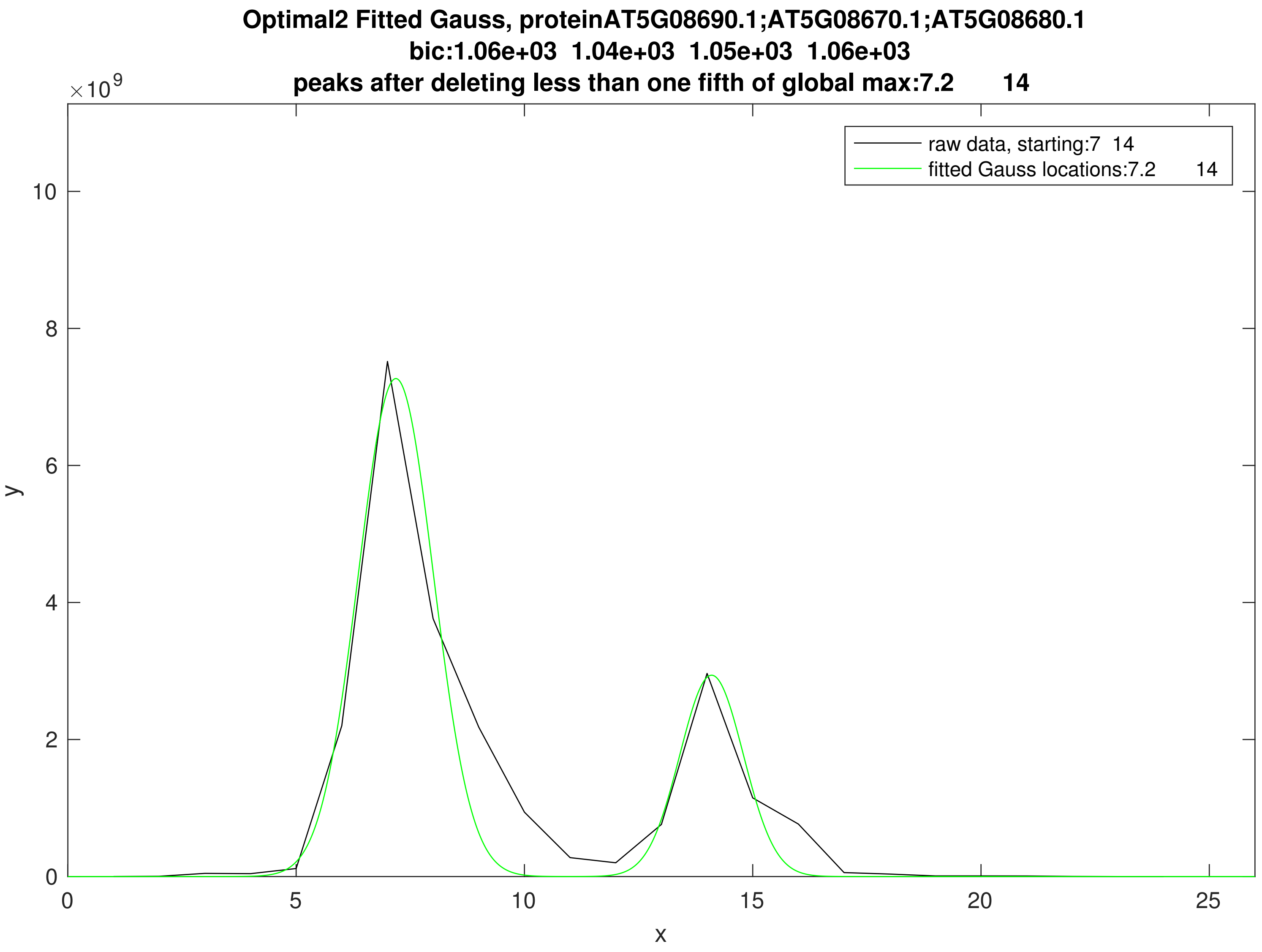

Optimal2 Fitted Gauss, proteinAT5G08690.1;AT5G08670.1;AT5G08680.1
bic:1.06e+03  1.04e+03  1.05e+03  1.06e+03
peaks after deleting less than one fifth of global max:7.2      14
×10^9
raw data, starting:7  14
fitted Gauss locations:7.2      14
y
x
0
2
4
6
8
10
0
5
10
15
20
25

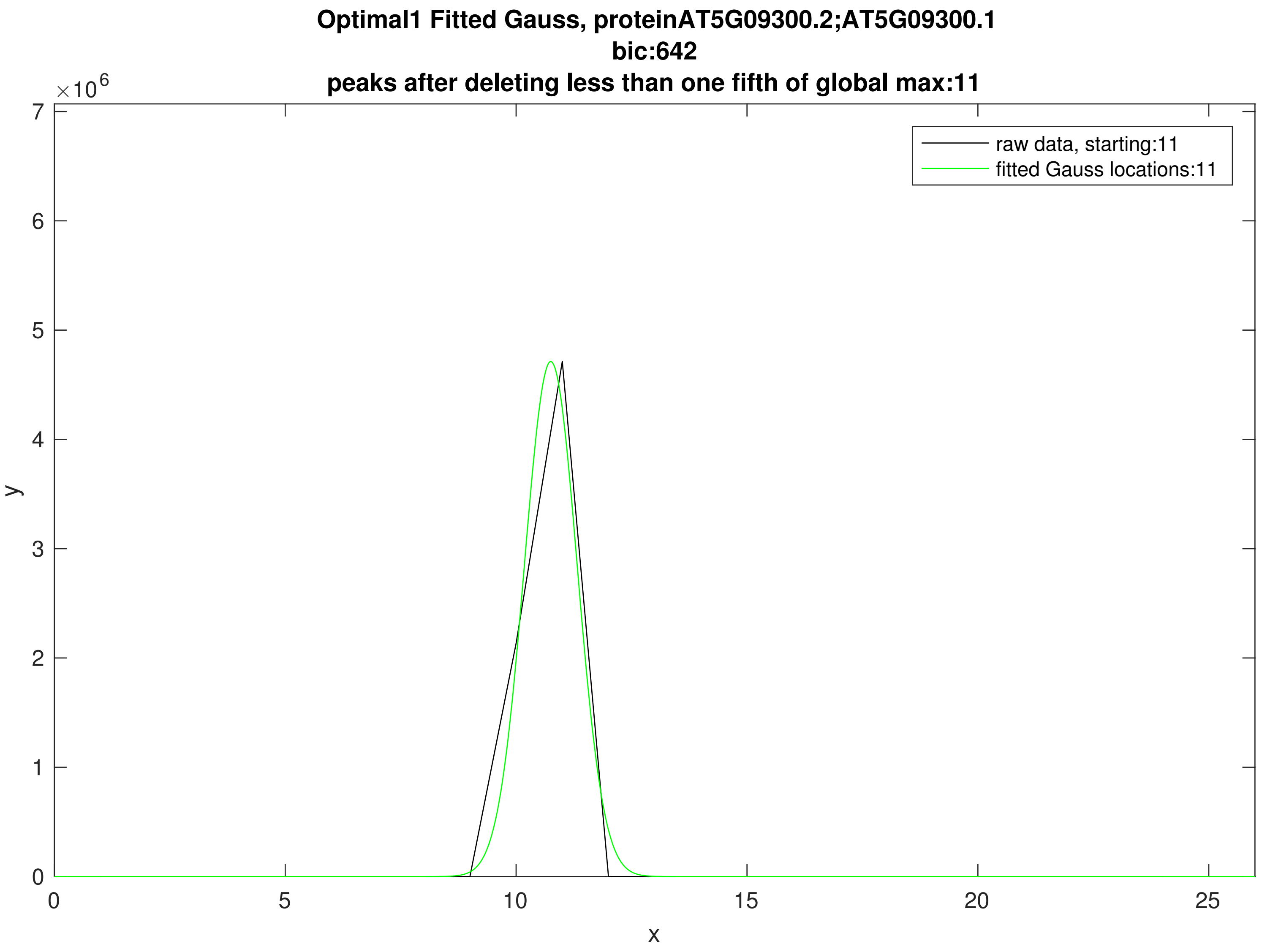
Optimal1 Fitted Gauss, proteinAT5G09300.2;AT5G09300.1
bic:642
peaks after deleting less than one fifth of global max:11
×10^6
raw data, starting:11
fitted Gauss locations:11
y
x
0
1
2
3
4
5
6
7
0
5
10
15
20
25

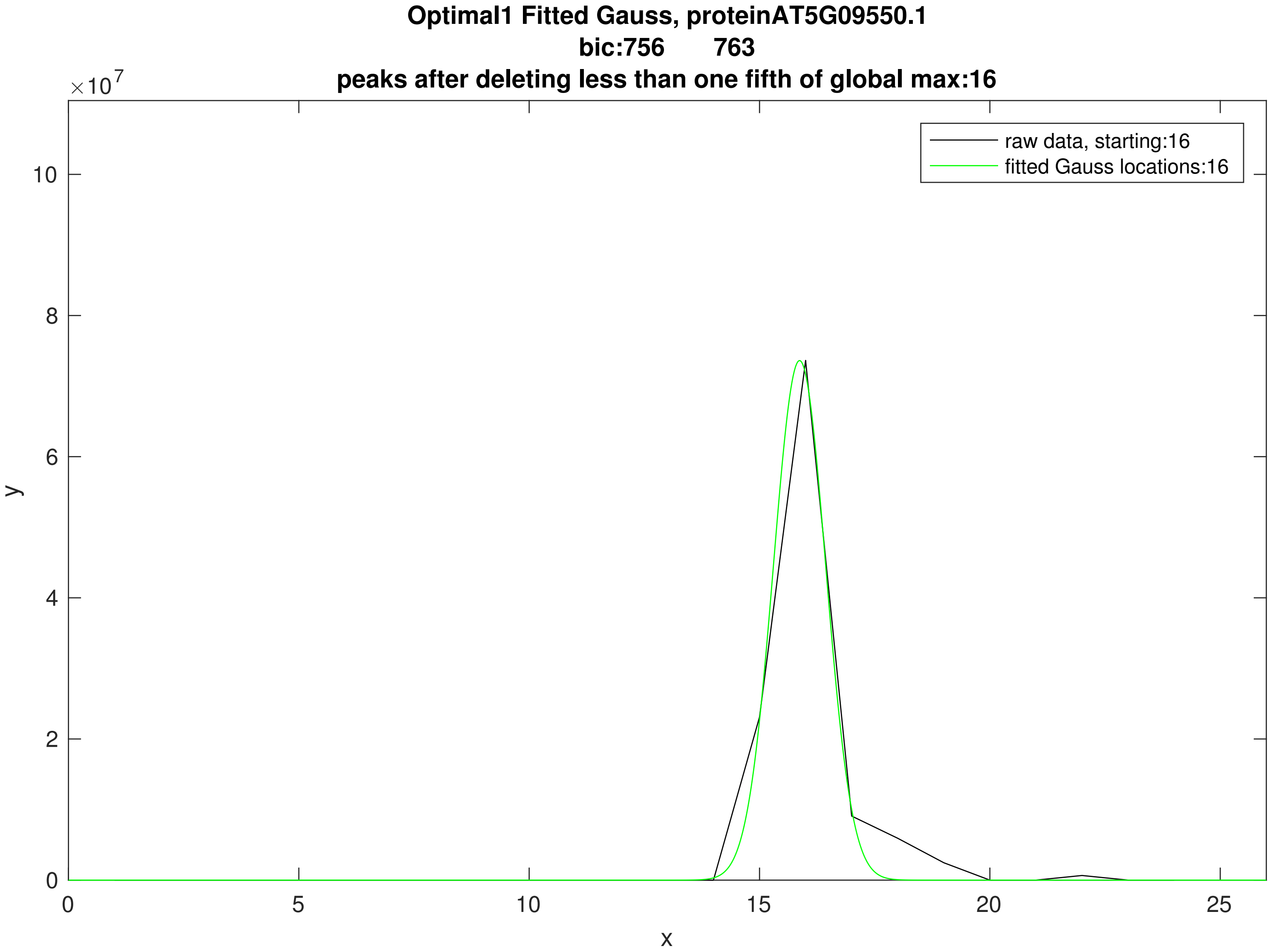
Optimal1 Fitted Gauss, proteinAT5G09550.1
bic:756 763
peaks after deleting less than one fifth of global max:16
×10^7
10
8
6
4
2
0
y
0
5
10
15
20
25
x
raw data, starting:16
fitted Gauss locations:16

**Optimal1 Fitted Gauss, proteinAT5G09590.1**
**bic:934 943**
**peaks after deleting less than one fifth of global max:12**

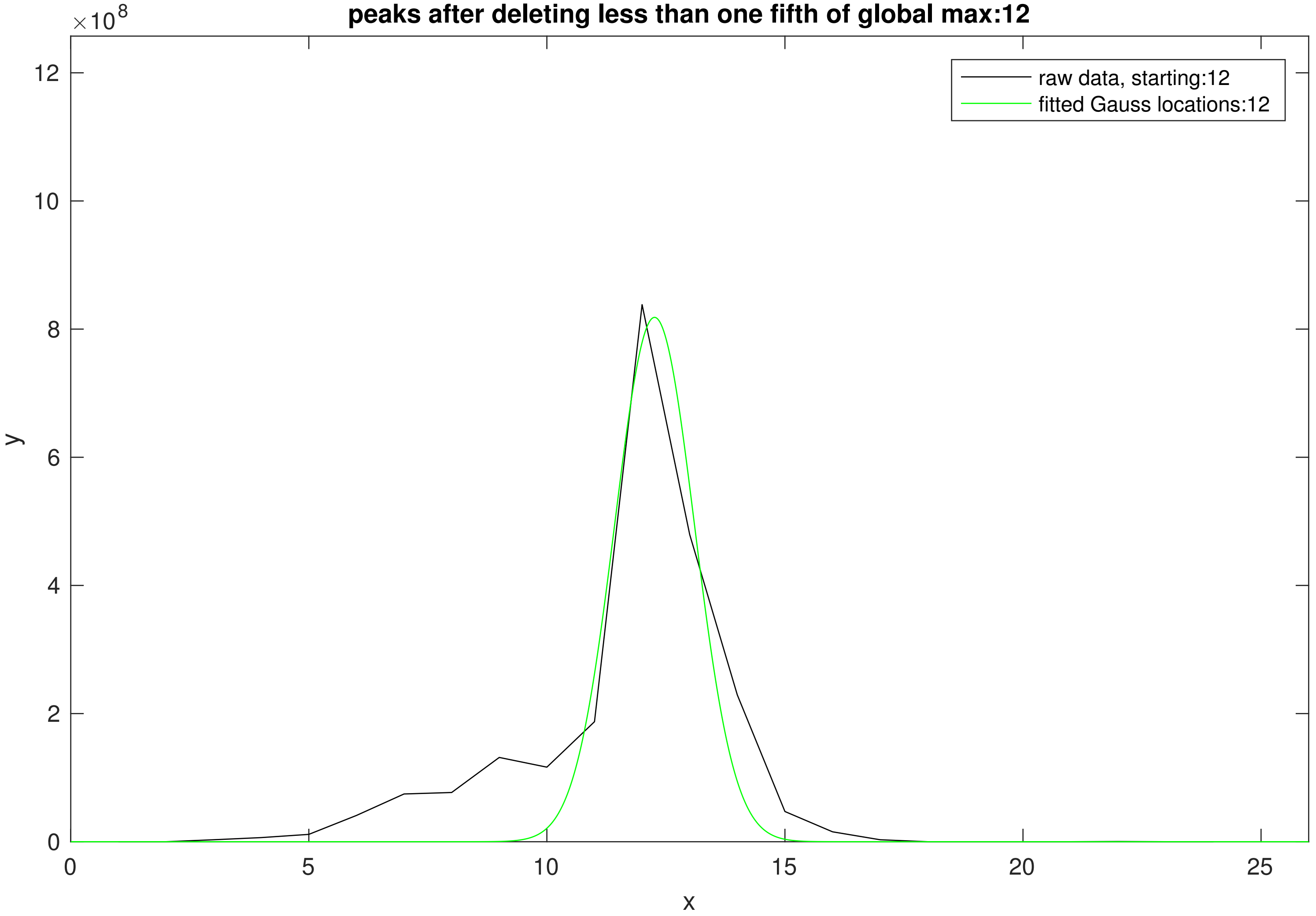

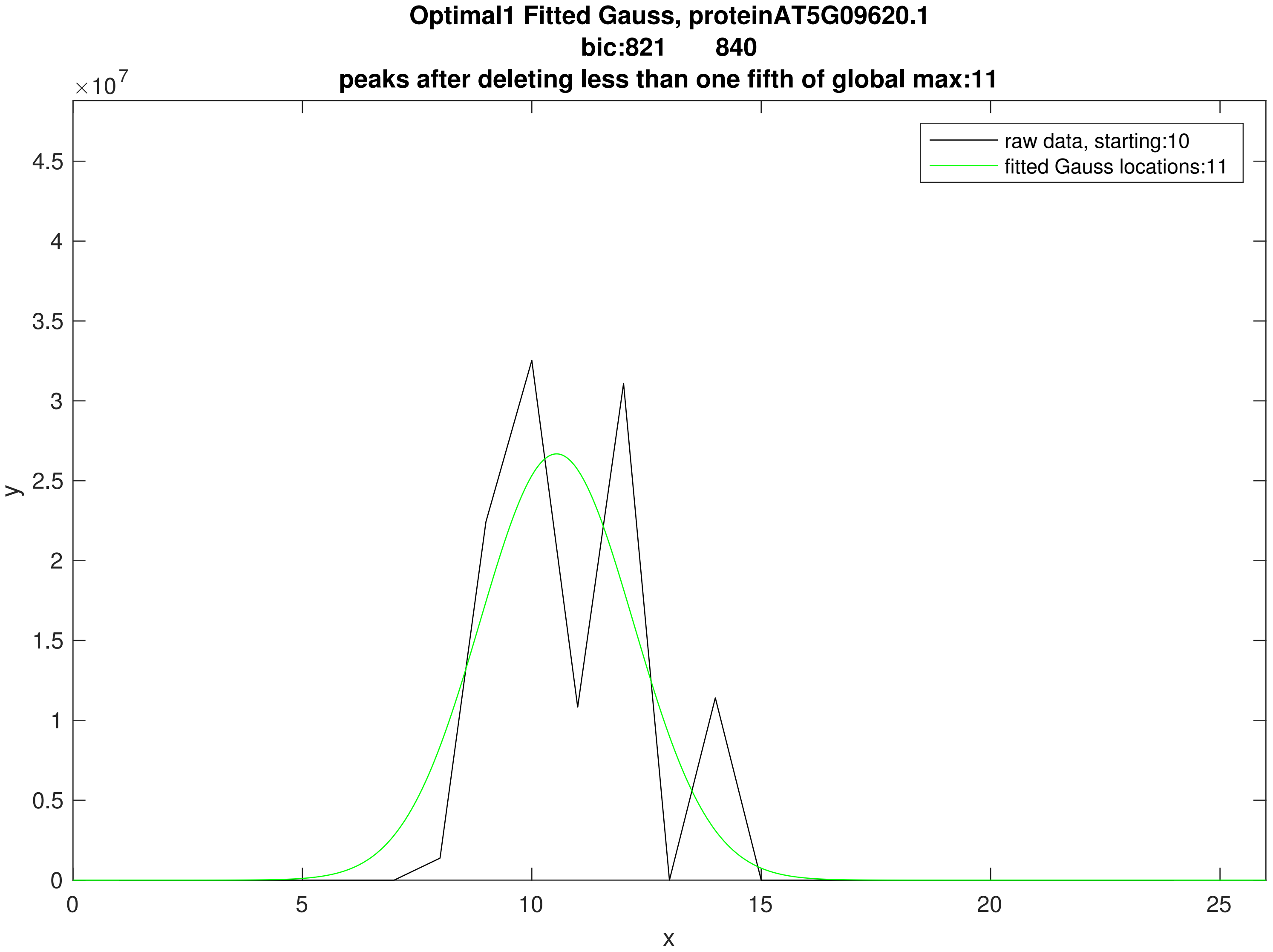
Optimal1 Fitted Gauss, proteinAT5G09620.1
bic:821 840
peaks after deleting less than one fifth of global max:11
×10^7
4.5
4
3.5
3
2.5
2
1.5
1
0.5
0
y
0
5
10
15
20
25
x
raw data, starting:10
fitted Gauss locations:11

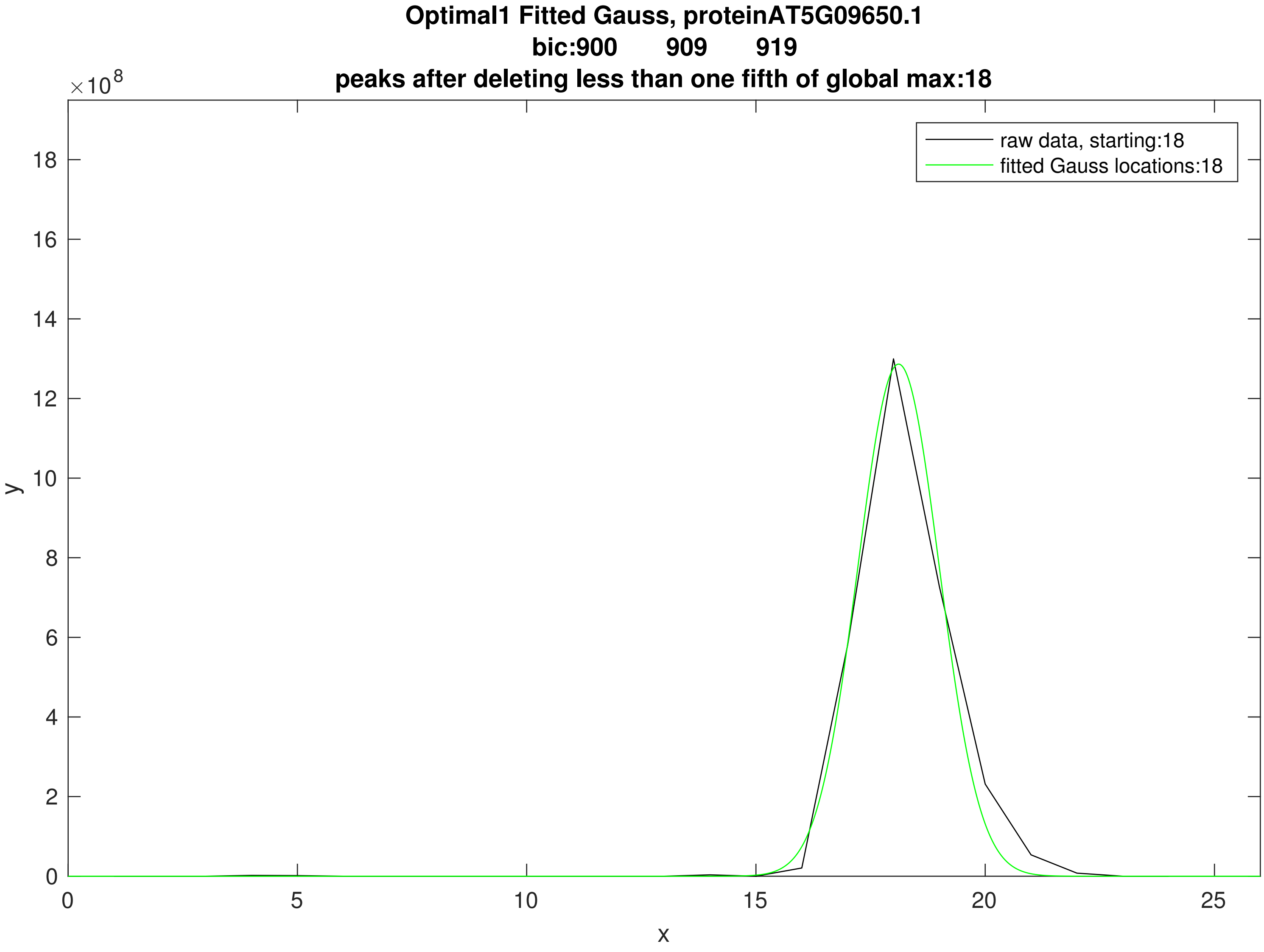
Optimal1 Fitted Gauss, proteinAT5G09650.1
bic:900 909 919
peaks after deleting less than one fifth of global max:18
×10^8
raw data, starting:18
fitted Gauss locations:18
y
x
0
2
4
6
8
10
12
14
16
18
0
5
10
15
20
25

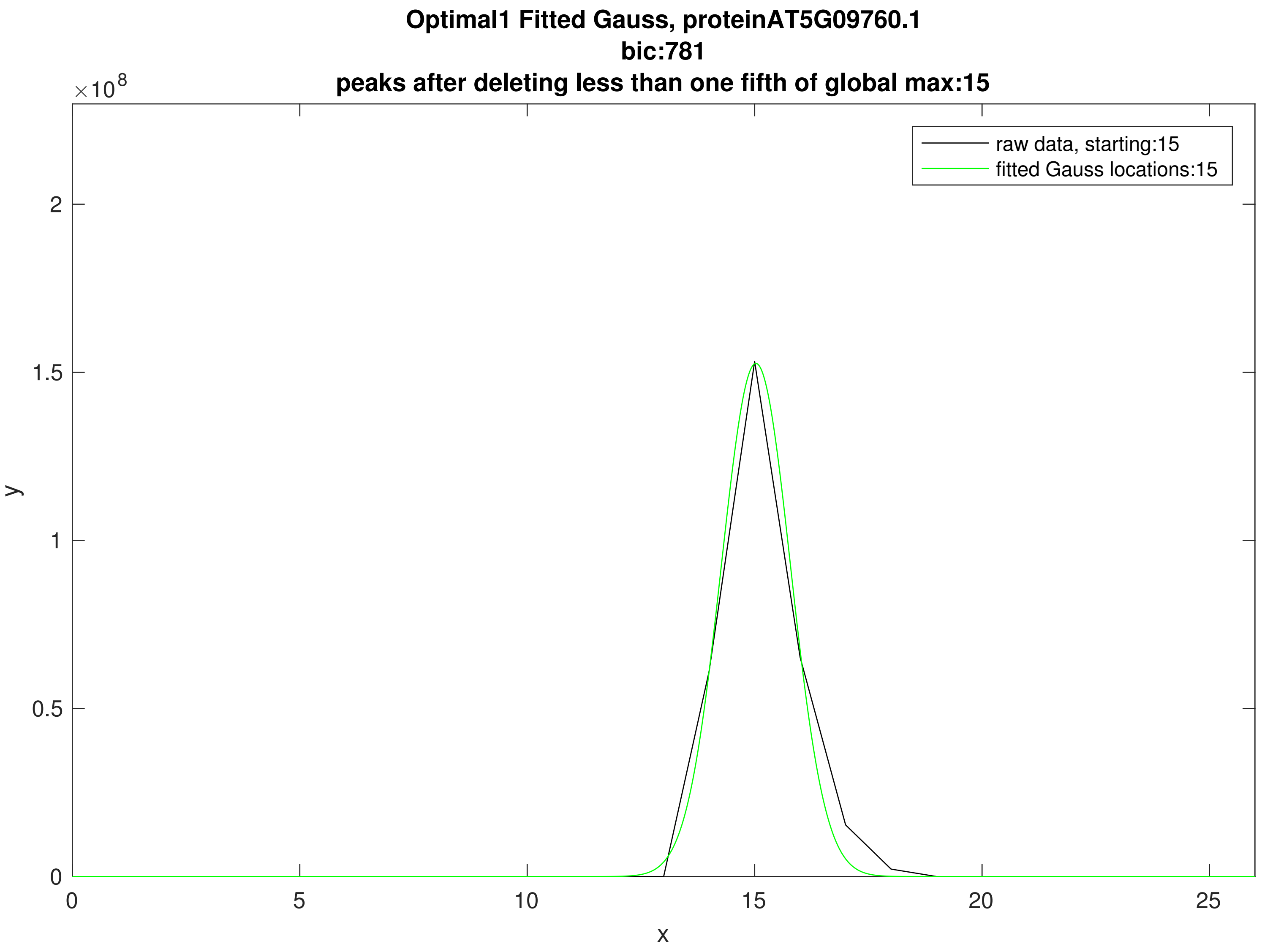

Optimal1 Fitted Gauss, proteinAT5G09760.1
bic:781
peaks after deleting less than one fifth of global max:15
×10^8
raw data, starting:15
fitted Gauss locations:15
2
1.5
1
0.5
0
y
0
5
10
15
20
25
x

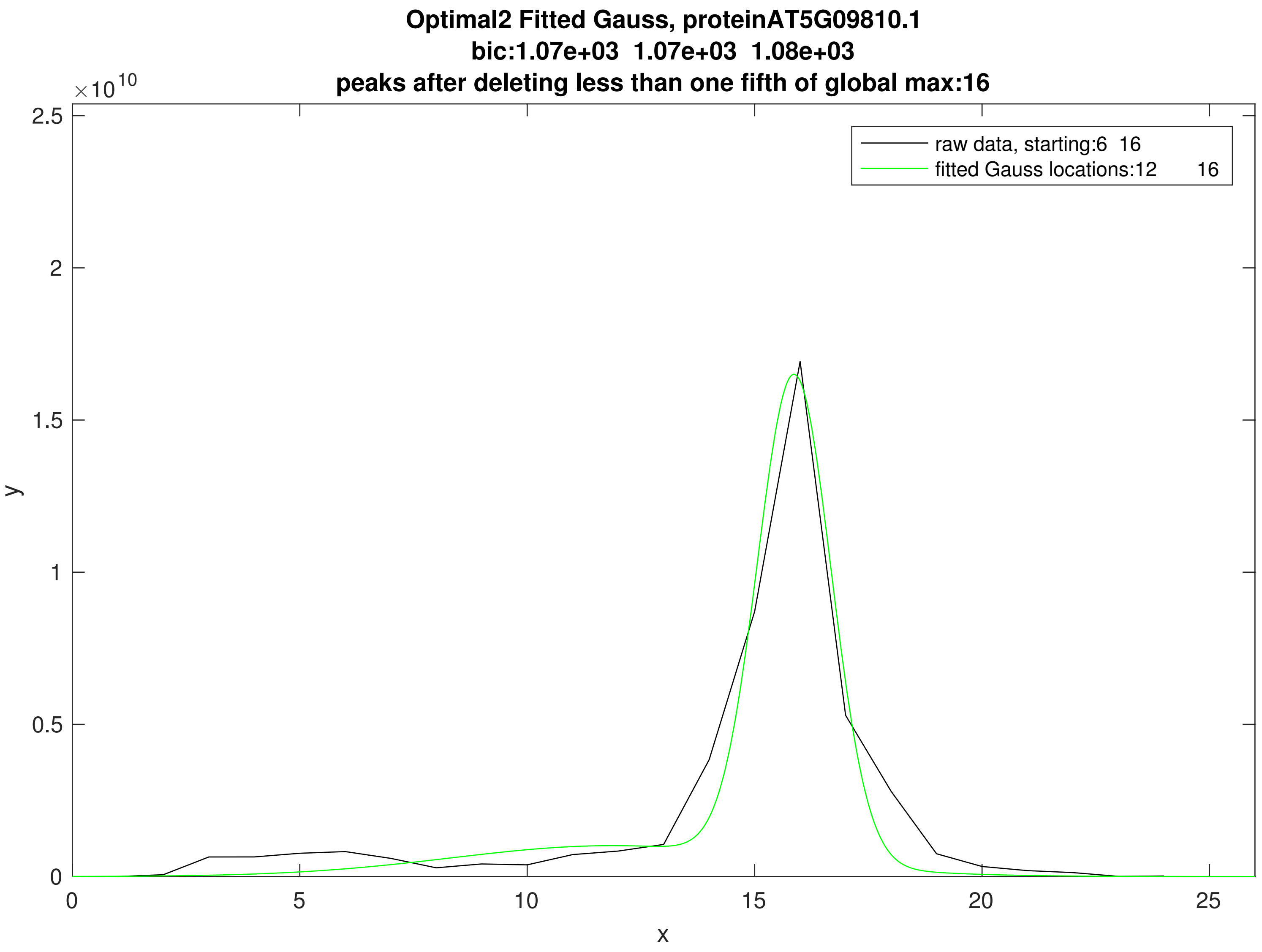
Optimal2 Fitted Gauss, proteinAT5G09810.1
bic:1.07e+03 1.07e+03 1.08e+03
peaks after deleting less than one fifth of global max:16
×10^10
raw data, starting:6 16
fitted Gauss locations:12 16
y
x
0
0.5
1
1.5
2
2.5
5
10
15
20
25

#### Optimal1 Fitted Gauss, proteinAT5G09830.1
#### bic:729
#### peaks after deleting less than one fifth of global max:20

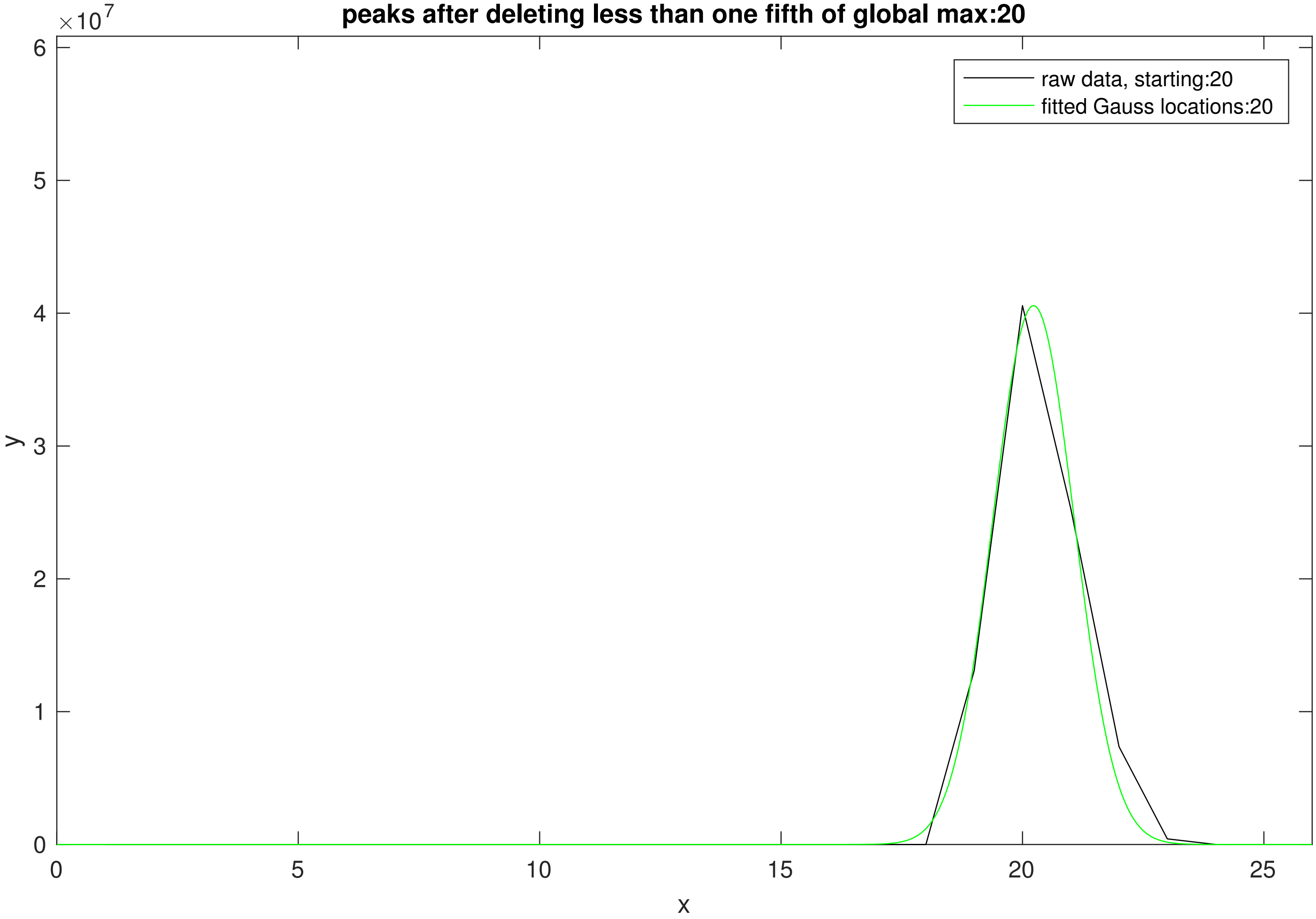

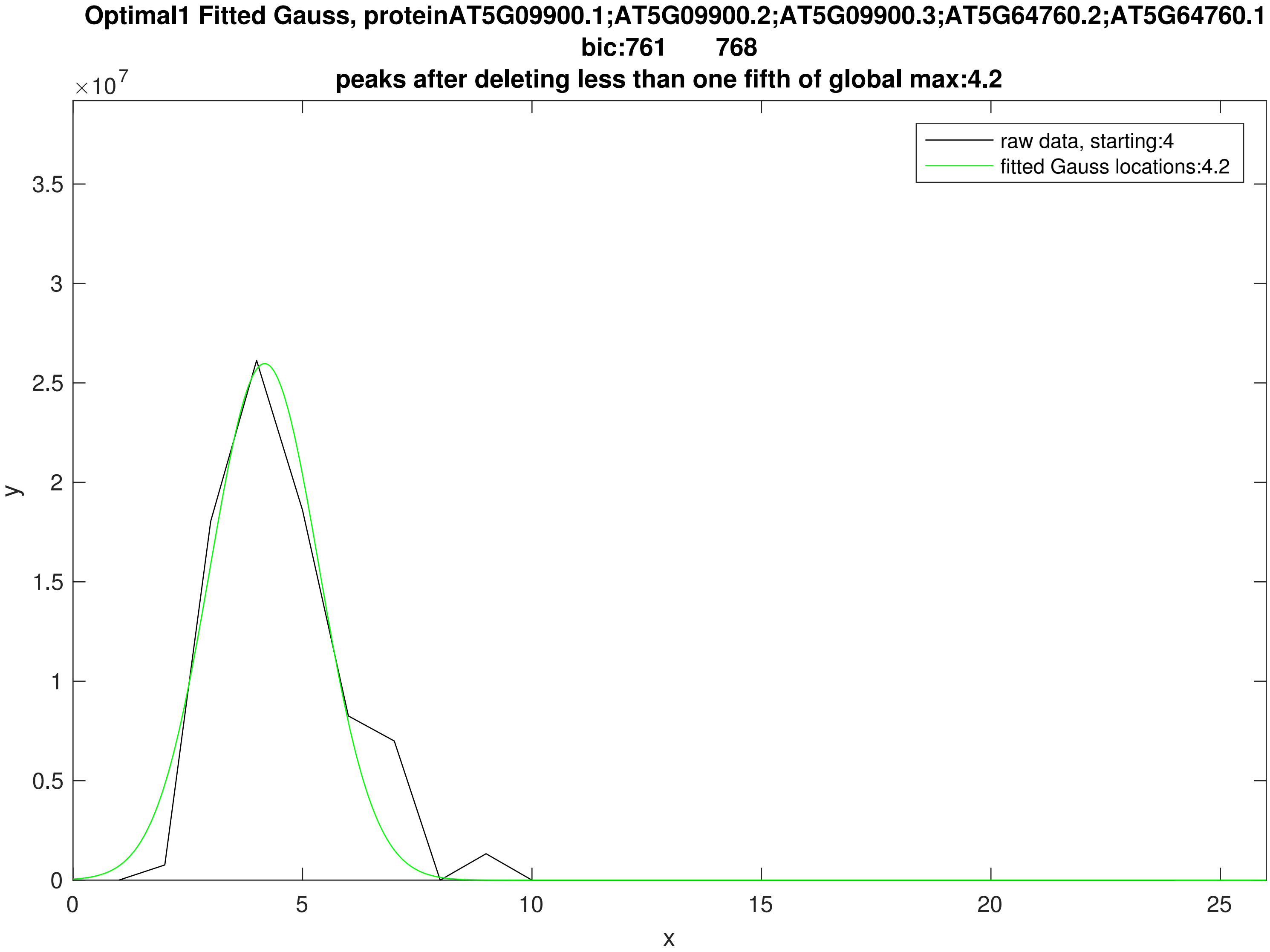

Optimal1 Fitted Gauss, proteinAT5G09900.1;AT5G09900.2;AT5G09900.3;AT5G64760.2;AT5G64760.1
bic:761 768
peaks after deleting less than one fifth of global max:4.2
×10^7
raw data, starting:4
fitted Gauss locations:4.2
3.5
3
2.5
2
1.5
1
0.5
0
y
0
5
10
15
20
25
x

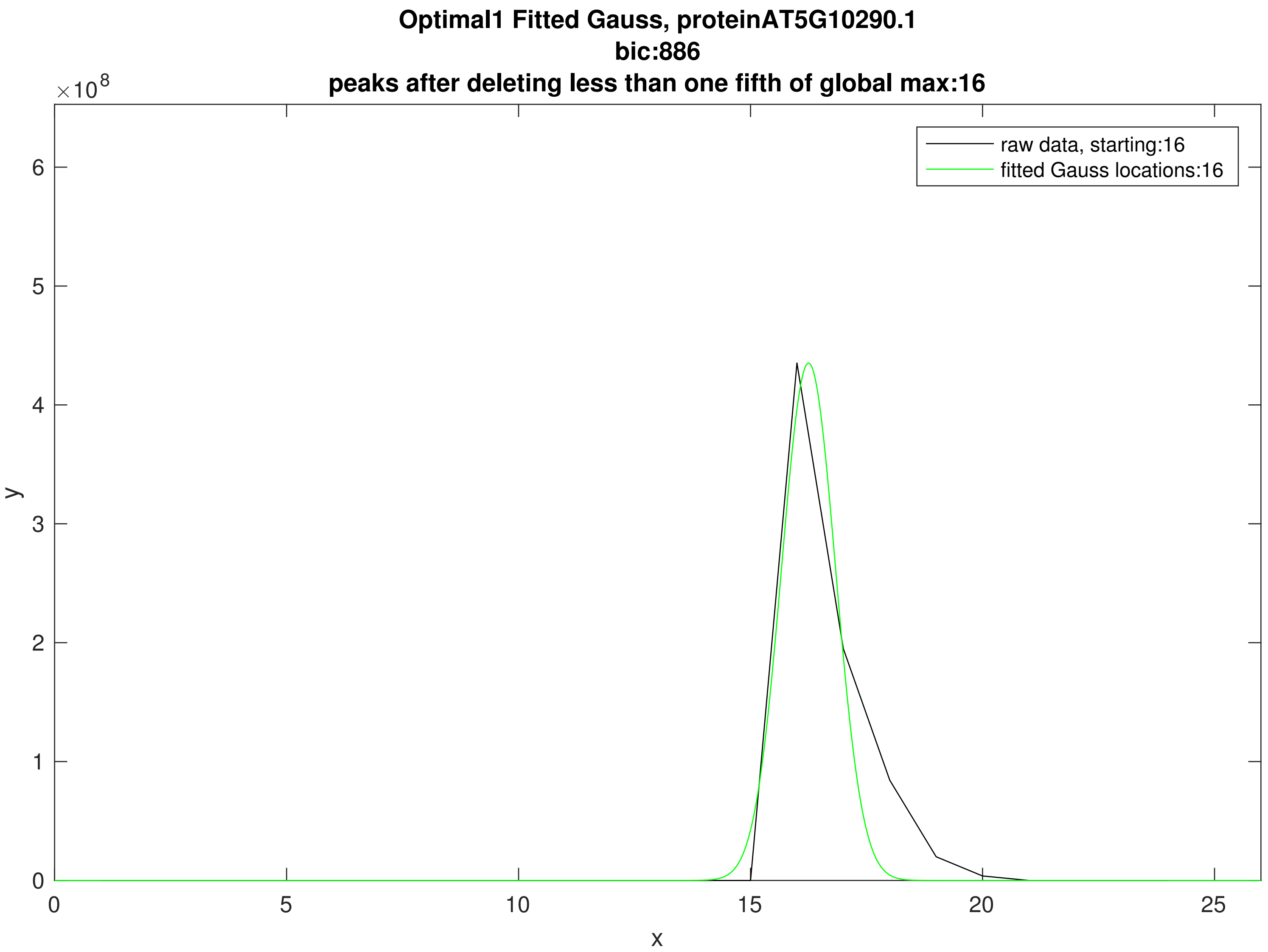

Optimal1 Fitted Gauss, proteinAT5G10290.1
bic:886
peaks after deleting less than one fifth of global max:16
×10^8
6
5
4
3
2
1
0
y
0
5
10
15
20
25
x
raw data, starting:16
fitted Gauss locations:16

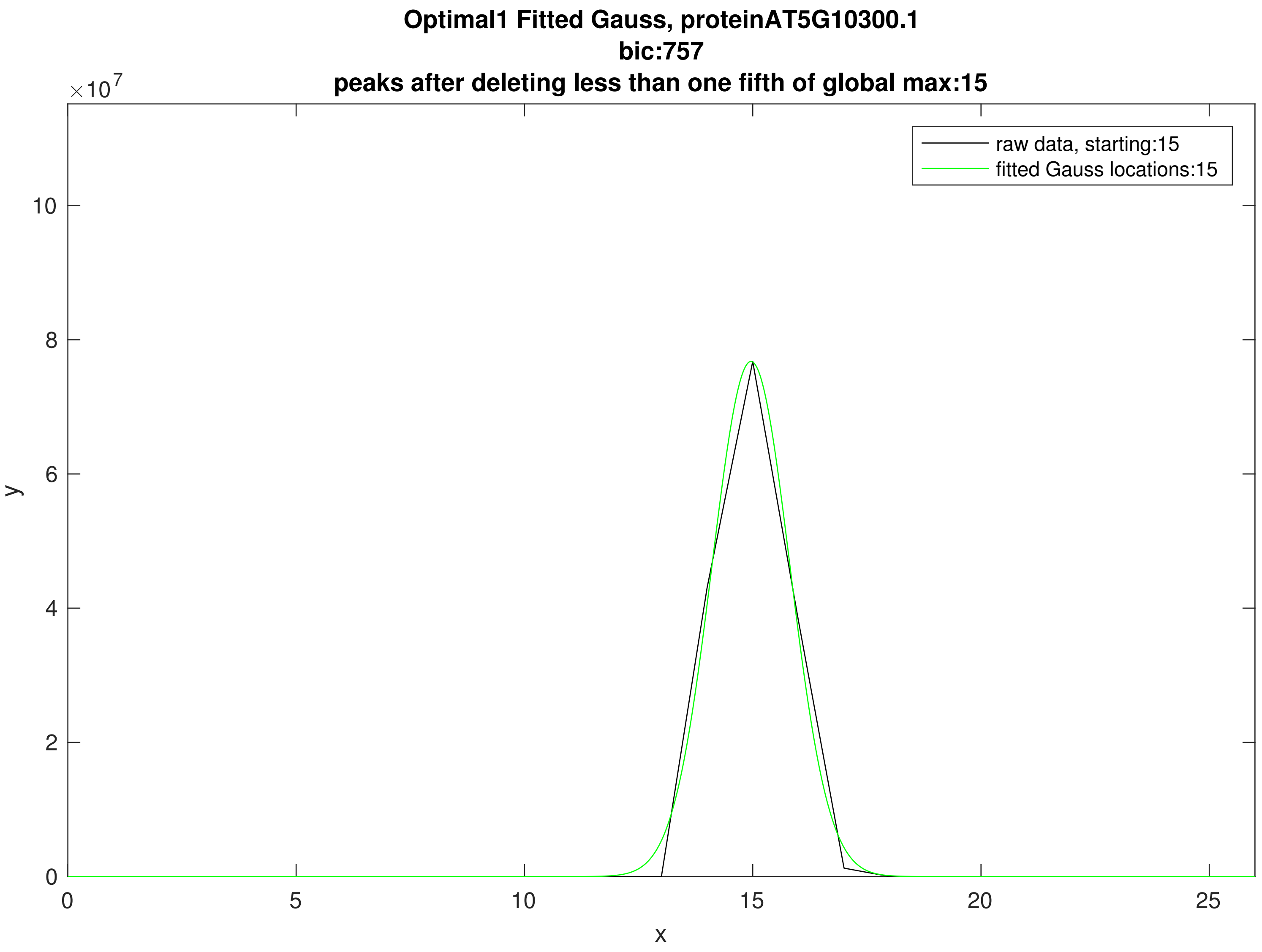
Optimal1 Fitted Gauss, proteinAT5G10300.1
bic:757
peaks after deleting less than one fifth of global max:15
×10^7
raw data, starting:15
fitted Gauss locations:15
y
x
0
2
4
6
8
10
0
5
10
15
20
25

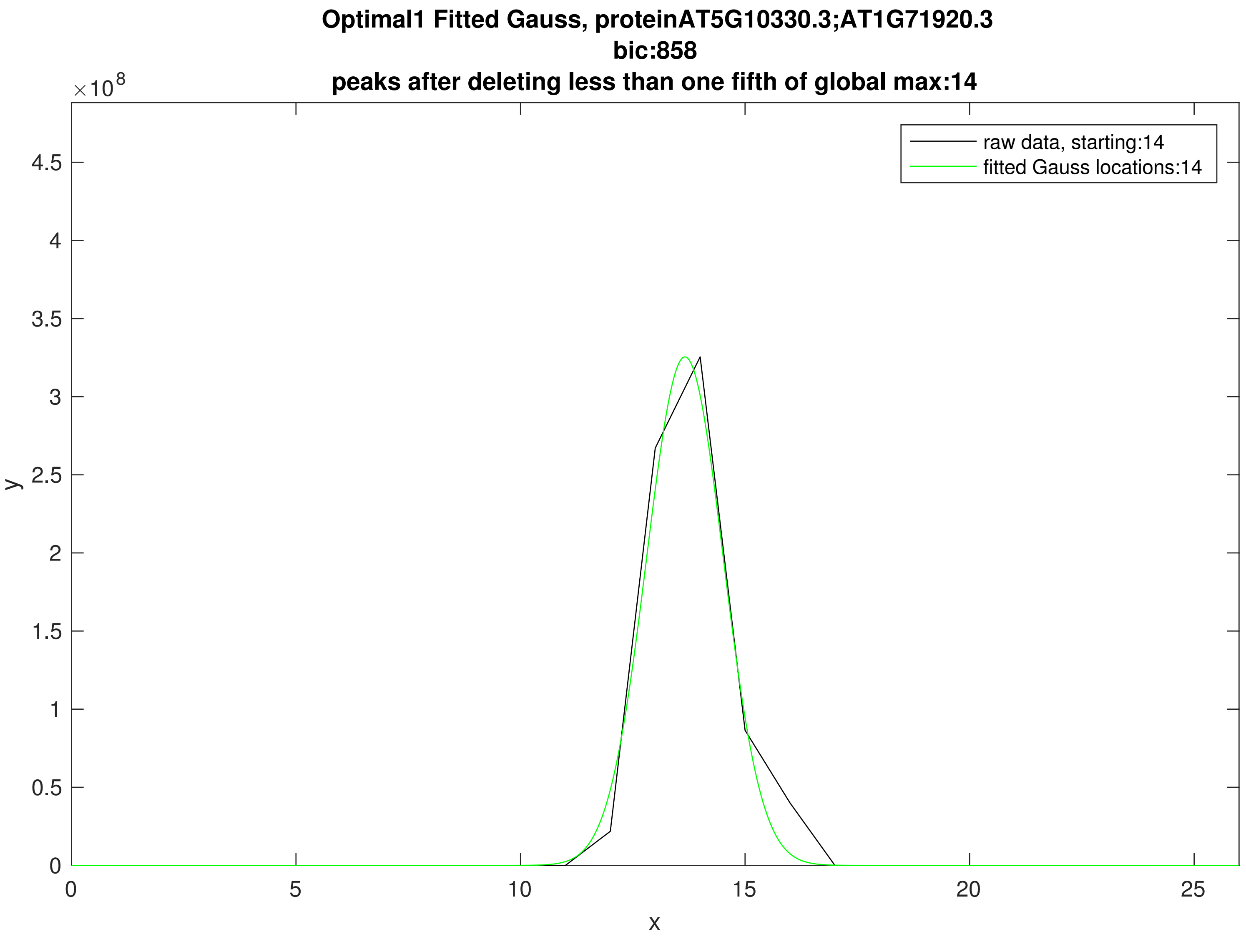
Optimal1 Fitted Gauss, proteinAT5G10330.3;AT1G71920.3
bic:858
peaks after deleting less than one fifth of global max:14
×10^8
raw data, starting:14
fitted Gauss locations:14
4.5
4
3.5
3
2.5
2
1.5
1
0.5
0
y
0
5
10
15
20
25
x

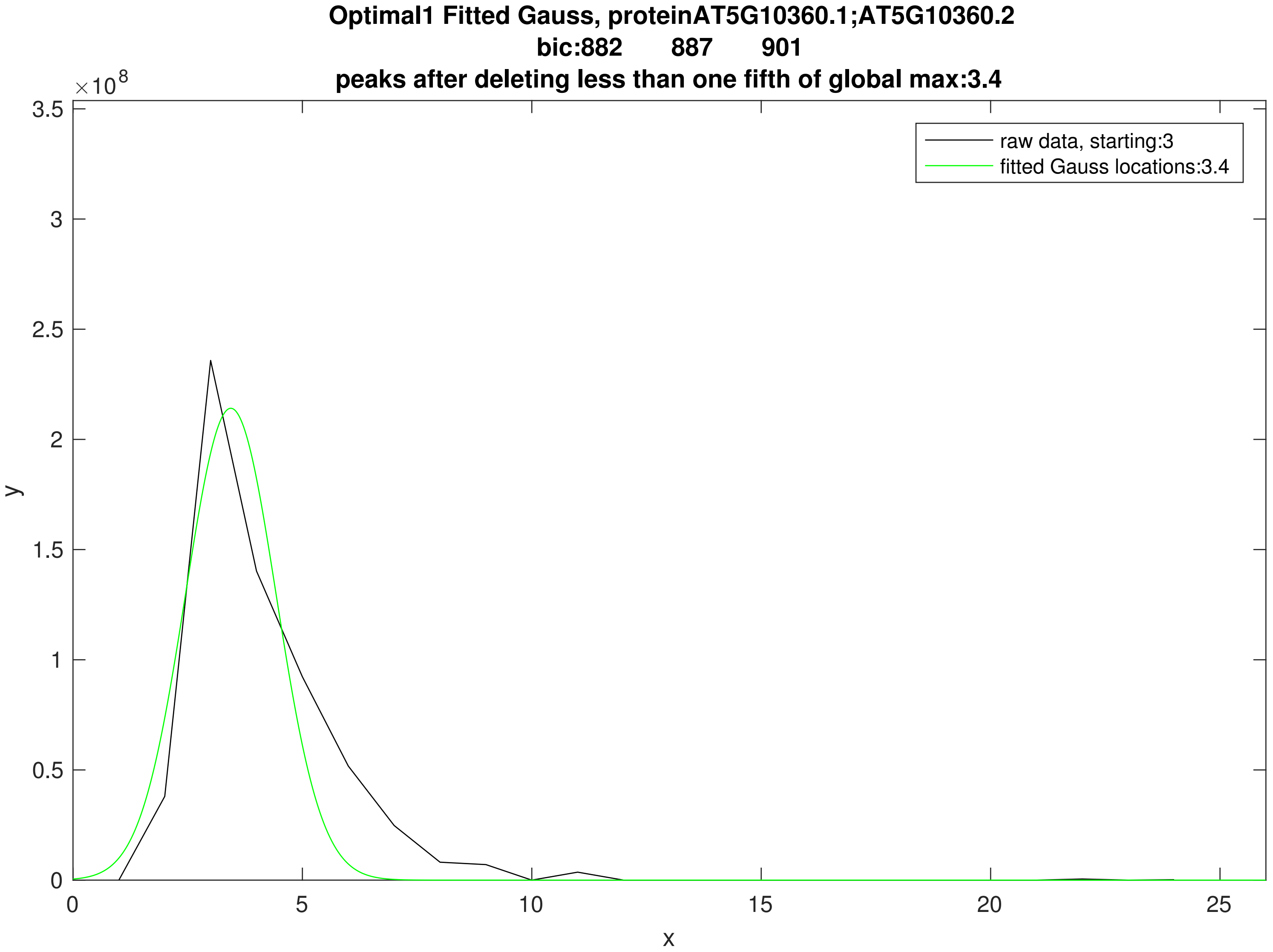
Optimal1 Fitted Gauss, proteinAT5G10360.1;AT5G10360.2
bic:882 887 901
peaks after deleting less than one fifth of global max:3.4
×10^8
raw data, starting:3
fitted Gauss locations:3.4
y
x
0
0.5
1
1.5
2
2.5
3
3.5
5
10
15
20
25

**Optimal1 Fitted Gauss, proteinAT5G10450.1;AT5G10450.2;AT5G10450.3;AT5G10450.4**
**bic:1.01e+03  1.02e+03  1.03e+03  1.04e+03**
**peaks after deleting less than one fifth of global max:13**

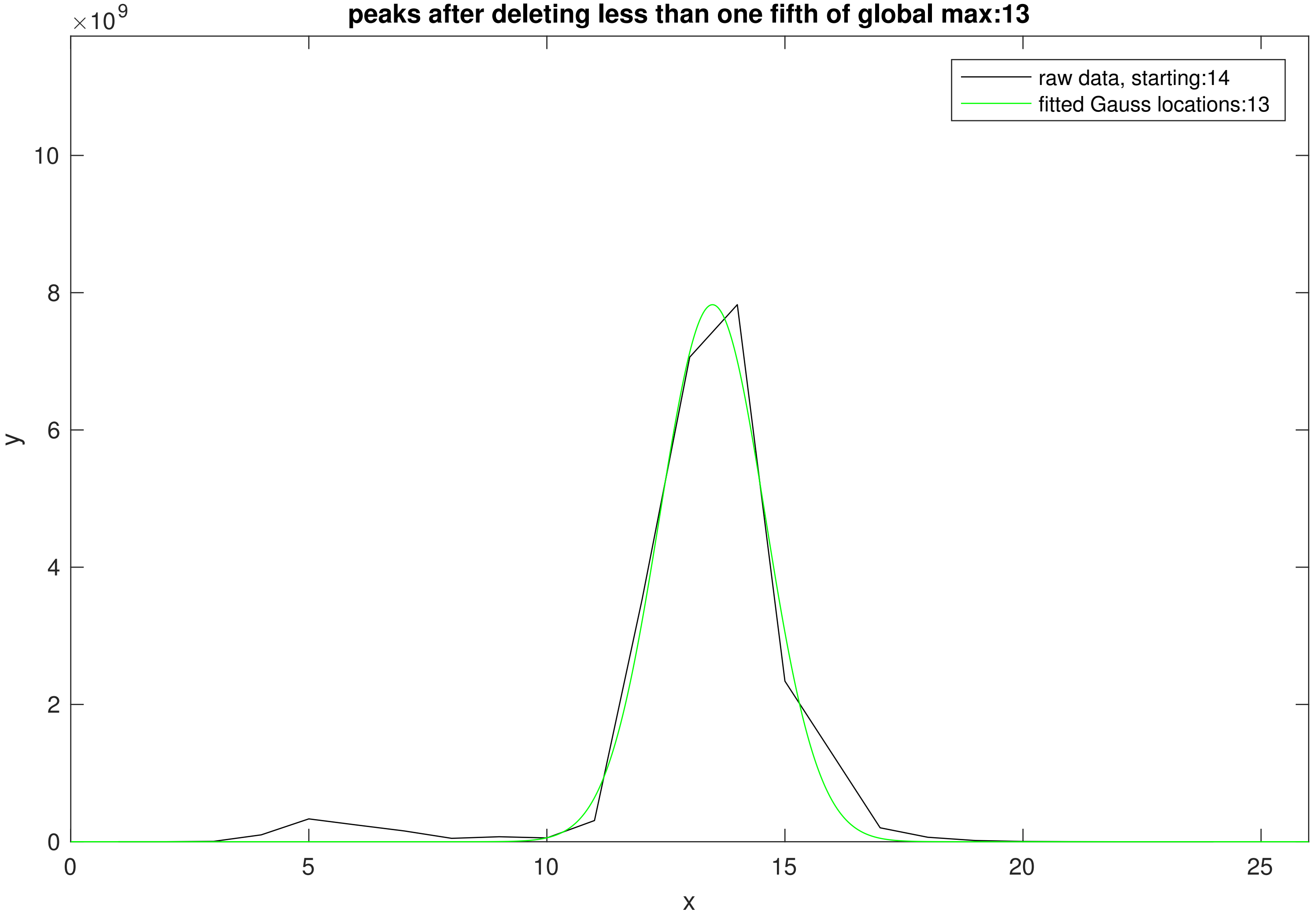

**Optimal1 Fitted Gauss, proteinAT5G10460.1**
**bic:691**
**peaks after deleting less than one fifth of global max:18**

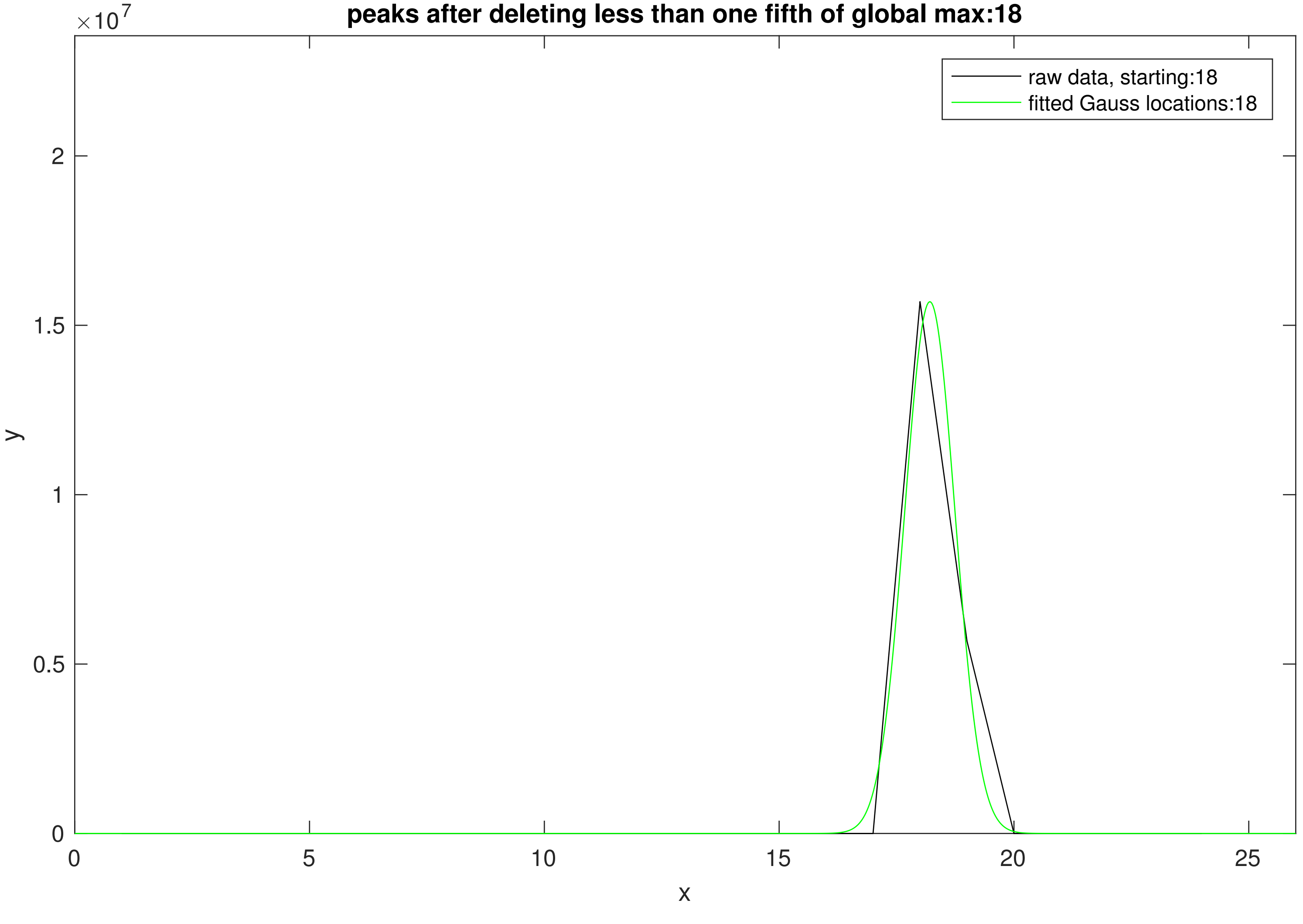

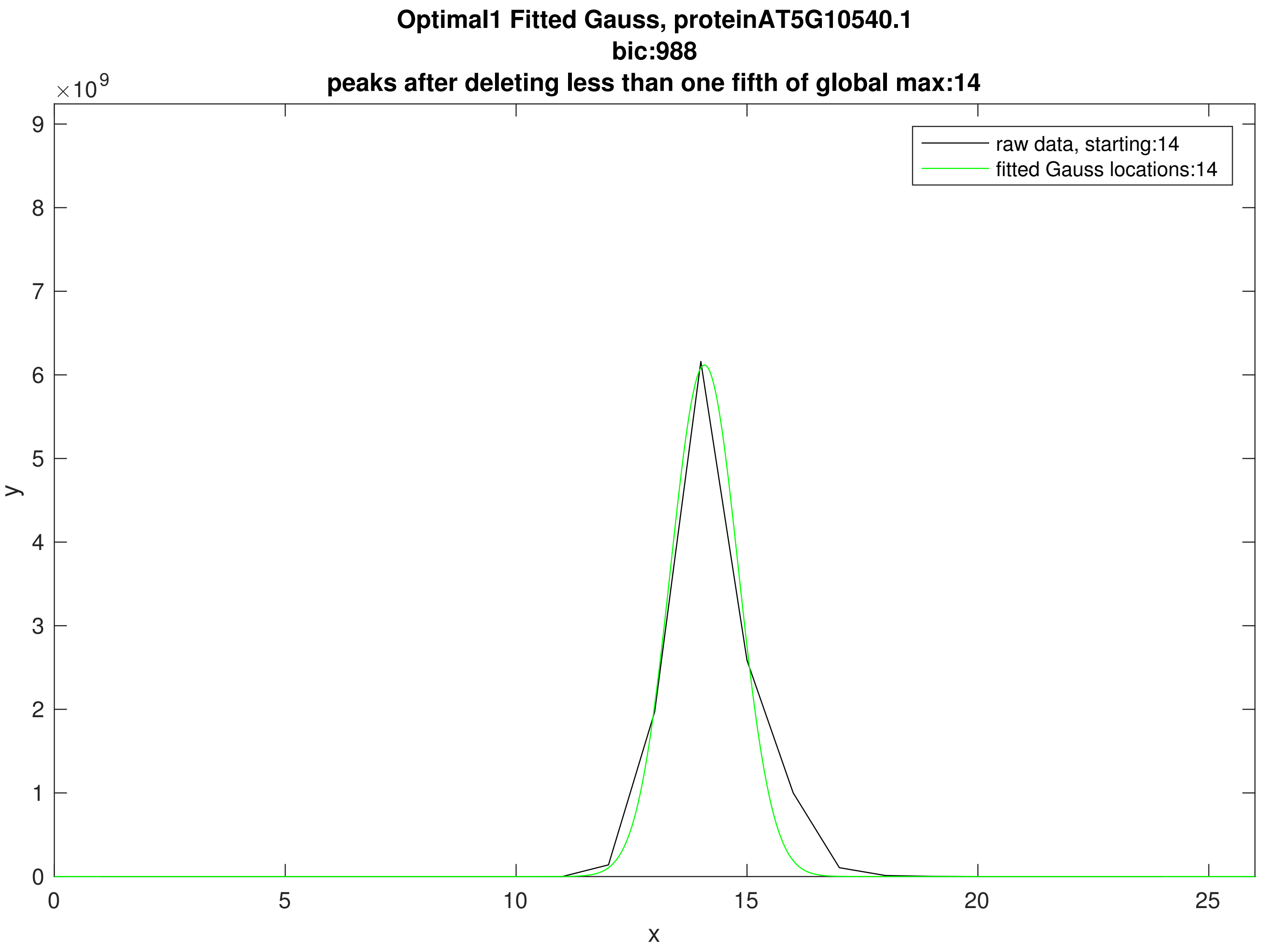
Optimal1 Fitted Gauss, proteinAT5G10540.1
bic:988
peaks after deleting less than one fifth of global max:14
×10^9
raw data, starting:14
fitted Gauss locations:14
y
x
0
1
2
3
4
5
6
7
8
9
0
5
10
15
20
25

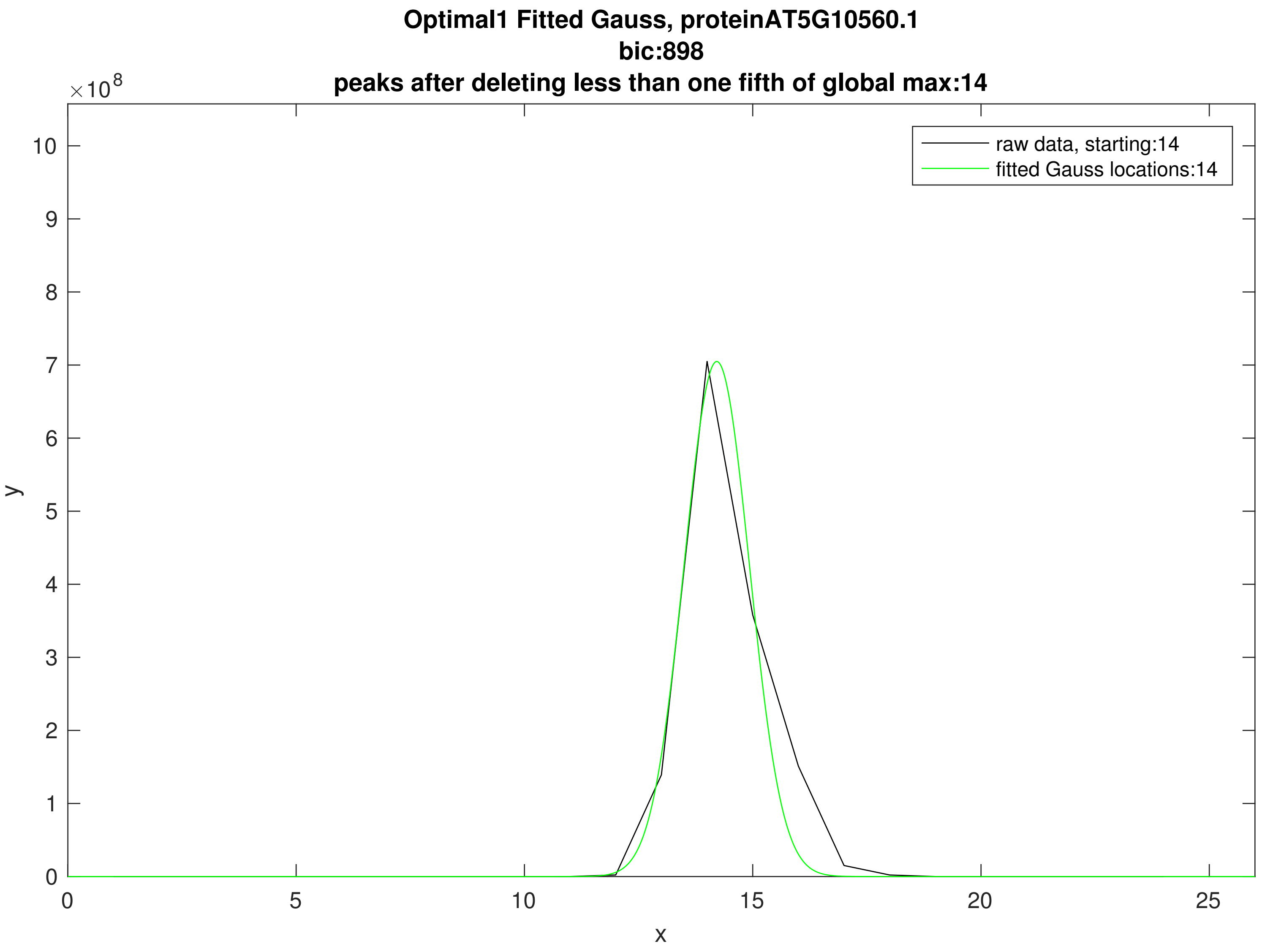
Optimal1 Fitted Gauss, proteinAT5G10560.1
bic:898
peaks after deleting less than one fifth of global max:14
×10^8
raw data, starting:14
fitted Gauss locations:14
y
x
0
1
2
3
4
5
6
7
8
9
10
0
5
10
15
20
25

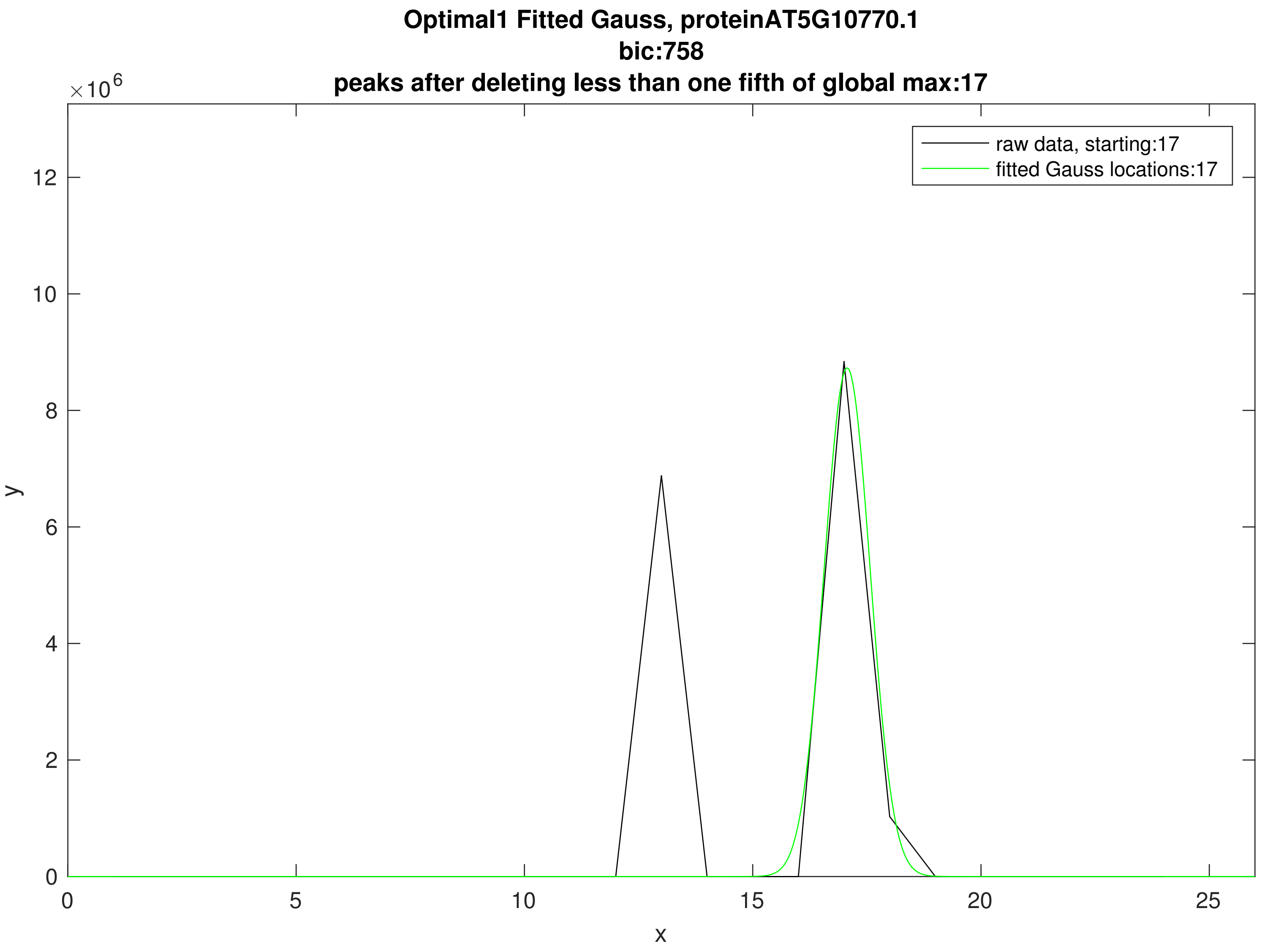
Optimal1 Fitted Gauss, proteinAT5G10770.1
bic:758
peaks after deleting less than one fifth of global max:17
$\times 10^6$
12
10
8
6
4
2
0
y
0
5
10
15
20
25
x
raw data, starting:17
fitted Gauss locations:17

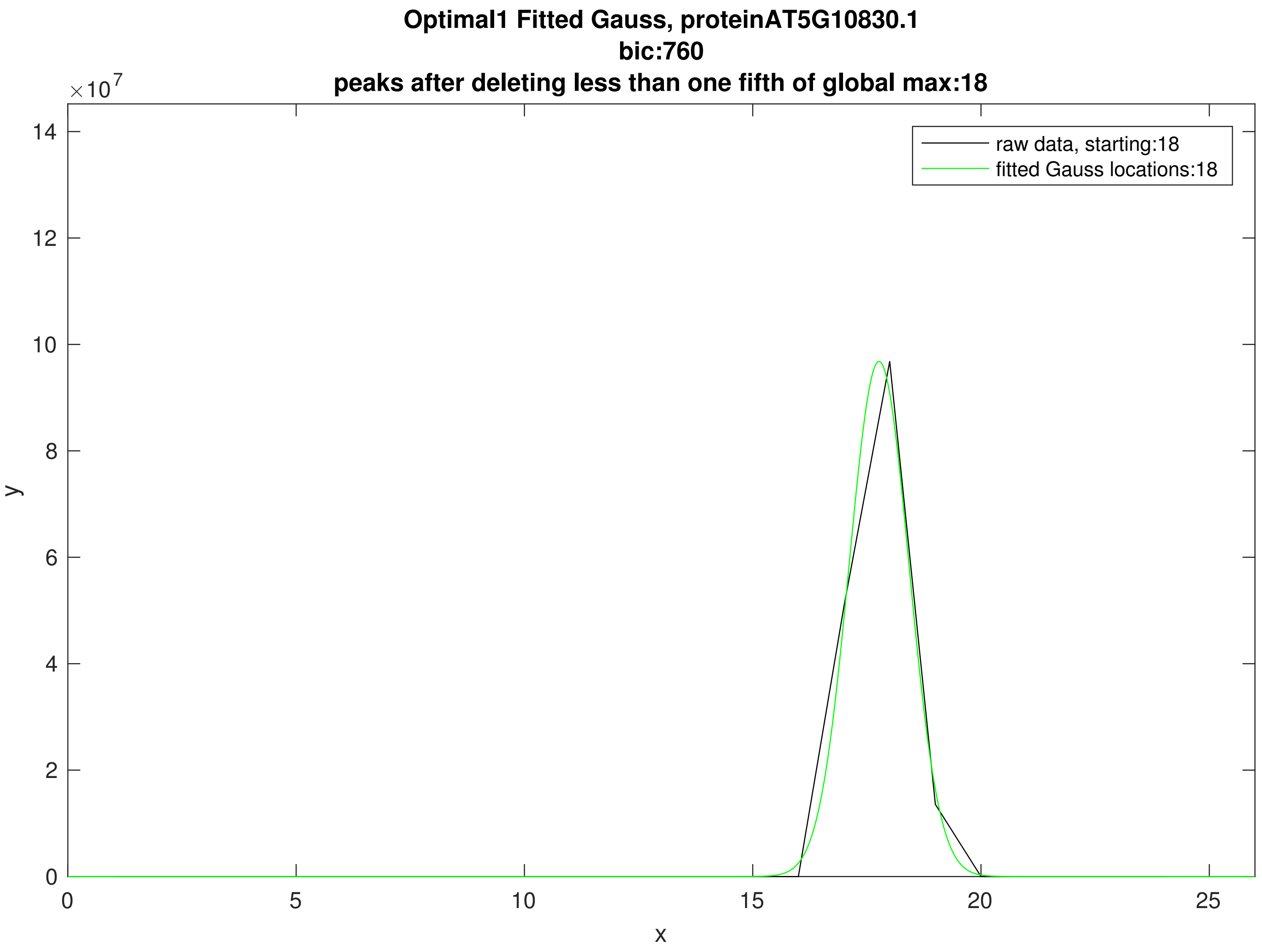
Optimal1 Fitted Gauss, proteinAT5G10830.1
bic:760
peaks after deleting less than one fifth of global max:18
×10^7
14
12
10
8
6
4
2
0
y
0
5
10
15
20
25
x
raw data, starting:18
fitted Gauss locations:18

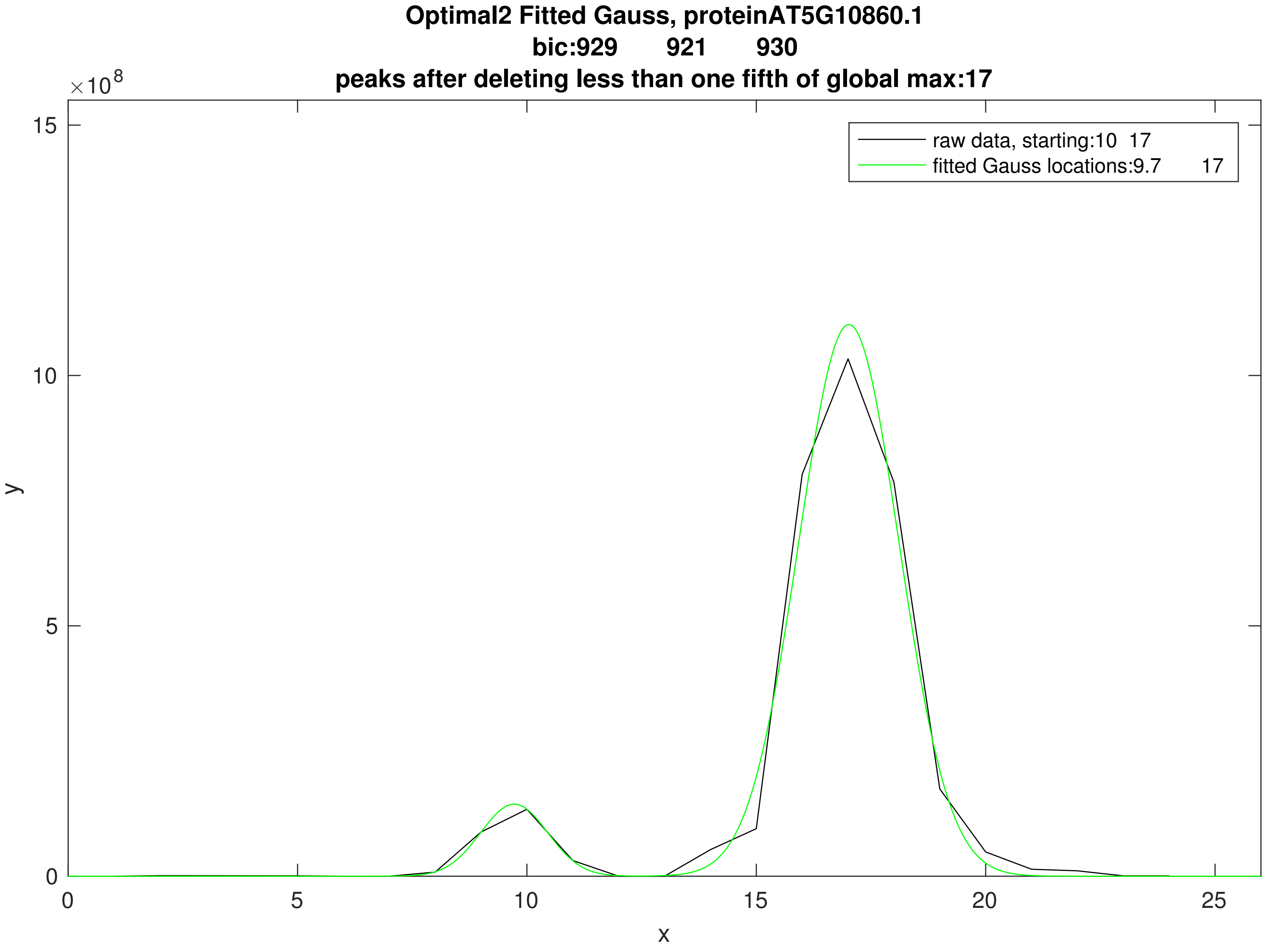
Optimal2 Fitted Gauss, proteinAT5G10860.1
bic:929 921 930
peaks after deleting less than one fifth of global max:17
×10^8
raw data, starting:10 17
fitted Gauss locations:9.7 17
y
x
0
5
10
15
20
25

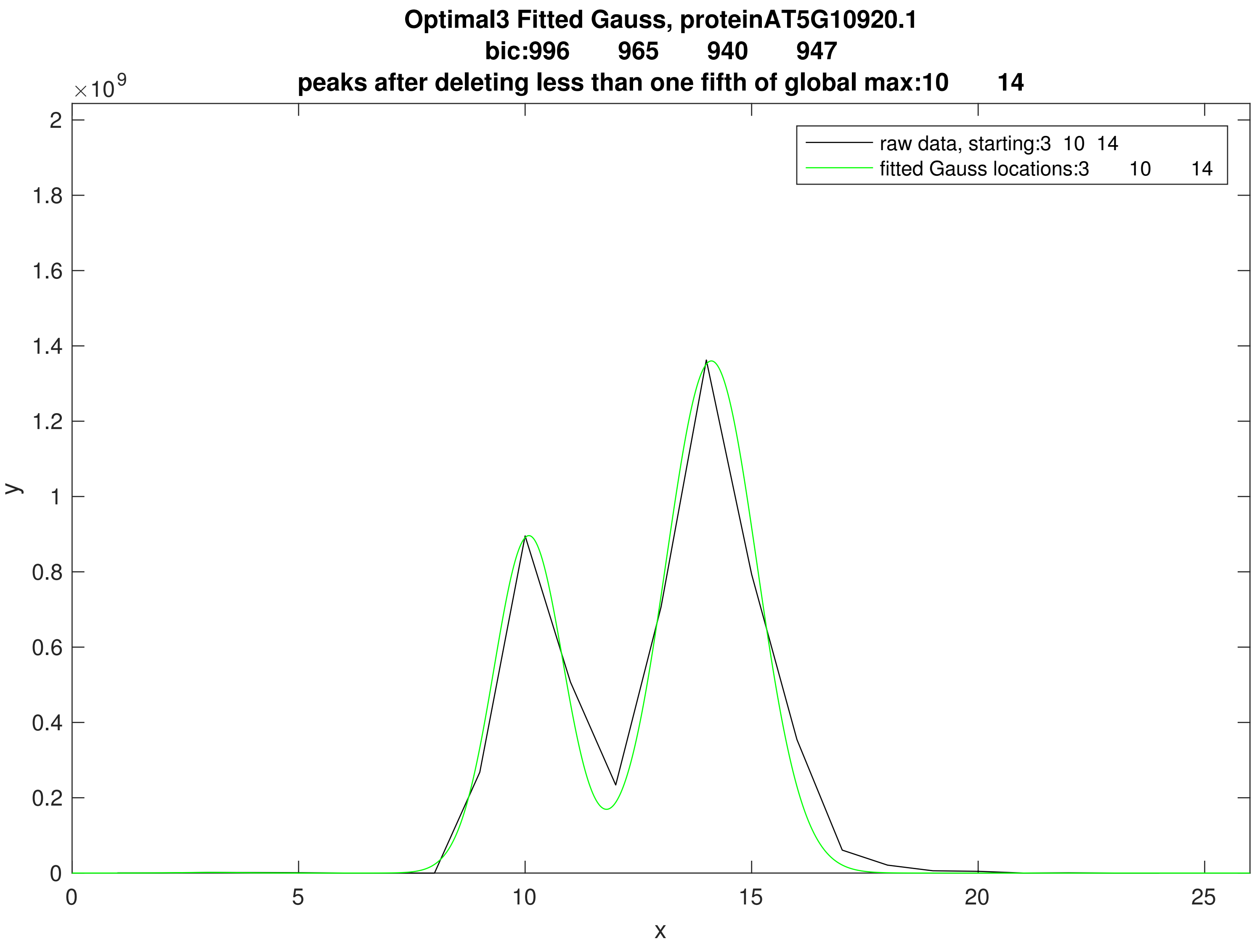
Optimal3 Fitted Gauss, proteinAT5G10920.1
bic:996 965 940 947
peaks after deleting less than one fifth of global max:10 14
×10^9
raw data, starting:3 10 14
fitted Gauss locations:3 10 14
y
x
0
0.2
0.4
0.6
0.8
1
1.2
1.4
1.6
1.8
2
5
10
15
20
25

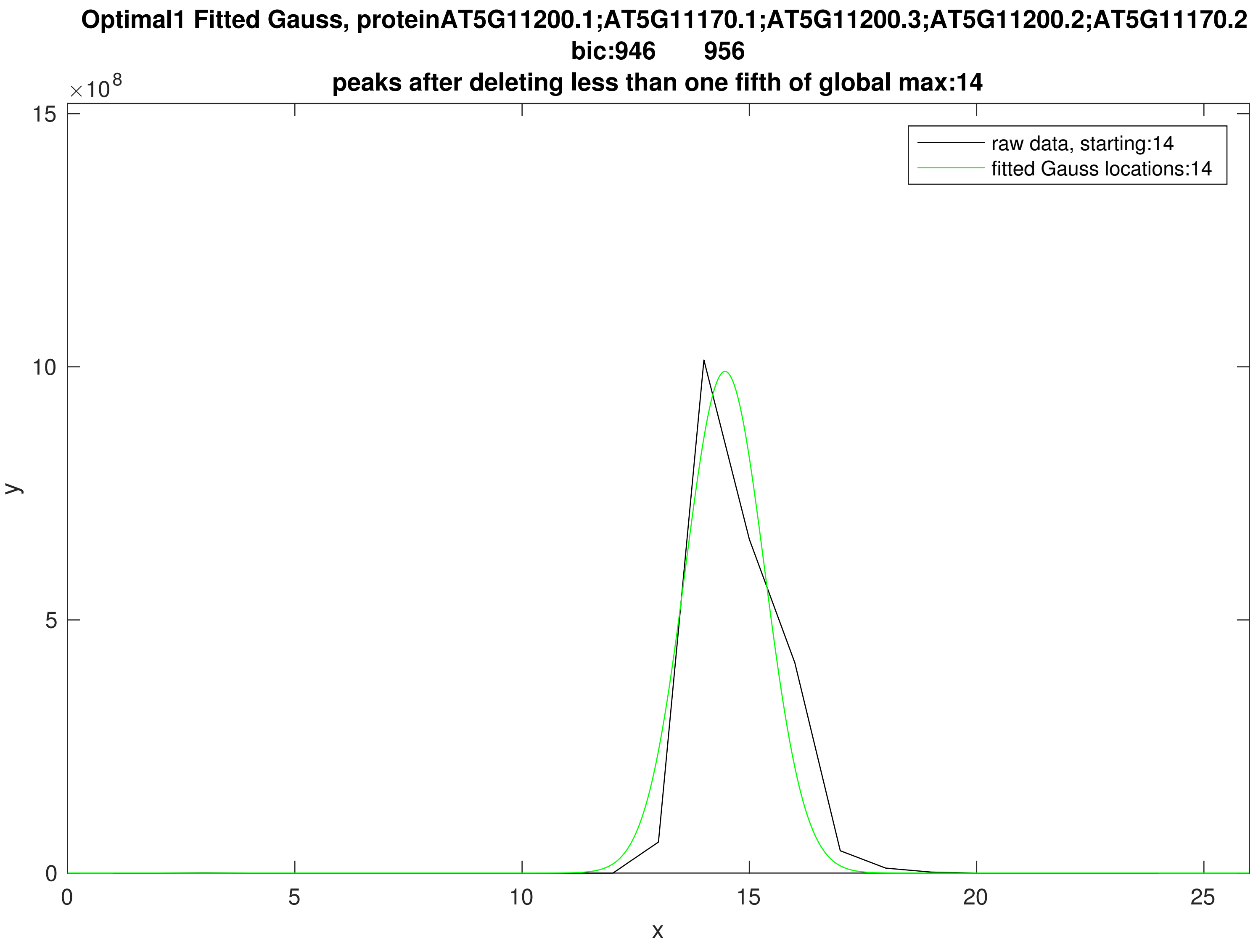
Optimal1 Fitted Gauss, proteinAT5G11200.1;AT5G11170.1;AT5G11200.3;AT5G11200.2;AT5G11170.2
bic:946 956
peaks after deleting less than one fifth of global max:14
×10^8
15
10
5
0
0
5
10
15
20
25
x
y
raw data, starting:14
fitted Gauss locations:14

**Optimal1 Fitted Gauss, proteinAT5G11330.1;AT5G11330.2**
**bic:747**
**peaks after deleting less than one fifth of global max:16**

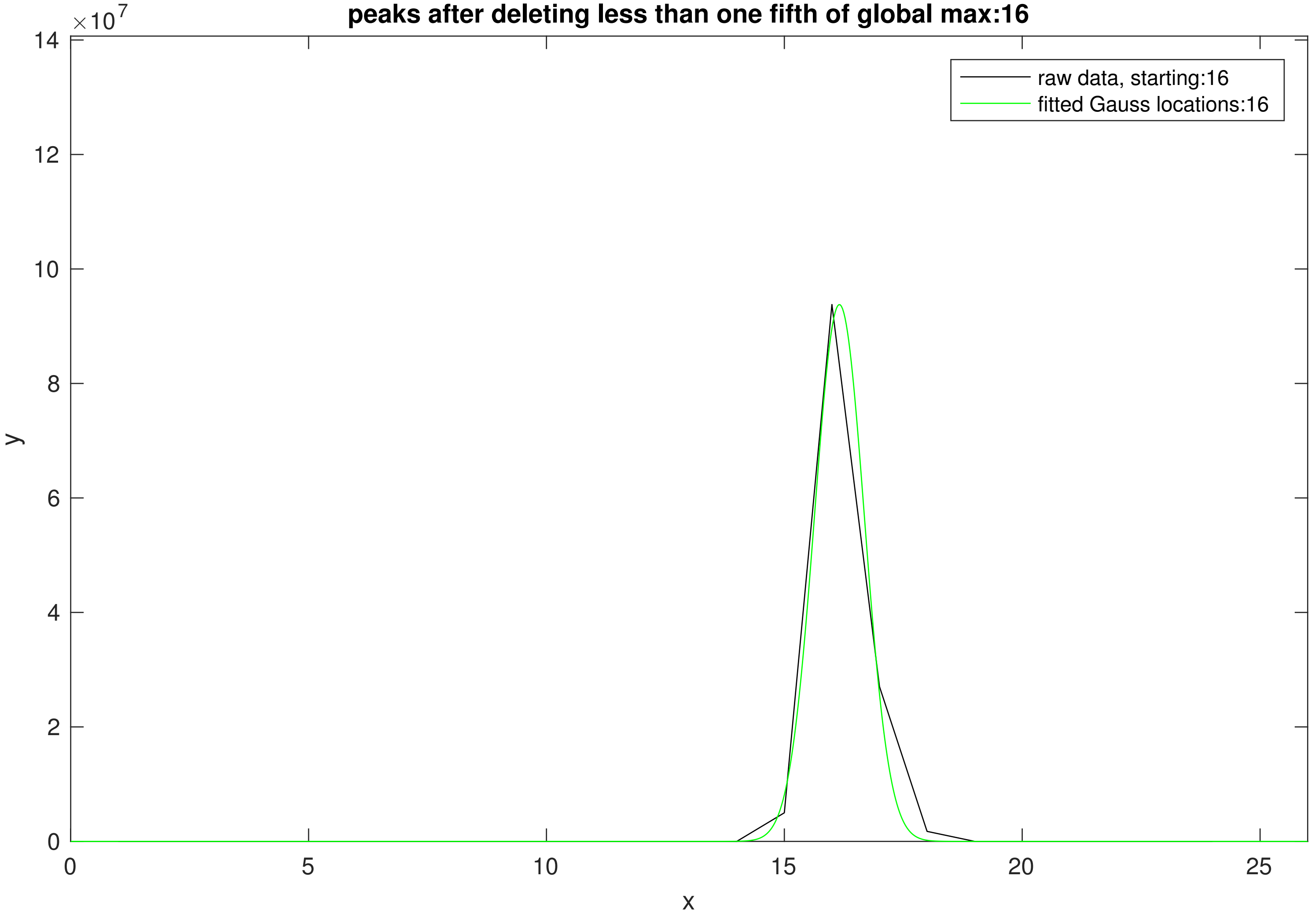

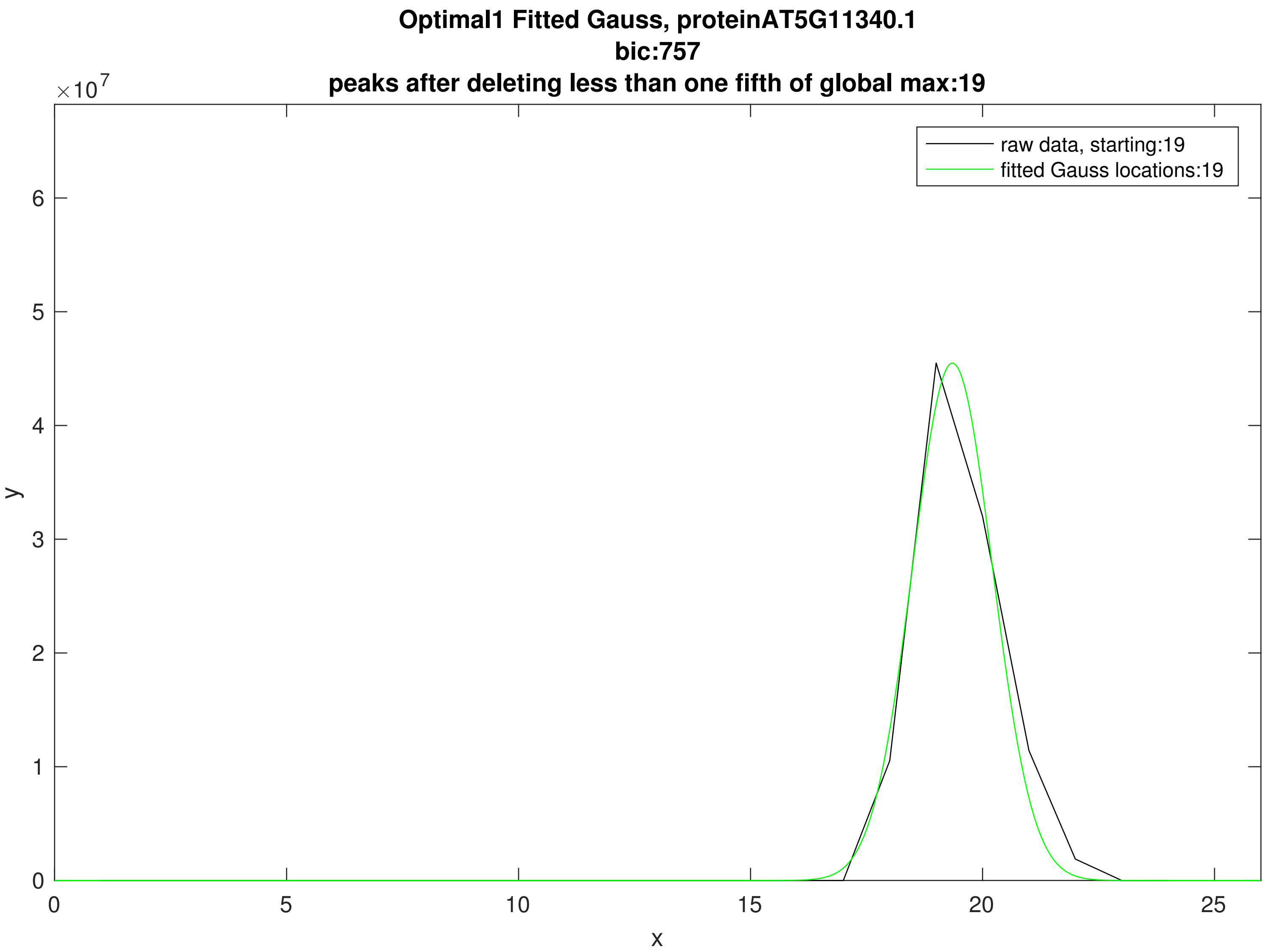
Optimal1 Fitted Gauss, proteinAT5G11340.1
bic:757
peaks after deleting less than one fifth of global max:19
×10^7
raw data, starting:19
fitted Gauss locations:19
y
x
0
1
2
3
4
5
6
0
5
10
15
20
25

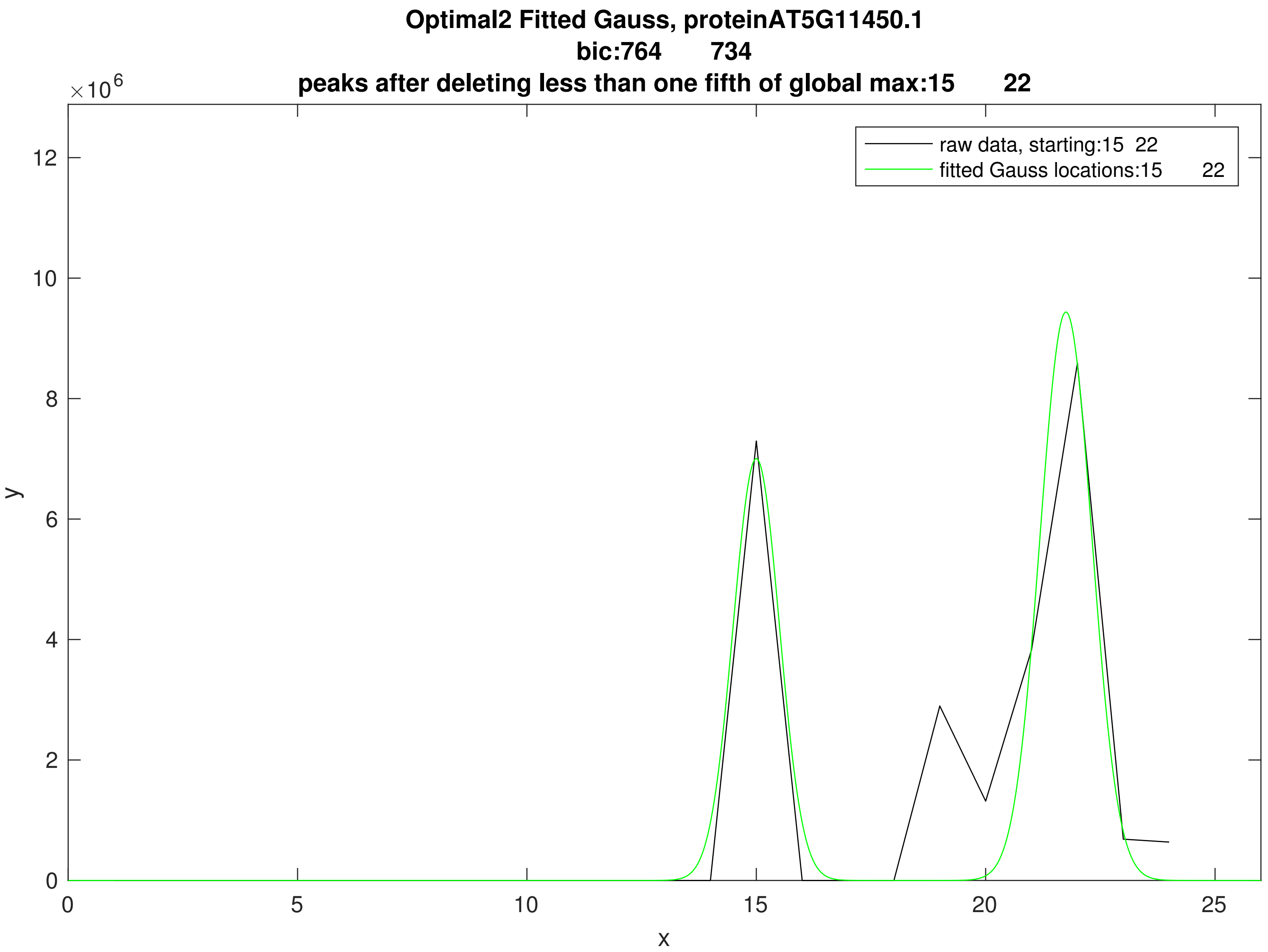

Optimal2 Fitted Gauss, proteinAT5G11450.1
bic:764 734
peaks after deleting less than one fifth of global max:15 22
×10^6
raw data, starting:15 22
fitted Gauss locations:15 22
12
10
8
6
4
2
0
y
0
5
10
15
20
25
x

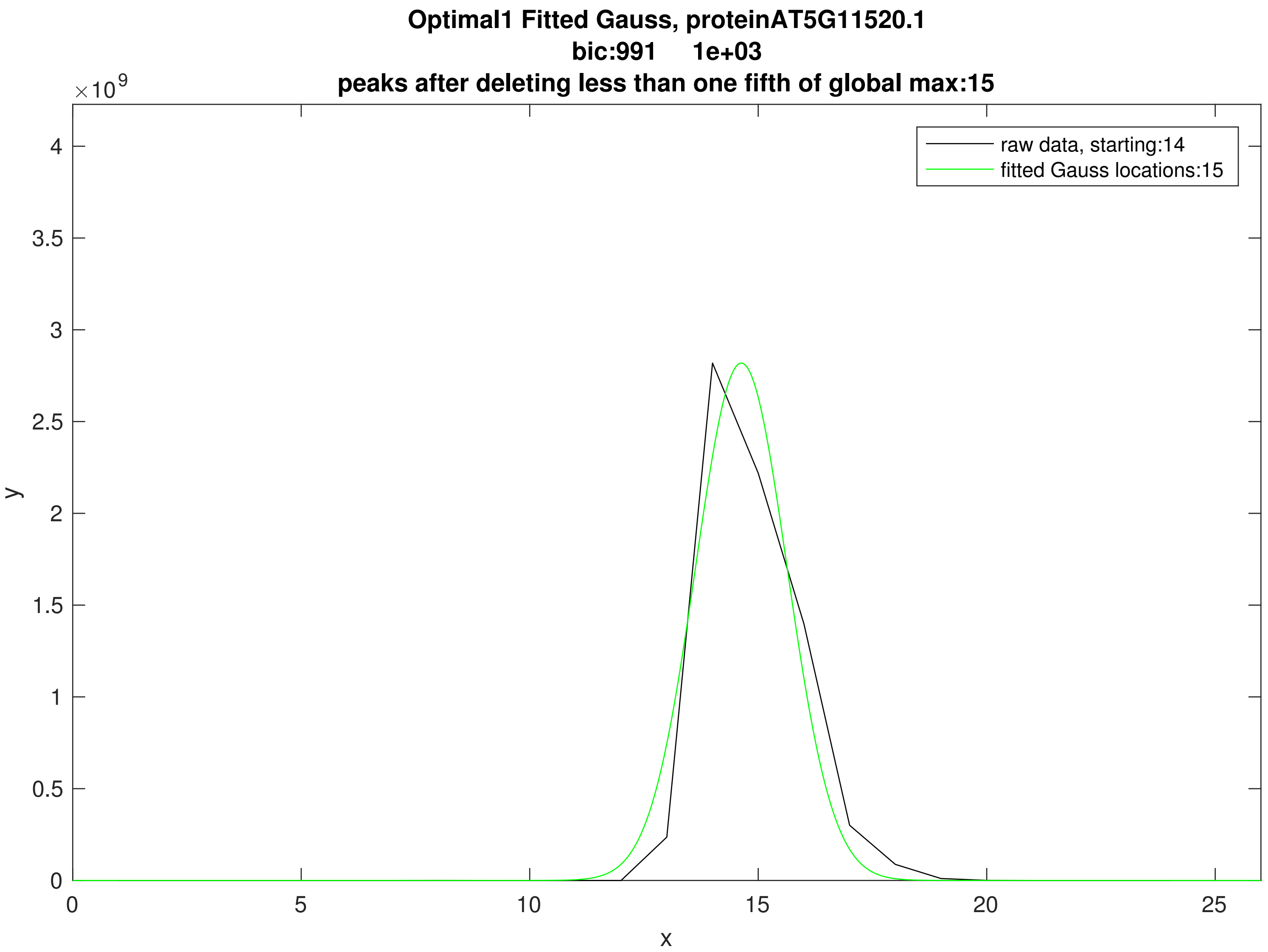
Optimal1 Fitted Gauss, proteinAT5G11520.1
bic:991 1e+03
peaks after deleting less than one fifth of global max:15
×10^9
raw data, starting:14
fitted Gauss locations:15
y
x
0
0.5
1
1.5
2
2.5
3
3.5
4
0
5
10
15
20
25

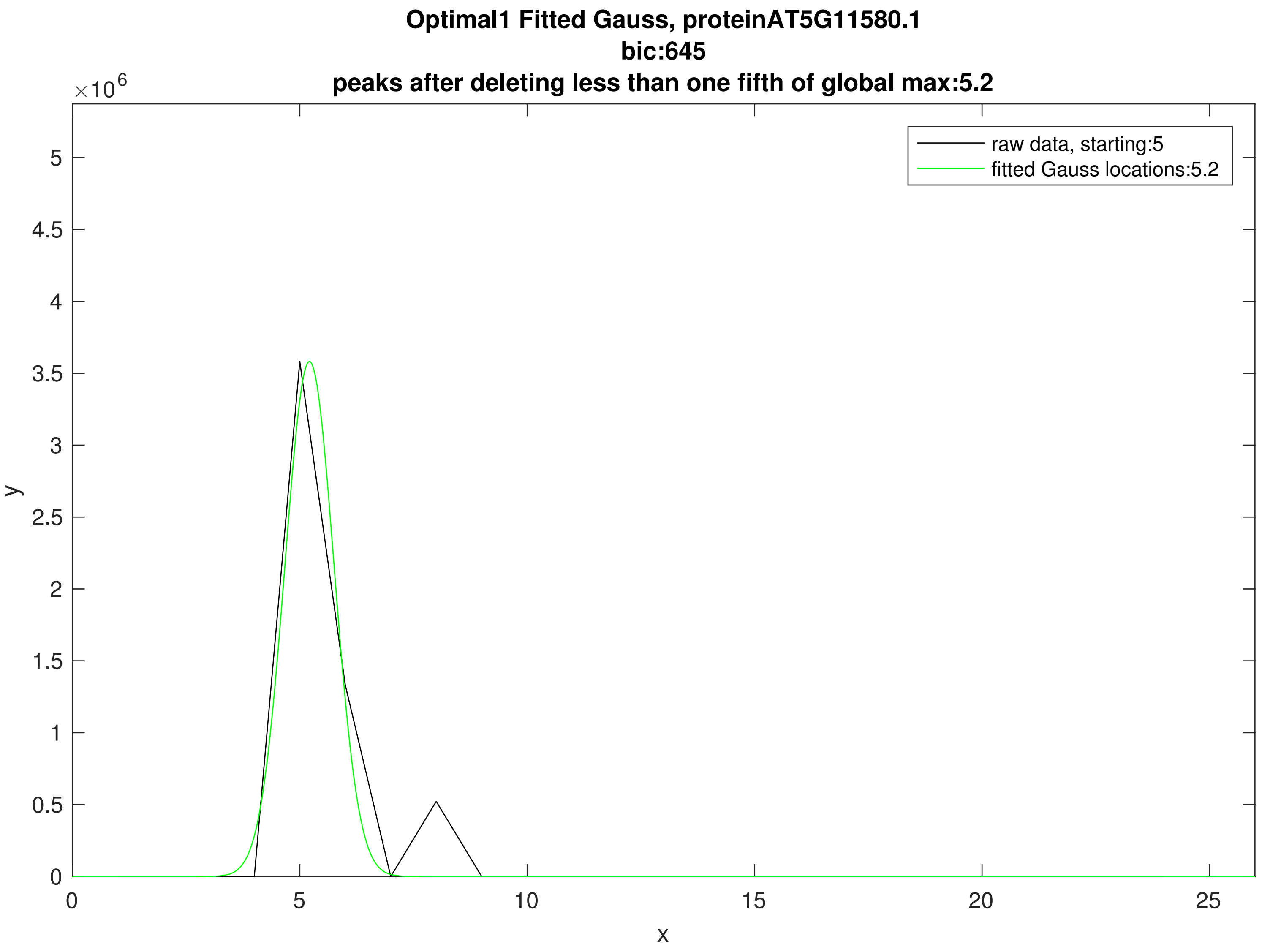

Optimal1 Fitted Gauss, proteinAT5G11580.1
bic:645
peaks after deleting less than one fifth of global max:5.2
×10^6
raw data, starting:5
fitted Gauss locations:5.2
y
x
0
0.5
1
1.5
2
2.5
3
3.5
4
4.5
5
0
5
10
15
20
25

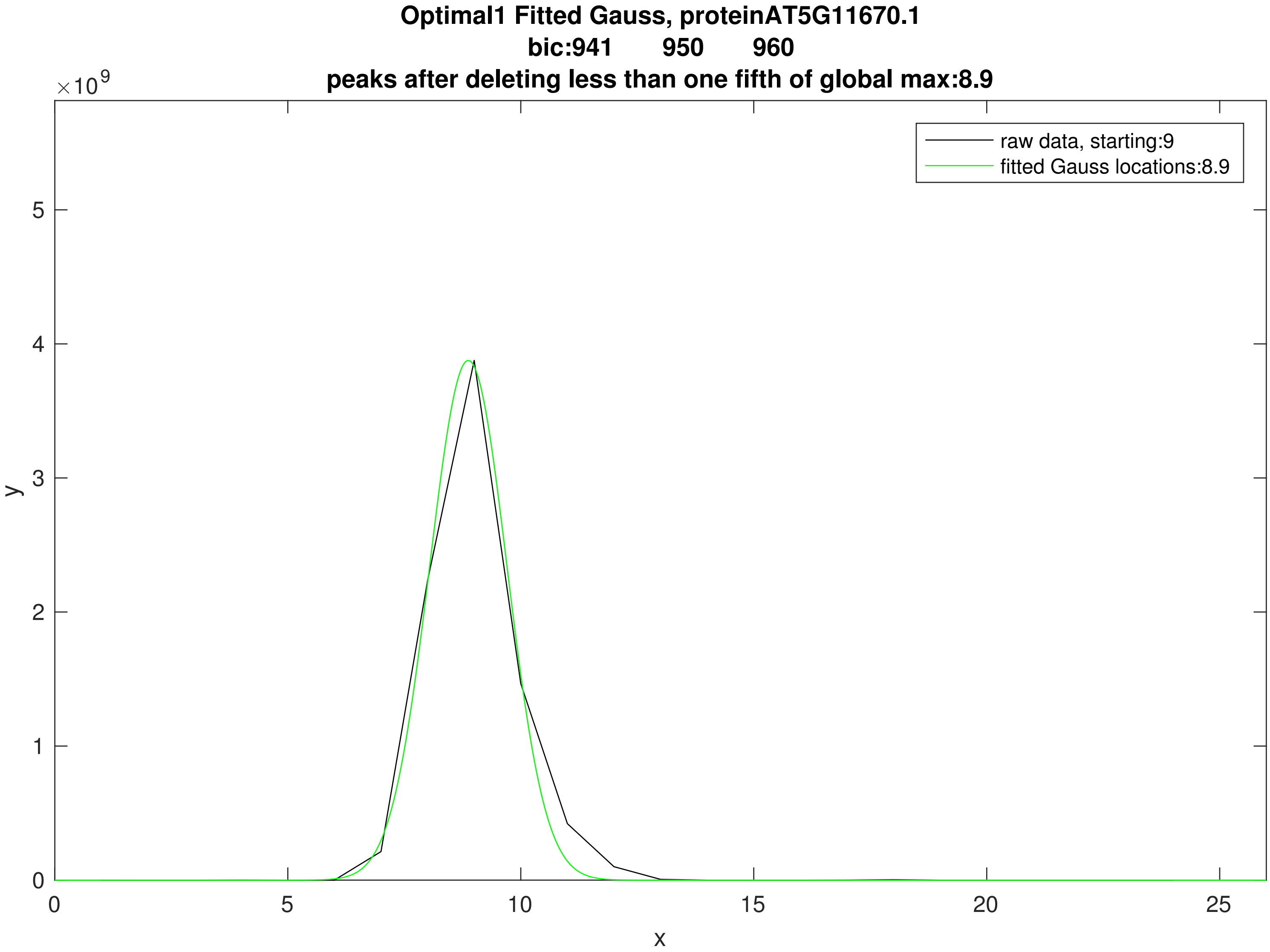
Optimal1 Fitted Gauss, proteinAT5G11670.1
bic:941 950 960
peaks after deleting less than one fifth of global max:8.9
×10^9
raw data, starting:9
fitted Gauss locations:8.9
y
x
0
1
2
3
4
5
0
5
10
15
20
25

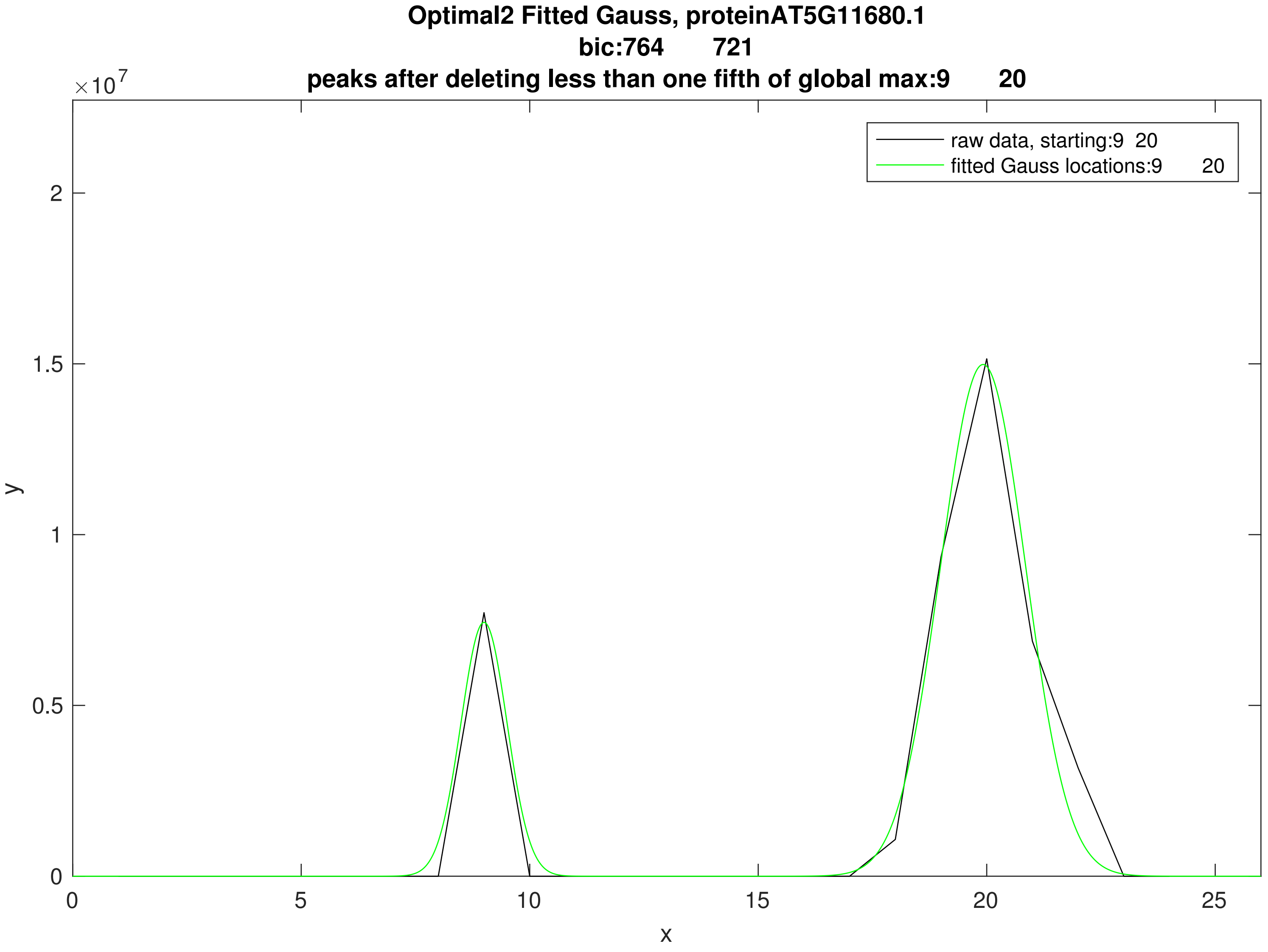
Optimal2 Fitted Gauss, proteinAT5G11680.1
bic:764 721
peaks after deleting less than one fifth of global max:9 20
×10^7
raw data, starting:9 20
fitted Gauss locations:9 20
2
1.5
1
0.5
0
y
0
5
10
15
20
25
x

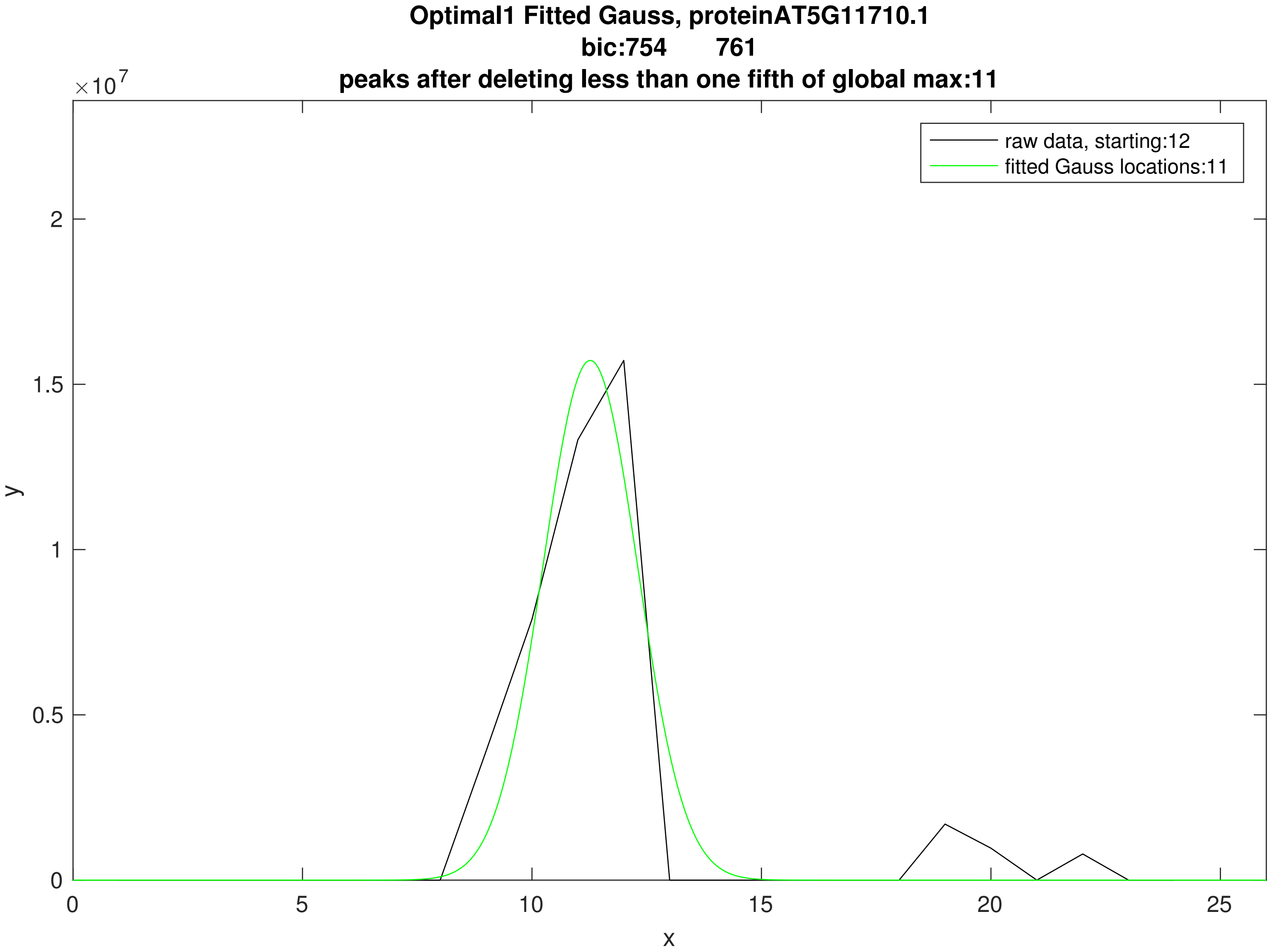
Optimal1 Fitted Gauss, proteinAT5G11710.1
bic:754 761
peaks after deleting less than one fifth of global max:11
×10^7
raw data, starting:12
fitted Gauss locations:11
2
1.5
1
0.5
0
y
0
5
10
15
20
25
x

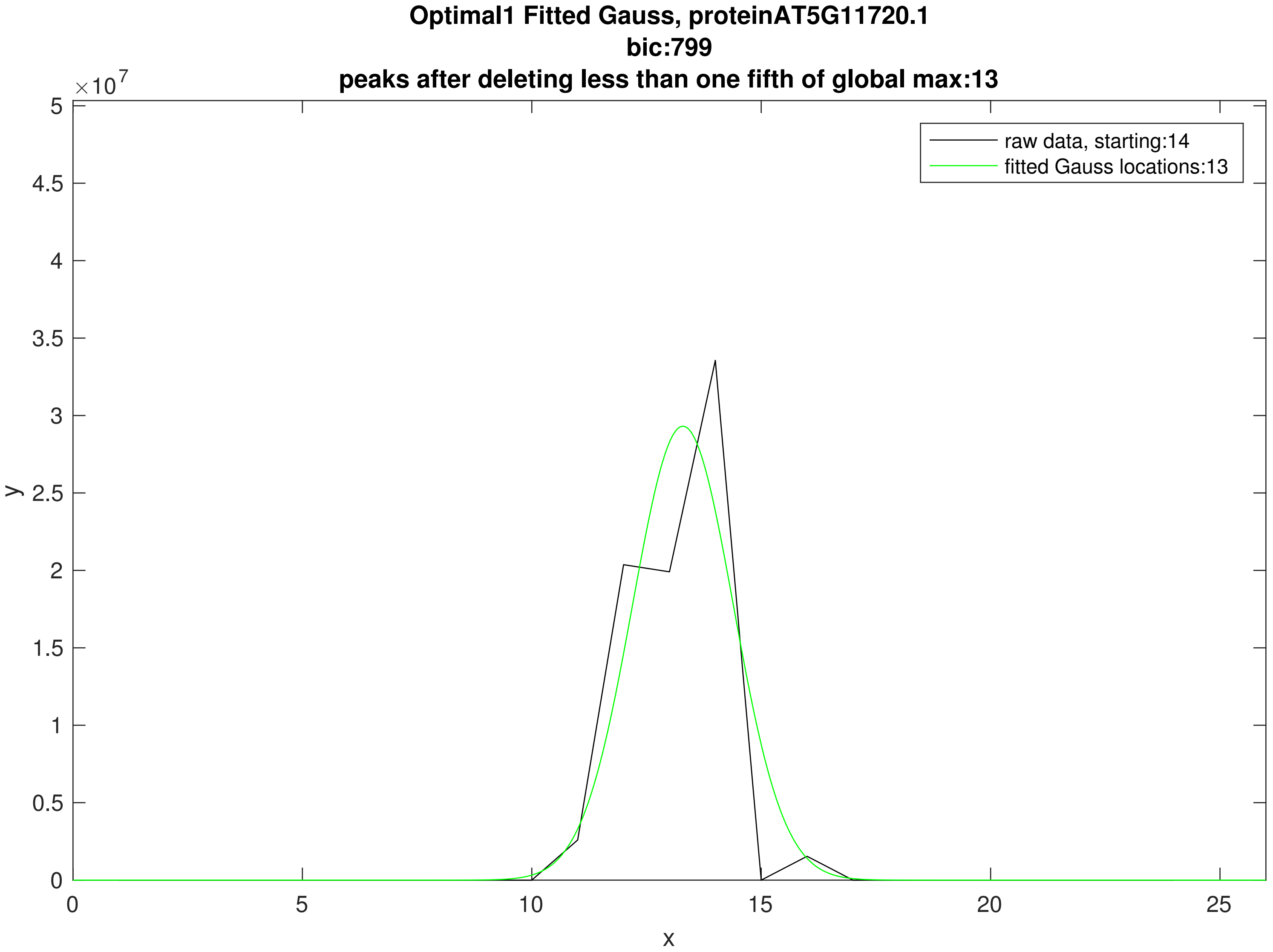

Optimal1 Fitted Gauss, proteinAT5G11720.1
bic:799
peaks after deleting less than one fifth of global max:13
×10^7
5
4.5
4
3.5
3
2.5
2
1.5
1
0.5
0
y
0
5
10
15
20
25
x
raw data, starting:14
fitted Gauss locations:13

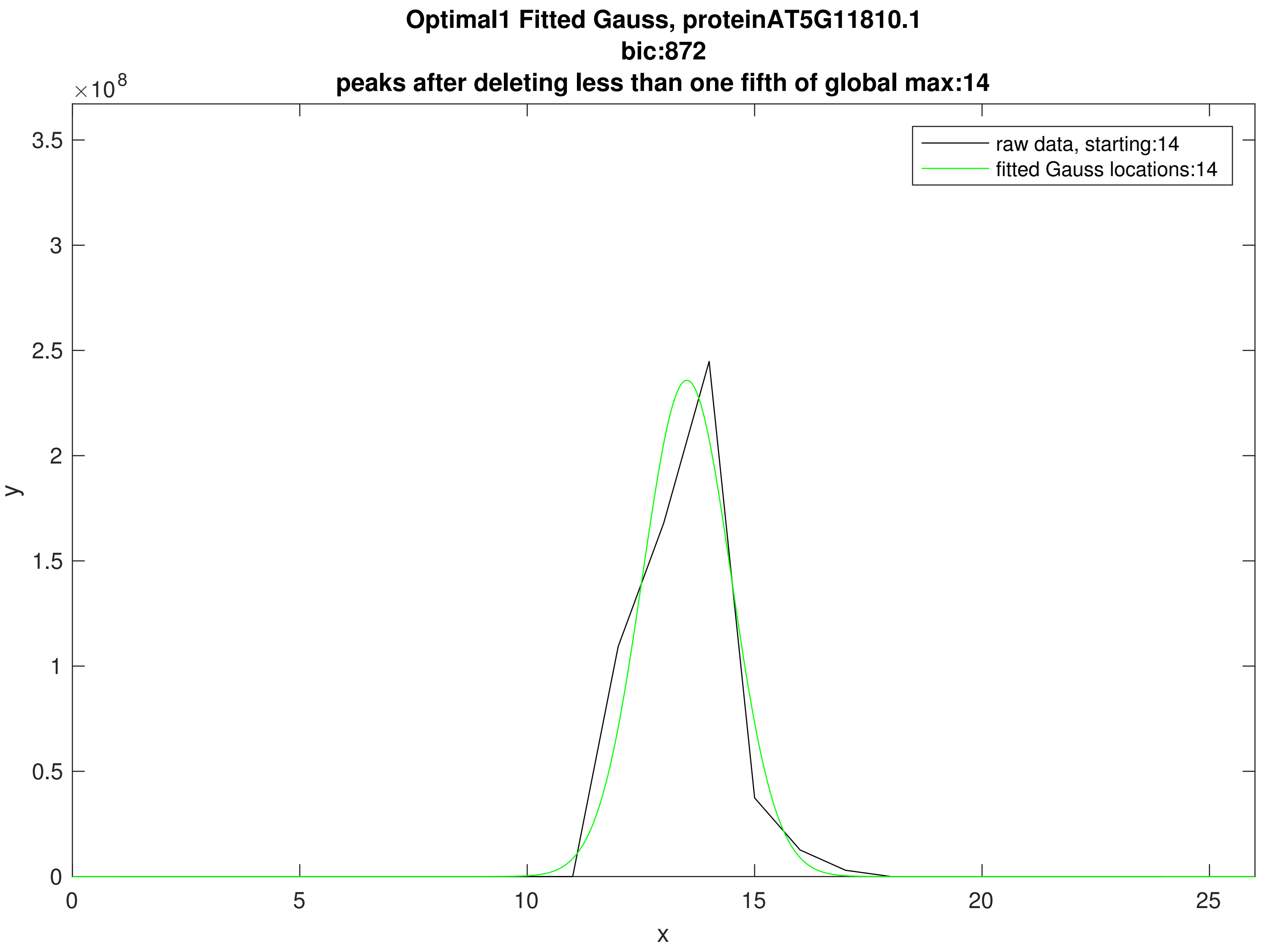

Optimal1 Fitted Gauss, proteinAT5G11810.1
bic:872
peaks after deleting less than one fifth of global max:14
×10^8
raw data, starting:14
fitted Gauss locations:14
3.5
3
2.5
2
1.5
1
0.5
0
y
0
5
10
15
20
25
x

**Optimal1 Fitted Gauss, proteinAT5G11880.1**
**bic:948**
**peaks after deleting less than one fifth of global max:13**

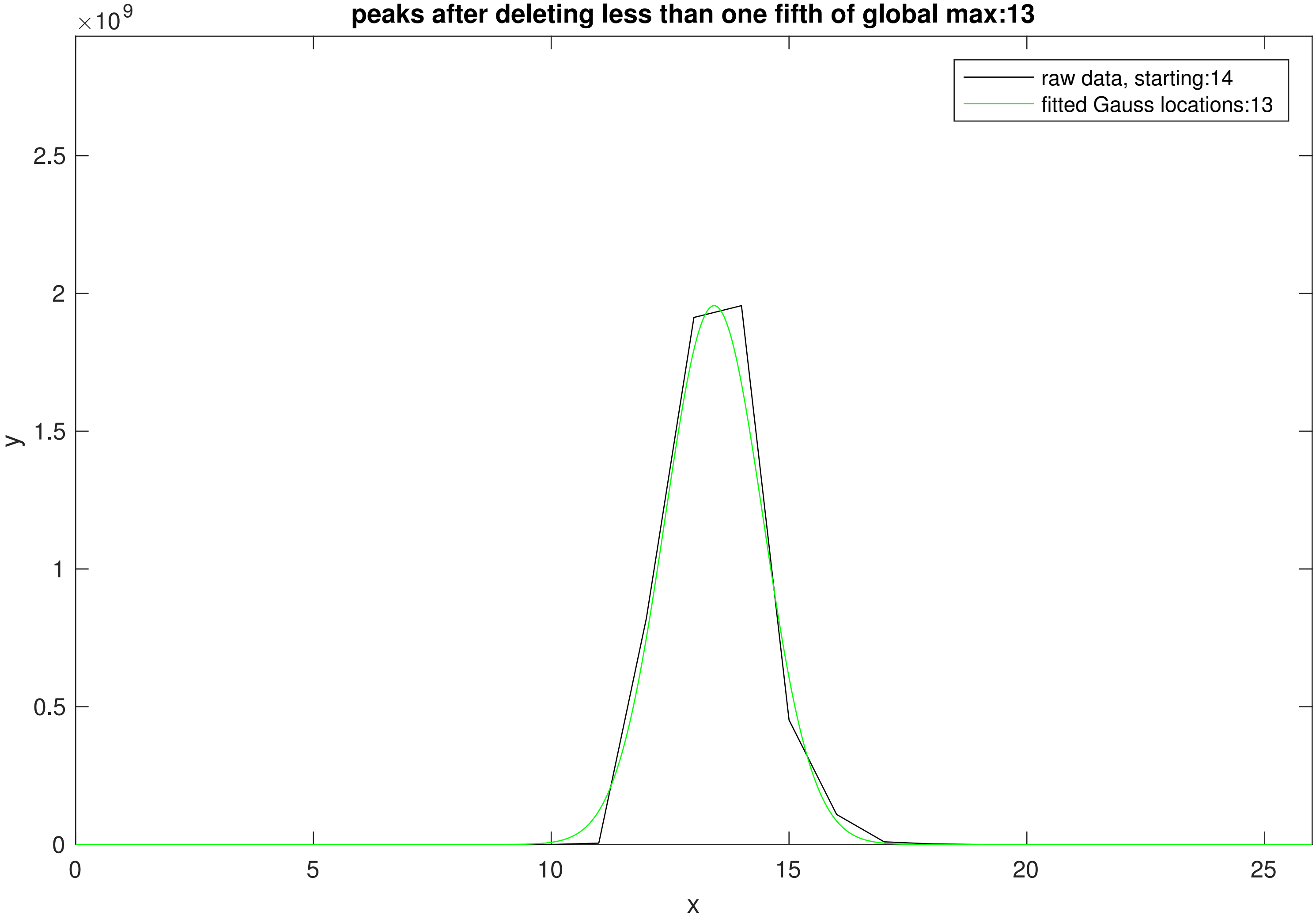

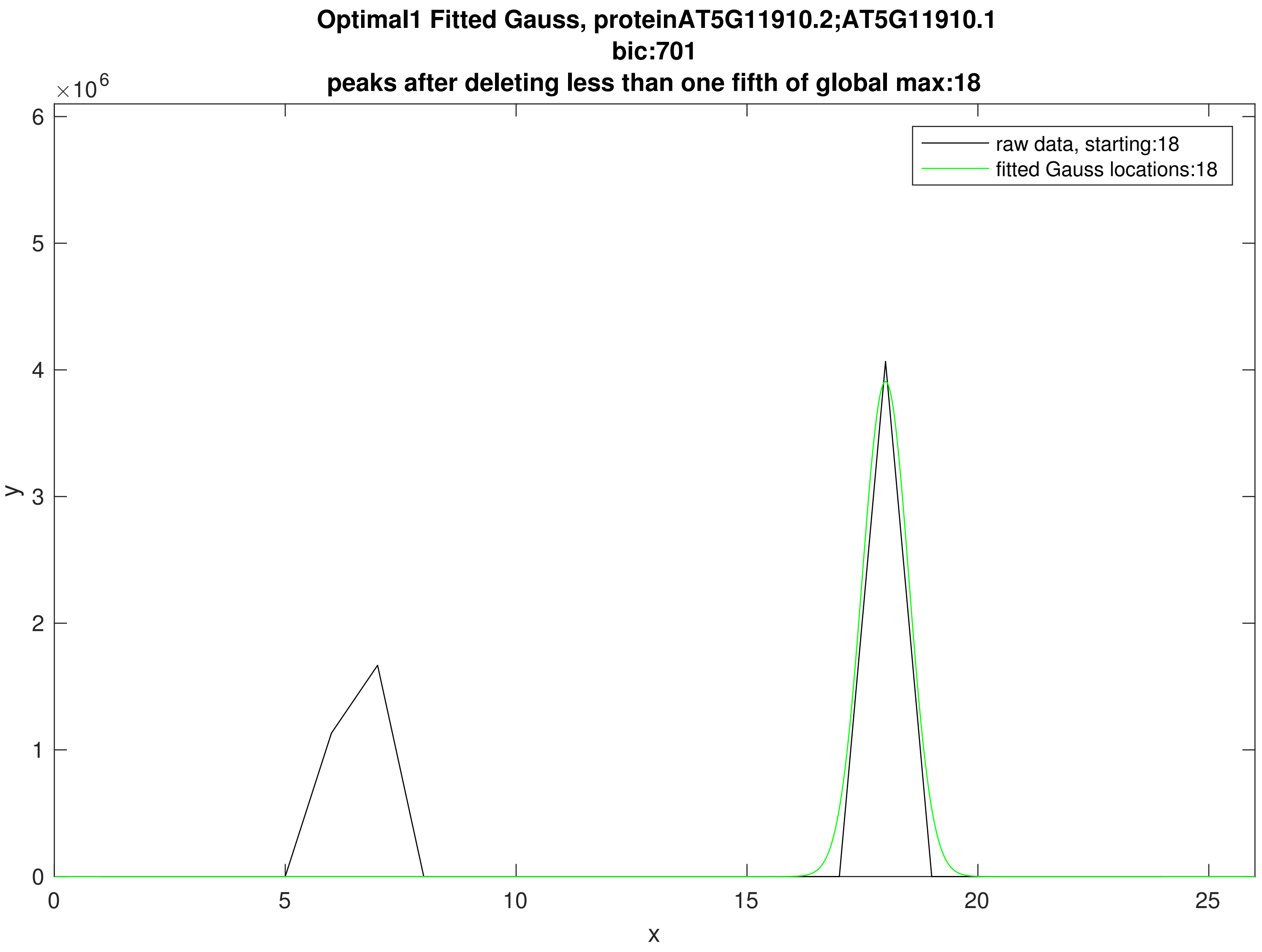
Optimal1 Fitted Gauss, proteinAT5G11910.2;AT5G11910.1
bic:701
peaks after deleting less than one fifth of global max:18
×10^6
y
x
raw data, starting:18
fitted Gauss locations:18

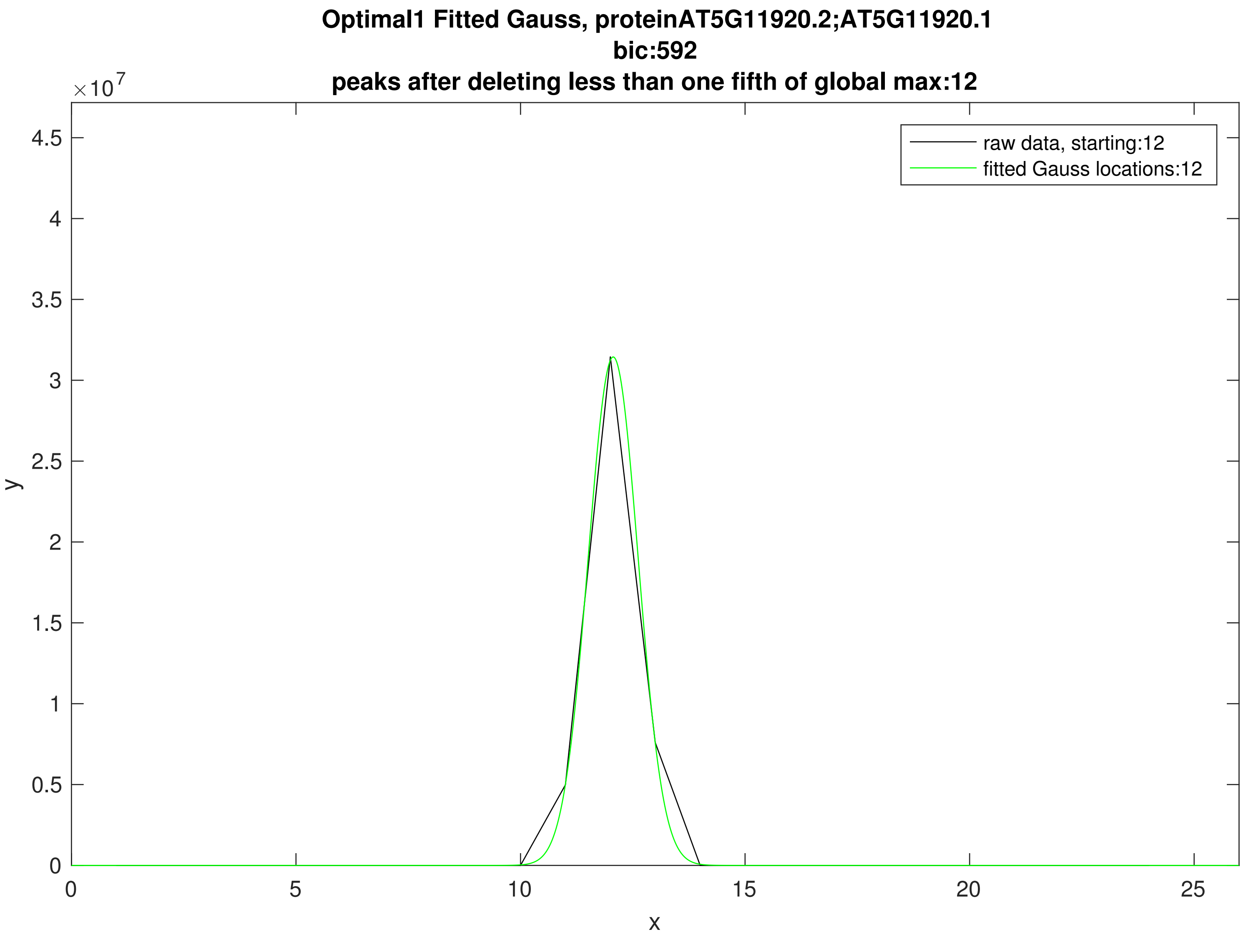

Optimal1 Fitted Gauss, proteinAT5G11920.2;AT5G11920.1
bic:592
peaks after deleting less than one fifth of global max:12
×10^7
4.5
4
3.5
3
2.5
2
1.5
1
0.5
0
y
0
5
10
15
20
25
x
raw data, starting:12
fitted Gauss locations:12

**Optimal1 Fitted Gauss, proteinAT5G11950.2;AT5G11950.1**
**bic:751**
**peaks after deleting less than one fifth of global max:17**

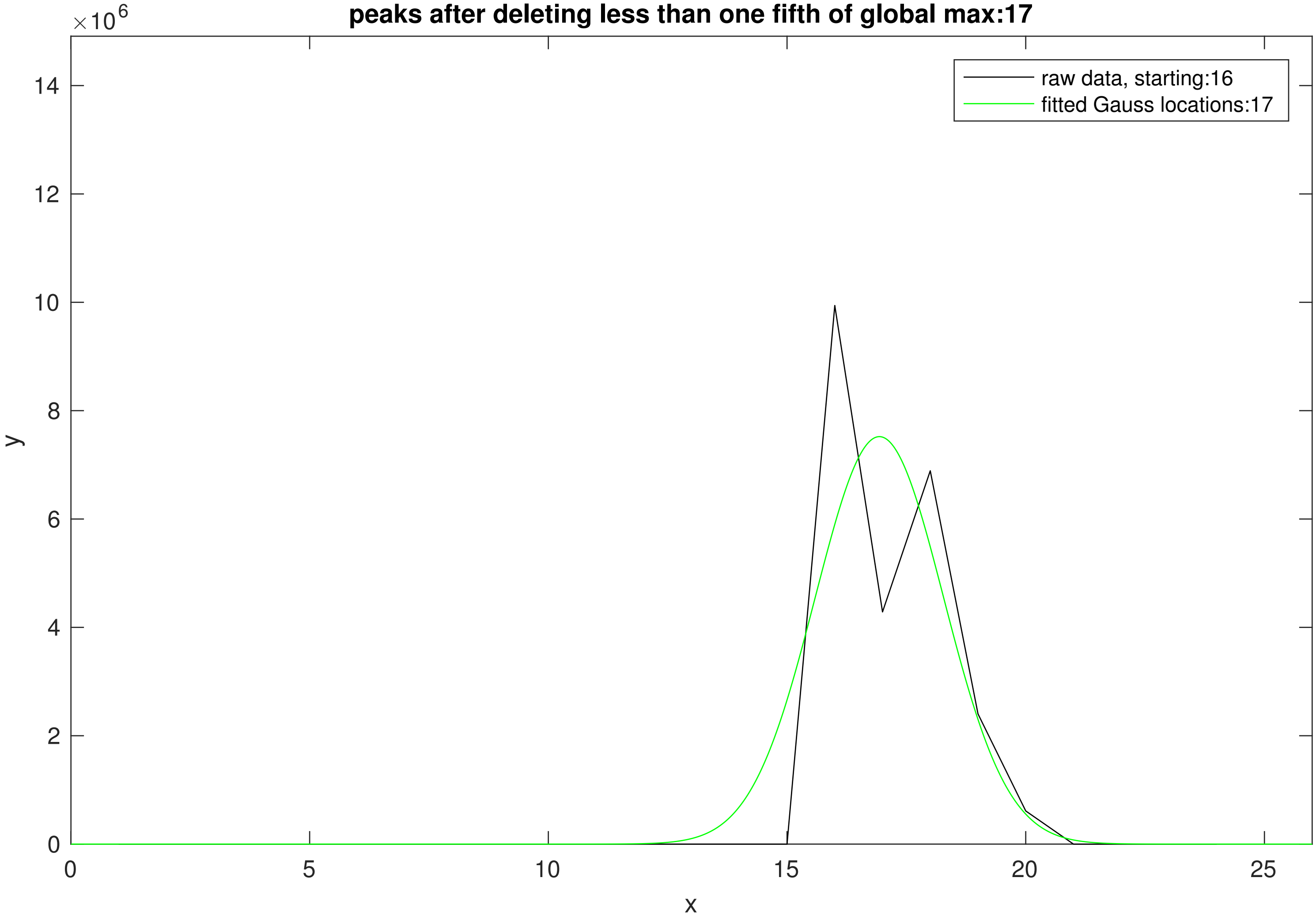

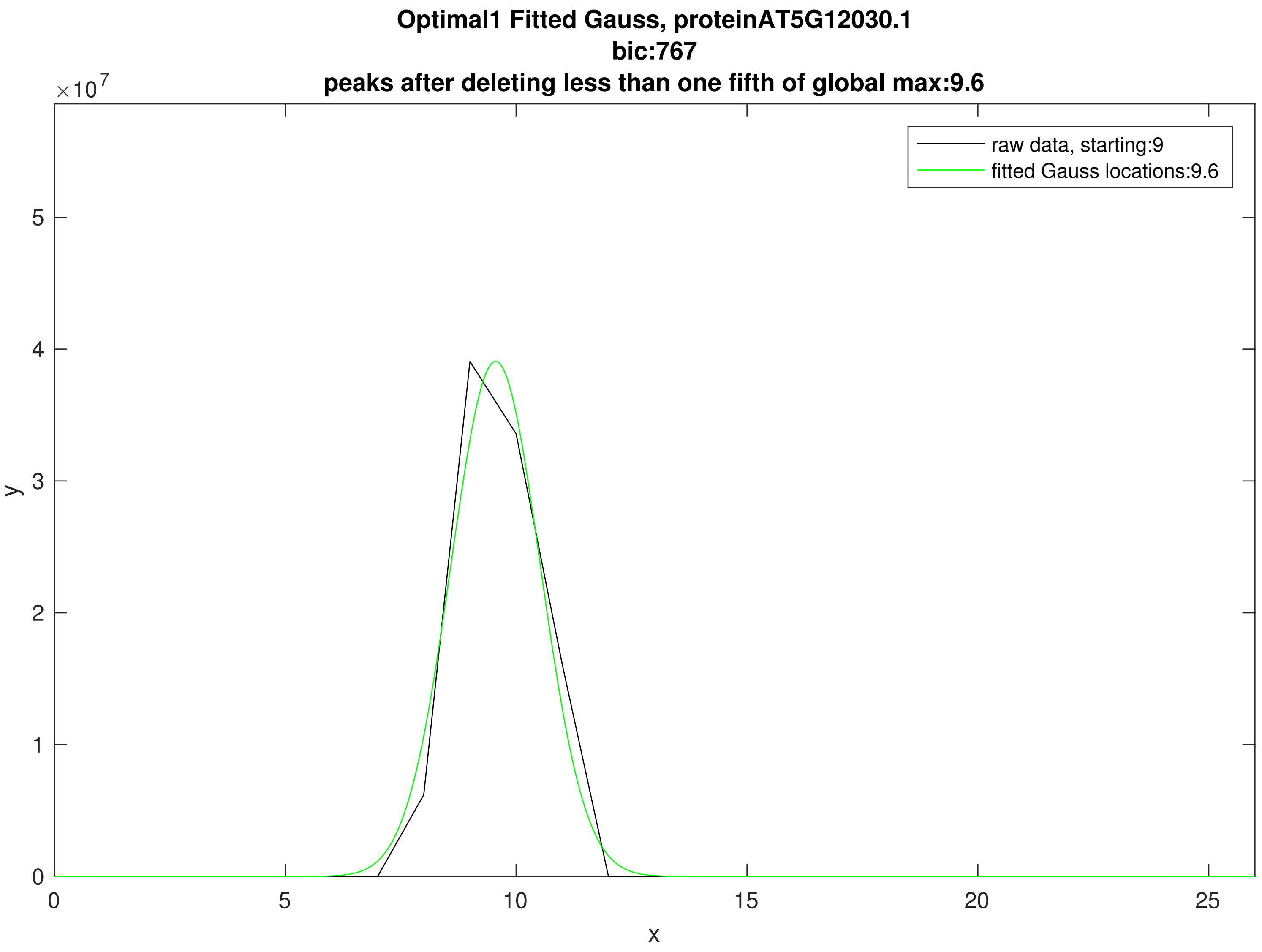

Optimal1 Fitted Gauss, proteinAT5G12030.1
bic:767
peaks after deleting less than one fifth of global max:9.6
×10^7
raw data, starting:9
fitted Gauss locations:9.6
y
0
1
2
3
4
5
0
5
10
15
20
25
x

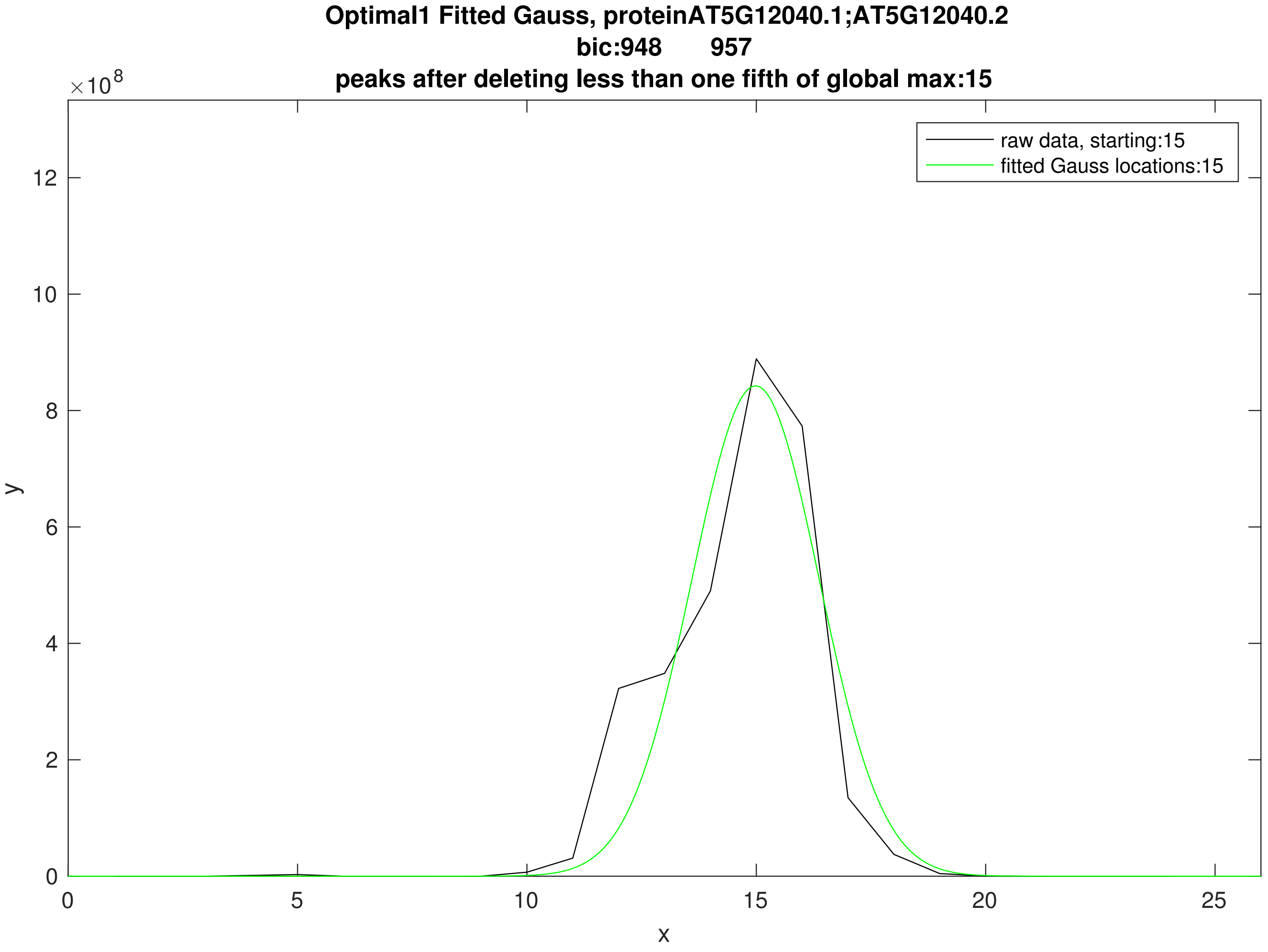

Optimal1 Fitted Gauss, proteinAT5G12040.1;AT5G12040.2
bic:948 957
peaks after deleting less than one fifth of global max:15
×10^8
raw data, starting:15
fitted Gauss locations:15
y
x
0
2
4
6
8
10
12
0
5
10
15
20
25

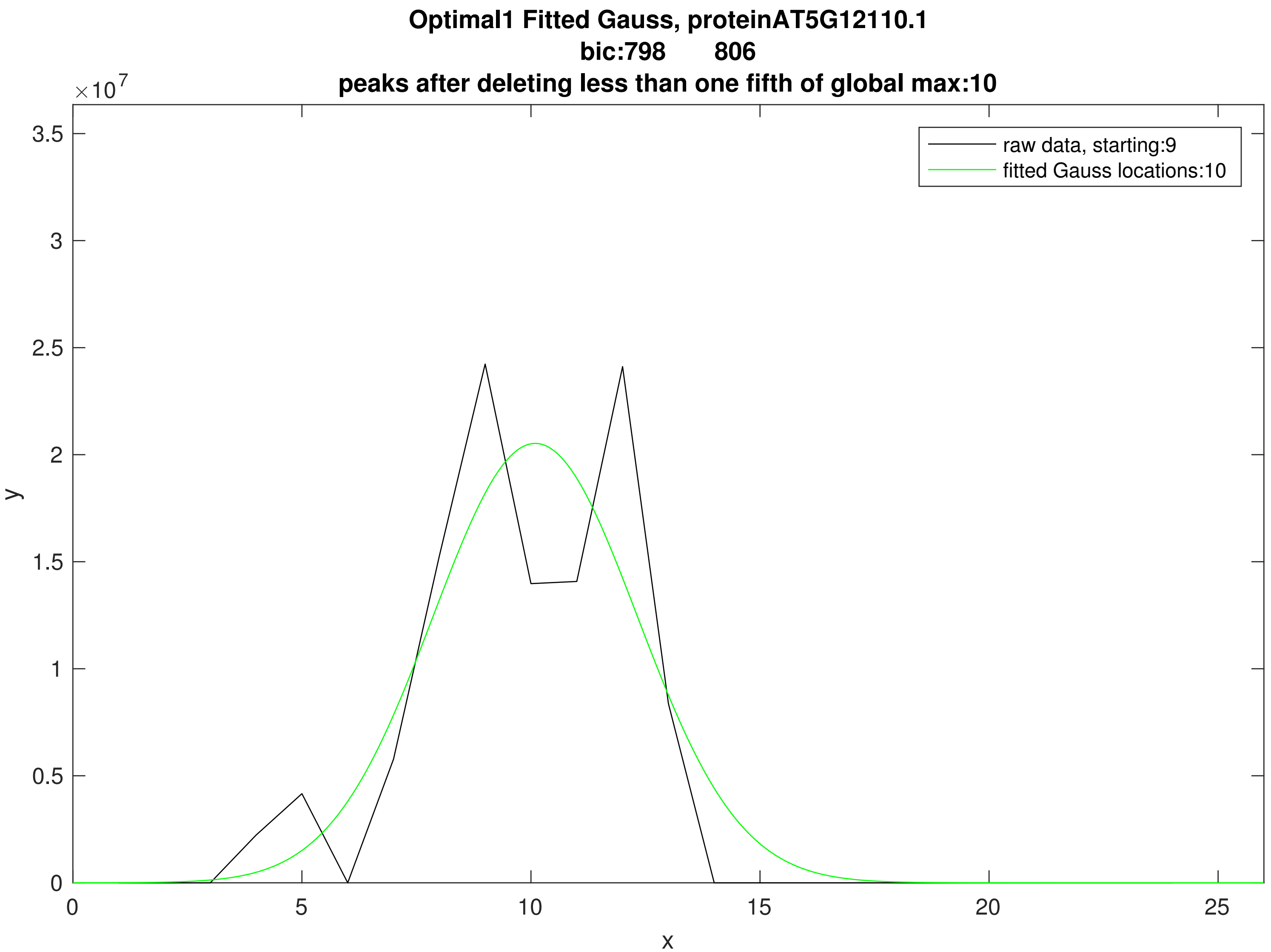
Optimal1 Fitted Gauss, proteinAT5G12110.1
bic:798 806
peaks after deleting less than one fifth of global max:10
×10^7
3.5
3
2.5
2
1.5
1
0.5
0
y
0
5
10
15
20
25
x
raw data, starting:9
fitted Gauss locations:10

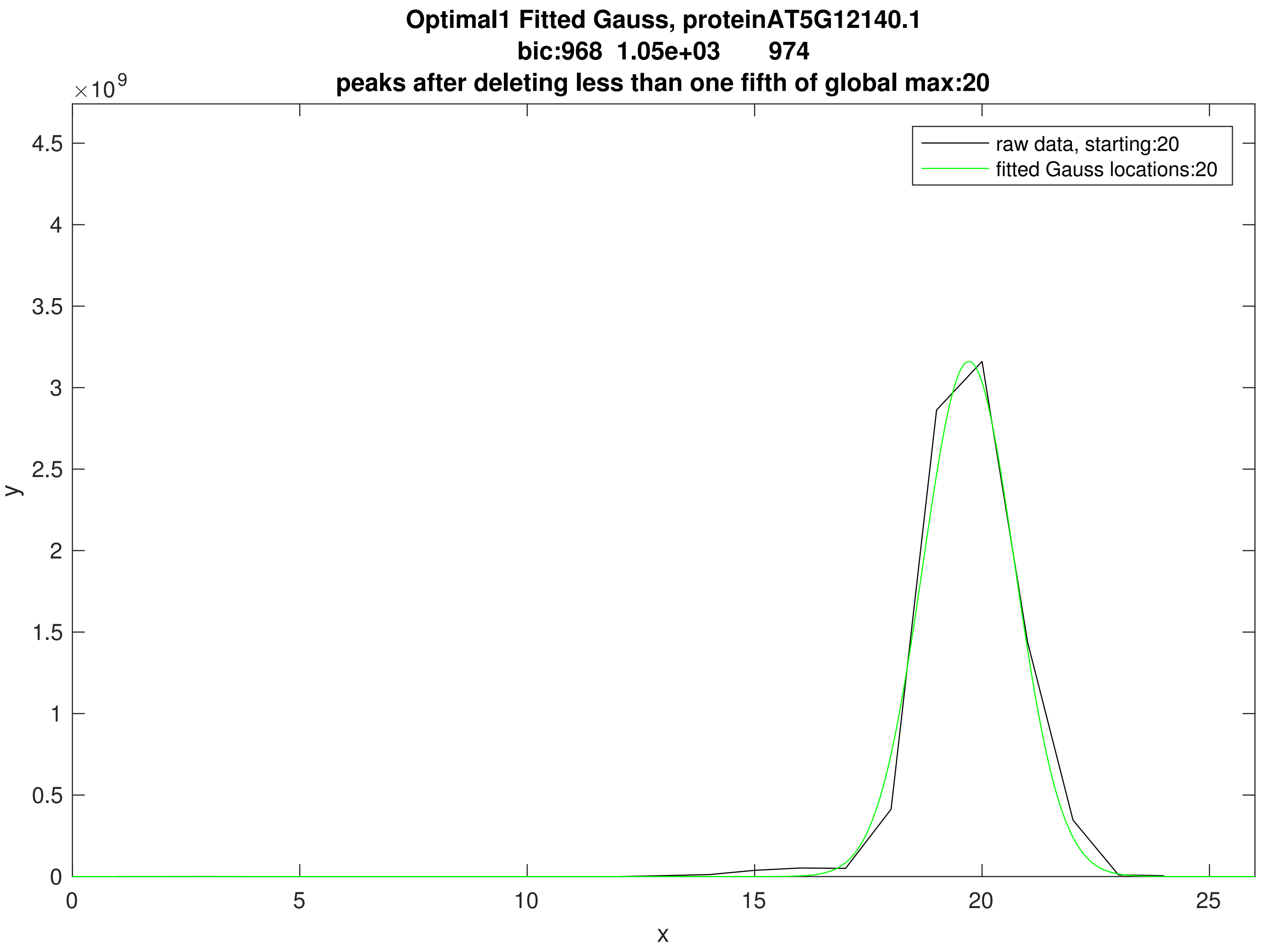

Optimal1 Fitted Gauss, proteinAT5G12140.1
bic:968 1.05e+03 974
peaks after deleting less than one fifth of global max:20
×10^9
4.5
4
3.5
3
2.5
2
1.5
1
0.5
0
y
0
5
10
15
20
25
x
raw data, starting:20
fitted Gauss locations:20

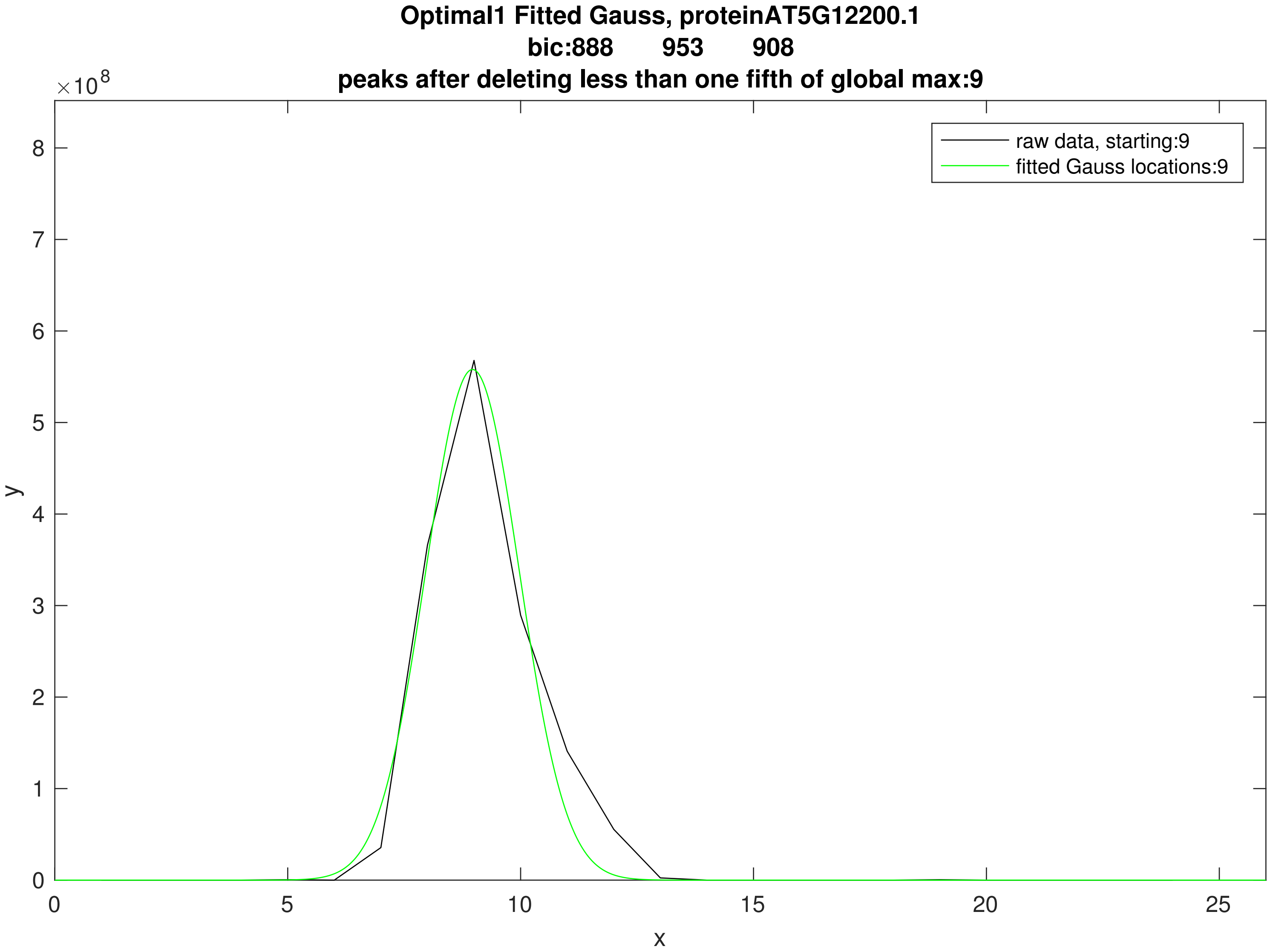
Optimal1 Fitted Gauss, proteinAT5G12200.1
bic:888 953 908
peaks after deleting less than one fifth of global max:9
×10^8
raw data, starting:9
fitted Gauss locations:9
y
x
0
1
2
3
4
5
6
7
8
0
5
10
15
20
25

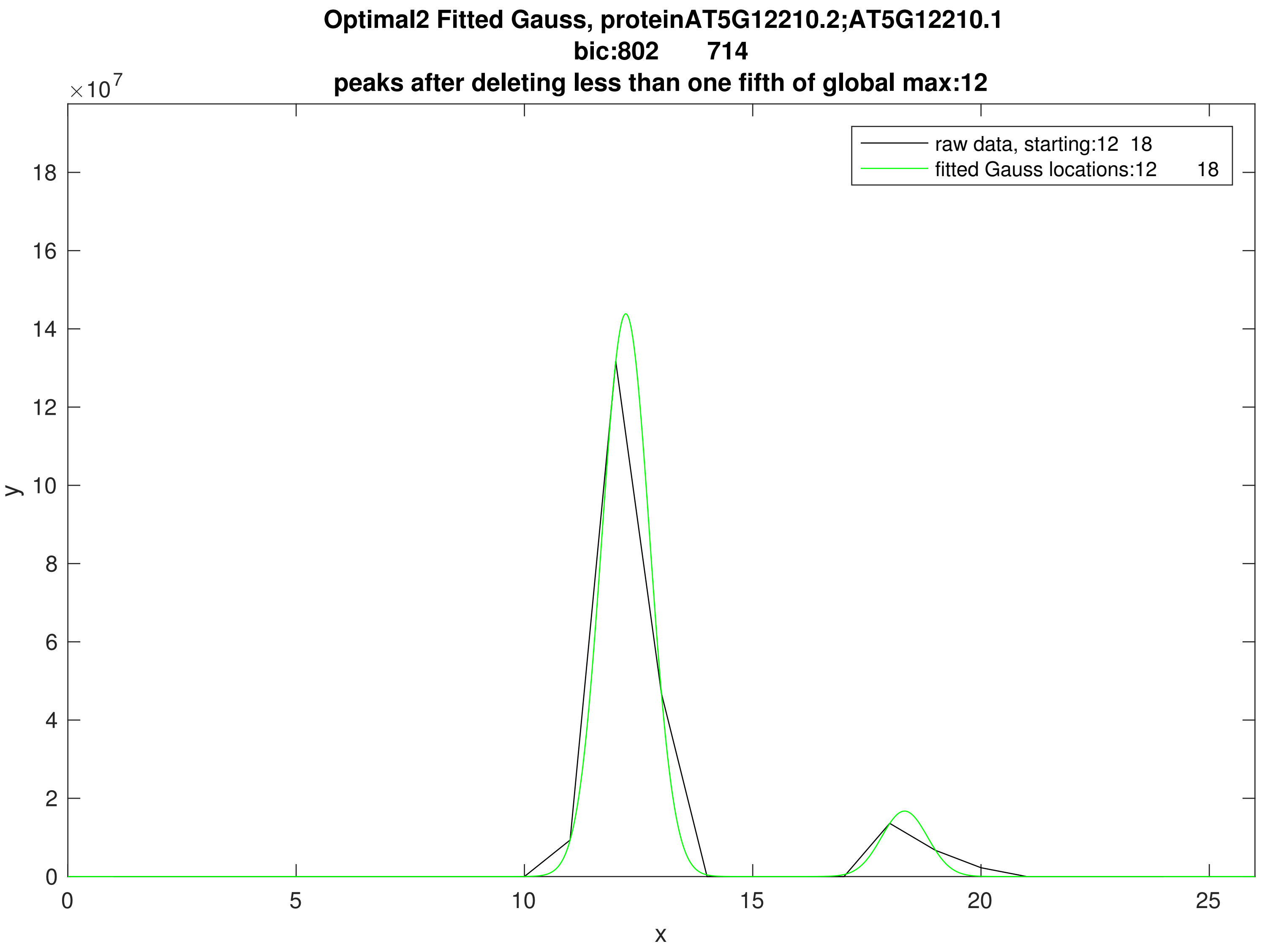
Optimal2 Fitted Gauss, proteinAT5G12210.2;AT5G12210.1
bic:802 714
peaks after deleting less than one fifth of global max:12
×10^7
raw data, starting:12 18
fitted Gauss locations:12 18
y
x
0
2
4
6
8
10
12
14
16
18
0
5
10
15
20
25

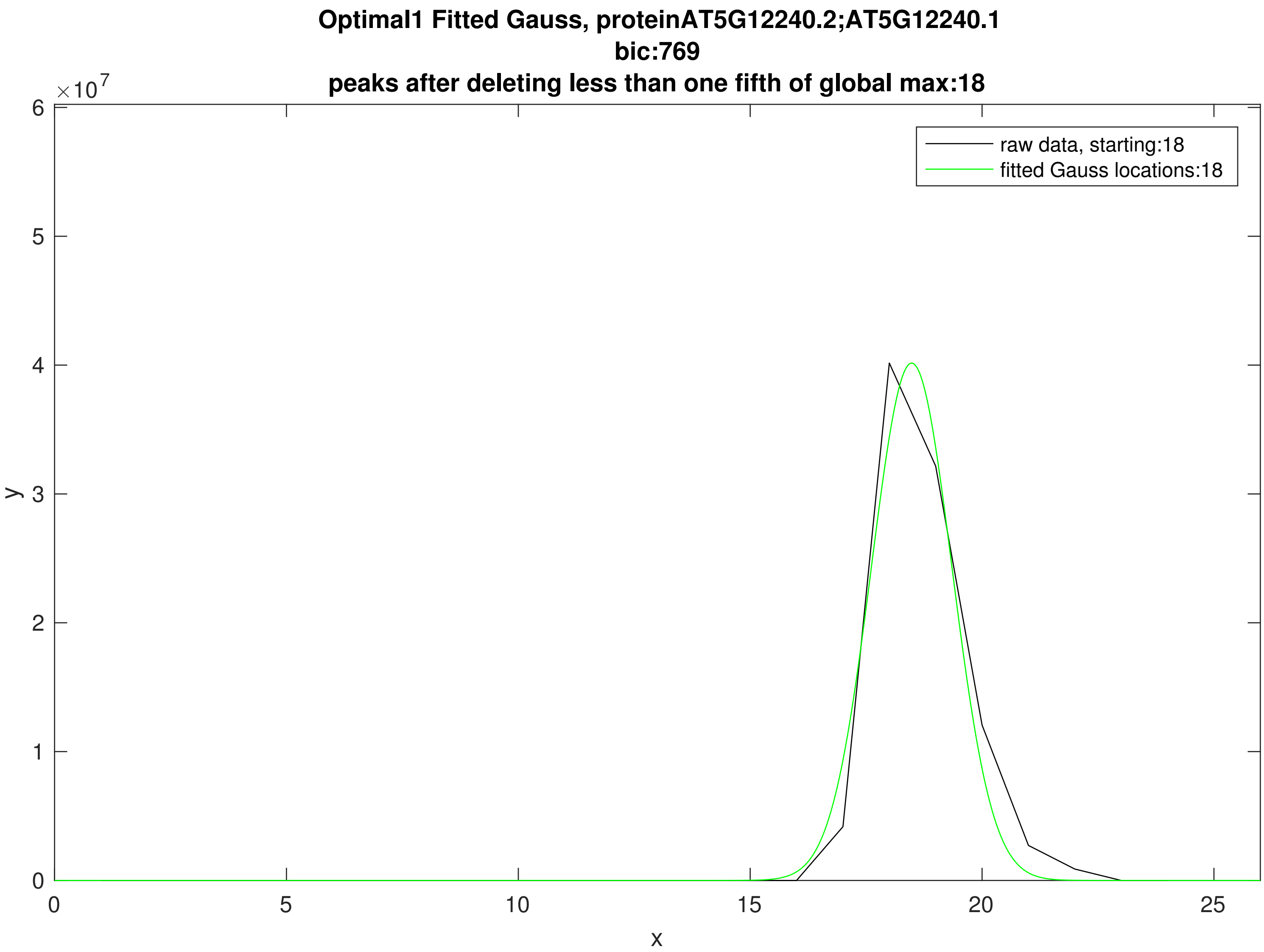

Optimal1 Fitted Gauss, proteinAT5G12240.2;AT5G12240.1
bic:769
peaks after deleting less than one fifth of global max:18
×10^7
raw data, starting:18
fitted Gauss locations:18
y
x
0
1
2
3
4
5
6
0
5
10
15
20
25

Optimal1 Fitted Gauss, proteinAT5G12250.1
bic:881 891
peaks after deleting less than one fifth of global max:14

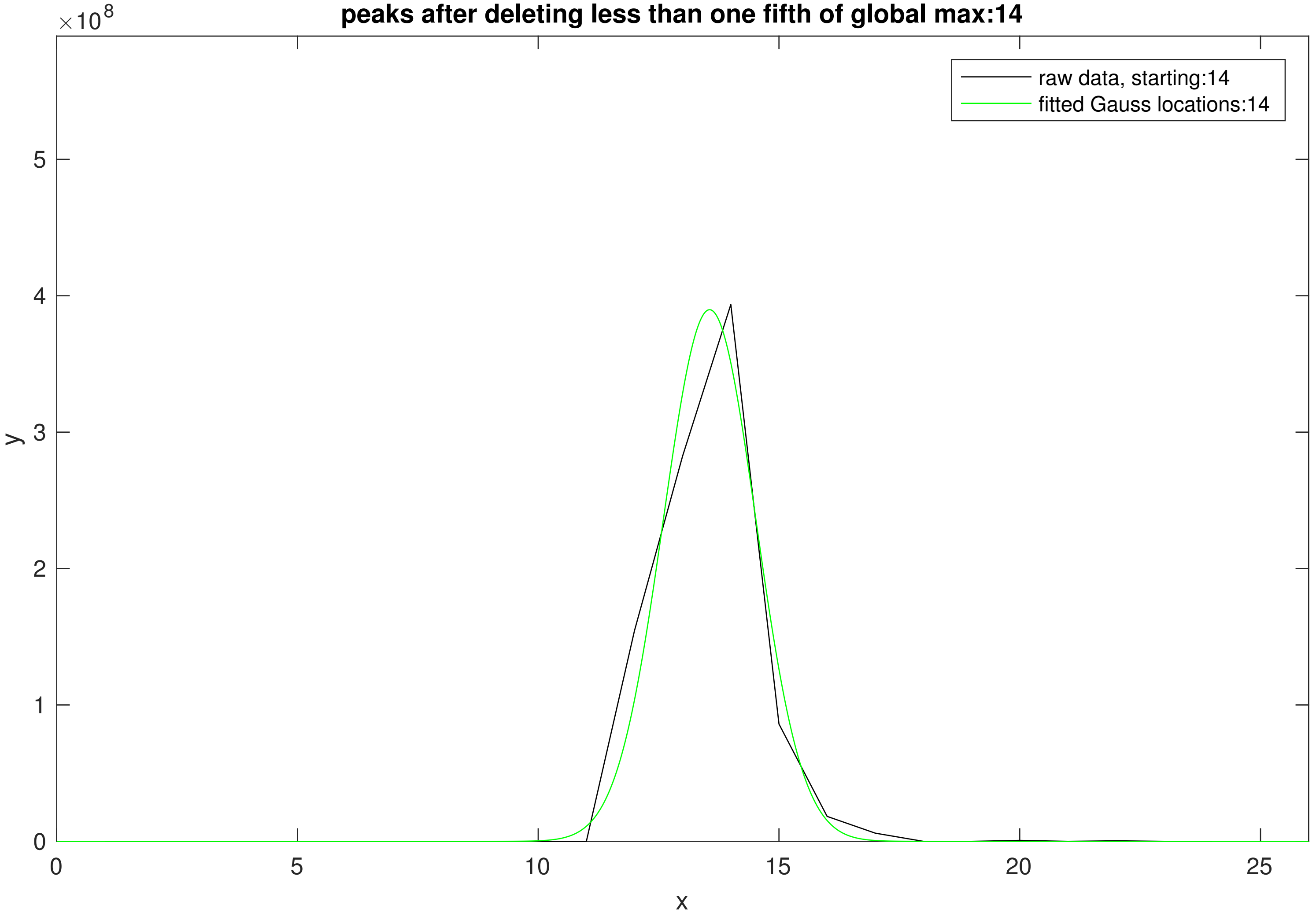

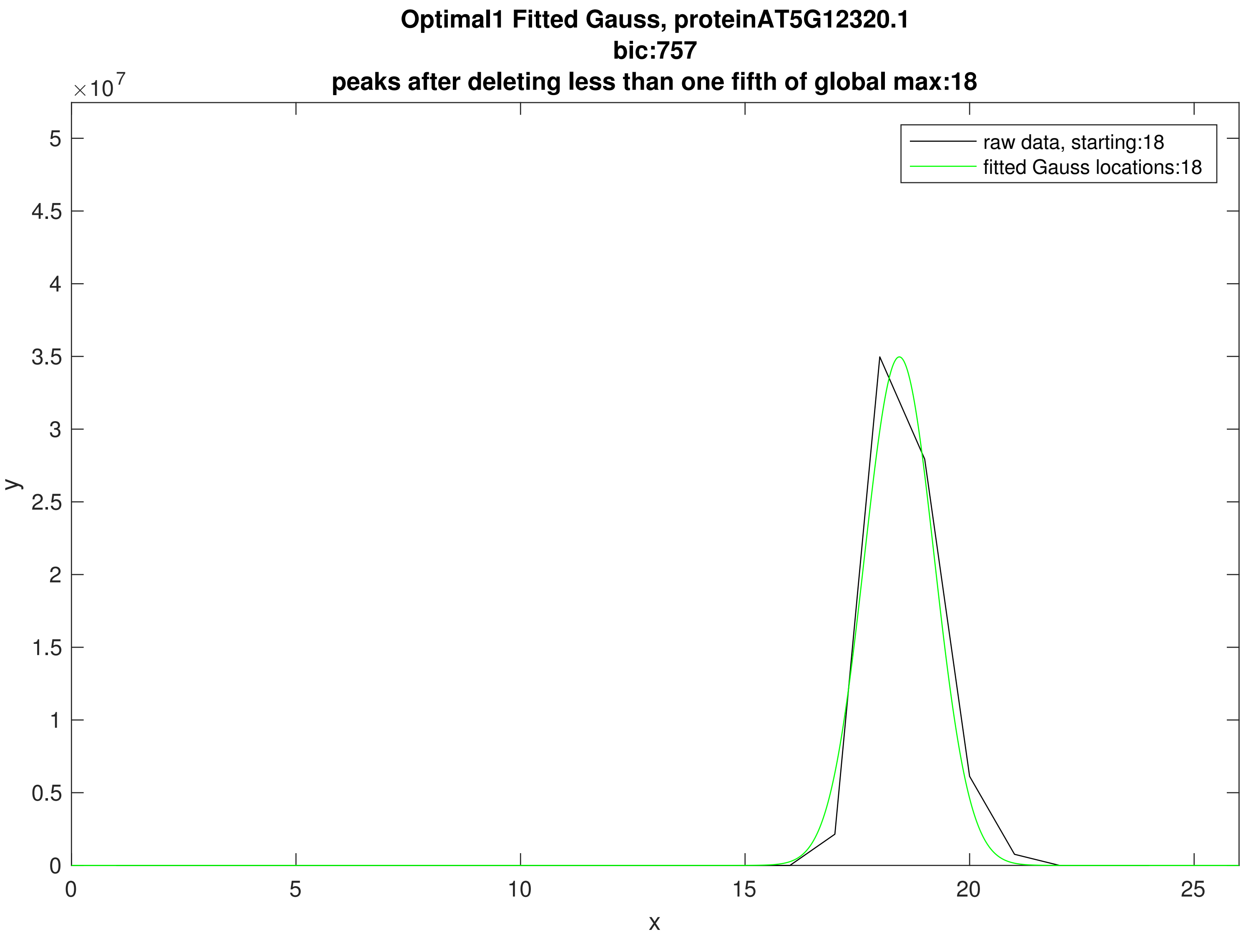
Optimal1 Fitted Gauss, proteinAT5G12320.1
bic:757
peaks after deleting less than one fifth of global max:18
×10^7
raw data, starting:18
fitted Gauss locations:18
y
x
0
0.5
1
1.5
2
2.5
3
3.5
4
4.5
5
0
5
10
15
20
25

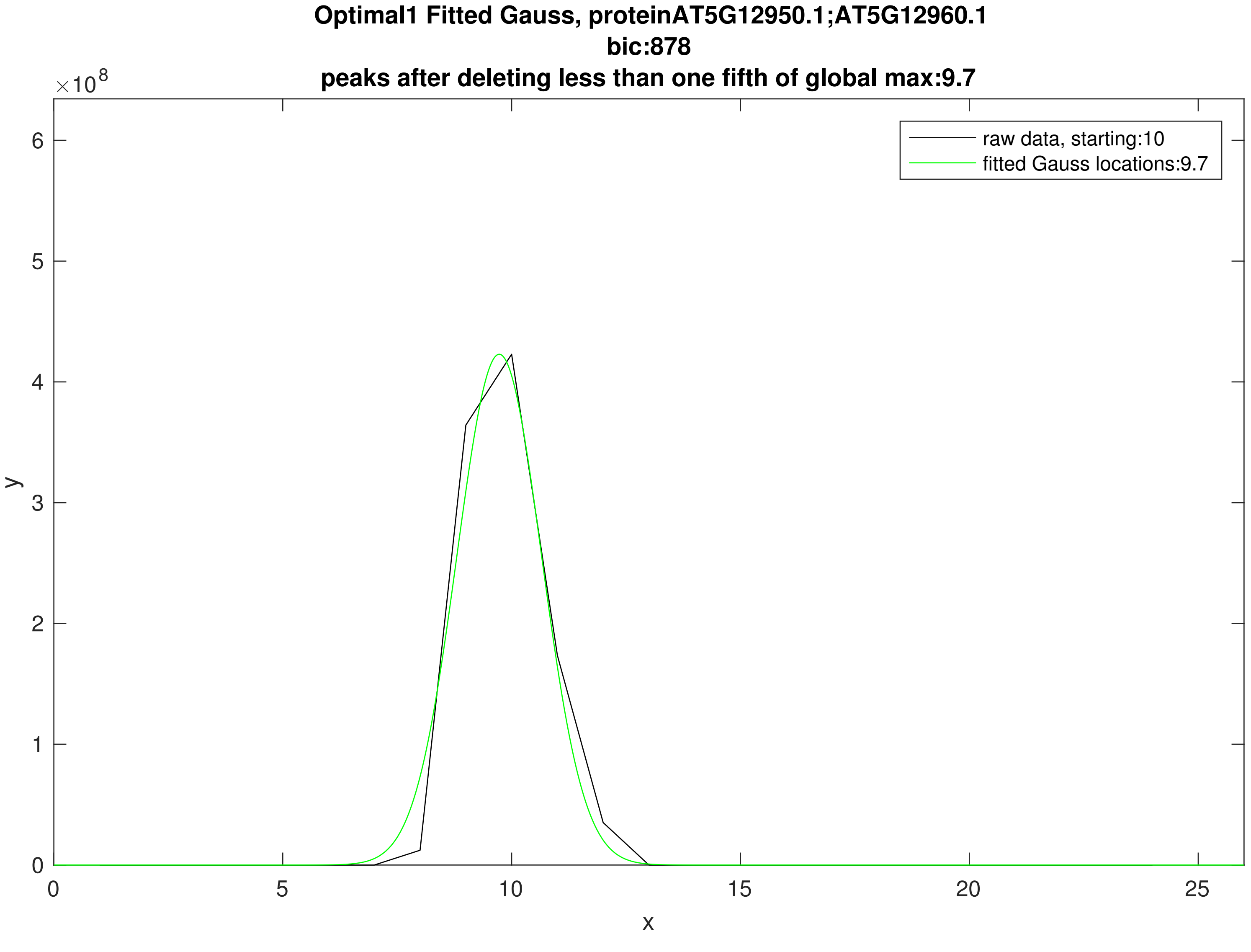
Optimal1 Fitted Gauss, proteinAT5G12950.1;AT5G12960.1
bic:878
peaks after deleting less than one fifth of global max:9.7
×10^8
6
5
4
3
2
1
0
y
0
5
10
15
20
25
x
raw data, starting:10
fitted Gauss locations:9.7

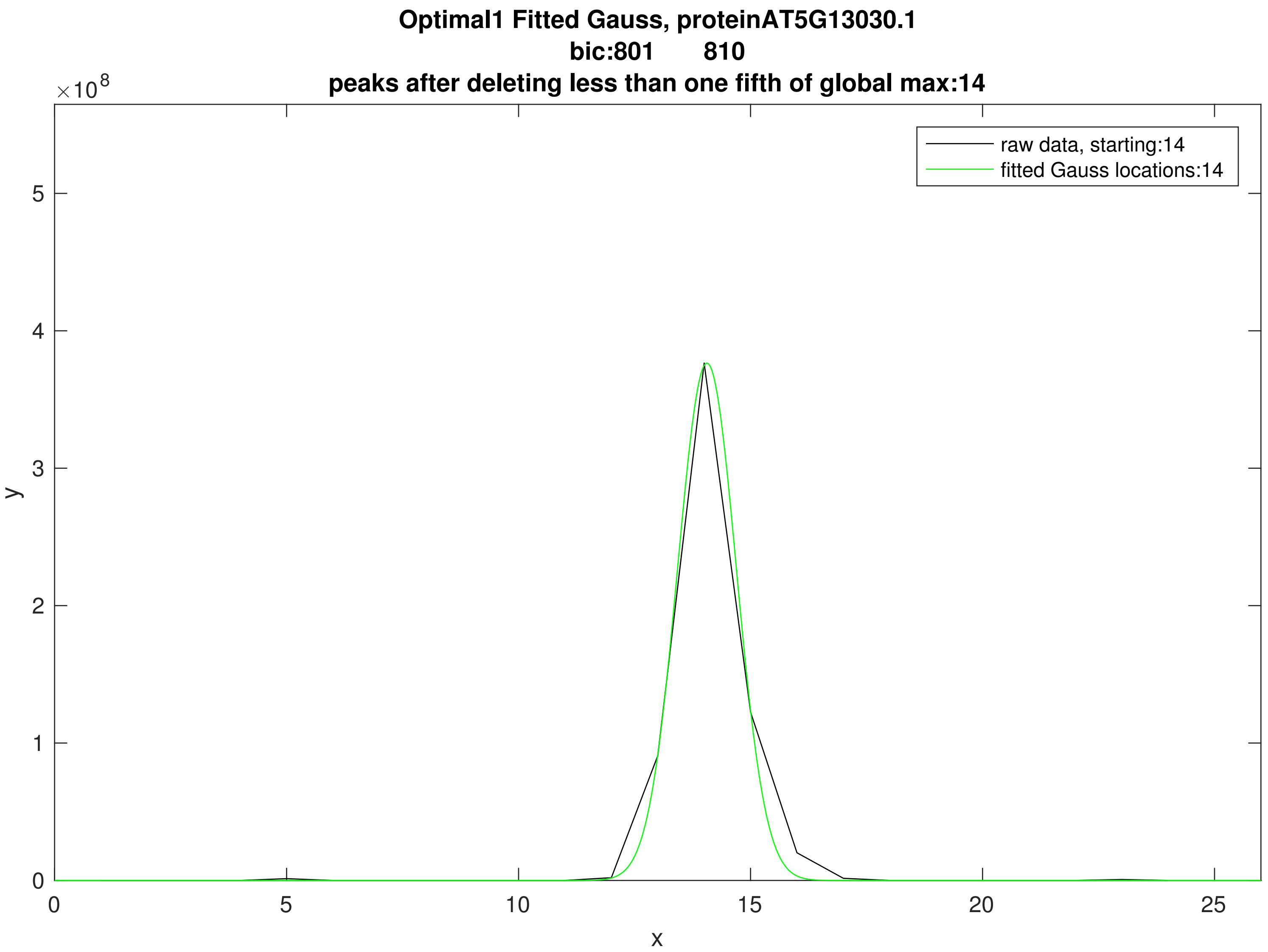
Optimal1 Fitted Gauss, proteinAT5G13030.1
bic:801 810
peaks after deleting less than one fifth of global max:14
×10^8
raw data, starting:14
fitted Gauss locations:14
y
x
0
1
2
3
4
5
0
5
10
15
20
25

Optimal1 Fitted Gauss, proteinAT5G13110.1
bic:759 765
peaks after deleting less than one fifth of global max:10

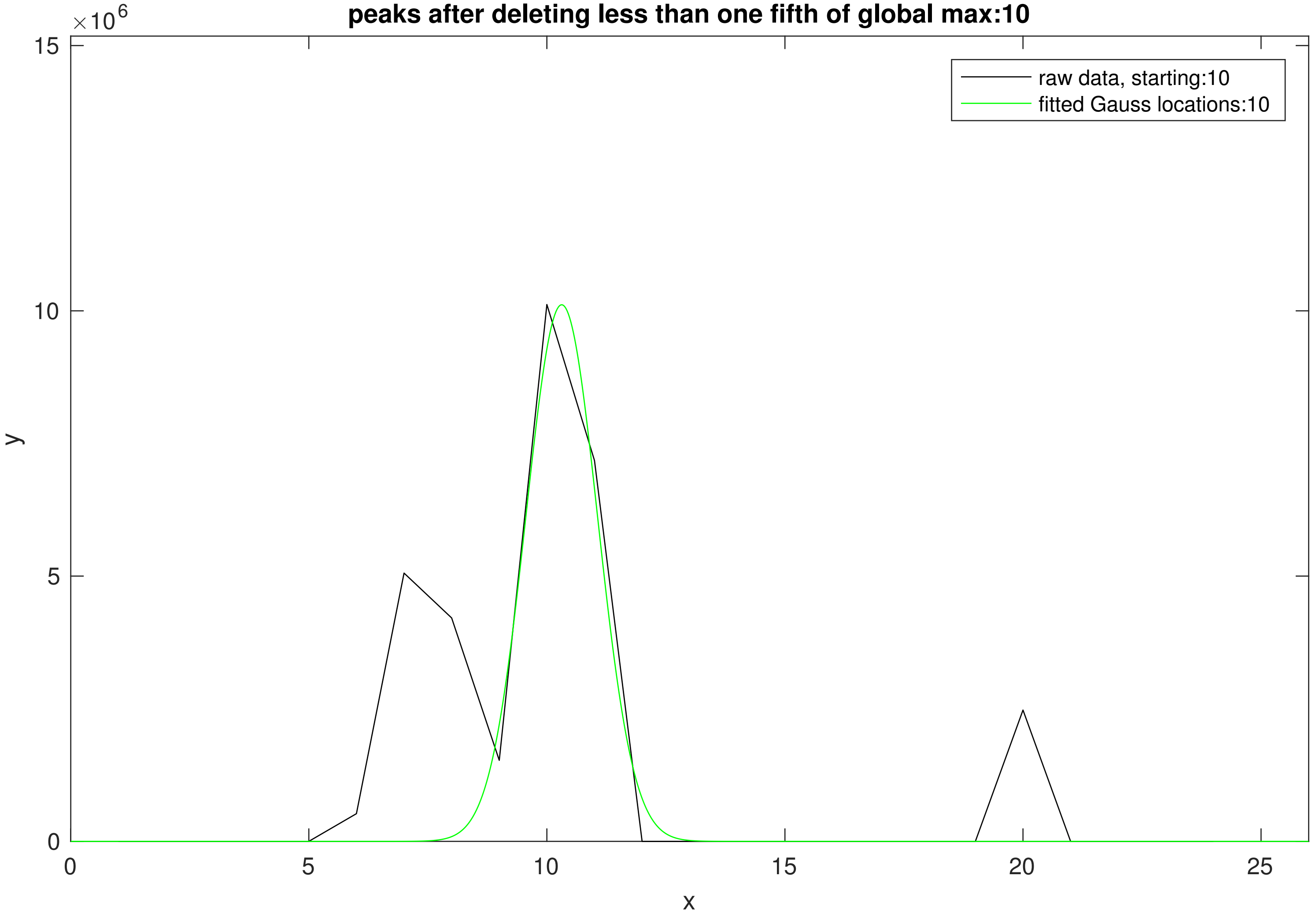

**Optimal1 Fitted Gauss, proteinAT5G13120.2;AT5G13120.1**
**bic:842 852 861**
**peaks after deleting less than one fifth of global max:20**

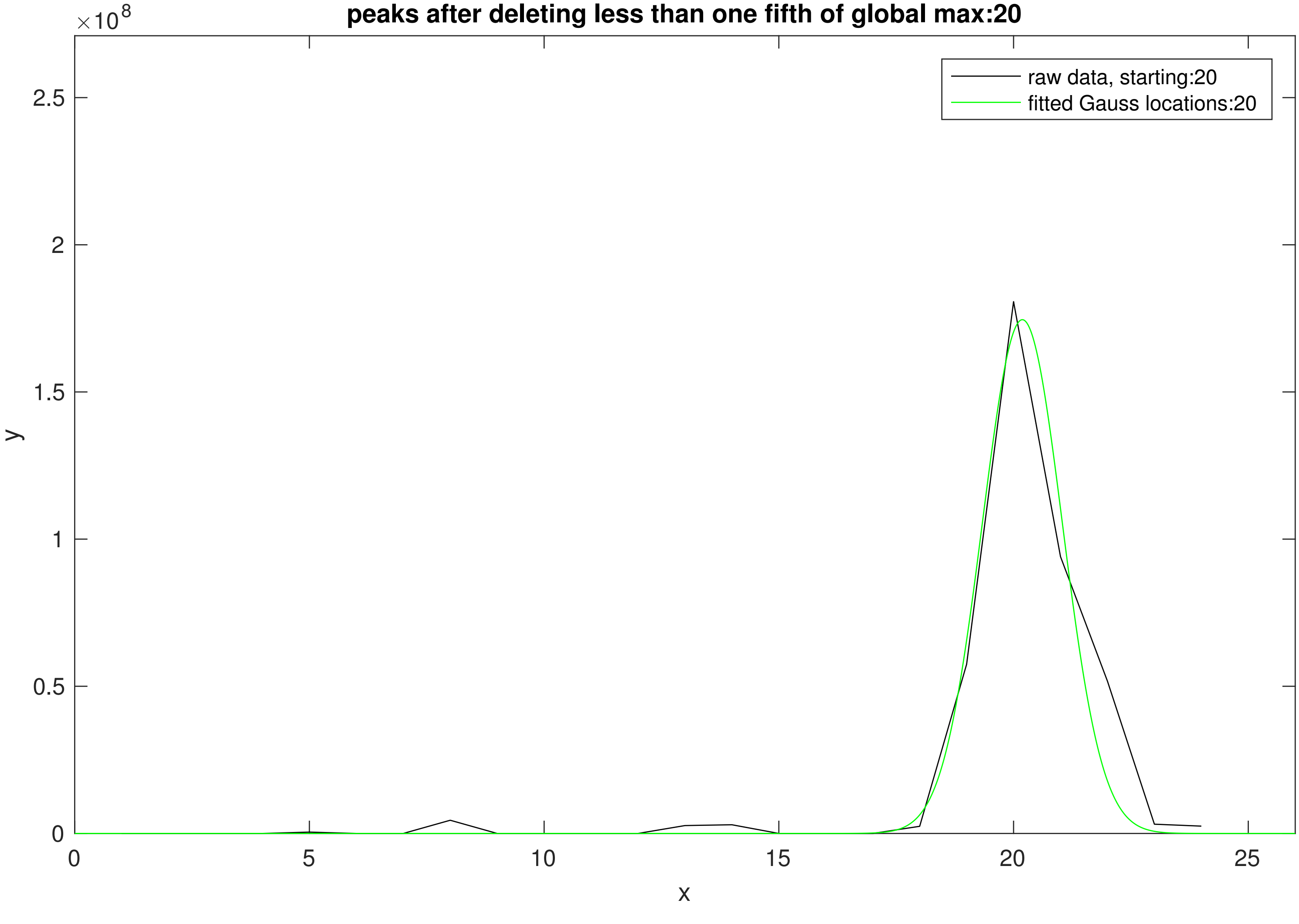

**Optimal1 Fitted Gauss, proteinAT5G13360.3;AT5G13360.2;AT5G13360.1**
**bic:906 907**
**peaks after deleting less than one fifth of global max:15**

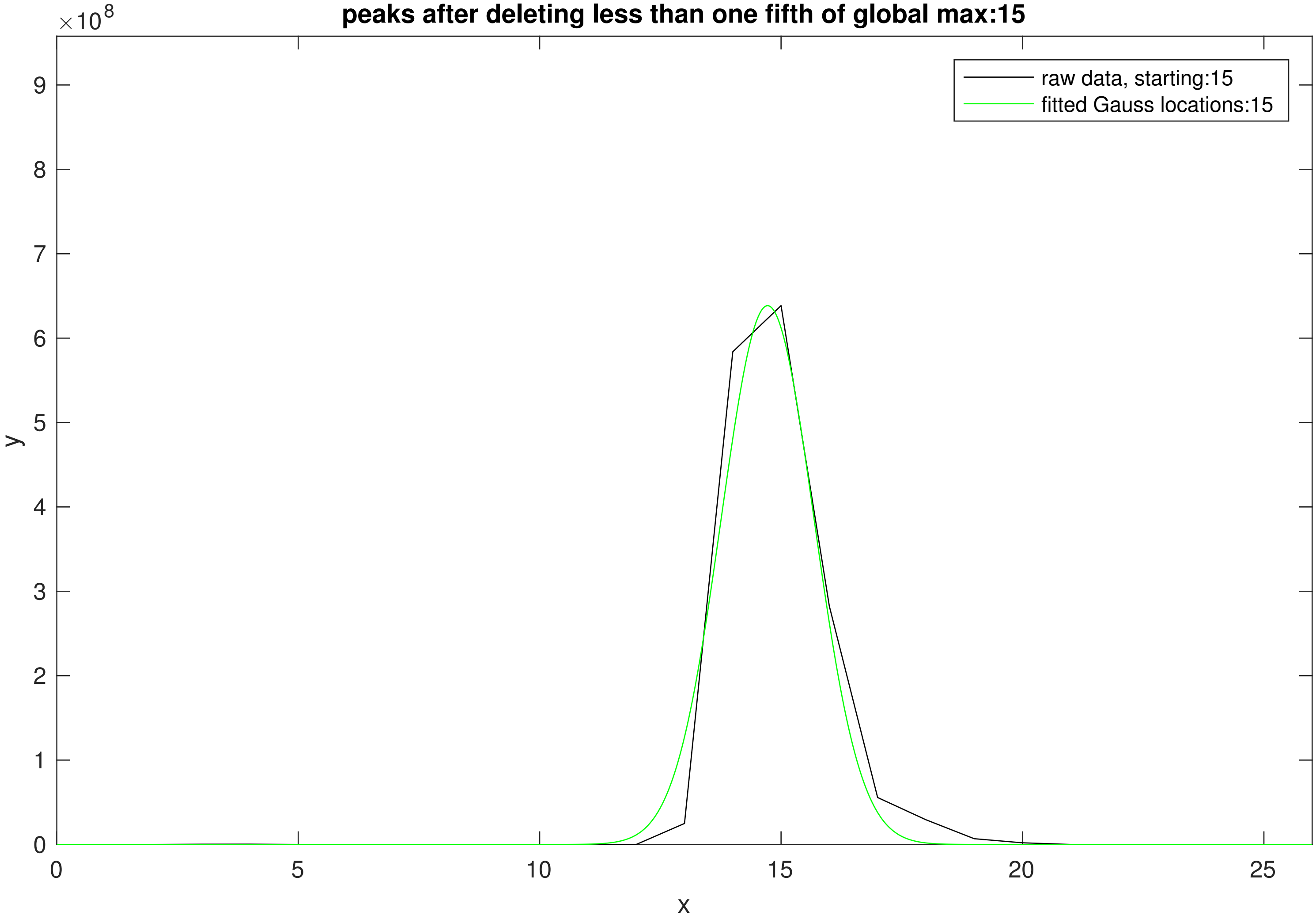

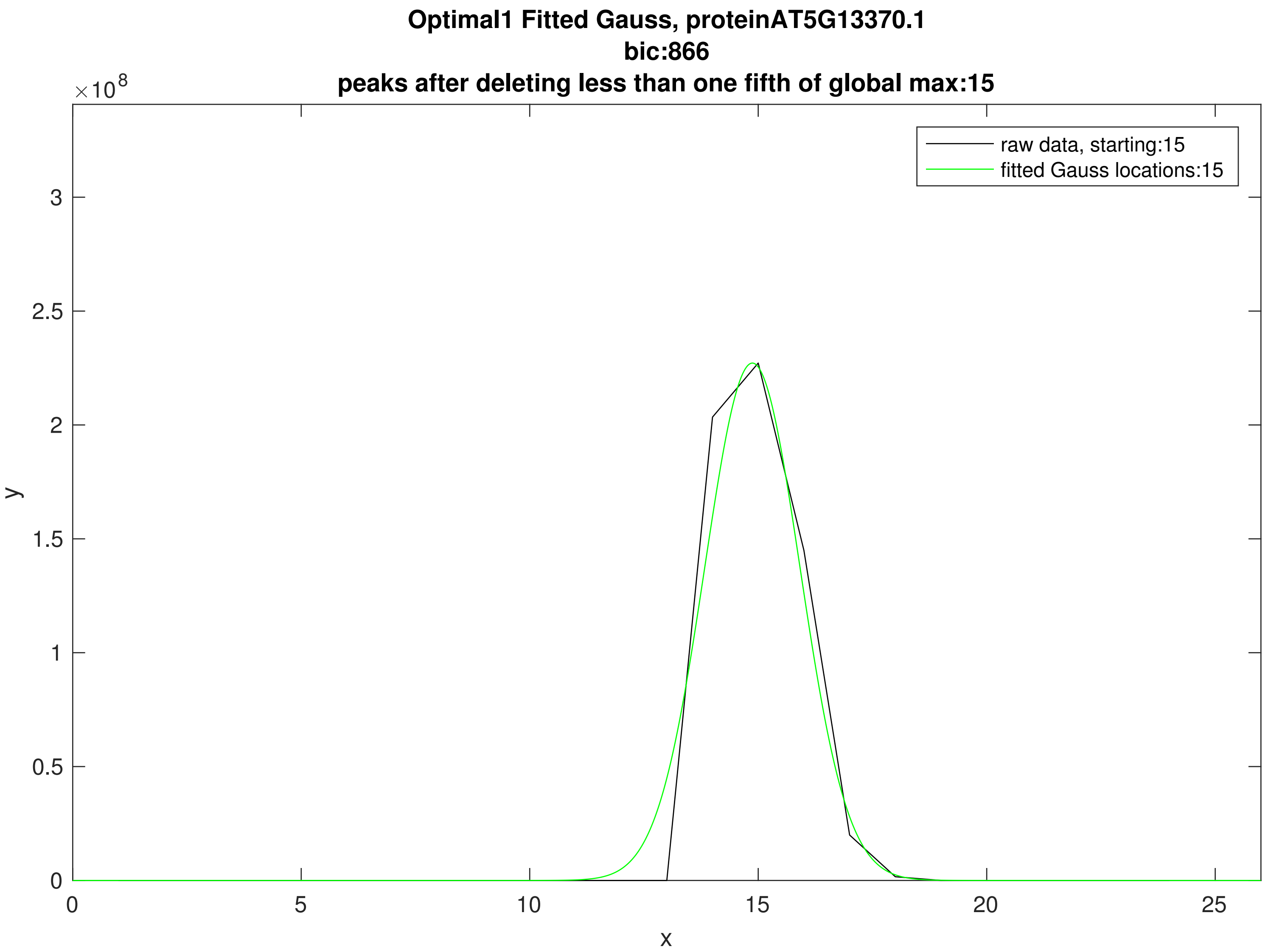

Optimal1 Fitted Gauss, proteinAT5G13370.1
bic:866
peaks after deleting less than one fifth of global max:15
×10^8
raw data, starting:15
fitted Gauss locations:15
y
x
0
0.5
1
1.5
2
2.5
3
0
5
10
15
20
25

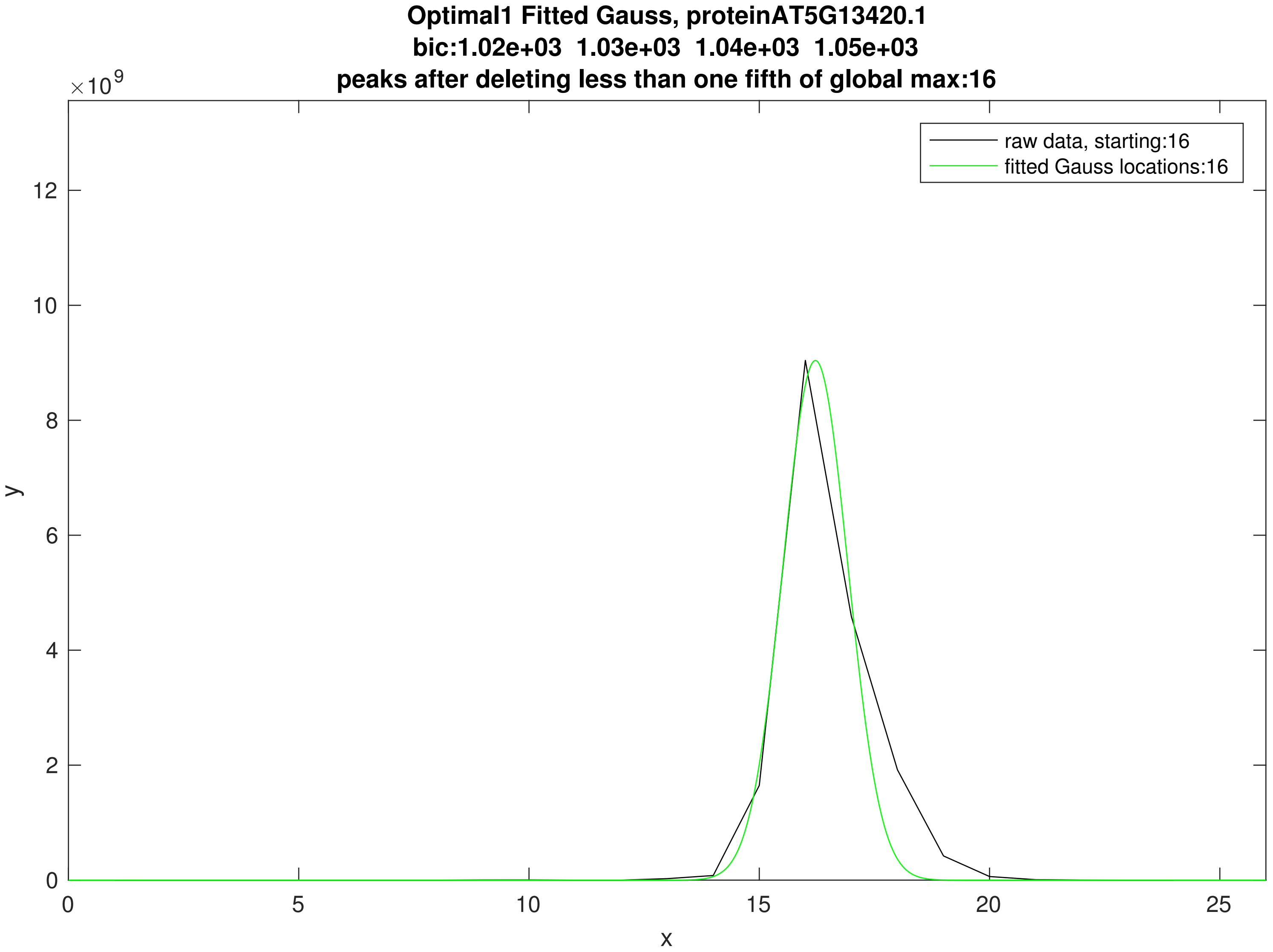

Optimal1 Fitted Gauss, proteinAT5G13420.1
bic:1.02e+03 1.03e+03 1.04e+03 1.05e+03
peaks after deleting less than one fifth of global max:16
×10^9
raw data, starting:16
fitted Gauss locations:16
y
x
0
2
4
6
8
10
12
0
5
10
15
20
25

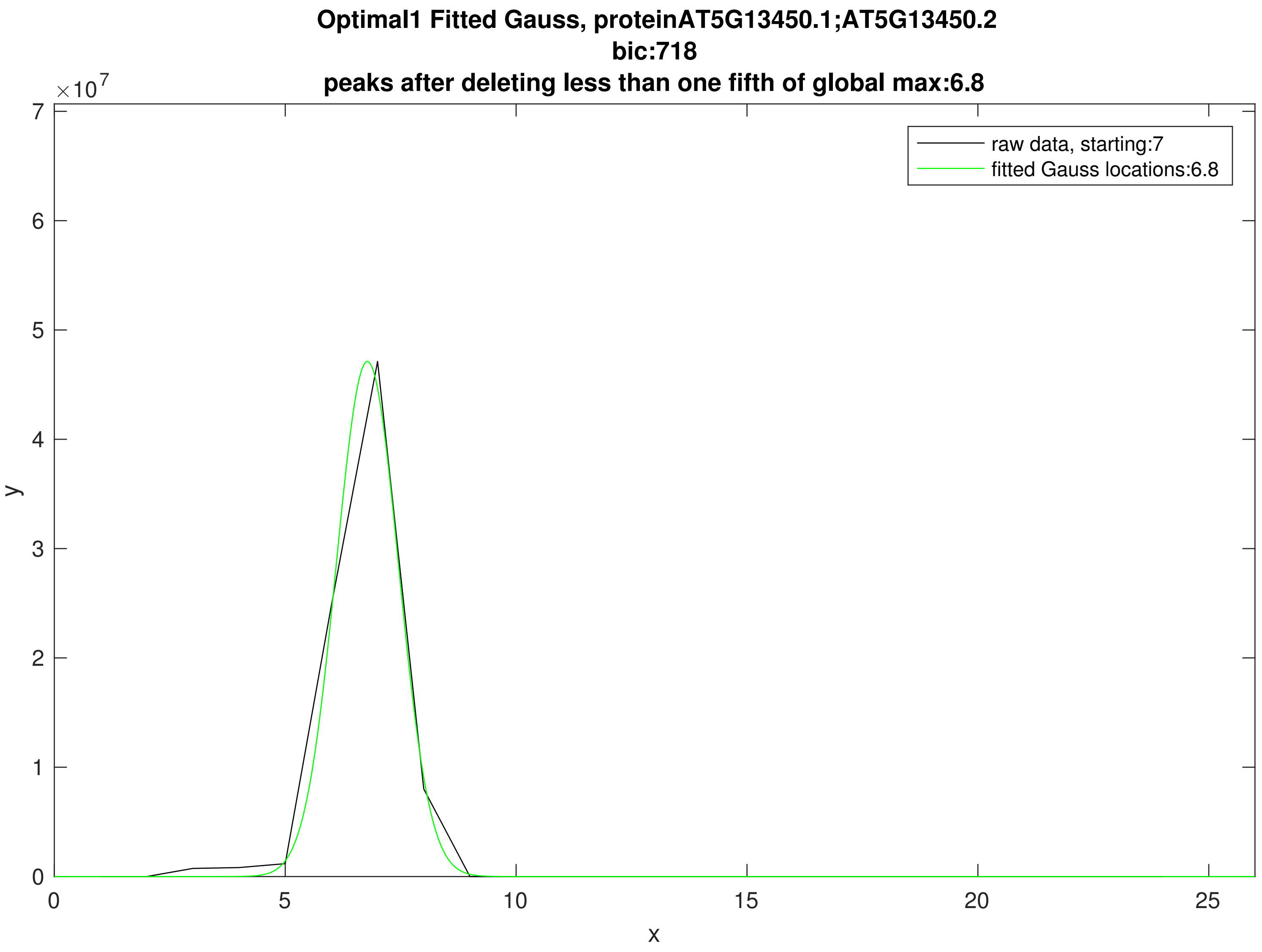
Optimal1 Fitted Gauss, proteinAT5G13450.1;AT5G13450.2
bic:718
peaks after deleting less than one fifth of global max:6.8
×10^7
y
x
raw data, starting:7
fitted Gauss locations:6.8

**Optimal1 Fitted Gauss, proteinAT5G13490.2;AT5G13490.1**
**bic:662**
**peaks after deleting less than one fifth of global max:3.8**

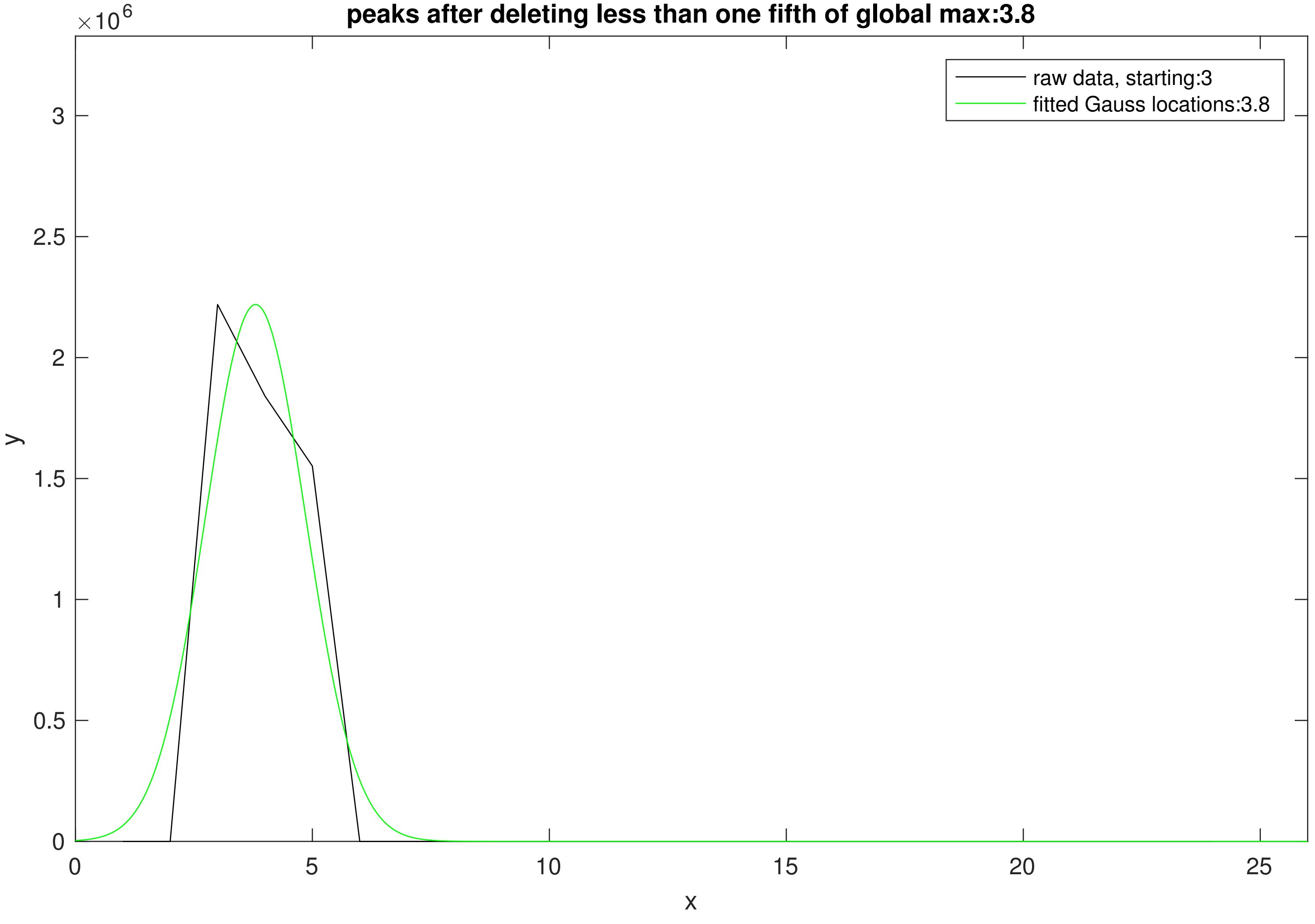

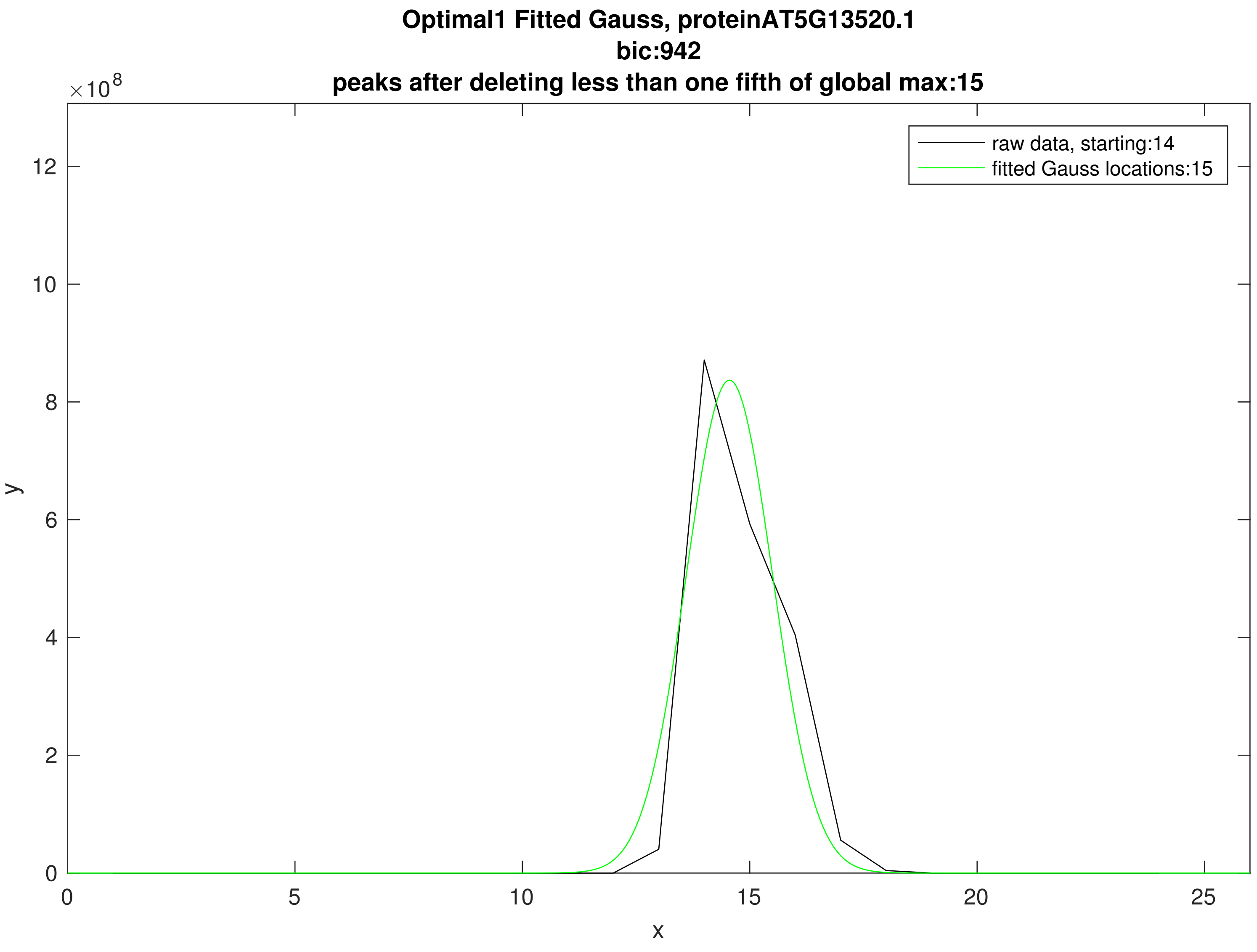

Optimal1 Fitted Gauss, proteinAT5G13520.1
bic:942
peaks after deleting less than one fifth of global max:15
×10^8
raw data, starting:14
fitted Gauss locations:15
y
x
0
2
4
6
8
10
12
0
5
10
15
20
25

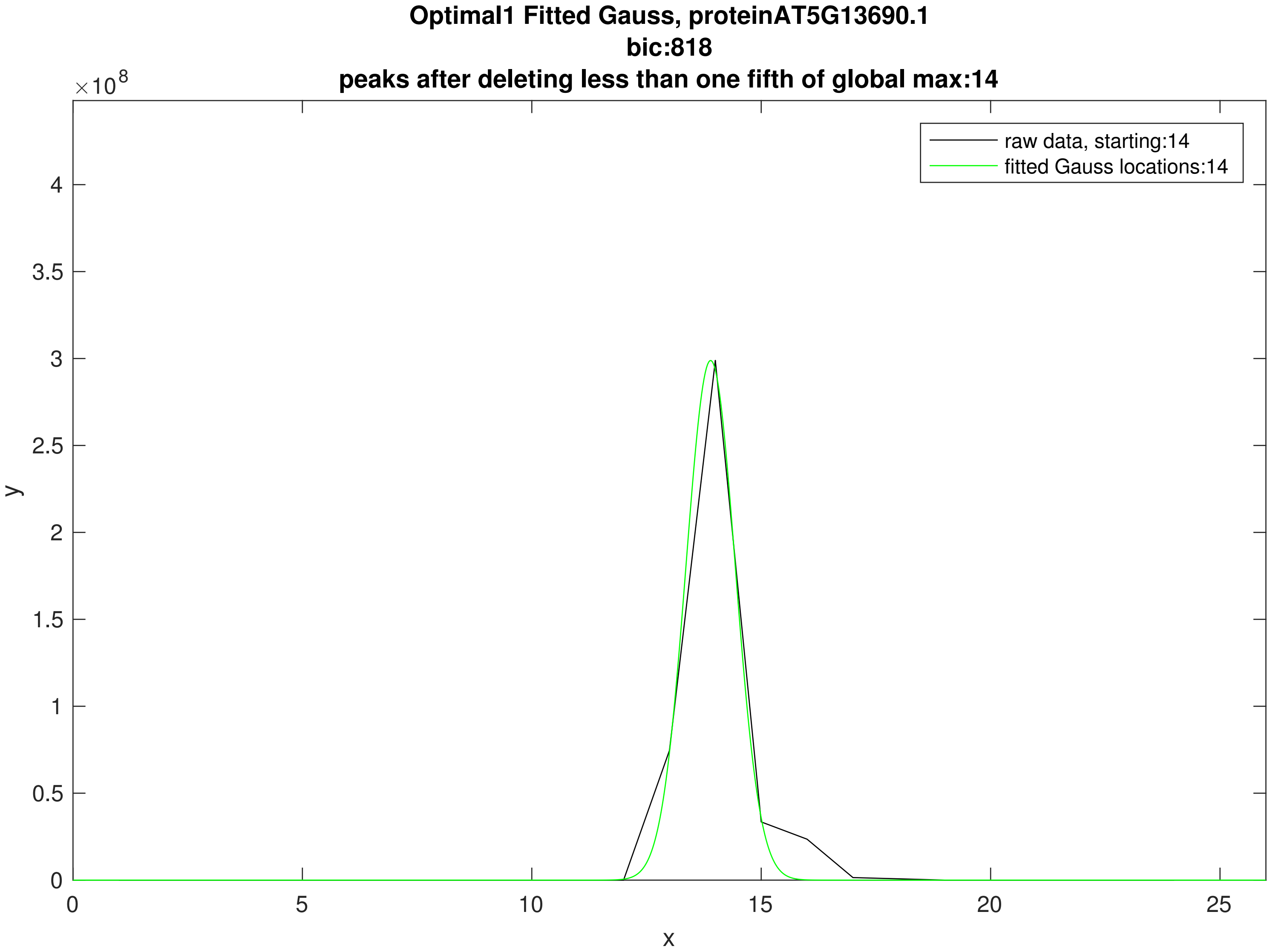
Optimal1 Fitted Gauss, proteinAT5G13690.1
bic:818
peaks after deleting less than one fifth of global max:14
×10^8
raw data, starting:14
fitted Gauss locations:14
4
3.5
3
2.5
2
1.5
1
0.5
0
y
0
5
10
15
20
25
x

#### Optimal1 Fitted Gauss, proteinAT5G13850.1
#### bic:839 842
#### peaks after deleting less than one fifth of global max:14

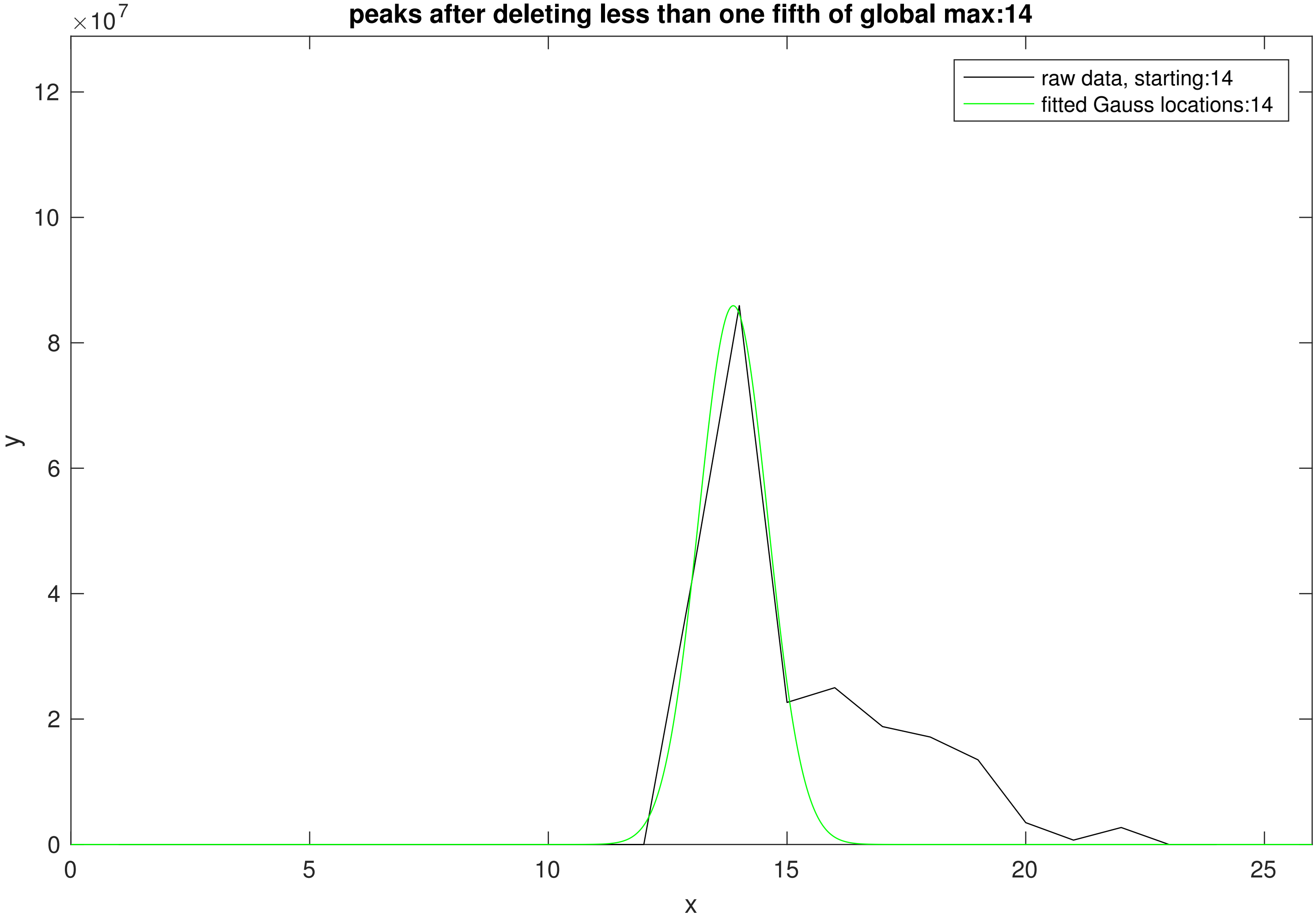

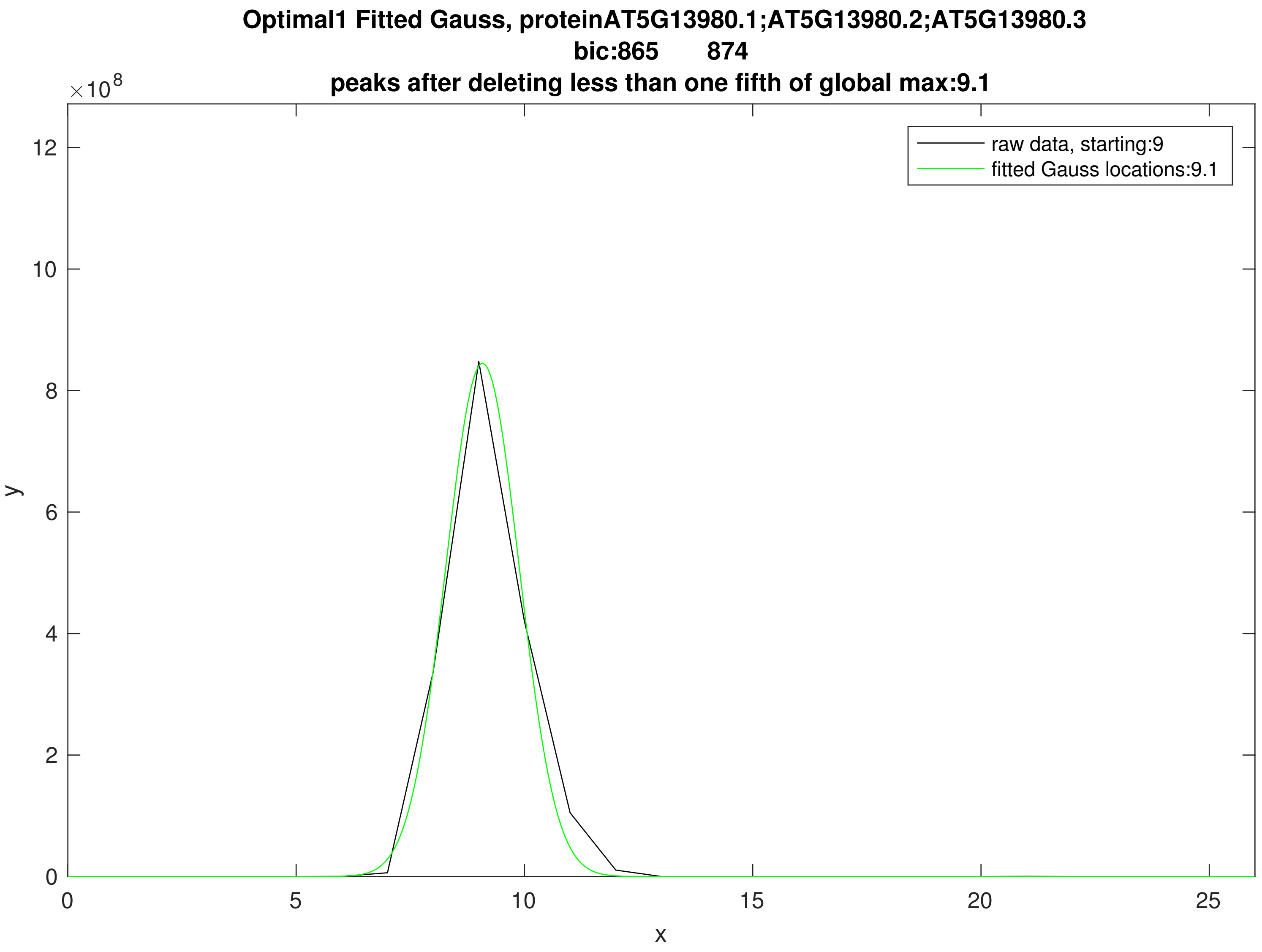
Optimal1 Fitted Gauss, proteinAT5G13980.1;AT5G13980.2;AT5G13980.3
bic:865 874
peaks after deleting less than one fifth of global max:9.1
×10^8
raw data, starting:9
fitted Gauss locations:9.1
y
x
0
2
4
6
8
10
12
5
15
20
25

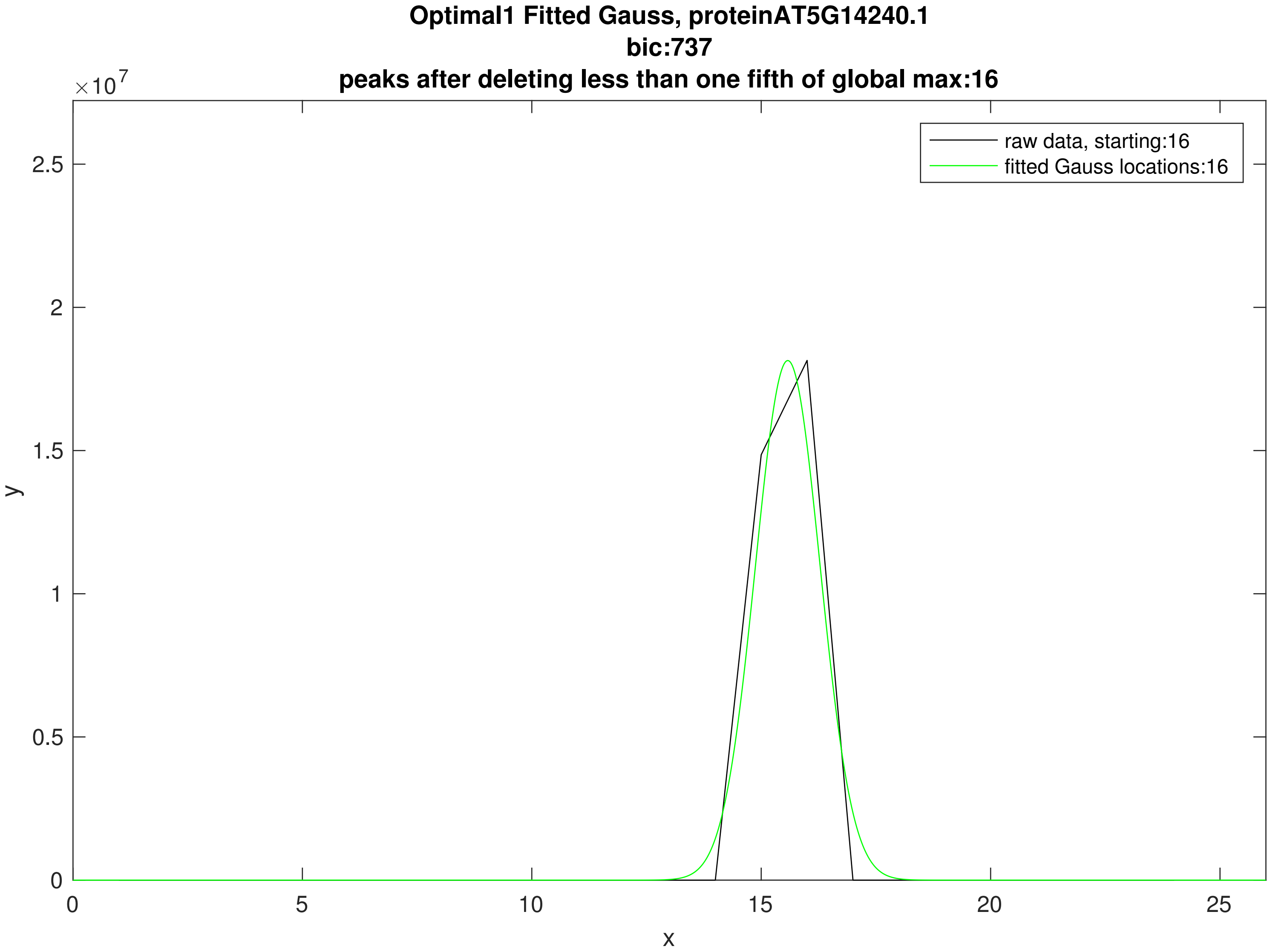
Optimal1 Fitted Gauss, proteinAT5G14240.1
bic:737
peaks after deleting less than one fifth of global max:16
×10^7
2.5
2
1.5
1
0.5
0
y
0
5
10
15
20
25
x
raw data, starting:16
fitted Gauss locations:16

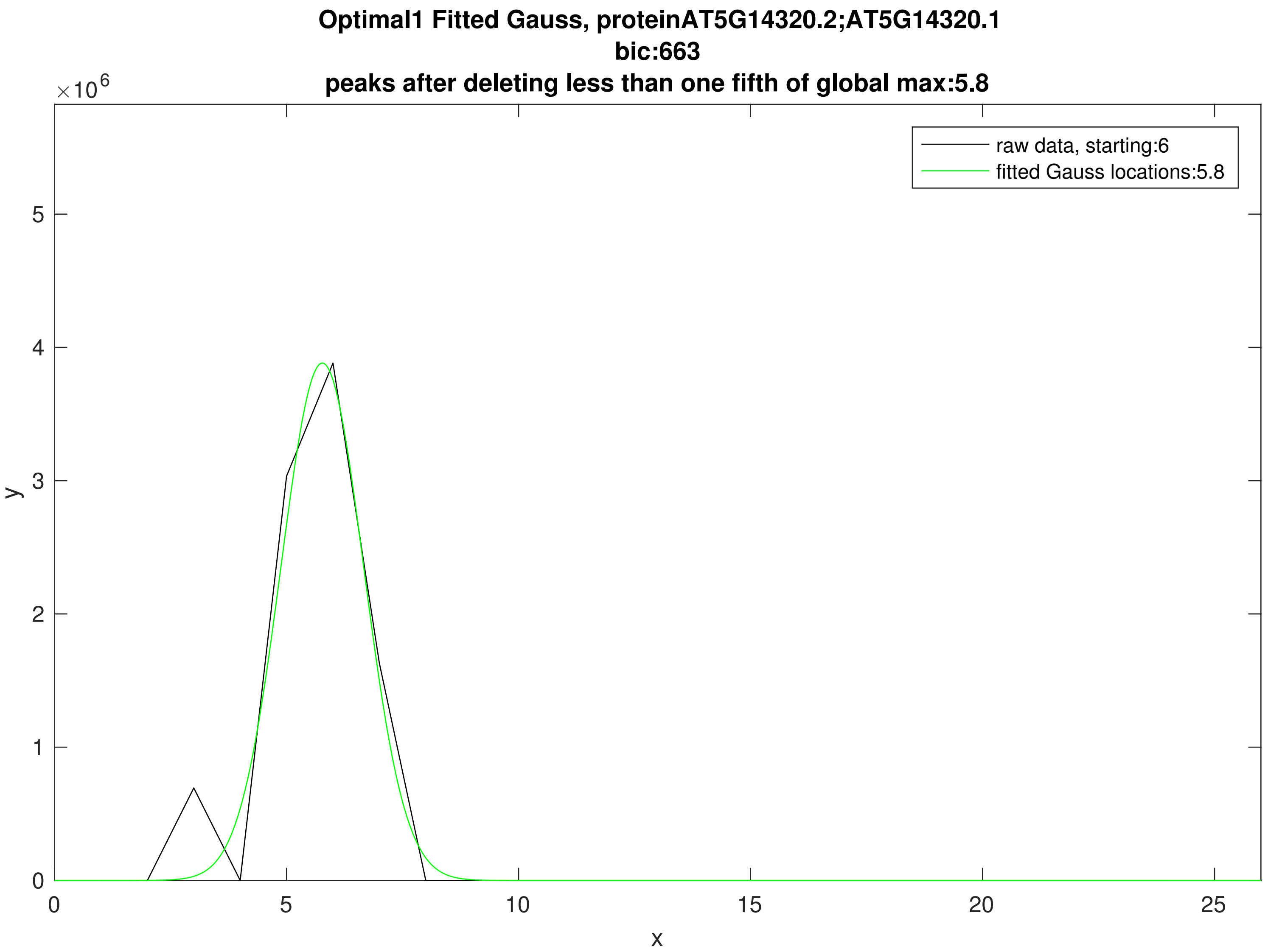
Optimal1 Fitted Gauss, proteinAT5G14320.2;AT5G14320.1
bic:663
peaks after deleting less than one fifth of global max:5.8
×10^6
raw data, starting:6
fitted Gauss locations:5.8
y
x
0
1
2
3
4
5
0
5
10
15
20
25

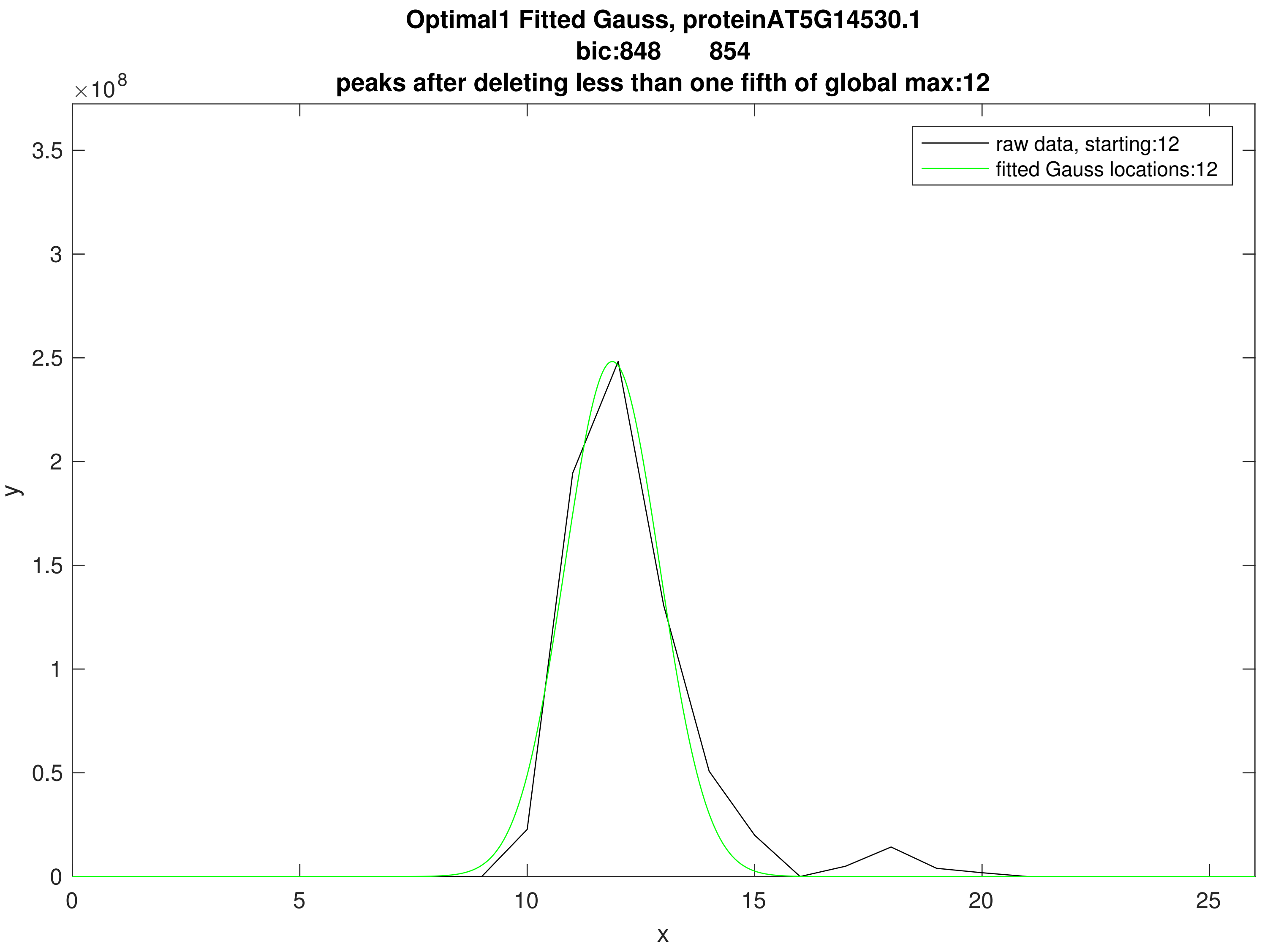
Optimal1 Fitted Gauss, proteinAT5G14530.1
bic:848 854
peaks after deleting less than one fifth of global max:12
×10^8
3.5
3
2.5
2
1.5
1
0.5
0
y
0
5
10
15
20
25
x
raw data, starting:12
fitted Gauss locations:12

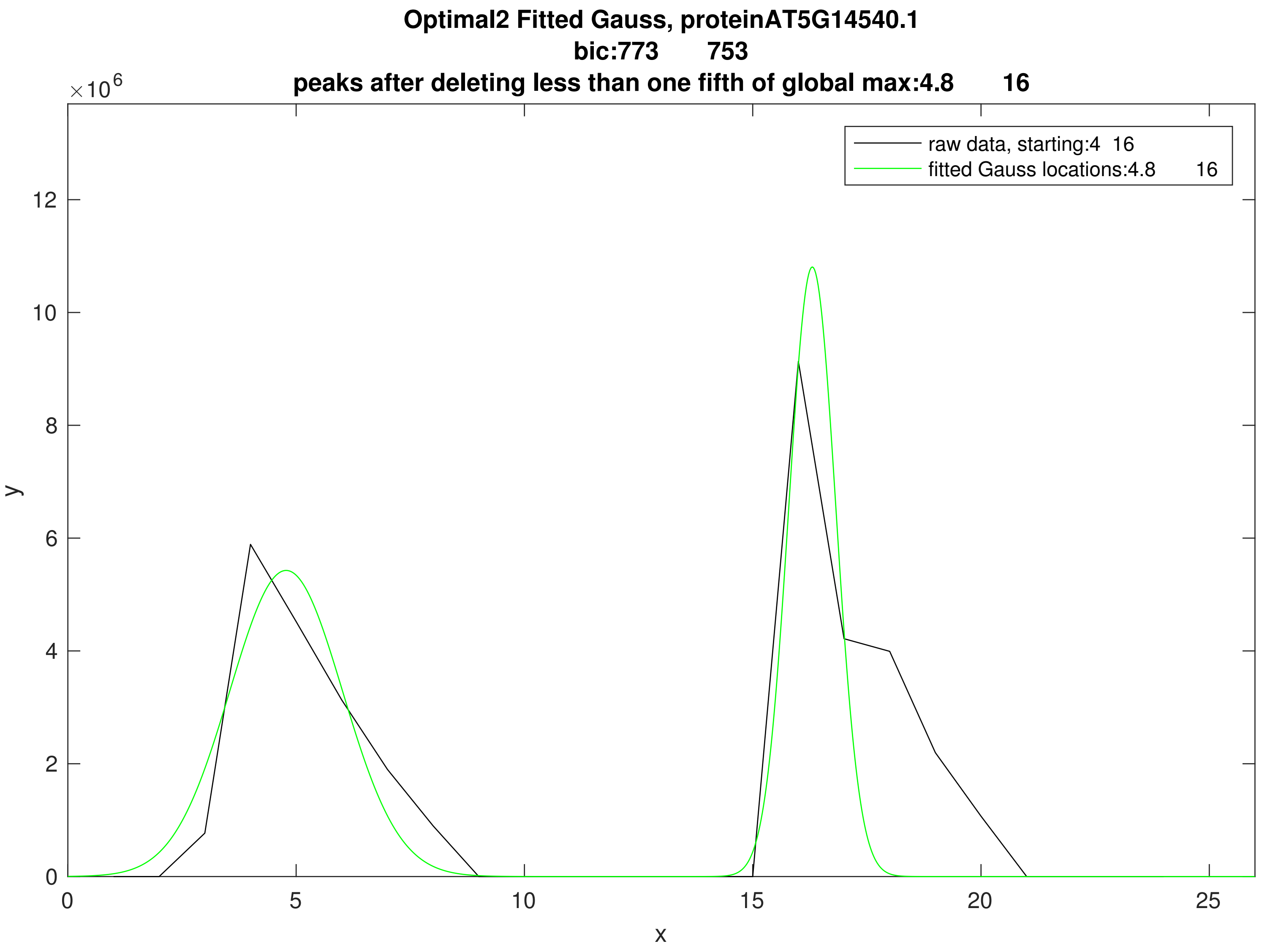
Optimal2 Fitted Gauss, proteinAT5G14540.1
bic:773 753
peaks after deleting less than one fifth of global max:4.8 16
×10^6
raw data, starting:4 16
fitted Gauss locations:4.8 16
y
x
0
2
4
6
8
10
12
0
5
10
15
20
25

Optimal1 Fitted Gauss, proteinAT5G14590.1
bic:895
peaks after deleting less than one fifth of global max:13

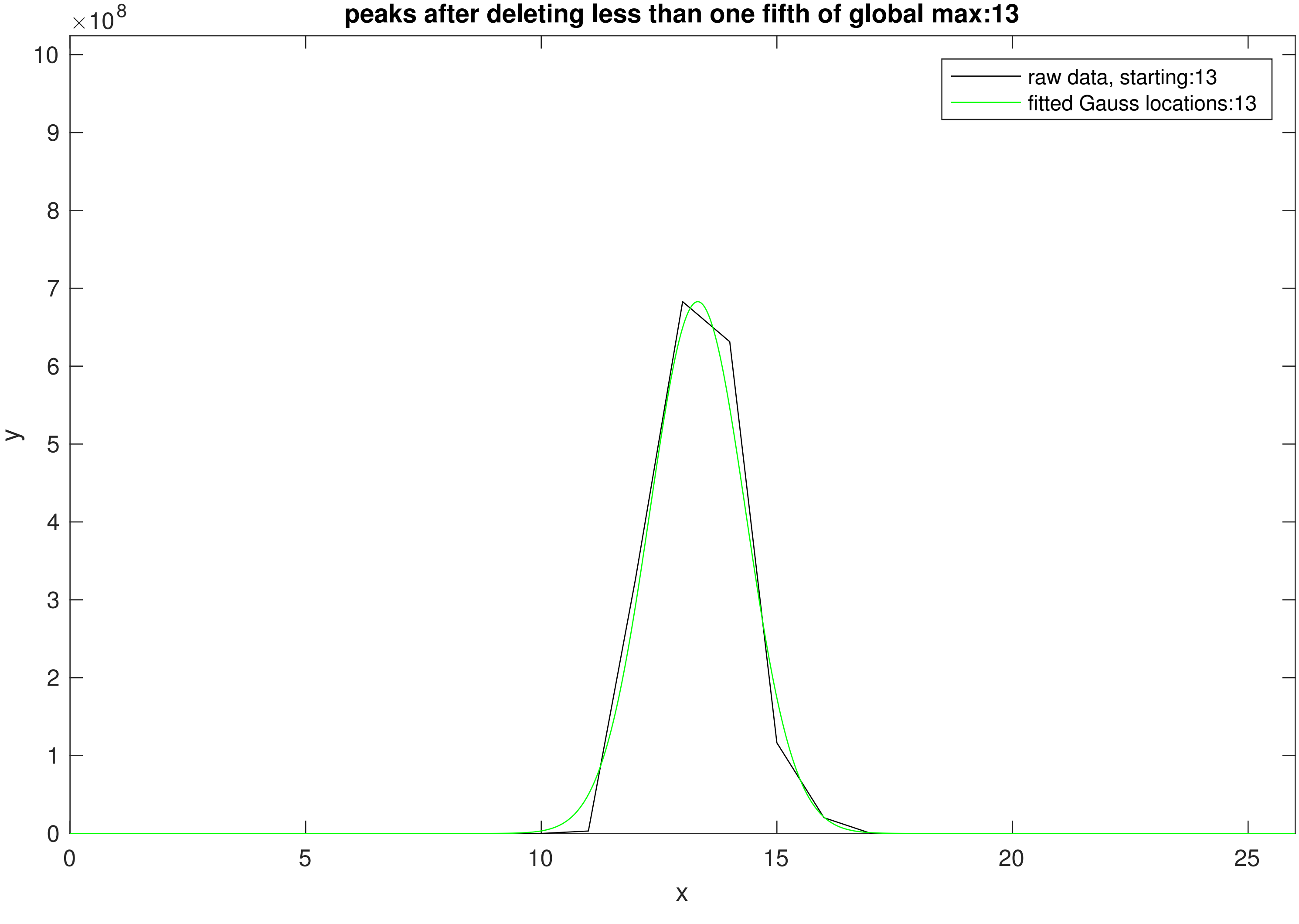

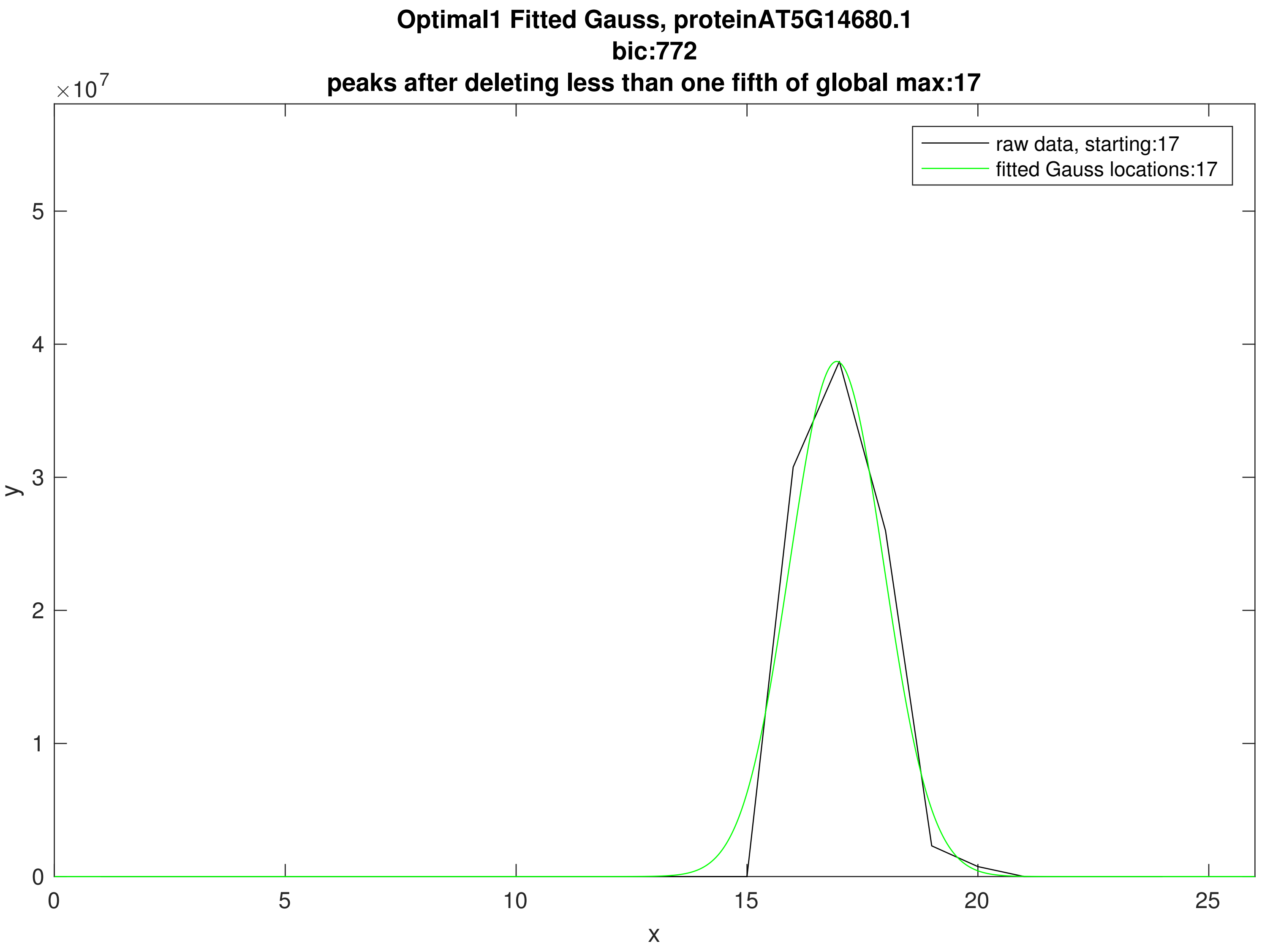

Optimal1 Fitted Gauss, proteinAT5G14680.1
bic:772
peaks after deleting less than one fifth of global max:17
×10^7
raw data, starting:17
fitted Gauss locations:17
y
x
0
1
2
3
4
5
0
5
10
15
20
25

**Optimal1 Fitted Gauss, proteinAT5G14740.2;AT5G14740.1;AT5G14740.5;AT5G14740.4;AT5G14740.3**
**bic:835 838 847 857**
**peaks after deleting less than one fifth of global max:8.8**

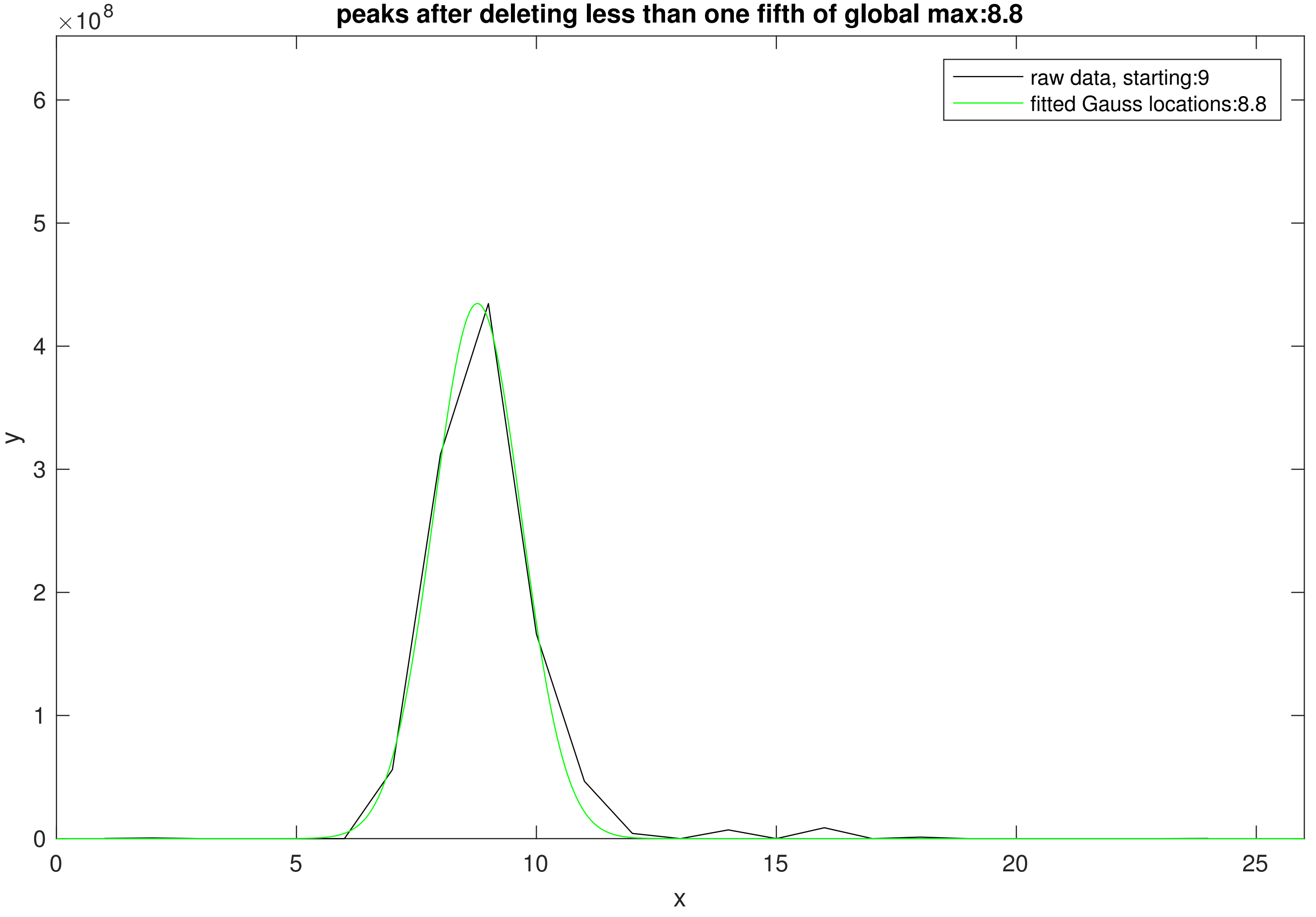

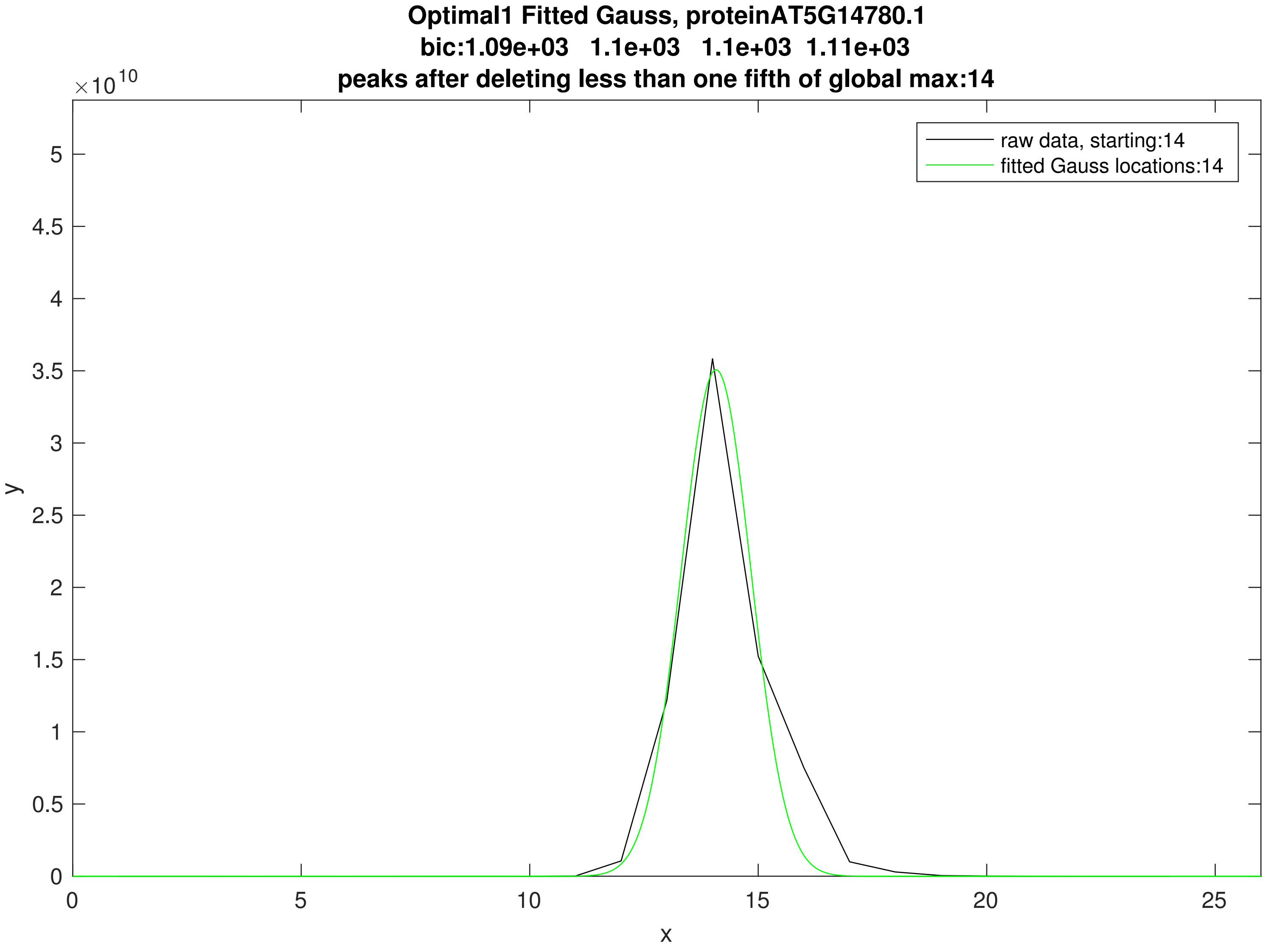

Optimal1 Fitted Gauss, proteinAT5G14780.1
bic:1.09e+03 1.1e+03 1.1e+03 1.11e+03
peaks after deleting less than one fifth of global max:14
×10^10
raw data, starting:14
fitted Gauss locations:14
y
x
0
0.5
1
1.5
2
2.5
3
3.5
4
4.5
5
0
5
10
15
20
25

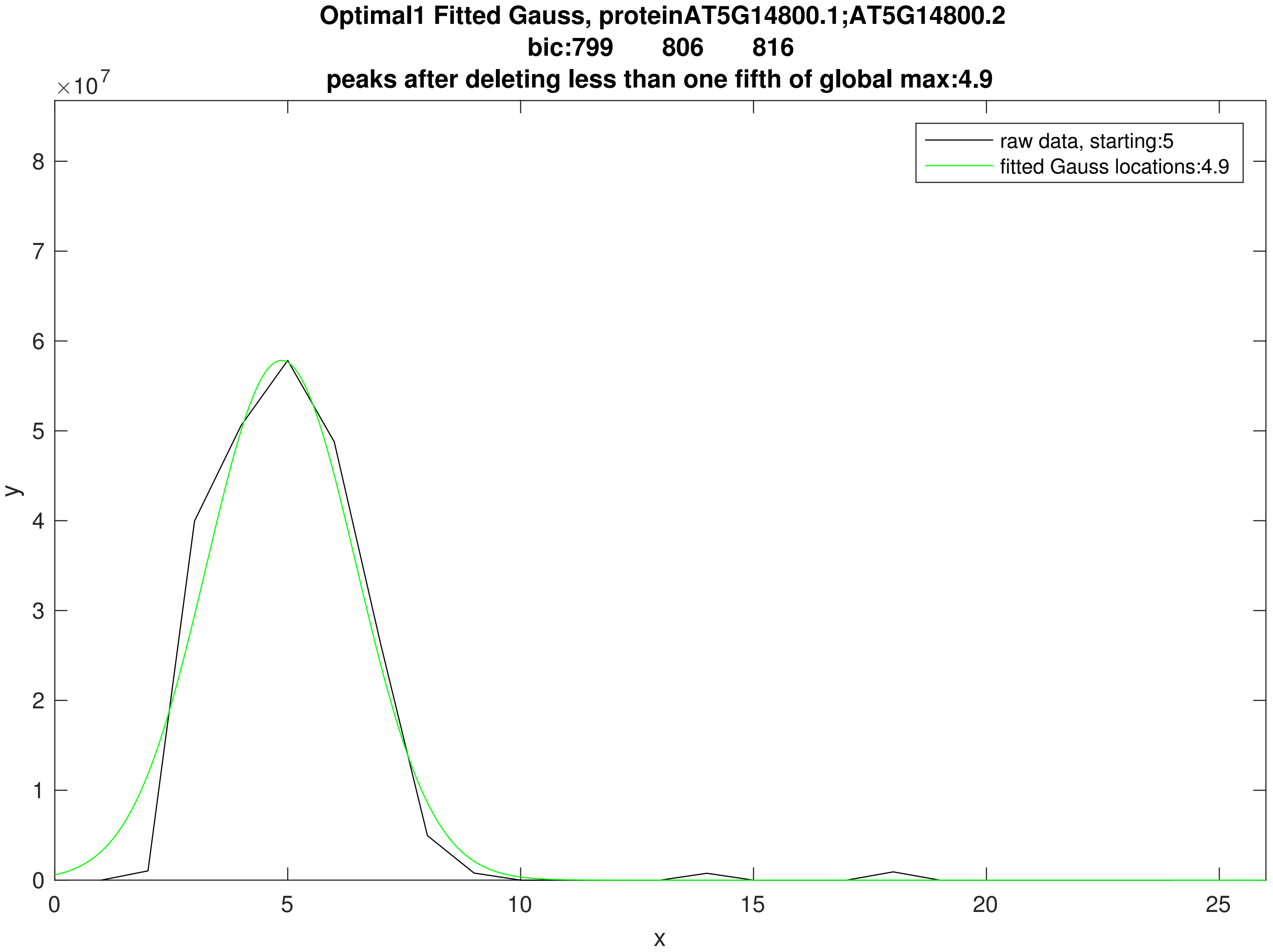
Optimal1 Fitted Gauss, proteinAT5G14800.1;AT5G14800.2
bic:799 806 816
peaks after deleting less than one fifth of global max:4.9
×10^7
raw data, starting:5
fitted Gauss locations:4.9
y
x
0
1
2
3
4
5
6
7
8
10
15
20
25

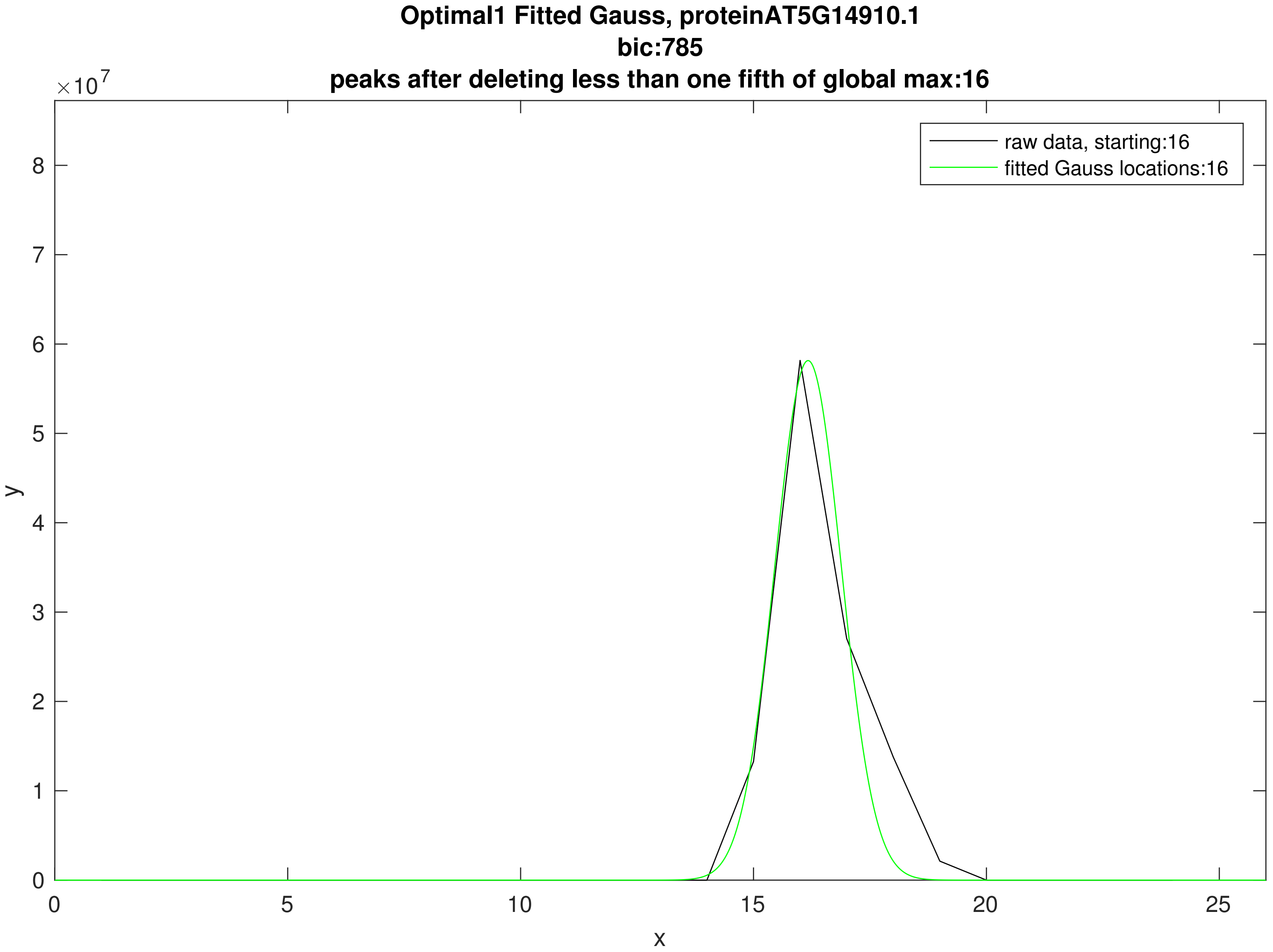
Optimal1 Fitted Gauss, proteinAT5G14910.1
bic:785
peaks after deleting less than one fifth of global max:16
×10^7
raw data, starting:16
fitted Gauss locations:16
y
x
0
1
2
3
4
5
6
7
8
0
5
10
15
20
25

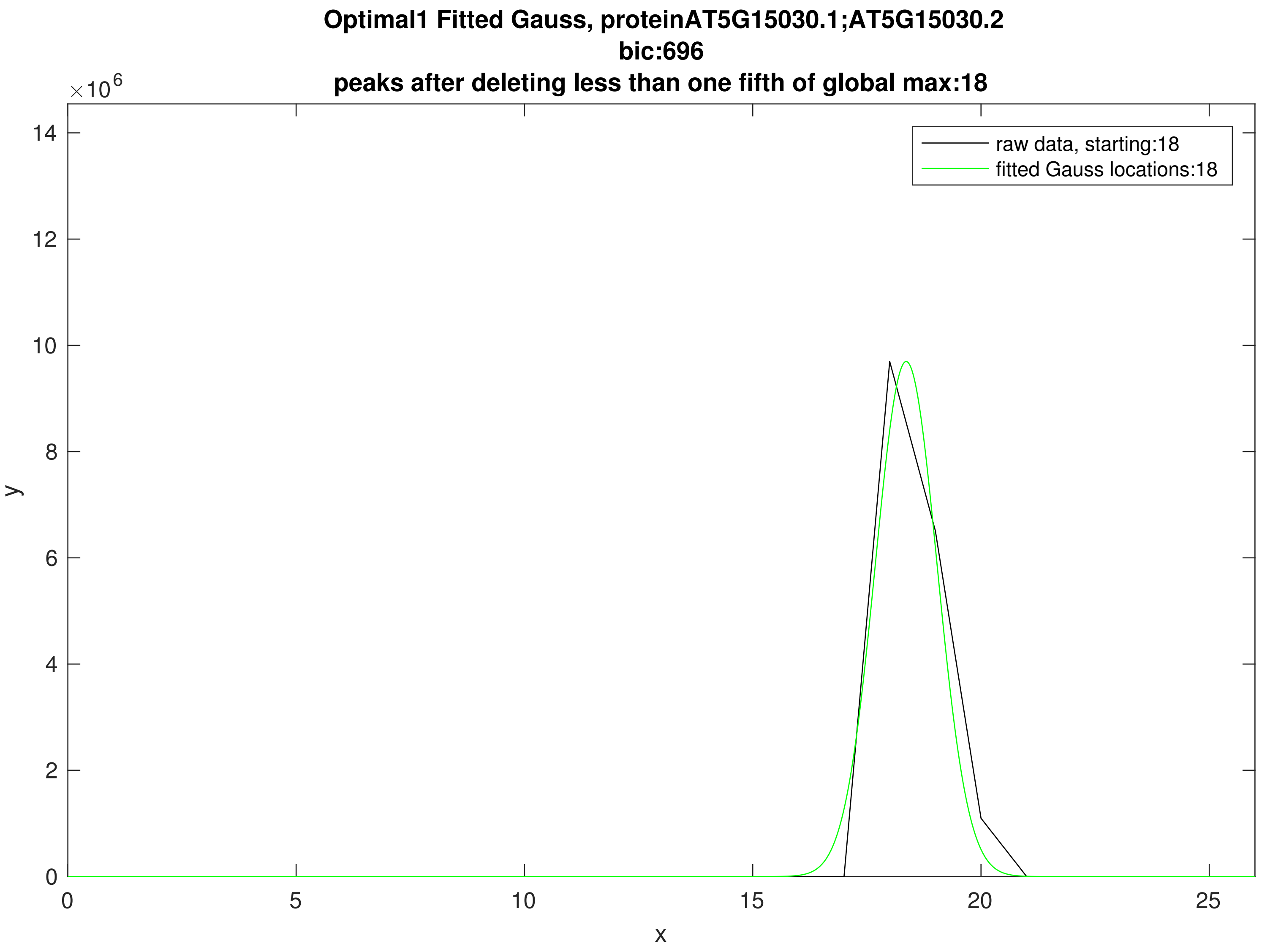
Optimal1 Fitted Gauss, proteinAT5G15030.1;AT5G15030.2
bic:696
peaks after deleting less than one fifth of global max:18
×10^6
raw data, starting:18
fitted Gauss locations:18
0
2
4
6
8
10
12
14
y
0
5
10
15
20
25
x

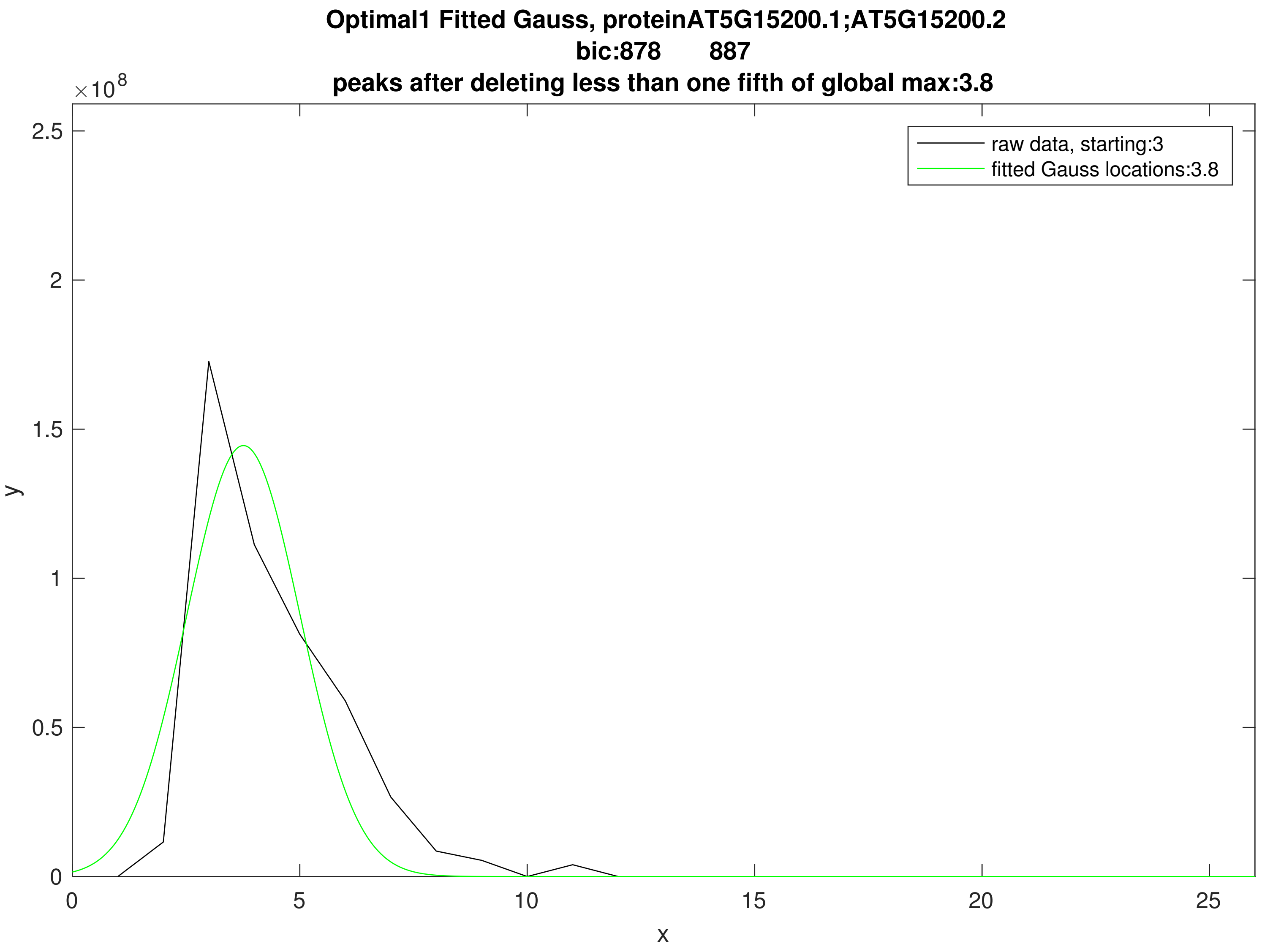
Optimal1 Fitted Gauss, proteinAT5G15200.1;AT5G15200.2
bic:878 887
peaks after deleting less than one fifth of global max:3.8
×10^8
2.5
2
1.5
1
0.5
0
y
0
5
10
15
20
25
x
raw data, starting:3
fitted Gauss locations:3.8

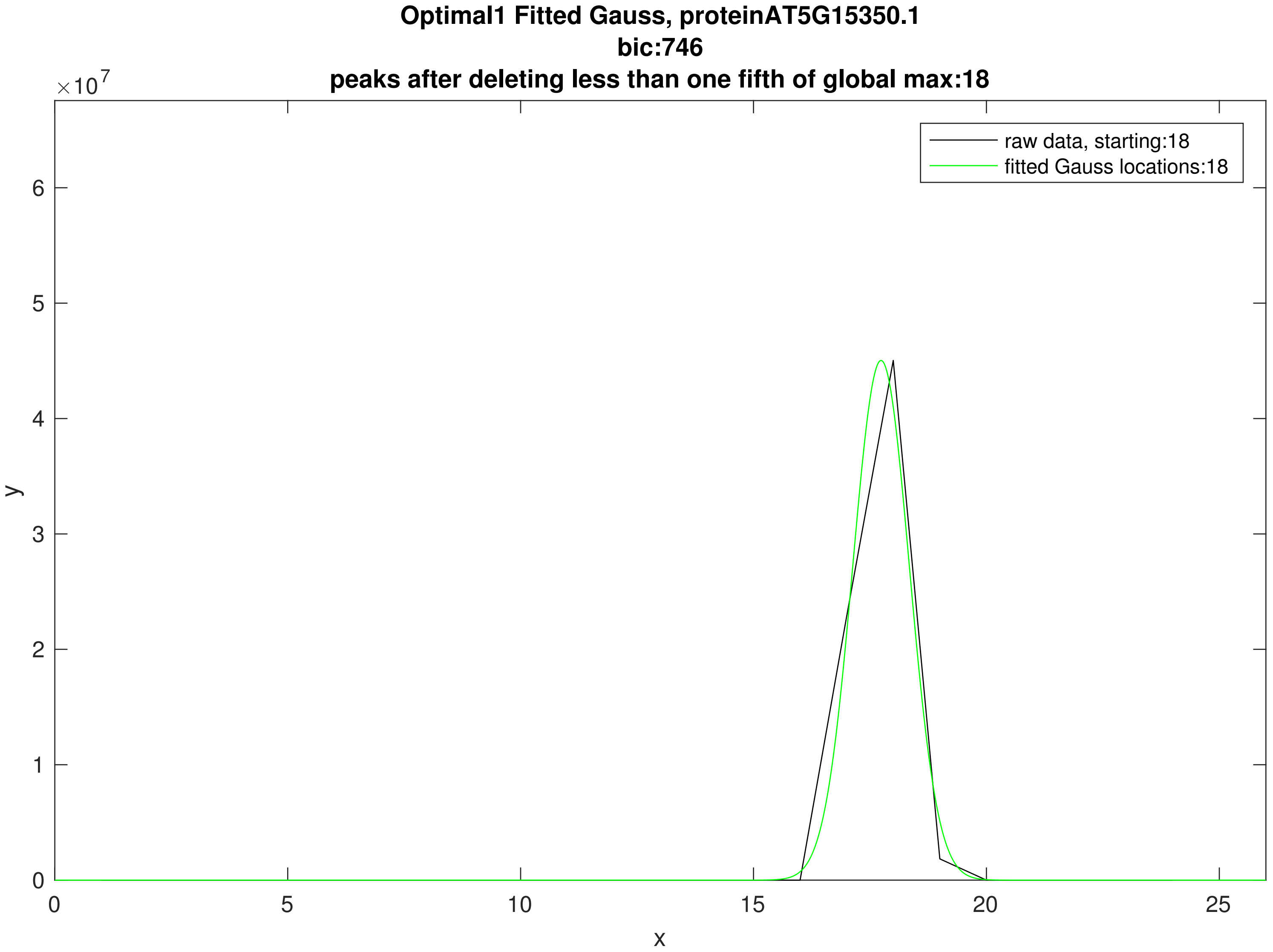

Optimal1 Fitted Gauss, proteinAT5G15350.1
bic:746
peaks after deleting less than one fifth of global max:18
×10^7
raw data, starting:18
fitted Gauss locations:18
y
x
0
1
2
3
4
5
6
0
5
10
15
20
25

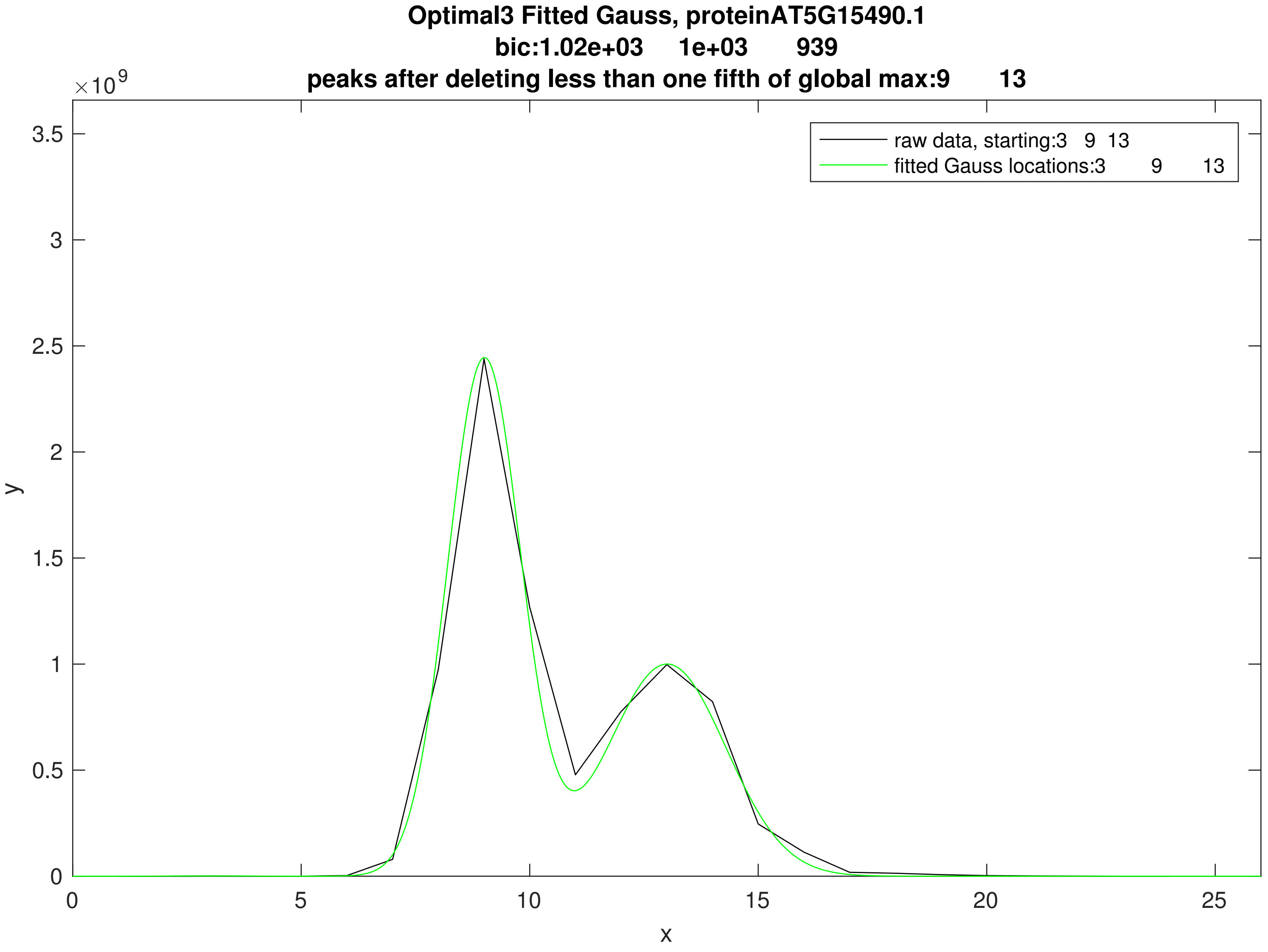
Optimal3 Fitted Gauss, proteinAT5G15490.1
bic:1.02e+03 1e+03 939
peaks after deleting less than one fifth of global max:9 13
×10^9
3.5
3
2.5
2
1.5
1
0.5
0
y
0
5
10
15
20
25
x
raw data, starting:3 9 13
fitted Gauss locations:3 9 13

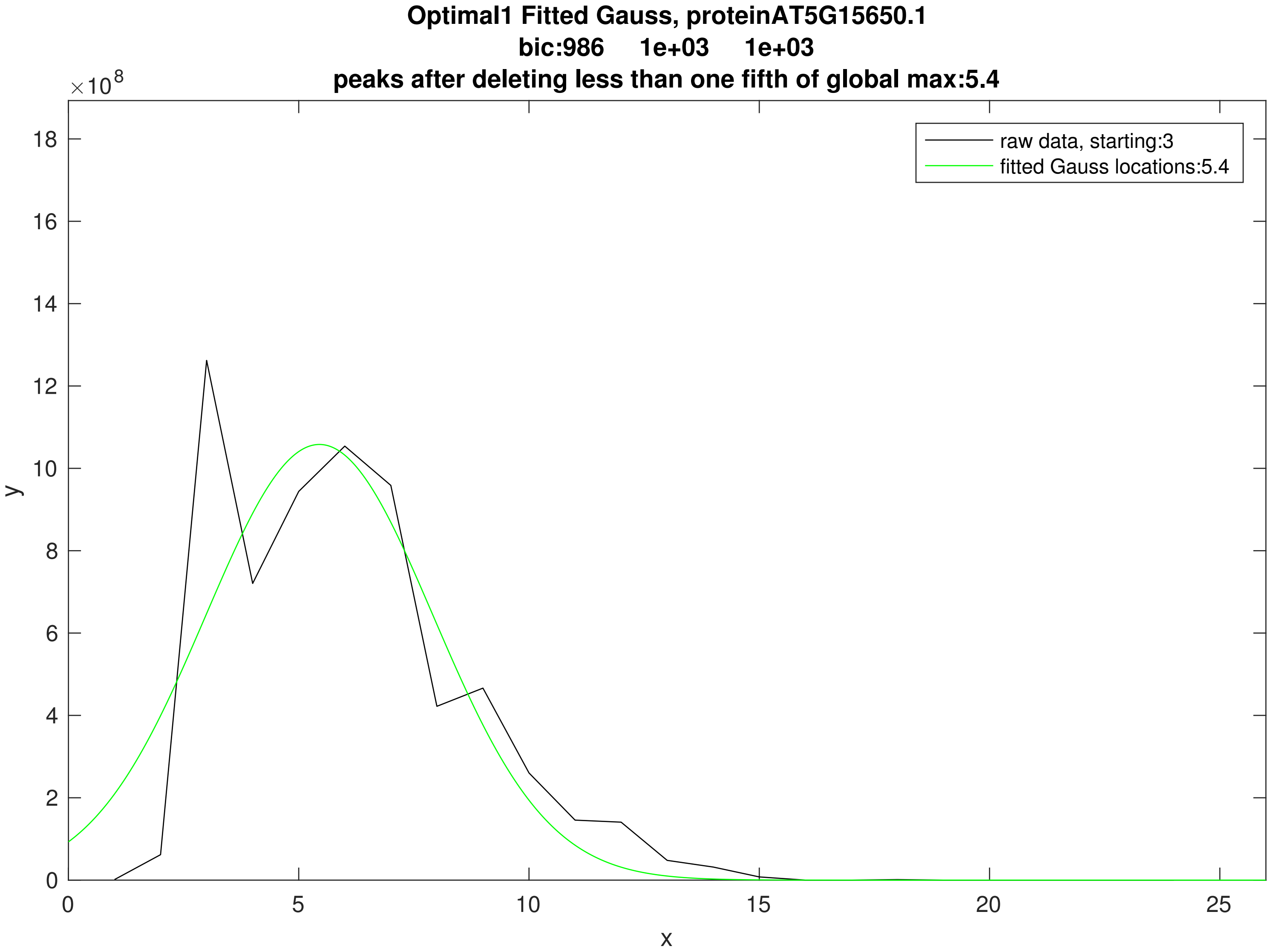
Optimal1 Fitted Gauss, proteinAT5G15650.1
bic:986 1e+03 1e+03
peaks after deleting less than one fifth of global max:5.4
×10^8
raw data, starting:3
fitted Gauss locations:5.4
y
x
0
2
4
6
8
10
12
14
16
18
0
5
10
15
20
25

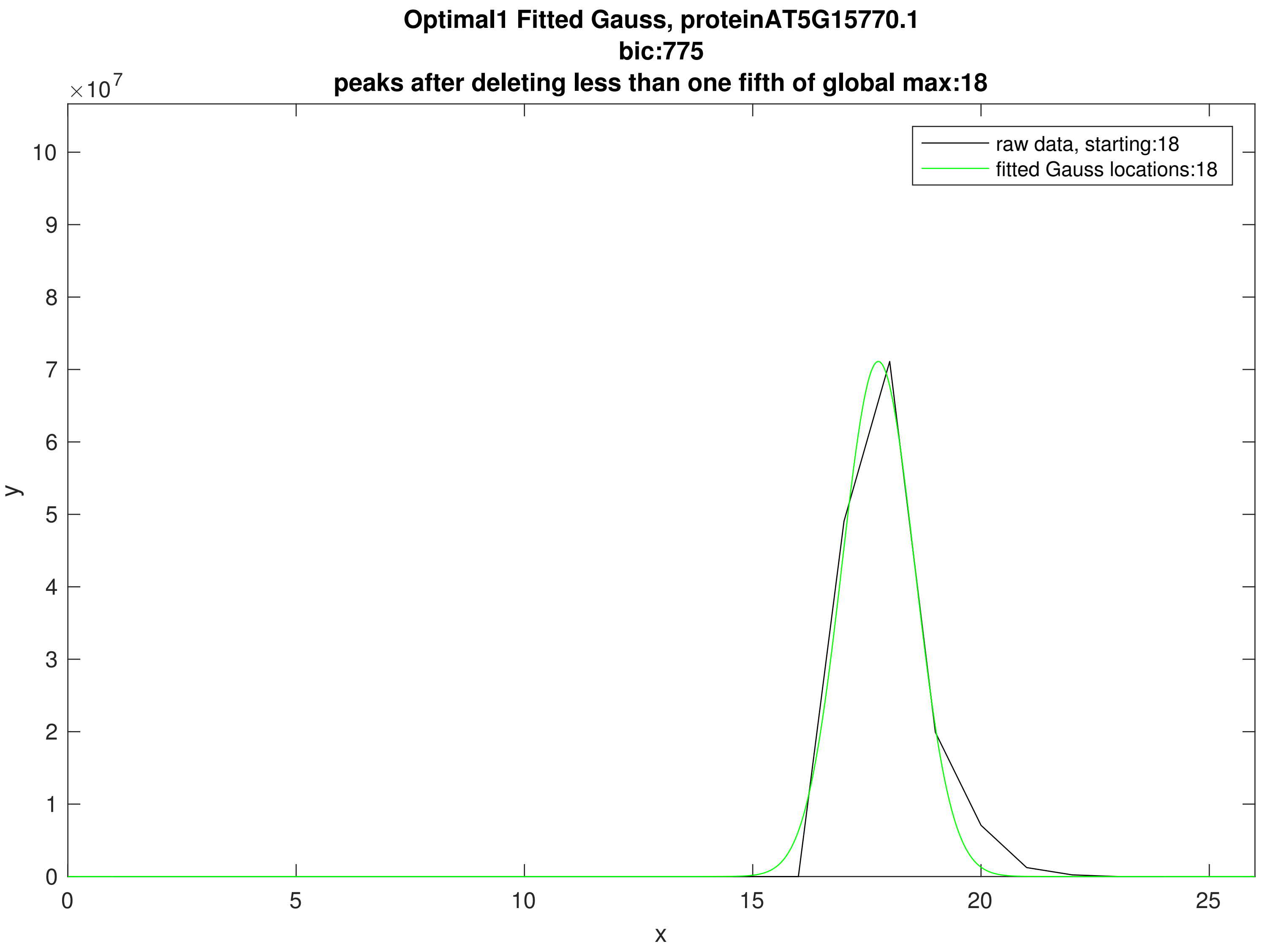

Optimal1 Fitted Gauss, proteinAT5G15770.1
bic:775
peaks after deleting less than one fifth of global max:18
×10^7
raw data, starting:18
fitted Gauss locations:18
y
x
0
1
2
3
4
5
6
7
8
9
10
0
5
10
15
20
25

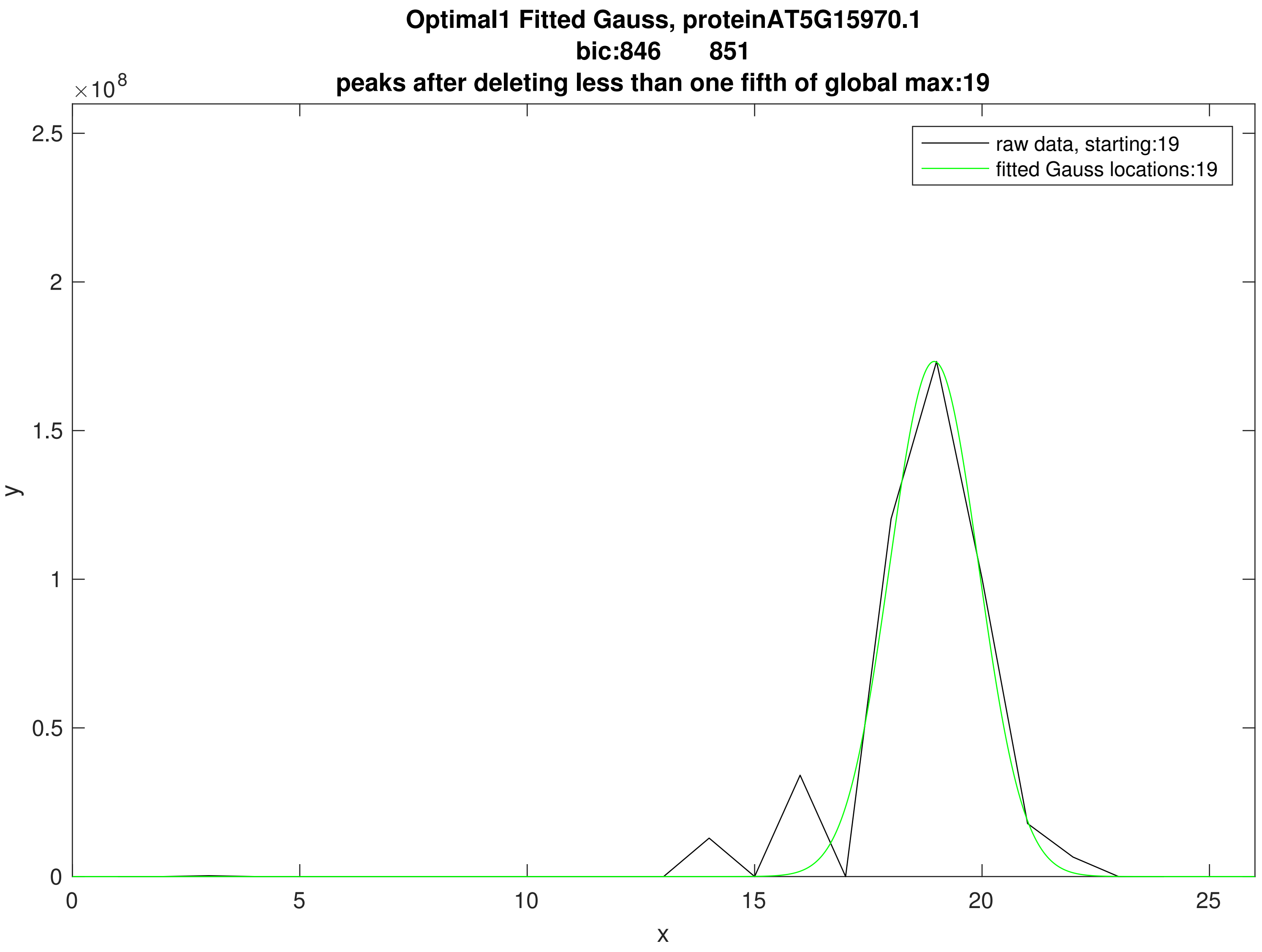
Optimal1 Fitted Gauss, proteinAT5G15970.1
bic:846 851
peaks after deleting less than one fifth of global max:19
×10^8
2.5
2
1.5
1
0.5
0
y
0
5
10
15
20
25
x
raw data, starting:19
fitted Gauss locations:19

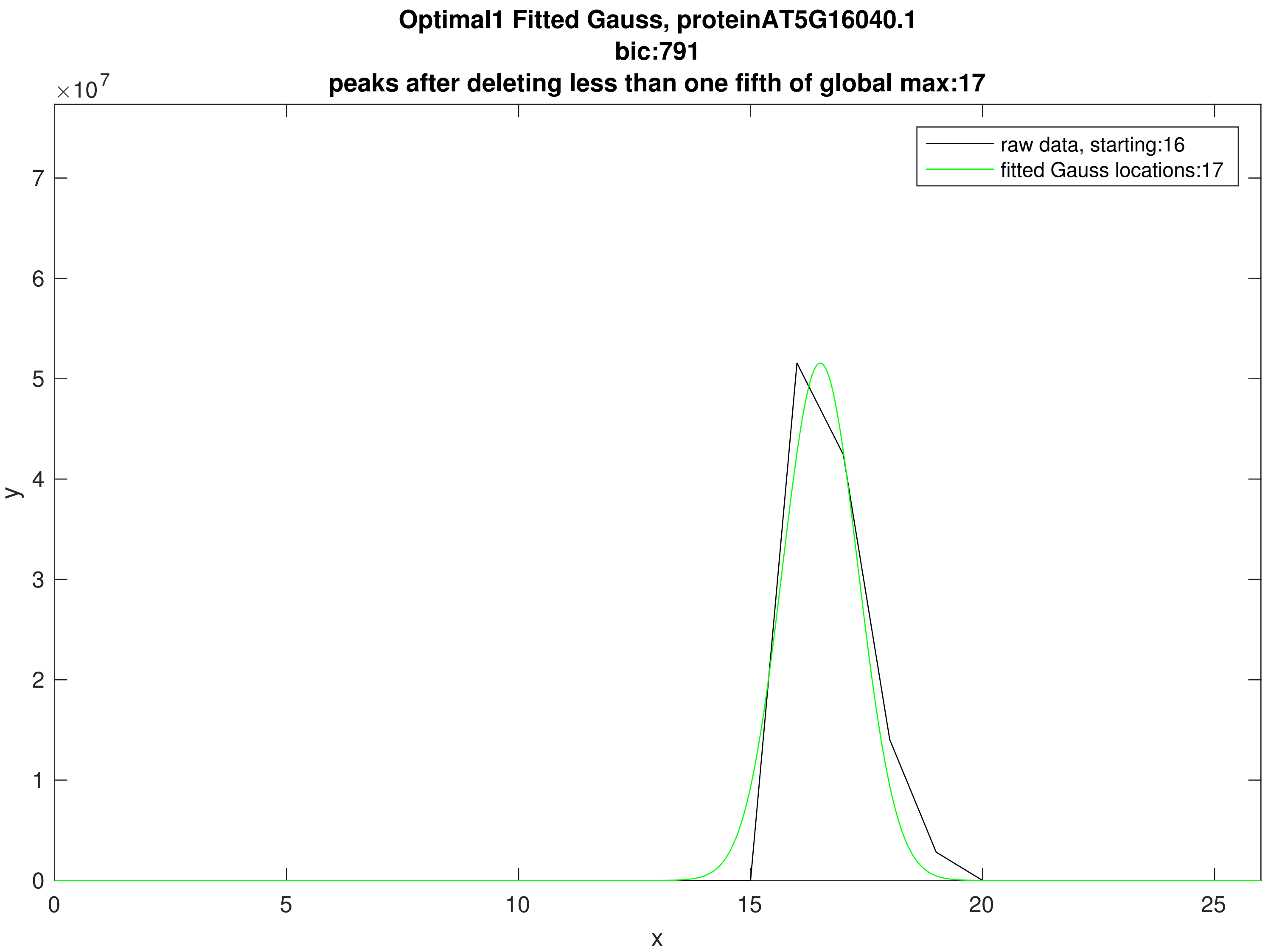

Optimal1 Fitted Gauss, proteinAT5G16040.1
bic:791
peaks after deleting less than one fifth of global max:17
×10^7
raw data, starting:16
fitted Gauss locations:17
y
x
0
1
2
3
4
5
6
7
0
5
10
15
20
25

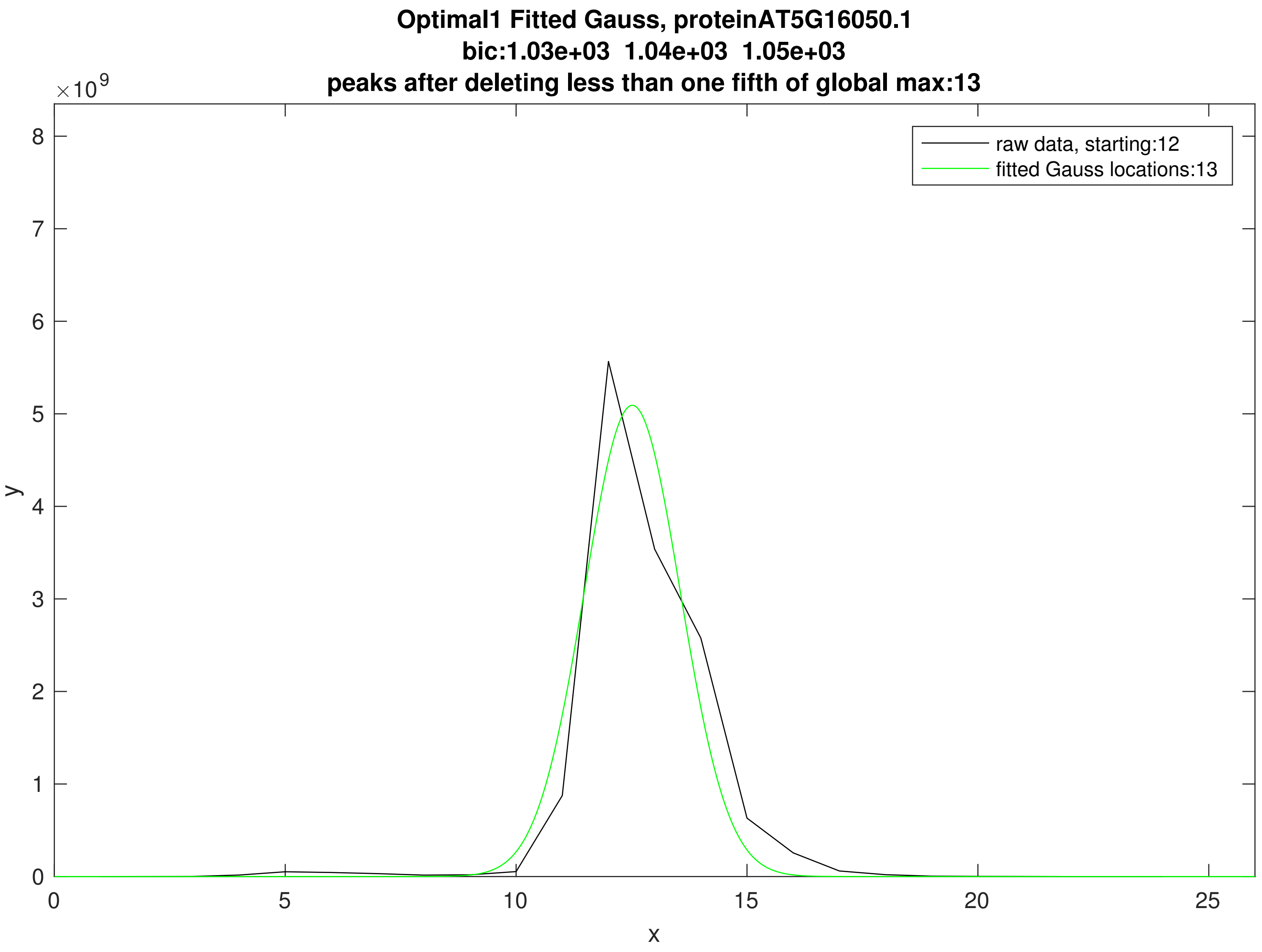

Optimal1 Fitted Gauss, proteinAT5G16050.1
bic:1.03e+03  1.04e+03  1.05e+03
peaks after deleting less than one fifth of global max:13
×10^9
raw data, starting:12
fitted Gauss locations:13
y
x
0
1
2
3
4
5
6
7
8
0
5
10
15
20
25

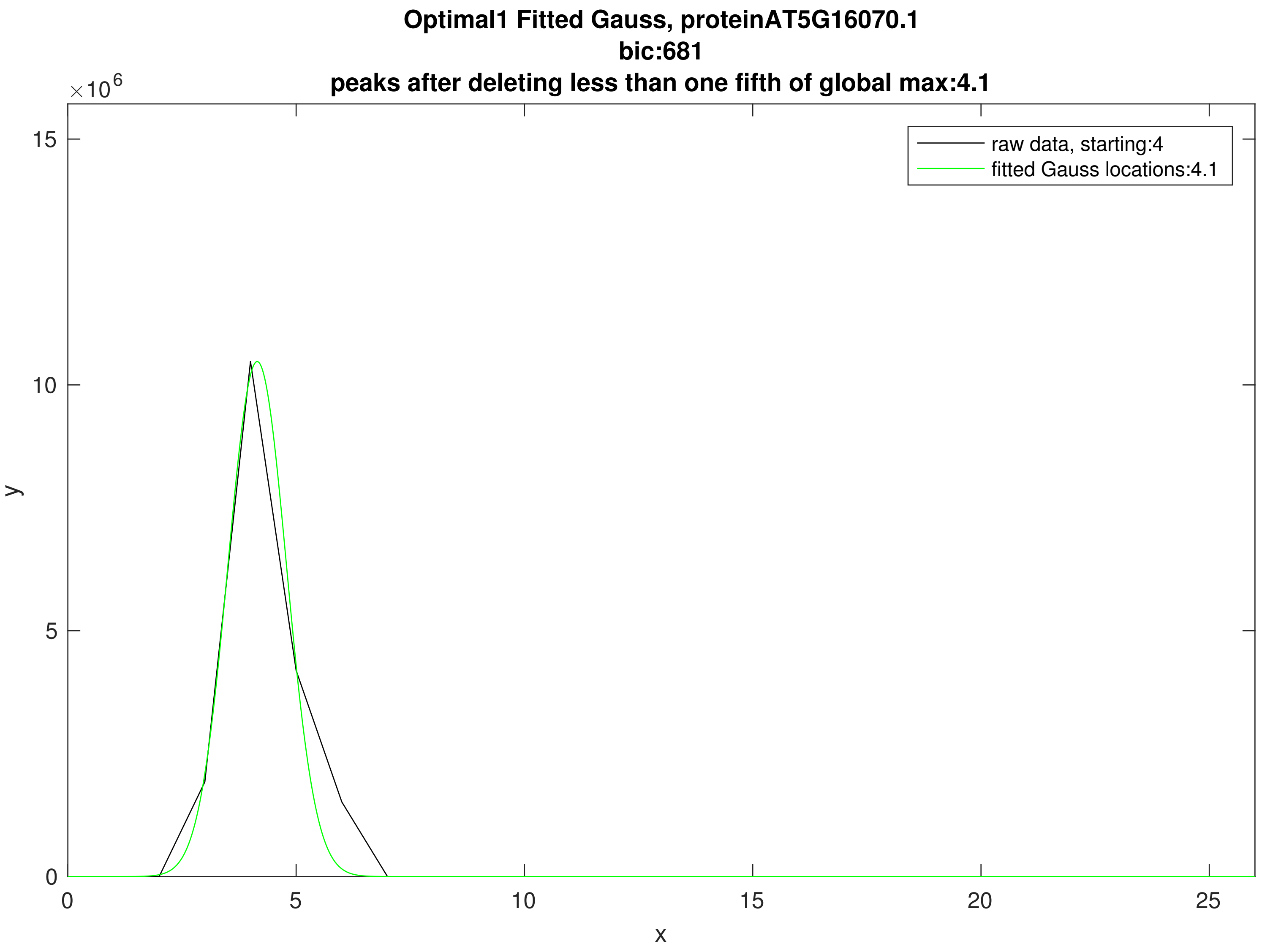
Optimal1 Fitted Gauss, proteinAT5G16070.1
bic:681
peaks after deleting less than one fifth of global max:4.1
×10^6
raw data, starting:4
fitted Gauss locations:4.1
y
x
0
5
10
15
20
25

Optimal1 Fitted Gauss, proteinAT5G16130.1
bic:790
peaks after deleting less than one fifth of global max:3.6

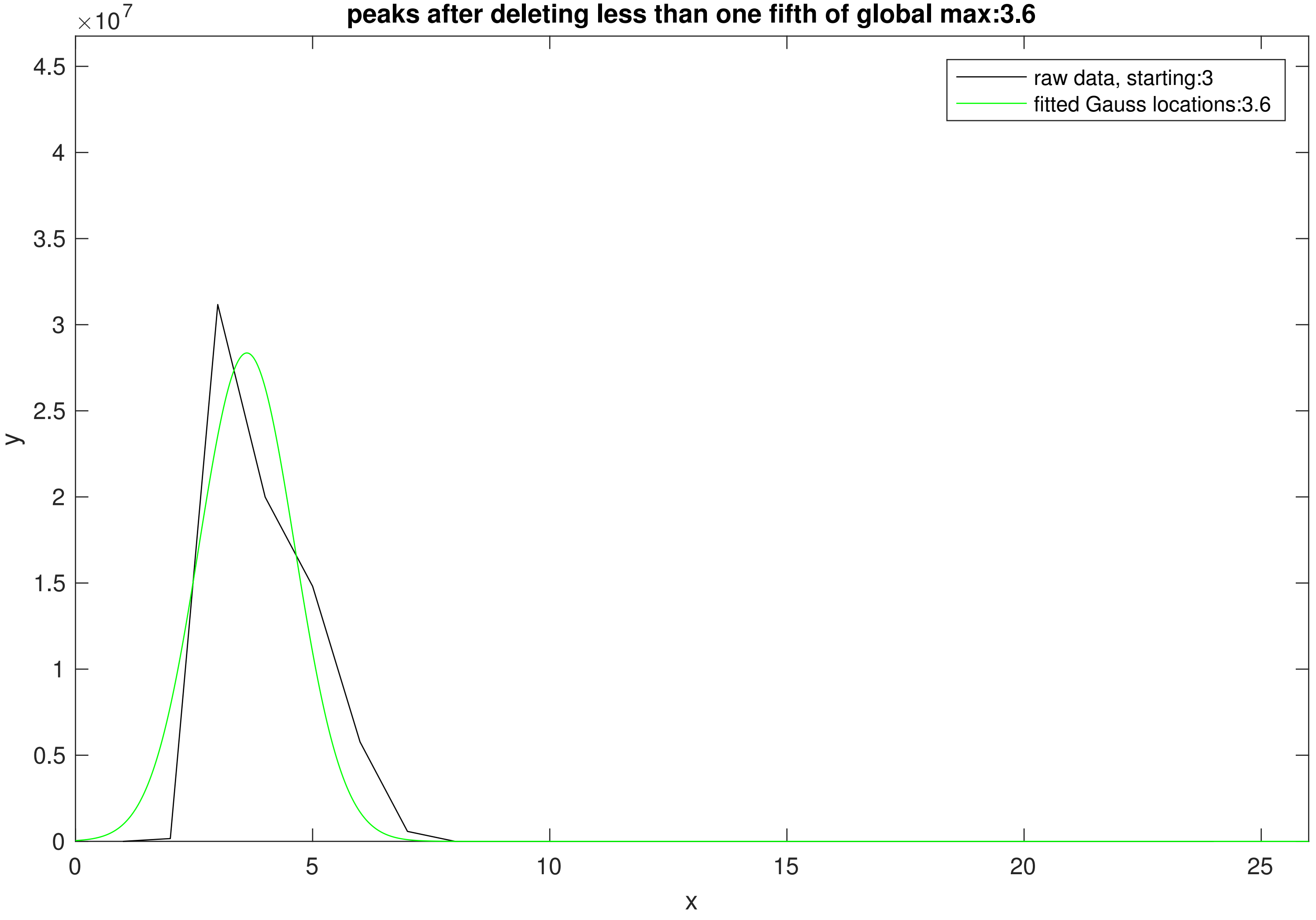

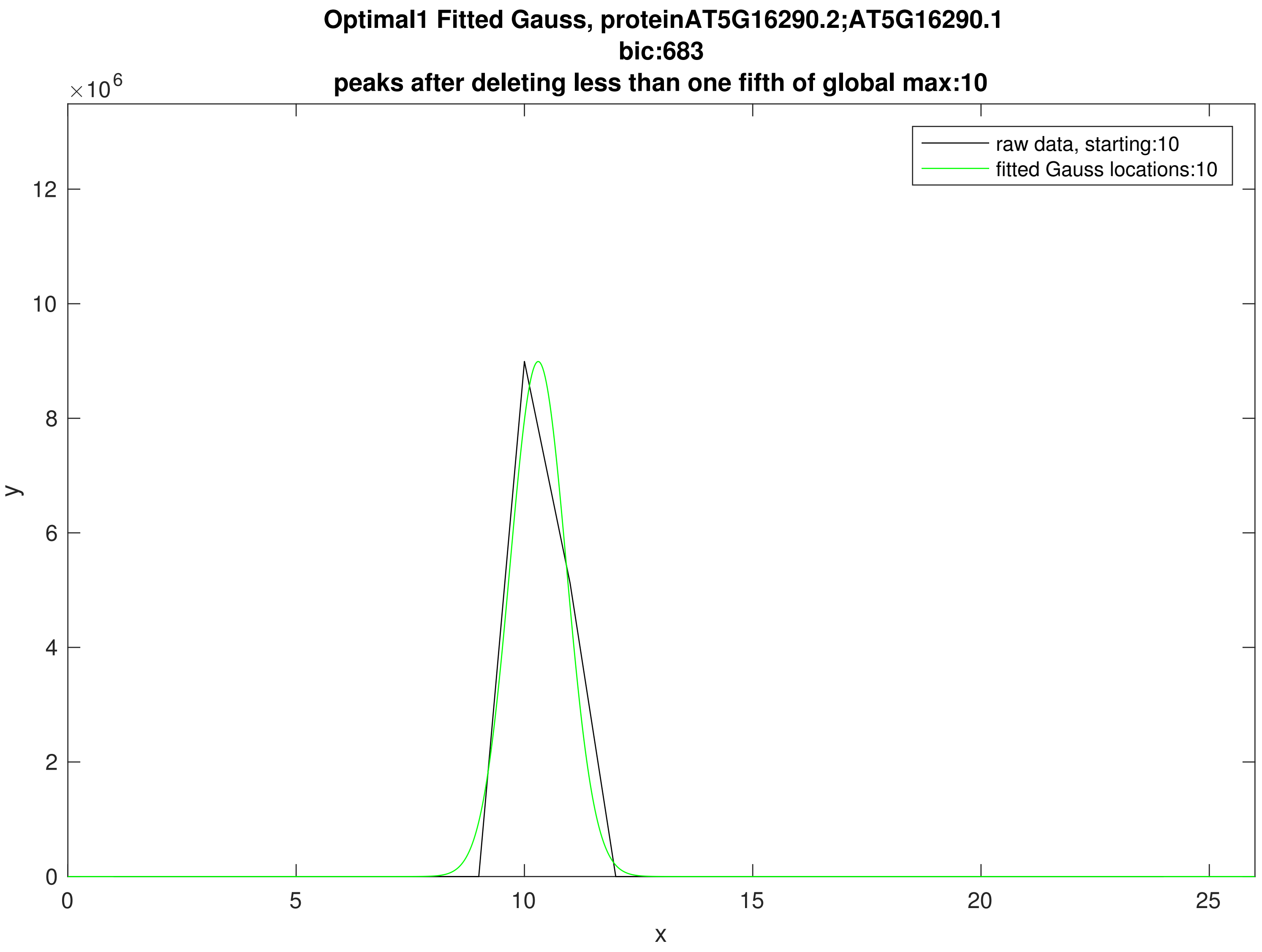

Optimal1 Fitted Gauss, proteinAT5G16290.2;AT5G16290.1
bic:683
peaks after deleting less than one fifth of global max:10
×10^6
raw data, starting:10
fitted Gauss locations:10
y
x
0
2
4
6
8
10
12
0
5
10
15
20
25

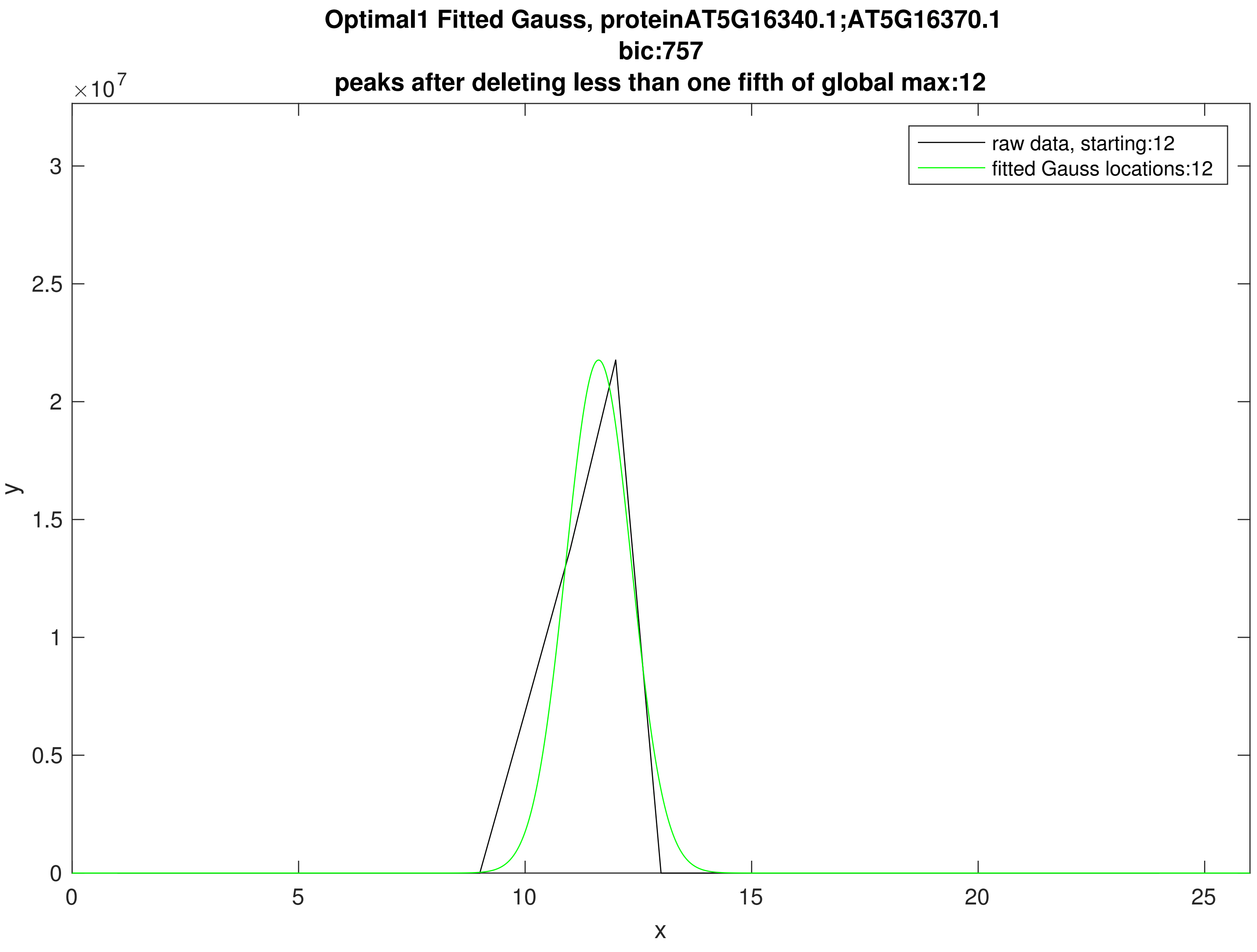
Optimal1 Fitted Gauss, proteinAT5G16340.1;AT5G16370.1
bic:757
peaks after deleting less than one fifth of global max:12
×10^7
raw data, starting:12
fitted Gauss locations:12
3
2.5
2
1.5
1
0.5
0
y
0
5
10
15
20
25
x

Optimal1 Fitted Gauss, proteinAT5G16390.2;AT5G16390.1
bic:889 899
peaks after deleting less than one fifth of global max:14

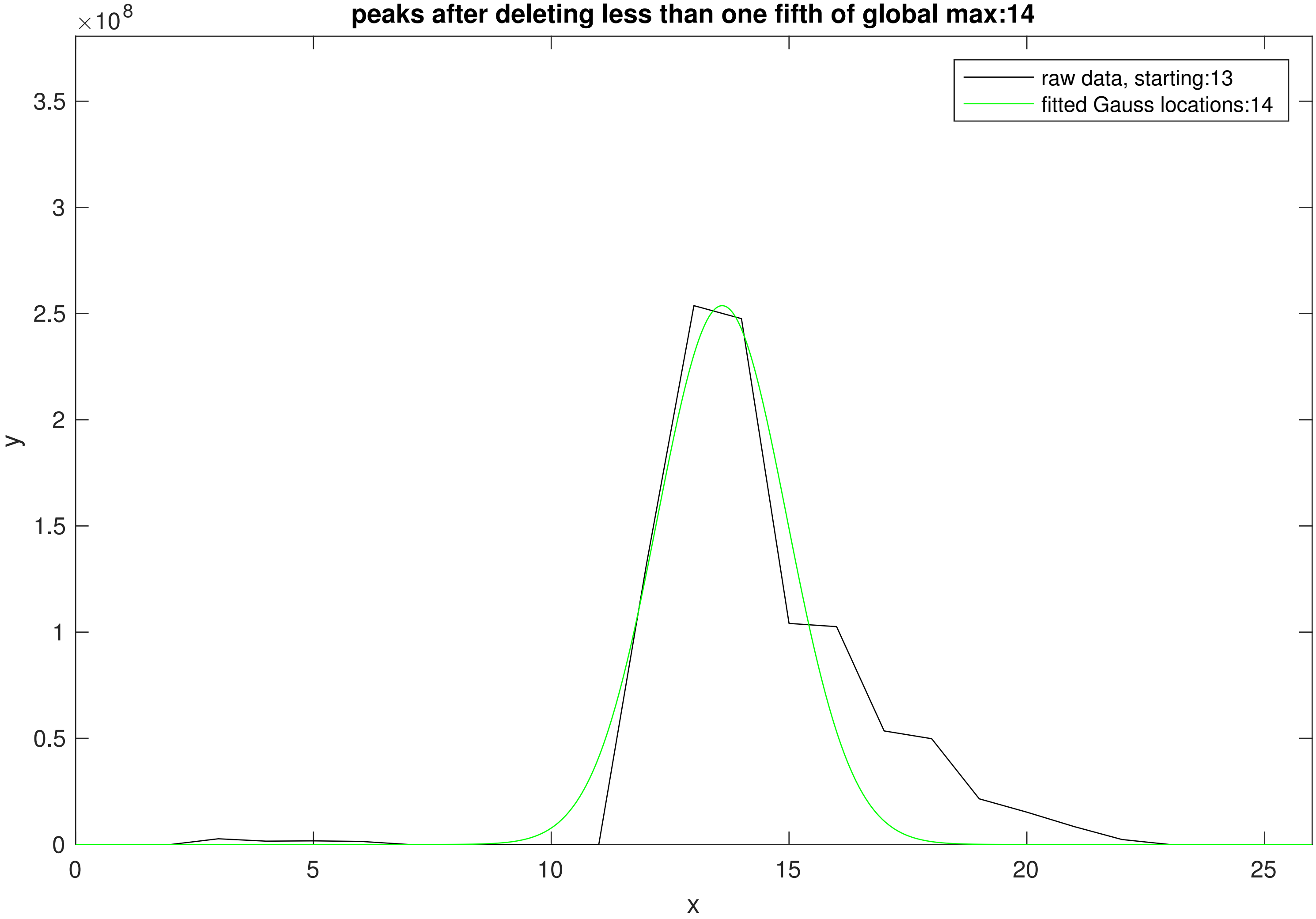

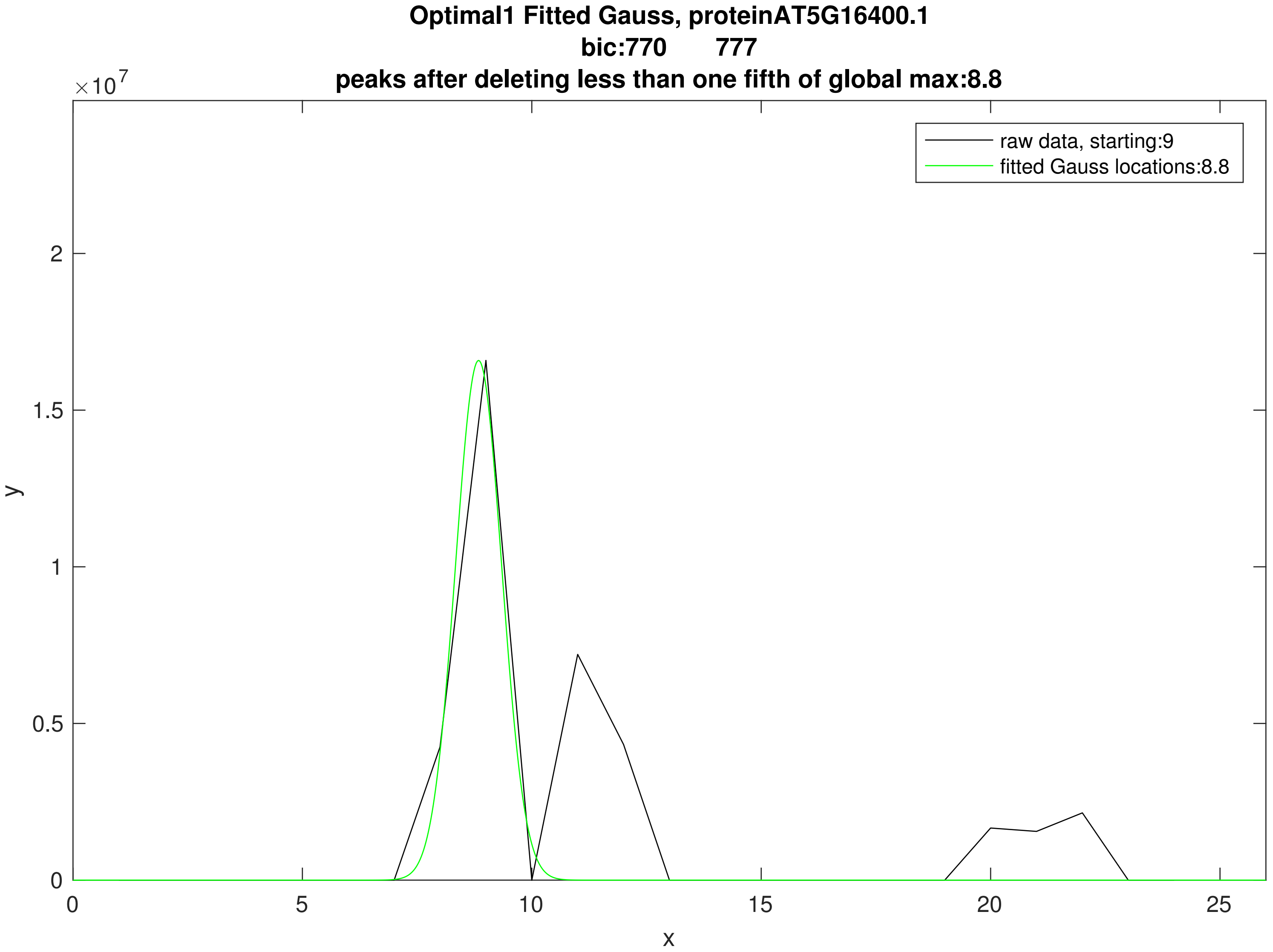
Optimal1 Fitted Gauss, proteinAT5G16400.1
bic:770 777
peaks after deleting less than one fifth of global max:8.8
×10^7
raw data, starting:9
fitted Gauss locations:8.8
y
x
0
0.5
1
1.5
2
5
10
15
20
25

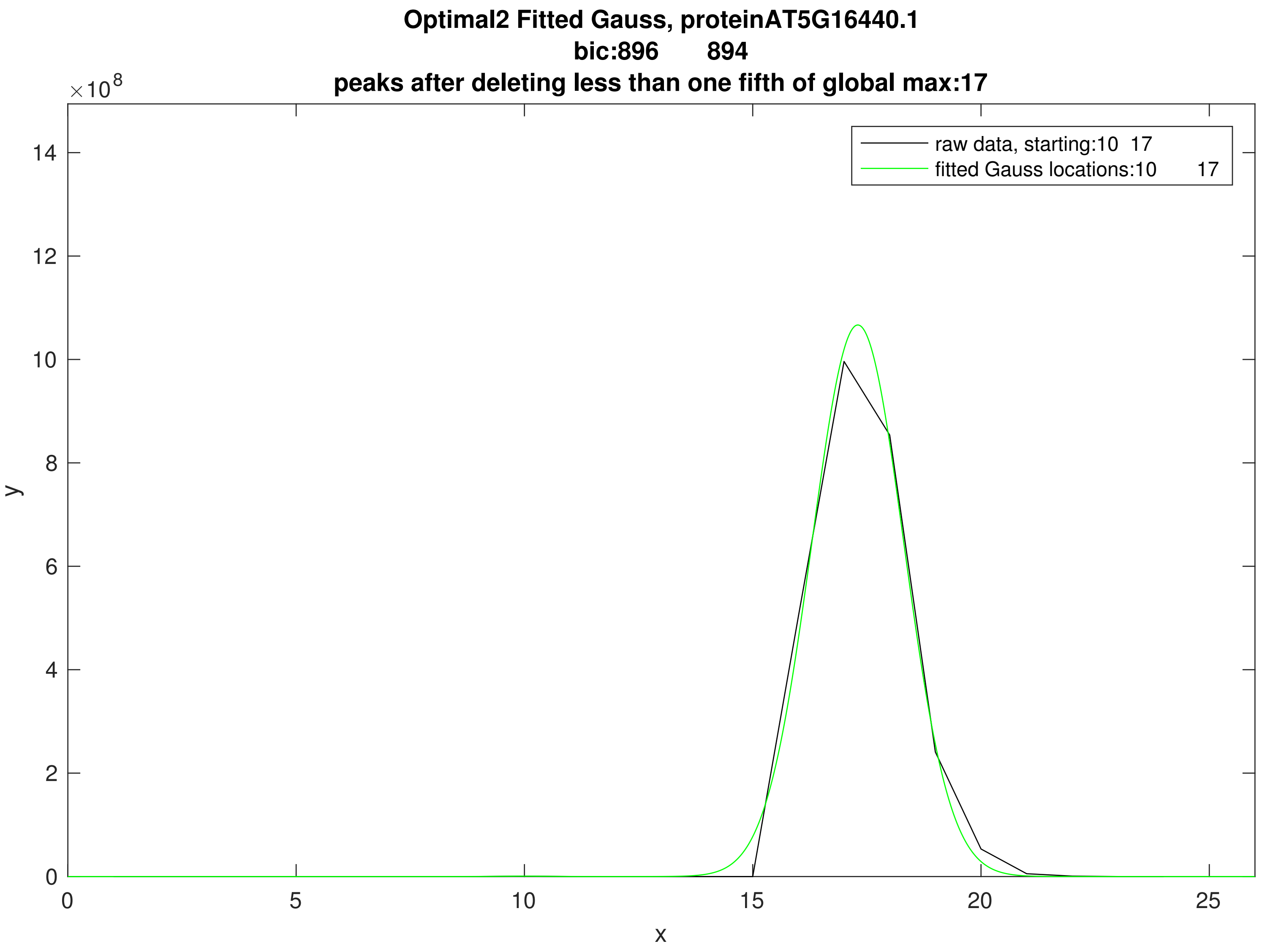
Optimal2 Fitted Gauss, proteinAT5G16440.1
bic:896 894
peaks after deleting less than one fifth of global max:17
×10^8
14
12
10
8
6
4
2
0
y
0
5
10
15
20
25
x
raw data, starting:10 17
fitted Gauss locations:10 17

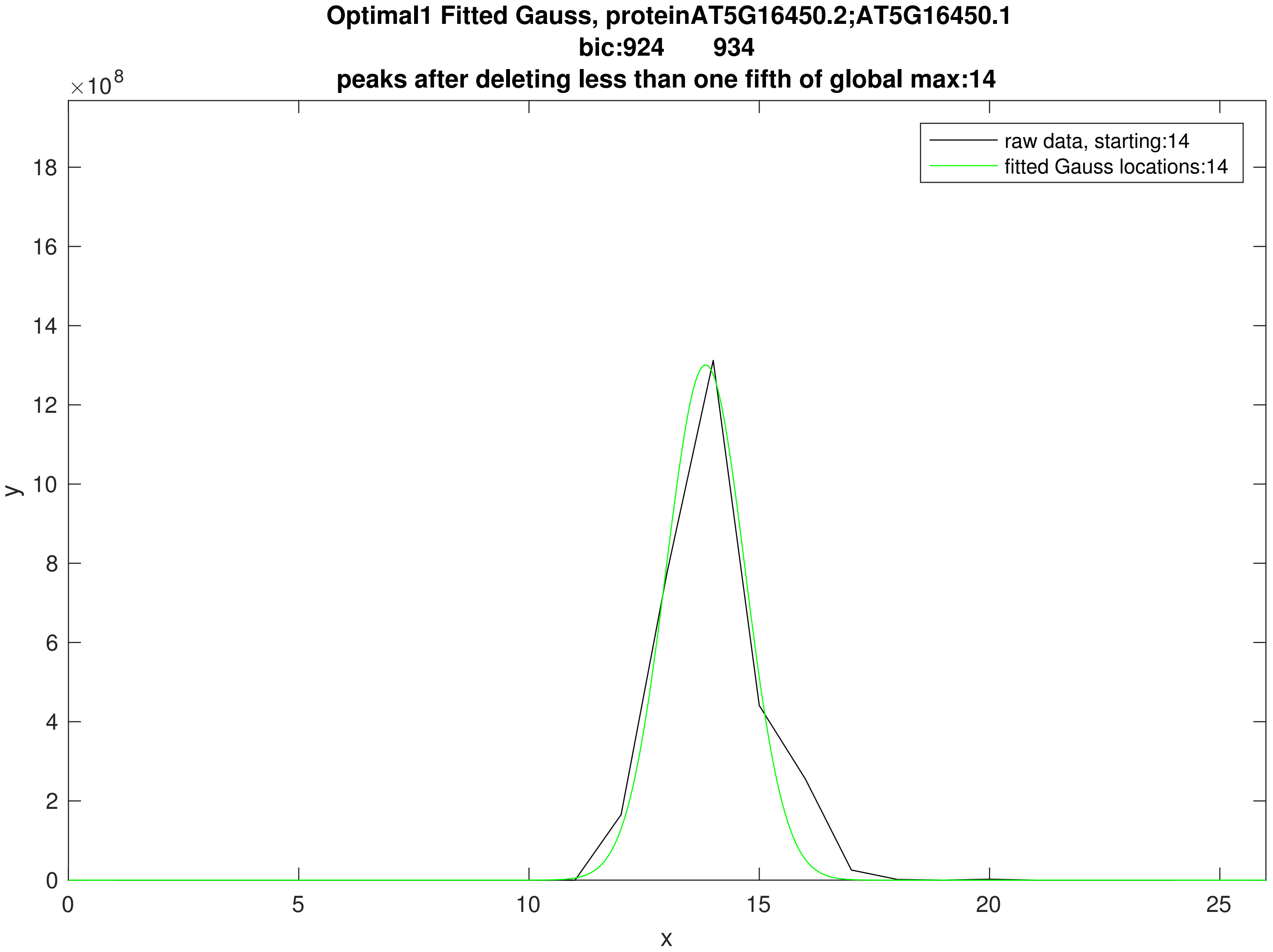
Optimal1 Fitted Gauss, proteinAT5G16450.2;AT5G16450.1
bic:924 934
peaks after deleting less than one fifth of global max:14
×10^8
raw data, starting:14
fitted Gauss locations:14
y
x
0
2
4
6
8
10
12
14
16
18
0
5
10
15
20
25

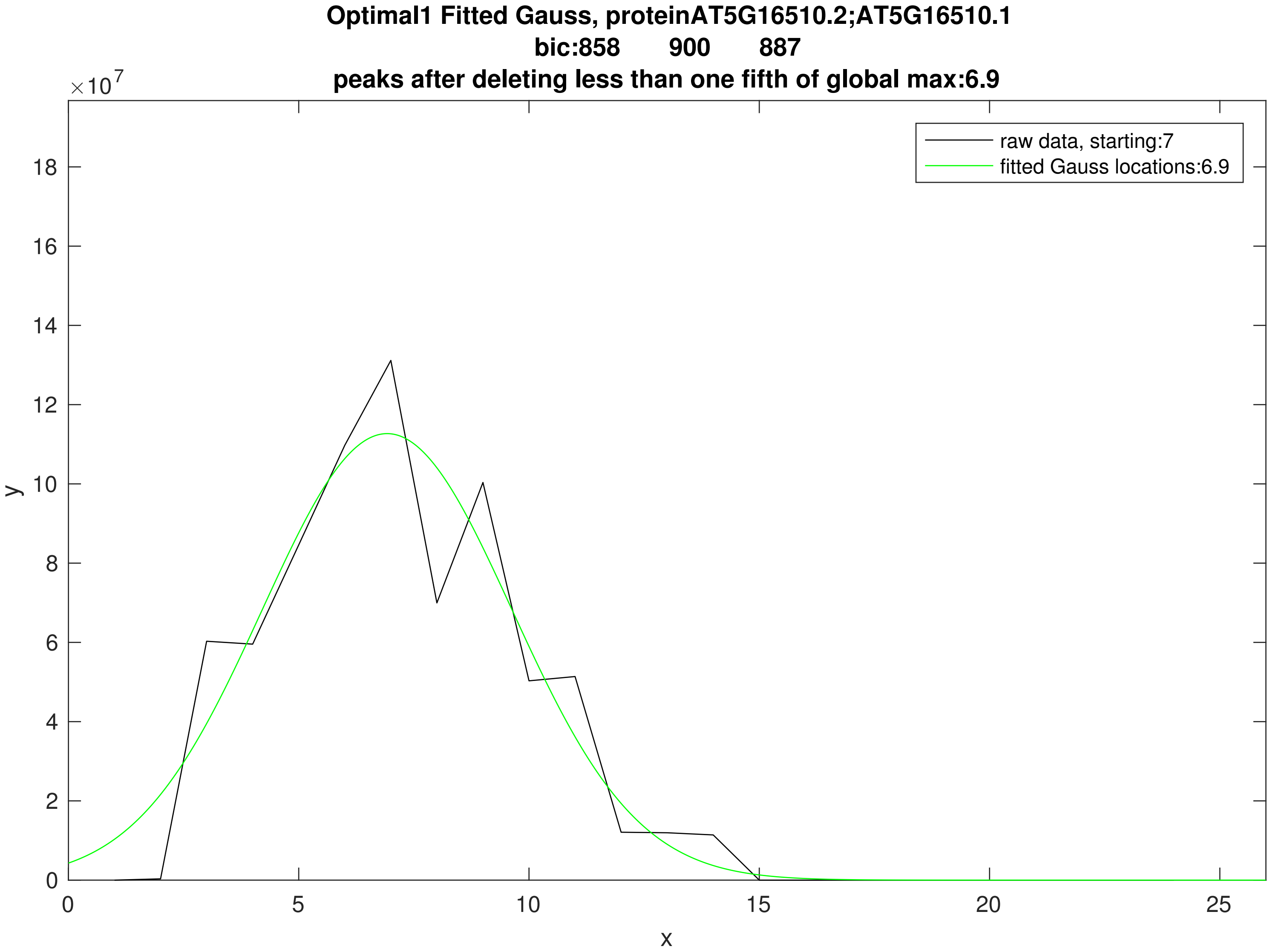
Optimal1 Fitted Gauss, proteinAT5G16510.2;AT5G16510.1
bic:858 900 887
peaks after deleting less than one fifth of global max:6.9
×10^7
raw data, starting:7
fitted Gauss locations:6.9
y
x
0
2
4
6
8
10
12
14
16
18
0
5
10
15
20
25

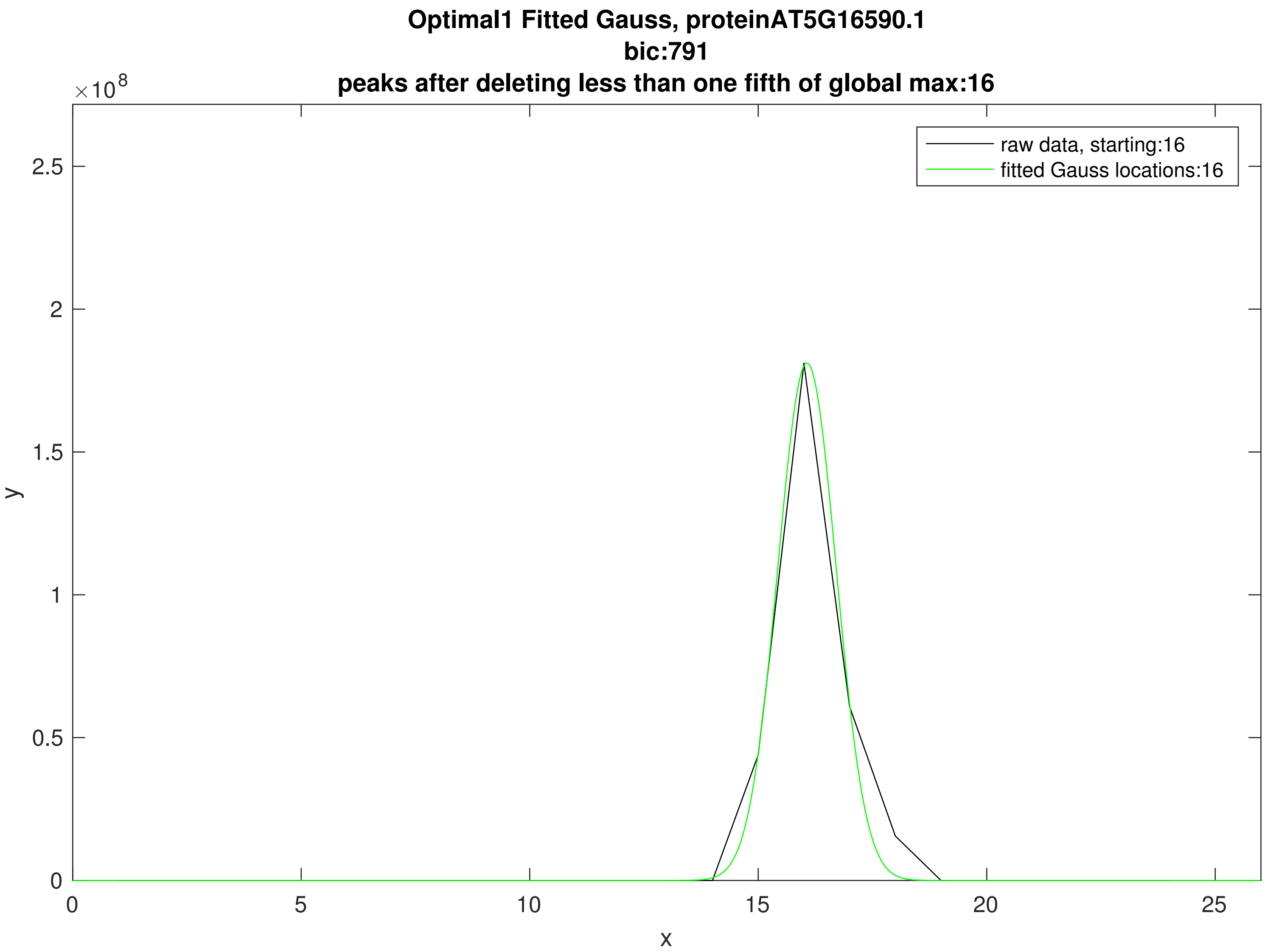
Optimal1 Fitted Gauss, proteinAT5G16590.1
bic:791
peaks after deleting less than one fifth of global max:16
×10^8
raw data, starting:16
fitted Gauss locations:16
2.5
2
1.5
1
0.5
0
y
0
5
10
15
20
25
x

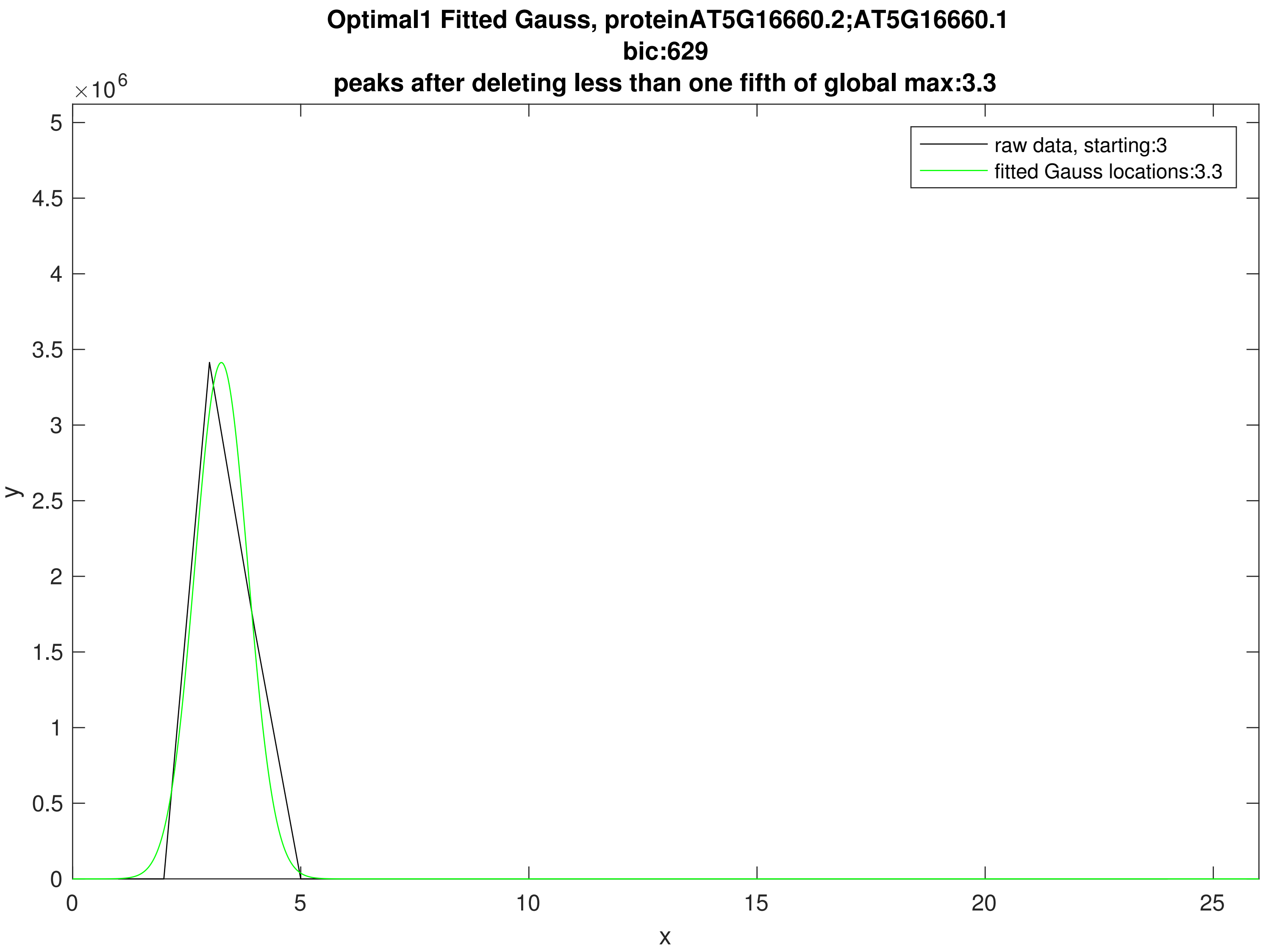
Optimal1 Fitted Gauss, proteinAT5G16660.2;AT5G16660.1
bic:629
peaks after deleting less than one fifth of global max:3.3
×10^6
5
4.5
4
3.5
3
2.5
2
1.5
1
0.5
0
y
0
5
10
15
20
25
x
raw data, starting:3
fitted Gauss locations:3.3

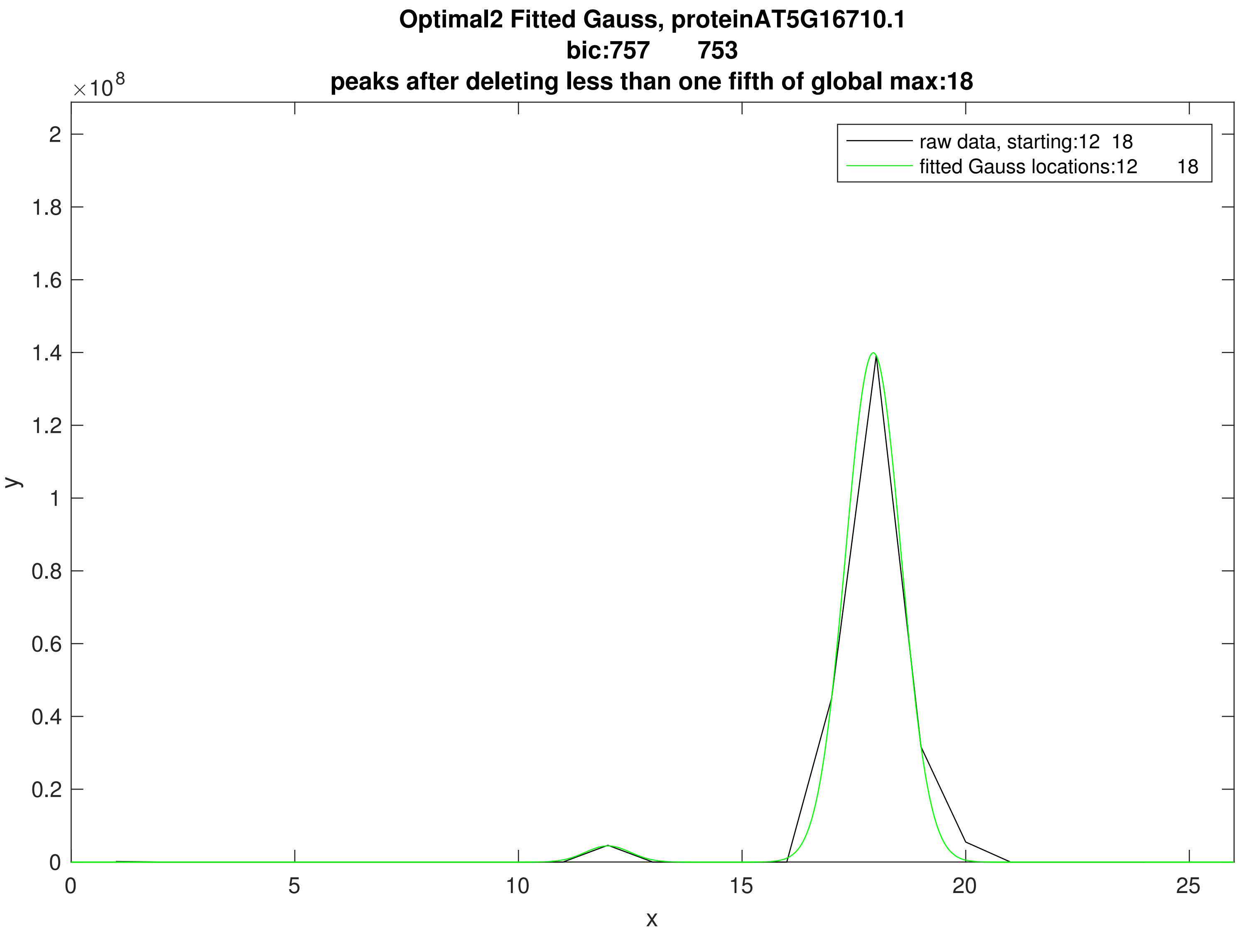
Optimal2 Fitted Gauss, proteinAT5G16710.1
bic:757 753
peaks after deleting less than one fifth of global max:18
raw data, starting:12 18
fitted Gauss locations:12 18
×10^8
2
1.8
1.6
1.4
1.2
1
0.8
0.6
0.4
0.2
0
y
0
5
10
15
20
25
x

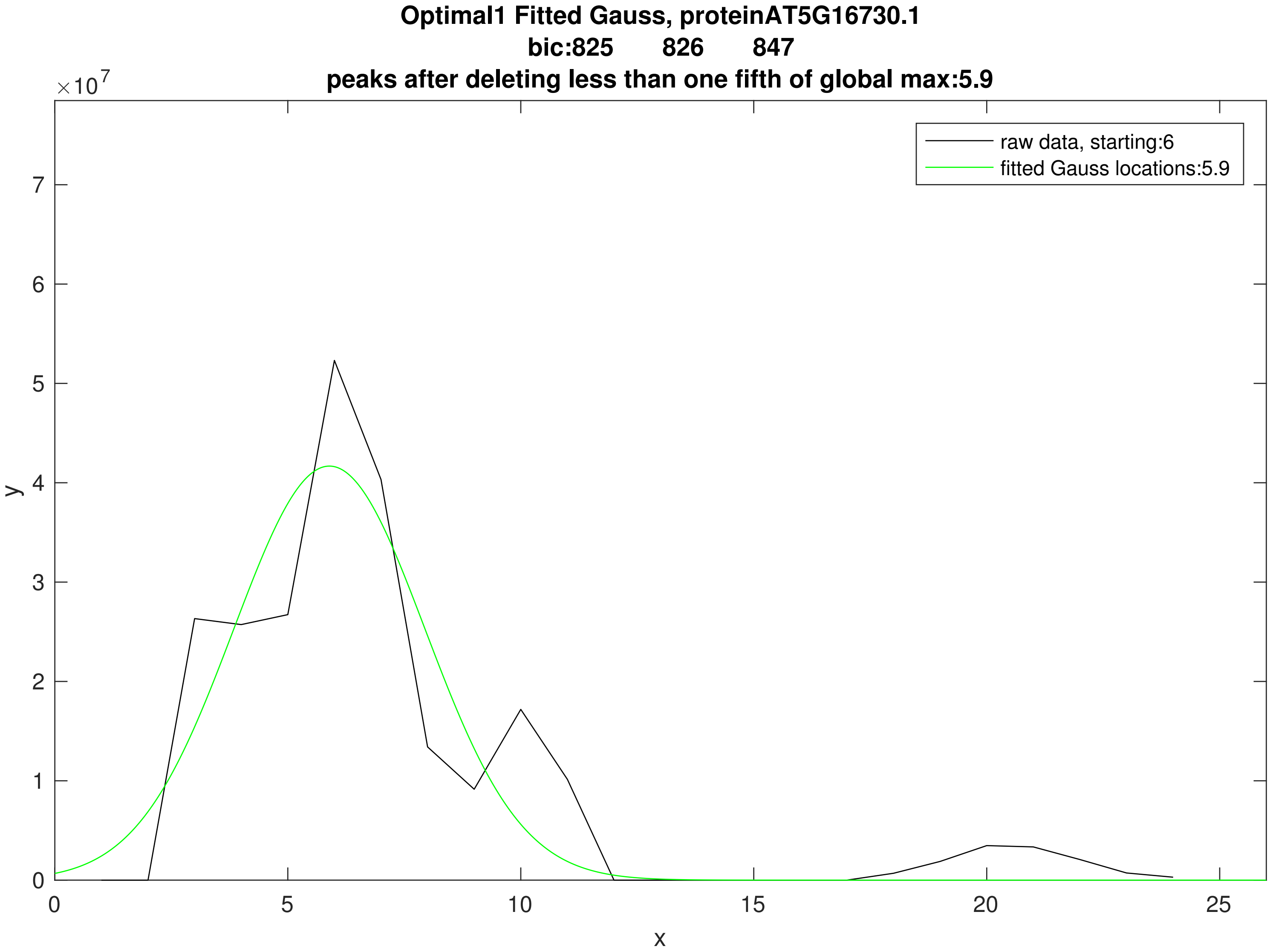
Optimal1 Fitted Gauss, proteinAT5G16730.1
bic:825 826 847
peaks after deleting less than one fifth of global max:5.9
×10^7
raw data, starting:6
fitted Gauss locations:5.9
y
x
0
1
2
3
4
5
6
7
0
5
10
15
20
25

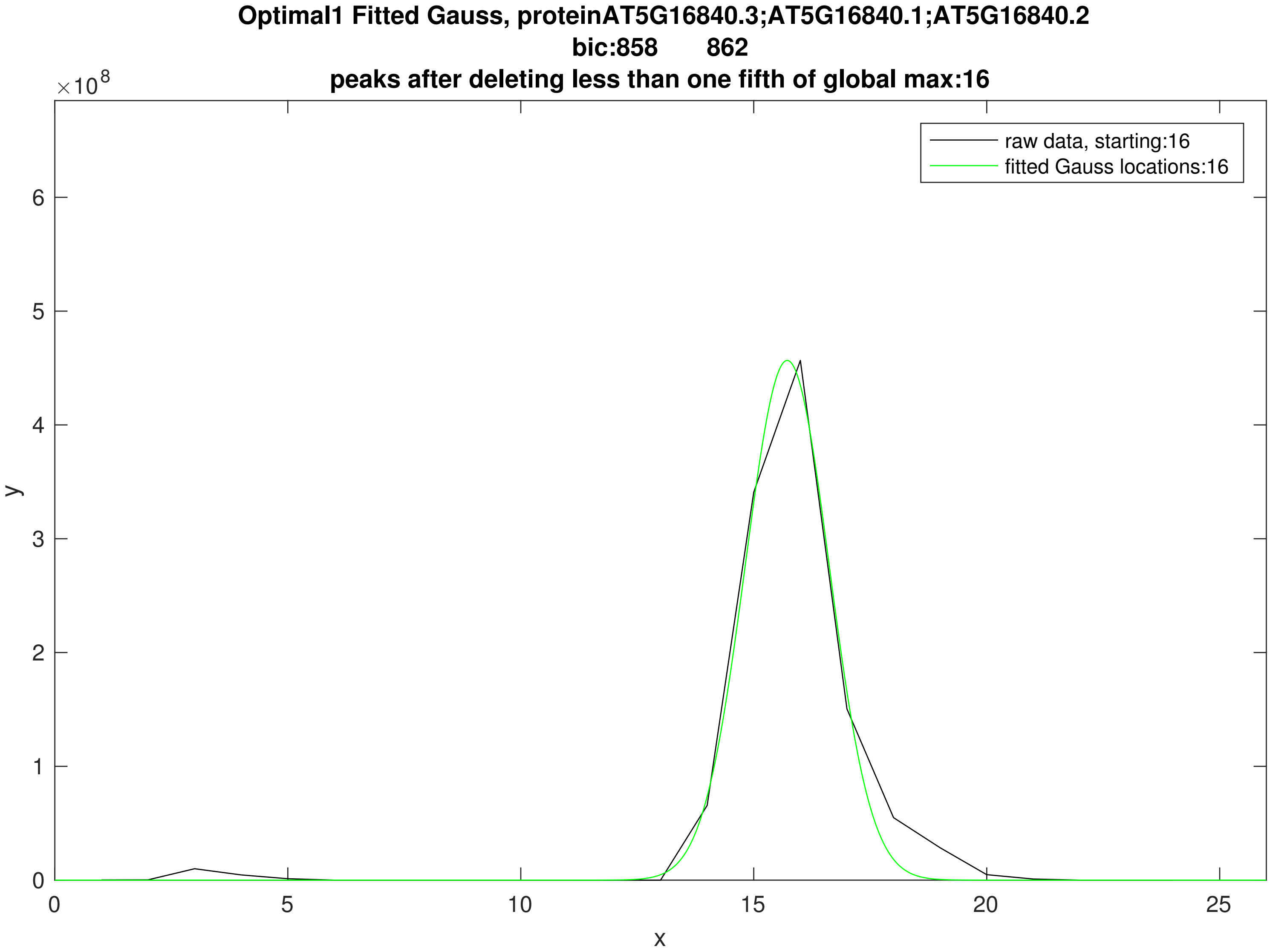
Optimal1 Fitted Gauss, proteinAT5G16840.3;AT5G16840.1;AT5G16840.2
bic:858 862
peaks after deleting less than one fifth of global max:16
×10^8
raw data, starting:16
fitted Gauss locations:16
y
x
0
1
2
3
4
5
6
0
5
10
15
20
25

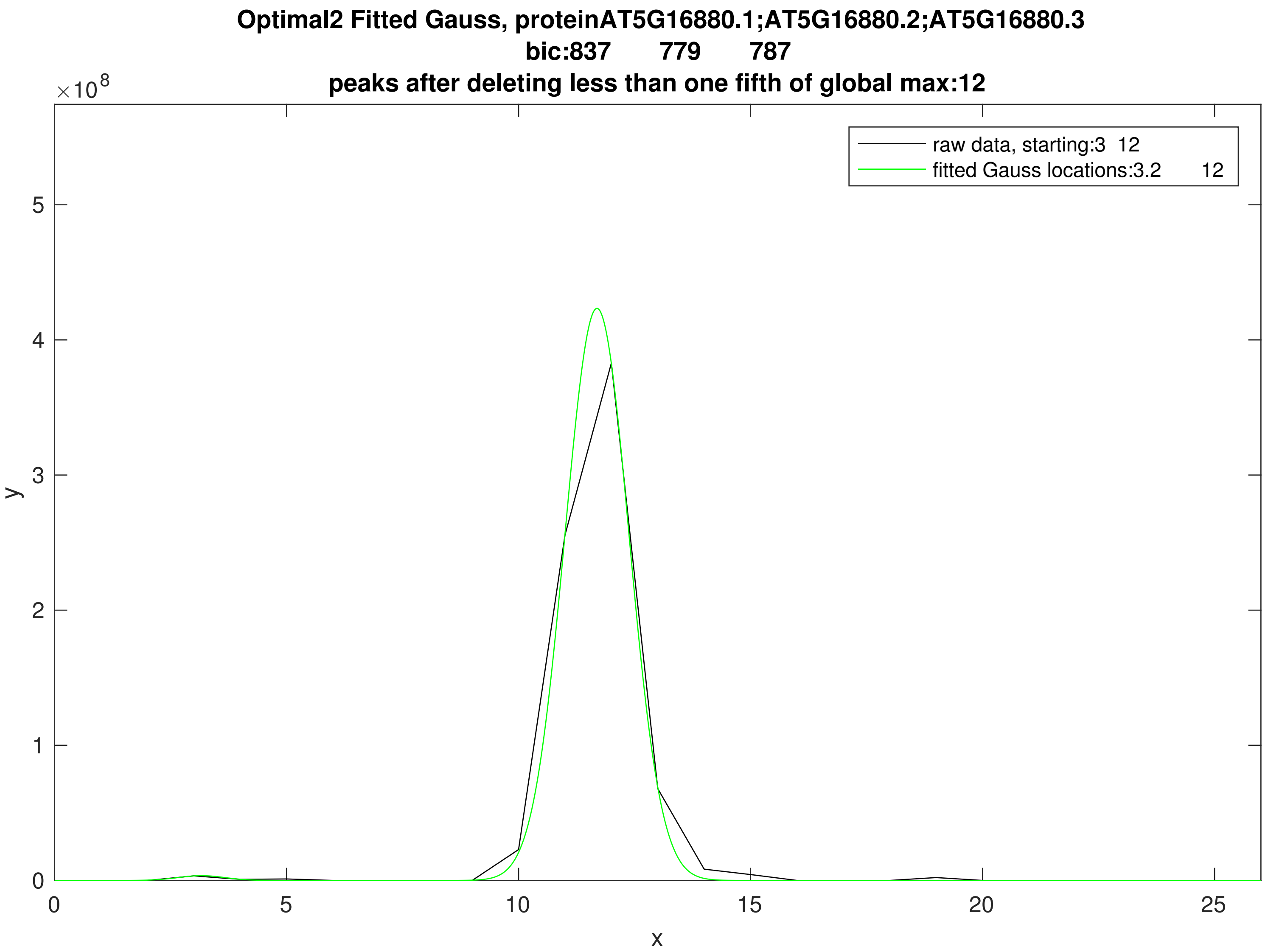
Optimal2 Fitted Gauss, proteinAT5G16880.1;AT5G16880.2;AT5G16880.3
bic:837 779 787
peaks after deleting less than one fifth of global max:12
×10^8
raw data, starting:3 12
fitted Gauss locations:3.2 12
5
4
3
2
1
0
y
0
5
10
15
20
25
x

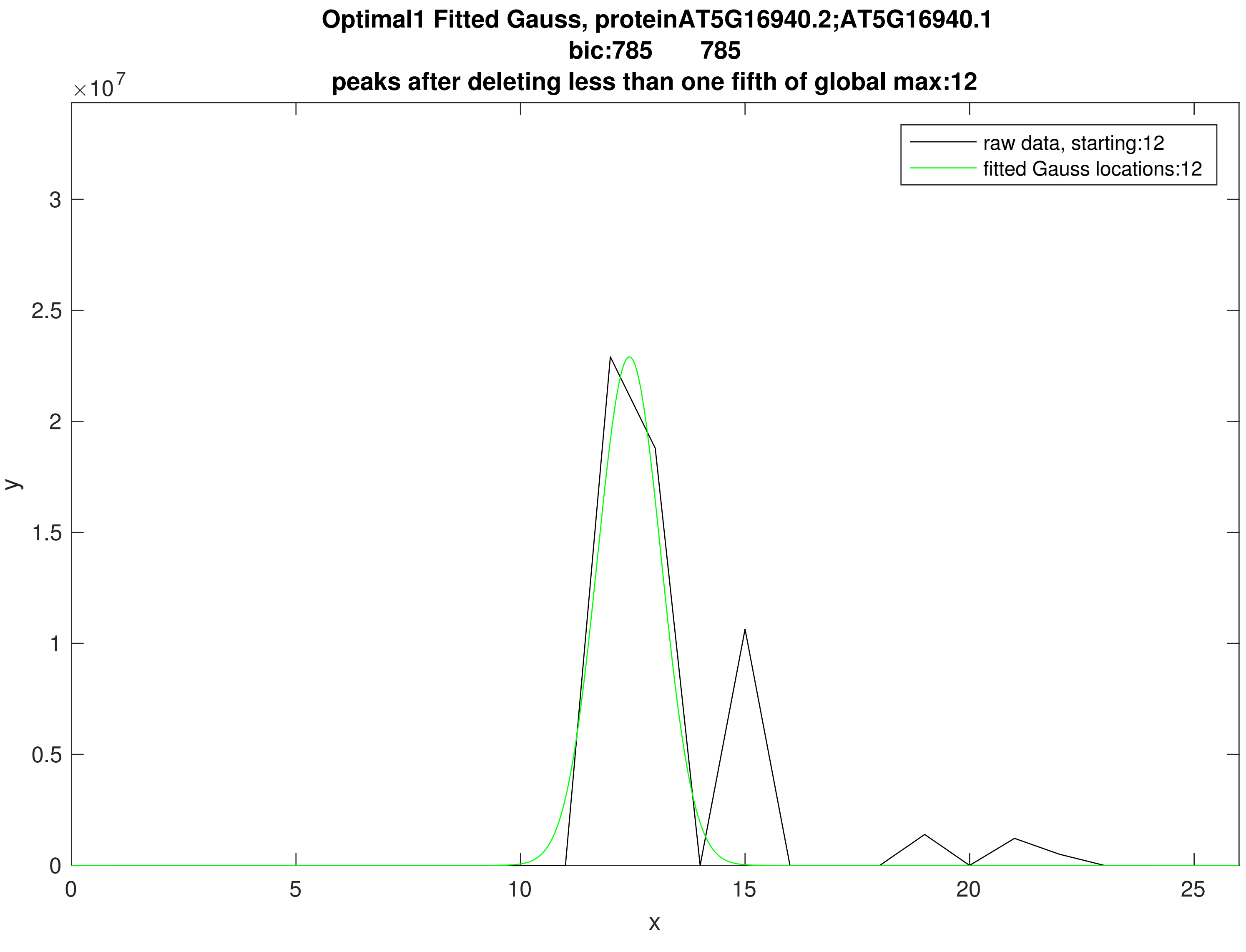

Optimal1 Fitted Gauss, proteinAT5G16940.2;AT5G16940.1
bic:785 785
peaks after deleting less than one fifth of global max:12
×10^7
raw data, starting:12
fitted Gauss locations:12
y
x
0
0.5
1
1.5
2
2.5
3
0
5
10
15
20
25

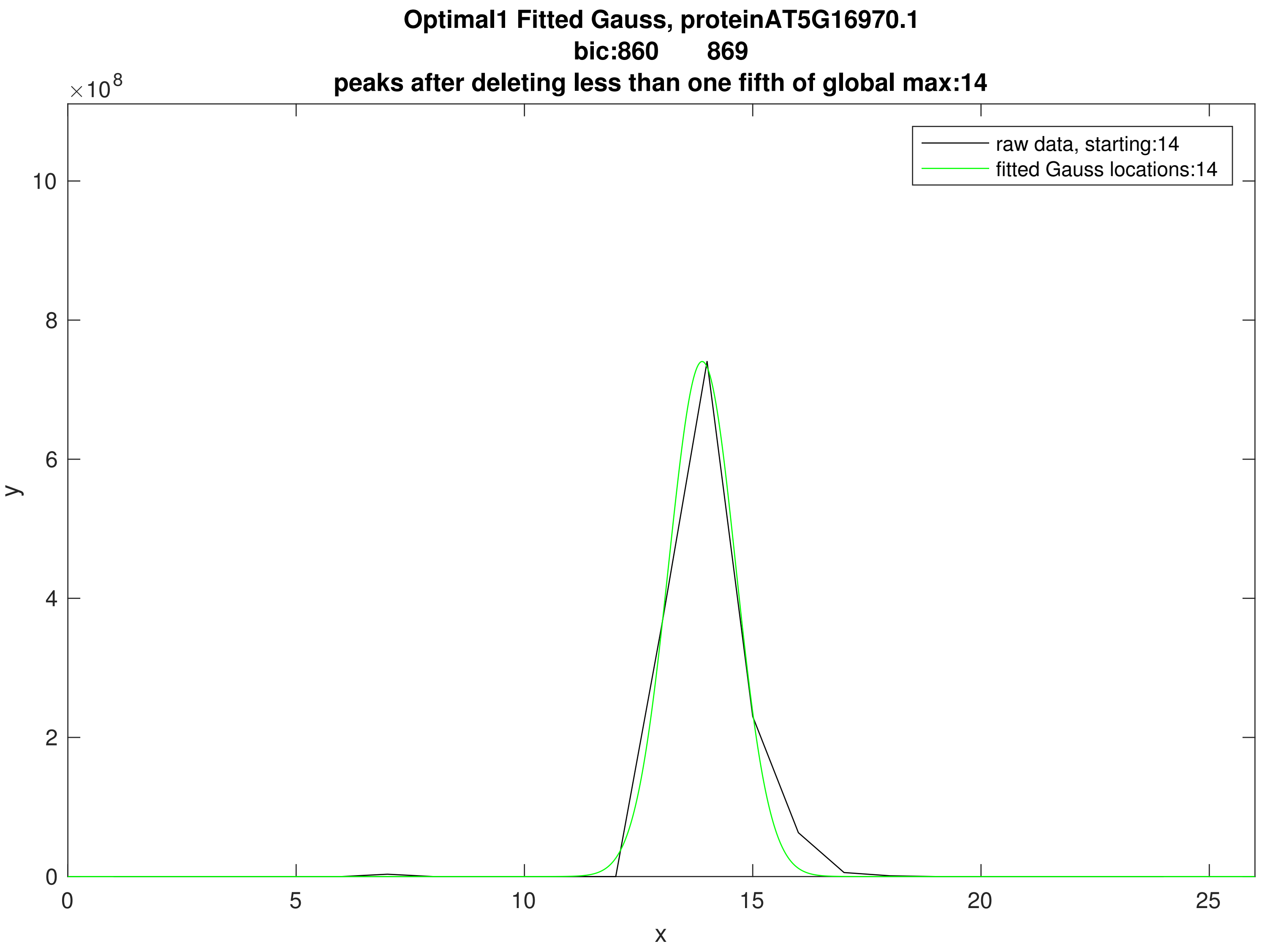

Optimal1 Fitted Gauss, proteinAT5G16970.1
bic:860 869
peaks after deleting less than one fifth of global max:14
×10^8
raw data, starting:14
fitted Gauss locations:14
y
x
0
2
4
6
8
10
0
5
10
15
20
25

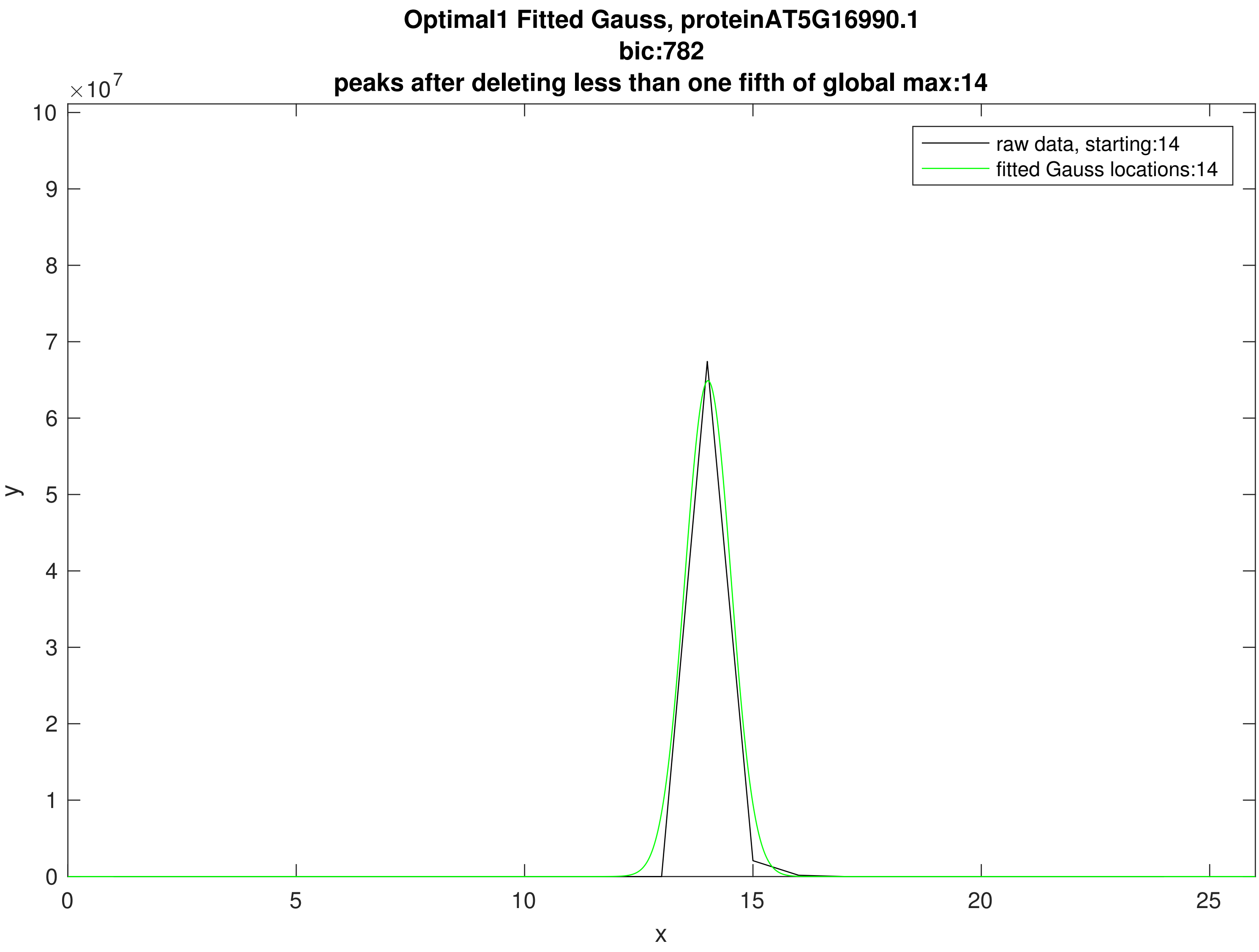
Optimal1 Fitted Gauss, proteinAT5G16990.1
bic:782
peaks after deleting less than one fifth of global max:14
×10^7
raw data, starting:14
fitted Gauss locations:14
y
x
0
1
2
3
4
5
6
7
8
9
10
0
5
10
15
20
25

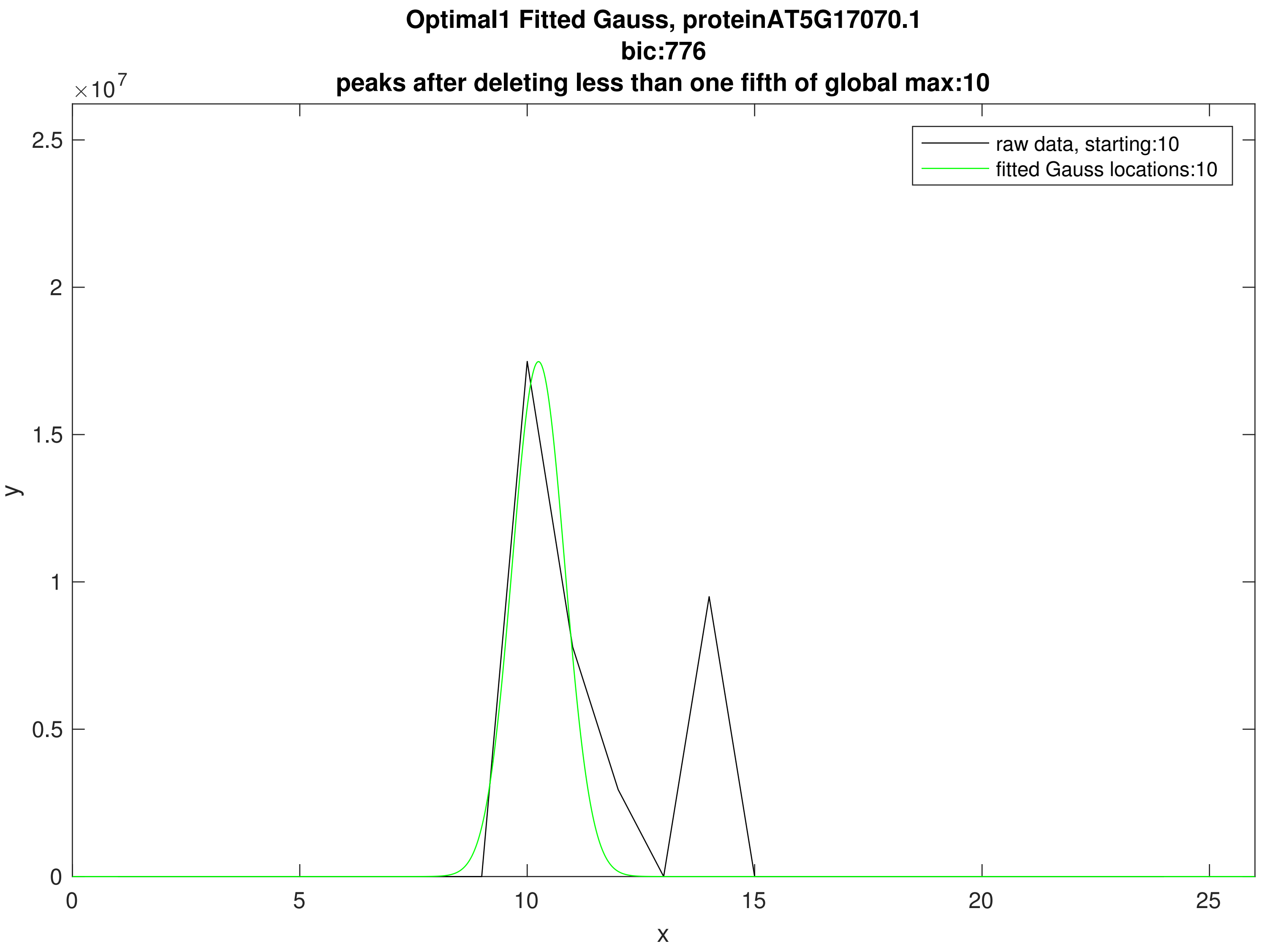
Optimal1 Fitted Gauss, proteinAT5G17070.1
bic:776
peaks after deleting less than one fifth of global max:10
×10^7
2.5
2
1.5
1
0.5
0
y
0
5
10
15
20
25
x
raw data, starting:10
fitted Gauss locations:10

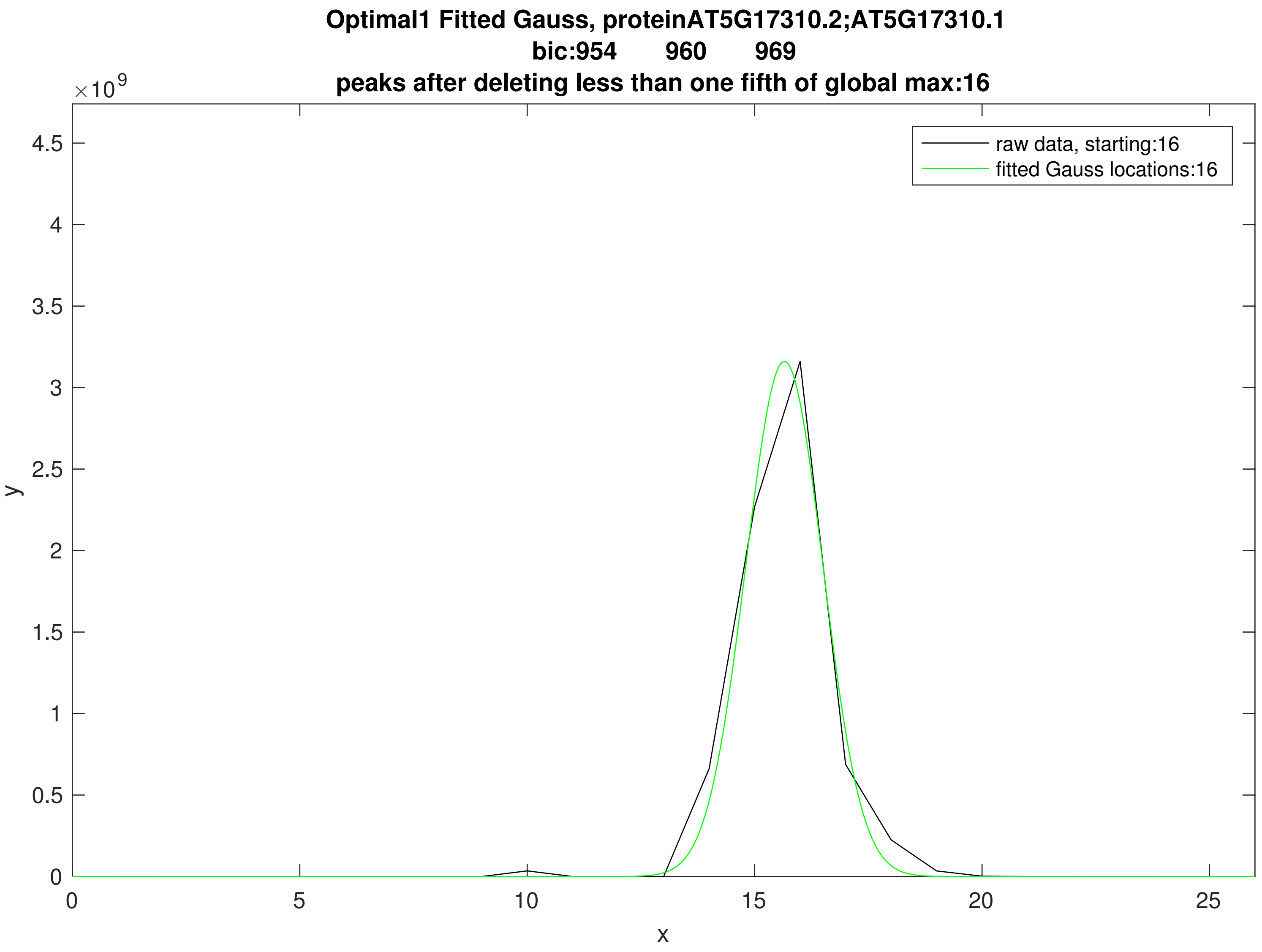
Optimal1 Fitted Gauss, proteinAT5G17310.2;AT5G17310.1
bic:954 960 969
peaks after deleting less than one fifth of global max:16
×10^9
raw data, starting:16
fitted Gauss locations:16
y
x
0
5
10
15
20
25
0.5
1
1.5
2
2.5
3
3.5
4
4.5

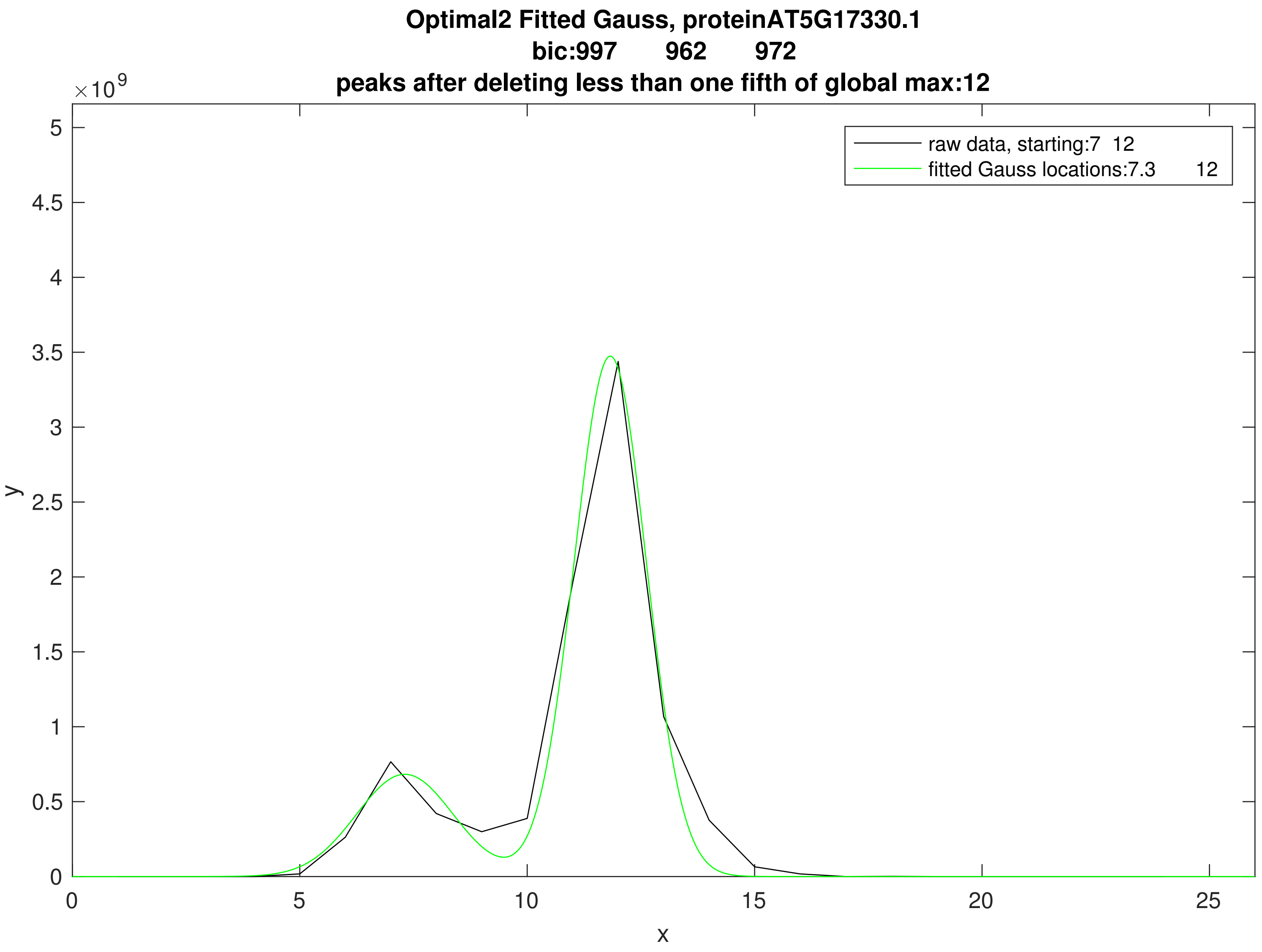
Optimal2 Fitted Gauss, proteinAT5G17330.1
bic:997 962 972
peaks after deleting less than one fifth of global max:12
×10^9
raw data, starting:7 12
fitted Gauss locations:7.3 12
y
x
0
0.5
1
1.5
2
2.5
3
3.5
4
4.5
5
10
15
20
25

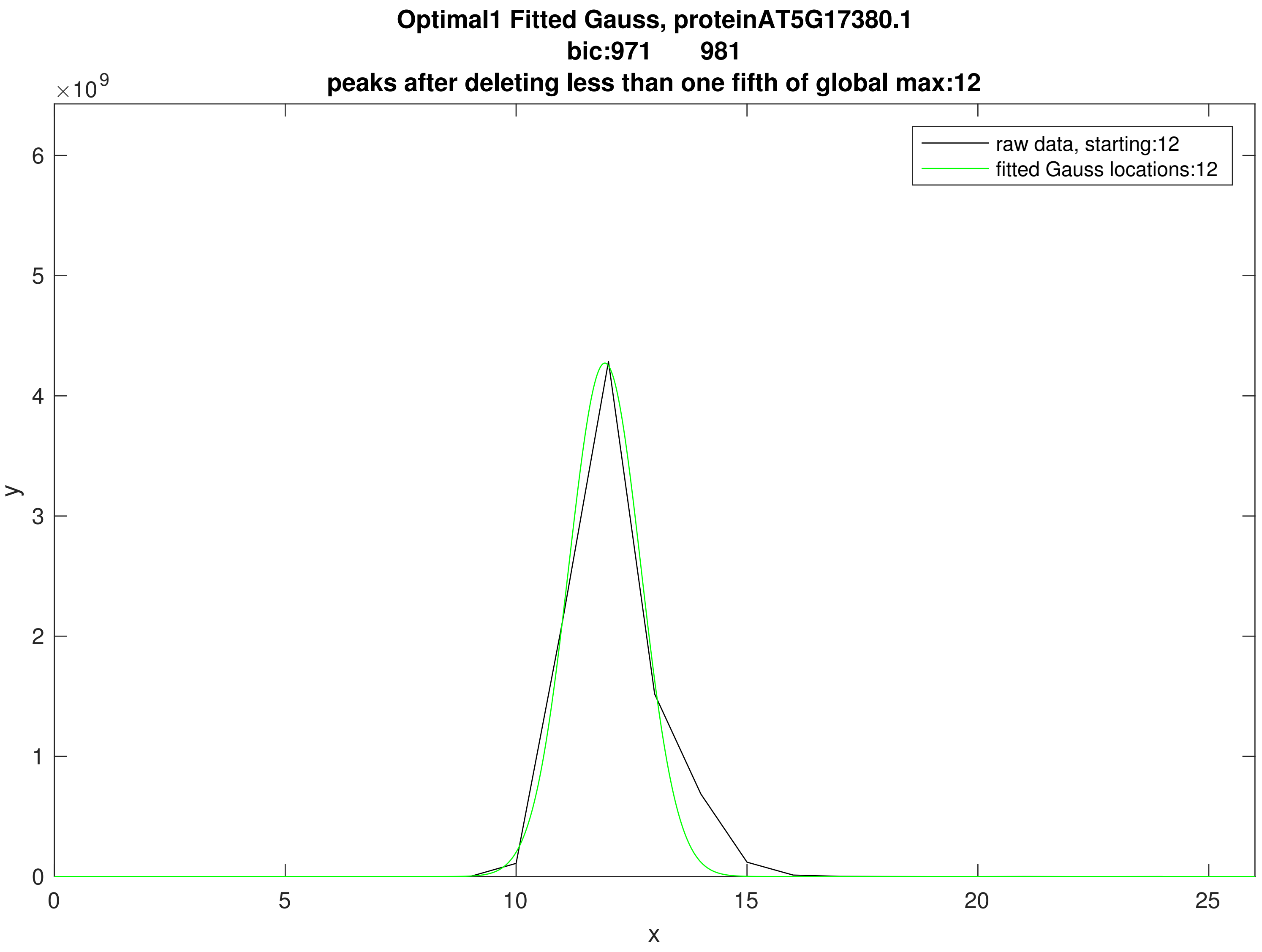

Optimal1 Fitted Gauss, proteinAT5G17380.1
bic:971 981
peaks after deleting less than one fifth of global max:12
×10^9
raw data, starting:12
fitted Gauss locations:12
y
x
0
1
2
3
4
5
6
0
5
10
15
20
25

**Optimal1 Fitted Gauss, proteinAT5G17530.1;AT5G17530.2;AT5G17530.3**
**bic:871 877**
**peaks after deleting less than one fifth of global max:15**

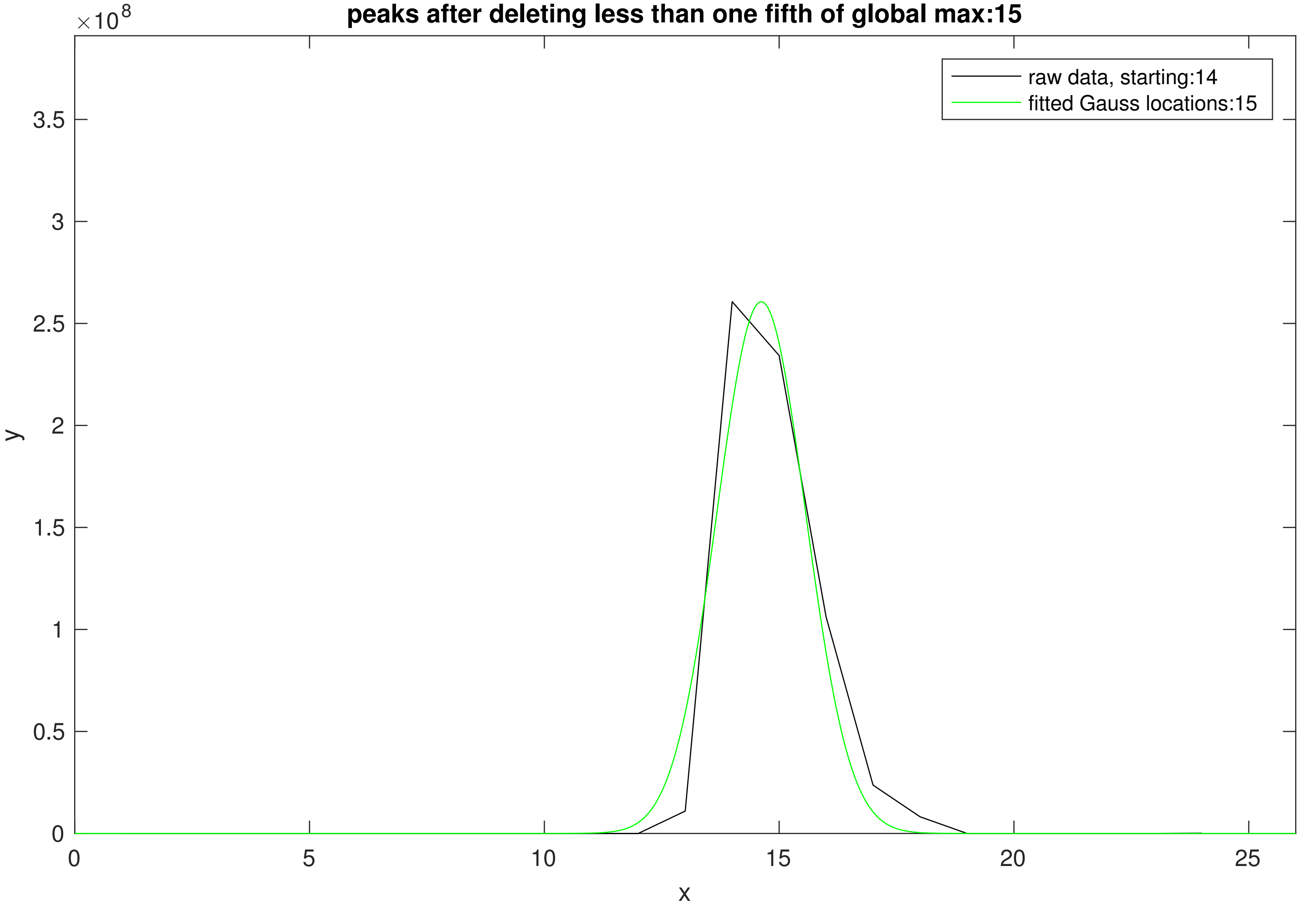

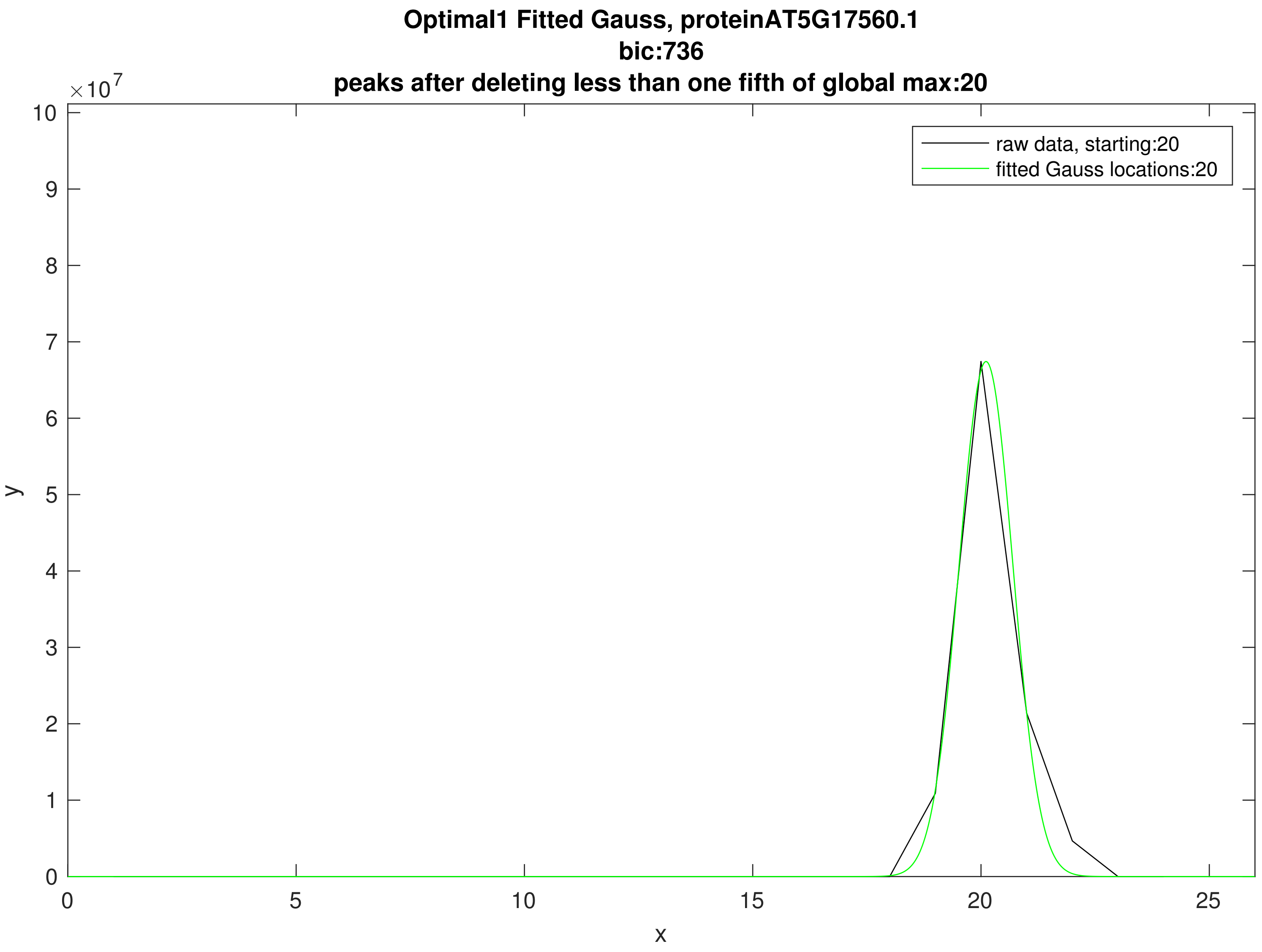

Optimal1 Fitted Gauss, proteinAT5G17560.1
bic:736
peaks after deleting less than one fifth of global max:20
×10^7
raw data, starting:20
fitted Gauss locations:20
y
x
0
1
2
3
4
5
6
7
8
9
10
0
5
10
15
20
25

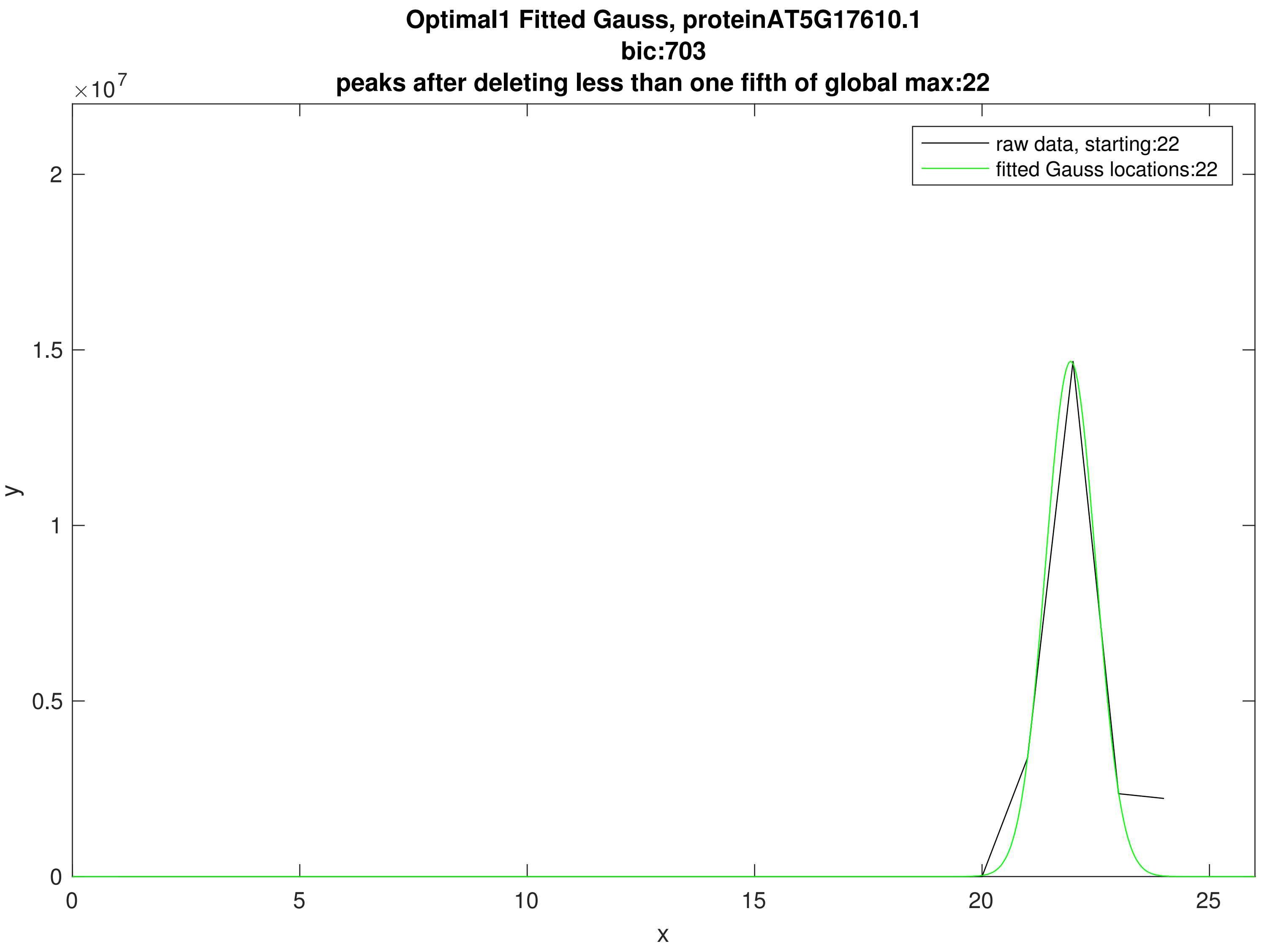

Optimal1 Fitted Gauss, proteinAT5G17610.1
bic:703
peaks after deleting less than one fifth of global max:22
×10^7
raw data, starting:22
fitted Gauss locations:22
y
x
0
0.5
1
1.5
2
0
5
10
15
20
25

Optimal1 Fitted Gauss, proteinAT5G17710.1;AT5G17710.2
bic:804
peaks after deleting less than one fifth of global max:8.7

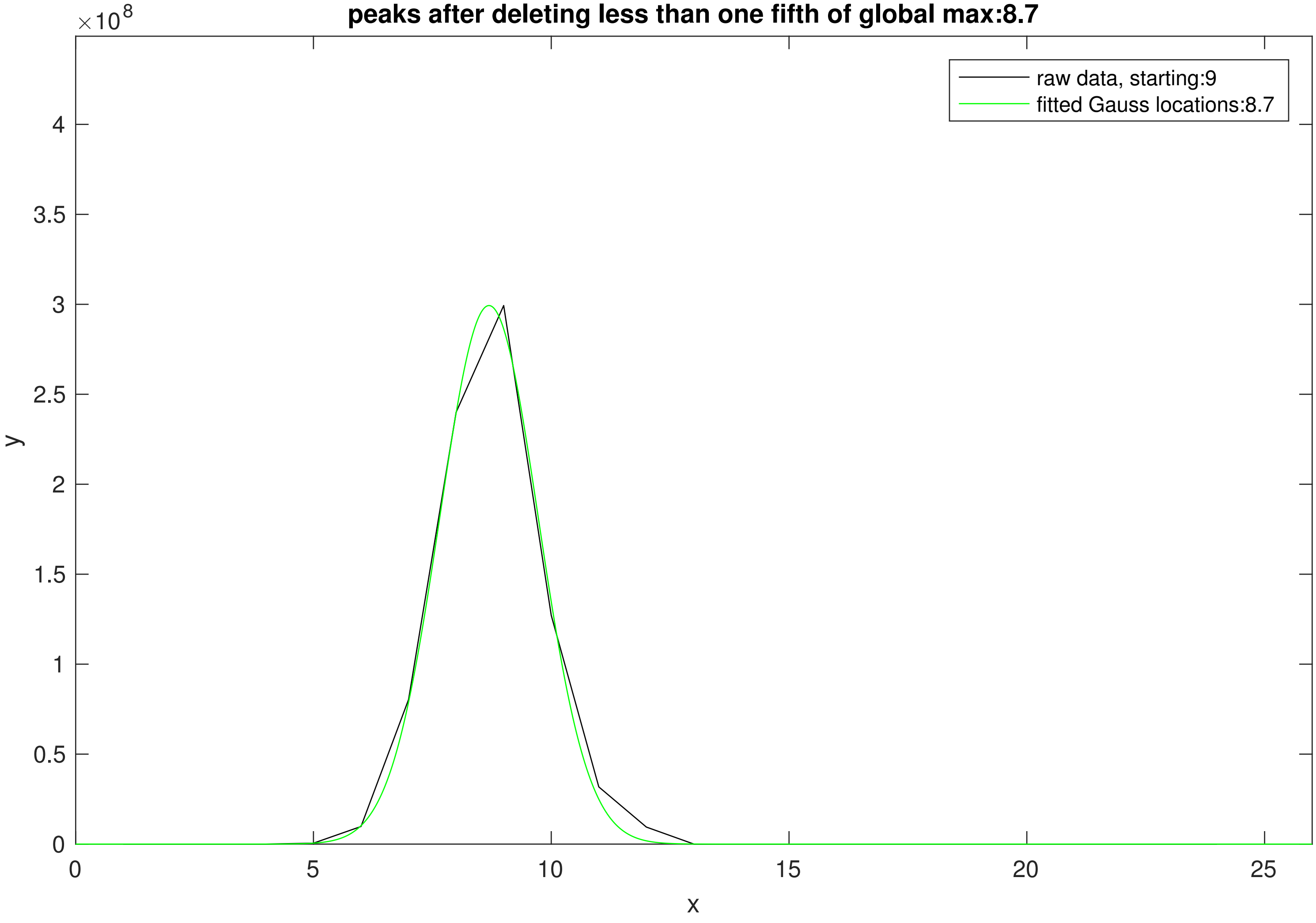

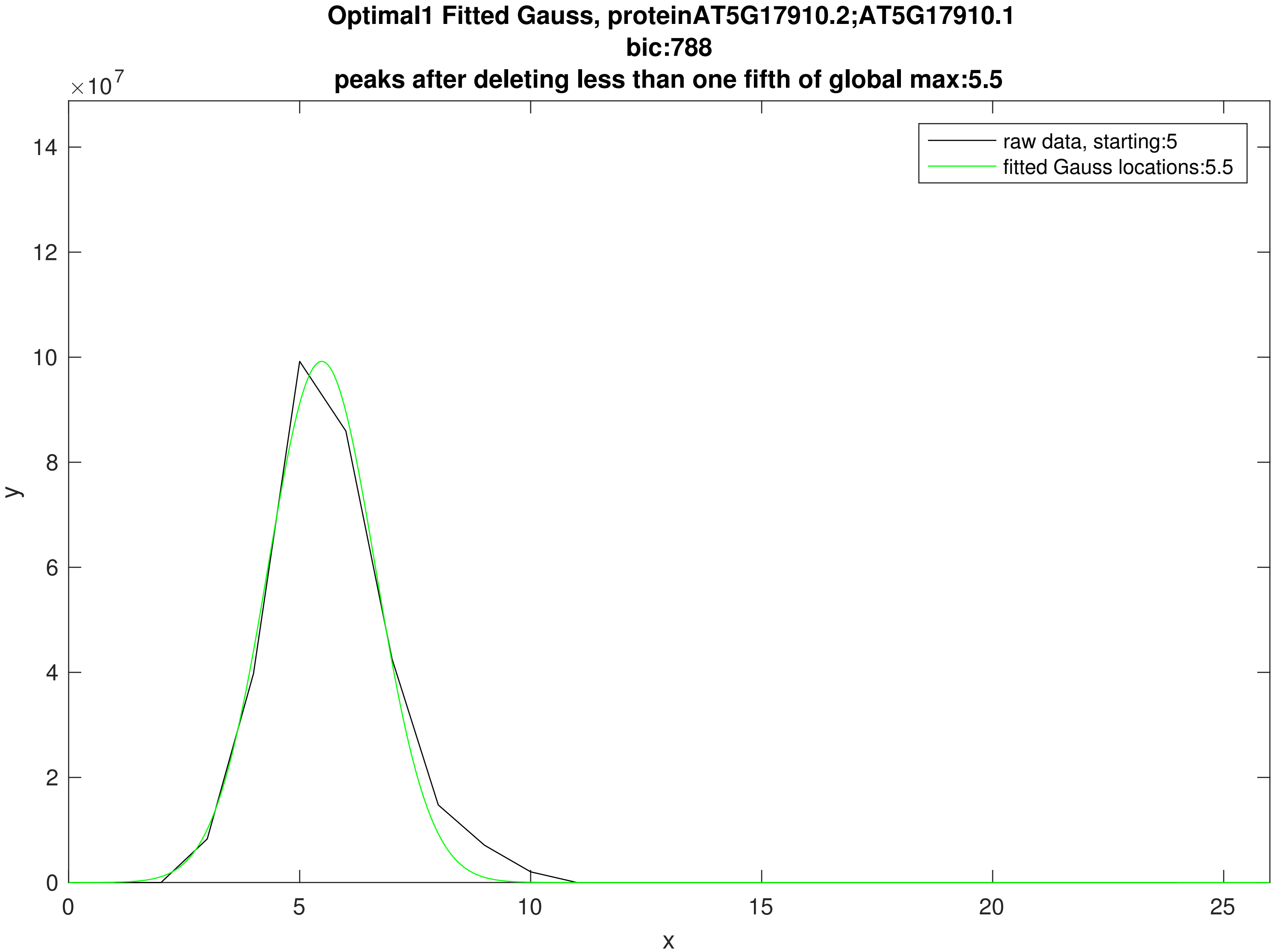
Optimal1 Fitted Gauss, proteinAT5G17910.2;AT5G17910.1
bic:788
peaks after deleting less than one fifth of global max:5.5
×10^7
raw data, starting:5
fitted Gauss locations:5.5
y
x
0
2
4
6
8
10
12
14
0
5
10
15
20
25

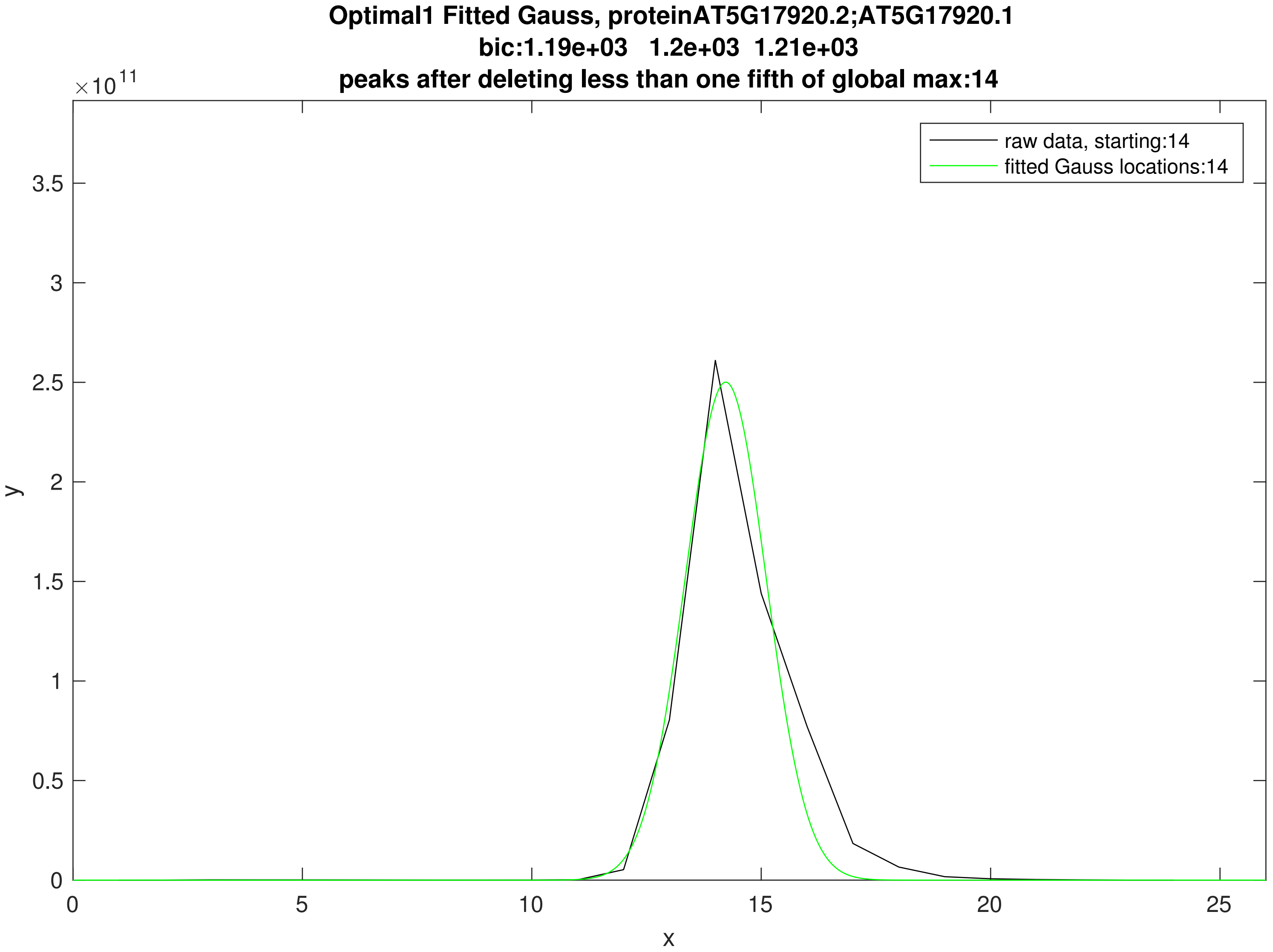
Optimal1 Fitted Gauss, proteinAT5G17920.2;AT5G17920.1
bic:1.19e+03 1.2e+03 1.21e+03
peaks after deleting less than one fifth of global max:14
×10^11
raw data, starting:14
fitted Gauss locations:14
y
x
0
0.5
1
1.5
2
2.5
3
3.5
0
5
10
15
20
25

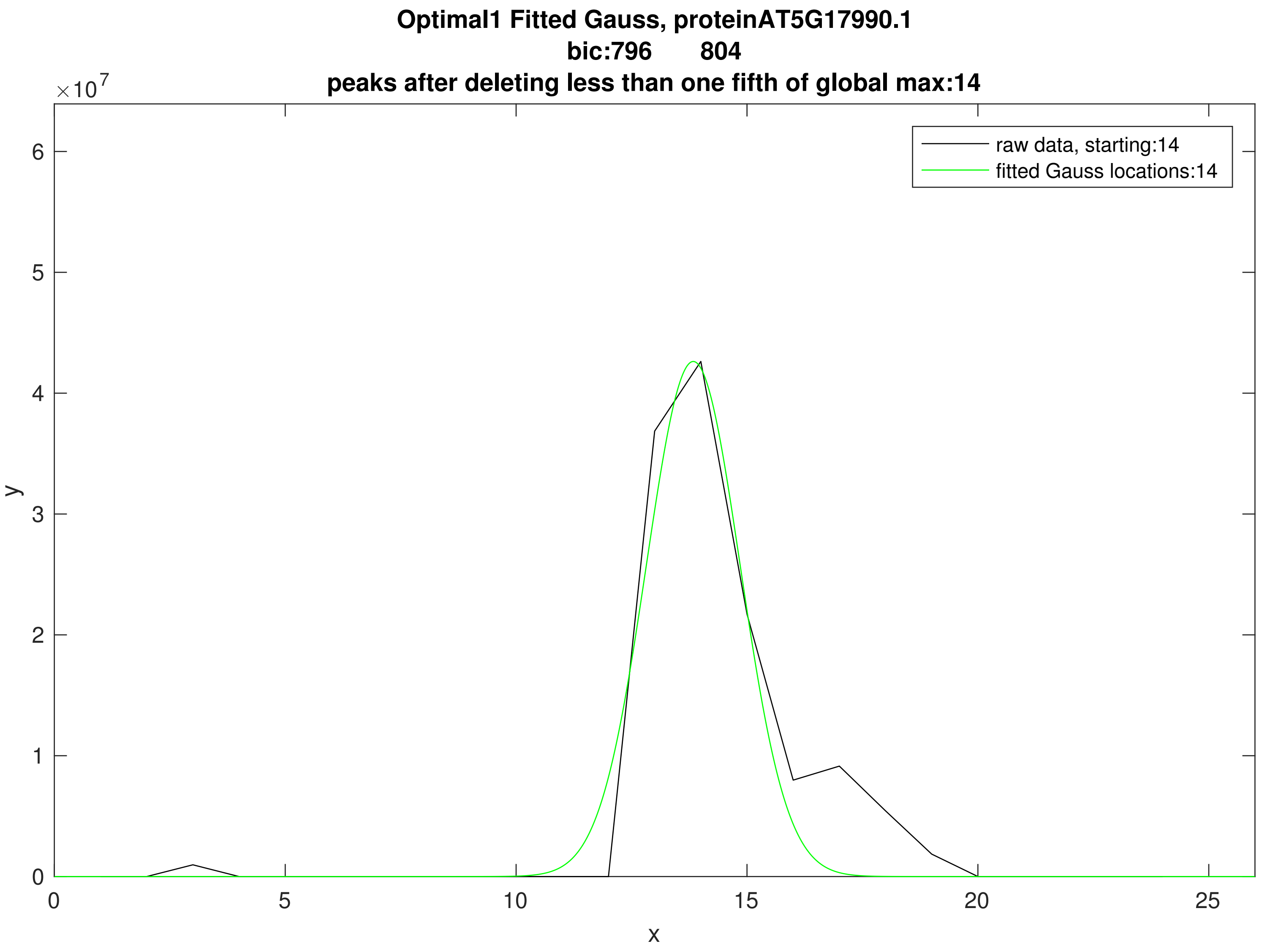

Optimal1 Fitted Gauss, proteinAT5G17990.1
bic:796 804
peaks after deleting less than one fifth of global max:14
×10^7
raw data, starting:14
fitted Gauss locations:14
y
x
0
1
2
3
4
5
6
0
5
10
15
20
25

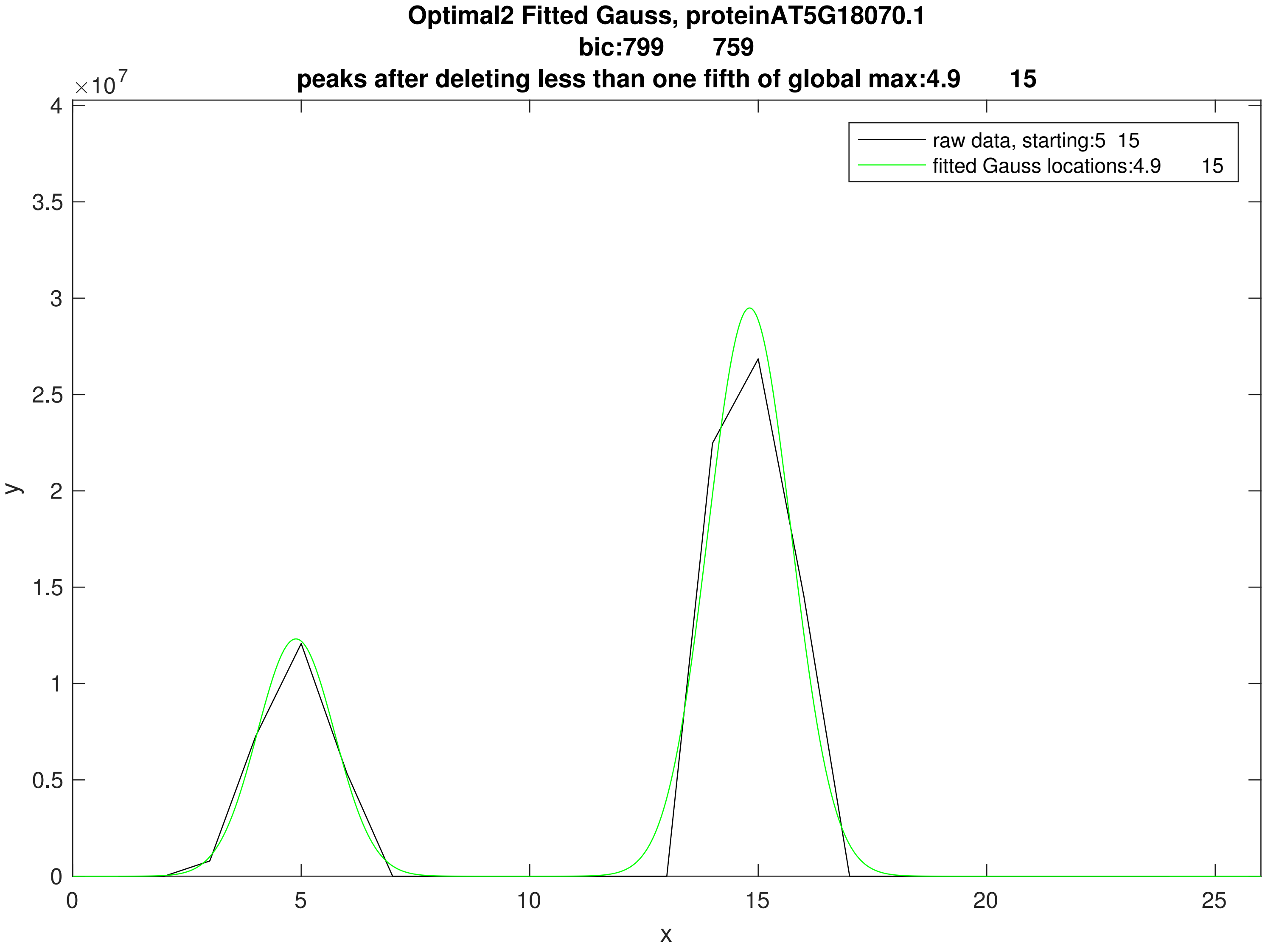

Optimal2 Fitted Gauss, proteinAT5G18070.1
bic:799 759
peaks after deleting less than one fifth of global max:4.9 15
×10^7
4
3.5
3
2.5
2
1.5
1
0.5
0
y
0
5
10
15
20
25
x
raw data, starting:5 15
fitted Gauss locations:4.9 15

Optimal1 Fitted Gauss, proteinAT5G18100.1;AT5G18100.2
bic:791
peaks after deleting less than one fifth of global max:17

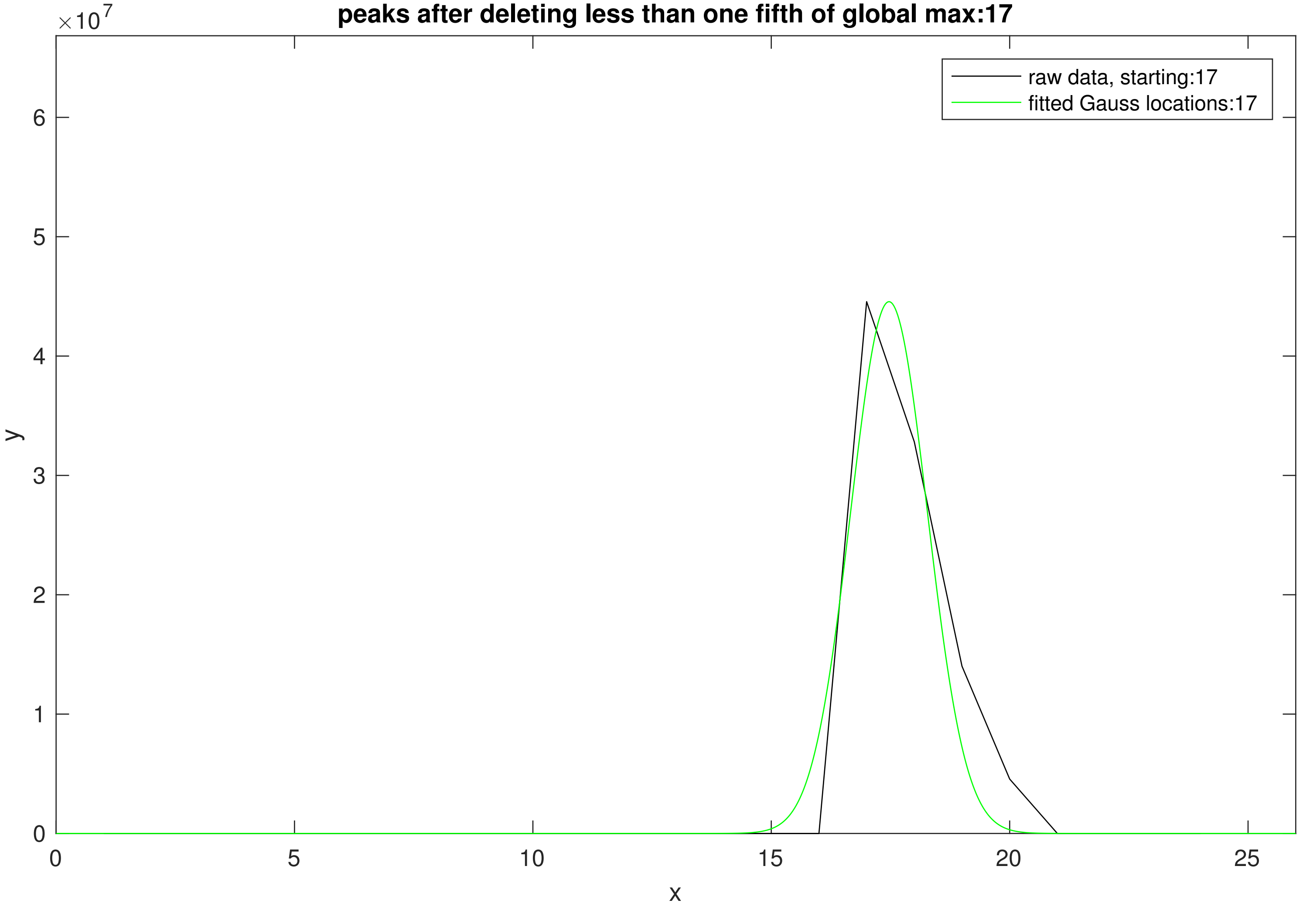

Optimal1 Fitted Gauss, proteinAT5G18170.1
bic:788
peaks after deleting less than one fifth of global max:9

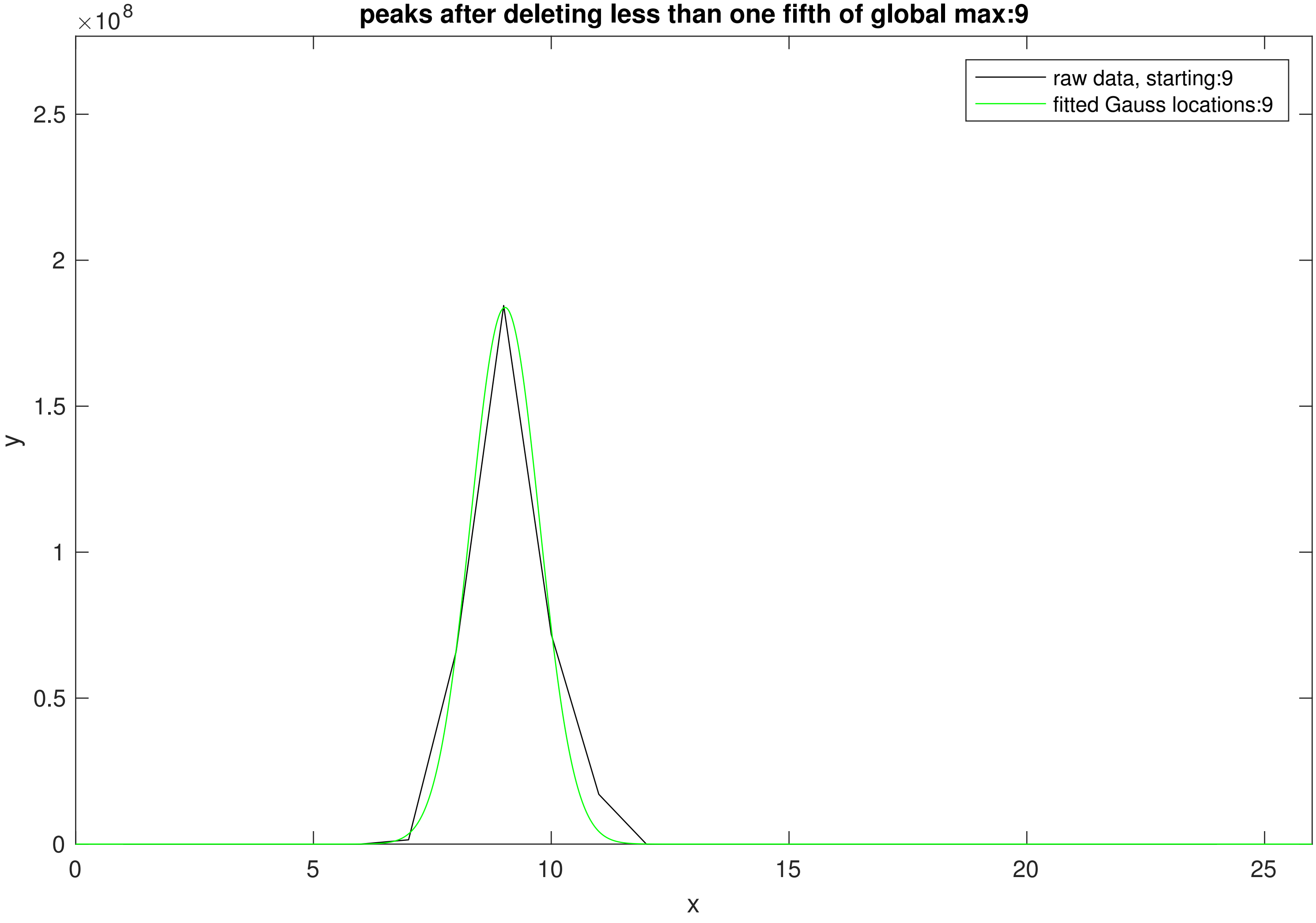

Optimal1 Fitted Gauss, proteinAT5G18200.1
bic:807
peaks after deleting less than one fifth of global max:14

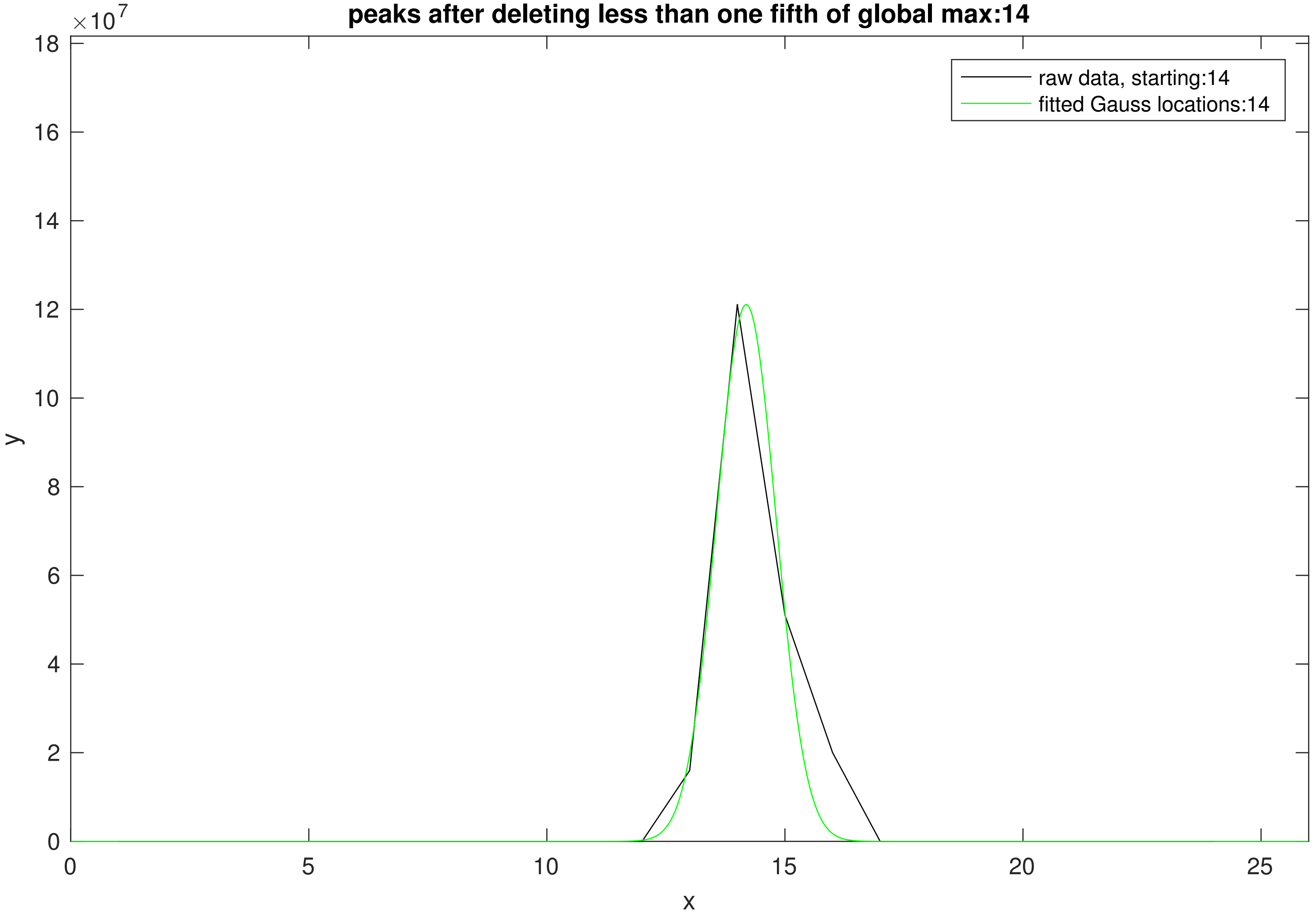

**Optimal1 Fitted Gauss, proteinAT5G18380.2;AT5G18380.1;AT2G09990.1;AT5G18380.3;AT3G04230.1**
**bic:864 874**
**peaks after deleting less than one fifth of global max:3.6**

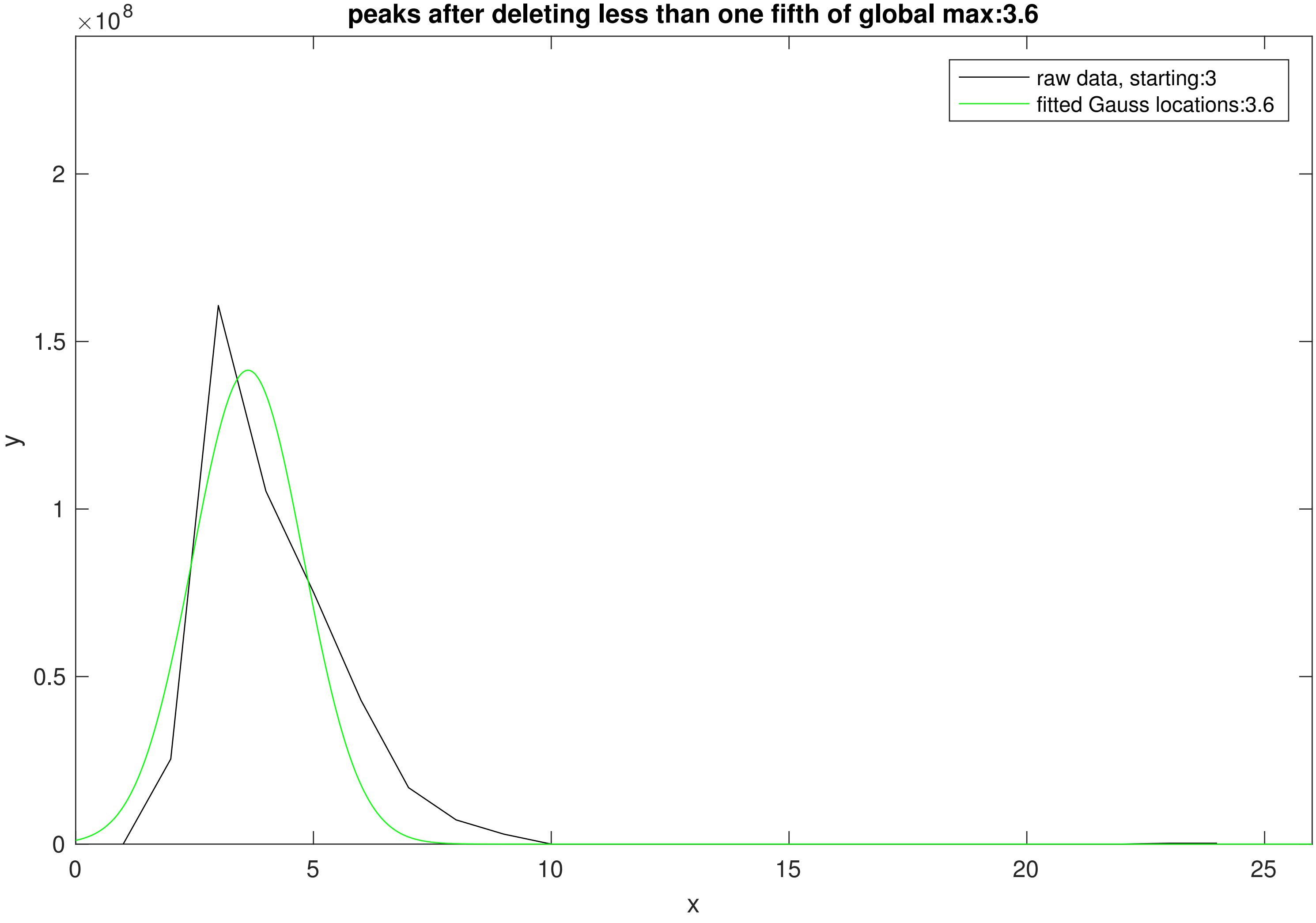

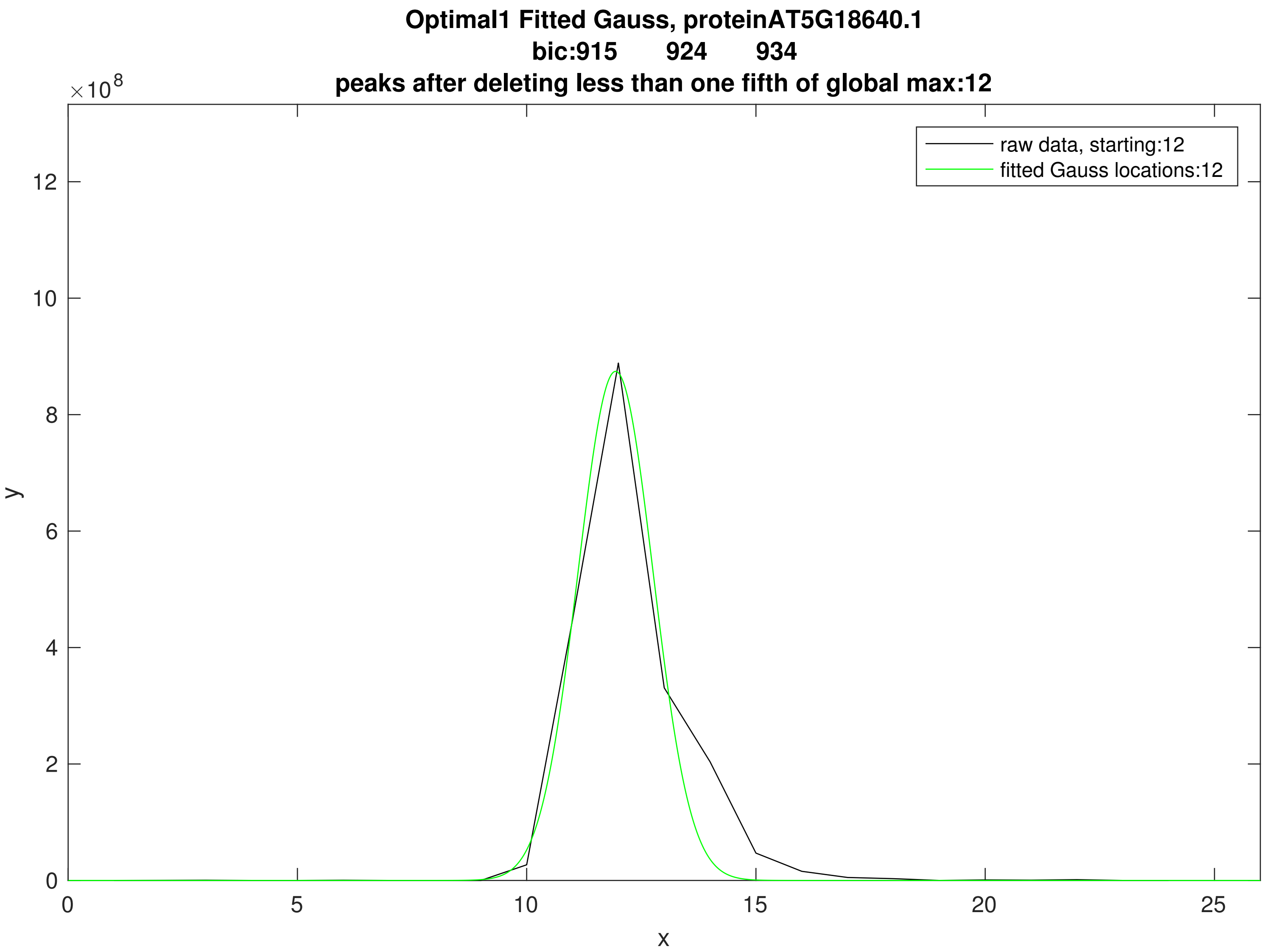

Optimal1 Fitted Gauss, proteinAT5G18640.1
bic:915 924 934
peaks after deleting less than one fifth of global max:12
×10^8
raw data, starting:12
fitted Gauss locations:12
y
x
0
2
4
6
8
10
12
0
5
10
15
20
25

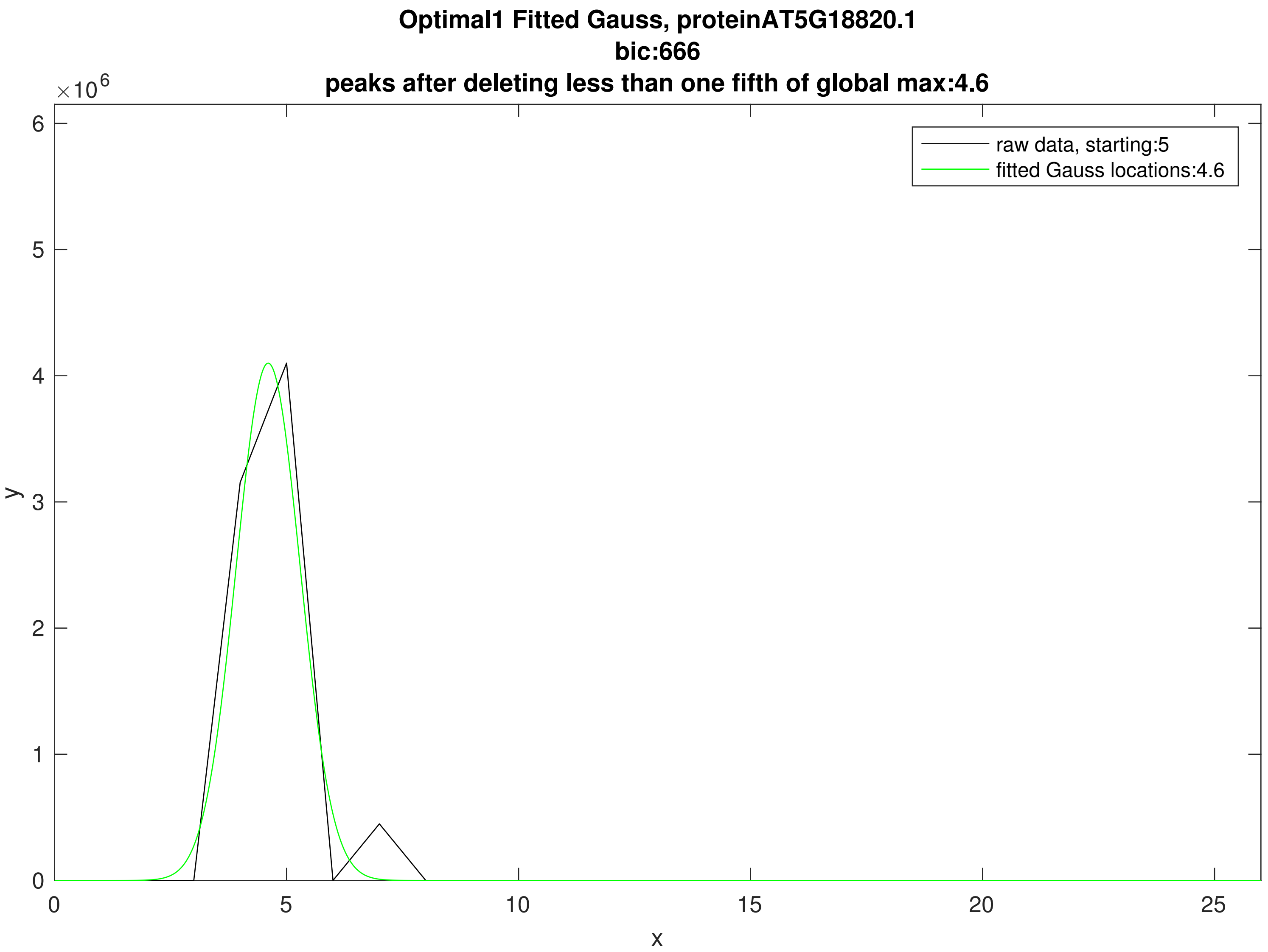
Optimal1 Fitted Gauss, proteinAT5G18820.1
bic:666
peaks after deleting less than one fifth of global max:4.6
×10^6
6
5
4
3
2
1
0
y
0
5
10
15
20
25
x
raw data, starting:5
fitted Gauss locations:4.6

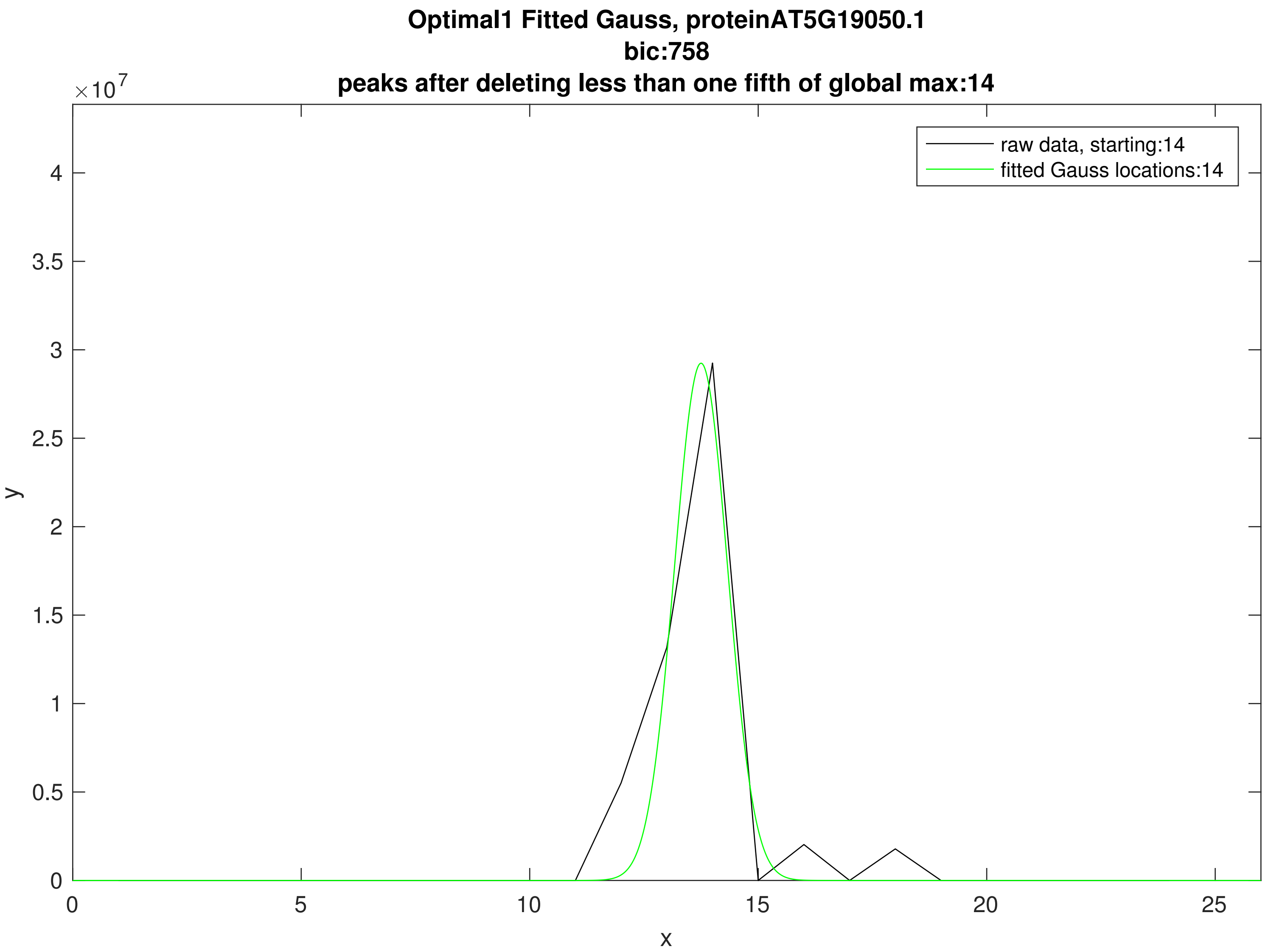

Optimal1 Fitted Gauss, proteinAT5G19050.1
bic:758
peaks after deleting less than one fifth of global max:14
×10^7
raw data, starting:14
fitted Gauss locations:14
y
x
0
0.5
1
1.5
2
2.5
3
3.5
4
0
5
10
15
20
25

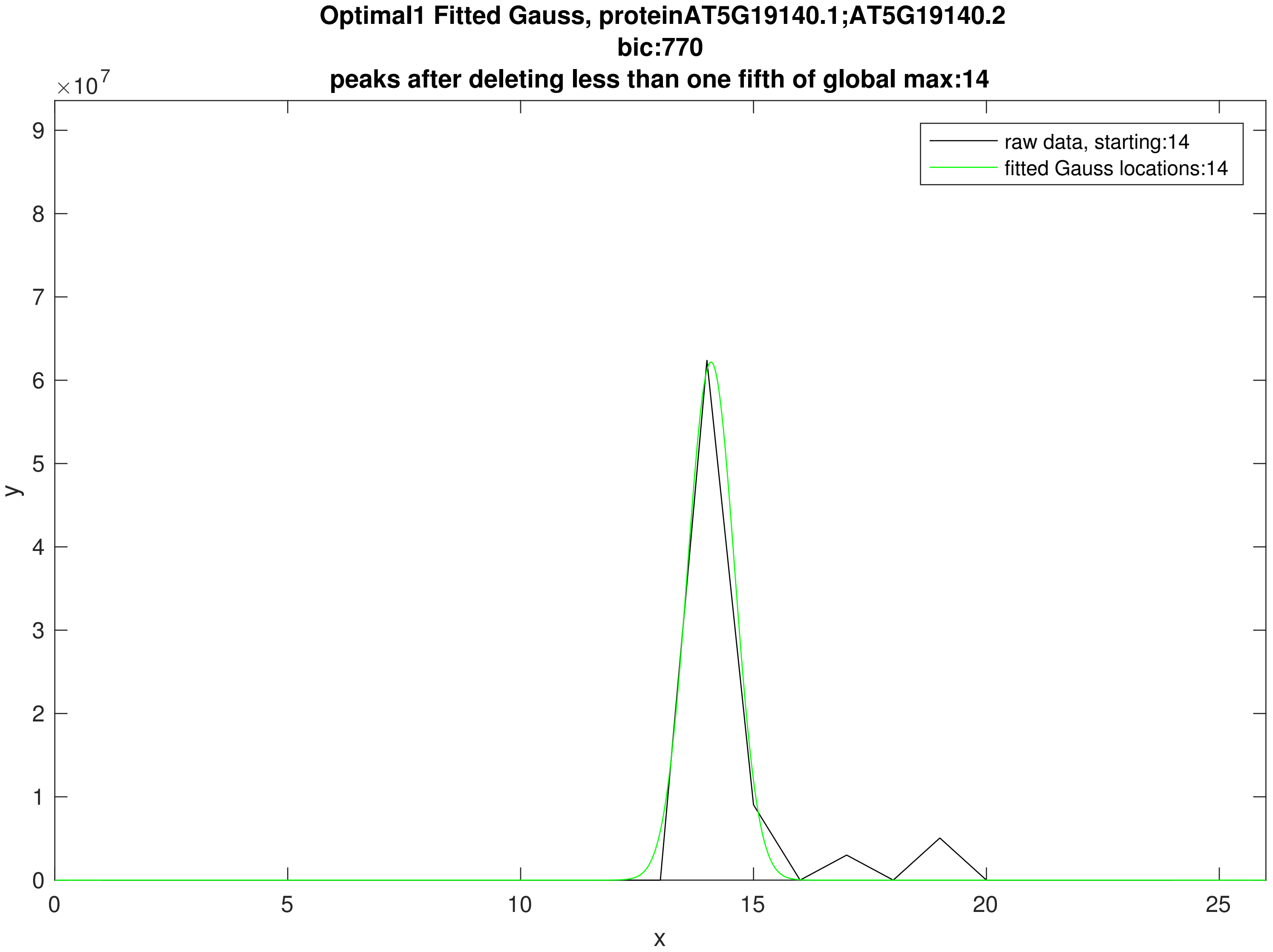
Optimal1 Fitted Gauss, proteinAT5G19140.1;AT5G19140.2
bic:770
peaks after deleting less than one fifth of global max:14
×10^7
9
8
7
6
5
4
3
2
1
0
y
0
5
10
15
20
25
x
raw data, starting:14
fitted Gauss locations:14

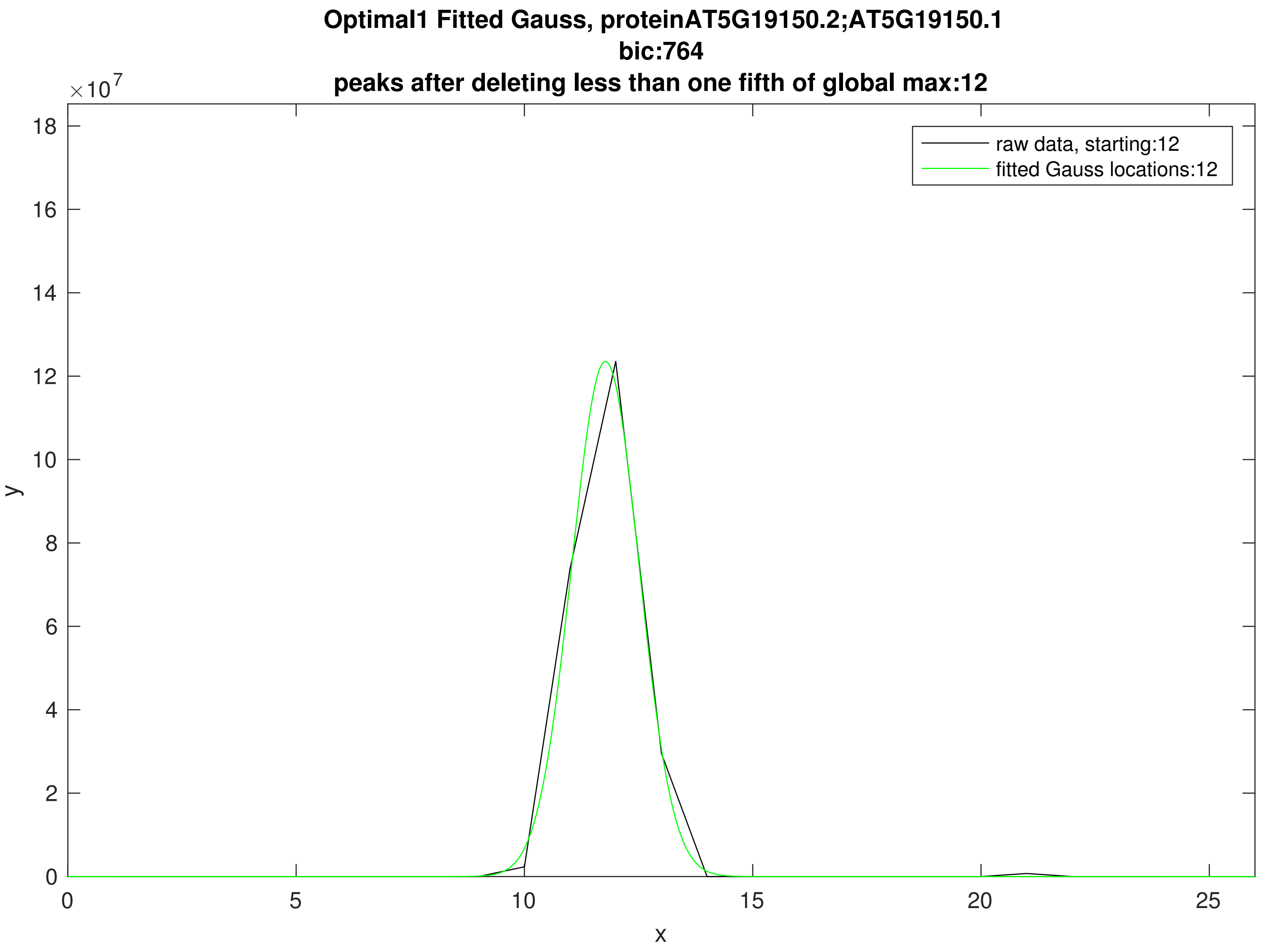
Optimal1 Fitted Gauss, proteinAT5G19150.2;AT5G19150.1
bic:764
peaks after deleting less than one fifth of global max:12
×10^7
raw data, starting:12
fitted Gauss locations:12
y
x
0
2
4
6
8
10
12
14
16
18
0
5
10
15
20
25

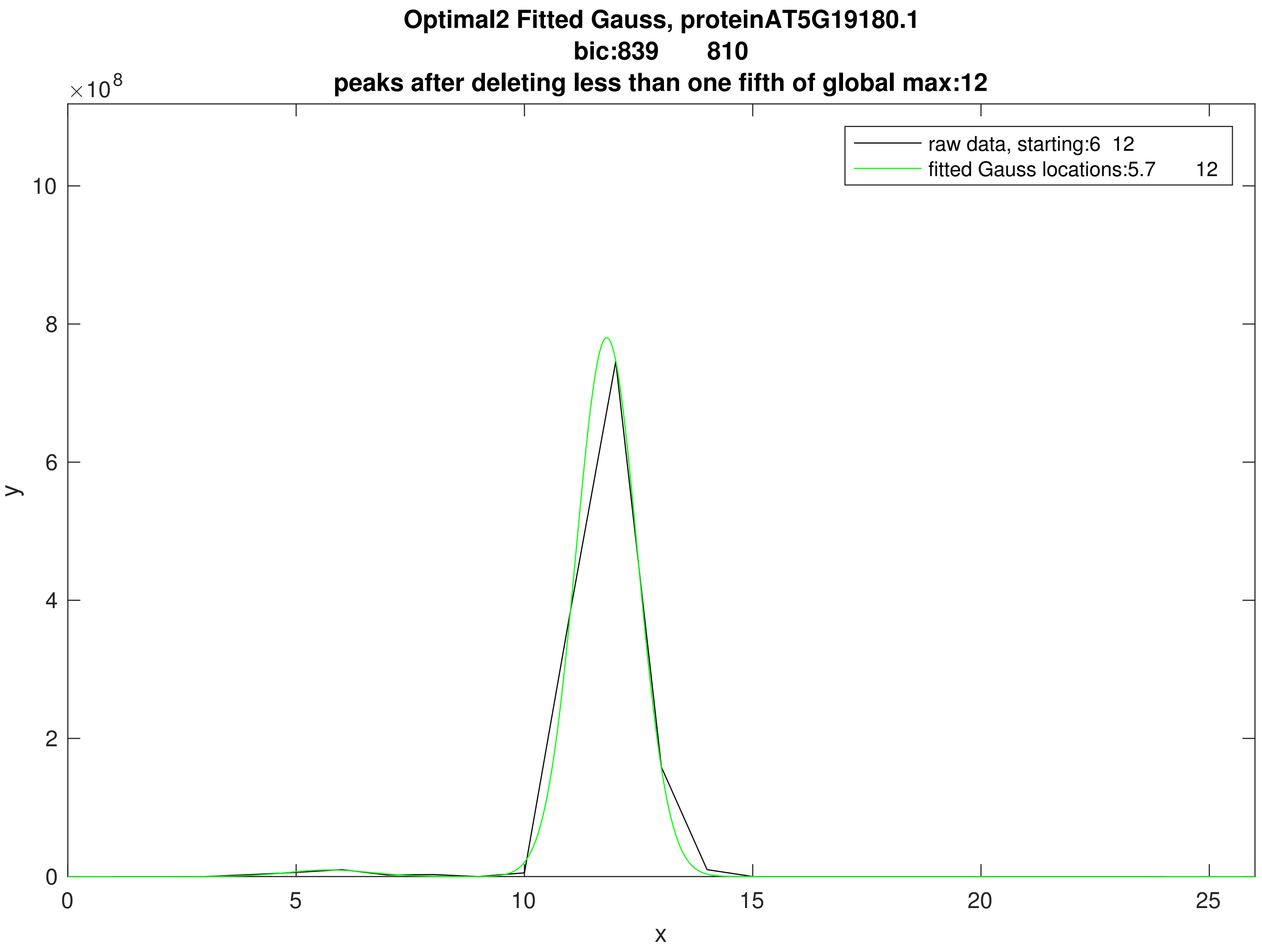
Optimal2 Fitted Gauss, proteinAT5G19180.1
bic:839 810
peaks after deleting less than one fifth of global max:12
raw data, starting:6 12
fitted Gauss locations:5.7 12
×10^8
y
x
0
2
4
6
8
10
5
15
20
25

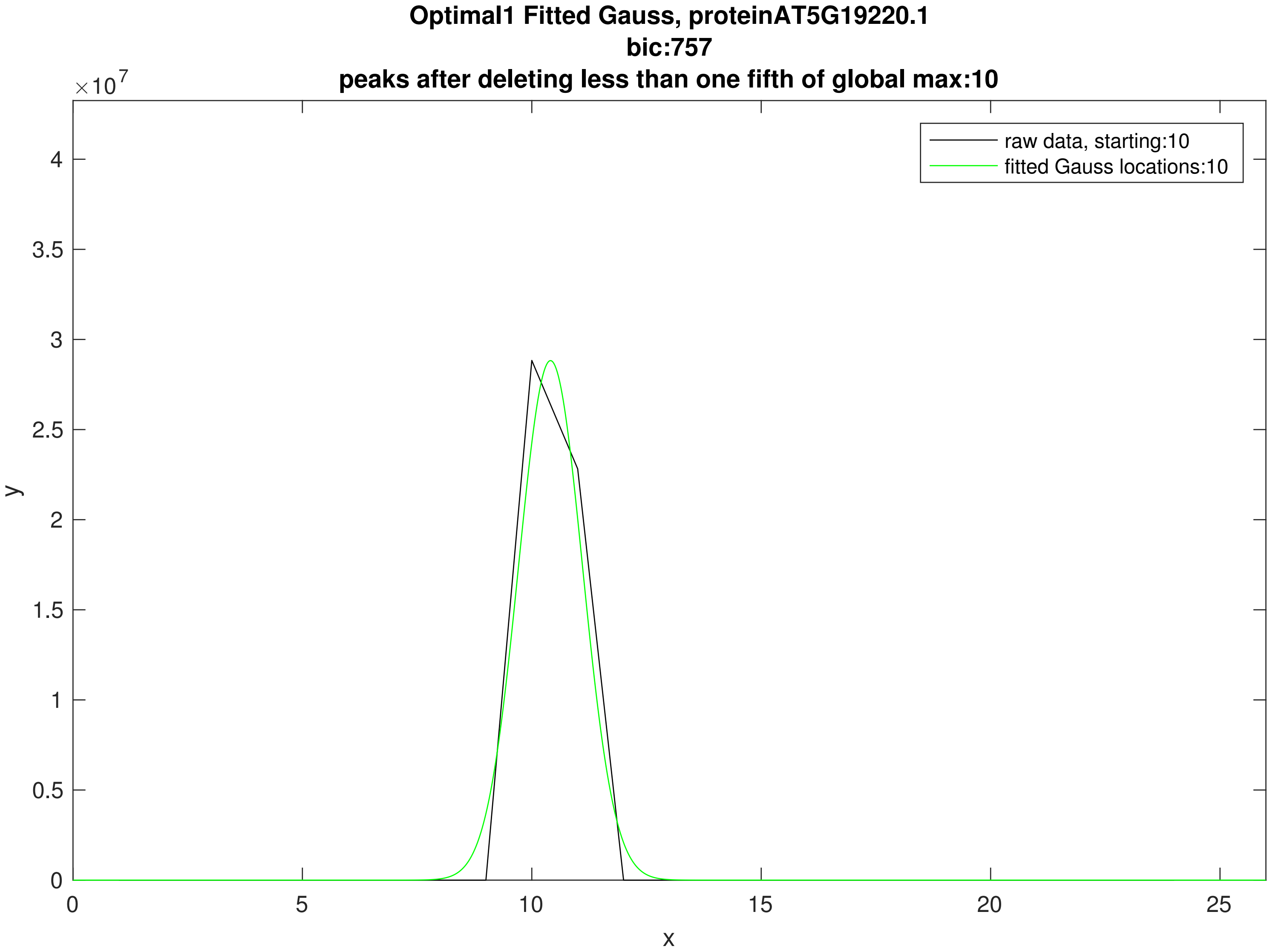
Optimal1 Fitted Gauss, proteinAT5G19220.1
bic:757
peaks after deleting less than one fifth of global max:10
×10^7
4
3.5
3
2.5
2
1.5
1
0.5
0
y
0
5
10
15
20
25
x
raw data, starting:10
fitted Gauss locations:10

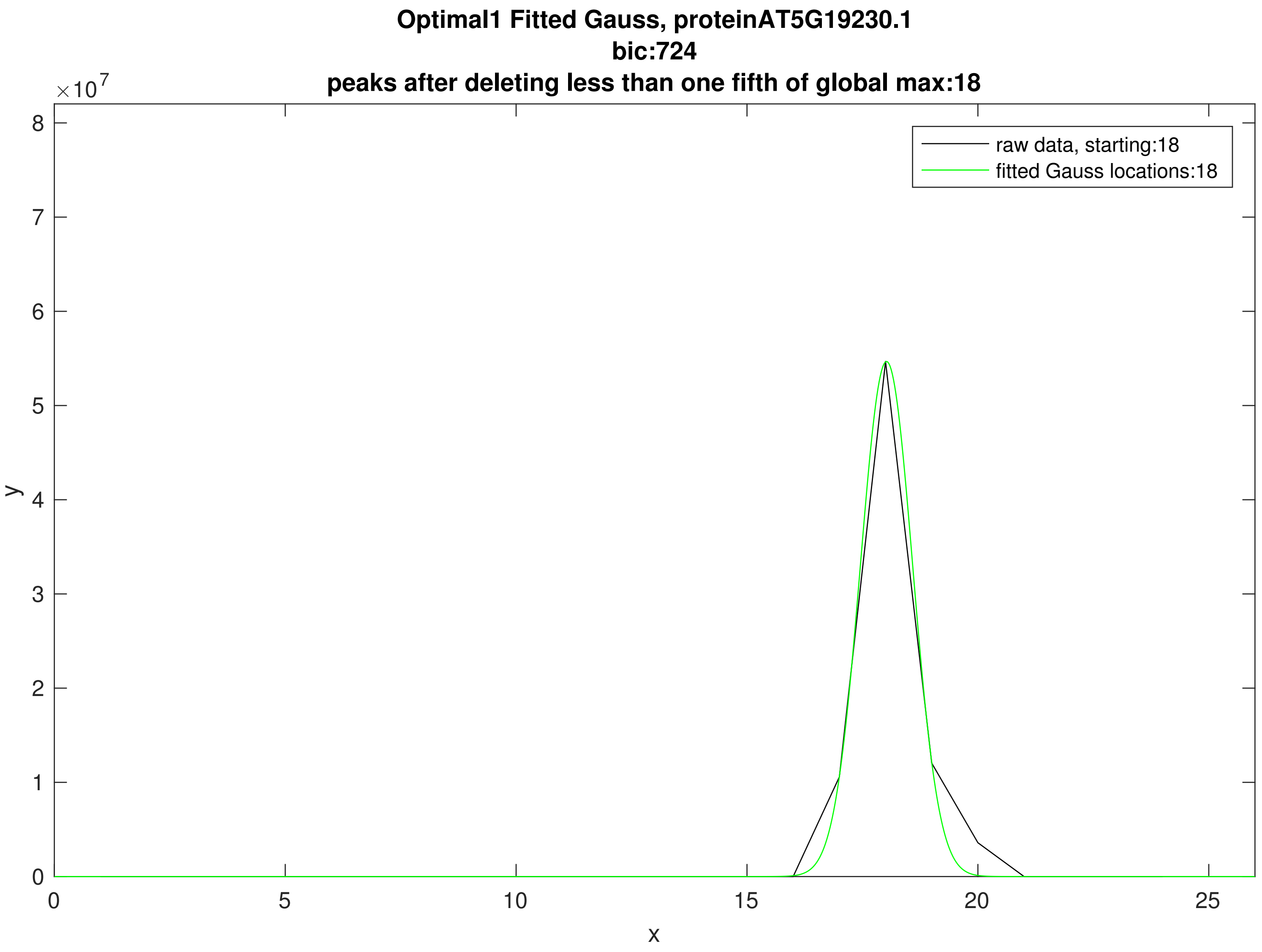
Optimal1 Fitted Gauss, proteinAT5G19230.1
bic:724
peaks after deleting less than one fifth of global max:18
$\times 10^7$
8
7
6
5
4
3
2
1
0
y
0
5
10
15
20
25
x
raw data, starting:18
fitted Gauss locations:18

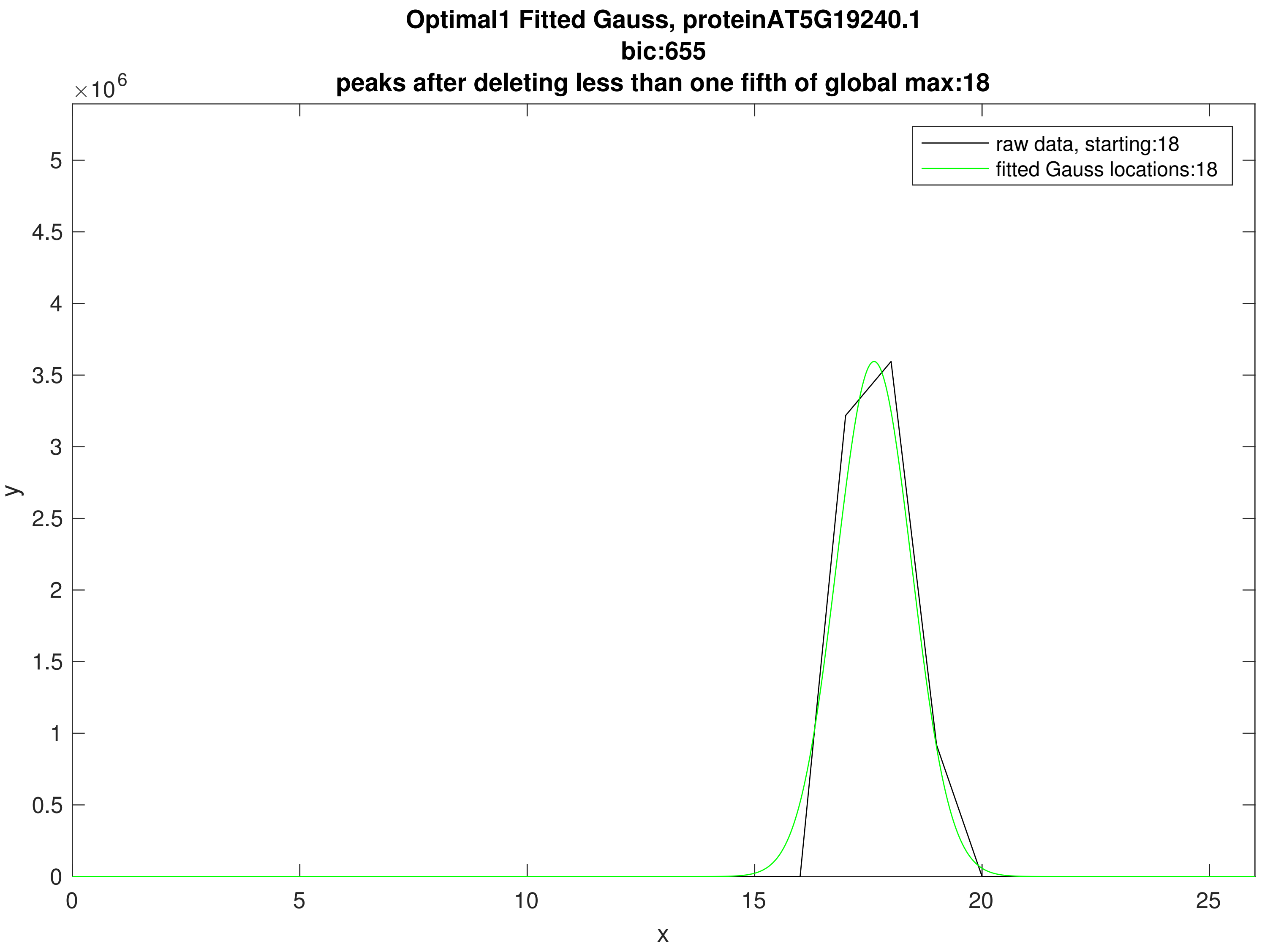

Optimal1 Fitted Gauss, proteinAT5G19240.1
bic:655
peaks after deleting less than one fifth of global max:18
×10^6
raw data, starting:18
fitted Gauss locations:18
y
x

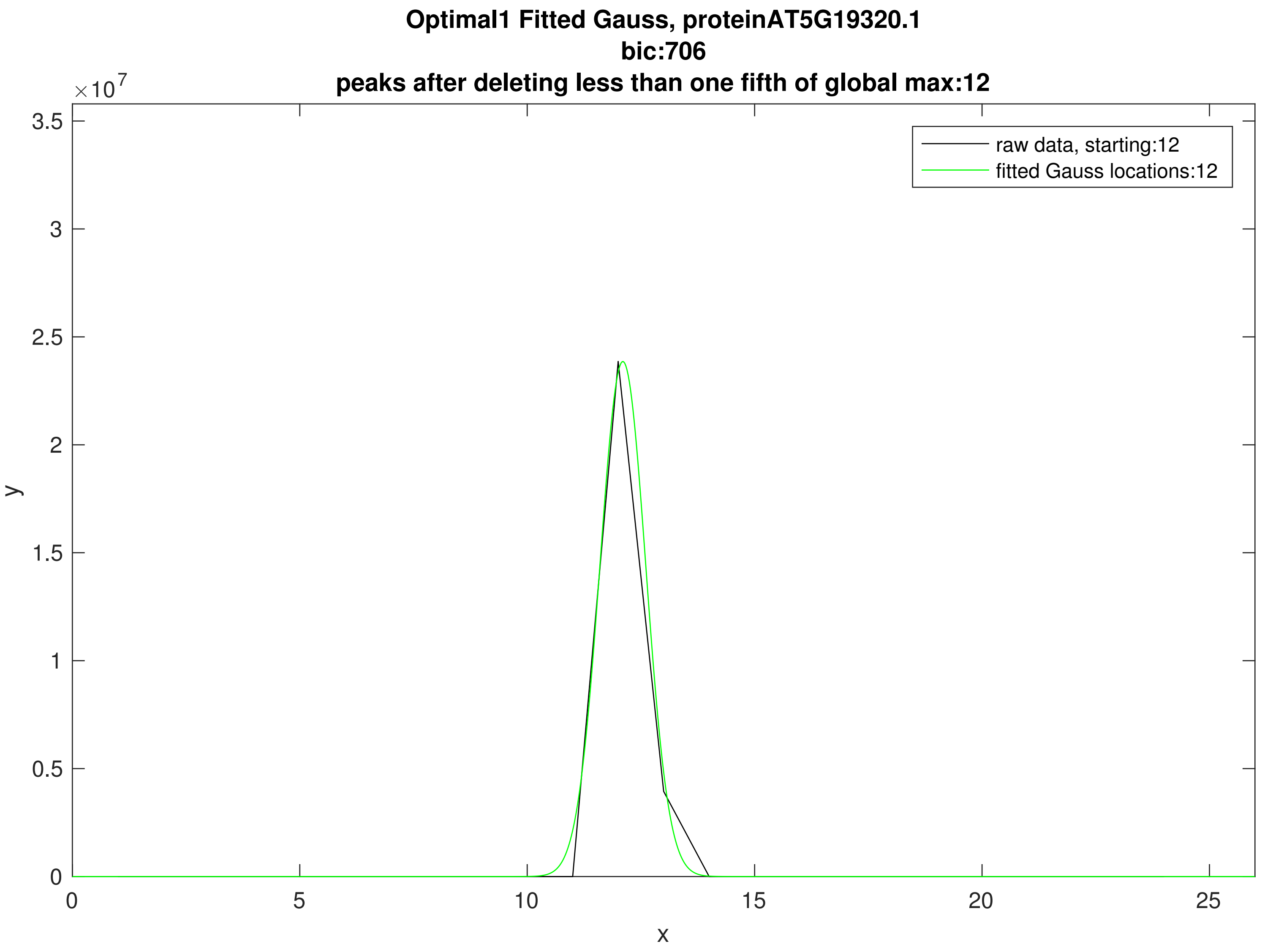
Optimal1 Fitted Gauss, proteinAT5G19320.1
bic:706
peaks after deleting less than one fifth of global max:12
×10^7
3.5
3
2.5
2
1.5
1
0.5
0
y
0
5
10
15
20
25
x
raw data, starting:12
fitted Gauss locations:12

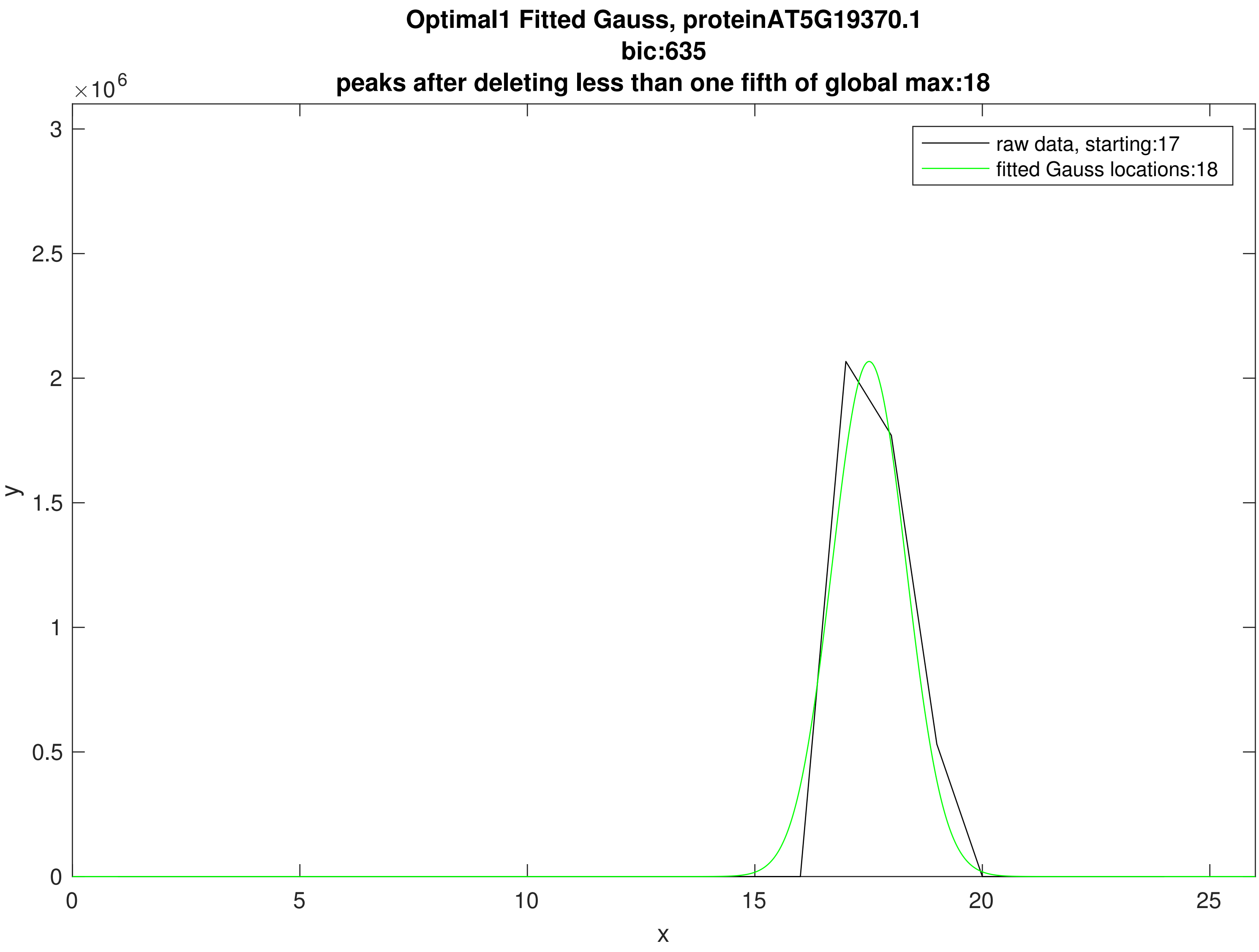

Optimal1 Fitted Gauss, proteinAT5G19370.1
bic:635
peaks after deleting less than one fifth of global max:18
×10^6
raw data, starting:17
fitted Gauss locations:18
3
2.5
2
1.5
1
0.5
0
y
0
5
10
15
20
25
x

Optimal1 Fitted Gauss, proteinAT5G19440.1
bic:883 887 897
peaks after deleting less than one fifth of global max:17

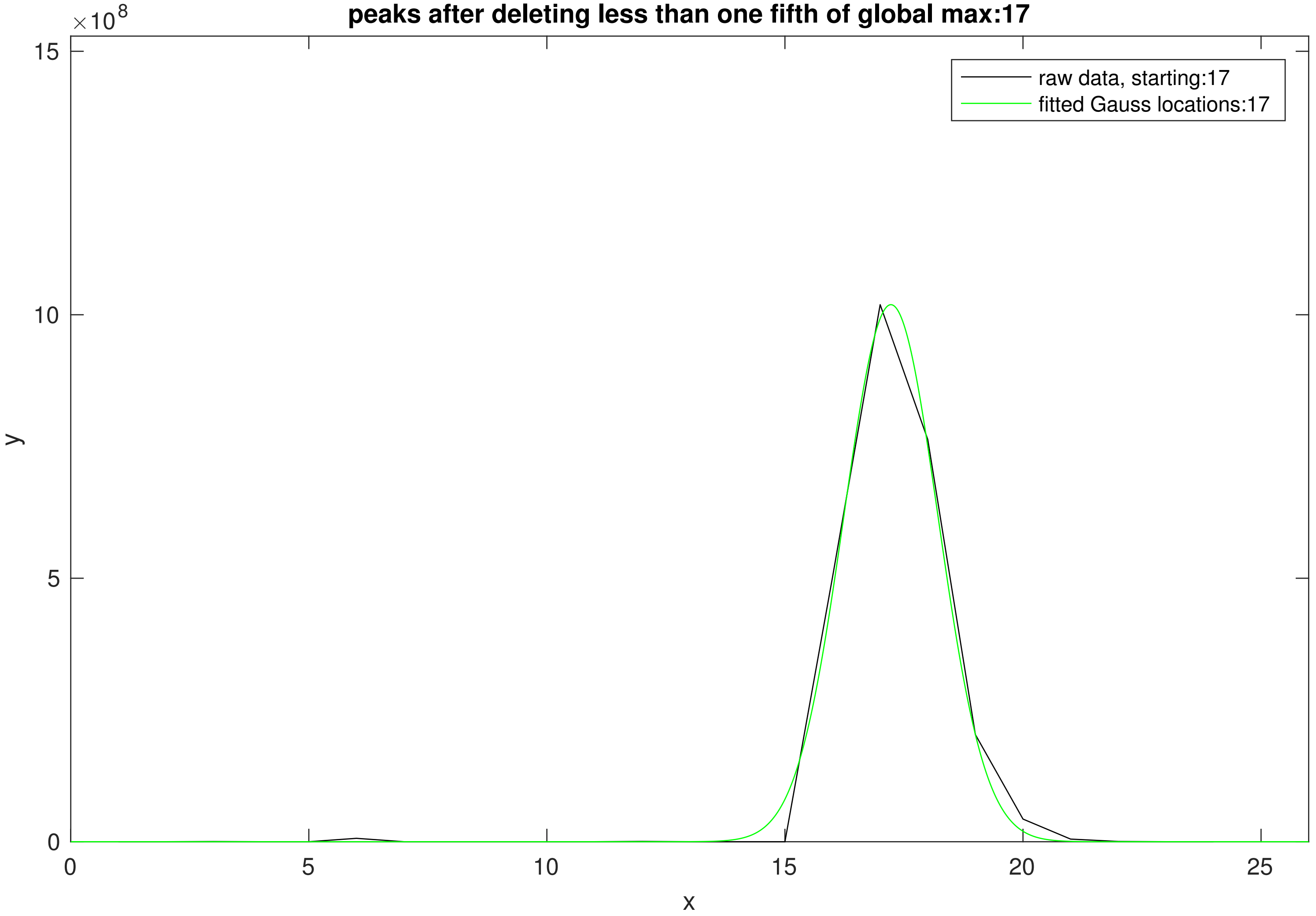

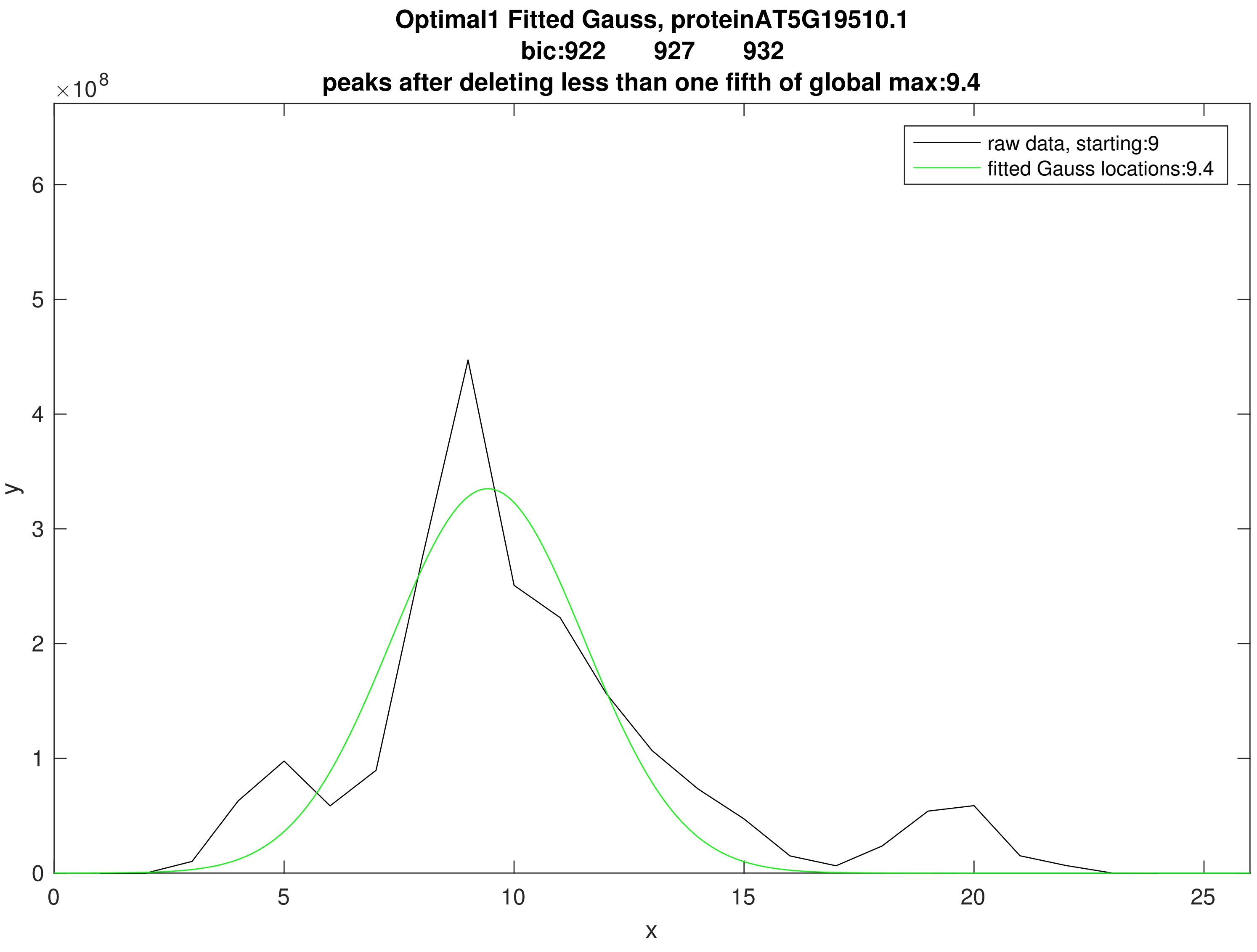
Optimal1 Fitted Gauss, proteinAT5G19510.1
bic:922 927 932
peaks after deleting less than one fifth of global max:9.4
×10^8
raw data, starting:9
fitted Gauss locations:9.4
y
x
0
1
2
3
4
5
6
0
5
10
15
20
25

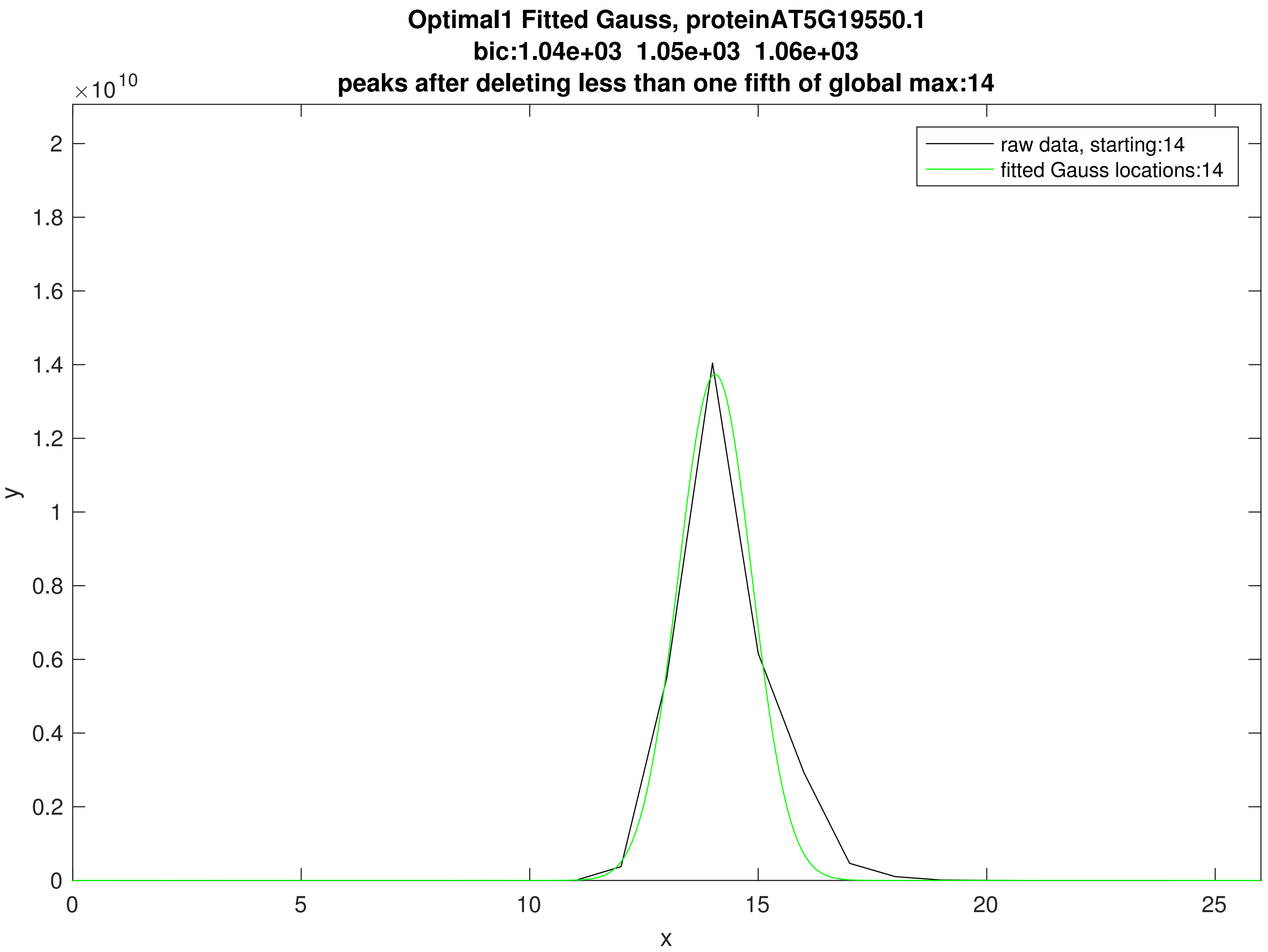
Optimal1 Fitted Gauss, proteinAT5G19550.1
bic:1.04e+03 1.05e+03 1.06e+03
peaks after deleting less than one fifth of global max:14
×10^10
2
1.8
1.6
1.4
1.2
1
0.8
0.6
0.4
0.2
0
y
0
5
10
15
20
25
x
raw data, starting:14
fitted Gauss locations:14

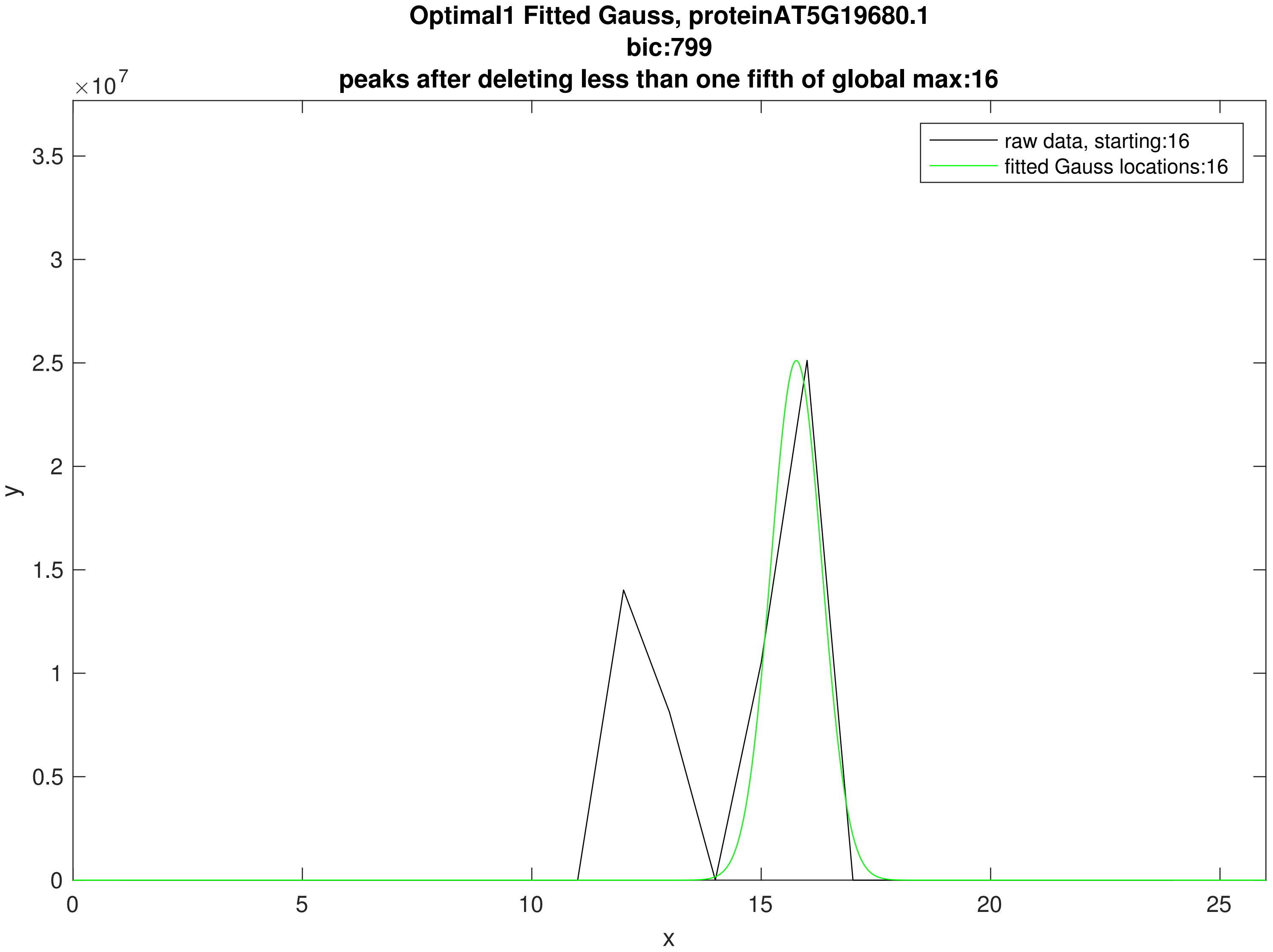
Optimal1 Fitted Gauss, proteinAT5G19680.1
bic:799
peaks after deleting less than one fifth of global max:16
×10^7
3.5
3
2.5
2
1.5
1
0.5
0
y
0
5
10
15
20
25
x
raw data, starting:16
fitted Gauss locations:16

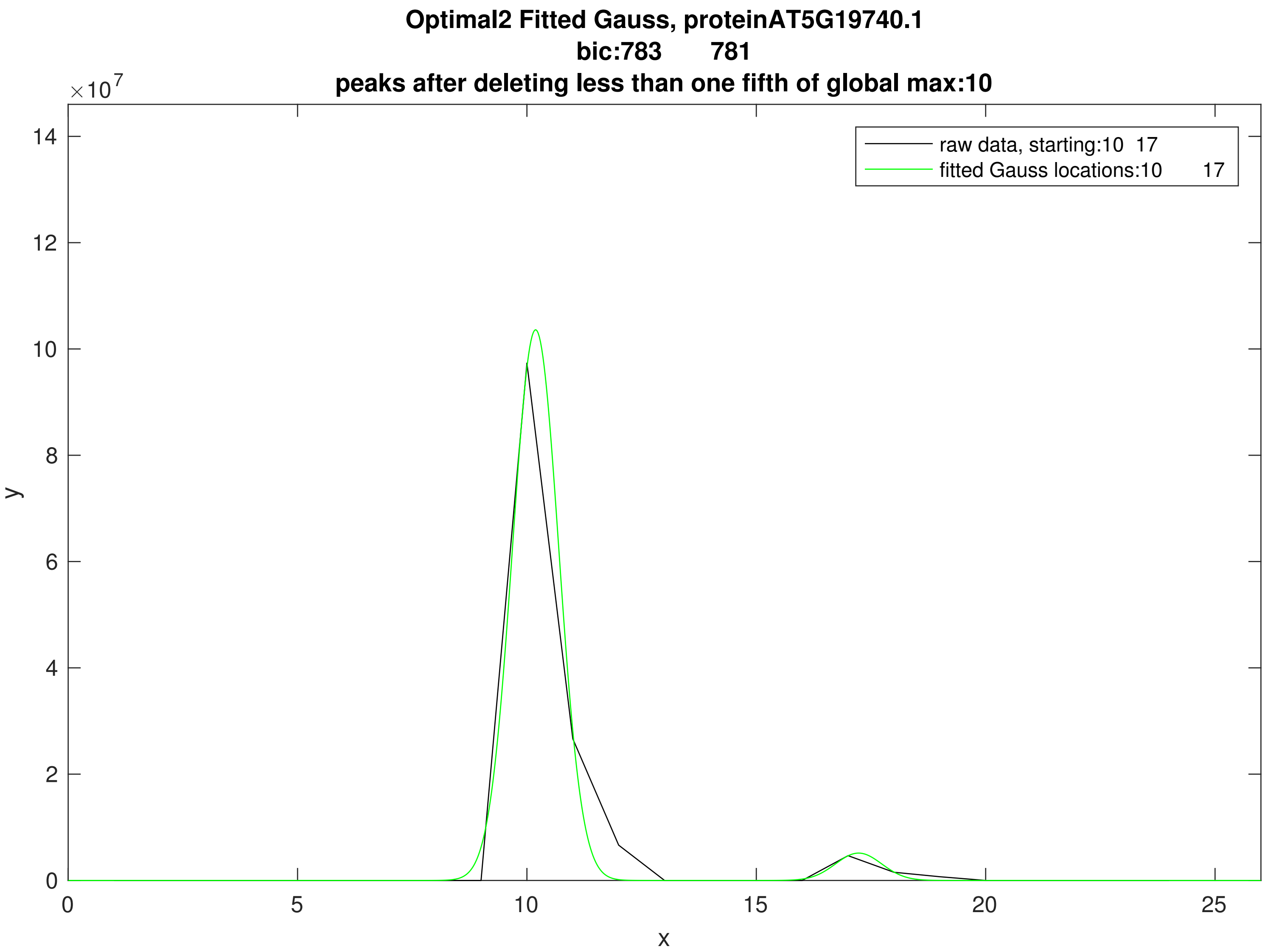
Optimal2 Fitted Gauss, proteinAT5G19740.1
bic:783 781
peaks after deleting less than one fifth of global max:10
×10^7
14
12
10
8
6
4
2
0
y
0
5
10
15
20
25
x
raw data, starting:10 17
fitted Gauss locations:10 17

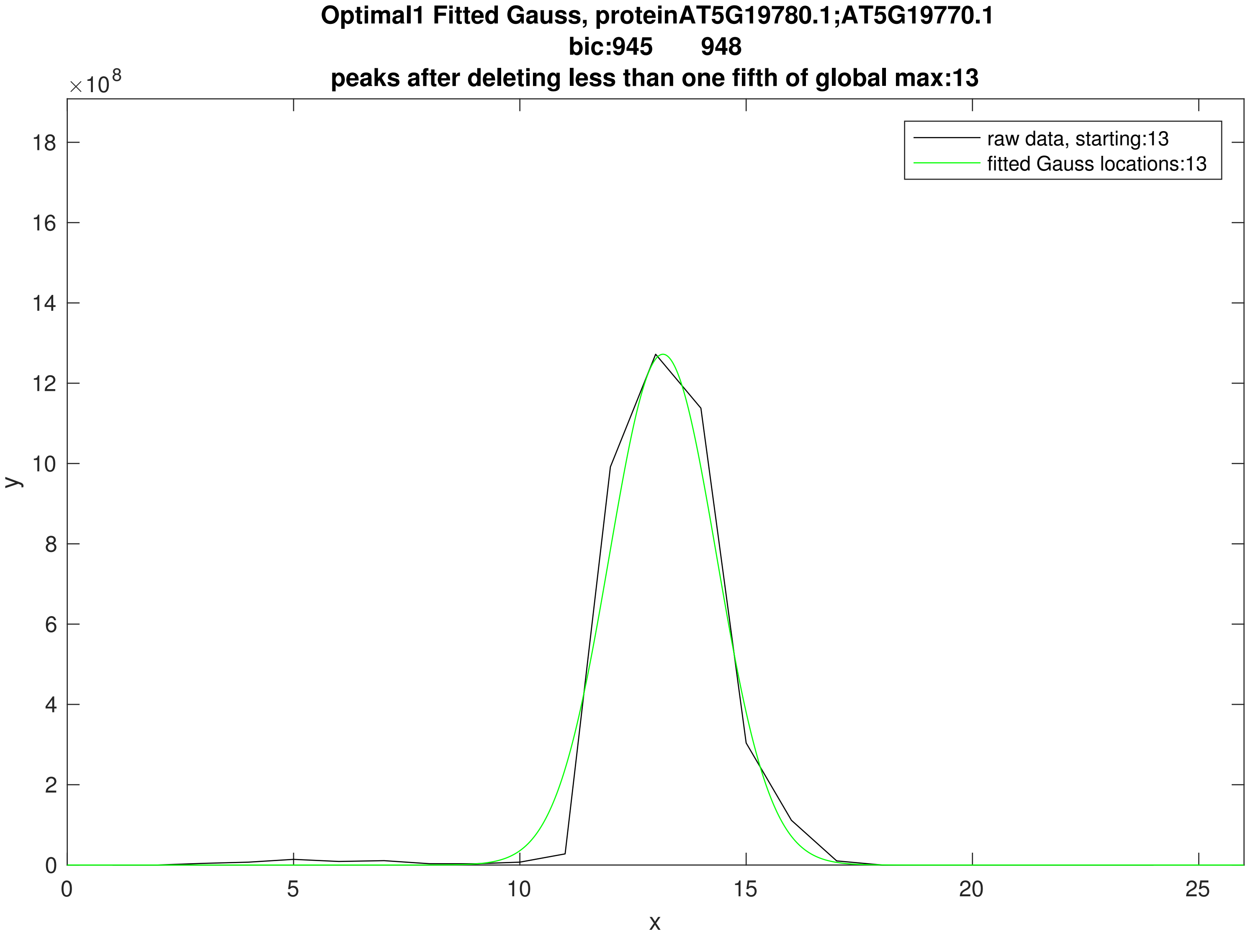
Optimal1 Fitted Gauss, proteinAT5G19780.1;AT5G19770.1
bic:945 948
peaks after deleting less than one fifth of global max:13
×10^8
raw data, starting:13
fitted Gauss locations:13
y
x
0
2
4
6
8
10
12
14
16
18
0
5
10
15
20
25

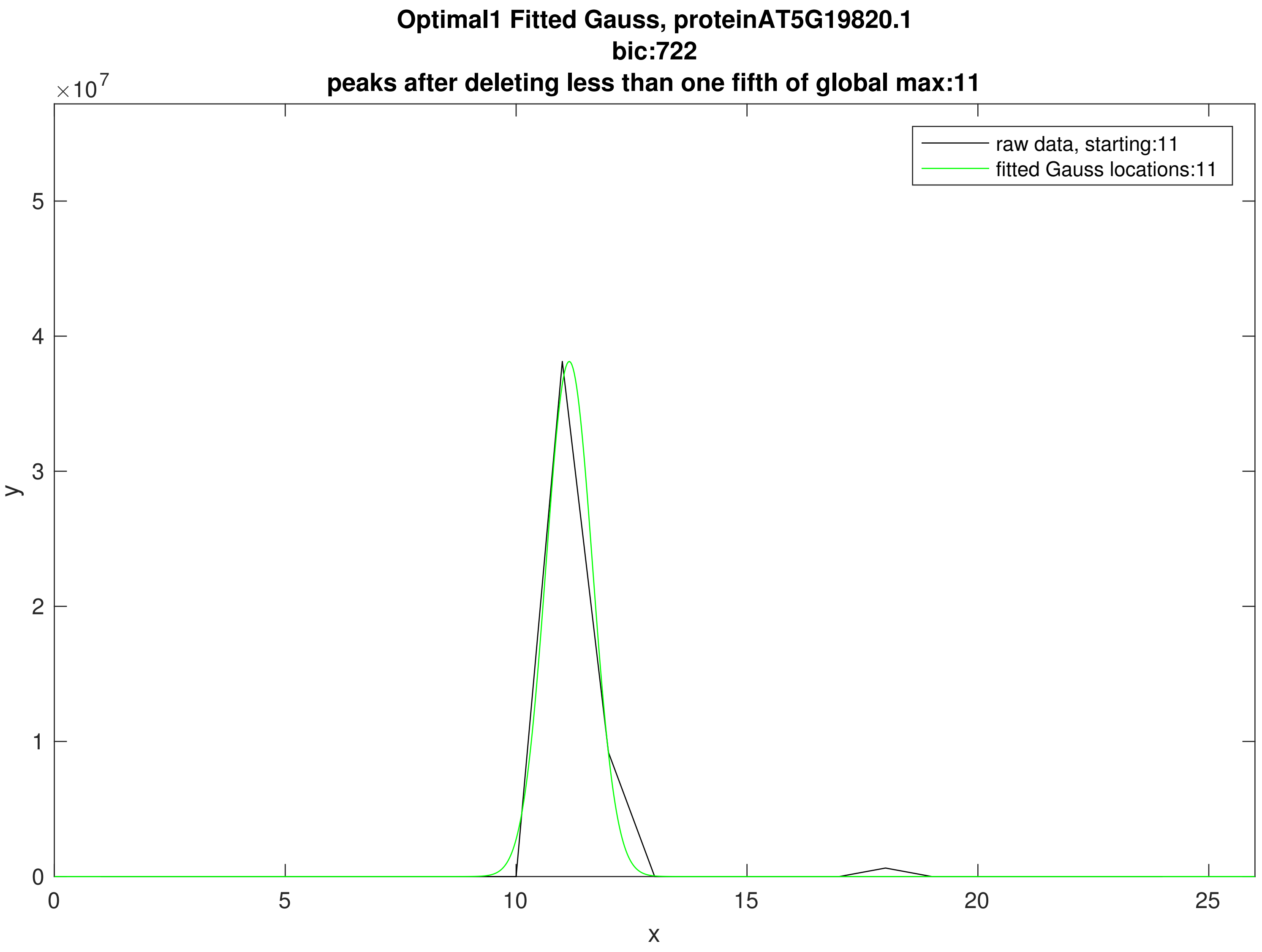
Optimal1 Fitted Gauss, proteinAT5G19820.1
bic:722
peaks after deleting less than one fifth of global max:11
×10^7
raw data, starting:11
fitted Gauss locations:11
y
x
0
1
2
3
4
5
0
5
10
15
20
25

**Optimal1 Fitted Gauss, proteinAT5G19860.1**
**bic:771**
**peaks after deleting less than one fifth of global max:20**

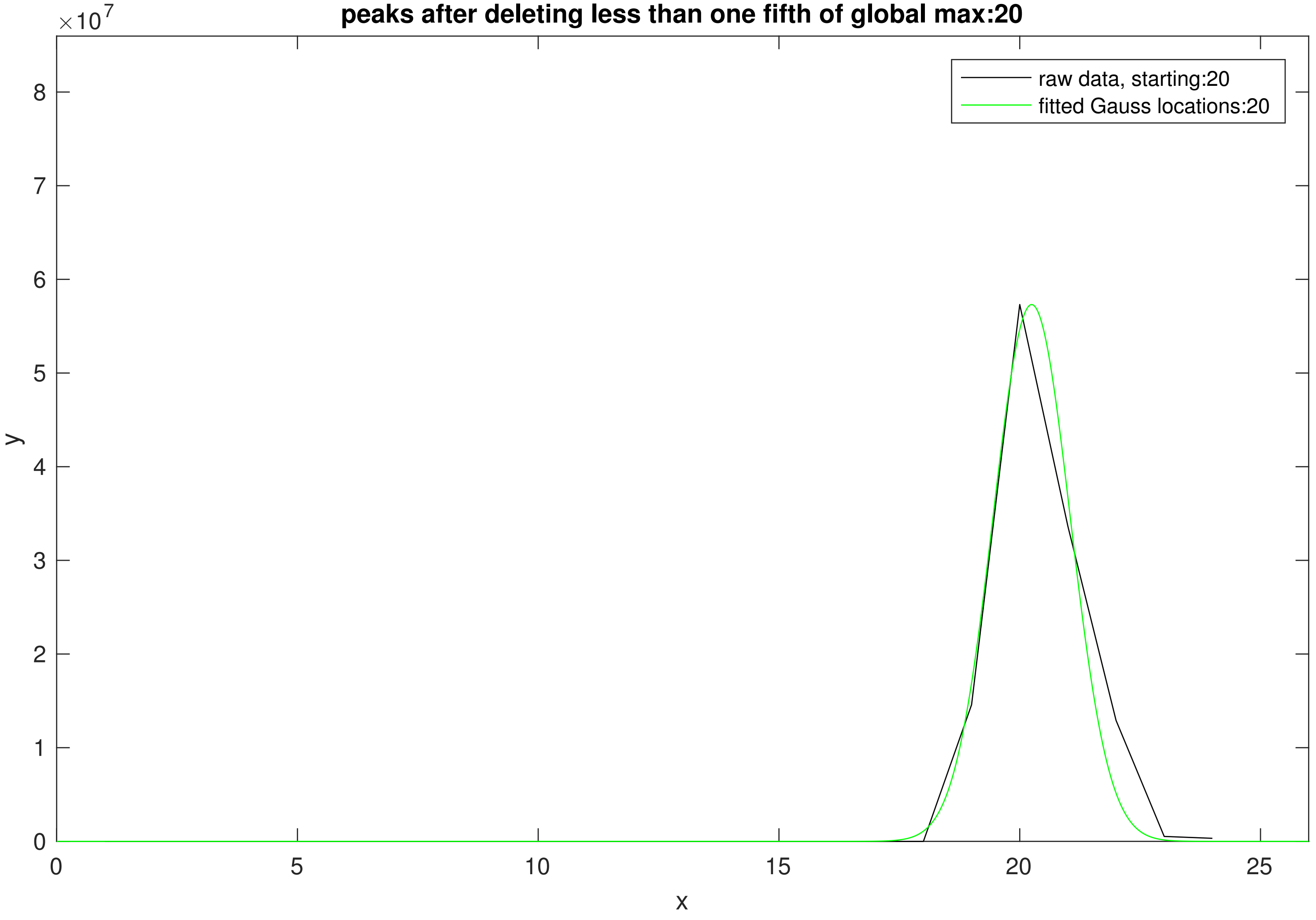

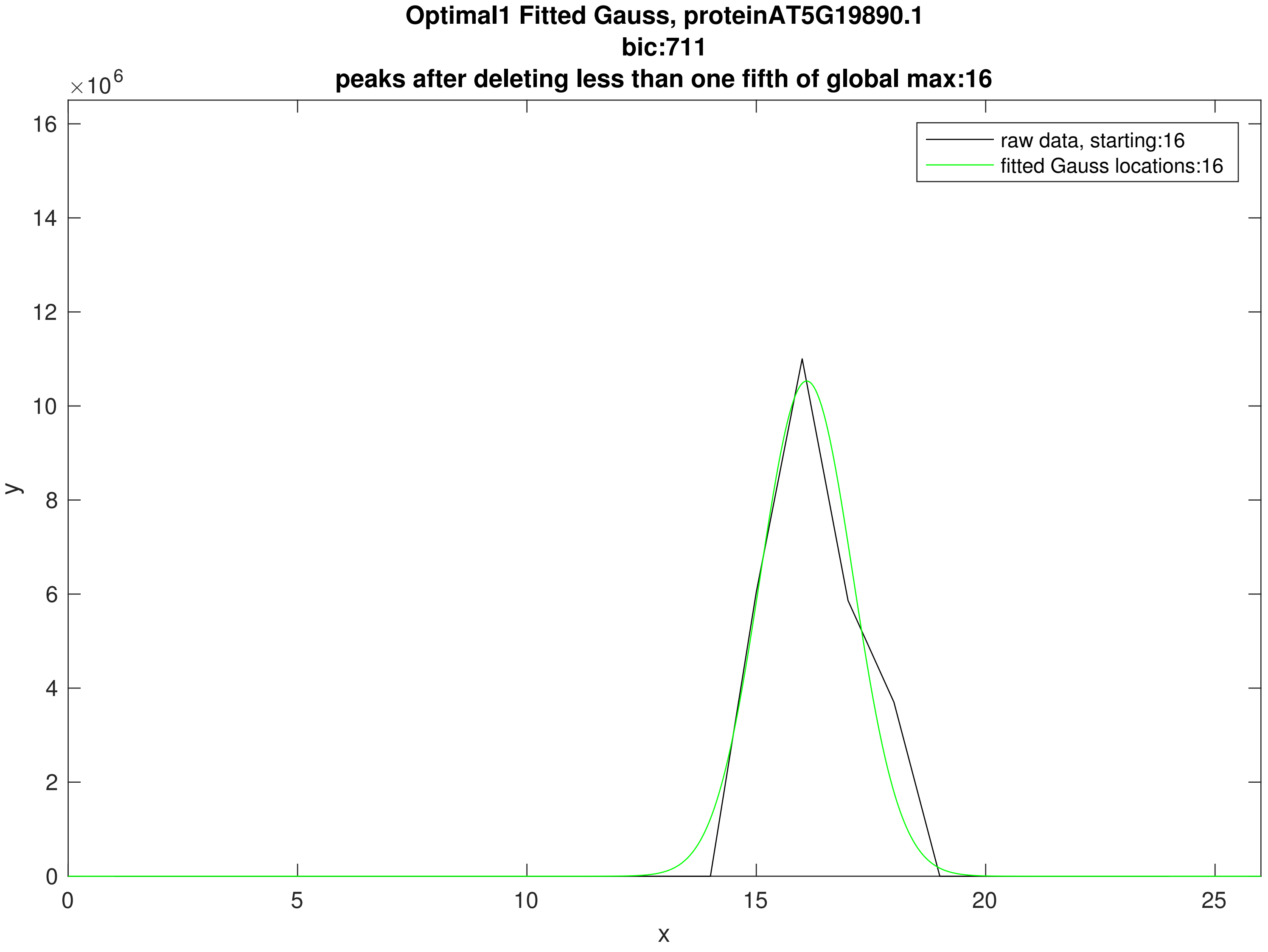

Optimal1 Fitted Gauss, proteinAT5G19890.1
bic:711
peaks after deleting less than one fifth of global max:16
×10^6
raw data, starting:16
fitted Gauss locations:16
y
x
0
2
4
6
8
10
12
14
16
0
5
10
15
20
25

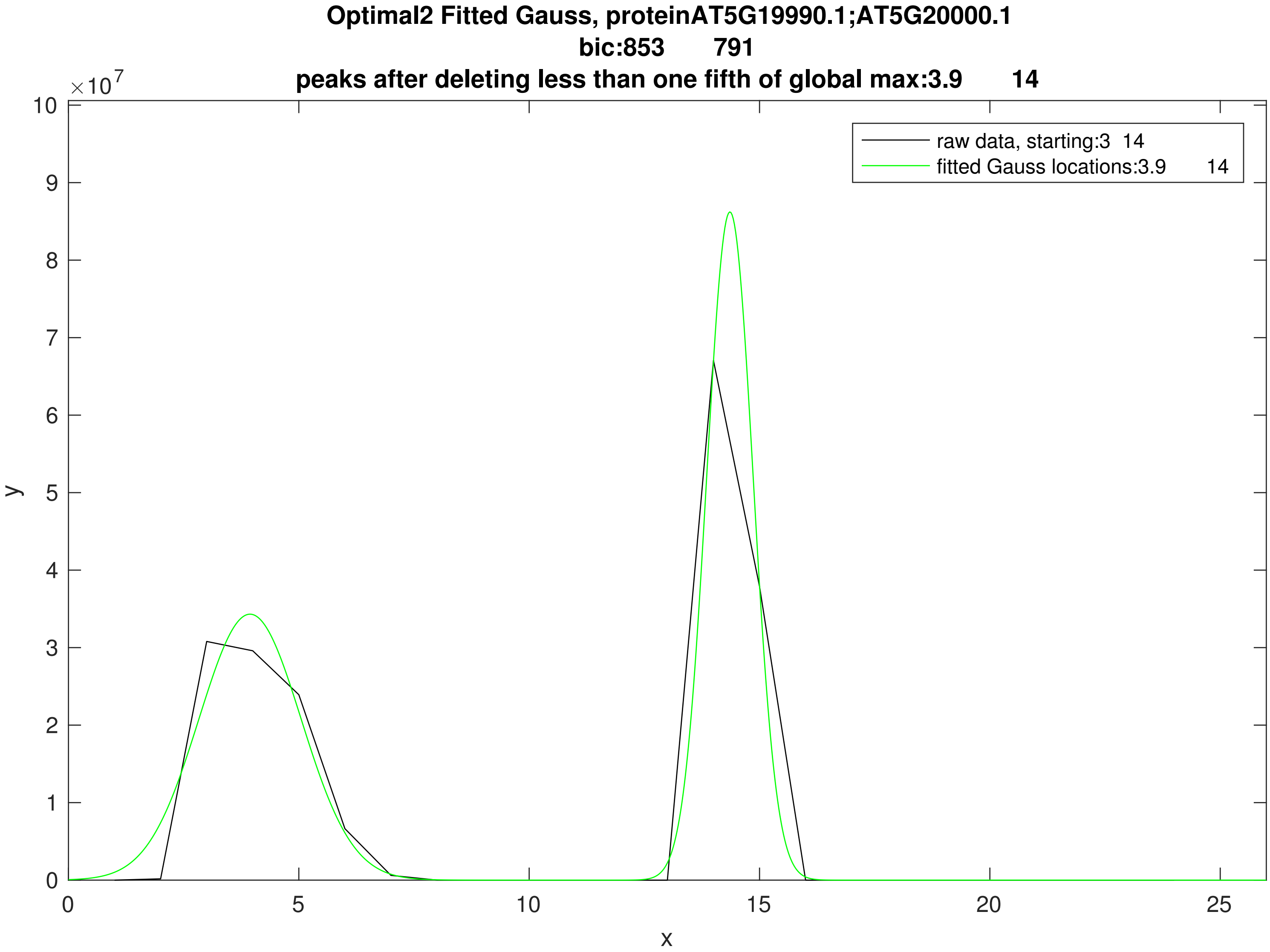

Optimal2 Fitted Gauss, proteinAT5G19990.1;AT5G20000.1
bic:853 791
peaks after deleting less than one fifth of global max:3.9 14
×10^7
raw data, starting:3 14
fitted Gauss locations:3.9 14
y
x
0
5
10
15
20
25
1
2
3
4
6
7
8
9

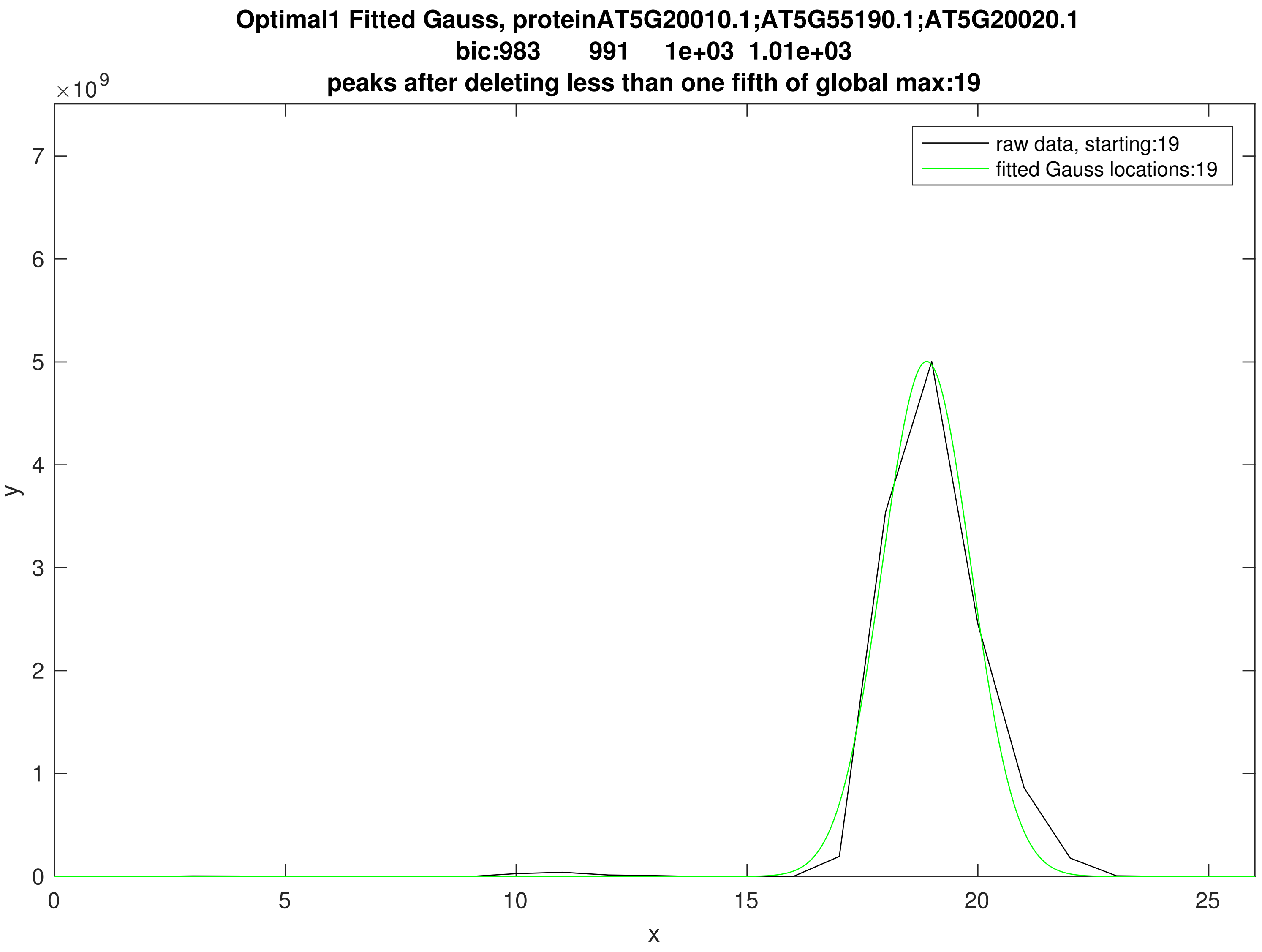

Optimal1 Fitted Gauss, proteinAT5G20010.1;AT5G55190.1;AT5G20020.1
bic:983 991 1e+03 1.01e+03
peaks after deleting less than one fifth of global max:19
×10^9
raw data, starting:19
fitted Gauss locations:19
y
x

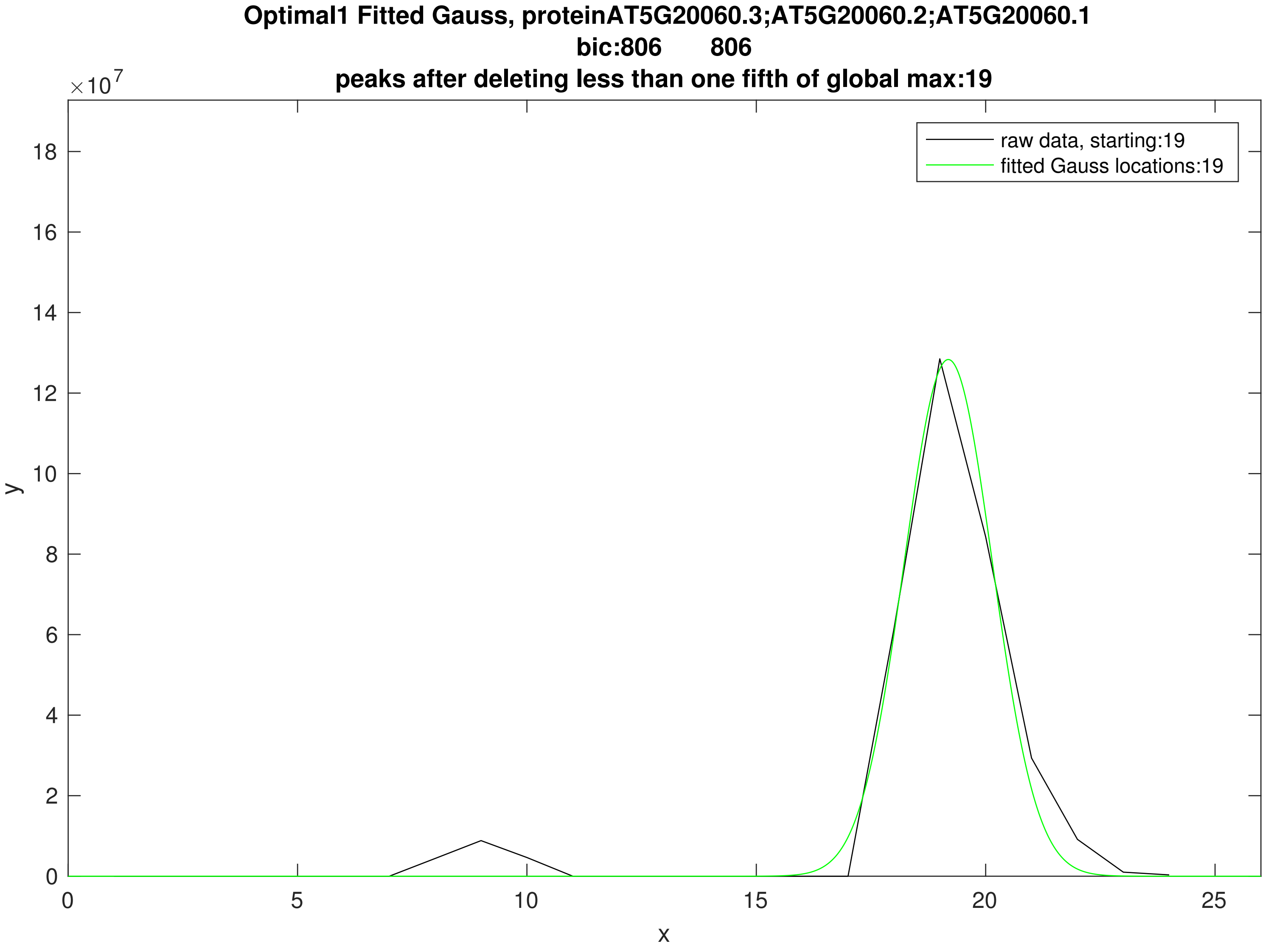
Optimal1 Fitted Gauss, proteinAT5G20060.3;AT5G20060.2;AT5G20060.1
bic:806 806
peaks after deleting less than one fifth of global max:19
×10^7
raw data, starting:19
fitted Gauss locations:19
y
x
0
2
4
6
8
10
12
14
16
18
0
5
10
15
20
25

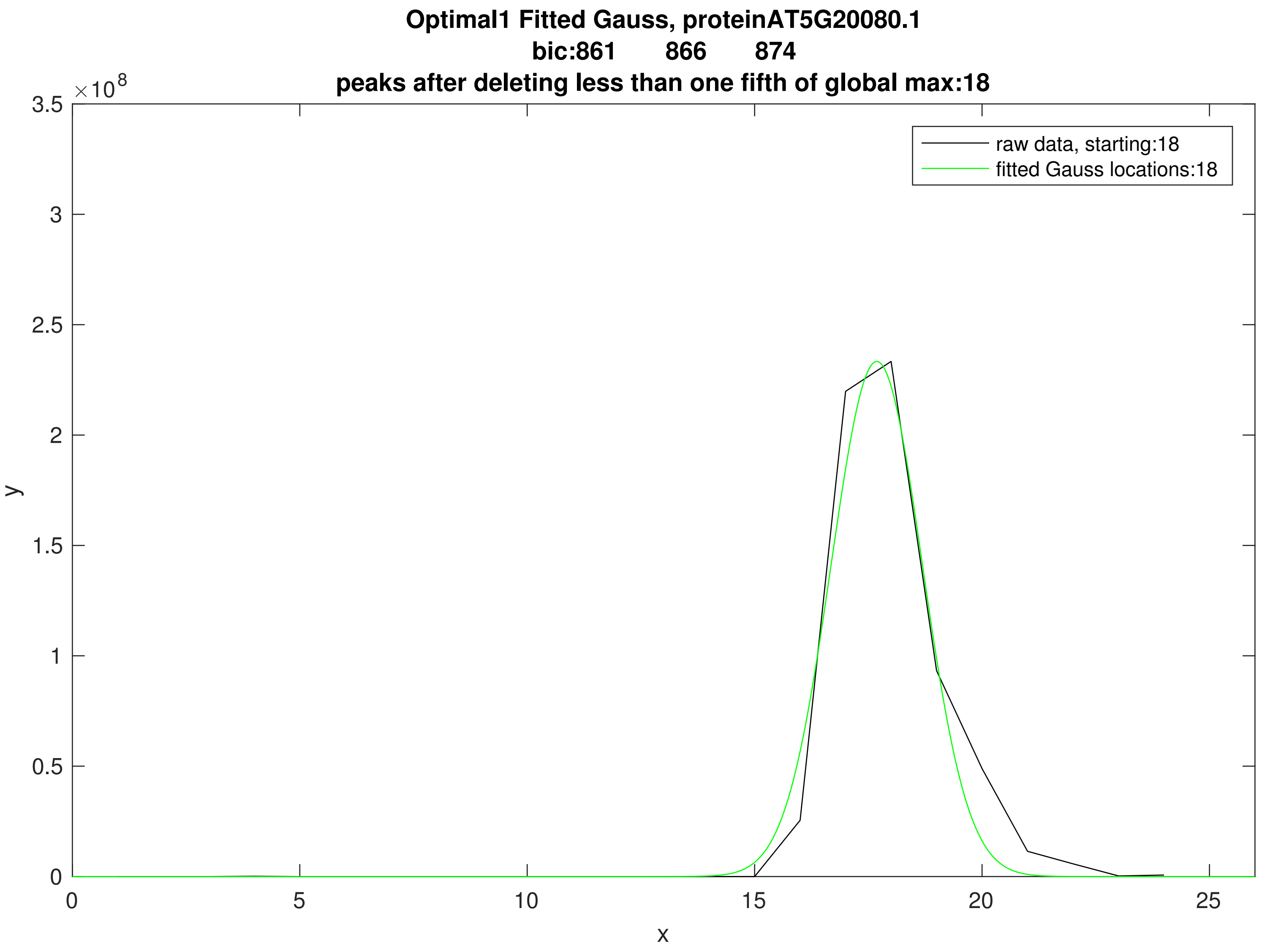
Optimal1 Fitted Gauss, proteinAT5G20080.1
bic:861 866 874
peaks after deleting less than one fifth of global max:18
raw data, starting:18
fitted Gauss locations:18
y
x

Optimal1 Fitted Gauss, proteinAT5G20250.4;AT5G20250.3;AT5G20250.2;AT5G20250.1
bic:1e+03 1.01e+03 1.02e+03 1.03e+03
peaks after deleting less than one fifth of global max:14

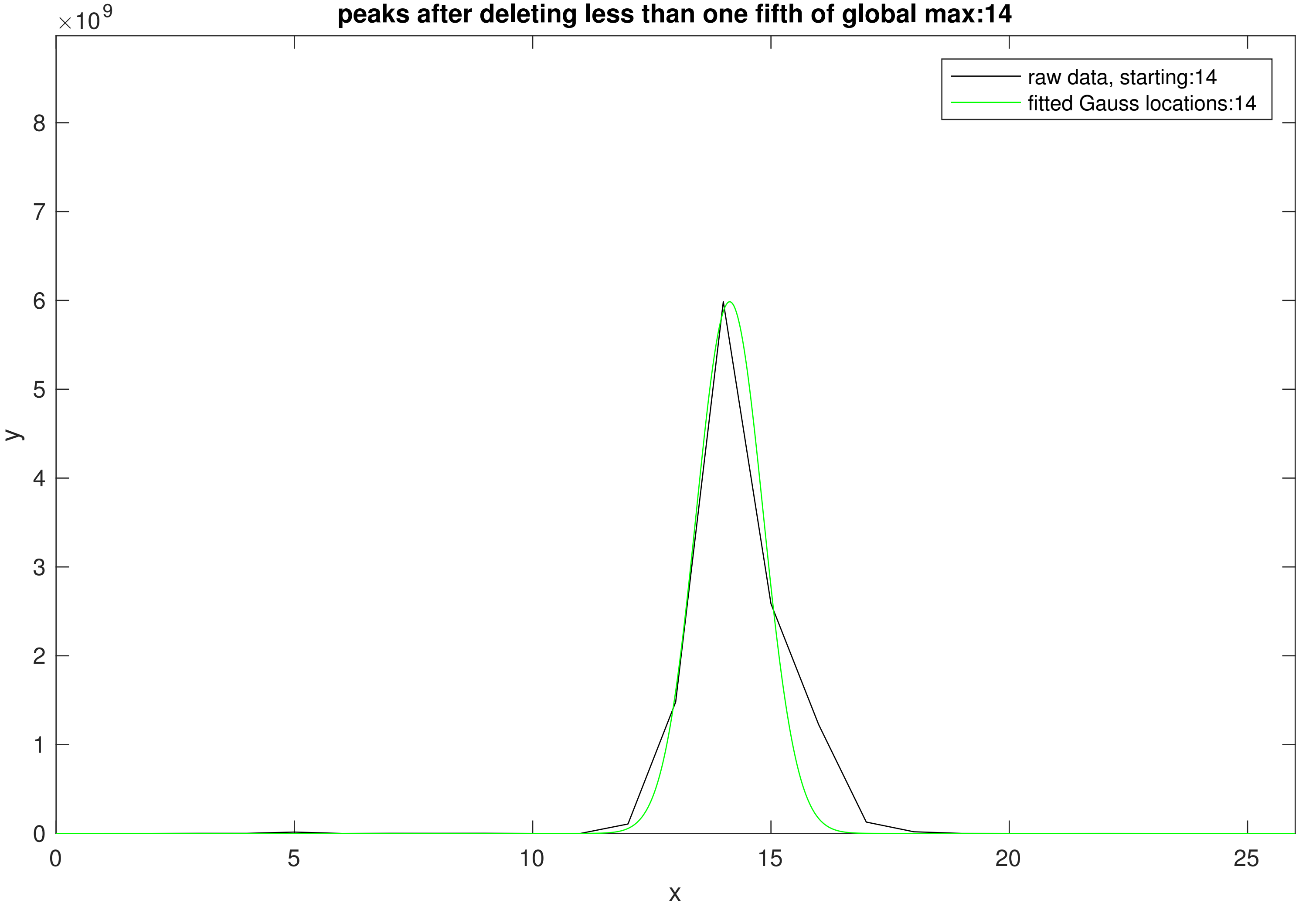

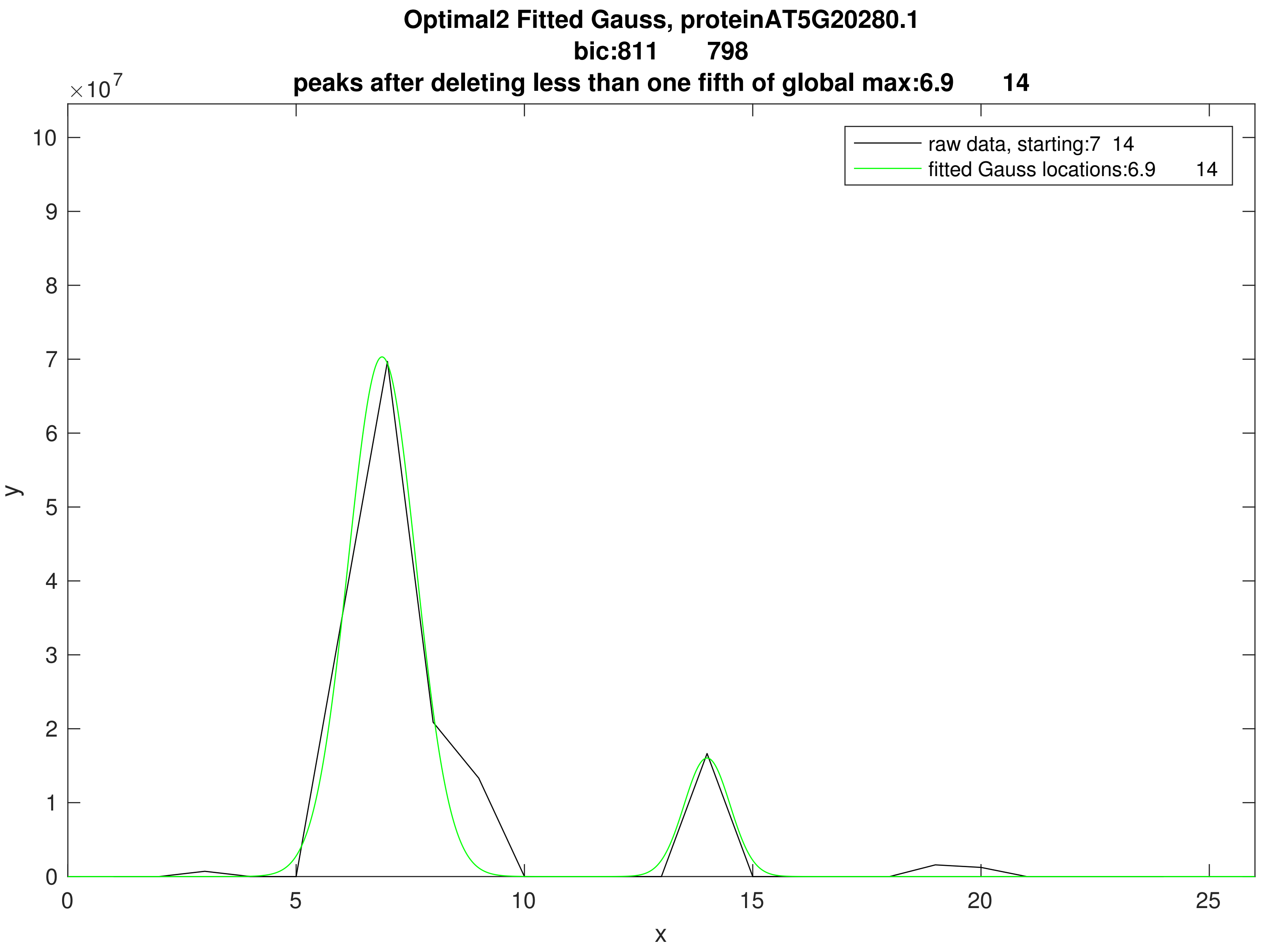

Optimal2 Fitted Gauss, proteinAT5G20280.1
bic:811 798
peaks after deleting less than one fifth of global max:6.9 14
×10^7
raw data, starting:7 14
fitted Gauss locations:6.9 14
y
x
0
1
2
3
4
5
6
7
8
9
10
0
5
10
15
20
25

Optimal1 Fitted Gauss, proteinAT5G20290.1
bic:896 905
peaks after deleting less than one fifth of global max:3.8

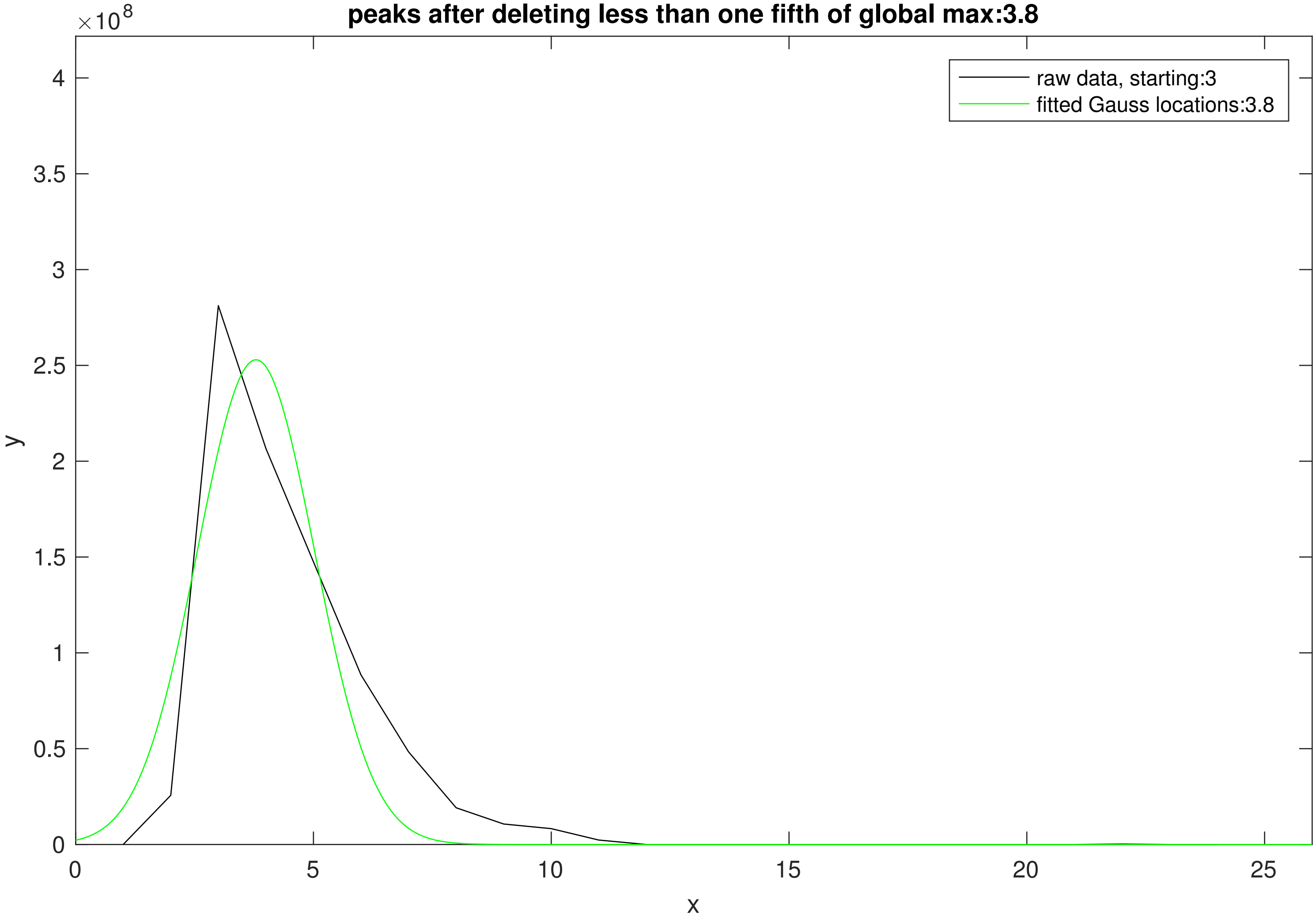

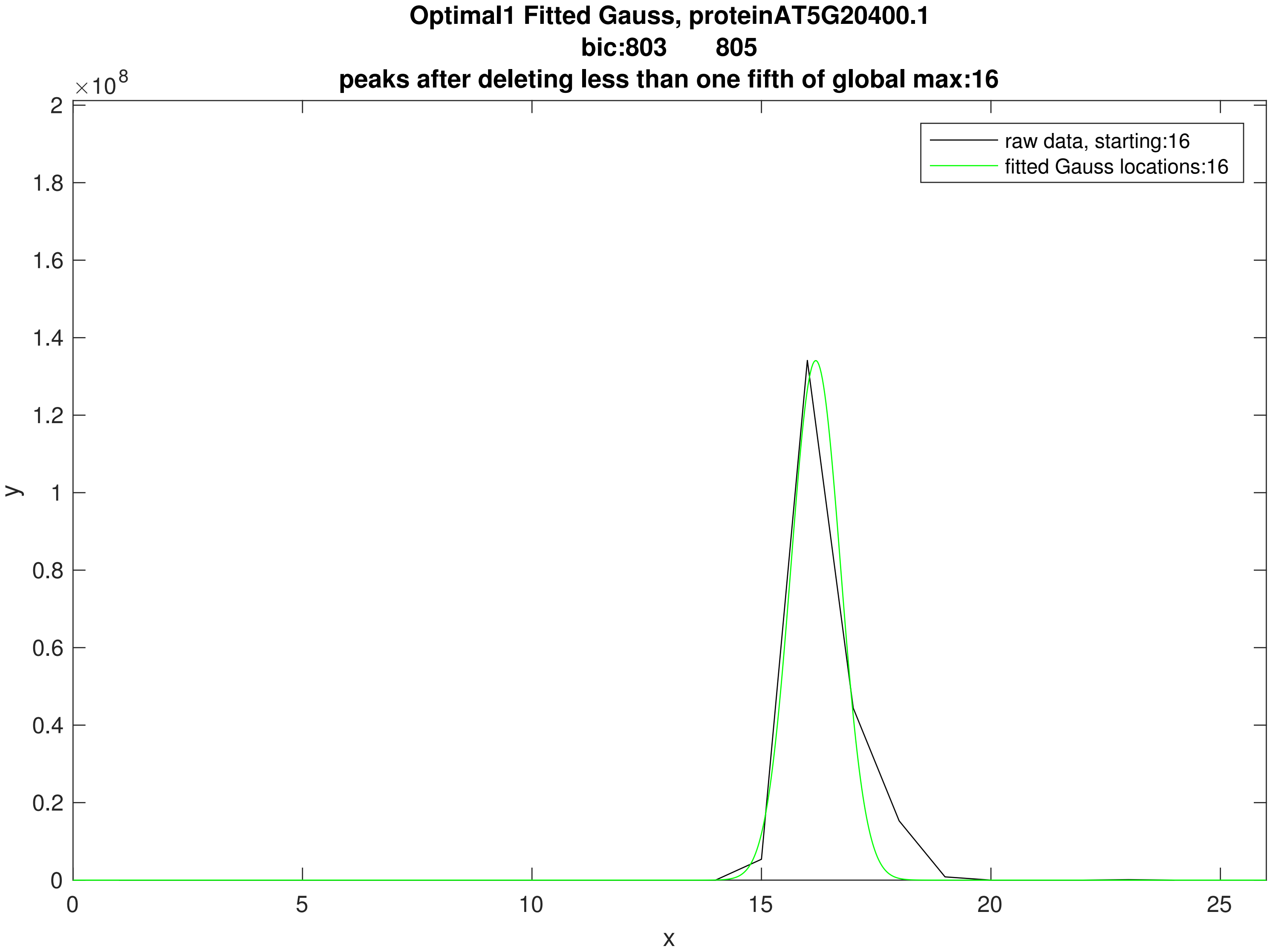
Optimal1 Fitted Gauss, proteinAT5G20400.1
bic:803 805
peaks after deleting less than one fifth of global max:16
×10^8
2
1.8
1.6
1.4
1.2
1
0.8
0.6
0.4
0.2
0
y
0
5
10
15
20
25
x
raw data, starting:16
fitted Gauss locations:16

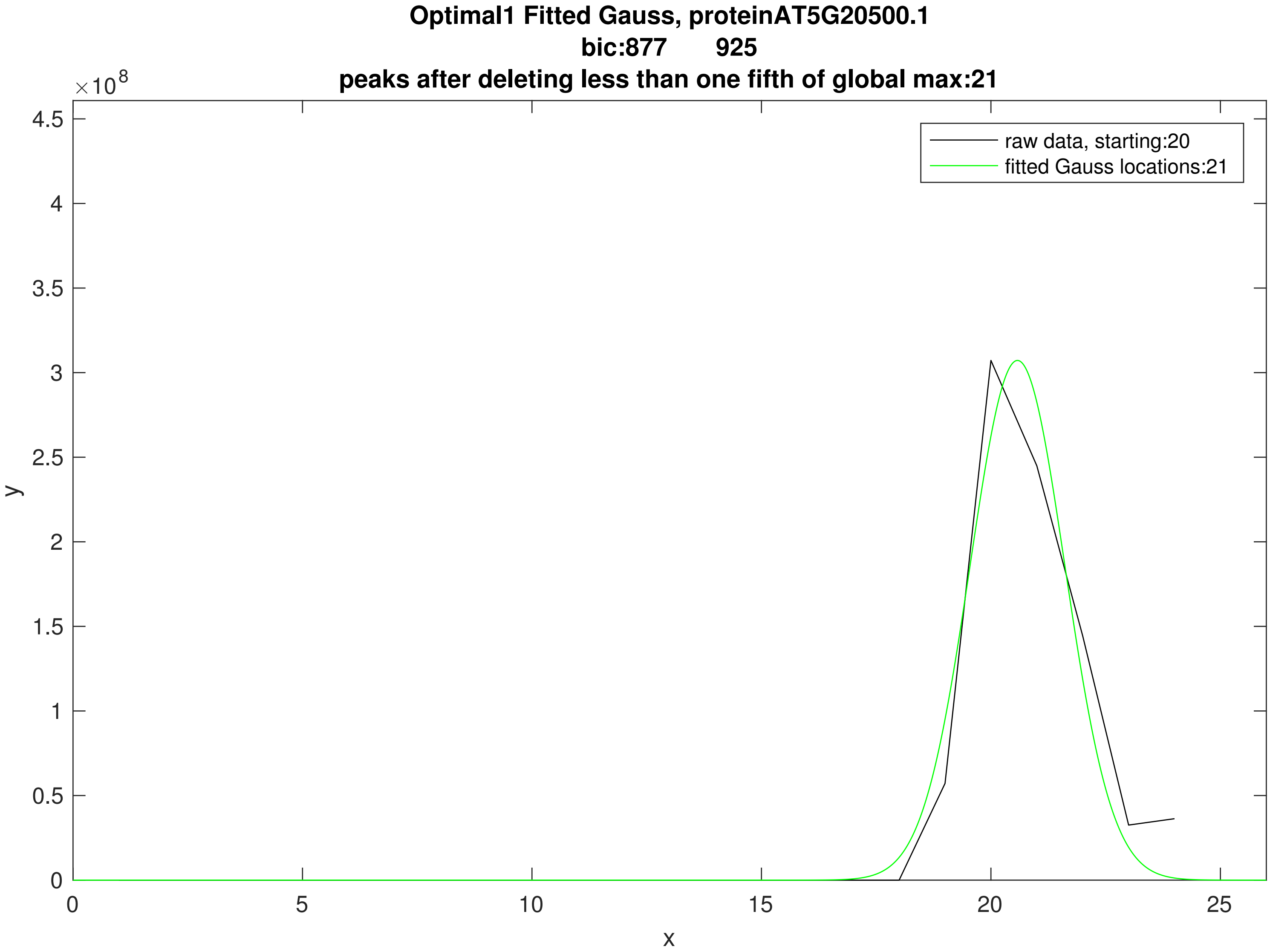
Optimal1 Fitted Gauss, proteinAT5G20500.1
bic:877 925
peaks after deleting less than one fifth of global max:21
×10^8
raw data, starting:20
fitted Gauss locations:21
y
x
0
0.5
1
1.5
2
2.5
3
3.5
4
4.5
0
5
10
15
20
25

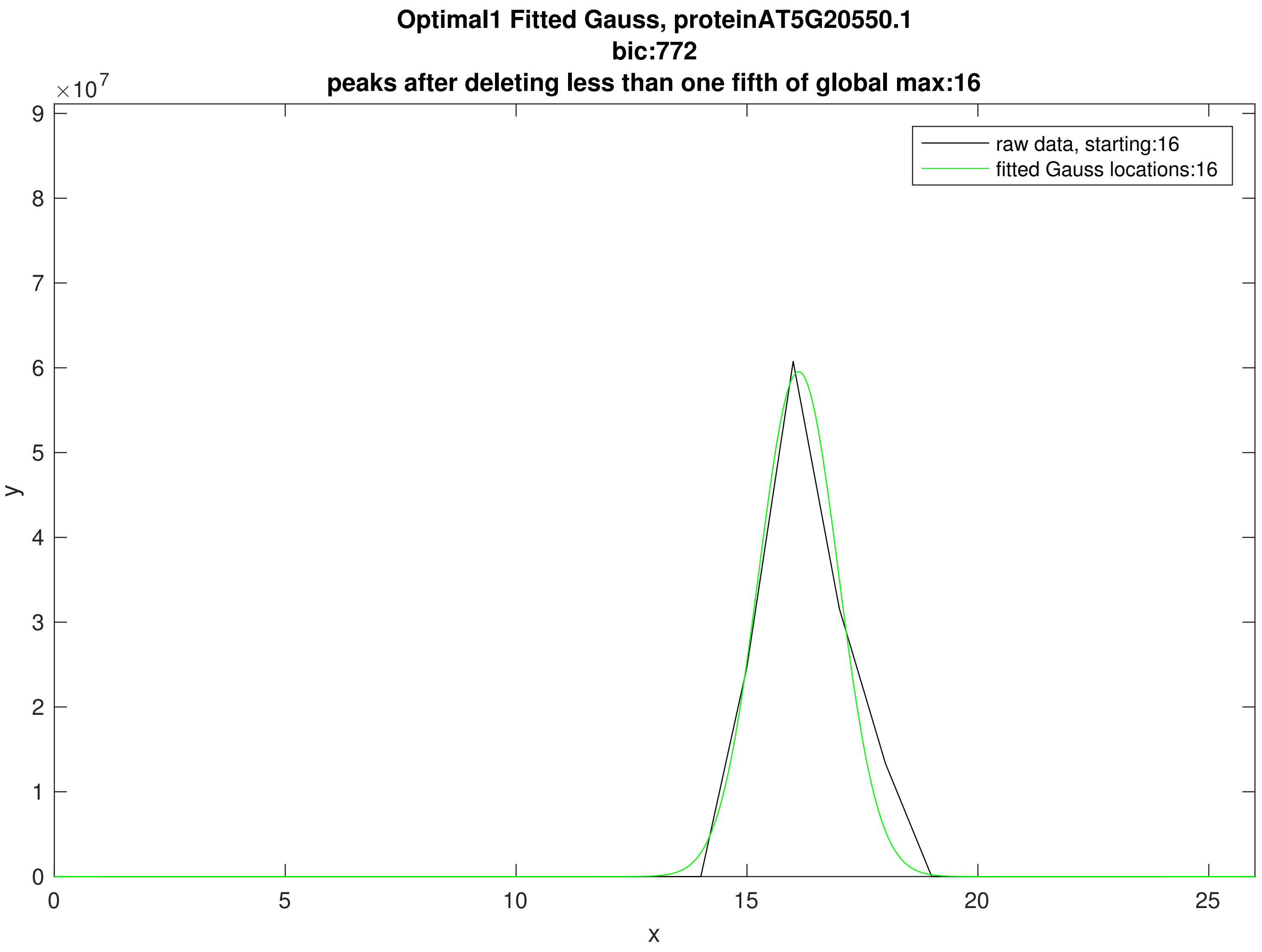

Optimal1 Fitted Gauss, proteinAT5G20550.1
bic:772
peaks after deleting less than one fifth of global max:16
×10^7
9
8
7
6
5
4
3
2
1
0
y
0
5
10
15
20
25
x
raw data, starting:16
fitted Gauss locations:16

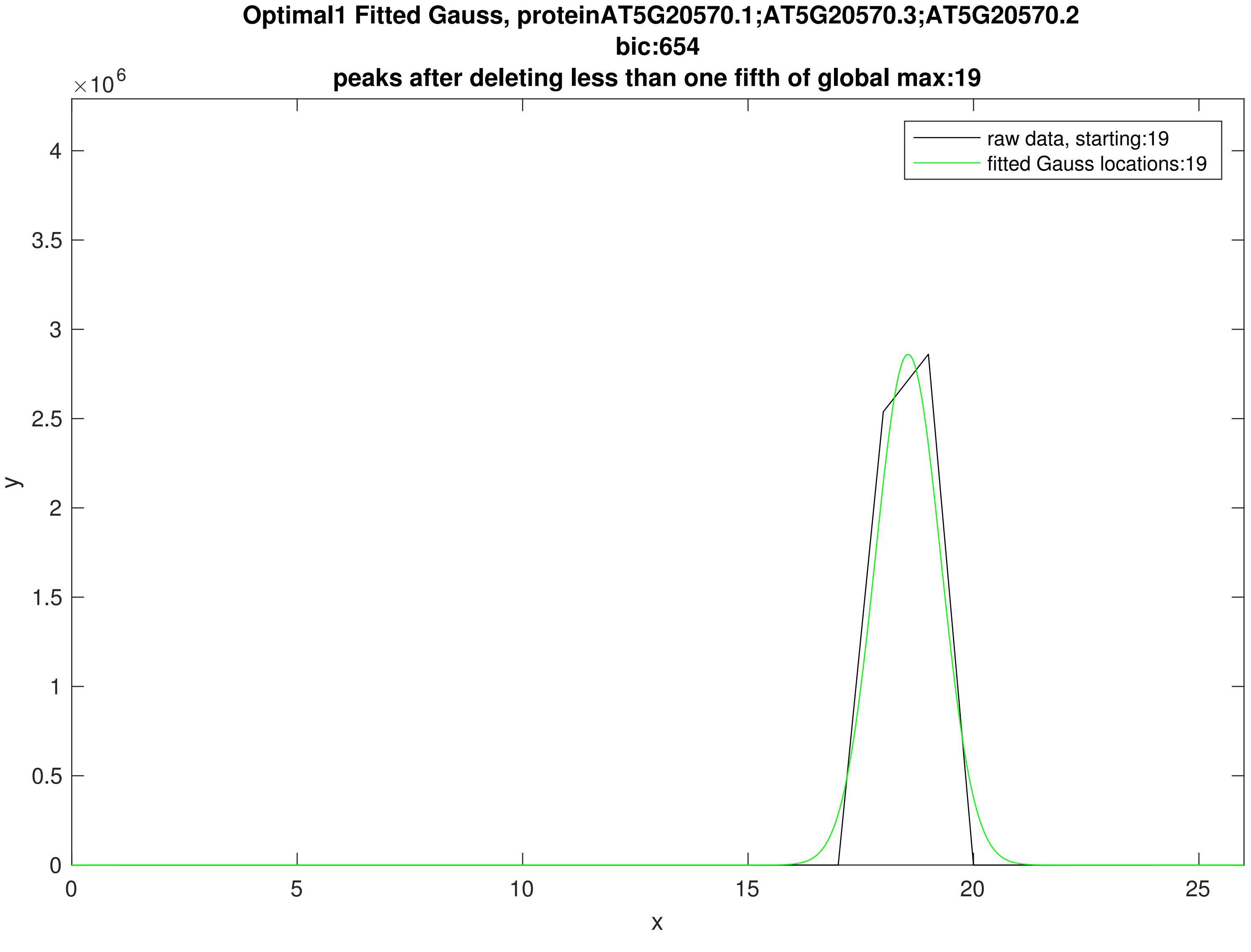
Optimal1 Fitted Gauss, proteinAT5G20570.1;AT5G20570.3;AT5G20570.2
bic:654
peaks after deleting less than one fifth of global max:19
×10^6
raw data, starting:19
fitted Gauss locations:19
y
x
0
0.5
1
1.5
2
2.5
3
3.5
4
5
10
15
20
25

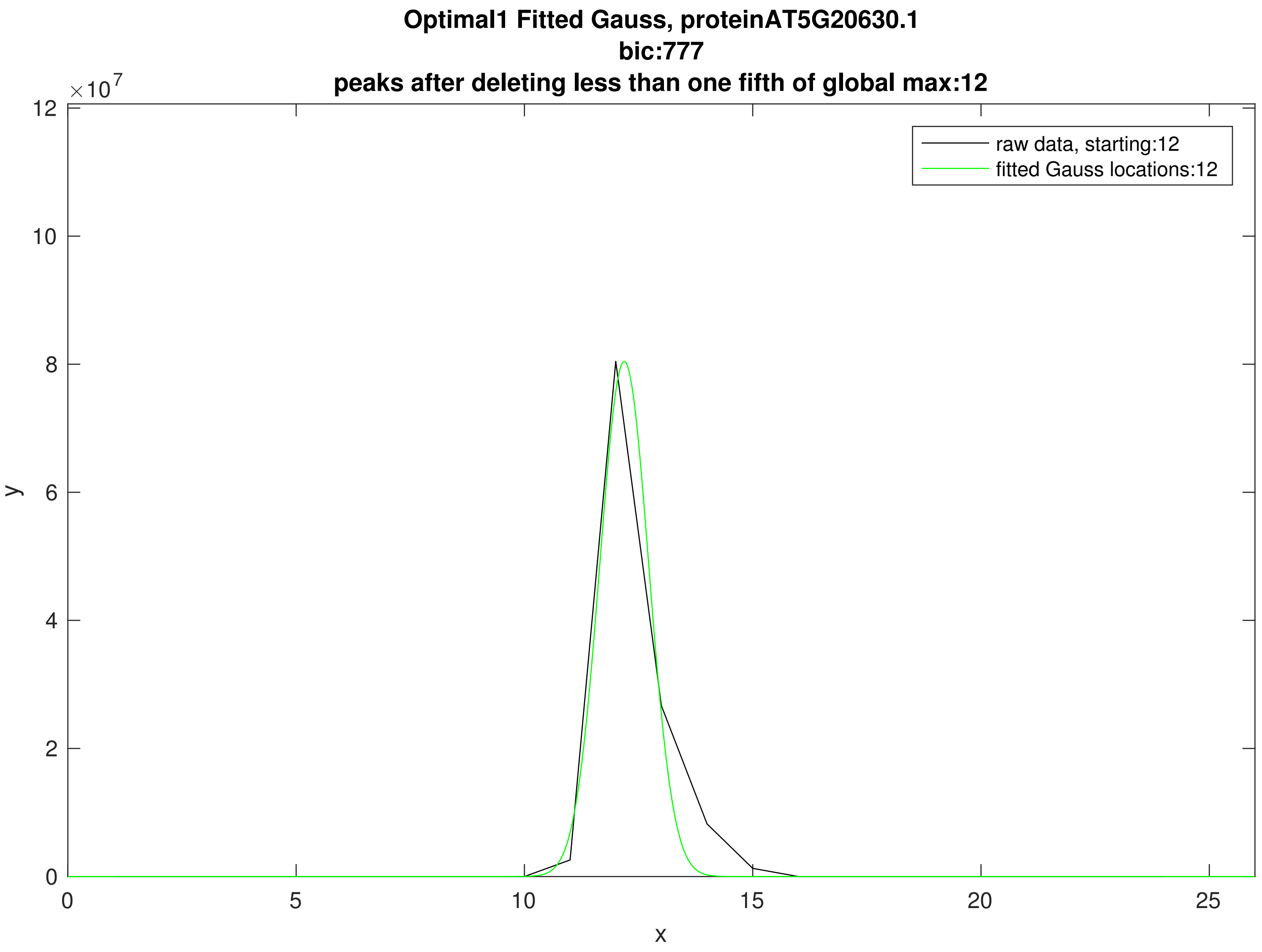
Optimal1 Fitted Gauss, proteinAT5G20630.1
bic:777
peaks after deleting less than one fifth of global max:12
×10^7
12
10
8
6
4
2
0
y
0
5
10
15
20
25
x
raw data, starting:12
fitted Gauss locations:12

#### Optimal1 Fitted Gauss, proteinAT5G20720.3;AT5G20720.2;AT5G20720.1
#### bic:986 989 1.01e+03 1.02e+03
#### peaks after deleting less than one fifth of global max:13

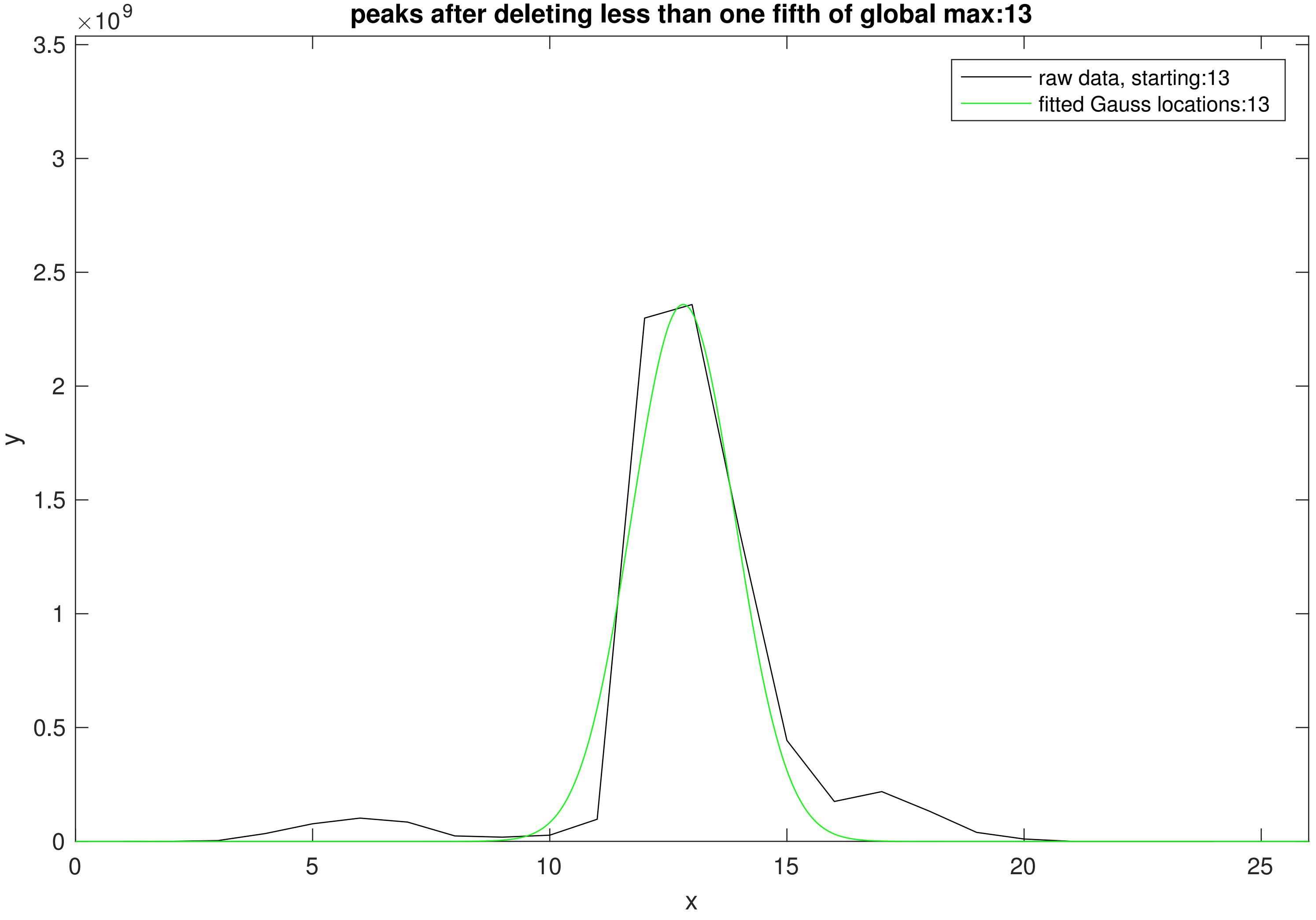

Optimal1 Fitted Gauss, proteinAT5G20890.1
bic:841 842 852
peaks after deleting less than one fifth of global max:4.5

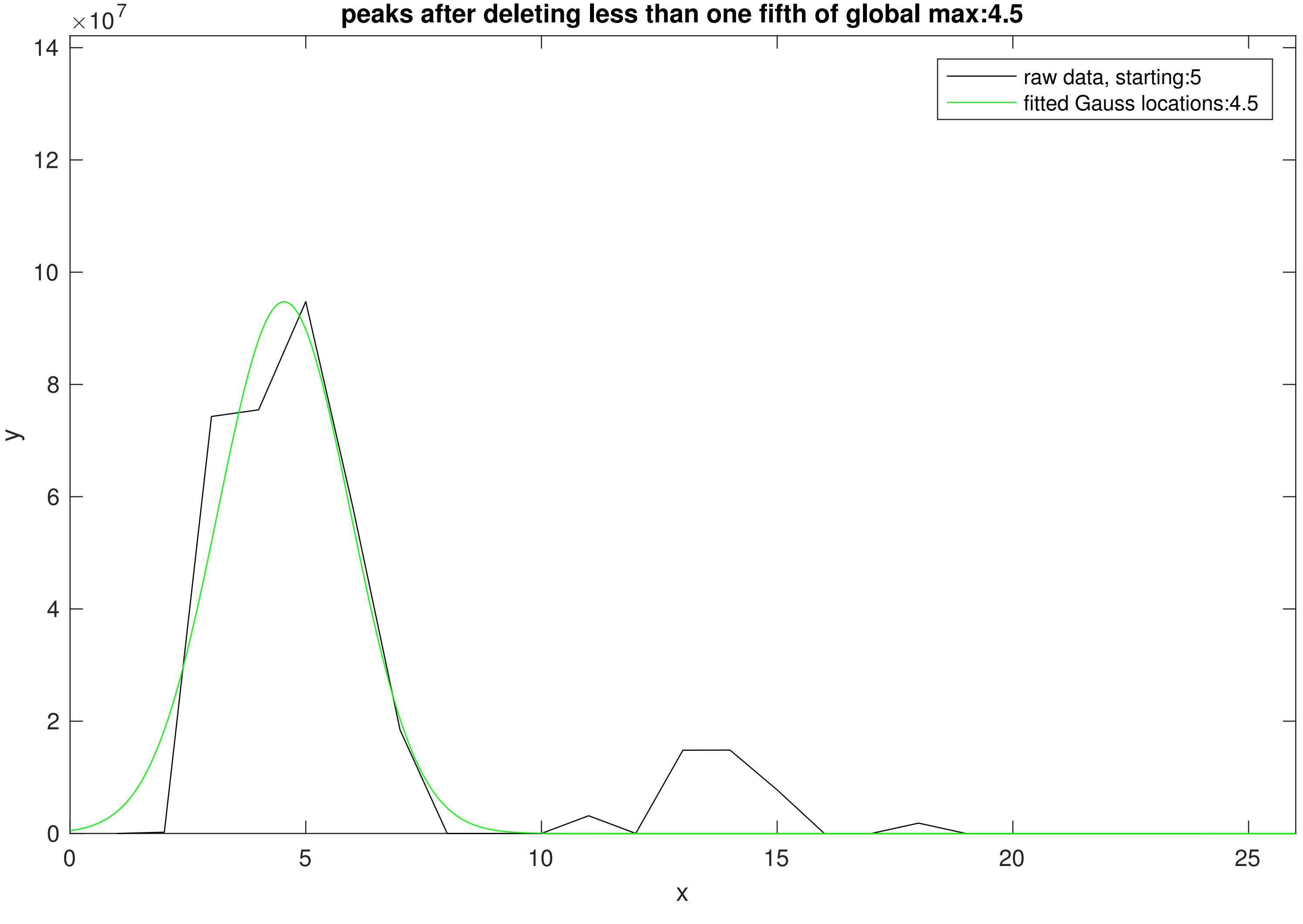

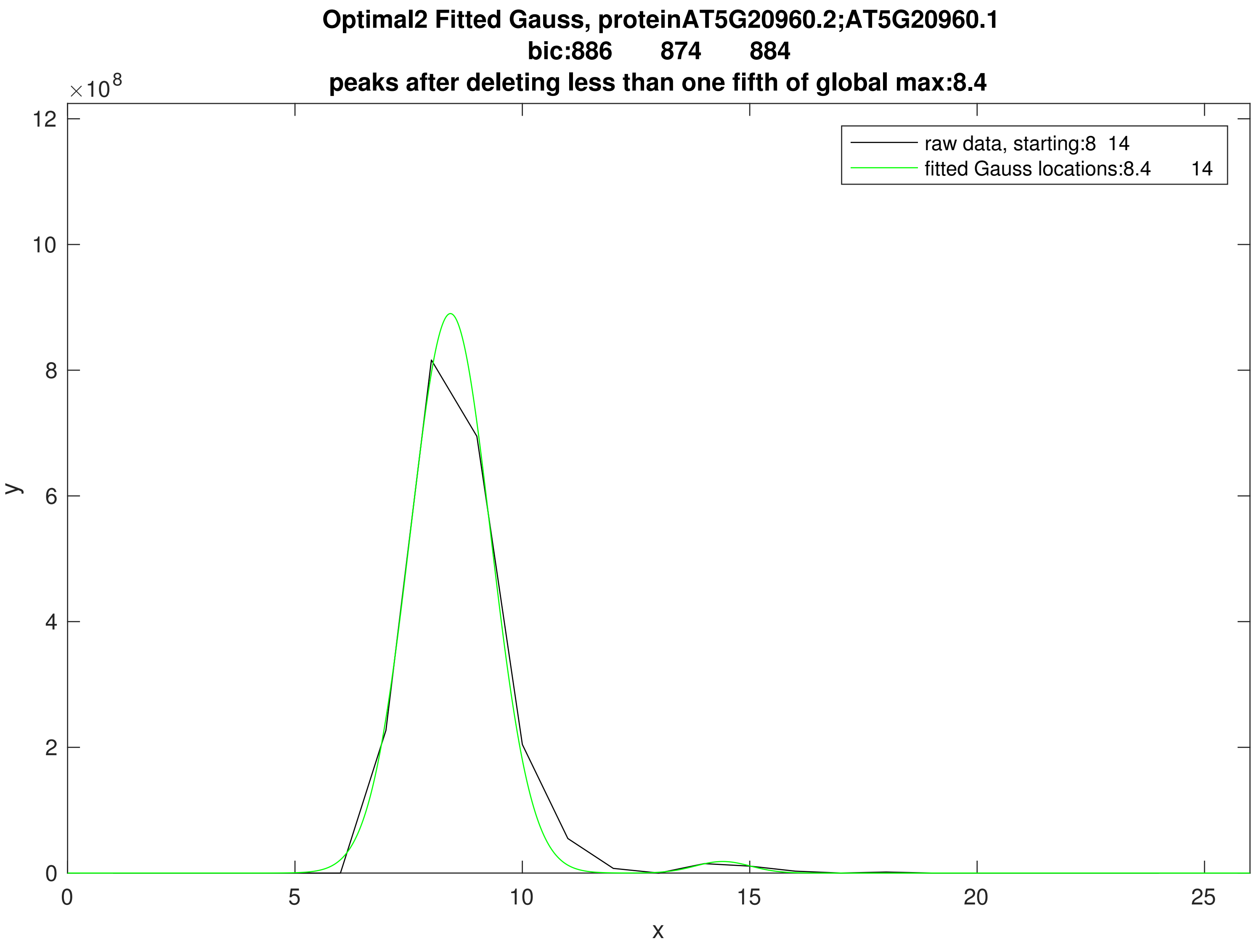

Optimal2 Fitted Gauss, proteinAT5G20960.2;AT5G20960.1
bic:886 874 884
peaks after deleting less than one fifth of global max:8.4
×10^8
12
10
8
6
4
2
0
y
raw data, starting:8 14
fitted Gauss locations:8.4 14
0
5
10
15
20
25
x

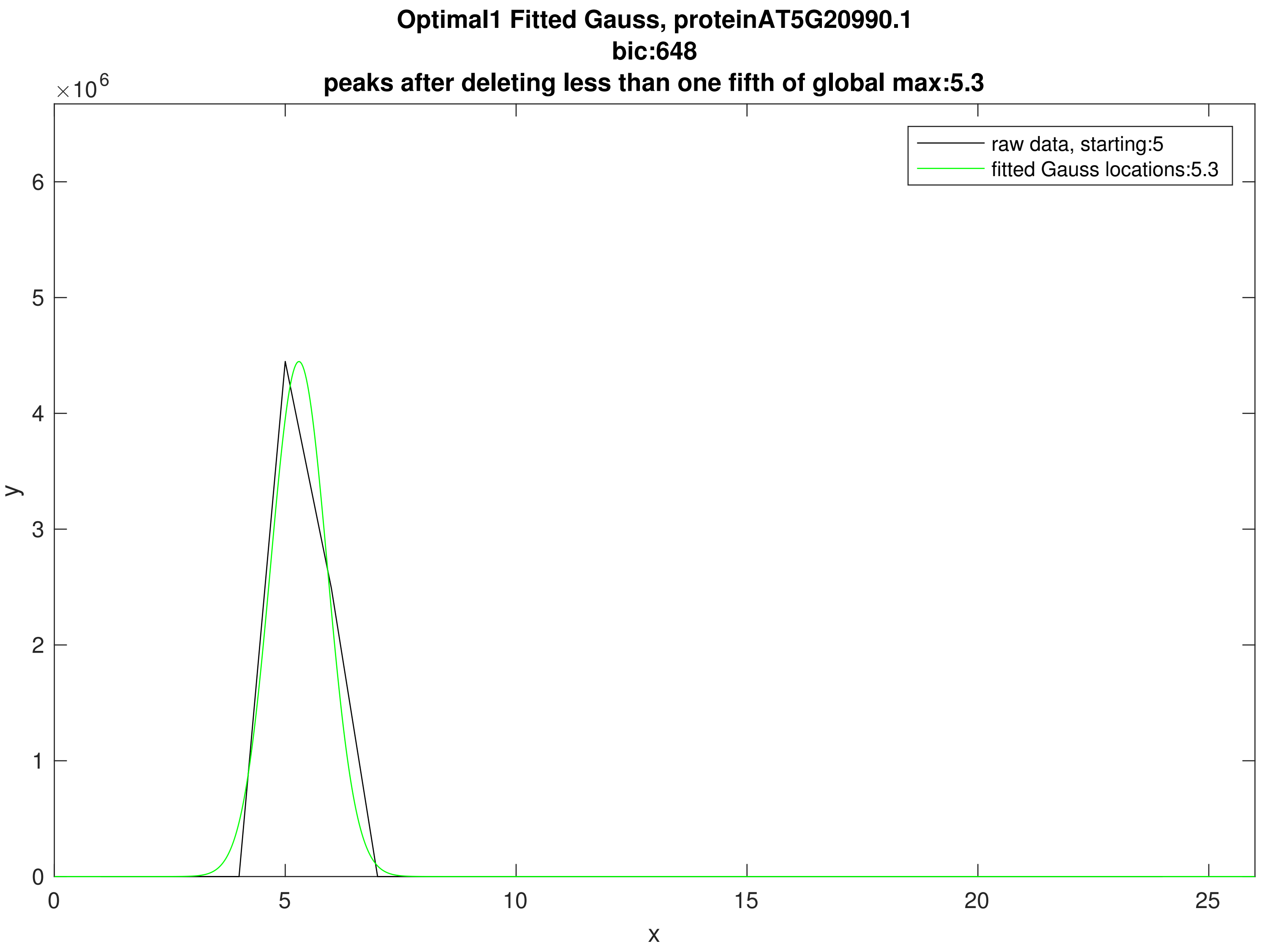

Optimal1 Fitted Gauss, proteinAT5G20990.1
bic:648
peaks after deleting less than one fifth of global max:5.3
×10^6
raw data, starting:5
fitted Gauss locations:5.3
y
x
0
1
2
3
4
5
6
0
5
10
15
20
25

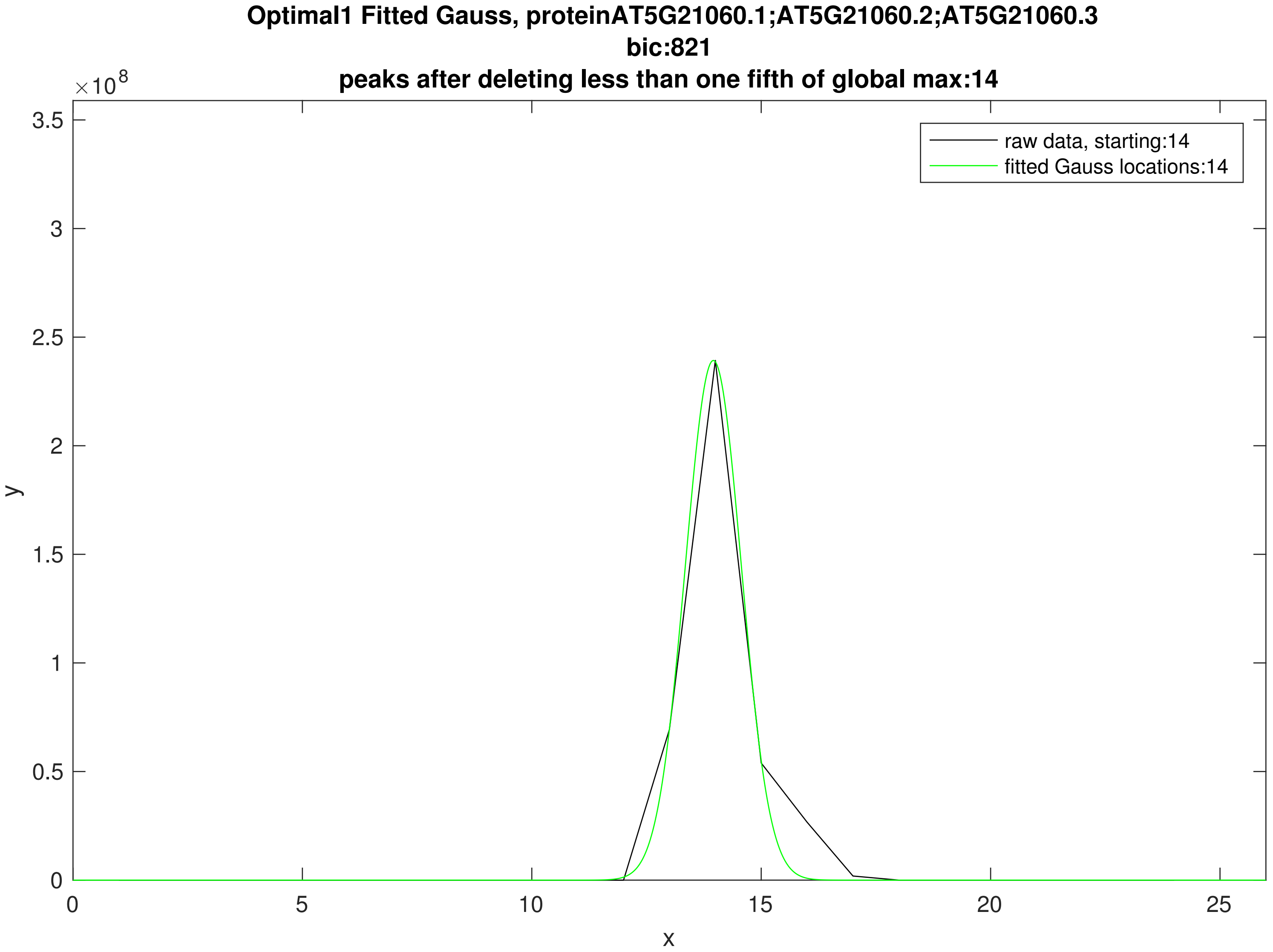

Optimal1 Fitted Gauss, proteinAT5G21060.1;AT5G21060.2;AT5G21060.3
bic:821
peaks after deleting less than one fifth of global max:14
×10^8
3.5
3
2.5
2
1.5
1
0.5
0
y
0
5
10
15
20
25
x
raw data, starting:14
fitted Gauss locations:14

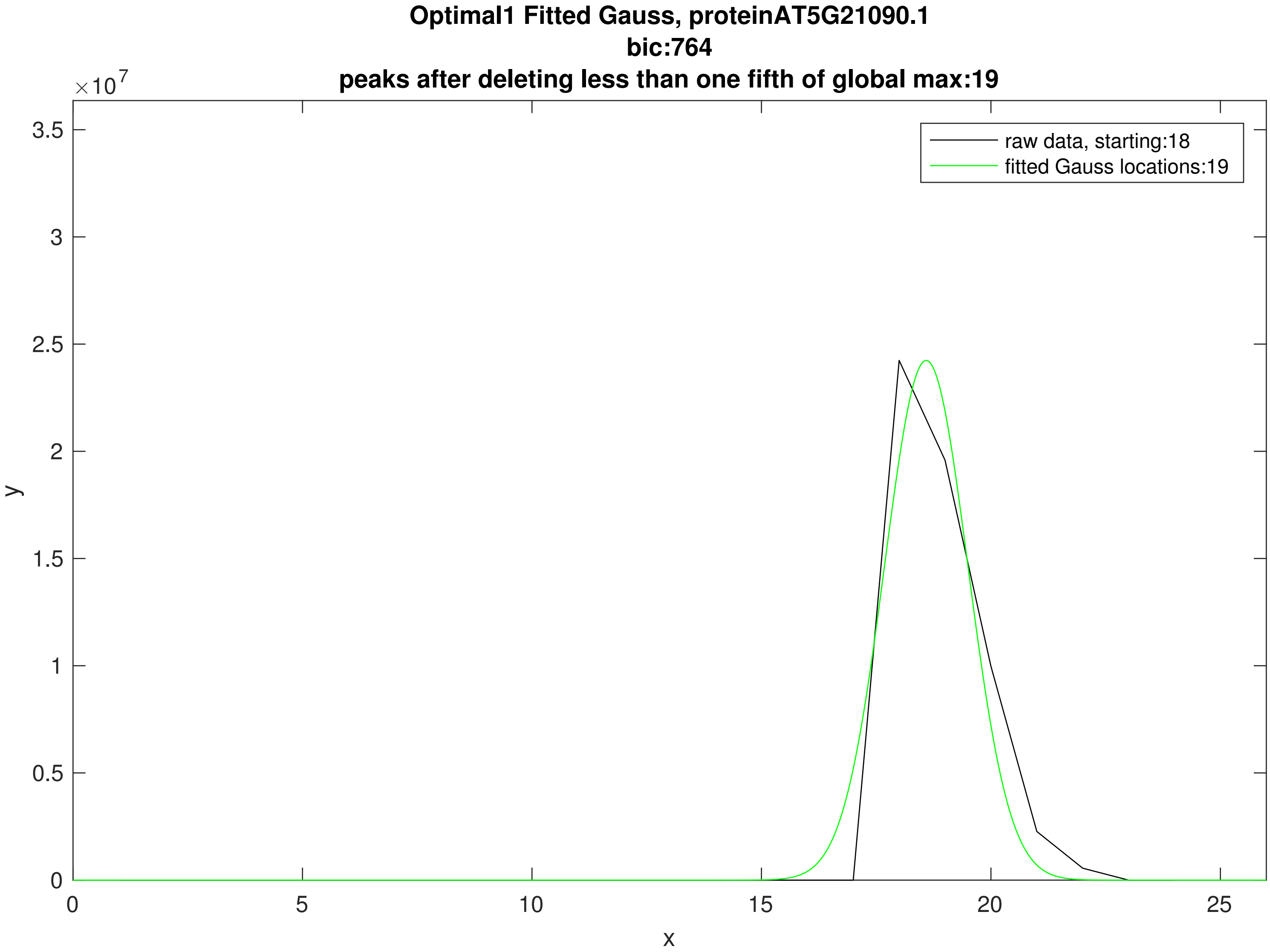
Optimal1 Fitted Gauss, proteinAT5G21090.1
bic:764
peaks after deleting less than one fifth of global max:19
×10^7
3.5
3
2.5
2
1.5
1
0.5
0
y
0
5
10
15
20
25
x
raw data, starting:18
fitted Gauss locations:19

Optimal1 Fitted Gauss, proteinAT5G22470.1
bic:726
peaks after deleting less than one fifth of global max:12

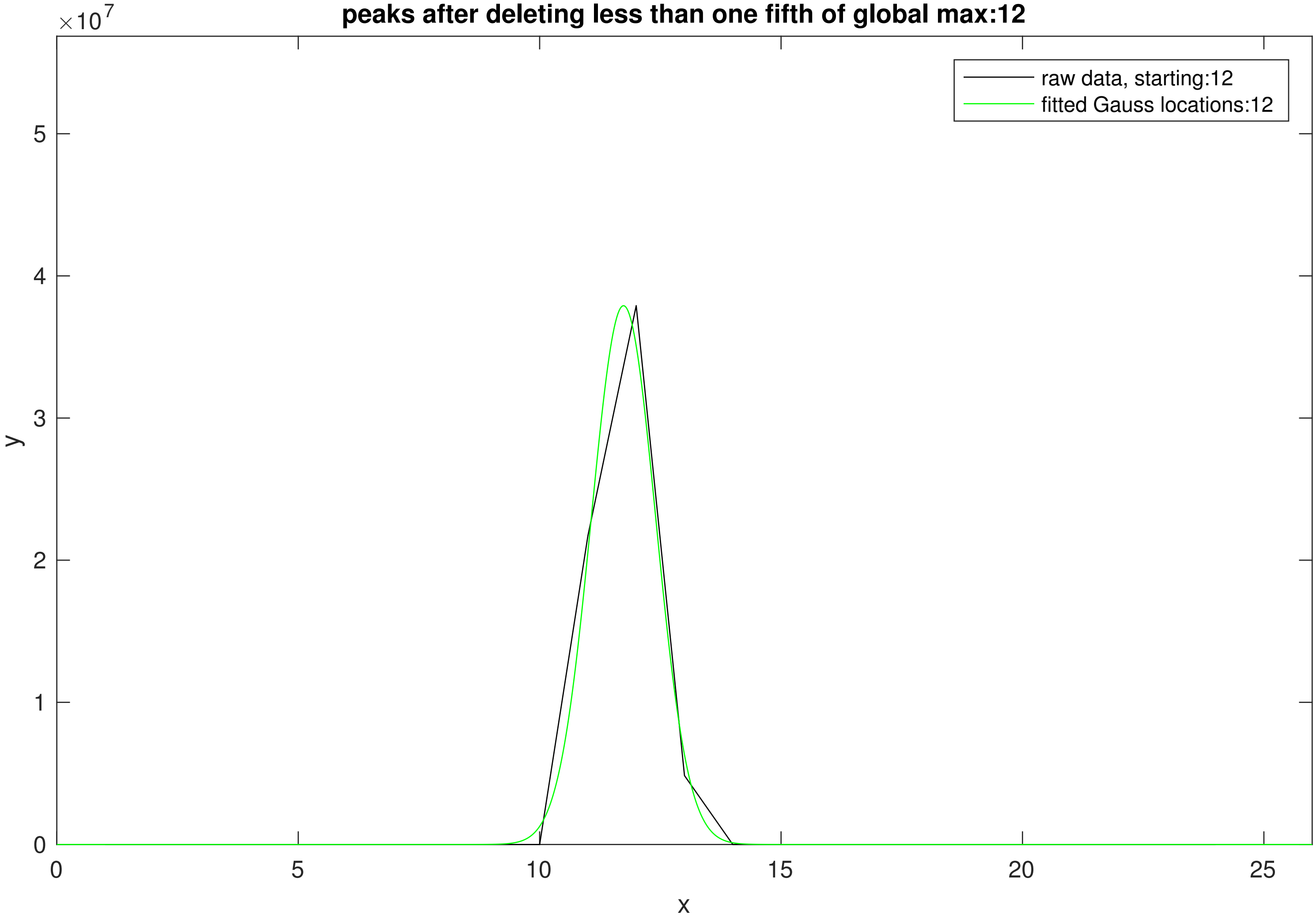

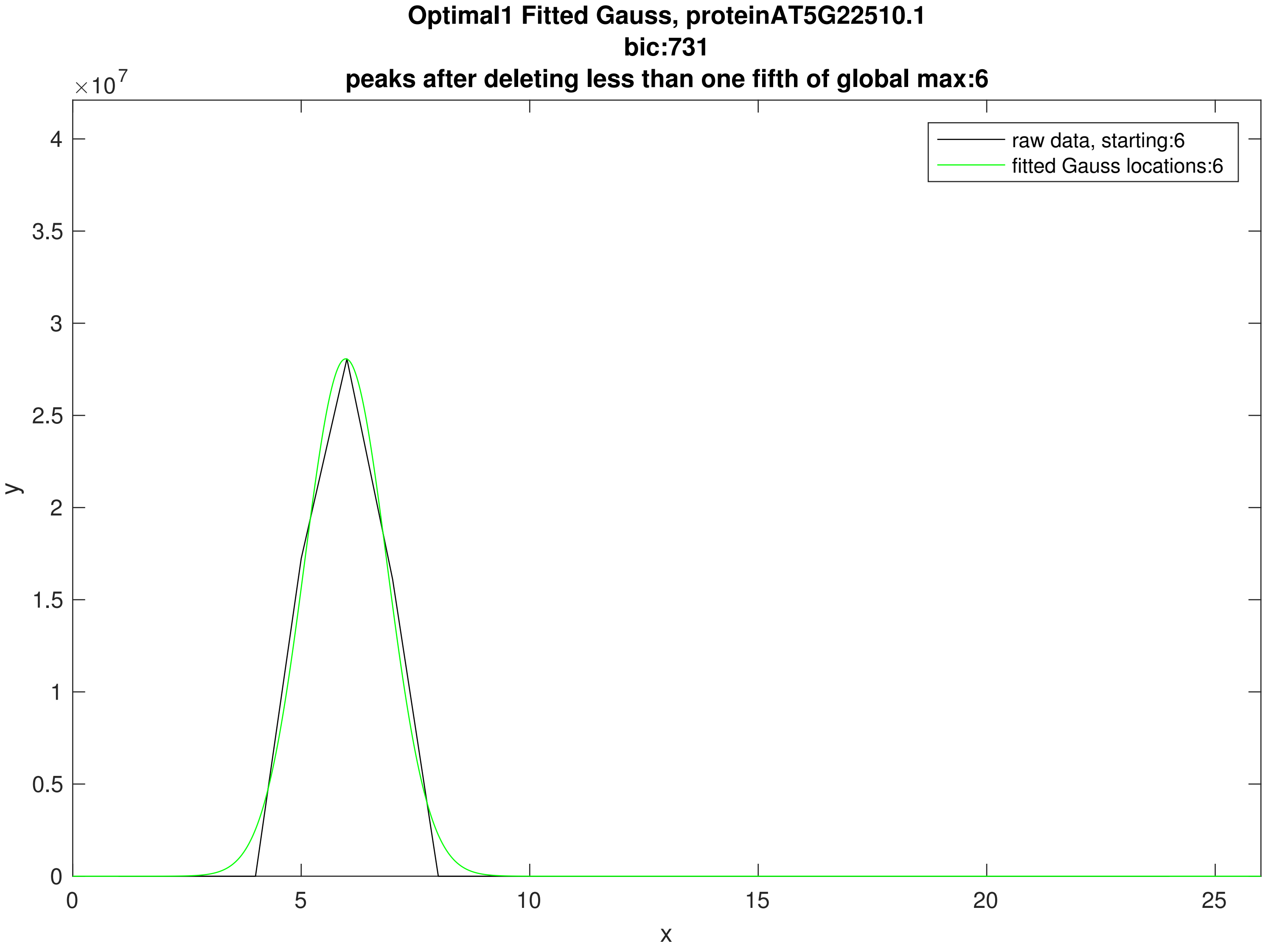

Optimal1 Fitted Gauss, proteinAT5G22510.1
bic:731
peaks after deleting less than one fifth of global max:6
×10^7
raw data, starting:6
fitted Gauss locations:6
y
x
0
0.5
1
1.5
2
2.5
3
3.5
4
5
10
15
20
25

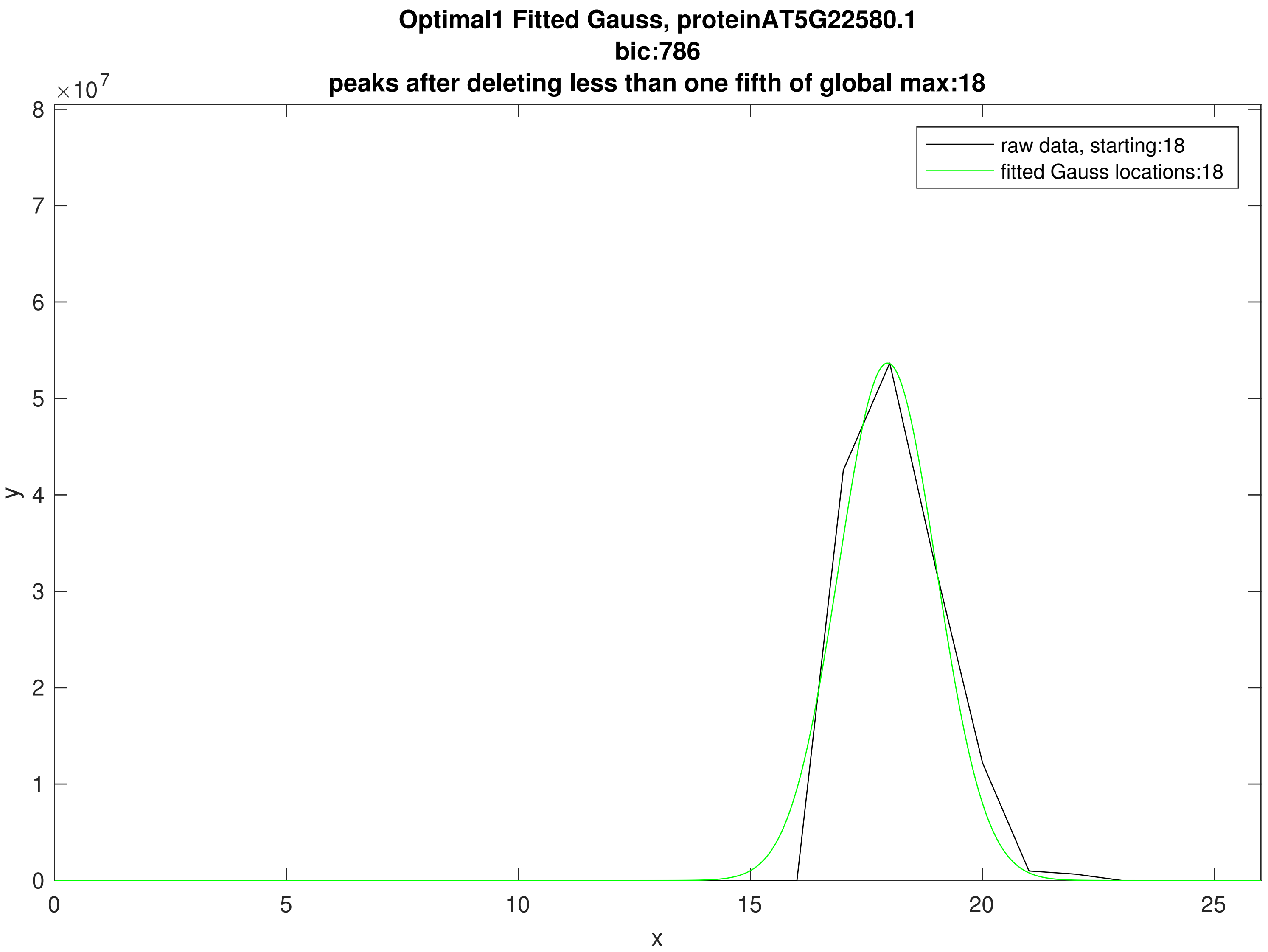

Optimal1 Fitted Gauss, proteinAT5G22580.1
bic:786
peaks after deleting less than one fifth of global max:18
×10^7
8
7
6
5
4
3
2
1
0
y
0
5
10
15
20
25
x
raw data, starting:18
fitted Gauss locations:18

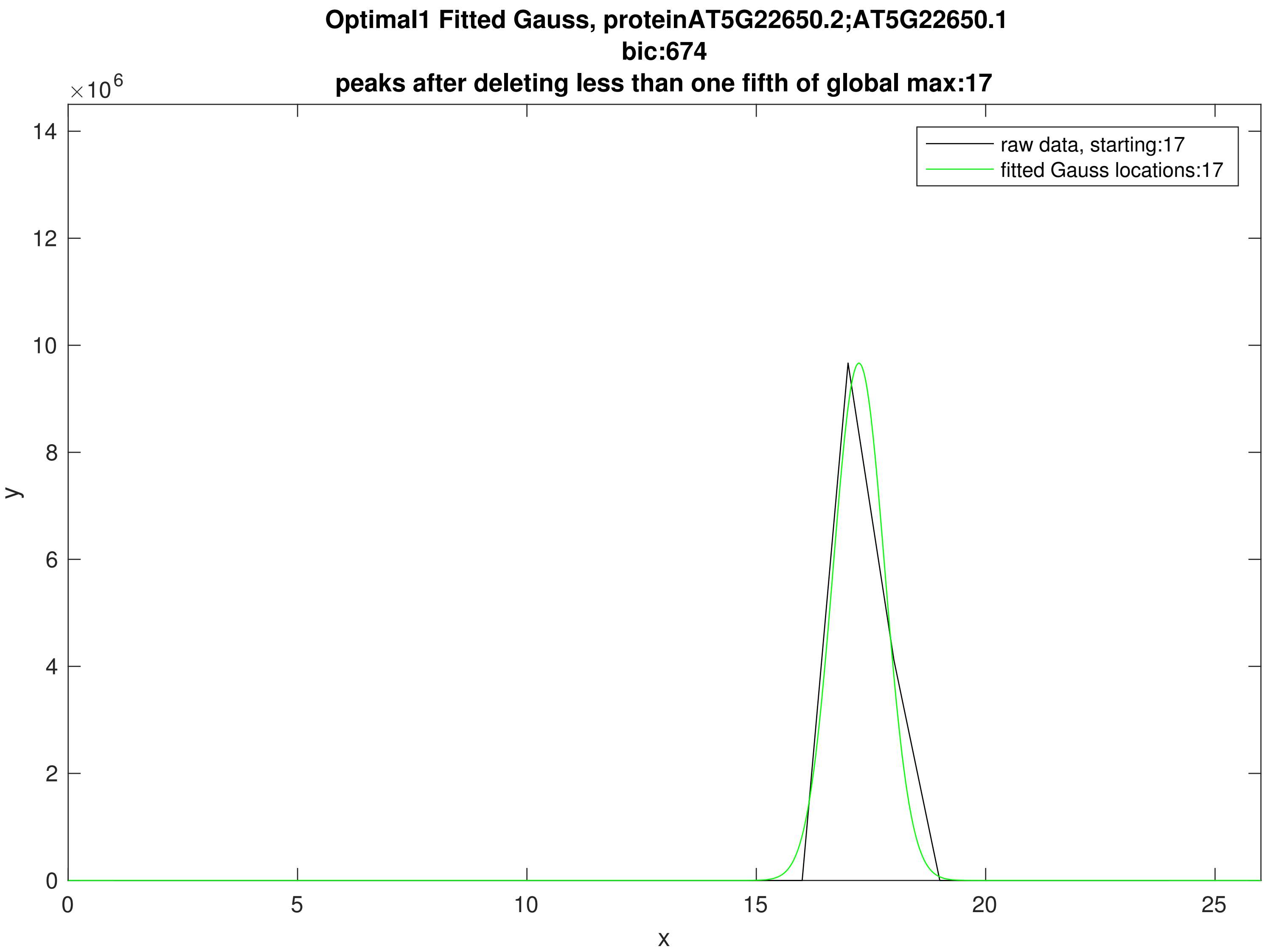
Optimal1 Fitted Gauss, proteinAT5G22650.2;AT5G22650.1
bic:674
peaks after deleting less than one fifth of global max:17
×10^6
14
12
10
8
6
4
2
0
y
0
5
10
15
20
25
x
raw data, starting:17
fitted Gauss locations:17

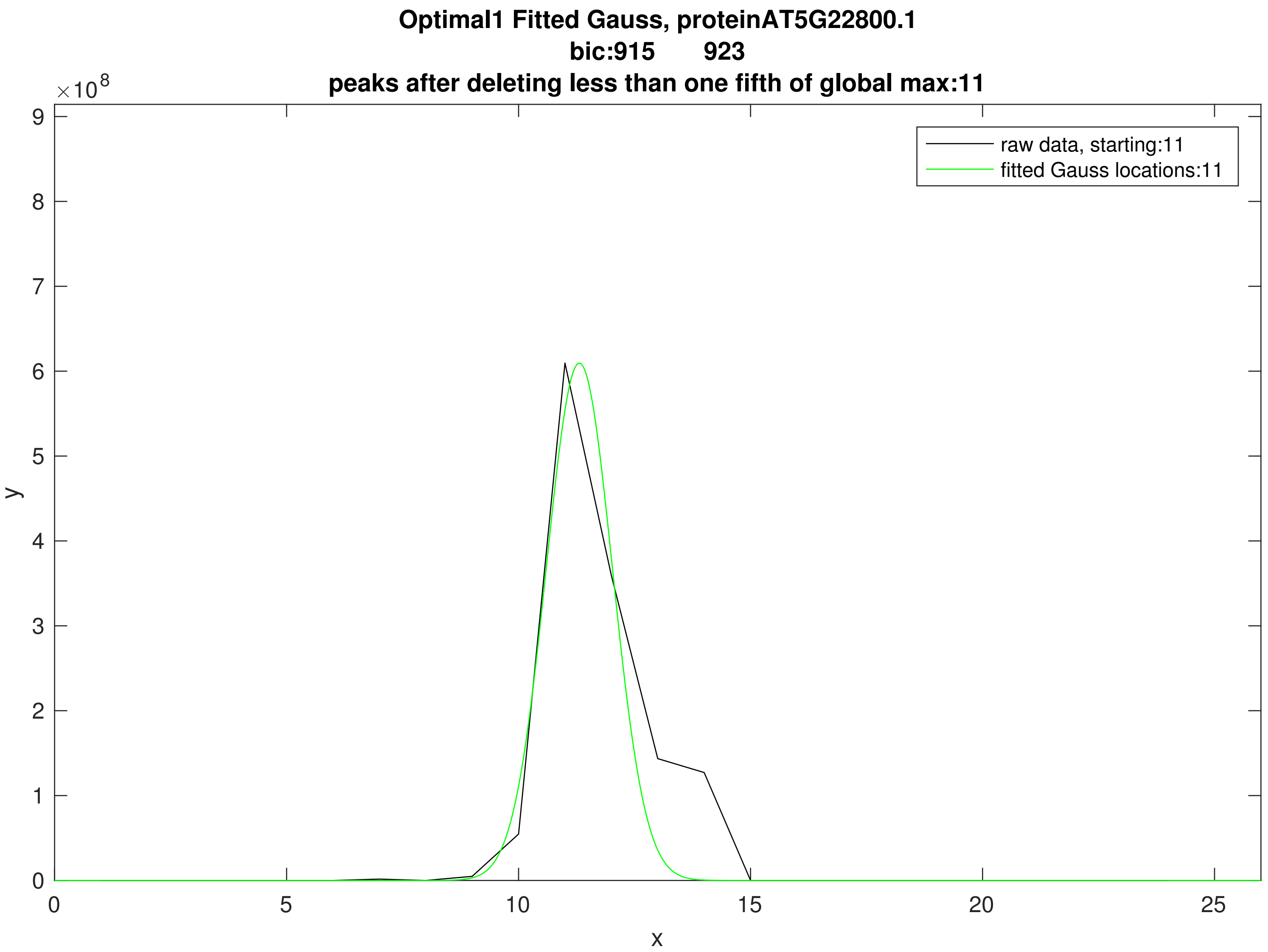
Optimal1 Fitted Gauss, proteinAT5G22800.1
bic:915 923
peaks after deleting less than one fifth of global max:11
raw data, starting:11
fitted Gauss locations:11
$\times 10^8$
y
x
0
1
2
3
4
5
6
7
8
9
0
5
10
15
20
25

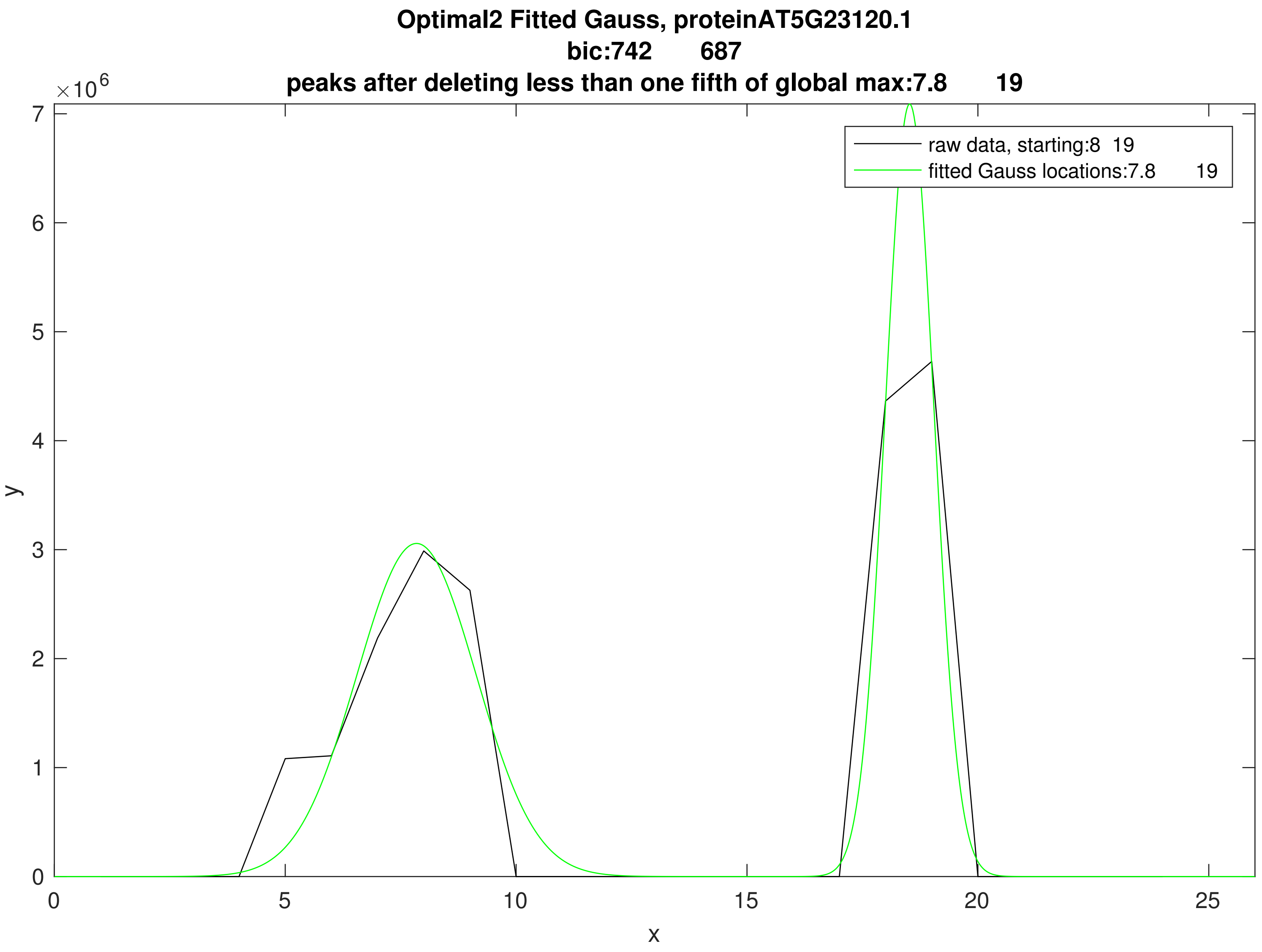
Optimal2 Fitted Gauss, proteinAT5G23120.1
bic:742 687
peaks after deleting less than one fifth of global max:7.8 19
×10^6
raw data, starting:8 19
fitted Gauss locations:7.8 19
y
x
0
1
2
3
4
5
6
7
0
5
10
15
20
25

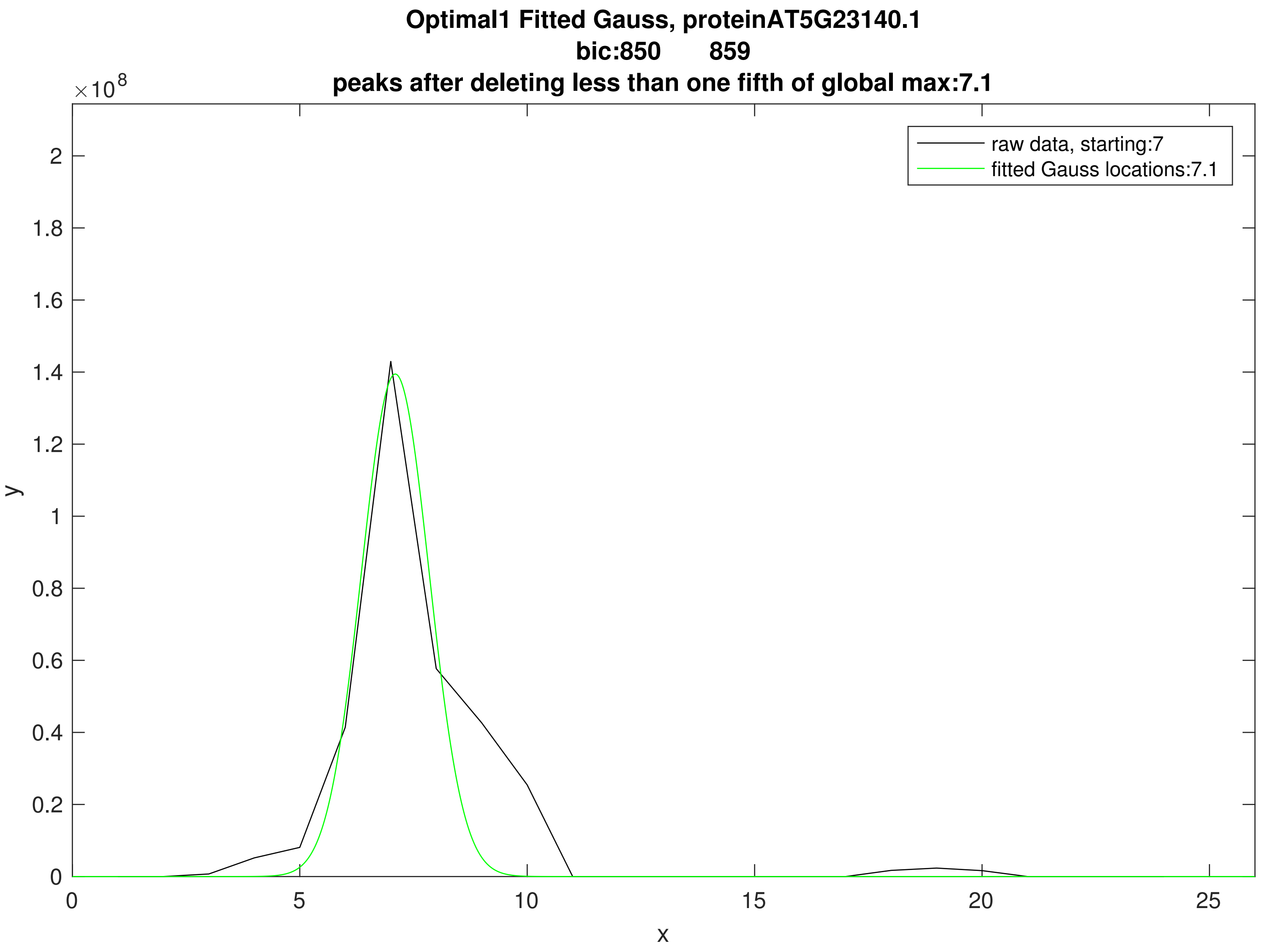

Optimal1 Fitted Gauss, proteinAT5G23140.1
bic:850 859
peaks after deleting less than one fifth of global max:7.1
×10^8
raw data, starting:7
fitted Gauss locations:7.1
y
x
0
0.2
0.4
0.6
0.8
1
1.2
1.4
1.6
1.8
2
0
5
10
15
20
25

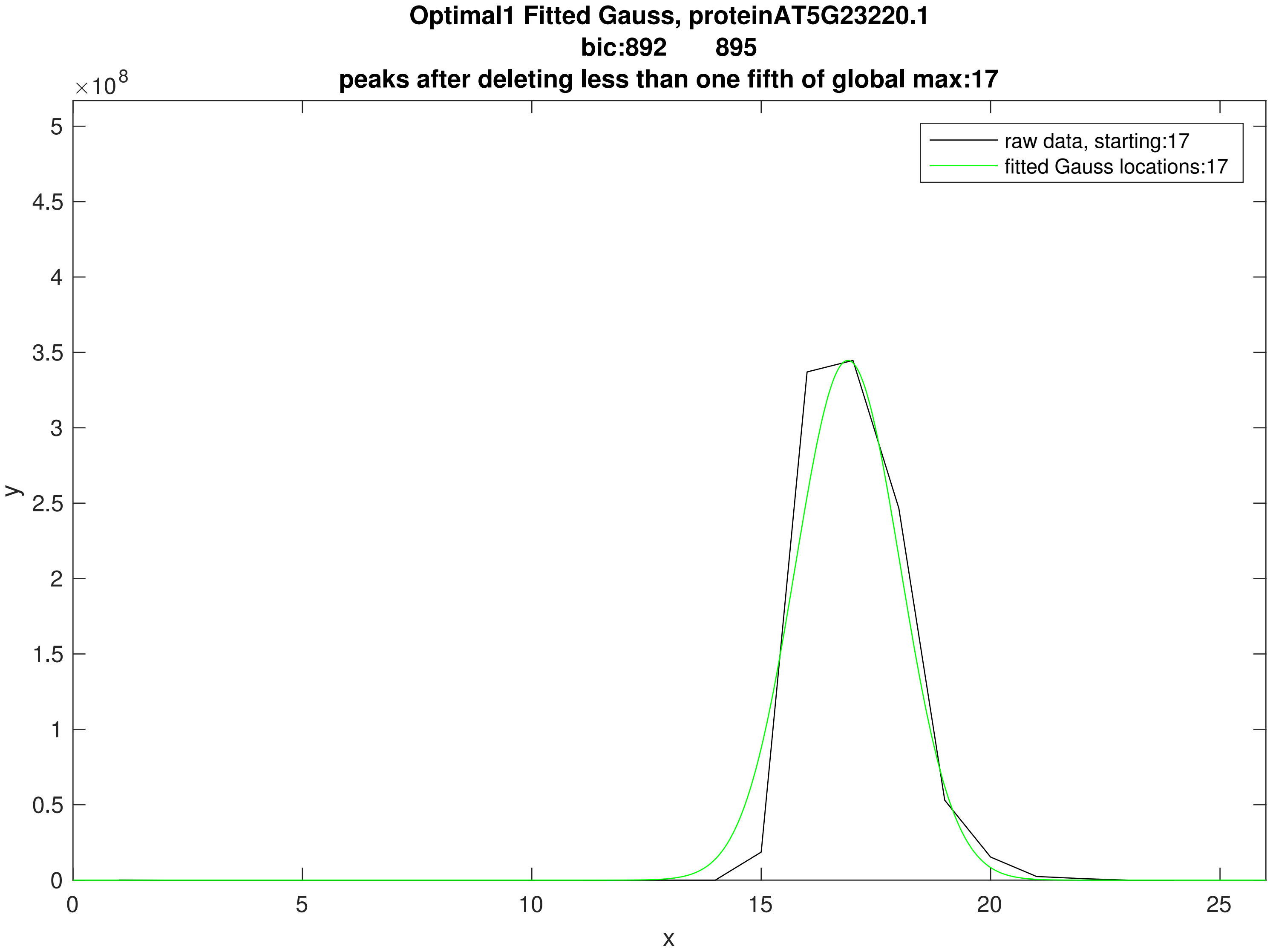
Optimal1 Fitted Gauss, proteinAT5G23220.1
bic:892 895
peaks after deleting less than one fifth of global max:17
×10^8
raw data, starting:17
fitted Gauss locations:17
y
x
0
0.5
1
1.5
2
2.5
3
3.5
4
4.5
5
0
5
10
15
20
25

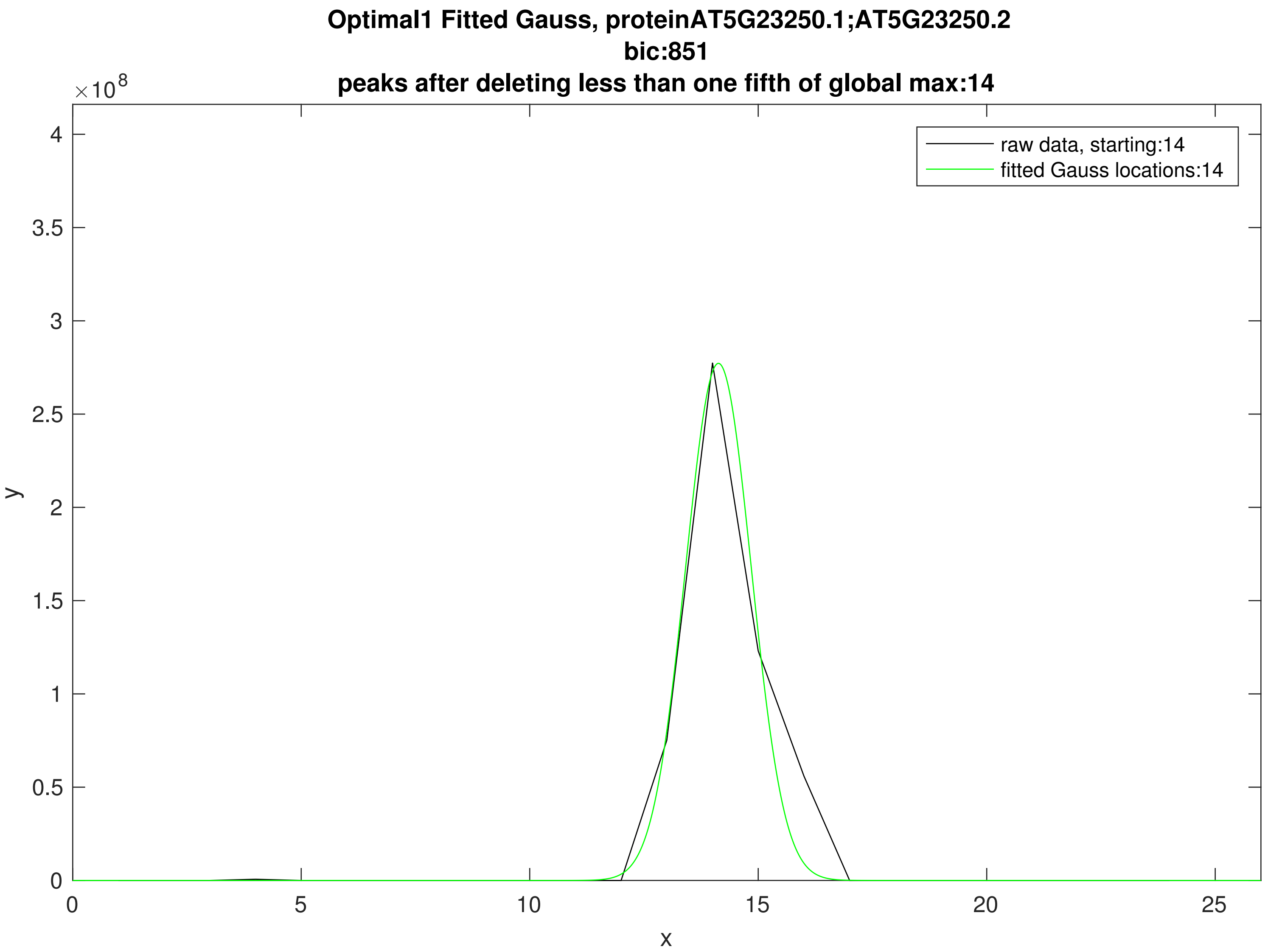
Optimal1 Fitted Gauss, proteinAT5G23250.1;AT5G23250.2
bic:851
peaks after deleting less than one fifth of global max:14
×10^8
raw data, starting:14
fitted Gauss locations:14
y
x
0
0.5
1
1.5
2
2.5
3
3.5
4
5
10
15
20
25

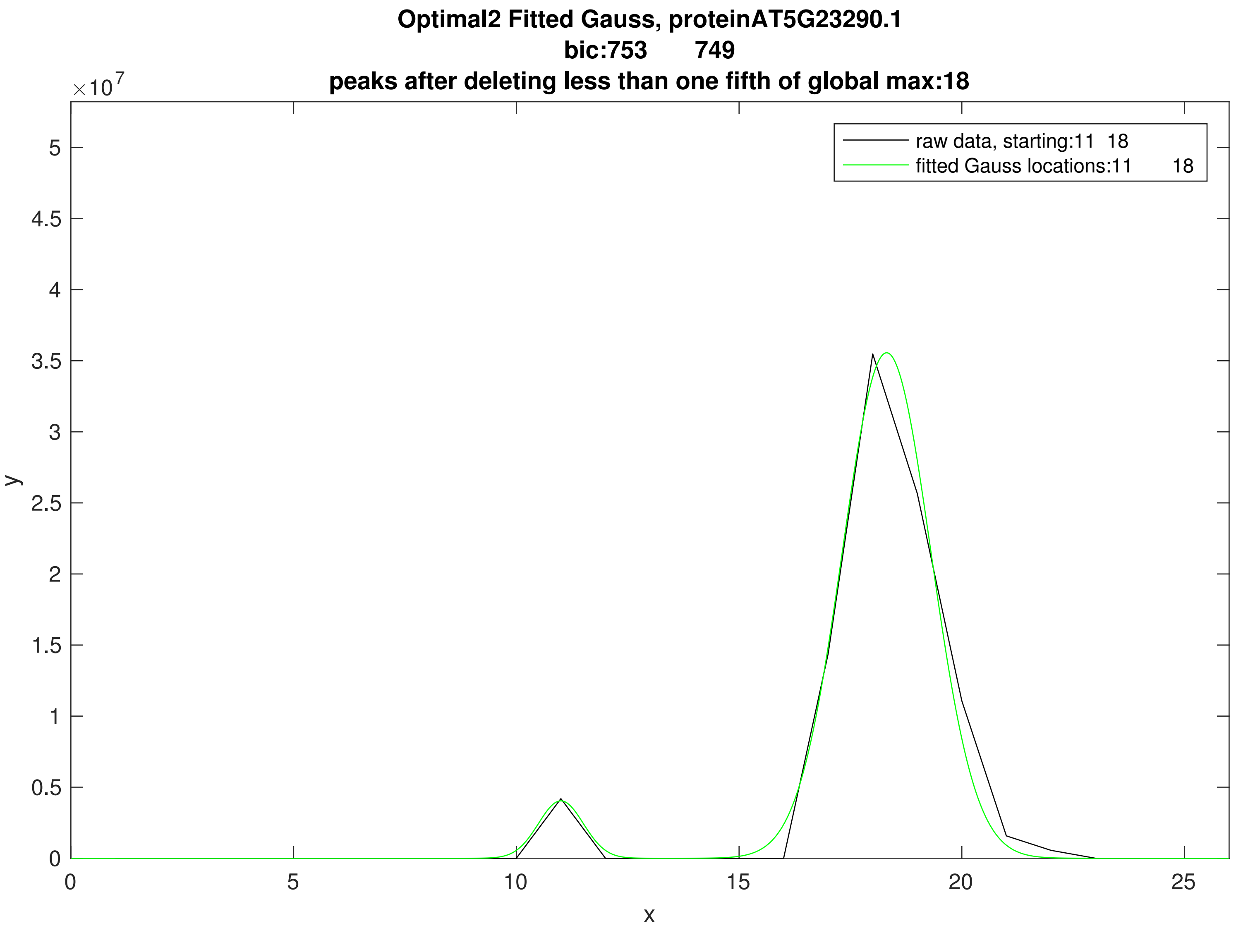
Optimal2 Fitted Gauss, proteinAT5G23290.1
bic:753 749
peaks after deleting less than one fifth of global max:18
×10^7
raw data, starting:11 18
fitted Gauss locations:11 18
y
x
0
0.5
1
1.5
2
2.5
3
3.5
4
4.5
5
0
5
10
15
20
25

Optimal1 Fitted Gauss, proteinAT5G23540.2;AT5G23540.1
bic:810
peaks after deleting less than one fifth of global max:3.8

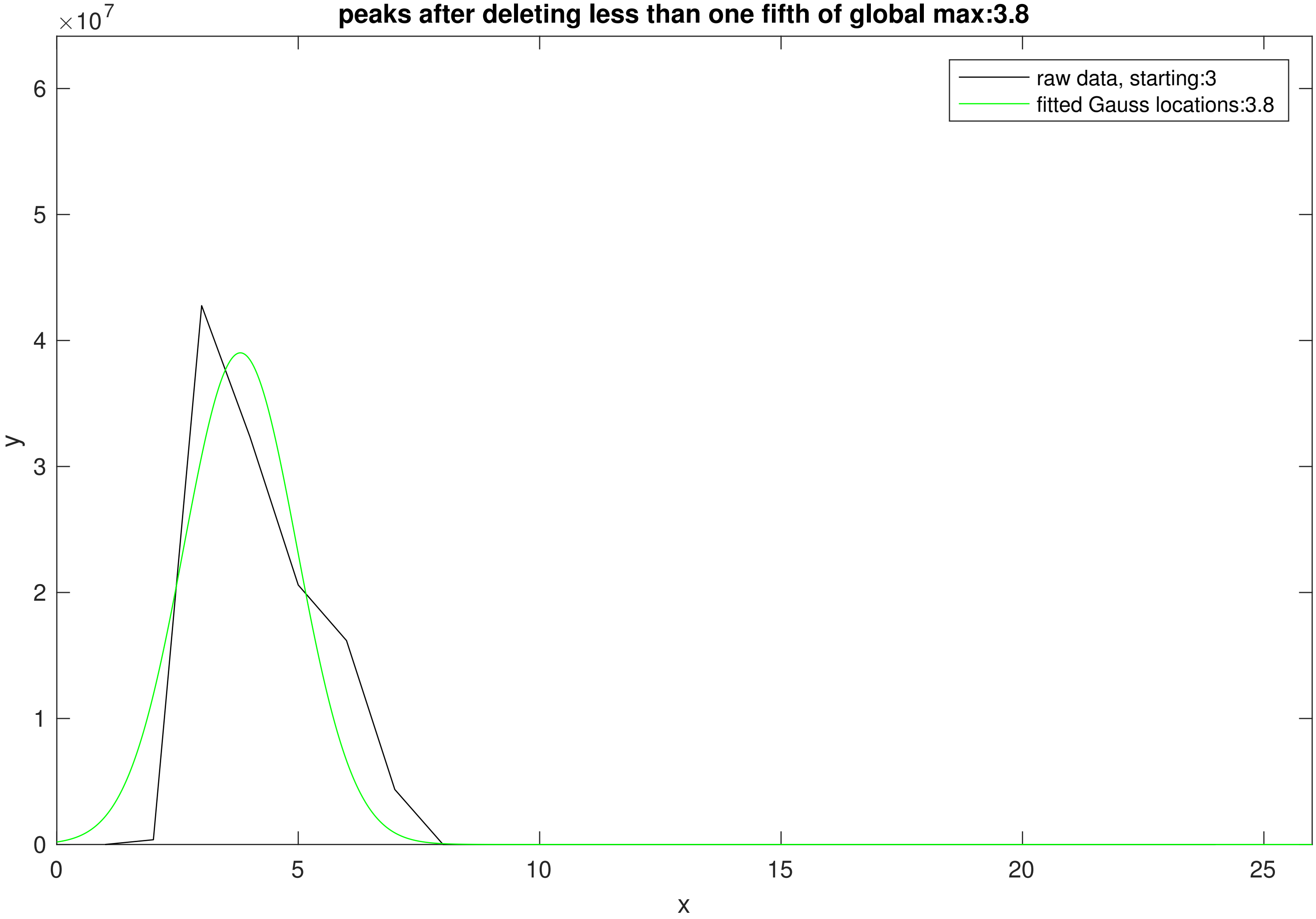

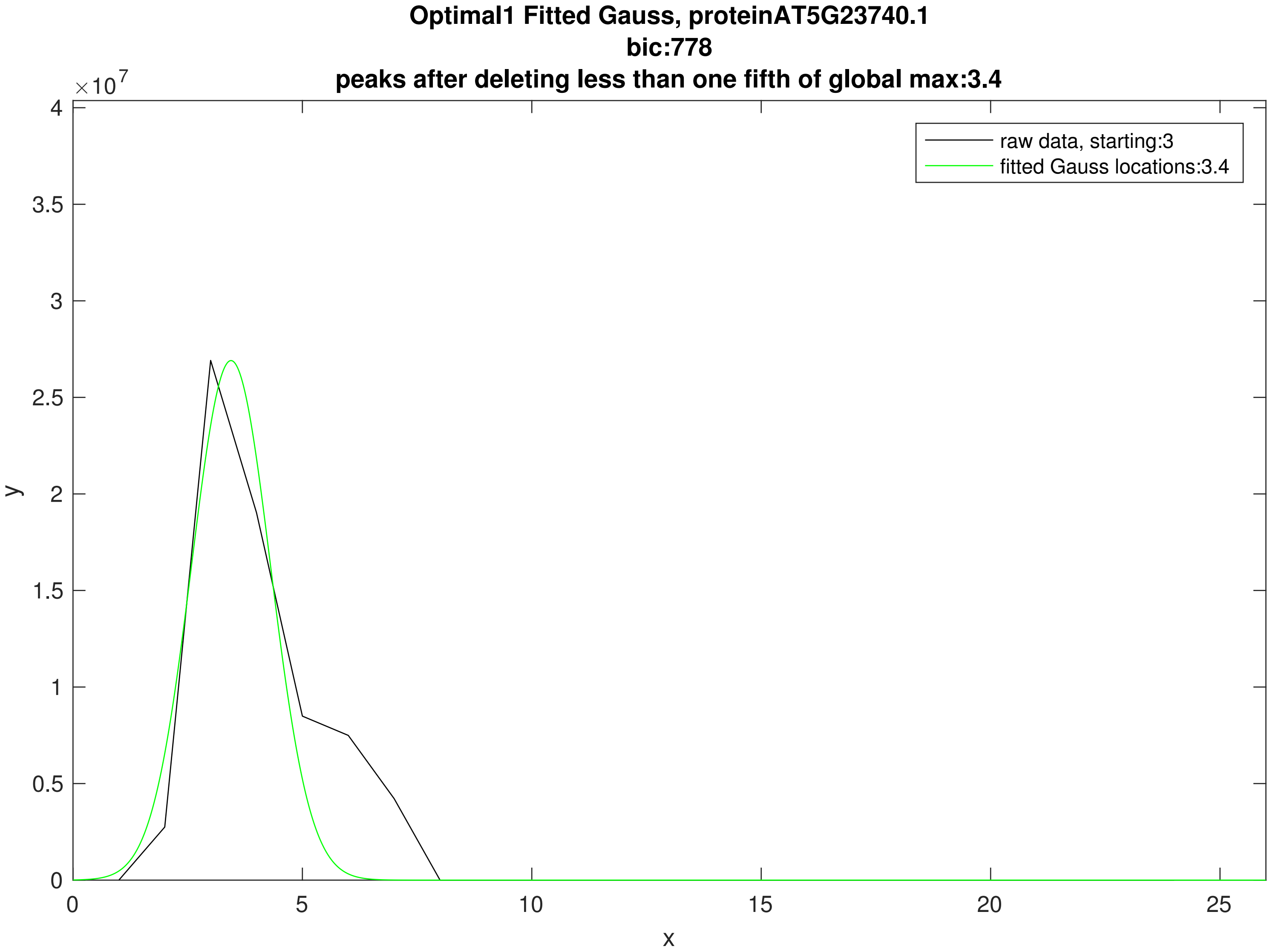

Optimal1 Fitted Gauss, proteinAT5G23740.1
bic:778
peaks after deleting less than one fifth of global max:3.4
×10^7
raw data, starting:3
fitted Gauss locations:3.4
y
x
0
0.5
1
1.5
2
2.5
3
3.5
4
5
10
15
20
25

**Optimal1 Fitted Gauss, proteinAT5G23820.1**
**bic:924**
**peaks after deleting less than one fifth of global max:18**

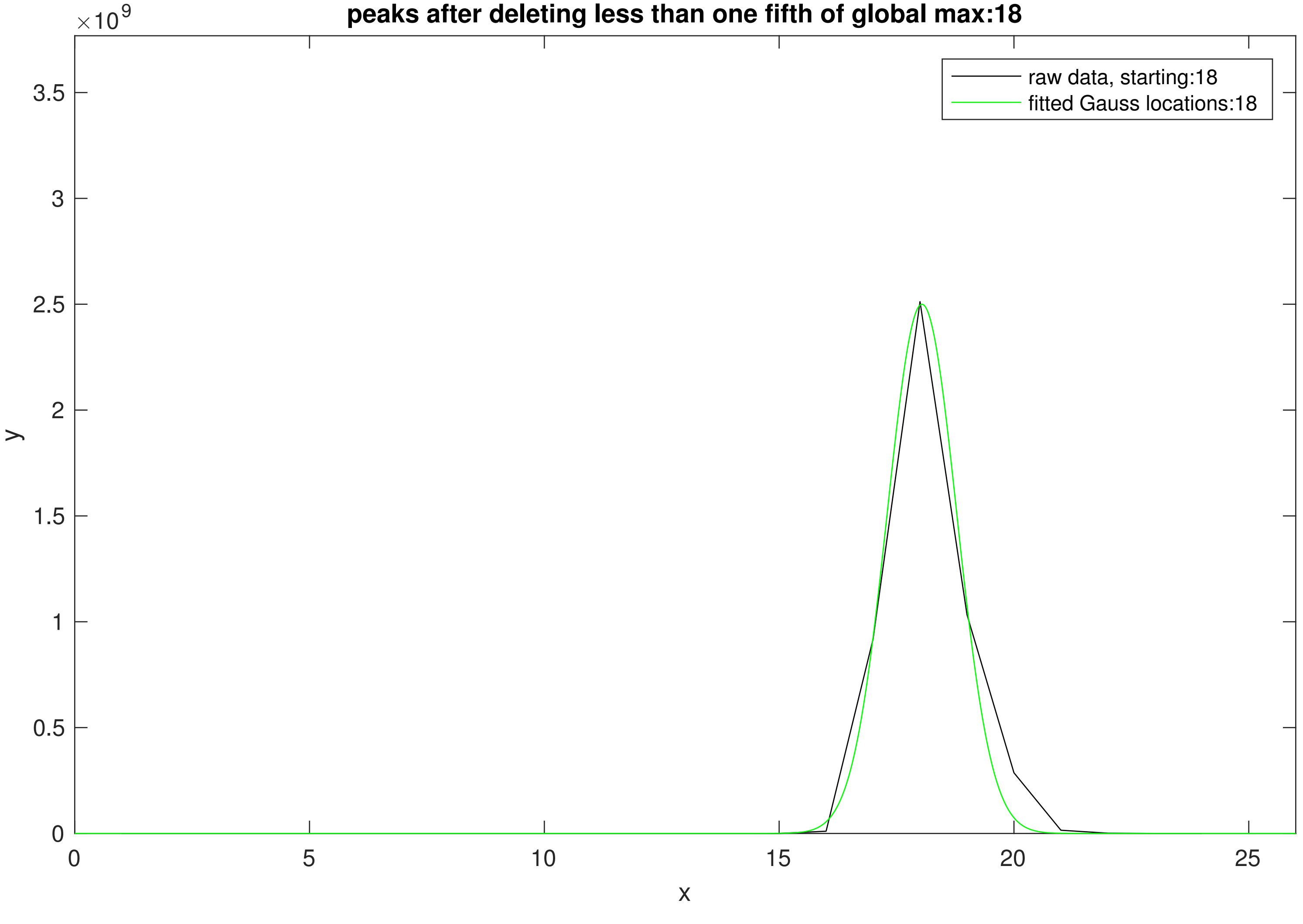

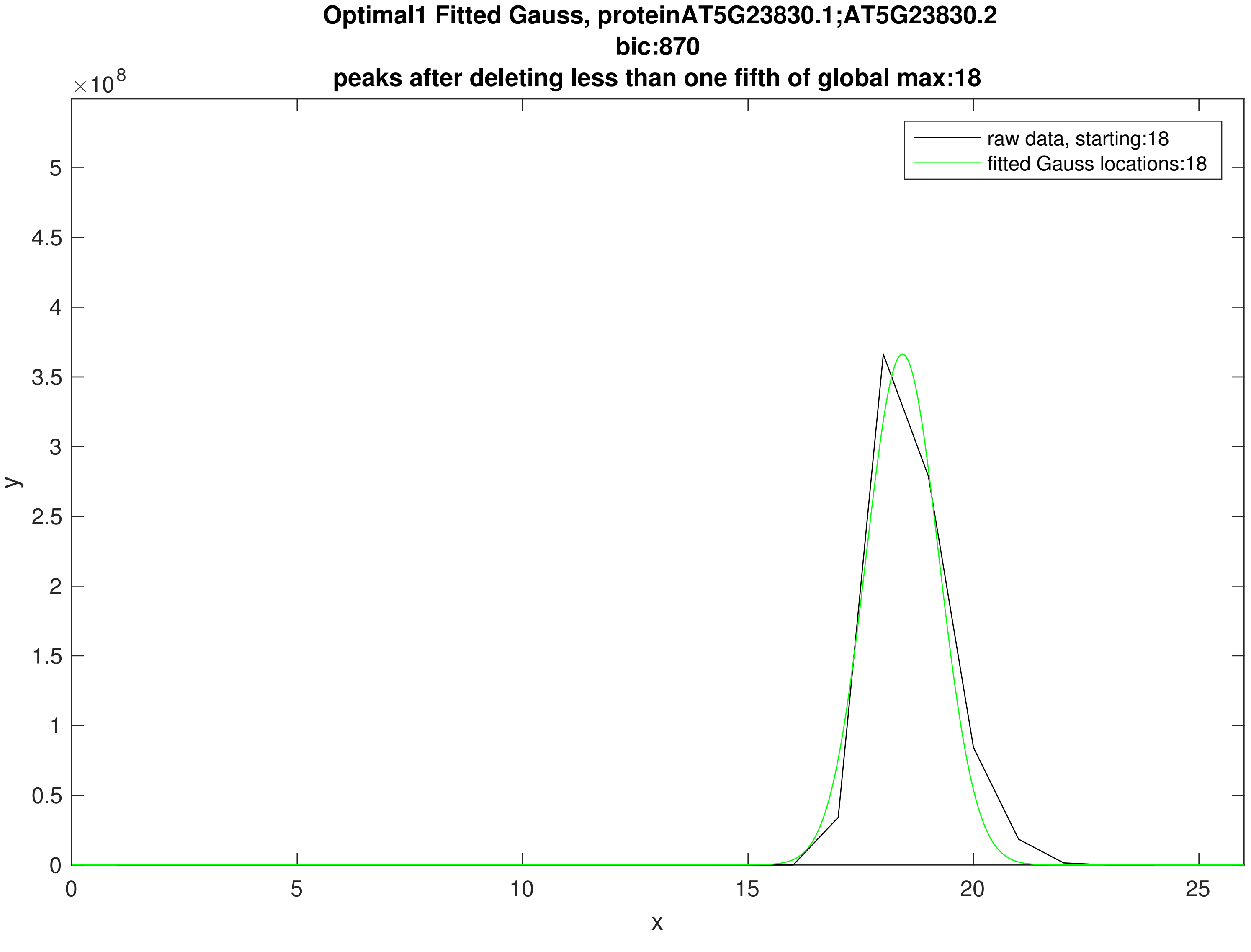
Optimal1 Fitted Gauss, proteinAT5G23830.1;AT5G23830.2
bic:870
peaks after deleting less than one fifth of global max:18
×10^8
raw data, starting:18
fitted Gauss locations:18
y
x
0
0.5
1
1.5
2
2.5
3
3.5
4
4.5
5
0
5
10
15
20
25

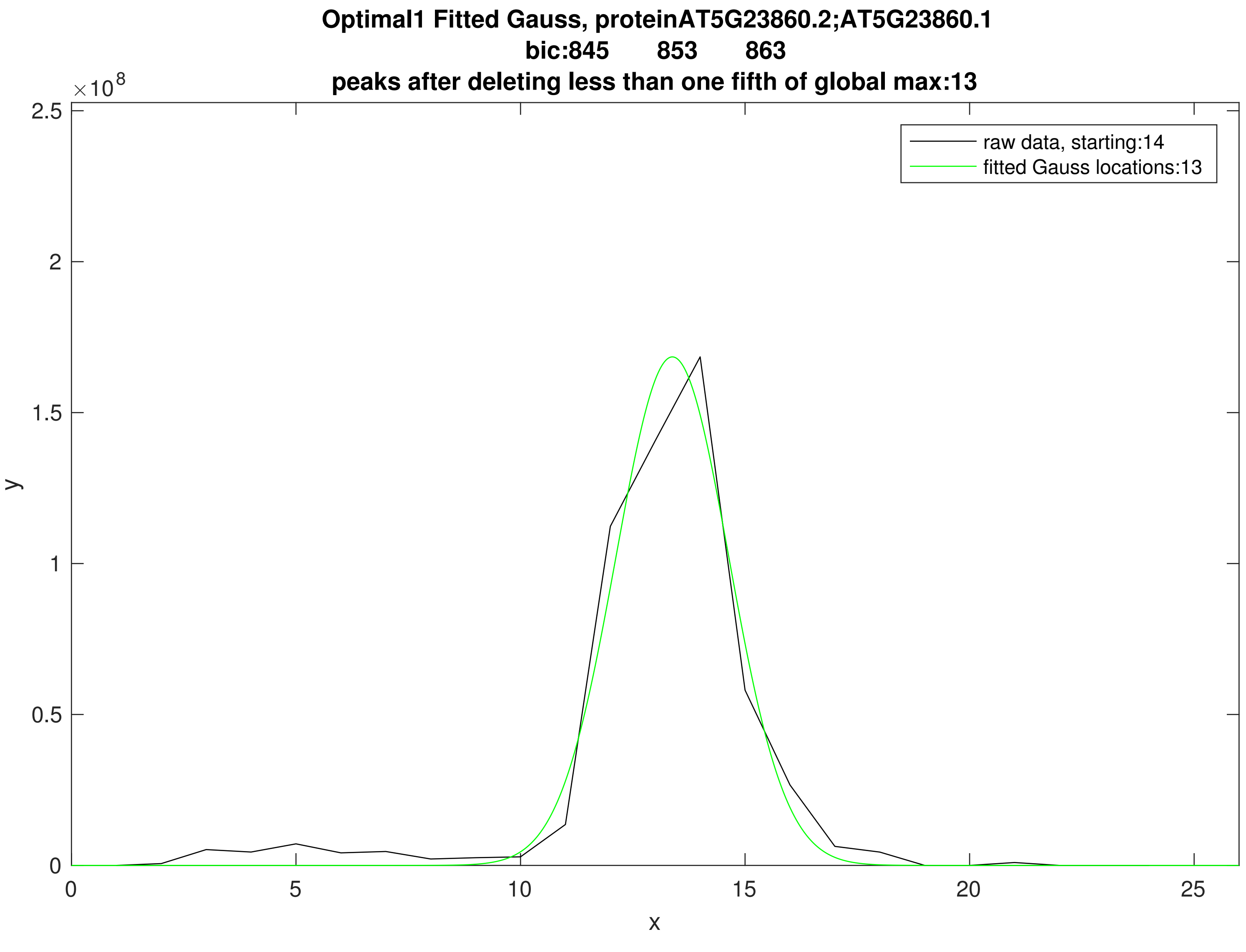
Optimal1 Fitted Gauss, proteinAT5G23860.2;AT5G23860.1
bic:845 853 863
peaks after deleting less than one fifth of global max:13
×10^8
2.5
2
1.5
1
0.5
0
y
0
5
10
15
20
25
x
raw data, starting:14
fitted Gauss locations:13

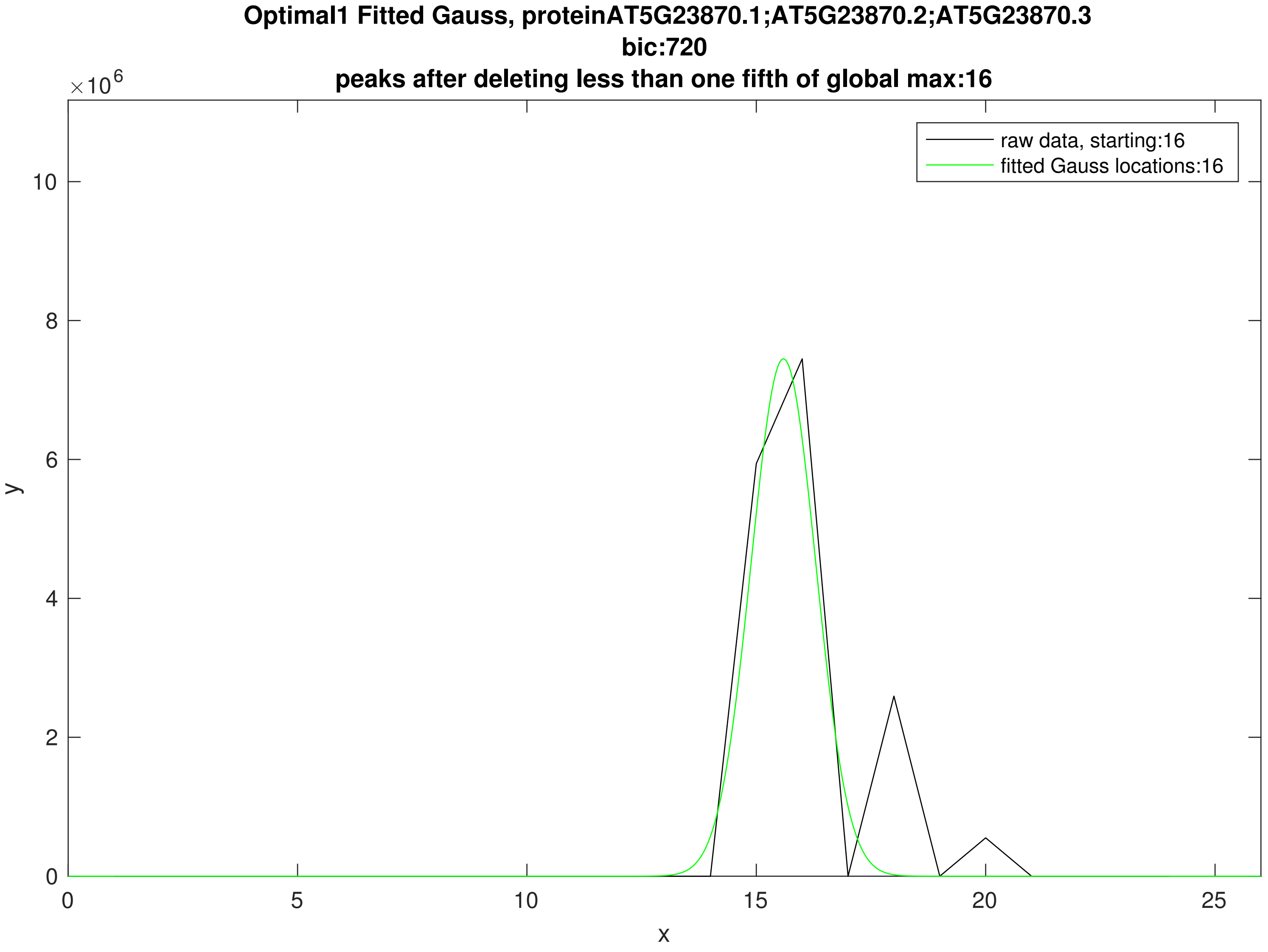

Optimal1 Fitted Gauss, proteinAT5G23870.1;AT5G23870.2;AT5G23870.3
bic:720
peaks after deleting less than one fifth of global max:16
×10^6
raw data, starting:16
fitted Gauss locations:16
y
x
0
2
4
6
8
10
0
5
10
15
20
25

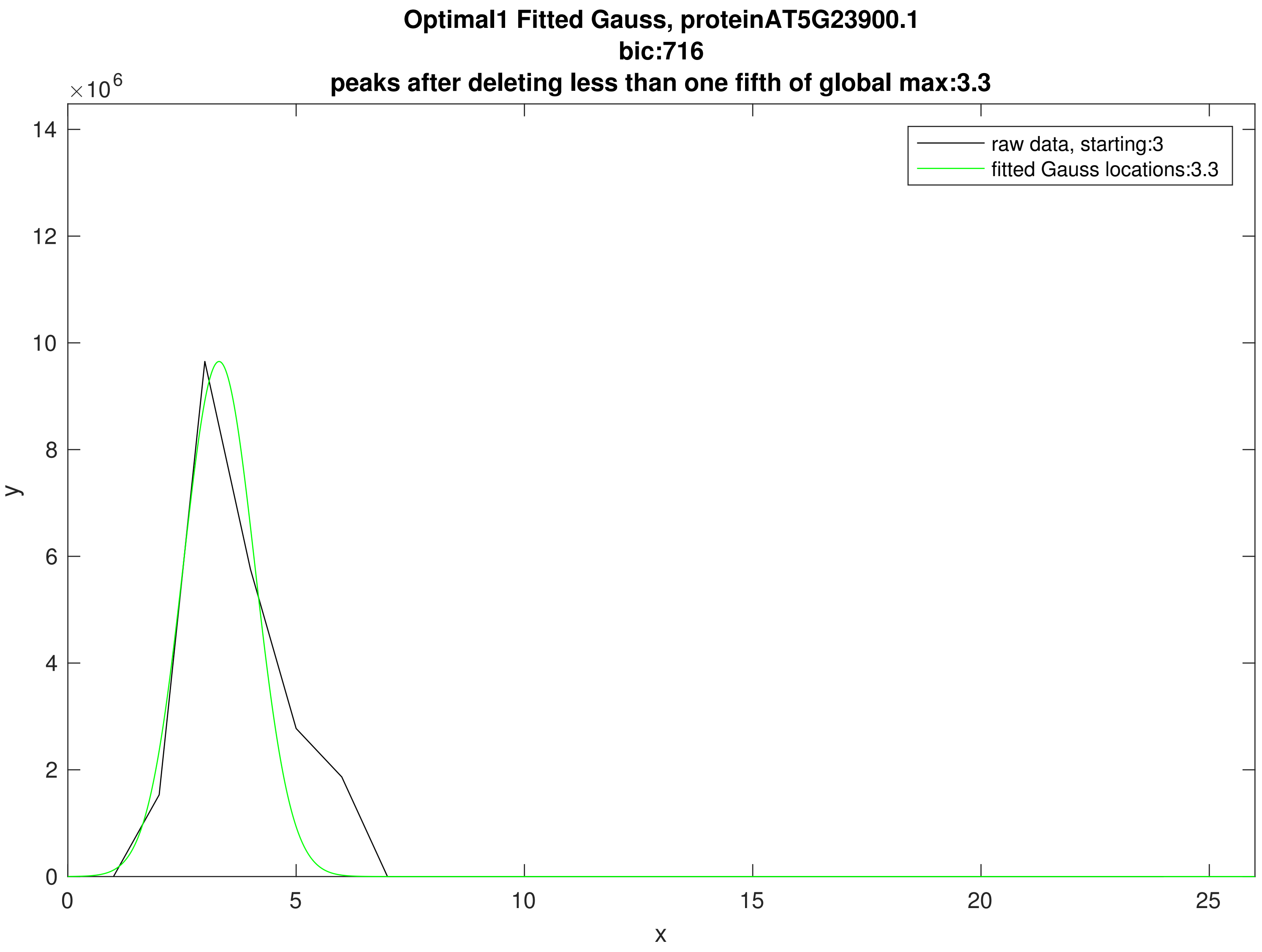

Optimal1 Fitted Gauss, proteinAT5G23900.1
bic:716
peaks after deleting less than one fifth of global max:3.3
×10^6
raw data, starting:3
fitted Gauss locations:3.3
14
12
10
8
6
4
2
0
y
0
5
10
15
20
25
x

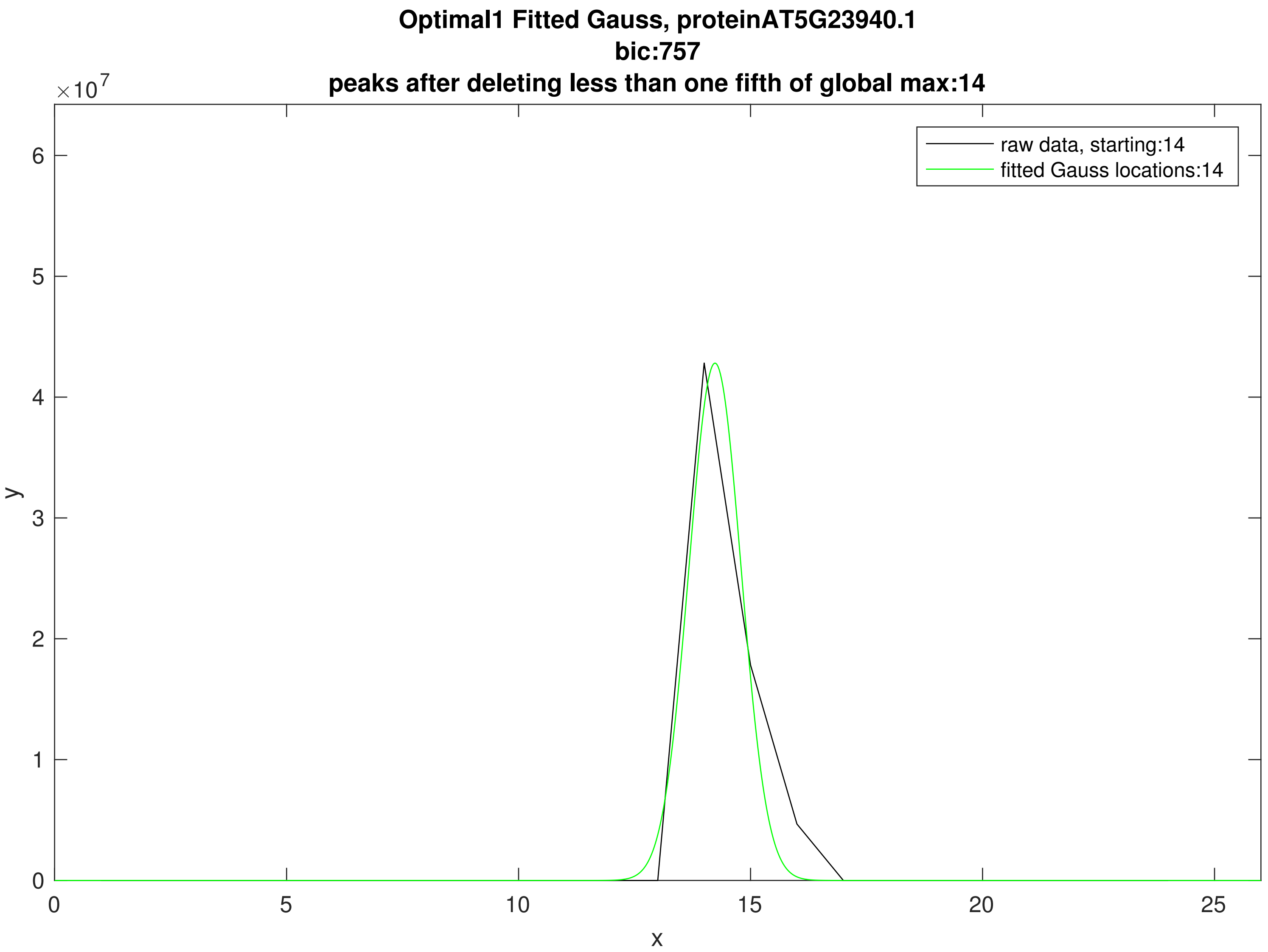

Optimal1 Fitted Gauss, proteinAT5G23940.1
bic:757
peaks after deleting less than one fifth of global max:14
×10^7
raw data, starting:14
fitted Gauss locations:14
0
1
2
3
4
5
6
y
0
5
10
15
20
25
x

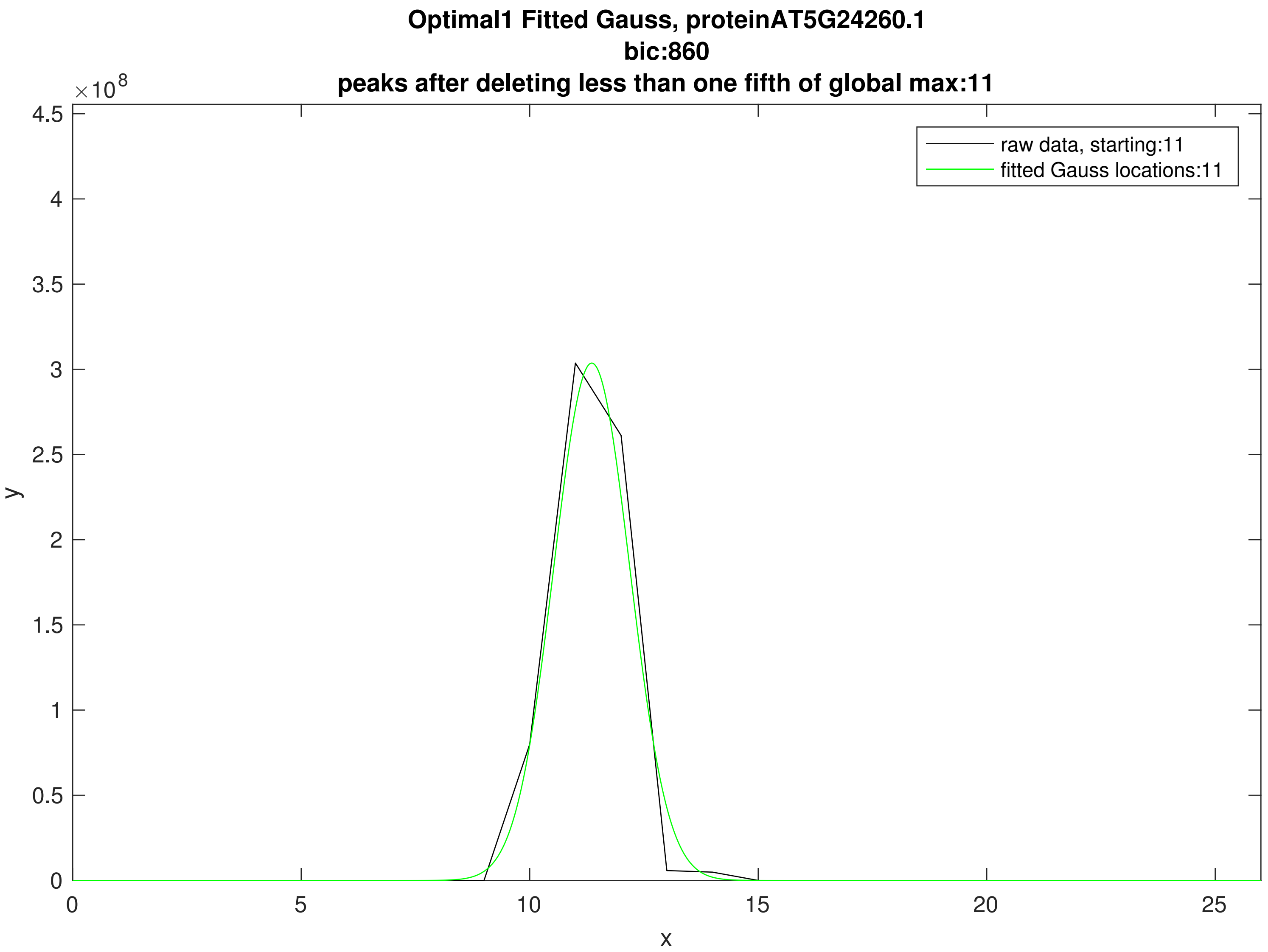

Optimal1 Fitted Gauss, proteinAT5G24260.1
bic:860
peaks after deleting less than one fifth of global max:11
×10^8
4.5
4
3.5
3
2.5
2
1.5
1
0.5
0
y
0
5
10
15
20
25
x
raw data, starting:11
fitted Gauss locations:11

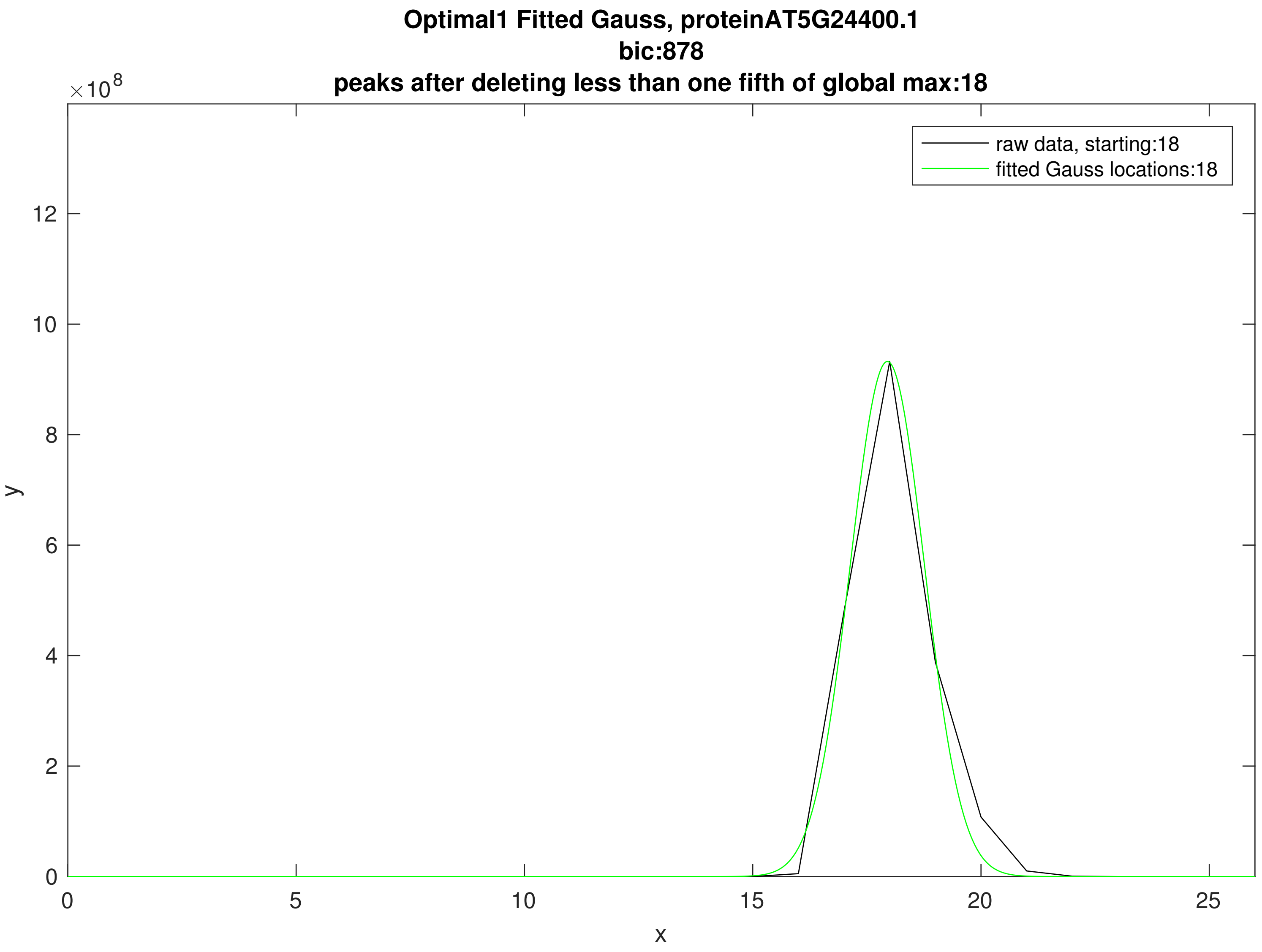

Optimal1 Fitted Gauss, proteinAT5G24400.1
bic:878
peaks after deleting less than one fifth of global max:18
×10^8
raw data, starting:18
fitted Gauss locations:18
12
10
8
6
4
2
0
y
0
5
10
15
20
25
x

#### Optimal1 Fitted Gauss, proteinAT5G24520.3;AT5G24520.2;AT5G24520.1
#### bic:756 761
#### peaks after deleting less than one fifth of global max:16

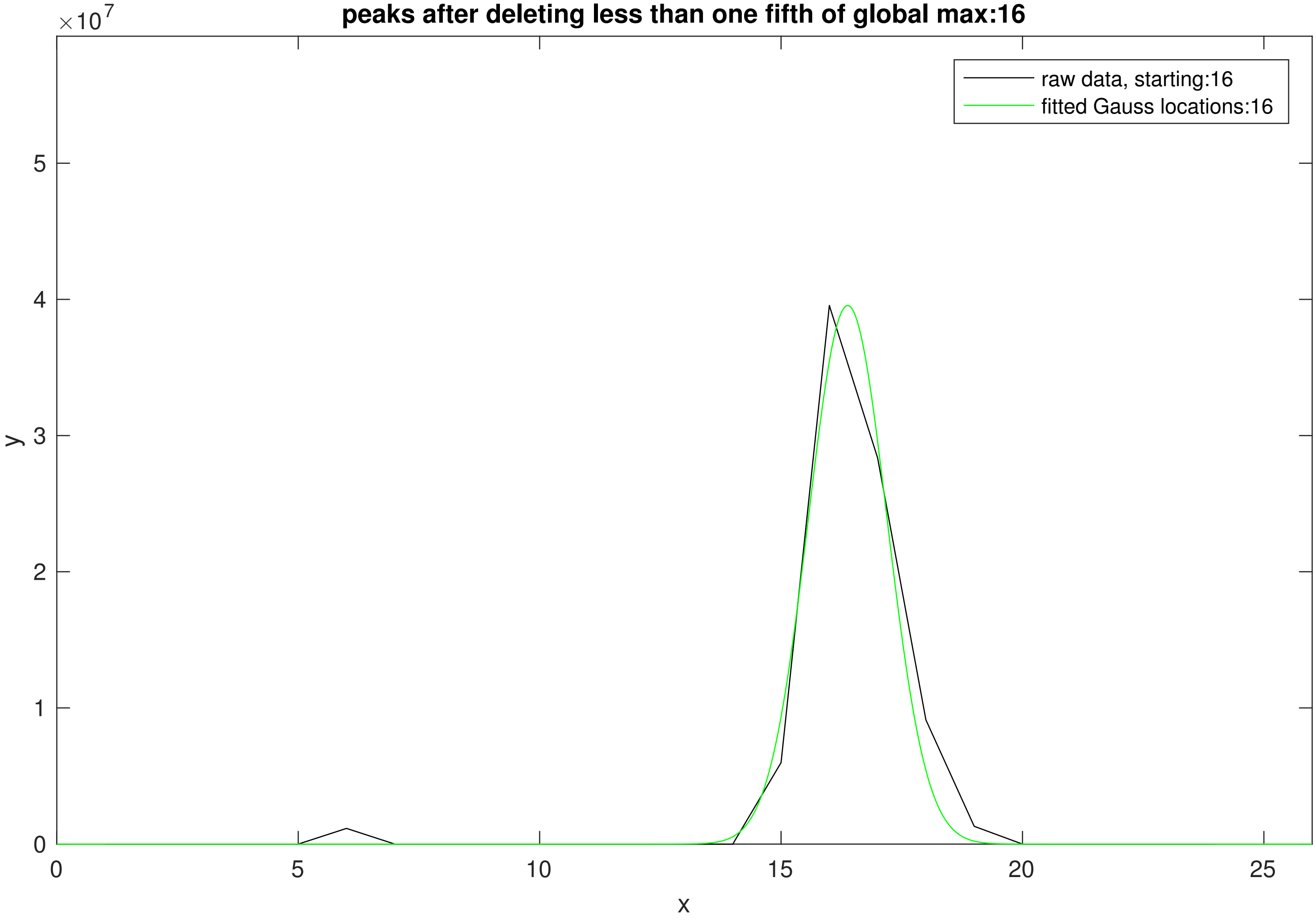

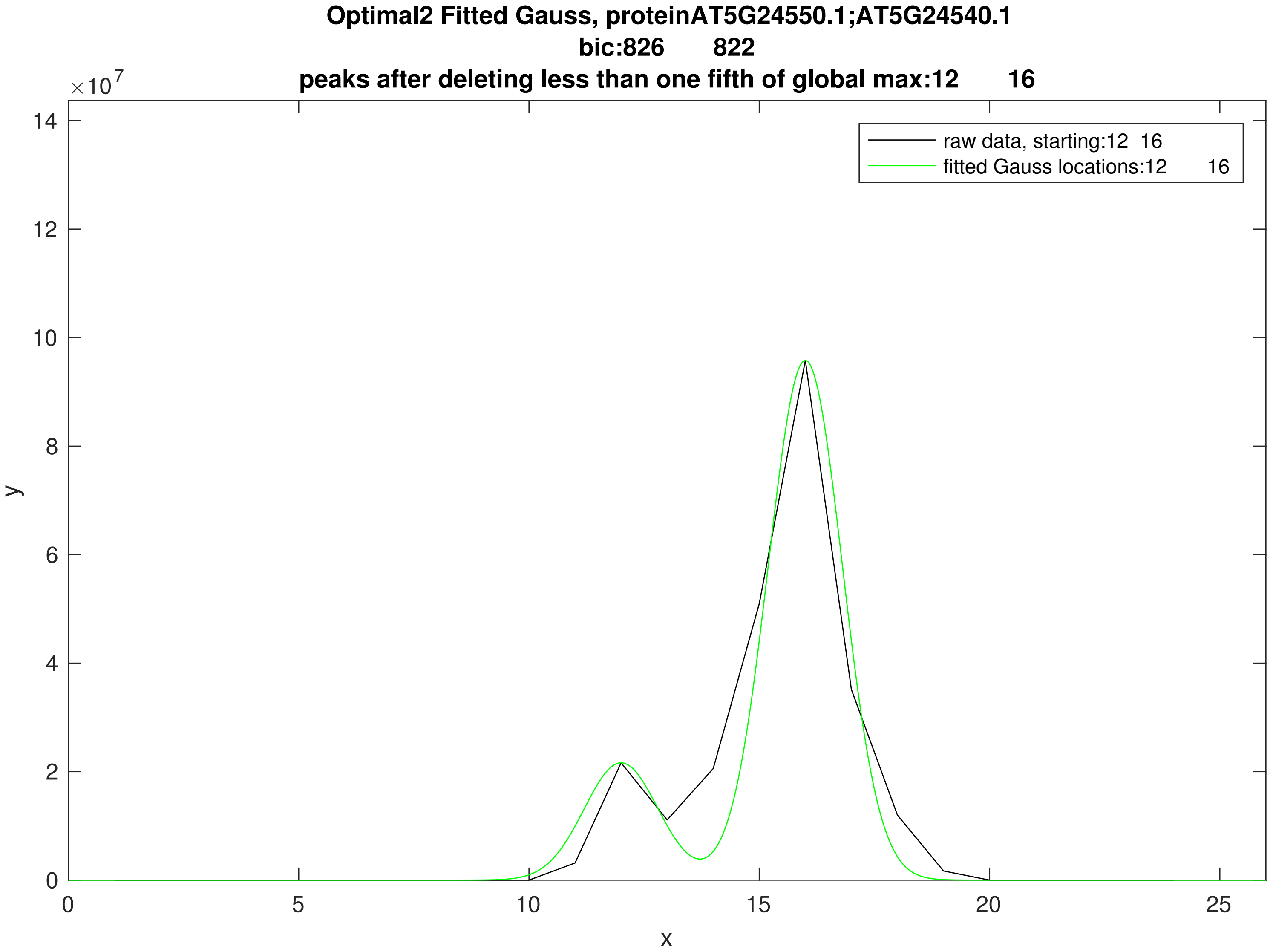
Optimal2 Fitted Gauss, proteinAT5G24550.1;AT5G24540.1
bic:826 822
peaks after deleting less than one fifth of global max:12 16
×10^7
raw data, starting:12 16
fitted Gauss locations:12 16
y
x
0
2
4
6
8
10
12
14
0
5
10
15
20
25

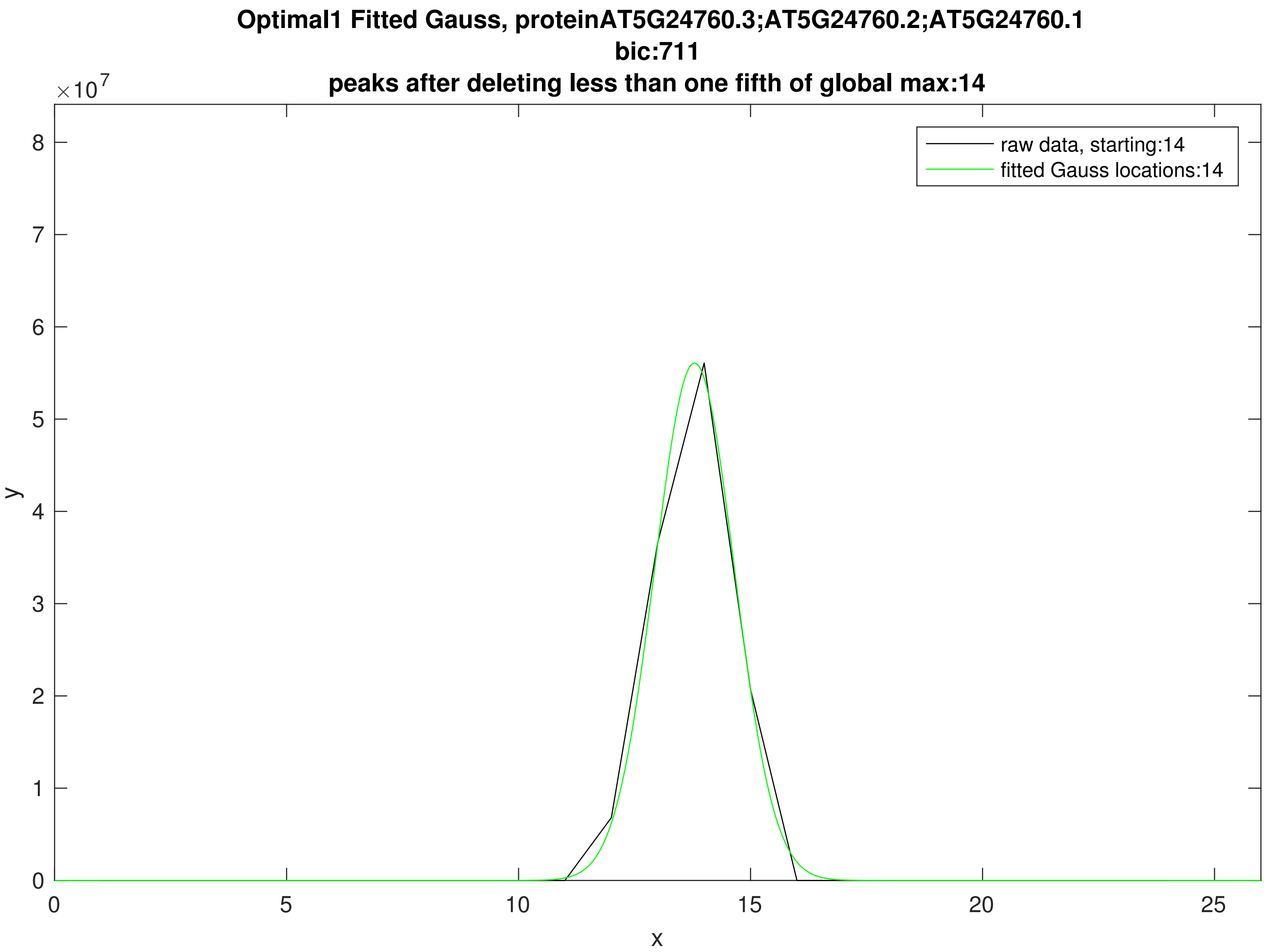

Optimal1 Fitted Gauss, proteinAT5G24760.3;AT5G24760.2;AT5G24760.1
bic:711
peaks after deleting less than one fifth of global max:14
×10^7
y
x
raw data, starting:14
fitted Gauss locations:14
0
1
2
3
4
5
6
7
8
0
5
10
15
20
25

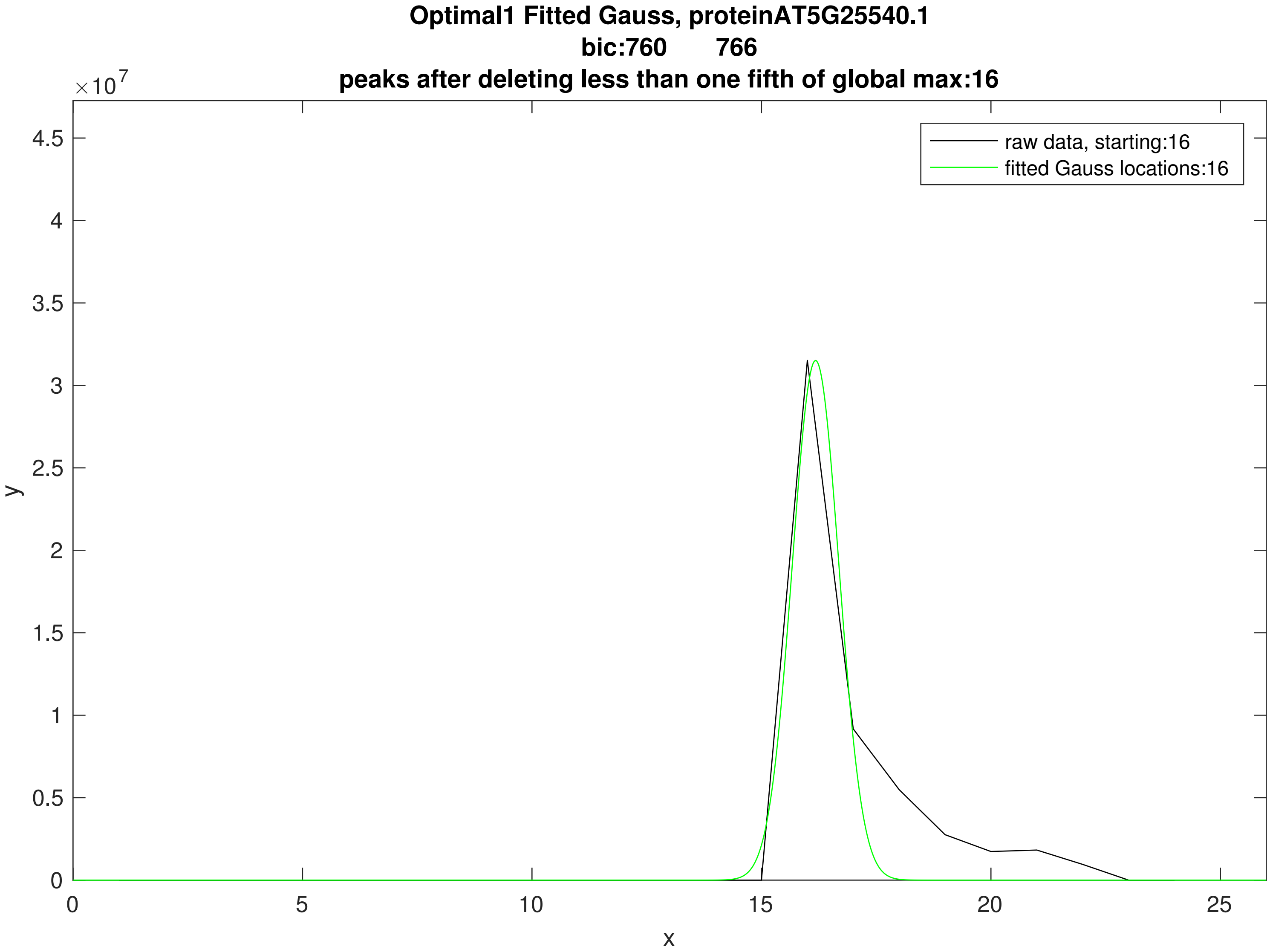
Optimal1 Fitted Gauss, proteinAT5G25540.1
bic:760 766
peaks after deleting less than one fifth of global max:16
×10^7
4.5
4
3.5
3
2.5
2
1.5
1
0.5
0
y
0
5
10
15
20
25
x
raw data, starting:16
fitted Gauss locations:16

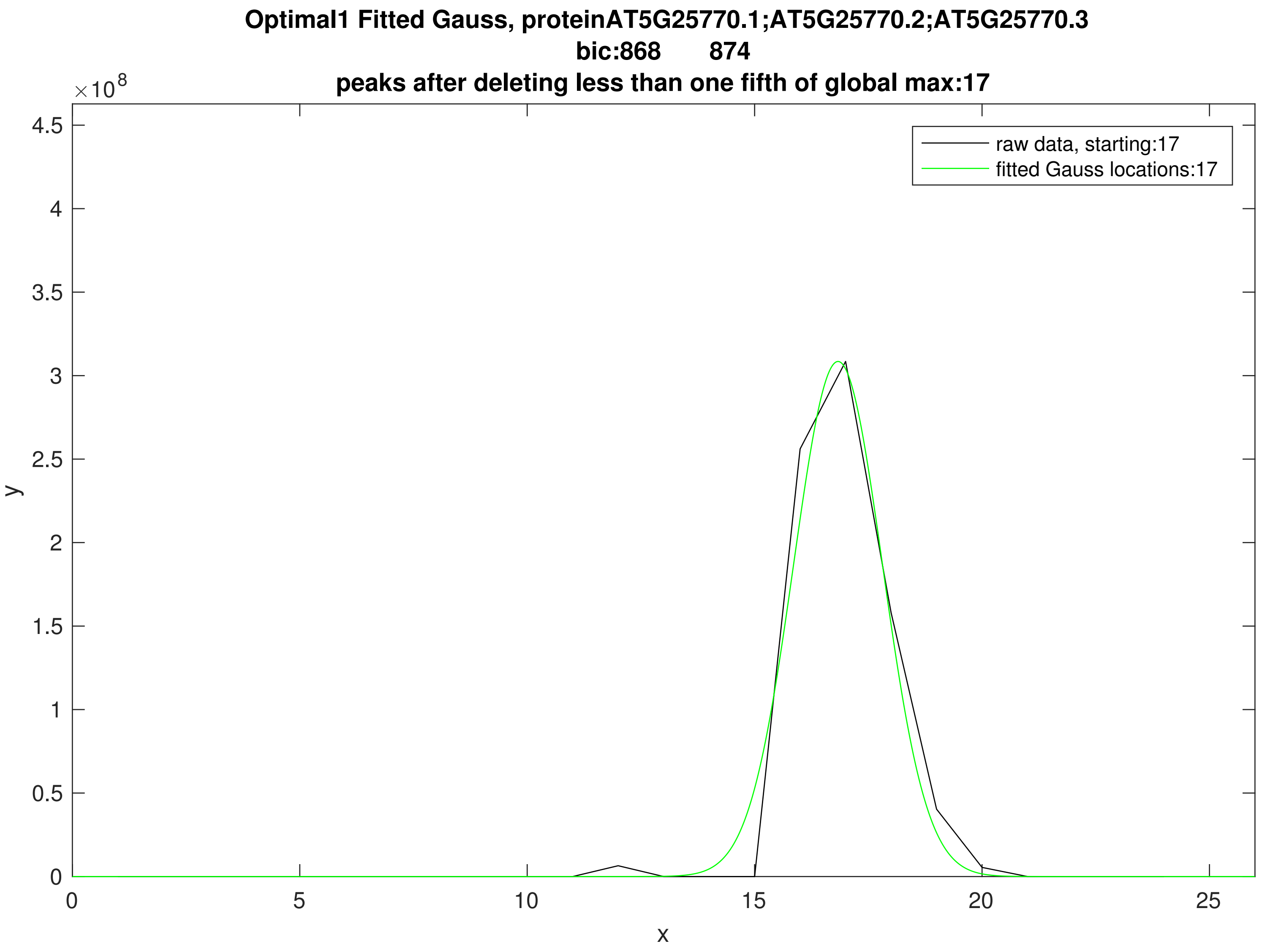
Optimal1 Fitted Gauss, proteinAT5G25770.1;AT5G25770.2;AT5G25770.3
bic:868 874
peaks after deleting less than one fifth of global max:17
×10^8
4.5
4
3.5
3
2.5
2
1.5
1
0.5
0
y
0
5
10
15
20
25
x
raw data, starting:17
fitted Gauss locations:17

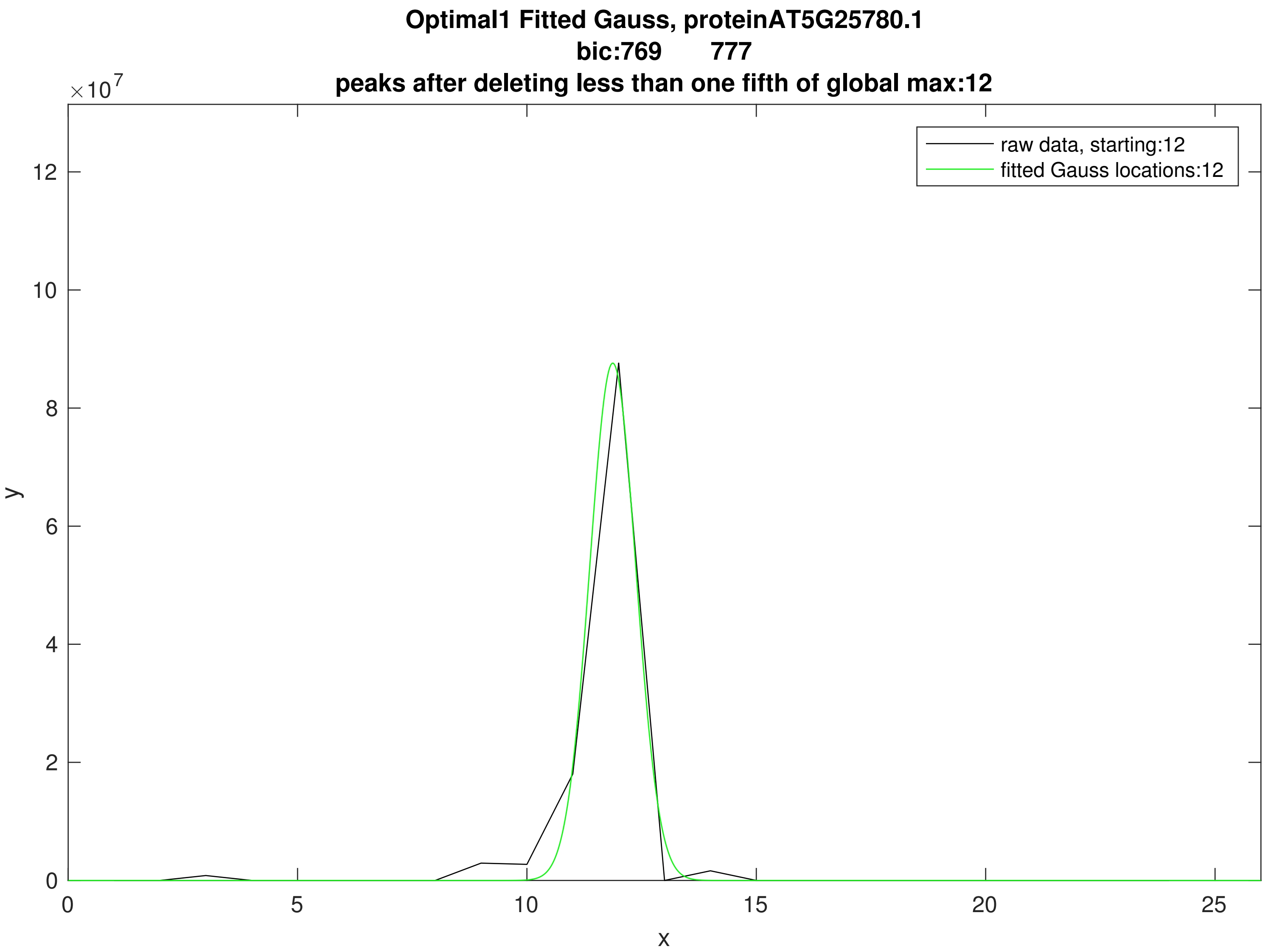
Optimal1 Fitted Gauss, proteinAT5G25780.1
bic:769 777
peaks after deleting less than one fifth of global max:12
×10^7
12
10
8
6
4
2
0
y
0
5
10
15
20
25
x
raw data, starting:12
fitted Gauss locations:12

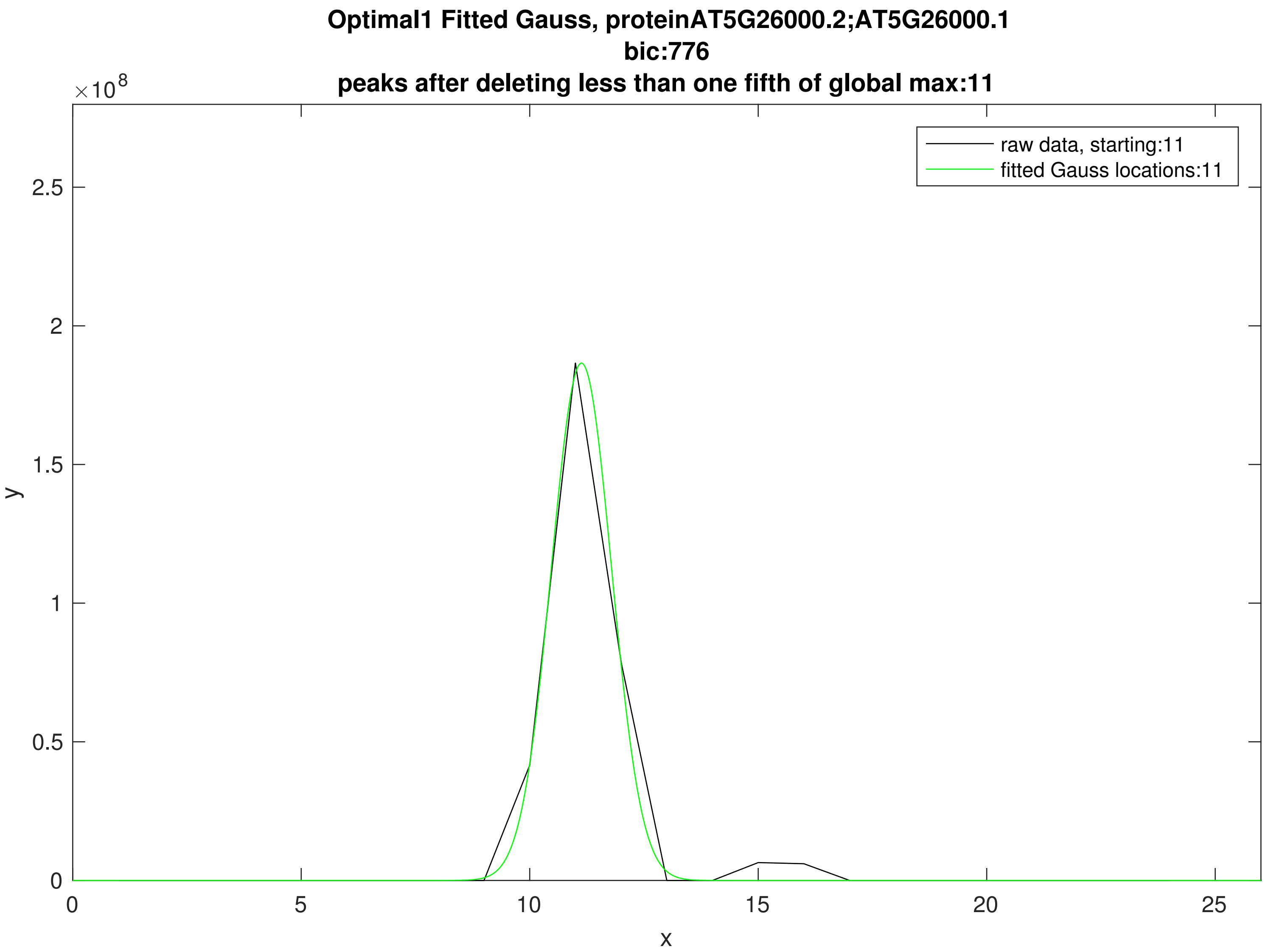

Optimal1 Fitted Gauss, proteinAT5G26000.2;AT5G26000.1
bic:776
peaks after deleting less than one fifth of global max:11
×10^8
2.5
2
1.5
1
0.5
0
y
0
5
10
15
20
25
x
raw data, starting:11
fitted Gauss locations:11

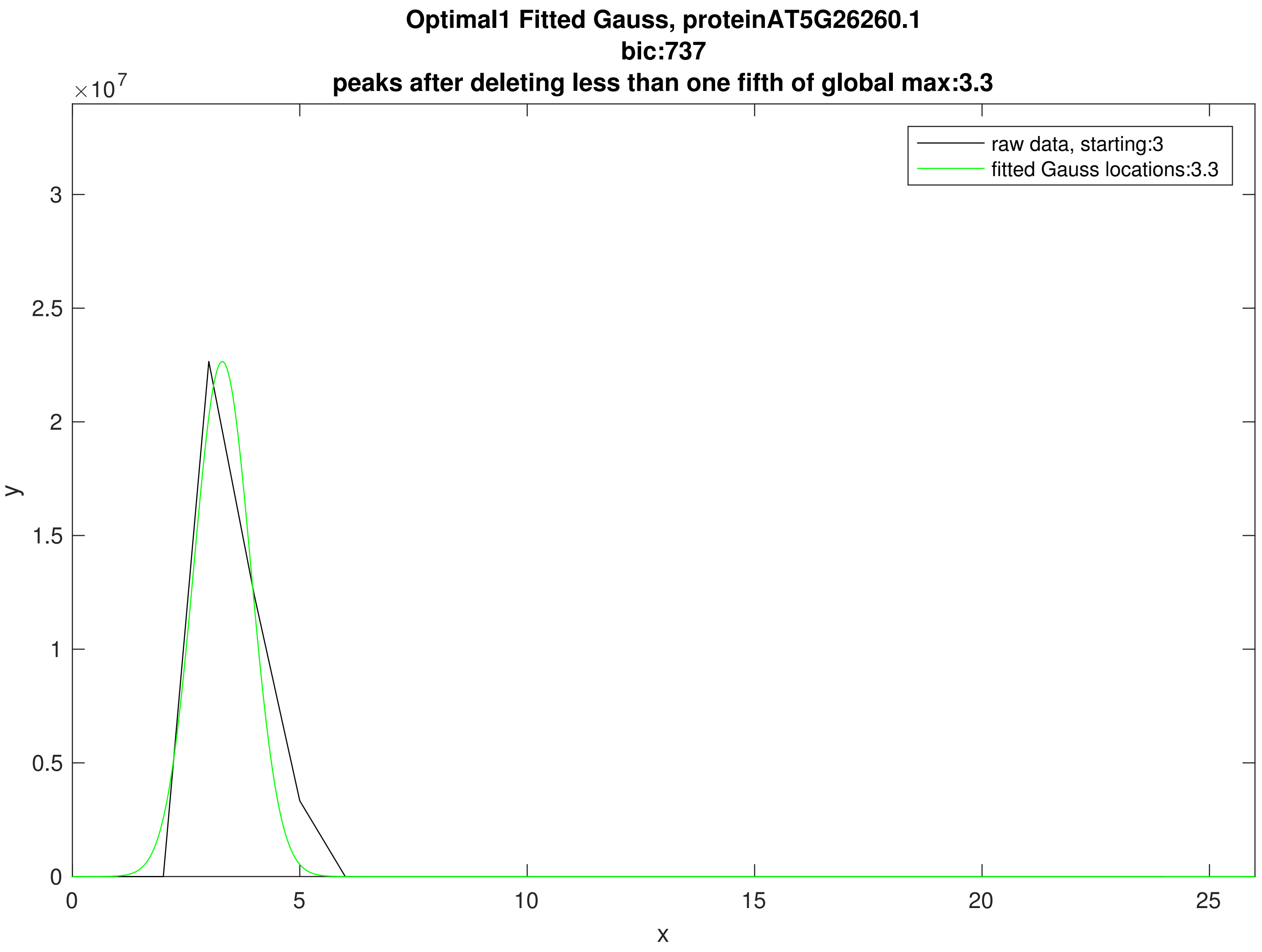

Optimal1 Fitted Gauss, proteinAT5G26260.1
bic:737
peaks after deleting less than one fifth of global max:3.3
×10^7
raw data, starting:3
fitted Gauss locations:3.3
y
x
0
0.5
1
1.5
2
2.5
3
0
5
10
15
20
25

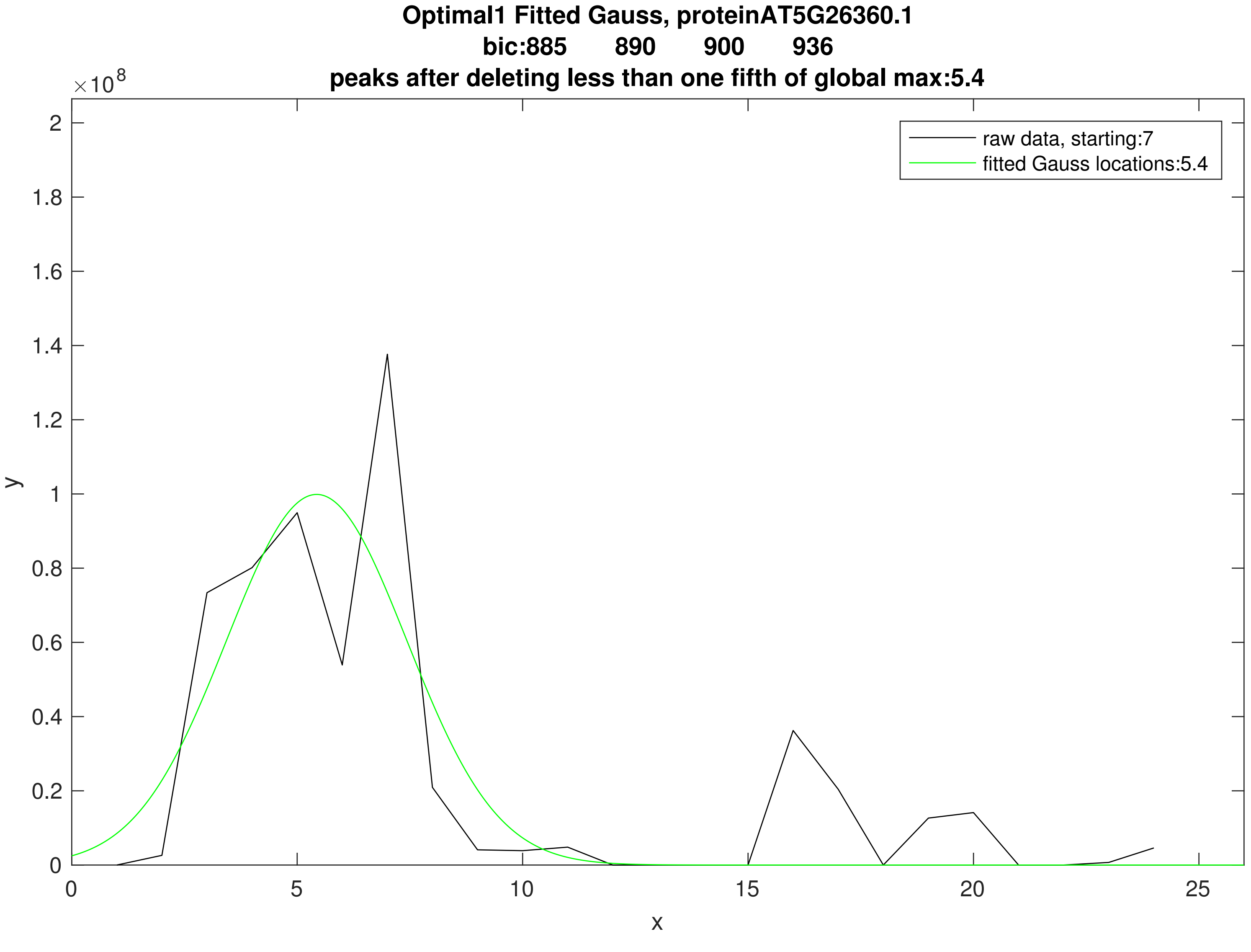

Optimal1 Fitted Gauss, proteinAT5G26360.1
bic:885 890 900 936
peaks after deleting less than one fifth of global max:5.4
×10^8
raw data, starting:7
fitted Gauss locations:5.4
y
x
0
0.2
0.4
0.6
0.8
1
1.2
1.4
1.6
1.8
2
5
10
15
20
25

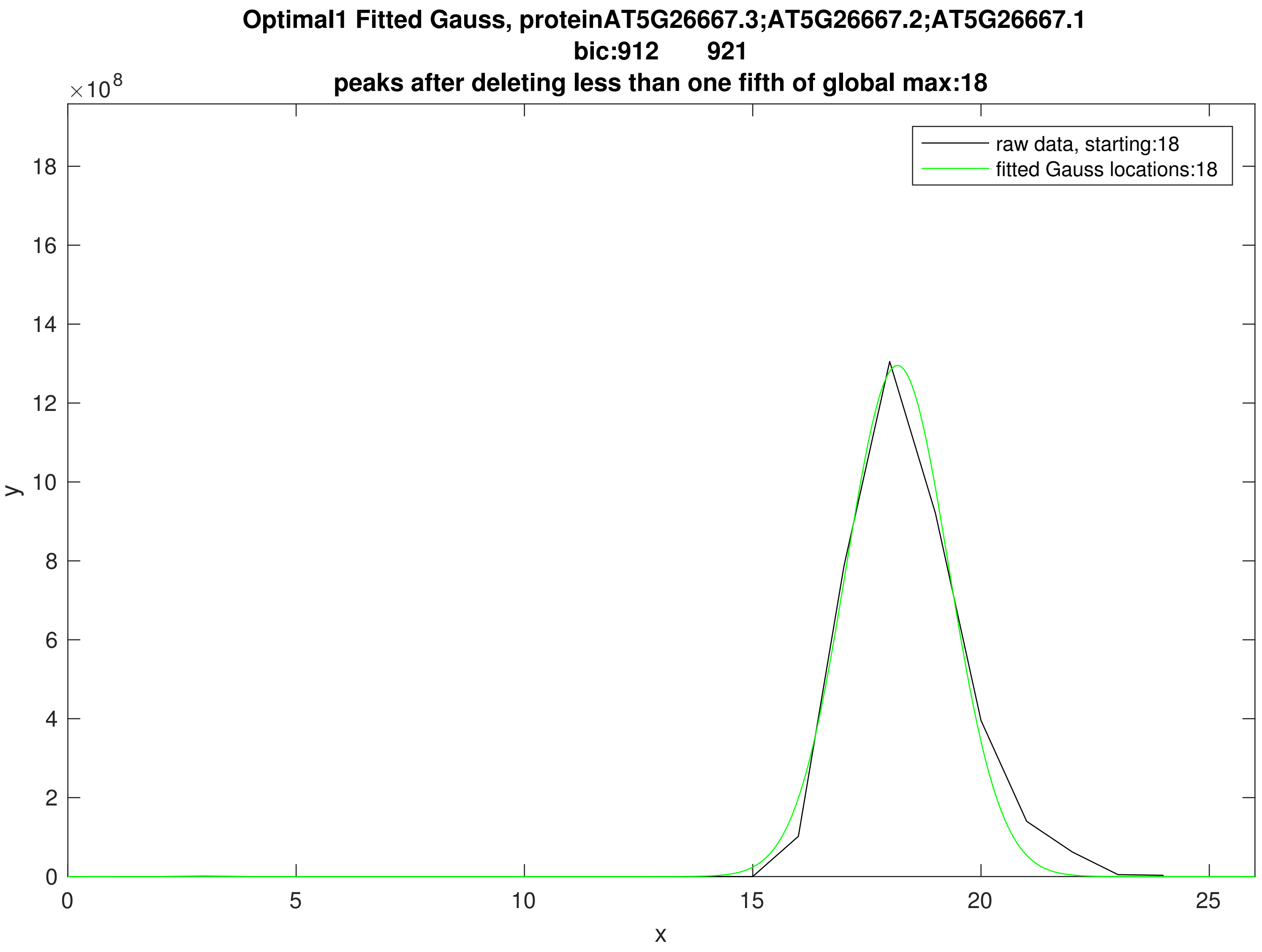

Optimal1 Fitted Gauss, proteinAT5G26667.3;AT5G26667.2;AT5G26667.1
bic:912 921
peaks after deleting less than one fifth of global max:18
×10^8
y
x
raw data, starting:18
fitted Gauss locations:18
0
2
4
6
8
10
12
14
16
18
5
15
20
25

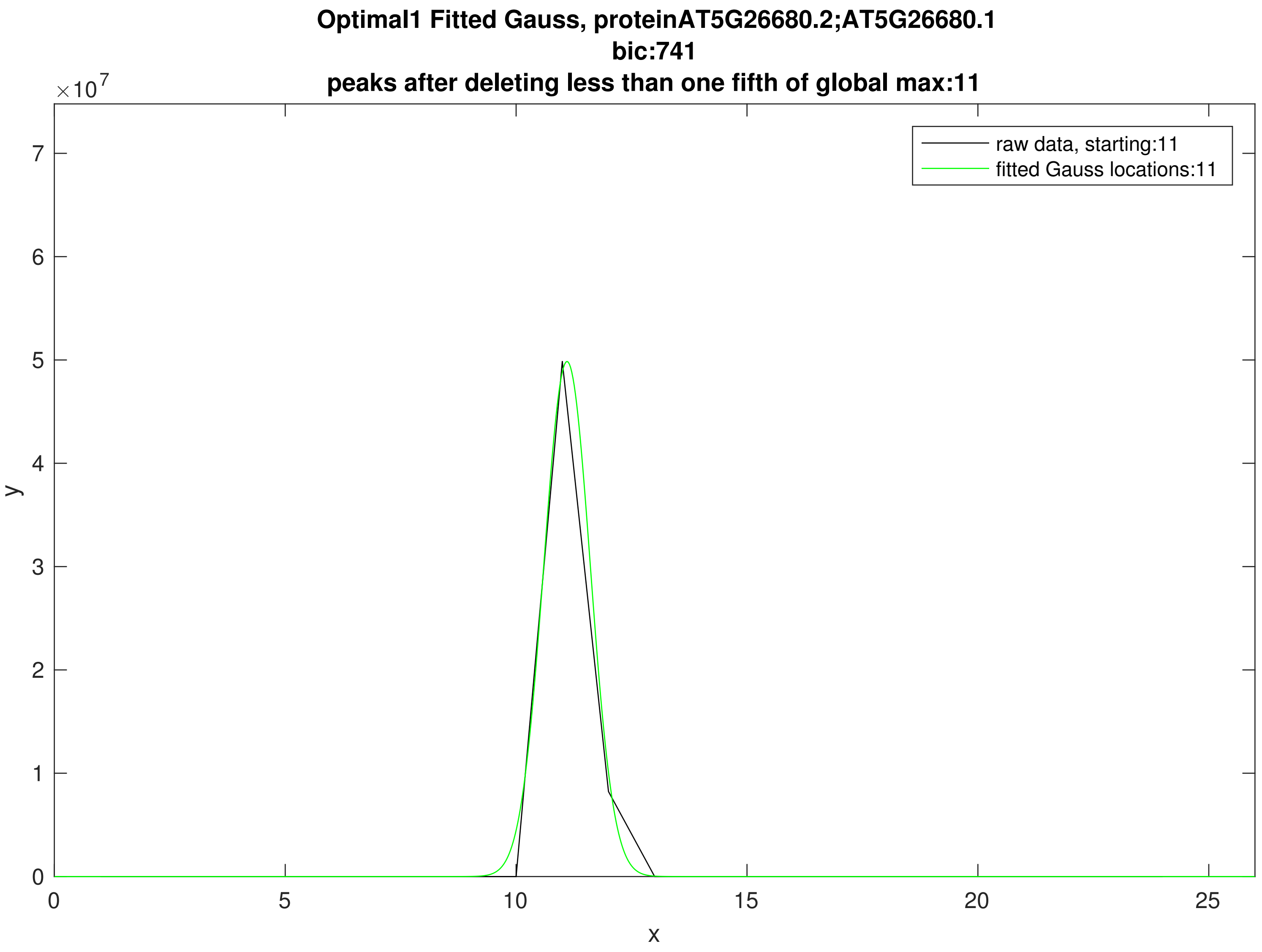
Optimal1 Fitted Gauss, proteinAT5G26680.2;AT5G26680.1
bic:741
peaks after deleting less than one fifth of global max:11
×10^7
raw data, starting:11
fitted Gauss locations:11
y
x
0
1
2
3
4
5
6
7
0
5
10
15
20
25

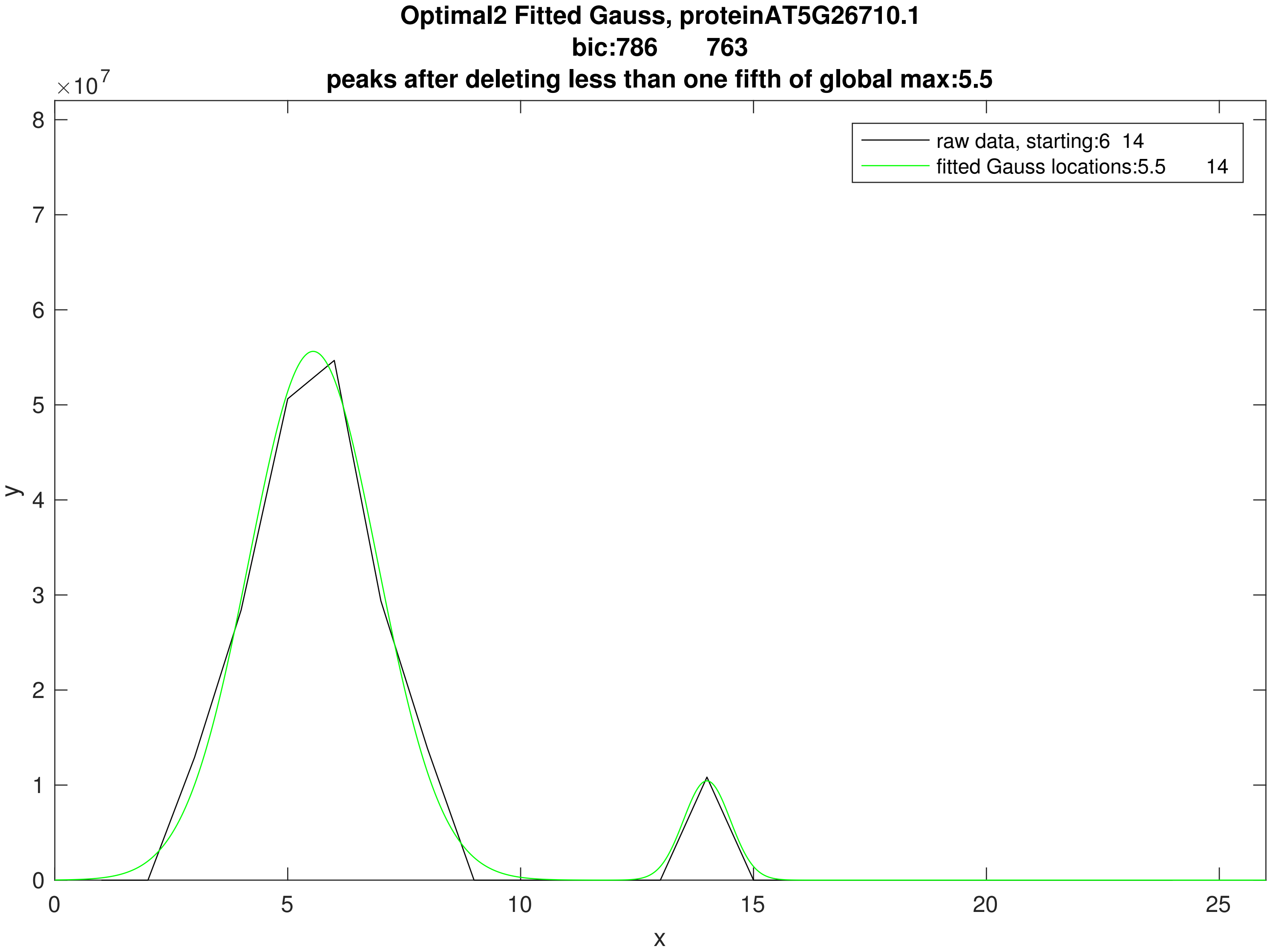

Optimal2 Fitted Gauss, proteinAT5G26710.1
bic:786 763
peaks after deleting less than one fifth of global max:5.5
×10^7
raw data, starting:6 14
fitted Gauss locations:5.5 14
y
x
0
1
2
3
4
5
6
7
8
0
5
10
15
20
25

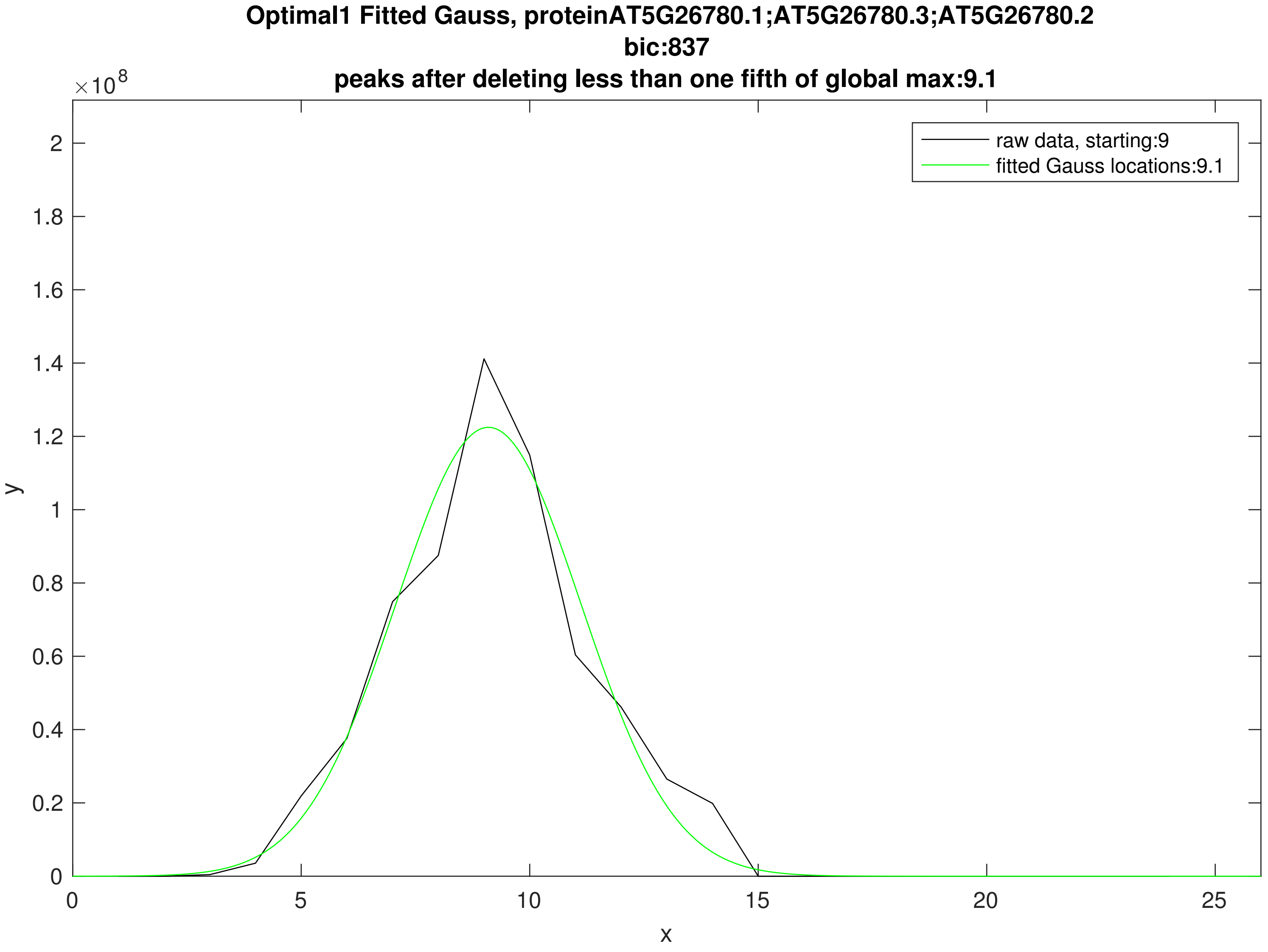
Optimal1 Fitted Gauss, proteinAT5G26780.1;AT5G26780.3;AT5G26780.2
bic:837
peaks after deleting less than one fifth of global max:9.1
×10^8
raw data, starting:9
fitted Gauss locations:9.1
y
x
0
0.2
0.4
0.6
0.8
1
1.2
1.4
1.6
1.8
2
0
5
10
15
20
25

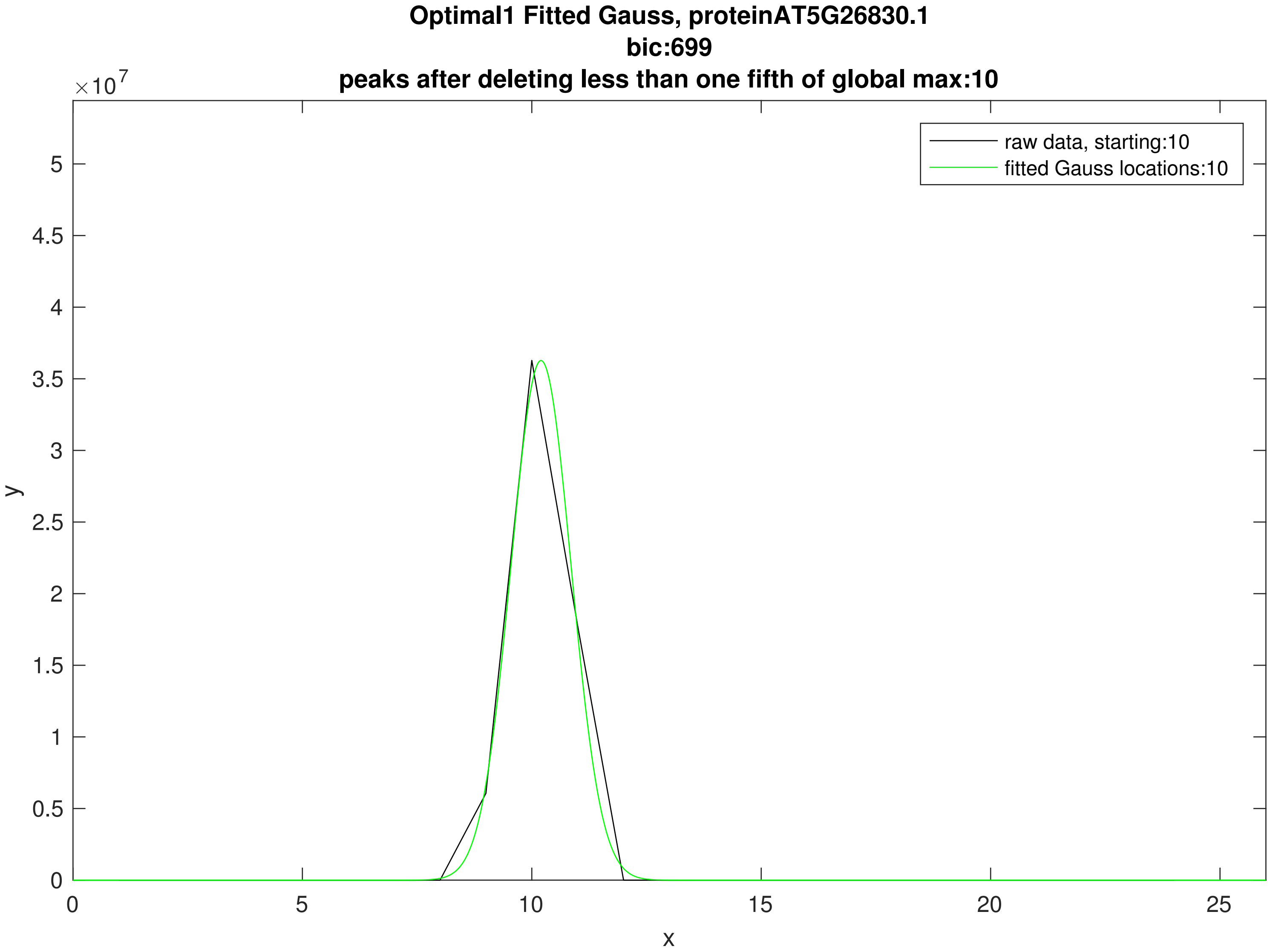
Optimal1 Fitted Gauss, proteinAT5G26830.1
bic:699
peaks after deleting less than one fifth of global max:10
×10^7
raw data, starting:10
fitted Gauss locations:10
5
4.5
4
3.5
3
2.5
2
1.5
1
0.5
0
y
0
5
10
15
20
25
x

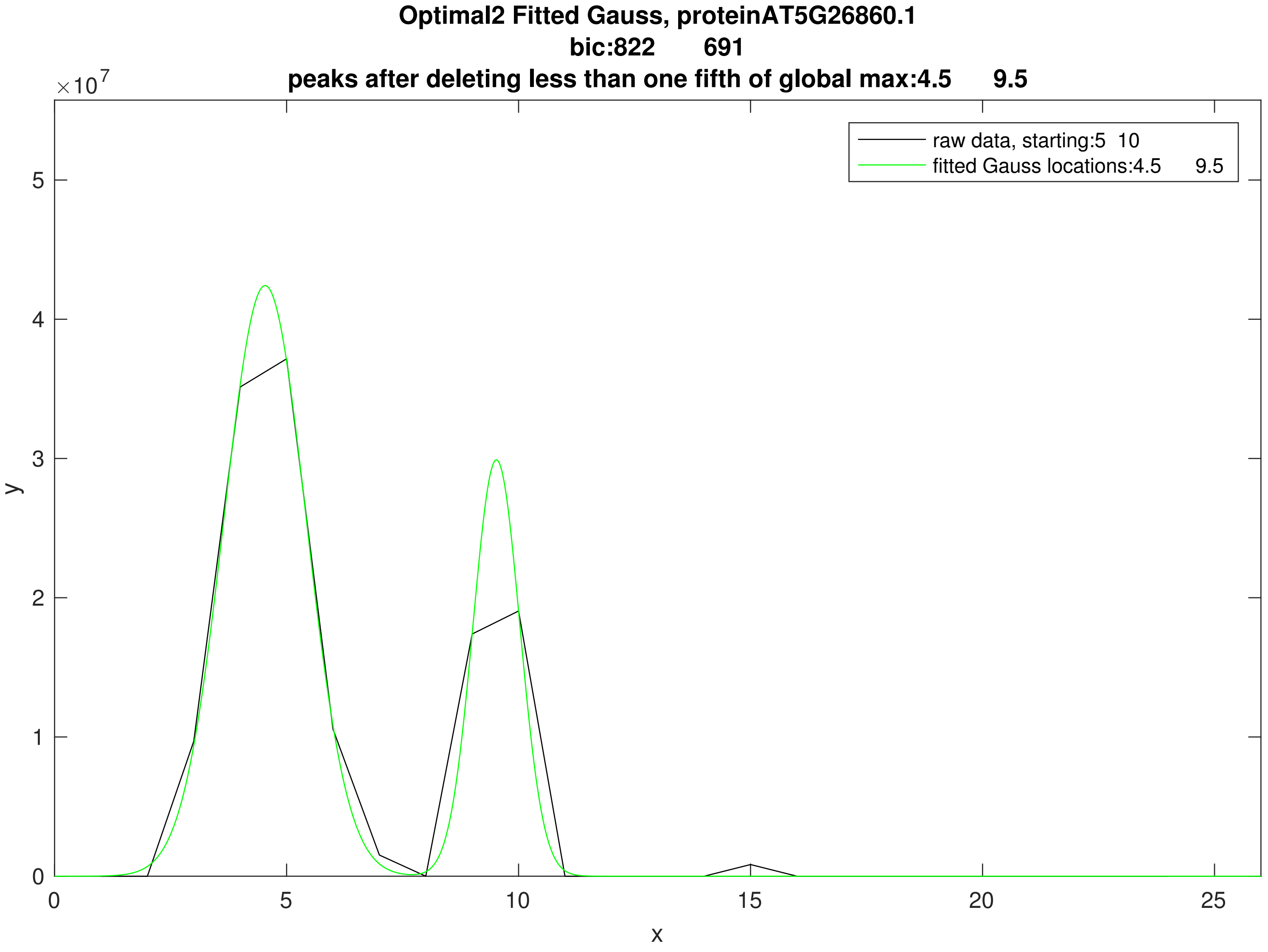

Optimal2 Fitted Gauss, proteinAT5G26860.1
bic:822 691
peaks after deleting less than one fifth of global max:4.5 9.5
×10^7
raw data, starting:5 10
fitted Gauss locations:4.5 9.5
y
x
0
1
2
3
4
5
0
5
10
15
20
25

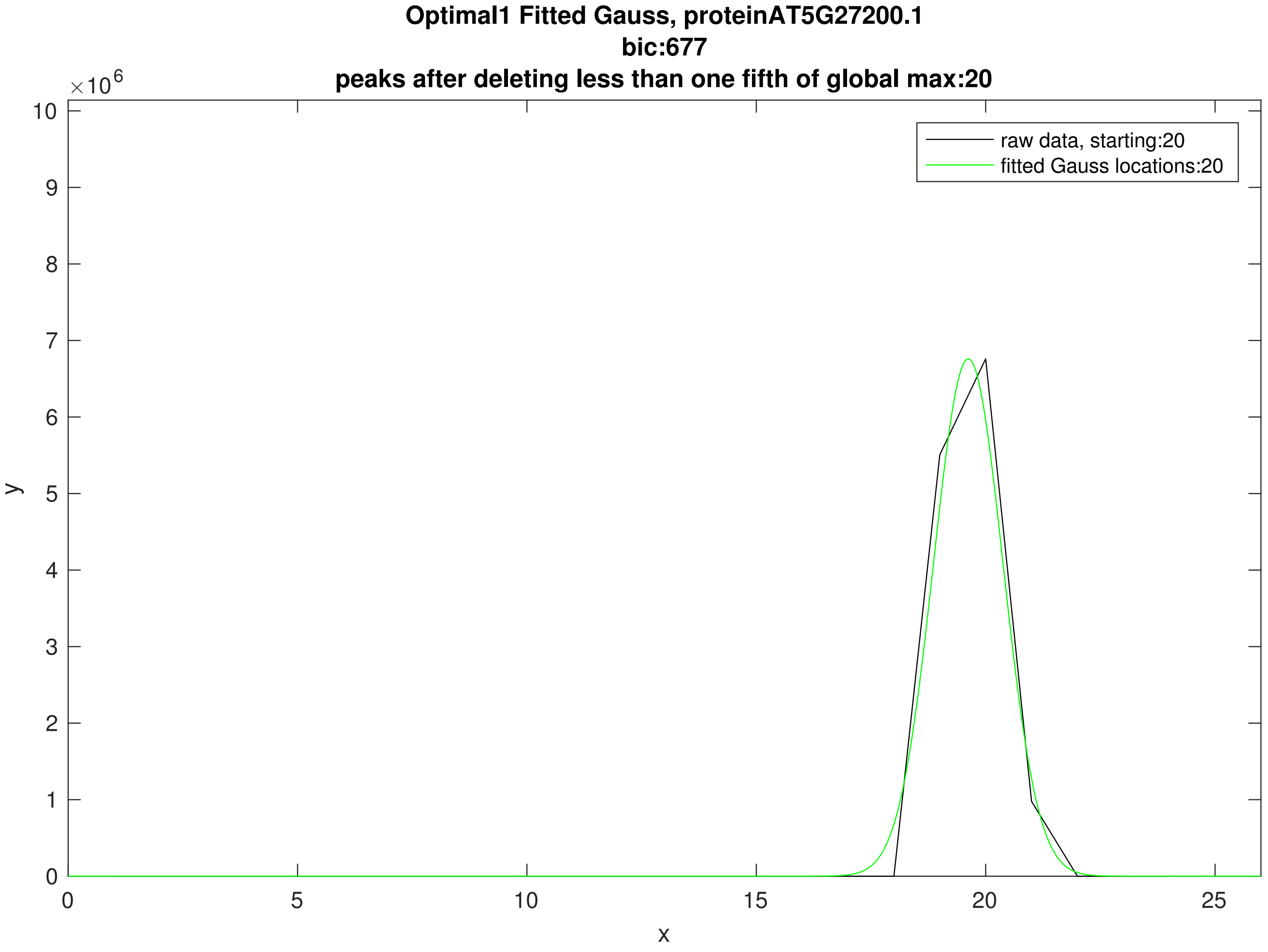
Optimal1 Fitted Gauss, proteinAT5G27200.1
bic:677
peaks after deleting less than one fifth of global max:20
×10^6
10
9
8
7
6
5
4
3
2
1
0
y
0
5
10
15
20
25
x
raw data, starting:20
fitted Gauss locations:20

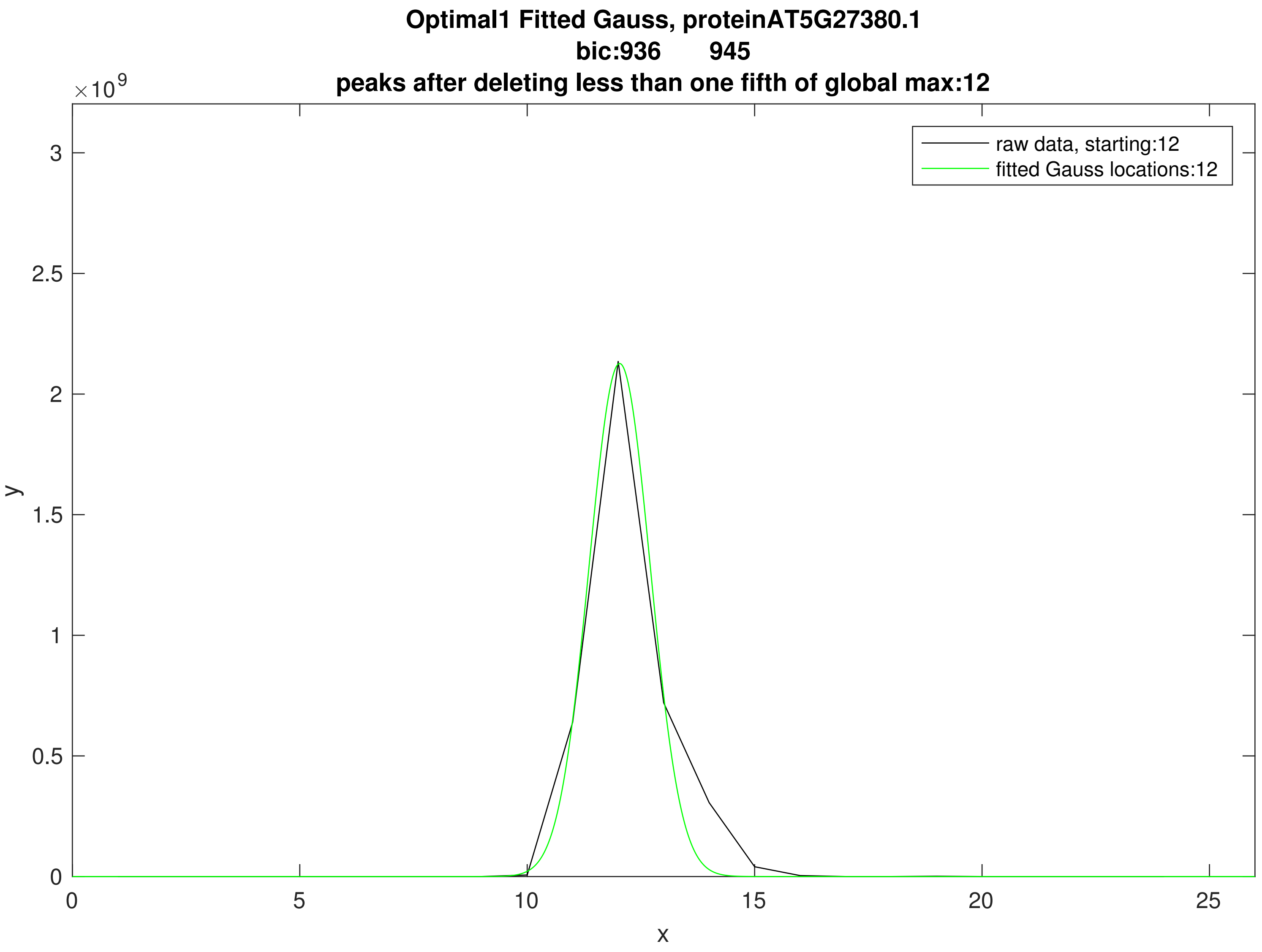

Optimal1 Fitted Gauss, proteinAT5G27380.1
bic:936 945
peaks after deleting less than one fifth of global max:12
×10^9
3
2.5
2
1.5
1
0.5
0
y
0
5
10
15
20
25
x
raw data, starting:12
fitted Gauss locations:12

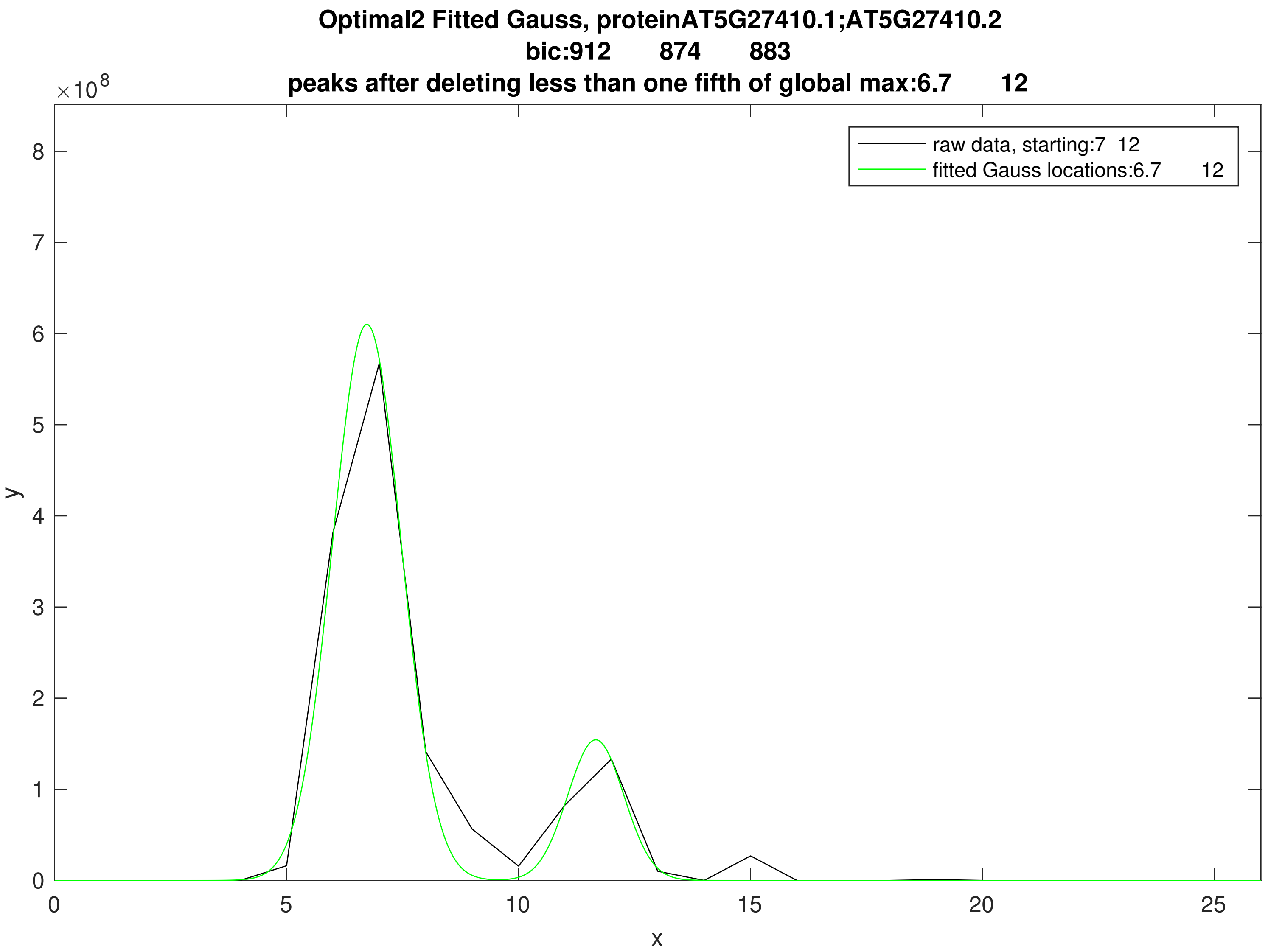

Optimal2 Fitted Gauss, proteinAT5G27410.1;AT5G27410.2
bic:912 874 883
peaks after deleting less than one fifth of global max:6.7 12
×10^8
raw data, starting:7 12
fitted Gauss locations:6.7 12
y
x
0
1
2
3
4
5
6
7
8
0
5
10
15
20
25

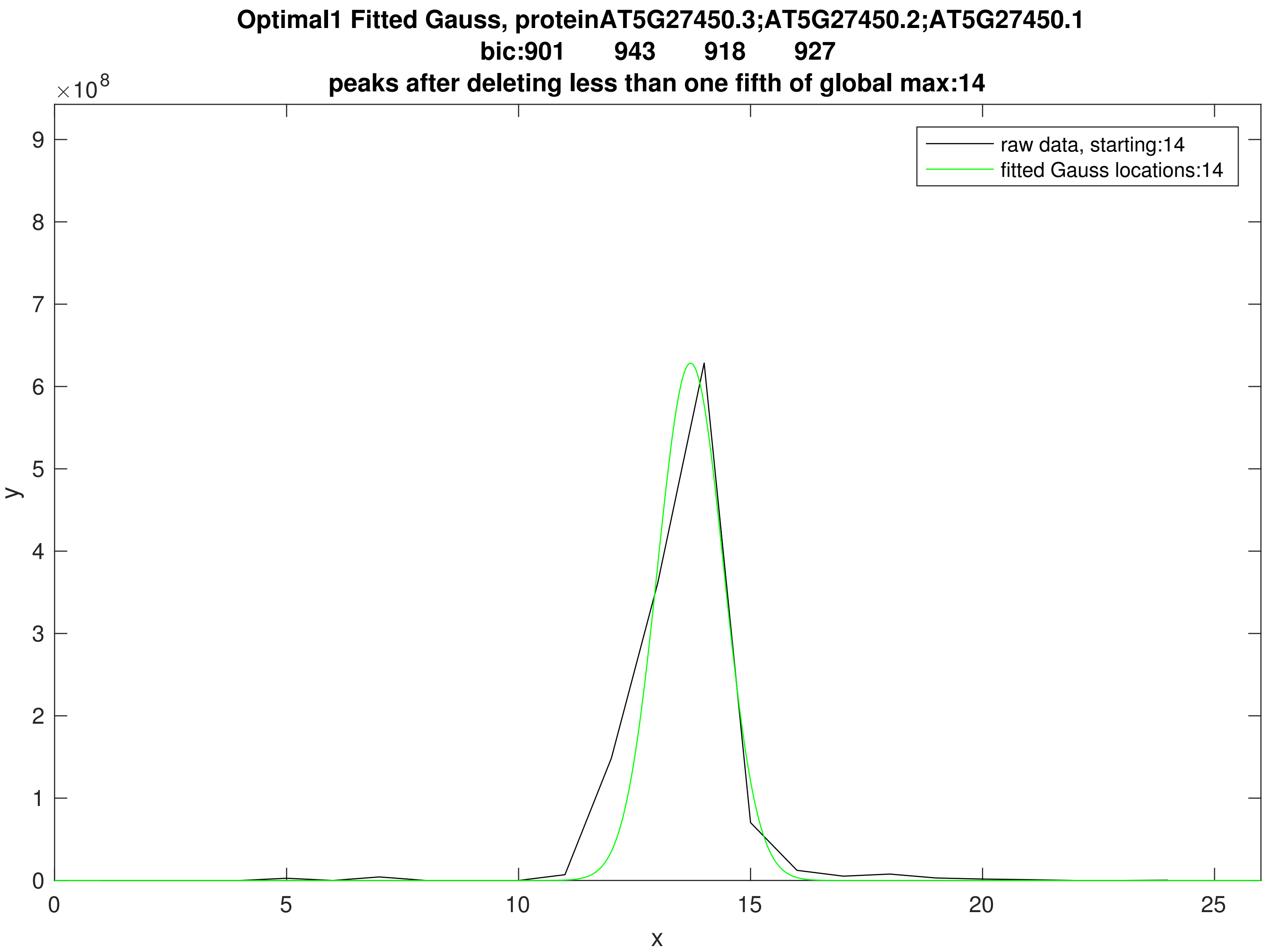
Optimal1 Fitted Gauss, proteinAT5G27450.3;AT5G27450.2;AT5G27450.1
bic:901 943 918 927
peaks after deleting less than one fifth of global max:14
×10^8
raw data, starting:14
fitted Gauss locations:14
y
x
0
1
2
3
4
5
6
7
8
9
0
5
10
15
20
25

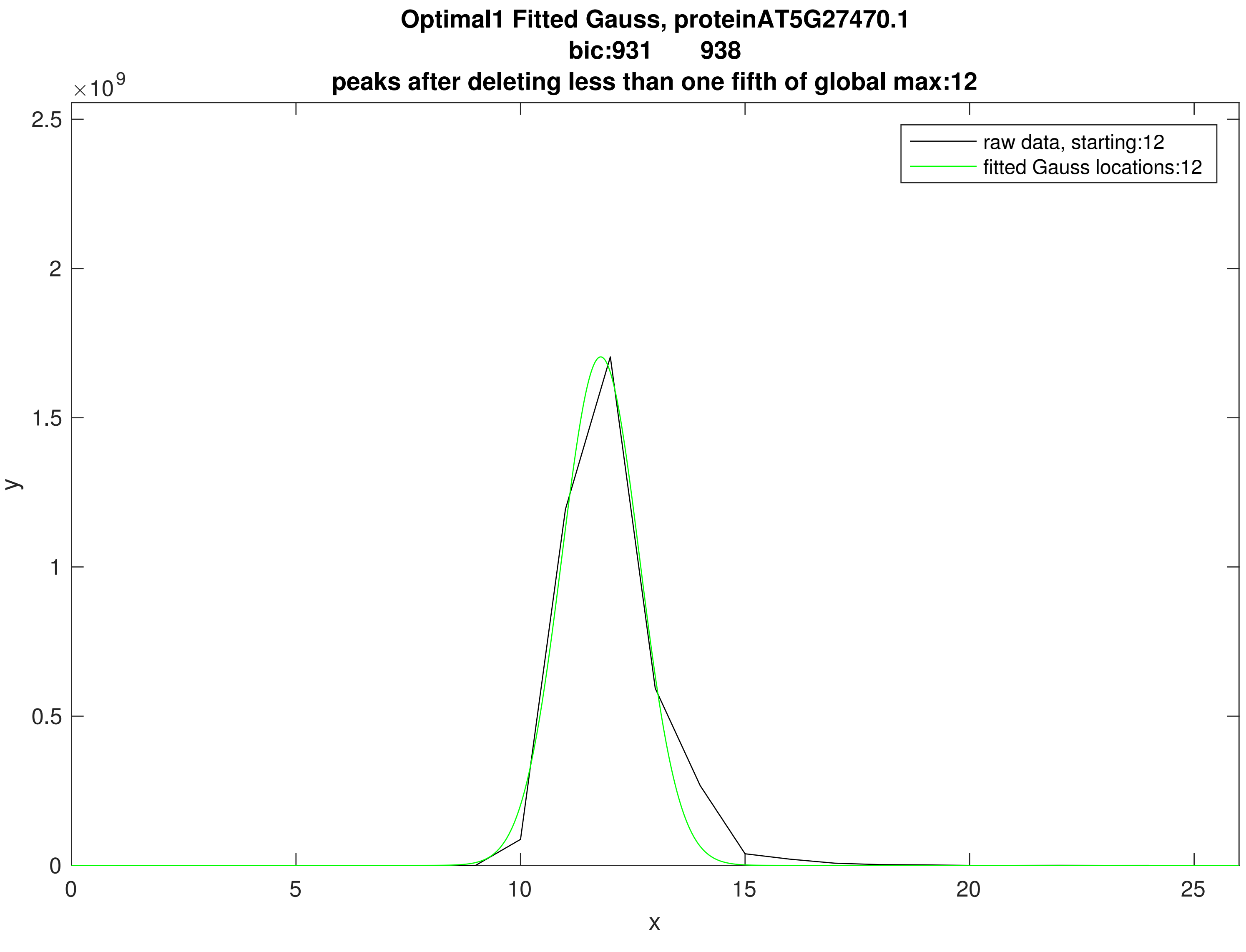
Optimal1 Fitted Gauss, proteinAT5G27470.1
bic:931 938
peaks after deleting less than one fifth of global max:12
×10^9
2.5
2
1.5
1
0.5
0
y
0
5
10
15
20
25
x
raw data, starting:12
fitted Gauss locations:12

**Optimal1 Fitted Gauss, proteinAT5G27640.1;AT5G27640.2**
**bic:693**
**peaks after deleting less than one fifth of global max:11**

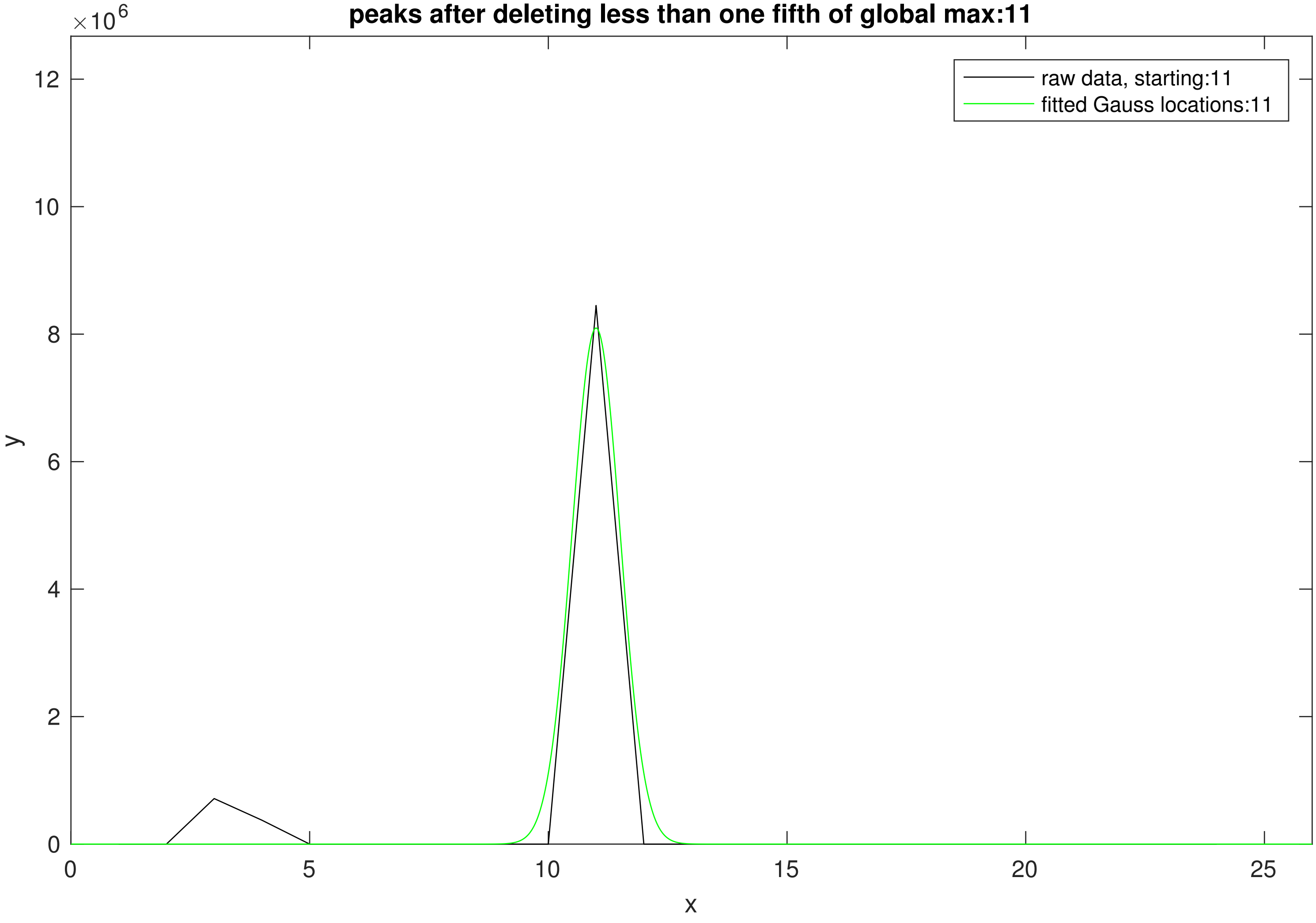

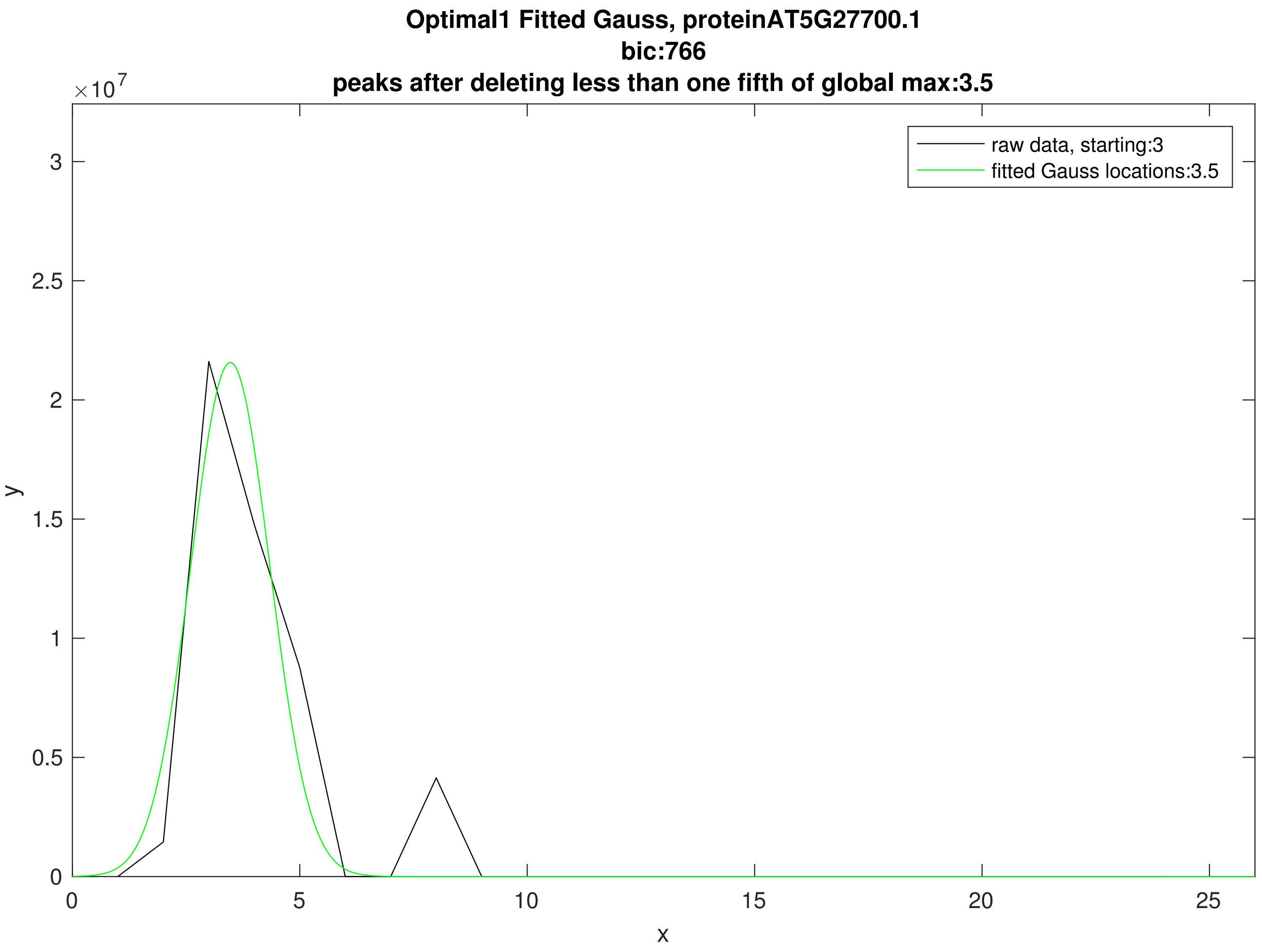
Optimal1 Fitted Gauss, proteinAT5G27700.1
bic:766
peaks after deleting less than one fifth of global max:3.5
×10^7
3
2.5
2
1.5
1
0.5
0
y
0
5
10
15
20
25
x
raw data, starting:3
fitted Gauss locations:3.5

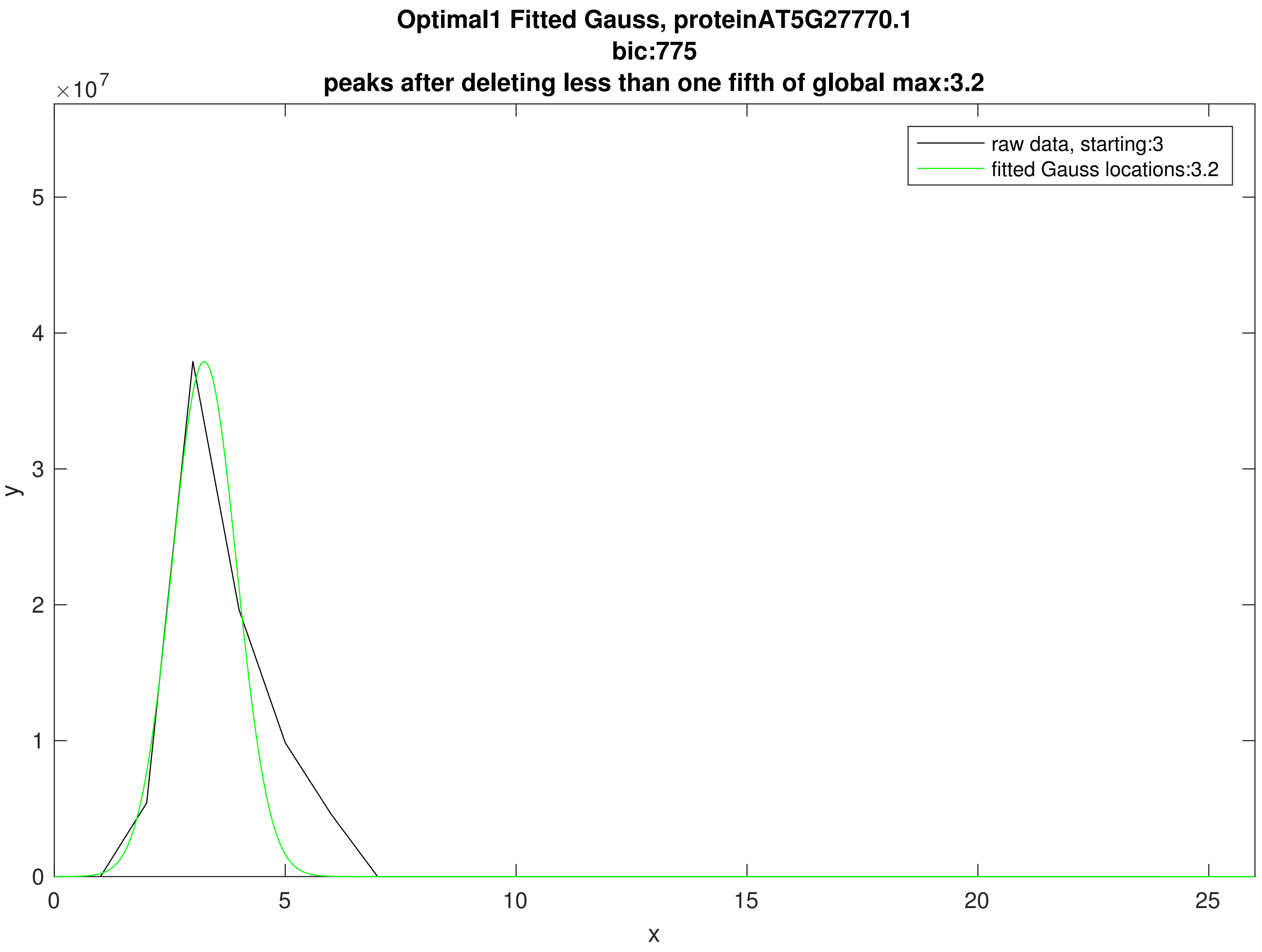
Optimal1 Fitted Gauss, proteinAT5G27770.1
bic:775
peaks after deleting less than one fifth of global max:3.2
×10^7
raw data, starting:3
fitted Gauss locations:3.2
y
x
0
1
2
3
4
5
0
5
10
15
20
25

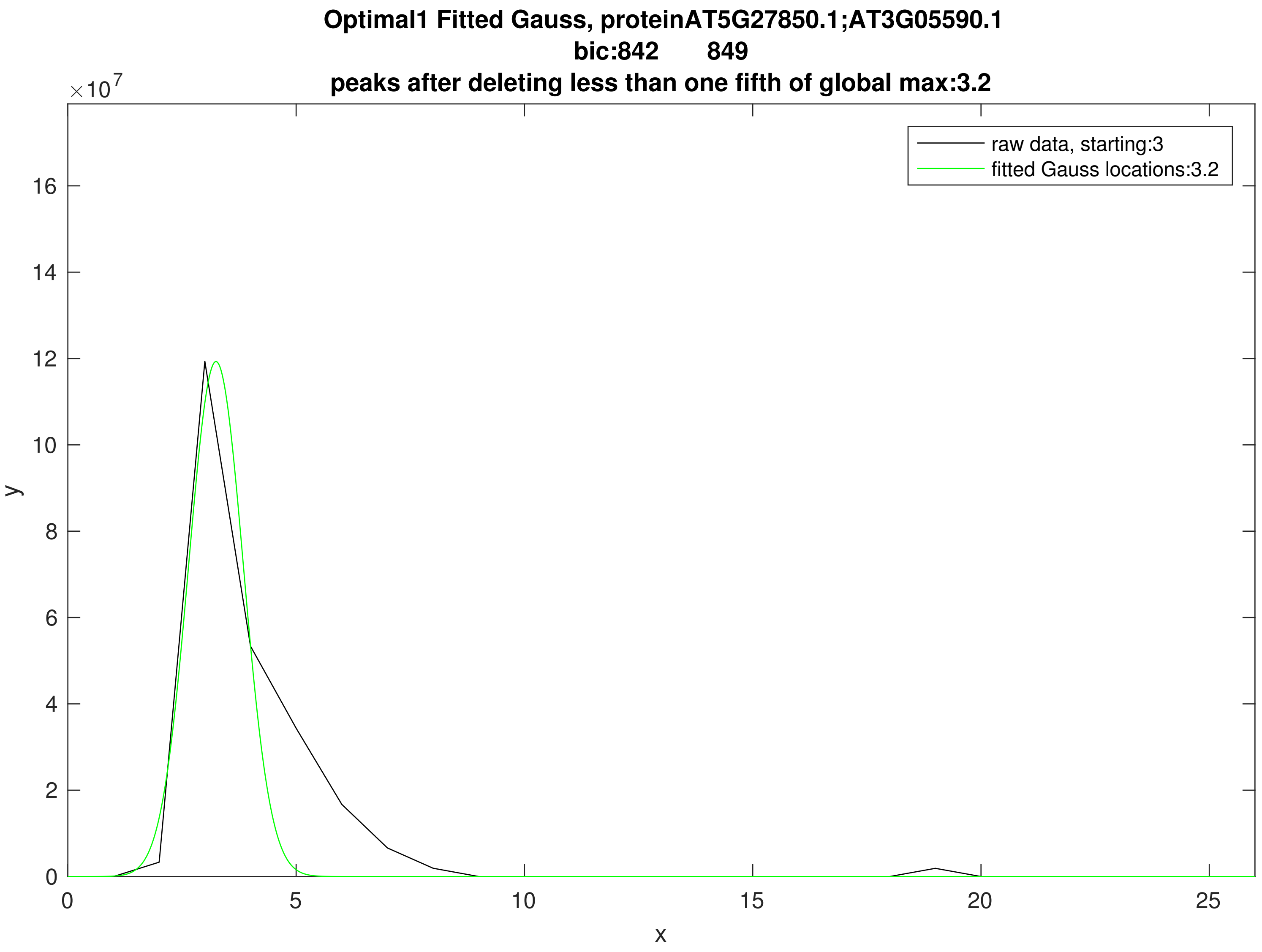
Optimal1 Fitted Gauss, proteinAT5G27850.1;AT3G05590.1
bic:842 849
peaks after deleting less than one fifth of global max:3.2
×10^7
raw data, starting:3
fitted Gauss locations:3.2
y
x
0
2
4
6
8
10
12
14
16
0
5
10
15
20
25

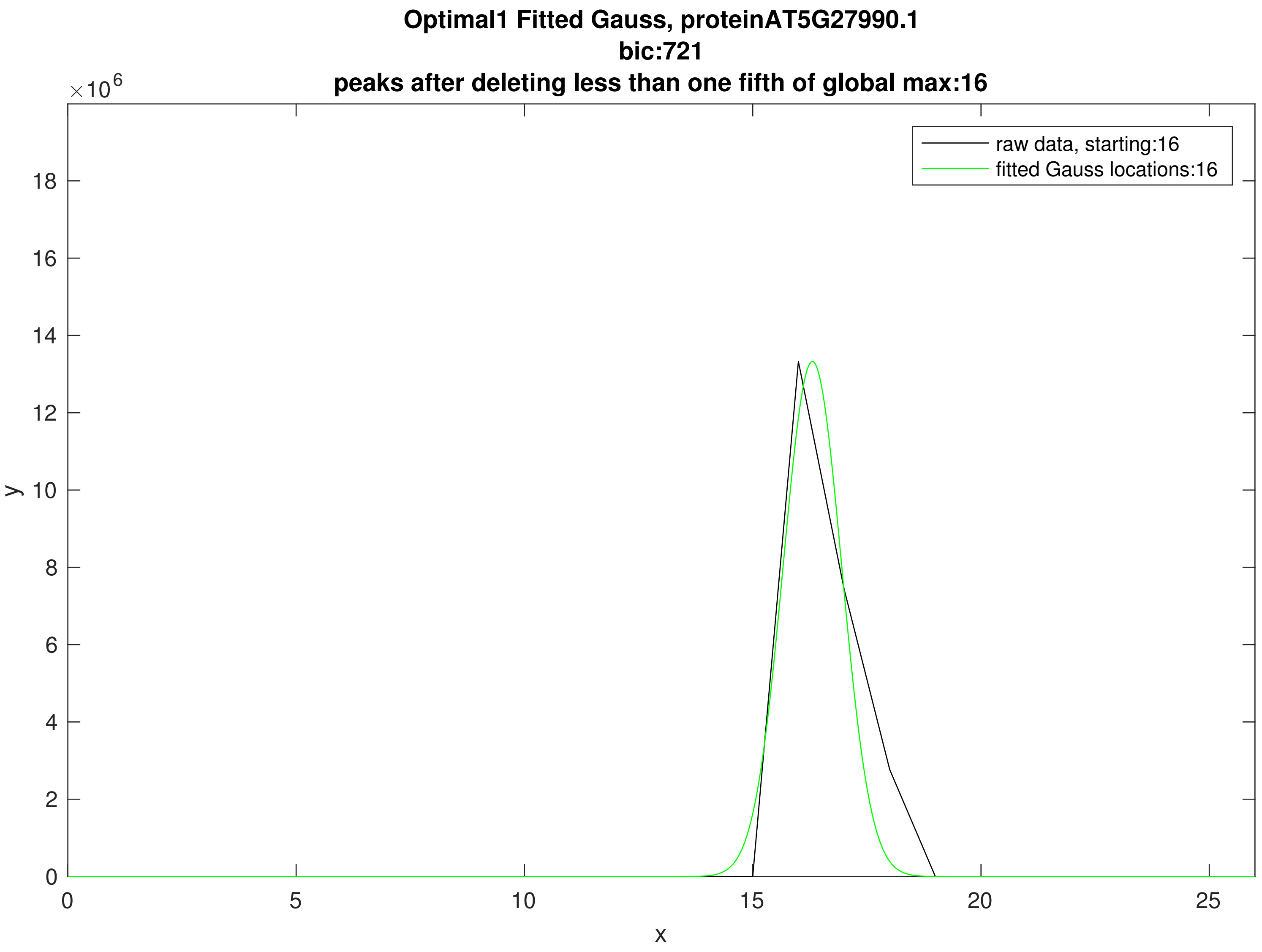

Optimal1 Fitted Gauss, proteinAT5G27990.1
bic:721
peaks after deleting less than one fifth of global max:16
×10^6
raw data, starting:16
fitted Gauss locations:16
y
x
0
2
4
6
8
10
12
14
16
18
0
5
10
15
20
25

**Optimal2 Fitted Gauss, proteinAT5G28020.6;AT5G28020.4;AT5G28020.3;AT5G28020.2;AT5G28020.1;AT5G28020**
**bic:945 937**
**peaks after deleting less than one fifth of global max:15**

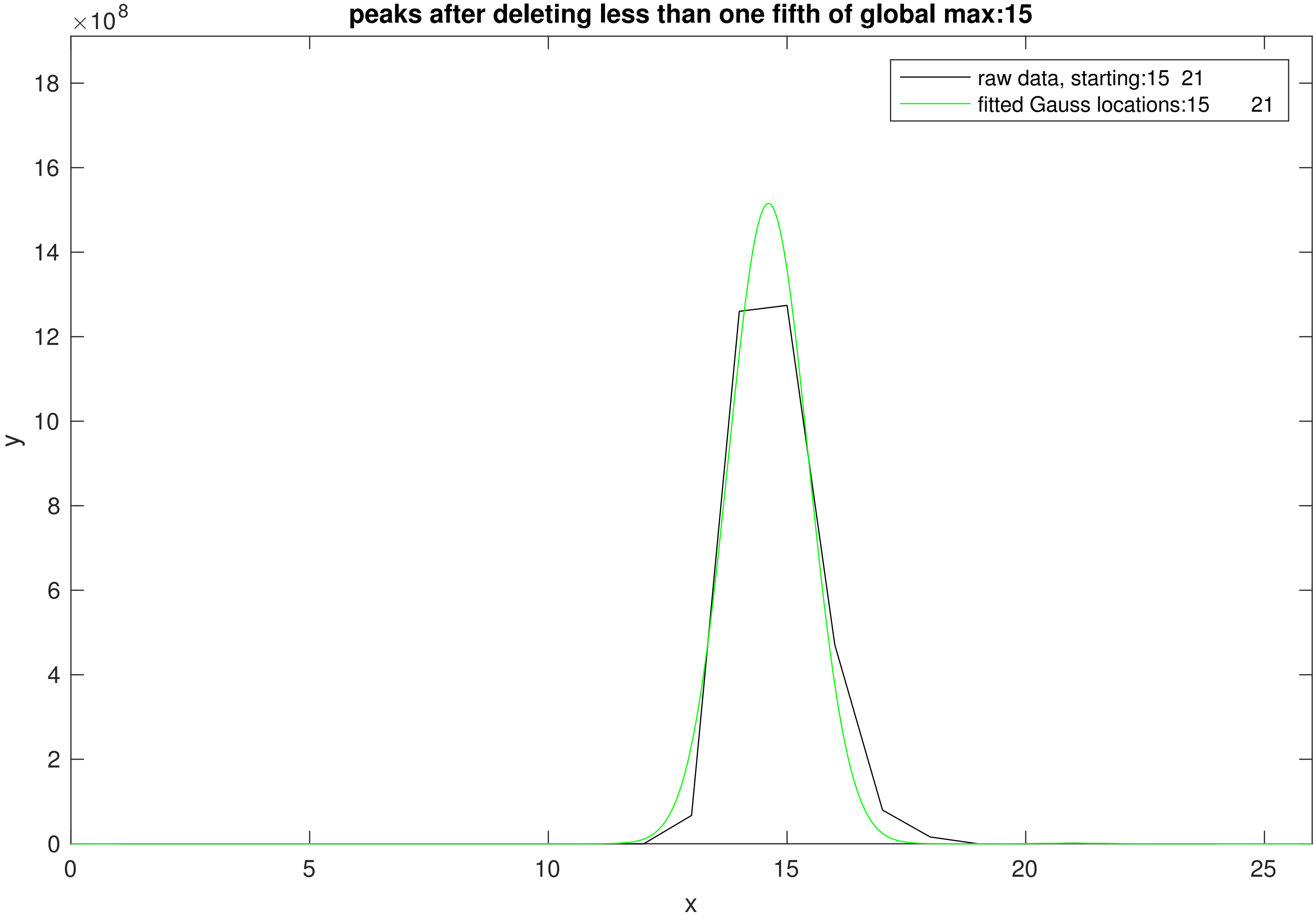

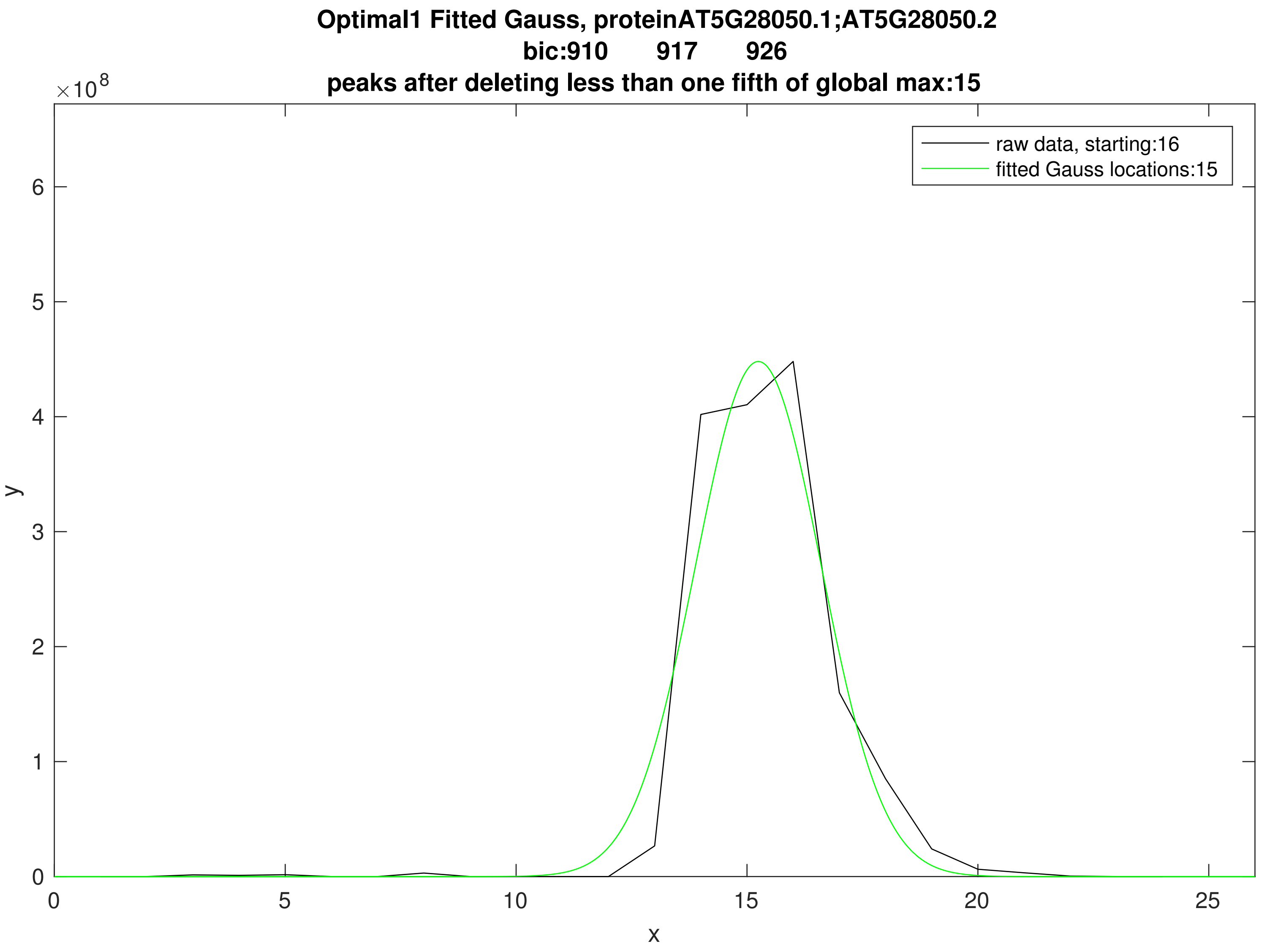
Optimal1 Fitted Gauss, proteinAT5G28050.1;AT5G28050.2
bic:910 917 926
peaks after deleting less than one fifth of global max:15
×10^8
raw data, starting:16
fitted Gauss locations:15
y
x
0
1
2
3
4
5
6
0
5
10
15
20
25

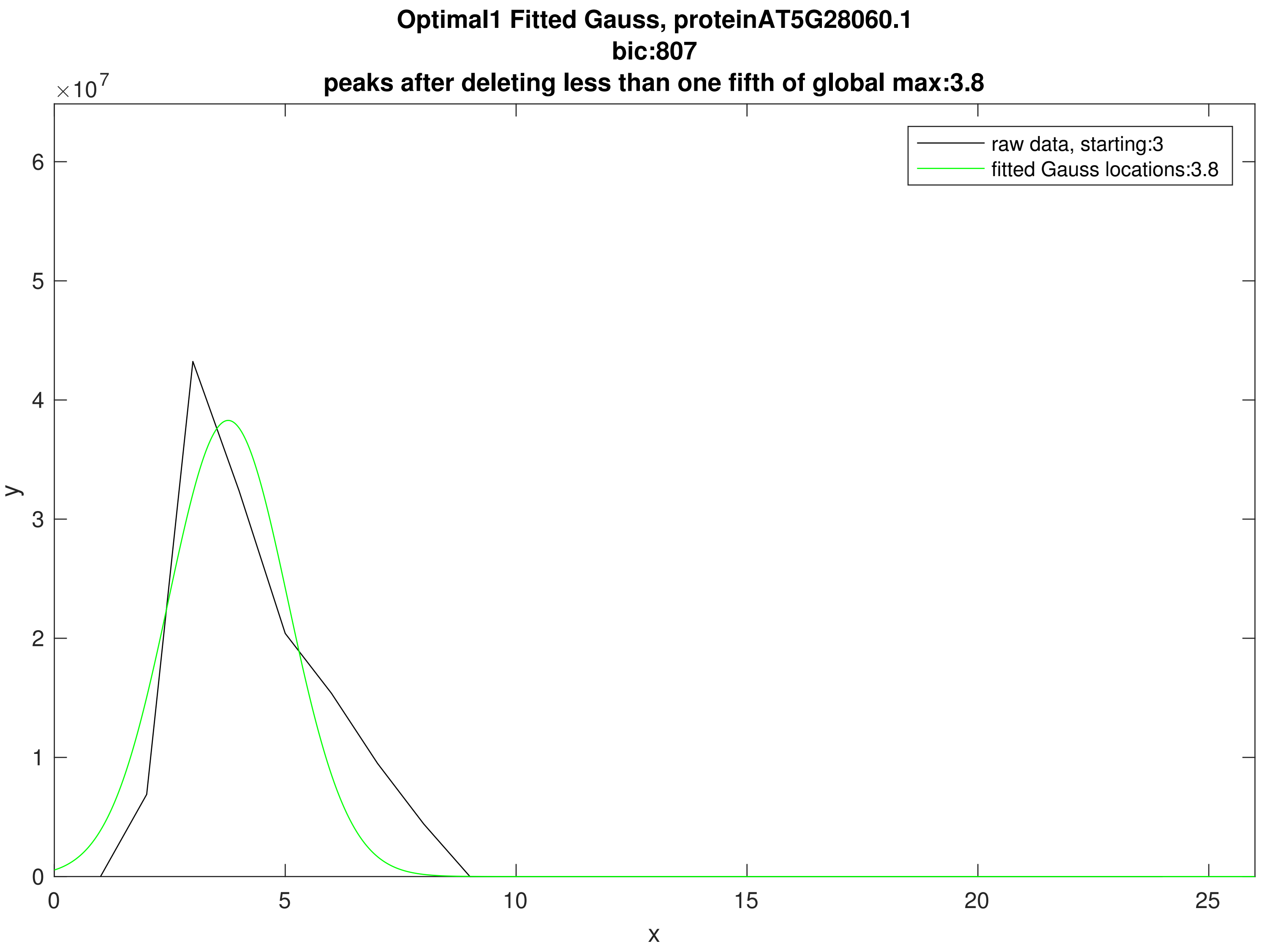
Optimal1 Fitted Gauss, proteinAT5G28060.1
bic:807
peaks after deleting less than one fifth of global max:3.8
×10^7
raw data, starting:3
fitted Gauss locations:3.8
y
x
0
1
2
3
4
5
6
0
5
10
15
20
25

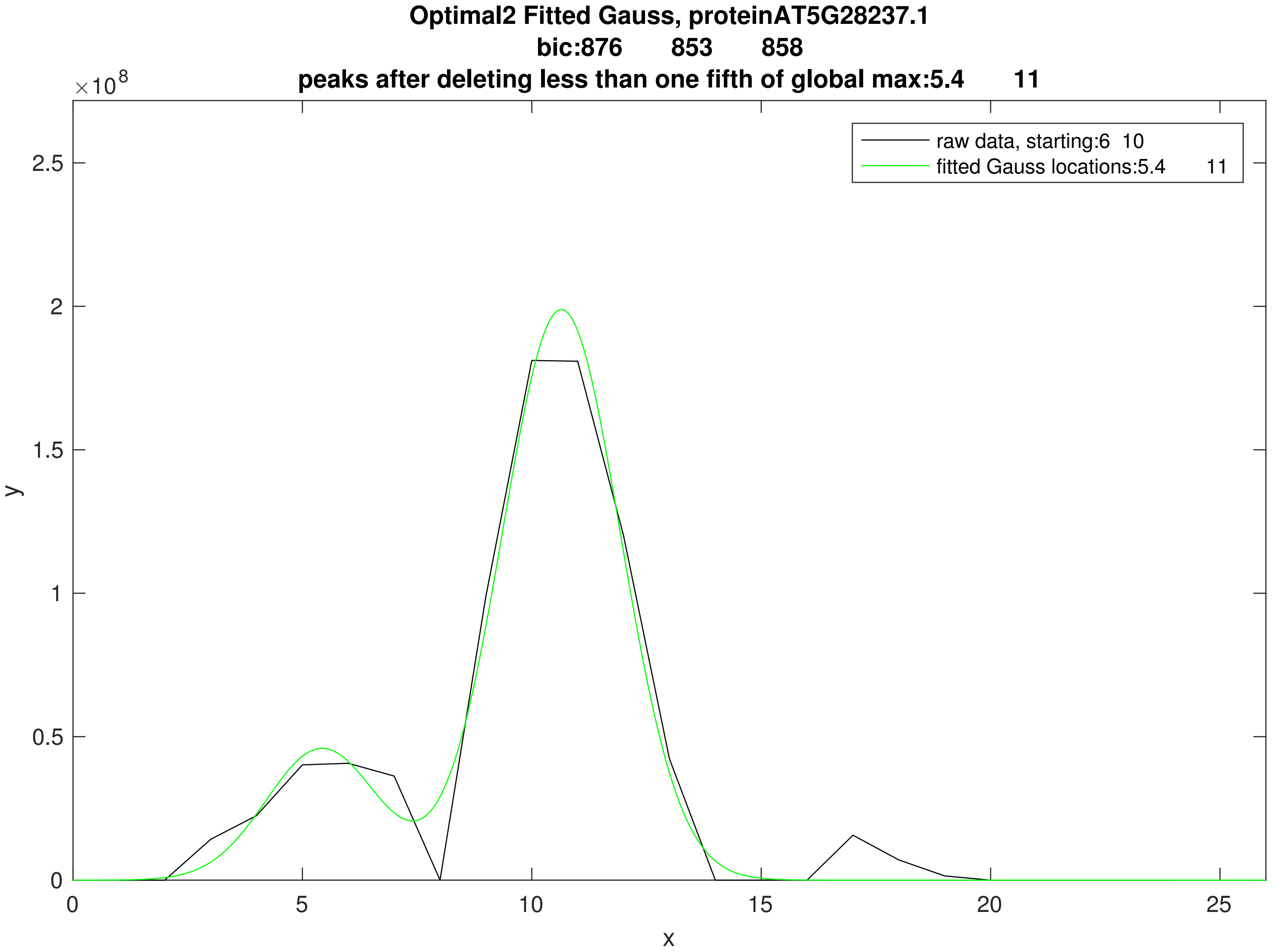

Optimal2 Fitted Gauss, proteinAT5G28237.1
bic:876 853 858
peaks after deleting less than one fifth of global max:5.4 11
×10^8
raw data, starting:6 10
fitted Gauss locations:5.4 11
y
x
0
0.5
1
1.5
2
2.5
0
5
10
15
20
25

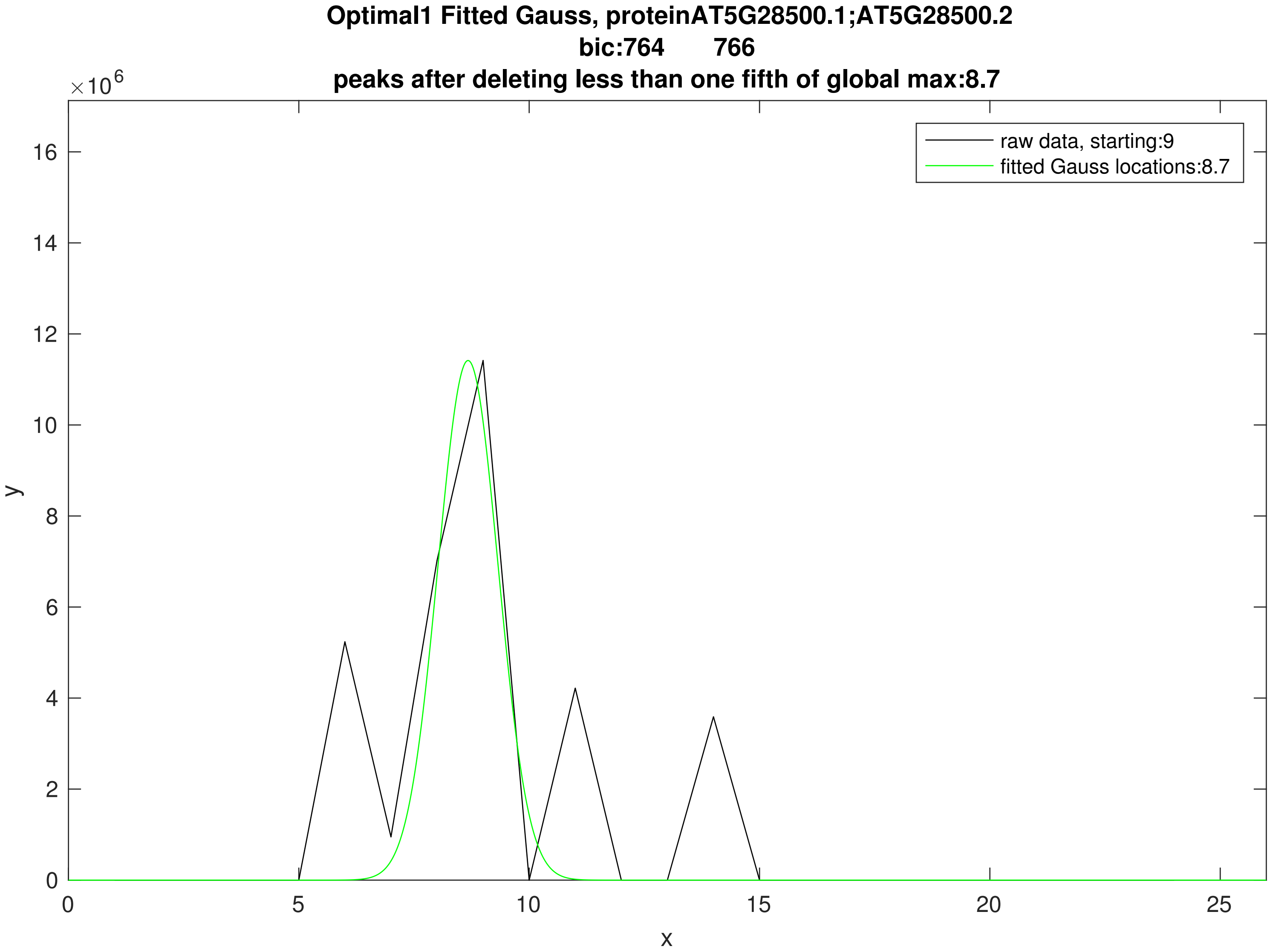
Optimal1 Fitted Gauss, proteinAT5G28500.1;AT5G28500.2
bic:764 766
peaks after deleting less than one fifth of global max:8.7
×10^6
16
14
12
10
8
6
4
2
0
y
0
5
10
15
20
25
x
raw data, starting:9
fitted Gauss locations:8.7

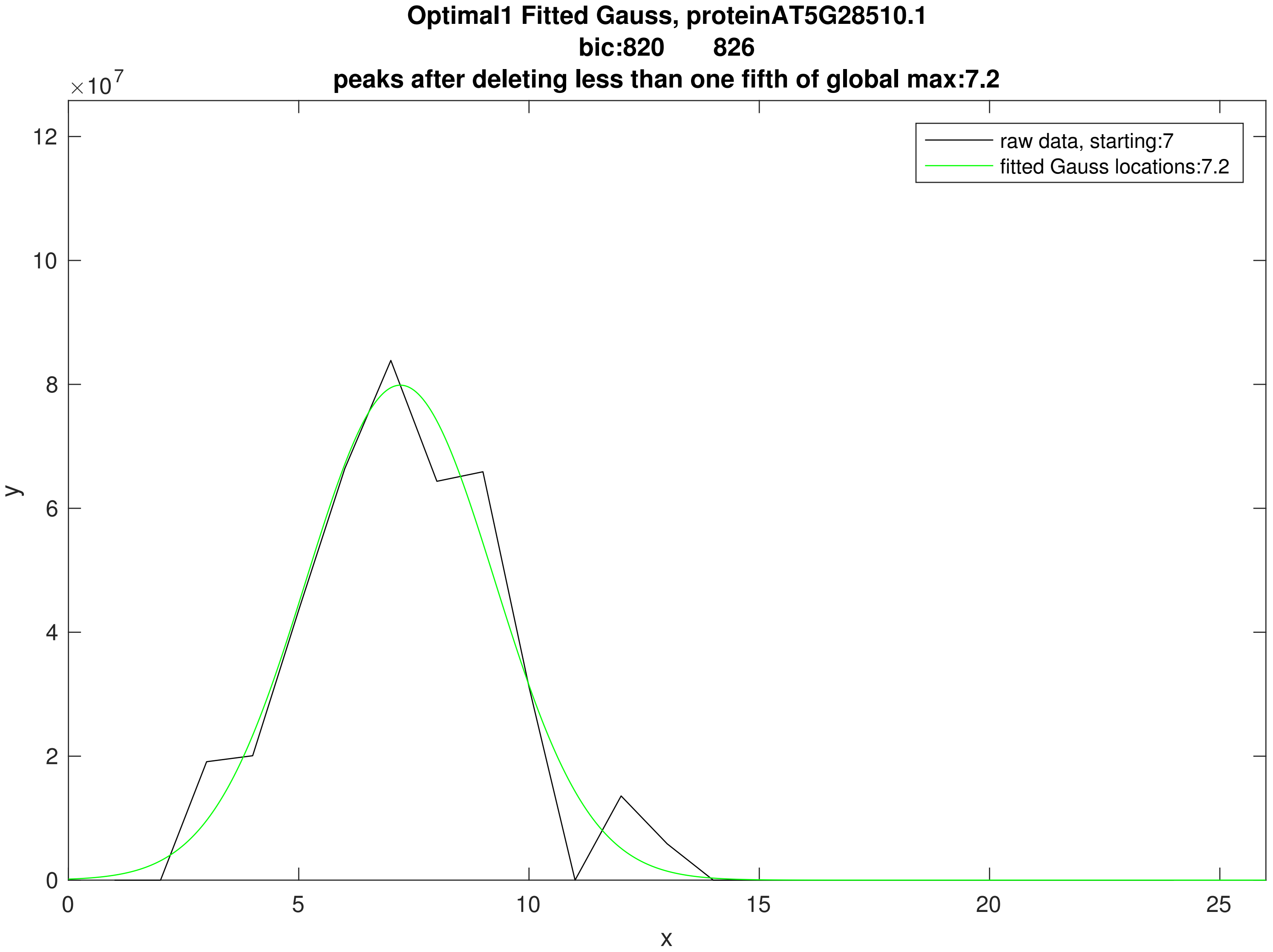

Optimal1 Fitted Gauss, proteinAT5G28510.1
bic:820 826
peaks after deleting less than one fifth of global max:7.2
×10^7
raw data, starting:7
fitted Gauss locations:7.2
y
x
0
2
4
6
8
10
12
0
5
10
15
20
25

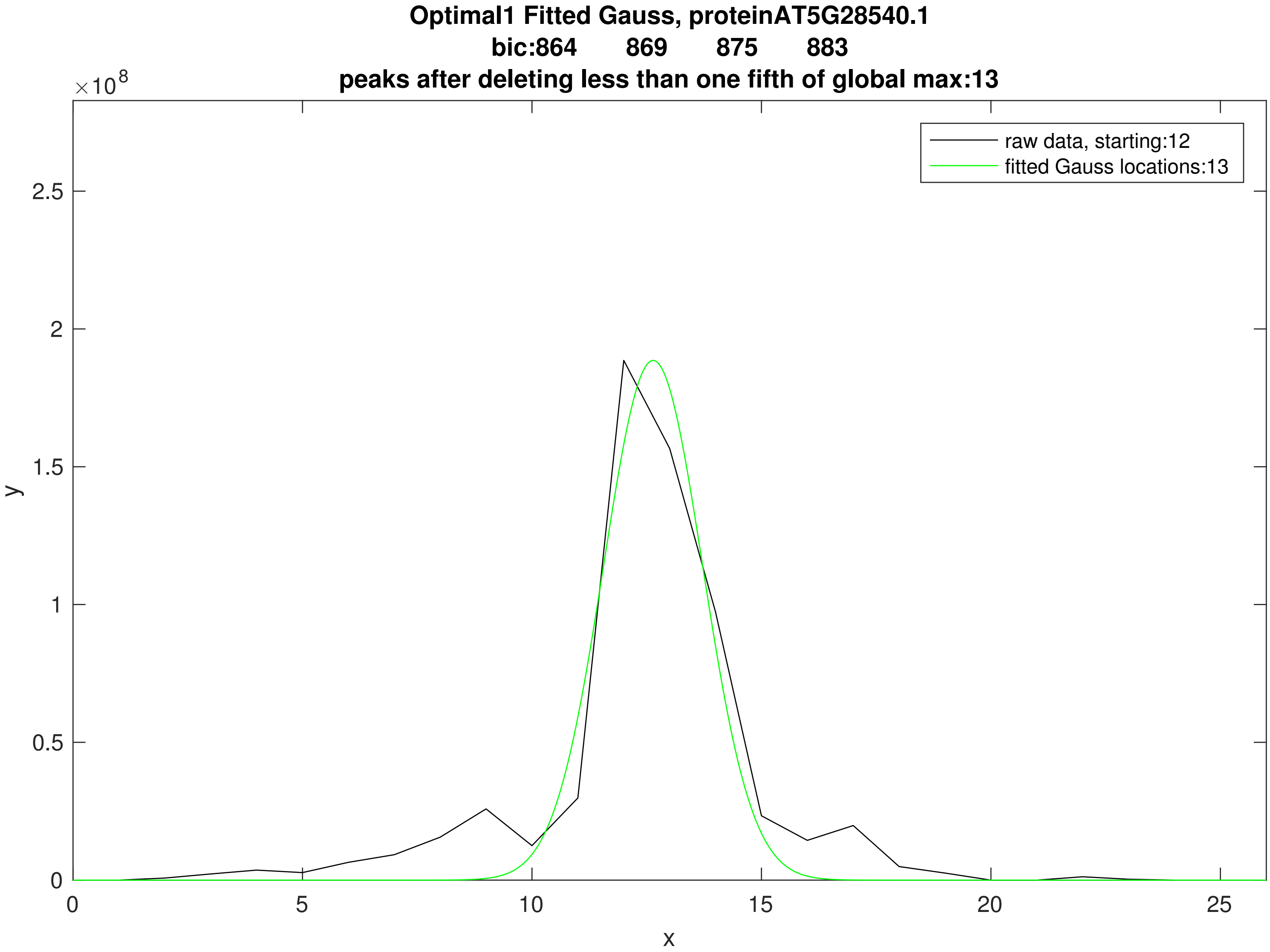

Optimal1 Fitted Gauss, proteinAT5G28540.1
bic:864 869 875 883
peaks after deleting less than one fifth of global max:13
×10^8
2.5
2
1.5
1
0.5
0
y
0
5
10
15
20
25
x
raw data, starting:12
fitted Gauss locations:13

Optimal1 Fitted Gauss, proteinAT5G28830.1
bic:500
peaks after deleting less than one fifth of global max:16

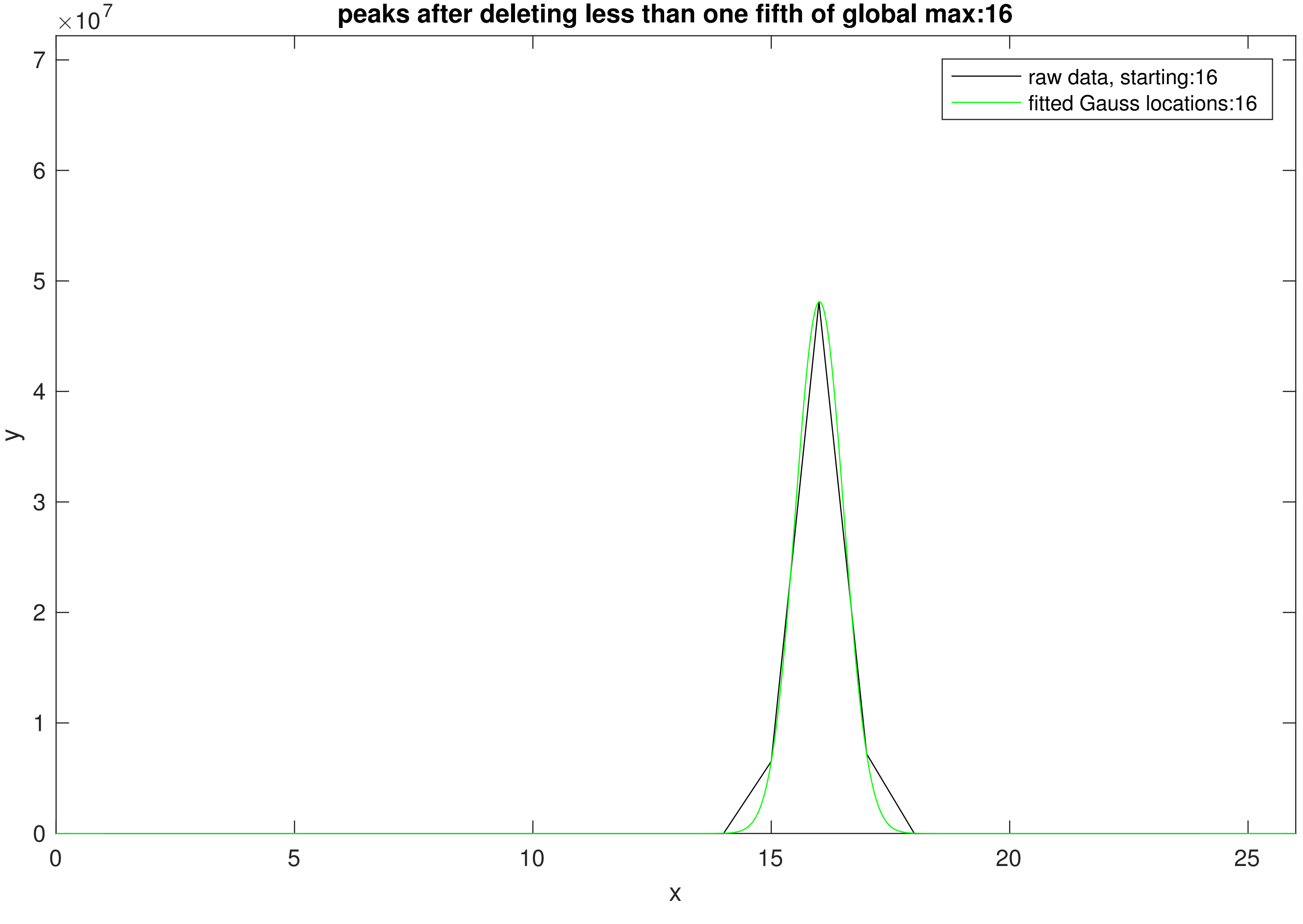

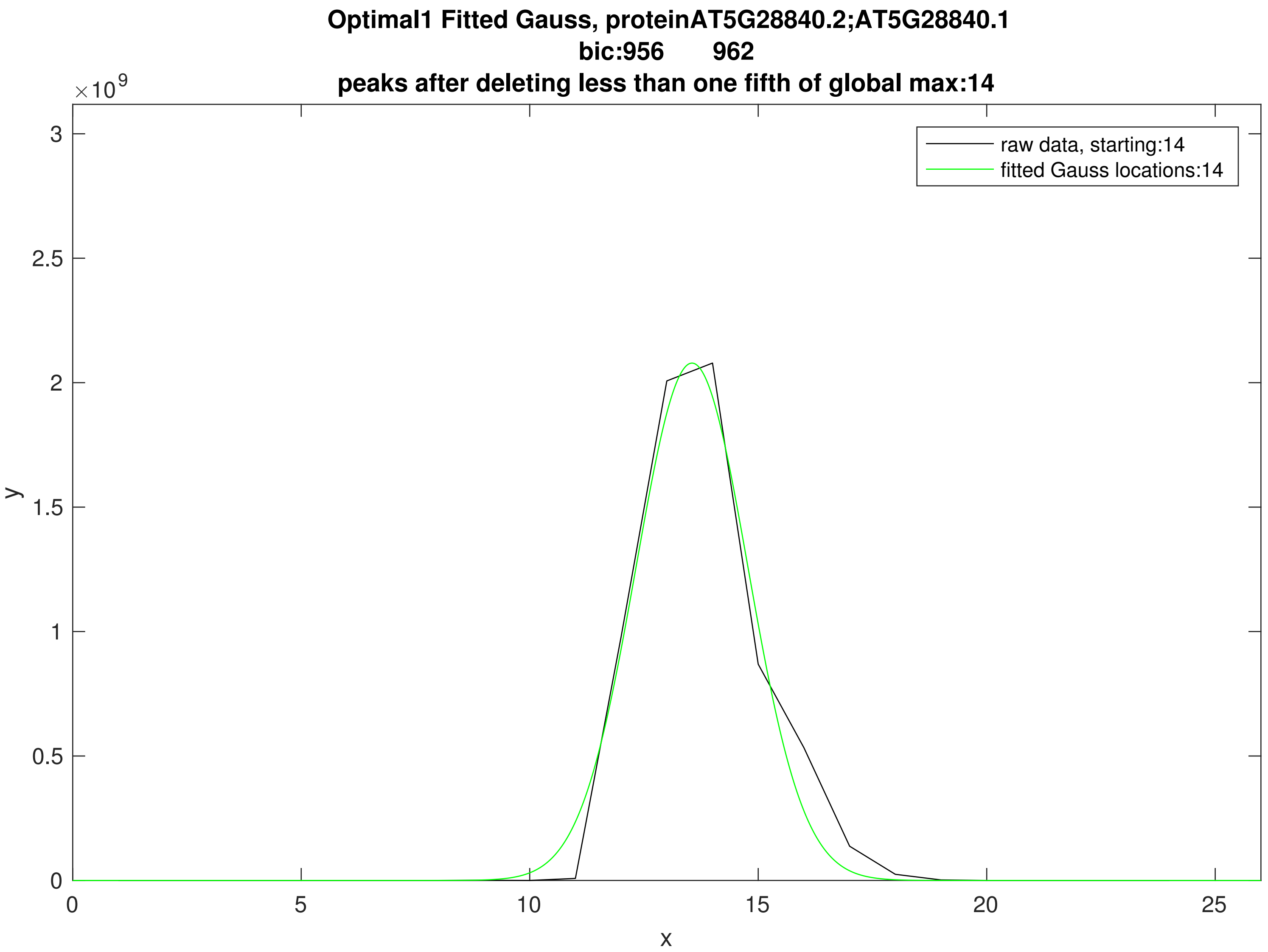
Optimal1 Fitted Gauss, proteinAT5G28840.2;AT5G28840.1
bic:956 962
peaks after deleting less than one fifth of global max:14
×10^9
raw data, starting:14
fitted Gauss locations:14
y
x
0
0.5
1
1.5
2
2.5
3
0
5
10
15
20
25

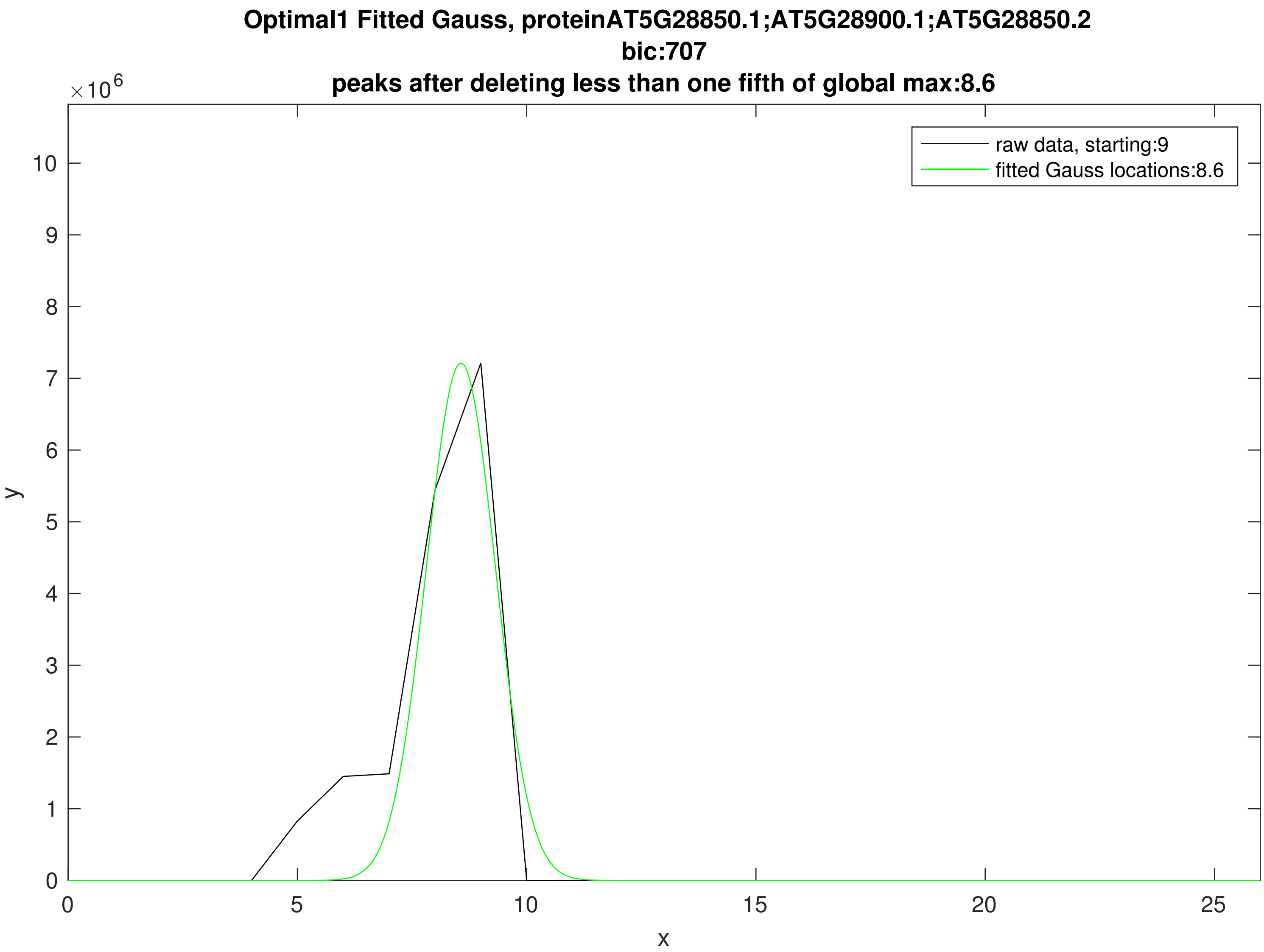
Optimal1 Fitted Gauss, proteinAT5G28850.1;AT5G28900.1;AT5G28850.2
bic:707
peaks after deleting less than one fifth of global max:8.6
×10^6
raw data, starting:9
fitted Gauss locations:8.6
y
x
0
1
2
3
4
5
6
7
8
9
10
15
20
25

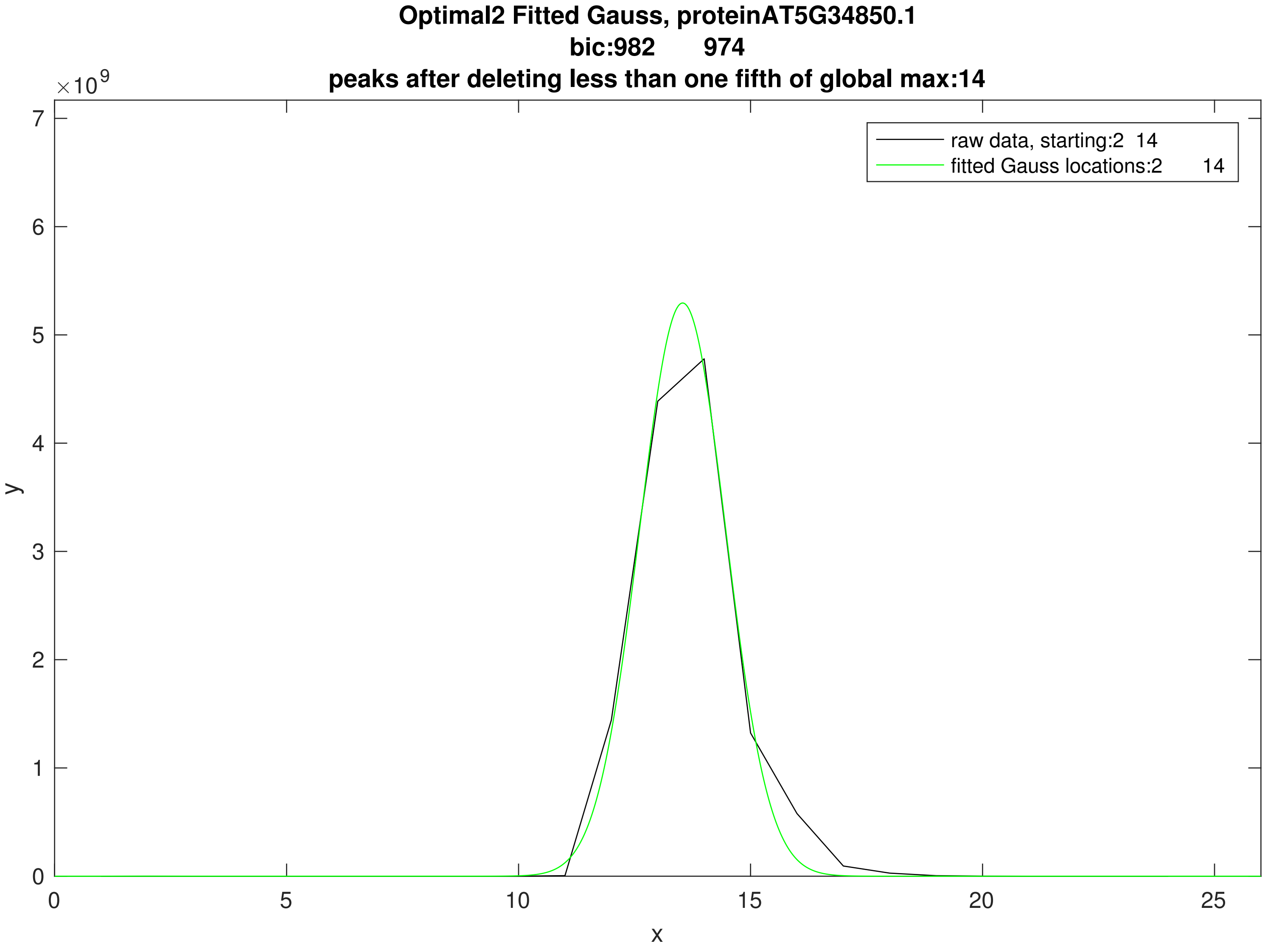
Optimal2 Fitted Gauss, proteinAT5G34850.1
bic:982 974
peaks after deleting less than one fifth of global max:14
×10^9
raw data, starting:2 14
fitted Gauss locations:2 14
y
x
0
1
2
3
4
5
6
7
0
5
10
15
20
25

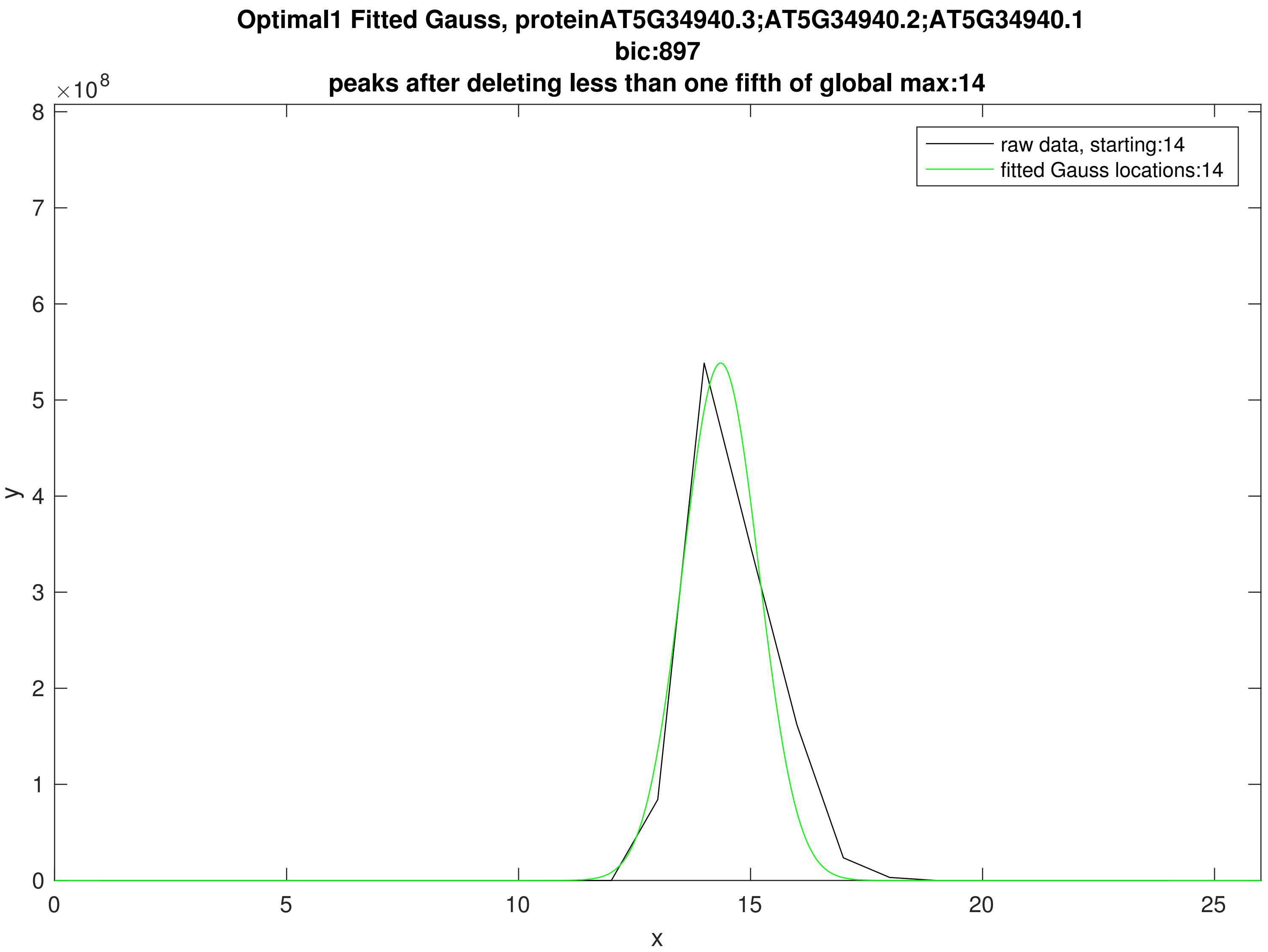

Optimal1 Fitted Gauss, proteinAT5G34940.3;AT5G34940.2;AT5G34940.1
bic:897
peaks after deleting less than one fifth of global max:14
×10^8
raw data, starting:14
fitted Gauss locations:14
y
x
0
1
2
3
4
5
6
7
8
0
5
10
15
20
25

#### Optimal1 Fitted Gauss, proteinAT5G35360.3;AT5G35360.1;AT5G35360.2
#### bic:880 928 939 948
#### peaks after deleting less than one fifth of global max:13

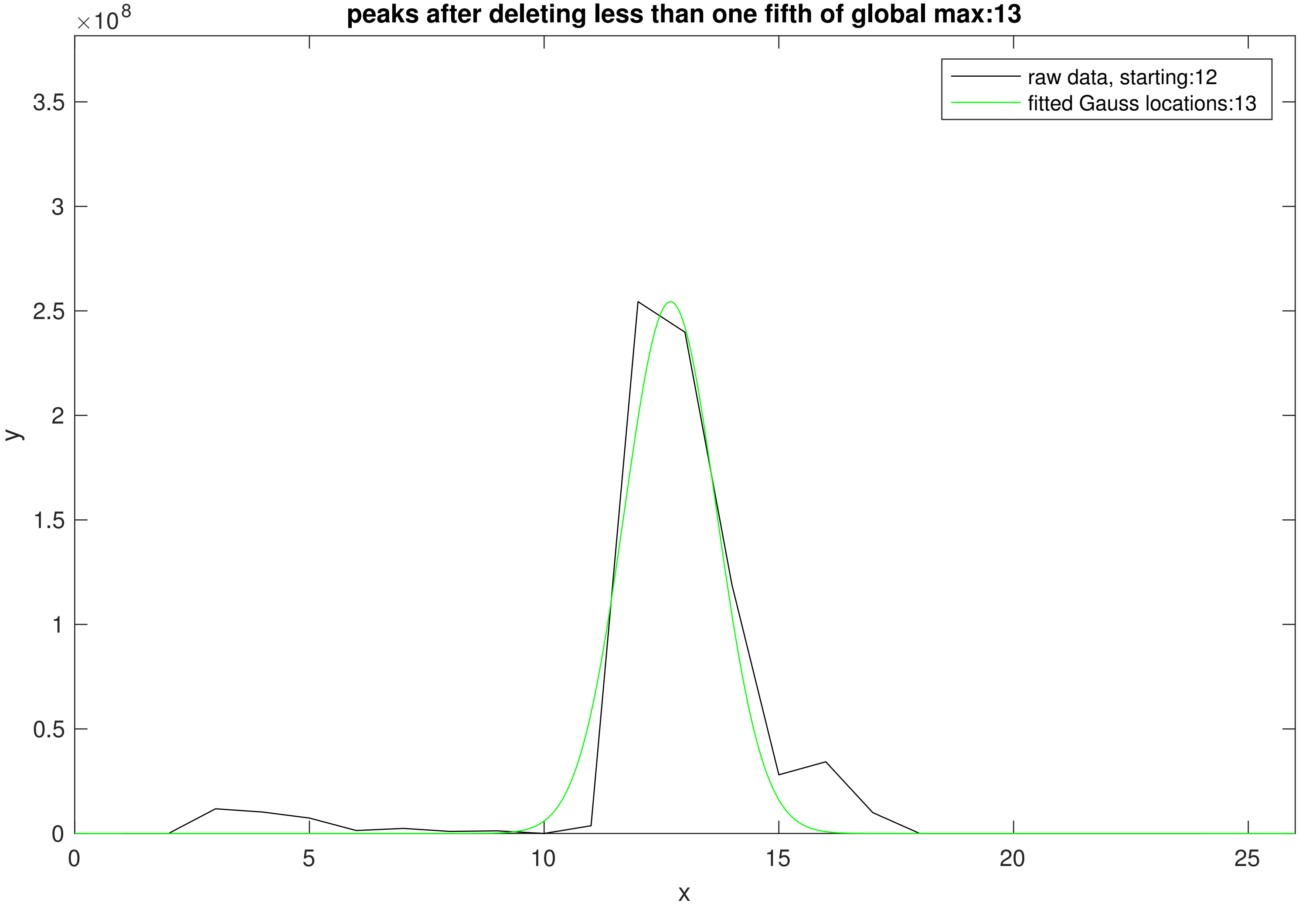

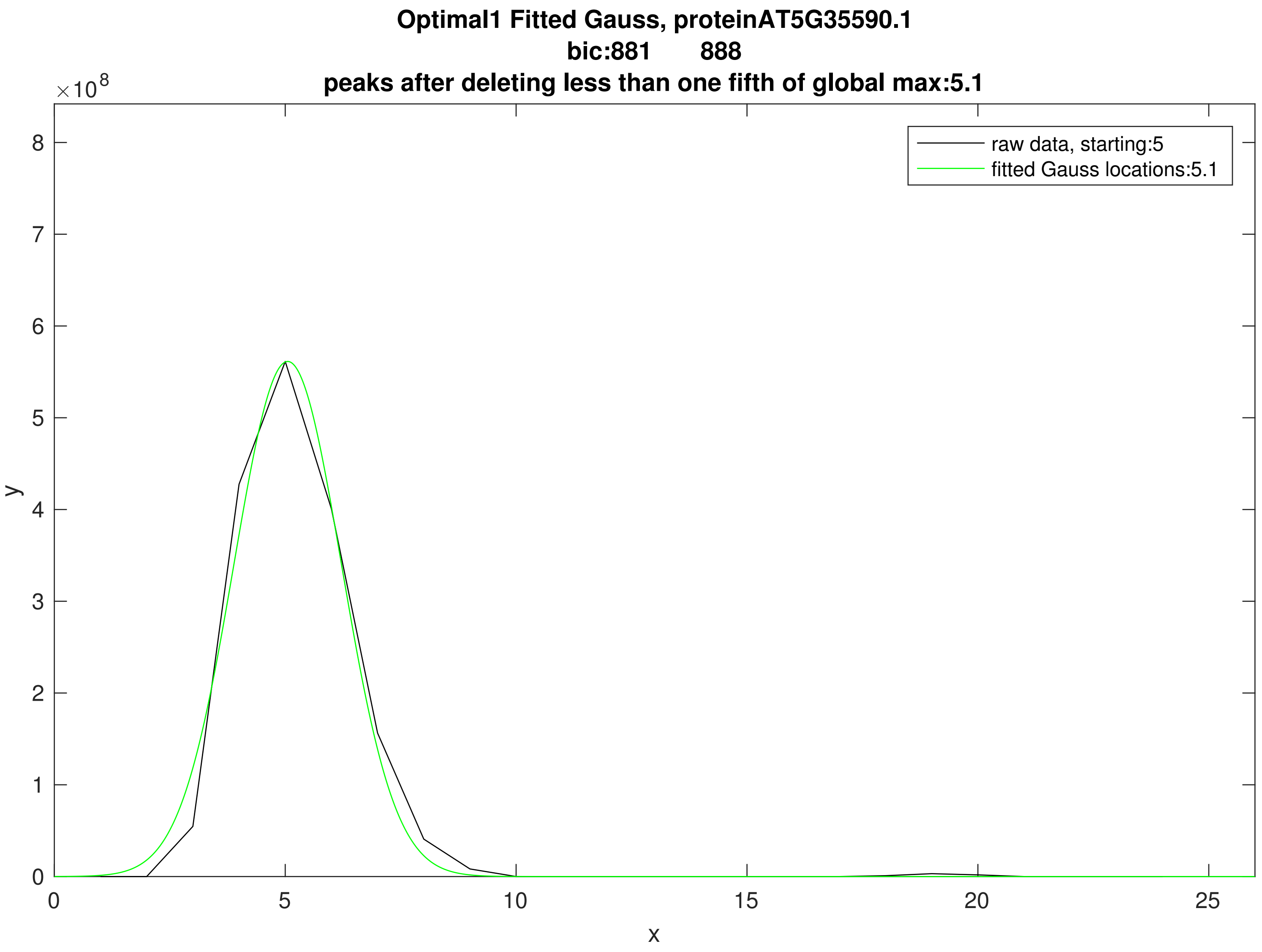
Optimal1 Fitted Gauss, proteinAT5G35590.1
bic:881 888
peaks after deleting less than one fifth of global max:5.1
×10^8
raw data, starting:5
fitted Gauss locations:5.1
y
x
0
1
2
3
4
5
6
7
8
0
5
10
15
20
25

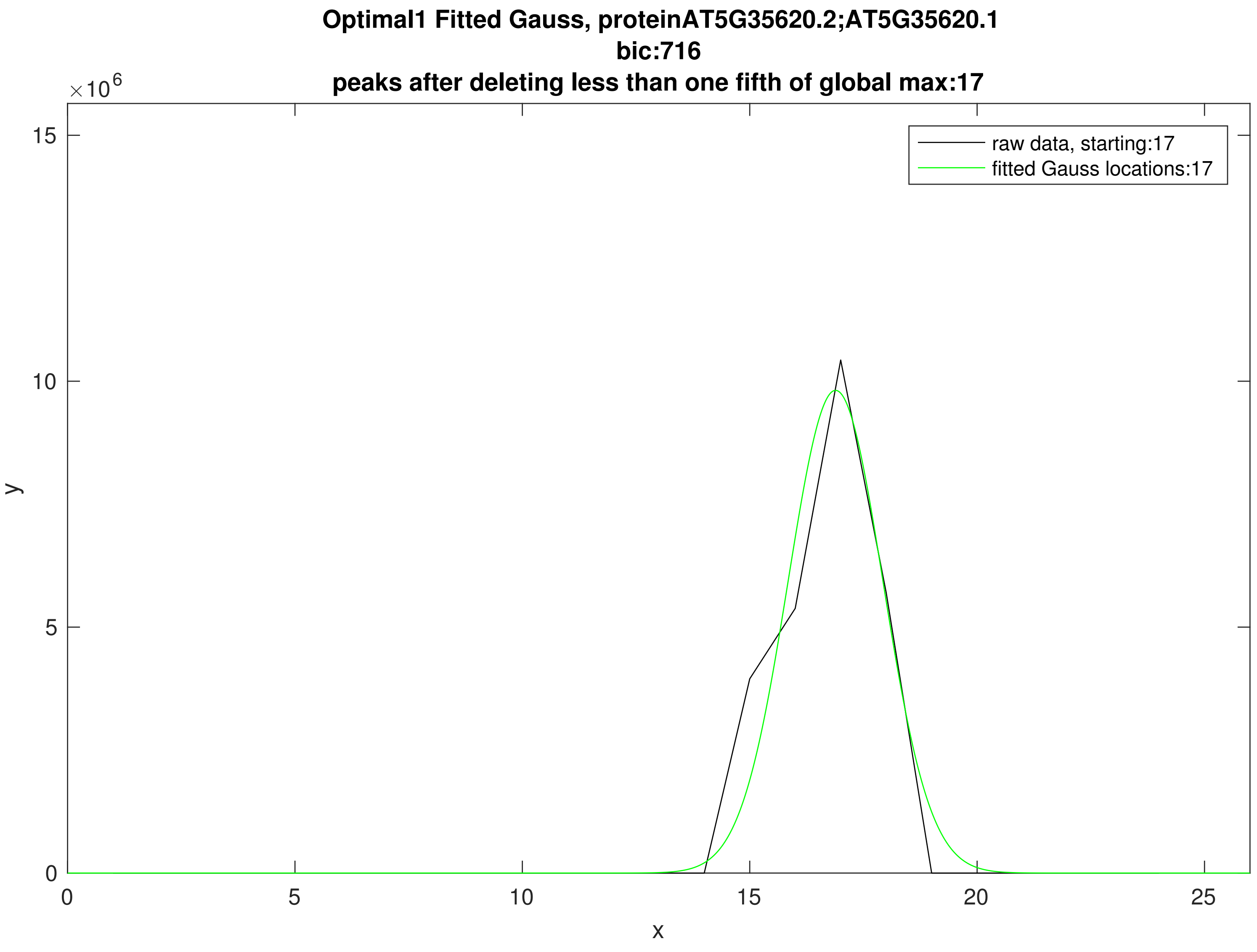

Optimal1 Fitted Gauss, proteinAT5G35620.2;AT5G35620.1
bic:716
peaks after deleting less than one fifth of global max:17
×10^6
raw data, starting:17
fitted Gauss locations:17
y
x
0
5
10
15
20
25

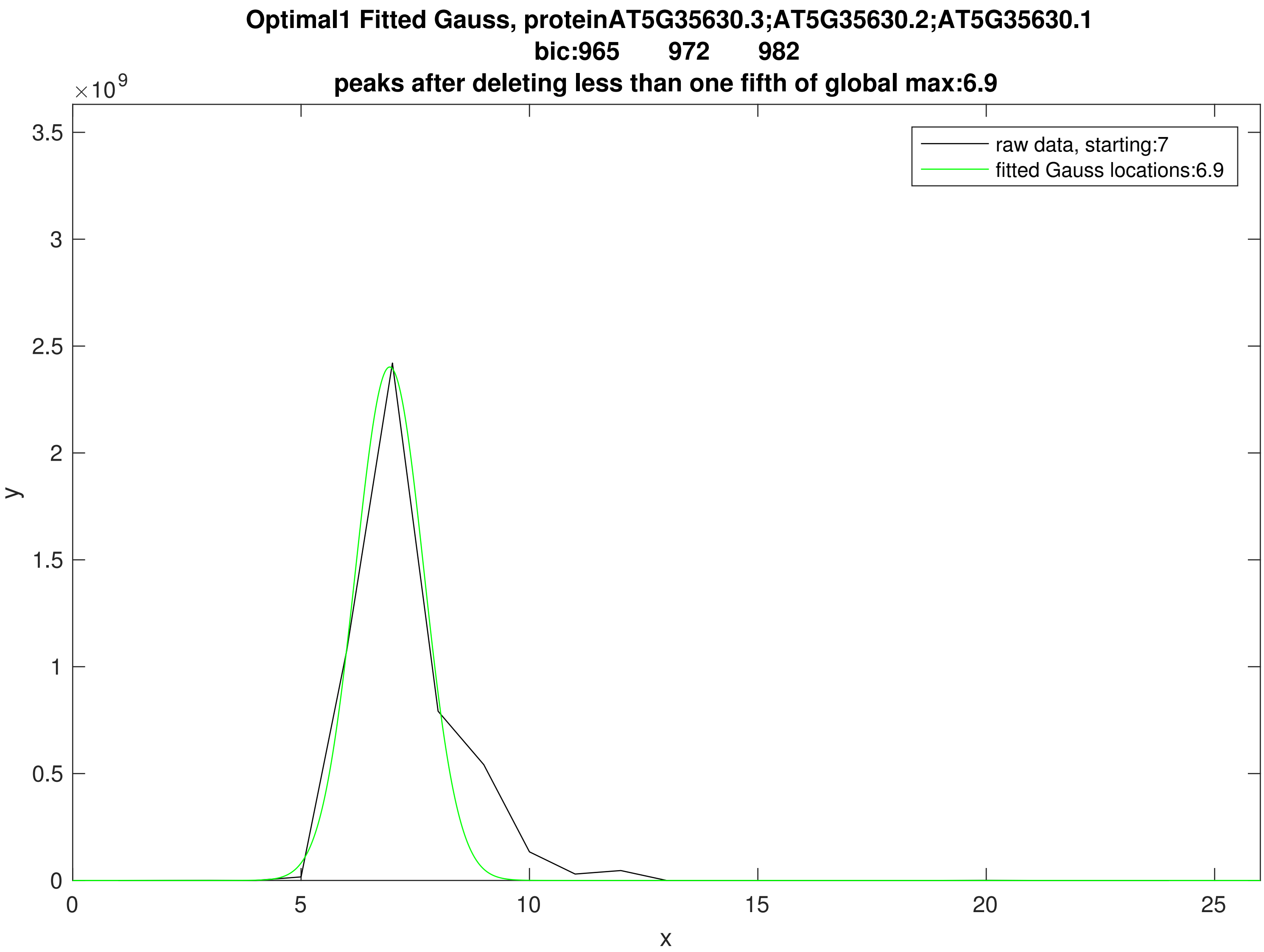
Optimal1 Fitted Gauss, proteinAT5G35630.3;AT5G35630.2;AT5G35630.1
bic:965 972 982
peaks after deleting less than one fifth of global max:6.9
×10^9
3.5
3
2.5
2
1.5
1
0.5
0
y
0
5
10
15
20
25
x
raw data, starting:7
fitted Gauss locations:6.9

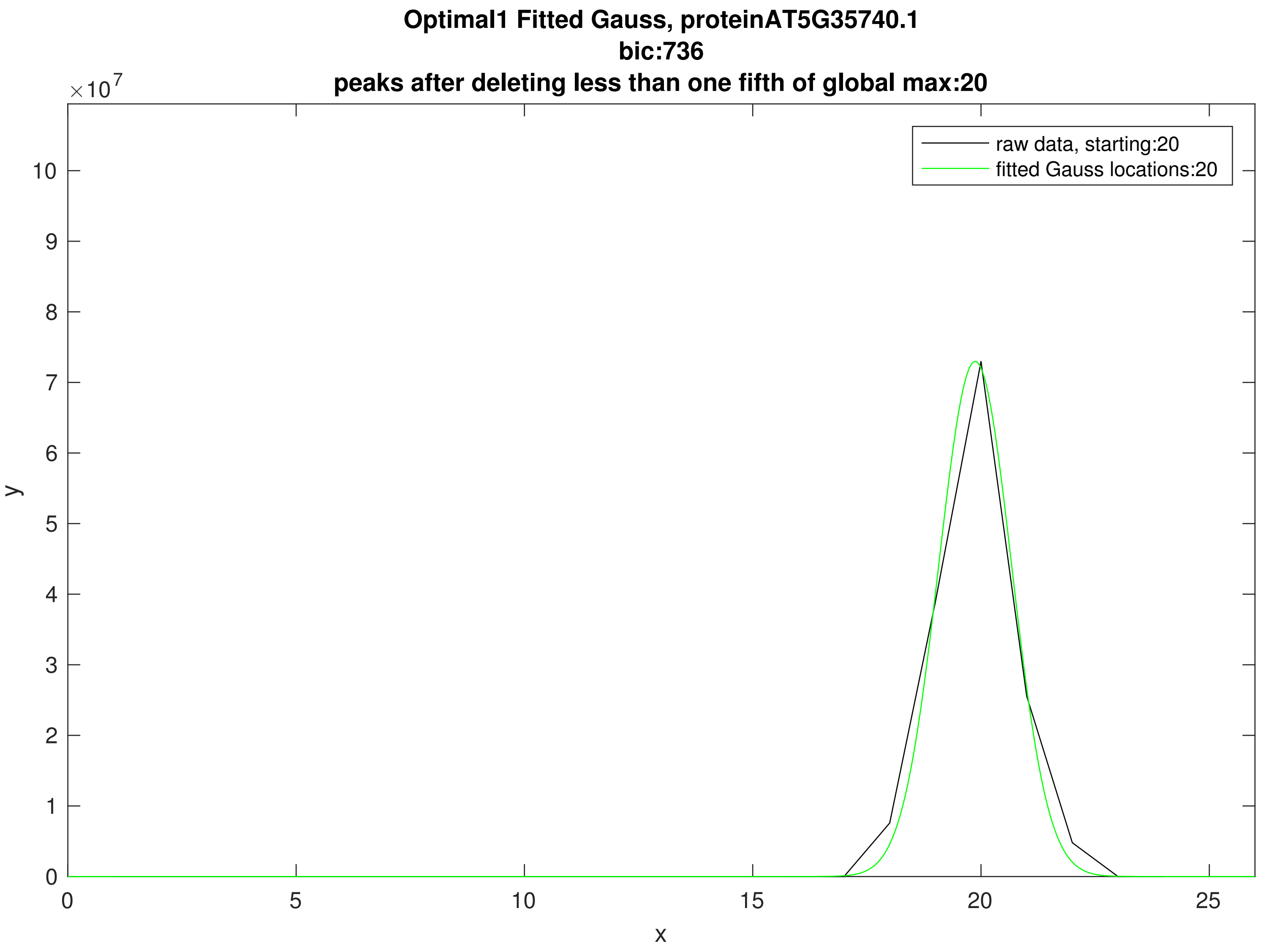

Optimal1 Fitted Gauss, proteinAT5G35740.1
bic:736
peaks after deleting less than one fifth of global max:20
×10^7
10
9
8
7
6
5
4
3
2
1
0
y
0
5
10
15
20
25
x
raw data, starting:20
fitted Gauss locations:20

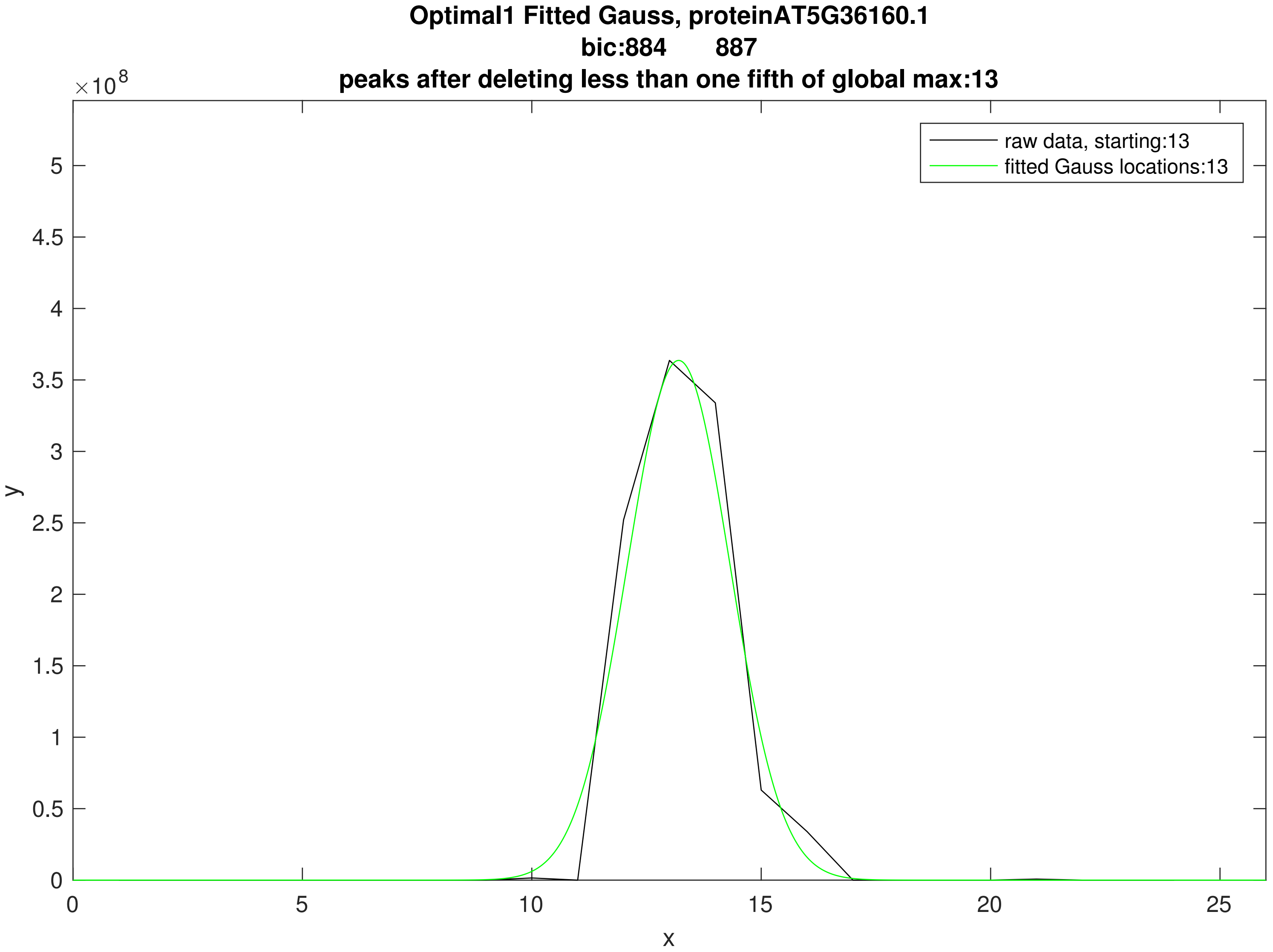
Optimal1 Fitted Gauss, proteinAT5G36160.1
bic:884 887
peaks after deleting less than one fifth of global max:13
×10^8
raw data, starting:13
fitted Gauss locations:13
y
x
0
0.5
1
1.5
2
2.5
3
3.5
4
4.5
5
0
5
10
15
20
25

Optimal1 Fitted Gauss, proteinAT5G36180.1;AT1G73300.1
bic:730
peaks after deleting less than one fifth of global max:12

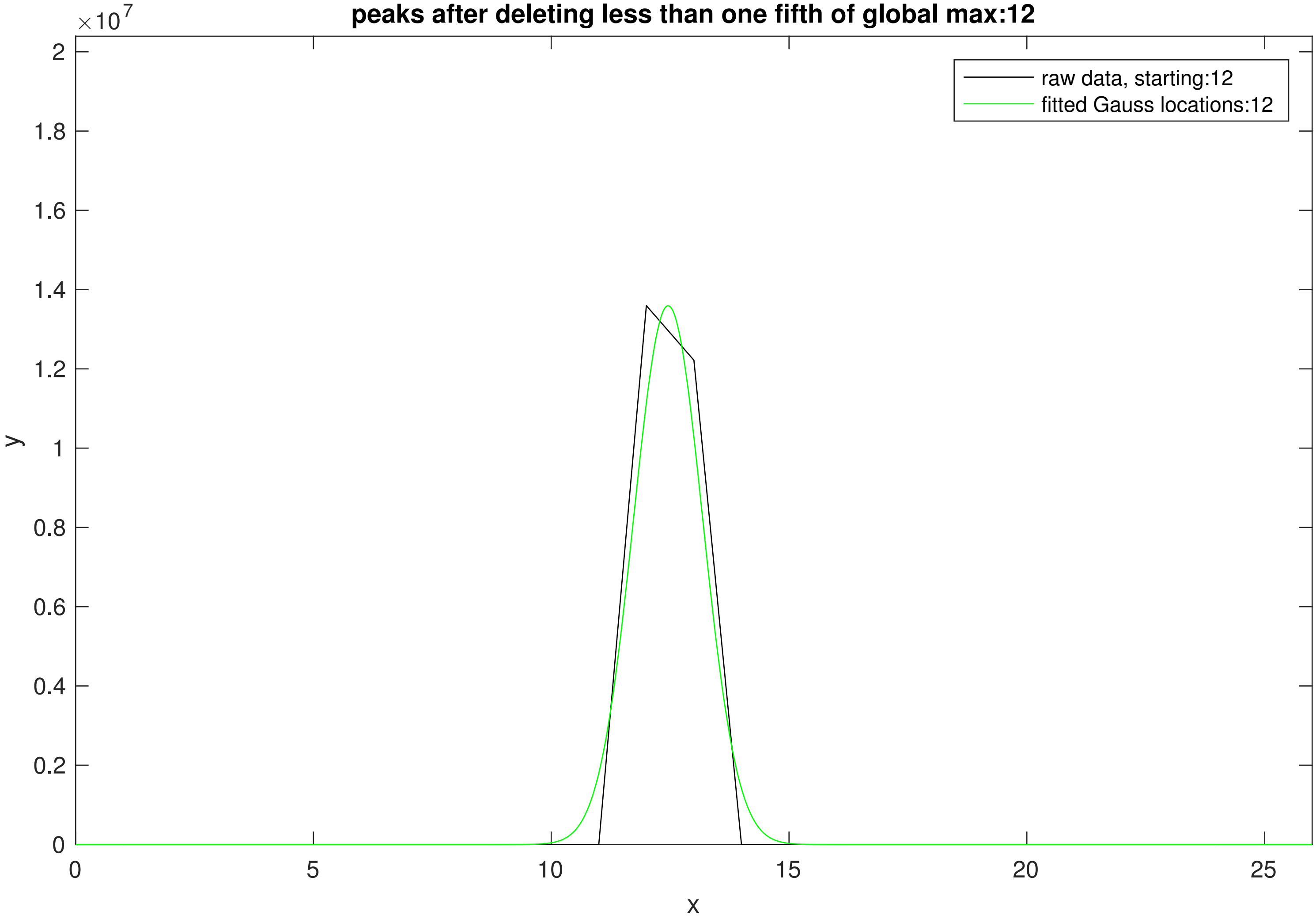

**Optimal1 Fitted Gauss, proteinAT5G36210.1**
**bic:935 943 953 962**
**peaks after deleting less than one fifth of global max:14**

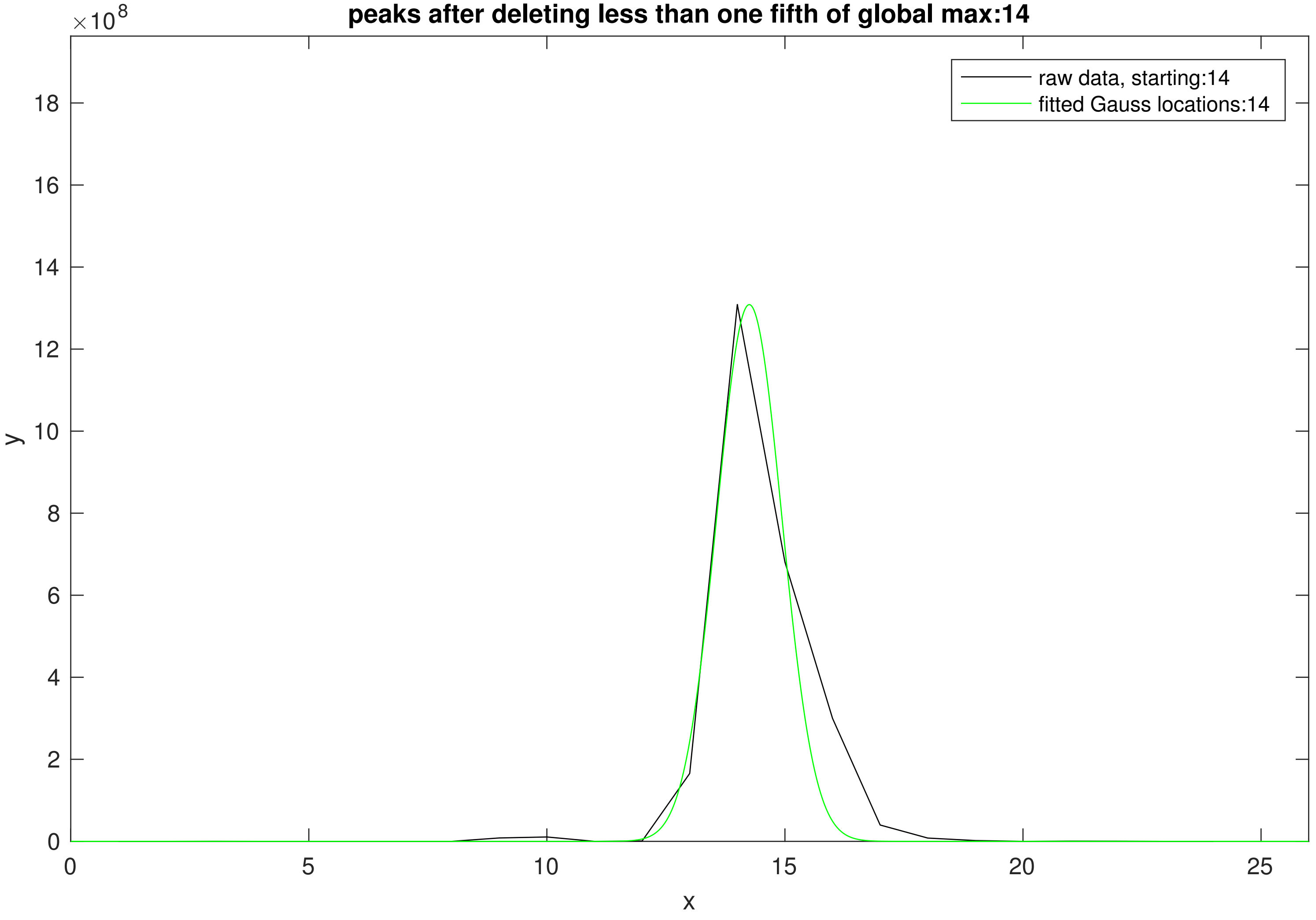

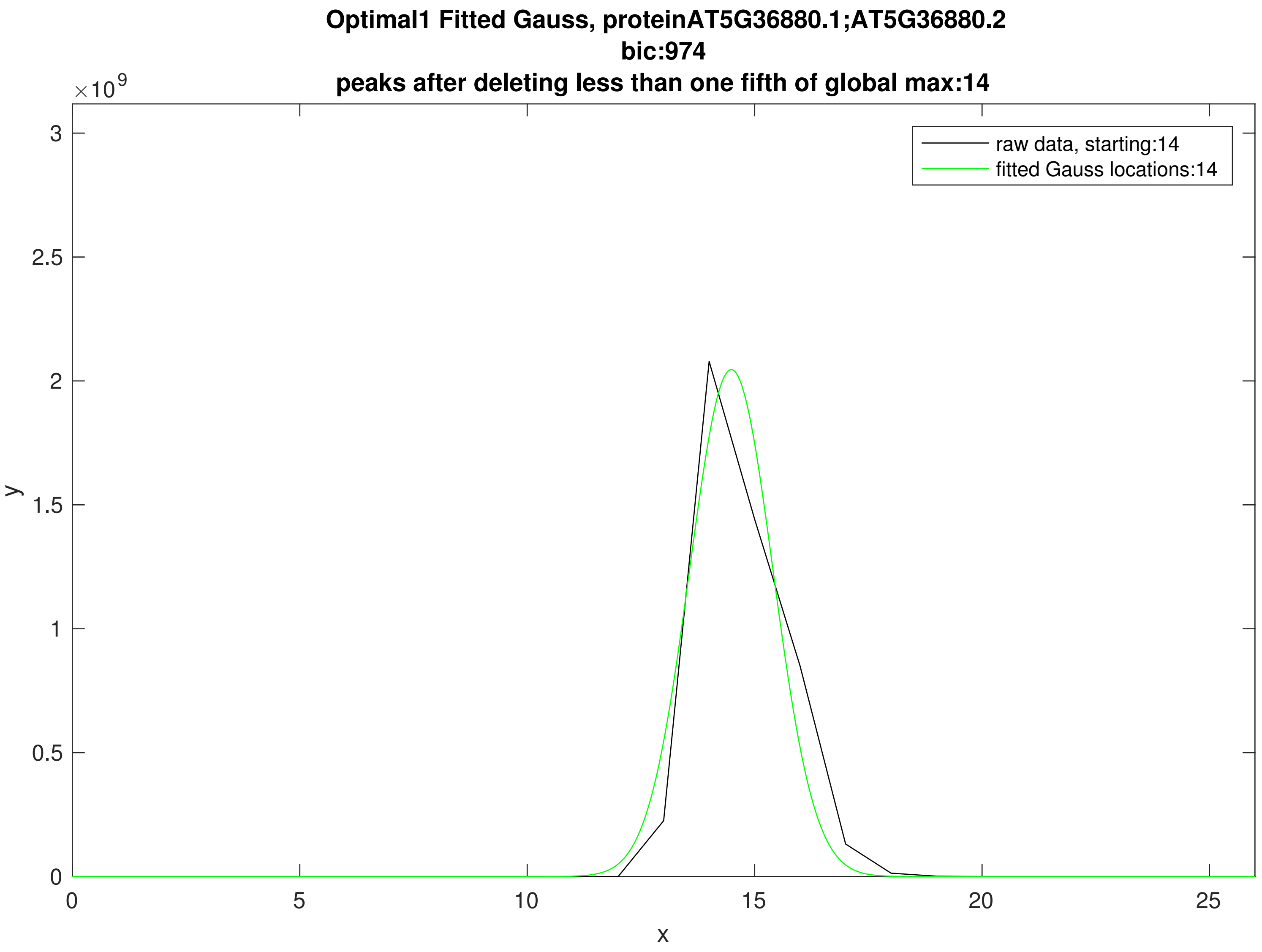

Optimal1 Fitted Gauss, proteinAT5G36880.1;AT5G36880.2
bic:974
peaks after deleting less than one fifth of global max:14
×10^9
raw data, starting:14
fitted Gauss locations:14
y
x
0
0.5
1
1.5
2
2.5
3
0
5
10
15
20
25

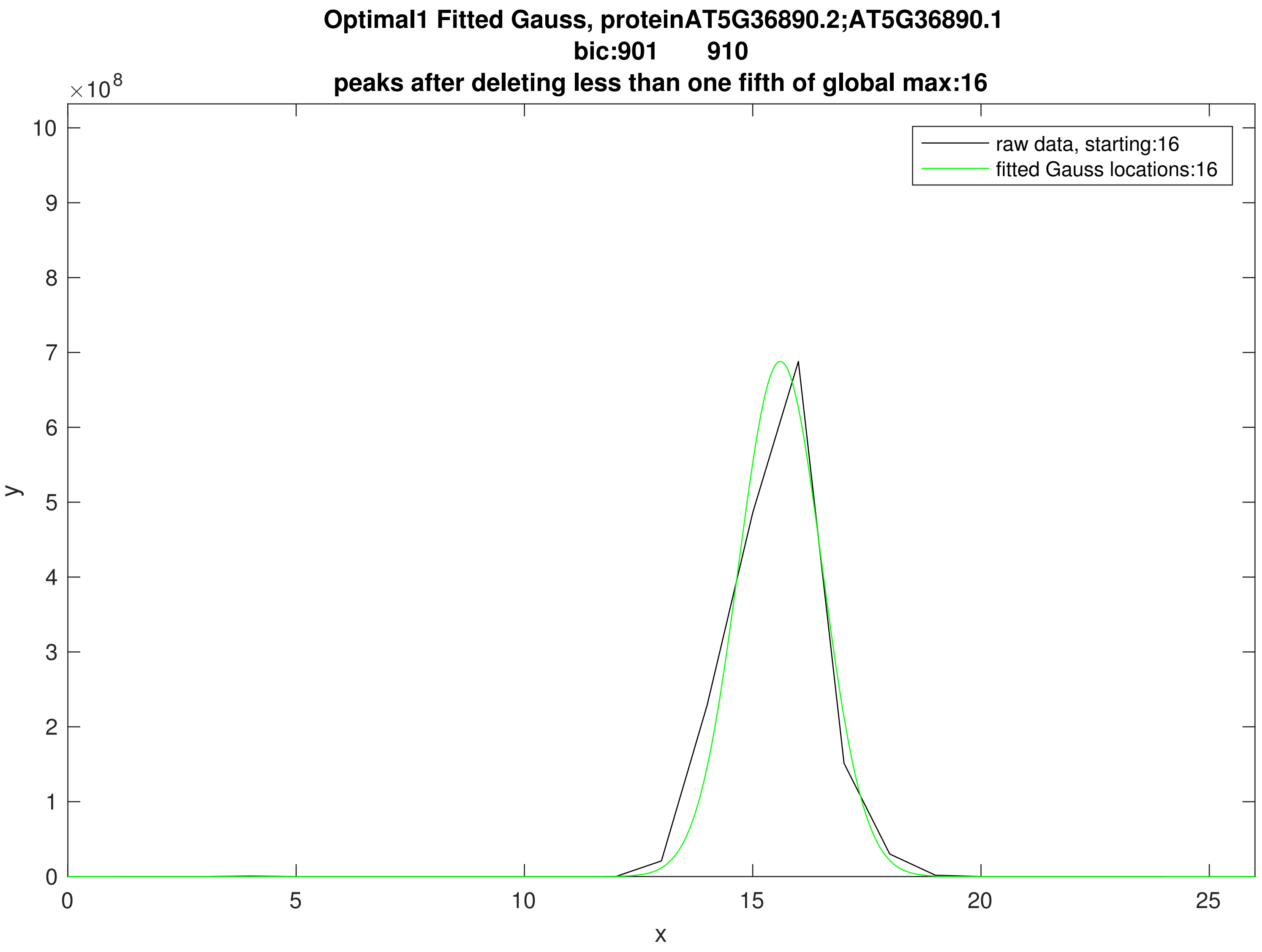
Optimal1 Fitted Gauss, proteinAT5G36890.2;AT5G36890.1
bic:901 910
peaks after deleting less than one fifth of global max:16
raw data, starting:16
fitted Gauss locations:16
y
x

#### Optimal1 Fitted Gauss, proteinAT5G37475.1
#### bic:722
#### peaks after deleting less than one fifth of global max:12

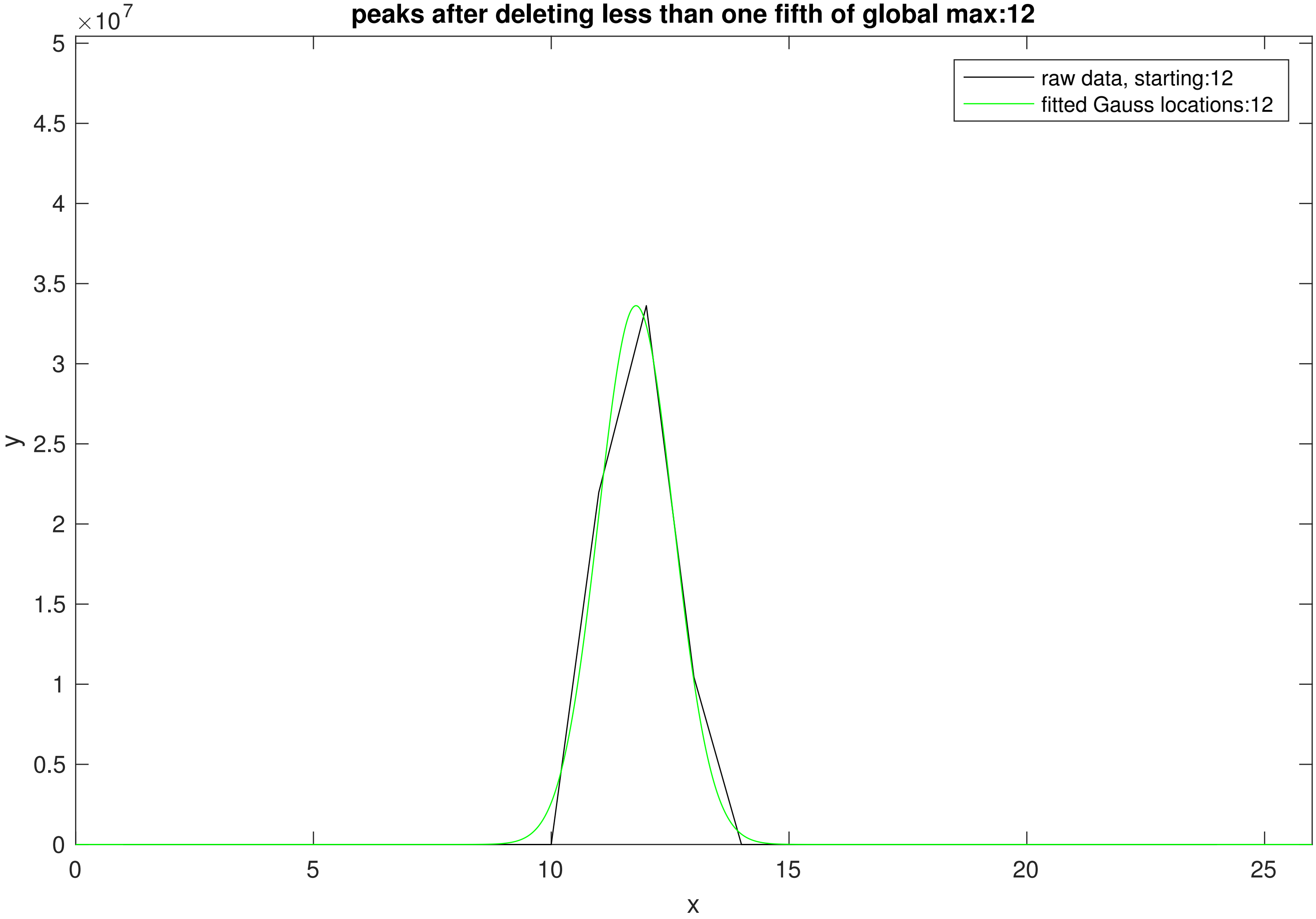

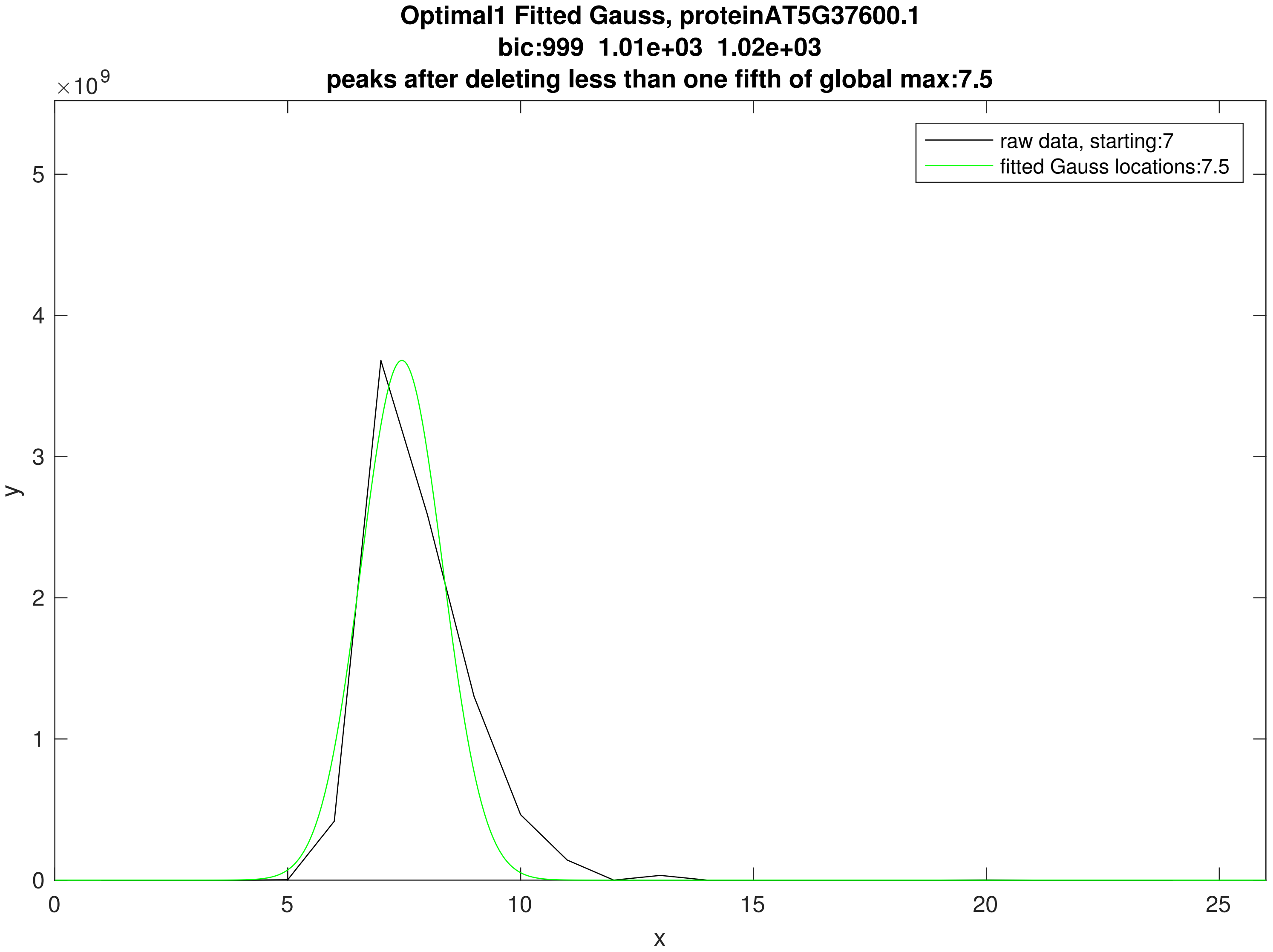

Optimal1 Fitted Gauss, proteinAT5G37600.1
bic:999 1.01e+03 1.02e+03
peaks after deleting less than one fifth of global max:7.5
×10^9
raw data, starting:7
fitted Gauss locations:7.5
y
x
0
1
2
3
4
5
0
5
10
15
20
25

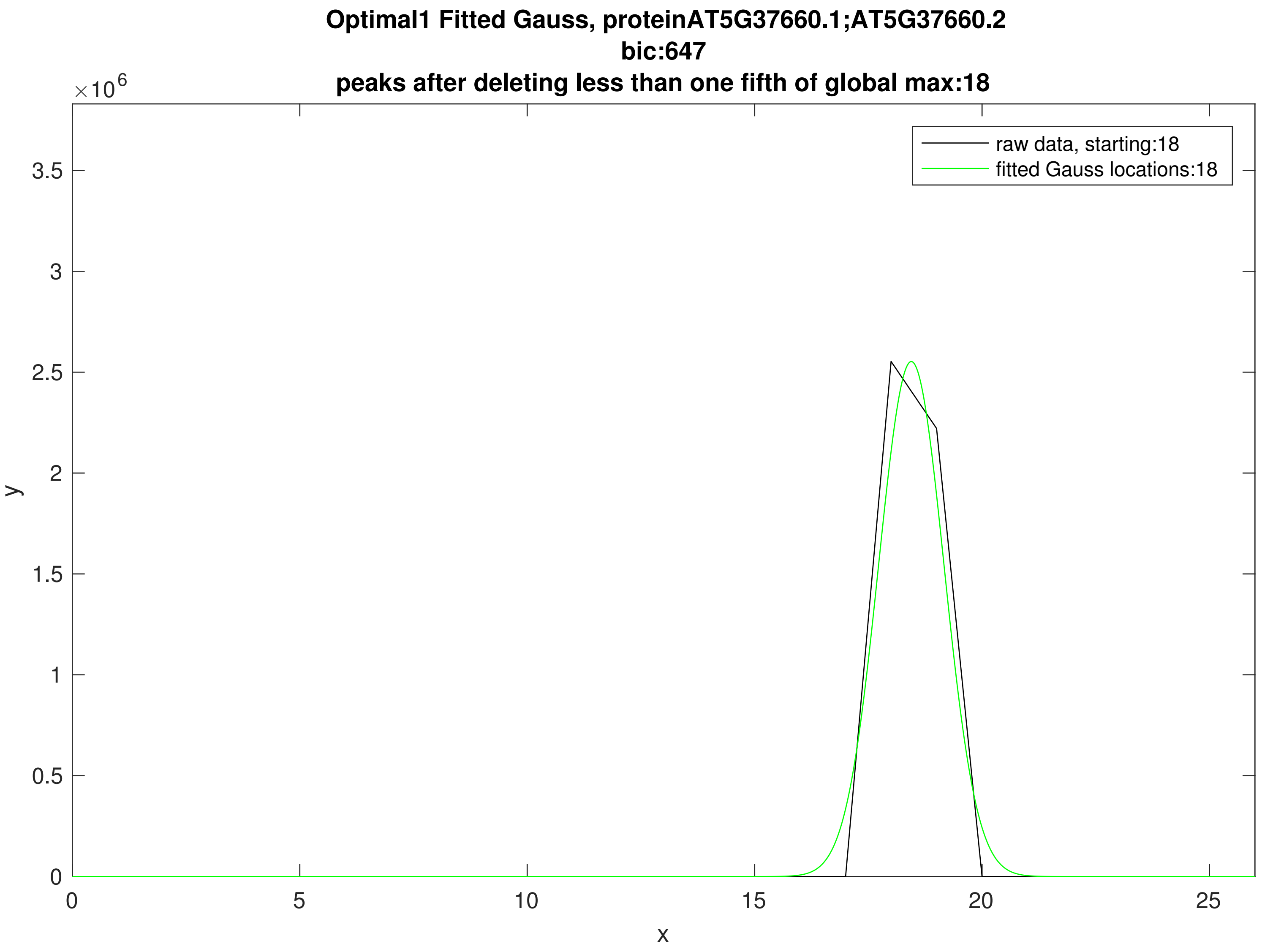
Optimal1 Fitted Gauss, proteinAT5G37660.1;AT5G37660.2
bic:647
peaks after deleting less than one fifth of global max:18
×10^6
raw data, starting:18
fitted Gauss locations:18
y
x
0
0.5
1
1.5
2
2.5
3
3.5
0
5
10
15
20
25

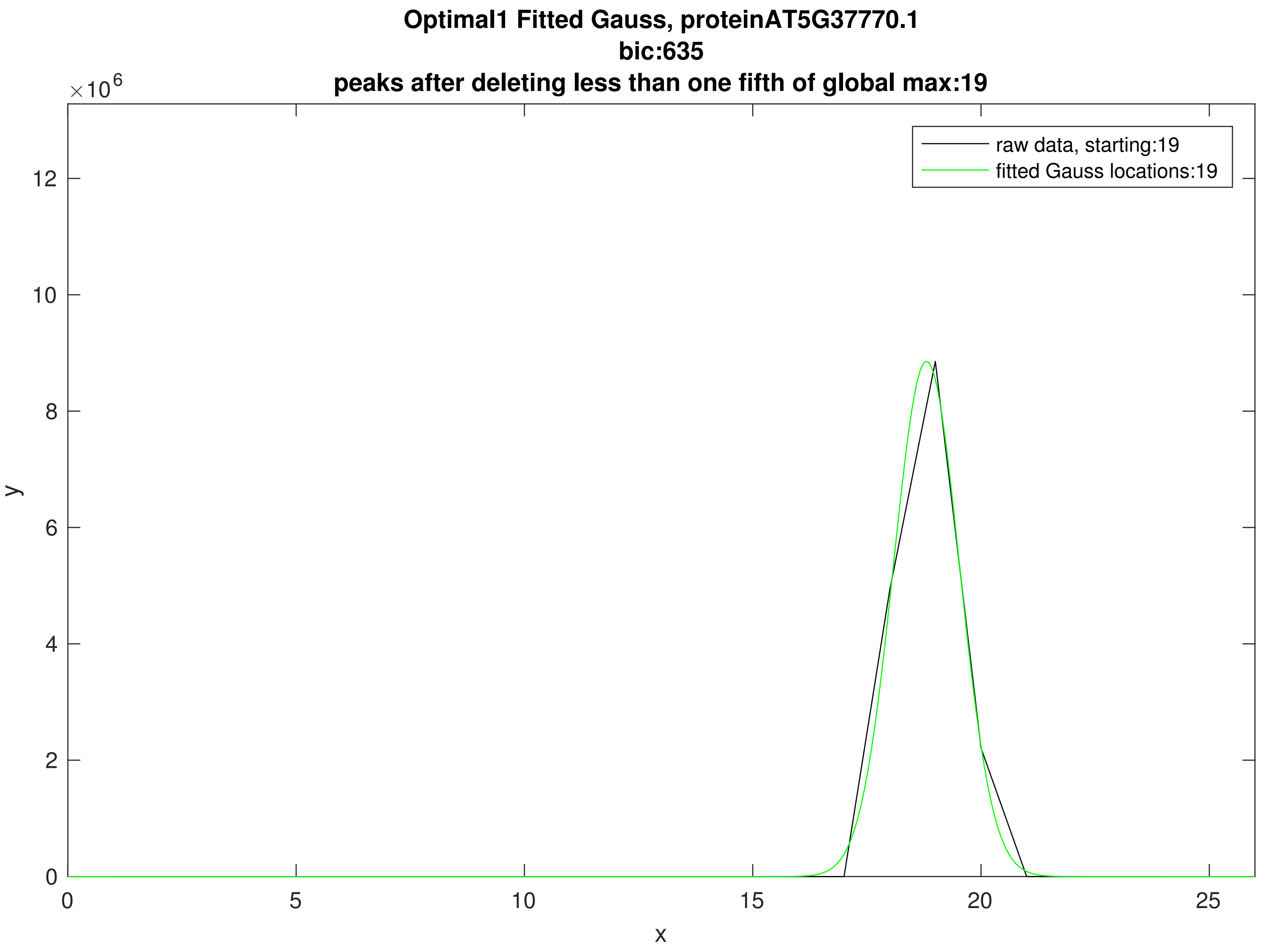
Optimal1 Fitted Gauss, proteinAT5G37770.1
bic:635
peaks after deleting less than one fifth of global max:19
×10^6
raw data, starting:19
fitted Gauss locations:19
12
10
8
6
4
2
0
y
0
5
10
15
20
25
x

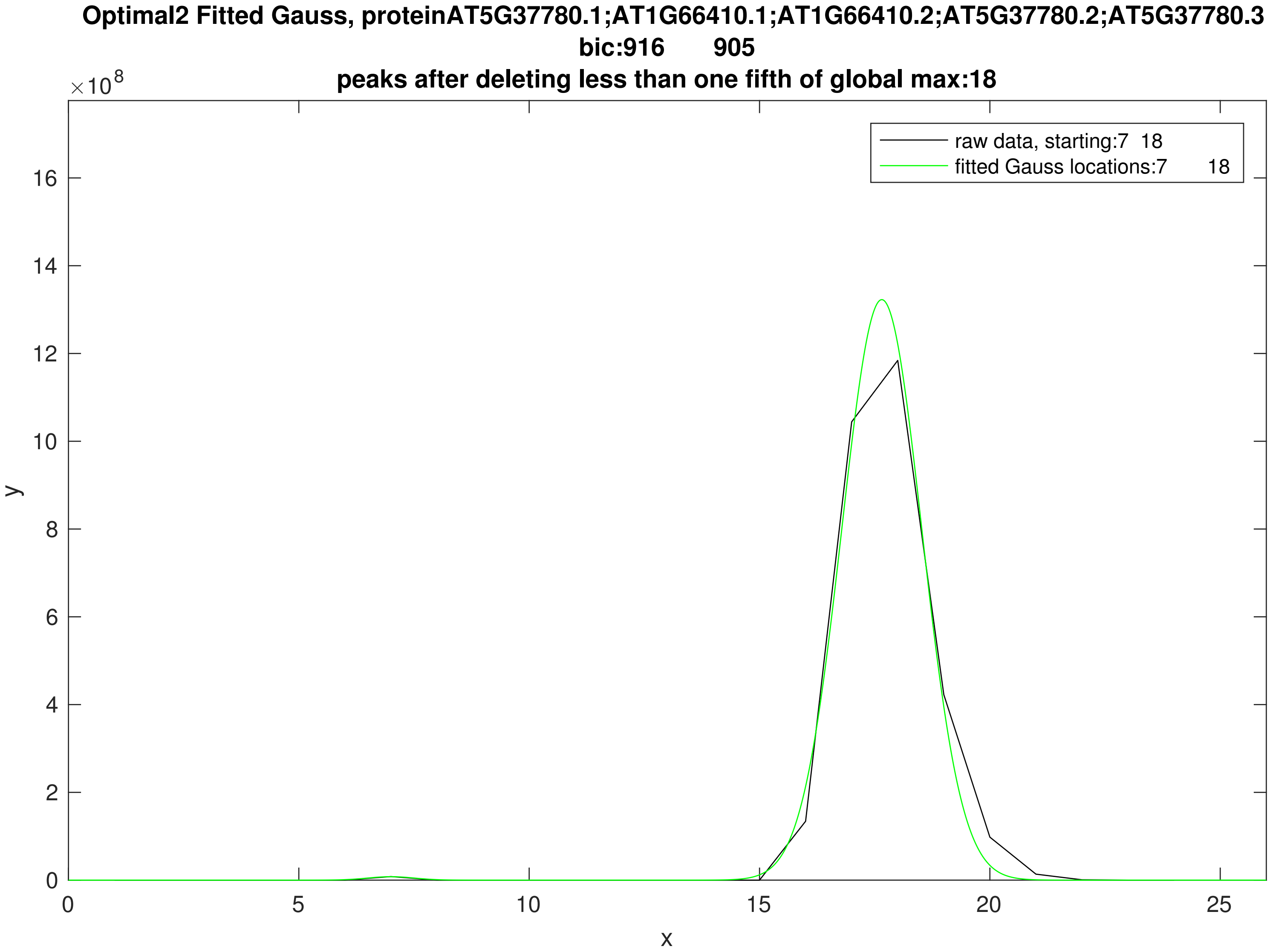

Optimal2 Fitted Gauss, proteinAT5G37780.1;AT1G66410.1;AT1G66410.2;AT5G37780.2;AT5G37780.3
bic:916 905
peaks after deleting less than one fifth of global max:18
×10^8
raw data, starting:7 18
fitted Gauss locations:7 18
y
x
0
2
4
6
8
10
12
14
16
5
15
20
25

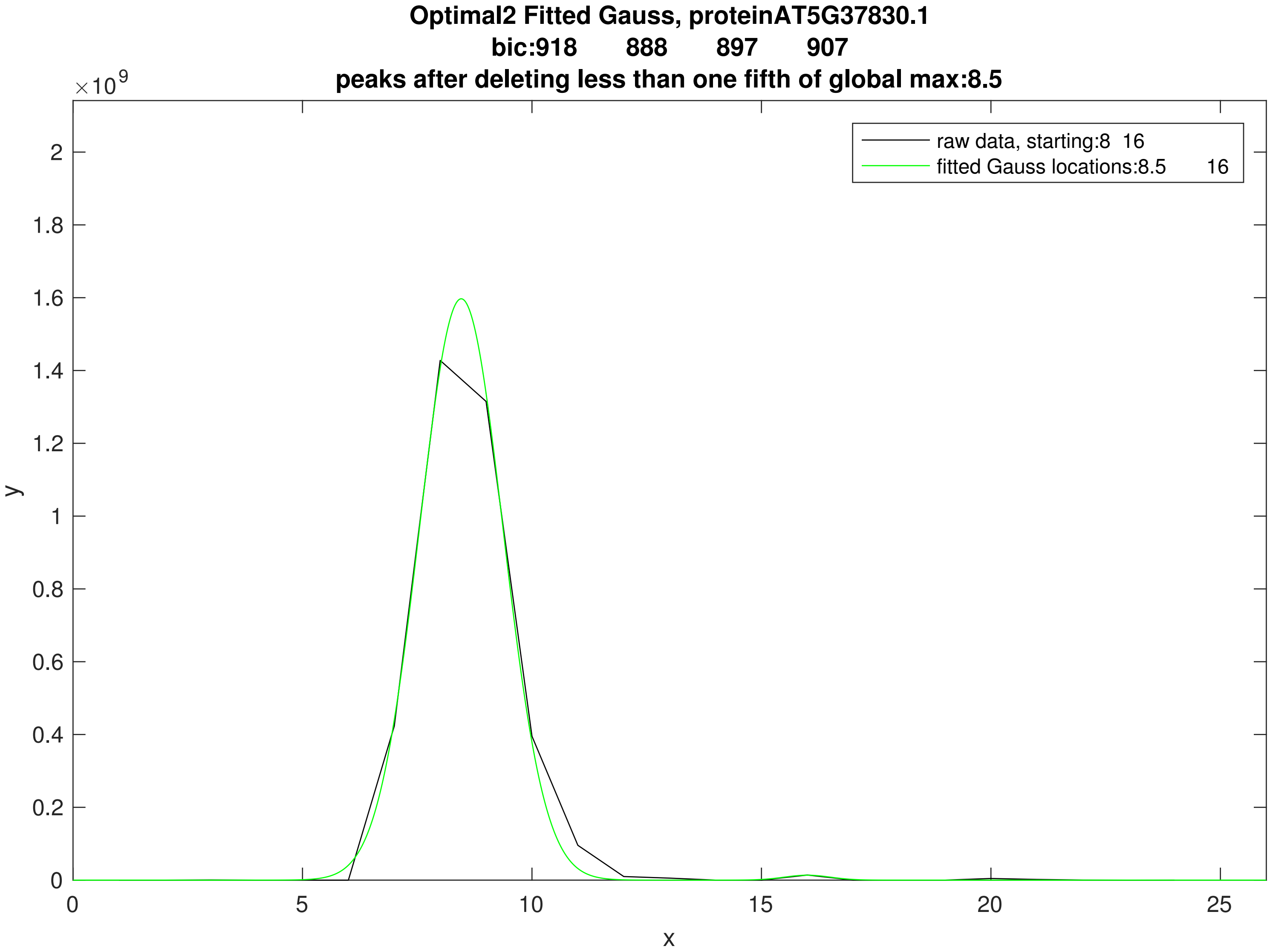

Optimal2 Fitted Gauss, proteinAT5G37830.1
bic:918 888 897 907
peaks after deleting less than one fifth of global max:8.5
×10^9
raw data, starting:8 16
fitted Gauss locations:8.5 16
y
x
0
0.2
0.4
0.6
0.8
1
1.2
1.4
1.6
1.8
2
0
5
10
15
20
25

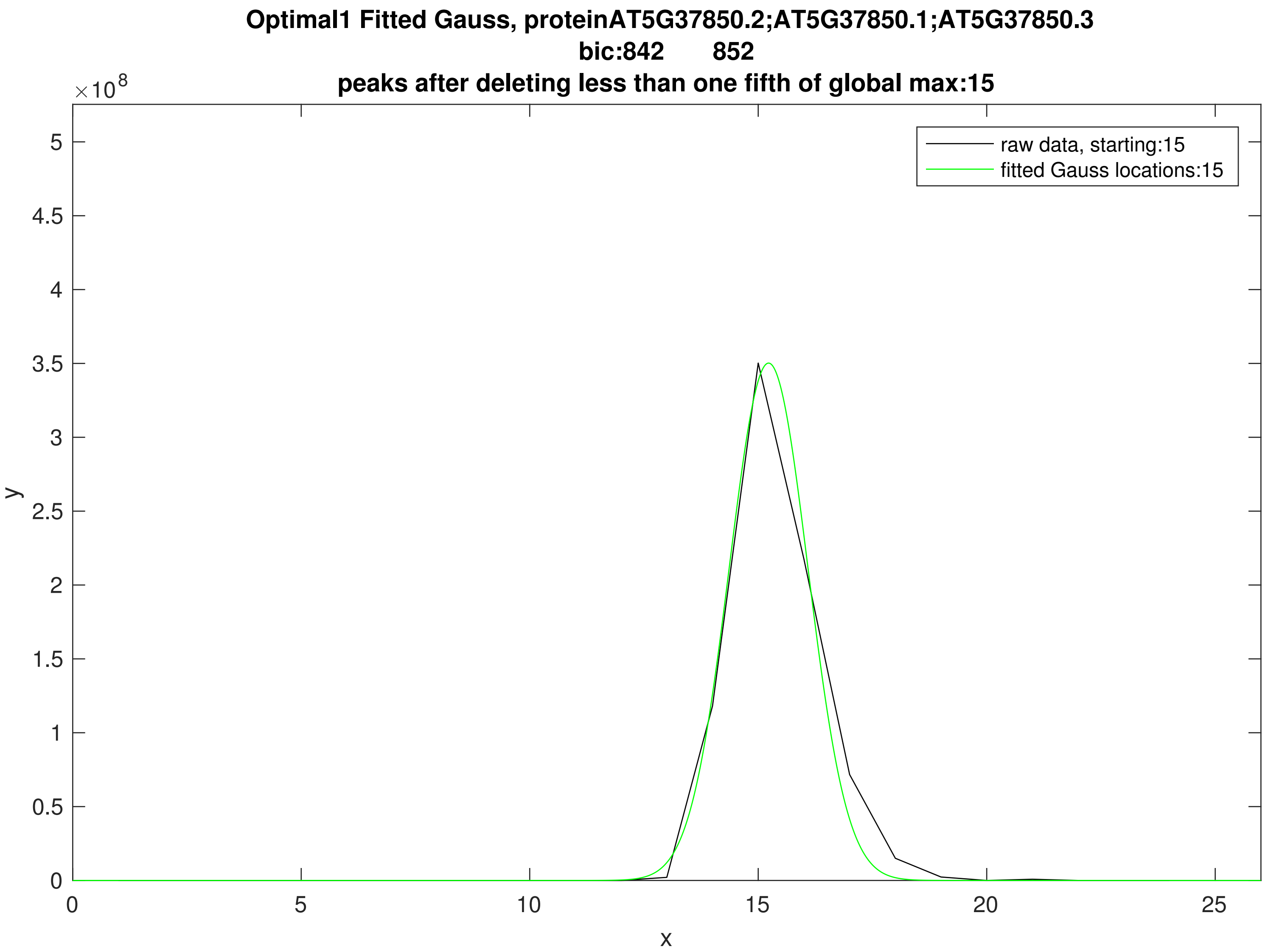

Optimal1 Fitted Gauss, proteinAT5G37850.2;AT5G37850.1;AT5G37850.3
bic:842 852
peaks after deleting less than one fifth of global max:15
×10^8
raw data, starting:15
fitted Gauss locations:15
y
x
0
0.5
1
1.5
2
2.5
3
3.5
4
4.5
5
0
5
10
15
20
25

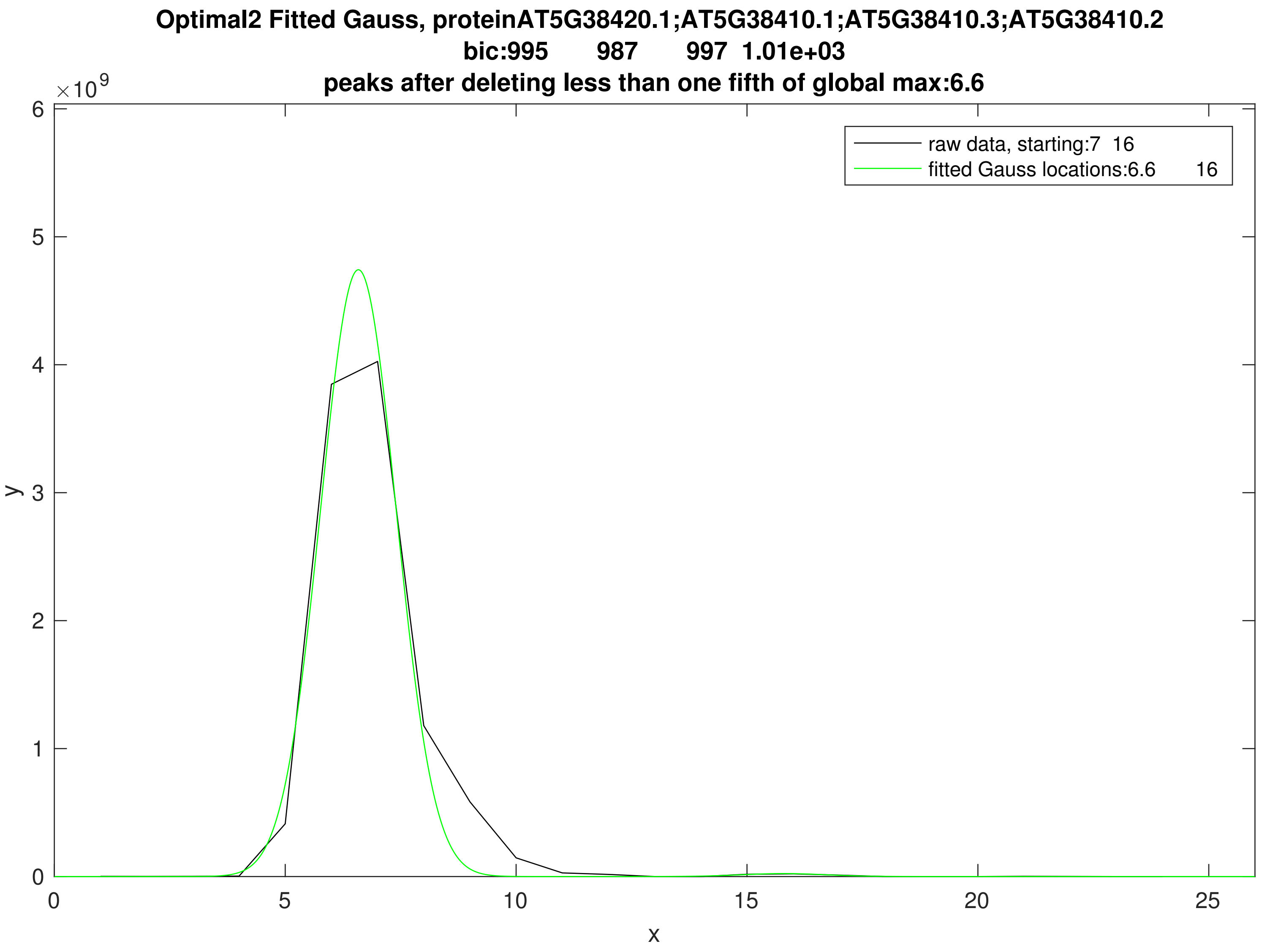

Optimal2 Fitted Gauss, proteinAT5G38420.1;AT5G38410.1;AT5G38410.3;AT5G38410.2
bic:995 987 997 1.01e+03
peaks after deleting less than one fifth of global max:6.6
×10^9
6
5
4
3
2
1
0
y
raw data, starting:7 16
fitted Gauss locations:6.6 16
0
5
10
15
20
25
x

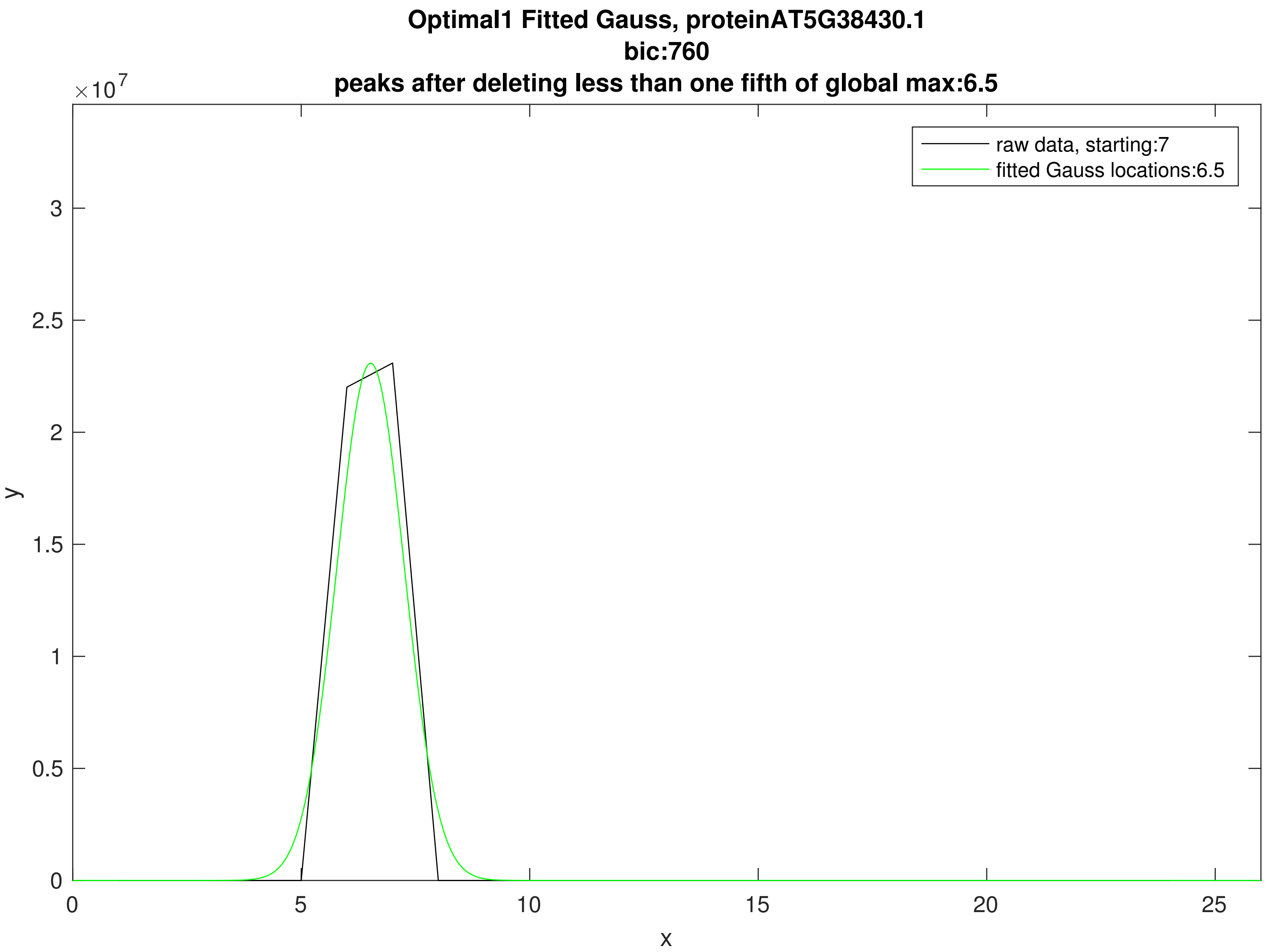

Optimal1 Fitted Gauss, proteinAT5G38430.1
bic:760
peaks after deleting less than one fifth of global max:6.5
×10^7
raw data, starting:7
fitted Gauss locations:6.5
3
2.5
2
1.5
1
0.5
0
y
0
5
10
15
20
25
x

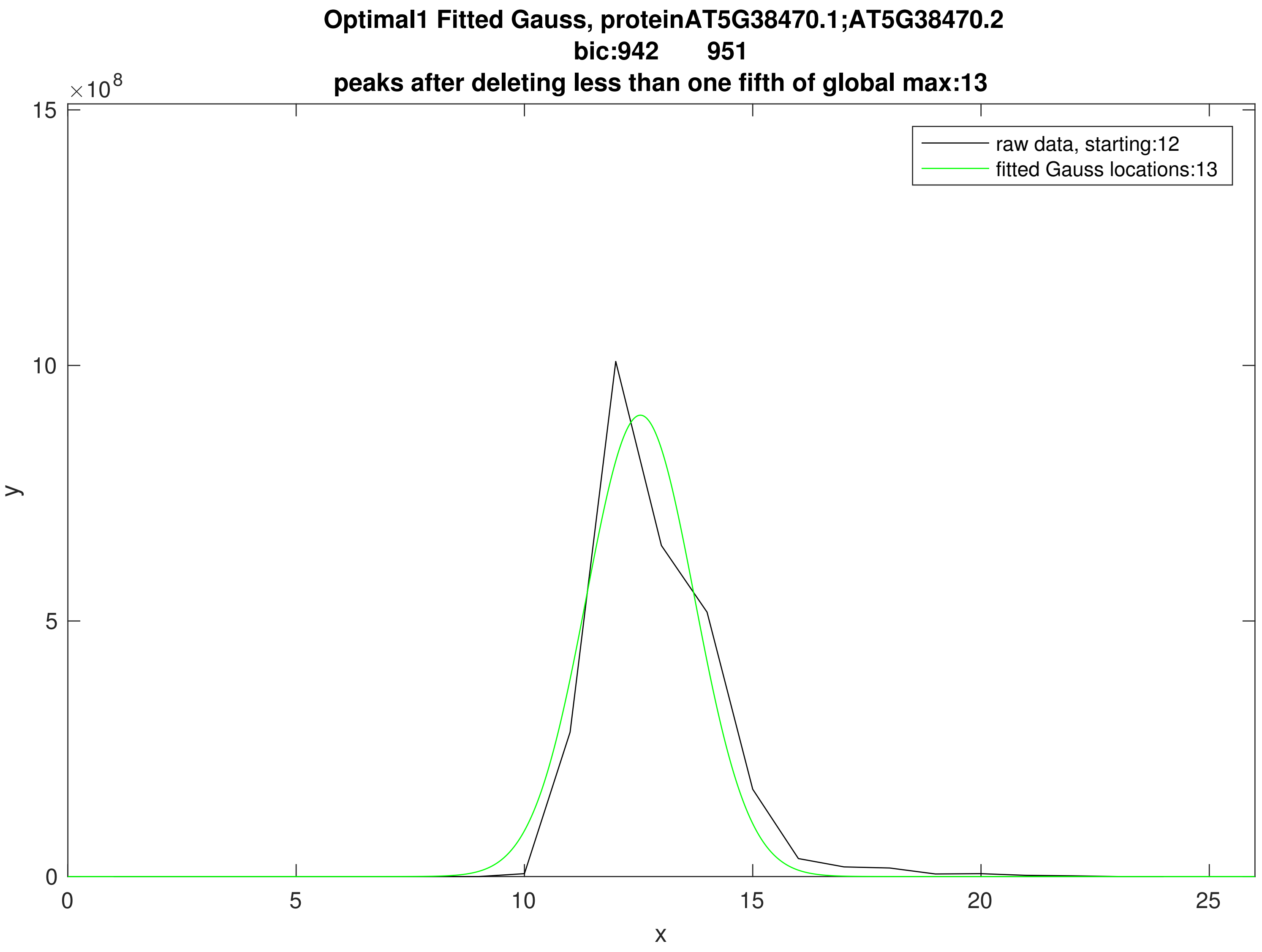

Optimal1 Fitted Gauss, proteinAT5G38470.1;AT5G38470.2
bic:942 951
peaks after deleting less than one fifth of global max:13
×10^8
15
10
5
0
y
0
5
10
15
20
25
x
raw data, starting:12
fitted Gauss locations:13

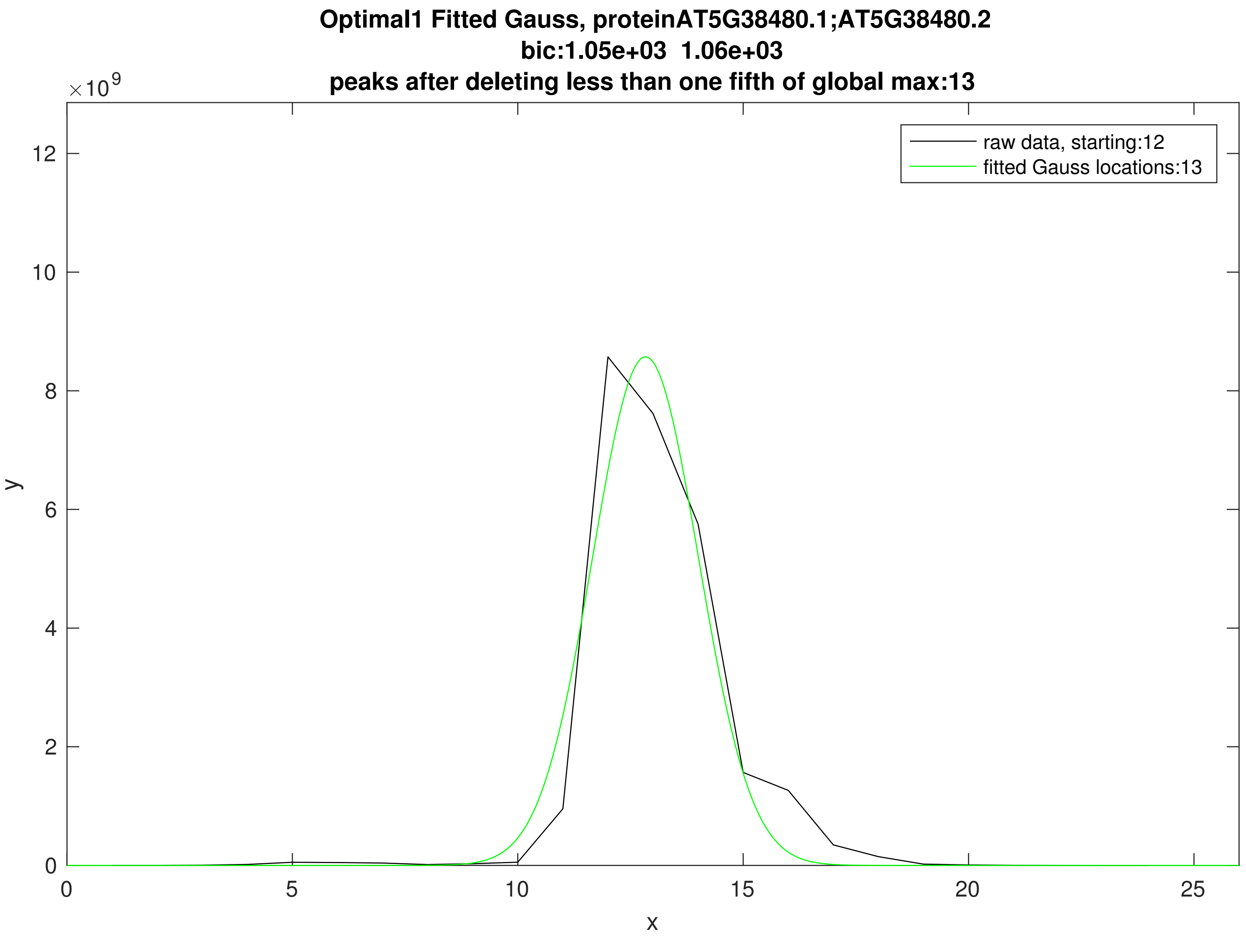
Optimal1 Fitted Gauss, proteinAT5G38480.1;AT5G38480.2
bic:1.05e+03 1.06e+03
peaks after deleting less than one fifth of global max:13
×10^9
raw data, starting:12
fitted Gauss locations:13
y
x
0
2
4
6
8
10
12
0
5
10
15
20
25

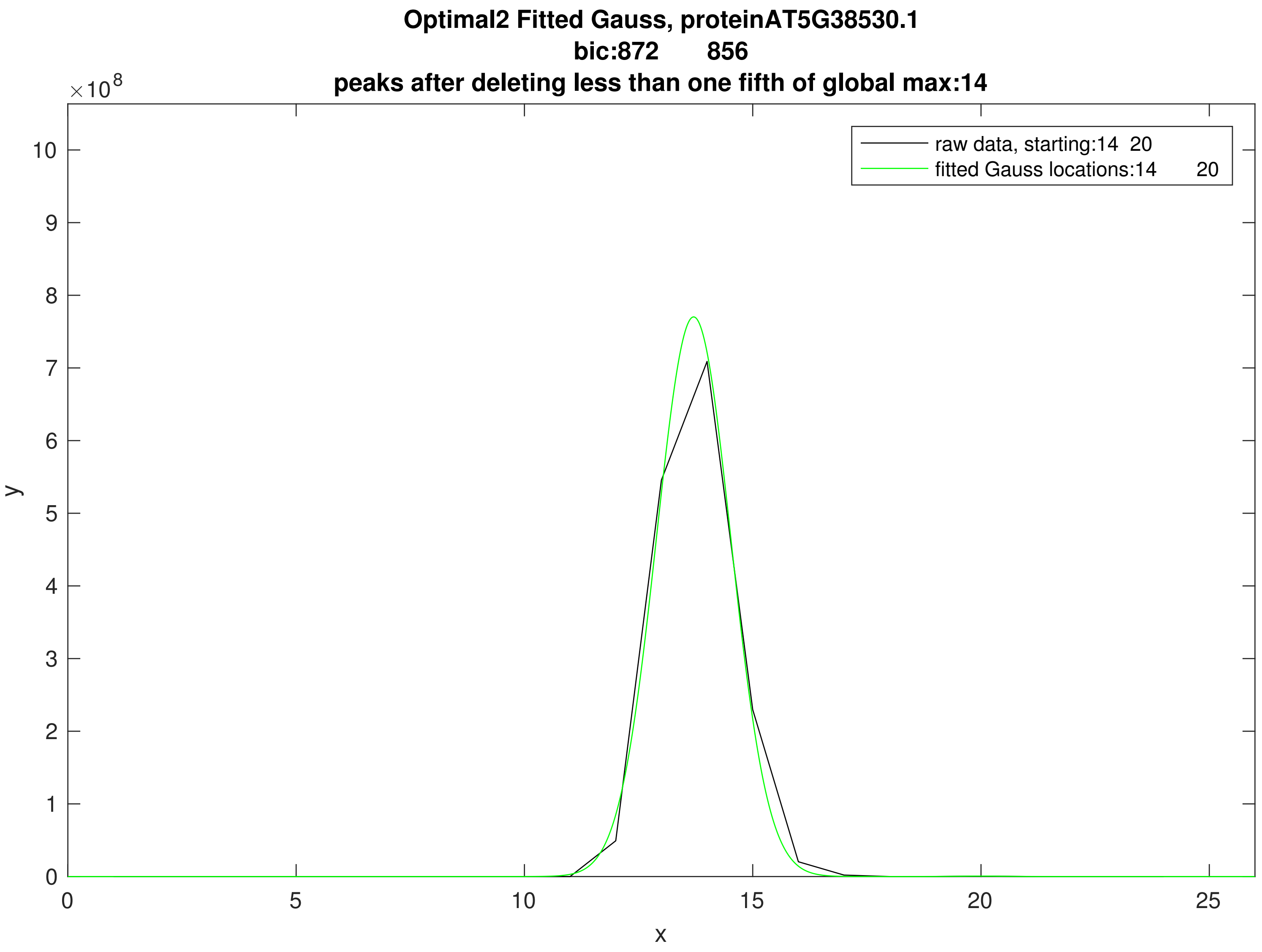
Optimal2 Fitted Gauss, proteinAT5G38530.1
bic:872 856
peaks after deleting less than one fifth of global max:14
$\times 10^8$
raw data, starting:14 20
fitted Gauss locations:14 20
y
x
0
1
2
3
4
5
6
7
8
9
10
0
5
10
15
20
25

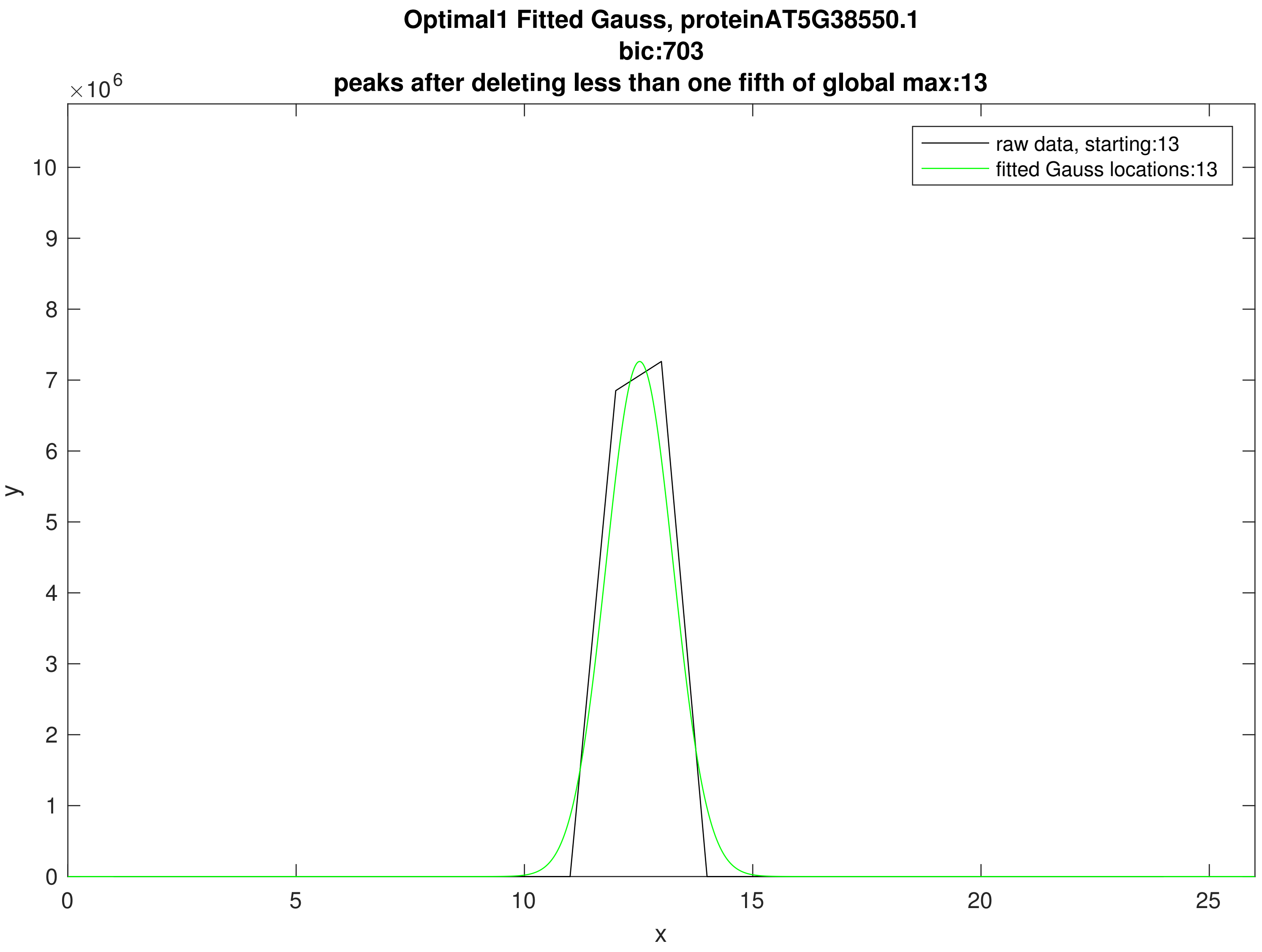

Optimal1 Fitted Gauss, proteinAT5G38550.1
bic:703
peaks after deleting less than one fifth of global max:13
×10^6
raw data, starting:13
fitted Gauss locations:13
y
x
0
1
2
3
4
5
6
7
8
9
10
0
5
10
15
20
25

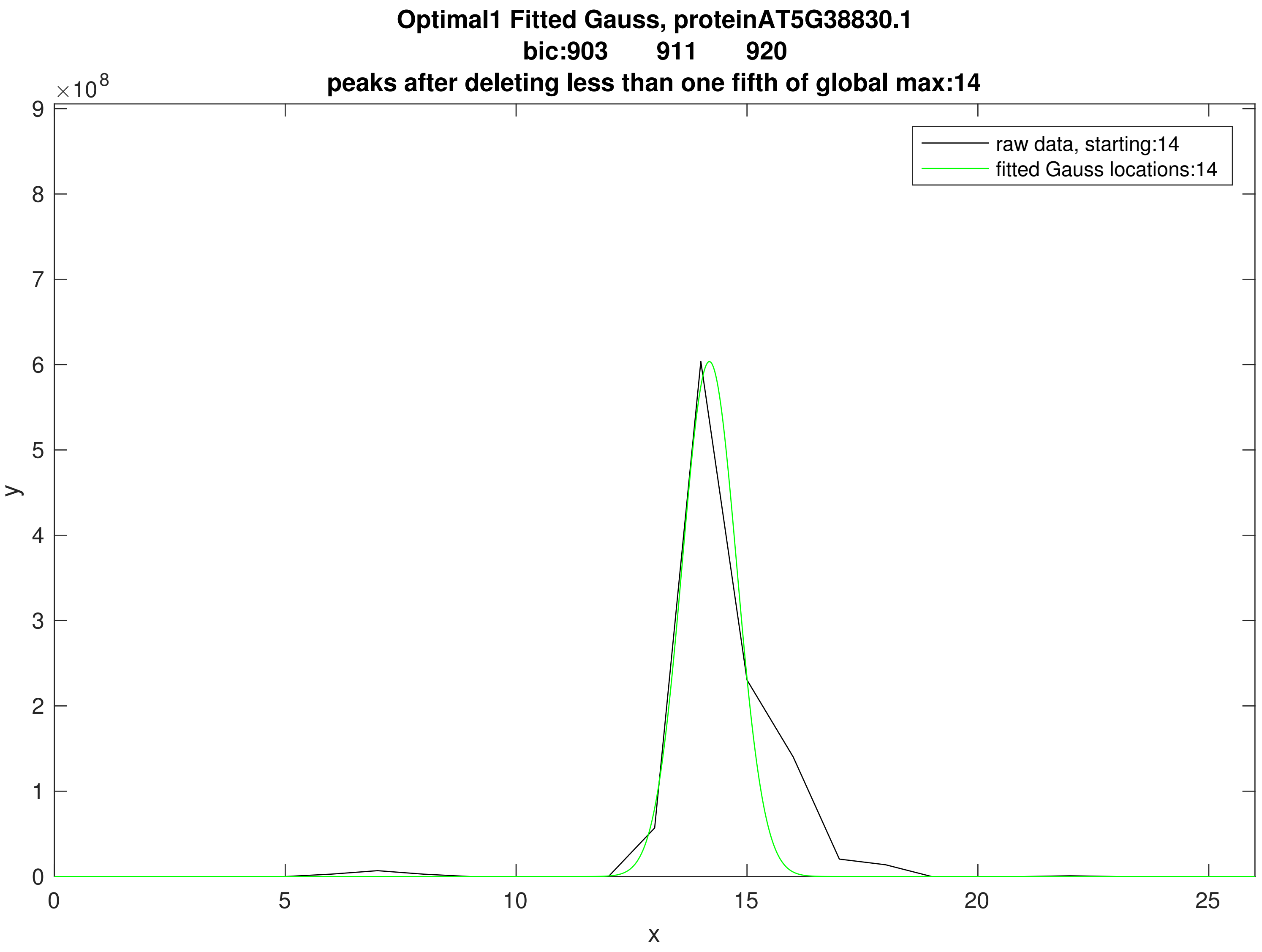
Optimal1 Fitted Gauss, proteinAT5G38830.1
bic:903 911 920
peaks after deleting less than one fifth of global max:14
×10^8
raw data, starting:14
fitted Gauss locations:14
y
x
0
1
2
3
4
5
6
7
8
9
0
5
10
15
20
25

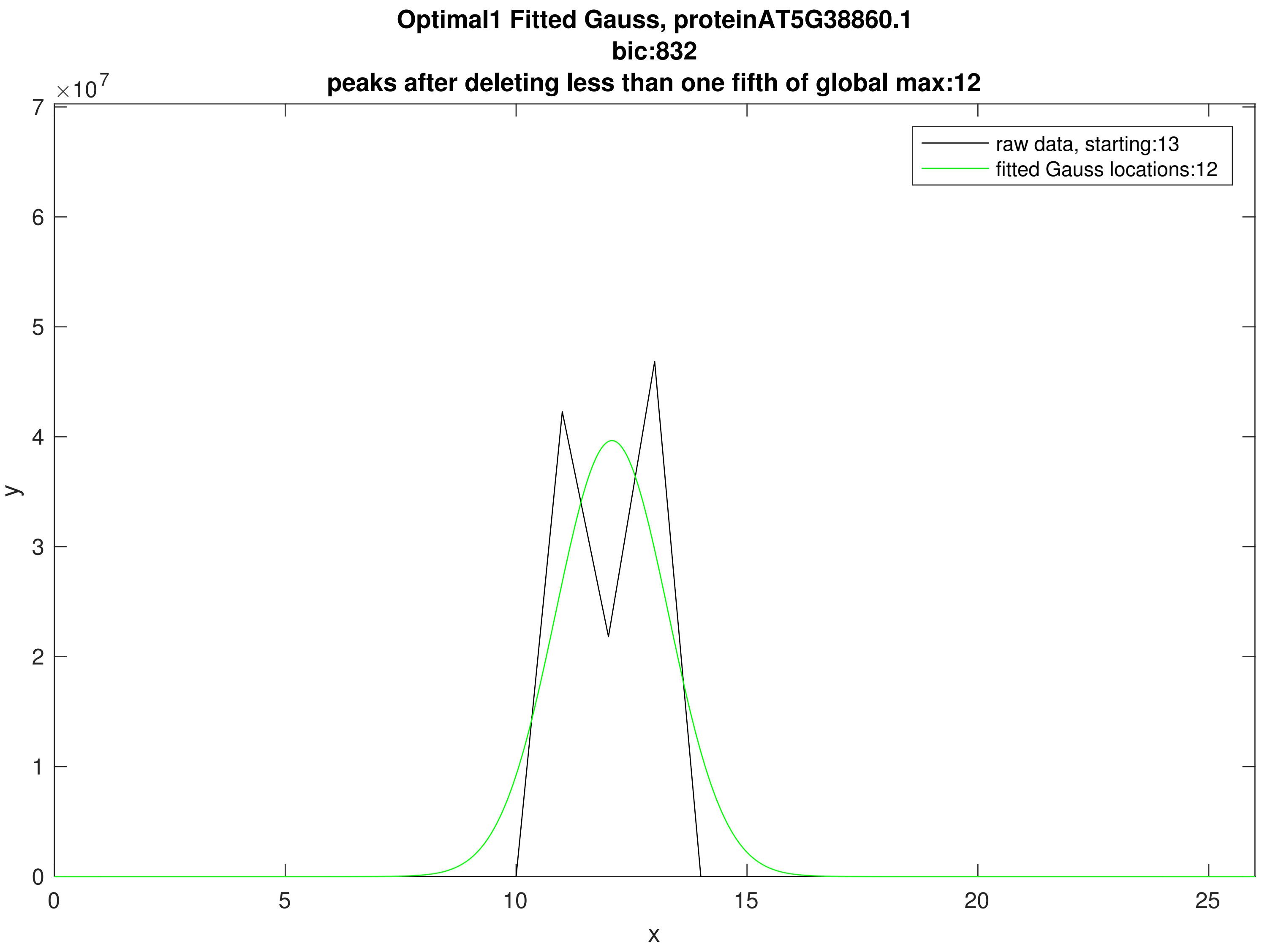

Optimal1 Fitted Gauss, proteinAT5G38860.1
bic:832
peaks after deleting less than one fifth of global max:12
×10^7
raw data, starting:13
fitted Gauss locations:12
y
x
0
1
2
3
4
5
6
7
0
5
10
15
20
25

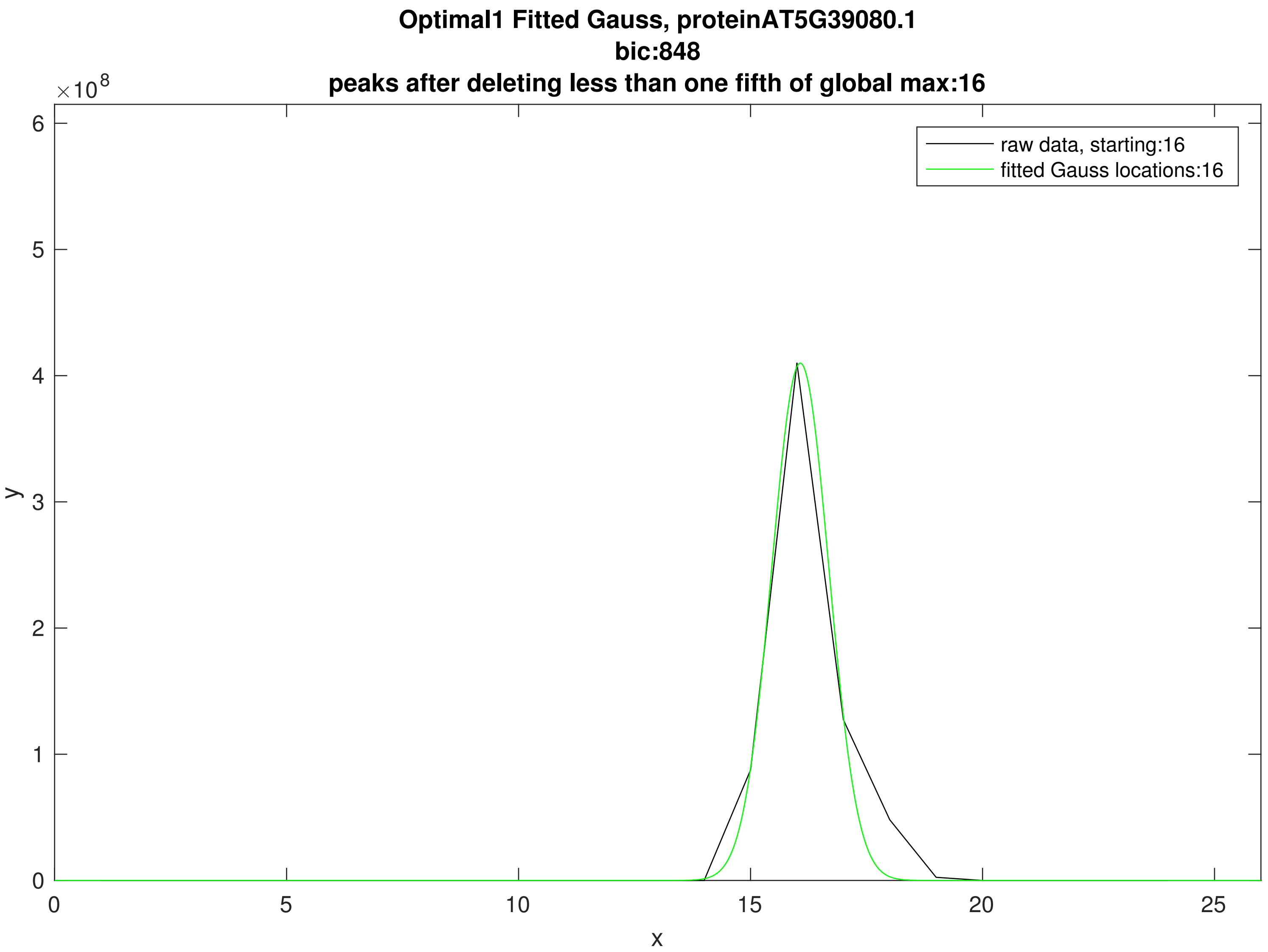
Optimal1 Fitted Gauss, proteinAT5G39080.1
bic:848
peaks after deleting less than one fifth of global max:16
×10^8
raw data, starting:16
fitted Gauss locations:16
y
x
0
1
2
3
4
5
6
0
5
10
15
20
25

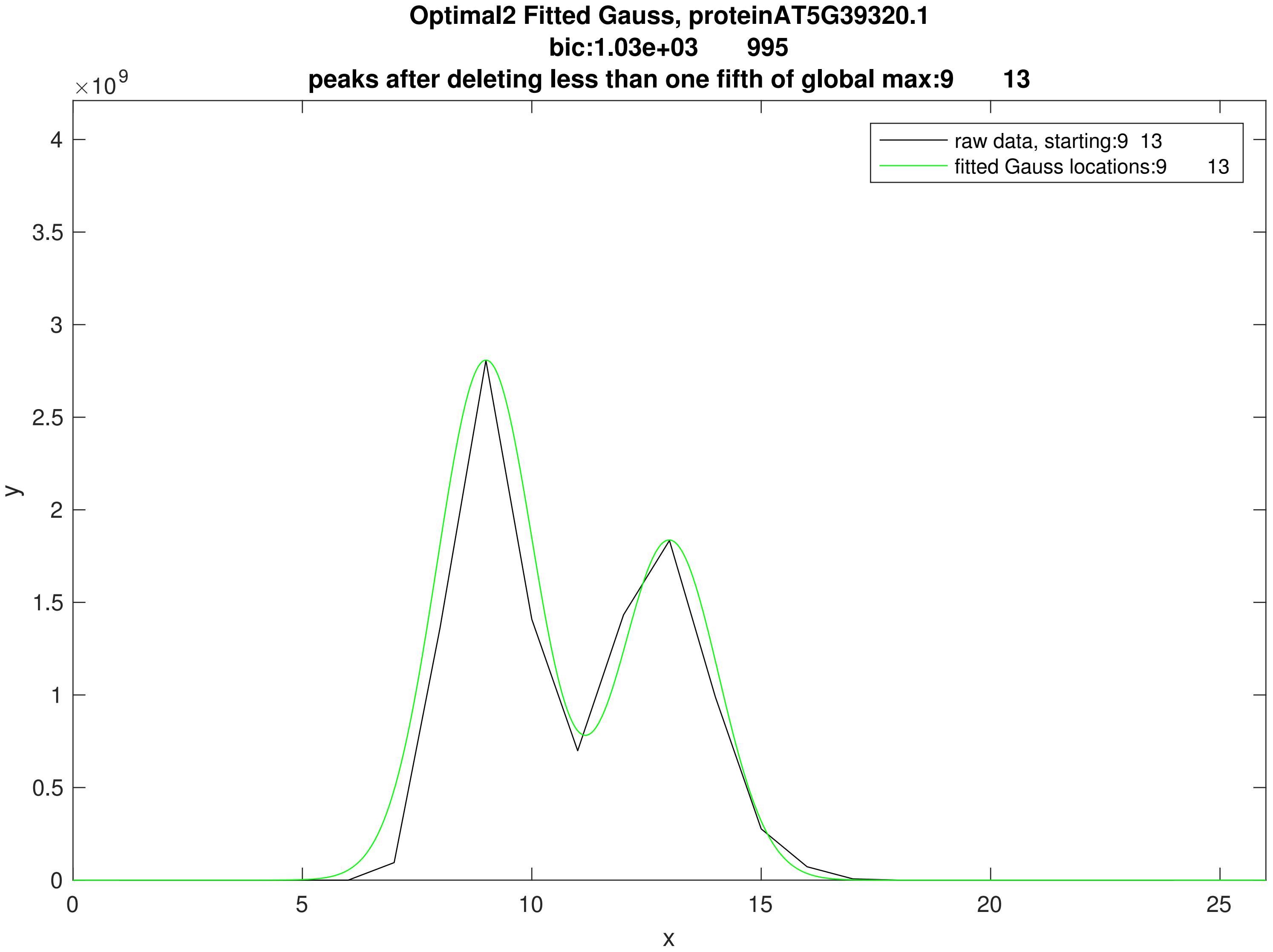

Optimal2 Fitted Gauss, proteinAT5G39320.1
bic:1.03e+03 995
peaks after deleting less than one fifth of global max:9 13
×10^9
raw data, starting:9 13
fitted Gauss locations:9 13
y
x
0
0.5
1
1.5
2
2.5
3
3.5
4
0
5
10
15
20
25

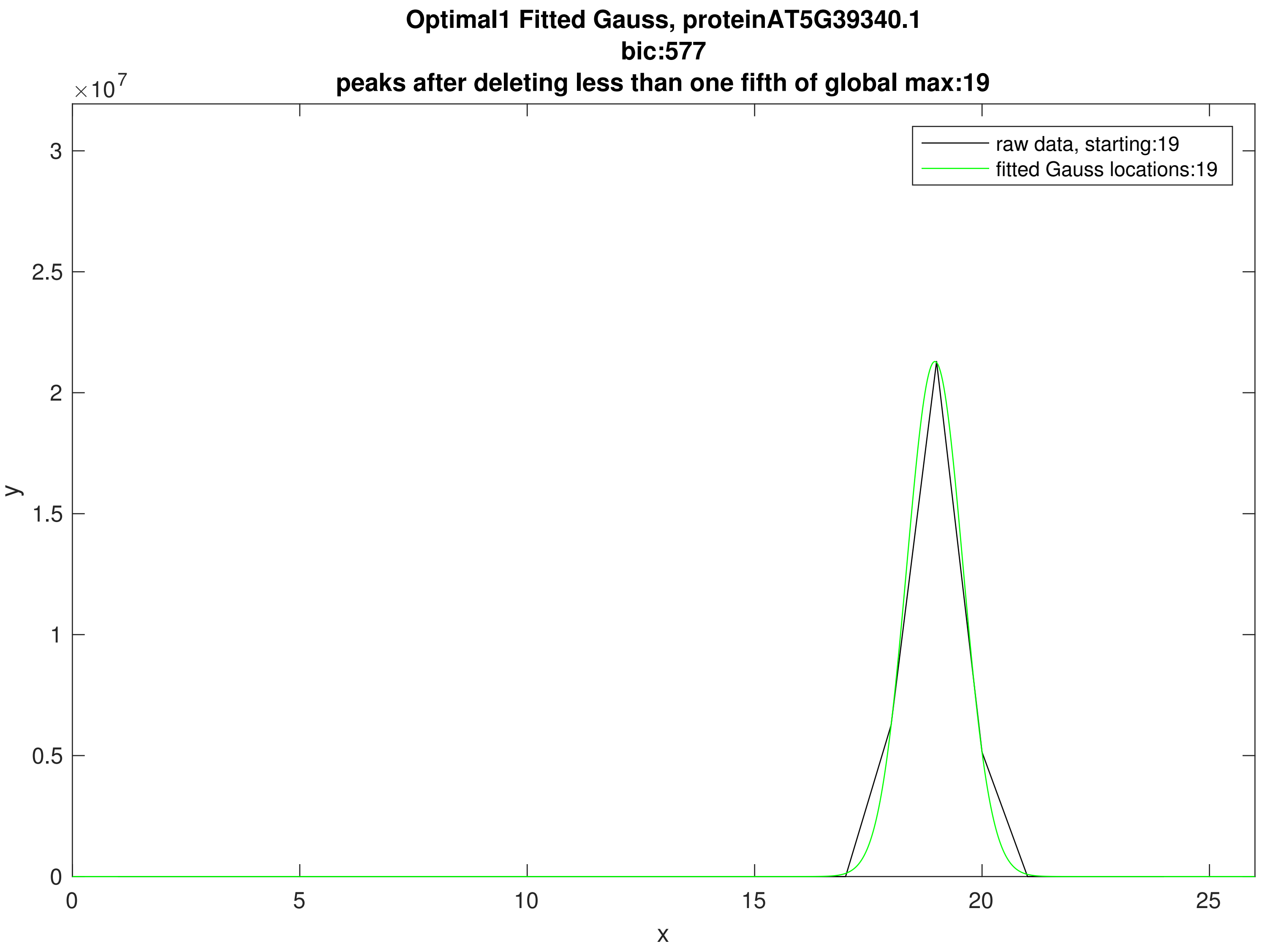

Optimal1 Fitted Gauss, proteinAT5G39340.1
bic:577
peaks after deleting less than one fifth of global max:19
×10^7
3
2.5
2
1.5
1
0.5
0
y
0
5
10
15
20
25
x
raw data, starting:19
fitted Gauss locations:19

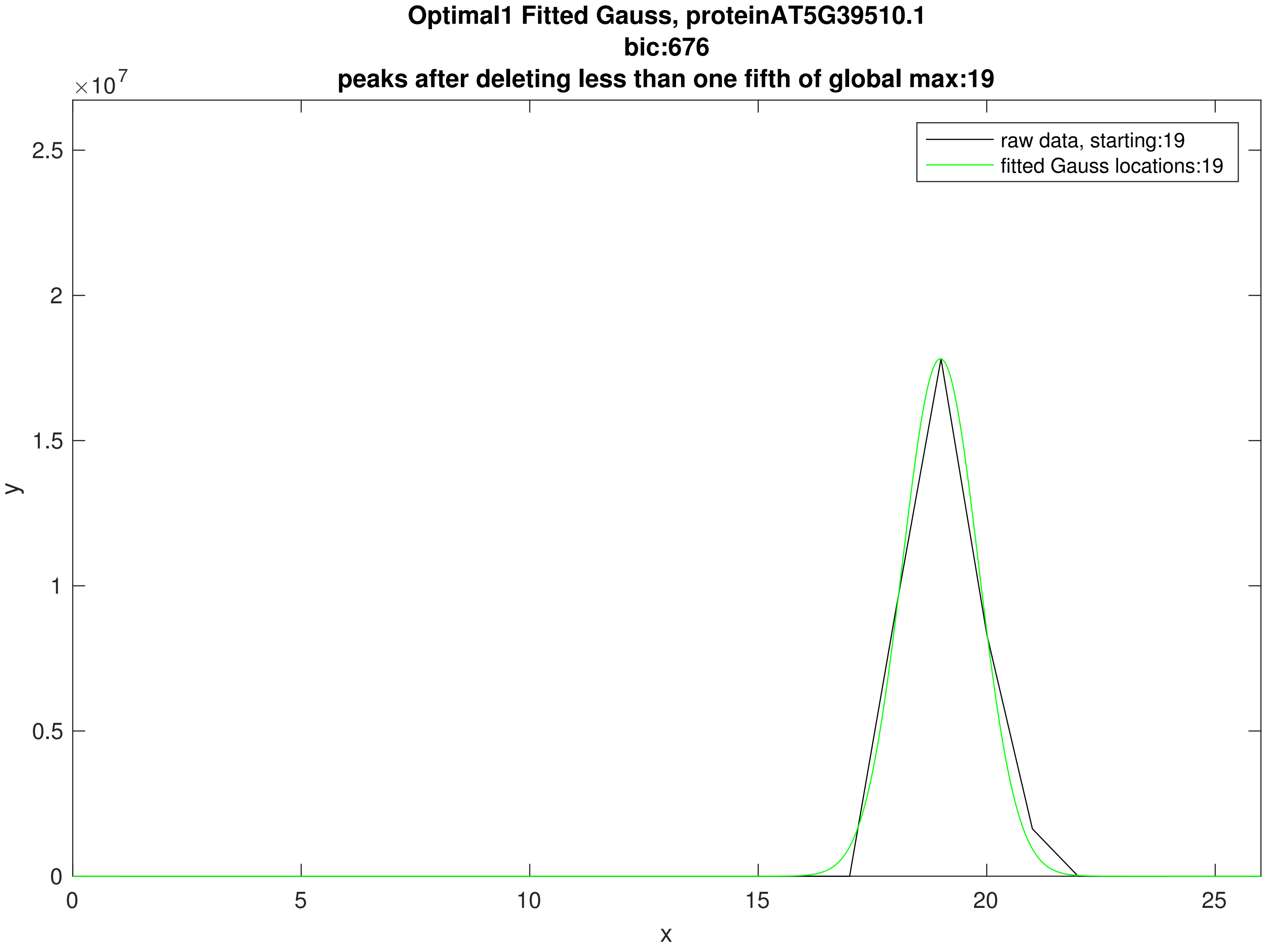
Optimal1 Fitted Gauss, proteinAT5G39510.1
bic:676
peaks after deleting less than one fifth of global max:19
×10^7
2.5
2
1.5
1
0.5
0
y
0
5
10
15
20
25
x
raw data, starting:19
fitted Gauss locations:19

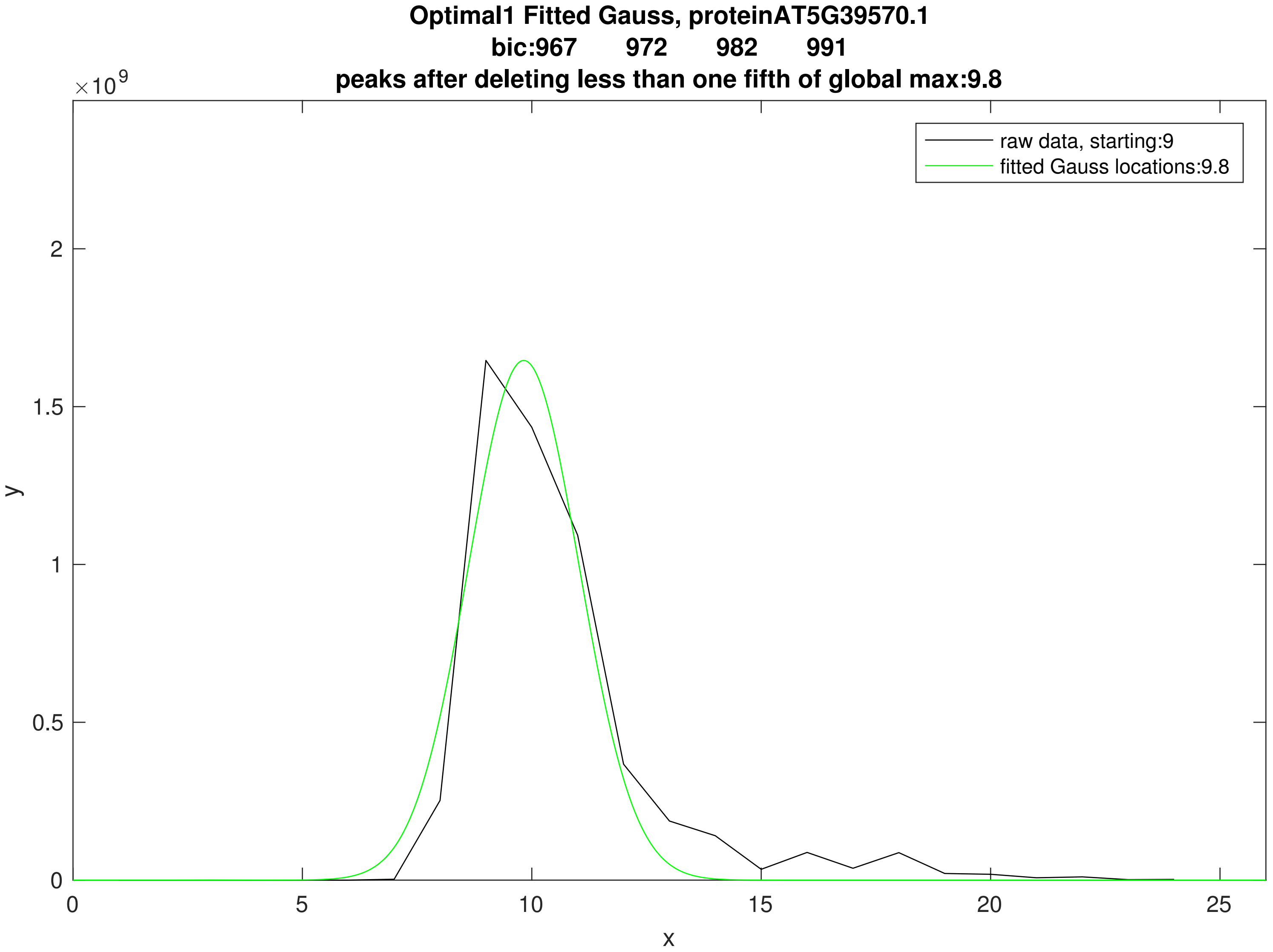
Optimal1 Fitted Gauss, proteinAT5G39570.1
bic:967 972 982 991
peaks after deleting less than one fifth of global max:9.8
×10^9
raw data, starting:9
fitted Gauss locations:9.8
y
x
0
0.5
1
1.5
2
0
5
10
15
20
25

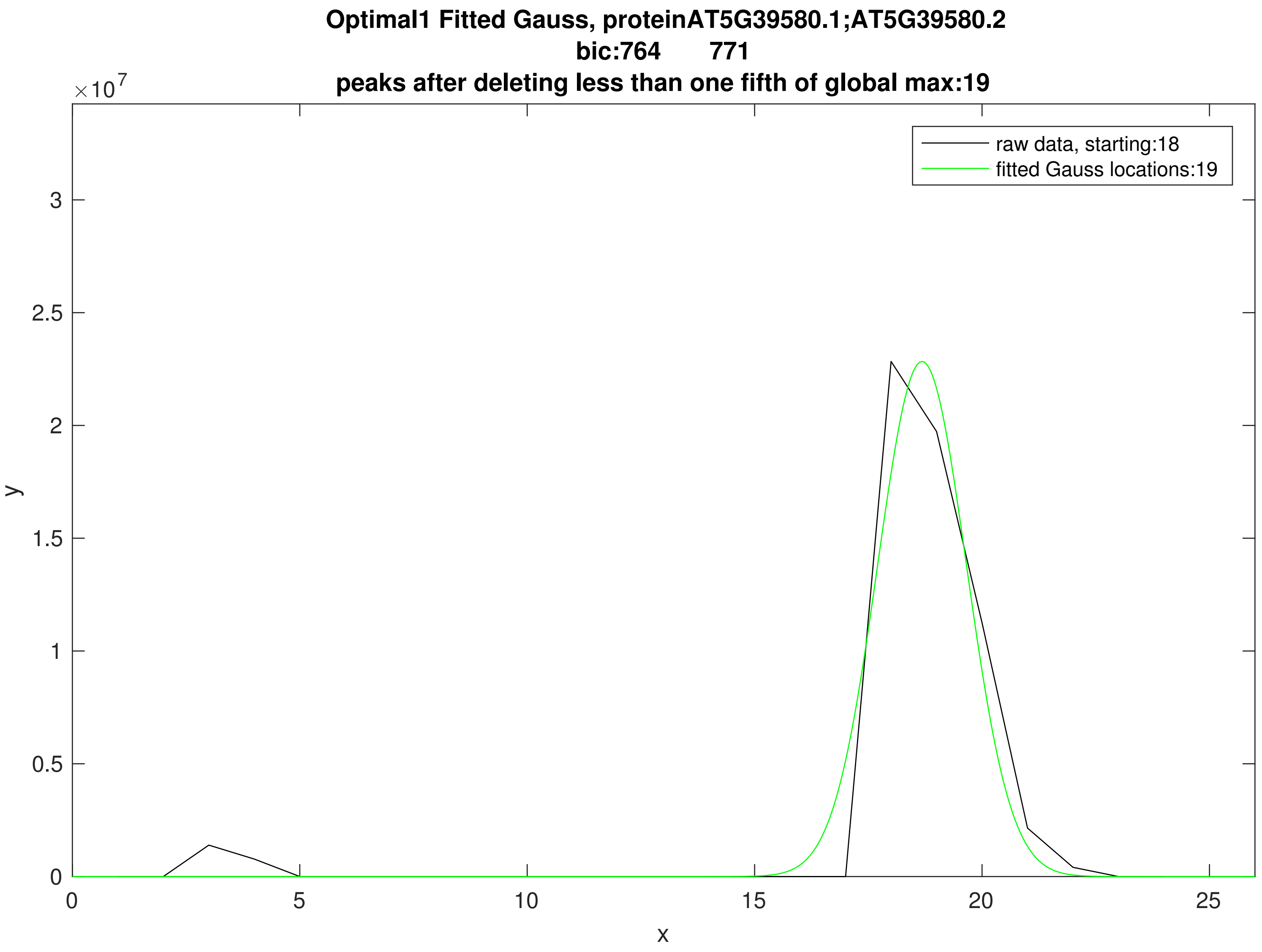
Optimal1 Fitted Gauss, proteinAT5G39580.1;AT5G39580.2
bic:764 771
peaks after deleting less than one fifth of global max:19
×10^7
raw data, starting:18
fitted Gauss locations:19
y
x
0
0.5
1
1.5
2
2.5
3
0
5
10
15
20
25

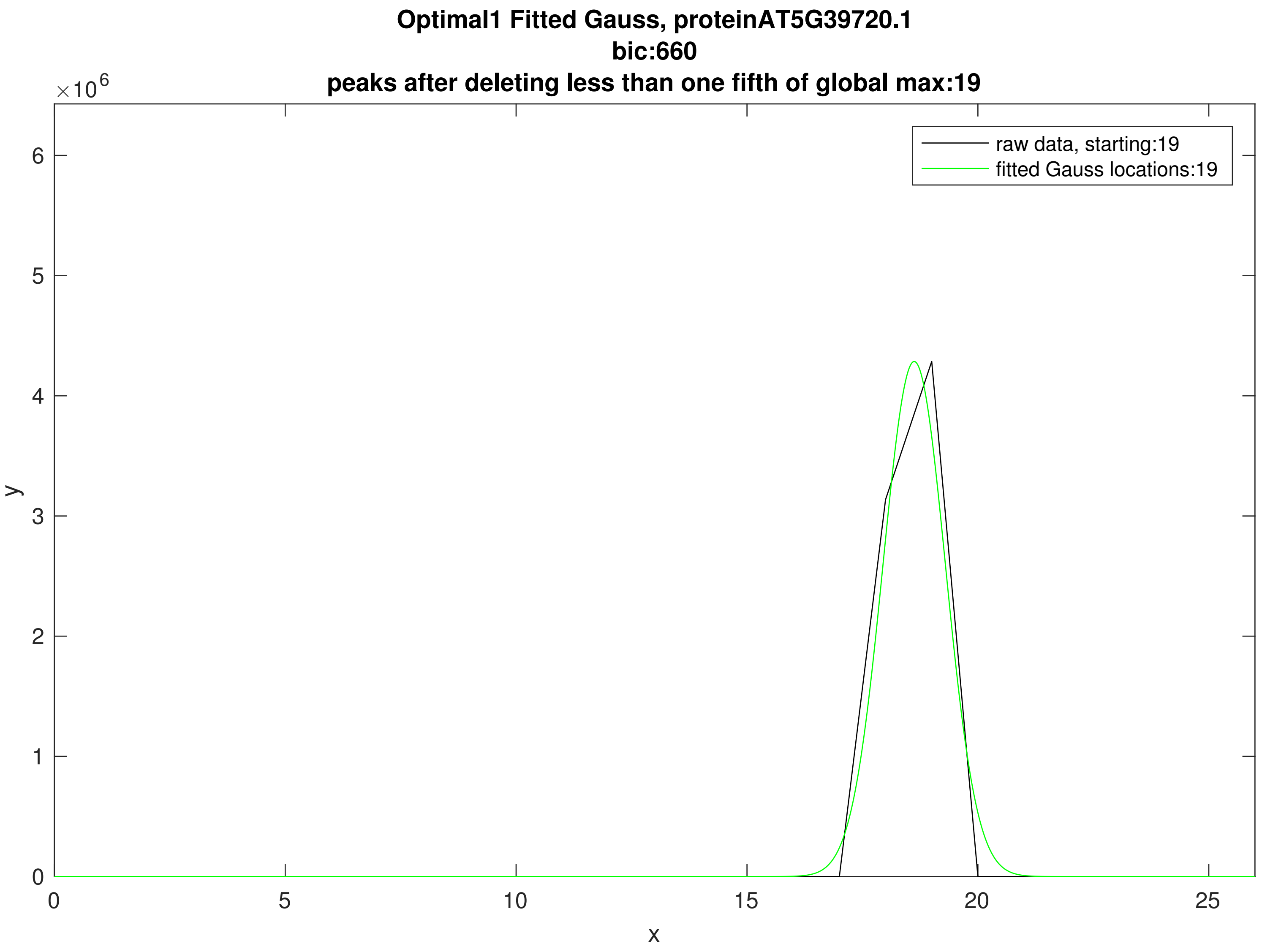
Optimal1 Fitted Gauss, proteinAT5G39720.1
bic:660
peaks after deleting less than one fifth of global max:19
×10^6
raw data, starting:19
fitted Gauss locations:19
y
x
0
1
2
3
4
5
6
0
5
10
15
20
25

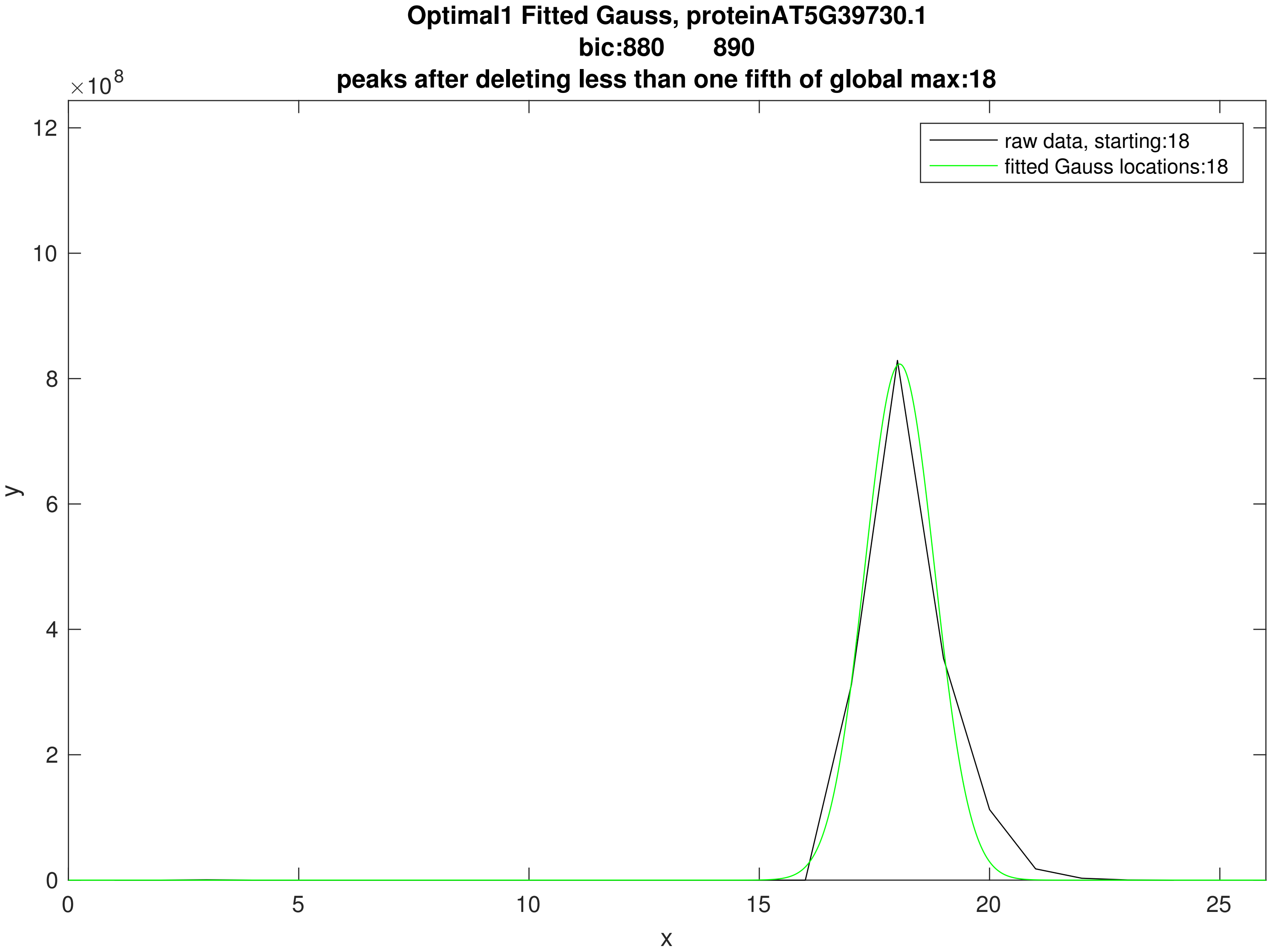

Optimal1 Fitted Gauss, proteinAT5G39730.1
bic:880 890
peaks after deleting less than one fifth of global max:18
×10^8
12
10
8
6
4
2
0
y
0
5
10
15
20
25
x
raw data, starting:18
fitted Gauss locations:18

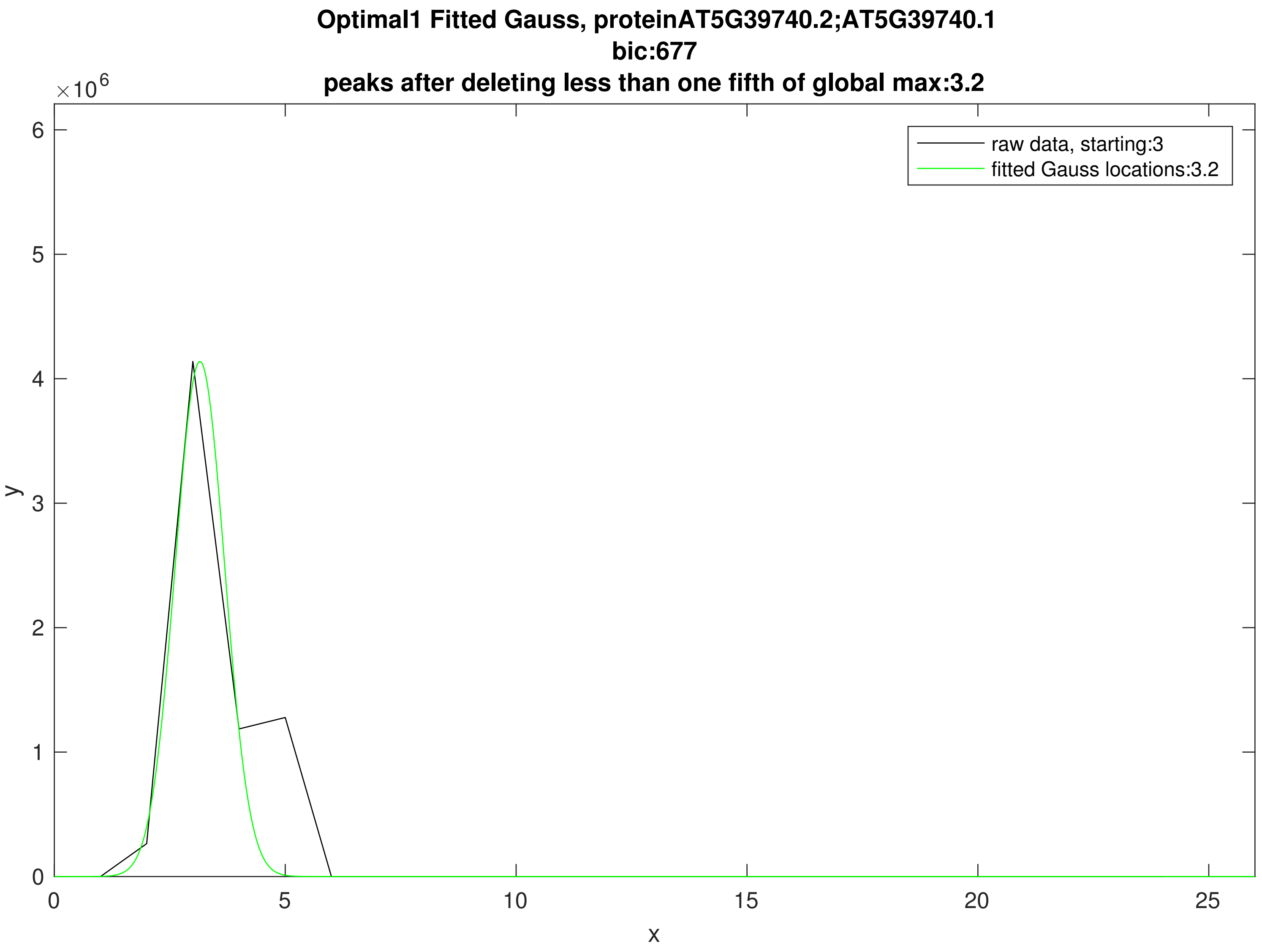

Optimal1 Fitted Gauss, proteinAT5G39740.2;AT5G39740.1
bic:677
peaks after deleting less than one fifth of global max:3.2
×10^6
raw data, starting:3
fitted Gauss locations:3.2
y
x
0
1
2
3
4
5
6
0
5
10
15
20
25

#### Optimal1 Fitted Gauss, proteinAT5G40060.1
#### bic:742
#### peaks after deleting less than one fifth of global max:14

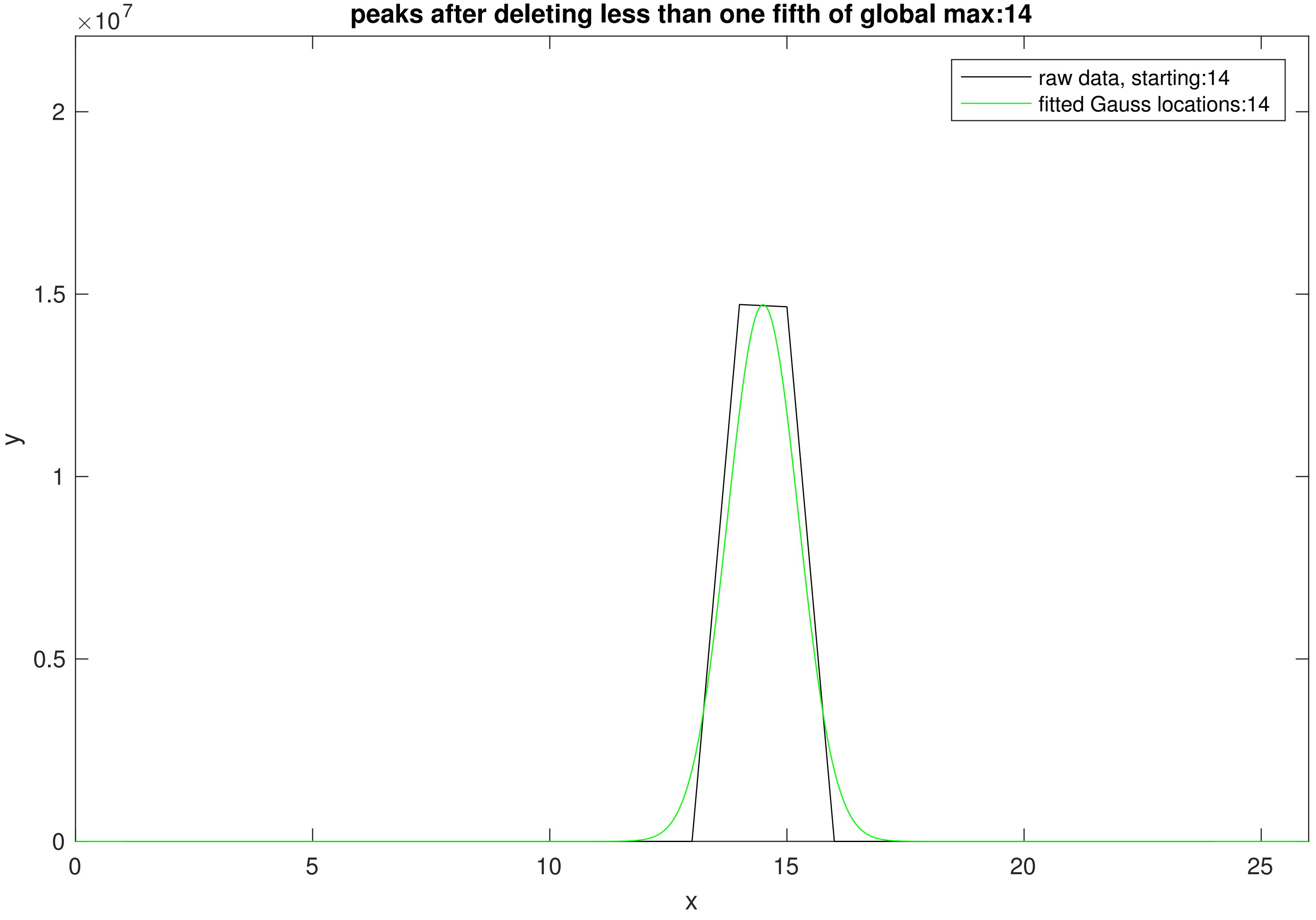

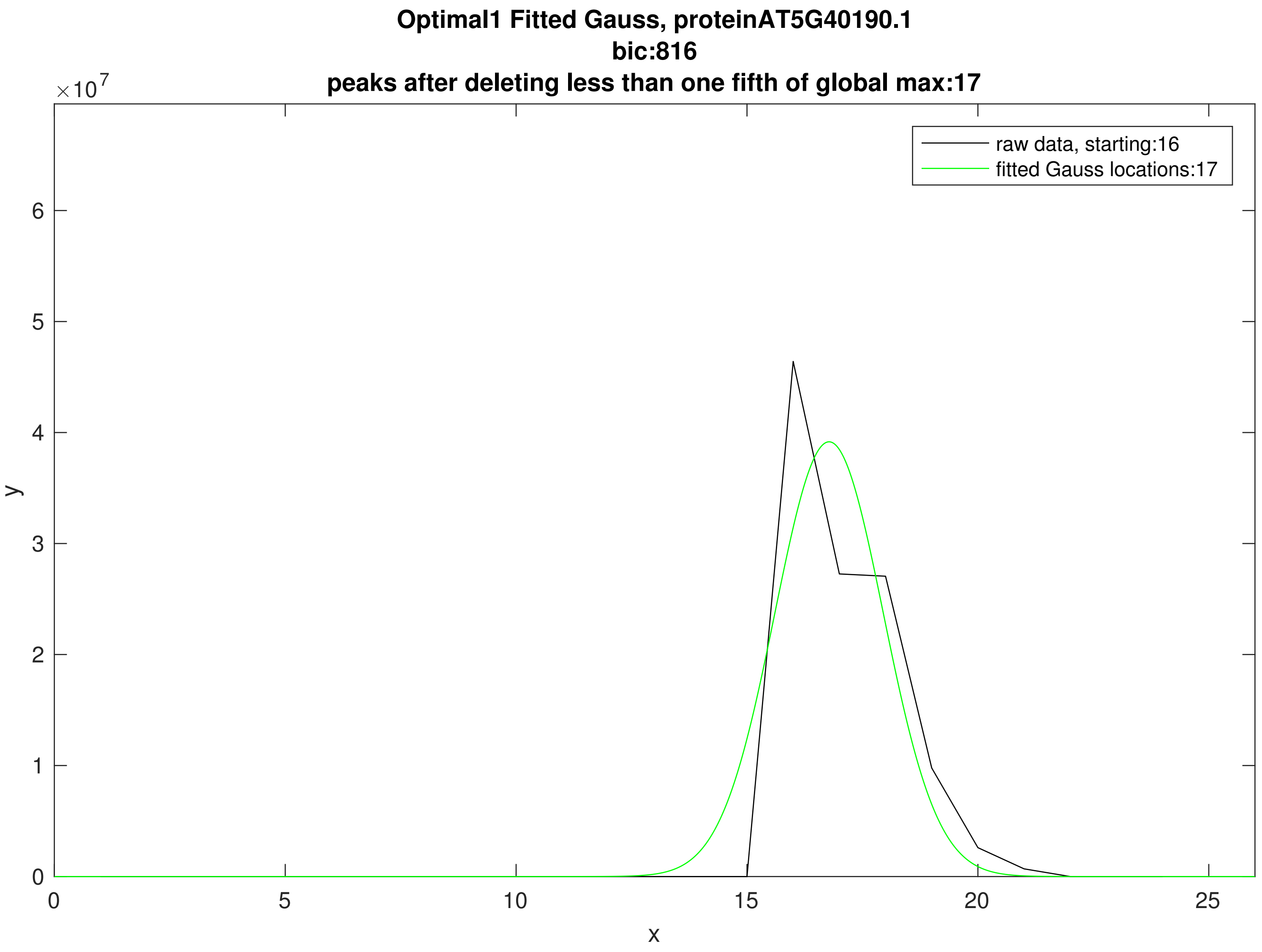
Optimal1 Fitted Gauss, proteinAT5G40190.1
bic:816
peaks after deleting less than one fifth of global max:17
×10^7
raw data, starting:16
fitted Gauss locations:17
y
x
0
1
2
3
4
5
6
0
5
10
15
20
25

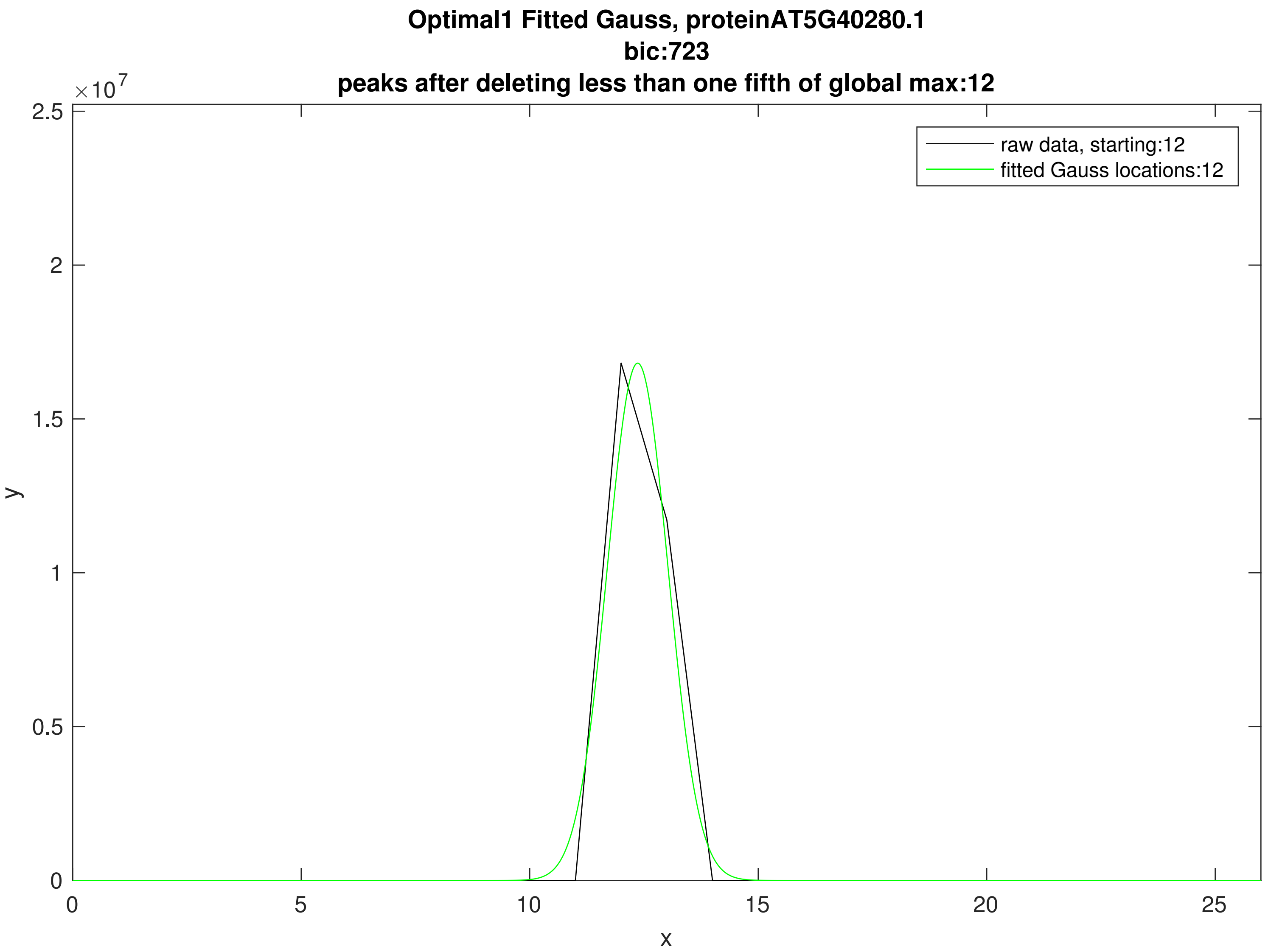

Optimal1 Fitted Gauss, proteinAT5G40280.1
bic:723
peaks after deleting less than one fifth of global max:12
×10^7
2.5
2
1.5
1
0.5
0
y
0
5
10
15
20
25
x
raw data, starting:12
fitted Gauss locations:12

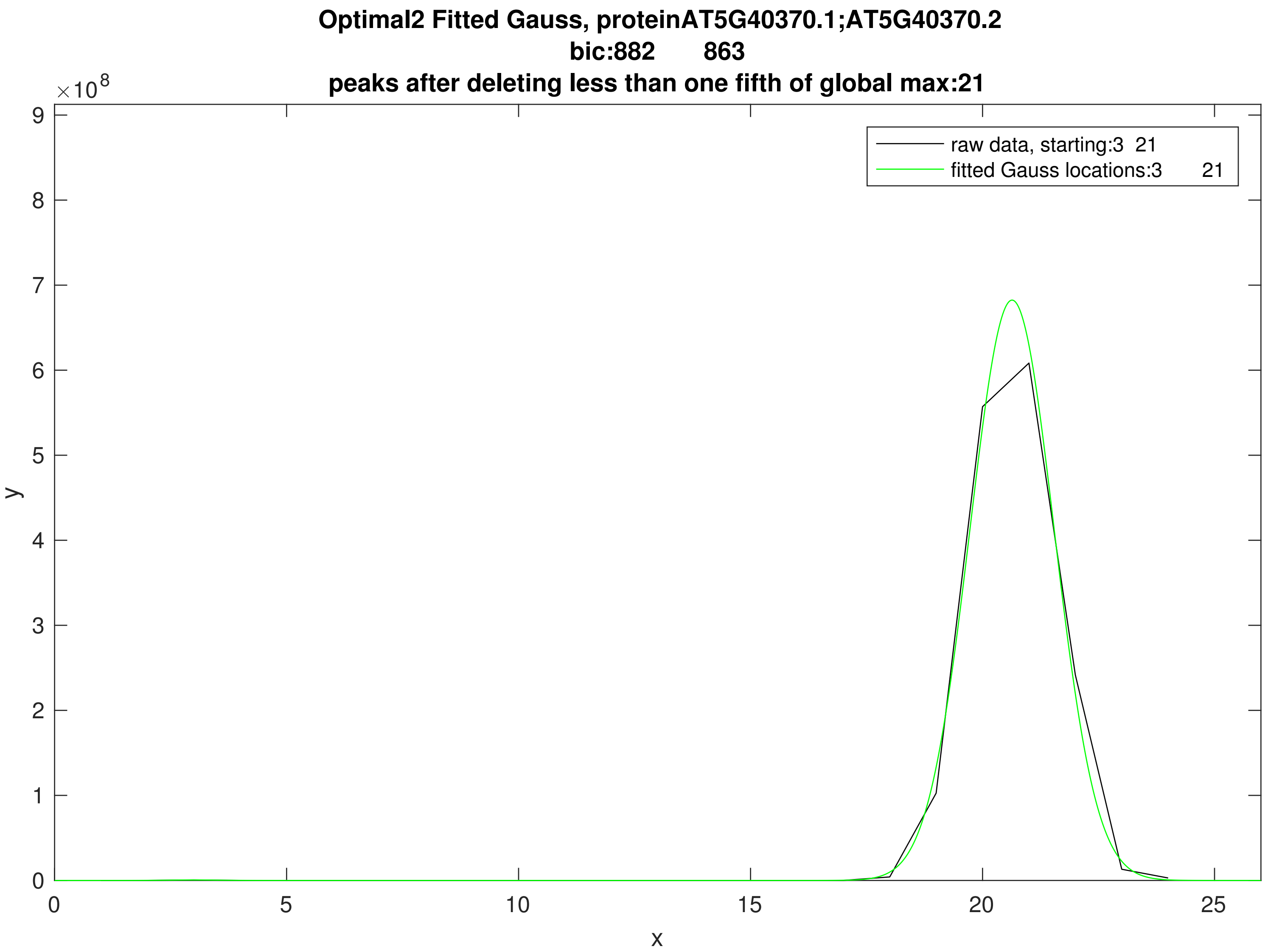

Optimal2 Fitted Gauss, proteinAT5G40370.1;AT5G40370.2
bic:882 863
peaks after deleting less than one fifth of global max:21
×10^8
raw data, starting:3 21
fitted Gauss locations:3 21
y
x
0
1
2
3
4
5
6
7
8
9
0
5
10
15
20
25

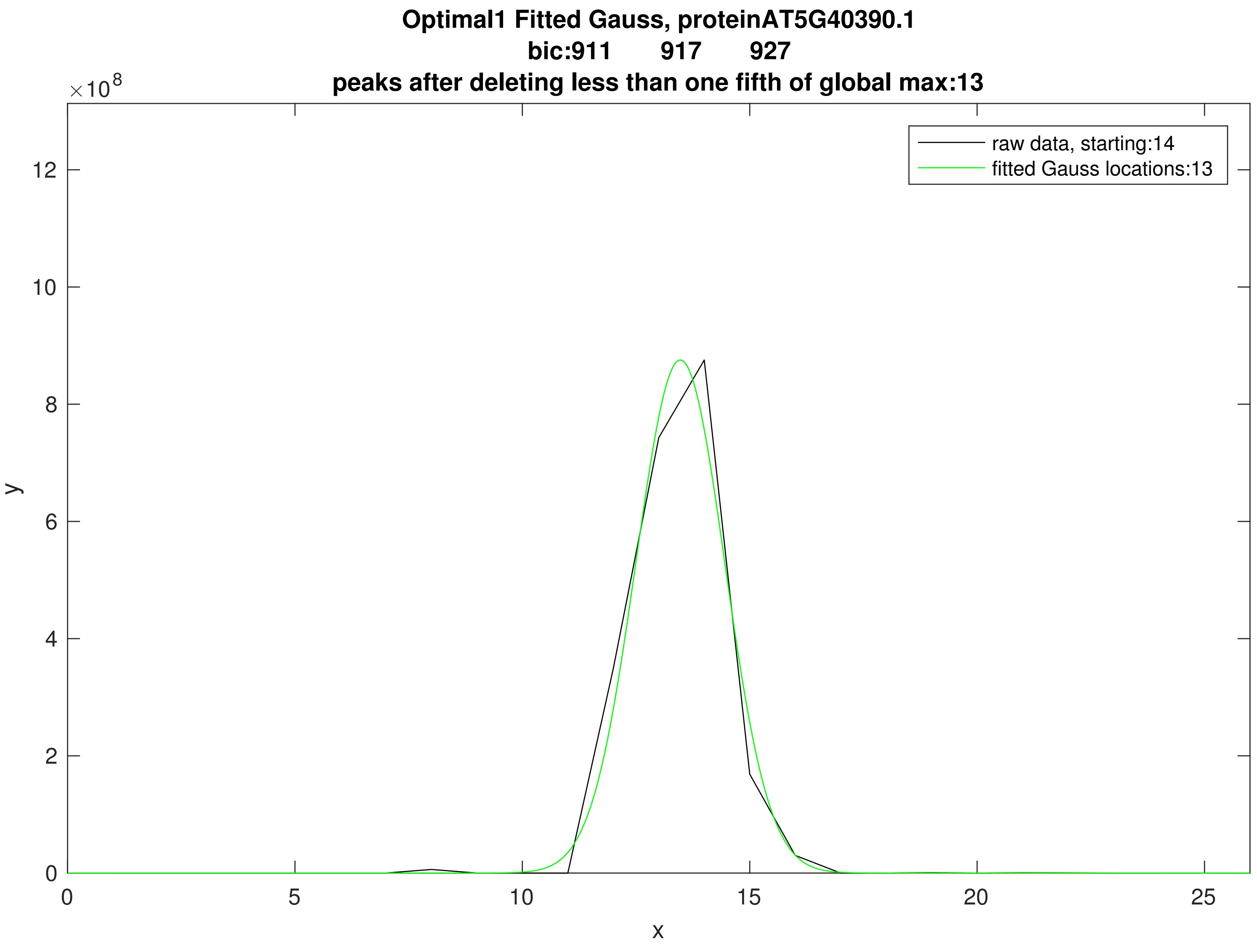
Optimal1 Fitted Gauss, proteinAT5G40390.1
bic:911 917 927
peaks after deleting less than one fifth of global max:13
×10^8
raw data, starting:14
fitted Gauss locations:13
12
10
8
6
4
2
0
y
0
5
10
15
20
25
x

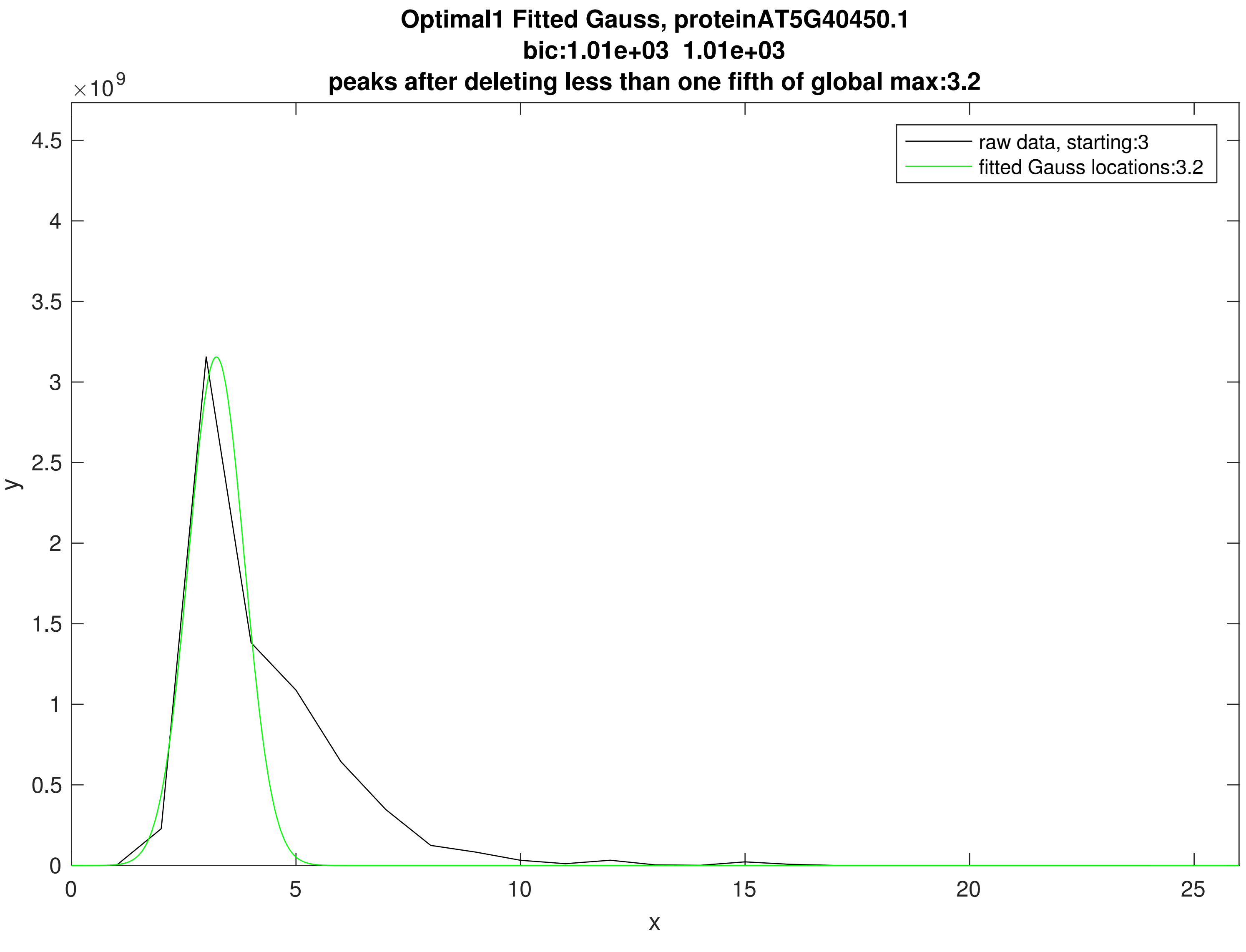
Optimal1 Fitted Gauss, proteinAT5G40450.1
bic:1.01e+03 1.01e+03
peaks after deleting less than one fifth of global max:3.2
×10^9
y
x
raw data, starting:3
fitted Gauss locations:3.2
0
0.5
1
1.5
2
2.5
3
3.5
4
4.5
5
10
15
20
25

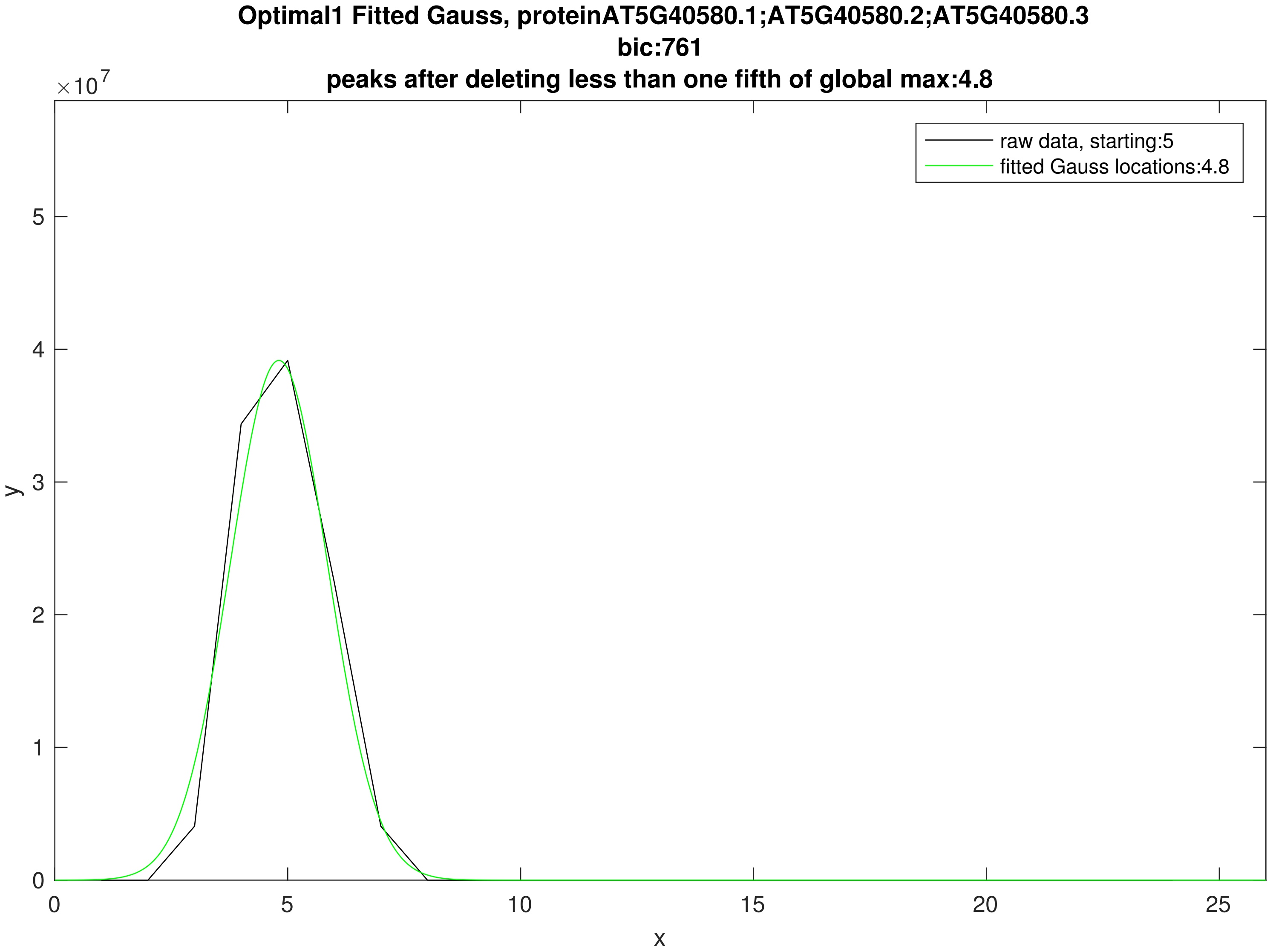
Optimal1 Fitted Gauss, proteinAT5G40580.1;AT5G40580.2;AT5G40580.3
bic:761
peaks after deleting less than one fifth of global max:4.8
×10^7
raw data, starting:5
fitted Gauss locations:4.8
y
x
0
1
2
3
4
5
10
15
20
25

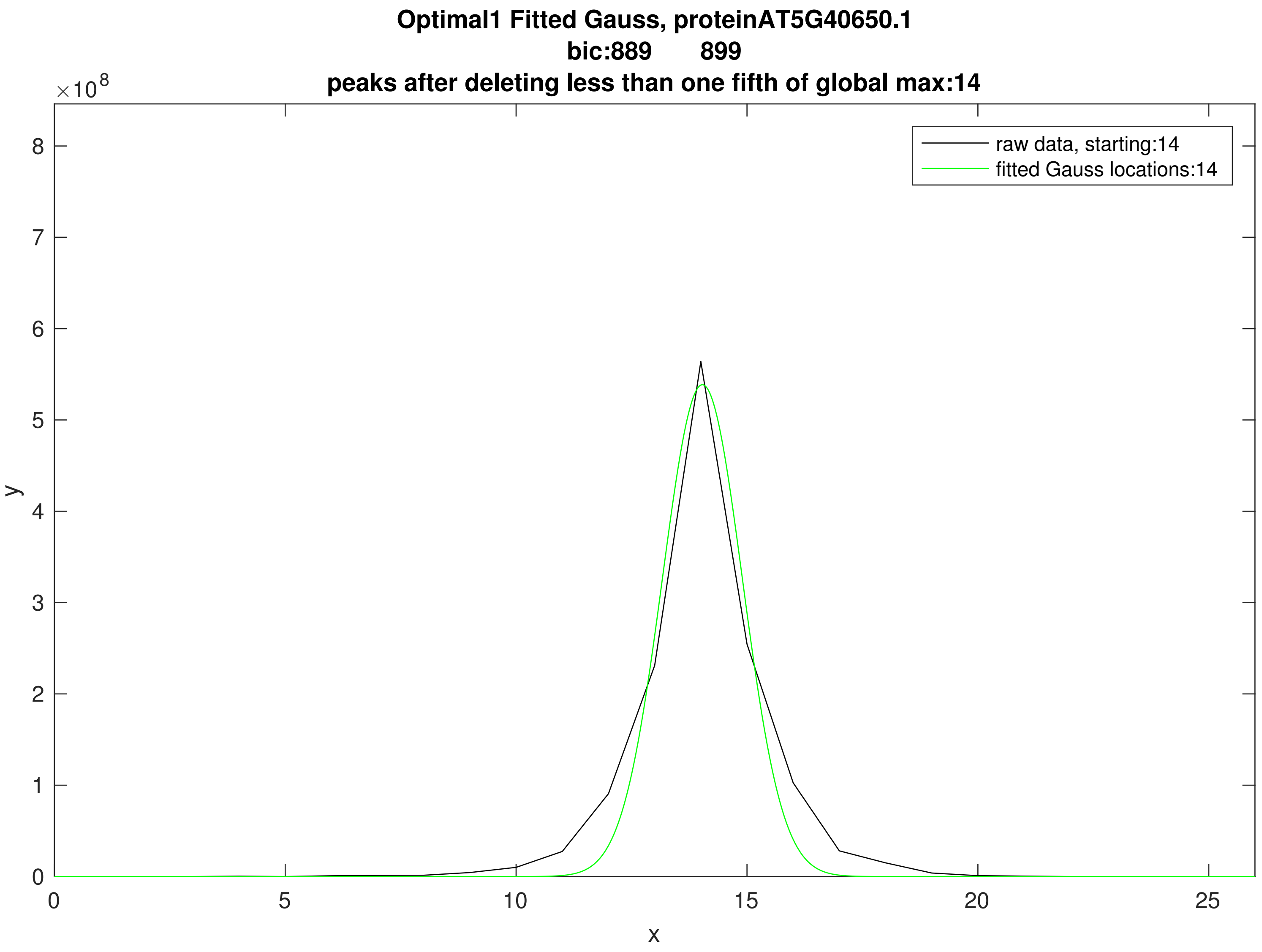
Optimal1 Fitted Gauss, proteinAT5G40650.1
bic:889 899
peaks after deleting less than one fifth of global max:14
×10^8
raw data, starting:14
fitted Gauss locations:14
y
x
0
1
2
3
4
5
6
7
8
0
5
10
15
20
25

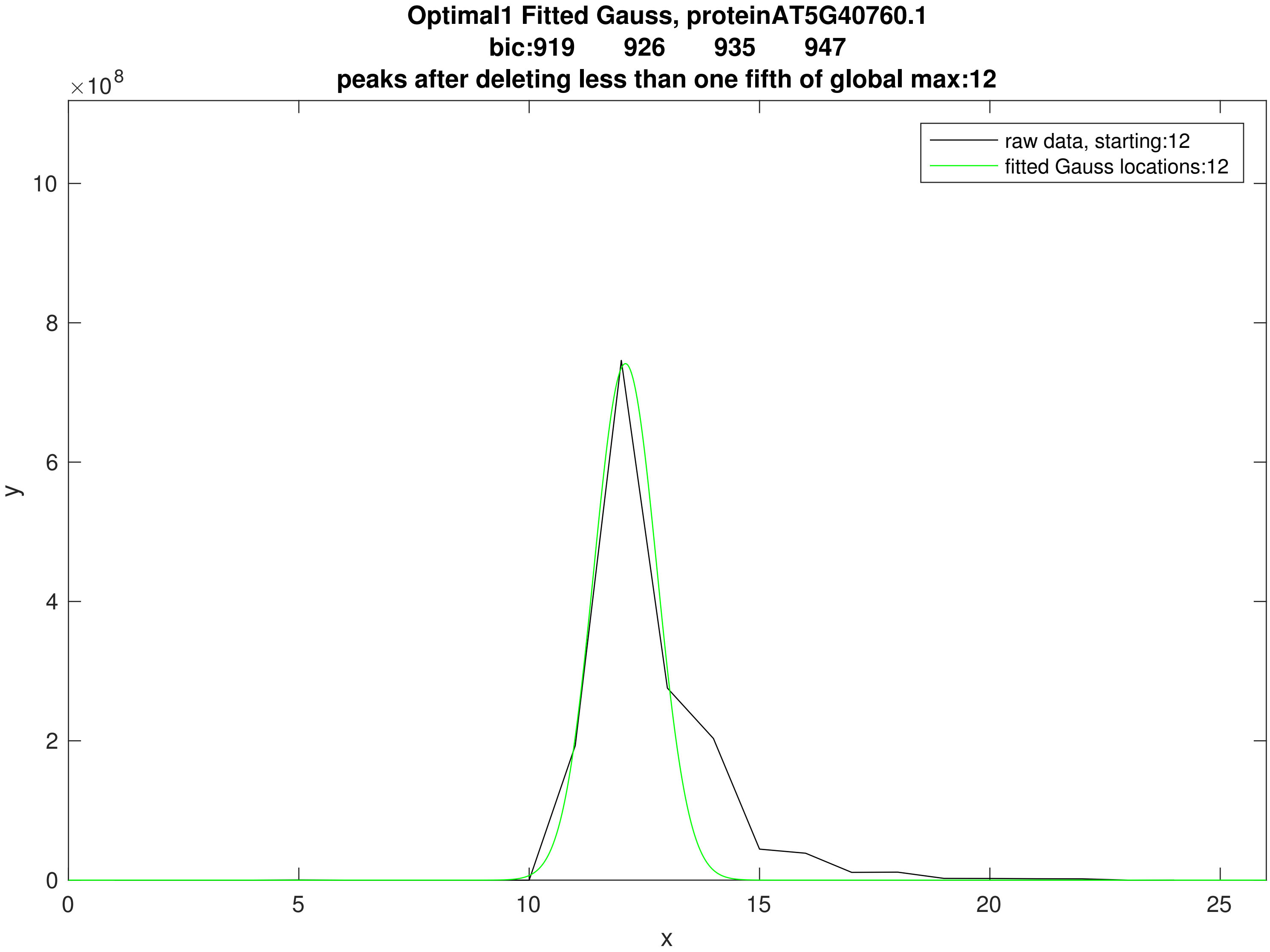

Optimal1 Fitted Gauss, proteinAT5G40760.1
bic:919 926 935 947
peaks after deleting less than one fifth of global max:12
$\times 10^8$
raw data, starting:12
fitted Gauss locations:12
10
8
6
4
2
0
y
0
5
10
15
20
25
x

Optimal1 Fitted Gauss, proteinAT5G40950.1
bic:702
peaks after deleting less than one fifth of global max:3.9

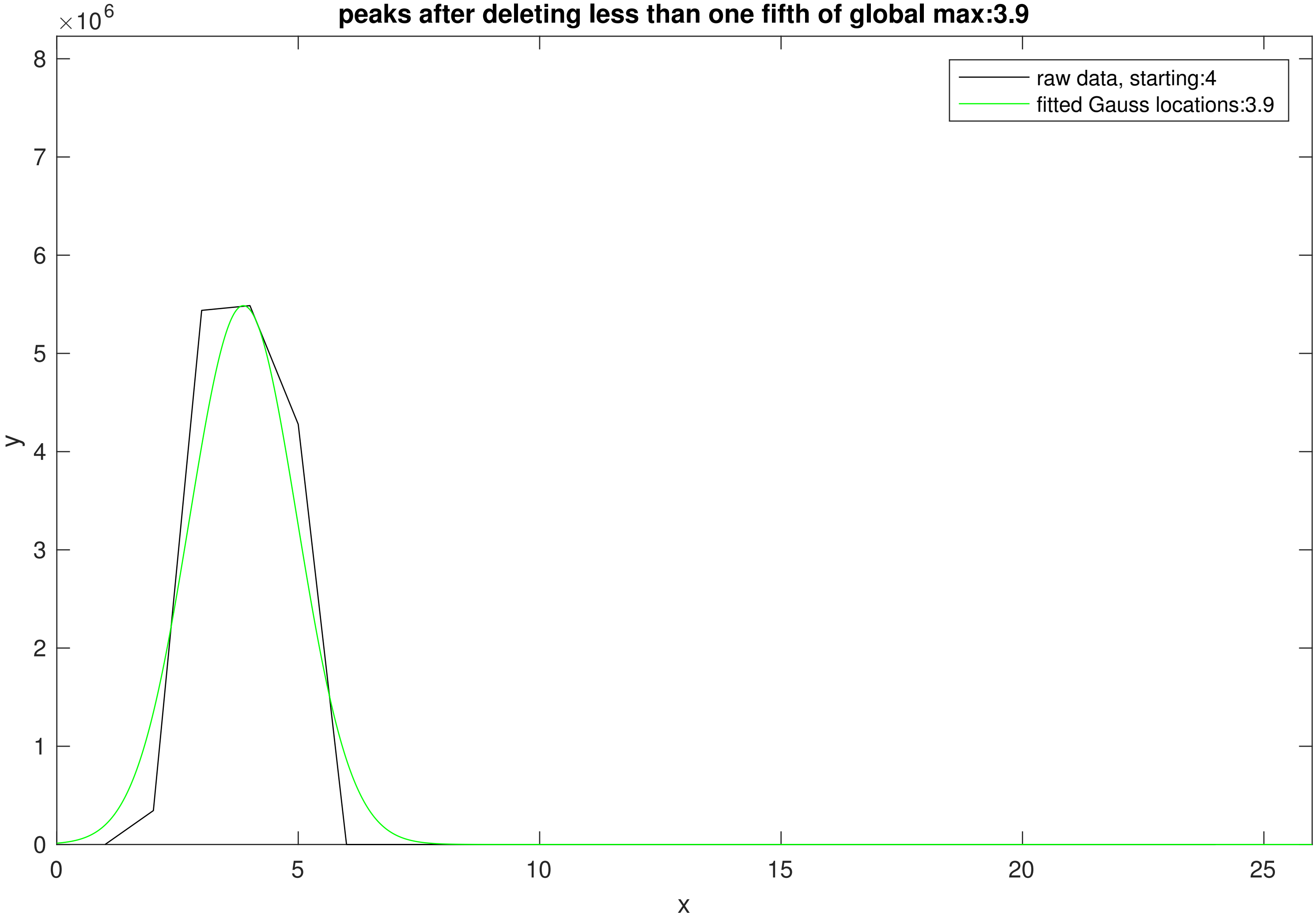

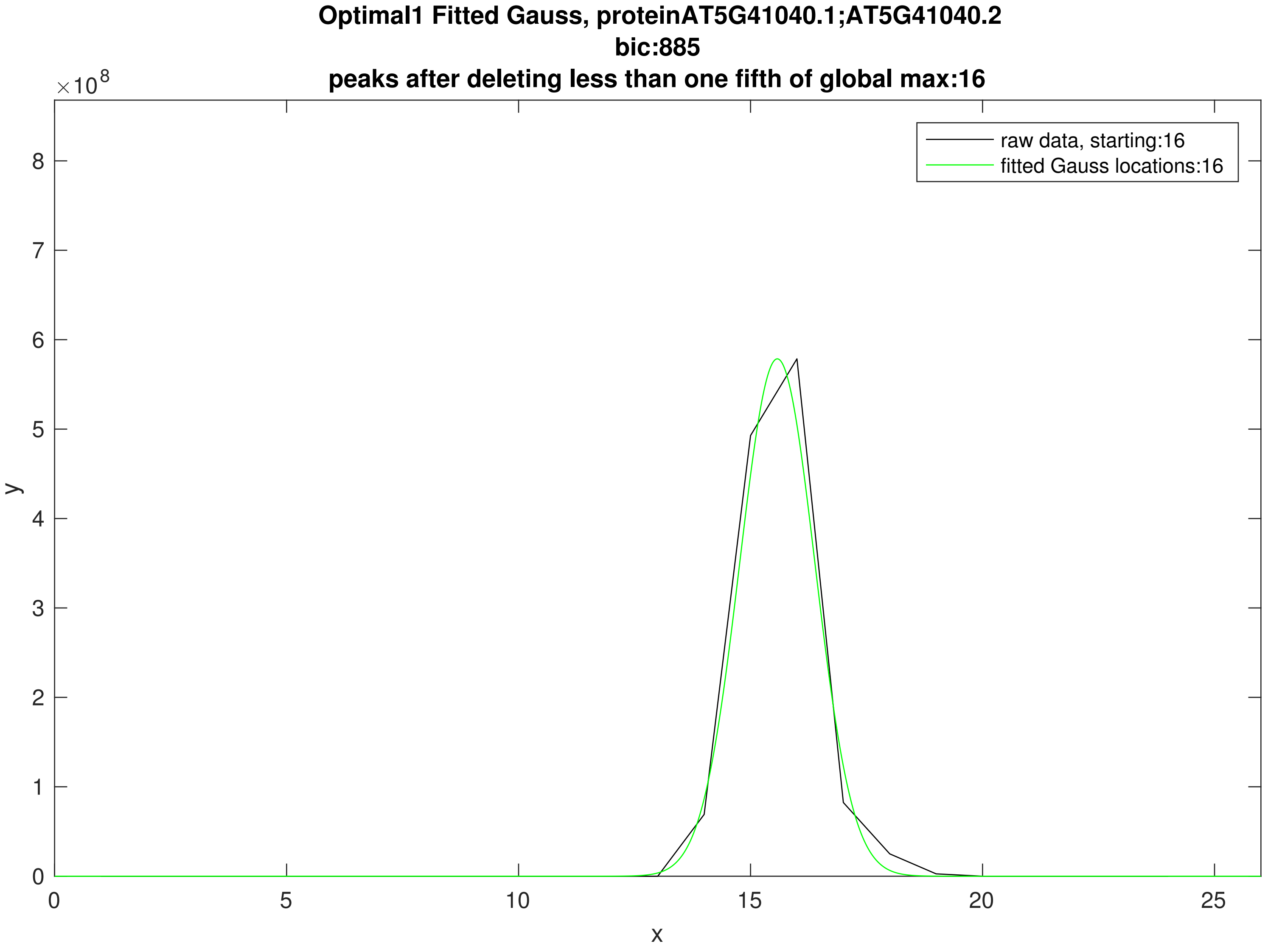
Optimal1 Fitted Gauss, proteinAT5G41040.1;AT5G41040.2
bic:885
peaks after deleting less than one fifth of global max:16
×10^8
raw data, starting:16
fitted Gauss locations:16
y
x
0
1
2
3
4
5
6
7
8
0
5
10
15
20
25

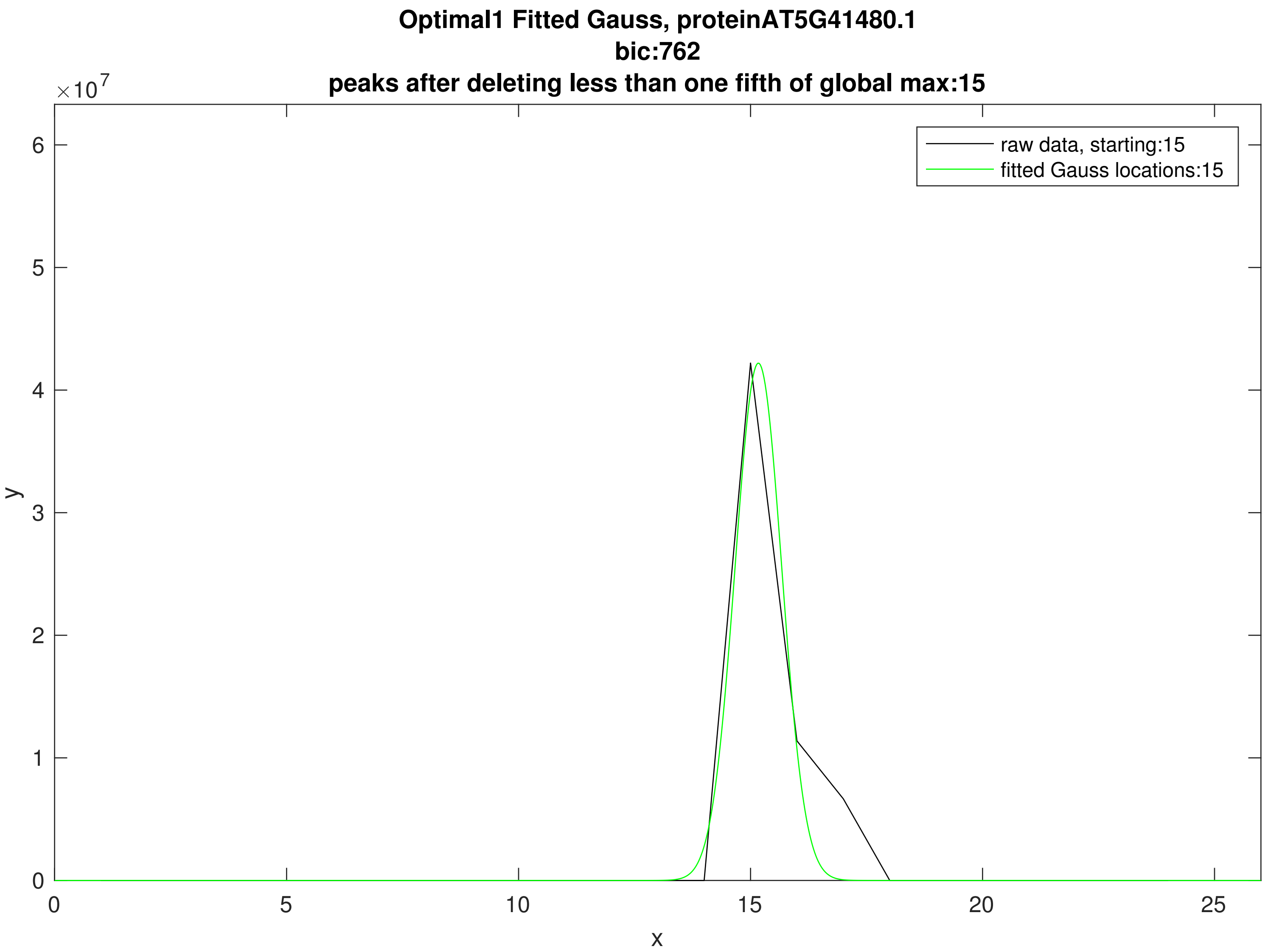

Optimal1 Fitted Gauss, proteinAT5G41480.1
bic:762
peaks after deleting less than one fifth of global max:15
×10^7
raw data, starting:15
fitted Gauss locations:15
y
x
0
1
2
3
4
5
6
0
5
10
15
20
25

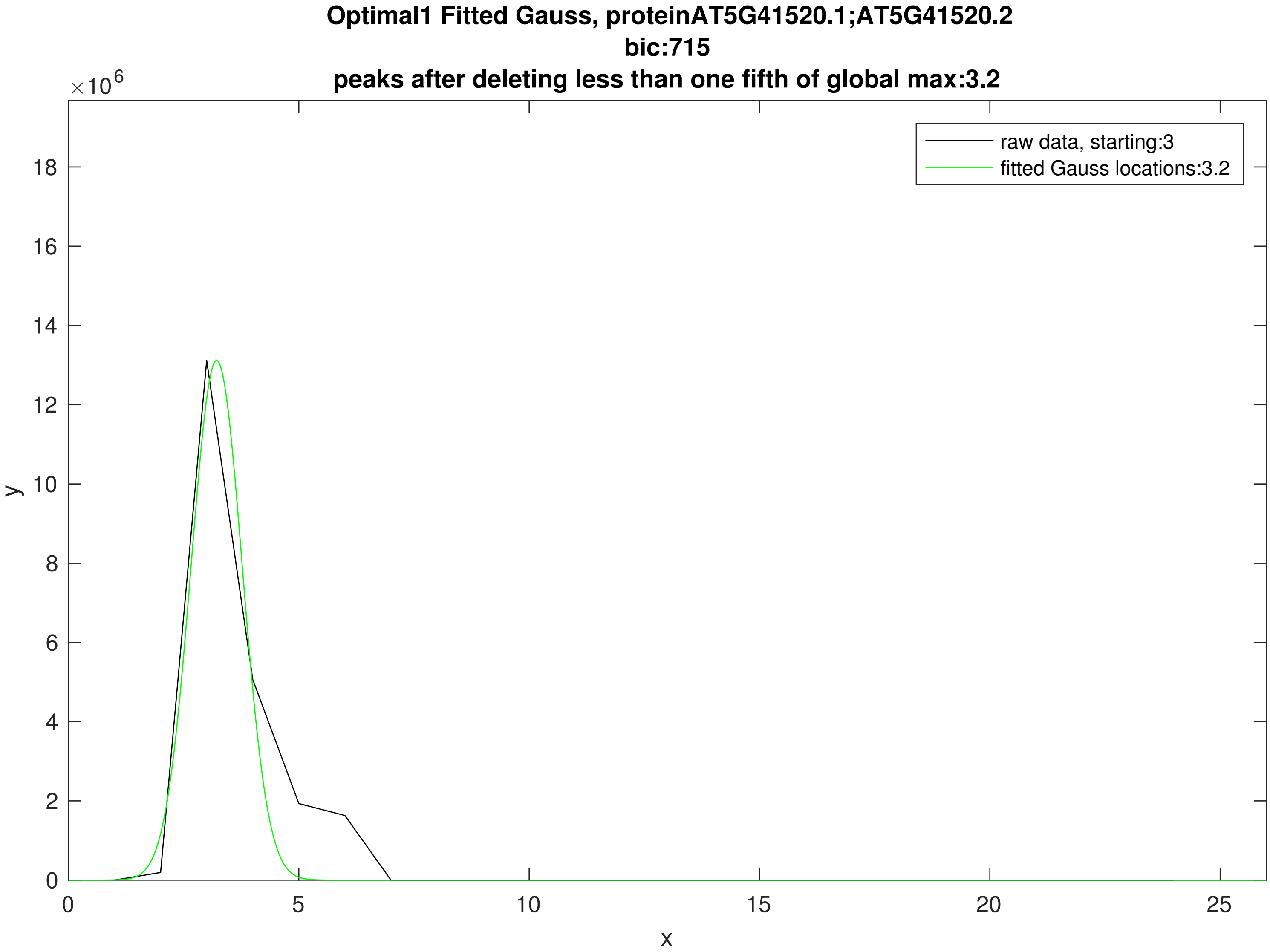
Optimal1 Fitted Gauss, proteinAT5G41520.1;AT5G41520.2
bic:715
peaks after deleting less than one fifth of global max:3.2
×10^6
raw data, starting:3
fitted Gauss locations:3.2
y
x
0
2
4
6
8
10
12
14
16
18
0
5
10
15
20
25

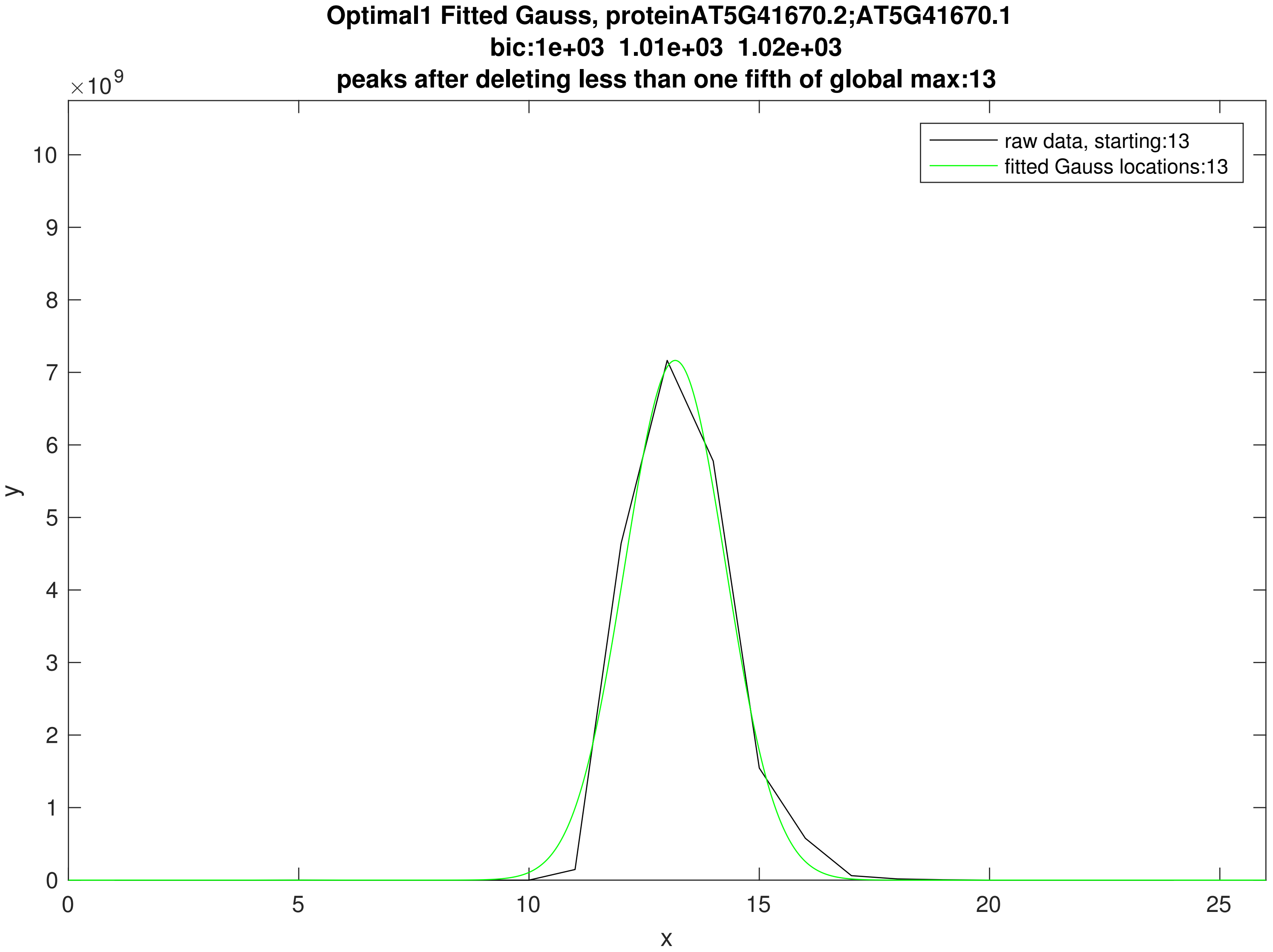

Optimal1 Fitted Gauss, proteinAT5G41670.2;AT5G41670.1
bic:1e+03 1.01e+03 1.02e+03
peaks after deleting less than one fifth of global max:13
×10^9
raw data, starting:13
fitted Gauss locations:13
y
x
0
1
2
3
4
5
6
7
8
9
10
0
5
10
15
20
25

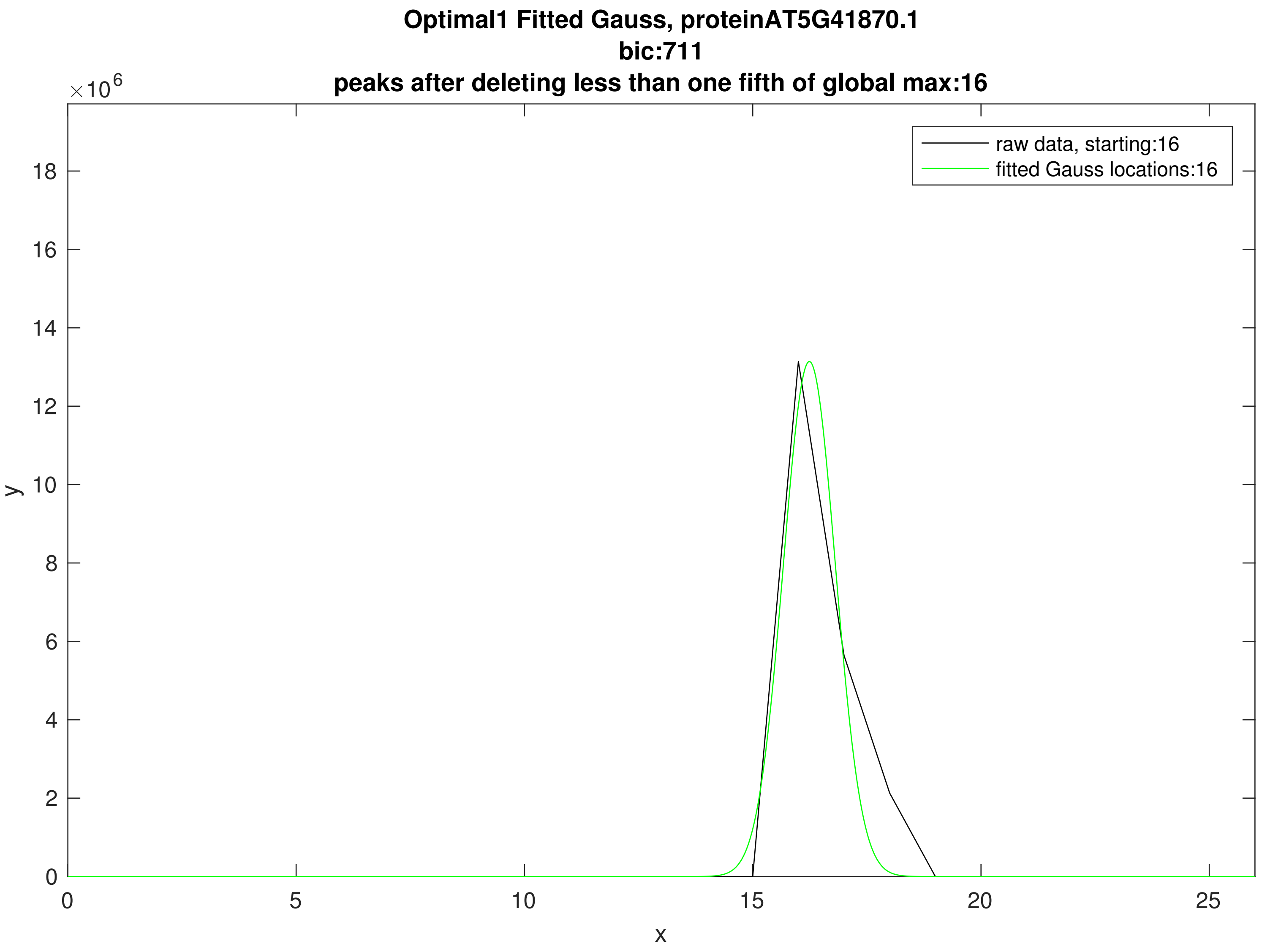

Optimal1 Fitted Gauss, proteinAT5G41870.1
bic:711
peaks after deleting less than one fifth of global max:16
×10^6
raw data, starting:16
fitted Gauss locations:16
y
x
0
2
4
6
8
10
12
14
16
18
0
5
10
15
20
25

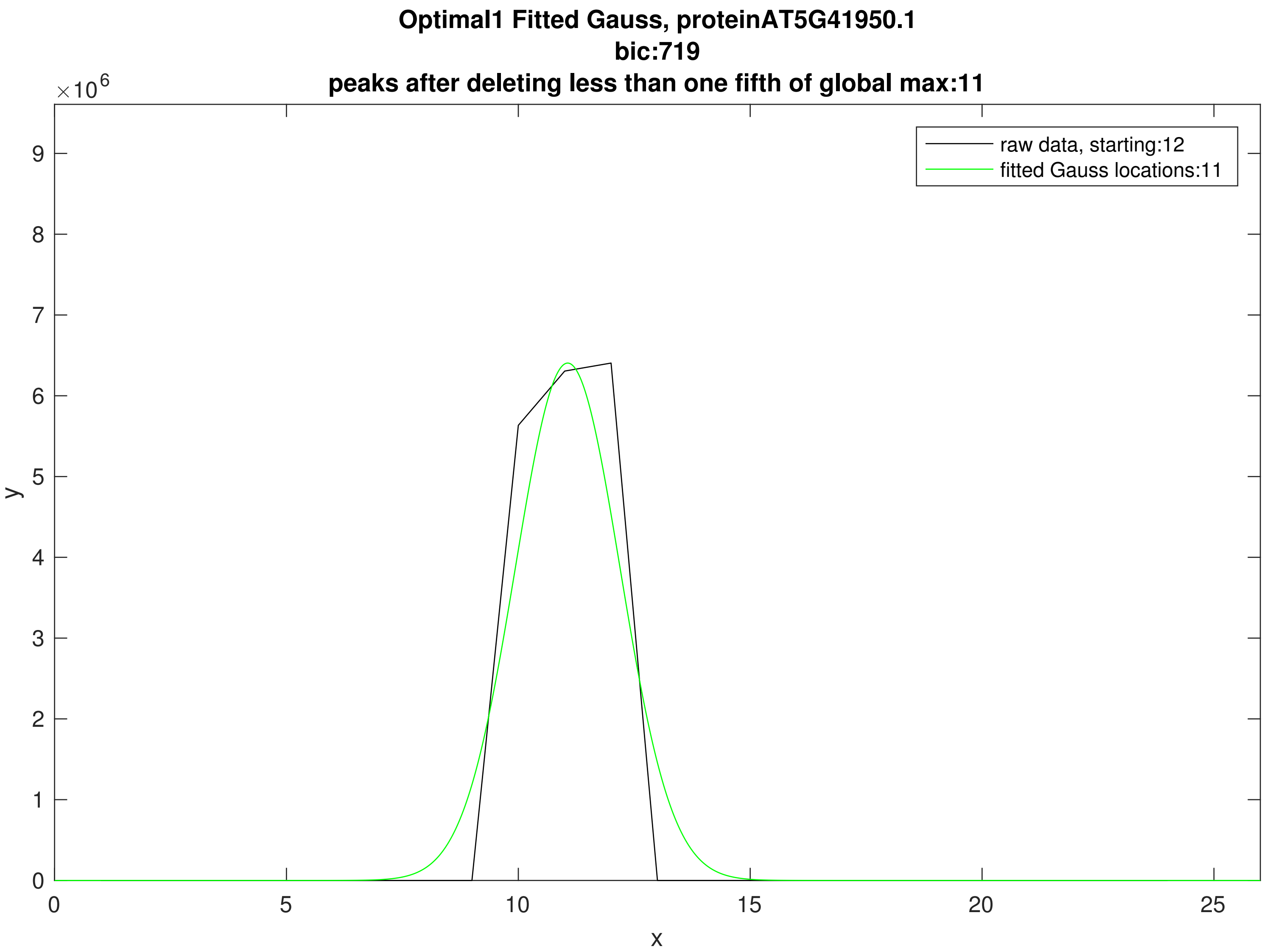

Optimal1 Fitted Gauss, proteinAT5G41950.1
bic:719
peaks after deleting less than one fifth of global max:11
×10^6
raw data, starting:12
fitted Gauss locations:11
y
x
0
1
2
3
4
5
6
7
8
9
0
5
10
15
20
25

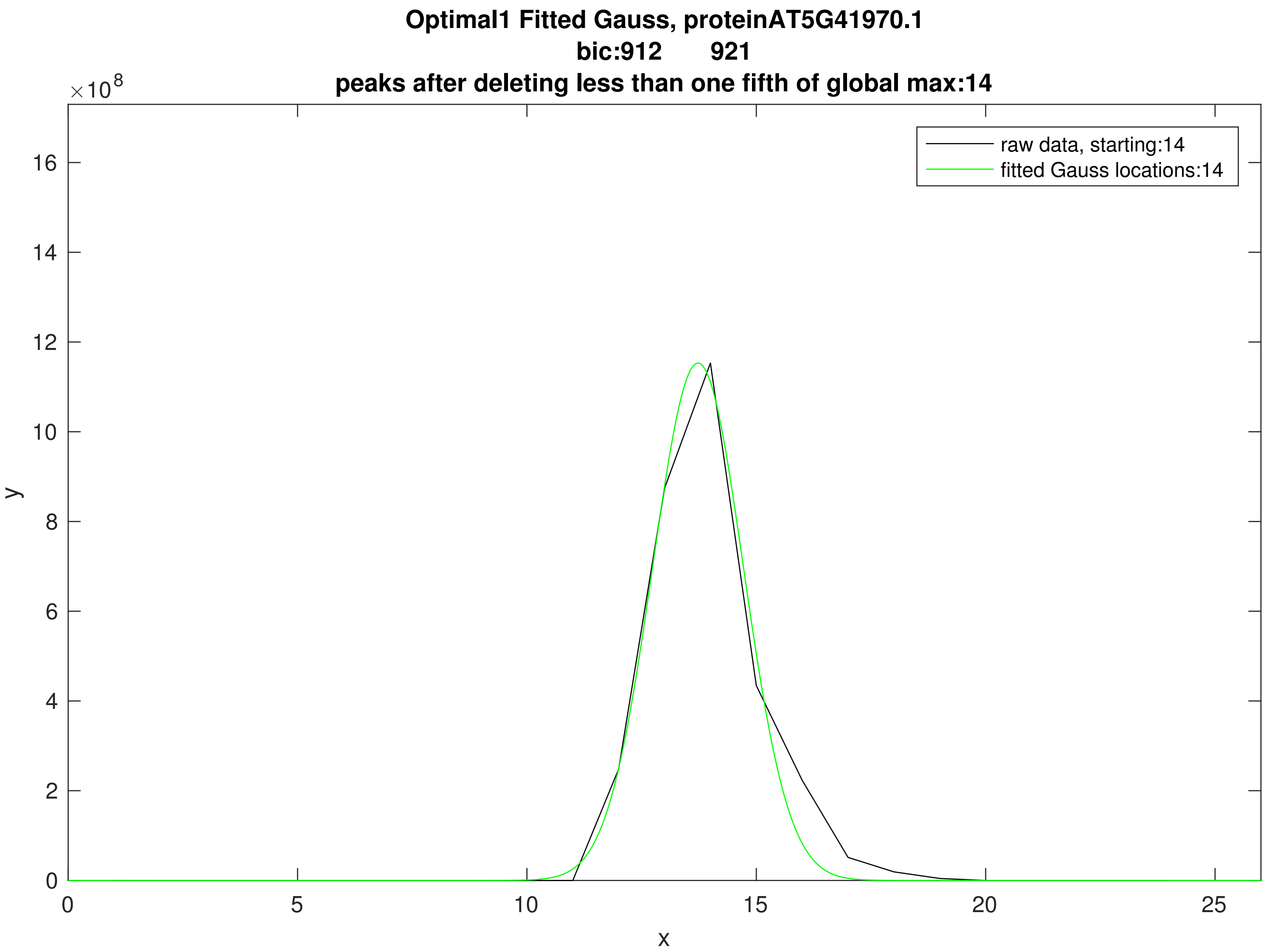

Optimal1 Fitted Gauss, proteinAT5G41970.1
bic:912 921
peaks after deleting less than one fifth of global max:14
×10^8
raw data, starting:14
fitted Gauss locations:14
16
14
12
10
8
6
4
2
0
y
0
5
10
15
20
25
x

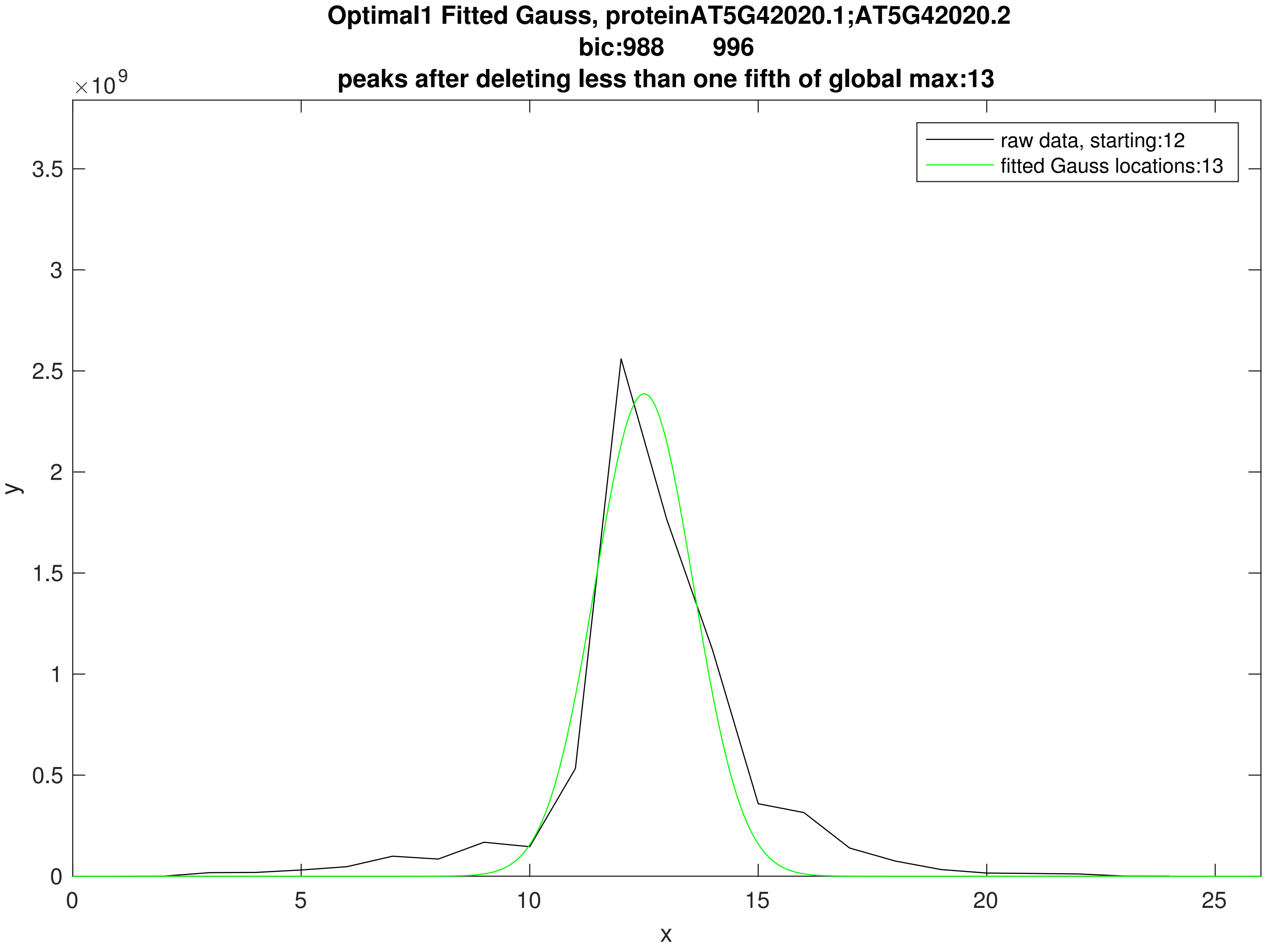
Optimal1 Fitted Gauss, proteinAT5G42020.1;AT5G42020.2
bic:988 996
peaks after deleting less than one fifth of global max:13
×10^9
raw data, starting:12
fitted Gauss locations:13
3.5
3
2.5
2
1.5
1
0.5
0
y
0
5
10
15
20
25
x

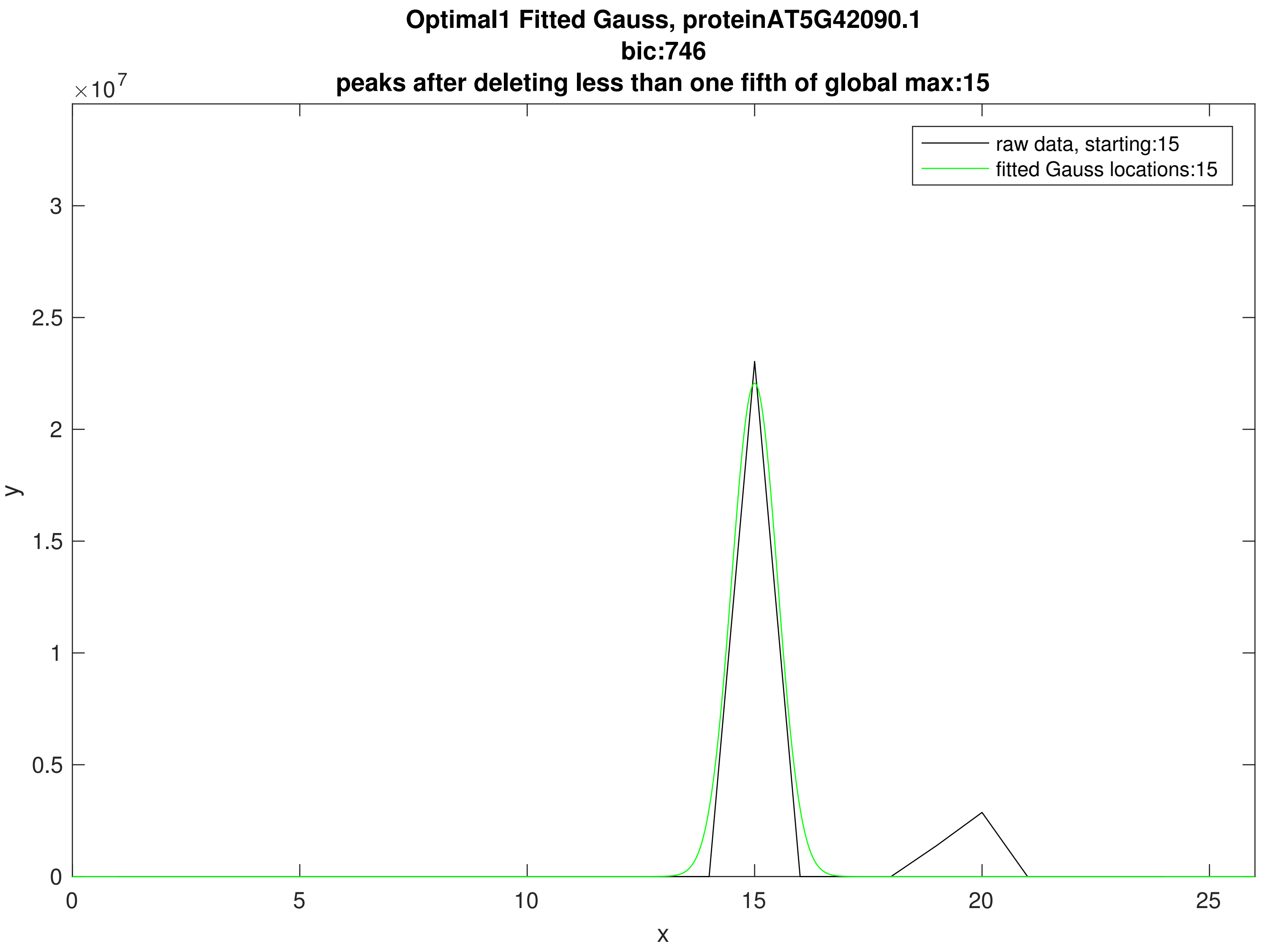
Optimal1 Fitted Gauss, proteinAT5G42090.1
bic:746
peaks after deleting less than one fifth of global max:15
×10^7
raw data, starting:15
fitted Gauss locations:15
y
x
0
0.5
1
1.5
2
2.5
3
5
10
15
20
25

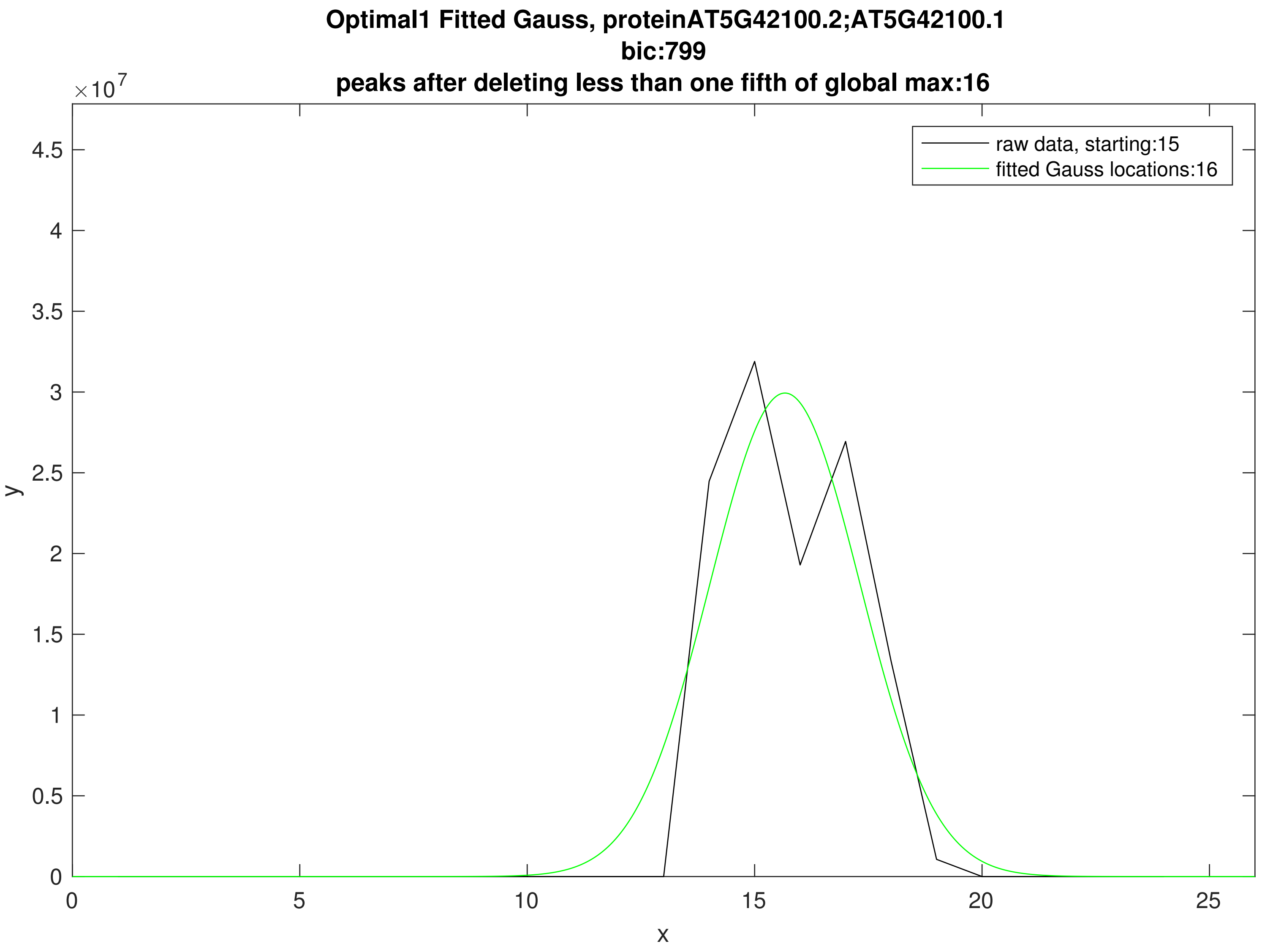
Optimal1 Fitted Gauss, proteinAT5G42100.2;AT5G42100.1
bic:799
peaks after deleting less than one fifth of global max:16
×10^7
raw data, starting:15
fitted Gauss locations:16
y
x
0
0.5
1
1.5
2
2.5
3
3.5
4
4.5
5
10
15
20
25

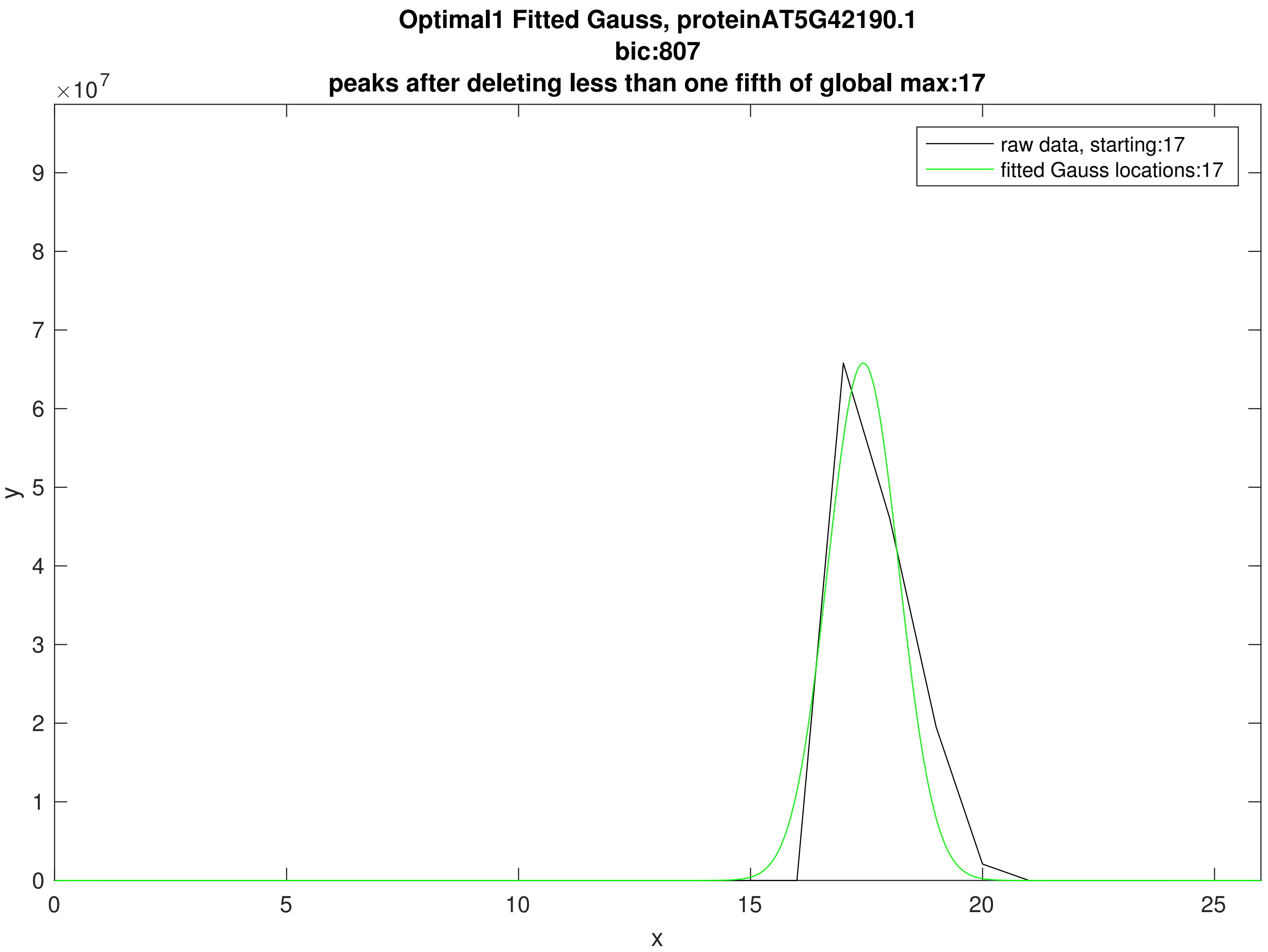
Optimal1 Fitted Gauss, proteinAT5G42190.1
bic:807
peaks after deleting less than one fifth of global max:17
×10^7
raw data, starting:17
fitted Gauss locations:17
y
x
0
1
2
3
4
5
6
7
8
9
0
5
10
15
20
25

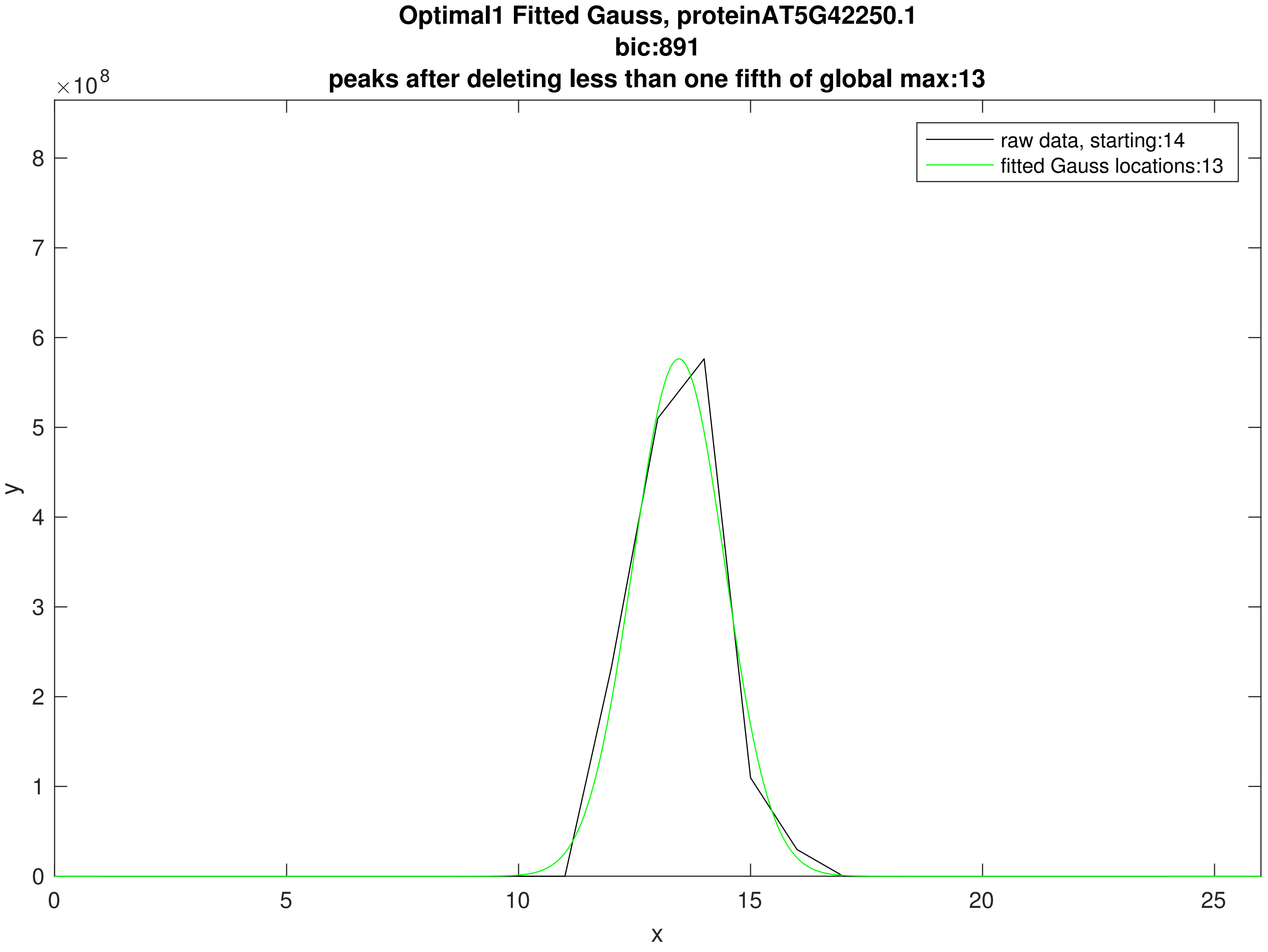

Optimal1 Fitted Gauss, proteinAT5G42250.1
bic:891
peaks after deleting less than one fifth of global max:13
×10^8
raw data, starting:14
fitted Gauss locations:13
y
x
0
1
2
3
4
5
6
7
8
0
5
10
15
20
25

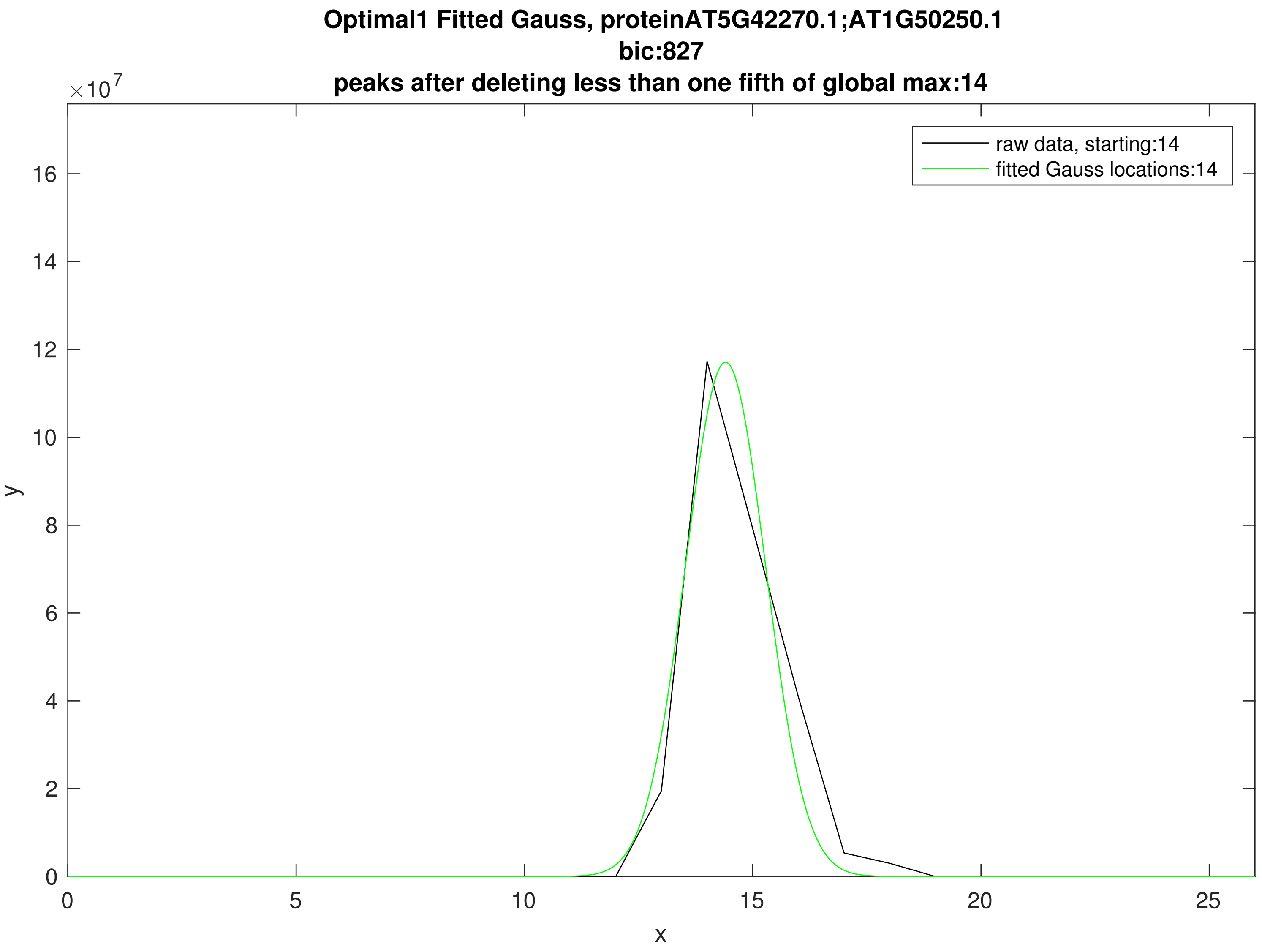

Optimal1 Fitted Gauss, proteinAT5G42270.1;AT1G50250.1
bic:827
peaks after deleting less than one fifth of global max:14
$\times 10^7$
raw data, starting:14
fitted Gauss locations:14
16
14
12
10
8
6
4
2
0
y
0
5
10
15
20
25
x

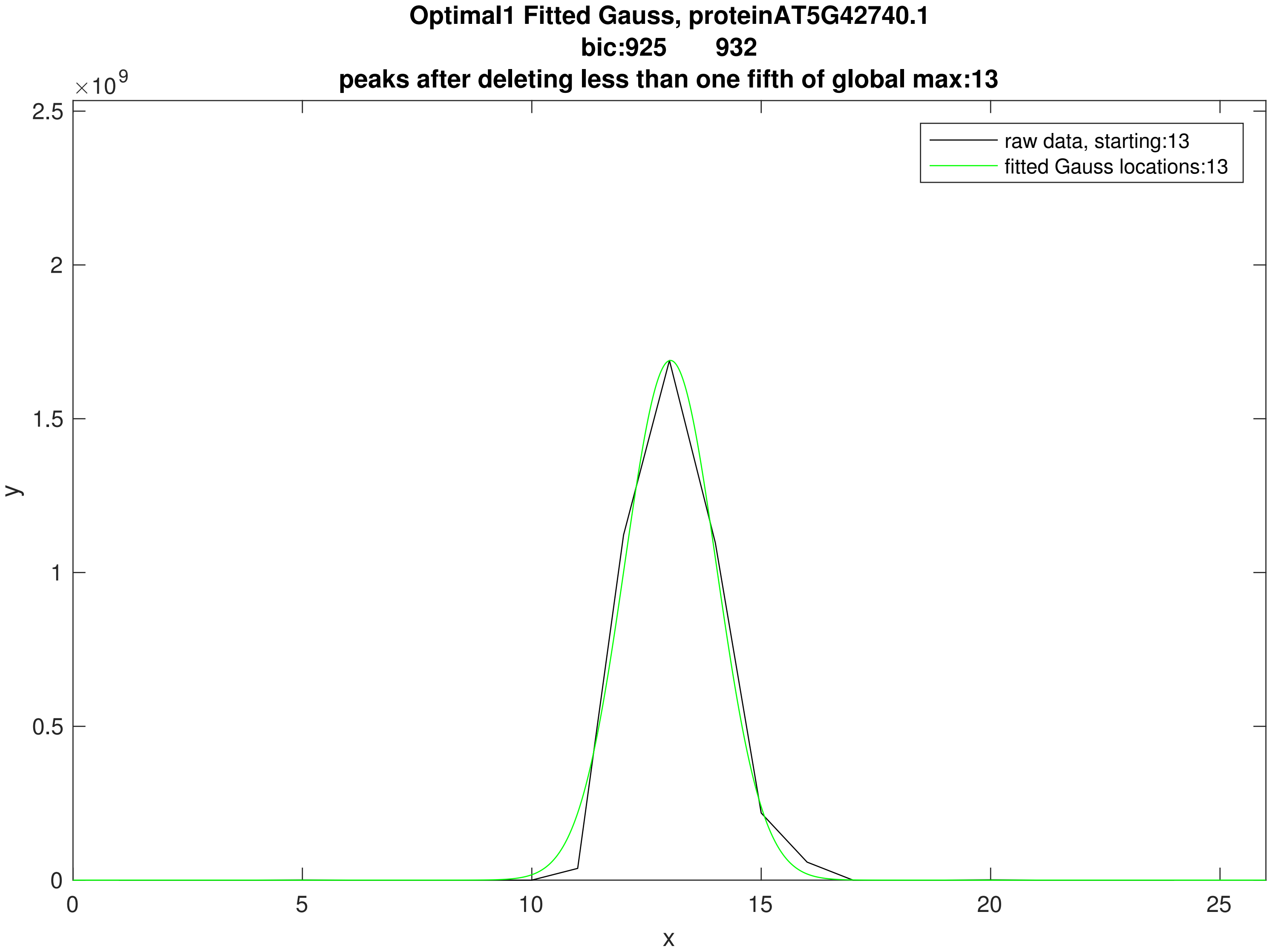
Optimal1 Fitted Gauss, proteinAT5G42740.1
bic:925 932
peaks after deleting less than one fifth of global max:13
×10^9
2.5
2
1.5
1
0.5
0
y
0
5
10
15
20
25
x
raw data, starting:13
fitted Gauss locations:13

**Optimal2 Fitted Gauss, proteinAT5G42770.2;AT5G42770.1**
**bic:801 800**
**peaks after deleting less than one fifth of global max:18**

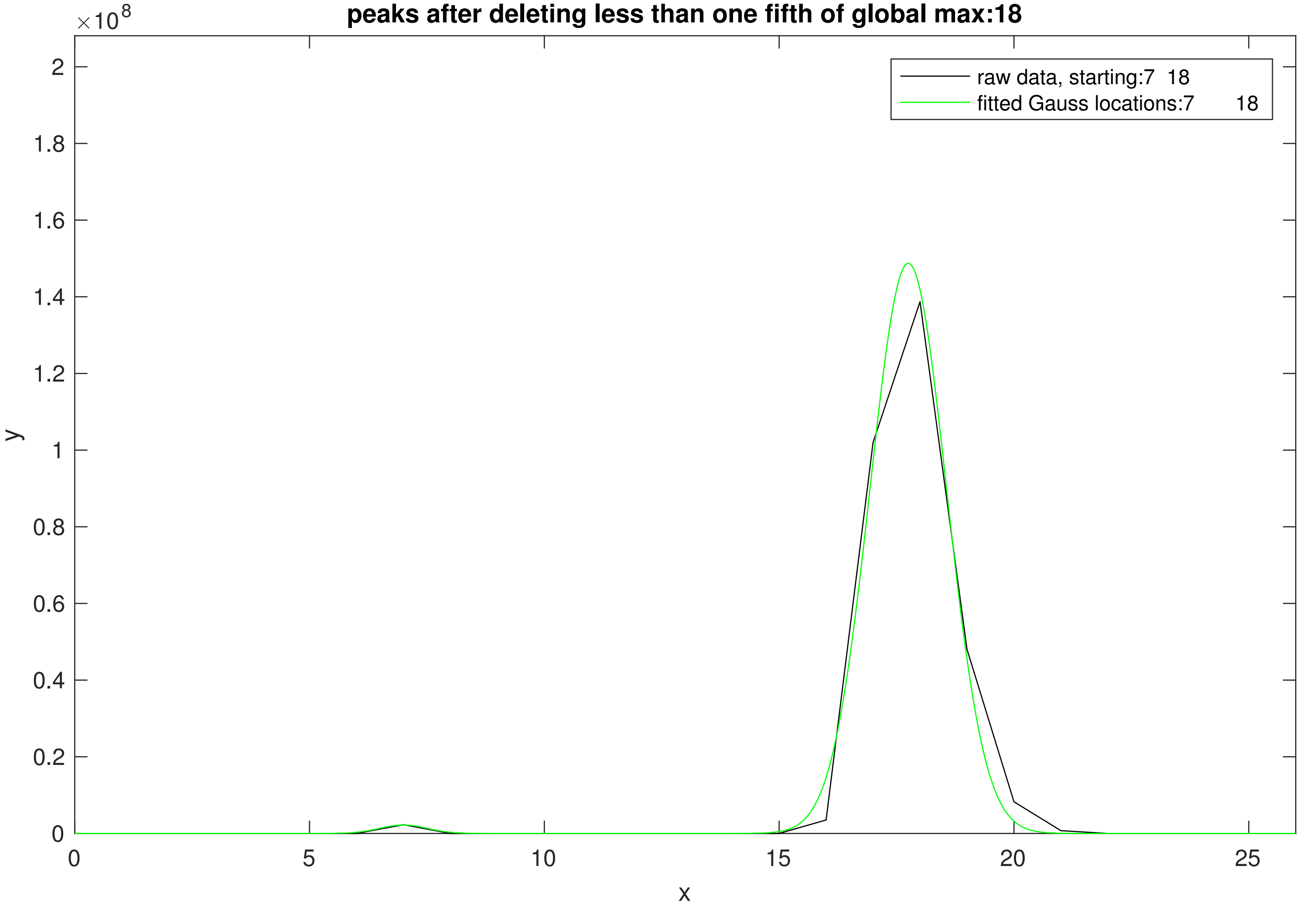

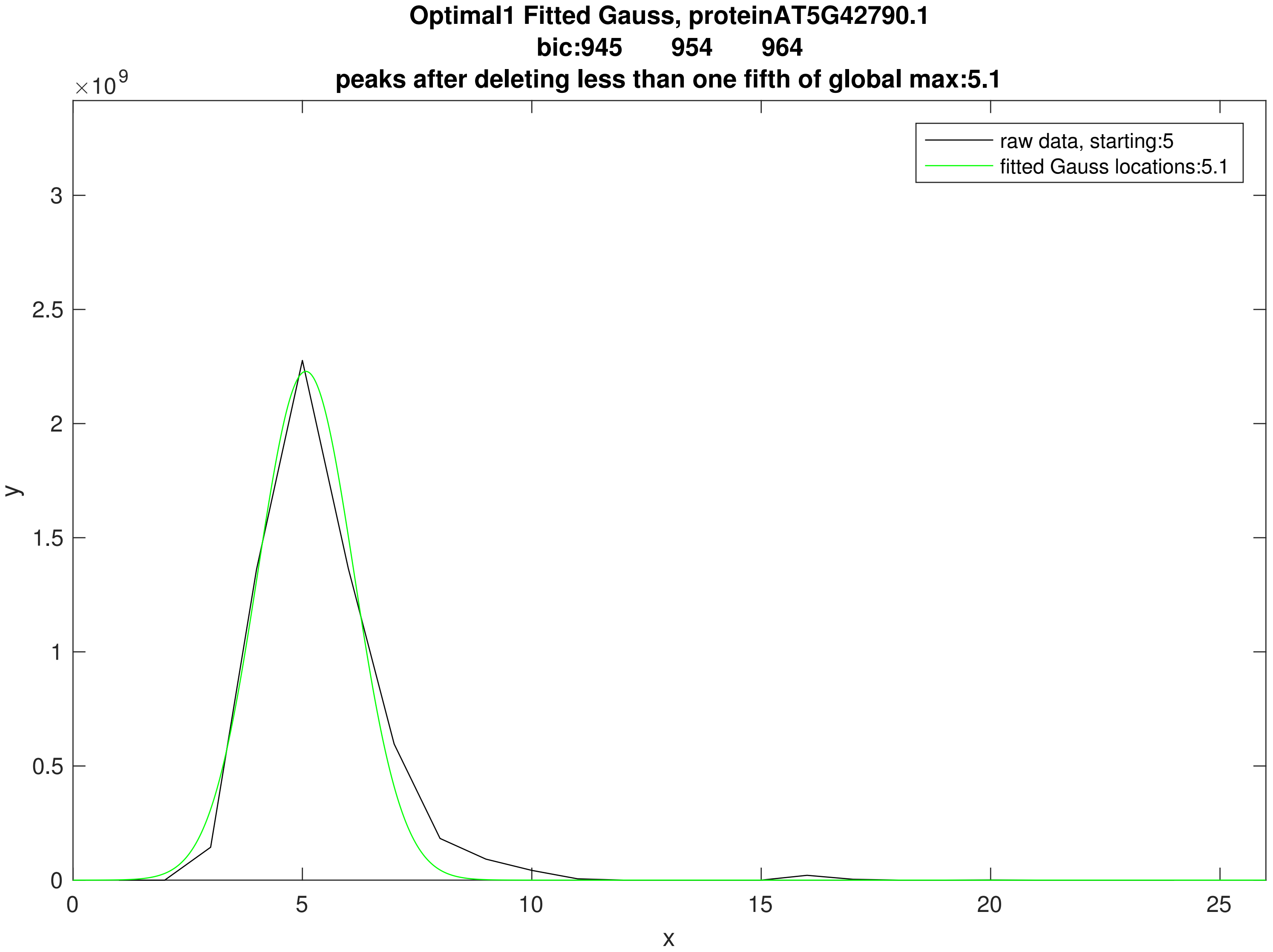
Optimal1 Fitted Gauss, proteinAT5G42790.1
bic:945 954 964
peaks after deleting less than one fifth of global max:5.1
×10^9
raw data, starting:5
fitted Gauss locations:5.1
y
x
0
0.5
1
1.5
2
2.5
3
0
5
10
15
20
25

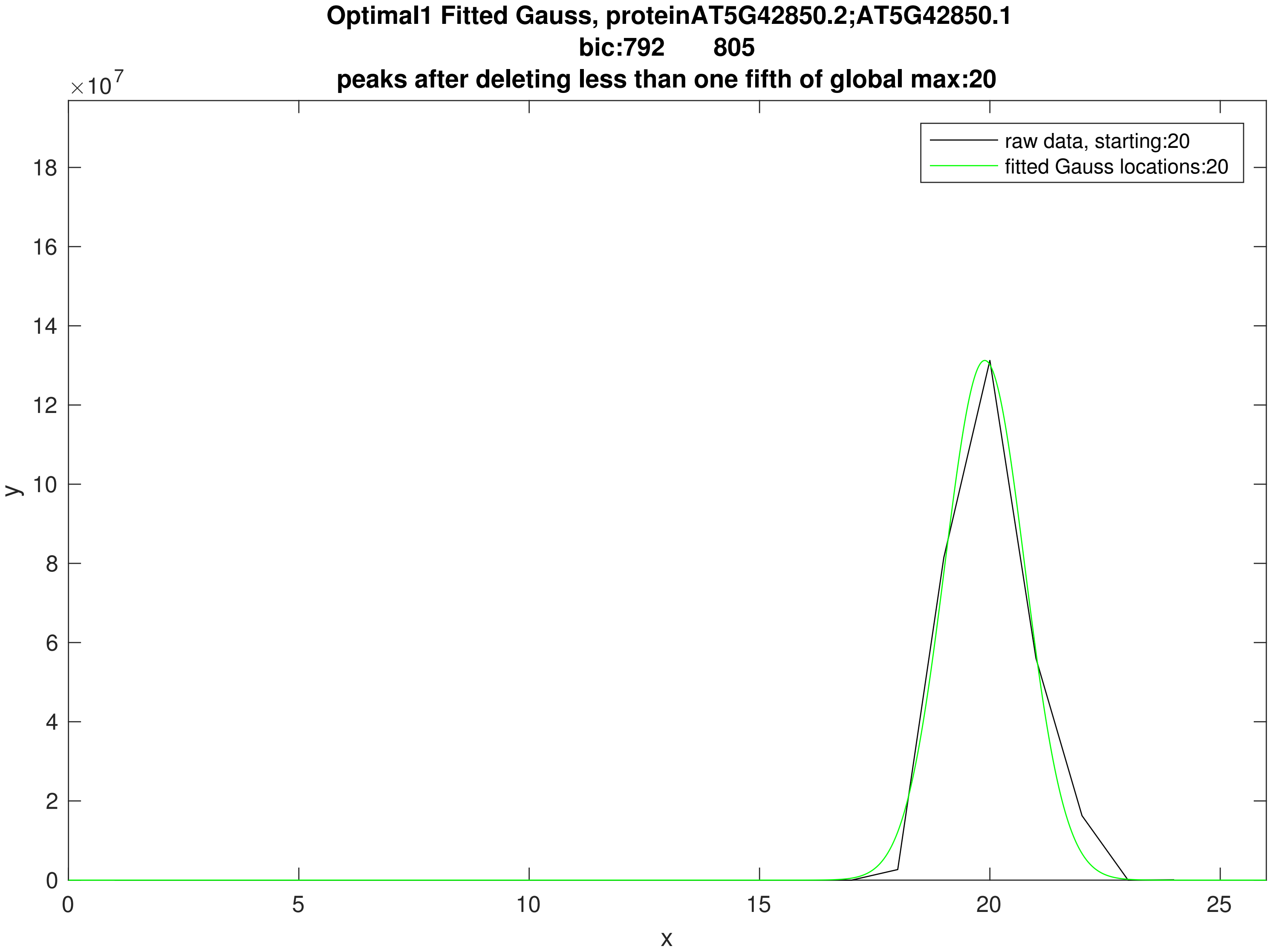

Optimal1 Fitted Gauss, proteinAT5G42850.2;AT5G42850.1
bic:792 805
peaks after deleting less than one fifth of global max:20
×10^7
raw data, starting:20
fitted Gauss locations:20
18
16
14
12
10
8
6
4
2
0
y
0
5
10
15
20
25
x

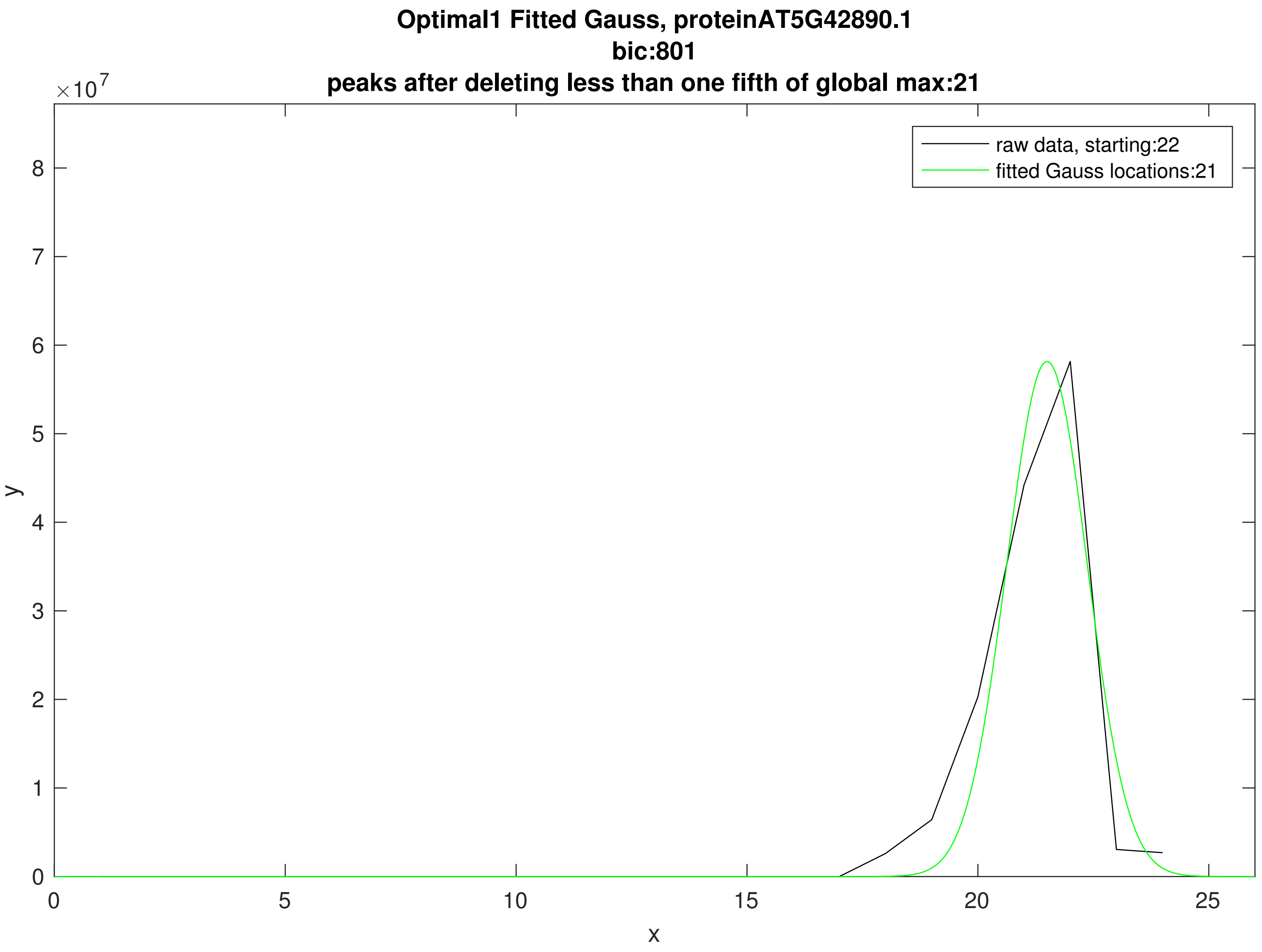
Optimal1 Fitted Gauss, proteinAT5G42890.1
bic:801
peaks after deleting less than one fifth of global max:21
×10^7
raw data, starting:22
fitted Gauss locations:21
y
x
0
1
2
3
4
5
6
7
8
0
5
10
15
20
25

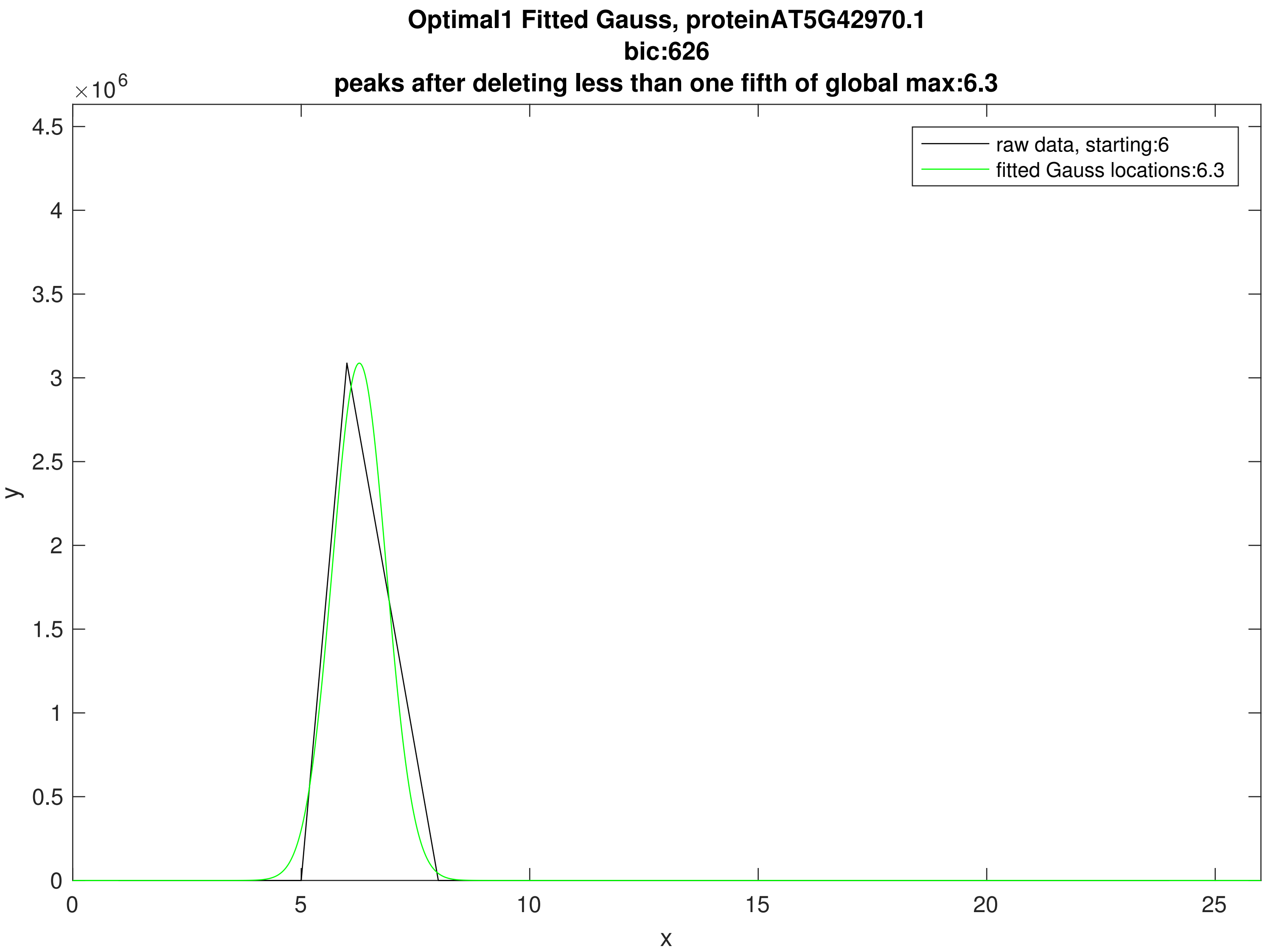

Optimal1 Fitted Gauss, proteinAT5G42970.1
bic:626
peaks after deleting less than one fifth of global max:6.3
×10^6
raw data, starting:6
fitted Gauss locations:6.3
y
x
0
0.5
1
1.5
2
2.5
3
3.5
4
4.5
5
10
15
20
25

**Optimal1 Fitted Gauss, proteinAT5G42980.1**
**bic:923 923**
**peaks after deleting less than one fifth of global max:20**

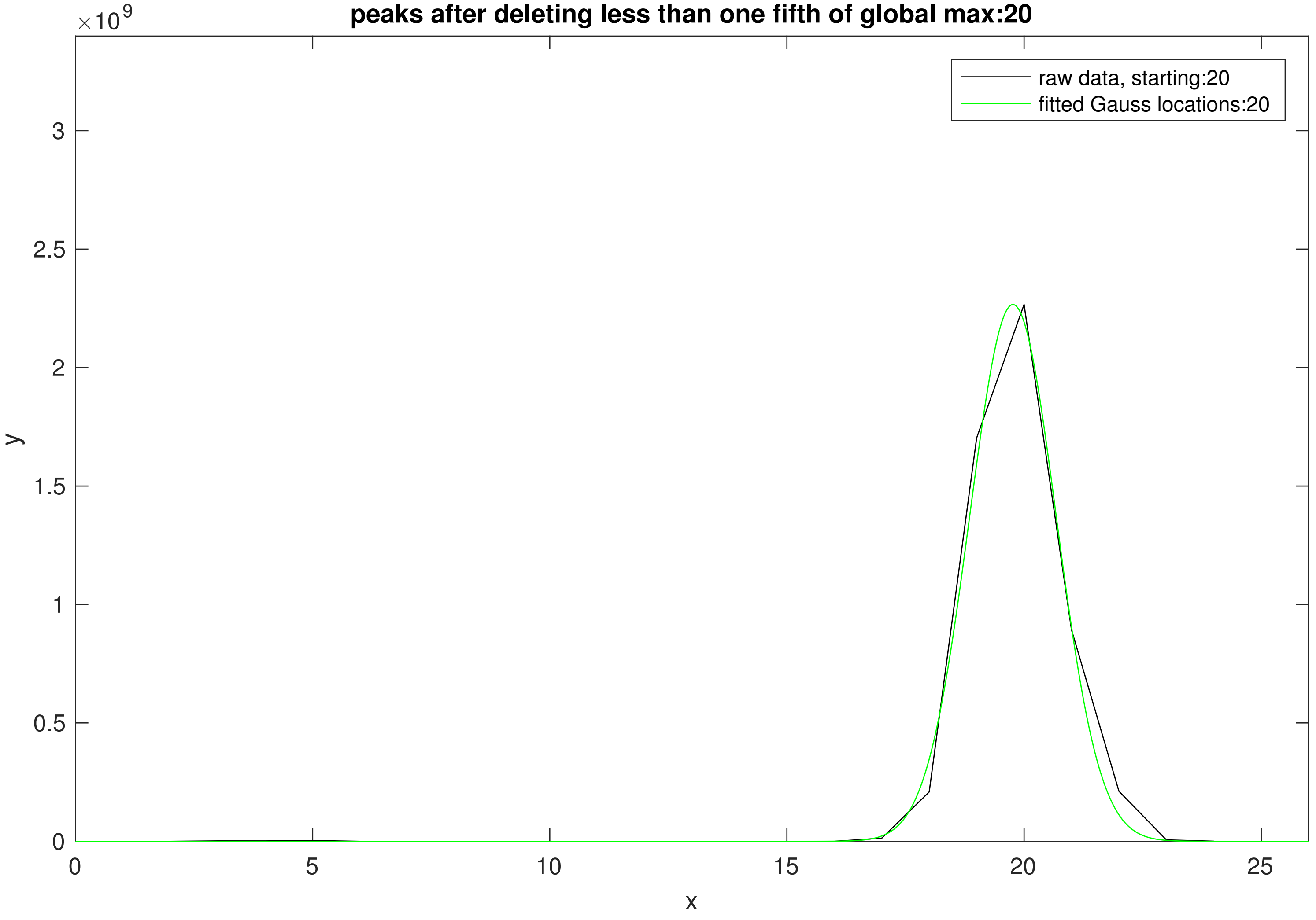

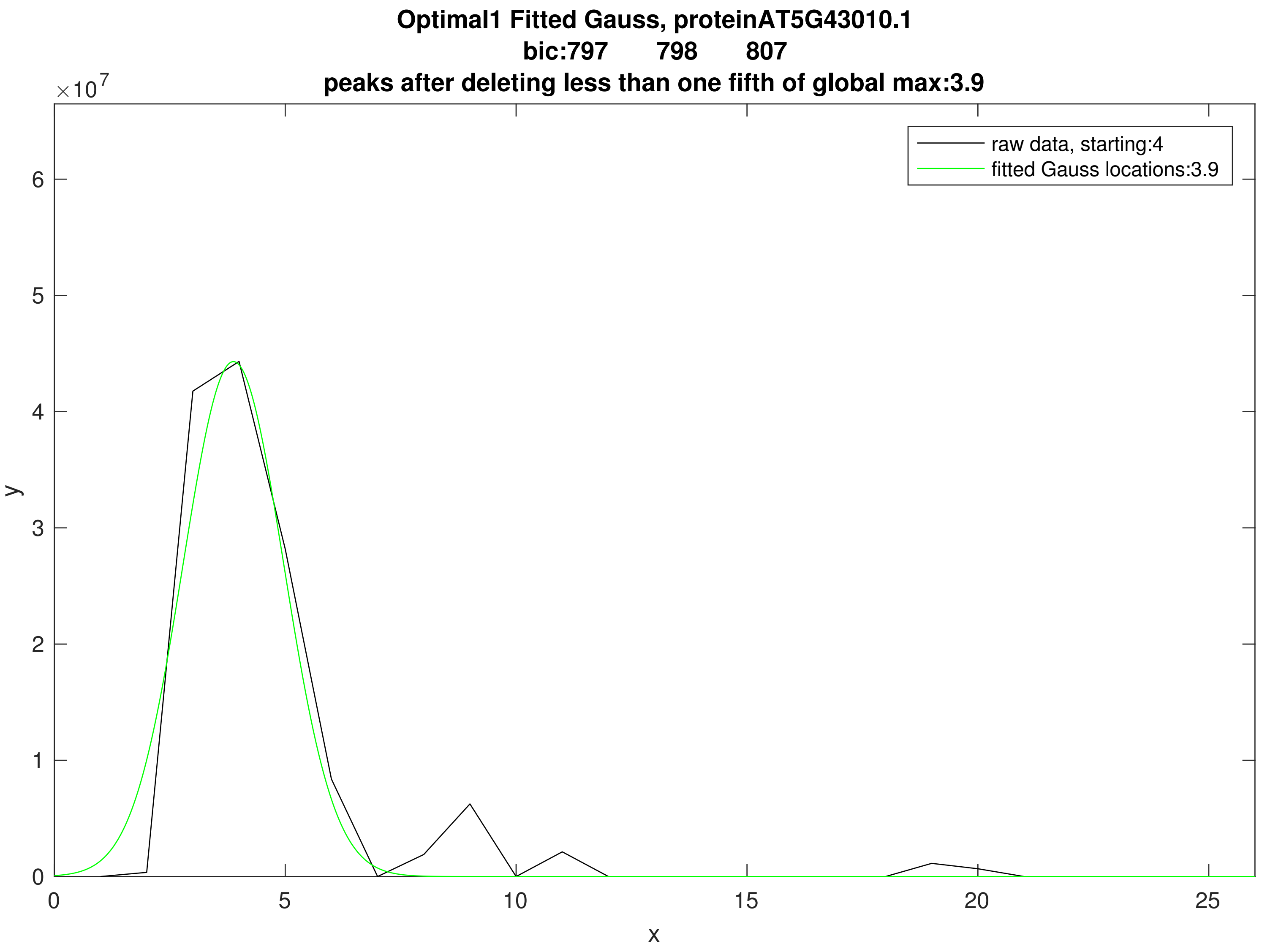
Optimal1 Fitted Gauss, proteinAT5G43010.1
bic:797 798 807
peaks after deleting less than one fifth of global max:3.9
×10^7
raw data, starting:4
fitted Gauss locations:3.9
y
x
0
1
2
3
4
5
6
0
5
10
15
20
25

**Optimal2 Fitted Gauss, proteinAT5G43060.1**
**bic:903 901 910 919**
**peaks after deleting less than one fifth of global max:13 17**

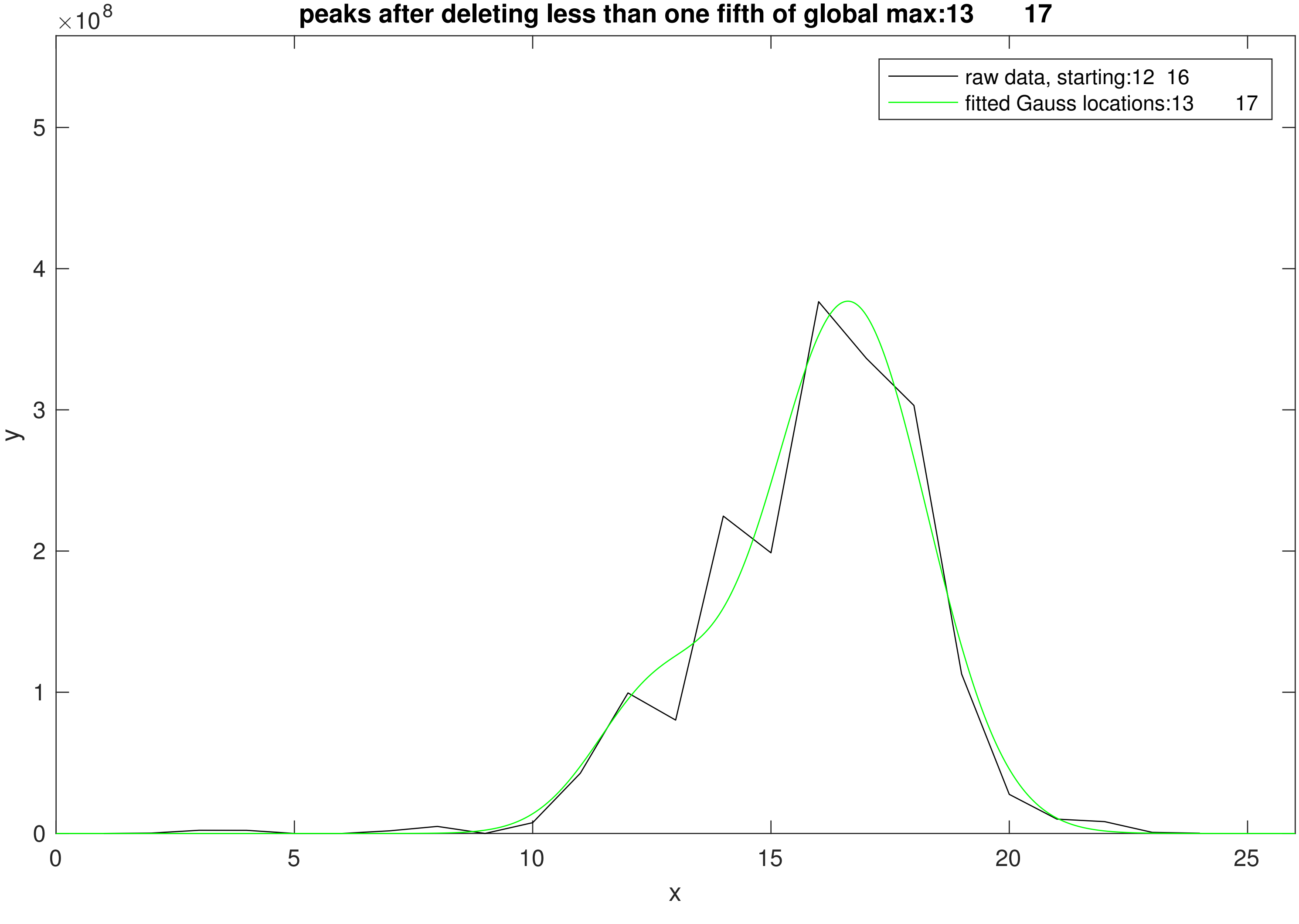

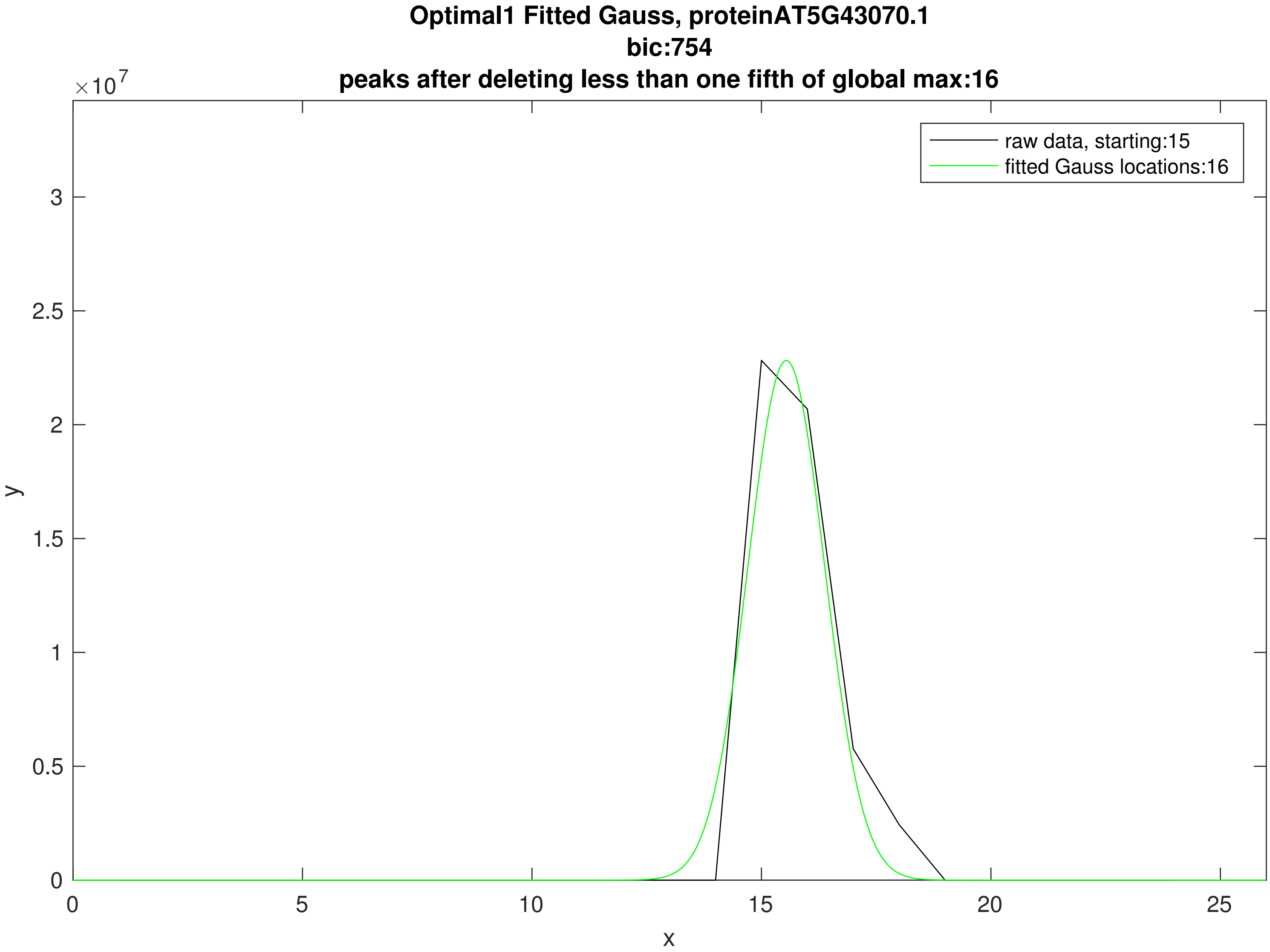
Optimal1 Fitted Gauss, proteinAT5G43070.1
bic:754
peaks after deleting less than one fifth of global max:16
×10^7
raw data, starting:15
fitted Gauss locations:16
y
x
0
0.5
1
1.5
2
2.5
3
0
5
10
15
20
25

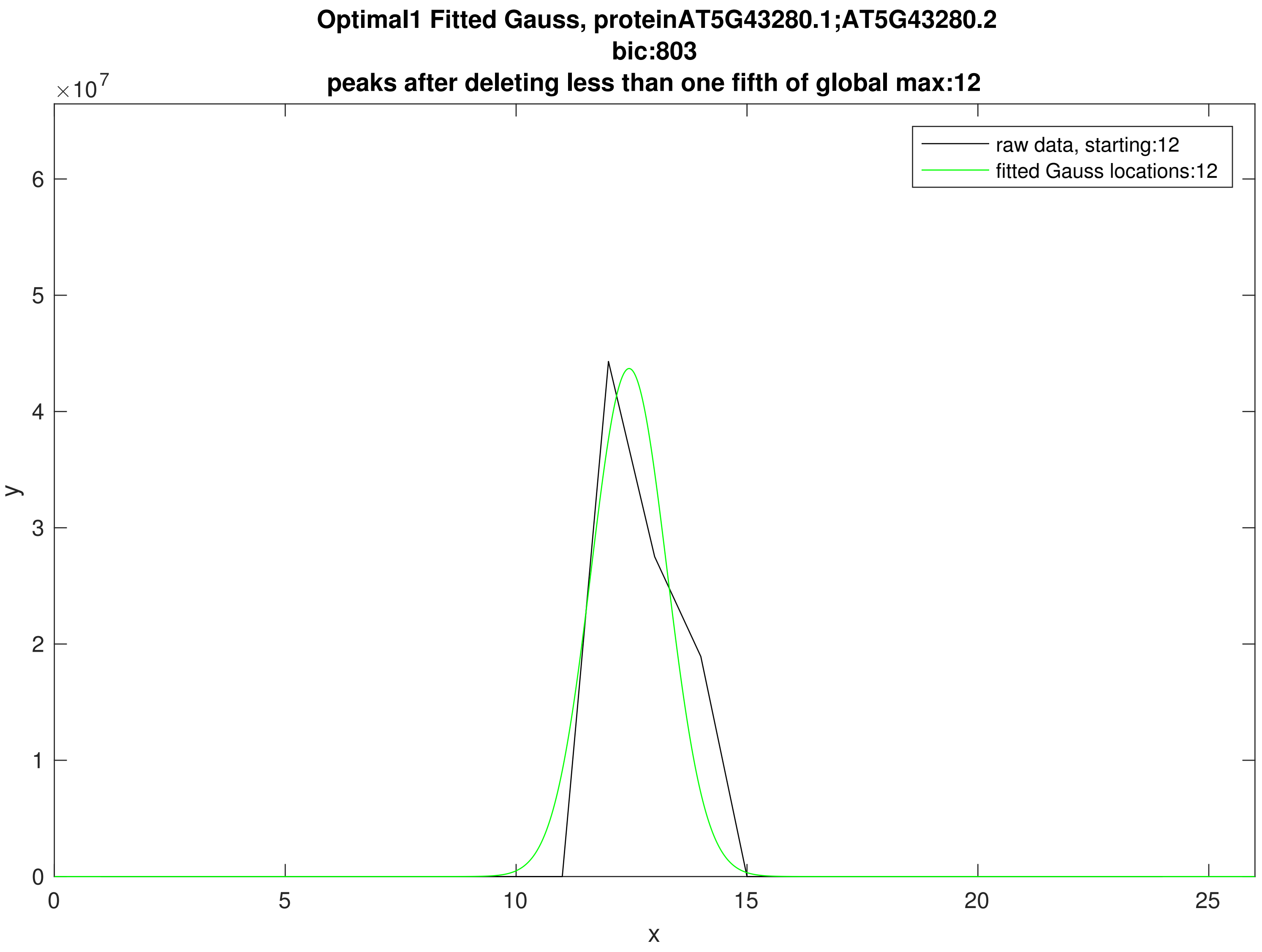
Optimal1 Fitted Gauss, proteinAT5G43280.1;AT5G43280.2
bic:803
peaks after deleting less than one fifth of global max:12
×10^7
raw data, starting:12
fitted Gauss locations:12
y
x
0
1
2
3
4
5
6
0
5
10
15
20
25

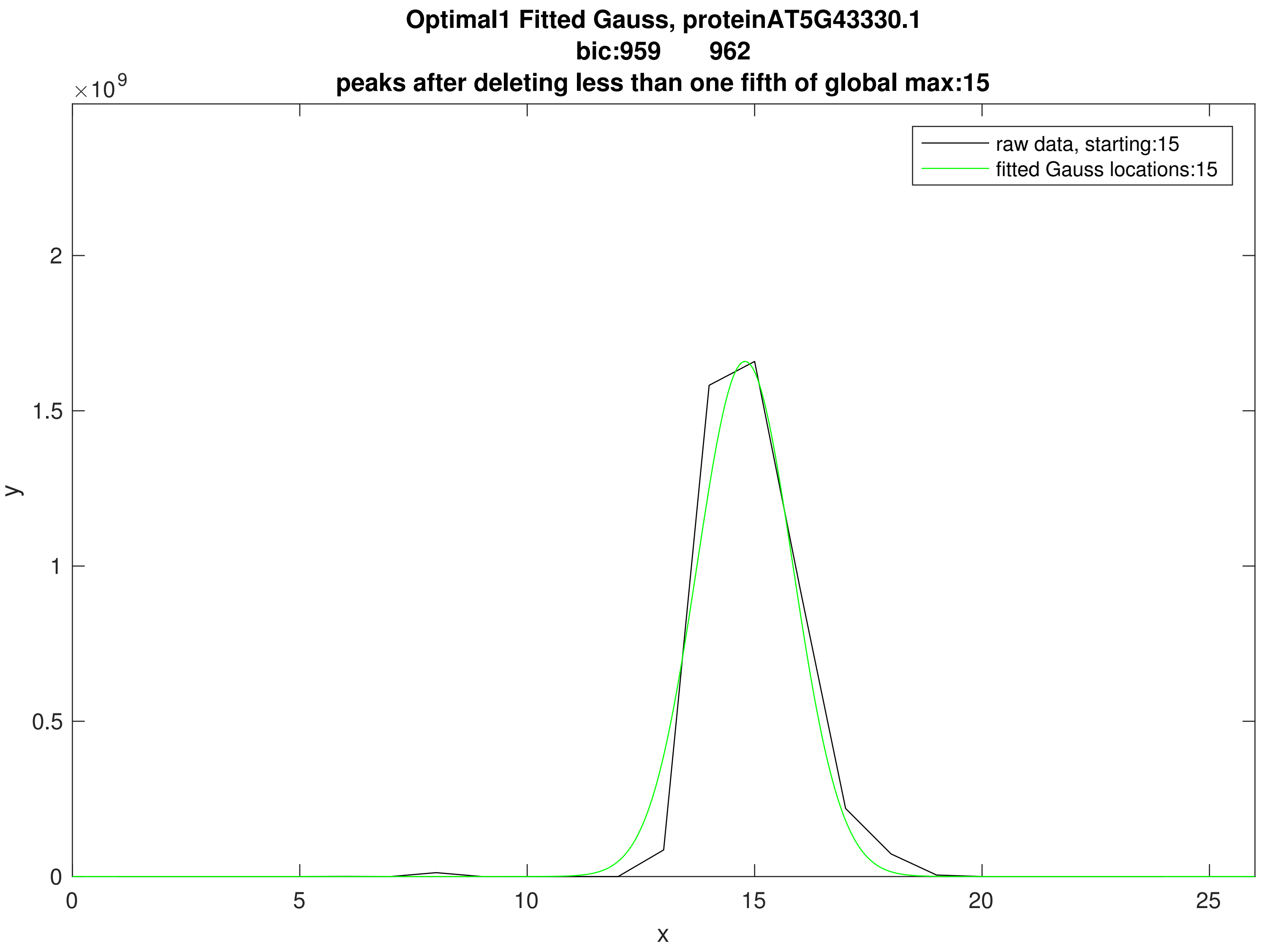
Optimal1 Fitted Gauss, proteinAT5G43330.1
bic:959 962
peaks after deleting less than one fifth of global max:15
×10^9
raw data, starting:15
fitted Gauss locations:15
y
x
0
0.5
1
1.5
2
0
5
10
15
20
25

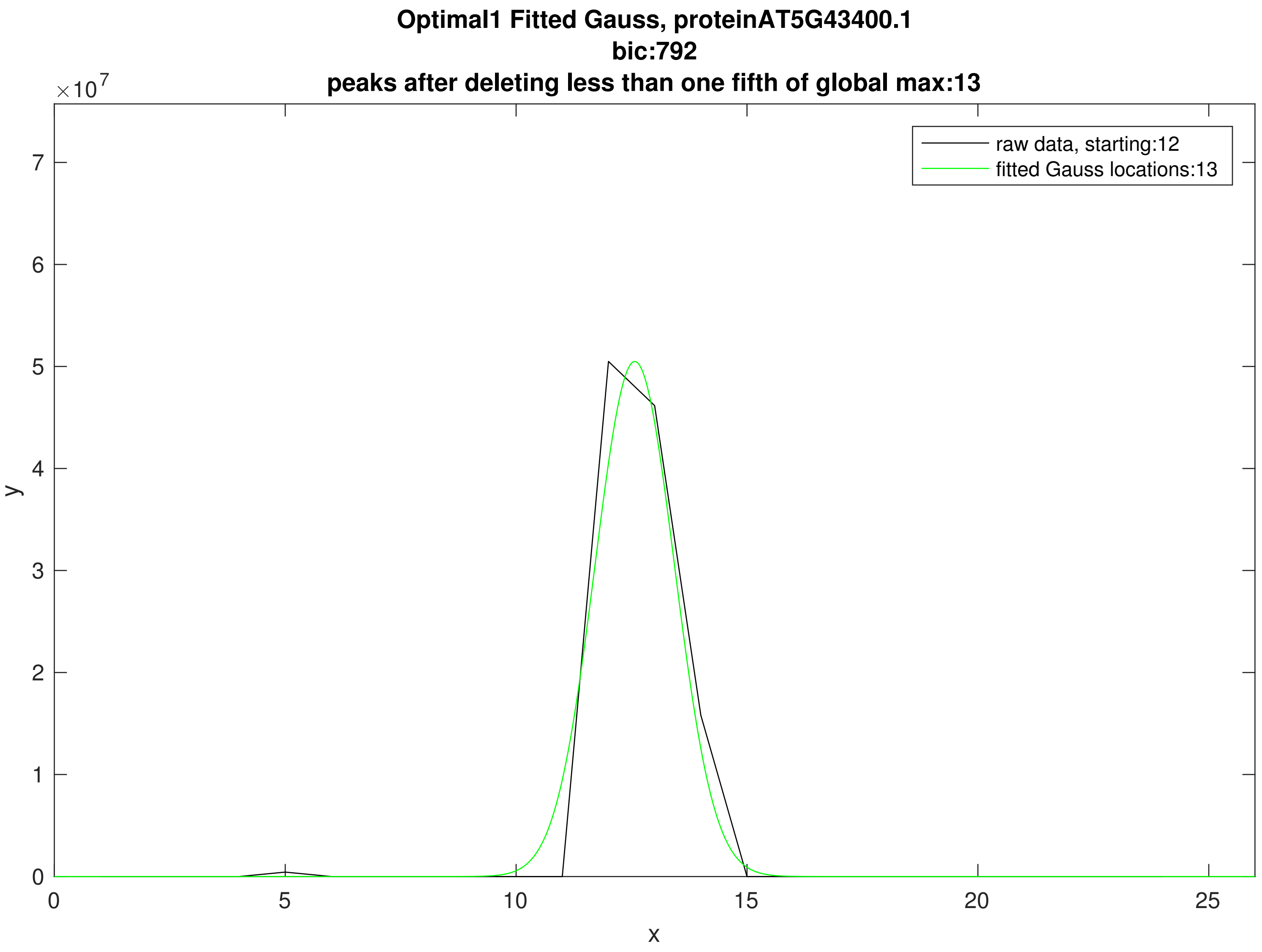
Optimal1 Fitted Gauss, proteinAT5G43400.1
bic:792
peaks after deleting less than one fifth of global max:13
×10^7
raw data, starting:12
fitted Gauss locations:13
y
x
0
1
2
3
4
5
6
7
0
5
10
15
20
25

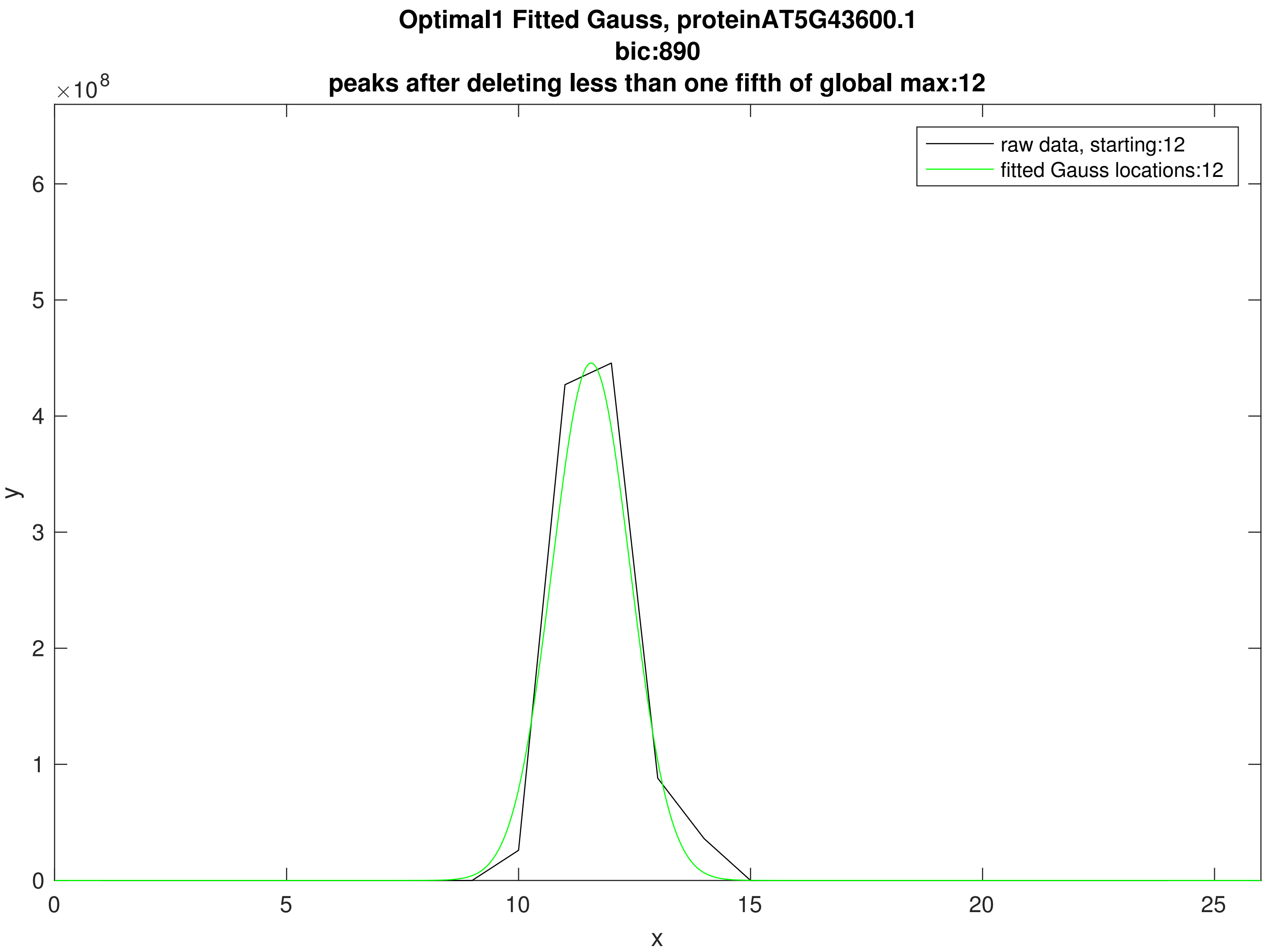
Optimal1 Fitted Gauss, proteinAT5G43600.1
bic:890
peaks after deleting less than one fifth of global max:12
×10^8
raw data, starting:12
fitted Gauss locations:12
y
x
0
1
2
3
4
5
6
0
5
10
15
20
25

#### Optimal1 Fitted Gauss, proteinAT5G43780.1
#### bic:886 905 918
#### peaks after deleting less than one fifth of global max:14

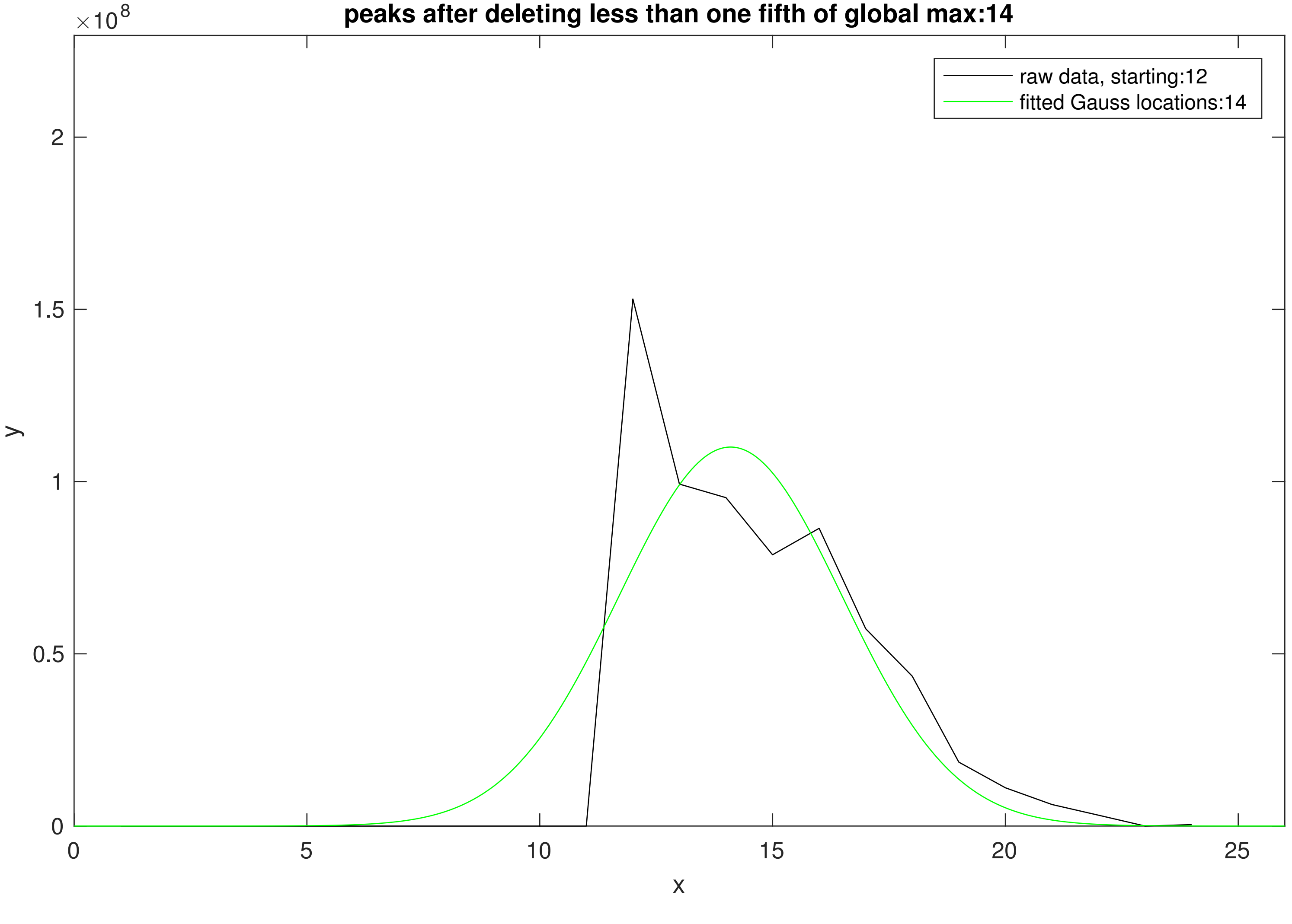

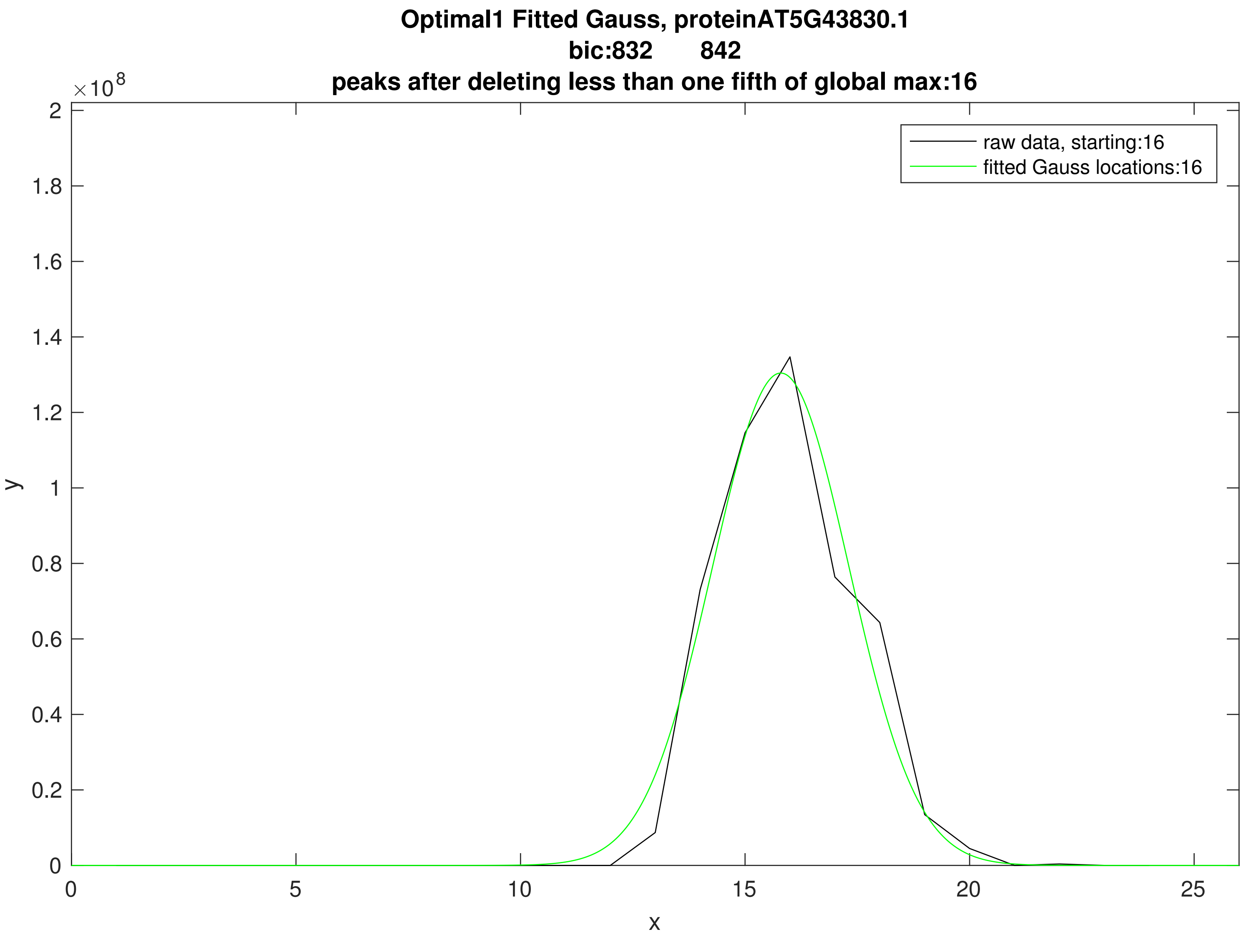
Optimal1 Fitted Gauss, proteinAT5G43830.1
bic:832 842
peaks after deleting less than one fifth of global max:16
×10^8
2
1.8
1.6
1.4
1.2
1
0.8
0.6
0.4
0.2
0
y
0
5
10
15
20
25
x
raw data, starting:16
fitted Gauss locations:16

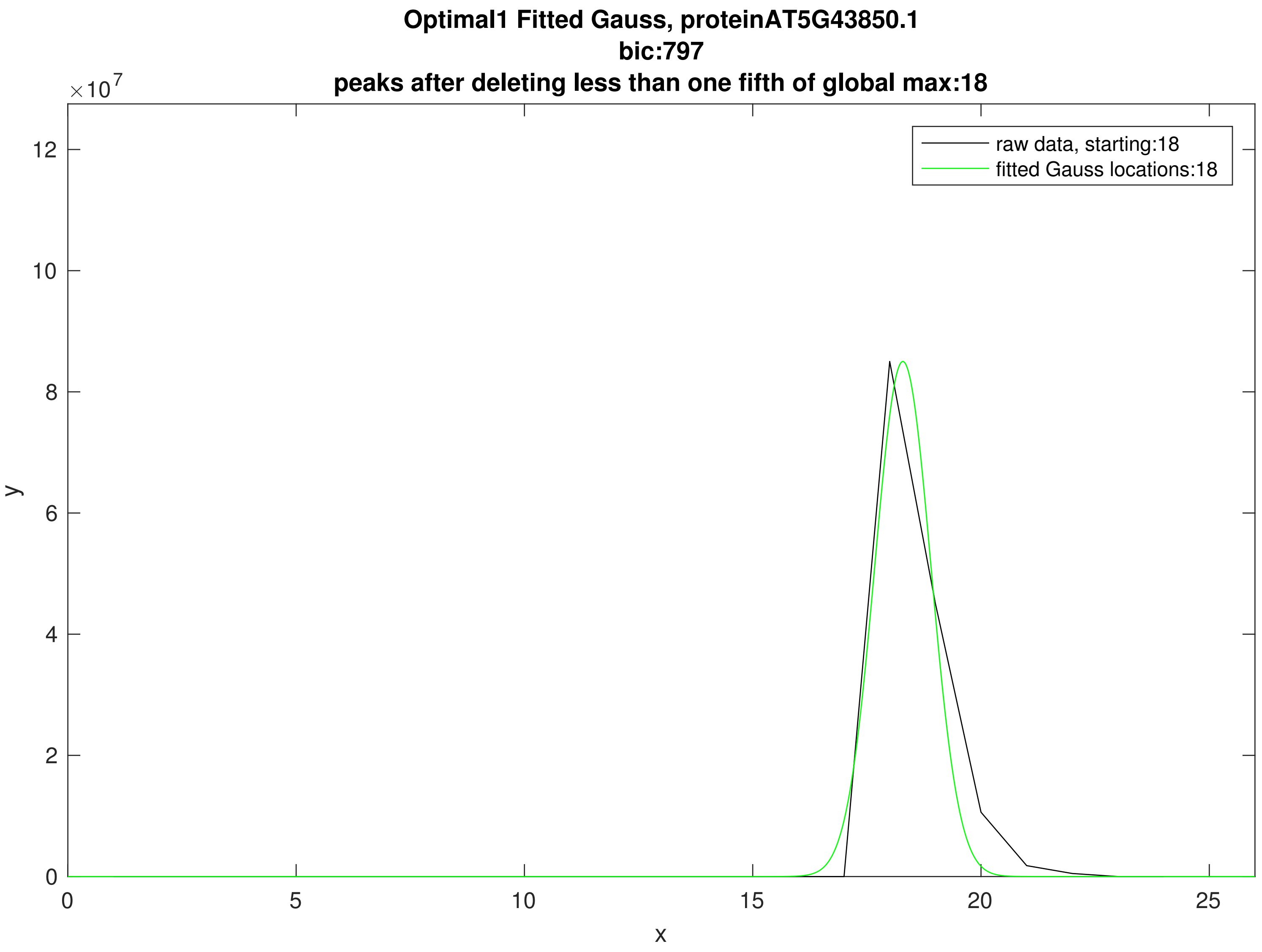

Optimal1 Fitted Gauss, proteinAT5G43850.1
bic:797
peaks after deleting less than one fifth of global max:18
×10^7
12
10
8
6
4
2
0
y
0
5
10
15
20
25
x
raw data, starting:18
fitted Gauss locations:18

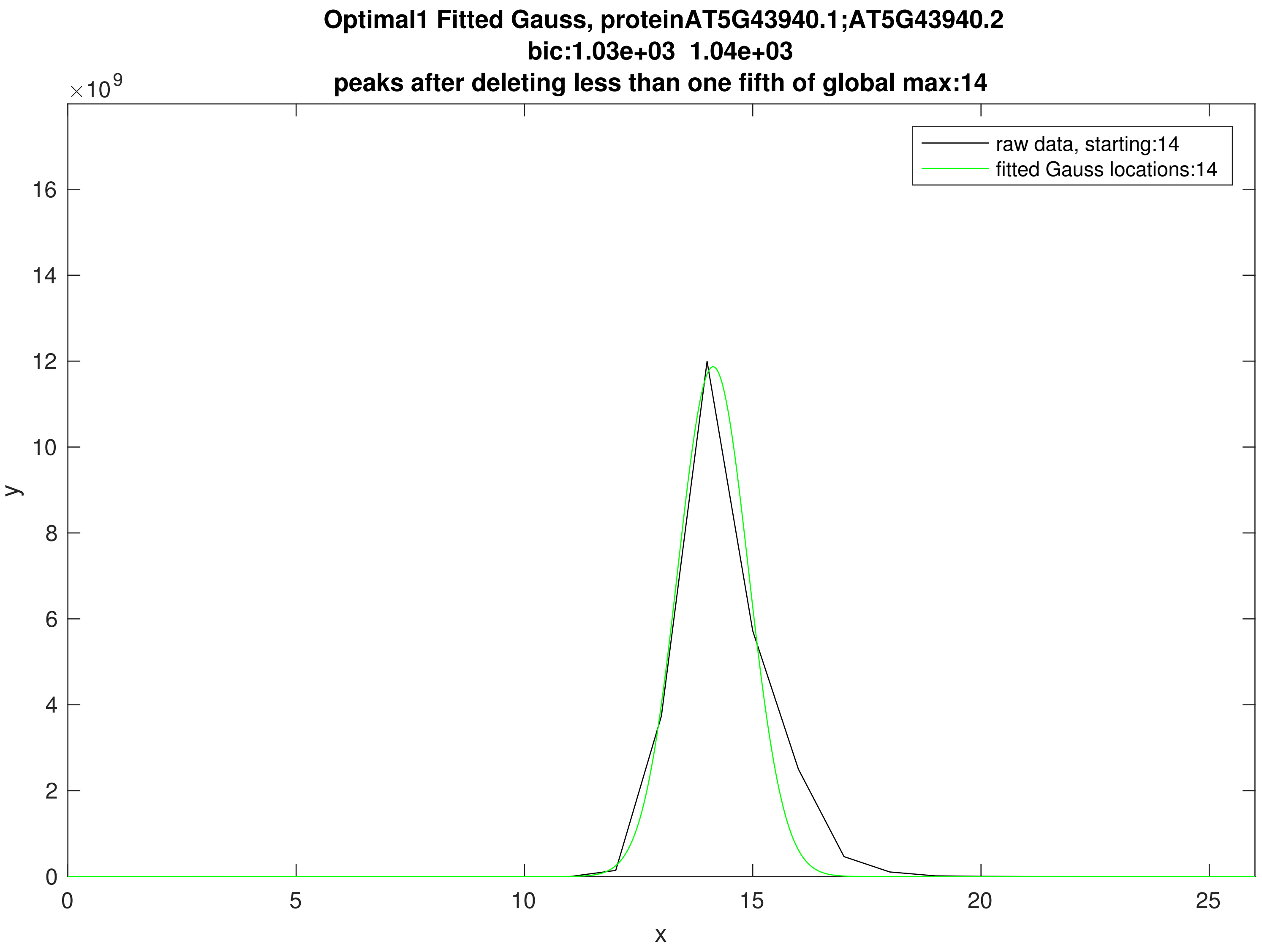
Optimal1 Fitted Gauss, proteinAT5G43940.1;AT5G43940.2
bic:1.03e+03  1.04e+03
peaks after deleting less than one fifth of global max:14
×10^9
raw data, starting:14
fitted Gauss locations:14
y
x
0
2
4
6
8
10
12
14
16
0
5
10
15
20
25

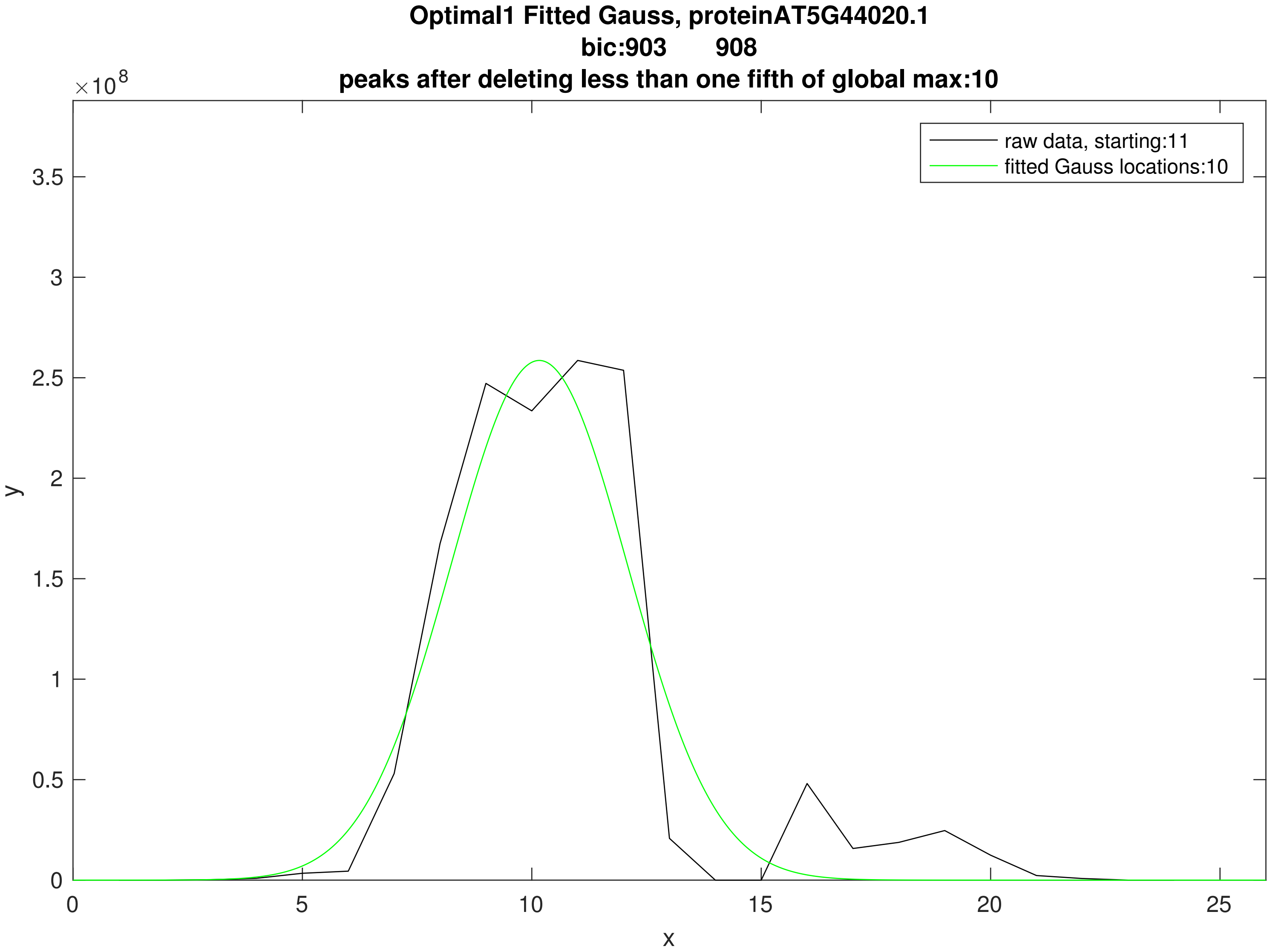
Optimal1 Fitted Gauss, proteinAT5G44020.1
bic:903 908
peaks after deleting less than one fifth of global max:10
×10^8
3.5
3
2.5
2
1.5
1
0.5
0
y
0
5
10
15
20
25
x
raw data, starting:11
fitted Gauss locations:10

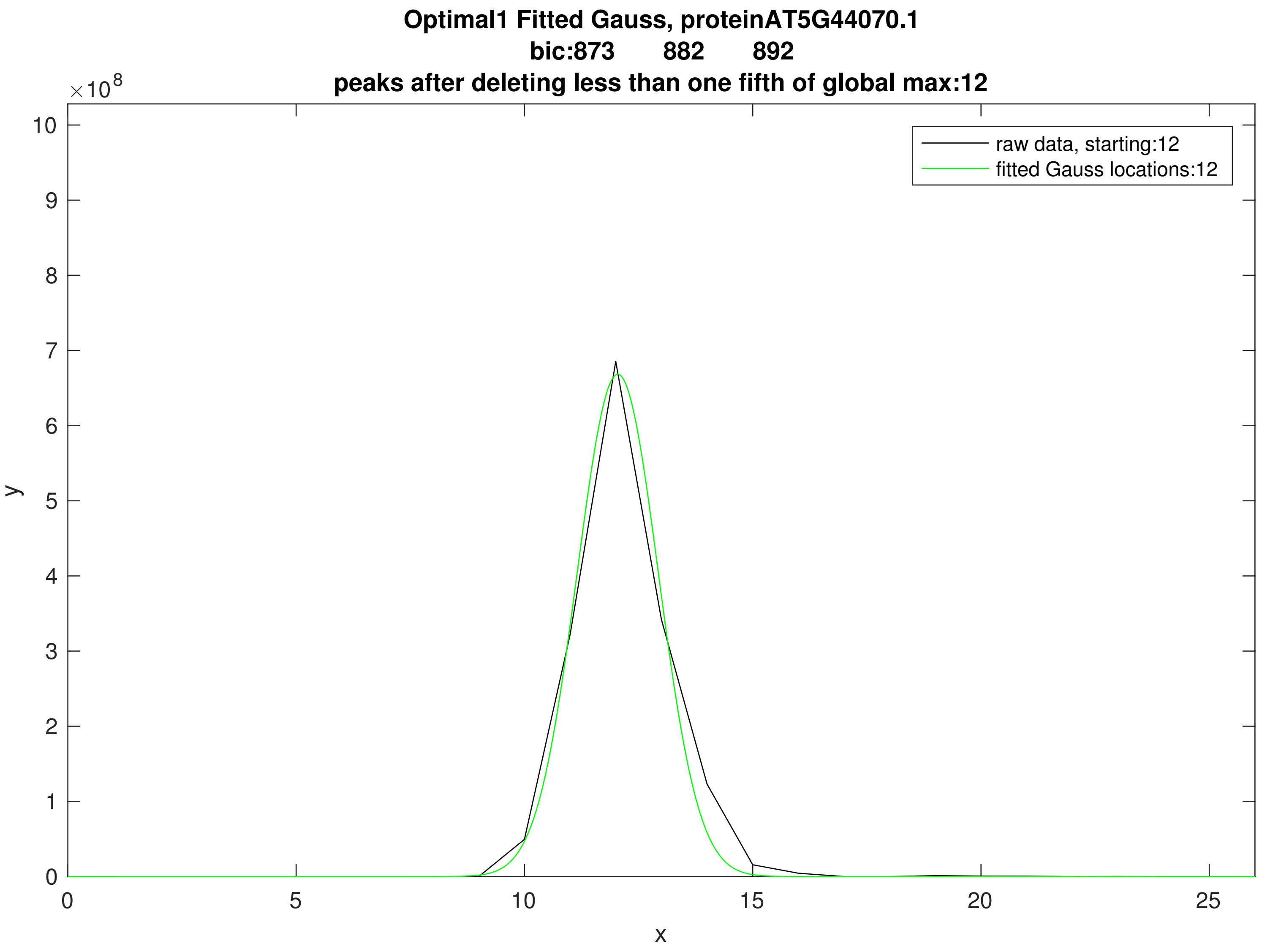
Optimal1 Fitted Gauss, proteinAT5G44070.1
bic:873 882 892
peaks after deleting less than one fifth of global max:12
×10^8
raw data, starting:12
fitted Gauss locations:12
y
x
0
1
2
3
4
5
6
7
8
9
10
0
5
10
15
20
25

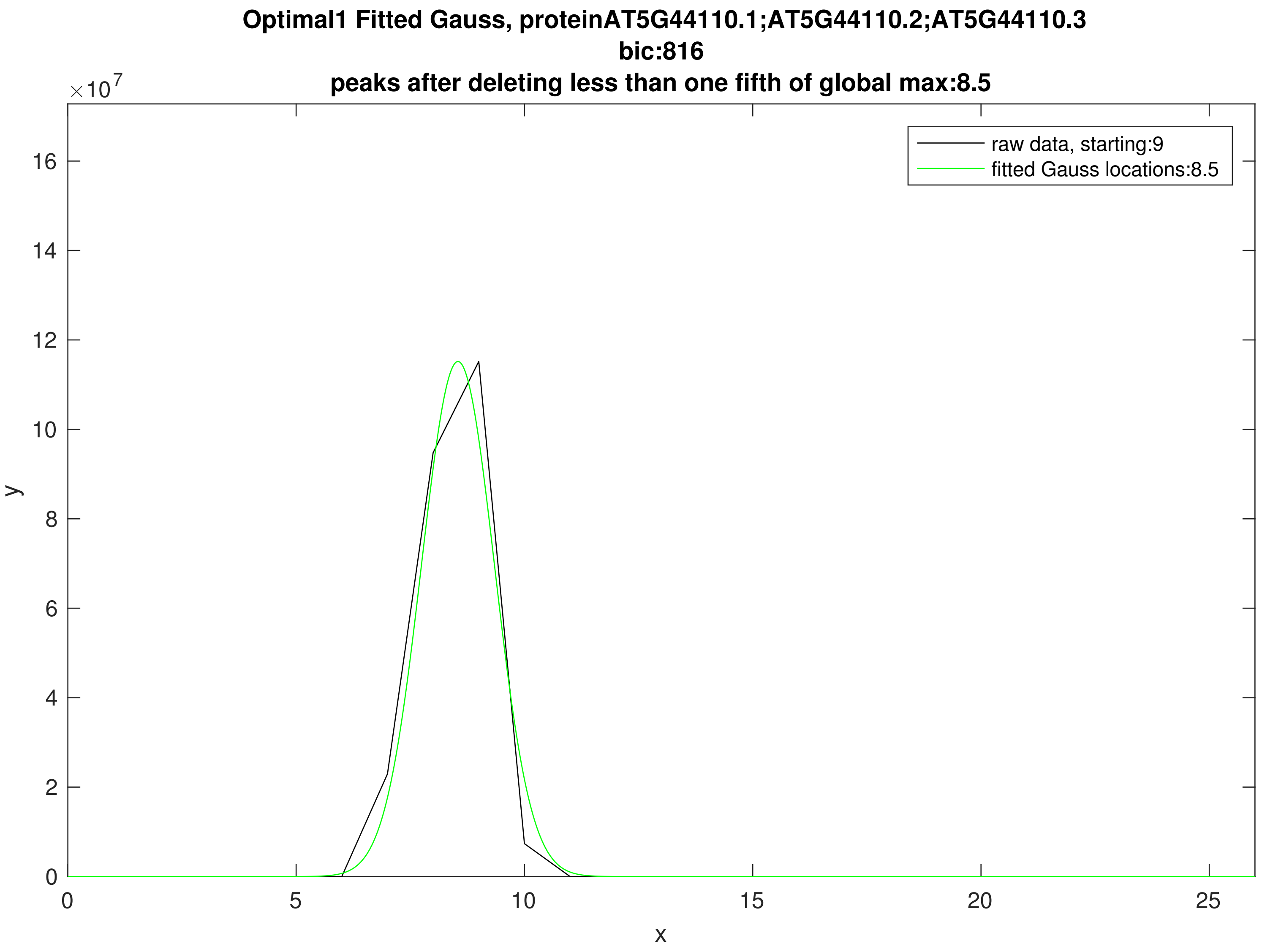
Optimal1 Fitted Gauss, proteinAT5G44110.1;AT5G44110.2;AT5G44110.3
bic:816
peaks after deleting less than one fifth of global max:8.5
×10^7
raw data, starting:9
fitted Gauss locations:8.5
y
x
0
2
4
6
8
10
12
14
16
0
5
10
15
20
25

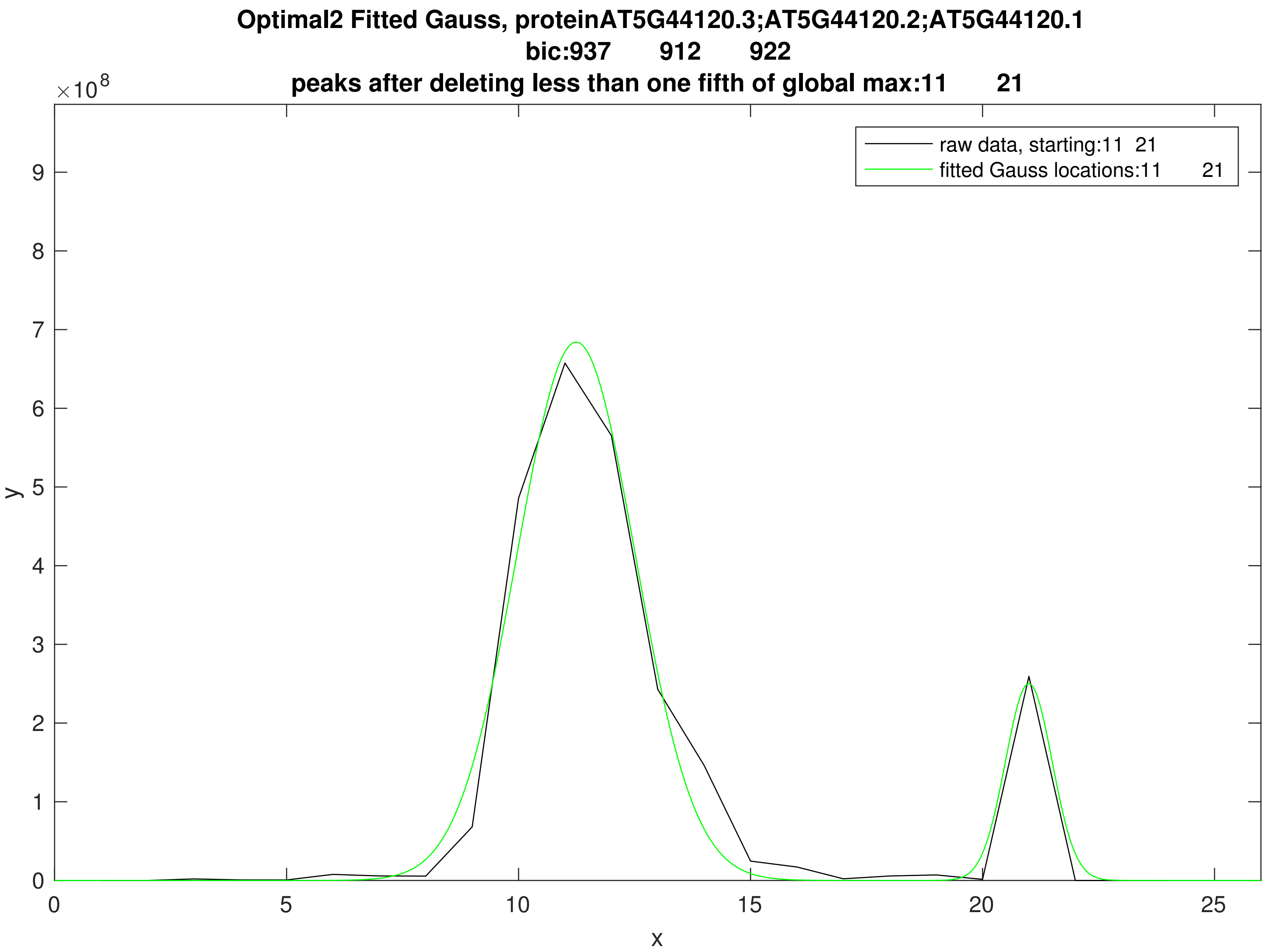

Optimal2 Fitted Gauss, proteinAT5G44120.3;AT5G44120.2;AT5G44120.1
bic:937 912 922
peaks after deleting less than one fifth of global max:11 21
×10^8
raw data, starting:11 21
fitted Gauss locations:11 21
y
x
0
1
2
3
4
5
6
7
8
9
10
15
20
25

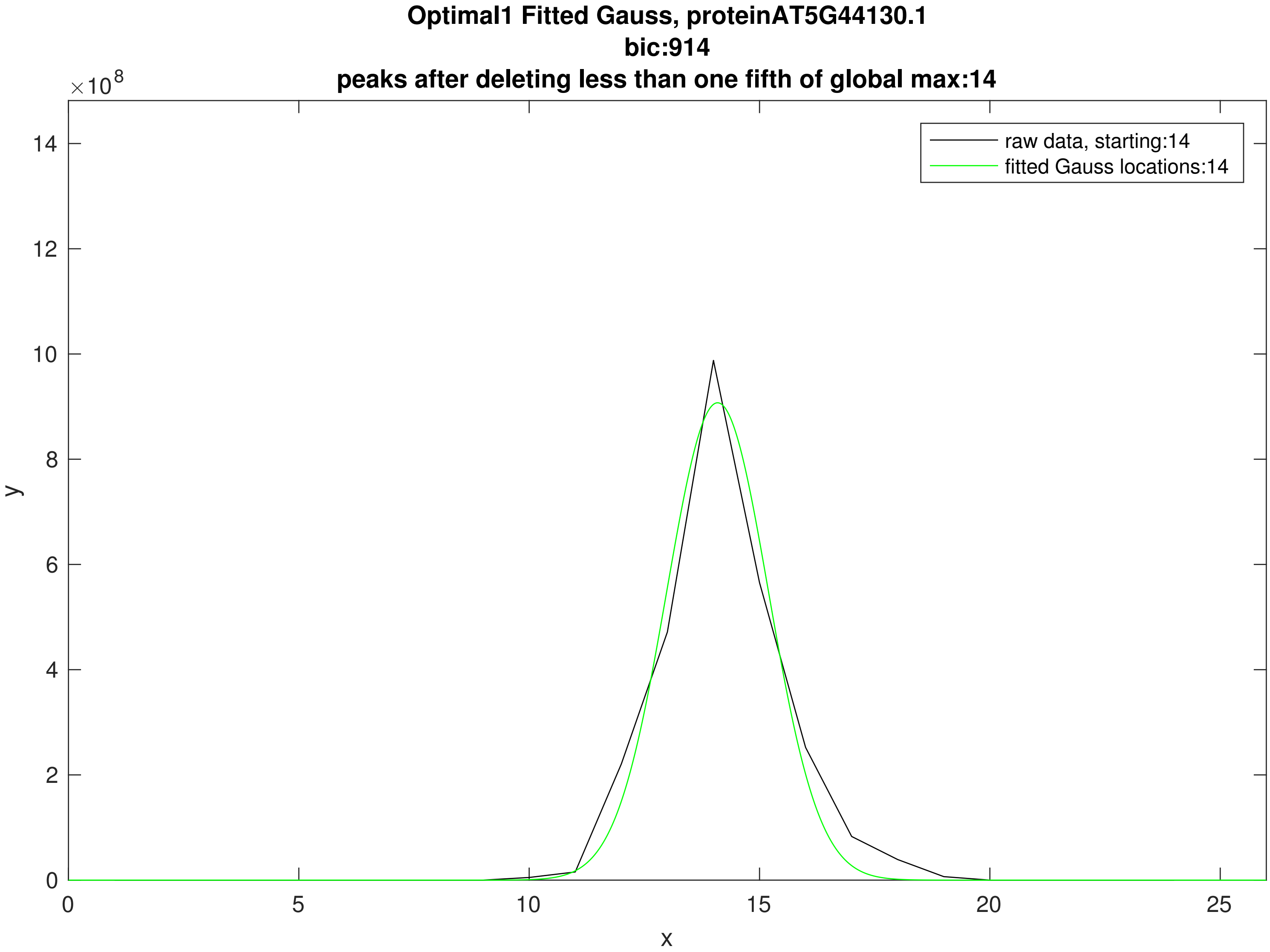
Optimal1 Fitted Gauss, proteinAT5G44130.1
bic:914
peaks after deleting less than one fifth of global max:14
×10^8
raw data, starting:14
fitted Gauss locations:14
y
x
0
2
4
6
8
10
12
14
0
5
10
15
20
25

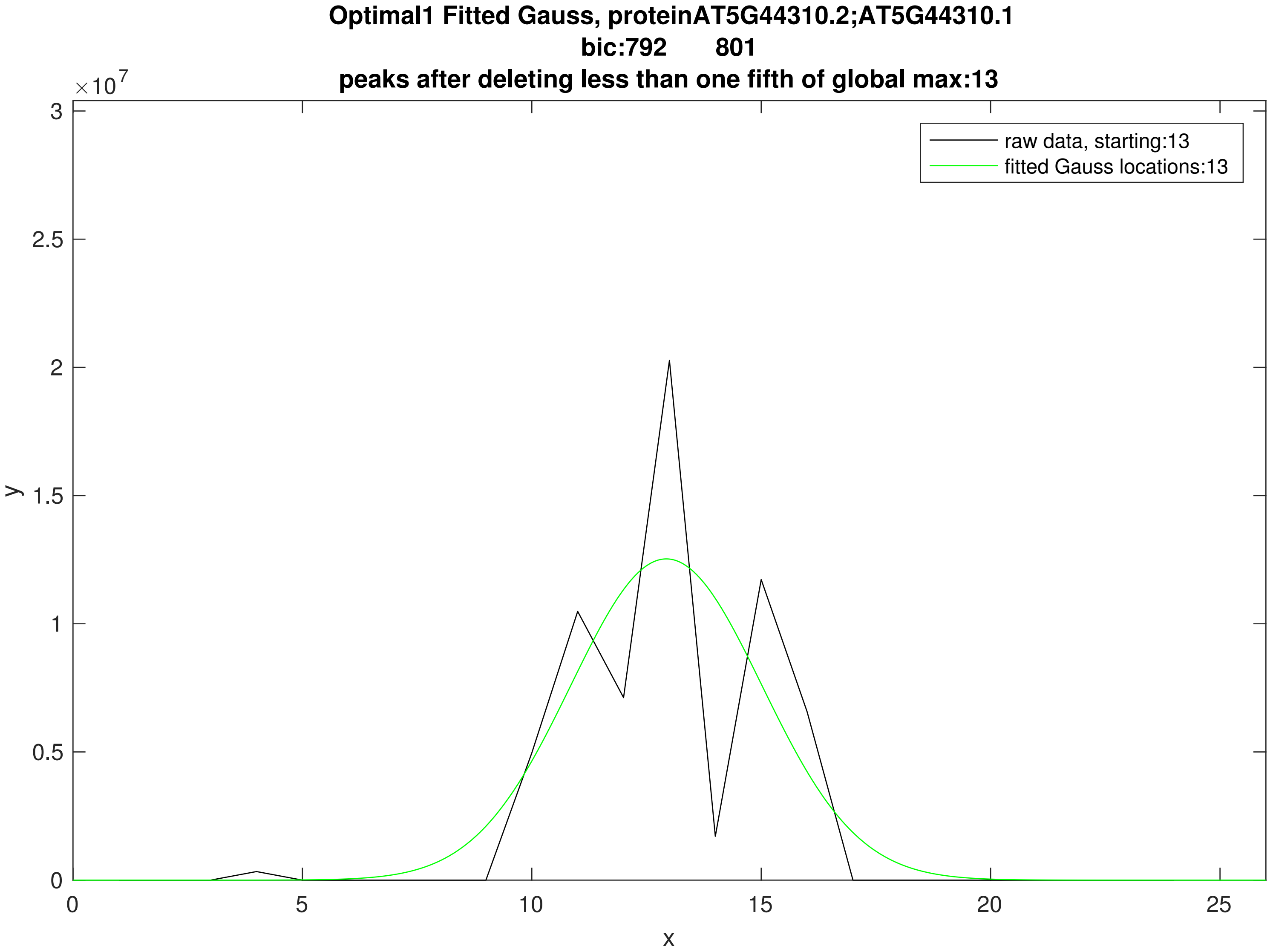

Optimal1 Fitted Gauss, proteinAT5G44310.2;AT5G44310.1
bic:792 801
peaks after deleting less than one fifth of global max:13
×10^7
raw data, starting:13
fitted Gauss locations:13
y
x
0
0.5
1
1.5
2
2.5
3
0
5
10
15
20
25

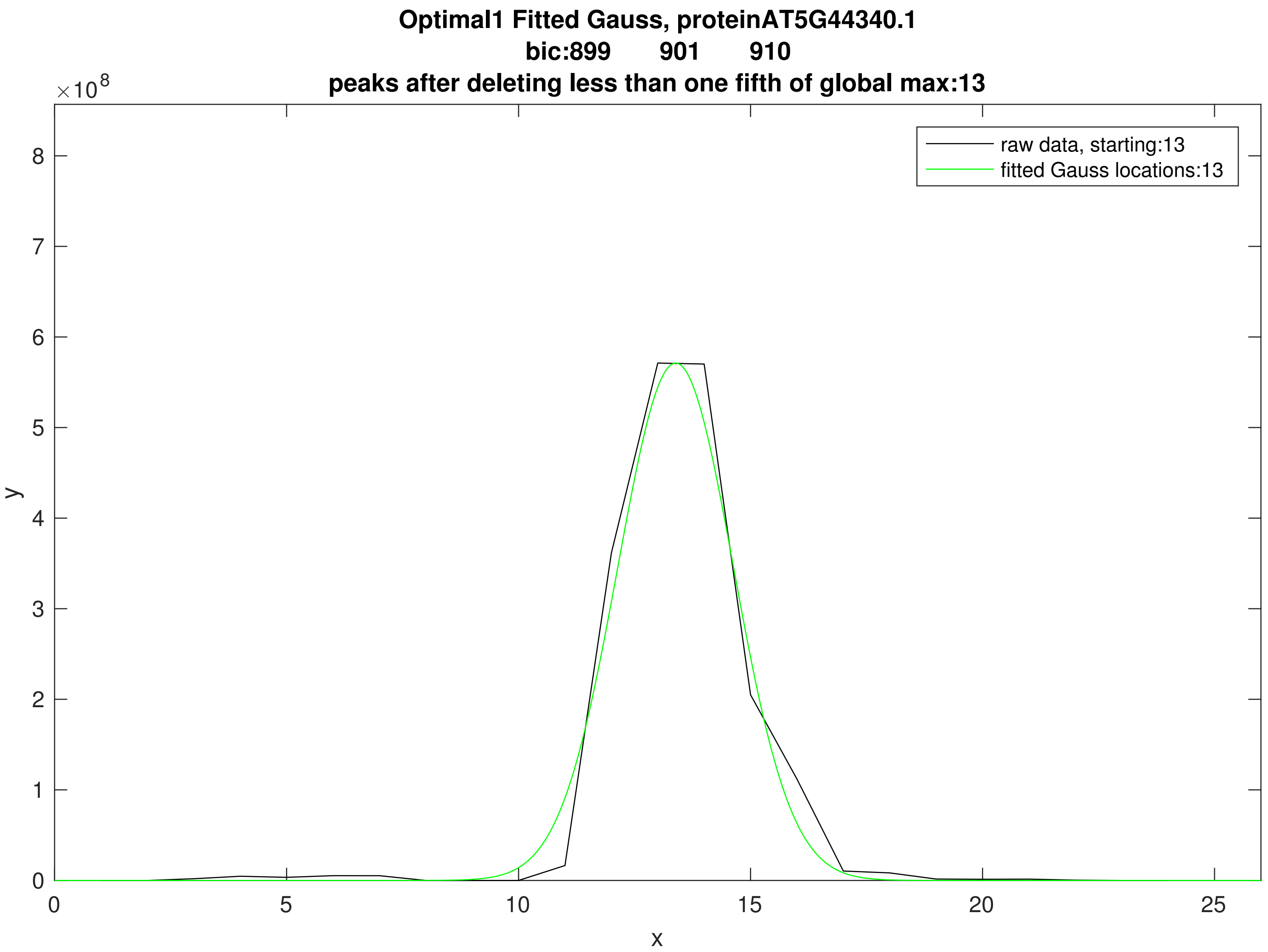
Optimal1 Fitted Gauss, proteinAT5G44340.1
bic:899 901 910
peaks after deleting less than one fifth of global max:13
×10^8
raw data, starting:13
fitted Gauss locations:13
y
x
0
1
2
3
4
5
6
7
8
0
5
10
15
20
25

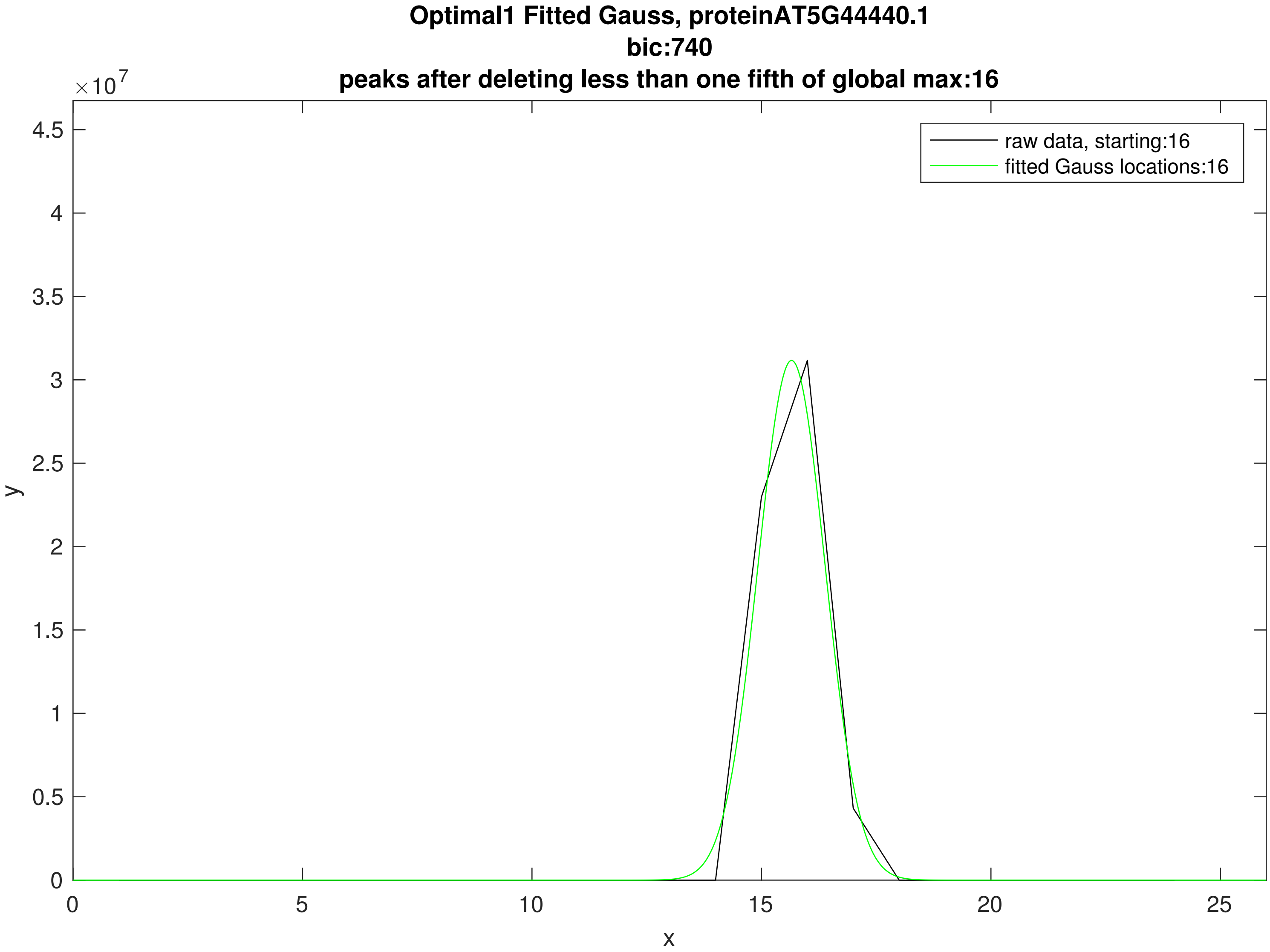
Optimal1 Fitted Gauss, proteinAT5G44440.1
bic:740
peaks after deleting less than one fifth of global max:16
×10^7
raw data, starting:16
fitted Gauss locations:16
y
x

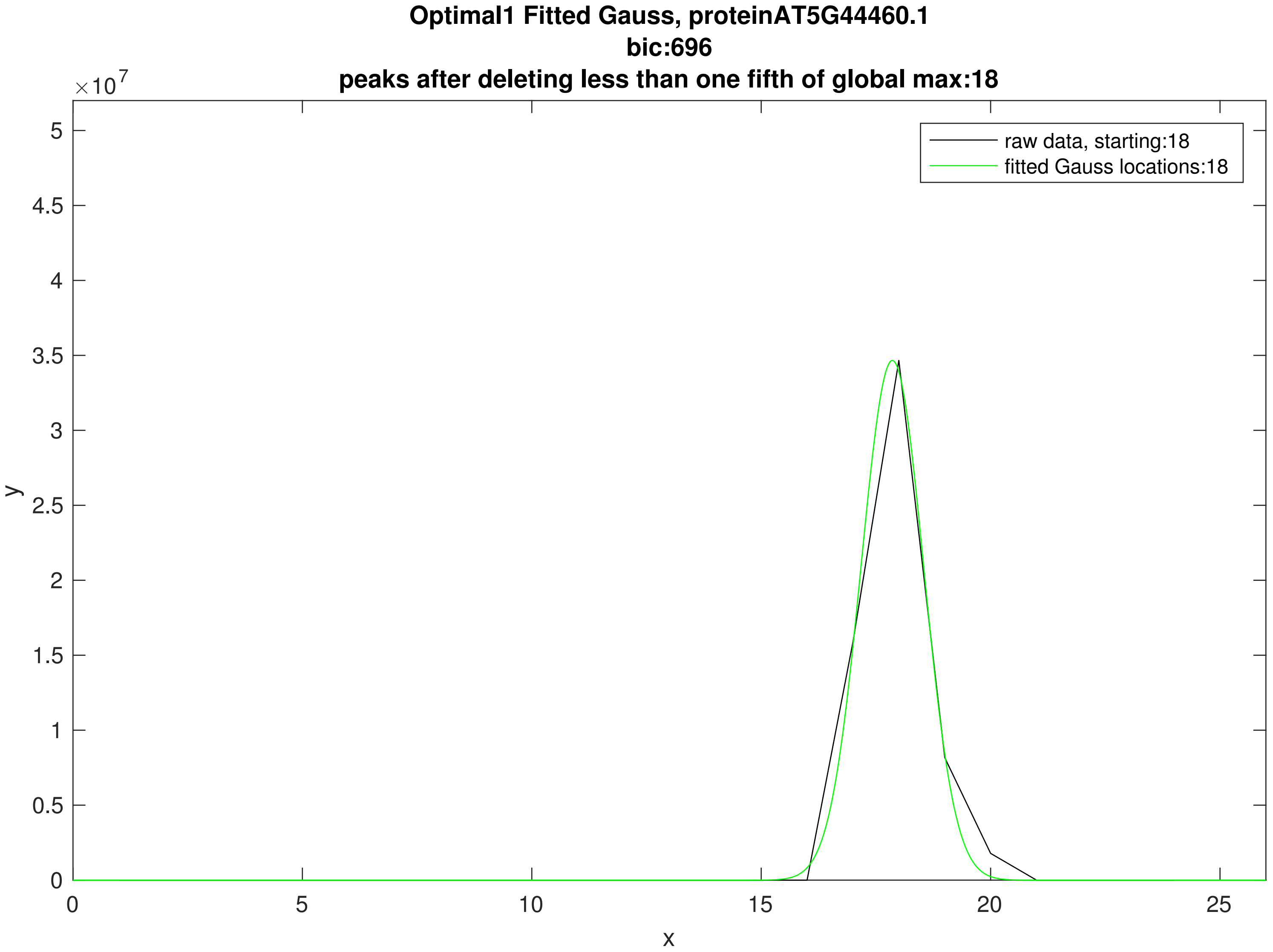

Optimal1 Fitted Gauss, proteinAT5G44460.1
bic:696
peaks after deleting less than one fifth of global max:18
×10^7
raw data, starting:18
fitted Gauss locations:18
y
x
0
0.5
1
1.5
2
2.5
3
3.5
4
4.5
5
0
5
10
15
20
25

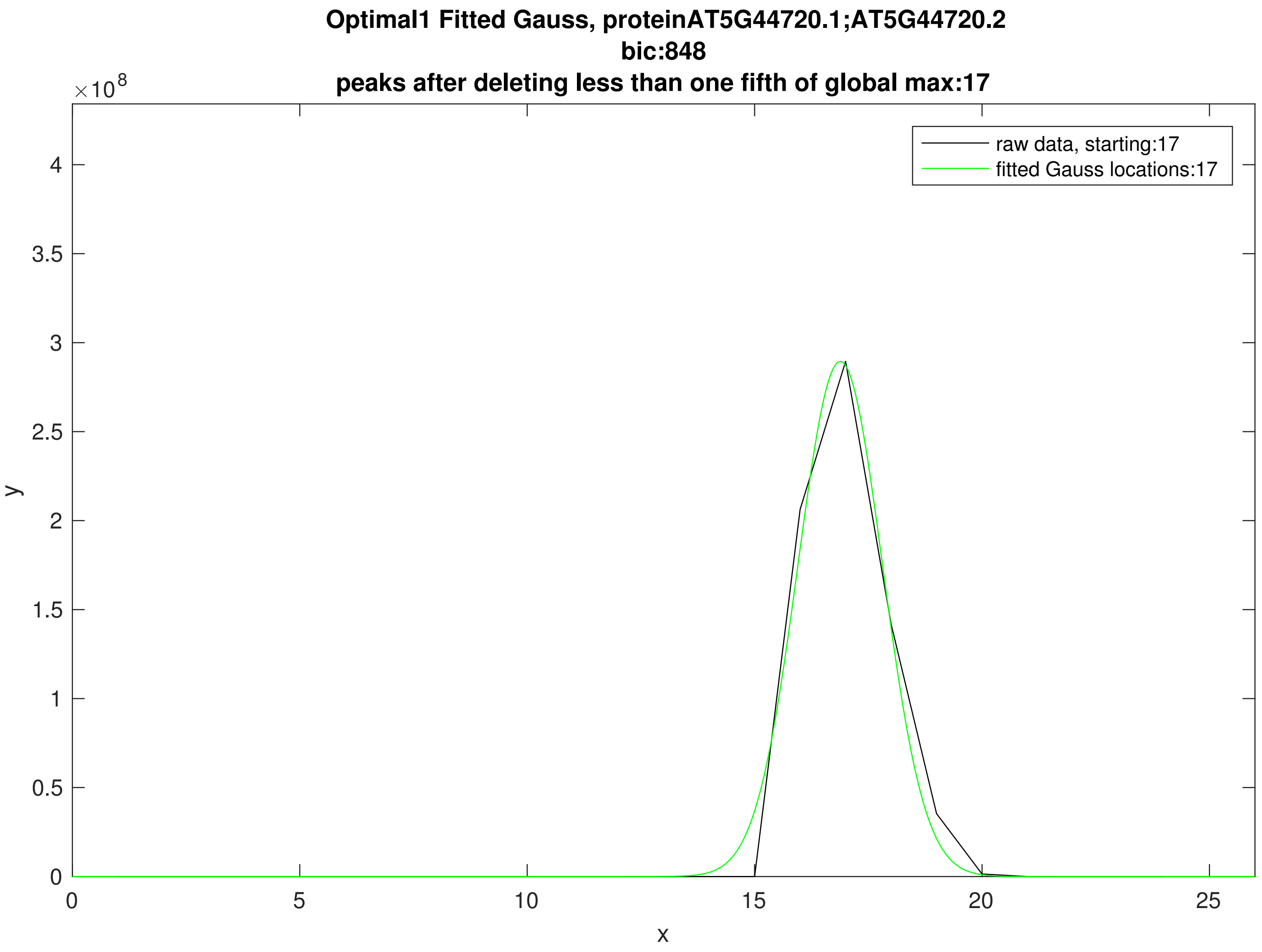
Optimal1 Fitted Gauss, proteinAT5G44720.1;AT5G44720.2
bic:848
peaks after deleting less than one fifth of global max:17
×10^8
raw data, starting:17
fitted Gauss locations:17
y
x
0
0.5
1
1.5
2
2.5
3
3.5
4
0
5
10
15
20
25

Optimal1 Fitted Gauss, proteinAT5G45020.2;AT5G45020.1
bic:719
peaks after deleting less than one fifth of global max:14

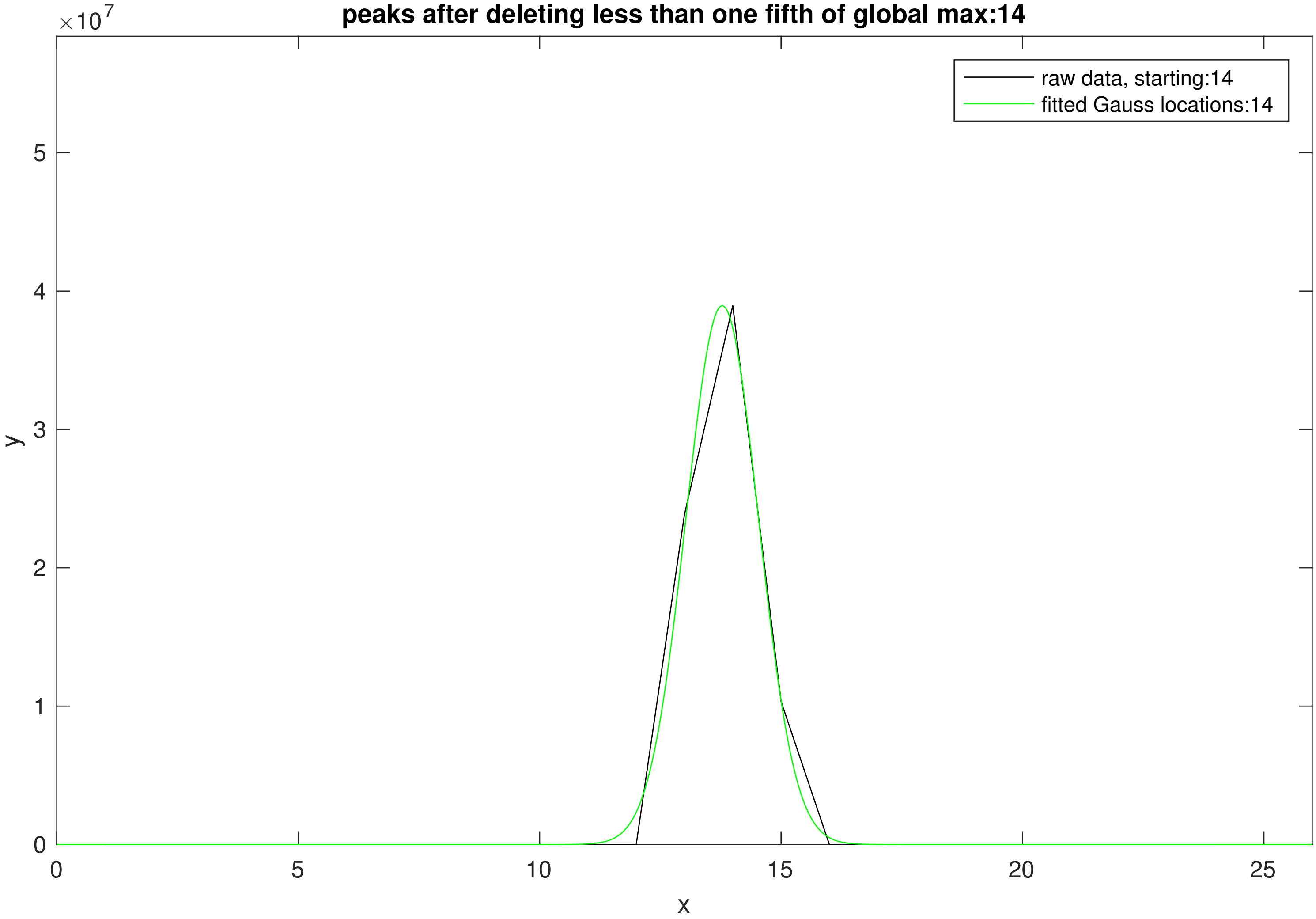

Optimal1 Fitted Gauss, proteinAT5G45130.1
bic:734
peaks after deleting less than one fifth of global max:19

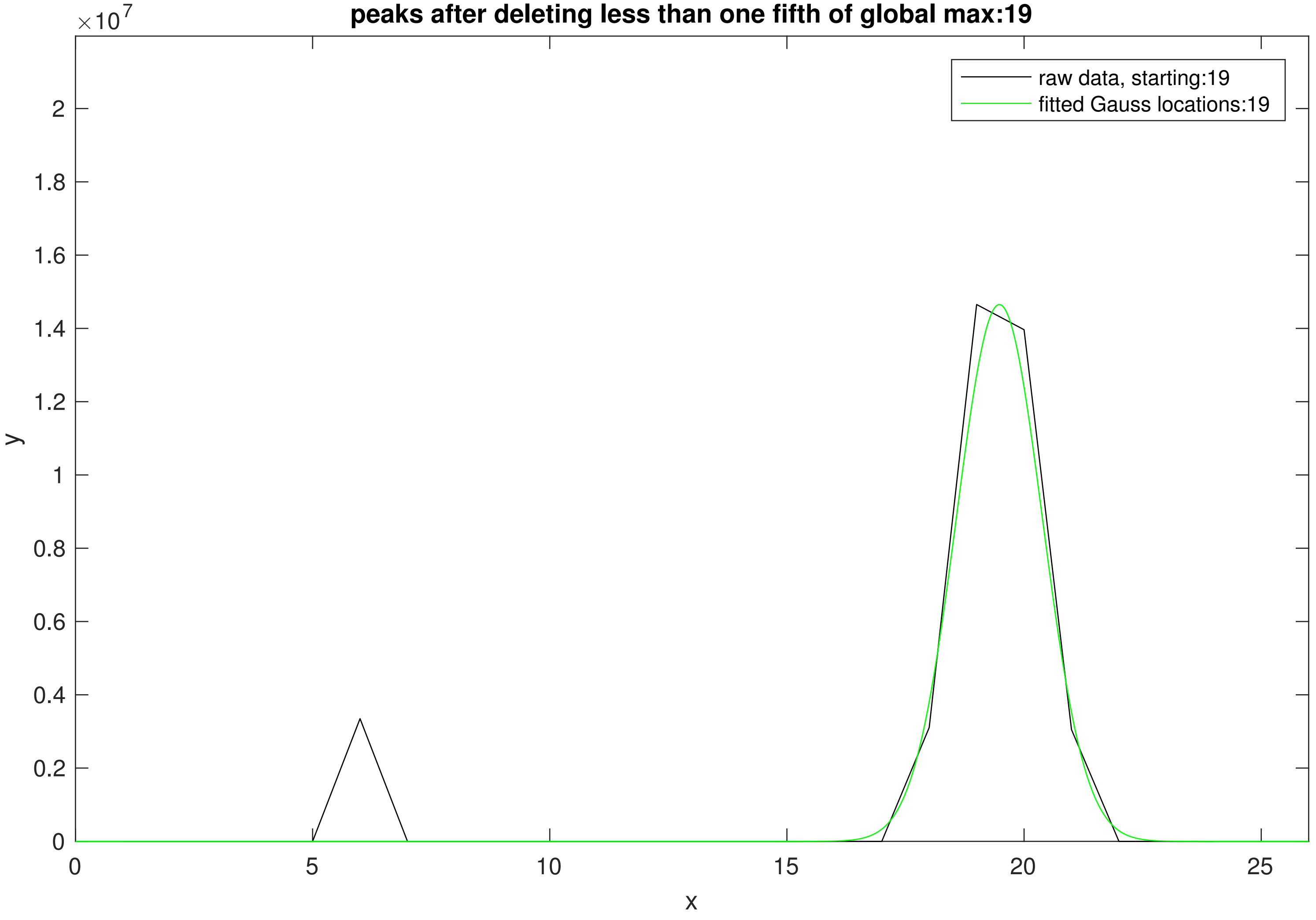

**Optimal1 Fitted Gauss, proteinAT5G45280.2;AT5G45280.1**
**bic:887 895**
**peaks after deleting less than one fifth of global max:18**

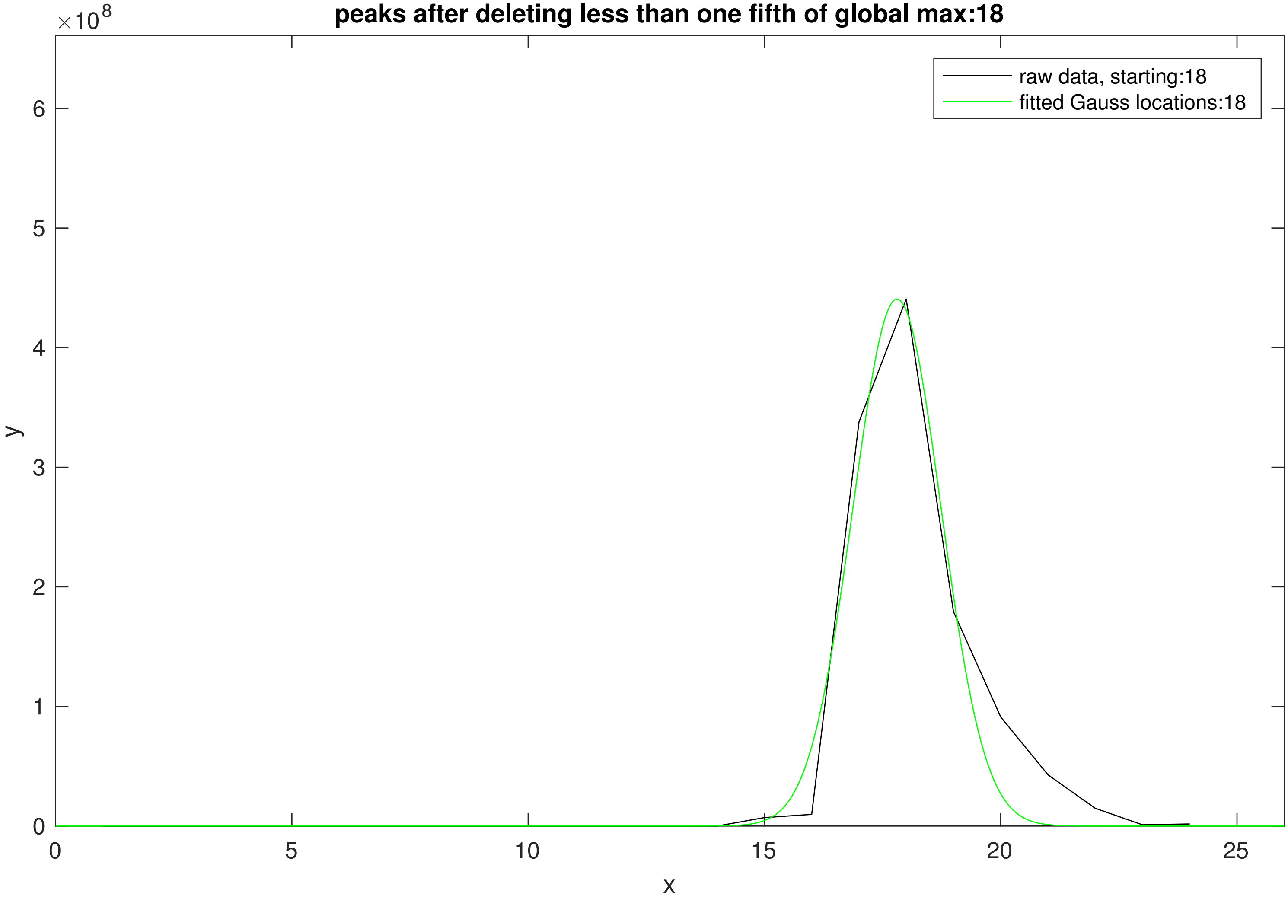

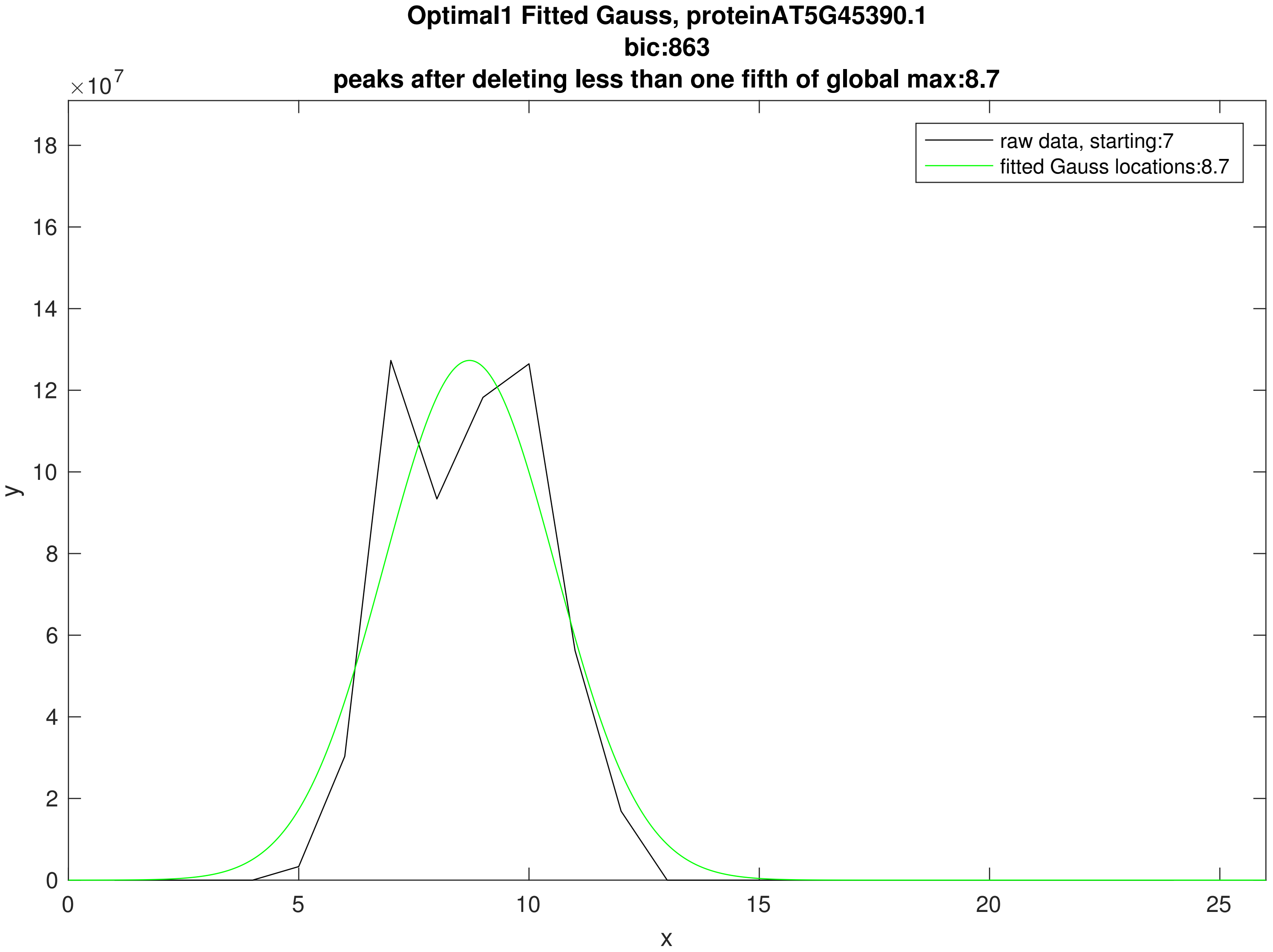
Optimal1 Fitted Gauss, proteinAT5G45390.1
bic:863
peaks after deleting less than one fifth of global max:8.7
×10^7
raw data, starting:7
fitted Gauss locations:8.7
y
x
0
2
4
6
8
10
12
14
16
18
0
5
10
15
20
25

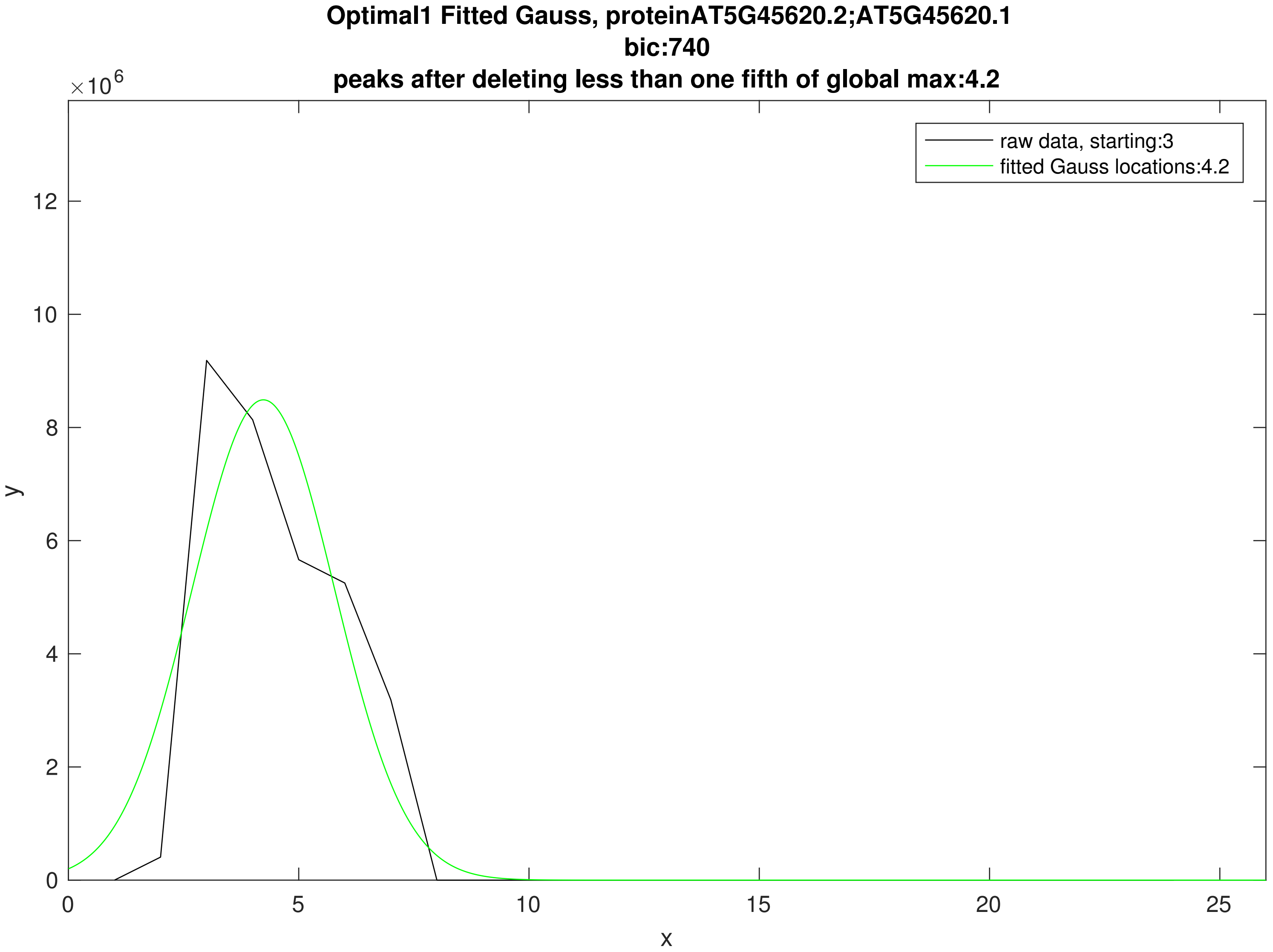
Optimal1 Fitted Gauss, proteinAT5G45620.2;AT5G45620.1
bic:740
peaks after deleting less than one fifth of global max:4.2
×10^6
raw data, starting:3
fitted Gauss locations:4.2
y
x
0
2
4
6
8
10
12
0
5
10
15
20
25

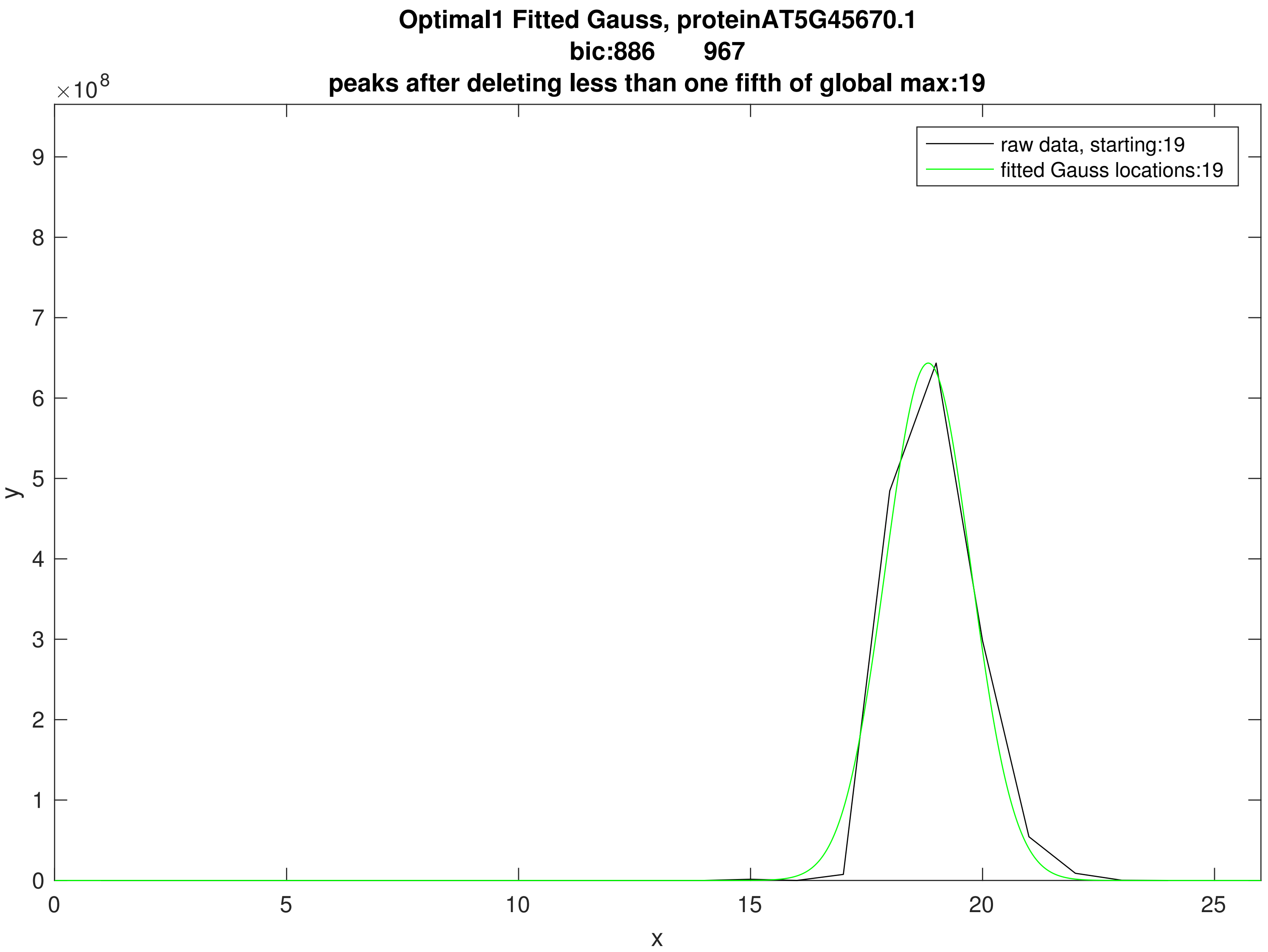
Optimal1 Fitted Gauss, proteinAT5G45670.1
bic:886 967
peaks after deleting less than one fifth of global max:19
×10^8
raw data, starting:19
fitted Gauss locations:19
y
x
0
1
2
3
4
5
6
7
8
9
0
5
10
15
20
25

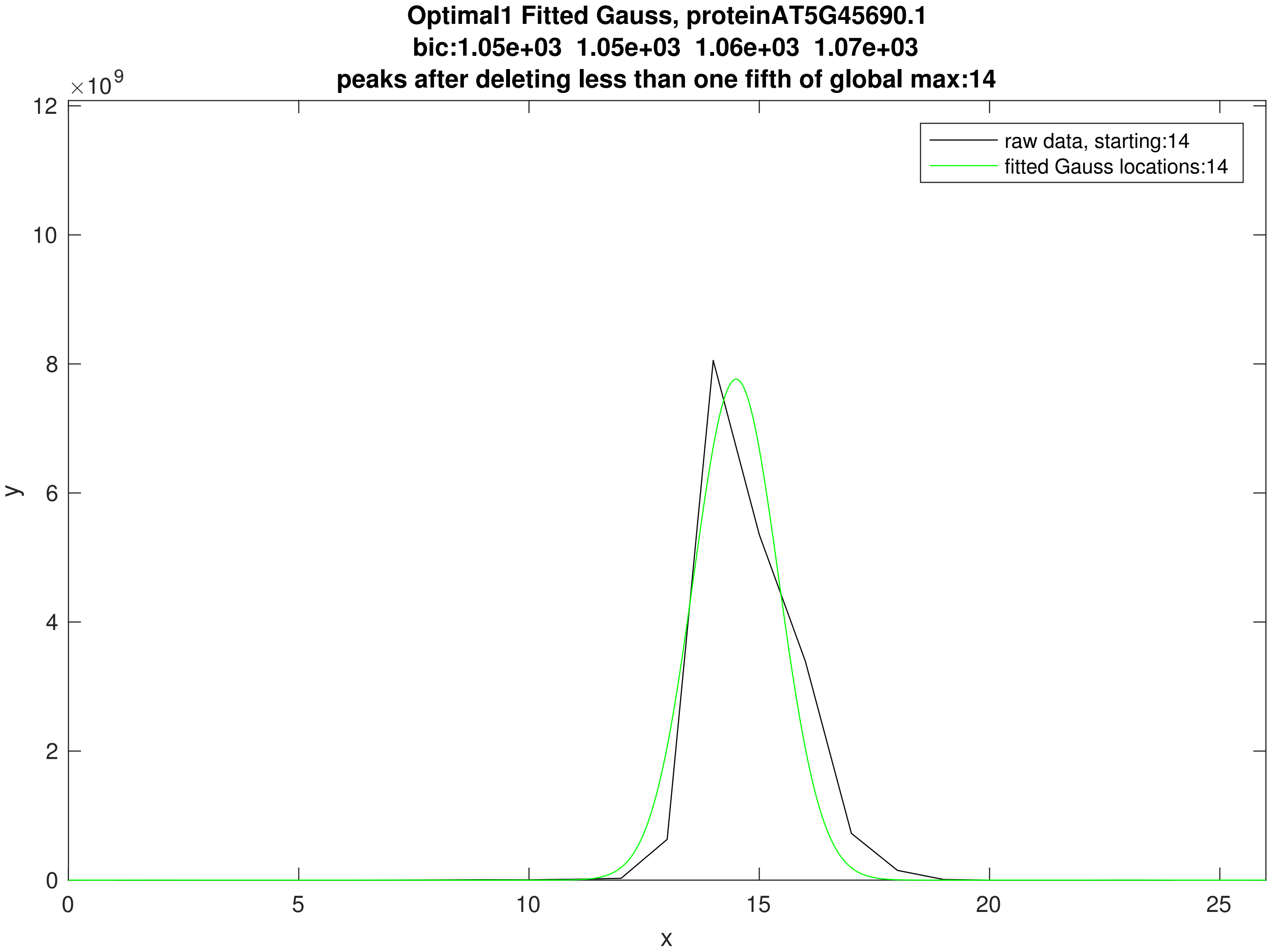

Optimal1 Fitted Gauss, proteinAT5G45690.1
bic:1.05e+03 1.05e+03 1.06e+03 1.07e+03
peaks after deleting less than one fifth of global max:14
×10^9
12
10
8
6
4
2
0
y
0
5
10
15
20
25
x
raw data, starting:14
fitted Gauss locations:14

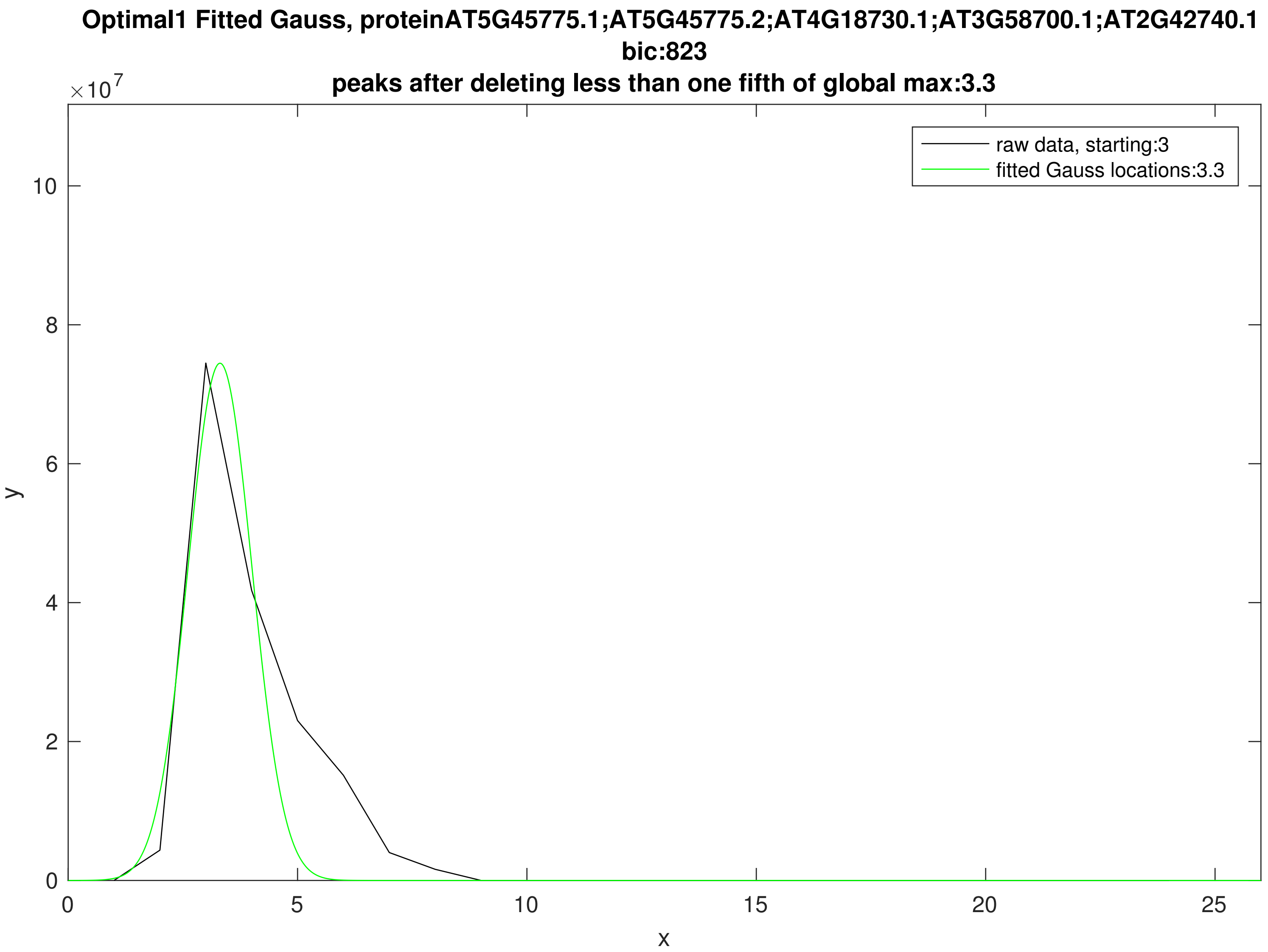
Optimal1 Fitted Gauss, proteinAT5G45775.1;AT5G45775.2;AT4G18730.1;AT3G58700.1;AT2G42740.1
bic:823
peaks after deleting less than one fifth of global max:3.3
×10^7
raw data, starting:3
fitted Gauss locations:3.3
y
x
0
2
4
6
8
10
0
5
10
15
20
25

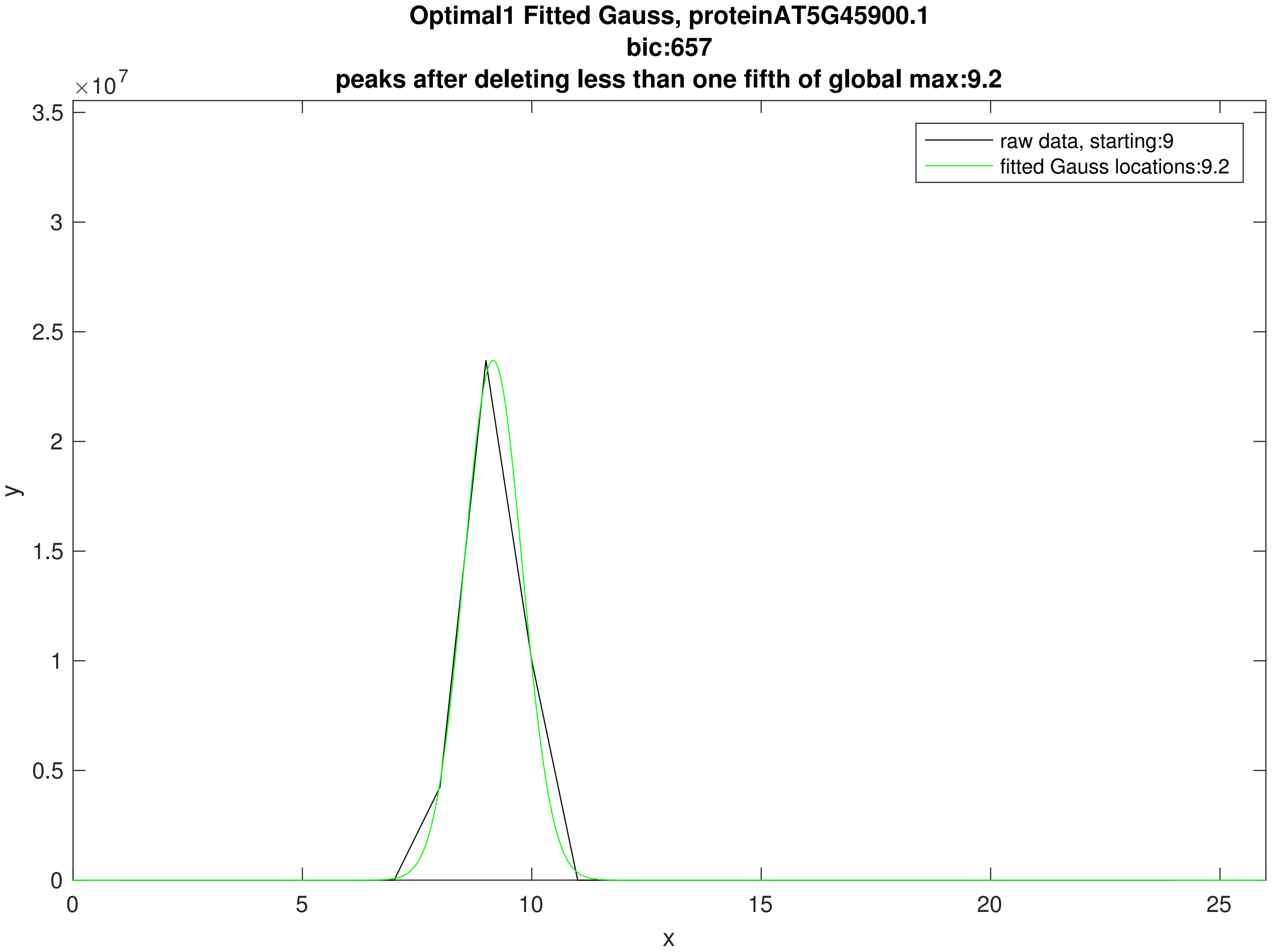
Optimal1 Fitted Gauss, proteinAT5G45900.1
bic:657
peaks after deleting less than one fifth of global max:9.2
×10^7
3.5
3
2.5
2
1.5
1
0.5
0
y
0
5
10
15
20
25
x
raw data, starting:9
fitted Gauss locations:9.2

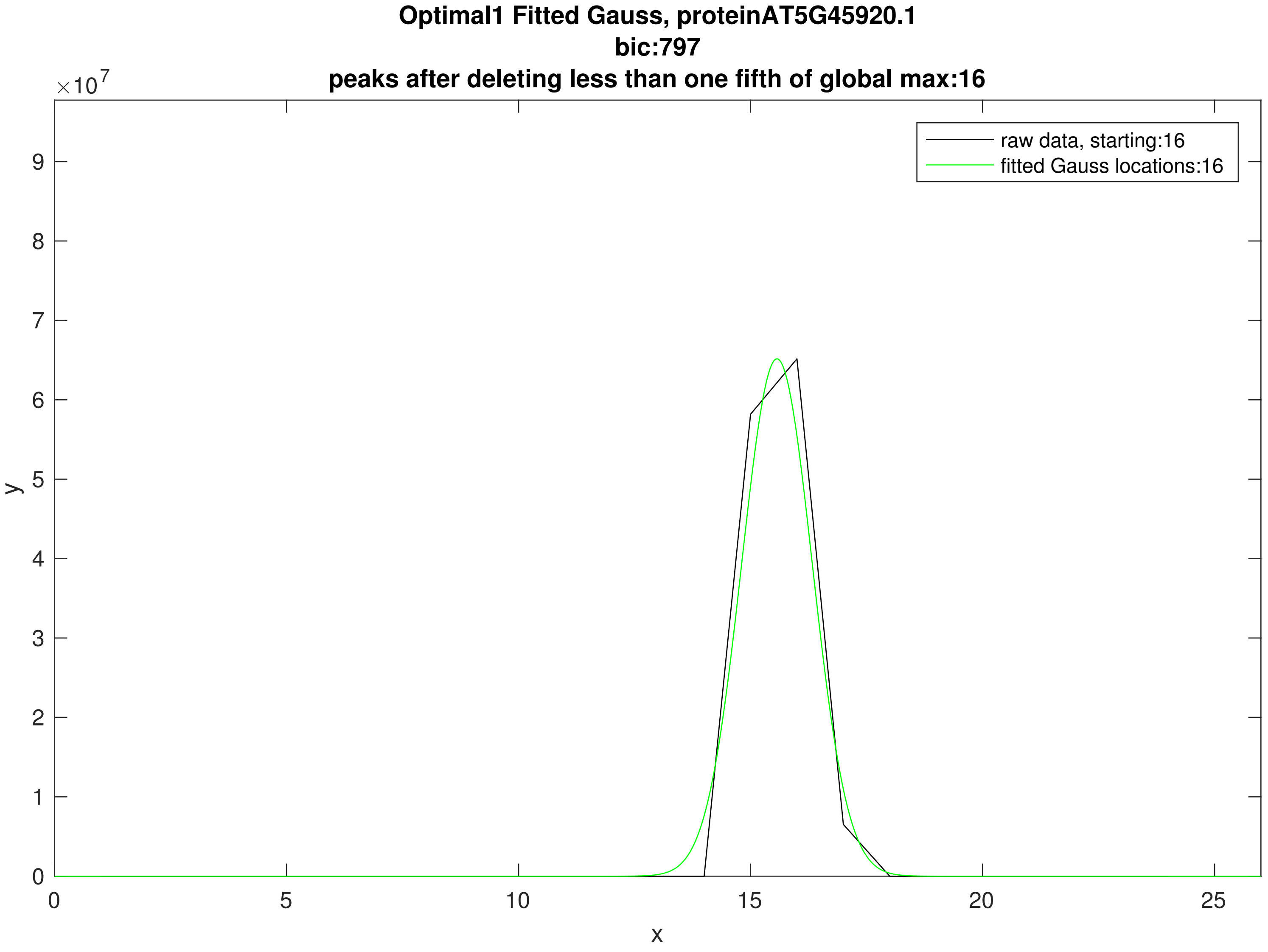

Optimal1 Fitted Gauss, proteinAT5G45920.1
bic:797
peaks after deleting less than one fifth of global max:16
×10^7
9
8
7
6
5
4
3
2
1
0
y
0
5
10
15
20
25
x
raw data, starting:16
fitted Gauss locations:16

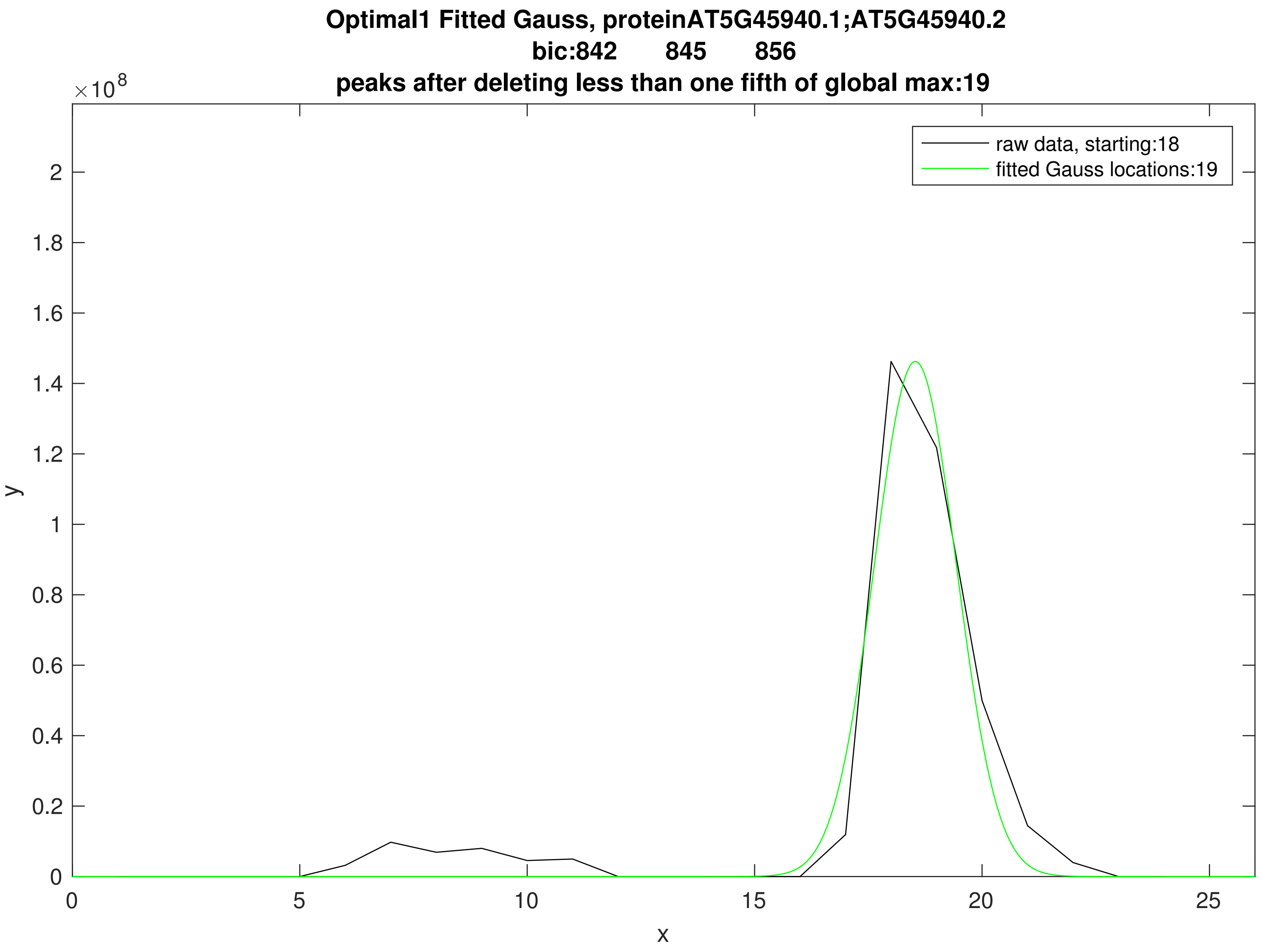

Optimal1 Fitted Gauss, proteinAT5G45940.1;AT5G45940.2
bic:842 845 856
peaks after deleting less than one fifth of global max:19
×10^8
raw data, starting:18
fitted Gauss locations:19
y
x
0
0.2
0.4
0.6
0.8
1
1.2
1.4
1.6
1.8
2
5
10
15
20
25

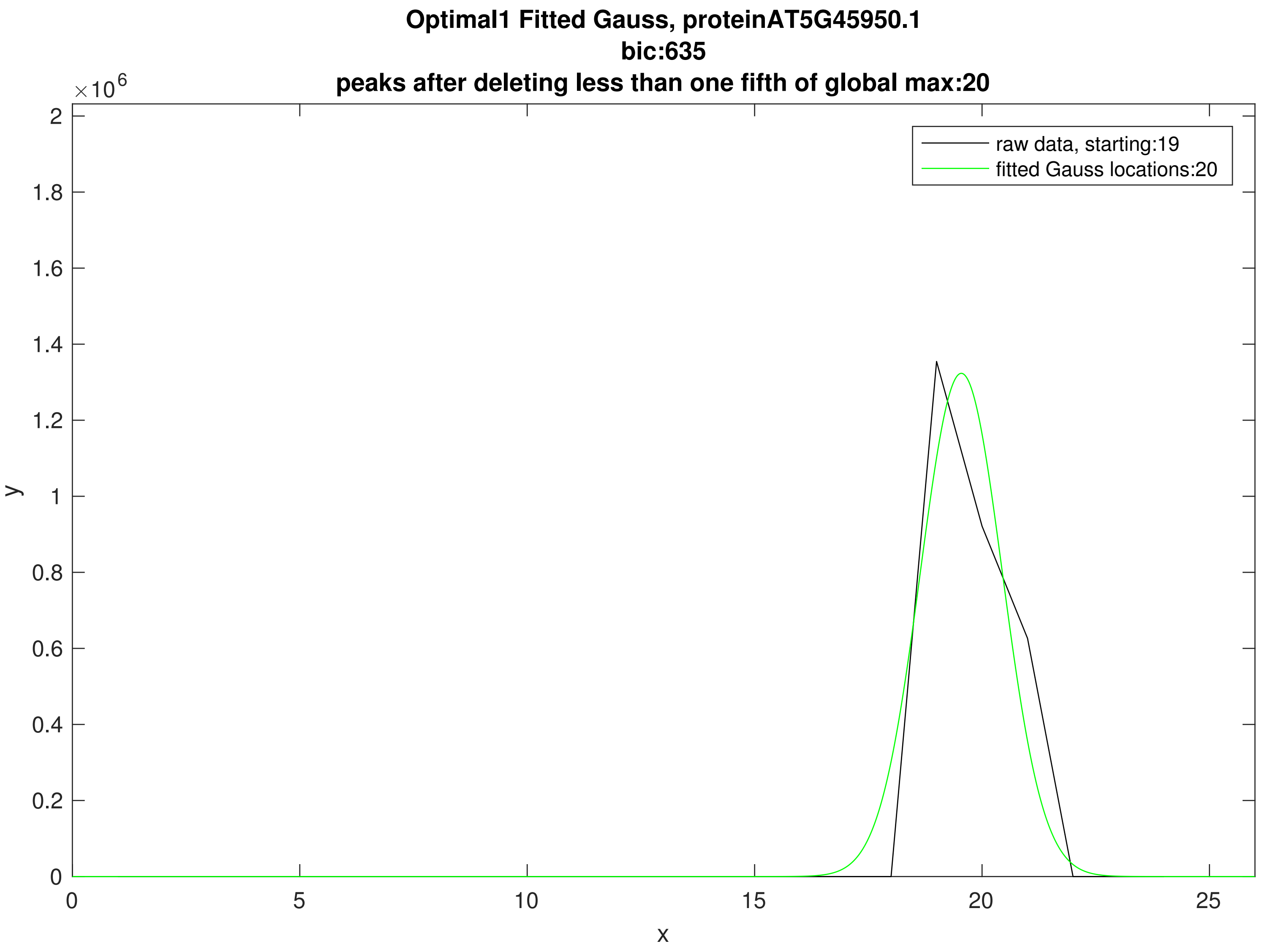
Optimal1 Fitted Gauss, proteinAT5G45950.1
bic:635
peaks after deleting less than one fifth of global max:20
×10^6
raw data, starting:19
fitted Gauss locations:20
y
x
0
0.2
0.4
0.6
0.8
1
1.2
1.4
1.6
1.8
2
0
5
10
15
20
25

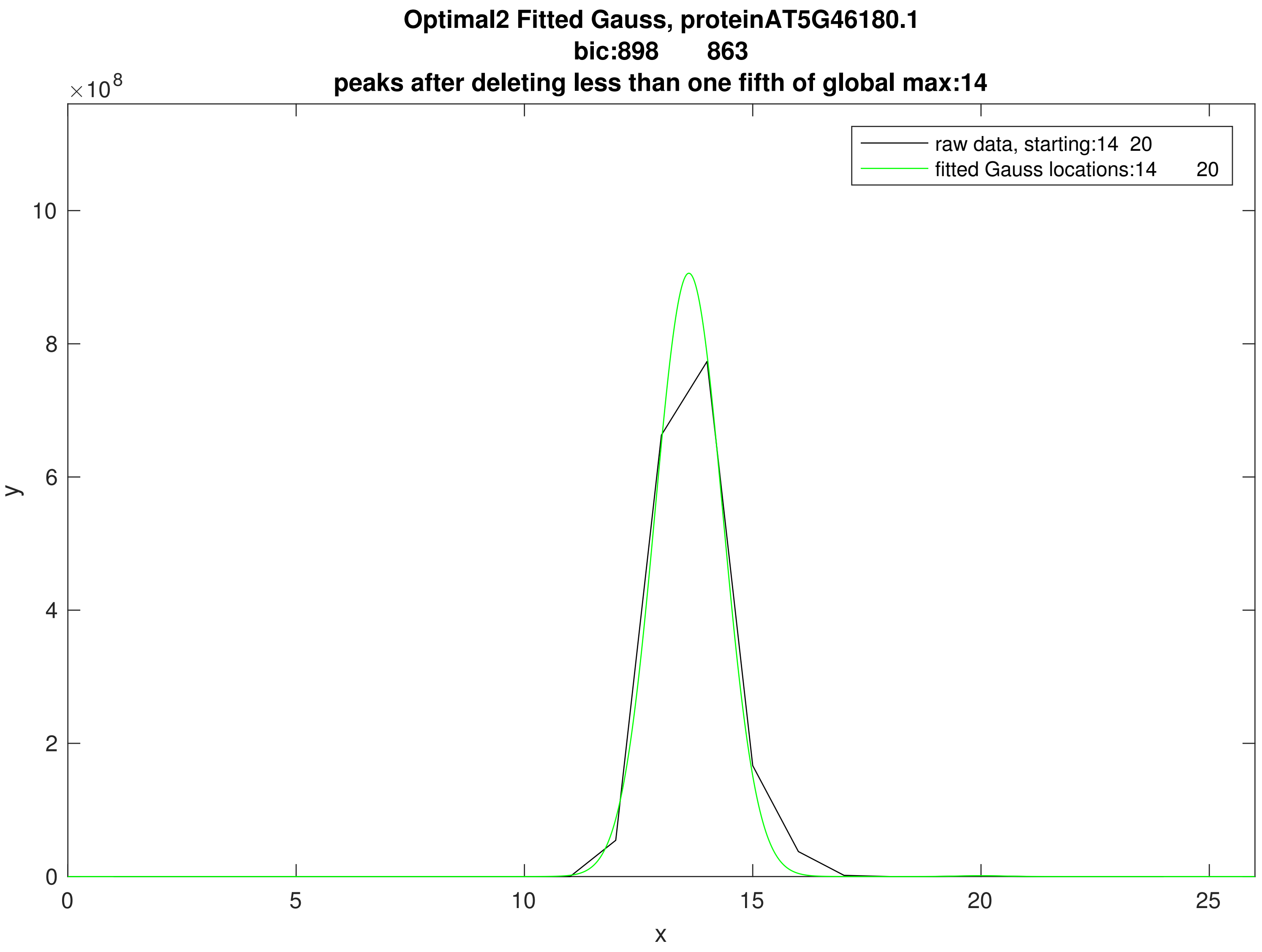

Optimal2 Fitted Gauss, proteinAT5G46180.1
bic:898 863
peaks after deleting less than one fifth of global max:14
×10^8
raw data, starting:14 20
fitted Gauss locations:14 20
10
8
6
4
2
0
y
0
5
10
15
20
25
x

**Optimal1 Fitted Gauss, proteinAT5G46230.1**
**bic:774 782**
**peaks after deleting less than one fifth of global max:20**

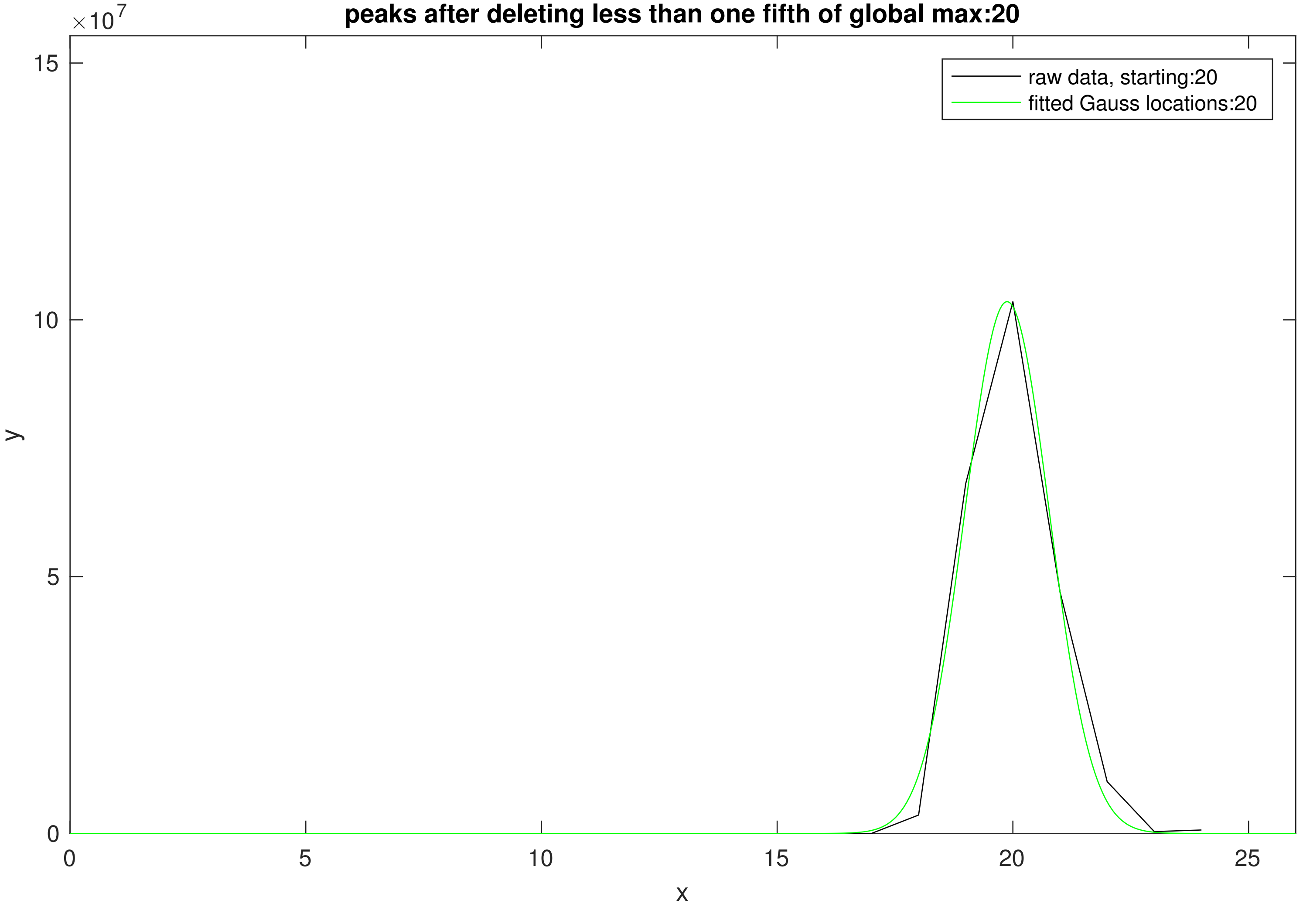

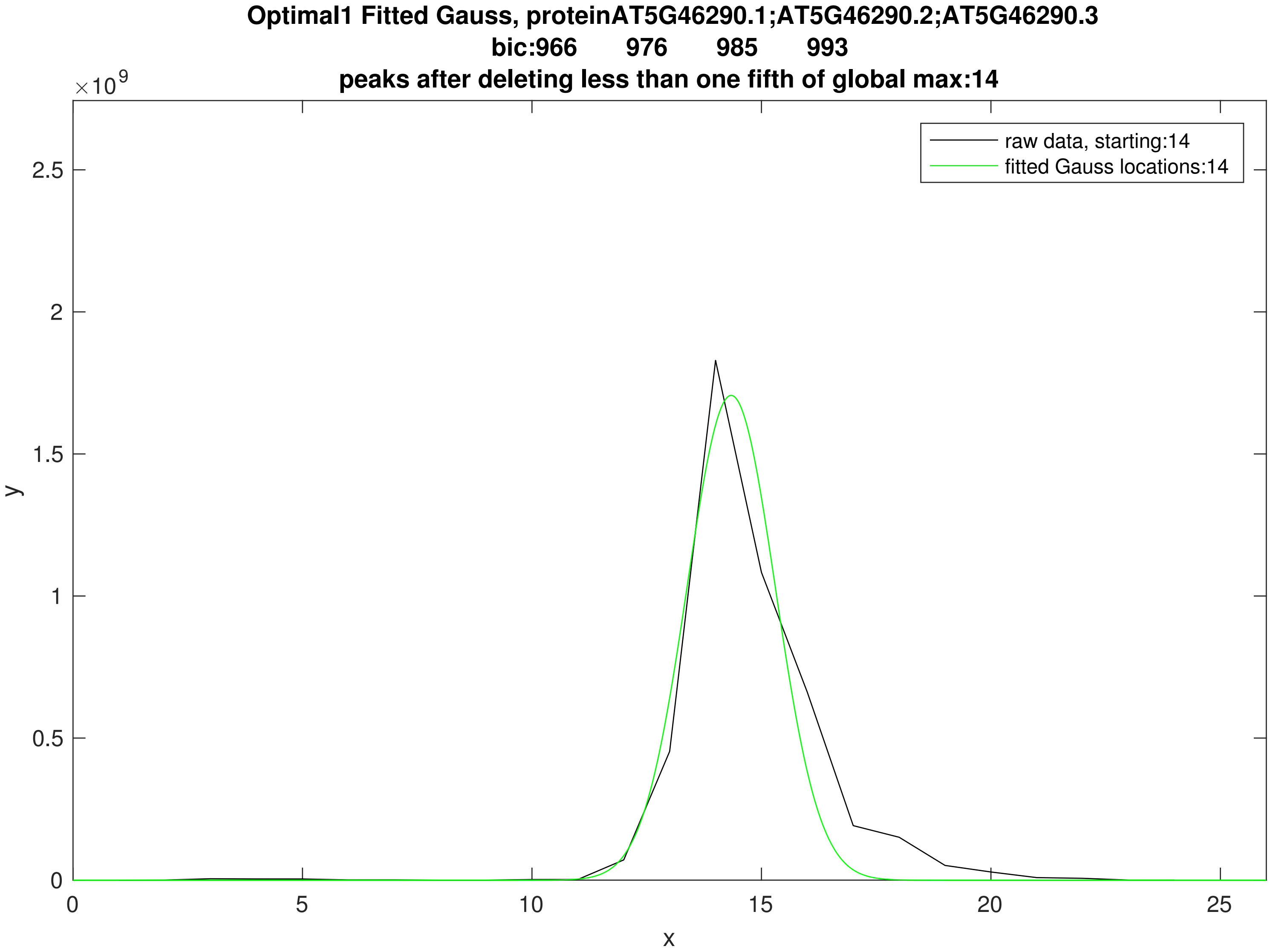
Optimal1 Fitted Gauss, proteinAT5G46290.1;AT5G46290.2;AT5G46290.3
bic:966 976 985 993
peaks after deleting less than one fifth of global max:14
×10^9
2.5
2
1.5
1
0.5
0
y
0
5
10
15
20
25
x
raw data, starting:14
fitted Gauss locations:14

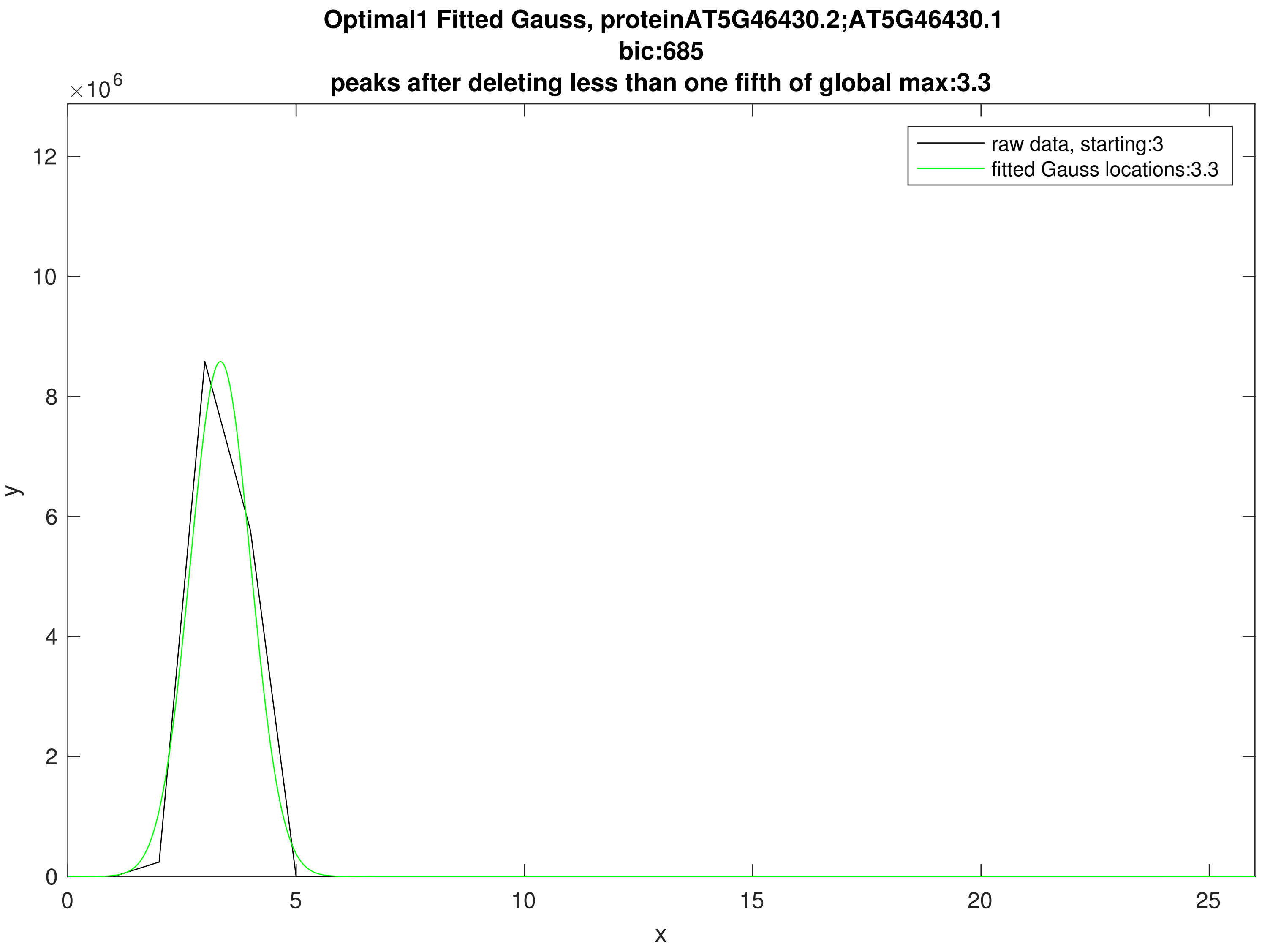
Optimal1 Fitted Gauss, proteinAT5G46430.2;AT5G46430.1
bic:685
peaks after deleting less than one fifth of global max:3.3
×10^6
raw data, starting:3
fitted Gauss locations:3.3
y
x
0
2
4
6
8
10
12
0
5
10
15
20
25

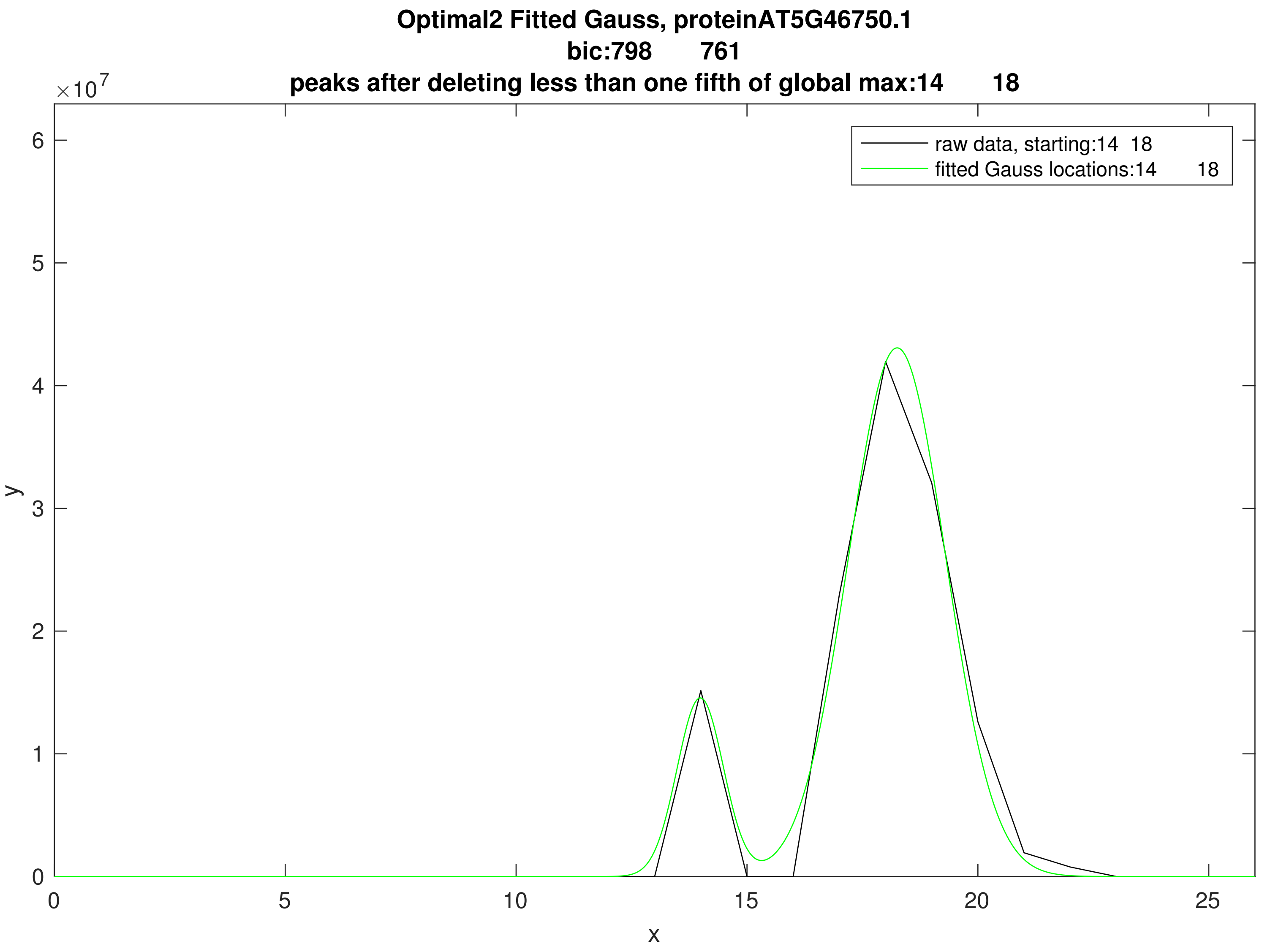
Optimal2 Fitted Gauss, proteinAT5G46750.1
bic:798 761
peaks after deleting less than one fifth of global max:14 18
×10^7
raw data, starting:14 18
fitted Gauss locations:14 18
y
x
0
1
2
3
4
5
6
0
5
10
15
20
25

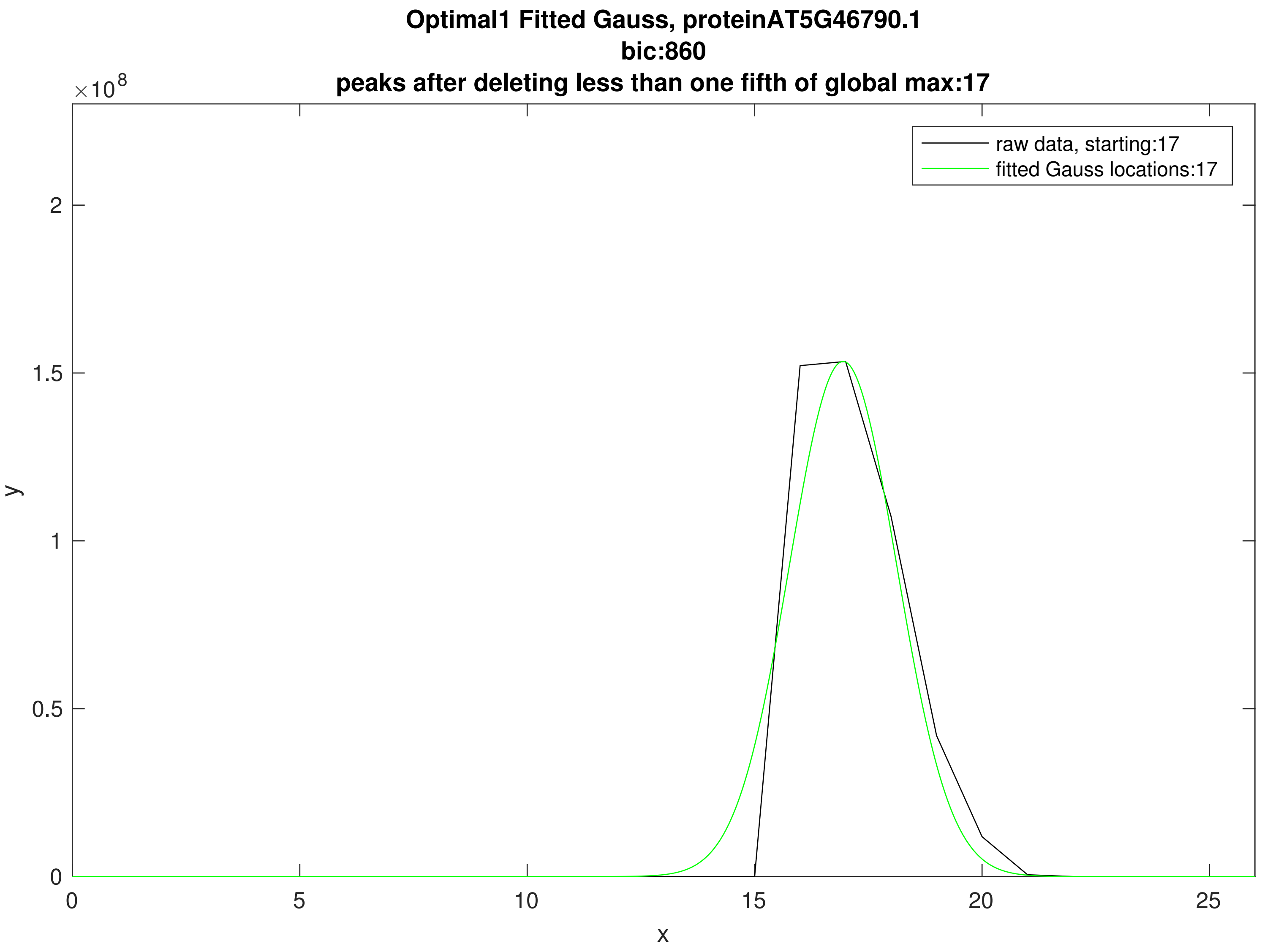
Optimal1 Fitted Gauss, proteinAT5G46790.1
bic:860
peaks after deleting less than one fifth of global max:17
×10^8
raw data, starting:17
fitted Gauss locations:17
2
1.5
1
0.5
0
y
0
5
10
15
20
25
x

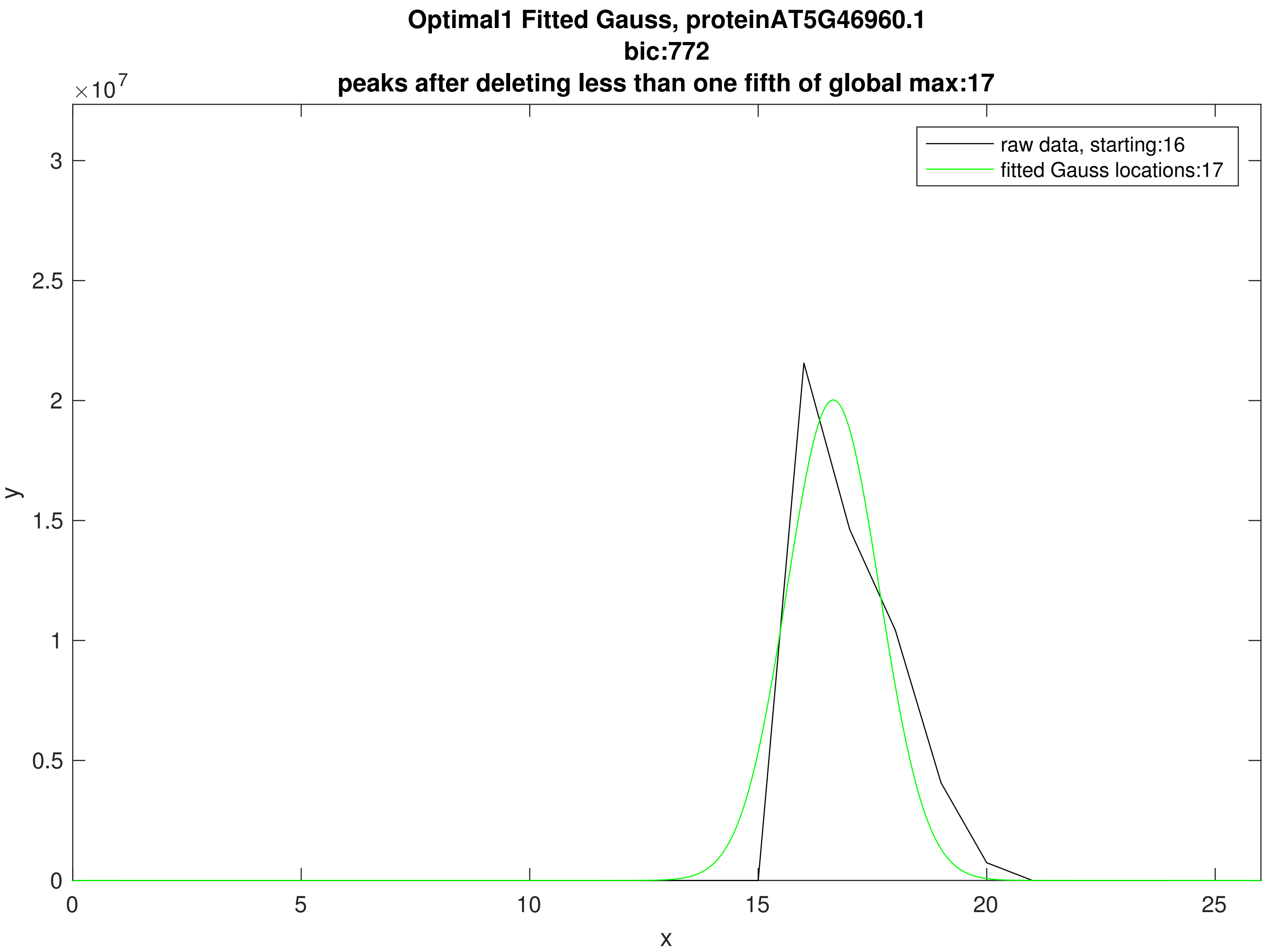
Optimal1 Fitted Gauss, proteinAT5G46960.1
bic:772
peaks after deleting less than one fifth of global max:17
×10^7
3
2.5
2
1.5
1
0.5
0
y
0
5
10
15
20
25
x
raw data, starting:16
fitted Gauss locations:17

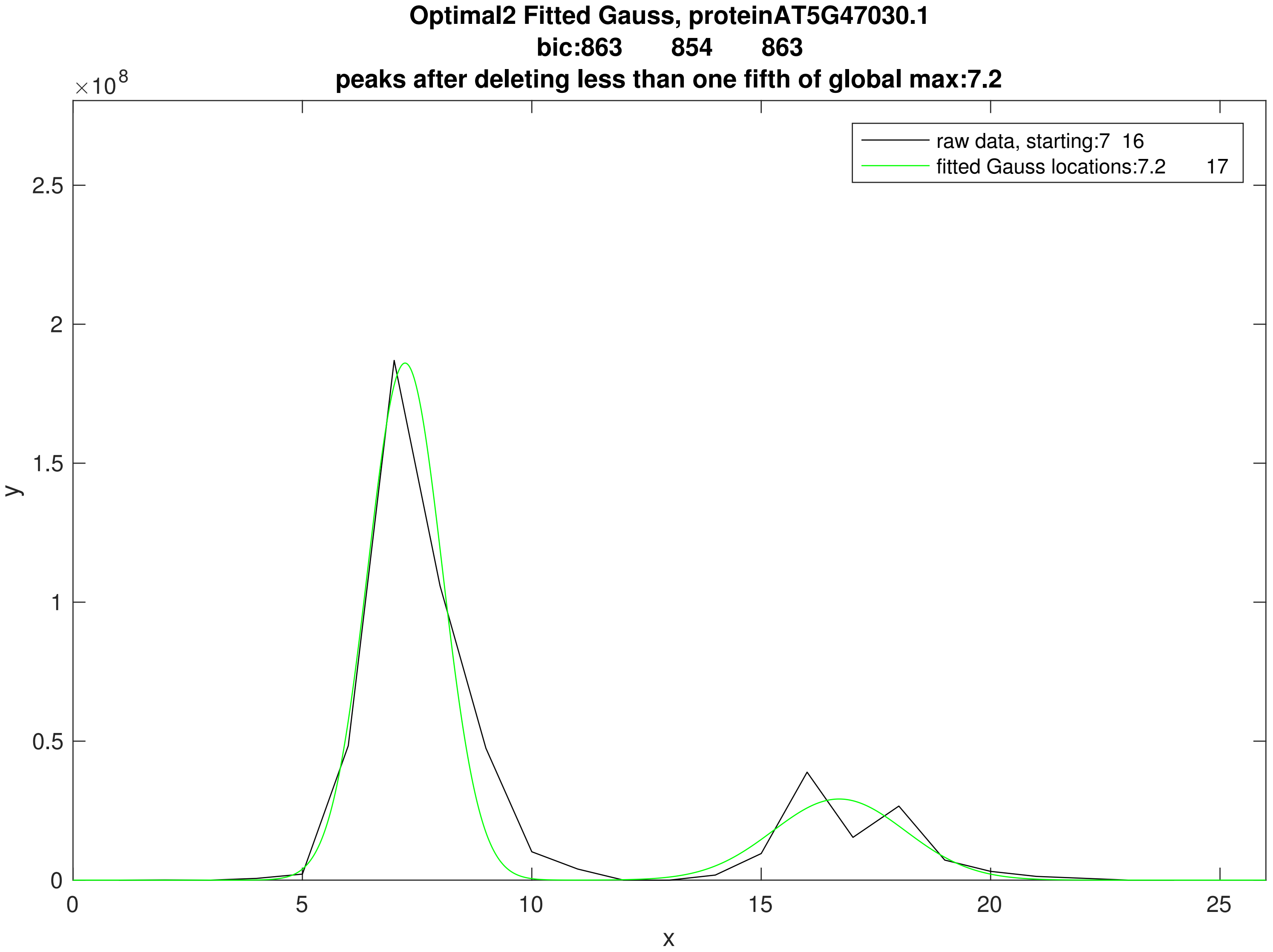
Optimal2 Fitted Gauss, proteinAT5G47030.1
bic:863 854 863
peaks after deleting less than one fifth of global max:7.2
×10^8
raw data, starting:7 16
fitted Gauss locations:7.2 17
y
x
0
0.5
1
1.5
2
2.5
0
5
10
15
20
25

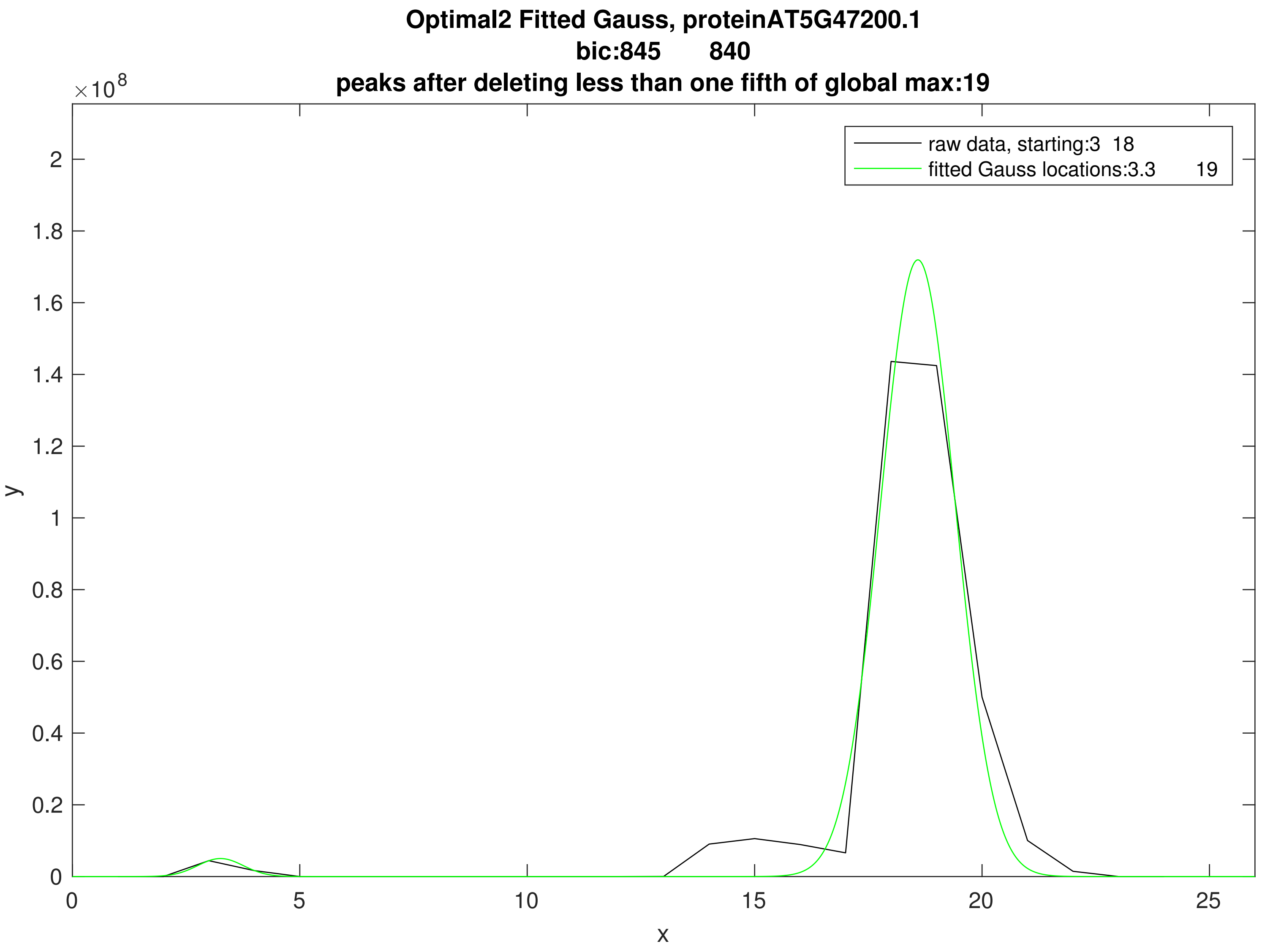
Optimal2 Fitted Gauss, proteinAT5G47200.1
bic:845 840
peaks after deleting less than one fifth of global max:19
×10^8
raw data, starting:3 18
fitted Gauss locations:3.3 19
y
x
0
0.2
0.4
0.6
0.8
1
1.2
1.4
1.6
1.8
2
5
10
15
20
25

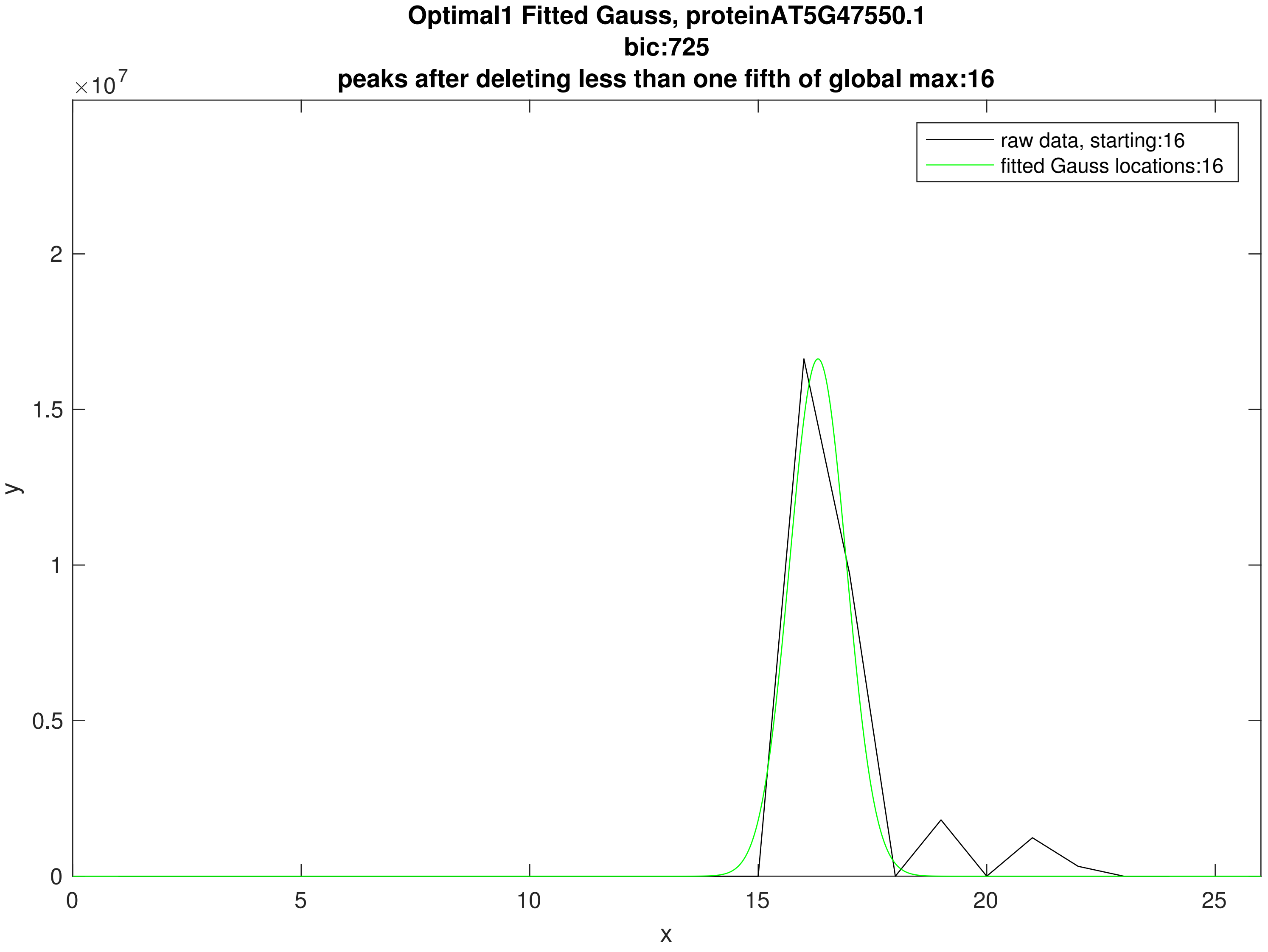

Optimal1 Fitted Gauss, proteinAT5G47550.1
bic:725
peaks after deleting less than one fifth of global max:16
×10^7
raw data, starting:16
fitted Gauss locations:16
y
x
0
0.5
1
1.5
2
0
5
10
15
20
25

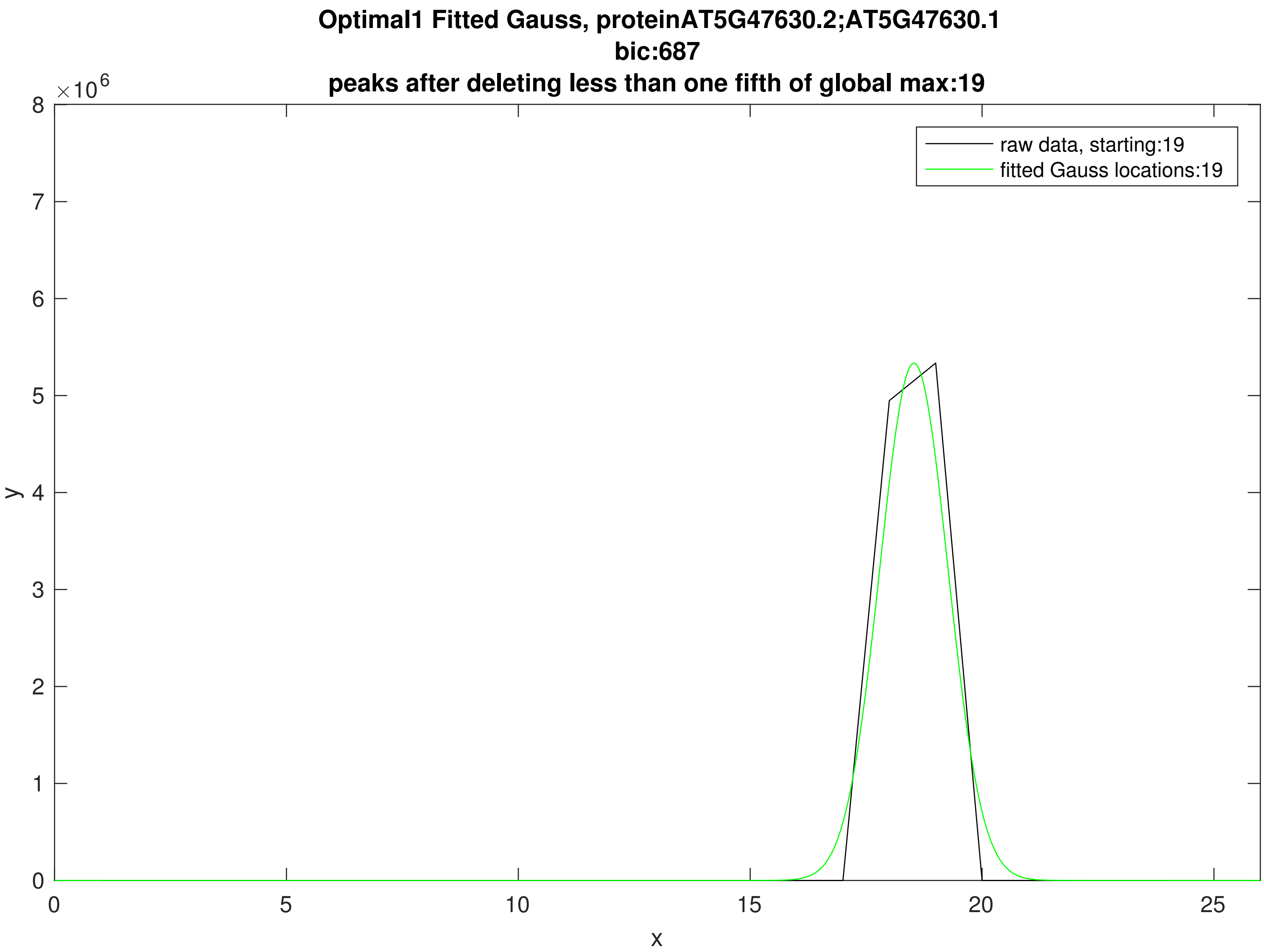
Optimal1 Fitted Gauss, proteinAT5G47630.2;AT5G47630.1
bic:687
peaks after deleting less than one fifth of global max:19
×10^6
8
7
6
5
4
3
2
1
0
y
raw data, starting:19
fitted Gauss locations:19
0
5
10
15
20
25
x

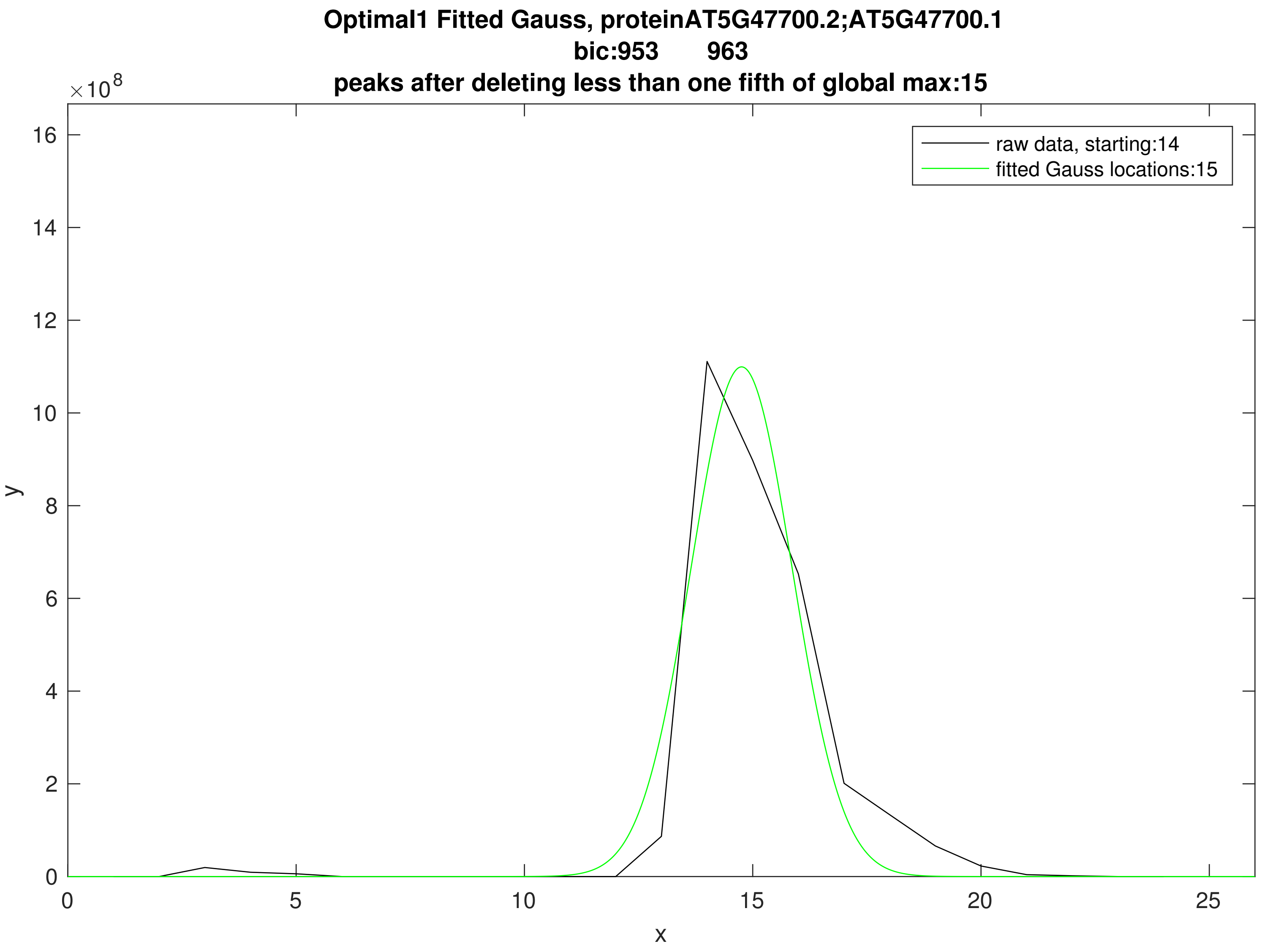
Optimal1 Fitted Gauss, proteinAT5G47700.2;AT5G47700.1
bic:953 963
peaks after deleting less than one fifth of global max:15
×10^8
raw data, starting:14
fitted Gauss locations:15
y
x
0
2
4
6
8
10
12
14
16
0
5
10
15
20
25

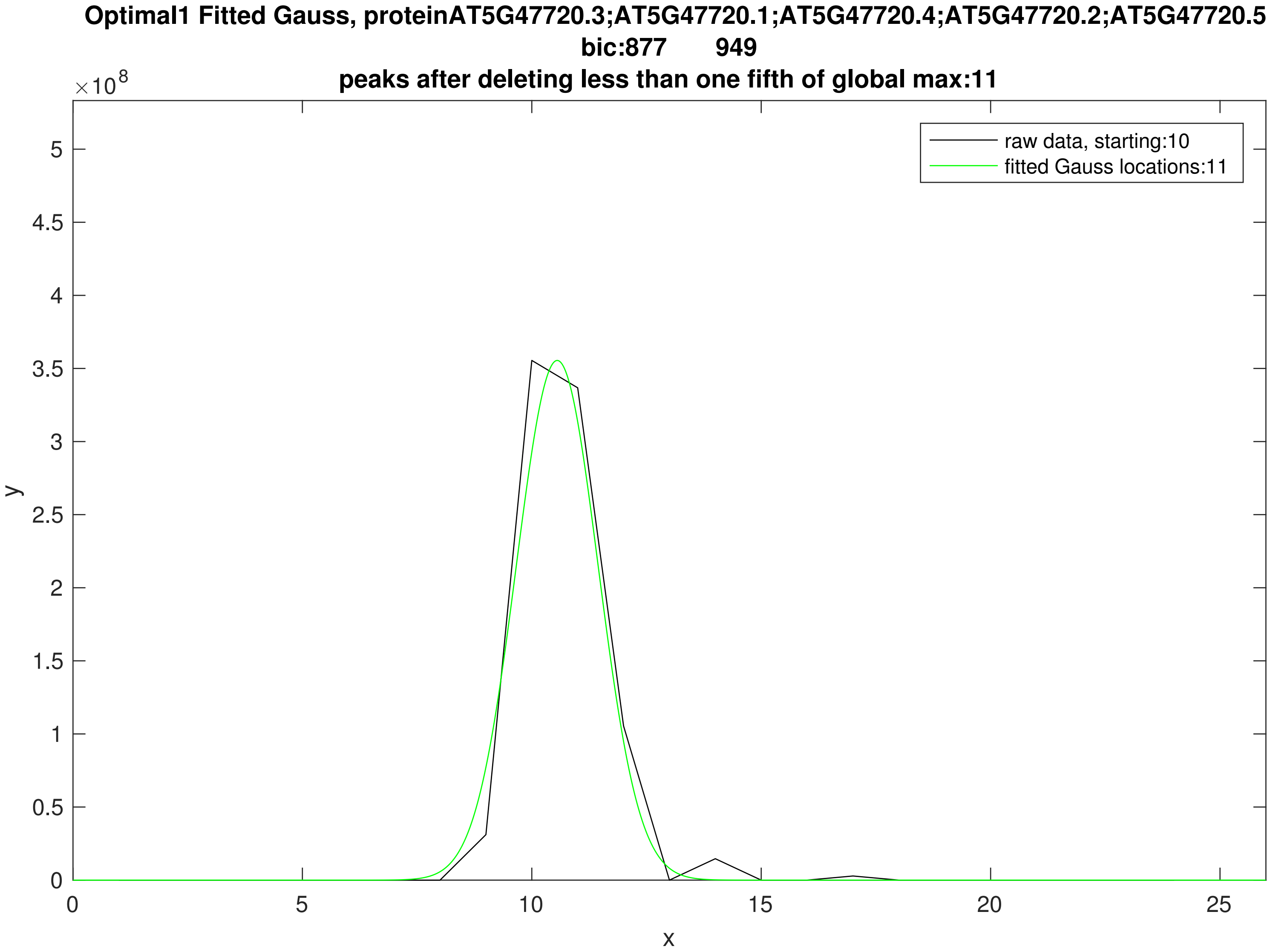

Optimal1 Fitted Gauss, proteinAT5G47720.3;AT5G47720.1;AT5G47720.4;AT5G47720.2;AT5G47720.5
bic:877 949
peaks after deleting less than one fifth of global max:11
×10^8
5
4.5
4
3.5
3
2.5
2
1.5
1
0.5
0
y
0
5
10
15
20
25
x
raw data, starting:10
fitted Gauss locations:11

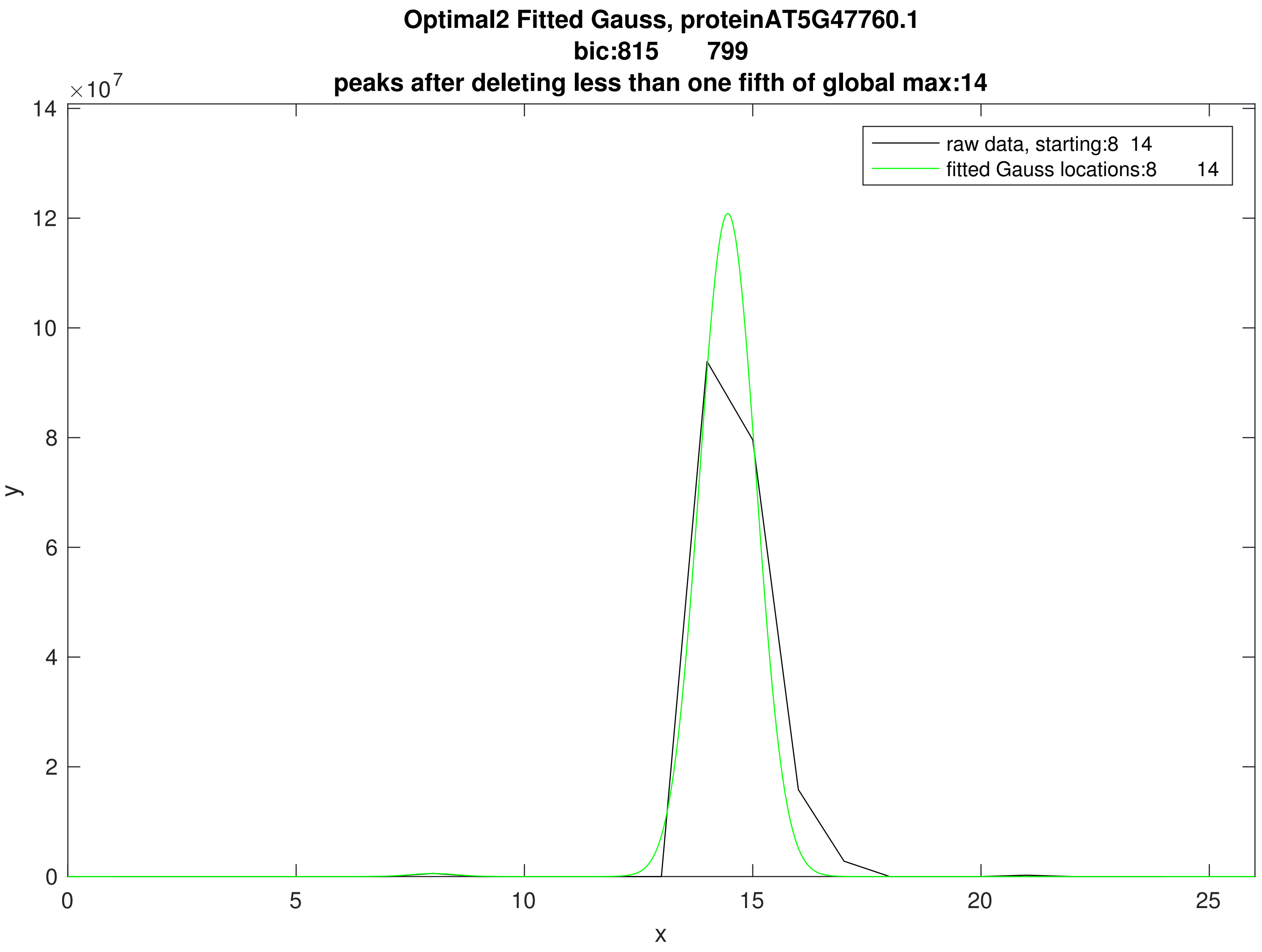
Optimal2 Fitted Gauss, proteinAT5G47760.1
bic:815 799
peaks after deleting less than one fifth of global max:14
×10^7
14
12
10
8
6
4
2
0
y
0
5
10
15
20
25
x
raw data, starting:8 14
fitted Gauss locations:8 14

**Optimal1 Fitted Gauss, proteinAT5G47770.1**
**bic:979 988**
**peaks after deleting less than one fifth of global max:14**

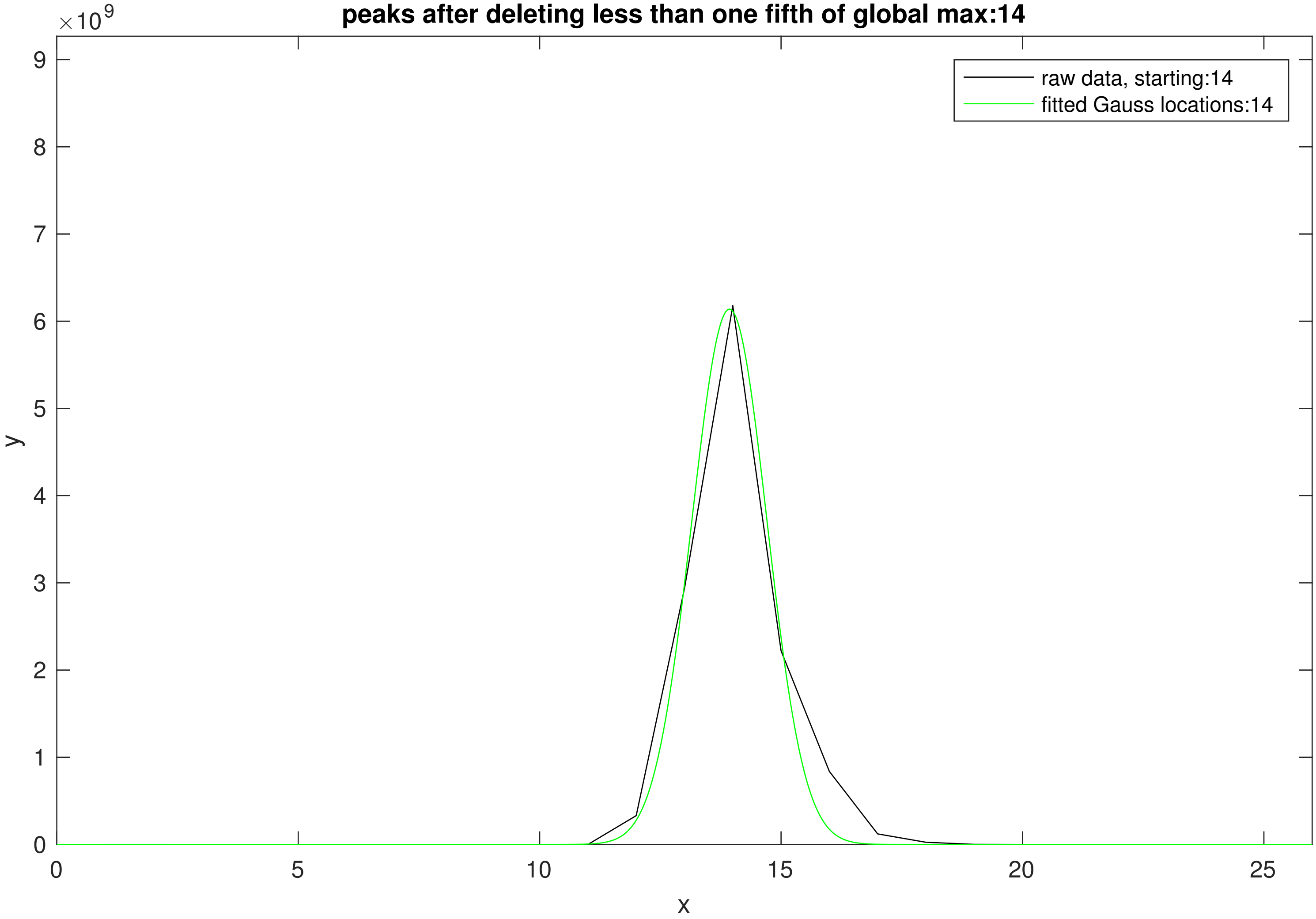

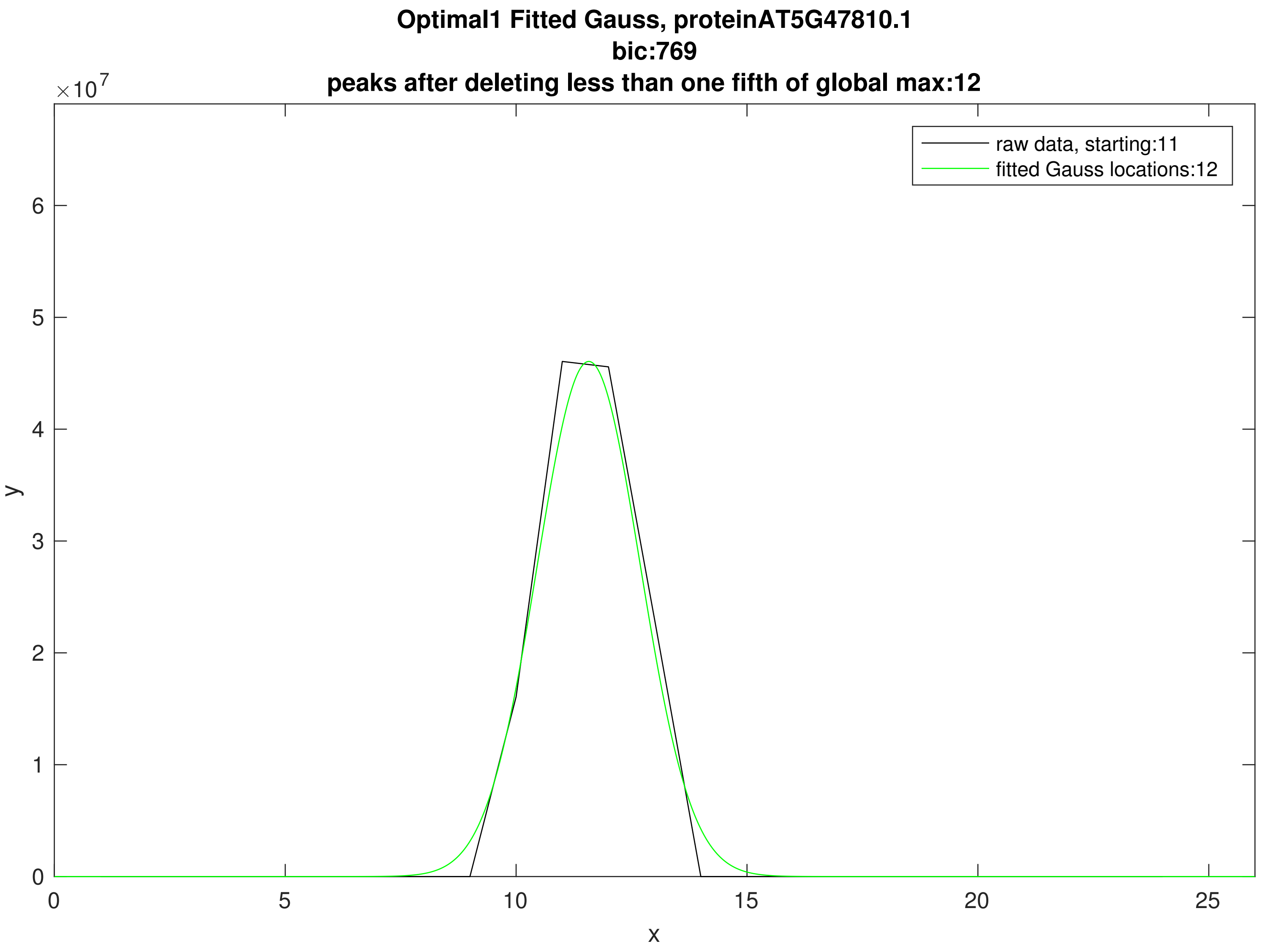
Optimal1 Fitted Gauss, proteinAT5G47810.1
bic:769
peaks after deleting less than one fifth of global max:12
×10^7
raw data, starting:11
fitted Gauss locations:12
y
x
0
1
2
3
4
5
6
0
5
10
15
20
25

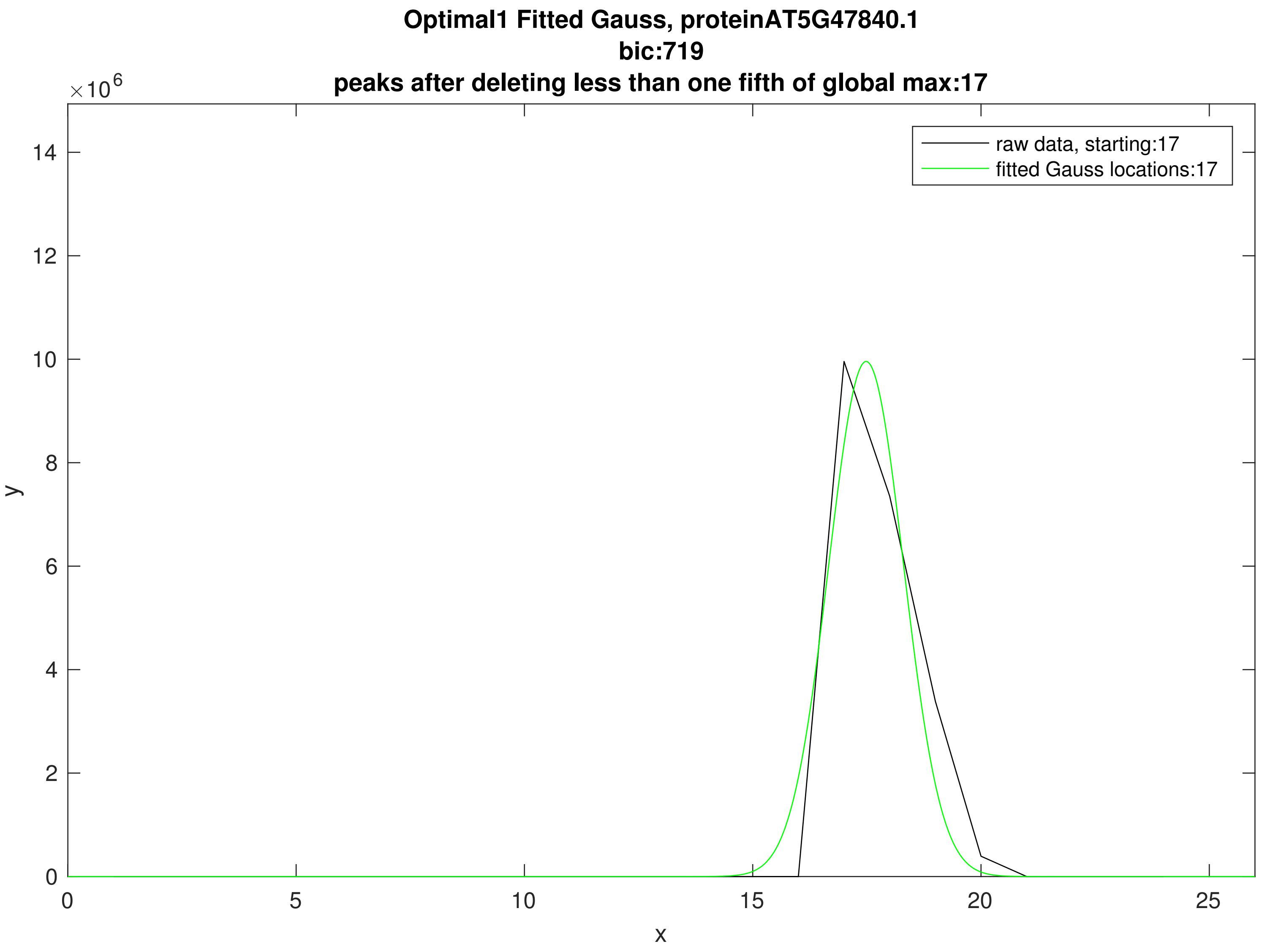

Optimal1 Fitted Gauss, proteinAT5G47840.1
bic:719
peaks after deleting less than one fifth of global max:17
×10^6
14
12
10
8
6
4
2
0
y
0
5
10
15
20
25
x
raw data, starting:17
fitted Gauss locations:17

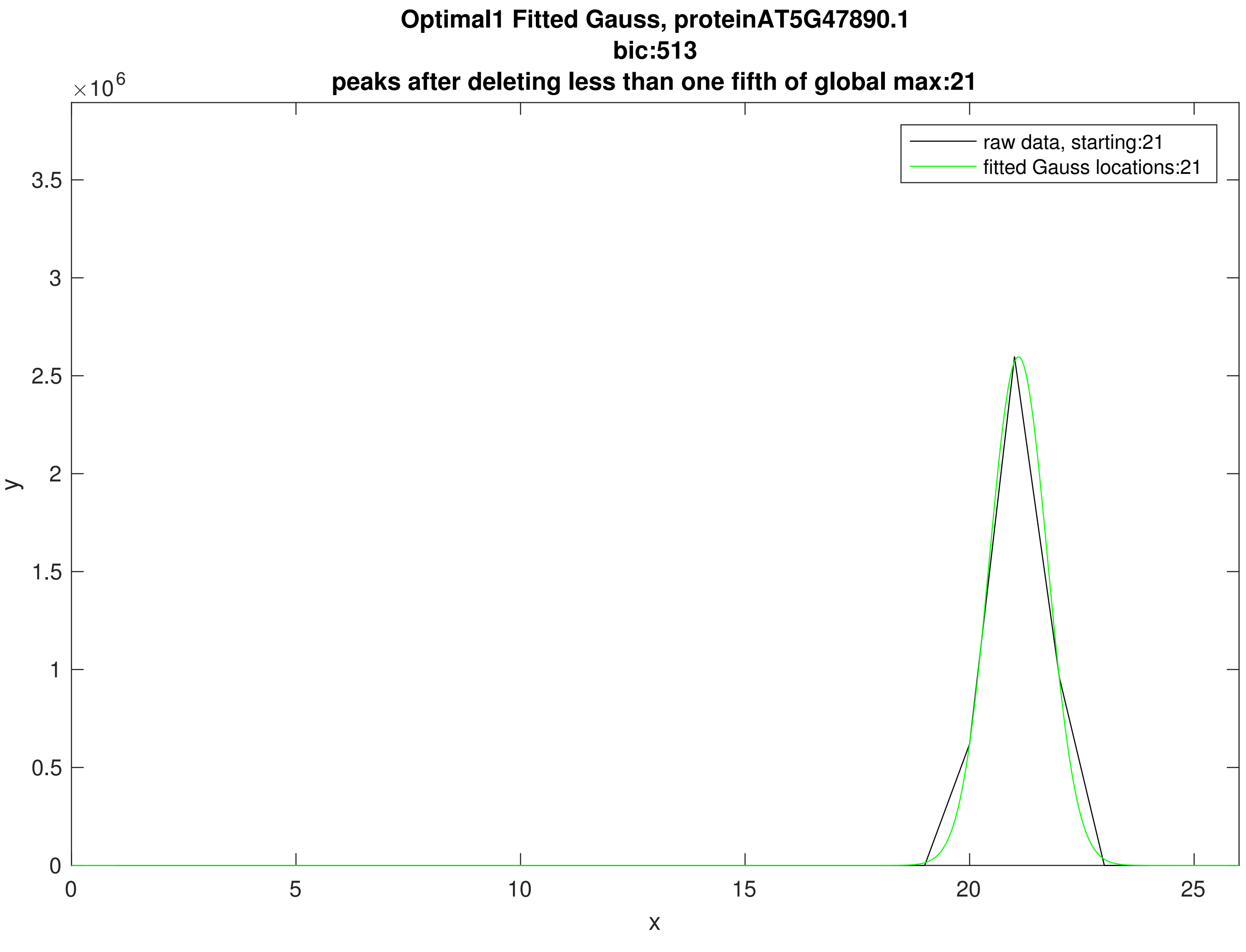
Optimal1 Fitted Gauss, proteinAT5G47890.1
bic:513
peaks after deleting less than one fifth of global max:21
×10^6
3.5
3
2.5
2
1.5
1
0.5
0
y
0
5
10
15
20
25
x
raw data, starting:21
fitted Gauss locations:21

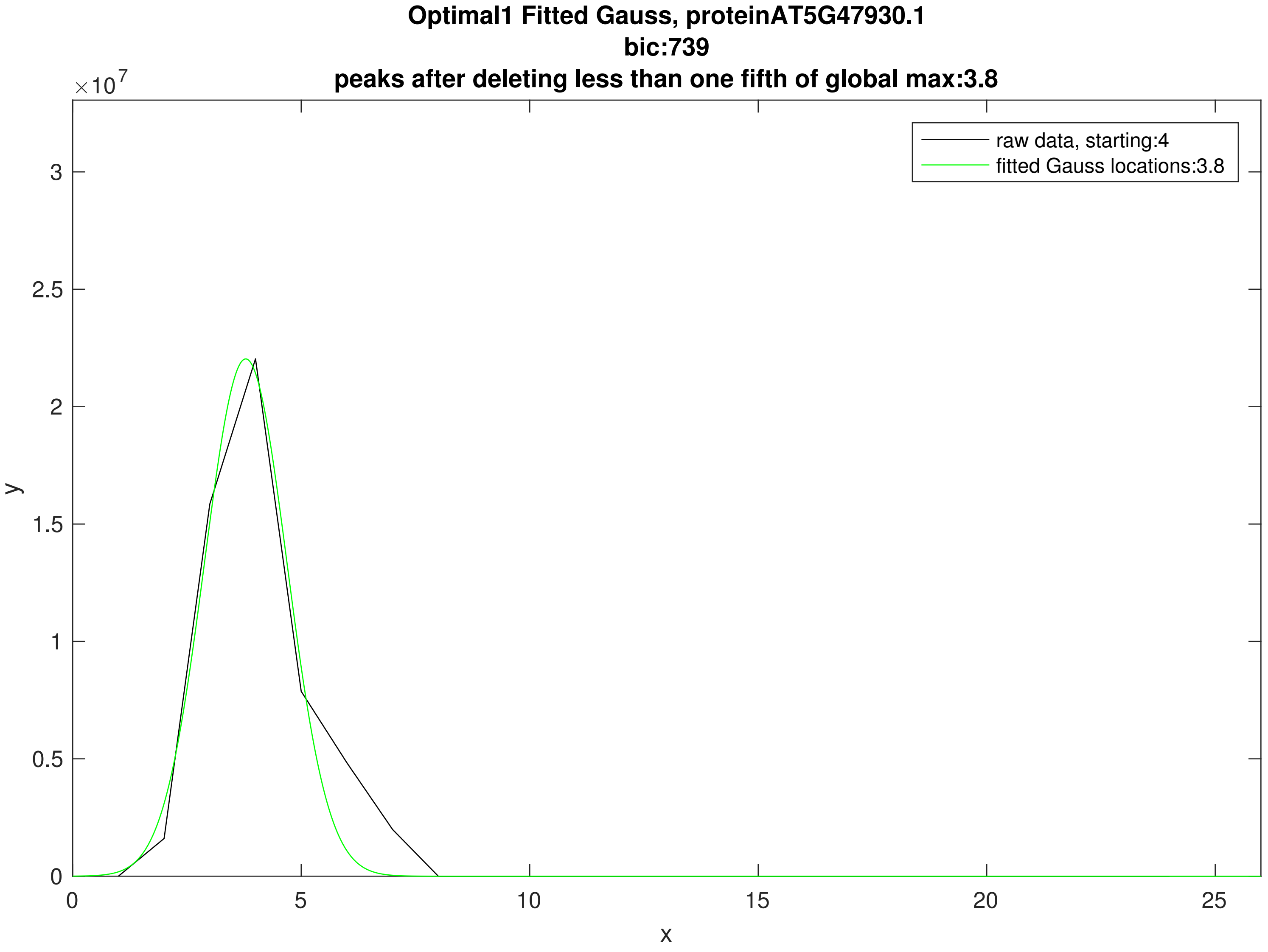
Optimal1 Fitted Gauss, proteinAT5G47930.1
bic:739
peaks after deleting less than one fifth of global max:3.8
×10^7
raw data, starting:4
fitted Gauss locations:3.8
y
x
0
0.5
1
1.5
2
2.5
3
0
5
10
15
20
25

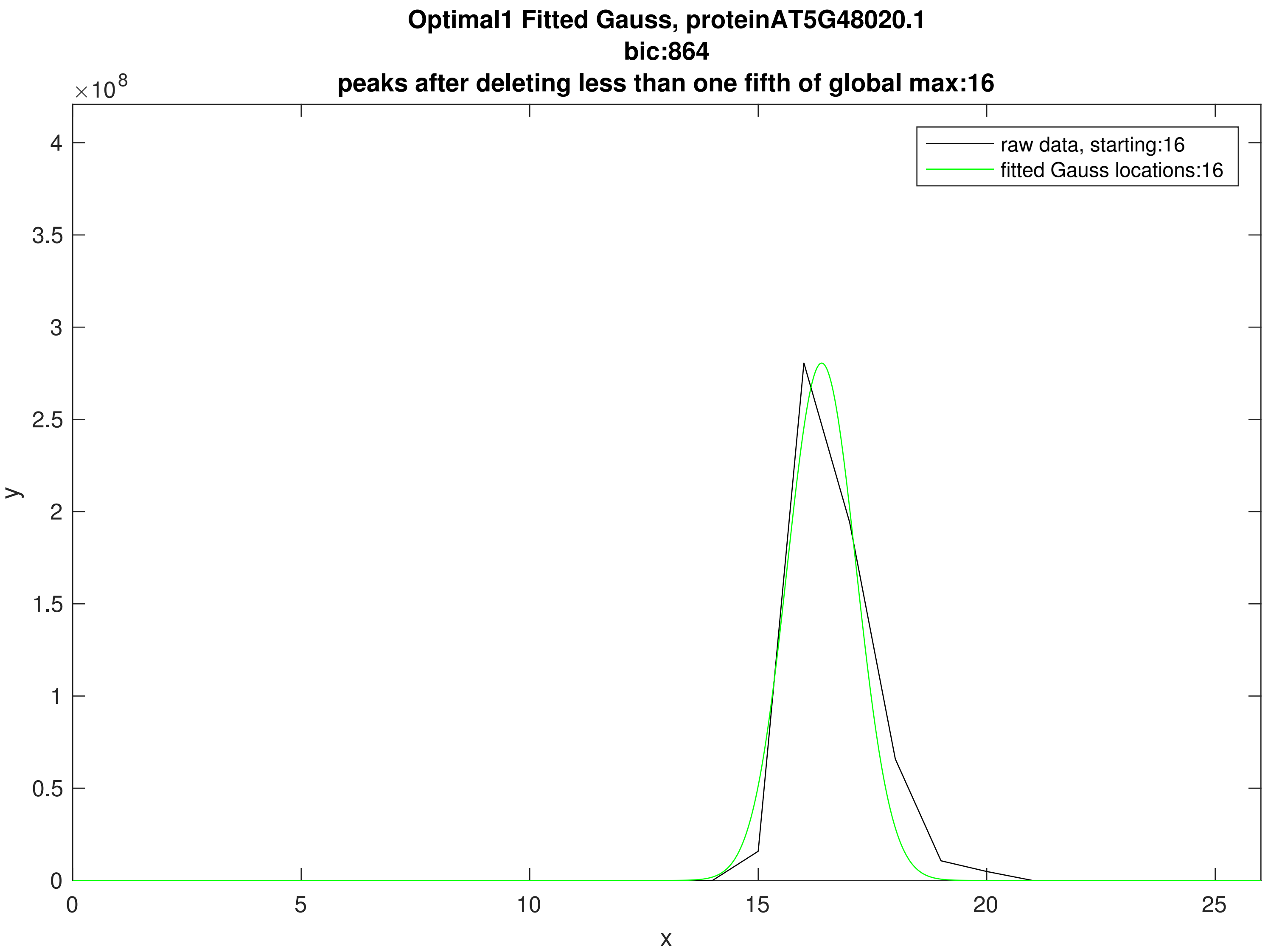
Optimal1 Fitted Gauss, proteinAT5G48020.1
bic:864
peaks after deleting less than one fifth of global max:16
×10^8
raw data, starting:16
fitted Gauss locations:16
y
x
0
0.5
1
1.5
2
2.5
3
3.5
4
0
5
10
15
20
25

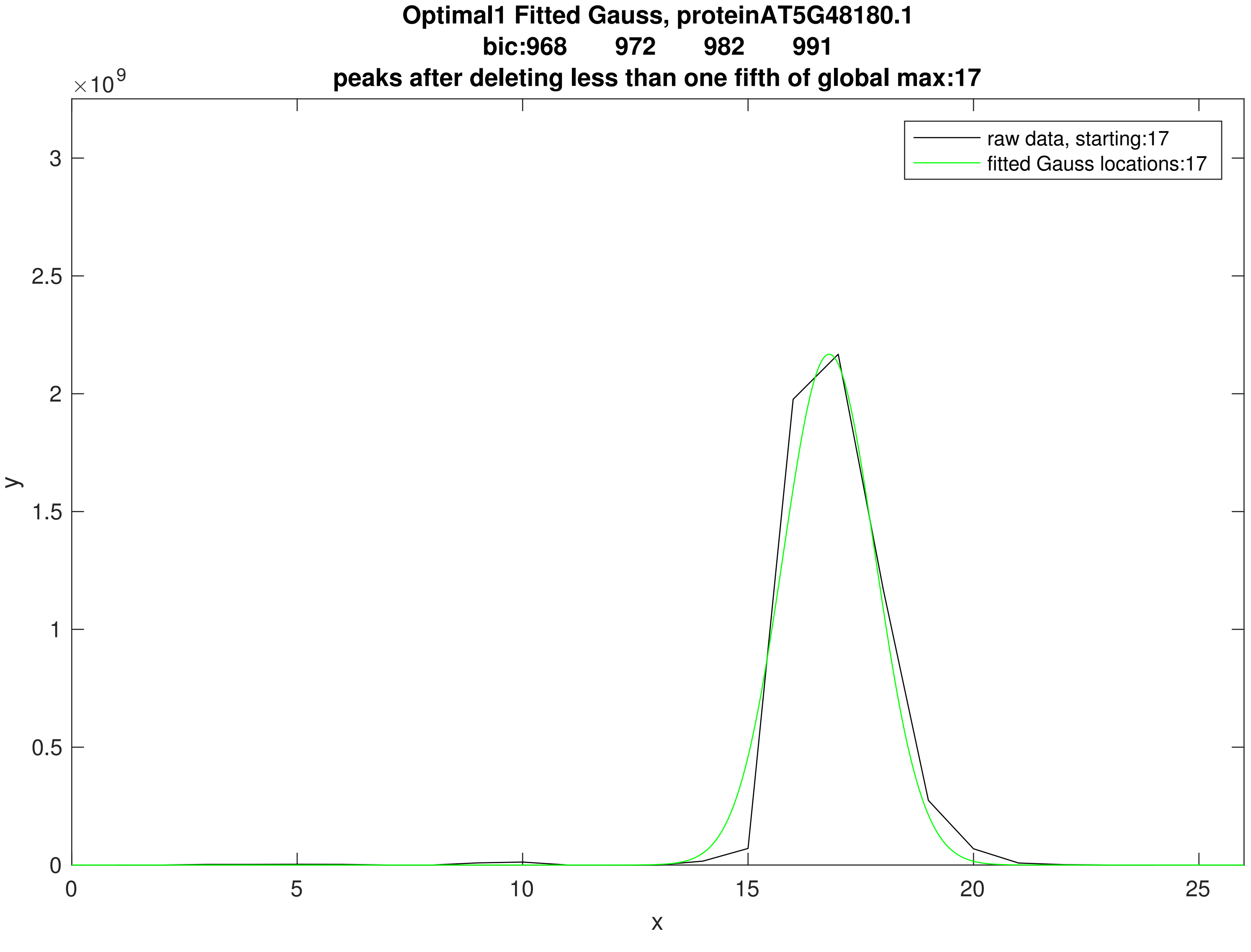
Optimal1 Fitted Gauss, proteinAT5G48180.1
bic:968 972 982 991
peaks after deleting less than one fifth of global max:17
×10^9
raw data, starting:17
fitted Gauss locations:17
y
x
0
0.5
1
1.5
2
2.5
3
5
10
15
20
25

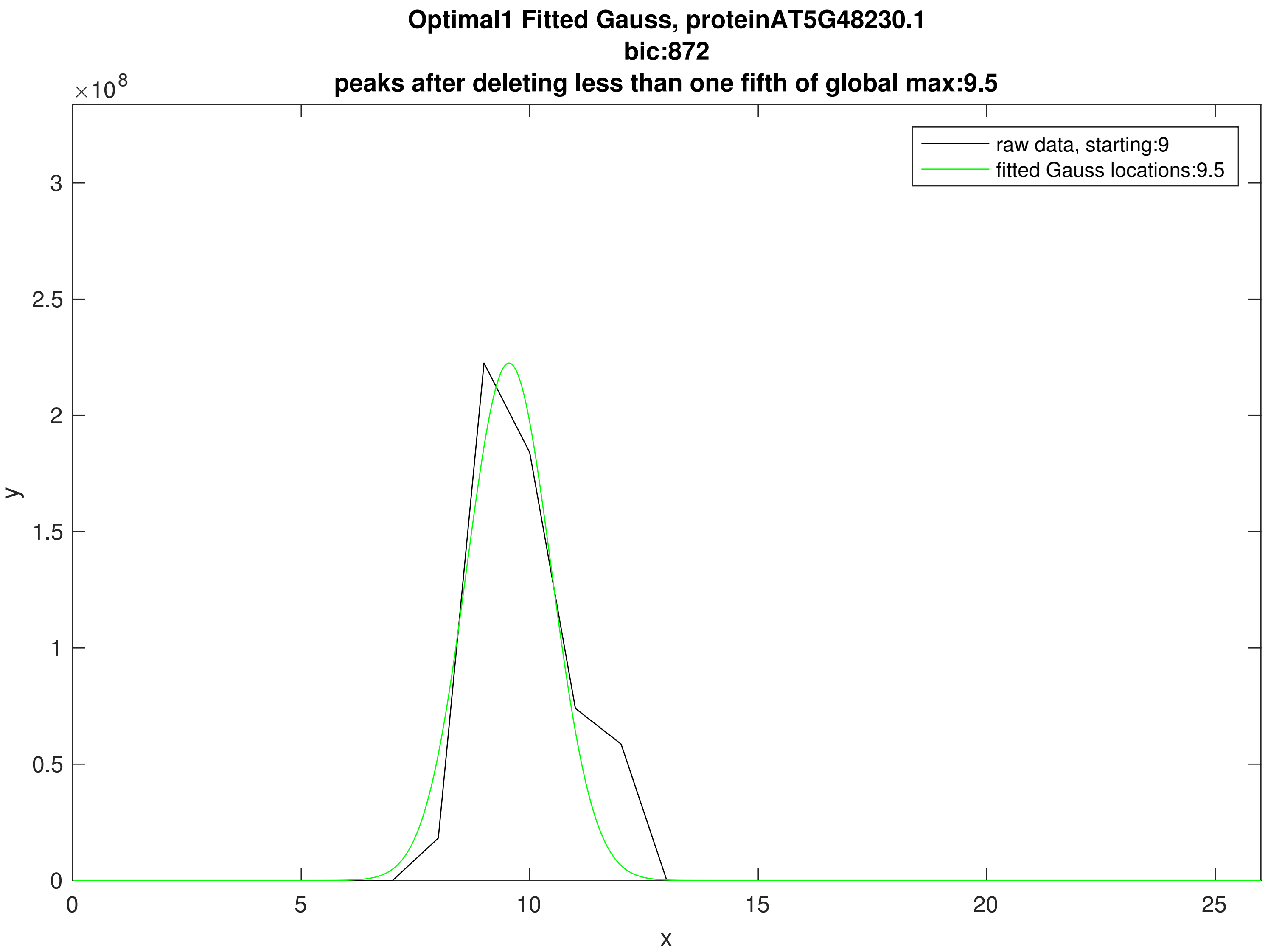
Optimal1 Fitted Gauss, proteinAT5G48230.1
bic:872
peaks after deleting less than one fifth of global max:9.5
×10^8
raw data, starting:9
fitted Gauss locations:9.5
0
0.5
1
1.5
2
2.5
3
y
0
5
10
15
20
25
x

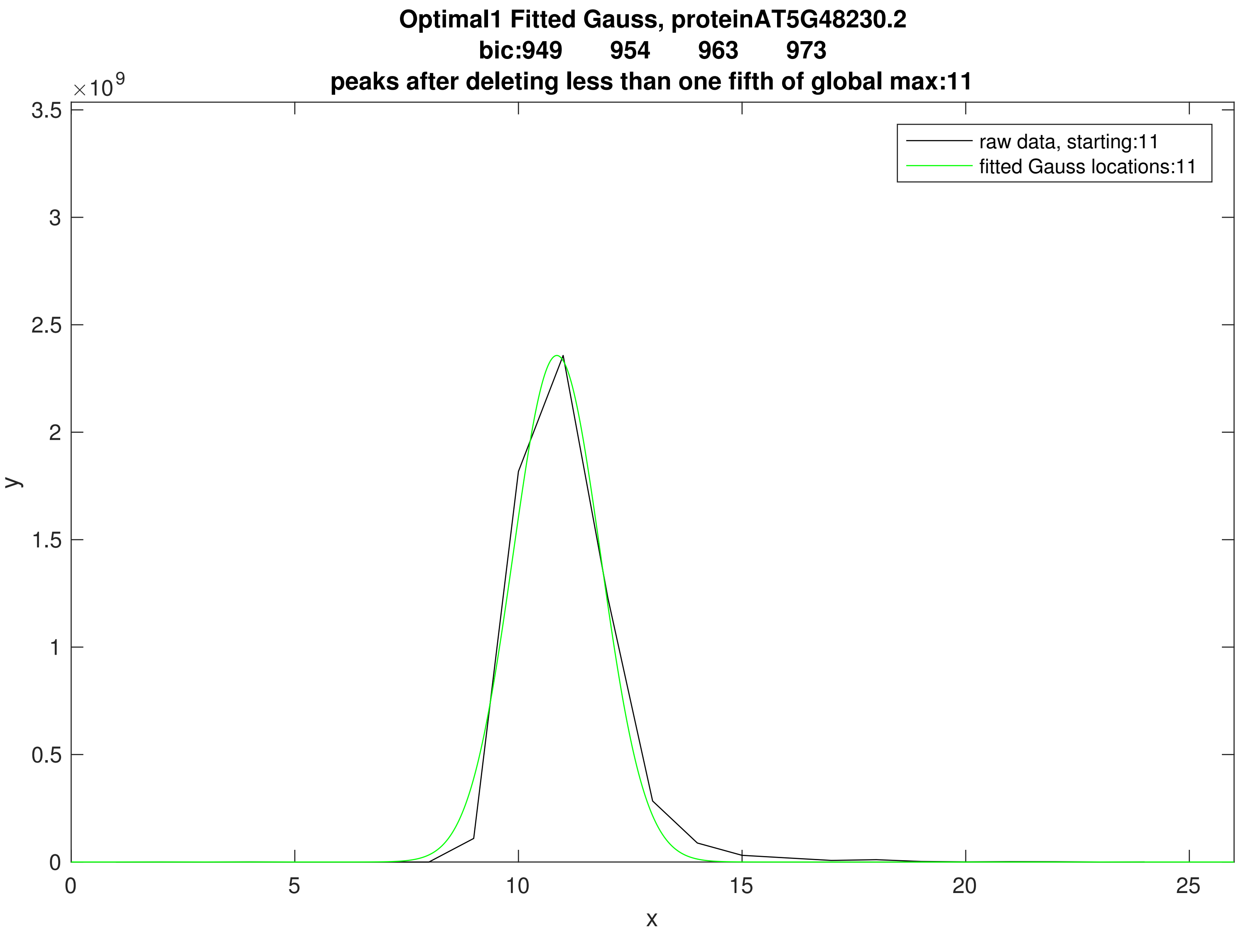

Optimal1 Fitted Gauss, proteinAT5G48230.2
bic:949 954 963 973
peaks after deleting less than one fifth of global max:11
×10^9
3.5
3
2.5
2
1.5
1
0.5
0
y
0
5
10
15
20
25
x
raw data, starting:11
fitted Gauss locations:11

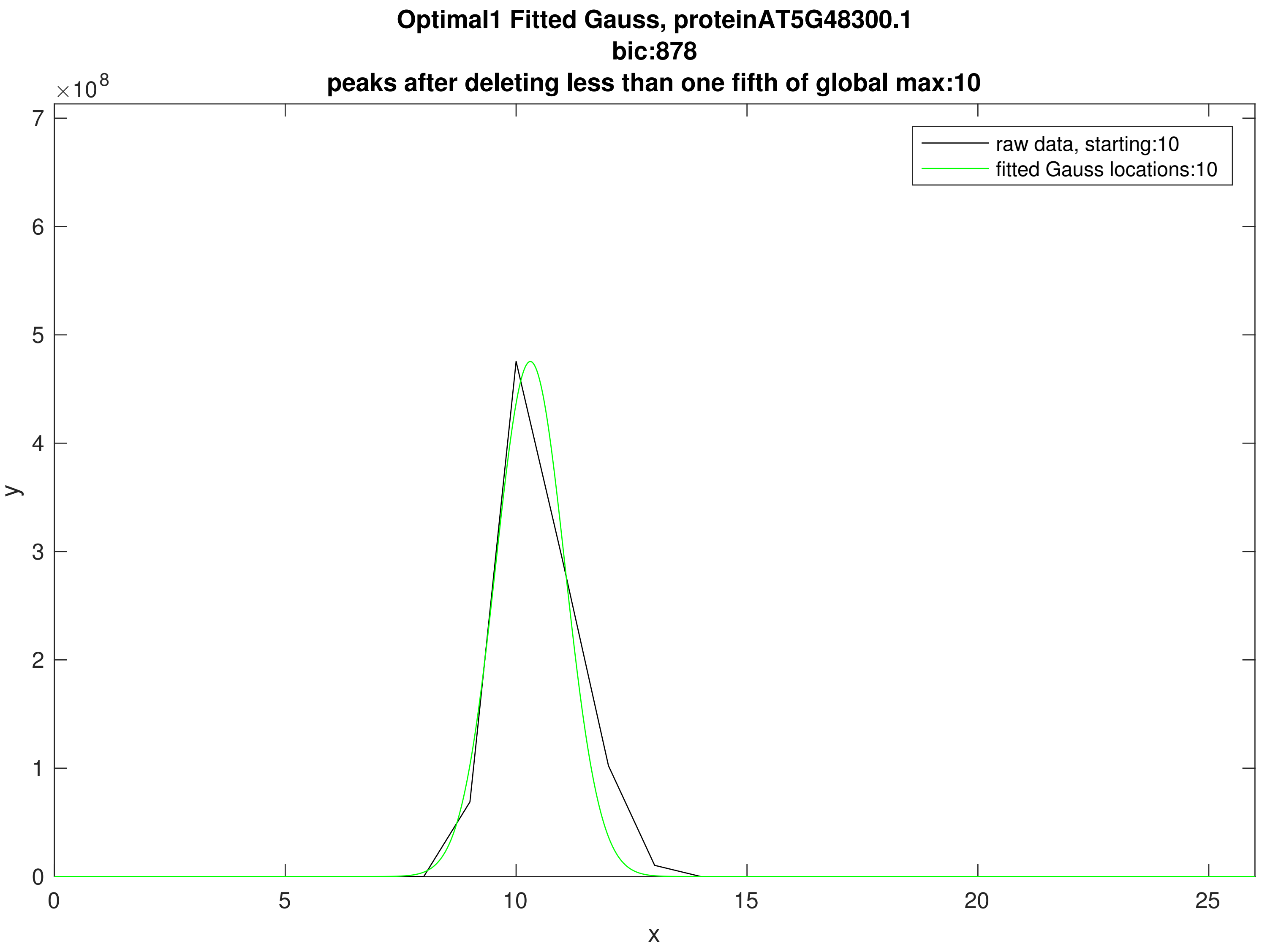
Optimal1 Fitted Gauss, proteinAT5G48300.1
bic:878
peaks after deleting less than one fifth of global max:10
×10^8
7
6
5
4
3
2
1
0
y
0
5
10
15
20
25
x
raw data, starting:10
fitted Gauss locations:10

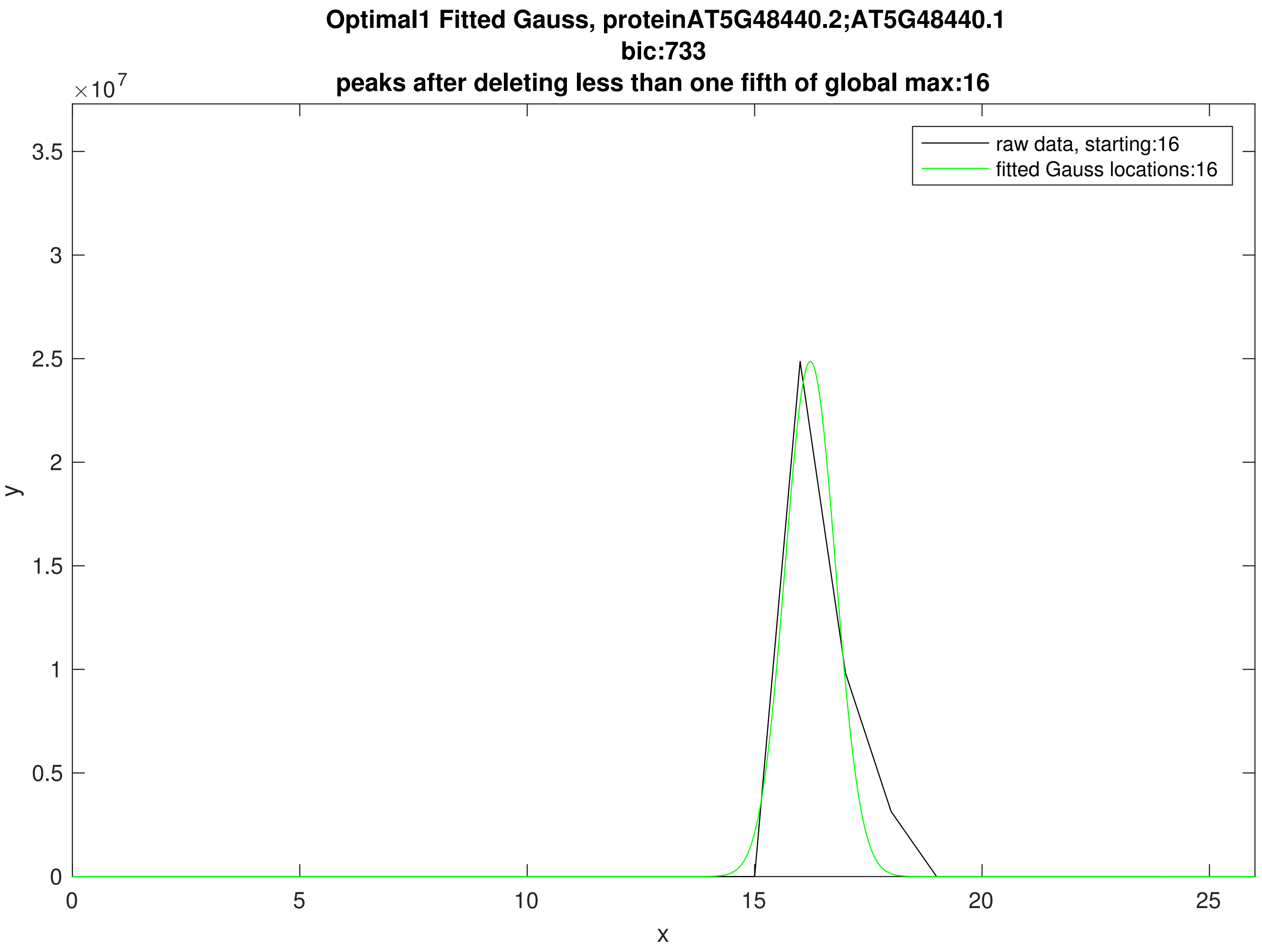

Optimal1 Fitted Gauss, proteinAT5G48440.2;AT5G48440.1
bic:733
peaks after deleting less than one fifth of global max:16
×10^7
3.5
3
2.5
2
1.5
1
0.5
0
y
0
5
10
15
20
25
x
raw data, starting:16
fitted Gauss locations:16

**Optimal2 Fitted Gauss, proteinAT5G48480.1**
**bic:816 813**
**peaks after deleting less than one fifth of global max:14 18**

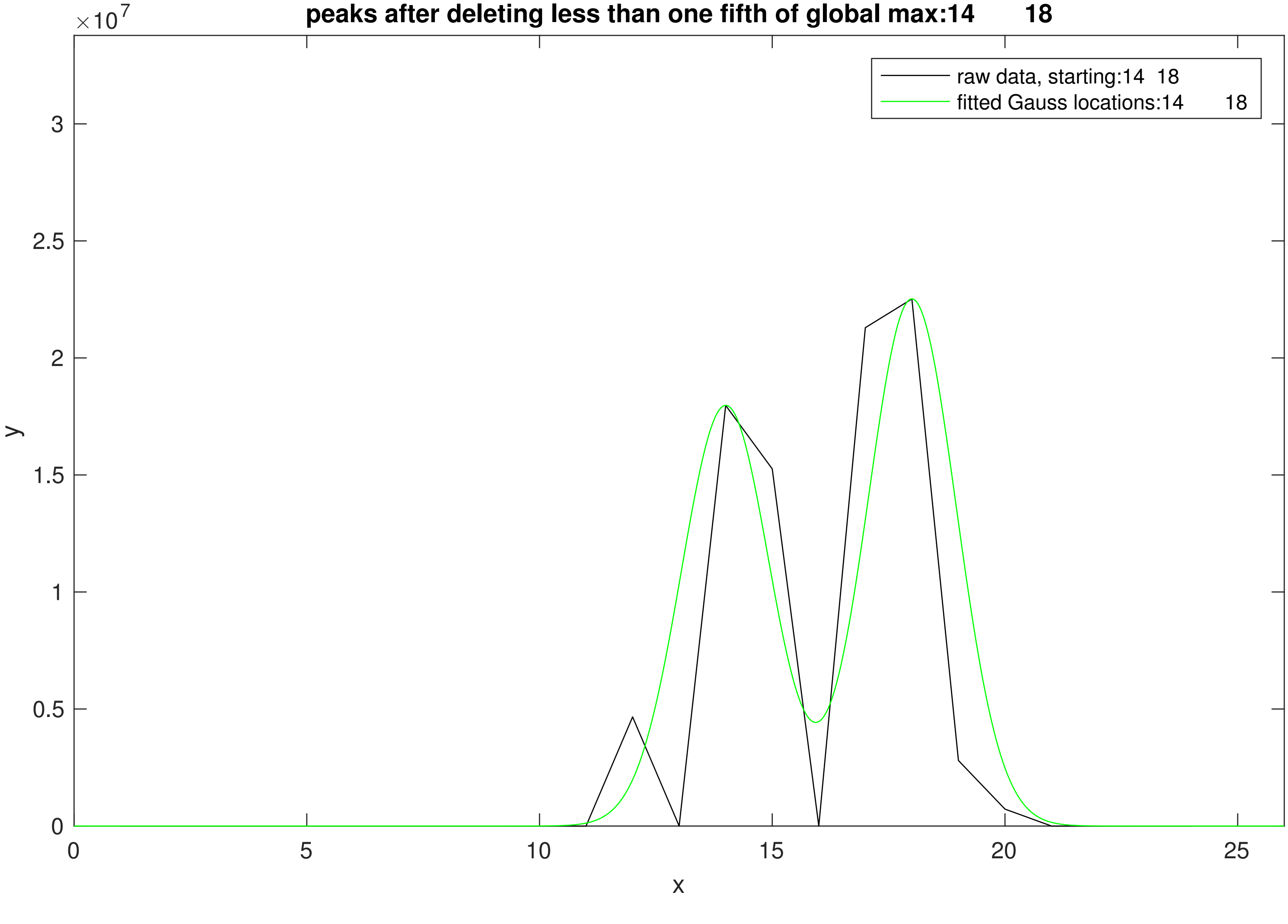

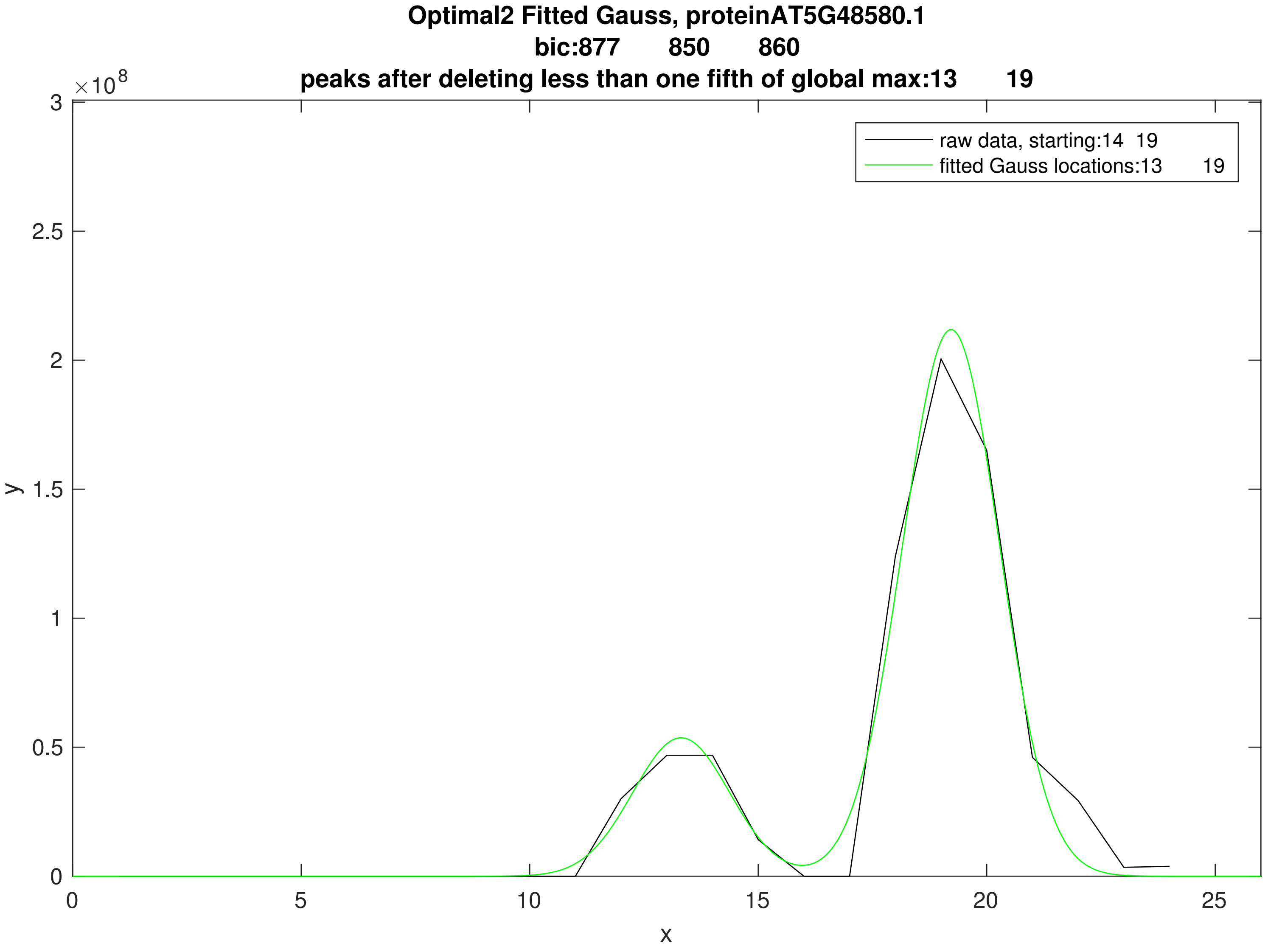
Optimal2 Fitted Gauss, proteinAT5G48580.1
bic:877 850 860
peaks after deleting less than one fifth of global max:13 19
raw data, starting:14 19
fitted Gauss locations:13 19
×10^8
3
2.5
2
1.5
1
0.5
0
y
0
5
10
15
20
25
x

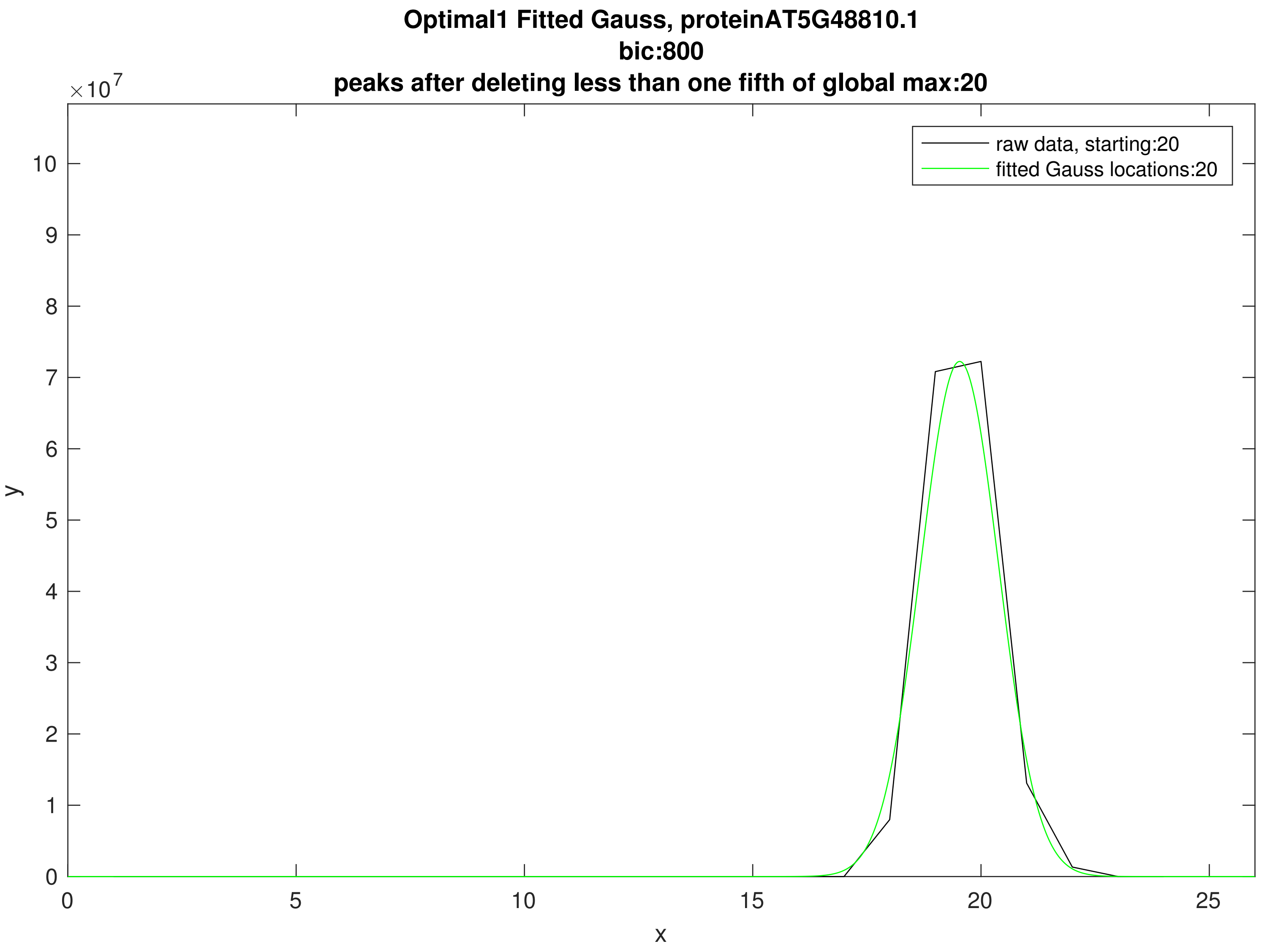

Optimal1 Fitted Gauss, proteinAT5G48810.1
bic:800
peaks after deleting less than one fifth of global max:20
×10^7
raw data, starting:20
fitted Gauss locations:20
y
x
0
1
2
3
4
5
6
7
8
9
10
15
20
25

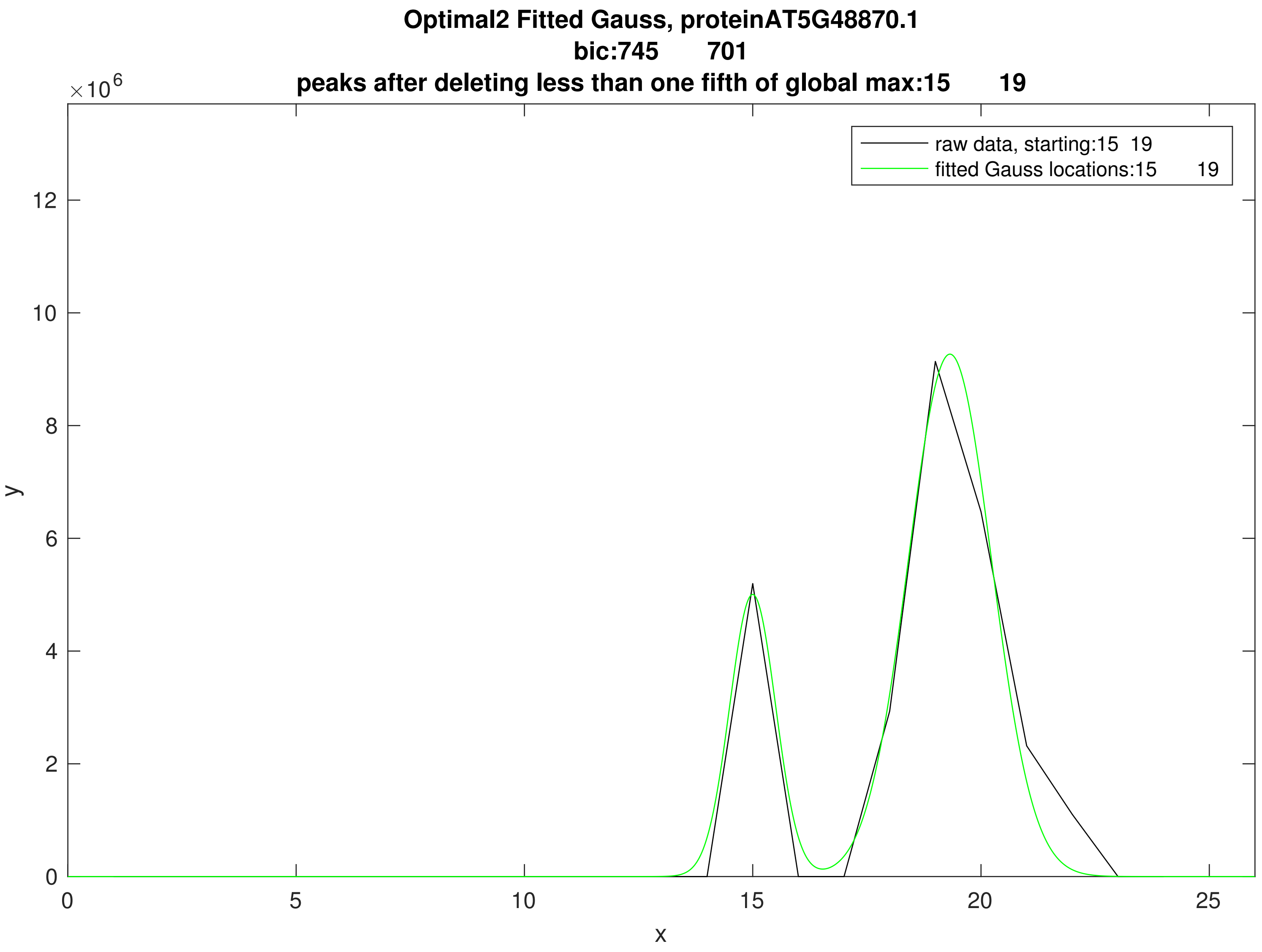

Optimal2 Fitted Gauss, proteinAT5G48870.1
bic:745 701
peaks after deleting less than one fifth of global max:15 19
×10^6
raw data, starting:15 19
fitted Gauss locations:15 19
12
10
8
6
4
2
0
y
0
5
10
15
20
25
x

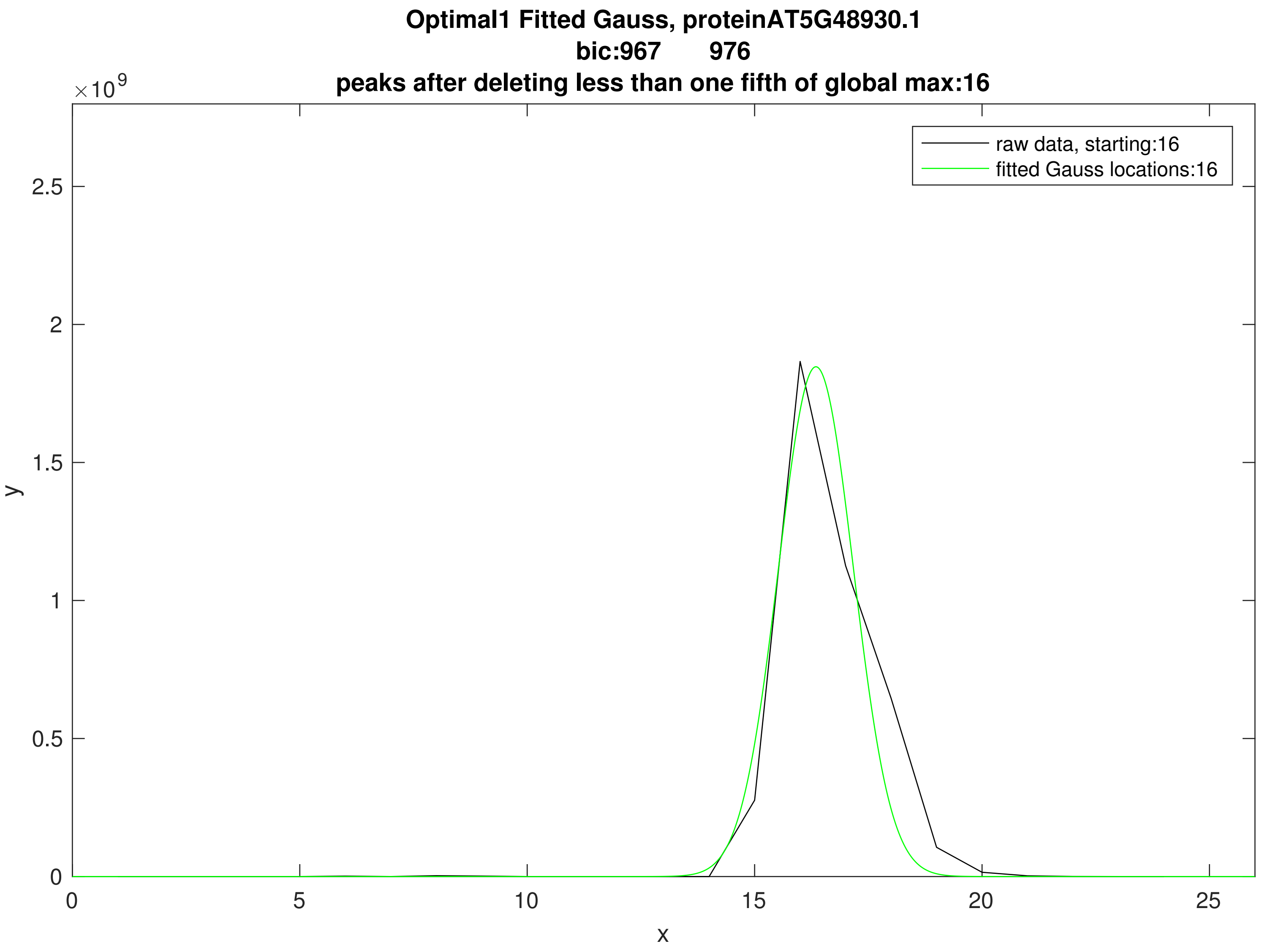
Optimal1 Fitted Gauss, proteinAT5G48930.1
bic:967 976
peaks after deleting less than one fifth of global max:16
×10^9
2.5
2
1.5
1
0.5
0
y
0
5
10
15
20
25
x
raw data, starting:16
fitted Gauss locations:16

**Optimal1 Fitted Gauss, proteinAT5G49020.1;AT5G49020.2**
**bic:758**
**peaks after deleting less than one fifth of global max:6.2**

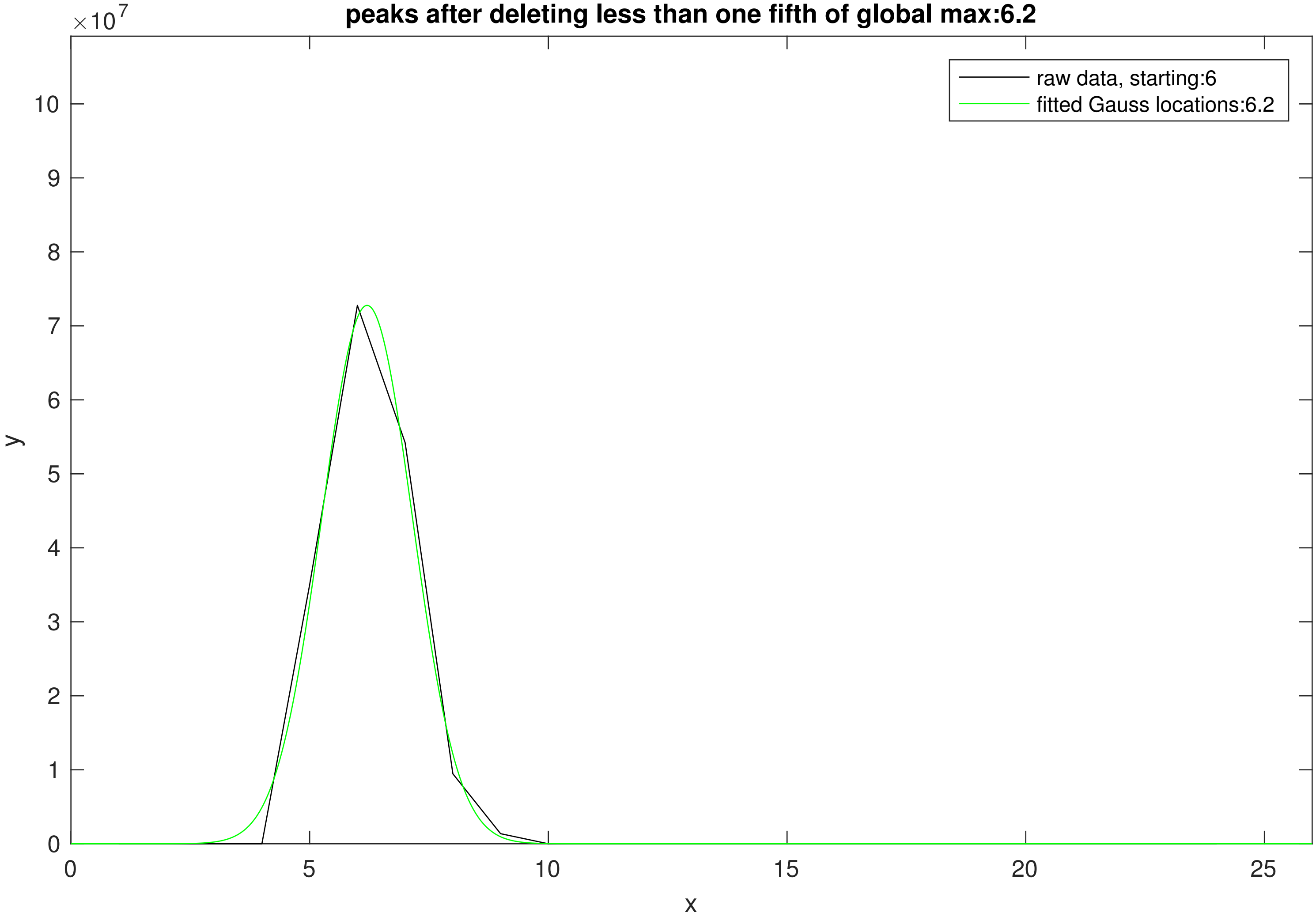

#### Optimal1 Fitted Gauss, proteinAT5G49030.3;AT5G49030.1;AT5G49030.2
#### bic:781
#### peaks after deleting less than one fifth of global max:12

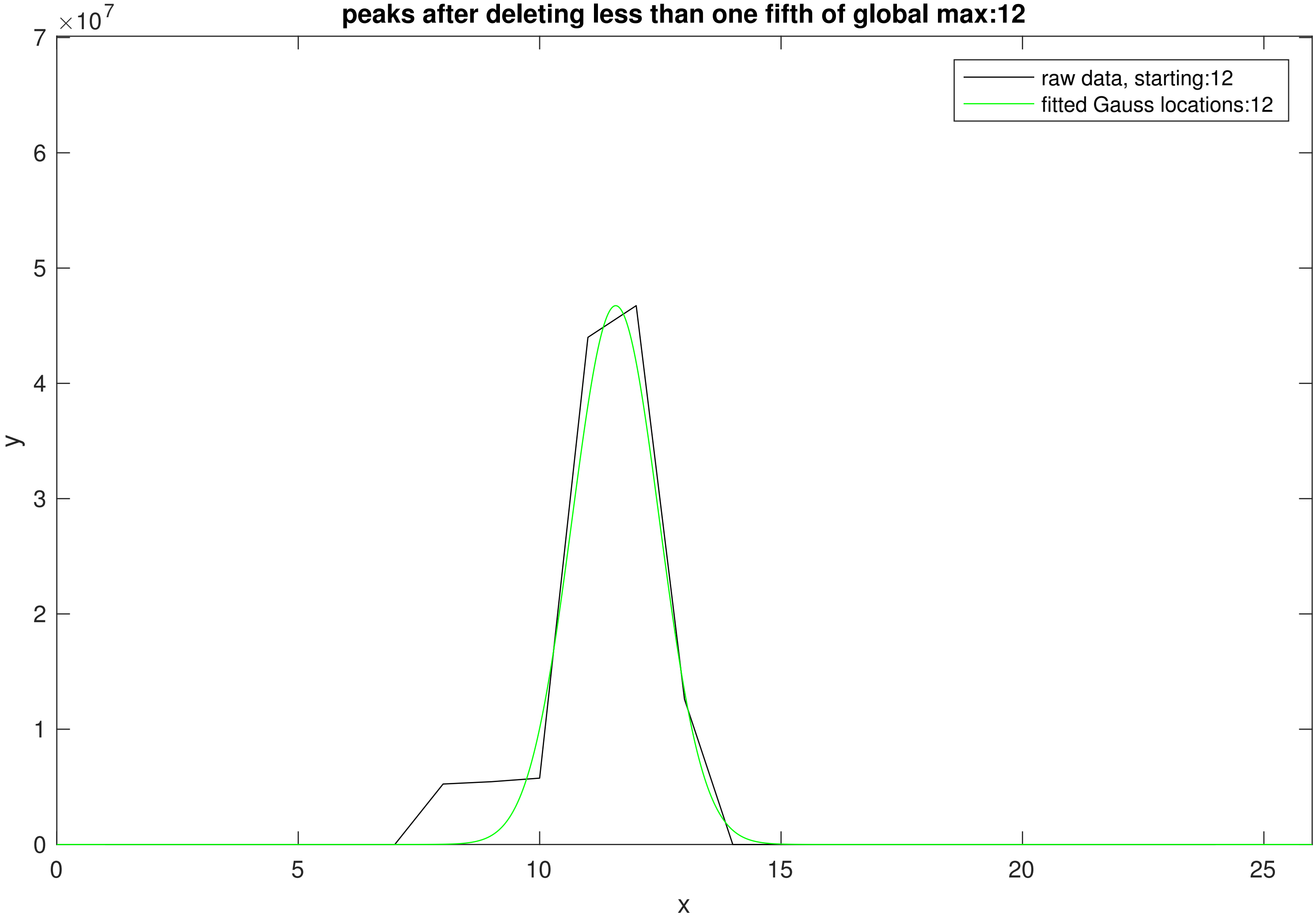

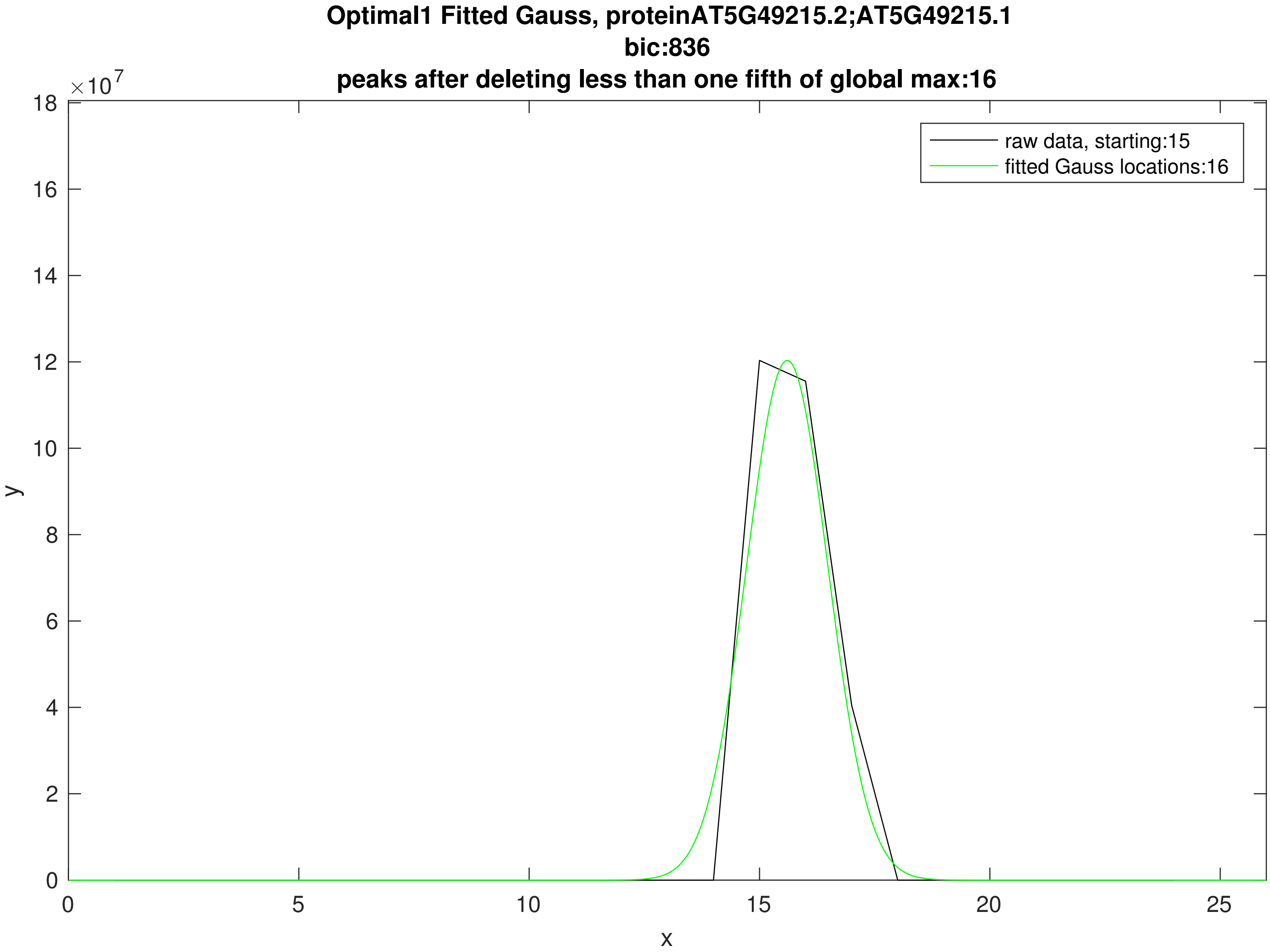
Optimal1 Fitted Gauss, proteinAT5G49215.2;AT5G49215.1
bic:836
peaks after deleting less than one fifth of global max:16
×10^7
18
16
14
12
10
8
6
4
2
0
y
0
5
10
15
20
25
x
raw data, starting:15
fitted Gauss locations:16

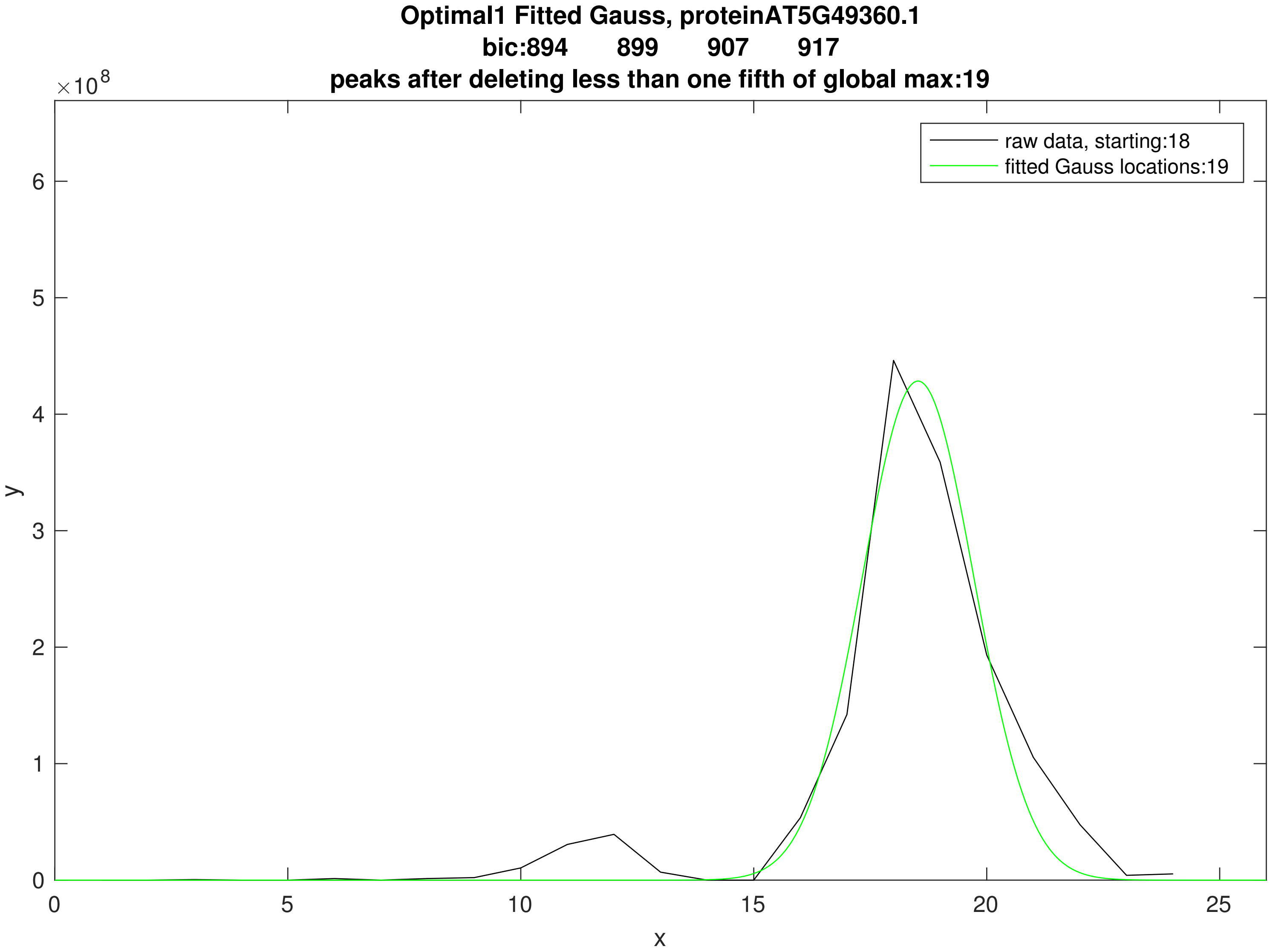

Optimal1 Fitted Gauss, proteinAT5G49360.1
bic:894 899 907 917
peaks after deleting less than one fifth of global max:19
×10^8
raw data, starting:18
fitted Gauss locations:19
y
x
0
1
2
3
4
5
6
0
5
10
15
20
25

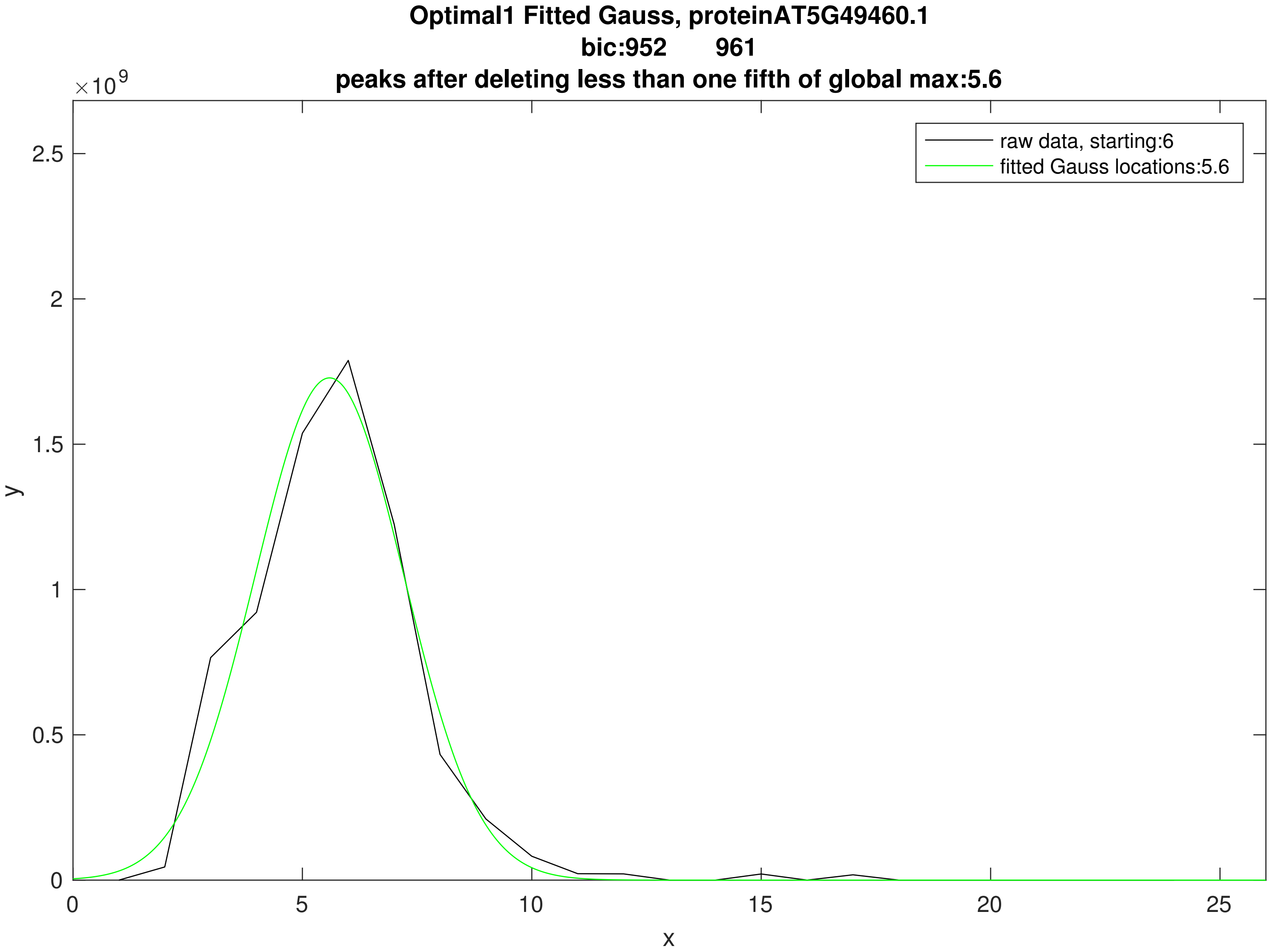

Optimal1 Fitted Gauss, proteinAT5G49460.1
bic:952 961
peaks after deleting less than one fifth of global max:5.6
×10^9
2.5
2
1.5
1
0.5
0
y
0
5
10
15
20
25
x
raw data, starting:6
fitted Gauss locations:5.6

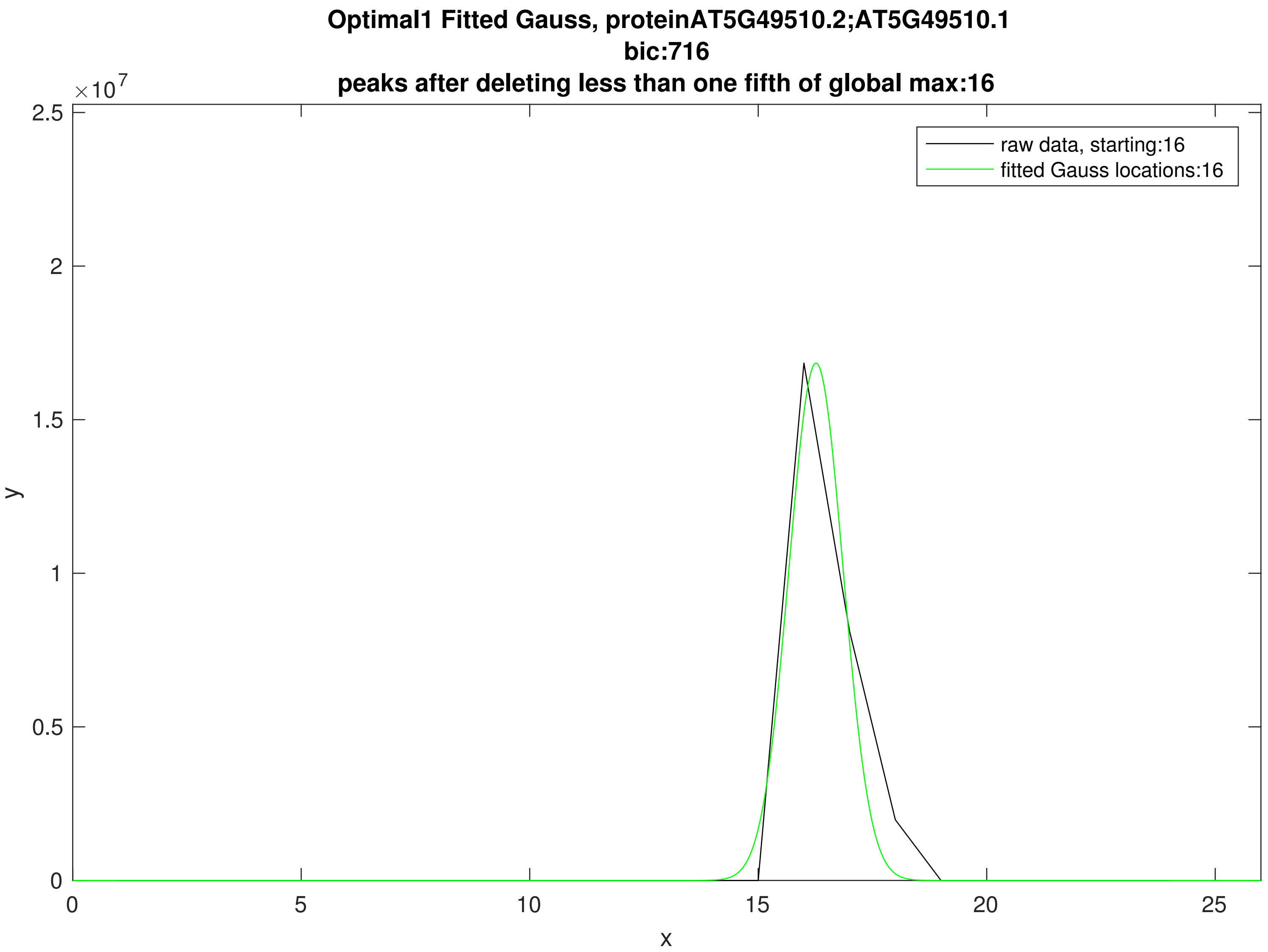
Optimal1 Fitted Gauss, proteinAT5G49510.2;AT5G49510.1
bic:716
peaks after deleting less than one fifth of global max:16
×10^7
2.5
2
1.5
1
0.5
0
y
0
5
10
15
20
25
x
raw data, starting:16
fitted Gauss locations:16

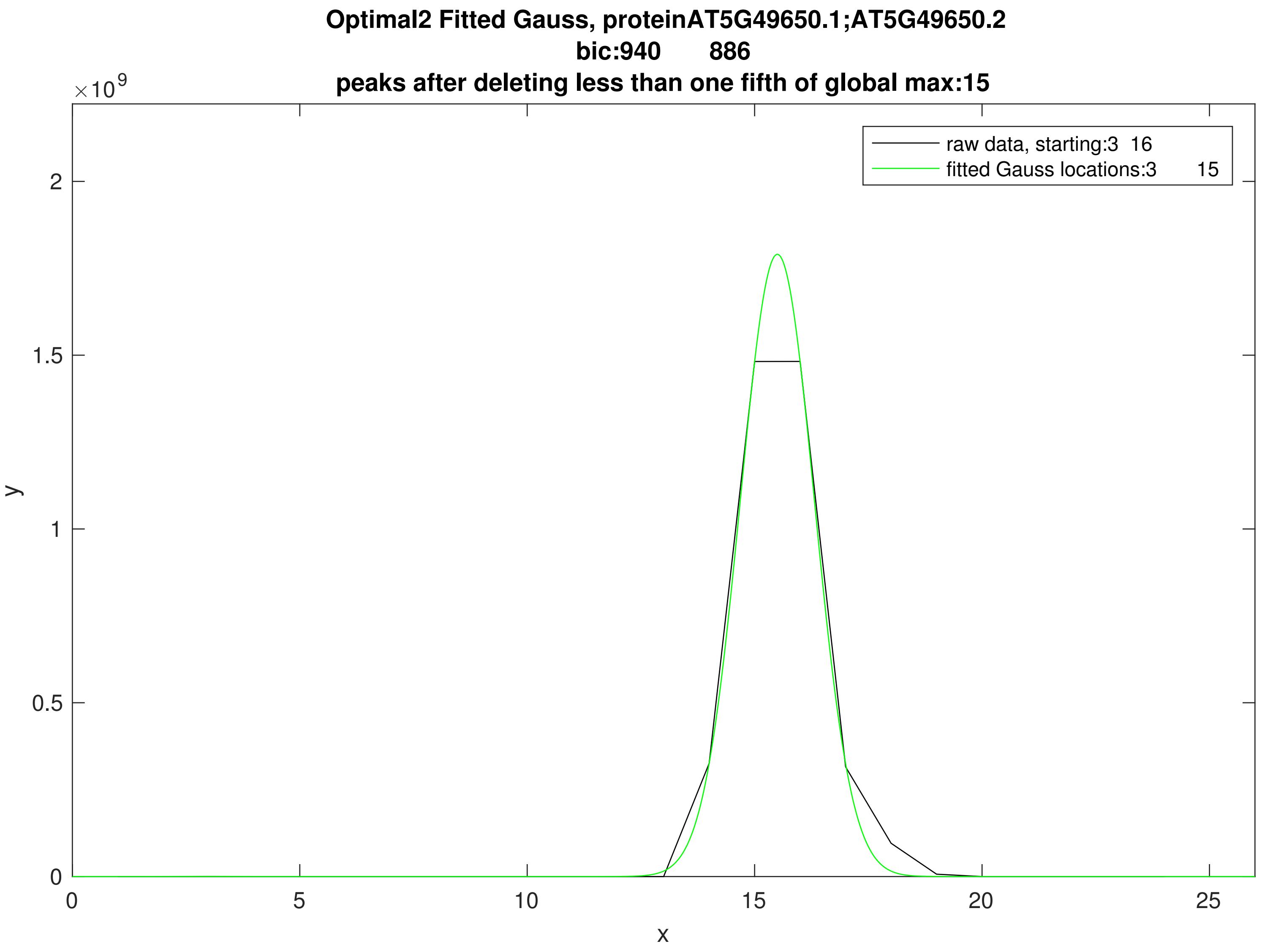

Optimal2 Fitted Gauss, proteinAT5G49650.1;AT5G49650.2
bic:940 886
peaks after deleting less than one fifth of global max:15
×10^9
raw data, starting:3 16
fitted Gauss locations:3 15
2
1.5
1
0.5
0
y
0
5
10
15
20
25
x

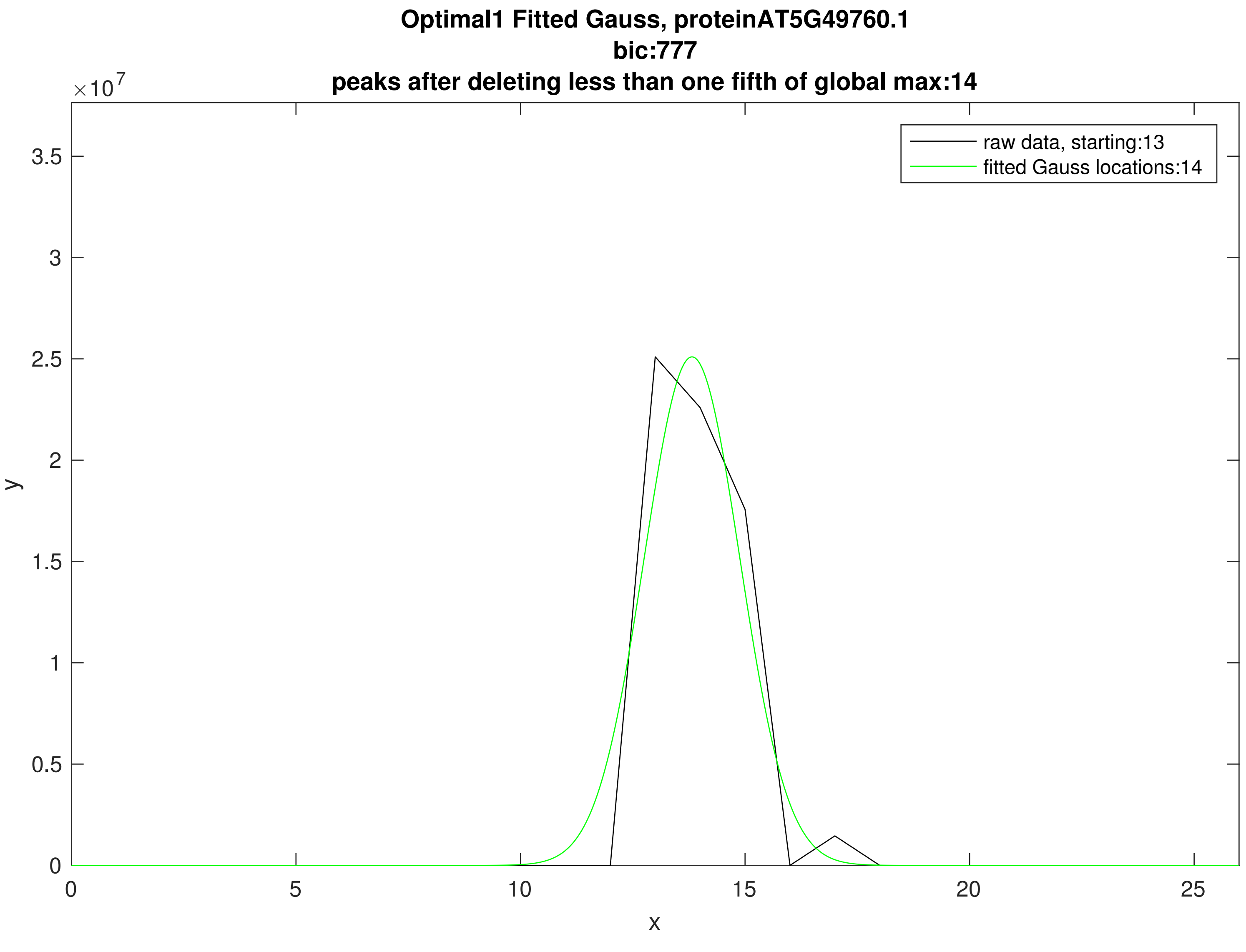

Optimal1 Fitted Gauss, proteinAT5G49760.1
bic:777
peaks after deleting less than one fifth of global max:14
×10^7
3.5
3
2.5
2
1.5
1
0.5
0
y
0
5
10
15
20
25
x
raw data, starting:13
fitted Gauss locations:14

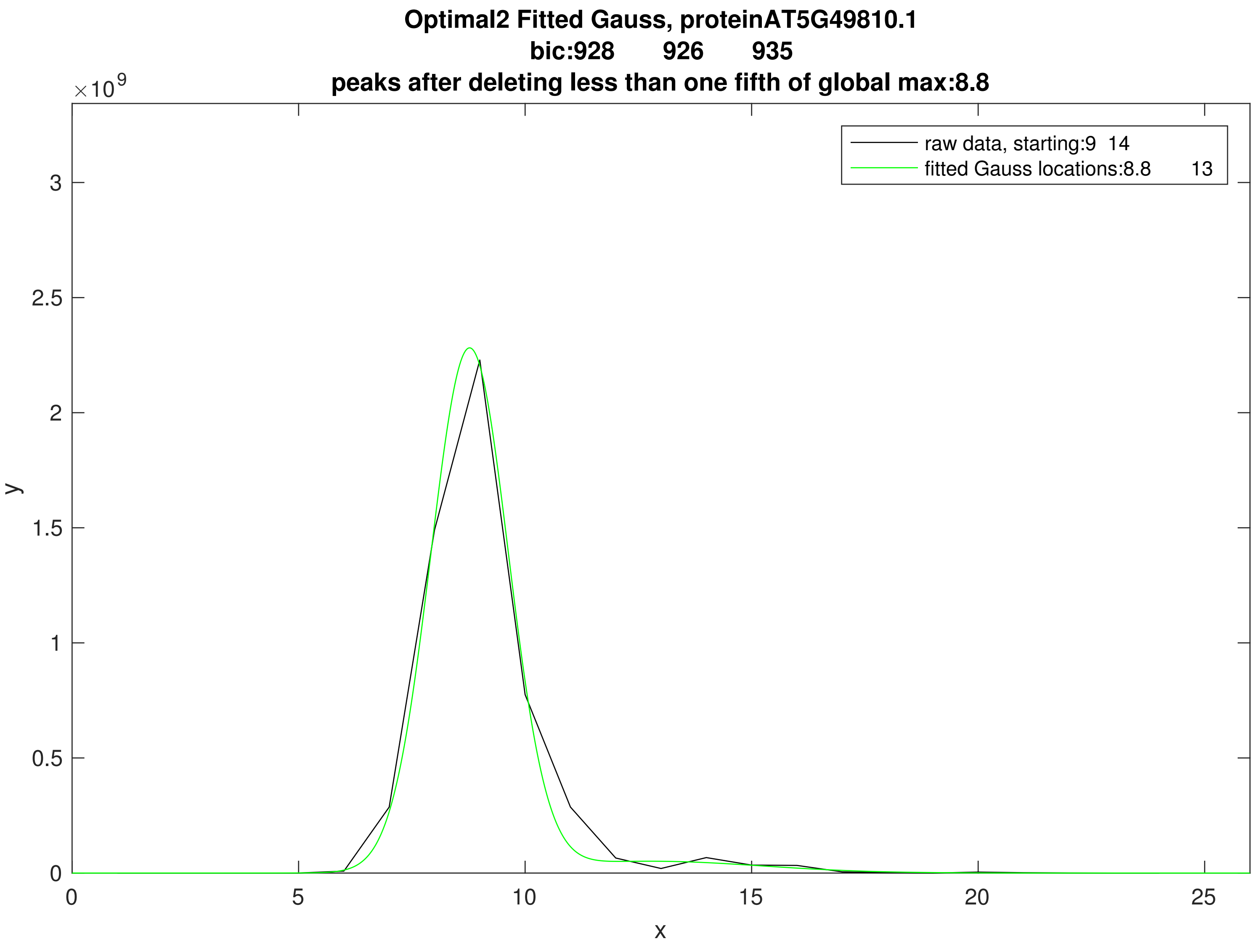
Optimal2 Fitted Gauss, proteinAT5G49810.1
bic:928 926 935
peaks after deleting less than one fifth of global max:8.8
raw data, starting:9 14
fitted Gauss locations:8.8 13
y
x
×10^9
0
0.5
1
1.5
2
2.5
3
0
5
10
15
20
25

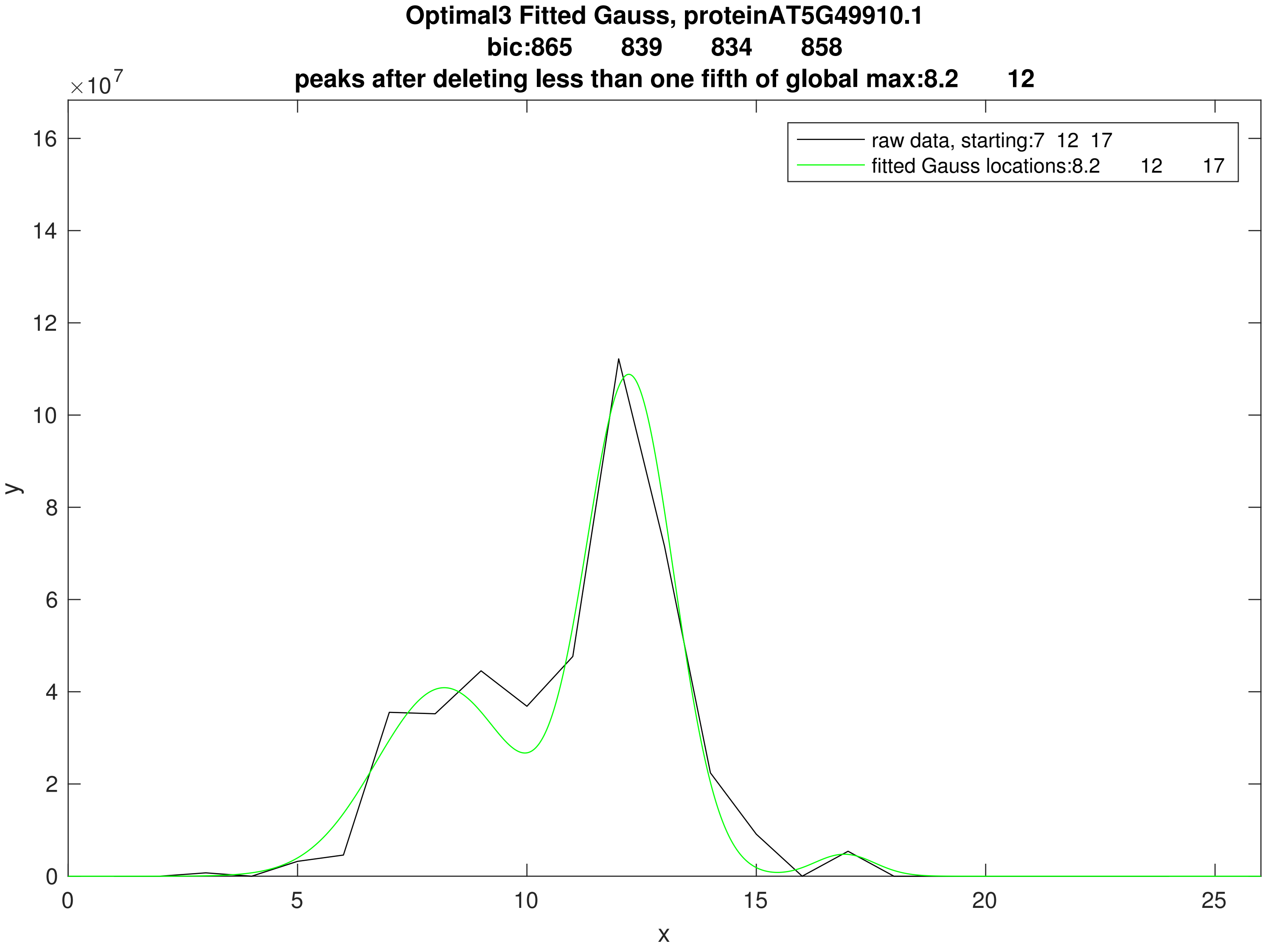
Optimal3 Fitted Gauss, proteinAT5G49910.1
bic:865 839 834 858
peaks after deleting less than one fifth of global max:8.2 12
×10^7
raw data, starting:7 12 17
fitted Gauss locations:8.2 12 17
y
x
0
2
4
6
8
10
12
14
16
5
15
20
25

Optimal1 Fitted Gauss, proteinAT5G49970.1;AT5G49970.2
bic:793
peaks after deleting less than one fifth of global max:12

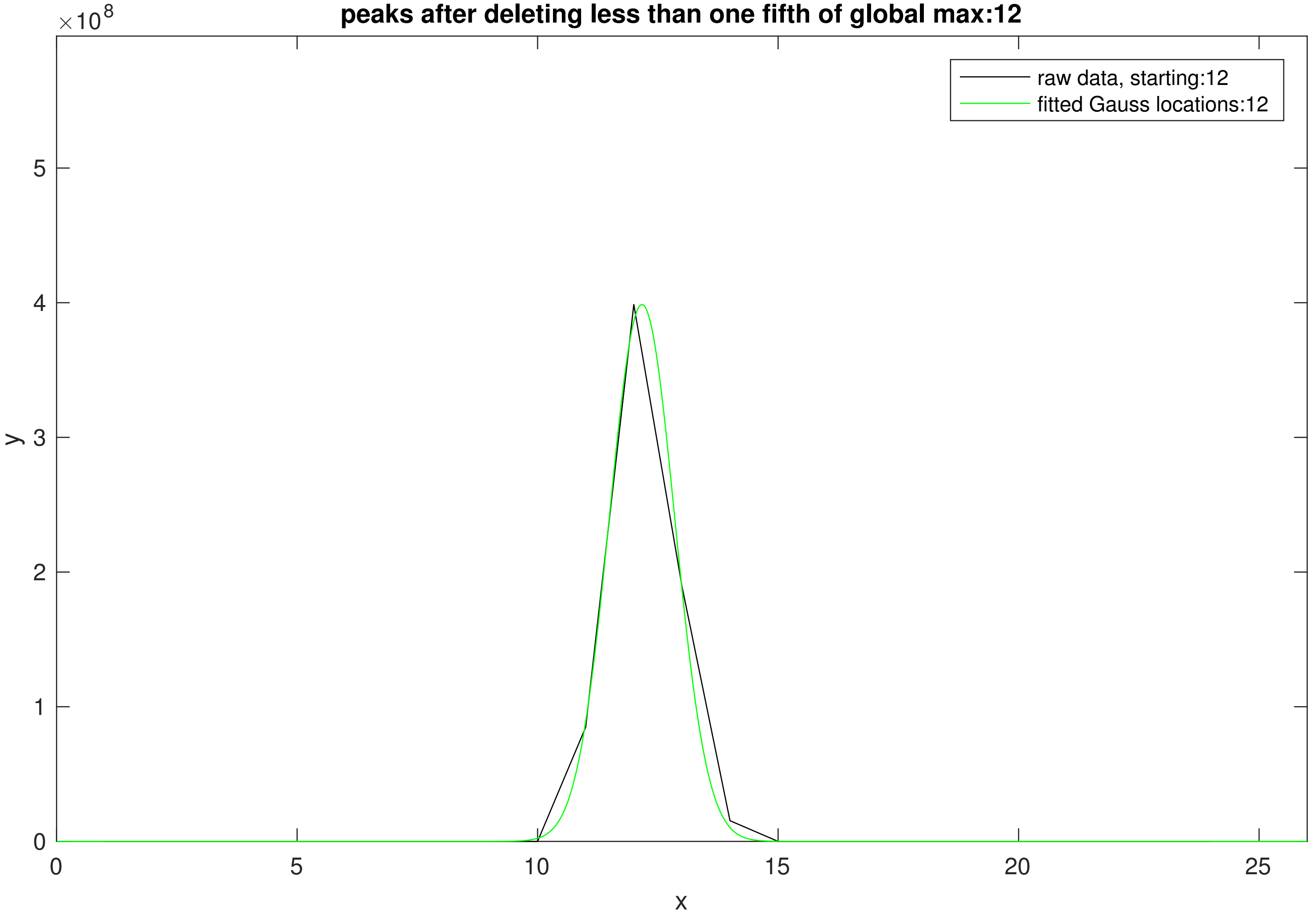

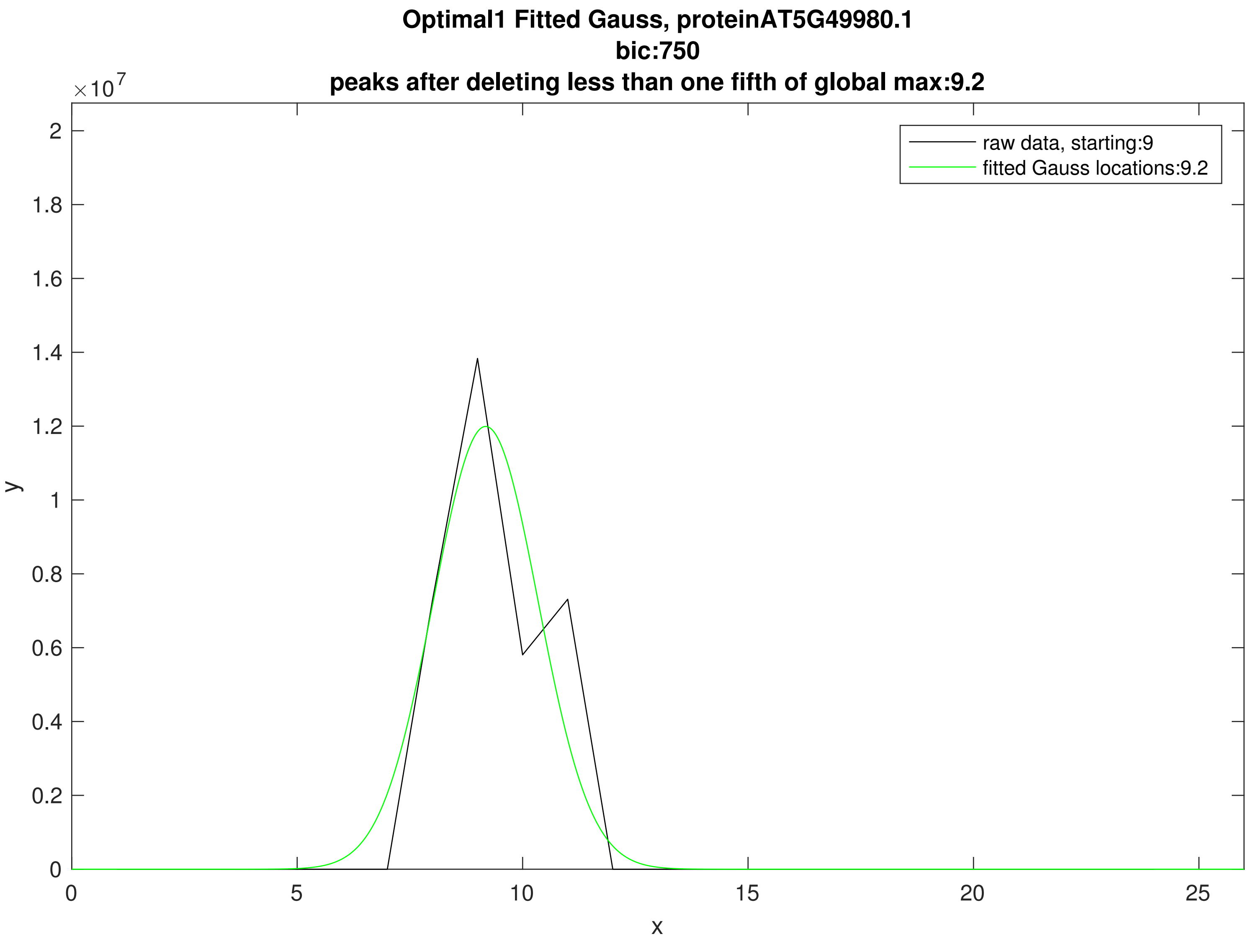

Optimal1 Fitted Gauss, proteinAT5G49980.1
bic:750
peaks after deleting less than one fifth of global max:9.2
×10^7
raw data, starting:9
fitted Gauss locations:9.2
y
x
0
0.2
0.4
0.6
0.8
1
1.2
1.4
1.6
1.8
2
0
5
10
15
20
25

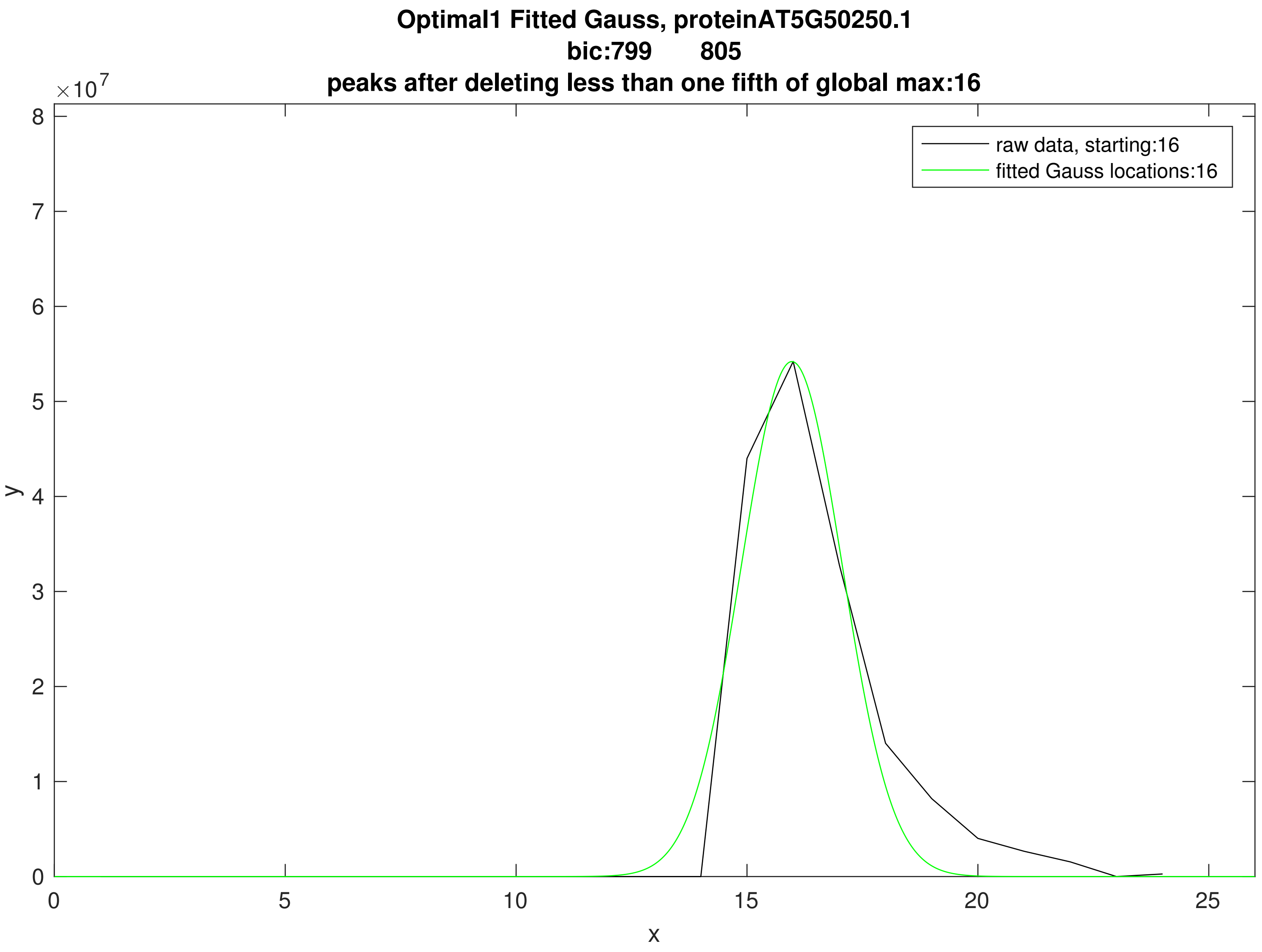

Optimal1 Fitted Gauss, proteinAT5G50250.1
bic:799 805
peaks after deleting less than one fifth of global max:16
×10^7
raw data, starting:16
fitted Gauss locations:16
y
x
0
1
2
3
4
5
6
7
8
0
5
10
15
20
25

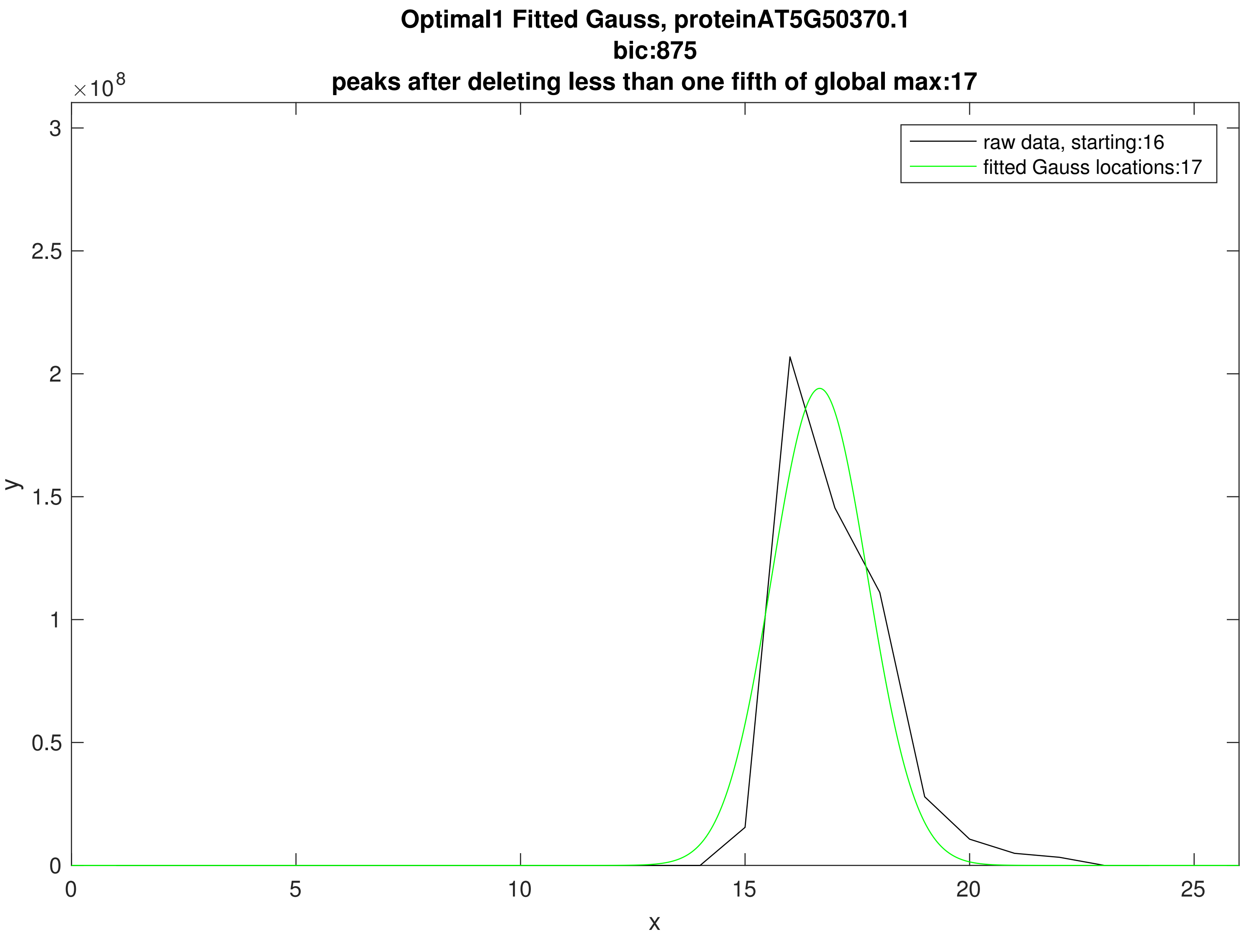

Optimal1 Fitted Gauss, proteinAT5G50370.1
bic:875
peaks after deleting less than one fifth of global max:17
×10^8
3
2.5
2
1.5
1
0.5
0
y
0
5
10
15
20
25
x
raw data, starting:16
fitted Gauss locations:17

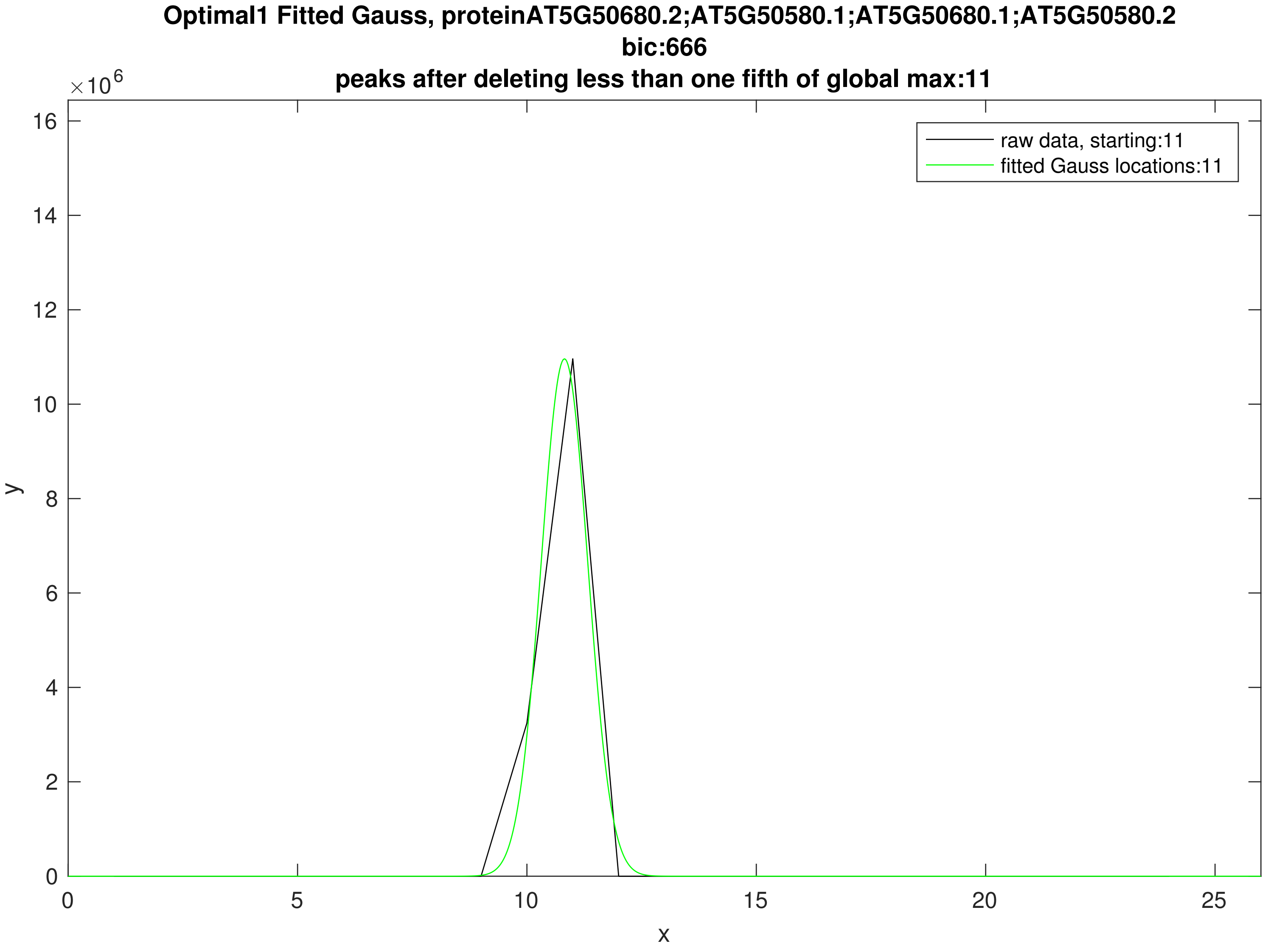

Optimal1 Fitted Gauss, proteinAT5G50680.2;AT5G50580.1;AT5G50680.1;AT5G50580.2
bic:666
peaks after deleting less than one fifth of global max:11
×10^6
16
14
12
10
8
6
4
2
0
y
0
5
10
15
20
25
x
raw data, starting:11
fitted Gauss locations:11

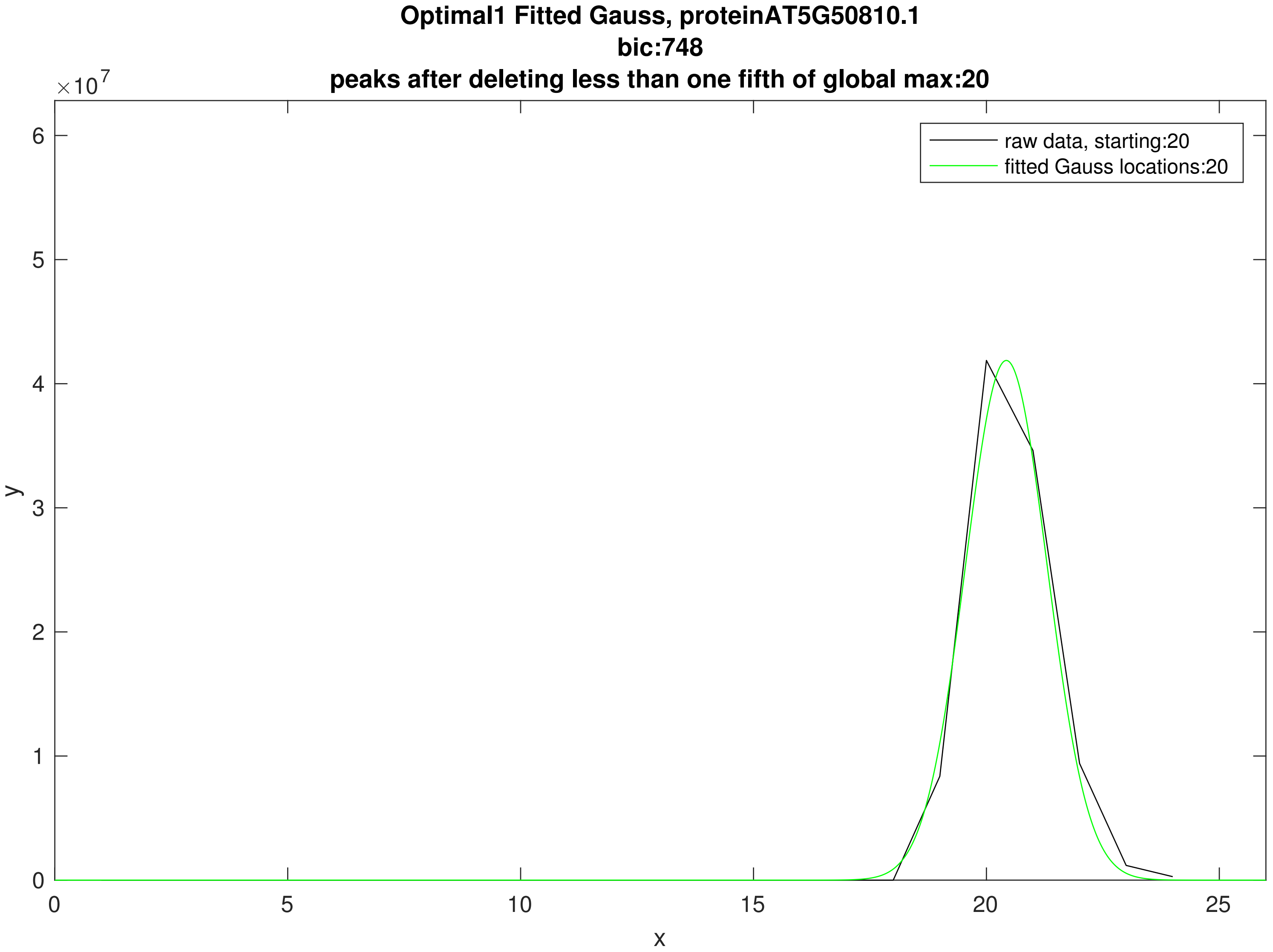
Optimal1 Fitted Gauss, proteinAT5G50810.1
bic:748
peaks after deleting less than one fifth of global max:20
×10^7
raw data, starting:20
fitted Gauss locations:20
y
x
0
1
2
3
4
5
6
0
5
10
15
20
25

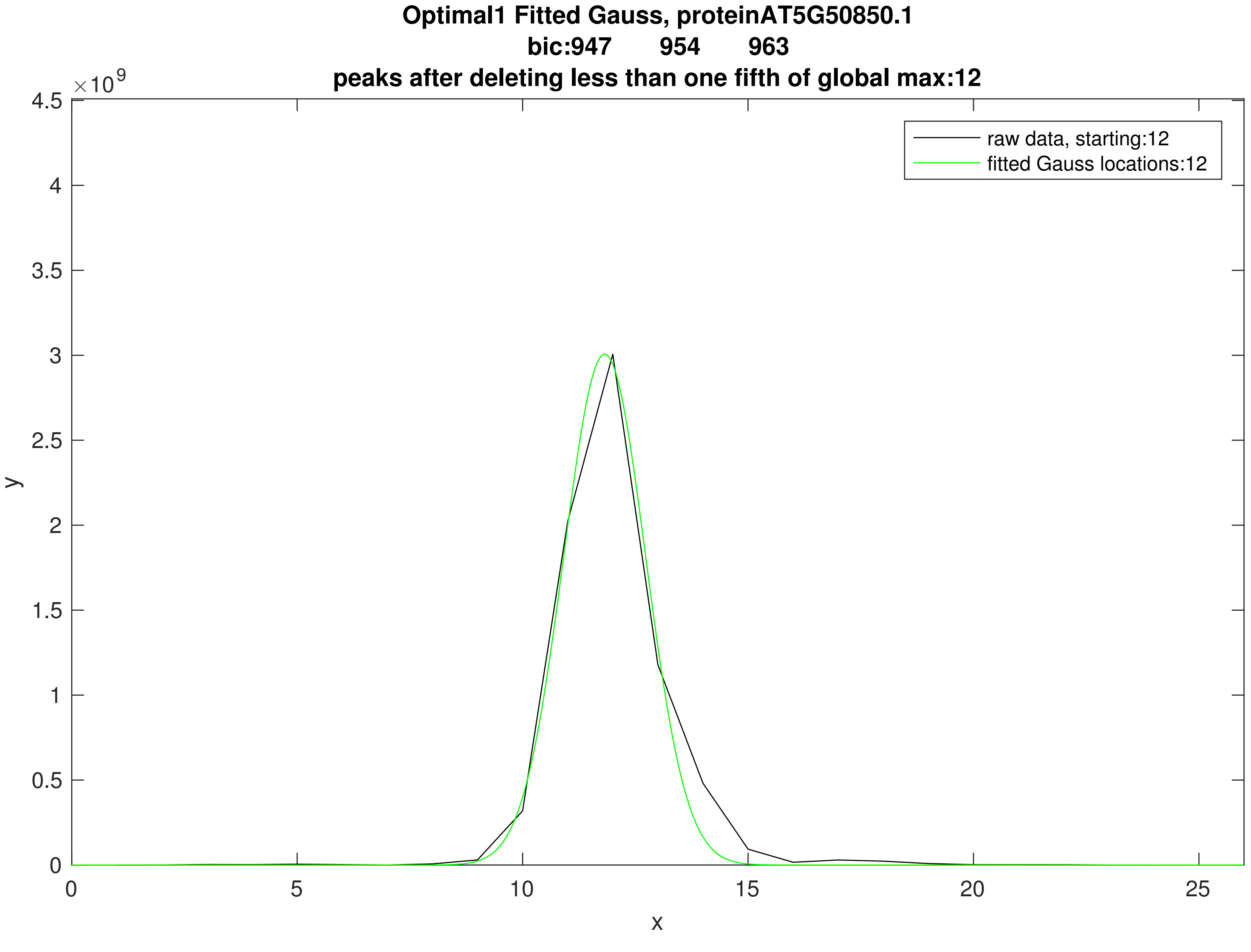
Optimal1 Fitted Gauss, proteinAT5G50850.1
bic:947 954 963
peaks after deleting less than one fifth of global max:12
raw data, starting:12
fitted Gauss locations:12
y
x
×10^9
0
0.5
1
1.5
2
2.5
3
3.5
4
4.5
5
10
15
20
25

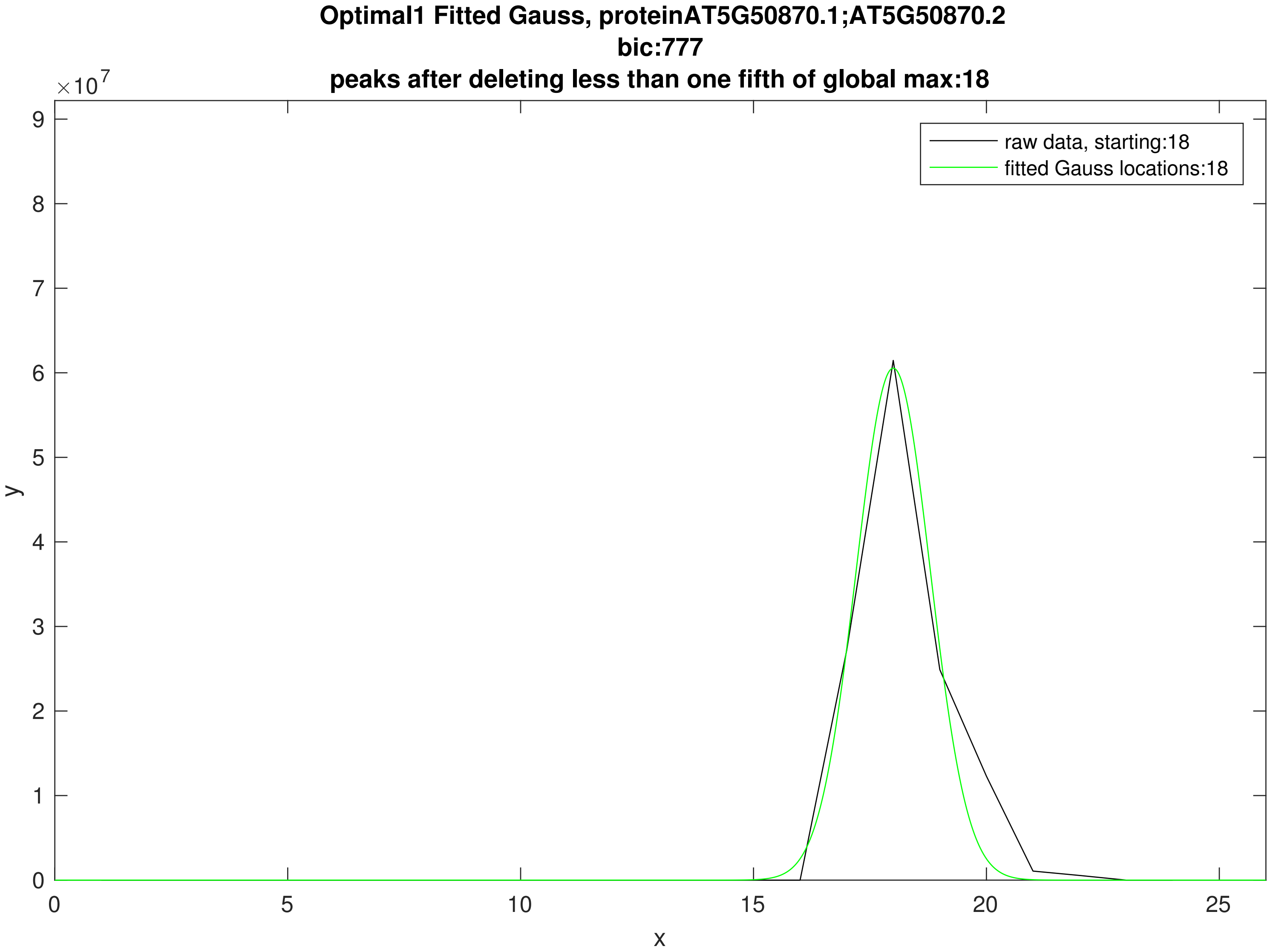
Optimal1 Fitted Gauss, proteinAT5G50870.1;AT5G50870.2
bic:777
peaks after deleting less than one fifth of global max:18
×10^7
raw data, starting:18
fitted Gauss locations:18
y
x

Optimal1 Fitted Gauss, proteinAT5G50920.1;AT3G48870.2;AT3G48870.1
bic:784
peaks after deleting less than one fifth of global max:11

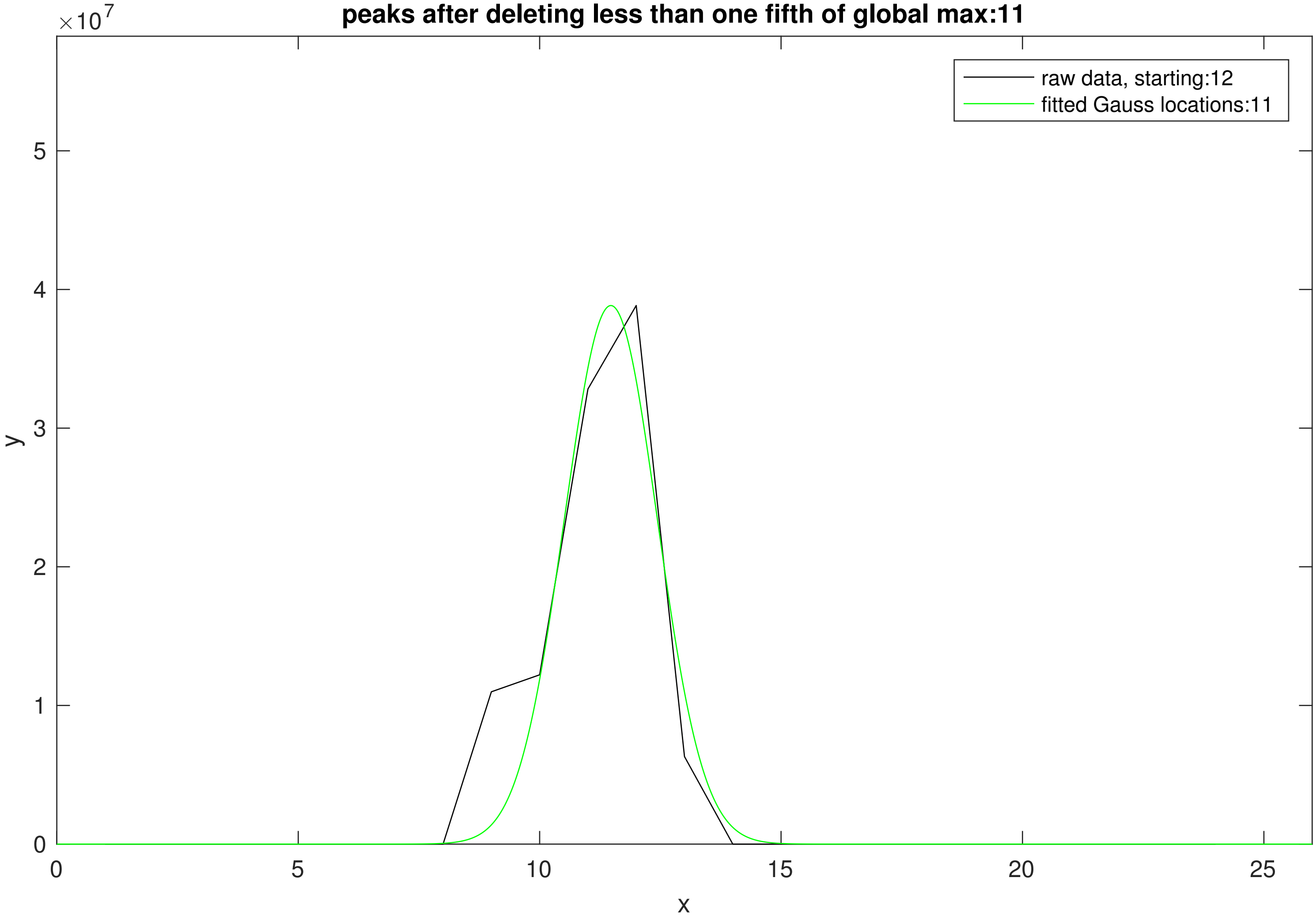

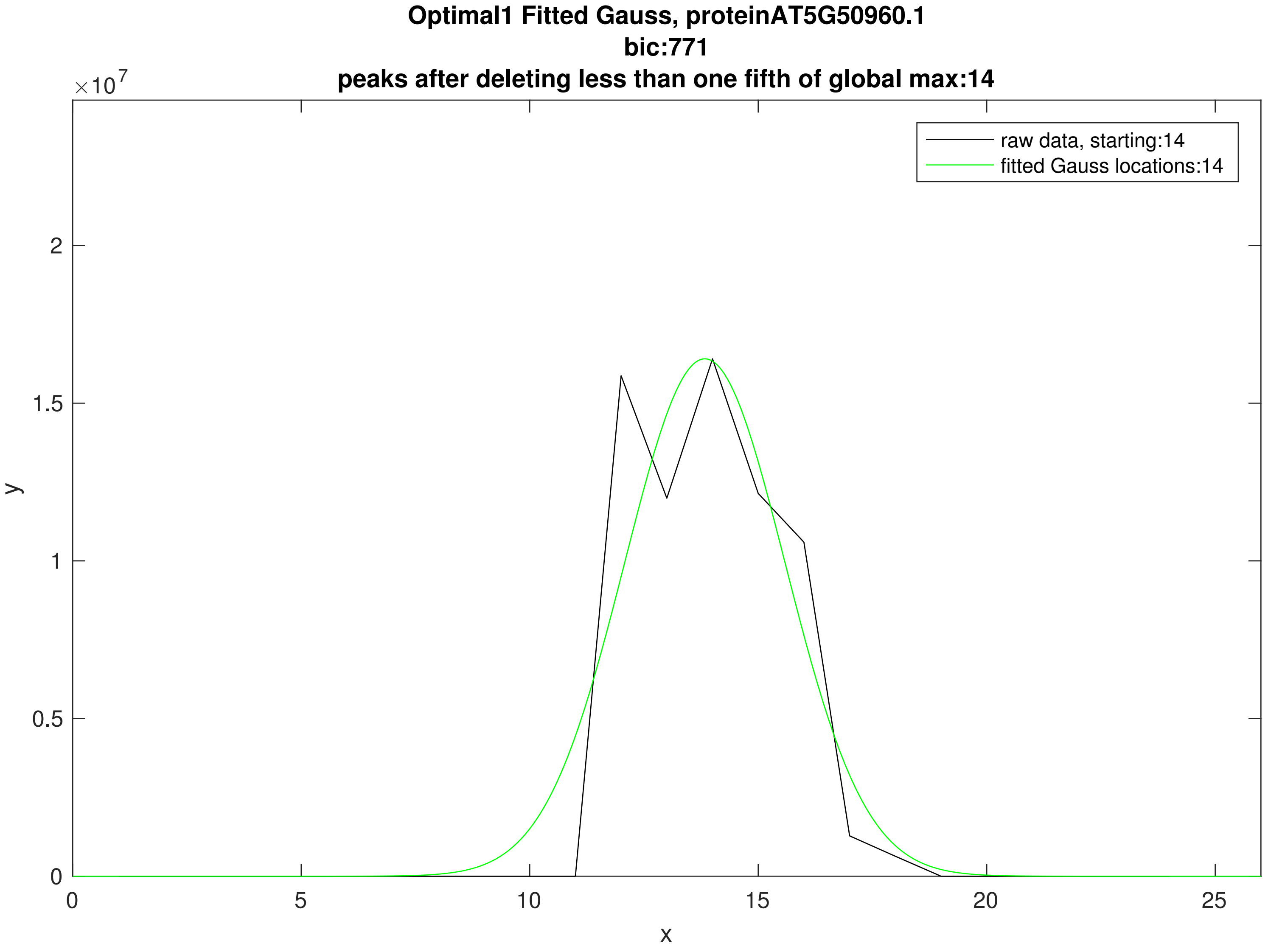
Optimal1 Fitted Gauss, proteinAT5G50960.1
bic:771
peaks after deleting less than one fifth of global max:14
×10^7
raw data, starting:14
fitted Gauss locations:14
2
1.5
1
0.5
0
y
0
5
10
15
20
25
x

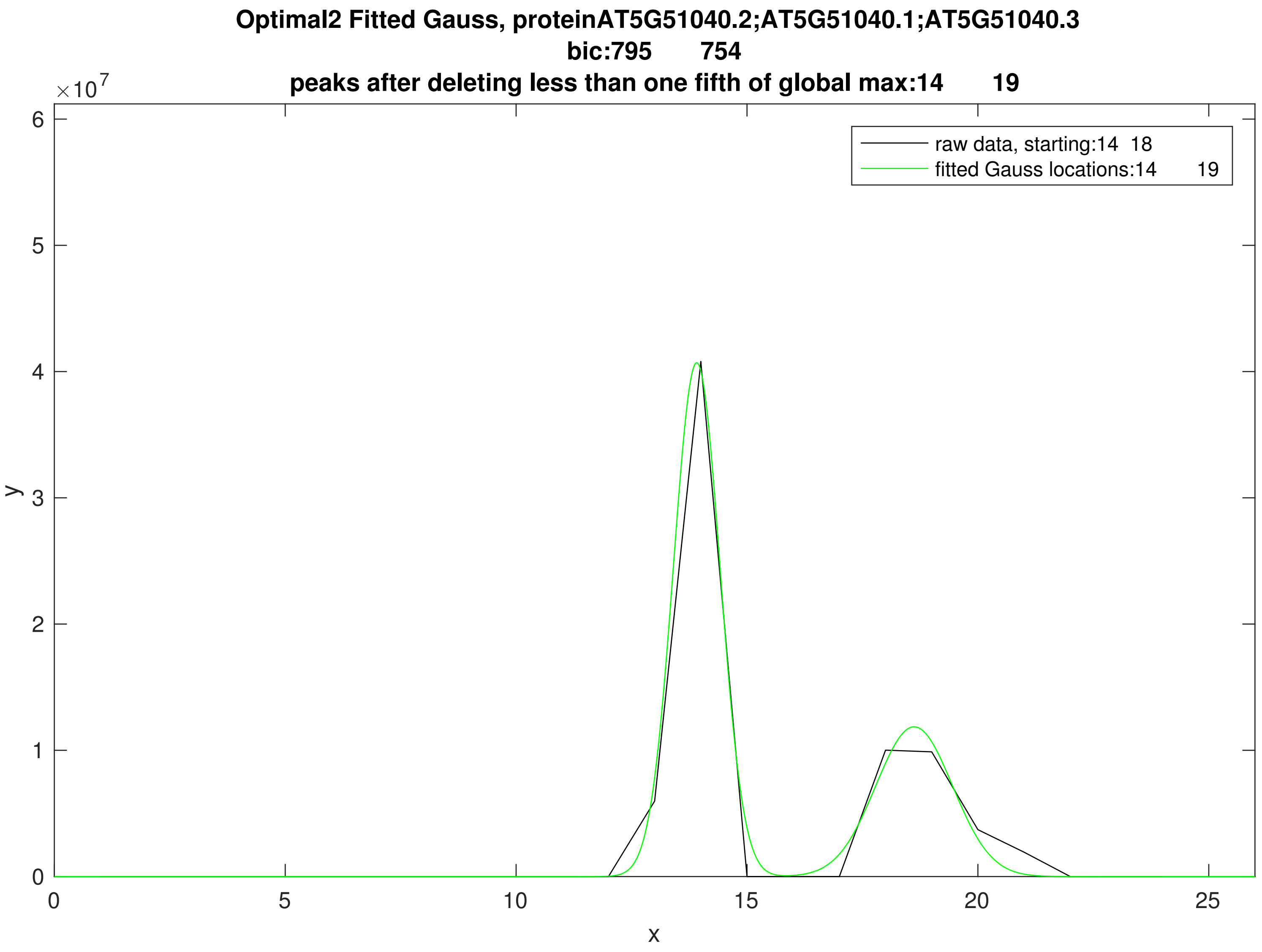

Optimal2 Fitted Gauss, proteinAT5G51040.2;AT5G51040.1;AT5G51040.3
bic:795 754
peaks after deleting less than one fifth of global max:14 19
×10^7
raw data, starting:14 18
fitted Gauss locations:14 19
y
x
0
1
2
3
4
5
6
0
5
10
15
20
25

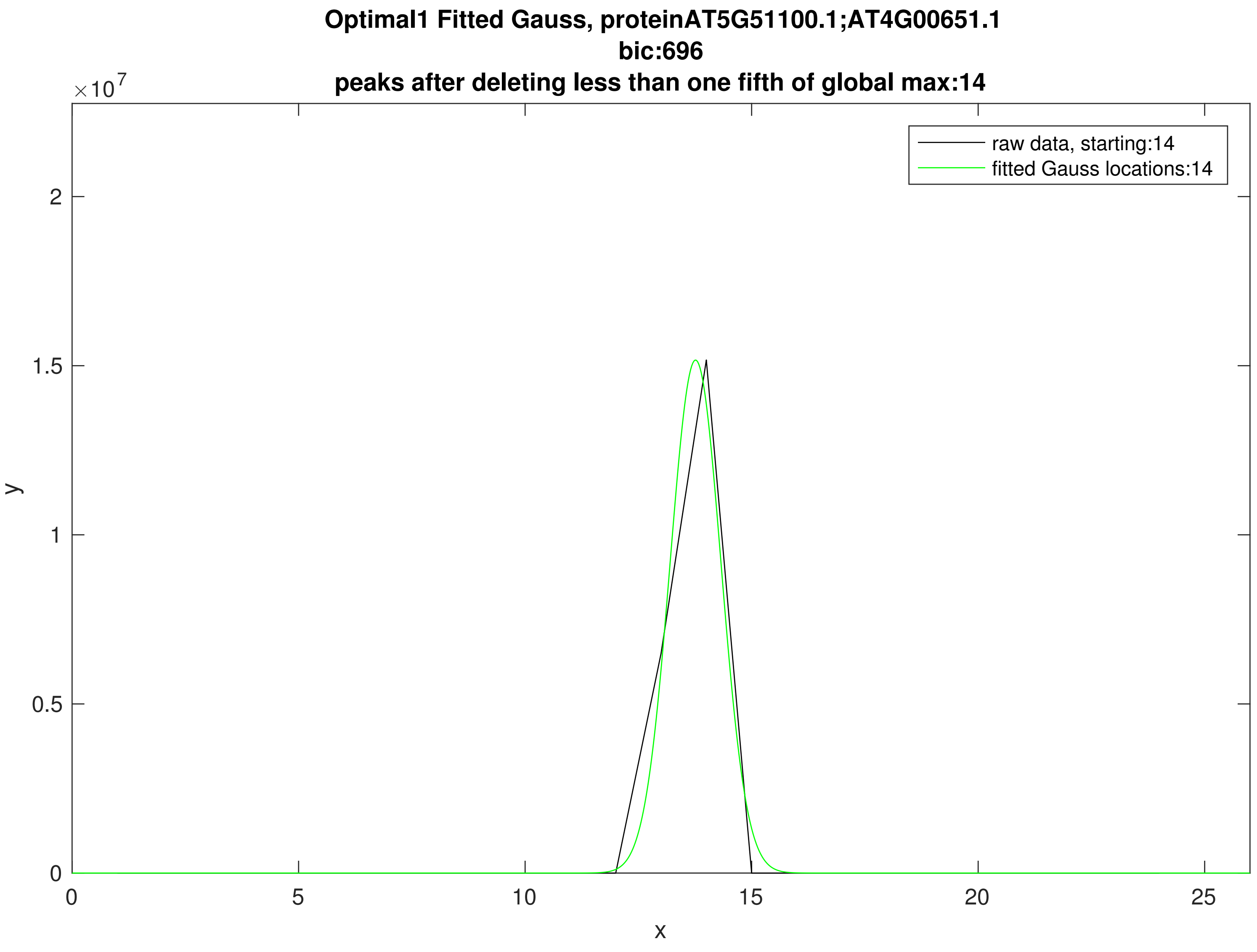
Optimal1 Fitted Gauss, proteinAT5G51100.1;AT4G00651.1
bic:696
peaks after deleting less than one fifth of global max:14
×10^7
raw data, starting:14
fitted Gauss locations:14
y
x
0
0.5
1
1.5
2
0
5
10
15
20
25

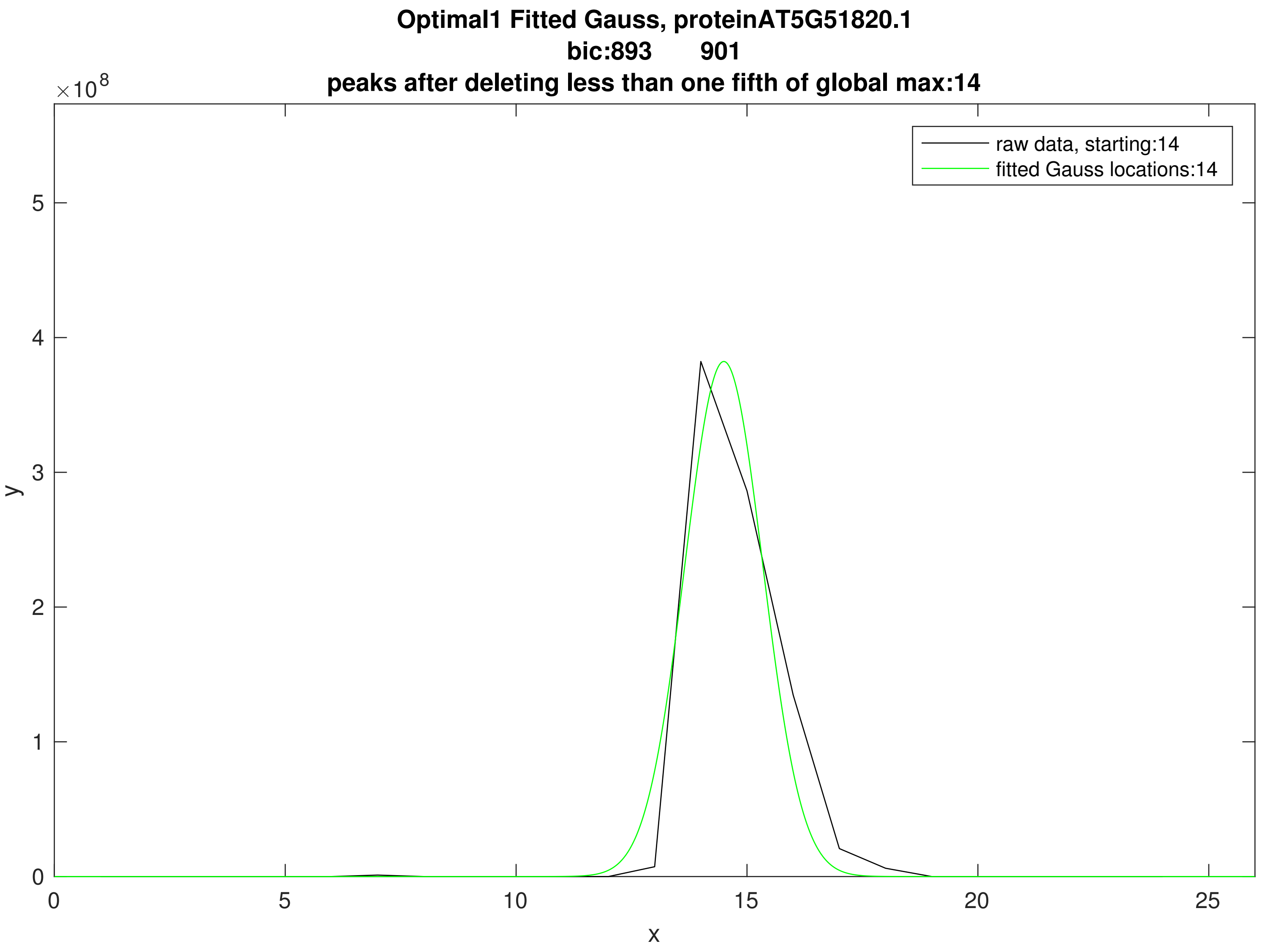

Optimal1 Fitted Gauss, proteinAT5G51820.1
bic:893 901
peaks after deleting less than one fifth of global max:14
×10^8
raw data, starting:14
fitted Gauss locations:14
0
1
2
3
4
5
y
0
5
10
15
20
25
x

Optimal1 Fitted Gauss, proteinAT5G51830.1
bic:742
peaks after deleting less than one fifth of global max:14

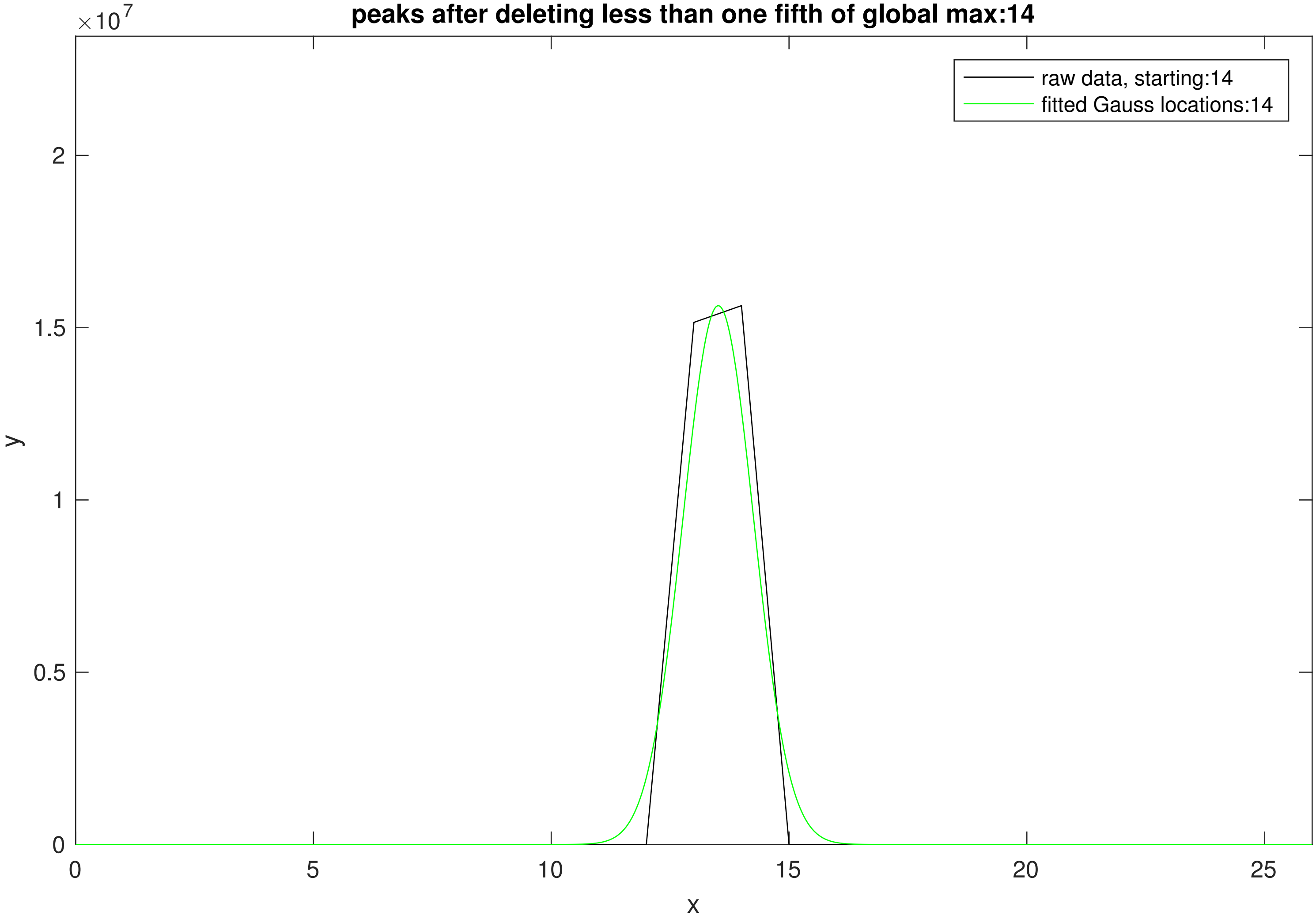

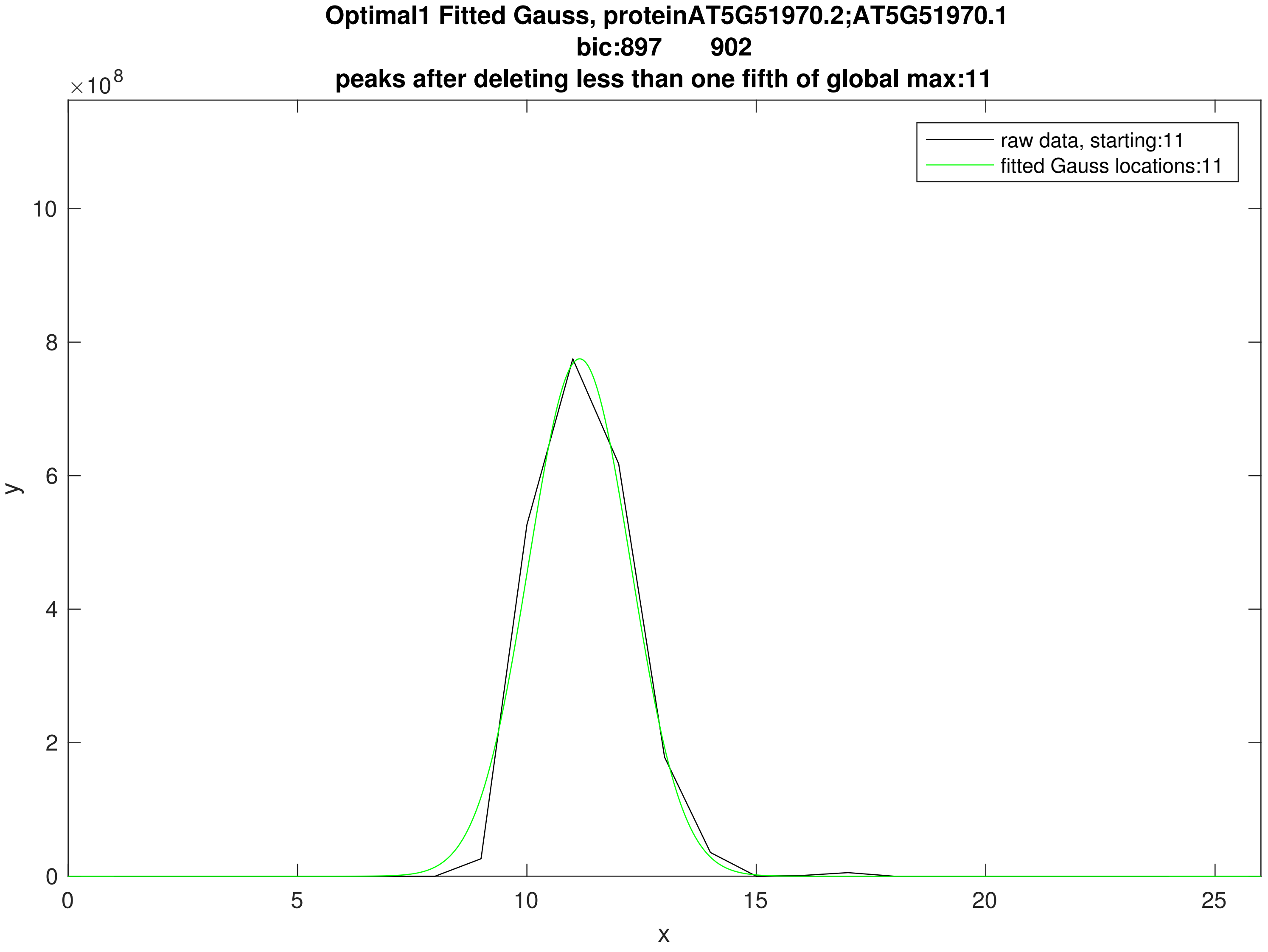

Optimal1 Fitted Gauss, proteinAT5G51970.2;AT5G51970.1
bic:897 902
peaks after deleting less than one fifth of global max:11
×10^8
raw data, starting:11
fitted Gauss locations:11
y
x
0
2
4
6
8
10
0
5
10
15
20
25

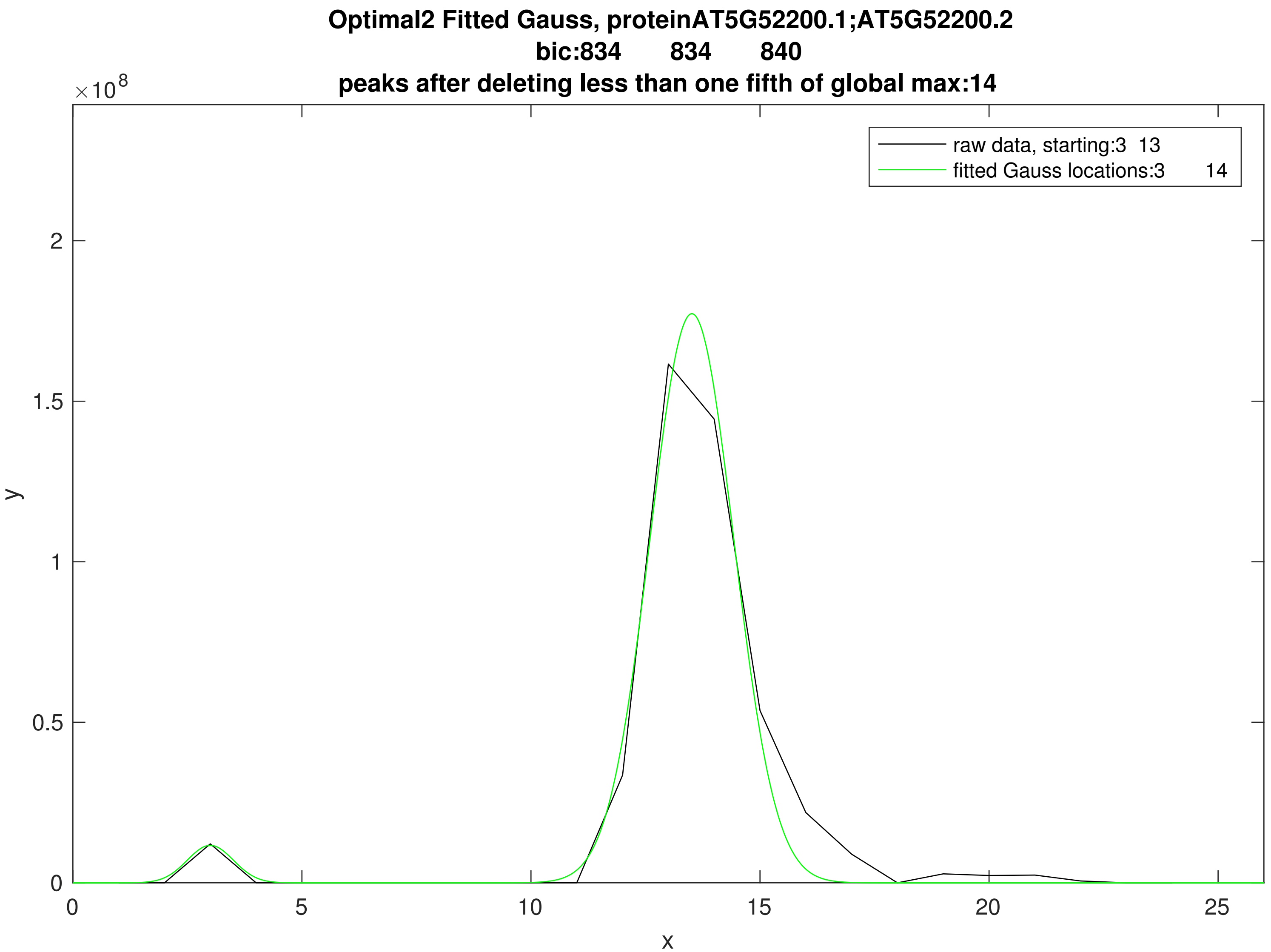
Optimal2 Fitted Gauss, proteinAT5G52200.1;AT5G52200.2
bic:834 834 840
peaks after deleting less than one fifth of global max:14
×10^8
raw data, starting:3 13
fitted Gauss locations:3 14
y
x
0
0.5
1
1.5
2
0
5
10
15
20
25

Optimal1 Fitted Gauss, proteinAT5G52210.2;AT5G52210.1
bic:747
peaks after deleting less than one fifth of global max:19

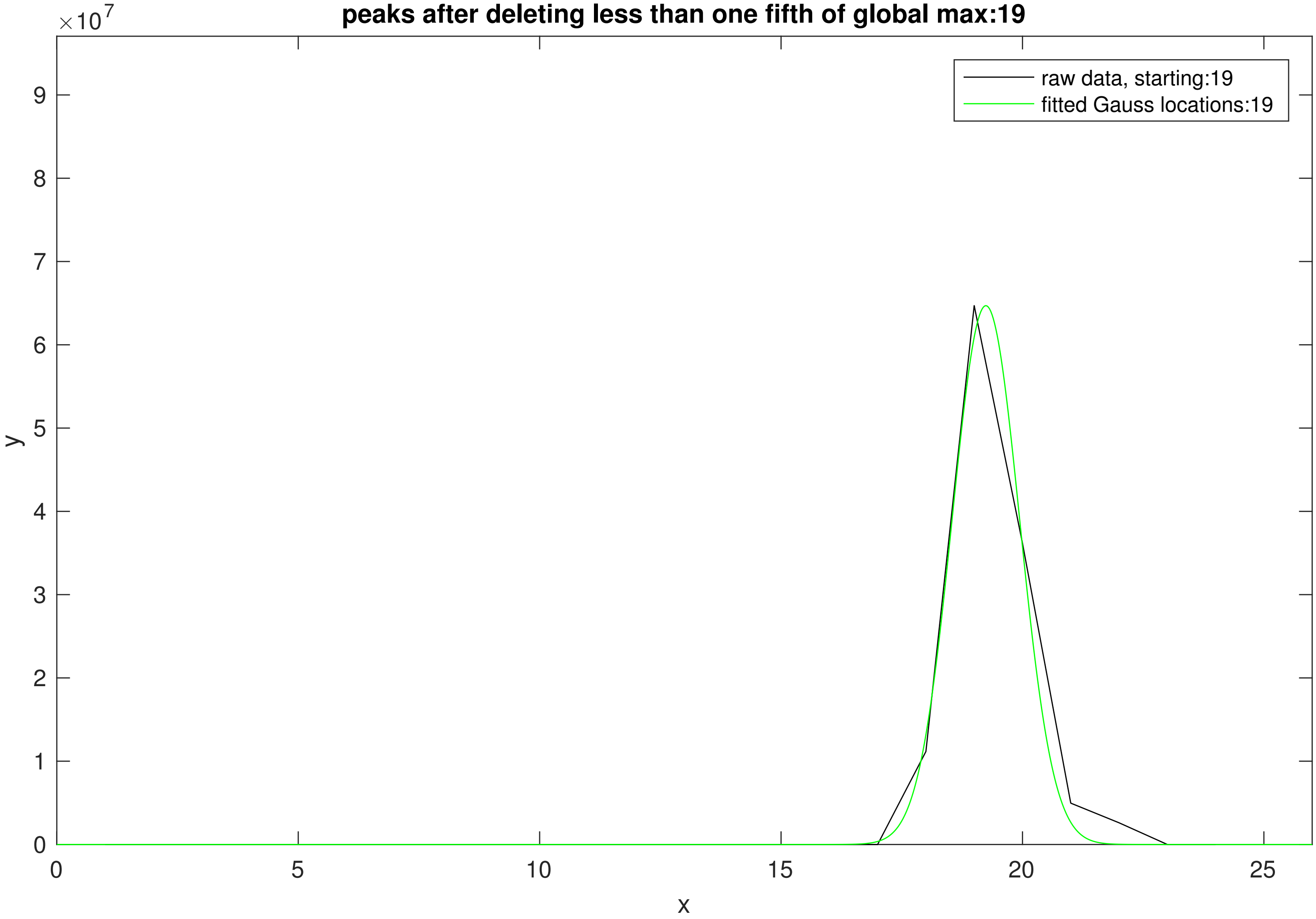

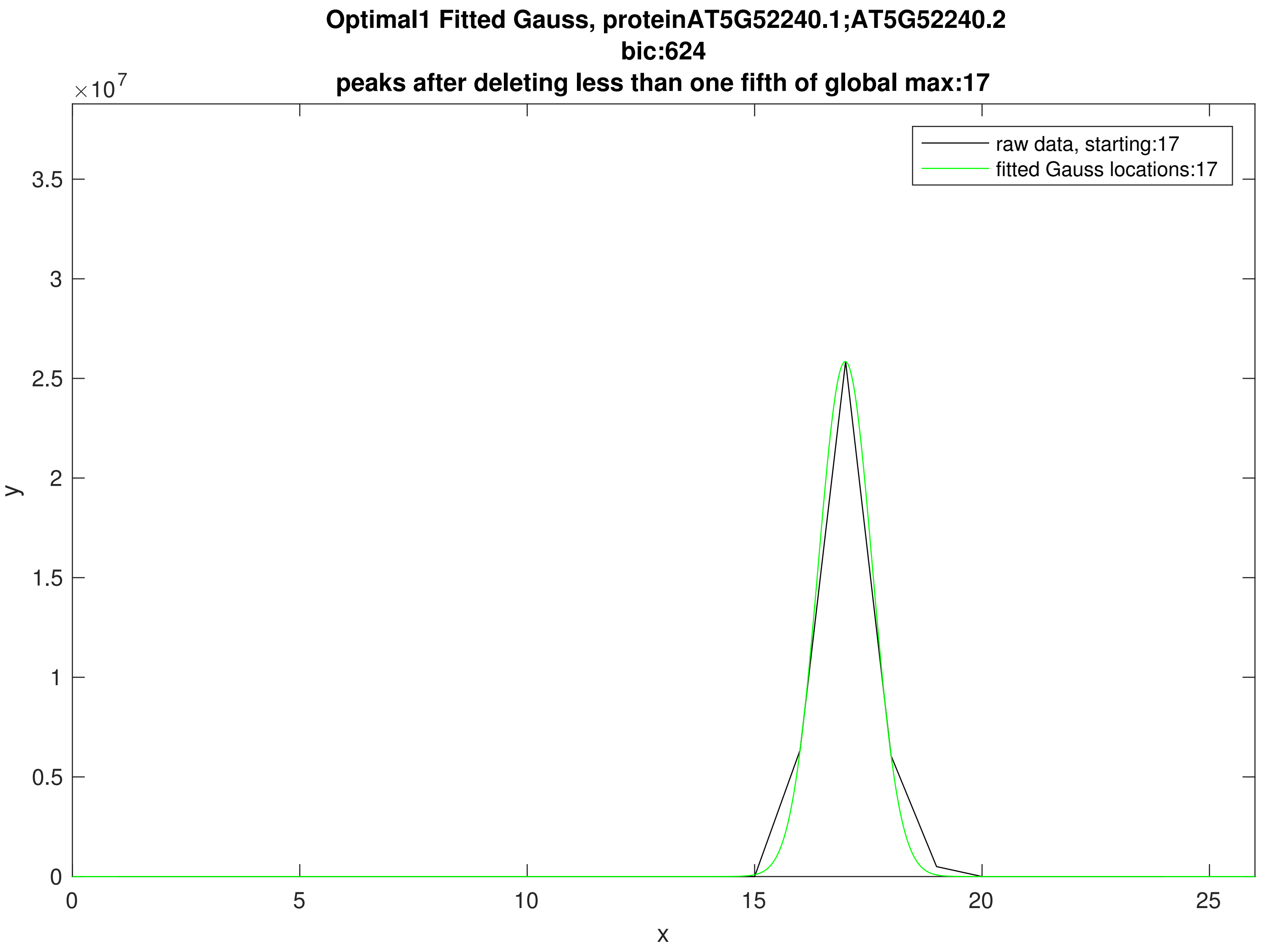
Optimal1 Fitted Gauss, proteinAT5G52240.1;AT5G52240.2
bic:624
peaks after deleting less than one fifth of global max:17
×10^7
3.5
3
2.5
2
1.5
1
0.5
0
y
0
5
10
15
20
25
x
raw data, starting:17
fitted Gauss locations:17

**Optimal1 Fitted Gauss, proteinAT5G52300.2;AT5G52300.1**
**bic:736**
**peaks after deleting less than one fifth of global max:7.6**

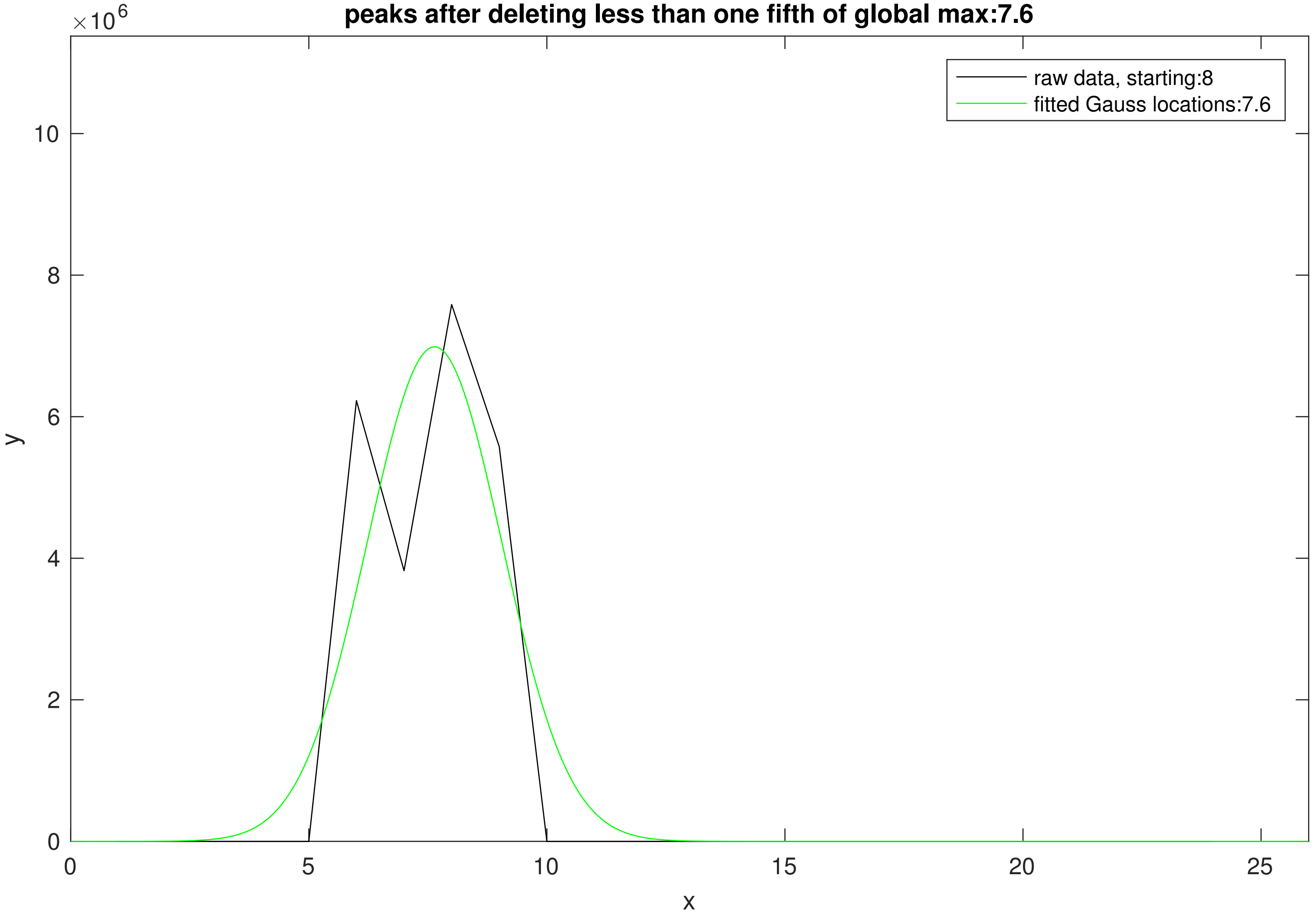

**Optimal1 Fitted Gauss, proteinAT5G52310.1**
**bic:894**
**peaks after deleting less than one fifth of global max:5.9**

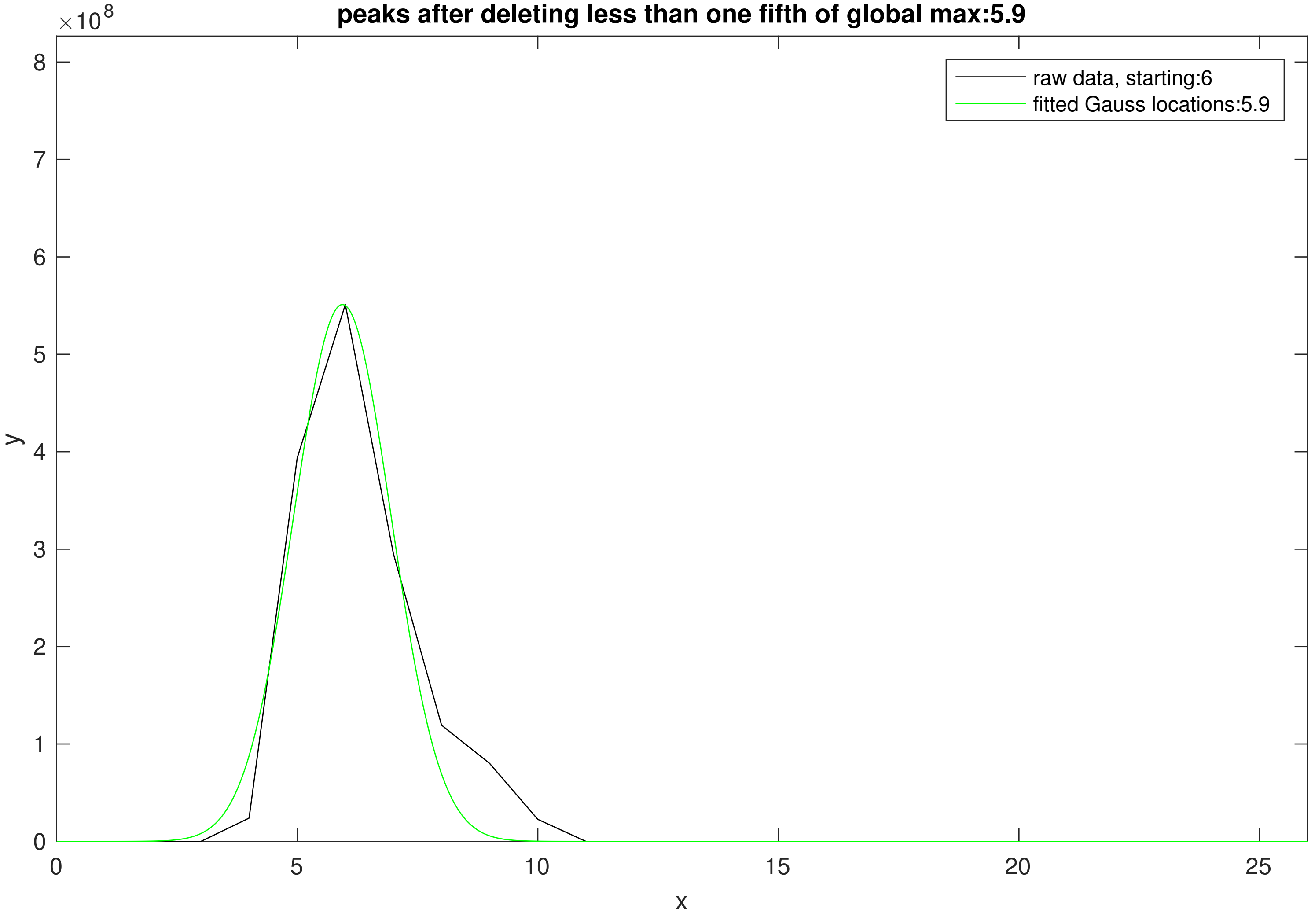

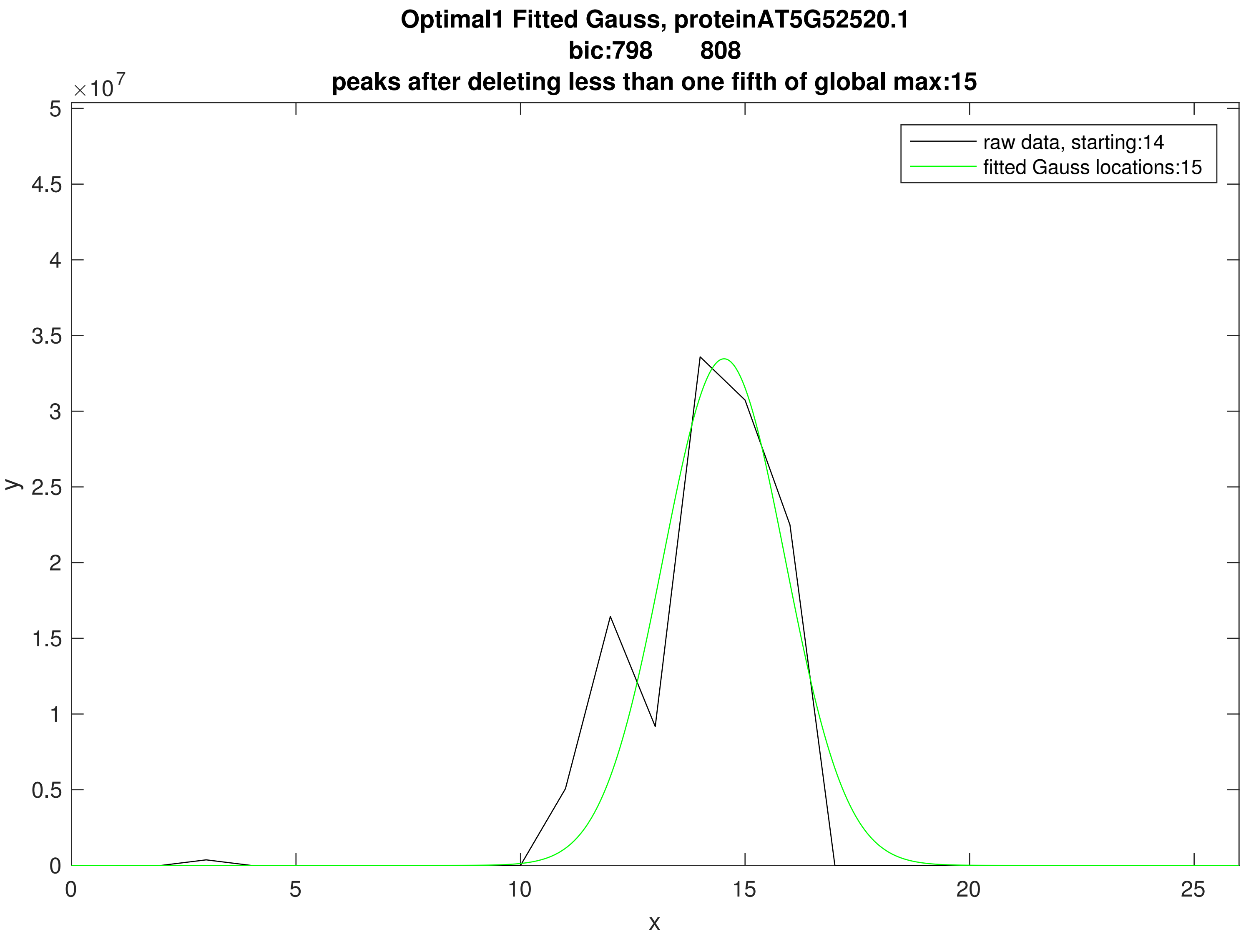

Optimal1 Fitted Gauss, proteinAT5G52520.1
bic:798 808
peaks after deleting less than one fifth of global max:15
×10^7
5
4.5
4
3.5
3
2.5
2
1.5
1
0.5
0
y
0
5
10
15
20
25
x
raw data, starting:14
fitted Gauss locations:15

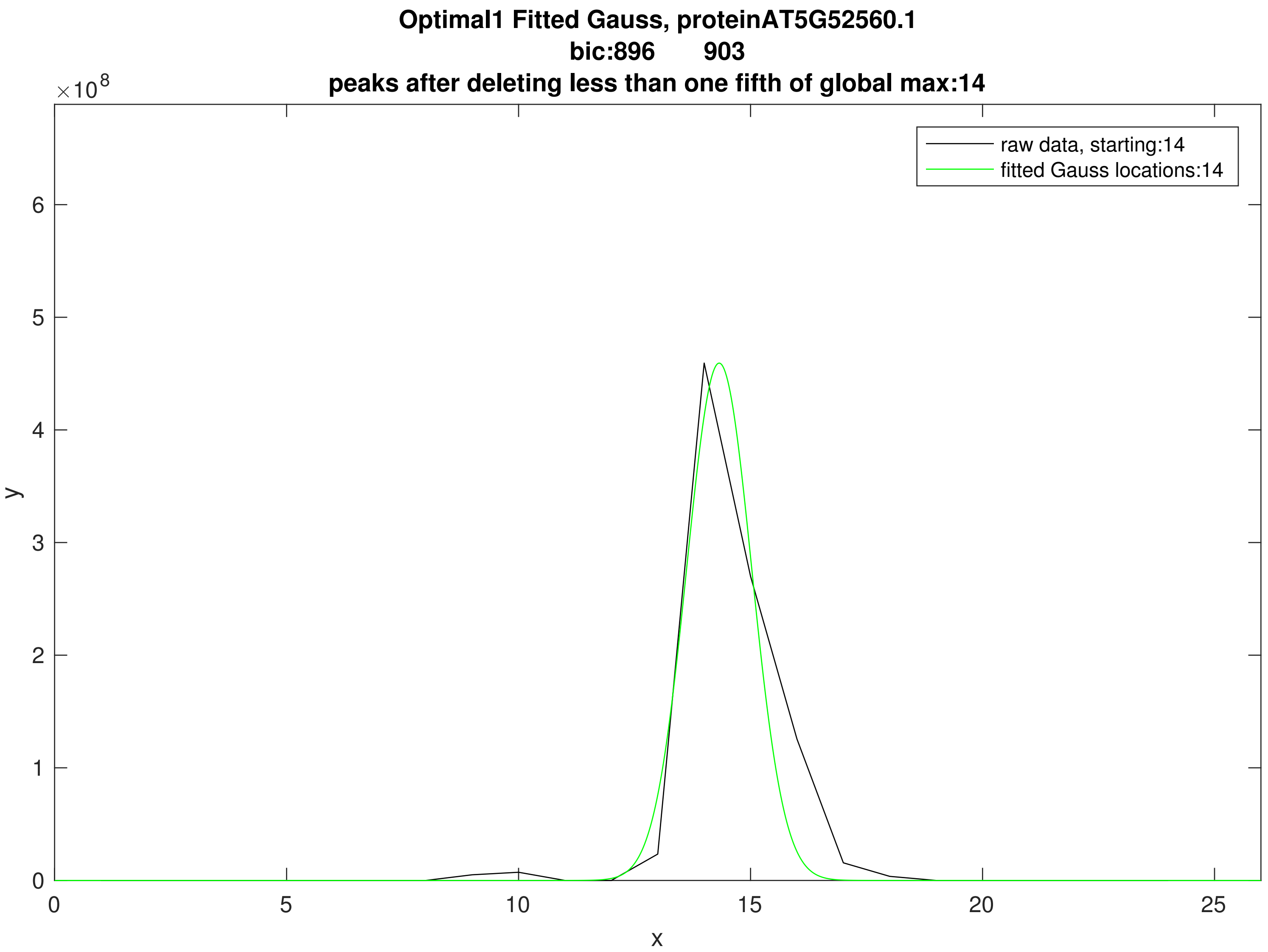
Optimal1 Fitted Gauss, proteinAT5G52560.1
bic:896 903
peaks after deleting less than one fifth of global max:14
×10^8
raw data, starting:14
fitted Gauss locations:14
y
x
0
1
2
3
4
5
6
0
5
10
15
20
25

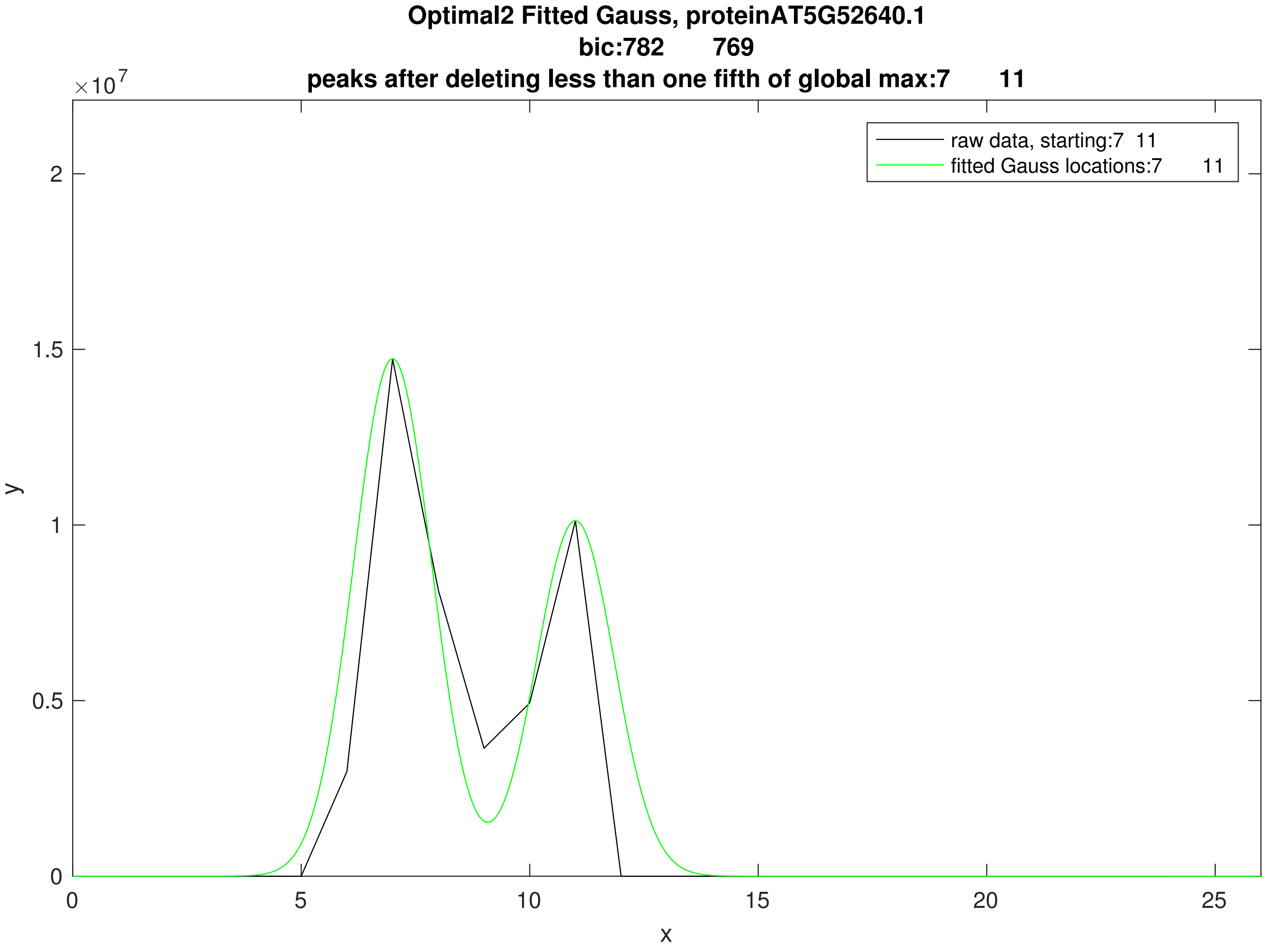

Optimal2 Fitted Gauss, proteinAT5G52640.1
bic:782 769
peaks after deleting less than one fifth of global max:7 11
×10^7
raw data, starting:7 11
fitted Gauss locations:7 11
y
x
0
0.5
1
1.5
2
0
5
10
15
20
25

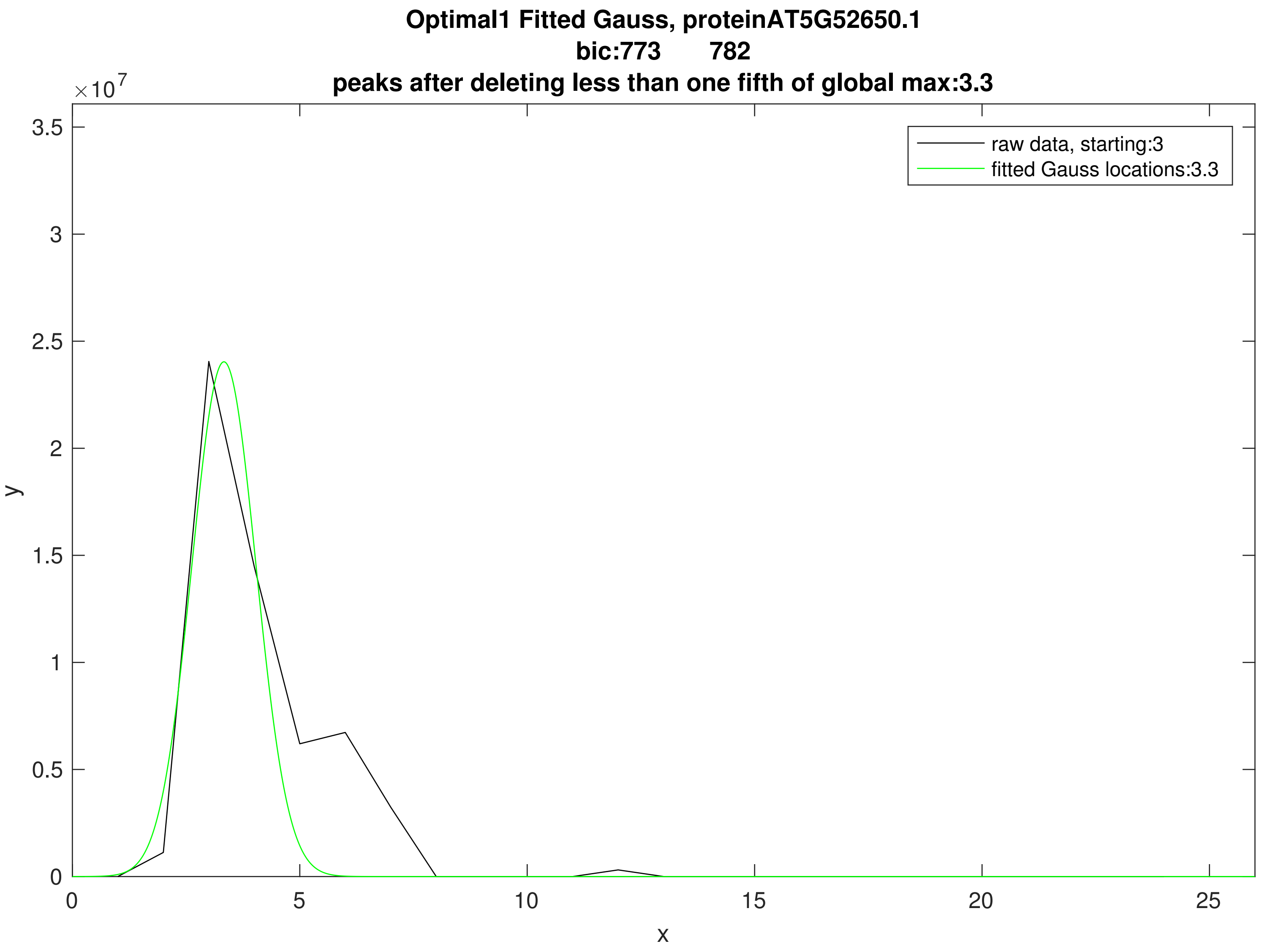

Optimal1 Fitted Gauss, proteinAT5G52650.1
bic:773 782
peaks after deleting less than one fifth of global max:3.3
×10^7
3.5
3
2.5
2
1.5
1
0.5
0
y
0
5
10
15
20
25
x
raw data, starting:3
fitted Gauss locations:3.3

**Optimal1 Fitted Gauss, proteinAT5G52840.1**
**bic:863 871**
**peaks after deleting less than one fifth of global max:16**

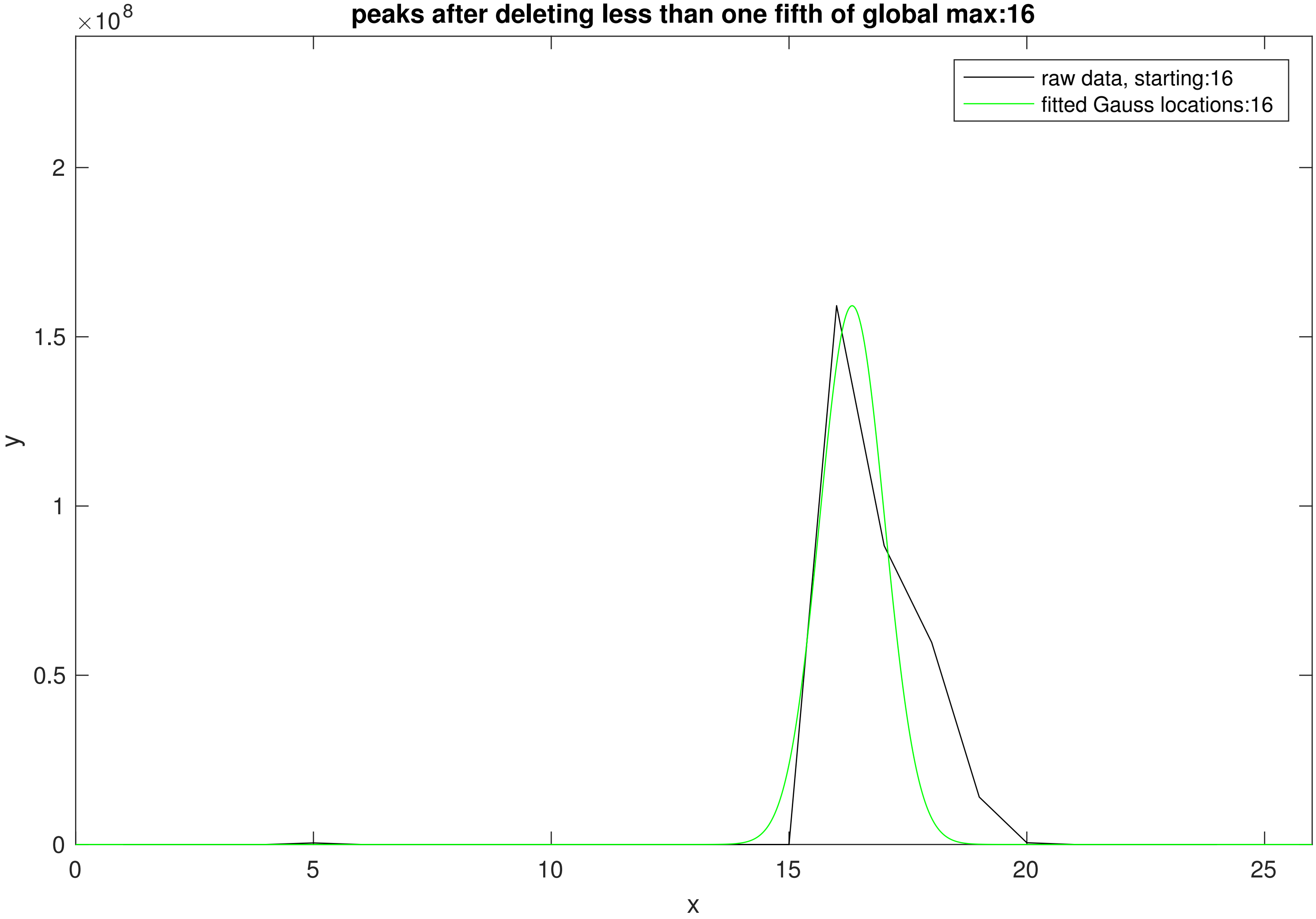

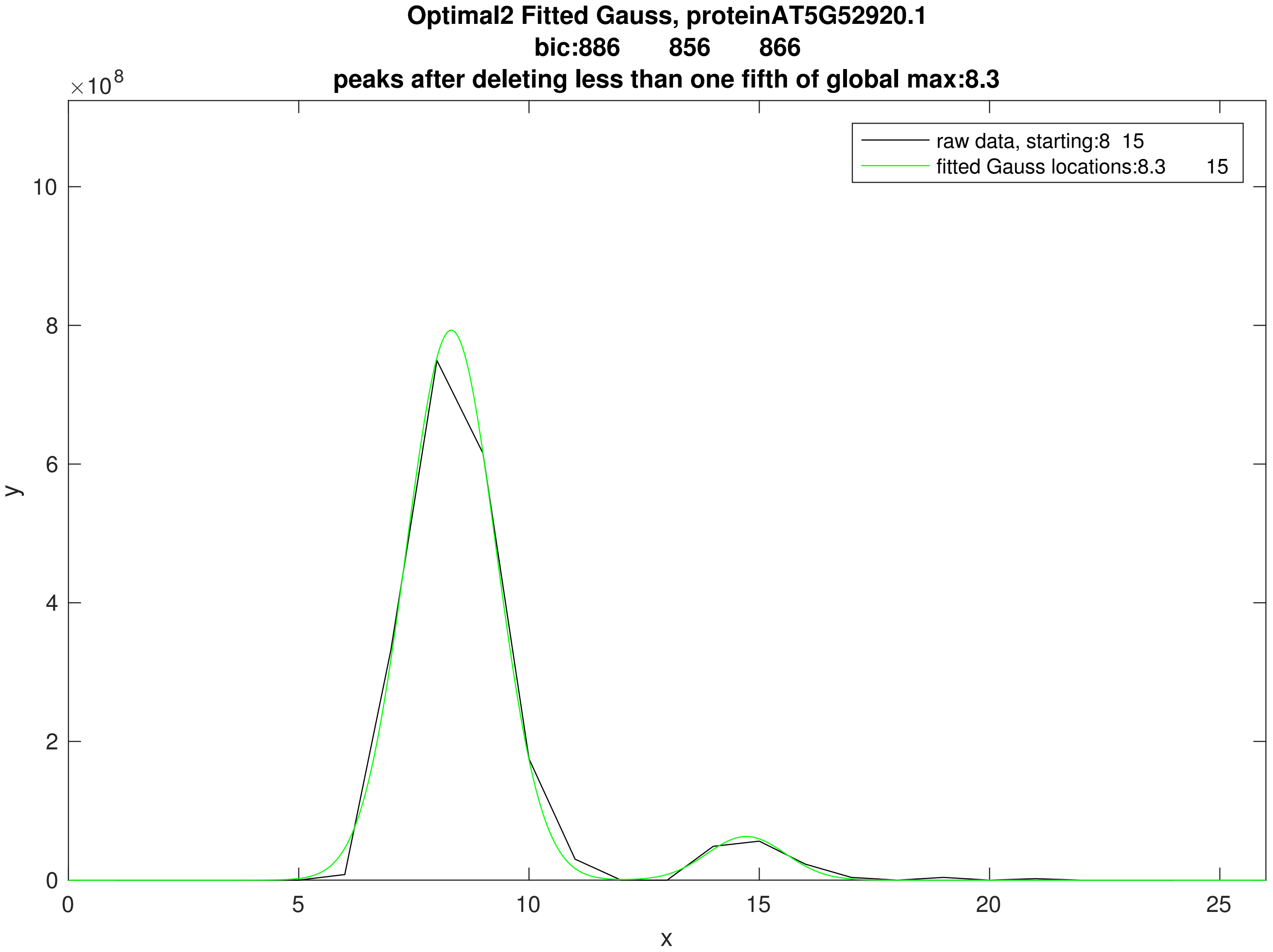

Optimal2 Fitted Gauss, proteinAT5G52920.1
bic:886 856 866
peaks after deleting less than one fifth of global max:8.3
×10^8
raw data, starting:8 15
fitted Gauss locations:8.3 15
10
8
6
4
2
0
y
0
5
10
15
20
25
x

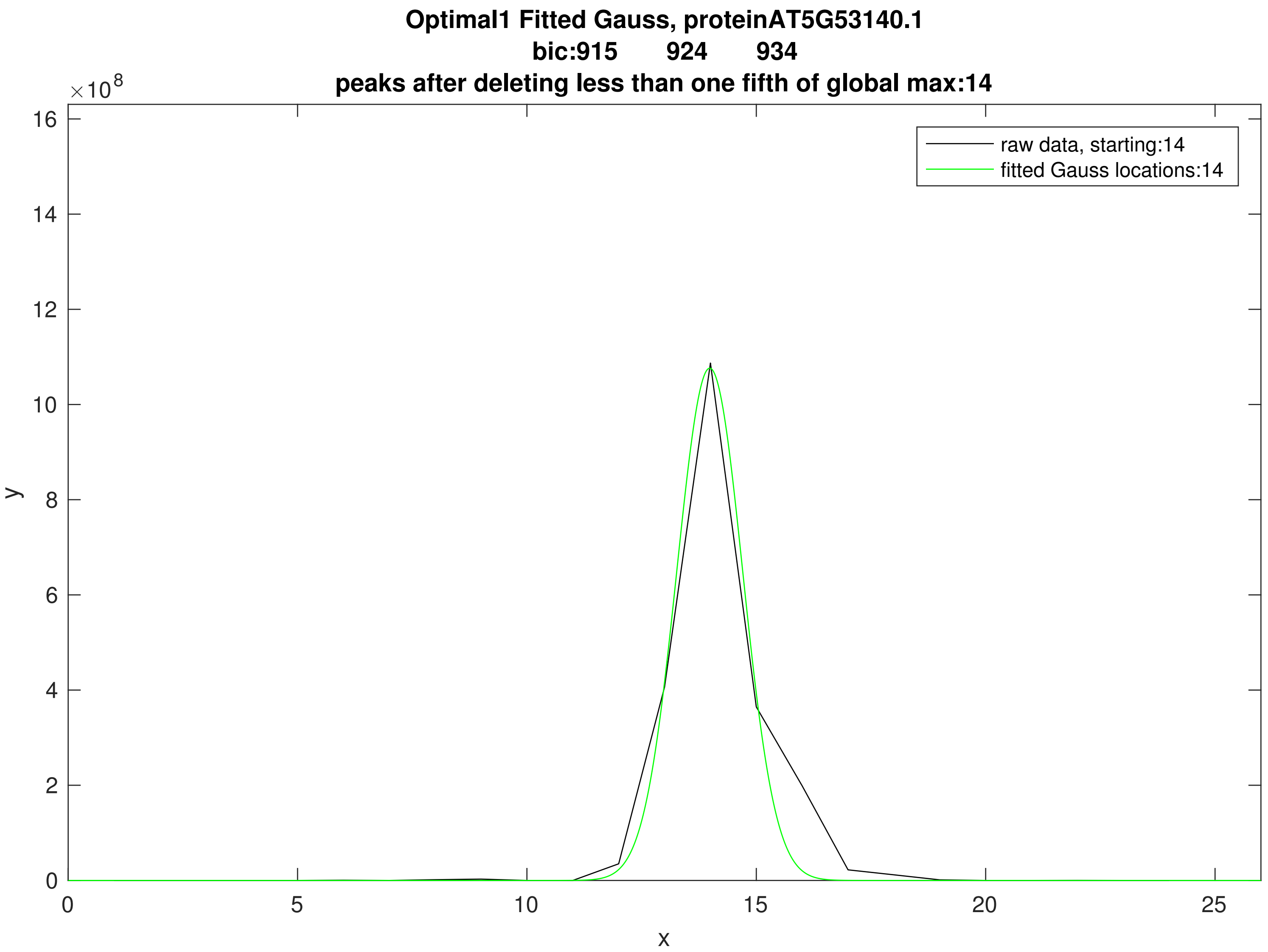

Optimal1 Fitted Gauss, proteinAT5G53140.1
bic:915 924 934
peaks after deleting less than one fifth of global max:14
×10^8
raw data, starting:14
fitted Gauss locations:14
y
x

G53300.4;AT4G27960.2;AT4G27960.1;AT1G64230.4;AT5G41700.2;AT1G64230.5;AT1G64230.3;AT5G41700.4;AT5G417[illegible]
bic:739
peaks after deleting less than one fifth of global max:20

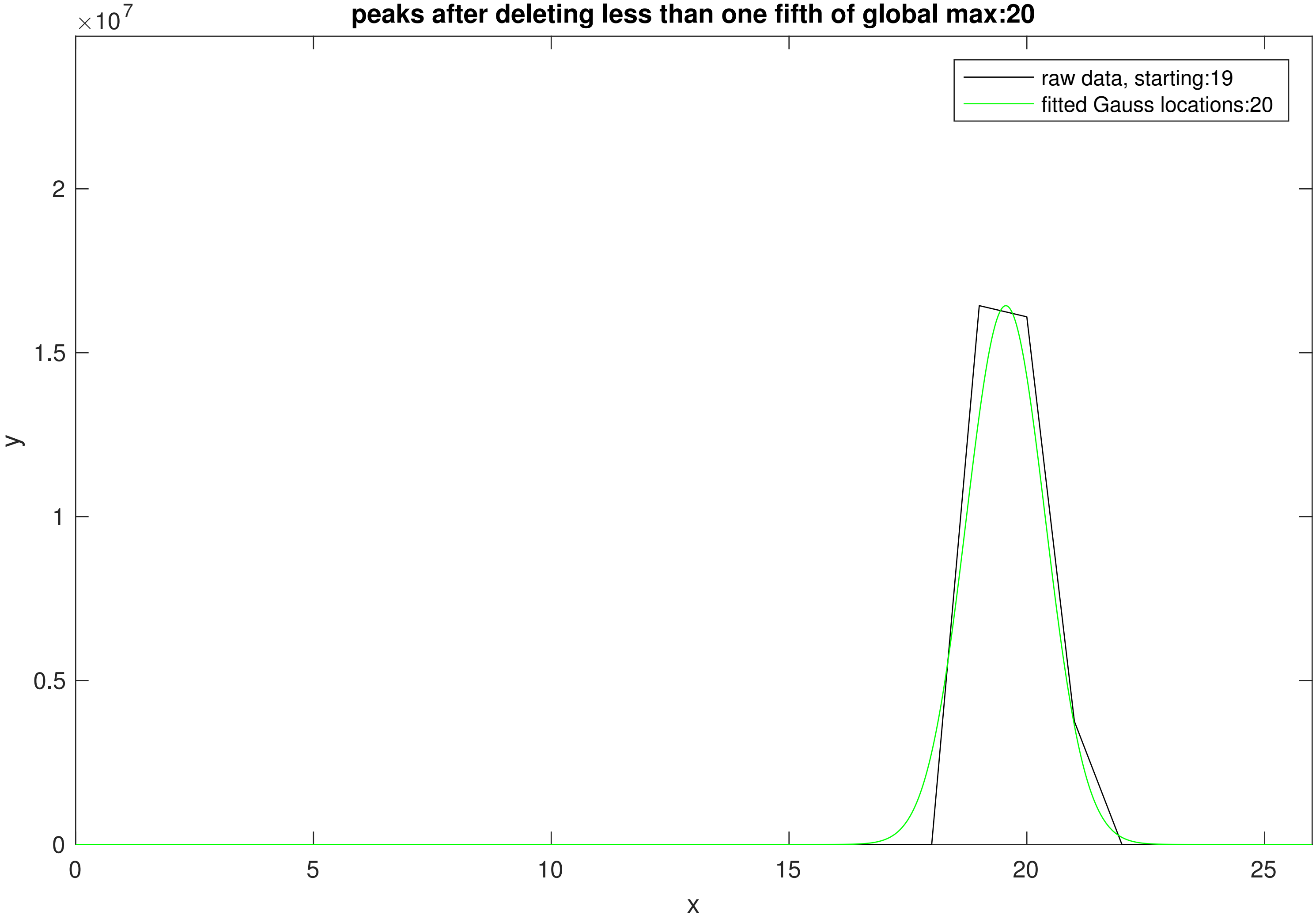

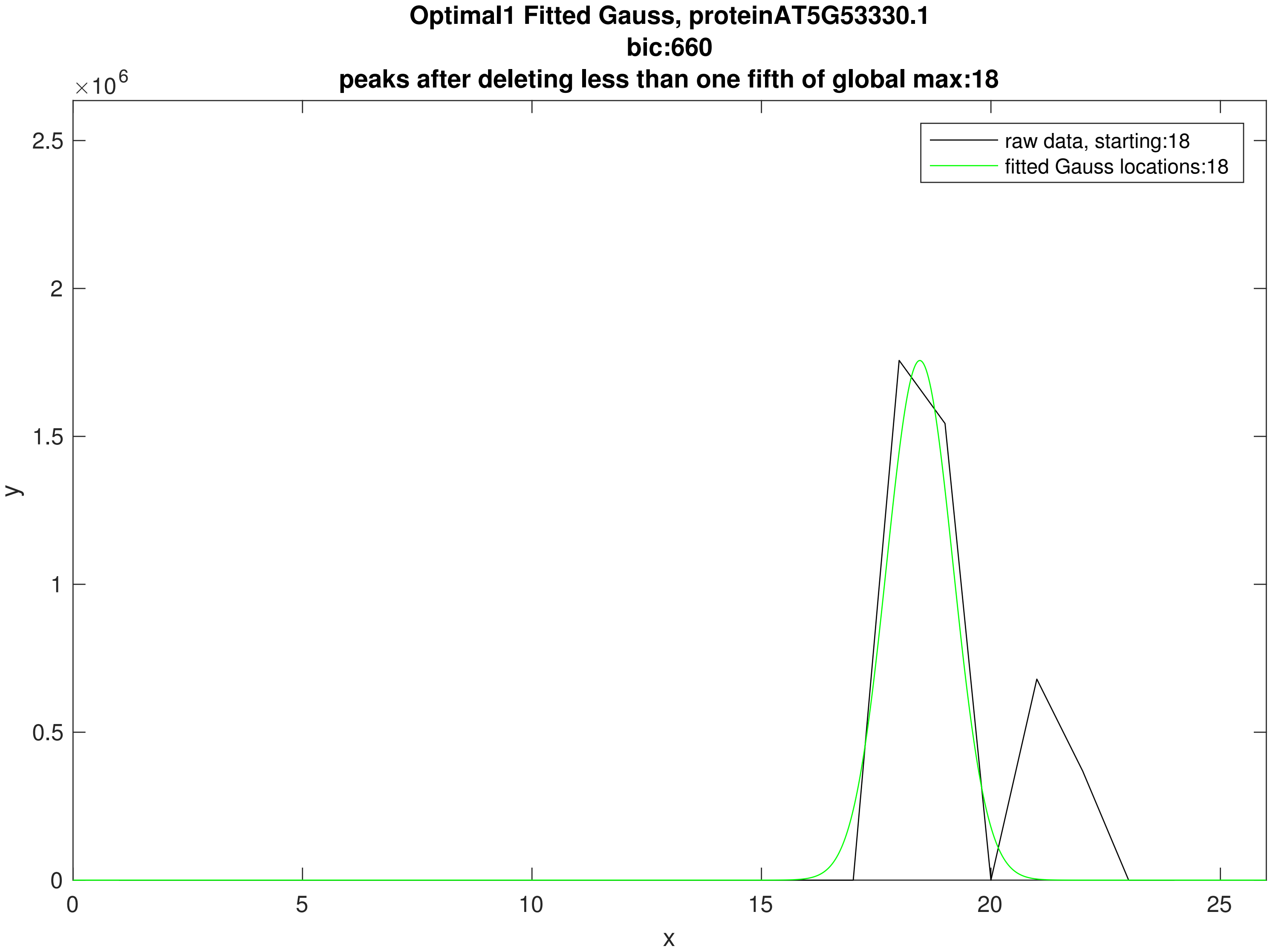
Optimal1 Fitted Gauss, proteinAT5G53330.1
bic:660
peaks after deleting less than one fifth of global max:18
×10^6
2.5
2
1.5
1
0.5
0
y
0
5
10
15
20
25
x
raw data, starting:18
fitted Gauss locations:18

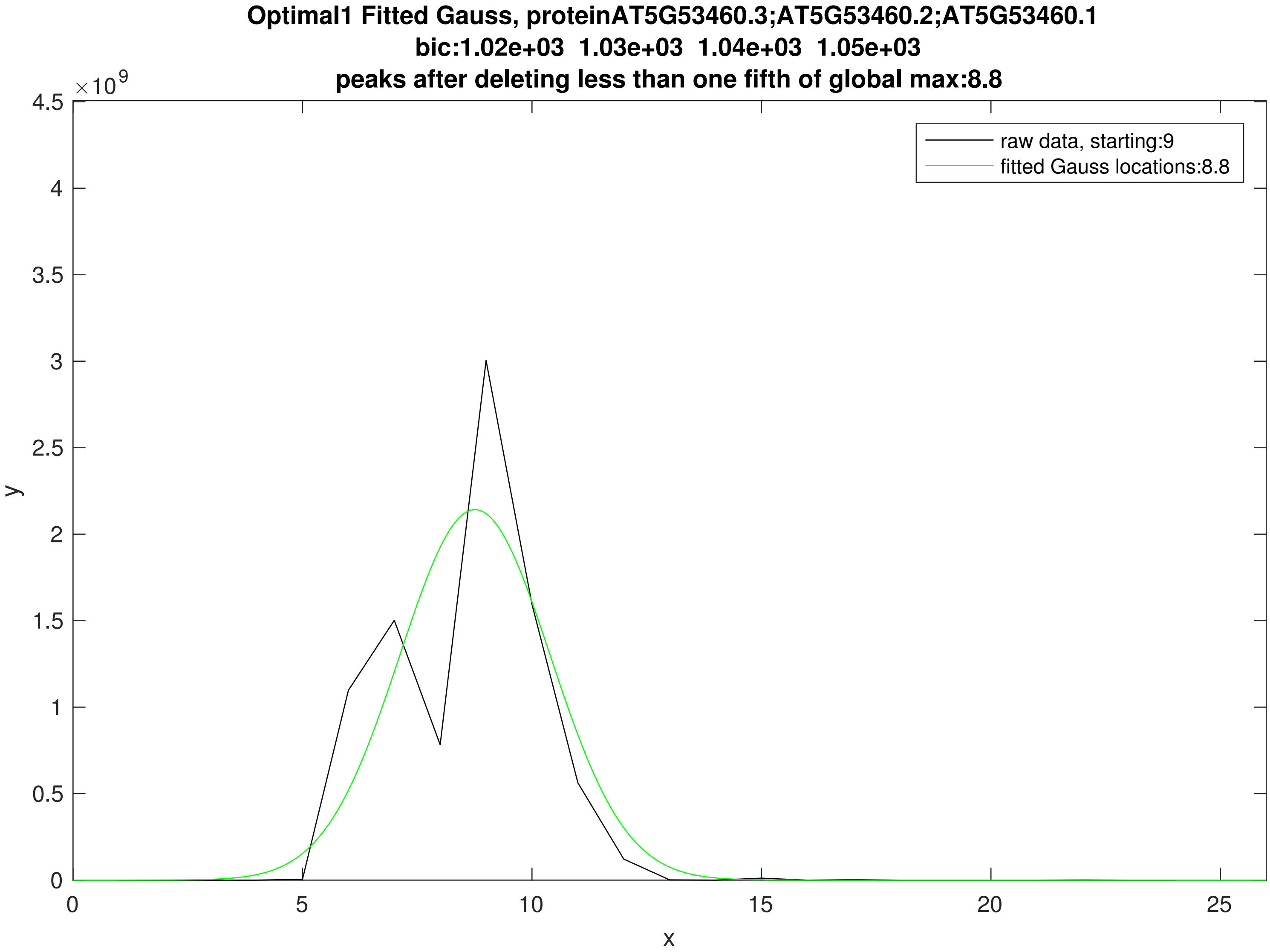
Optimal1 Fitted Gauss, proteinAT5G53460.3;AT5G53460.2;AT5G53460.1
bic:1.02e+03  1.03e+03  1.04e+03  1.05e+03
peaks after deleting less than one fifth of global max:8.8
×10^9
4.5
4
3.5
3
2.5
2
1.5
1
0.5
0
y
0
5
10
15
20
25
x
raw data, starting:9
fitted Gauss locations:8.8

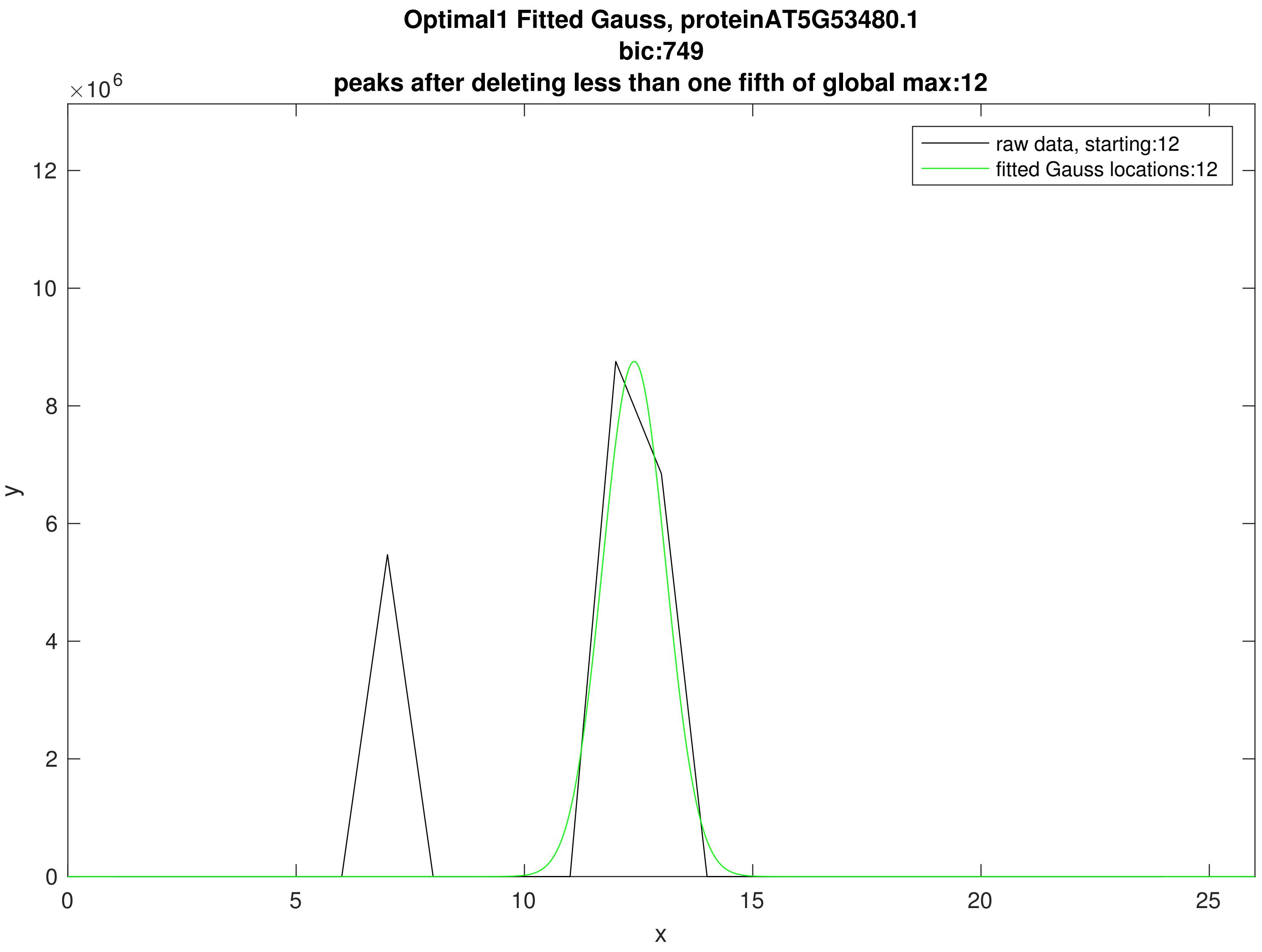

Optimal1 Fitted Gauss, proteinAT5G53480.1
bic:749
peaks after deleting less than one fifth of global max:12
×10^6
raw data, starting:12
fitted Gauss locations:12
12
10
8
6
4
2
0
y
0
5
10
15
20
25
x

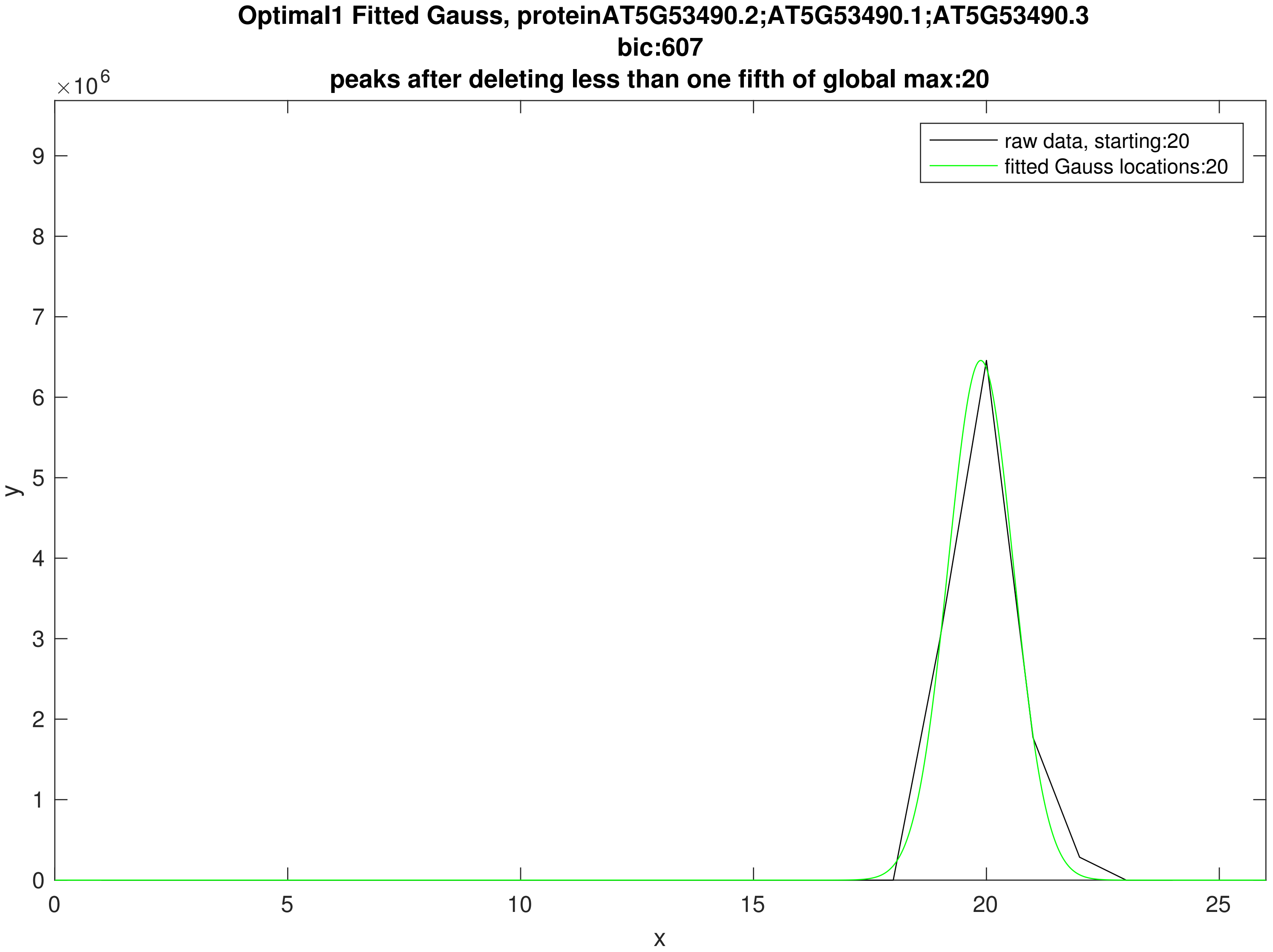

Optimal1 Fitted Gauss, proteinAT5G53490.2;AT5G53490.1;AT5G53490.3
bic:607
peaks after deleting less than one fifth of global max:20
×10^6
raw data, starting:20
fitted Gauss locations:20
y
x
0
1
2
3
4
5
6
7
8
9
0
5
10
15
20
25

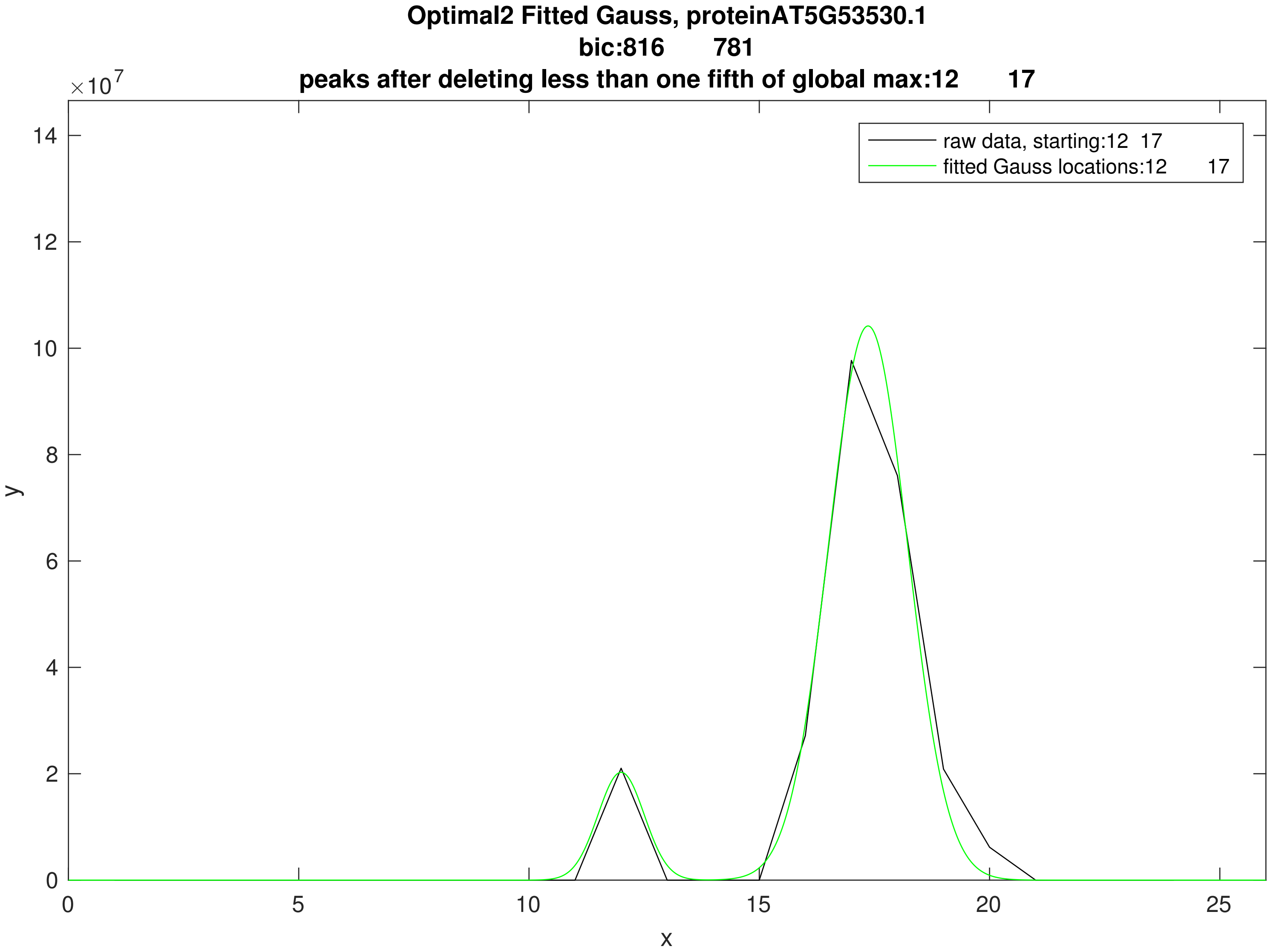
Optimal2 Fitted Gauss, proteinAT5G53530.1
bic:816 781
peaks after deleting less than one fifth of global max:12 17
raw data, starting:12 17
fitted Gauss locations:12 17
$\times 10^7$
y
x
0
2
4
6
8
10
12
14
0
5
10
15
20
25

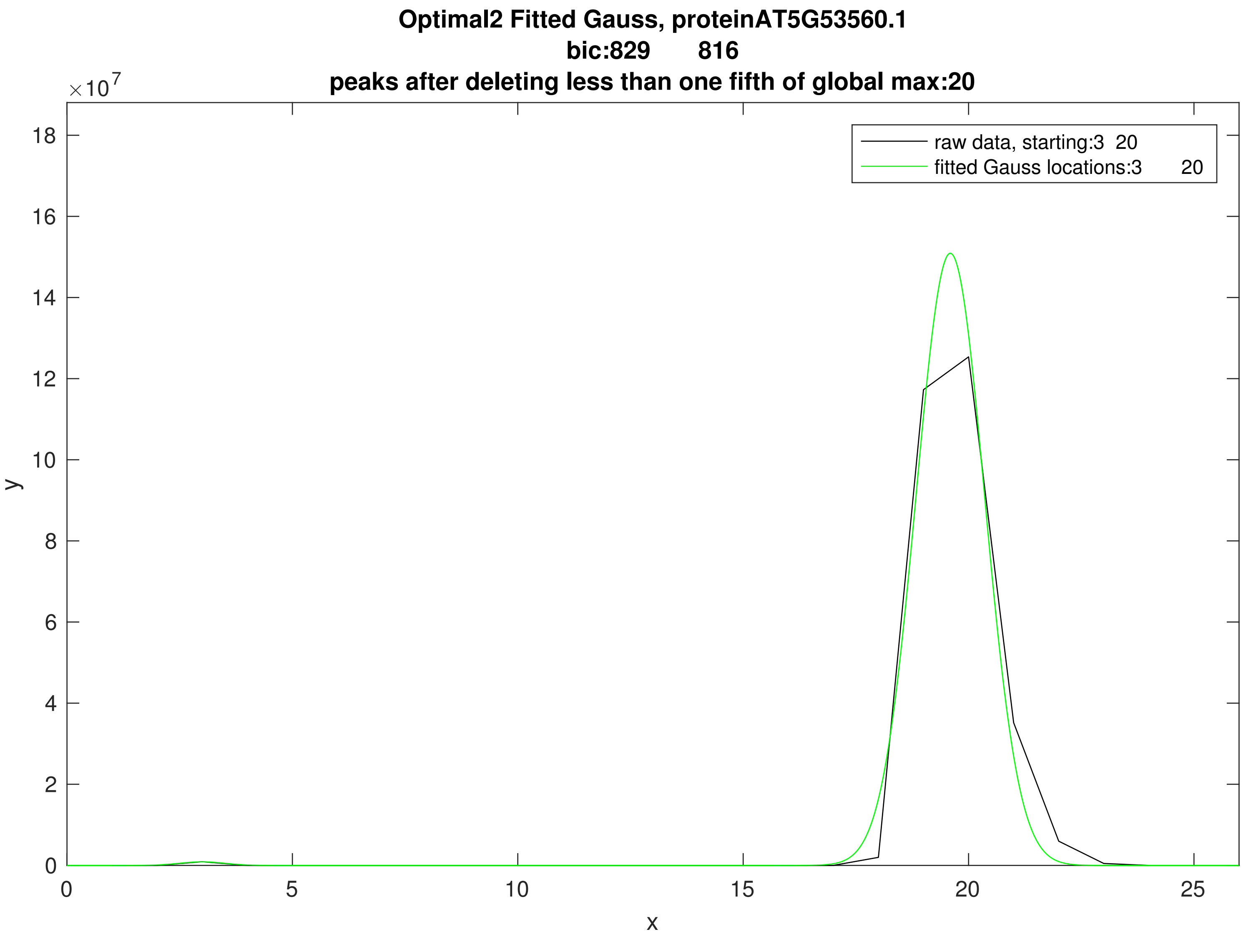
Optimal2 Fitted Gauss, proteinAT5G53560.1
bic:829 816
peaks after deleting less than one fifth of global max:20
$\times 10^7$
18
16
14
12
10
8
6
4
2
0
y
0
5
10
15
20
25
x
raw data, starting:3 20
fitted Gauss locations:3 20

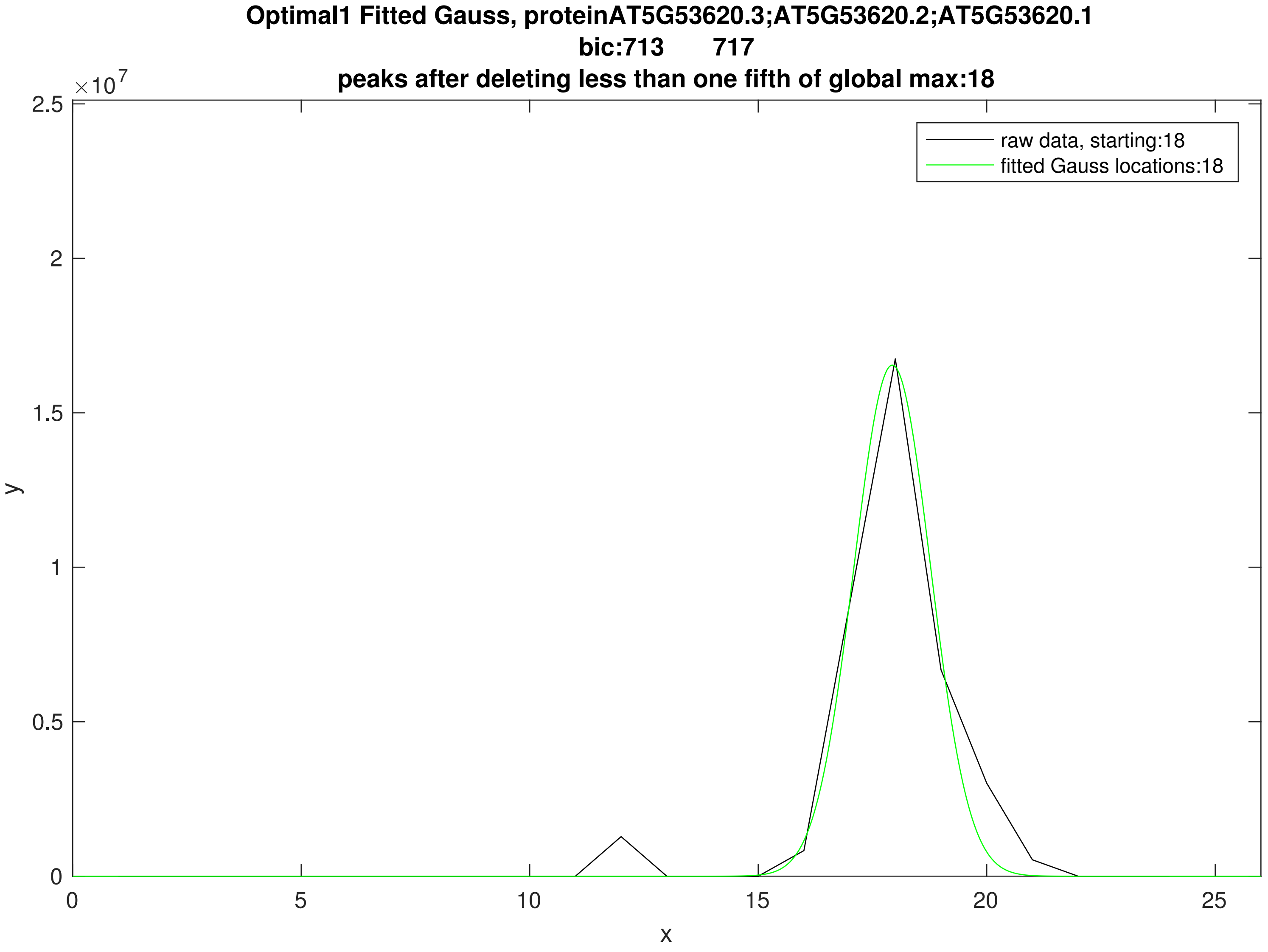
Optimal1 Fitted Gauss, proteinAT5G53620.3;AT5G53620.2;AT5G53620.1
bic:713 717
peaks after deleting less than one fifth of global max:18
×10^7
2.5
2
1.5
1
0.5
0
y
0
5
10
15
20
25
x
raw data, starting:18
fitted Gauss locations:18

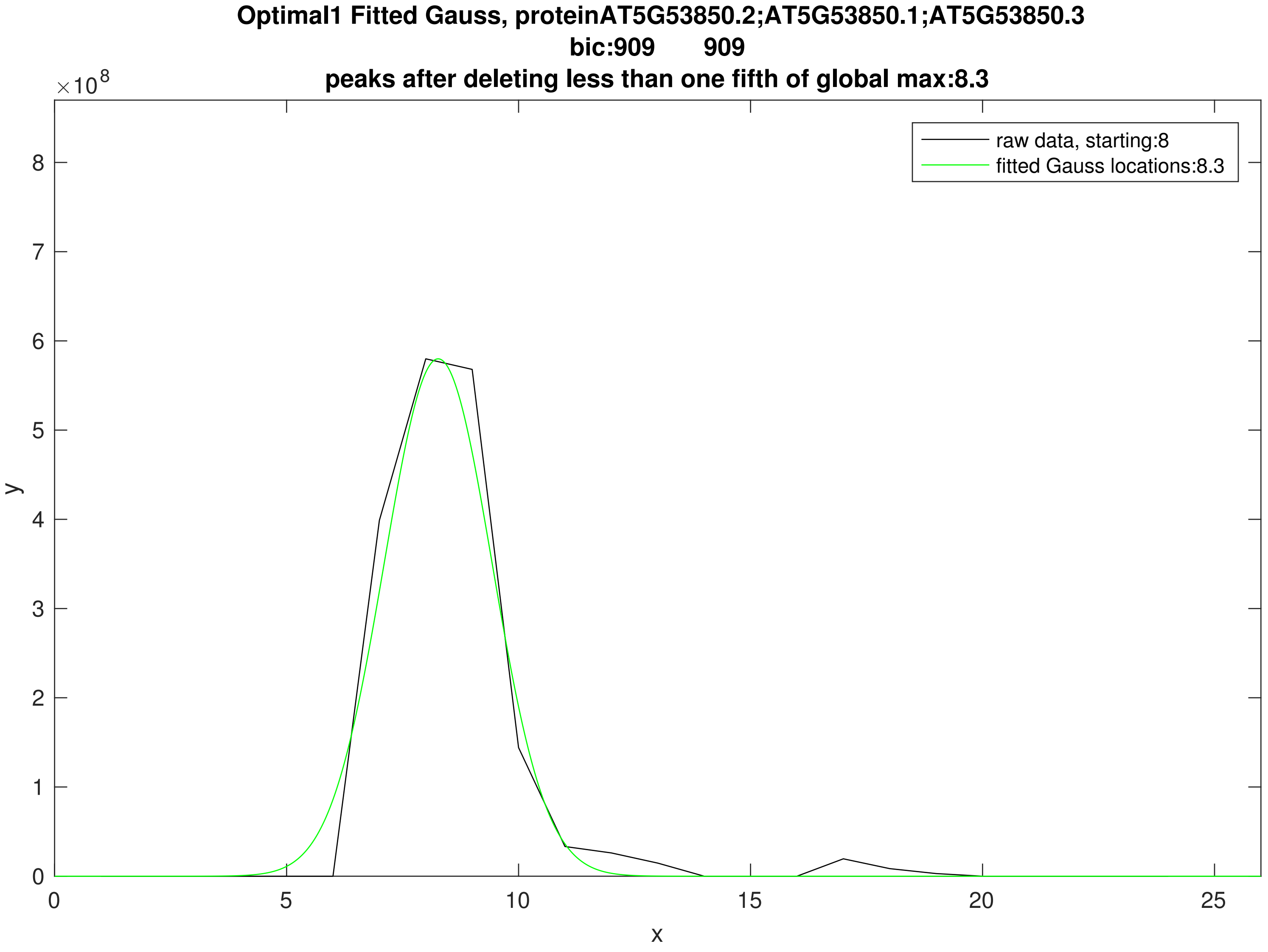

Optimal1 Fitted Gauss, proteinAT5G53850.2;AT5G53850.1;AT5G53850.3
bic:909 909
peaks after deleting less than one fifth of global max:8.3
×10^8
raw data, starting:8
fitted Gauss locations:8.3
y
x
0
1
2
3
4
5
6
7
8
0
5
10
15
20
25

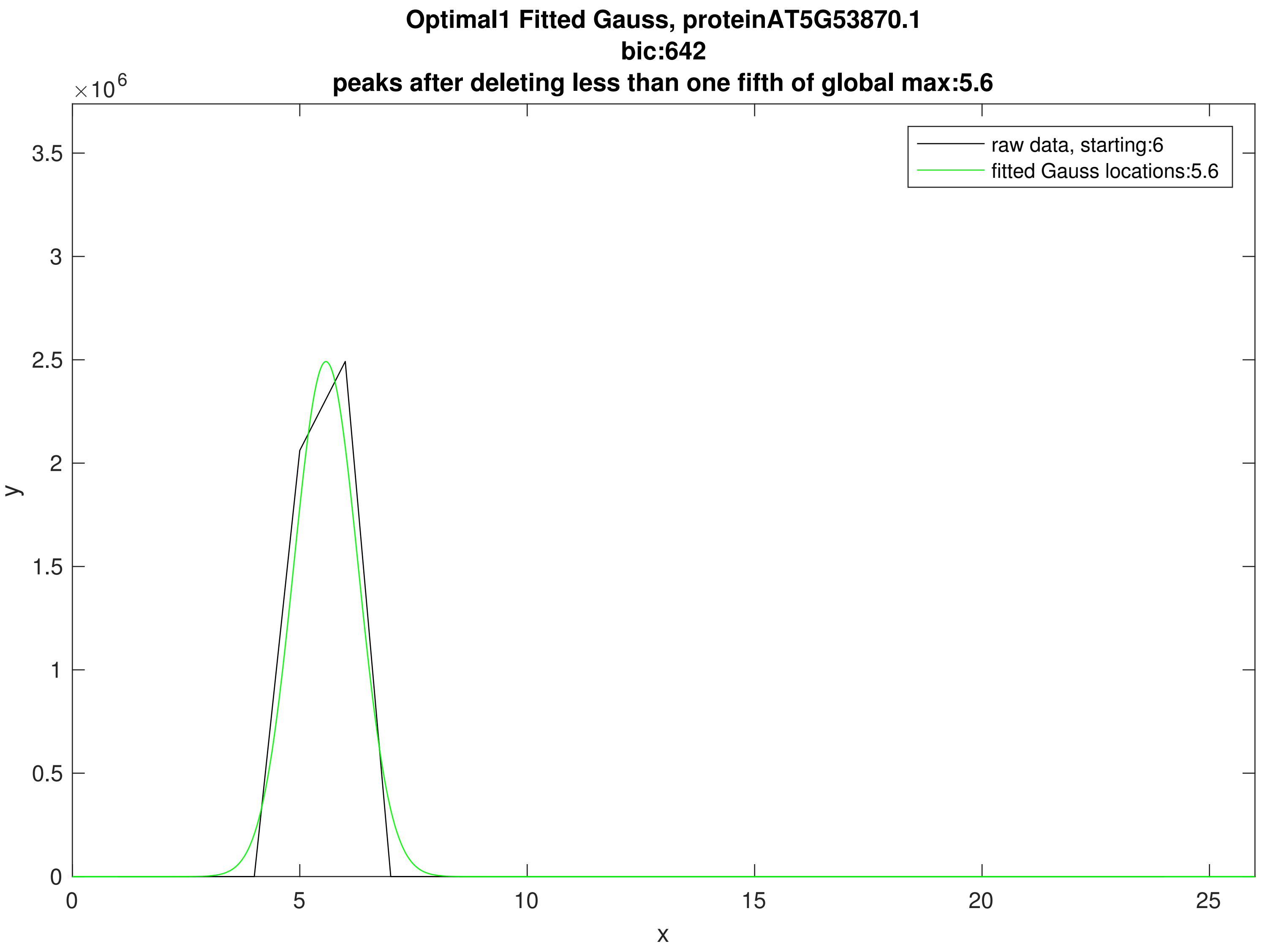
Optimal1 Fitted Gauss, proteinAT5G53870.1
bic:642
peaks after deleting less than one fifth of global max:5.6
×10^6
raw data, starting:6
fitted Gauss locations:5.6
y
x
0
0.5
1
1.5
2
2.5
3
3.5
0
5
10
15
20
25

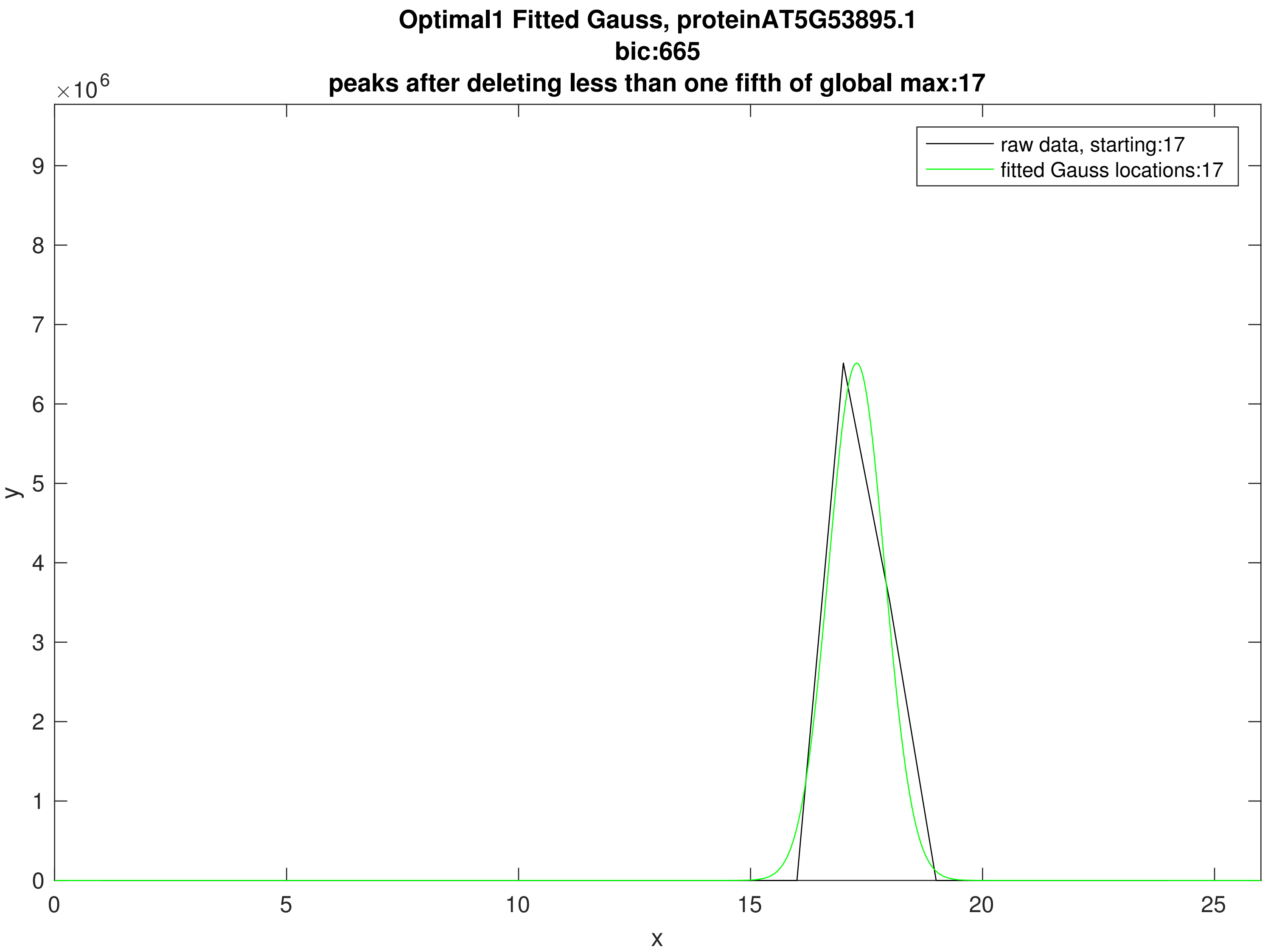

Optimal1 Fitted Gauss, proteinAT5G53895.1
bic:665
peaks after deleting less than one fifth of global max:17
×10^6
raw data, starting:17
fitted Gauss locations:17
y
x
0
1
2
3
4
5
6
7
8
9
0
5
10
15
20
25

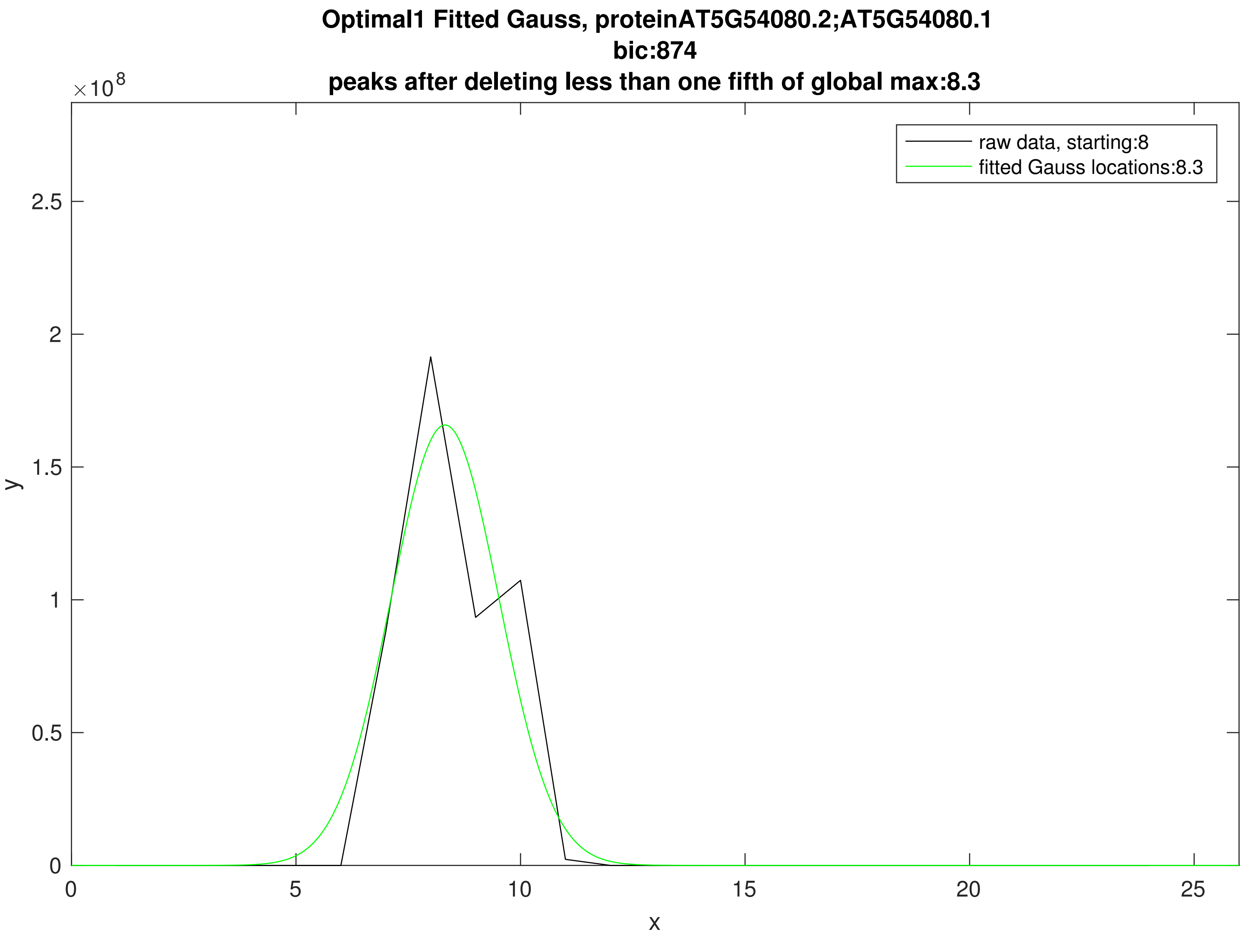
Optimal1 Fitted Gauss, proteinAT5G54080.2;AT5G54080.1
bic:874
peaks after deleting less than one fifth of global max:8.3
raw data, starting:8
fitted Gauss locations:8.3
y
x

**Optimal1 Fitted Gauss, proteinAT5G54160.1**
**bic:994 1e+03 1.01e+03**
**peaks after deleting less than one fifth of global max:14**

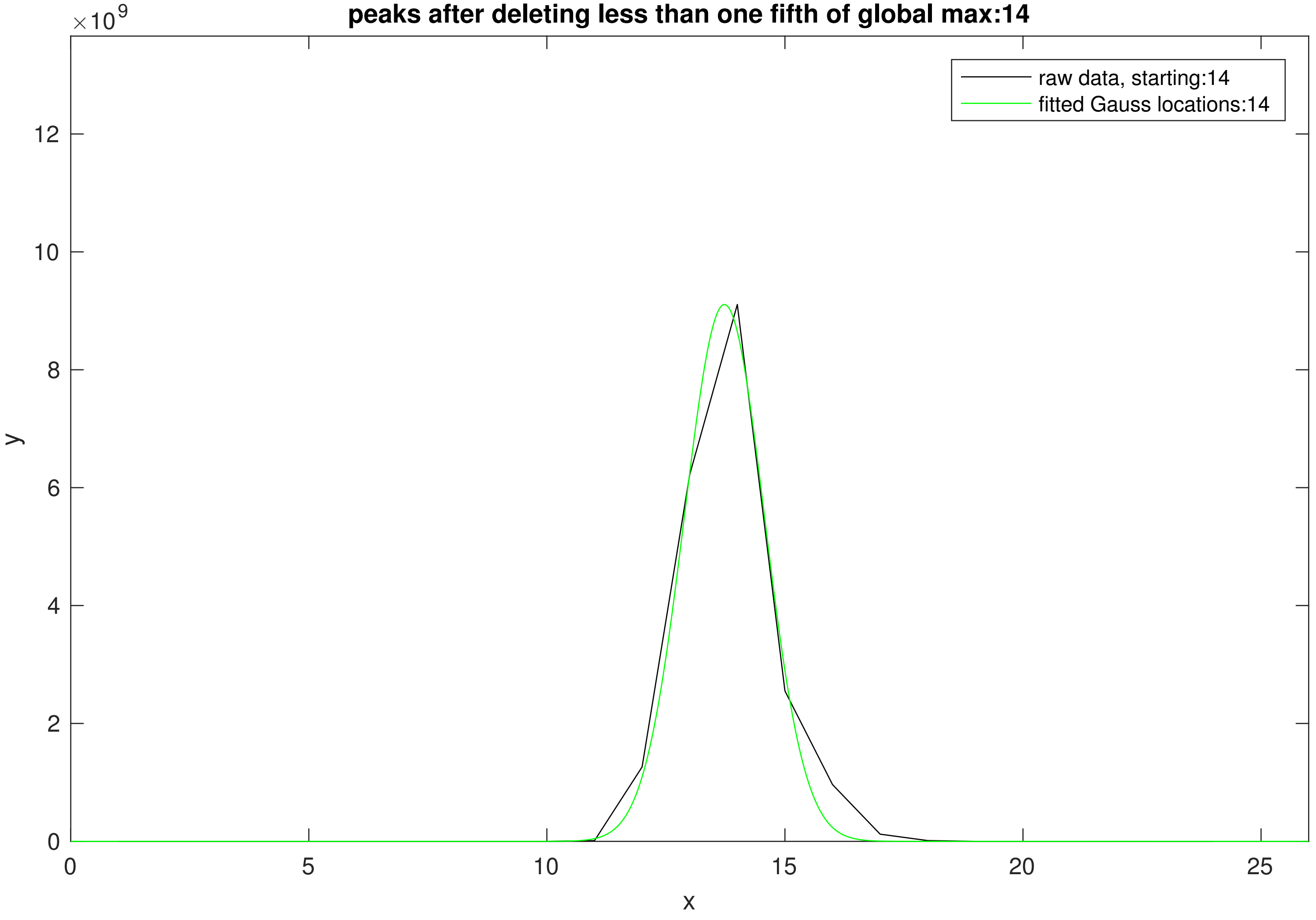

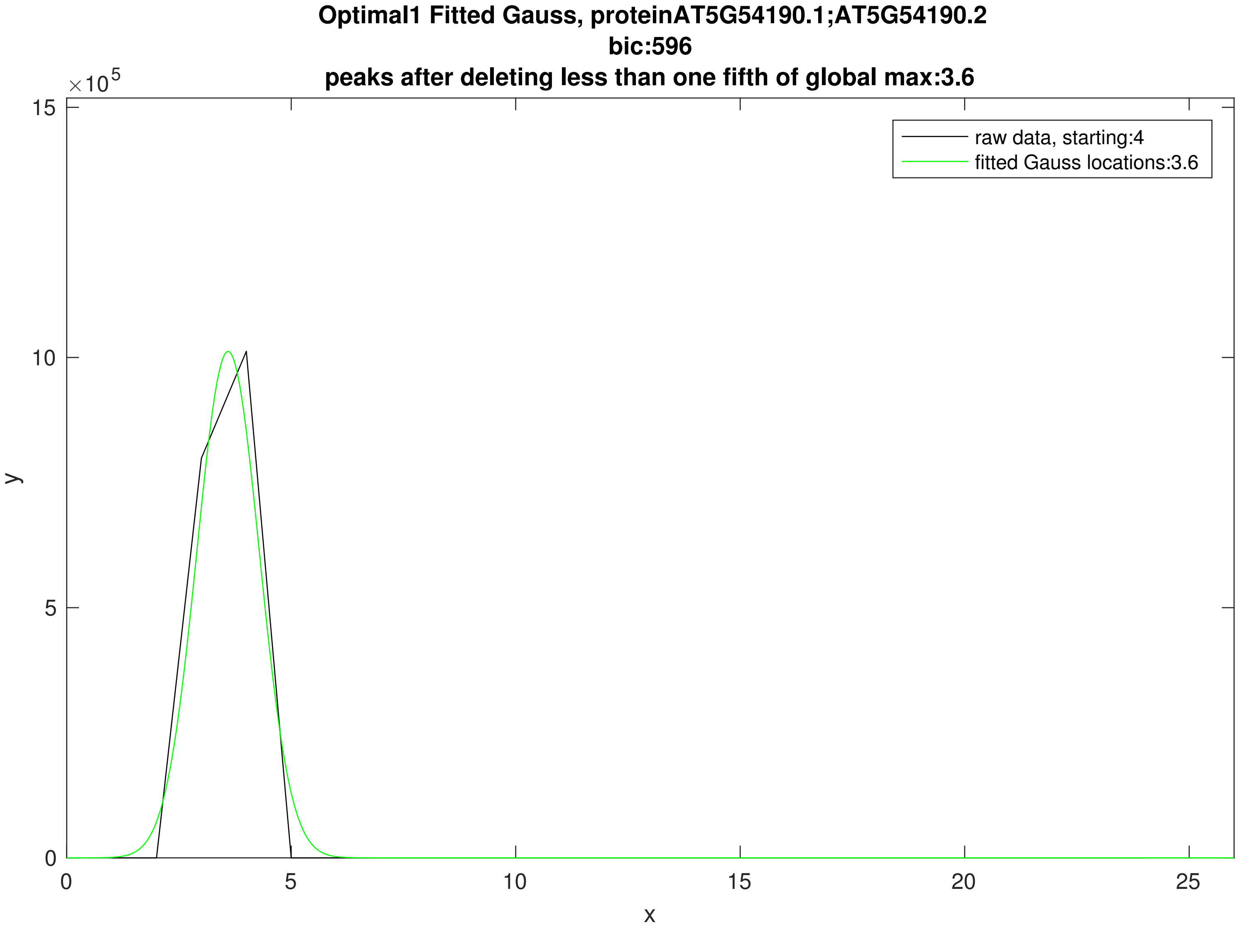
Optimal1 Fitted Gauss, proteinAT5G54190.1;AT5G54190.2
bic:596
peaks after deleting less than one fifth of global max:3.6
×10^5
15
10
5
0
y
0
5
10
15
20
25
x
raw data, starting:4
fitted Gauss locations:3.6

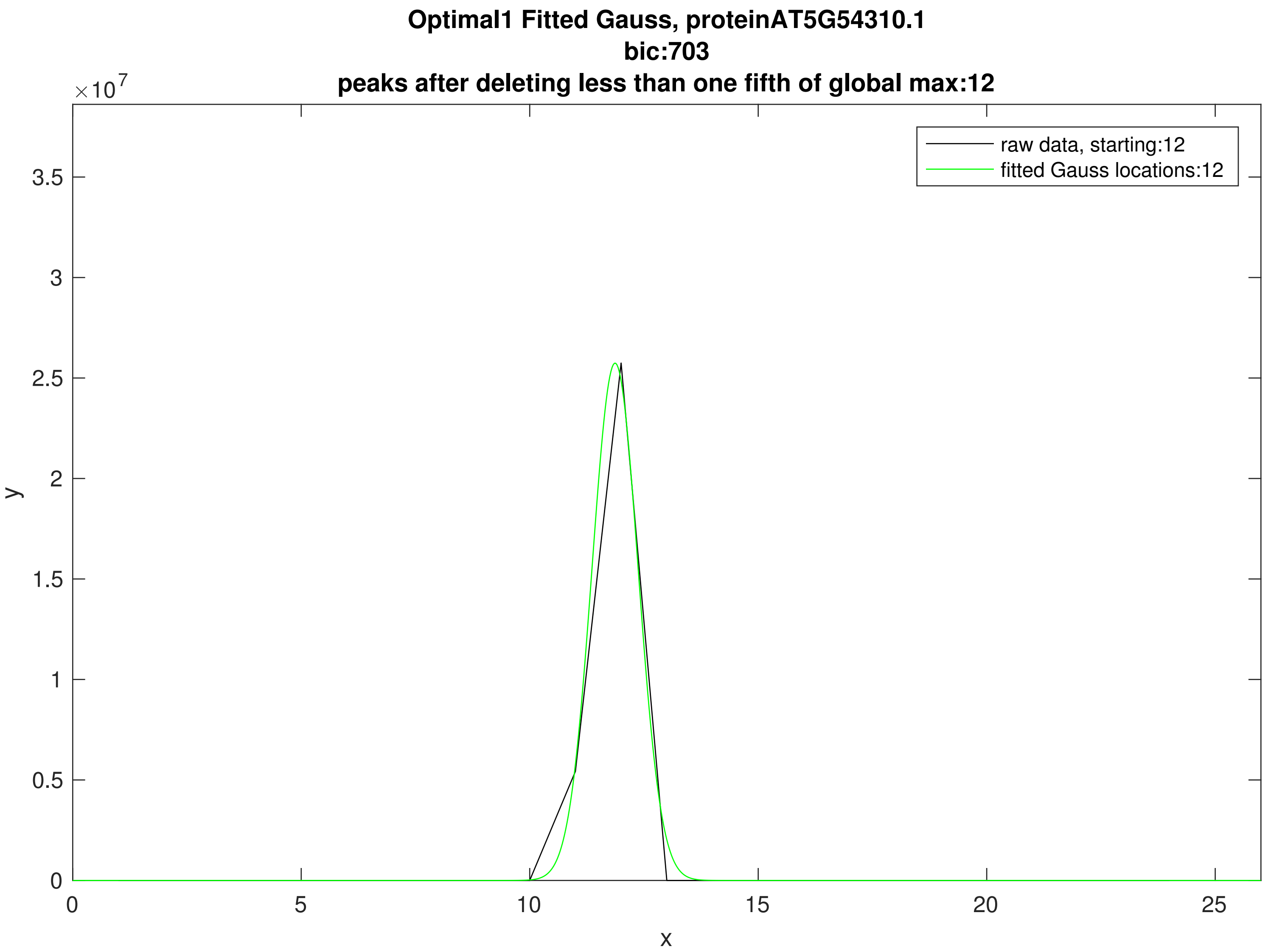
Optimal1 Fitted Gauss, proteinAT5G54310.1
bic:703
peaks after deleting less than one fifth of global max:12
×10^7
raw data, starting:12
fitted Gauss locations:12
3.5
3
2.5
2
1.5
1
0.5
0
y
0
5
10
15
20
25
x

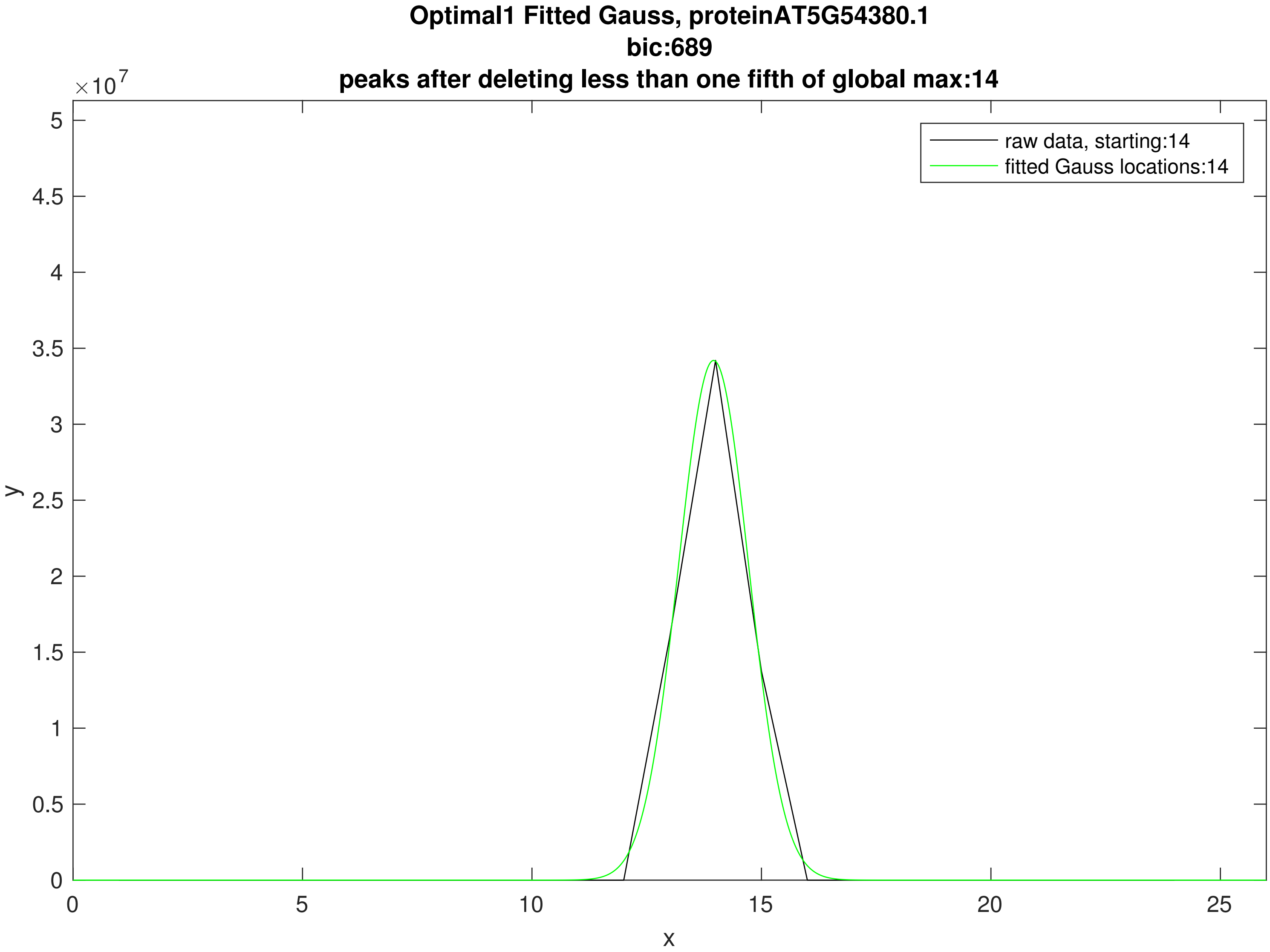
Optimal1 Fitted Gauss, proteinAT5G54380.1
bic:689
peaks after deleting less than one fifth of global max:14
×10^7
raw data, starting:14
fitted Gauss locations:14
5
4.5
4
3.5
3
2.5
2
1.5
1
0.5
0
y
0
5
10
15
20
25
x

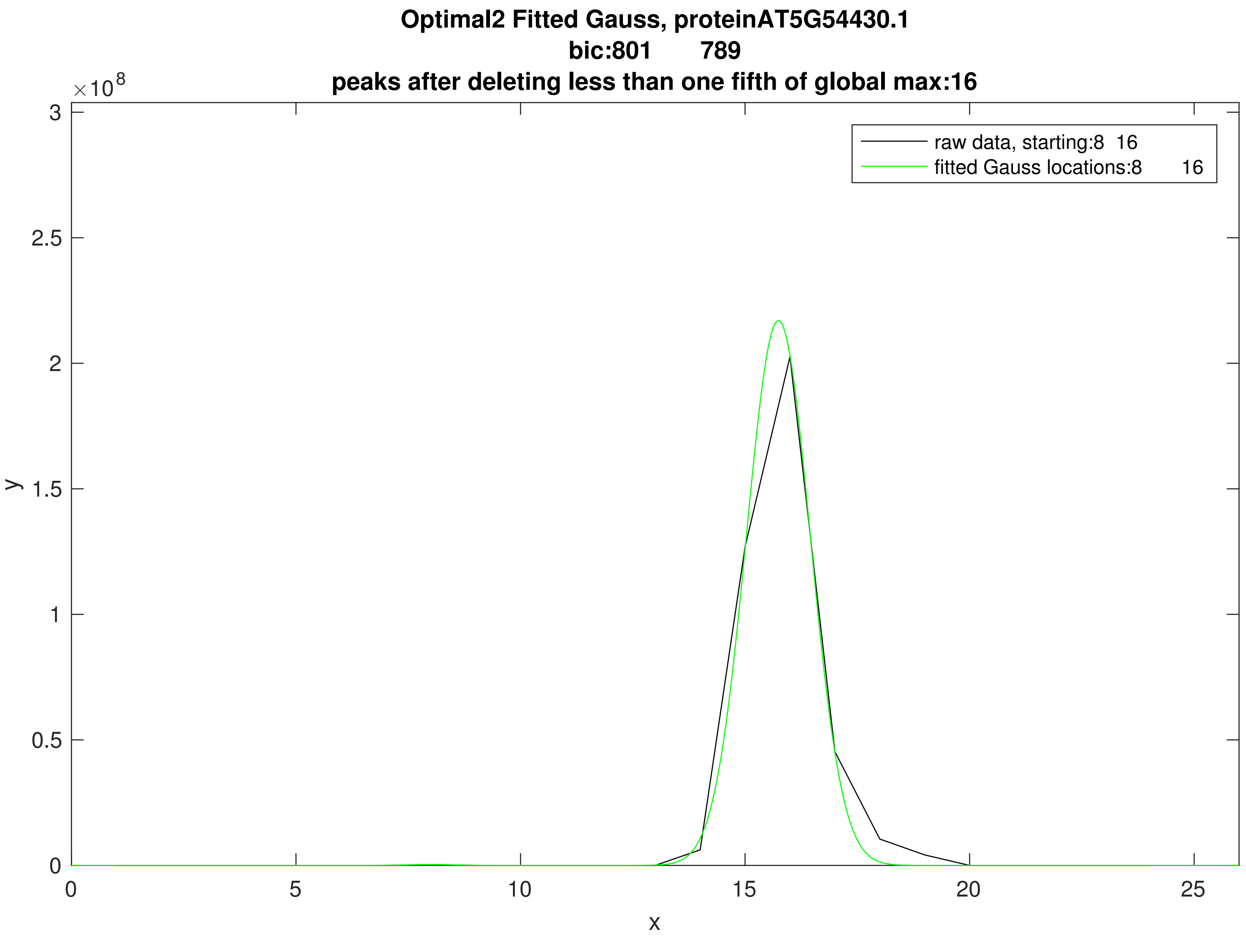

Optimal2 Fitted Gauss, proteinAT5G54430.1
bic:801 789
peaks after deleting less than one fifth of global max:16
×10^8
3
2.5
2
1.5
1
0.5
0
y
0
5
10
15
20
25
x
raw data, starting:8 16
fitted Gauss locations:8 16

**Optimal1 Fitted Gauss, proteinAT5G54500.1;AT5G54500.2**
**bic:976 985 994**
**peaks after deleting less than one fifth of global max:14**

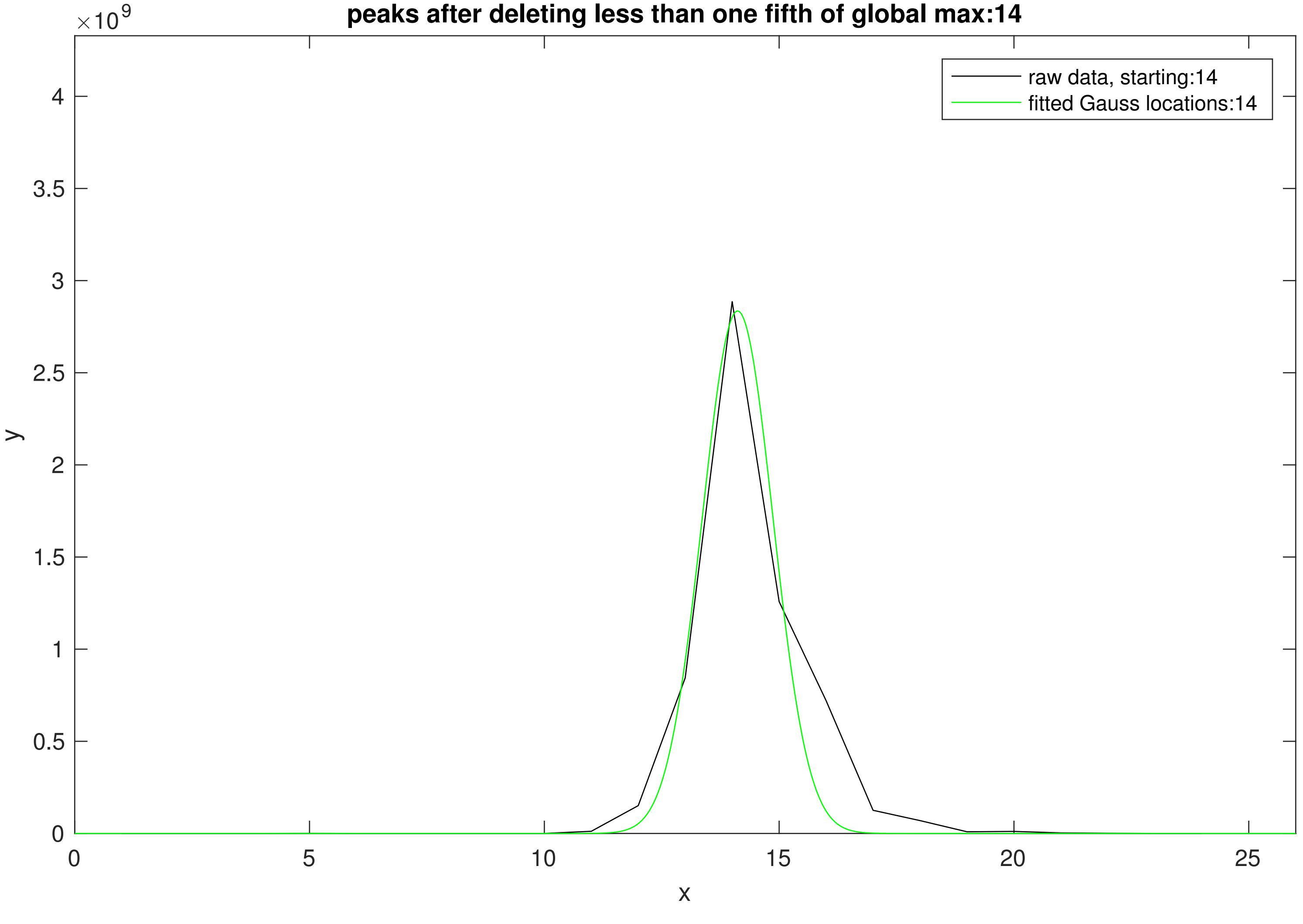

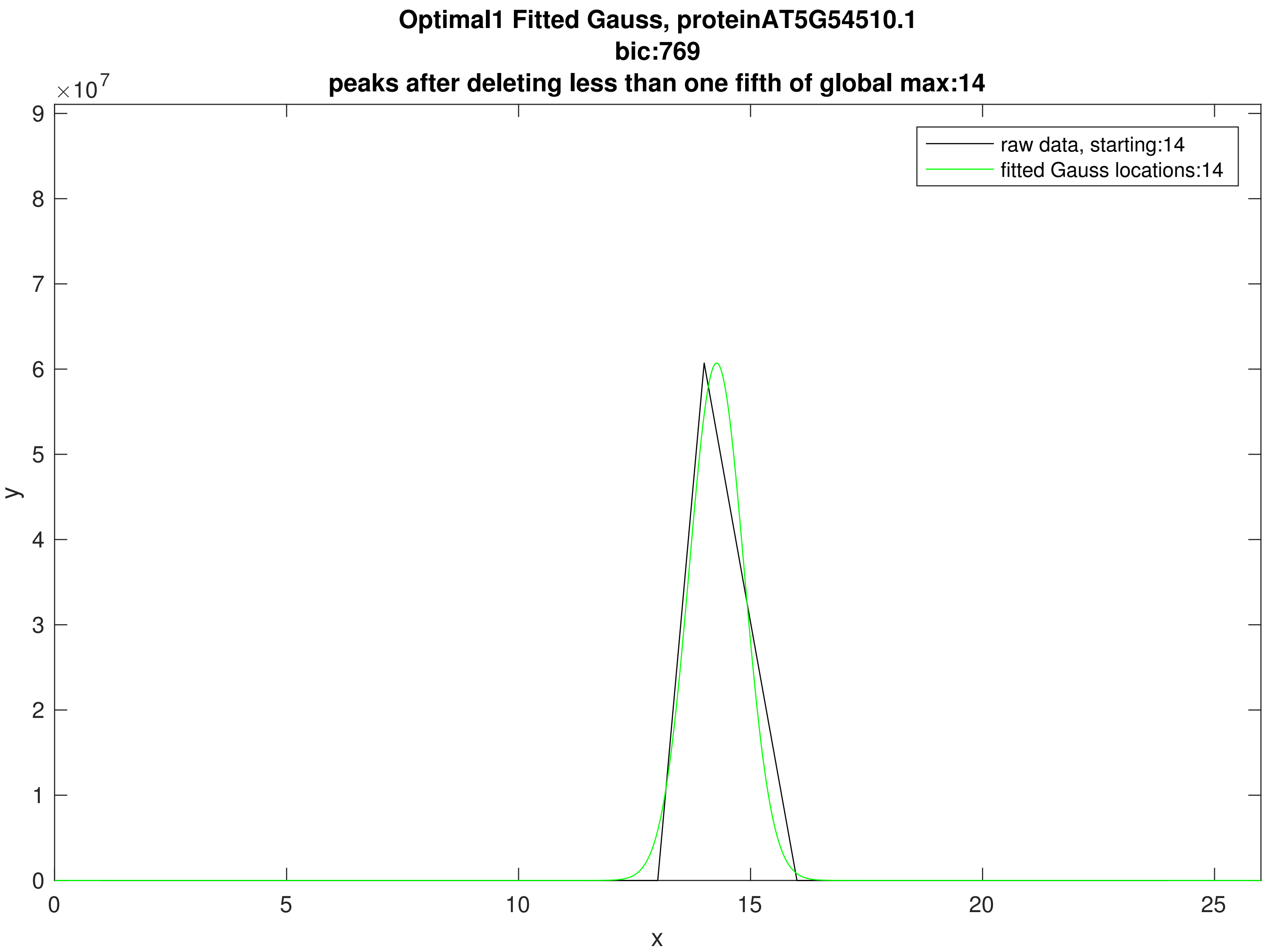
Optimal1 Fitted Gauss, proteinAT5G54510.1
bic:769
peaks after deleting less than one fifth of global max:14
×10^7
9
8
7
6
5
4
3
2
1
0
y
0
5
10
15
20
25
x
raw data, starting:14
fitted Gauss locations:14

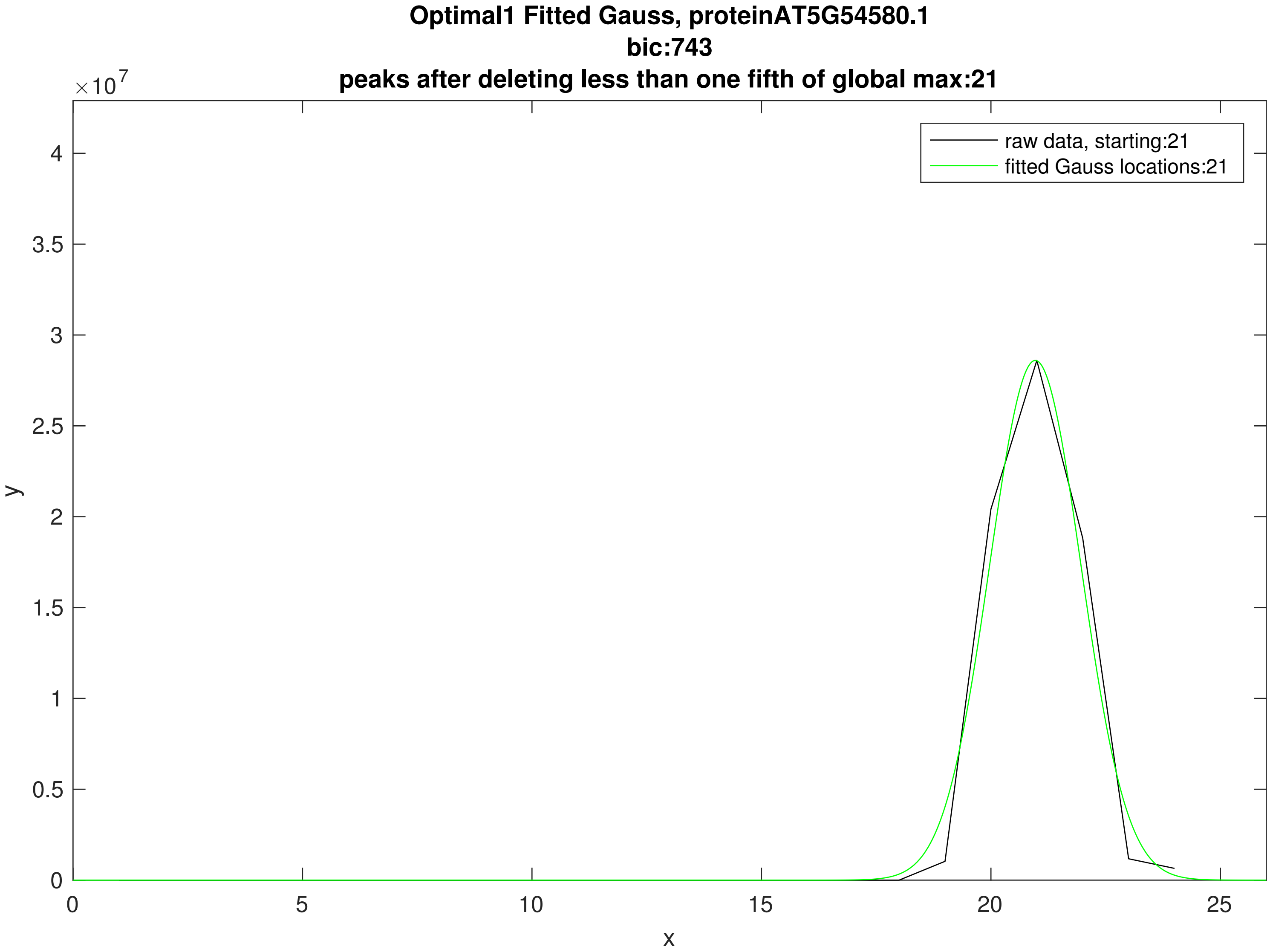

Optimal1 Fitted Gauss, proteinAT5G54580.1
bic:743
peaks after deleting less than one fifth of global max:21
$\times 10^7$
raw data, starting:21
fitted Gauss locations:21
4
3.5
3
2.5
2
1.5
1
0.5
0
y
0
5
10
15
20
25
x

Optimal1 Fitted Gauss, proteinAT5G54600.2;AT5G54600.1
bic:591
peaks after deleting less than one fifth of global max:3.6

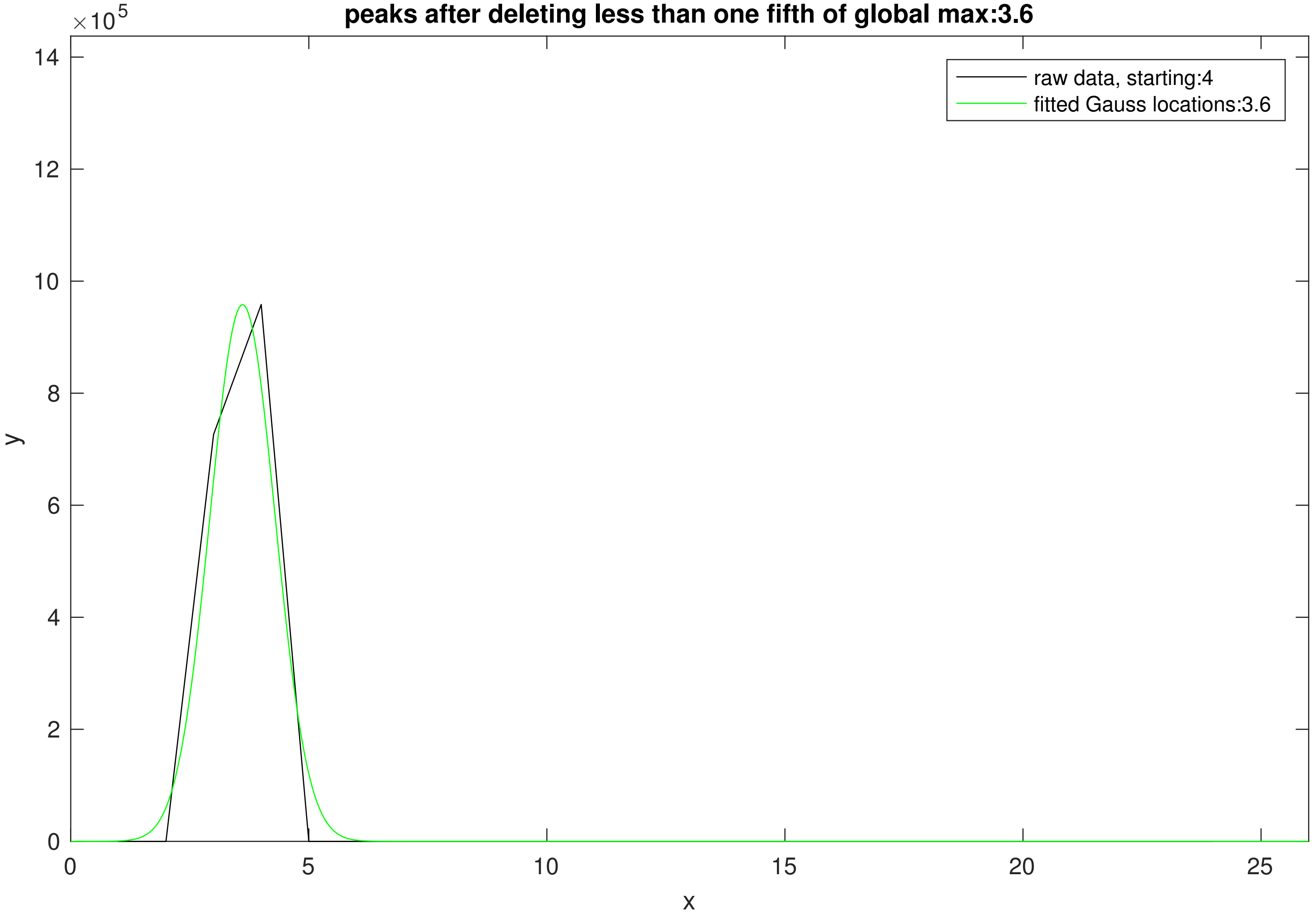

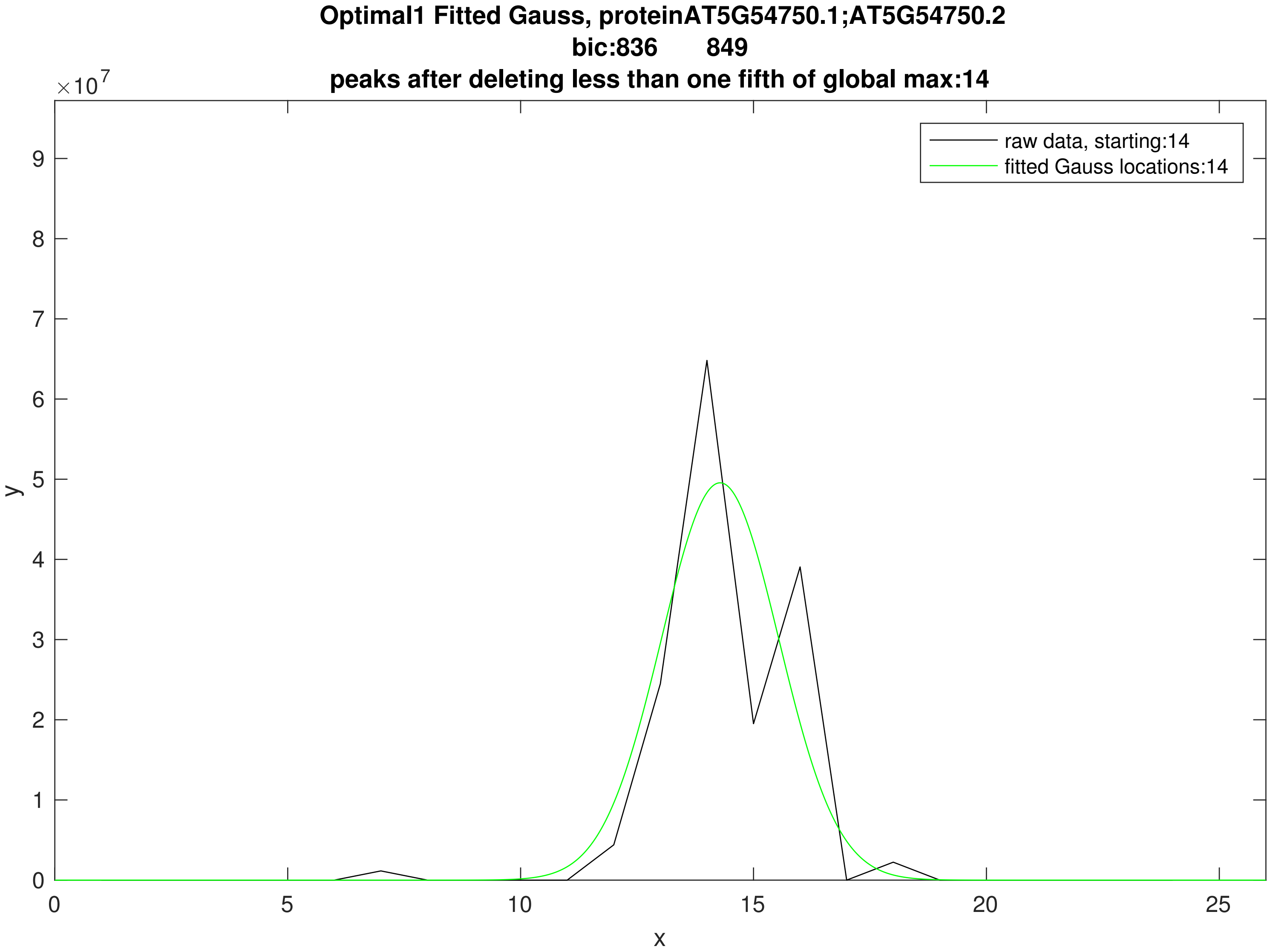

Optimal1 Fitted Gauss, proteinAT5G54750.1;AT5G54750.2
bic:836 849
peaks after deleting less than one fifth of global max:14
×10^7
raw data, starting:14
fitted Gauss locations:14
y
x

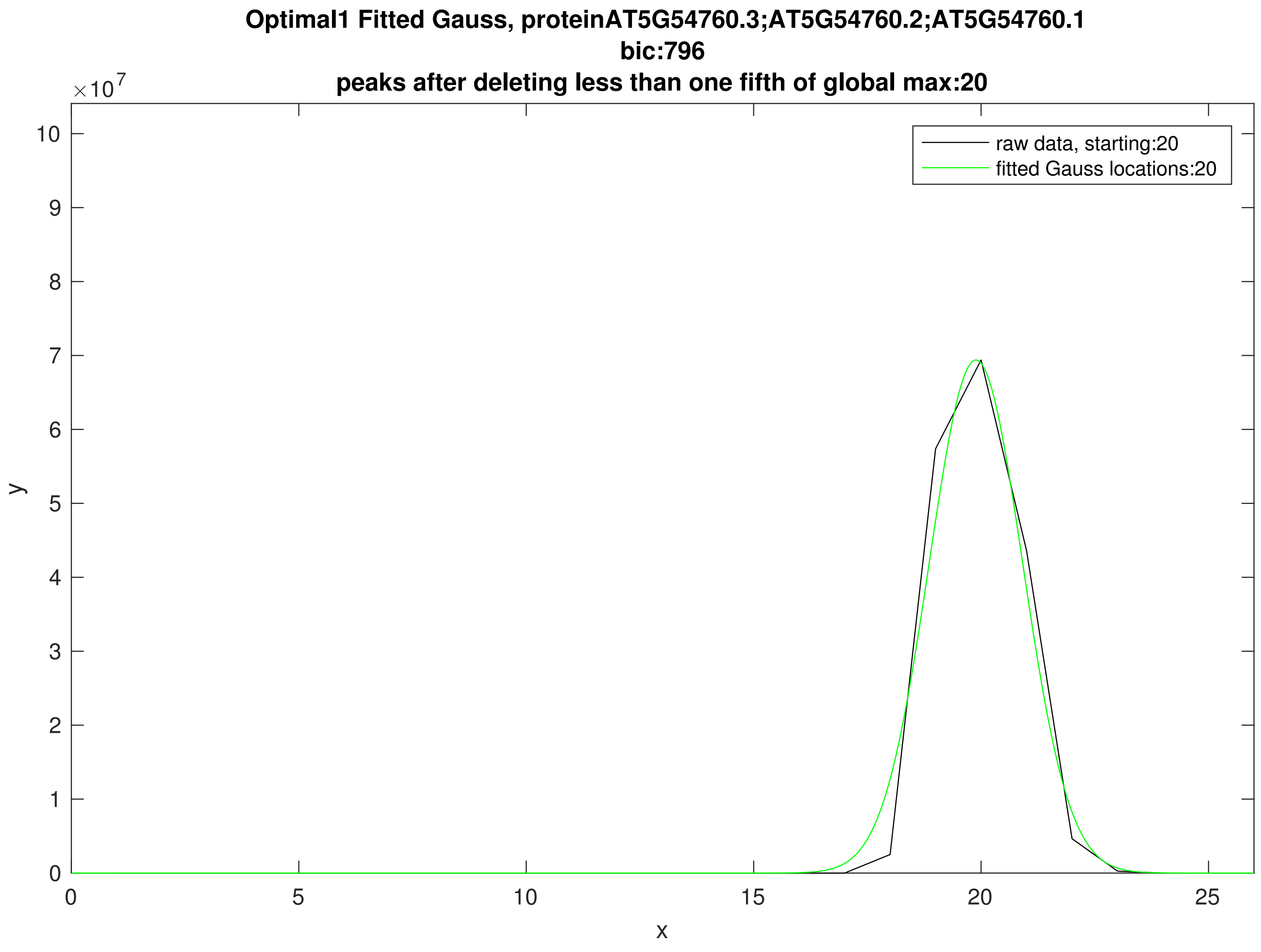
Optimal1 Fitted Gauss, proteinAT5G54760.3;AT5G54760.2;AT5G54760.1
bic:796
peaks after deleting less than one fifth of global max:20
×10^7
raw data, starting:20
fitted Gauss locations:20
y
x

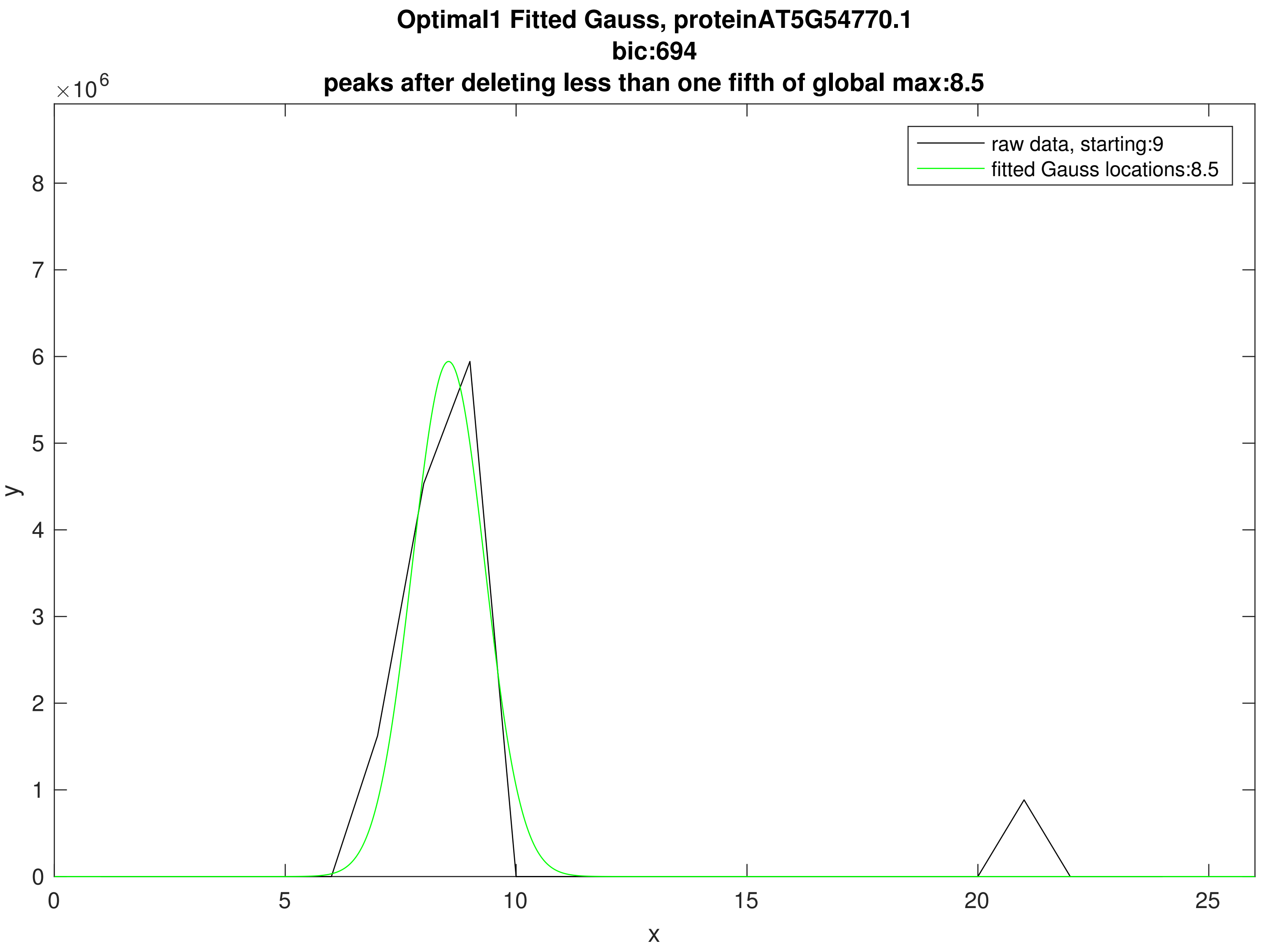

Optimal1 Fitted Gauss, proteinAT5G54770.1
bic:694
peaks after deleting less than one fifth of global max:8.5
×10^6
raw data, starting:9
fitted Gauss locations:8.5
y
x
0
1
2
3
4
5
6
7
8
0
5
10
15
20
25

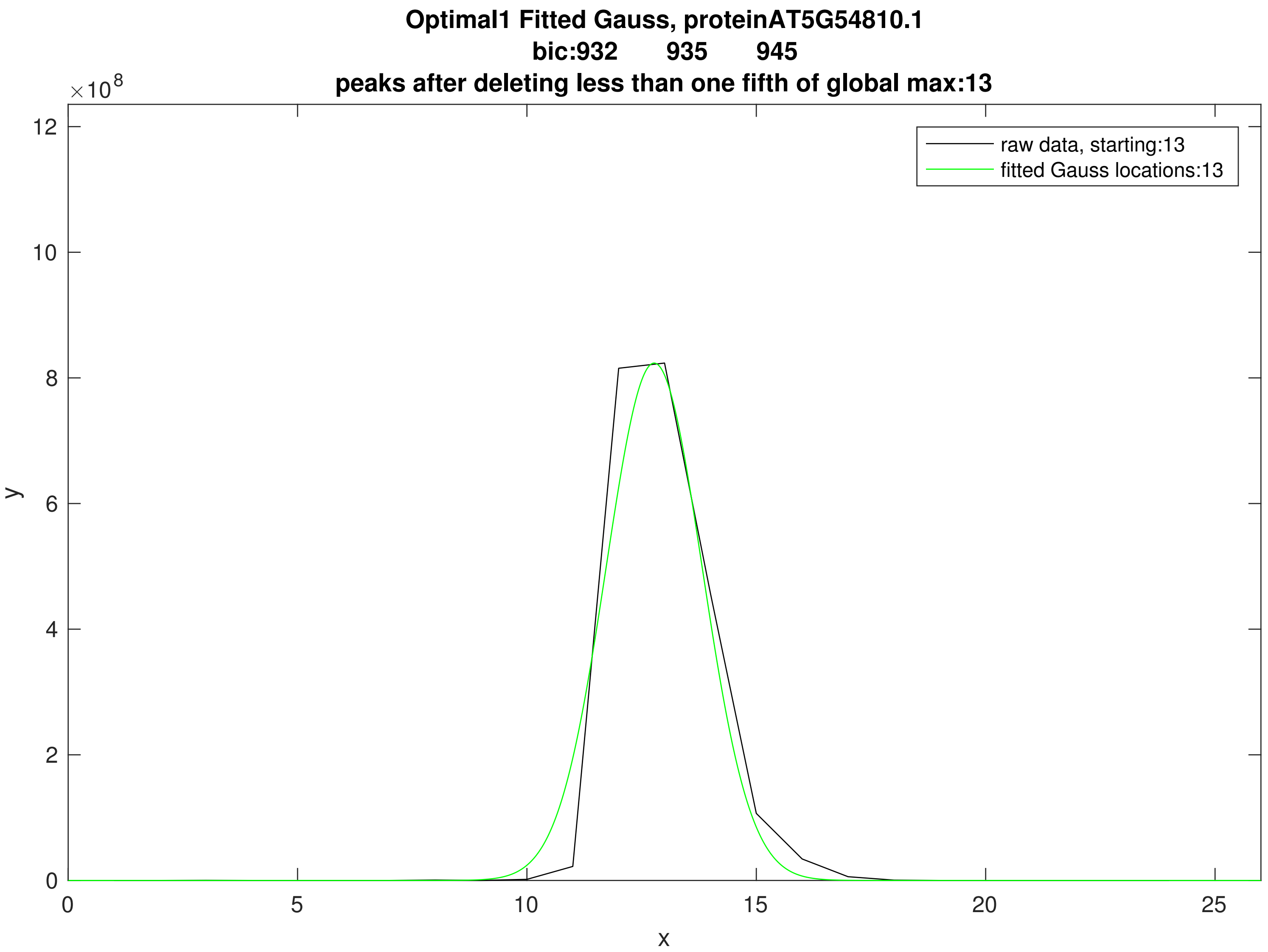
Optimal1 Fitted Gauss, proteinAT5G54810.1
bic:932 935 945
peaks after deleting less than one fifth of global max:13
×10^8
12
10
8
6
4
2
0
y
0
5
10
15
20
25
x
raw data, starting:13
fitted Gauss locations:13

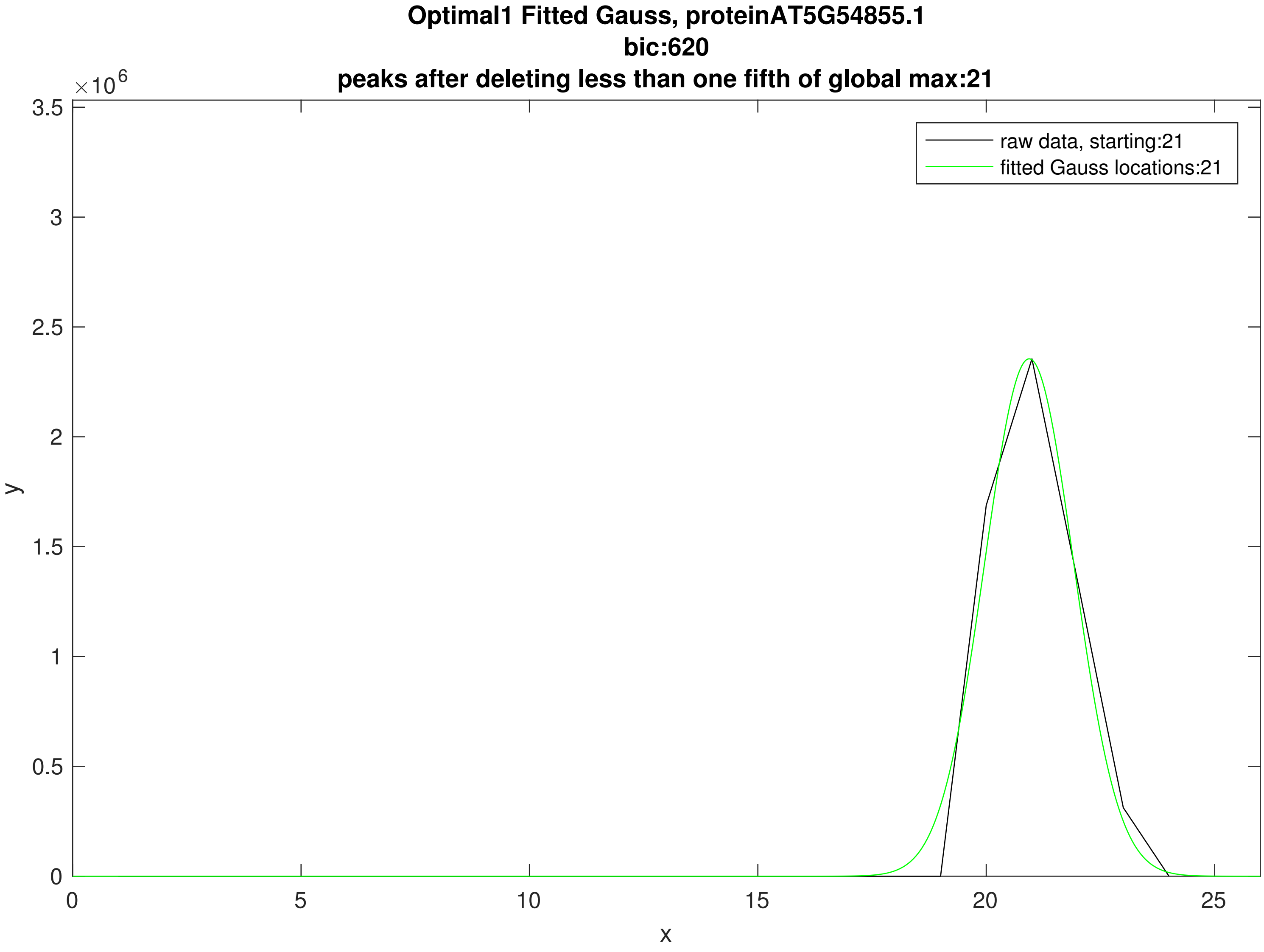
Optimal1 Fitted Gauss, proteinAT5G54855.1
bic:620
peaks after deleting less than one fifth of global max:21
$\times 10^6$
3.5
3
2.5
2
1.5
1
0.5
0
y
0
5
10
15
20
25
x
raw data, starting:21
fitted Gauss locations:21

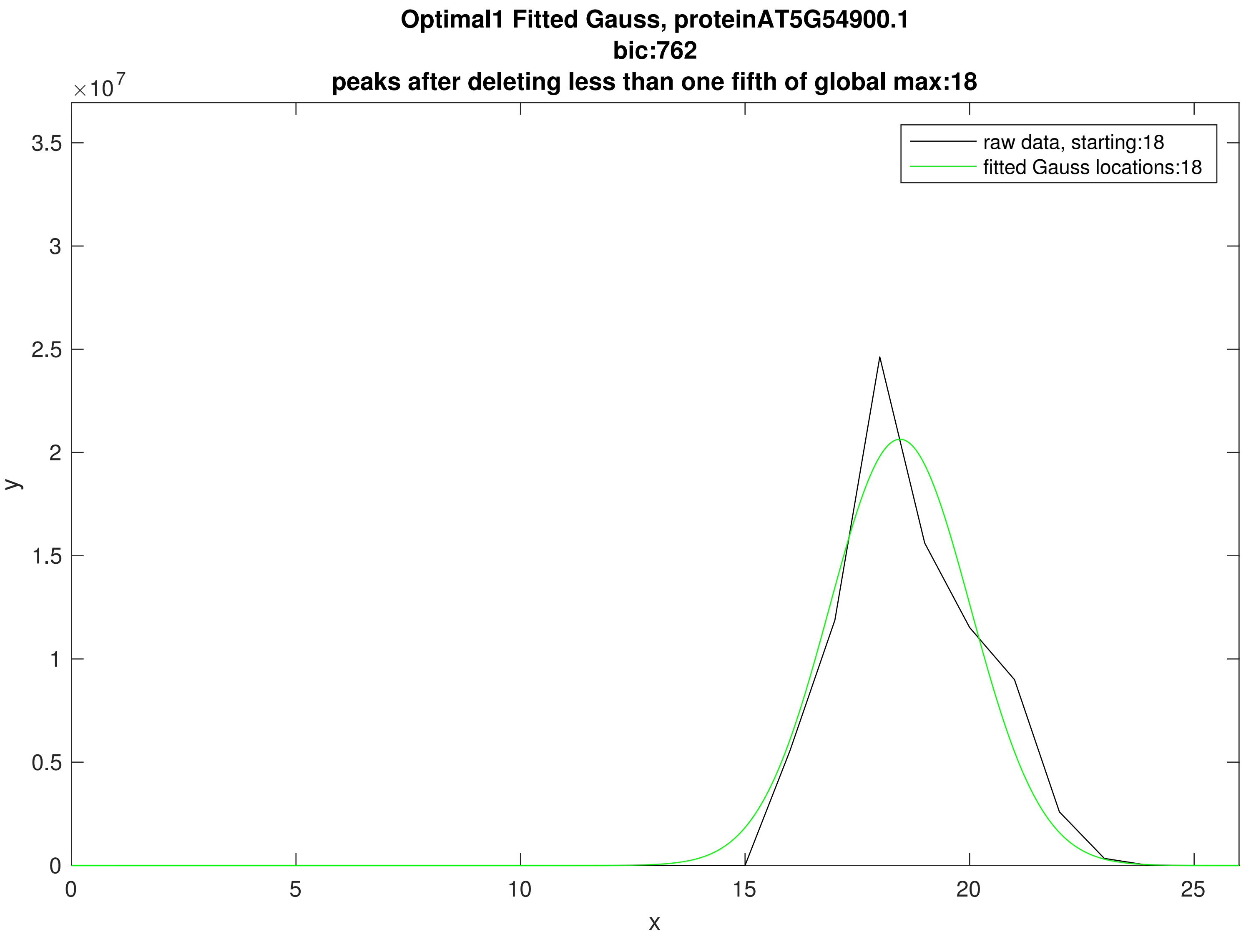

Optimal1 Fitted Gauss, proteinAT5G54900.1
bic:762
peaks after deleting less than one fifth of global max:18
×10^7
3.5
3
2.5
2
1.5
1
0.5
0
y
0
5
10
15
20
25
x
raw data, starting:18
fitted Gauss locations:18

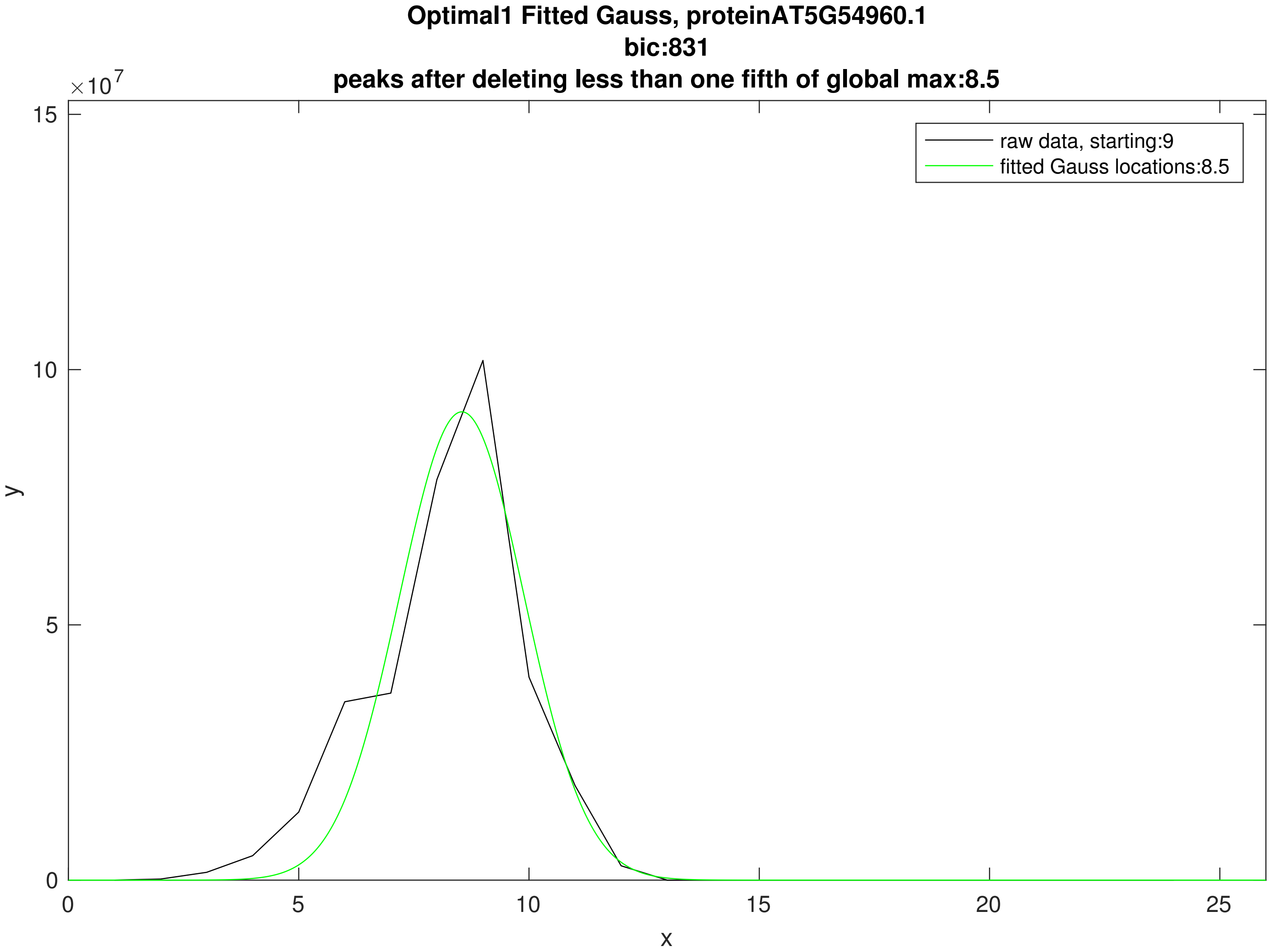
Optimal1 Fitted Gauss, proteinAT5G54960.1
bic:831
peaks after deleting less than one fifth of global max:8.5
×10^7
raw data, starting:9
fitted Gauss locations:8.5
y
x
0
5
10
15
20
25

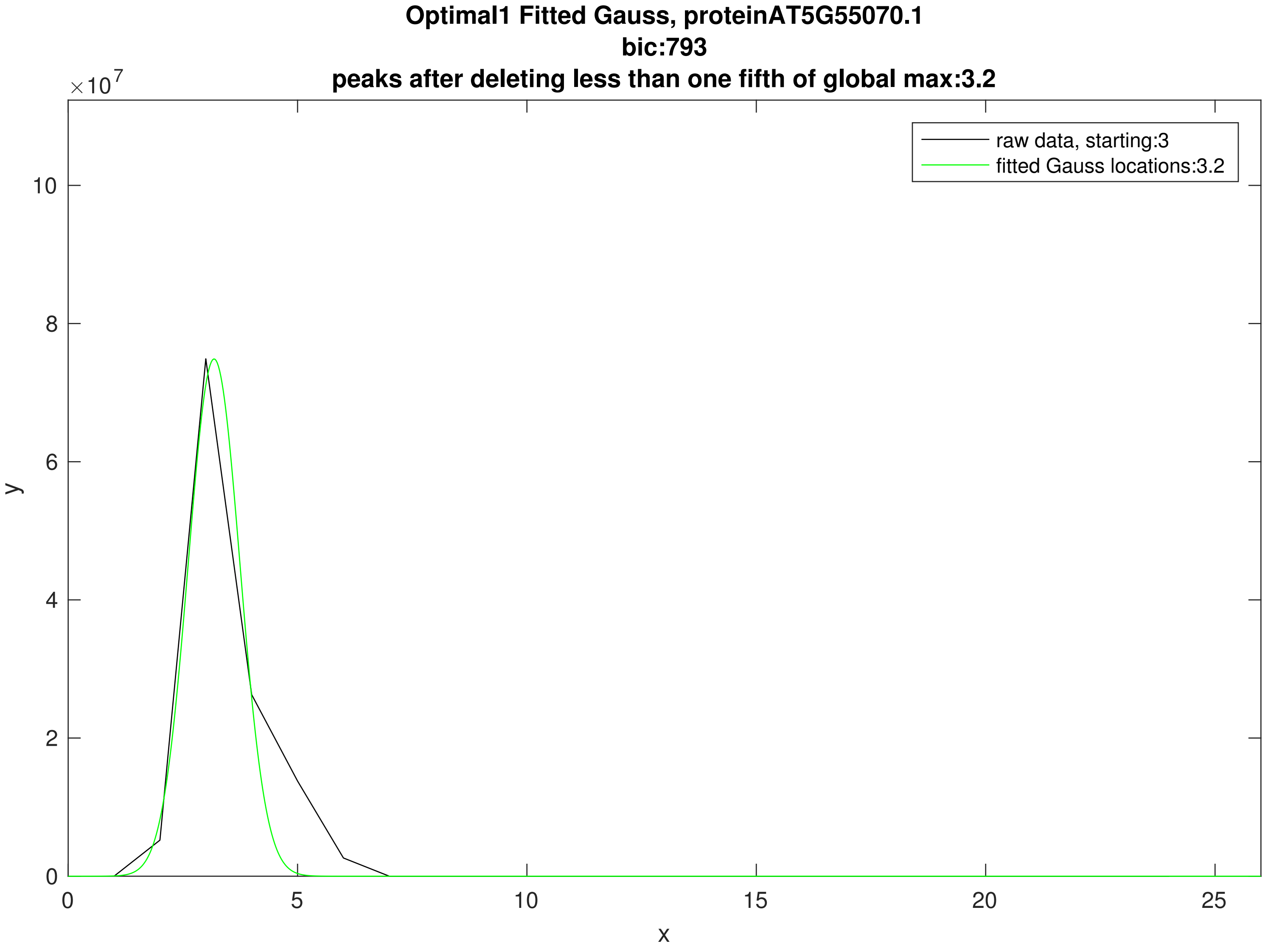
Optimal1 Fitted Gauss, proteinAT5G55070.1
bic:793
peaks after deleting less than one fifth of global max:3.2
×10^7
raw data, starting:3
fitted Gauss locations:3.2
10
8
6
4
2
0
y
0
5
10
15
20
25
x

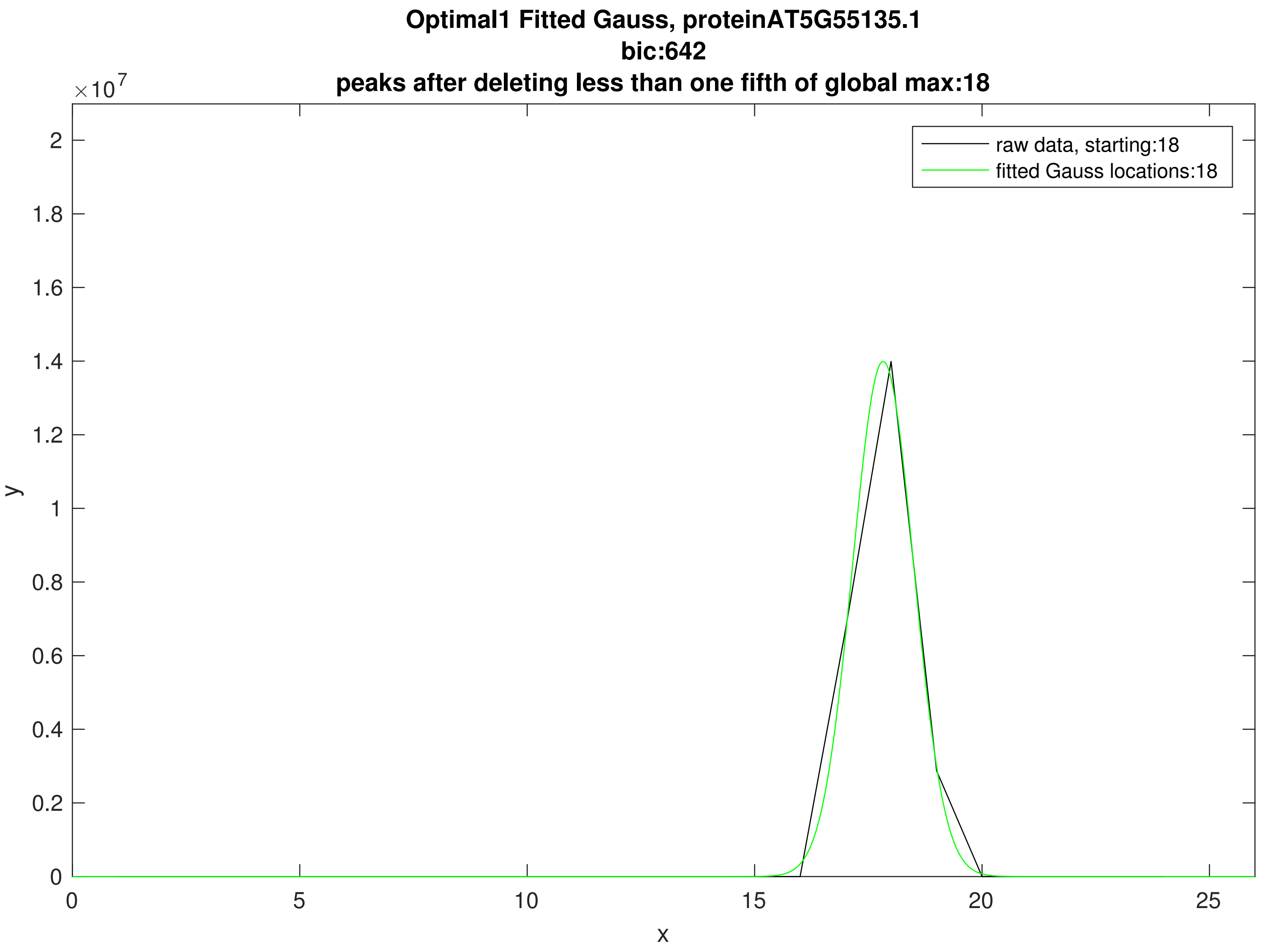

Optimal1 Fitted Gauss, proteinAT5G55135.1
bic:642
peaks after deleting less than one fifth of global max:18
×10^7
raw data, starting:18
fitted Gauss locations:18
y
x
0
0.2
0.4
0.6
0.8
1
1.2
1.4
1.6
1.8
2
0
5
10
15
20
25

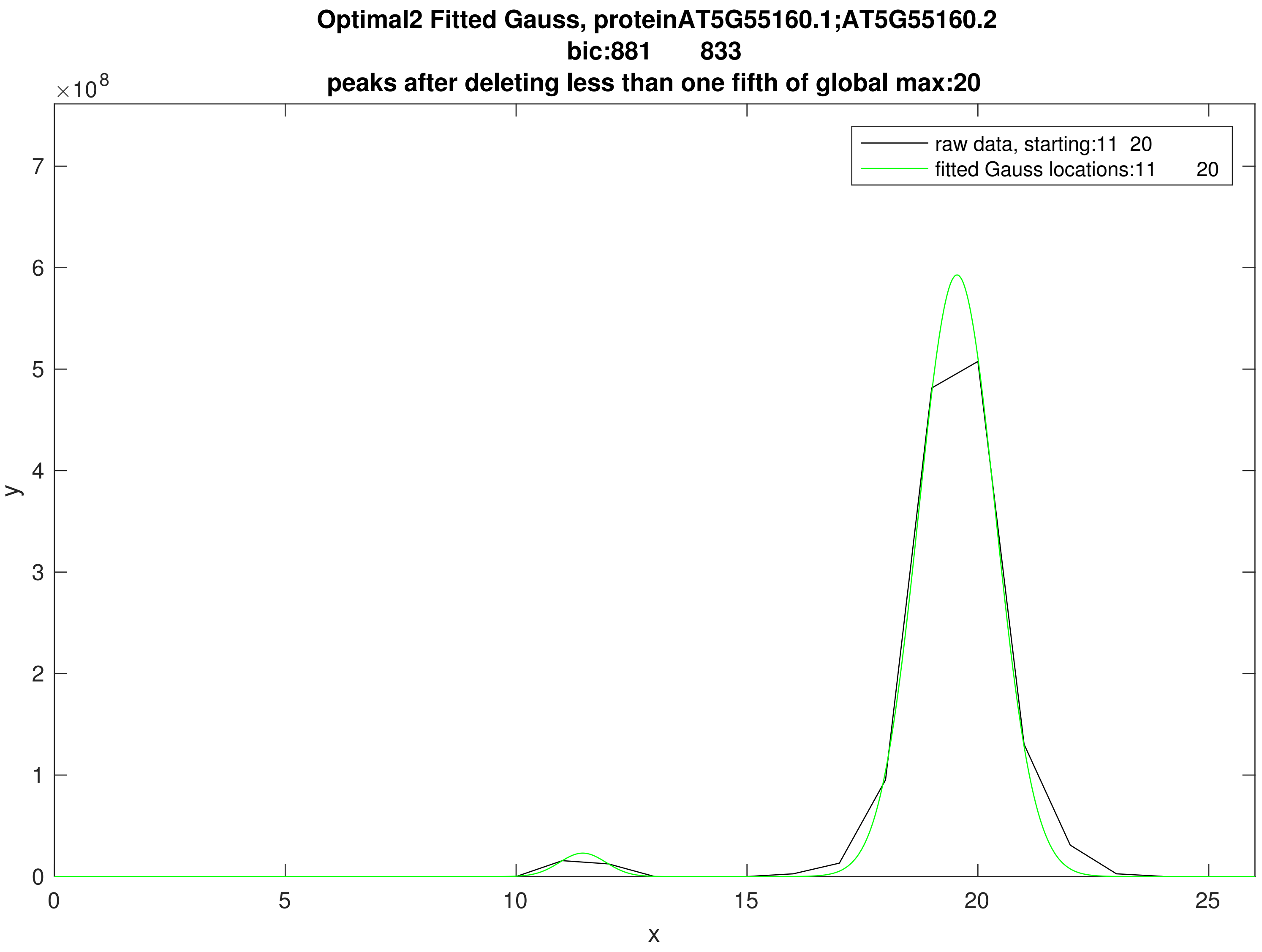
Optimal2 Fitted Gauss, proteinAT5G55160.1;AT5G55160.2
bic:881 833
peaks after deleting less than one fifth of global max:20
×10^8
raw data, starting:11 20
fitted Gauss locations:11 20
y
x
0
1
2
3
4
5
6
7
0
5
10
15
20
25

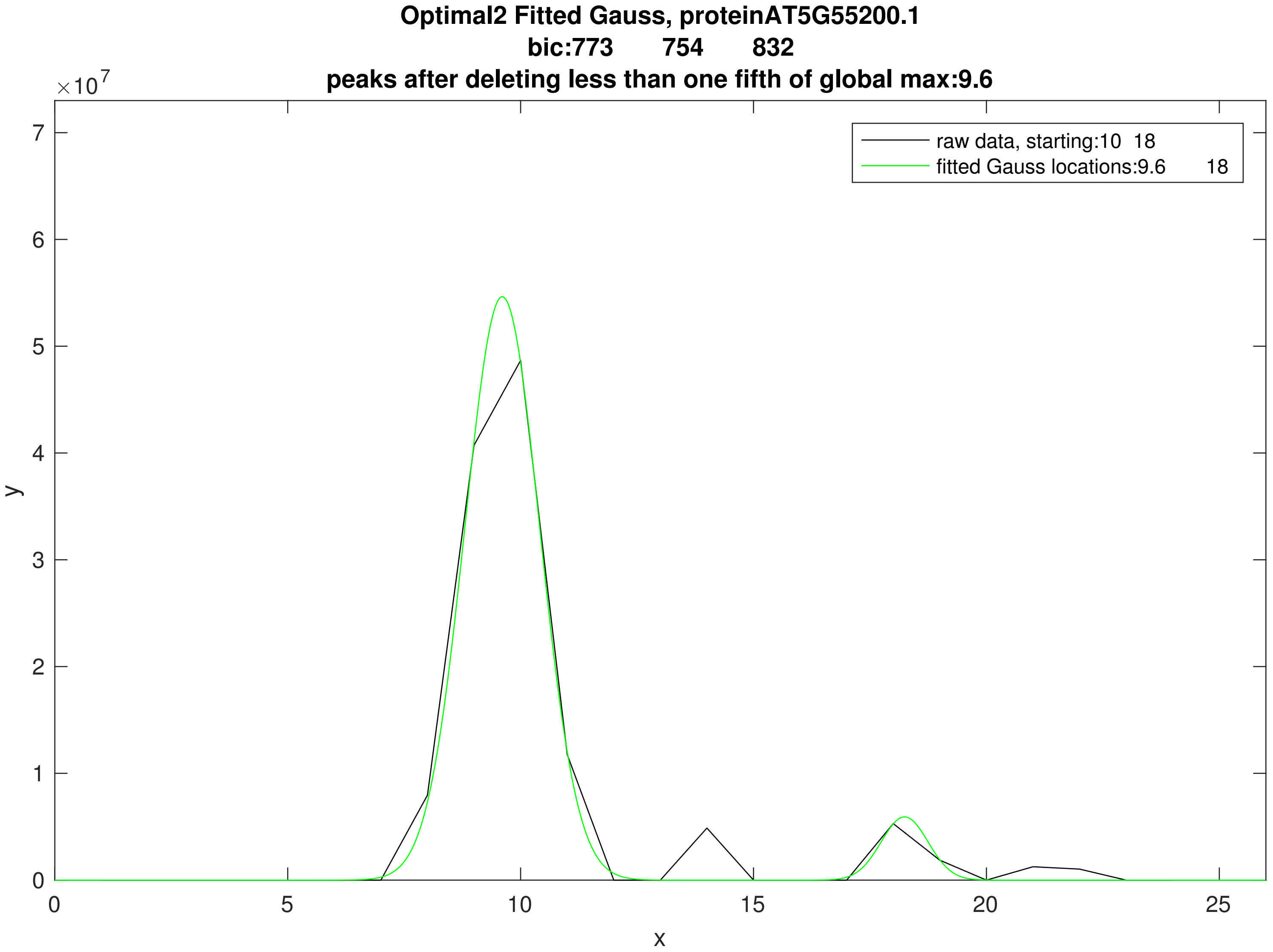

Optimal2 Fitted Gauss, proteinAT5G55200.1
bic:773 754 832
peaks after deleting less than one fifth of global max:9.6
×10^7
raw data, starting:10 18
fitted Gauss locations:9.6 18
y
x
0
1
2
3
4
5
6
7
0
5
10
15
20
25

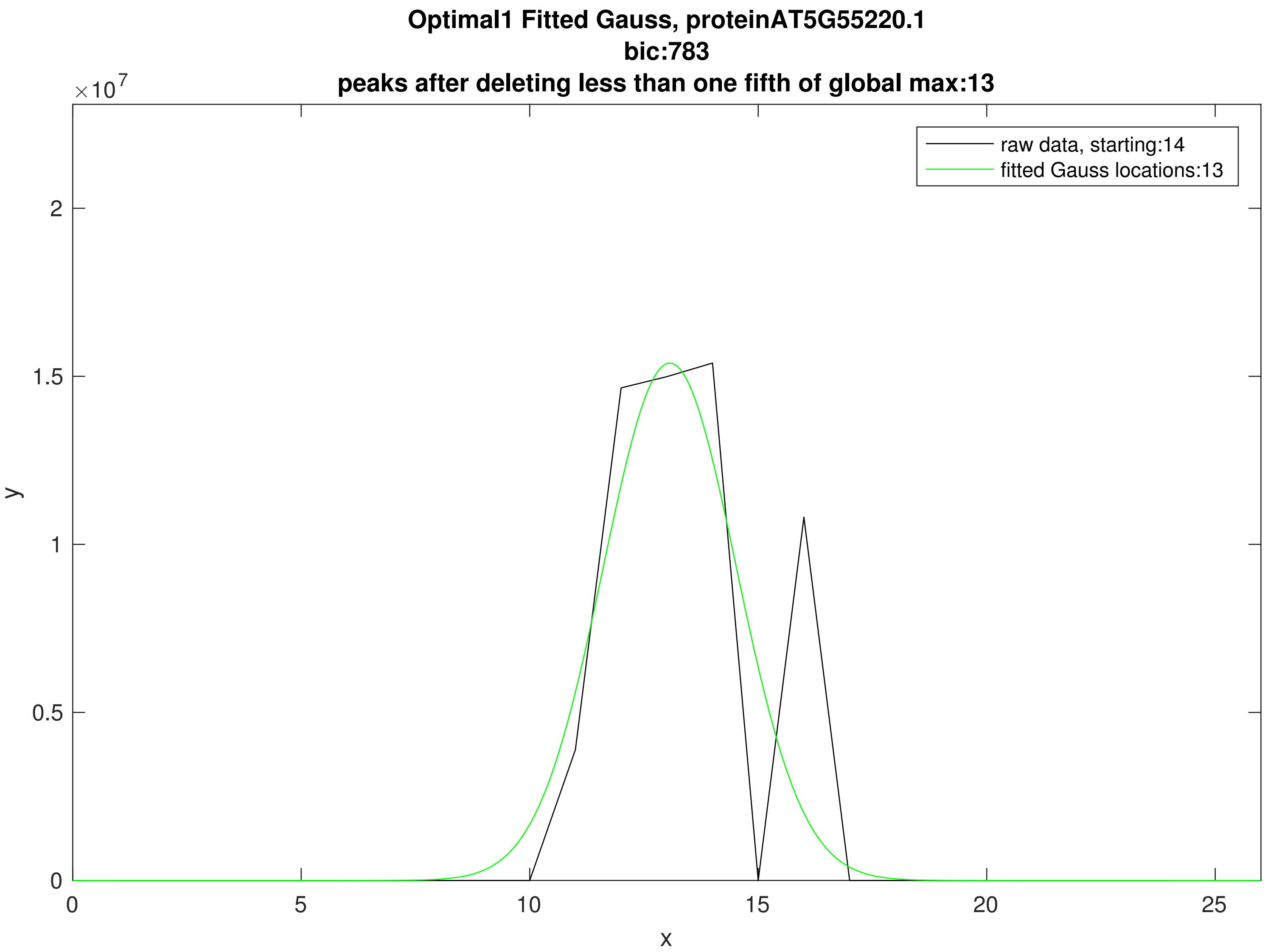
Optimal1 Fitted Gauss, proteinAT5G55220.1
bic:783
peaks after deleting less than one fifth of global max:13
×10^7
raw data, starting:14
fitted Gauss locations:13
y
x
0
0.5
1
1.5
2
0
5
10
15
20
25

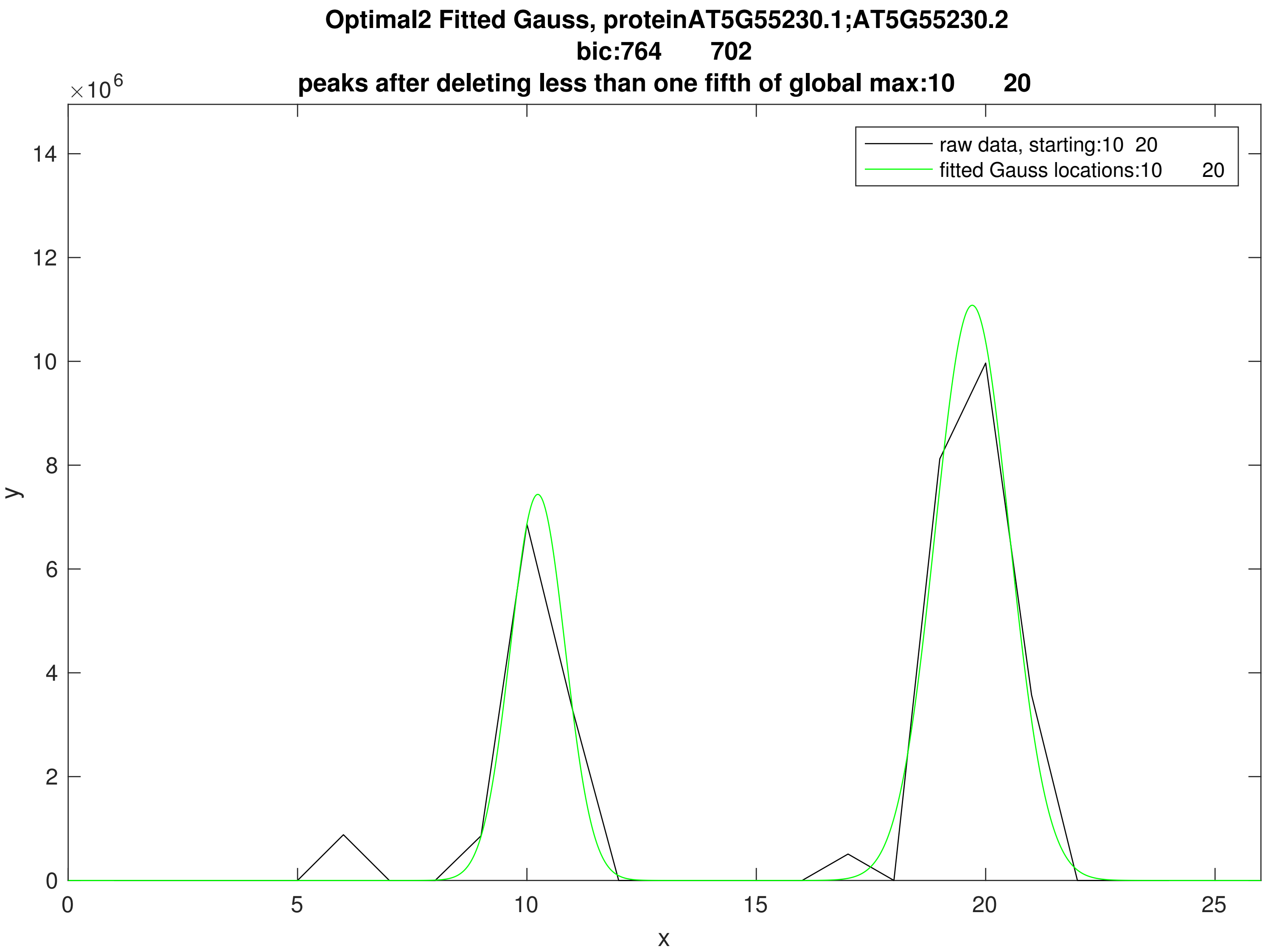
Optimal2 Fitted Gauss, proteinAT5G55230.1;AT5G55230.2
bic:764 702
peaks after deleting less than one fifth of global max:10 20
×10^6
raw data, starting:10 20
fitted Gauss locations:10 20
y
x
0
2
4
6
8
10
12
14
0
5
10
15
20
25

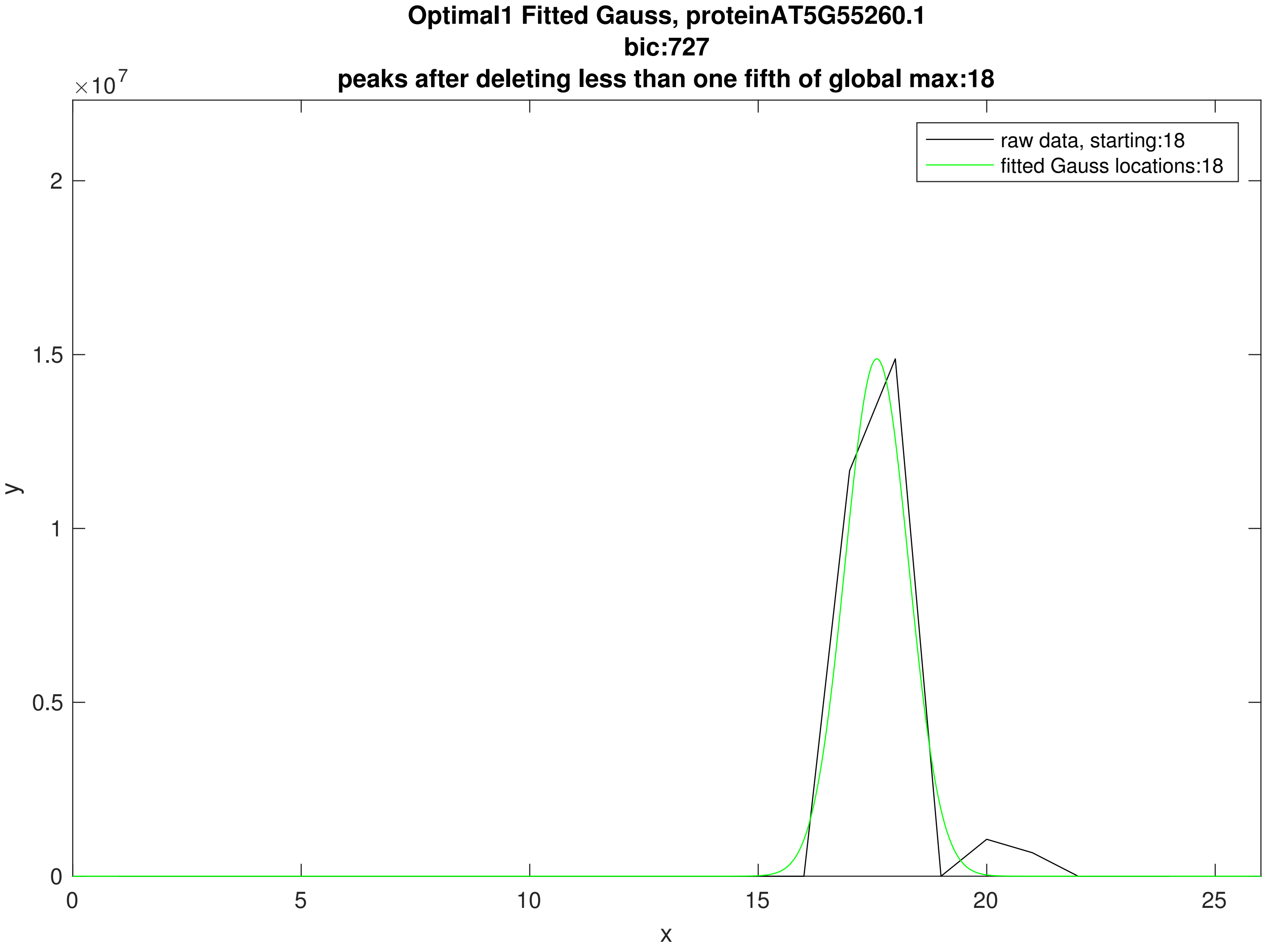
Optimal1 Fitted Gauss, proteinAT5G55260.1
bic:727
peaks after deleting less than one fifth of global max:18
×10^7
raw data, starting:18
fitted Gauss locations:18
y
x
0
0.5
1
1.5
2
0
5
10
15
20
25

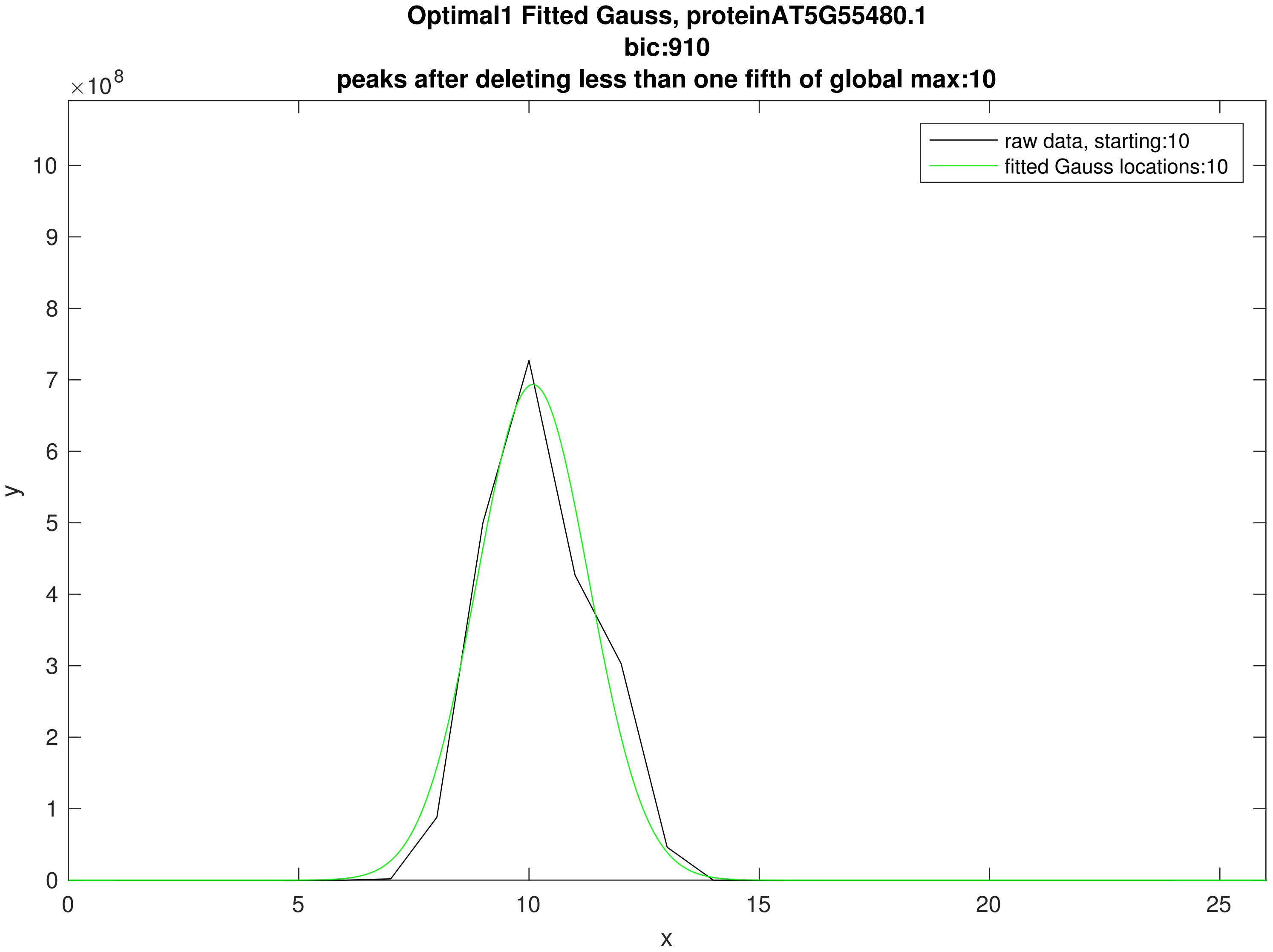

Optimal1 Fitted Gauss, proteinAT5G55480.1
bic:910
peaks after deleting less than one fifth of global max:10
×10^8
raw data, starting:10
fitted Gauss locations:10
10
9
8
7
6
5
4
3
2
1
0
y
0
5
10
15
20
25
x

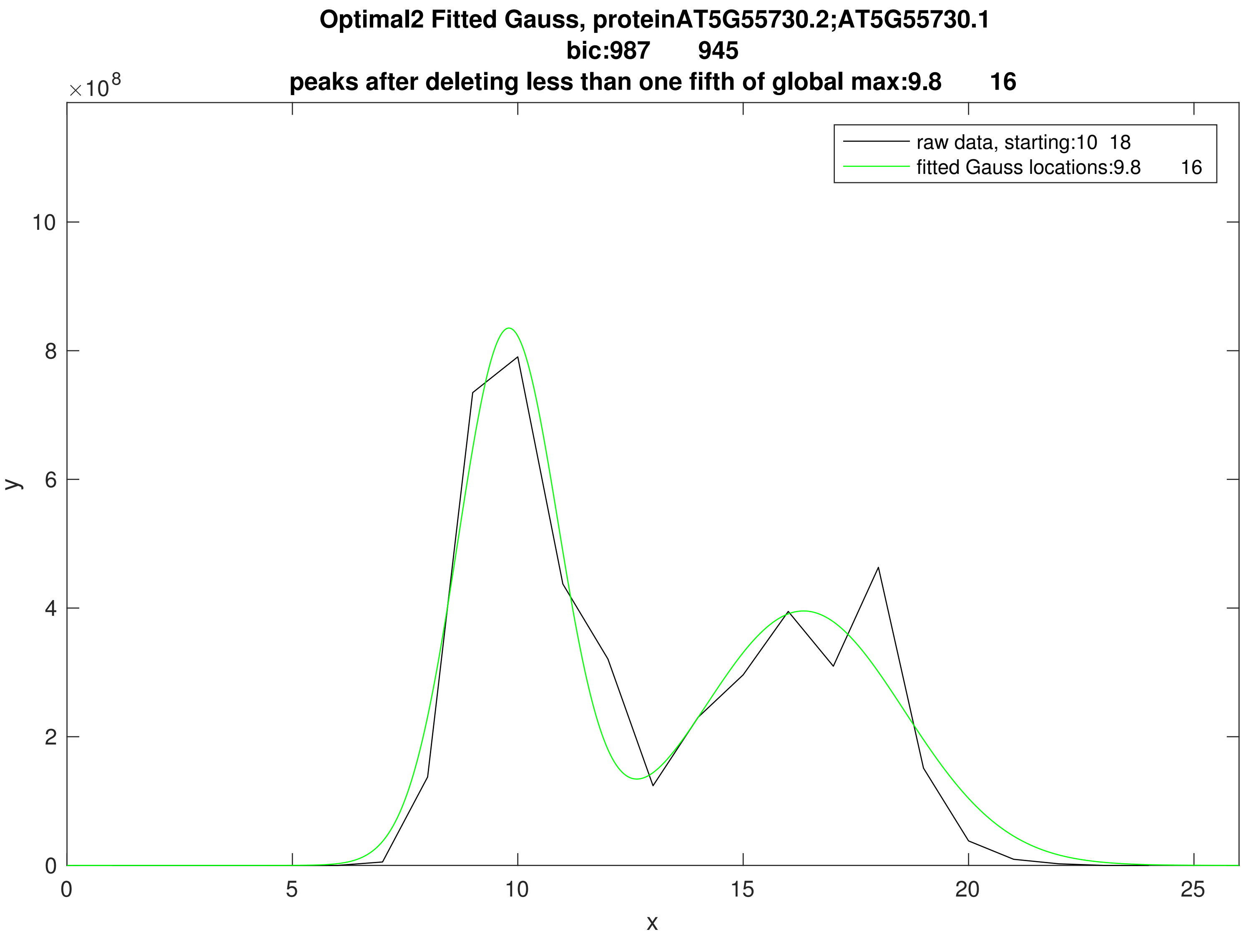
Optimal2 Fitted Gauss, proteinAT5G55730.2;AT5G55730.1
bic:987 945
peaks after deleting less than one fifth of global max:9.8 16
raw data, starting:10 18
fitted Gauss locations:9.8 16
×10^8
y
x
0
2
4
6
8
10
5
15
20
25

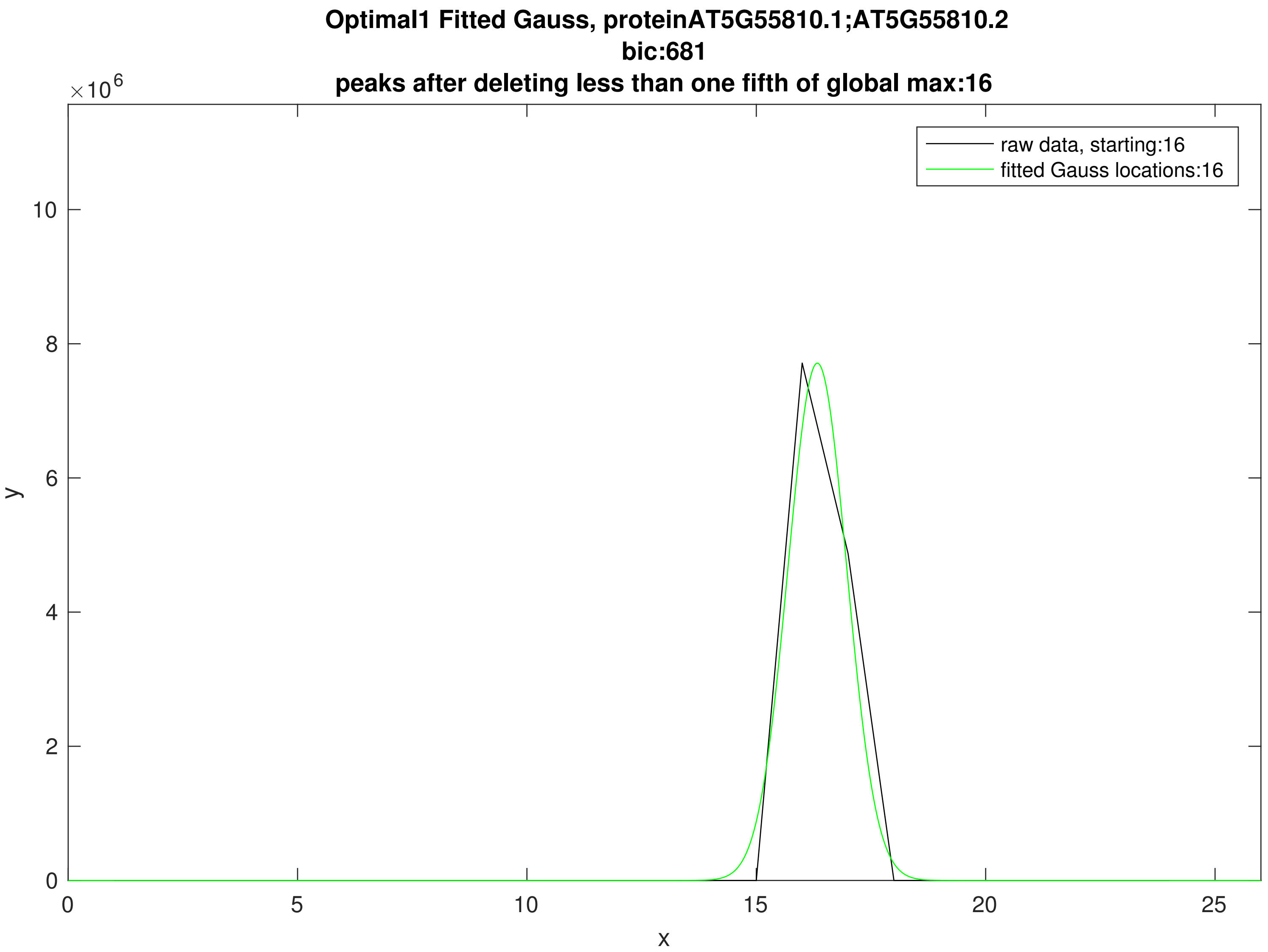

Optimal1 Fitted Gauss, proteinAT5G55810.1;AT5G55810.2
bic:681
peaks after deleting less than one fifth of global max:16
×10^6
raw data, starting:16
fitted Gauss locations:16
y
x
0
2
4
6
8
10
0
5
10
15
20
25

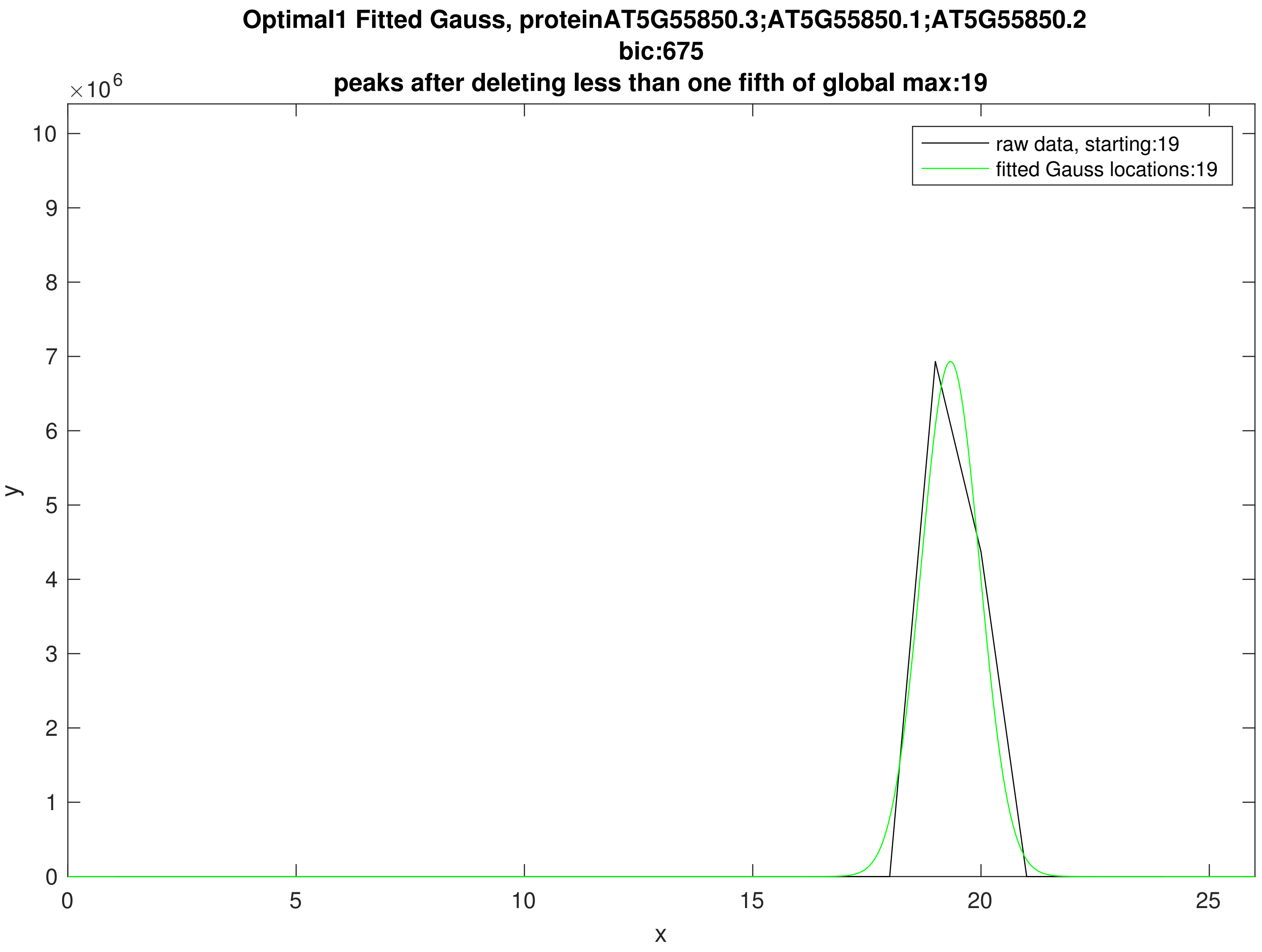

Optimal1 Fitted Gauss, proteinAT5G55850.3;AT5G55850.1;AT5G55850.2
bic:675
peaks after deleting less than one fifth of global max:19
×10^6
raw data, starting:19
fitted Gauss locations:19
y
x

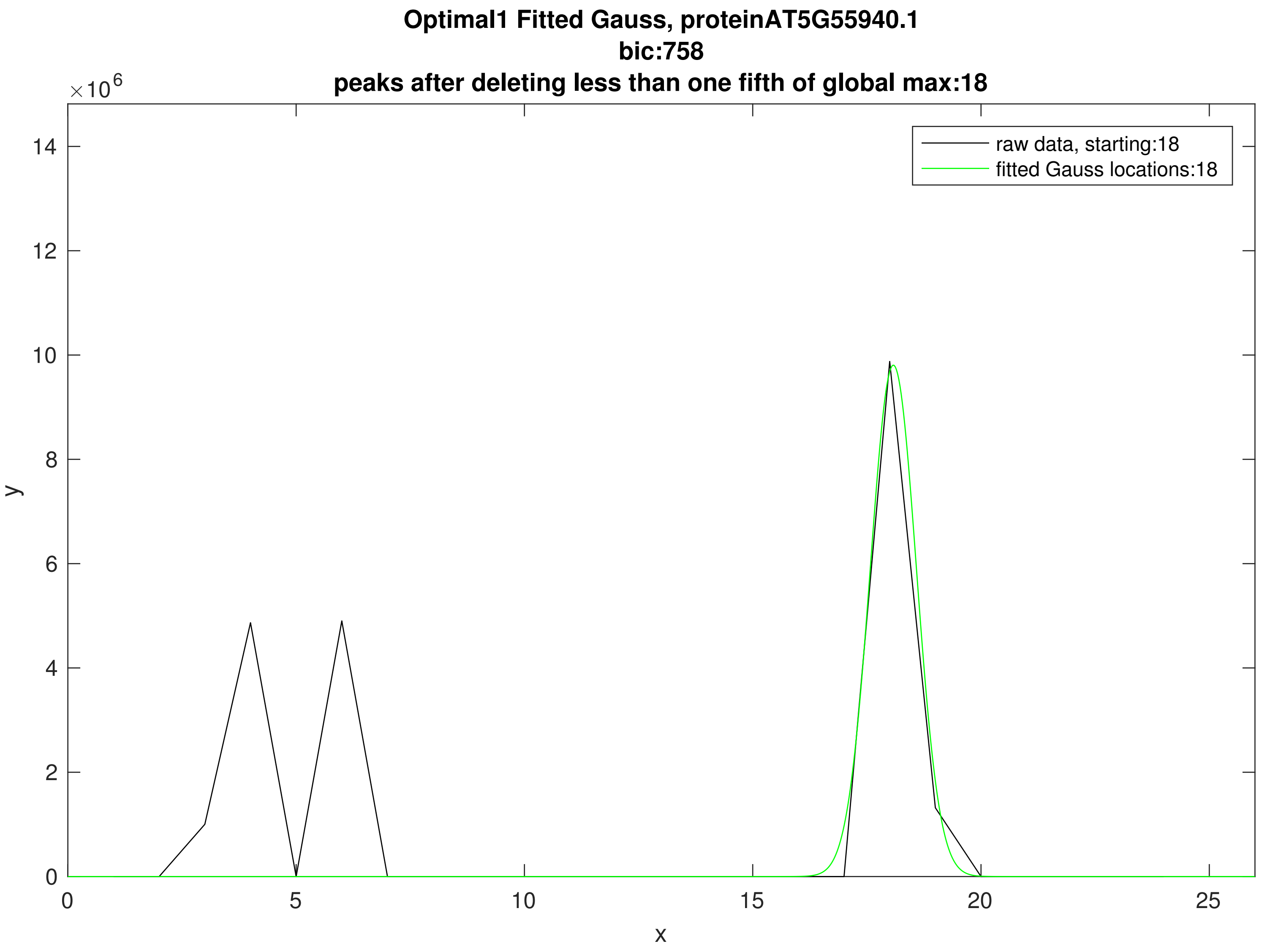
Optimal1 Fitted Gauss, proteinAT5G55940.1
bic:758
peaks after deleting less than one fifth of global max:18
×10^6
raw data, starting:18
fitted Gauss locations:18
y
x
0
2
4
6
8
10
12
14
0
5
10
15
20
25

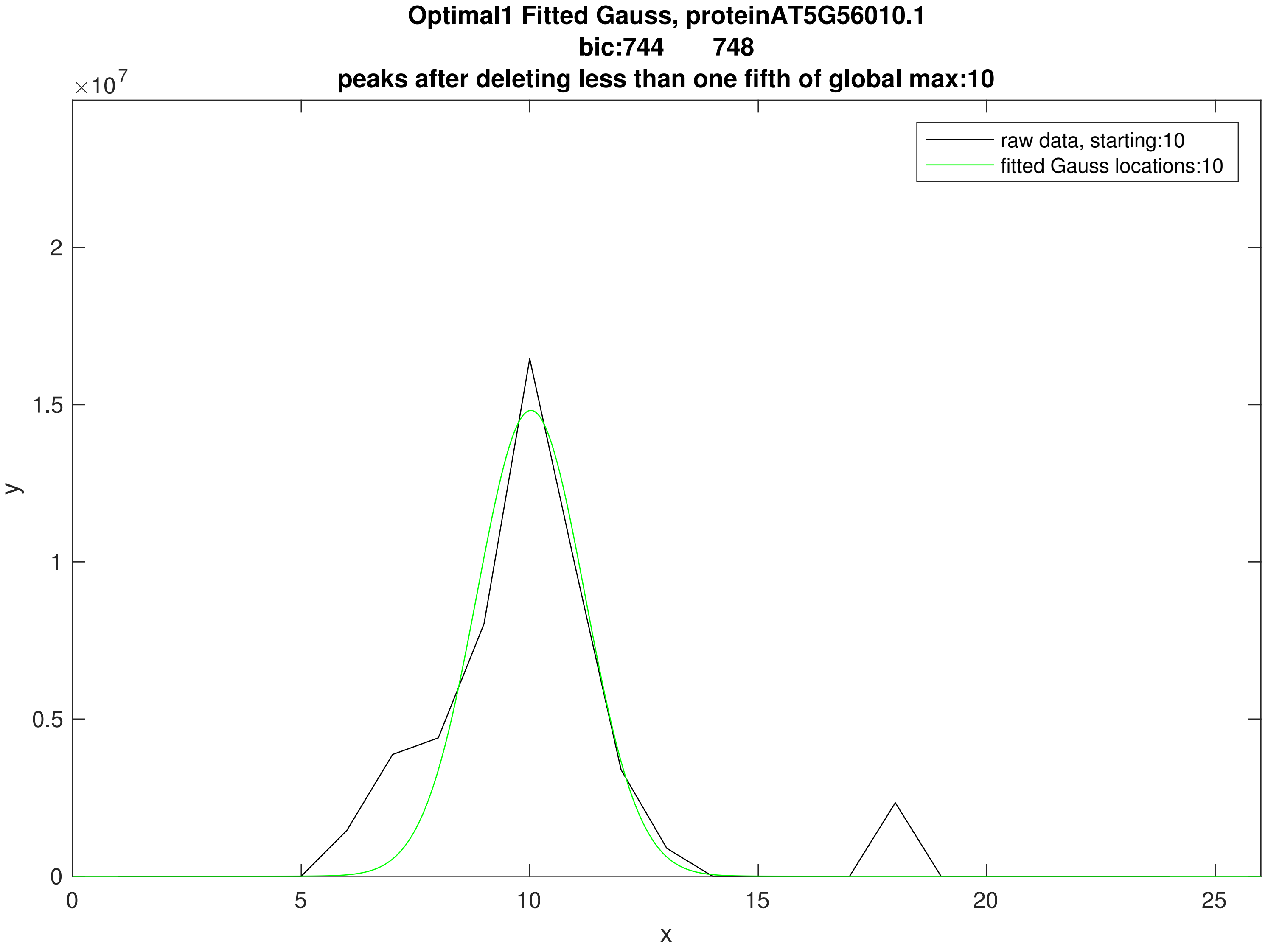
Optimal1 Fitted Gauss, proteinAT5G56010.1
bic:744 748
peaks after deleting less than one fifth of global max:10
×10^7
raw data, starting:10
fitted Gauss locations:10
y
x
0
0.5
1
1.5
2
5
10
15
20
25

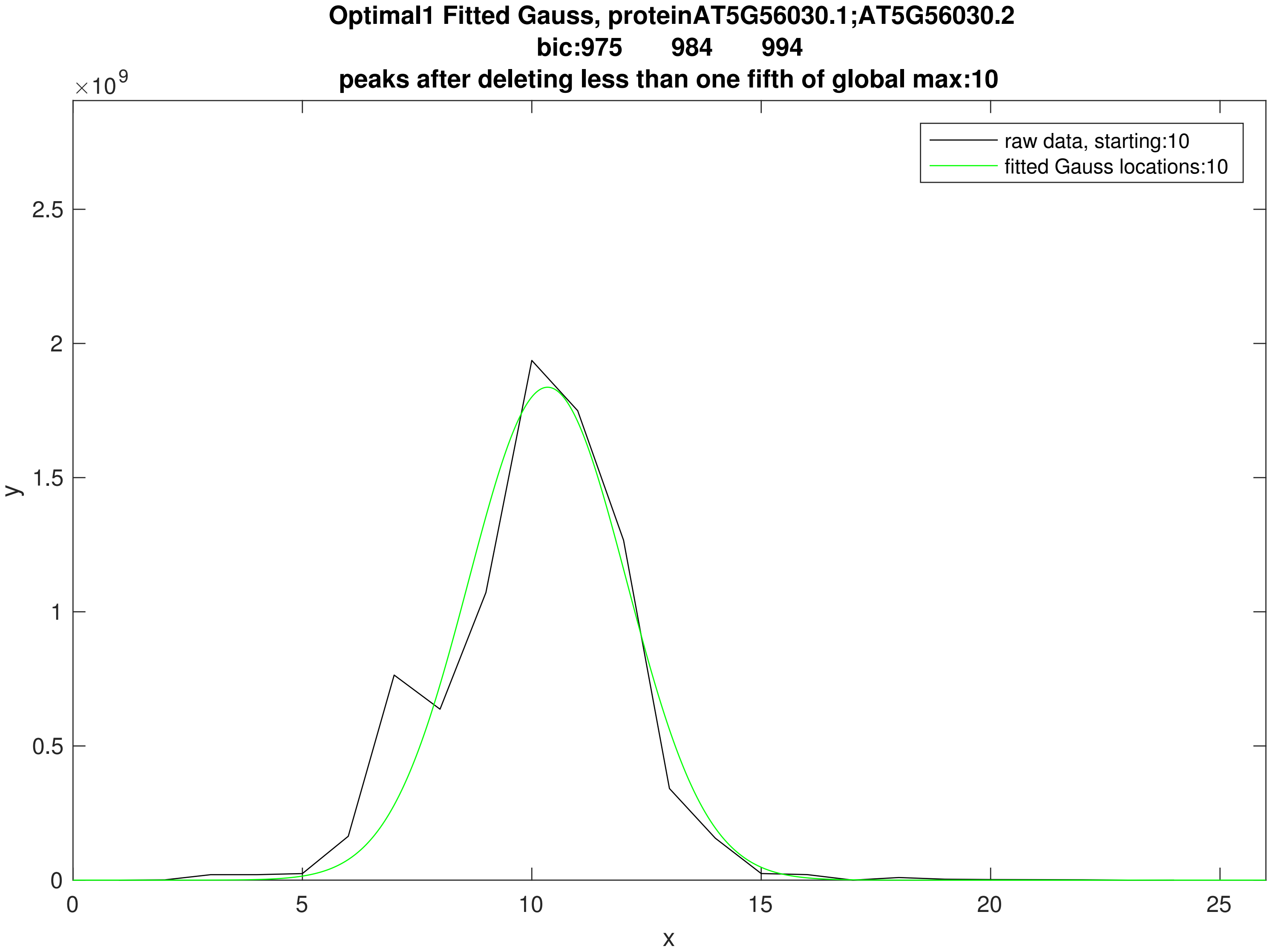
Optimal1 Fitted Gauss, proteinAT5G56030.1;AT5G56030.2
bic:975 984 994
peaks after deleting less than one fifth of global max:10
×10^9
raw data, starting:10
fitted Gauss locations:10
2.5
2
1.5
1
0.5
0
y
0
5
10
15
20
25
x

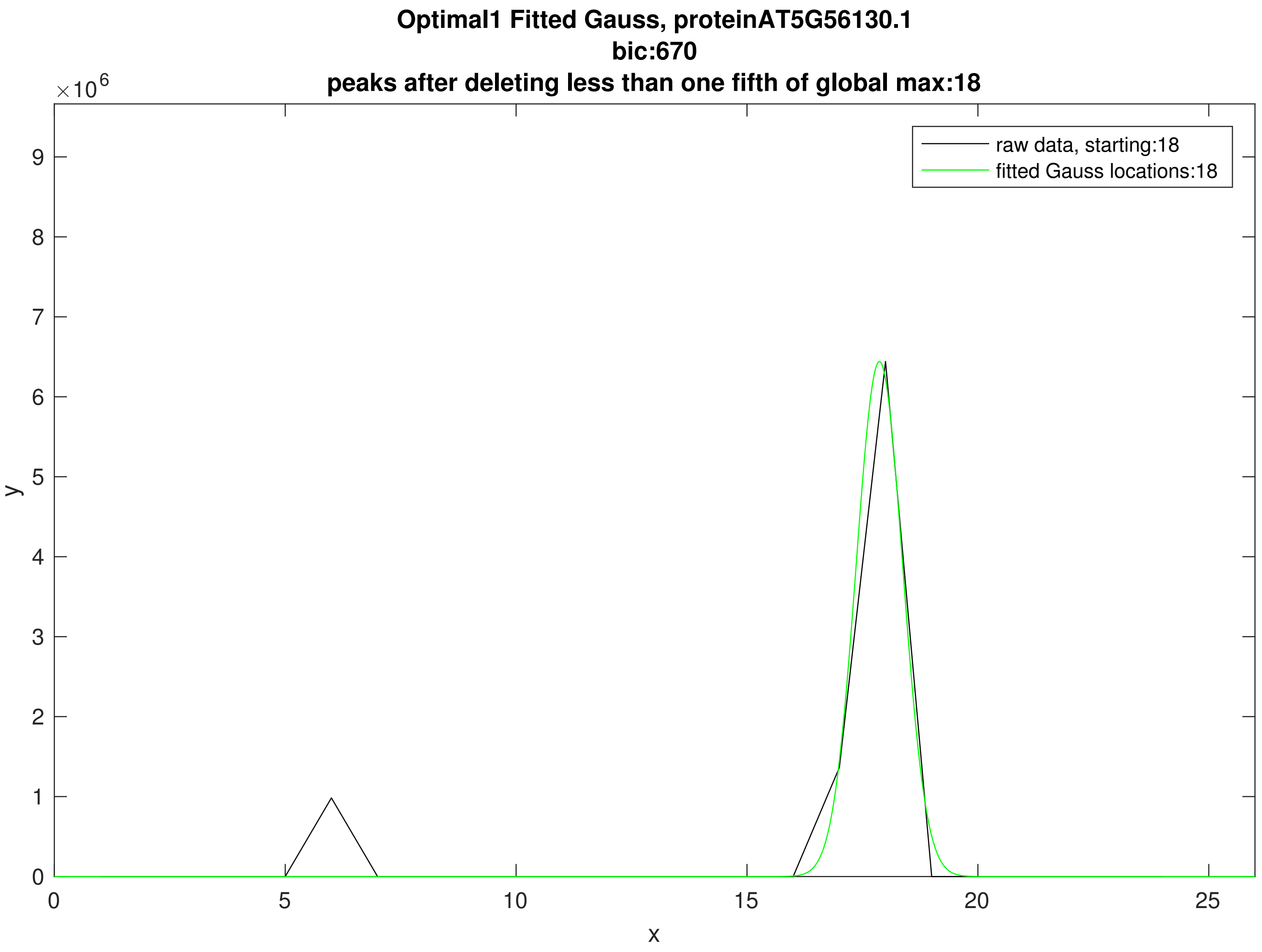
Optimal1 Fitted Gauss, proteinAT5G56130.1
bic:670
peaks after deleting less than one fifth of global max:18
×10^6
raw data, starting:18
fitted Gauss locations:18
y
x
0
1
2
3
4
5
6
7
8
9
0
5
10
15
20
25

#### Optimal1 Fitted Gauss, proteinAT5G56260.1
#### bic:881
#### peaks after deleting less than one fifth of global max:13

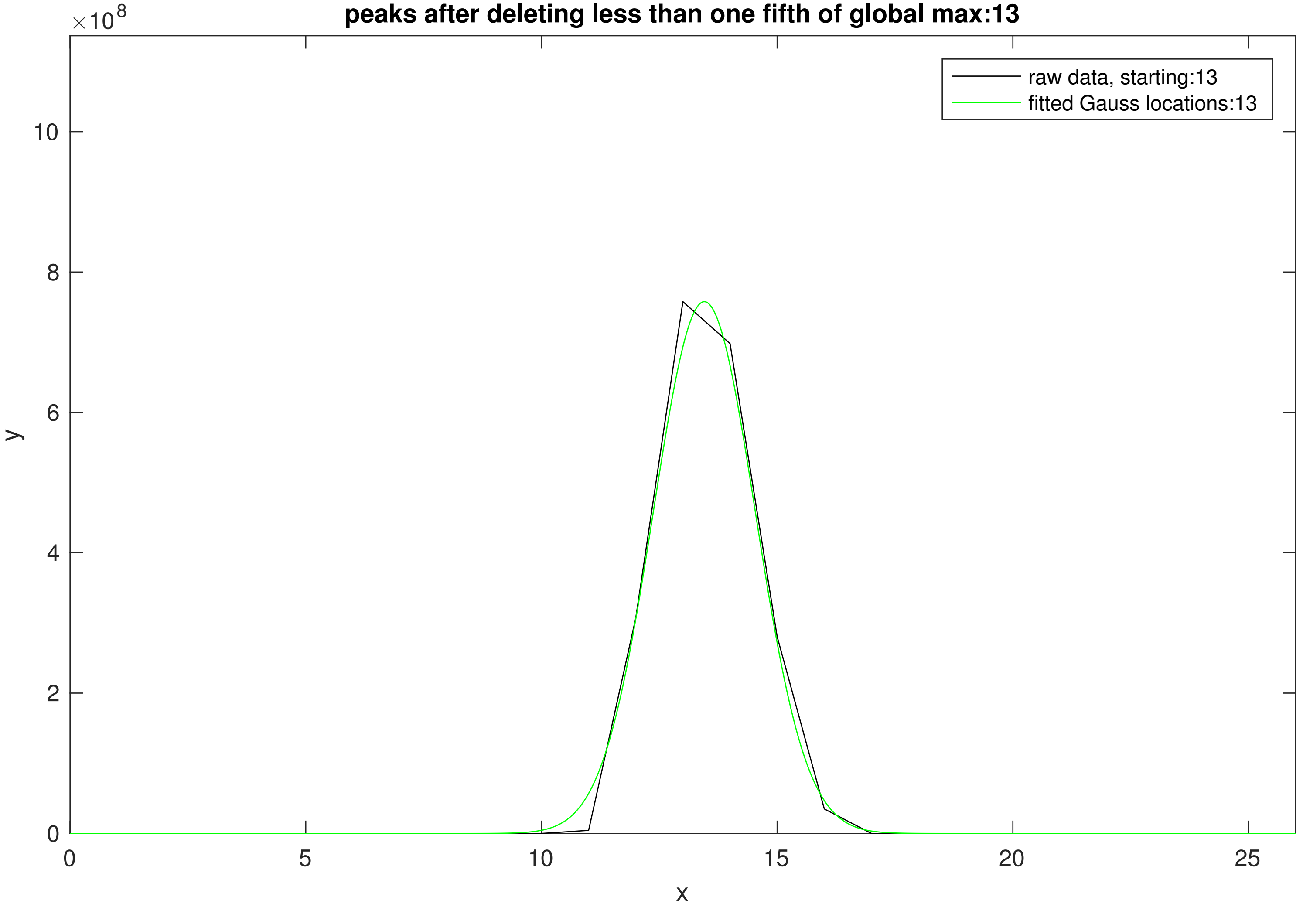

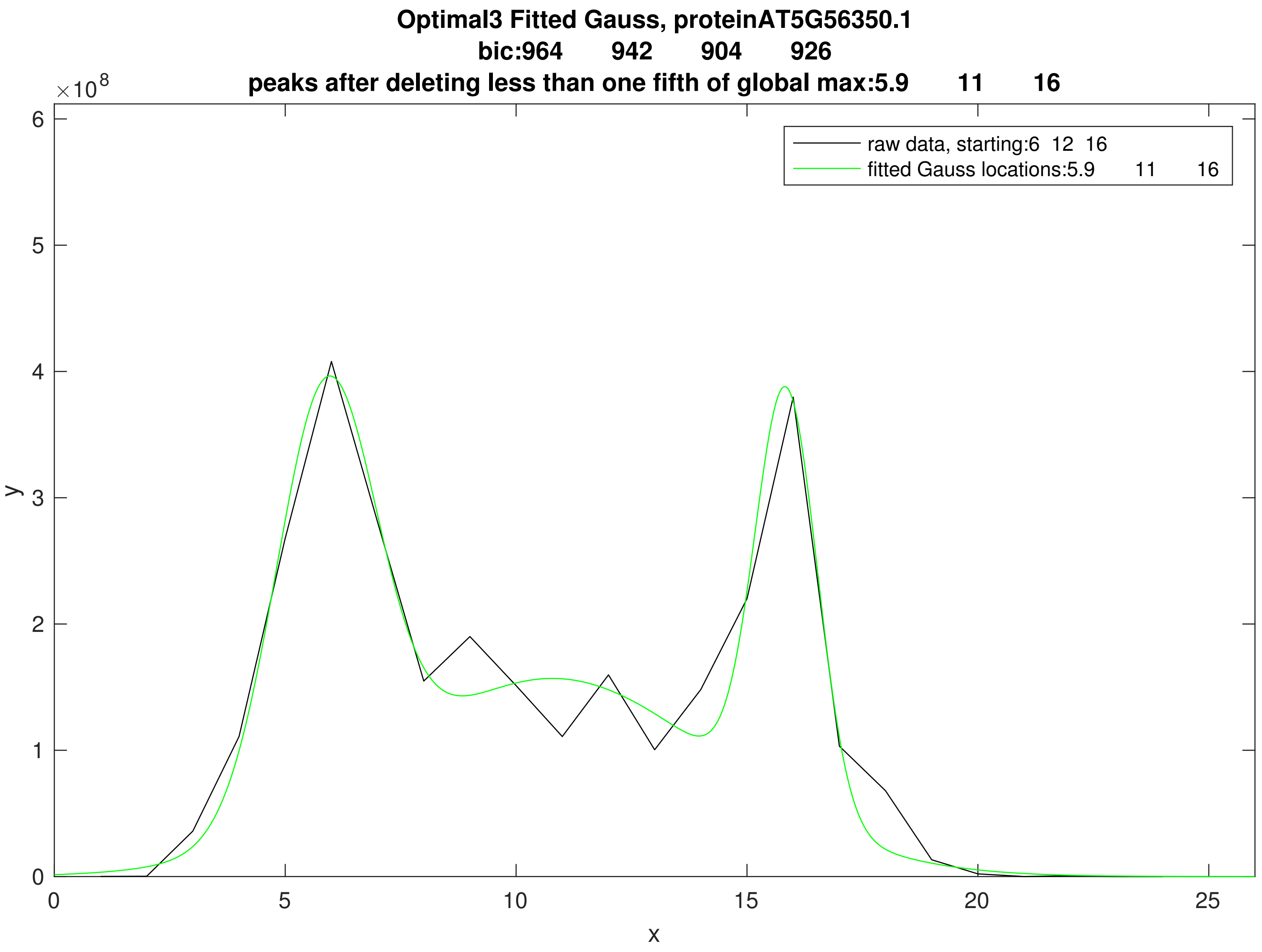
Optimal3 Fitted Gauss, proteinAT5G56350.1
bic:964 942 904 926
peaks after deleting less than one fifth of global max:5.9 11 16
×10^8
raw data, starting:6 12 16
fitted Gauss locations:5.9 11 16
y
x
0
1
2
3
4
5
6
0
5
10
15
20
25

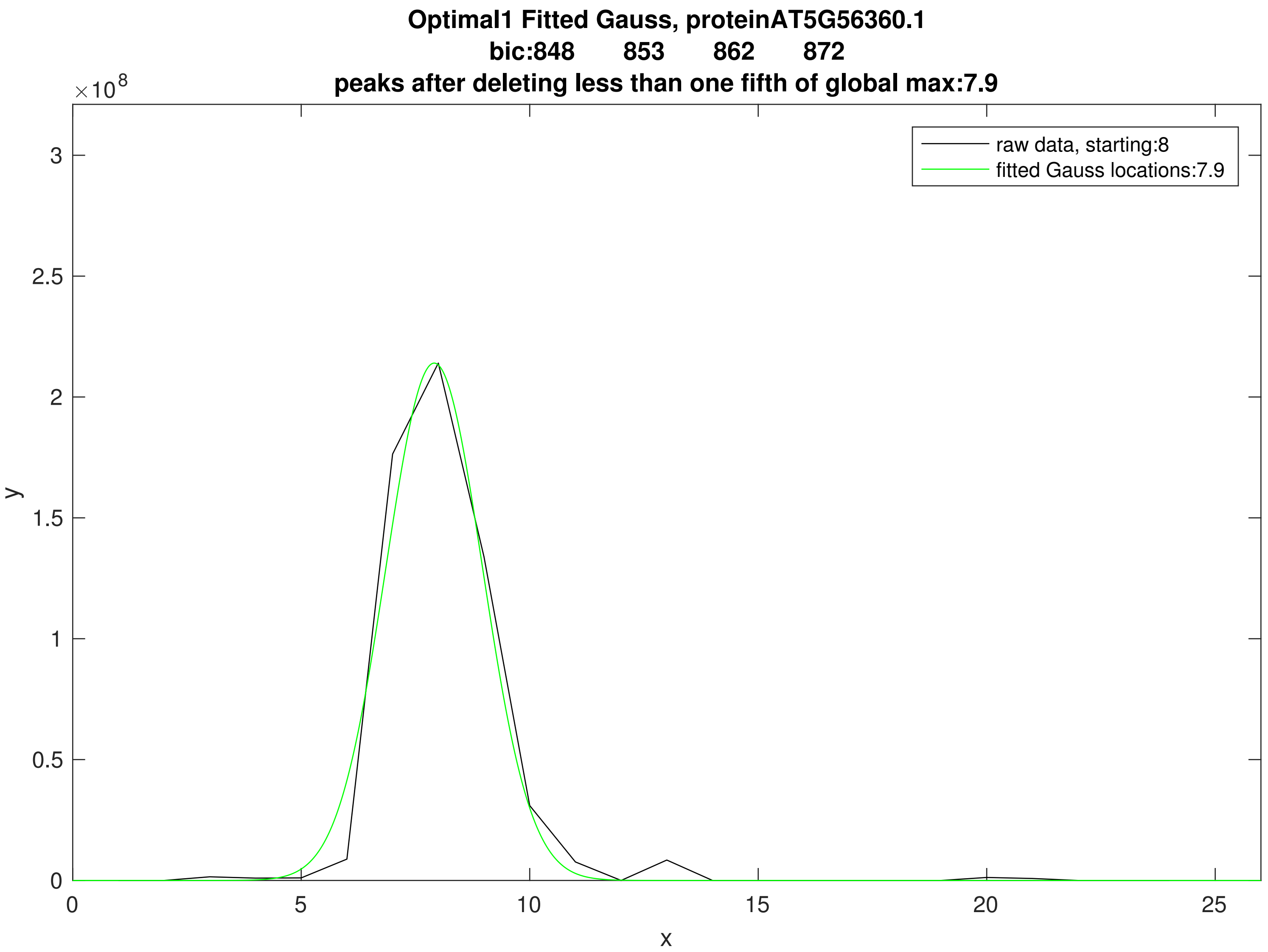

Optimal1 Fitted Gauss, proteinAT5G56360.1
bic:848 853 862 872
peaks after deleting less than one fifth of global max:7.9
×10^8
3
2.5
2
1.5
1
0.5
0
y
0
5
10
15
20
25
x
raw data, starting:8
fitted Gauss locations:7.9

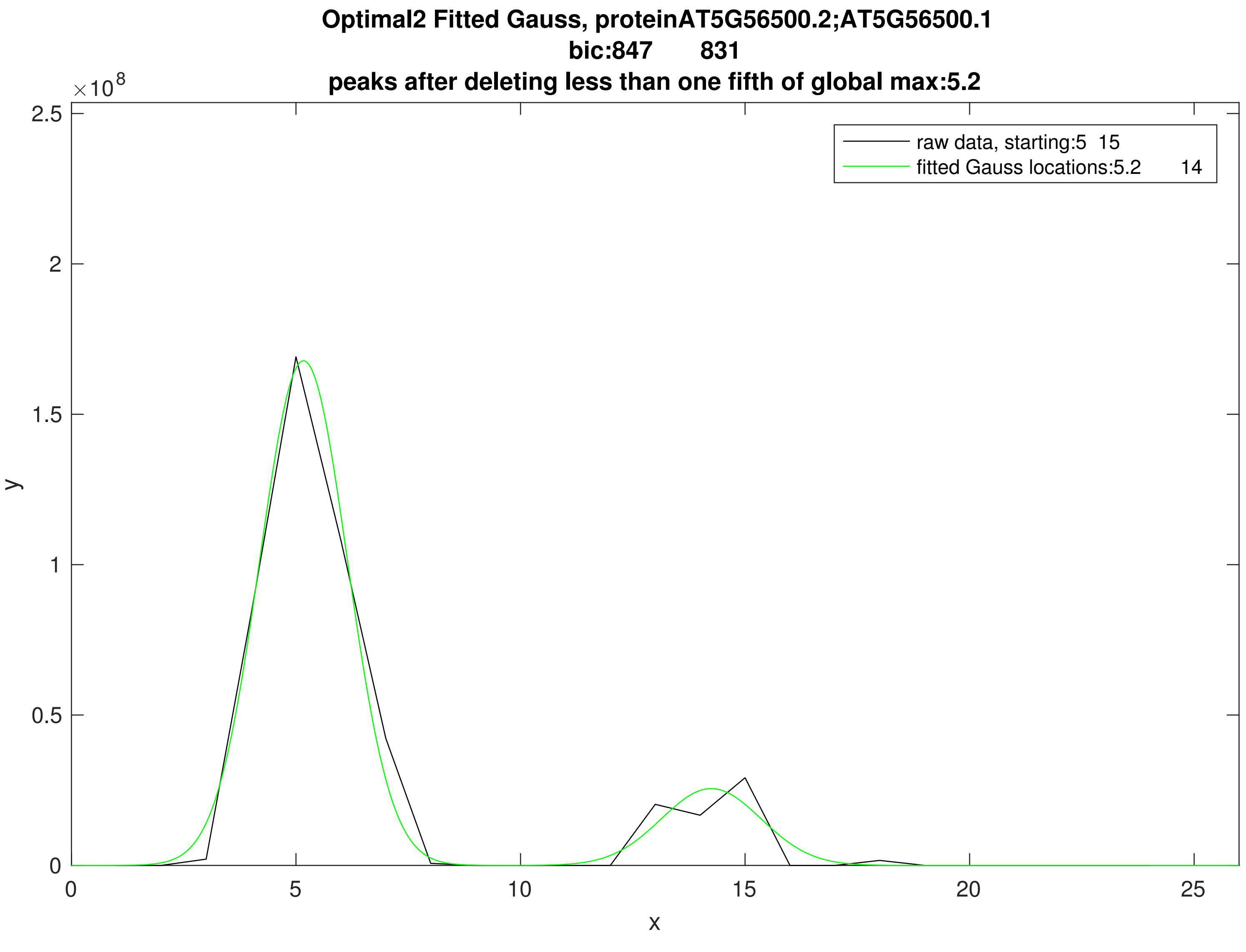
Optimal2 Fitted Gauss, proteinAT5G56500.2;AT5G56500.1
bic:847 831
peaks after deleting less than one fifth of global max:5.2
raw data, starting:5 15
fitted Gauss locations:5.2 14
y
x
×10^8
2.5
2
1.5
1
0.5
0
0
5
10
15
20
25

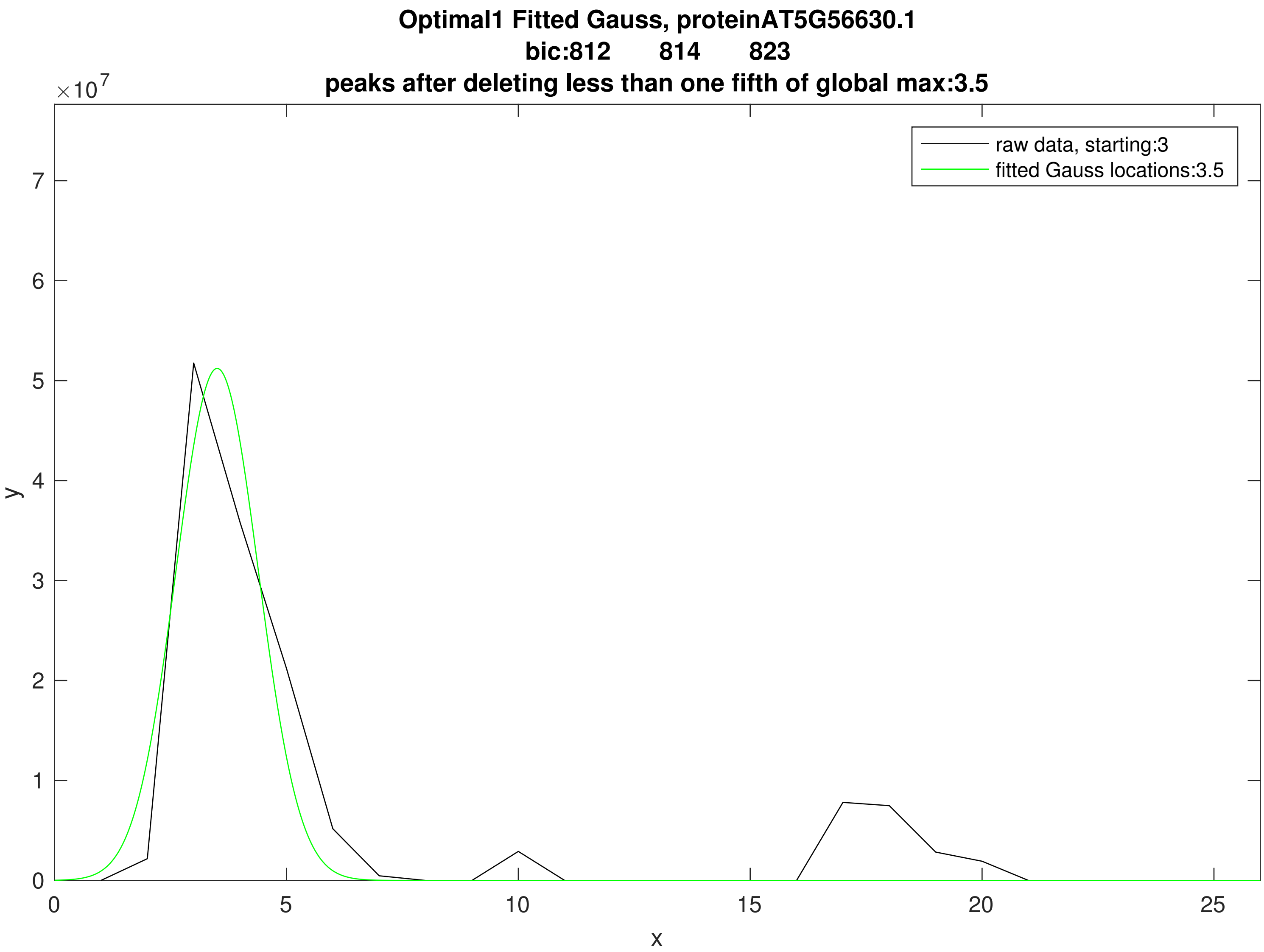

Optimal1 Fitted Gauss, proteinAT5G56630.1
bic:812 814 823
peaks after deleting less than one fifth of global max:3.5
×10^7
raw data, starting:3
fitted Gauss locations:3.5
y
x
0
1
2
3
4
5
6
7
0
5
10
15
20
25

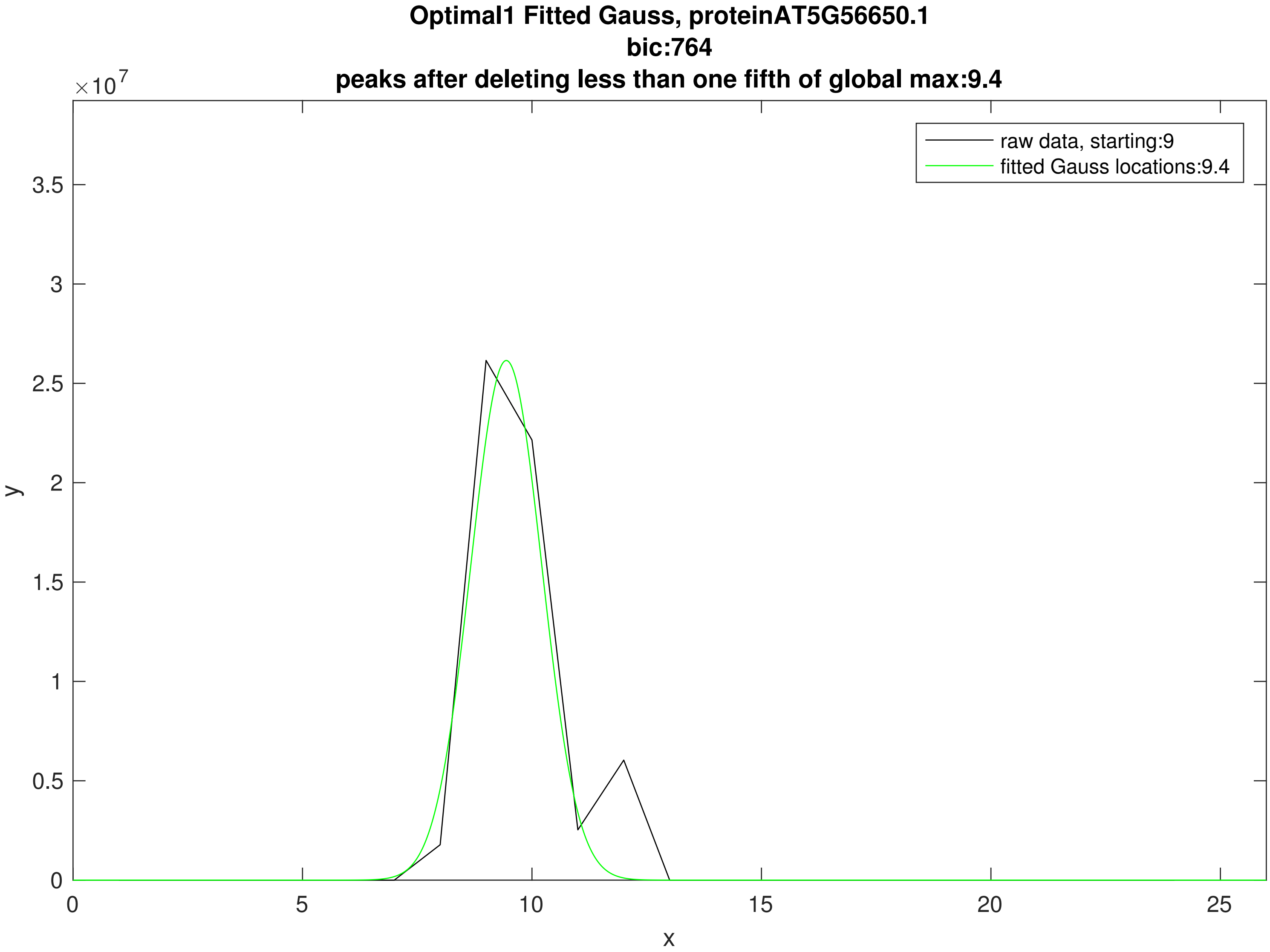
Optimal1 Fitted Gauss, proteinAT5G56650.1
bic:764
peaks after deleting less than one fifth of global max:9.4
×10^7
raw data, starting:9
fitted Gauss locations:9.4
0
0.5
1
1.5
2
2.5
3
3.5
y
0
5
10
15
20
25
x

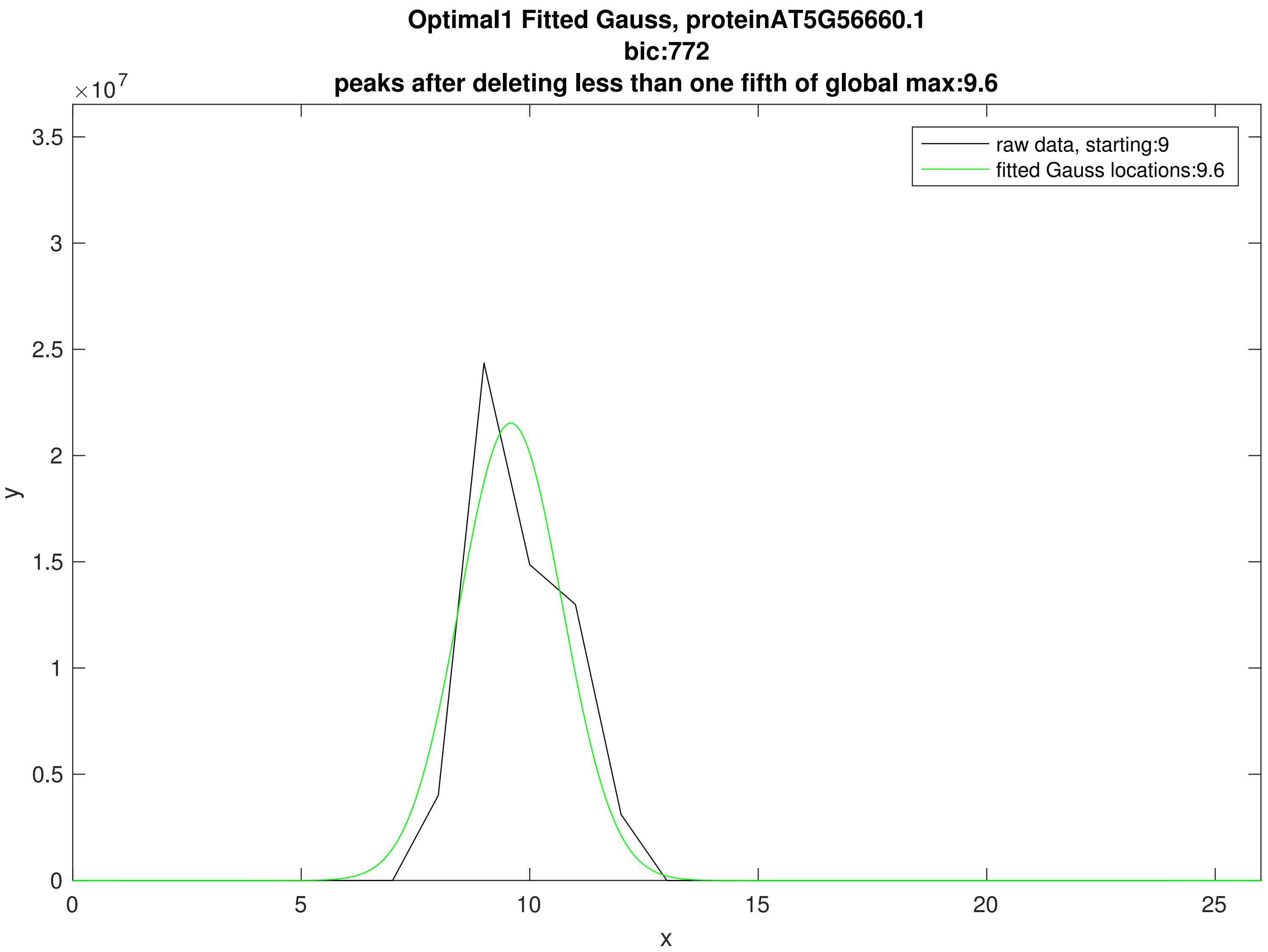
Optimal1 Fitted Gauss, proteinAT5G56660.1
bic:772
peaks after deleting less than one fifth of global max:9.6
×10^7
3.5
3
2.5
2
1.5
1
0.5
0
y
0
5
10
15
20
25
x
raw data, starting:9
fitted Gauss locations:9.6

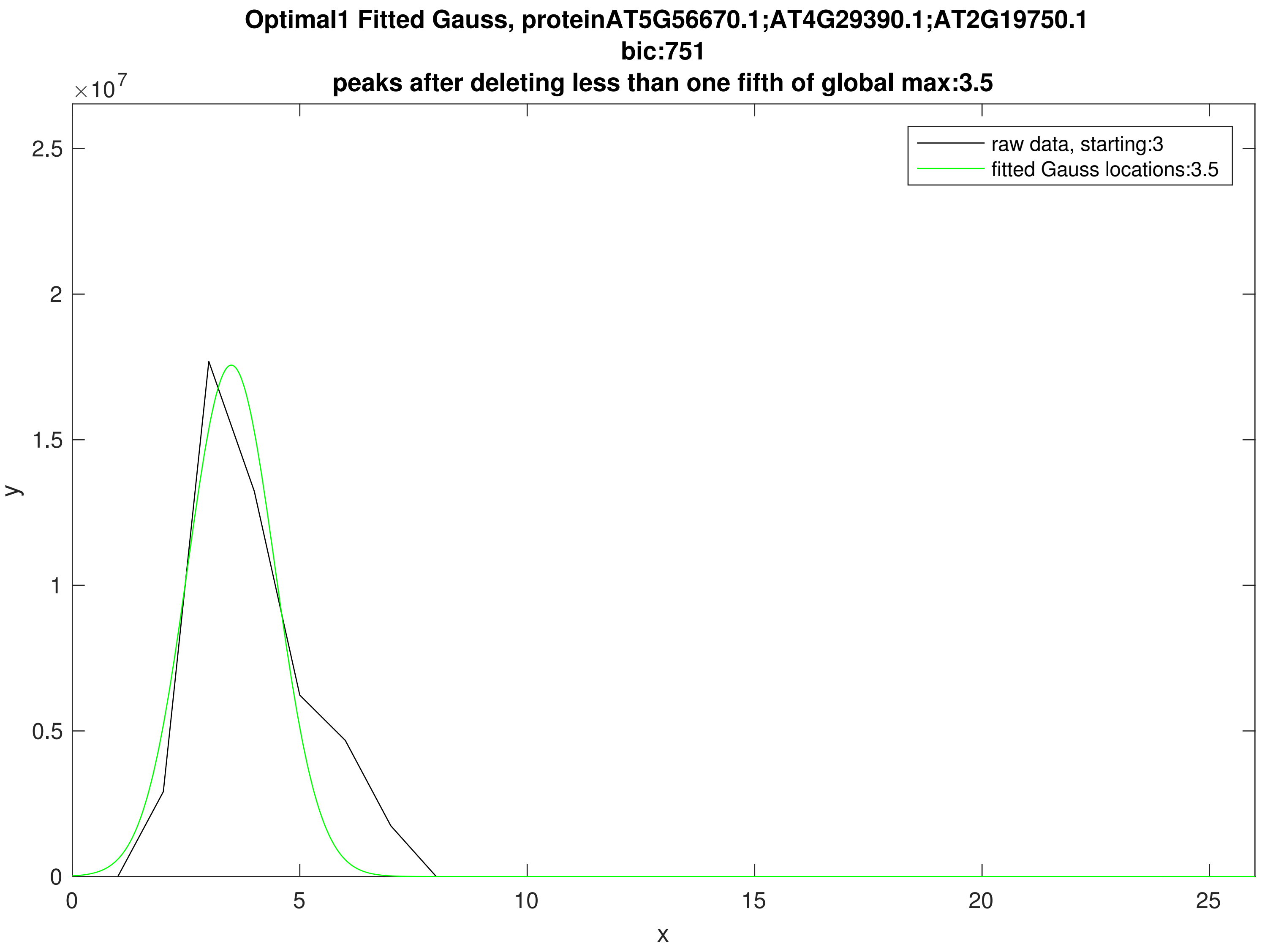
Optimal1 Fitted Gauss, proteinAT5G56670.1;AT4G29390.1;AT2G19750.1
bic:751
peaks after deleting less than one fifth of global max:3.5
×10^7
2.5
2
1.5
1
0.5
0
y
0
5
10
15
20
25
x
raw data, starting:3
fitted Gauss locations:3.5

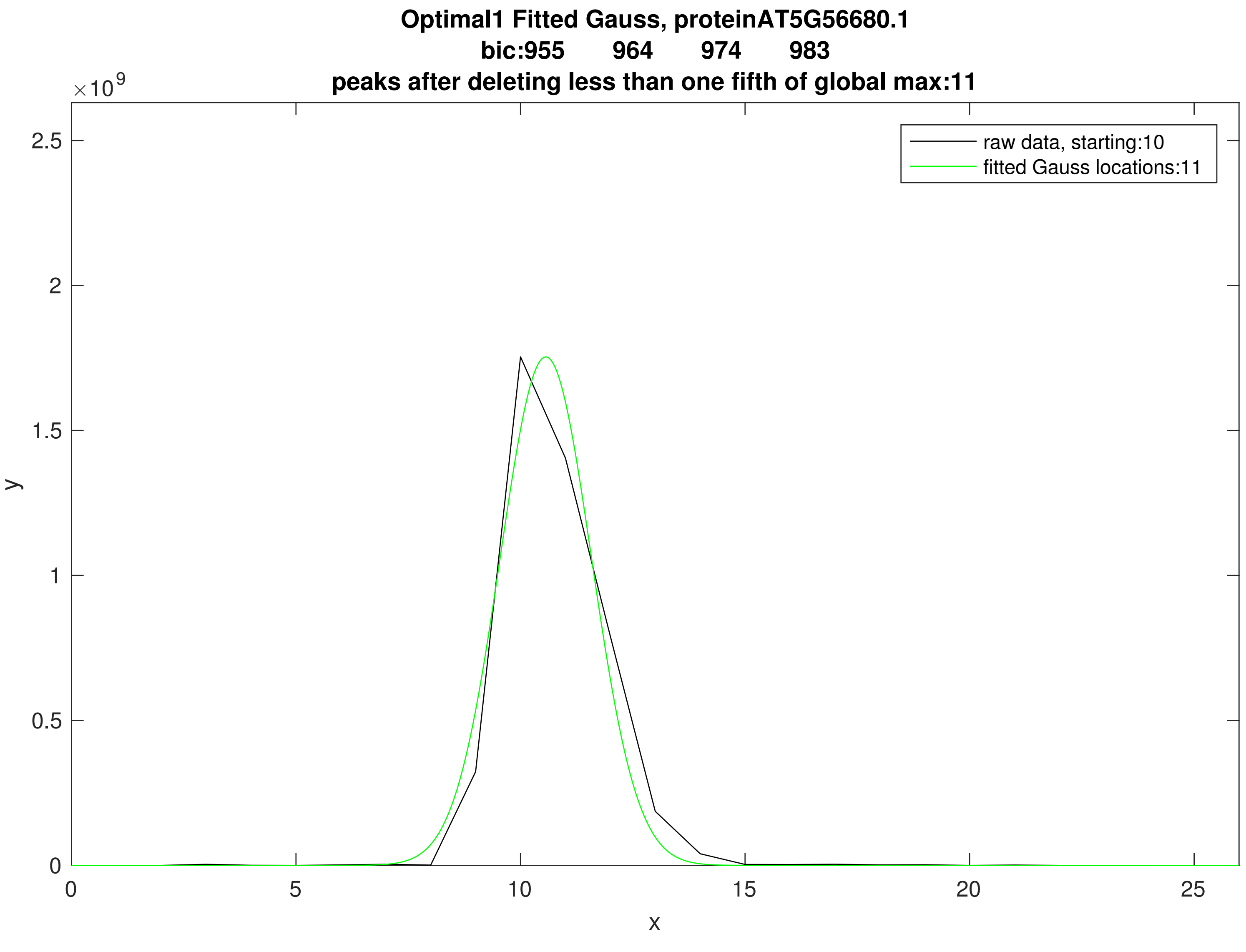
Optimal1 Fitted Gauss, proteinAT5G56680.1
bic:955 964 974 983
peaks after deleting less than one fifth of global max:11
×10^9
2.5
2
1.5
1
0.5
0
y
0
5
10
15
20
25
x
raw data, starting:10
fitted Gauss locations:11

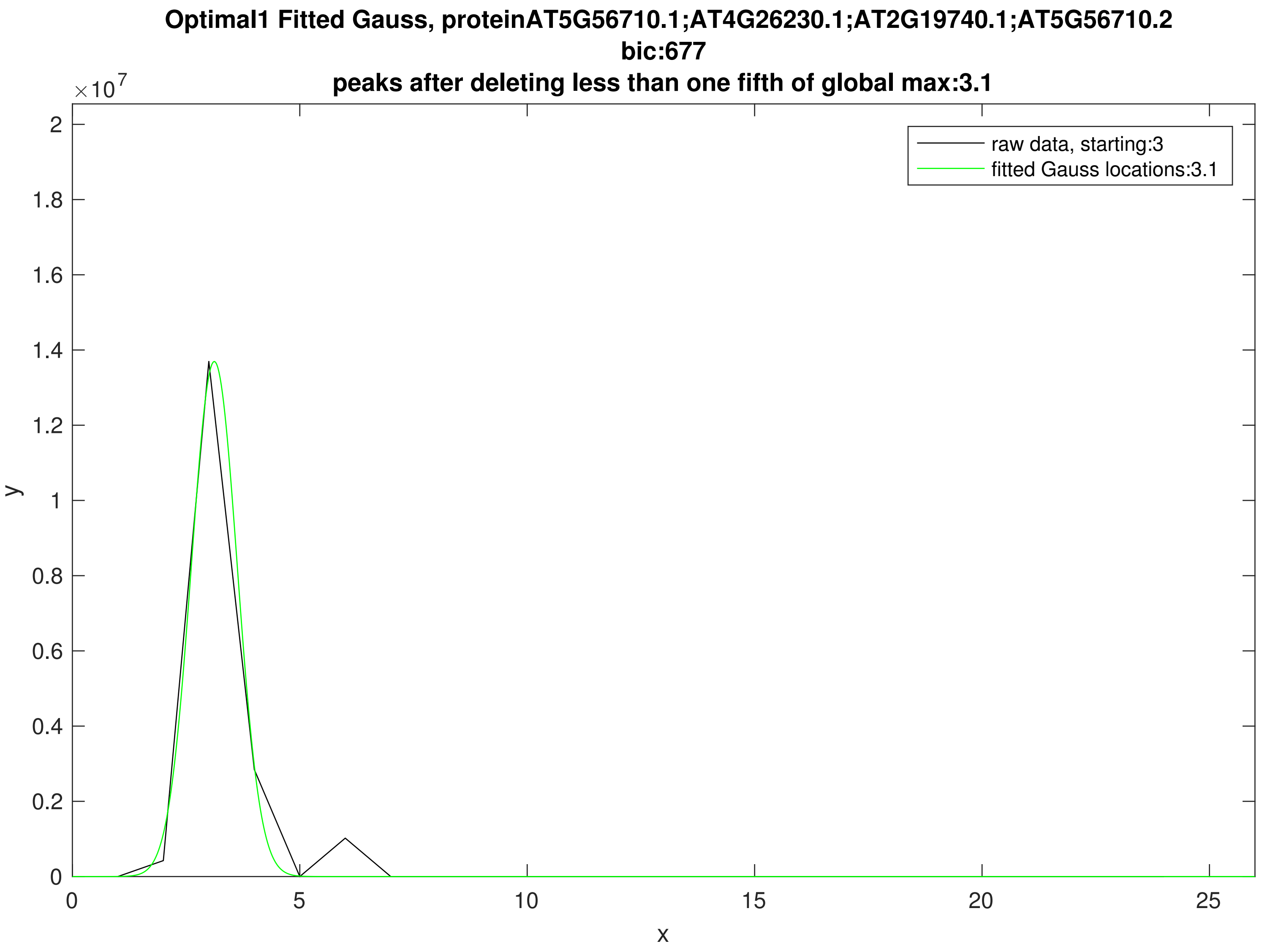
Optimal1 Fitted Gauss, proteinAT5G56710.1;AT4G26230.1;AT2G19740.1;AT5G56710.2
bic:677
peaks after deleting less than one fifth of global max:3.1
×10^7
raw data, starting:3
fitted Gauss locations:3.1
y
x
0
0.2
0.4
0.6
0.8
1
1.2
1.4
1.6
1.8
2
5
10
15
20
25

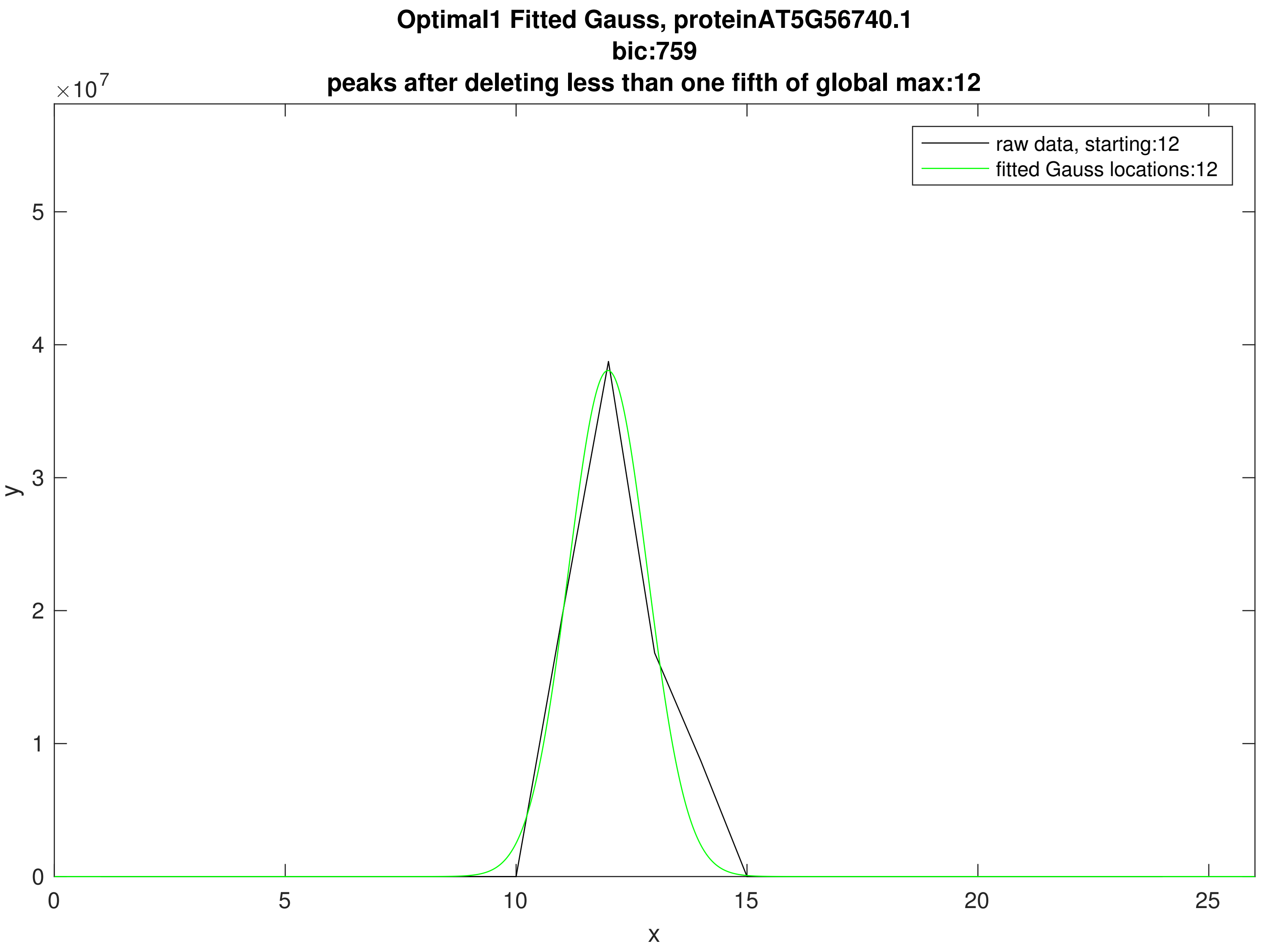
Optimal1 Fitted Gauss, proteinAT5G56740.1
bic:759
peaks after deleting less than one fifth of global max:12
×10^7
raw data, starting:12
fitted Gauss locations:12
y
x
0
1
2
3
4
5
0
5
10
15
20
25

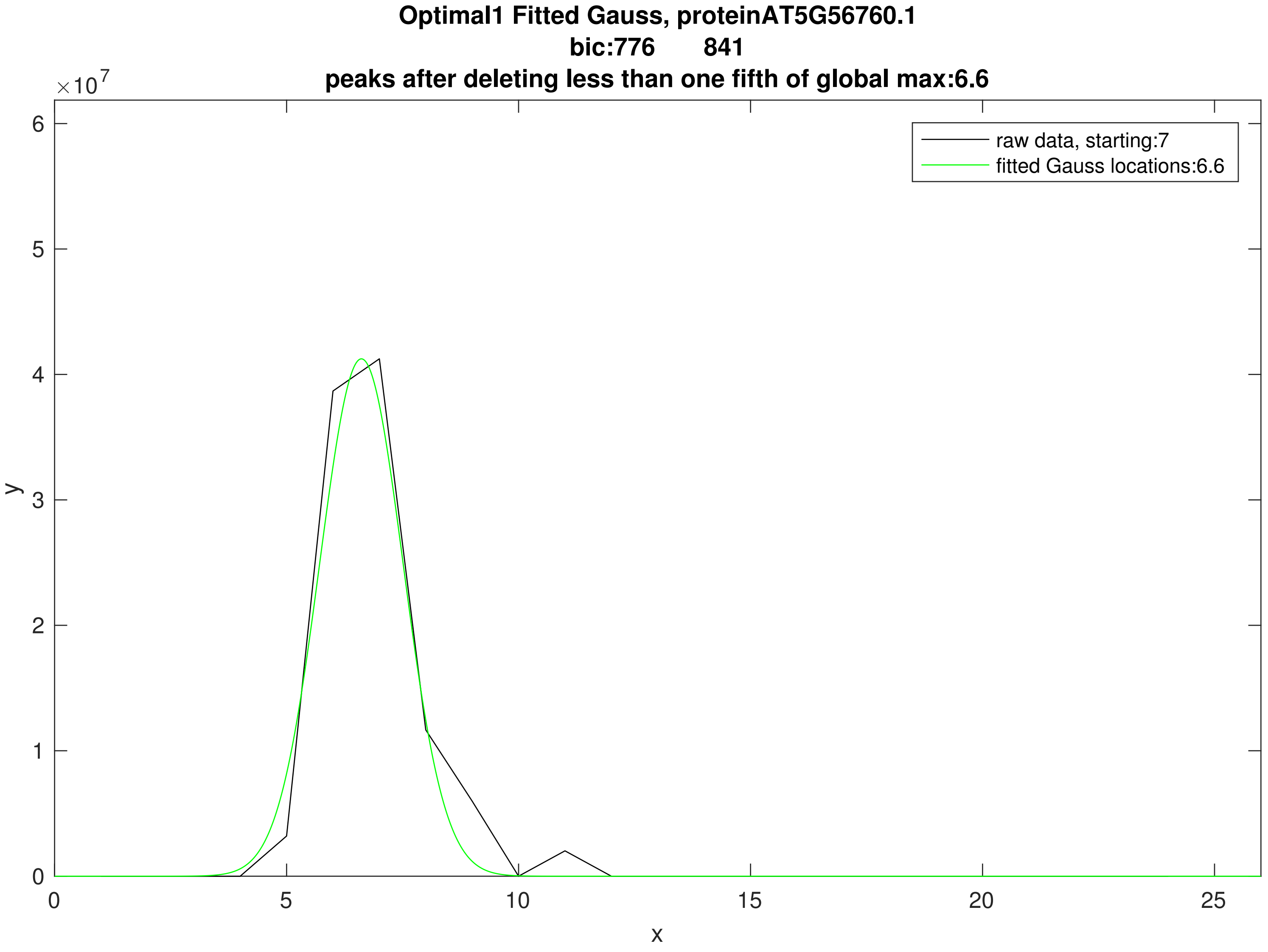
Optimal1 Fitted Gauss, proteinAT5G56760.1
bic:776 841
peaks after deleting less than one fifth of global max:6.6
×10^7
raw data, starting:7
fitted Gauss locations:6.6
y
x
0
1
2
3
4
5
6
0
5
10
15
20
25

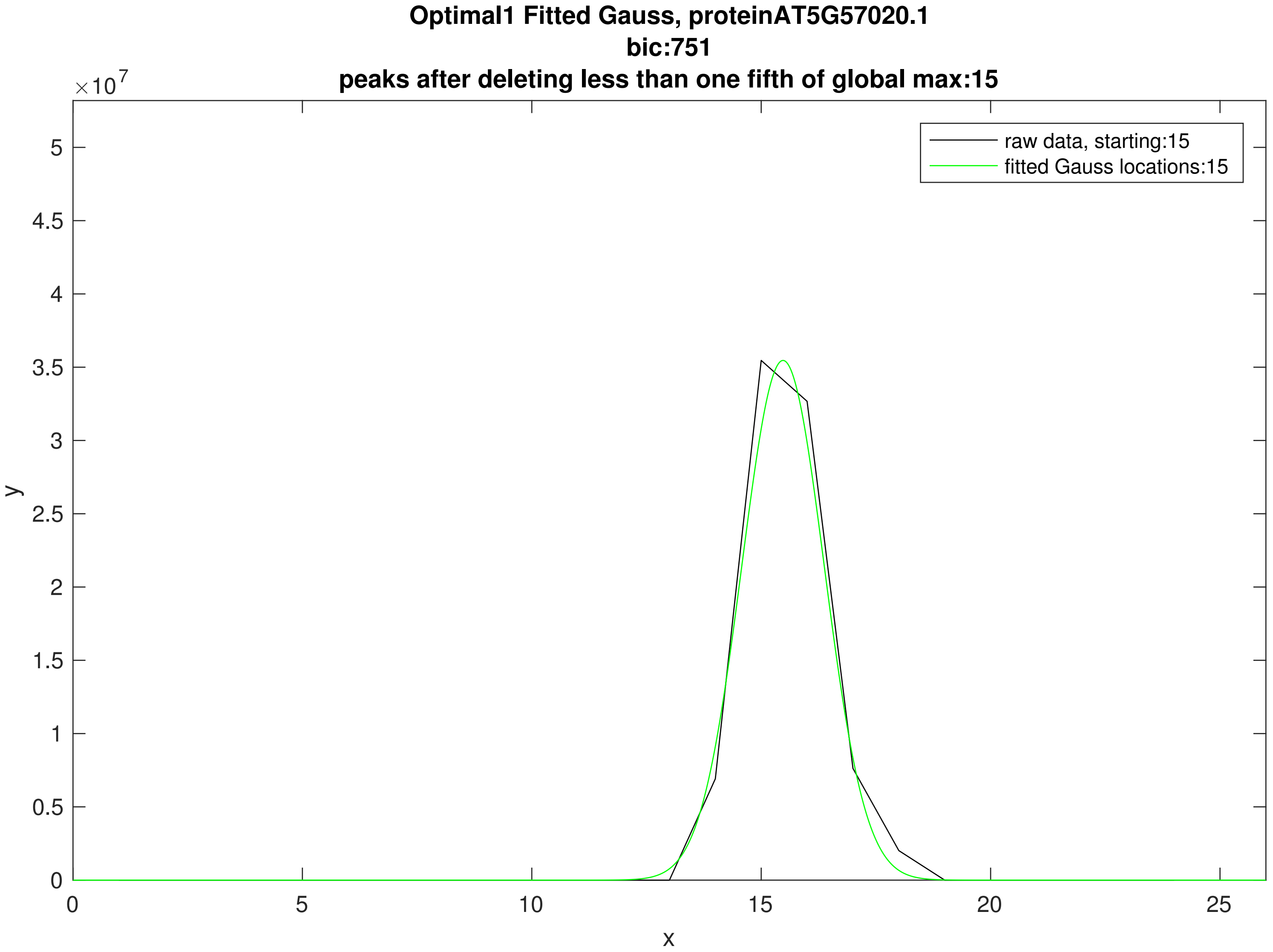
Optimal1 Fitted Gauss, proteinAT5G57020.1
bic:751
peaks after deleting less than one fifth of global max:15
×10^7
raw data, starting:15
fitted Gauss locations:15
y
x
0
0.5
1
1.5
2
2.5
3
3.5
4
4.5
5
0
5
10
15
20
25

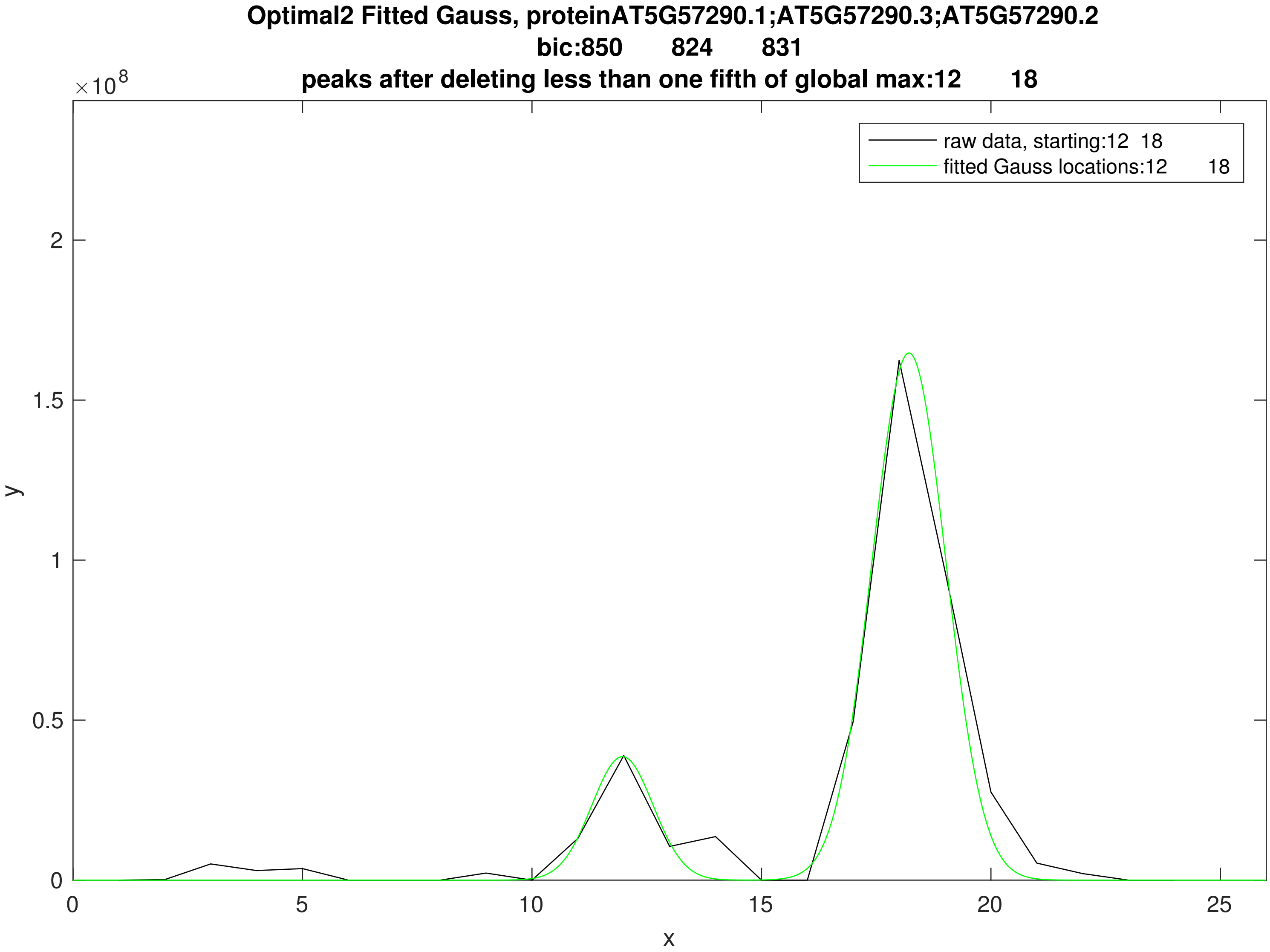

Optimal2 Fitted Gauss, proteinAT5G57290.1;AT5G57290.3;AT5G57290.2
bic:850 824 831
peaks after deleting less than one fifth of global max:12 18
×10^8
raw data, starting:12 18
fitted Gauss locations:12 18
y
x
0
0.5
1
1.5
2
5
10
15
20
25

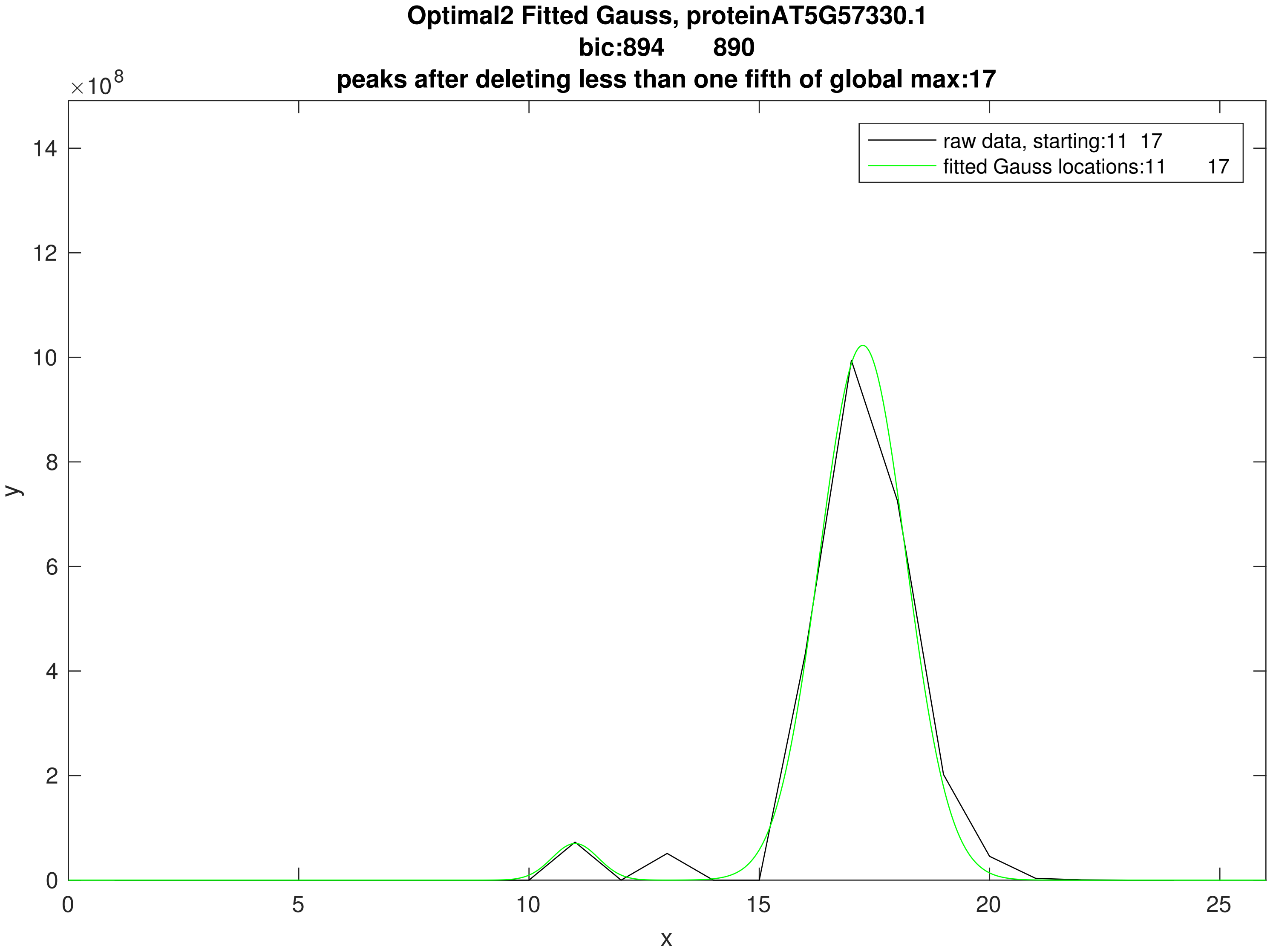

Optimal2 Fitted Gauss, proteinAT5G57330.1
bic:894 890
peaks after deleting less than one fifth of global max:17
×10^8
raw data, starting:11 17
fitted Gauss locations:11 17
y
x
0
2
4
6
8
10
12
14
5
15
20
25

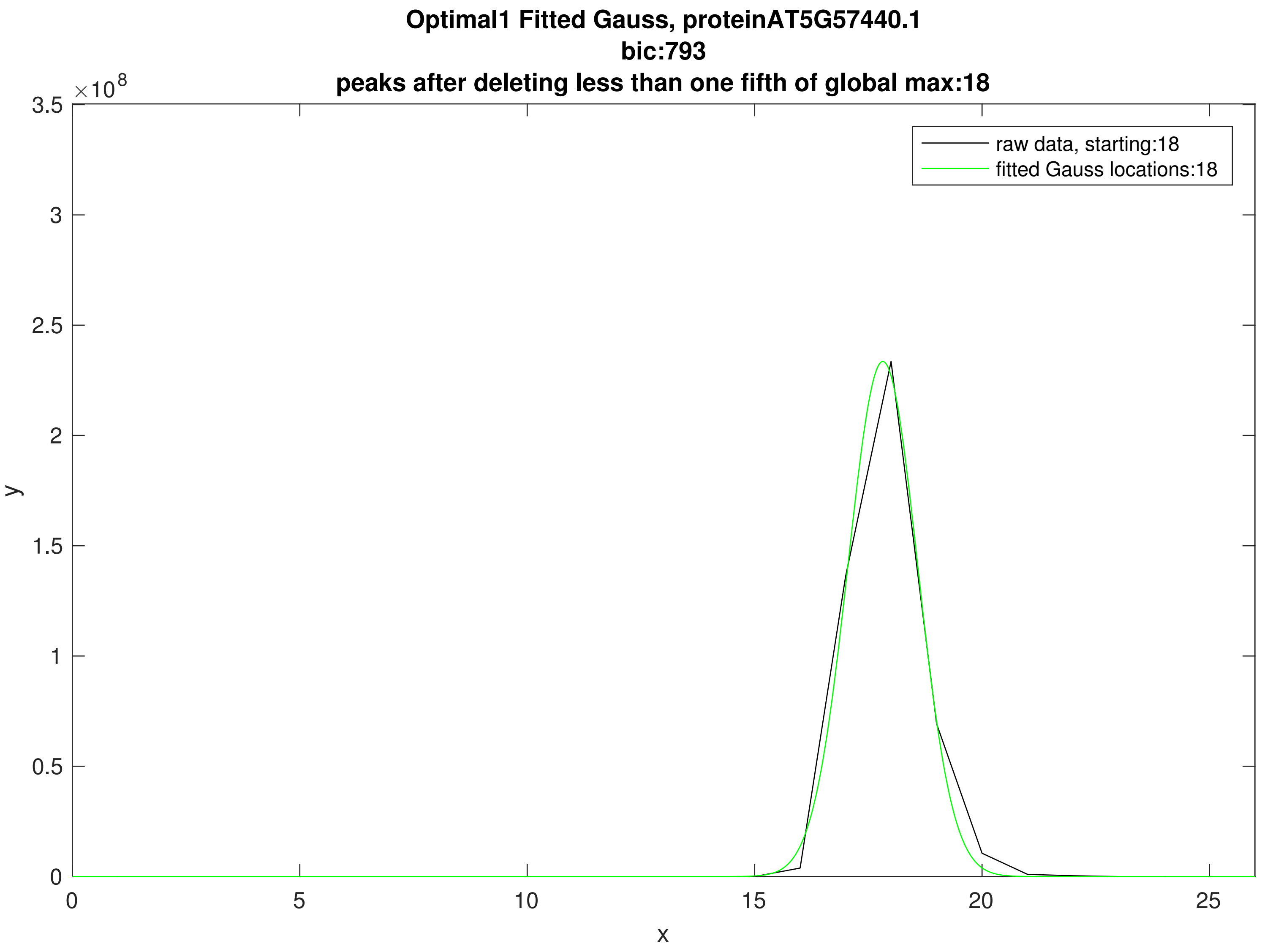

Optimal1 Fitted Gauss, proteinAT5G57440.1
bic:793
peaks after deleting less than one fifth of global max:18
×10^8
3.5
3
2.5
2
1.5
1
0.5
0
y
0
5
10
15
20
25
x
raw data, starting:18
fitted Gauss locations:18

Optimal1 Fitted Gauss, proteinAT5G57580.1
bic:778
peaks after deleting less than one fifth of global max:15

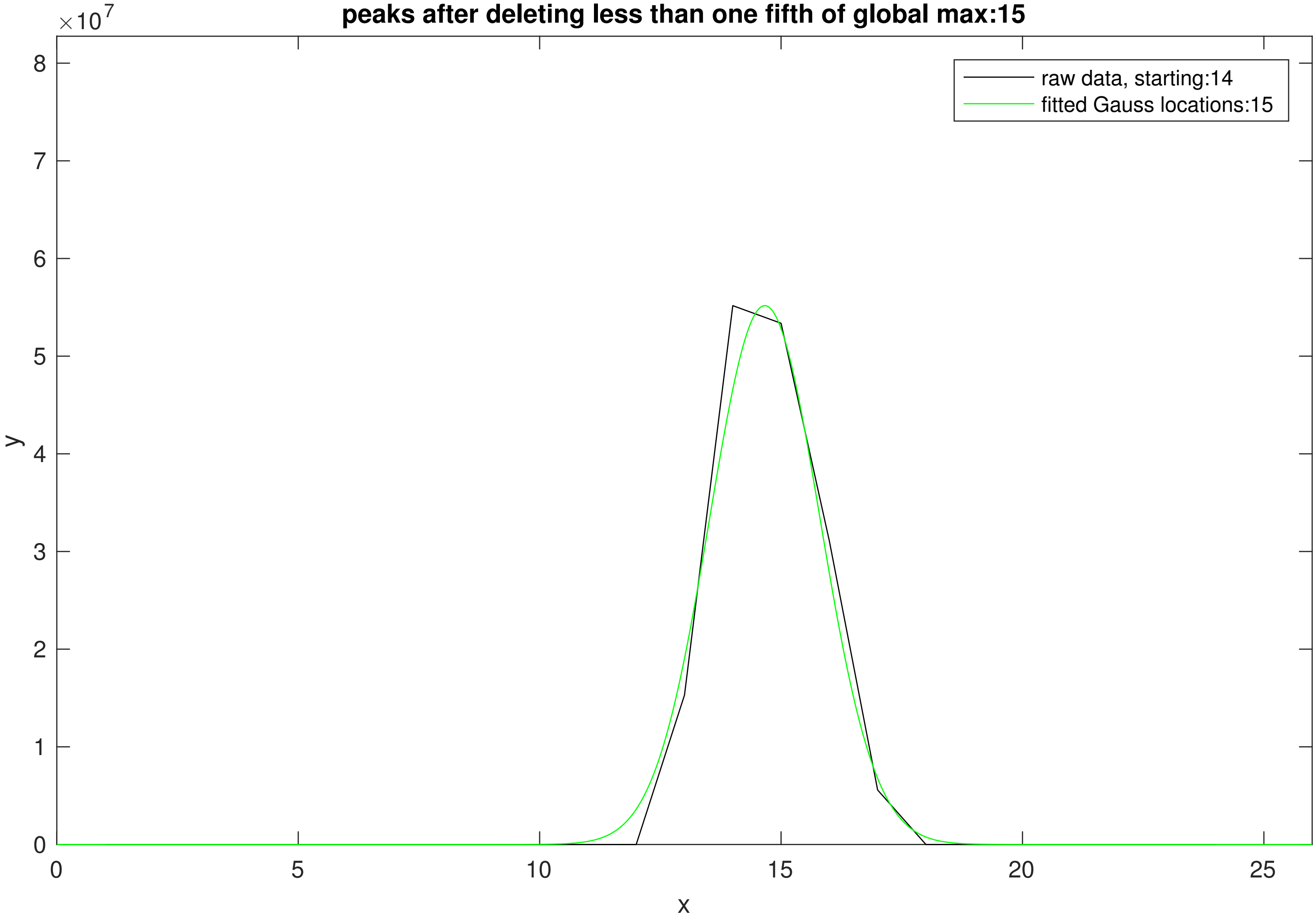

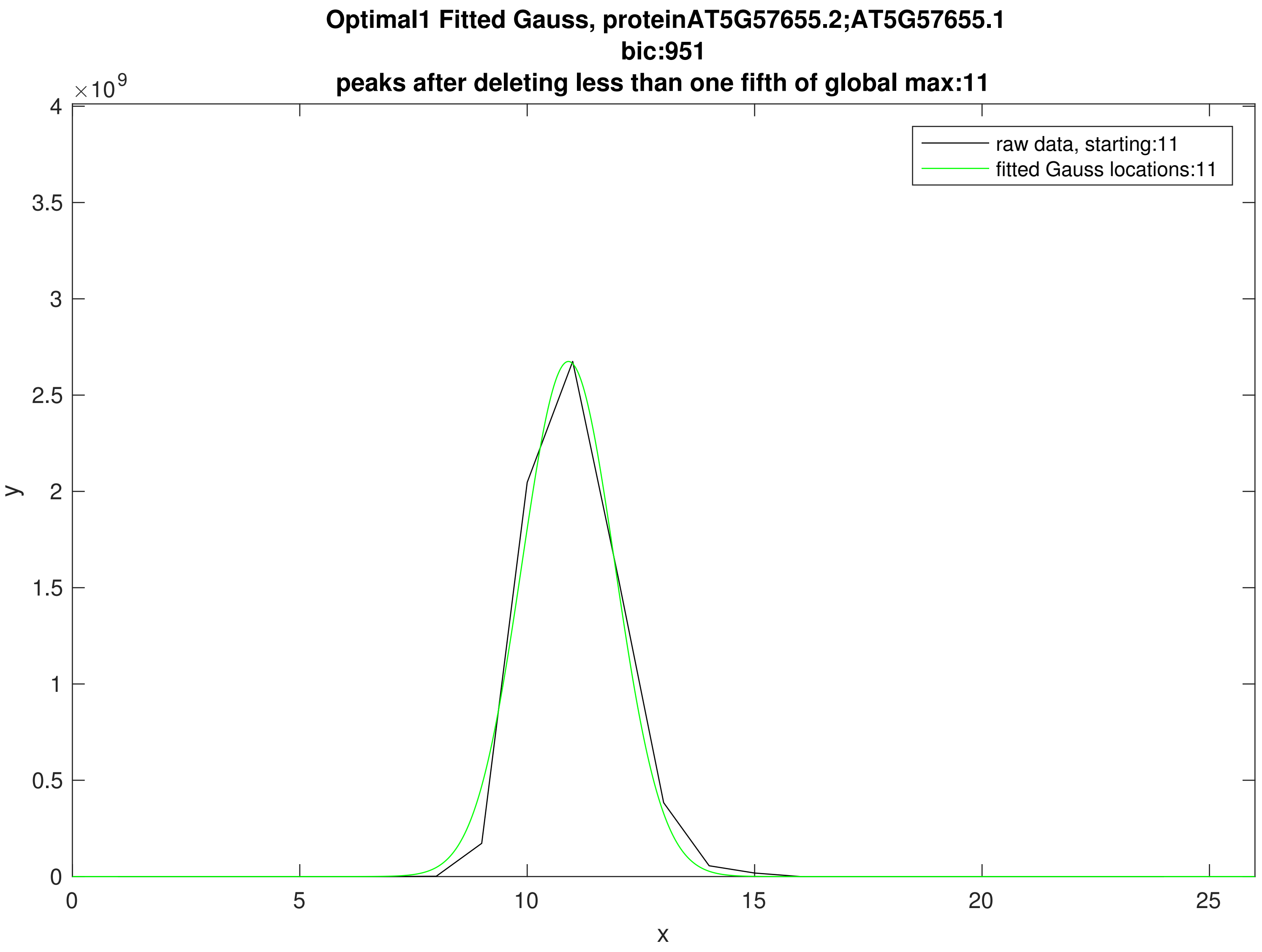
Optimal1 Fitted Gauss, proteinAT5G57655.2;AT5G57655.1
bic:951
peaks after deleting less than one fifth of global max:11
×10^9
raw data, starting:11
fitted Gauss locations:11
y
x
0
0.5
1
1.5
2
2.5
3
3.5
4
0
5
10
15
20
25

**Optimal1 Fitted Gauss, proteinAT5G57700.2;AT5G57700.4;AT5G57700.1;AT5G57700.3**
**bic:872 874**
**peaks after deleting less than one fifth of global max:13**

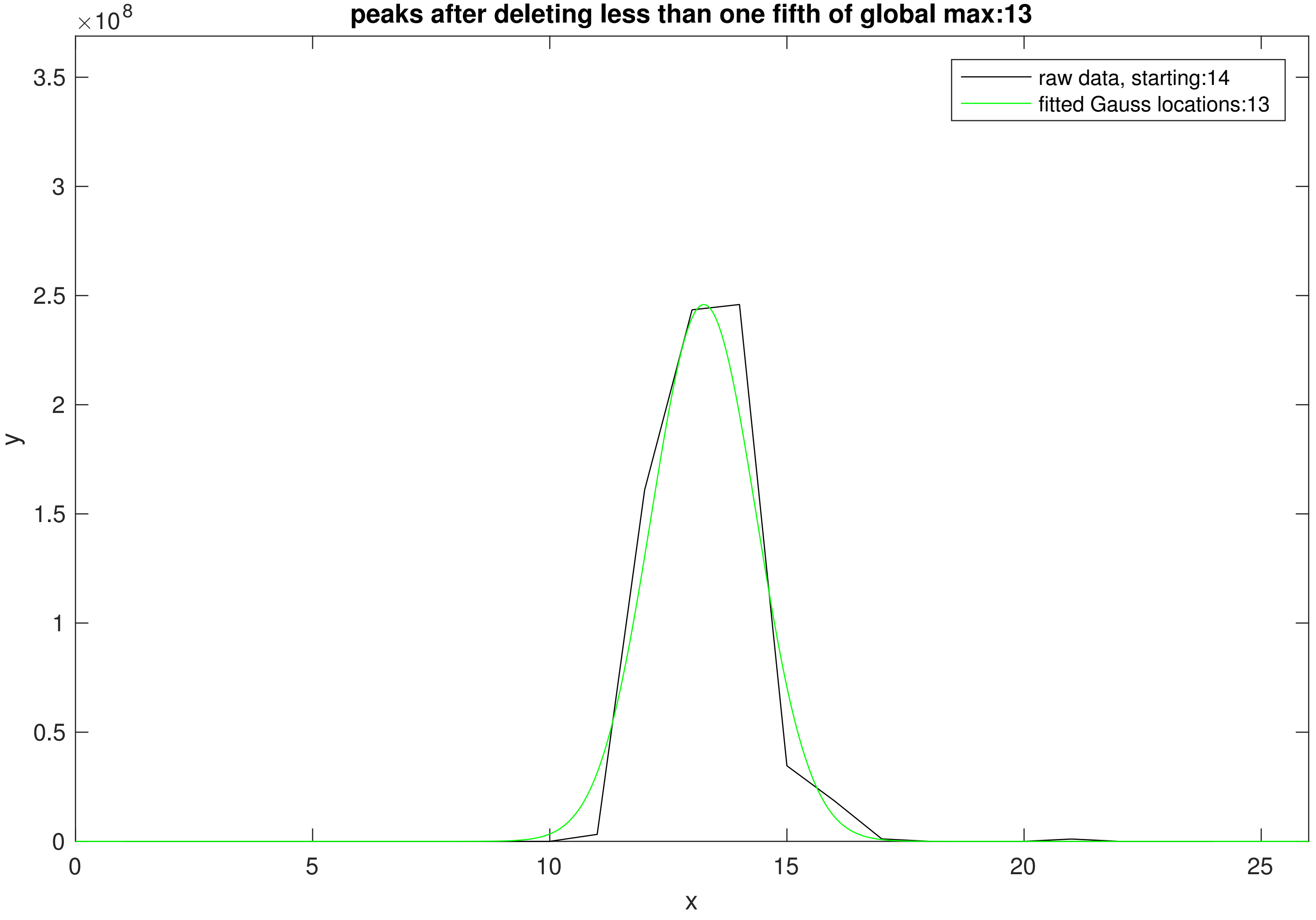

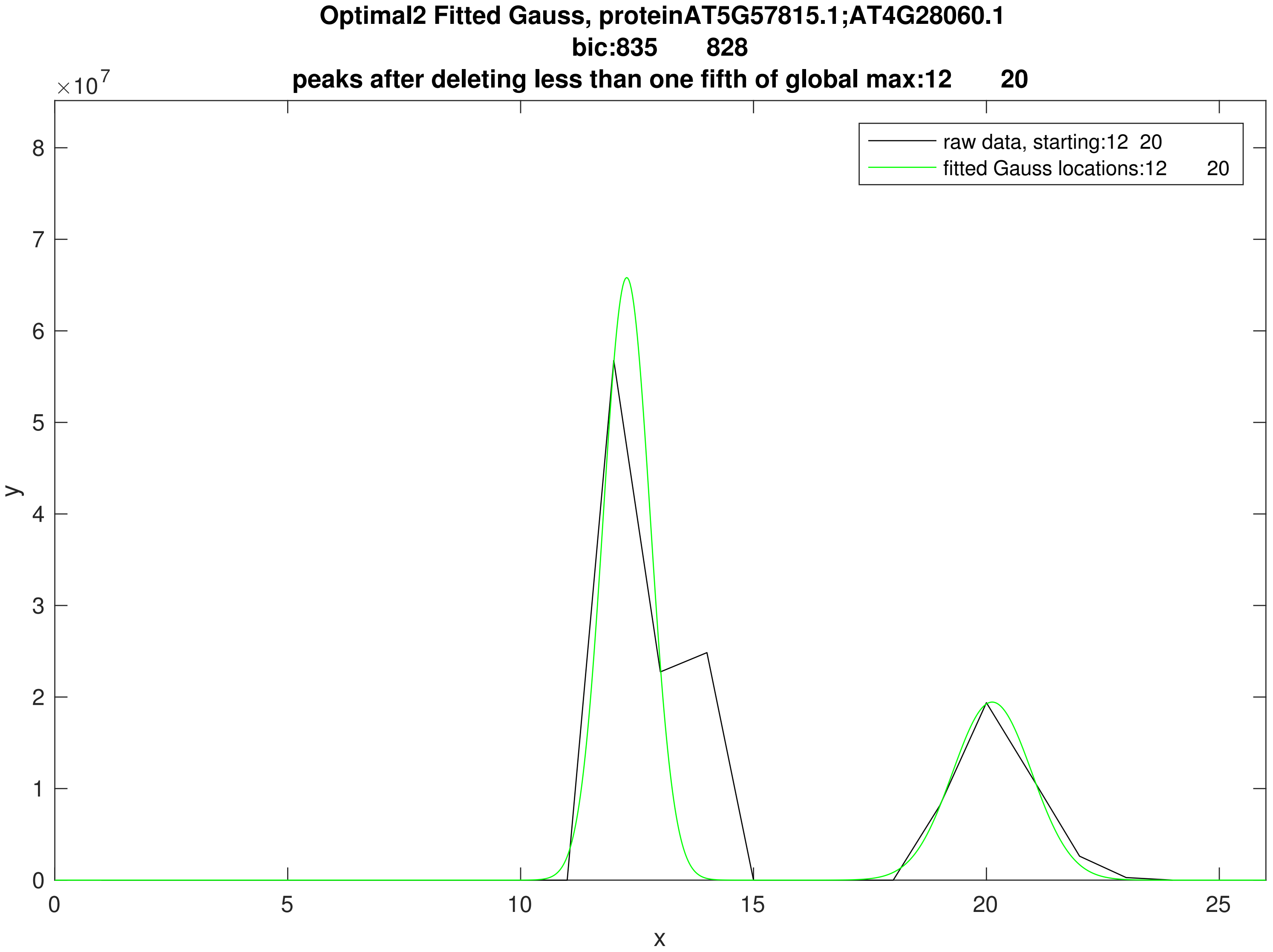
Optimal2 Fitted Gauss, proteinAT5G57815.1;AT4G28060.1
bic:835 828
peaks after deleting less than one fifth of global max:12 20
raw data, starting:12 20
fitted Gauss locations:12 20
y
x
×10^7

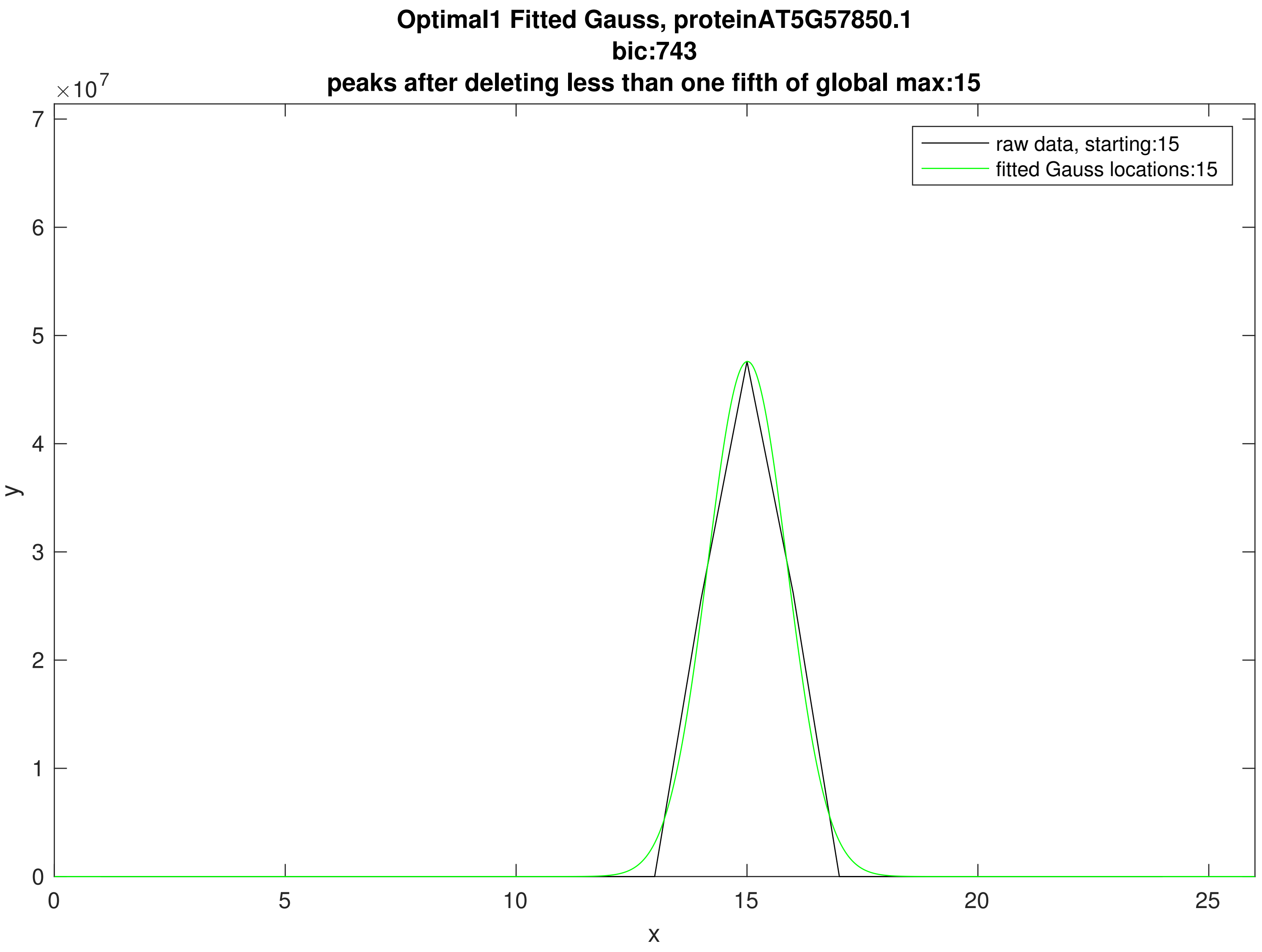
Optimal1 Fitted Gauss, proteinAT5G57850.1
bic:743
peaks after deleting less than one fifth of global max:15
×10^7
7
6
5
4
3
2
1
0
y
0
5
10
15
20
25
x
raw data, starting:15
fitted Gauss locations:15

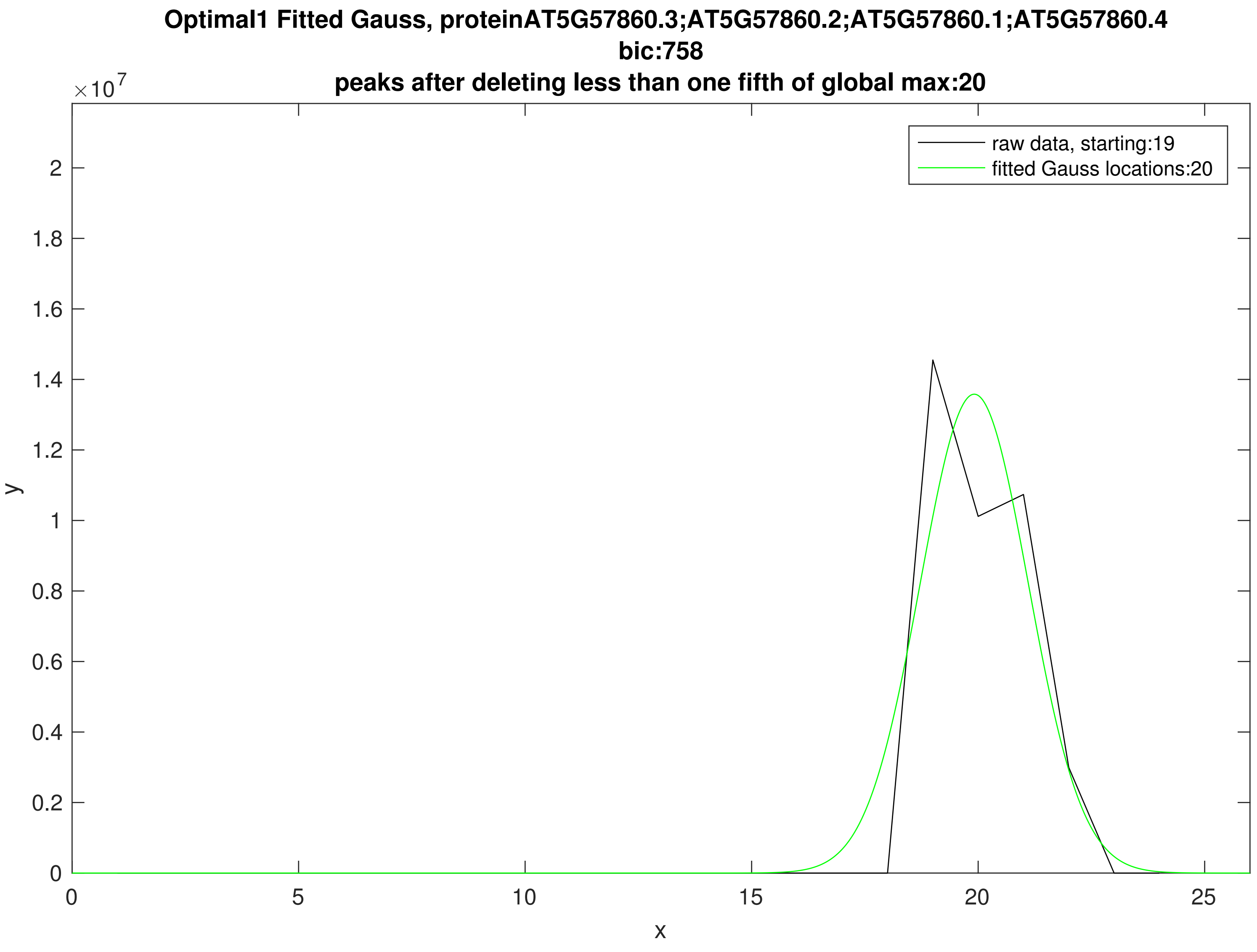
Optimal1 Fitted Gauss, proteinAT5G57860.3;AT5G57860.2;AT5G57860.1;AT5G57860.4
bic:758
peaks after deleting less than one fifth of global max:20
×10^7
raw data, starting:19
fitted Gauss locations:20
y
x
0
0.2
0.4
0.6
0.8
1
1.2
1.4
1.6
1.8
2
0
5
10
15
20
25

Optimal1 Fitted Gauss, proteinAT5G57870.2;AT5G57870.1
bic:774
peaks after deleting less than one fifth of global max:16

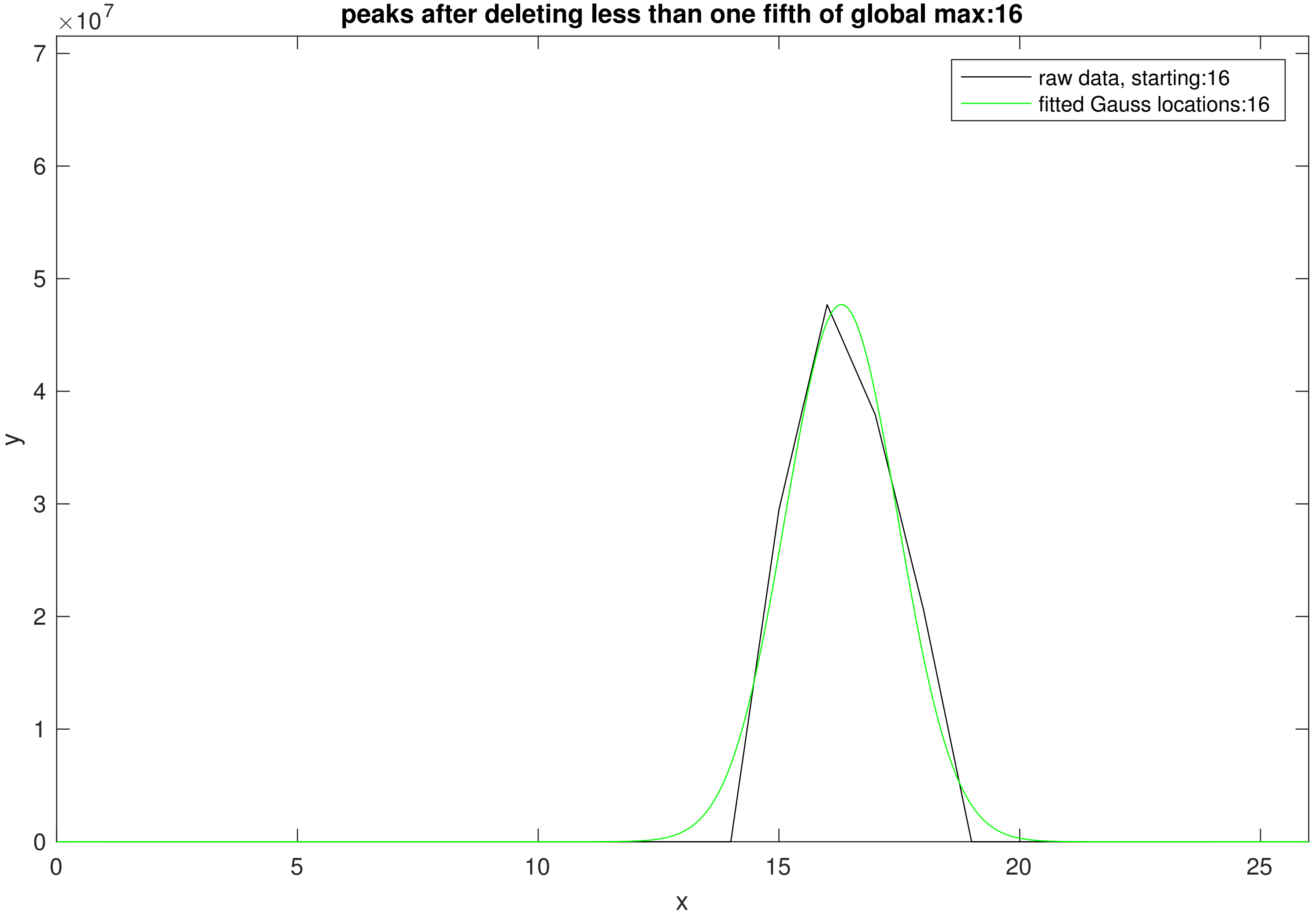

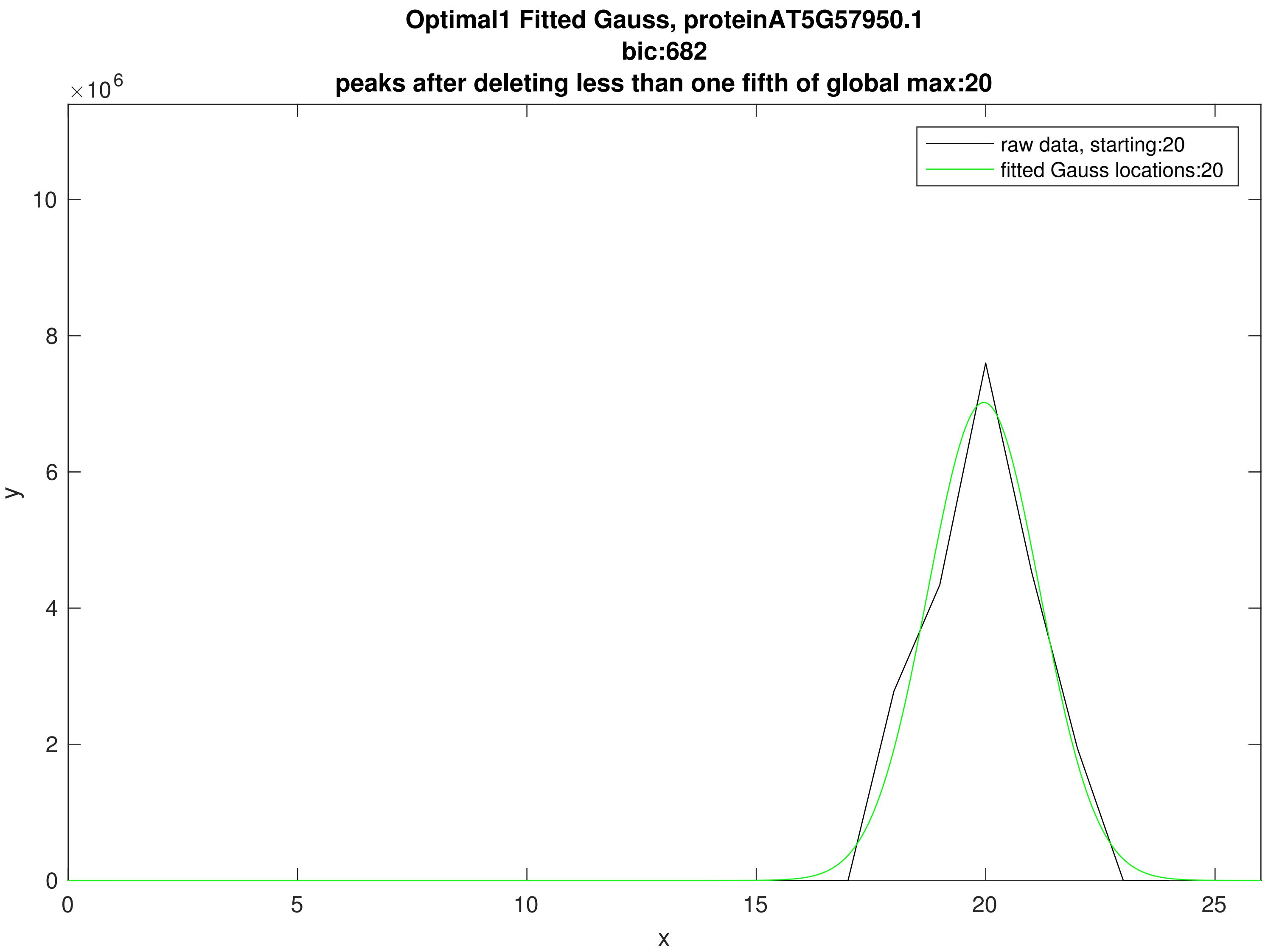

Optimal1 Fitted Gauss, proteinAT5G57950.1
bic:682
peaks after deleting less than one fifth of global max:20
×10^6
raw data, starting:20
fitted Gauss locations:20
10
8
6
4
2
0
y
0
5
10
15
20
25
x

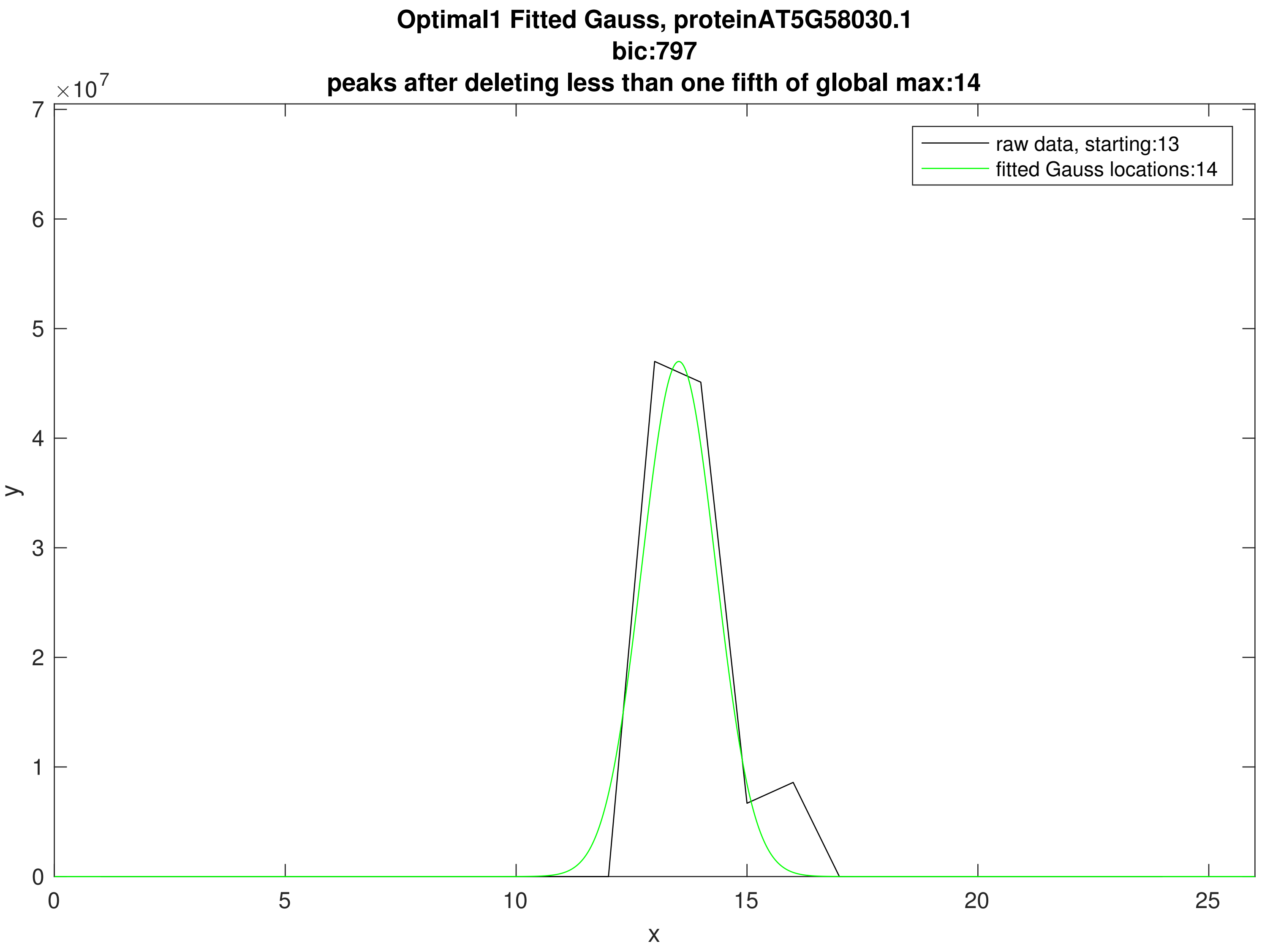

Optimal1 Fitted Gauss, proteinAT5G58030.1
bic:797
peaks after deleting less than one fifth of global max:14
×10^7
raw data, starting:13
fitted Gauss locations:14
y
x
0
1
2
3
4
5
6
7
0
5
10
15
20
25

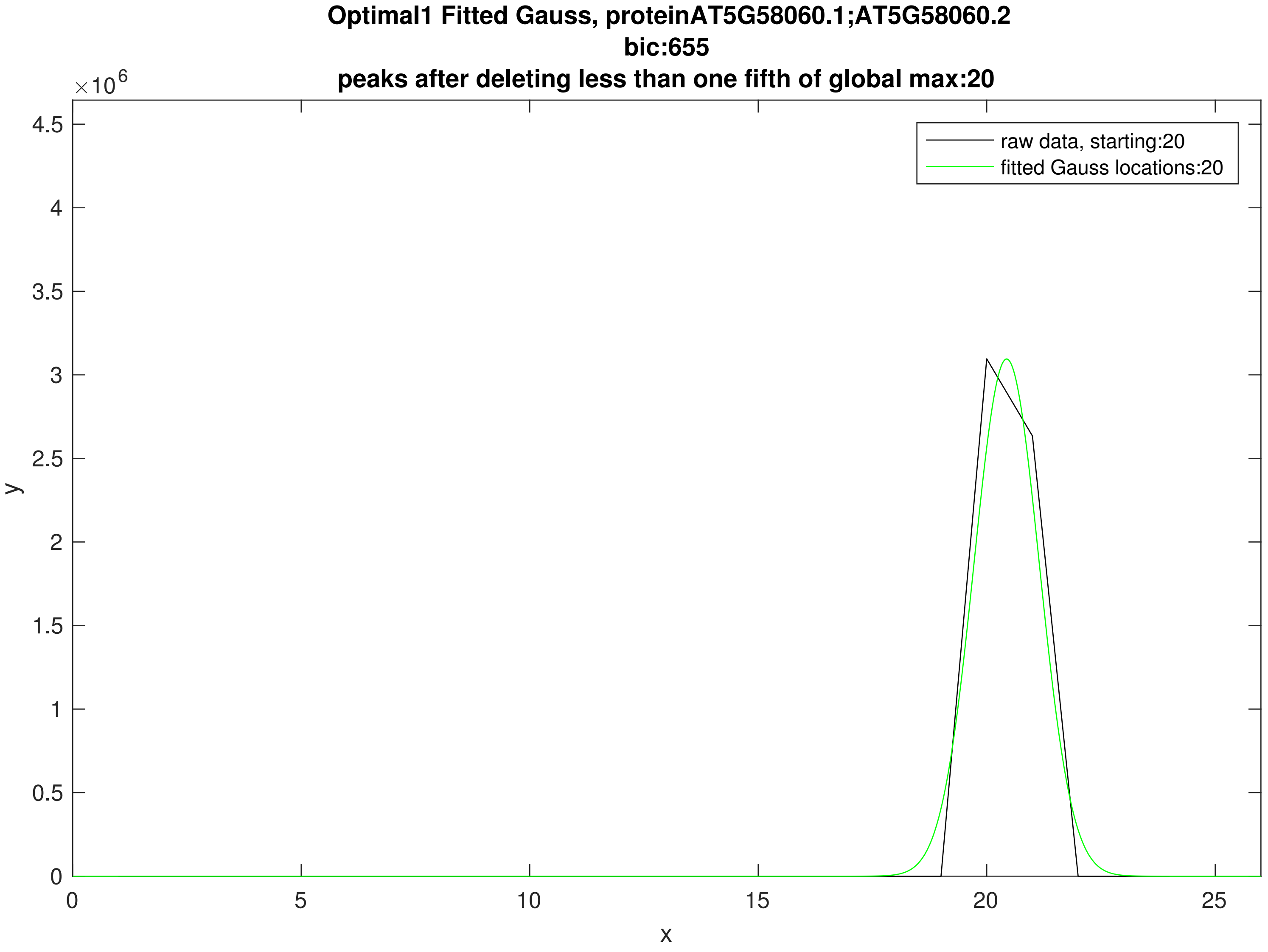
Optimal1 Fitted Gauss, proteinAT5G58060.1;AT5G58060.2
bic:655
peaks after deleting less than one fifth of global max:20
×10^6
raw data, starting:20
fitted Gauss locations:20
y
x
0
0.5
1
1.5
2
2.5
3
3.5
4
4.5
0
5
10
15
20
25

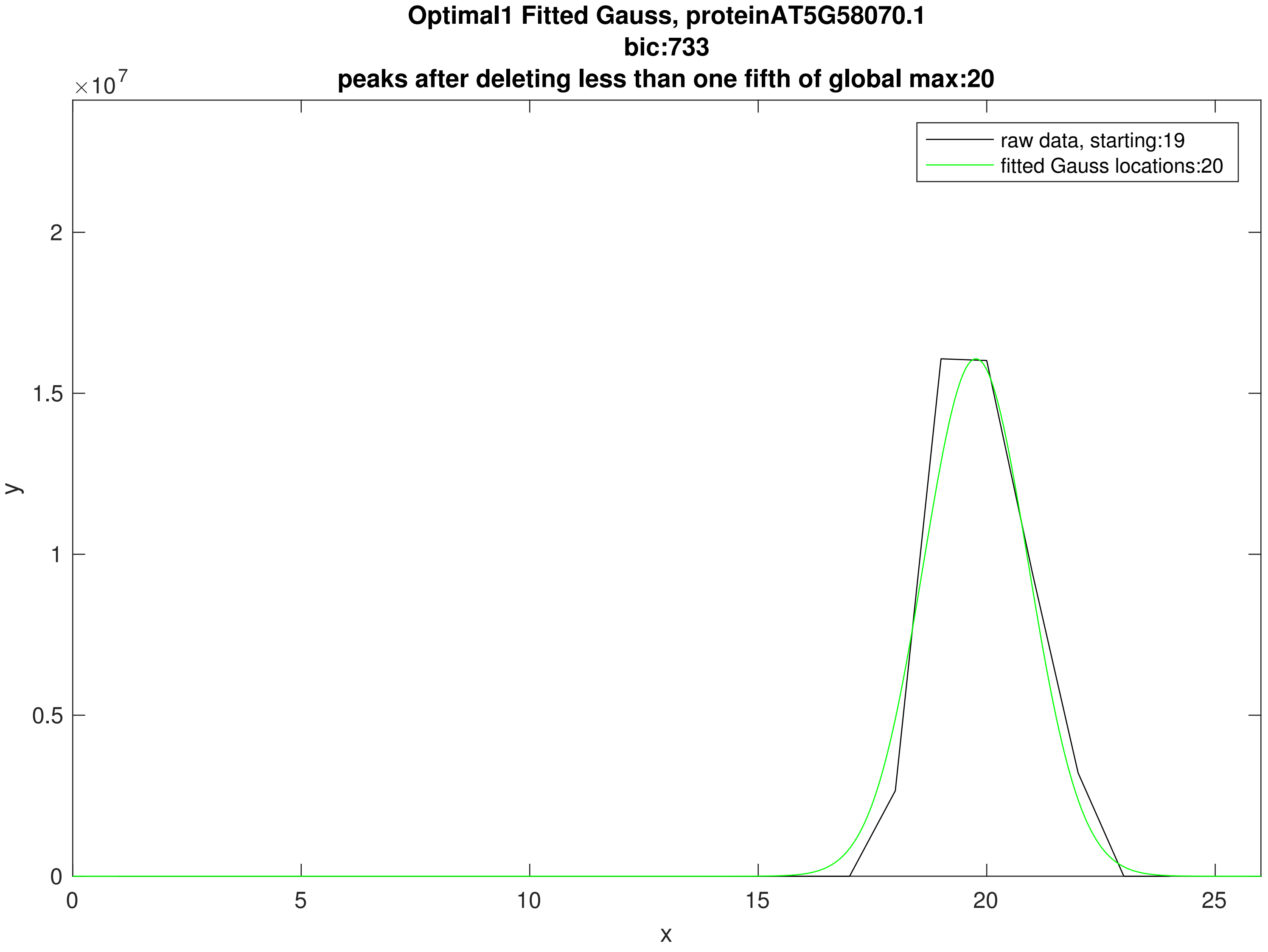
Optimal1 Fitted Gauss, proteinAT5G58070.1
bic:733
peaks after deleting less than one fifth of global max:20
×10^7
raw data, starting:19
fitted Gauss locations:20
2
1.5
1
0.5
0
y
0
5
10
15
20
25
x

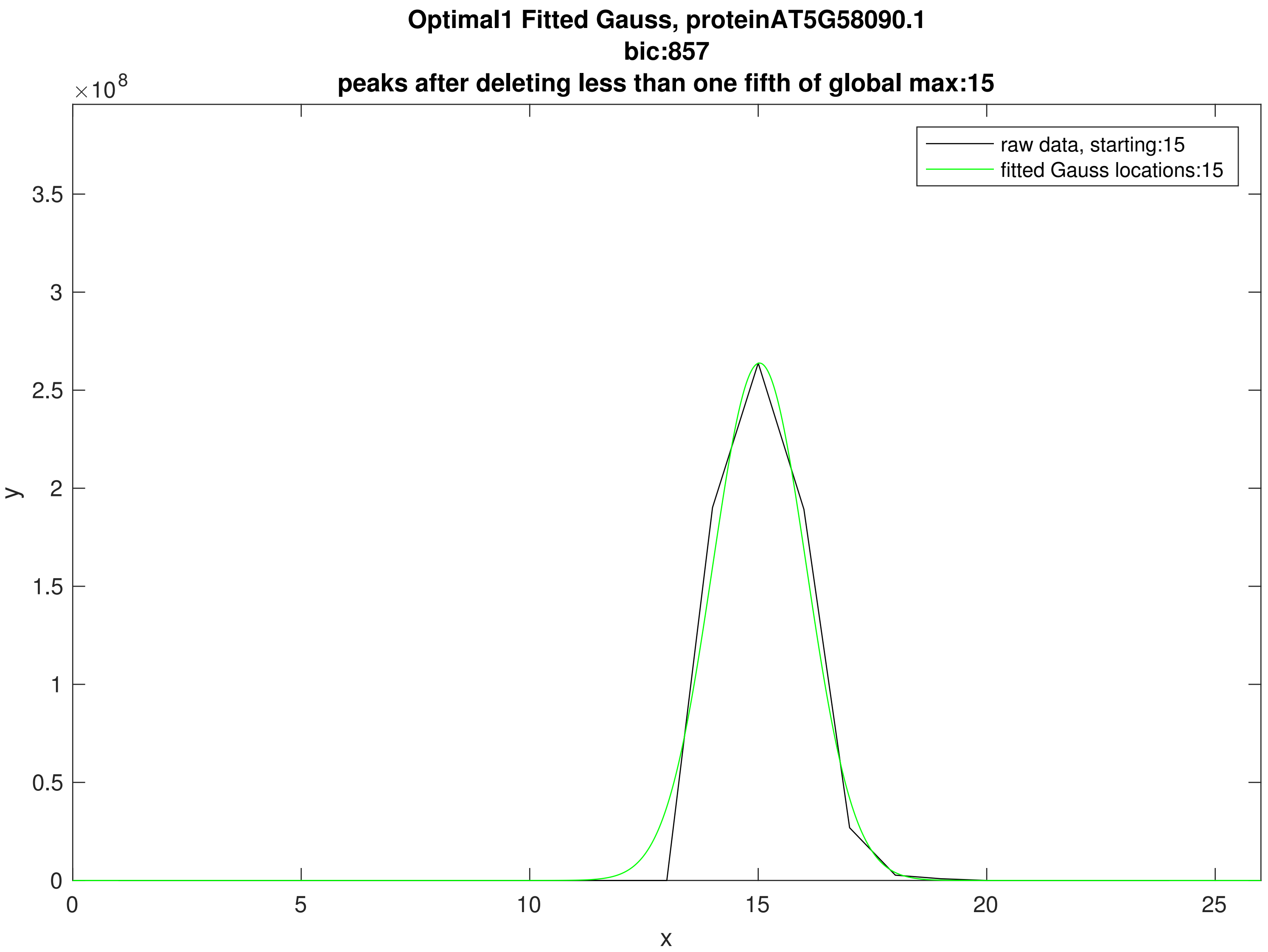
Optimal1 Fitted Gauss, proteinAT5G58090.1
bic:857
peaks after deleting less than one fifth of global max:15
×10^8
3.5
3
2.5
2
1.5
1
0.5
0
y
0
5
10
15
20
25
x
raw data, starting:15
fitted Gauss locations:15

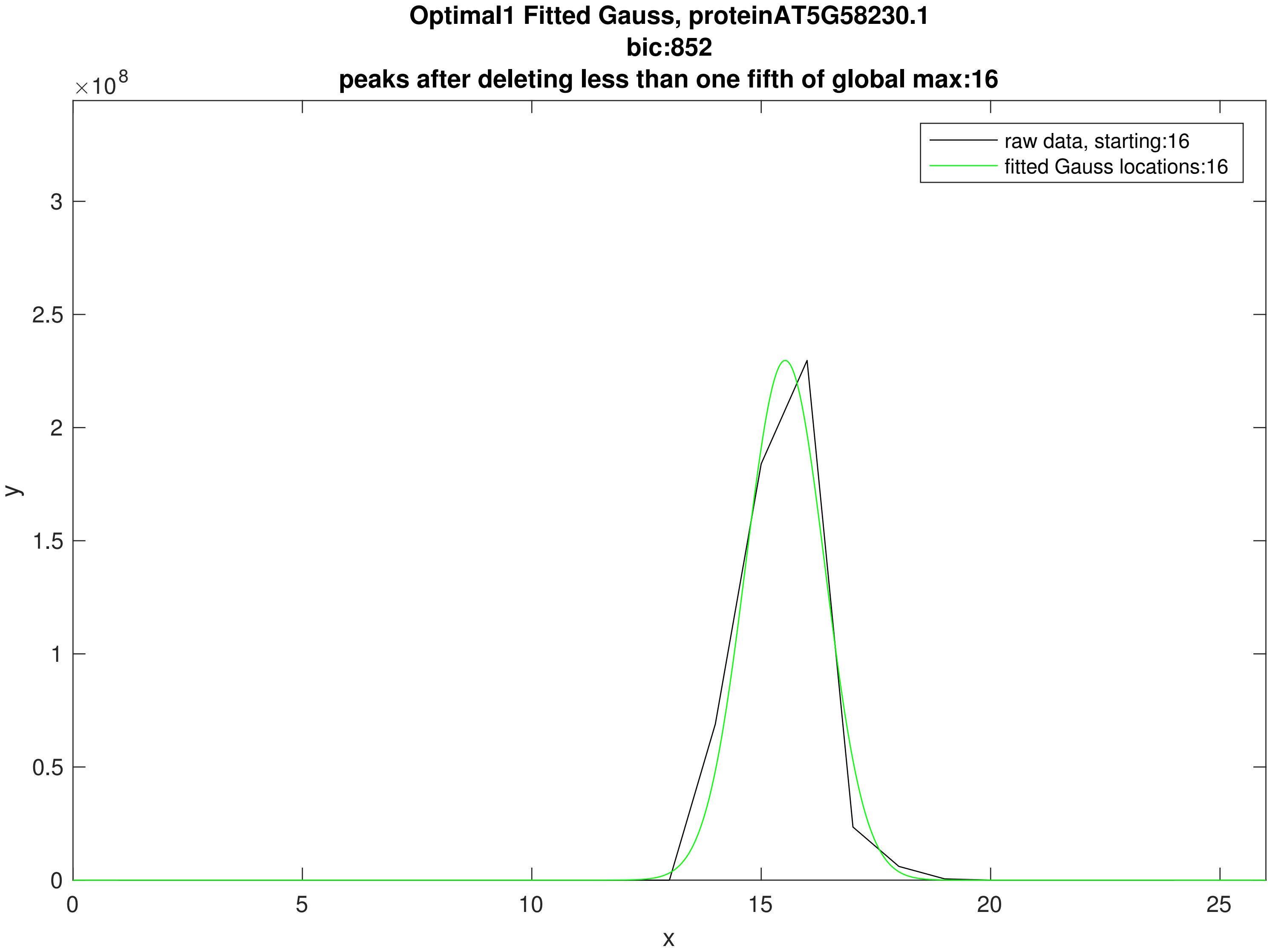

Optimal1 Fitted Gauss, proteinAT5G58230.1
bic:852
peaks after deleting less than one fifth of global max:16
×10^8
raw data, starting:16
fitted Gauss locations:16
y
x
0
0.5
1
1.5
2
2.5
3
0
5
10
15
20
25

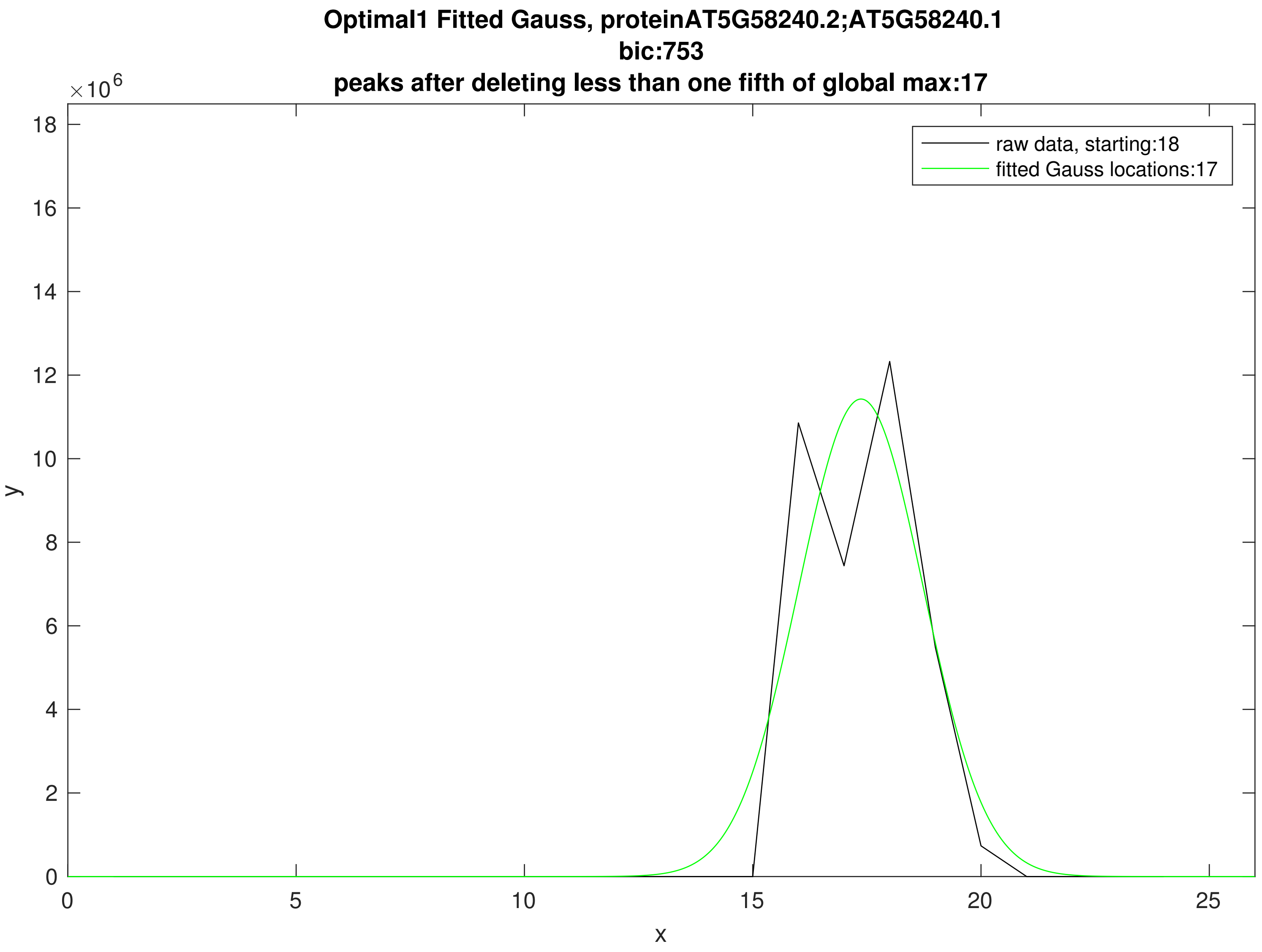
Optimal1 Fitted Gauss, proteinAT5G58240.2;AT5G58240.1
bic:753
peaks after deleting less than one fifth of global max:17
×10^6
raw data, starting:18
fitted Gauss locations:17
y
x
0
2
4
6
8
10
12
14
16
18
0
5
10
15
20
25

**Optimal1 Fitted Gauss, proteinAT5G58290.1**
**bic:830 839**
**peaks after deleting less than one fifth of global max:3.8**

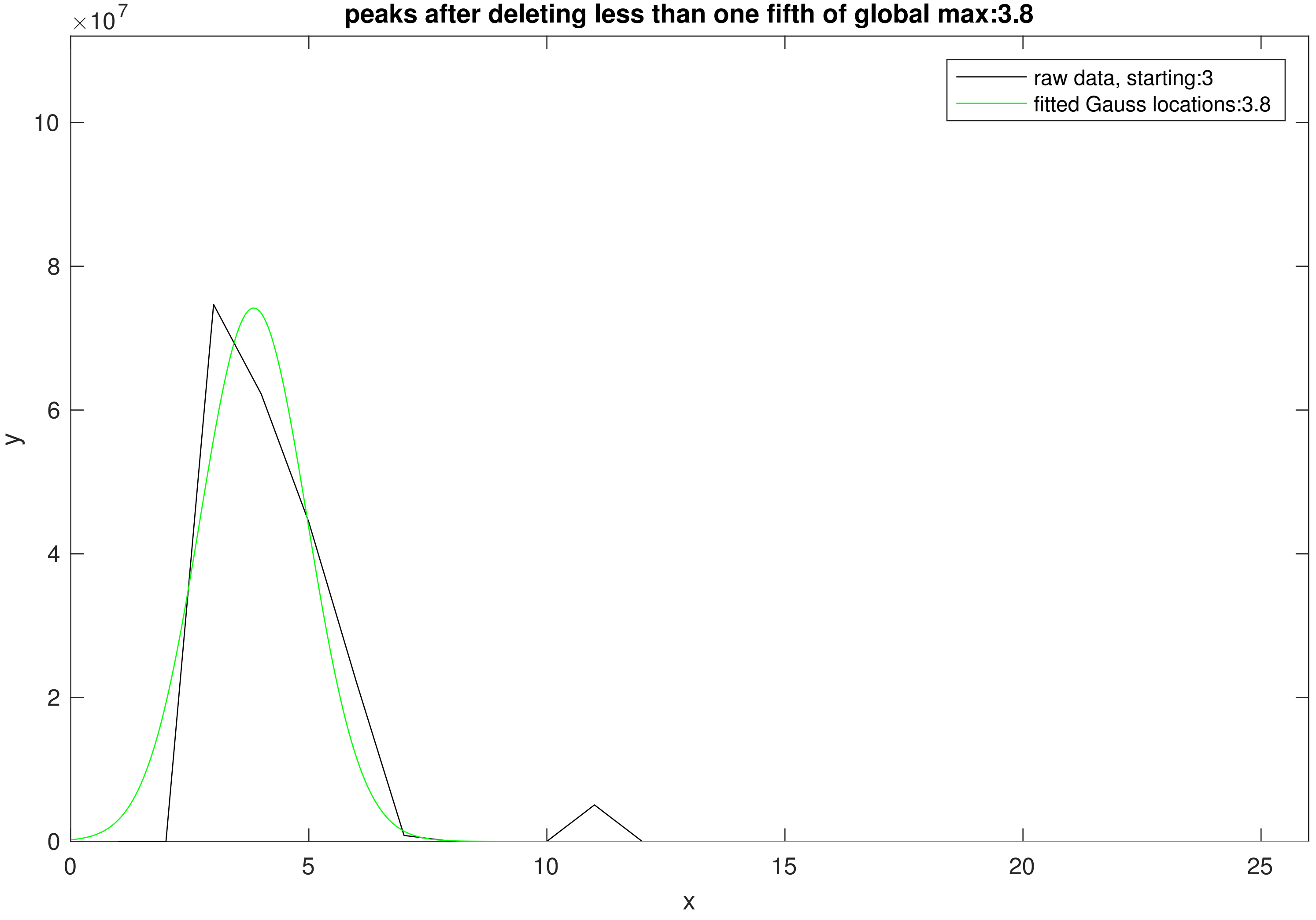

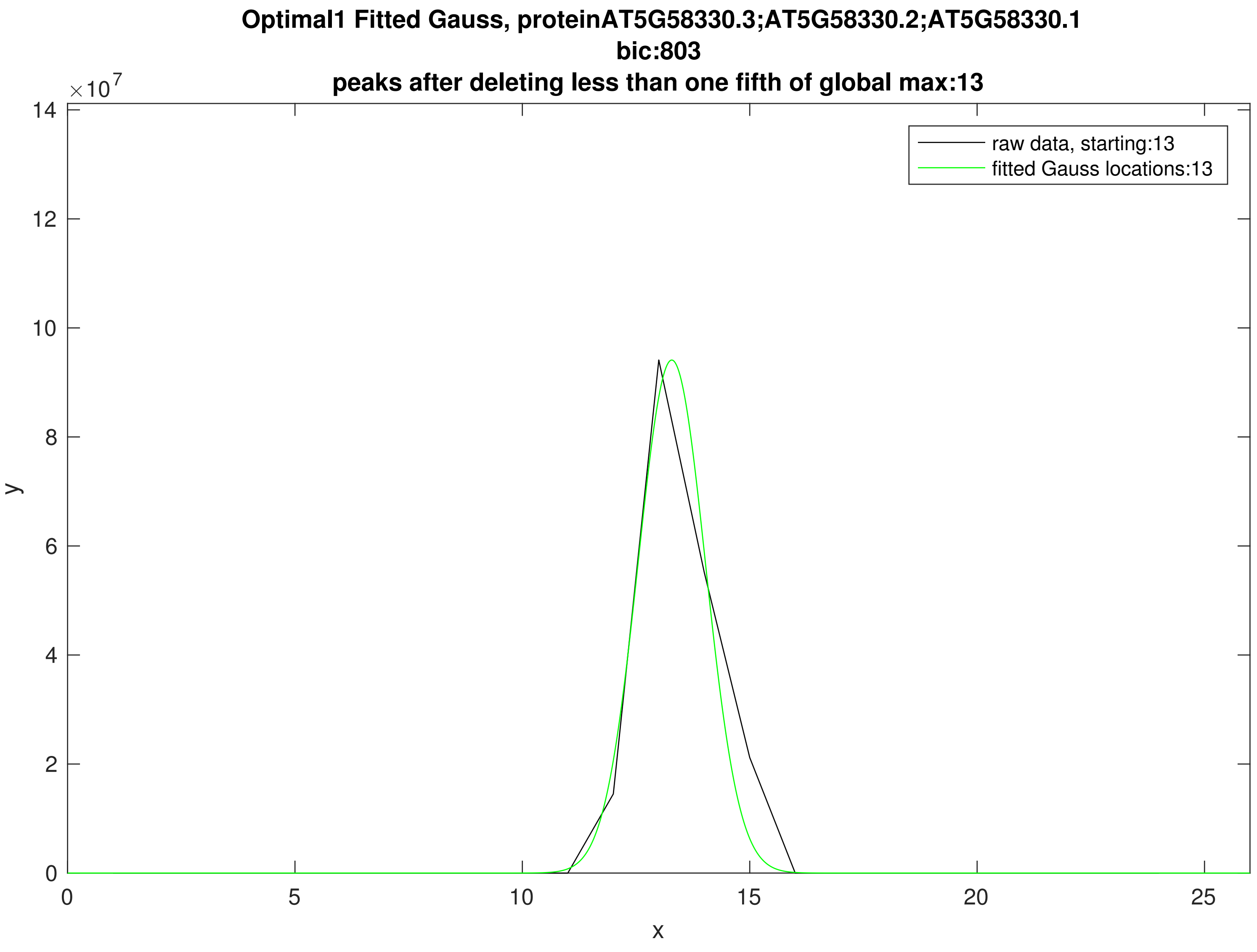

Optimal1 Fitted Gauss, proteinAT5G58330.3;AT5G58330.2;AT5G58330.1
bic:803
peaks after deleting less than one fifth of global max:13
×10^7
raw data, starting:13
fitted Gauss locations:13
y
x
0
2
4
6
8
10
12
14
5
15
20
25

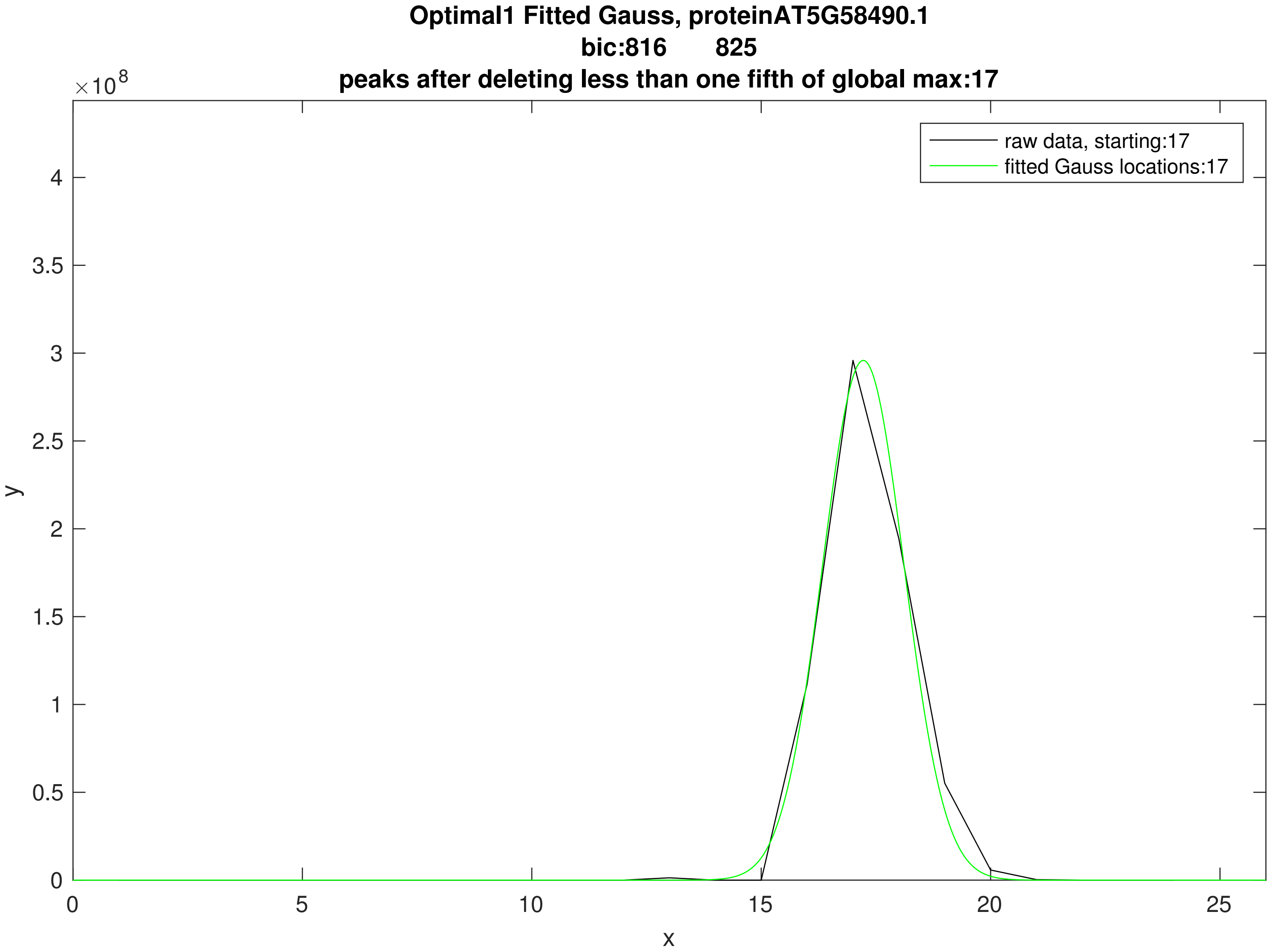
Optimal1 Fitted Gauss, proteinAT5G58490.1
bic:816 825
peaks after deleting less than one fifth of global max:17
×10^8
raw data, starting:17
fitted Gauss locations:17
y
x
0
0.5
1
1.5
2
2.5
3
3.5
4
0
5
10
15
20
25

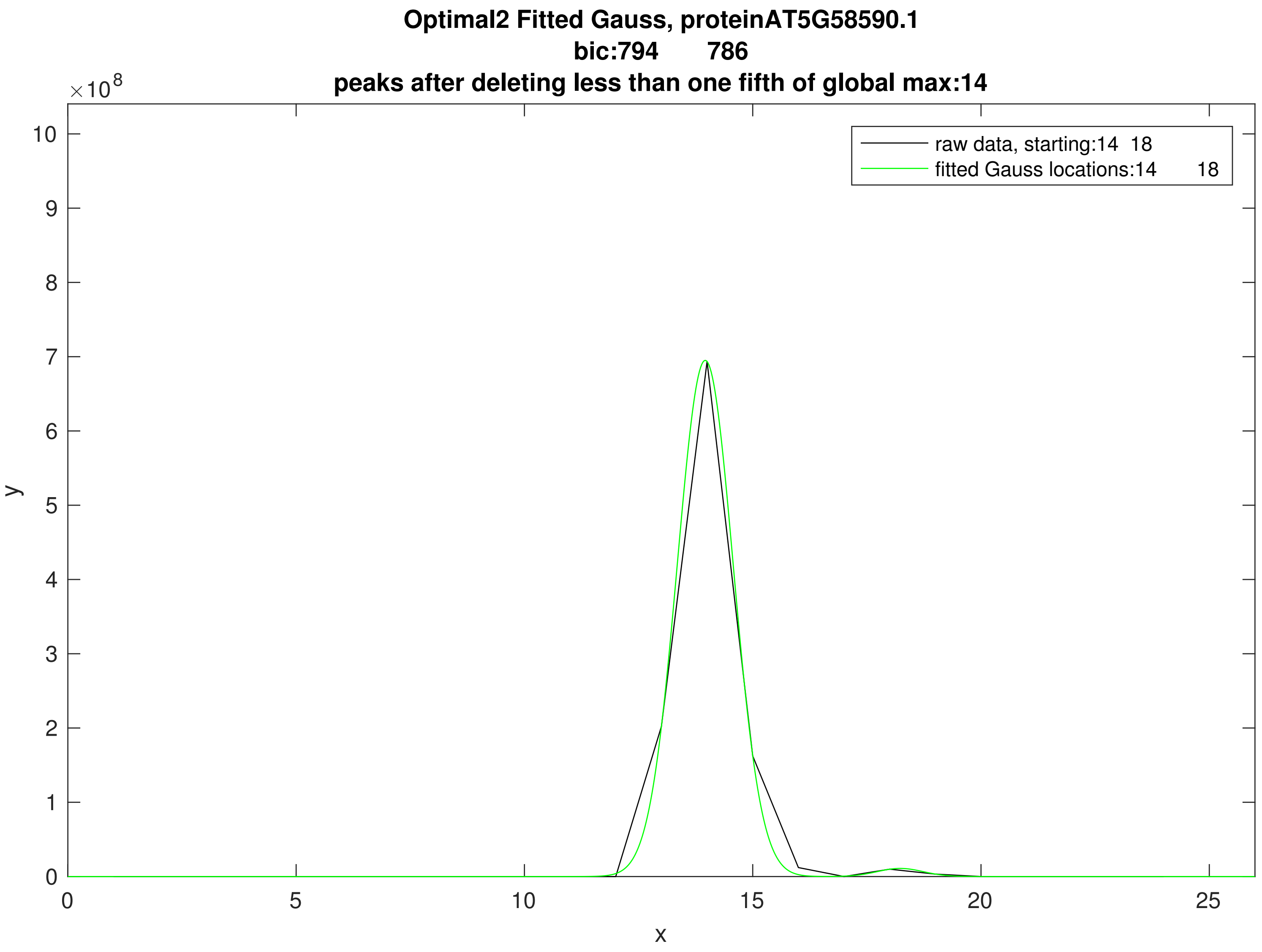
Optimal2 Fitted Gauss, proteinAT5G58590.1
bic:794 786
peaks after deleting less than one fifth of global max:14
×10^8
raw data, starting:14 18
fitted Gauss locations:14 18
y
x
0
1
2
3
4
5
6
7
8
9
10
0
5
10
15
20
25

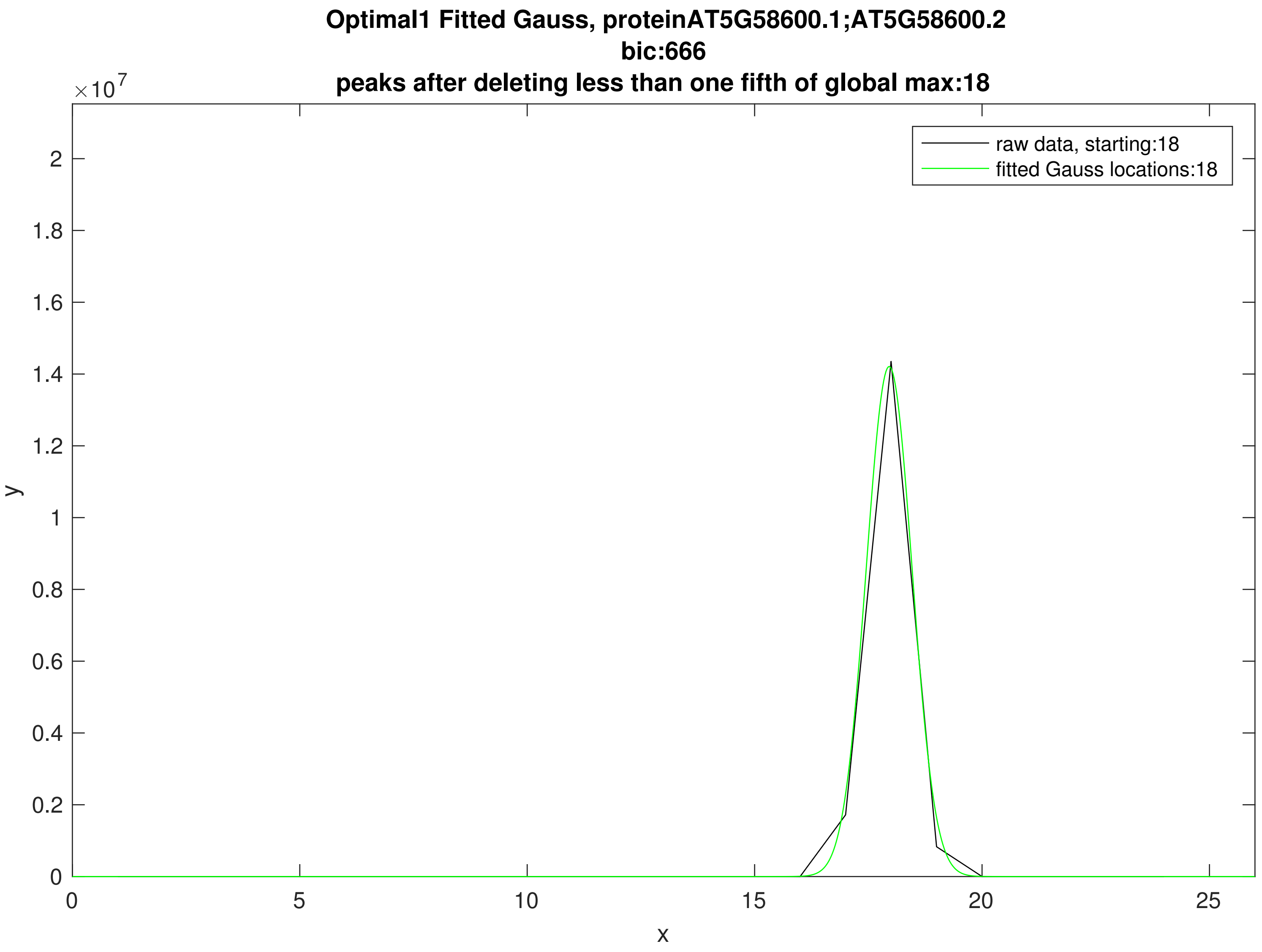
Optimal1 Fitted Gauss, proteinAT5G58600.1;AT5G58600.2
bic:666
peaks after deleting less than one fifth of global max:18
raw data, starting:18
fitted Gauss locations:18
y
x

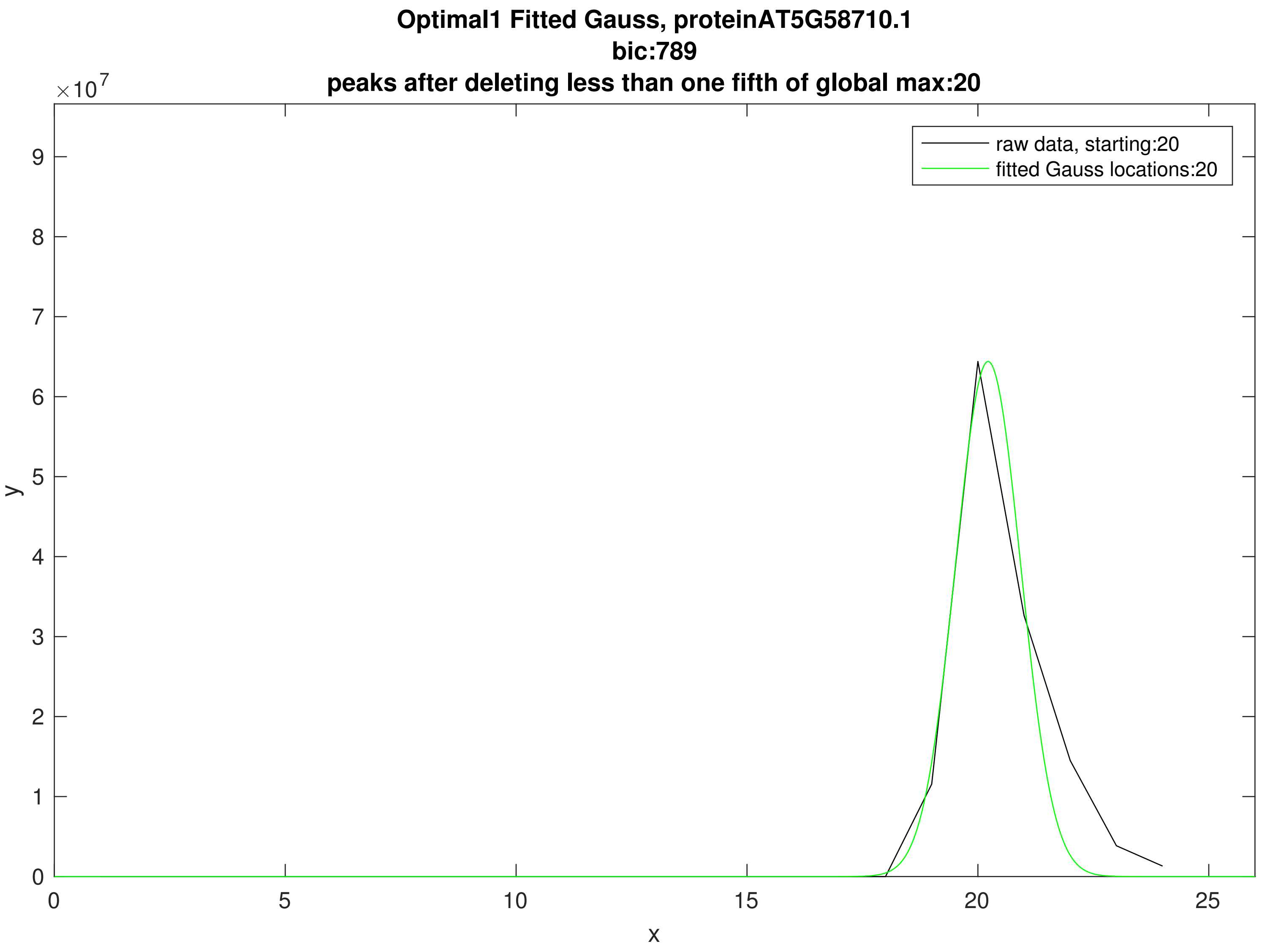
Optimal1 Fitted Gauss, proteinAT5G58710.1
bic:789
peaks after deleting less than one fifth of global max:20
×10^7
raw data, starting:20
fitted Gauss locations:20
y
x
0
1
2
3
4
5
6
7
8
9
0
5
10
15
20
25

**Optimal1 Fitted Gauss, proteinAT5G58750.1**
**bic:649**
**peaks after deleting less than one fifth of global max:17**

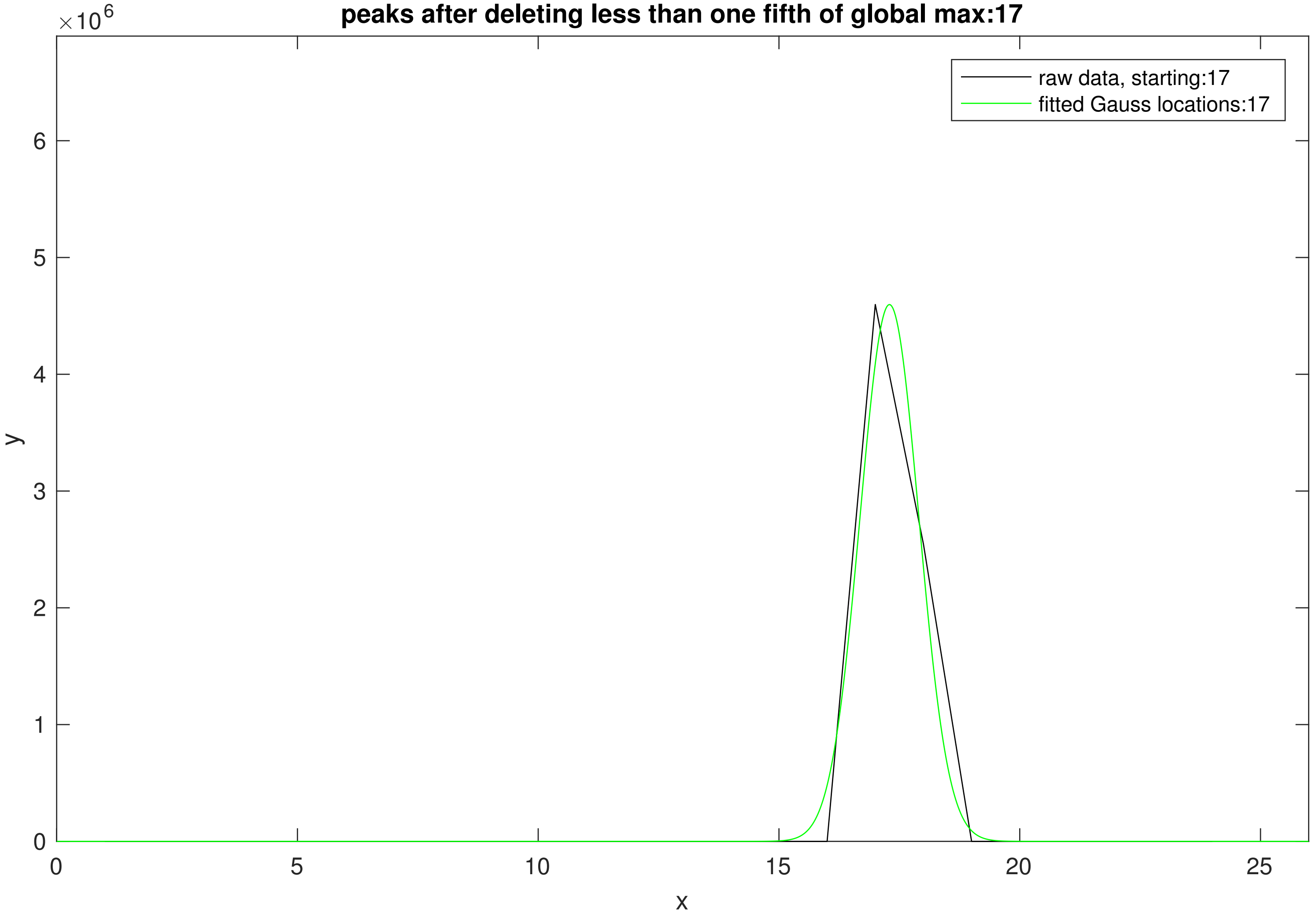

**Optimal2 Fitted Gauss, proteinAT5G59160.3;AT5G59160.2;AT5G59160.1;AT3G46820.1;AT2G39840.1**
**bic:880 811**
**peaks after deleting less than one fifth of global max:13 17**

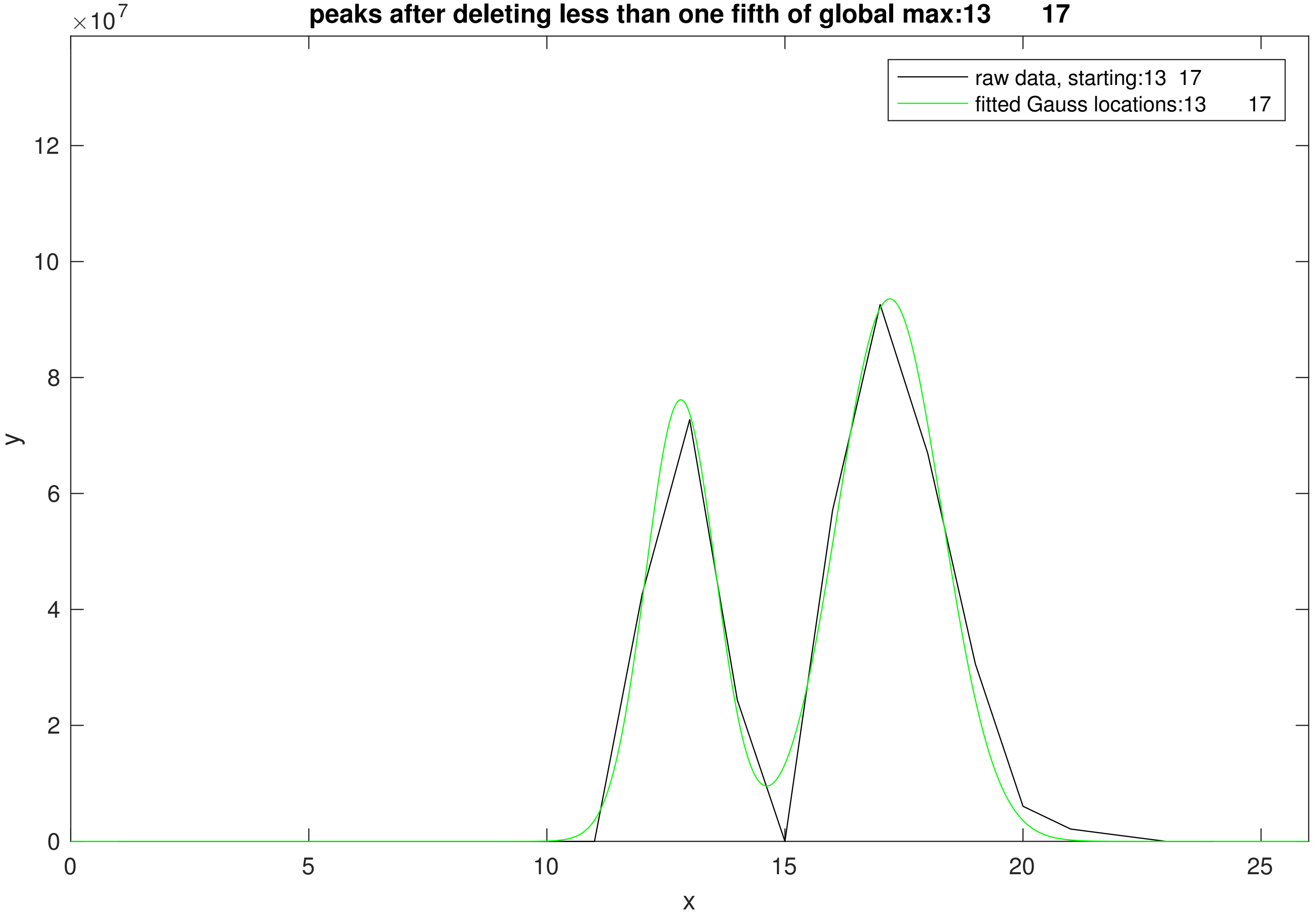

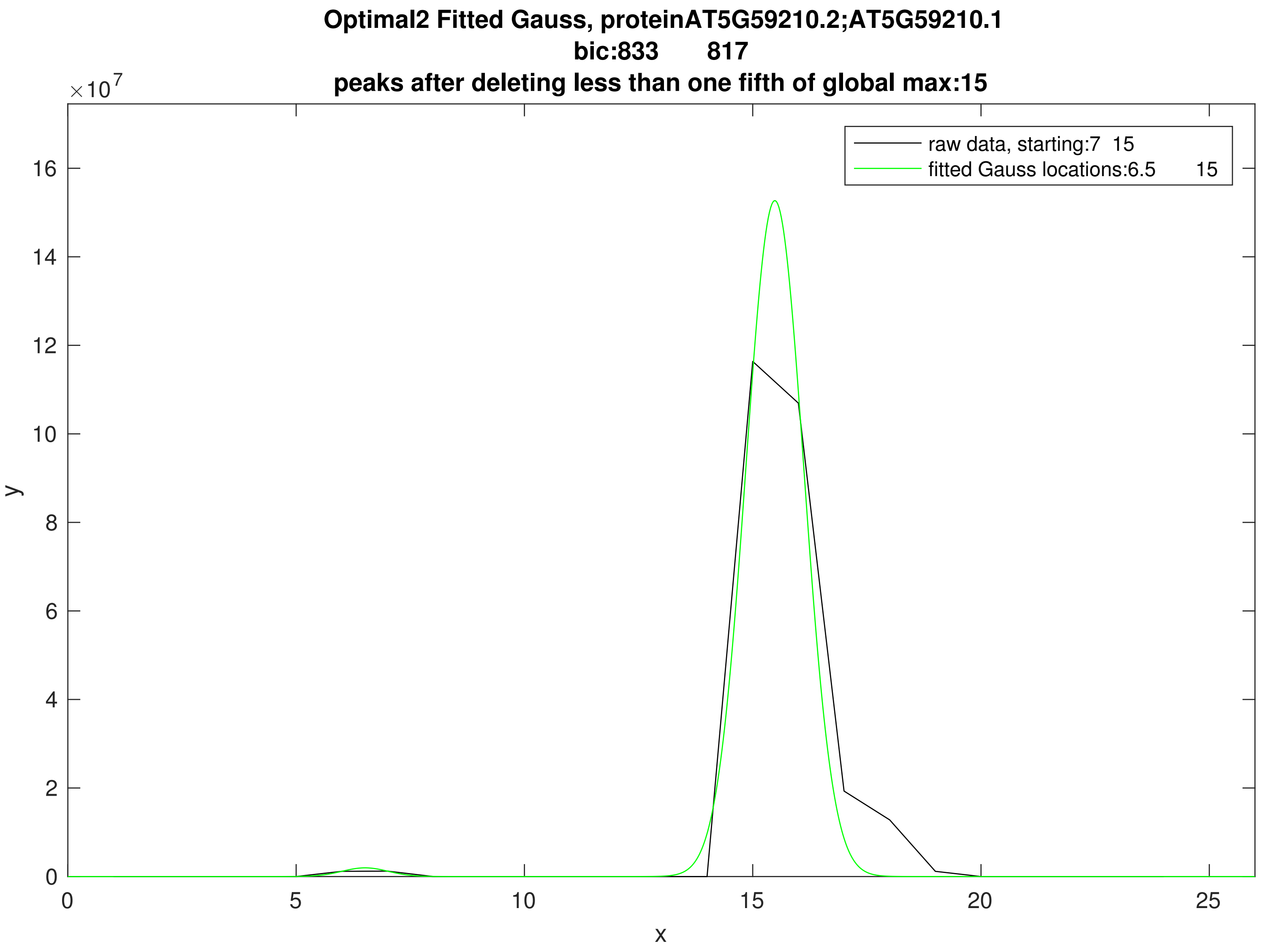

Optimal2 Fitted Gauss, proteinAT5G59210.2;AT5G59210.1
bic:833 817
peaks after deleting less than one fifth of global max:15
×10^7
raw data, starting:7 15
fitted Gauss locations:6.5 15
16
14
12
10
8
6
4
2
0
y
0
5
10
15
20
25
x

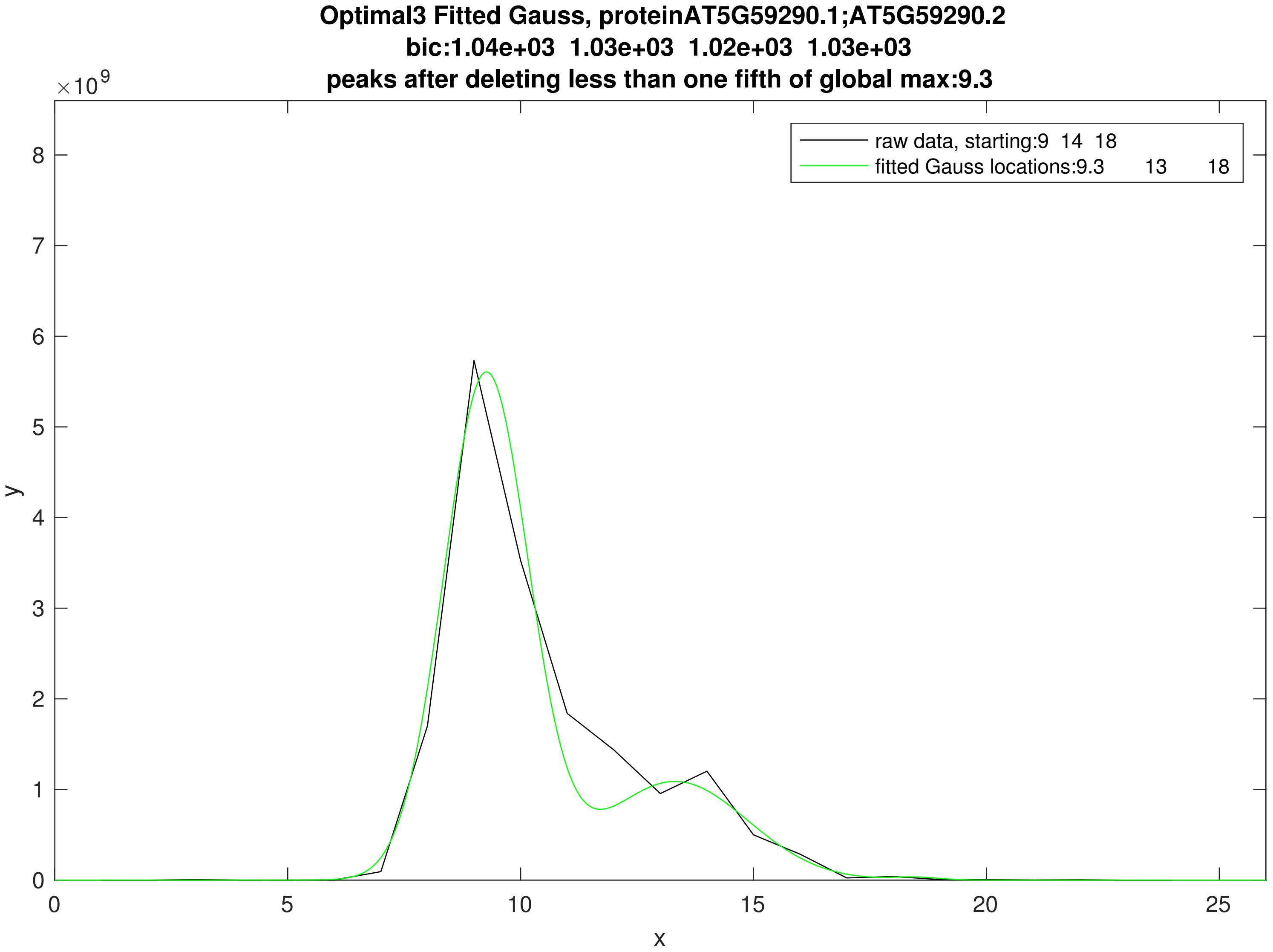

Optimal3 Fitted Gauss, proteinAT5G59290.1;AT5G59290.2
bic:1.04e+03 1.03e+03 1.02e+03 1.03e+03
peaks after deleting less than one fifth of global max:9.3
×10^9
raw data, starting:9 14 18
fitted Gauss locations:9.3 13 18
y
x
0
1
2
3
4
5
6
7
8
0
5
10
15
20
25

**Optimal1 Fitted Gauss, proteinAT5G59300.1**
**bic:746**
**peaks after deleting less than one fifth of global max:18**

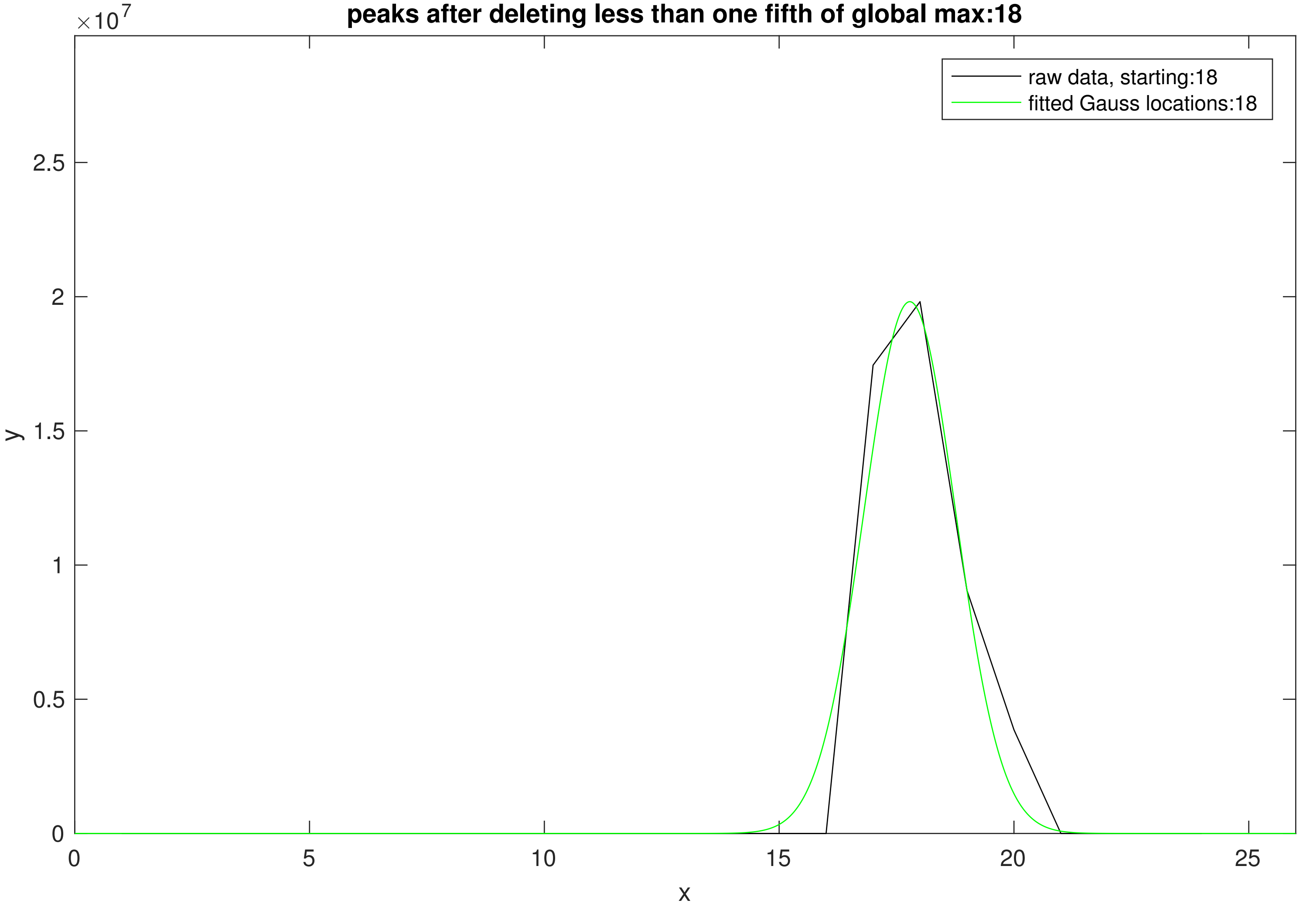

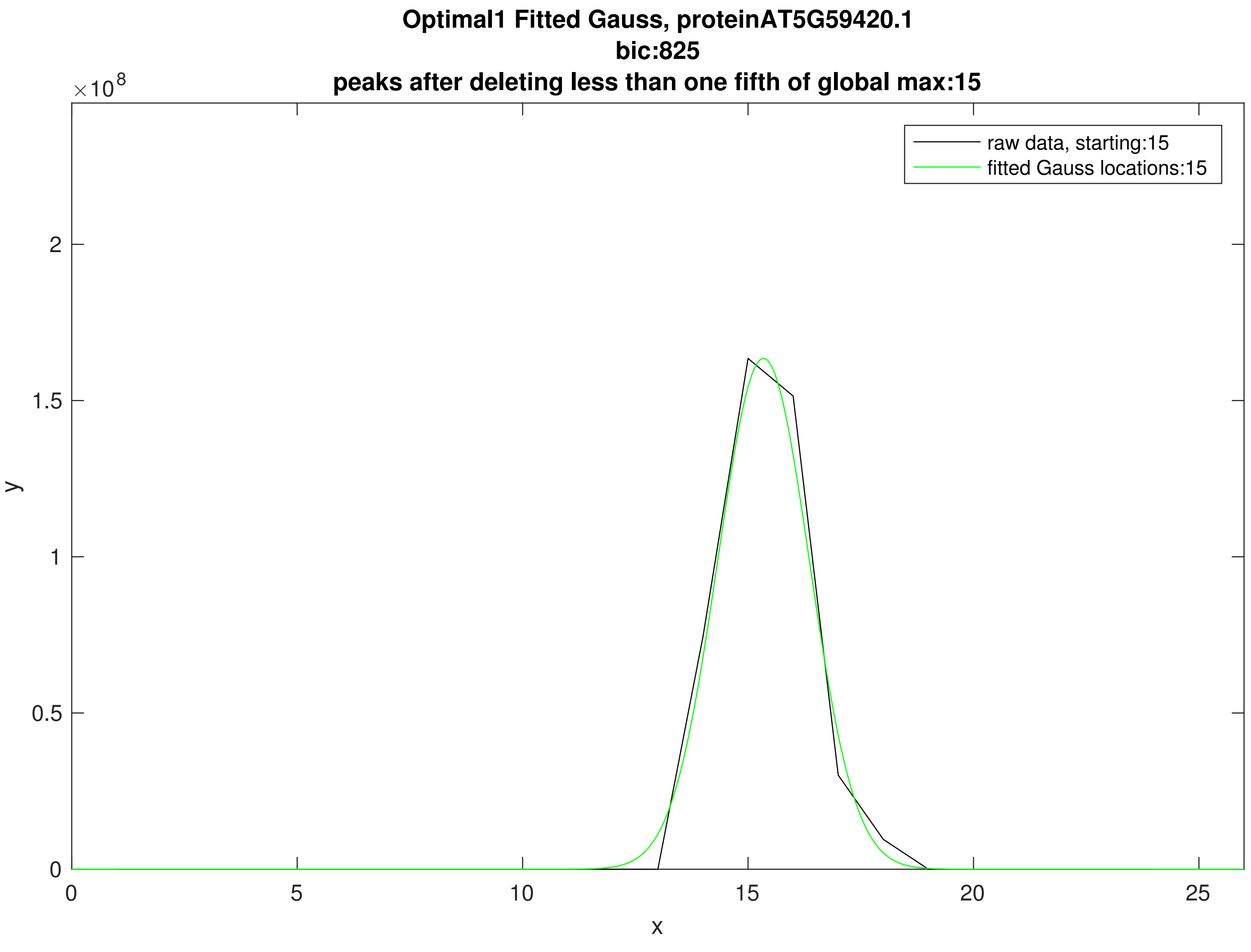

Optimal1 Fitted Gauss, proteinAT5G59420.1
bic:825
peaks after deleting less than one fifth of global max:15
×10^8
raw data, starting:15
fitted Gauss locations:15
y
x
0
0.5
1
1.5
2
0
5
10
15
20
25

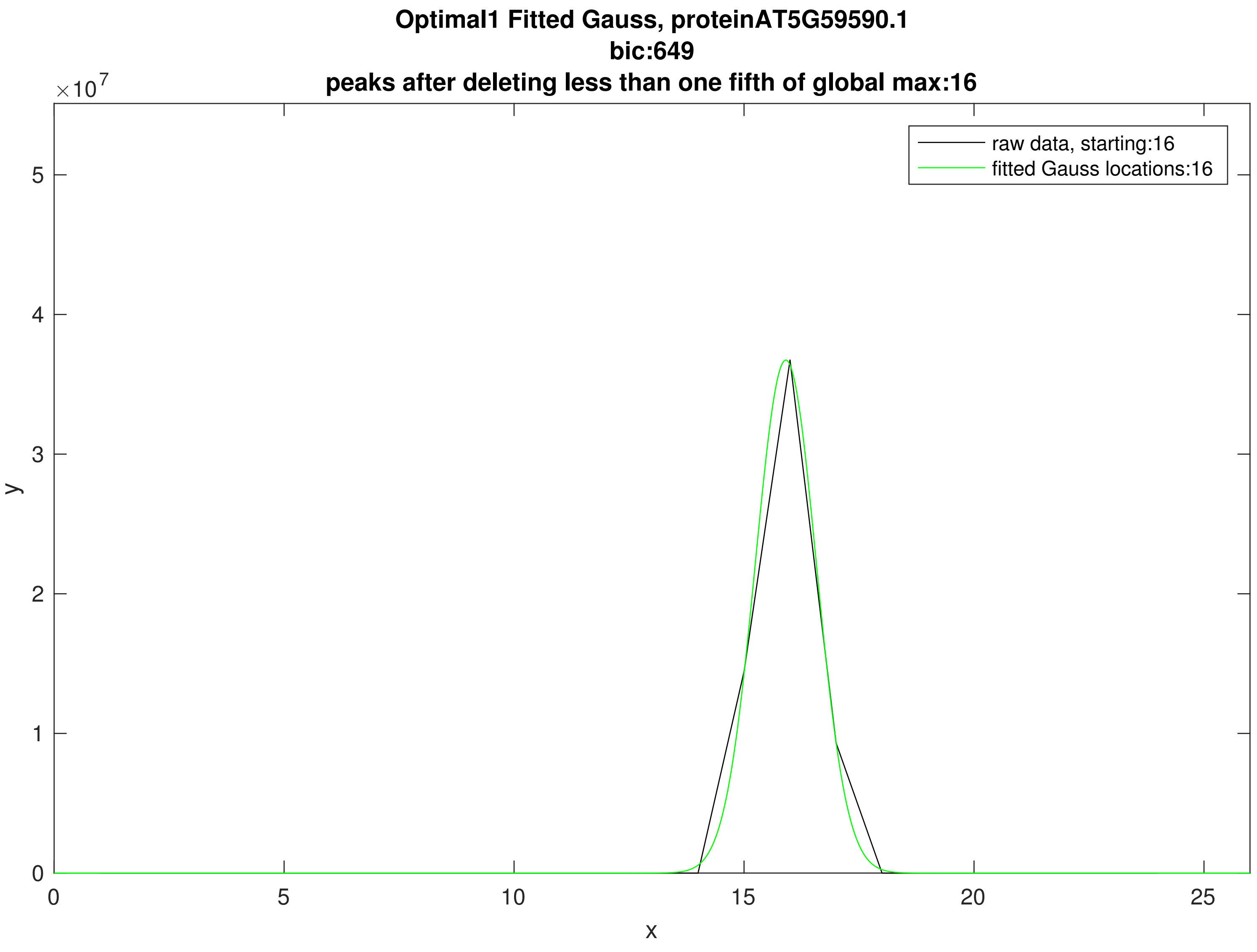
Optimal1 Fitted Gauss, proteinAT5G59590.1
bic:649
peaks after deleting less than one fifth of global max:16
×10^7
raw data, starting:16
fitted Gauss locations:16
y
x
0
1
2
3
4
5
0
5
10
15
20
25

**Optimal1 Fitted Gauss, proteinAT5G59850.1;AT1G07770.2;AT1G07770.1**
**bic:799**
**peaks after deleting less than one fifth of global max:3.8**

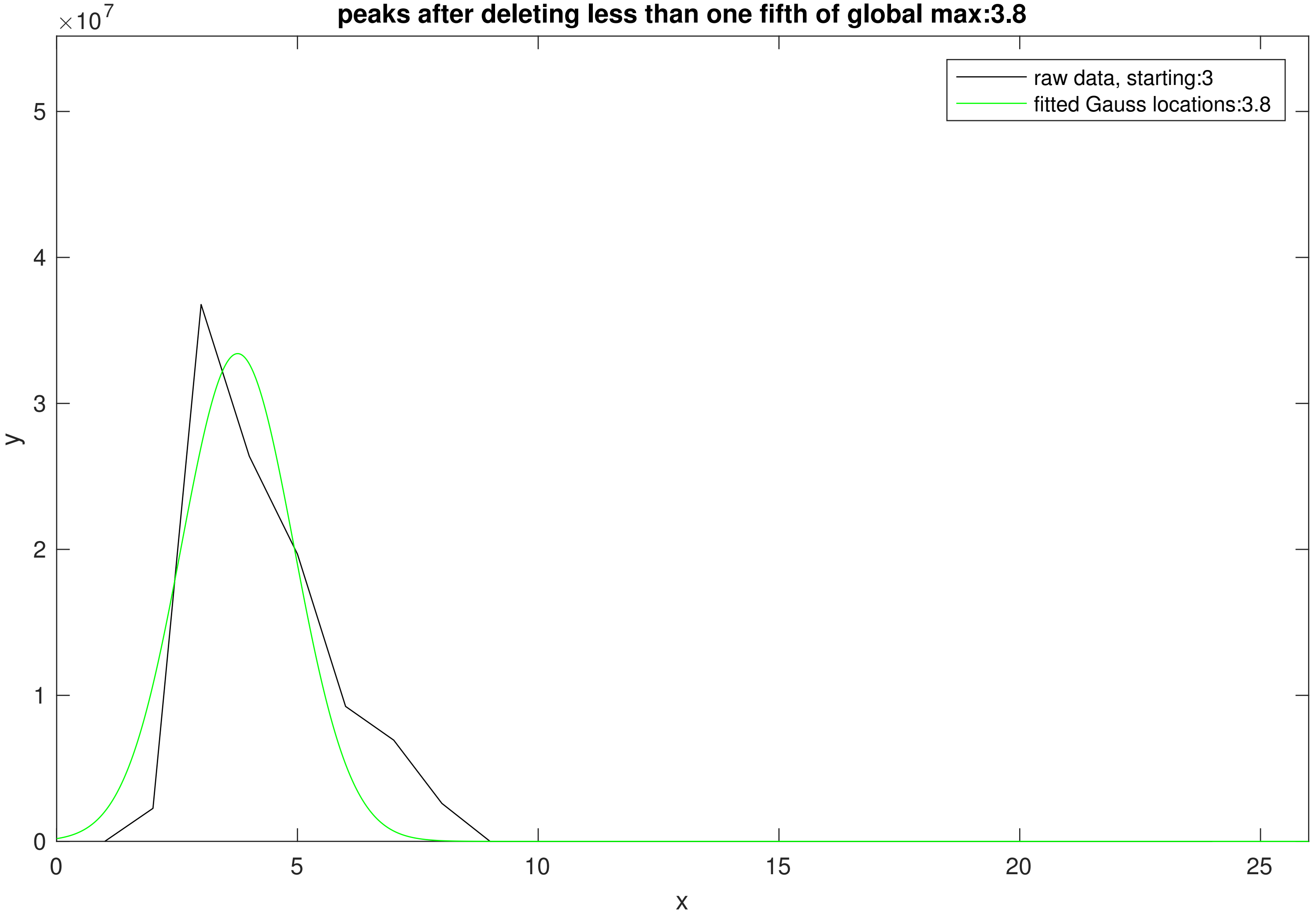

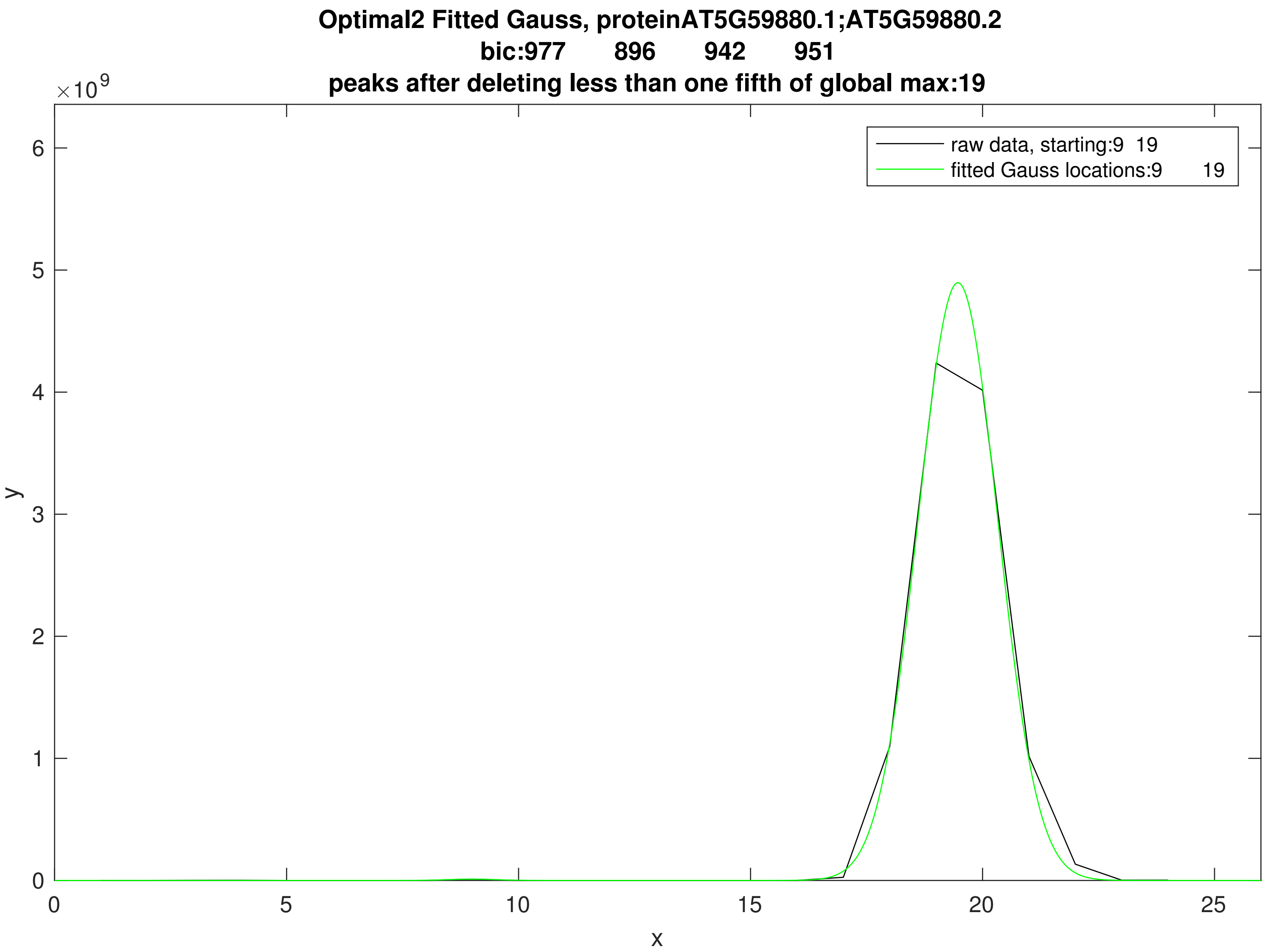

Optimal2 Fitted Gauss, proteinAT5G59880.1;AT5G59880.2
bic:977 896 942 951
peaks after deleting less than one fifth of global max:19
×10^9
raw data, starting:9 19
fitted Gauss locations:9 19
y
x
0
1
2
3
4
5
6
0
5
10
15
20
25

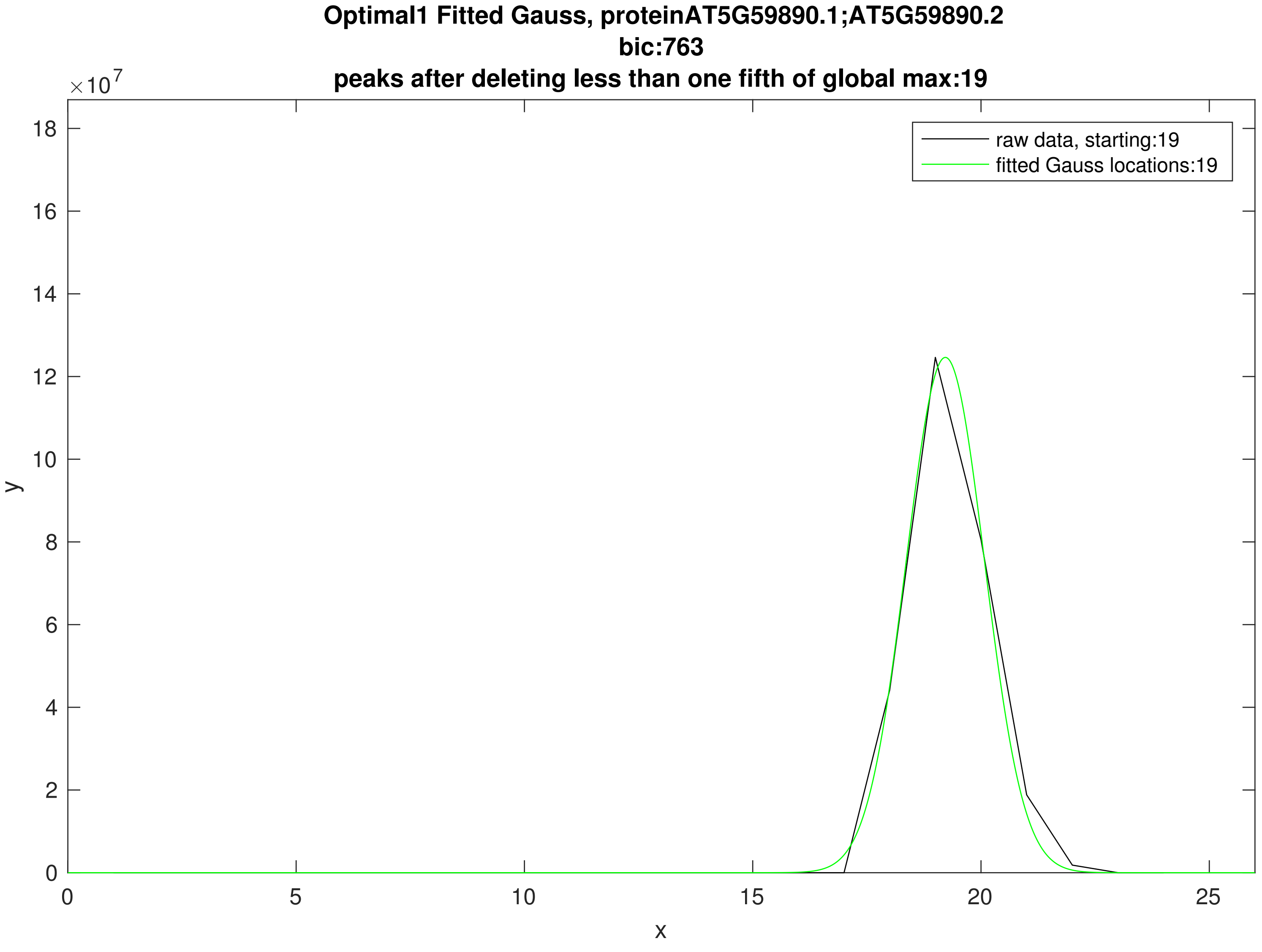

Optimal1 Fitted Gauss, proteinAT5G59890.1;AT5G59890.2
bic:763
peaks after deleting less than one fifth of global max:19
×10^7
raw data, starting:19
fitted Gauss locations:19
y
x
0
2
4
6
8
10
12
14
16
18
0
5
10
15
20
25

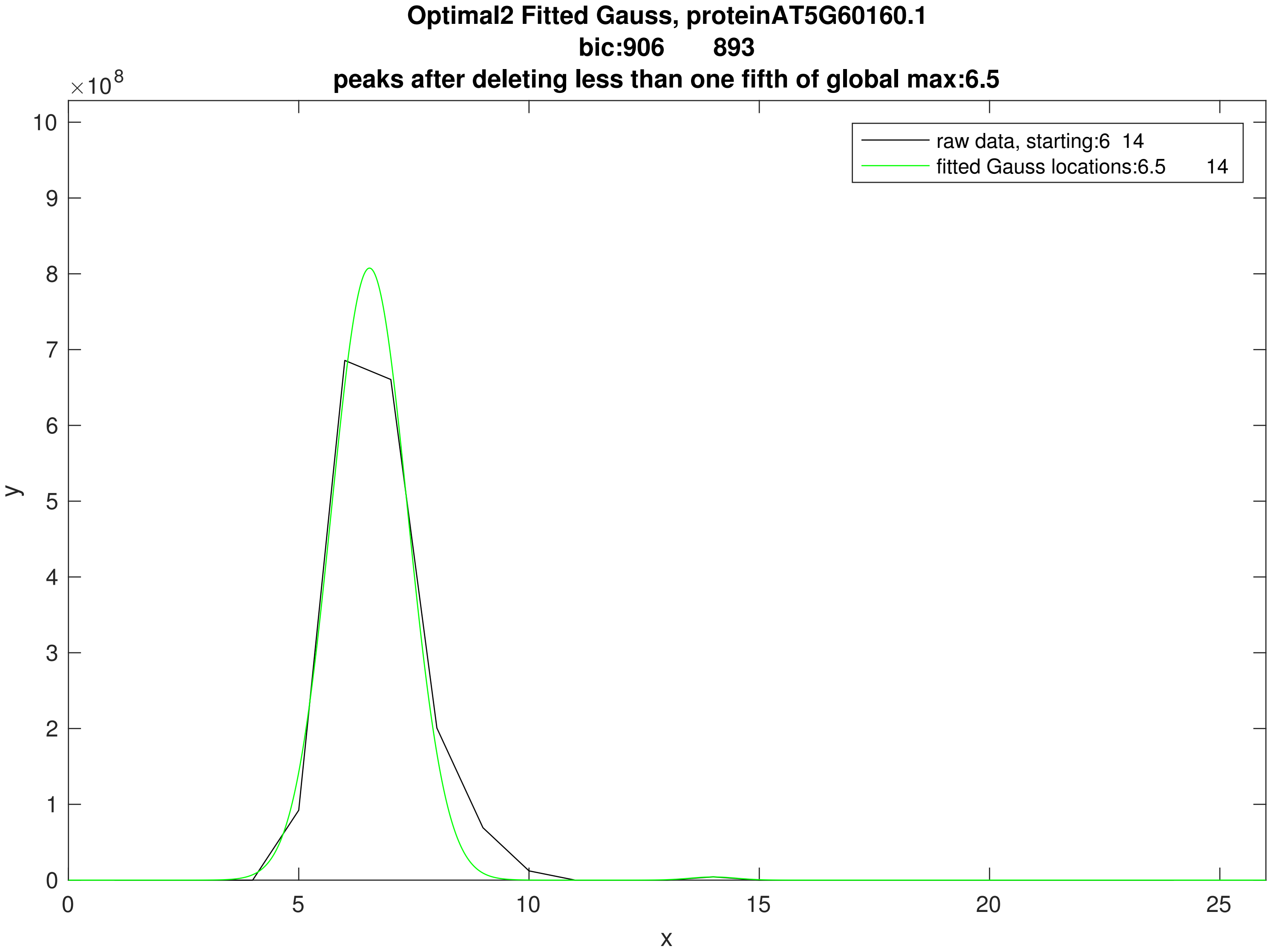

Optimal2 Fitted Gauss, proteinAT5G60160.1
bic:906 893
peaks after deleting less than one fifth of global max:6.5
×10^8
raw data, starting:6 14
fitted Gauss locations:6.5 14
y
x
0
1
2
3
4
5
6
7
8
9
10
0
5
10
15
20
25

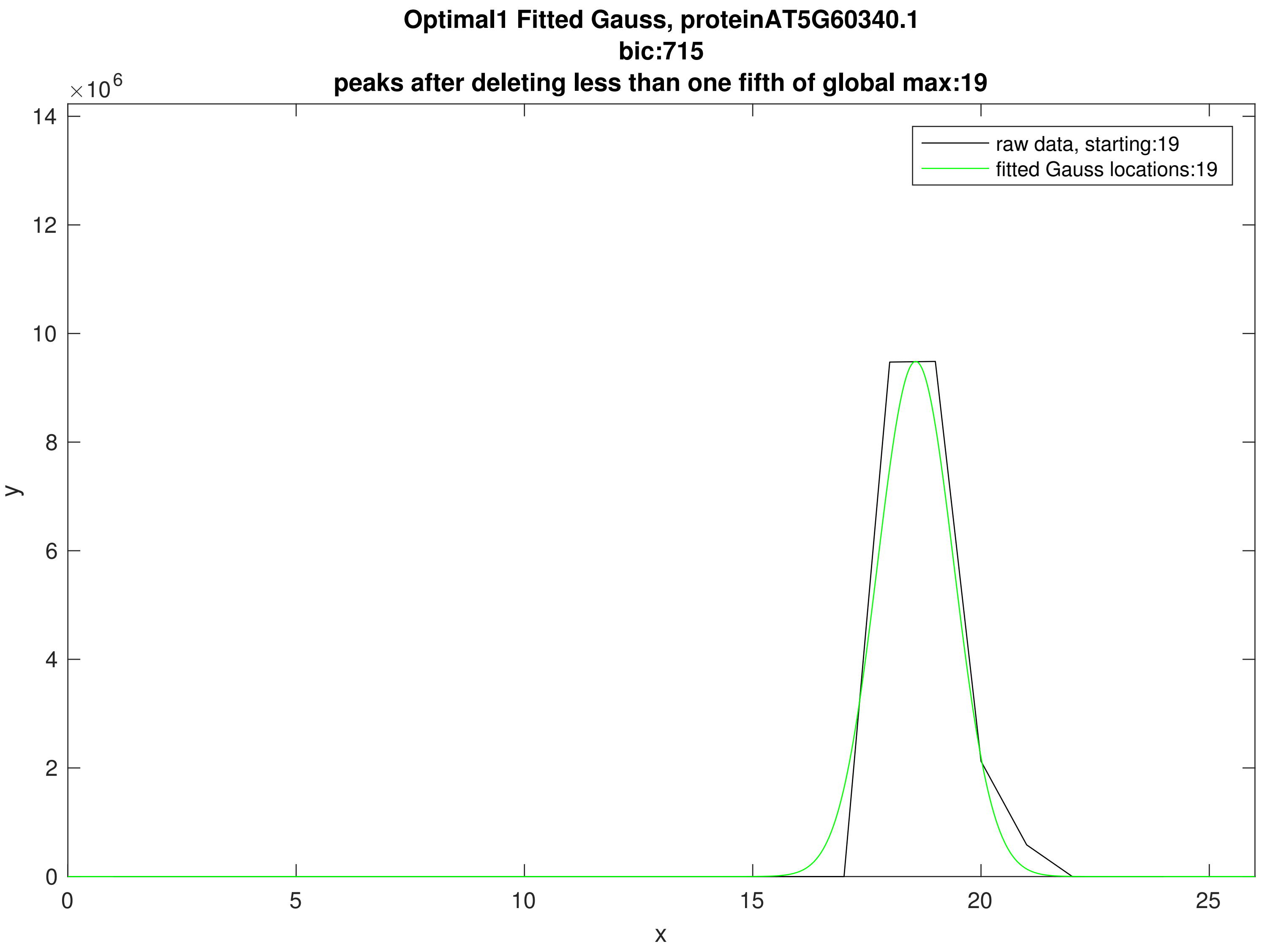

Optimal1 Fitted Gauss, proteinAT5G60340.1
bic:715
peaks after deleting less than one fifth of global max:19
×10^6
14
12
10
8
6
4
2
0
y
0
5
10
15
20
25
x
raw data, starting:19
fitted Gauss locations:19

**Optimal1 Fitted Gauss, proteinAT5G60360.1;AT5G60360.2;AT5G60360.3**
**bic:922 932**
**peaks after deleting less than one fifth of global max:18**

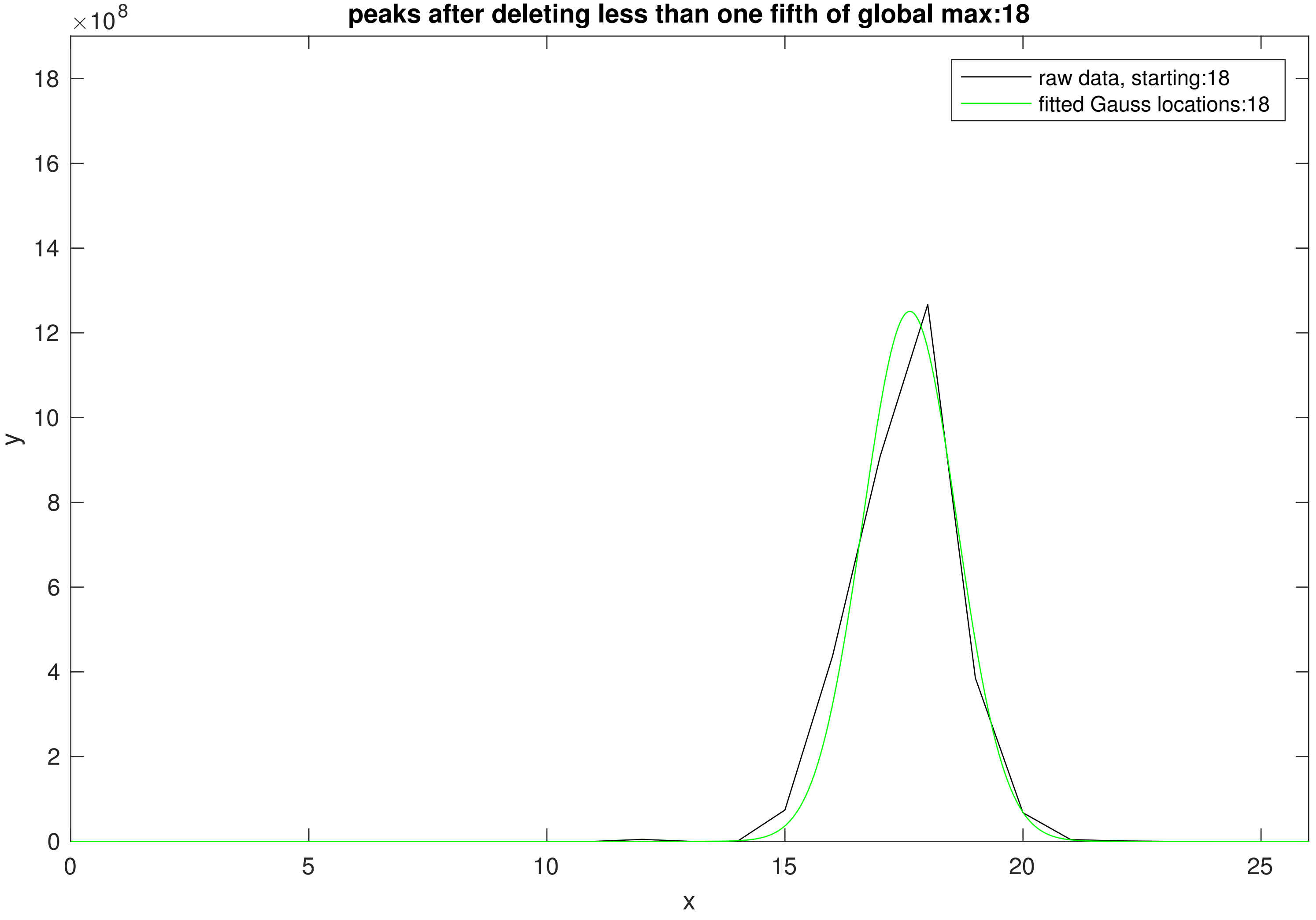

**2 Fitted Gauss, proteinAT5G60390.3;AT5G60390.1;AT1G07940.2;AT1G07940.1;AT1G07930.1;AT1G07920.1;AT5G6039[illegible]**

**bic:1.04e+03  1.04e+03  1.05e+03  1.05e+03**

**peaks after deleting less than one fifth of global max:15**

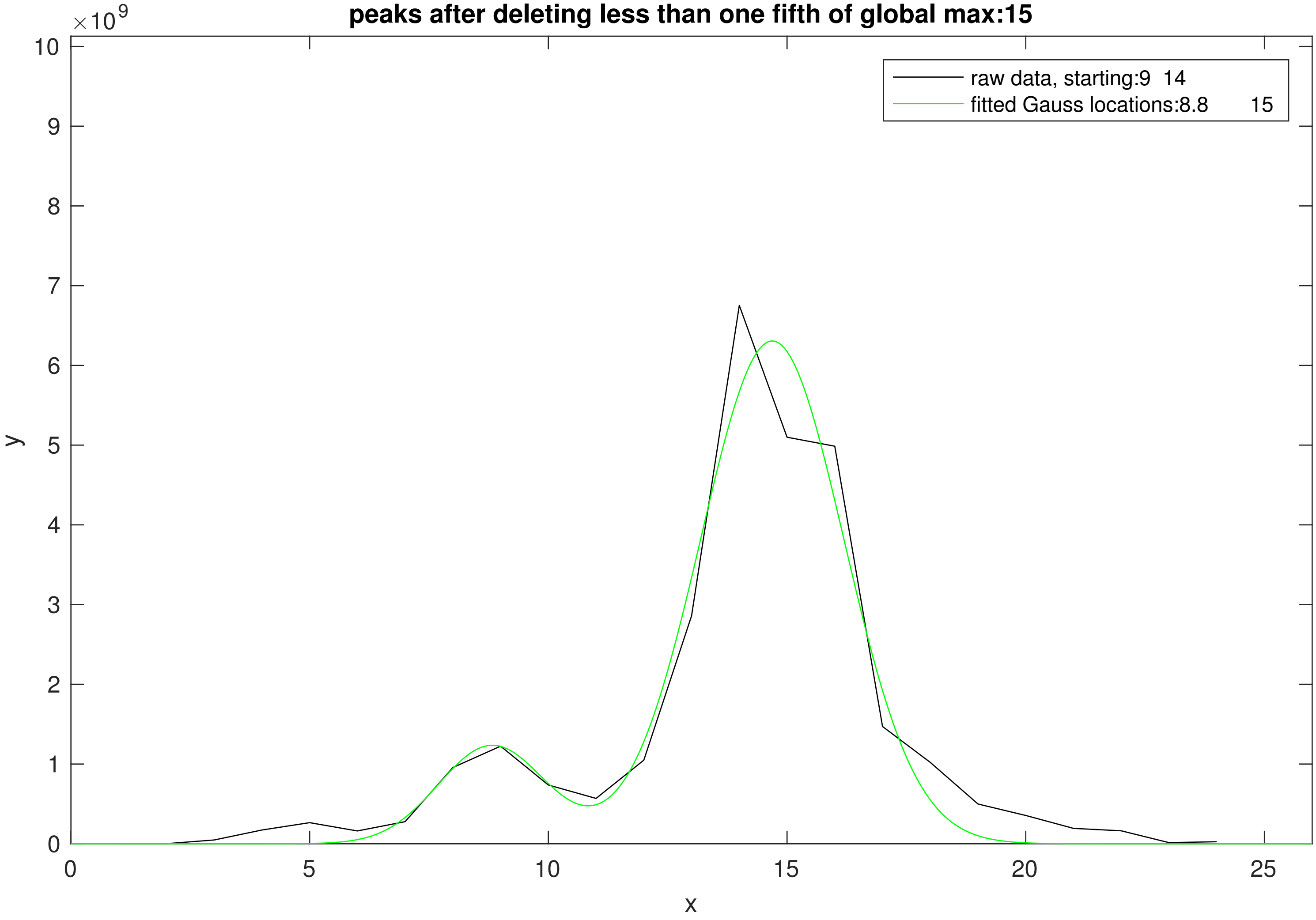

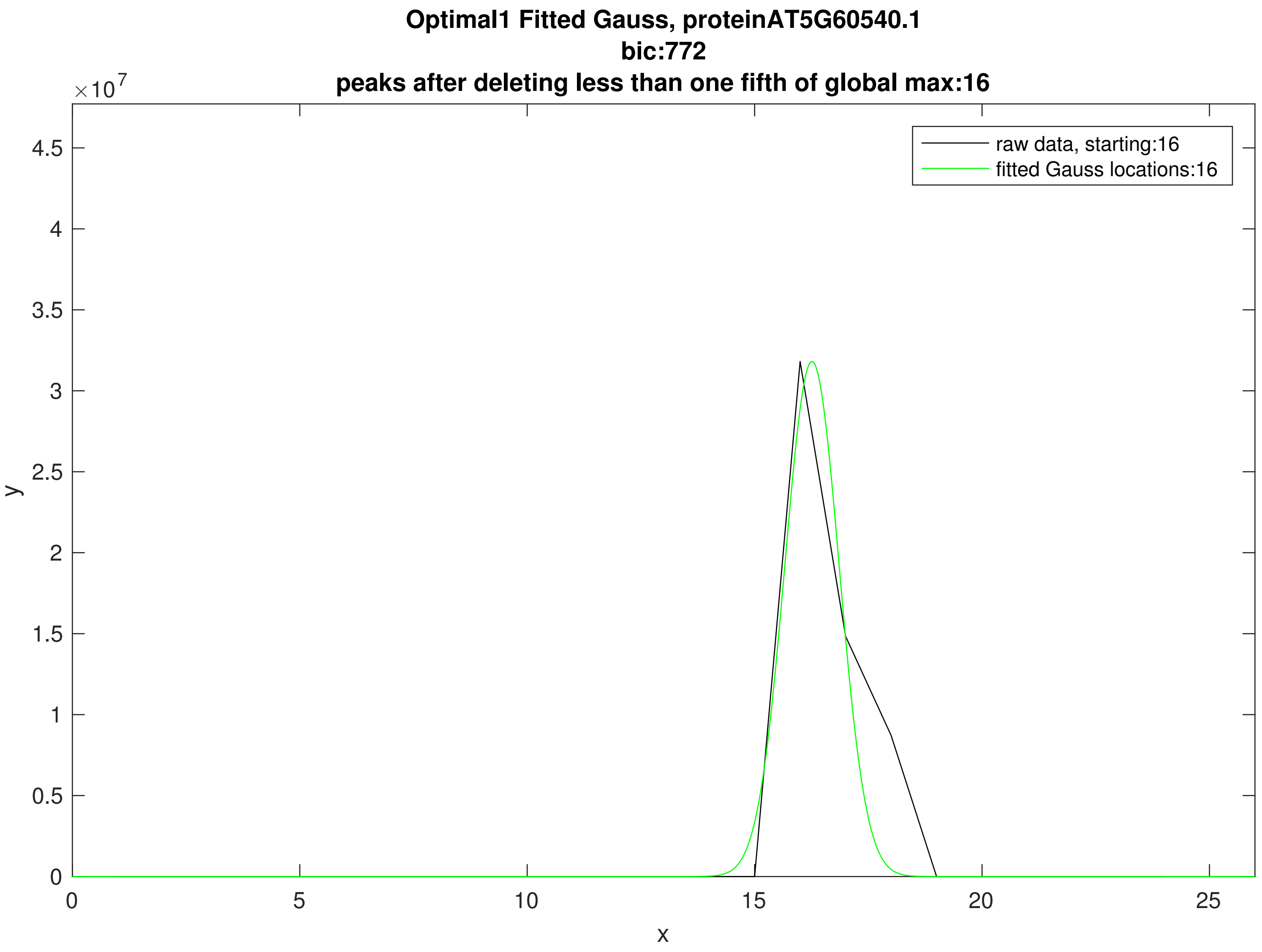

Optimal1 Fitted Gauss, proteinAT5G60540.1
bic:772
peaks after deleting less than one fifth of global max:16
×10^7
4.5
4
3.5
3
2.5
2
1.5
1
0.5
0
y
0
5
10
15
20
25
x
raw data, starting:16
fitted Gauss locations:16

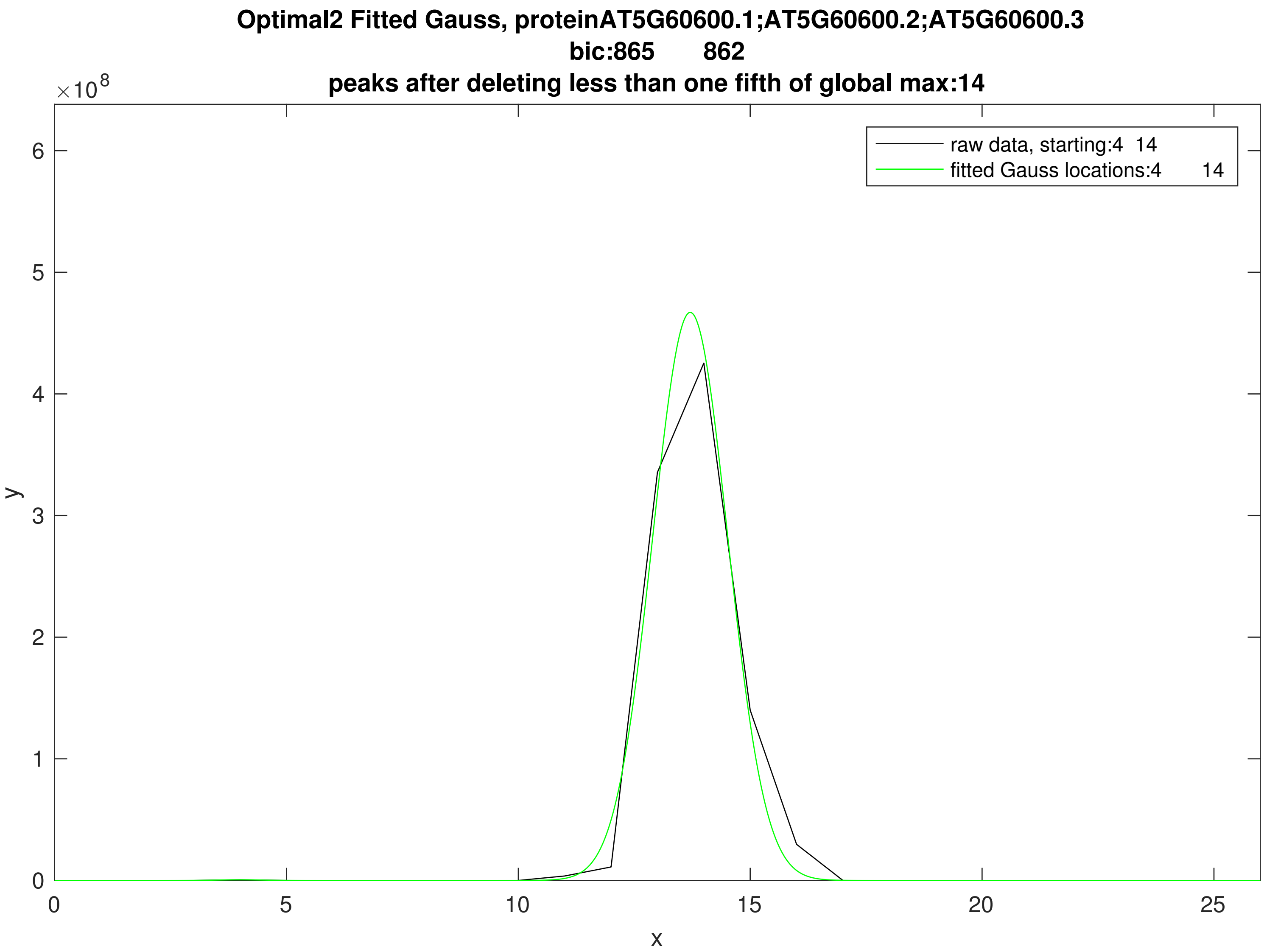
Optimal2 Fitted Gauss, proteinAT5G60600.1;AT5G60600.2;AT5G60600.3
bic:865 862
peaks after deleting less than one fifth of global max:14
×10^8
raw data, starting:4 14
fitted Gauss locations:4 14
y
x
0
1
2
3
4
5
6
0
5
10
15
20
25

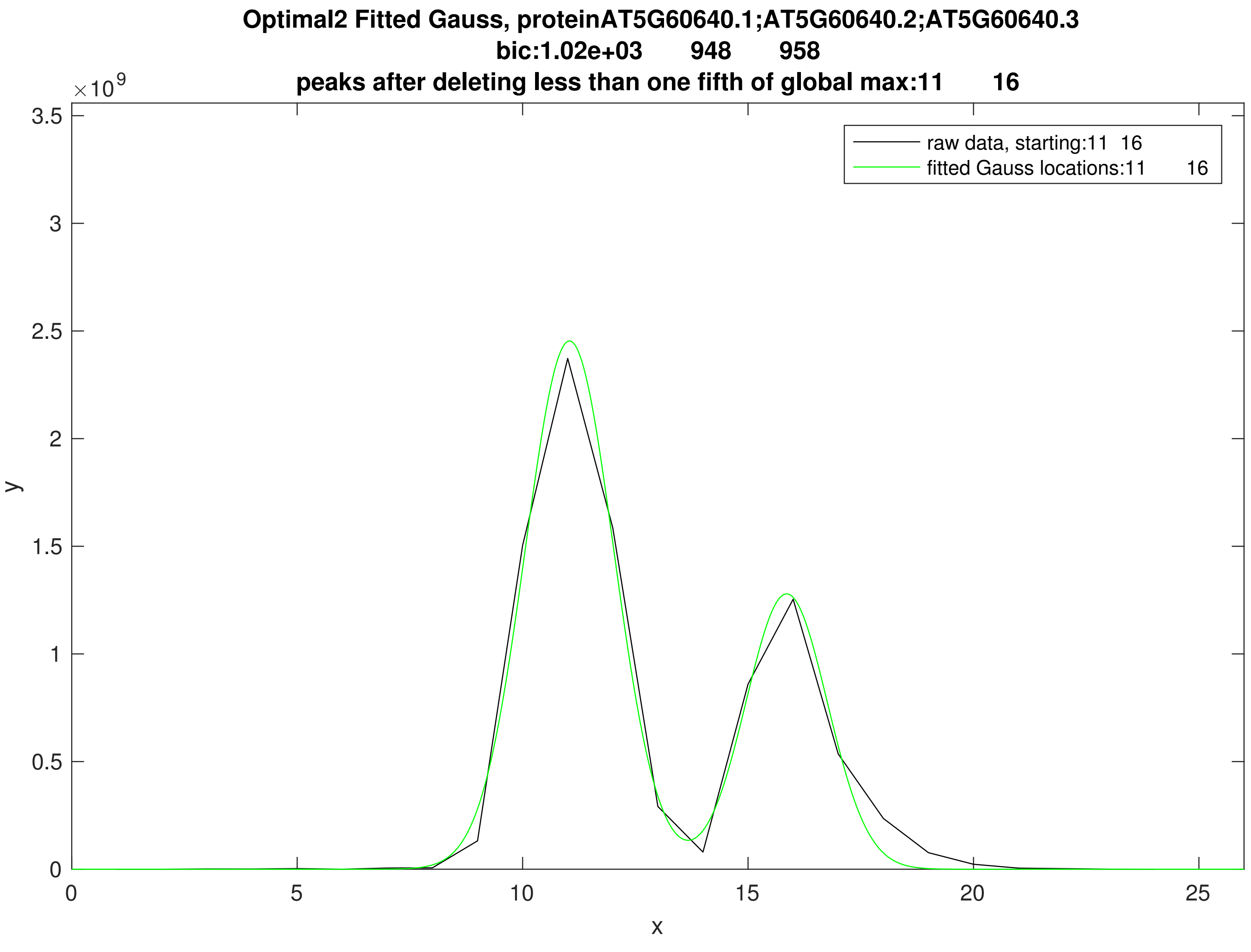
Optimal2 Fitted Gauss, proteinAT5G60640.1;AT5G60640.2;AT5G60640.3
bic:1.02e+03 948 958
peaks after deleting less than one fifth of global max:11 16
×10^9
raw data, starting:11 16
fitted Gauss locations:11 16
y
x
0
0.5
1
1.5
2
2.5
3
3.5
0
5
10
15
20
25

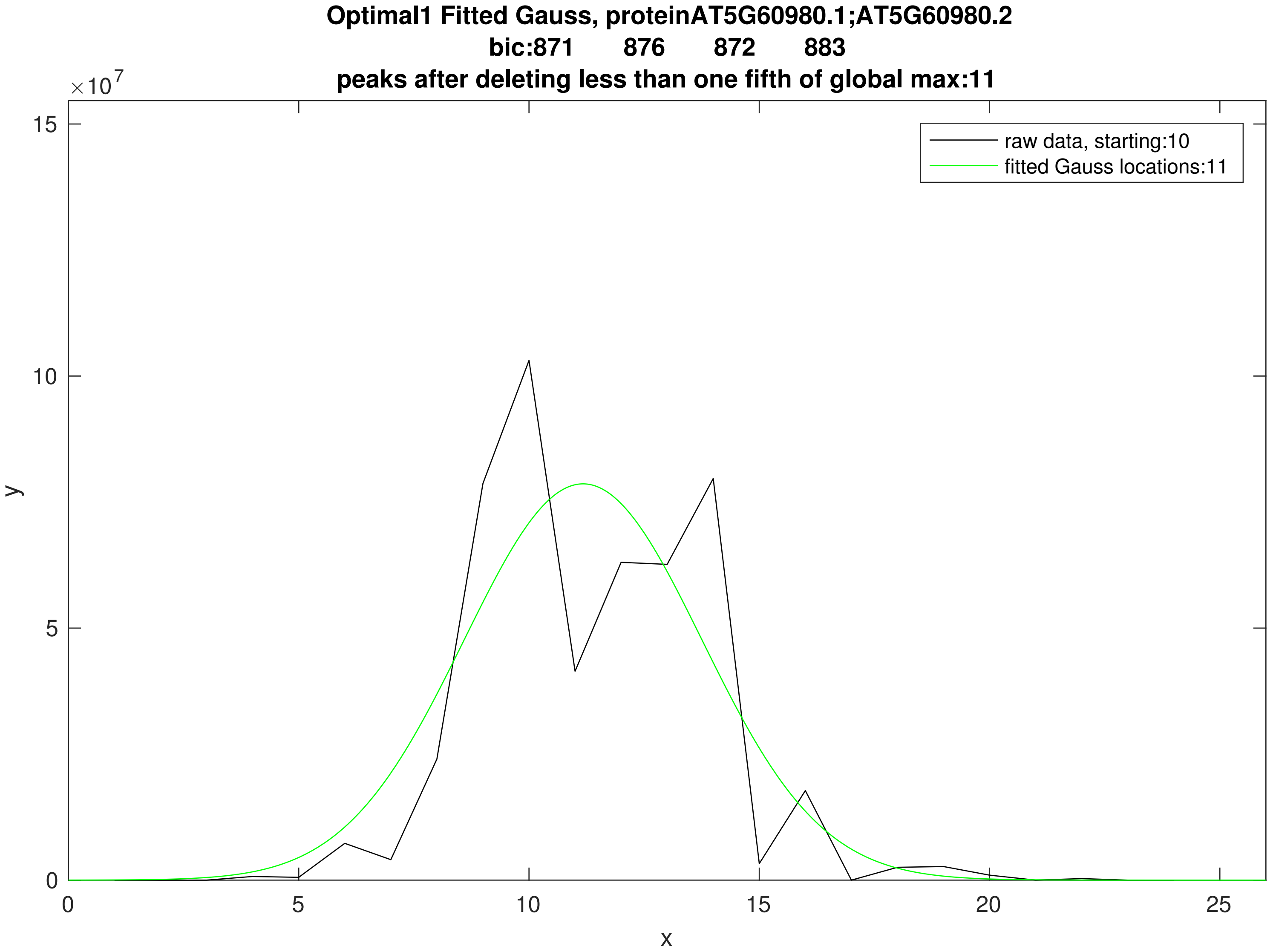
Optimal1 Fitted Gauss, proteinAT5G60980.1;AT5G60980.2
bic:871 876 872 883
peaks after deleting less than one fifth of global max:11
×10^7
raw data, starting:10
fitted Gauss locations:11
y
x
0
5
10
15
20
25

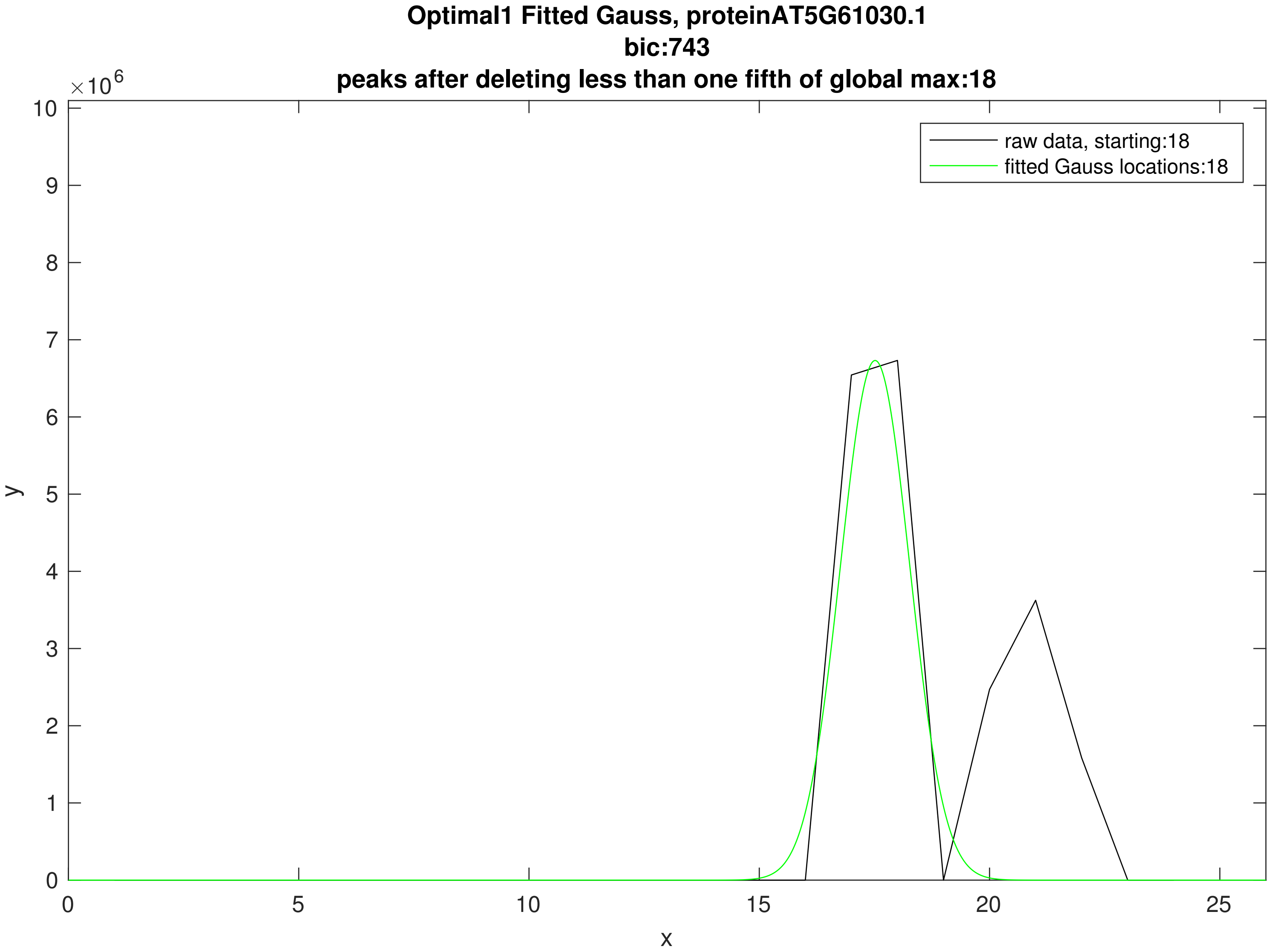

Optimal1 Fitted Gauss, proteinAT5G61030.1
bic:743
peaks after deleting less than one fifth of global max:18
×10^6
10
9
8
7
6
5
4
3
2
1
0
y
0
5
10
15
20
25
x
raw data, starting:18
fitted Gauss locations:18

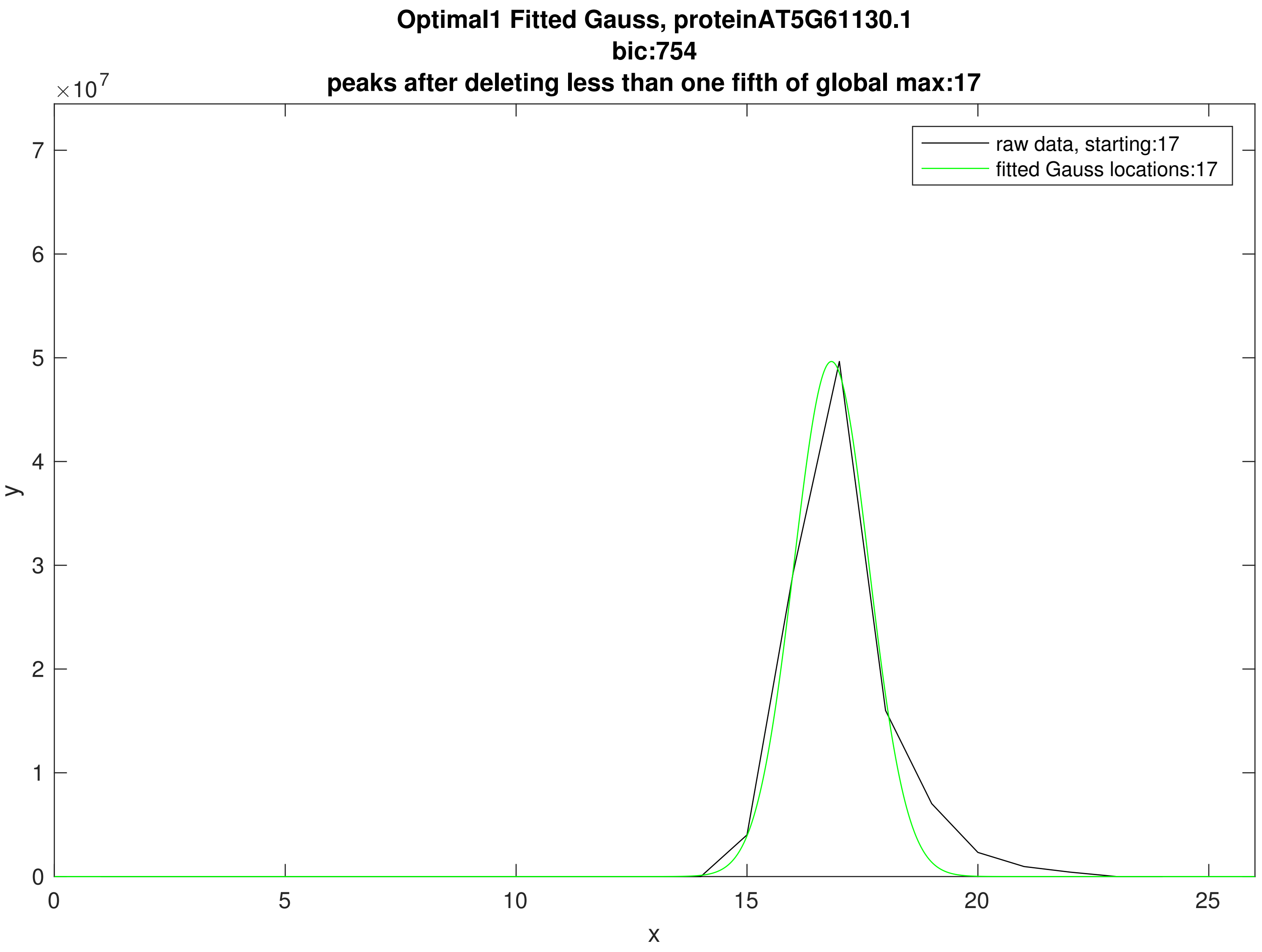

Optimal1 Fitted Gauss, proteinAT5G61130.1
bic:754
peaks after deleting less than one fifth of global max:17
×10^7
raw data, starting:17
fitted Gauss locations:17
y
x
0
1
2
3
4
5
6
7
0
5
10
15
20
25

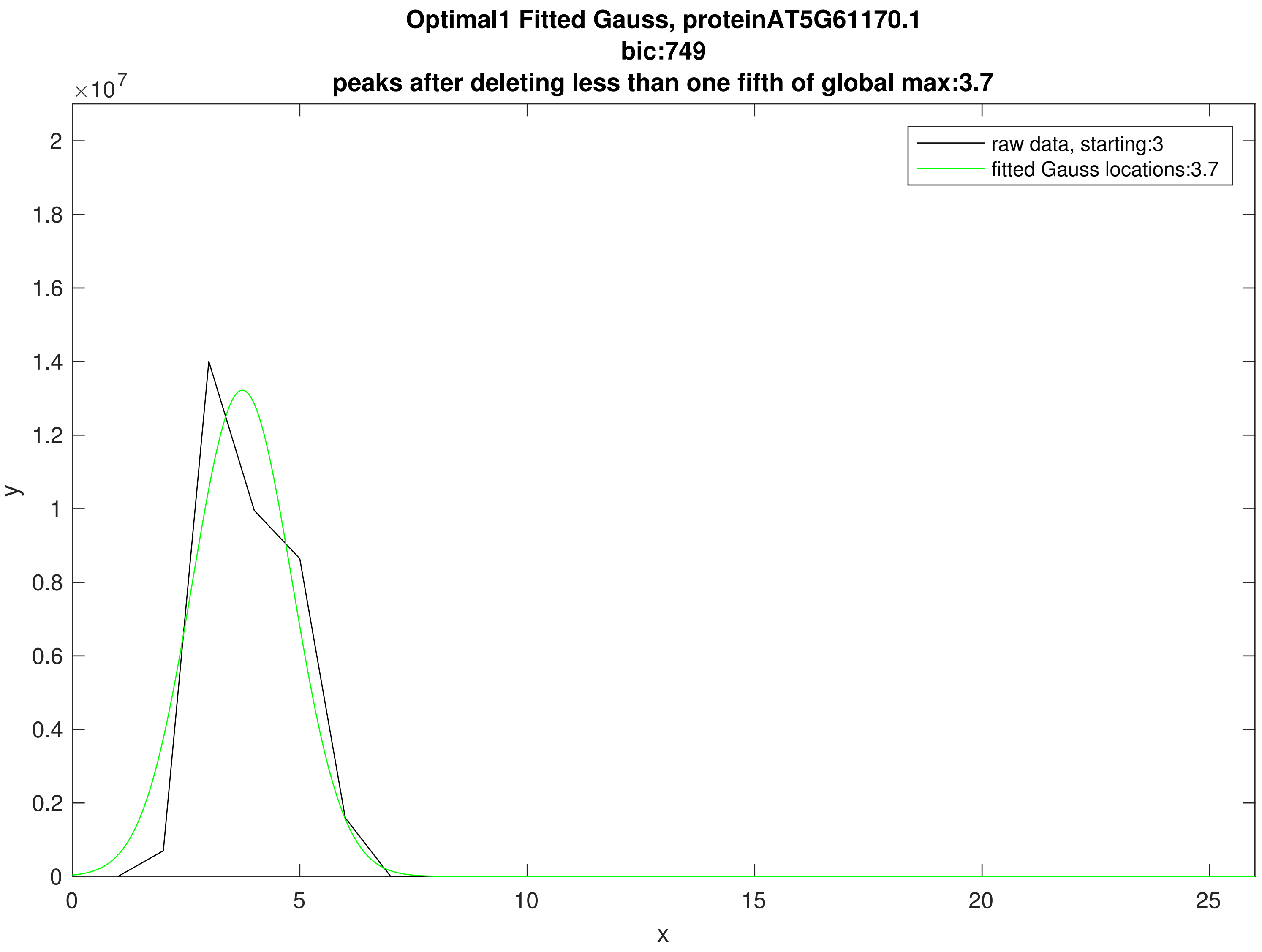

Optimal1 Fitted Gauss, proteinAT5G61170.1
bic:749
peaks after deleting less than one fifth of global max:3.7
×10^7
raw data, starting:3
fitted Gauss locations:3.7
y
x
0
0.2
0.4
0.6
0.8
1
1.2
1.4
1.6
1.8
2
5
10
15
20
25

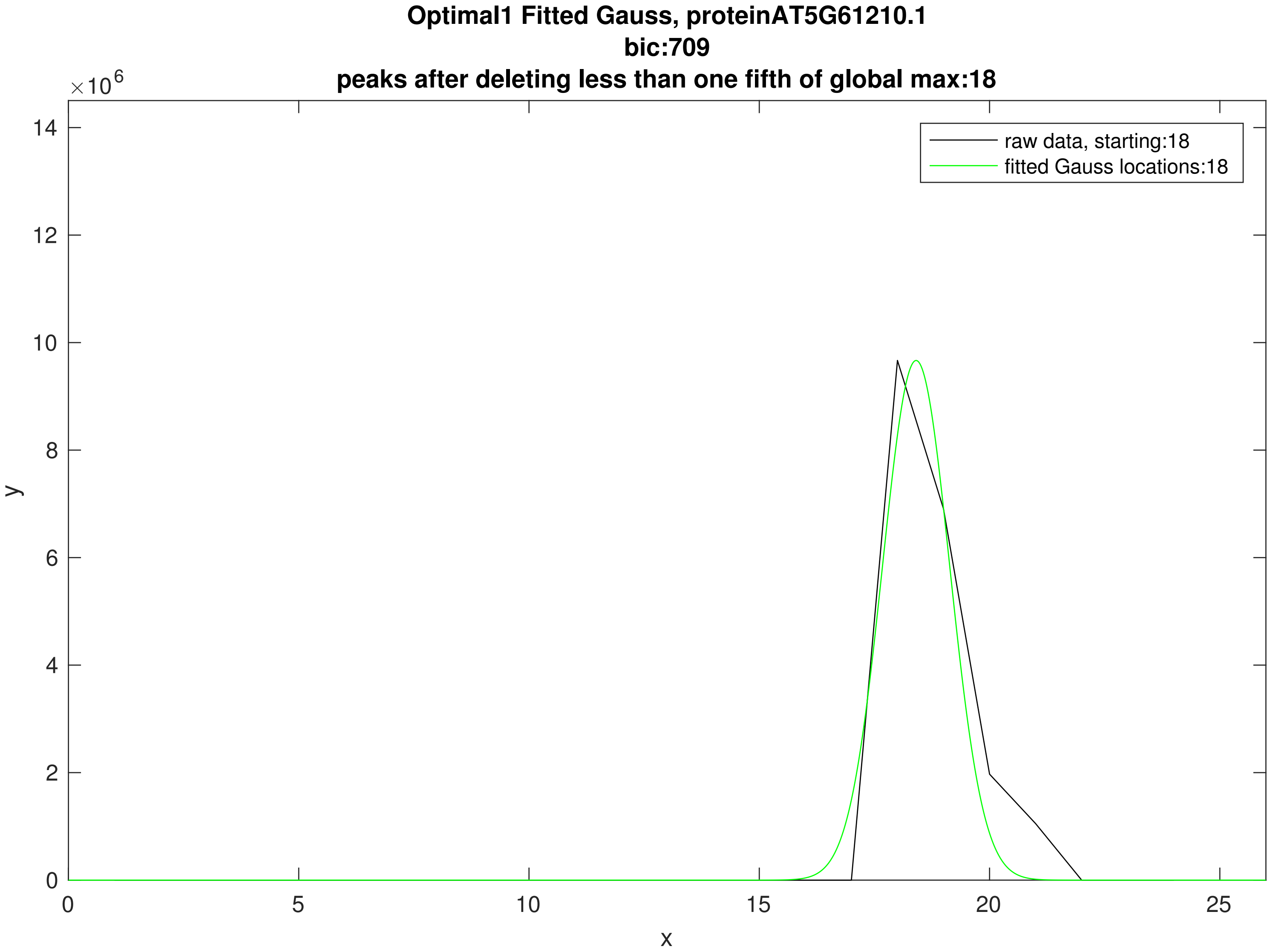
Optimal1 Fitted Gauss, proteinAT5G61210.1
bic:709
peaks after deleting less than one fifth of global max:18
×10^6
14
12
10
8
6
4
2
0
y
0
5
10
15
20
25
x
raw data, starting:18
fitted Gauss locations:18

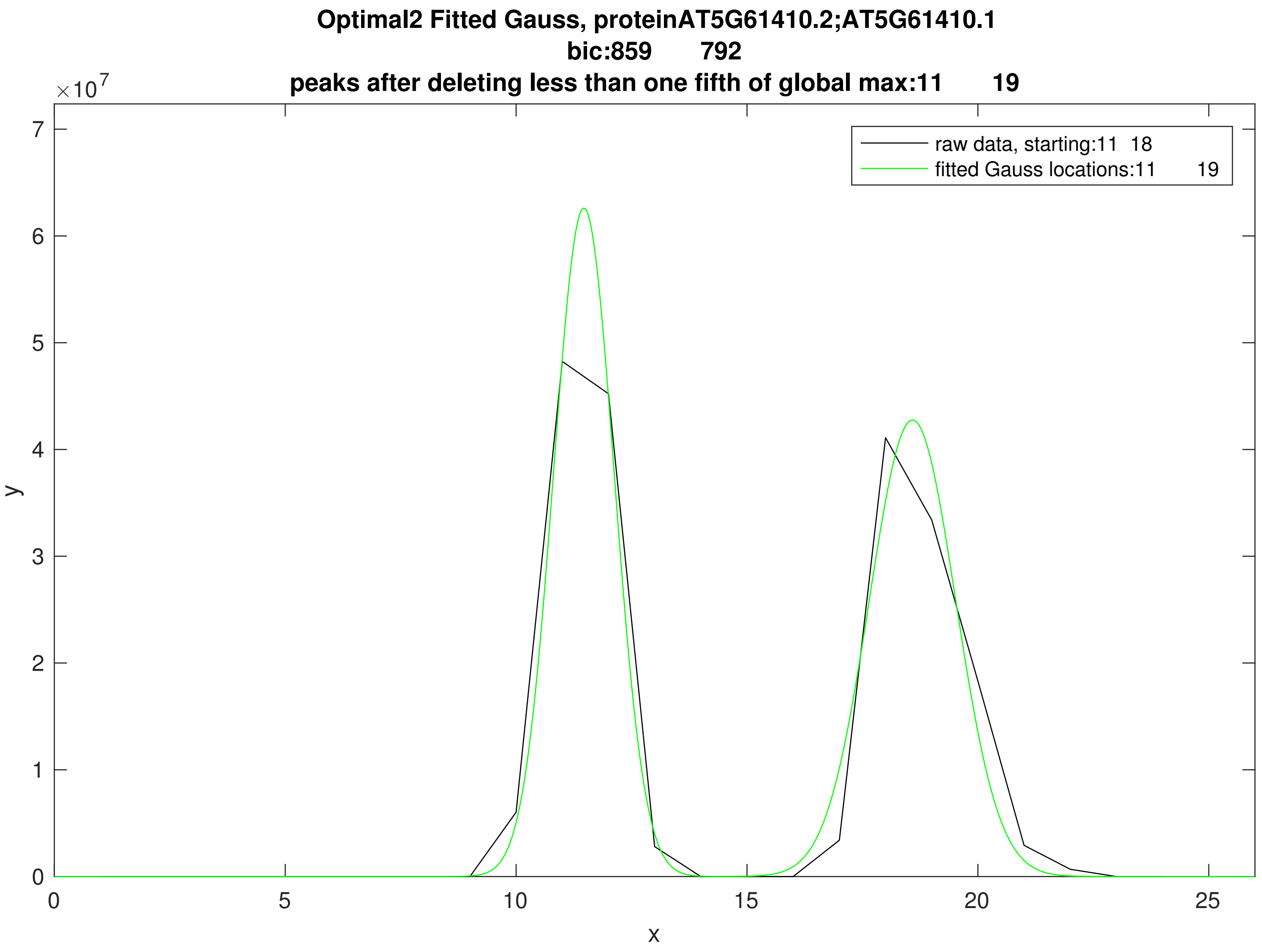

Optimal2 Fitted Gauss, proteinAT5G61410.2;AT5G61410.1
bic:859 792
peaks after deleting less than one fifth of global max:11 19
×10^7
raw data, starting:11 18
fitted Gauss locations:11 19
y
x
0
1
2
3
4
5
6
7
0
5
10
15
20
25

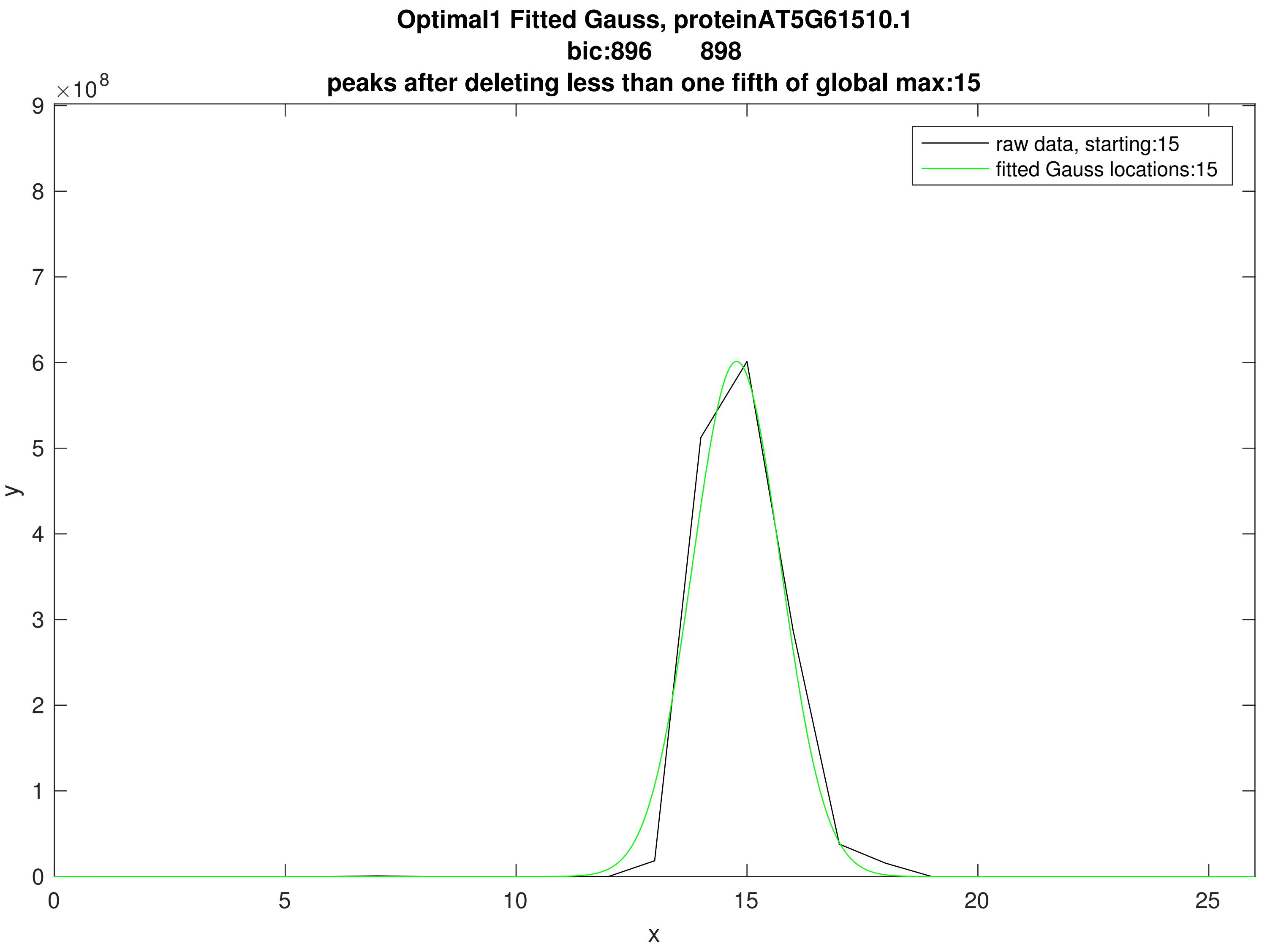
Optimal1 Fitted Gauss, proteinAT5G61510.1
bic:896 898
peaks after deleting less than one fifth of global max:15
×10^8
y
x
raw data, starting:15
fitted Gauss locations:15

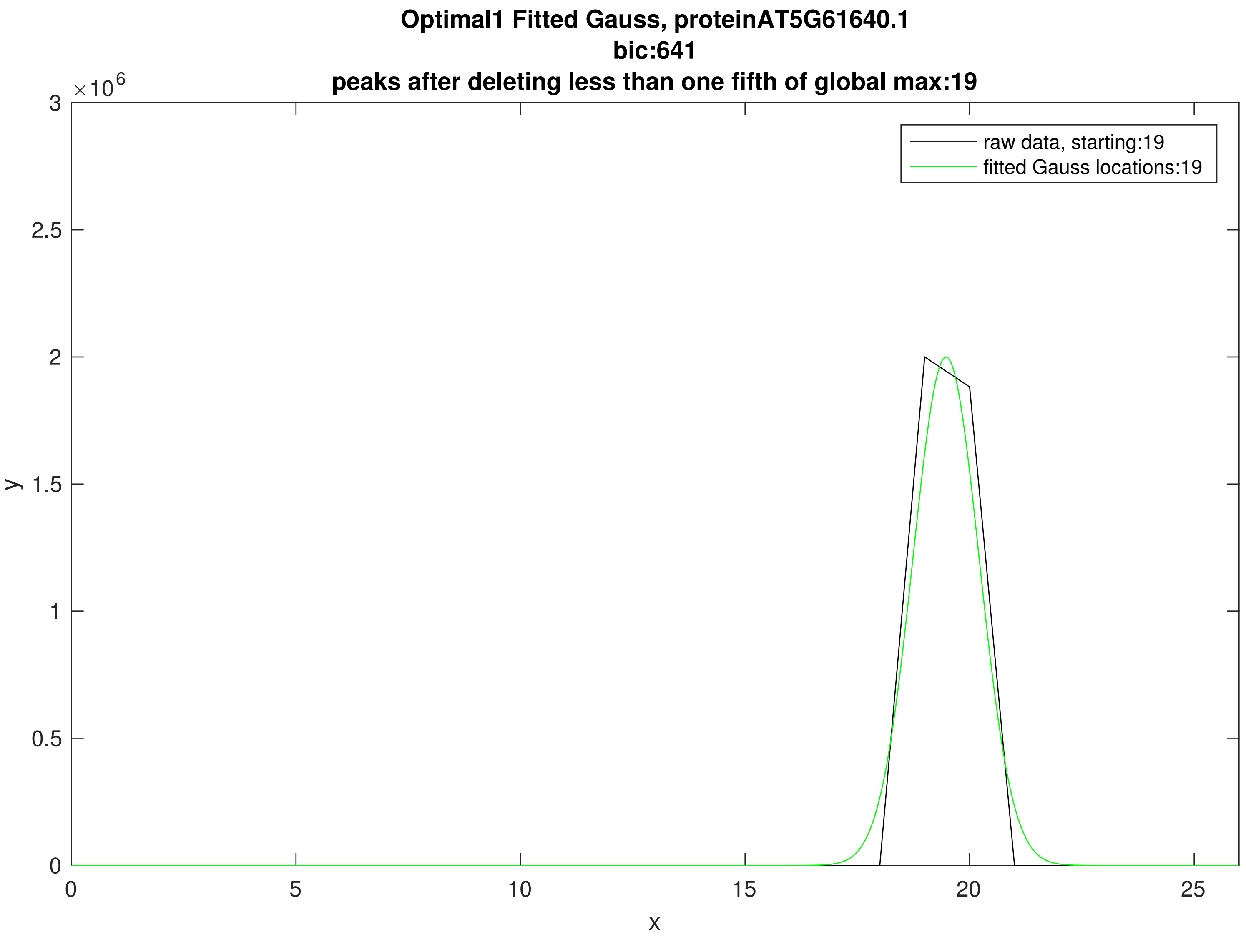

Optimal1 Fitted Gauss, proteinAT5G61640.1
bic:641
peaks after deleting less than one fifth of global max:19
×10^6
3
2.5
2
1.5
1
0.5
0
y
0
5
10
15
20
25
x
raw data, starting:19
fitted Gauss locations:19

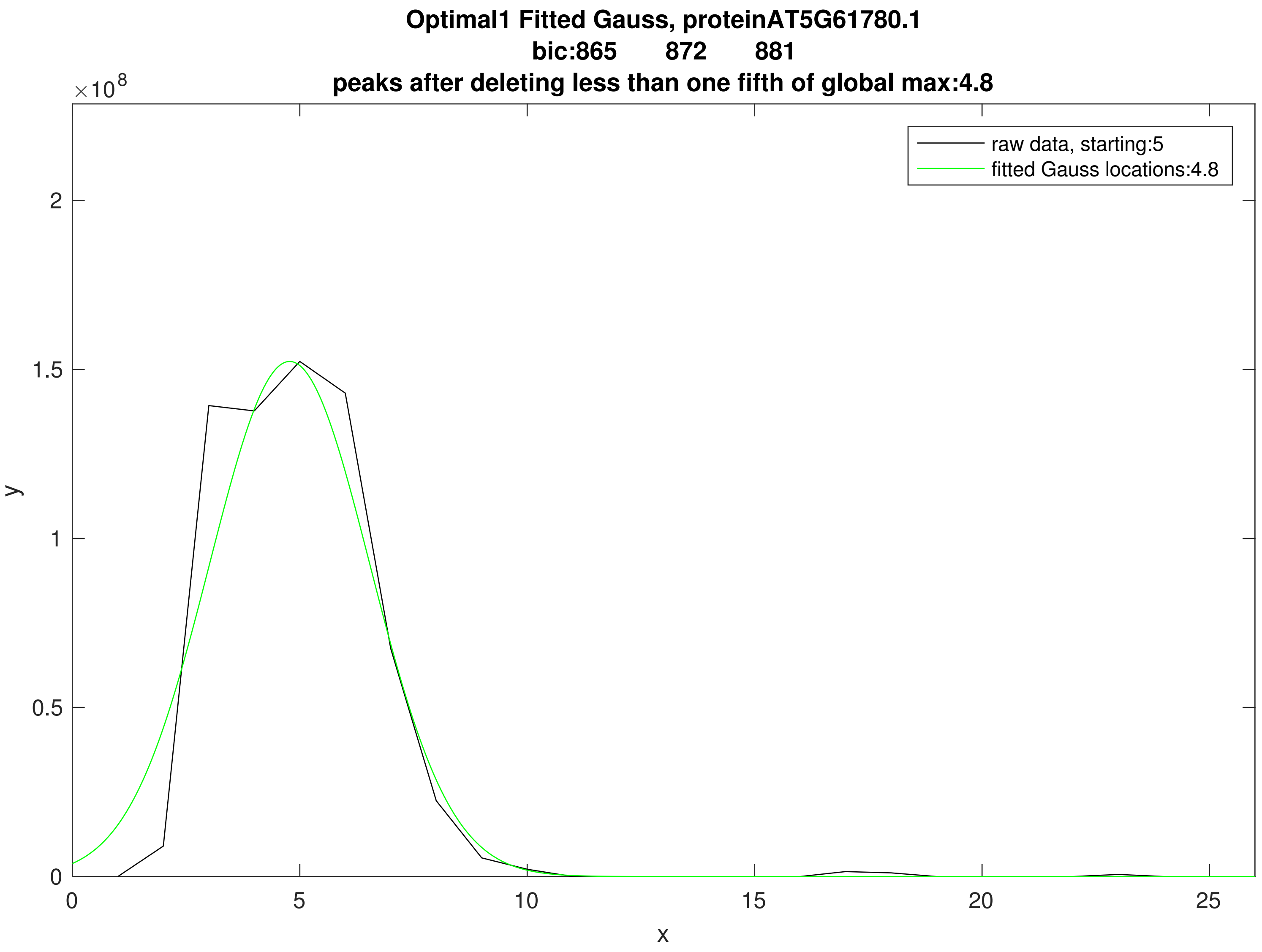
Optimal1 Fitted Gauss, proteinAT5G61780.1
bic:865 872 881
peaks after deleting less than one fifth of global max:4.8
×10^8
raw data, starting:5
fitted Gauss locations:4.8
y
x
0
0.5
1
1.5
2
5
10
15
20
25

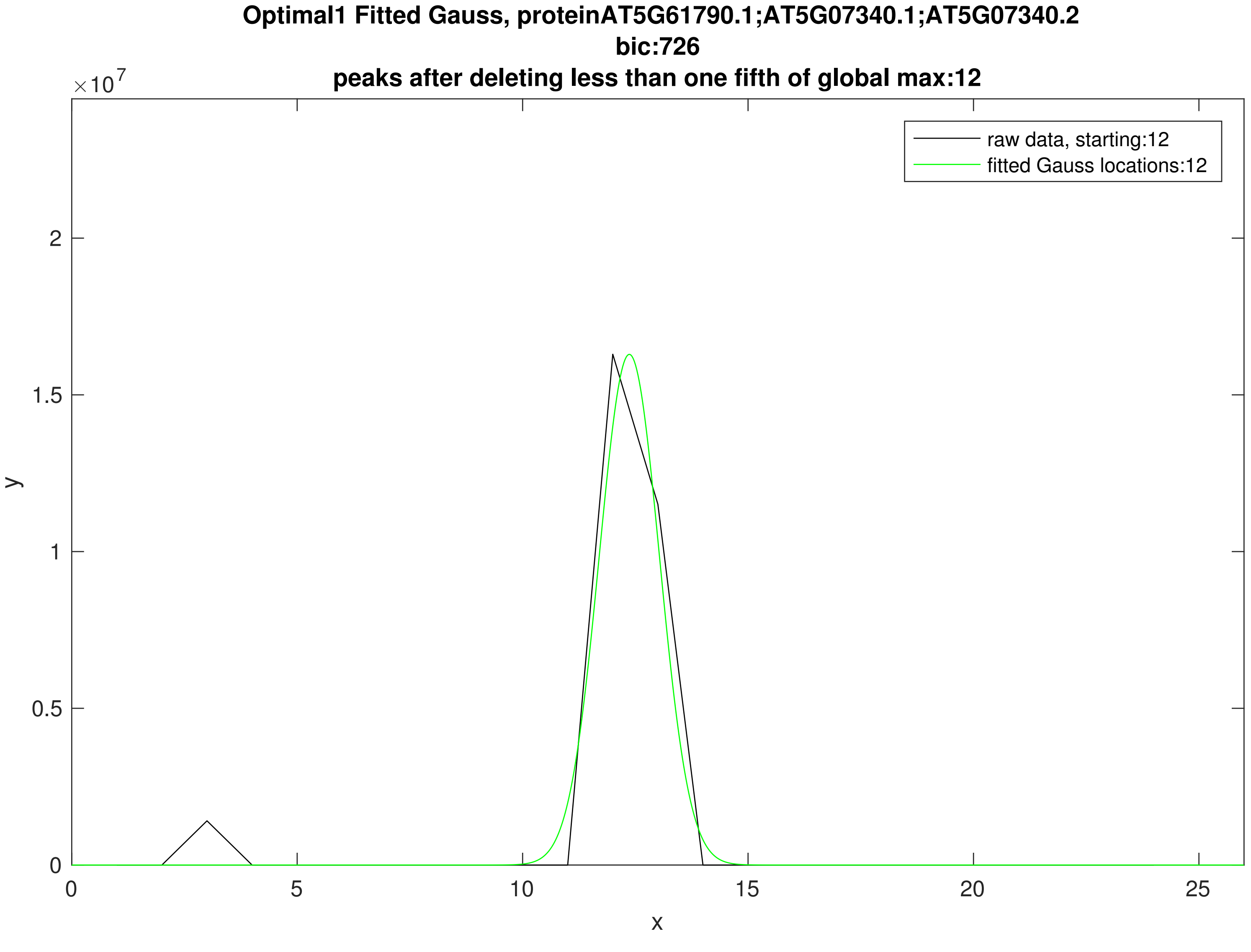
Optimal1 Fitted Gauss, proteinAT5G61790.1;AT5G07340.1;AT5G07340.2
bic:726
peaks after deleting less than one fifth of global max:12
×10^7
raw data, starting:12
fitted Gauss locations:12
y
x
0
0.5
1
1.5
2
5
10
15
20
25

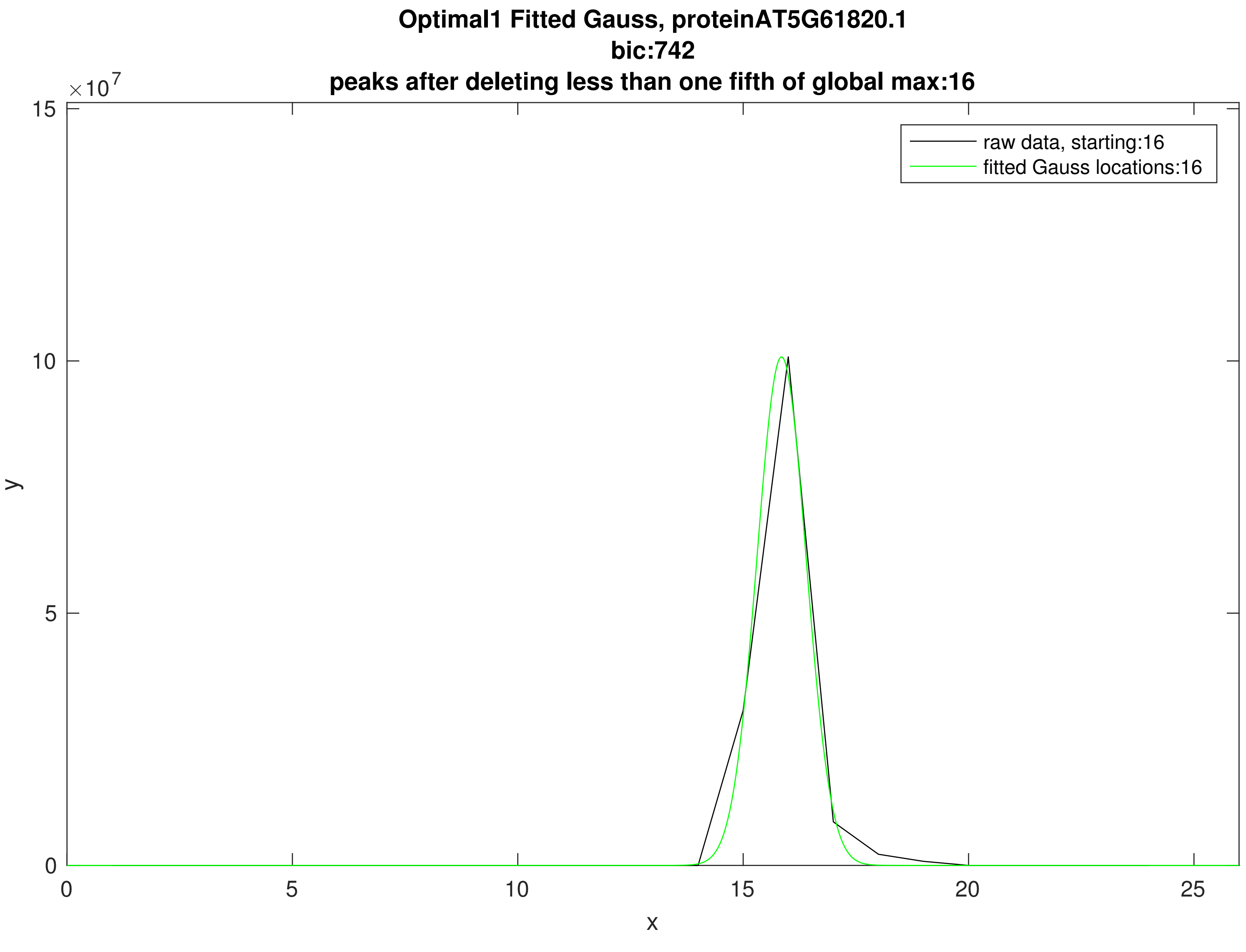
Optimal1 Fitted Gauss, proteinAT5G61820.1
bic:742
peaks after deleting less than one fifth of global max:16
×10^7
15
10
5
0
y
0
5
10
15
20
25
x
raw data, starting:16
fitted Gauss locations:16

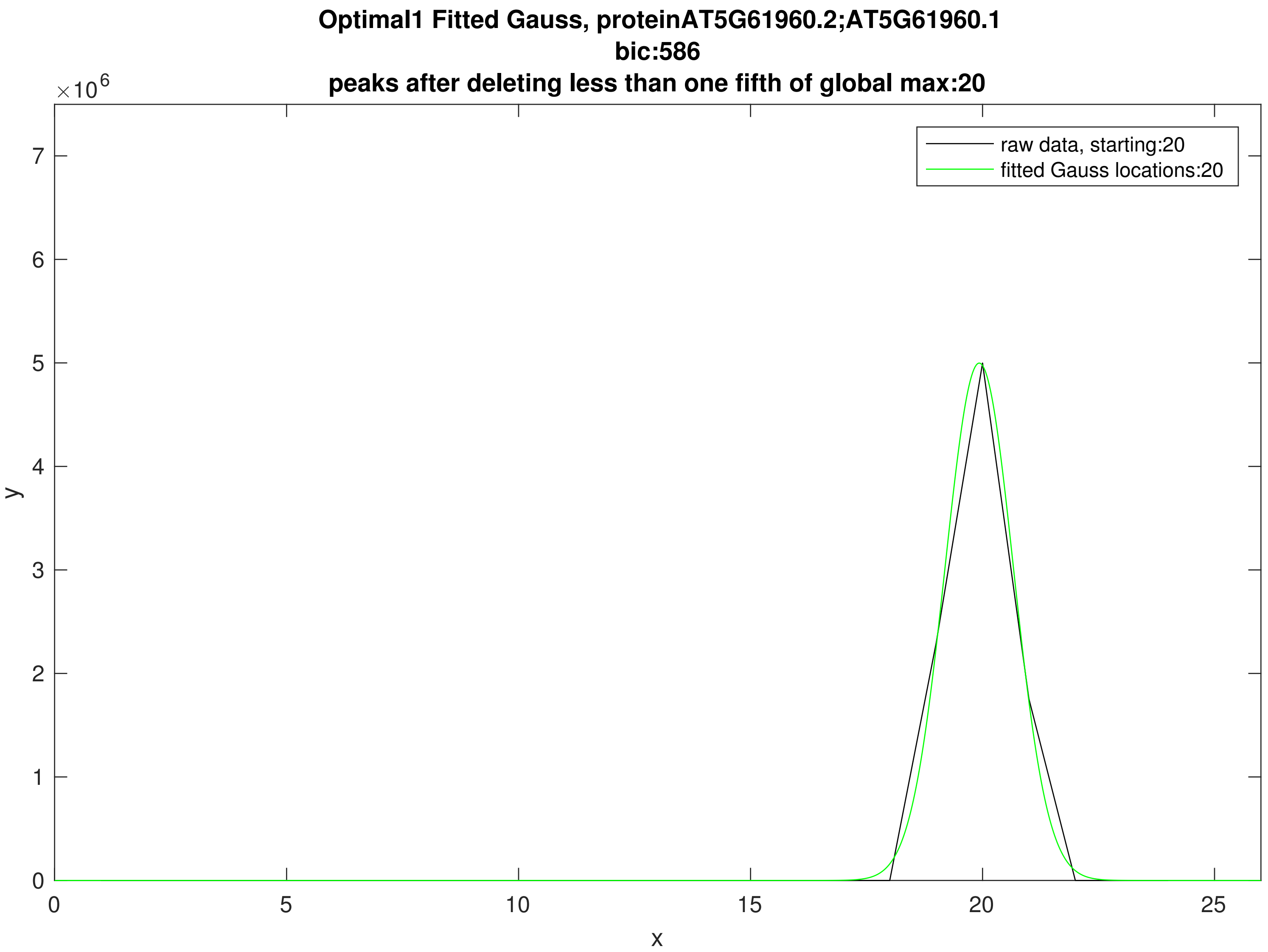
Optimal1 Fitted Gauss, proteinAT5G61960.2;AT5G61960.1
bic:586
peaks after deleting less than one fifth of global max:20
×10^6
raw data, starting:20
fitted Gauss locations:20
y
x
0
1
2
3
4
5
6
7
0
5
10
15
20
25

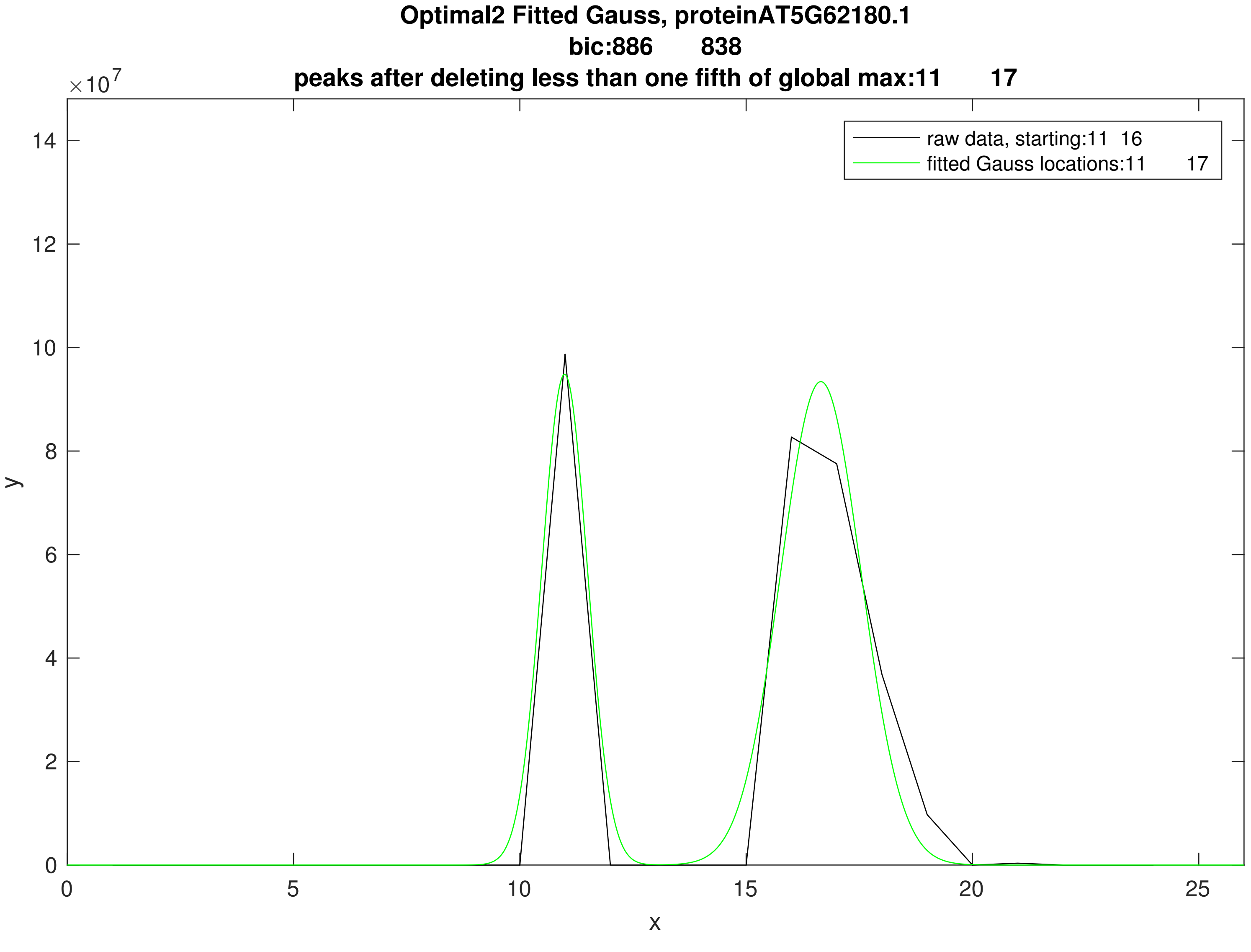

Optimal2 Fitted Gauss, proteinAT5G62180.1
bic:886 838
peaks after deleting less than one fifth of global max:11 17
×10^7
raw data, starting:11 16
fitted Gauss locations:11 17
y
x
0
2
4
6
8
10
12
14
0
5
10
15
20
25

#### Optimal1 Fitted Gauss, proteinAT5G62300.2;AT5G62300.1;AT3G45030.1
#### bic:842
#### peaks after deleting less than one fifth of global max:3.8

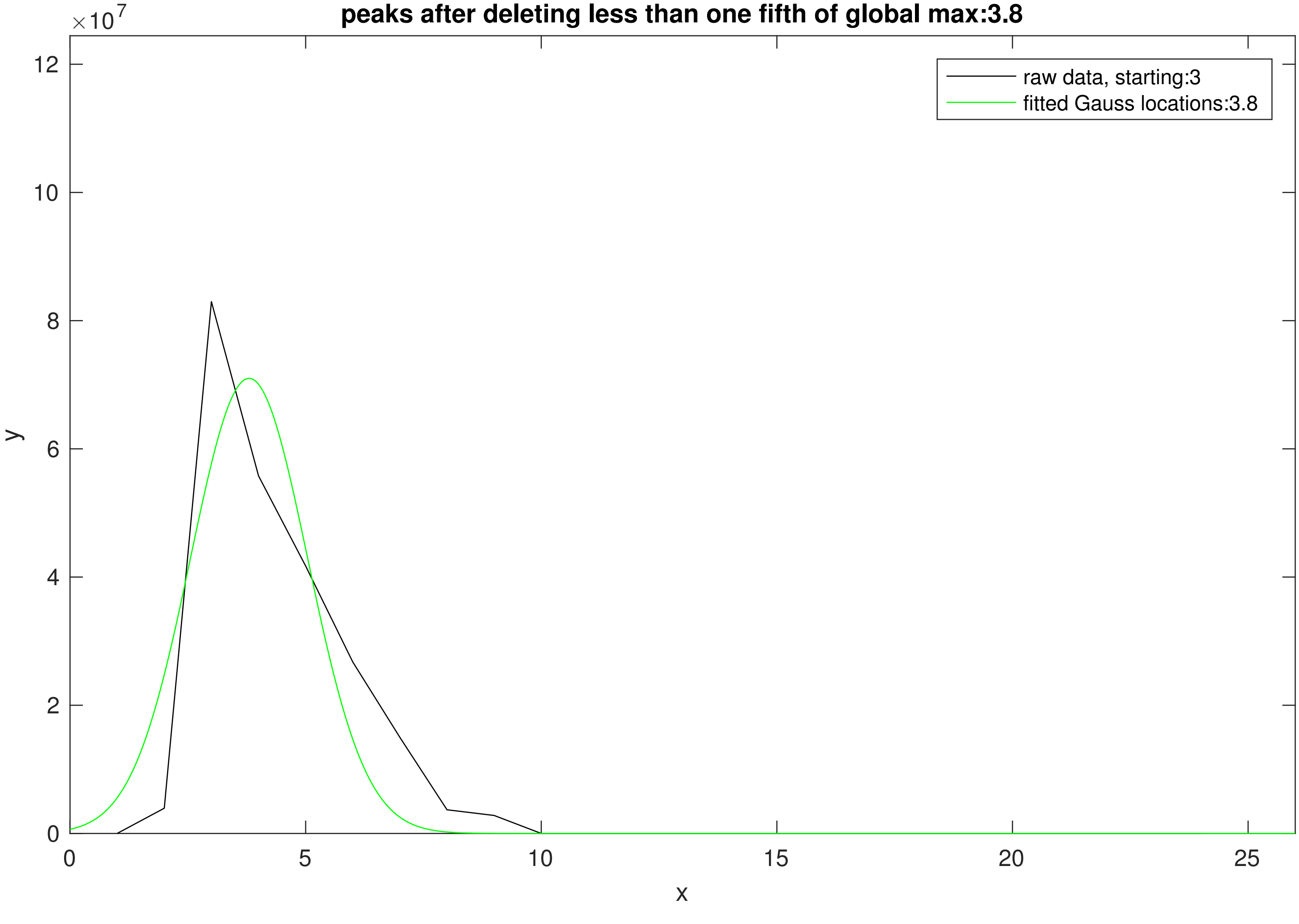

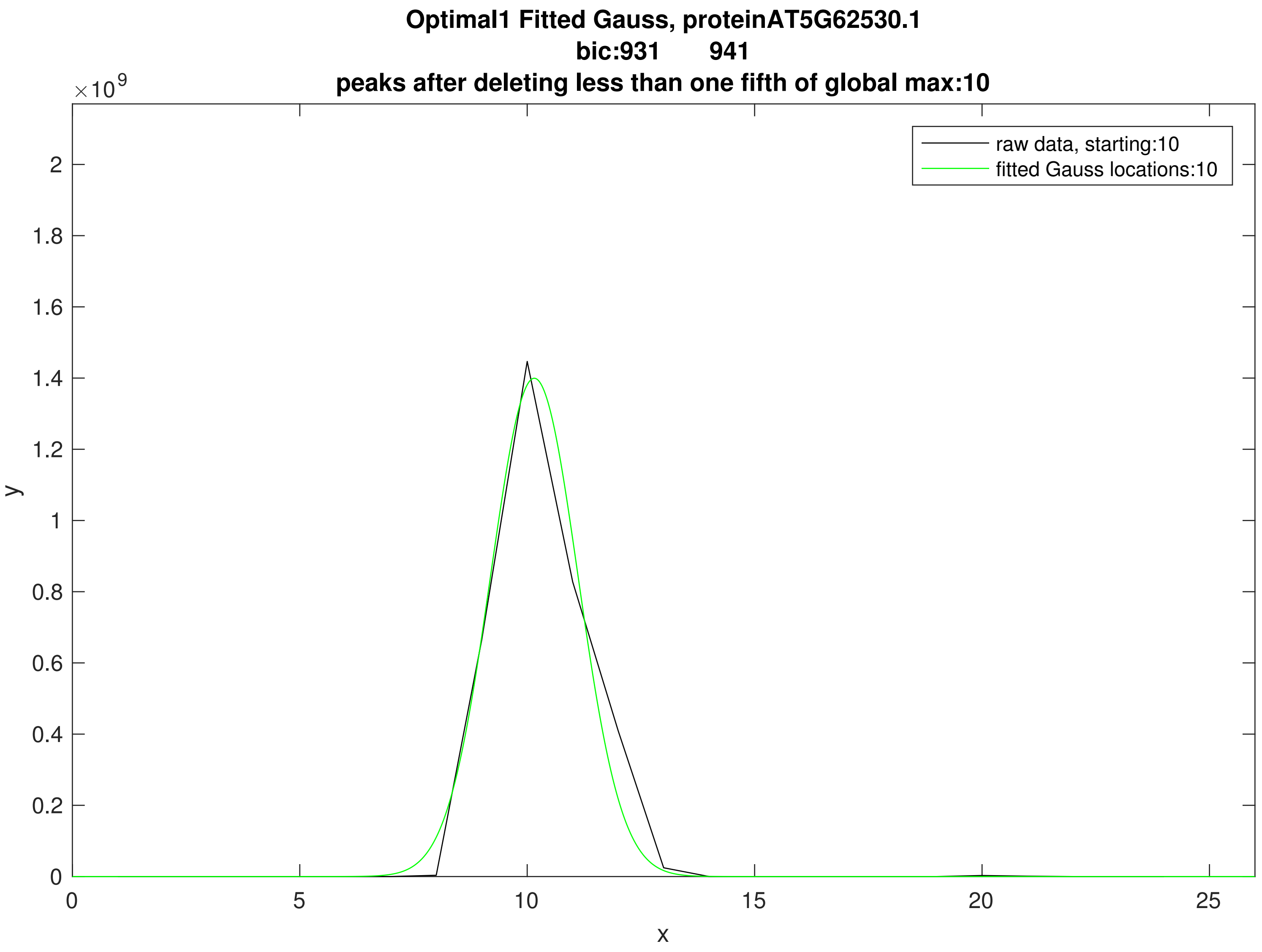
Optimal1 Fitted Gauss, proteinAT5G62530.1
bic:931 941
peaks after deleting less than one fifth of global max:10
×10^9
raw data, starting:10
fitted Gauss locations:10
2
1.8
1.6
1.4
1.2
1
0.8
0.6
0.4
0.2
0
y
0
5
10
15
20
25
x

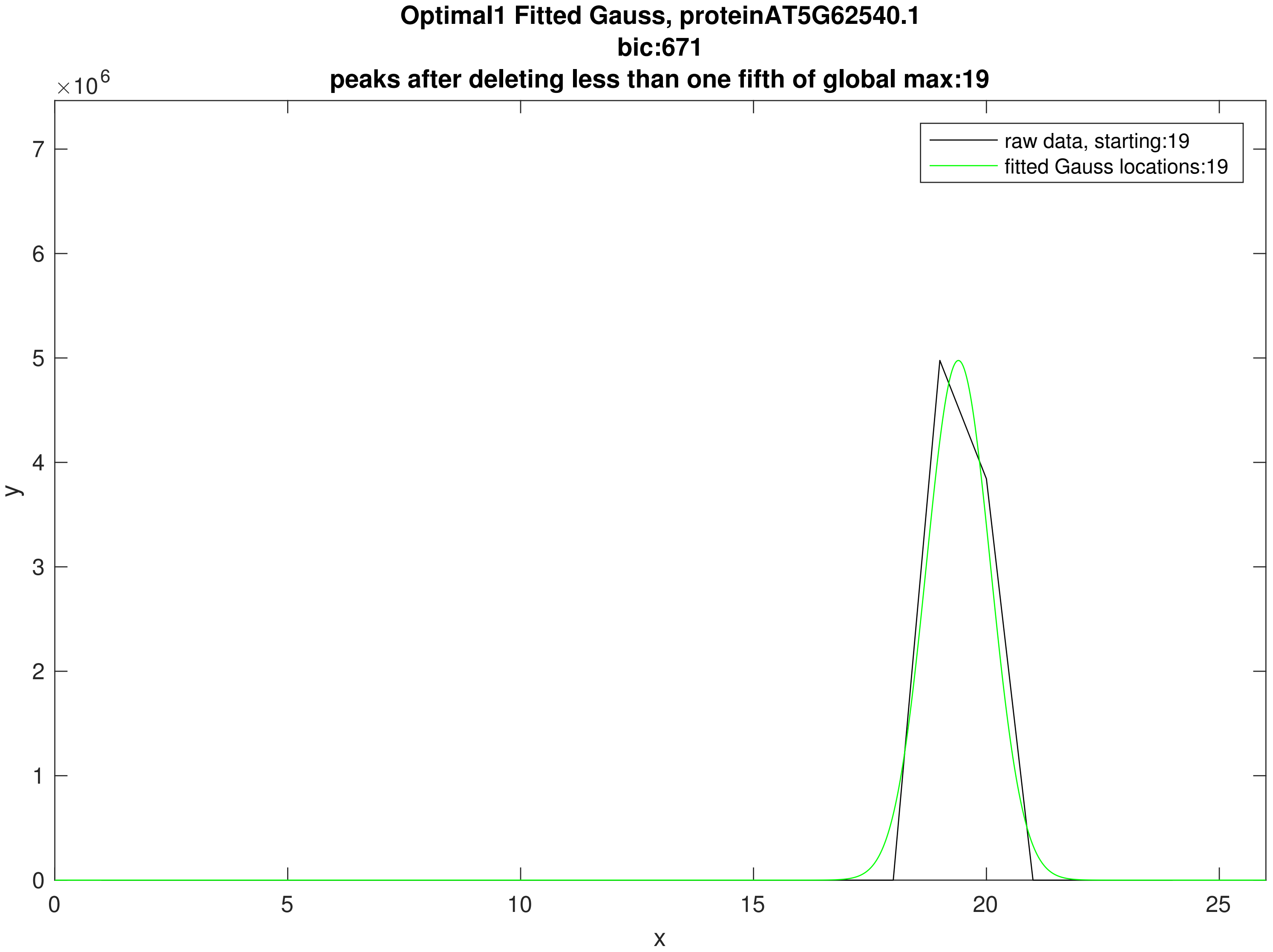

Optimal1 Fitted Gauss, proteinAT5G62540.1
bic:671
peaks after deleting less than one fifth of global max:19
×10^6
raw data, starting:19
fitted Gauss locations:19
y
x
0
1
2
3
4
5
6
7
0
5
10
15
20
25

**Optimal1 Fitted Gauss, proteinAT5G62700.1;AT5G62690.1**
**bic:853 855 865 875**
**peaks after deleting less than one fifth of global max:14**

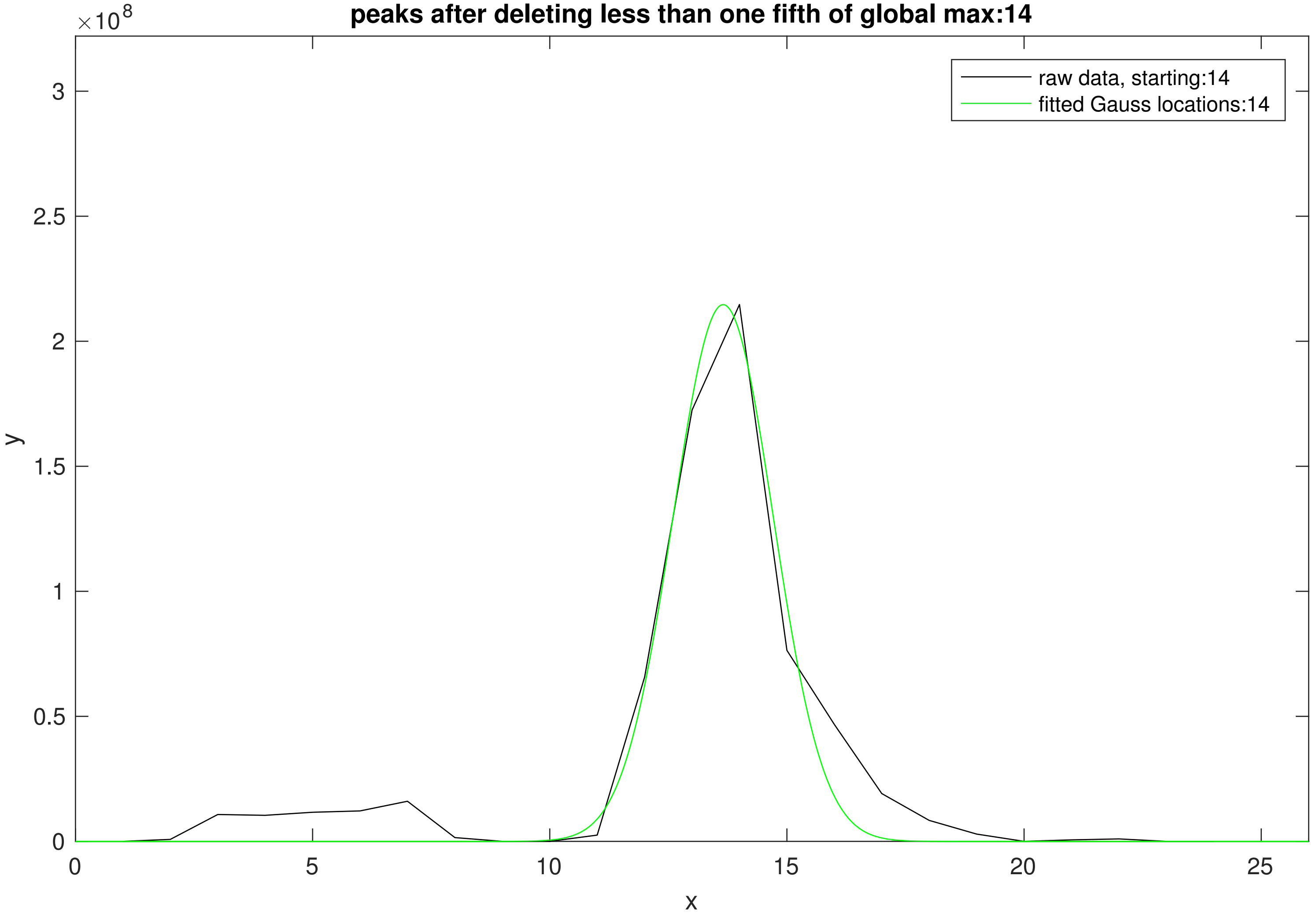

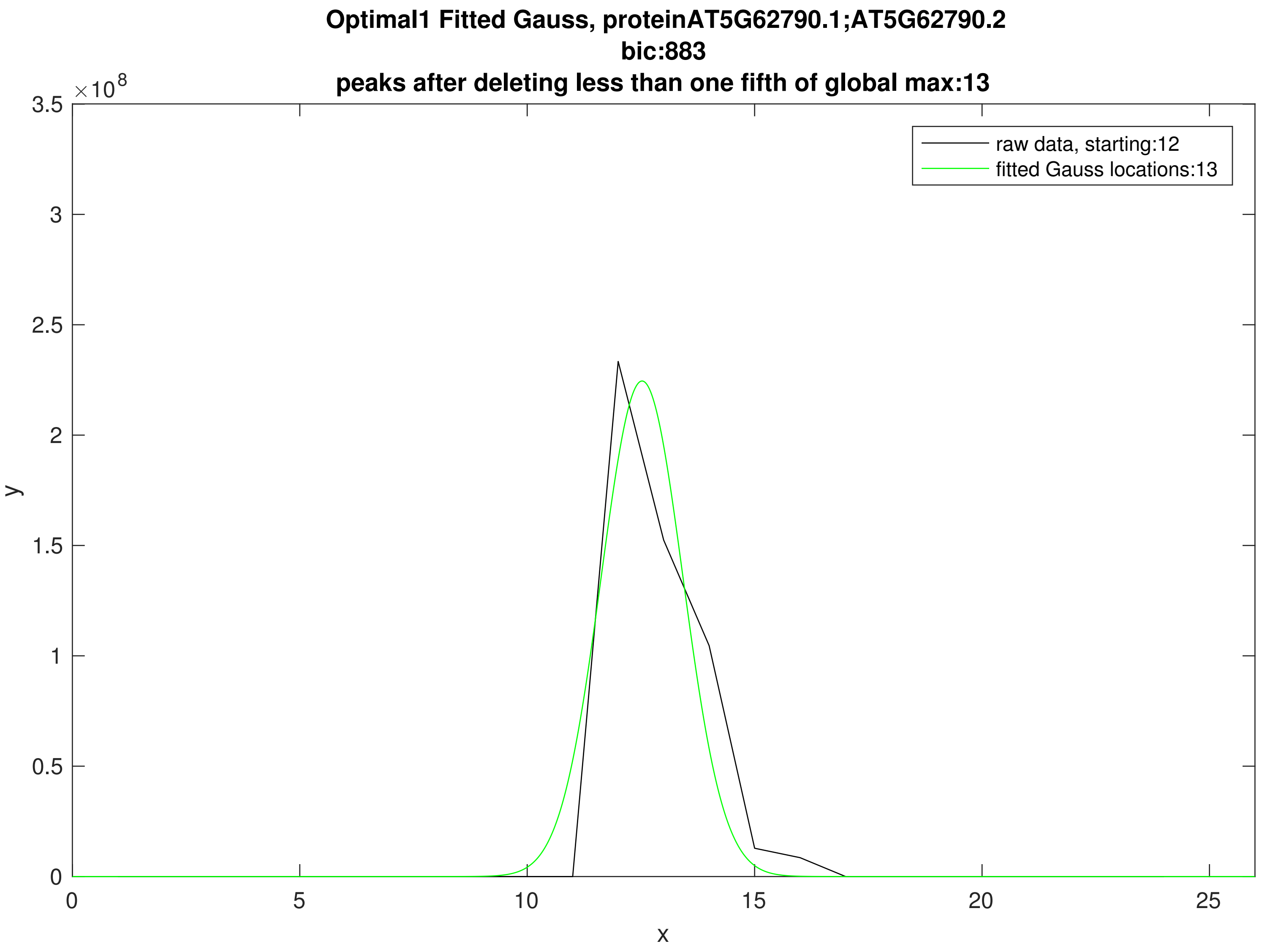
Optimal1 Fitted Gauss, proteinAT5G62790.1;AT5G62790.2
bic:883
peaks after deleting less than one fifth of global max:13
×10^8
3.5
3
2.5
2
1.5
1
0.5
0
y
0
5
10
15
20
25
x
raw data, starting:12
fitted Gauss locations:13

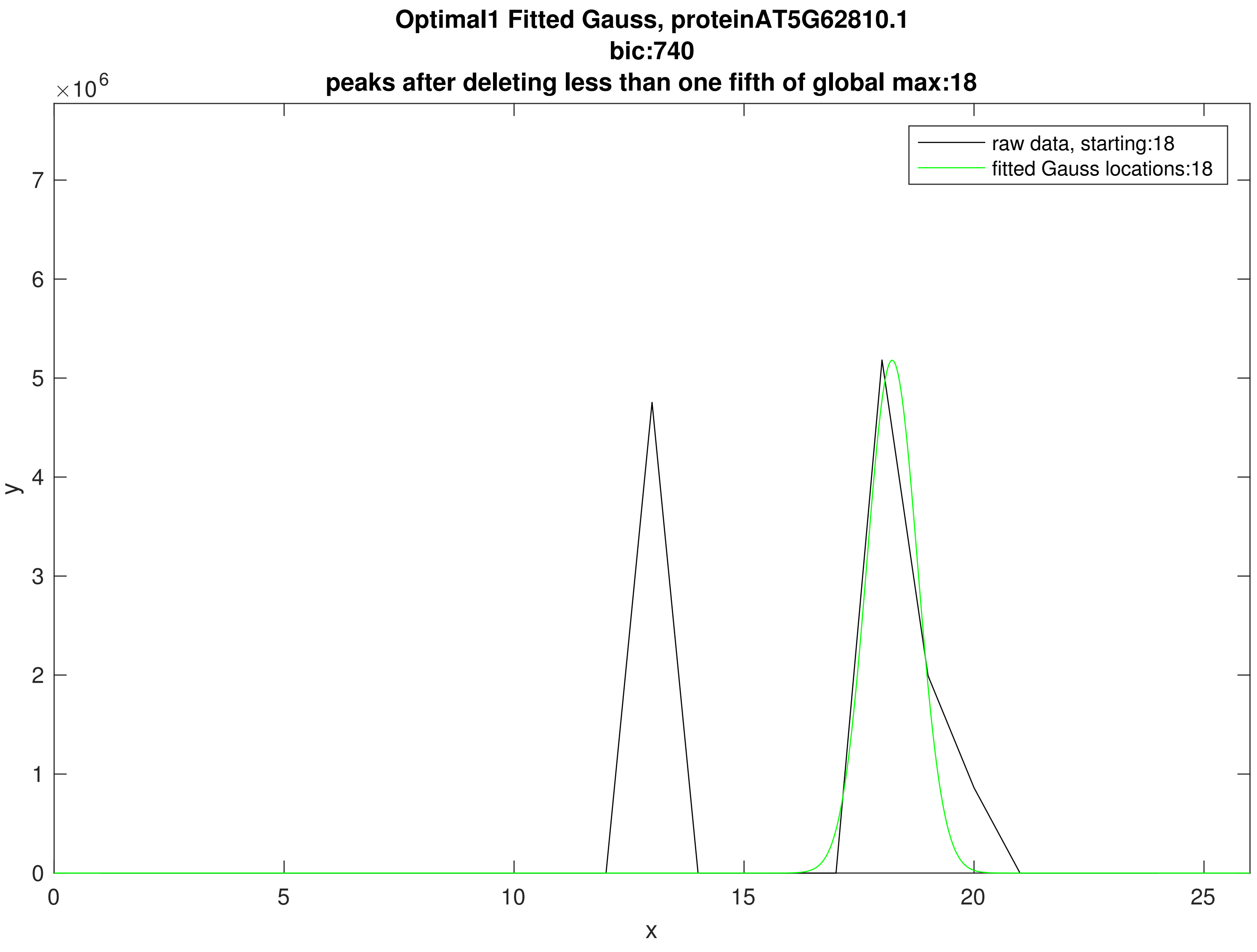
Optimal1 Fitted Gauss, proteinAT5G62810.1
bic:740
peaks after deleting less than one fifth of global max:18
×10^6
raw data, starting:18
fitted Gauss locations:18
y
x
0
1
2
3
4
5
6
7
0
5
10
15
20
25

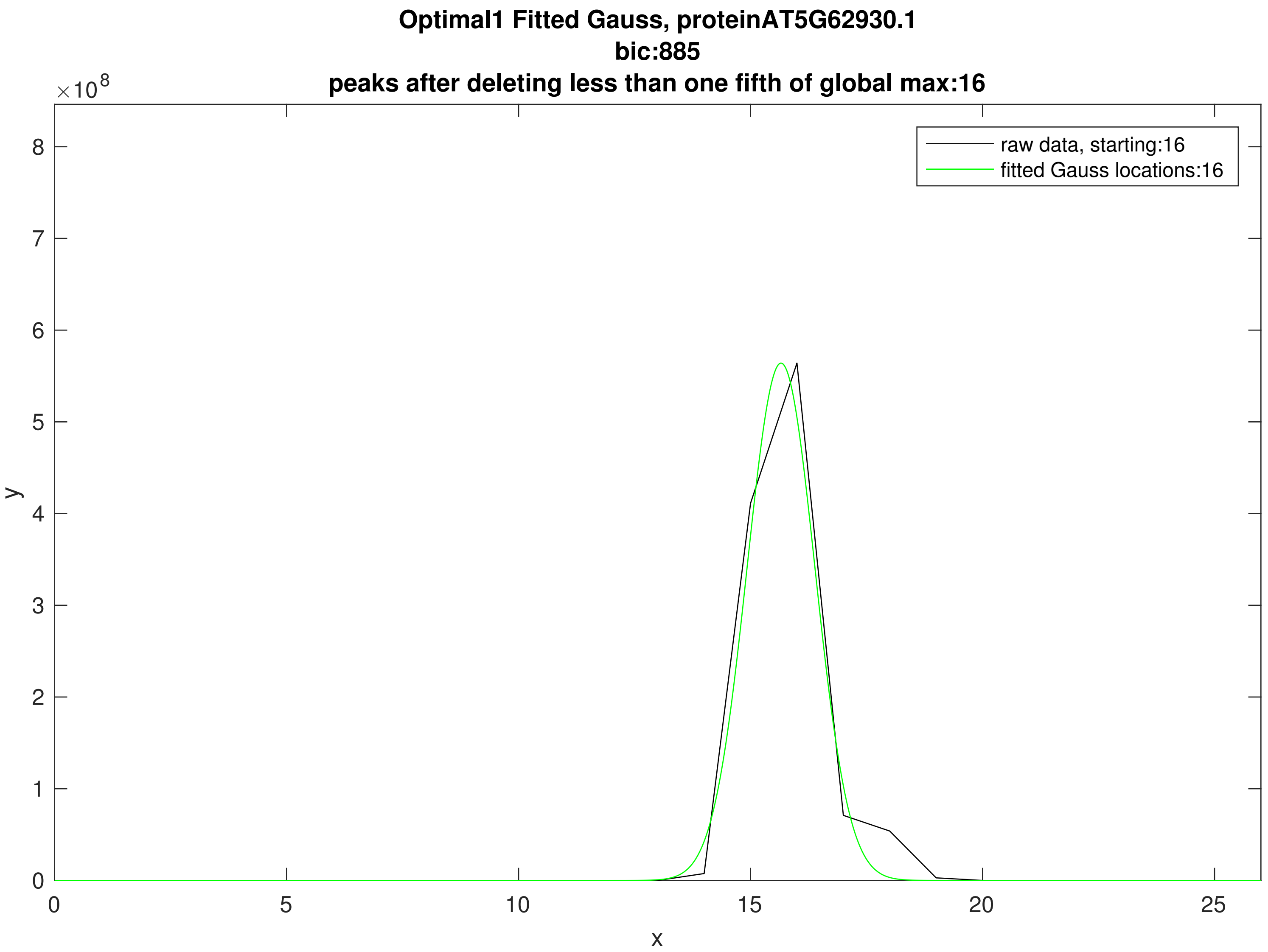
Optimal1 Fitted Gauss, proteinAT5G62930.1
bic:885
peaks after deleting less than one fifth of global max:16
×10^8
raw data, starting:16
fitted Gauss locations:16
y
x
0
1
2
3
4
5
6
7
8
0
5
10
15
20
25

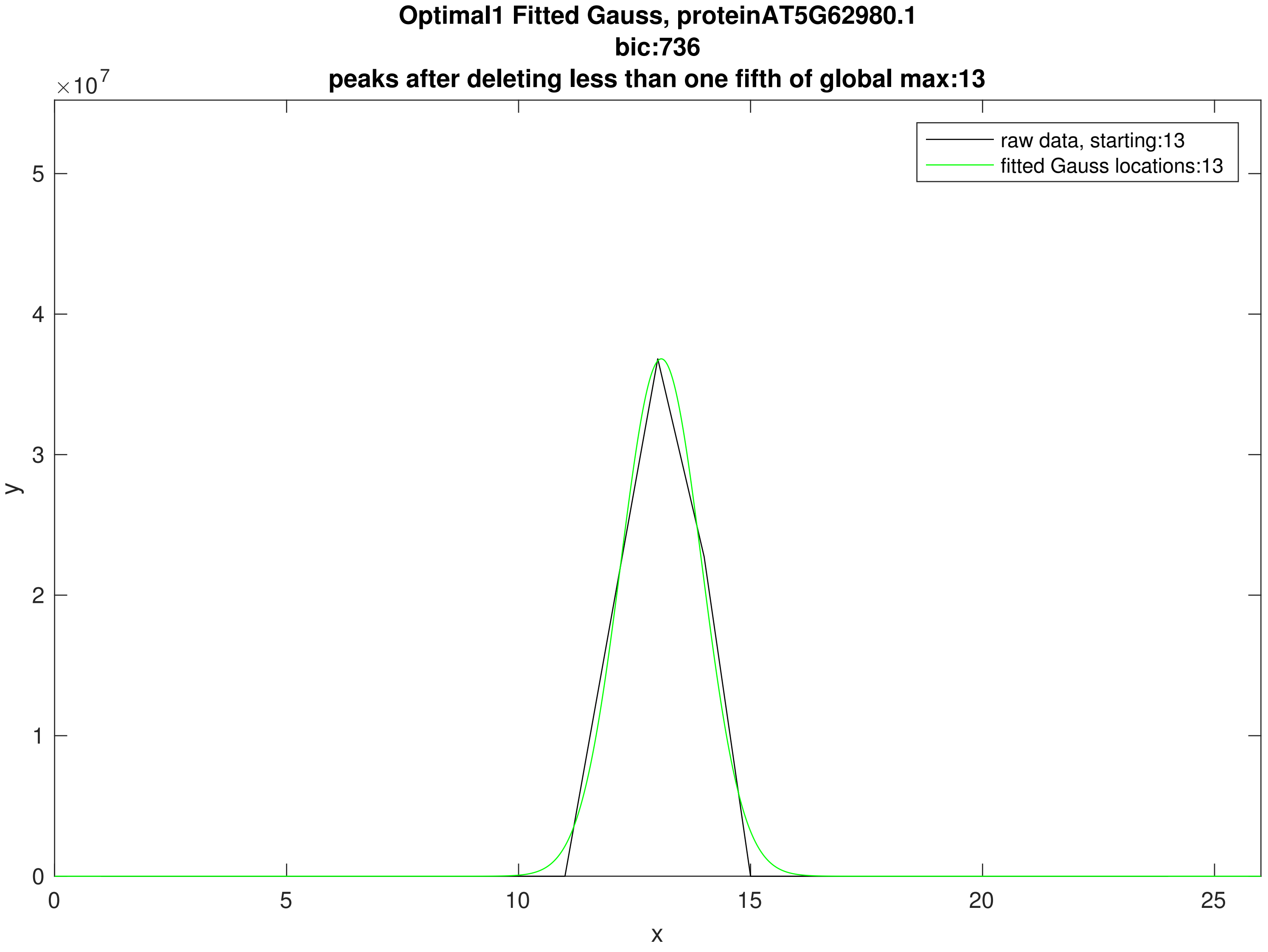
Optimal1 Fitted Gauss, proteinAT5G62980.1
bic:736
peaks after deleting less than one fifth of global max:13
×10^7
raw data, starting:13
fitted Gauss locations:13
y
x
0
1
2
3
4
5
0
5
10
15
20
25

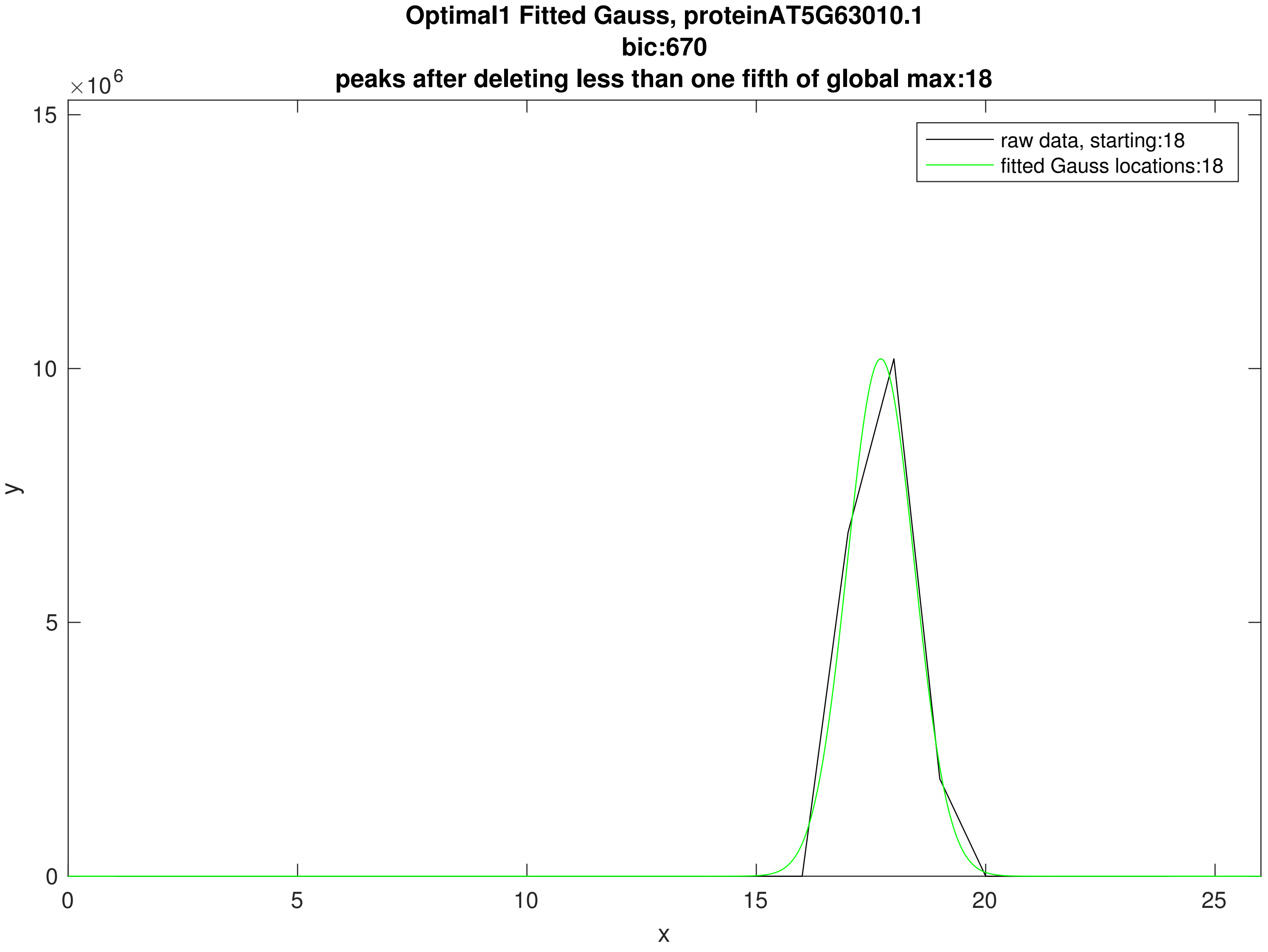

Optimal1 Fitted Gauss, proteinAT5G63010.1
bic:670
peaks after deleting less than one fifth of global max:18
×10^6
15
10
5
0
y
0
5
10
15
20
25
x
raw data, starting:18
fitted Gauss locations:18

**Optimal1 Fitted Gauss, proteinAT5G63030.1**
**bic:812 821**
**peaks after deleting less than one fifth of global max:20**

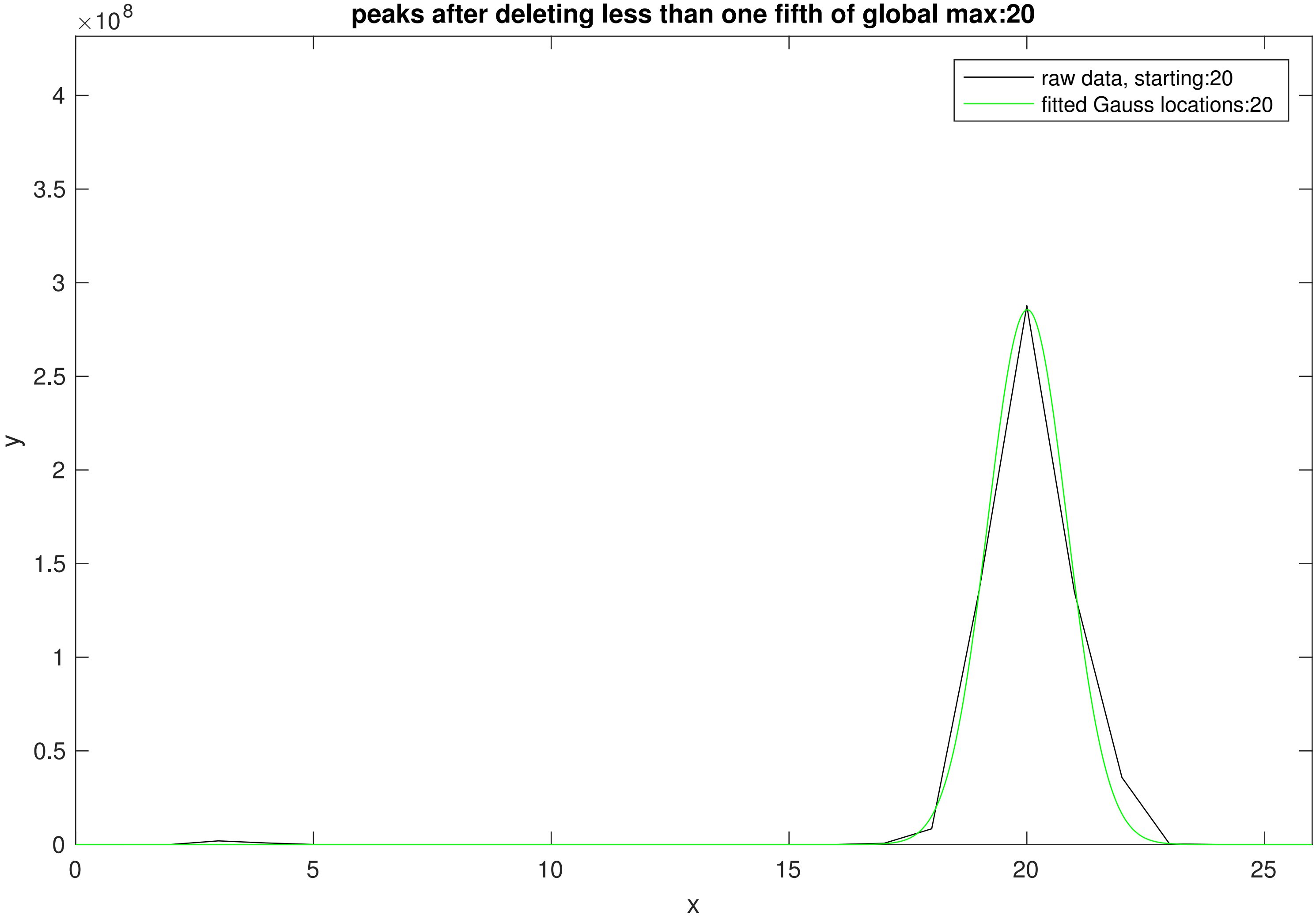

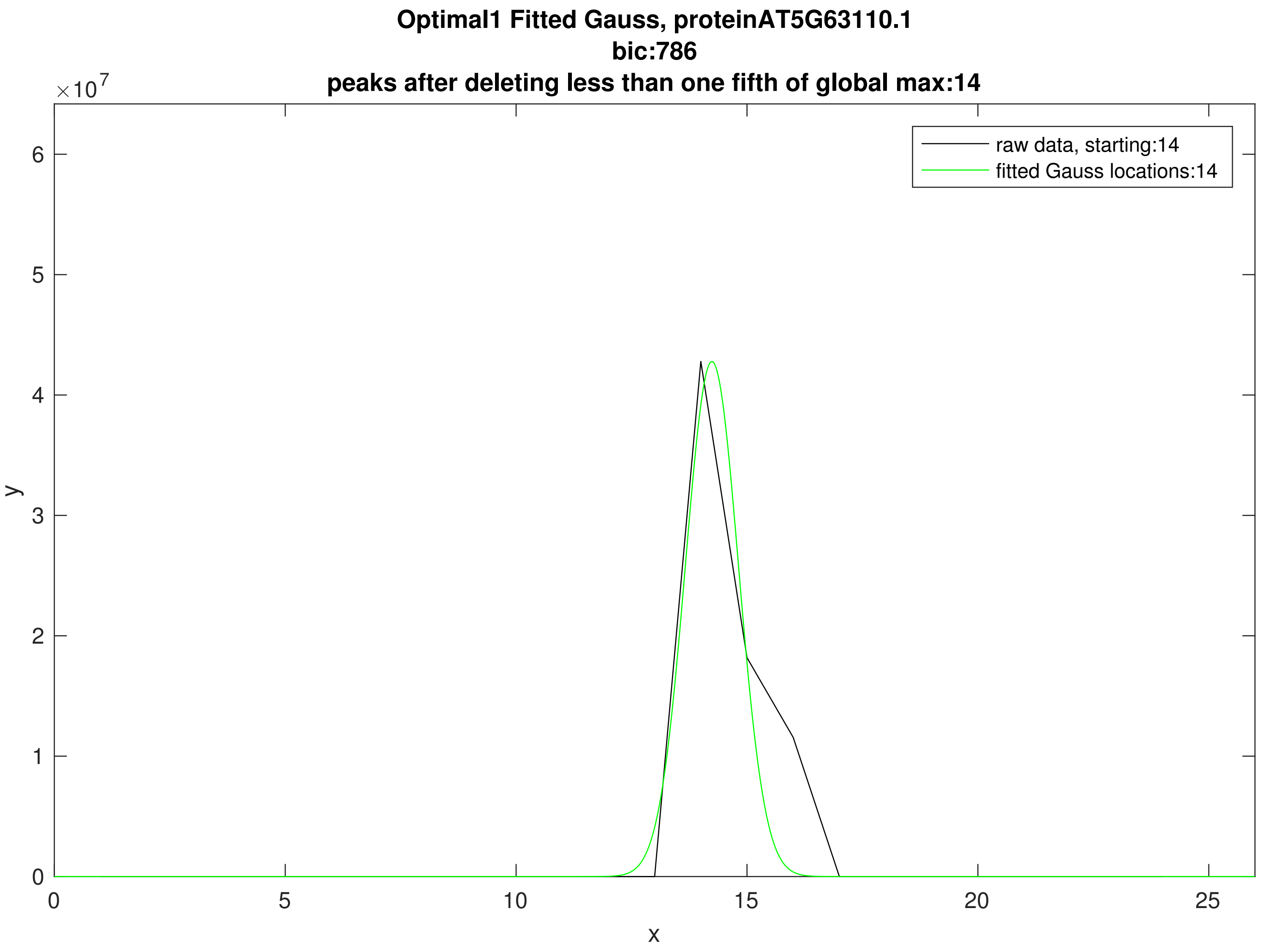
Optimal1 Fitted Gauss, proteinAT5G63110.1
bic:786
peaks after deleting less than one fifth of global max:14
×10^7
raw data, starting:14
fitted Gauss locations:14
y
x
0
1
2
3
4
5
6
0
5
10
15
20
25

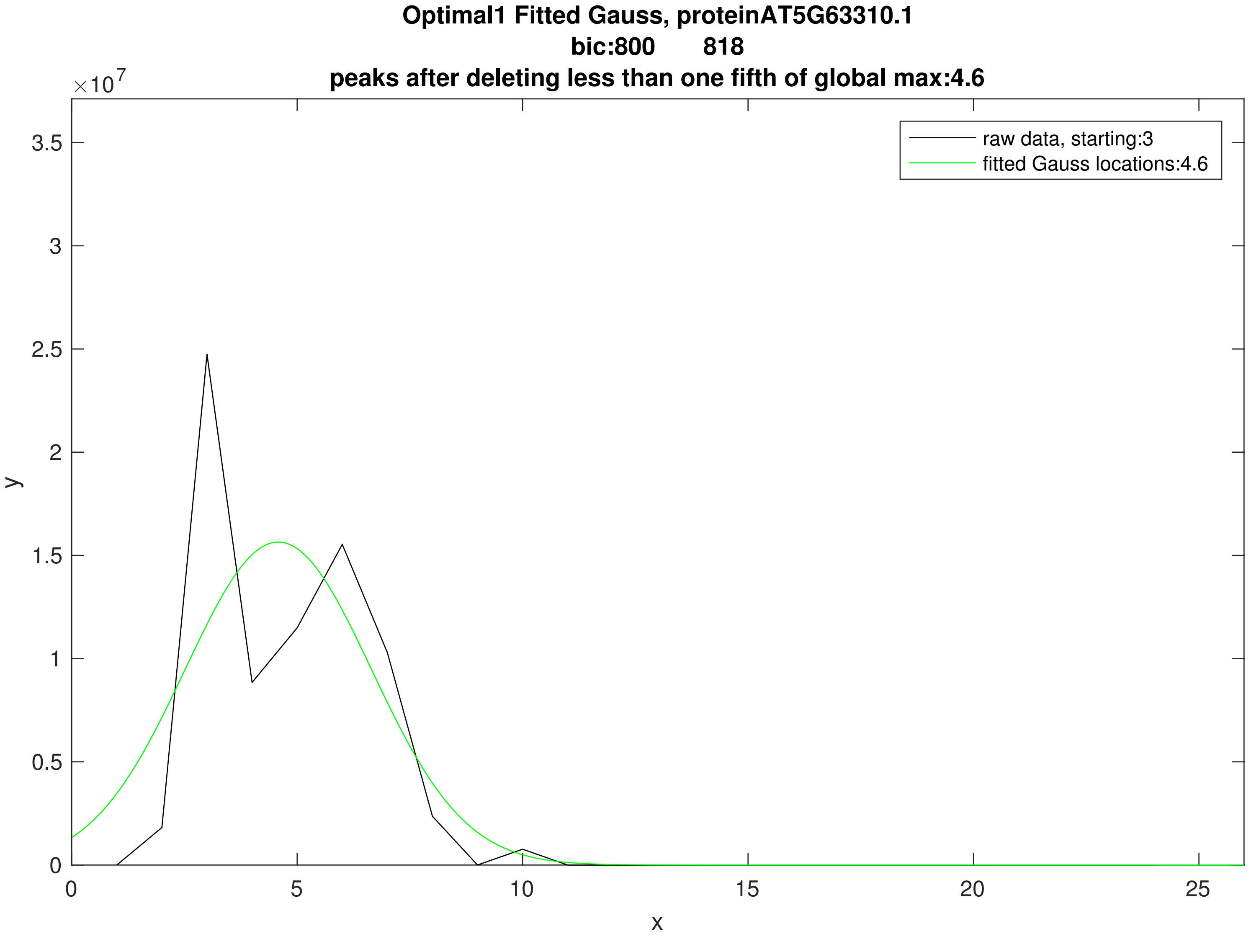
Optimal1 Fitted Gauss, proteinAT5G63310.1
bic:800 818
peaks after deleting less than one fifth of global max:4.6
raw data, starting:3
fitted Gauss locations:4.6
×10^7
y
x

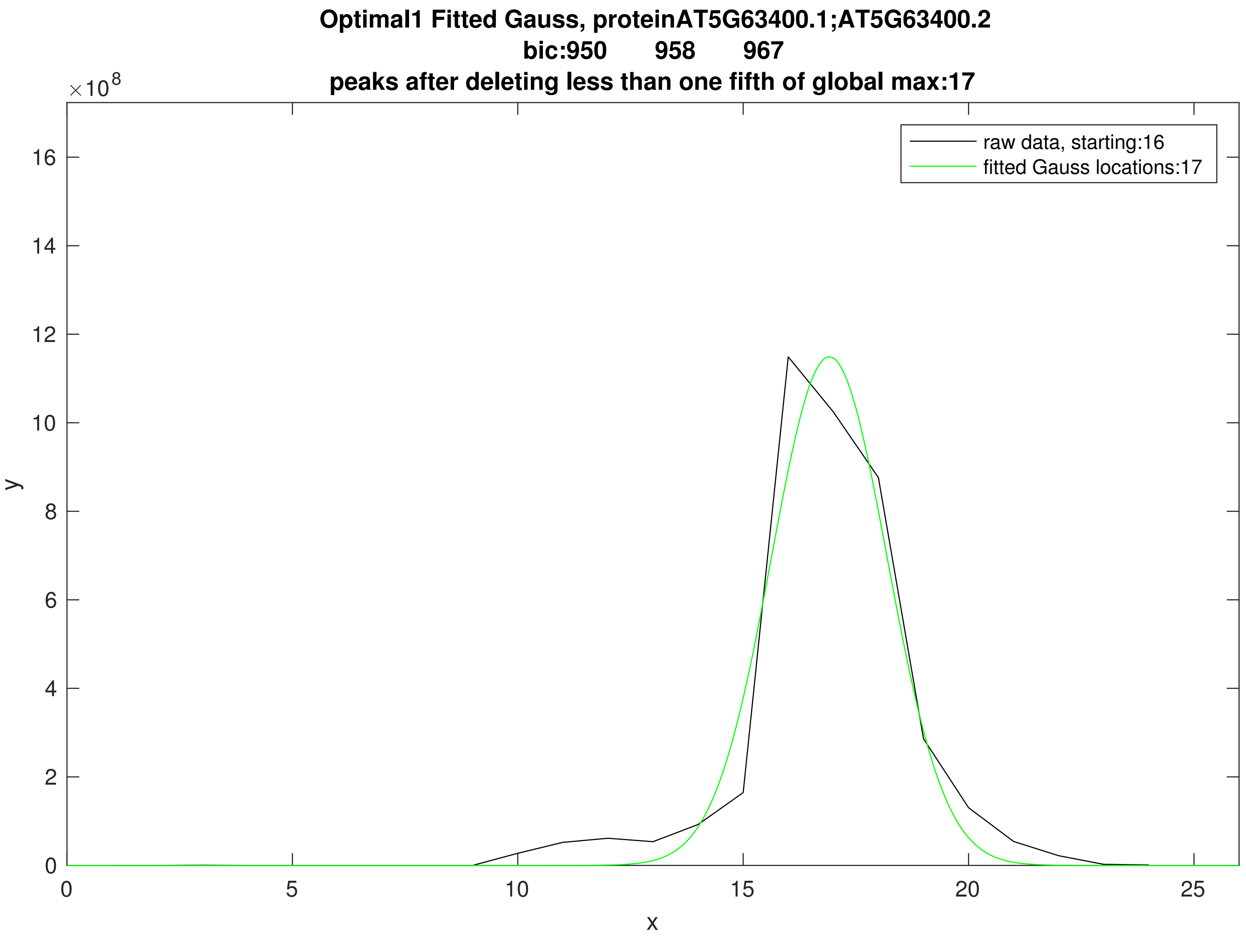
Optimal1 Fitted Gauss, proteinAT5G63400.1;AT5G63400.2
bic:950 958 967
peaks after deleting less than one fifth of global max:17
×10^8
raw data, starting:16
fitted Gauss locations:17
y
x
0
2
4
6
8
10
12
14
16
0
5
10
15
20
25

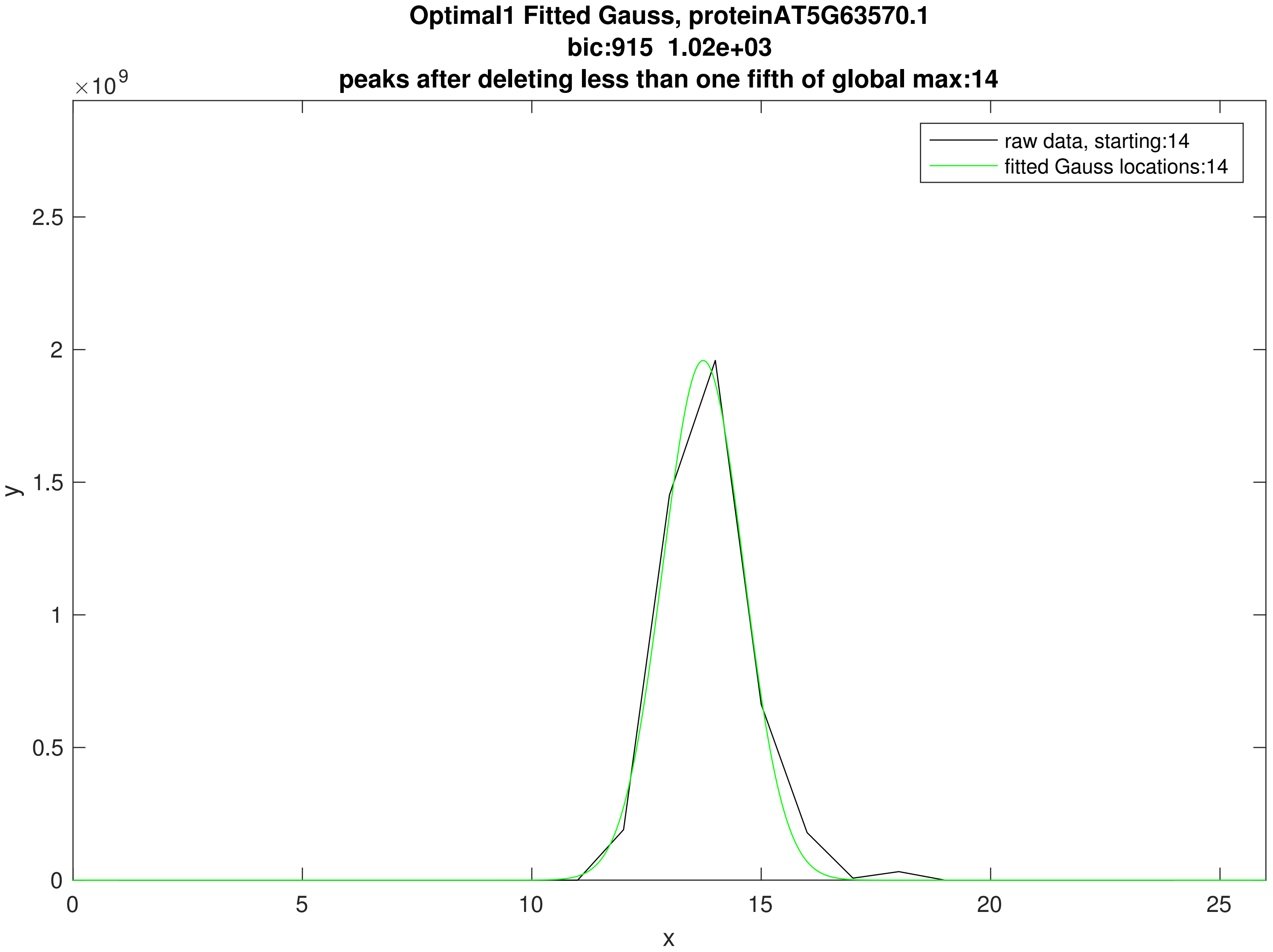
Optimal1 Fitted Gauss, proteinAT5G63570.1
bic:915 1.02e+03
peaks after deleting less than one fifth of global max:14
×10^9
2.5
2
1.5
1
0.5
0
y
0
5
10
15
20
25
x
raw data, starting:14
fitted Gauss locations:14

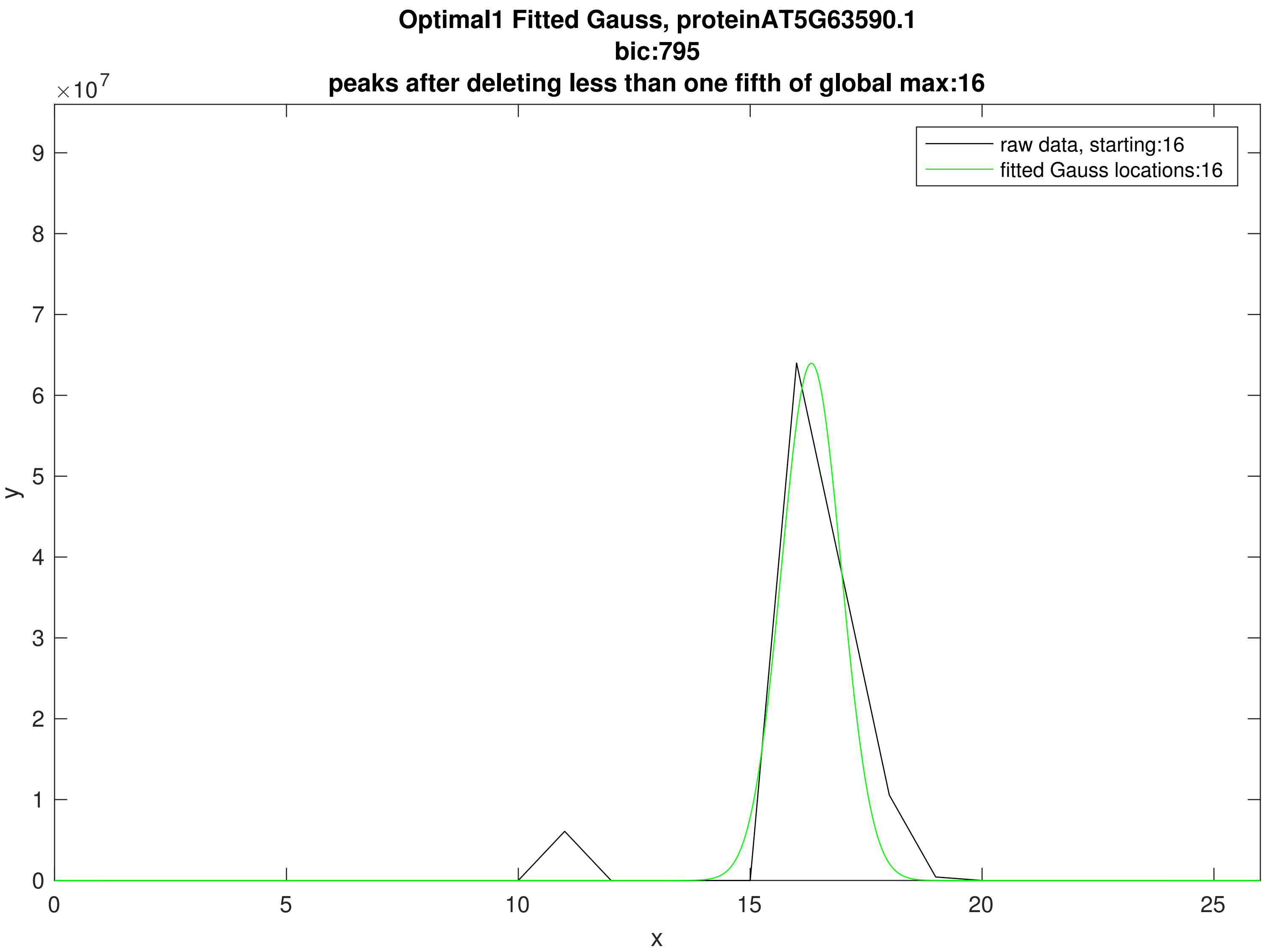

Optimal1 Fitted Gauss, proteinAT5G63590.1
bic:795
peaks after deleting less than one fifth of global max:16
×10^7
raw data, starting:16
fitted Gauss locations:16
y
x
0
1
2
3
4
5
6
7
8
9
0
5
10
15
20
25

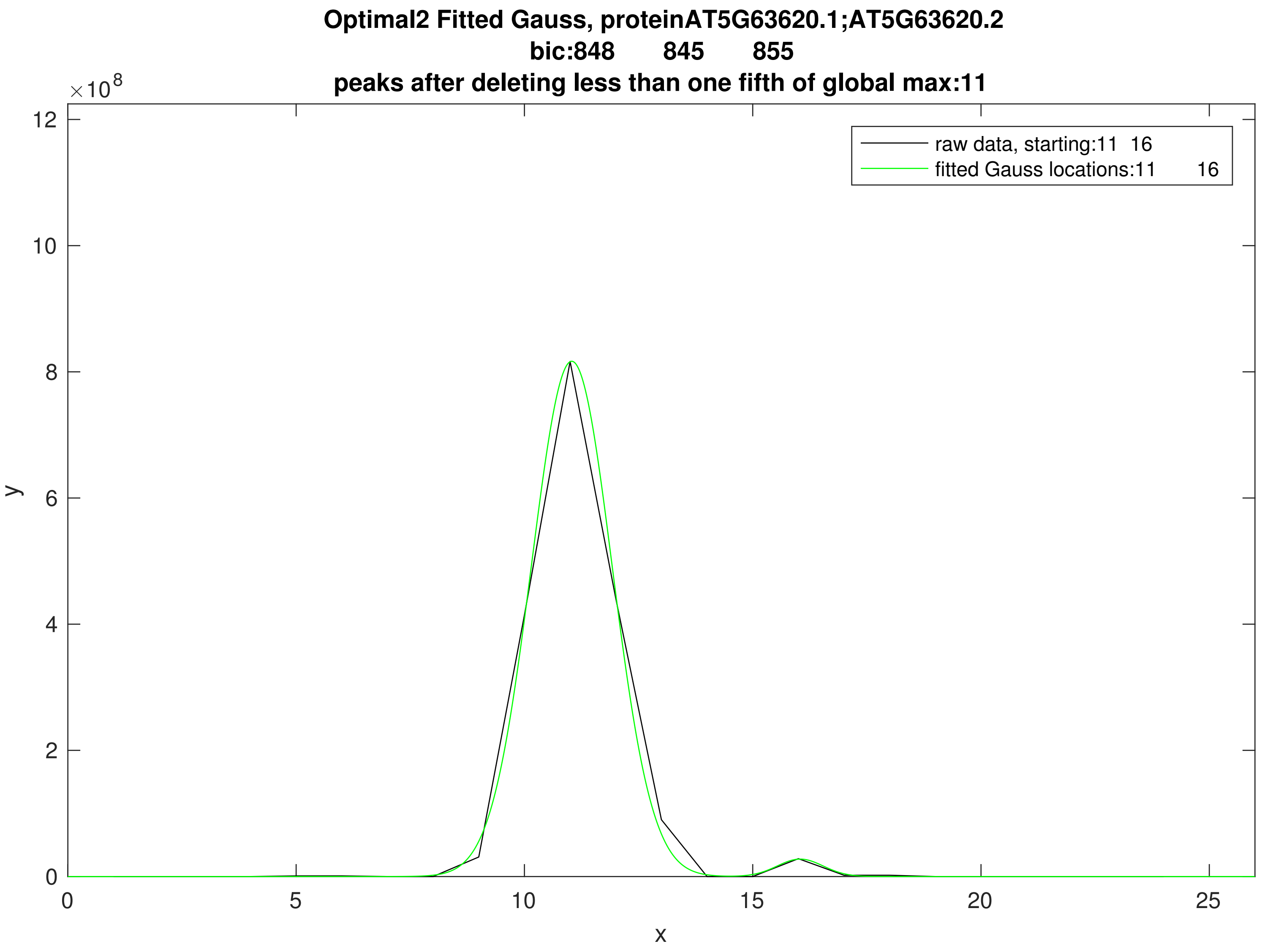
Optimal2 Fitted Gauss, proteinAT5G63620.1;AT5G63620.2
bic:848 845 855
peaks after deleting less than one fifth of global max:11
×10^8
y
x
raw data, starting:11 16
fitted Gauss locations:11 16
0
2
4
6
8
10
12
0
5
10
15
20
25

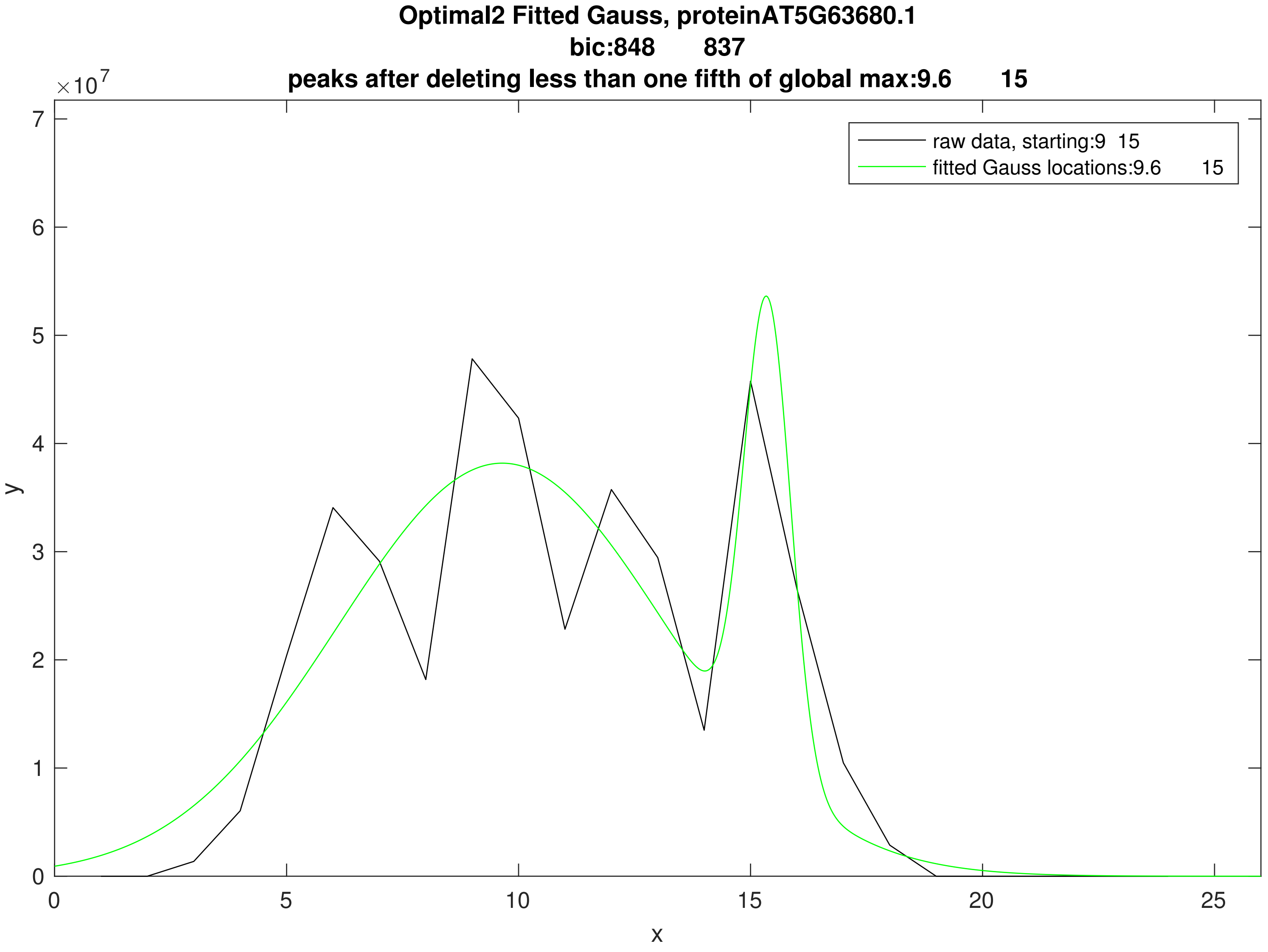
Optimal2 Fitted Gauss, proteinAT5G63680.1
bic:848 837
peaks after deleting less than one fifth of global max:9.6 15
×10^7
raw data, starting:9 15
fitted Gauss locations:9.6 15
y
x
0
1
2
3
4
5
6
7
0
5
10
15
20
25

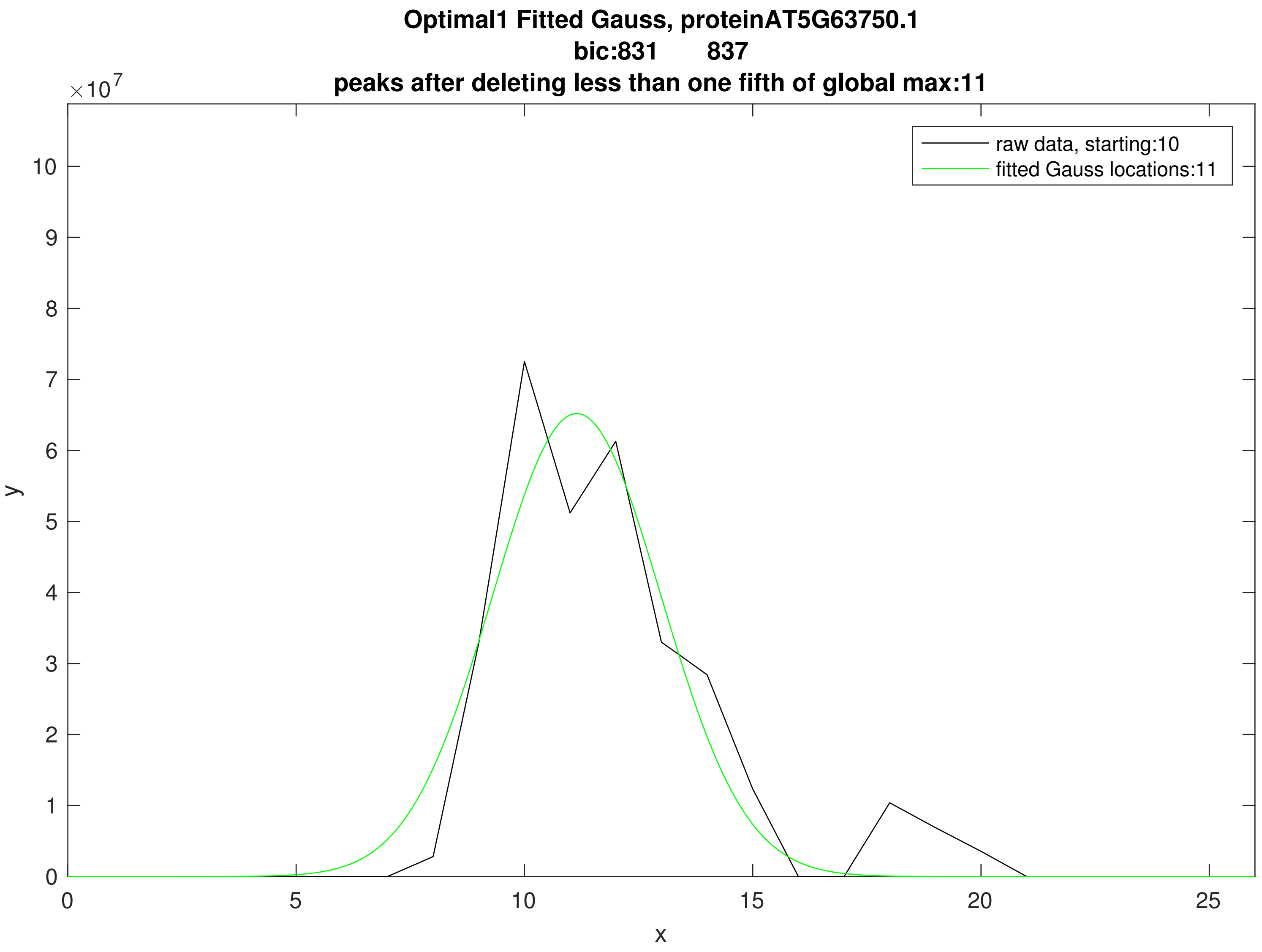
Optimal1 Fitted Gauss, proteinAT5G63750.1
bic:831 837
peaks after deleting less than one fifth of global max:11
raw data, starting:10
fitted Gauss locations:11
×10^7
y
x

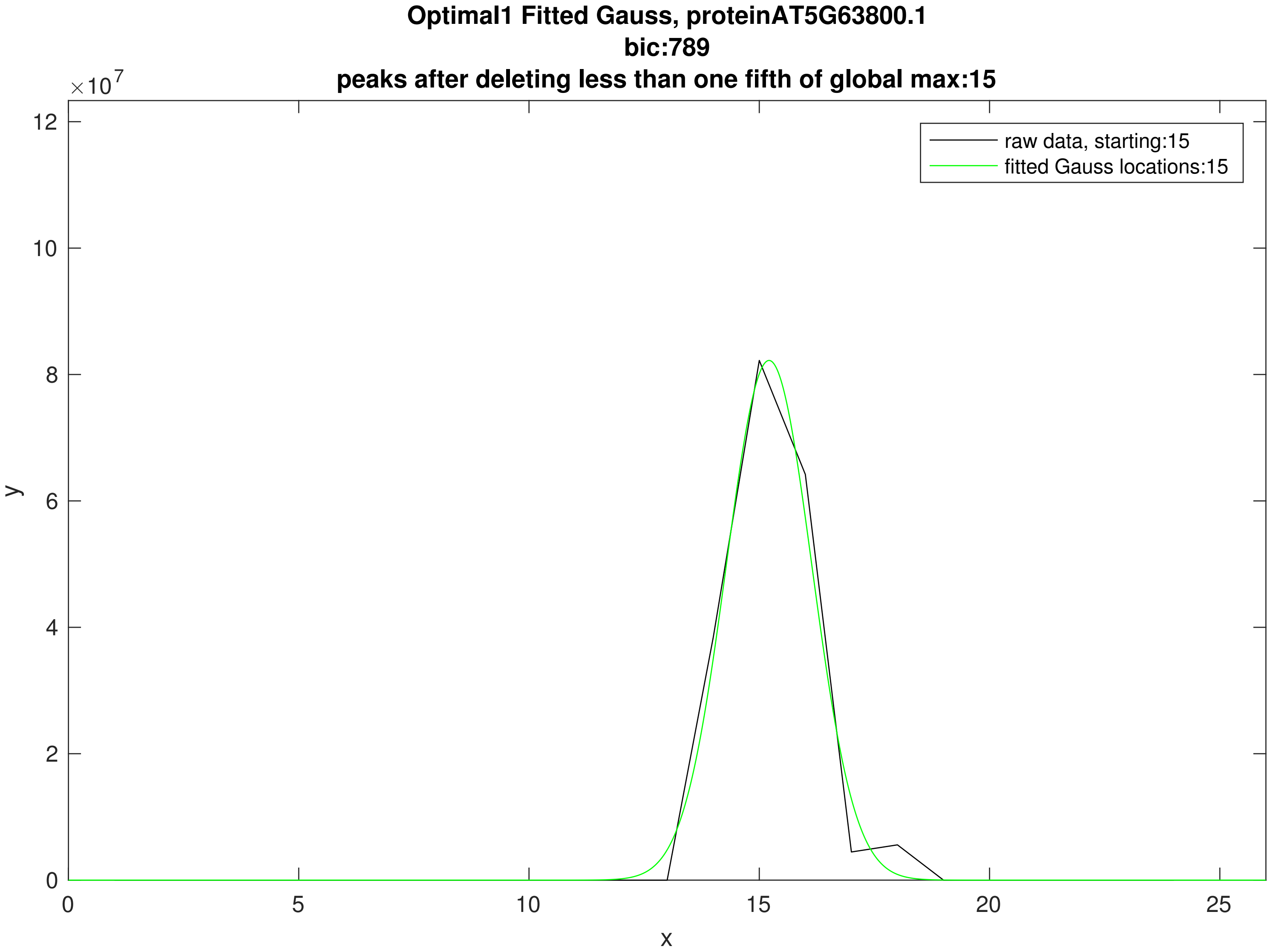

Optimal1 Fitted Gauss, proteinAT5G63800.1
bic:789
peaks after deleting less than one fifth of global max:15
×10^7
raw data, starting:15
fitted Gauss locations:15
y
x
0
2
4
6
8
10
12
5
15
20
25

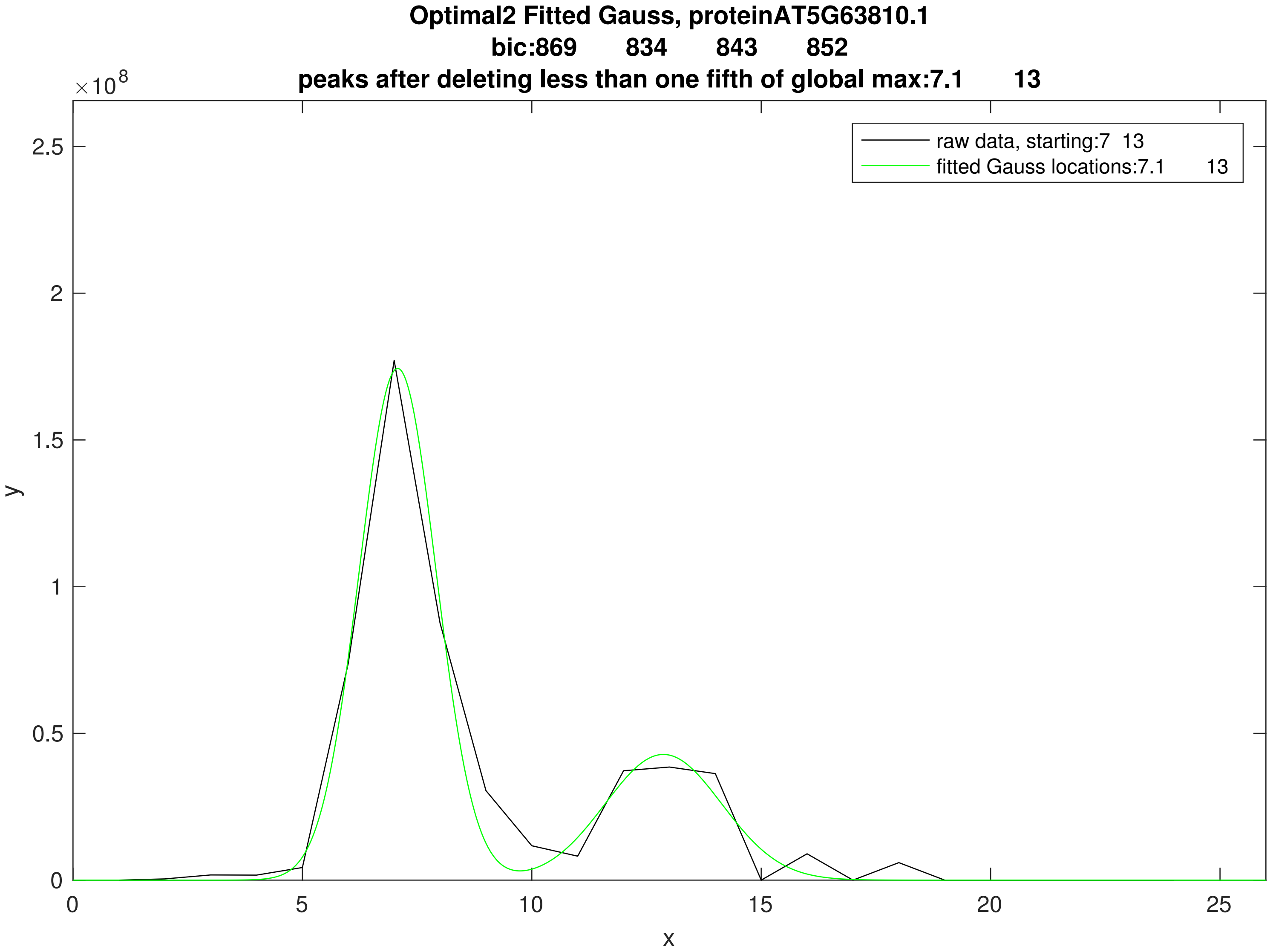
Optimal2 Fitted Gauss, proteinAT5G63810.1
bic:869 834 843 852
peaks after deleting less than one fifth of global max:7.1 13
×10^8
raw data, starting:7 13
fitted Gauss locations:7.1 13
y
x
0
0.5
1
1.5
2
2.5
5
10
15
20
25

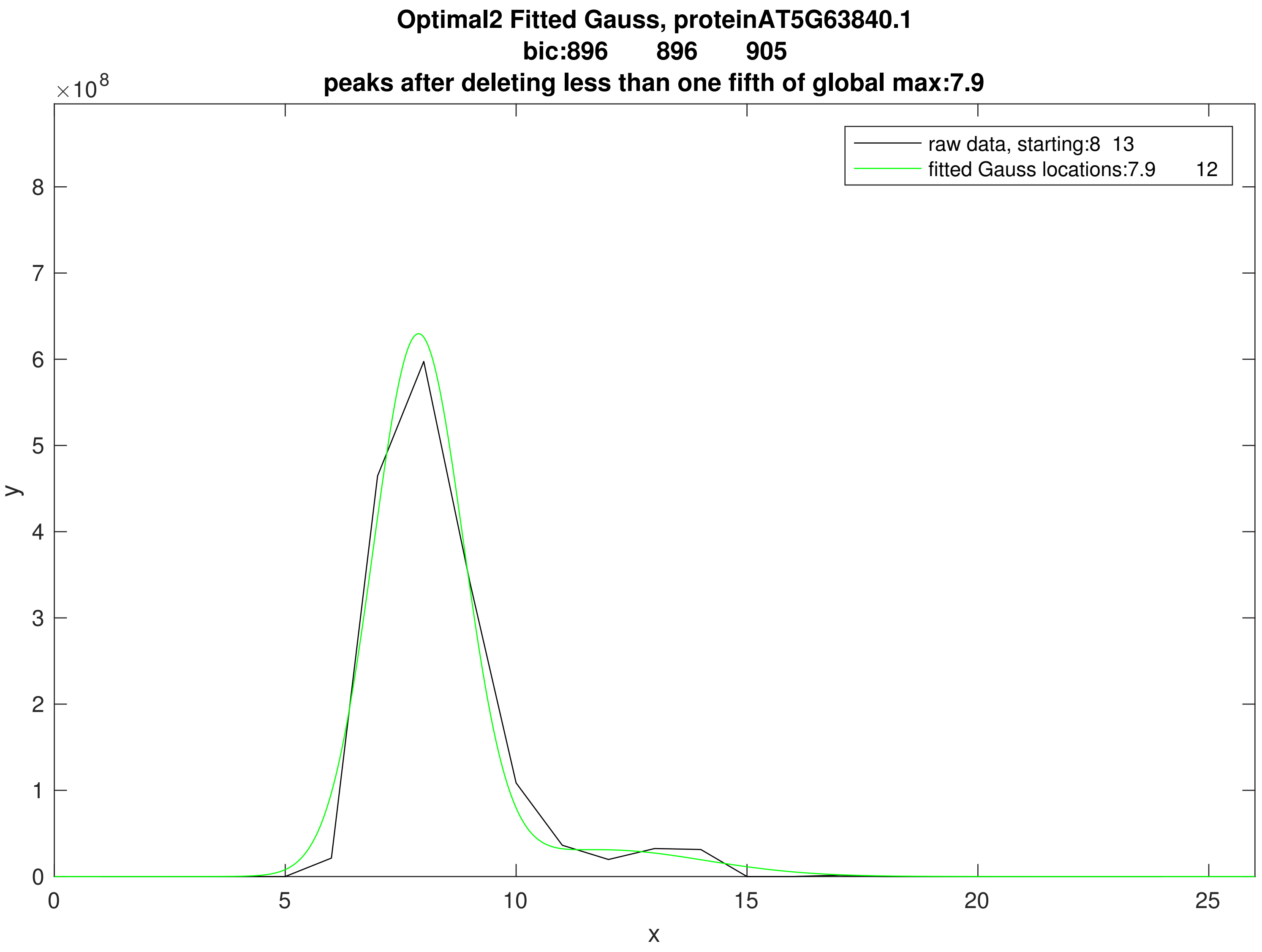
Optimal2 Fitted Gauss, proteinAT5G63840.1
bic:896 896 905
peaks after deleting less than one fifth of global max:7.9
×10^8
raw data, starting:8 13
fitted Gauss locations:7.9 12
y
x
0
1
2
3
4
5
6
7
8
0
5
10
15
20
25

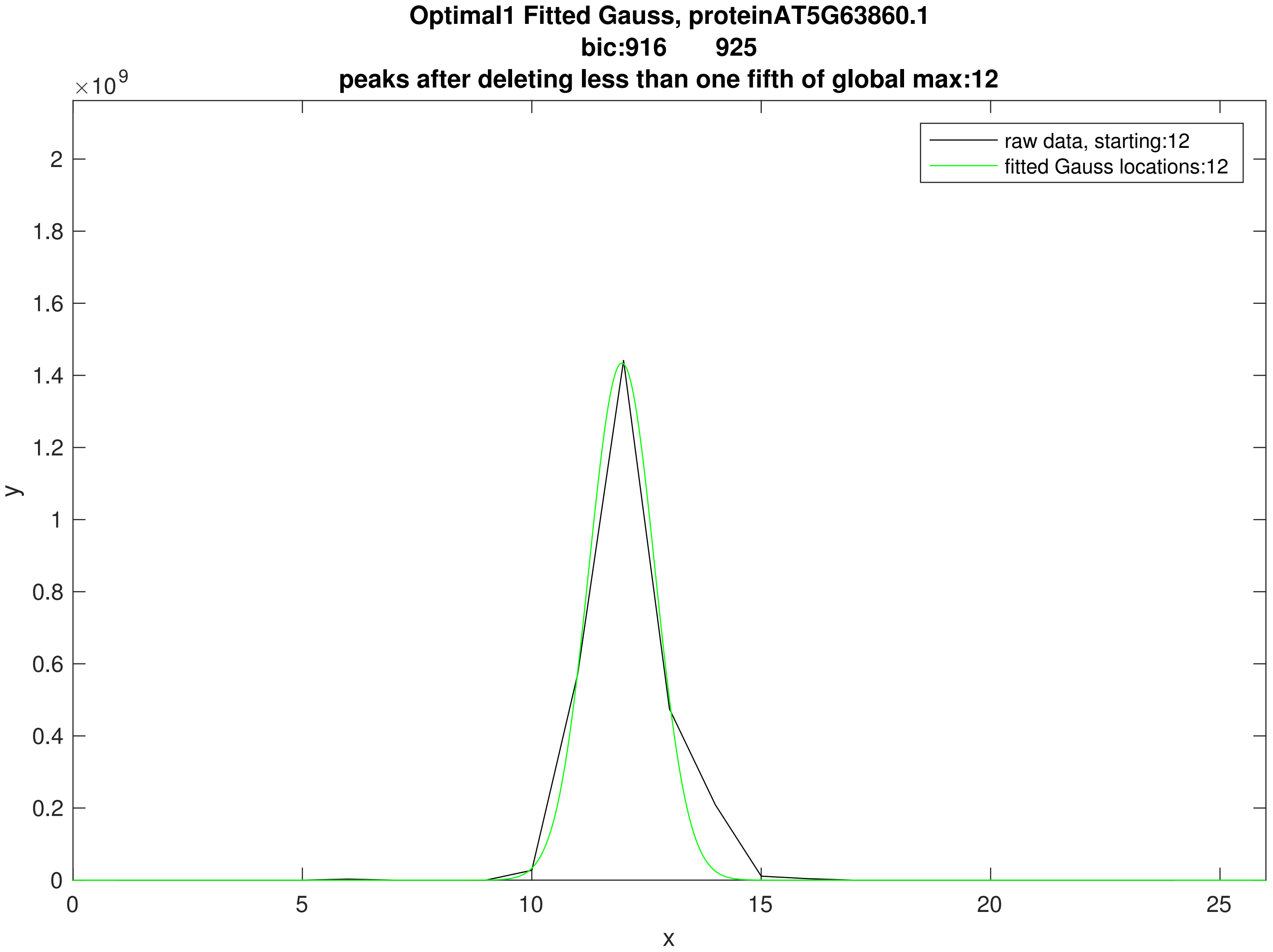

Optimal1 Fitted Gauss, proteinAT5G63860.1
bic:916 925
peaks after deleting less than one fifth of global max:12
×10^9
2
1.8
1.6
1.4
1.2
1
0.8
0.6
0.4
0.2
0
y
0
5
10
15
20
25
x
raw data, starting:12
fitted Gauss locations:12

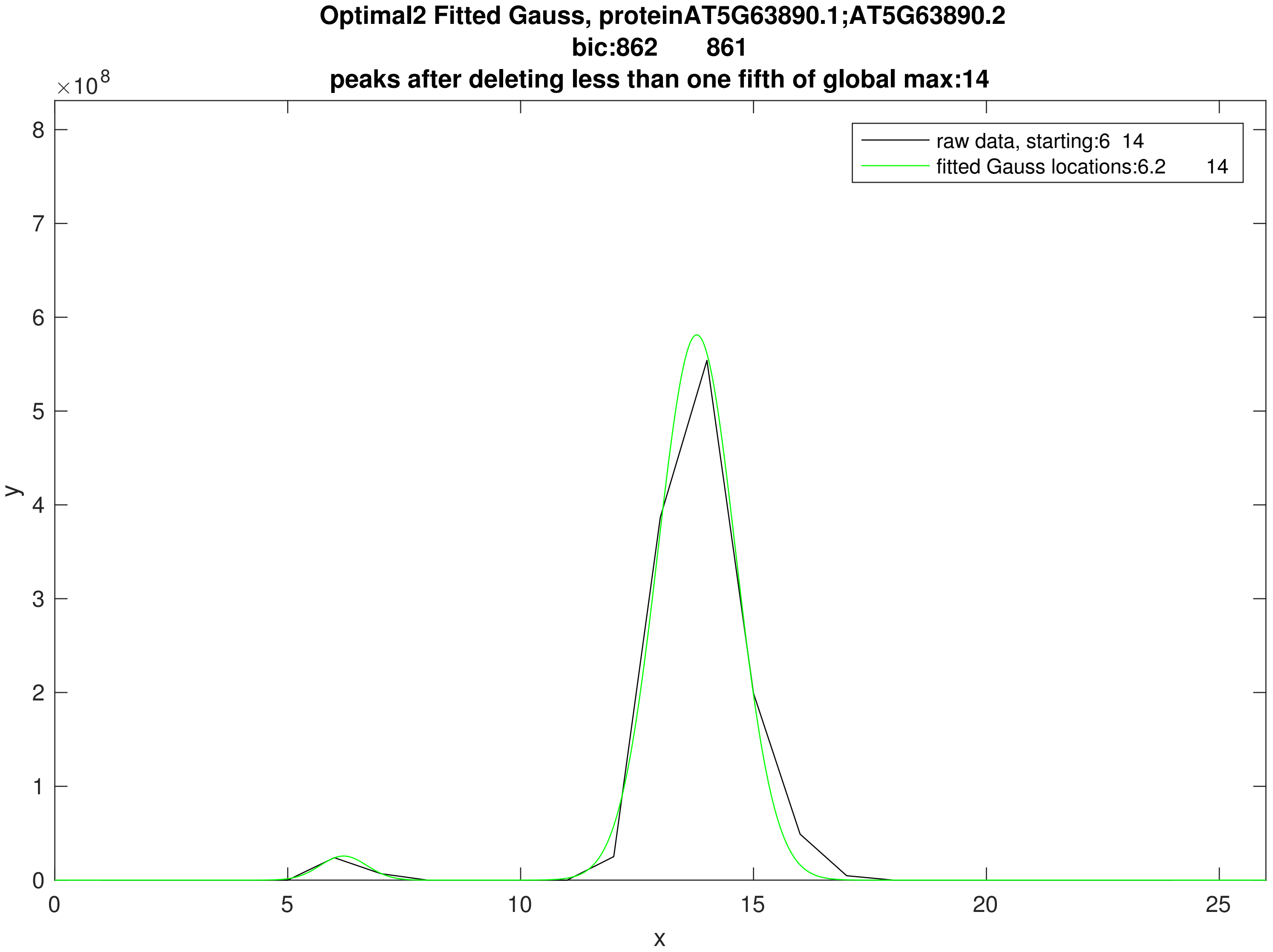

Optimal2 Fitted Gauss, proteinAT5G63890.1;AT5G63890.2
bic:862 861
peaks after deleting less than one fifth of global max:14
×10^8
raw data, starting:6 14
fitted Gauss locations:6.2 14
y
x
0
1
2
3
4
5
6
7
8
0
5
10
15
20
25

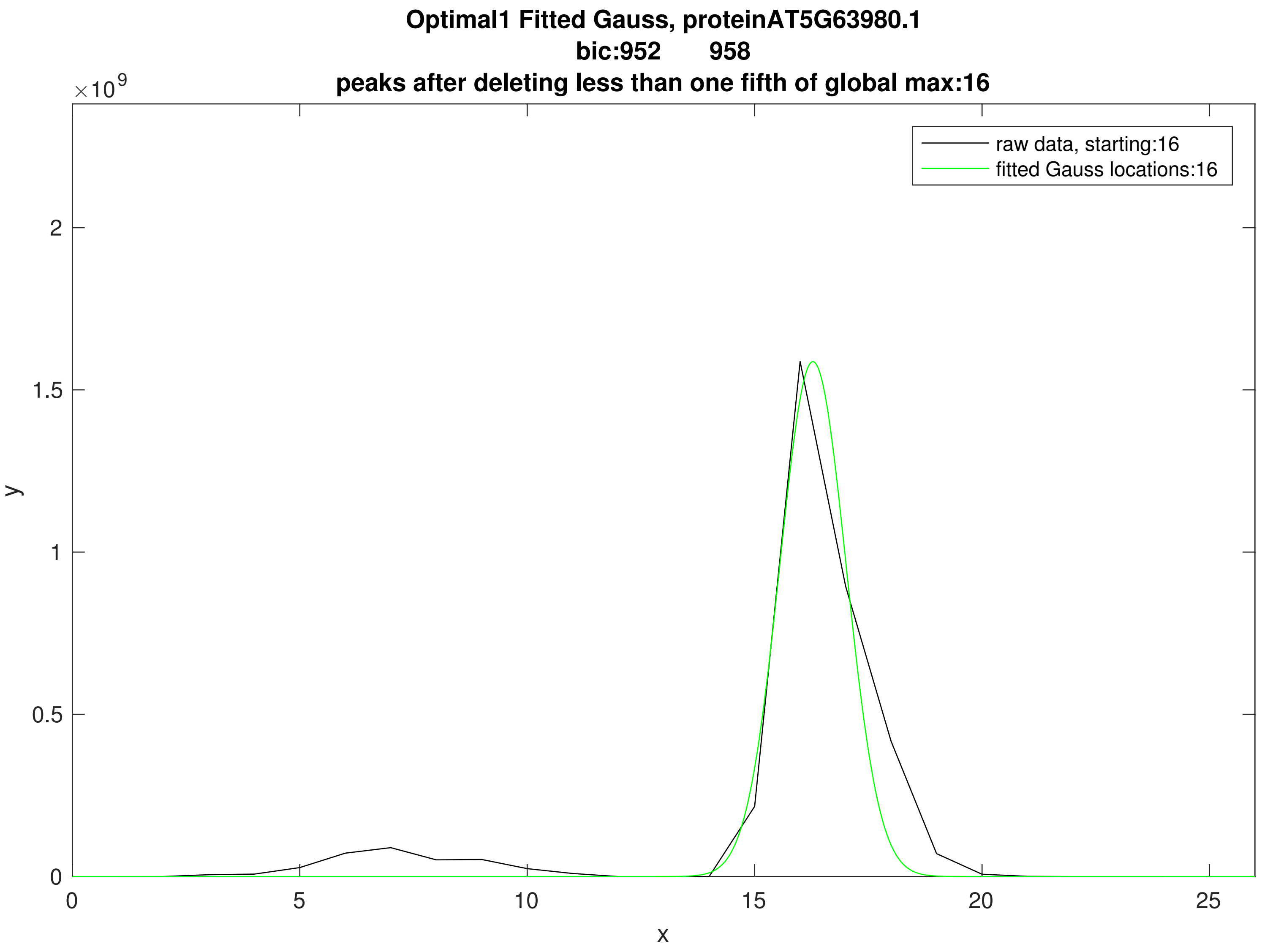
Optimal1 Fitted Gauss, proteinAT5G63980.1
bic:952 958
peaks after deleting less than one fifth of global max:16
×10^9
raw data, starting:16
fitted Gauss locations:16
y
x
0
0.5
1
1.5
2
0
5
10
15
20
25

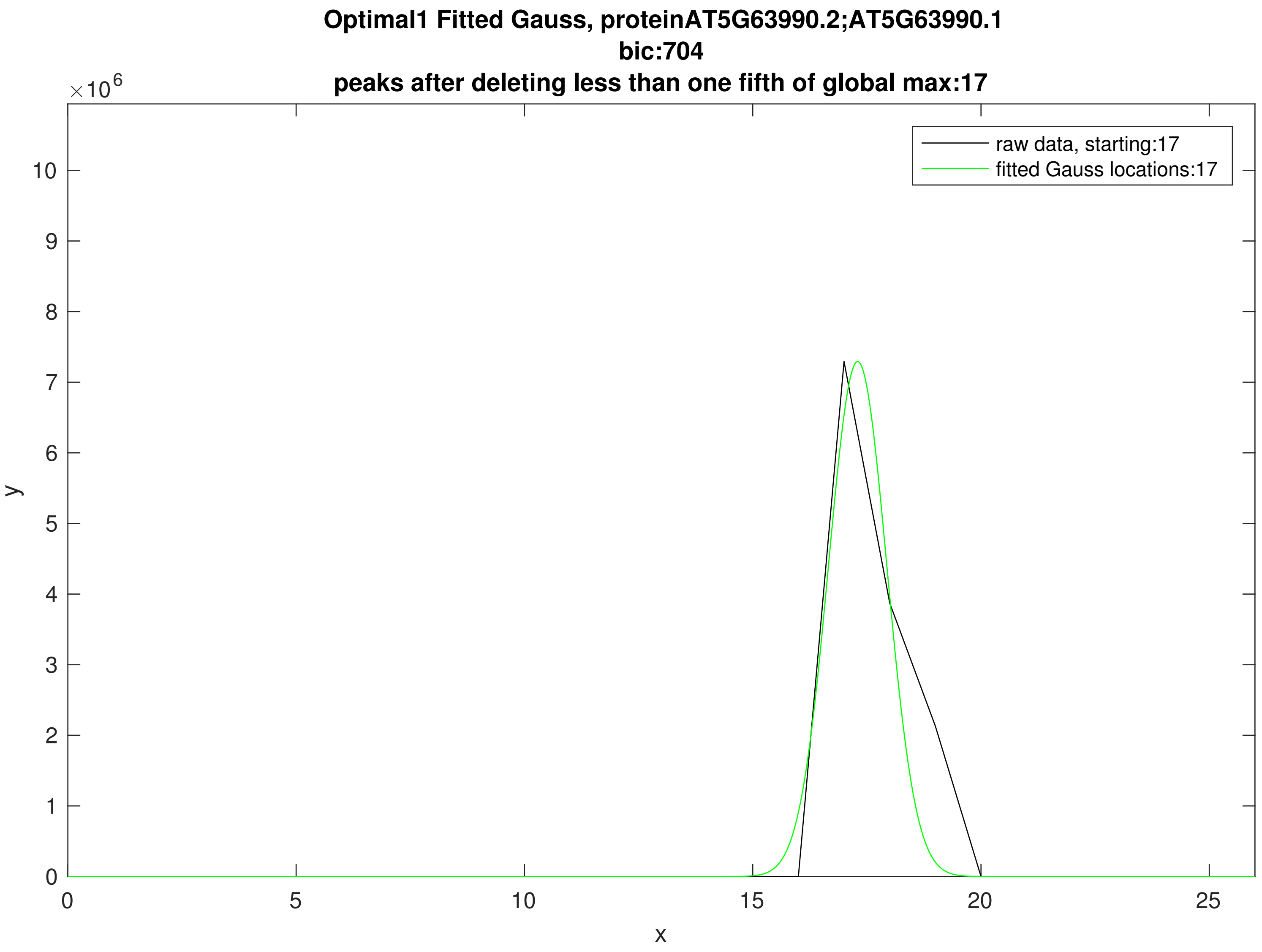

Optimal1 Fitted Gauss, proteinAT5G63990.2;AT5G63990.1
bic:704
peaks after deleting less than one fifth of global max:17
×10^6
raw data, starting:17
fitted Gauss locations:17
y
x
0
1
2
3
4
5
6
7
8
9
10
0
5
10
15
20
25

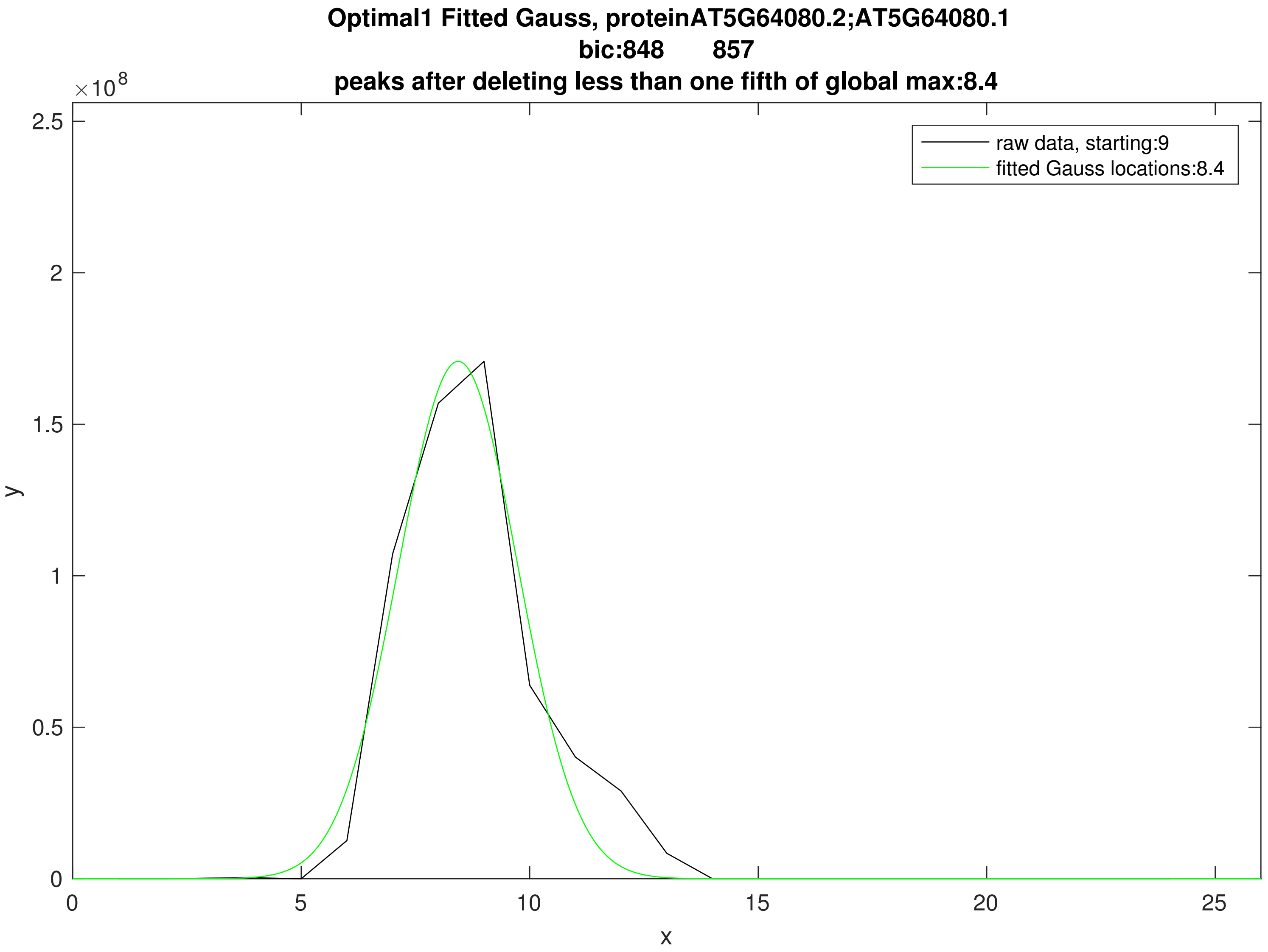
Optimal1 Fitted Gauss, proteinAT5G64080.2;AT5G64080.1
bic:848 857
peaks after deleting less than one fifth of global max:8.4
×10^8
2.5
2
1.5
1
0.5
0
y
0
5
10
15
20
25
x
raw data, starting:9
fitted Gauss locations:8.4

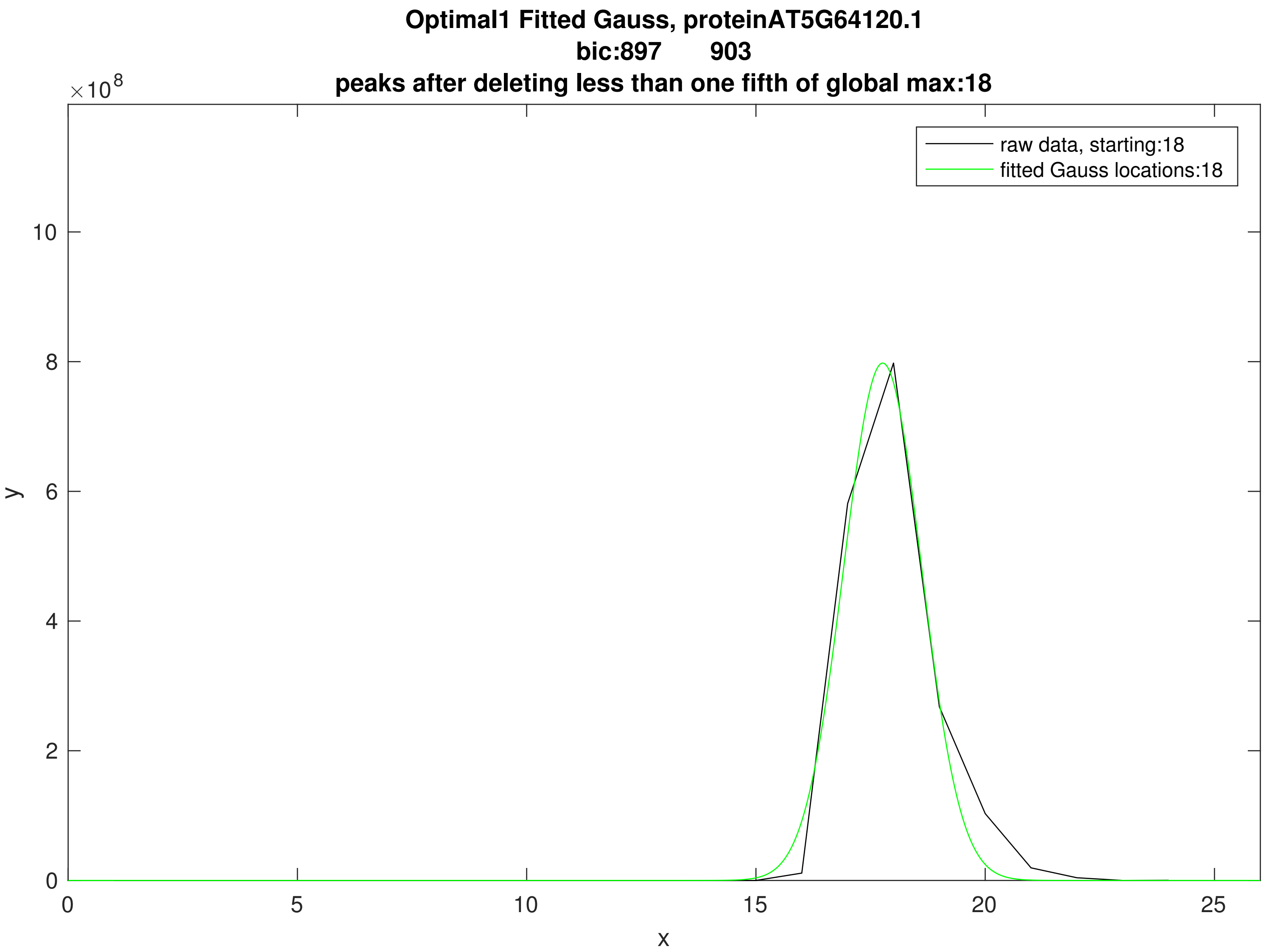

Optimal1 Fitted Gauss, proteinAT5G64120.1
bic:897 903
peaks after deleting less than one fifth of global max:18
×10^8
raw data, starting:18
fitted Gauss locations:18
10
8
6
4
2
0
y
0
5
10
15
20
25
x

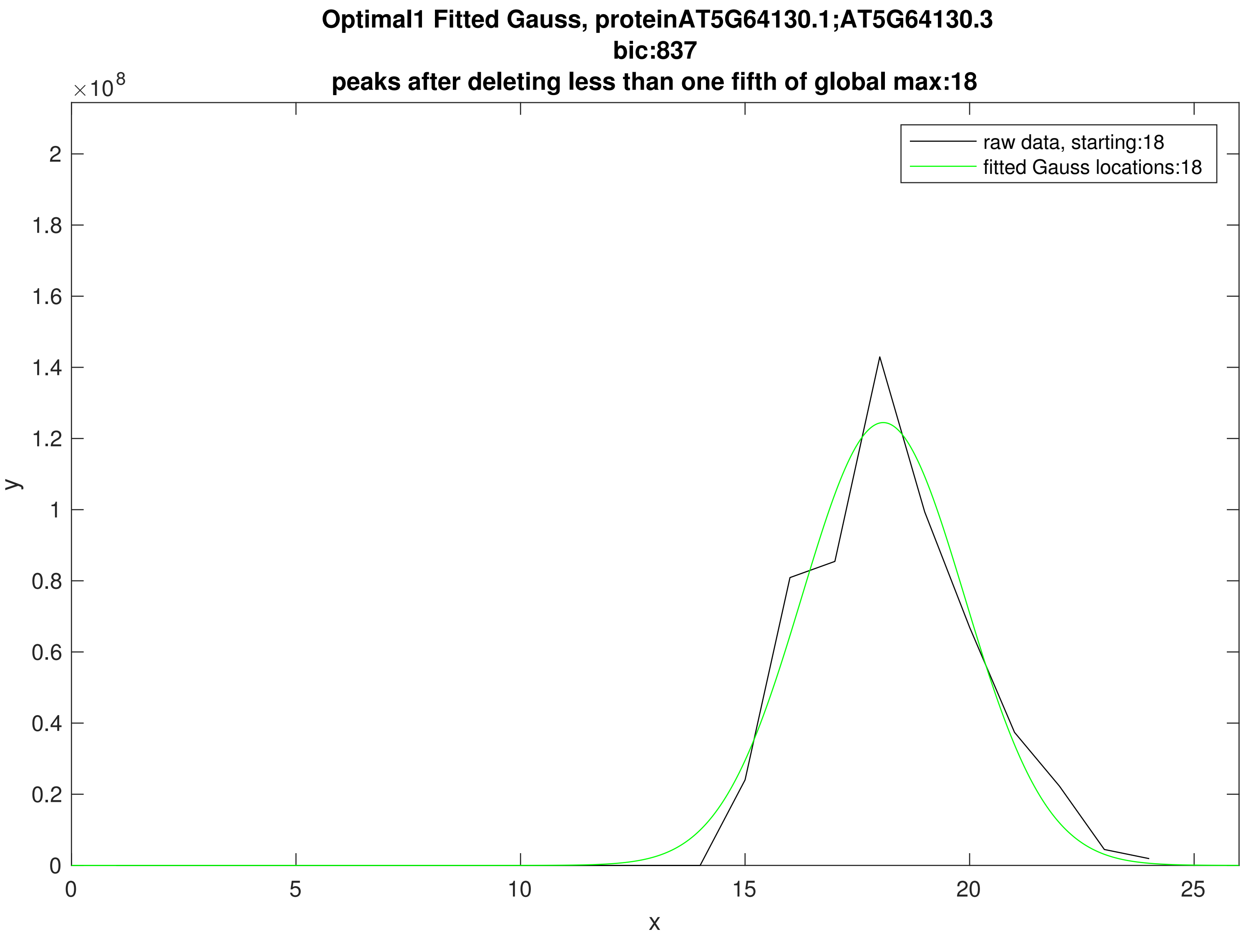

Optimal1 Fitted Gauss, proteinAT5G64130.1;AT5G64130.3
bic:837
peaks after deleting less than one fifth of global max:18
$\times 10^8$
raw data, starting:18
fitted Gauss locations:18
y
x
0
0.2
0.4
0.6
0.8
1
1.2
1.4
1.6
1.8
2
0
5
10
15
20
25

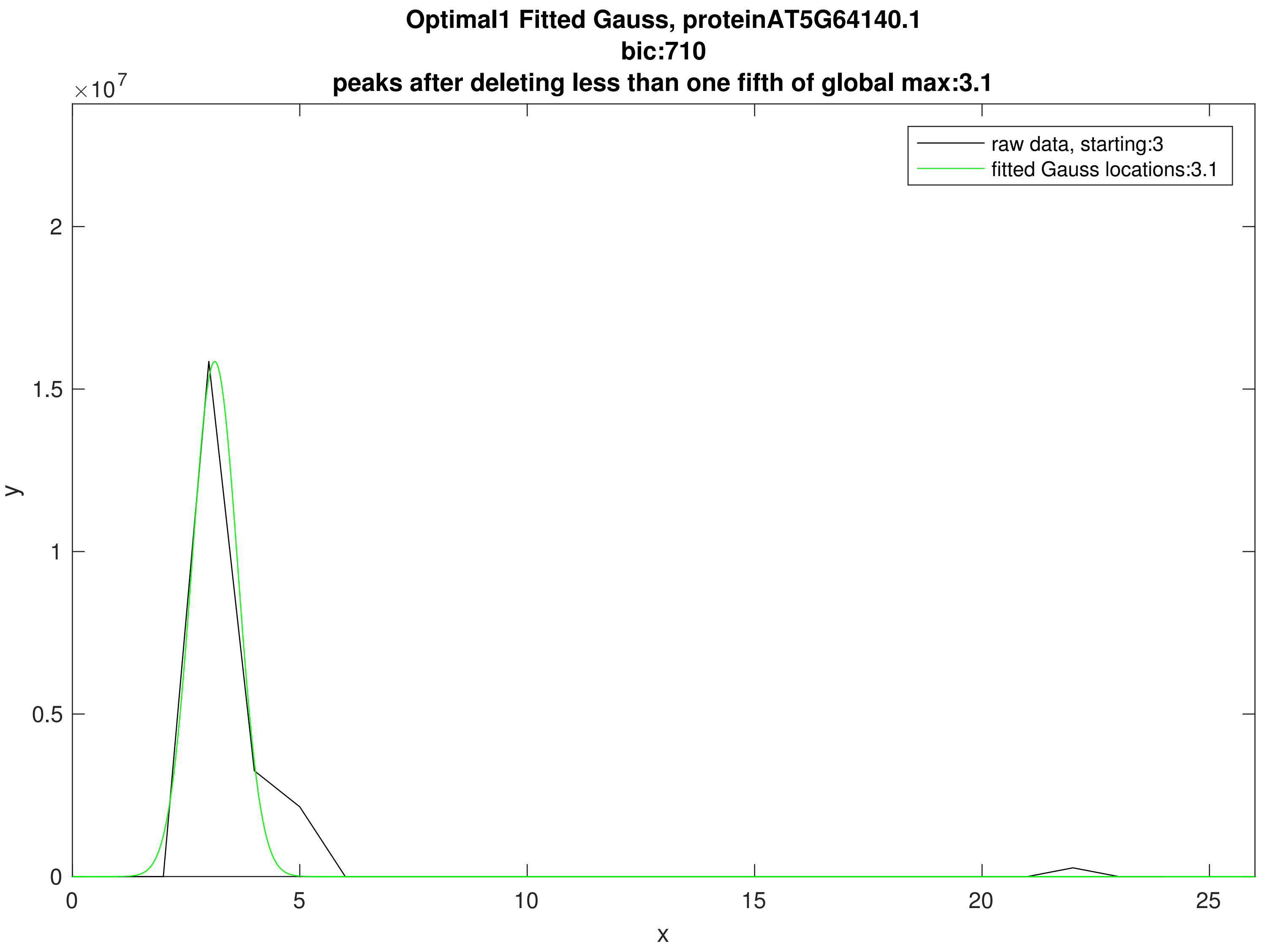
Optimal1 Fitted Gauss, proteinAT5G64140.1
bic:710
peaks after deleting less than one fifth of global max:3.1
×10^7
raw data, starting:3
fitted Gauss locations:3.1
y
x
0
0.5
1
1.5
2
5
10
15
20
25

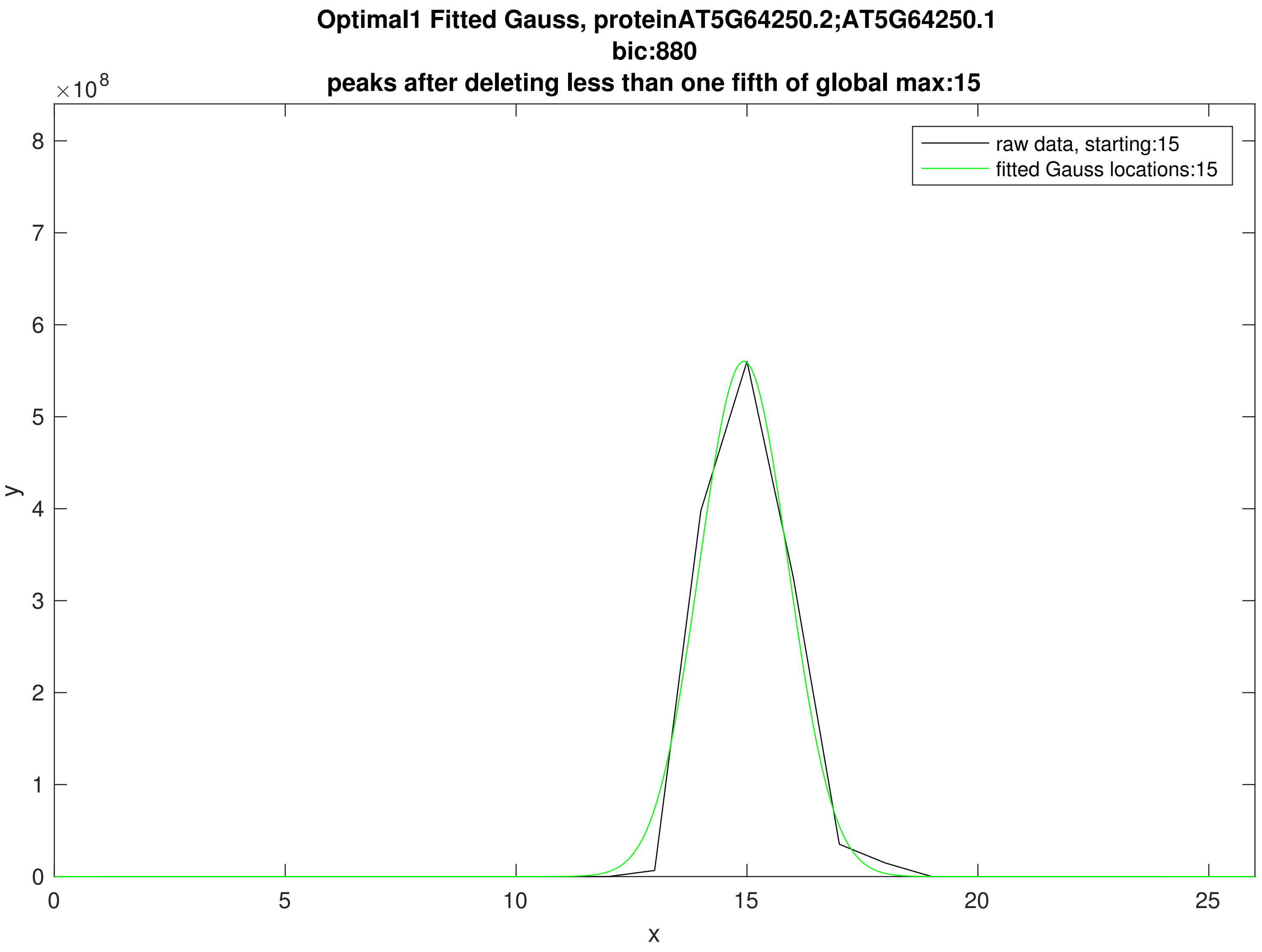
Optimal1 Fitted Gauss, proteinAT5G64250.2;AT5G64250.1
bic:880
peaks after deleting less than one fifth of global max:15
×10^8
raw data, starting:15
fitted Gauss locations:15
y
x
0
1
2
3
4
5
6
7
8
0
5
10
15
20
25

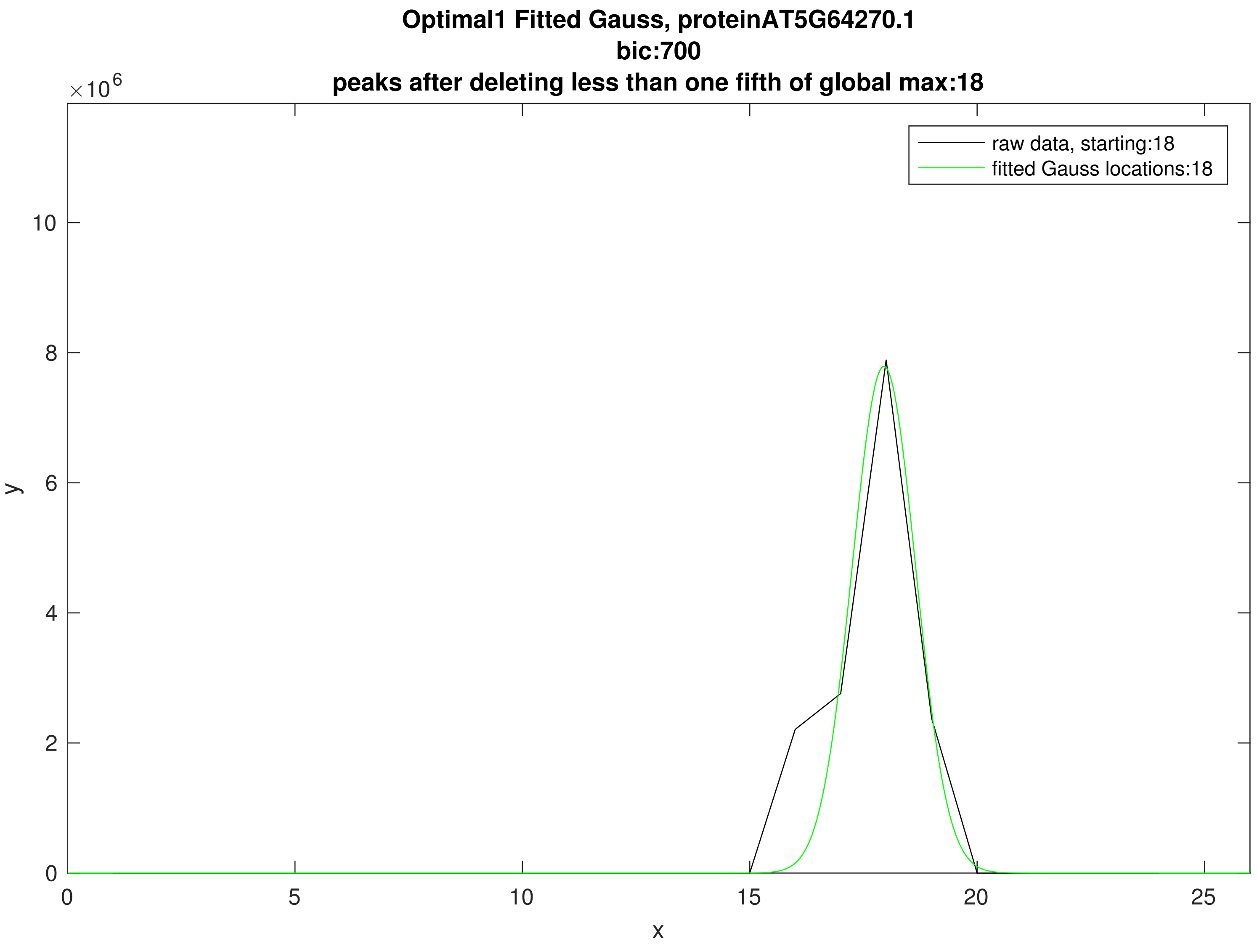
Optimal1 Fitted Gauss, proteinAT5G64270.1
bic:700
peaks after deleting less than one fifth of global max:18
$\times 10^6$
raw data, starting:18
fitted Gauss locations:18
10
8
6
4
2
0
y
0
5
10
15
20
25
x

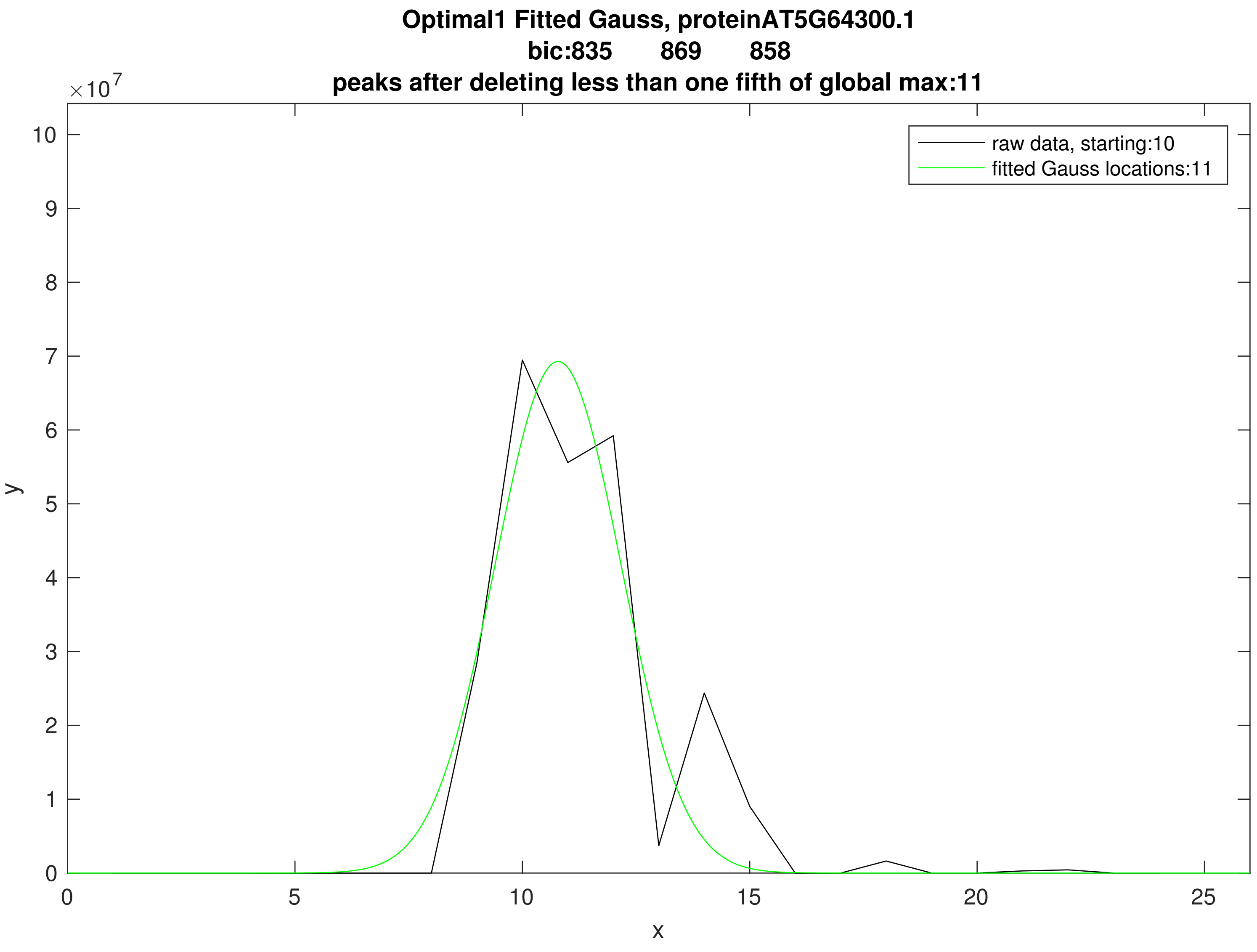
Optimal1 Fitted Gauss, proteinAT5G64300.1
bic:835 869 858
peaks after deleting less than one fifth of global max:11
×10^7
raw data, starting:10
fitted Gauss locations:11
y
x
0
1
2
3
4
5
6
7
8
9
10
0
5
10
15
20
25

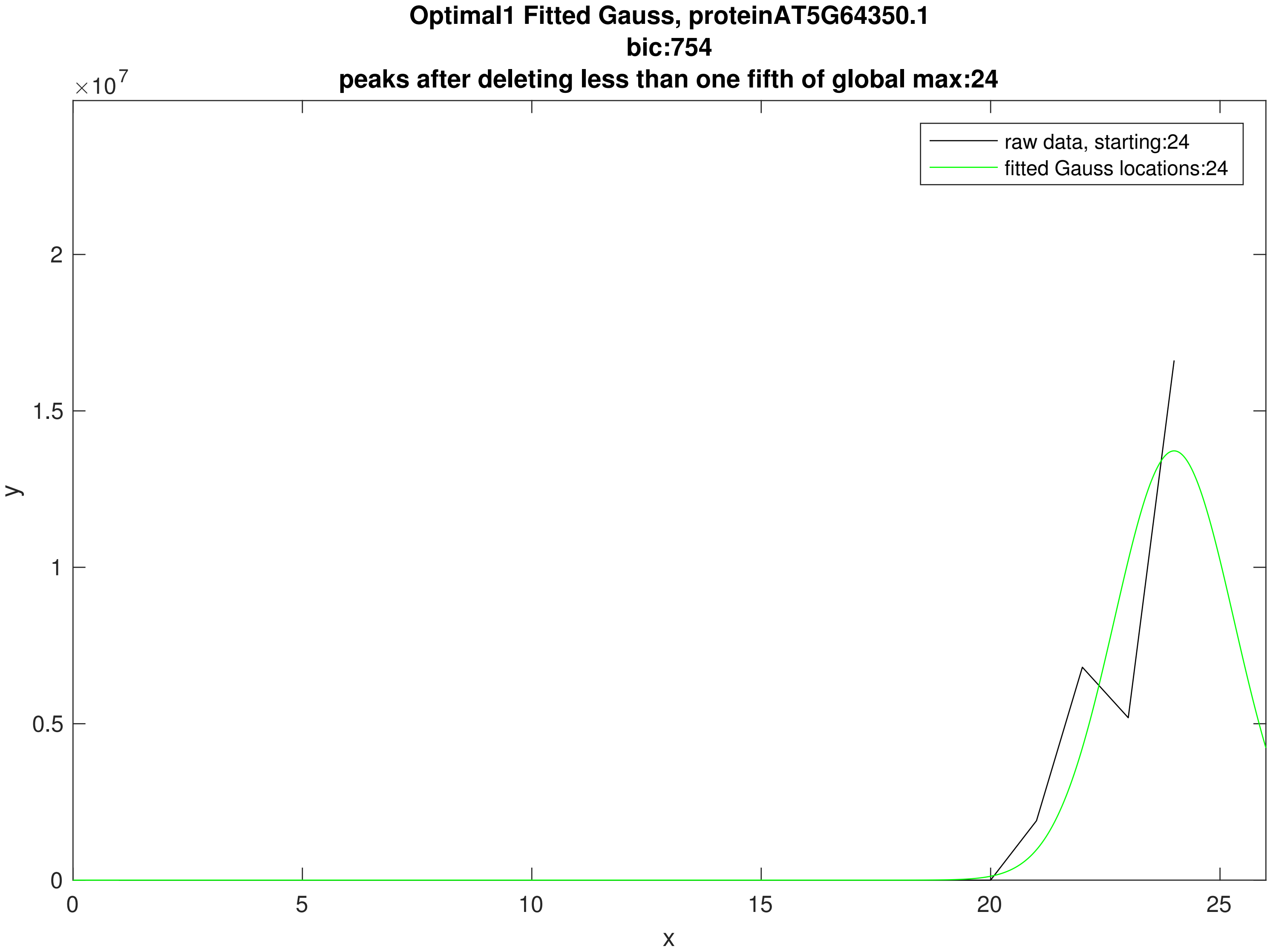
Optimal1 Fitted Gauss, proteinAT5G64350.1
bic:754
peaks after deleting less than one fifth of global max:24
×10^7
raw data, starting:24
fitted Gauss locations:24
y
x
0
0.5
1
1.5
2
0
5
10
15
20
25

Optimal1 Fitted Gauss, proteinAT5G64370.1
bic:876
peaks after deleting less than one fifth of global max:5

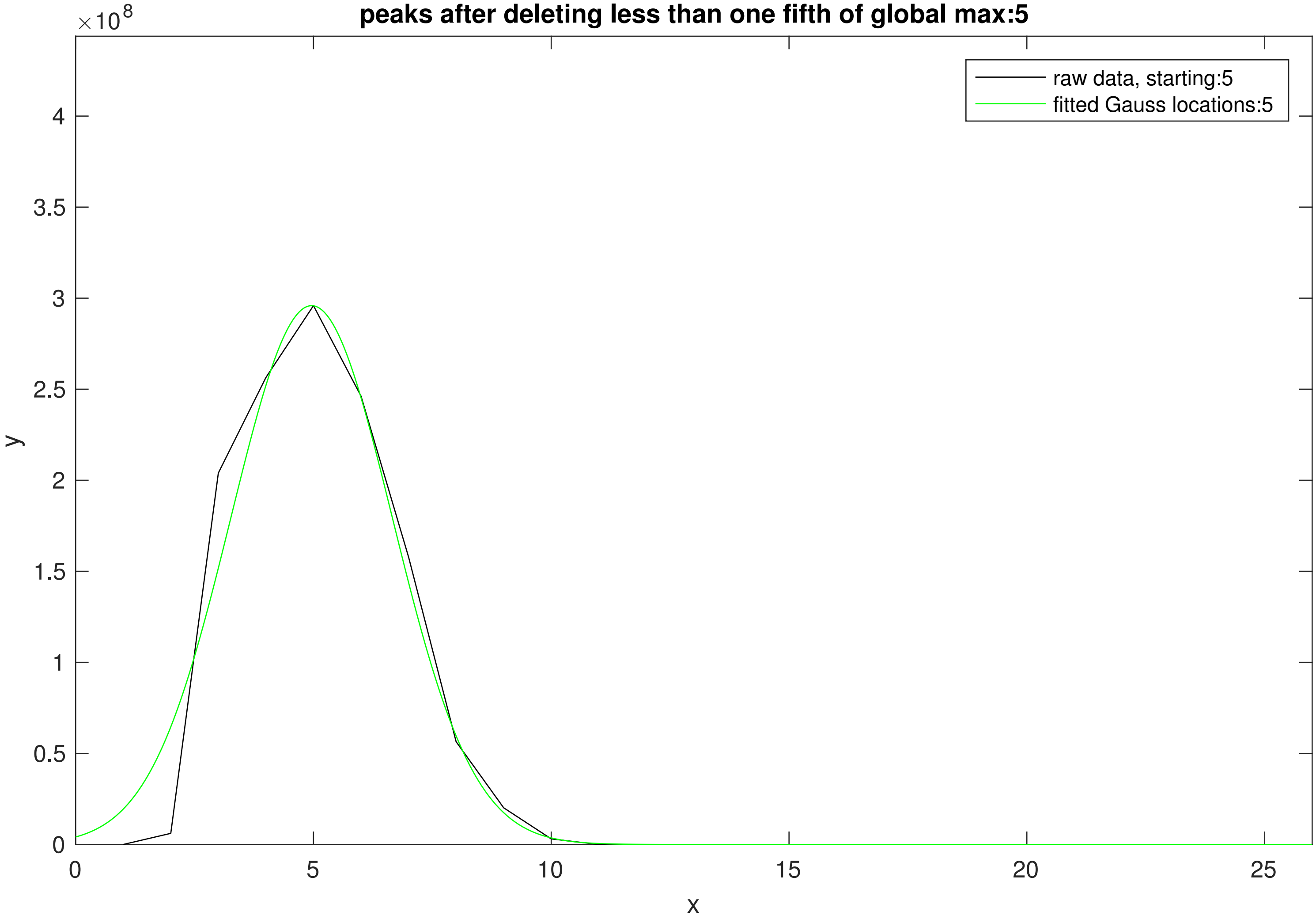

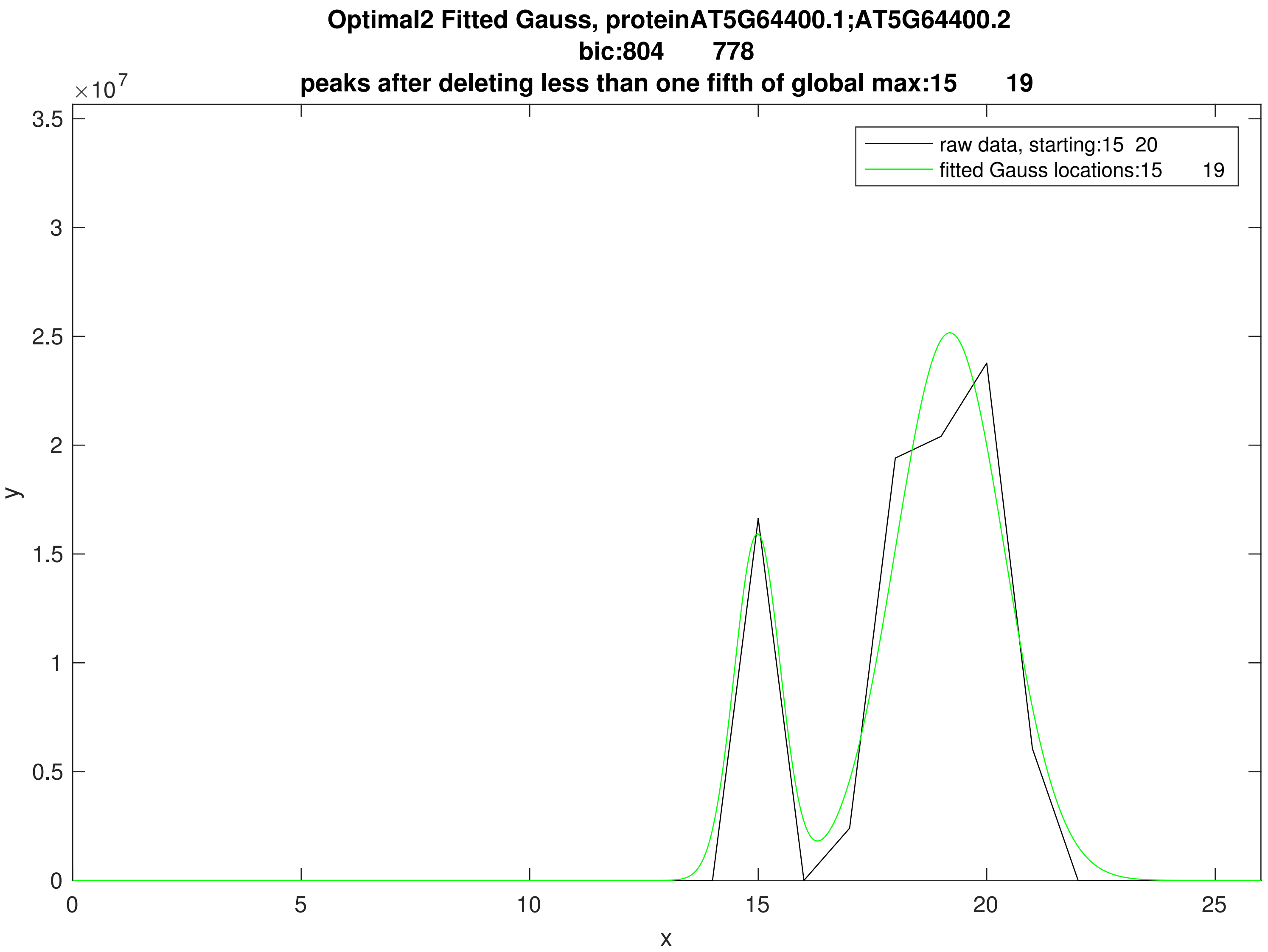
Optimal2 Fitted Gauss, proteinAT5G64400.1;AT5G64400.2
bic:804 778
peaks after deleting less than one fifth of global max:15 19
×10^7
raw data, starting:15 20
fitted Gauss locations:15 19
y
x
0
0.5
1
1.5
2
2.5
3
3.5
0
5
10
15
20
25

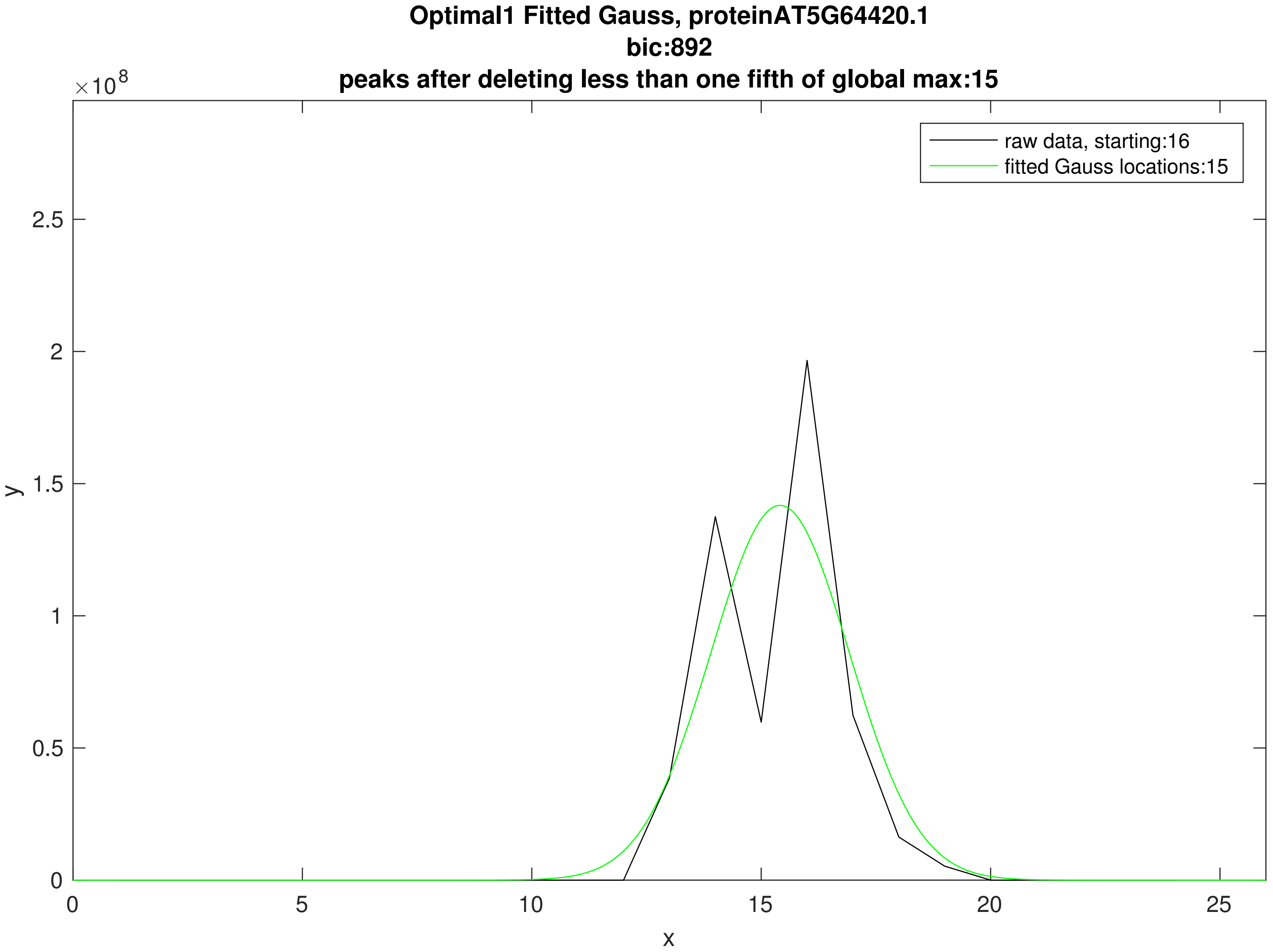
Optimal1 Fitted Gauss, proteinAT5G64420.1
bic:892
peaks after deleting less than one fifth of global max:15
×10^8
raw data, starting:16
fitted Gauss locations:15
y
x
0
0.5
1
1.5
2
2.5
0
5
10
15
20
25

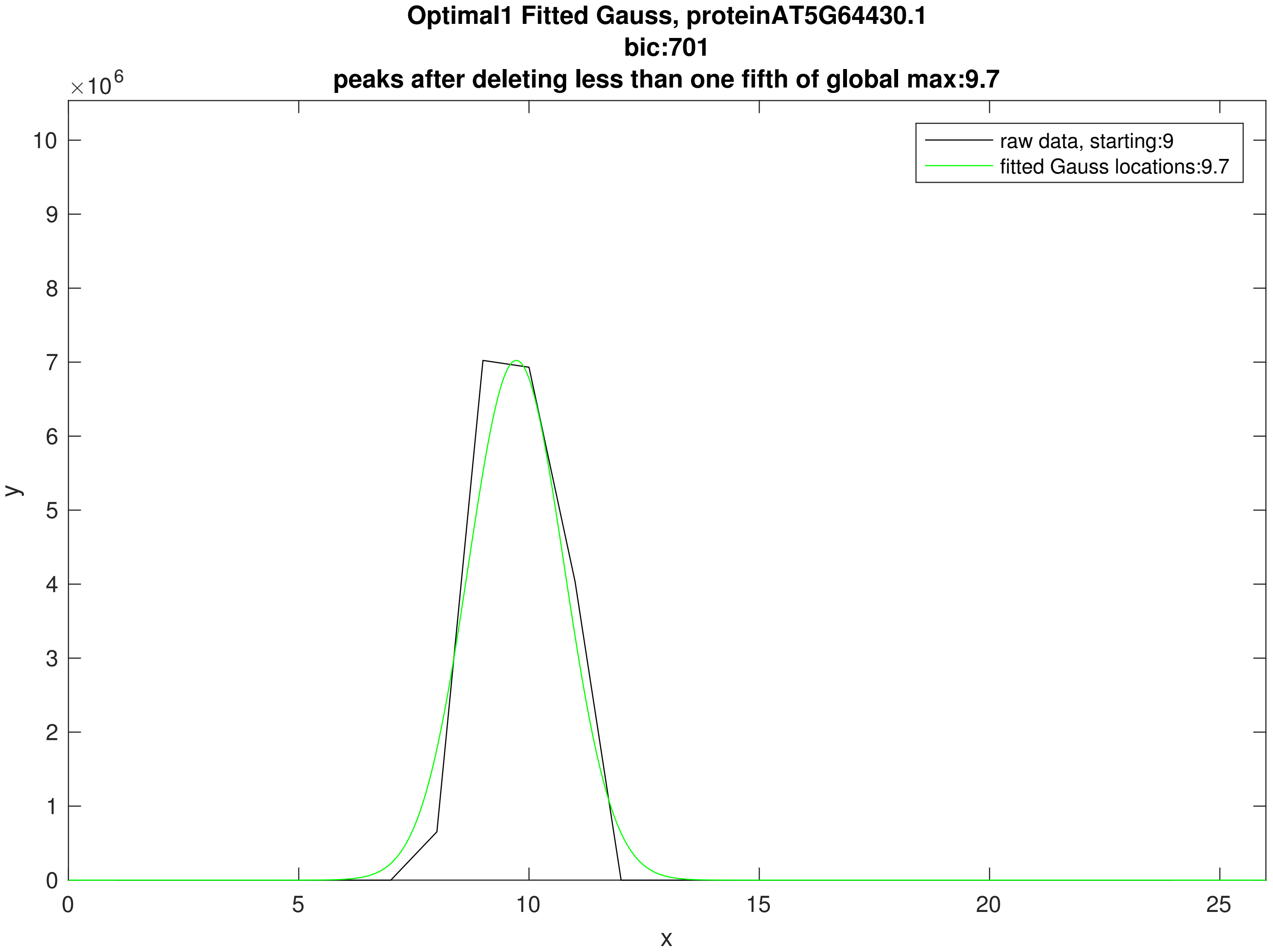
Optimal1 Fitted Gauss, proteinAT5G64430.1
bic:701
peaks after deleting less than one fifth of global max:9.7
×10^6
raw data, starting:9
fitted Gauss locations:9.7
y
x
0
1
2
3
4
5
6
7
8
9
10
0
5
10
15
20
25

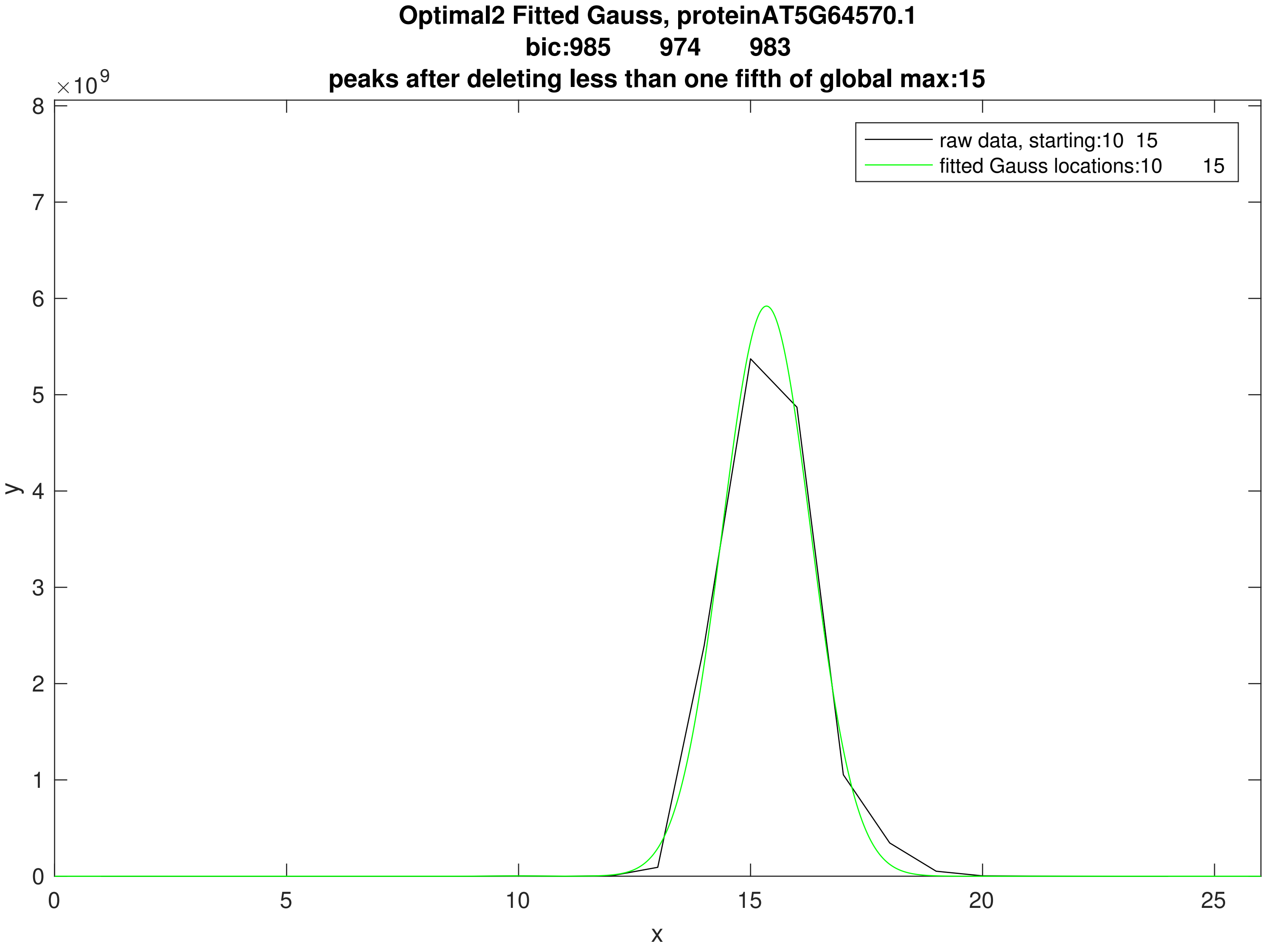
Optimal2 Fitted Gauss, proteinAT5G64570.1
bic:985 974 983
peaks after deleting less than one fifth of global max:15
raw data, starting:10 15
fitted Gauss locations:10 15
×10^9
y
x
0
1
2
3
4
5
6
7
8
0
5
10
15
20
25

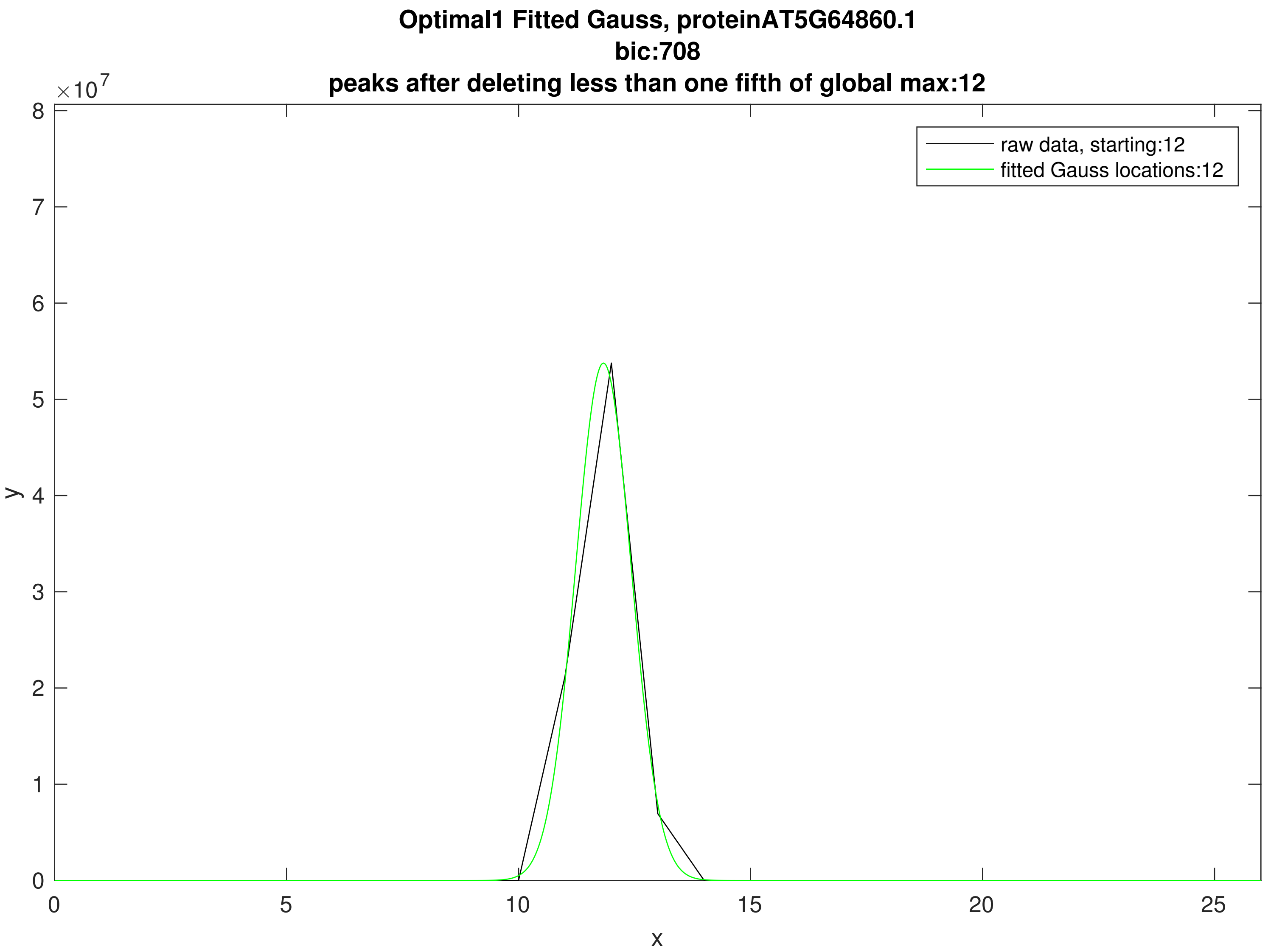
Optimal1 Fitted Gauss, proteinAT5G64860.1
bic:708
peaks after deleting less than one fifth of global max:12
×10^7
raw data, starting:12
fitted Gauss locations:12
y
x
0
1
2
3
4
5
6
7
8
0
5
10
15
20
25

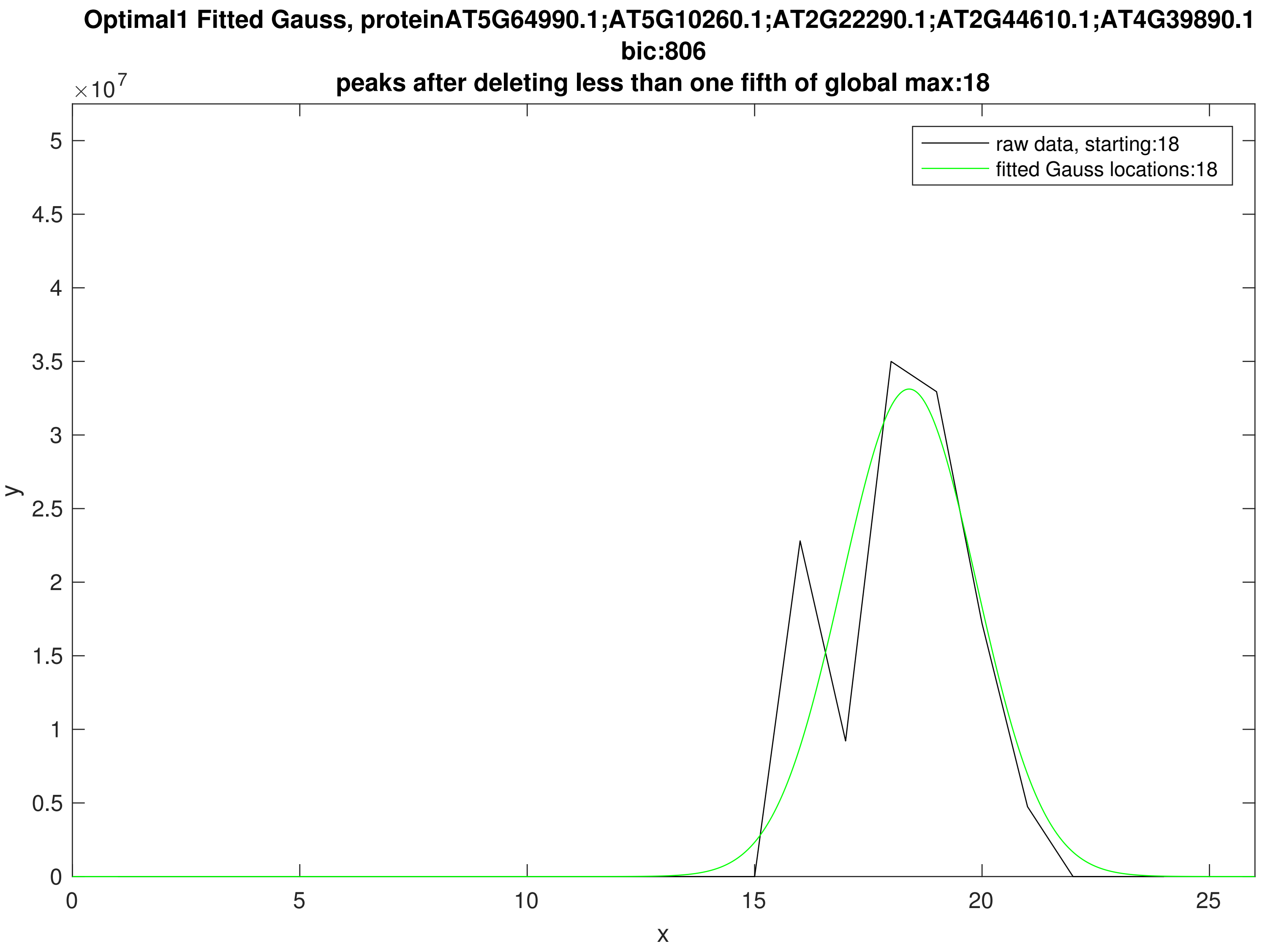
Optimal1 Fitted Gauss, proteinAT5G64990.1;AT5G10260.1;AT2G22290.1;AT2G44610.1;AT4G39890.1
bic:806
peaks after deleting less than one fifth of global max:18
×10^7
5
4.5
4
3.5
3
2.5
2
1.5
1
0.5
0
y
0
5
10
15
20
25
x
raw data, starting:18
fitted Gauss locations:18

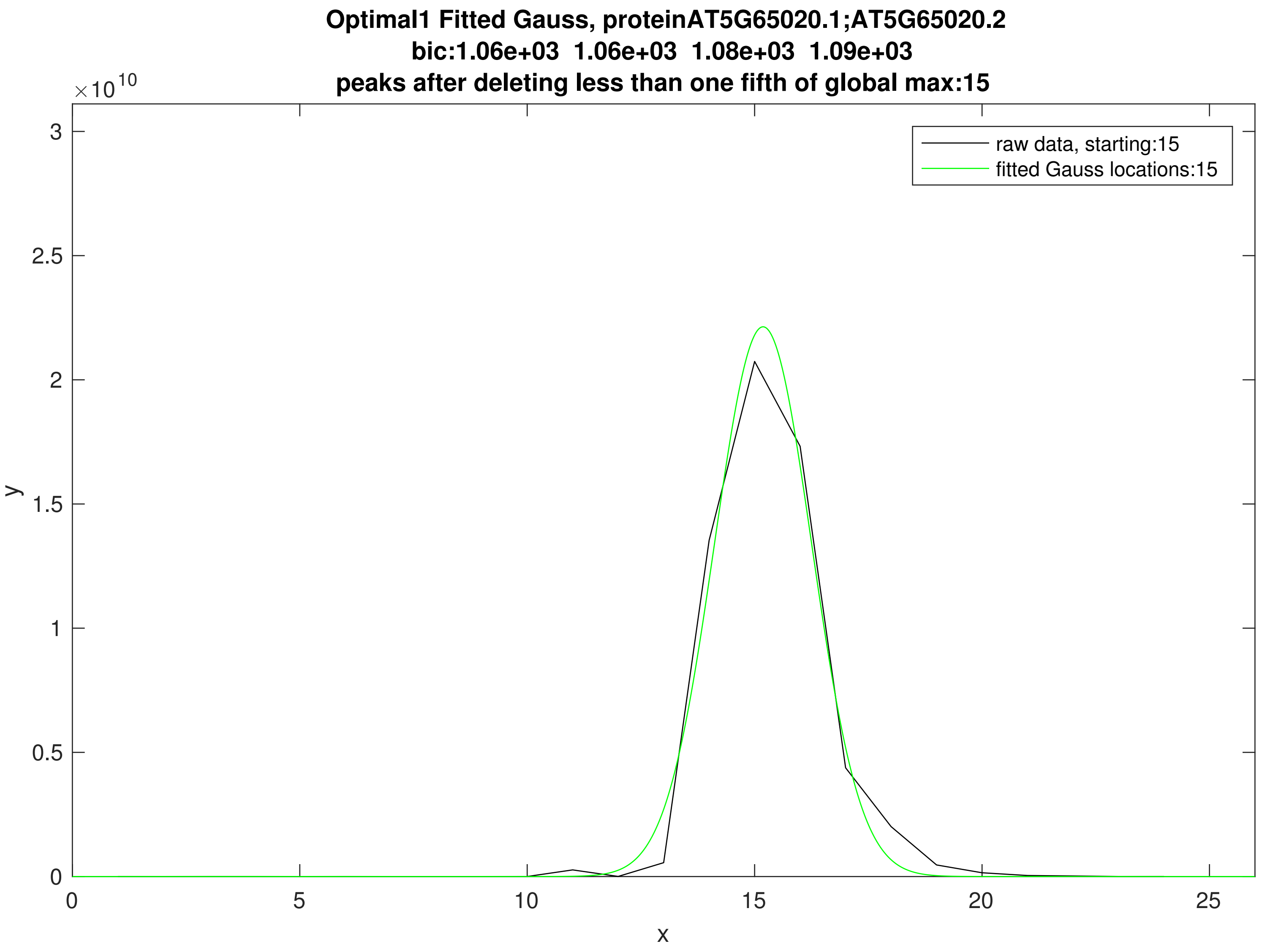
Optimal1 Fitted Gauss, proteinAT5G65020.1;AT5G65020.2
bic:1.06e+03 1.06e+03 1.08e+03 1.09e+03
peaks after deleting less than one fifth of global max:15
raw data, starting:15
fitted Gauss locations:15
×10^10
y
x
0
0.5
1
1.5
2
2.5
3
5
10
15
20
25

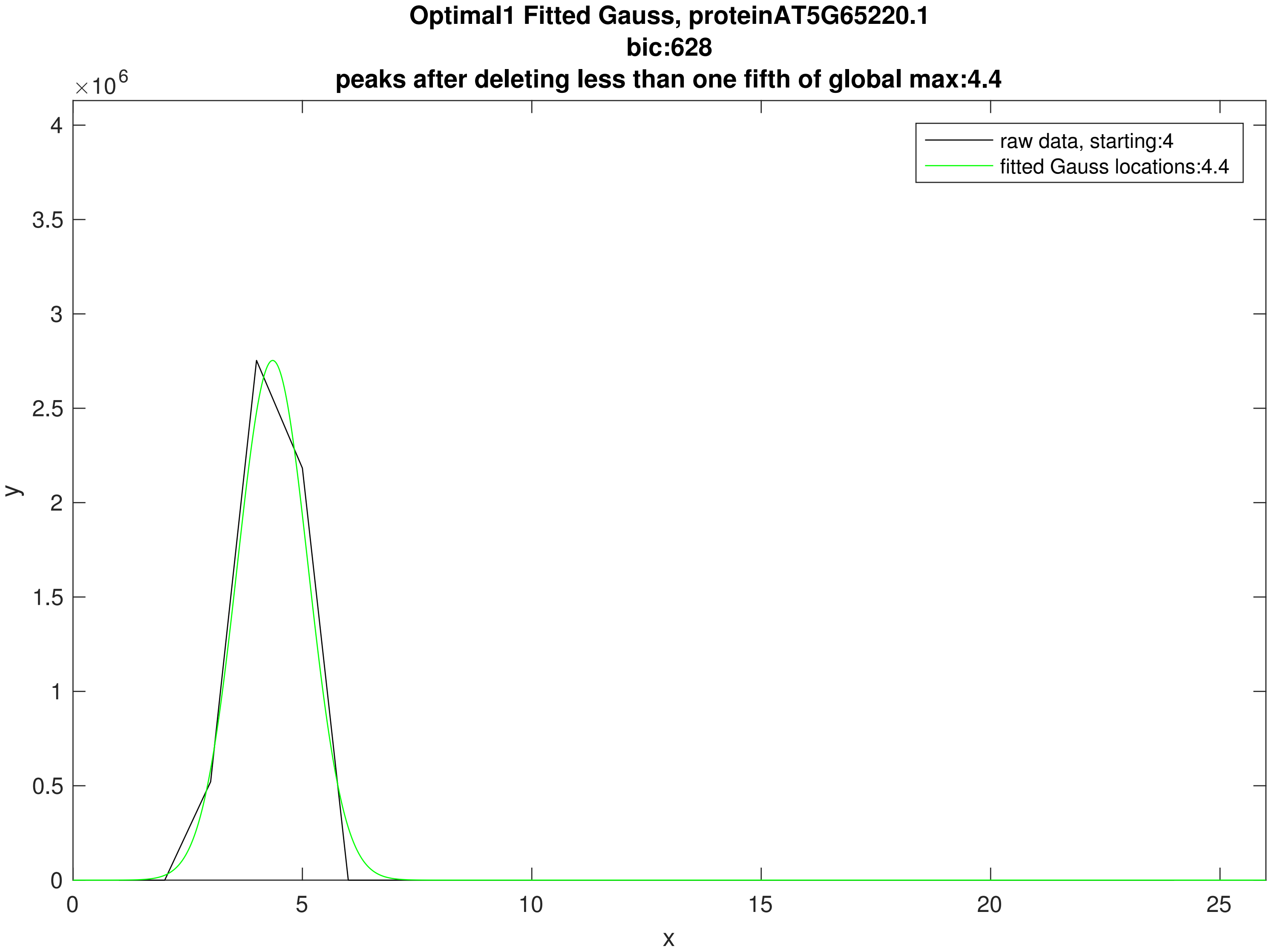
Optimal1 Fitted Gauss, proteinAT5G65220.1
bic:628
peaks after deleting less than one fifth of global max:4.4
×10^6
4
3.5
3
2.5
2
1.5
1
0.5
0
y
0
5
10
15
20
25
x
raw data, starting:4
fitted Gauss locations:4.4

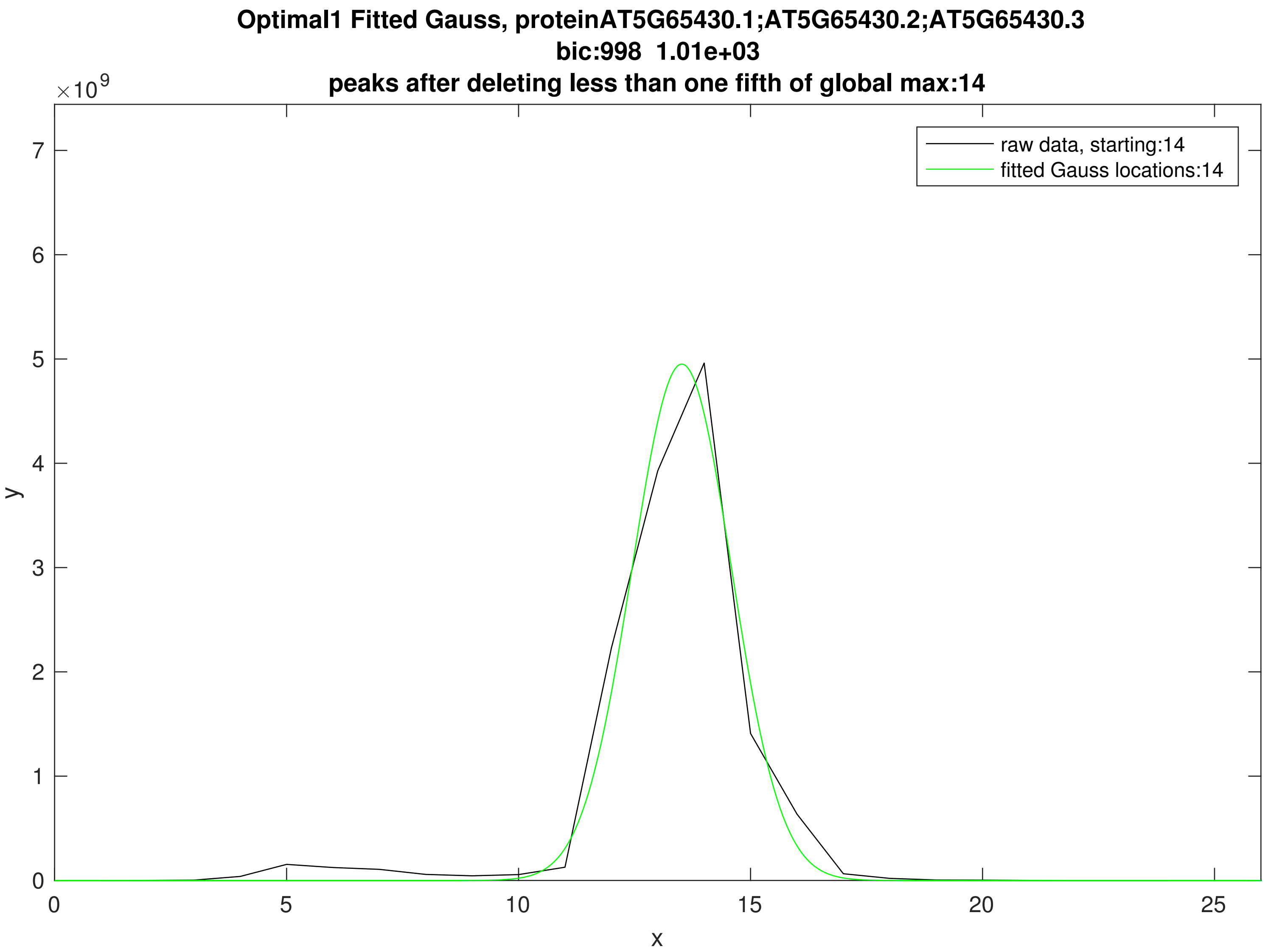
Optimal1 Fitted Gauss, proteinAT5G65430.1;AT5G65430.2;AT5G65430.3
bic:998  1.01e+03
peaks after deleting less than one fifth of global max:14
×10^9
raw data, starting:14
fitted Gauss locations:14
y
x
0
1
2
3
4
5
6
7
0
5
10
15
20
25

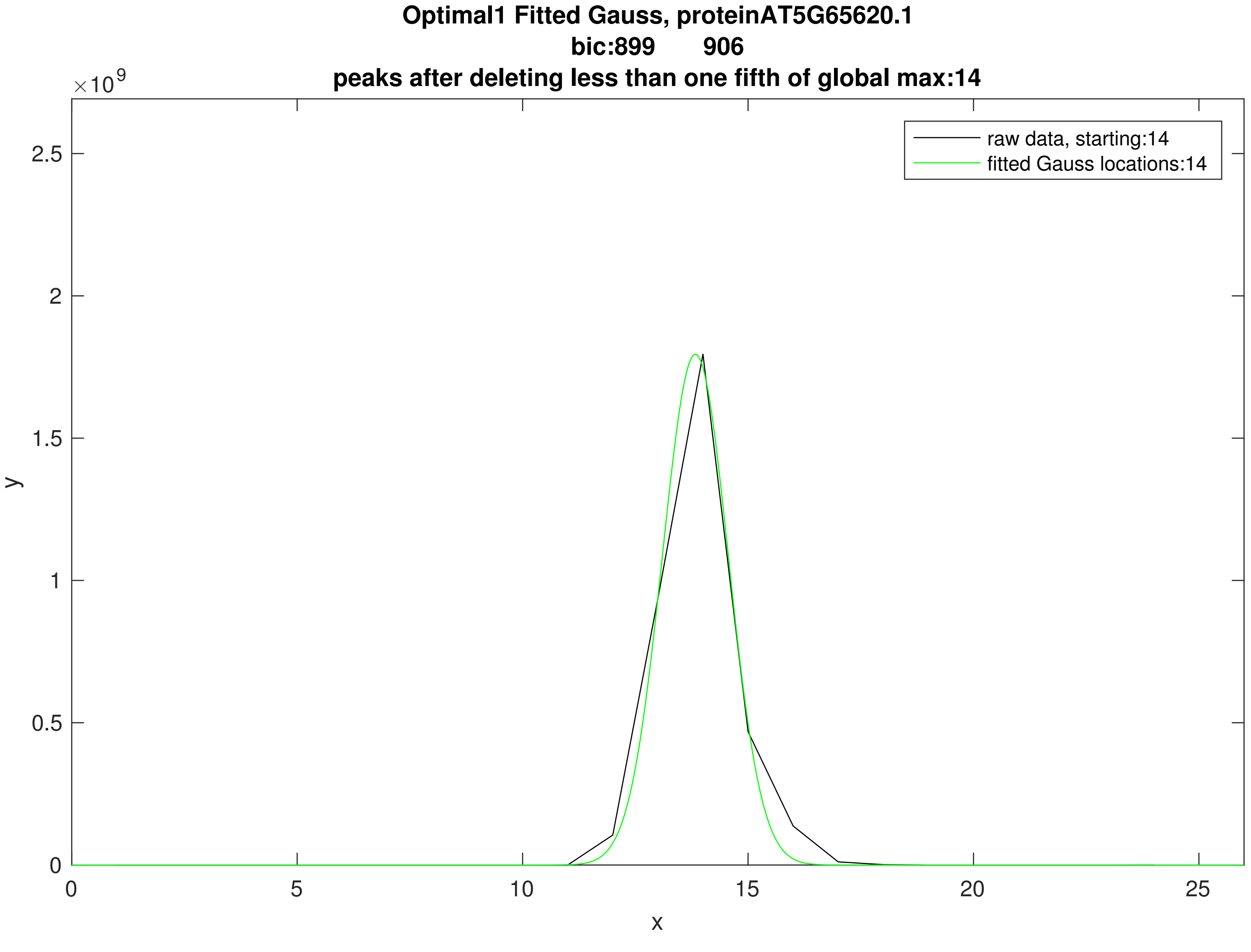

Optimal1 Fitted Gauss, proteinAT5G65620.1
bic:899 906
peaks after deleting less than one fifth of global max:14
×10^9
raw data, starting:14
fitted Gauss locations:14
2.5
2
1.5
1
0.5
0
y
0
5
10
15
20
25
x

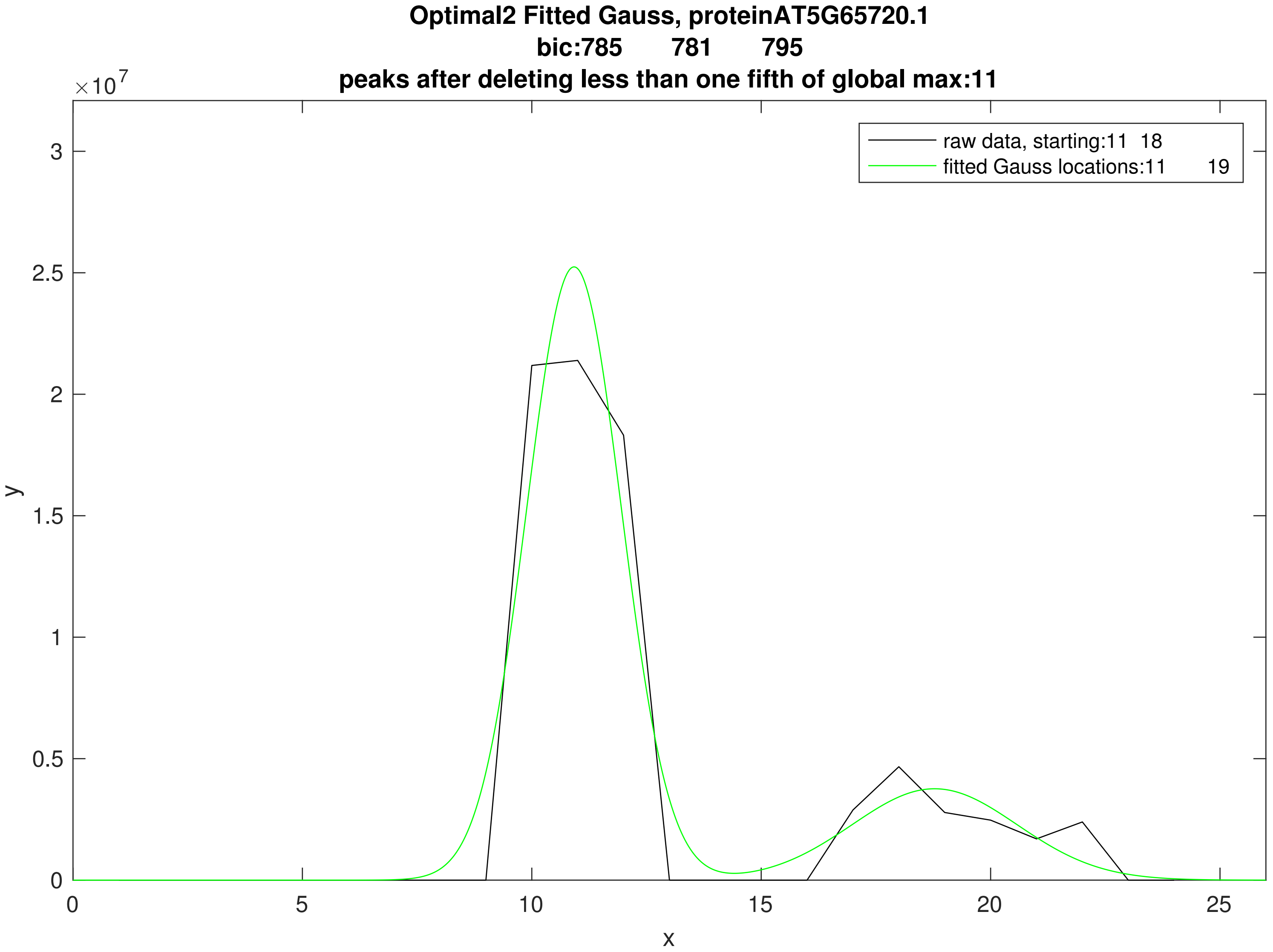

Optimal2 Fitted Gauss, proteinAT5G65720.1
bic:785 781 795
peaks after deleting less than one fifth of global max:11
×10^7
raw data, starting:11 18
fitted Gauss locations:11 19
3
2.5
2
1.5
1
0.5
0
y
0
5
10
15
20
25
x

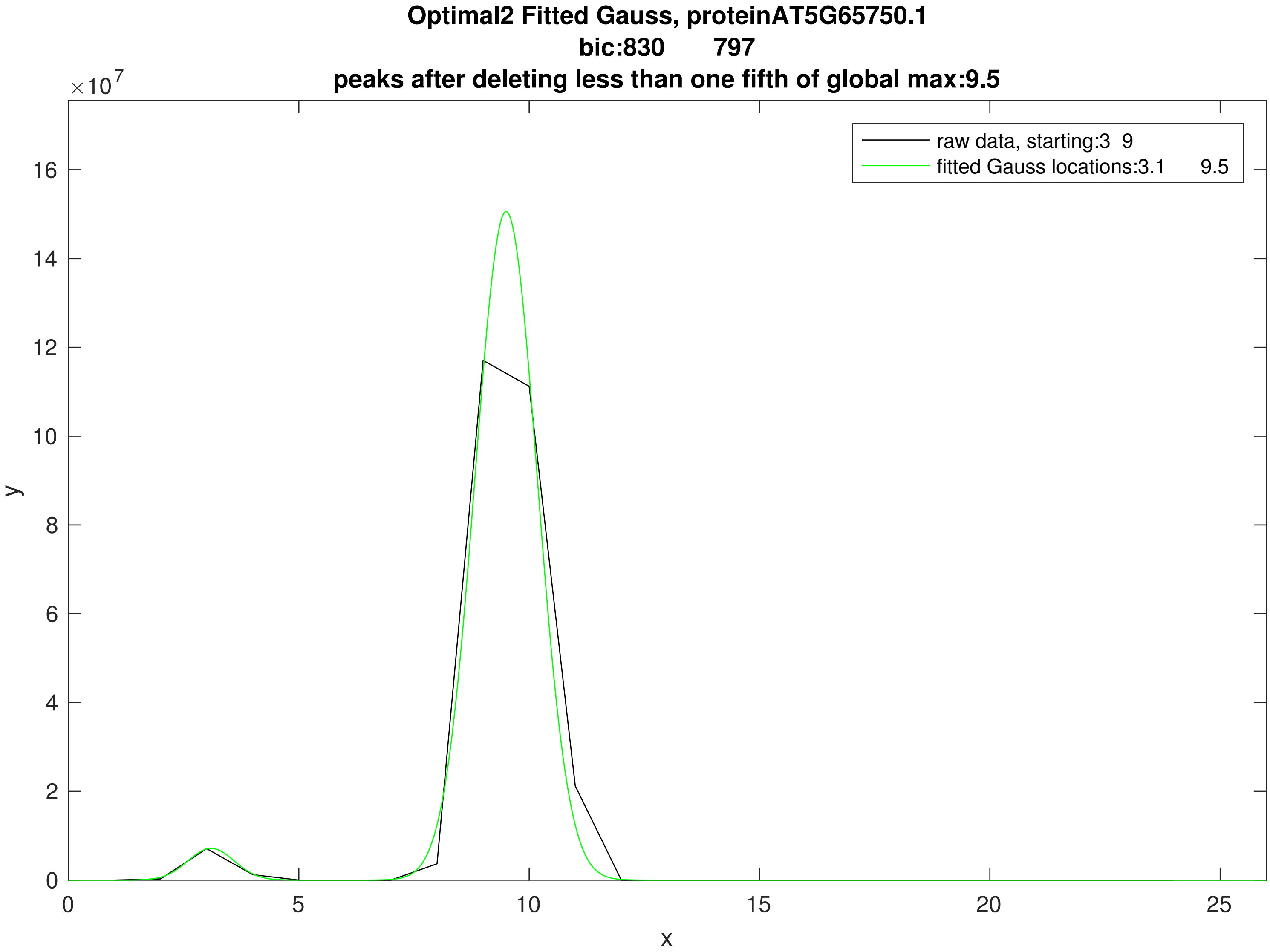

Optimal2 Fitted Gauss, proteinAT5G65750.1
bic:830 797
peaks after deleting less than one fifth of global max:9.5
×10^7
raw data, starting:3 9
fitted Gauss locations:3.1 9.5
16
14
12
10
8
6
4
2
0
y
0
5
10
15
20
25
x

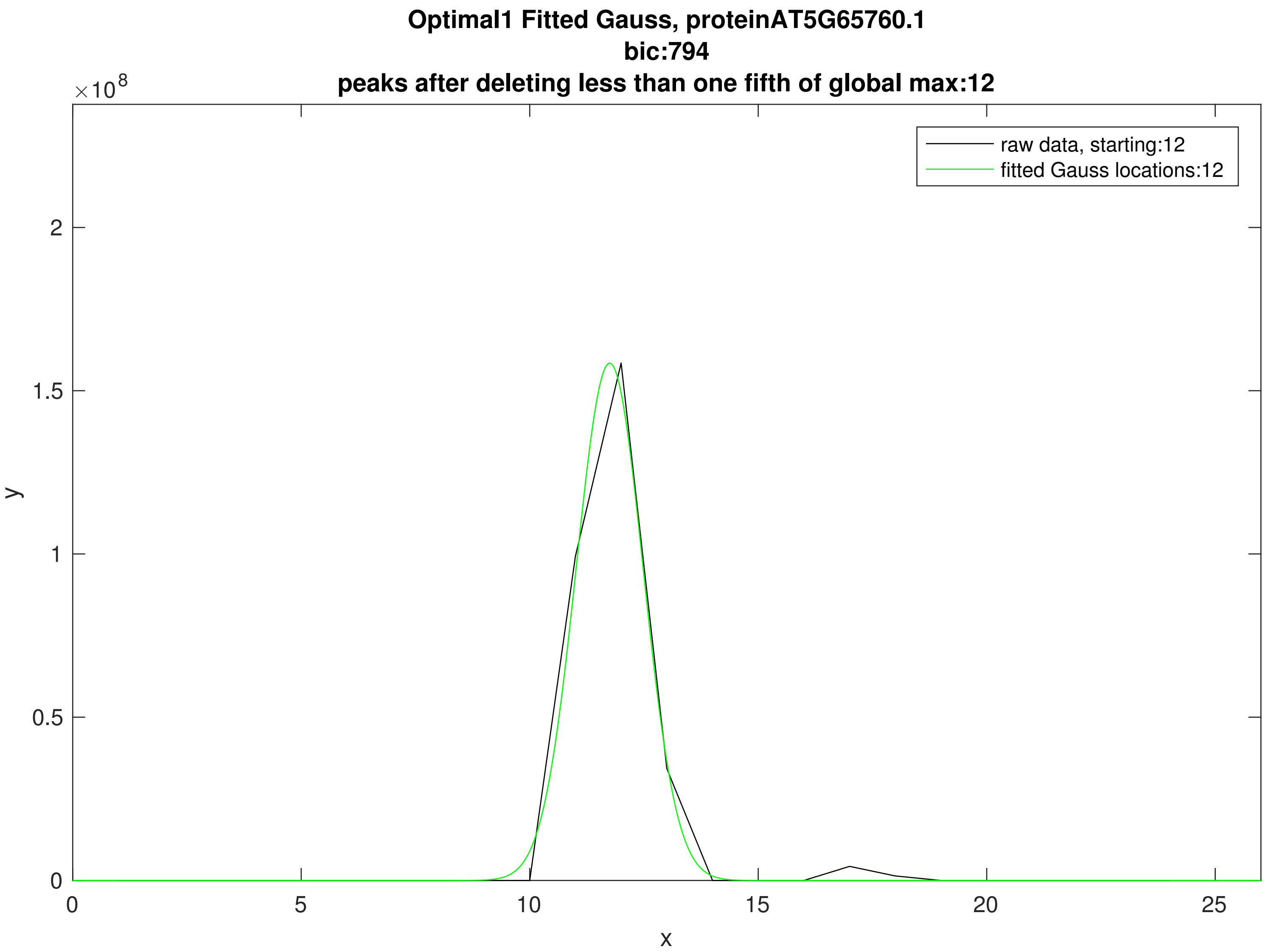

Optimal1 Fitted Gauss, proteinAT5G65760.1
bic:794
peaks after deleting less than one fifth of global max:12
×10^8
raw data, starting:12
fitted Gauss locations:12
y
x
0
0.5
1
1.5
2
0
5
10
15
20
25

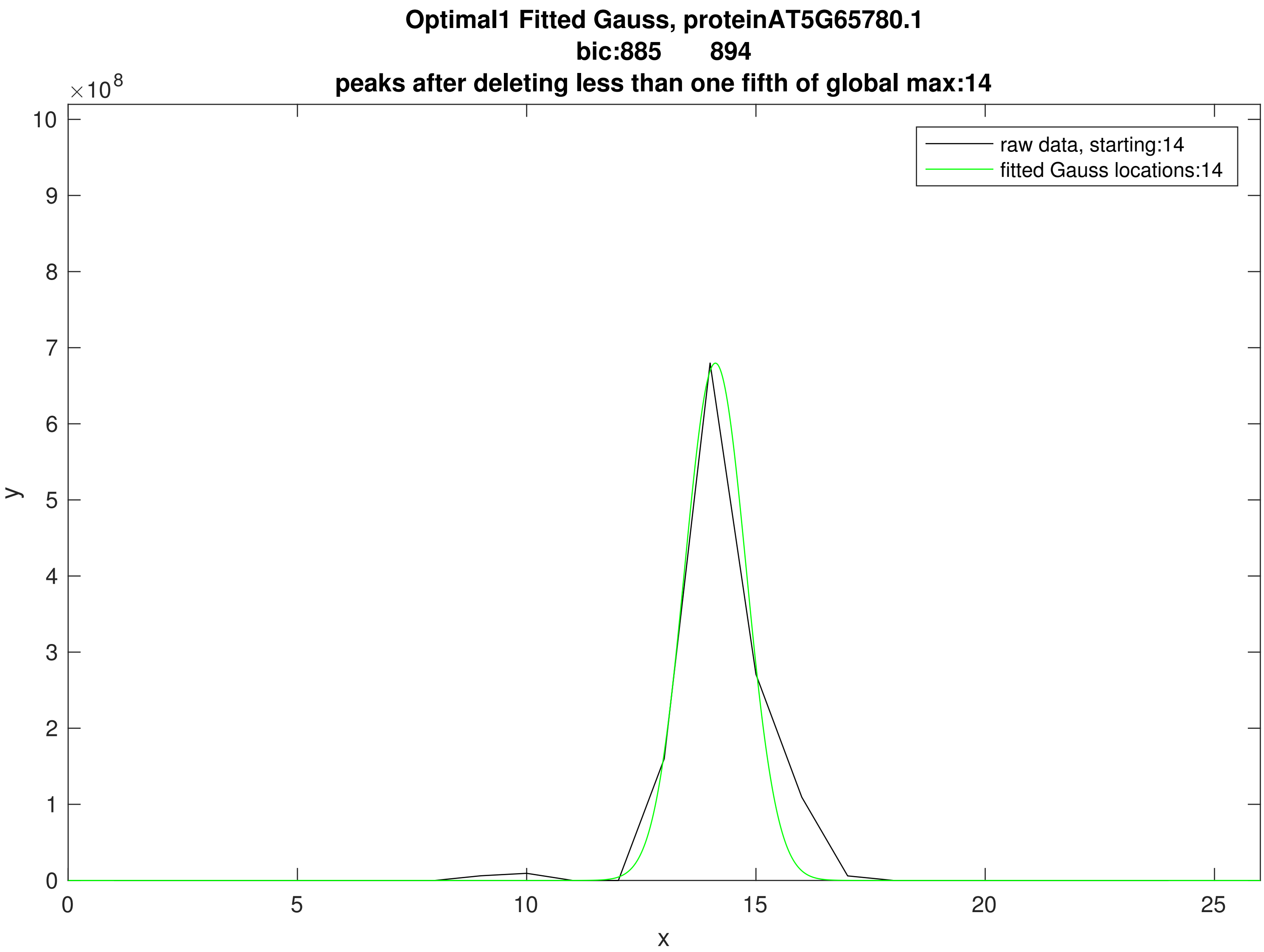
Optimal1 Fitted Gauss, proteinAT5G65780.1
bic:885 894
peaks after deleting less than one fifth of global max:14
×10^8
raw data, starting:14
fitted Gauss locations:14
y
x
0
1
2
3
4
5
6
7
8
9
10
0
5
10
15
20
25

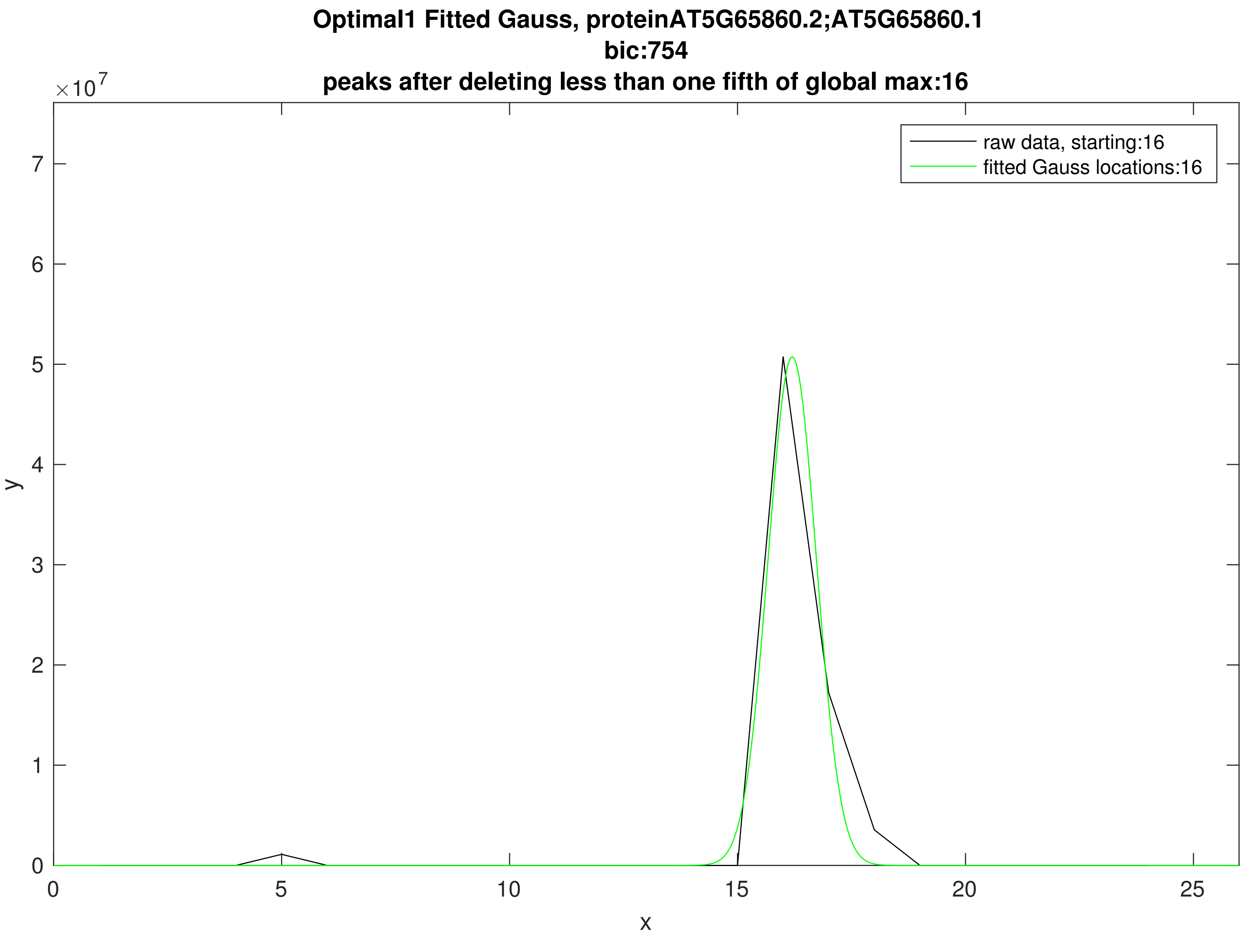

Optimal1 Fitted Gauss, proteinAT5G65860.2;AT5G65860.1
bic:754
peaks after deleting less than one fifth of global max:16
×10^7
raw data, starting:16
fitted Gauss locations:16
y
x

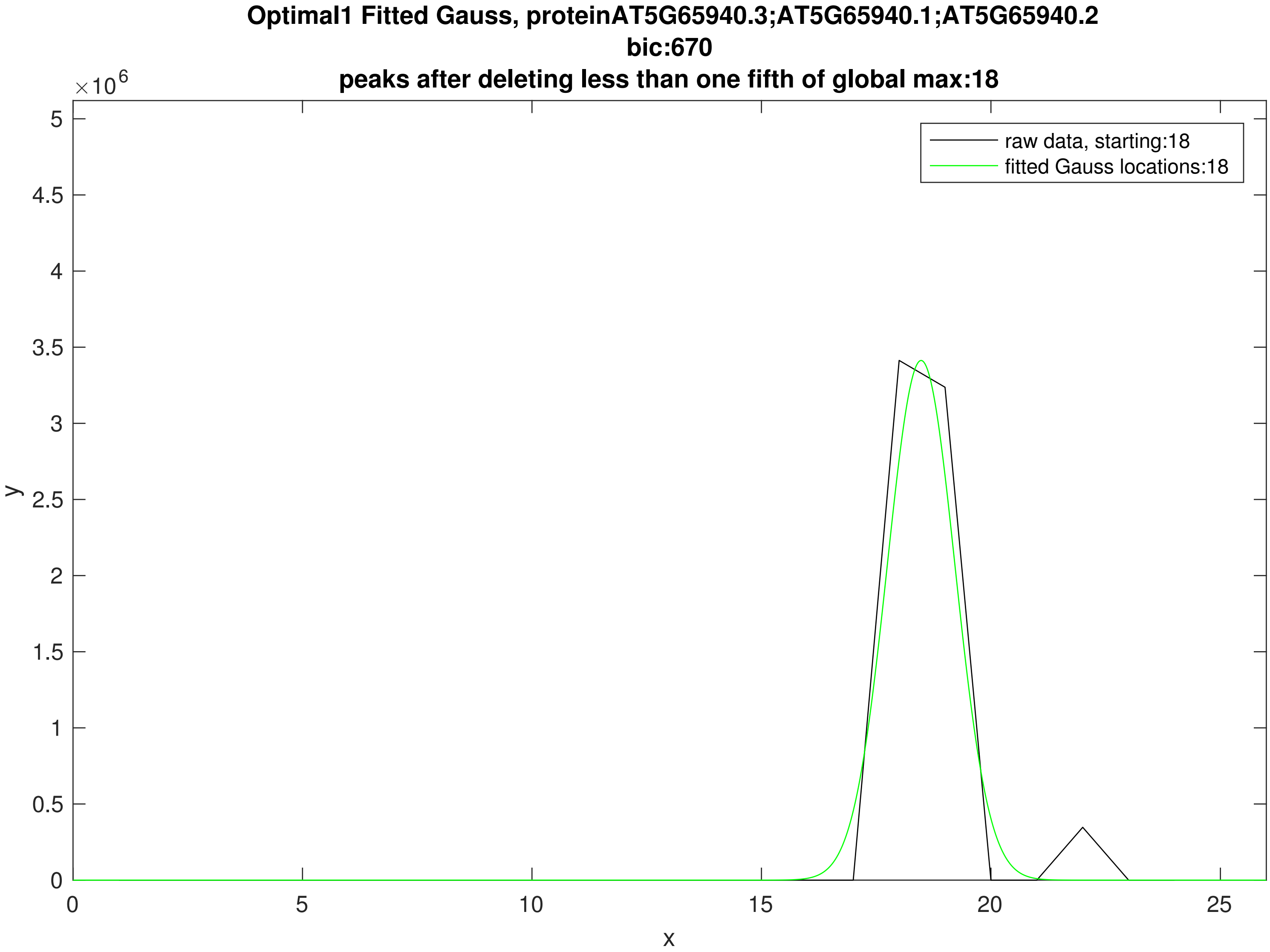

Optimal1 Fitted Gauss, proteinAT5G65940.3;AT5G65940.1;AT5G65940.2
bic:670
peaks after deleting less than one fifth of global max:18
×10^6
5
4.5
4
3.5
3
2.5
2
1.5
1
0.5
0
y
0
5
10
15
20
25
x
raw data, starting:18
fitted Gauss locations:18

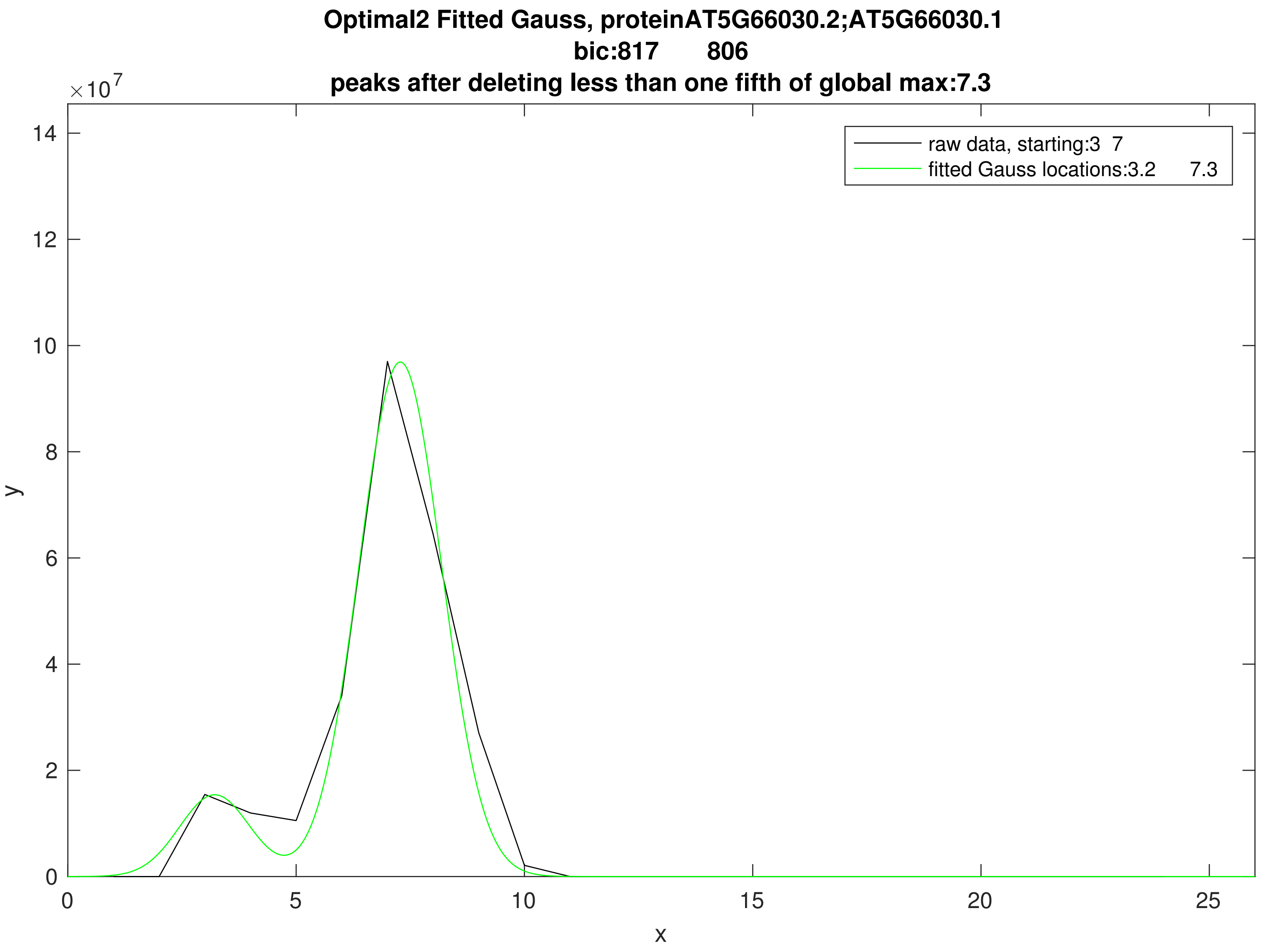
Optimal2 Fitted Gauss, proteinAT5G66030.2;AT5G66030.1
bic:817 806
peaks after deleting less than one fifth of global max:7.3
×10^7
raw data, starting:3 7
fitted Gauss locations:3.2 7.3
y
x
0
2
4
6
8
10
12
14
0
5
10
15
20
25

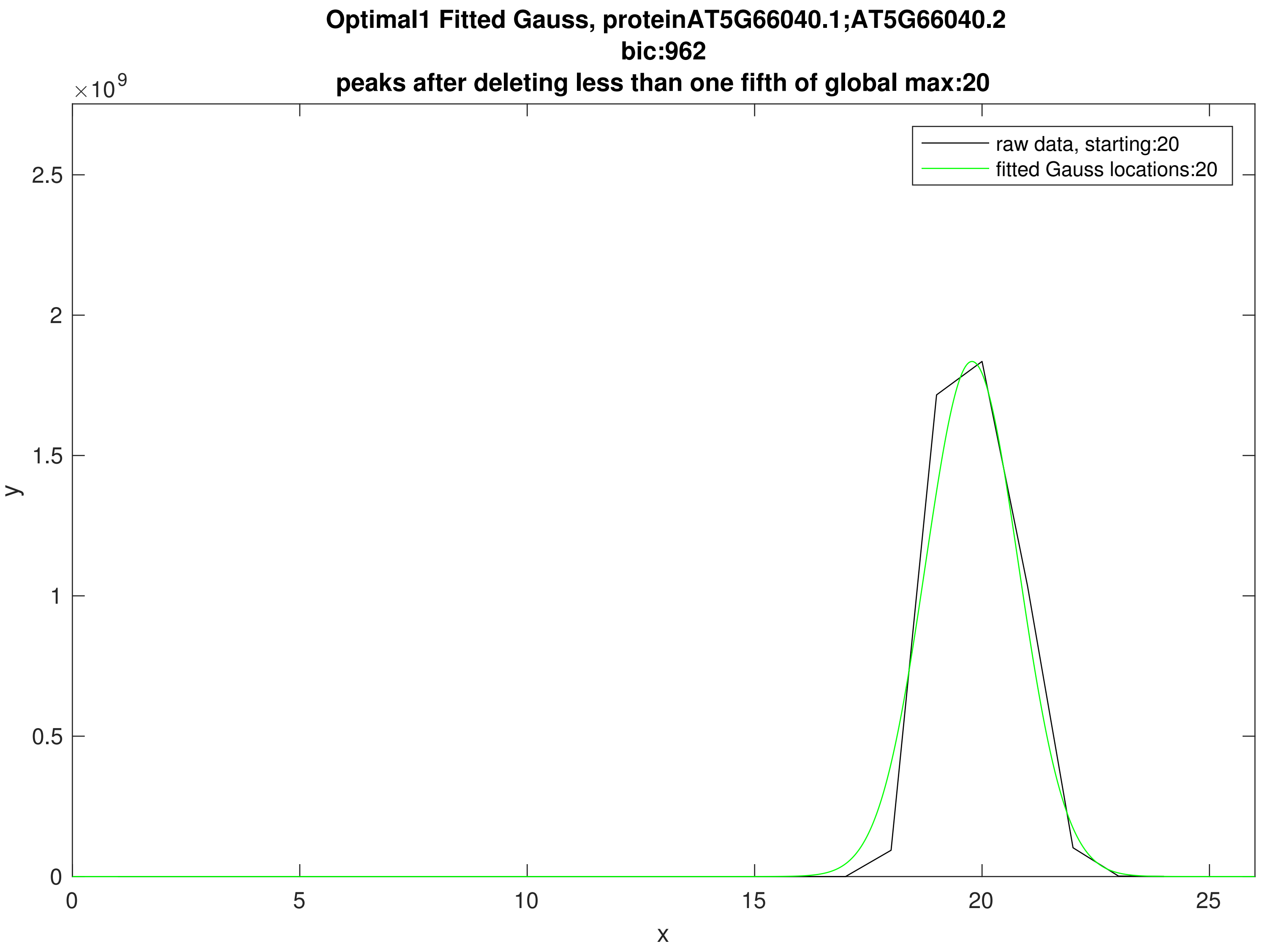
Optimal1 Fitted Gauss, proteinAT5G66040.1;AT5G66040.2
bic:962
peaks after deleting less than one fifth of global max:20
×10^9
2.5
2
1.5
1
0.5
0
y
0
5
10
15
20
25
x
raw data, starting:20
fitted Gauss locations:20

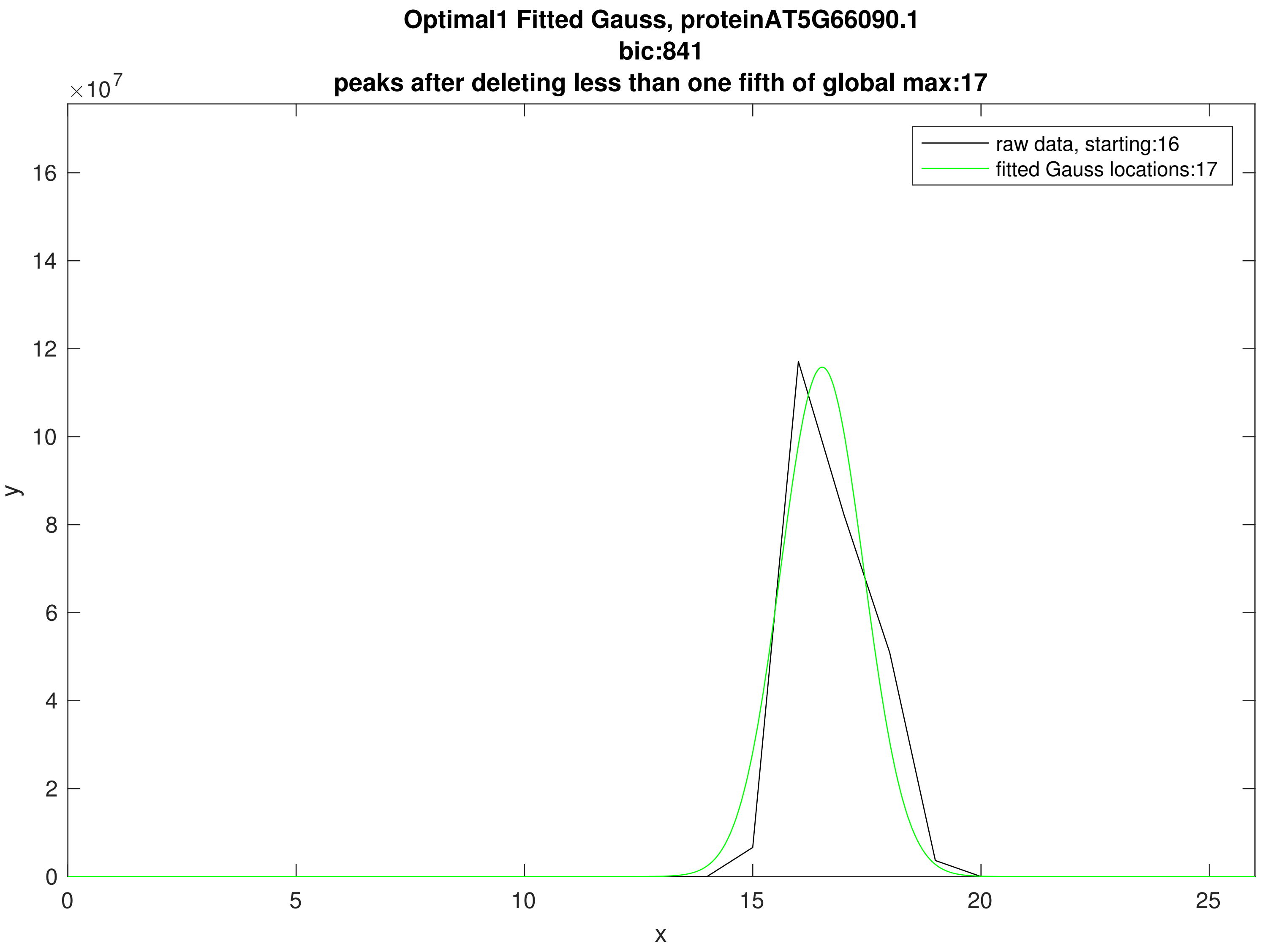
Optimal1 Fitted Gauss, proteinAT5G66090.1
bic:841
peaks after deleting less than one fifth of global max:17
×10^7
raw data, starting:16
fitted Gauss locations:17
y
x
0
2
4
6
8
10
12
14
16
0
5
10
15
20
25

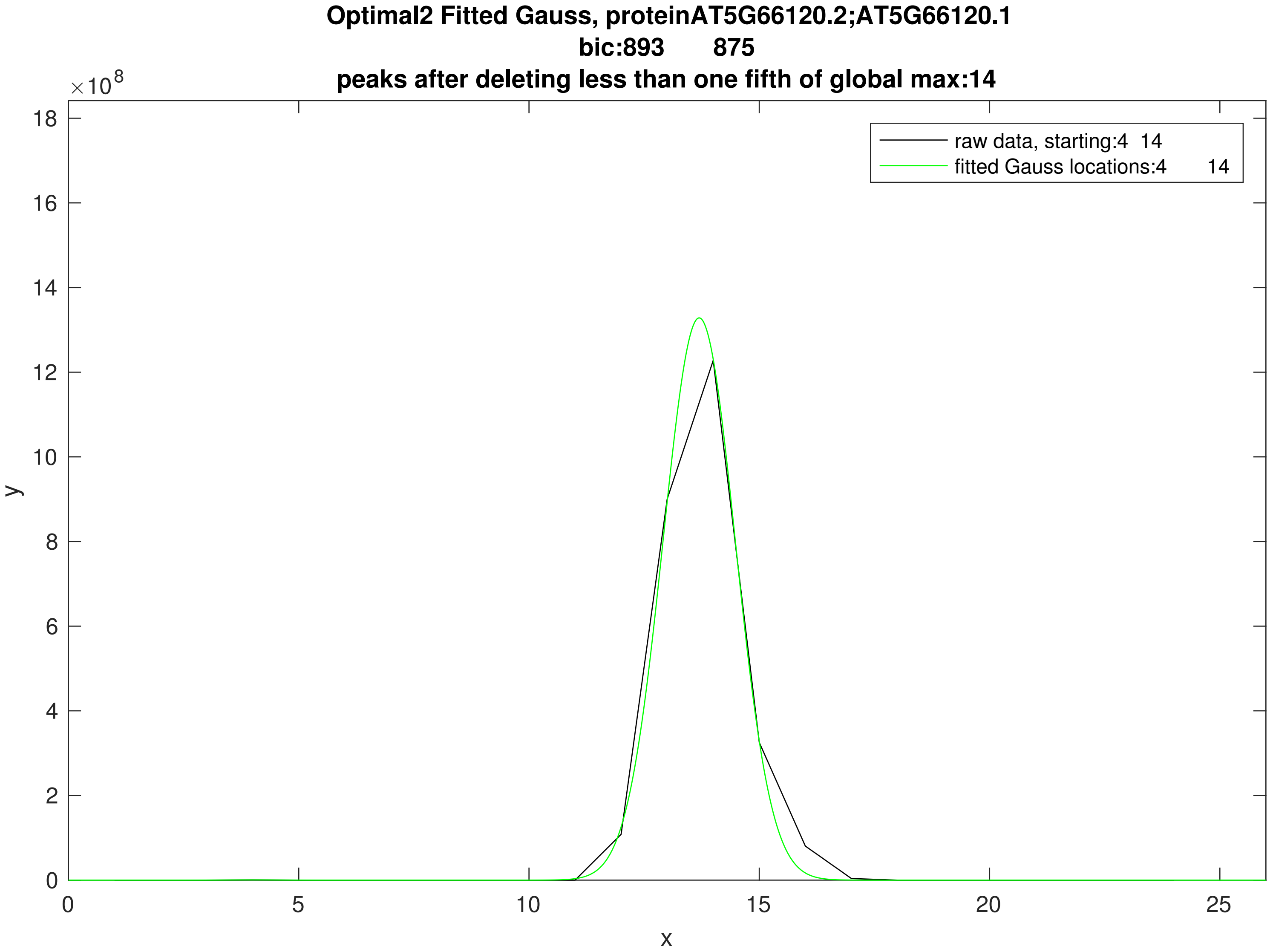
Optimal2 Fitted Gauss, proteinAT5G66120.2;AT5G66120.1
bic:893 875
peaks after deleting less than one fifth of global max:14
×10^8
raw data, starting:4 14
fitted Gauss locations:4 14
y
x

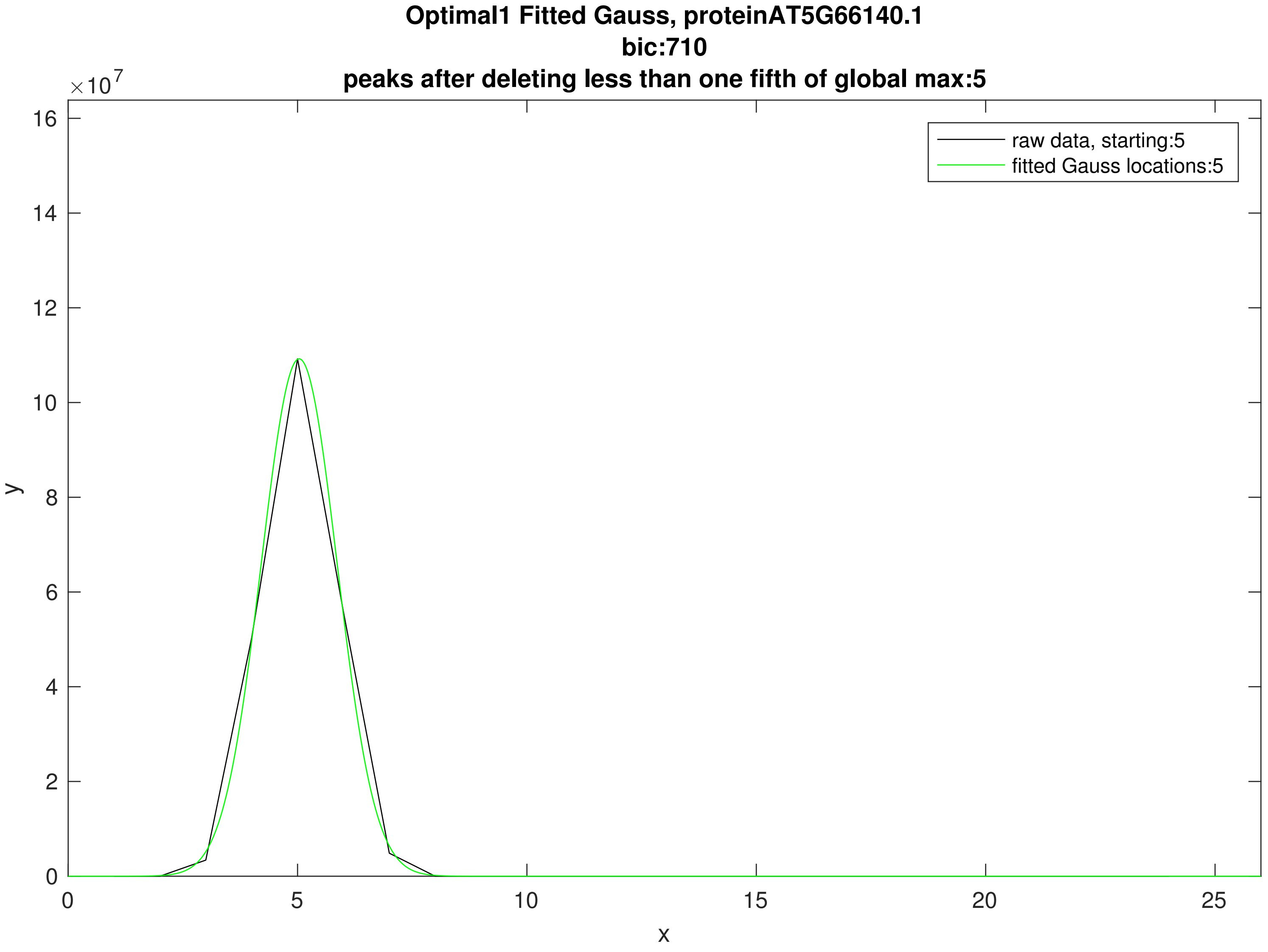
Optimal1 Fitted Gauss, proteinAT5G66140.1
bic:710
peaks after deleting less than one fifth of global max:5
×10^7
16
14
12
10
8
6
4
2
0
y
0
5
10
15
20
25
x
raw data, starting:5
fitted Gauss locations:5

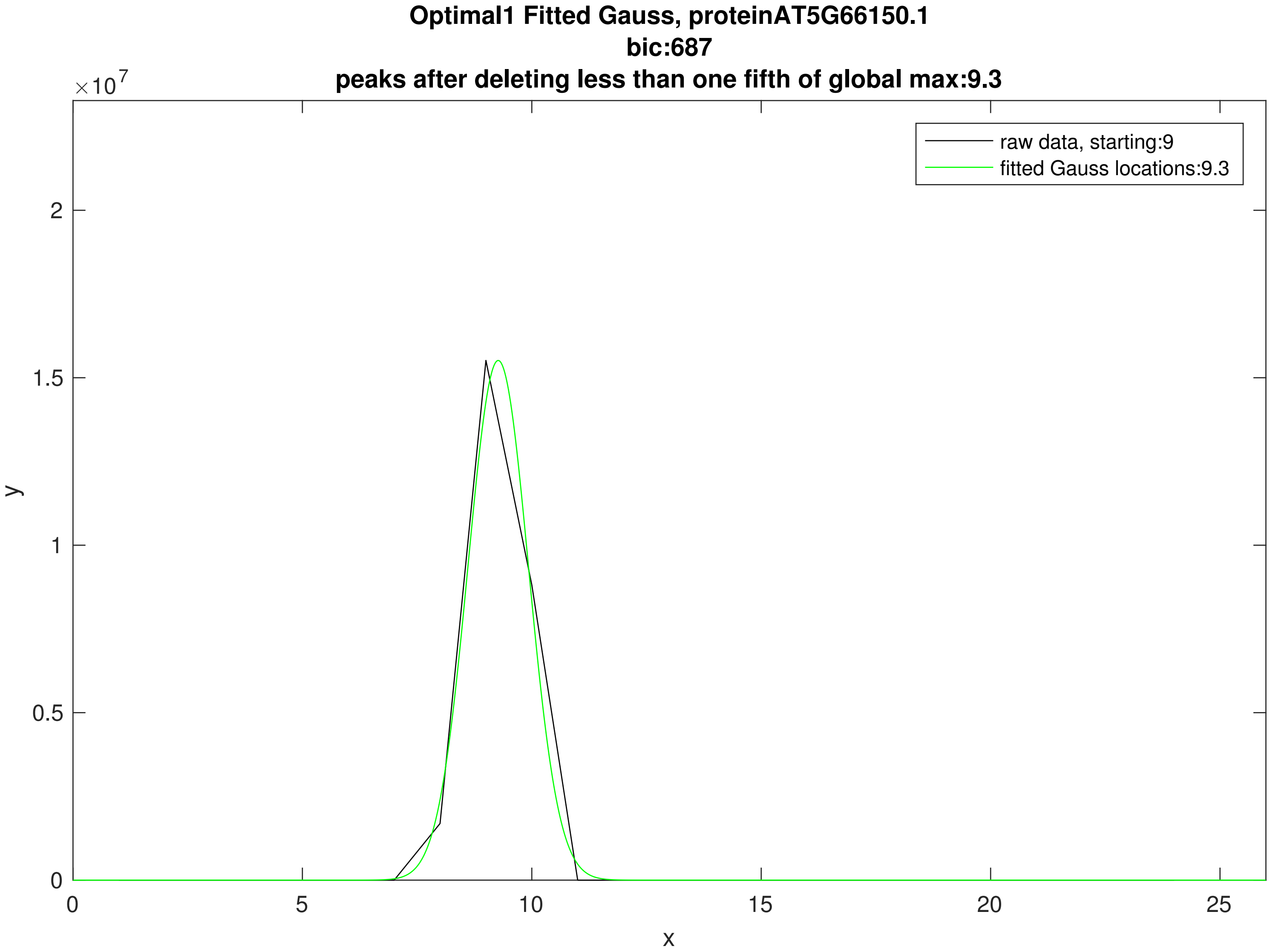
Optimal1 Fitted Gauss, proteinAT5G66150.1
bic:687
peaks after deleting less than one fifth of global max:9.3
×10^7
raw data, starting:9
fitted Gauss locations:9.3
2
1.5
1
0.5
0
y
0
5
10
15
20
25
x

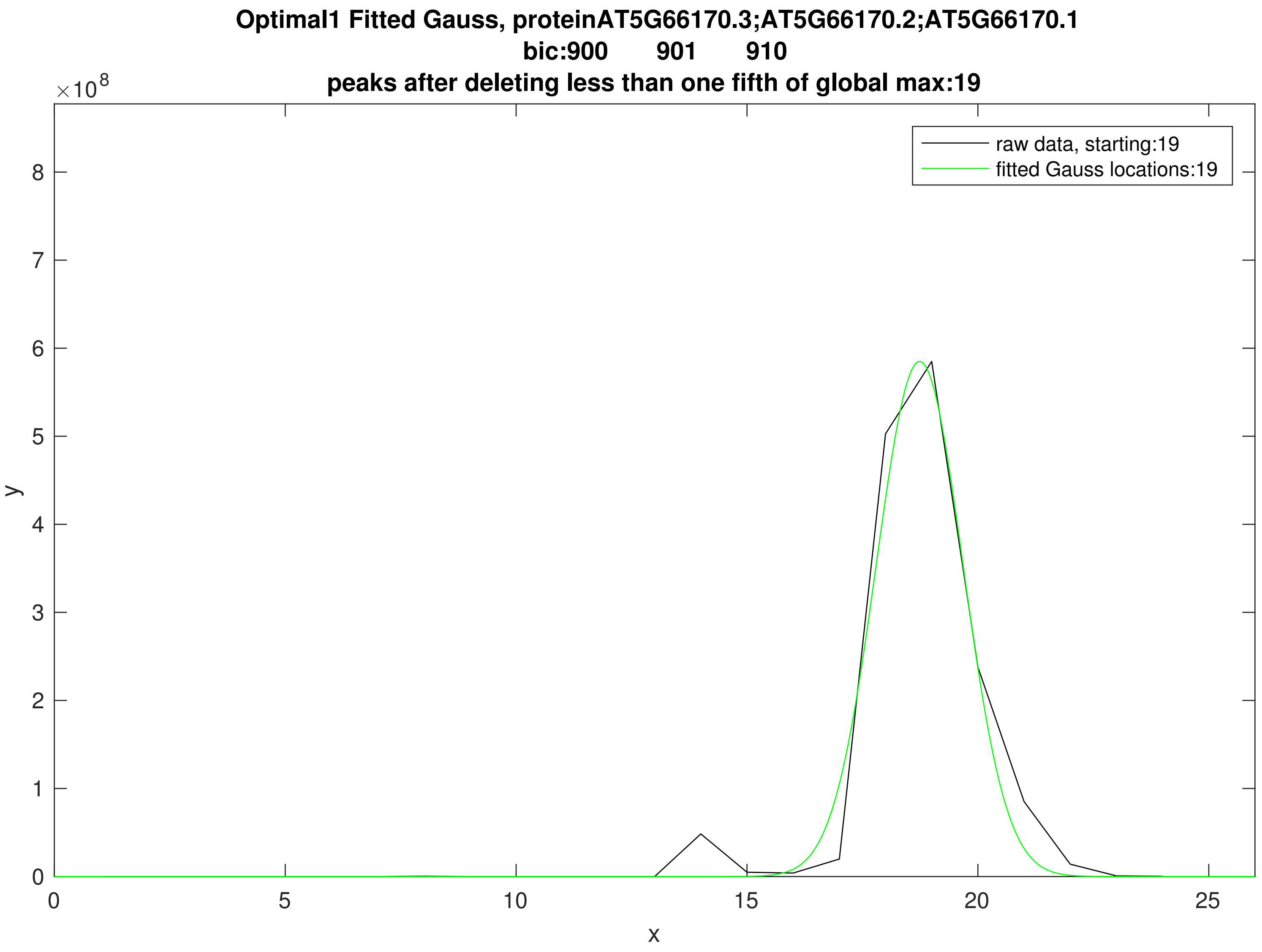
Optimal1 Fitted Gauss, proteinAT5G66170.3;AT5G66170.2;AT5G66170.1
bic:900 901 910
peaks after deleting less than one fifth of global max:19
×10^8
8
7
6
5
4
3
2
1
0
y
0
5
10
15
20
25
x
raw data, starting:19
fitted Gauss locations:19

Optimal2 Fitted Gauss, proteinAT5G66190.1;AT5G66190.2
bic:722 698
peaks after deleting less than one fifth of global max:16

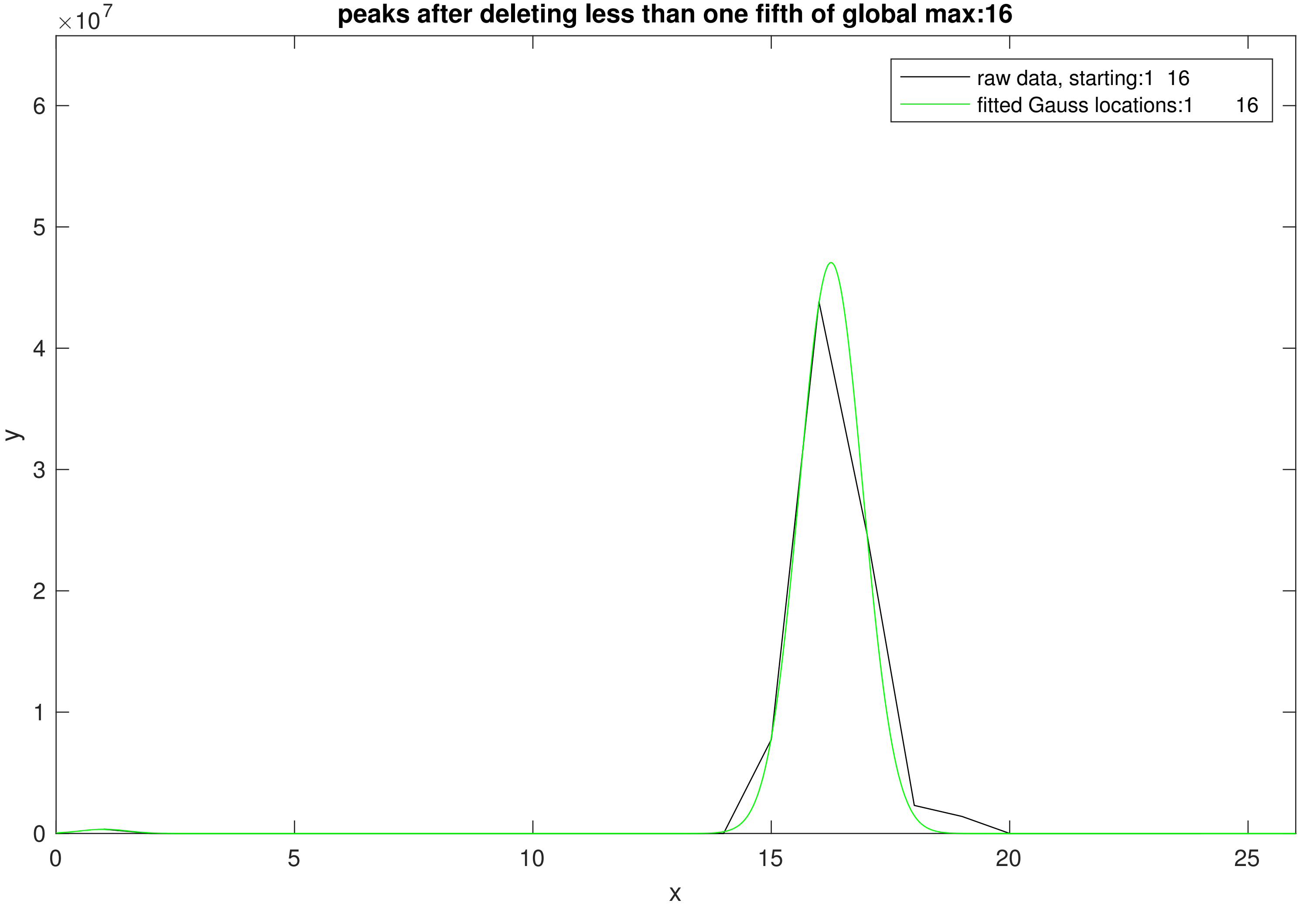

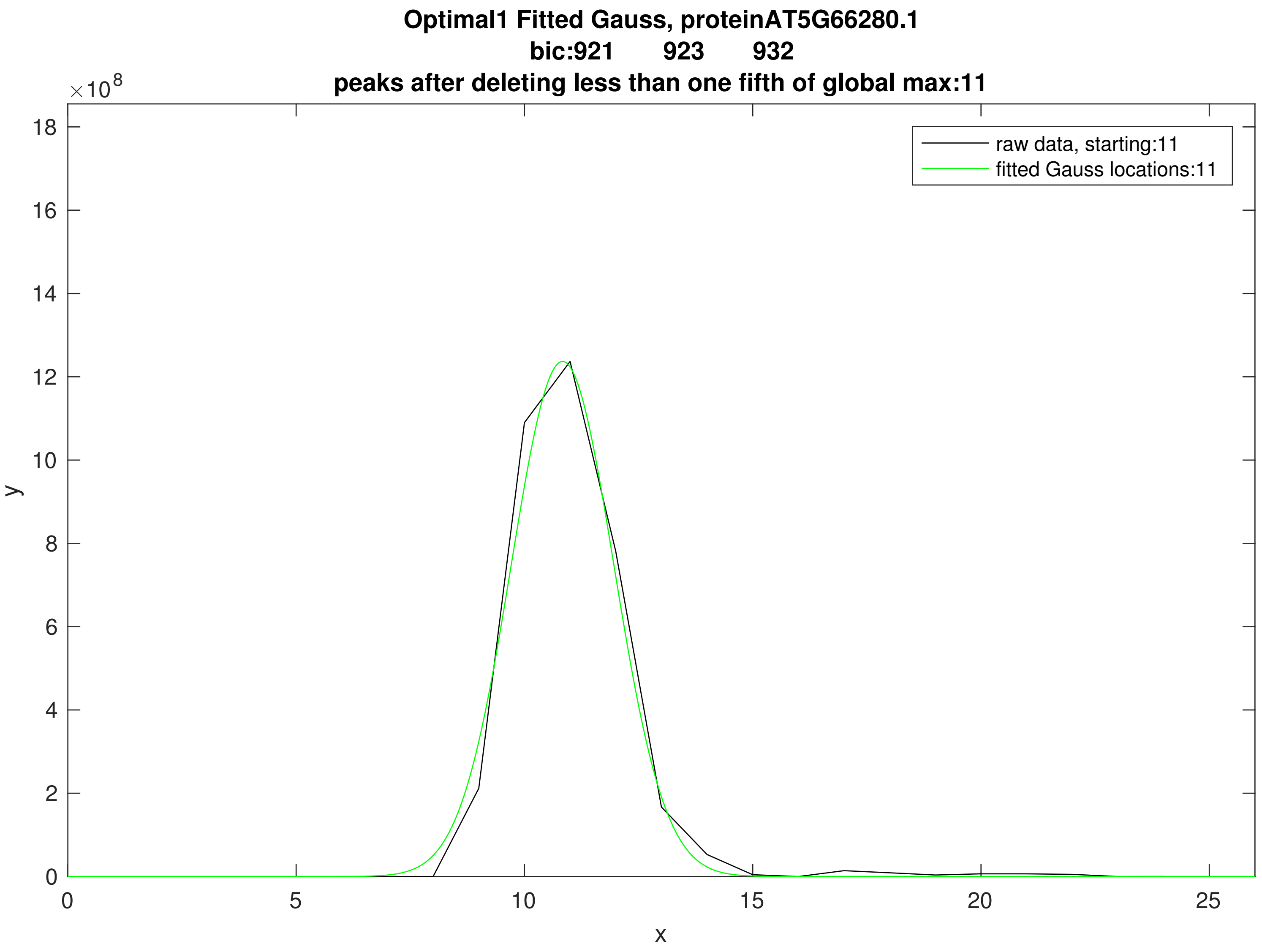

Optimal1 Fitted Gauss, proteinAT5G66280.1
bic:921 923 932
peaks after deleting less than one fifth of global max:11
×10^8
raw data, starting:11
fitted Gauss locations:11
y
x
0
2
4
6
8
10
12
14
16
18
0
5
10
15
20
25

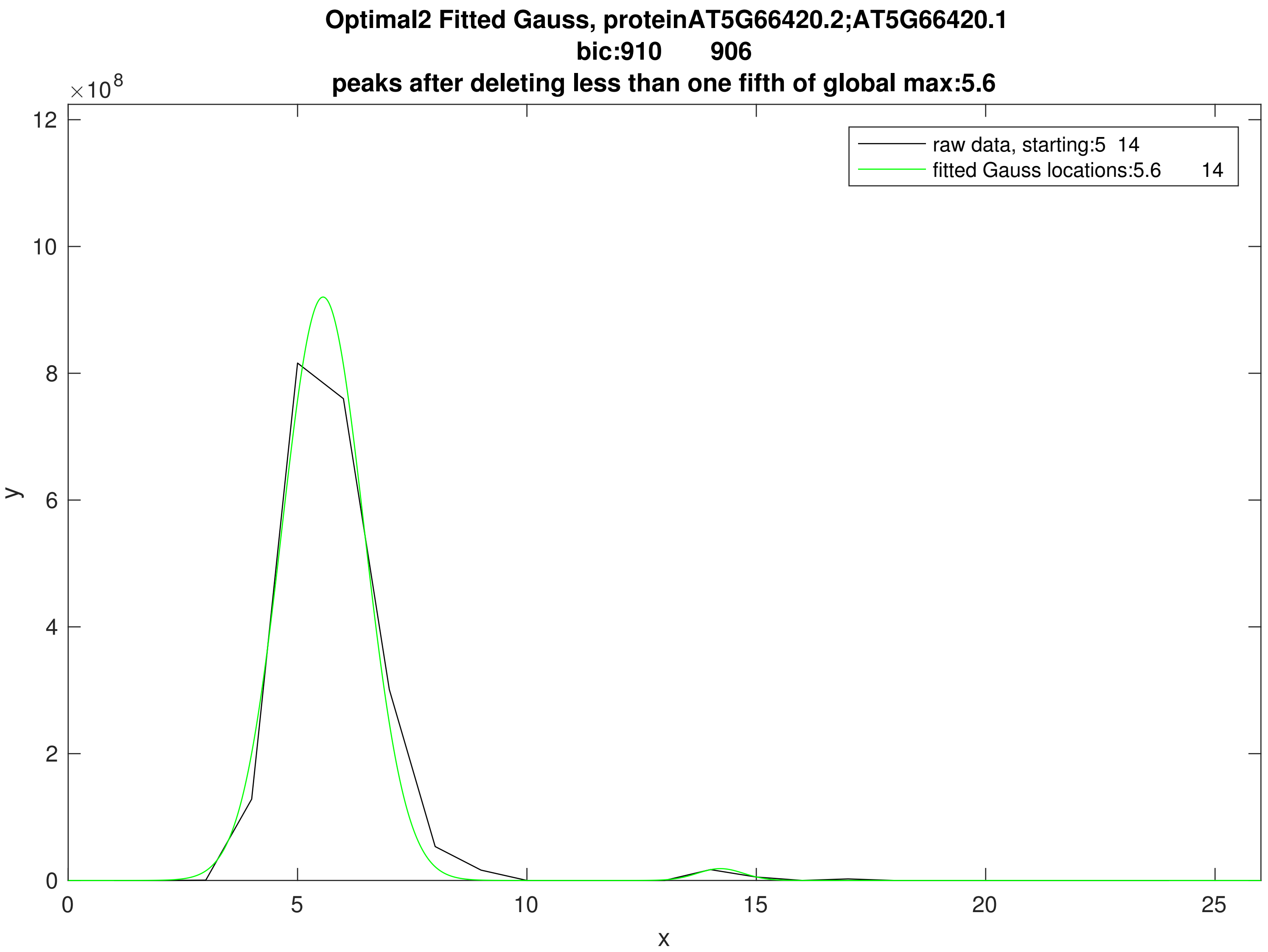
Optimal2 Fitted Gauss, proteinAT5G66420.2;AT5G66420.1
bic:910 906
peaks after deleting less than one fifth of global max:5.6
×10^8
raw data, starting:5 14
fitted Gauss locations:5.6 14
y
x
0
2
4
6
8
10
12
0
5
10
15
20
25

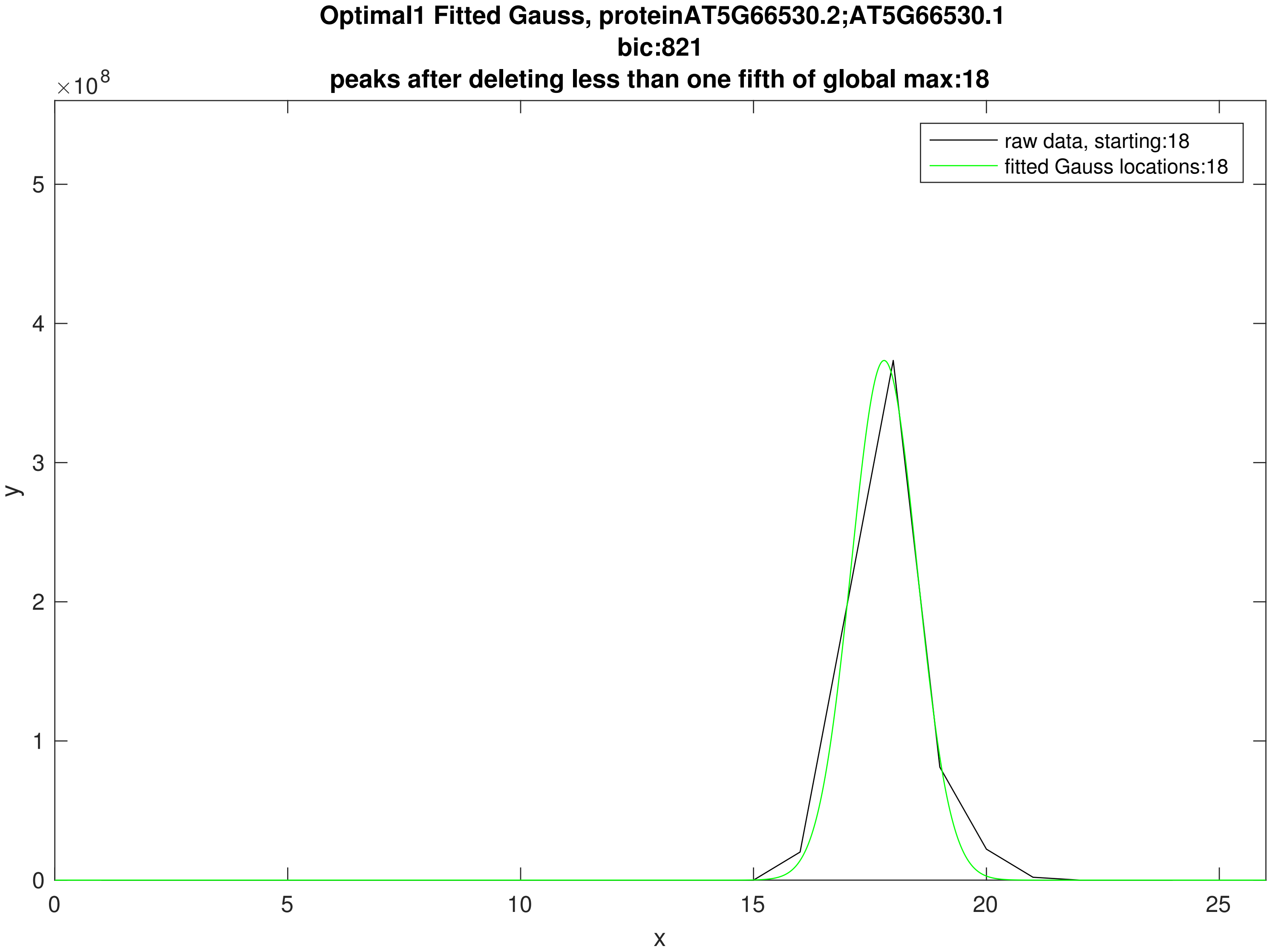
Optimal1 Fitted Gauss, proteinAT5G66530.2;AT5G66530.1
bic:821
peaks after deleting less than one fifth of global max:18
×10^8
raw data, starting:18
fitted Gauss locations:18
5
4
3
2
1
0
y
0
5
10
15
20
25
x

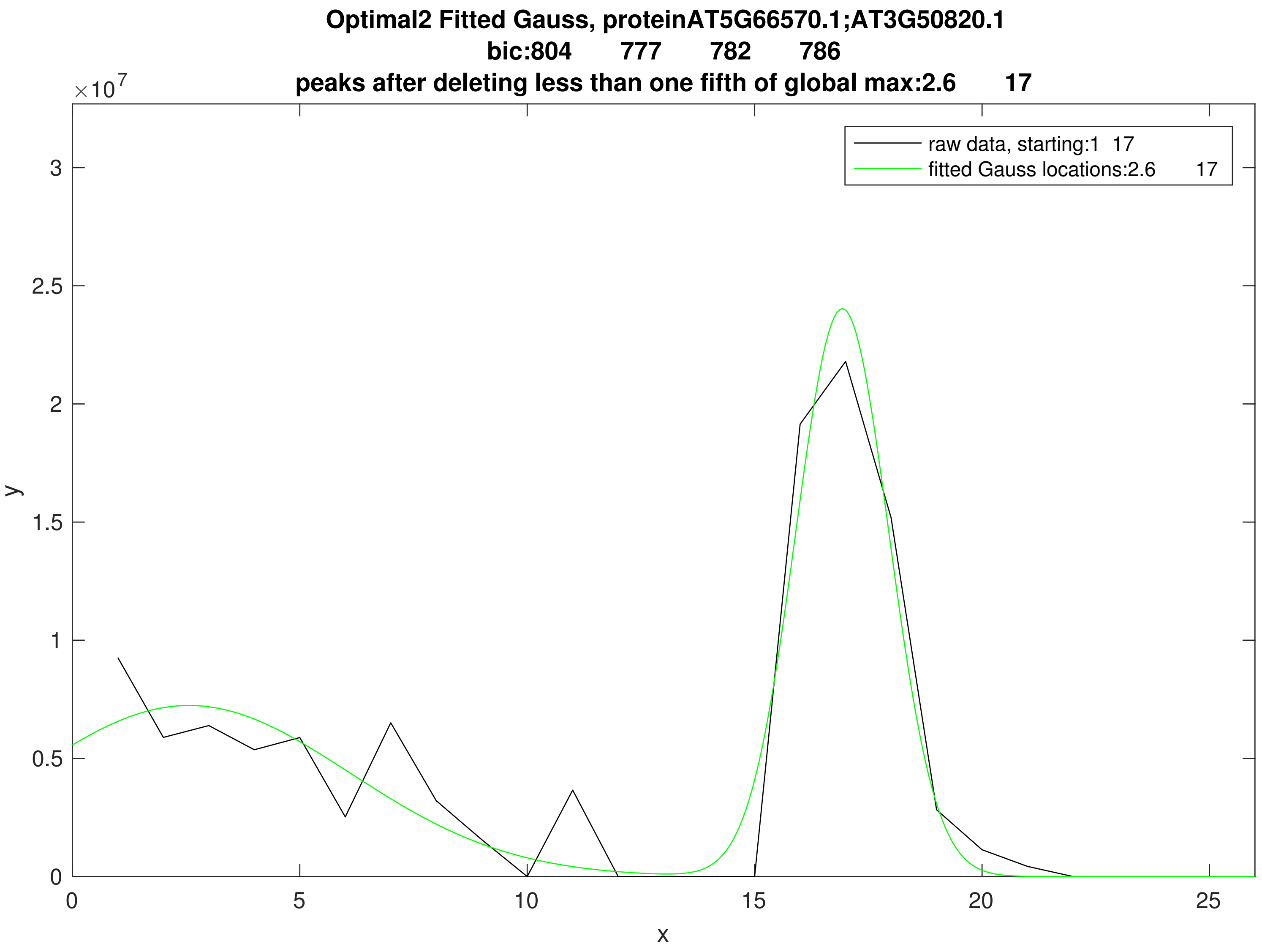
Optimal2 Fitted Gauss, proteinAT5G66570.1;AT3G50820.1
bic:804 777 782 786
peaks after deleting less than one fifth of global max:2.6 17
×10^7
raw data, starting:1 17
fitted Gauss locations:2.6 17
3
2.5
2
1.5
1
0.5
0
y
0
5
10
15
20
25
x

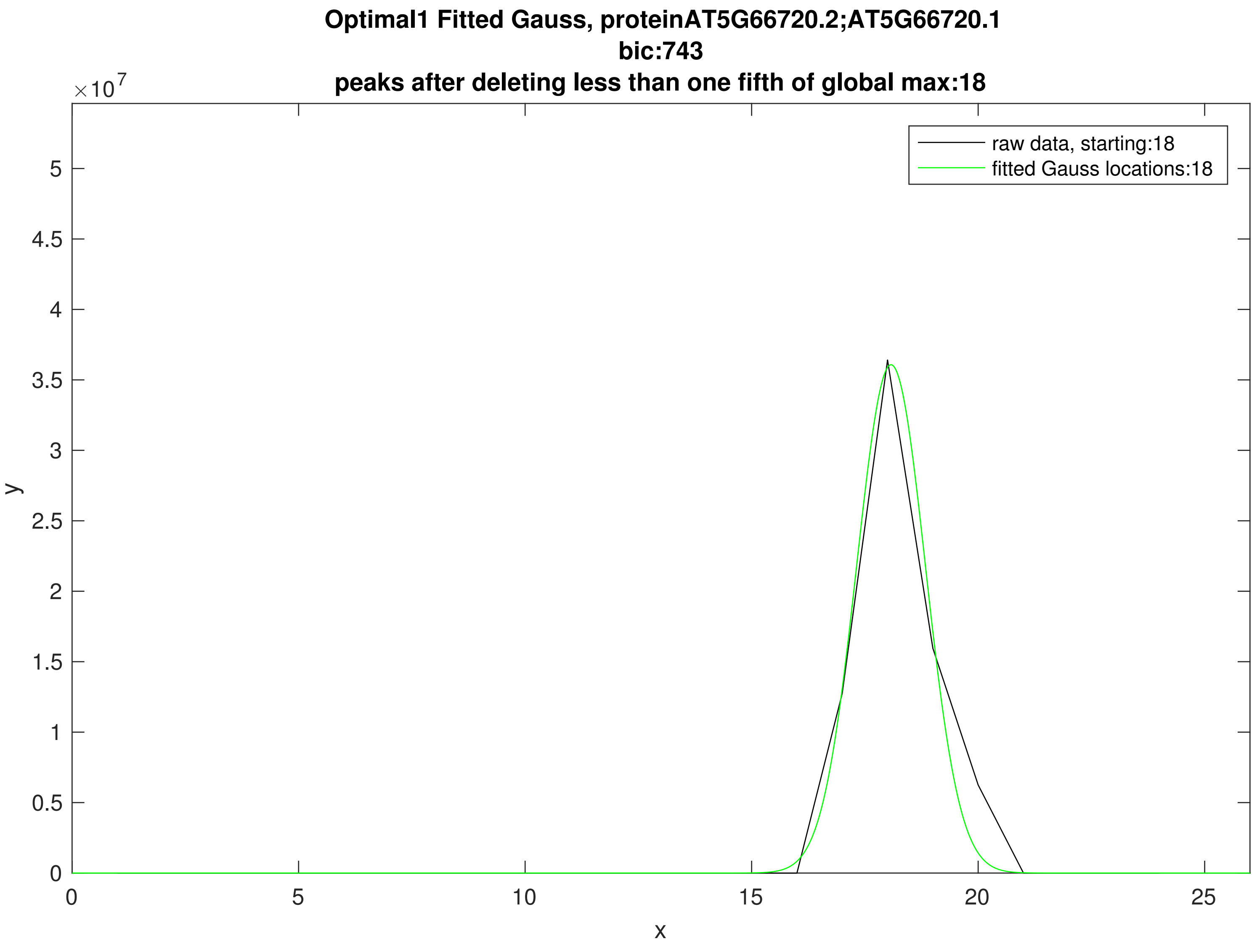
Optimal1 Fitted Gauss, proteinAT5G66720.2;AT5G66720.1
bic:743
peaks after deleting less than one fifth of global max:18
×10^7
raw data, starting:18
fitted Gauss locations:18
y
x
0
0.5
1
1.5
2
2.5
3
3.5
4
4.5
5
0
5
10
15
20
25

**Optimal1 Fitted Gauss, proteinAT5G66760.1**
**bic:978 988 997 1.01e+03**
**peaks after deleting less than one fifth of global max:14**

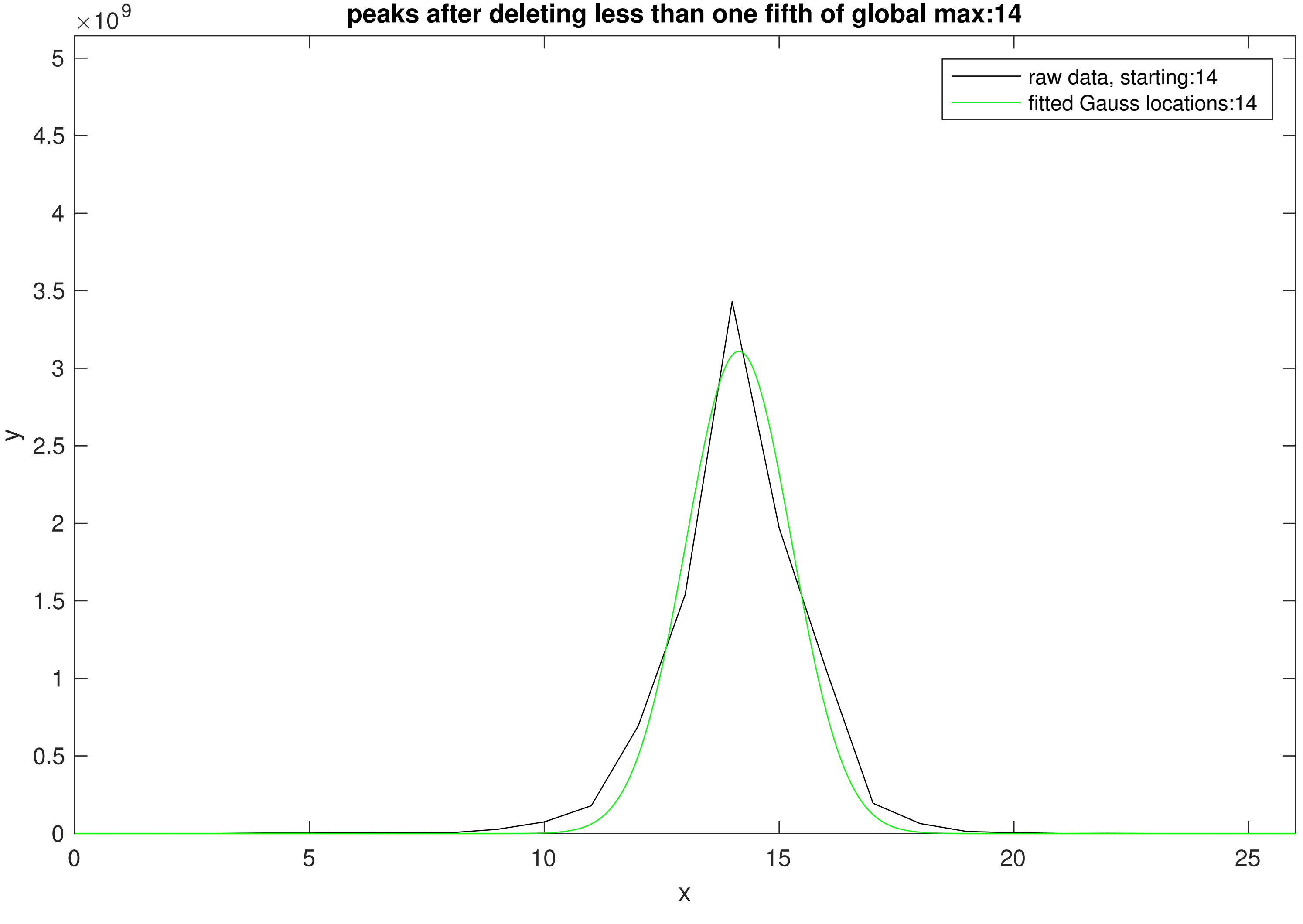

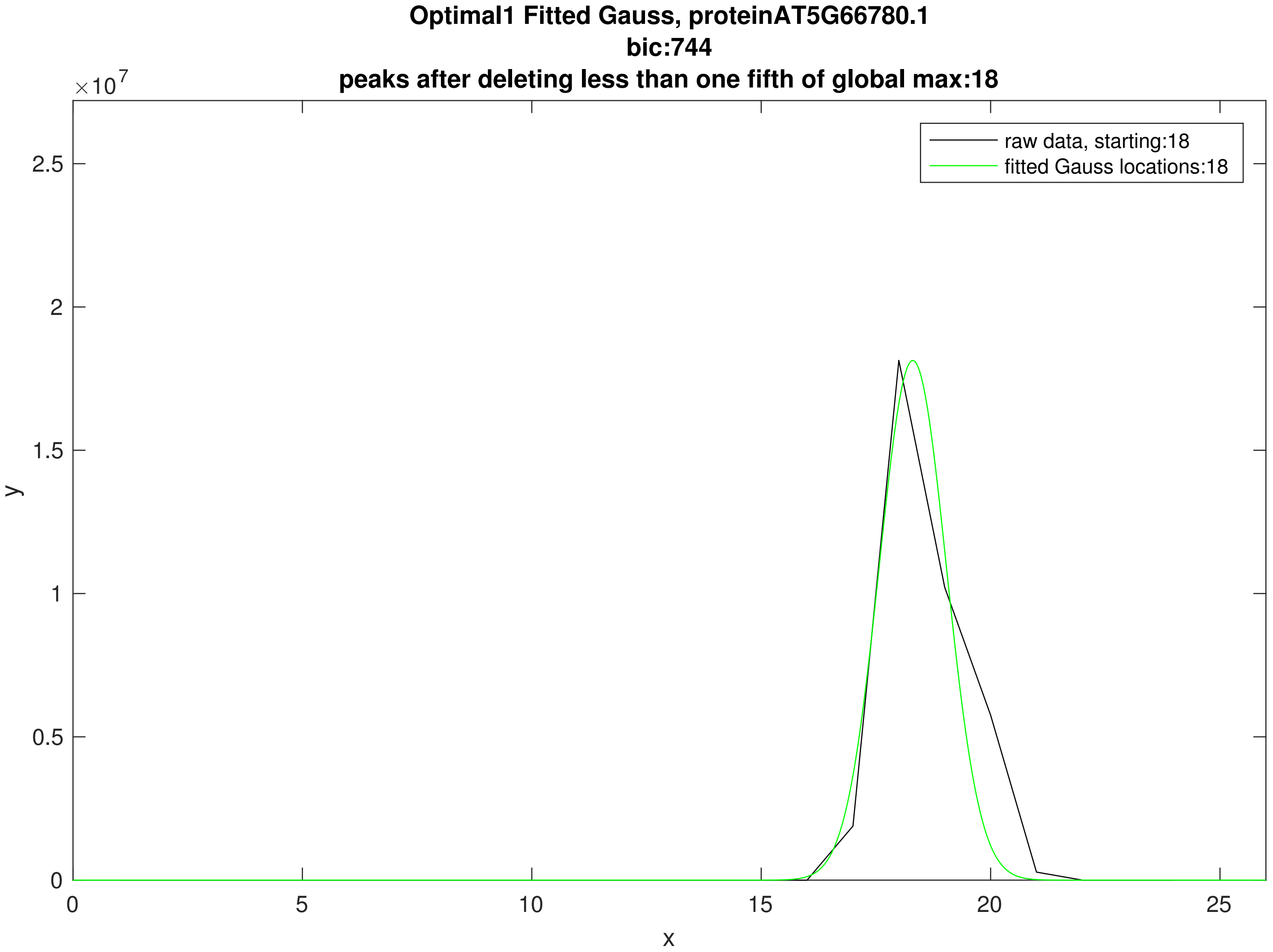
Optimal1 Fitted Gauss, proteinAT5G66780.1
bic:744
peaks after deleting less than one fifth of global max:18
×10^7
raw data, starting:18
fitted Gauss locations:18
2.5
2
1.5
1
0.5
0
y
0
5
10
15
20
25
x

#### Optimal1 Fitted Gauss, proteinAT5G66920.1
#### bic:813
#### peaks after deleting less than one fifth of global max:17

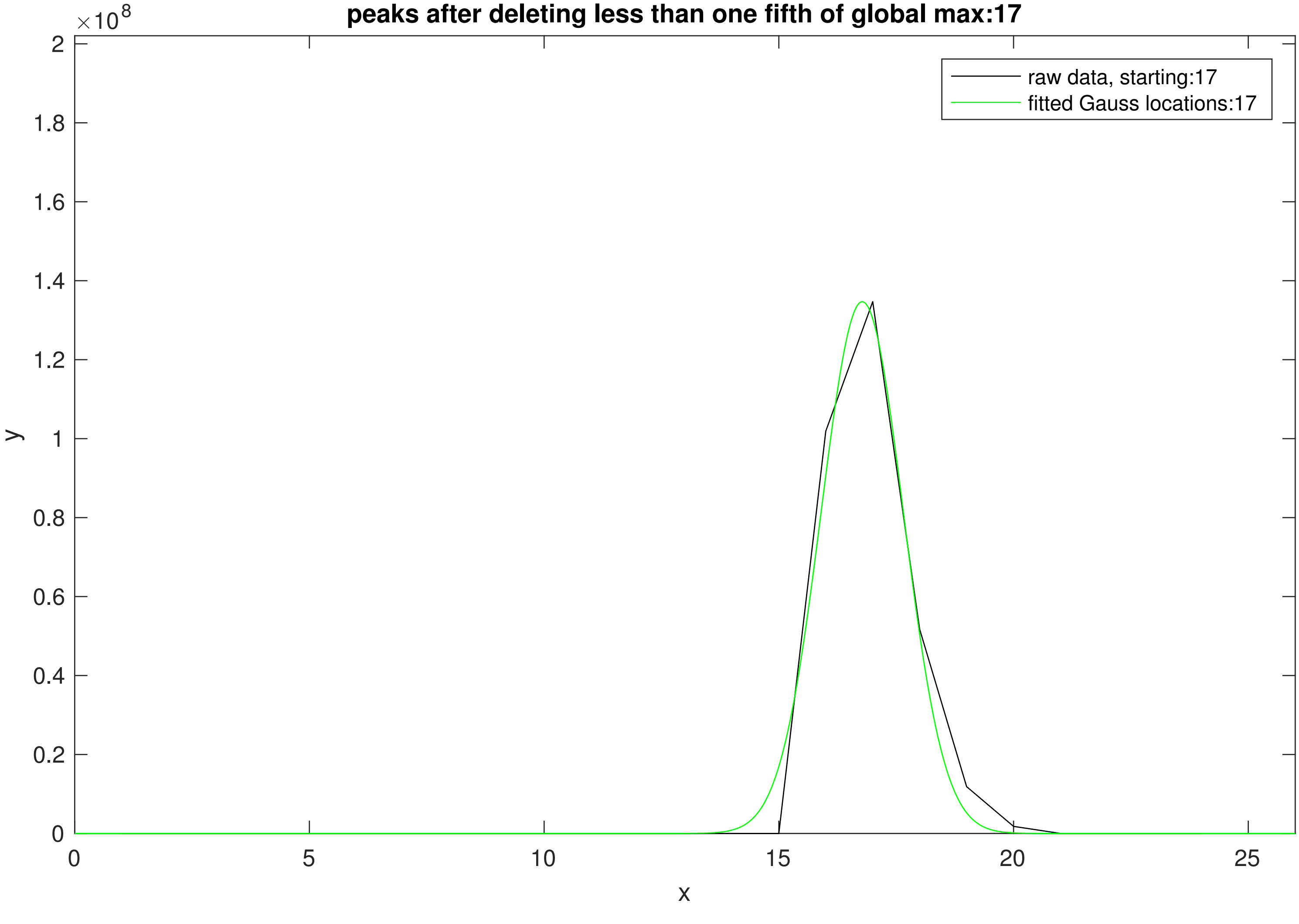

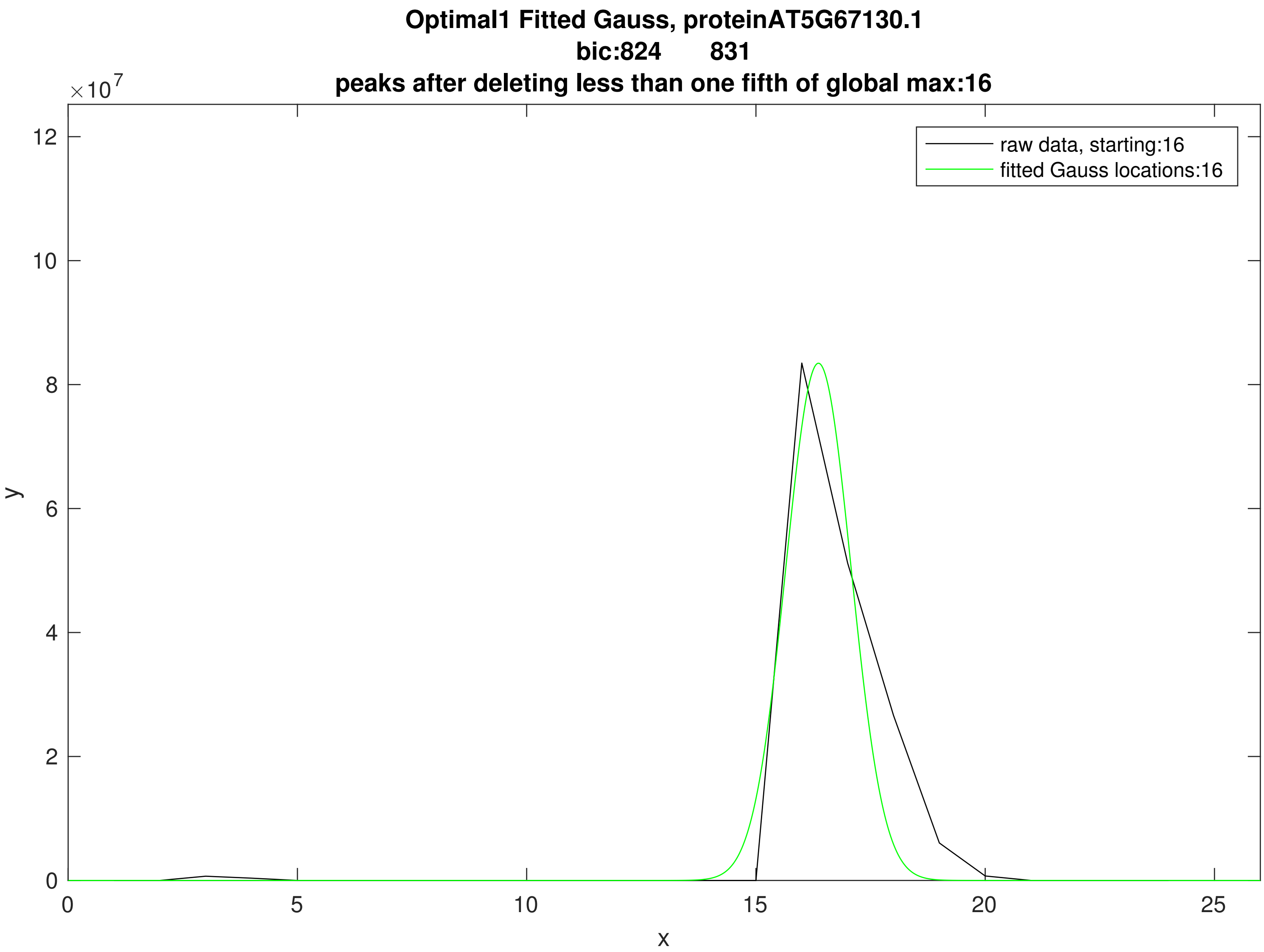
Optimal1 Fitted Gauss, proteinAT5G67130.1
bic:824 831
peaks after deleting less than one fifth of global max:16
×10^7
12
10
8
6
4
2
0
y
0
5
10
15
20
25
x
raw data, starting:16
fitted Gauss locations:16

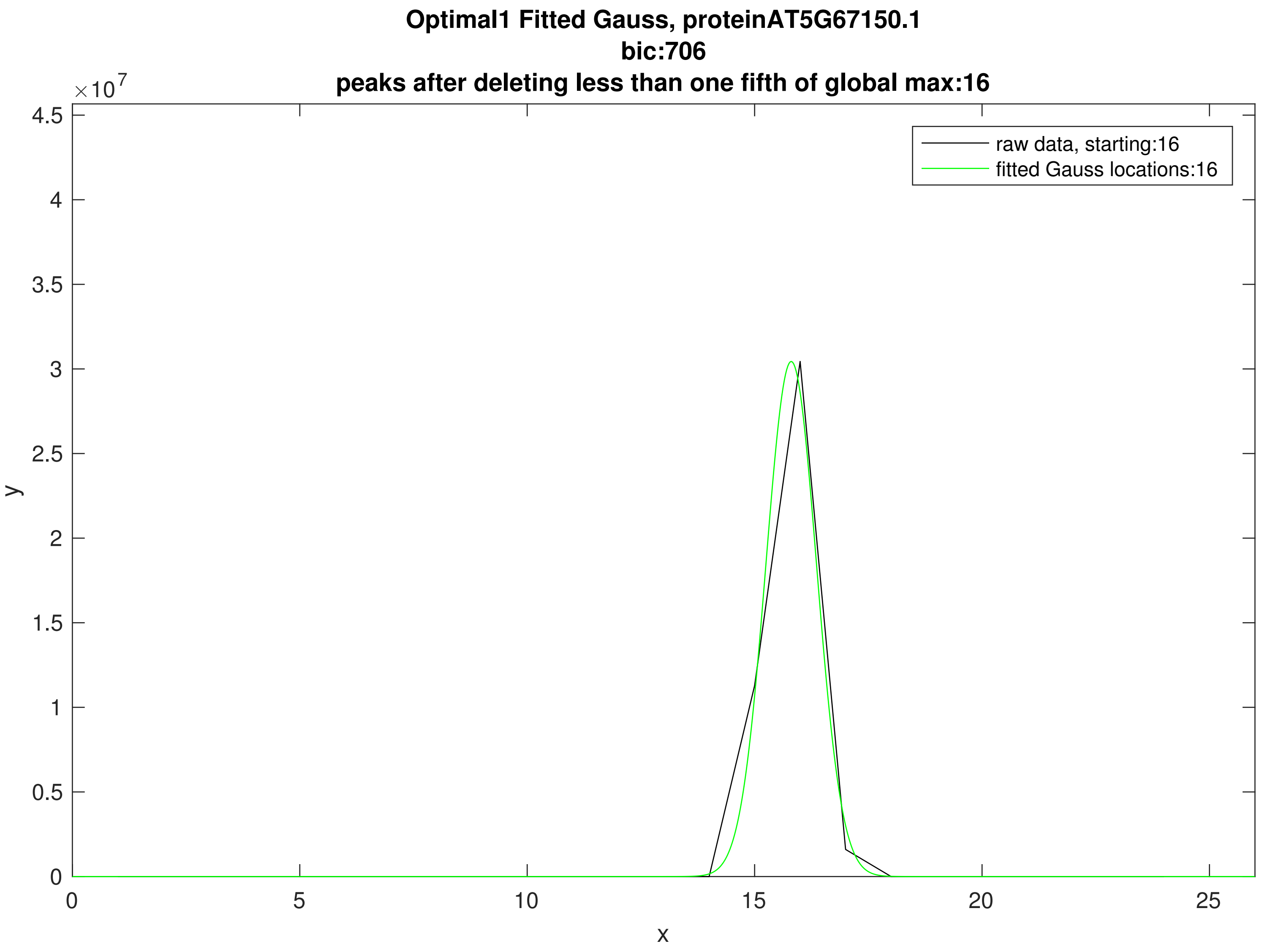
Optimal1 Fitted Gauss, proteinAT5G67150.1
bic:706
peaks after deleting less than one fifth of global max:16
×10^7
y
x
raw data, starting:16
fitted Gauss locations:16

**Optimal1 Fitted Gauss, proteinAT5G67290.1**
**bic:745**
**peaks after deleting less than one fifth of global max:17**

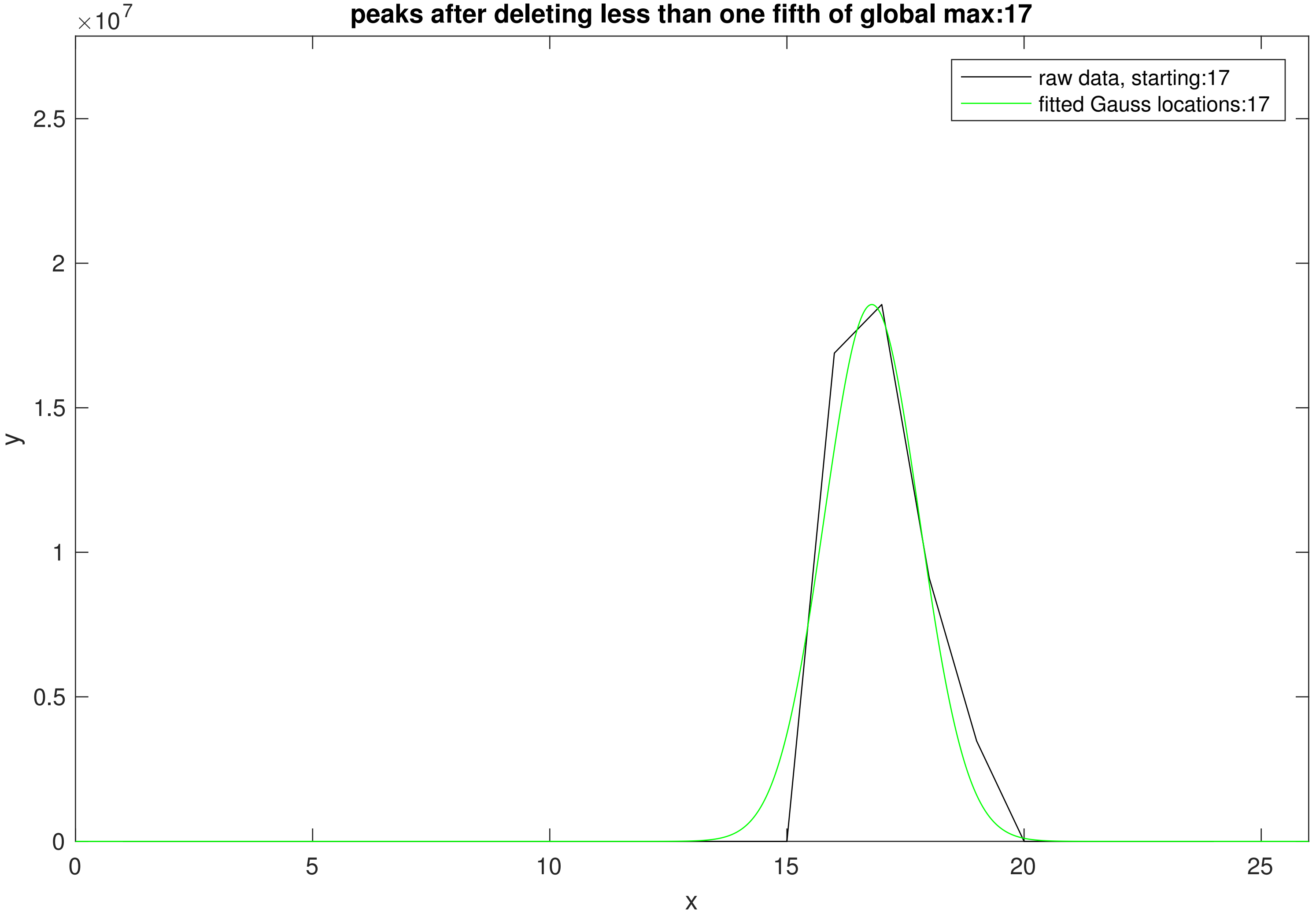

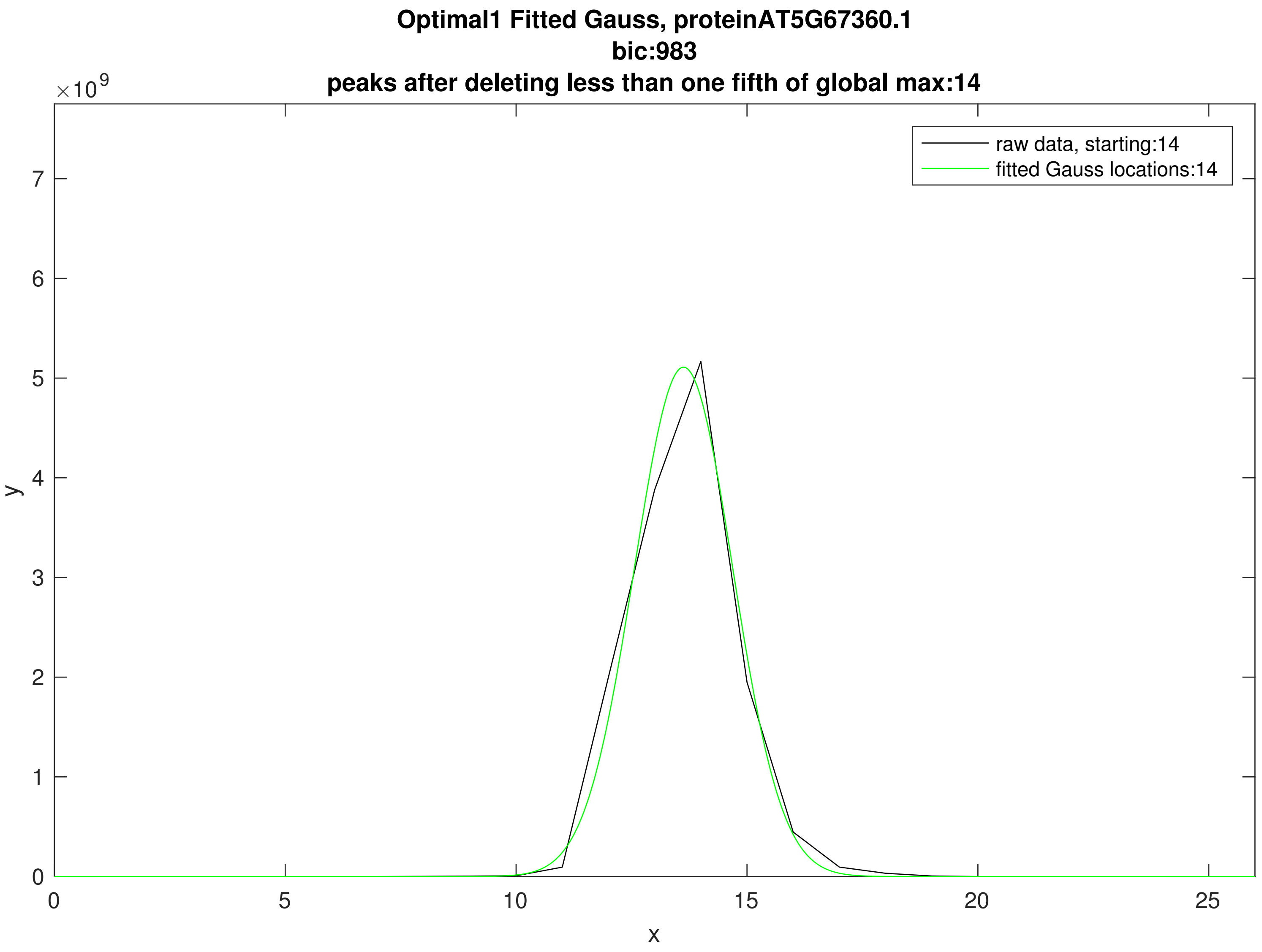
Optimal1 Fitted Gauss, proteinAT5G67360.1
bic:983
peaks after deleting less than one fifth of global max:14
×10^9
raw data, starting:14
fitted Gauss locations:14
y
x
0
1
2
3
4
5
6
7
0
5
10
15
20
25

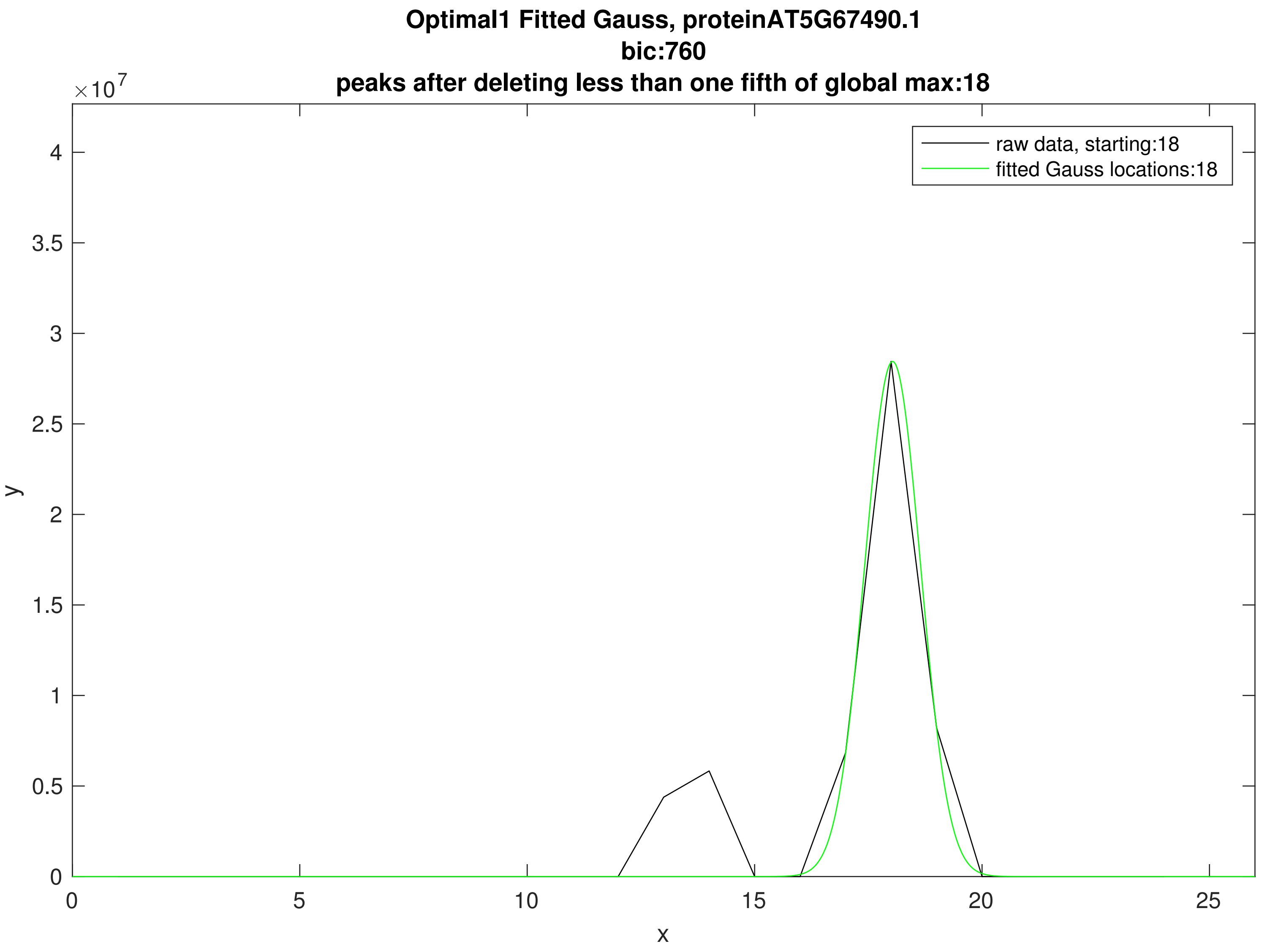

Optimal1 Fitted Gauss, proteinAT5G67490.1
bic:760
peaks after deleting less than one fifth of global max:18
×10^7
raw data, starting:18
fitted Gauss locations:18
y
x
0
0.5
1
1.5
2
2.5
3
3.5
4
0
5
10
15
20
25

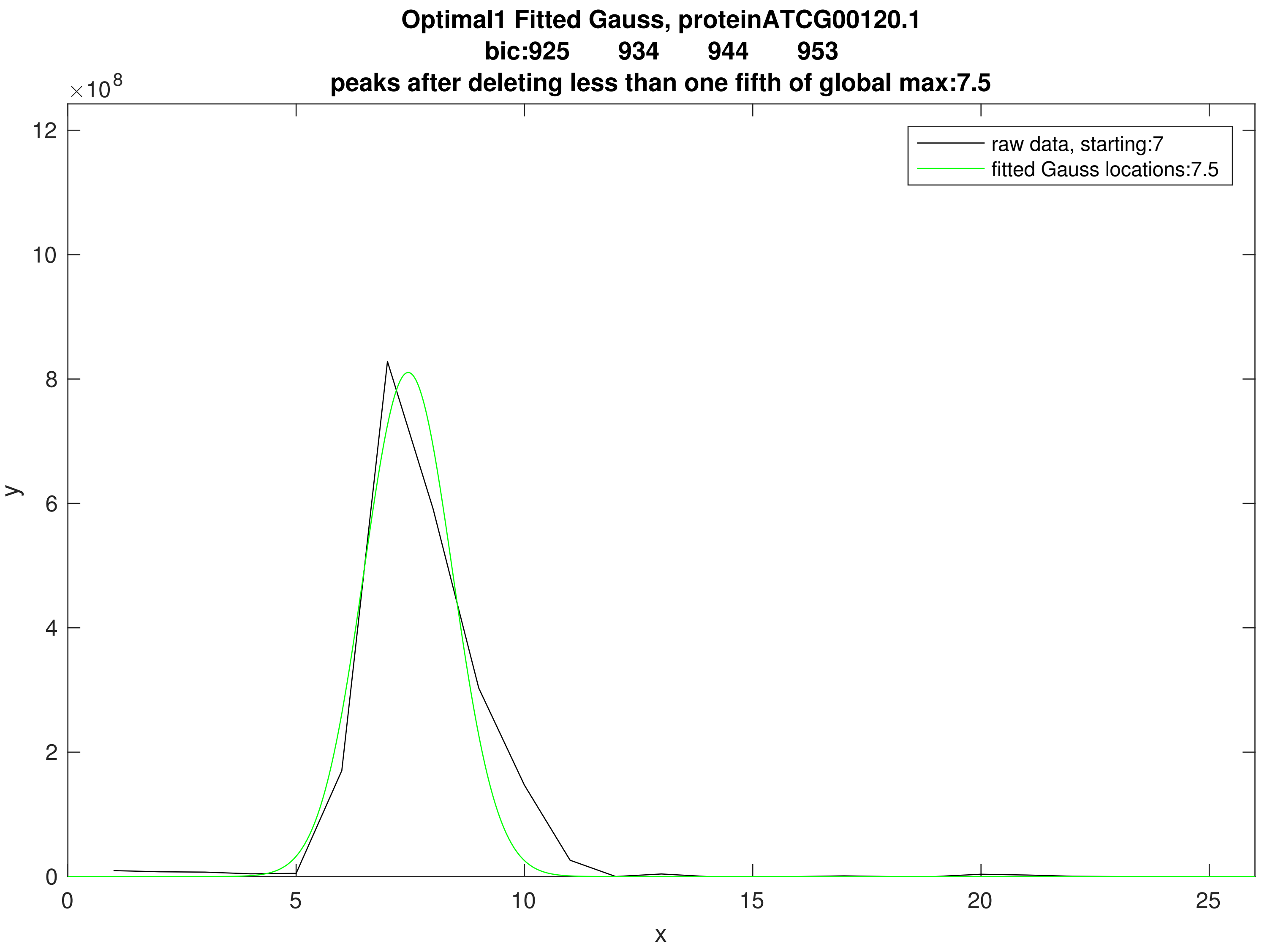
Optimal1 Fitted Gauss, proteinATCG00120.1
bic:925 934 944 953
peaks after deleting less than one fifth of global max:7.5
×10^8
raw data, starting:7
fitted Gauss locations:7.5
0
2
4
6
8
10
12
y
0
5
10
15
20
25
x

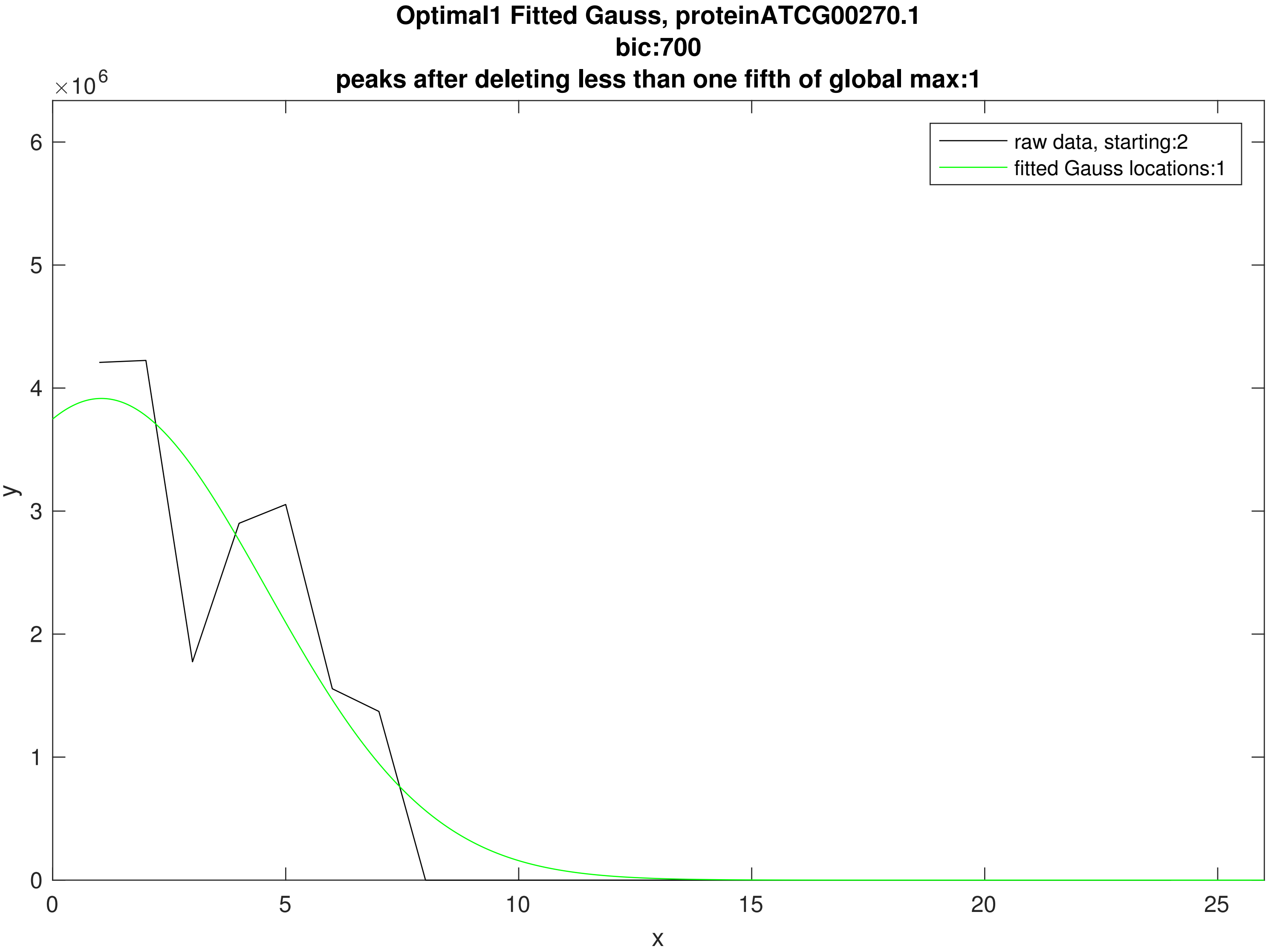

Optimal1 Fitted Gauss, proteinATCG00270.1
bic:700
peaks after deleting less than one fifth of global max:1
×10^6
raw data, starting:2
fitted Gauss locations:1
y
x
0
1
2
3
4
5
6
0
5
10
15
20
25

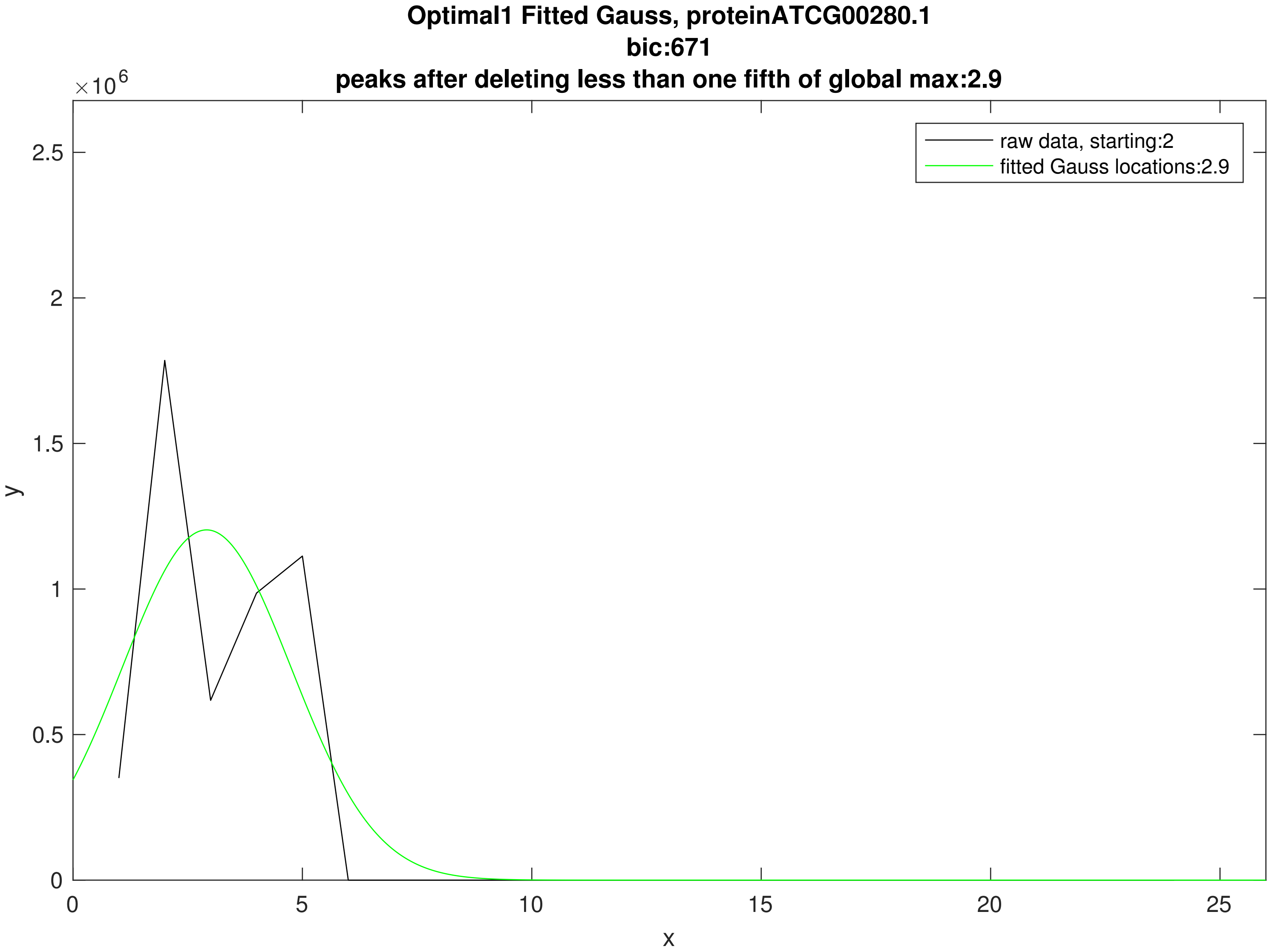

Optimal1 Fitted Gauss, proteinATCG00280.1
bic:671
peaks after deleting less than one fifth of global max:2.9
×10^6
raw data, starting:2
fitted Gauss locations:2.9
2.5
2
1.5
1
0.5
0
y
0
5
10
15
20
25
x

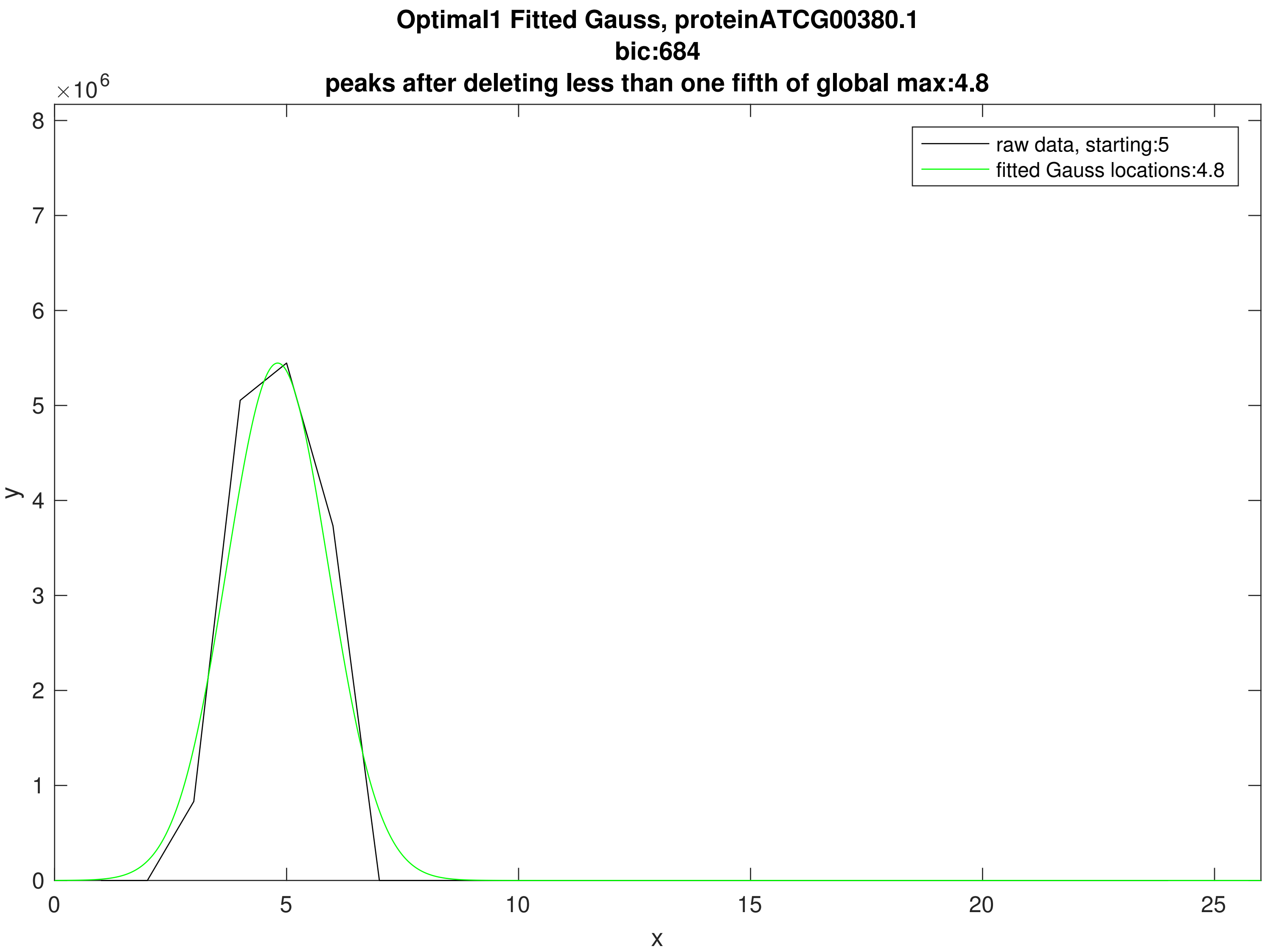
Optimal1 Fitted Gauss, proteinATCG00380.1
bic:684
peaks after deleting less than one fifth of global max:4.8
×10^6
8
7
6
5
4
3
2
1
0
y
0
5
10
15
20
25
x
raw data, starting:5
fitted Gauss locations:4.8

**Optimal1 Fitted Gauss, proteinATCG00470.1**
**bic:804**
**peaks after deleting less than one fifth of global max:7.5**

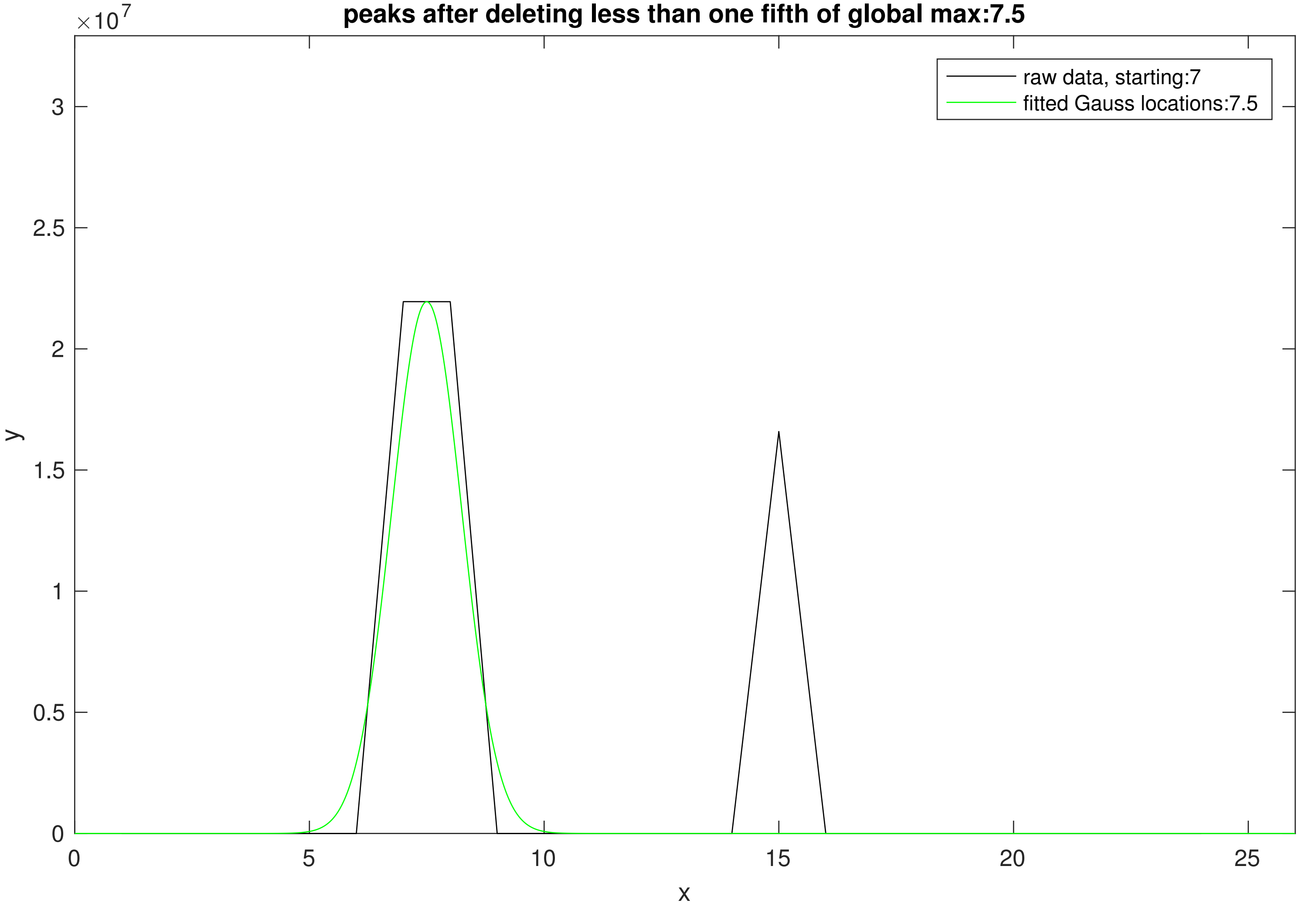

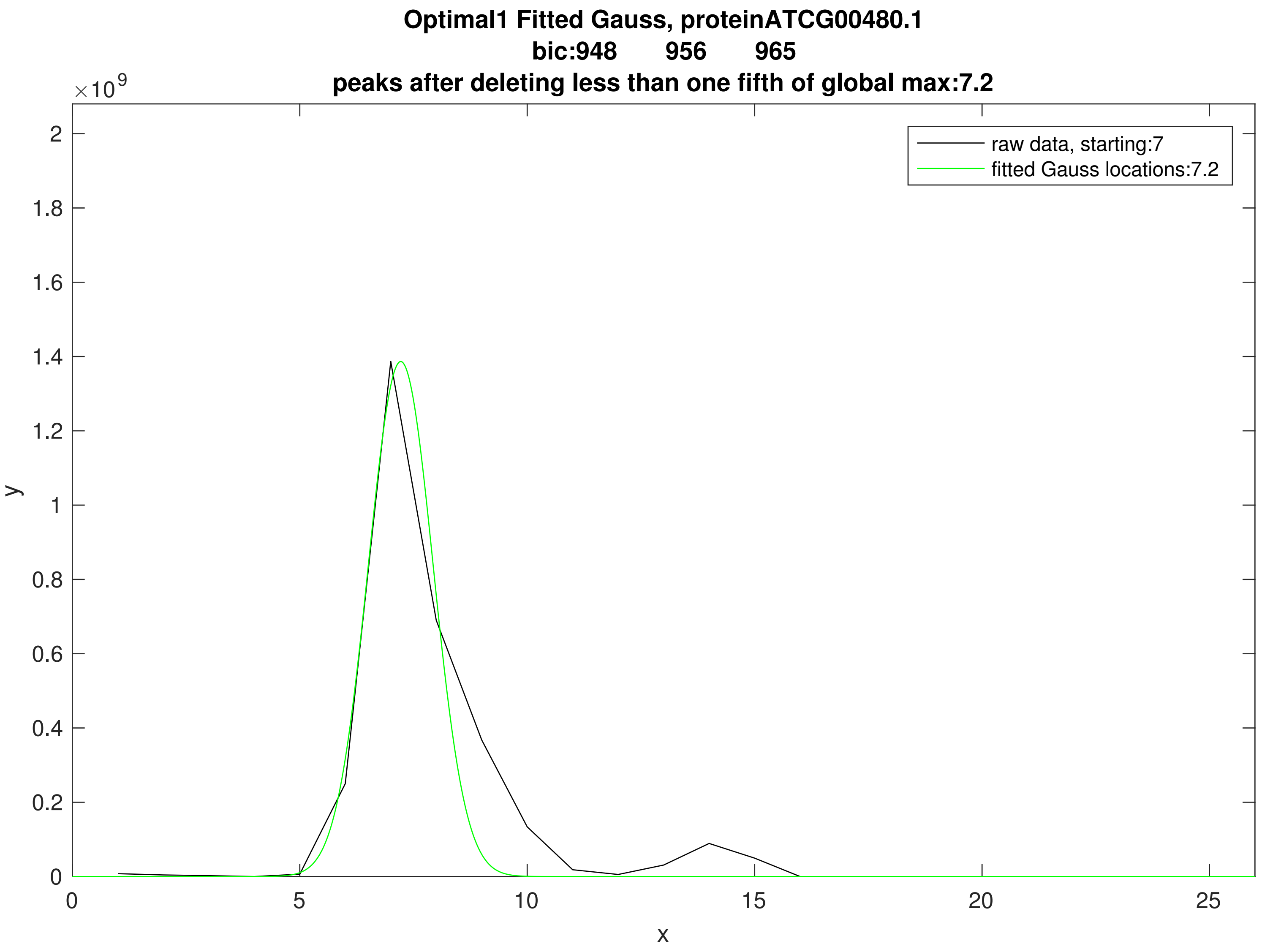
Optimal1 Fitted Gauss, proteinATCG00480.1
bic:948 956 965
peaks after deleting less than one fifth of global max:7.2
raw data, starting:7
fitted Gauss locations:7.2
×10^9
y
x
0
0.2
0.4
0.6
0.8
1
1.2
1.4
1.6
1.8
2
5
10
15
20
25

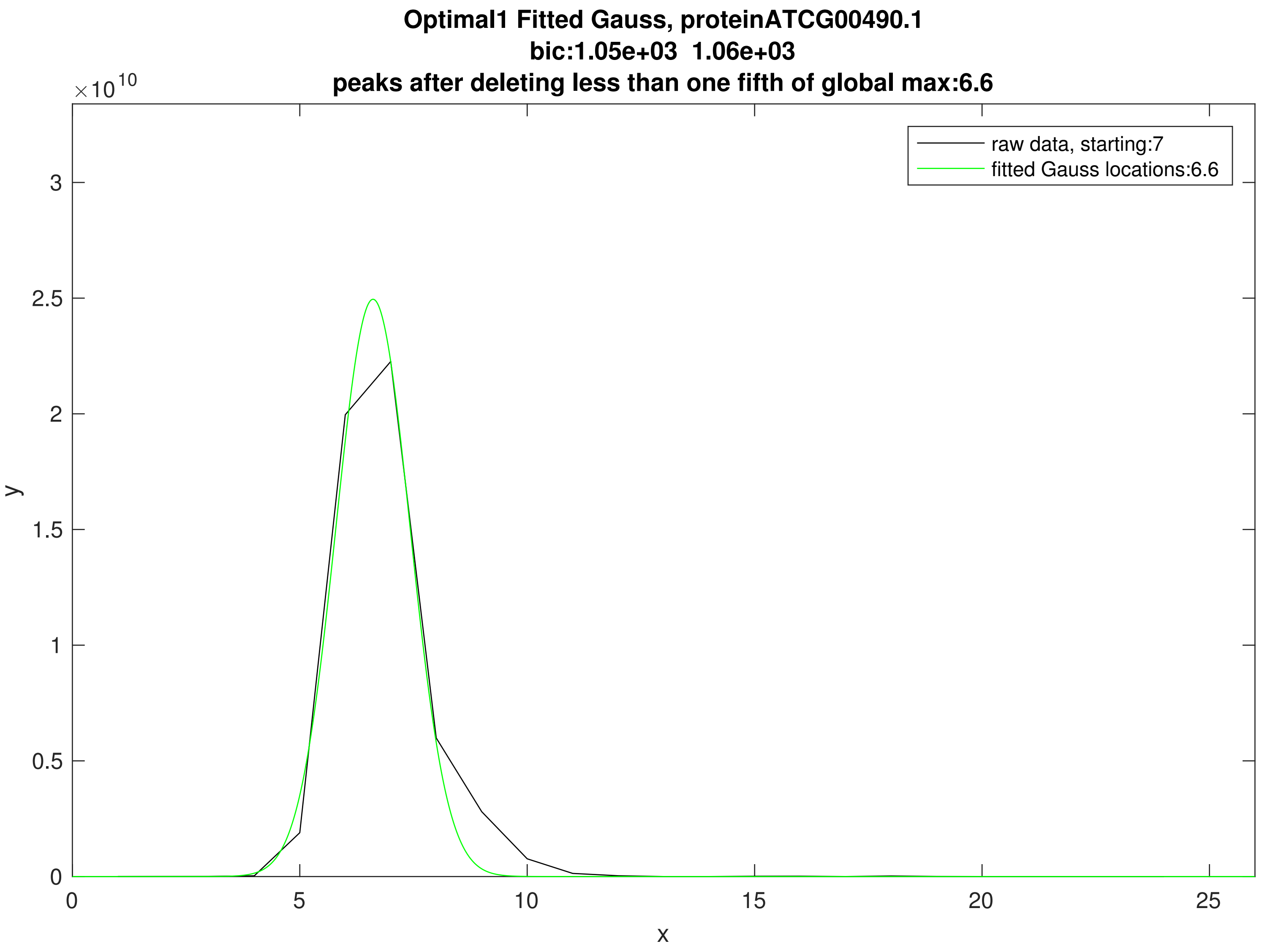
Optimal1 Fitted Gauss, proteinATCG00490.1
bic:1.05e+03  1.06e+03
peaks after deleting less than one fifth of global max:6.6
×10^10
raw data, starting:7
fitted Gauss locations:6.6
0
0.5
1
1.5
2
2.5
3
y
0
5
10
15
20
25
x

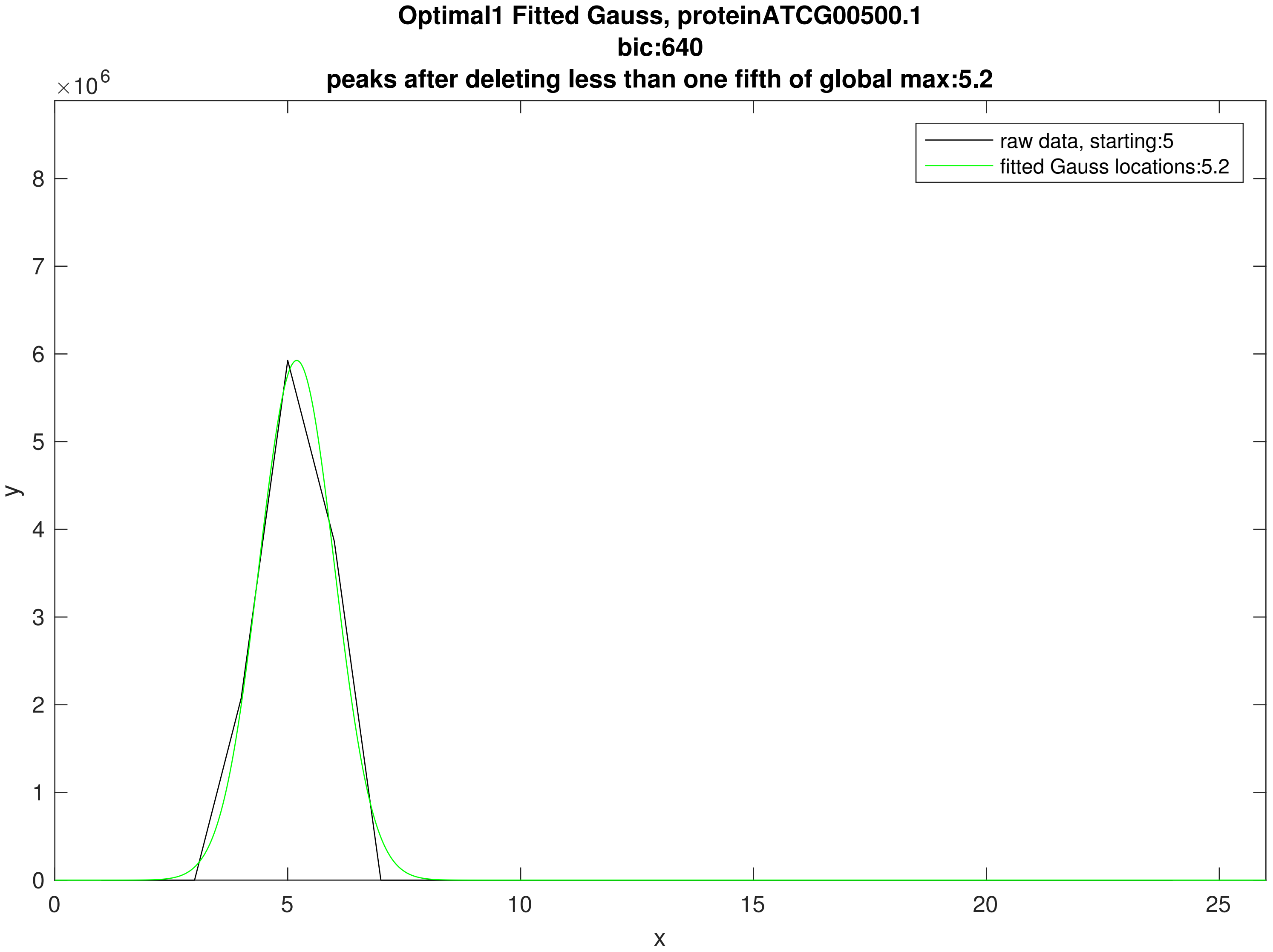

Optimal1 Fitted Gauss, proteinATCG00500.1
bic:640
peaks after deleting less than one fifth of global max:5.2
×10^6
raw data, starting:5
fitted Gauss locations:5.2
8
7
6
5
4
3
2
1
0
y
0
5
10
15
20
25
x

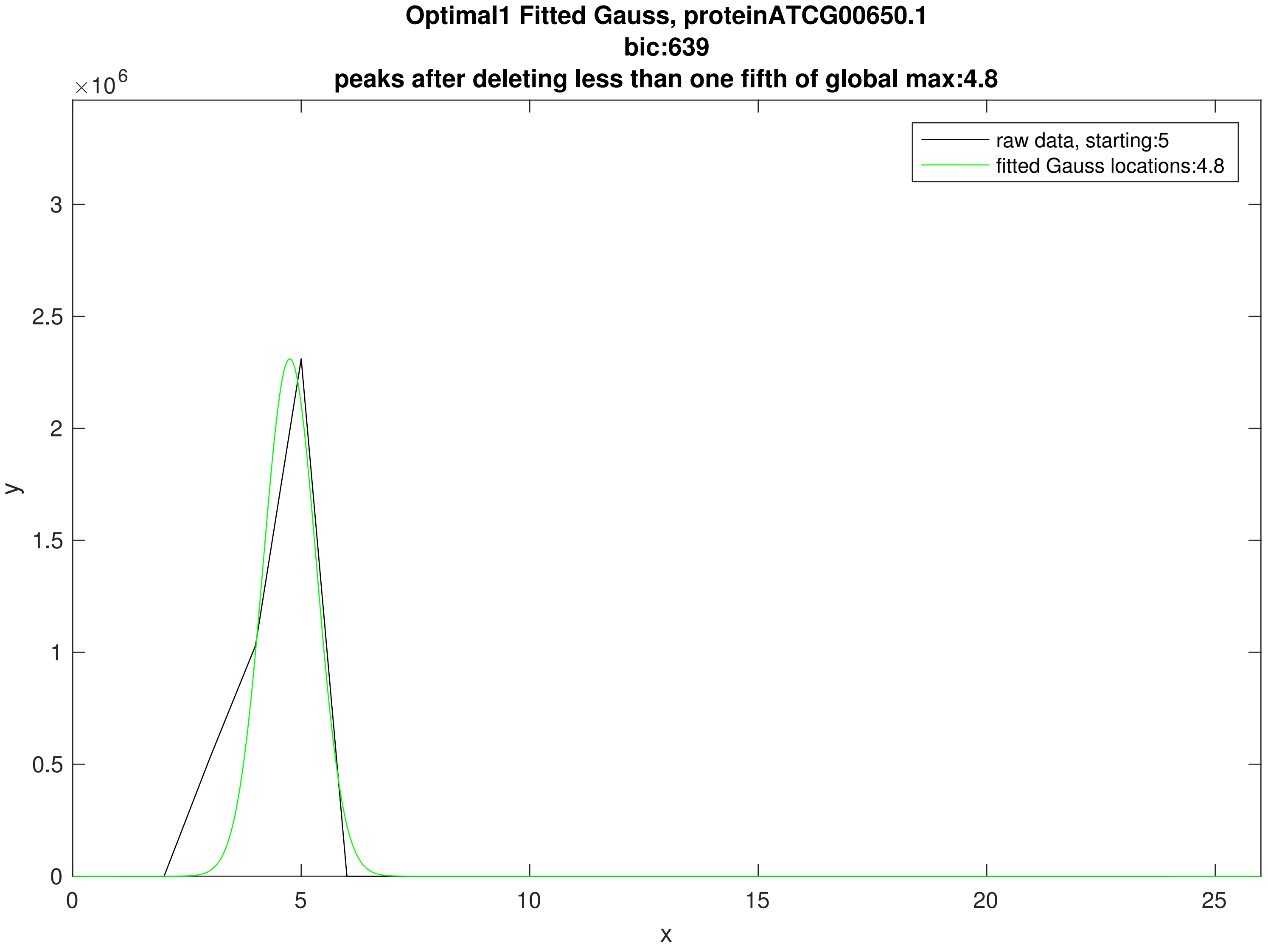
Optimal1 Fitted Gauss, proteinATCG00650.1
bic:639
peaks after deleting less than one fifth of global max:4.8
×10^6
raw data, starting:5
fitted Gauss locations:4.8
y
x
0
0.5
1
1.5
2
2.5
3
5
10
15
20
25

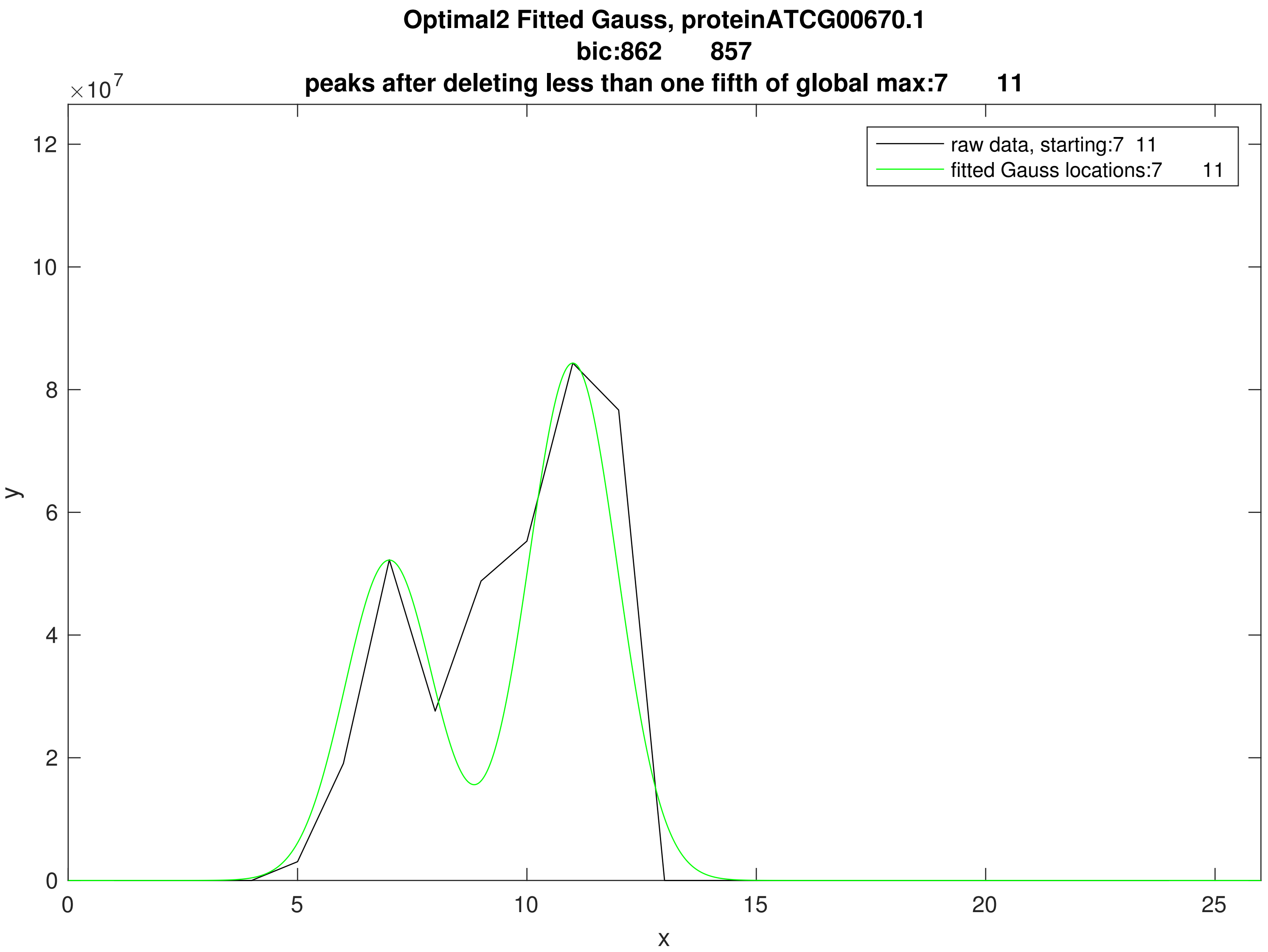
Optimal2 Fitted Gauss, proteinATCG00670.1
bic:862 857
peaks after deleting less than one fifth of global max:7 11
×10^7
raw data, starting:7 11
fitted Gauss locations:7 11
y
x
0
2
4
6
8
10
12
0
5
10
15
20
25

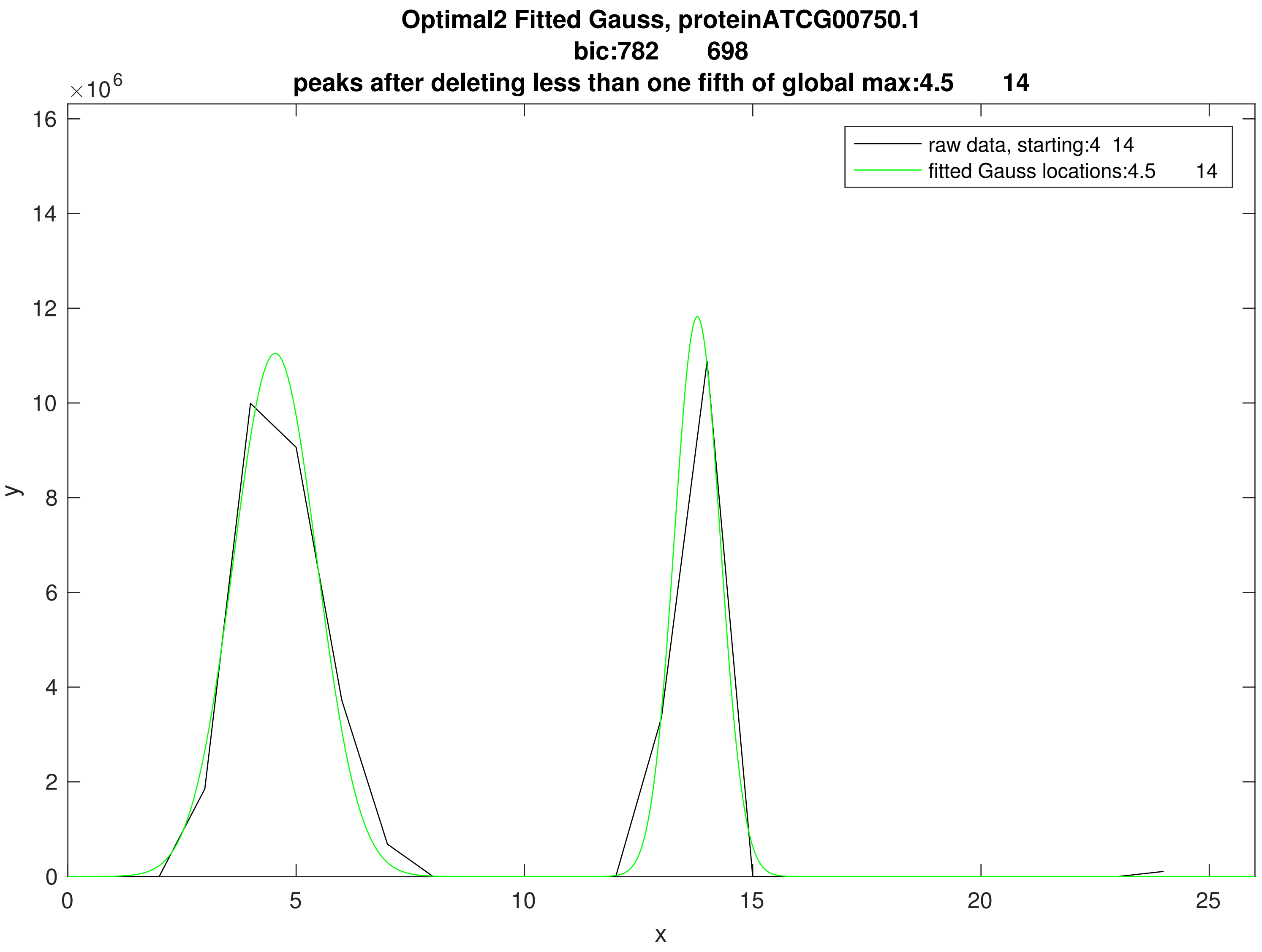

Optimal2 Fitted Gauss, proteinATCG00750.1
bic:782 698
peaks after deleting less than one fifth of global max:4.5 14
×10^6
raw data, starting:4 14
fitted Gauss locations:4.5 14
y
x
0
2
4
6
8
10
12
14
16
0
5
10
15
20
25

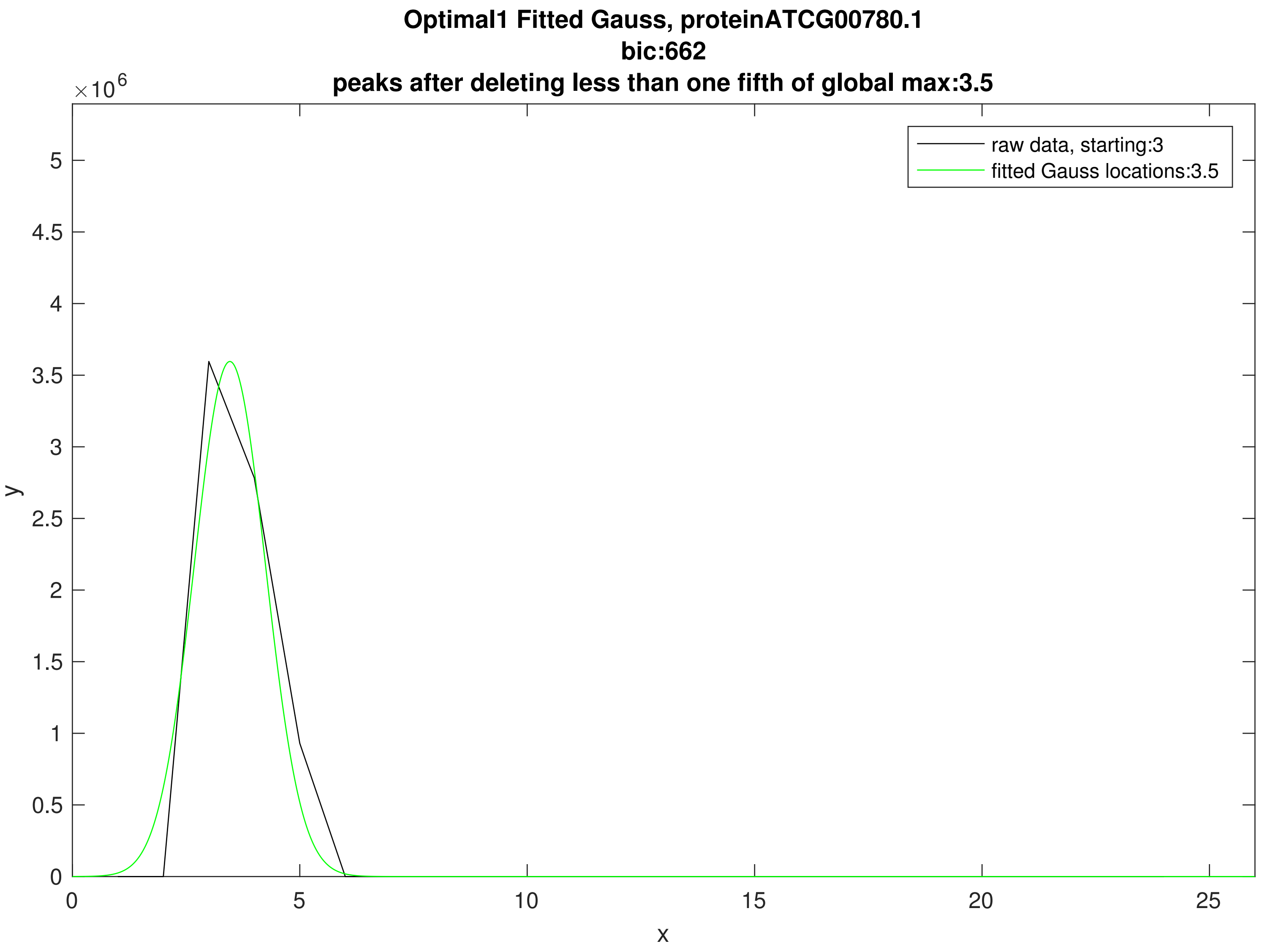
Optimal1 Fitted Gauss, proteinATCG00780.1
bic:662
peaks after deleting less than one fifth of global max:3.5
×10^6
raw data, starting:3
fitted Gauss locations:3.5
y
x
0
0.5
1
1.5
2
2.5
3
3.5
4
4.5
5
0
5
10
15
20
25

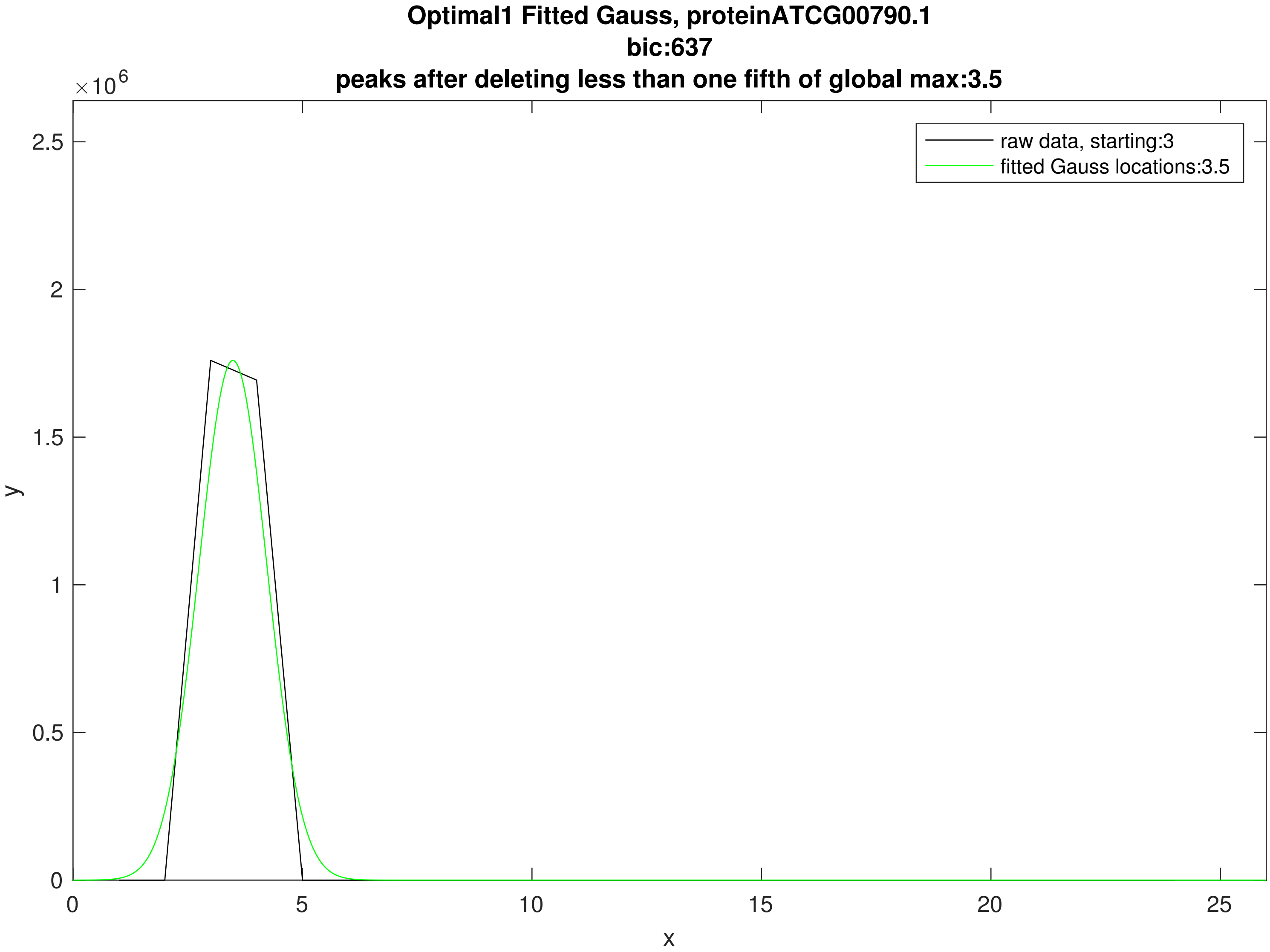
Optimal1 Fitted Gauss, proteinATCG00790.1
bic:637
peaks after deleting less than one fifth of global max:3.5
×10^6
2.5
2
1.5
1
0.5
0
y
0
5
10
15
20
25
x
raw data, starting:3
fitted Gauss locations:3.5

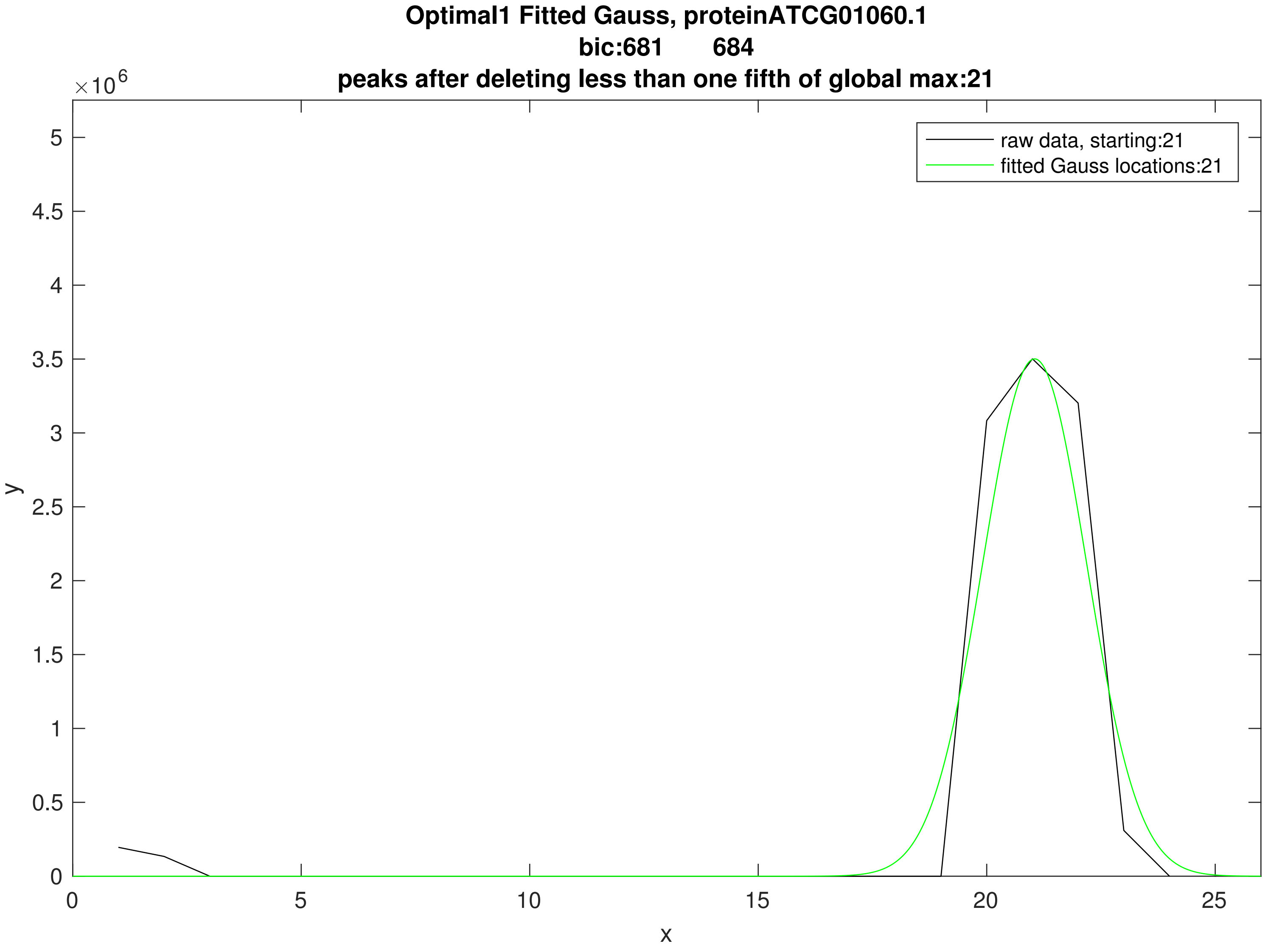
Optimal1 Fitted Gauss, proteinATCG01060.1
bic:681 684
peaks after deleting less than one fifth of global max:21
×10^6
raw data, starting:21
fitted Gauss locations:21
y
x
0
0.5
1
1.5
2
2.5
3
3.5
4
4.5
5
0
5
10
15
20
25

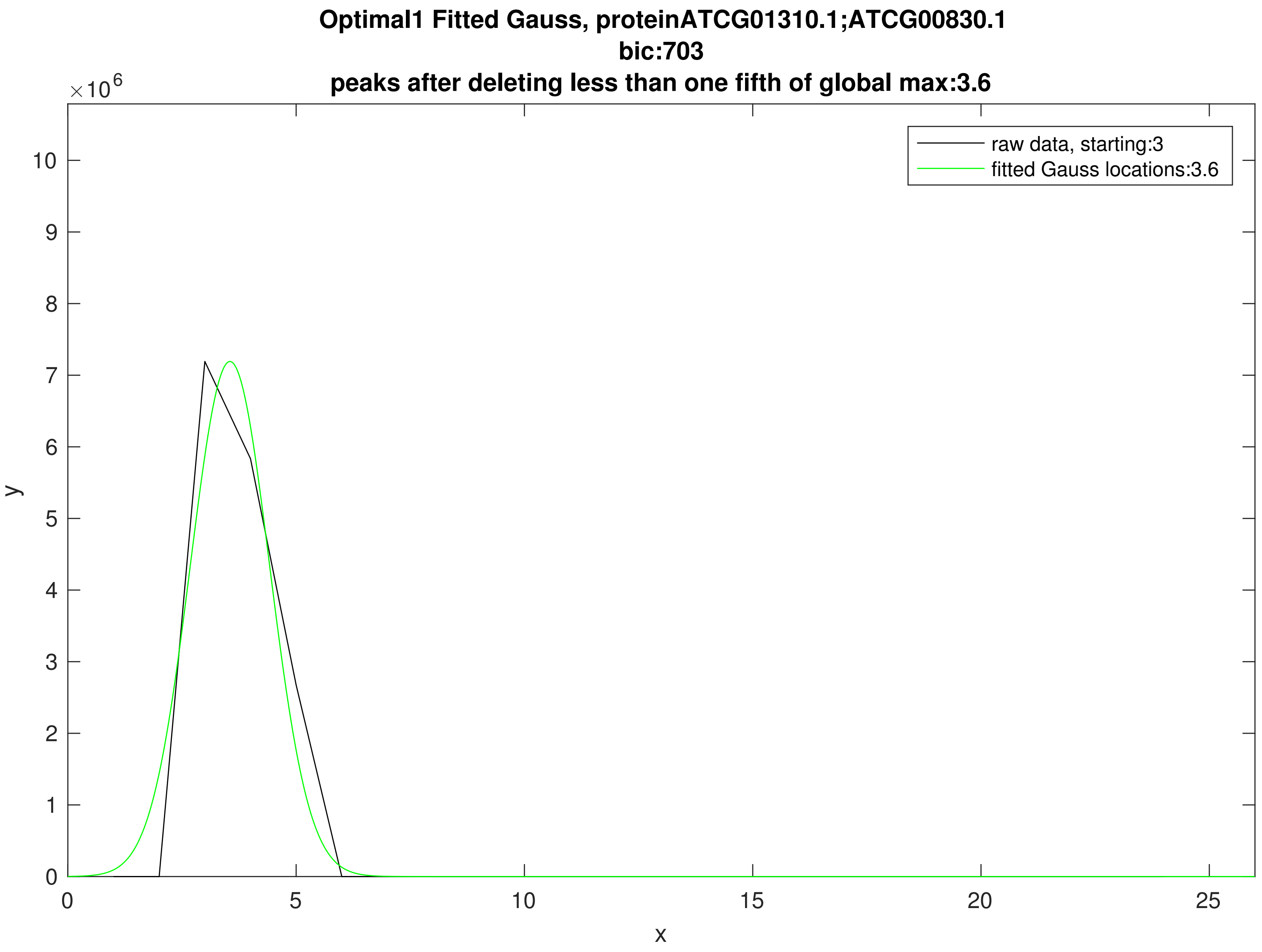
Optimal1 Fitted Gauss, proteinATCG01310.1;ATCG00830.1
bic:703
peaks after deleting less than one fifth of global max:3.6
×10^6
raw data, starting:3
fitted Gauss locations:3.6
y
x
0
5
10
15
20
25
0
1
2
3
4
5
6
7
8
9
10

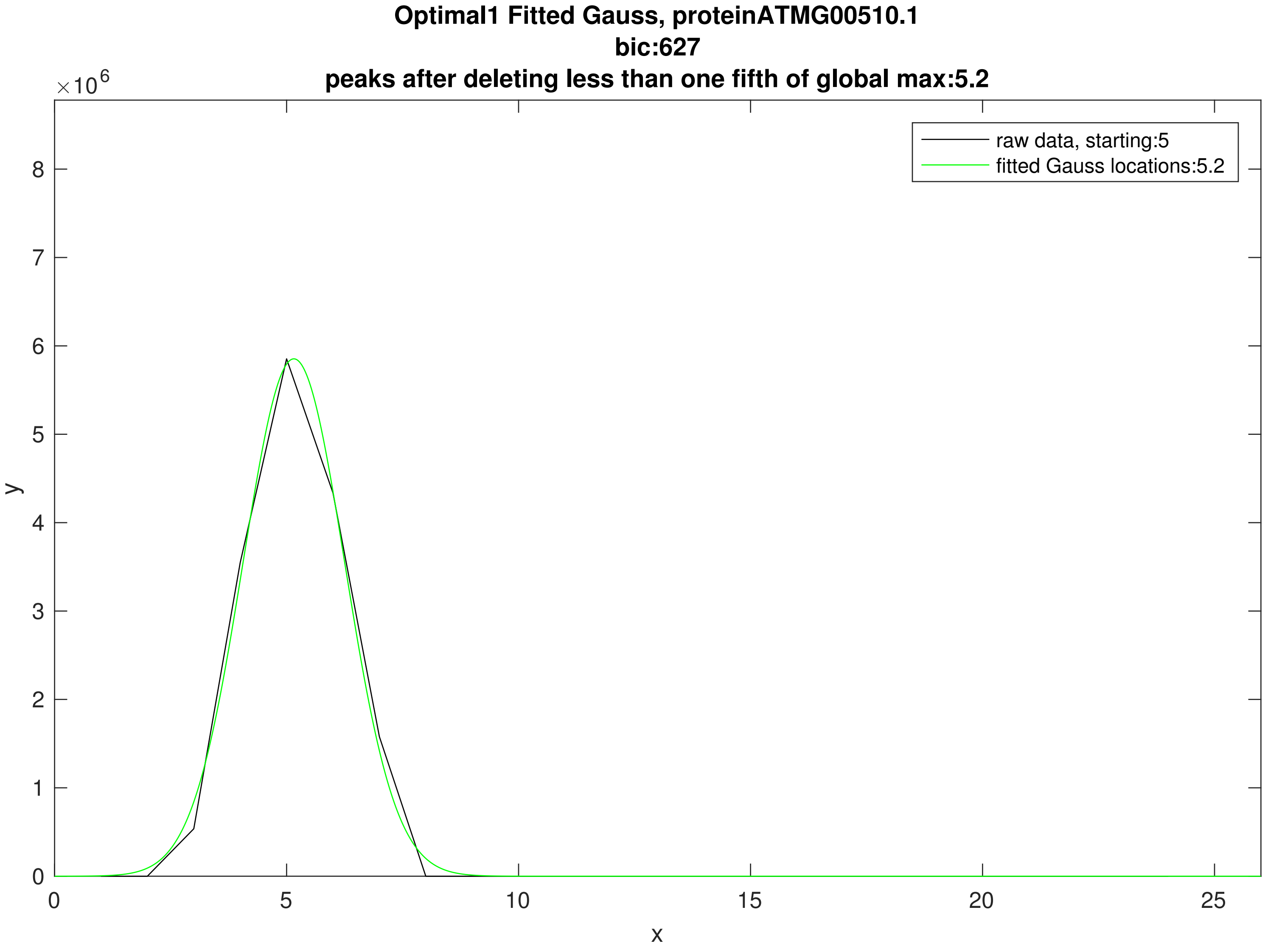
Optimal1 Fitted Gauss, proteinATMG00510.1
bic:627
peaks after deleting less than one fifth of global max:5.2
×10^6
raw data, starting:5
fitted Gauss locations:5.2
y
x
0
1
2
3
4
5
6
7
8
0
5
10
15
20
25
