## Supplemental Table S1 for "Rapid ethylene-triggered protein complex remodeling in dark grown hypocotyls"

**Supplemental Table S1. Quantitative RT-PCR Primers**

| **Group1 (Statistically differentially expressed proteins)** | |
| --- | --- |
| Putatively ethylene down-regulated | |
| AT1G03130-F  AT1G03130-R  AT4G28750-F  AT4G28750-R  AT2G20260-F  AT2G20260-R  AT5G66570-F  AT5G66570-R  AT1G06680-F  AT1G06680-R  AT4G05180-F  AT4G05180-R  AT4G21280-F  AT4G21280-R  AT4G27520-F  AT4G27520-R  AT3G62410-F  AT3G62410-R | GGTCTCCTCCGTAAAGCACAA  ACTGTTCTTTCCTCGCCAGTT  TGGATCAGTTGTTGCCGTTG  AAGCTGCAACTTCTTCGACCT  ATAGTTCTTCCACCACCGCT  GCCACAACTGATCCAACGTT  CAGGATATGACAACGCCGTG  GATCACCTCTCCTGTCTCCG  TGAAGACGATAACTCCGCCG  CCCAAACACGTTTGCAGCTT  CCGTTACGATCTCAACACCG  TACTCTTTGATCTCGCCGCA  CCTTCCAGCTGGGACAGAT  TTCCTGTCGATCAATGGCTTCA  CCGTCAGATTCTCCGTCAGG  AACGGTCATACCATTCGCCG  GGAAGGAGGGATATCGGACG  TCCGAACCATCAGCCTTCTT |
| Putatively ethylene up-regulated | |
| AT4G27170-F  AT4G27170-R | GGCCGTTAGATTCCAGGGAC  GGATCTGGAAGGGGCAAACA |
| **Group2 (Differentially expressed proteins based on MS/MS counts)** | |
| Putatively ethylene down-regulated | |
| AT5G09830-F  AT5G09830-R  AT4G11210-F  AT4G11210-R  AT4G26110-F  AT4G26110-R  AT1G02640-F  AT1G02640-R  AT1G02305-F  AT1G02305-R  AT1G33680-F  AT1G33680-R  AT3G02320-F  AT3G02320-R  AT1G04530-F  AT1G04530-R  AT2G37250-F  AT2G37250-R | AGATTGCTGGAGAGGCATCG  CAGAGTCTTGTGATGGTGGCT  CAACACAACACAAAGGGGCA  CGTCAATGTGATCGGTCGTT  TGGTGCAAGATTGAAGAGCCT  TAGCCTTCTCAAGCAGTGGC  TCTCTCTCTCGGAGTCCACG  GCAACCAGCTGTTTCTTCGG  CGACAATACTGGTTGCTCGC  ACCTTGTACGCACTAACACCAT  CTATCCATCGGCAGGTGGTC  TGGTAAGGCACATCACCAGG  GTCCAAAGCTTAGGGCTGGT  TCTGGCGTTTTGGATCAACCT  AACGCTATGCAGATACGCGA  CCGAGCGTACTCTCCAAGAA  GGGTGAGAATGGAAGACCTGG  TCAAGAGGCTGGCTCGTTTC |
| Putatively ethylene up-regulated | |
| AT2G19590-F  AT2G19590-R  AT3G18680-F  AT3G18680-R | CAGAATGCCCACGTCCTGAG  TGGCGGTATAGGAACCCACT  AAACGCTCGTCTGCTTGACT  GCATGTGGTCAAGGTTTGAGAG |
| ERF1-RT-F  ERF1-RT-R  ACTIN2-RT-F  ACTIN2-RT-R | ACGTTCTCAACCGCCTACAG  CGGACTCGCTCTCTGGTG  ATTCAGATGCCCAGAAGTCTTGTT  ACGGTCAGCGATACCTGAGAAC |
